# A comprehensive mechanistic model of adipocyte signaling with layers of confidence

**DOI:** 10.1101/2022.03.11.483974

**Authors:** William Lövfors, Cecilia Jönsson, Charlotta S. Olofsson, Gunnar Cedersund, Elin Nyman

## Abstract

Adipocyte cellular signaling, normally and in type 2 diabetes, is far from fully studied. We have earlier developed detailed dynamic mathematical models for some well-studied, and partially overlapping, signaling pathways in adipocytes. Still, these models only cover a fraction of the total cellular response. For a broader coverage of the response, large-scale phosphoproteomic data is key. There exists such data for the insulin response of adipocytes, as well as prior knowledge on possible protein-protein interactions associated with a confidence level. However, methods to combine detailed dynamic models with large-scale data, using information about the confidence of included interactions, are lacking. In our new method, we first establish a core model by connecting our partially overlapping models of adipocyte cellular signaling with focus on: 1) lipolysis and fatty acid release, 2) glucose uptake, and 3) the release of adiponectin. We use the phosphoproteome data and prior knowledge to identify phosphosites adjacent to the core model, and then try to add the adjacent phosphosites to the model. The additions of the adjacent phosphosites is tested in a parallel, pairwise approach with low computation time. We then iteratively collect the accepted additions into a *layer*, and use the newly added layer to find new adjacent phosphosites. We find that the first 15 layers (60 added phosphosites) with the highest confidence can correctly predict independent inhibitor-data (70-90 % correct), and that this ability decrease when we add layers of decreasing confidence. In total, 60 layers (3926 phosphosites) can be added to the model and still keep predictive ability. Finally, we use the comprehensive adipocyte model to simulate systems-wide alterations in adipocytes in type 2 diabetes. This new method provide a tool to create large models that keeps track of varying confidence.

## Introduction

A major challenge within biomedical research, from clinical studies to drug development, is how to handle and make optimal use of the increasing availability of omics data, e.g. proteomics, phosphoproteomics and transcriptomics. The handling of large data is today mostly done within the fields of bioinformatics, where data are mapped to static networks. The information in these networks are found in databases with known interactions of different level of confidence. Methods within bioinformatics can typically not be used to simulate time-resolved scenarios, e.g. to predict the clinical effect of a treatment or the change in signaling inside cells in response to new drugs. Such methods also lack the ability to integrate data from several sources into a consistent body of knowledge. This is instead the strength of systems biology models based on ordinary differential equations (ODEs).

Systems biology ODE-based models are built on mechanistic details known about the system at study, such as intracellular signaling pathways. Furthermore, ODE-based modelling is the most common framework to model biological systems when time-resolved data is available [1]. This is evident in e.g. the BioModels database [2] which contain over 1000 manually curated models, where over 80% of the models are ODE-based. Also, there exists ODE-based models for most biological systems, which are often partially overlapping and not interconnected.

The ODE-based models need to be fed with numerical values for the kinetic rate constants corresponding to the included mechanistic details. These rate constants are model parameters that are usually unknown since they are often impossible to measure explicitly, and must instead be estimated based on indirect time-resolved measurements. A central challenge with large ODE-based models is thus how to handle the large amount of unknown parameters and how to estimate the parameter values.

This challenge has been approached using adjoint sensitivities [3] which improves the scaling of the parameter estimation problem with the number of parameters, together with the use of a sparse linear solver. The authors [3] speed up the optimization process ~40,000 times for a model with ~4000 free parameters. This development is important to go from smaller to larger ODE-based models. However, methods developed to estimate parameters for large ODE-based models, still cannot handle the whole omics scale.

Other methods to handle semi-large protein data using ODE-based models have been developed [4–7]. These approaches use information rich multi-perturbation data to create data-driven ODE-based models. These methods have been used to predict the response of new drug candidates and rank their ability to overcome drug resistance. Even though these methods can handle a large number of parameters, for all possible interactions between states, they are limited to ~200 proteins since all possible interactions are tested. Also, the information-rich perturbation data used in these methods is rare and costly to obtain experimentally.

One approach to avoid the estimation problem is to use parameter-free Boolean logical models instead of ODE-based models. Such models have been used to study large protein-activity datasets [8, 9]. Boolean logical models are useful in the search for biologically relevant interactions between proteins, and can thus be used for hypothesis generation when it comes to new interactions. However, Boolean logical models cannot be used to predict the effect of new drugs/perturbations or to simulate scenarios, another important aspect in biomedical research.

Another solution to the large parameter estimation problem is to divide the problem into smaller sub-problems, adding a single protein/gene (and corresponding data) at a time, preferably starting from an established core model. The additions of new proteins/genes can then be done in parallel, substantially reducing the computation time. We have earlier developed a method to do such an iterative model expansion [10]. However, this method does not take the confidence of included interactions into account and can only add a single layer of new interactions.

One final approach is to develop smaller, partially overlapping, models and to interconnect them to yield a large model. In this way, the development of each submodel corresponds to a manageable parameter estimation problem, each resulting in a submodel of high confidence. Unfortunately, this process is both time-consuming and there is no guarantee that the submodels can be combined.

In summary, it is an increasingly common situation to have large-scale omics data that need to be analyzed, and to have partially overlapping small ODE-based models, which are of high confidence, but which cannot be used to analyze the omics data. One challenge is to use the existing data to establish a core model of high confidence, another challenge is to have a differentiated view of the variables that are not in core model with different degrees of confidence. One example of such a situation is adipocyte signalling.

Within adipocyte signaling, we have developed several partially overlapping models for different aspects of the adipocyte, such as glucose uptake, lipolysis, and adiponectin secretion [11–16]. These models have not been combined into a single high confidence core model. Furthermore, there is large-scale phosphoproteomics data available from 3T3-L1 adipocytes, which has not been analyzed using scalable mechanistic approaches. Adipocyte signalling is therefore a good use-case to solve an increasingly common problem.

Here, we present a method to create comprehensive mechanistic models with layers of confidence, starting with the interconnection of three partially overlapping models into a core model containing detailed interactions of high confidence. From there, we add large-scale data in layers starting from high confidence interactions to purely data-driven interactions (Fig. 1). When adding these layers, we use a parallel, pairwise approach with a manageable computation time. The biological system we focus on is adipocyte cellular signaling, due to the relevance for type 2 diabetes.

**Figure 1:**
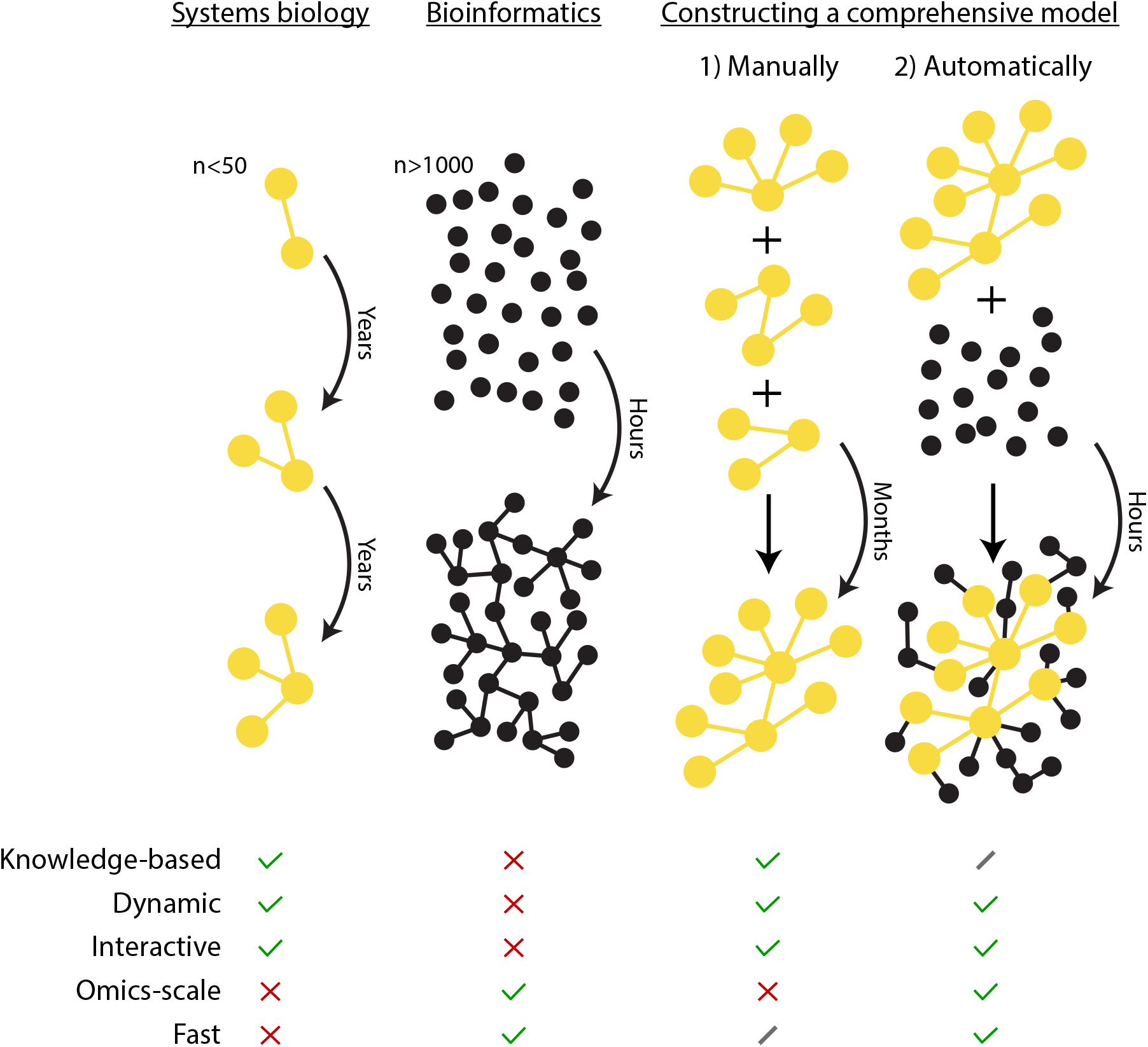
Overview of traditional approaches and the developed approach. In the field of systems biology, typically small mechanistic models of high confidence are developed. These dynamic models can be used to simulate new experiments, such as new drug interventions. However, they are slow to develop and are thus not suitable for analyzing omics data. Conversely, such large omics data are commonly analyzed using bioinformatics methods. However, the networks generated from such methods are typically static, and the individual interactions are often of lower confidence. In our new approach, we first manually combine three separate mechanistic models into a connected core model. This core model is then expanded automatically using omics data and lists of protein-protein interactions. During this expansion, we gradually introduce lower confidence additions. Thus, we end up with an expanded model with a highly confident core with layers of gradually decreasing confidence, with the possibility to simulate experiments on a large portion of the phosphoproteome.

## Results

We first established a core model of adipocyte signaling (Fig. 2D), based on our earlier work [12, 15, 16]. The included earlier works are the following three models: 1) a glucose uptake model [12] that includes major insulin signaling pathways of adipocytes, based on data from both non-diabetic and type 2 diabetic patients (Fig. 2A). 2) a lipolysis model [16] that describes the release of fatty acids and glycerol in response to *α*- and *β*_2_-adrenergic receptor agonists and inhibitors, as well as the anti-lipolytic effect of insulin (Fig. 2B). The lipolysis model includes intracellular signaling intermediaries and is also based on data from both non-diabetic and type 2 diabetic patients. 3) an adiponectin release model [15] that unravel the mechanisms of adiponectin release in response to epinephrine, and a *β*_3_-adrenergic receptor agonist, as well as intracellular signaling intermediaries. This adiponectin release model includes patch-clamp experiments in the 3T3-L1 adipocyte cell line, as well as confirming studies in adipocytes from non-diabetic patients (Fig. 2C).

**Figure 2:**
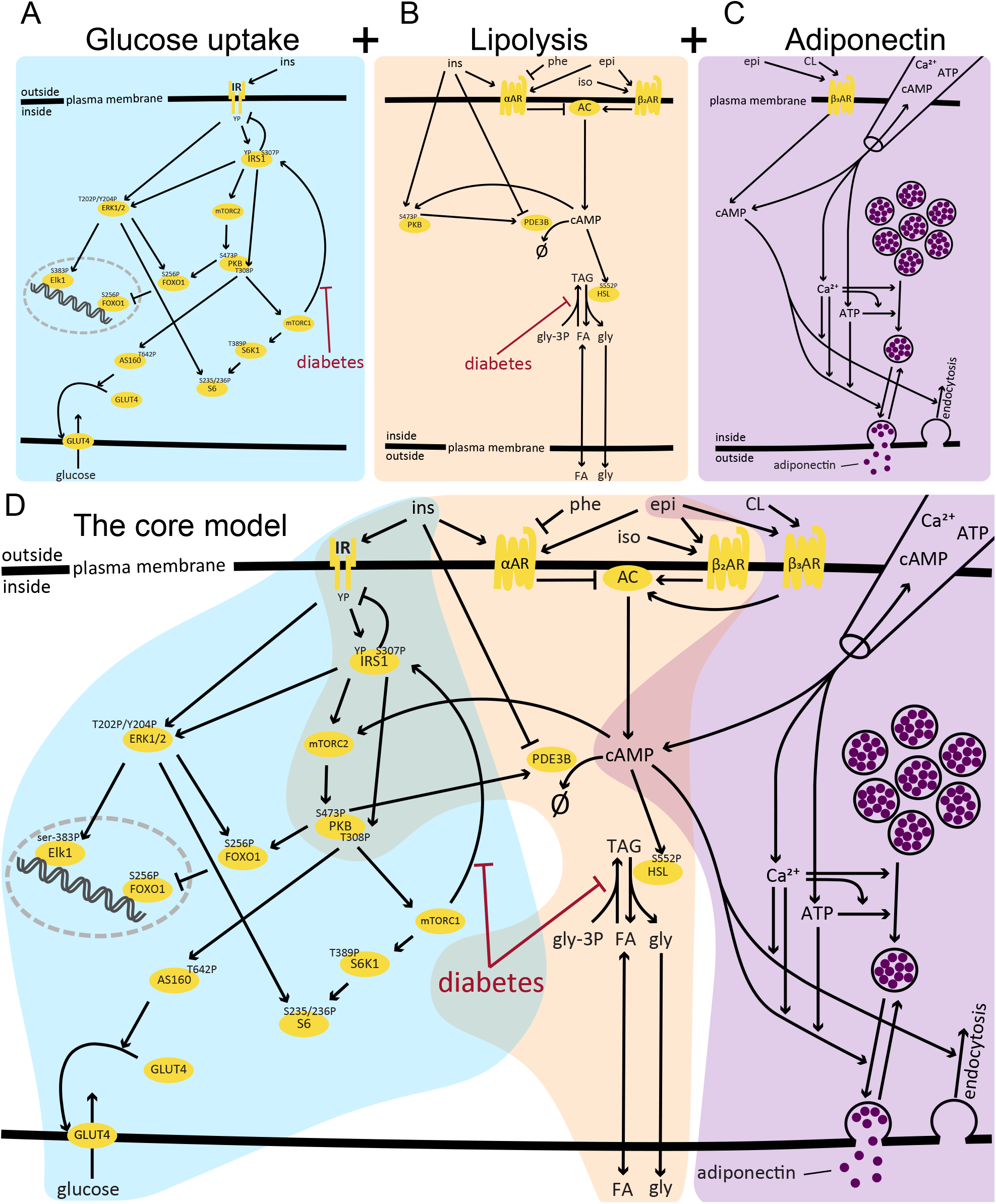
The submodels and the connected core model of adipocyte signaling. The core model includes crosstalk between glucose uptake in response to insulin (ins), fatty acid (FA) and glycerol release in response to *α*- and *β*_2_-adrenergic receptor signaling, and adiponectin secretion in response to cAMP, ATP and Ca^2+^(added inside the cell with a pipette) and extracellularly stimulated *β*_2_-adrenergic receptor signaling. The model includes key differences between signaling normally and in type 2 diabetes. phe - phentolamine, iso - isoprenaline, epi - epinephrine, CL - CL 316243.

### The crosstalk between different pathways in adipocyte signaling

To connect the models, we looked for crosstalk in between the included signaling pathways (Fig. 2). There are clear crosstalk between glucose uptake and lipolysis, through the signaling pathways of both insulin and *α*- and *β*-adrenergic receptors. A central node for this crosstalk is cAMP. cAMP is stimulated by *β*-adrenergic receptors and inhibited by *α*-adrenergic receptors, and in turn affects the phosphoinositide 3-kinase inhibitor (PI3K*α*) [17]. PI3K*α* is also involved in the insulin – glucose uptake signaling network. Furthermore, stimulation with insulin (at physiological concentrations) will lead to an activation of phosphodiesterase 3B (PDE3B), which directly breaks down cAMP. cAMP is also a central mediator for adiponectin secretion [14].

### The creation of a core model from previous models and data

The three separate models [12, 15, 16] were connected in two steps. Firstly, we connected the lipolysis model [16] with the glucose uptake model [12]. This was done by replacing the insulin action (*Ins*_1_) and the protein kinase B (PKB) equations from the lipolysis model with the insulin receptor and PKB from the glucose uptake model, and by having cAMP from the lipolysis model activate the mammalian target of rapamycin complex 2 (mTORC2) in the glucose uptake model. This interaction of cAMP to mTORC2 is a simplified mechanism of the activation of PI3K with subsequent activation of mTORC2. Secondly, we connected the newly combined lipolysis–glucose uptake model with the adiponectin model [15]. In essence, we combined the model by letting the *β*_3_-adrenergic receptors from the adiponectin model activate adenylyl cyclase (AC) from the lipolysis model instead of directly leading to a production of cAMP. All details regarding the connection of the model and the full connected core-model is found in all details in *S2 Model equations and parameter values*.

### The core model can reproduce all previously used data

We tested the core model by separating the experimental data used in the previous works [12, 15, 16] into a training set and a validation set. To be able to compare the combined core model to the ingoing individual models, we divided the data in the same way as done in the previous works. In practice, this meant that we used the data for the *β*_3_-adrenergic receptor agonist CL 316243 and ATP stimulated adiponectin release [15], and the data for the phosphorylation of HSL in response to stimulation with insulin and isoprenaline [16], for model validation and thus removed this validation data from the set of training data. After estimating the model parameters using the training data, we found the best agreement to be acceptable, supported by a χ^2^-test 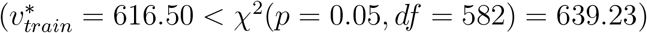. We then estimated the model uncertainty by finding the maximal and minimal simulation values in each time point for each experiment, while simultaneously requiring the agreement with the training data to be sufficiently good 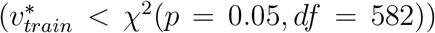, described in detail in the *uncertainty estimation* section. While estimating the model uncertainty, we also collected the parameter set giving the best agreement with the validation data set. We found that the best model agreement to the validation data was statistically acceptable 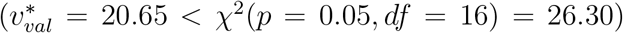. The full description and graphical representations of the model agreement to data is given in the supplemental file *S1 Recreation of the model agreement to data and validations*.

After the model validation, we included both the training and validation data into an extended set of data. We tested if the model could explain the extended set of data sufficiently well. We found the model agreement to the extended data to be acceptable 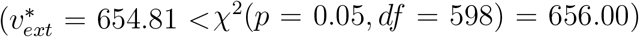, and thus decided to use the connected model as a core for further extensions. The model agreement to the extended data is shown in Figs. 3 to 5. We again estimated the model uncertainty by finding the maximal and minimal simulation values in each time point for each experiment, while simultaneously requiring the agreement with the extended set of data to be sufficiently good 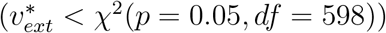, described in detail in the *uncertainty estimation* section.

**Figure 3:**
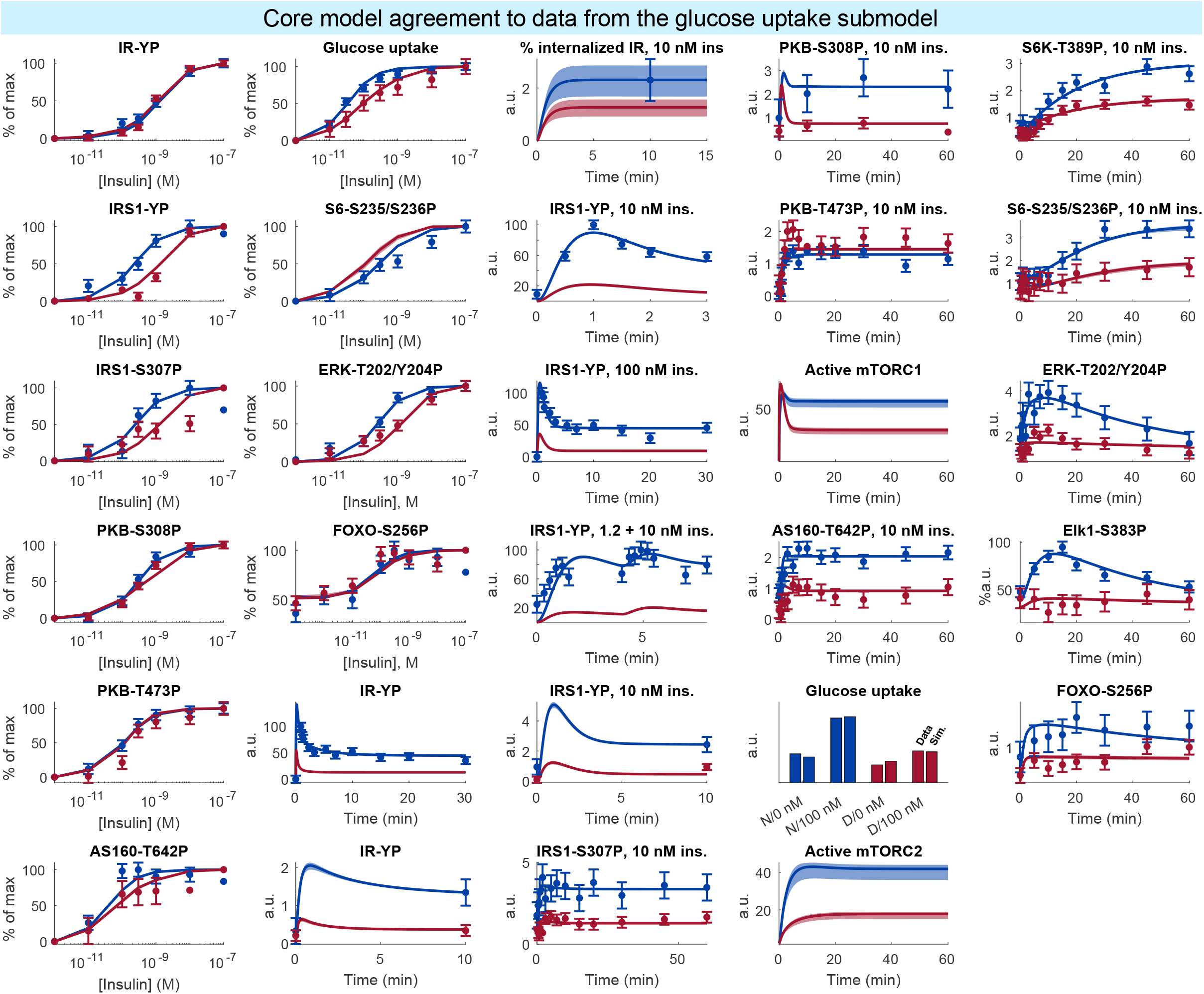
Model agreement to the data from the previous insulin signaling work. Data comes from isolated human adipocytes stimulated with insulin in different doses and for different times [12]. In all panels, lines represent the model simulation with the best agreement to data, the shaded areas represent the model uncertainty, and experimental data points are represented as mean values with error bars (SEM). Simulations and experimental data in red corresponds to experiments under type 2 diabetic conditions, and in blue under non-diabetic conditions.

**Figure 4:**
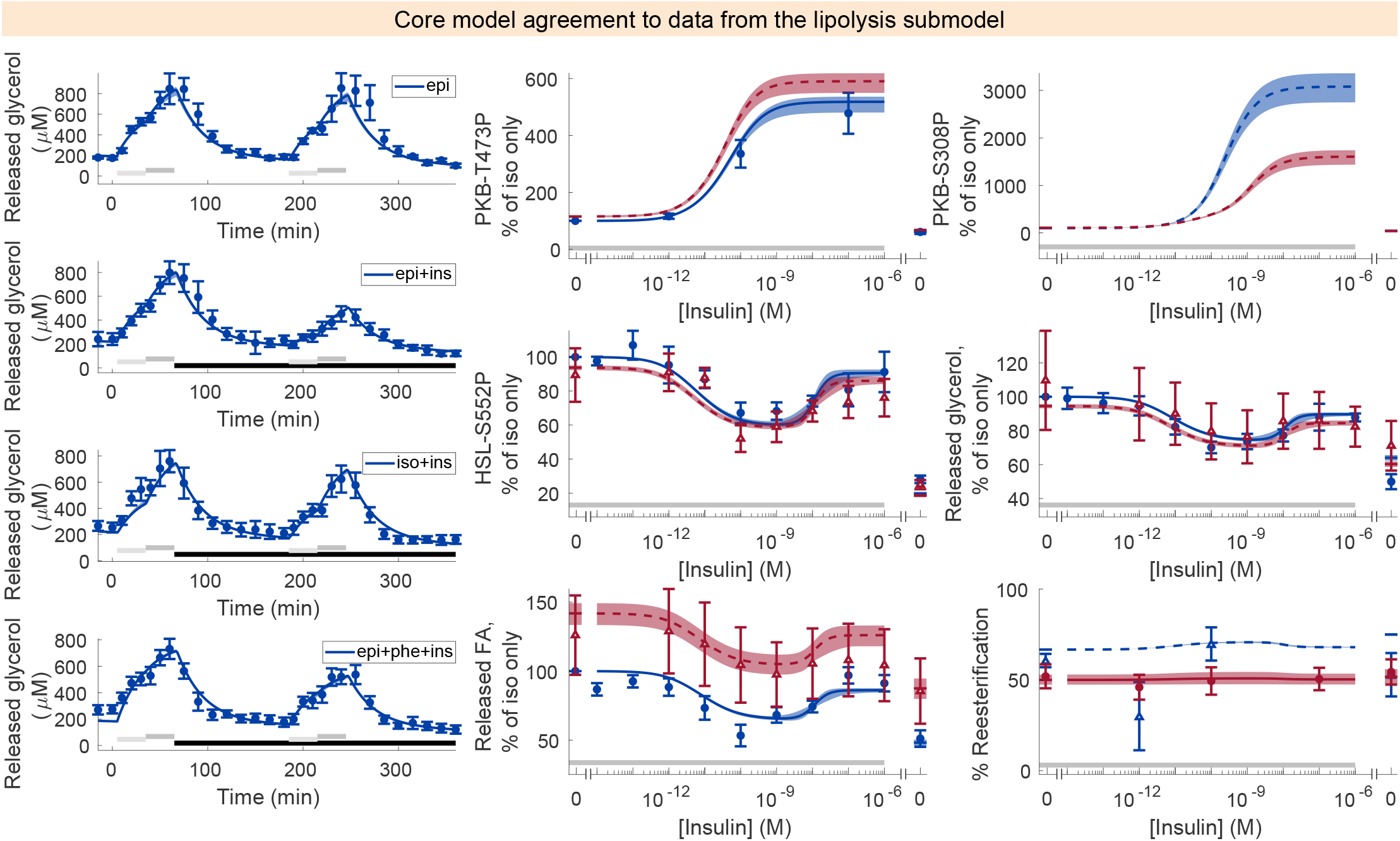
Model agreement with the data from the previous lipolysis work. Released glycerol (left) was measured using microdialysis in the adipose tissue in situ [18]. All other data (middle, right) was measured in isolated human adipocytes [17]. In all panels, lines represent the model simulation with the best agreement to data, the shaded areas represent the model uncertainty, and experimental data points are represented as mean values with error bars (SEM). Simulations and experimental data in red corresponds to experiments under type 2 diabetic conditions, and in blue under non-diabetic conditions. Dashed lines and experimental data with open triangles were not used to estimate the model parameters. Light/dark gray horizontal bars indicate adrenergic stimulation, and black horizontal bars in the left figures indicate insulin stimulation. epi – epinephrine, ins – insulin, iso – isoprenaline, FA – released fatty acids

**Figure 5:**
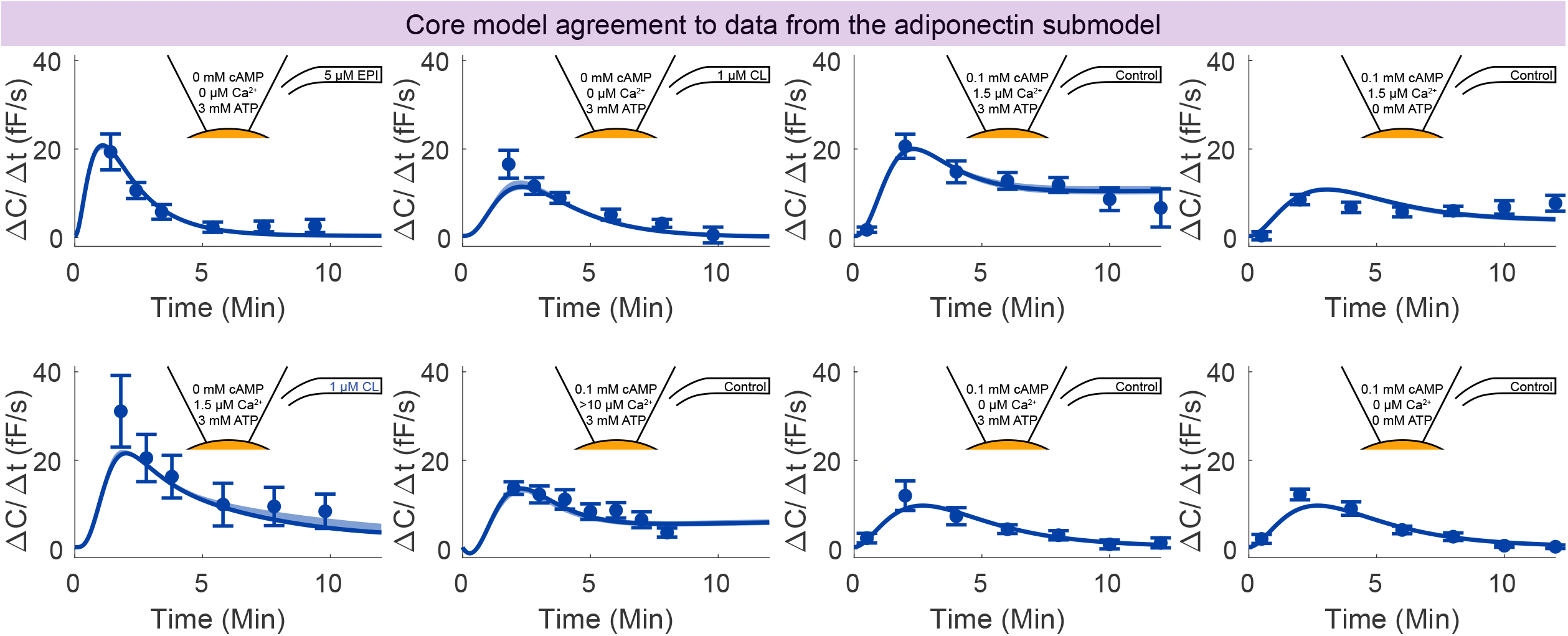
Model agreement with the data from previous adiponectin release modeling work. Data represent patch-clamp capacitance recordings in 3T3-L1 adipocytes [16]. In all panels, lines represent the model simulation with the best agreement to data, the shaded areas represent the model uncertainty, and experimental data points are represented as mean values with error bars (SEM). EPI - epinephrine, CL - *β*_3_-adrenergic receptor agonist CL 316243, Ca^2+^- Calcium.

### From the core model to the comprehensive mechanistic model of adipocyte signaling

To go from the core model to the comprehensive adipocyte, we use phosphoproteomic data for the time-resolved insulin response in 3T3-L1 adipocytes [19]. Prior knowledge on possible protein–protein interactions was collected using the python package OmniPathDB [20], which compiles a list of interactions from several sources, such as Reactome and PhosphoSitePlus. The list includes a confidence level in the sense of number of primary sources that have studied each interaction.

We first decided whether to employ a top-down or a bottom-up approach. By filtering the list of interactions to only include interactions where phosphoproteomic data were available, we could estimate the size of the maximal model if constructed using all available interactions with data. Such a model would contain 6642 states and 59169 parameters, assuming each phosphorylation site can switch between a phosphorylated and an unphosphorylated state, returning to the unphosphorylated state using a single parameter, and that each input in the list of interactions would contribute with one parameter. Clearly, the amount of unknown parameters would be too many to optimize using even the most state-of-art methods. We thus conceived of a bottom-up approach instead.

We divided the phosphoproteomic data into two sets as done by the authors in [19]: one set with sites responding to insulin stimulation and another set with sites not responding to insulin stimulation, based on if the sites are effected by inhibitors of PI3K and PKB. We refer to these data sets as *responders* or *nonresponders*. For our model expansion we started with the responder data.

Using a pairwise approach, we expand the core model towards the comprehensive mechanistic model in multiple layers (Fig. 6). We refer to the additions of the adjacent sites as the addition of a *layer* of sites to the model. For the first layer, we start with the core model and add phosphorylation sites by identifying the sites that are adjacent to the core model given prior knowledge about interactions. We start with interactions of the highest confidence, i.e. with the most (20) primary sources. From that level of confidence, we test in a random order if the phosphorylation sites can be added using interaction from the core model as input. Once no more adjacent proteins could be added to the layer, we used the last added layer as inputs to find new adjacent sites. This layered addition of sites was continued until no more sites could be added to the model.

**Figure 6:**
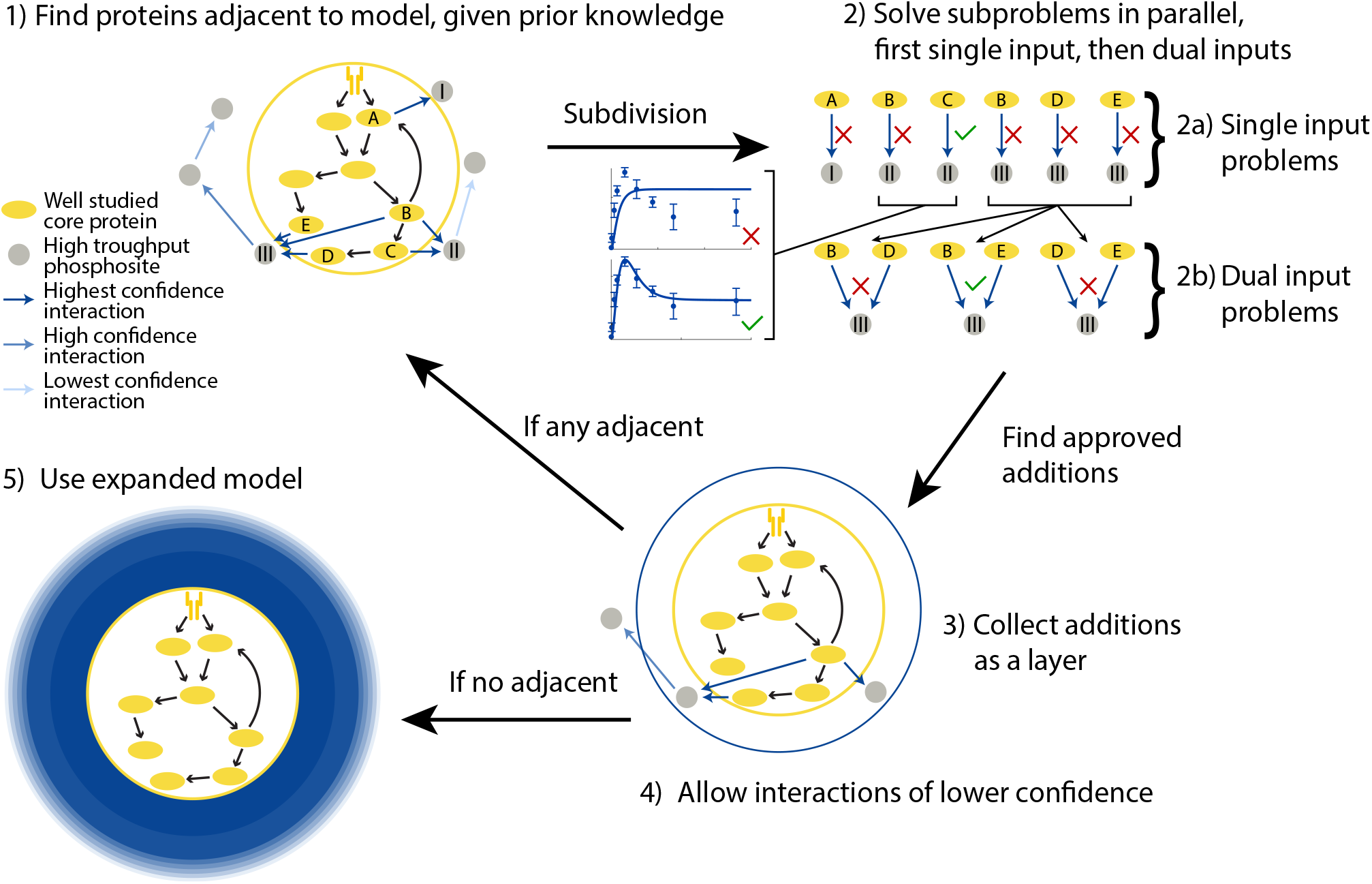
Overview of the developed approach. At the core of the method is our connected model. Outside the core connected model, there exists phosphoproteomic data (gray circles) not covered by the model. Using lists of interactions and our automatic model expansion algorithm we are able to add parts of this phosphoproteomic data, for proteins adjacent to the core model, to an expanded model. These additions are then subsequently used to further add proteins to the expanded model. The core model has the highest degree of confidence, but has the lowest coverage of the phosphoproteome. For each added layer, a larger portion of the phosphoproteome is covered but with a lower confidence.

**Figure 7:**
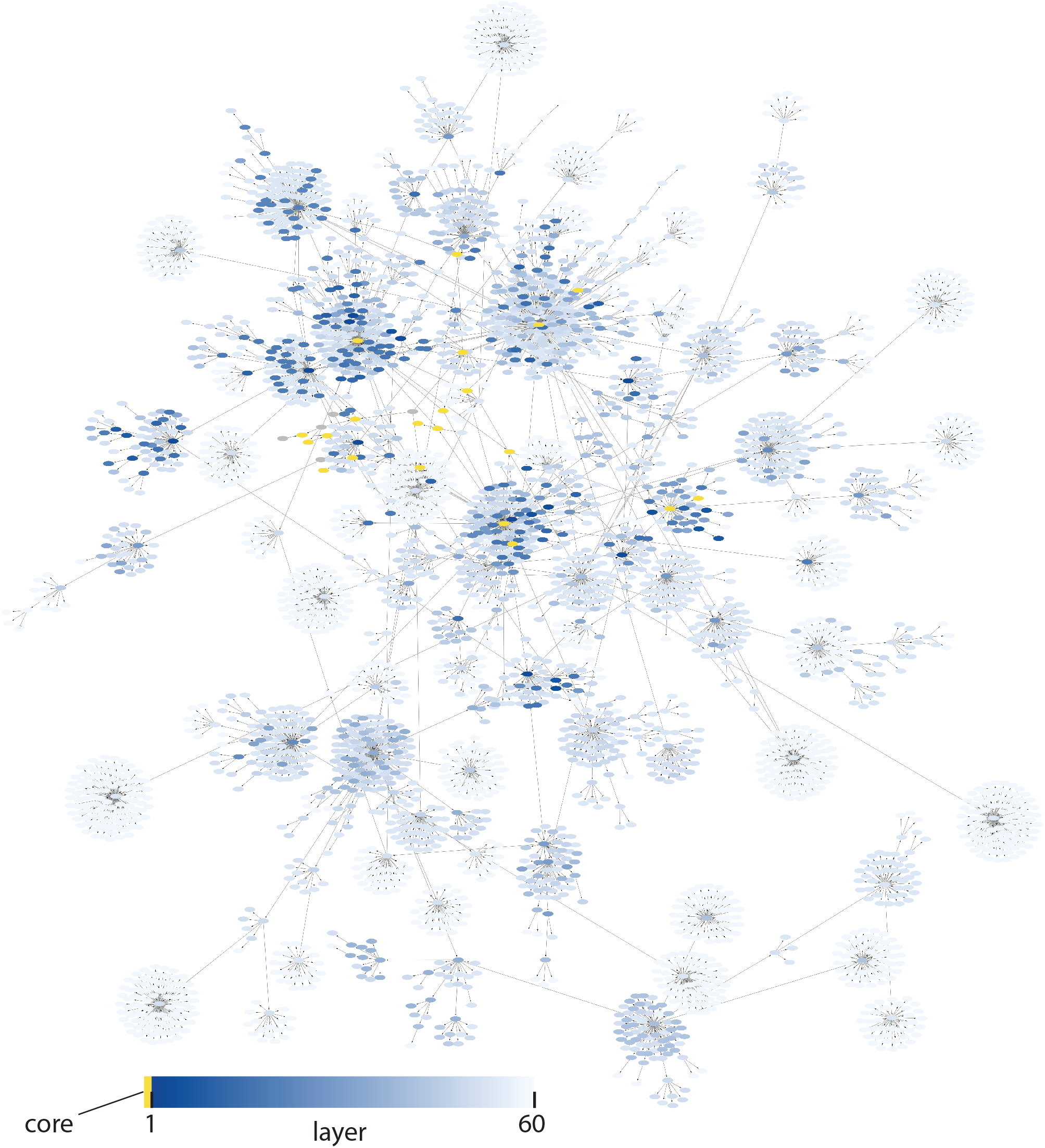
The comprehensive model of adipocyte signaling. The network structure of the expanded model, color coded by the order addition for the 60 added layers in blue. The core model is shown in yellow. Additions with high confidence is shown in dark blue, and additions with lower confidence in light blue. The full network is also available in *S4 Cytoscape file of the expanded model*. The simulations and experimental data for all phosphosites in the model is available in *S3 Time series of all sites added to the model*.

Once no more sites could be added to the model, we also included the interactions with the second-highest number of primary sources and attempted to add adjacent sites. We continued to include interactions with a lower confidence in a stepwise manner until no more sites could be added even with the interactions with the lowest number of primary sources. Using the responder data and all interactions, we added 189 sites to the model.

We then used all of the data (including the nonresponders), and again started with the interactions with the highest number of primary sources. Again, we step-wise included interactions with a lower number of primary sources. Using all data and all interactions we could add 2146 additional sites, to a total of 2335 sites. When no more interactions could be added using all the data and all the interactions, we continued to add sites not yet included in the model using a data-driven approach. Here, we created a new list of potential interactions, based on the agreement between the simulations of the sites in the expanded model with the experimental data of the sites that had not been added. Using the data-driven approach we could add 1591 additional sites, to a total of 3926 sites. The final expanded model contains 7795 states and 7877 parameters.

#### Predicting inhibitions

To test the validity of the expanded model, we made predictions by simulating the change in the insulin response with a PKB inhibitor and with a PI3K inhibitor, and compared the predictions to experimental data for the insulin response in the presence of the PKB inhibitor MK2206 and the PI3K inhibitor LY294002, gathered in the same study as the large-scale time series data [19]. Since the model was expanded using both interactions and data with different levels of confidence, we evaluate the model’s predictive capabilities on a layer-to-layer basis. The simulated time series of all phosphorylation sites, and the effect of the inhibitors, can be found in *S3 Time series of all sites added to the model*. The model is able to correctly predict the direction of the two separate inhibition experiments in a majority of all sites, in all confidence layers (Fig. 8). Note that the model predictions are the most accurate when the most confident interactions and data is used, and that the accuracy of the predictions fall as the confidence is decreased (when going from left to right in Fig. 8A,B). Since the model expansion is done using the most confident interactions and data first - and then gradually decrease in confidence - it is clear that additions with higher confidence give the most accurate model predictions (left to right in Fig. 8A,B).

**Figure 8:**
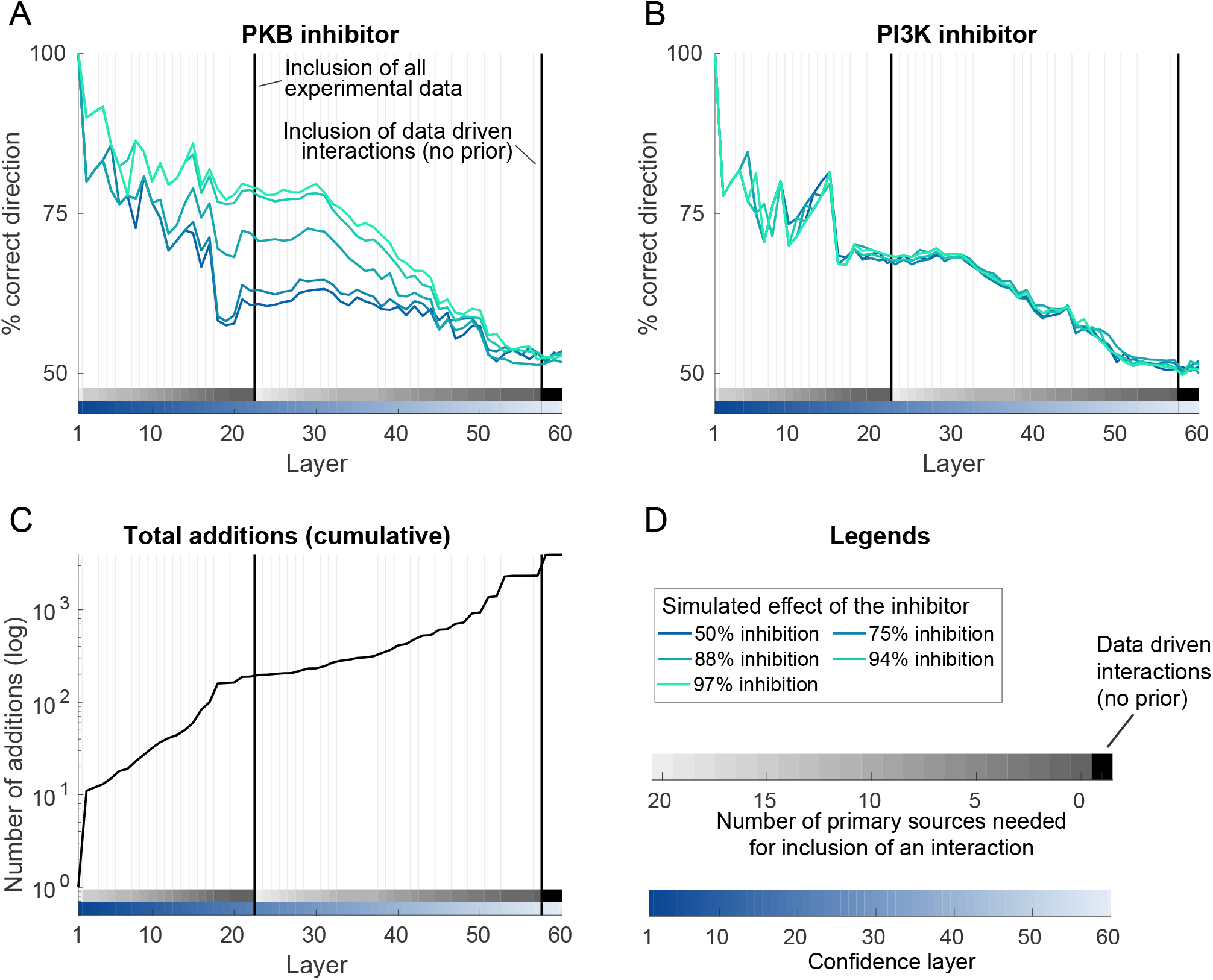
The predictive ability of the comprehensive model. For each added layer, we compute the predictive ability of the model using independent inhibitor data. (A) shows the ability of the model to correctly predict the direction (inhibitory or stimulatory) for the effect of PKB inhibition at 20 minutes after insulin stimulation, for the automatically added phosphosites. (B) shows the ability of the model to correctly predict the direction (inhibitory or stimulatory) for the effect of PI3K inhibition at 20 minutes after insulin stimulation, for the automatically added phosphosites. (C) shows the total number of added phosphosites. (D) shows the legends to the lines and gradients in (A-C). Vertical black lines correspond to when additional (nonresponder) data and purely data-driven interactions were allowed.

#### Simulating type 2 diabetes

Since the connected model used as a core for the automatic model expansion could simulate type 2 diabetes, we can now simulate how to the effect of type 2 diabetes would spread to a large portion of the phosphoproteome. We evaluated the effect of type 2 diabetes by simulating the same insulin stimulated experiment as done in the phosphoproteomic data under both normal and type 2 diabetic conditions. The time-resolved comparison between normal and type 2 diabetic conditions are shown for the first 5 layers in Fig. 9, and for all phosphorylation sites in *S3 Time series of all sites added to the model*. Furthermore, we quantified the change between normal and type 2 diabetic conditions by finding the fold change relative to the normal condition at *t* = 20 minutes (Fig. 10).

**Figure 9:**
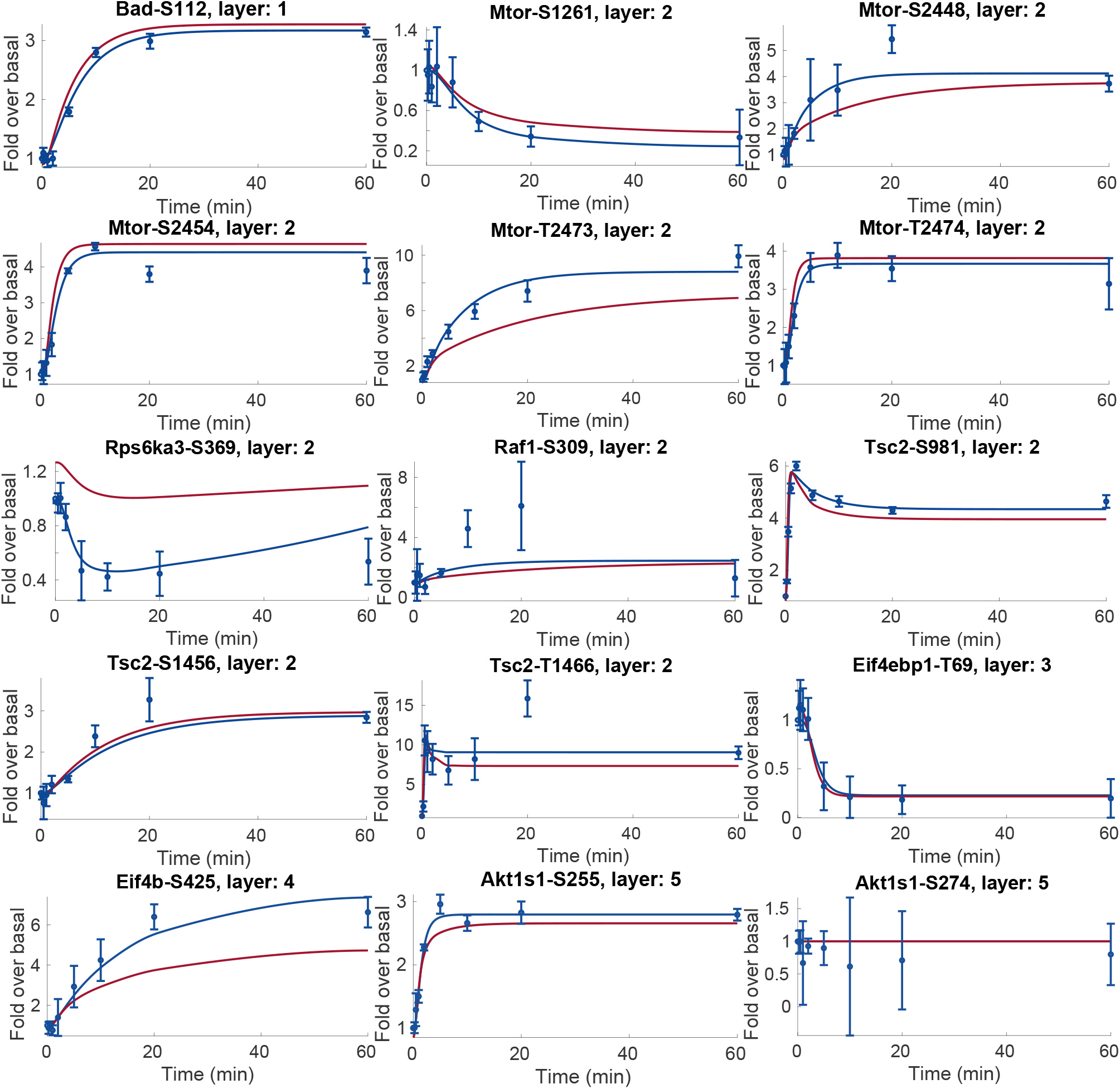
Simulations and data for the first 5 layers of the comprehensive model. The first 5 layers contain 15 added sites. In all panels, blue lines represent model simulation with the best agreement to data and experimental data points are represented as mean values with error bars (SEM). Red lines are simulations of the type 2 diabetic conditions, and these simulations are model predictions without corresponding data. All model simulations of the effect of type 2 diabetes are available in *S3 Time series of all sites added to the model*.

**Figure 10:**
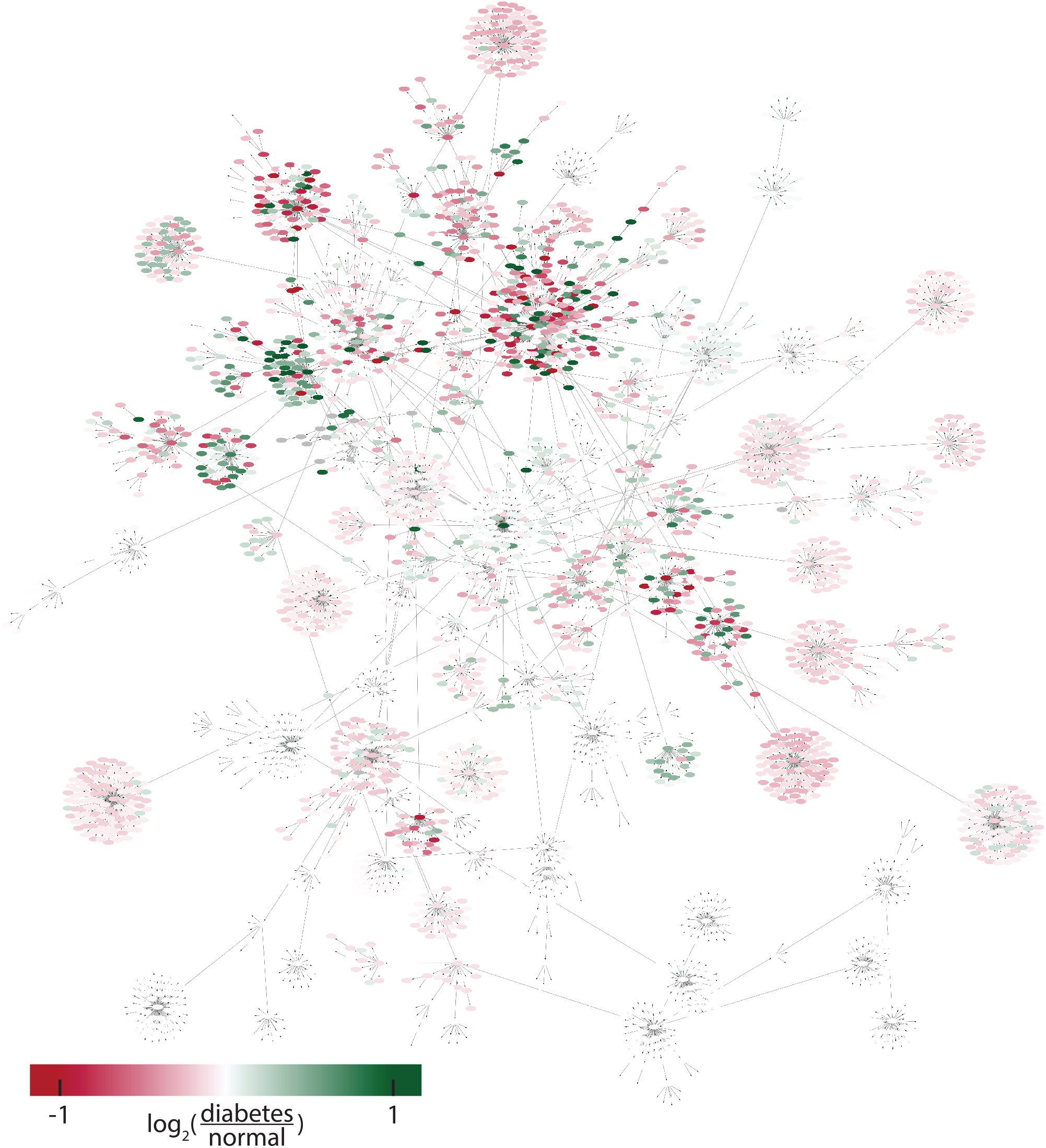
The effect of type 2 diabetes propagated to all sites in the expanded model. The color code goes from red corresponding to decreased signaling, to green corresponding to increased signaling in type 2 diabetes. White corresponds to no, or low, effect. The full network is also available in *S4 Cytoscape file of the expanded model*. The simulations and experimental data for all phosphosites in the model is available in *S3 Time series of all sites added to the model*.

## Discussion

In this work, we present a new methodology for automatic model expansion from a dynamic core model to include large-scale data and keep track of the level of confidence of both data and interactions (Fig. 6). In the methodology, the new data is added in parallel in layers, to reduce computation time and allow for thousands of added time-resolved measurements. To create a connected high-confidence core model of adipocyte signaling, we have connected major events related to lipolysis, glucose uptake, and adiponectin release into a connected model (Fig. 2). This connected model, developed herein, agrees with numerous observations from adipocytes (Figs. 3 to 5). From the core model, we use our new methodology to add layers of experimentally measured phosphosites [19] based on a list of possible interactions, including level of confidence, compiled using OmniPathDB [20]. The developed methodology allows for combining detailed mechanistic models, common in the field of systems biology, with large-scale data, traditionally analyzed in the field of bioinformatics, thus acting as a bridge between such methods.

The created comprehensive model has a layered level of confidence based on both interactions and data (Fig. 7). Most of the data from the insulin signaling models [12, 16] comes from isolated primary adipocytes from non-diabetic and type 2 diabetic patients, where protein levels and activities have been measured using antibodies and the western blot technique. The adiponectin release and phosphoproteome data instead comes from the 3T3-L1 adipocyte cell line, where the release of adiponectin was measured using a patch-clamp technique, and the protein phosphorylations was measured with mass-spectrometry. Furthermore, the small-scale data (western blot and patch-clamp) are typically more reliable since there are more repeats (typically *n* = 5 – 15) compared to the large-scale phosphoproteomic data (*n* = 0 – 3) per measurement. The additions in the expanded model is based on interactions with varying degrees of confidence. During the expansion, we first start with the interactions with the highest confidence, and then incrementally allow interactions with lower confidence (Fig. 8).

The development of new mechanistic ODE-based models for biological systems is a long and iterative process. First, you start with some data and hypothesis, then you refine the model in many steps, while using the model to make predictions for new data to gather. The development of the models included in the core model here, as well as the gathering of corresponding data, have taken us and our collaborators decades of work. Thus, it is not feasible to cover the entire phosphoproteome with consistent high quality data and mechanistic ODE-based models. The developed automated methodology therefore provides a reasonable trade-off between confidence in the model and the coverage of the phosphoproteome (Fig. 6).

The most confident part of the model is the connected core model, which agrees with the full set of original experimental data sufficiently well (Figs. 3 to 5), supported by a χ^2^-test: 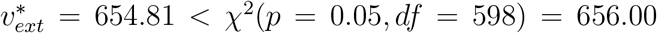. However, at the same time the optimal cost 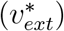 is close to the threshold of rejection (from 654.81 to 656.00). This means that only changes in the parameter values resulting in a maximum cost increase by 1.19 is accepted when estimating the model uncertainty. In other words, a cost increase by around 0.2 %. This narrow range of acceptable cost increase, results in a model uncertainty that is also narrow. We do not believe this narrow uncertainty to be the result of failed optimizations, but rather due to the small allowed increase in cost. Perhaps alternative thresholds of rejection should be considered, such as allowing a cost increase from the optimal cost corresponding to one degree of freedom (χ^2^(*p* = 0.05, *df* =1) = 3.84).

Alternative attempts at modelling the phosphoproteome have been made, such as the machine learning clustering analysis in the original work that collected the phosphoproteome data used herein [19], or the PhosR method for identifying “signalomes” from the same research group [21]. Other attempts include the multi-omics analysis method MEPHISTO [22], or the employment of protein-protein interaction databases to infer time-dependent kinase-substrate relationships [23]. However, to the best of our knowledge, none of these methods are able to explain the time dynamics of the phosphoproteome using a mechanistic approach. Furthermore, they are not able to simulate new situations and make predictions, something our expanded model is able to do.

The biological relevance of a comprehensive model of adipocytes signaling is related to type 2 diabetes. Insulin resistance is a hallmark of type 2 diabetes, and the core of the model describe in detail the intracellular insulin resistance of adipocytes (Fig. 3). A major function of adipocytes is to act as energy reserve for other organs, i.e. to store and release fatty acids. The release of fatty acids is increased in type 2 diabetes, due to a decrease in the reesterification (Fig. 4). Also, circulating adiponectin has been associated with a reduced risk of type 2 diabetes [24], and an increased insulin sensitivity [25]. All of these processes have already been established in detail in models, and here we provide a connection including crosstalk that allow for their simultaneous simulations. Even more important, we provide a link between these highly established processes to the whole phosphoproteome, where the model allow anyone to simulate, e.g. the effect of a drug. We provide a type 2 diabetes version of the model (Figs. 9 and 10), therefore the model can be used to simulate the effect of anti-diabetic drugs throughout the adipocyte. In several efforts (e.g. [26, 27]), plasma omics measurements of diabetic patients have been related to clinical parameters using statistical models. Such models provide important insight to relations between clinical parameters and specific proteins and/or genes, and can be used for classifications of patient groups. Such purely statistical models cannot, however, go beyond the data used for training to e.g. simulate the effect of new treatments.

Other approaches to model human adipocytes on a large scale is the use of genome-scale metabolic models [28]. Such models provide another view on adipocyte changes in obesity and type 2 diabetes. These models integrate genomics, transcriptomics, and proteomics data and apply to a fixed metabolic map based on known human metabolic reactions. Such models are complementary to the work herein since they do not consider the signaling pathways, and our approach does so far not include metabolic reactions. We are currently gathering labelled metabolite data for metabolic flux analysis from human adipocytes. These data will be integrated in the next version of the comprehensive adipocyte.

In summary, we here present a method that can scale small-scale dynamic models to the omics level, while still preserving the dynamic qualities of the model. Our method can integrate data from different sources into a cohesive framework, and can be further extended when additional data becomes available in the future. The expanded model can be used to predict events, such as perturbations by an inhibitor or a drug, and the changes to the signaling network as a response to a disease. Furthermore, the method should be applicable to other biological systems where detailed small-scale models, and omics-level data, is available.

## Method

### Data collection and filtration

We used publicly available data, both data previously used to develop our models of glucose uptake, lipolysis, and adiponectin release[11, 12, 15, 16] and from other sources [19].

For the newly used phosphoproteomic data [19], we excluded sites where any time point had less than two repeats. For the inhibition data, sites with less than two repeats, or with an unclear effect of the inhibition (such that the uncertainty covered both an increase and a decrease) were ignored when the prediction accuracy was calculated. Furthermore, we divided the data into two groups (*responders* and *nonresponders* to insulin) based on the original classification done by the authors of the phosphoproteome work [19].

### Quantifying the model agreement to experimental data

In order to evaluate the model’s performance, we quantified the model agreement to data using a function typically referred to as a cost function. In this case, we used the normalized sum of squared residual as cost function (Eq. (1)).

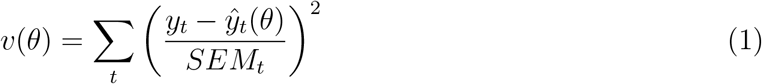

Here, *v*(*θ*) is the cost for the set of parameter values *θ*, equal to the sum of the normalized residual over all measured time points, *t*; *p* is the parameters; *y_t_* is the measured data and 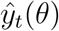 is the model simulations; *SEM_t_* is the standard error of the mean for the measured data.

### Statistical analysis

To reject models, we used the χ^2^-test with a significance level of 0.05. We used 582 degrees of freedom for the original training data leading to a threshold for rejection of χ^2^(*p* = 0.05, *df* = 582) ≈ 639.23. For the extended set of experimental data including all the experimental data from the connected model we used 598 degrees of freedom, resulting in a threshold for rejection of χ^2^(*p* = 0.05, *df* = 598) ≈ 656.00. Any combination of model and parameter set that results in a cost (Eq. (1)) above the threshold must be rejected. If no parameter set exists for a model that results in a sufficiently low cost, the model structure must be rejected.

For the automatic model expansion we allowed additions to the model where the cost for the tested phosphorylation site was below χ^2^(0.05, 8) ≈ 15.5.

### Optimization and software

We used MATLAB R2021a (MathWorks, Natick, MA) and IQM tools (IntiQuan GmbH, Basel, Switzerland), a continuation of [29], for modelling. IQM tools uses CVODES [30] to numerically integrate the ODEs. The parameter values were estimated using the enhanced scatter search (eSS) algorithm from the MEIGO toolbox [31]. We allowed the parameter estimation to freely find the best possible combinations of parameter values, within boundaries. Both the optimal parameter values and the bounds for the parameter values are given in *S2 Model equations and parameter values*.

### Model and code availability

The model as well as the complete code for data analysis and modelling will be freely available upon submission for peer review. The model equations and parameter values are available in *S2 Model equations and parameter values*.

### Uncertainty estimation

To estimate the model uncertainty, we employed the formulation of the uncertainty estimation problem as described in [32] and implemented in [15, 16]. In short, we estimated the model uncertainty by maximizing or minimizing a specific model prediction 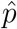 (such as the simulation of an experiment at a specific time point), while requiring that the model agreement with data is sufficiently good (the cost being less than the χ^2^-limit) (Eq. (2)).

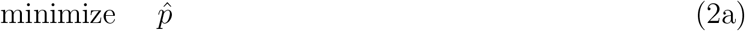

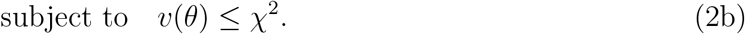

Here, some prediction 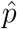 is minimized to find the lower bound on the value of the prediction, while requiring the cost *v*(*θ*) to be below the χ^2^-threshold. To get the upper bound of the prediction, the problem in Eq. (2) can be solved as a maximization problem. In practice, we solved the maximization problem as a minimization problem by changing the sign of the objective function (Eq. (2)) to 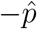. Furthermore, we relaxed the constraint (Eq. (2b)) into the objective function as a L1 penalty term with an offset if *v*(*θ*) > χ^2^ (Eq. (3)).

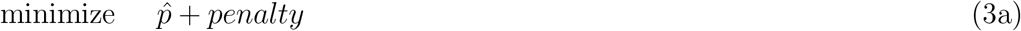

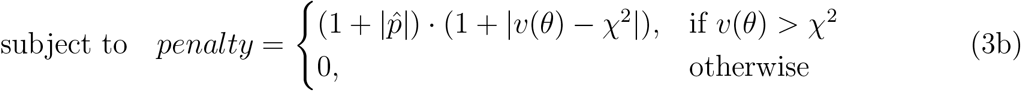

### Automatic model expansion

In essence, we create an expanded model by iteratively adding proteins to the core model by identifying the proteins that are closest to, but not yet in, the model, and test if they can be added to the model in a pairwise fashion. This was done using phosphoproteomic data collected using mass-spectrometry [19], and a list of protein-protein interactions compiled using the python package OmniPathDB [20]. In detail, we find all proteins that are adjacent to the model, i.e. having at least one direct interaction in the list of interactions going from the model to the proteins. Using the list of adjacent proteins and corresponding prior knowledge interactions, we test if the proteins can be added to the model one by one (in parallel). In practice, each addition of an adjacent protein is tested using three different types of interactions. Firstly, we test if the interaction can be a single pairwise interaction corresponding to a phosphorylation, a dephosphorylation, or a saturated phosphorylation of the adjacent protein. Secondly, we test if the interaction must be a double interaction with two different inputs (double phosphorylation, double dephosphorylation or one phosphorylation with one dephosphorylation). Lastly, we test if the interaction results in a phosphorylation leading to a subsequent secondary state (such as an internalization) before returning to the unphosphorylated state. The adjacent proteins and interactions are tested by comparing the model simulation of the adjacent protein with the available experimental data. All adjacent proteins where the model simulations agree with the data sufficiently well are added to the model. We refer to the addition of the adjacent proteins that agree with data sufficiently well as the addition of a *layer* to the model. Once one such layer have been added to the model, we repeat the process of finding and adding another layer of adjacent proteins. For subsequent addition of layers, proteins in the previous layers are also used to find the new adjacent proteins. The algorithm for the model expansions is given as pseudocode in Algorithm 1.

## Supporting information

S1 Recreation of the model agreement to data and validations

S2 Model equations and parameter values

S3 Time series of all sites added to the model

S4 Cytoscape file of the expanded model

S5 Excel spreadsheet with the impact of type 2 diabetes

## Funding information

GC acknowledges support from the Swedish Research Council (2018-05418, 2018-03319), CENIIT (15.09), the Swedish Foundation for Strategic Research (ITM17-0245), SciLifeLab National COVID-19 Research Program financed by the Knut and Alice Wallenberg Foundation (2020.0182), the H2020 project PRECISE4Q (777107), the Swedish Fund for Research without Animal Experiments (F2019-0010), ELLIIT (2020-A12), and VINNOVA (VisualSweden, 2020-04711). EN acknowledges support from the Swedish Research Council (Dnr 2019-03767), the Heart and Lung Foundation, CENIIT (20.08), Åke Wibergs Stiftelse (M19-0449, M21-0030), and the Swedish Fund for Research without Animal Experiments (S2021-0008). The funders had no role in study design, data collection and analysis, decision to publish, or preparation of the manuscript.

### Algorithm 1 Automatic model expansion

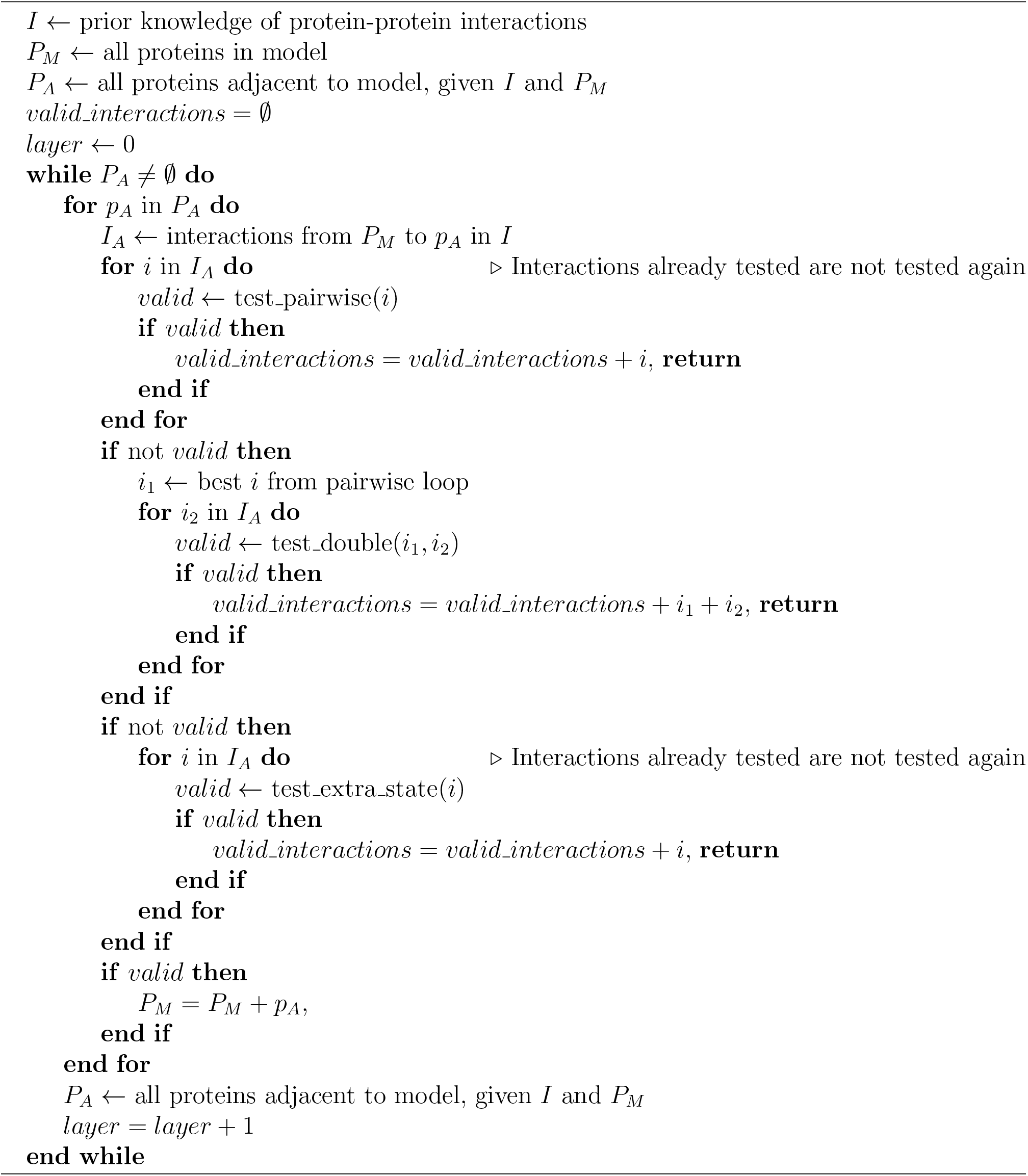

The following files are available as supplemental files.

**S1 Recreation of the model agreement to data and validations**

**S2 Model equations and parameter values**

**S3 Time series of all sites added to the model**

**S4 Cytoscape file of the expanded model**

**S5 Excel spreadsheet with the impact of type 2 diabetes**

