## Supplementary material for "A comprehensive mechanistic model of adipocyte signaling with layers of confidence": S1 Recreation of the model agreement to data and validations

### Recreating the model agreement to data and validations from previous works.

May 2, 2022

This document shows that the combined model can recreate model agreement with data and the validations from the previous modelling works. We first trained the model to the dataset without the subsets previously used for validations. In practice, this meant that we divided the full data set into a training data set and a validation data set. The validation data set consisted of the data for adiponectin release stimulate by CL and ATP from citeadiponectin2, and the phosphorylation of HSL in response to stimulation with insulin and isoproterenol from [1]. The training data set consisted of the rest of the data, i.e. all data not in the validation data set. We first trained the model on the training data set and tested if the model agreement to data was sufficiently good. We then tested the models predictive power by simulating the experiments corresponding to the data in the validation data set, and again tested if the agreement to data was sufficiently good. For the validation from the insulin signaling – glucose uptake work [2, 3], we recreated the inhibitions with rapamycin, torin and PD184352.

### 1 The combined model can explain all previous data

We first tested if the model could explain the set of training data sufficiently well. This was done by optimizing the model parameters such that the model agreement to data was as good possible. The model agreement to data was estimated using the *objective function* defined in Eq. (1).

$$v(\theta) = \sum_t \left( \frac{y_t - \hat{y}_t(\theta)}{SEM_t} \right)^2 \tag{1}$$

Here, $v$ is typically referred to as the cost, and it is equal to the sum of the normalized residual over all measured time points, $t$; $\theta$ is the parameters; $y_t$ is the measured data at time $t$ and $\hat{y}_t(\theta)$ is the model simulations at time $t$; $SEM_t$ is the standard error of the mean for the measured data at time $t$.

The better the agreement between the model simulations and the data becomes, the lower the cost ($v(\theta)$) becomes. To test if the agreement is sufficiently good, a $\chi^2$-test is used. If the cost is greater than the $\chi^2$-statistic ($v(\theta) > \chi^2$) the model given the parameter values $\theta$ must be rejected. If the cost given the *optimal* parameters $\theta^*$ is greater than the $\chi^2$-statistic ($v(\theta^*) > \chi^2$) then the model structure must be rejected. Conversely, if we can find a parameter set resulting in a cost lower than the $\chi^2$-statistic ($v(\theta) < \chi^2$) we deem the model sufficiently good.

Here, we found the best cost ($v^*$) for the combined model when trained to the set of training data to be below the threshold of rejection ($v^*_{train} = 616.50 < \chi^2(p = 0.05, df = 582) = 639.23$). The model agreement to the training data is shown in Figs. 1 to 3.

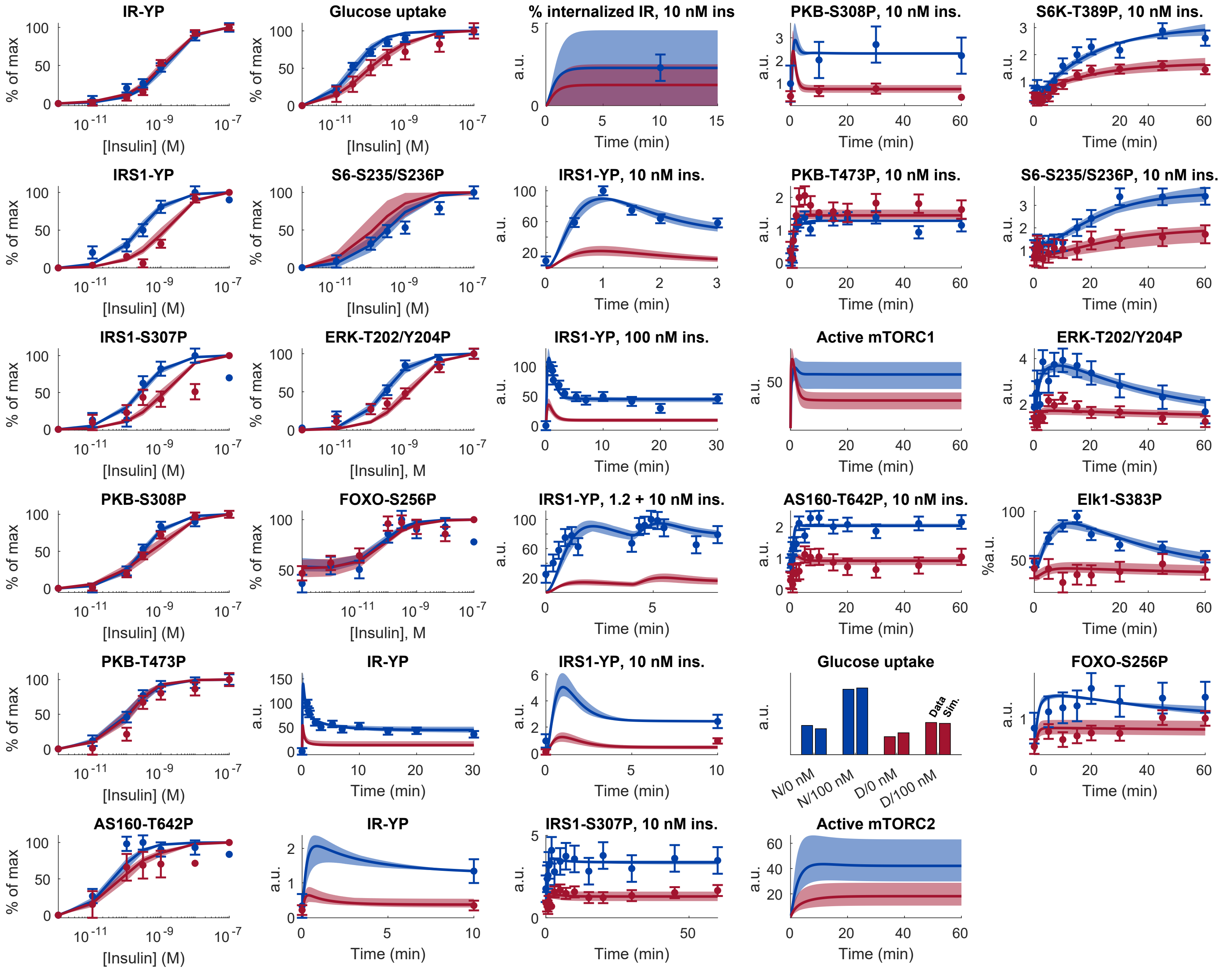


Figure 1: Model agreement with data from [2]. Data comes from isolated human adipocytes stimulated with insulin in different doses and for different times [2]. In all panels, lines represent the model simulation with the best agreement to data, the shaded areas represent the model uncertainty, and experimental data points are represented as mean values with error bars (SEM). Simulations and experimental data in red corresponds to experiments under diabetic conditions, and in blue under non-diabetic conditions.

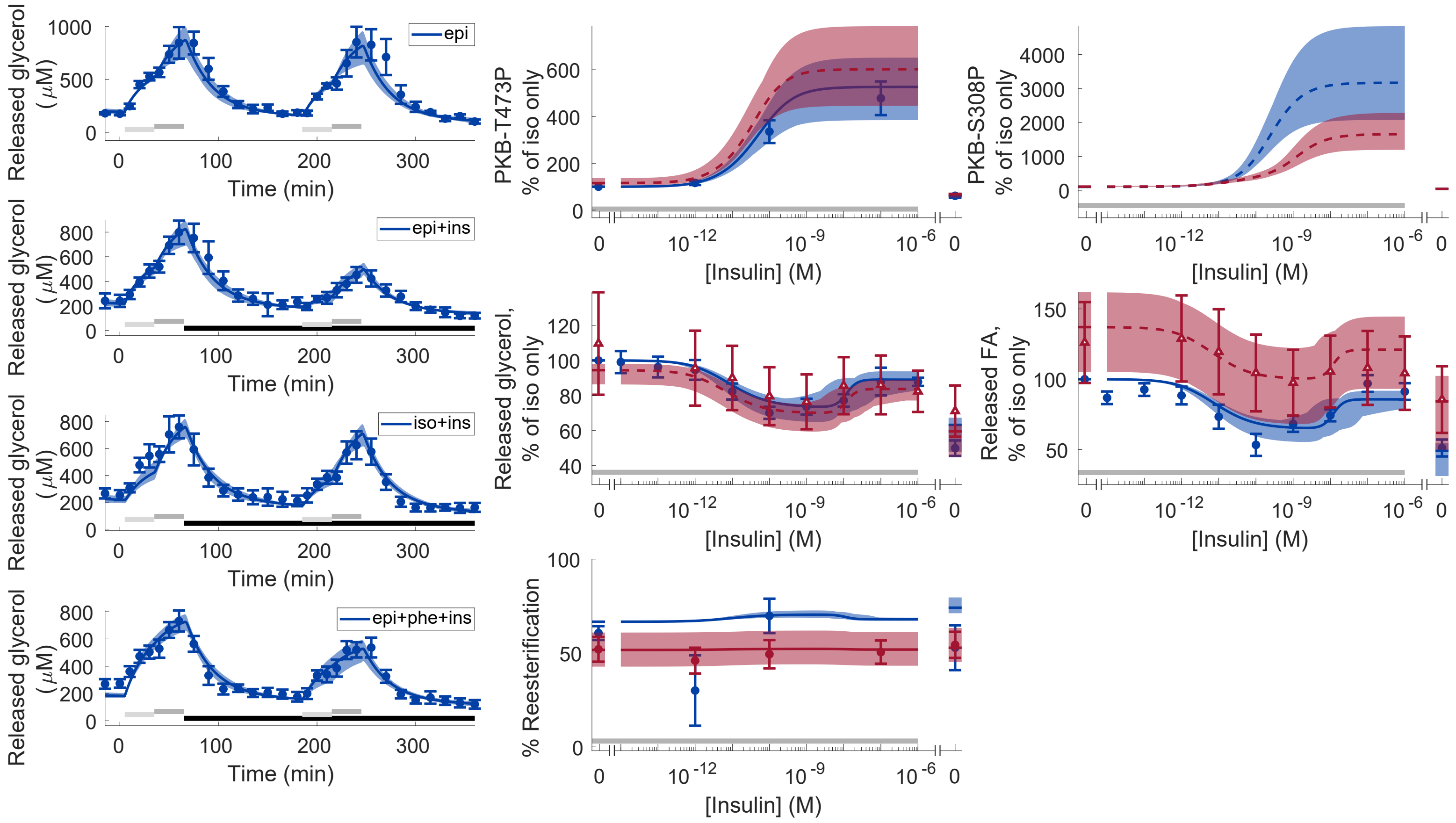


Figure 2: Model agreement with lipolysis data from [4]. Released glycerol (left) was measured using microdialysis in the adipose tissue in situ [5]. All other data (middle, right) was measured in isolated human adipocytes [4]. In all panels, lines represent the model simulation with the best agreement to data, the shaded areas represent the model uncertainty, and experimental data points are represented as mean values with error bars (SEM). Simulations and experimental data in red corresponds to experiments under type 2 diabetic conditions, and in blue under non-diabetic conditions. Dashed lines and experimental data with open triangles were not used to estimate the model parameters. Light/dark gray horizontal bars indicate adrenergic stimulation, and black horizontal bars in the left figures indicate insulin stimulation. epi - epinephrine, ins - insulin, iso - isoprenaline, FA - released fatty acids

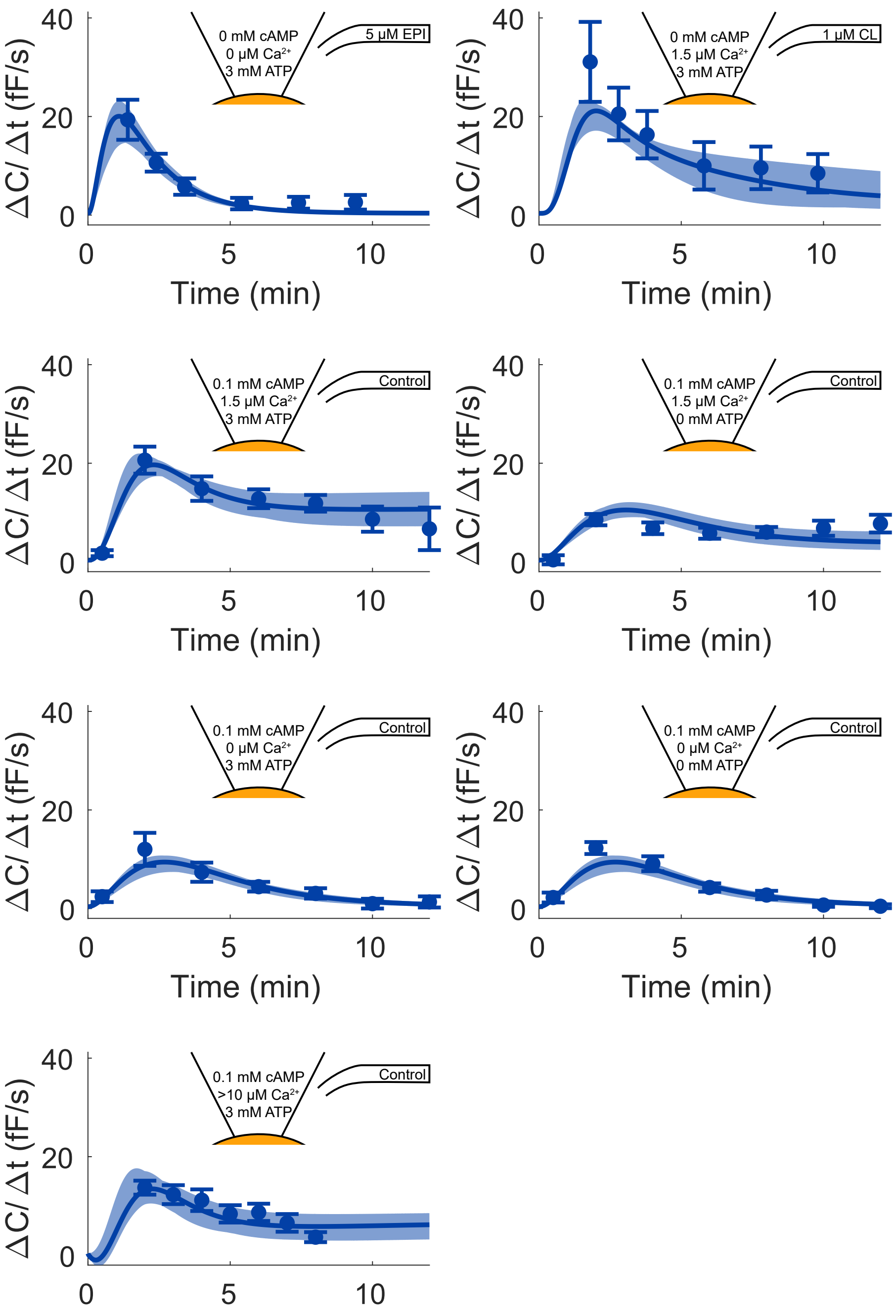


Figure 3: Model agreement with the adiponectin release data used in [1]. Data represent patch-clamp capacitance recordings in 3T3-L1 adipocytes [1]. In all panels, lines represent the model simulation with the best agreement to data and experimental data points are represented as mean values with error bars (SEM). EPI - epinephrine, CL - β3-adrenergic receptor agonist CL 316243, $Ca^{2+}$- Calcium.

### 2 The combined model can accurately predict all previous validation data

The validation data set consisted of the data for adiponectin release stimulate by CL and ATP from citeadiponectin2, and the phosphorylation of HSL in response to stimulation with insulin and isoproterenol from [1].

#### 2.1 Lipolysis

We used the dose response for HSL phosphorylation (HSLp) as validation data in the original lipolysis work [1]. Here, we have also excluded the HSLp data from the pool of training data and used the HSLp data as validation data. We retrained the model to the limited training data and predicted the dose response of HSLp. The model prediction and experimental data is shown in Fig. 4A.

#### 2.2 Adiponectin

We used the exocytosis time series data for stimulation with 1 µM CL and 3 mM ATP (CL+ATP dataset) as validation data in the work of the adiponectin submodel [6]. Here, we also excluded the CL+ATP dataset from the pool of training data and used the CL+ATP dataset for model validation. We retrained the model to the limited pool of training data and predicted the exocytosis when stimulated with CL+ATP. The model prediction and experimental data is shown in Fig. 4B.

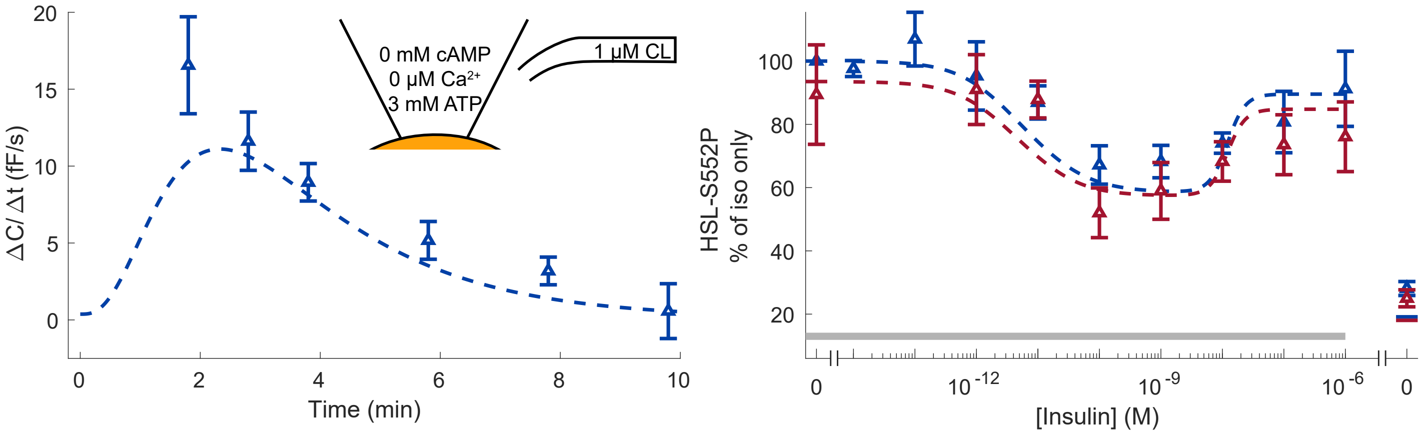


Figure 4: Recreation of the model validation from the previous lipolysis work ([1, Fig. 4 and S2 Fig.]) and adiponectin work ([6, Fig. 4]). In both panels, lines represent the model simulation with the best agreement to data, and experimental data points are represented as mean values with error bars (SEM). Simulations and experimental data in red corresponds to experiments under diabetic conditions, and in blue under non-diabetic conditions. Dashed lines and experimental data with open triangles were not used to estimate the model parameters.

#### 2.3 Glucose uptake

As in the original work, we simulated the effect of the inhibitions ranging from 50% to 93% inhibition. First, we recreated the validations from [3]. Here, we predicted the effect of the mTORC1 inhibitor rapamycin on the phosphorylation of of S6-S235/236P, IR-YP, IRS1-307P, AS160-T642P, PKB-S473P and FOXO1-S256P. Both model predictions and experimental data are shown in Fig. 5. In short, rapamycin should have an inhibiting effect on mTORC1, and thus on IRS1-307P, IR-YP and S6-S235/236P. At the same time, rapamycin should not have an inhibiting effect on mTORC2 and downstream phosphorylations. Thus, PKB-473P, AS160-T642P and FOXO1-S256P should not be inhibited.

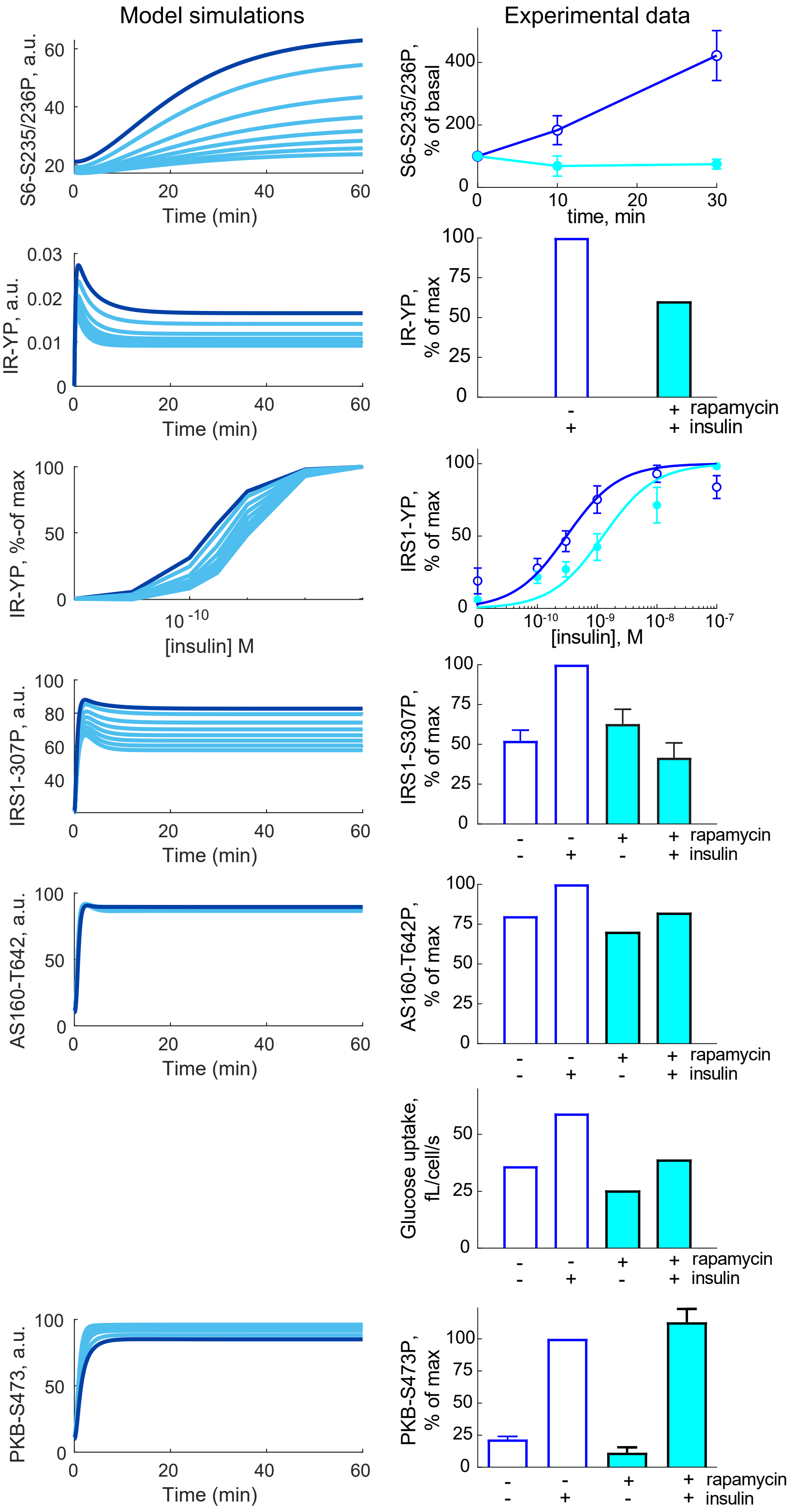


Figure 5: Recreation of the model validation in [3, Fig. 7].

We then recreated the validations from [7]. Here, the mTORC1 inhibitor rapamycin was again used, as well as the mTORC1 and mTORC2 inhibitor torin, and the MEK inhibitor akti.

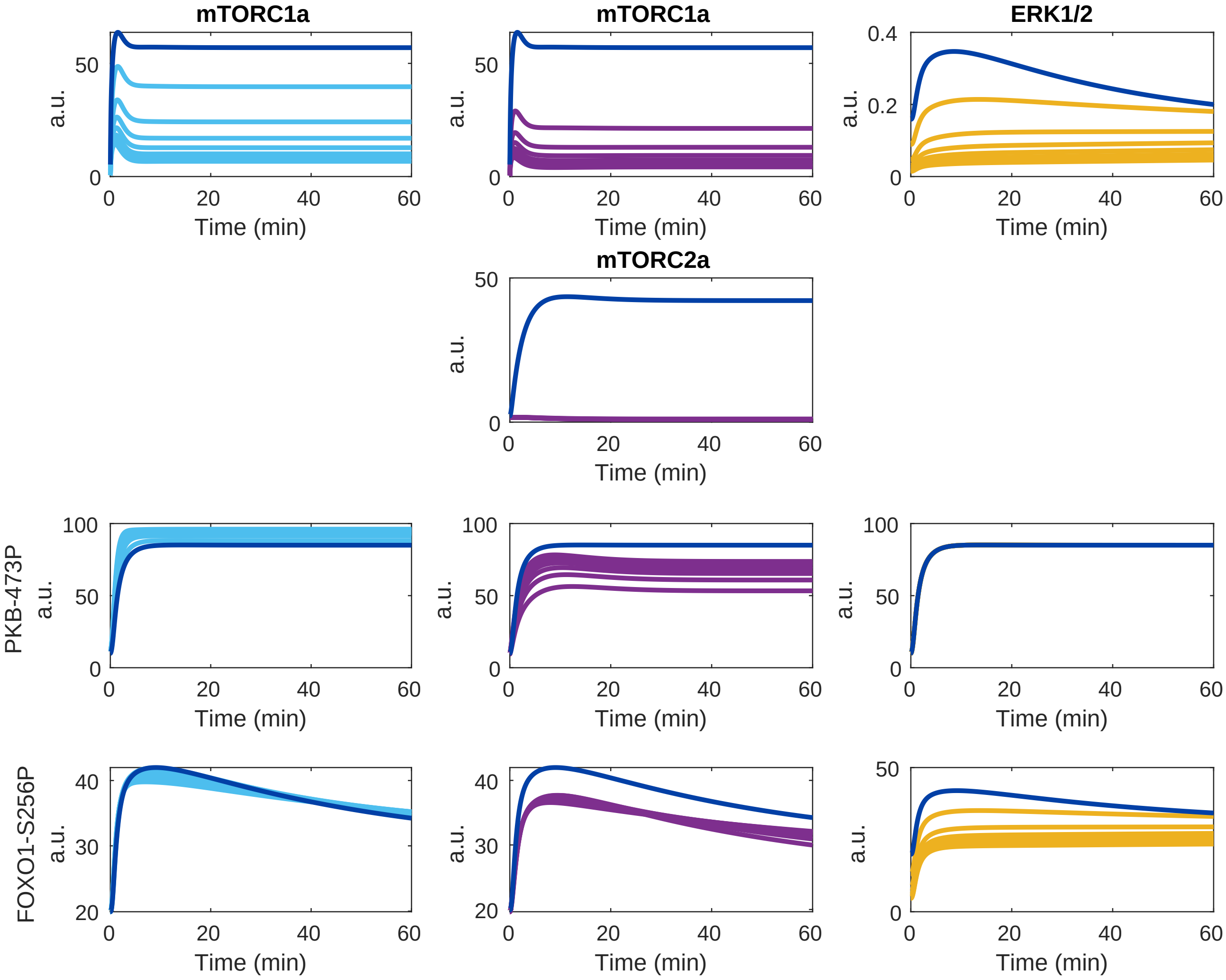


Figure 6: Recreation of the model validation in [2, Fig. 11].
