## Supplementary material for "A comprehensive mechanistic model of adipocyte signaling with layers of confidence": S2 Model equations and parameter values

May 2, 2022

The connected model is based on ordinary differential equations (ODEs), and is a combination of three different previously published models for glucose uptake [1], adiponectin secretion [2] and lipolysis [3]. A typical ODE used in this work looks similar to Eq. (1).

$$\begin{aligned} d/dt(x) &= -va + vb \\ va &= ka \cdot x \\ vb &= kb \cdot input \end{aligned} \tag{1}$$

Here, $x$ is a state in model, $va$ and $vb$ are reaction rates, $ka$ and $kb$ are rate determining parameters, and $input$ is some input to the state. In other words, the amount of the state $x$ is decreased by the reaction $va$ with the speed $ka$, and increased by the reaction $vb$ with the speed $kb$ depending on some input $input$.

In all ODEs in this document, $vGx$ and $kGx$ corresponds to reaction rates and rate parameters for the glucose uptake submodel, $vLx$ and $kLx$ to the lipolysis submodel and $vAx$ and $kAx$ to the adiponectin submodel. The final connected model is shown in Fig. 3, and the equation are given in TODO, however we will first highlight the changes made to the original submodel equations due to the connection of the submodels before we go through the equations for the connected model.

### 1 Connecting the submodels

For these three submodels, there are important points of crosstalk that had not previously been addressed. The glucose uptake submodel [1] and lipolysis submodel [3] have an overlap in the insulin receptor and subsequent signaling intermediaries leading up to the phosphorylation of PKB-S473 (Fig. 1). In the lipolysis model, we had previously simplified the signalling pathway from the insulin receptor to PKB as a direct interaction (action-1 in Fig. 1).

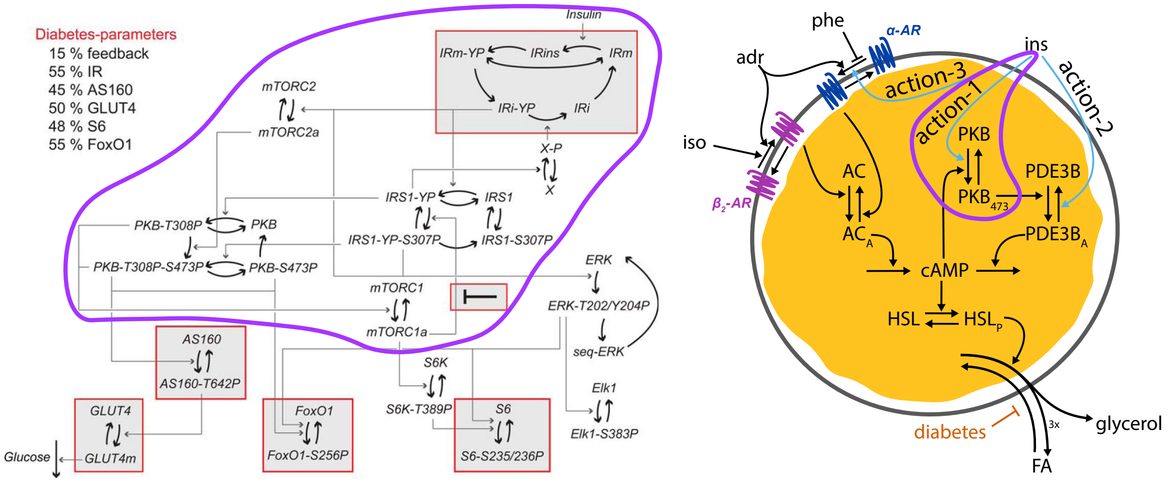


Figure 1: The glucose uptake model [1] and the lipolysis model [3]. The overlap between the models is highlighted in purple.

To connect these two submodels, we updated the lipolysis submodel to use the insulin signalling pathway from the glucose uptake submodel instead of insulin action-1. In practice, we removed the equations for $PKB$ in the lipolysis model, and changed the input to PDE3B from $PKB_{473}$ as in the lipolysis model to $(PKB\_T473P + PKB\_S308P\_T473P)$ from the glucose uptake model. The equations for PDE3B used in the original lipolysis work is given in Eq. (2).

$$\begin{aligned} d/dt(PDE3B) &= -vL2a + vL2b \\ d/dt(PDE3Ba) &= vL2a - vL2b \\ vL2a &= kL2a \cdot \boldsymbol{PKB473} \cdot PDE3B \\ vL2b &= kL2b \cdot PDE3Ba \cdot Ins\_2 \end{aligned} \tag{2}$$

The updated reaction rate equations ($vL2a$, $VL2b$) for PDE3B after the models had been connected is given in Eq. (3).

$$\begin{aligned} vL2a &= kL2a \cdot \boldsymbol{(PKB\_T473P + PKB\_S308P\_T473P)} \cdot PDE3B \\ vL2b &= kL2b \cdot PDE3Ba \cdot Ins\_2 \end{aligned} \tag{3}$$

Furthermore, we added the effect of cAMP on PKB from the lipolysis model as an activation of mTORC2 in the pathway from the glucose uptake model, shown in bold in Eq. (4).

$$\begin{aligned} d/dt(mTORC2) &= -\, vG5c + vG5d \\ d/dt(mTORC2a) &= vG5c - vG5d \\ vG5c &= mTORC2 \cdot (kG5c \cdot IRi\_YP + \boldsymbol{kL1a \cdot cAMP}) \\ vG5d &= kG5d \cdot mTORC2a \end{aligned} \tag{4}$$

In addition to the overlap between the glucose uptake and lipolysis submodels, there is also an overlap between the lipolysis submodel and the adiponectin secretion submodel(Fig. 2). This overlap is cAMP.

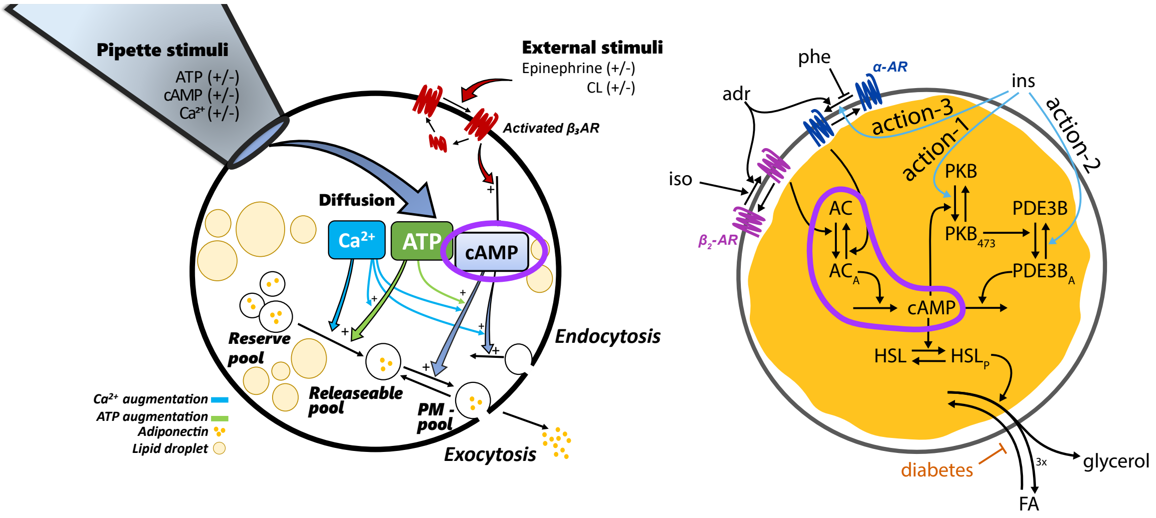


Figure 2: The adiponectin model [2] and the lipolysis model [3]. The overlap between the model is highlighted in purple.

To connect the lipolysis and adiponectin submodels, we combined the reactions relating to cAMP. Furthermore, the simplified effect from the $\beta_3$ receptors to directly lead to cAMP production was instead directed to the activation of the adenylyl complex (AC) in the lipolysis submodel. The equations for cAMP from the adiponectin submodel is given in Eq. (5).

$$\begin{aligned} d/dt(cAMP) &= vApip\_cAMP + vA6a - vAdegcAMP \\ vApip\_cAMP &= kADiffcAMP \cdot (pipcAMP - cAMP) \cdot pip \\ vA6a &= kA6a \cdot BETA3a \\ vA6b &= kAdegcAMP \cdot cAMP \end{aligned} \tag{5}$$

The equations for cAMP from the lipolysis submodel is given in Eq. (6).

$$\begin{aligned} d/dt(cAMP) &= vL6a - vL6b \\ vL6a &= kL6a \cdot ACa \\ vL6b &= kL6b \cdot PDE3Ba \cdot cAMP \end{aligned} \tag{6}$$

The equations for cAMP for the connected model is given in Eq. (7).

$$\begin{aligned}
d/dt(cAMP) &= vApip_cAMP - vA6b + vL6a - vL6b \\
vApip_cAMP &= kADiffcAMP \cdot (pipcAMP - cAMP) \cdot pip \\
vA6b &= kAdegcAMP \cdot cAMP \\
vL6a &= kL6a \cdot ACa \\
vL6b &= kL6b \cdot PDE3Ba \cdot cAMP
\end{aligned} \tag{7}$$

Here it should be noted that $ACa$ is now activated by both the $\beta_2$ receptor from the lipolysis submodel and $\beta_3$ from the adiponectin submodel. The equations for the combined AC activation is given in Eq. (8).

$$\begin{aligned}
d/dt(AC) &= -vL5a + vL5b \\
d/dt(ACa) &= vL5a - vL5b + vA5a \\
vL5a &= kL5a \cdot BETA2a \cdot AC \\
vL5b &= kL5b \cdot ALPHAa \cdot ACa \\
vA5a &= kA5a \cdot BETA3a
\end{aligned} \tag{8}$$

### 2 Description of the equations for the connected model

Here we will go through all the equations for the fully connected model Fig. 3. For readability, we will, as much as possible, go through the signalling pathways in the glucose uptake submodel first, then the lipolysis submodel and lastly the adiponectin submodel. Note that all model parameters with names starting with $k$ are rate determining parameters, for which the parameter values are estimated based on the experimental data. These rate parameters will not be mentioned in detail below.

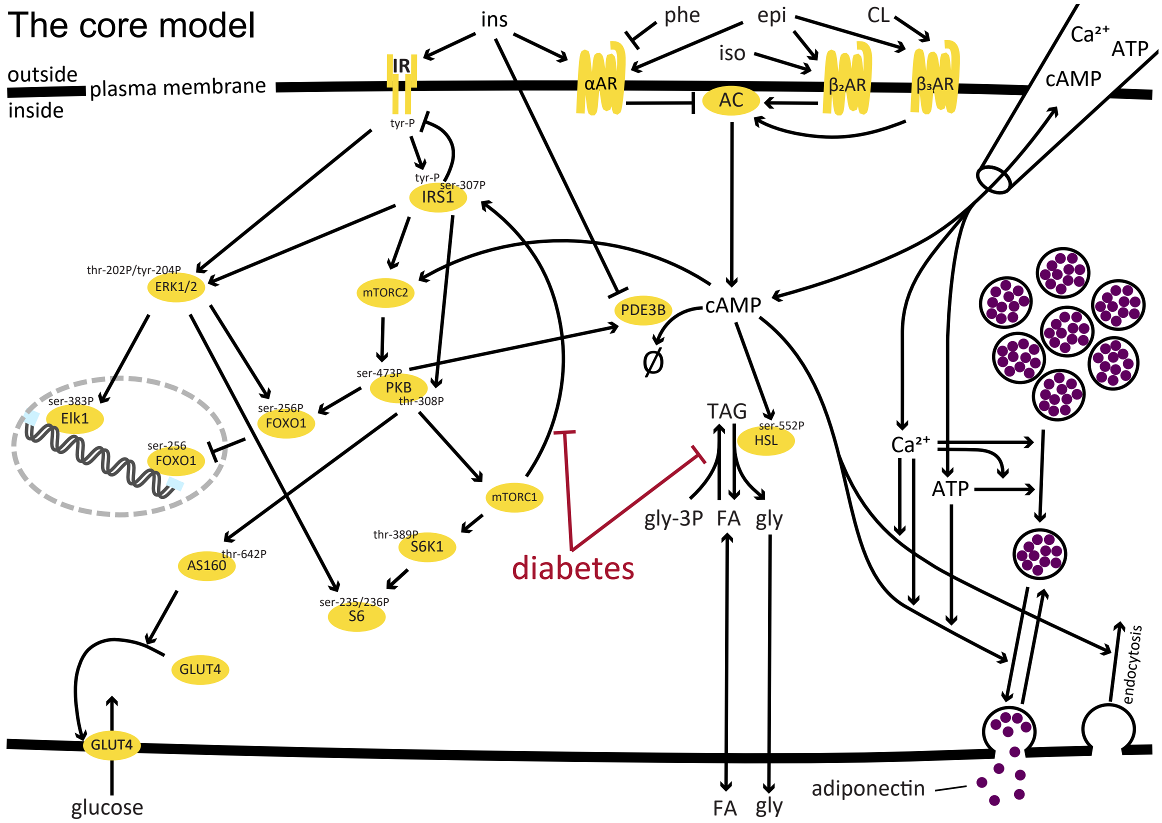


Figure 3: **The core model of adipocyte signaling**. The core model includes crosstalk between glucose uptake in response to insulin, fatty acid and glycerol release in response to alpha- and beta-adrenergic receptor signaling and adiponectin secretion in response to cAMP, ATP and Ca2+ (added into the cell with a pipette).

#### 2.1 The glucose uptake submodel

This section describes the equations of the insulin signalling submodel, based on the model developed in [1]. Note that all reactions and parameters below are renamed from $vx$ and $kx$ to $v\boldsymbol{G}x$ and $k\boldsymbol{G}x$ relative to the original model in [1].

The equations for the insulin receptor is described in Eq. (9).

$$\begin{aligned}
d/dt(IR) &= -vG1a - vG1basal + vG1r + vG1g \\
d/dt(IR_YP) &= vG1basal + vG1c - vG1d - vG1g \\
d/dt(IRins) &= vG1a - vG1c \\
d/dt(IRi_YP) &= vG1d - vG1e \\
d/dt(IRi) &= vG1e - vG1r \\
\\
vG1a &= kG1a \cdot IR \cdot ins \\
vG1basal &= kG1basal \cdot IR \\
vG1c &= IRins \cdot kG1c \\
vG1d &= IR_YP \cdot kG1d \\
vG1e &= IRi_YP \cdot kG1f \cdot X_P \\
vG1g &= IR_YP \cdot kG1g \\
vG1r &= IRi \cdot kG1r
\end{aligned} \tag{9}$$

The equations for IRS1 is given in Eq. (10)

$$\begin{aligned}
d/dt(IRS1) &= vG2b + vG2g - vG2a - vG2basal \\
d/dt(IRS1_YP) &= vG2a + vG2d - vG2b - vG2c \\
d/dt(IRS1_YP_S307P) &= vG2c - vG2d - vG2f \\
d/dt(IRS1_S307P) &= vG2basal + vG2f - vG2g \\
\\
vG2a &= IRS1 \cdot kG2a \cdot IRi_YP \\
vG2b &= IRS1_YP \cdot kG2b \\
vG2c &= IRS1_YP \cdot kG2c \cdot mTORC1a \cdot diabetes \\
vG2d &= IRS1_YP_S307P \cdot kG2d \\
vG2f &= IRS1_YP_S307P \cdot kG2f \\
vG2basal &= IRS1 \cdot kG2basal \\
vG2g &= IRS1_S307P \cdot kG2g
\end{aligned} \tag{10}$$

The equation for the unknown protein X is given in Eq. (11)

$$\begin{aligned}
d/dt(X) &= vG3b - vG3a \\
d/dt(X_P) &= vG3a - vG3b \\
\\
vG3a &= X \cdot kG3a \cdot IRS1_YP \\
vG3b &= X_P \cdot kG3b
\end{aligned} \tag{11}$$

The equations for PKB is given in Eq. (12)

$$
\begin{aligned}
d/dt(PKB) &= -vG4a + vG4b + vG4h \\
d/dt(PKB\_S308P) &= vG4a - vG4b - vG4c \\
d/dt(PKB\_T473P) &= -vG4e + vG4f - vG4h \\
d/dt(PKB\_S308P\_T473P) &= vG4c + vG4e - vG4f \\
\\
vG4a &= kG4a \cdot PKB \cdot IRS1\_YP \\
vG4b &= kG4b \cdot PKB\_S308P \\
vG4c &= kG4c \cdot PKB\_S308P \cdot mTORC2a \\
vG4e &= kG4e \cdot PKB\_T473P \cdot IRS1\_YP\_S307P \\
vG4f &= kG4f \cdot PKB\_S308P\_T473P \\
vG4h &= kG4h \cdot PKB\_T473P
\end{aligned}
\tag{12}
$$

The equations for mTORC1 is given in Eq. (13)

$$
\begin{aligned}
d/dt(mTORC1) &= vG5b - vG5a \\
d/dt(mTORC1a) &= vG5a - vG5b \\
\\
vG5a &= mTORC1 \cdot (kG5a1 \cdot PKB\_S308P\_T473P + kG5a2 \cdot PKB\_S308P) \\
vG5b &= mTORC1a \cdot kG5b
\end{aligned}
\tag{13}
$$

The equations for mTORC2 is given in Eq. (14)

$$
\begin{aligned}
d/dt(mTORC2) &= -vG5c + vG5d \\
d/dt(mTORC2a) &= vG5c - vG5d \\
\\
vG5c &= mTORC2 \cdot (kG5c \cdot IRi\_YP + kL1a \cdot cAMP) \\
vG5d &= kG5d \cdot mTORC2a
\end{aligned}
\tag{14}
$$

The equations for AS160 is given in Eq. (15)

$$
\begin{aligned}
d/dt(AS160) &= vG6b - vG6a \\
d/dt(AS160\_T642P) &= vG6a - vG6b \\
\\
vG6a &= AS160 \cdot (kG6a1 \cdot PKB\_S308P\_T473P + kG6a2 \cdot PKB\_T473P) \\
vG6b &= AS160\_T642P \cdot kG6b
\end{aligned}
\tag{15}
$$

The equations for GLUT4 is given in Eq. (16)

$$\begin{aligned}
d/dt(GLUT4m) &= vG7a - vG7b \\
d/dt(GLUT4) &= -vG7a + vG7b \\
\\
vG7a &= GLUT4 \cdot kG7a \cdot AS160_T642P \\
vG7b &= GLUT4m \cdot kG7b
\end{aligned} \tag{16}$$

The equations for the glucose uptake is given in Eq. (17)

$$\begin{aligned}
d/dt(GLUCOSE) &= vG8 \\
\\
vG8 &= kG8 \cdot GLUT4m \cdot gluc/(kGmG4 + gluc) + kGglut1 \cdot gluc/(kGmG1 + gluc)
\end{aligned} \tag{17}$$

The equations for S6K is given in Eq. (18)

$$\begin{aligned}
d/dt(S6K) &= vG9b - vG9a \\
d/dt(S6K_T389P) &= vG9a - vG9b \\
\\
vG9a &= S6K \cdot kG9a \cdot mTORC1a \\
vG9b &= S6K_T389P \cdot kG9b
\end{aligned} \tag{18}$$

The equations for S6 is given in Eq. (19)

$$\begin{aligned}
d/dt(S6) &= vG9d - vG9c \\
d/dt(S6_S235_S236P) &= vG9c - vG9d \\
\\
vG9c &= S6 \cdot kG9c1 \cdot S6K_T389P + kG9c2 \cdot S6 \cdot ERK_T202_Y204P \\
vG9d &= S6_S235_S236P \cdot kG9d
\end{aligned} \tag{19}$$

The equations for ERK is given in Eq. (20)

$$\begin{aligned}
d/dt(ERK) &= -vG10a - vG10basal + vG10c \\
d/dt(ERK_T202_Y204P) &= vG10a + vG10basal - vG10b \\
d/dt(seqERK) &= vG10b - vG10c \\
\\
vG10basal &= kG10basal \cdot ERK \\
vG10a &= kG10a1 \cdot ERK \cdot IRi_YP + kG10a2 \cdot ERK \cdot IRS1_YP_S307P \\
vG10b &= kG10b \cdot ERK_T202_Y204P \\
vG10c &= kG10c \cdot seqERK
\end{aligned} \tag{20}$$

The equations for Elk1 is given in Eq. (21)

$$
\begin{aligned}
d/dt(Elk1) &= -vG11a + vG11b \\
d/dt(Elk1\_S383P) &= vG11a - vG11b \\
\\
vG11a &= kG11a \cdot Elk1 \cdot ERK\_T202\_Y204P \\
vG11b &= kG11b \cdot Elk1\_S383P
\end{aligned}
\tag{21}
$$

The equations for FOXO is given in Eq. (22)

$$
\begin{aligned}
d/dt(FOXO) &= -vG12a + vG12b \\
d/dt(FOXO\_S256P) &= vG12a - vG12b \\
\\
vG12a &= (kG12a1 \cdot (PKB\_T473P + PKB\_S308P\_T473P) + kG12a2 \cdot ERK\_T202\_Y204P)/ \\
vG12b &= kG12b \cdot FOXO\_S256P
\end{aligned}
\tag{22}
$$

#### 2.2 The lipolysis submodel

This section describes the equations of the lipolysis submodel, based on the model developed in [3]. Note that all reactions and parameters below are renamed from $vx$ and $kx$ to $v\boldsymbol{L}x$ and $k\boldsymbol{L}x$ relative to the original model.

The equations for PDE3B is given in Eq. (23)

$$
\begin{aligned}
d/dt(PDE3B) &= -vL2a + vL2b \\
d/dt(PDE3Ba) &= vL2a - vL2b \\
\\
vL2a &= kL2a \cdot (PKB\_T473P + PKB\_S308P\_T473P) \cdot PDE3B \\
vL2b &= kL2b \cdot PDE3Ba \cdot Ins\_2
\end{aligned}
\tag{23}
$$

The equations for the $\alpha$ adrenergic receptor is given in Eq. (24)

$$
\begin{aligned}
d/dt(ALPHA) &= -vL3a + vL3b \\
d/dt(ALPHAa) &= vL3a - vL3b \\
\\
vL3a &= (kL3a \cdot Ins\_3 \cdot epi + kL3a2) \cdot (1 - phe\_effect \cdot phe) \cdot ALPHA \\
vL3b &= kL3b \cdot ALPHAa
\end{aligned}
\tag{24}
$$

The equations for the $\beta_2$ adrenergic receptor is given in Eq. (25)

$$
\begin{aligned}
d/dt(BETA2) &= -vL4a + vL4b \\
d/dt(BETA2a) &= vL4a - vL4b \\
\\
vL4a &= (kL4a \cdot (iso \cdot isoscale + epi) + kL4a2) \cdot BETA2 \\
vL4b &= kL4b \cdot BETA2a
\end{aligned}
\tag{25}
$$

The equations for the adenylyl cyclase is given in Eq. (26)

$$
\begin{aligned}
d/dt(AC) &= -vL5a + vL5b \\
d/dt(ACa) &= vL5a - vL5b + vA5a \\
\\
vL5a &= kL5a \cdot BETA2a \cdot AC \\
vL5b &= kL5b \cdot ALPHAa \cdot ACa \\
vA5a &= kA5a \cdot BETA3a
\end{aligned}
\tag{26}
$$

The equations for cAMP is given in Eq. (27)

$$
\begin{aligned}
d/dt(cAMP) &= vApip\_cAMP - vA6b + vL6a - vL6b \\
\\
vL6a &= kL6a \cdot ACa \\
vL6b &= kL6b \cdot PDE3Ba \cdot cAMP \\
vA6b &= kAdegcAMP \cdot cAMP
\end{aligned}
\tag{27}
$$

The equations for HSL is given in Eq. (28)

$$
\begin{aligned}
d/dt(HSL) &= -vL7a + vL7b \\
d/dt(HSLp) &= vL7a - vL7b \\
\\
vL7a &= kL7a \cdot cAMP \cdot HSL \\
vL7b &= kL7b \cdot HSLp
\end{aligned}
\tag{28}
$$

The equations for the release of glycerol and fatty acids is given in Eq. (29)

$$
\begin{aligned}
d/dt(Gly) &= vL8a - vL8b \\
d/dt(FFA) &= 3 \cdot vL8a - vL8c \\
\\
vL8a &= kL8a \cdot HSLp \\
vL8b &= kLclear \cdot Gly \\
vL8c &= (kLclear + kL8c \cdot diab\_reest) \cdot FFA
\end{aligned}
\tag{29}
$$

The dose dependent insulin actions are given in Eq. (30).

$$\begin{aligned} Ins_2 &= 100 + \frac{min_2 - 100}{1 + (ins/EC50_2)^{n_2}} \\ Ins_3 &= 100 + \frac{min_3 - 100}{1 + (ins/EC50_3)^{n_3}} \end{aligned} \tag{30}$$

#### 2.3 The adiponectin submodel

This section describes the equations of the lipolysis submodel, based on the model developed in [2]. Note that all reactions and parameters below are renamed from $vx$ and $kx$ to $v\boldsymbol{A}x$ and $k\boldsymbol{A}x$ relative to the original model.

The equations for the $\beta_3$ adrenergic receptor is given in Eq. (31)

$$\begin{aligned} d/dt(BETA3) &= vA4c - vA4a \\ d/dt(BETA3a) &= vA4a - vA4b \\ d/dt(BETA3de) &= vA4b - vA4c \\ \\ vA4a &= kA4a \cdot (epi + kACL \cdot CL) \cdot BETA3 \\ vA4b &= kA4b \cdot BETA3a \\ vA4c &= kA4c \cdot BETA3de \end{aligned} \tag{31}$$

The equations for the intracellular mediators Ca2+ and ATP is given in Eq. (32)

$$\begin{aligned} d/dt(Ca) &= vApip\_Ca - vAremCa \\ d/dt(ATP) &= vApip\_ATP - vAdegATP \\ \\ vAremCa &= kAremCa \cdot Ca \\ vAdegATP &= kAdegATP \cdot ATP \\ vApip\_Ca &= kADiffCa \cdot (pipCa - Ca) \cdot pip \\ vApip\_ATP &= kADiffATP \cdot (pipATP - ATP) \cdot pip \end{aligned} \tag{32}$$

The equations for the reserve pool is given in Eq. (33)

$$d/dt(Res) = 0 \tag{33}$$

The equations for releasable pool is given in Eq. (34)

$$\begin{aligned} d/dt(Rel) &= vARes\_Rel - vARel\_PM + vAPM\_Rel \\ \\ vARes\_Rel &= ((Ca/(kAm + Ca)) \cdot (kACa2 + kAATP2 \cdot ATP) + kARelBasal) \cdot Res \\ vARel\_PM &= cAMP \cdot (kAcAMP + (Ca/(kAm + Ca)) \cdot ATP \cdot kACaATP) \cdot Rel \\ vAPM\_Rel &= kArel \cdot PM \end{aligned} \tag{34}$$

The equations for vesicles at the plasma membrane is given in Eq. (35)

$$\begin{aligned} d/dt(PM) &= vARel\_PM - vAPM\_Rel - vAPM \\ vARel\_PM &= cAMP \cdot (kAcAMP + (Ca/(kAm + Ca)) \cdot ATP \cdot kACaATP) \cdot Rel \\ vAPM\_Rel &= kArel \cdot PM \\ vAPM &= kAexo \cdot PM \end{aligned} \tag{35}$$

The equations for the endocytosis is given in Eq. (36)

$$d/dt(Endo) = vACa\_Endo - vAEndo \tag{36}$$

The equations for the released adiponectin is given in Eq. (37)

$$\begin{aligned} d/dt(Adiponectin) &= vAPM \\ \\ vACa\_Endo &= kACacAMP \cdot cAMP \cdot Ca \\ vAEndo &= kAEndo \cdot Endo \end{aligned} \tag{37}$$

The equations for the mediators in the pipette is given in Eq. (38). The equations for the mediator cAMP have previously been described in Eq. (27).

$$\begin{aligned} d/dt(pipCa) &= -vApip\_Ca \cdot kAVcell/kAVpip \\ d/dt(pipATP) &= -vApip\_ATP \cdot kAVcell/kAVpip \\ d/dt(pipcAMP) &= -vApip\_cAMP \cdot kAVcell/kAVpip \\ \\ vApip\_cAMP &= kADiffcAMP \cdot (pipcAMP - cAMP) \cdot pip \\ vApip\_Ca &= kADiffCa \cdot (pipCa - Ca) \cdot pip \\ vApip\_ATP &= kADiffATP \cdot (pipATP - ATP) \cdot pip \end{aligned} \tag{38}$$

### 3 Parameter values

This section gives the optimal parameter values for the connected model both when estimated to the estimation data set (columns $\theta^*_{est}$), and to the total data set (columns $\theta^*_{tot}$). Furthermore, the bounds used in the optimization for all parameters are also given (columns *lowerbound* and *upperbound*). The parameter values and optimization bounds for the glucose uptake submodel are given in Table 1 and Table 2. The parameter values and optimization bounds for the lipolysis submodel are given in Table 3. The parameter values and optimization bounds for the adiponectin submodel are given in Table 4.

| Parameter | $\theta^*_{est}$ | $\theta^*_{tot}$ | lower bound | upper bound |
|---|---|---|---|---|
| $kG1a$ | $1.2696 \cdot 10^{-1}$ | $1.2501 \cdot 10^{-1}$ | $1.0000 \cdot 10^{-8}$ | $1.0000 \cdot 10^{6}$ |
| $kG1basal$ | $2.2526 \cdot 10^{-4}$ | $2.2526 \cdot 10^{-4}$ | $1.0000 \cdot 10^{-8}$ | $1.0000 \cdot 10^{6}$ |
| $kG1c$ | $1.7234 \cdot 10^{-1}$ | $1.7224 \cdot 10^{-1}$ | $1.0000 \cdot 10^{-8}$ | $1.0000 \cdot 10^{6}$ |
| $kG1d$ | $9.9995 \cdot 10^{5}$ | $9.9997 \cdot 10^{5}$ | $1.0000 \cdot 10^{-8}$ | $1.0000 \cdot 10^{6}$ |
| $kG1f$ | $2.0184 \cdot 10^{1}$ | $2.0267 \cdot 10^{1}$ | $1.0000 \cdot 10^{-8}$ | $1.0000 \cdot 10^{6}$ |
| $kG1g$ | $5.3850 \cdot 10^{-8}$ | $3.1332 \cdot 10^{-8}$ | $1.0000 \cdot 10^{-8}$ | $1.0000 \cdot 10^{6}$ |
| $kG1r$ | $6.5382$ | $6.4662$ | $1.0000 \cdot 10^{-8}$ | $1.0000 \cdot 10^{6}$ |
| $kG2a$ | $7.2702 \cdot 10^{1}$ | $7.2938 \cdot 10^{1}$ | $1.0000 \cdot 10^{-8}$ | $1.0000 \cdot 10^{6}$ |
| $kG2c$ | $2.0619 \cdot 10^{3}$ | $2.0024 \cdot 10^{3}$ | $1.0000 \cdot 10^{-8}$ | $1.0000 \cdot 10^{6}$ |
| $kG2basal$ | $7.2956 \cdot 10^{-2}$ | $7.1708 \cdot 10^{-2}$ | $1.0000 \cdot 10^{-8}$ | $1.0000 \cdot 10^{6}$ |
| $kG2b$ | $3.7448 \cdot 10^{3}$ | $3.7448 \cdot 10^{3}$ | $1.0000 \cdot 10^{-8}$ | $1.0000 \cdot 10^{6}$ |
| $kG2d$ | $4.8589$ | $4.8512$ | $1.0000 \cdot 10^{-8}$ | $1.0000 \cdot 10^{6}$ |
| $kG2f$ | $9.1342 \cdot 10^{-1}$ | $9.1327 \cdot 10^{-1}$ | $1.0000 \cdot 10^{-8}$ | $1.0000 \cdot 10^{6}$ |
| $kG2g$ | $2.9109 \cdot 10^{-1}$ | $2.8790 \cdot 10^{-1}$ | $1.0000 \cdot 10^{-8}$ | $1.0000 \cdot 10^{6}$ |
| $kG3a$ | $8.6395 \cdot 10^{1}$ | $8.5992 \cdot 10^{1}$ | $1.0000 \cdot 10^{-8}$ | $1.0000 \cdot 10^{6}$ |
| $kG3b$ | $1.0026 \cdot 10^{-1}$ | $1.0452 \cdot 10^{-1}$ | $1.0000 \cdot 10^{-8}$ | $1.0000 \cdot 10^{6}$ |
| $kG4a$ | $6.7328 \cdot 10^{2}$ | $6.7328 \cdot 10^{2}$ | $1.0000 \cdot 10^{-8}$ | $1.0000 \cdot 10^{6}$ |
| $kG4b$ | $9.6161 \cdot 10^{4}$ | $9.6157 \cdot 10^{4}$ | $1.0000 \cdot 10^{-8}$ | $1.0000 \cdot 10^{6}$ |
| $kG4c$ | $6.2629 \cdot 10^{3}$ | $6.1511 \cdot 10^{3}$ | $1.0000 \cdot 10^{-8}$ | $1.0000 \cdot 10^{6}$ |
| $kG4e$ | $4.8452 \cdot 10^{-2}$ | $4.8419 \cdot 10^{-2}$ | $1.0000 \cdot 10^{-8}$ | $1.0000 \cdot 10^{6}$ |
| $kG4f$ | $7.7143$ | $7.3032$ | $1.0000 \cdot 10^{-8}$ | $1.0000 \cdot 10^{6}$ |
| $kG4h$ | $9.1781 \cdot 10^{-2}$ | $9.1778 \cdot 10^{-2}$ | $1.0000 \cdot 10^{-8}$ | $1.0000 \cdot 10^{6}$ |
| $kG5a1$ | $3.0828$ | $2.9201$ | $1.0000 \cdot 10^{-8}$ | $1.0000 \cdot 10^{6}$ |
| $kG5a2$ | $1.3173 \cdot 10^{4}$ | $1.3540 \cdot 10^{4}$ | $1.0000 \cdot 10^{-8}$ | $1.0000 \cdot 10^{6}$ |
| $kG5b$ | $2.3183 \cdot 10^{1}$ | $2.3184 \cdot 10^{1}$ | $1.0000 \cdot 10^{-8}$ | $1.0000 \cdot 10^{6}$ |
| $kG5d$ | $1.4851 \cdot 10^{-1}$ | $1.4851 \cdot 10^{-1}$ | $1.0000 \cdot 10^{-8}$ | $1.0000 \cdot 10^{6}$ |
| $kG5c$ | $6.4703$ | $6.3330$ | $1.0000 \cdot 10^{-8}$ | $1.0000 \cdot 10^{6}$ |
| $kG6a1$ | $1.9667 \cdot 10^{-1}$ | $1.9198 \cdot 10^{-1}$ | $1.0000 \cdot 10^{-8}$ | $1.0000 \cdot 10^{6}$ |
| $kG6a2$ | $1.1012 \cdot 10^{-7}$ | $1.0513 \cdot 10^{-7}$ | $1.0000 \cdot 10^{-8}$ | $1.0000 \cdot 10^{6}$ |
| $kG6b$ | $2.2643 \cdot 10^{-1}$ | $2.3368 \cdot 10^{-1}$ | $1.0000 \cdot 10^{-8}$ | $1.0000 \cdot 10^{6}$ |
| $kG7a$ | $2.5980 \cdot 10^{-6}$ | $2.6309 \cdot 10^{-6}$ | $1.0000 \cdot 10^{-8}$ | $1.0000 \cdot 10^{6}$ |
| $kG7b$ | $1.6018 \cdot 10^{3}$ | $1.6330 \cdot 10^{3}$ | $1.0000 \cdot 10^{-8}$ | $1.0000 \cdot 10^{6}$ |
| $kG8$ | $3.7913 \cdot 10^{4}$ | $3.8195 \cdot 10^{4}$ | $1.0000 \cdot 10^{-8}$ | $1.0000 \cdot 10^{6}$ |
| $kGglut1$ | $2.0747 \cdot 10^{-2}$ | $2.0805 \cdot 10^{-2}$ | $1.0000 \cdot 10^{-8}$ | $1.0000 \cdot 10^{6}$ |
| $kGmG4$ | $1.1159 \cdot 10^{1}$ | $1.1071 \cdot 10^{1}$ | $1.0000 \cdot 10^{-8}$ | $1.0000 \cdot 10^{6}$ |
| $kGmG1$ | $8.8508 \cdot 10^{-1}$ | $8.8402 \cdot 10^{-1}$ | $1.0000 \cdot 10^{-8}$ | $1.0000 \cdot 10^{6}$ |
| $kG9a$ | $1.1660 \cdot 10^{-6}$ | $1.1786 \cdot 10^{-6}$ | $1.0000 \cdot 10^{-8}$ | $1.0000 \cdot 10^{6}$ |
| $kG9b$ | $4.9870 \cdot 10^{-2}$ | $4.9403 \cdot 10^{-2}$ | $1.0000 \cdot 10^{-8}$ | $1.0000 \cdot 10^{6}$ |
| $kG9c1$ | $4.2332 \cdot 10^{-1}$ | $4.1739 \cdot 10^{-1}$ | $1.0000 \cdot 10^{-8}$ | $1.0000 \cdot 10^{6}$ |
| $kG9c2$ | $1.9737 \cdot 10^{-2}$ | $1.9038 \cdot 10^{-2}$ | $1.0000 \cdot 10^{-8}$ | $1.0000 \cdot 10^{6}$ |
| $kG9d$ | $3.2024 \cdot 10^{-2}$ | $3.2662 \cdot 10^{-2}$ | $1.0000 \cdot 10^{-8}$ | $1.0000 \cdot 10^{6}$ |

Table 1: **Parameter values related to the glucose uptake submodel.**

| | | | | |
|---|---|---|---|---|
| $kG10a1$ | $9.4534 \cdot 10^{-7}$ | $2.4530 \cdot 10^{-7}$ | $1.0000 \cdot 10^{-8}$ | $1.0000 \cdot 10^{6}$ |
| $kG10a2$ | $8.2693 \cdot 10^{-4}$ | $8.3407 \cdot 10^{-4}$ | $1.0000 \cdot 10^{-8}$ | $1.0000 \cdot 10^{6}$ |
| $kG10basal$ | $9.5726 \cdot 10^{-3}$ | $9.5430 \cdot 10^{-3}$ | $1.0000 \cdot 10^{-8}$ | $1.0000 \cdot 10^{6}$ |
| $kG10b$ | $1.8650 \cdot 10^{-1}$ | $1.8514 \cdot 10^{-1}$ | $1.0000 \cdot 10^{-8}$ | $1.0000 \cdot 10^{6}$ |
| $kG10c$ | $2.5402 \cdot 10^{-4}$ | $2.5572 \cdot 10^{-4}$ | $1.0000 \cdot 10^{-8}$ | $1.0000 \cdot 10^{6}$ |
| $kG11a$ | $9.2964 \cdot 10^{-8}$ | $9.2595 \cdot 10^{-8}$ | $1.0000 \cdot 10^{-8}$ | $1.0000 \cdot 10^{6}$ |
| $kG11b$ | $3.9138 \cdot 10^{-1}$ | $3.9123 \cdot 10^{-1}$ | $1.0000 \cdot 10^{-8}$ | $1.0000 \cdot 10^{6}$ |
| $kG12a1$ | $1.0188 \cdot 10^{3}$ | $1.0328 \cdot 10^{3}$ | $1.0000 \cdot 10^{-8}$ | $1.0000 \cdot 10^{6}$ |
| $kG12a2$ | $4.9313 \cdot 10^{5}$ | $4.9310 \cdot 10^{5}$ | $1.0000 \cdot 10^{-8}$ | $1.0000 \cdot 10^{6}$ |
| $kG12b$ | $1.9751 \cdot 10^{1}$ | $1.9997 \cdot 10^{1}$ | $1.0000 \cdot 10^{-8}$ | $1.0000 \cdot 10^{6}$ |
| $kGm12$ | $1.7943 \cdot 10^{4}$ | $1.7943 \cdot 10^{4}$ | $1.0000 \cdot 10^{-8}$ | $1.0000 \cdot 10^{6}$ |
| $diabetes$ | $5.5870 \cdot 10^{-2}$ | $5.6831 \cdot 10^{-2}$ | 0 | 1.0000 |

Table 2: **Parameter values related to the glucose uptake submodel.**

| | | | | |
|---|---|---|---|---|
| $kLdrift$ | $2.3544 \cdot 10^{2}$ | $2.3859 \cdot 10^{2}$ | $1.0000 \cdot 10^{-8}$ | $1.0000 \cdot 10^{6}$ |
| $kL1a$ | 6.3543 | 6.5066 | $1.0000 \cdot 10^{-8}$ | $1.0000 \cdot 10^{6}$ |
| $kL2a$ | $2.0167 \cdot 10^{-2}$ | $2.0173 \cdot 10^{-2}$ | $1.0000 \cdot 10^{-8}$ | $1.0000 \cdot 10^{6}$ |
| $kL2b$ | $1.5134 \cdot 10^{-2}$ | $1.5133 \cdot 10^{-2}$ | $1.0000 \cdot 10^{-8}$ | $1.0000 \cdot 10^{6}$ |
| $kL3b$ | $2.6030 \cdot 10^{3}$ | $2.6139 \cdot 10^{3}$ | $1.0000 \cdot 10^{-8}$ | $1.0000 \cdot 10^{6}$ |
| $kL3a$ | $1.3271 \cdot 10^{2}$ | $1.4695 \cdot 10^{2}$ | $1.0000 \cdot 10^{-8}$ | $1.0000 \cdot 10^{6}$ |
| $kL3a2$ | $2.5765 \cdot 10^{2}$ | $2.5765 \cdot 10^{2}$ | $1.0000 \cdot 10^{-8}$ | $1.0000 \cdot 10^{6}$ |
| $kL4a$ | $1.1421 \cdot 10^{-2}$ | $1.1415 \cdot 10^{-2}$ | $1.0000 \cdot 10^{-8}$ | $1.0000 \cdot 10^{6}$ |
| $kL4a2$ | $2.3029 \cdot 10^{-4}$ | $2.3122 \cdot 10^{-4}$ | $1.0000 \cdot 10^{-8}$ | $1.0000 \cdot 10^{6}$ |
| $kL4b$ | 2.1055 | 2.1060 | $1.0000 \cdot 10^{-8}$ | $1.0000 \cdot 10^{6}$ |
| $kL5a$ | $2.5570 \cdot 10^{-1}$ | $2.5654 \cdot 10^{-1}$ | $1.0000 \cdot 10^{-8}$ | $1.0000 \cdot 10^{6}$ |
| $kL5b$ | 7.5766 | 7.5763 | $1.0000 \cdot 10^{-8}$ | $1.0000 \cdot 10^{6}$ |
| $kL6a$ | $8.4296 \cdot 10^{-2}$ | $8.4395 \cdot 10^{-2}$ | $1.0000 \cdot 10^{-8}$ | $1.0000 \cdot 10^{6}$ |
| $kL6b$ | $2.3022 \cdot 10^{-2}$ | $2.3084 \cdot 10^{-2}$ | $1.0000 \cdot 10^{-8}$ | $1.0000 \cdot 10^{6}$ |
| $kL7a$ | $7.2582 \cdot 10^{2}$ | $9.0592 \cdot 10^{2}$ | $1.0000 \cdot 10^{-8}$ | $1.0000 \cdot 10^{6}$ |
| $kL7b$ | $1.3458 \cdot 10^{1}$ | $1.2935 \cdot 10^{1}$ | $1.0000 \cdot 10^{-8}$ | $1.0000 \cdot 10^{6}$ |
| $kL8a$ | $2.9542 \cdot 10^{2}$ | $2.6349 \cdot 10^{2}$ | $1.0000 \cdot 10^{-8}$ | $1.0000 \cdot 10^{6}$ |
| $kL8c$ | $2.2077 \cdot 10^{-2}$ | $2.5507 \cdot 10^{-2}$ | $1.0000 \cdot 10^{-8}$ | $1.0000 \cdot 10^{6}$ |
| $kLclear$ | $3.7677 \cdot 10^{-2}$ | $3.6746 \cdot 10^{-2}$ | $1.0000 \cdot 10^{-8}$ | $1.0000 \cdot 10^{6}$ |
| $isoscale$ | $1.2000 \cdot 10^{1}$ | $1.2000 \cdot 10^{1}$ | 8.0000 | $1.2000 \cdot 10^{1}$ |
| $phe\_effect$ | $8.9227 \cdot 10^{-1}$ | $8.8912 \cdot 10^{-1}$ | $6.0000 \cdot 10^{-1}$ | 1.0000 |
| $min2$ | $2.0854 \cdot 10^{1}$ | $2.0854 \cdot 10^{1}$ | 0 | $1.0000 \cdot 10^{2}$ |
| $min3$ | $9.8712 \cdot 10^{-3}$ | $9.8712 \cdot 10^{-3}$ | 0 | $1.0000 \cdot 10^{2}$ |
| $EC502$ | 1.0882 | 1.0720 | −5.0000 | 3.0000 |
| $EC503$ | $8.2934 \cdot 10^{-2}$ | $6.6394 \cdot 10^{-2}$ | −5.0000 | 3.0000 |
| $n2$ | 3.0000 | 2.1674 | $5.0000 \cdot 10^{-1}$ | 3.0000 |
| $n3$ | 3.0000 | 2.9723 | $5.0000 \cdot 10^{-1}$ | 3.0000 |
| $diab\_reest$ | $9.6659 \cdot 10^{-2}$ | $1.2111 \cdot 10^{-1}$ | 0 | 1.0000 |

Table 3: **Parameter values related to the lipolysis submodel.**

| | | | | |
|---|---|---|---|---|
| $kArel$ | $2.0500 \cdot 10^{-7}$ | $3.2440 \cdot 10^{-7}$ | $1.0000 \cdot 10^{-8}$ | $1.0000 \cdot 10^{6}$ |
| $kAexo$ | $6.6957 \cdot 10^{-1}$ | $6.6957 \cdot 10^{-1}$ | $1.0000 \cdot 10^{-8}$ | $1.0000 \cdot 10^{6}$ |
| $kACaATP$ | $2.9179 \cdot 10^{1}$ | $2.9226 \cdot 10^{1}$ | $1.0000 \cdot 10^{-8}$ | $1.0000 \cdot 10^{6}$ |
| $kAcAMP$ | $1.8864 \cdot 10^{1}$ | $1.8864 \cdot 10^{1}$ | $1.0000 \cdot 10^{-8}$ | $1.0000 \cdot 10^{6}$ |
| $kACa2$ | $1.1695 \cdot 10^{-3}$ | $1.1638 \cdot 10^{-3}$ | $1.0000 \cdot 10^{-8}$ | $1.0000 \cdot 10^{6}$ |
| $kAATP2$ | $9.2660 \cdot 10^{-4}$ | $9.2651 \cdot 10^{-4}$ | $1.0000 \cdot 10^{-8}$ | $1.0000 \cdot 10^{6}$ |
| $kAEndo$ | $8.0992 \cdot 10^{5}$ | $9.9826 \cdot 10^{5}$ | $1.0000 \cdot 10^{-8}$ | $1.0000 \cdot 10^{6}$ |
| $kACacAMP$ | $6.0089 \cdot 10^{2}$ | $6.0768 \cdot 10^{2}$ | $1.0000 \cdot 10^{-8}$ | $1.0000 \cdot 10^{6}$ |
| $kAm$ | $1.2535 \cdot 10^{-8}$ | $1.2535 \cdot 10^{-8}$ | $1.0000 \cdot 10^{-8}$ | $1.0000 \cdot 10^{6}$ |
| $kARelBasal$ | $1.0534 \cdot 10^{-4}$ | $1.0749 \cdot 10^{-4}$ | $1.0000 \cdot 10^{-8}$ | $1.0000 \cdot 10^{6}$ |
| $kAdegcAMP$ | $2.0201 \cdot 10^{-6}$ | $1.8269 \cdot 10^{-6}$ | $1.0000 \cdot 10^{-8}$ | $1.0000 \cdot 10^{6}$ |
| $kAremCa$ | $5.5039 \cdot 10^{-5}$ | $4.8329 \cdot 10^{-5}$ | $1.0000 \cdot 10^{-8}$ | $1.0000 \cdot 10^{6}$ |
| $kAdegATP$ | $2.2781 \cdot 10^{-1}$ | $2.4347 \cdot 10^{-1}$ | $1.0000 \cdot 10^{-8}$ | $1.0000 \cdot 10^{6}$ |
| $kADiffcAMP$ | $2.7110 \cdot 10^{-1}$ | $2.7104 \cdot 10^{-1}$ | $1.0000 \cdot 10^{-8}$ | $1.0000 \cdot 10^{6}$ |
| $kADiffCa$ | $1.6417 \cdot 10^{4}$ | $1.6417 \cdot 10^{4}$ | $1.0000 \cdot 10^{-8}$ | $1.0000 \cdot 10^{6}$ |
| $kADiffATP$ | $5.3773 \cdot 10^{-1}$ | $4.9675 \cdot 10^{-1}$ | $1.0000 \cdot 10^{-8}$ | $1.0000 \cdot 10^{6}$ |
| $kA4a$ | $1.7383 \cdot 10^{4}$ | $1.7384 \cdot 10^{4}$ | $1.0000 \cdot 10^{-8}$ | $1.0000 \cdot 10^{6}$ |
| $kACL$ | $2.4186 \cdot 10^{-5}$ | $2.4874 \cdot 10^{-5}$ | $1.0000 \cdot 10^{-8}$ | $1.0000 \cdot 10^{6}$ |
| $kA4c$ | $1.0638 \cdot 10^{-8}$ | $1.0001 \cdot 10^{-8}$ | $1.0000 \cdot 10^{-8}$ | $1.0000 \cdot 10^{6}$ |
| $kA4b$ | 2.8284 | 2.8344 | $1.0000 \cdot 10^{-8}$ | $1.0000 \cdot 10^{6}$ |
| $kA5a$ | 3.2559 | 3.2559 | $1.0000 \cdot 10^{-8}$ | $1.0000 \cdot 10^{6}$ |
| $kAVpip$ | $2.2330 \cdot 10^{-5}$ | $2.3223 \cdot 10^{-5}$ | $2.0000 \cdot 10^{-5}$ | $6.0000 \cdot 10^{-5}$ |
| $kAVcell$ | $1.6689 \cdot 10^{-13}$ | $1.6613 \cdot 10^{-13}$ | $8.6400 \cdot 10^{-15}$ | $1.7168 \cdot 10^{-13}$ |

Table 4: **Parameter values related to the adiponectin submodel.**

### 4 Initial values

The model was given arbitrary initial values and then simulated without any inputs until steady state was reached to get a stable set of inti before any experiment was started. In most cases, the values are expressed as percentages. In other words, all state of a component in the model sum to 100. Note that the initial values for the mediators in the pipett ($pipCa$, $pipcAMP$, and $pipATP$) was set to the corresponding experimental concentrations during the start of the simulation of the adiponectin experiments (after steady state), and was set to 0 during all other experiments. The initial values used - before simulation to steady state - are given in Table 5, Table 6, and Table 7 for the glucose uptake, lipolysis, and adiponectin submodels respectively.

| State | Initial values |
|---|---|
| $IR$ | 100 |
| $IR\_YP$ | 0 |
| $IRins$ | 0 |
| $IRi\_YP$ | 0 |
| $IRi$ | 0 |
| $IRS1$ | 100 |
| $IRS1\_YP$ | 0 |
| $IRS1\_YP\_S307P$ | 0 |
| $IRS1\_S307P$ | 0 |
| $X$ | 100 |
| $X\_P$ | 0 |
| $PKB$ | 100 |
| $PKB\_S308P$ | 0 |
| $PKB\_T473P$ | 0 |
| $PKB\_S308P\_T473P$ | 0 |
| $mTORC1$ | 100 |
| $mTORC1a$ | 0 |
| $mTORC2$ | 100 |
| $mTORC2a$ | 0 |
| $AS160$ | 100 |
| $AS160\_T642P$ | 0 |
| $GLUT4m$ | 0 |
| $GLUT4$ | 100 |
| $GLUCOSE$ | 0 |
| $S6K$ | 100 |
| $S6K\_T389P$ | 0 |
| $S6$ | 100 |
| $S6\_S235\_S236P$ | 0 |
| $ERK$ | 100 |
| $ERK\_T202\_Y204P$ | 0 |
| $seqERK$ | 0 |
| $Elk1$ | 100 |
| $Elk1\_S383P$ | 0 |
| $FOXO$ | 100 |
| $FOXO\_S256P$ | 0 |

Table 5: **Initial values for the glucose uptake submodel.**

| | |
|---|---|
| $BETA2$ | 100 |
| $BETA2a$ | 0 |
| $ALPHA$ | 80 |
| $ALPHAa$ | 20 |
| $AC$ | 80 |
| $ACa$ | 20 |
| $PDE3B$ | 80 |
| $PDE3Ba$ | 20 |
| $HSL$ | 80 |
| $HSLp$ | 20 |
| $Gly$ | 0 |
| $FFA$ | 0 |

Table 6: **Initial values for the lipolysis submodel.**

| | |
|---|---|
| $BETA3$ | 100 |
| $BETA3a$ | 0 |
| $BETA3de$ | 0 |
| $Ca$ | 0 |
| $ATP$ | 0 |
| $cAMP$ | 0 |
| $Res$ | 99 |
| $Rel$ | 1 |
| $PM$ | 0 |
| $Endo$ | 0 |
| $Adiponectin$ | 0 |
| $pipCa$ | 0 |
| $pipATP$ | 0 |
| $pipcAMP$ | 0 |

Table 7: **Initial values for the adiponectin submodel.**
