## Supplementary material for "A comprehensive mechanistic model of adipocyte signaling with layers of confidence": S3 Time series of all sites added to the model

### Time series of all sites added to the model.
#### A supplement to "A comprehensive mechanistic model of adipocyte signaling with varying layers of confidence"

William Lövfors[1,2], Cecilia Jönsson[1,3], Charlotta S. Olofsson[4], Elin Nyman[1,*,**], and Gunnar Cedersund[1,5, *,**]

[1]Department of Biomedical Engineering, Linköping University, Linköping, Sweden
[2]Department of Mathematics, Linköping University, Linköping, Sweden
[3]Department of Biomedical and Clinical Sciences, Linköping University, Linköping, Sweden
[4]Department of Physiology/Metabolic Physiology, Institute of Neuroscience and Physiology, The Sahlgrenska Academy at University of Gothenburg, Gothenburg, Sweden
[5]Center for Medical Image Science and Visualization (CMIV), Linköping University, Linköping, Sweden
[*]Shared last author
[**]Corresponding author

May 2, 2022

### 1 How to interpret the figures.

All figures show the time series of a phosphosite added to the model during the model expansion. In all figures, dots and error bars corresponds to the experimental data (mean and SEM), and lines correspond to model simulations. The different simulation and data are colored coded based on what conditions are being applied. For normal conditions, the simulations and data are blue. Simulations under type 2 diabetic conditions are red. An example is shown in Fig. 1. For simulations of experiments with the PKB inhibitor MK2206 (MK) and the PI3K inhibitor LY294002 (LY), the simulations are shifting from the "normal blue" to turquoise corresponding to the amount of inhibition exerted by the inhibitor. En example is shown in Fig. 2. The color blue for normal simulations shift from a dark blue to a light blue the further away from the core model a site is added, corresponding to the confidence of the addition. Compare the color for the normal simulations in Fig. 2 and Fig. 3.

The title of a figure tells which phosphosite that was used an input to the added phosphosite (input -> added). Furthermore, the title also gives the cost, corresponding to the model agreement to data (cost: value). The cutoff for acceptable agreement was set to $cost = \chi^2(p = 0.05, df = 8) \approx 15.5$. Finally, the title also gives which layer the phosphosite was added to.

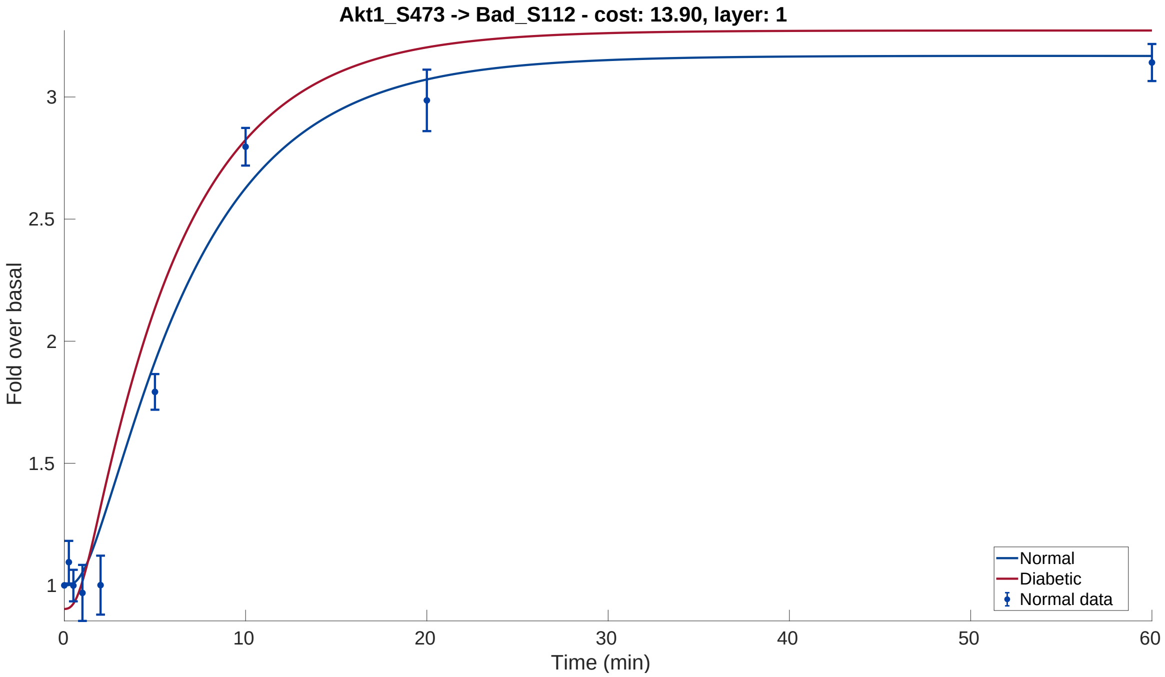


Figure 1: **Simulation of Bad-S112, in response to insulin stimulation normally and in type 2 diabetes.**

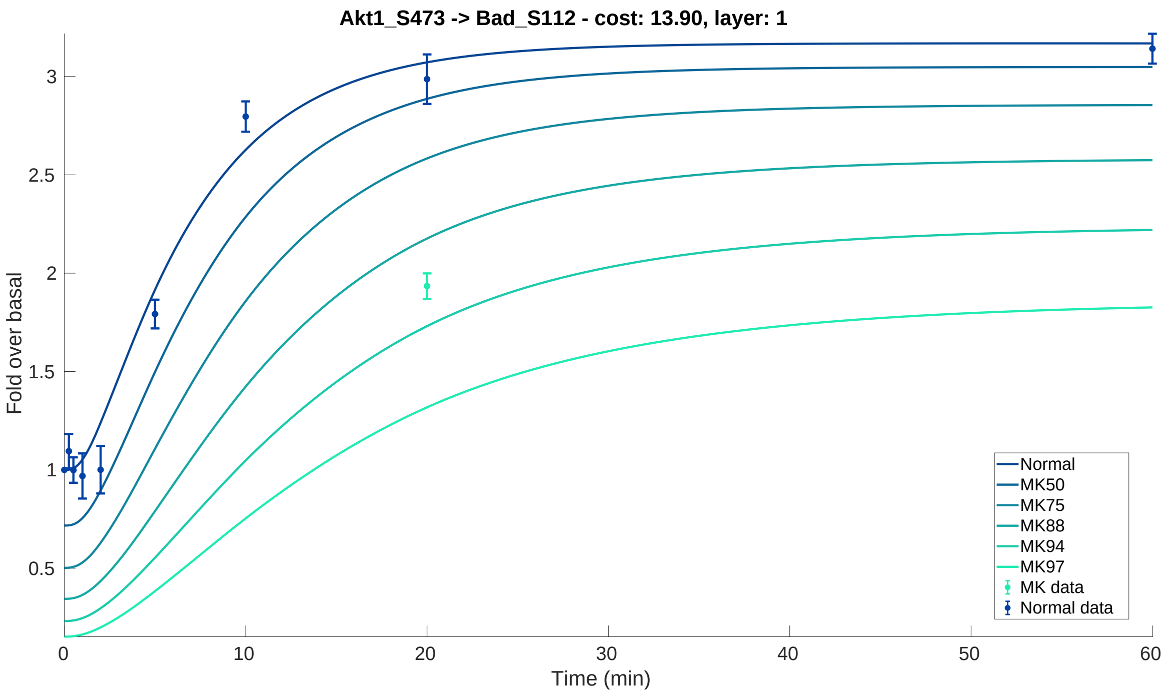


Figure 2: **Simulation of Bad-S112, in response to insulin stimulation normally and with the inhibitor MK2206.**

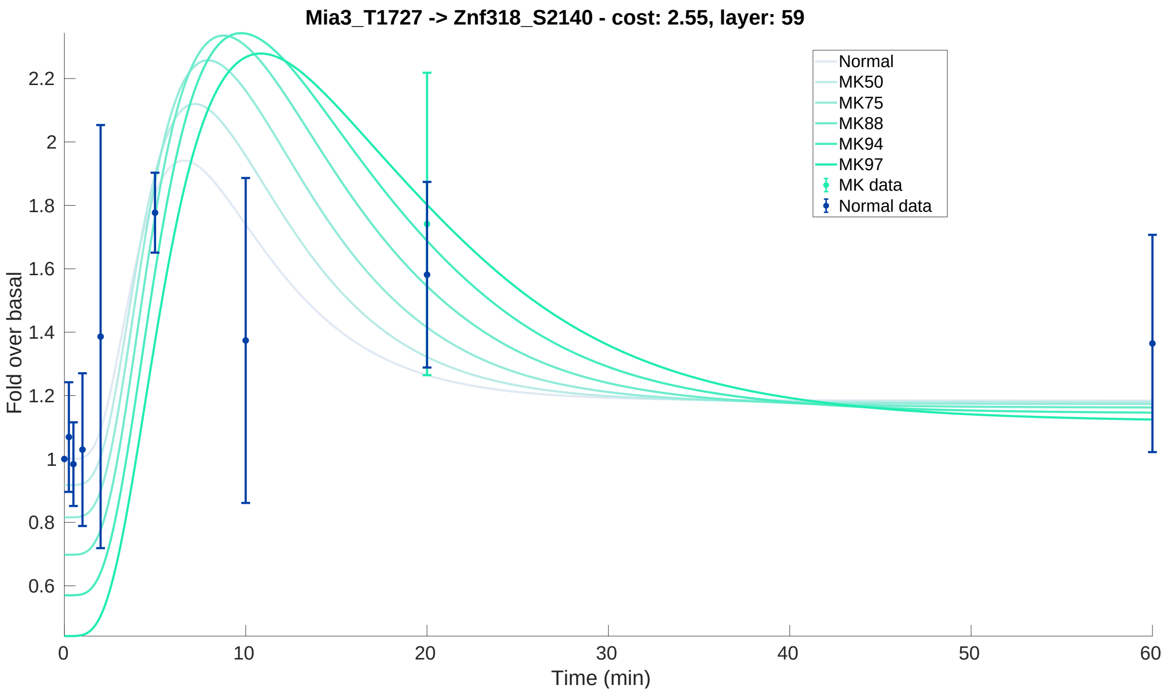


Figure 3: **Simulation of Znf318-S214 - a phosphosite added in a distant layer - in response to insulin stimulation normally and with the inhibitor MK2206.**

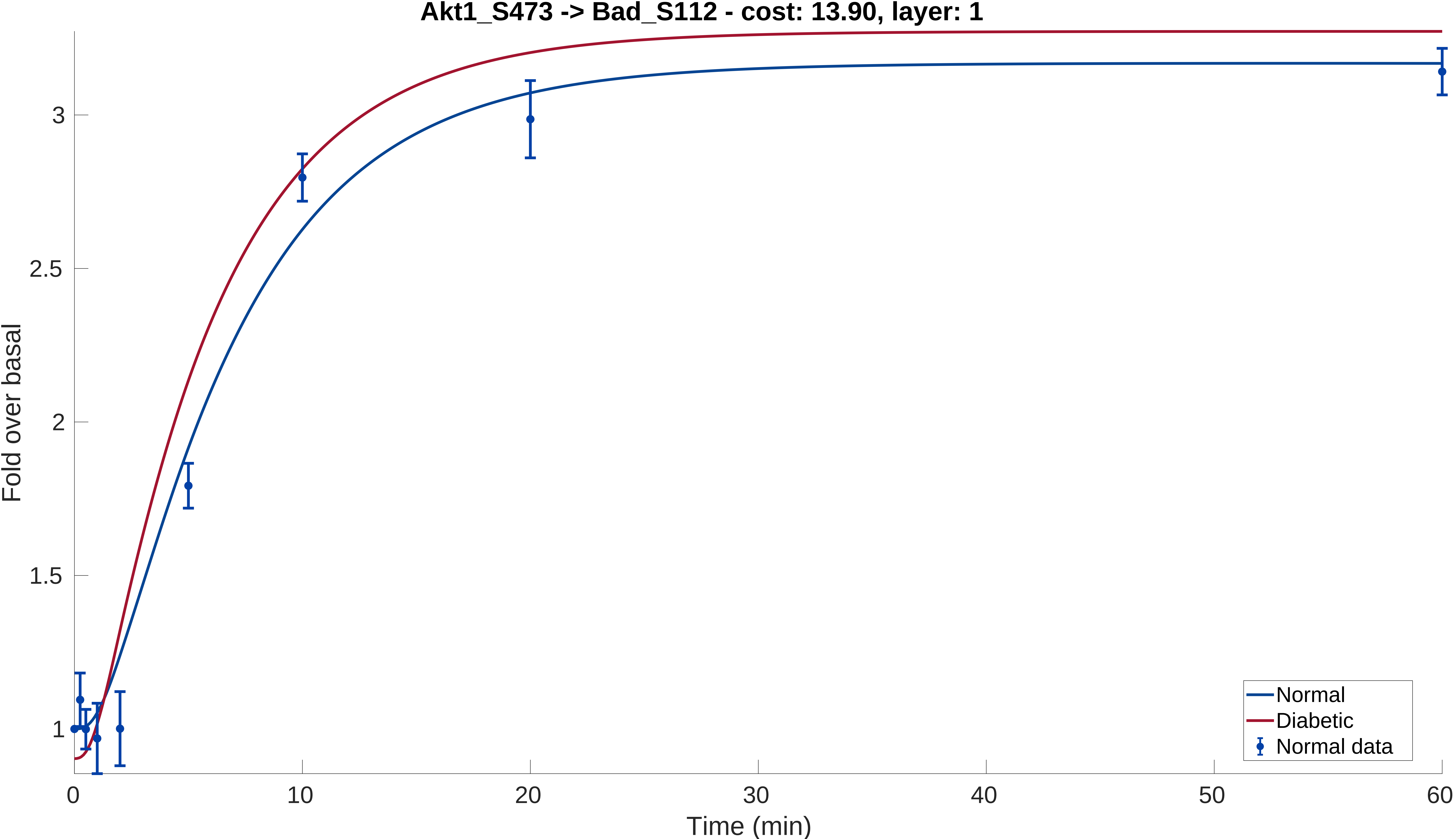
Akt1_S473 -> Bad_S112 - cost: 13.90, layer: 1
Fold over basal
Time (min)
3
2.5
2
1.5
1
0
10
20
30
40
50
60
Normal
Diabetic
Normal data

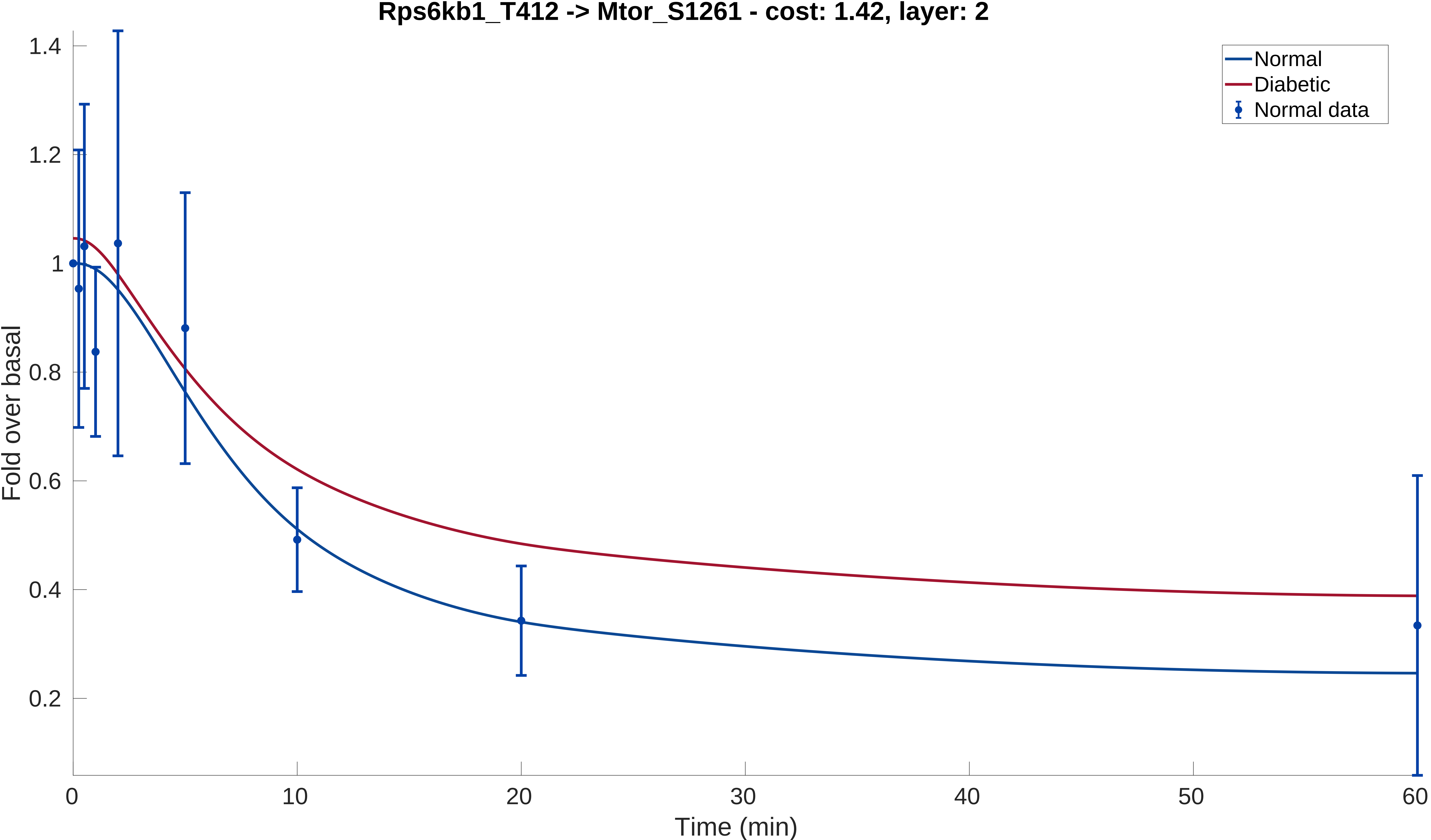

Rps6kb1_T412 -> Mtor_S1261 - cost: 1.42, layer: 2
Normal
Diabetic
Normal data
Fold over basal
Time (min)
0.2
0.4
0.6
0.8
1
1.2
1.4
0
10
20
30
40
50
60

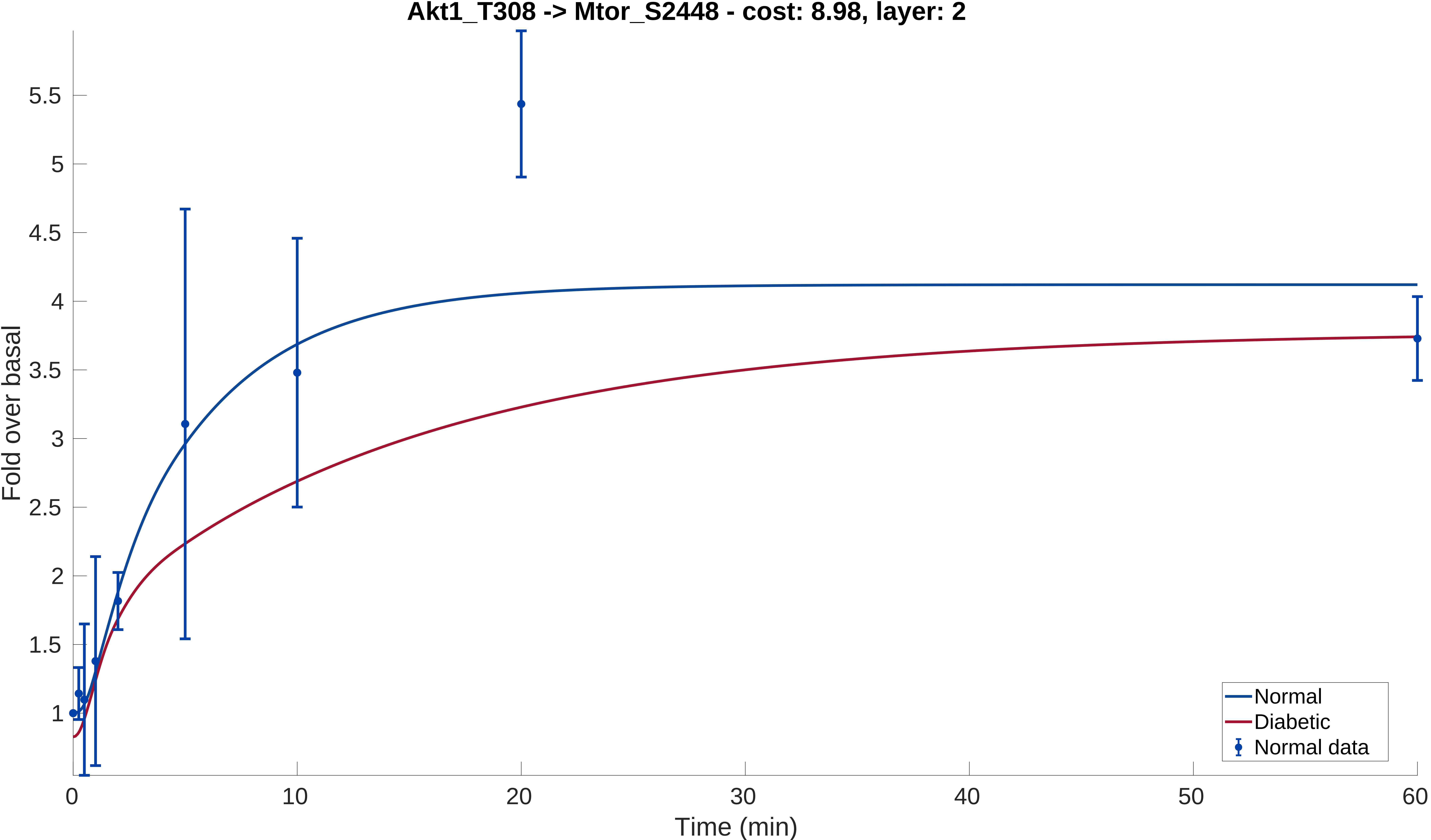

Akt1_T308 -> Mtor_S2448 - cost: 8.98, layer: 2
Fold over basal
5.5
5
4.5
4
3.5
3
2.5
2
1.5
1
0
10
20
30
40
50
60
Time (min)
Normal
Diabetic
Normal data

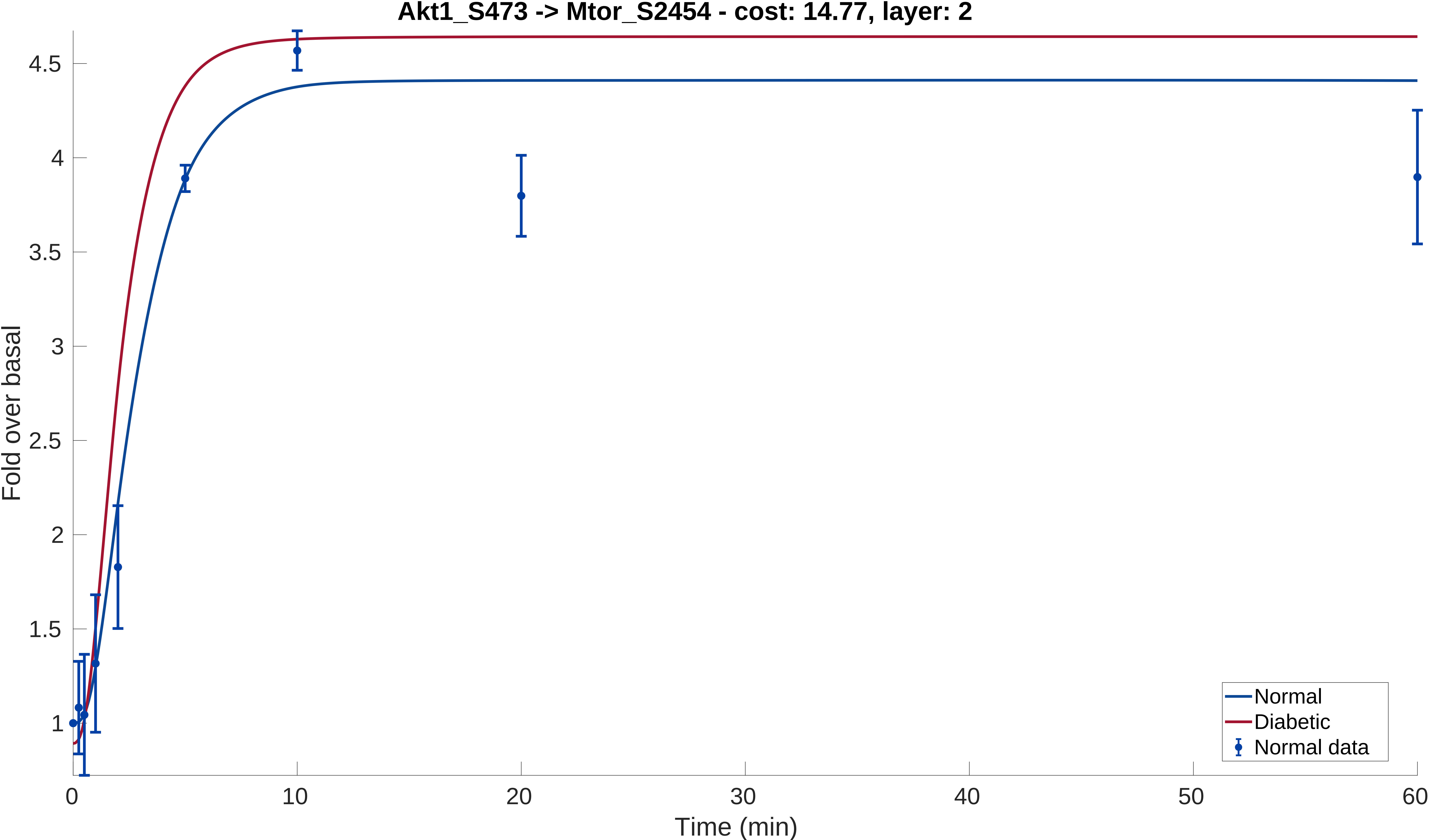

Akt1_S473 -> Mtor_S2454 - cost: 14.77, layer: 2
Fold over basal
4.5
4
3.5
3
2.5
2
1.5
1
0
10
20
30
40
50
60
Time (min)
Normal
Diabetic
Normal data

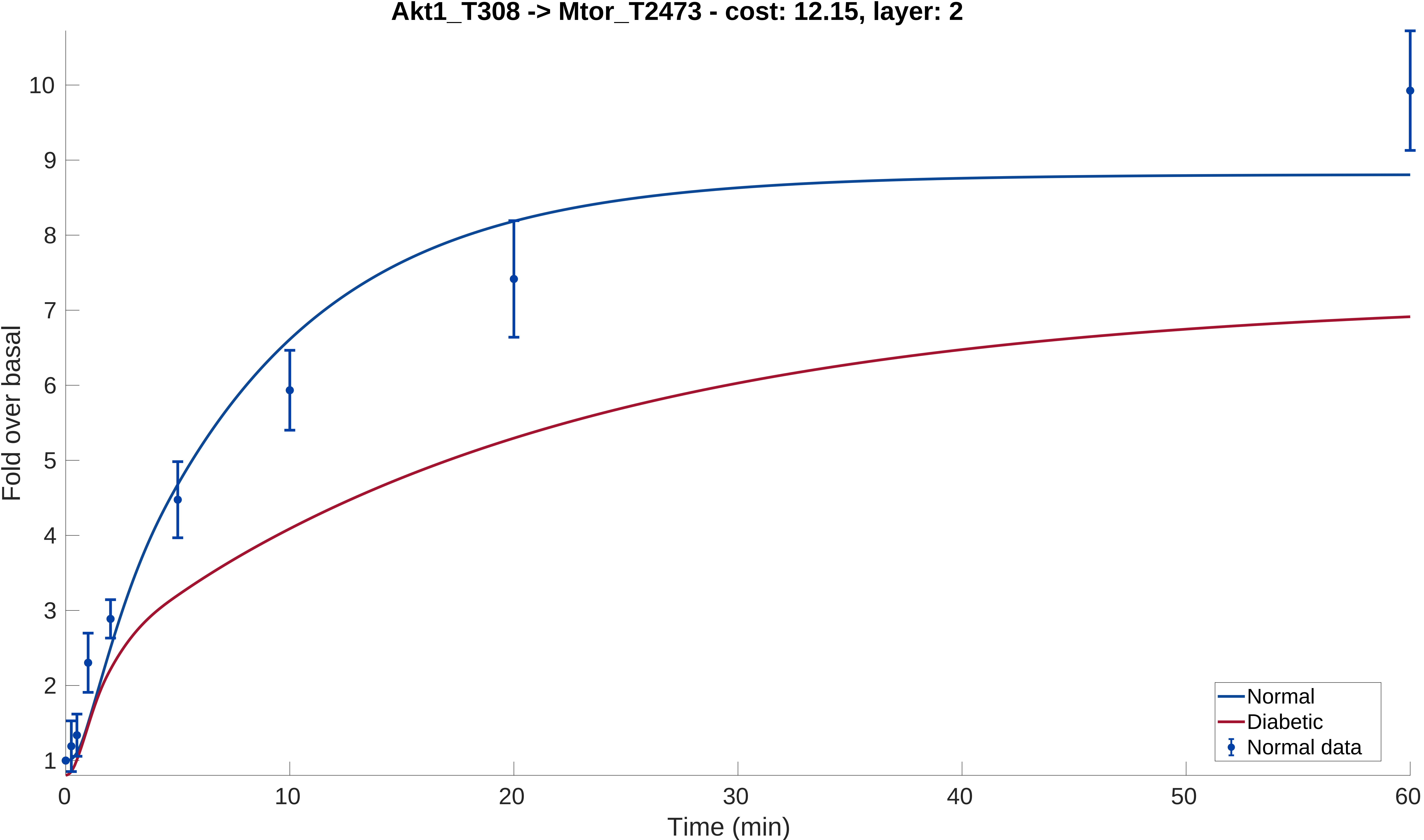
Akt1_T308 -> Mtor_T2473 - cost: 12.15, layer: 2
Fold over basal
Time (min)
Normal
Diabetic
Normal data
1
2
3
4
5
6
7
8
9
10
0
10
20
30
40
50
60

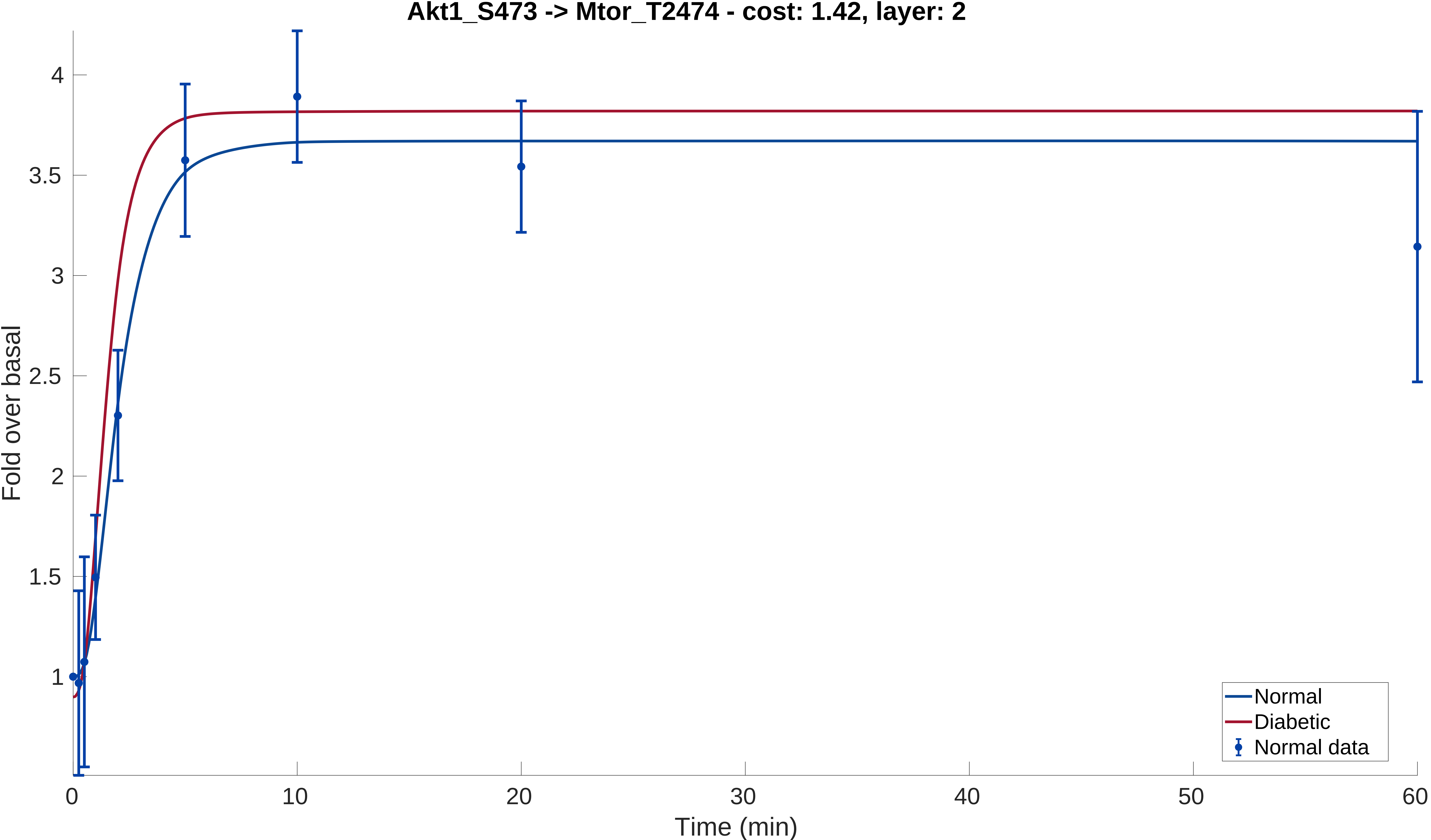

Akt1_S473 -> Mtor_T2474 - cost: 1.42, layer: 2
Fold over basal
Time (min)
4
3.5
3
2.5
2
1.5
1
0
10
20
30
40
50
60
Normal
Diabetic
Normal data

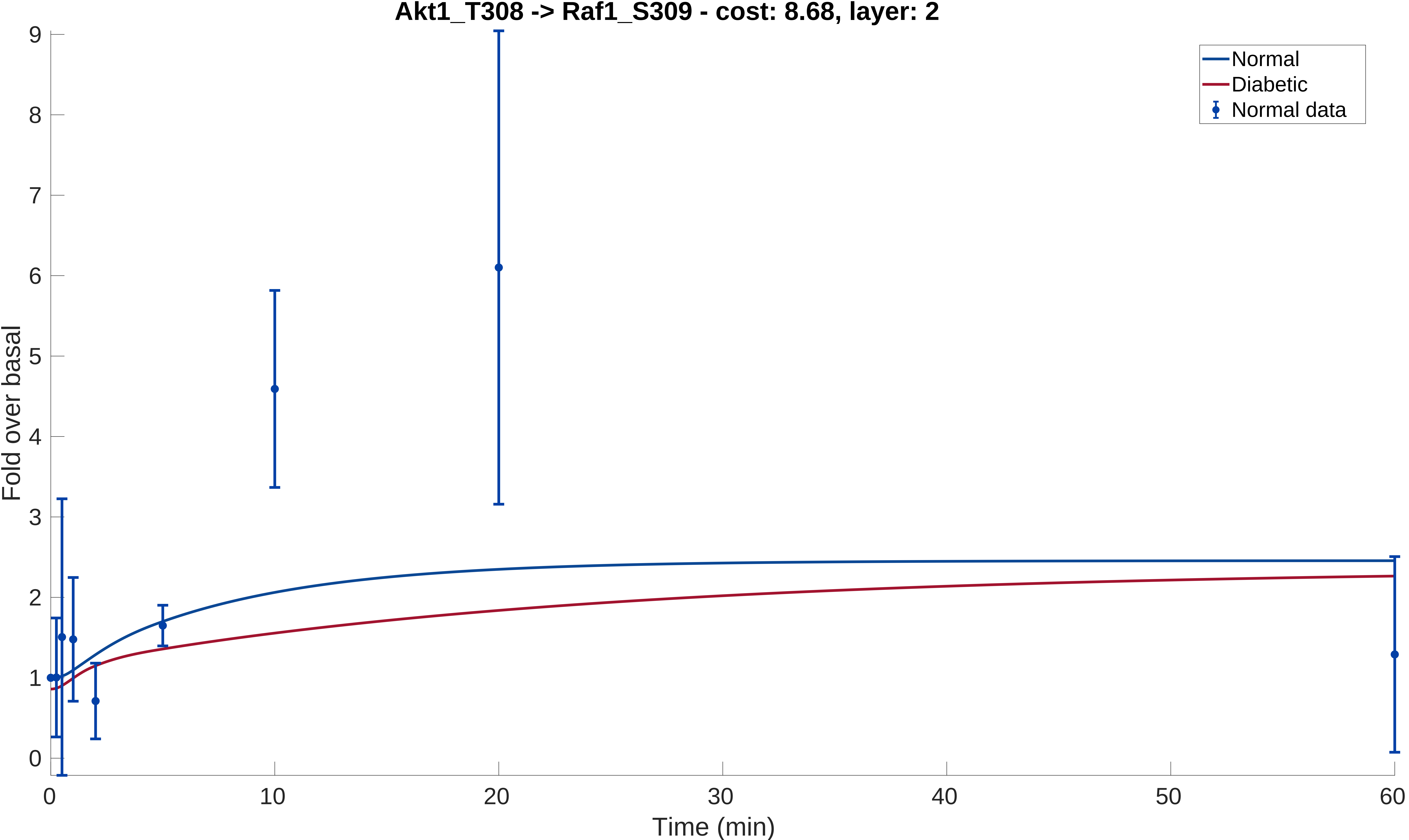

Akt1_T308 -> Raf1_S309 - cost: 8.68, layer: 2
Fold over basal
Time (min)
Normal
Diabetic
Normal data
0
1
2
3
4
5
6
7
8
9
0
10
20
30
40
50
60

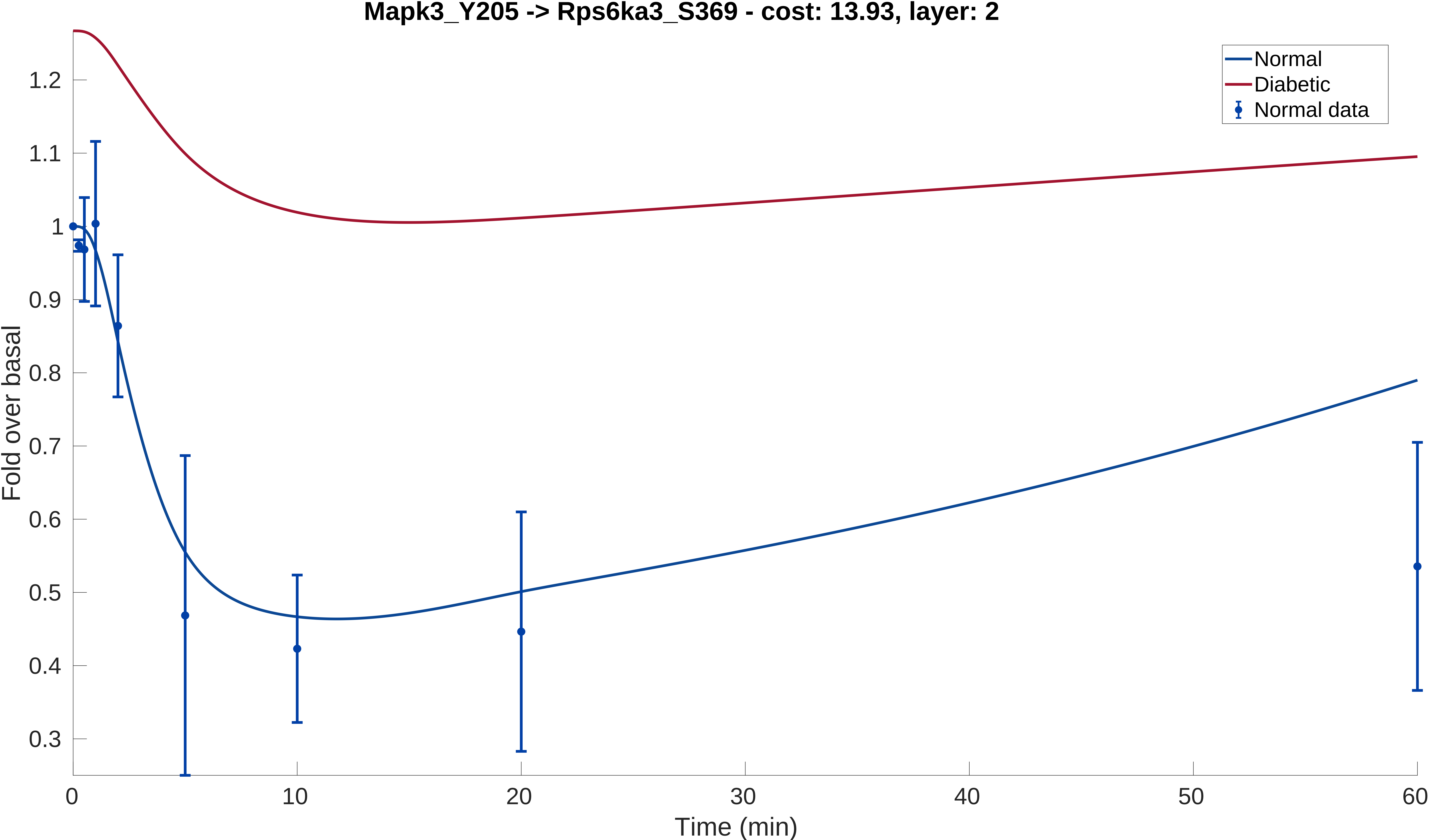

Mapk3_Y205 -> Rps6ka3_S369 - cost: 13.93, layer: 2
Normal
Diabetic
Normal data
Fold over basal
Time (min)
0.3
0.4
0.5
0.6
0.7
0.8
0.9
1
1.1
1.2
0
10
20
30
40
50
60

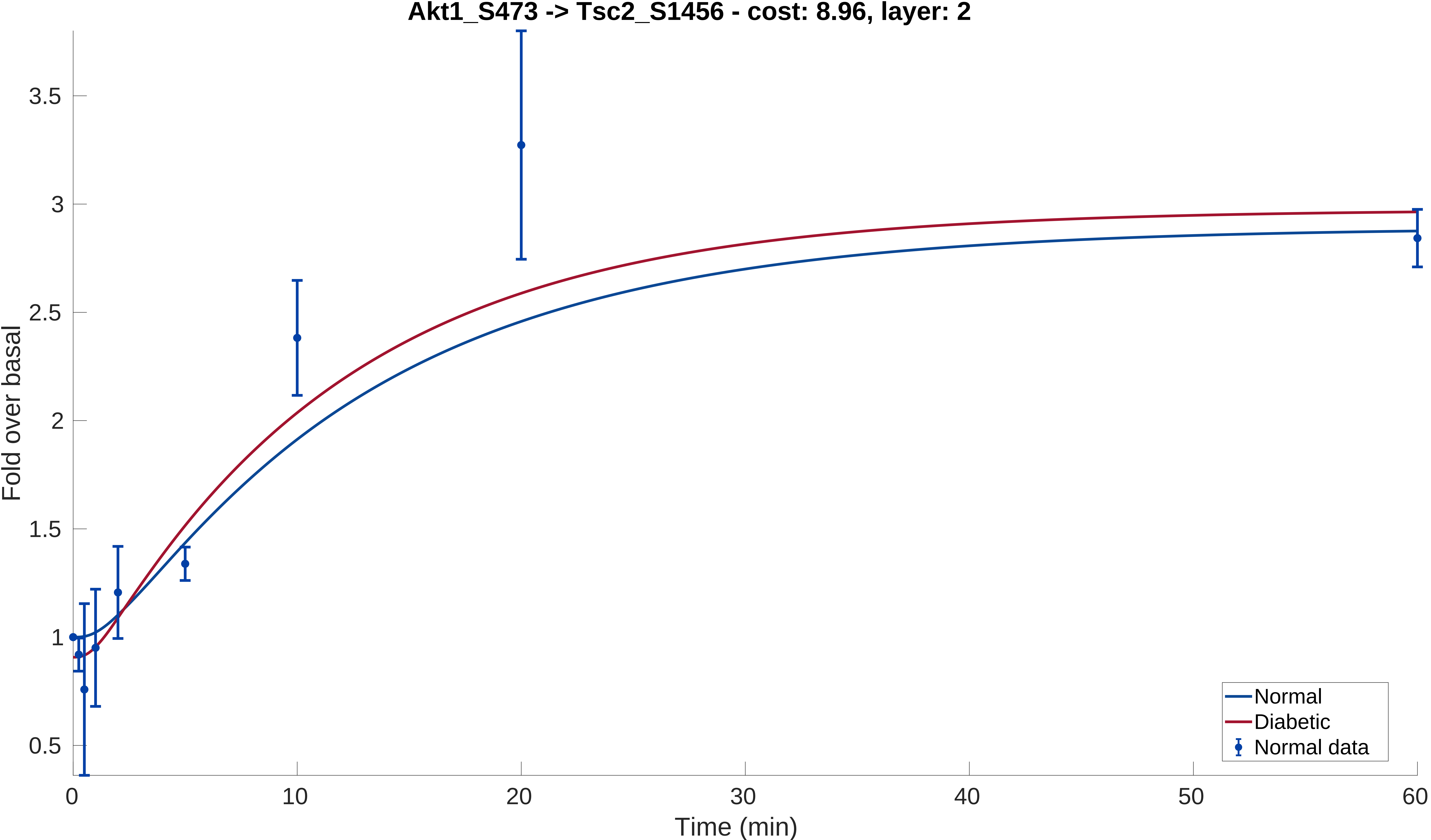

Akt1_S473 -> Tsc2_S1456 - cost: 8.96, layer: 2
3.5
3
2.5
2
1.5
1
0.5
Fold over basal
0
10
20
30
40
50
60
Time (min)
Normal
Diabetic
Normal data

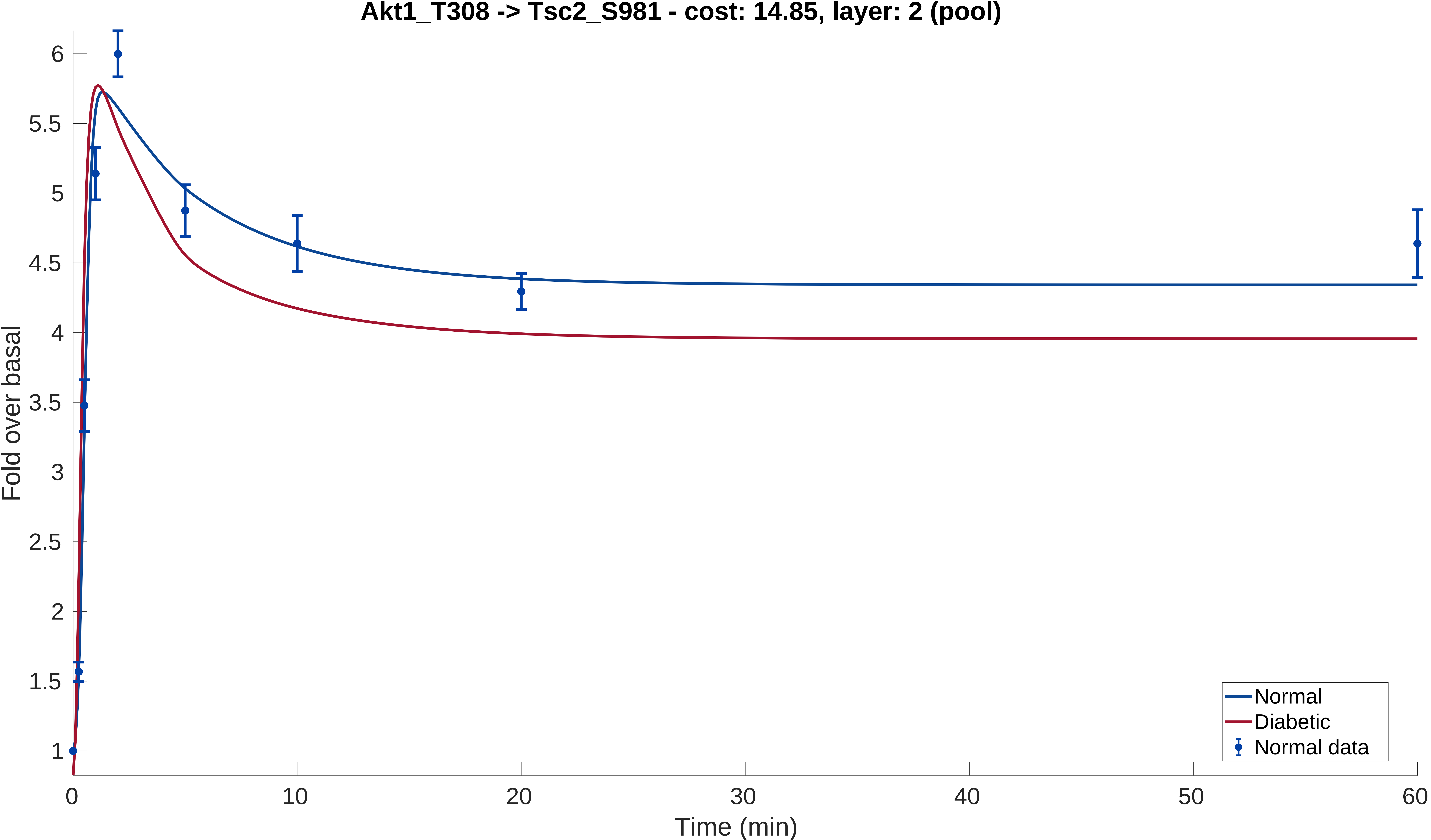

Akt1_T308 -> Tsc2_S981 - cost: 14.85, layer: 2 (pool)
Fold over basal
6
5.5
5
4.5
4
3.5
3
2.5
2
1.5
1
0
10
20
30
40
50
60
Time (min)
Normal
Diabetic
Normal data

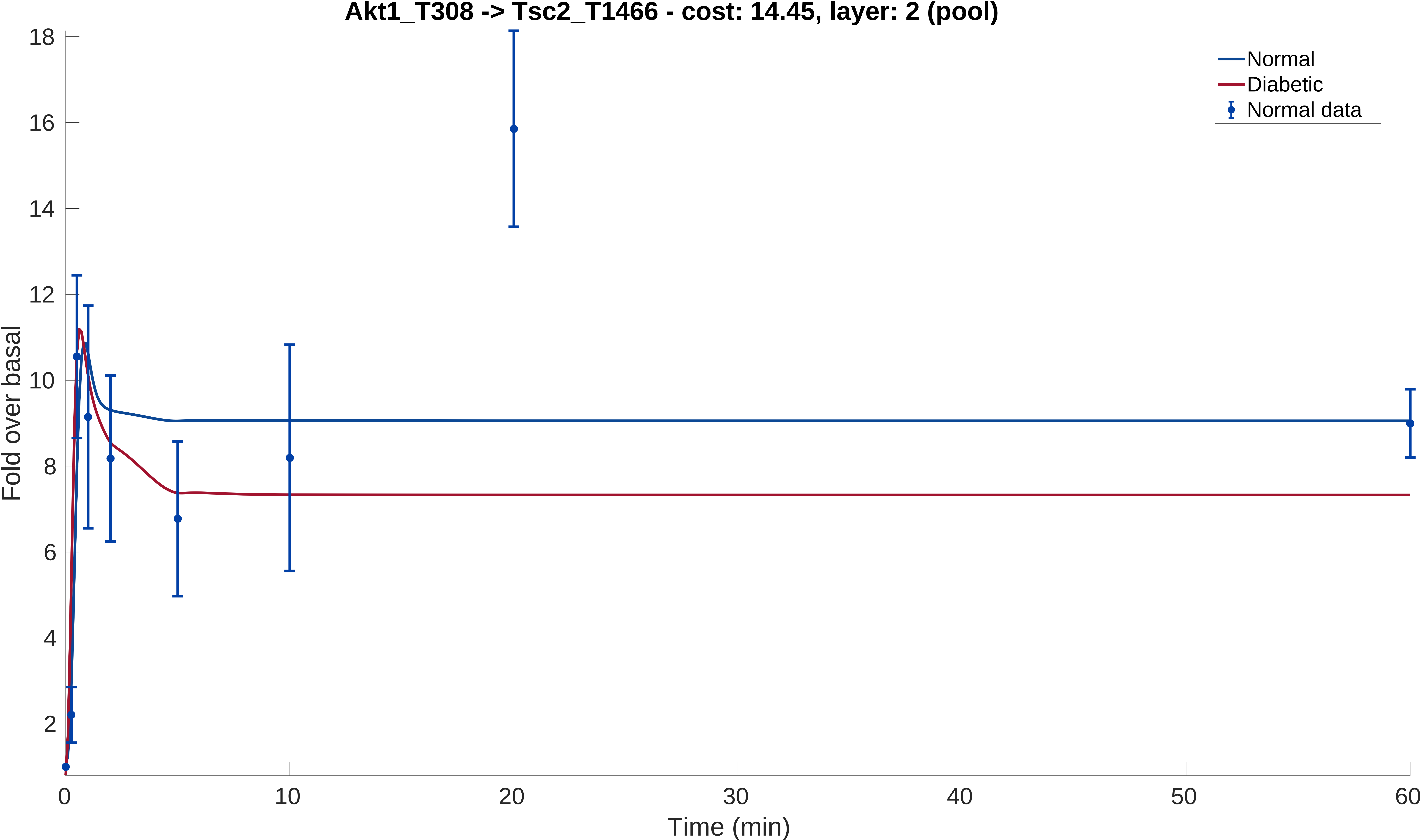

Akt1_T308 -> Tsc2_T1466 - cost: 14.45, layer: 2 (pool)
Normal
Diabetic
Normal data
Fold over basal
Time (min)
18
16
14
12
10
8
6
4
2
0
10
20
30
40
50
60

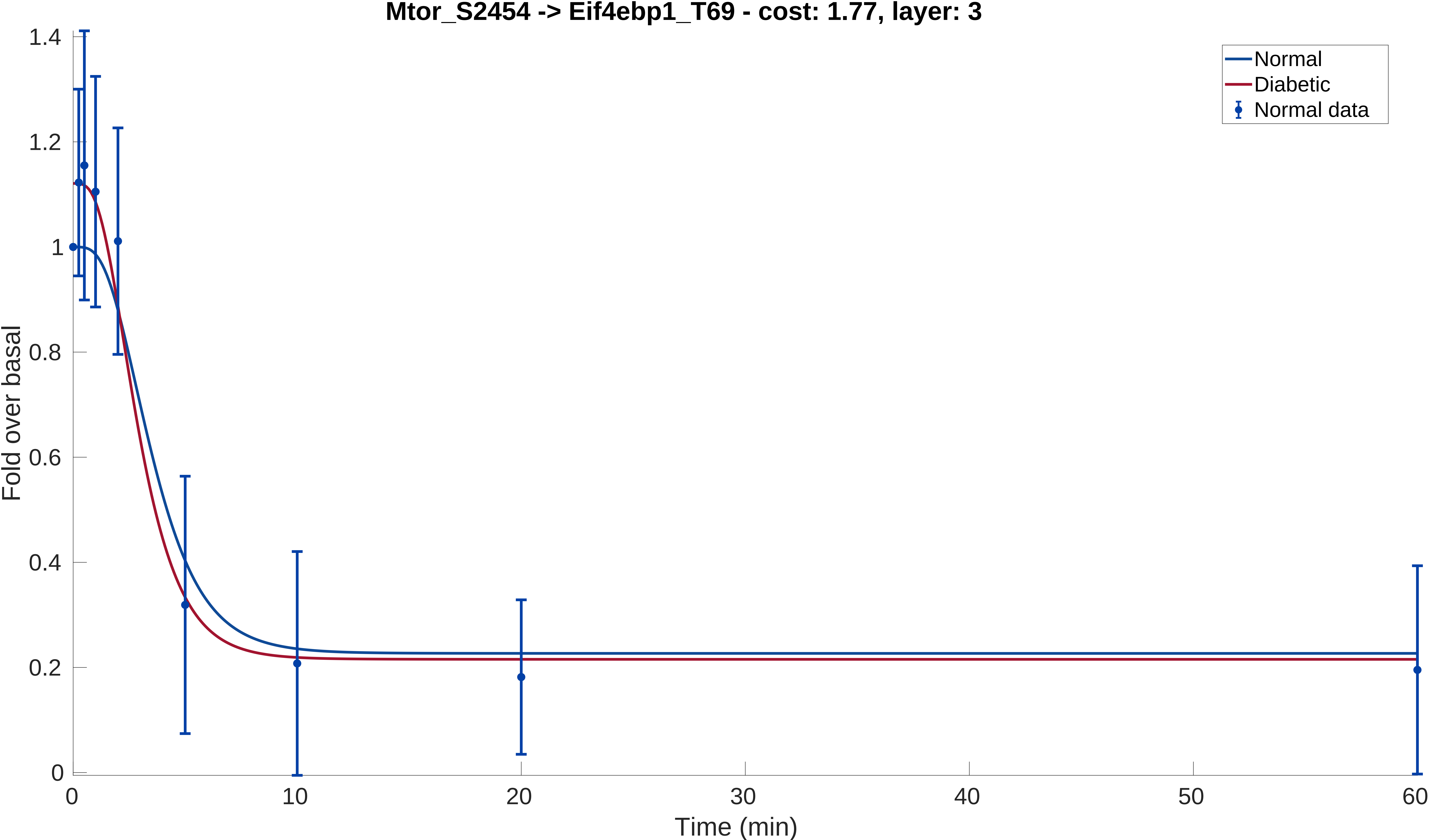

Mtor_S2454 -> Eif4ebp1_T69 - cost: 1.77, layer: 3
Normal
Diabetic
Normal data
Fold over basal
Time (min)
0
0.2
0.4
0.6
0.8
1
1.2
1.4
0
10
20
30
40
50
60

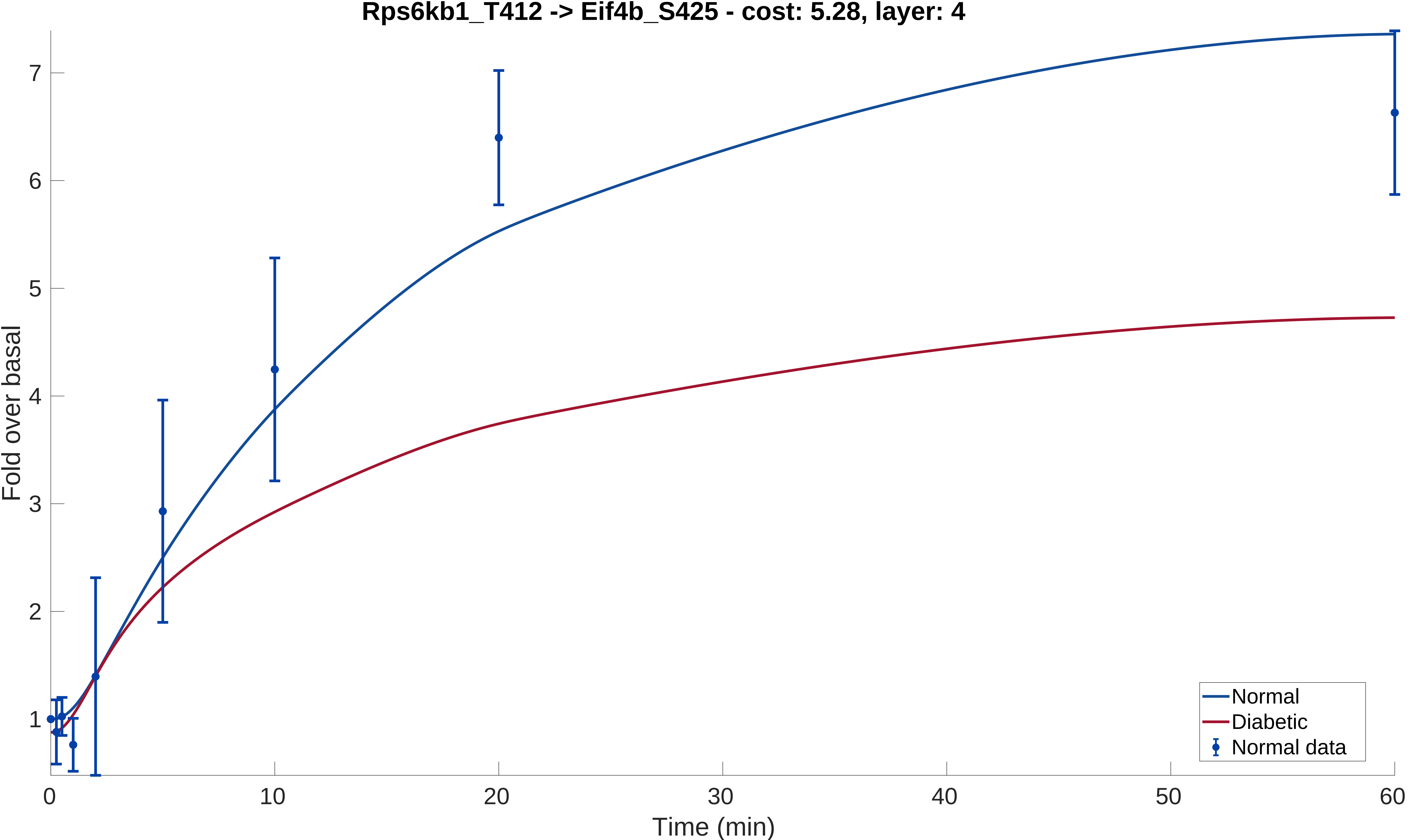

Rps6kb1_T412 -> Eif4b_S425 - cost: 5.28, layer: 4
Fold over basal
1
2
3
4
5
6
7
0
10
20
30
40
50
60
Time (min)
Normal
Diabetic
Normal data

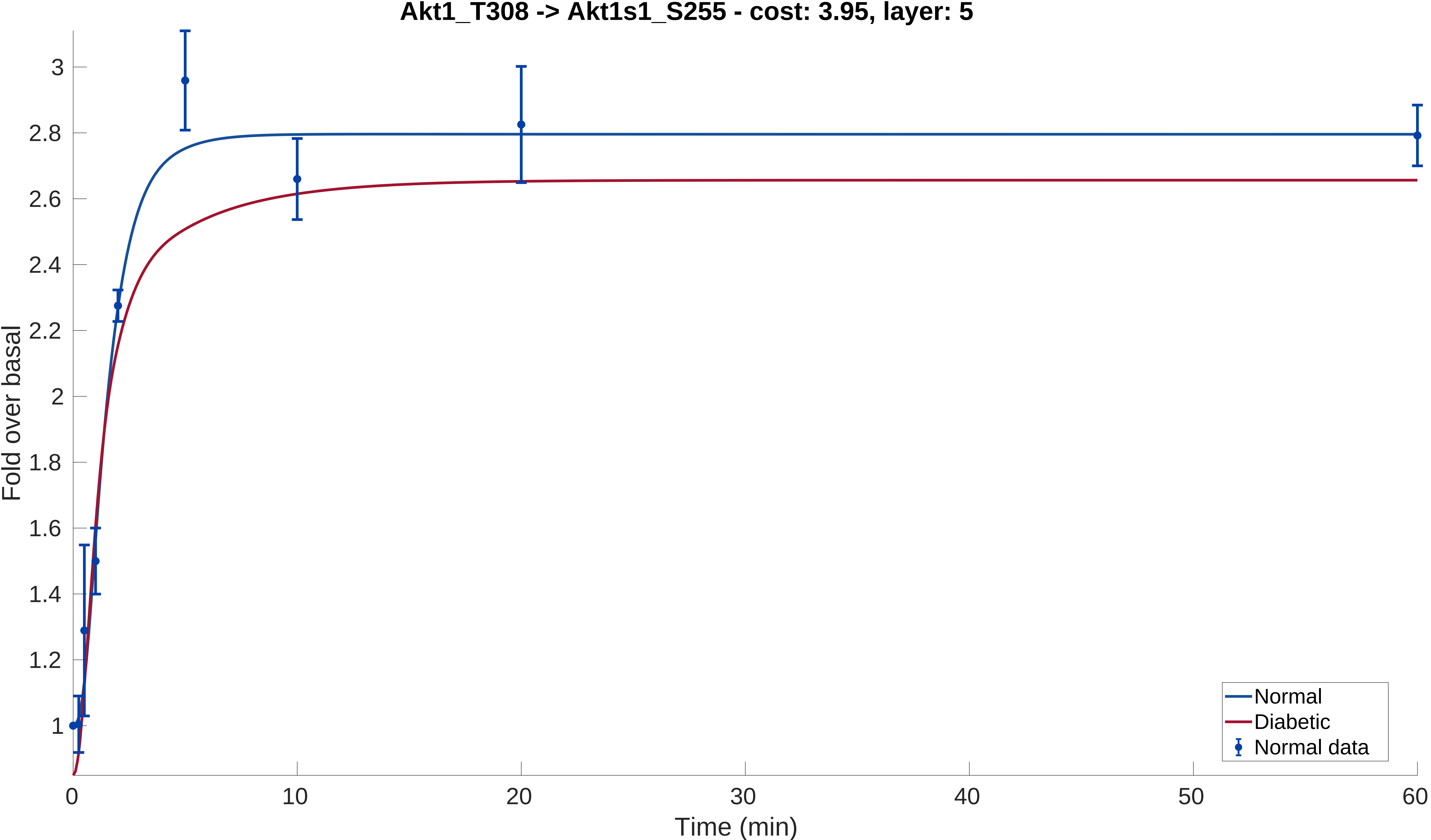

Akt1_T308 -> Akt1s1_S255 - cost: 3.95, layer: 5
Fold over basal
Time (min)
3
2.8
2.6
2.4
2.2
2
1.8
1.6
1.4
1.2
1
0
10
20
30
40
50
60
Normal
Diabetic
Normal data

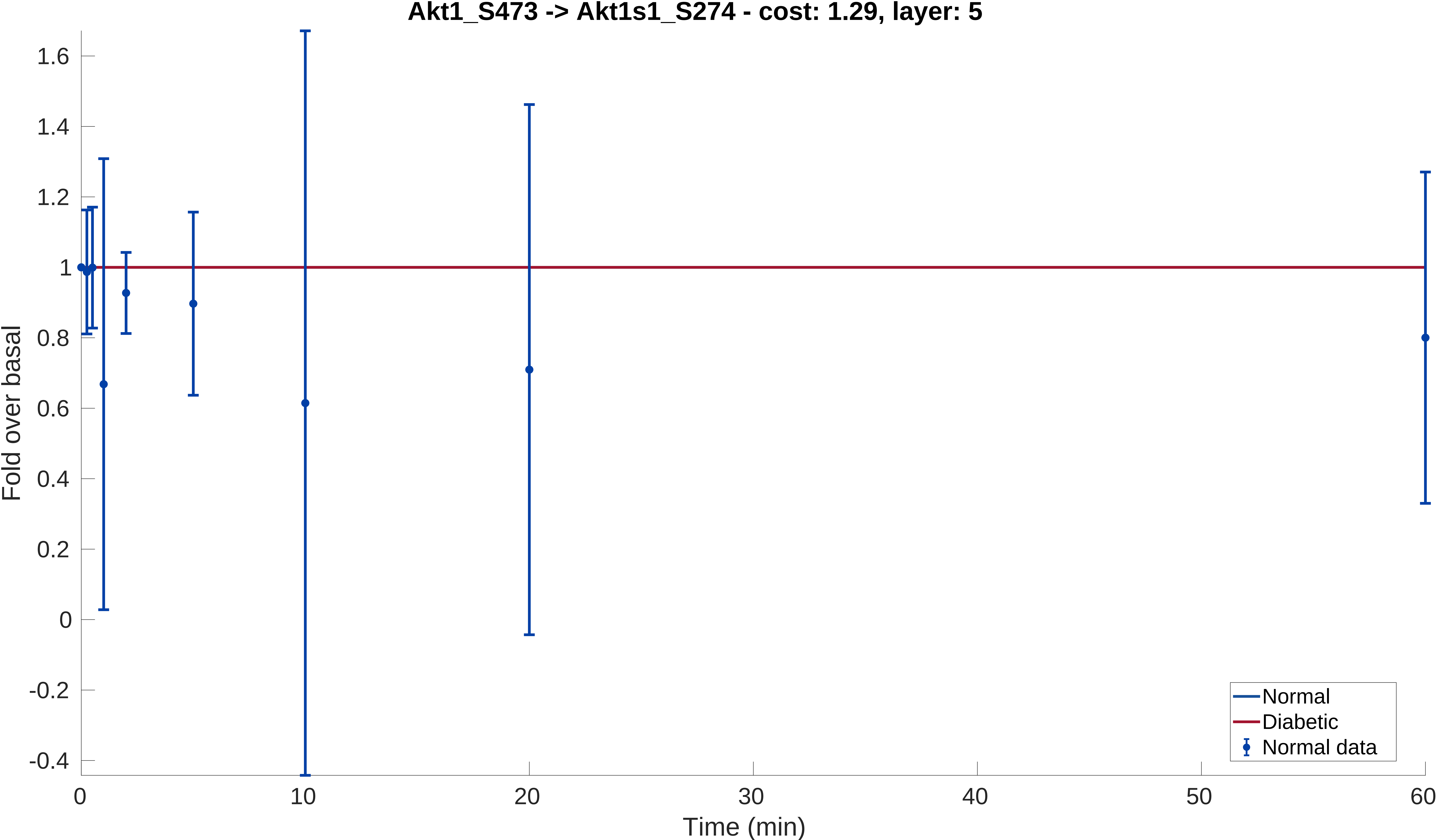

Akt1_S473 -> Akt1s1_S274 - cost: 1.29, layer: 5
Fold over basal
1.6
1.4
1.2
1
0.8
0.6
0.4
0.2
0
-0.2
-0.4
0
10
20
30
40
50
60
Time (min)
Normal
Diabetic
Normal data

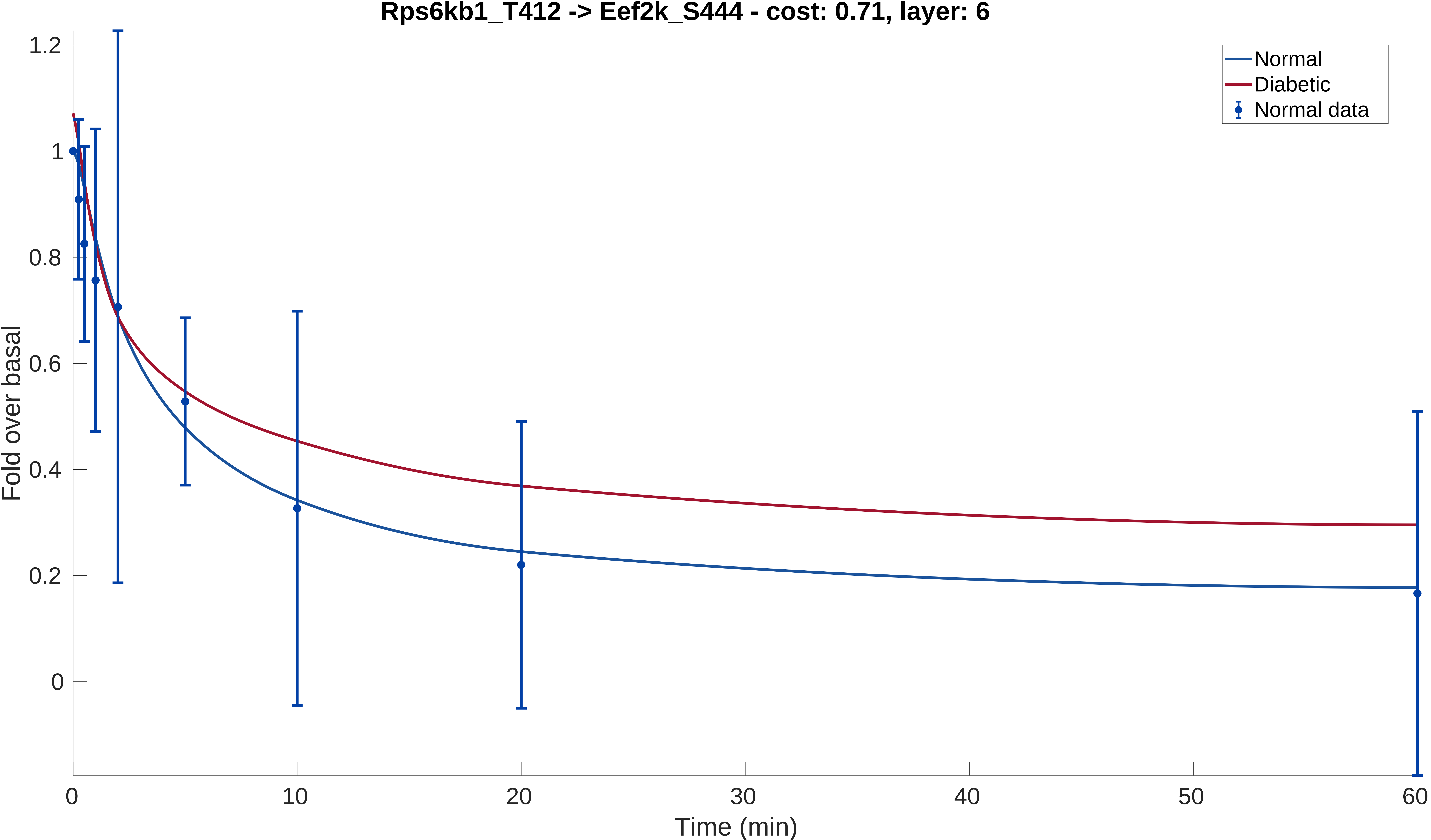

Rps6kb1_T412 -> Eef2k_S444 - cost: 0.71, layer: 6
Normal
Diabetic
Normal data
Fold over basal
1.2
1
0.8
0.6
0.4
0.2
0
Time (min)
0
10
20
30
40
50
60

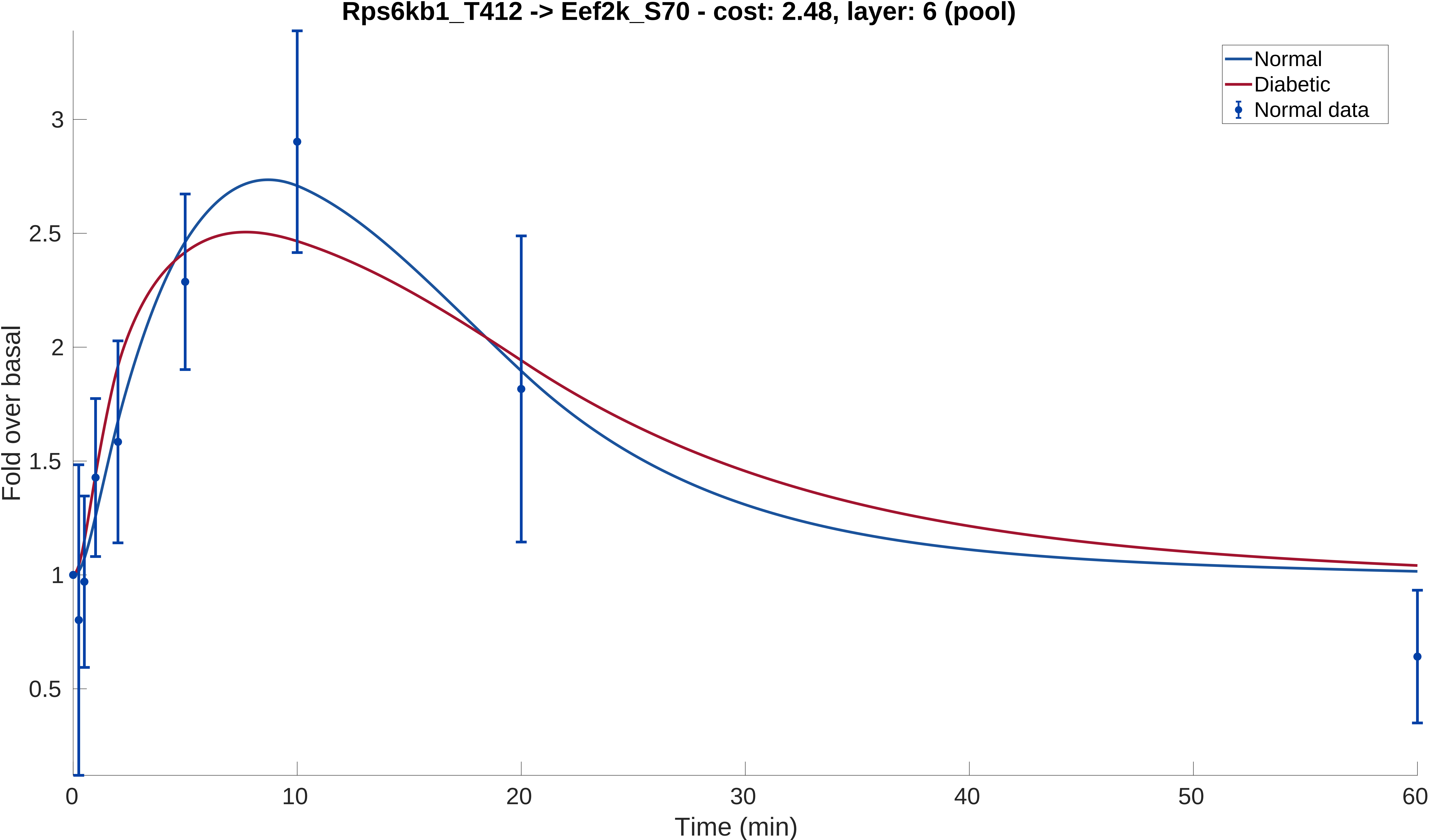
Rps6kb1_T412 -> Eef2k_S70 - cost: 2.48, layer: 6 (pool)
Normal
Diabetic
Normal data
Fold over basal
Time (min)
0
10
20
30
40
50
60
0.5
1
1.5
2
2.5
3

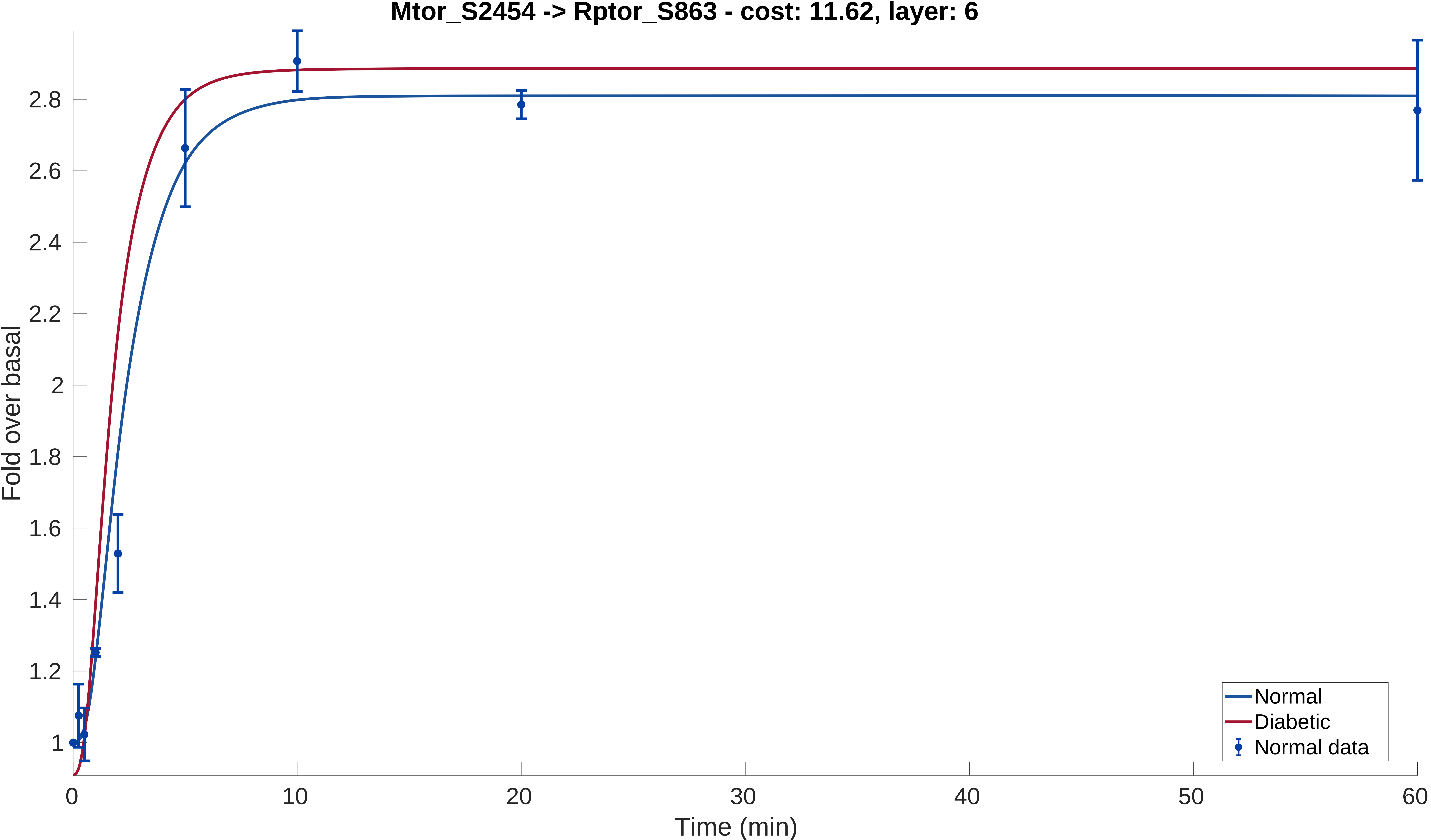

Mtor_S2454 -> Rptor_S863 - cost: 11.62, layer: 6
Fold over basal
Time (min)
2.8
2.6
2.4
2.2
2
1.8
1.6
1.4
1.2
1
0
10
20
30
40
50
60
Normal
Diabetic
Normal data

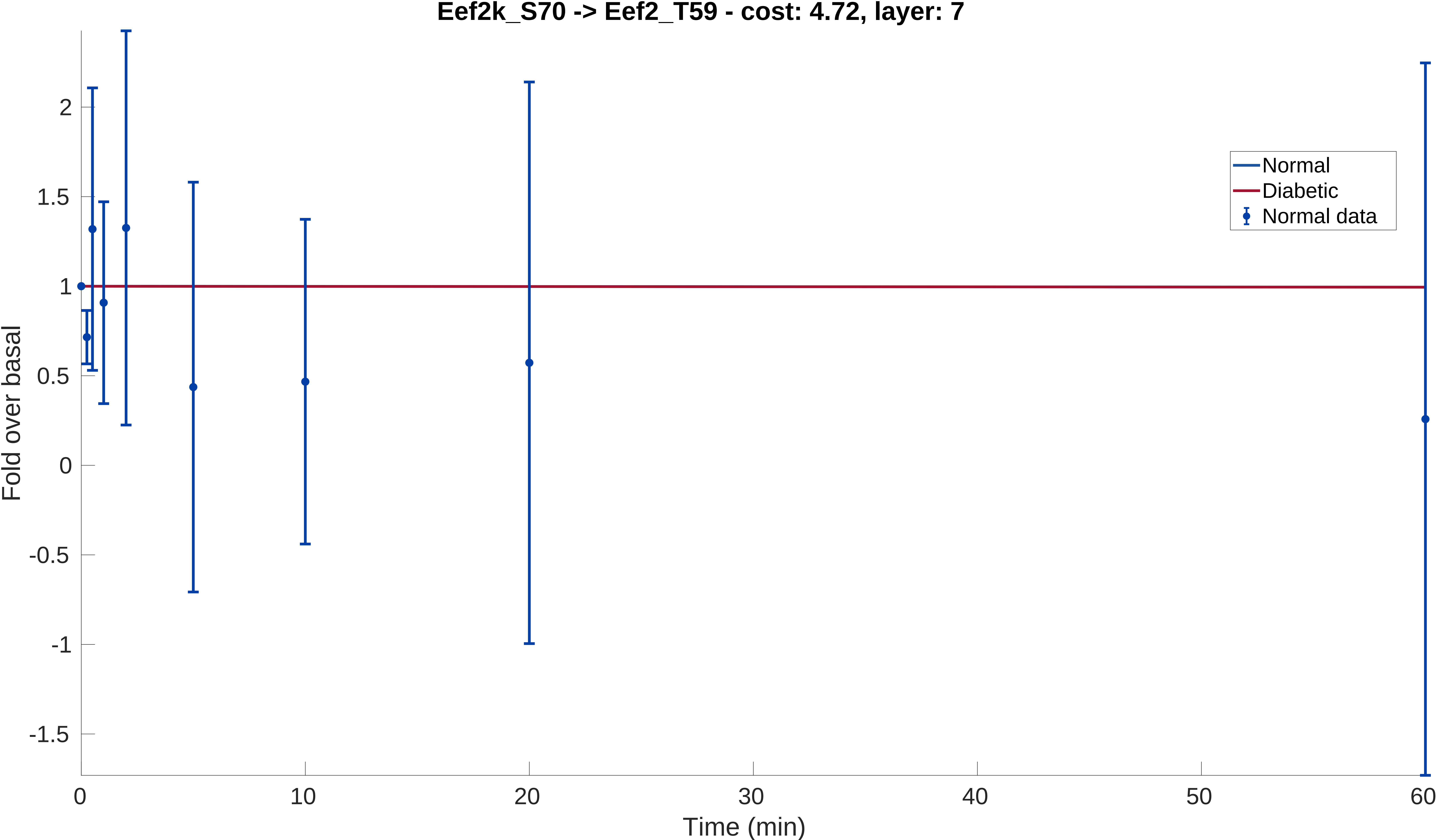

Eef2k_S70 -> Eef2_T59 - cost: 4.72, layer: 7
Fold over basal
Time (min)
Normal
Diabetic
Normal data
2
1.5
1
0.5
0
-0.5
-1
-1.5
0
10
20
30
40
50
60

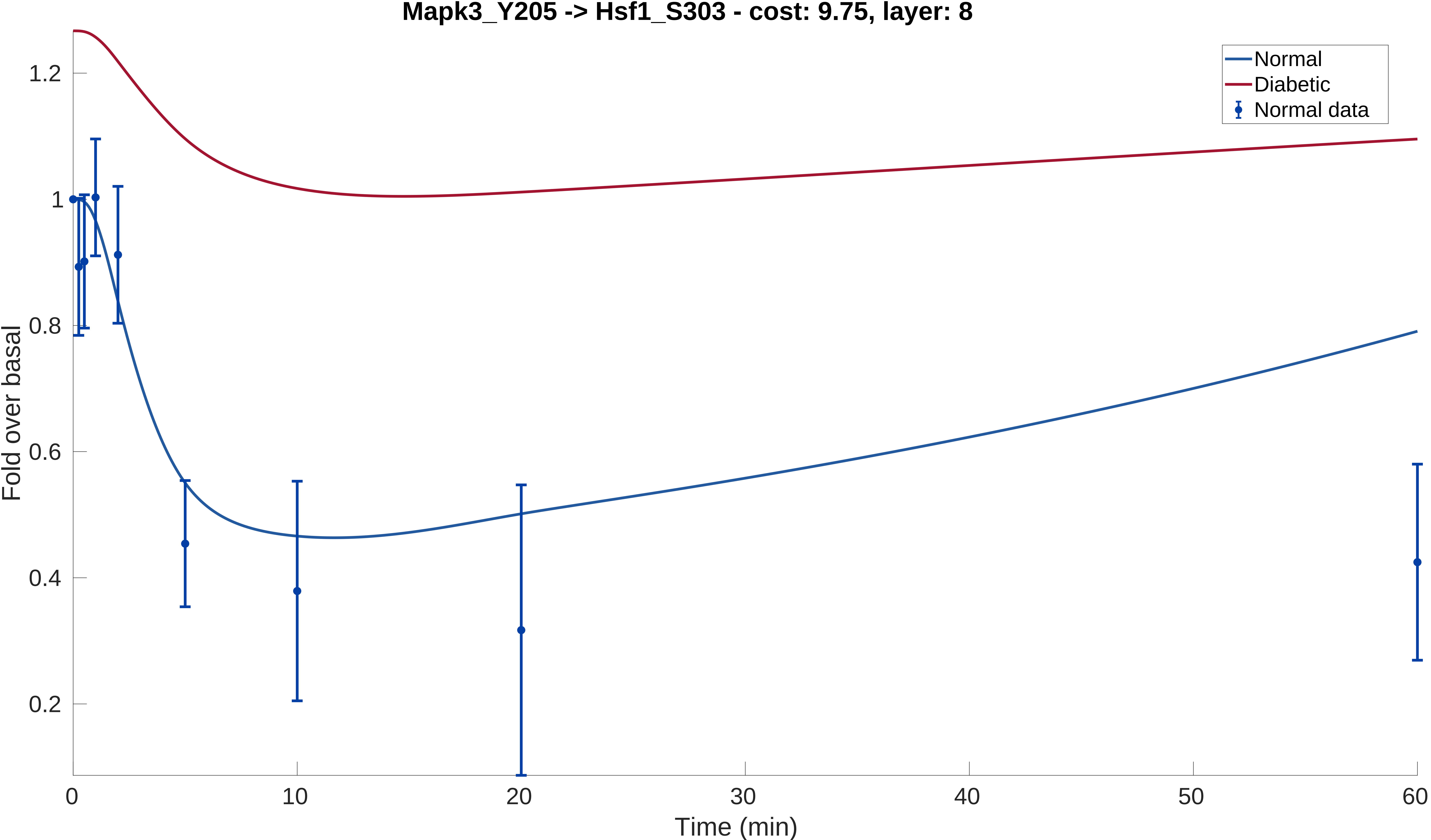

Mapk3_Y205 -> Hsf1_S303 - cost: 9.75, layer: 8
Normal
Diabetic
Normal data
Fold over basal
Time (min)
1.2
1
0.8
0.6
0.4
0.2
0
10
20
30
40
50
60

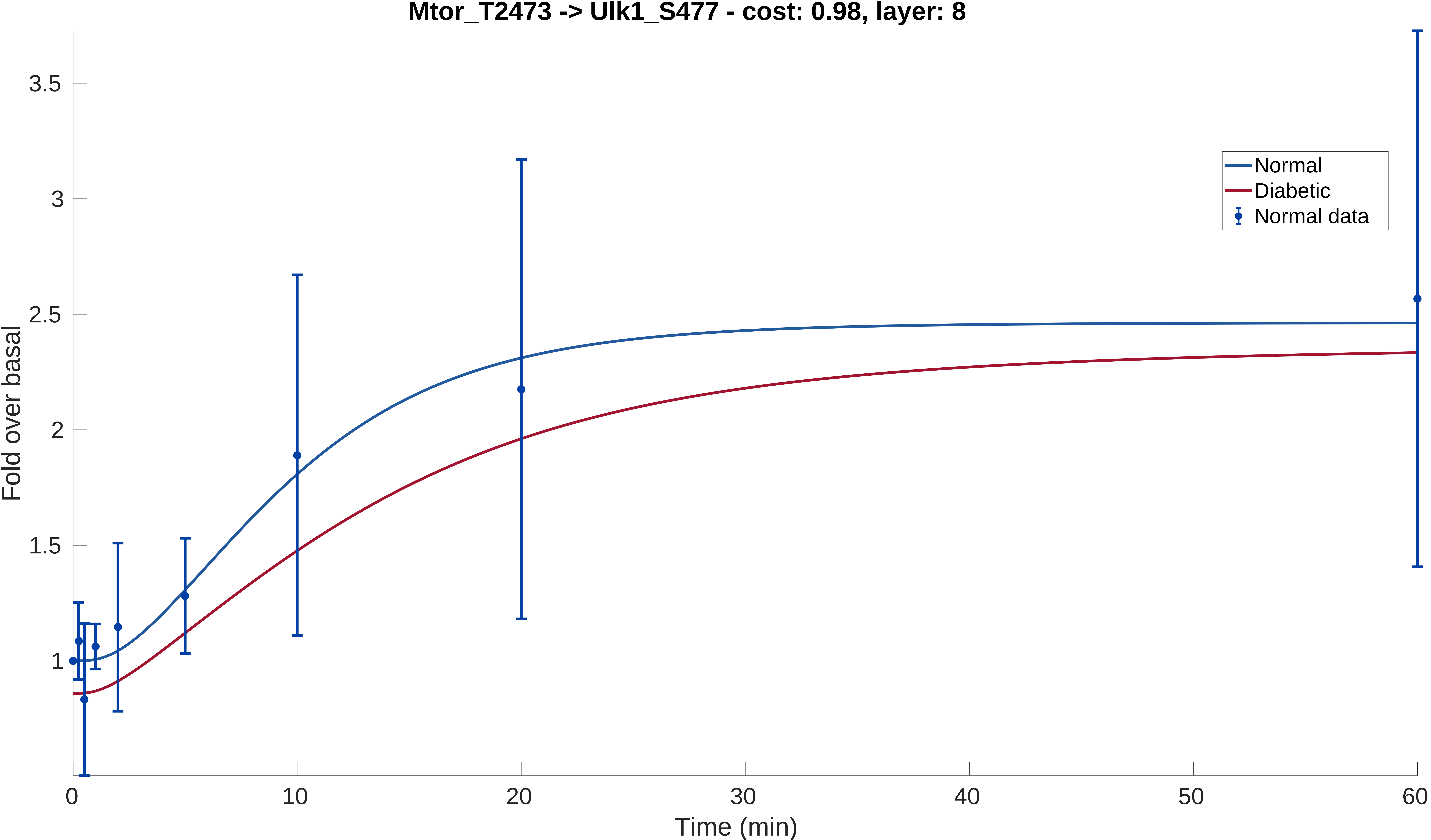

Mtor_T2473 -> Ulk1_S477 - cost: 0.98, layer: 8
Normal
Diabetic
Normal data
Fold over basal
3.5
3
2.5
2
1.5
1
0
10
20
30
40
50
60
Time (min)

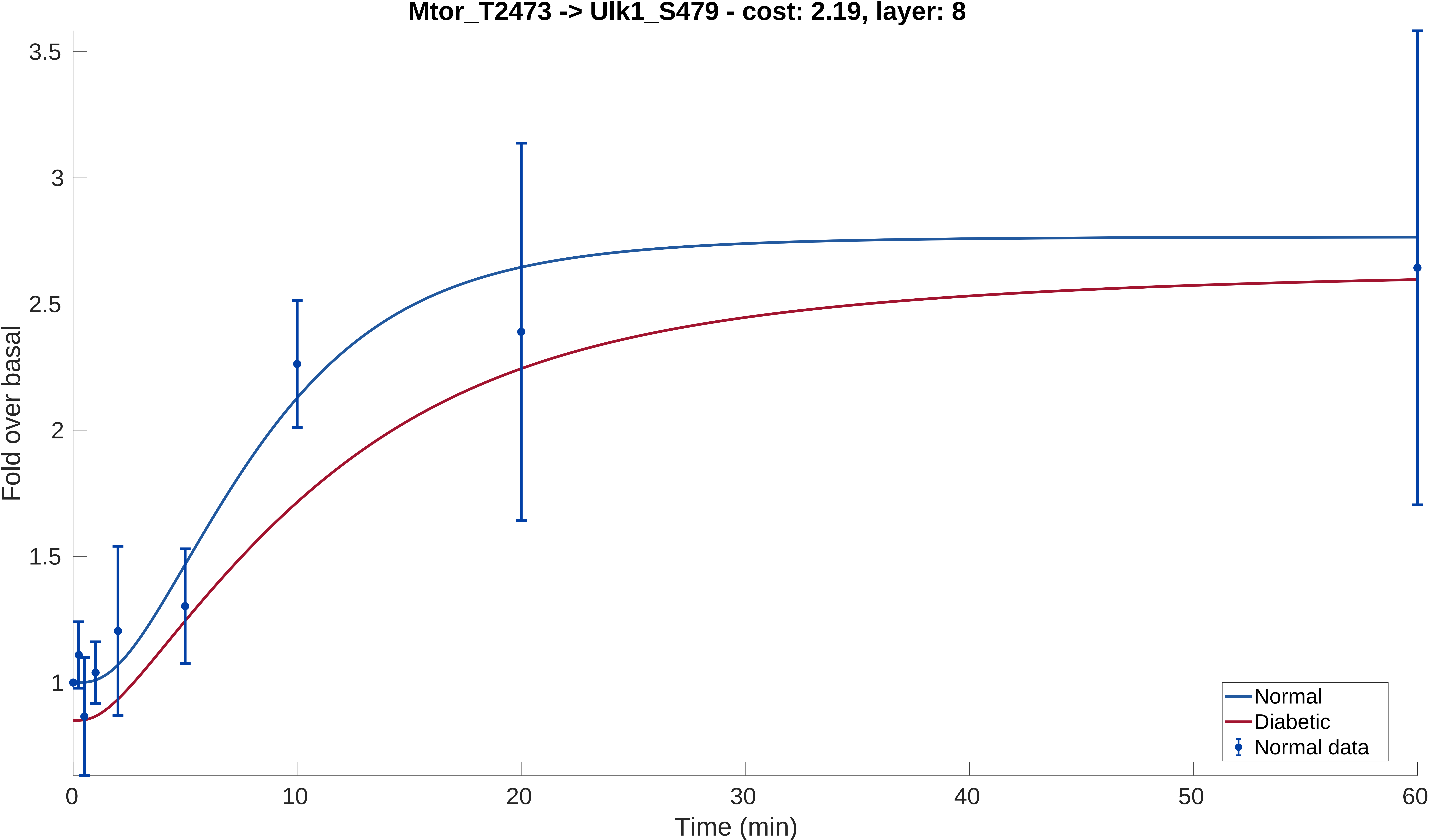

Mtor_T2473 -> Ulk1_S479 - cost: 2.19, layer: 8
Fold over basal
3.5
3
2.5
2
1.5
1
0
10
20
30
40
50
60
Time (min)
Normal
Diabetic
Normal data

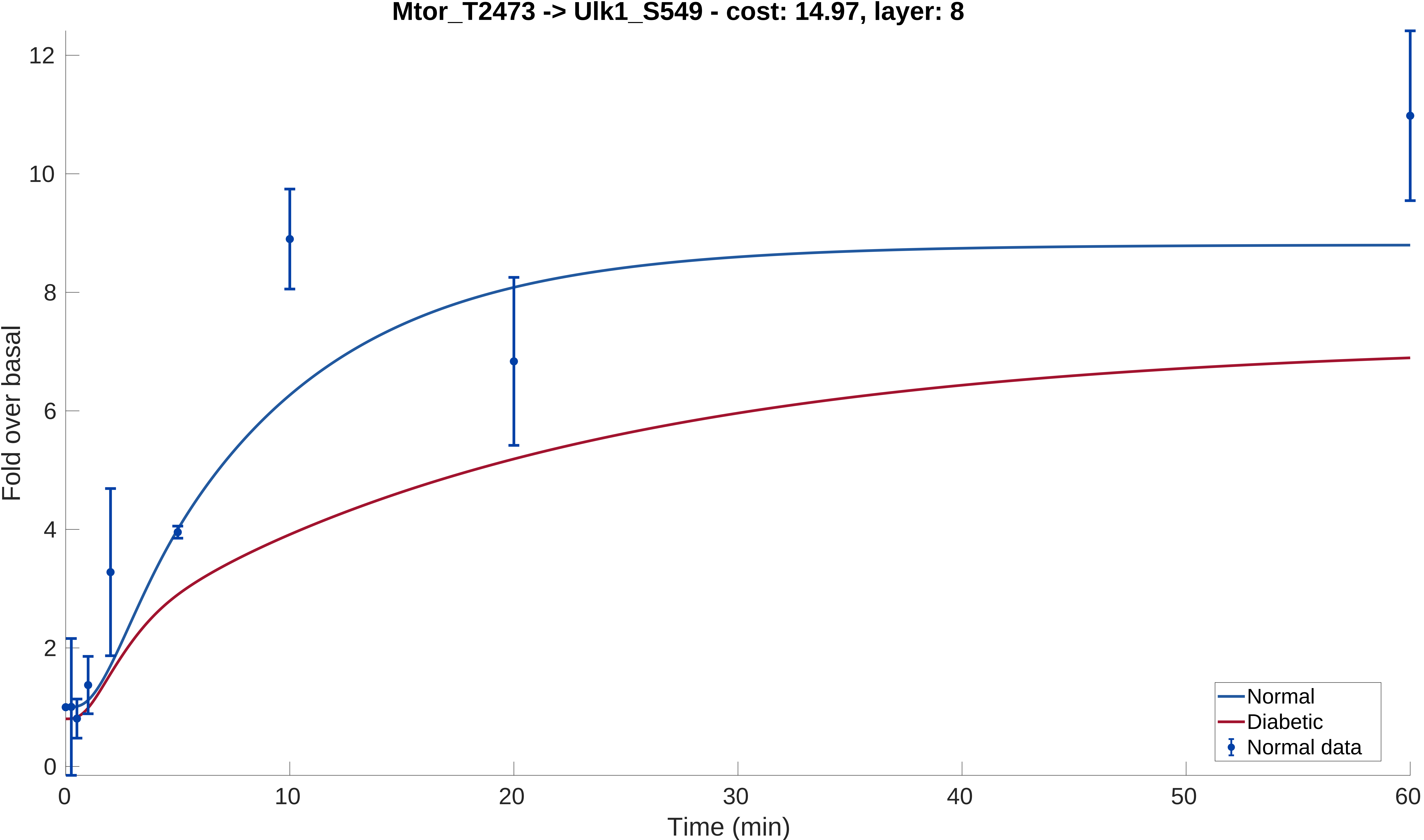

Mtor_T2473 -> Ulk1_S549 - cost: 14.97, layer: 8
Fold over basal
Time (min)
0
2
4
6
8
10
12
0
10
20
30
40
50
60
Normal
Diabetic
Normal data

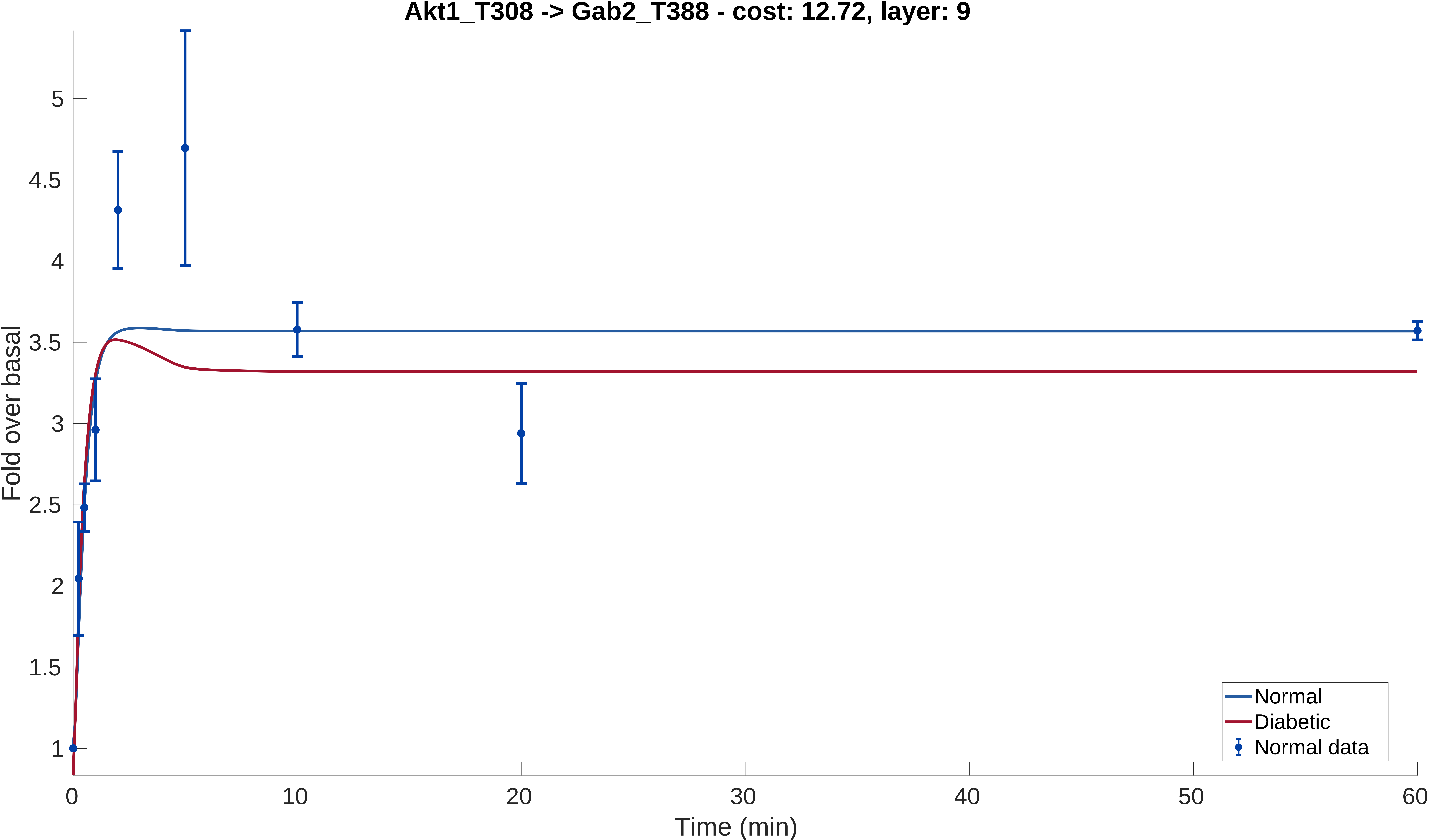

Akt1_T308 -> Gab2_T388 - cost: 12.72, layer: 9
Fold over basal
Time (min)
5
4.5
4
3.5
3
2.5
2
1.5
1
0
10
20
30
40
50
60
Normal
Diabetic
Normal data

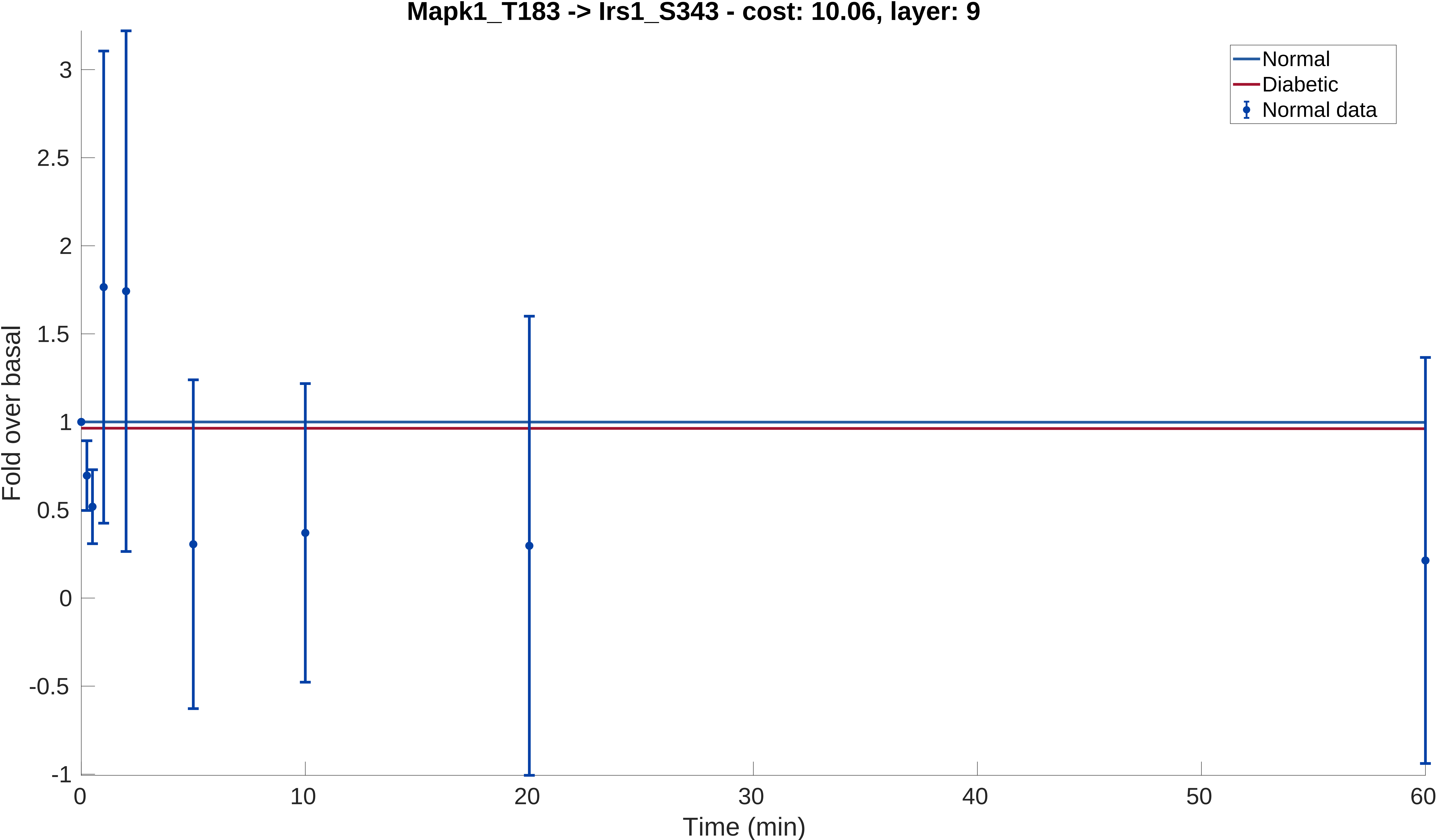
Mapk1_T183 -> Irs1_S343 - cost: 10.06, layer: 9
Fold over basal
Time (min)
Normal
Diabetic
Normal data
3
2.5
2
1.5
1
0.5
0
-0.5
-1
0
10
20
30
40
50
60

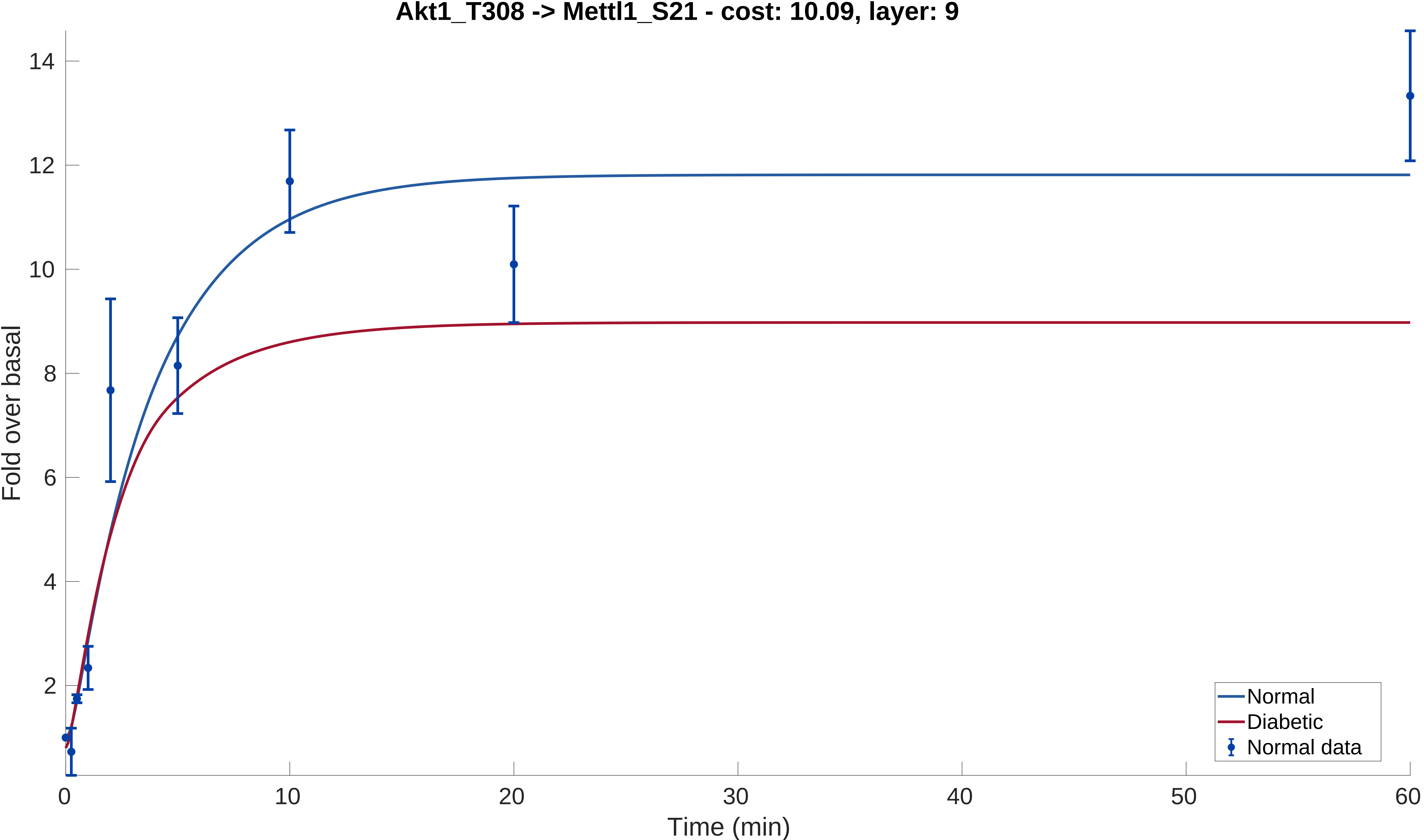

Akt1_T308 -> Mettl1_S21 - cost: 10.09, layer: 9
Fold over basal
Time (min)
0
10
20
30
40
50
60
2
4
6
8
10
12
14
Normal
Diabetic
Normal data

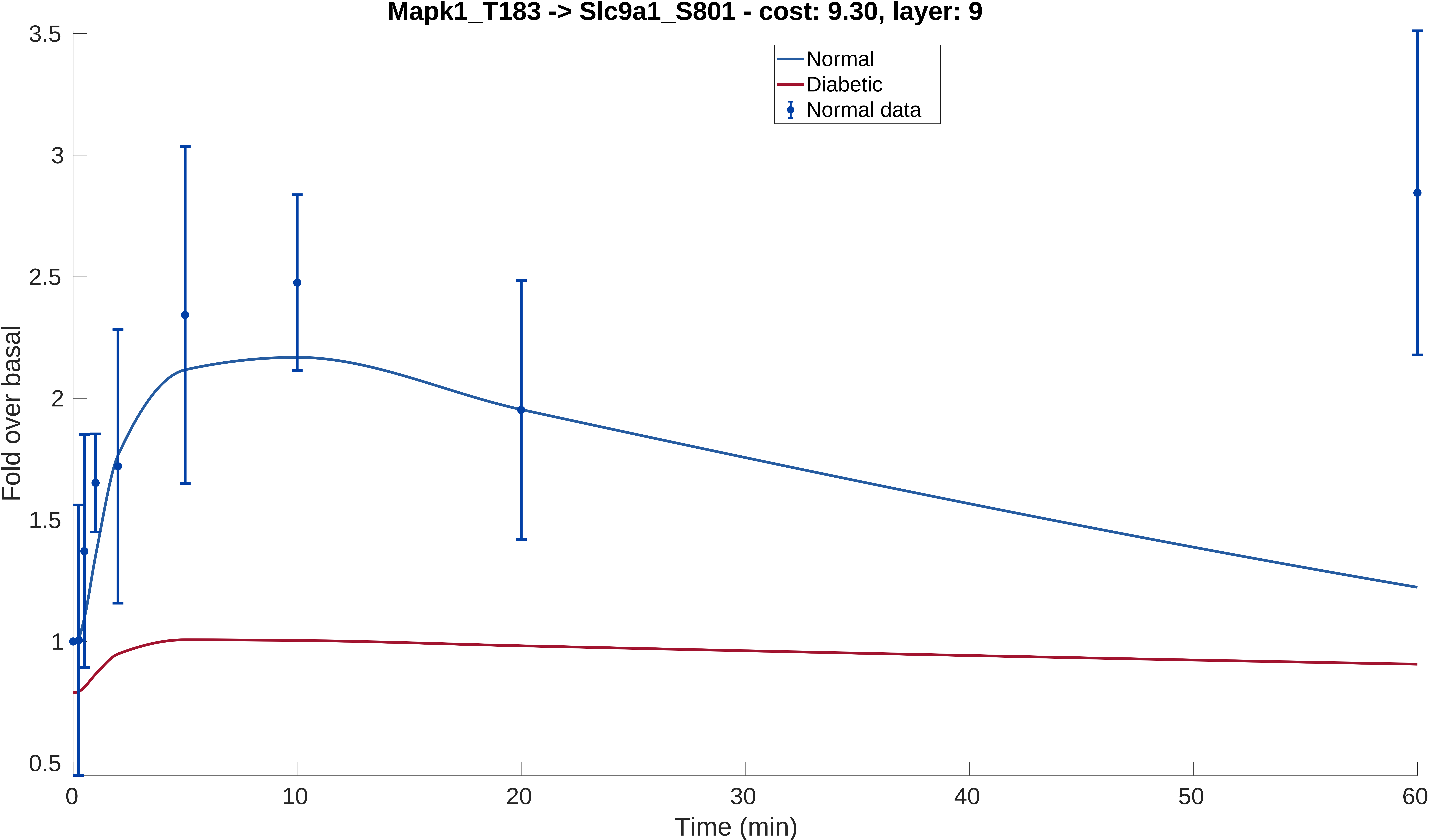
Mapk1_T183 -> Slc9a1_S801 - cost: 9.30, layer: 9
Normal
Diabetic
Normal data
Fold over basal
Time (min)
0.5
1
1.5
2
2.5
3
3.5
0
10
20
30
40
50
60

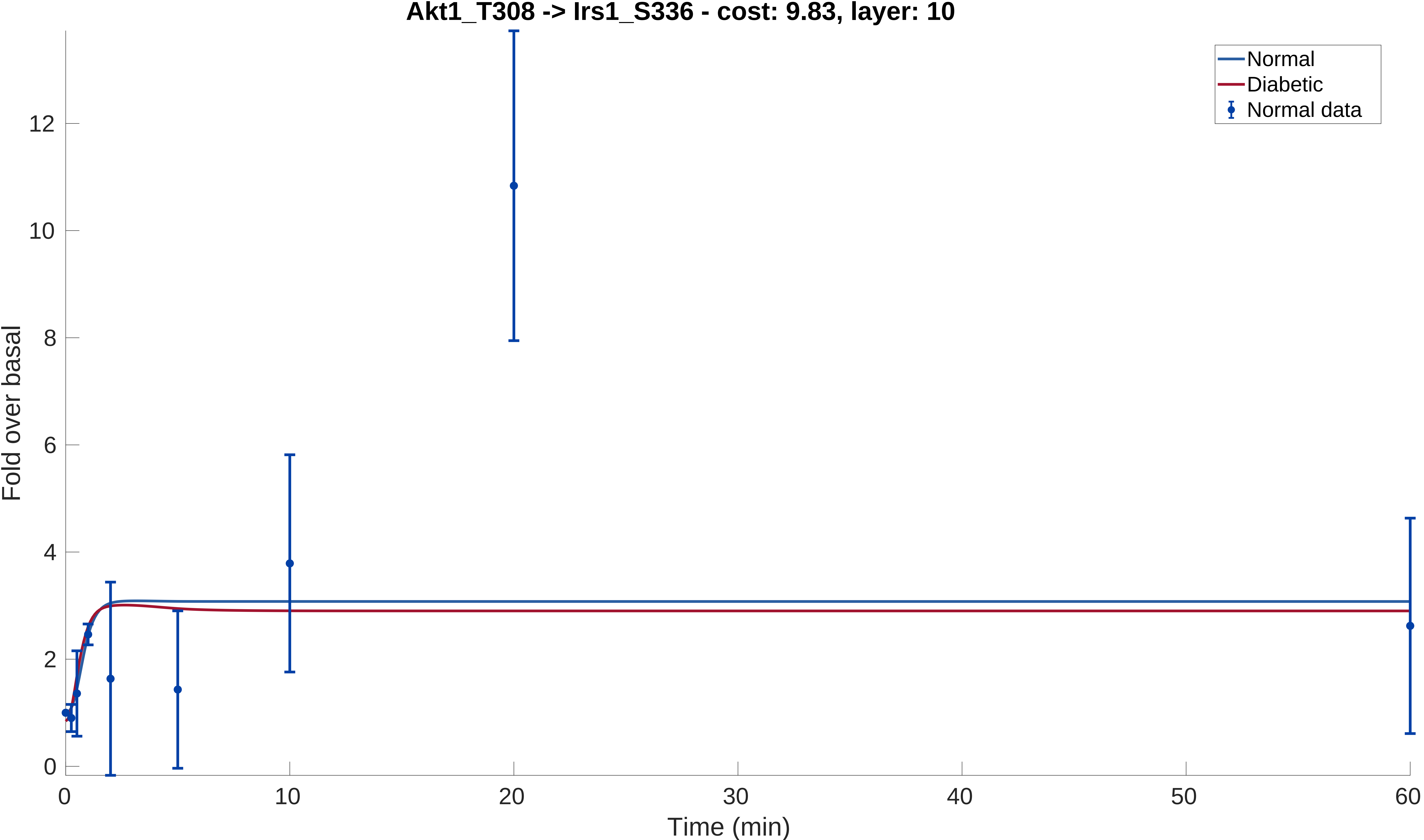
Akt1_T308 -> Irs1_S336 - cost: 9.83, layer: 10
Normal
Diabetic
Normal data
Fold over basal
0
2
4
6
8
10
12
0
10
20
30
40
50
60
Time (min)

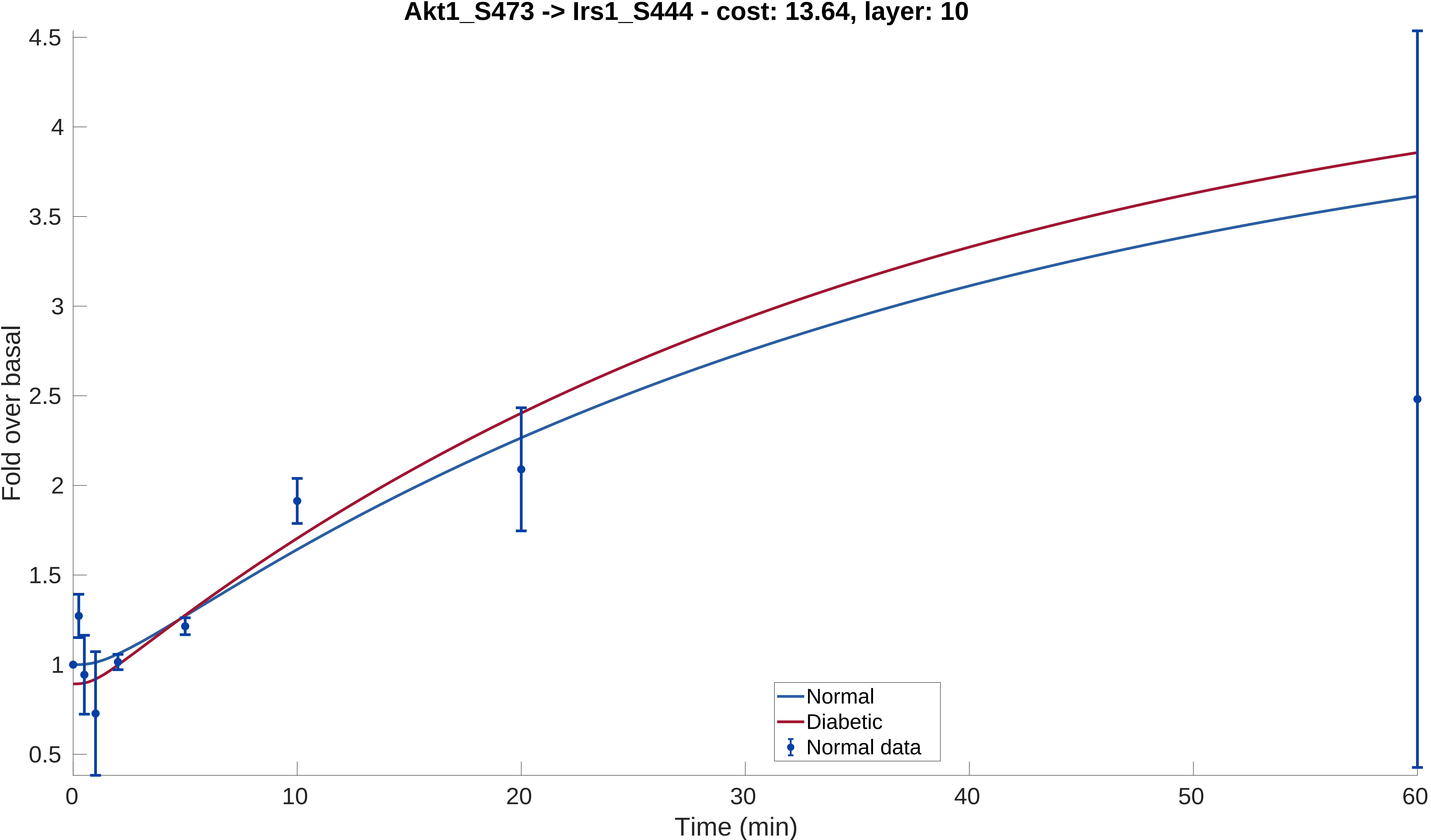

Akt1_S473 -> Irs1_S444 - cost: 13.64, layer: 10
Fold over basal
4.5
4
3.5
3
2.5
2
1.5
1
0.5
0
10
20
30
40
50
60
Time (min)
Normal
Diabetic
Normal data

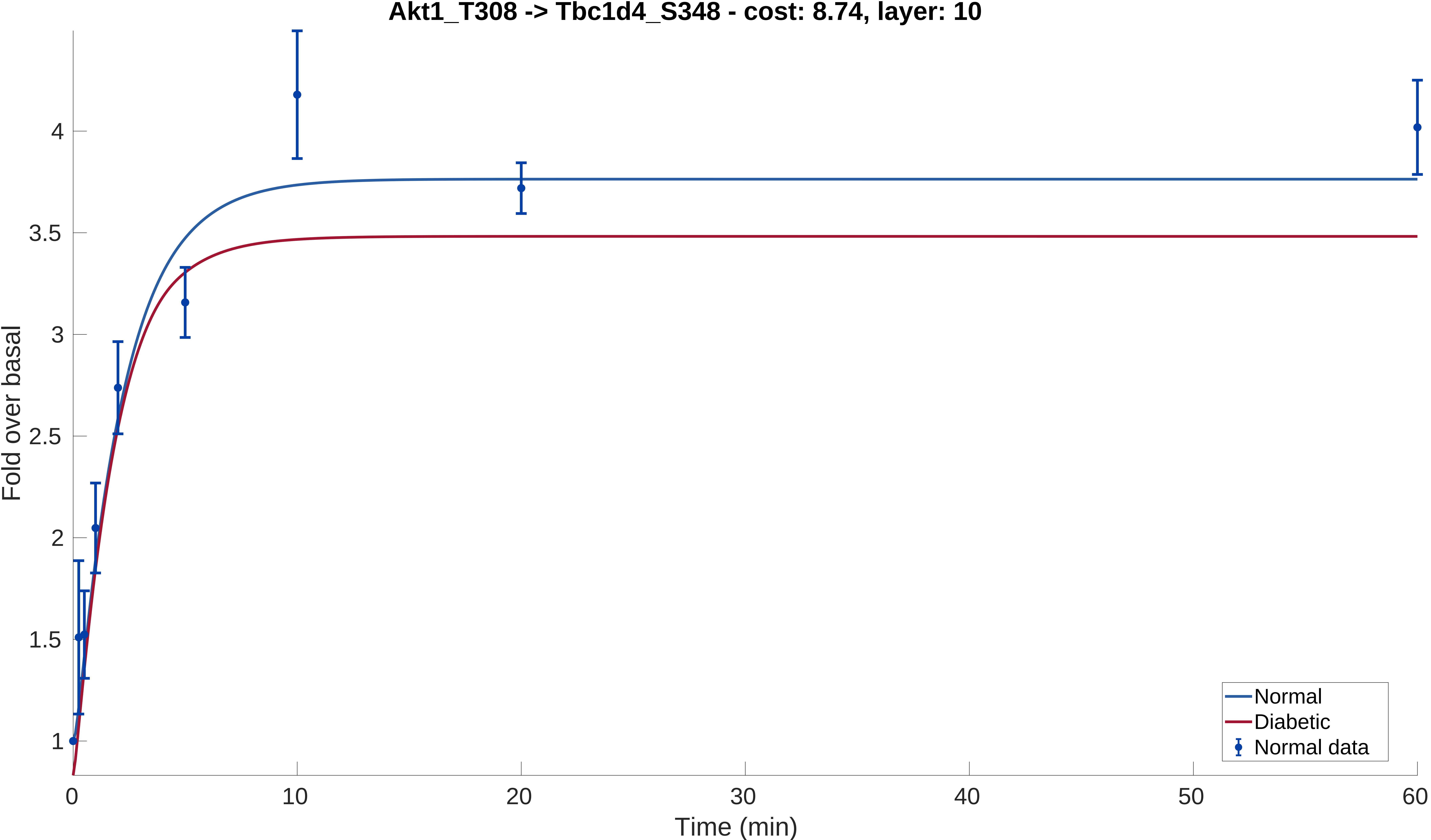
Akt1_T308 -> Tbc1d4_S348 - cost: 8.74, layer: 10
Fold over basal
Time (min)
0
10
20
30
40
50
60
1
1.5
2
2.5
3
3.5
4
Normal
Diabetic
Normal data

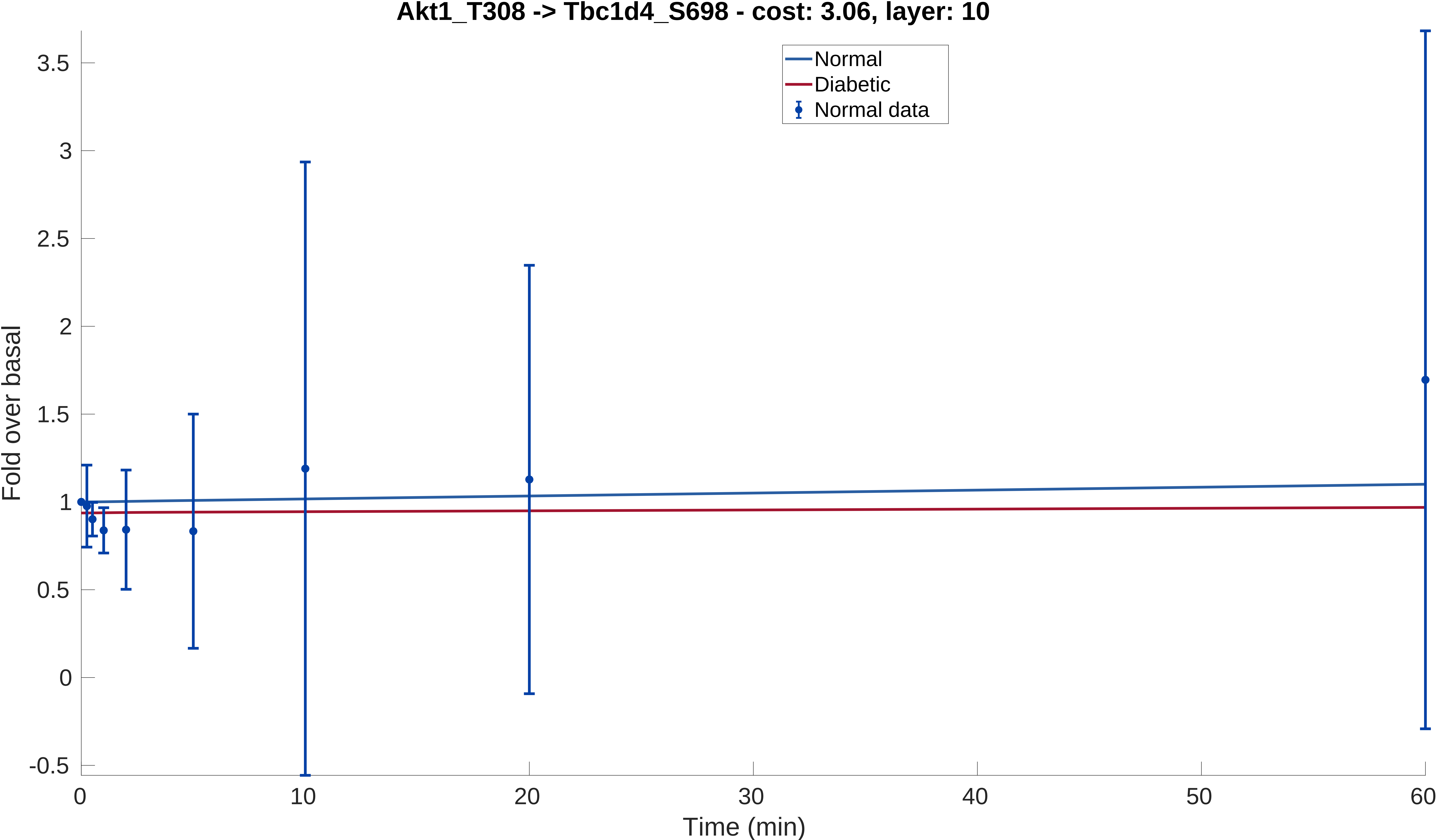

Akt1_T308 -> Tbc1d4_S698 - cost: 3.06, layer: 10
Normal
Diabetic
Normal data
Fold over basal
3.5
3
2.5
2
1.5
1
0.5
0
-0.5
0
10
20
30
40
50
60
Time (min)

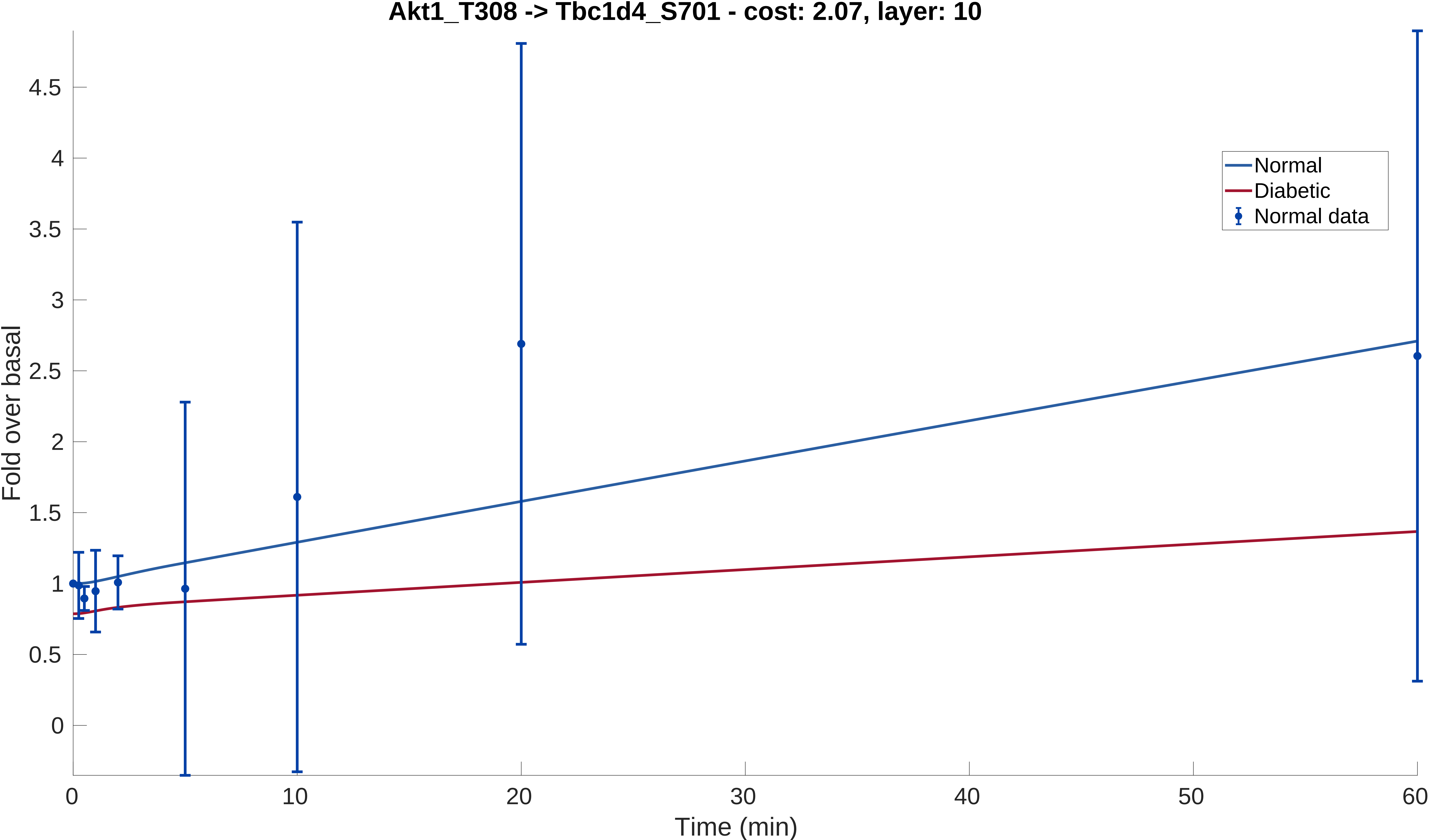

Akt1_T308 -> Tbc1d4_S701 - cost: 2.07, layer: 10
Fold over basal
Time (min)
Normal
Diabetic
Normal data
0
0.5
1
1.5
2
2.5
3
3.5
4
4.5
0
10
20
30
40
50
60

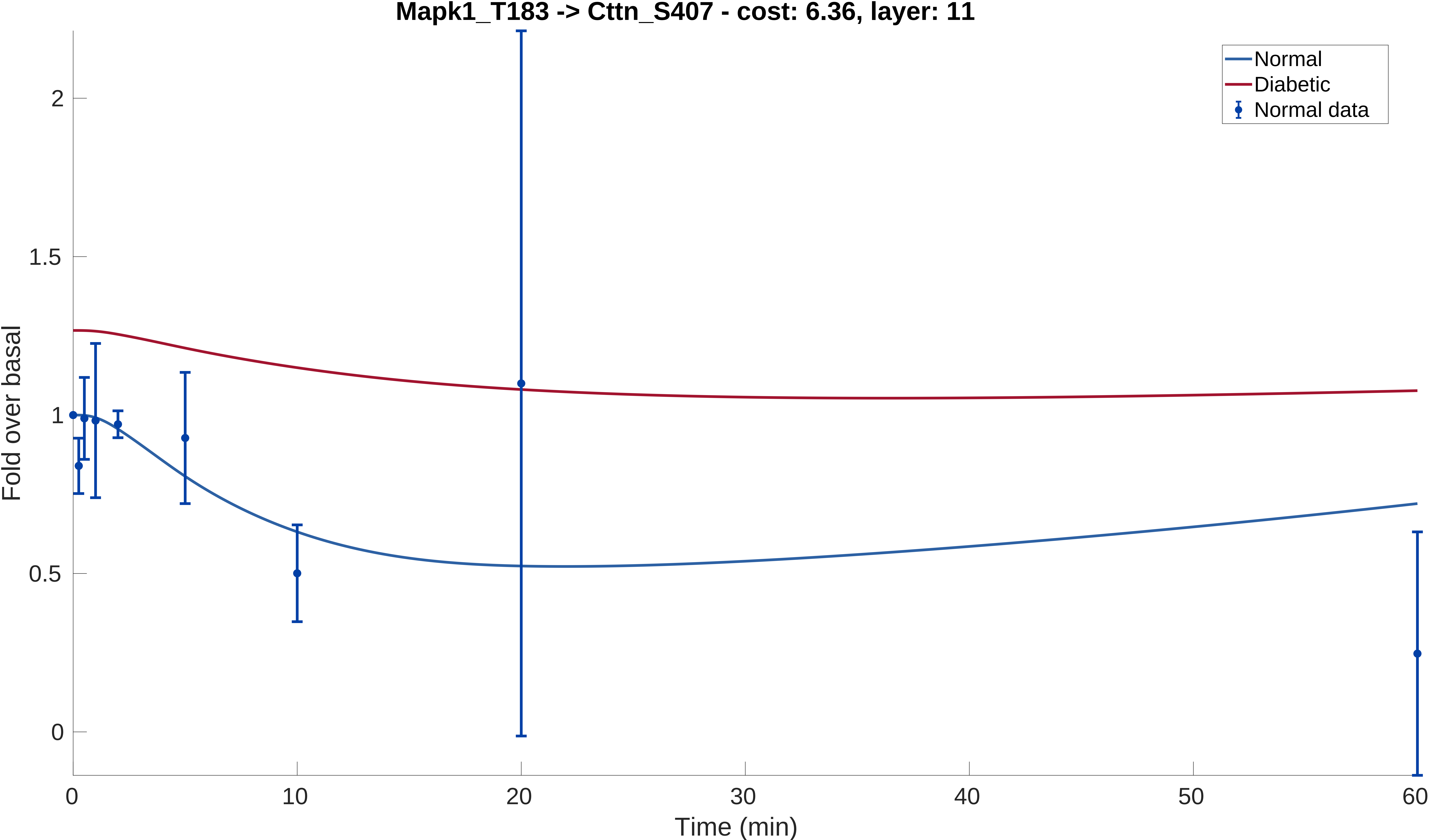

Mapk1_T183 -> Cttn_S407 - cost: 6.36, layer: 11
Normal
Diabetic
Normal data
Fold over basal
2
1.5
1
0.5
0
0
10
20
30
40
50
60
Time (min)

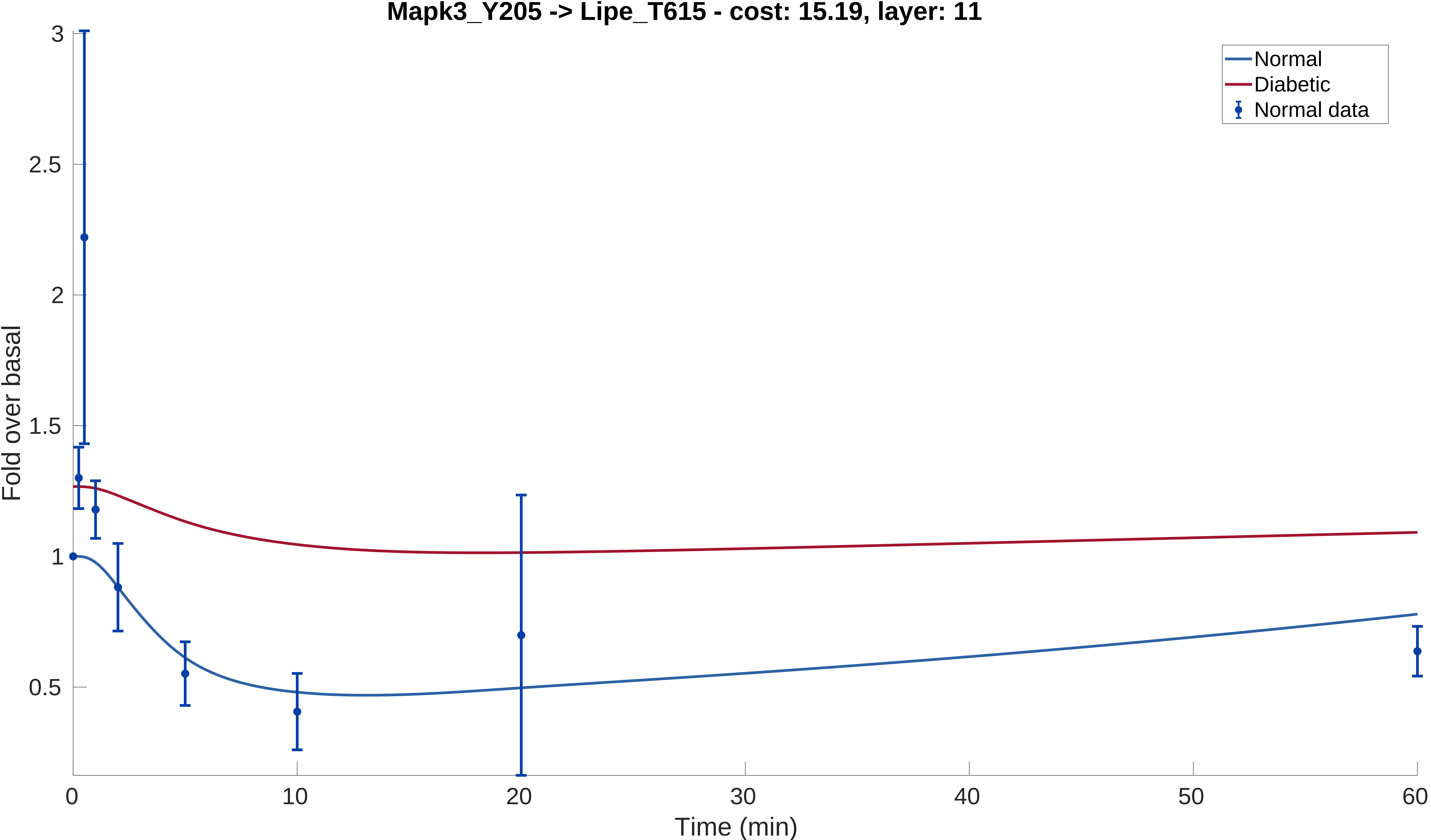

Mapk3_Y205 -> Lipe_T615 - cost: 15.19, layer: 11
Normal
Diabetic
Normal data
Fold over basal
Time (min)
0
10
20
30
40
50
60
0.5
1
1.5
2
2.5
3

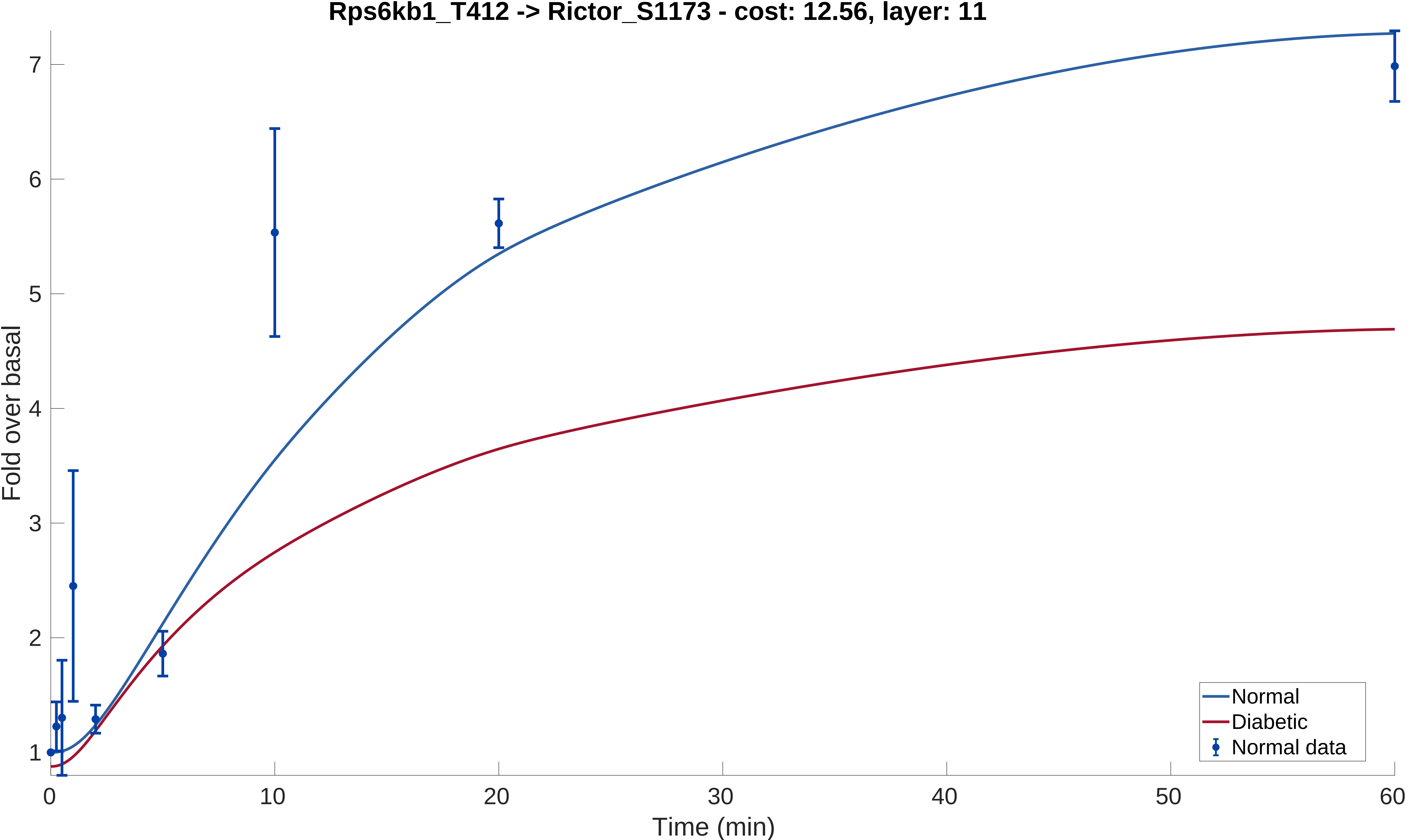

Rps6kb1_T412 -> Rictor_S1173 - cost: 12.56, layer: 11
Fold over basal
Time (min)
Normal
Diabetic
Normal data
0
10
20
30
40
50
60
1
2
3
4
5
6
7

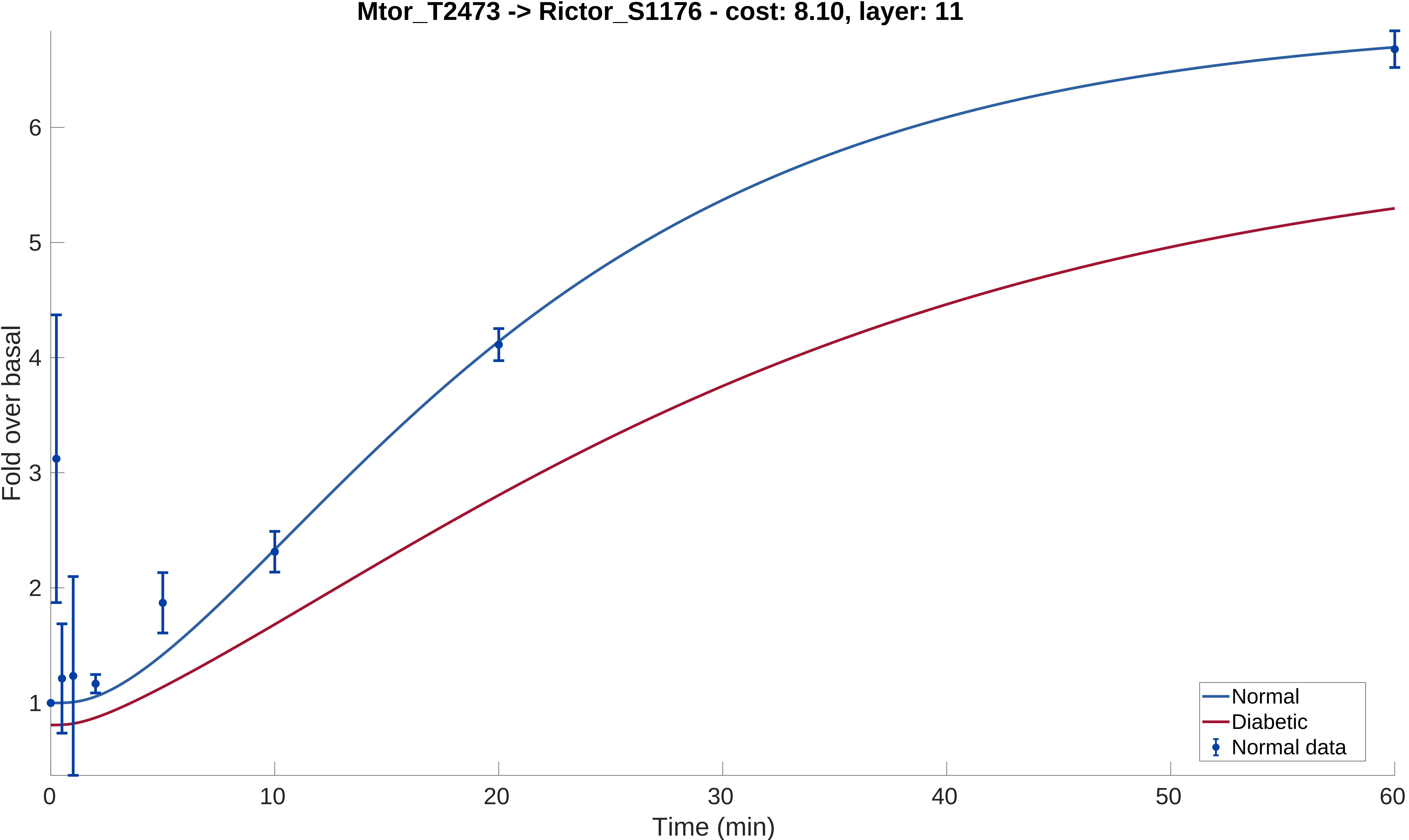
Mtor_T2473 -> Rictor_S1176 - cost: 8.10, layer: 11
Fold over basal
Time (min)
1
2
3
4
5
6
0
10
20
30
40
50
60
Normal
Diabetic
Normal data

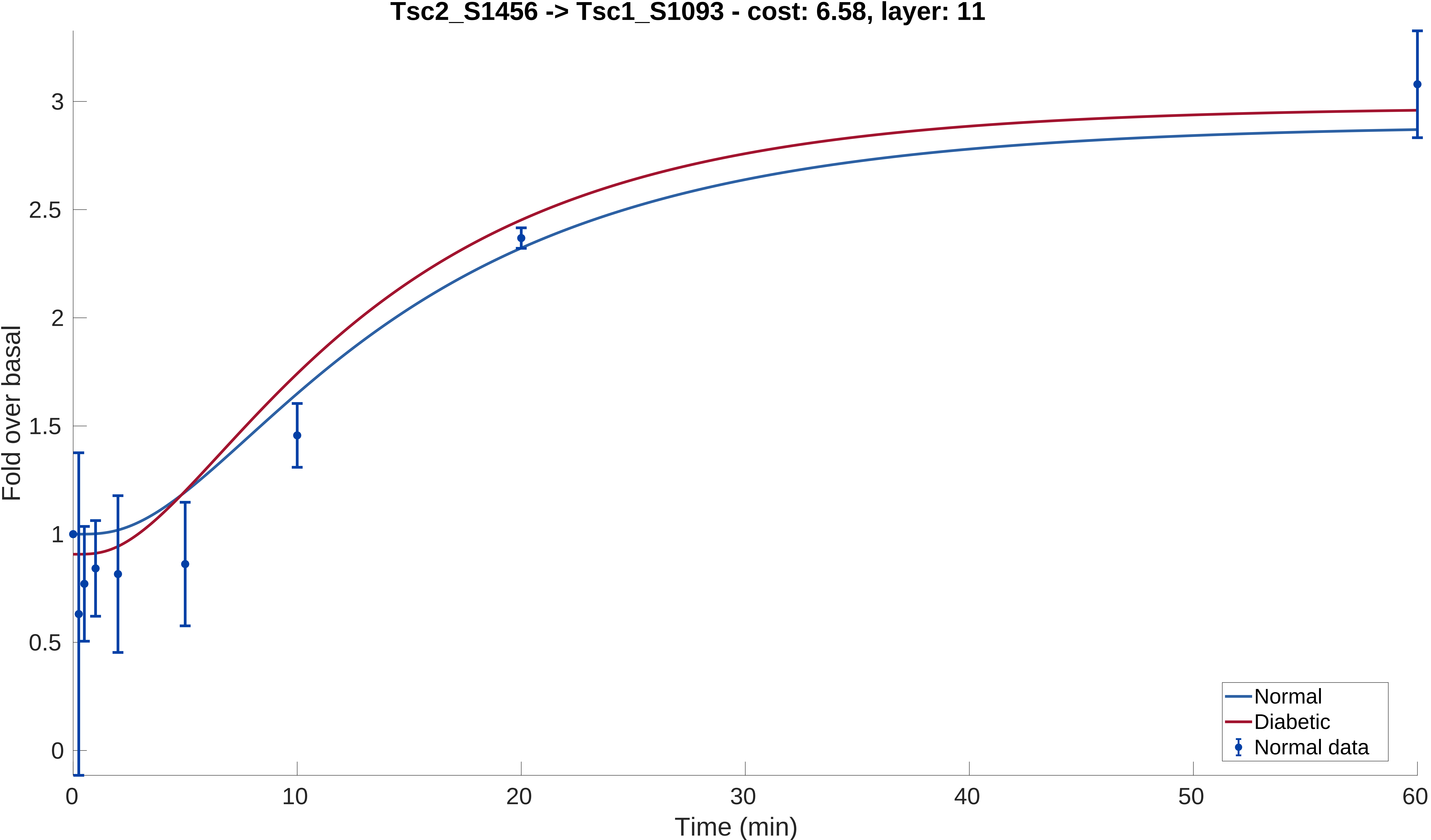

Tsc2_S1456 -> Tsc1_S1093 - cost: 6.58, layer: 11
Fold over basal
3
2.5
2
1.5
1
0.5
0
0
10
20
30
40
50
60
Time (min)
Normal
Diabetic
Normal data

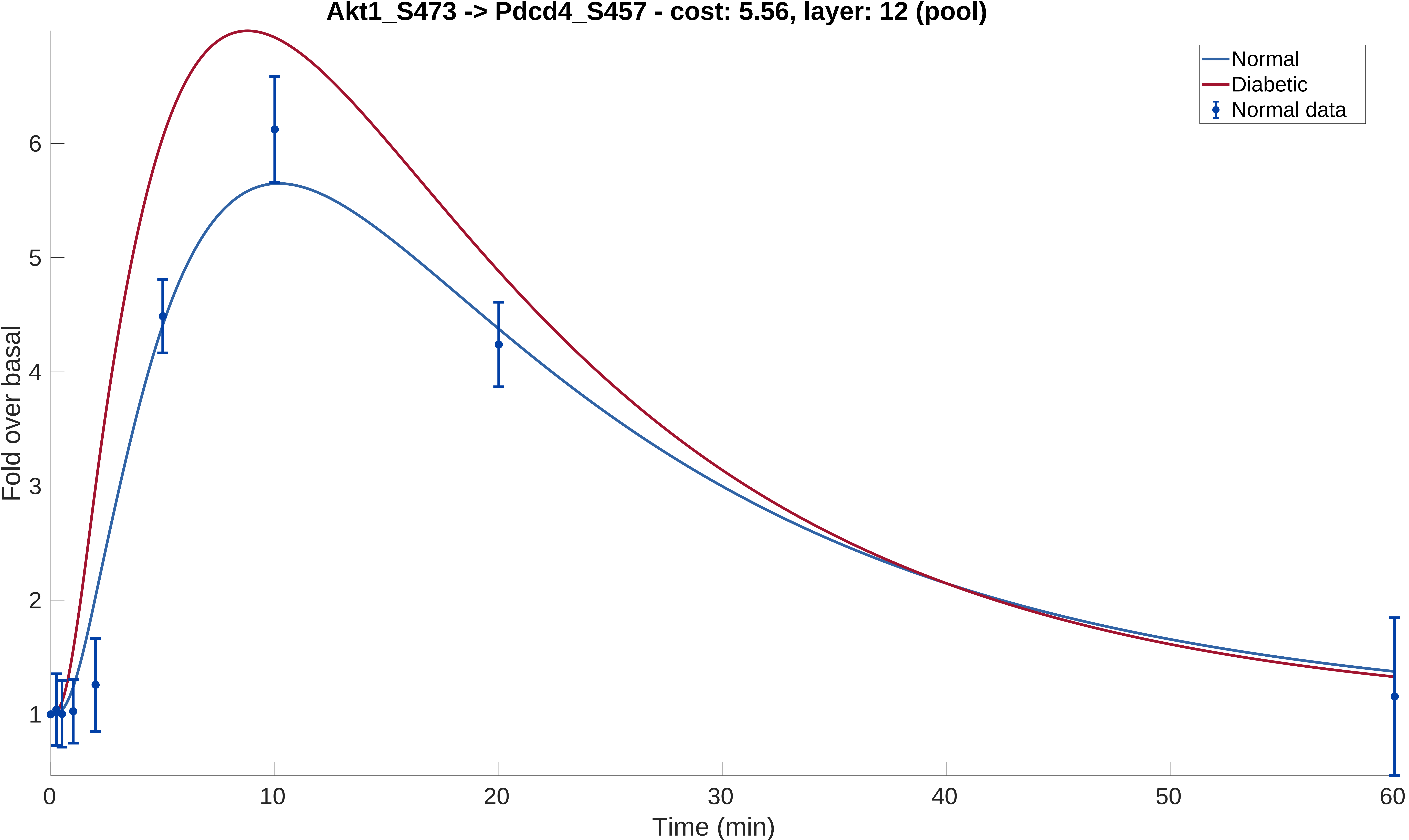
Akt1_S473 -> Pdcd4_S457 - cost: 5.56, layer: 12 (pool)
Normal
Diabetic
Normal data
Fold over basal
1
2
3
4
5
6
Time (min)
0
10
20
30
40
50
60

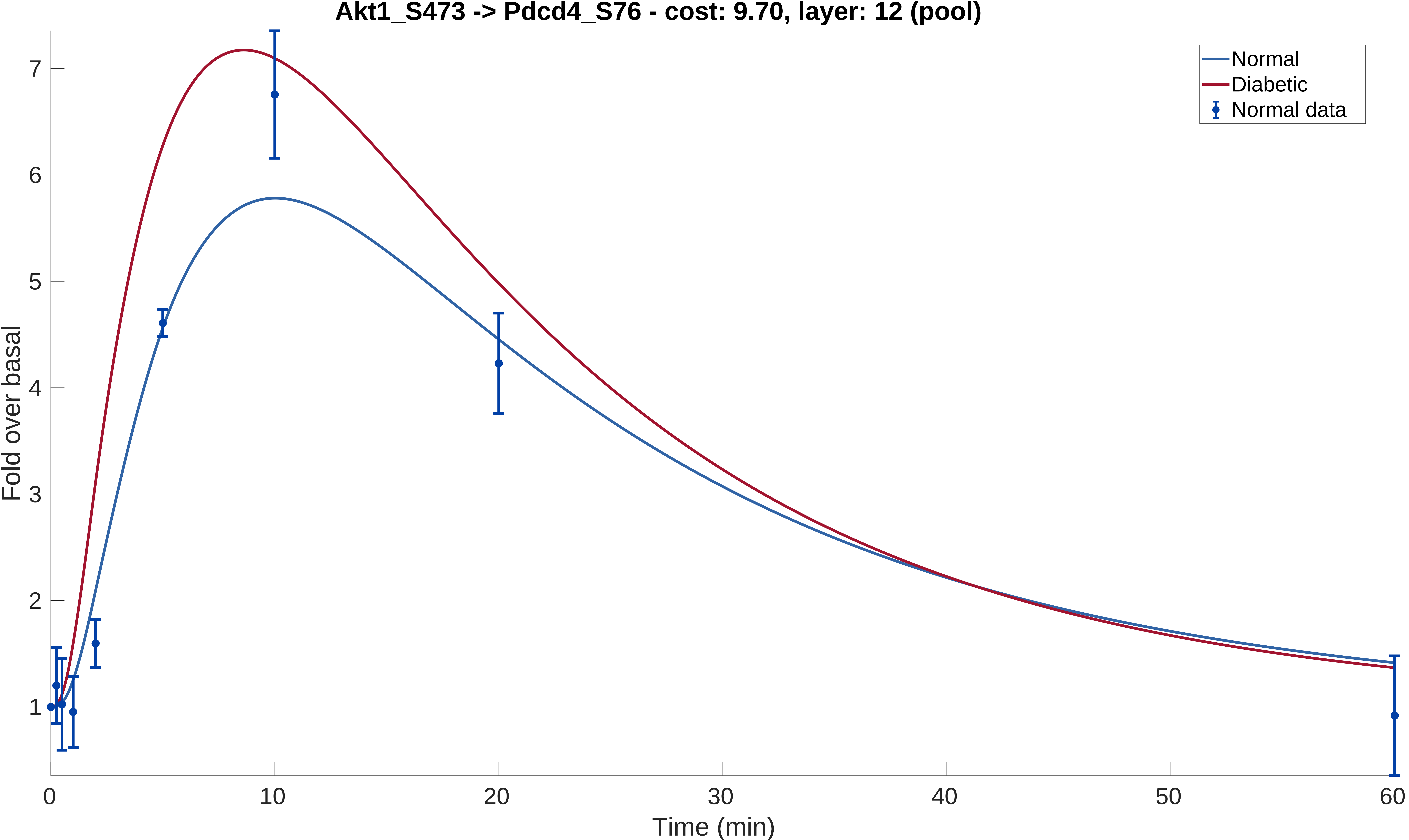

Akt1_S473 -> Pdcd4_S76 - cost: 9.70, layer: 12 (pool)
Normal
Diabetic
Normal data
Fold over basal
Time (min)
1
2
3
4
5
6
7
0
10
20
30
40
50
60

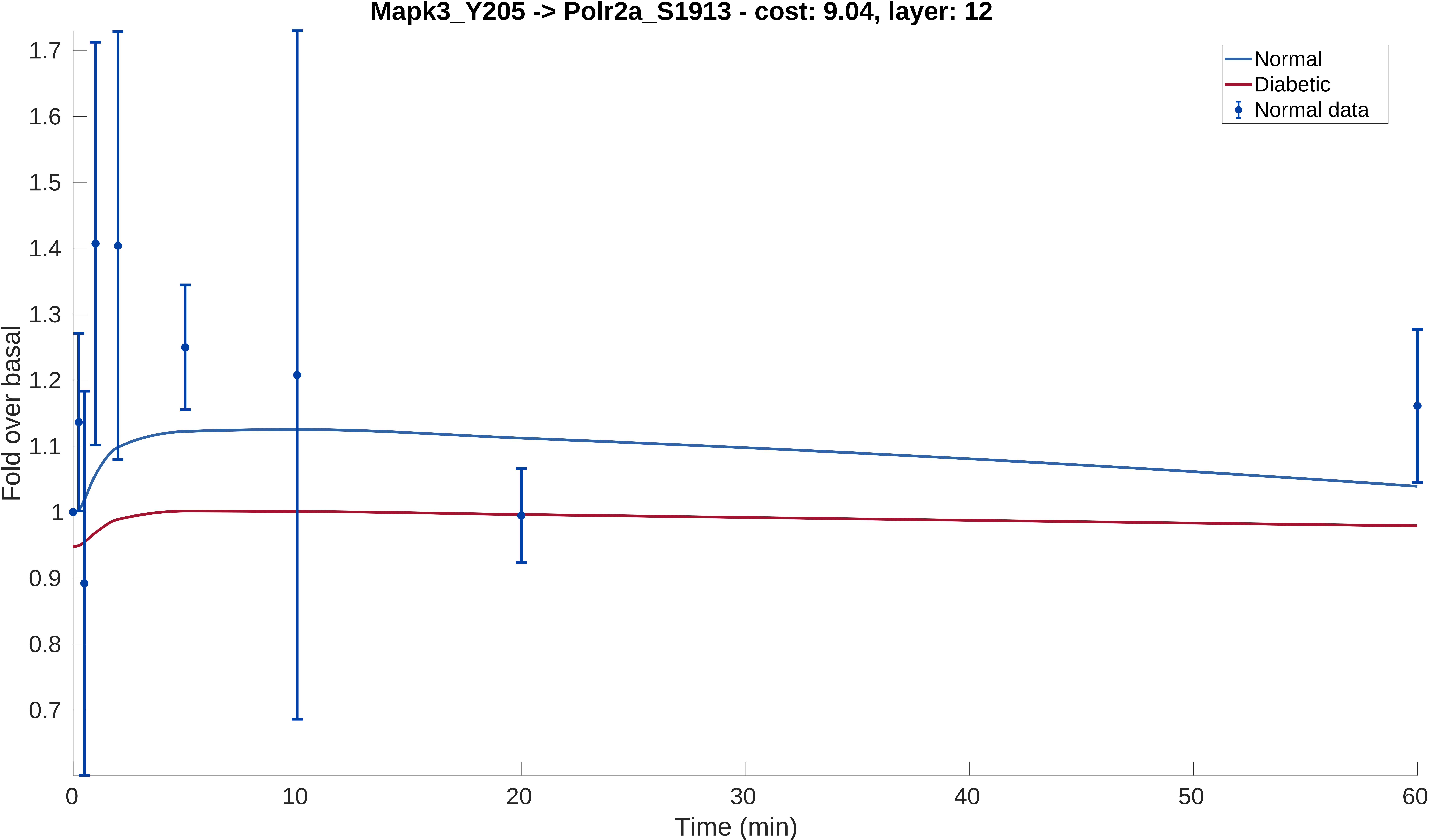

Mapk3_Y205 -> Polr2a_S1913 - cost: 9.04, layer: 12
Normal
Diabetic
Normal data
Fold over basal
Time (min)
0.7
0.8
0.9
1
1.1
1.2
1.3
1.4
1.5
1.6
1.7
0
10
20
30
40
50
60

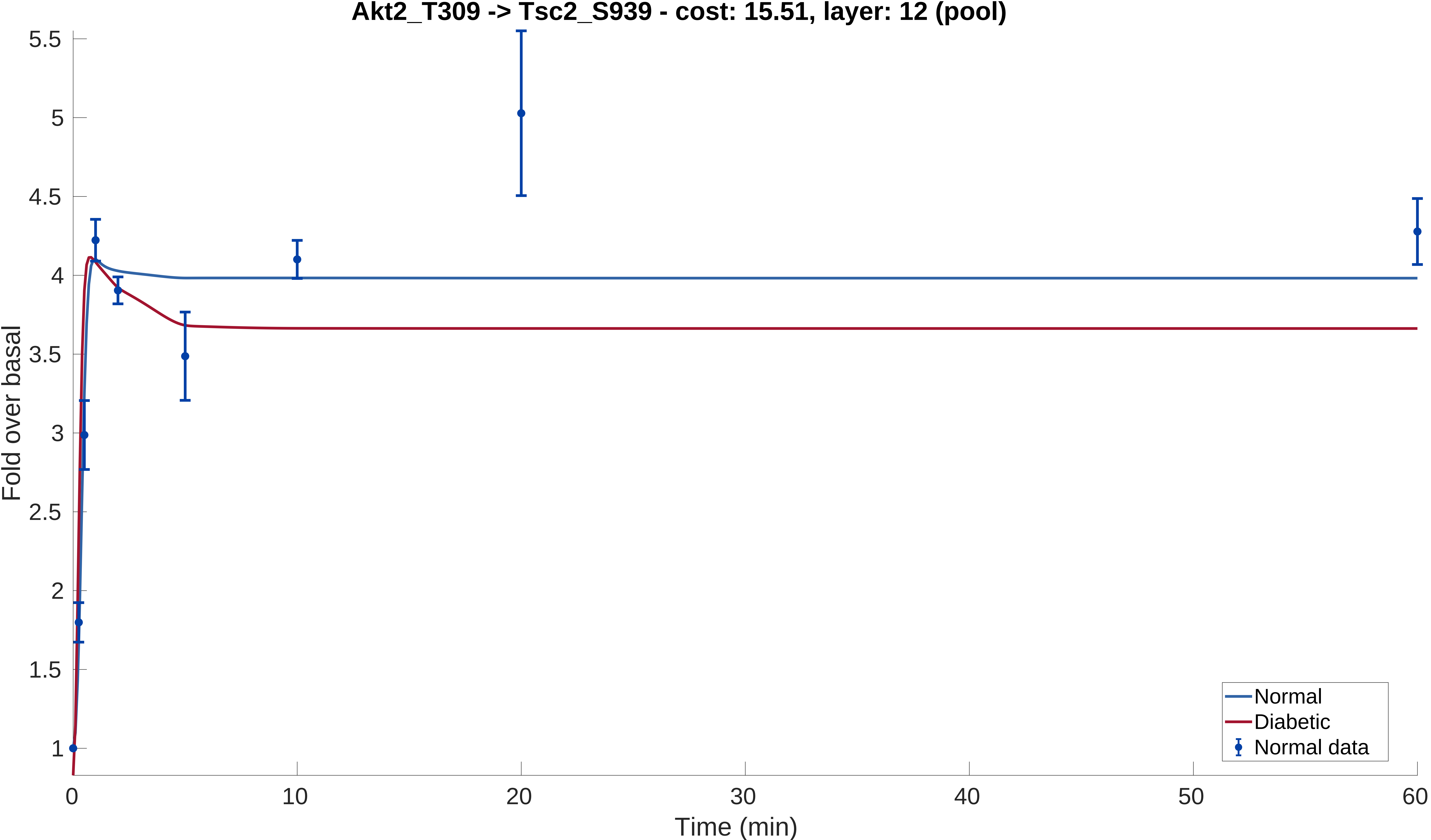

Akt2_T309 -> Tsc2_S939 - cost: 15.51, layer: 12 (pool)
Fold over basal
Time (min)
Normal
Diabetic
Normal data

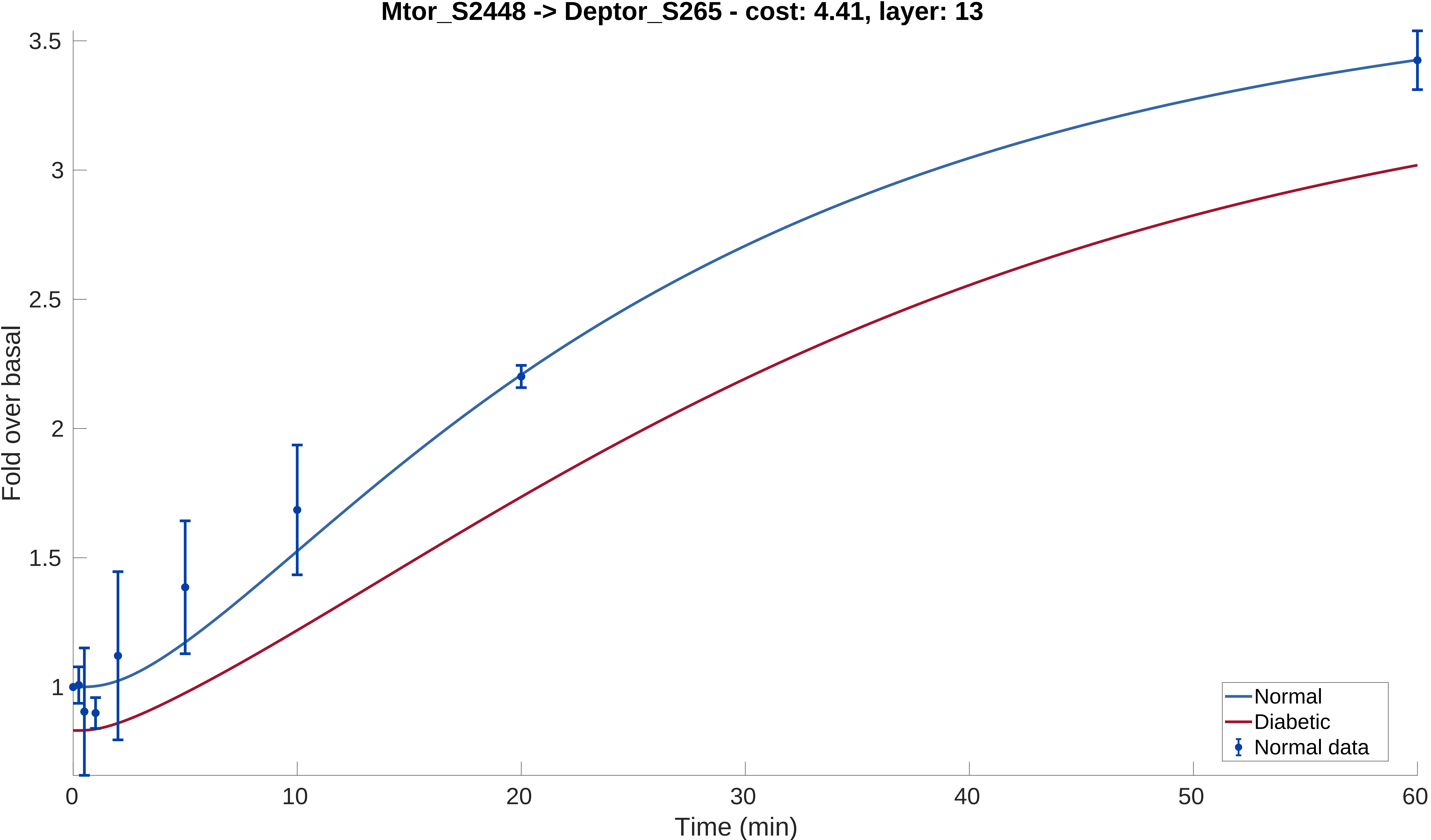
Mtor_S2448 -> Deptor_S265 - cost: 4.41, layer: 13
Fold over basal
3.5
3
2.5
2
1.5
1
0
10
20
30
40
50
60
Time (min)
Normal
Diabetic
Normal data

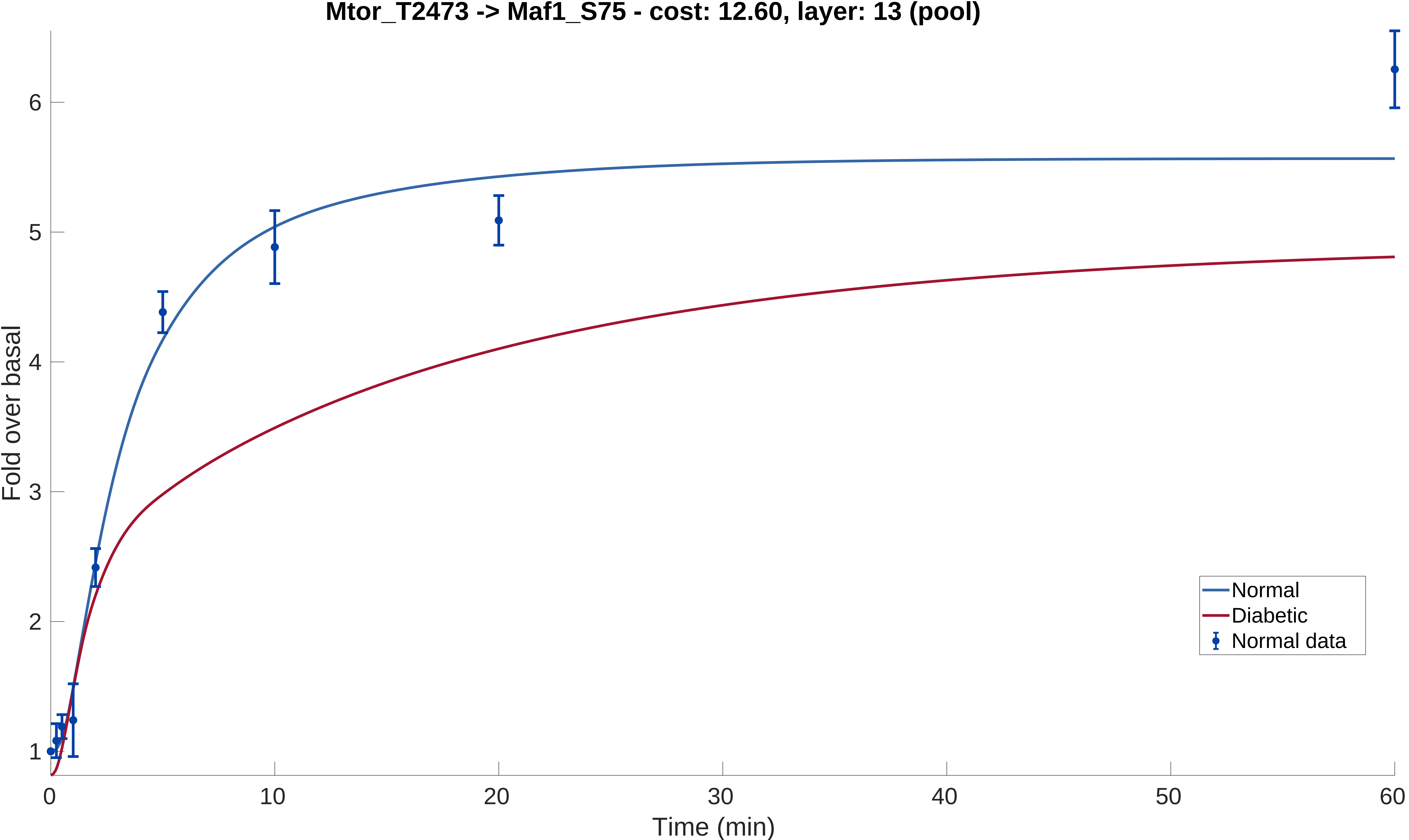

Mtor_T2473 -> Maf1_S75 - cost: 12.60, layer: 13 (pool)
Fold over basal
Time (min)
1
2
3
4
5
6
0
10
20
30
40
50
60
Normal
Diabetic
Normal data

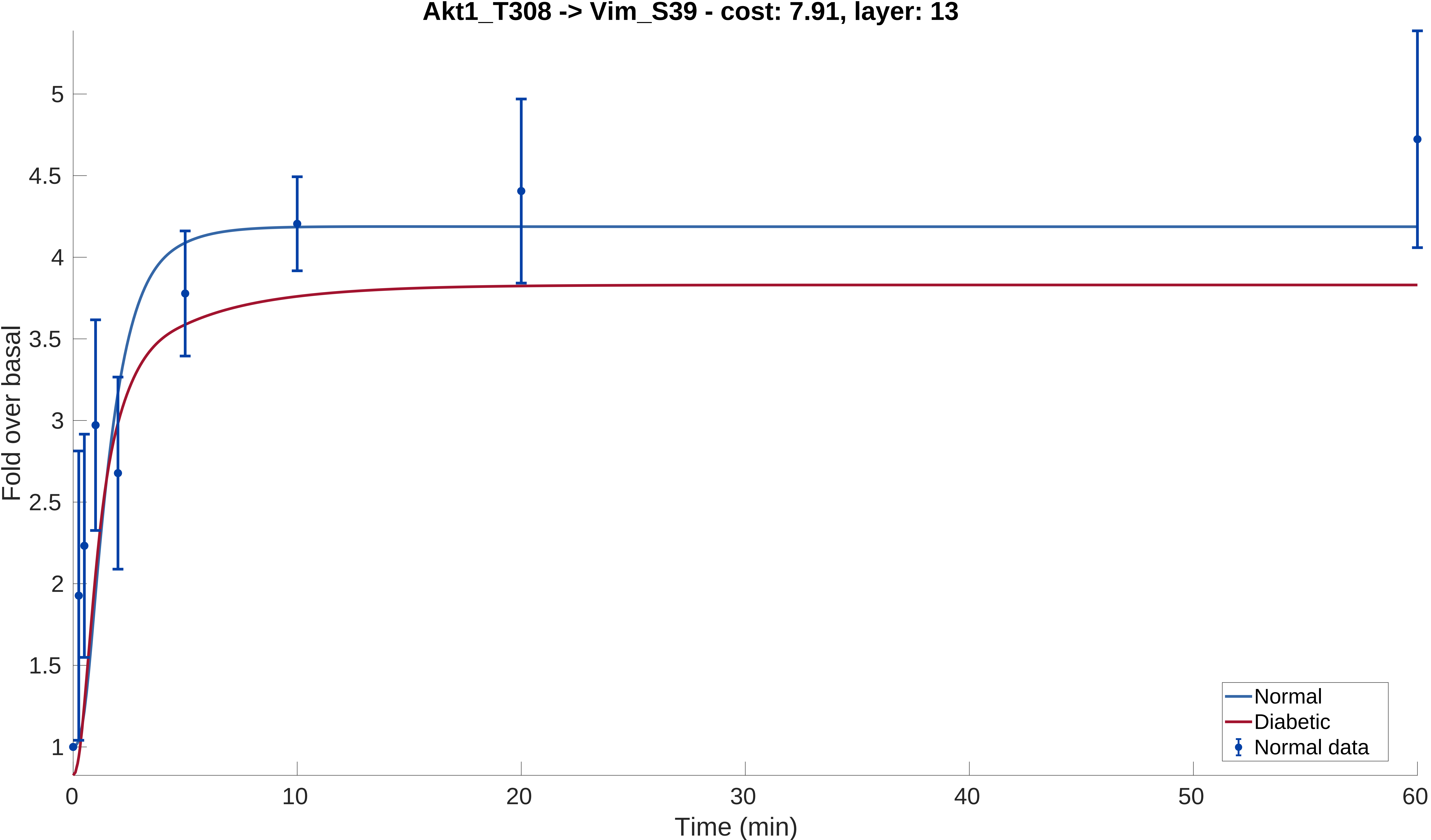
Akt1_T308 -> Vim_S39 - cost: 7.91, layer: 13
Fold over basal
5
4.5
4
3.5
3
2.5
2
1.5
1
0
10
20
30
40
50
60
Time (min)
Normal
Diabetic
Normal data

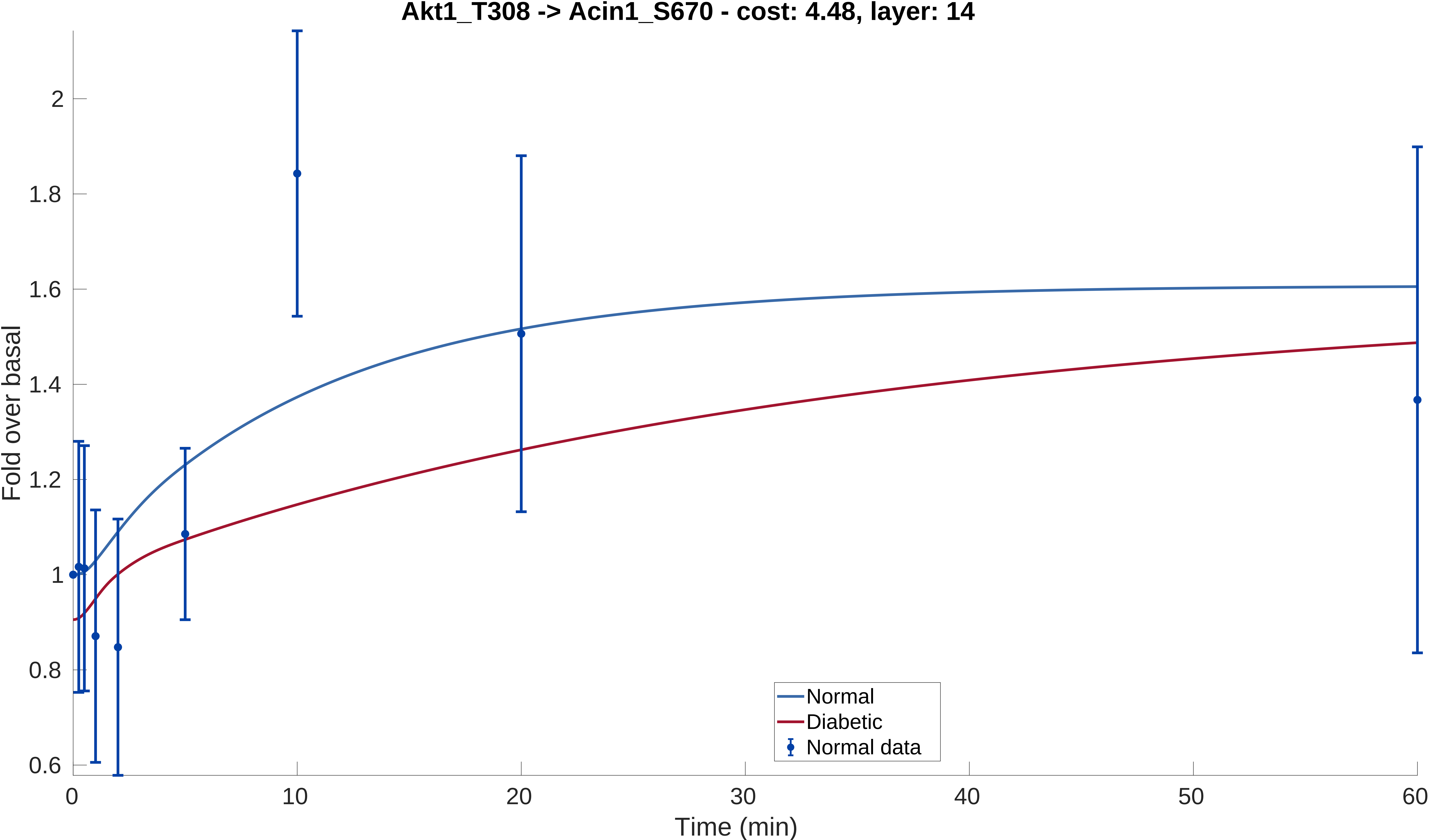

Akt1_T308 -> Acin1_S670 - cost: 4.48, layer: 14
Fold over basal
2
1.8
1.6
1.4
1.2
1
0.8
0.6
0
10
20
30
40
50
60
Time (min)
Normal
Diabetic
Normal data

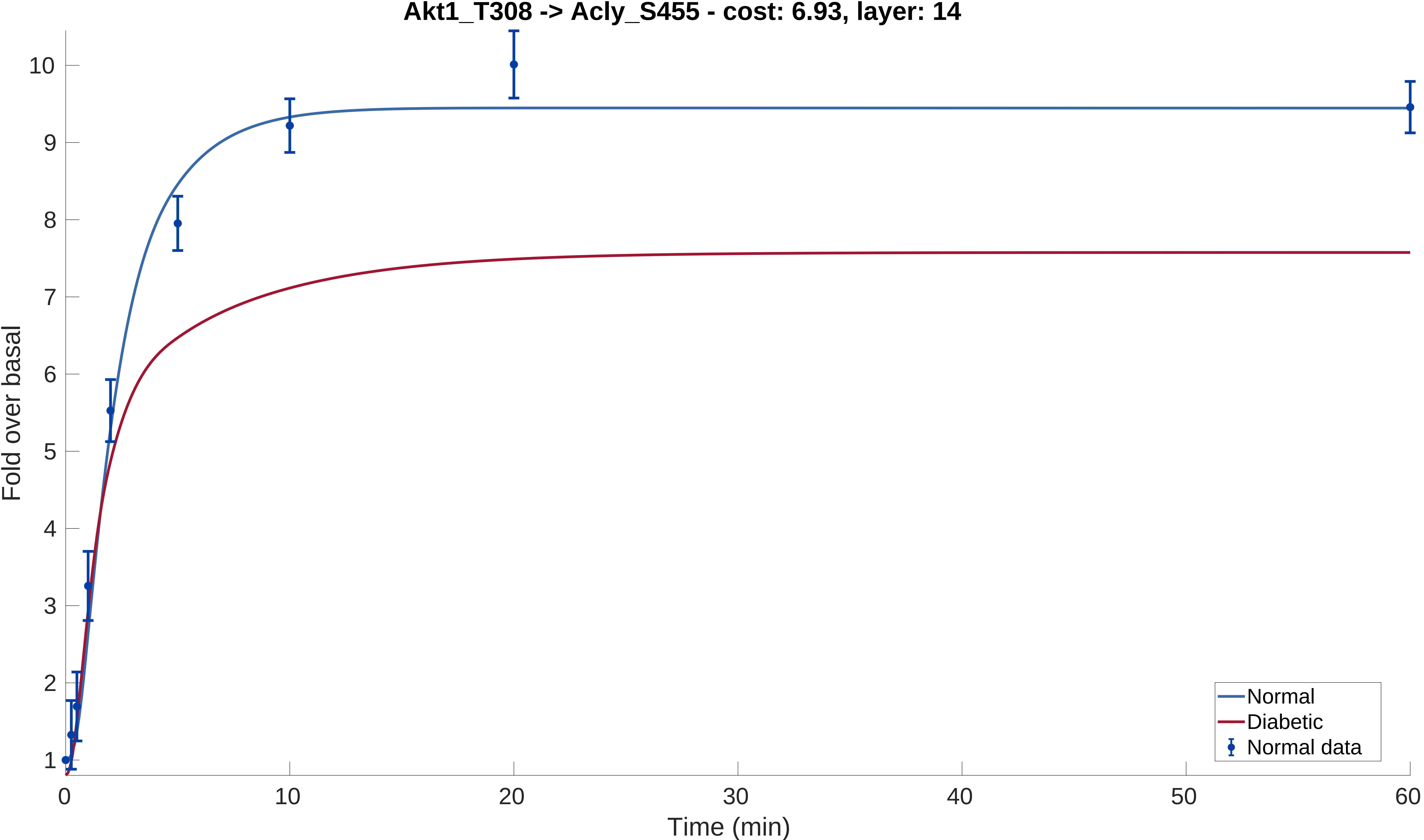

Akt1_T308 -> Acly_S455 - cost: 6.93, layer: 14
Fold over basal
Time (min)
Normal
Diabetic
Normal data

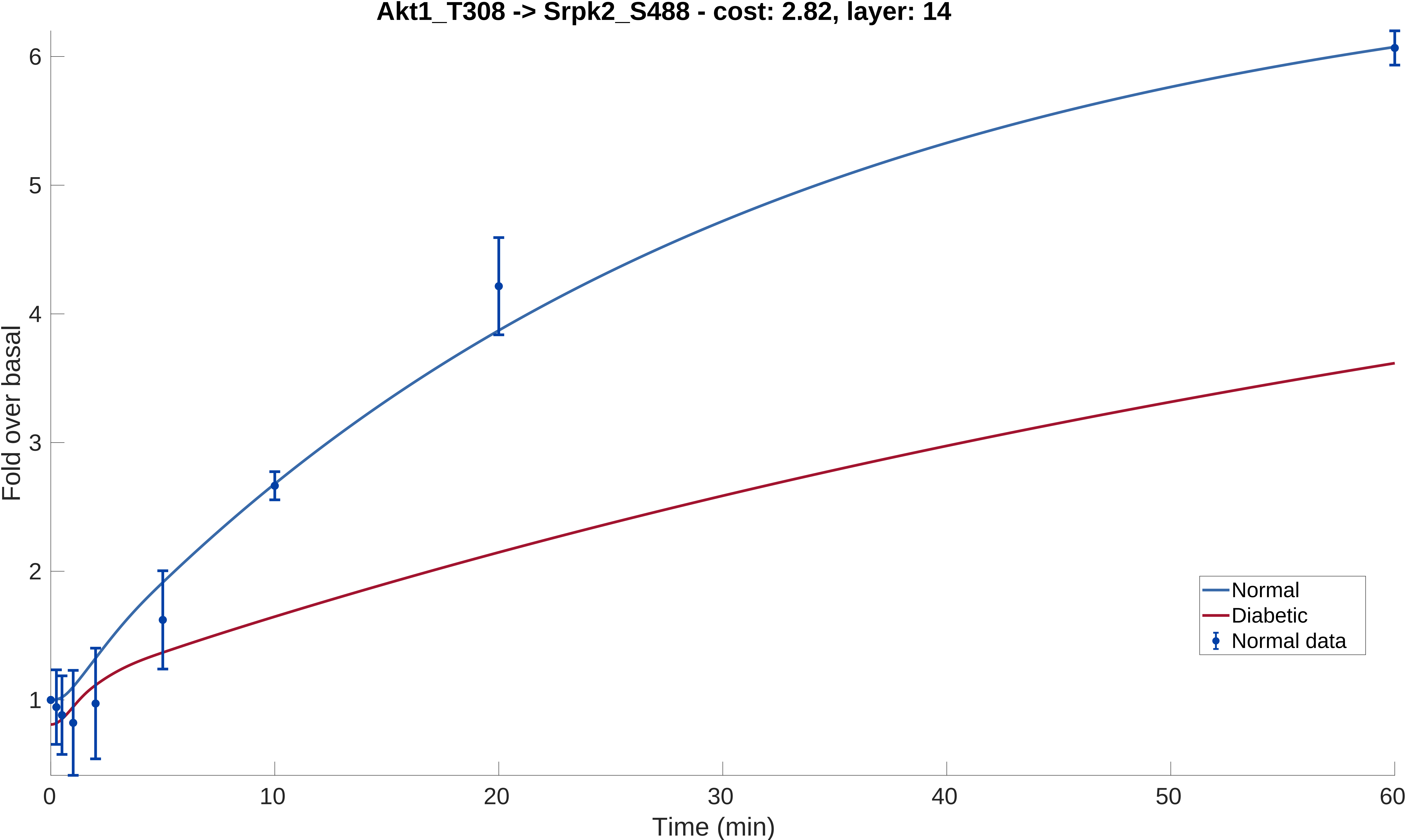
Akt1_T308 -> Srpk2_S488 - cost: 2.82, layer: 14
Fold over basal
6
5
4
3
2
1
0
10
20
30
40
50
60
Time (min)
Normal
Diabetic
Normal data

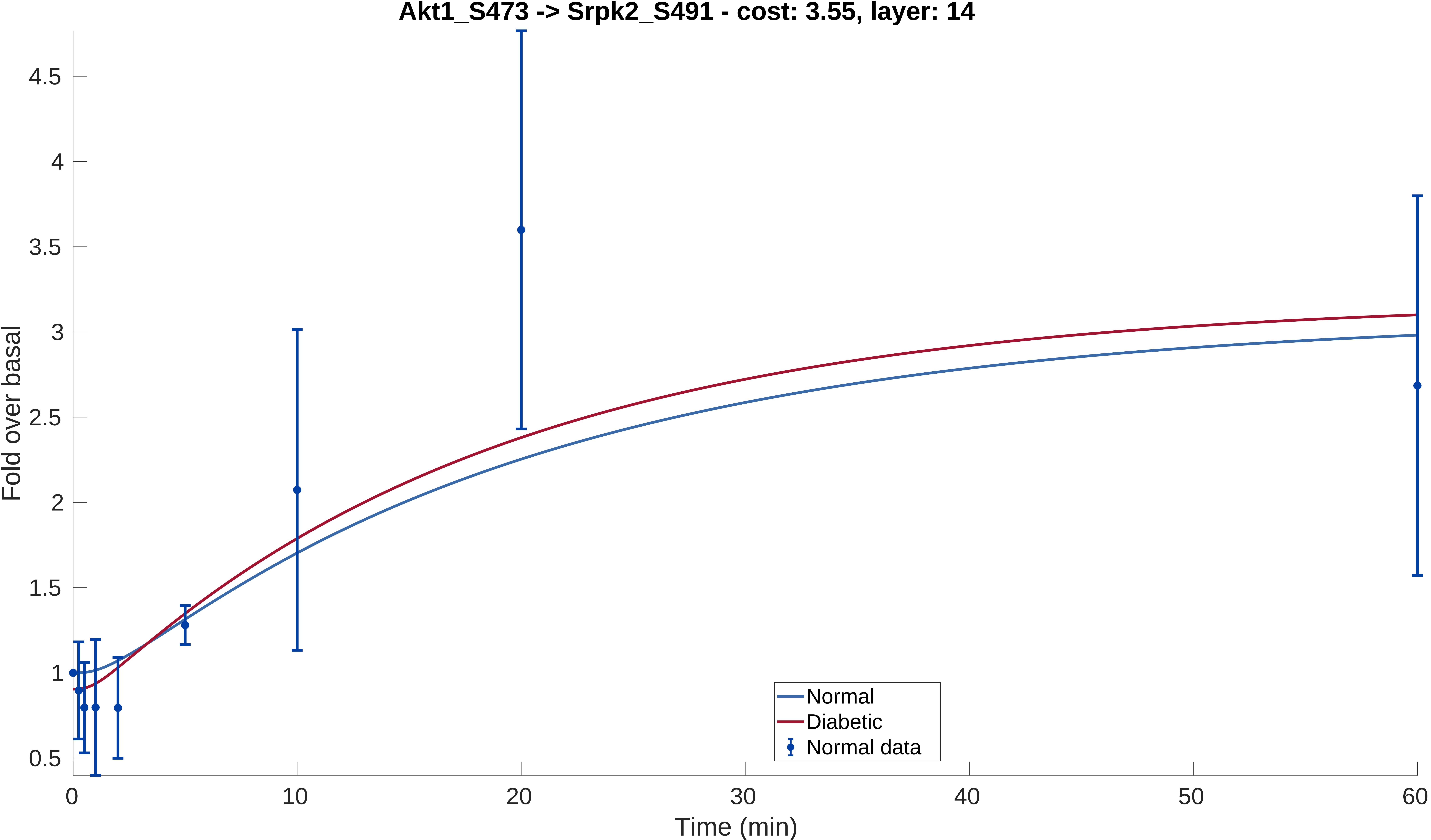

Akt1_S473 -> Srpk2_S491 - cost: 3.55, layer: 14
Fold over basal
4.5
4
3.5
3
2.5
2
1.5
1
0.5
0
10
20
30
40
50
60
Time (min)
Normal
Diabetic
Normal data

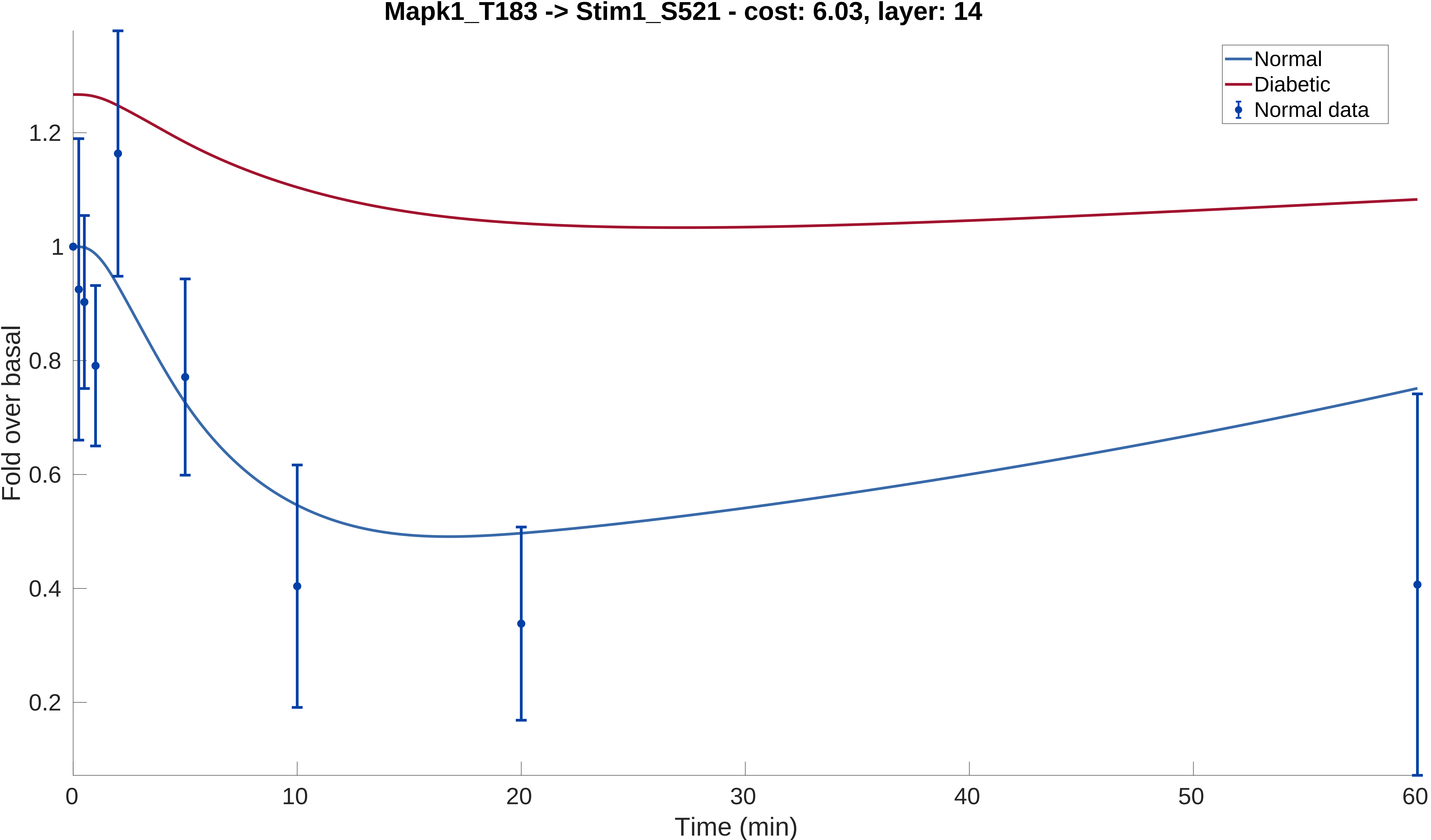

Mapk1_T183 -> Stim1_S521 - cost: 6.03, layer: 14
Normal
Diabetic
Normal data
1.2
1
0.8
0.6
0.4
0.2
Fold over basal
0
10
20
30
40
50
60
Time (min)

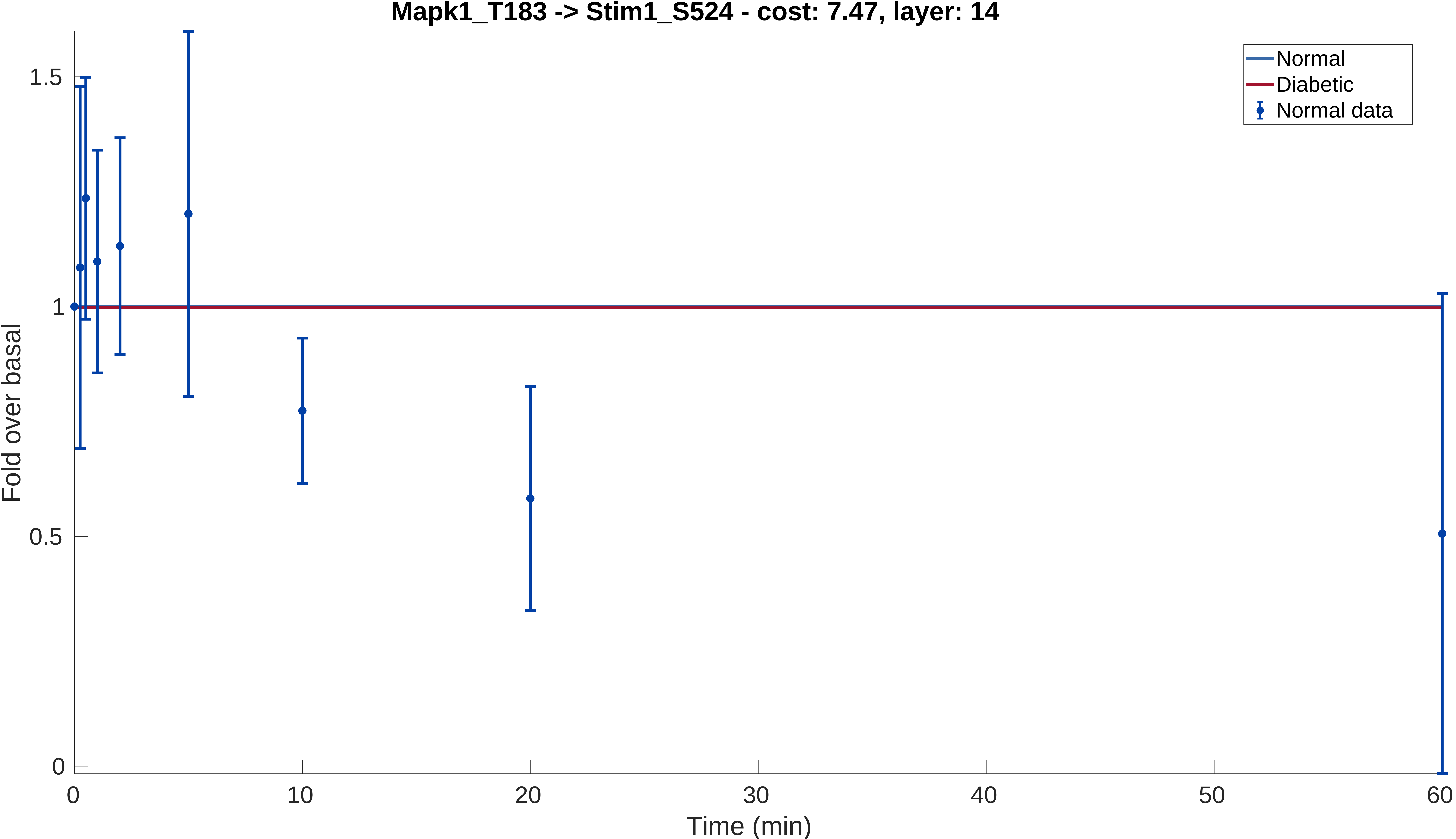
Mapk1_T183 -> Stim1_S524 - cost: 7.47, layer: 14
Normal
Diabetic
Normal data
Fold over basal
Time (min)
0
0.5
1
1.5
0
10
20
30
40
50
60

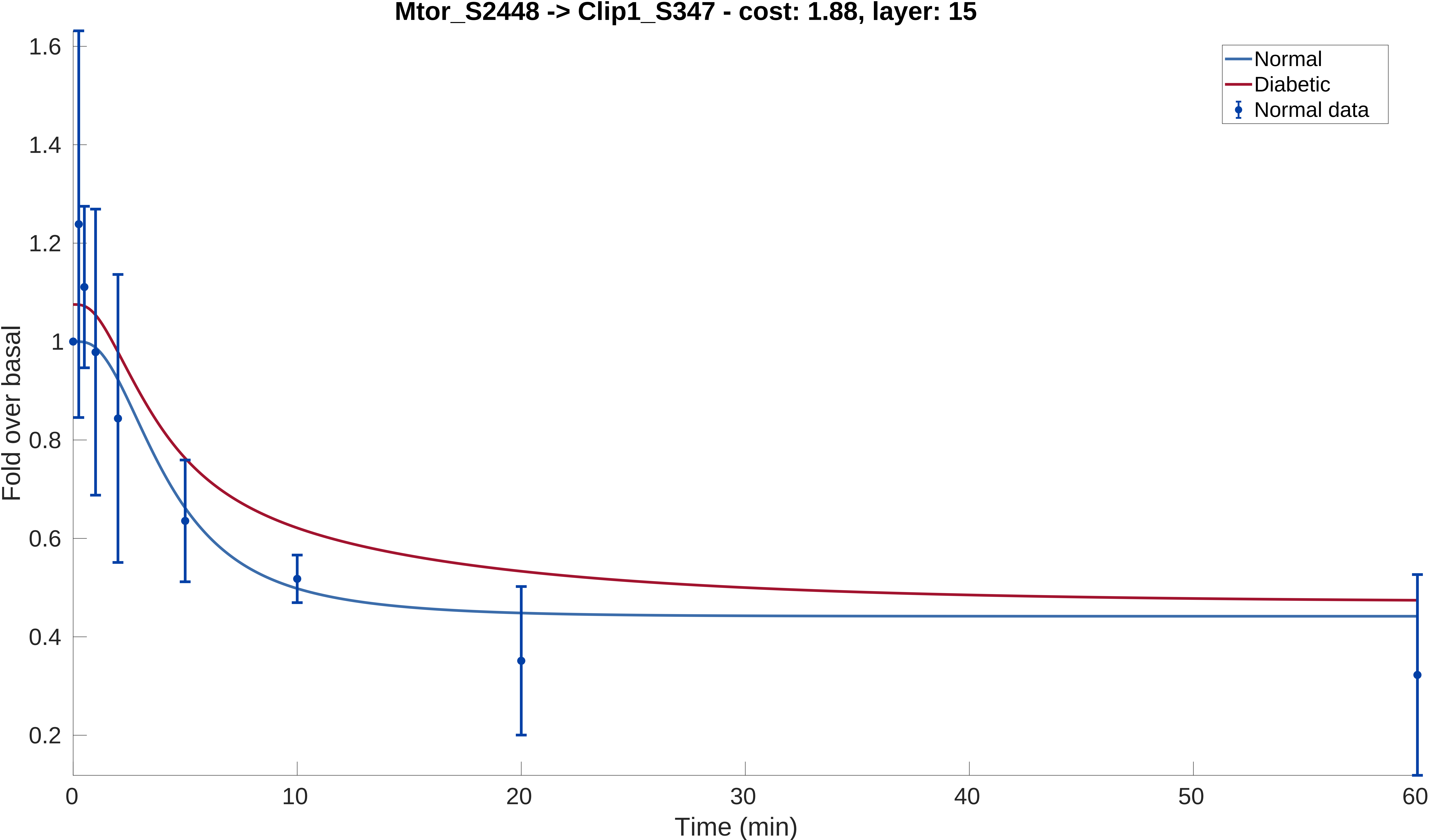

Mtor_S2448 -> Clip1_S347 - cost: 1.88, layer: 15
Normal
Diabetic
Normal data
Fold over basal
Time (min)
0.2
0.4
0.6
0.8
1
1.2
1.4
1.6
0
10
20
30
40
50
60

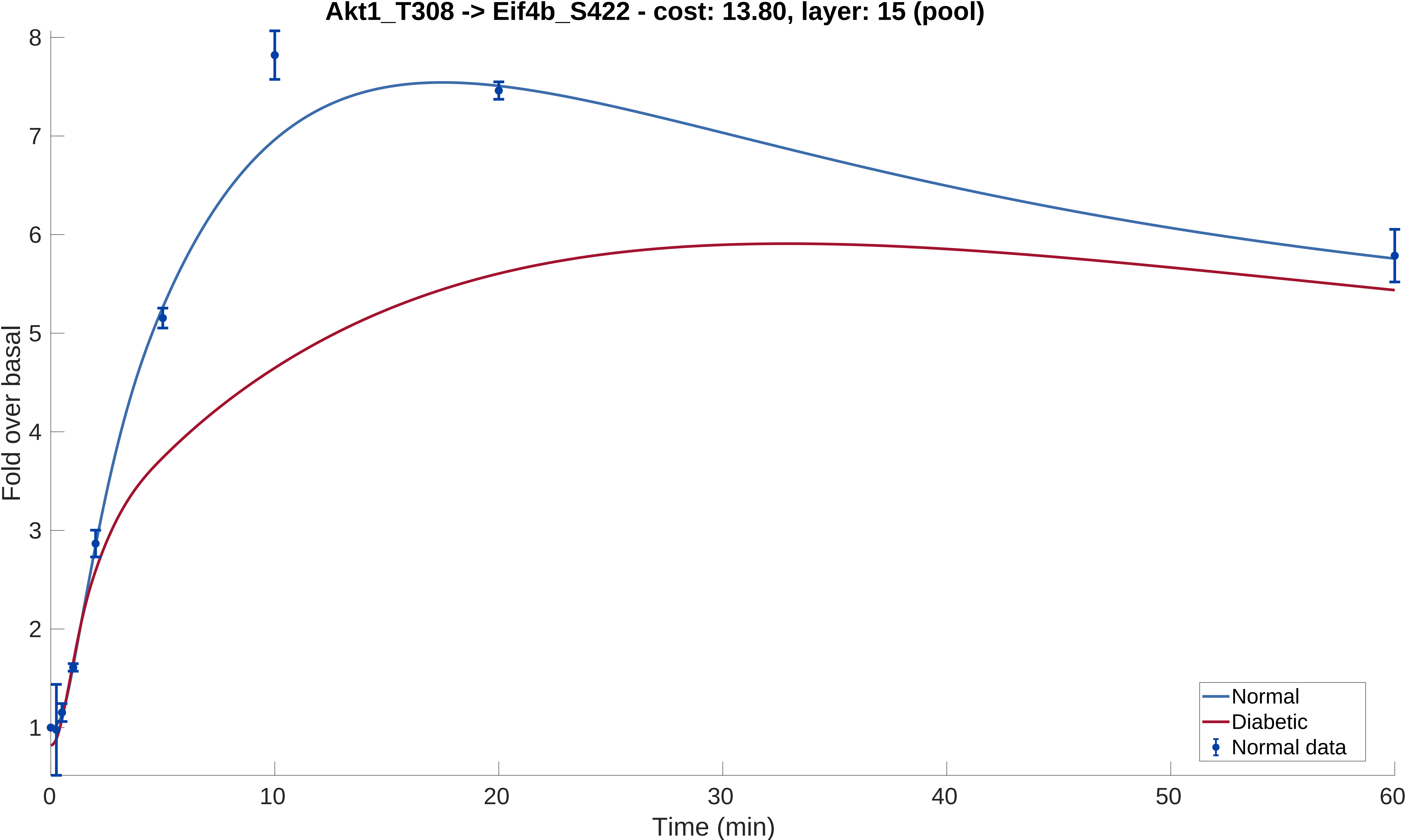

Akt1_T308 -> Eif4b_S422 - cost: 13.80, layer: 15 (pool)
Fold over basal
Time (min)
0
10
20
30
40
50
60
1
2
3
4
5
6
7
8
Normal
Diabetic
Normal data

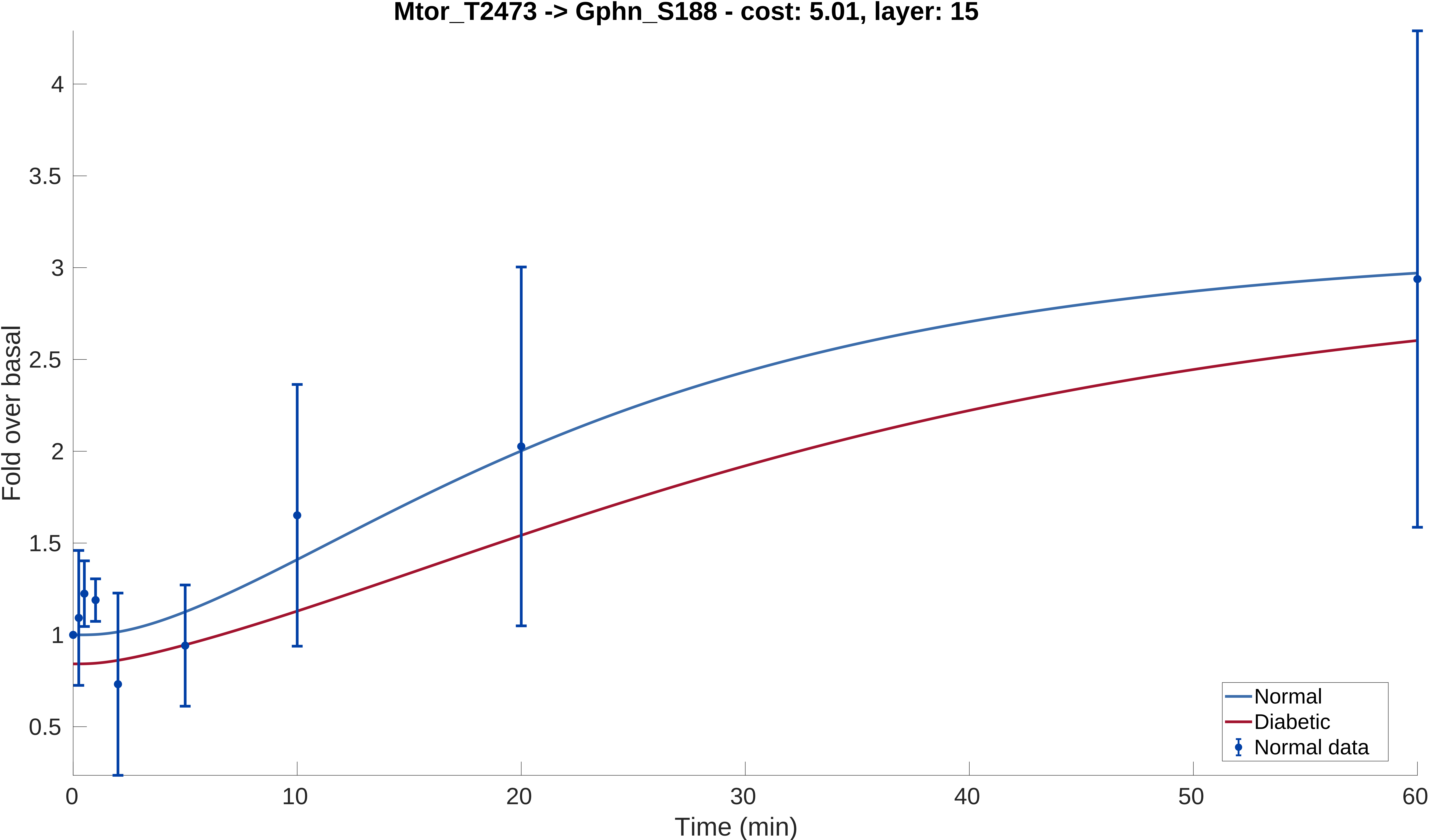
Mtor_T2473 -> Gphn_S188 - cost: 5.01, layer: 15
Fold over basal
Time (min)
0
10
20
30
40
50
60
0.5
1
1.5
2
2.5
3
3.5
4
Normal
Diabetic
Normal data

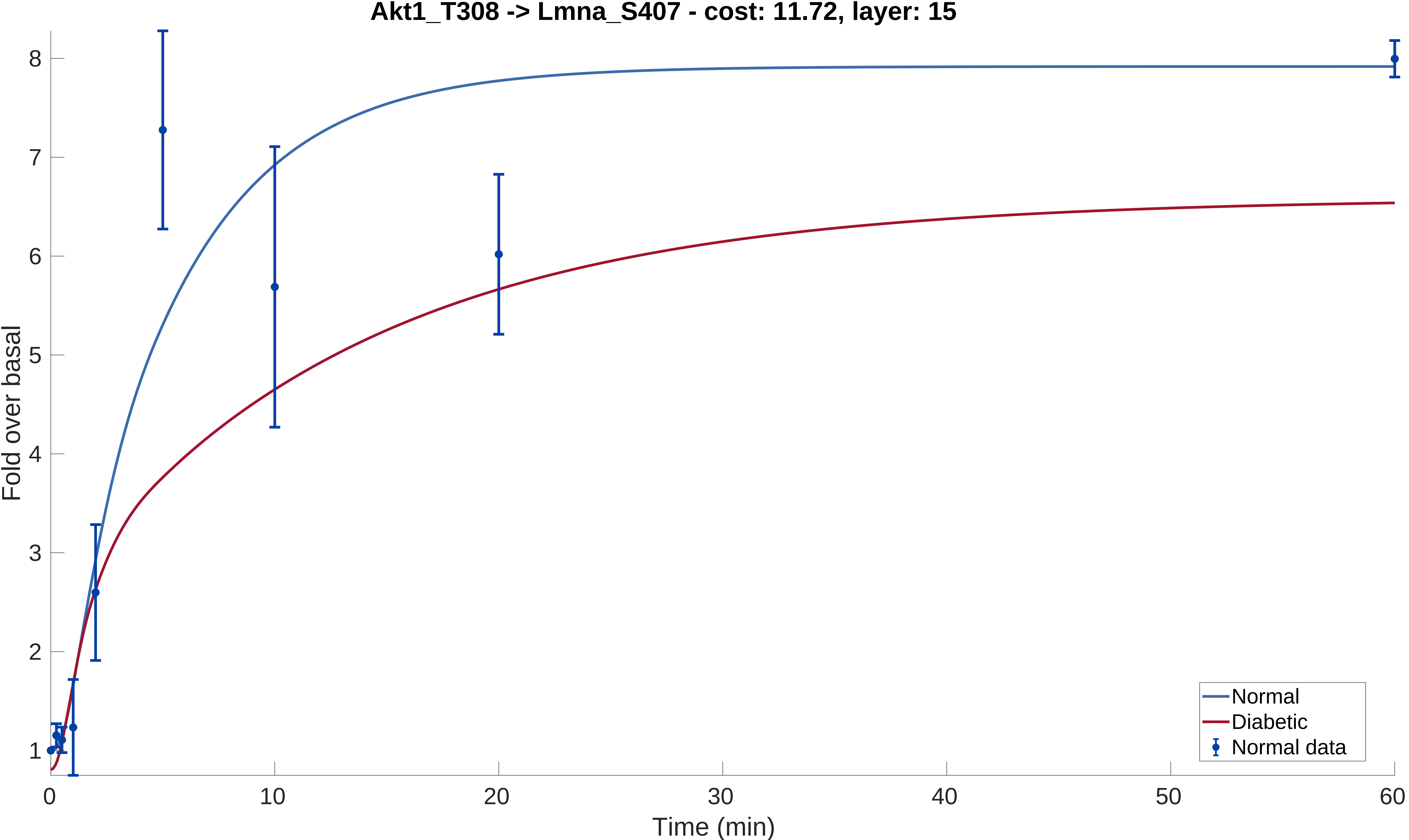
Akt1_T308 -> Lmna_S407 - cost: 11.72, layer: 15
Fold over basal
Time (min)
0
10
20
30
40
50
60
1
2
3
4
5
6
7
8
Normal
Diabetic
Normal data

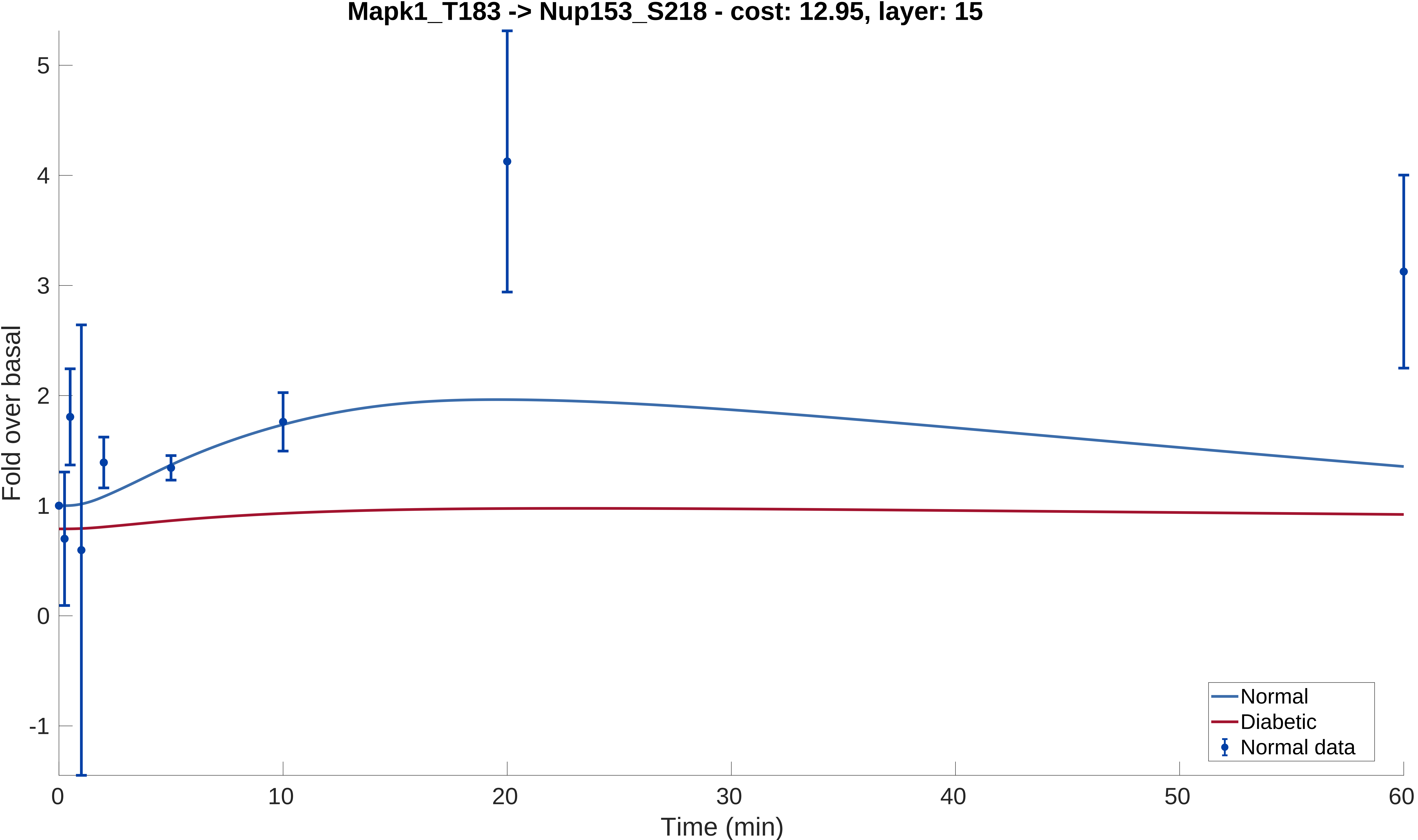

Mapk1_T183 -> Nup153_S218 - cost: 12.95, layer: 15
Fold over basal
Time (min)
5
4
3
2
1
0
-1
0
10
20
30
40
50
60
Normal
Diabetic
Normal data

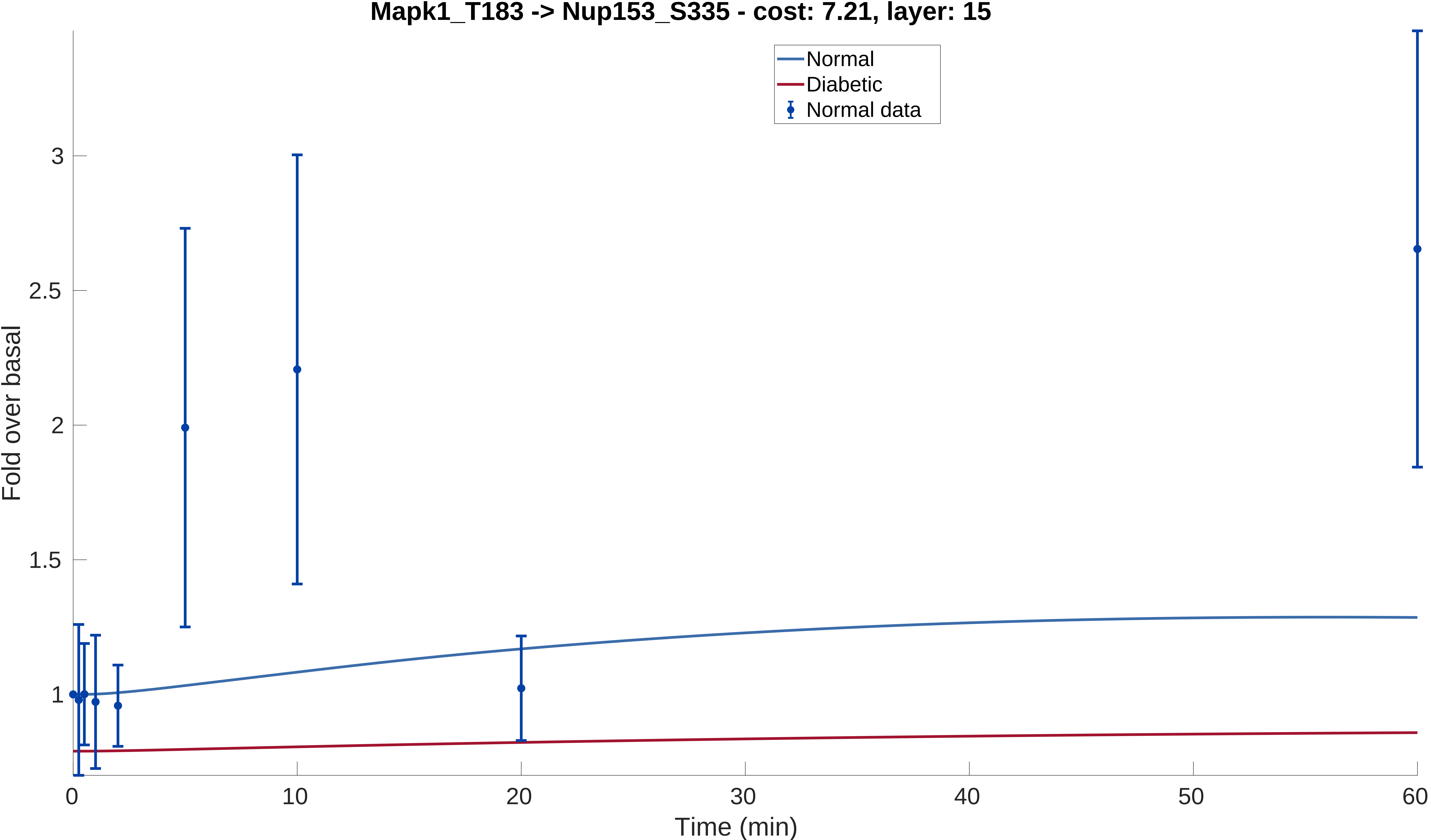

Mapk1_T183 -> Nup153_S335 - cost: 7.21, layer: 15
Normal
Diabetic
Normal data
Fold over basal
1
1.5
2
2.5
3
0
10
20
30
40
50
60
Time (min)

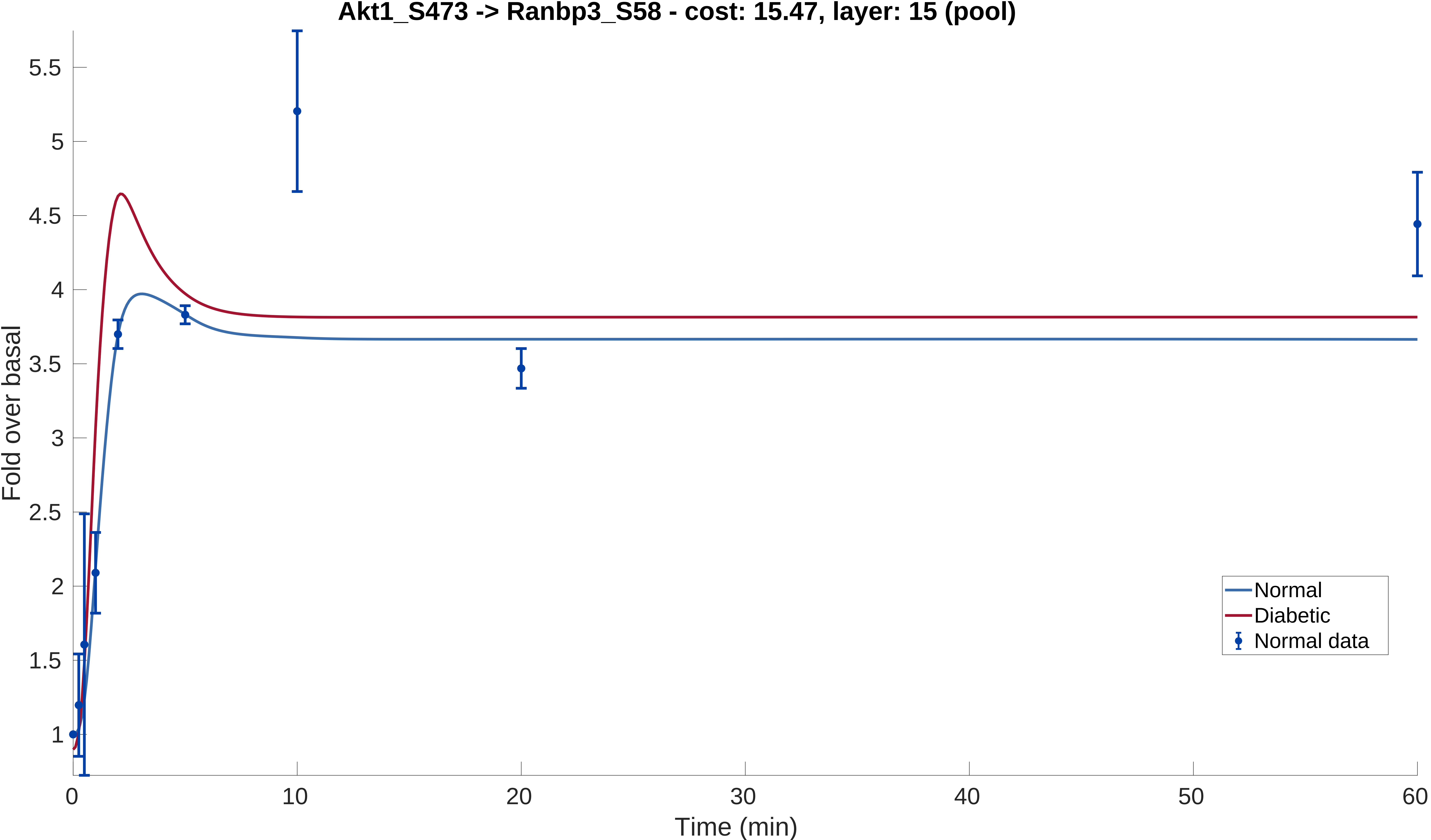

Akt1_S473 -> Ranbp3_S58 - cost: 15.47, layer: 15 (pool)
Fold over basal
5.5
5
4.5
4
3.5
3
2.5
2
1.5
1
0
10
20
30
40
50
60
Time (min)
Normal
Diabetic
Normal data

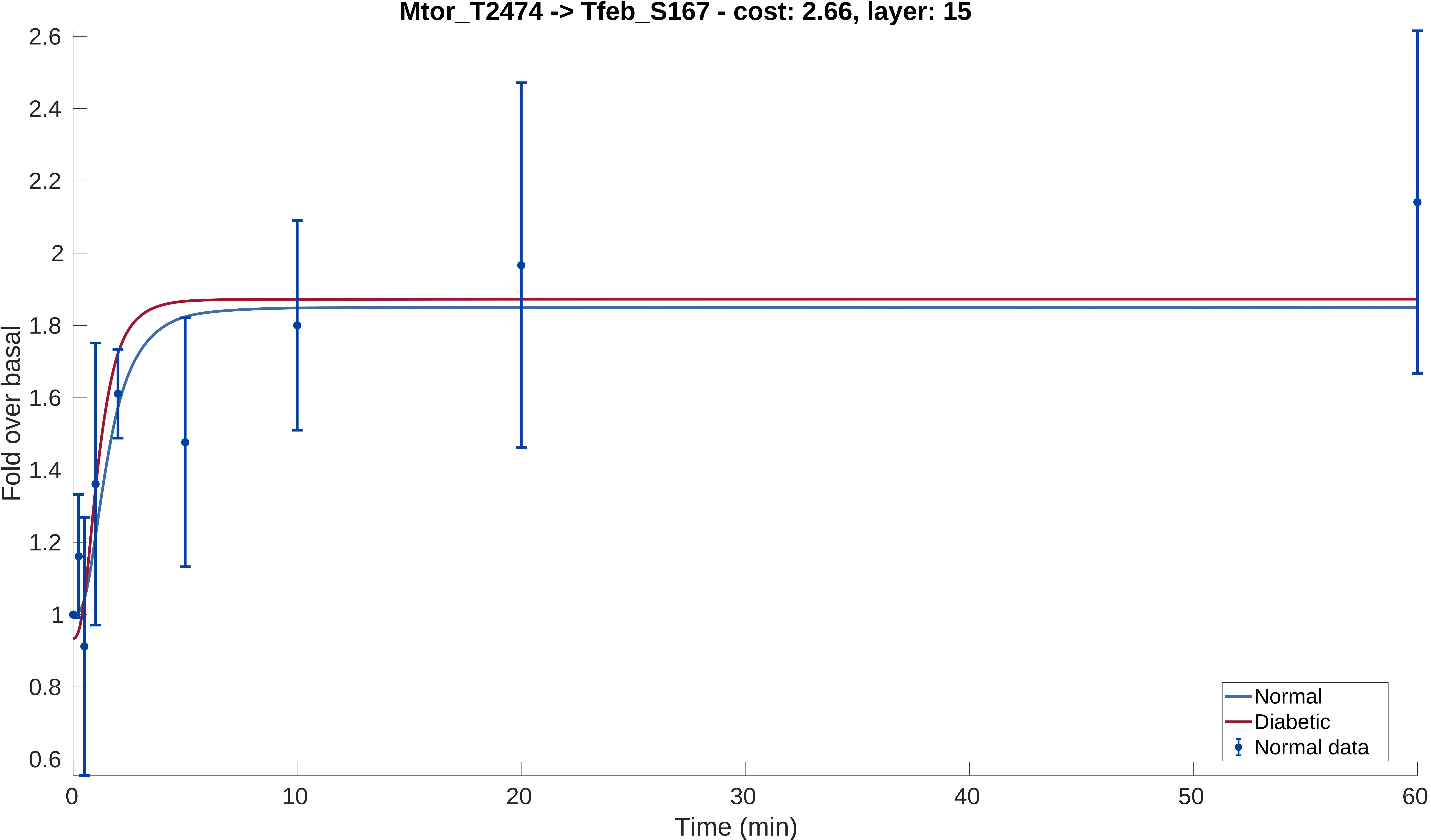

Mtor_T2474 -> Tfeb_S167 - cost: 2.66, layer: 15
Fold over basal
Time (min)
0.6
0.8
1
1.2
1.4
1.6
1.8
2
2.2
2.4
2.6
0
10
20
30
40
50
60
Normal
Diabetic
Normal data

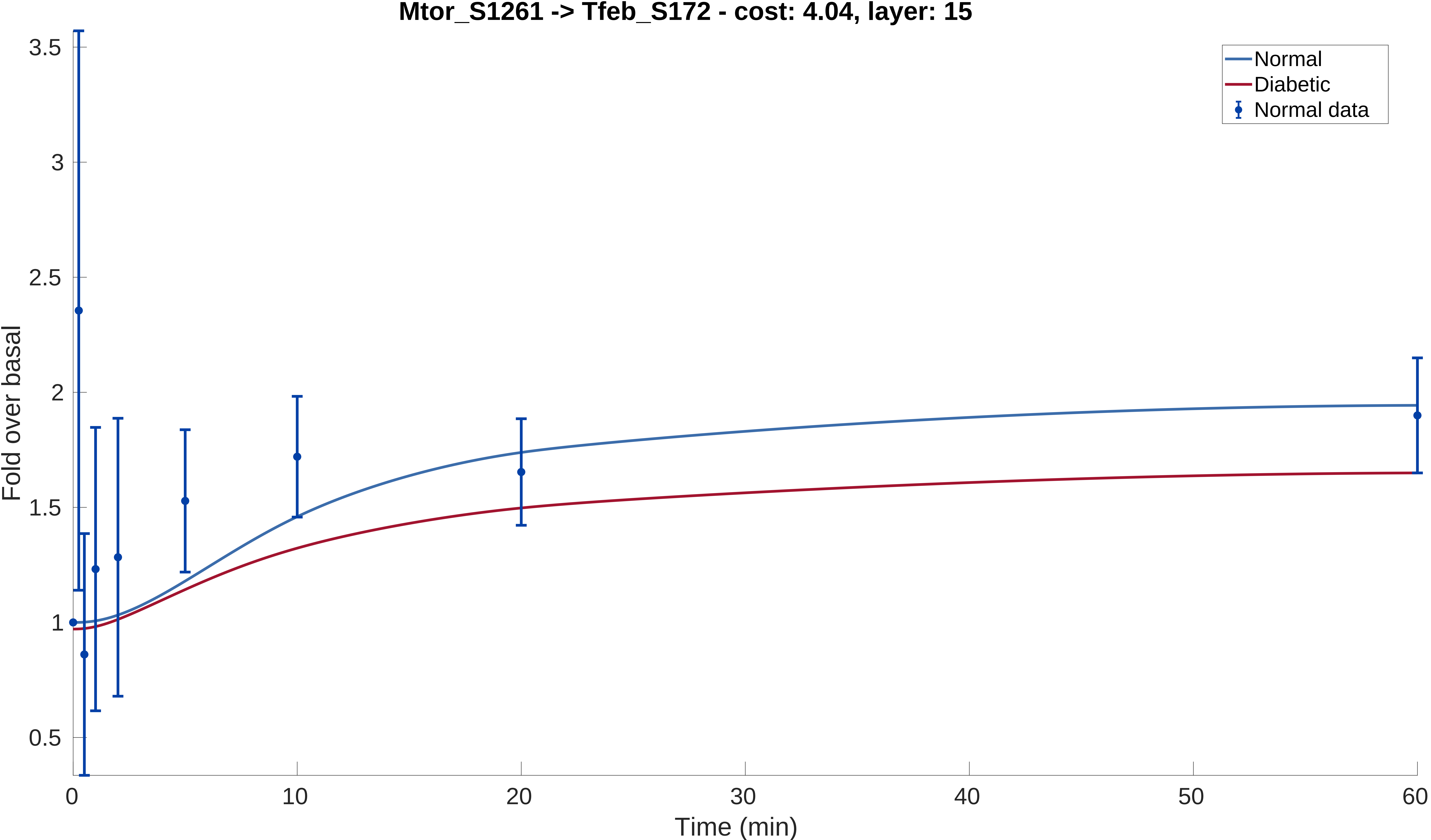

Mtor_S1261 -> Tfeb_S172 - cost: 4.04, layer: 15
Normal
Diabetic
Normal data
Fold over basal
3.5
3
2.5
2
1.5
1
0.5
0
10
20
30
40
50
60
Time (min)

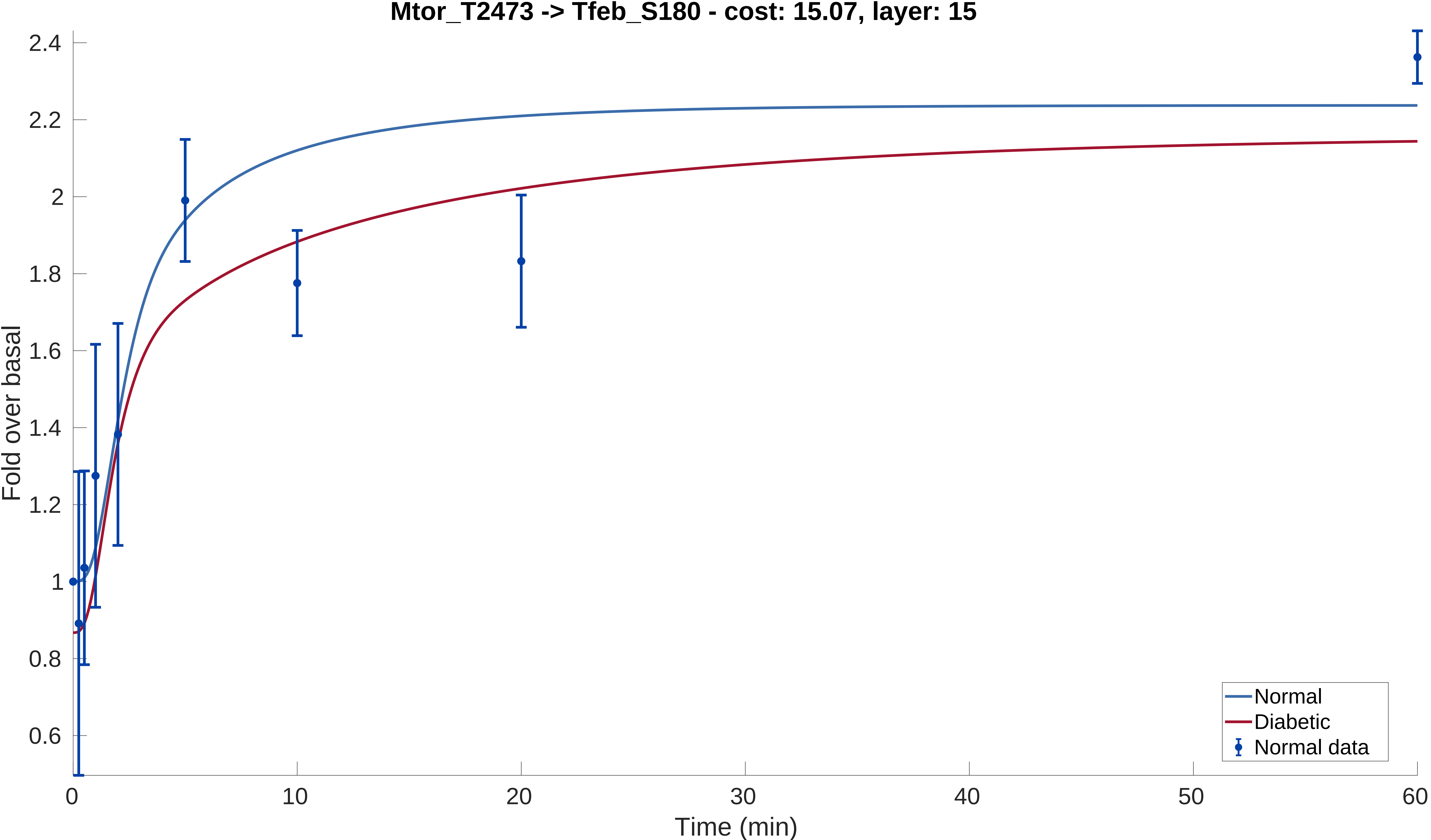

Mtor_T2473 -> Tfeb_S180 - cost: 15.07, layer: 15
Fold over basal
Time (min)
Normal
Diabetic
Normal data
0.6
0.8
1
1.2
1.4
1.6
1.8
2
2.2
2.4
0
10
20
30
40
50
60

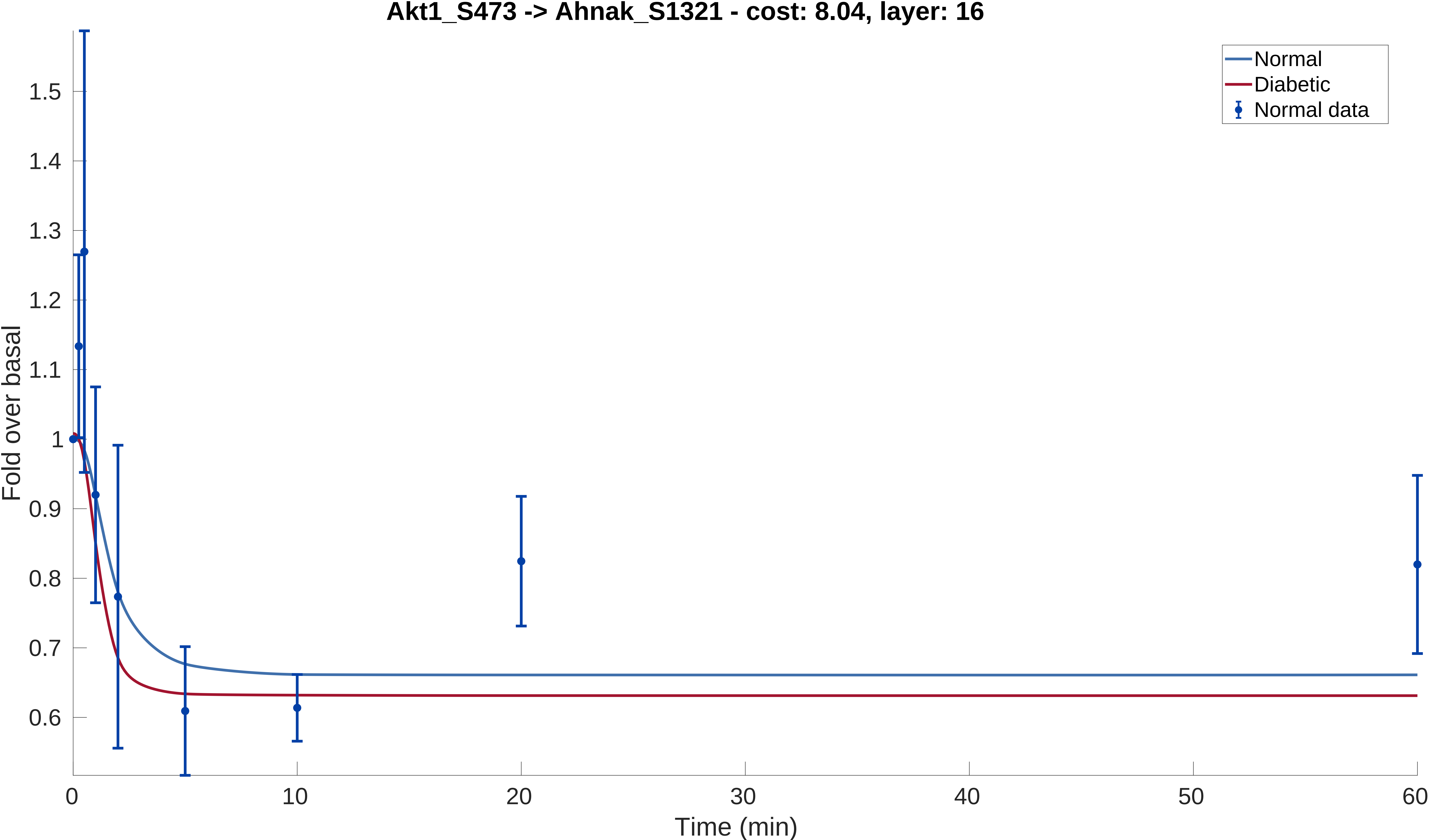

Akt1_S473 -> Ahnak_S1321 - cost: 8.04, layer: 16
Normal
Diabetic
Normal data
Fold over basal
1.5
1.4
1.3
1.2
1.1
1
0.9
0.8
0.7
0.6
0
10
20
30
40
50
60
Time (min)

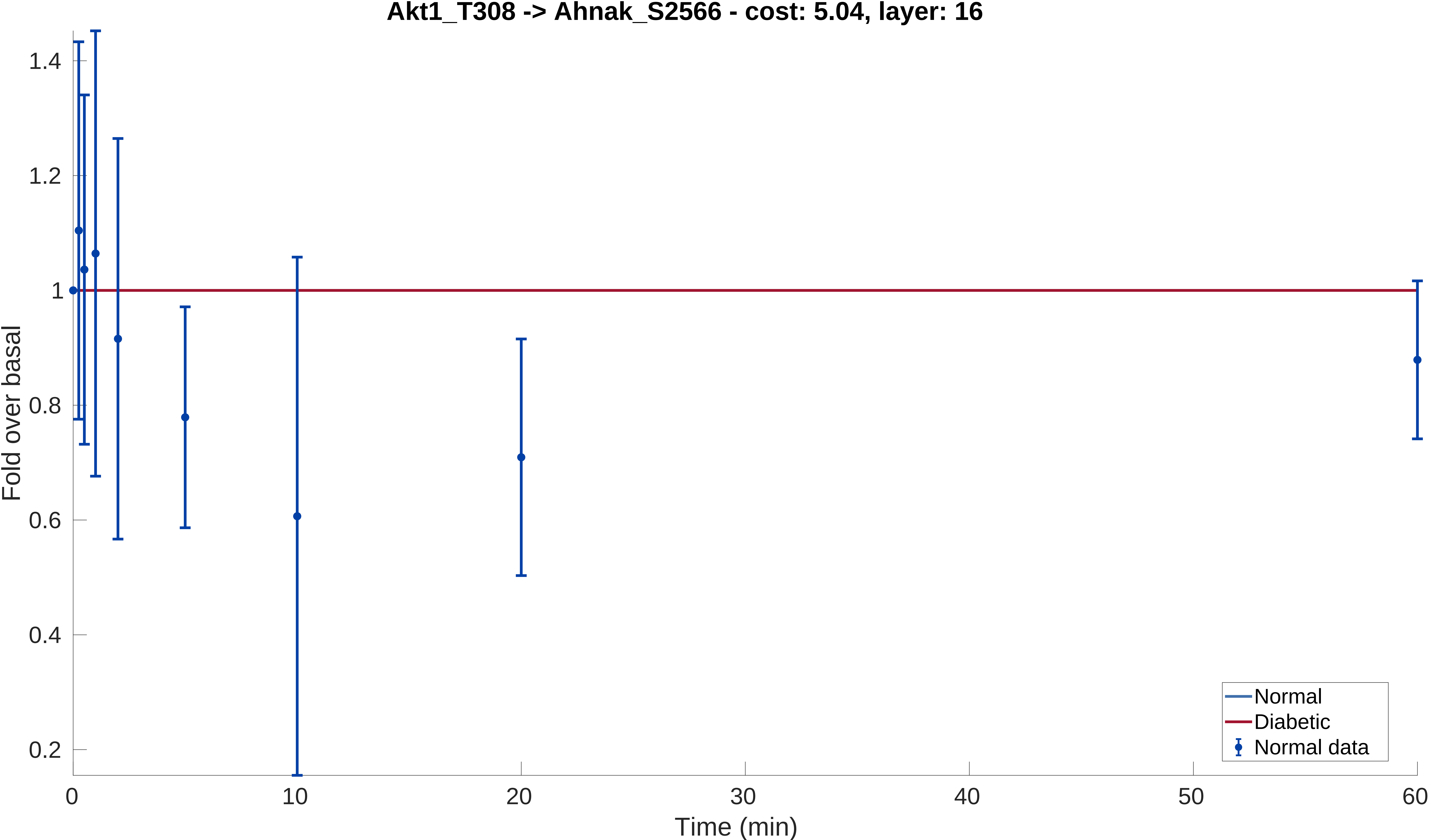

Akt1_T308 -> Ahnak_S2566 - cost: 5.04, layer: 16
Fold over basal
1.4
1.2
1
0.8
0.6
0.4
0.2
0
10
20
30
40
50
60
Time (min)
Normal
Diabetic
Normal data

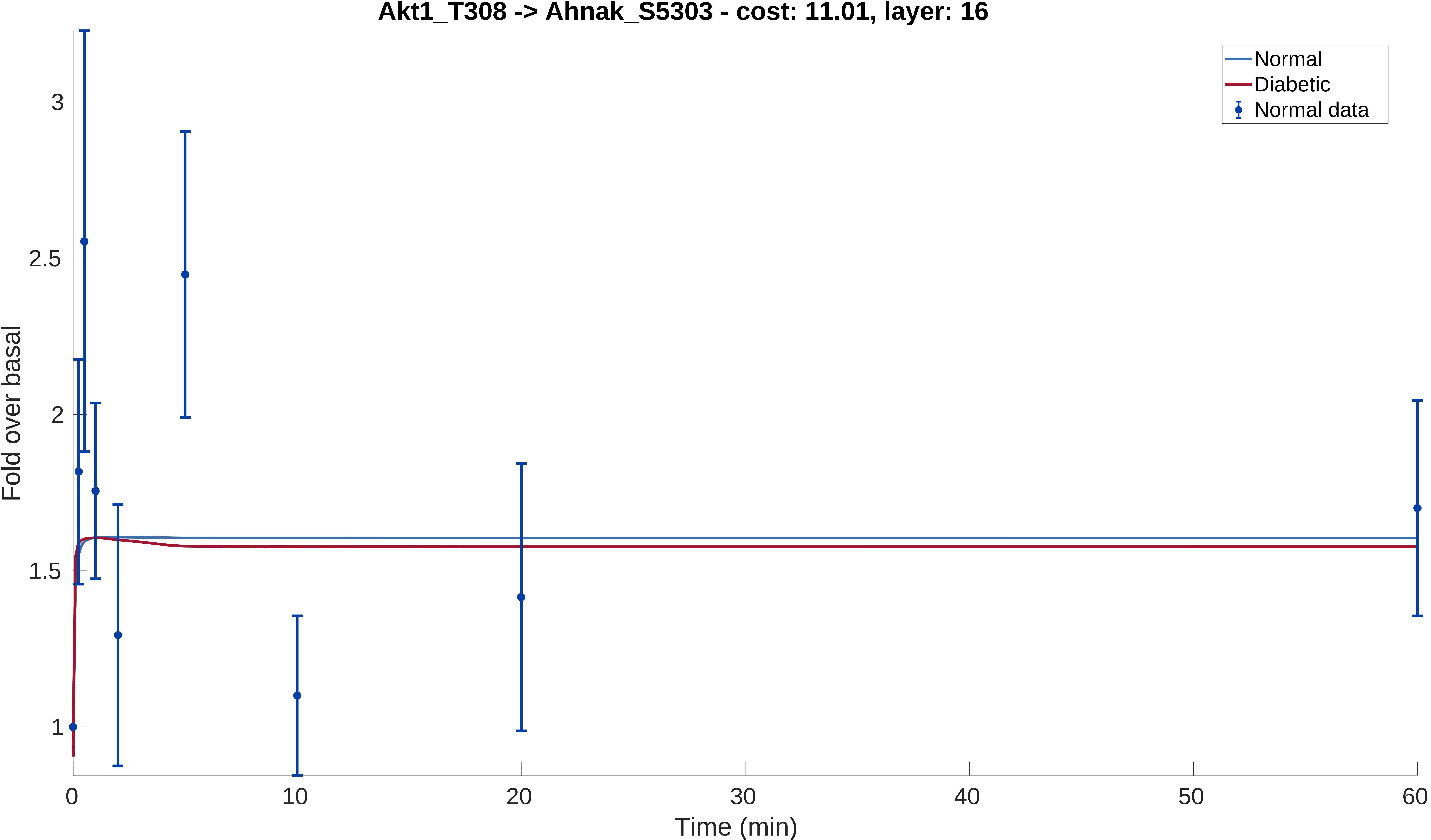

Akt1_T308 -> Ahnak_S5303 - cost: 11.01, layer: 16
Normal
Diabetic
Normal data
Fold over basal
Time (min)
3
2.5
2
1.5
1
0
10
20
30
40
50
60

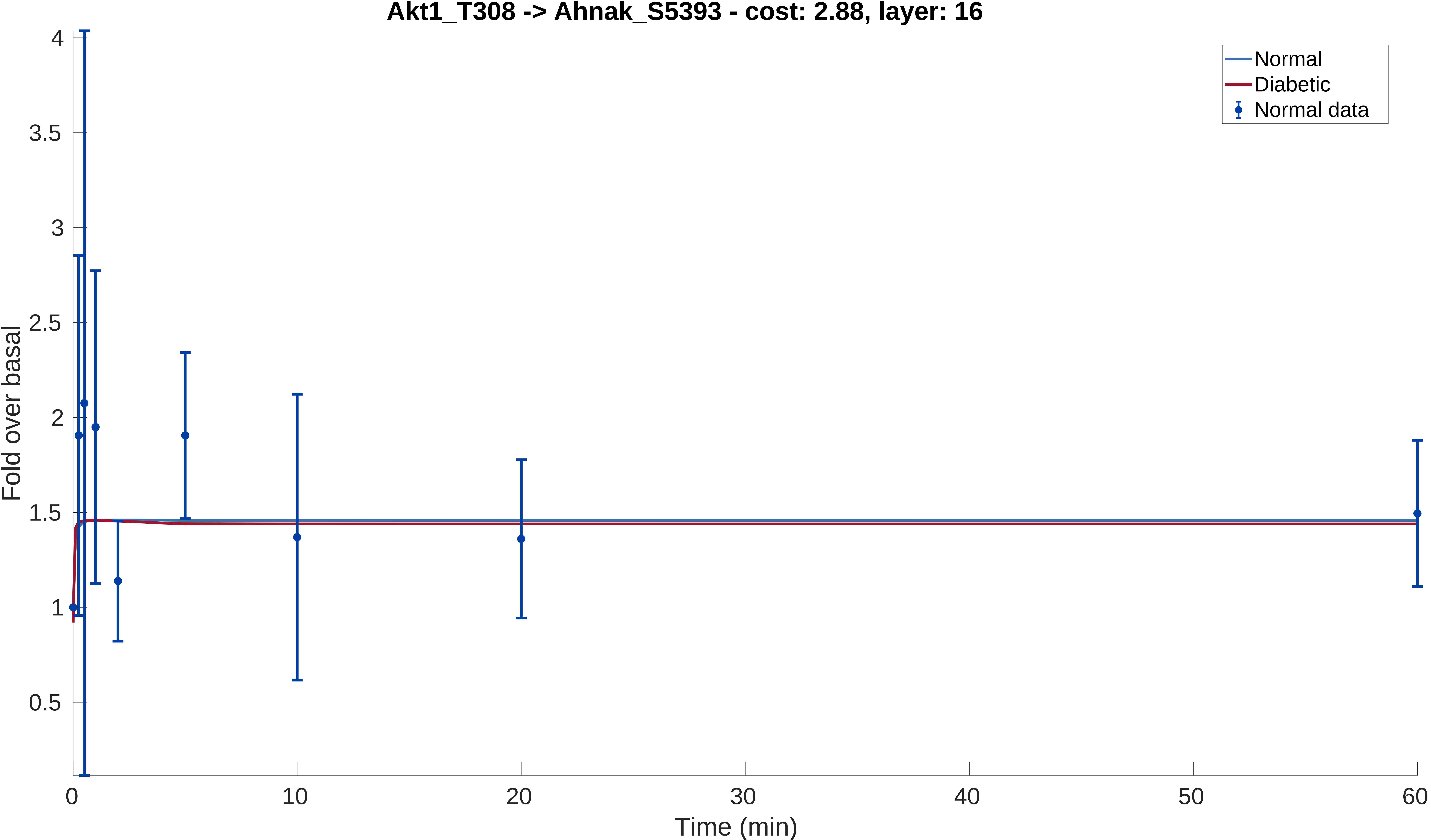
Akt1_T308 -> Ahnak_S5393 - cost: 2.88, layer: 16
Normal
Diabetic
Normal data
Fold over basal
Time (min)
4
3.5
3
2.5
2
1.5
1
0.5
0
10
20
30
40
50
60

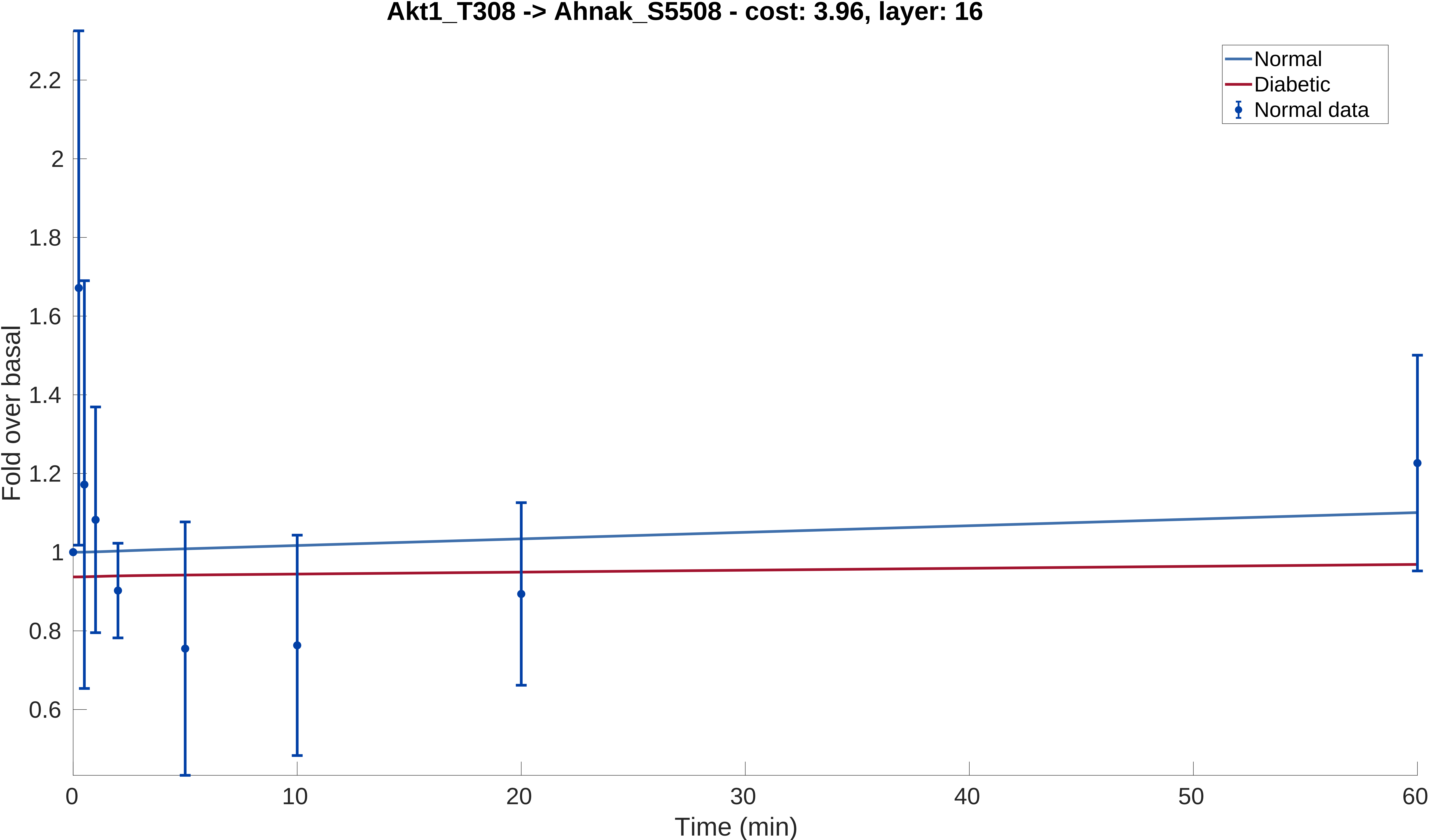

Akt1_T308 -> Ahnak_S5508 - cost: 3.96, layer: 16
Normal
Diabetic
Normal data
Fold over basal
Time (min)
0.6
0.8
1
1.2
1.4
1.6
1.8
2
2.2
0
10
20
30
40
50
60

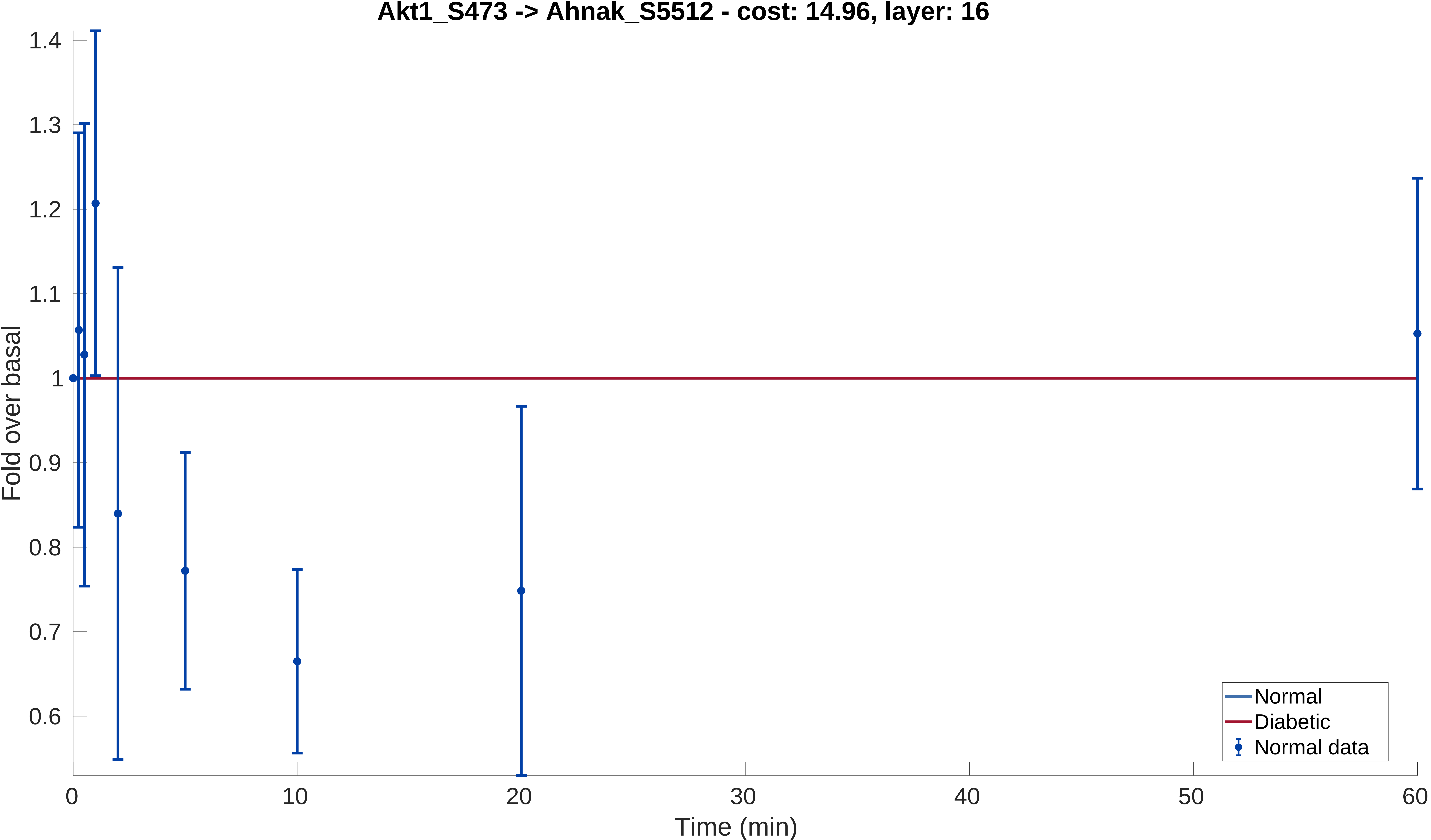

Akt1_S473 -> Ahnak_S5512 - cost: 14.96, layer: 16
Fold over basal
1.4
1.3
1.2
1.1
1
0.9
0.8
0.7
0.6
0
10
20
30
40
50
60
Time (min)
Normal
Diabetic
Normal data

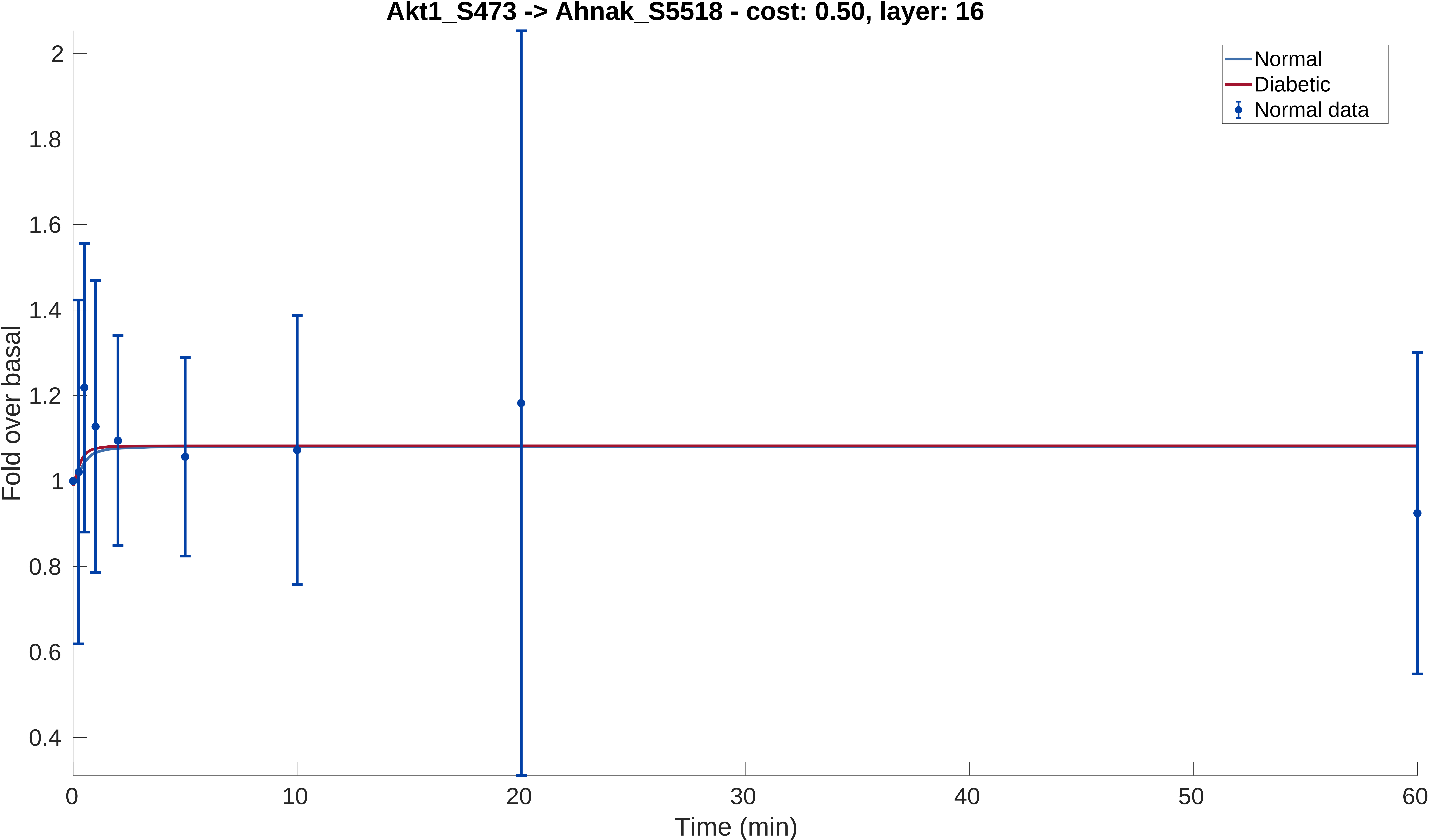
Akt1_S473 -> Ahnak_S5518 - cost: 0.50, layer: 16
Normal
Diabetic
Normal data
Fold over basal
Time (min)
2
1.8
1.6
1.4
1.2
1
0.8
0.6
0.4
0
10
20
30
40
50
60

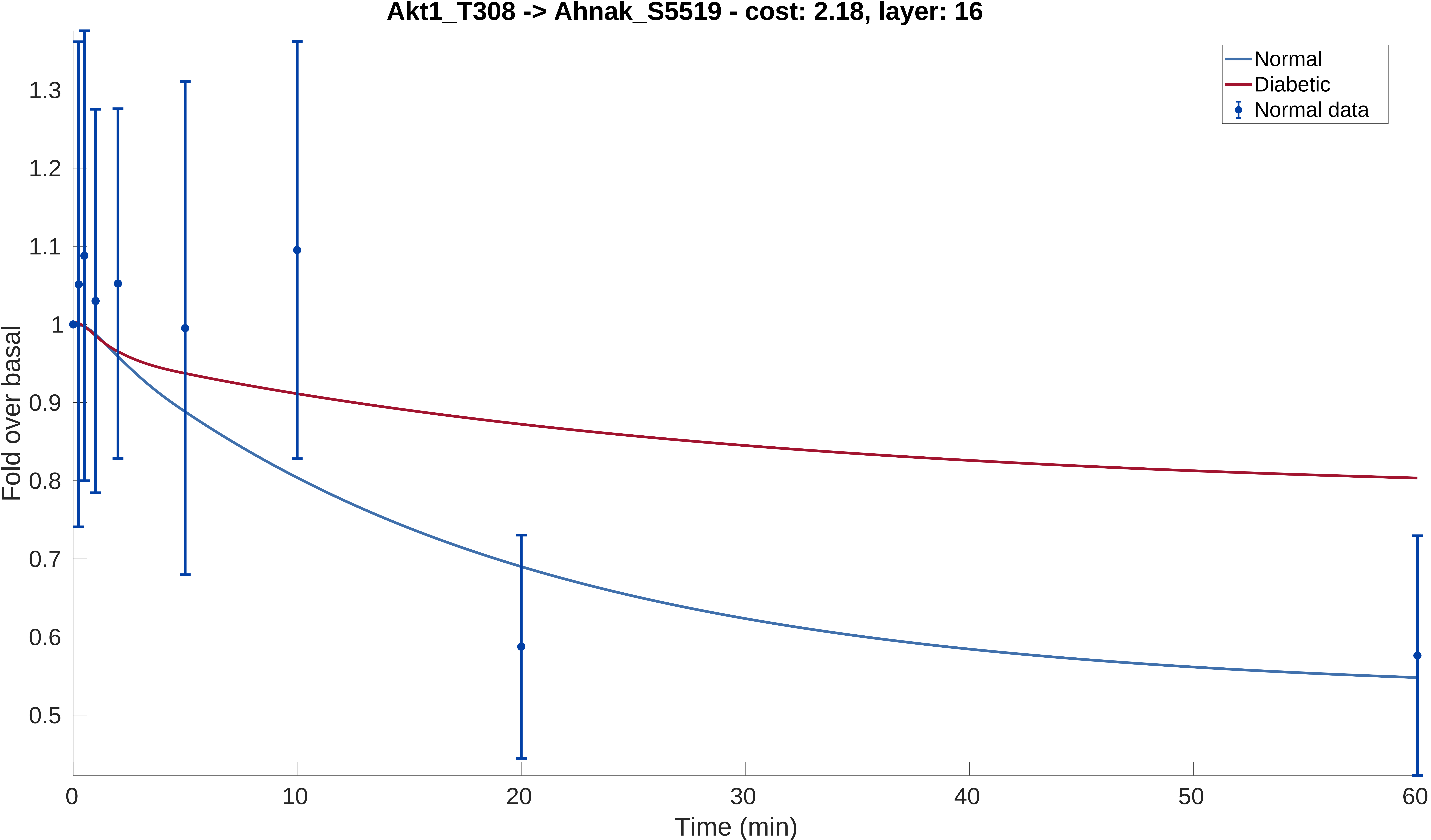

Akt1_T308 -> Ahnak_S5519 - cost: 2.18, layer: 16
Normal
Diabetic
Normal data
Fold over basal
1.3
1.2
1.1
1
0.9
0.8
0.7
0.6
0.5
0
10
20
30
40
50
60
Time (min)

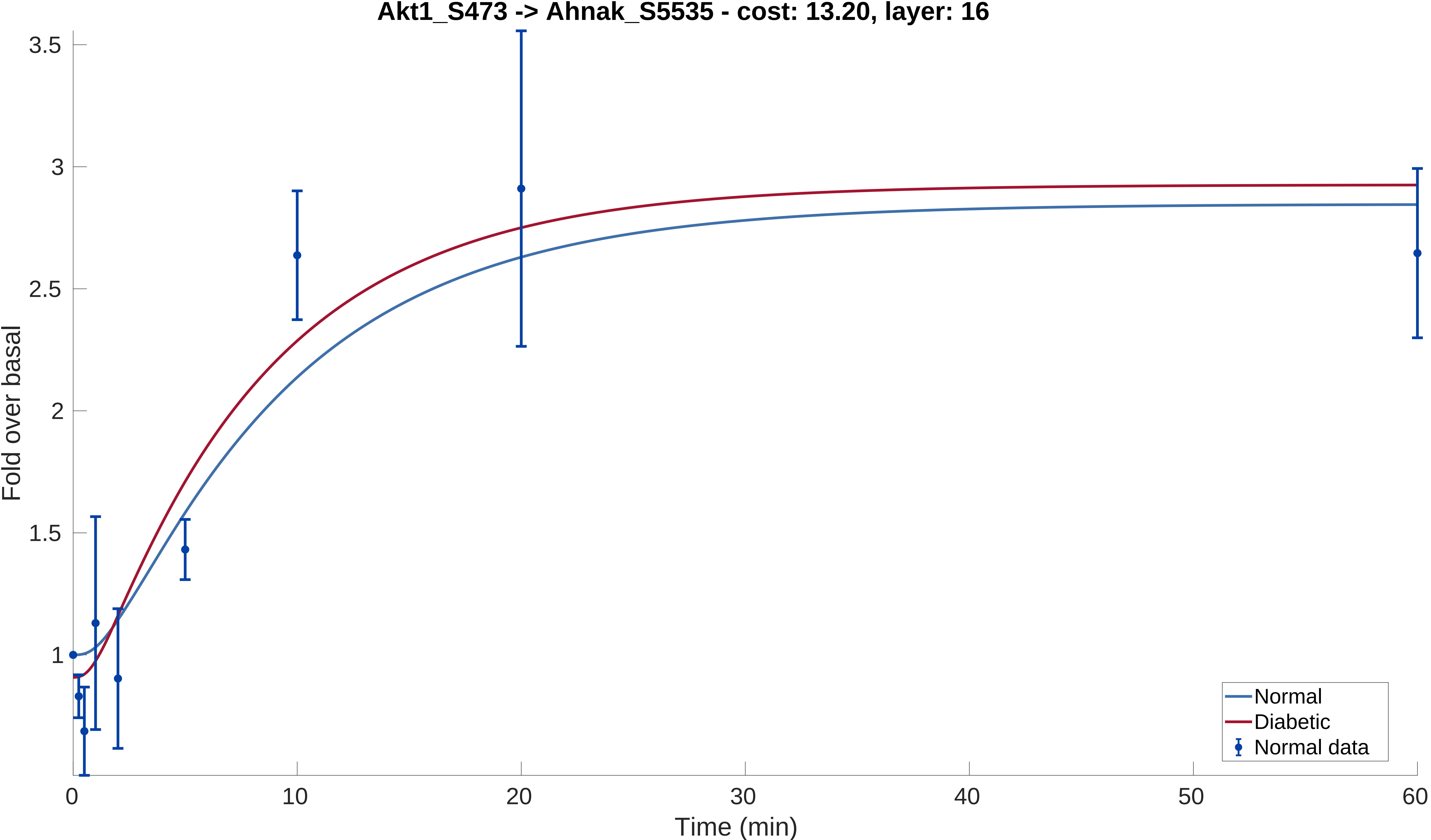

Akt1_S473 -> Ahnak_S5535 - cost: 13.20, layer: 16
Fold over basal
3.5
3
2.5
2
1.5
1
0
10
20
30
40
50
60
Time (min)
Normal
Diabetic
Normal data

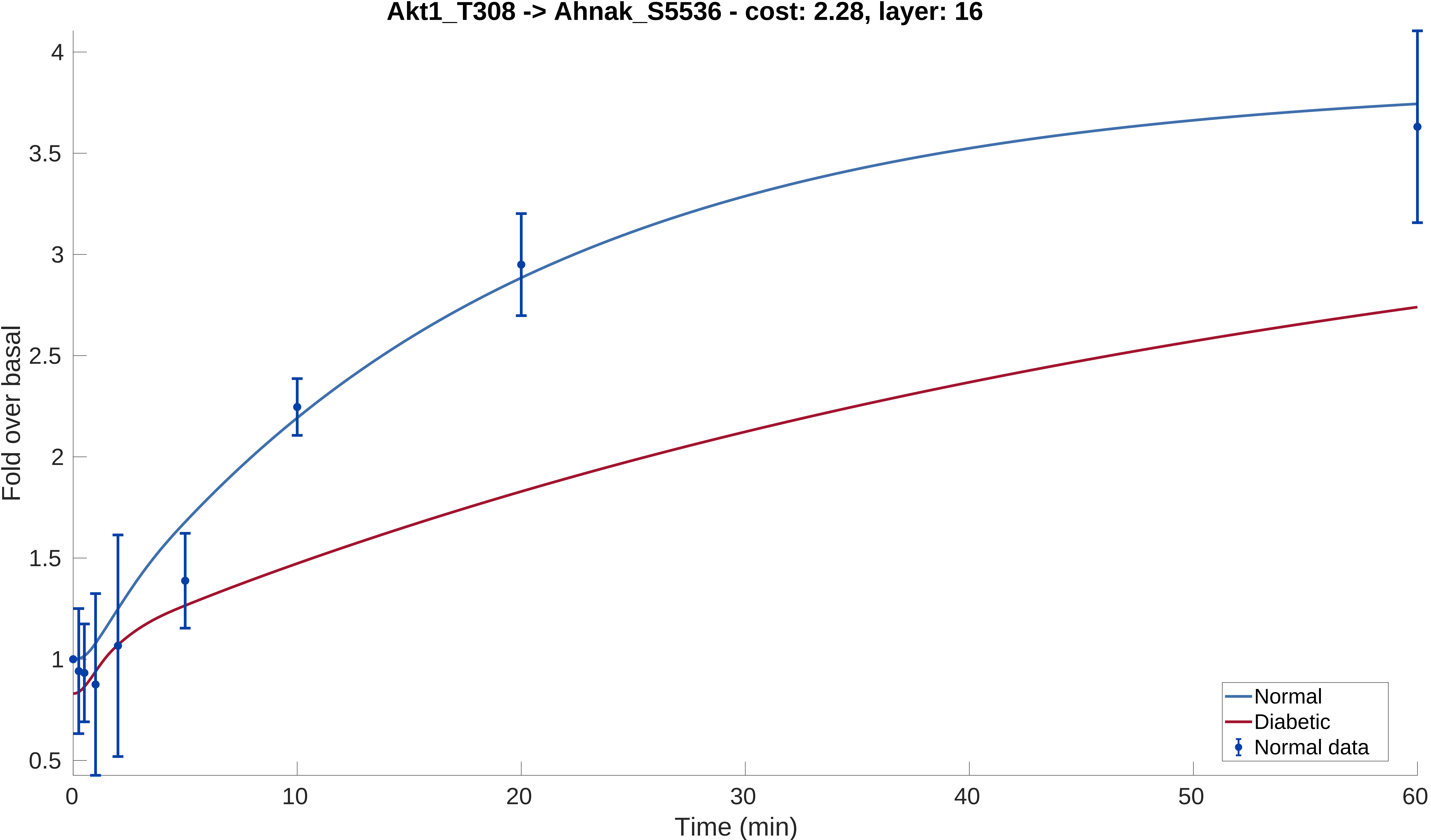
Akt1_T308 -> Ahnak_S5536 - cost: 2.28, layer: 16
Fold over basal
Time (min)
0.5
1
1.5
2
2.5
3
3.5
4
0
10
20
30
40
50
60
Normal
Diabetic
Normal data

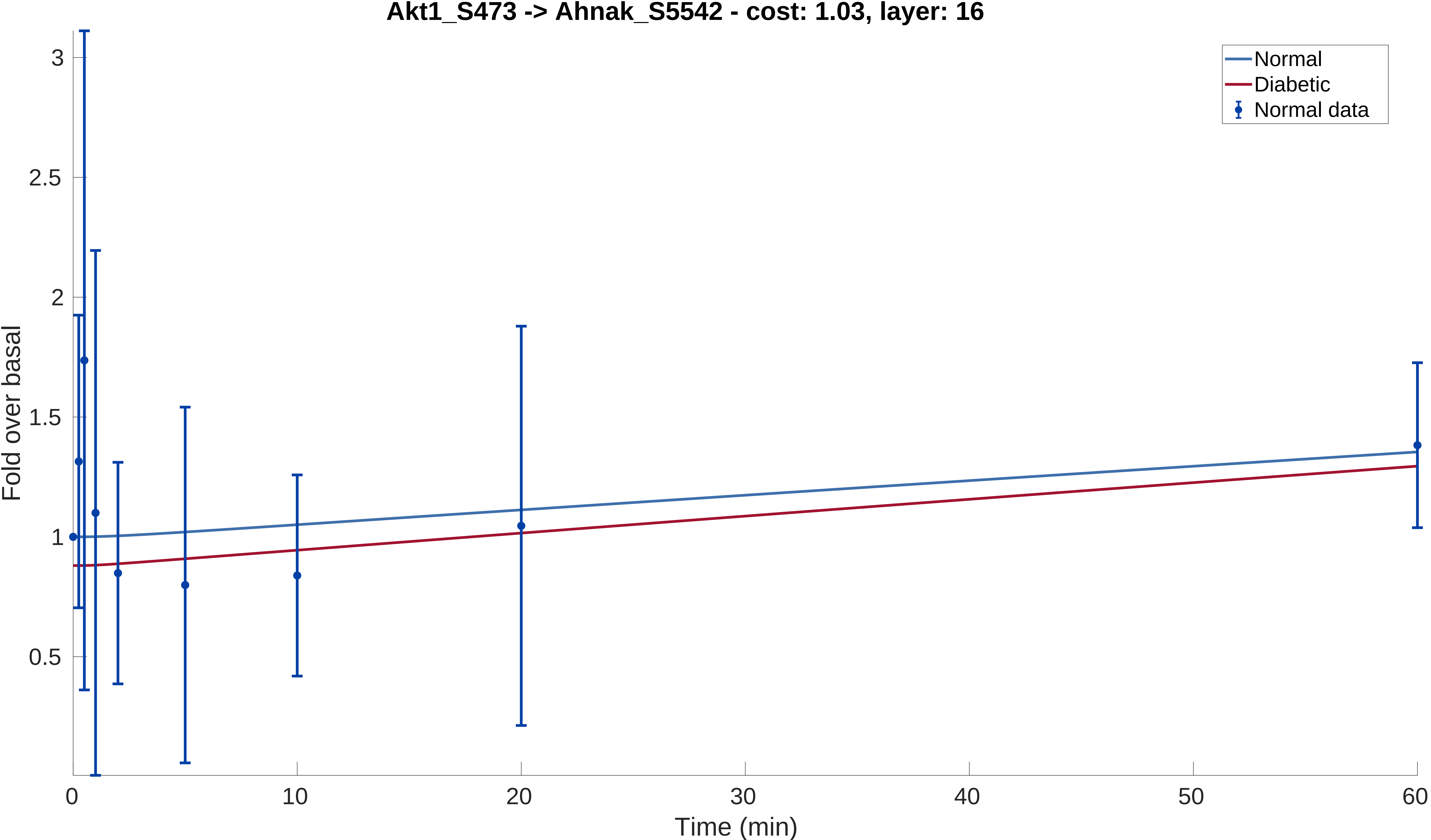

Akt1_S473 -> Ahnak_S5542 - cost: 1.03, layer: 16
Normal
Diabetic
Normal data
Fold over basal
3
2.5
2
1.5
1
0.5
0
10
20
30
40
50
60
Time (min)

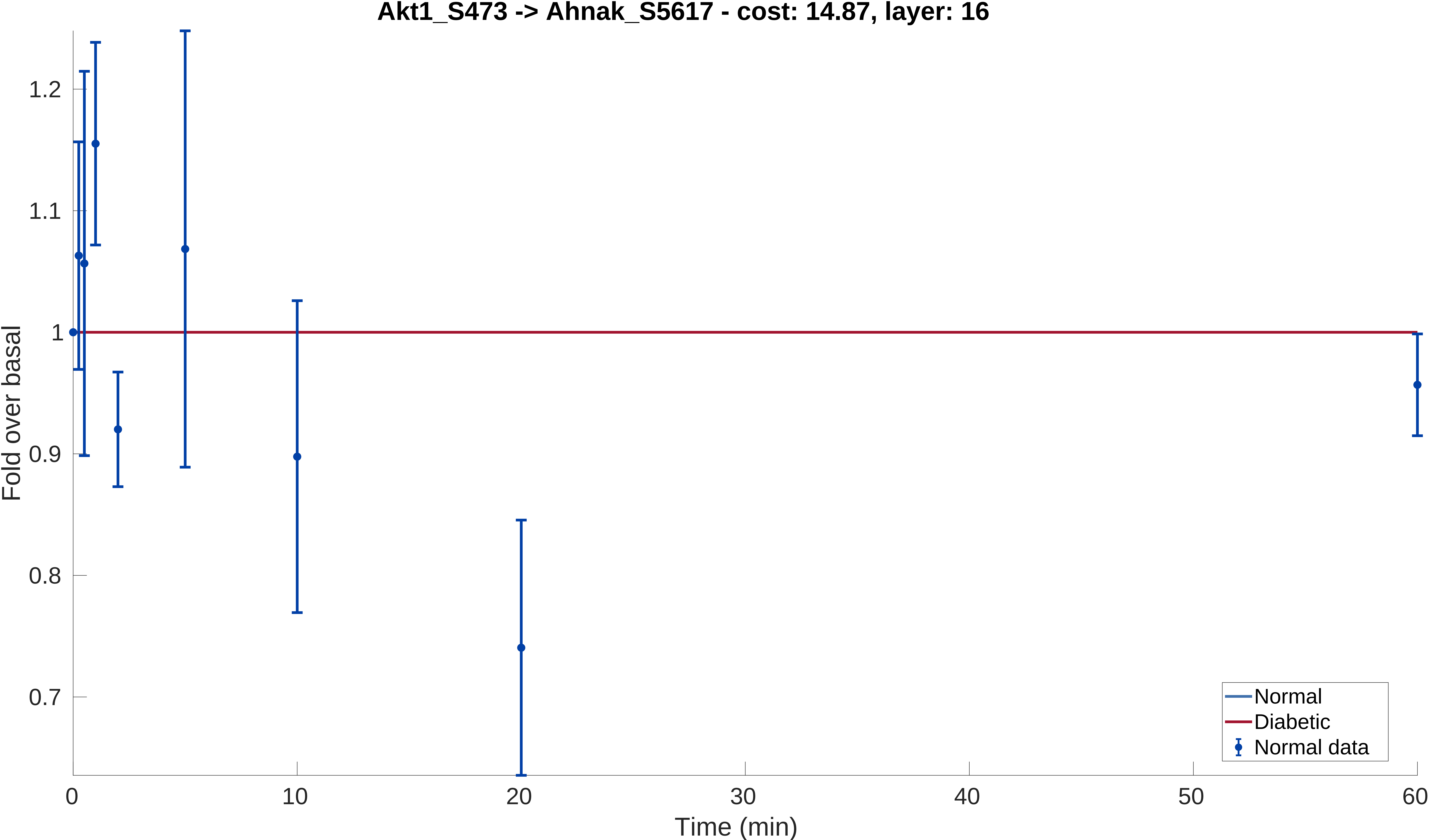

Akt1_S473 -> Ahnak_S5617 - cost: 14.87, layer: 16
Fold over basal
1.2
1.1
1
0.9
0.8
0.7
0
10
20
30
40
50
60
Time (min)
Normal
Diabetic
Normal data

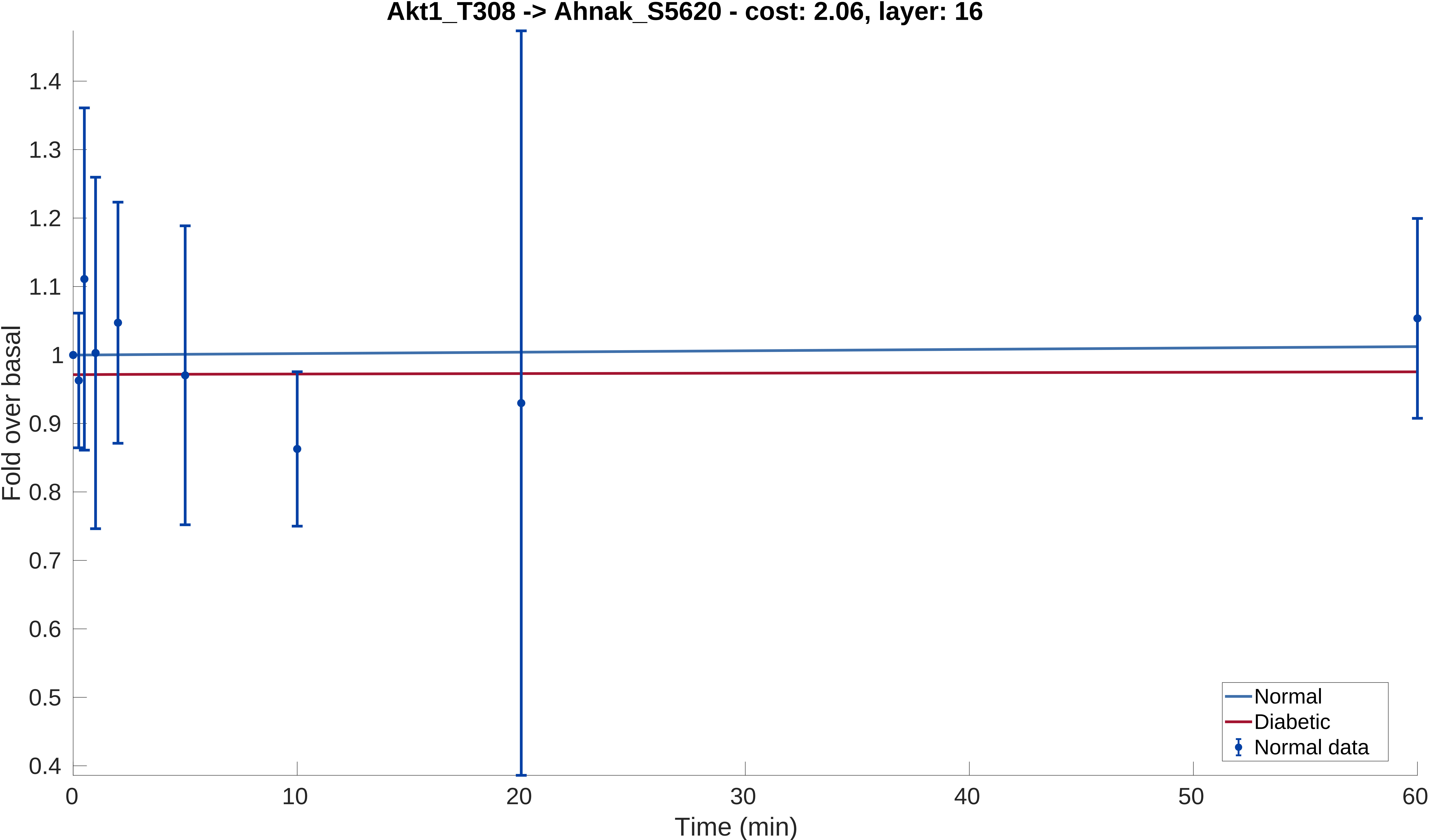

Akt1_T308 -> Ahnak_S5620 - cost: 2.06, layer: 16
Fold over basal
1.4
1.3
1.2
1.1
1
0.9
0.8
0.7
0.6
0.5
0.4
0
10
20
30
40
50
60
Time (min)
Normal
Diabetic
Normal data

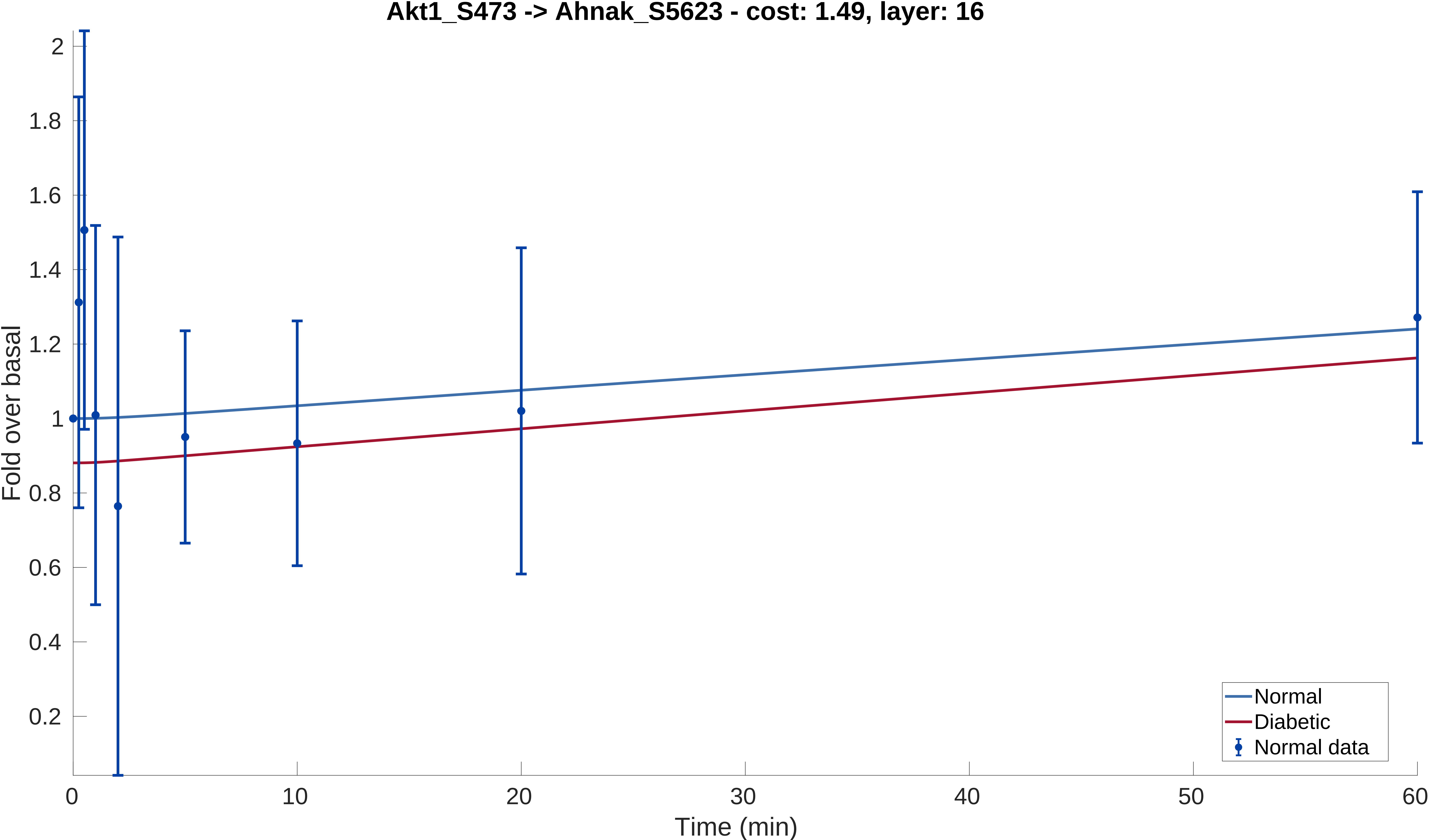

Akt1_S473 -> Ahnak_S5623 - cost: 1.49, layer: 16
Fold over basal
2
1.8
1.6
1.4
1.2
1
0.8
0.6
0.4
0.2
0
10
20
30
40
50
60
Time (min)
Normal
Diabetic
Normal data

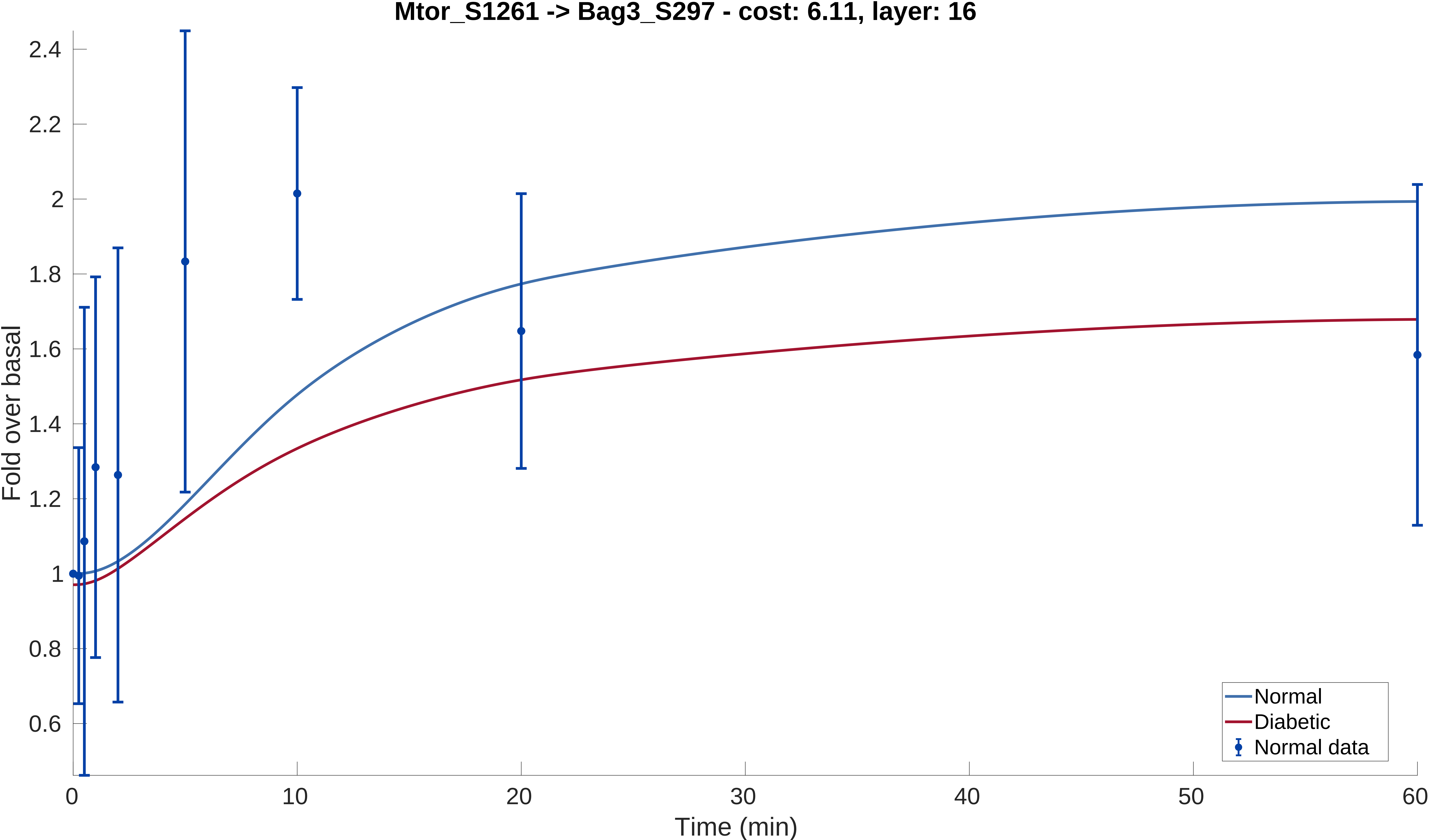

Mtor_S1261 -> Bag3_S297 - cost: 6.11, layer: 16
Fold over basal
2.4
2.2
2
1.8
1.6
1.4
1.2
1
0.8
0.6
0
10
20
30
40
50
60
Time (min)
Normal
Diabetic
Normal data

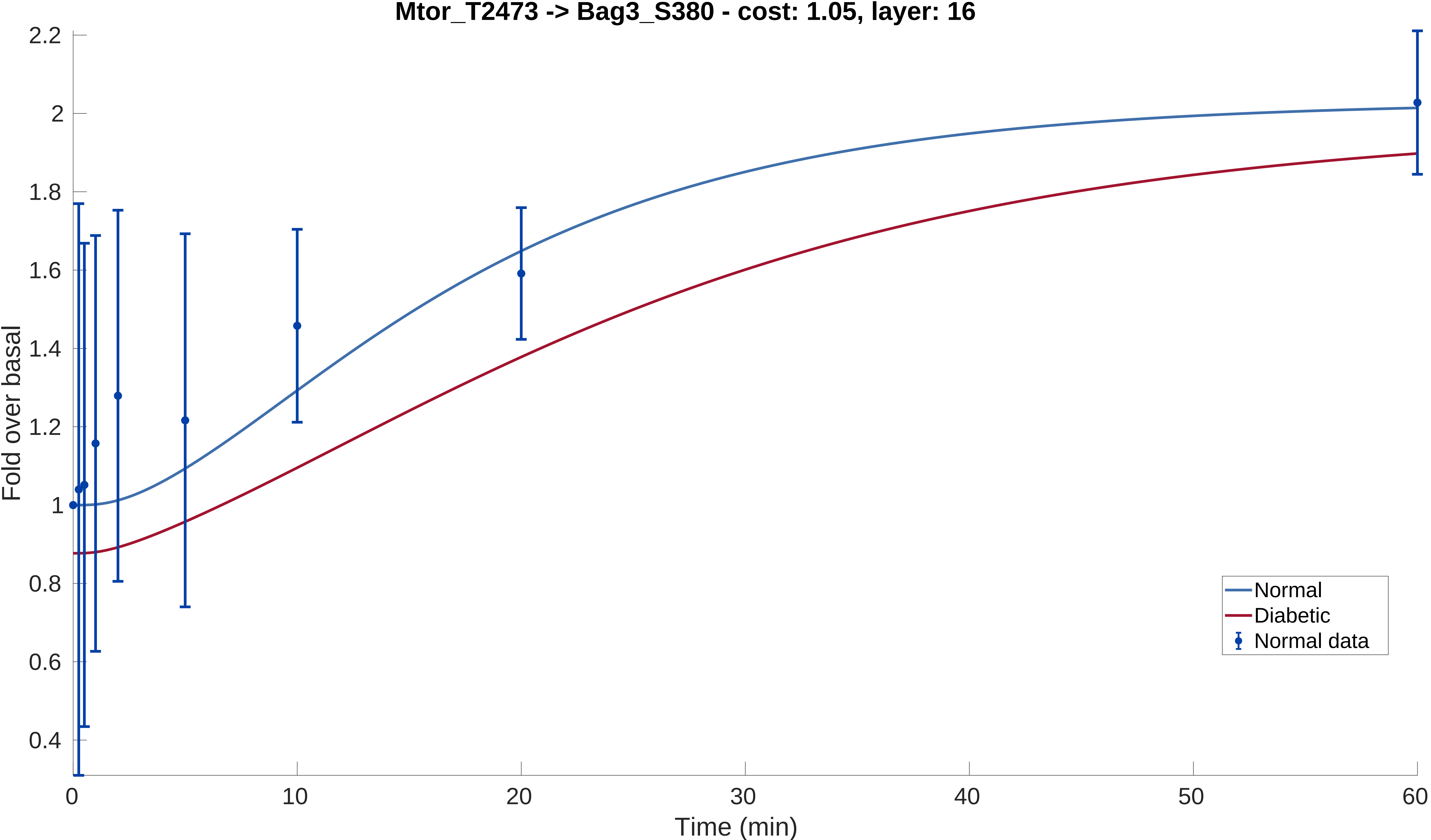
Mtor_T2473 -> Bag3_S380 - cost: 1.05, layer: 16
Fold over basal
Time (min)
2.2
2
1.8
1.6
1.4
1.2
1
0.8
0.6
0.4
0
10
20
30
40
50
60
Normal
Diabetic
Normal data

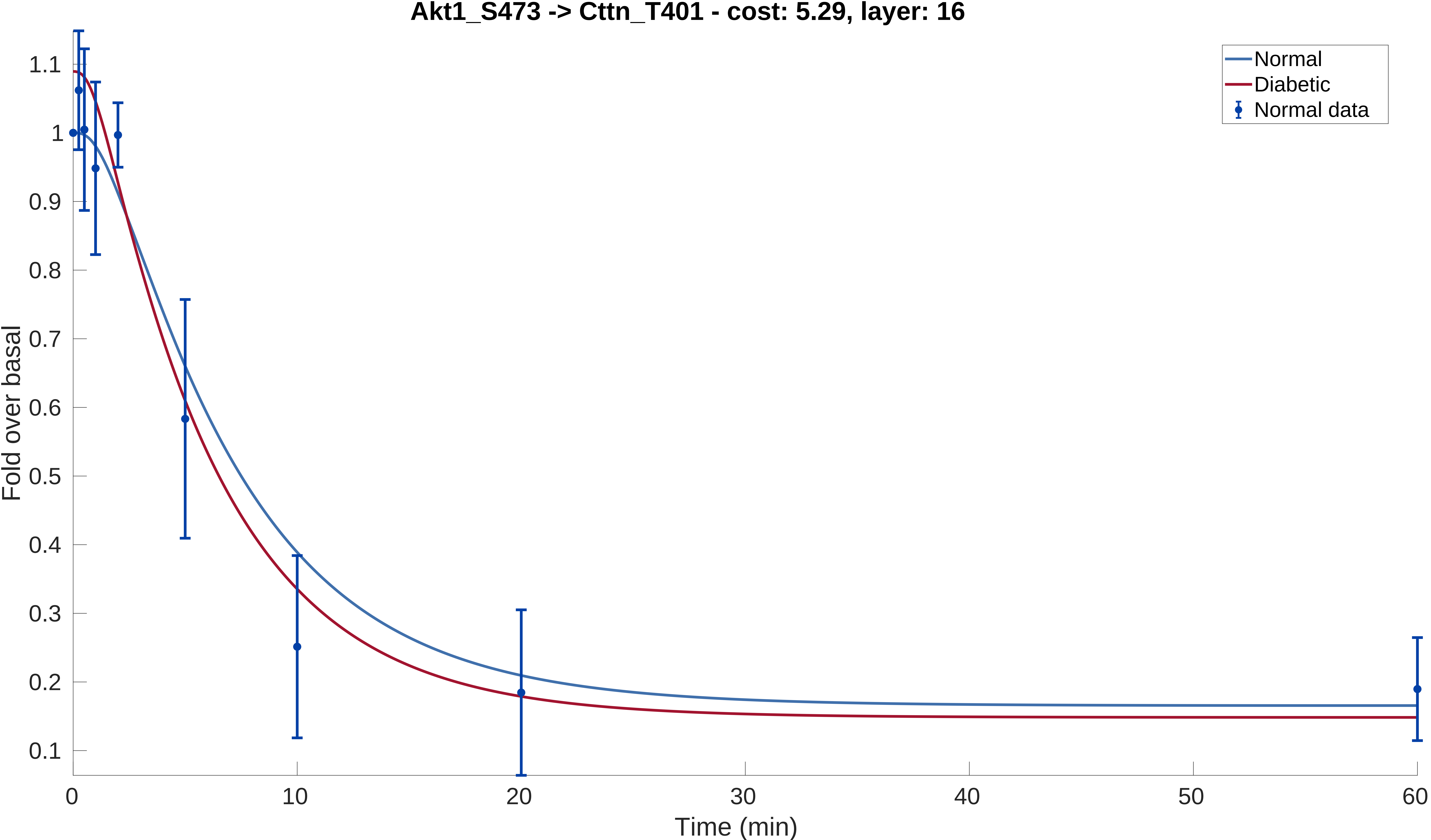

Akt1_S473 -> Cttn_T401 - cost: 5.29, layer: 16
Normal
Diabetic
Normal data
Fold over basal
Time (min)
0
10
20
30
40
50
60
0.1
0.2
0.3
0.4
0.5
0.6
0.7
0.8
0.9
1
1.1

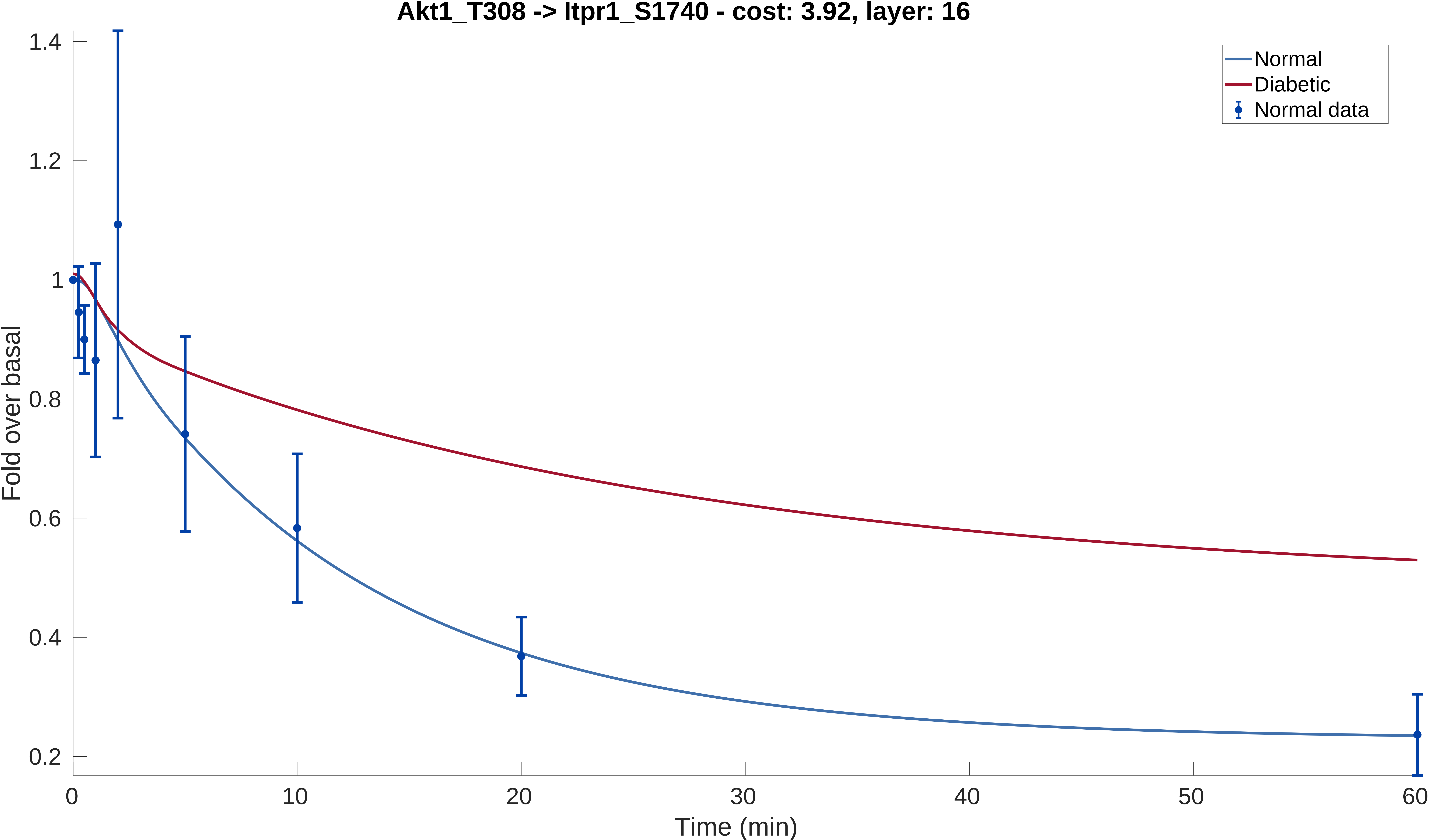
Akt1_T308 -> Itpr1_S1740 - cost: 3.92, layer: 16
Normal
Diabetic
Normal data
Fold over basal
1.4
1.2
1
0.8
0.6
0.4
0.2
0
10
20
30
40
50
60
Time (min)

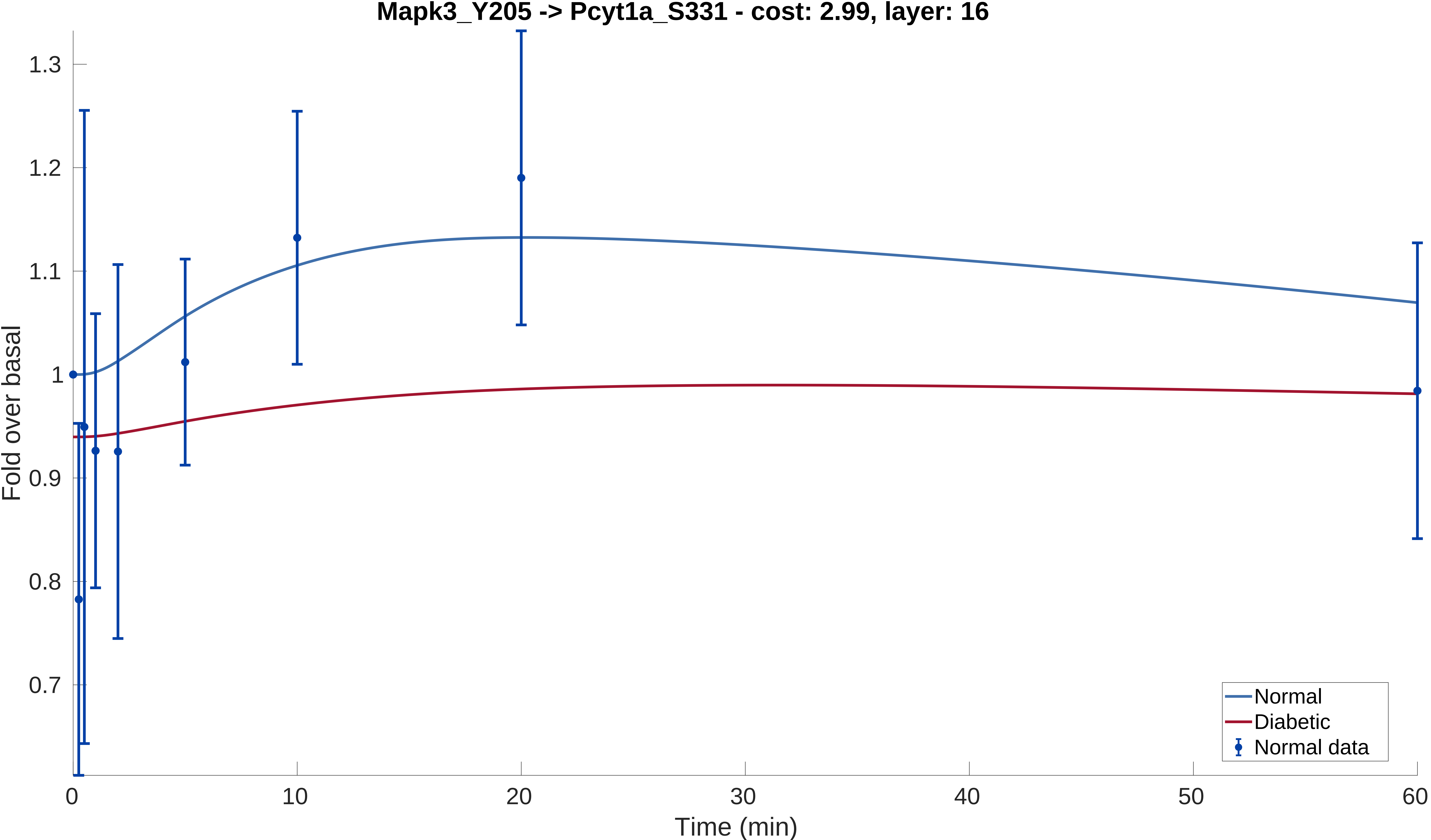
Mapk3_Y205 -> Pcyt1a_S331 - cost: 2.99, layer: 16
Fold over basal
1.3
1.2
1.1
1
0.9
0.8
0.7
0
10
20
30
40
50
60
Time (min)
Normal
Diabetic
Normal data

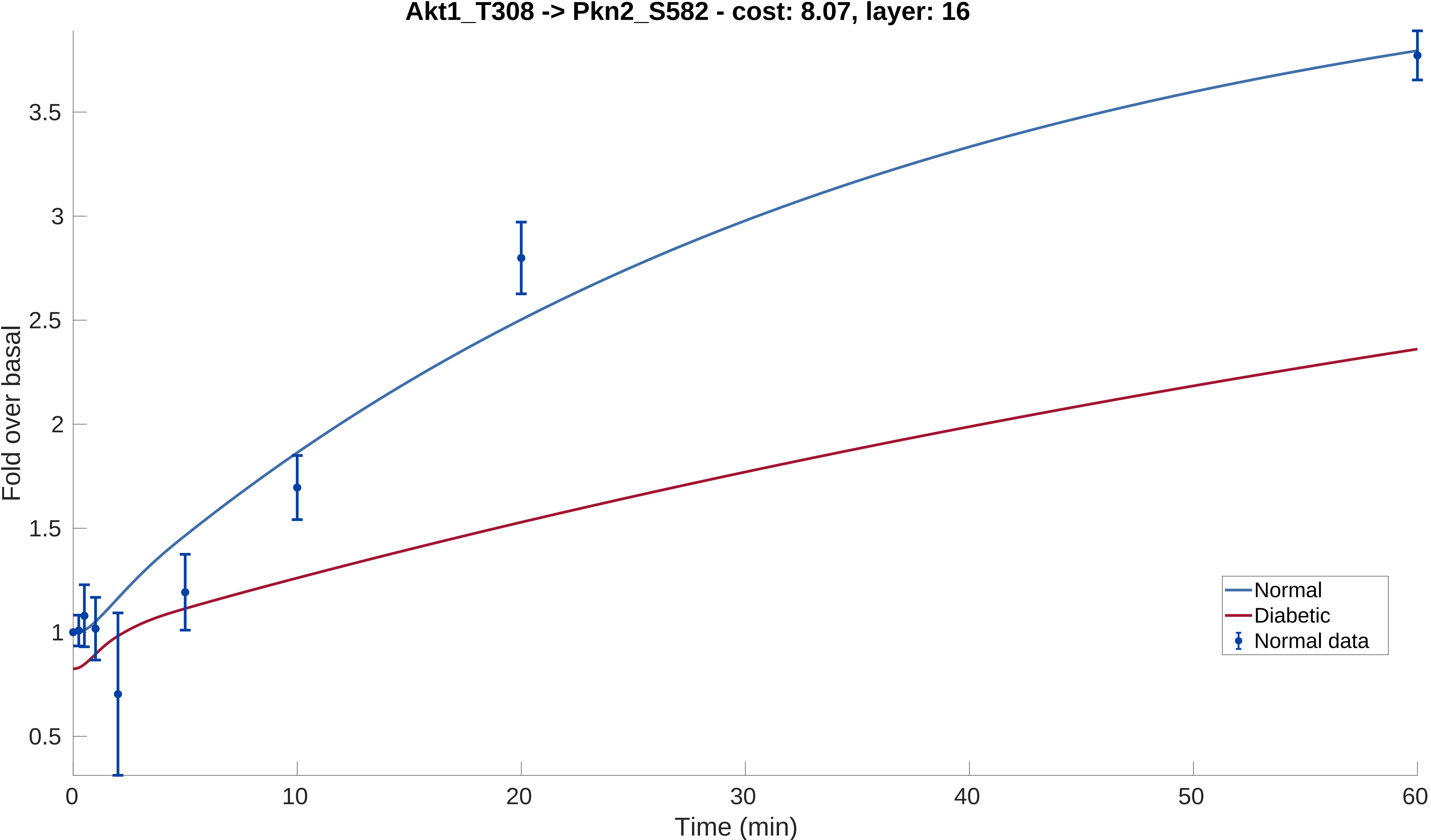

Akt1_T308 -> Pkn2_S582 - cost: 8.07, layer: 16
Fold over basal
3.5
3
2.5
2
1.5
1
0.5
0
10
20
30
40
50
60
Time (min)
Normal
Diabetic
Normal data

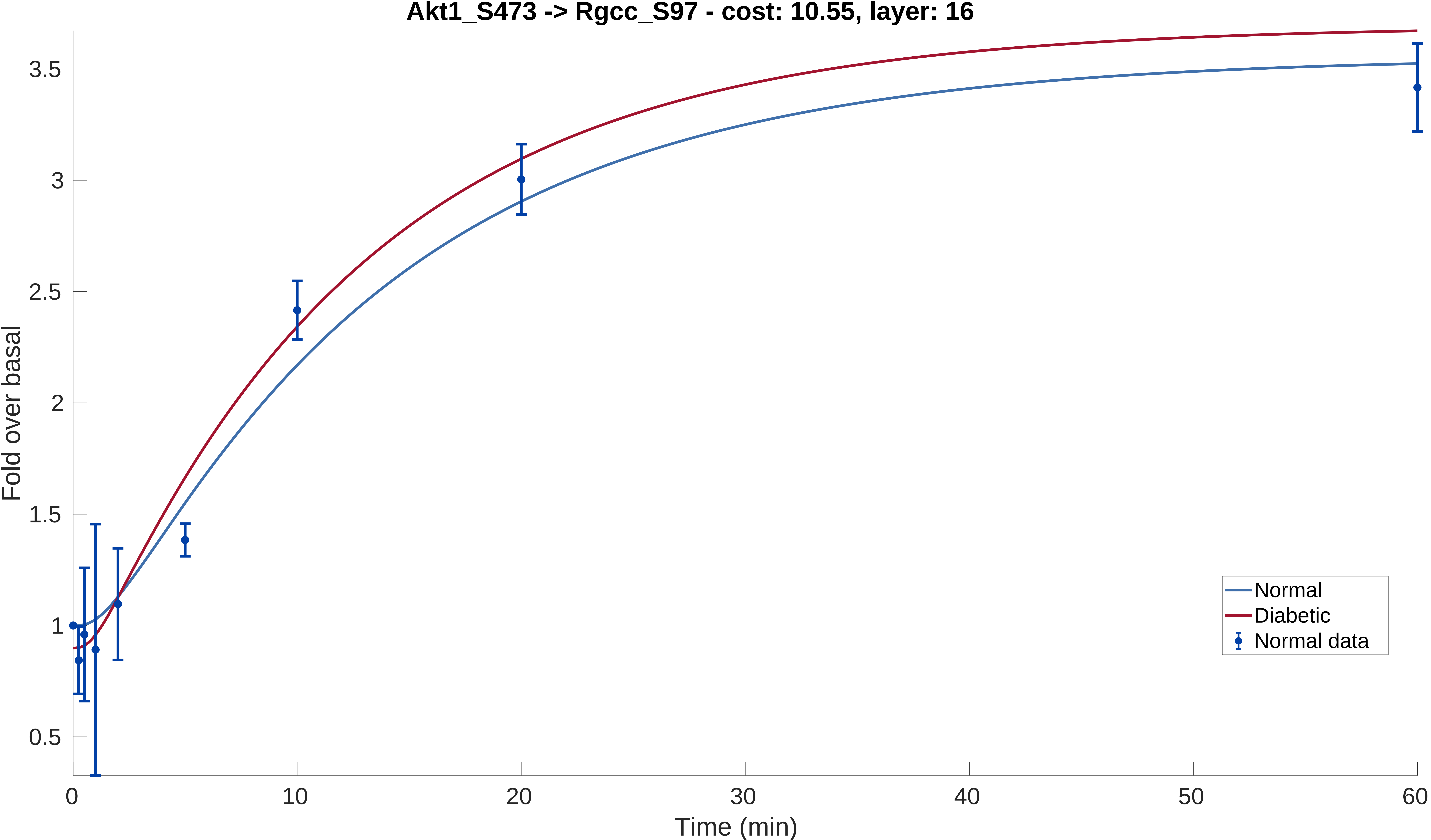

Akt1_S473 -> Rgcc_S97 - cost: 10.55, layer: 16
Fold over basal
Time (min)
0.5
1
1.5
2
2.5
3
3.5
0
10
20
30
40
50
60
Normal
Diabetic
Normal data

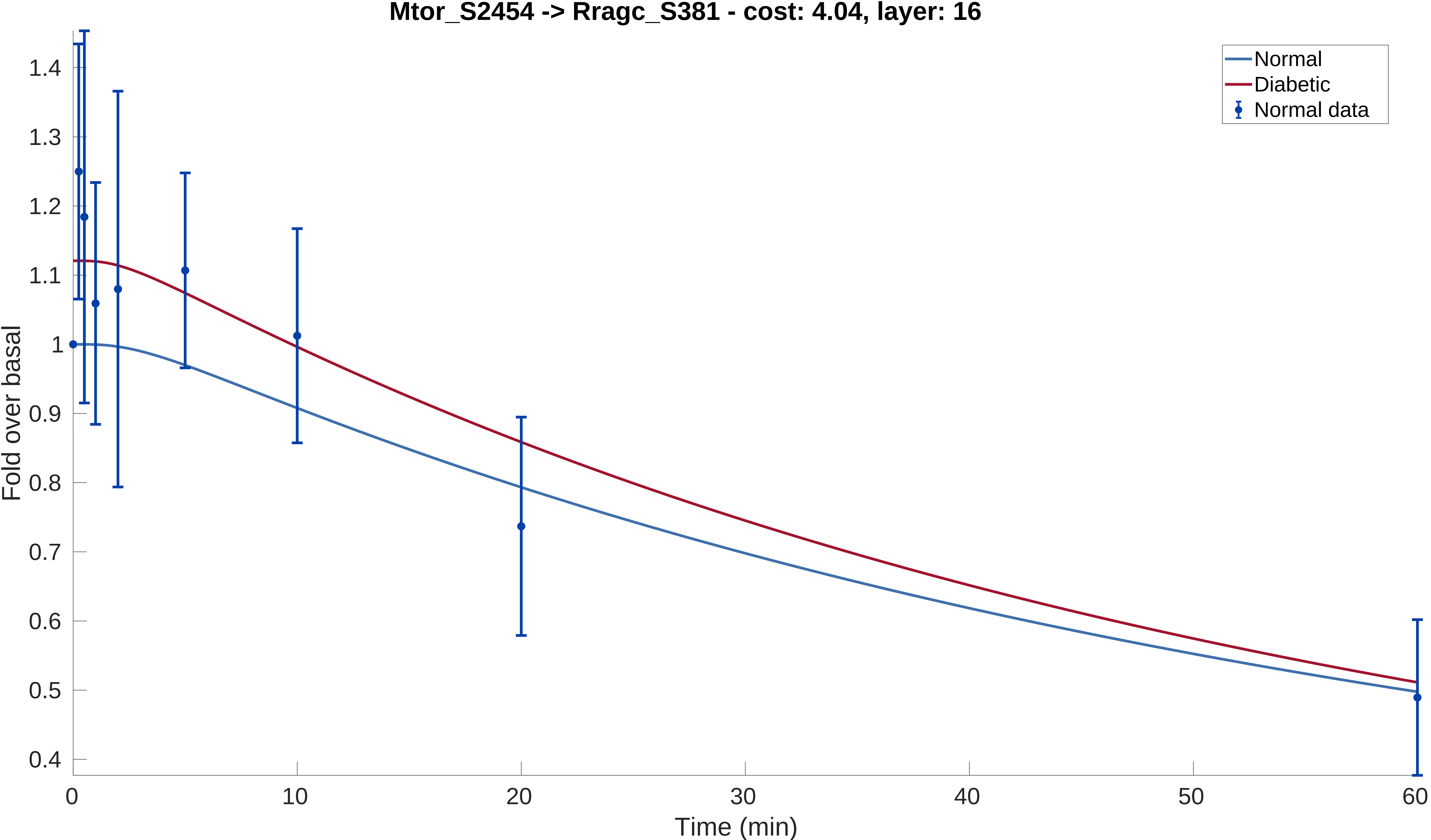

Mtor_S2454 -> Rragc_S381 - cost: 4.04, layer: 16
Normal
Diabetic
Normal data
Fold over basal
Time (min)
0.4
0.5
0.6
0.7
0.8
0.9
1
1.1
1.2
1.3
1.4
0
10
20
30
40
50
60

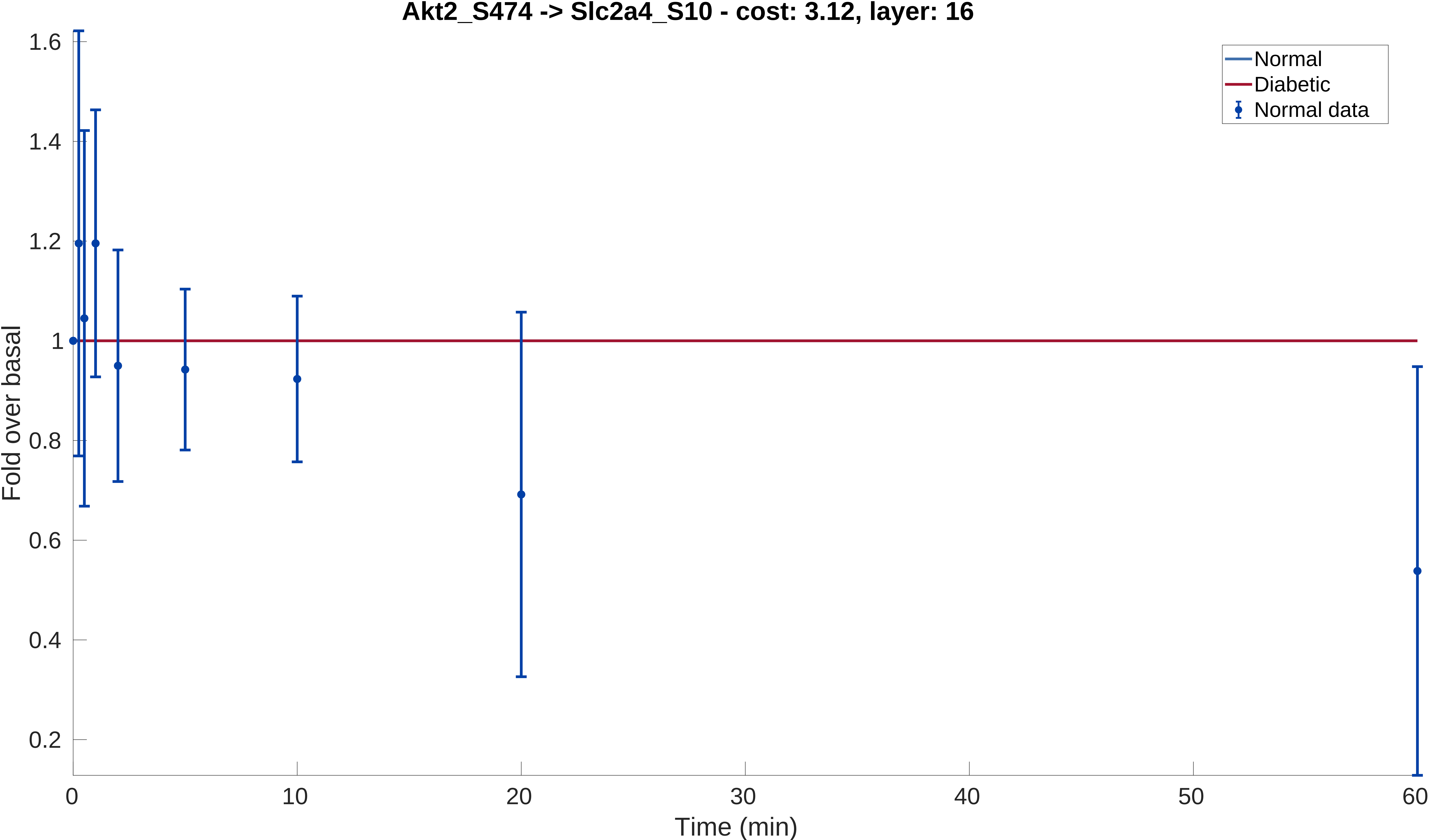

Akt2_S474 -> Slc2a4_S10 - cost: 3.12, layer: 16
Normal
Diabetic
Normal data
Fold over basal
1.6
1.4
1.2
1
0.8
0.6
0.4
0.2
0
10
20
30
40
50
60
Time (min)

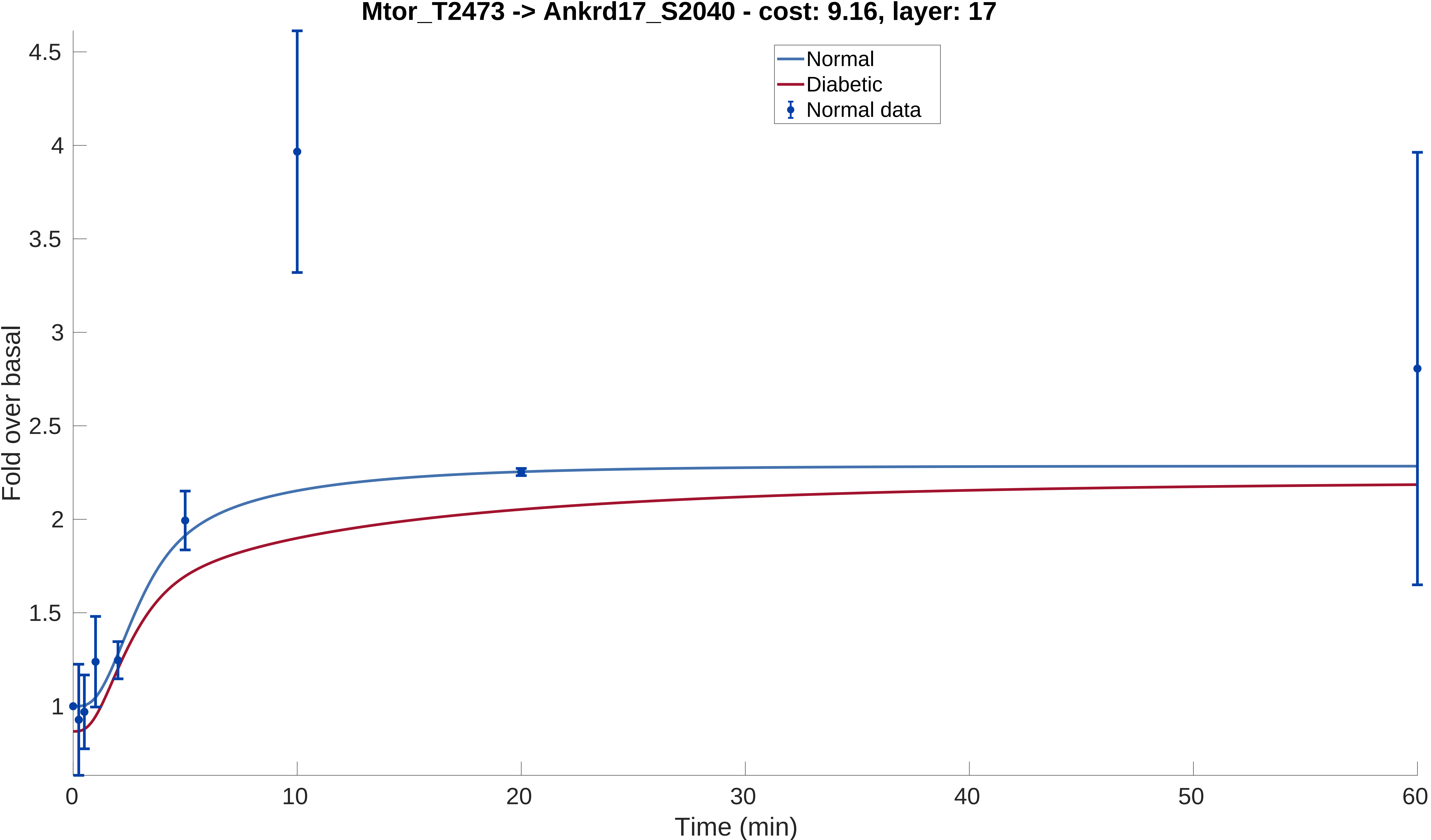

Mtor_T2473 -> Ankrd17_S2040 - cost: 9.16, layer: 17
Normal
Diabetic
Normal data
Fold over basal
Time (min)
4.5
4
3.5
3
2.5
2
1.5
1
0
10
20
30
40
50
60

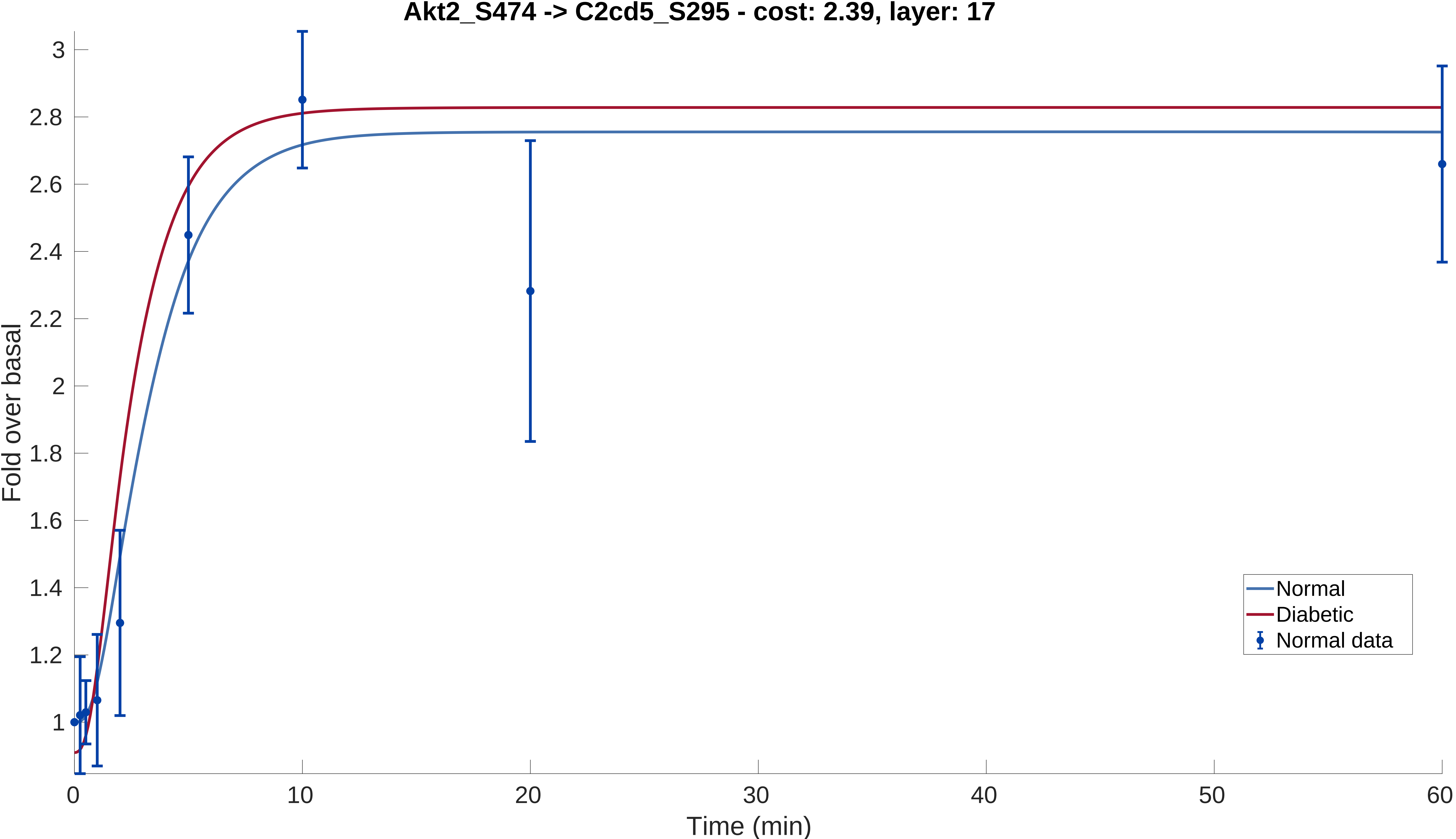
Akt2_S474 -> C2cd5_S295 - cost: 2.39, layer: 17
Fold over basal
3
2.8
2.6
2.4
2.2
2
1.8
1.6
1.4
1.2
1
0
10
20
30
40
50
60
Time (min)
Normal
Diabetic
Normal data

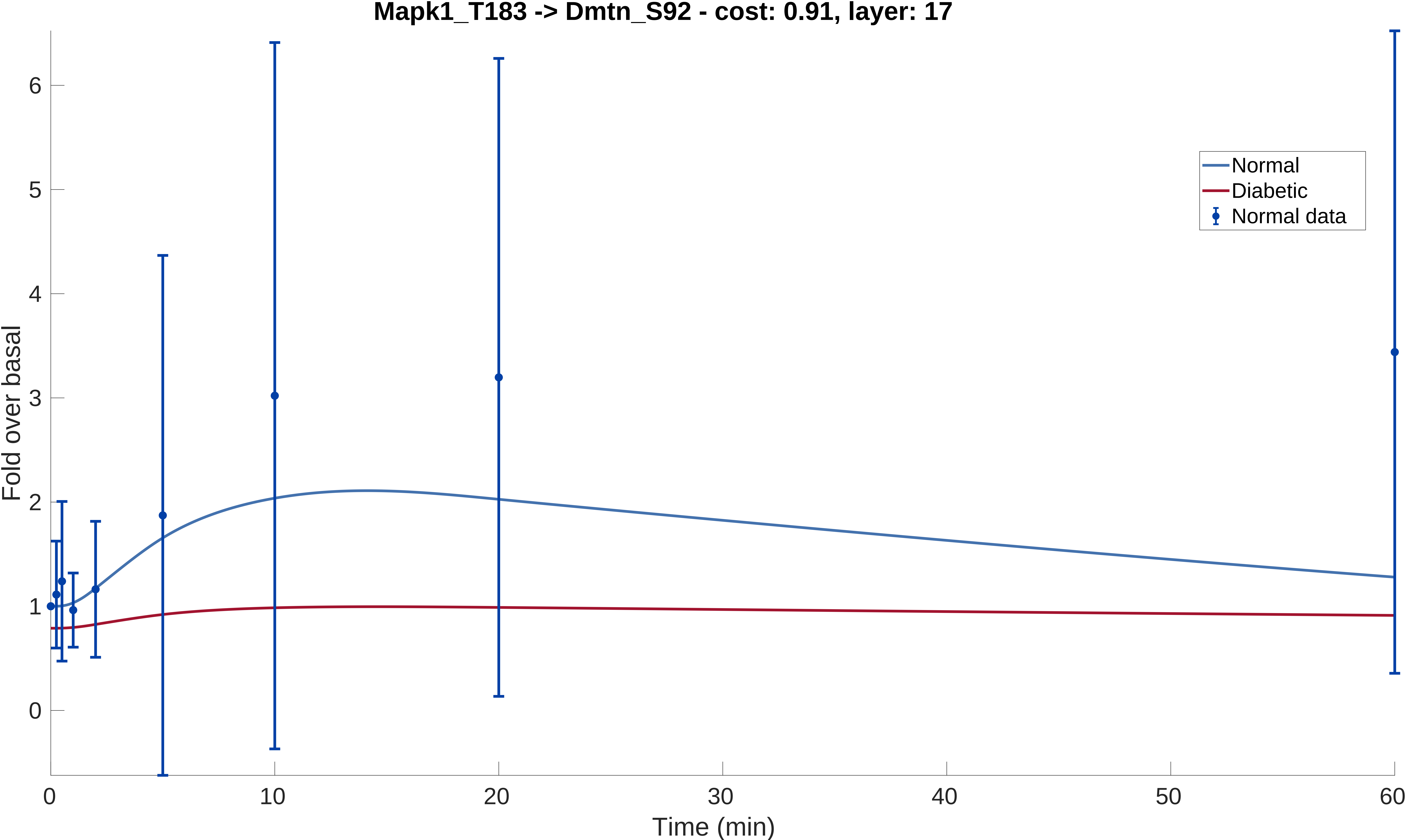
Mapk1_T183 -> Dmtn_S92 - cost: 0.91, layer: 17
Fold over basal
Time (min)
Normal
Diabetic
Normal data
0
1
2
3
4
5
6
0
10
20
30
40
50
60

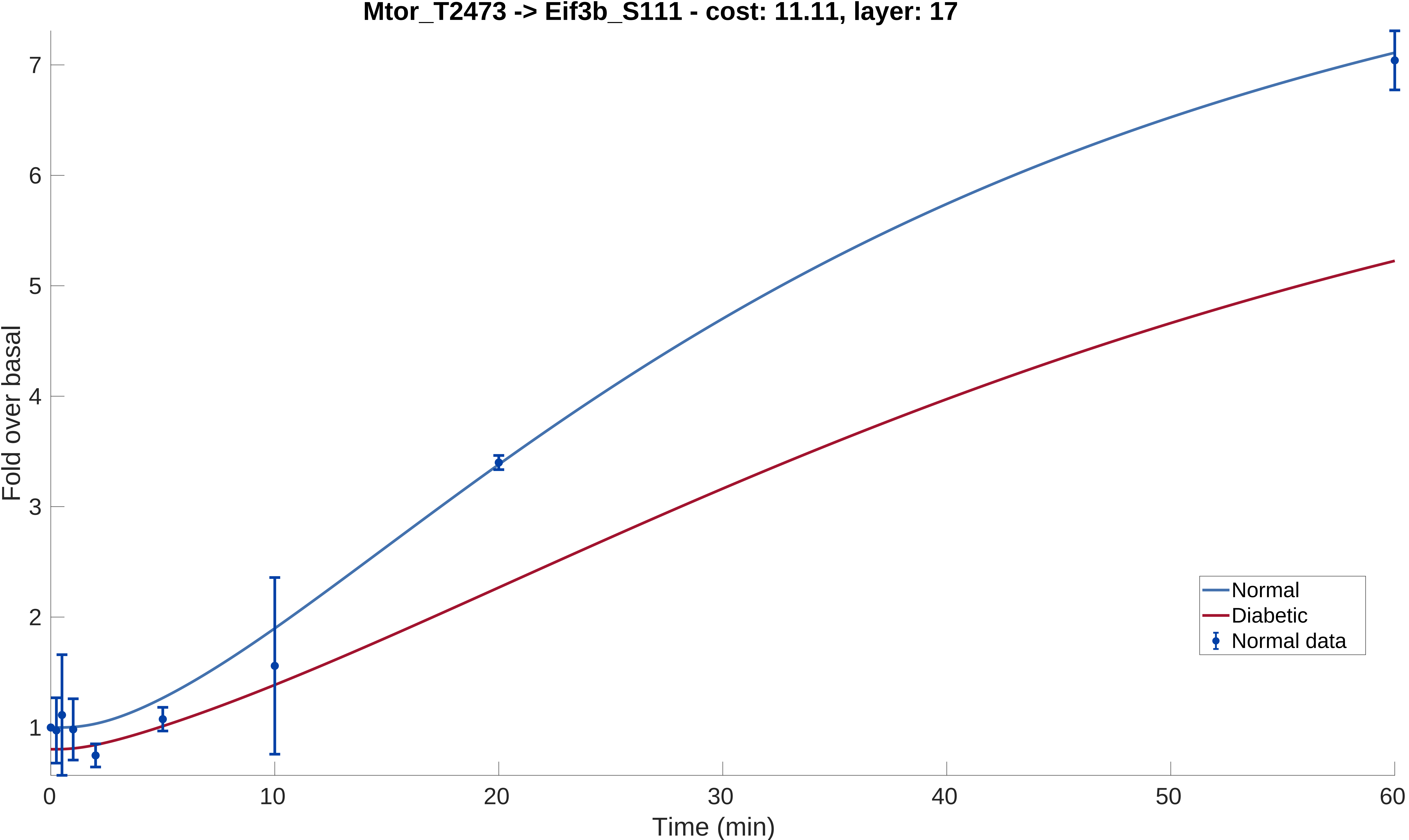

Mtor_T2473 -> Eif3b_S111 - cost: 11.11, layer: 17
Fold over basal
Time (min)
0
10
20
30
40
50
60
1
2
3
4
5
6
7
Normal
Diabetic
Normal data

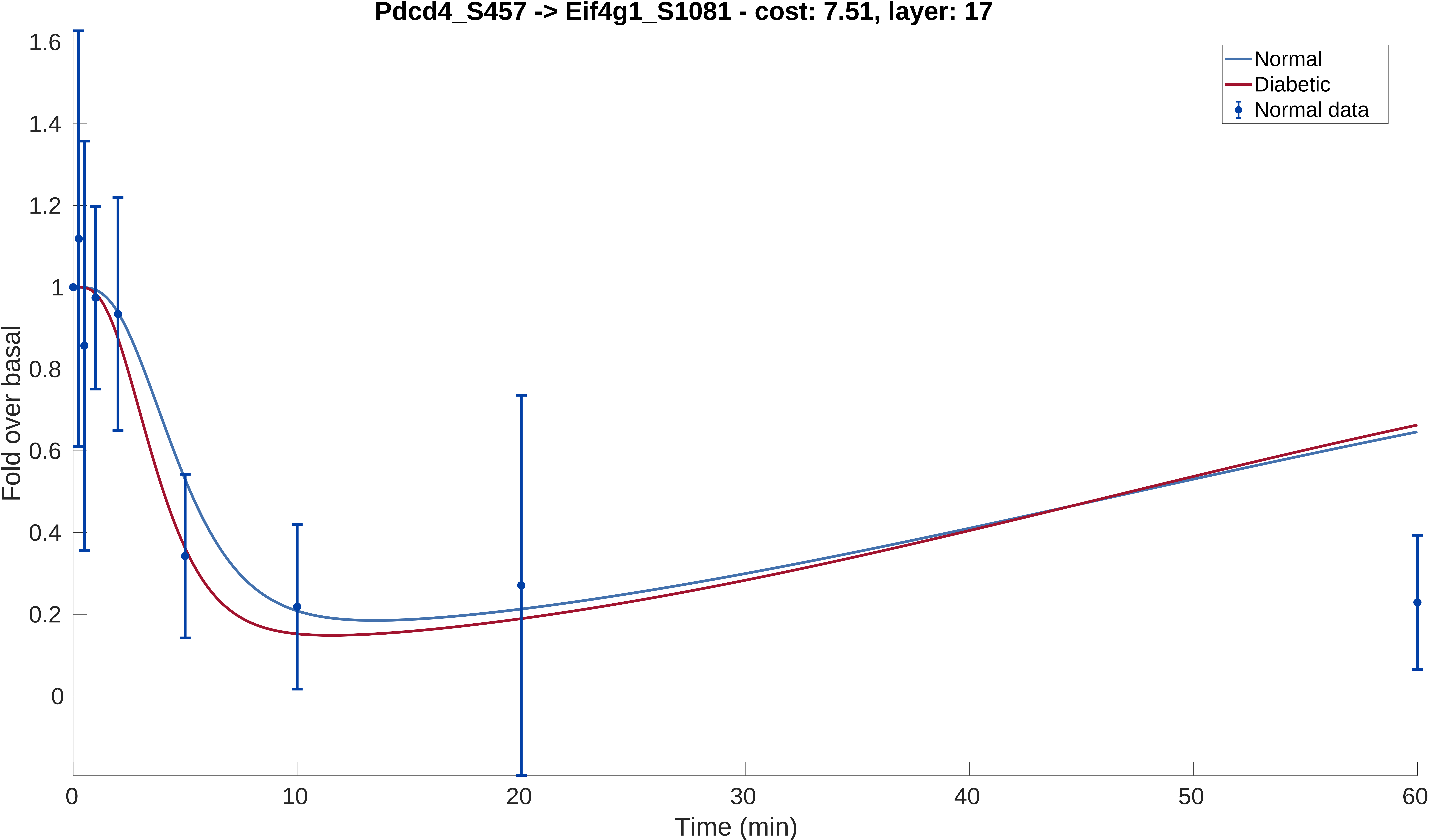

Pdcd4_S457 -> Eif4g1_S1081 - cost: 7.51, layer: 17
Normal
Diabetic
Normal data
Fold over basal
Time (min)
1.6
1.4
1.2
1
0.8
0.6
0.4
0.2
0
0
10
20
30
40
50
60

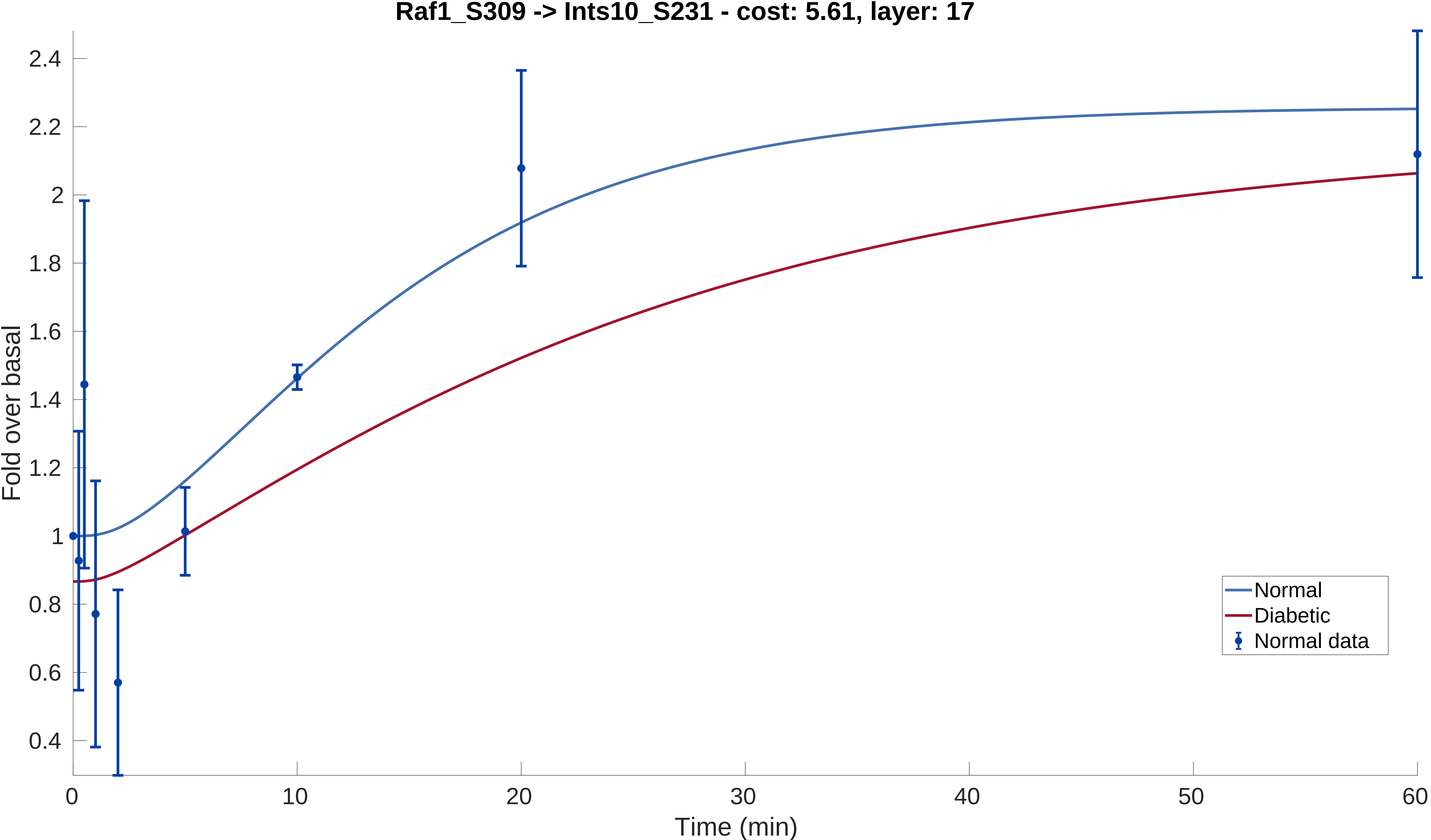

Raf1_S309 -> Ints10_S231 - cost: 5.61, layer: 17
Fold over basal
Time (min)
0
10
20
30
40
50
60
0.4
0.6
0.8
1
1.2
1.4
1.6
1.8
2
2.2
2.4
Normal
Diabetic
Normal data

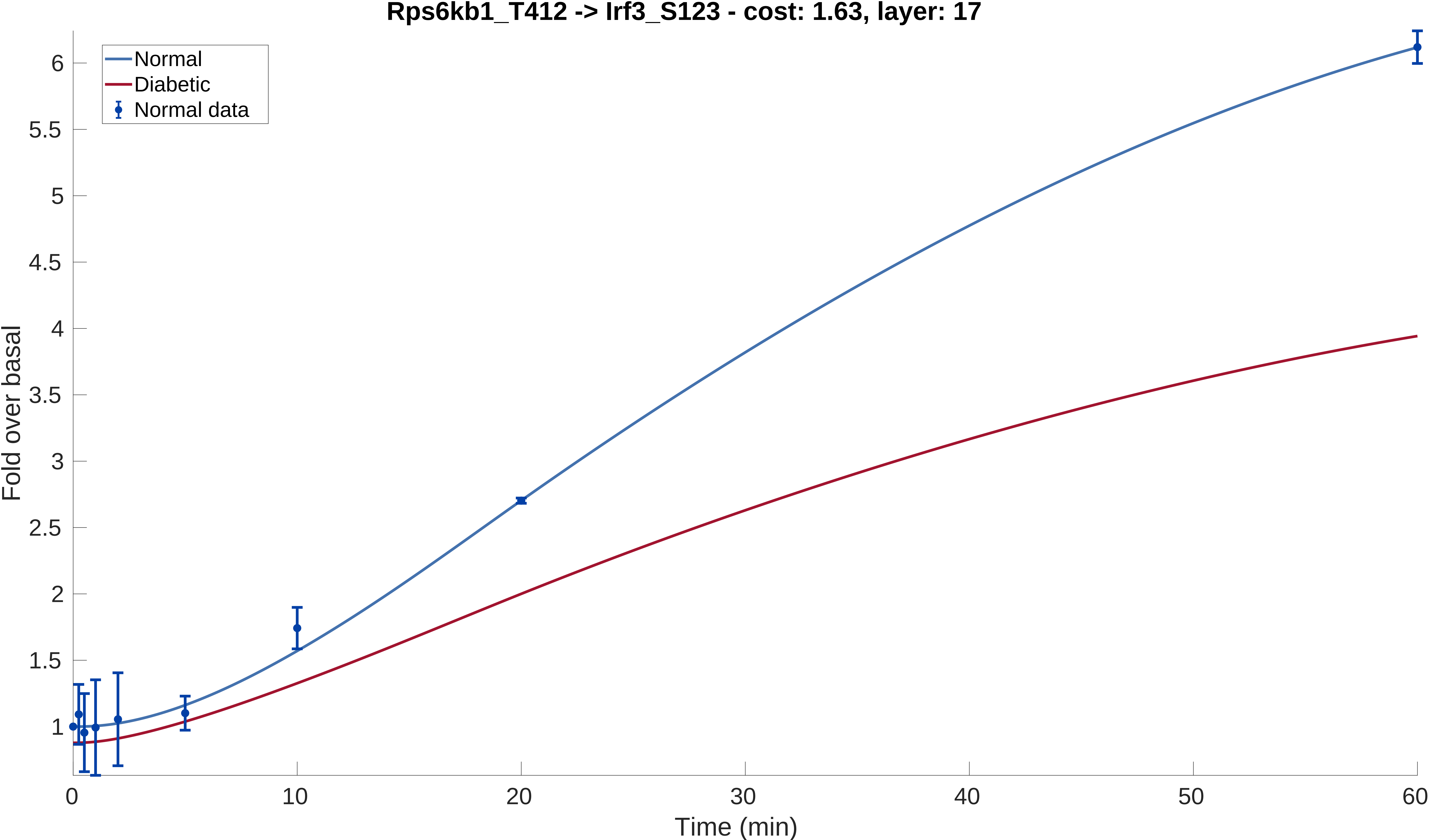

Rps6kb1_T412 -> Irf3_S123 - cost: 1.63, layer: 17
Normal
Diabetic
Normal data
Fold over basal
6
5.5
5
4.5
4
3.5
3
2.5
2
1.5
1
0
10
20
30
40
50
60
Time (min)

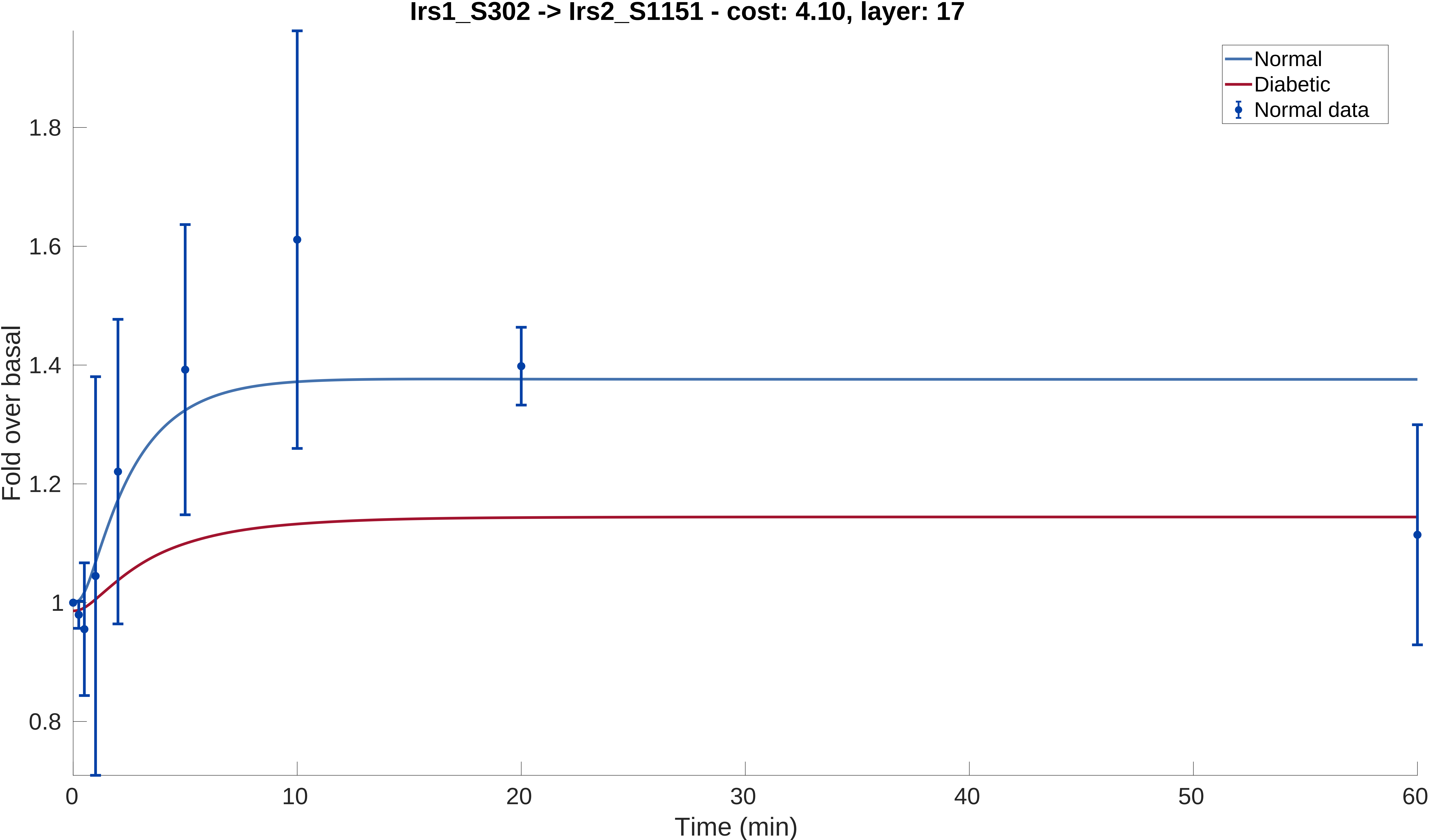
Irs1_S302 -> Irs2_S1151 - cost: 4.10, layer: 17
Normal
Diabetic
Normal data
Fold over basal
1.8
1.6
1.4
1.2
1
0.8
0
10
20
30
40
50
60
Time (min)

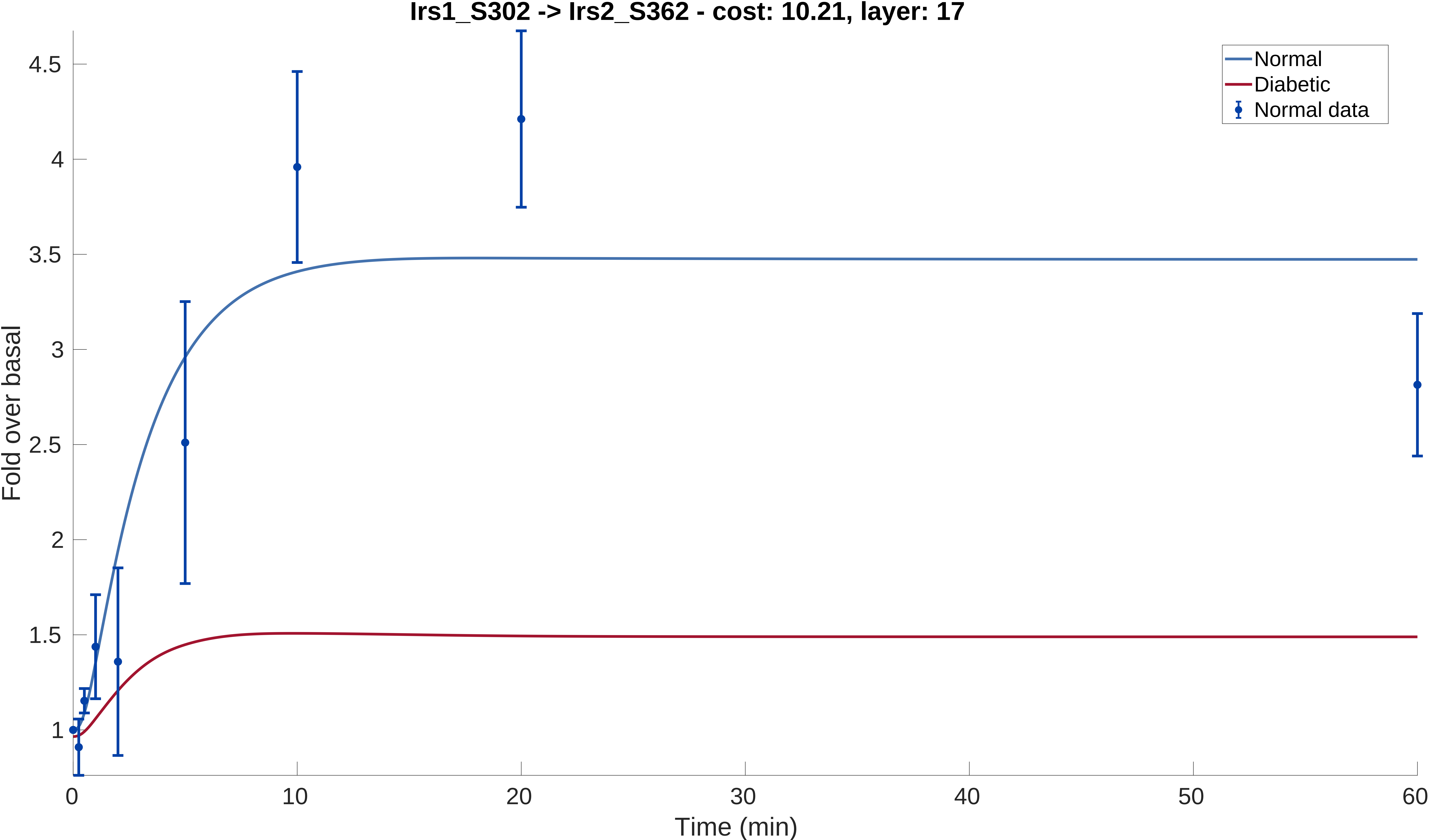

Irs1_S302 -> Irs2_S362 - cost: 10.21, layer: 17
Normal
Diabetic
Normal data
Fold over basal
Time (min)
4.5
4
3.5
3
2.5
2
1.5
1
0
10
20
30
40
50
60

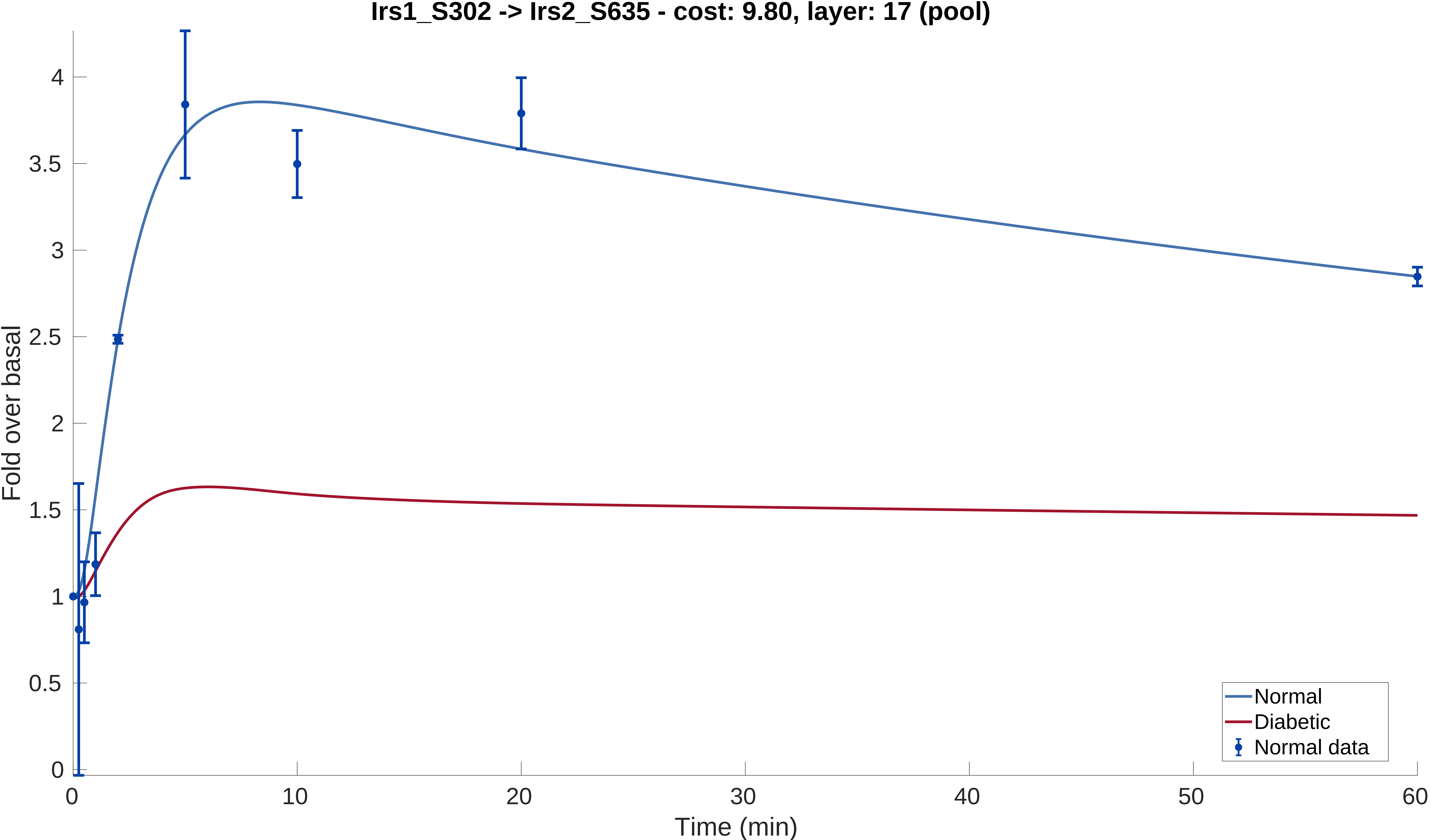

Irs1_S302 -> Irs2_S635 - cost: 9.80, layer: 17 (pool)
Fold over basal
Time (min)
0
0.5
1
1.5
2
2.5
3
3.5
4
0
10
20
30
40
50
60
Normal
Diabetic
Normal data

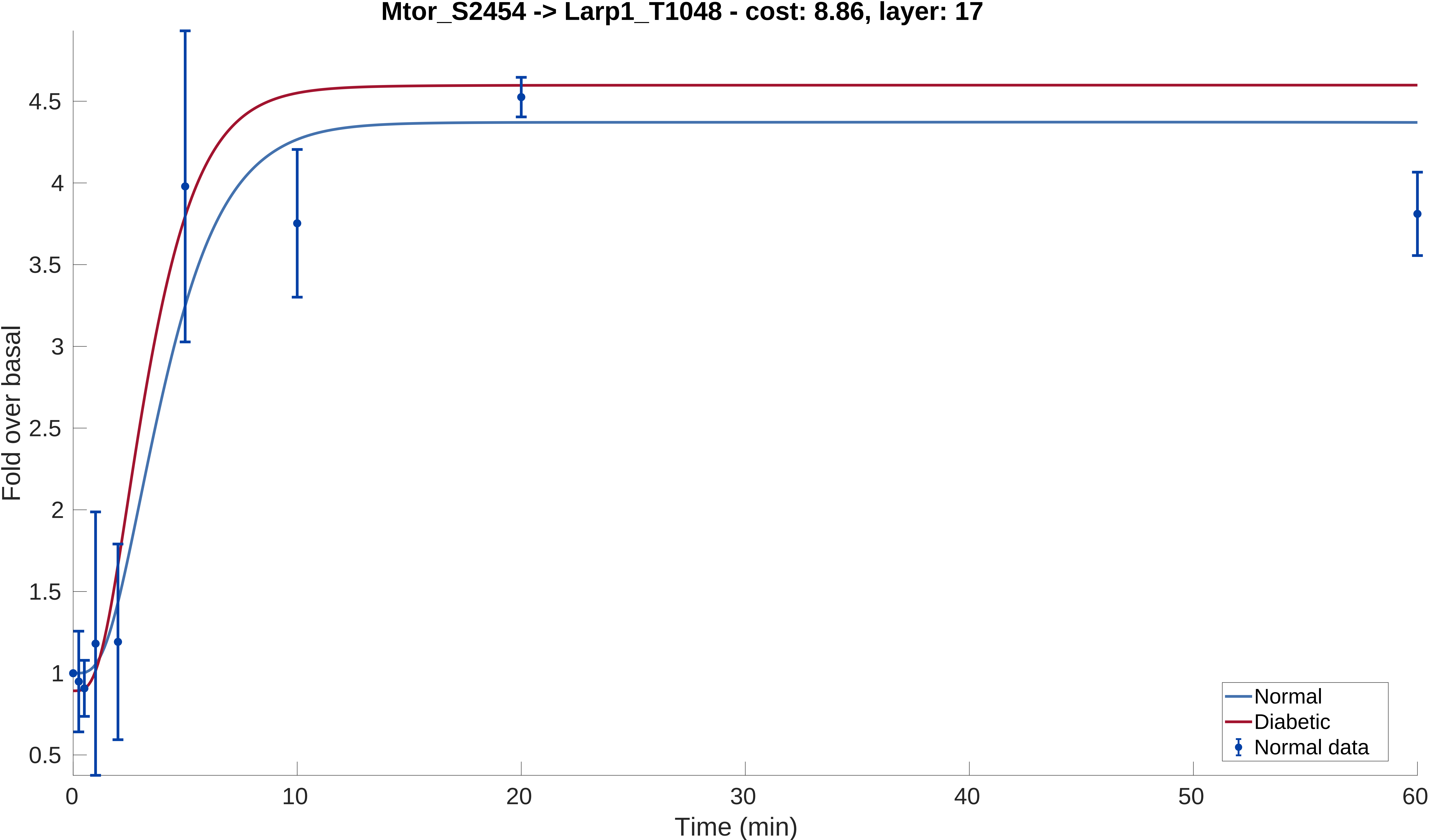

Mtor_S2454 -> Larp1_T1048 - cost: 8.86, layer: 17
Fold over basal
Time (min)
Normal
Diabetic
Normal data

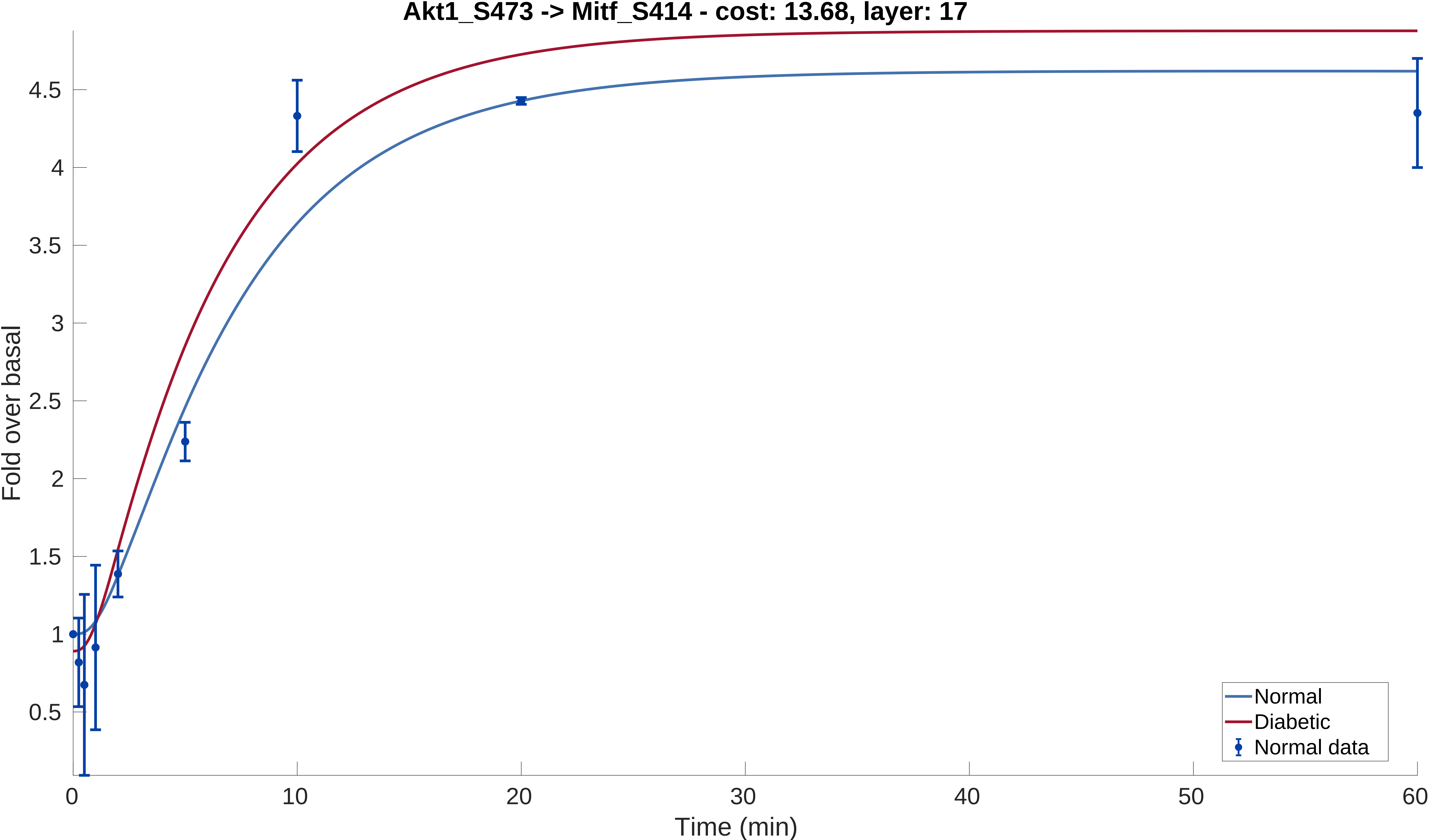

Akt1_S473 -> Mitf_S414 - cost: 13.68, layer: 17
Fold over basal
Time (min)
0.5
1
1.5
2
2.5
3
3.5
4
4.5
0
10
20
30
40
50
60
Normal
Diabetic
Normal data

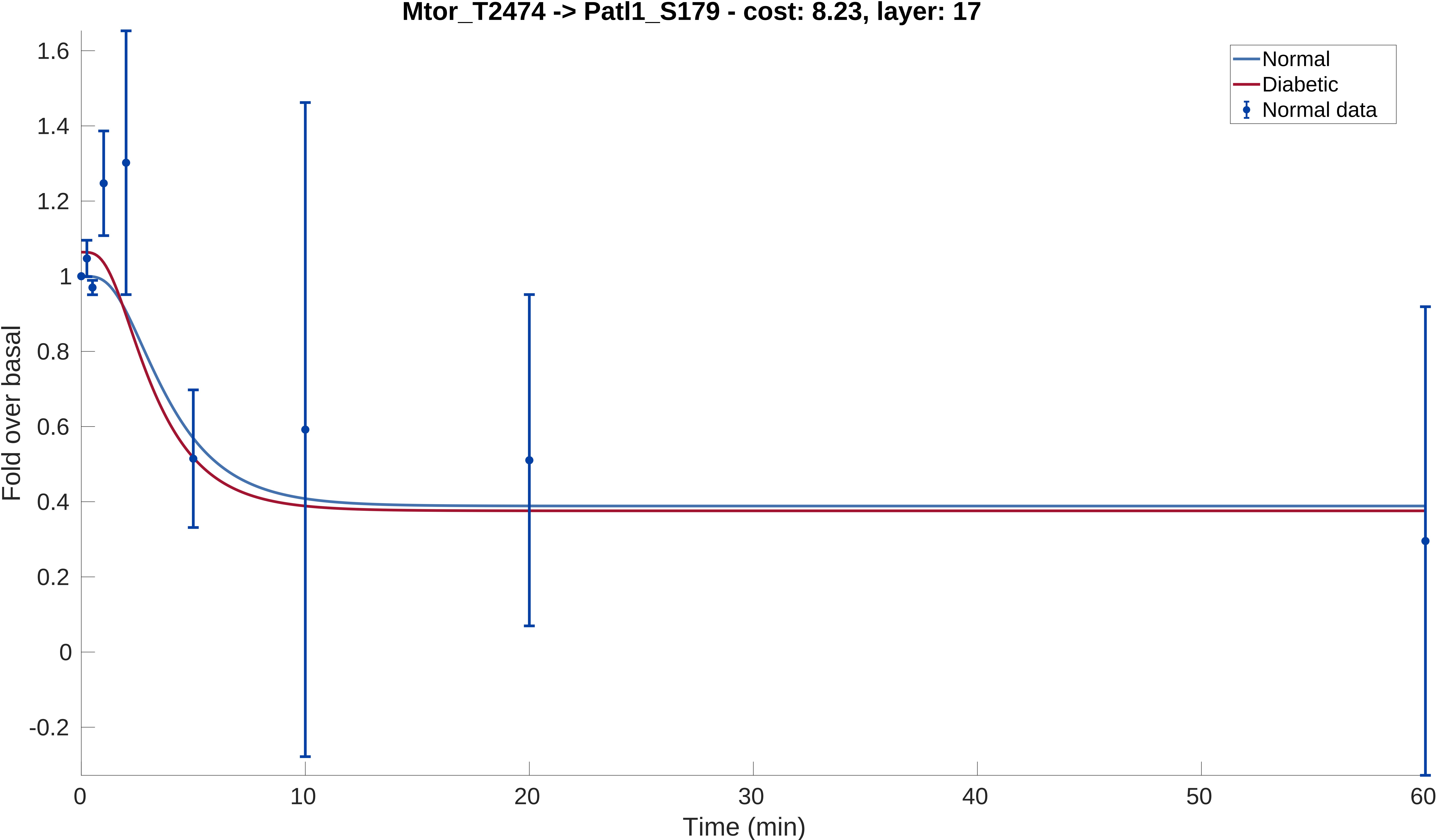
Mtor_T2474 -> Patl1_S179 - cost: 8.23, layer: 17
Normal
Diabetic
Normal data
Fold over basal
Time (min)
1.6
1.4
1.2
1
0.8
0.6
0.4
0.2
0
-0.2
0
10
20
30
40
50
60

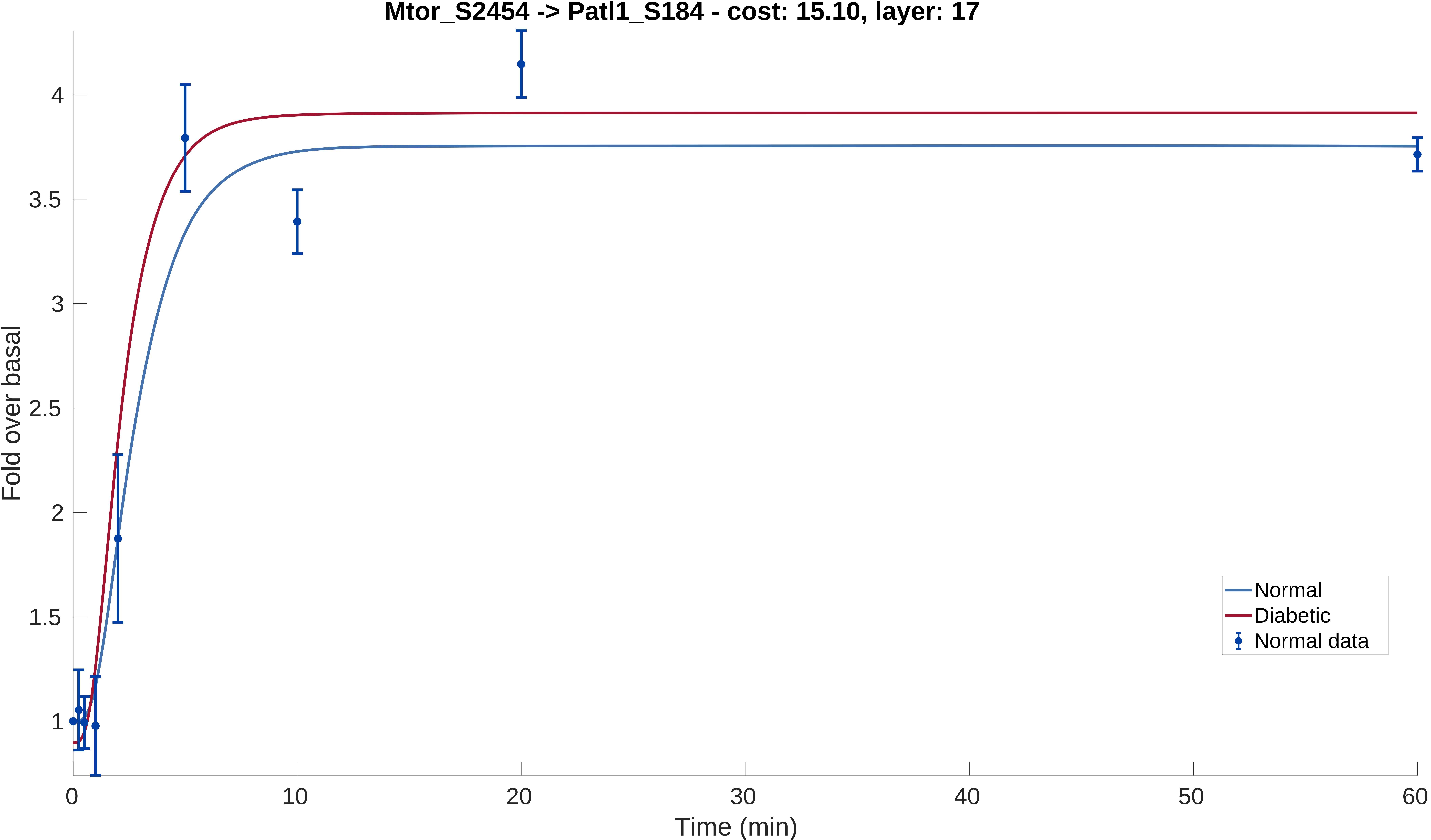

Mtor_S2454 -> Patl1_S184 - cost: 15.10, layer: 17
Fold over basal
4
3.5
3
2.5
2
1.5
1
0
10
20
30
40
50
60
Time (min)
Normal
Diabetic
Normal data

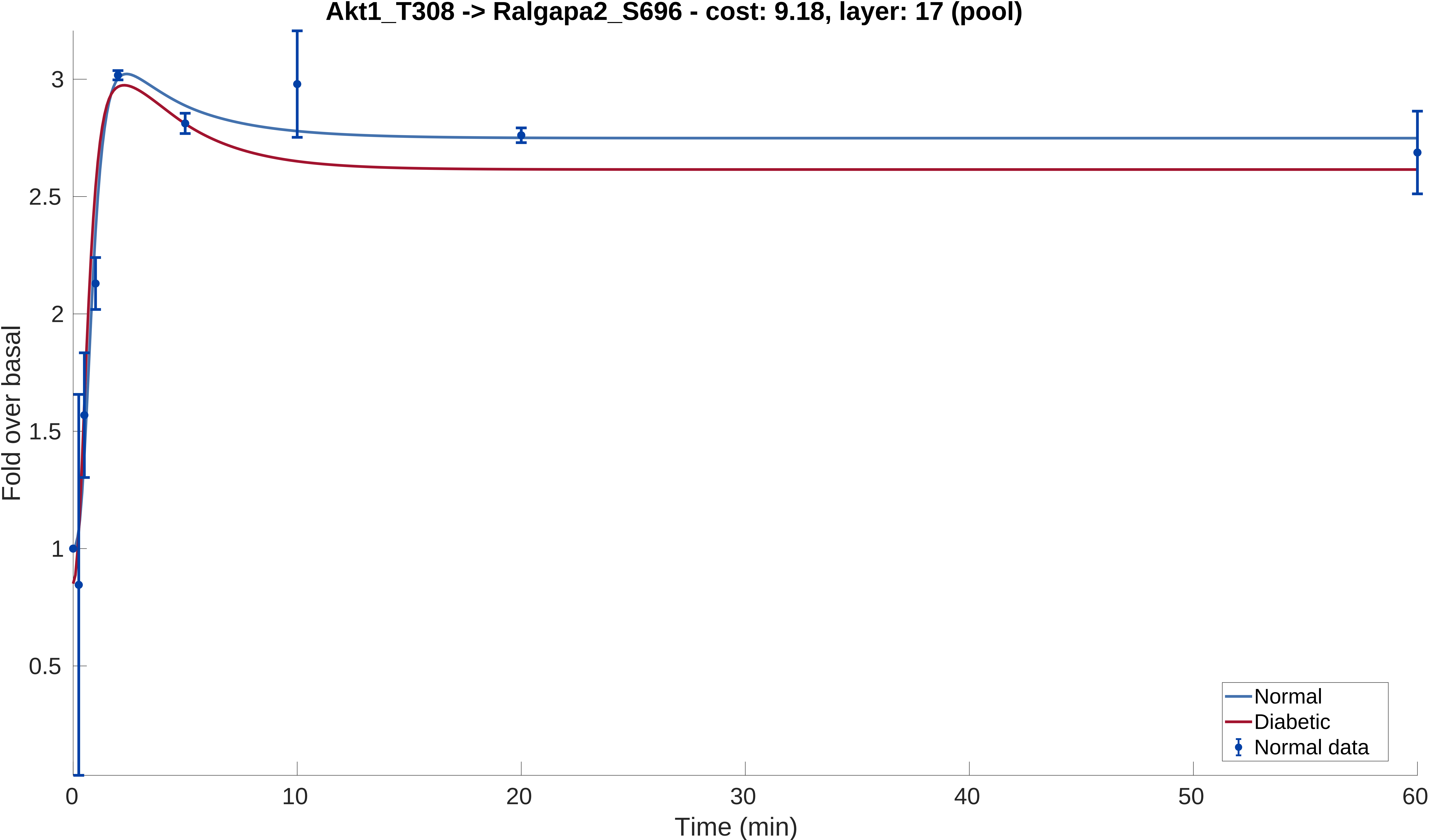
Akt1_T308 -> Ralgapa2_S696 - cost: 9.18, layer: 17 (pool)
Fold over basal
0.5
1
1.5
2
2.5
3
0
10
20
30
40
50
60
Time (min)
Normal
Diabetic
Normal data

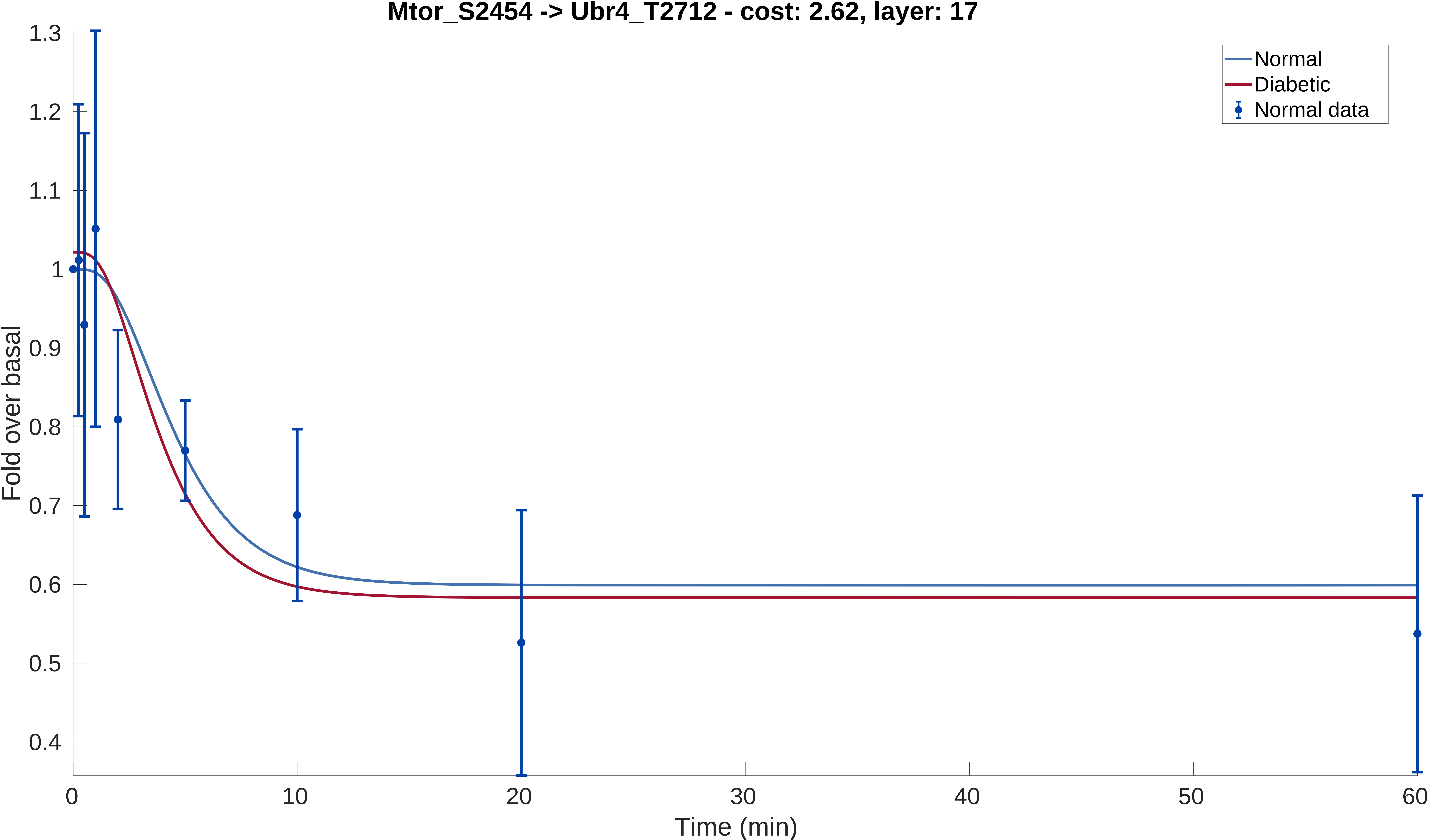
Mtor_S2454 -> Ubr4_T2712 - cost: 2.62, layer: 17
Normal
Diabetic
Normal data
Fold over basal
1.3
1.2
1.1
1
0.9
0.8
0.7
0.6
0.5
0.4
0
10
20
30
40
50
60
Time (min)

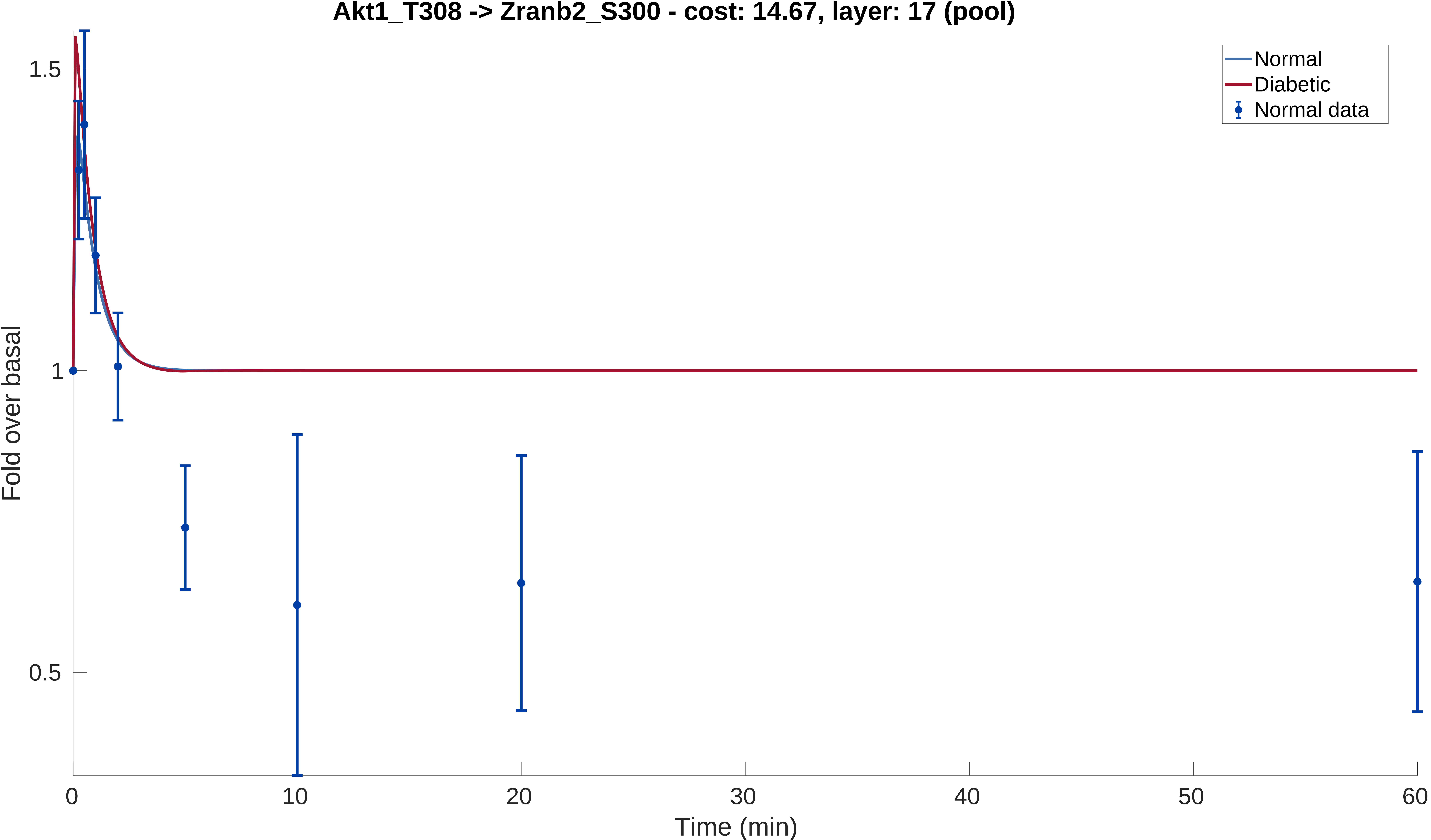

Akt1_T308 -> Zranb2_S300 - cost: 14.67, layer: 17 (pool)
Normal
Diabetic
Normal data
Fold over basal
1.5
1
0.5
0
10
20
30
40
50
60
Time (min)

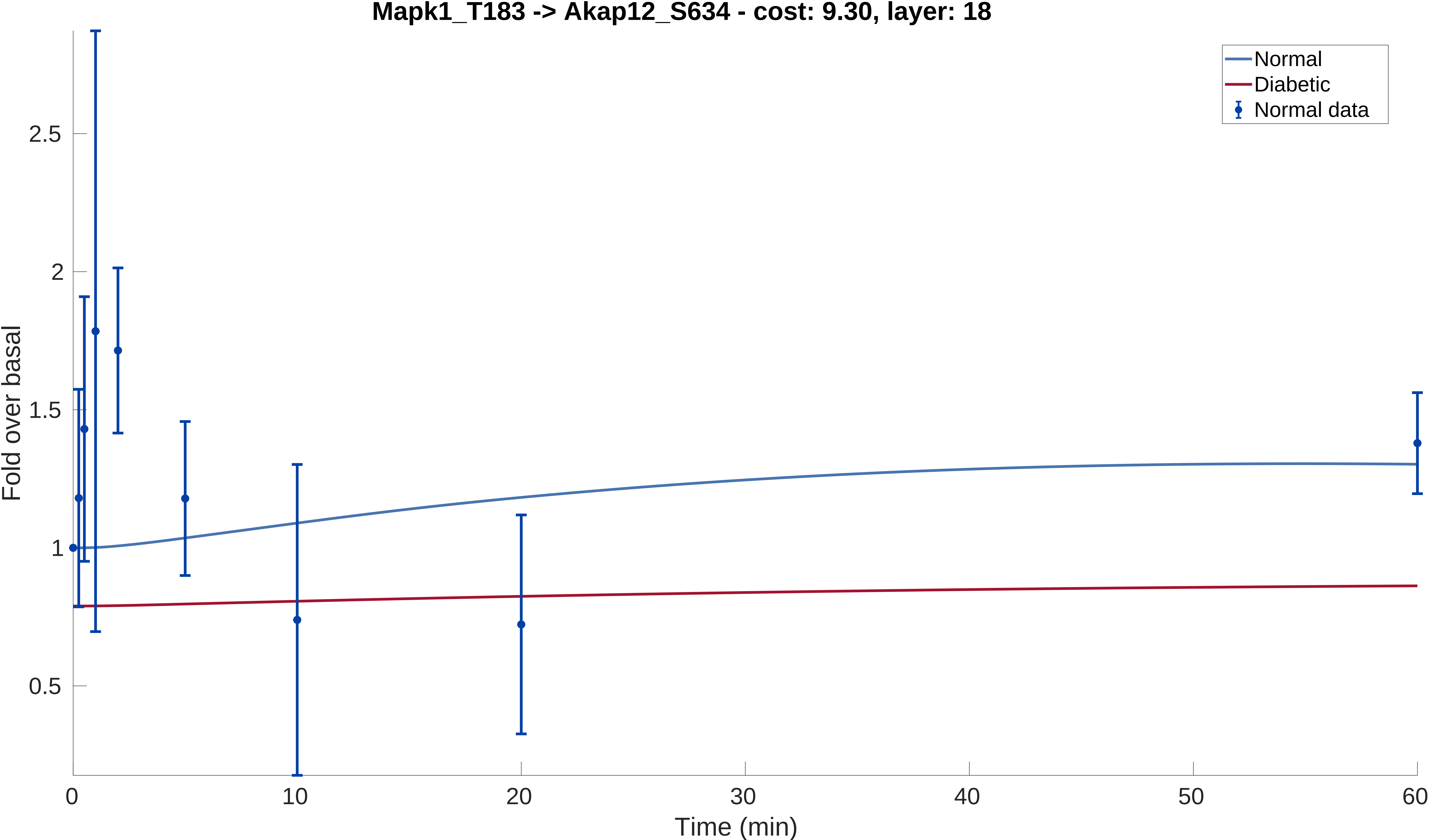
Mapk1_T183 -> Akap12_S634 - cost: 9.30, layer: 18
Normal
Diabetic
Normal data
Fold over basal
2.5
2
1.5
1
0.5
0
10
20
30
40
50
60
Time (min)

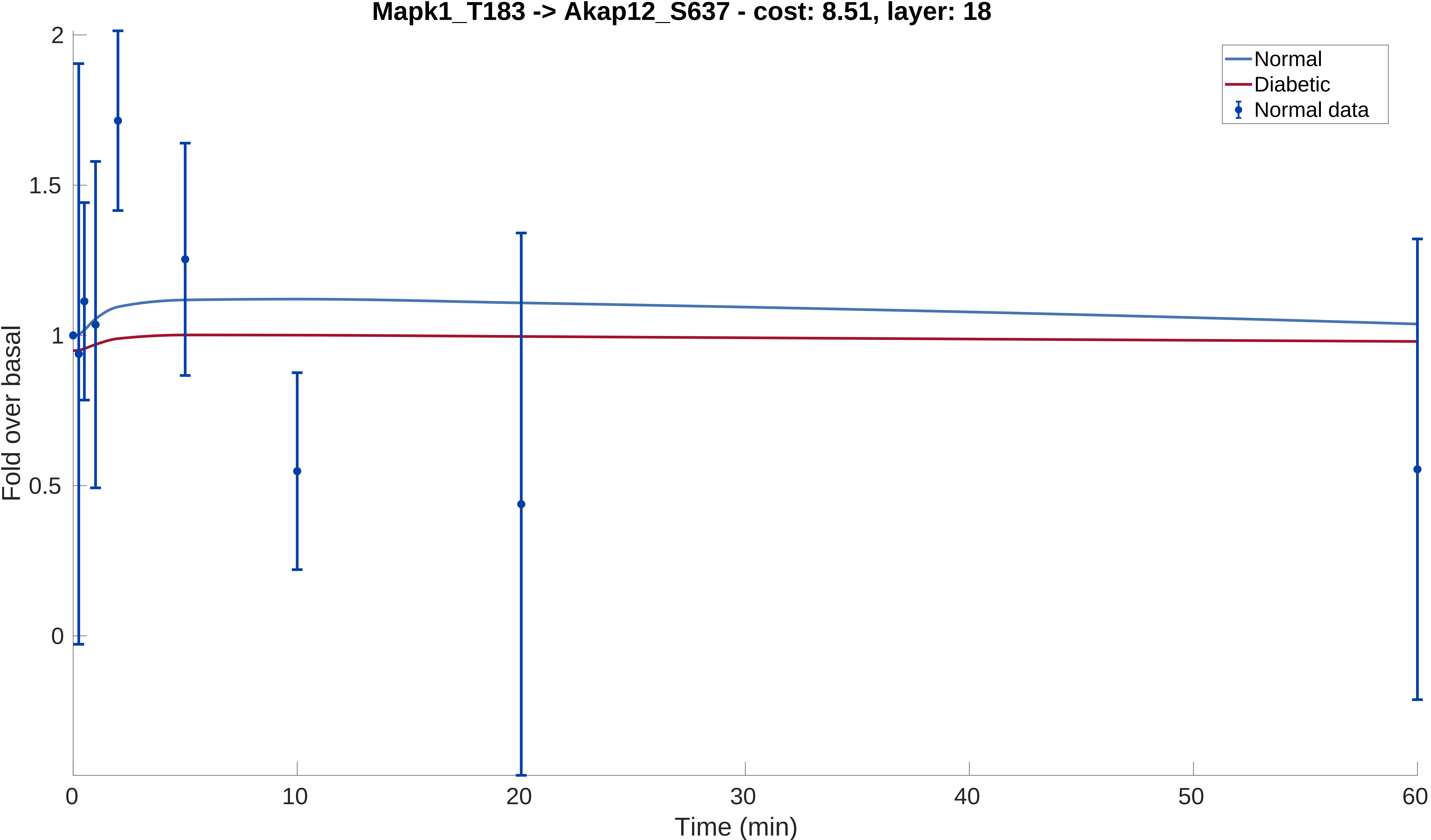

Mapk1_T183 -> Akap12_S637 - cost: 8.51, layer: 18
Normal
Diabetic
Normal data
Fold over basal
2
1.5
1
0.5
0
0
10
20
30
40
50
60
Time (min)

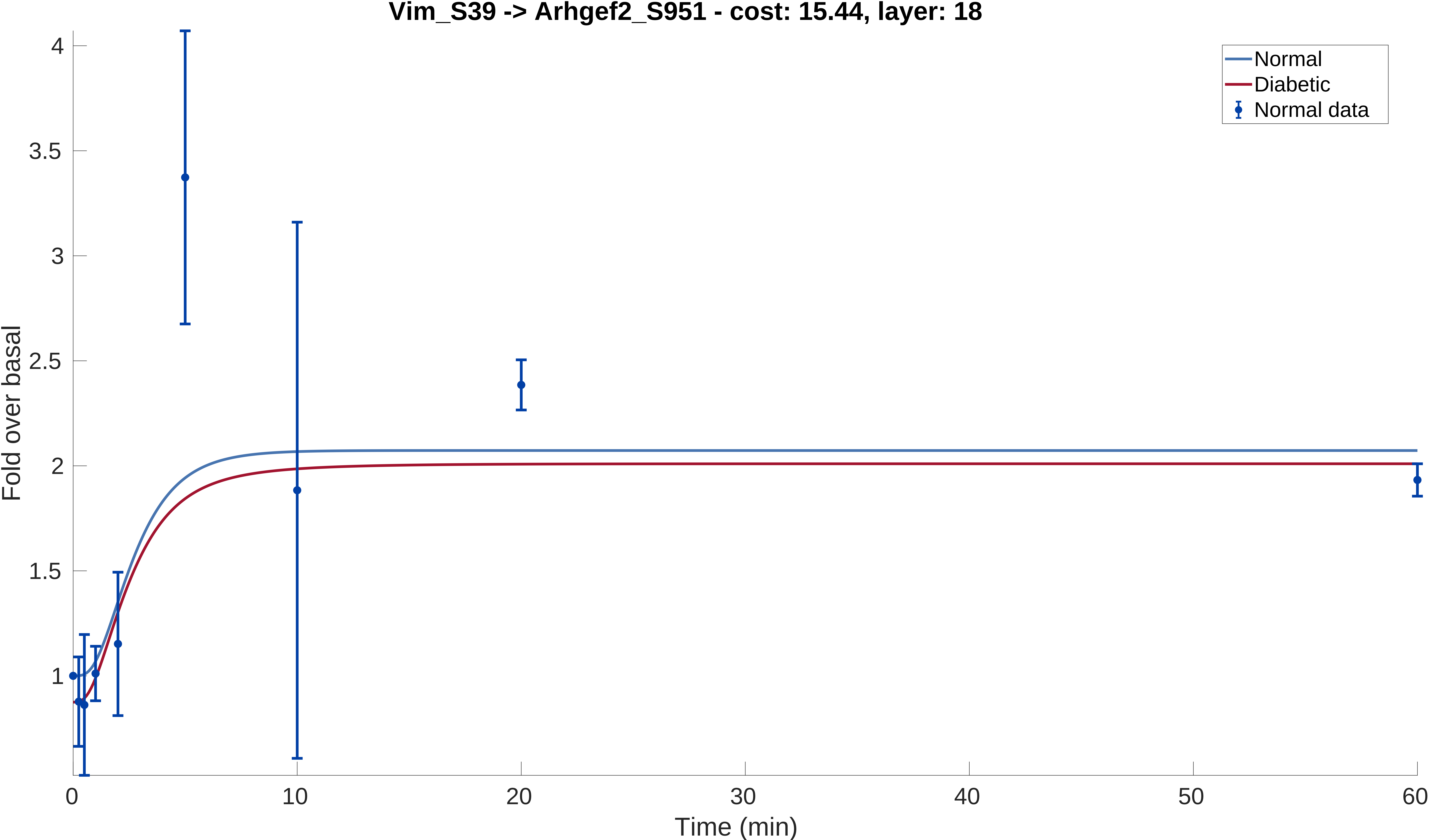

Vim_S39 -> Arhgef2_S951 - cost: 15.44, layer: 18
Normal
Diabetic
Normal data
Fold over basal
4
3.5
3
2.5
2
1.5
1
0
10
20
30
40
50
60
Time (min)

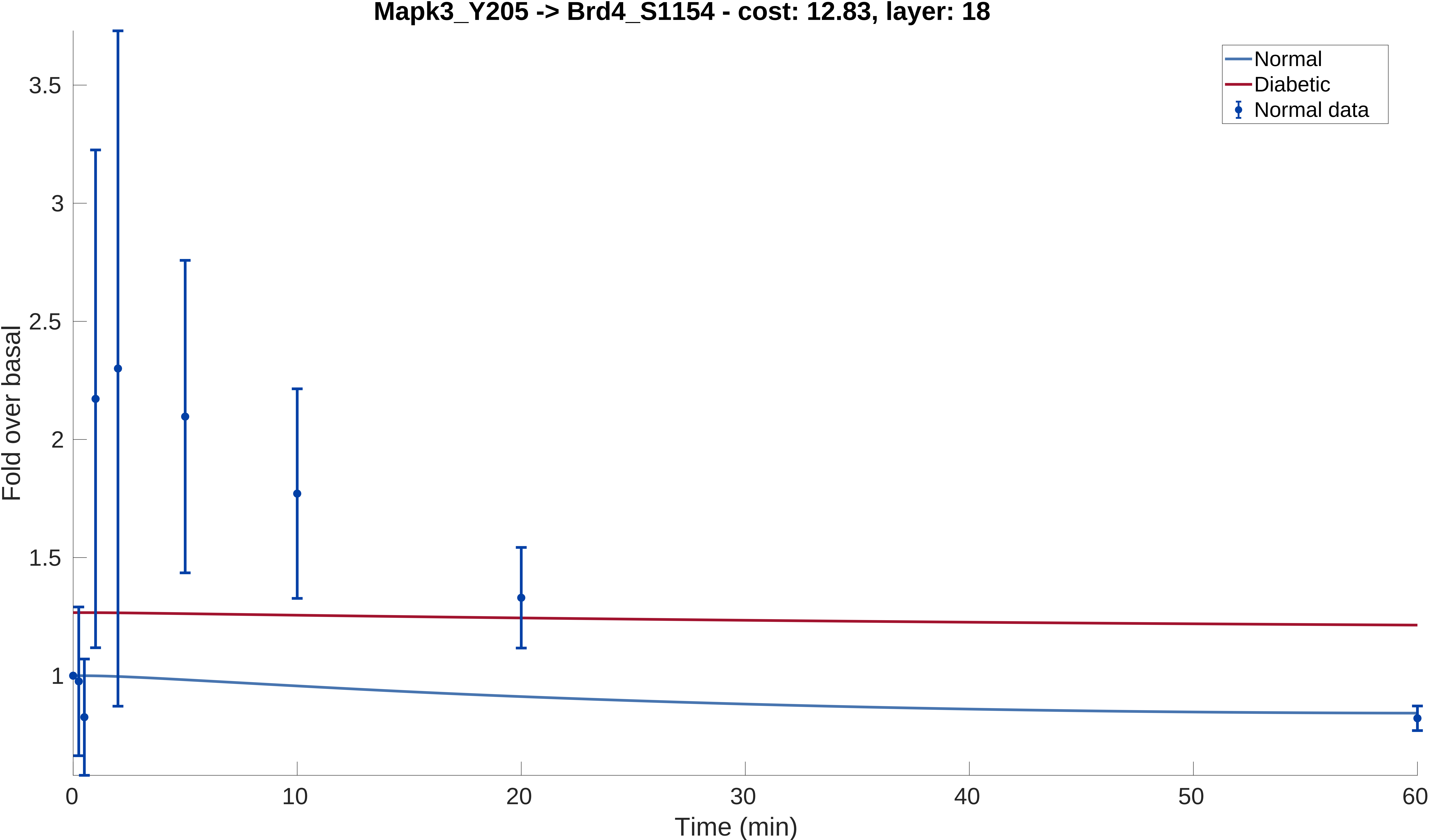

Mapk3_Y205 -> Brd4_S1154 - cost: 12.83, layer: 18
Normal
Diabetic
Normal data
Fold over basal
3.5
3
2.5
2
1.5
1
0
10
20
30
40
50
60
Time (min)

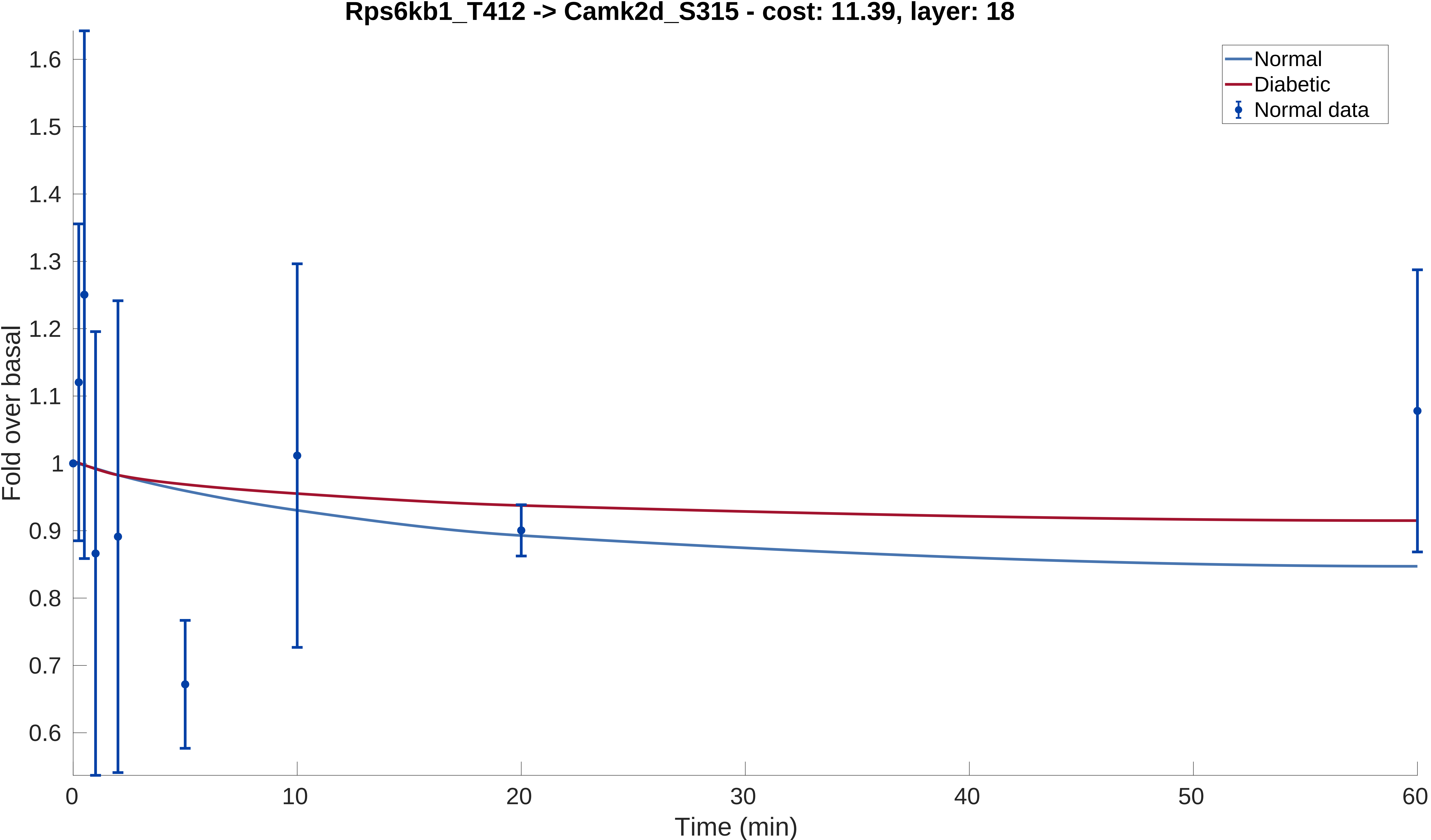

Rps6kb1_T412 -> Camk2d_S315 - cost: 11.39, layer: 18
Normal
Diabetic
Normal data
Fold over basal
1.6
1.5
1.4
1.3
1.2
1.1
1
0.9
0.8
0.7
0.6
0
10
20
30
40
50
60
Time (min)

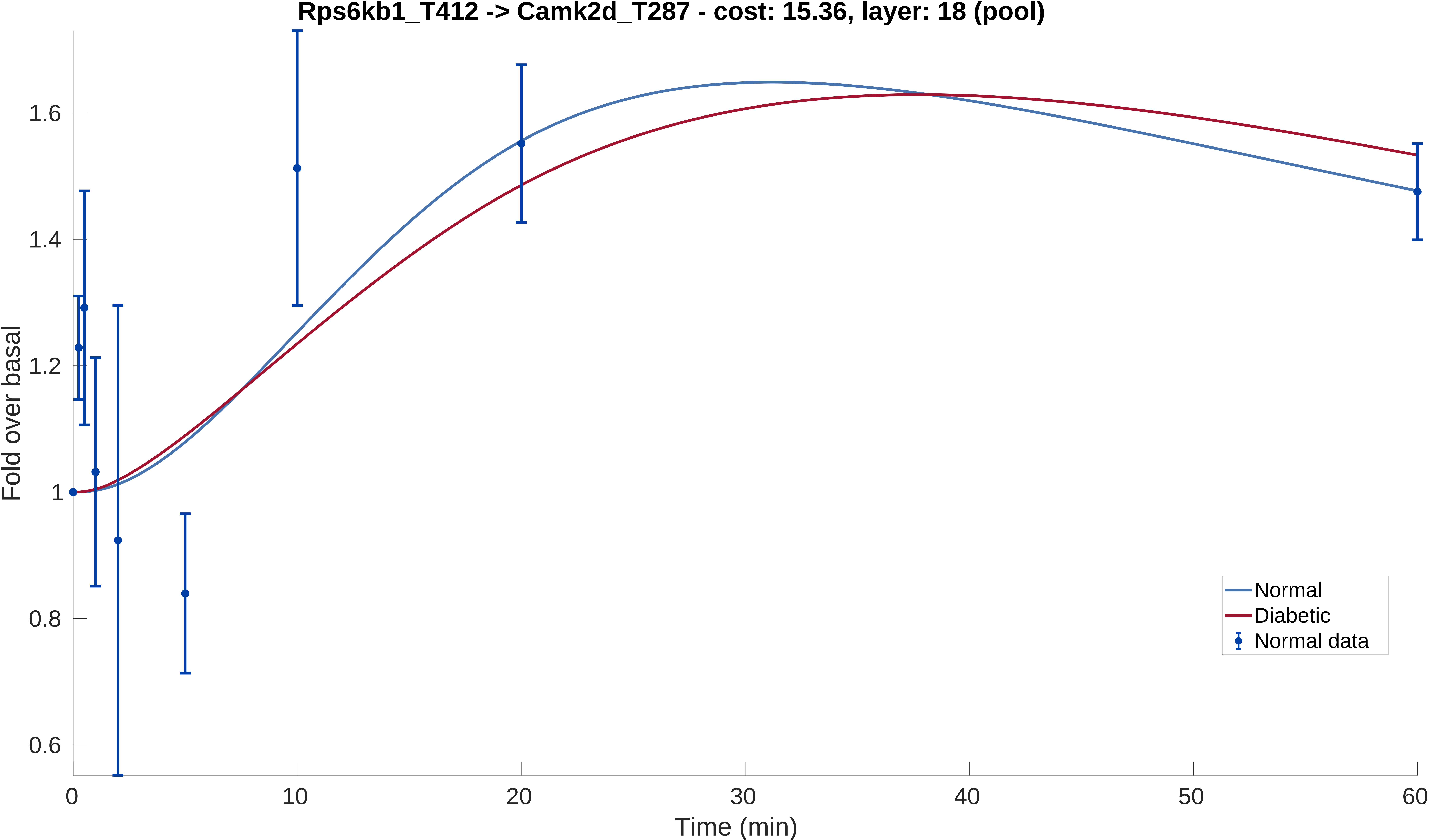

Rps6kb1_T412 -> Camk2d_T287 - cost: 15.36, layer: 18 (pool)
Fold over basal
1.6
1.4
1.2
1
0.8
0.6
0
10
20
30
40
50
60
Time (min)
Normal
Diabetic
Normal data

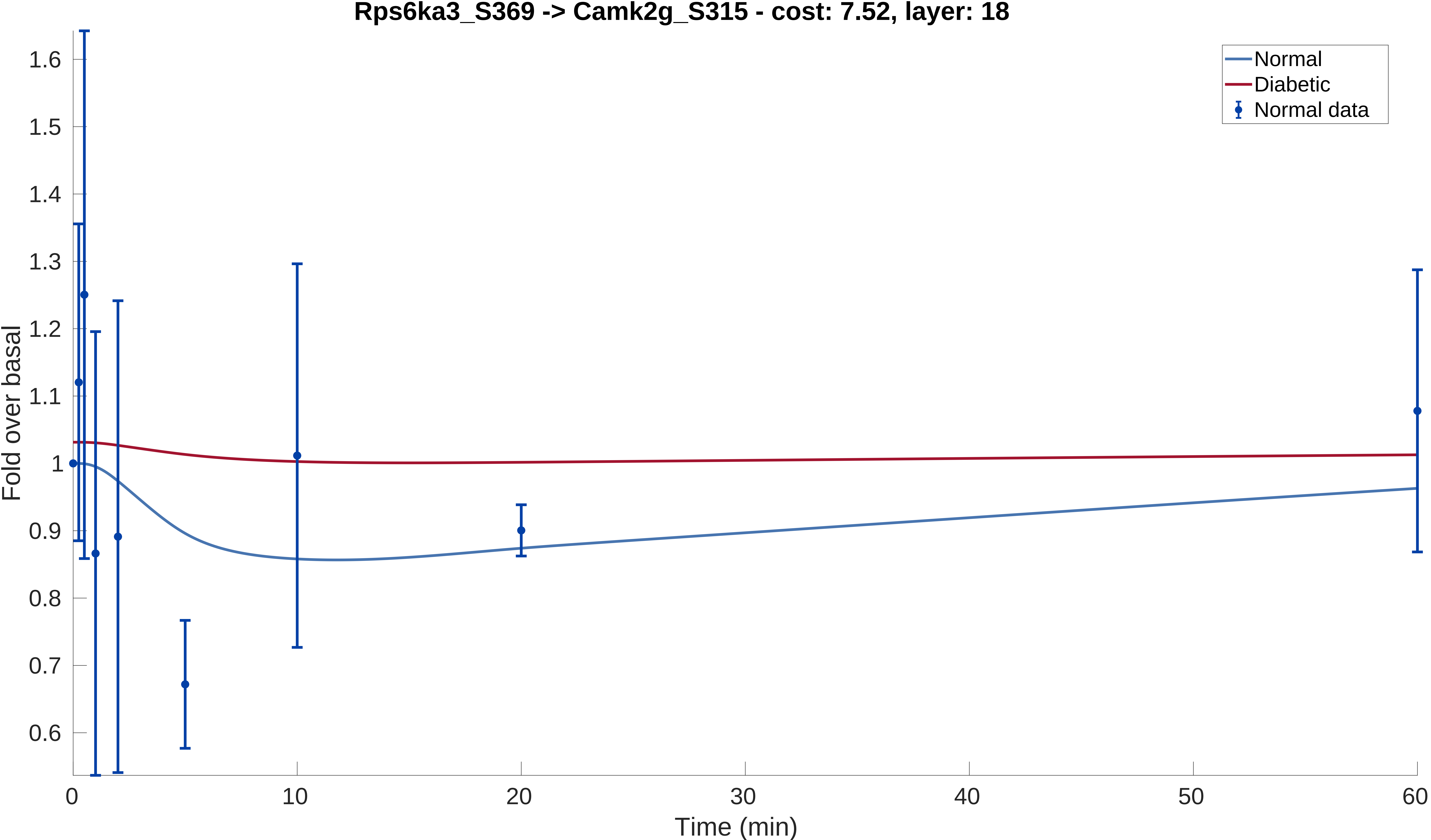

Rps6ka3_S369 -> Camk2g_S315 - cost: 7.52, layer: 18
Normal
Diabetic
Normal data
Fold over basal
Time (min)
1.6
1.5
1.4
1.3
1.2
1.1
1
0.9
0.8
0.7
0.6
0
10
20
30
40
50
60

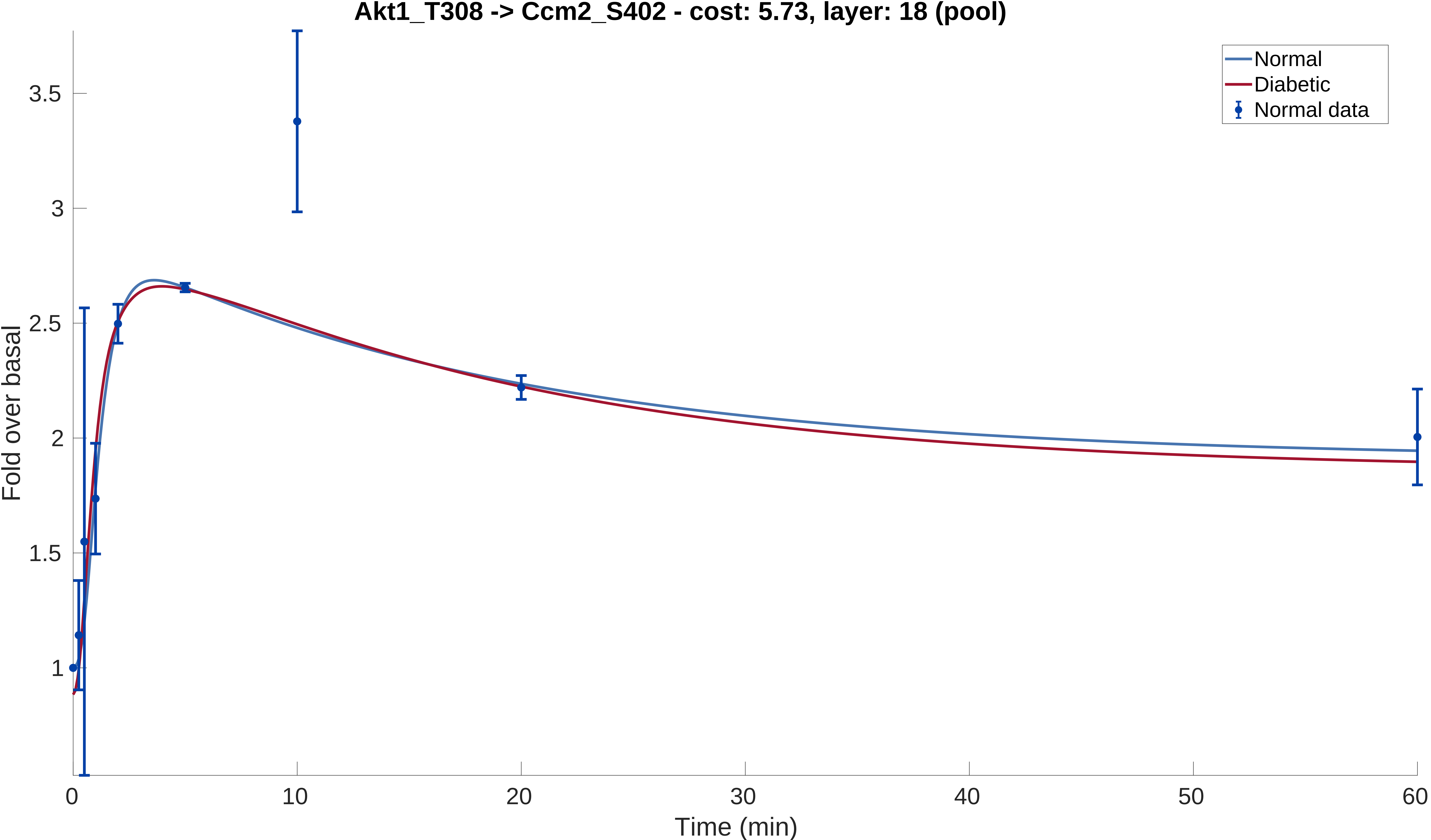

Akt1_T308 -> Ccm2_S402 - cost: 5.73, layer: 18 (pool)
Normal
Diabetic
Normal data
Fold over basal
Time (min)
3.5
3
2.5
2
1.5
1
0
10
20
30
40
50
60

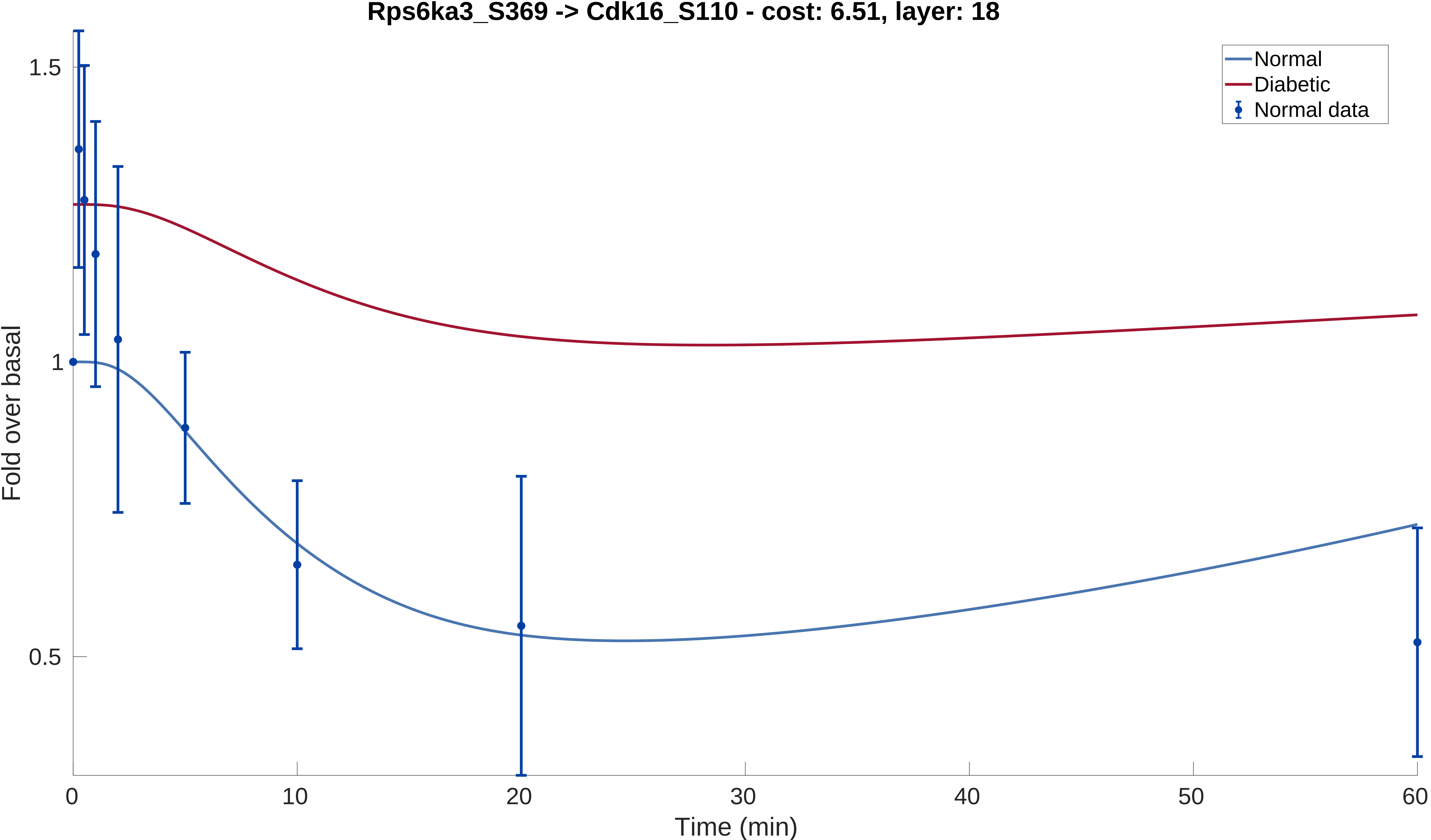

Rps6ka3_S369 -> Cdk16_S110 - cost: 6.51, layer: 18
Normal
Diabetic
Normal data
1.5
1
0.5
Fold over basal
0
10
20
30
40
50
60
Time (min)

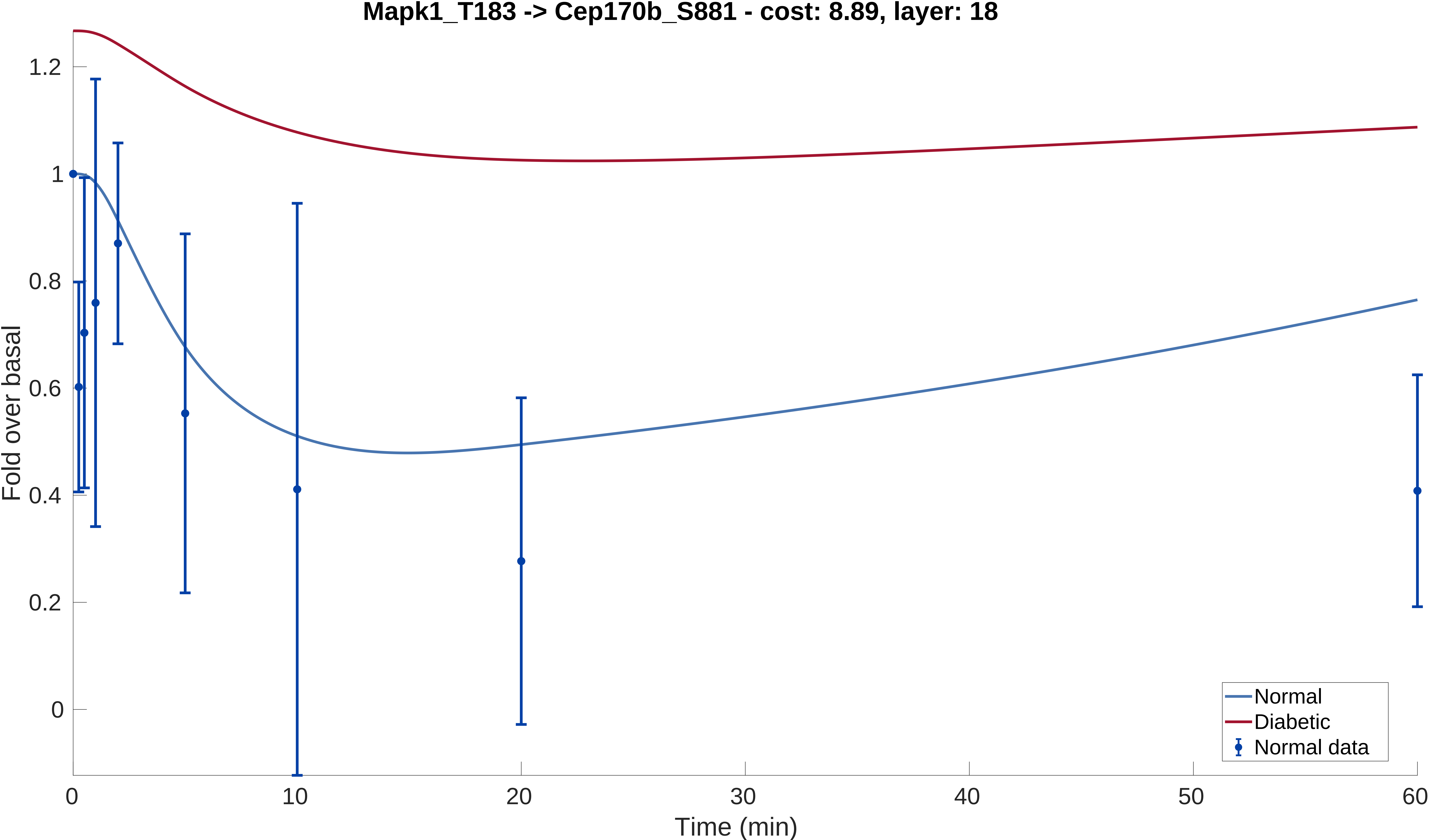

Mapk1_T183 -> Cep170b_S881 - cost: 8.89, layer: 18
Fold over basal
1.2
1
0.8
0.6
0.4
0.2
0
0
10
20
30
40
50
60
Time (min)
Normal
Diabetic
Normal data

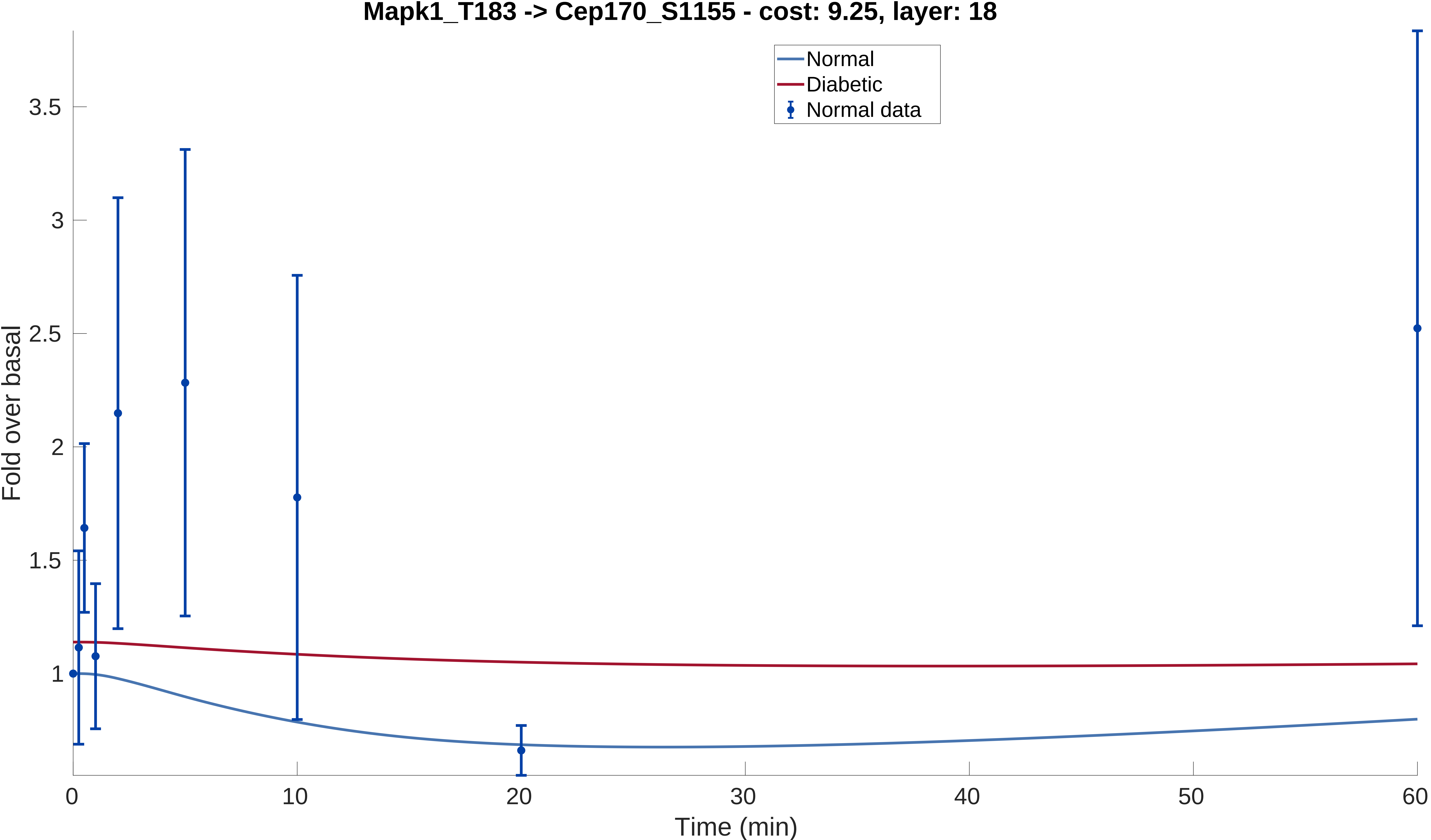
Mapk1_T183 -> Cep170_S1155 - cost: 9.25, layer: 18
Normal
Diabetic
Normal data
Fold over basal
Time (min)
3.5
3
2.5
2
1.5
1
0
10
20
30
40
50
60

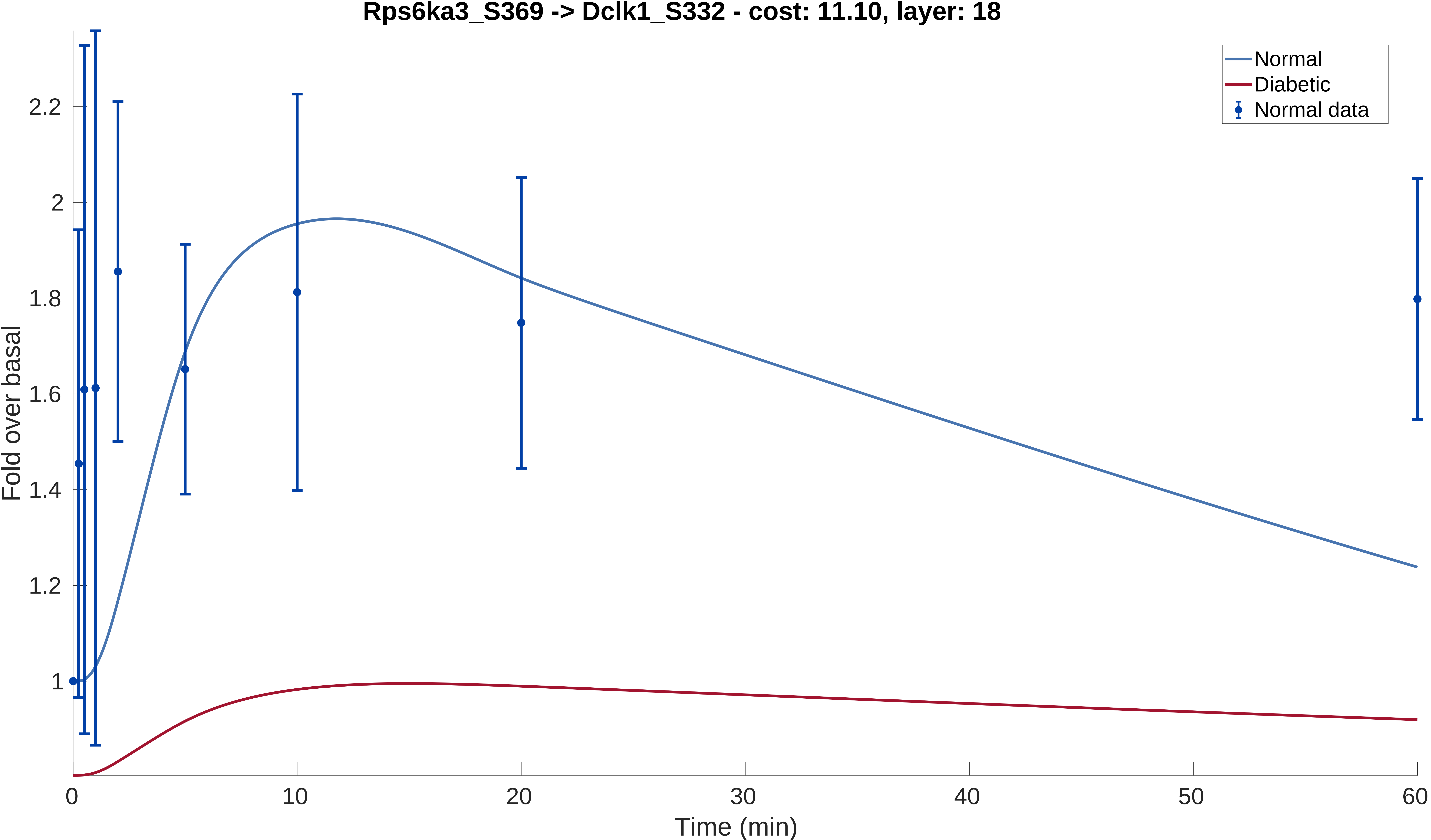
Rps6ka3_S369 -> Dclk1_S332 - cost: 11.10, layer: 18
Normal
Diabetic
Normal data
Fold over basal
Time (min)
0
10
20
30
40
50
60
1
1.2
1.4
1.6
1.8
2
2.2

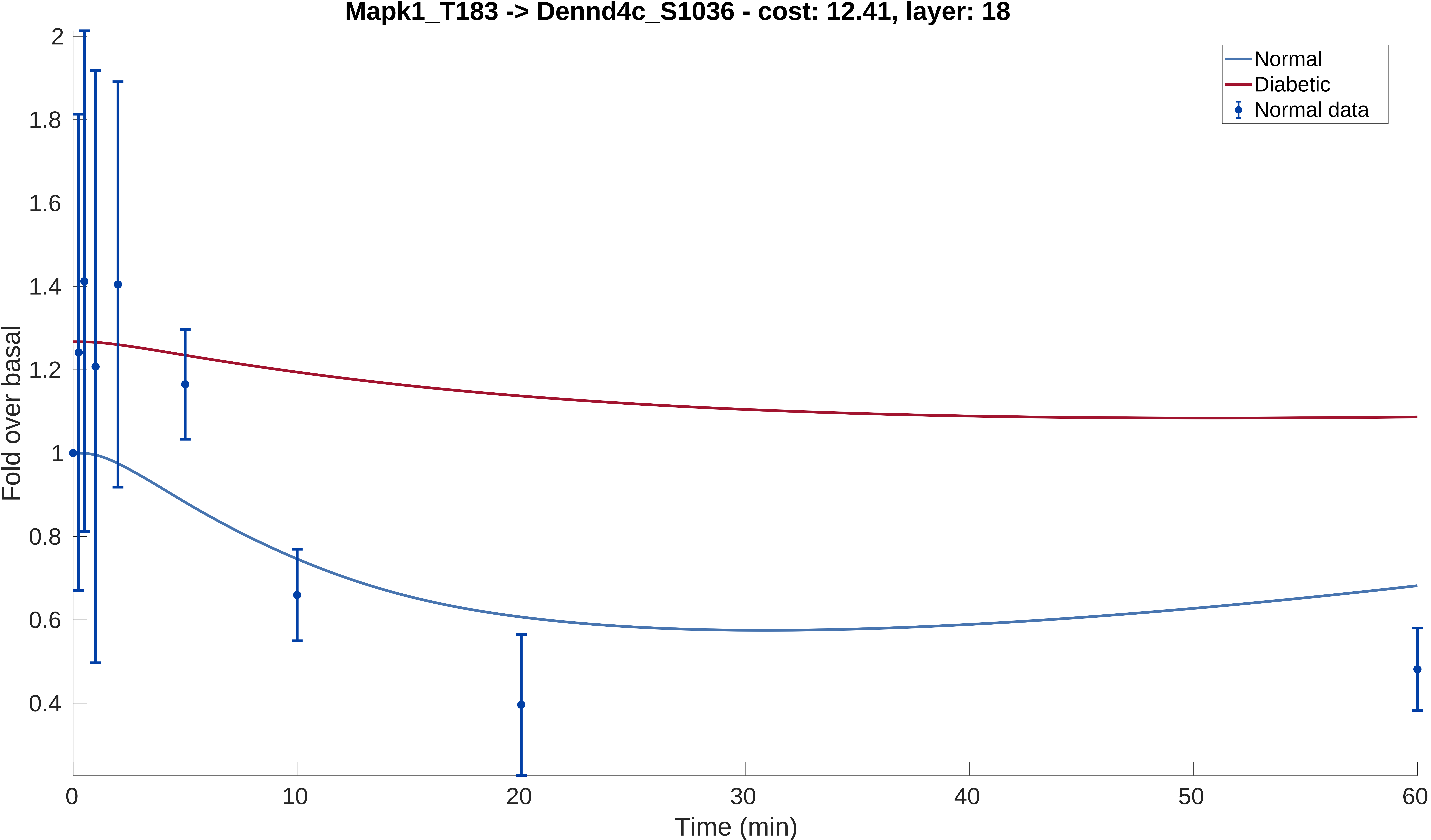

Mapk1_T183 -> Dennd4c_S1036 - cost: 12.41, layer: 18
Normal
Diabetic
Normal data
Fold over basal
Time (min)
2
1.8
1.6
1.4
1.2
1
0.8
0.6
0.4
0
10
20
30
40
50
60

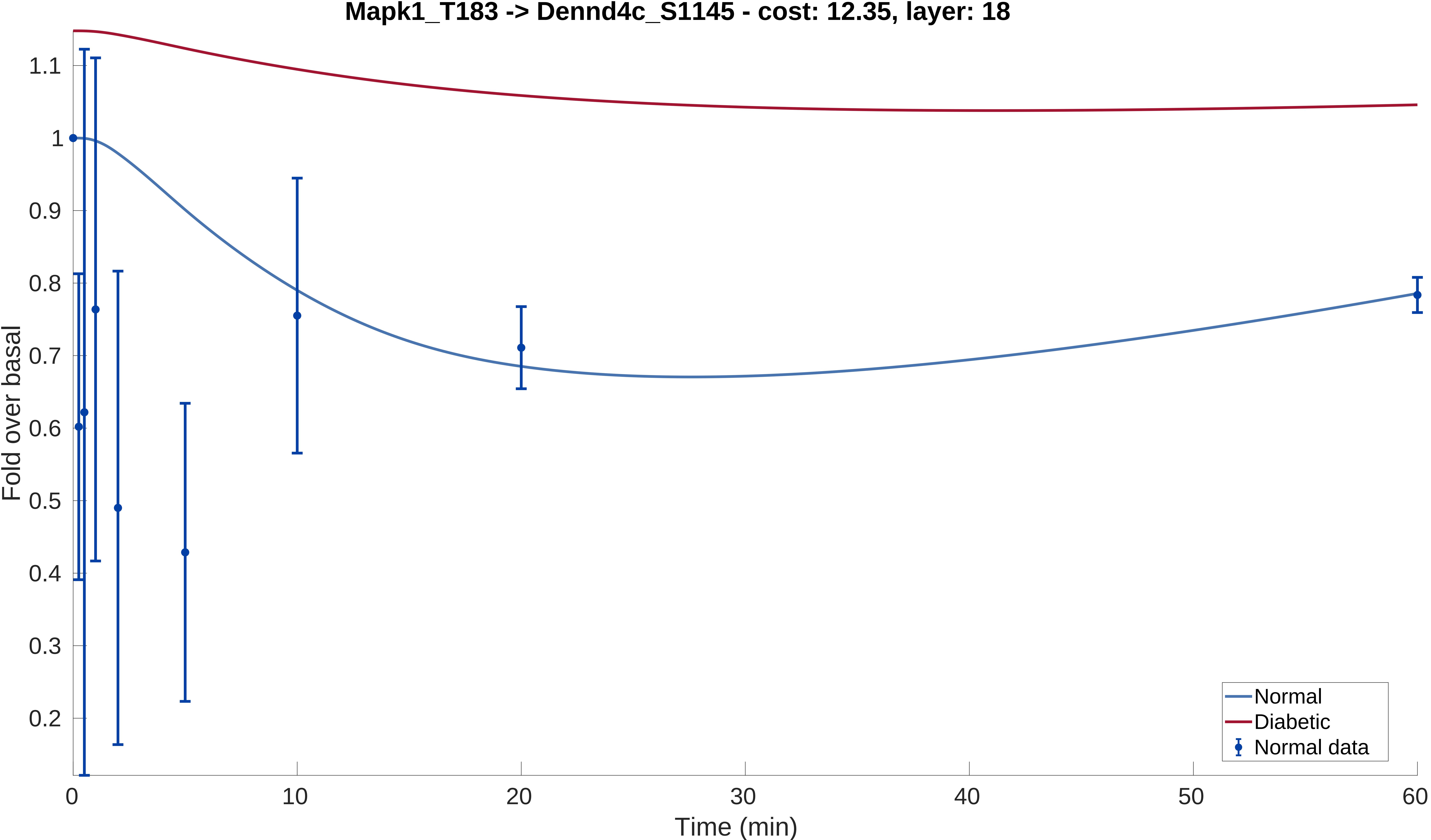
Mapk1_T183 -> Dennd4c_S1145 - cost: 12.35, layer: 18
Fold over basal
Time (min)
0
10
20
30
40
50
60
0.2
0.3
0.4
0.5
0.6
0.7
0.8
0.9
1
1.1
Normal
Diabetic
Normal data

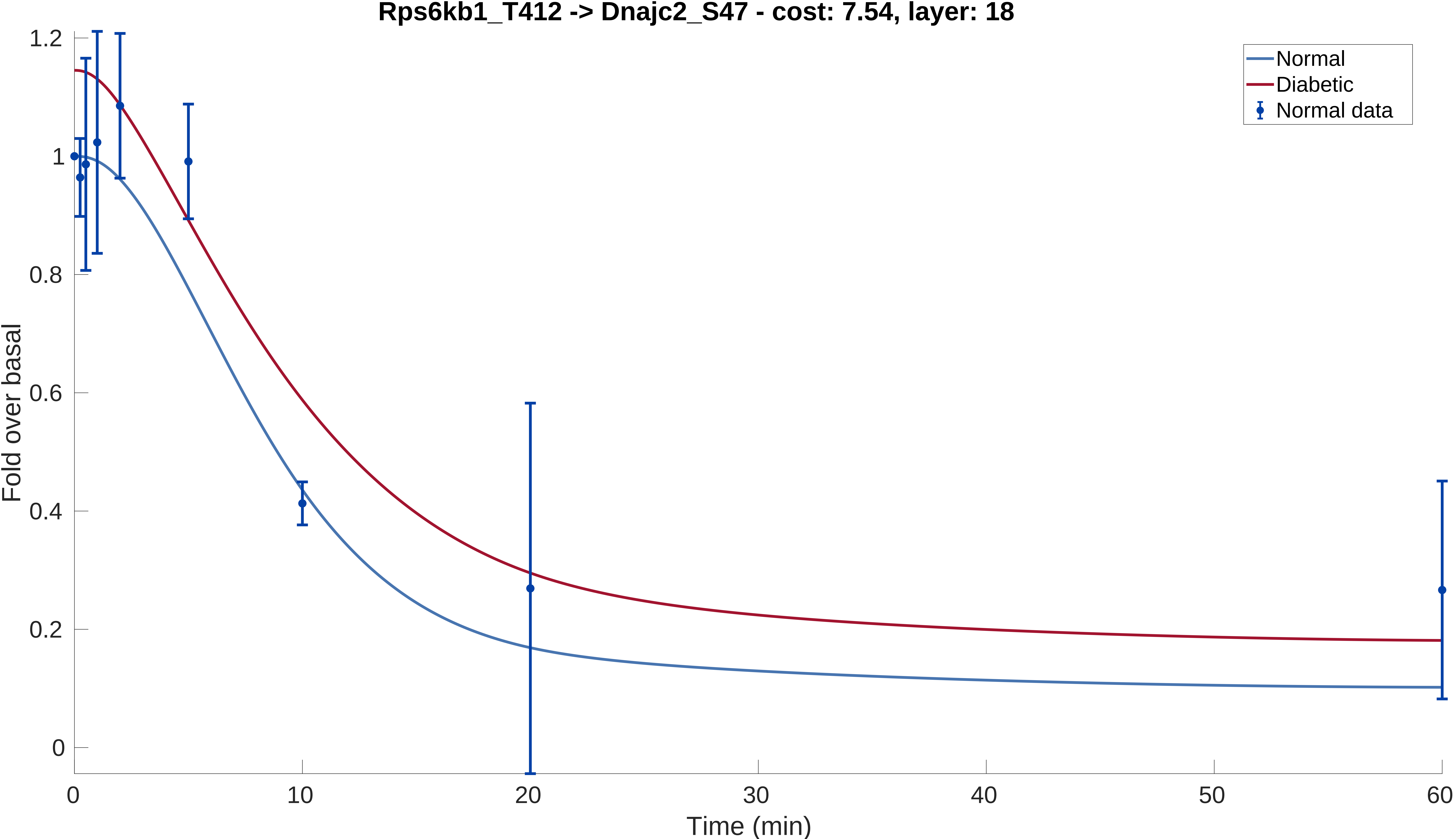

Rps6kb1_T412 -> Dnajc2_S47 - cost: 7.54, layer: 18
Normal
Diabetic
Normal data
Fold over basal
Time (min)
0
0.2
0.4
0.6
0.8
1
1.2
0
10
20
30
40
50
60

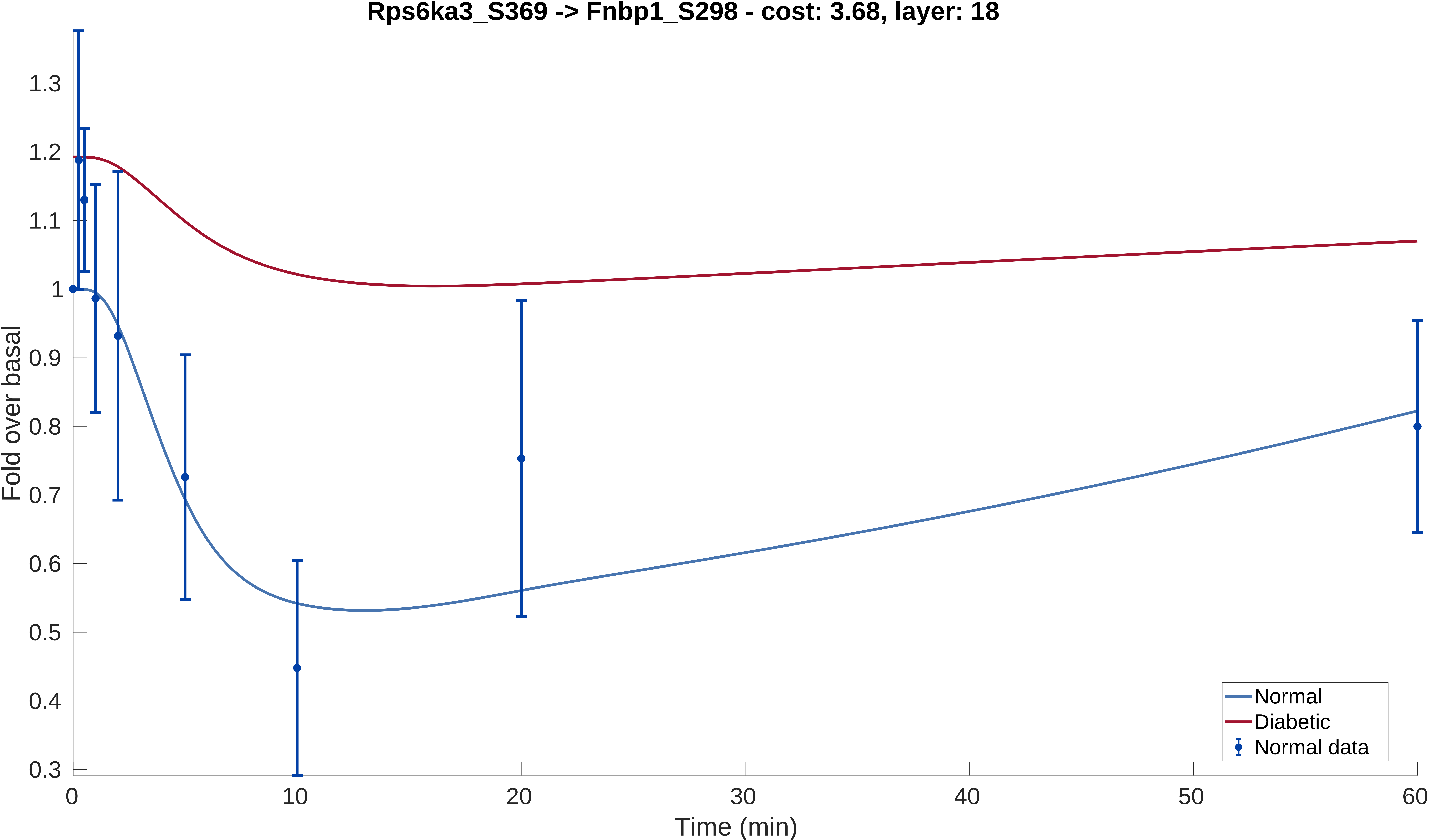

Rps6ka3_S369 -> Fnbp1_S298 - cost: 3.68, layer: 18
Fold over basal
1.3
1.2
1.1
1
0.9
0.8
0.7
0.6
0.5
0.4
0.3
0
10
20
30
40
50
60
Time (min)
Normal
Diabetic
Normal data

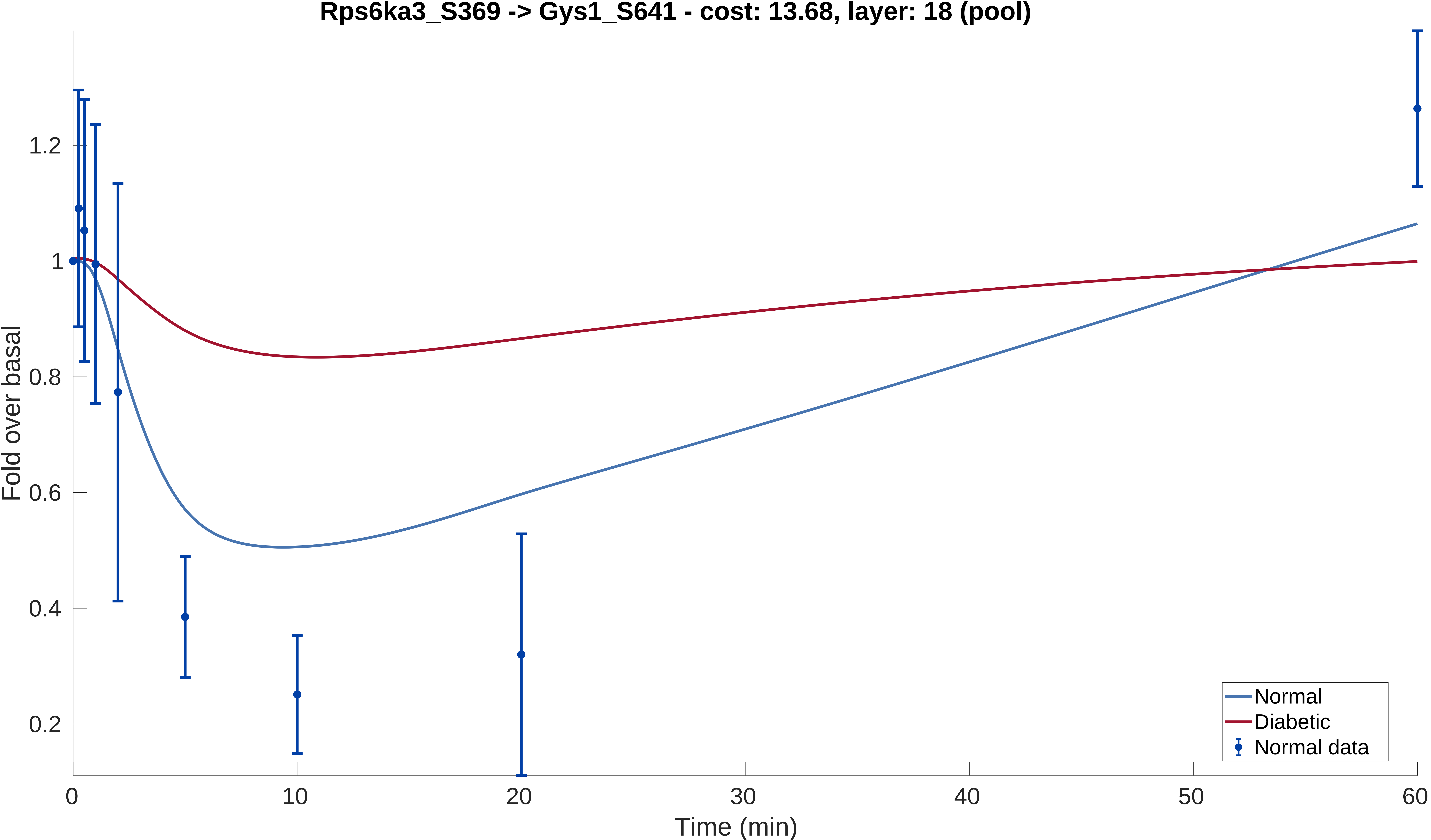

Rps6ka3_S369 -> Gys1_S641 - cost: 13.68, layer: 18 (pool)
Fold over basal
Time (min)
0
10
20
30
40
50
60
0.2
0.4
0.6
0.8
1
1.2
Normal
Diabetic
Normal data

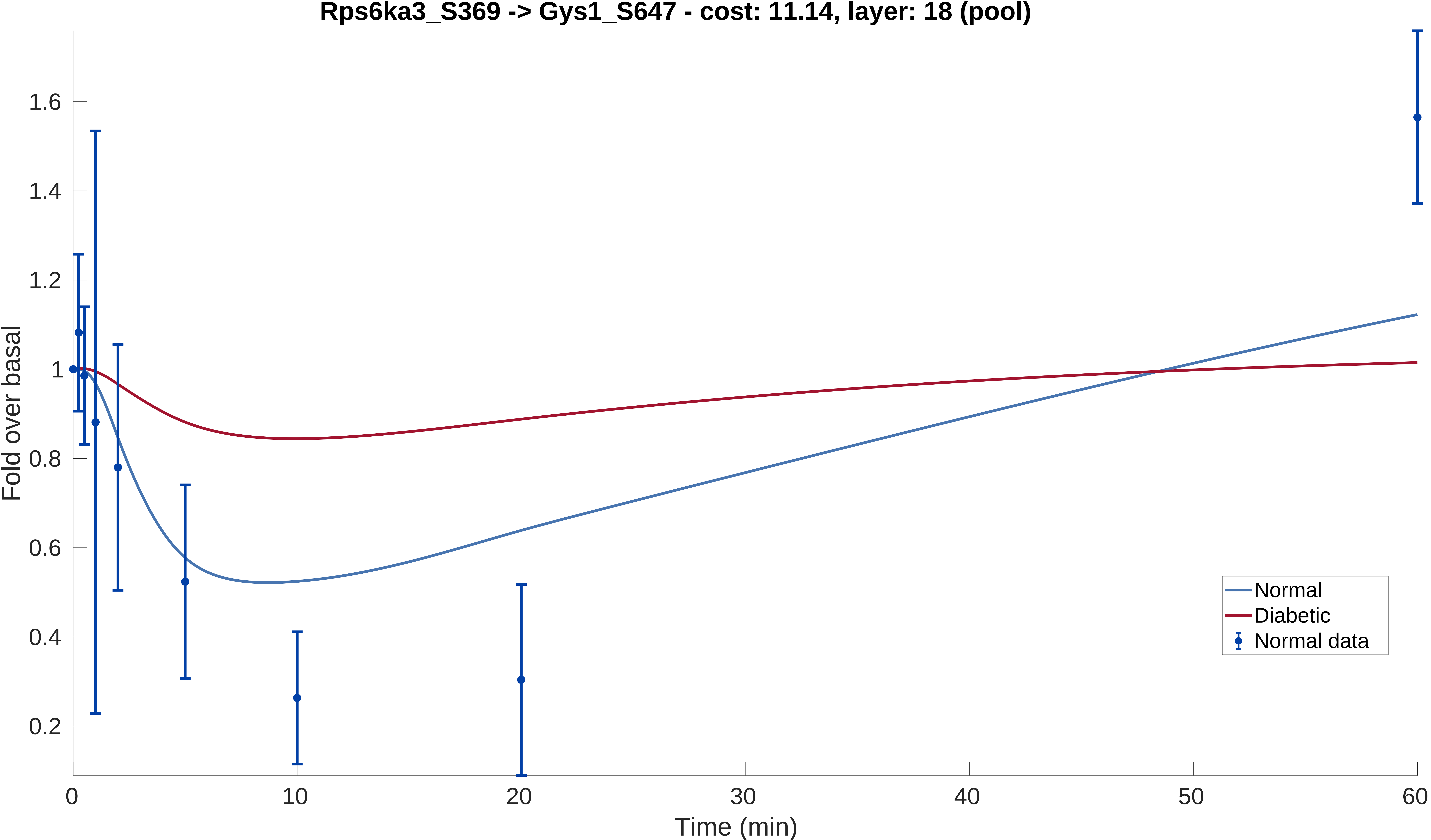

Rps6ka3_S369 -> Gys1_S647 - cost: 11.14, layer: 18 (pool)
1.6
1.4
1.2
1
0.8
0.6
0.4
0.2
Fold over basal
0
10
20
30
40
50
60
Time (min)
Normal
Diabetic
Normal data

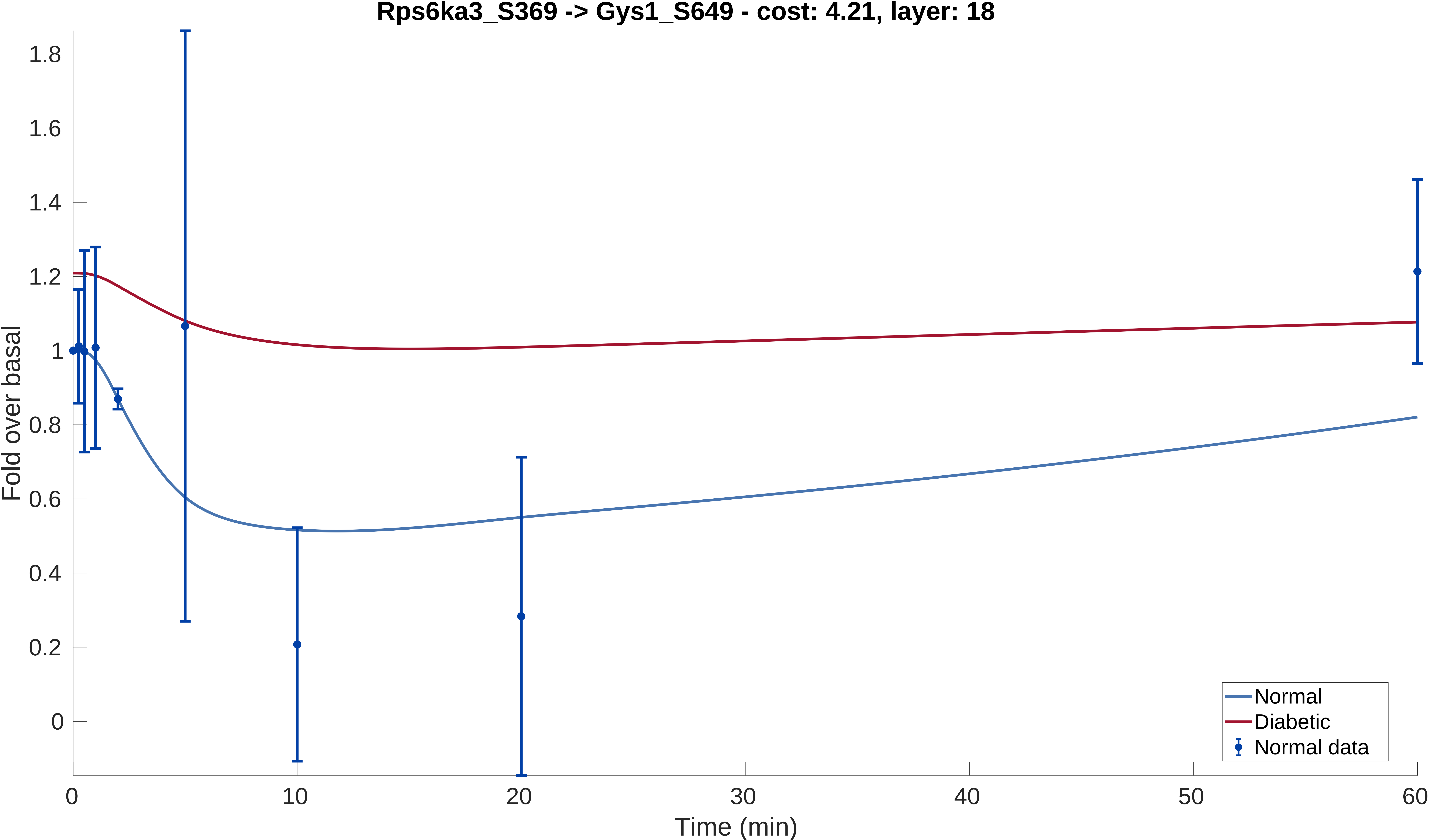

Rps6ka3_S369 -> Gys1_S649 - cost: 4.21, layer: 18
Fold over basal
Time (min)
1.8
1.6
1.4
1.2
1
0.8
0.6
0.4
0.2
0
10
20
30
40
50
60
Normal
Diabetic
Normal data

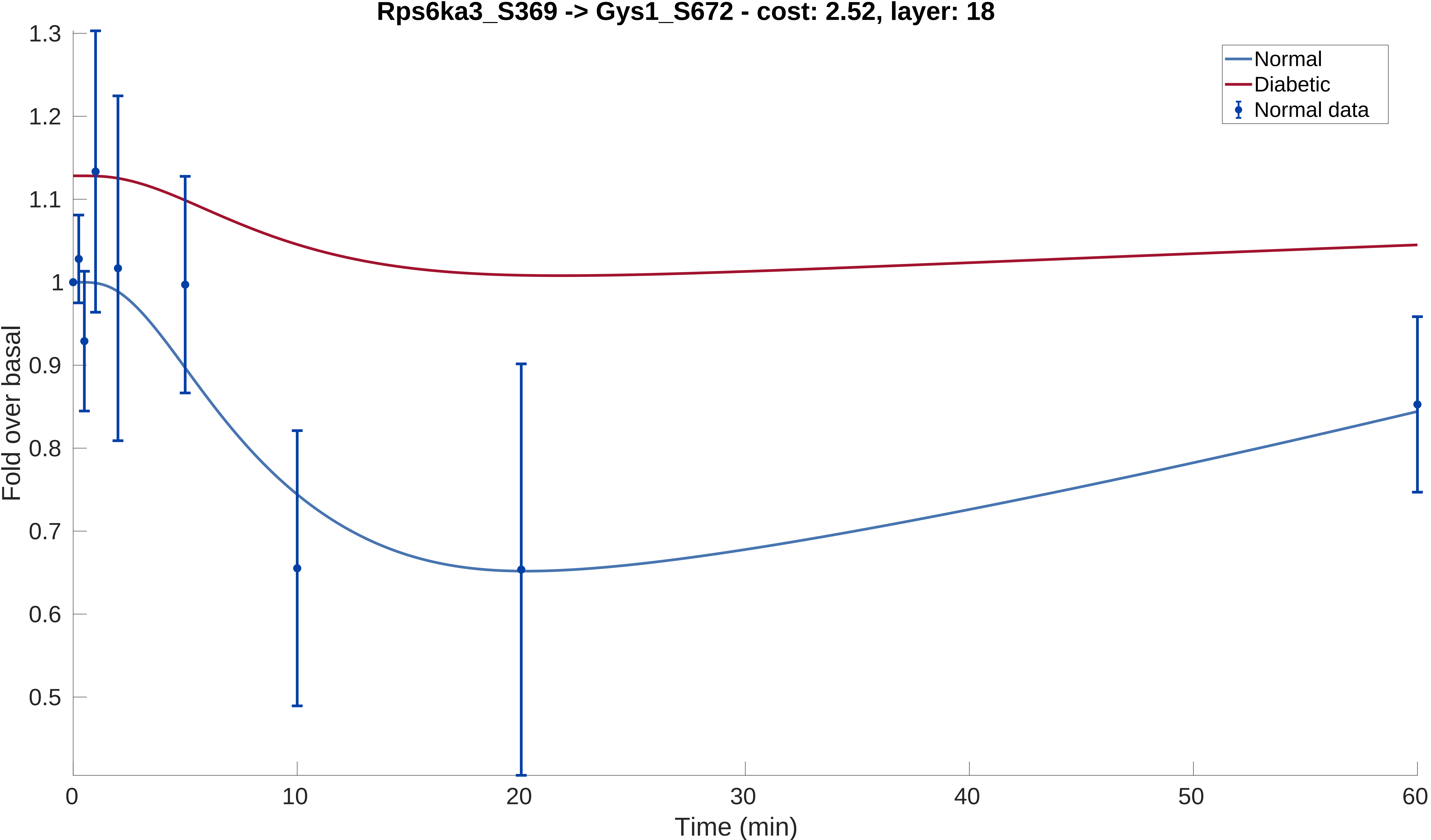

Rps6ka3_S369 -> Gys1_S672 - cost: 2.52, layer: 18
Normal
Diabetic
Normal data
Fold over basal
1.3
1.2
1.1
1
0.9
0.8
0.7
0.6
0.5
0
10
20
30
40
50
60
Time (min)

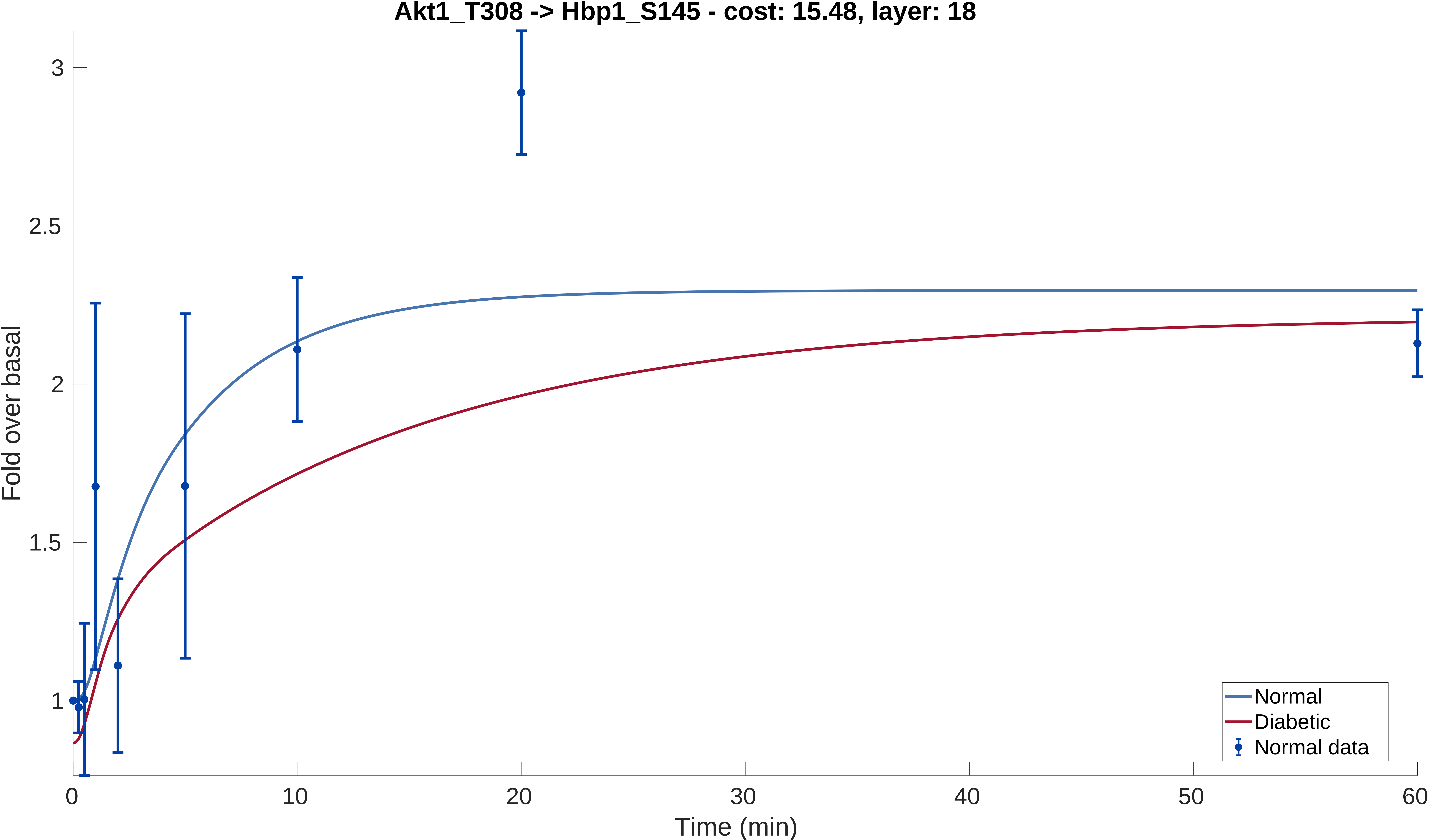

Akt1_T308 -> Hbp1_S145 - cost: 15.48, layer: 18
Fold over basal
3
2.5
2
1.5
1
0
10
20
30
40
50
60
Time (min)
Normal
Diabetic
Normal data

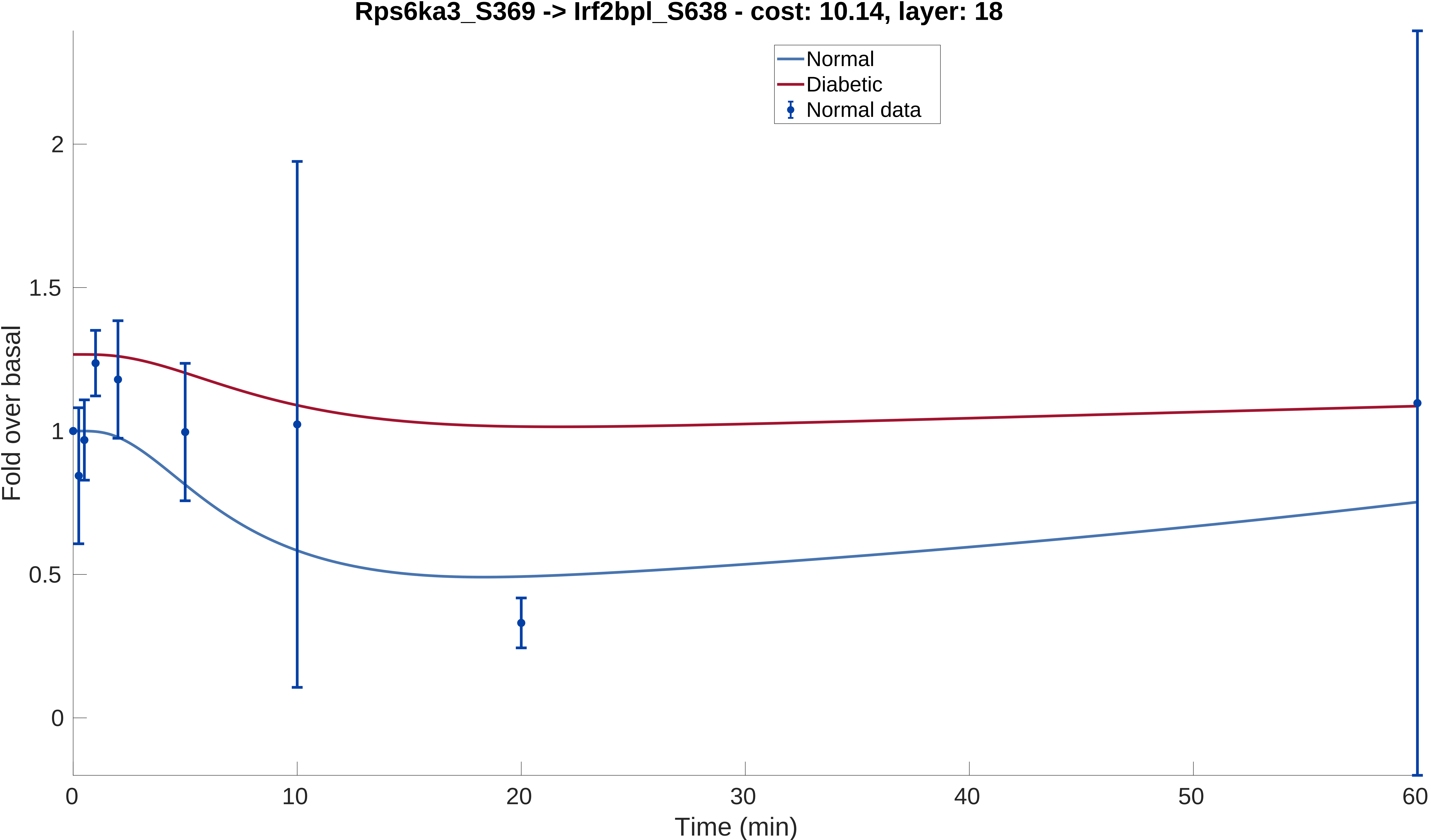

Rps6ka3_S369 -> Irf2bpl_S638 - cost: 10.14, layer: 18
Normal
Diabetic
Normal data
Fold over basal
0
0.5
1
1.5
2
0
10
20
30
40
50
60
Time (min)

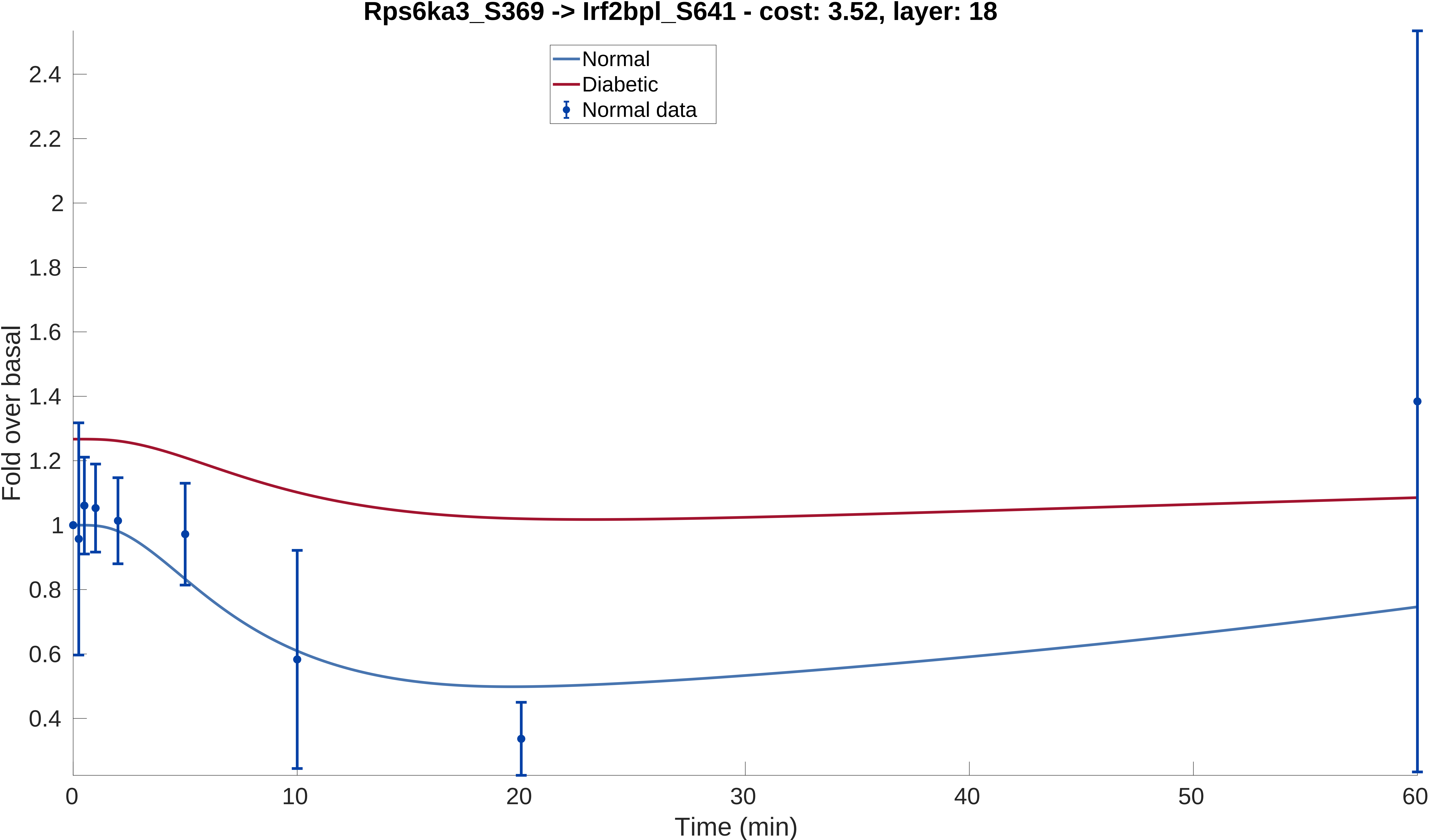

Rps6ka3_S369 -> Irf2bpl_S641 - cost: 3.52, layer: 18
Normal
Diabetic
Normal data
Fold over basal
Time (min)
2.4
2.2
2
1.8
1.6
1.4
1.2
1
0.8
0.6
0.4
0
10
20
30
40
50
60

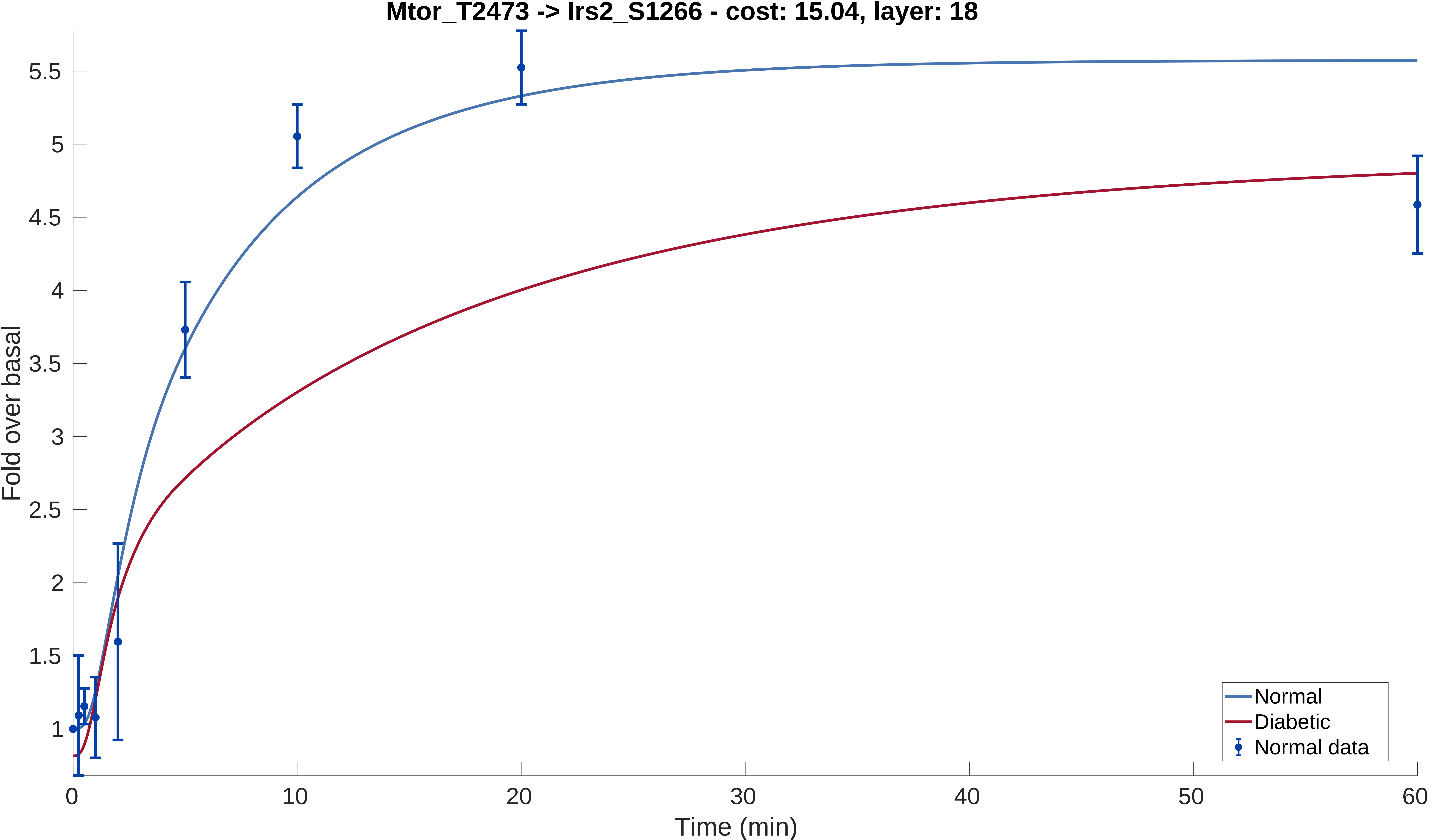

Mtor_T2473 -> Irs2_S1266 - cost: 15.04, layer: 18
Fold over basal
5.5
5
4.5
4
3.5
3
2.5
2
1.5
1
0
10
20
30
40
50
60
Time (min)
Normal
Diabetic
Normal data

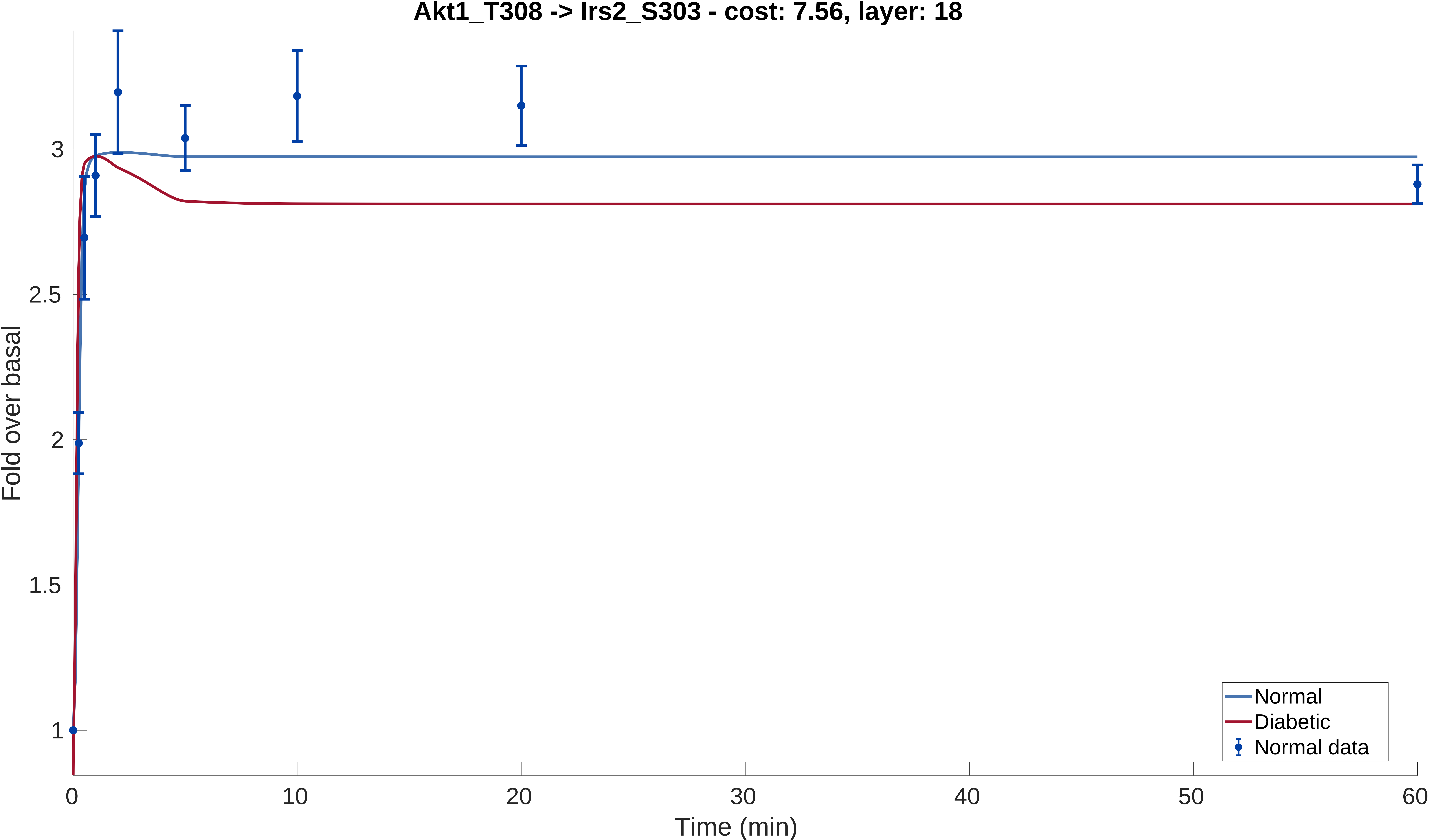
Akt1_T308 -> Irs2_S303 - cost: 7.56, layer: 18
Fold over basal
1
1.5
2
2.5
3
0
10
20
30
40
50
60
Time (min)
Normal
Diabetic
Normal data

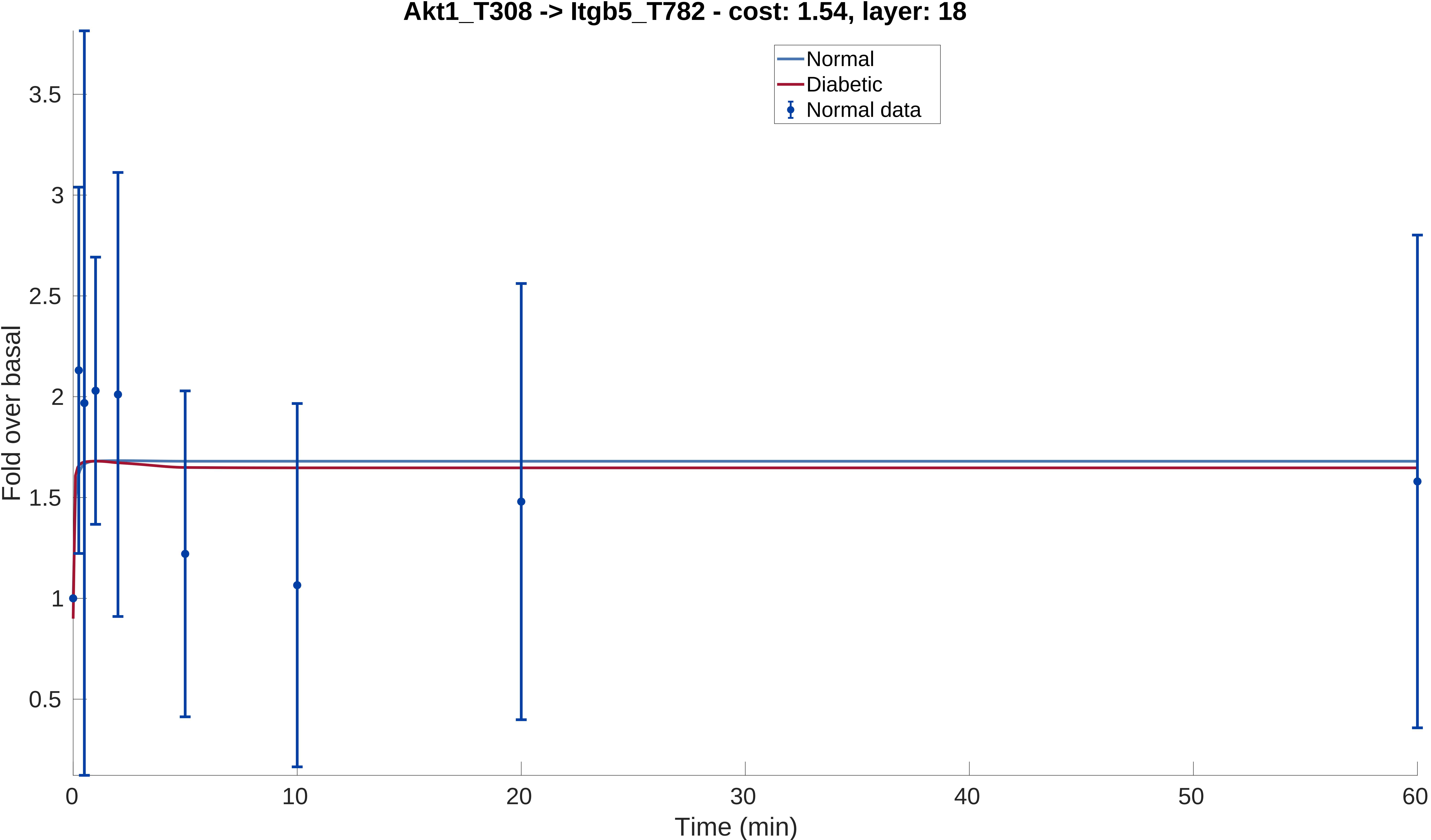

Akt1_T308 -> Itgb5_T782 - cost: 1.54, layer: 18
Normal
Diabetic
Normal data
Fold over basal
3.5
3
2.5
2
1.5
1
0.5
0
10
20
30
40
50
60
Time (min)

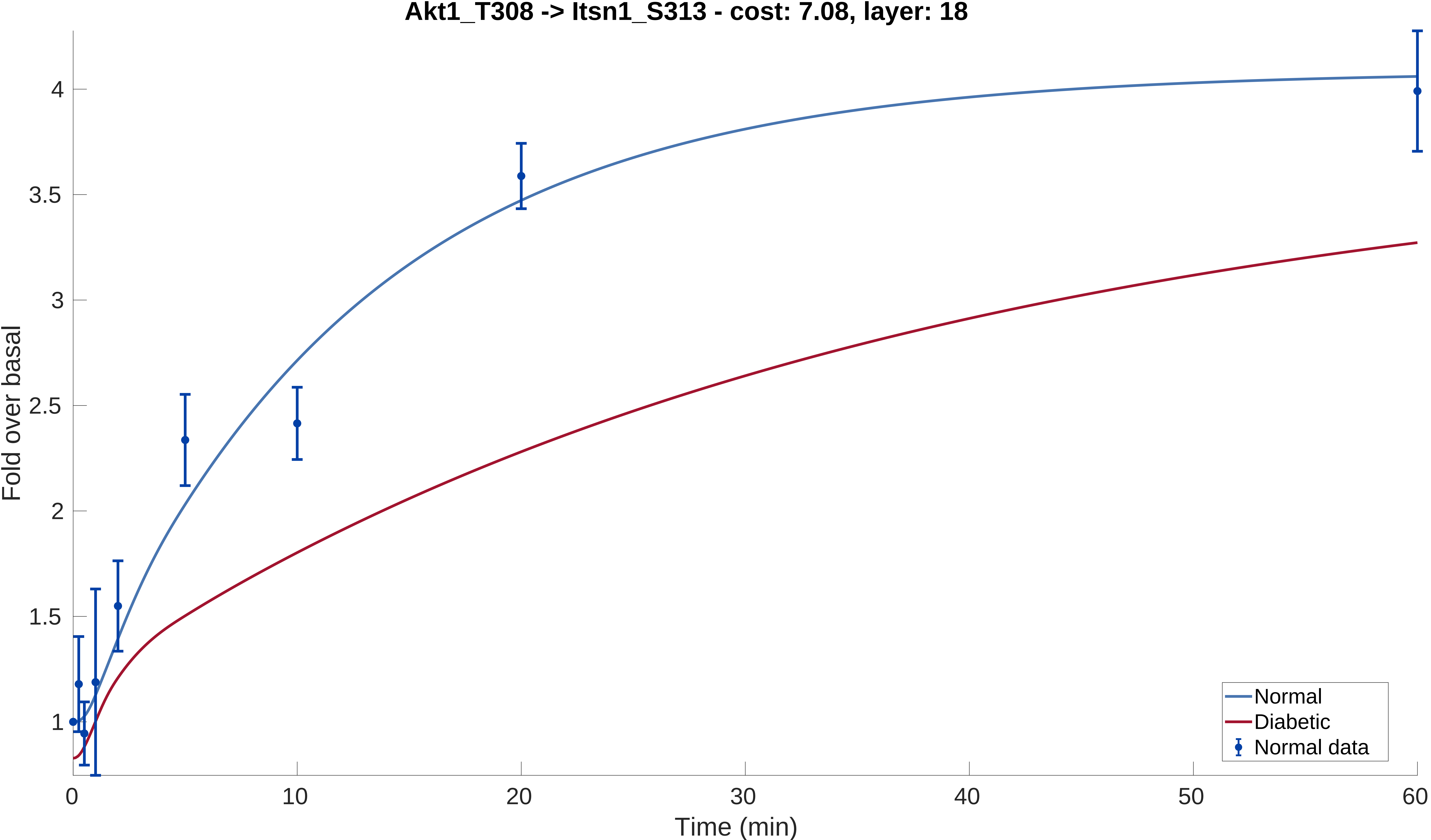

Akt1_T308 -> Itsn1_S313 - cost: 7.08, layer: 18
Fold over basal
4
3.5
3
2.5
2
1.5
1
0
10
20
30
40
50
60
Time (min)
Normal
Diabetic
Normal data

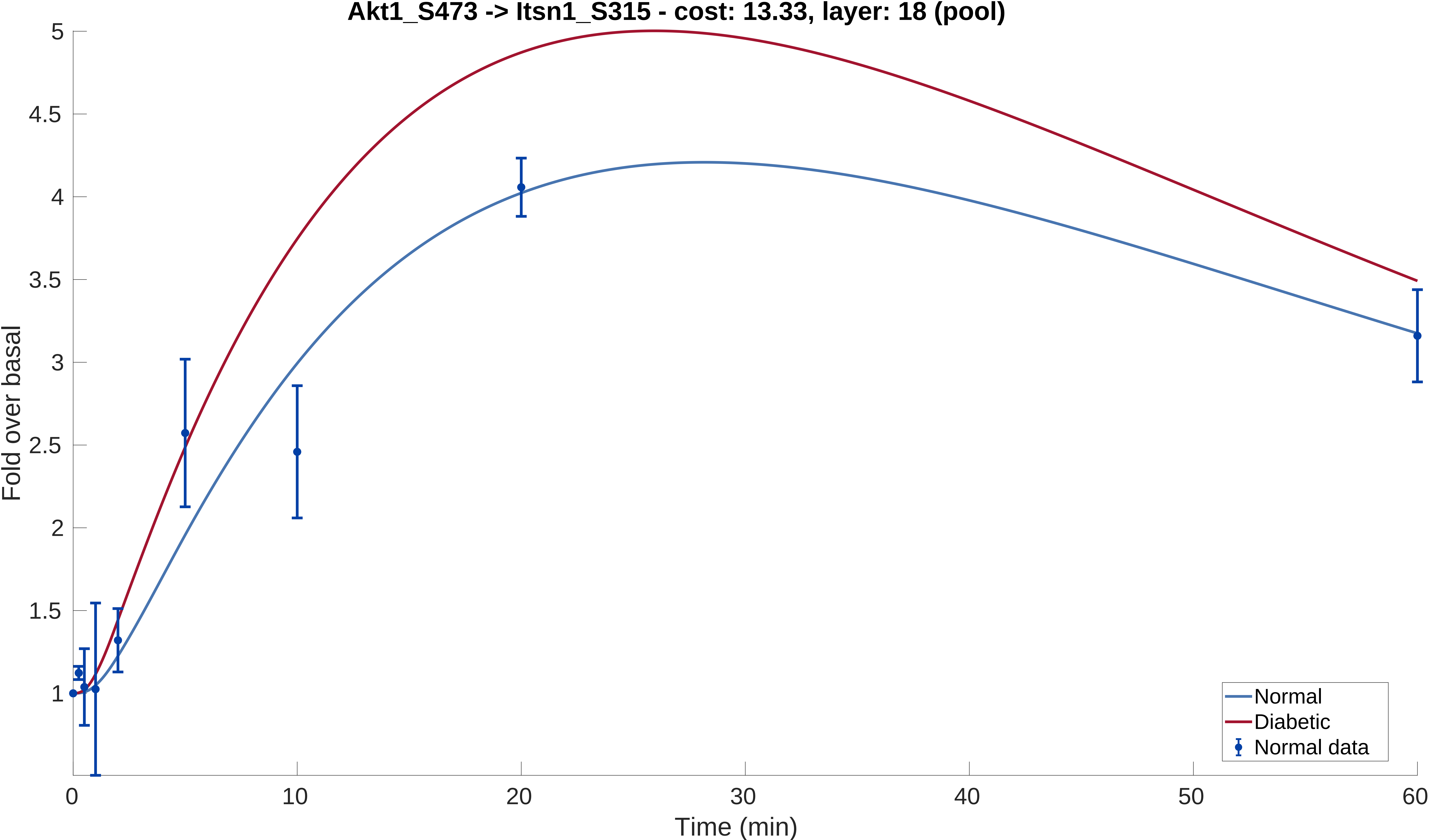

Akt1_S473 -> Itsn1_S315 - cost: 13.33, layer: 18 (pool)
Fold over basal
5
4.5
4
3.5
3
2.5
2
1.5
1
0
10
20
30
40
50
60
Time (min)
Normal
Diabetic
Normal data

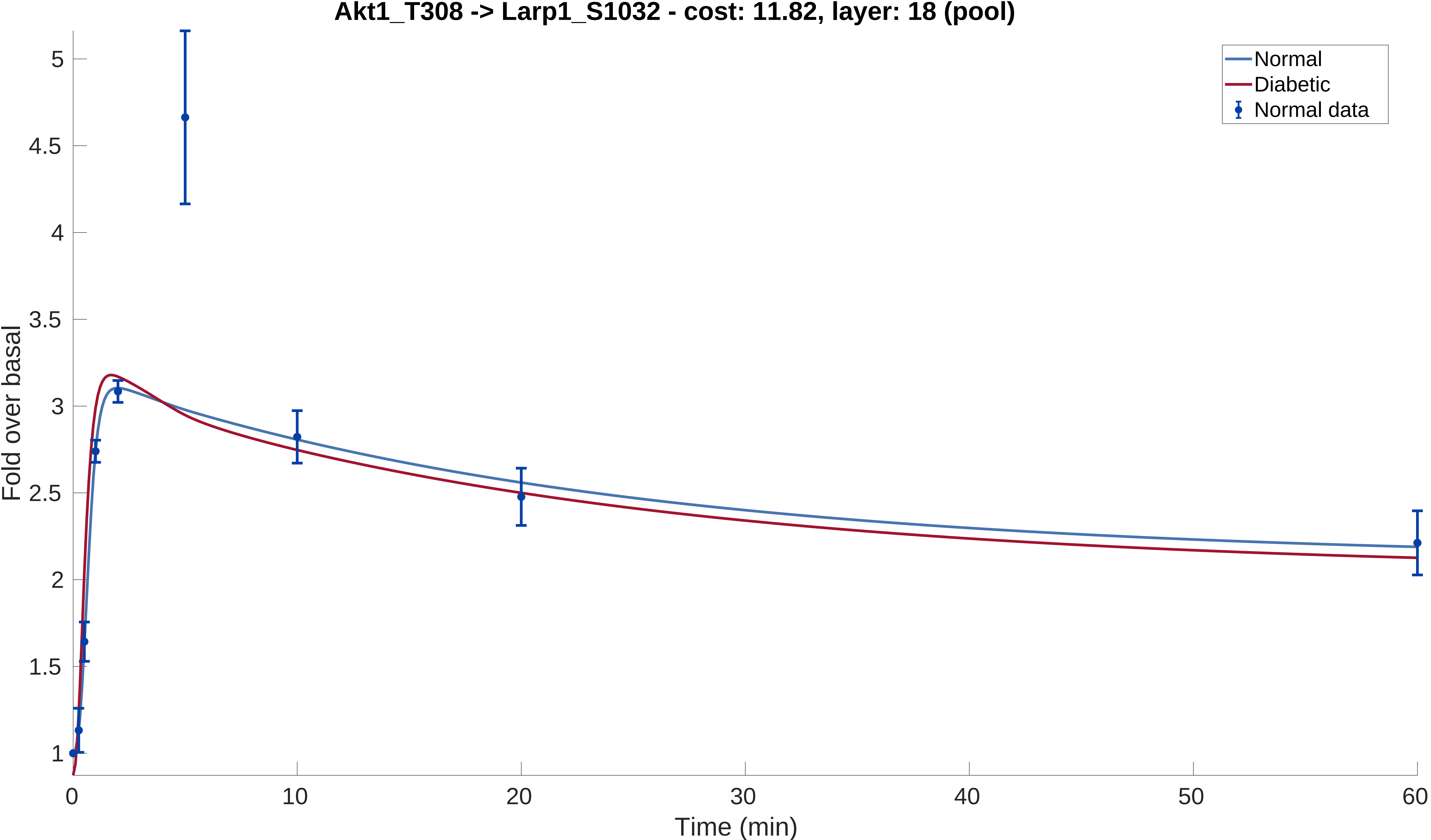

Akt1_T308 -> Larp1_S1032 - cost: 11.82, layer: 18 (pool)
Fold over basal
Time (min)
Normal
Diabetic
Normal data
5
4.5
4
3.5
3
2.5
2
1.5
1
0
10
20
30
40
50
60

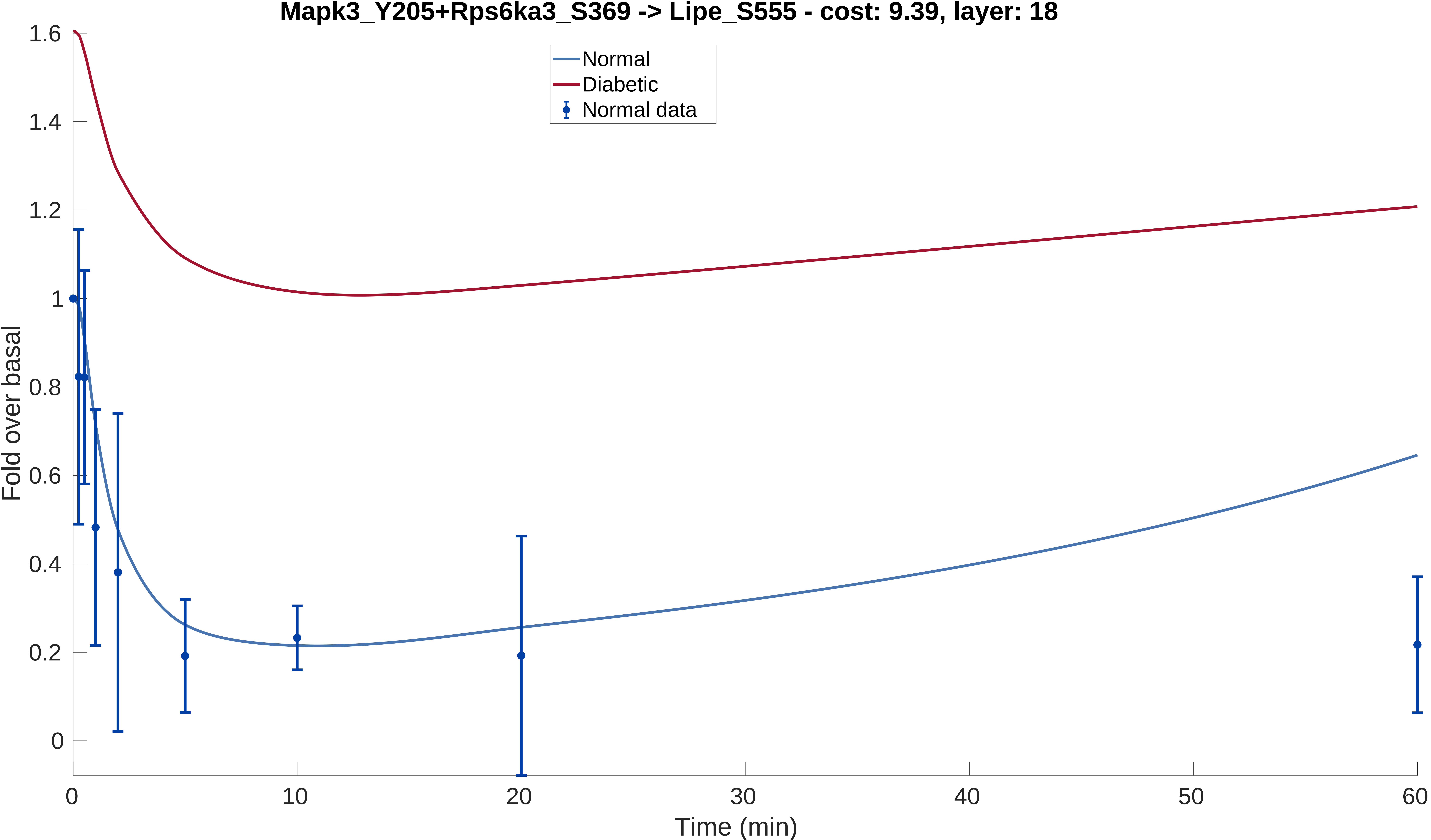

Mapk3_Y205+Rps6ka3_S369 -> Lipe_S555 - cost: 9.39, layer: 18
Normal
Diabetic
Normal data
Fold over basal
Time (min)
0
0.2
0.4
0.6
0.8
1
1.2
1.4
1.6
0
10
20
30
40
50
60

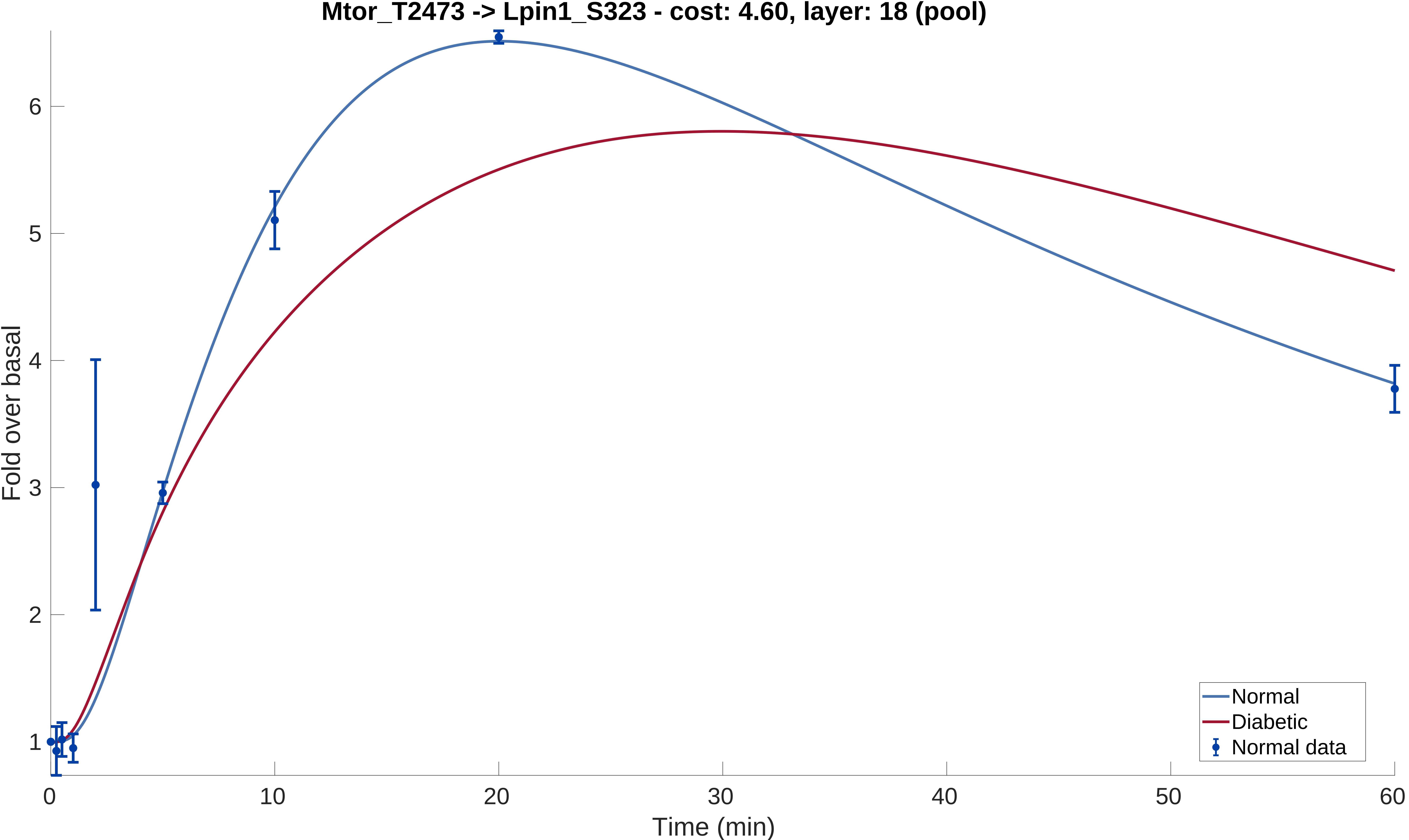

Mtor_T2473 -> Lpin1_S323 - cost: 4.60, layer: 18 (pool)
Fold over basal
Time (min)
0
10
20
30
40
50
60
1
2
3
4
5
6
Normal
Diabetic
Normal data

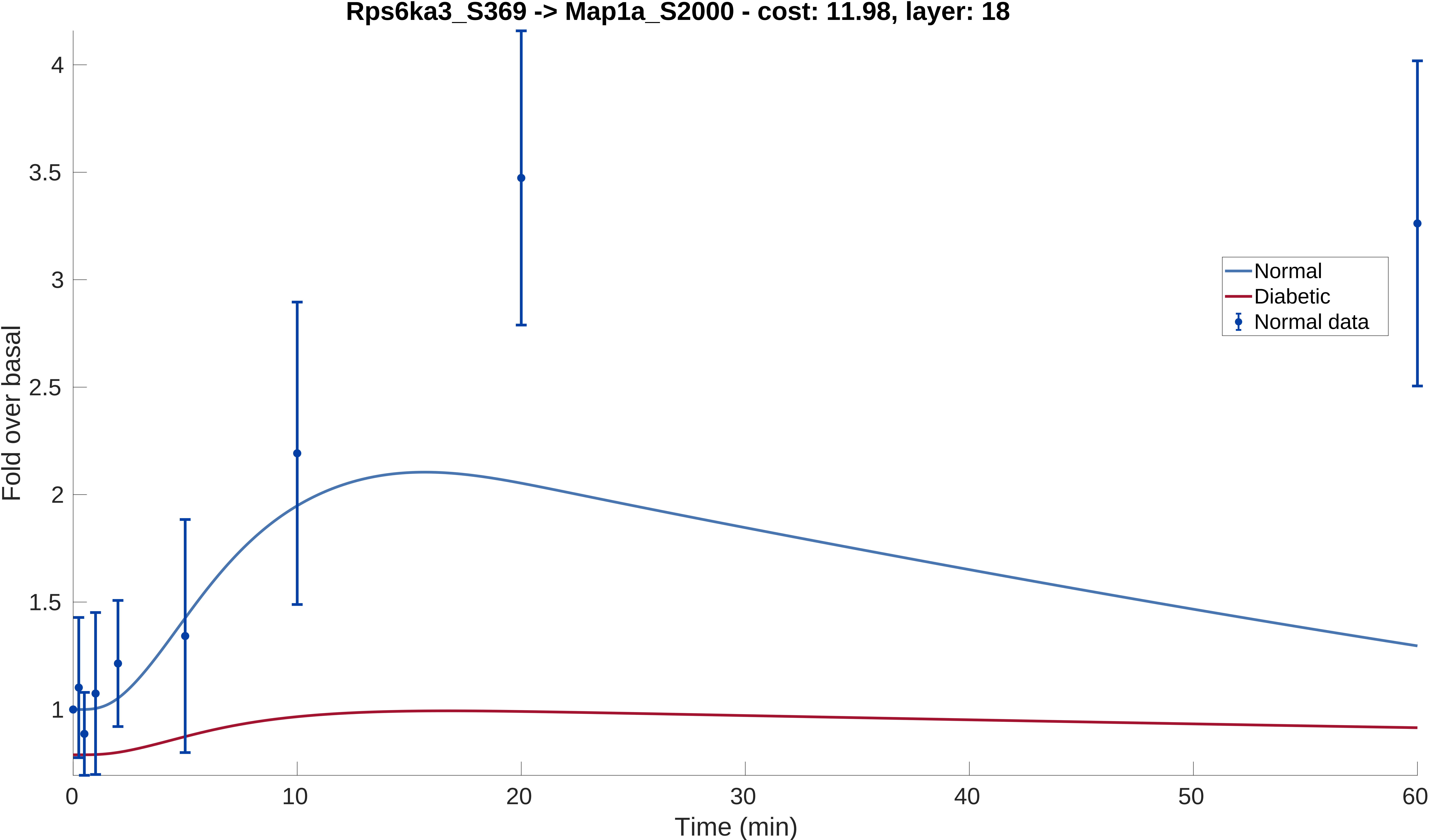

Rps6ka3_S369 -> Map1a_S2000 - cost: 11.98, layer: 18
Fold over basal
Time (min)
Normal
Diabetic
Normal data
4
3.5
3
2.5
2
1.5
1
0
10
20
30
40
50
60

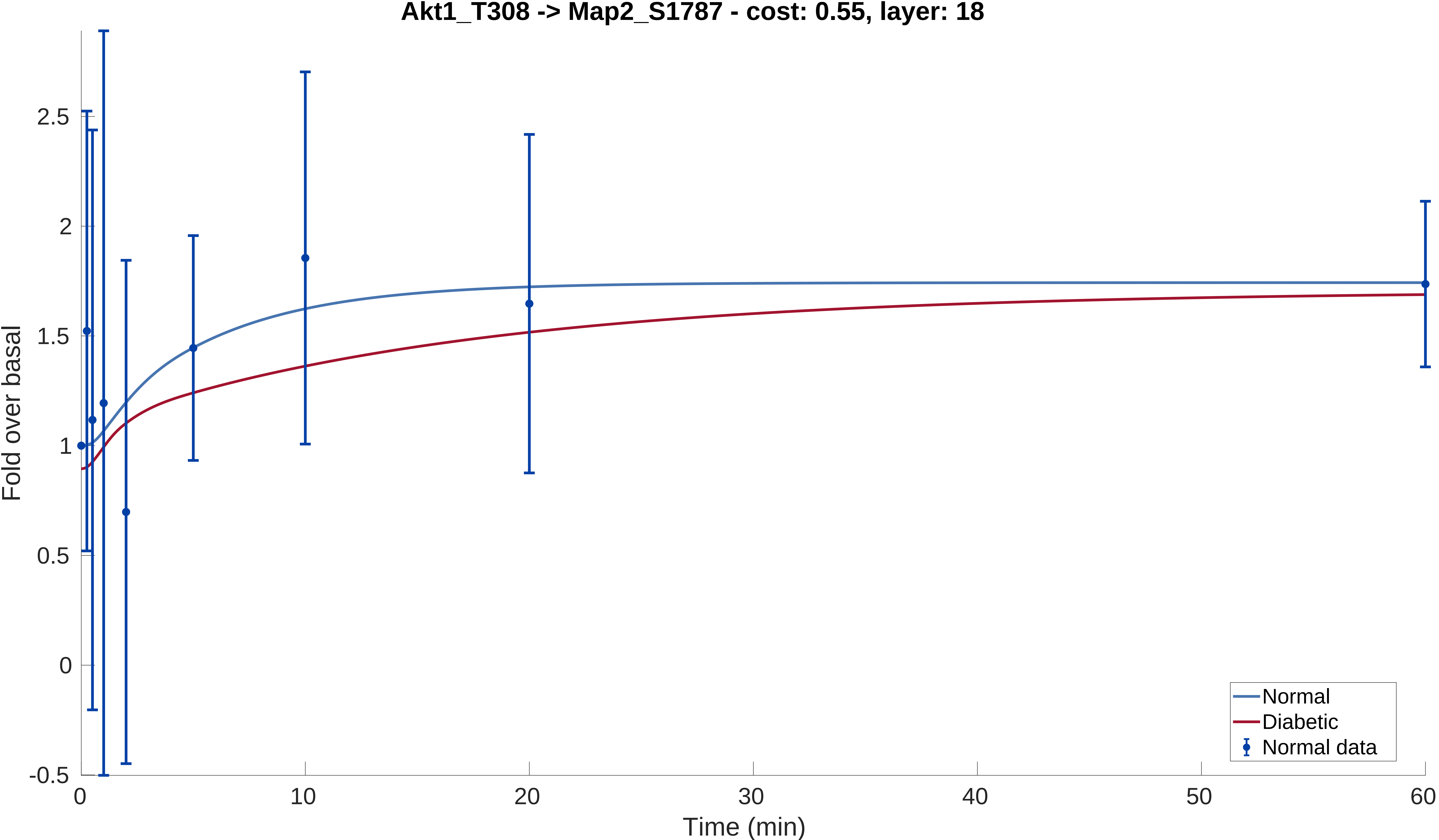

Akt1_T308 -> Map2_S1787 - cost: 0.55, layer: 18
Fold over basal
Time (min)
2.5
2
1.5
1
0.5
0
-0.5
0
10
20
30
40
50
60
Normal
Diabetic
Normal data

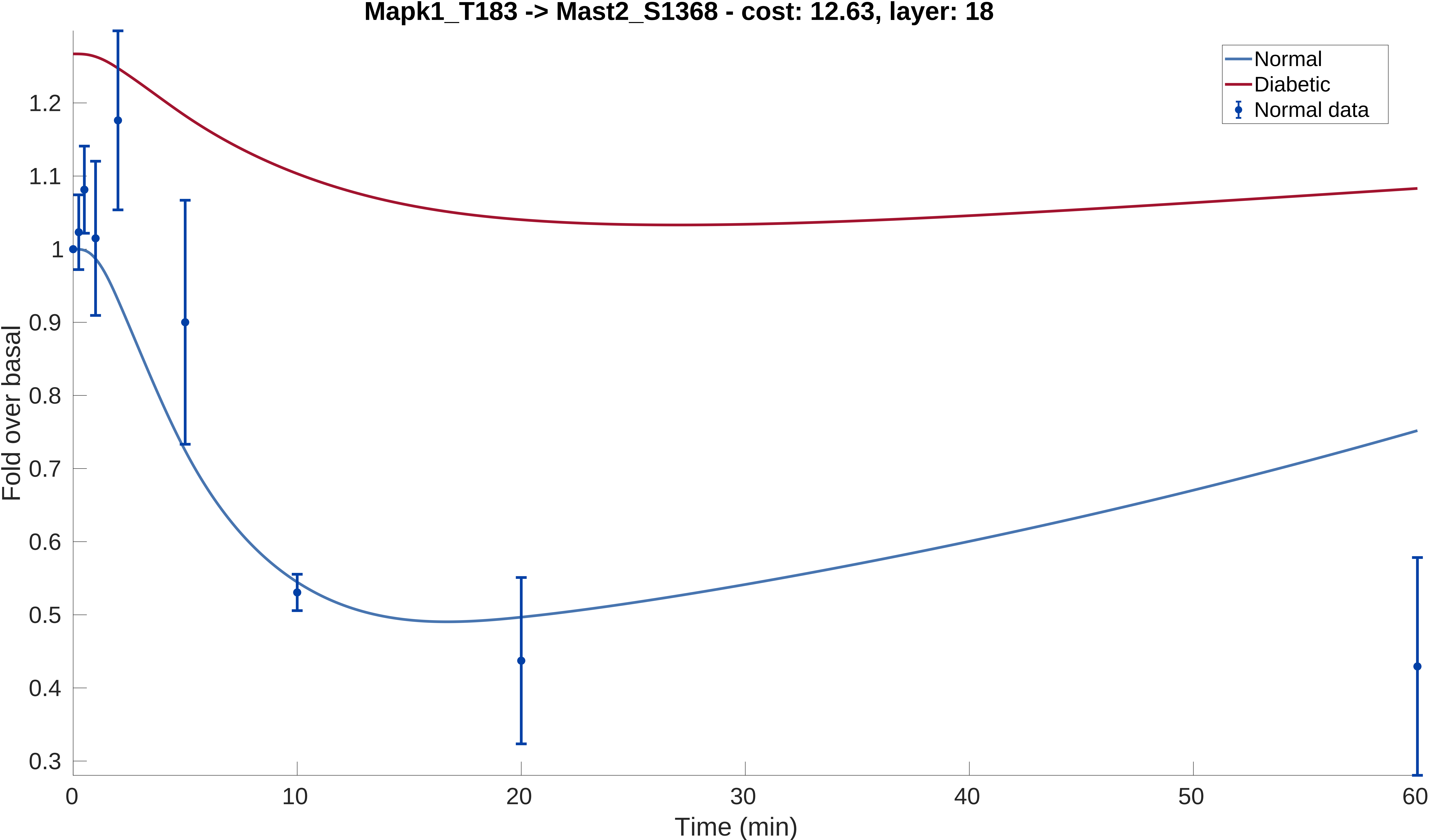

Mapk1_T183 -> Mast2_S1368 - cost: 12.63, layer: 18
Normal
Diabetic
Normal data
Fold over basal
Time (min)
0
10
20
30
40
50
60
0.3
0.4
0.5
0.6
0.7
0.8
0.9
1
1.1
1.2

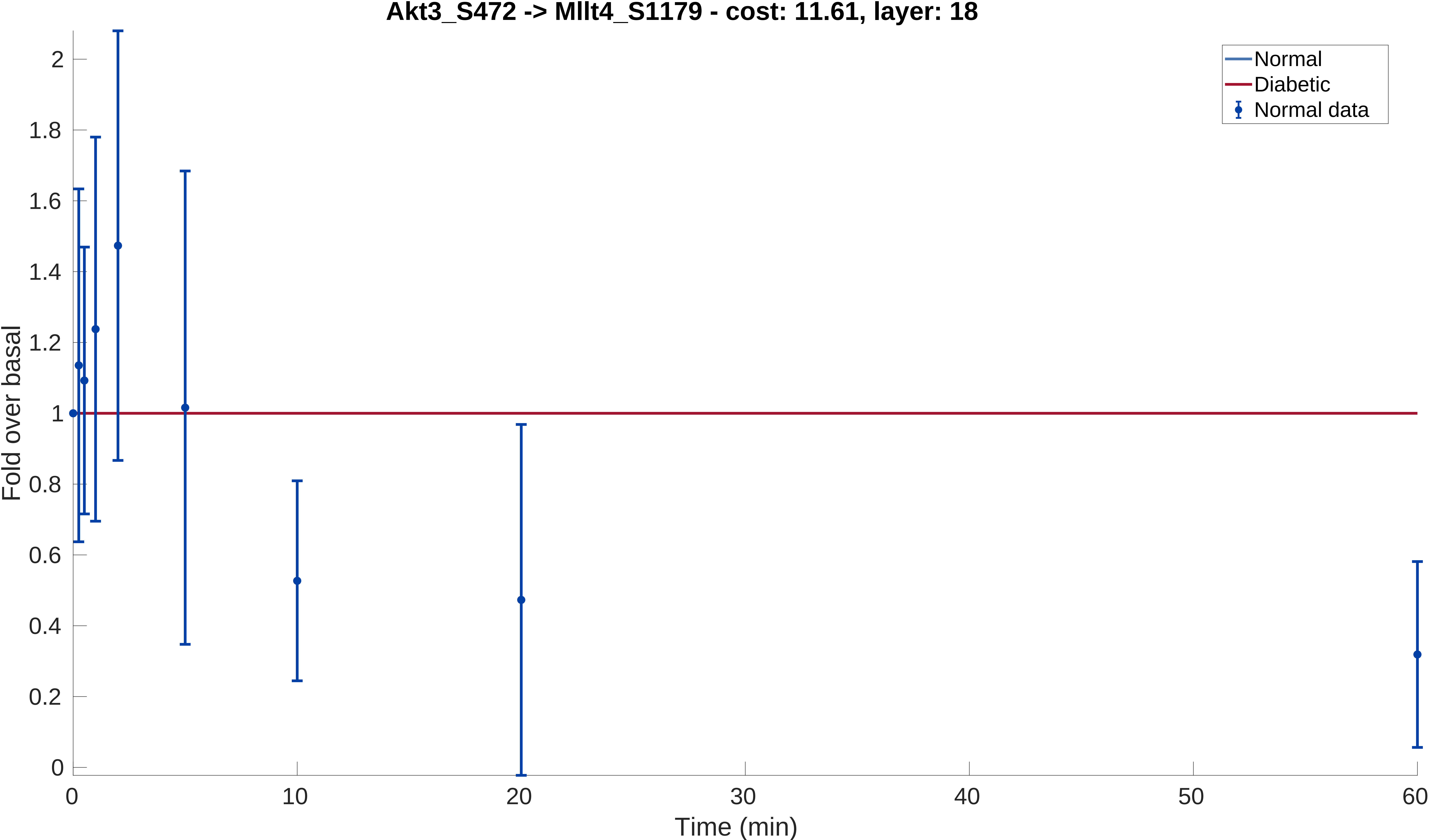
Akt3_S472 -> Mllt4_S1179 - cost: 11.61, layer: 18
Normal
Diabetic
Normal data
Fold over basal
Time (min)
0
0.2
0.4
0.6
0.8
1
1.2
1.4
1.6
1.8
2
0
10
20
30
40
50
60

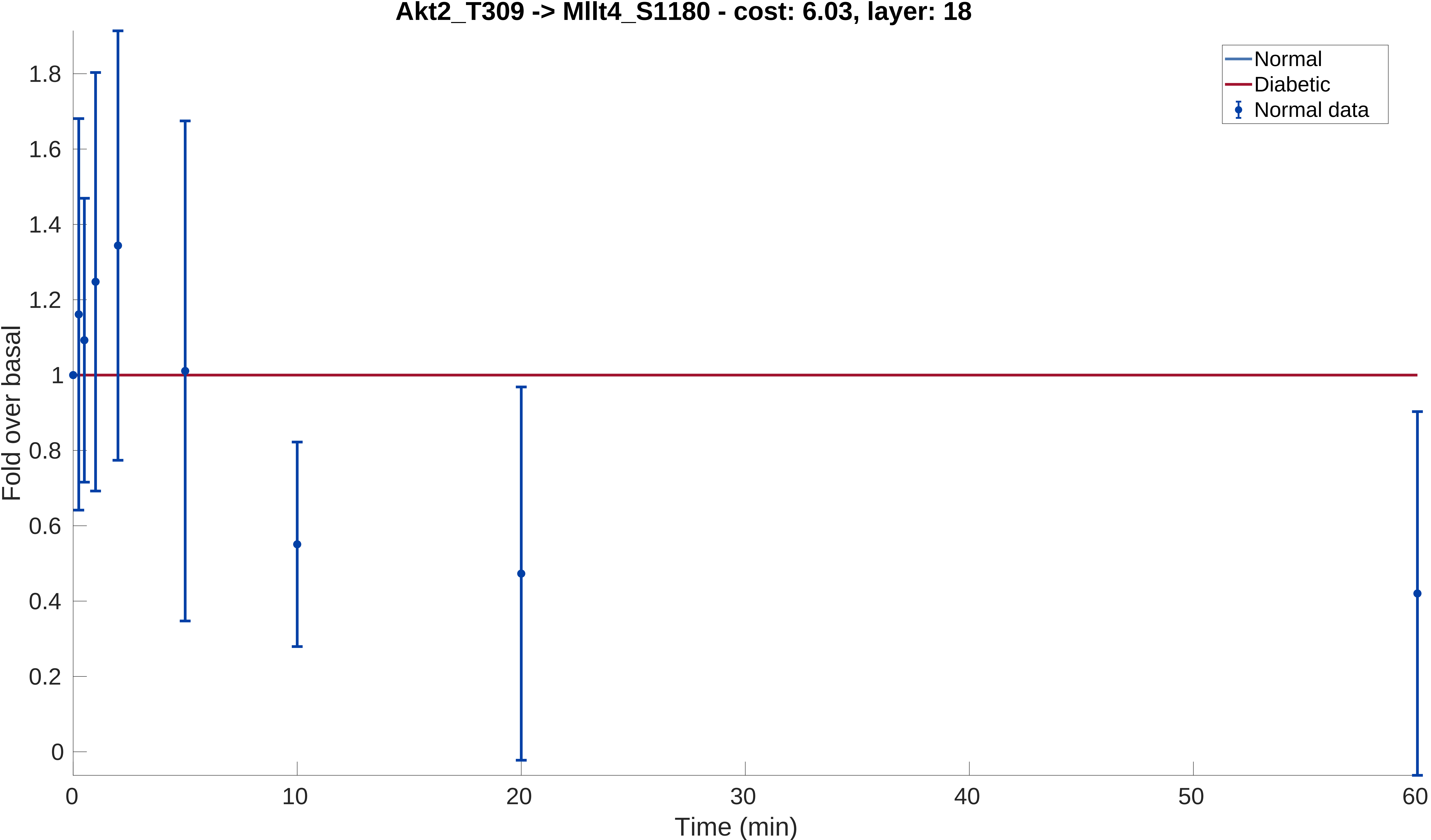

Akt2_T309 -> Mllt4_S1180 - cost: 6.03, layer: 18
Normal
Diabetic
Normal data
Fold over basal
Time (min)
0
0.2
0.4
0.6
0.8
1
1.2
1.4
1.6
1.8
0
10
20
30
40
50
60

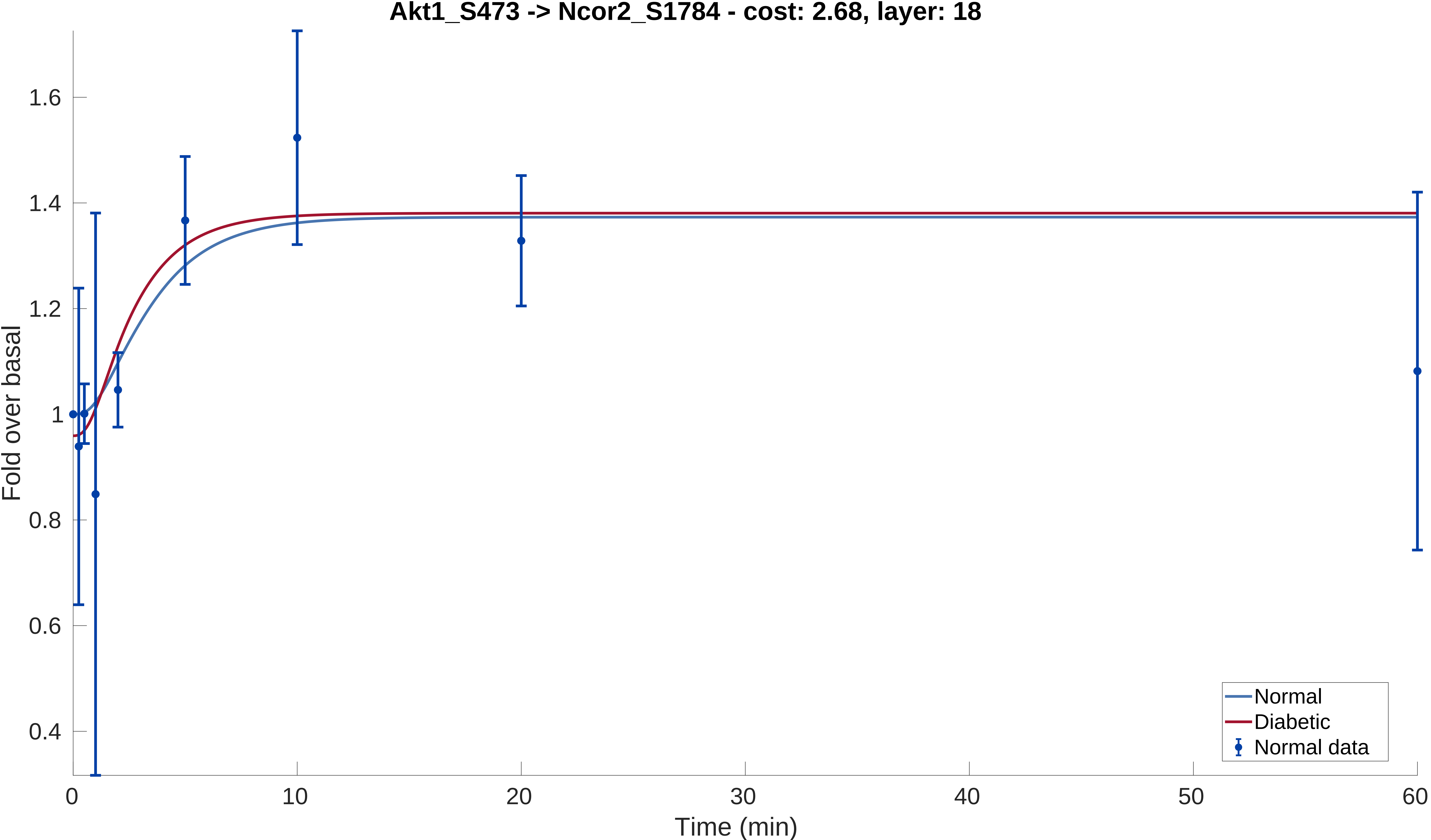
Akt1_S473 -> Ncor2_S1784 - cost: 2.68, layer: 18
Fold over basal
1.6
1.4
1.2
1
0.8
0.6
0.4
0
10
20
30
40
50
60
Time (min)
Normal
Diabetic
Normal data

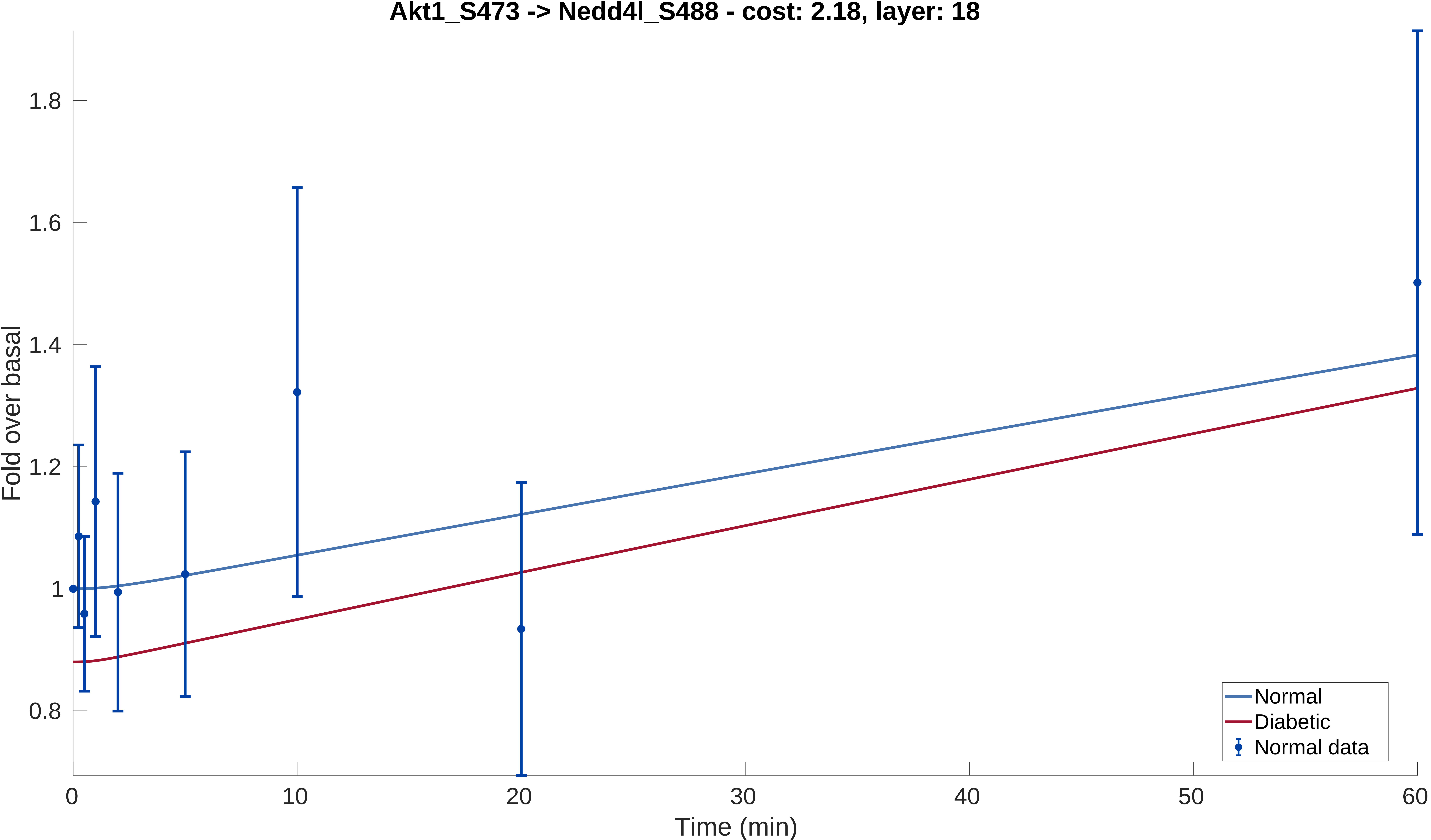

Akt1_S473 -> Nedd4l_S488 - cost: 2.18, layer: 18
Fold over basal
Time (min)
0.8
1
1.2
1.4
1.6
1.8
0
10
20
30
40
50
60
Normal
Diabetic
Normal data

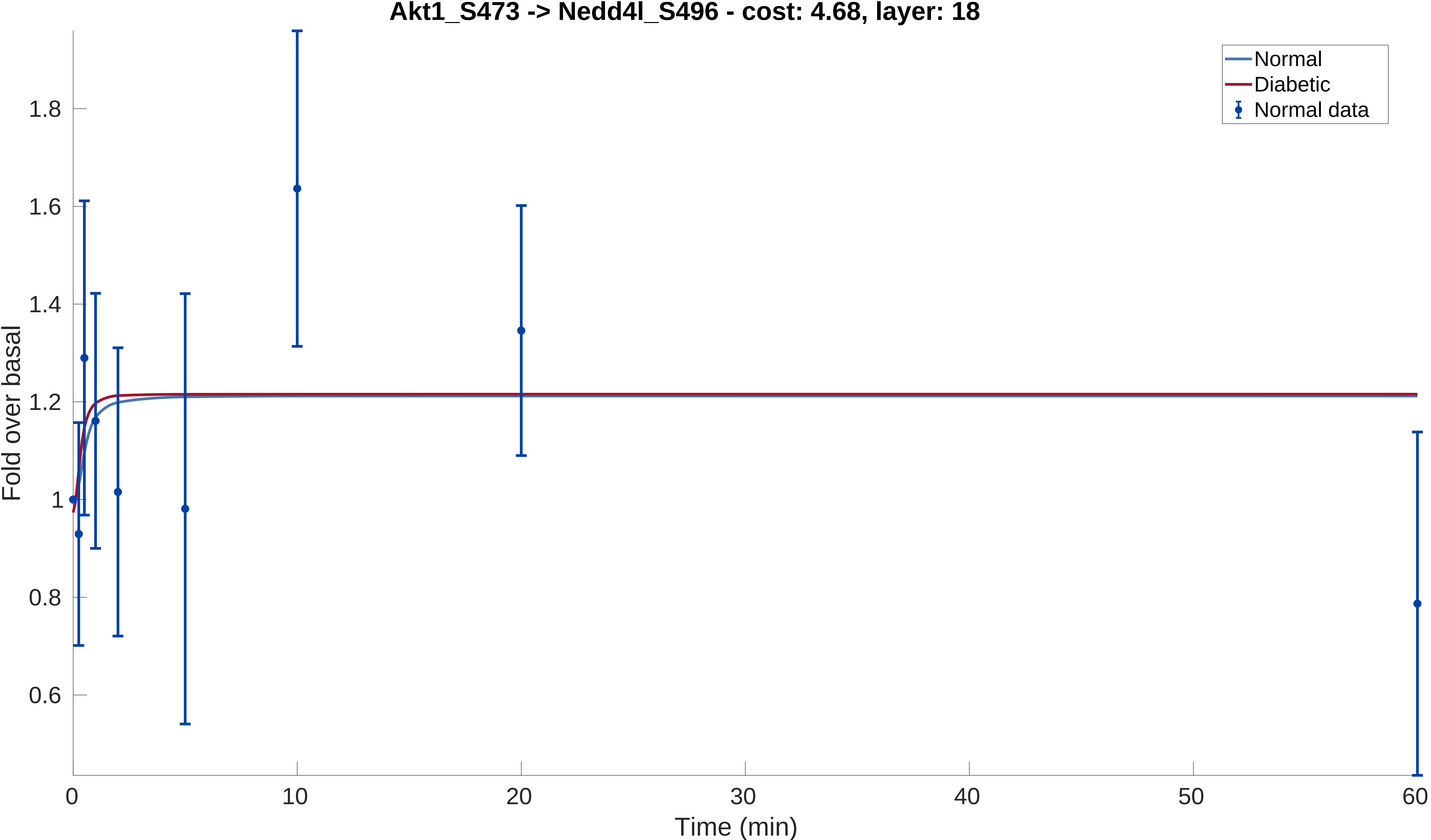

Akt1_S473 -> Nedd4l_S496 - cost: 4.68, layer: 18
Normal
Diabetic
Normal data
Fold over basal
1.8
1.6
1.4
1.2
1
0.8
0.6
0
10
20
30
40
50
60
Time (min)

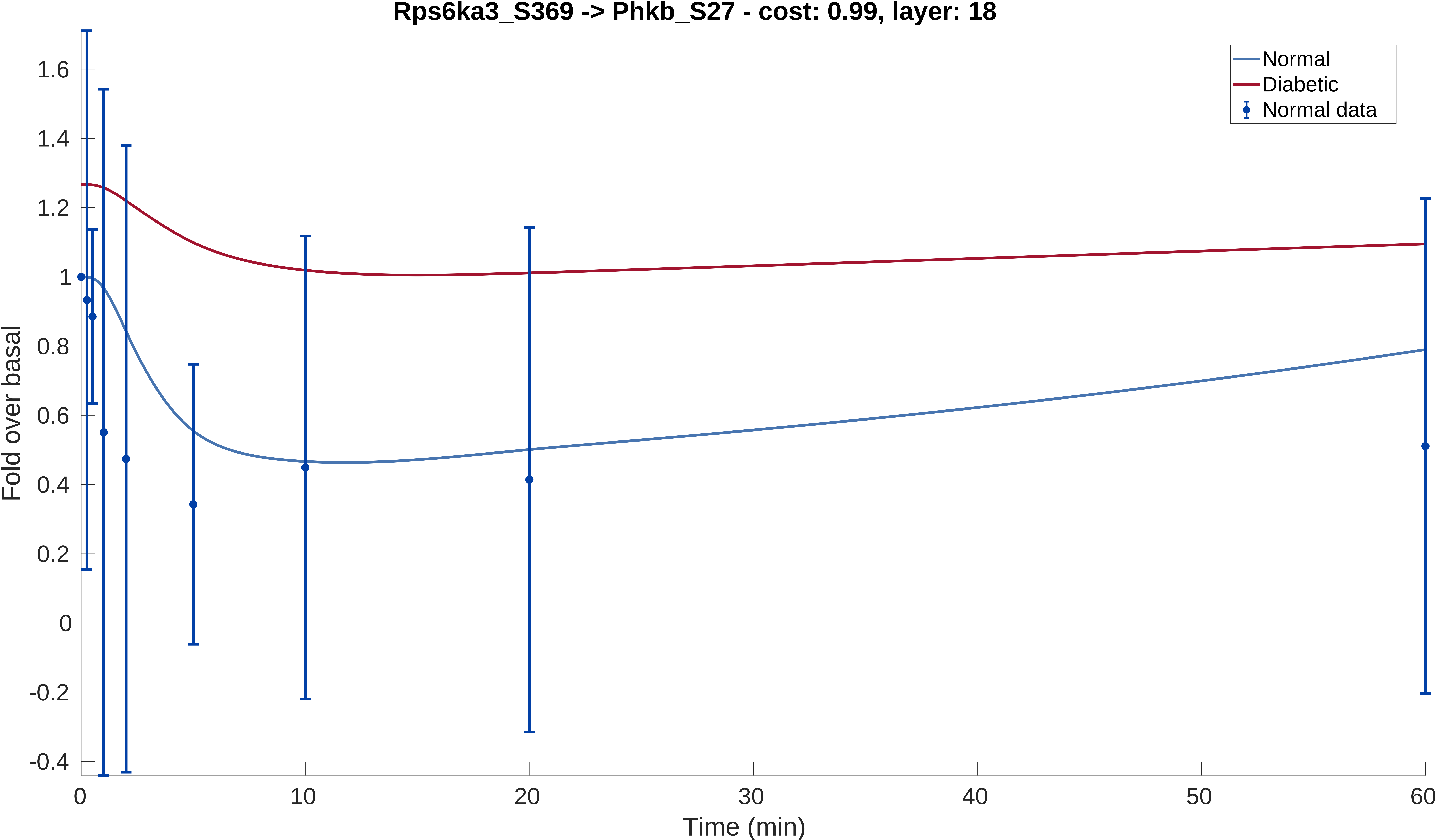
Rps6ka3_S369 -> Phkb_S27 - cost: 0.99, layer: 18
Normal
Diabetic
Normal data
Fold over basal
Time (min)
1.6
1.4
1.2
1
0.8
0.6
0.4
0.2
0
-0.2
-0.4
0
10
20
30
40
50
60

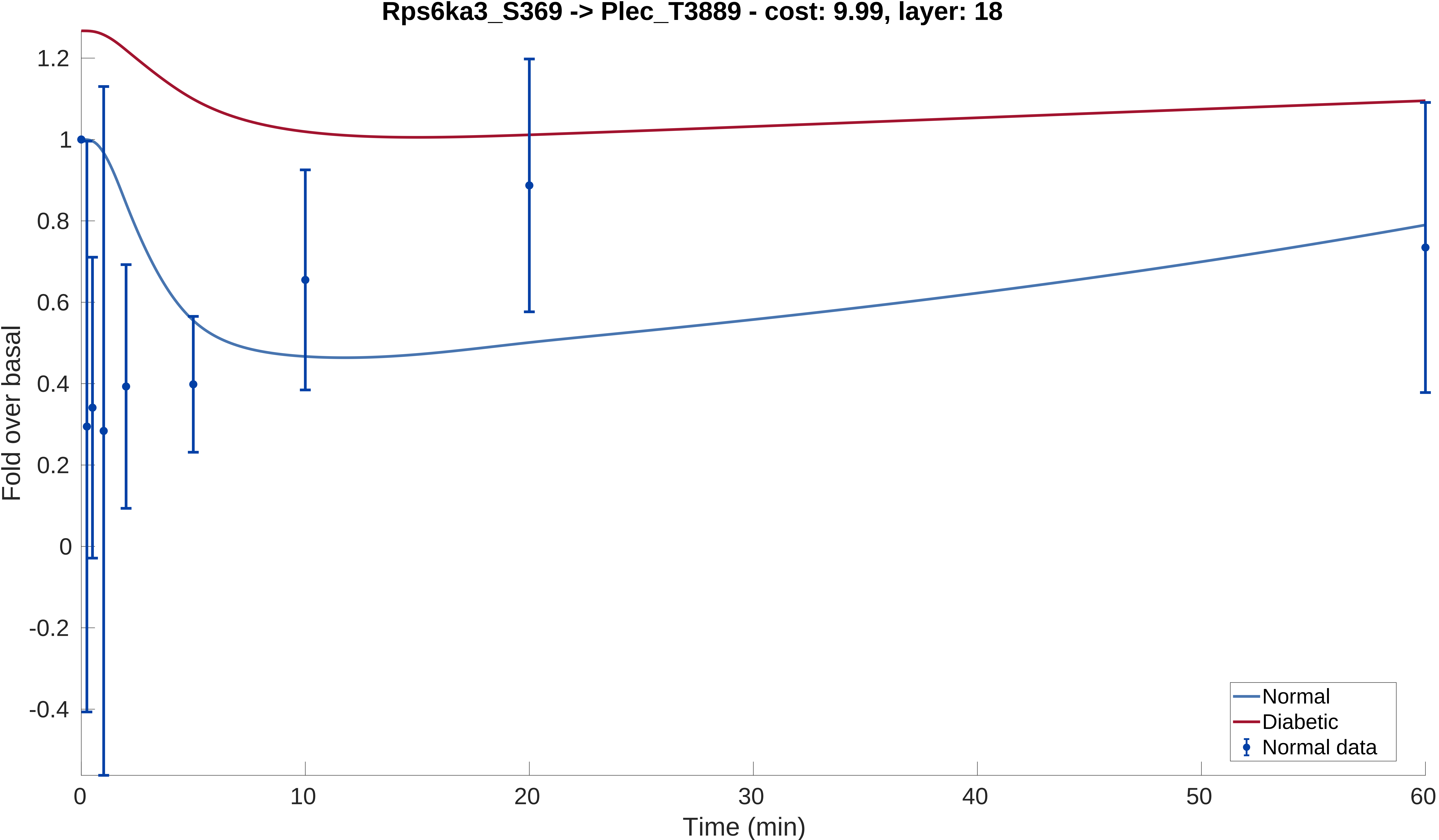

Rps6ka3_S369 -> Plec_T3889 - cost: 9.99, layer: 18
Fold over basal
1.2
1
0.8
0.6
0.4
0.2
0
-0.2
-0.4
0
10
20
30
40
50
60
Time (min)
Normal
Diabetic
Normal data

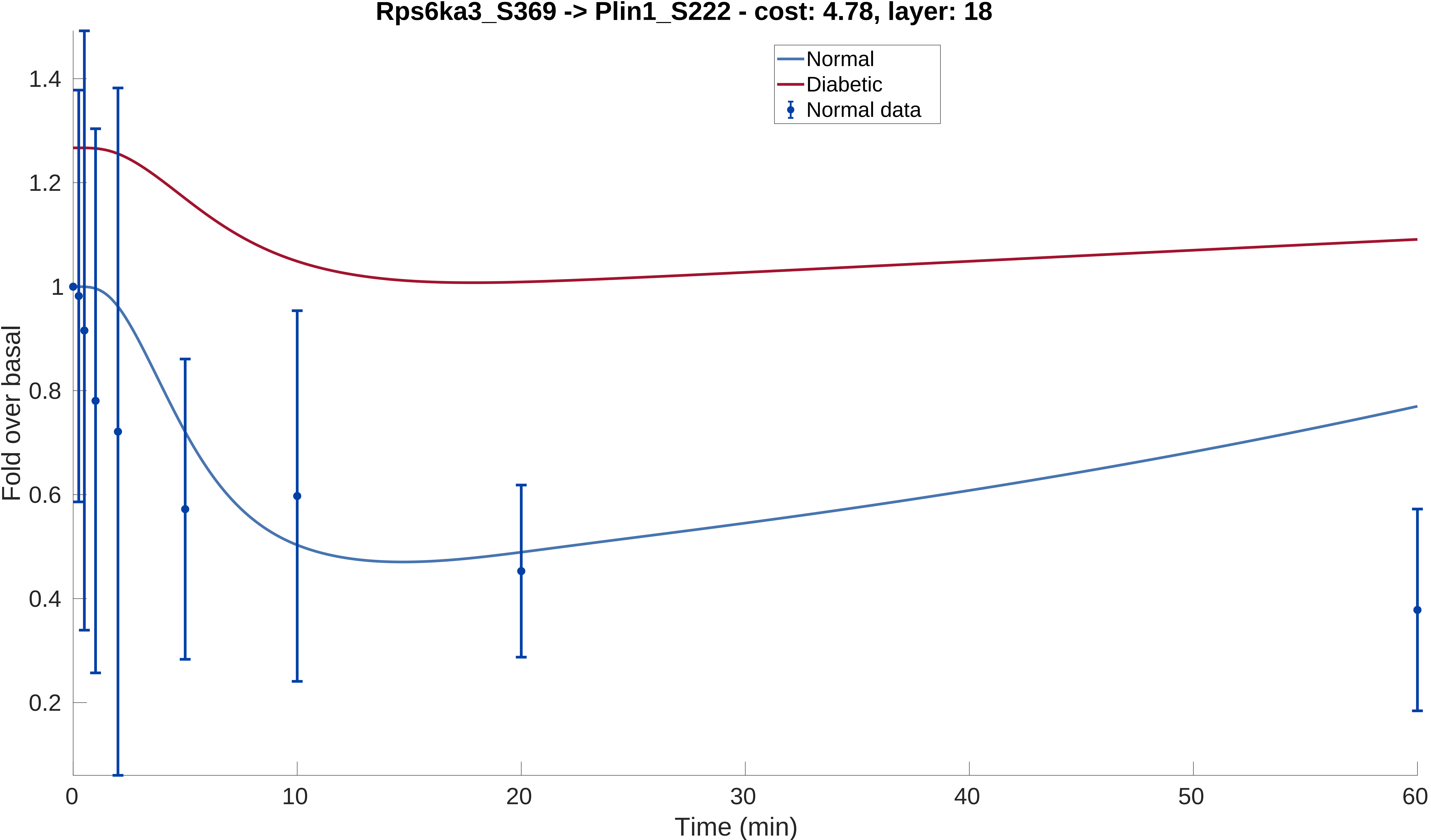

Rps6ka3_S369 -> Plin1_S222 - cost: 4.78, layer: 18
Normal
Diabetic
Normal data
Fold over basal
1.4
1.2
1
0.8
0.6
0.4
0.2
0
10
20
30
40
50
60
Time (min)

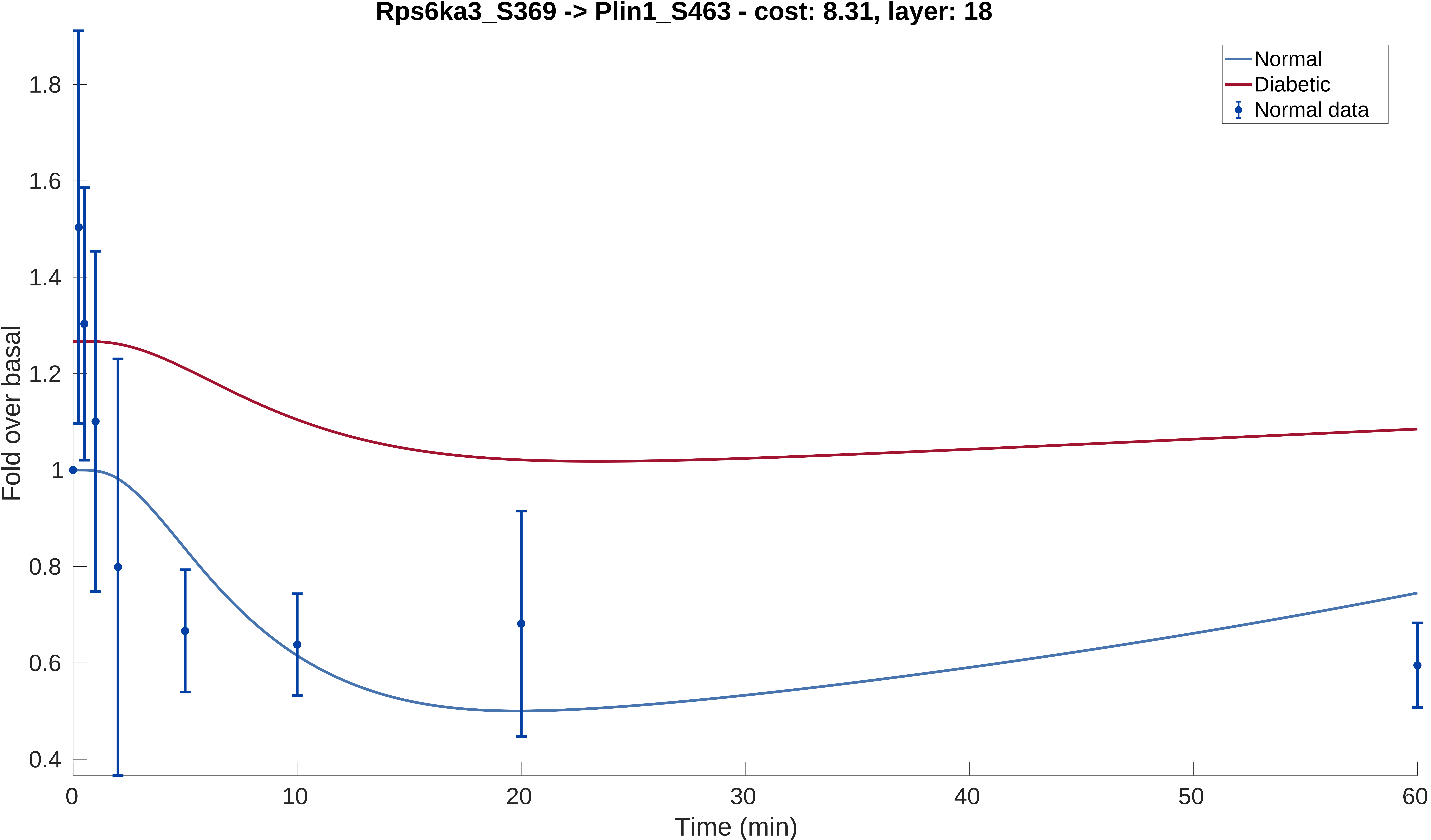

Rps6ka3_S369 -> Plin1_S463 - cost: 8.31, layer: 18
Normal
Diabetic
Normal data
Fold over basal
1.8
1.6
1.4
1.2
1
0.8
0.6
0.4
0
10
20
30
40
50
60
Time (min)

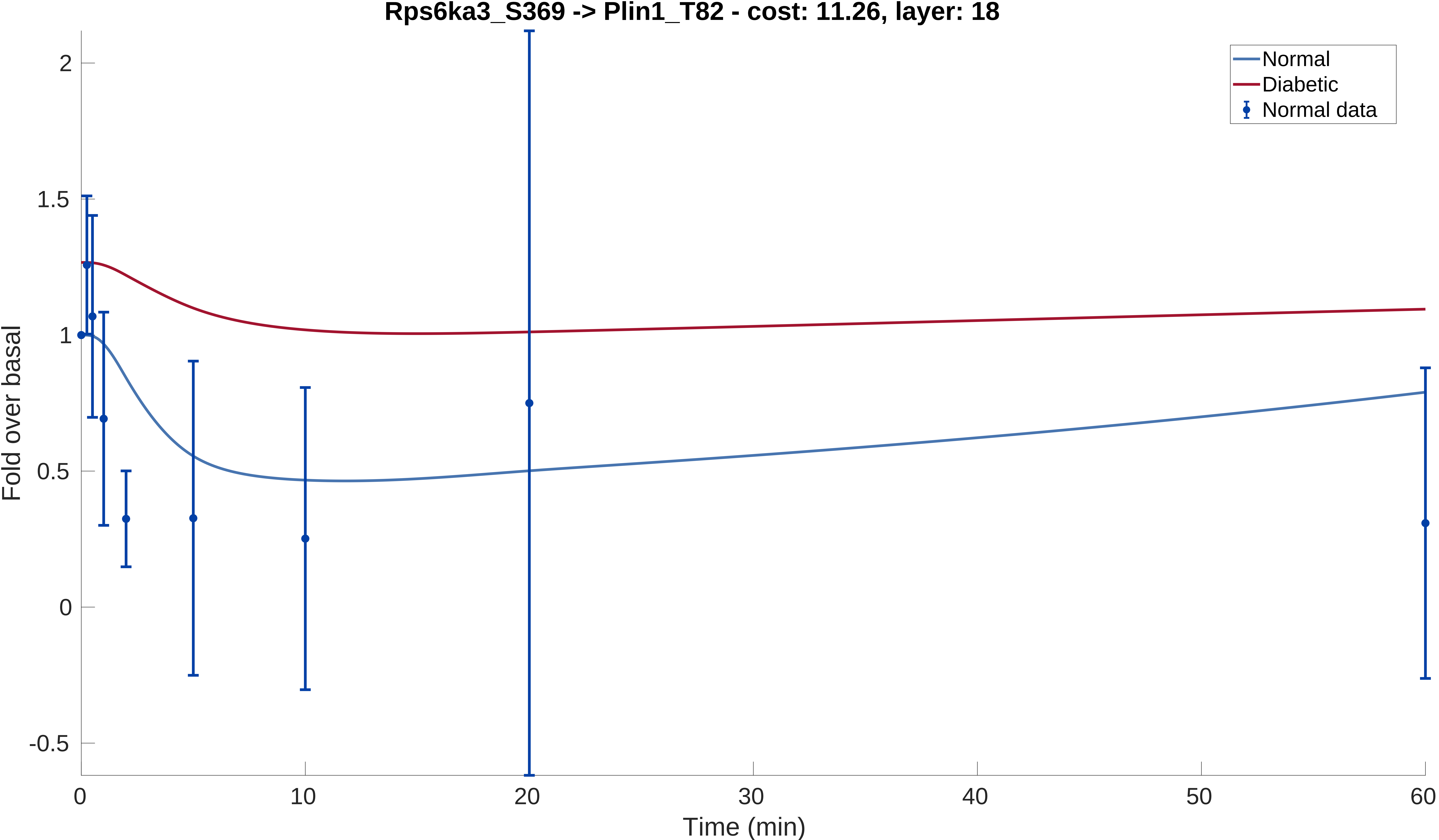

Rps6ka3_S369 -> Plin1_T82 - cost: 11.26, layer: 18
Normal
Diabetic
Normal data
Fold over basal
Time (min)
2
1.5
1
0.5
0
-0.5
0
10
20
30
40
50
60

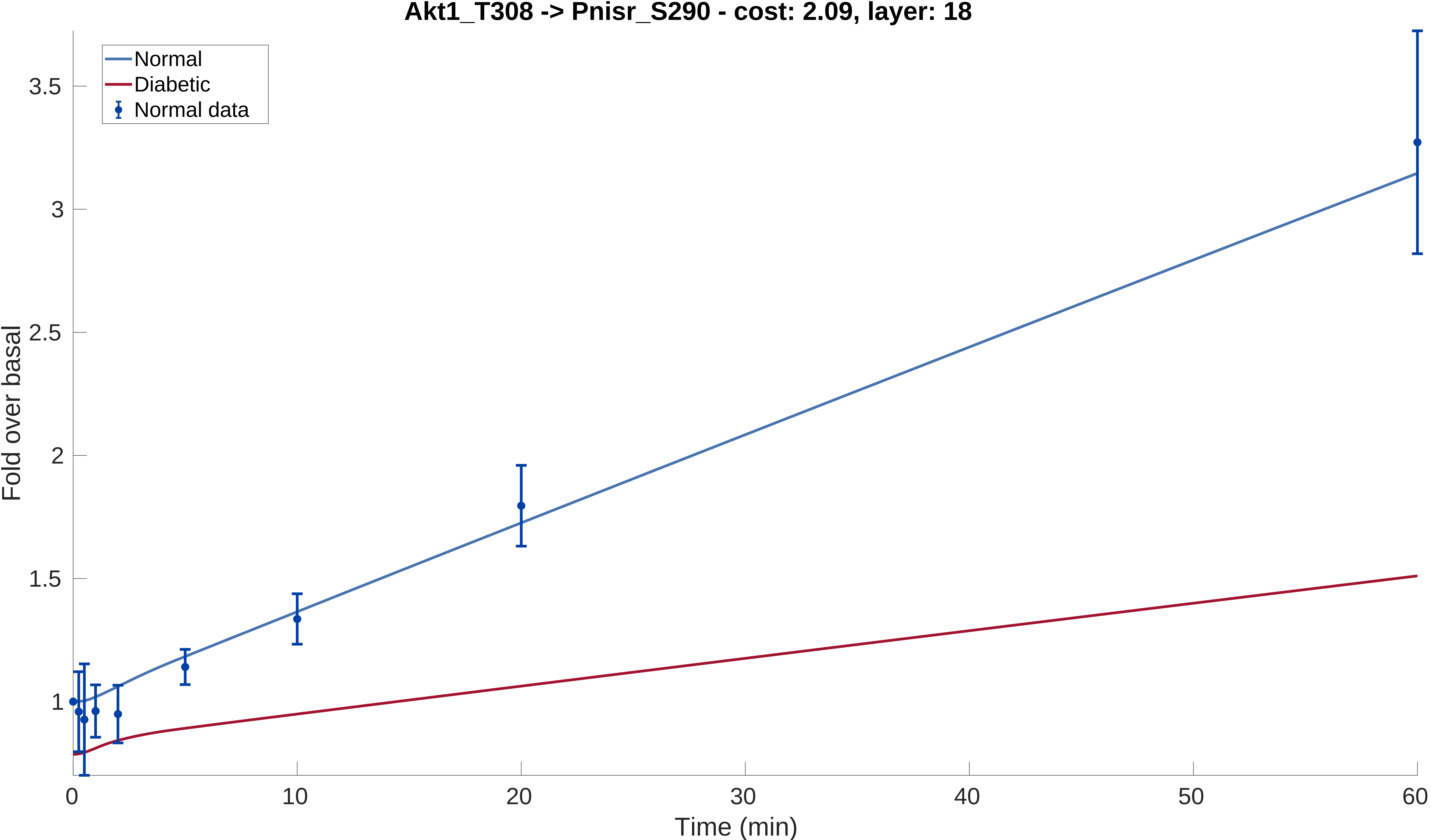
Akt1_T308 -> Pnisr_S290 - cost: 2.09, layer: 18
Normal
Diabetic
Normal data
Fold over basal
3.5
3
2.5
2
1.5
1
0
10
20
30
40
50
60
Time (min)

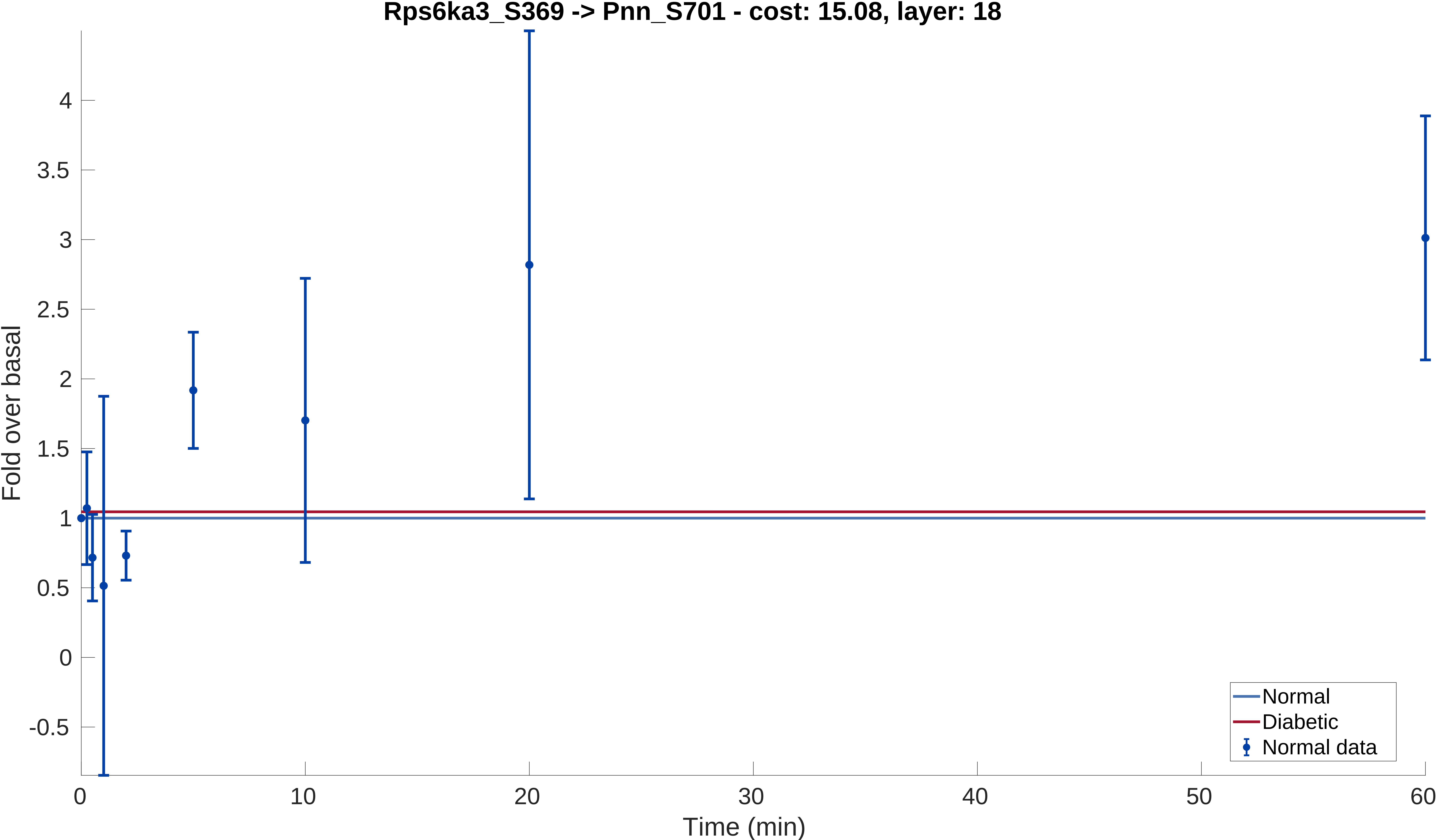

Rps6ka3_S369 -> Pnn_S701 - cost: 15.08, layer: 18
Fold over basal
Time (min)
Normal
Diabetic
Normal data
4
3.5
3
2.5
2
1.5
1
0.5
0
-0.5
0
10
20
30
40
50
60

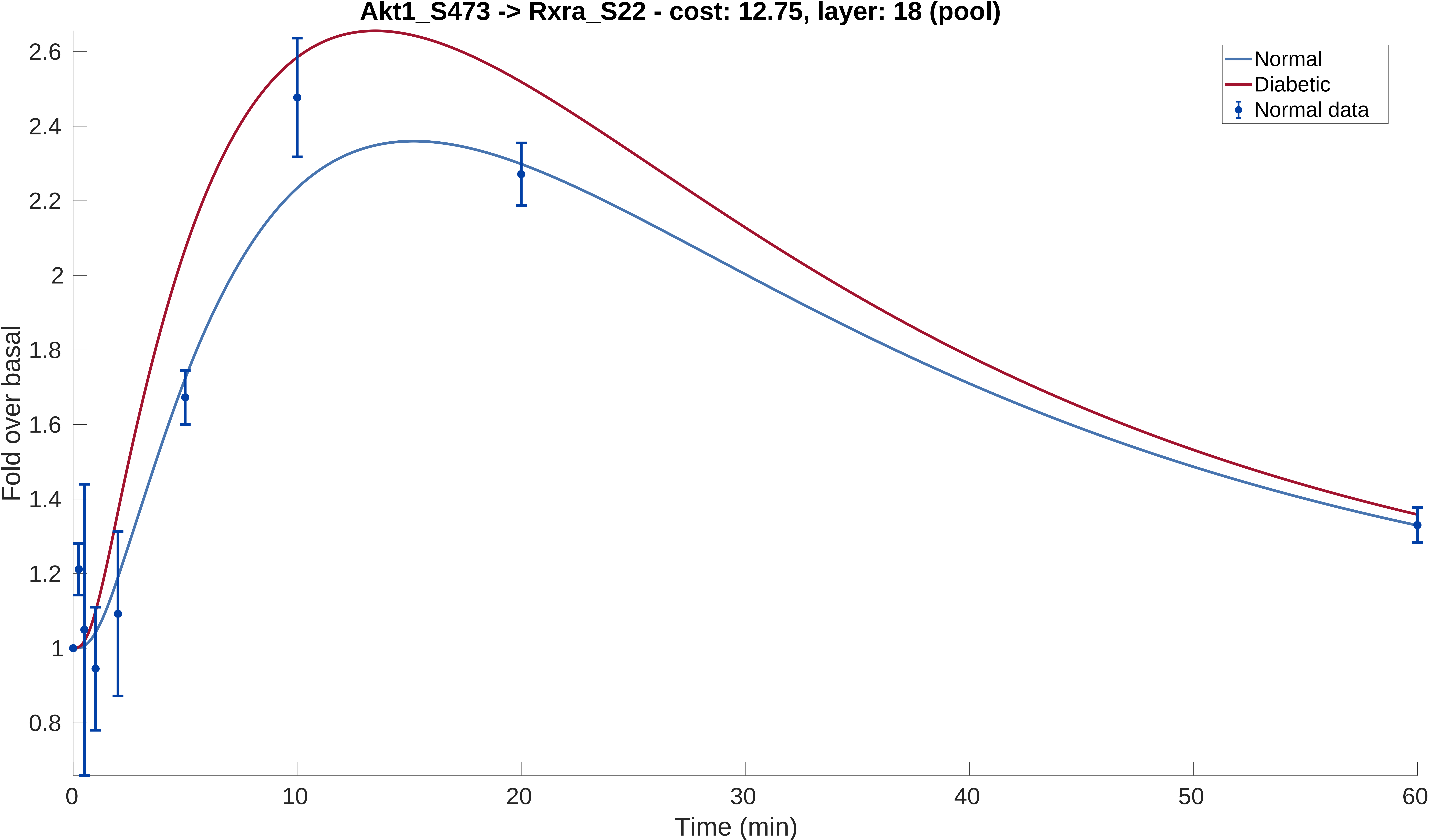

Akt1_S473 -> Rxra_S22 - cost: 12.75, layer: 18 (pool)
Normal
Diabetic
Normal data
Fold over basal
Time (min)
2.6
2.4
2.2
2
1.8
1.6
1.4
1.2
1
0.8
0
10
20
30
40
50
60

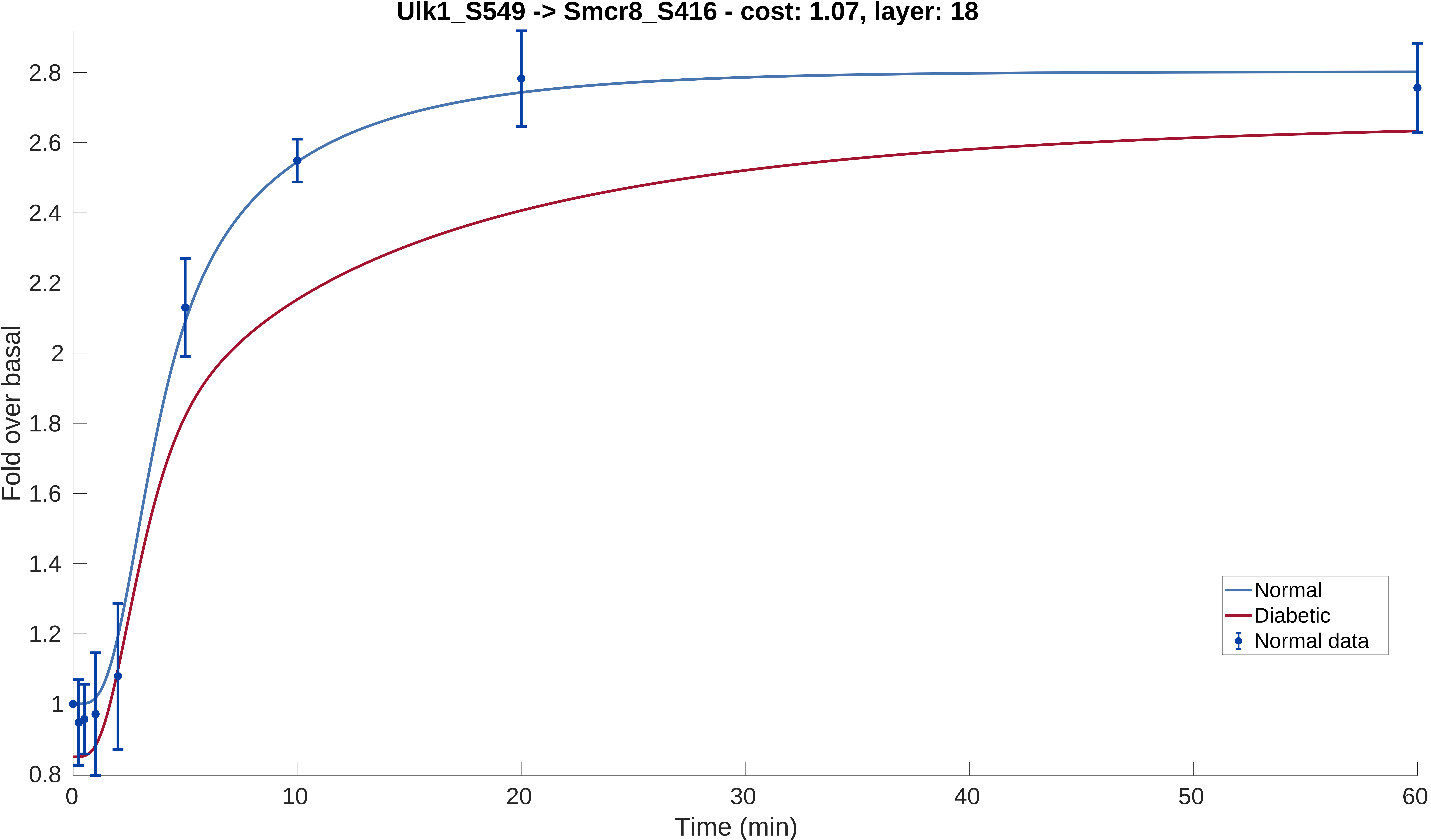

Ulk1_S549 -> Smcr8_S416 - cost: 1.07, layer: 18
Fold over basal
Time (min)
0.8
1
1.2
1.4
1.6
1.8
2
2.2
2.4
2.6
2.8
0
10
20
30
40
50
60
Normal
Diabetic
Normal data

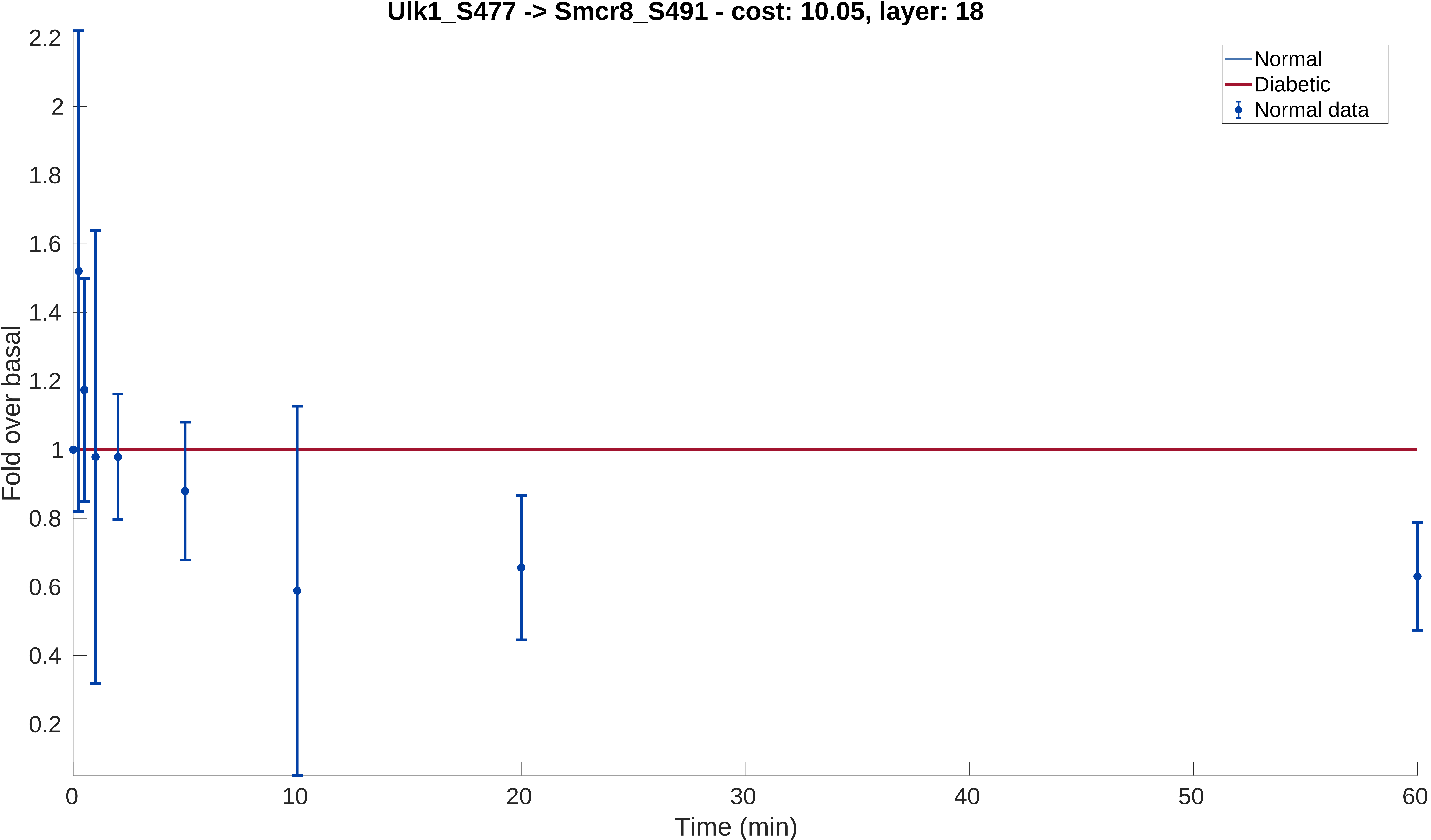
Ulk1_S477 -> Smcr8_S491 - cost: 10.05, layer: 18
Normal
Diabetic
Normal data
Fold over basal
Time (min)
2.2
2
1.8
1.6
1.4
1.2
1
0.8
0.6
0.4
0.2
0
10
20
30
40
50
60

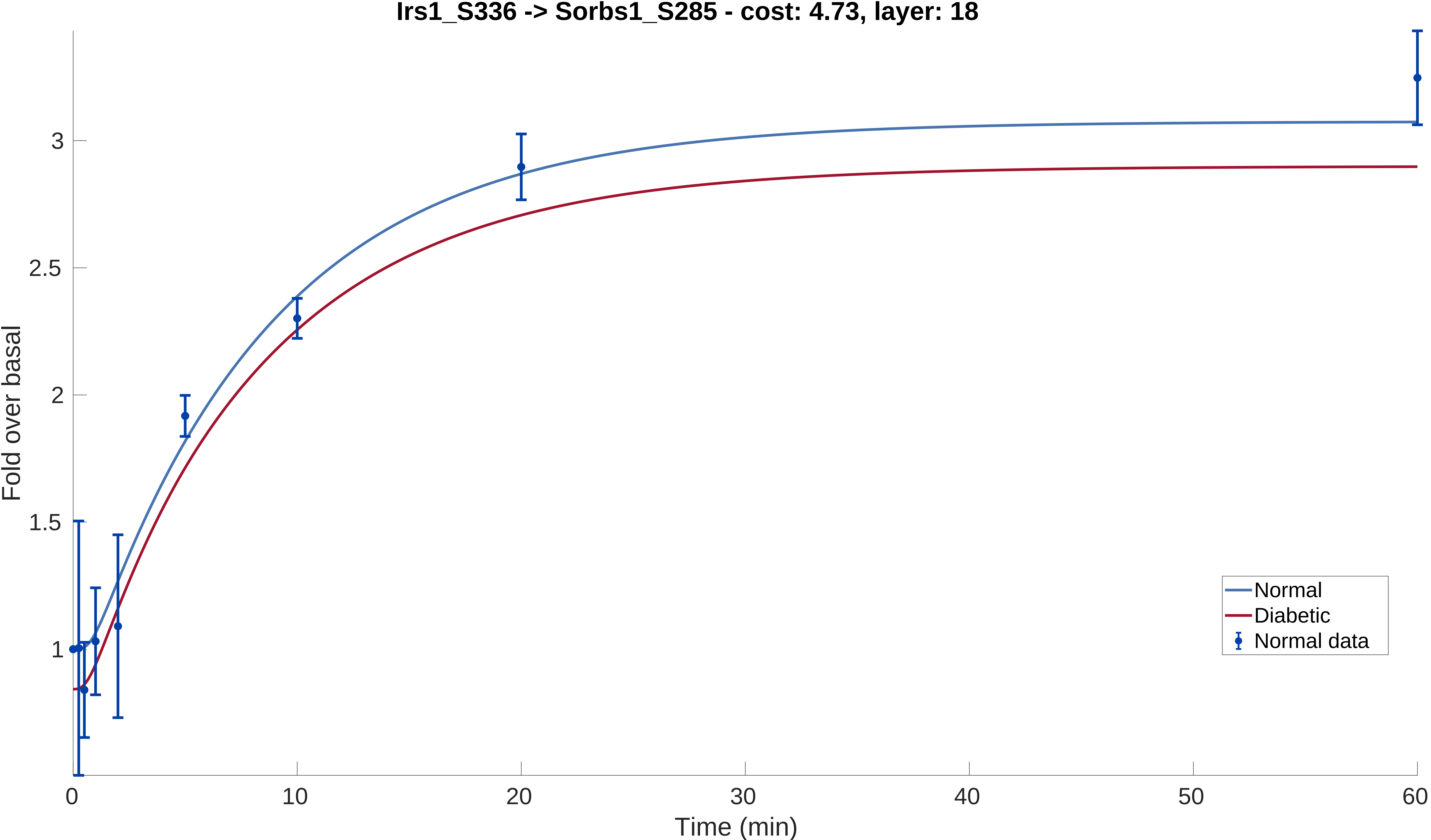

Irs1_S336 -> Sorbs1_S285 - cost: 4.73, layer: 18
Fold over basal
3
2.5
2
1.5
1
Time (min)
0
10
20
30
40
50
60
Normal
Diabetic
Normal data

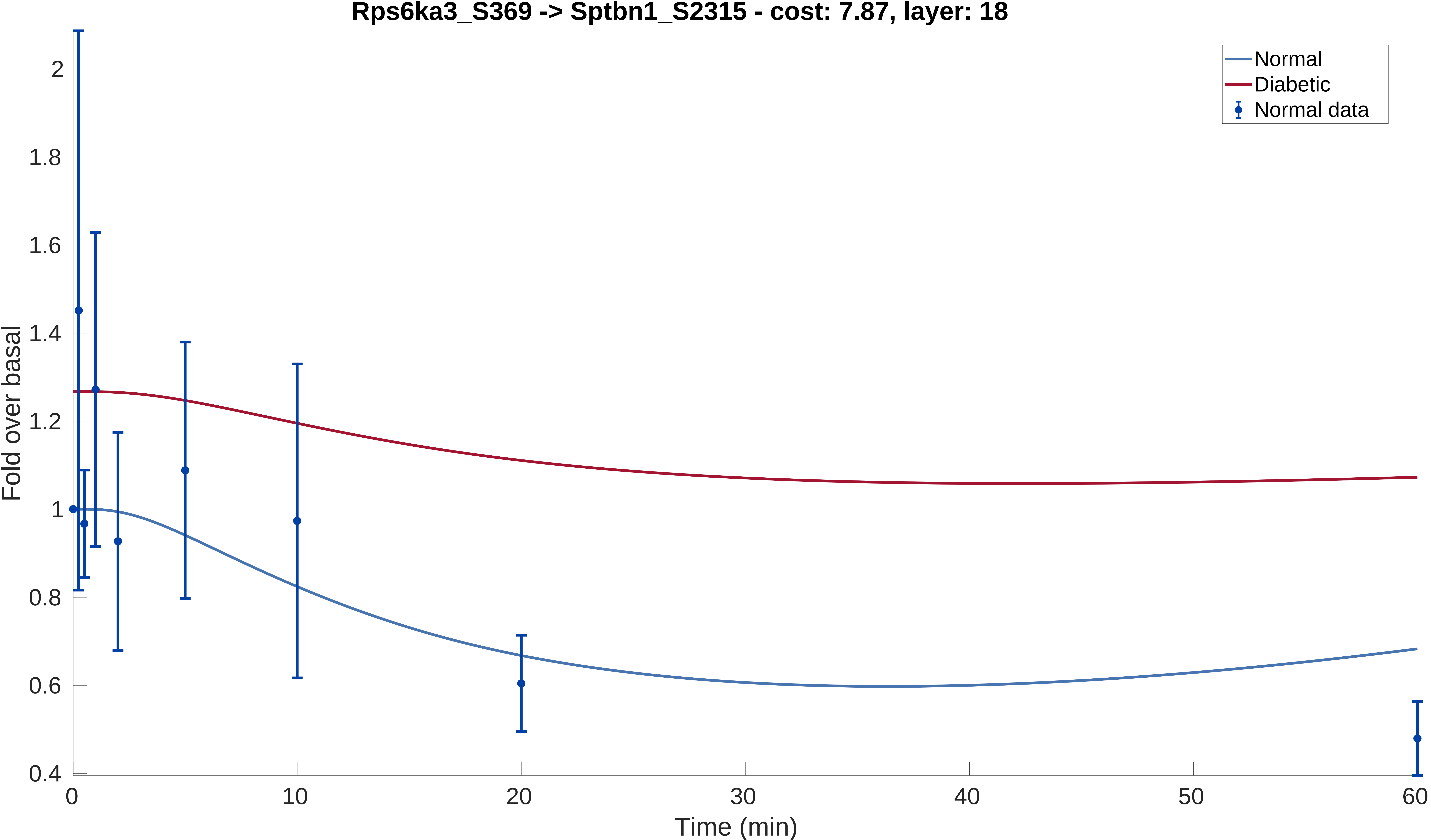

Rps6ka3_S369 -> Sptbn1_S2315 - cost: 7.87, layer: 18
Normal
Diabetic
Normal data
Fold over basal
2
1.8
1.6
1.4
1.2
1
0.8
0.6
0.4
0
10
20
30
40
50
60
Time (min)

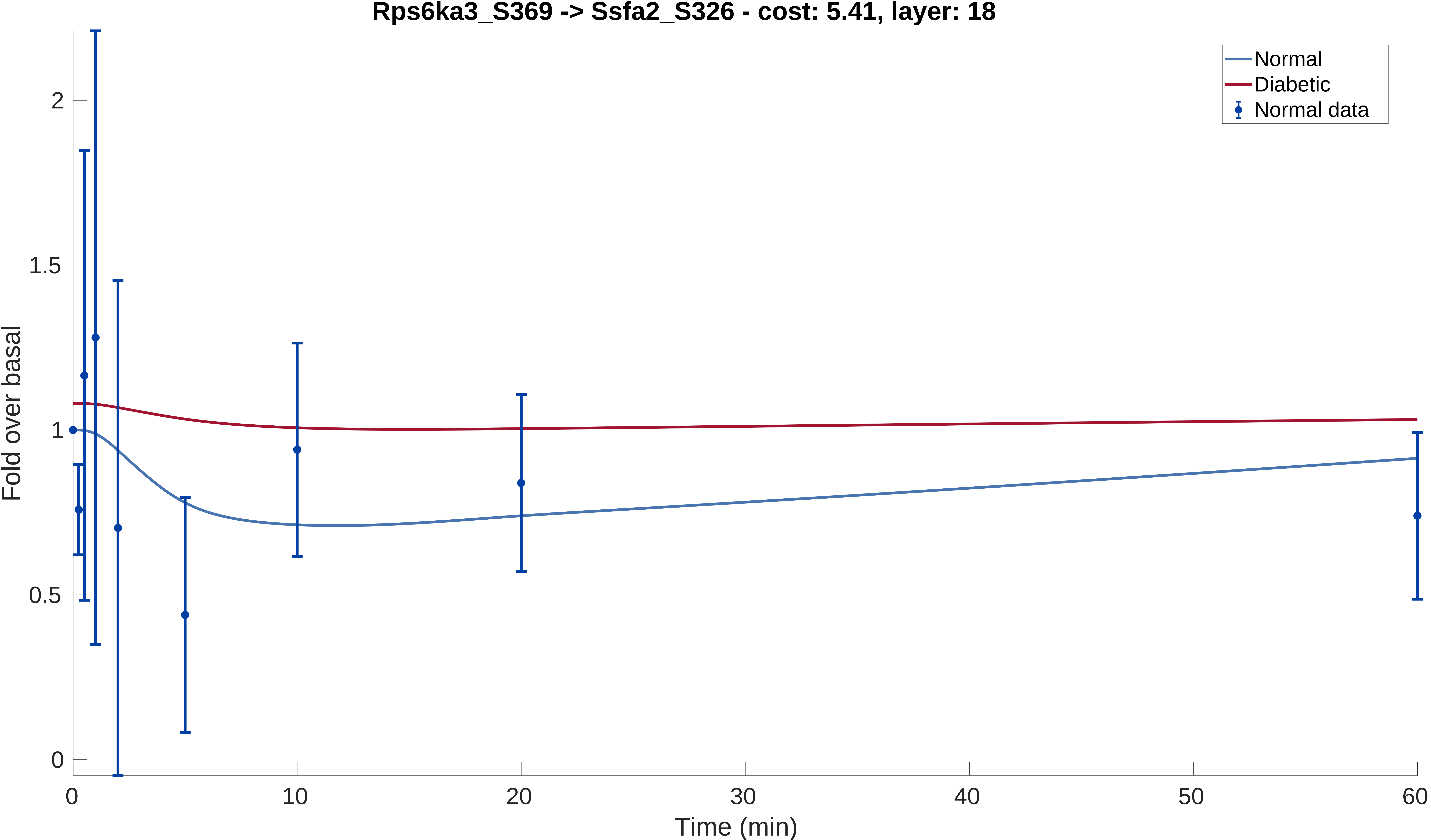
Rps6ka3_S369 -> Ssfa2_S326 - cost: 5.41, layer: 18
Normal
Diabetic
Normal data
Fold over basal
0
0.5
1
1.5
2
0
10
20
30
40
50
60
Time (min)

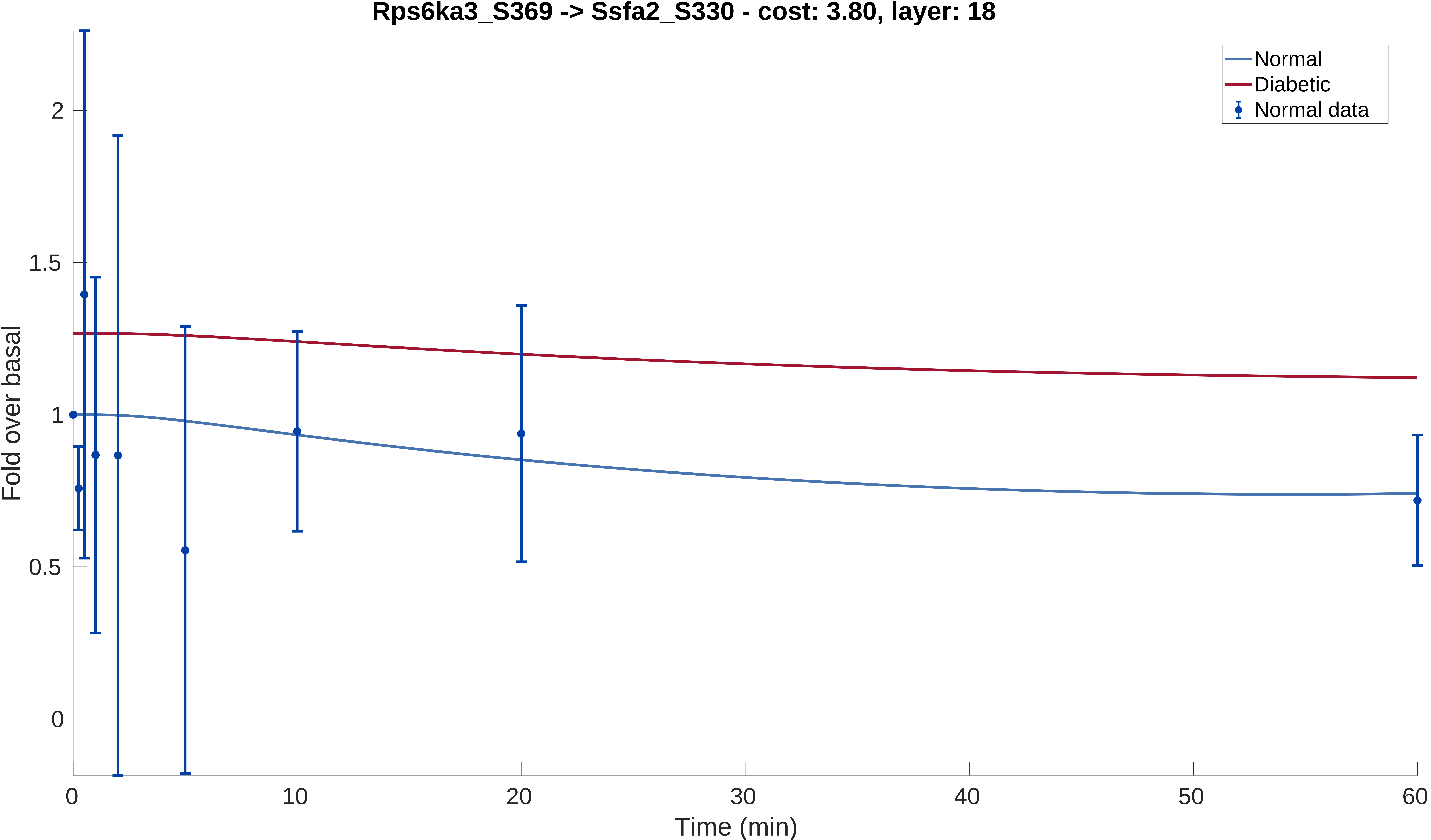

Rps6ka3_S369 -> Ssfa2_S330 - cost: 3.80, layer: 18
Normal
Diabetic
Normal data
Fold over basal
Time (min)
0
0.5
1
1.5
2
0
10
20
30
40
50
60

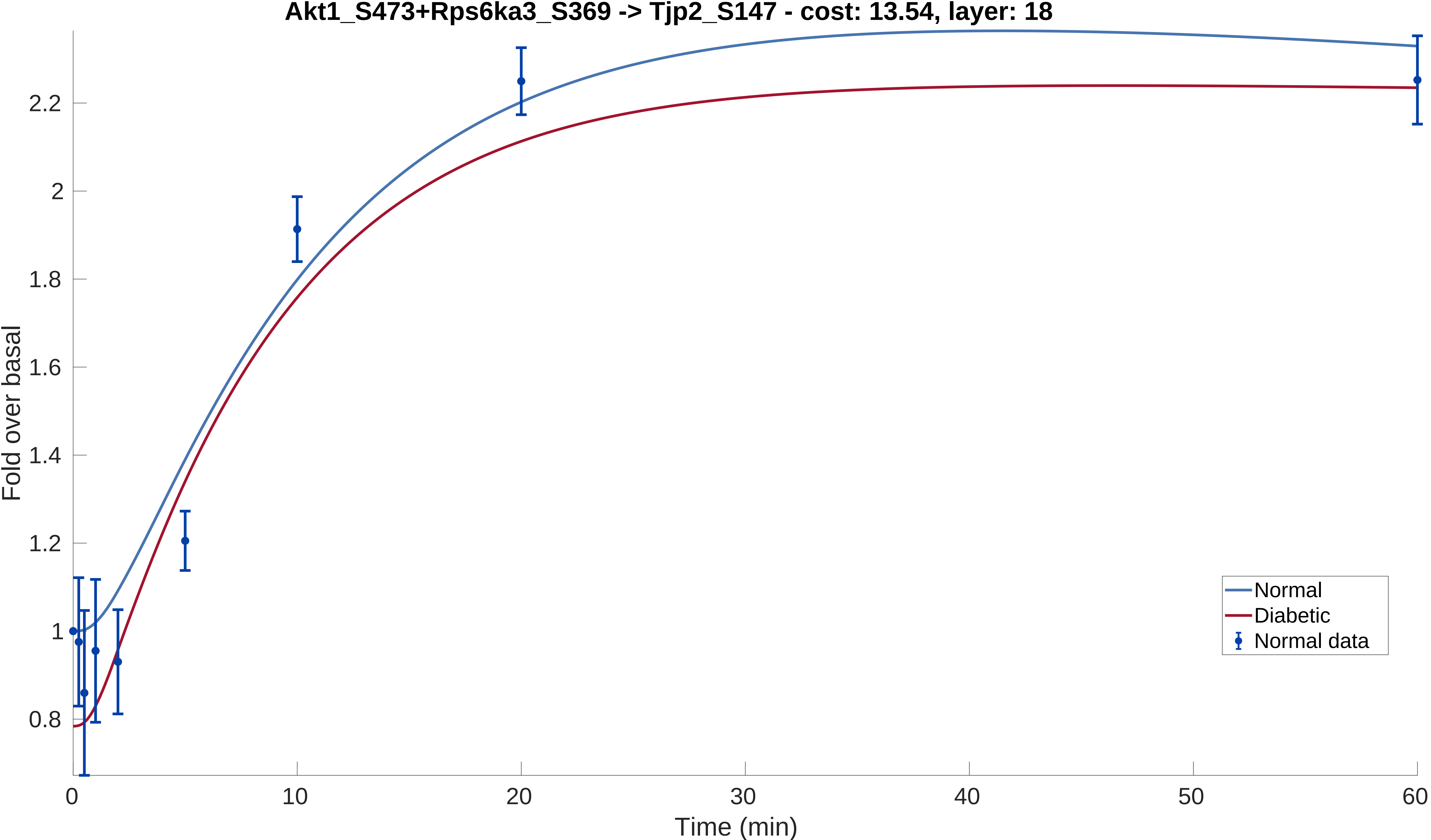
Akt1_S473+Rps6ka3_S369 -> Tjp2_S147 - cost: 13.54, layer: 18
Fold over basal
Time (min)
Normal
Diabetic
Normal data
0.8
1
1.2
1.4
1.6
1.8
2
2.2
0
10
20
30
40
50
60

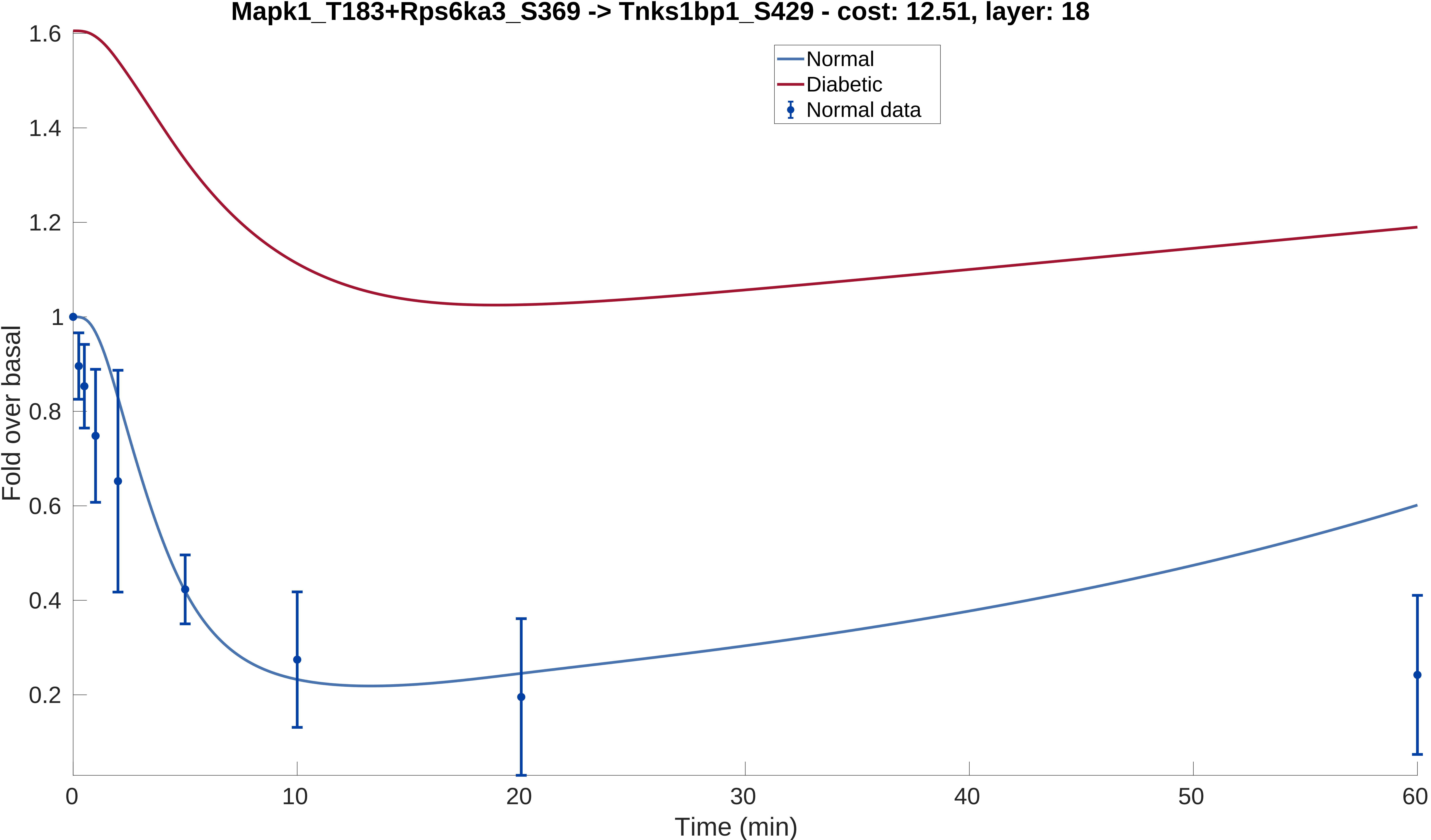

Mapk1_T183+Rps6ka3_S369 -> Tnks1bp1_S429 - cost: 12.51, layer: 18
Normal
Diabetic
Normal data
Fold over basal
Time (min)
1.6
1.4
1.2
1
0.8
0.6
0.4
0.2
0
10
20
30
40
50
60

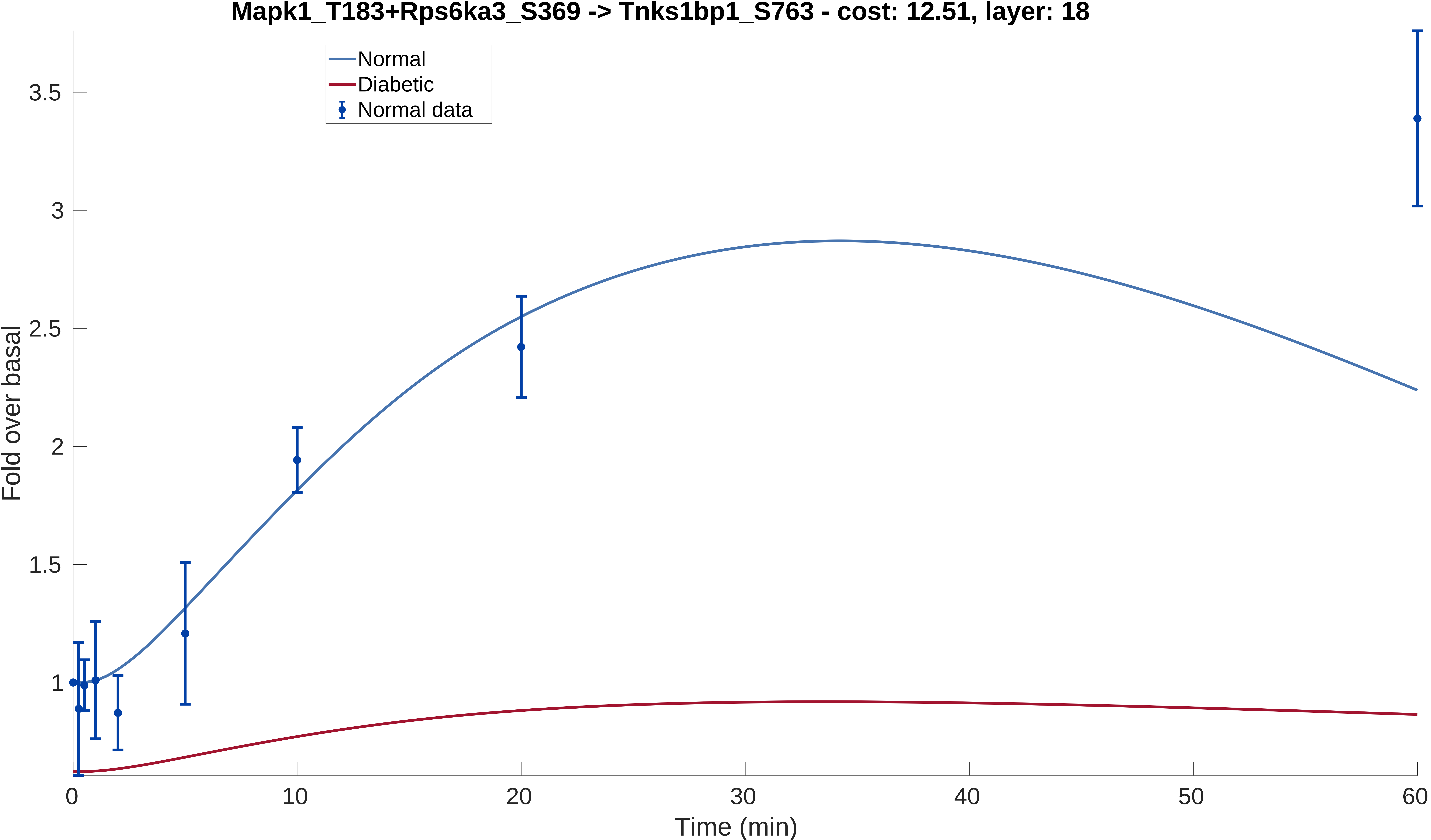
Mapk1_T183+Rps6ka3_S369 -> Tnks1bp1_S763 - cost: 12.51, layer: 18
Normal
Diabetic
Normal data
Fold over basal
3.5
3
2.5
2
1.5
1
0
10
20
30
40
50
60
Time (min)

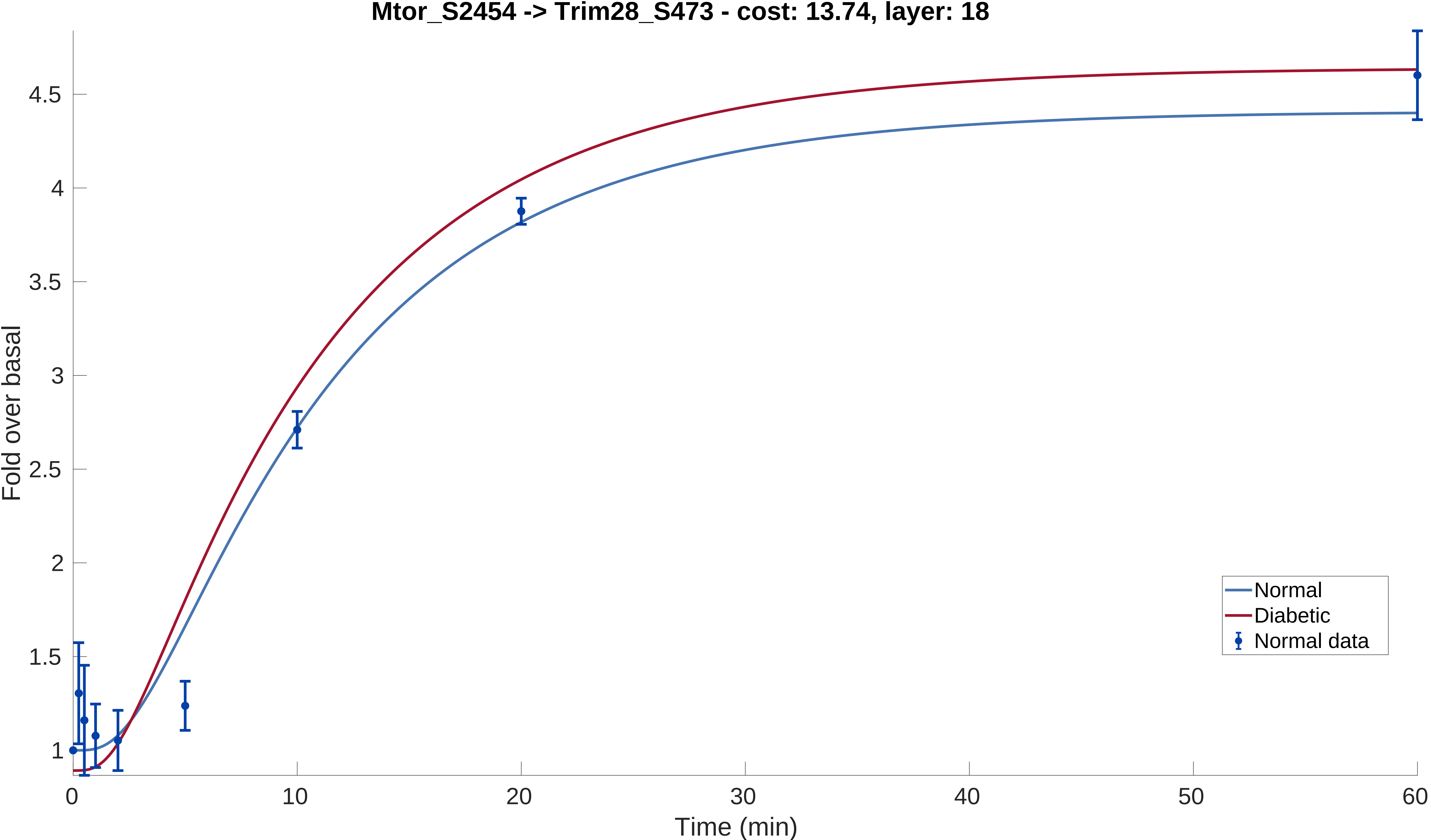
Mtor_S2454 -> Trim28_S473 - cost: 13.74, layer: 18
Fold over basal
Time (min)
Normal
Diabetic
Normal data
4.5
4
3.5
3
2.5
2
1.5
1
0
10
20
30
40
50
60

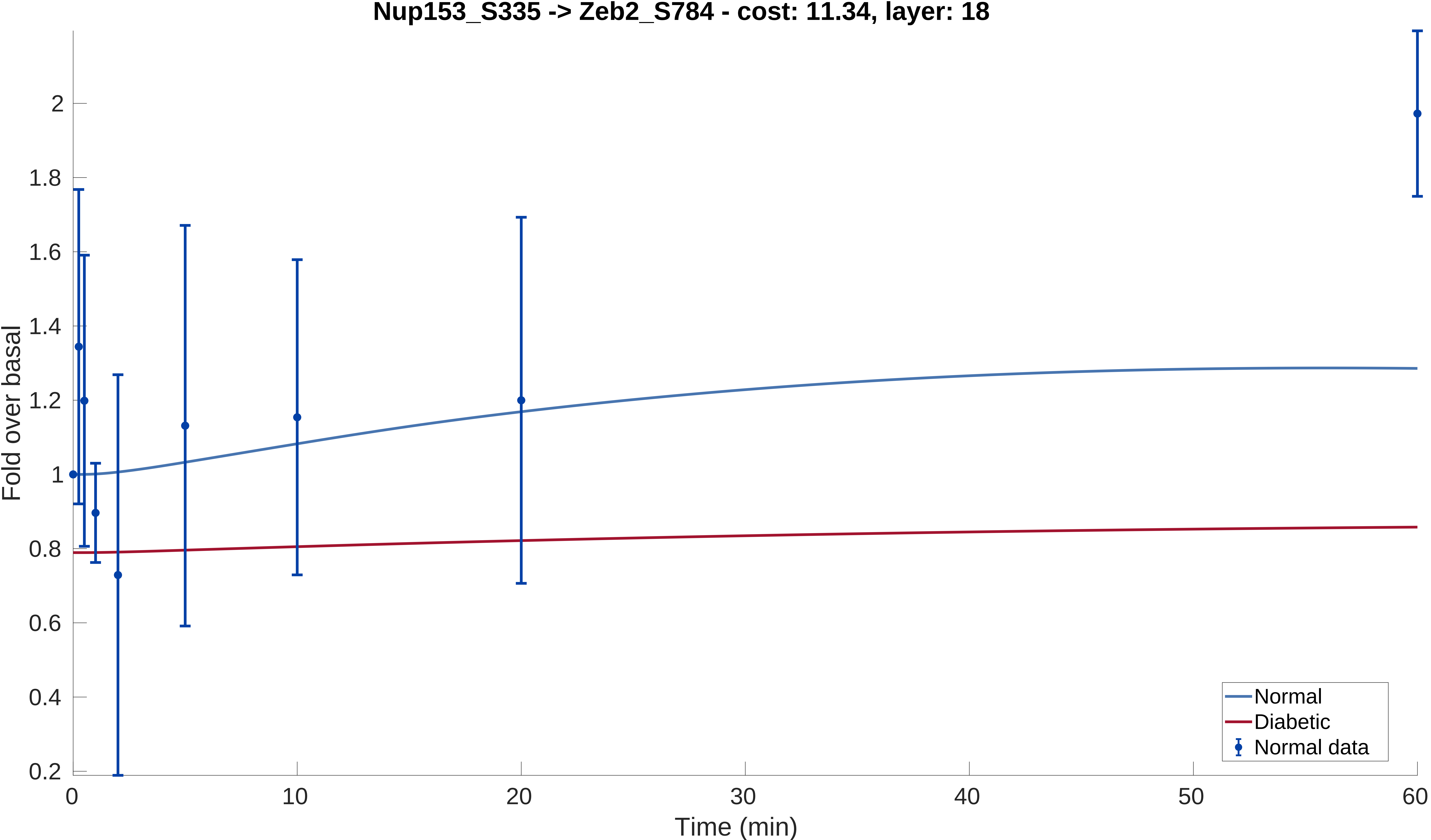
Nup153_S335 -> Zeb2_S784 - cost: 11.34, layer: 18
Fold over basal
Time (min)
0.2
0.4
0.6
0.8
1
1.2
1.4
1.6
1.8
2
0
10
20
30
40
50
60
Normal
Diabetic
Normal data

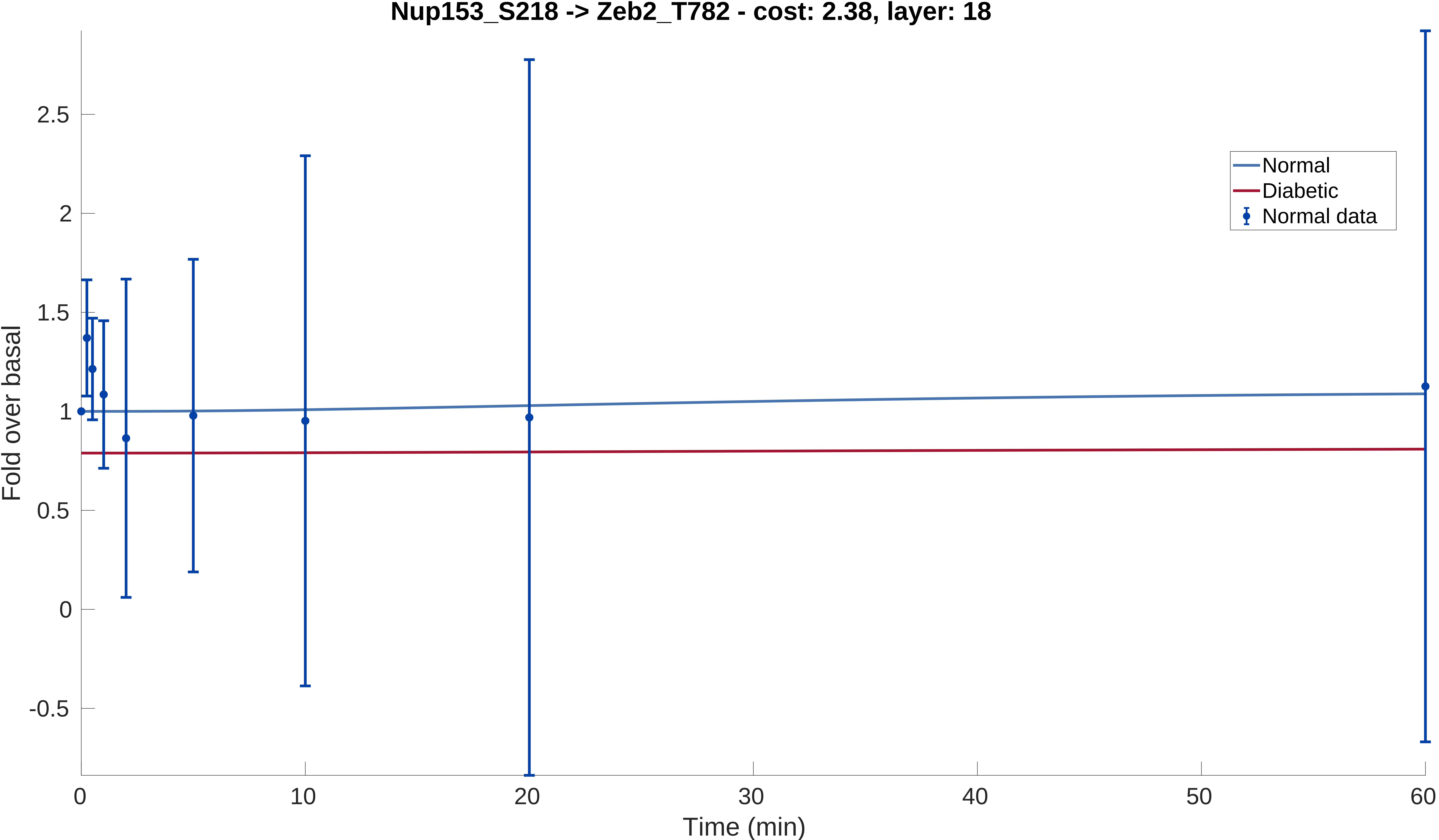
Nup153_S218 -> Zeb2_T782 - cost: 2.38, layer: 18
Fold over basal
Time (min)
Normal
Diabetic
Normal data
2.5
2
1.5
1
0.5
0
-0.5
0
10
20
30
40
50
60

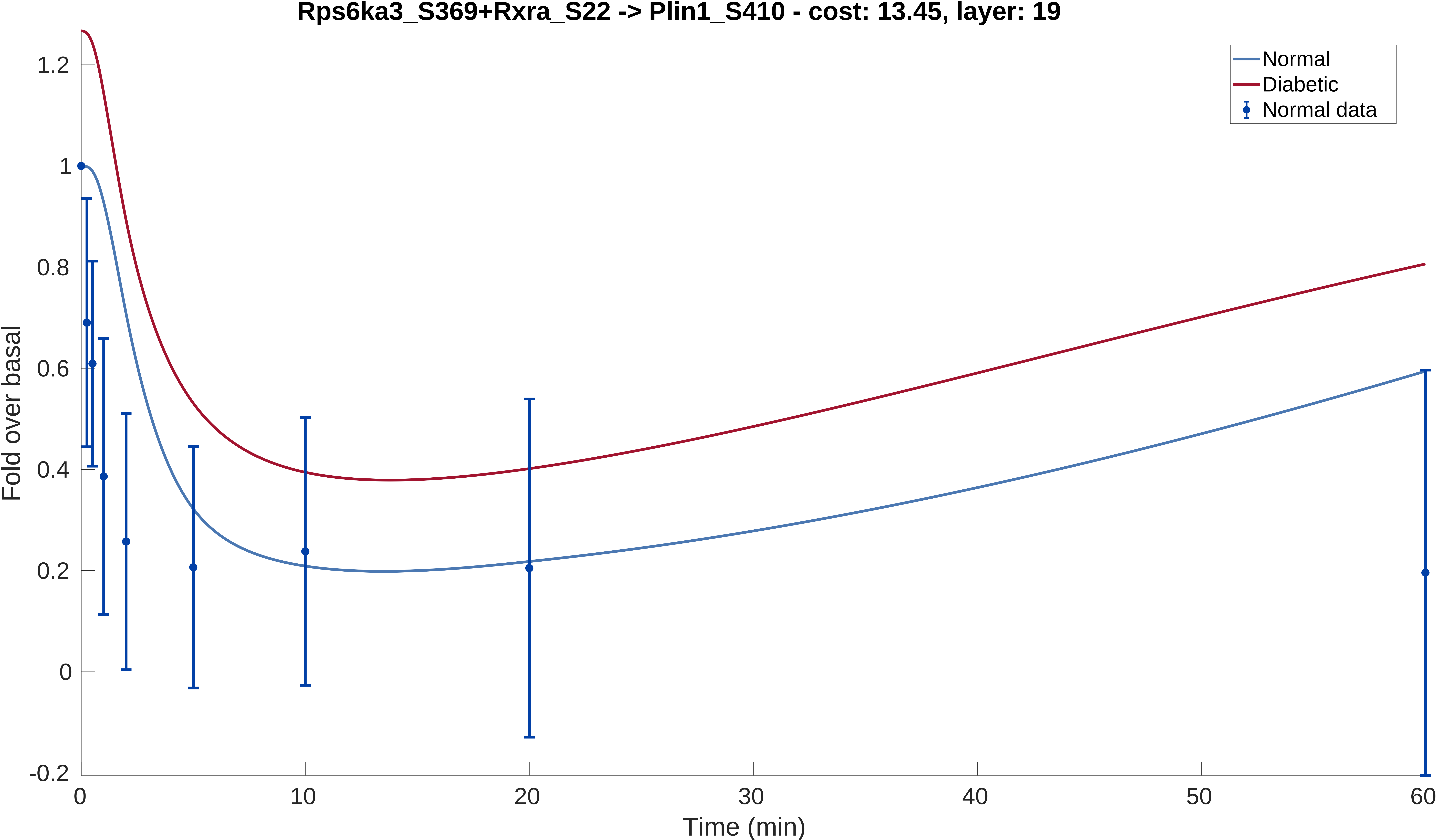

Rps6ka3_S369+Rxra_S22 -> Plin1_S410 - cost: 13.45, layer: 19
Normal
Diabetic
Normal data
Fold over basal
Time (min)
1.2
1
0.8
0.6
0.4
0.2
0
-0.2
0
10
20
30
40
50
60

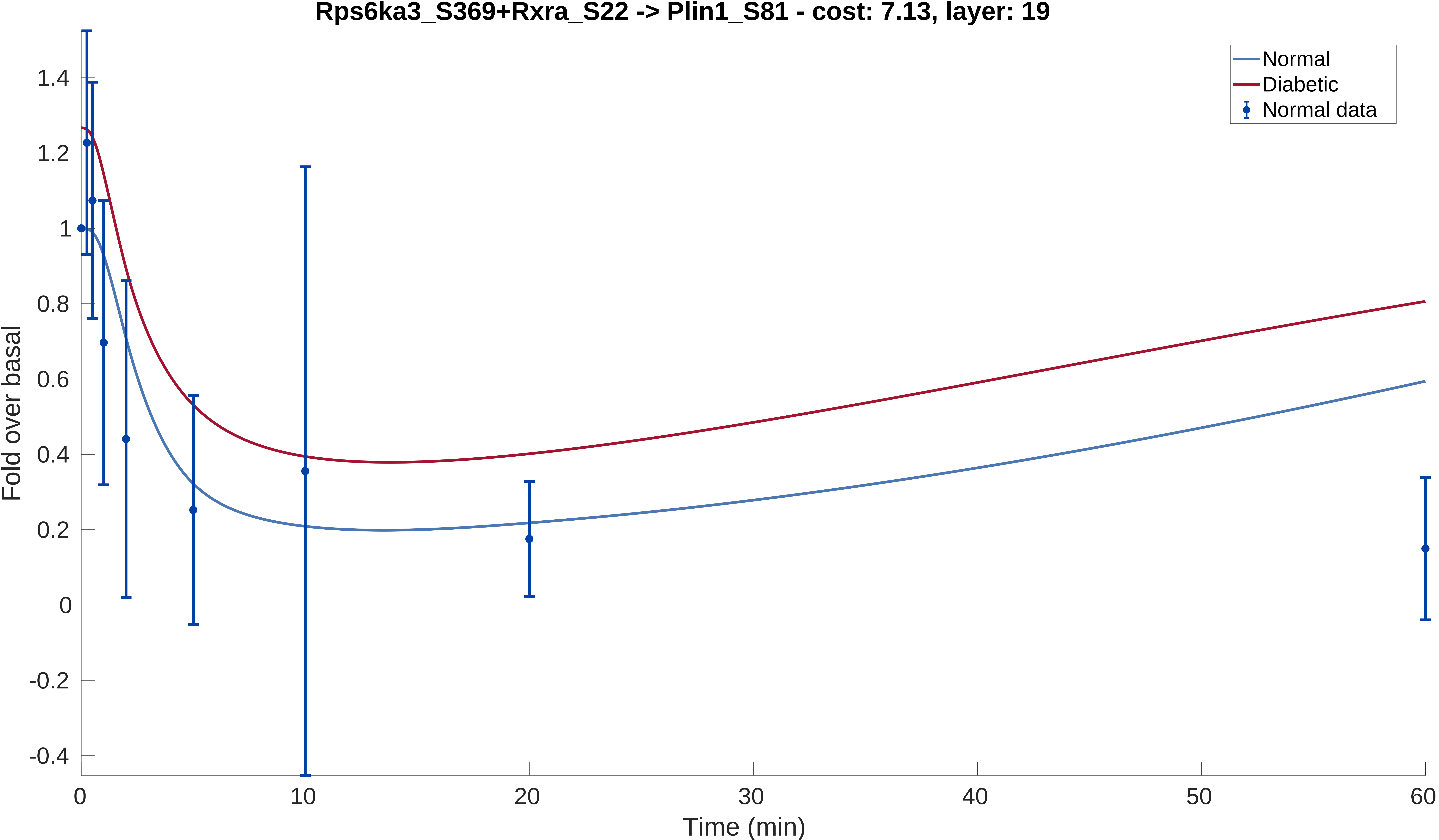

Rps6ka3_S369+Rxra_S22 -> Plin1_S81 - cost: 7.13, layer: 19
Normal
Diabetic
Normal data
Fold over basal
Time (min)
1.4
1.2
1
0.8
0.6
0.4
0.2
0
-0.2
-0.4
0
10
20
30
40
50
60

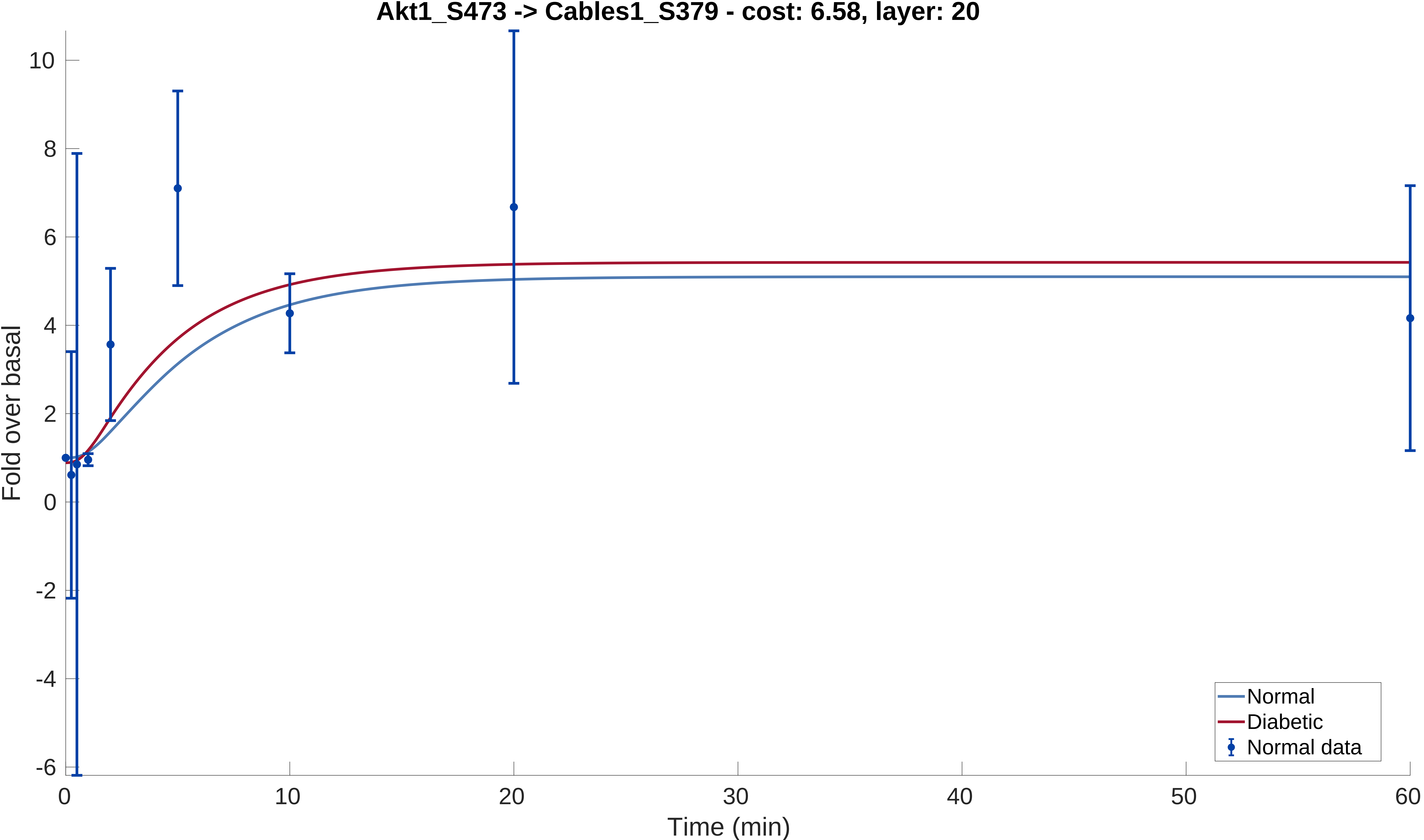

Akt1_S473 -> Cables1_S379 - cost: 6.58, layer: 20
Fold over basal
Time (min)
10
8
6
4
2
0
-2
-4
-6
0
10
20
30
40
50
60
Normal
Diabetic
Normal data

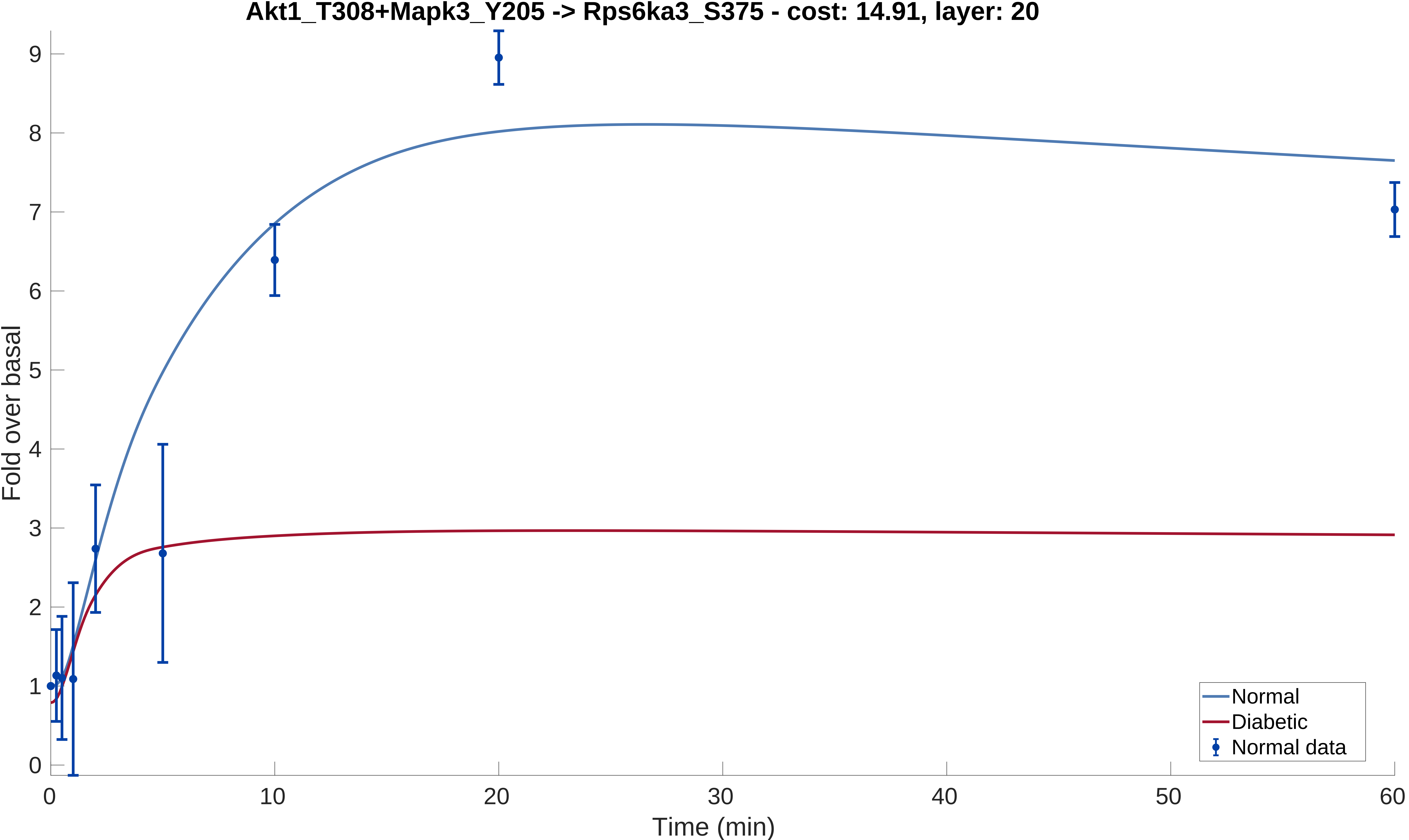

Akt1_T308+Mapk3_Y205 -> Rps6ka3_S375 - cost: 14.91, layer: 20
Fold over basal
Time (min)
0
1
2
3
4
5
6
7
8
9
0
10
20
30
40
50
60
Normal
Diabetic
Normal data

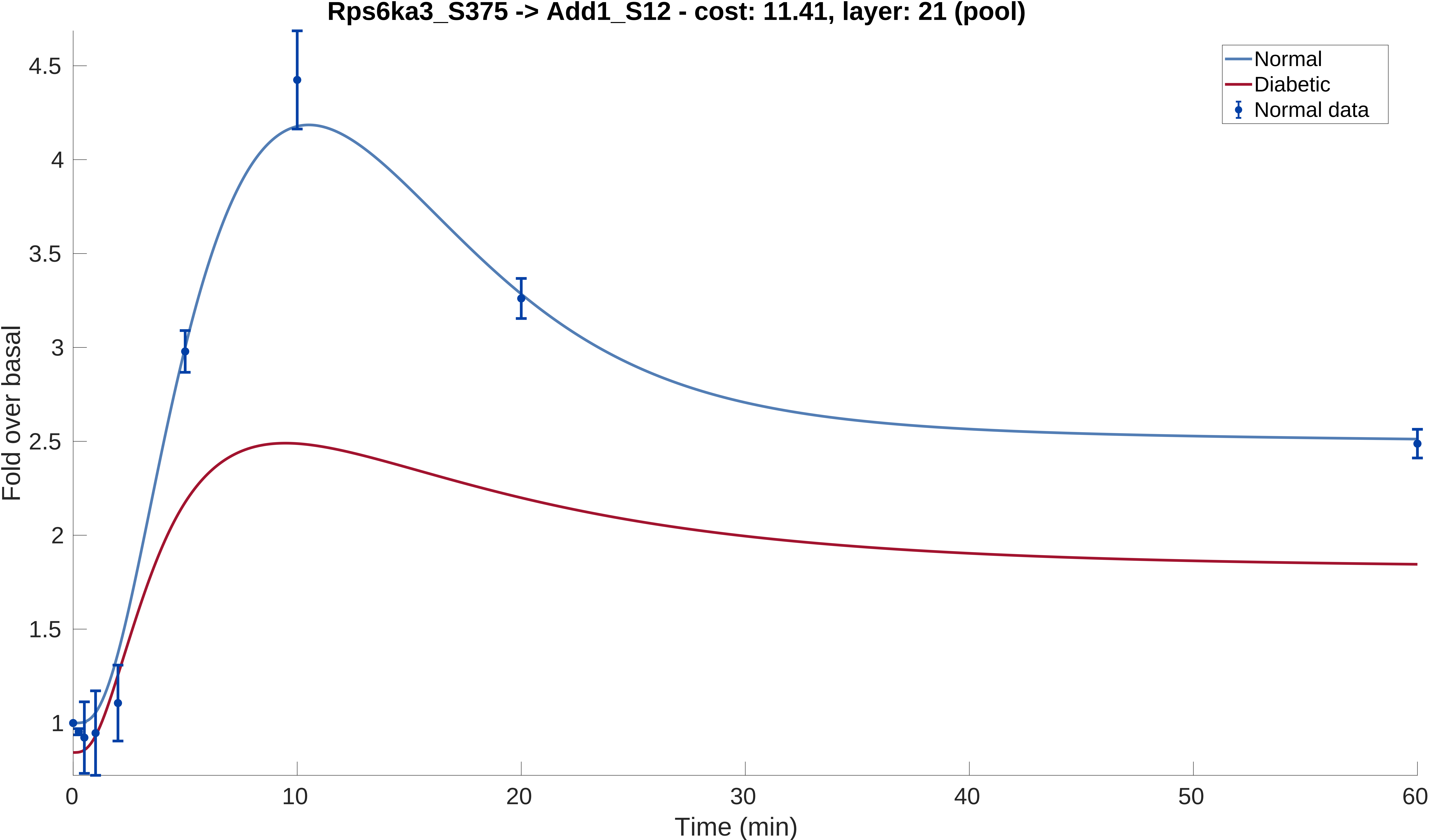

Rps6ka3_S375 -> Add1_S12 - cost: 11.41, layer: 21 (pool)
Normal
Diabetic
Normal data
Fold over basal
4.5
4
3.5
3
2.5
2
1.5
1
0
10
20
30
40
50
60
Time (min)

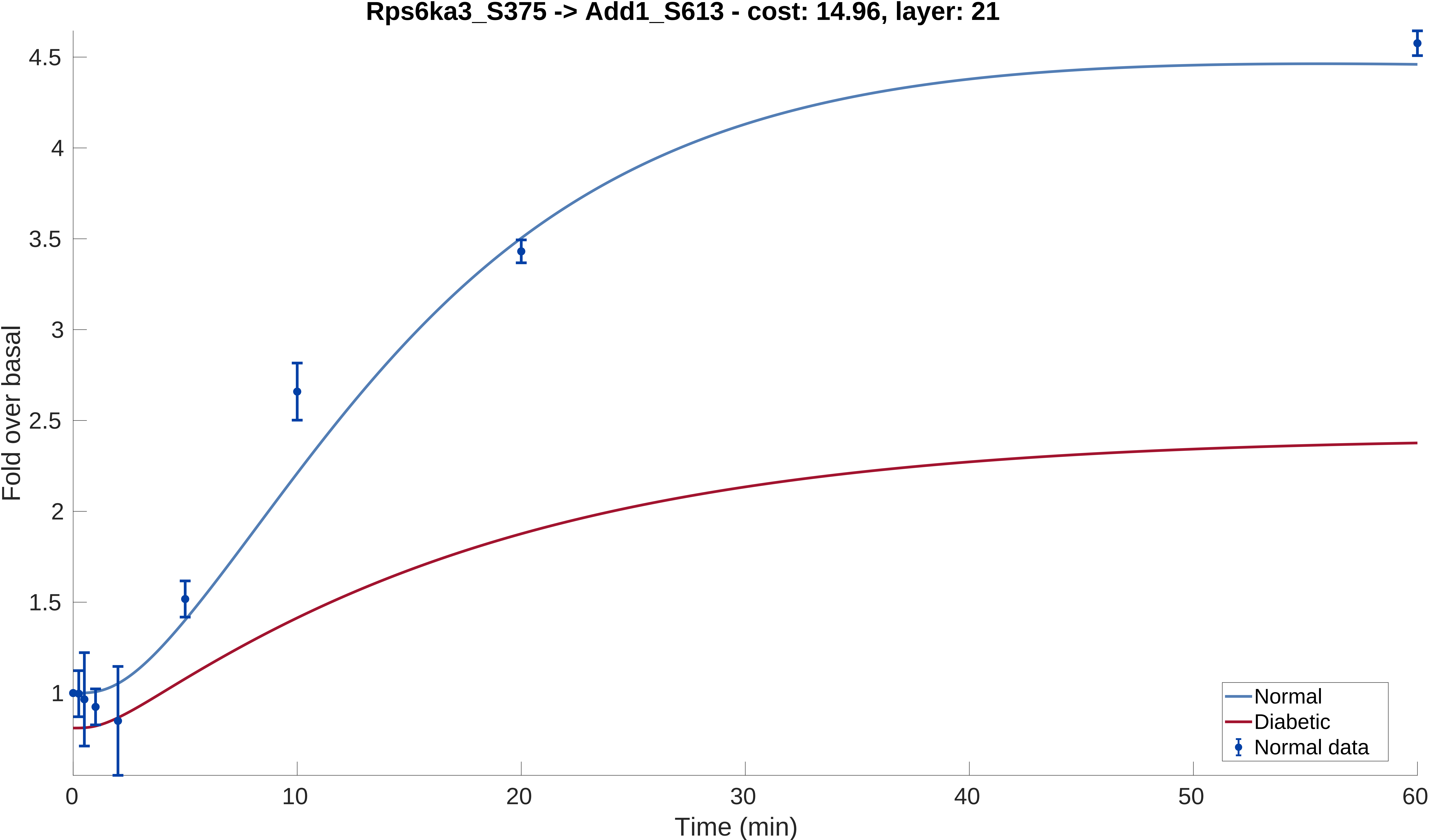

Rps6ka3_S375 -> Add1_S613 - cost: 14.96, layer: 21
Fold over basal
Time (min)
4.5
4
3.5
3
2.5
2
1.5
1
0
10
20
30
40
50
60
Normal
Diabetic
Normal data

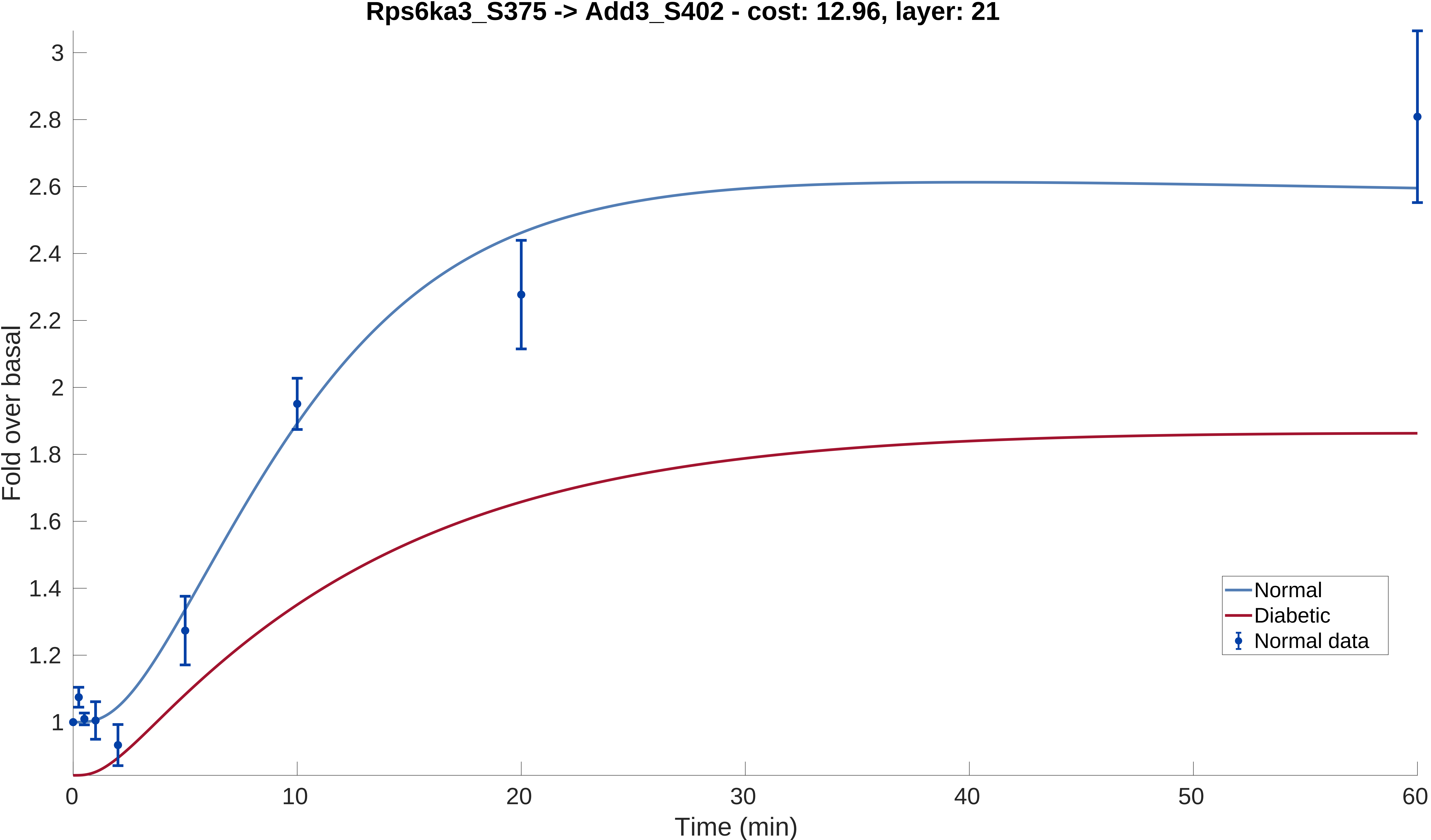
Rps6ka3_S375 -> Add3_S402 - cost: 12.96, layer: 21
Fold over basal
Time (min)
Normal
Diabetic
Normal data
3
2.8
2.6
2.4
2.2
2
1.8
1.6
1.4
1.2
1
0
10
20
30
40
50
60

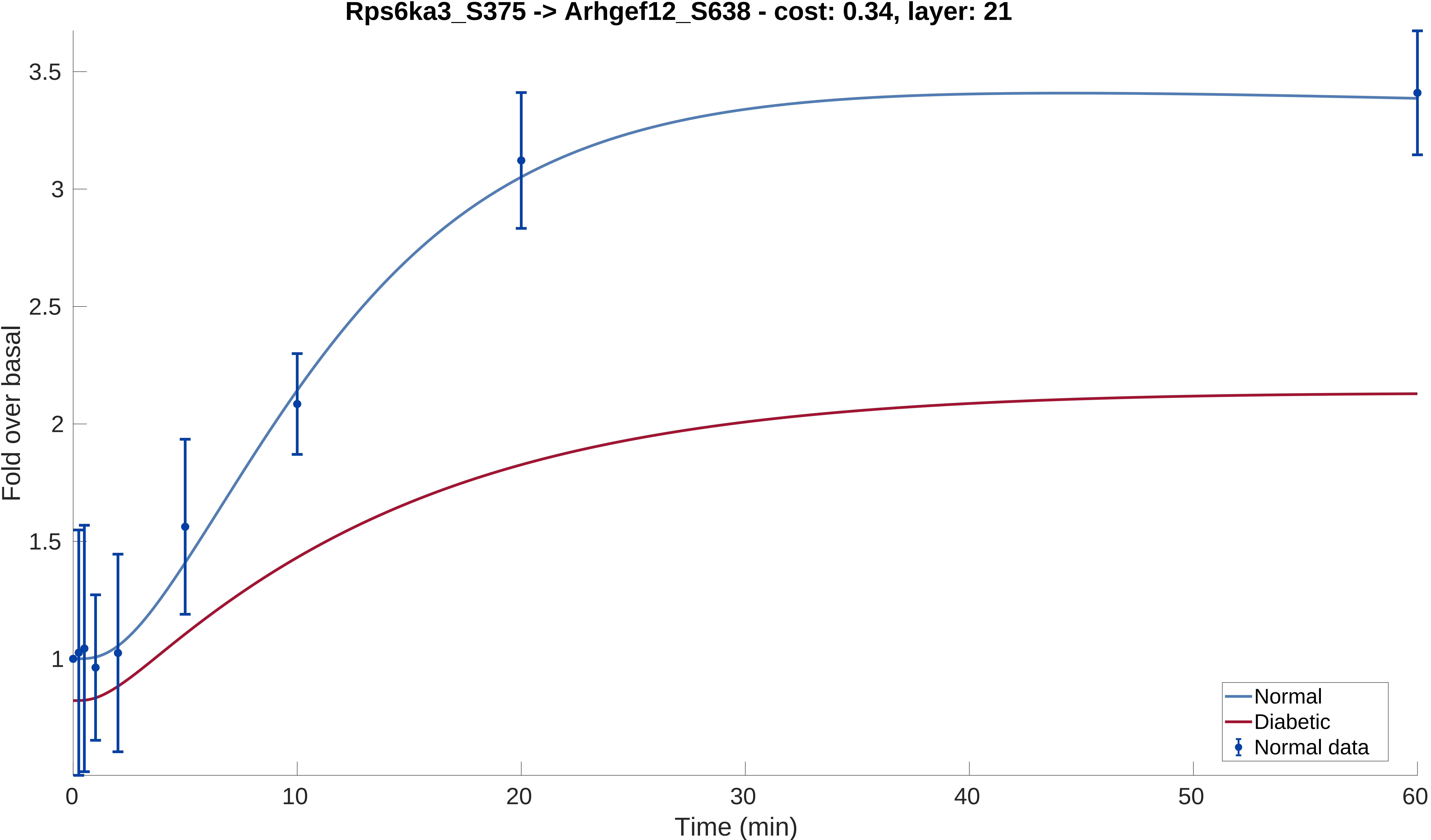

Rps6ka3_S375 -> Arhgef12_S638 - cost: 0.34, layer: 21
Fold over basal
3.5
3
2.5
2
1.5
1
0
10
20
30
40
50
60
Time (min)
Normal
Diabetic
Normal data

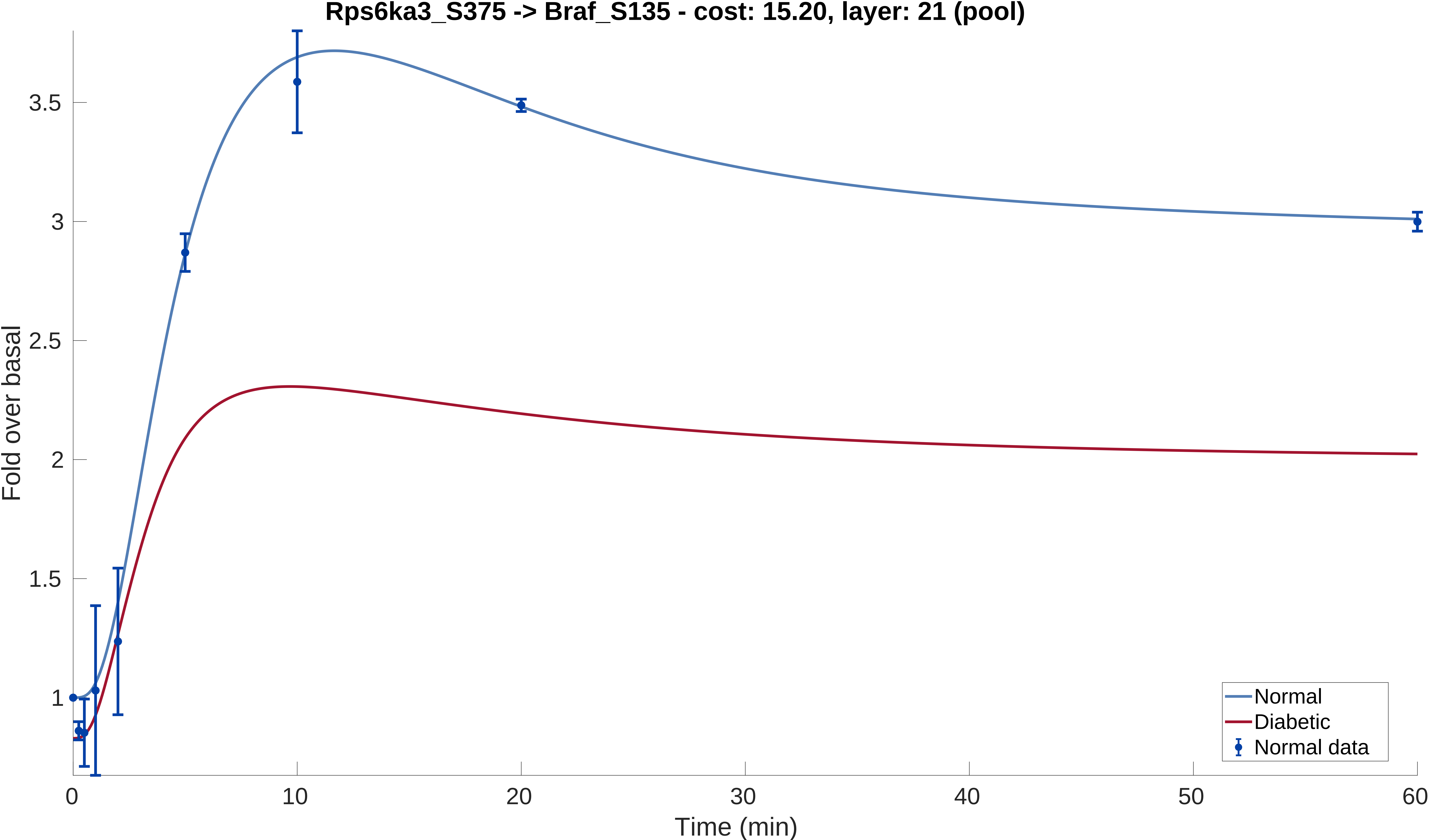

Rps6ka3_S375 -> Braf_S135 - cost: 15.20, layer: 21 (pool)
Fold over basal
3.5
3
2.5
2
1.5
1
0
10
20
30
40
50
60
Time (min)
Normal
Diabetic
Normal data

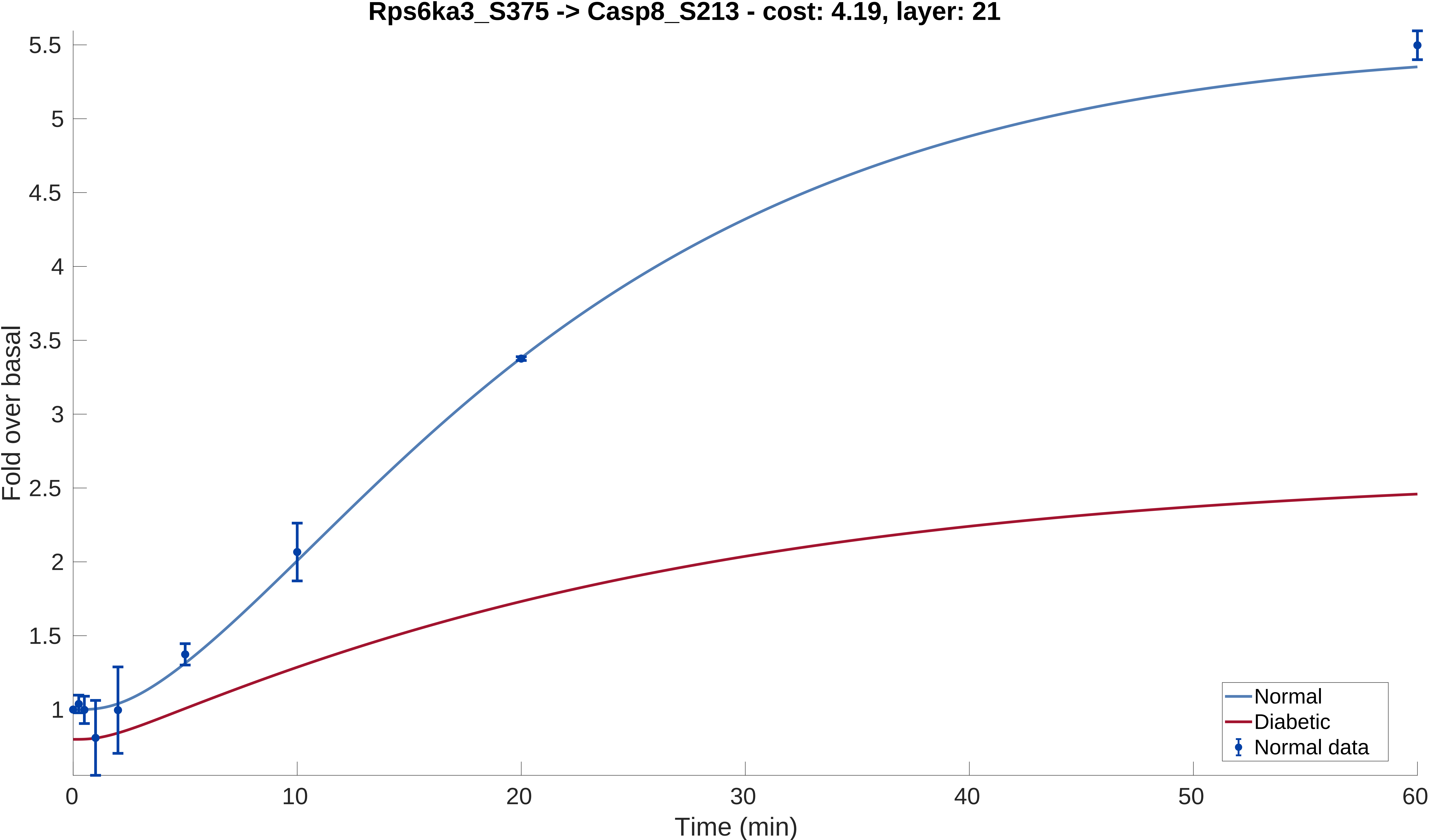

Rps6ka3_S375 -> Casp8_S213 - cost: 4.19, layer: 21
Fold over basal
5.5
5
4.5
4
3.5
3
2.5
2
1.5
1
0
10
20
30
40
50
60
Time (min)
Normal
Diabetic
Normal data

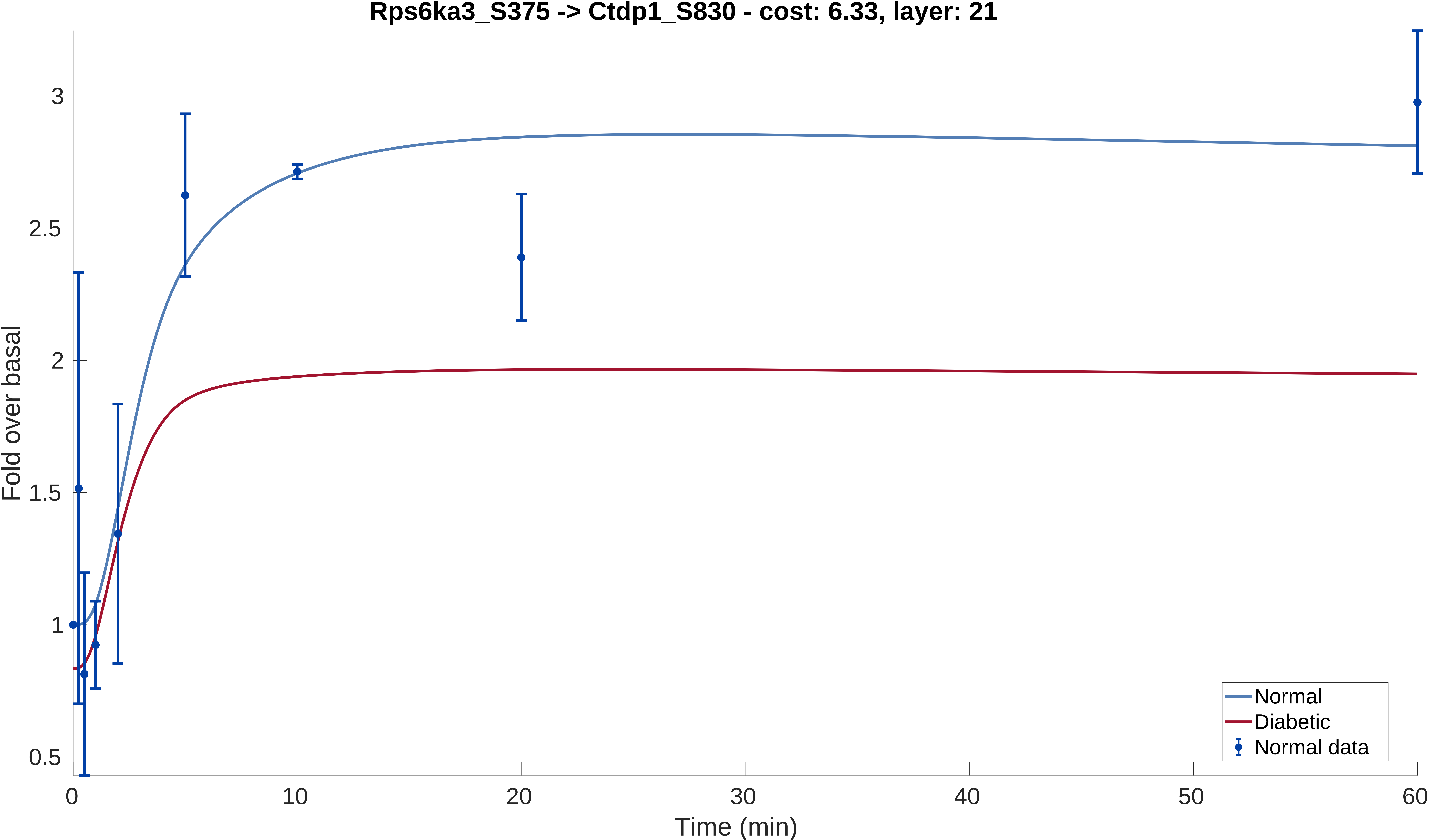
Rps6ka3_S375 -> Ctdp1_S830 - cost: 6.33, layer: 21
Fold over basal
Time (min)
0.5
1
1.5
2
2.5
3
0
10
20
30
40
50
60
Normal
Diabetic
Normal data

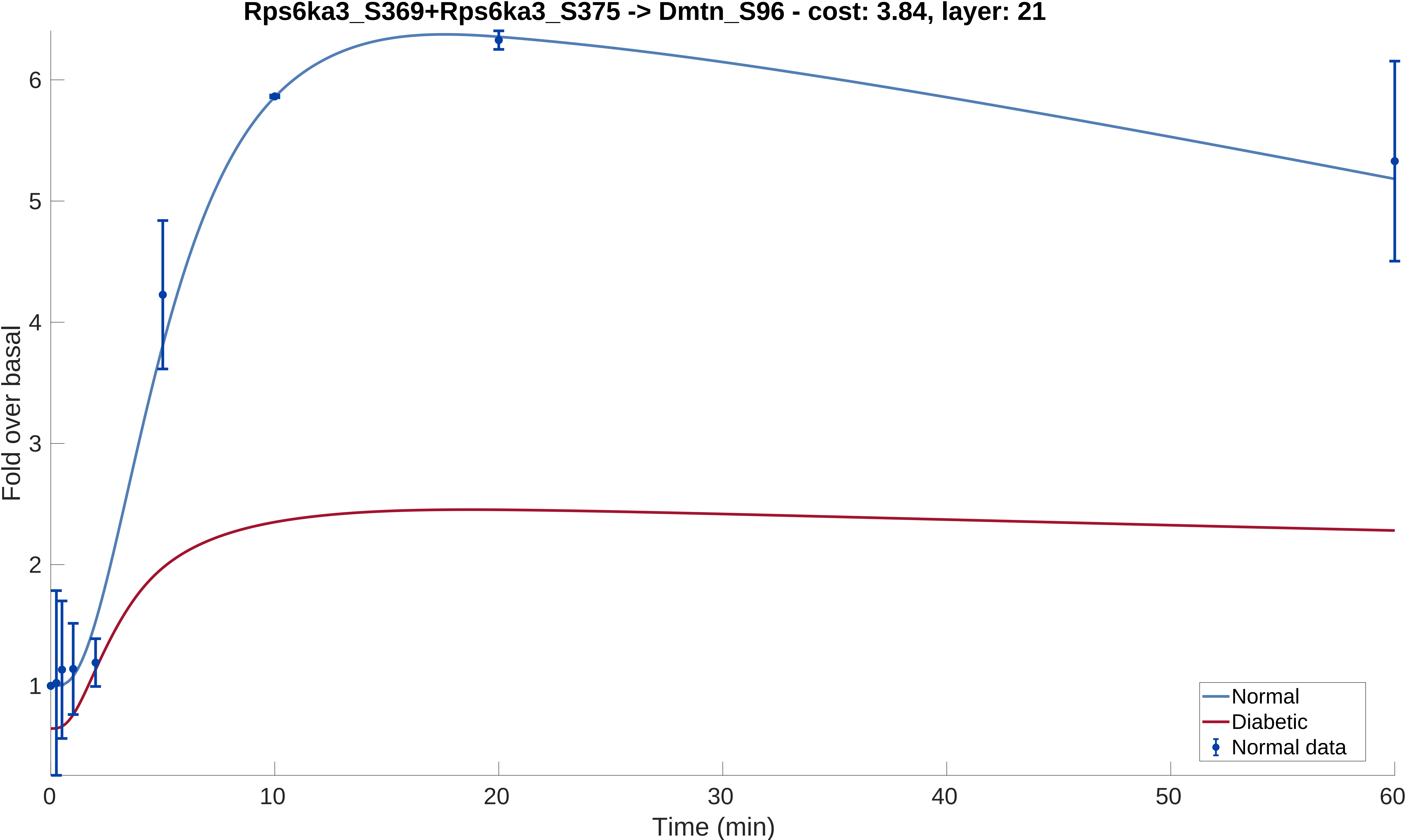
Rps6ka3_S369+Rps6ka3_S375 -> Dmtn_S96 - cost: 3.84, layer: 21
Fold over basal
Time (min)
0
10
20
30
40
50
60
1
2
3
4
5
6
Normal
Diabetic
Normal data

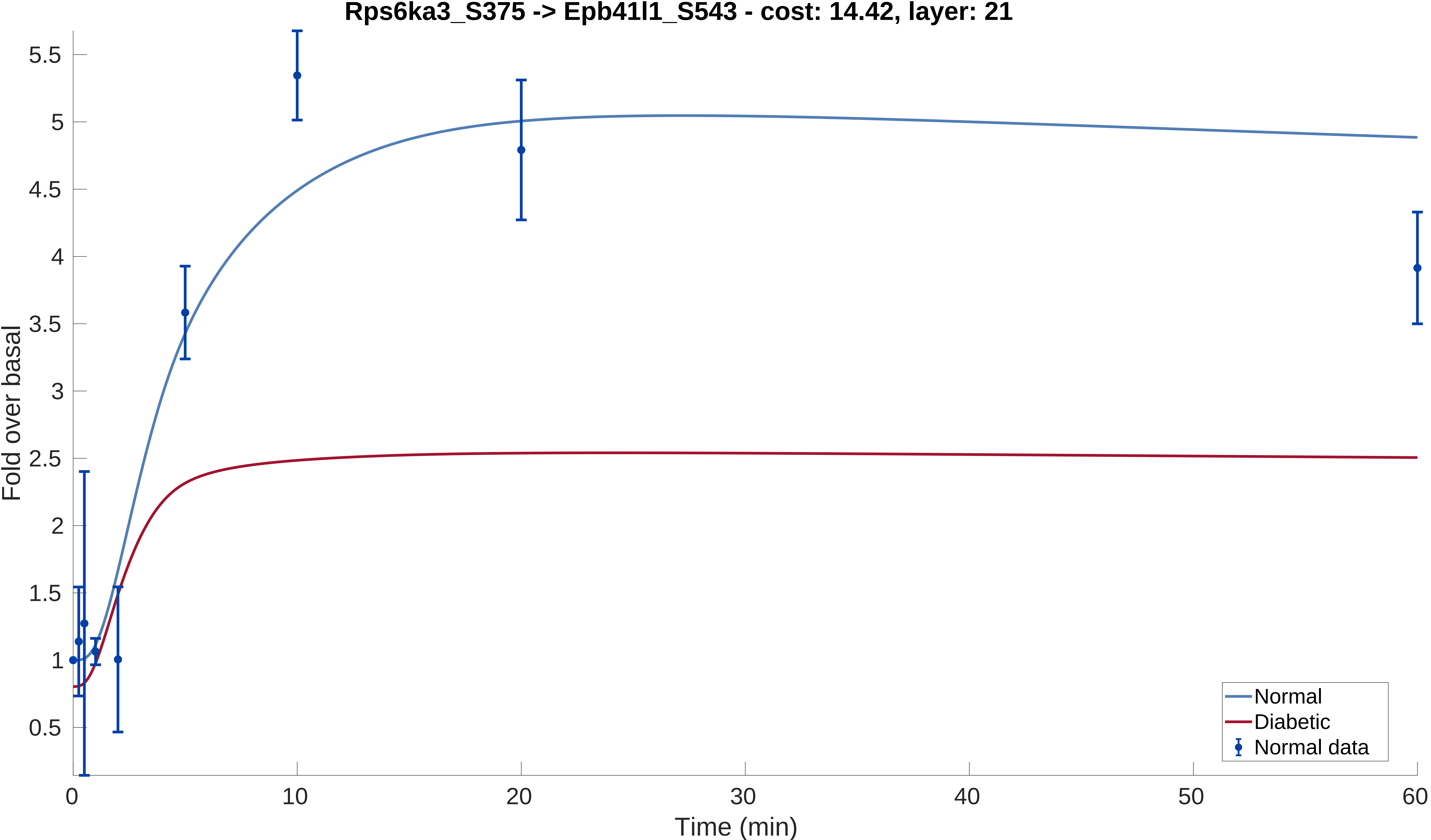
Rps6ka3_S375 -> Epb41l1_S543 - cost: 14.42, layer: 21
Fold over basal
Time (min)
Normal
Diabetic
Normal data
0.5
1
1.5
2
2.5
3
3.5
4
4.5
5
5.5
0
10
20
30
40
50
60

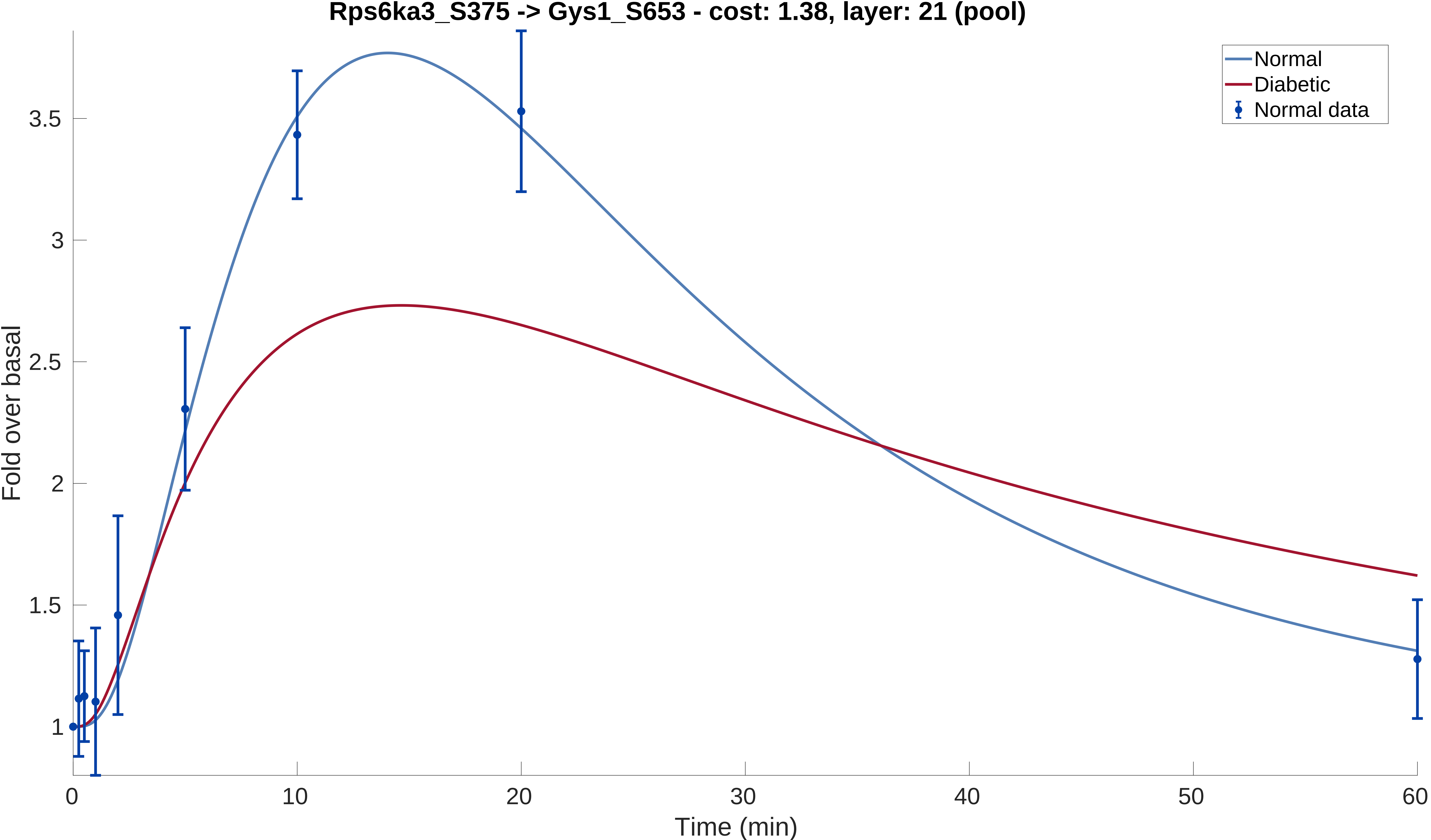
Rps6ka3_S375 -> Gys1_S653 - cost: 1.38, layer: 21 (pool)
Normal
Diabetic
Normal data
Fold over basal
Time (min)
3.5
3
2.5
2
1.5
1
0
10
20
30
40
50
60

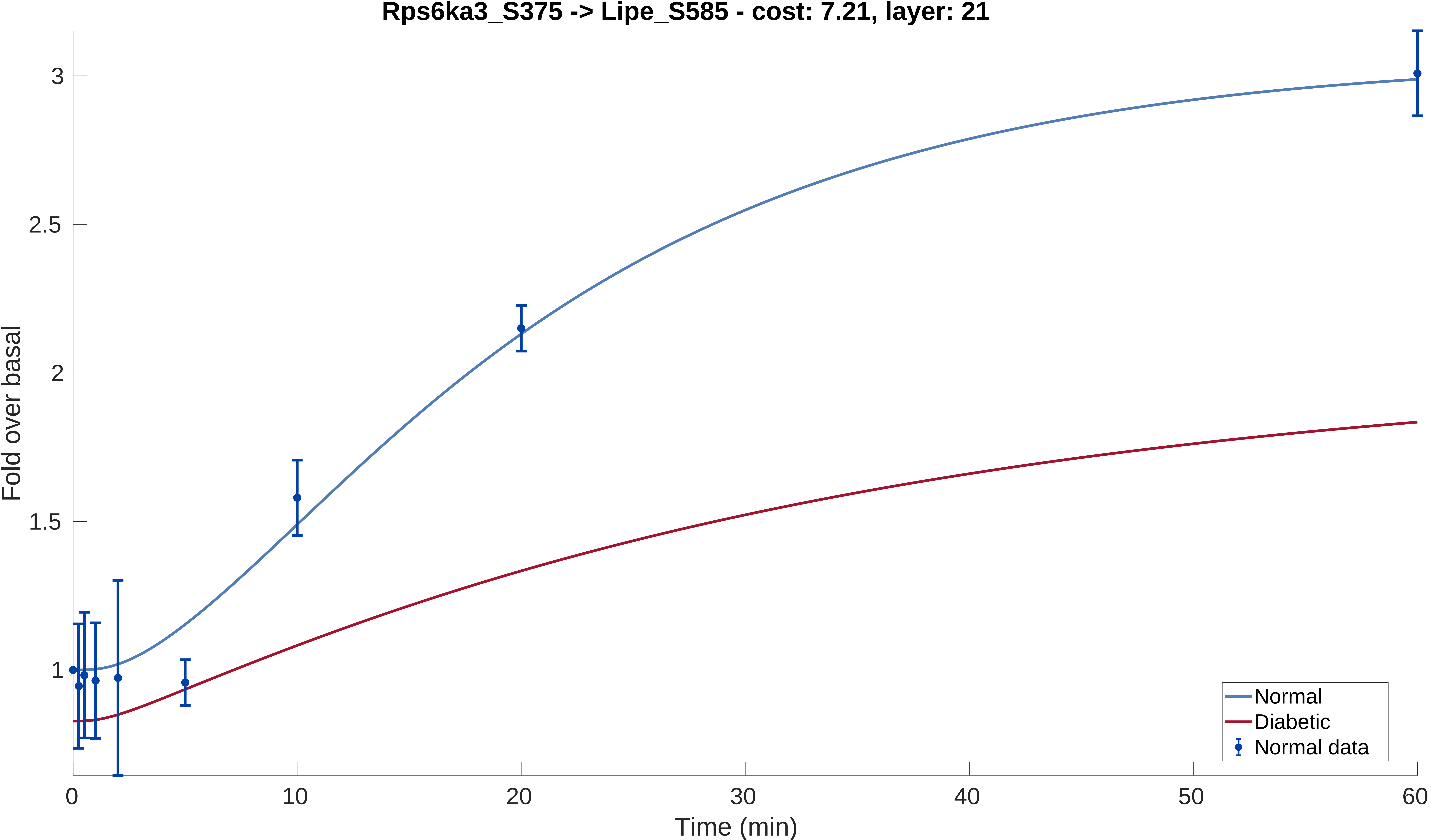

Rps6ka3_S375 -> Lipe_S585 - cost: 7.21, layer: 21
Fold over basal
3
2.5
2
1.5
1
0
10
20
30
40
50
60
Time (min)
Normal
Diabetic
Normal data

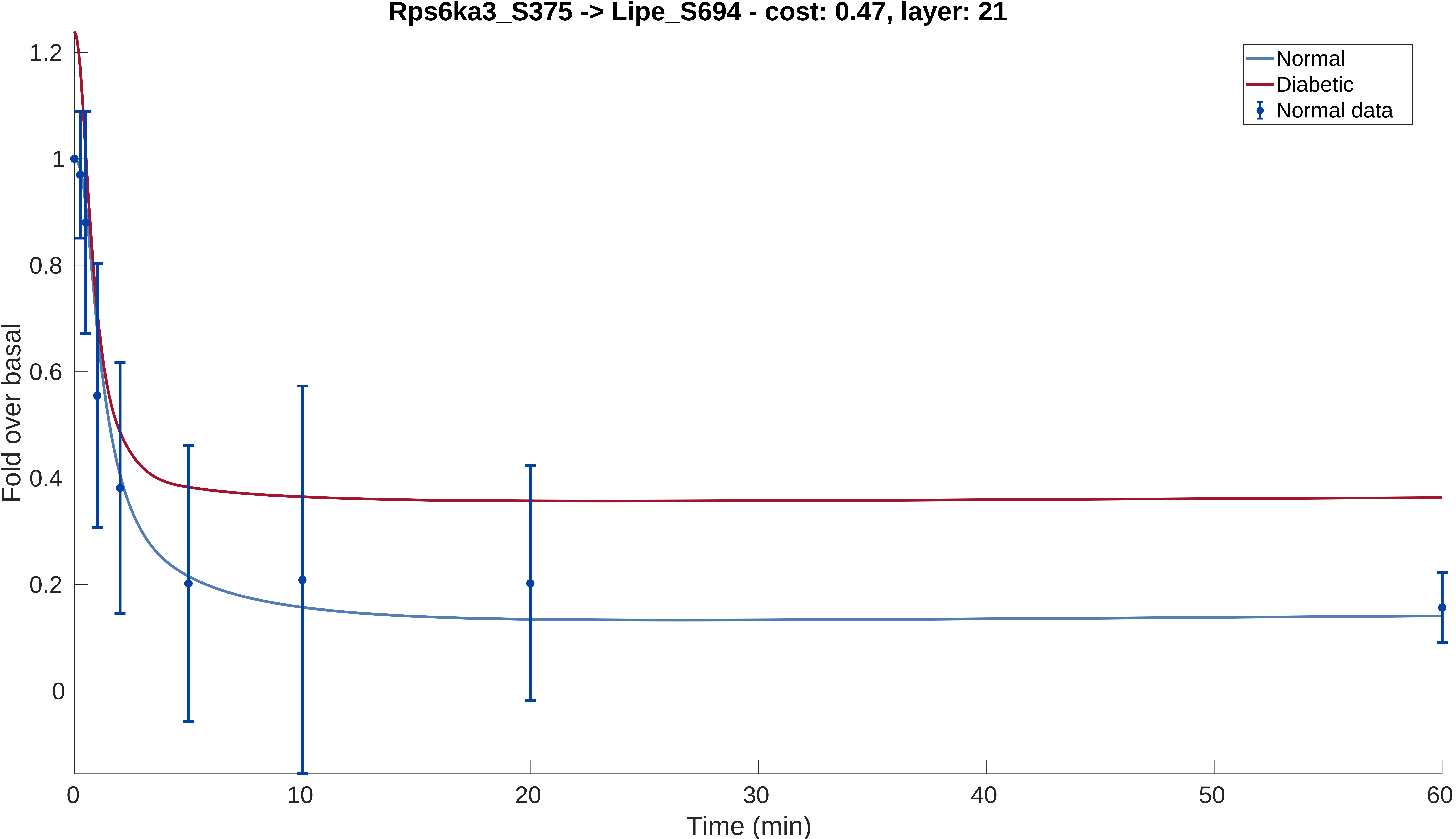

Rps6ka3_S375 -> Lipe_S694 - cost: 0.47, layer: 21
Normal
Diabetic
Normal data
Fold over basal
1.2
1
0.8
0.6
0.4
0.2
0
0
10
20
30
40
50
60
Time (min)

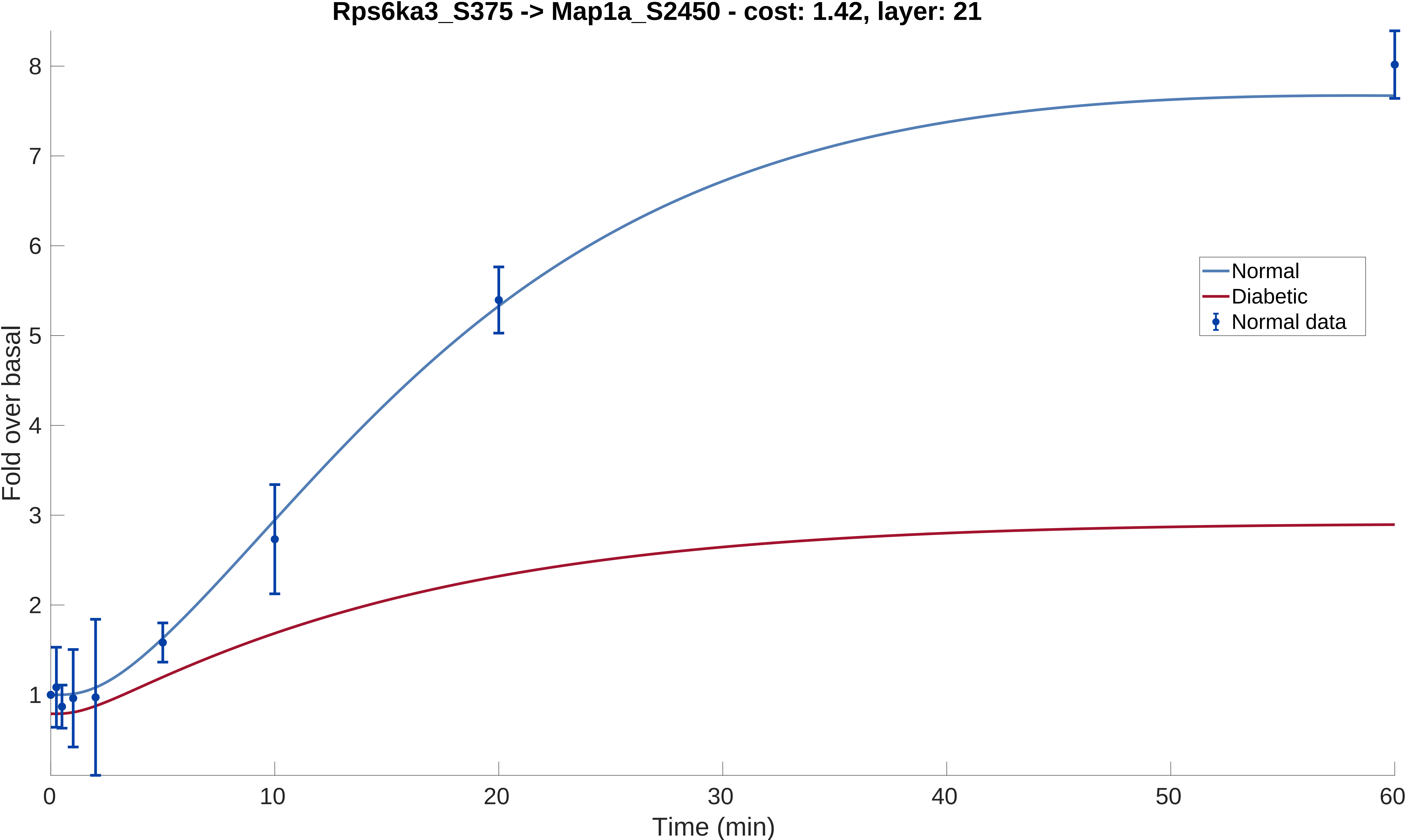

Rps6ka3_S375 -> Map1a_S2450 - cost: 1.42, layer: 21
Fold over basal
Time (min)
Normal
Diabetic
Normal data
0
10
20
30
40
50
60
1
2
3
4
5
6
7
8

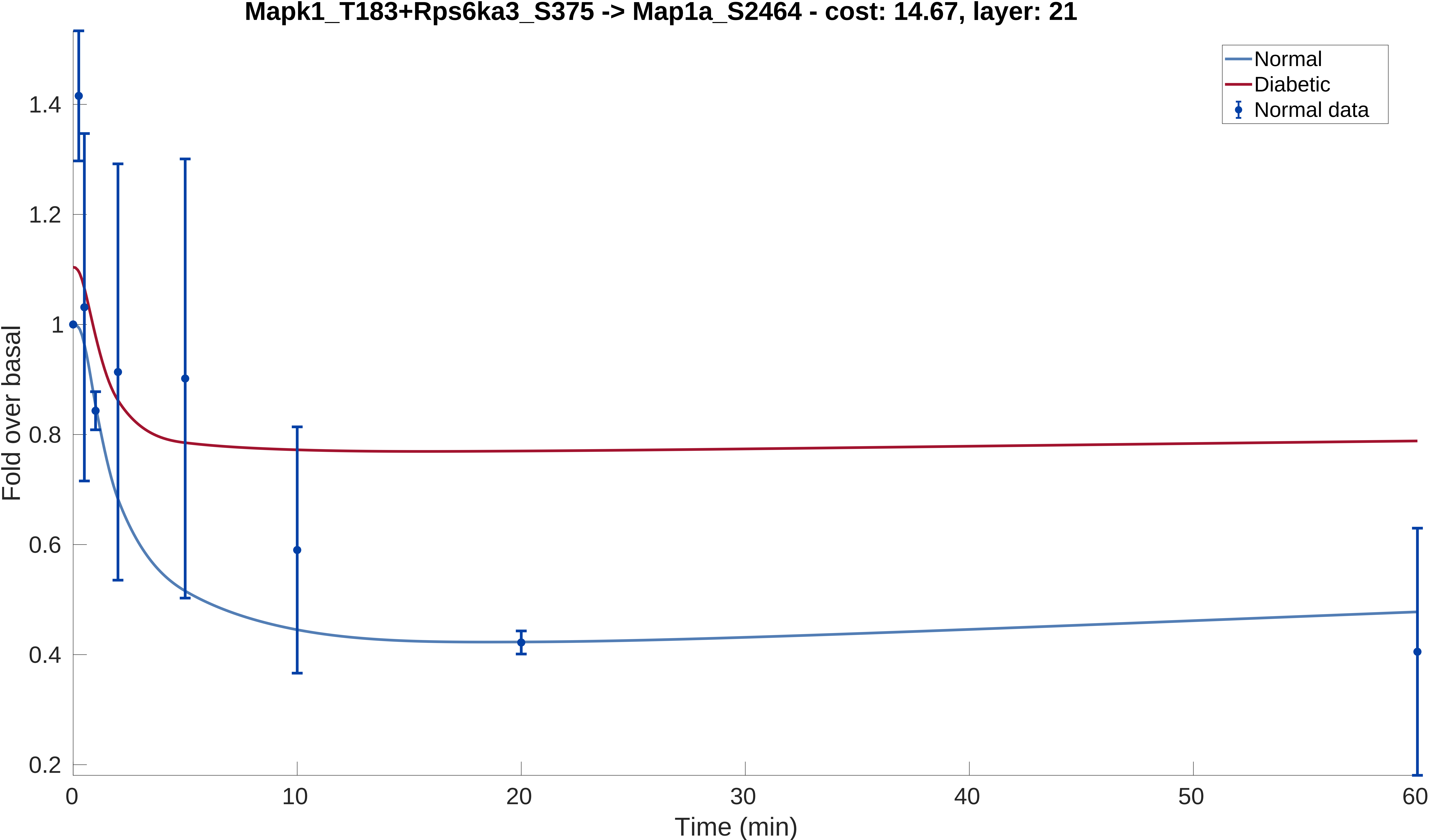

Mapk1_T183+Rps6ka3_S375 -> Map1a_S2464 - cost: 14.67, layer: 21
Normal
Diabetic
Normal data
Fold over basal
Time (min)
0.2
0.4
0.6
0.8
1
1.2
1.4
0
10
20
30
40
50
60

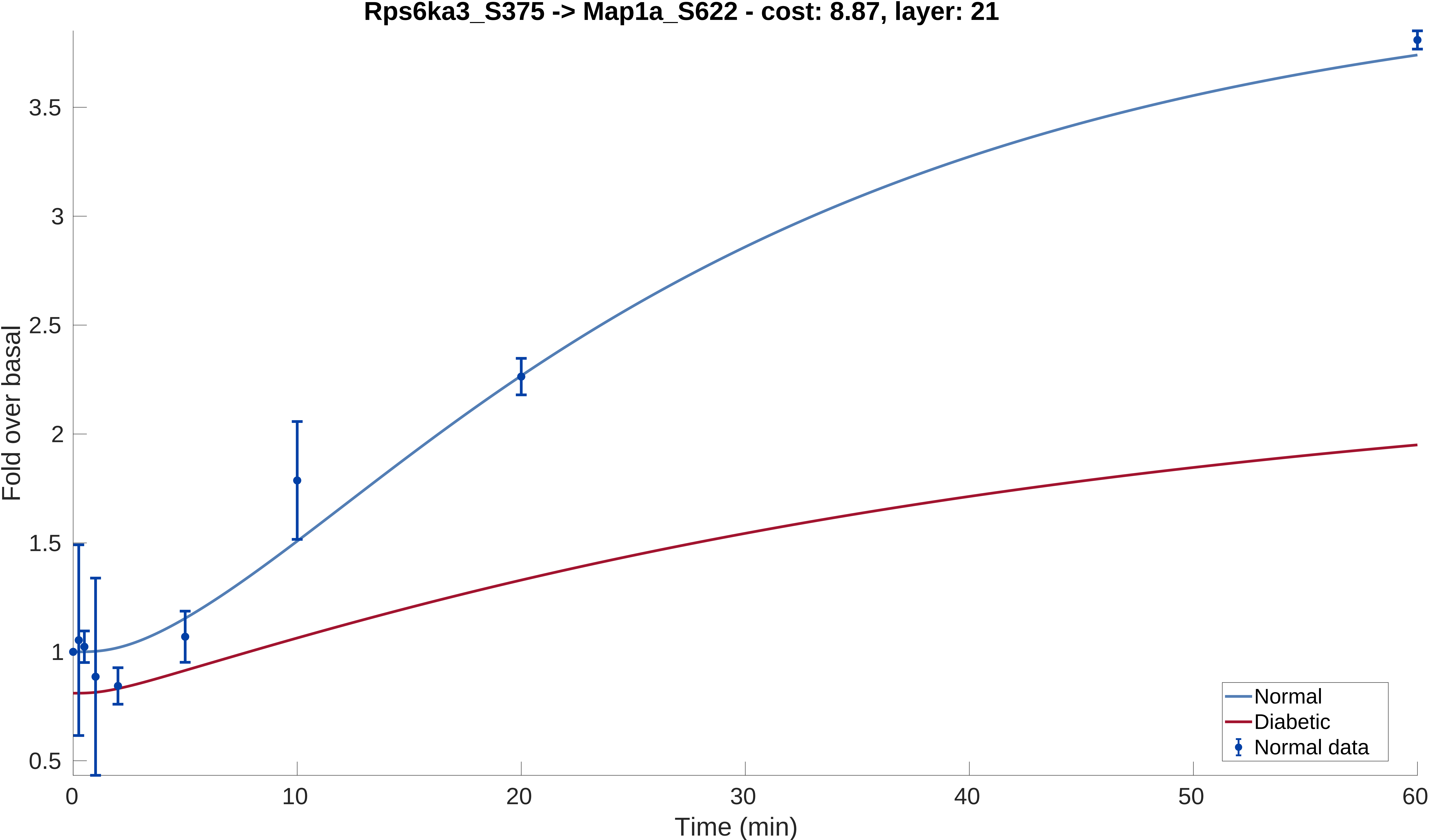

Rps6ka3_S375 -> Map1a_S622 - cost: 8.87, layer: 21
Fold over basal
Time (min)
0.5
1
1.5
2
2.5
3
3.5
0
10
20
30
40
50
60
Normal
Diabetic
Normal data

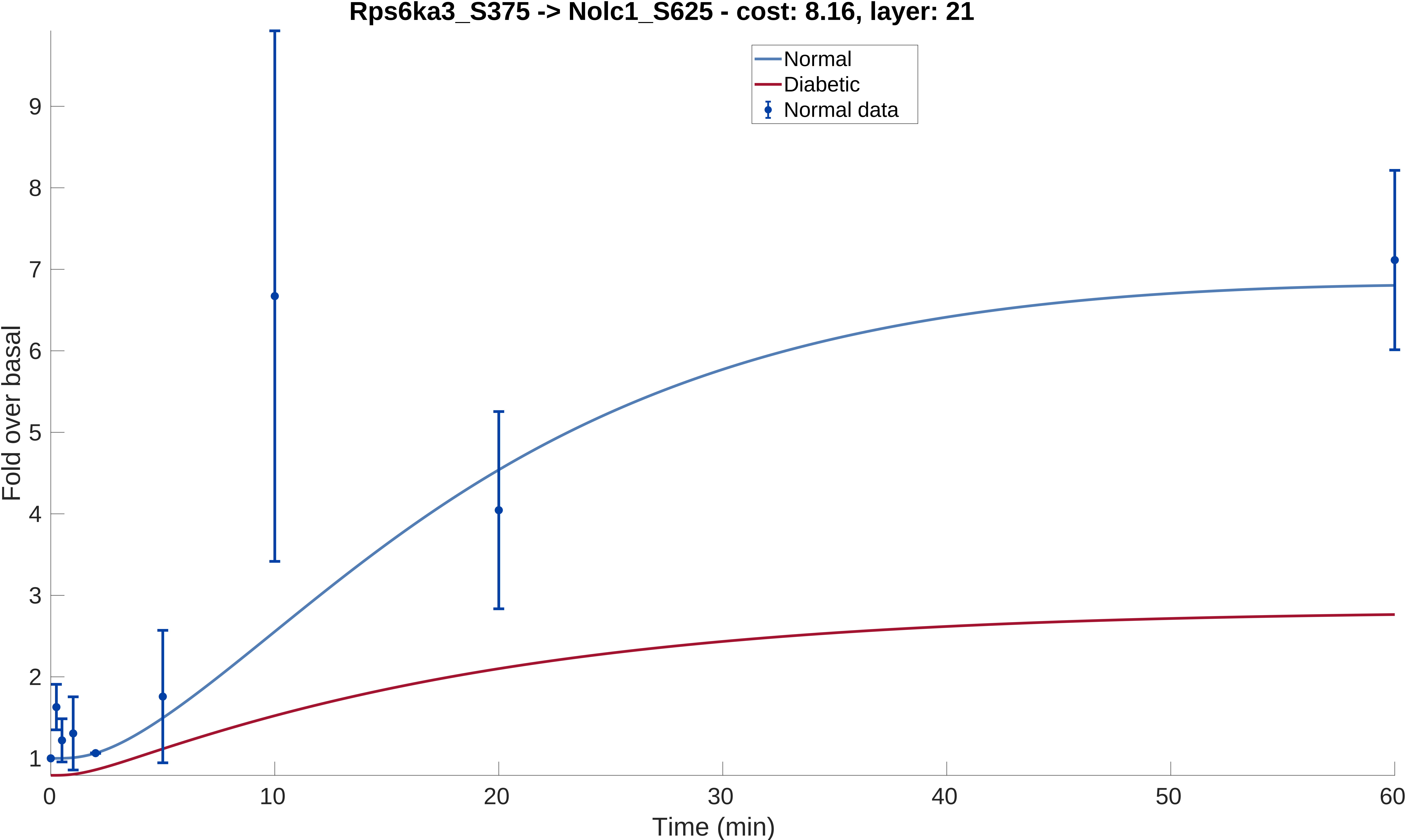

Rps6ka3_S375 -> Nolc1_S625 - cost: 8.16, layer: 21
Normal
Diabetic
Normal data
Fold over basal
Time (min)
1
2
3
4
5
6
7
8
9
0
10
20
30
40
50
60

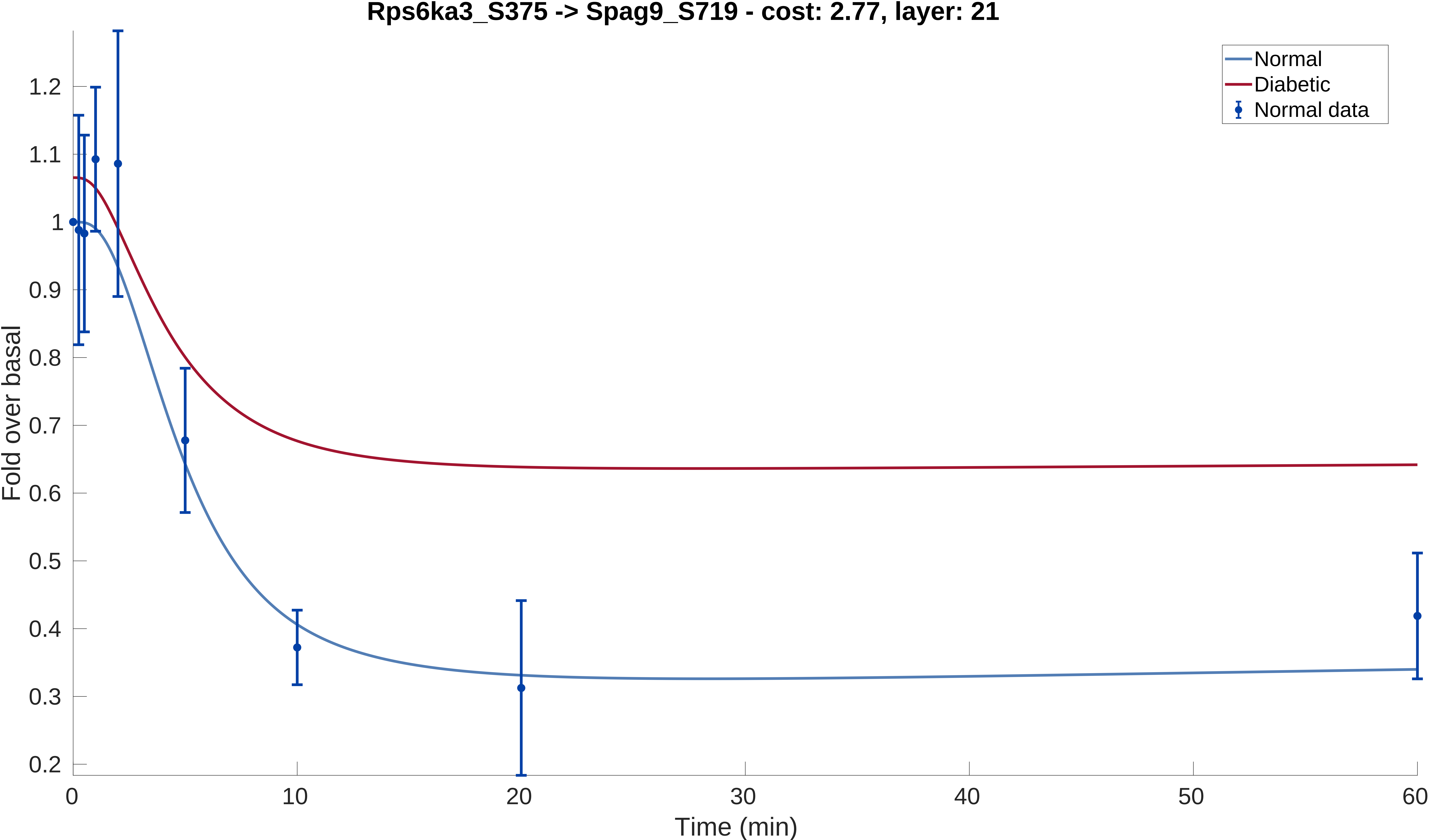

Rps6ka3_S375 -> Spag9_S719 - cost: 2.77, layer: 21
Normal
Diabetic
Normal data
Fold over basal
Time (min)
1.2
1.1
1
0.9
0.8
0.7
0.6
0.5
0.4
0.3
0.2
0
10
20
30
40
50
60

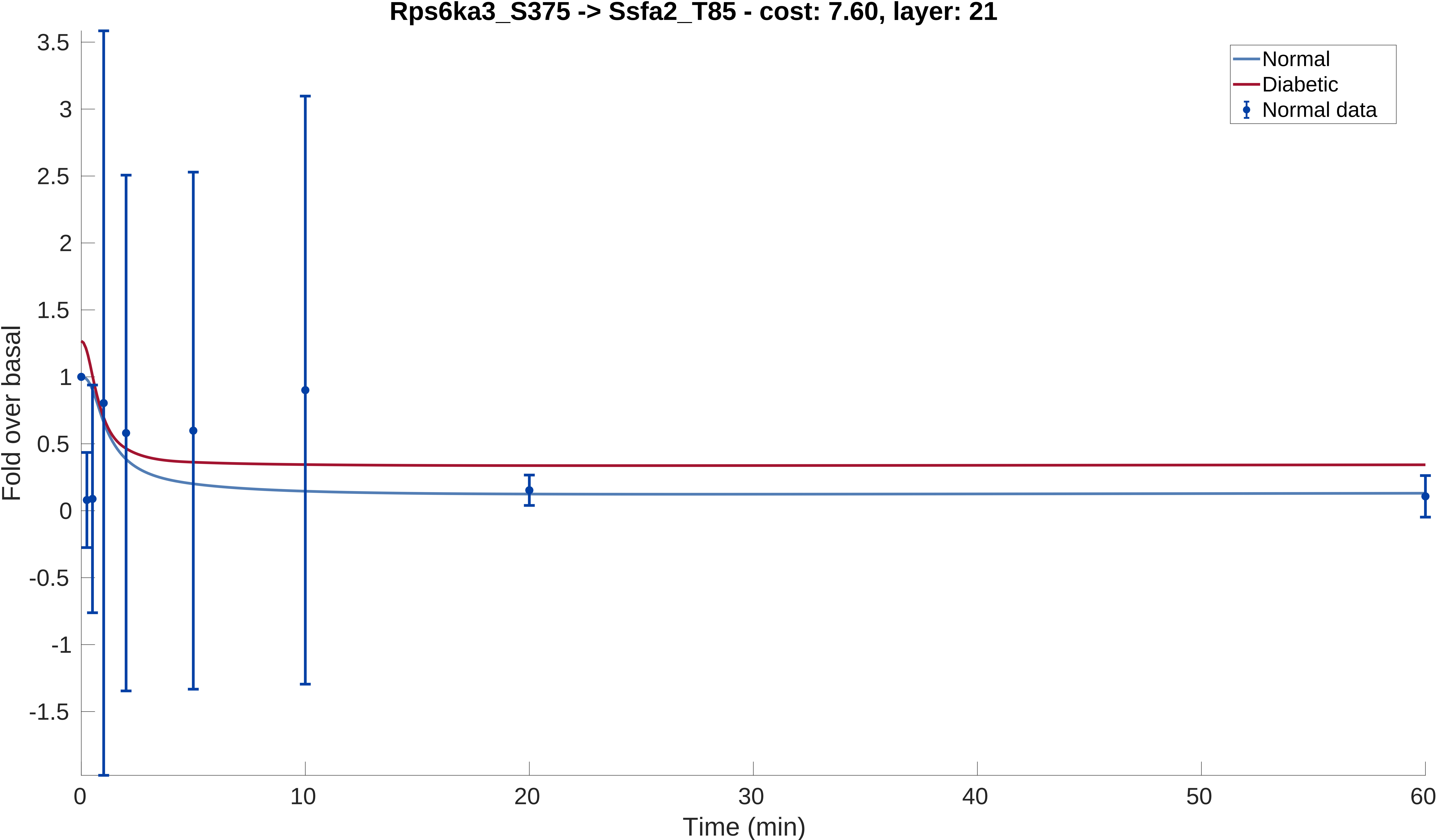

Rps6ka3_S375 -> Ssfa2_T85 - cost: 7.60, layer: 21
Normal
Diabetic
Normal data
Fold over basal
Time (min)
3.5
3
2.5
2
1.5
1
0.5
0
-0.5
-1
-1.5
0
10
20
30
40
50
60

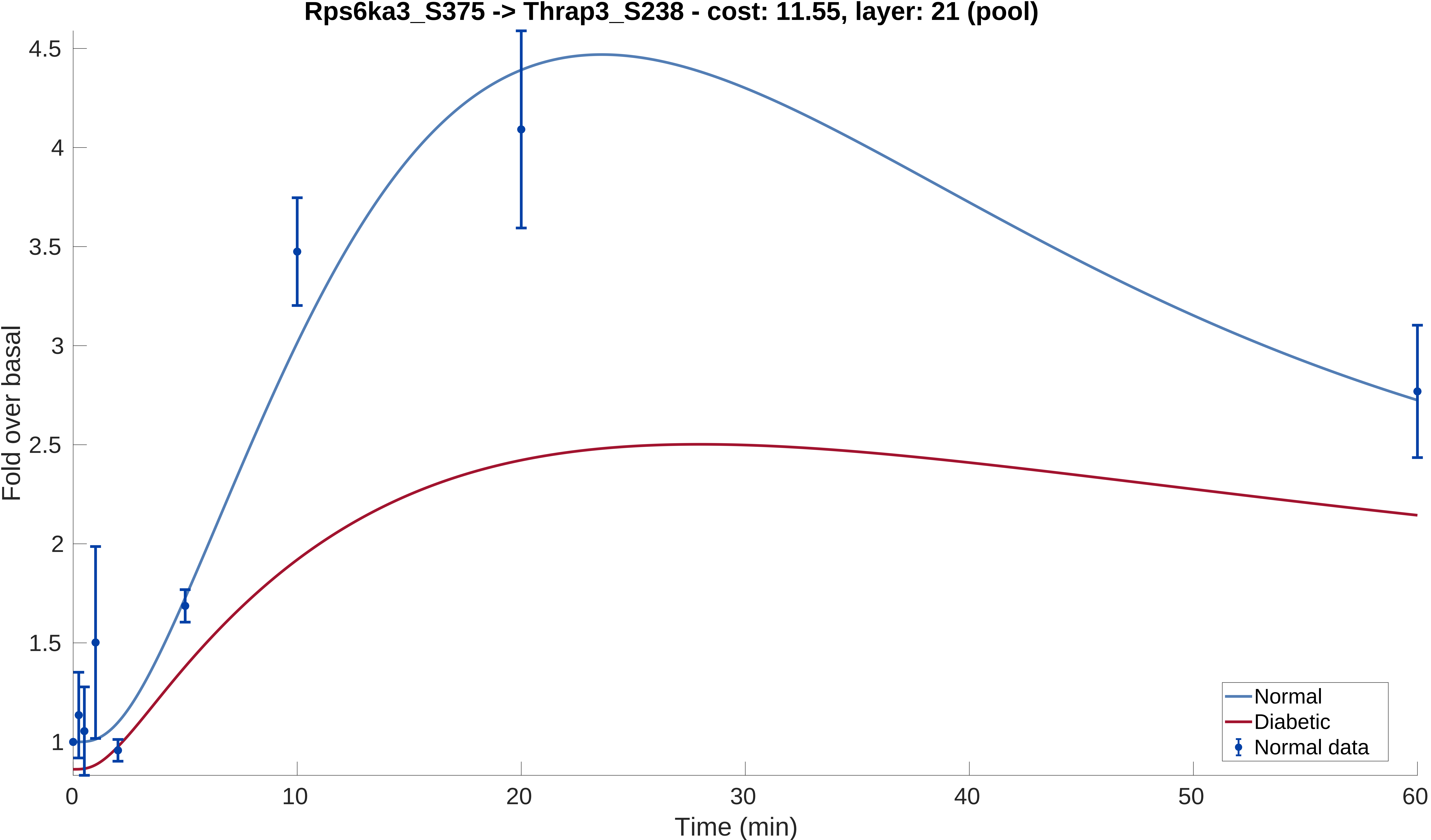

Rps6ka3_S375 -> Thrap3_S238 - cost: 11.55, layer: 21 (pool)
Fold over basal
4.5
4
3.5
3
2.5
2
1.5
1
0
10
20
30
40
50
60
Time (min)
Normal
Diabetic
Normal data

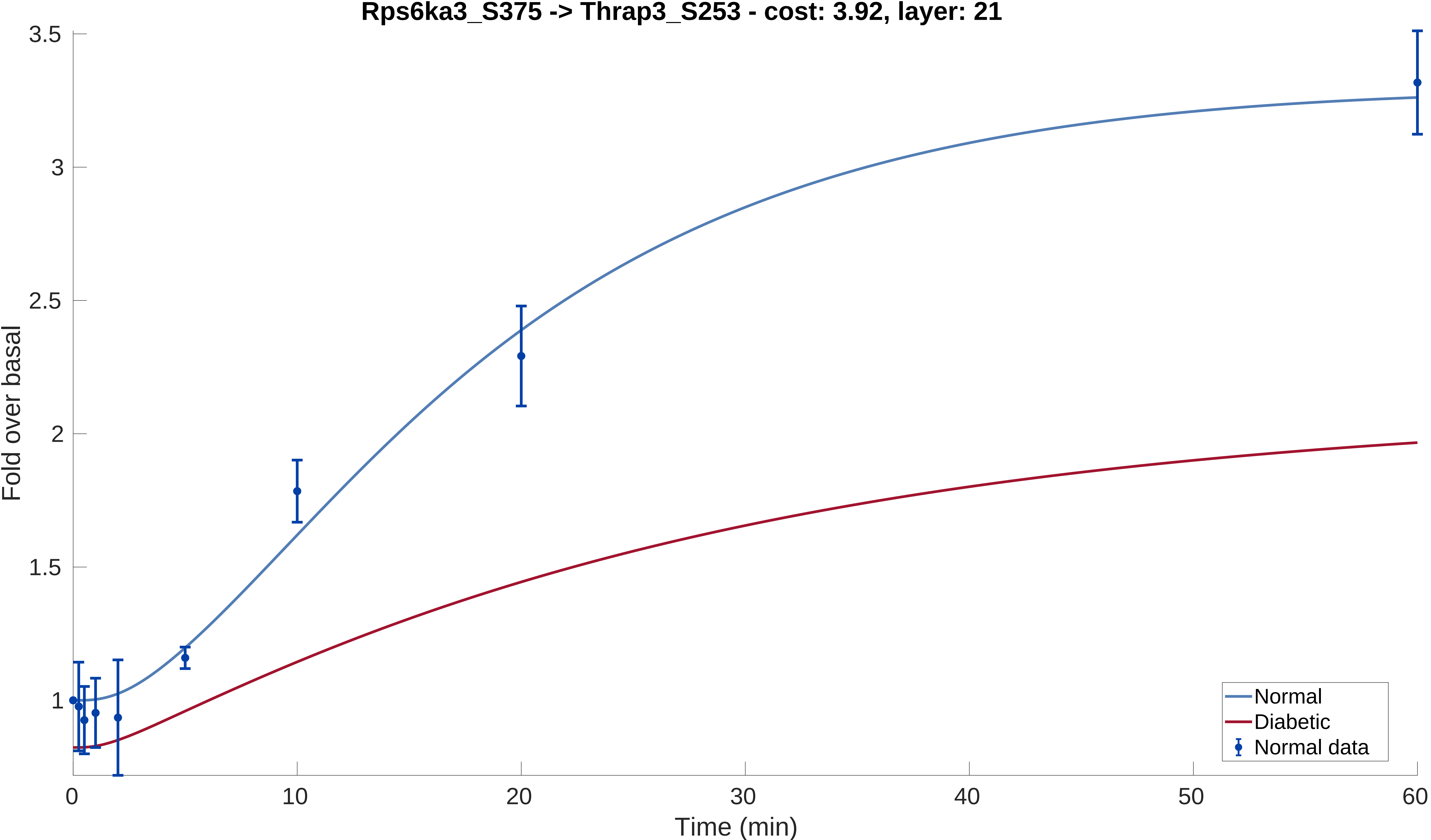

Rps6ka3_S375 -> Thrap3_S253 - cost: 3.92, layer: 21
Fold over basal
3.5
3
2.5
2
1.5
1
0
10
20
30
40
50
60
Time (min)
Normal
Diabetic
Normal data

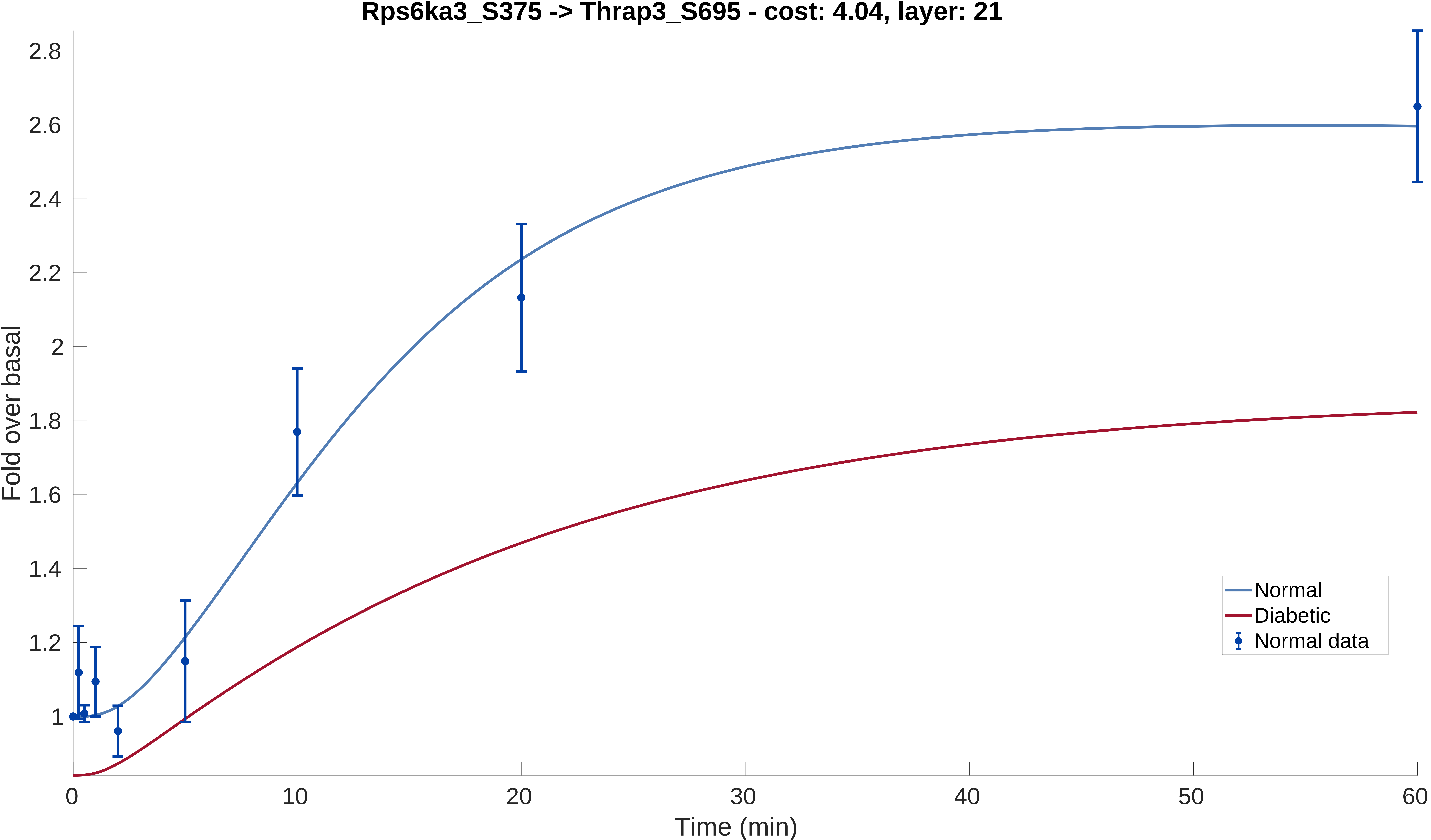
Rps6ka3_S375 -> Thrap3_S695 - cost: 4.04, layer: 21
Fold over basal
Time (min)
2.8
2.6
2.4
2.2
2
1.8
1.6
1.4
1.2
1
0
10
20
30
40
50
60
Normal
Diabetic
Normal data

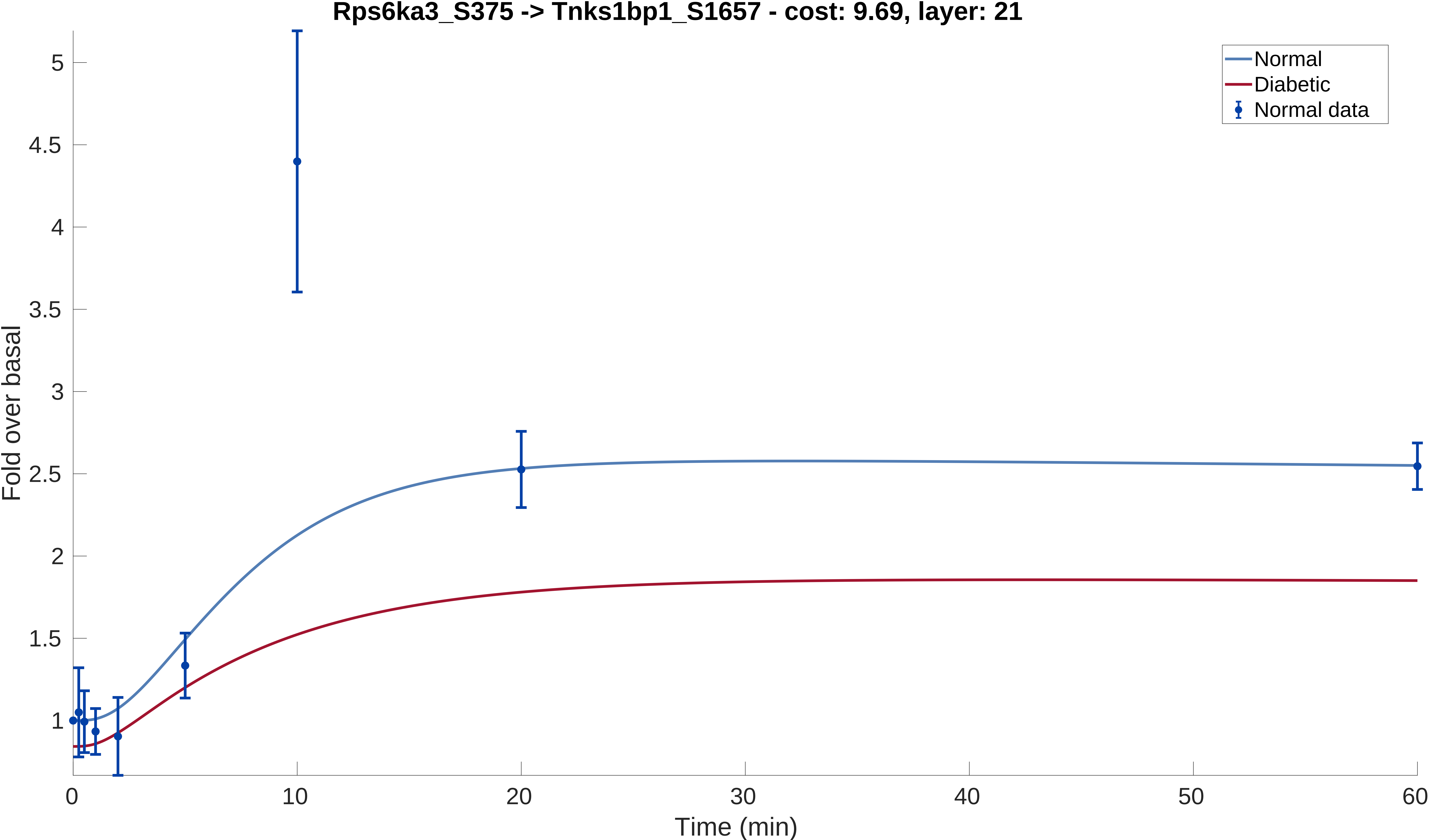

Rps6ka3_S375 -> Tnks1bp1_S1657 - cost: 9.69, layer: 21
Normal
Diabetic
Normal data
Fold over basal
Time (min)
5
4.5
4
3.5
3
2.5
2
1.5
1
0
10
20
30
40
50
60

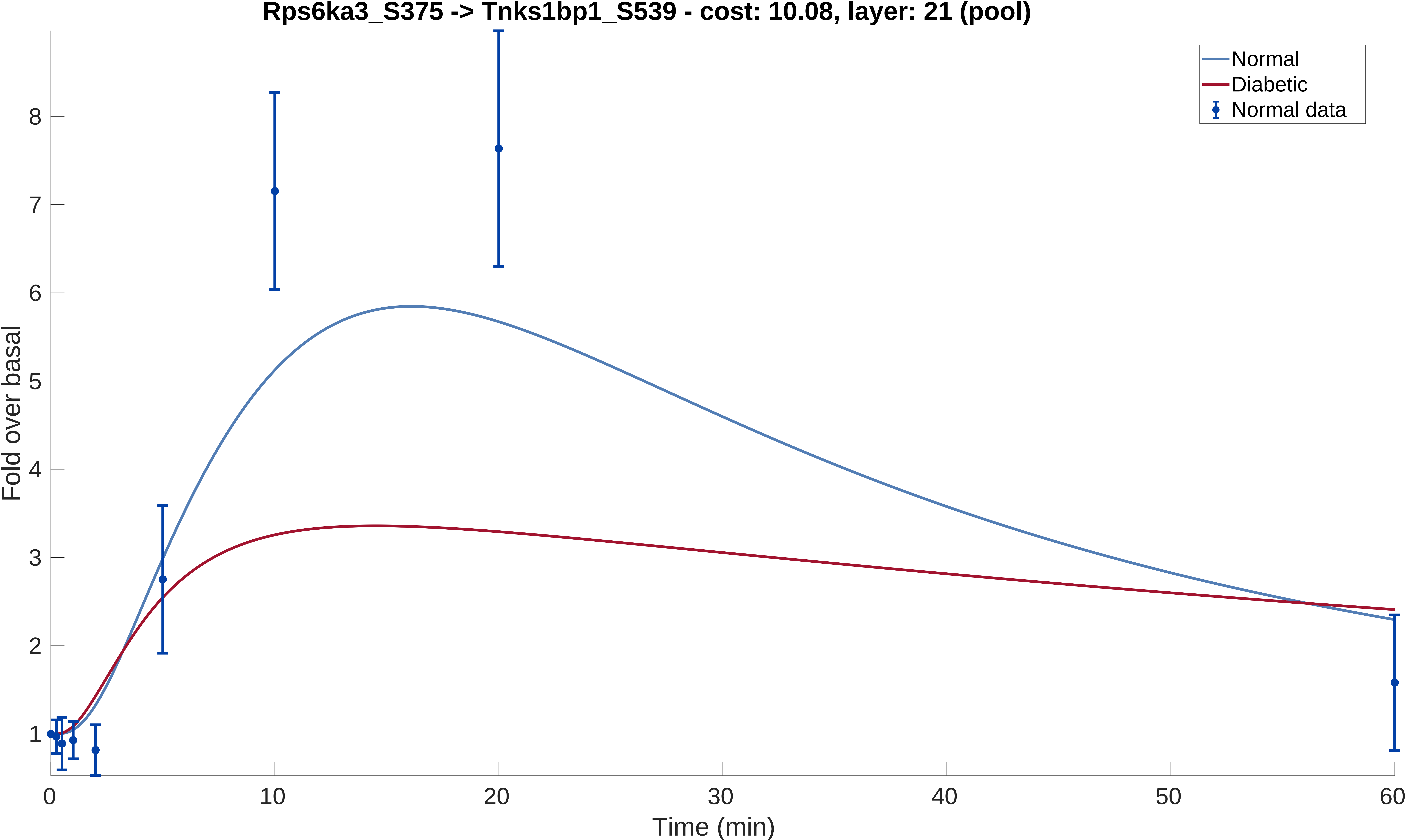

Rps6ka3_S375 -> Tnks1bp1_S539 - cost: 10.08, layer: 21 (pool)
Normal
Diabetic
Normal data
Fold over basal
Time (min)
1
2
3
4
5
6
7
8
0
10
20
30
40
50
60

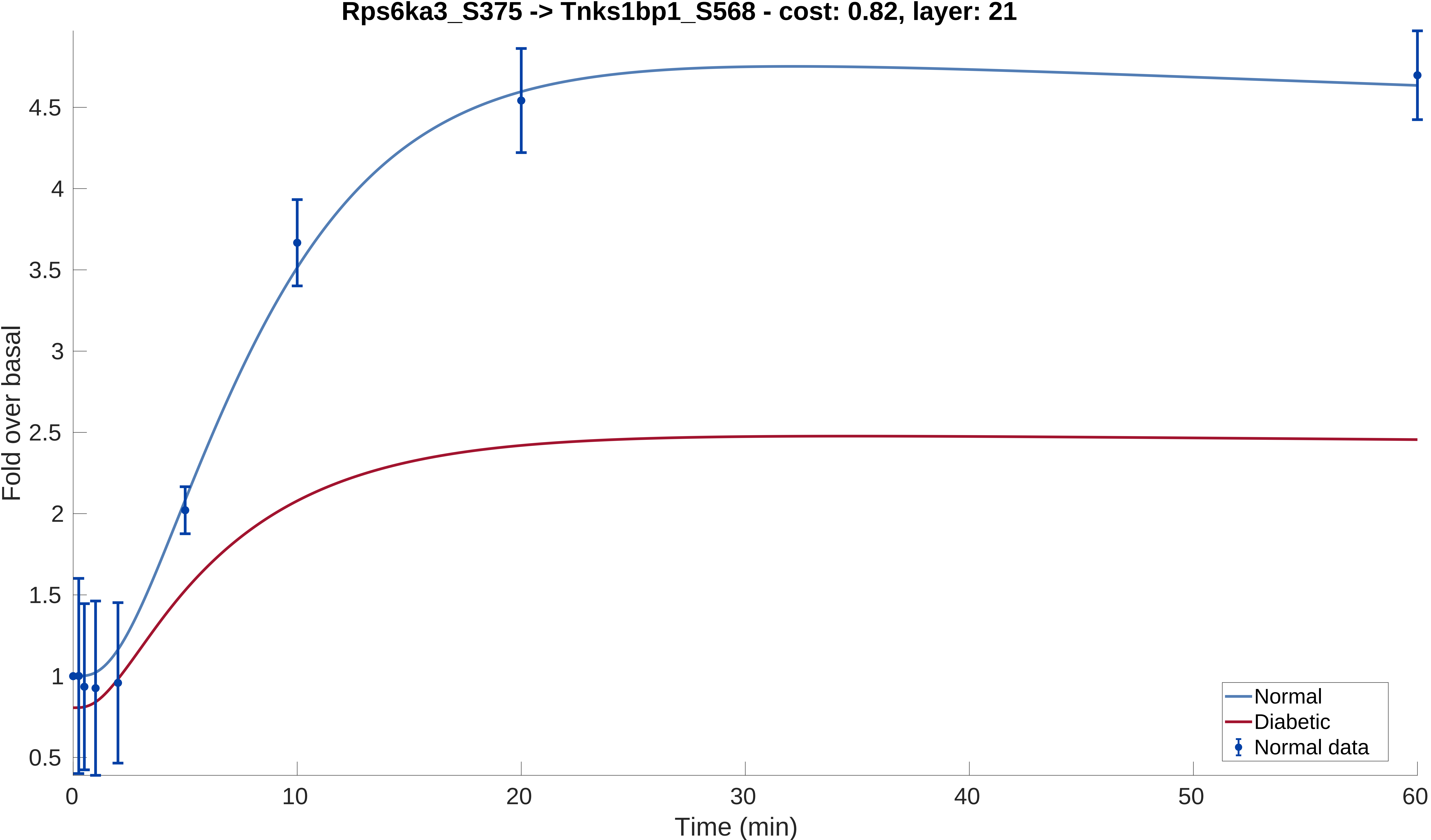

Rps6ka3_S375 -> Tnks1bp1_S568 - cost: 0.82, layer: 21
Fold over basal
Time (min)
0.5
1
1.5
2
2.5
3
3.5
4
4.5
0
10
20
30
40
50
60
Normal
Diabetic
Normal data

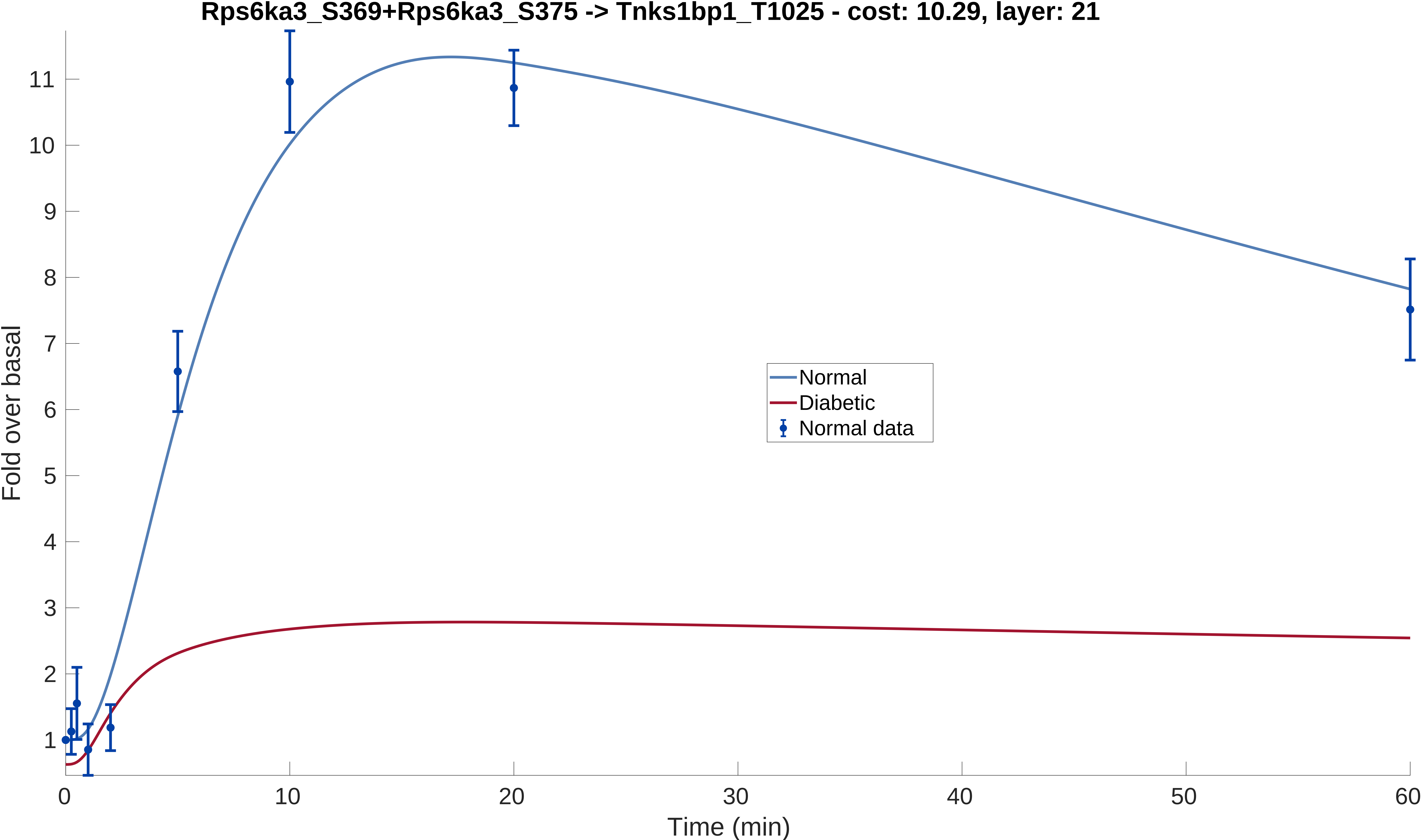

Rps6ka3_S369+Rps6ka3_S375 -> Tnks1bp1_T1025 - cost: 10.29, layer: 21
Fold over basal
Time (min)
Normal
Diabetic
Normal data
0
10
20
30
40
50
60
1
2
3
4
5
6
7
8
9
10
11

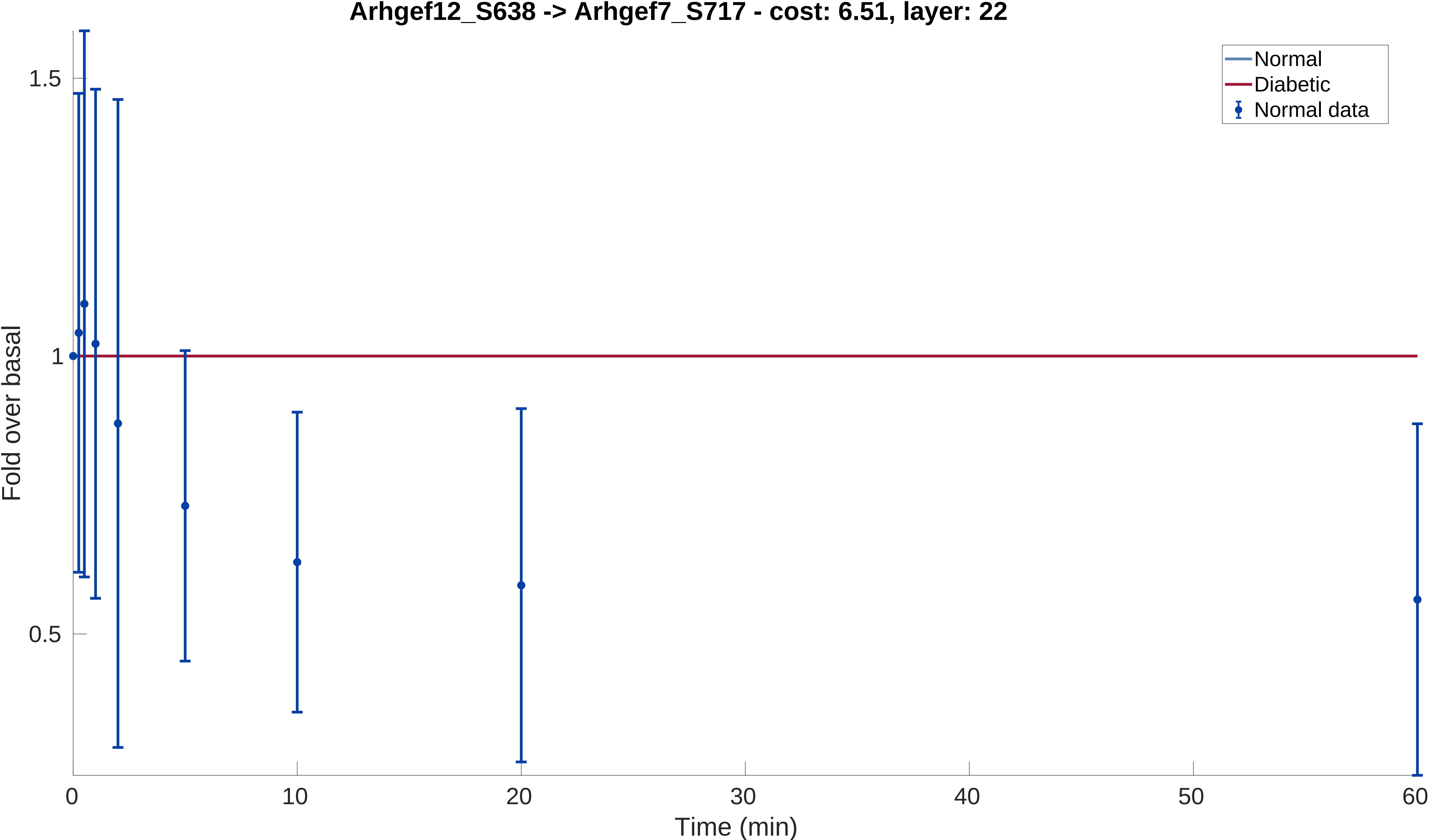

Arhgef12_S638 -> Arhgef7_S717 - cost: 6.51, layer: 22
Normal
Diabetic
Normal data
Fold over basal
1.5
1
0.5
0
10
20
30
40
50
60
Time (min)

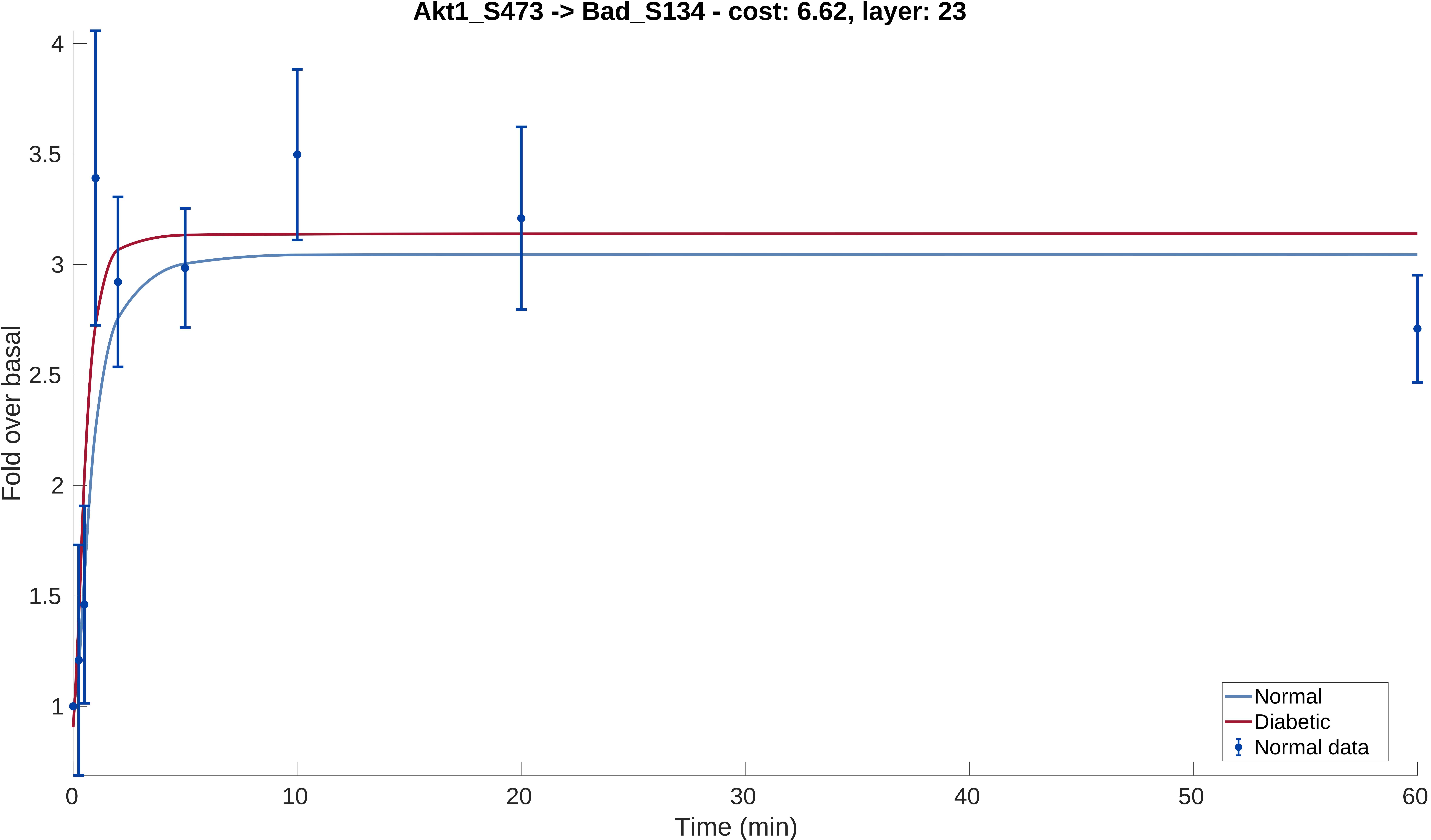
Akt1_S473 -> Bad_S134 - cost: 6.62, layer: 23
Fold over basal
4
3.5
3
2.5
2
1.5
1
0
10
20
30
40
50
60
Time (min)
Normal
Diabetic
Normal data

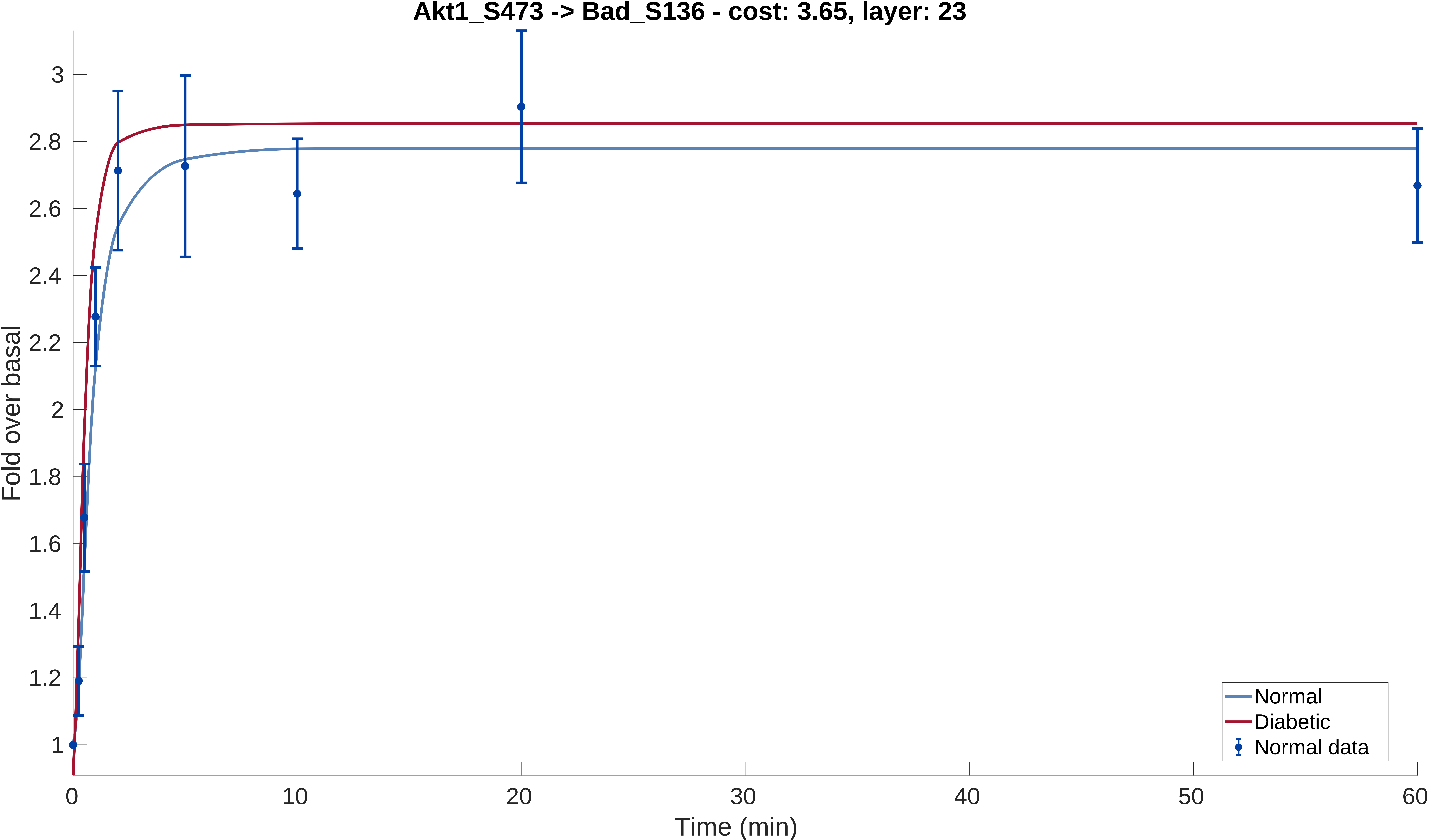

Akt1_S473 -> Bad_S136 - cost: 3.65, layer: 23
Fold over basal
3
2.8
2.6
2.4
2.2
2
1.8
1.6
1.4
1.2
1
0
10
20
30
40
50
60
Time (min)
Normal
Diabetic
Normal data

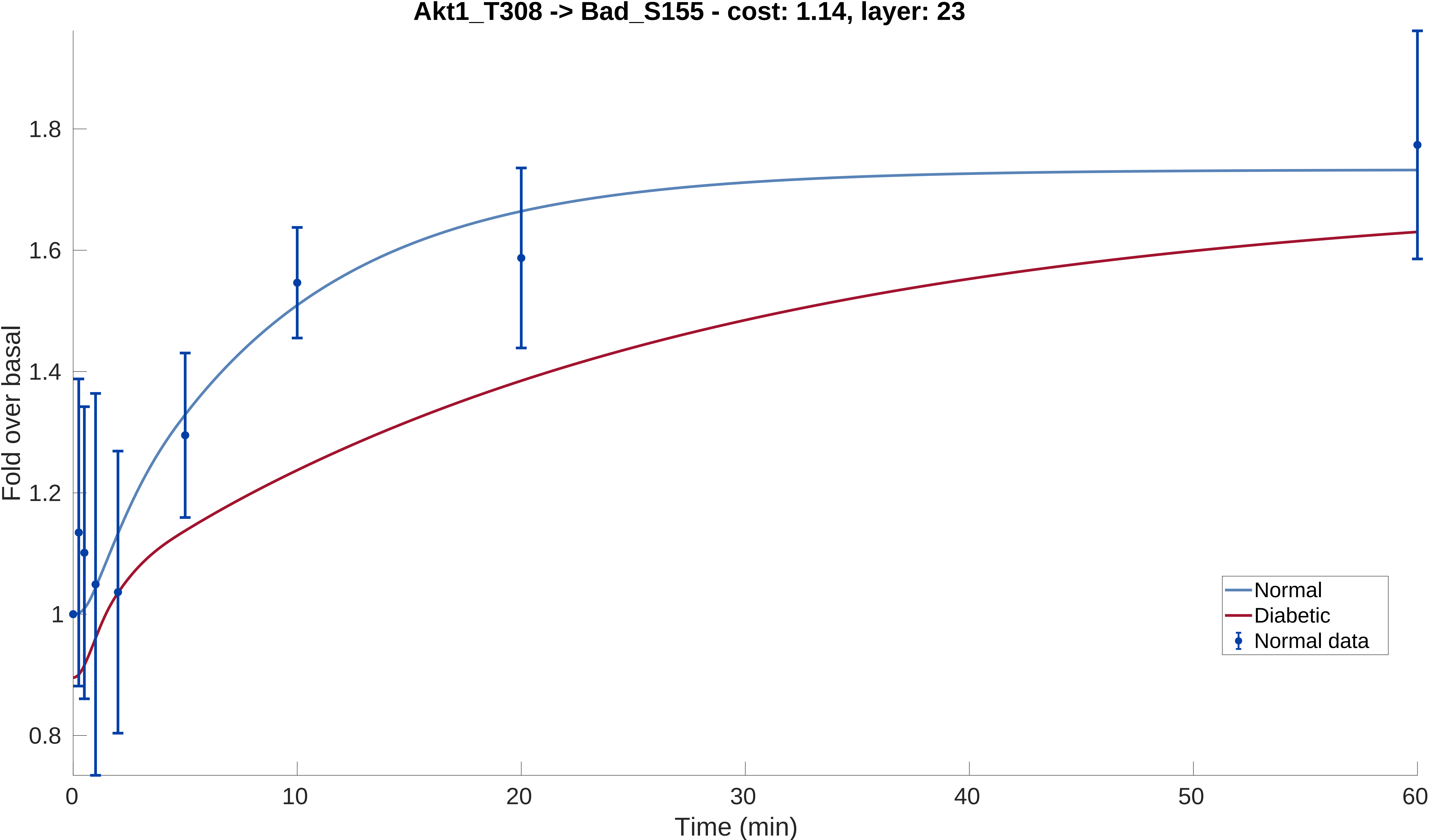
Akt1_T308 -> Bad_S155 - cost: 1.14, layer: 23
Fold over basal
1.8
1.6
1.4
1.2
1
0.8
0
10
20
30
40
50
60
Time (min)
Normal
Diabetic
Normal data

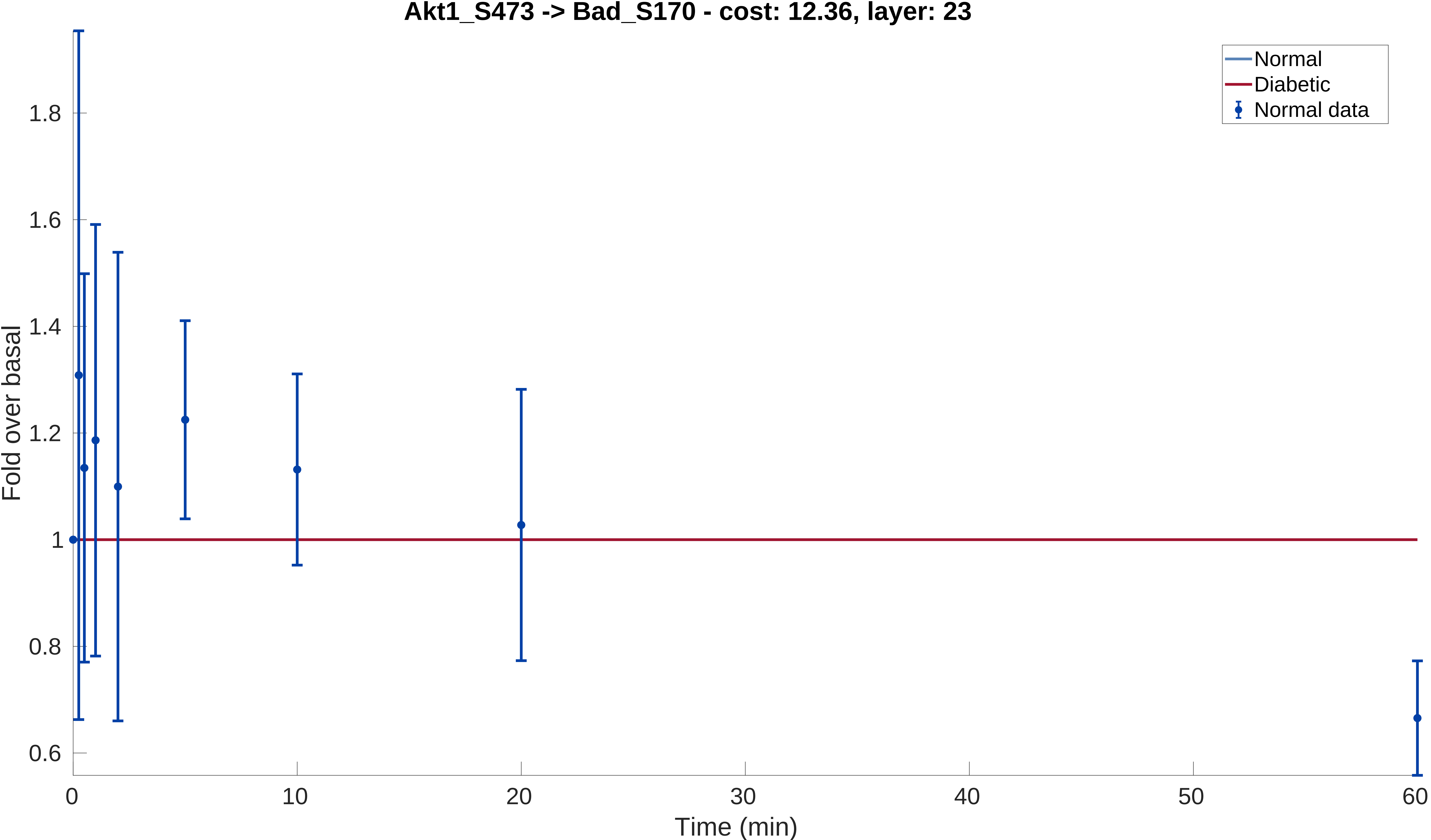
Akt1_S473 -> Bad_S170 - cost: 12.36, layer: 23
Normal
Diabetic
Normal data
Fold over basal
1.8
1.6
1.4
1.2
1
0.8
0.6
0
10
20
30
40
50
60
Time (min)

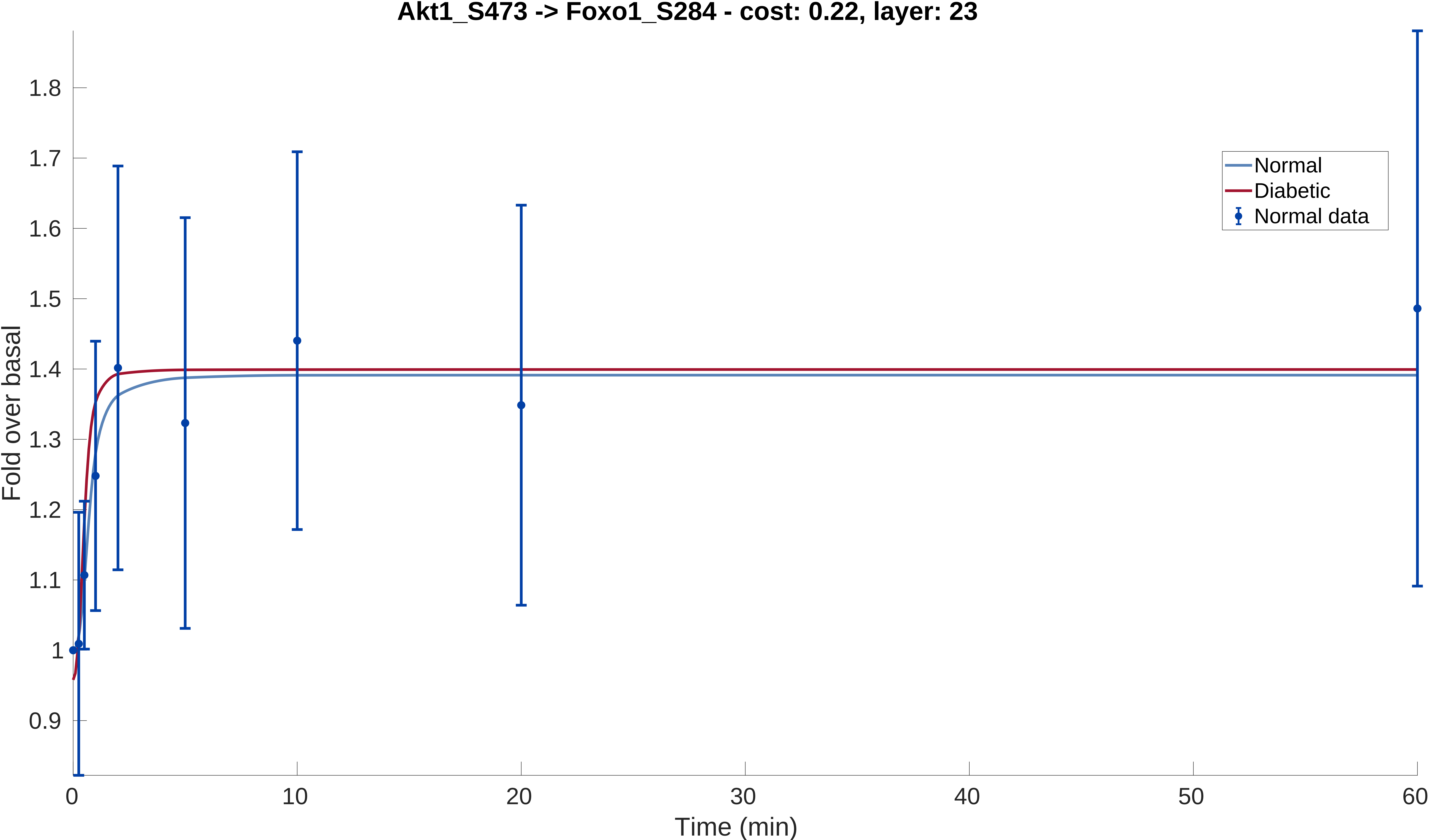

Akt1_S473 -> Foxo1_S284 - cost: 0.22, layer: 23
Normal
Diabetic
Normal data
Fold over basal
Time (min)
0.9
1
1.1
1.2
1.3
1.4
1.5
1.6
1.7
1.8
0
10
20
30
40
50
60

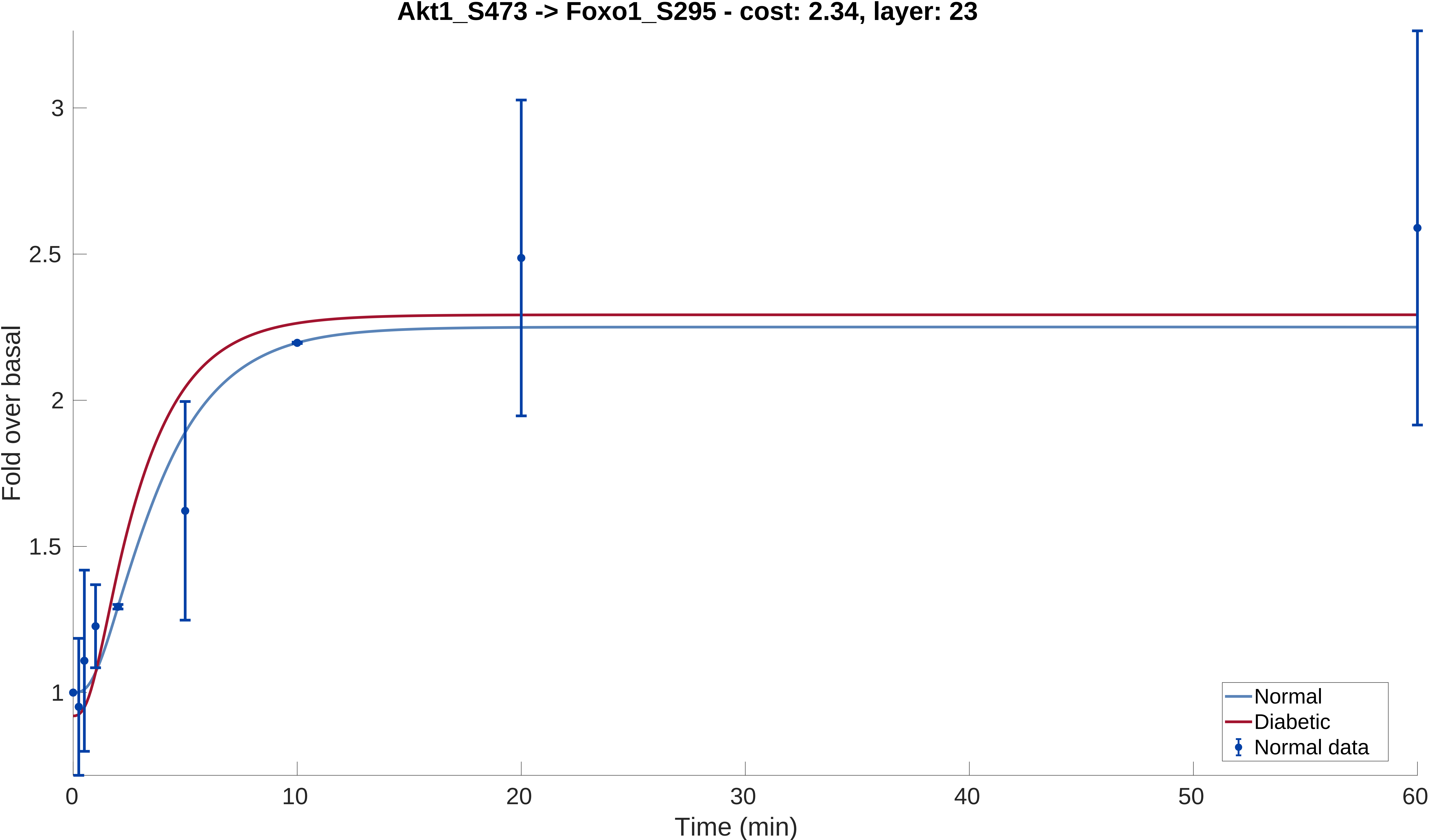

Akt1_S473 -> Foxo1_S295 - cost: 2.34, layer: 23
Fold over basal
3
2.5
2
1.5
1
0
10
20
30
40
50
60
Time (min)
Normal
Diabetic
Normal data

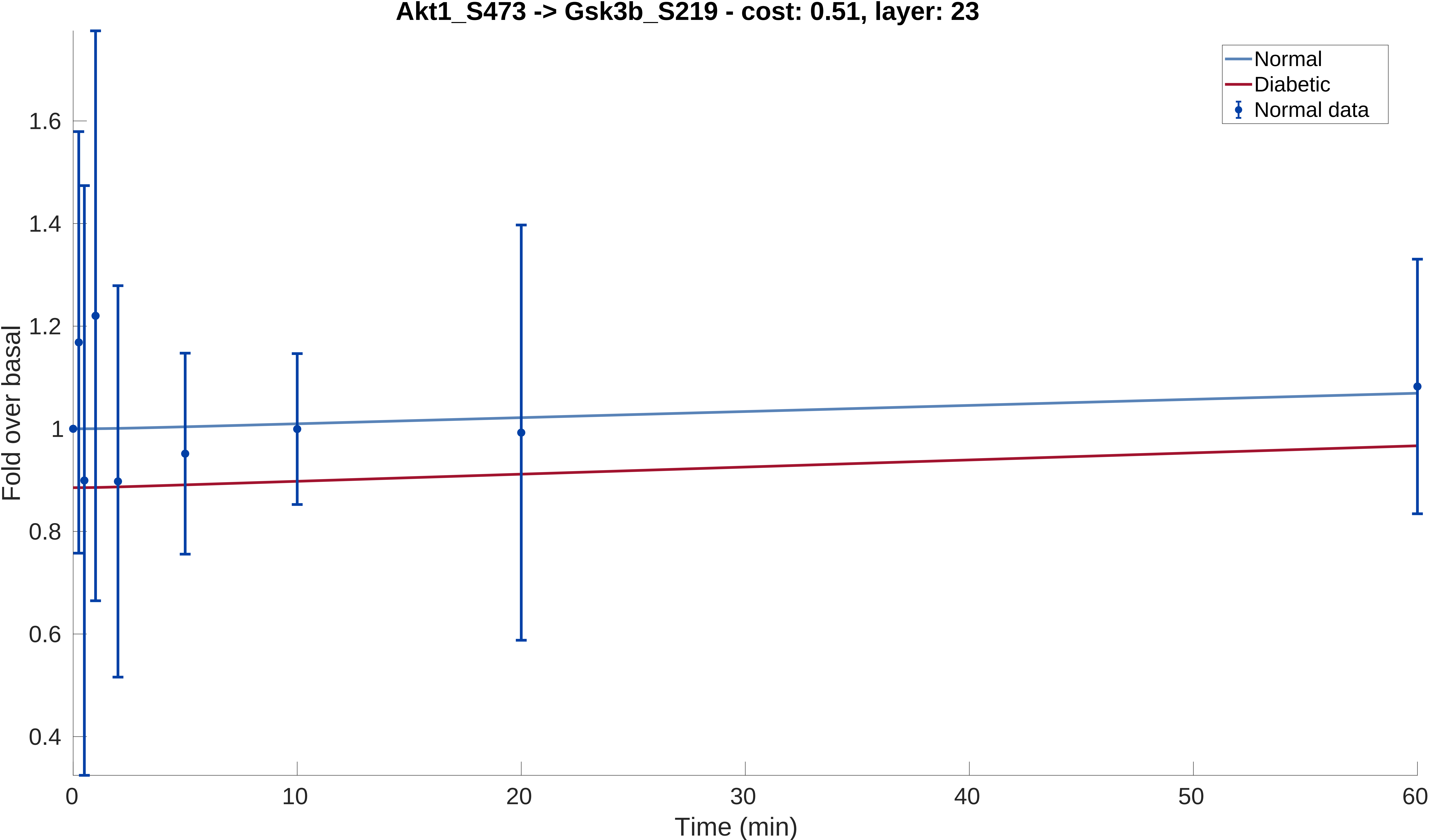
Akt1_S473 -> Gsk3b_S219 - cost: 0.51, layer: 23
Normal
Diabetic
Normal data
Fold over basal
1.6
1.4
1.2
1
0.8
0.6
0.4
0
10
20
30
40
50
60
Time (min)

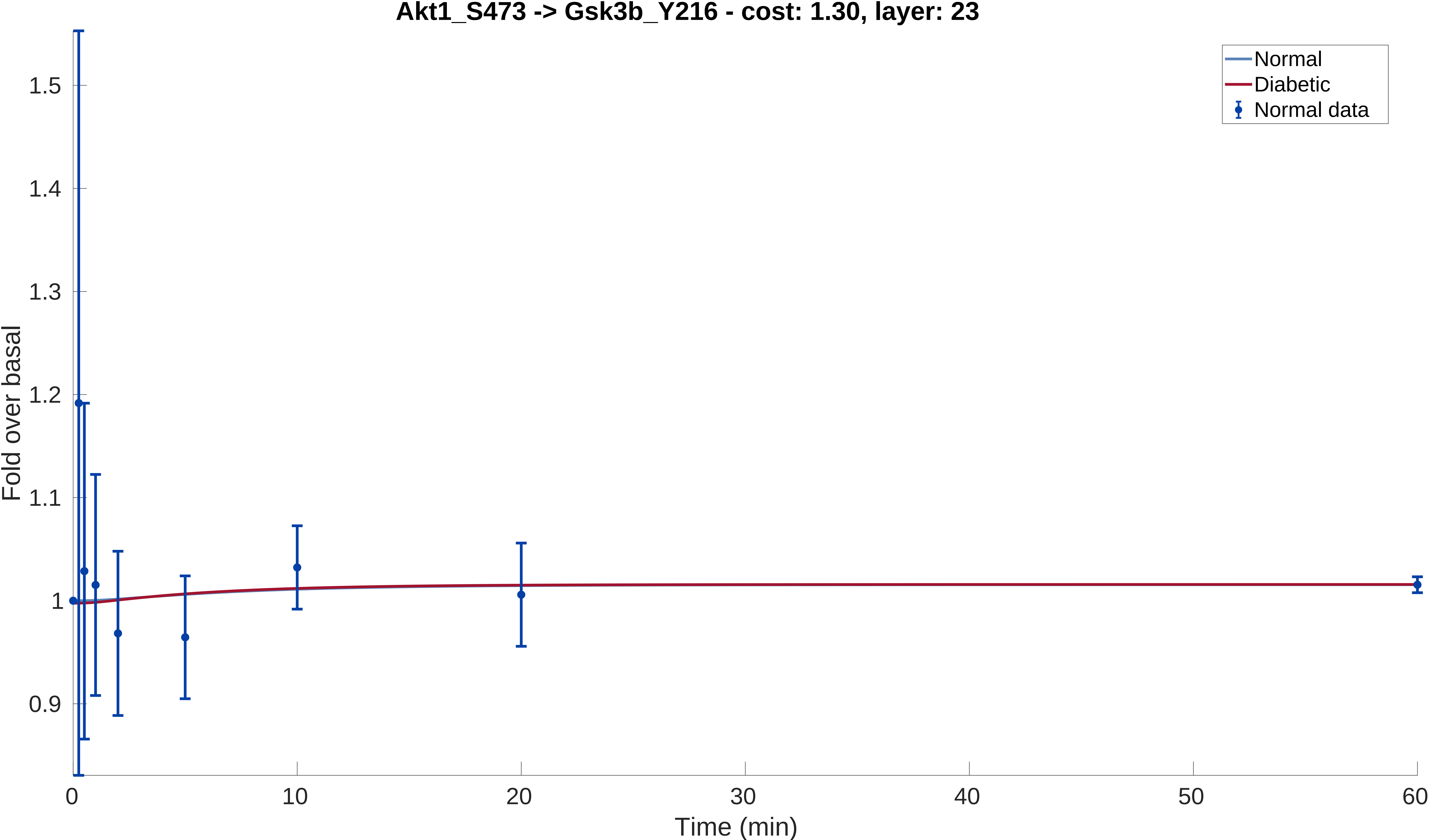

Akt1_S473 -> Gsk3b_Y216 - cost: 1.30, layer: 23
Normal
Diabetic
Normal data
Fold over basal
1.5
1.4
1.3
1.2
1.1
1
0.9
0
10
20
30
40
50
60
Time (min)

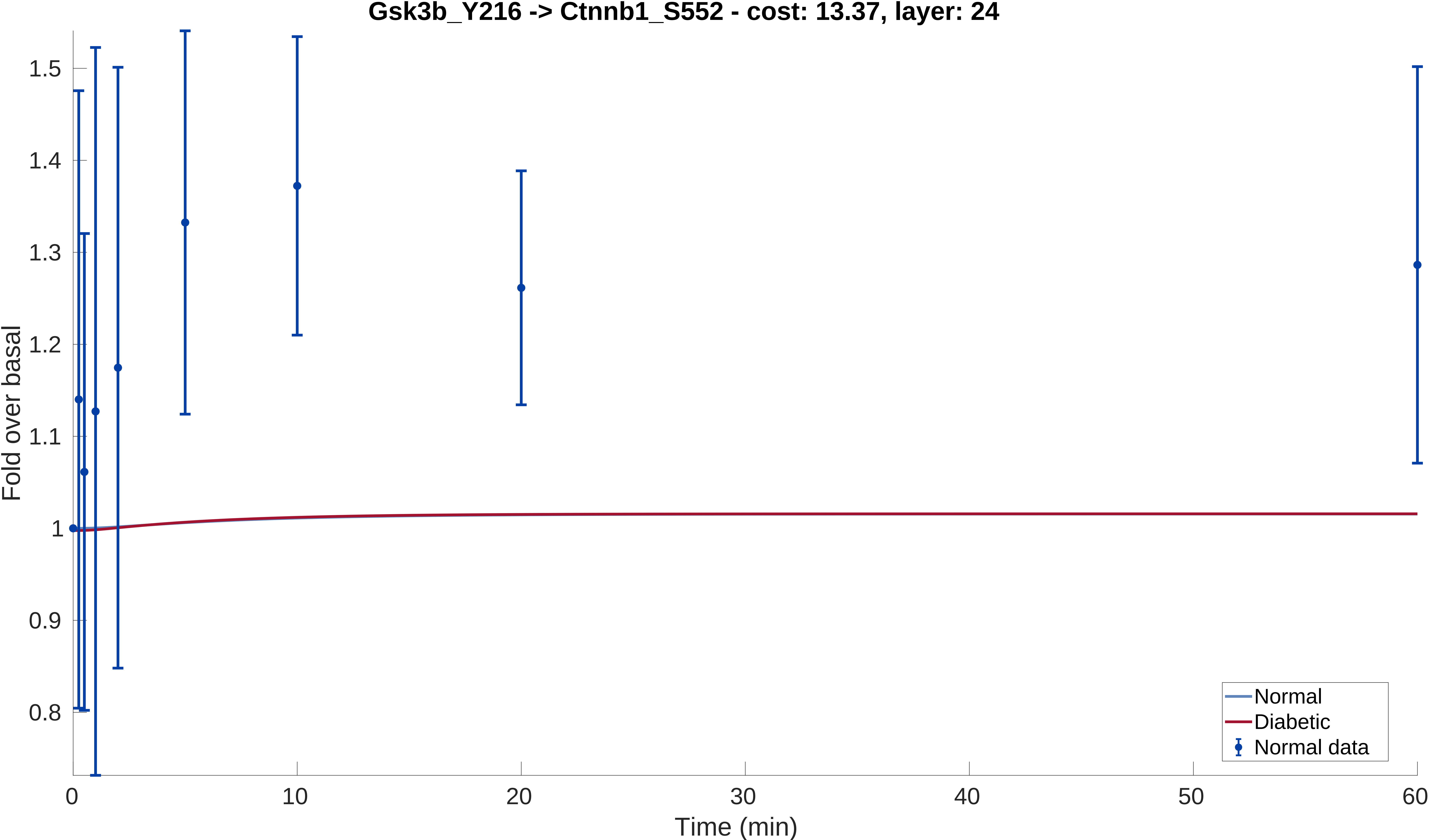

Gsk3b_Y216 -> Ctnnb1_S552 - cost: 13.37, layer: 24
1.5
1.4
1.3
1.2
1.1
1
0.9
0.8
Fold over basal
0
10
20
30
40
50
60
Time (min)
Normal
Diabetic
Normal data

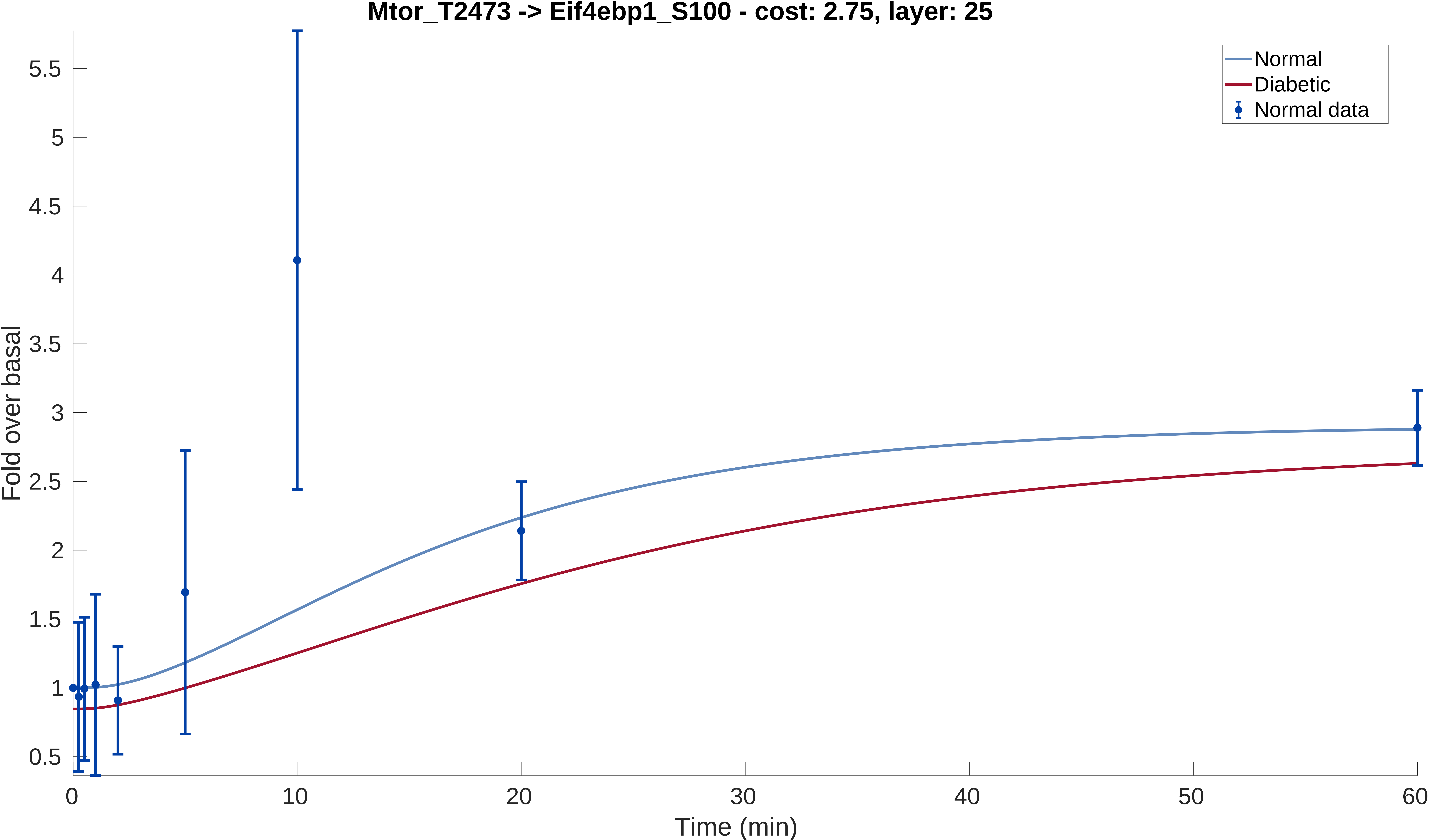

Mtor_T2473 -> Eif4ebp1_S100 - cost: 2.75, layer: 25
Normal
Diabetic
Normal data
Fold over basal
Time (min)
0.5
1
1.5
2
2.5
3
3.5
4
4.5
5
5.5
0
10
20
30
40
50
60

#### Mtor_S2448 -> Eif4ebp1_S64 - cost: 3.16, layer: 25

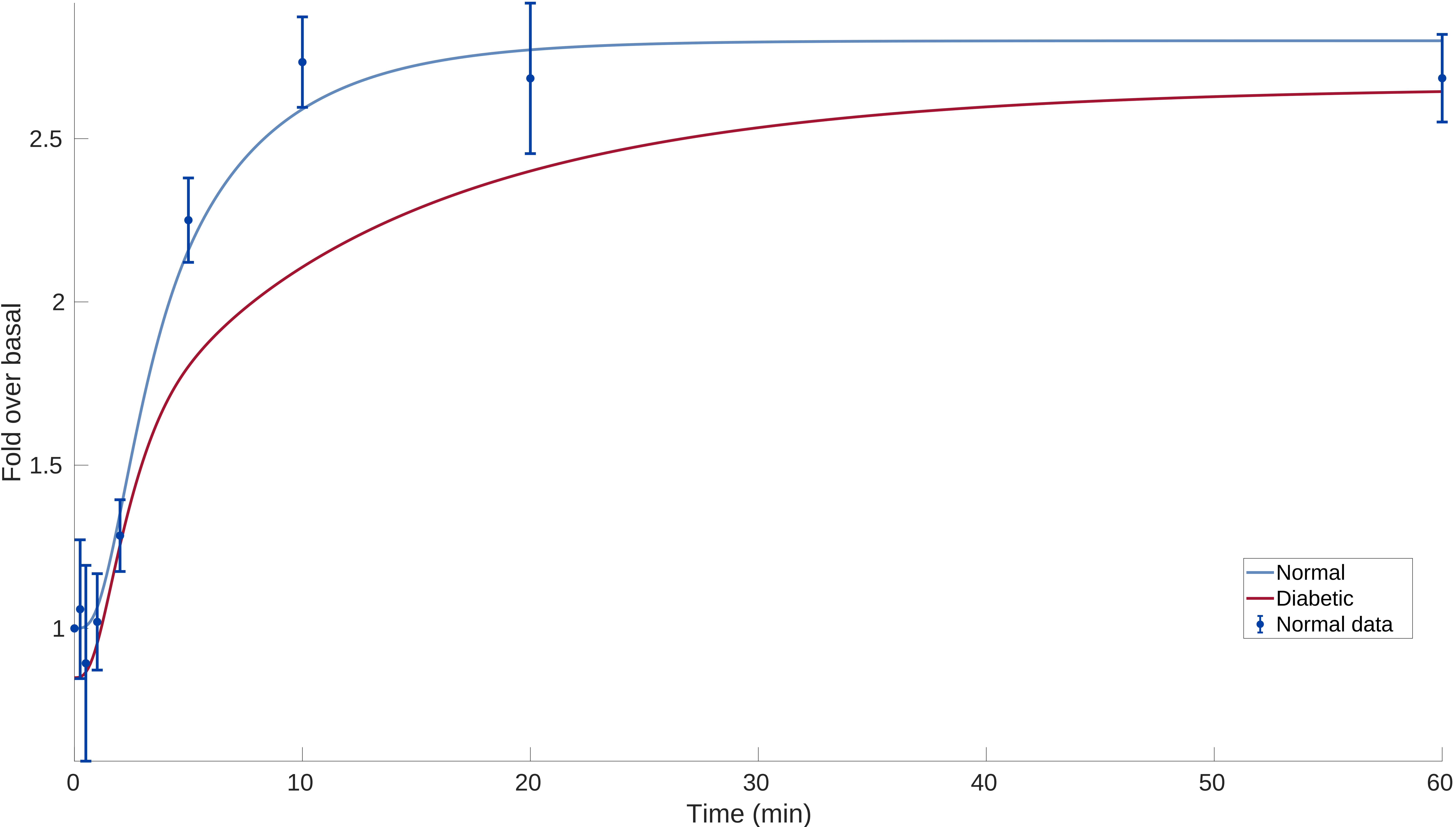

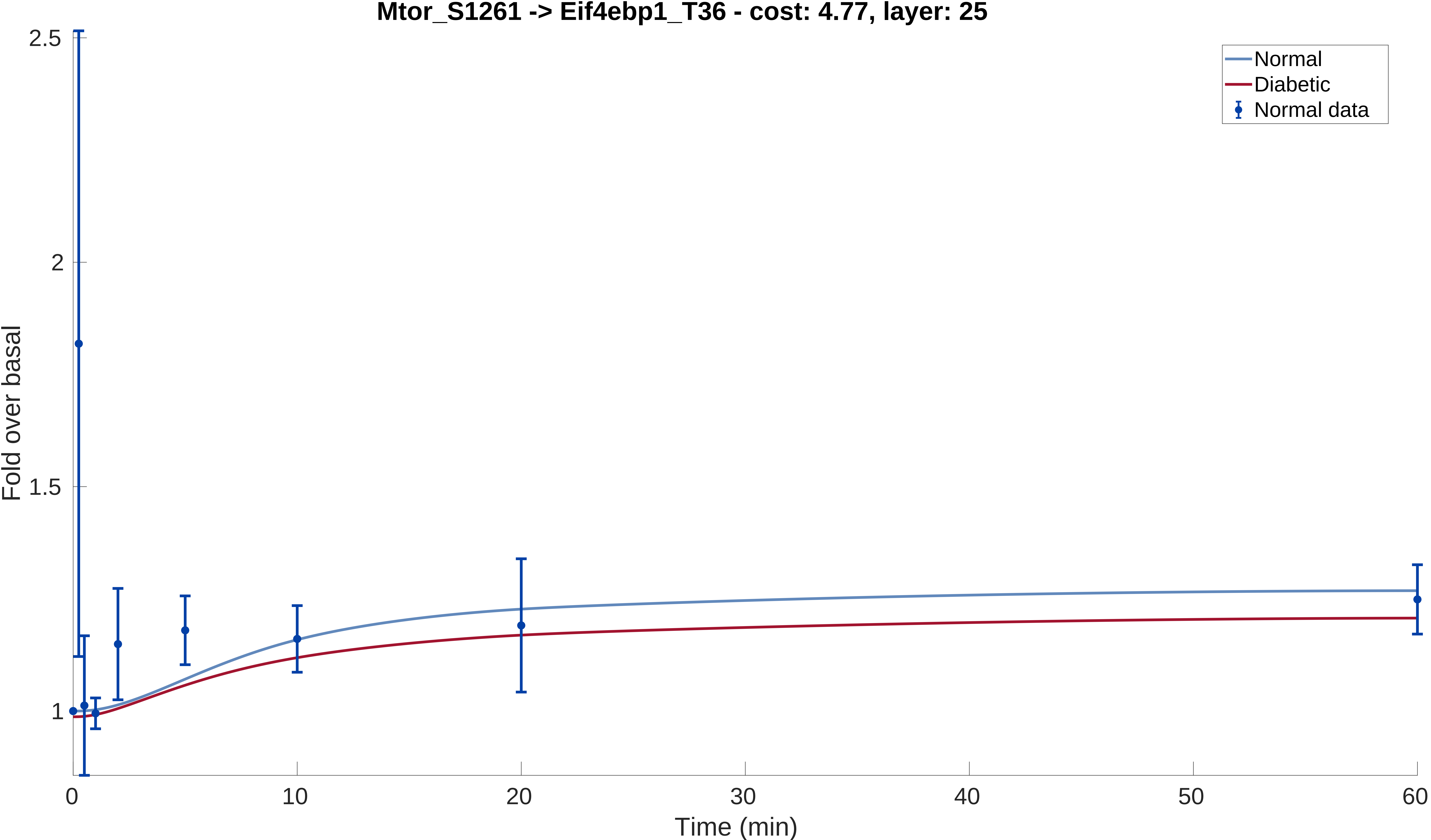

Mtor_S1261 -> Eif4ebp1_T36 - cost: 4.77, layer: 25
Normal
Diabetic
Normal data
Fold over basal
2.5
2
1.5
1
0
10
20
30
40
50
60
Time (min)

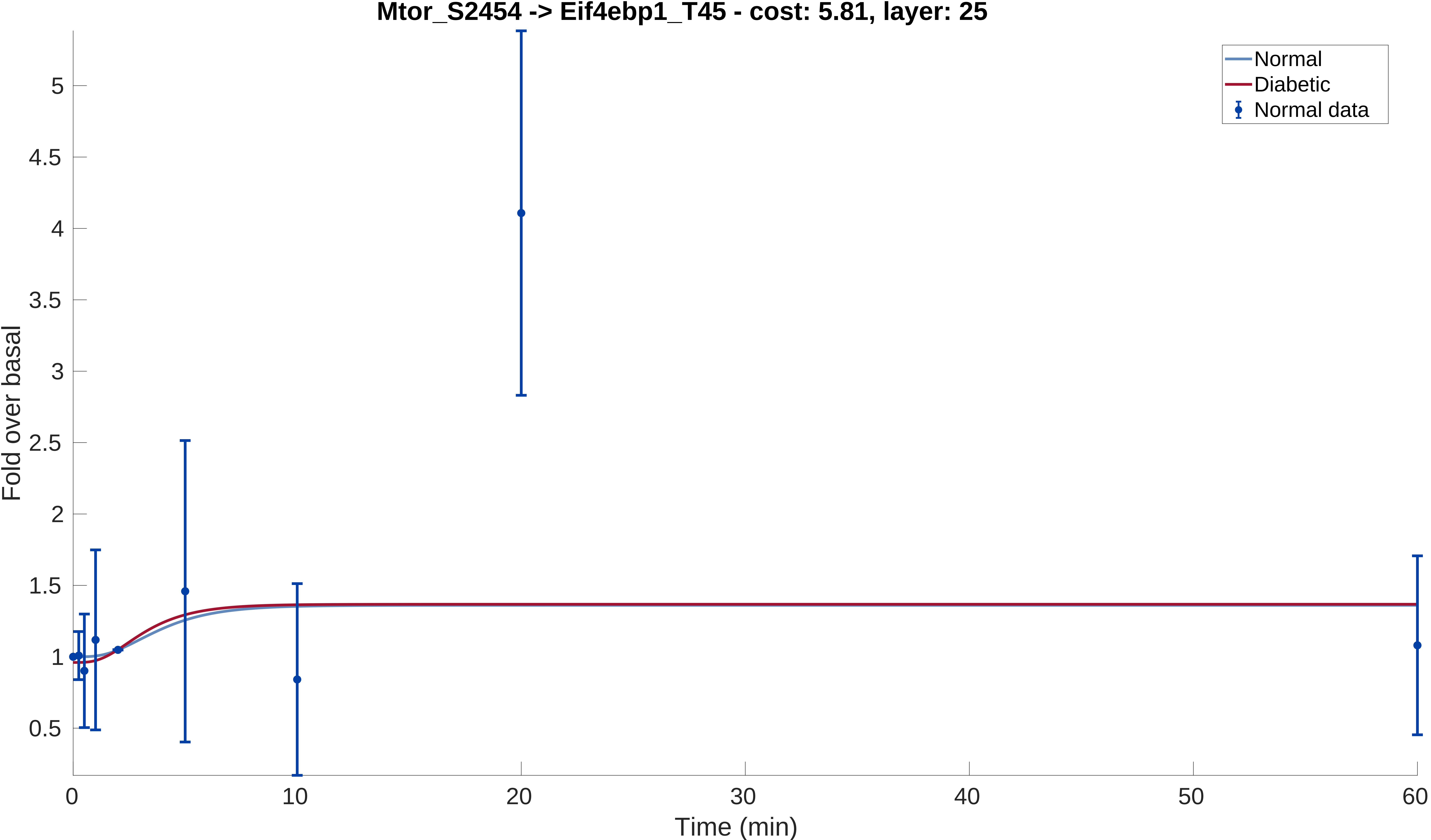

Mtor_S2454 -> Eif4ebp1_T45 - cost: 5.81, layer: 25
Normal
Diabetic
Normal data
Fold over basal
Time (min)
0.5
1
1.5
2
2.5
3
3.5
4
4.5
5
0
10
20
30
40
50
60

### Akt1_S473 -> Foxo4_S230 - cost: 8.77, layer: 26

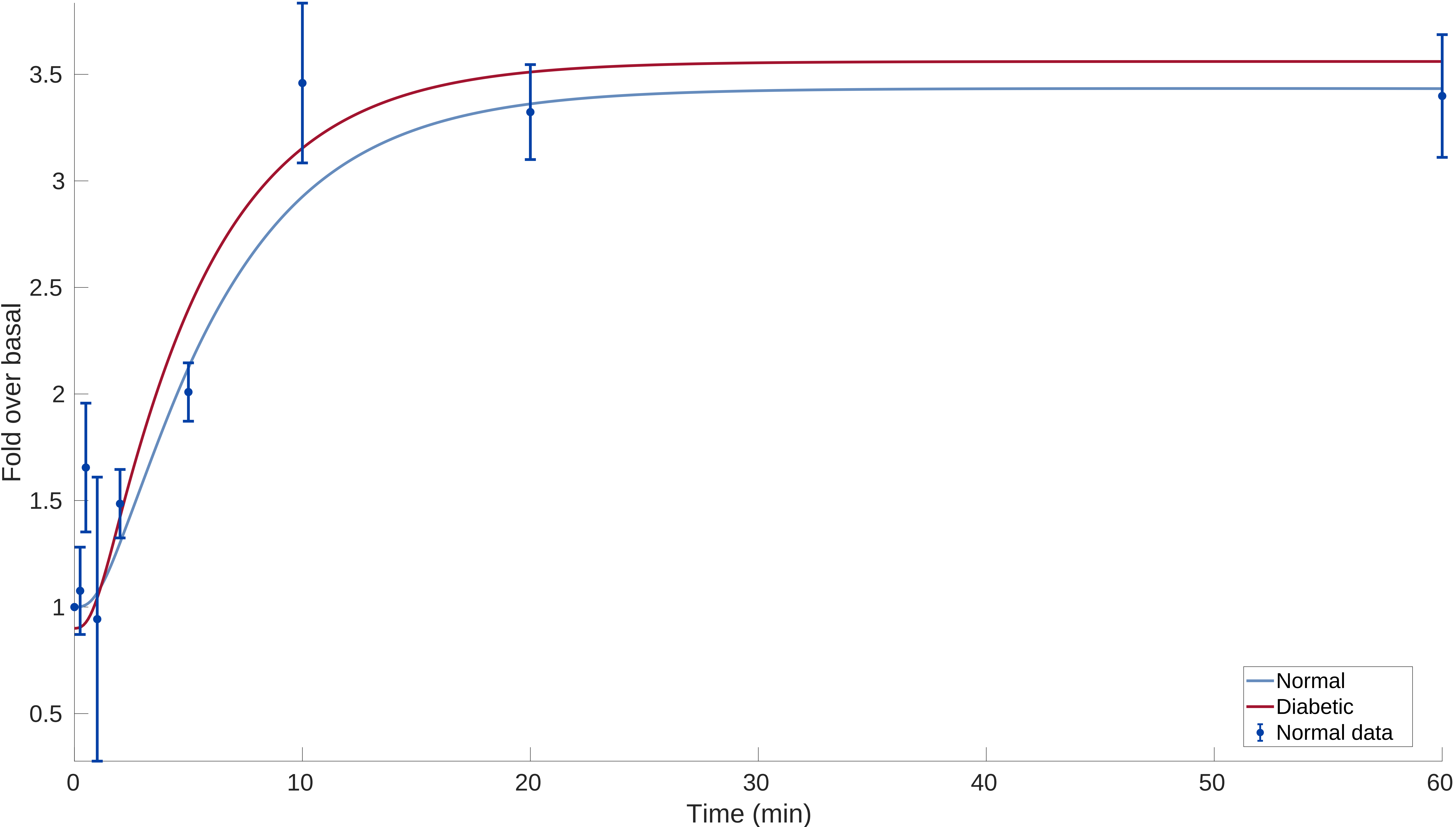

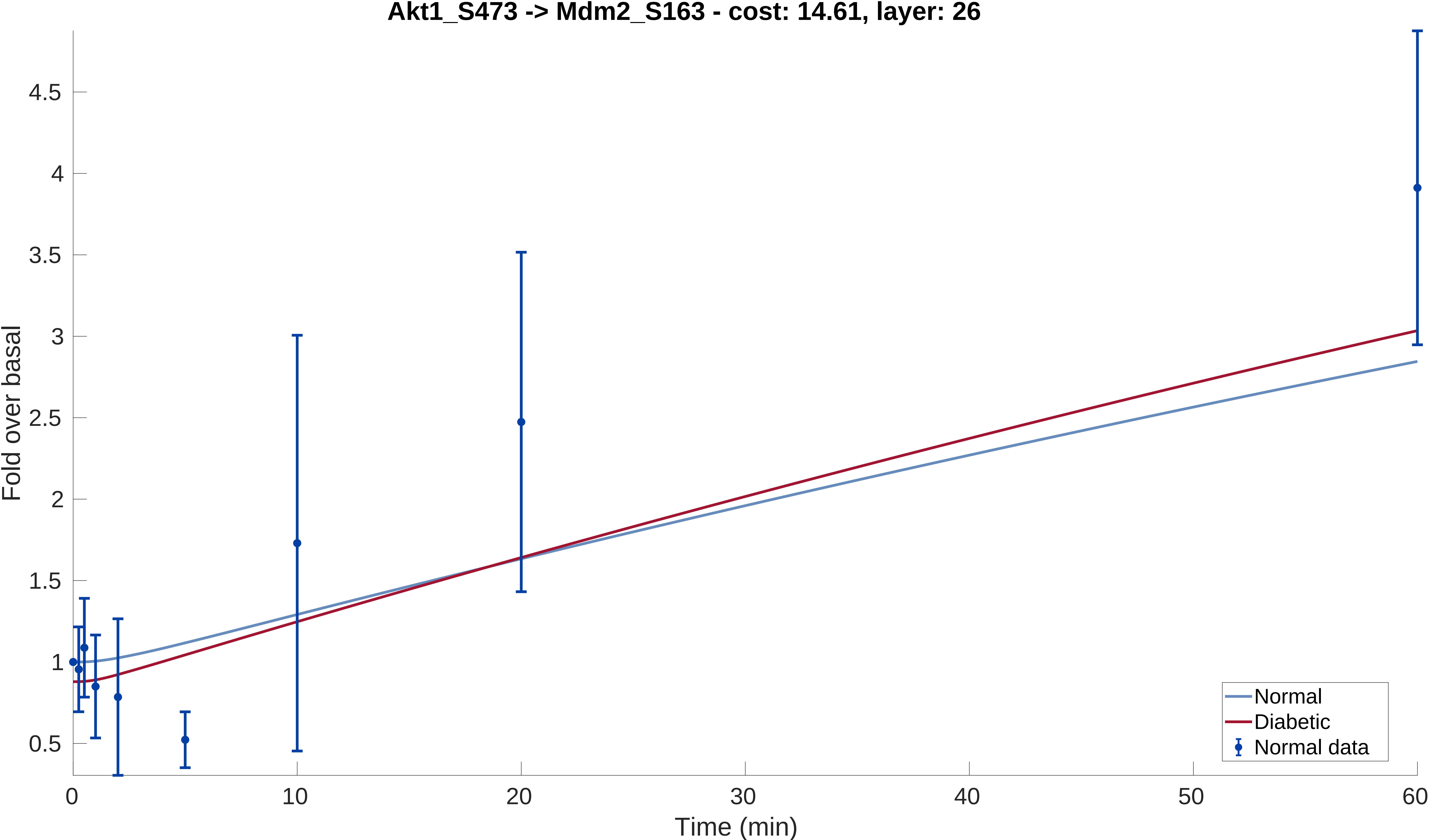

Akt1_S473 -> Mdm2_S163 - cost: 14.61, layer: 26
Fold over basal
Time (min)
0
10
20
30
40
50
60
0.5
1
1.5
2
2.5
3
3.5
4
4.5
Normal
Diabetic
Normal data

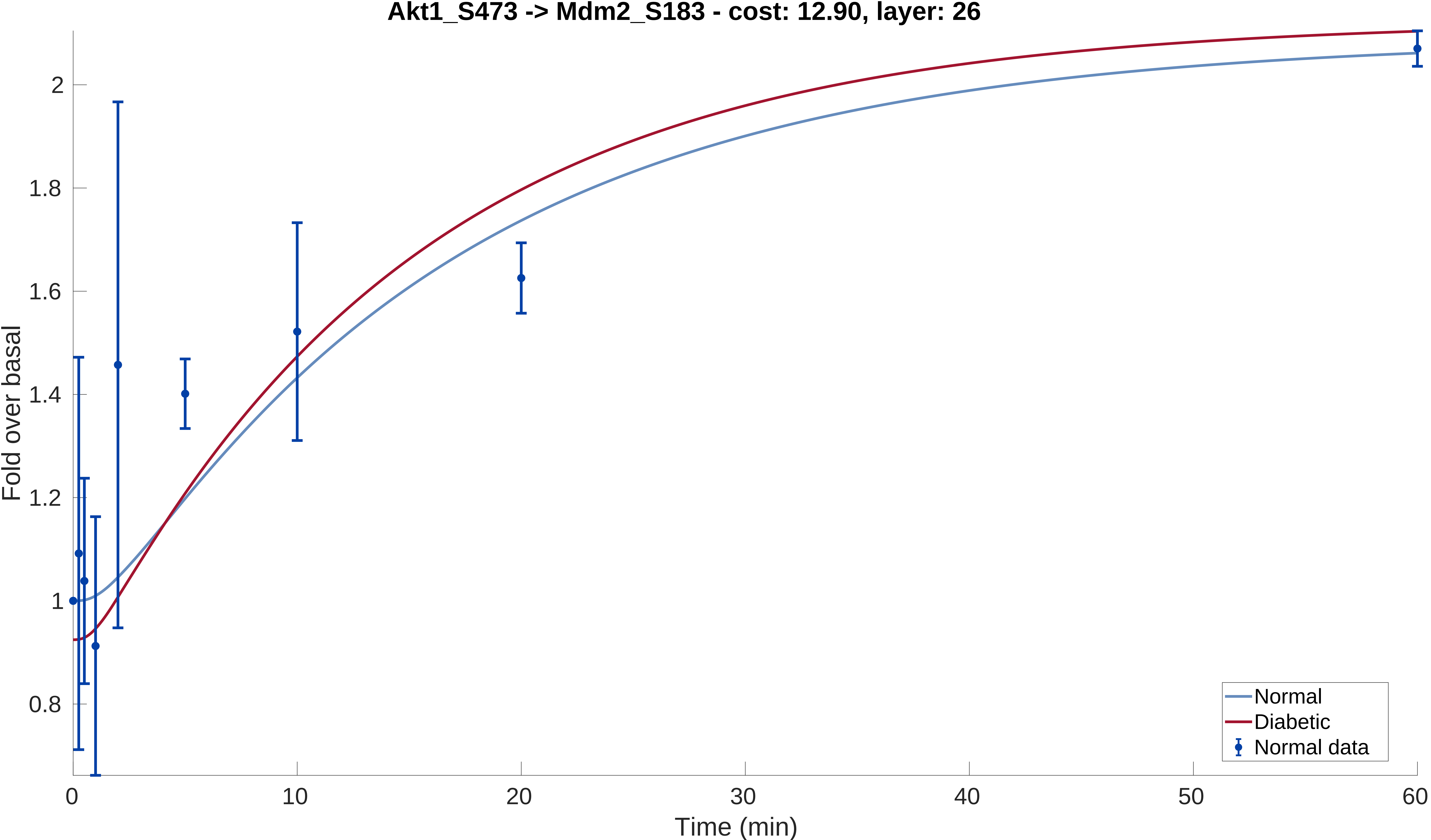
Akt1_S473 -> Mdm2_S183 - cost: 12.90, layer: 26
Fold over basal
0.8
1
1.2
1.4
1.6
1.8
2
0
10
20
30
40
50
60
Time (min)
Normal
Diabetic
Normal data

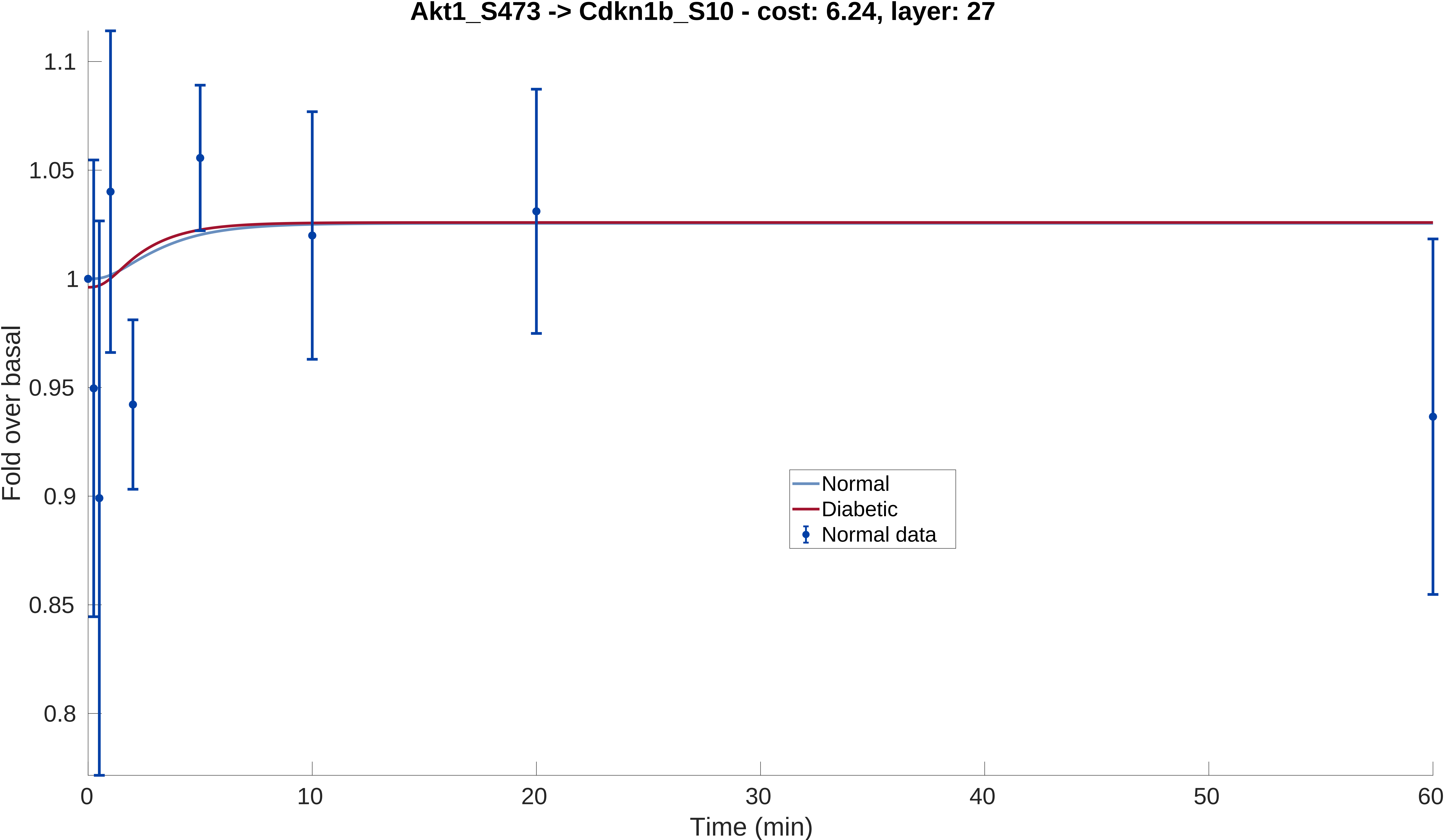

Akt1_S473 -> Cdkn1b_S10 - cost: 6.24, layer: 27
Fold over basal
Time (min)
1.1
1.05
1
0.95
0.9
0.85
0.8
0
10
20
30
40
50
60
Normal
Diabetic
Normal data

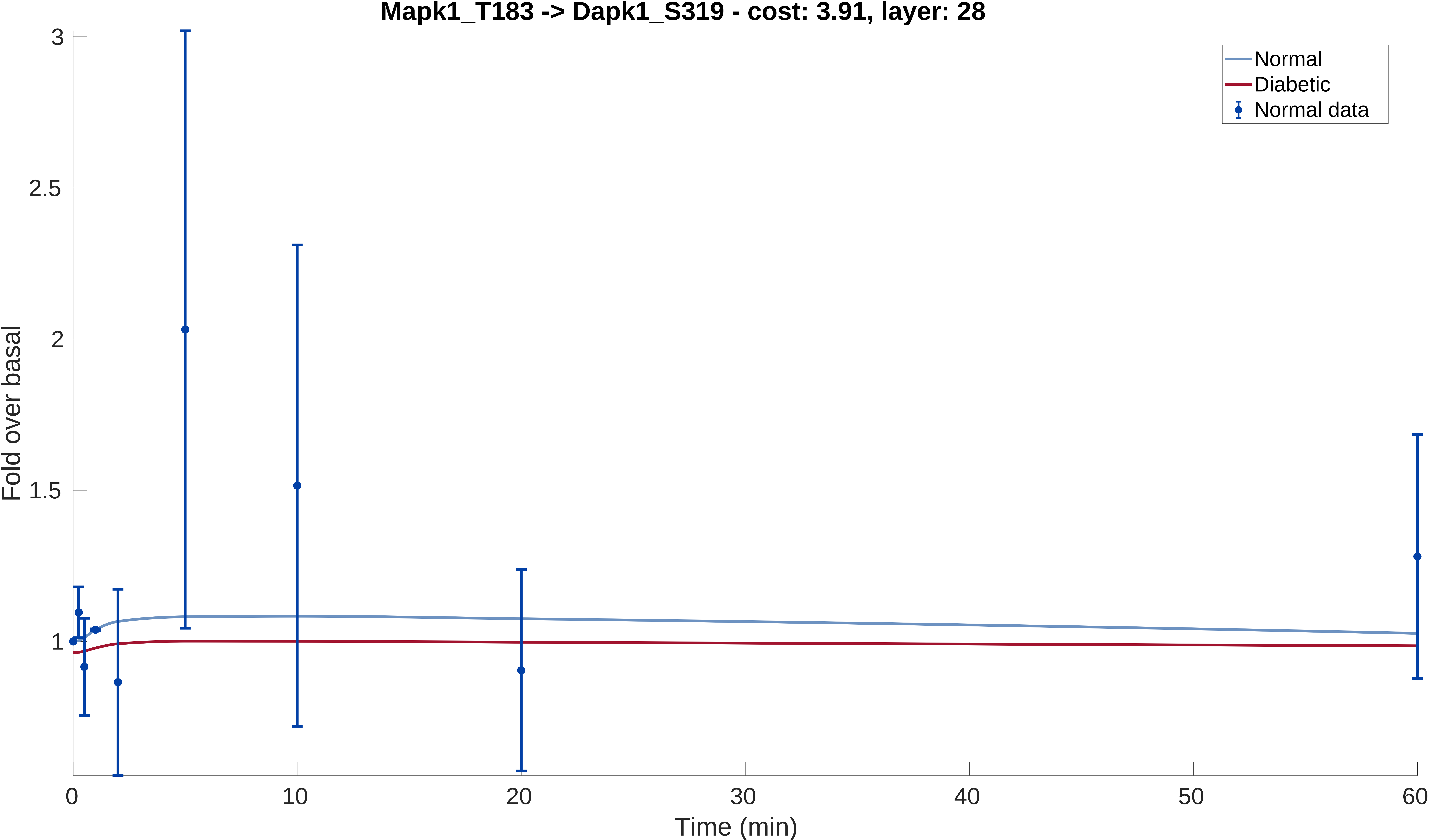
Mapk1_T183 -> Dapk1_S319 - cost: 3.91, layer: 28
Normal
Diabetic
Normal data
Fold over basal
3
2.5
2
1.5
1
0
10
20
30
40
50
60
Time (min)

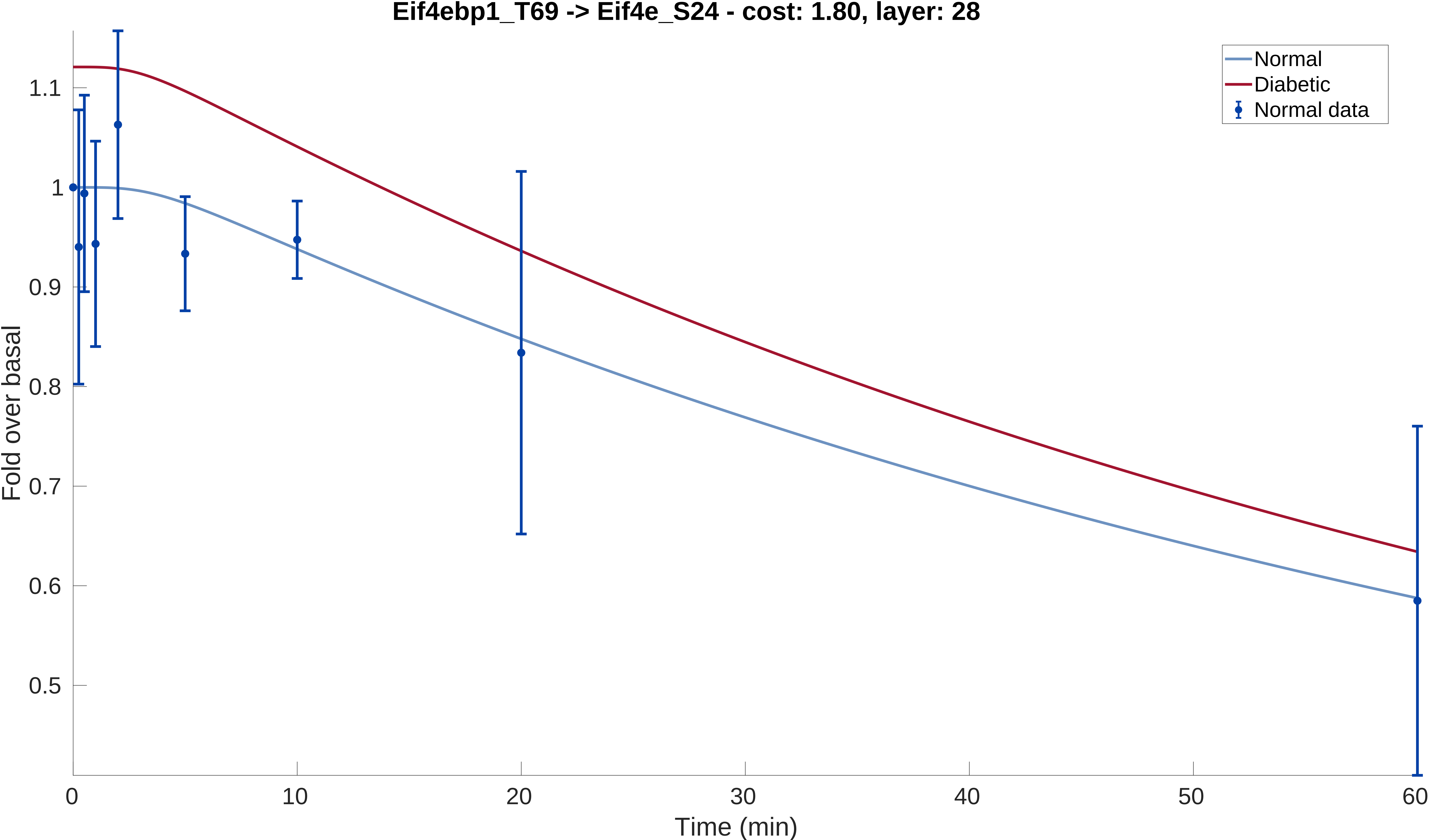

Eif4ebp1_T69 -> Eif4e_S24 - cost: 1.80, layer: 28
Normal
Diabetic
Normal data
1.1
1
0.9
0.8
0.7
0.6
0.5
Fold over basal
0
10
20
30
40
50
60
Time (min)

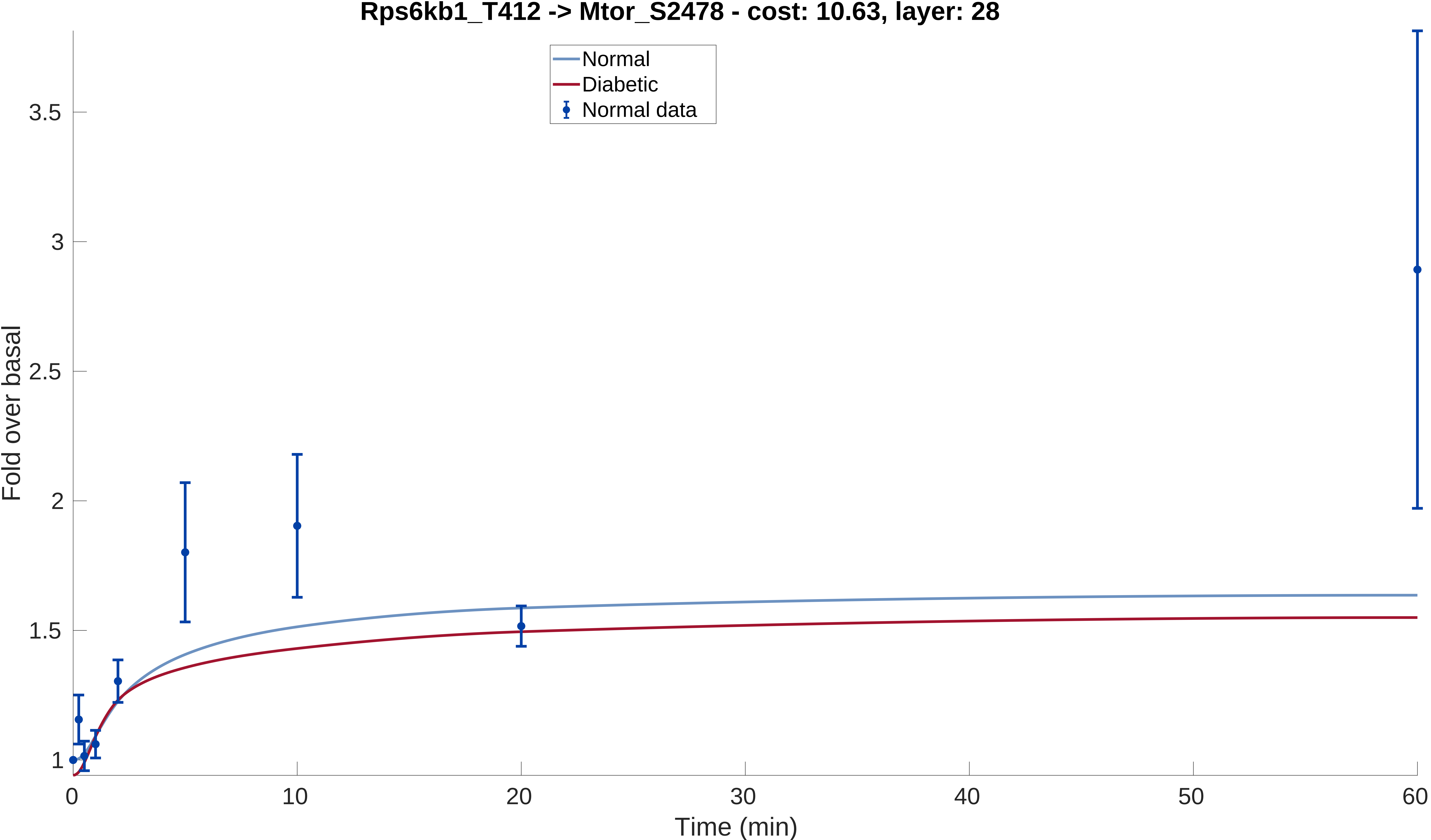
Rps6kb1_T412 -> Mtor_S2478 - cost: 10.63, layer: 28
Normal
Diabetic
Normal data
Fold over basal
Time (min)
1
1.5
2
2.5
3
3.5
0
10
20
30
40
50
60

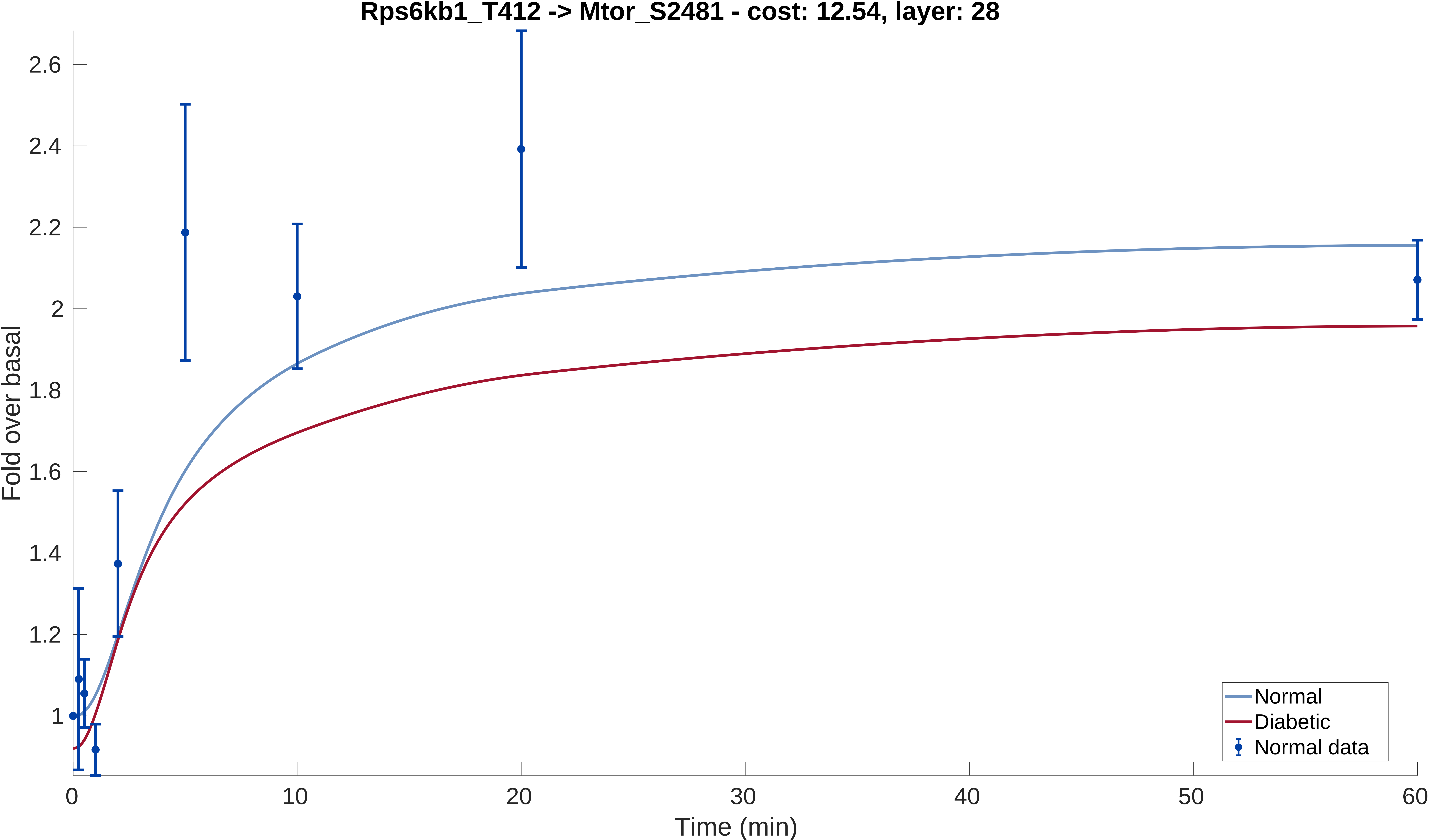
Rps6kb1_T412 -> Mtor_S2481 - cost: 12.54, layer: 28
Fold over basal
2.6
2.4
2.2
2
1.8
1.6
1.4
1.2
1
0
10
20
30
40
50
60
Time (min)
Normal
Diabetic
Normal data

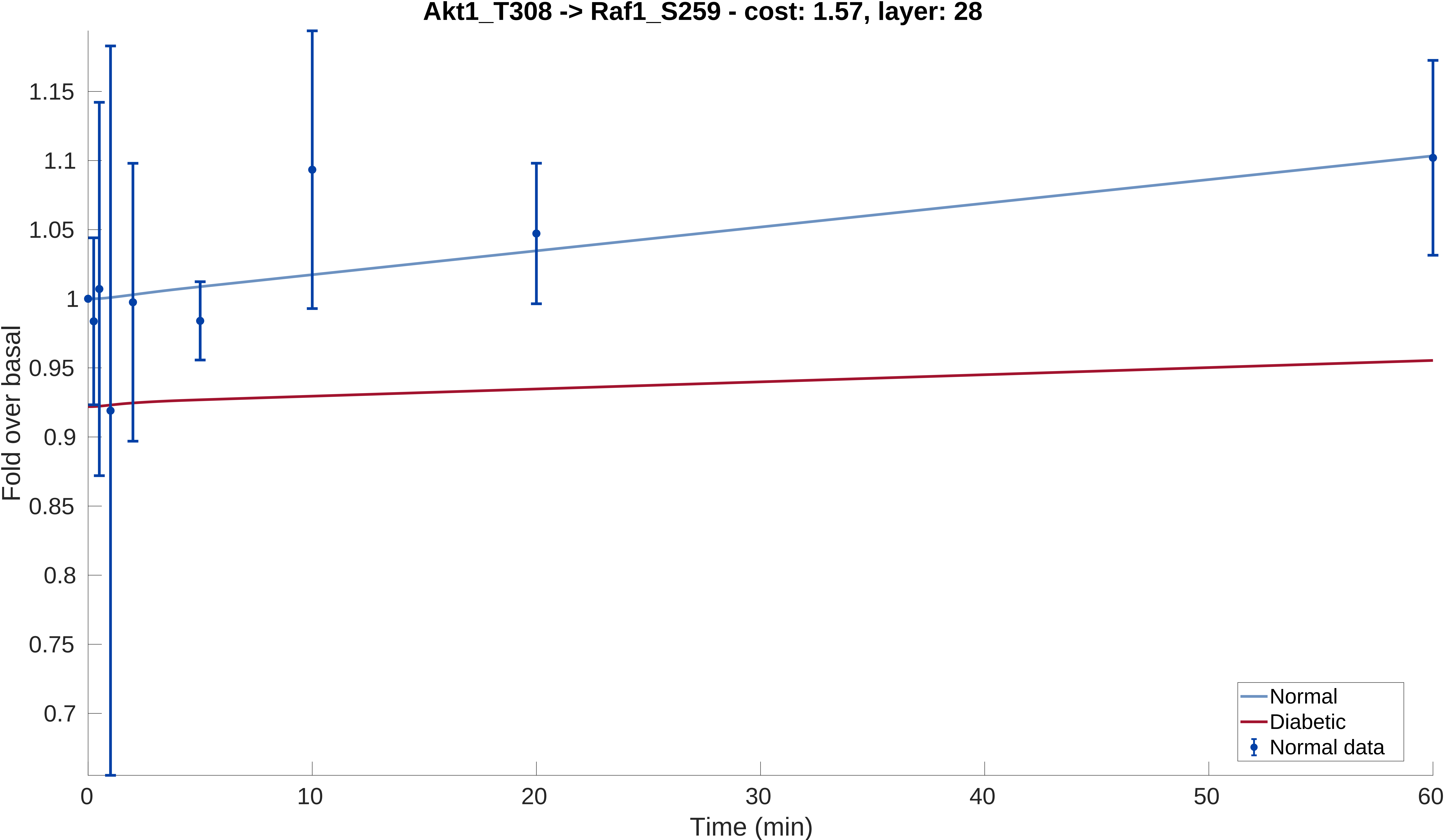

Akt1_T308 -> Raf1_S259 - cost: 1.57, layer: 28
Fold over basal
1.15
1.1
1.05
1
0.95
0.9
0.85
0.8
0.75
0.7
0
10
20
30
40
50
60
Time (min)
Normal
Diabetic
Normal data

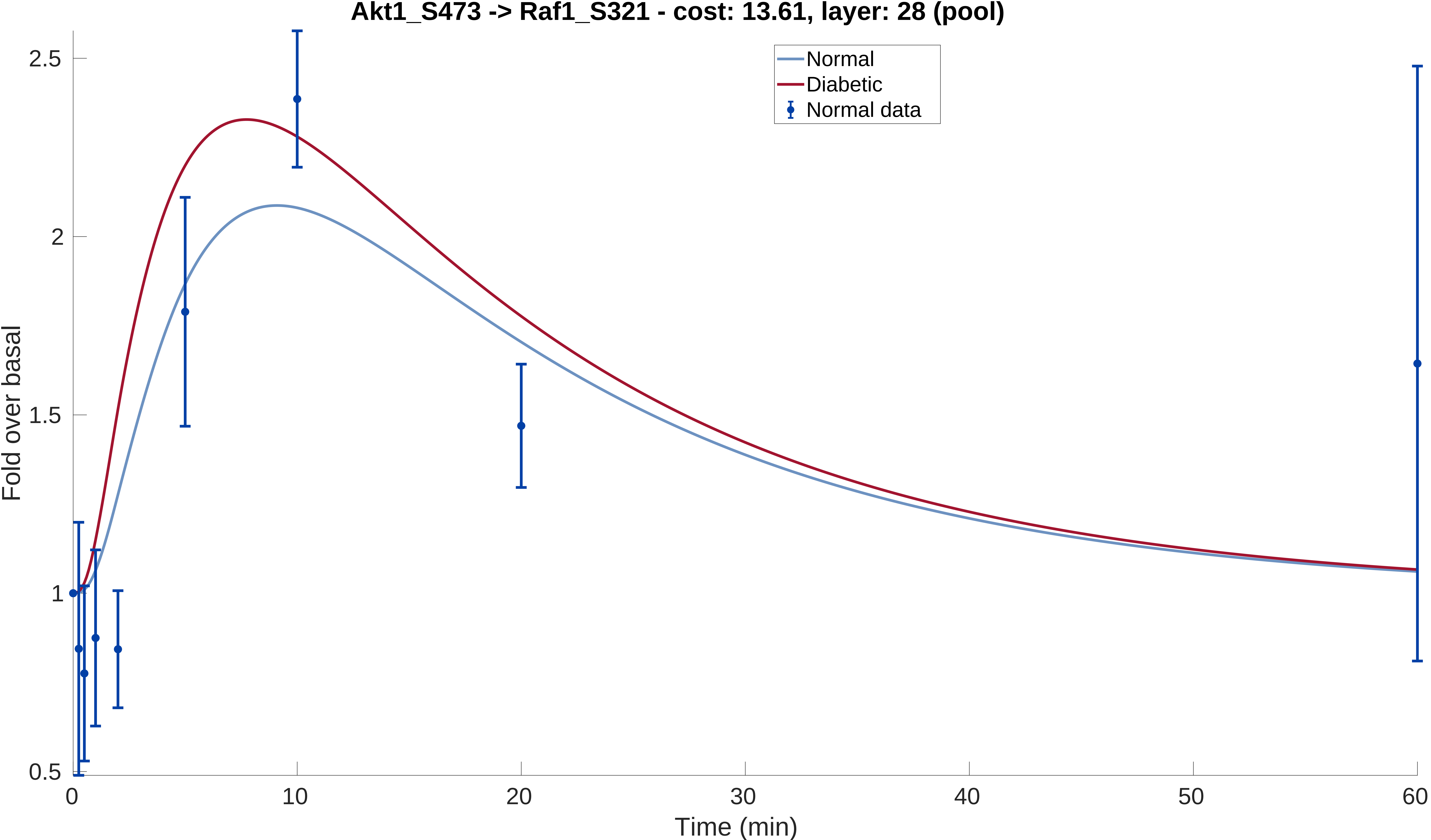

Akt1_S473 -> Raf1_S321 - cost: 13.61, layer: 28 (pool)
Normal
Diabetic
Normal data
Fold over basal
2.5
2
1.5
1
0.5
0
10
20
30
40
50
60
Time (min)

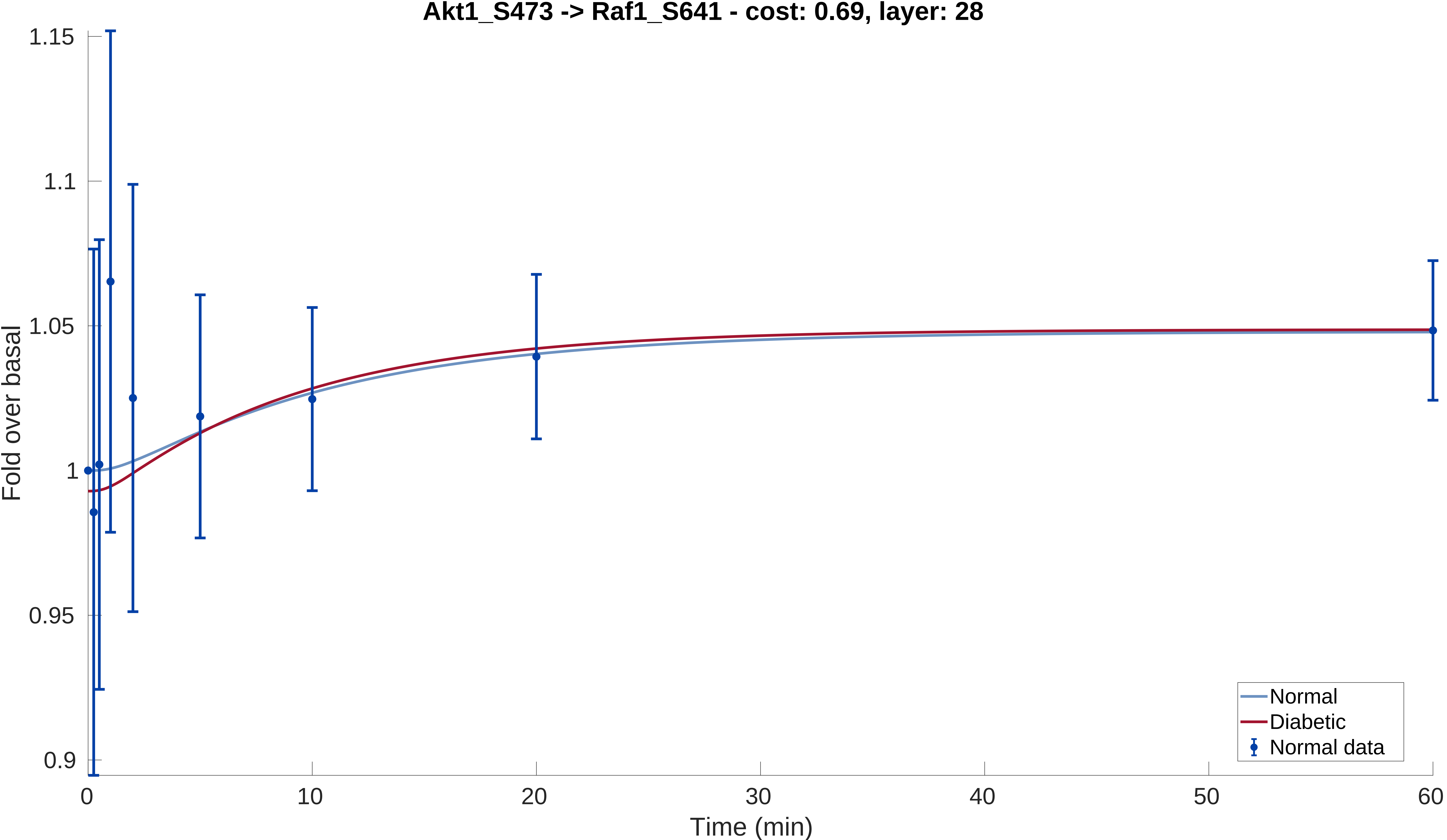

Akt1_S473 -> Raf1_S641 - cost: 0.69, layer: 28
1.15
1.1
1.05
1
0.95
0.9
Fold over basal
0
10
20
30
40
50
60
Time (min)
Normal
Diabetic
Normal data

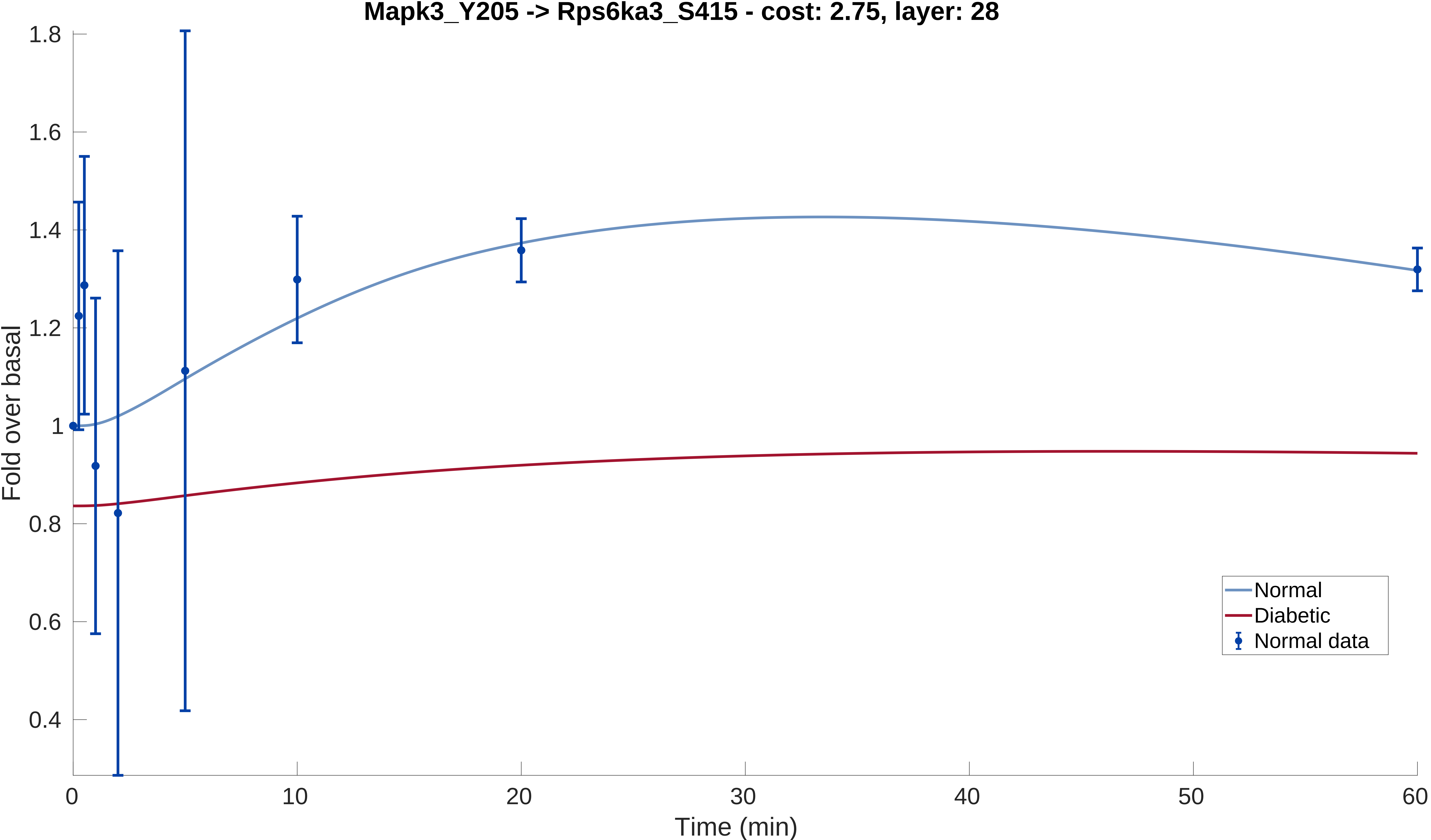

Mapk3_Y205 -> Rps6ka3_S415 - cost: 2.75, layer: 28
Fold over basal
Time (min)
0.4
0.6
0.8
1
1.2
1.4
1.6
1.8
0
10
20
30
40
50
60
Normal
Diabetic
Normal data

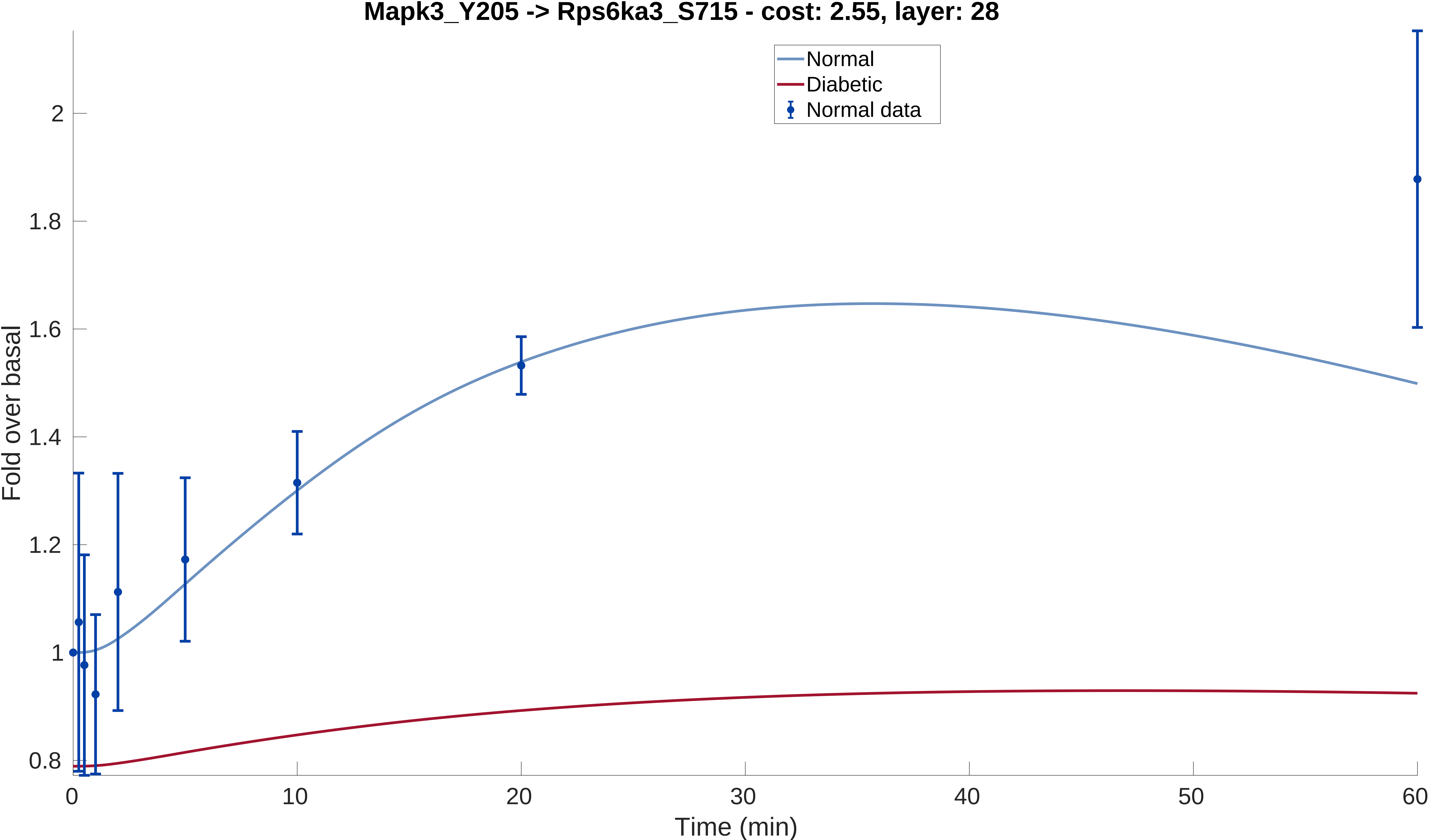

Mapk3_Y205 -> Rps6ka3_S715 - cost: 2.55, layer: 28
Normal
Diabetic
Normal data
Fold over basal
0.8
1
1.2
1.4
1.6
1.8
2
0
10
20
30
40
50
60
Time (min)

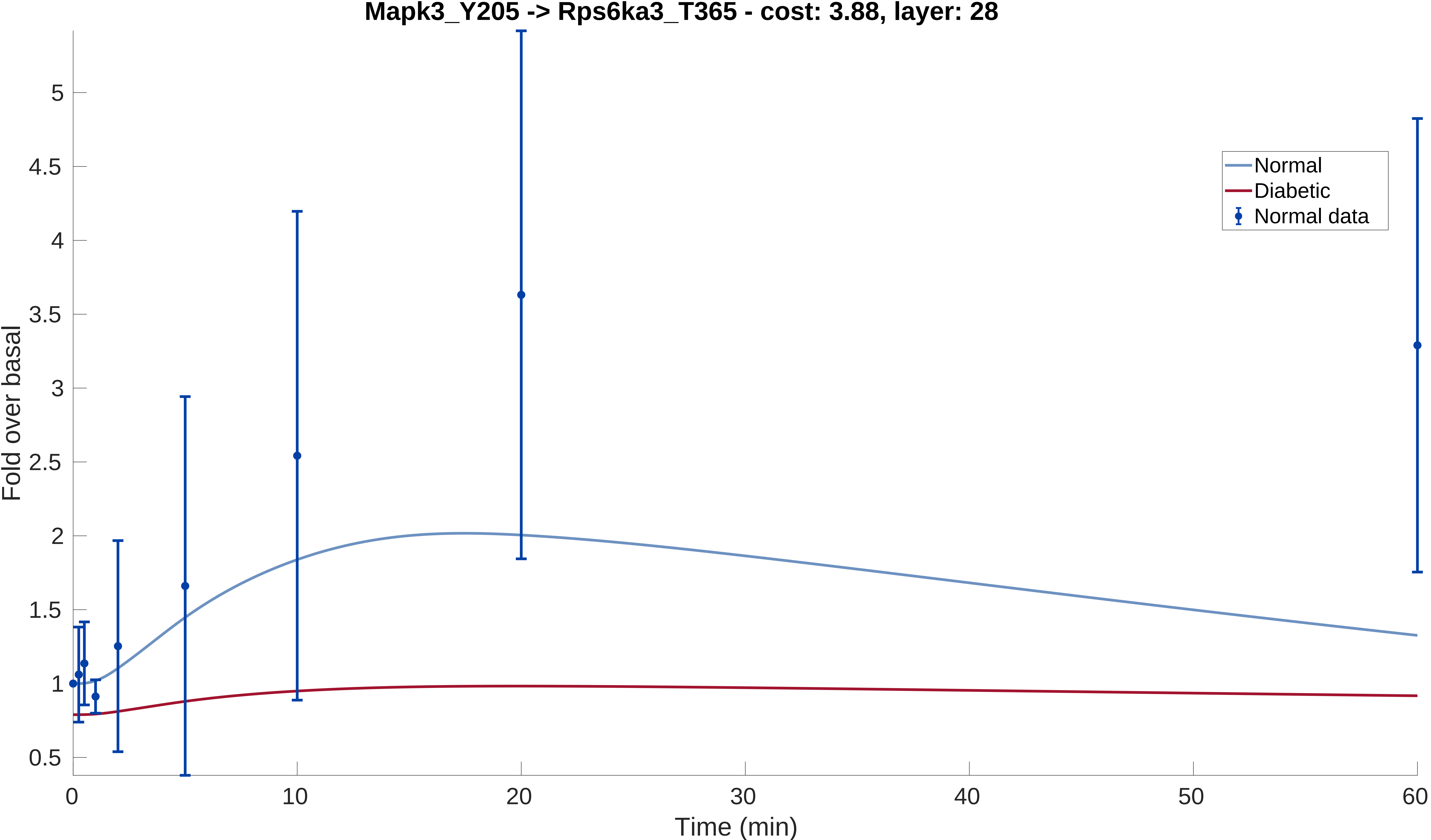

Mapk3_Y205 -> Rps6ka3_T365 - cost: 3.88, layer: 28
Fold over basal
Time (min)
Normal
Diabetic
Normal data
0.5
1
1.5
2
2.5
3
3.5
4
4.5
5
0
10
20
30
40
50
60

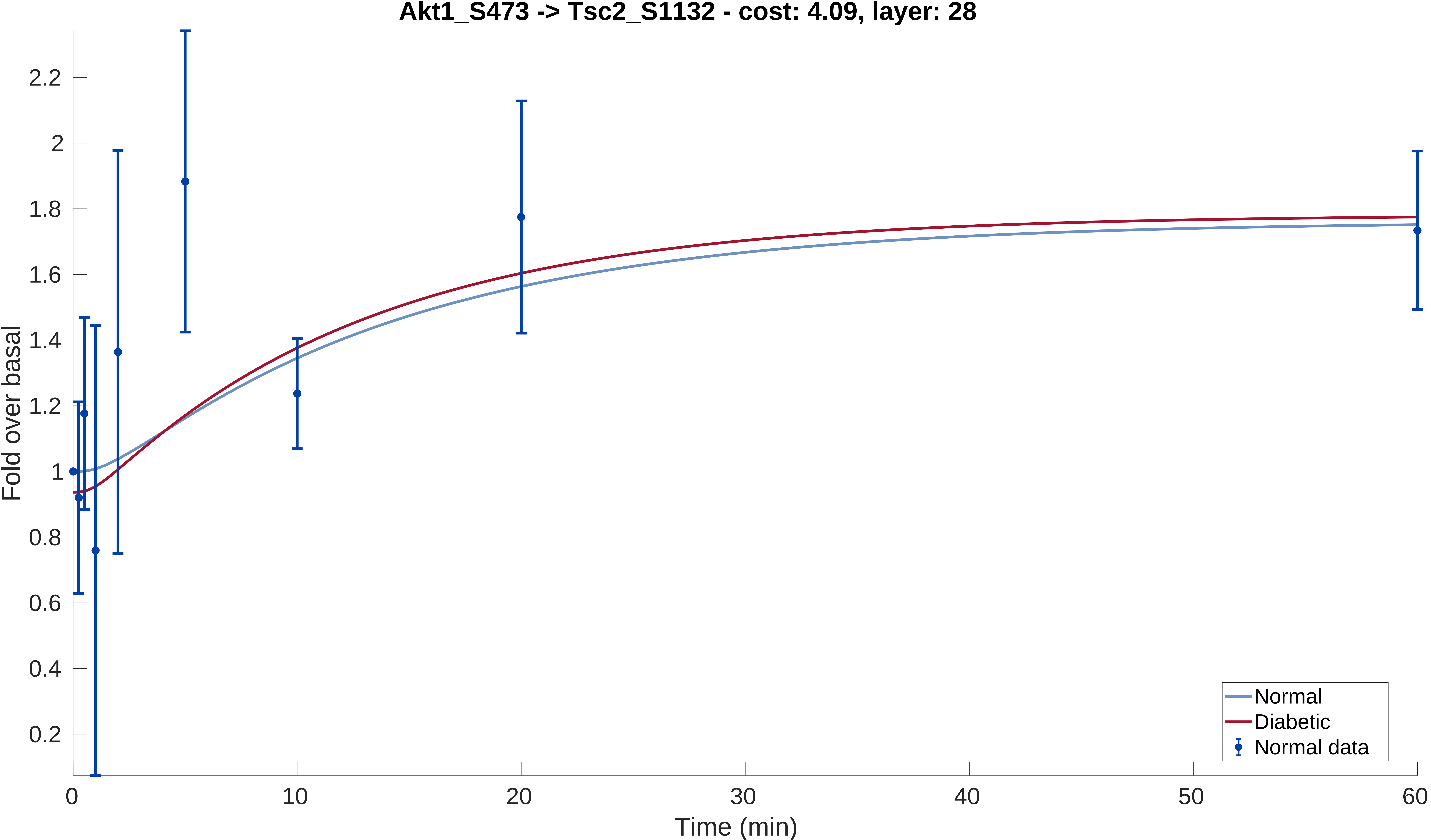

Akt1_S473 -> Tsc2_S1132 - cost: 4.09, layer: 28
Fold over basal
Time (min)
2.2
2
1.8
1.6
1.4
1.2
1
0.8
0.6
0.4
0.2
0
10
20
30
40
50
60
Normal
Diabetic
Normal data

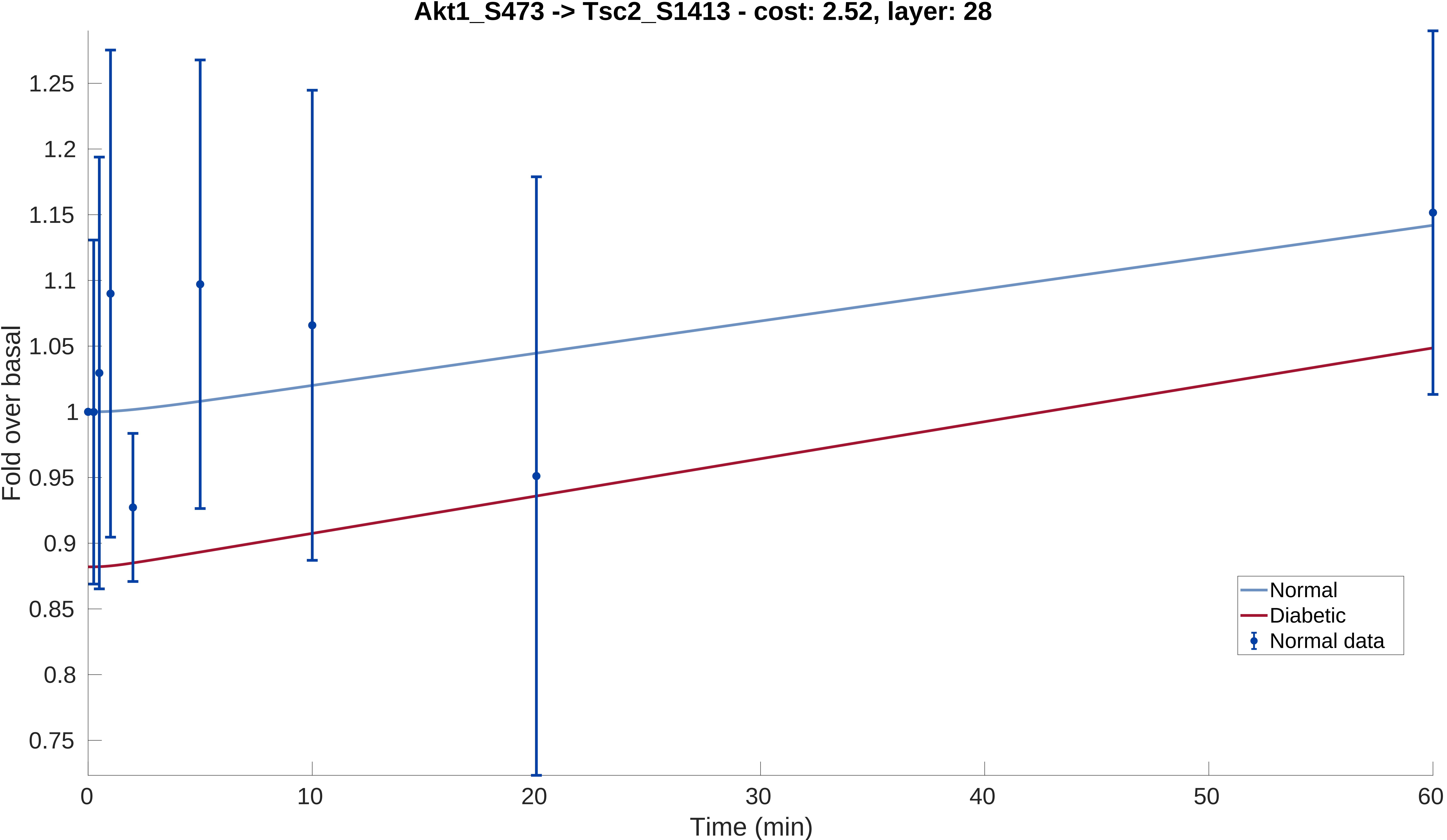

Akt1_S473 -> Tsc2_S1413 - cost: 2.52, layer: 28
Fold over basal
1.25
1.2
1.15
1.1
1.05
1
0.95
0.9
0.85
0.8
0.75
0
10
20
30
40
50
60
Time (min)
Normal
Diabetic
Normal data

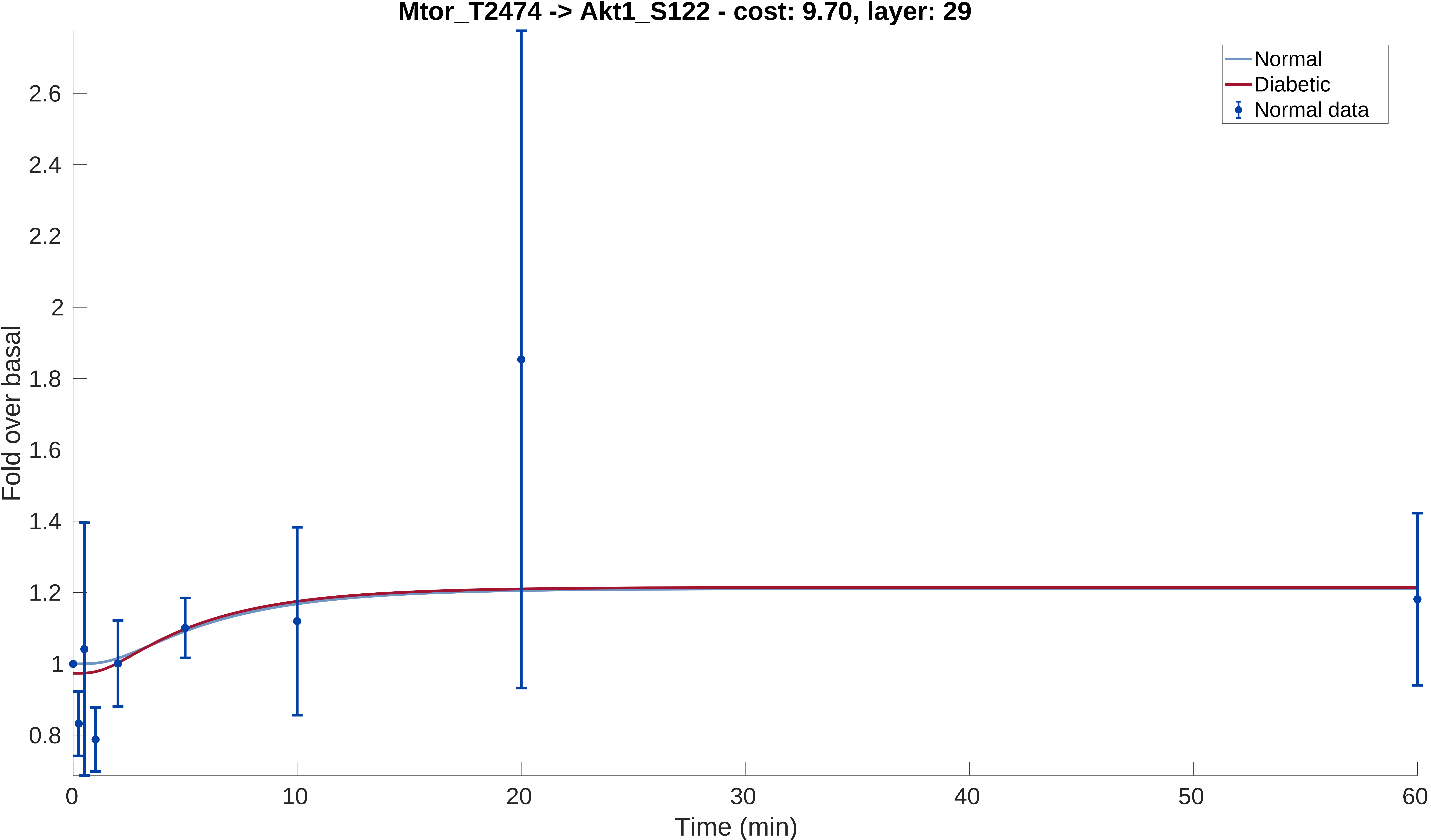

Mtor_T2474 -> Akt1_S122 - cost: 9.70, layer: 29
Normal
Diabetic
Normal data
Fold over basal
2.6
2.4
2.2
2
1.8
1.6
1.4
1.2
1
0.8
0
10
20
30
40
50
60
Time (min)

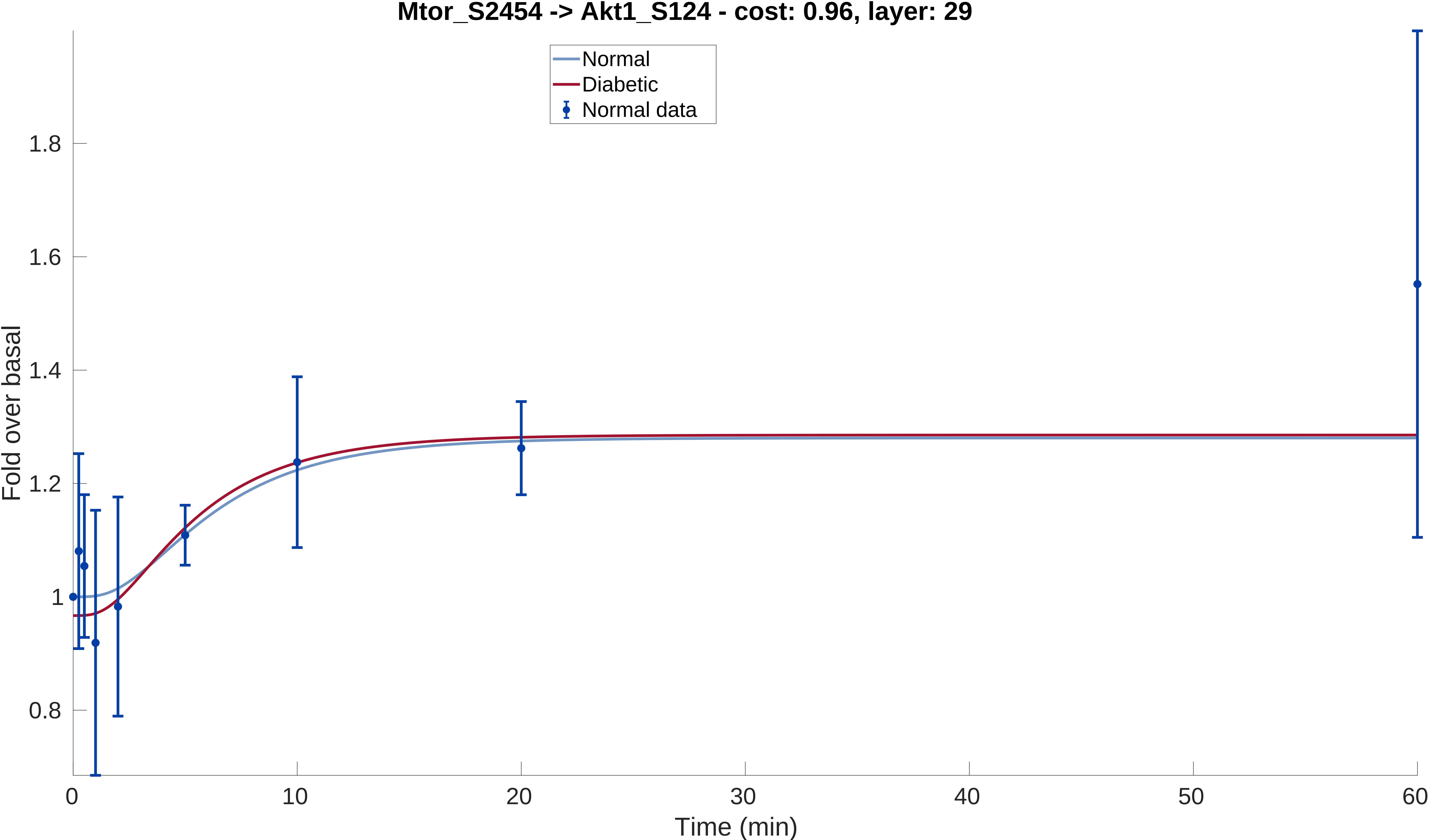
Mtor_S2454 -> Akt1_S124 - cost: 0.96, layer: 29
Normal
Diabetic
Normal data
Fold over basal
0.8
1
1.2
1.4
1.6
1.8
0
10
20
30
40
50
60
Time (min)

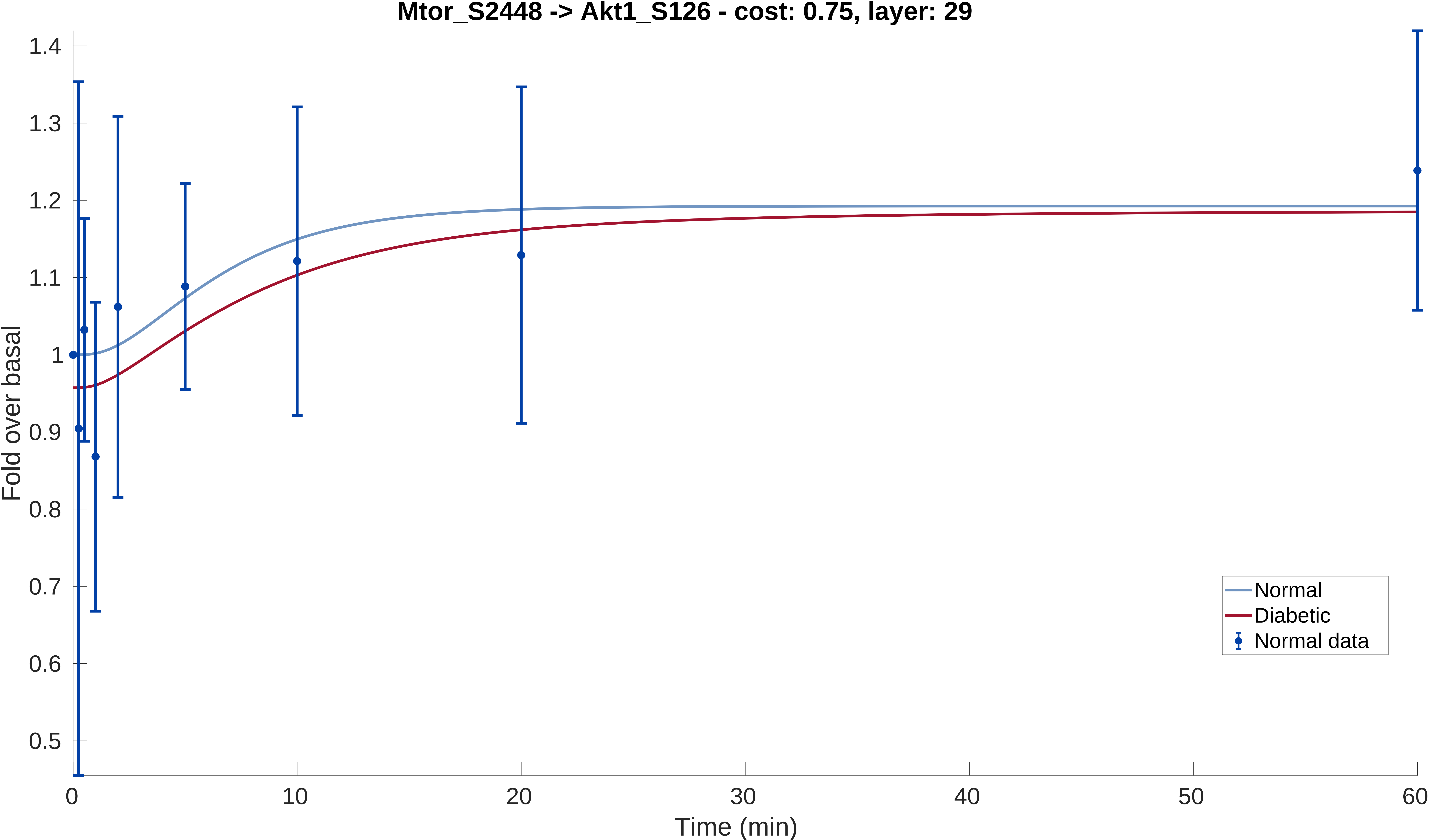
Mtor_S2448 -> Akt1_S126 - cost: 0.75, layer: 29
Fold over basal
1.4
1.3
1.2
1.1
1
0.9
0.8
0.7
0.6
0.5
0
10
20
30
40
50
60
Time (min)
Normal
Diabetic
Normal data

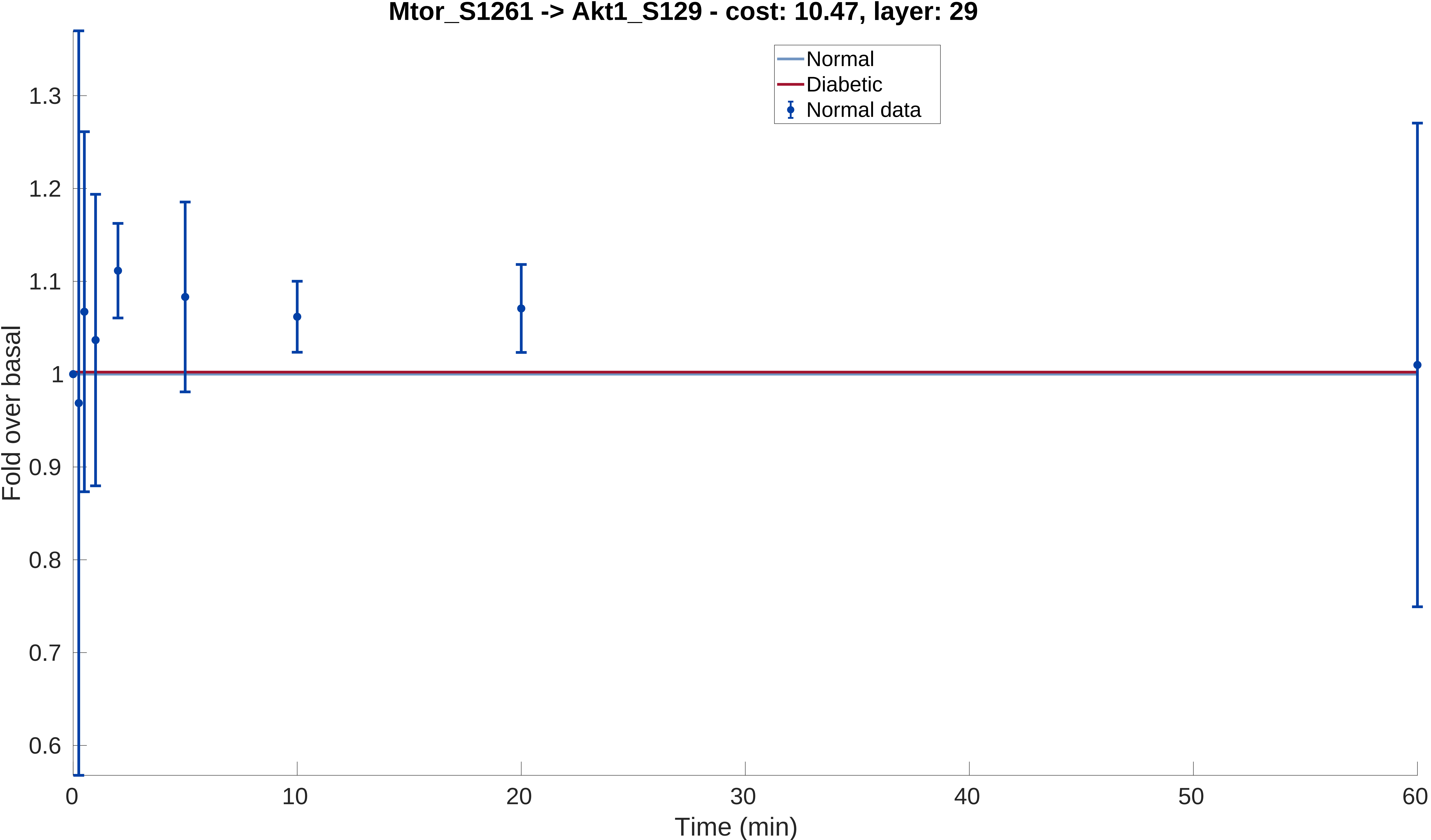

Mtor_S1261 -> Akt1_S129 - cost: 10.47, layer: 29
Normal
Diabetic
Normal data
1.3
1.2
1.1
1
0.9
0.8
0.7
0.6
Fold over basal
0
10
20
30
40
50
60
Time (min)

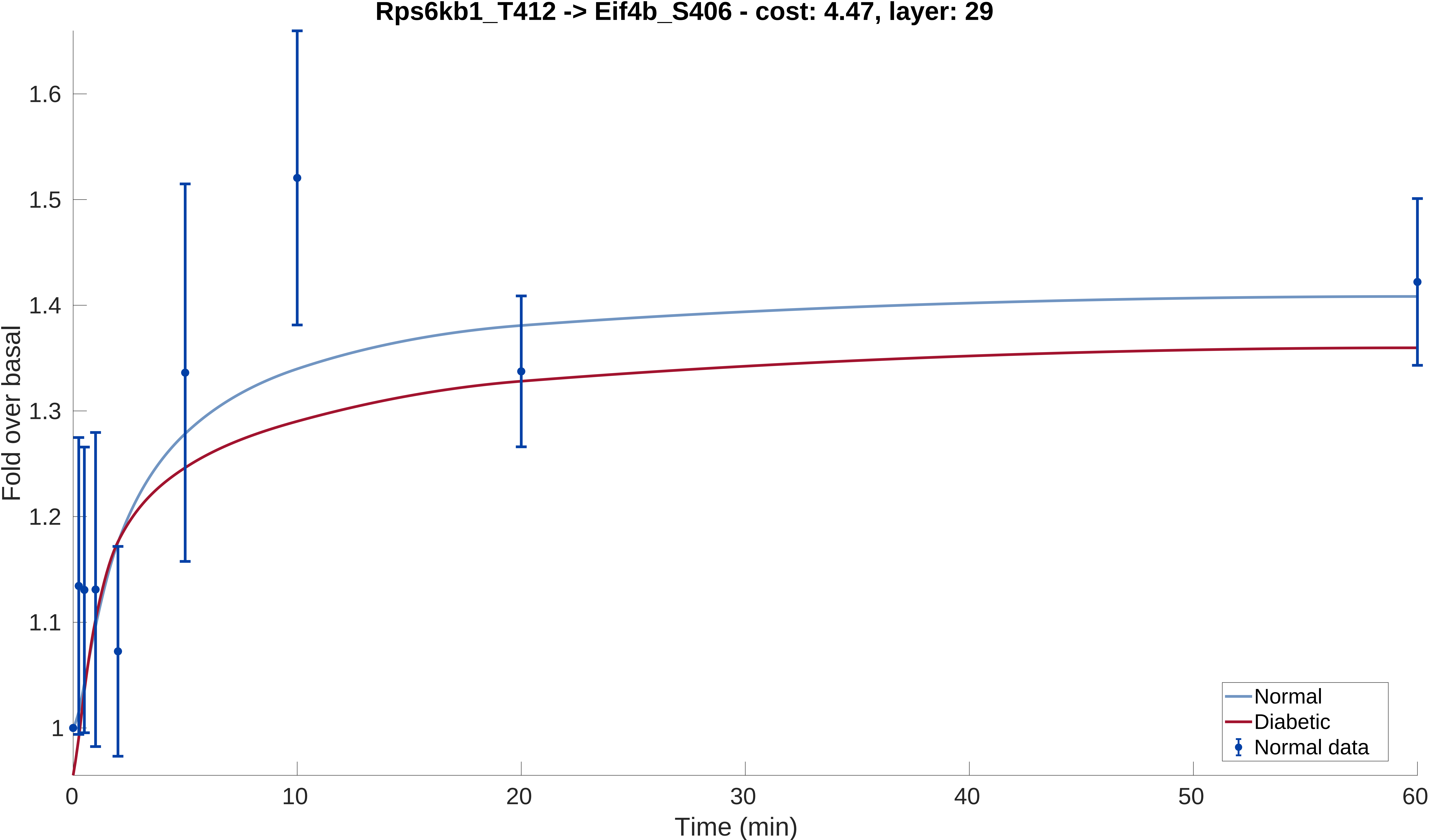

Rps6kb1_T412 -> Eif4b_S406 - cost: 4.47, layer: 29
1.6
1.5
1.4
1.3
1.2
1.1
1
Fold over basal
0
10
20
30
40
50
60
Time (min)
Normal
Diabetic
Normal data

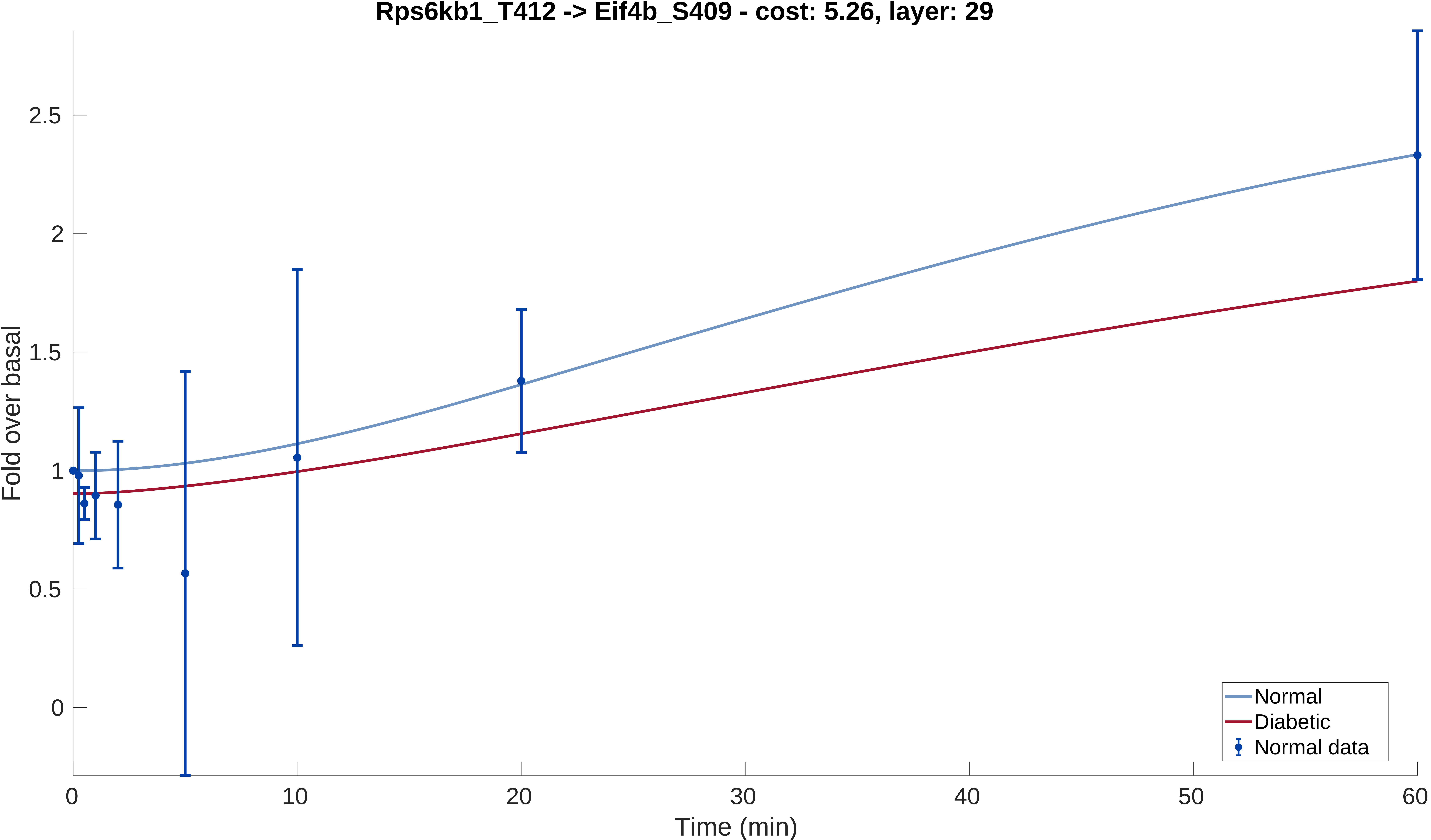

Rps6kb1_T412 -> Eif4b_S409 - cost: 5.26, layer: 29
Fold over basal
2.5
2
1.5
1
0.5
0
0
10
20
30
40
50
60
Time (min)
Normal
Diabetic
Normal data

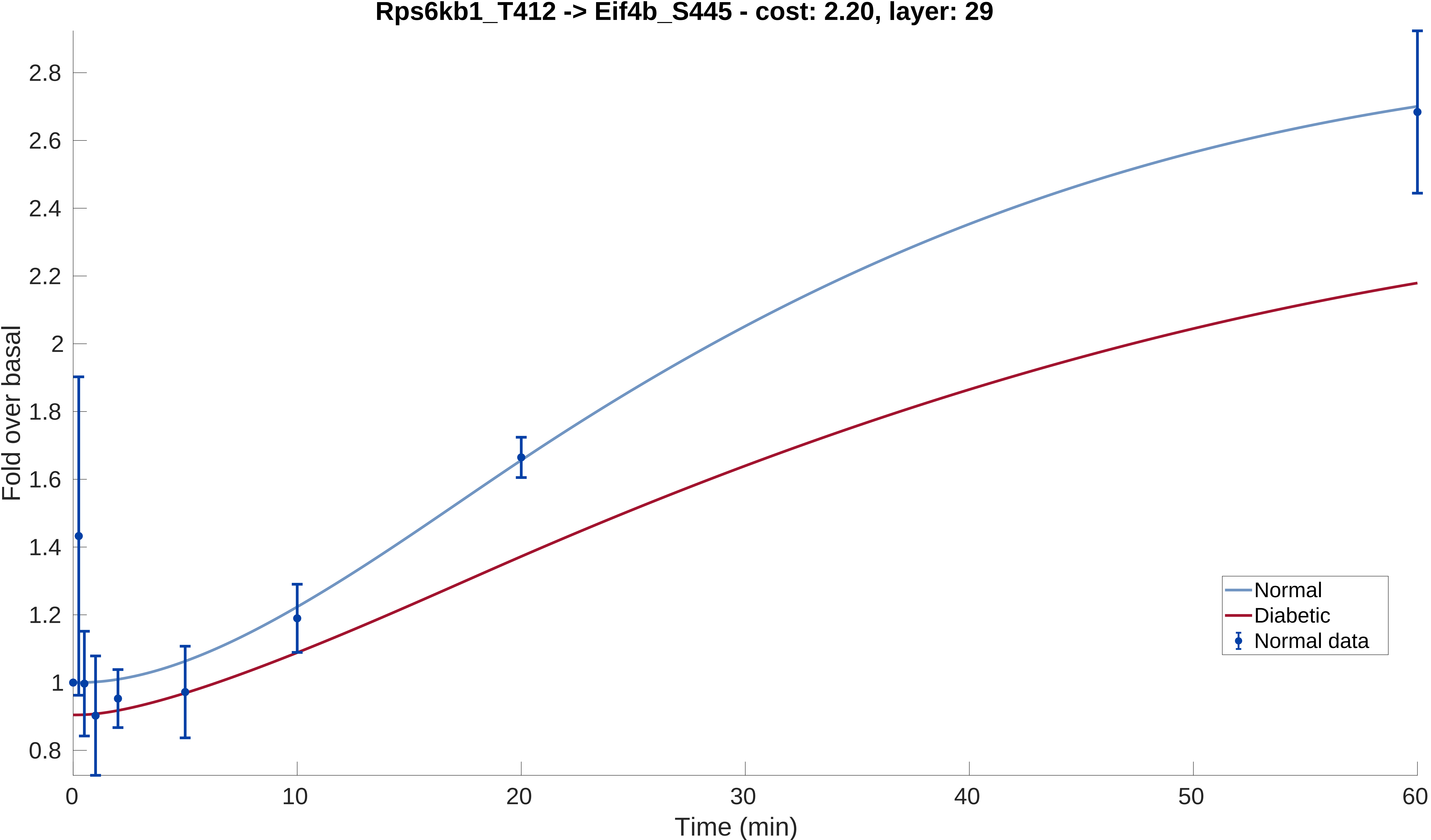

Rps6kb1_T412 -> Eif4b_S445 - cost: 2.20, layer: 29
Fold over basal
Time (min)
0.8
1
1.2
1.4
1.6
1.8
2
2.2
2.4
2.6
2.8
0
10
20
30
40
50
60
Normal
Diabetic
Normal data

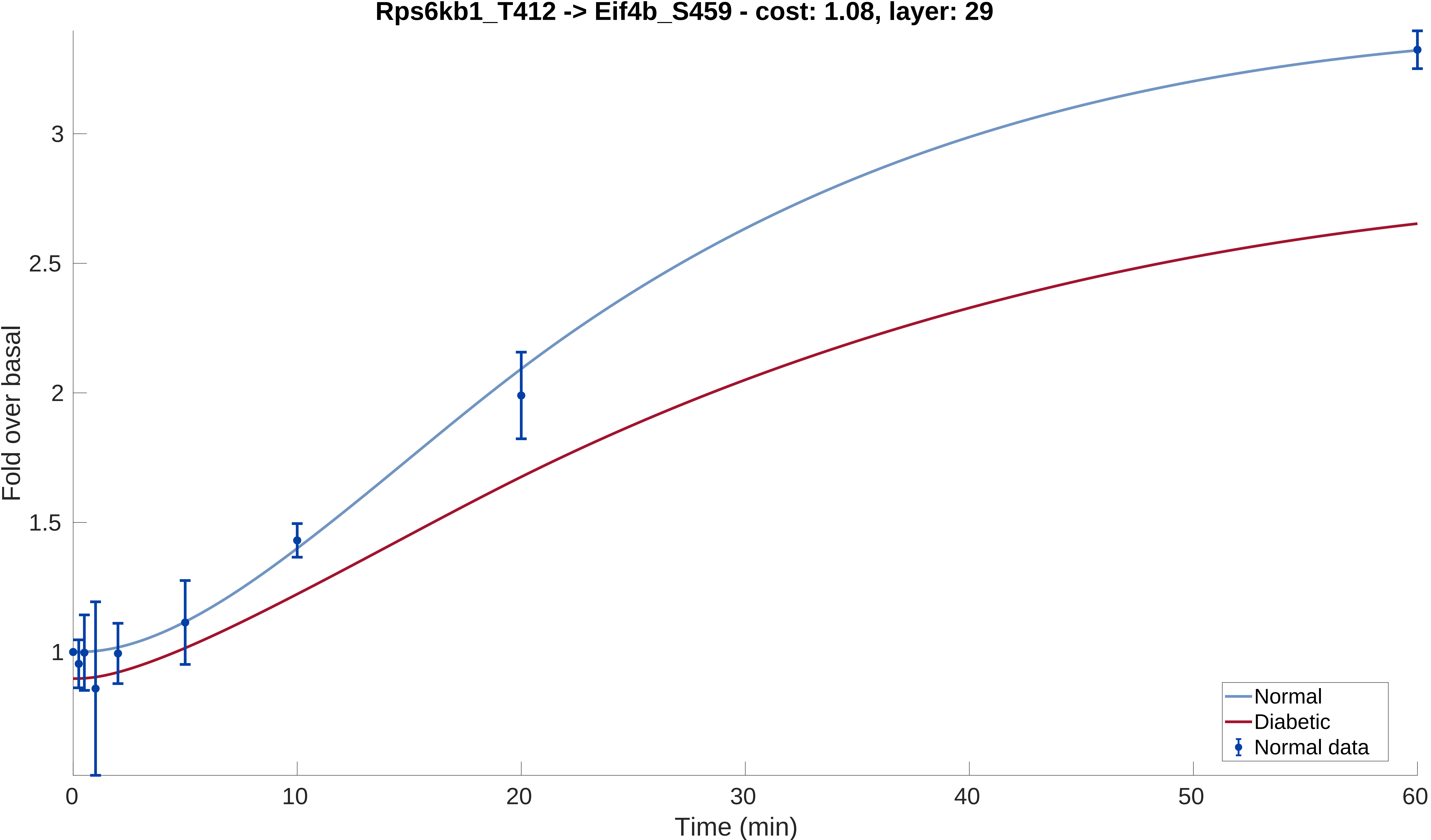
Rps6kb1_T412 -> Eif4b_S459 - cost: 1.08, layer: 29
Fold over basal
Time (min)
3
2.5
2
1.5
1
0
10
20
30
40
50
60
Normal
Diabetic
Normal data

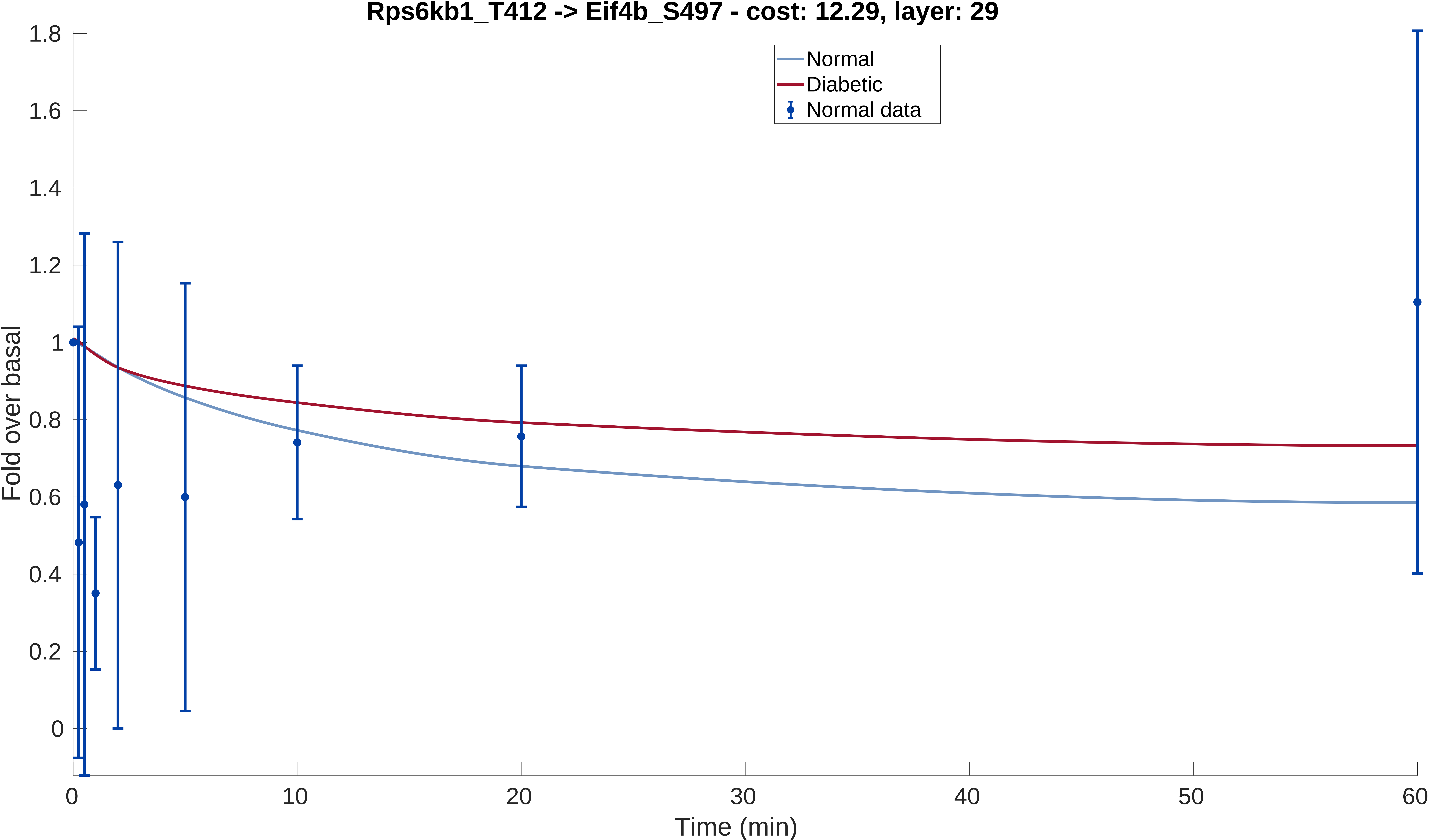
Rps6kb1_T412 -> Eif4b_S497 - cost: 12.29, layer: 29
Normal
Diabetic
Normal data
Fold over basal
Time (min)
1.8
1.6
1.4
1.2
1
0.8
0.6
0.4
0.2
0
0
10
20
30
40
50
60

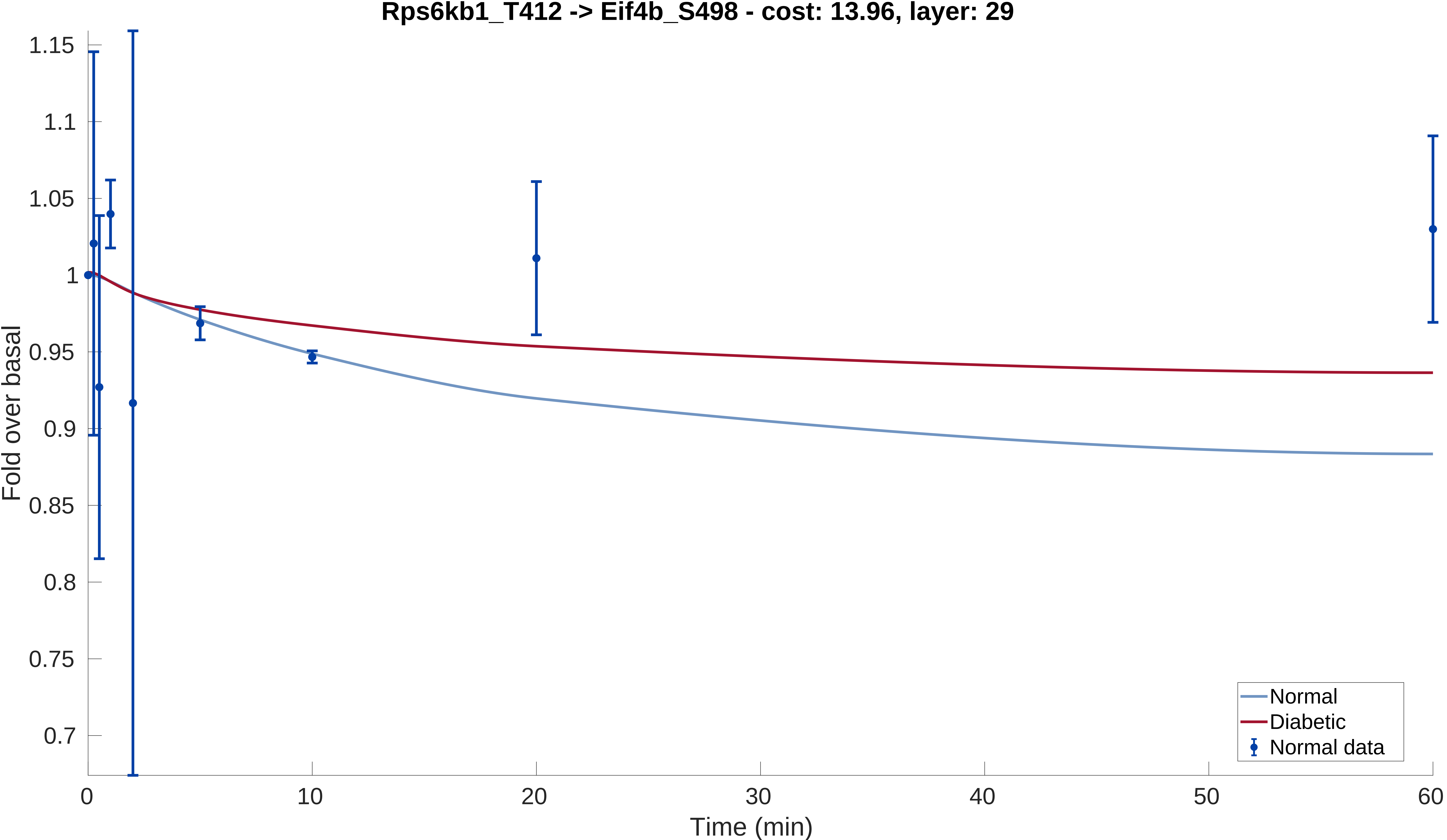

Rps6kb1_T412 -> Eif4b_S498 - cost: 13.96, layer: 29
1.15
1.1
1.05
1
0.95
0.9
0.85
0.8
0.75
0.7
Fold over basal
0
10
20
30
40
50
60
Time (min)
Normal
Diabetic
Normal data

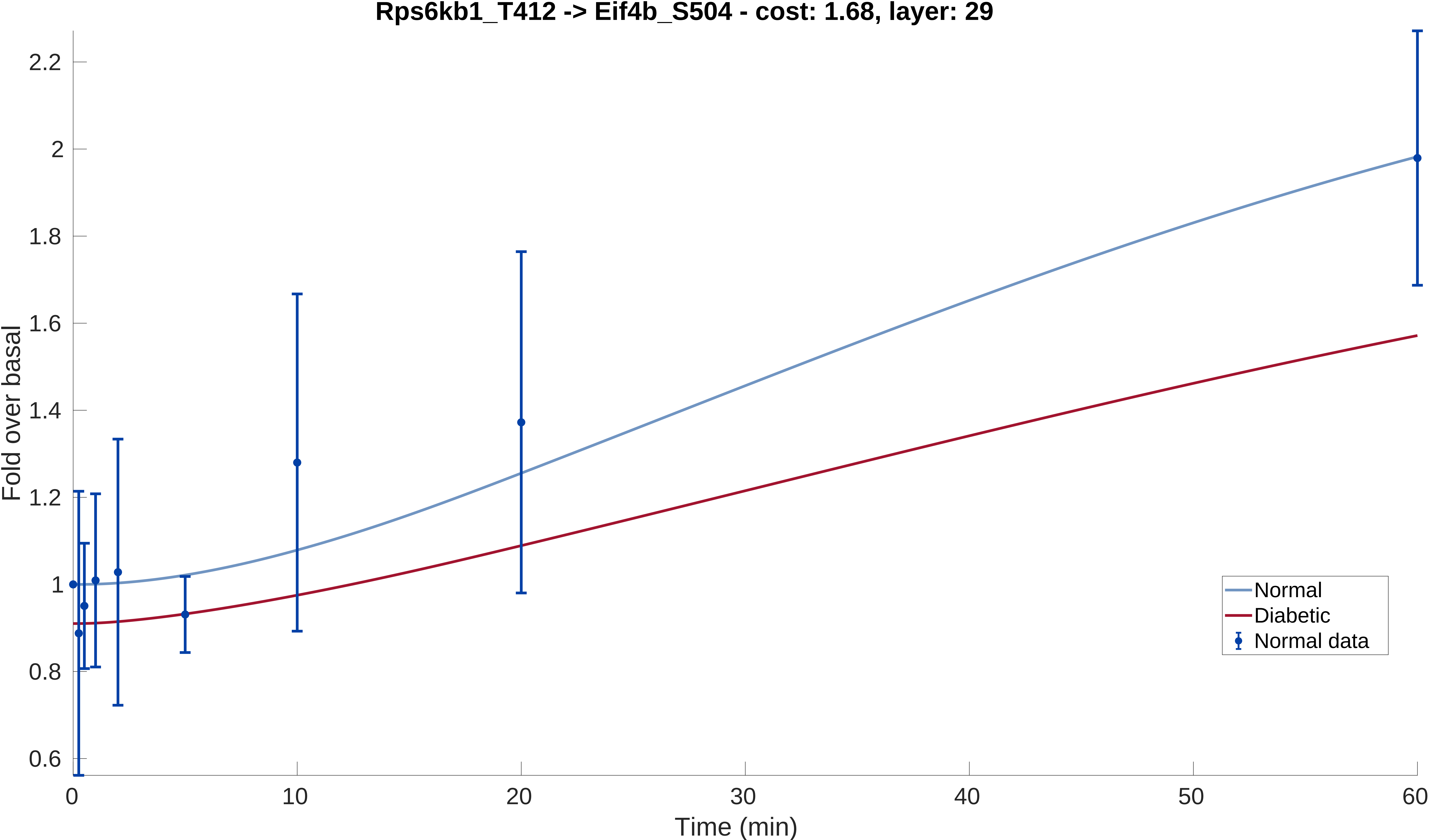

Rps6kb1_T412 -> Eif4b_S504 - cost: 1.68, layer: 29
Fold over basal
Time (min)
0.6
0.8
1
1.2
1.4
1.6
1.8
2
2.2
0
10
20
30
40
50
60
Normal
Diabetic
Normal data

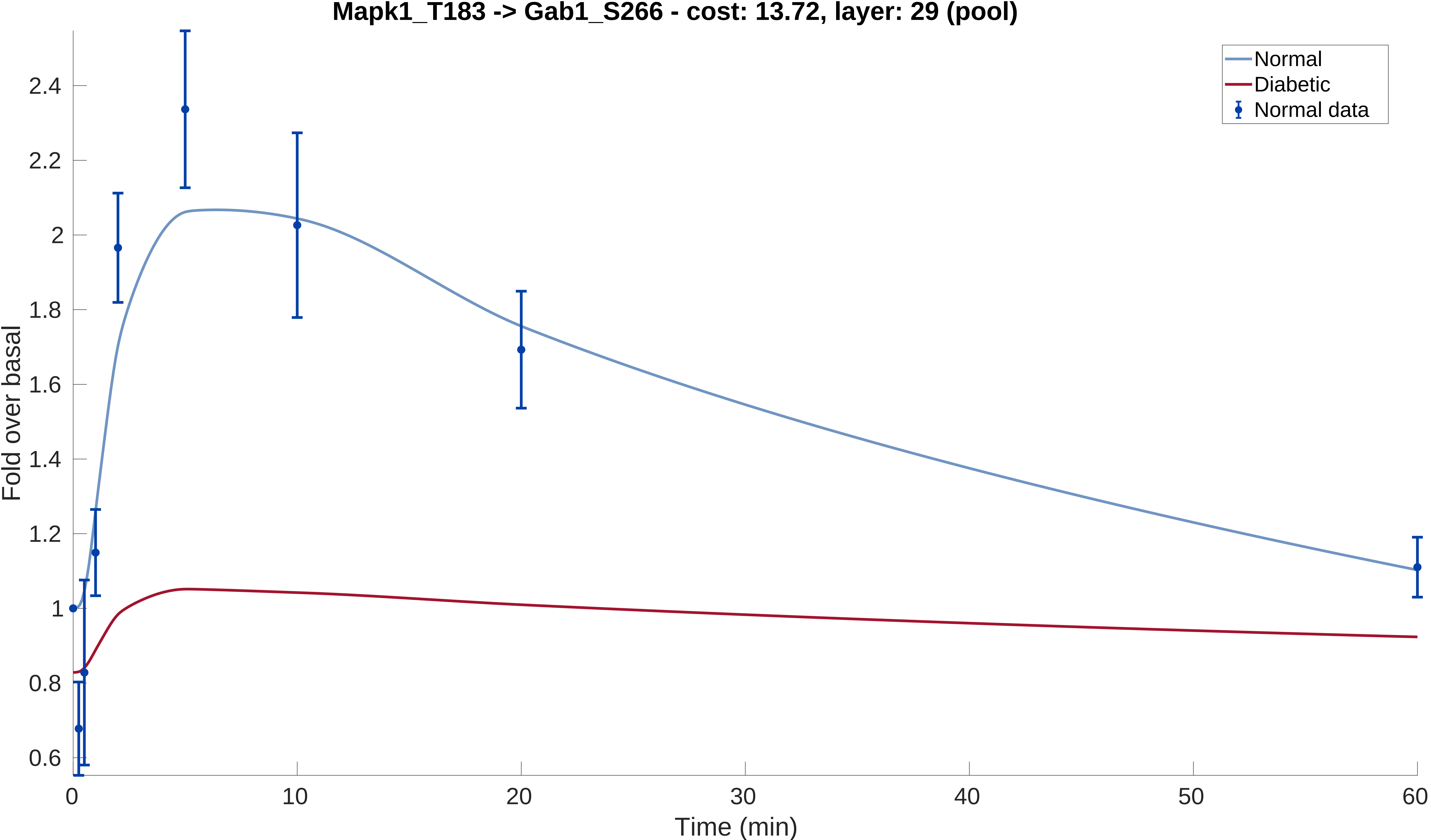

Mapk1_T183 -> Gab1_S266 - cost: 13.72, layer: 29 (pool)
Normal
Diabetic
Normal data
Fold over basal
Time (min)
0.6
0.8
1
1.2
1.4
1.6
1.8
2
2.2
2.4
0
10
20
30
40
50
60

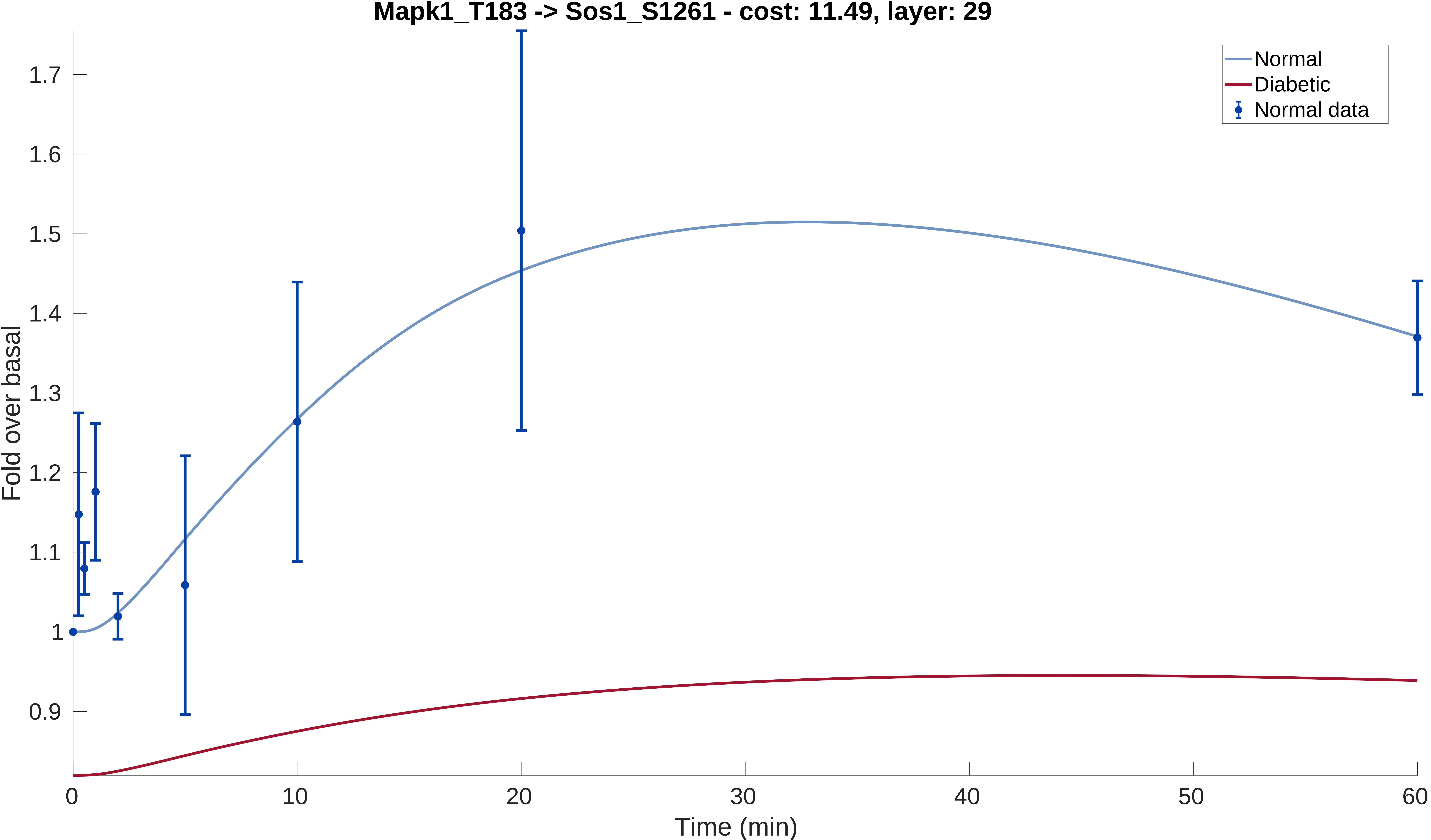

Mapk1_T183 -> Sos1_S1261 - cost: 11.49, layer: 29
Normal
Diabetic
Normal data
Fold over basal
1.7
1.6
1.5
1.4
1.3
1.2
1.1
1
0.9
0
10
20
30
40
50
60
Time (min)

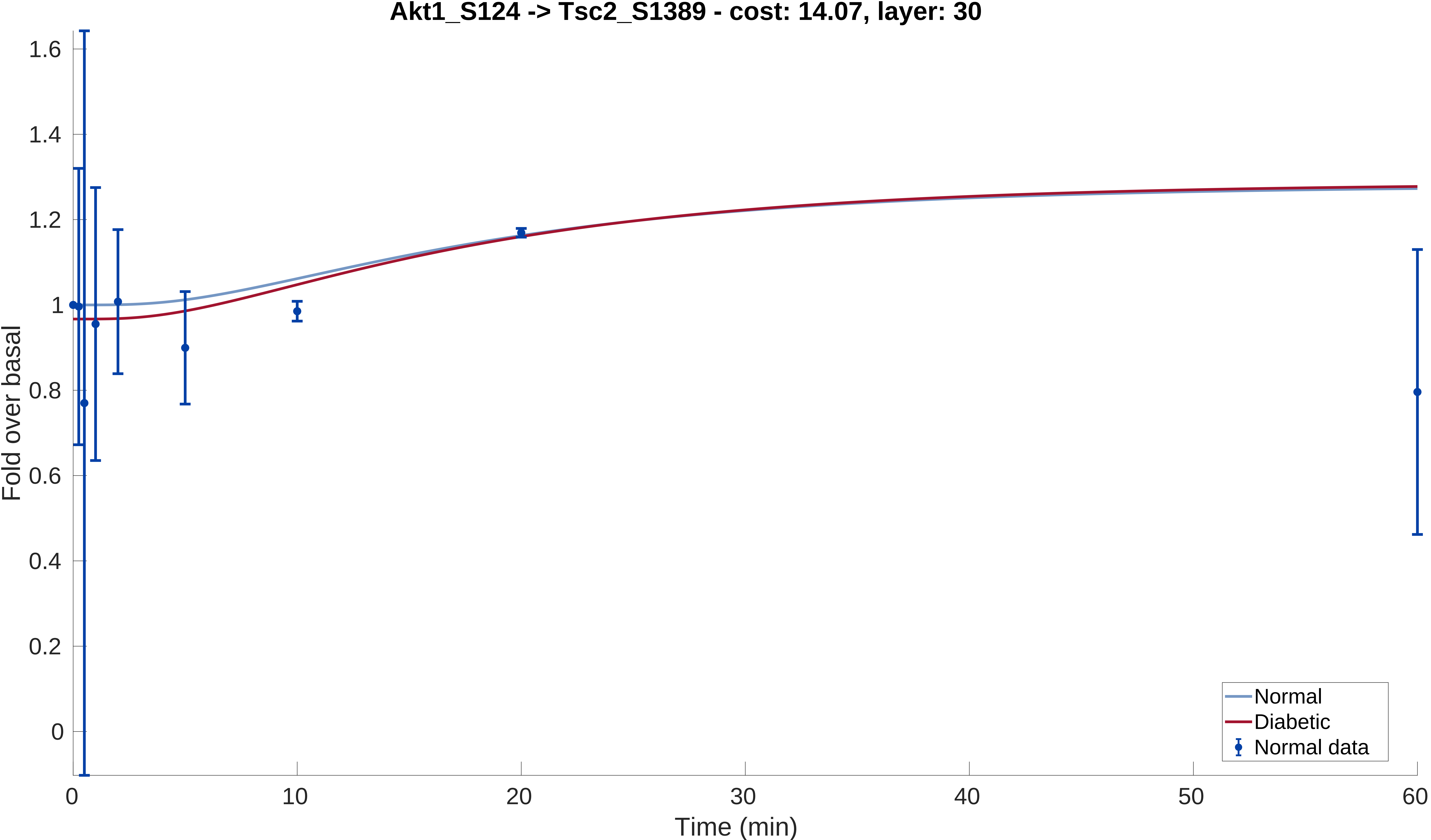

Akt1_S124 -> Tsc2_S1389 - cost: 14.07, layer: 30
Fold over basal
1.6
1.4
1.2
1
0.8
0.6
0.4
0.2
0
0
10
20
30
40
50
60
Time (min)
Normal
Diabetic
Normal data

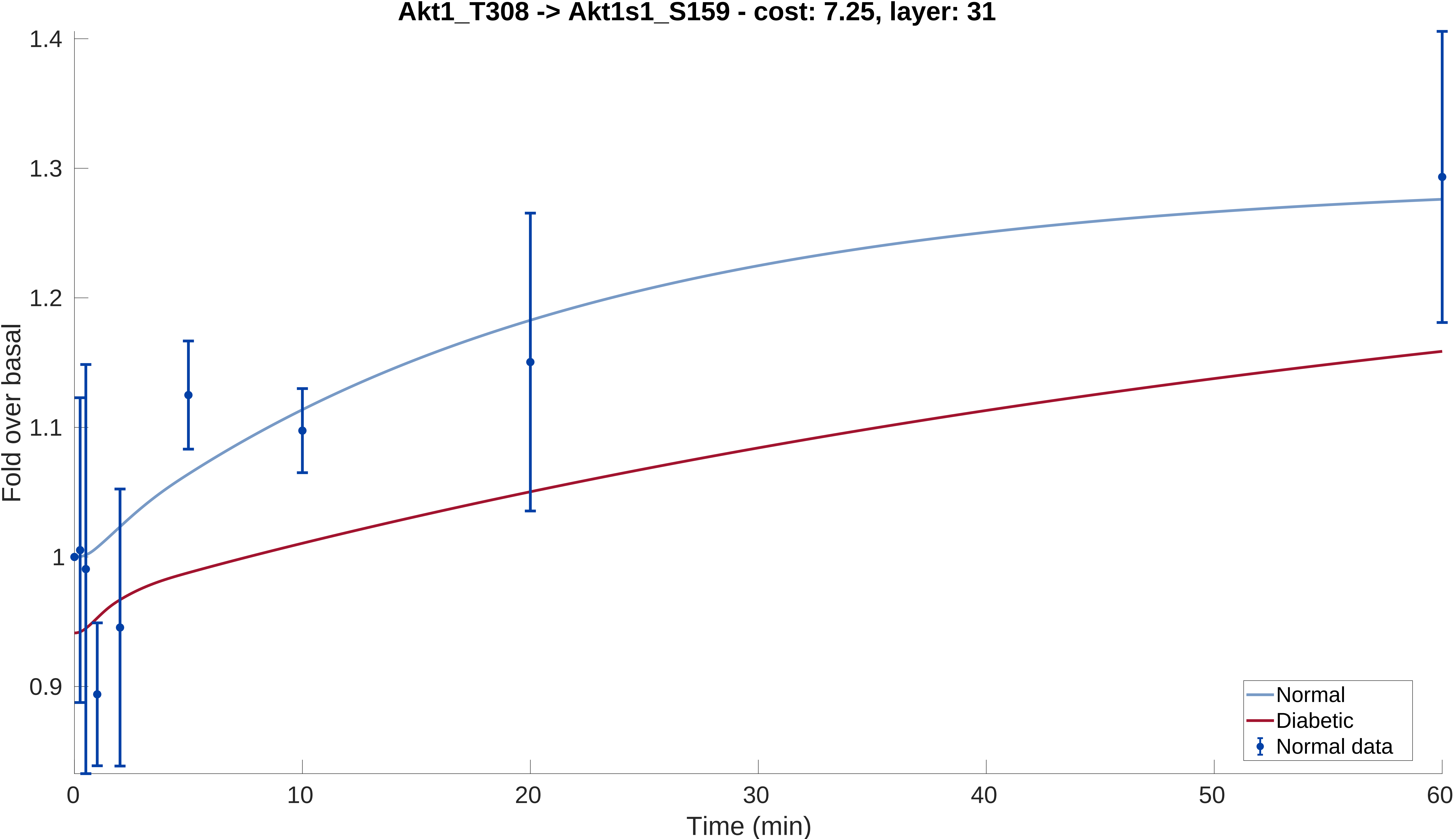

Akt1_T308 -> Akt1s1_S159 - cost: 7.25, layer: 31
Fold over basal
1.4
1.3
1.2
1.1
1
0.9
0
10
20
30
40
50
60
Time (min)
Normal
Diabetic
Normal data

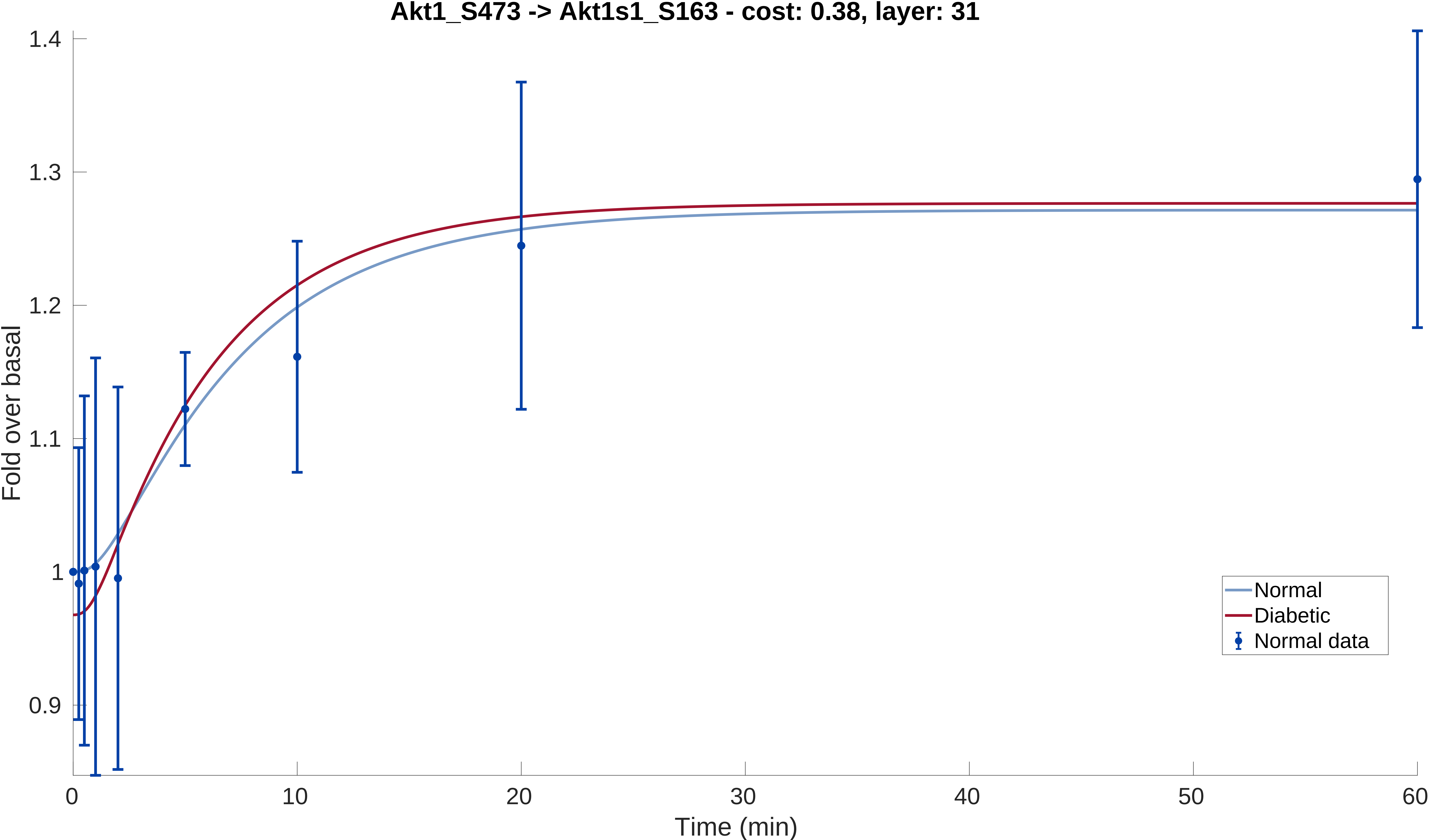

Akt1_S473 -> Akt1s1_S163 - cost: 0.38, layer: 31
Fold over basal
Time (min)
1.4
1.3
1.2
1.1
1
0.9
0
10
20
30
40
50
60
Normal
Diabetic
Normal data

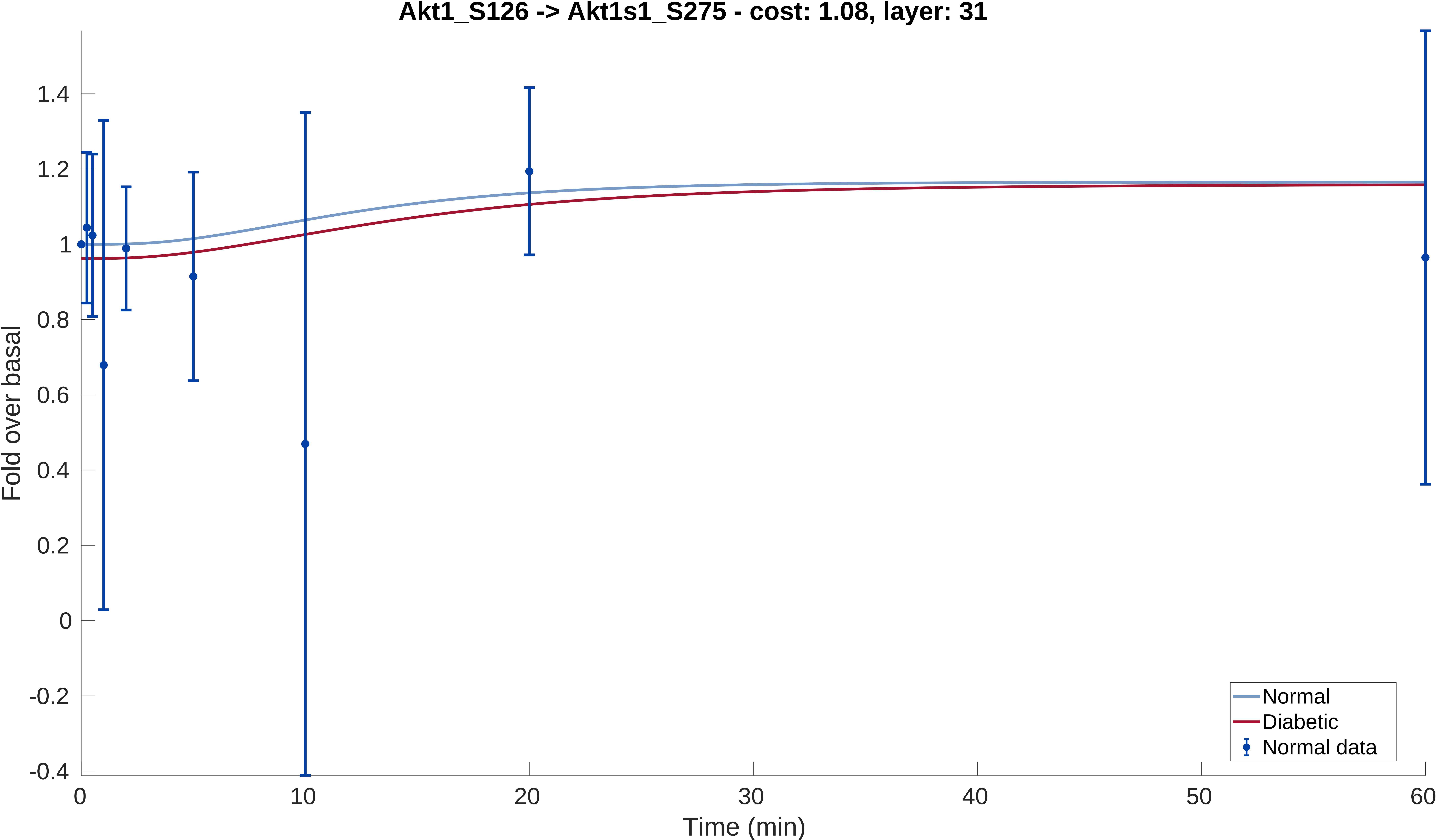

Akt1_S126 -> Akt1s1_S275 - cost: 1.08, layer: 31
Fold over basal
1.4
1.2
1
0.8
0.6
0.4
0.2
0
-0.2
-0.4
0
10
20
30
40
50
60
Time (min)
Normal
Diabetic
Normal data

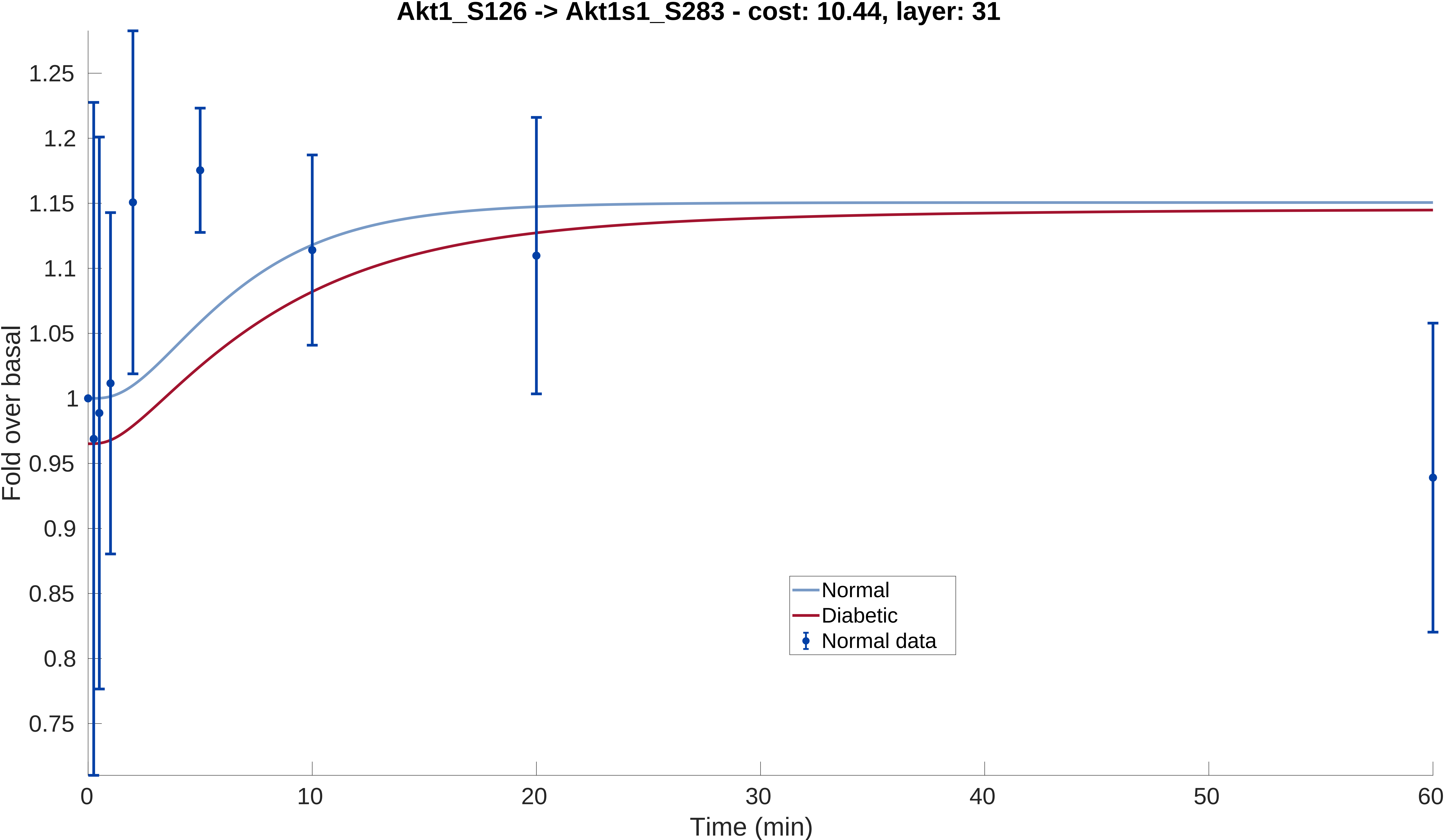
Akt1_S126 -> Akt1s1_S283 - cost: 10.44, layer: 31
Fold over basal
1.25
1.2
1.15
1.1
1.05
1
0.95
0.9
0.85
0.8
0.75
0
10
20
30
40
50
60
Time (min)
Normal
Diabetic
Normal data

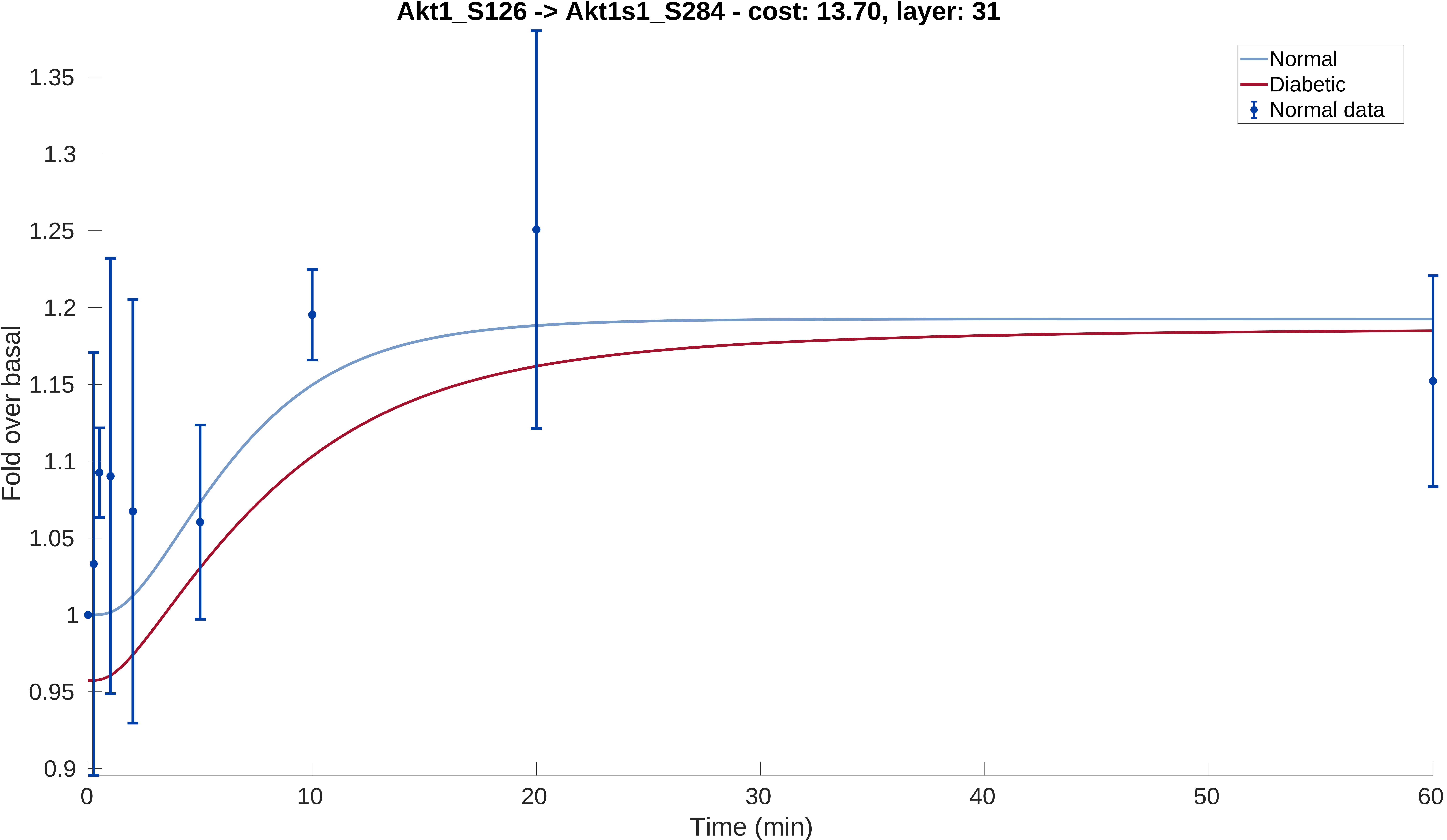

Akt1_S126 -> Akt1s1_S284 - cost: 13.70, layer: 31
Normal
Diabetic
Normal data
Fold over basal
Time (min)
0.9
0.95
1
1.05
1.1
1.15
1.2
1.25
1.3
1.35
0
10
20
30
40
50
60

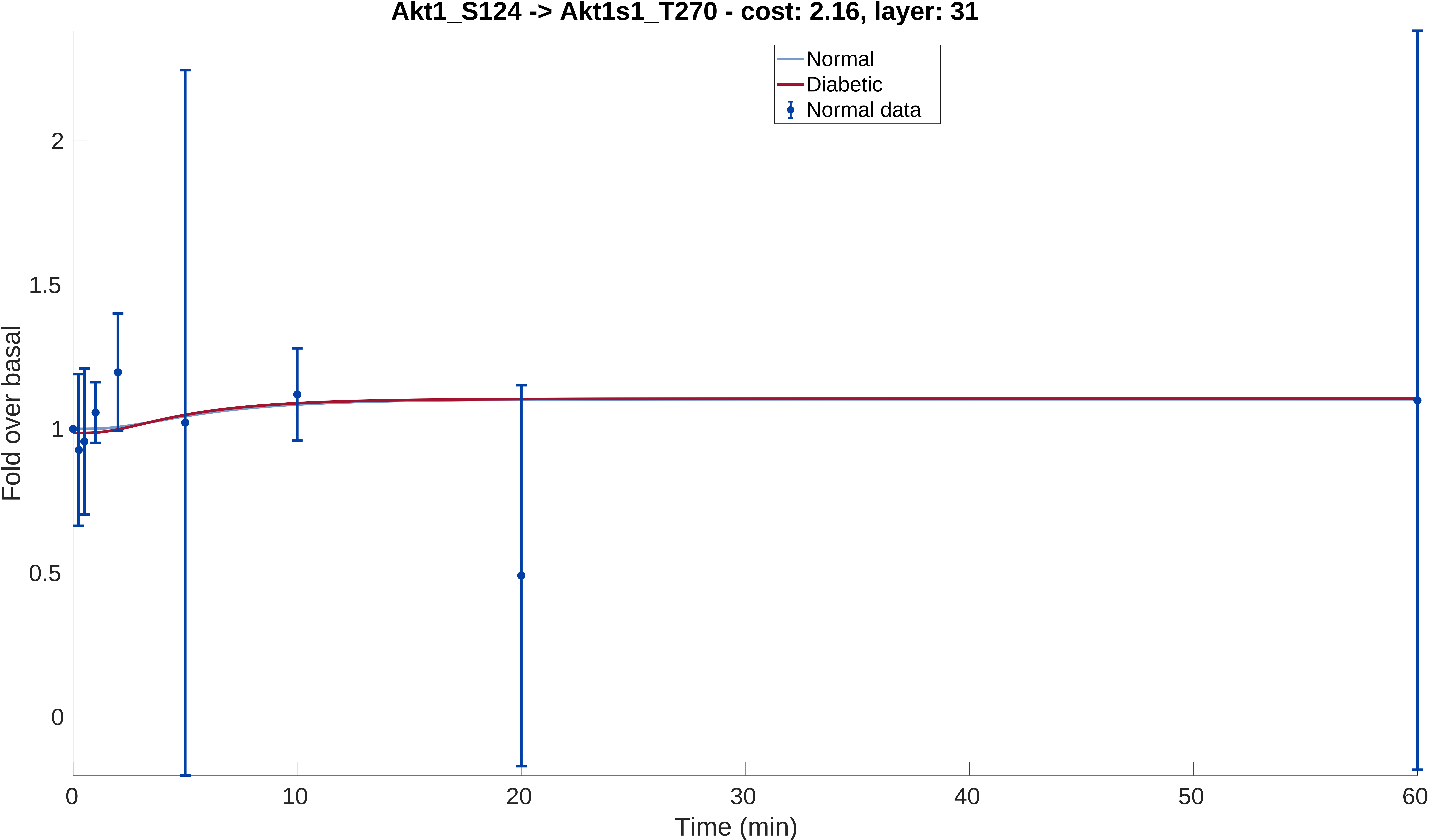

Akt1_S124 -> Akt1s1_T270 - cost: 2.16, layer: 31
Normal
Diabetic
Normal data
Fold over basal
Time (min)
0
0.5
1
1.5
2
0
10
20
30
40
50
60

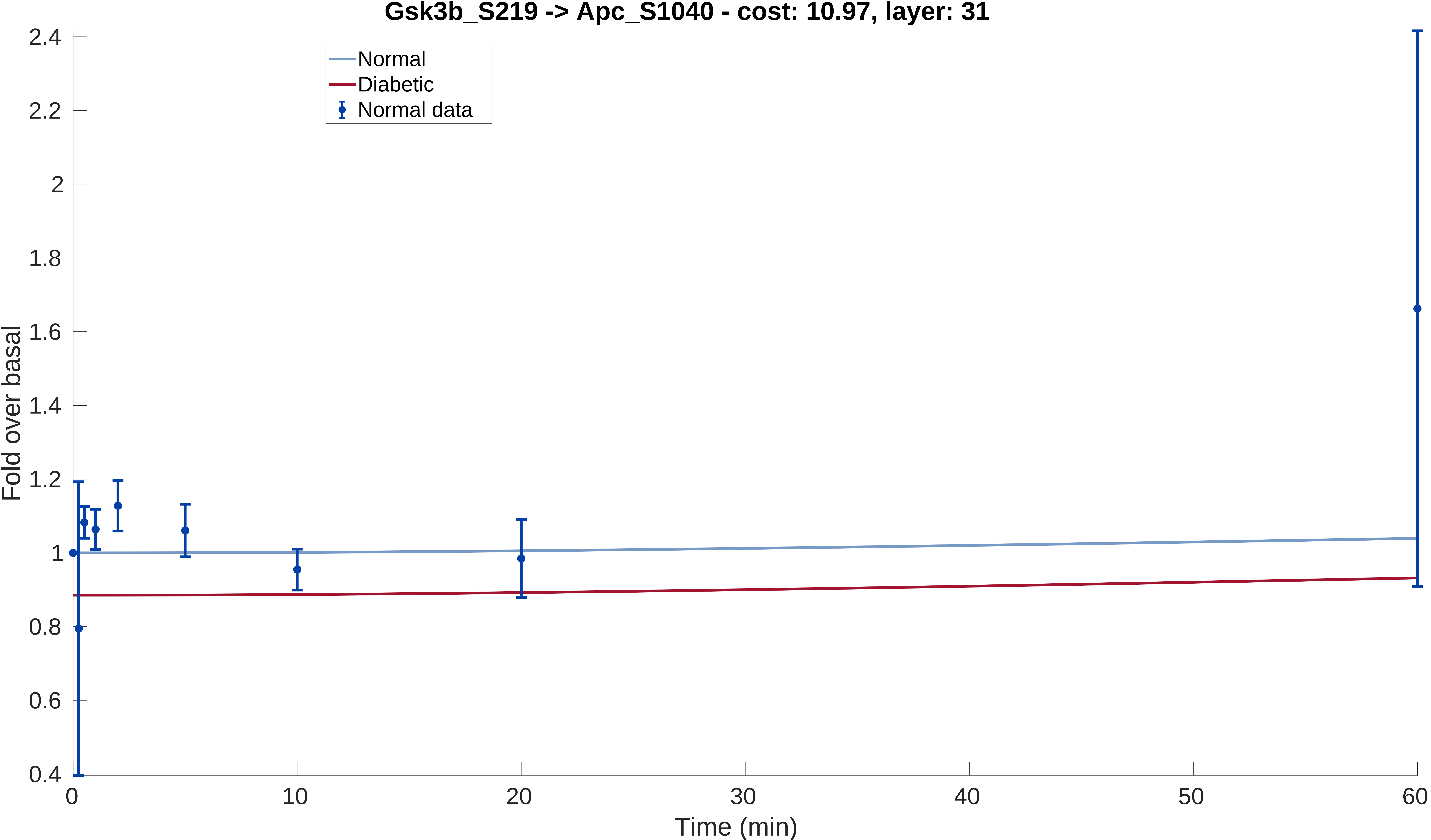

Gsk3b_S219 -> Apc_S1040 - cost: 10.97, layer: 31
Normal
Diabetic
Normal data
Fold over basal
Time (min)
0.4
0.6
0.8
1
1.2
1.4
1.6
1.8
2
2.2
2.4
0
10
20
30
40
50
60

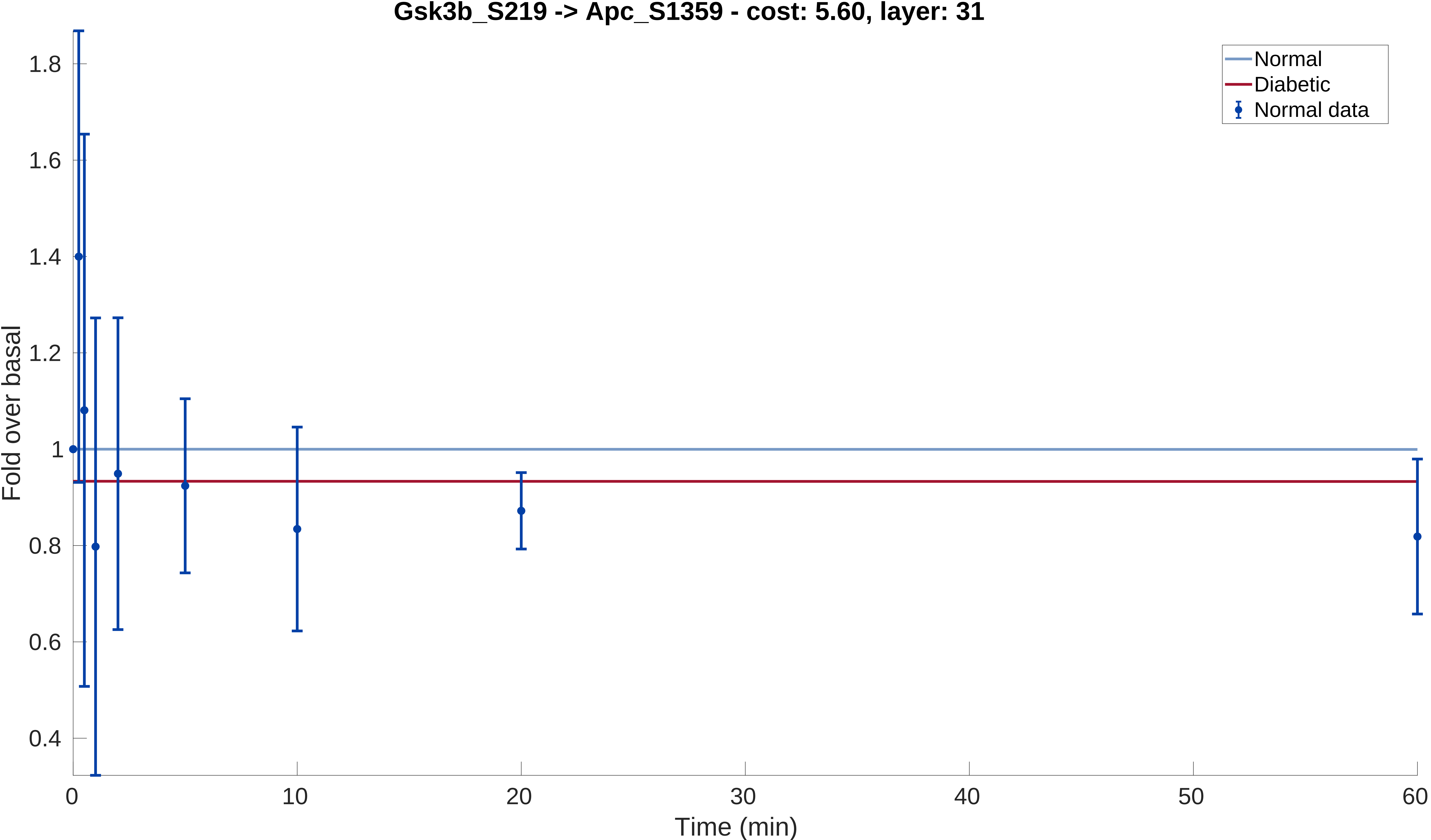

Gsk3b_S219 -> Apc_S1359 - cost: 5.60, layer: 31
Normal
Diabetic
Normal data
Fold over basal
1.8
1.6
1.4
1.2
1
0.8
0.6
0.4
0
10
20
30
40
50
60
Time (min)

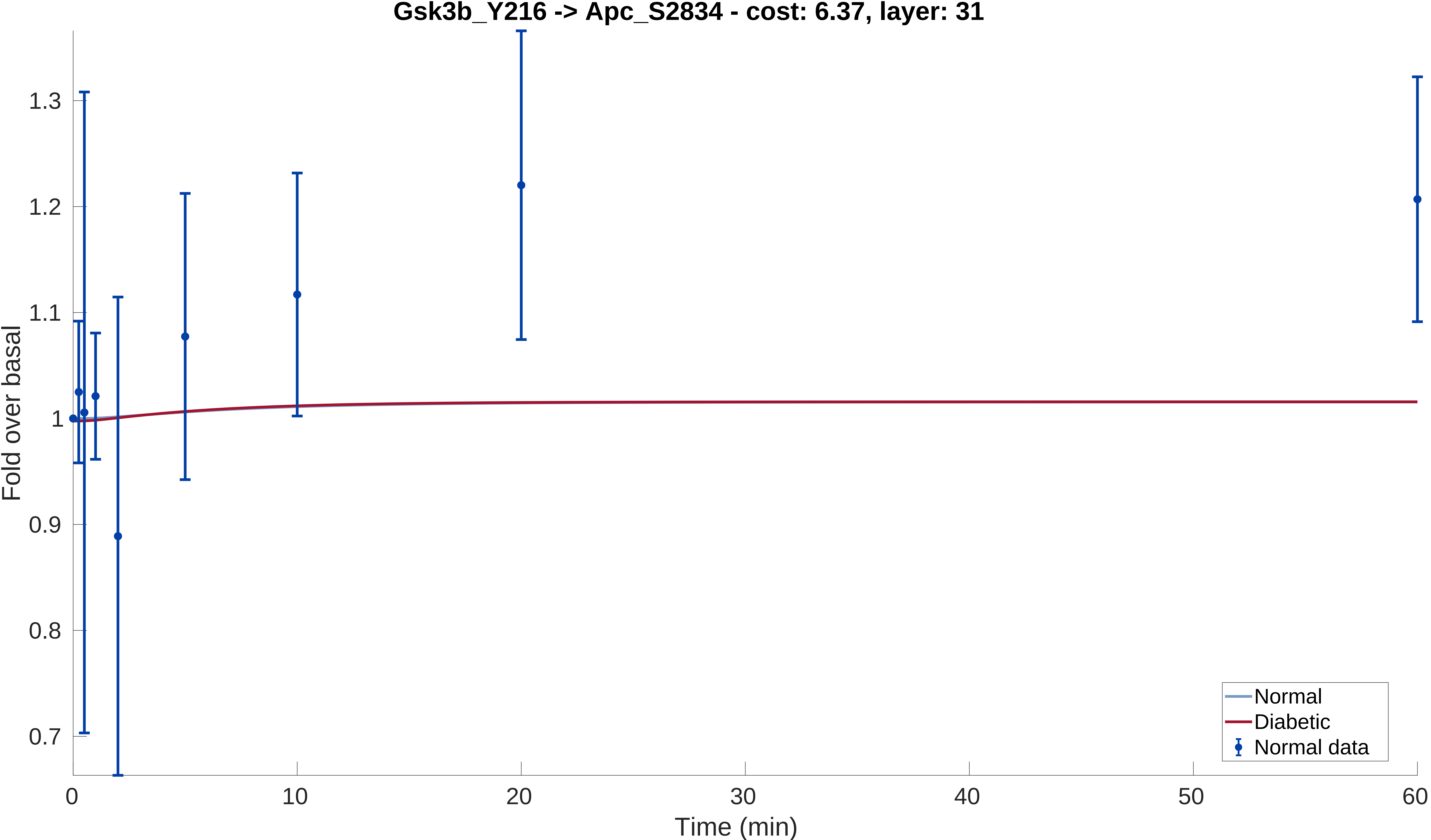

Gsk3b_Y216 -> Apc_S2834 - cost: 6.37, layer: 31
Fold over basal
1.3
1.2
1.1
1
0.9
0.8
0.7
0
10
20
30
40
50
60
Time (min)
Normal
Diabetic
Normal data

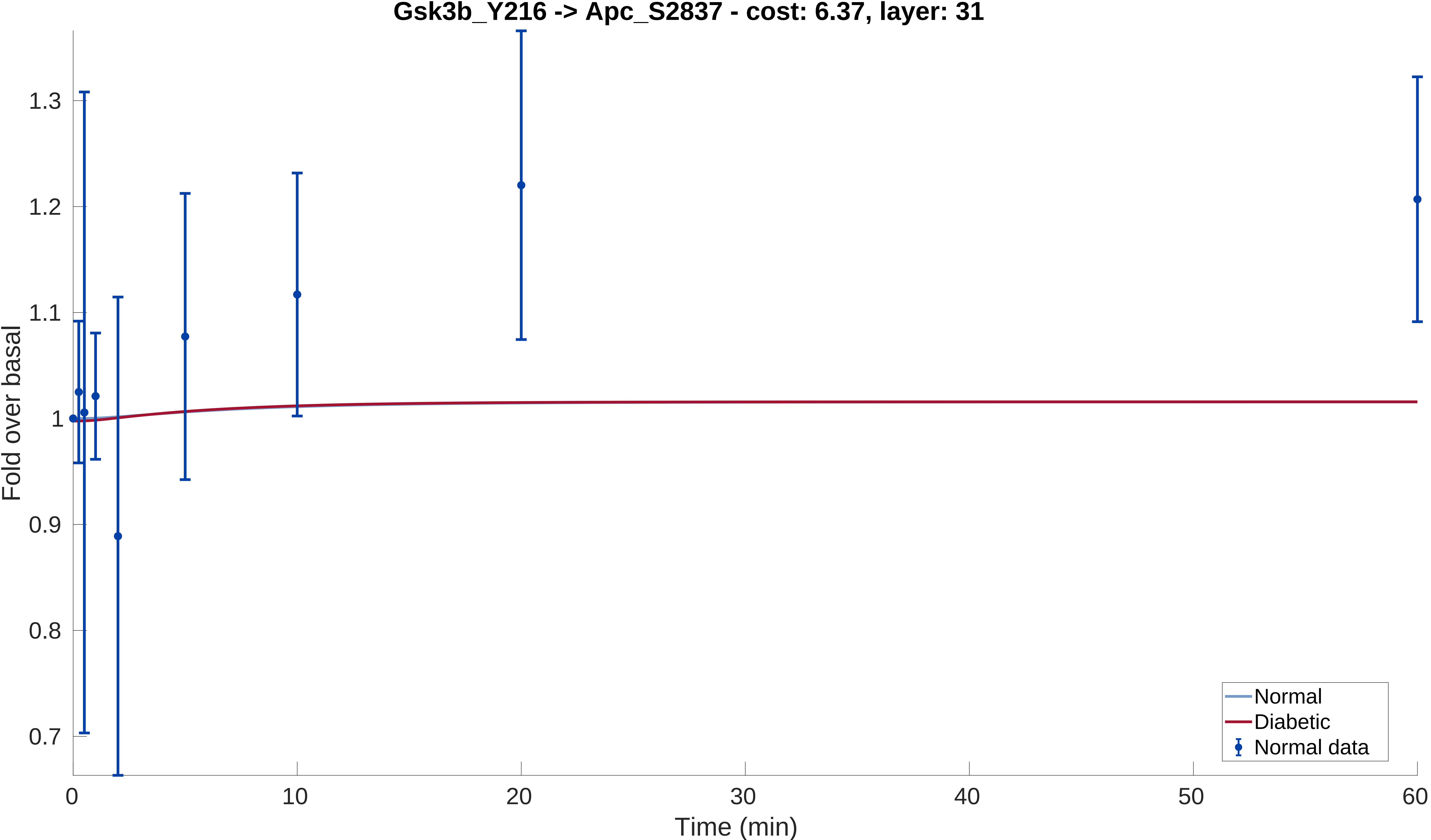

Gsk3b_Y216 -> Apc_S2837 - cost: 6.37, layer: 31
Fold over basal
1.3
1.2
1.1
1
0.9
0.8
0.7
0
10
20
30
40
50
60
Time (min)
Normal
Diabetic
Normal data

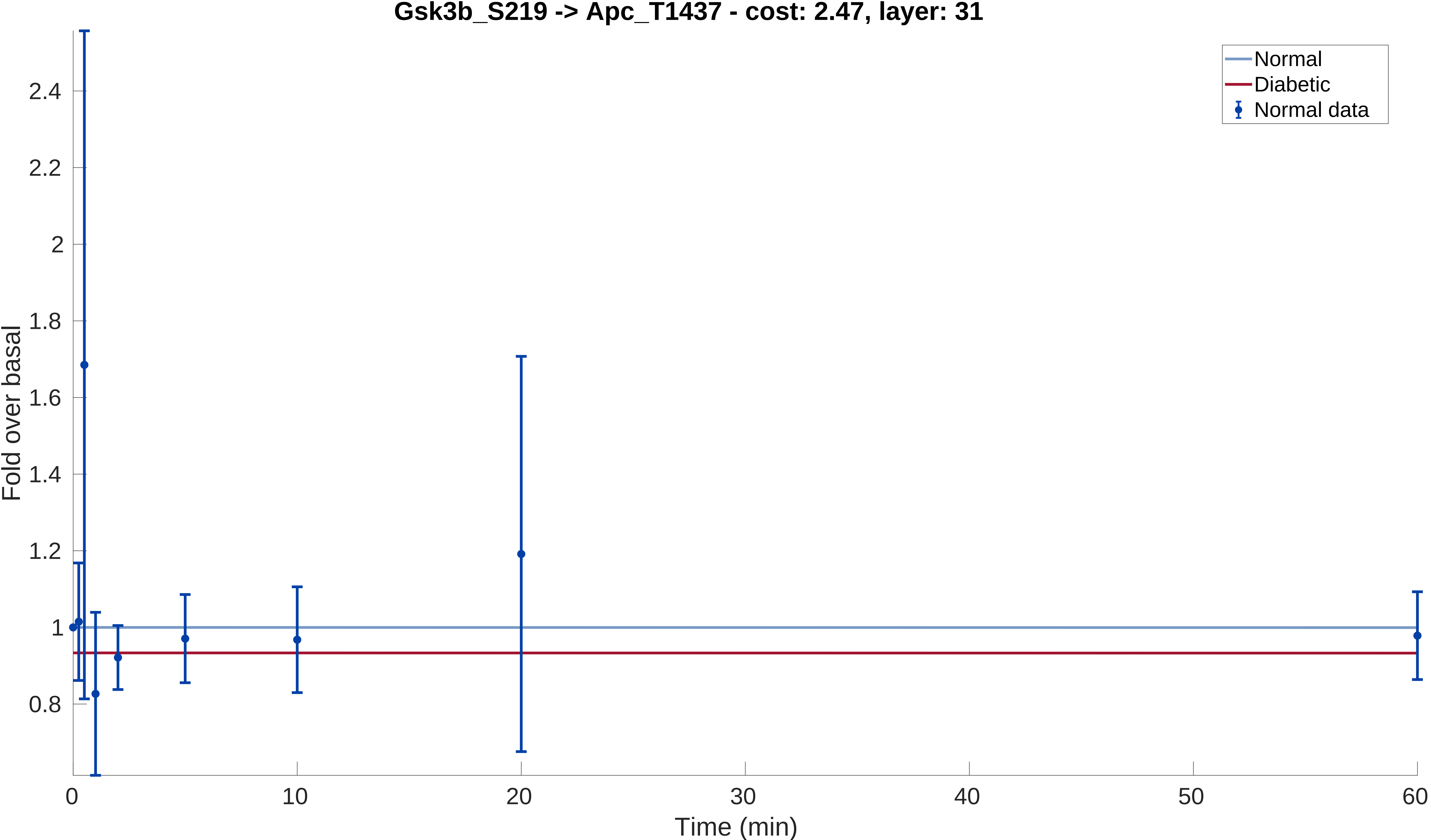

Gsk3b_S219 -> Apc_T1437 - cost: 2.47, layer: 31
Normal
Diabetic
Normal data
Fold over basal
2.4
2.2
2
1.8
1.6
1.4
1.2
1
0.8
0
10
20
30
40
50
60
Time (min)

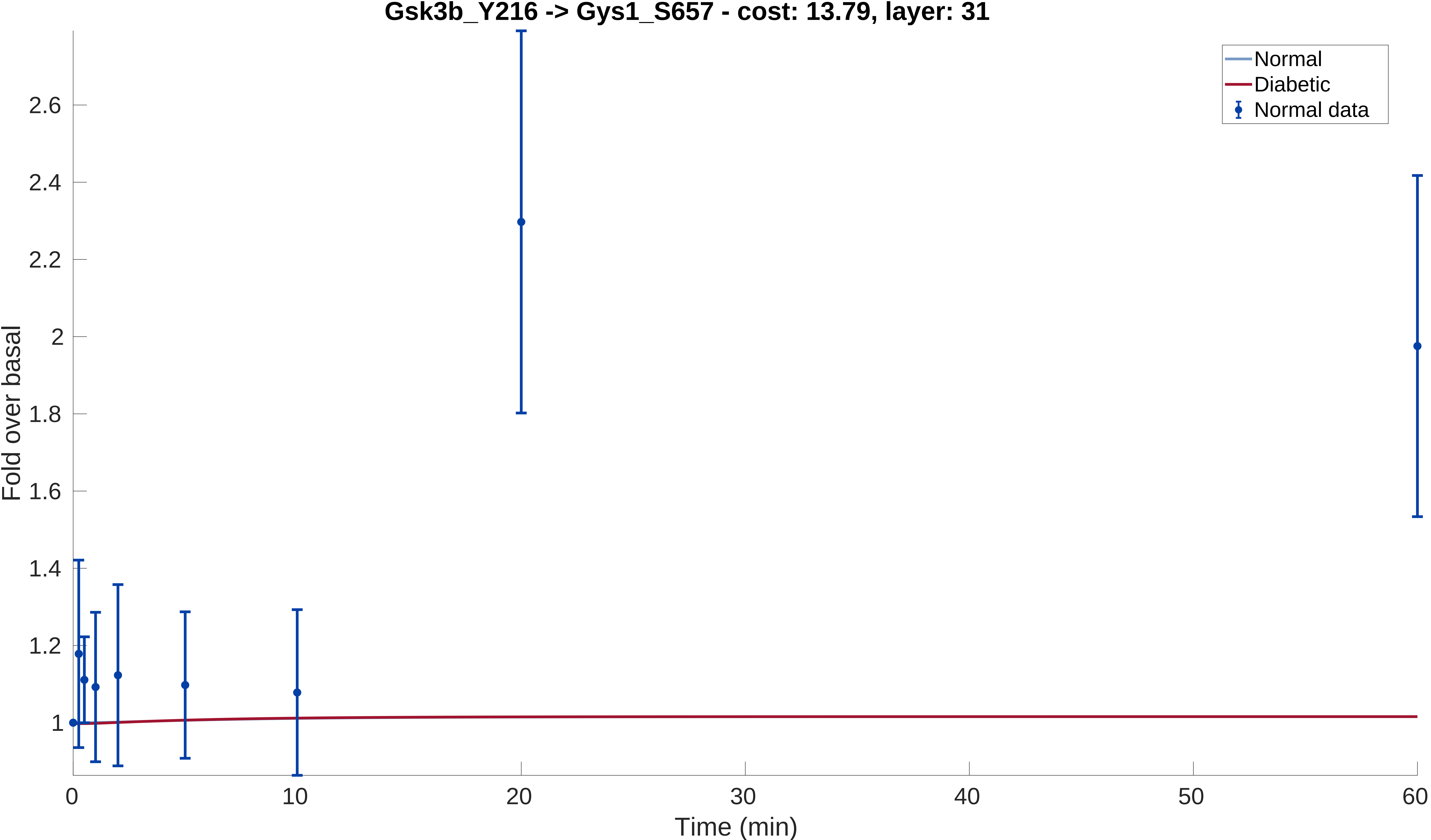

Gsk3b_Y216 -> Gys1_S657 - cost: 13.79, layer: 31
Normal
Diabetic
Normal data
Fold over basal
2.6
2.4
2.2
2
1.8
1.6
1.4
1.2
1
0
10
20
30
40
50
60
Time (min)

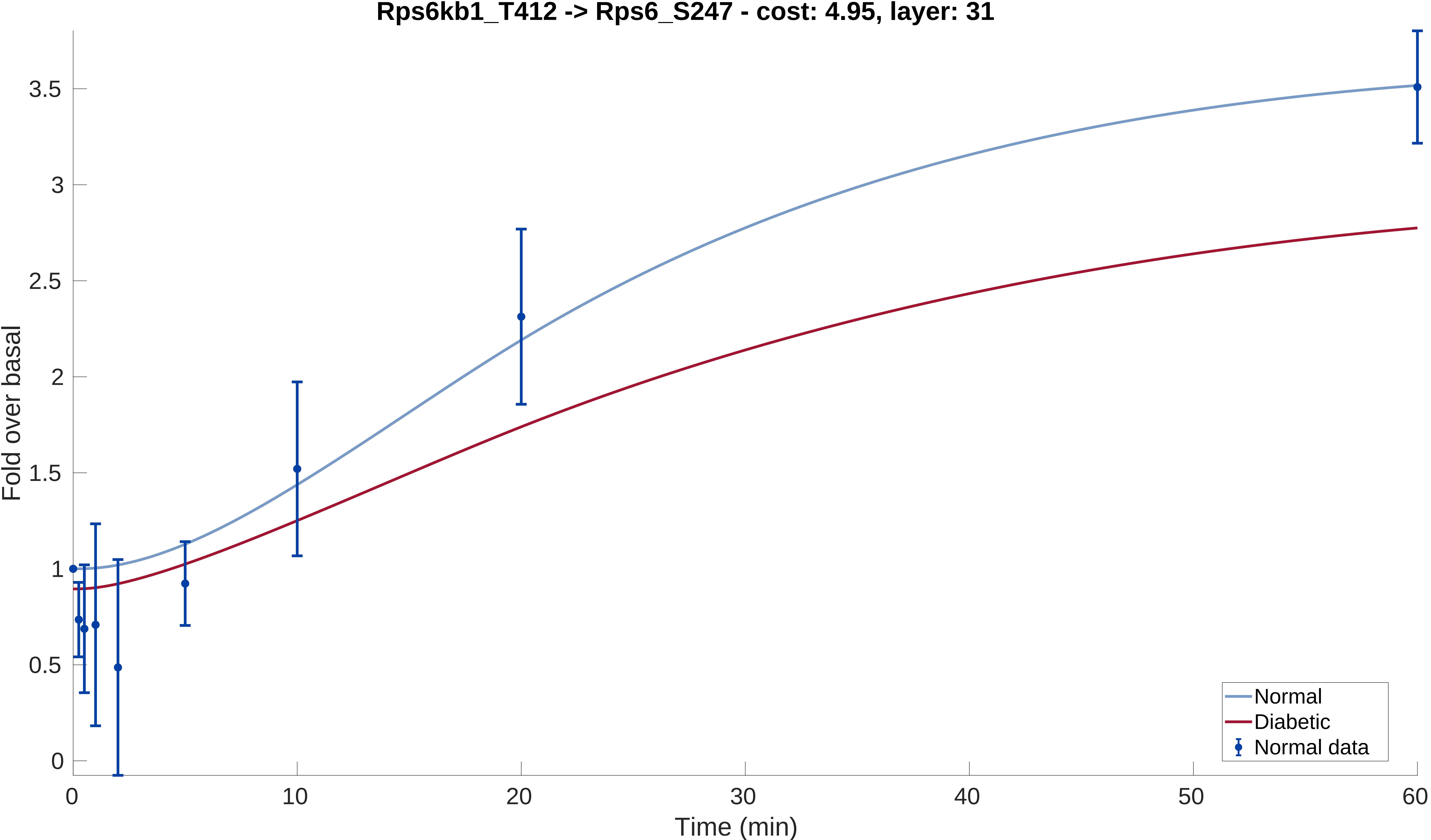
Rps6kb1_T412 -> Rps6_S247 - cost: 4.95, layer: 31
Fold over basal
3.5
3
2.5
2
1.5
1
0.5
0
0
10
20
30
40
50
60
Time (min)
Normal
Diabetic
Normal data

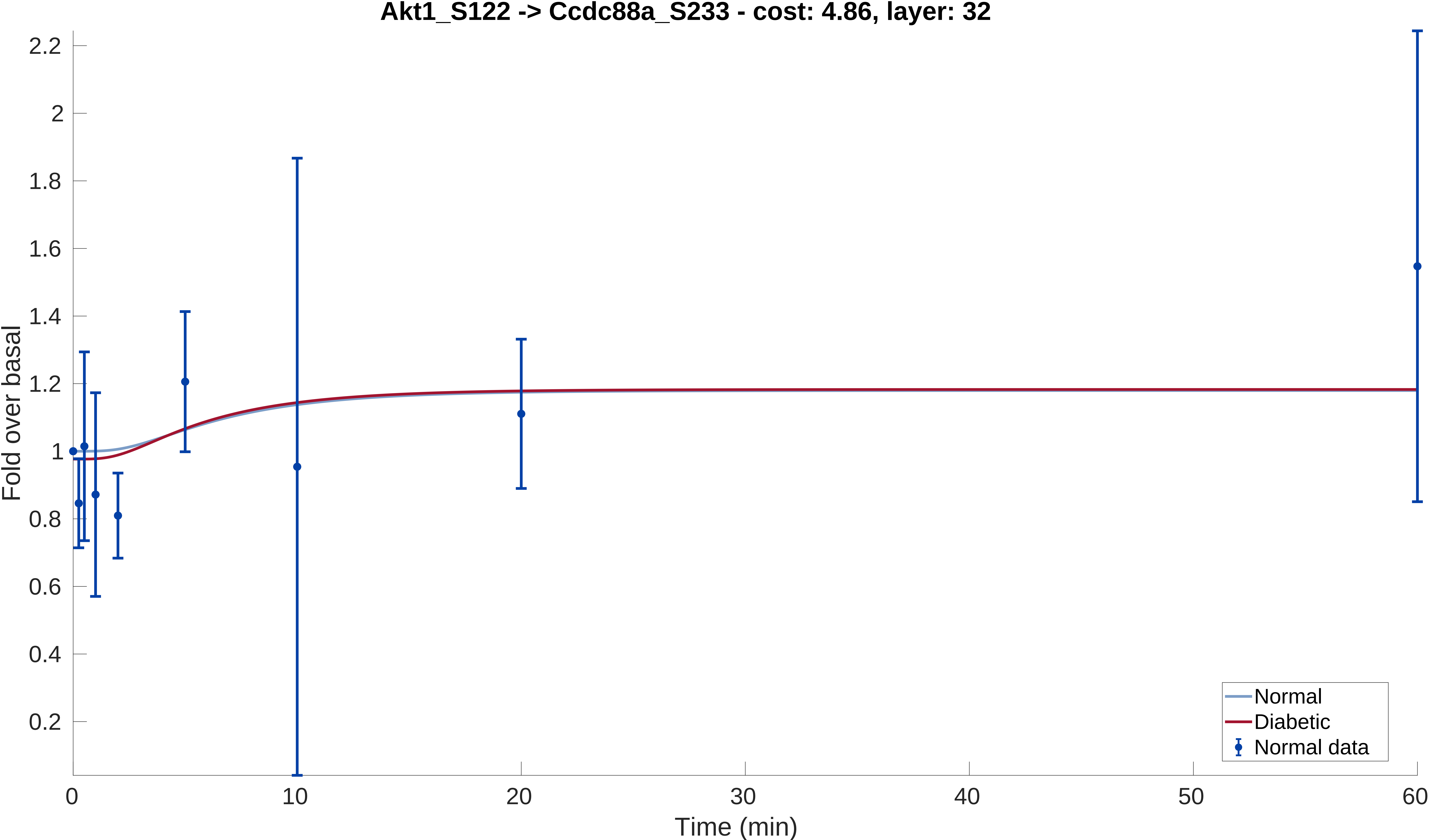
Akt1_S122 -> Ccdc88a_S233 - cost: 4.86, layer: 32
Fold over basal
Time (min)
2.2
2
1.8
1.6
1.4
1.2
1
0.8
0.6
0.4
0.2
0
10
20
30
40
50
60
Normal
Diabetic
Normal data

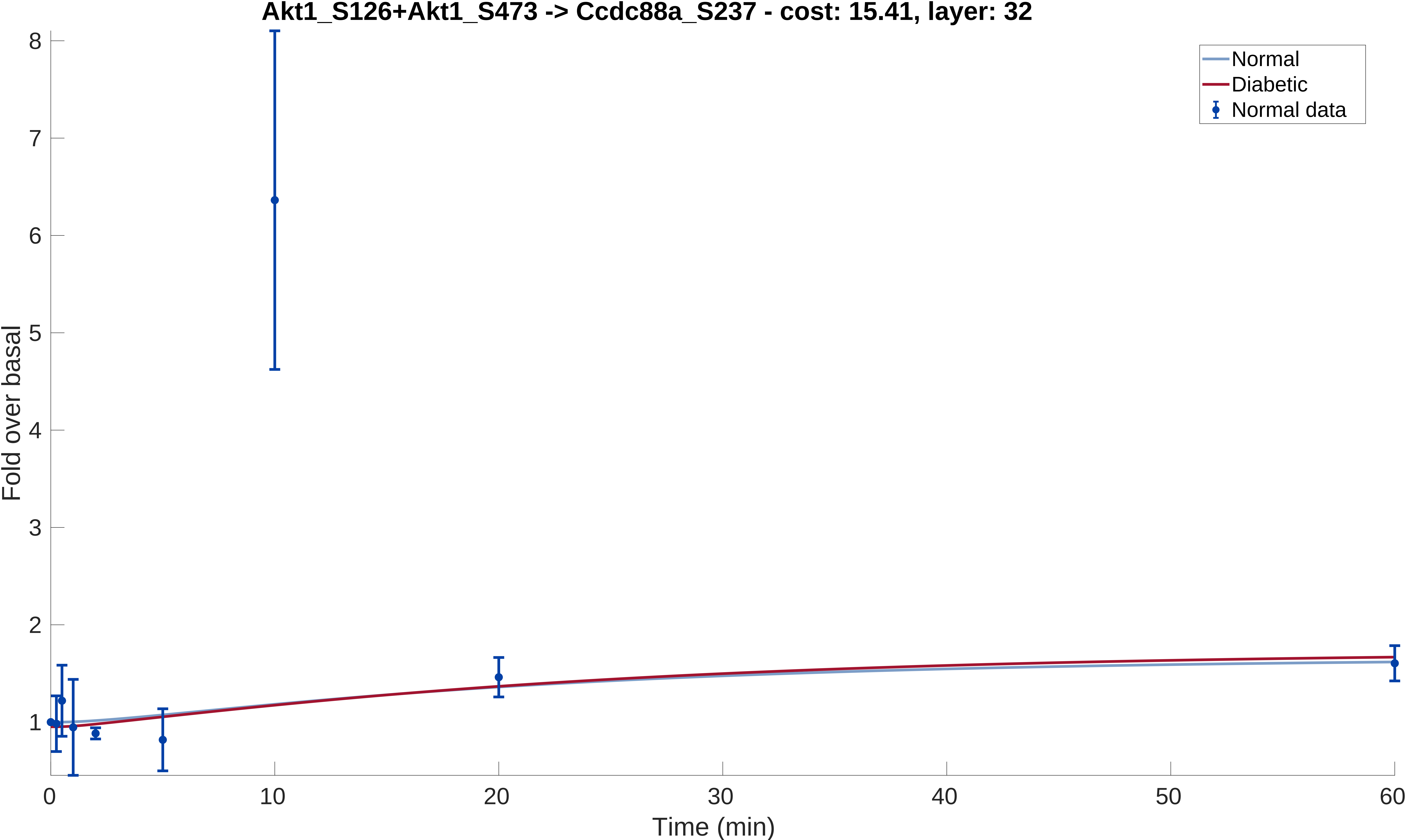
Akt1_S126+Akt1_S473 -> Ccdc88a_S237 - cost: 15.41, layer: 32
Fold over basal
Time (min)
Normal
Diabetic
Normal data
0
10
20
30
40
50
60
1
2
3
4
5
6
7
8

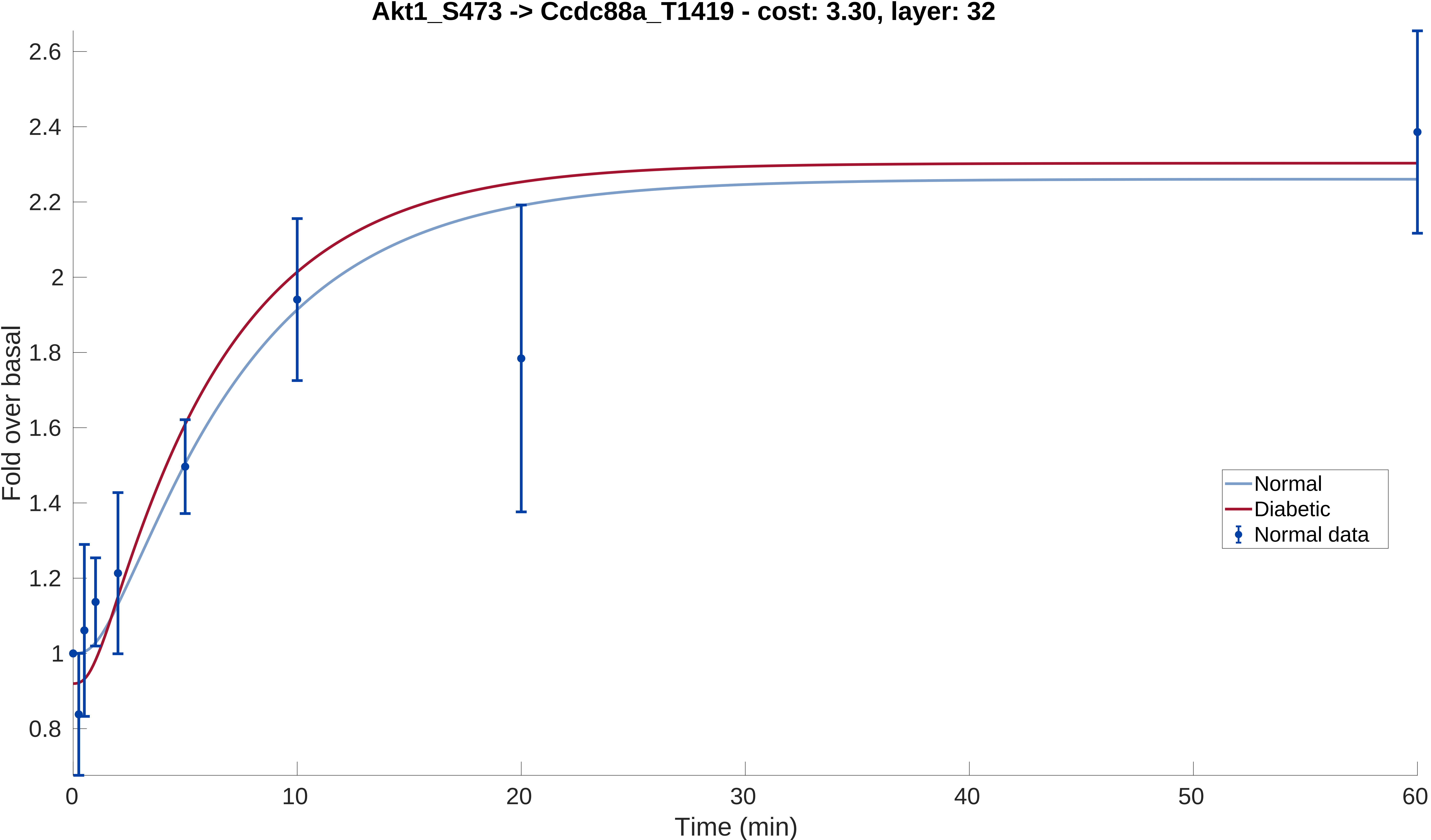
Akt1_S473 -> Ccdc88a_T1419 - cost: 3.30, layer: 32
Fold over basal
2.6
2.4
2.2
2
1.8
1.6
1.4
1.2
1
0.8
0
10
20
30
40
50
60
Time (min)
Normal
Diabetic
Normal data

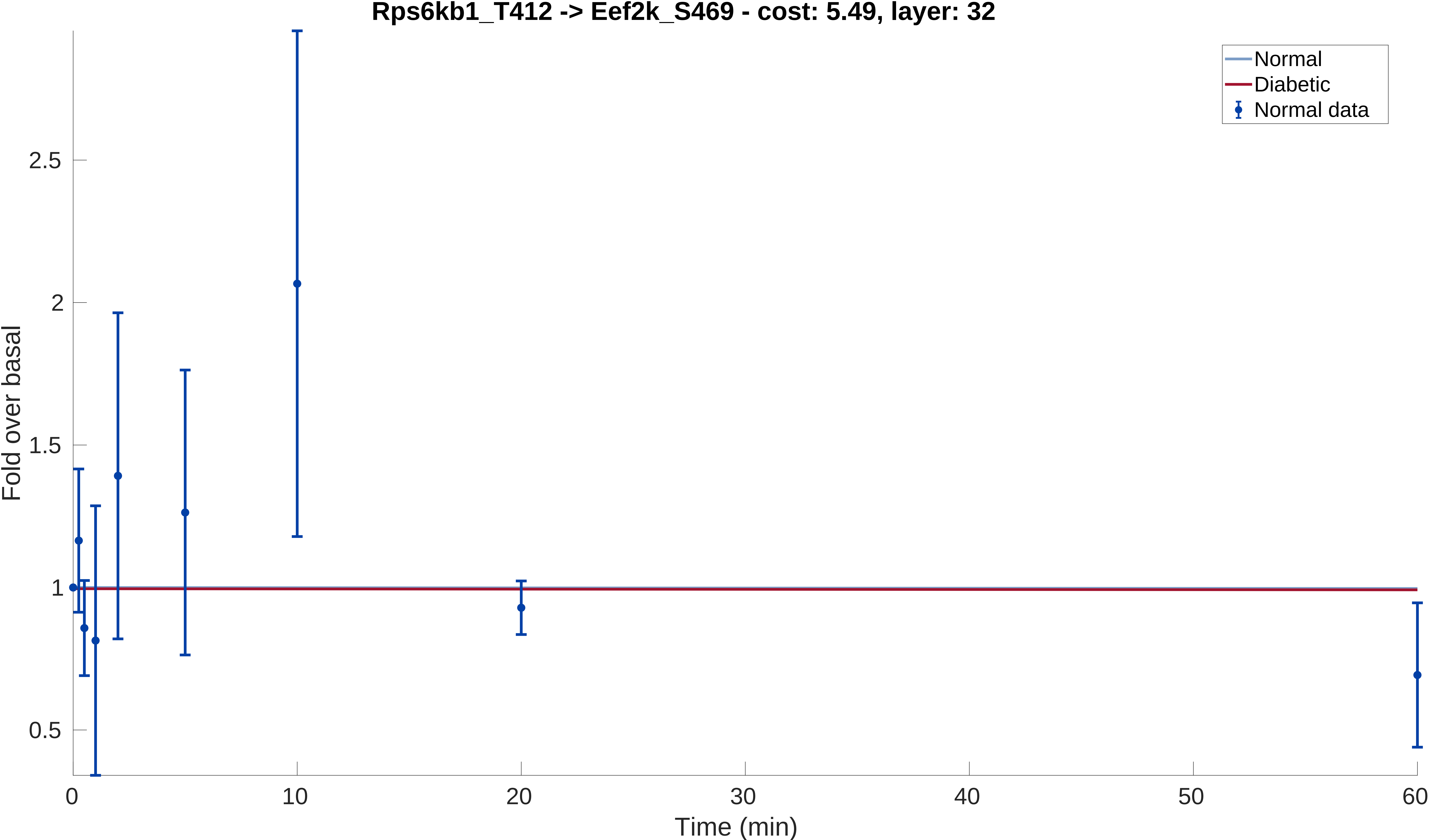

Rps6kb1_T412 -> Eef2k_S469 - cost: 5.49, layer: 32
Normal
Diabetic
Normal data
Fold over basal
0.5
1
1.5
2
2.5
0
10
20
30
40
50
60
Time (min)

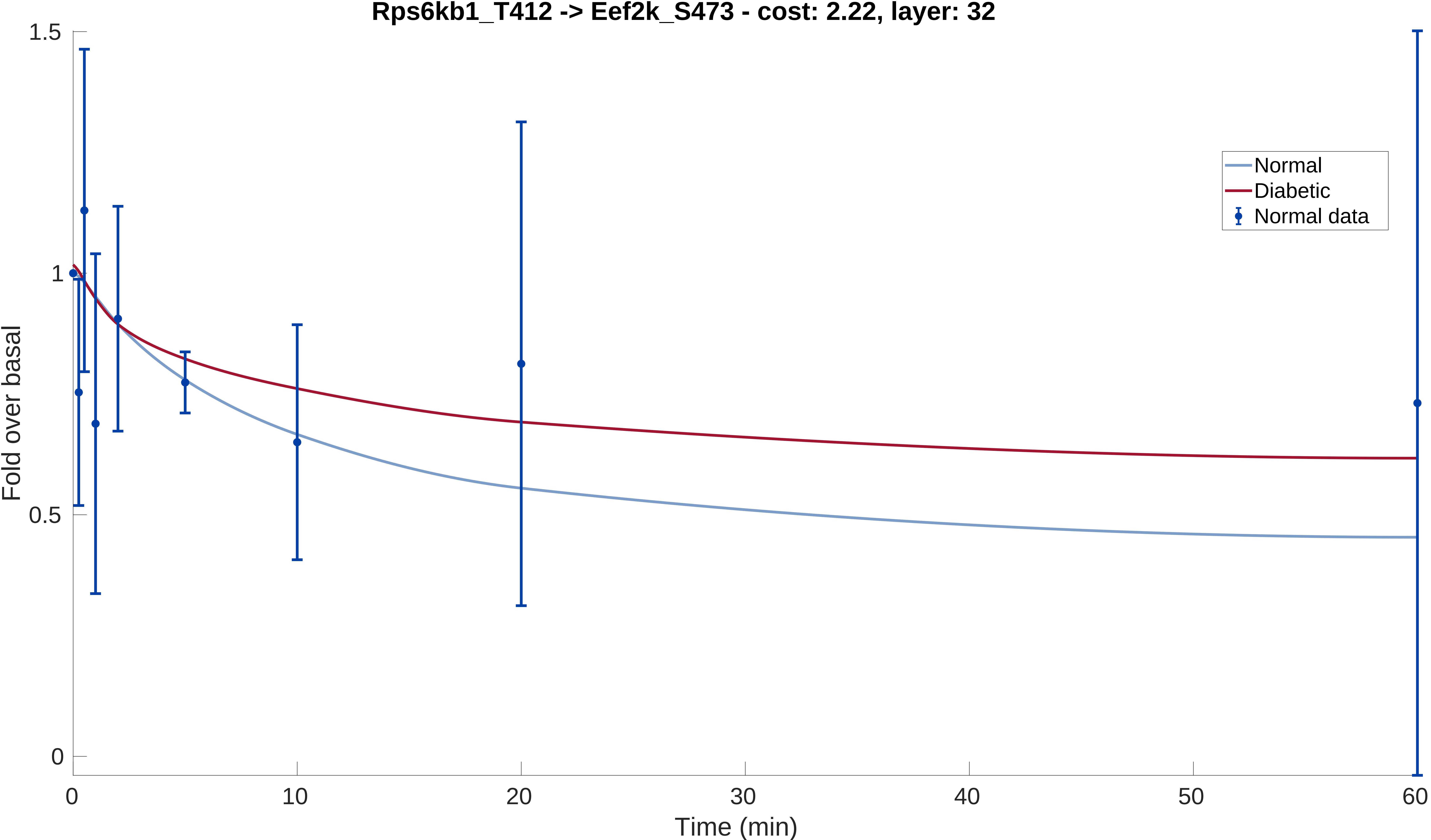

Rps6kb1_T412 -> Eef2k_S473 - cost: 2.22, layer: 32
Fold over basal
Time (min)
Normal
Diabetic
Normal data
0
0.5
1
1.5
0
10
20
30
40
50
60

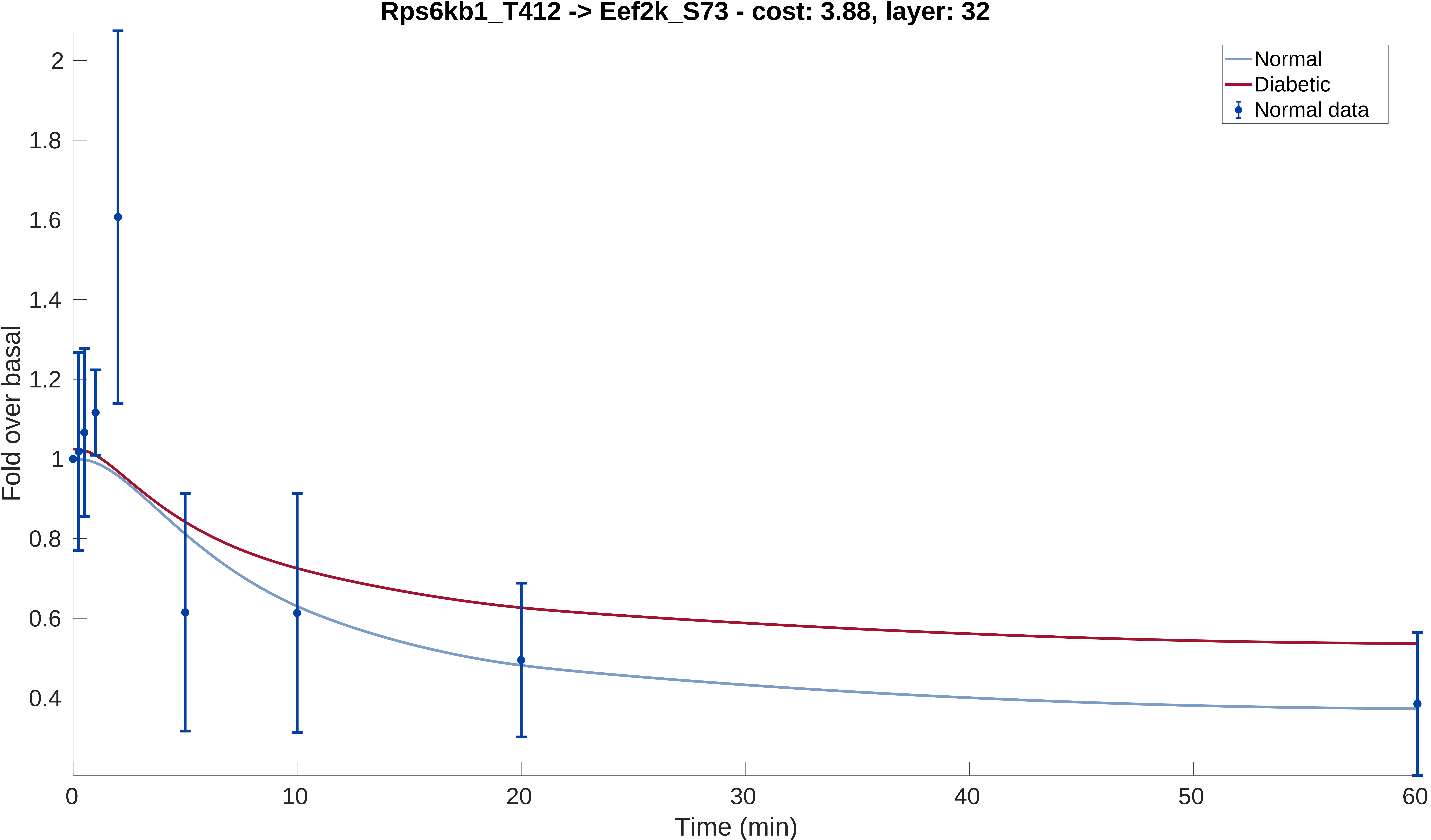

Rps6kb1_T412 -> Eef2k_S73 - cost: 3.88, layer: 32
Normal
Diabetic
Normal data
Fold over basal
2
1.8
1.6
1.4
1.2
1
0.8
0.6
0.4
0
10
20
30
40
50
60
Time (min)

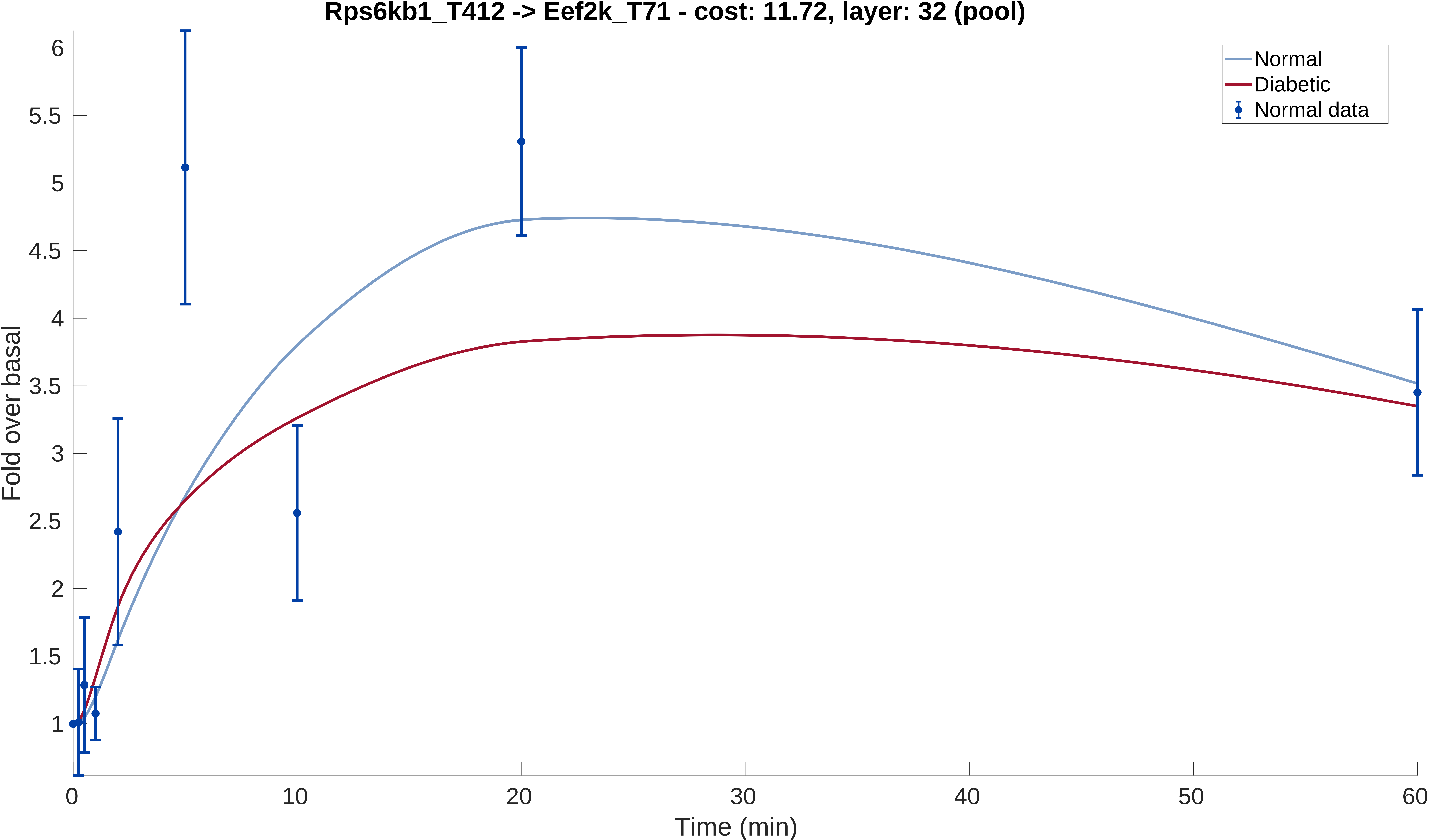

Rps6kb1_T412 -> Eef2k_T71 - cost: 11.72, layer: 32 (pool)
Fold over basal
Time (min)
Normal
Diabetic
Normal data
6
5.5
5
4.5
4
3.5
3
2.5
2
1.5
1
0
10
20
30
40
50
60

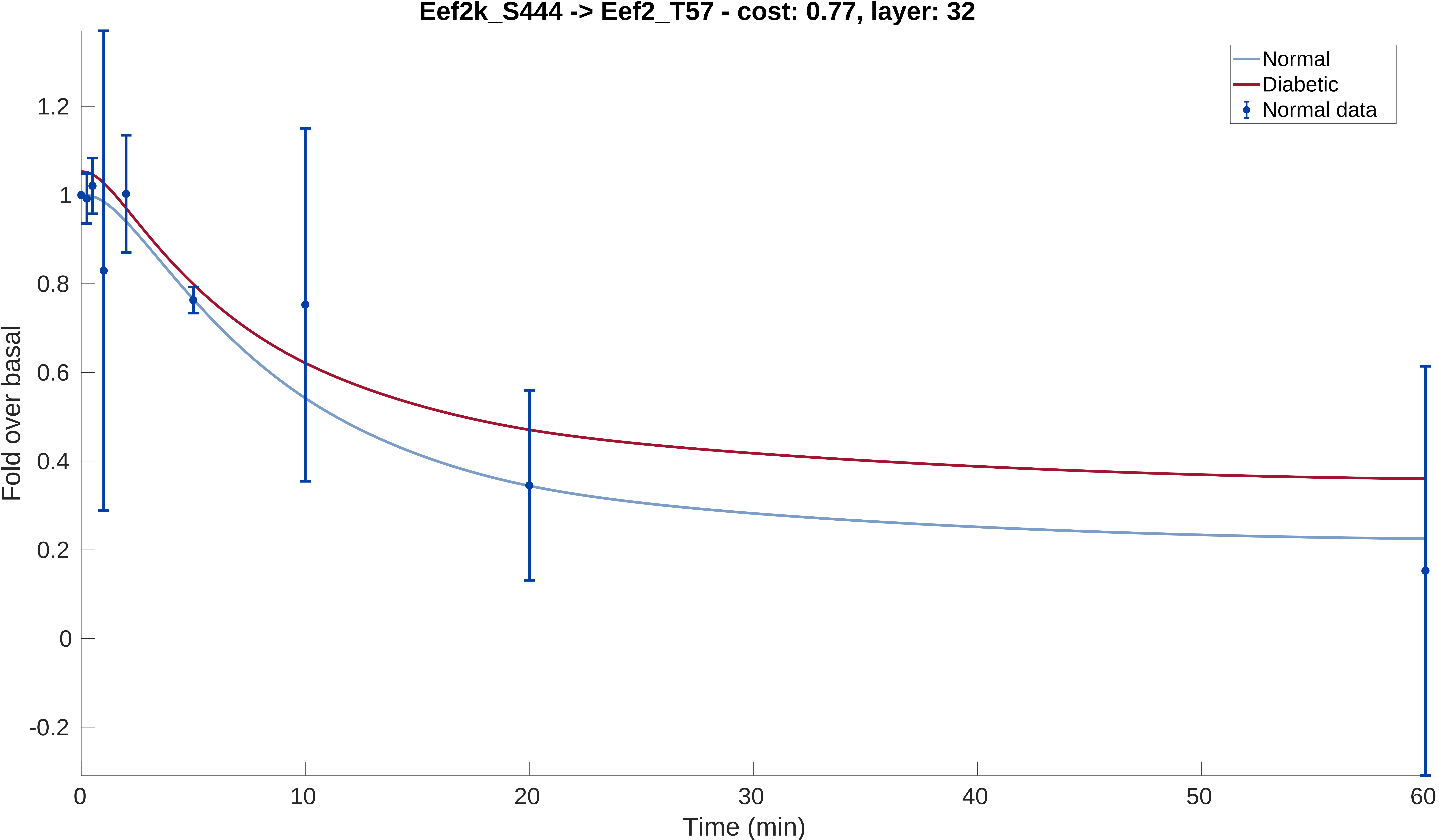

Eef2k_S444 -> Eef2_T57 - cost: 0.77, layer: 32
Normal
Diabetic
Normal data
Fold over basal
Time (min)
1.2
1
0.8
0.6
0.4
0.2
0
-0.2
0
10
20
30
40
50
60

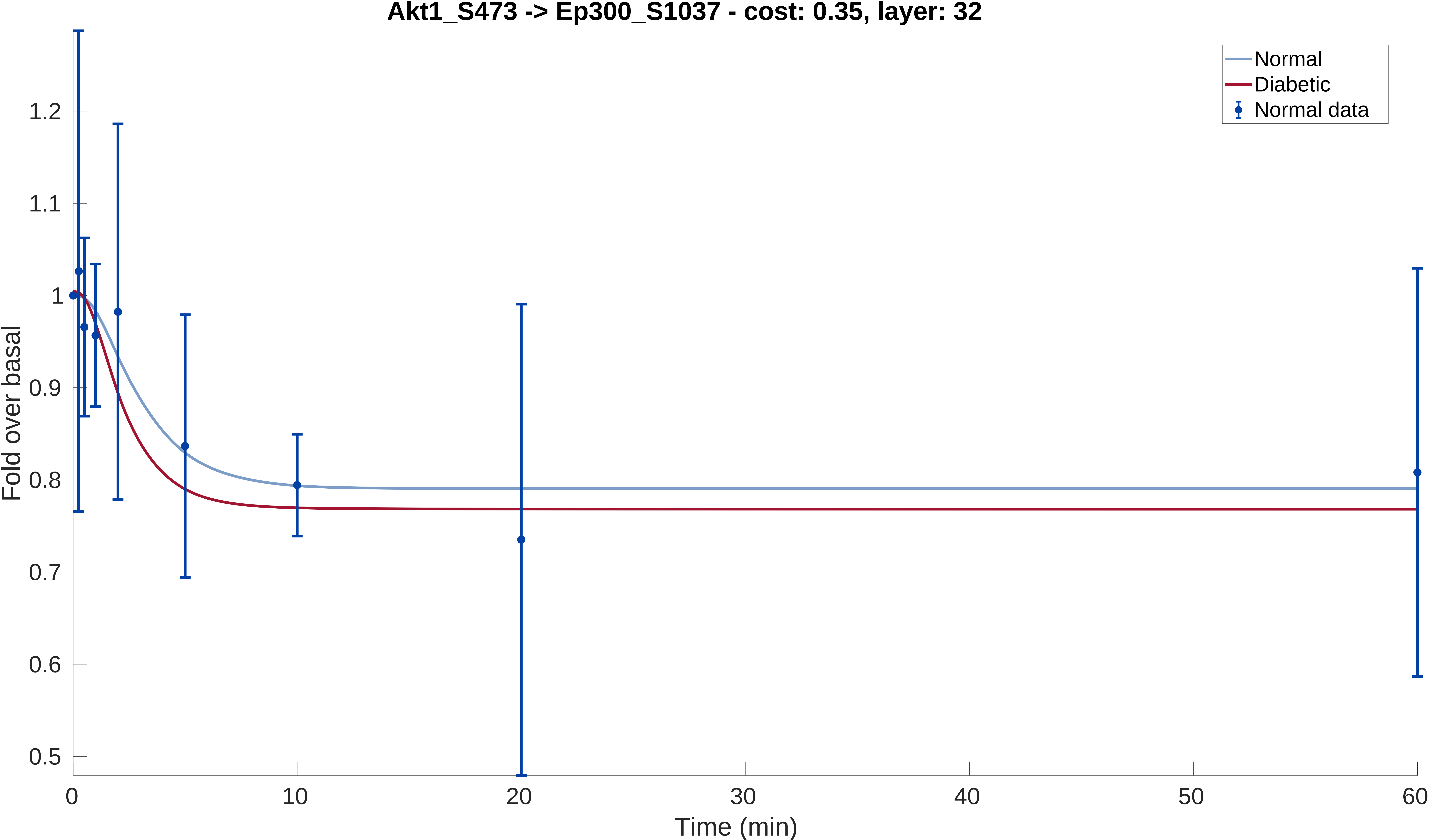

Akt1_S473 -> Ep300_S1037 - cost: 0.35, layer: 32
Normal
Diabetic
Normal data
Fold over basal
1.2
1.1
1
0.9
0.8
0.7
0.6
0.5
0
10
20
30
40
50
60
Time (min)

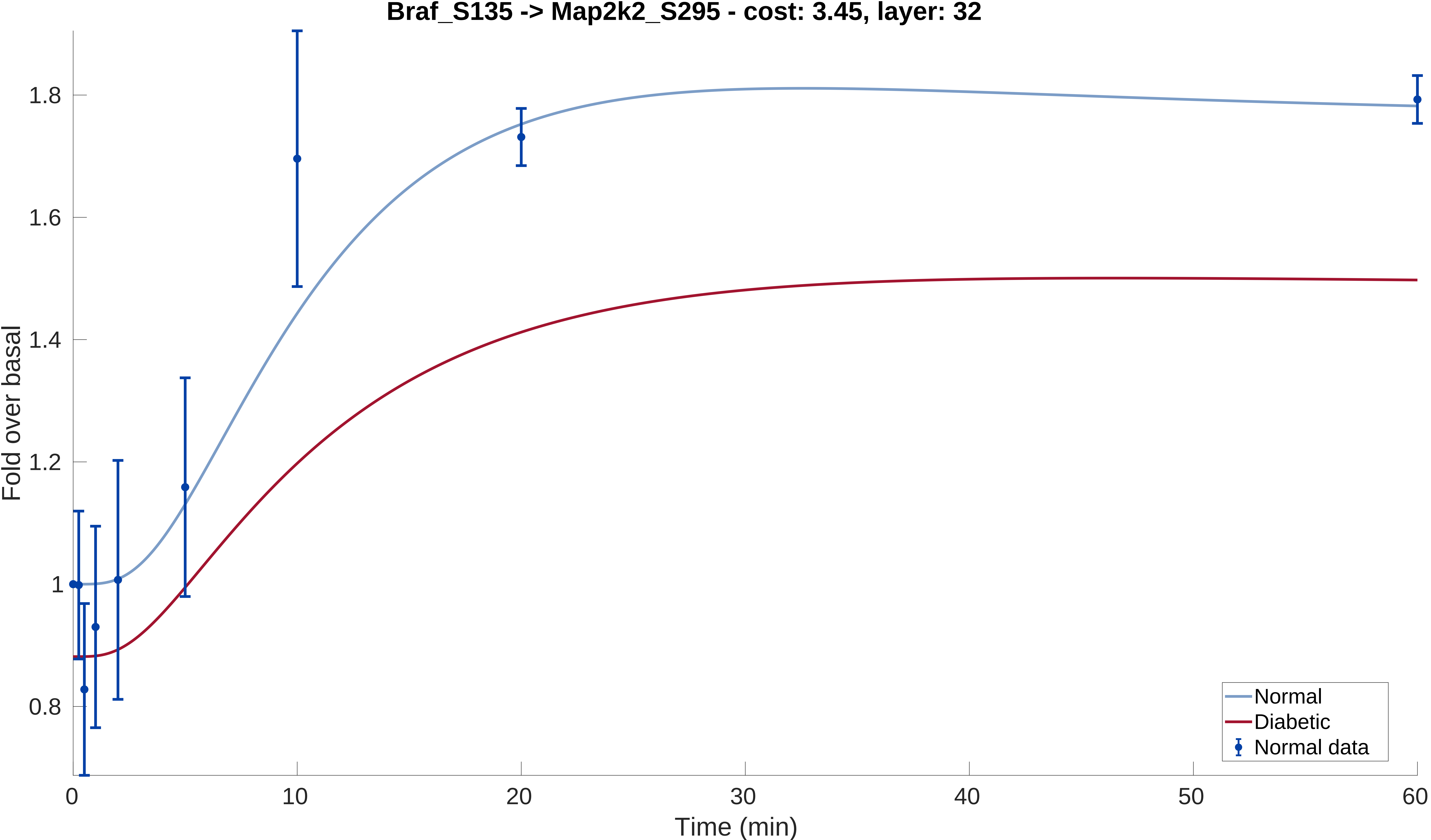
Braf_S135 -> Map2k2_S295 - cost: 3.45, layer: 32
Fold over basal
1.8
1.6
1.4
1.2
1
0.8
0
10
20
30
40
50
60
Time (min)
Normal
Diabetic
Normal data

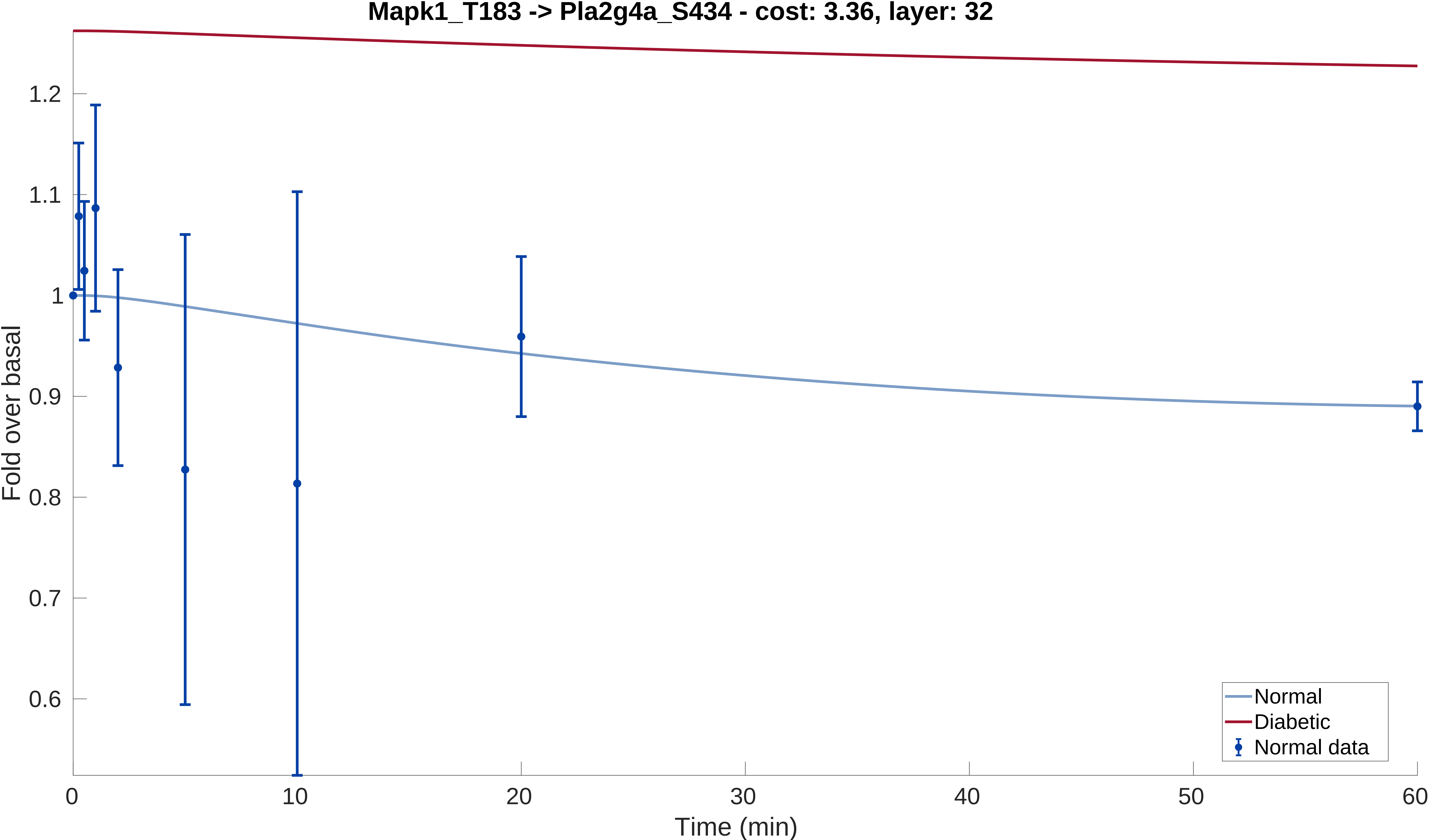

Mapk1_T183 -> Pla2g4a_S434 - cost: 3.36, layer: 32
Fold over basal
1.2
1.1
1
0.9
0.8
0.7
0.6
0
10
20
30
40
50
60
Time (min)
Normal
Diabetic
Normal data

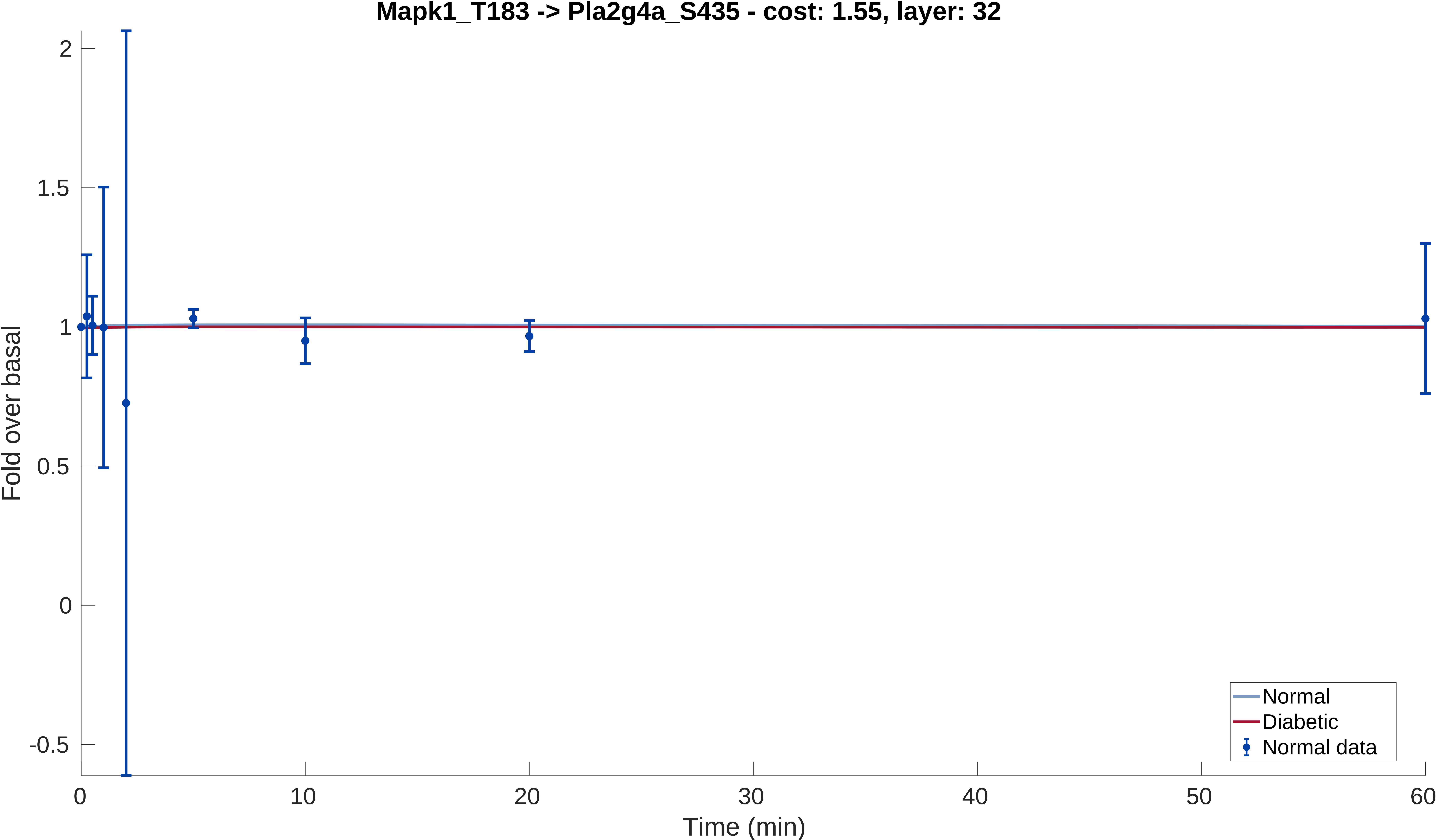

Mapk1_T183 -> Pla2g4a_S435 - cost: 1.55, layer: 32
Fold over basal
Time (min)
2
1.5
1
0.5
0
-0.5
0
10
20
30
40
50
60
Normal
Diabetic
Normal data

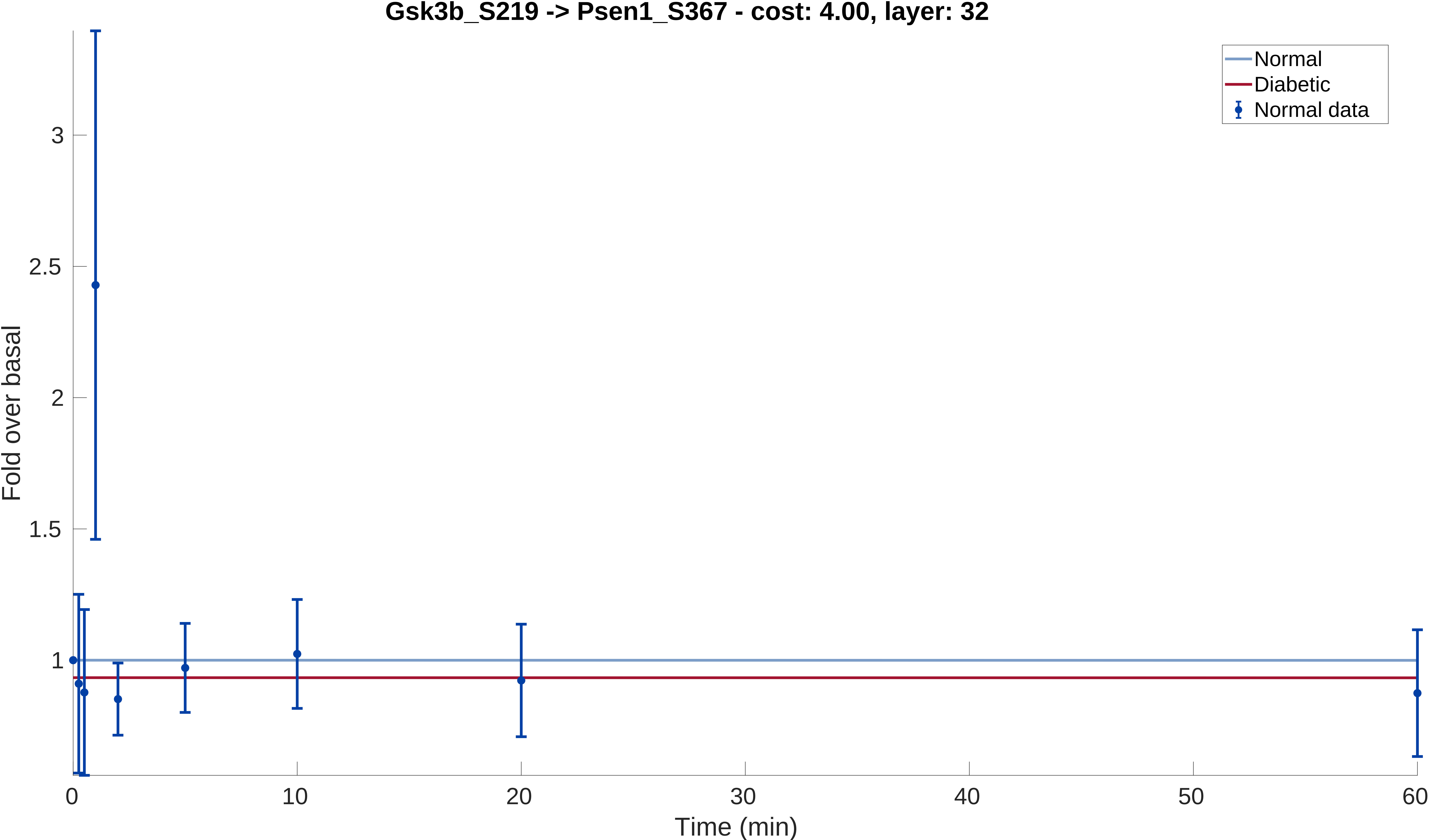

Gsk3b_S219 -> Psen1_S367 - cost: 4.00, layer: 32
Normal
Diabetic
Normal data
Fold over basal
3
2.5
2
1.5
1
0
10
20
30
40
50
60
Time (min)

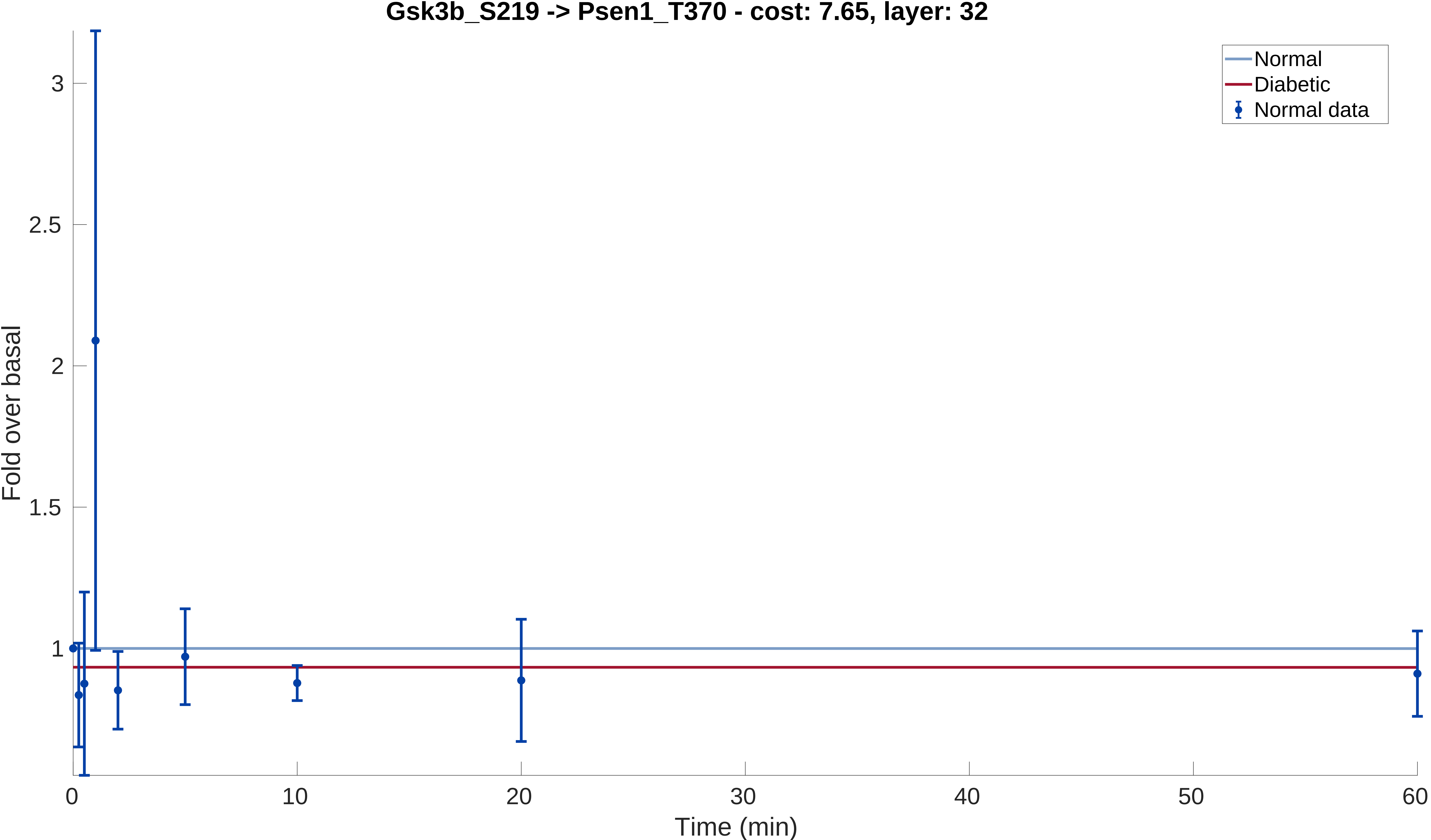

Gsk3b_S219 -> Psen1_T370 - cost: 7.65, layer: 32
Normal
Diabetic
Normal data
Fold over basal
3
2.5
2
1.5
1
0
10
20
30
40
50
60
Time (min)

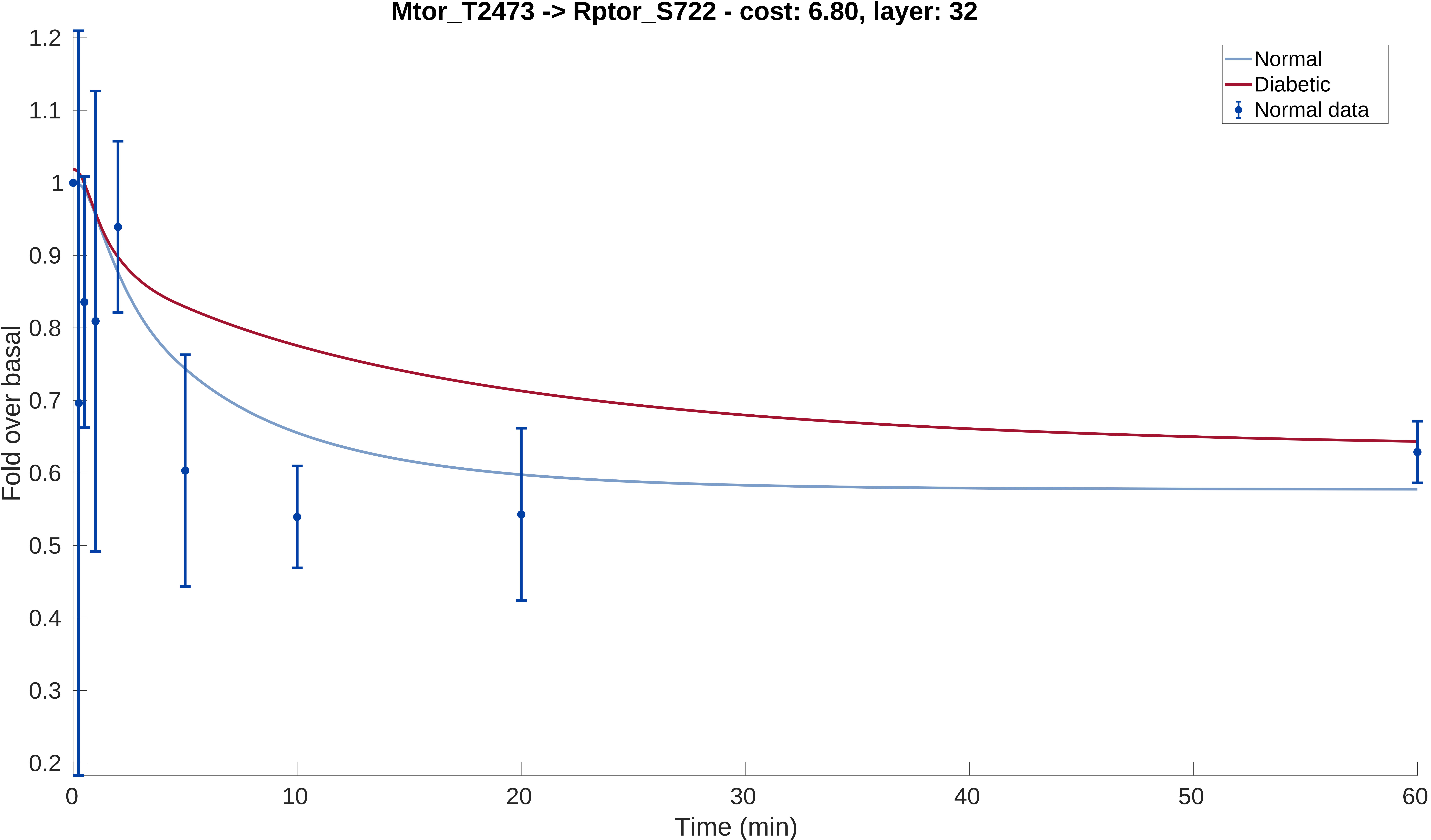

Mtor_T2473 -> Rptor_S722 - cost: 6.80, layer: 32
Normal
Diabetic
Normal data
Fold over basal
Time (min)
1.2
1.1
1
0.9
0.8
0.7
0.6
0.5
0.4
0.3
0.2
0
10
20
30
40
50
60

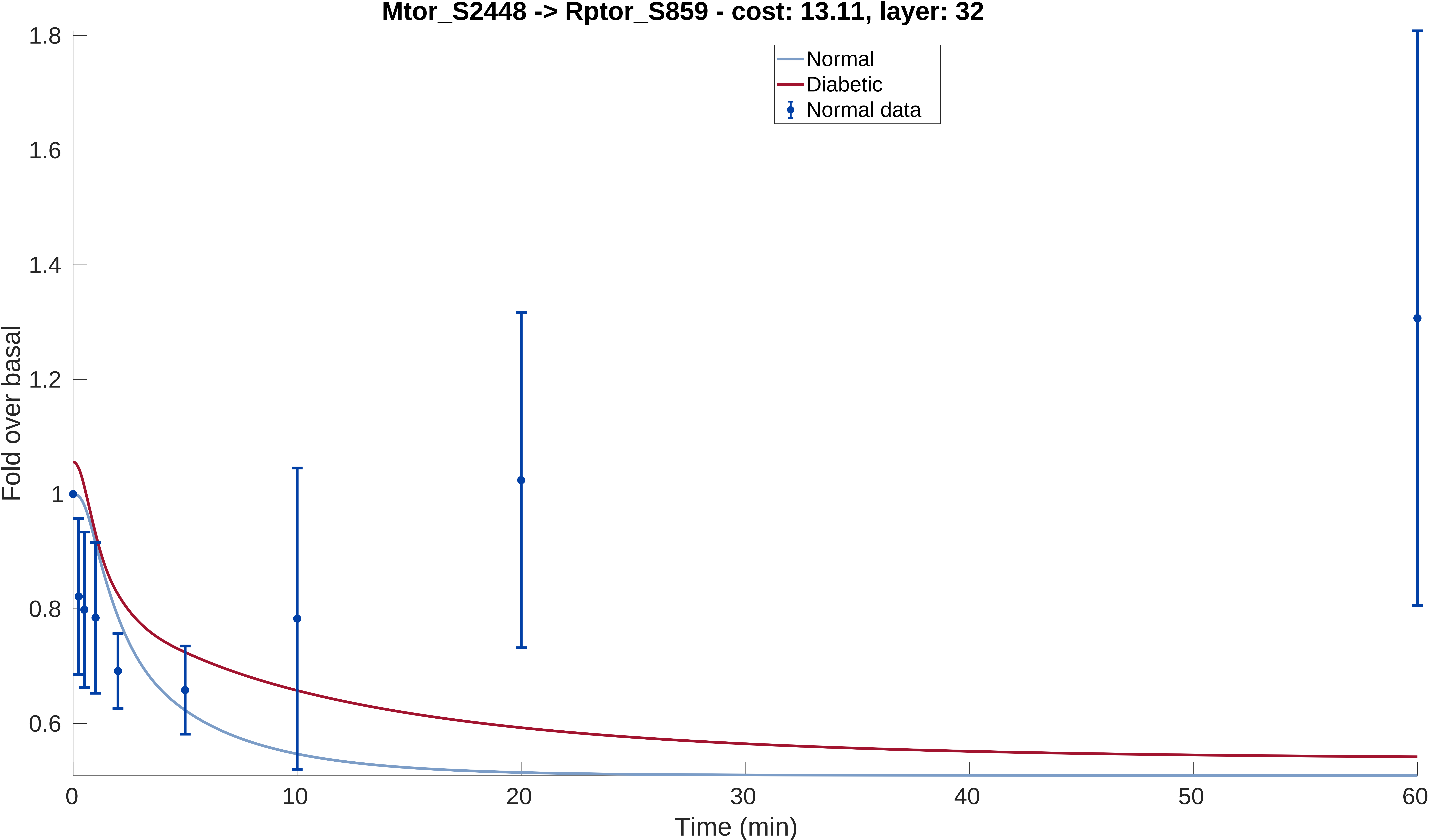
Mtor_S2448 -> Rptor_S859 - cost: 13.11, layer: 32
Normal
Diabetic
Normal data
Fold over basal
Time (min)
0.6
0.8
1
1.2
1.4
1.6
1.8
0
10
20
30
40
50
60

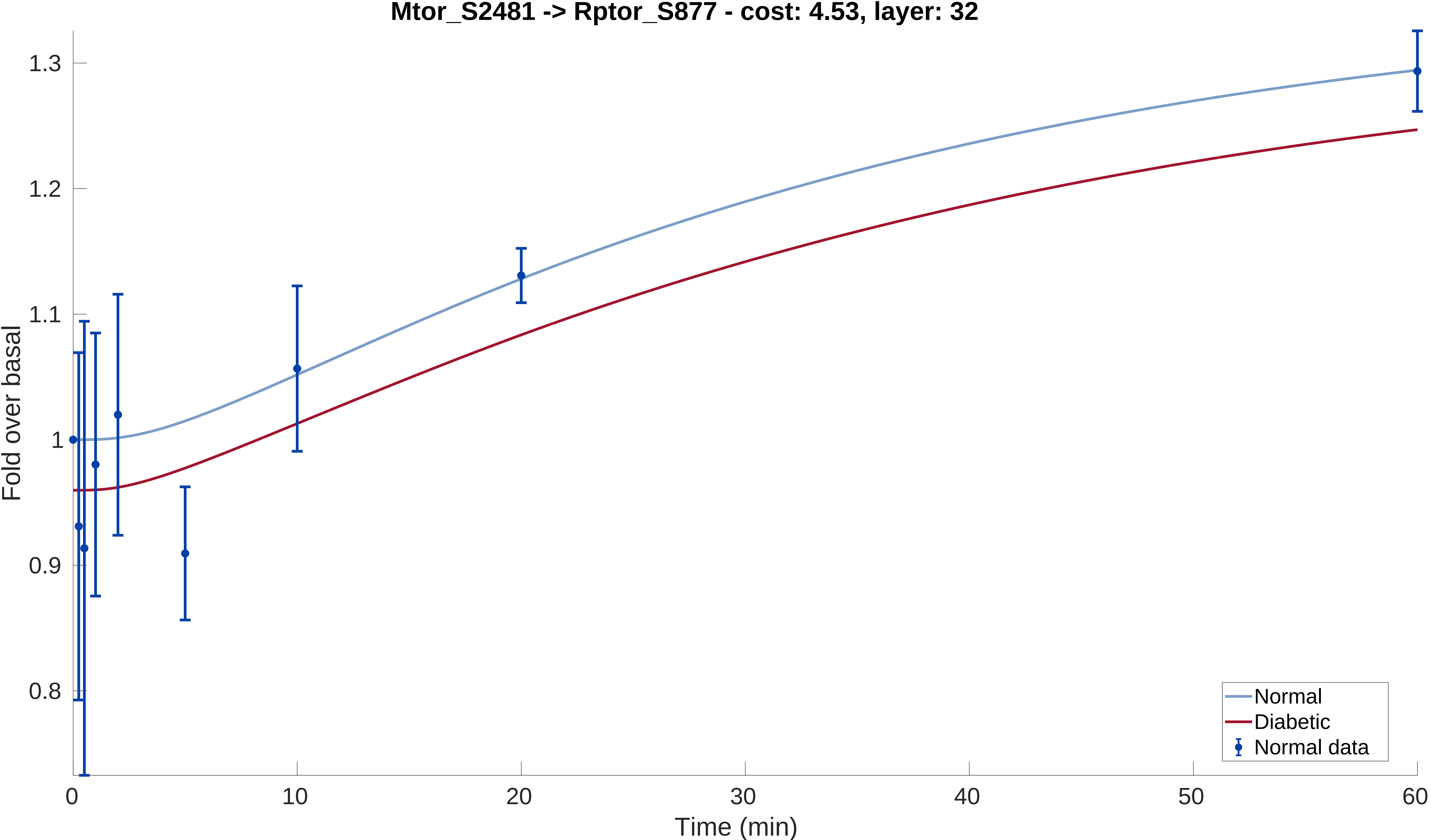

Mtor_S2481 -> Rptor_S877 - cost: 4.53, layer: 32
Fold over basal
1.3
1.2
1.1
1
0.9
0.8
0
10
20
30
40
50
60
Time (min)
Normal
Diabetic
Normal data

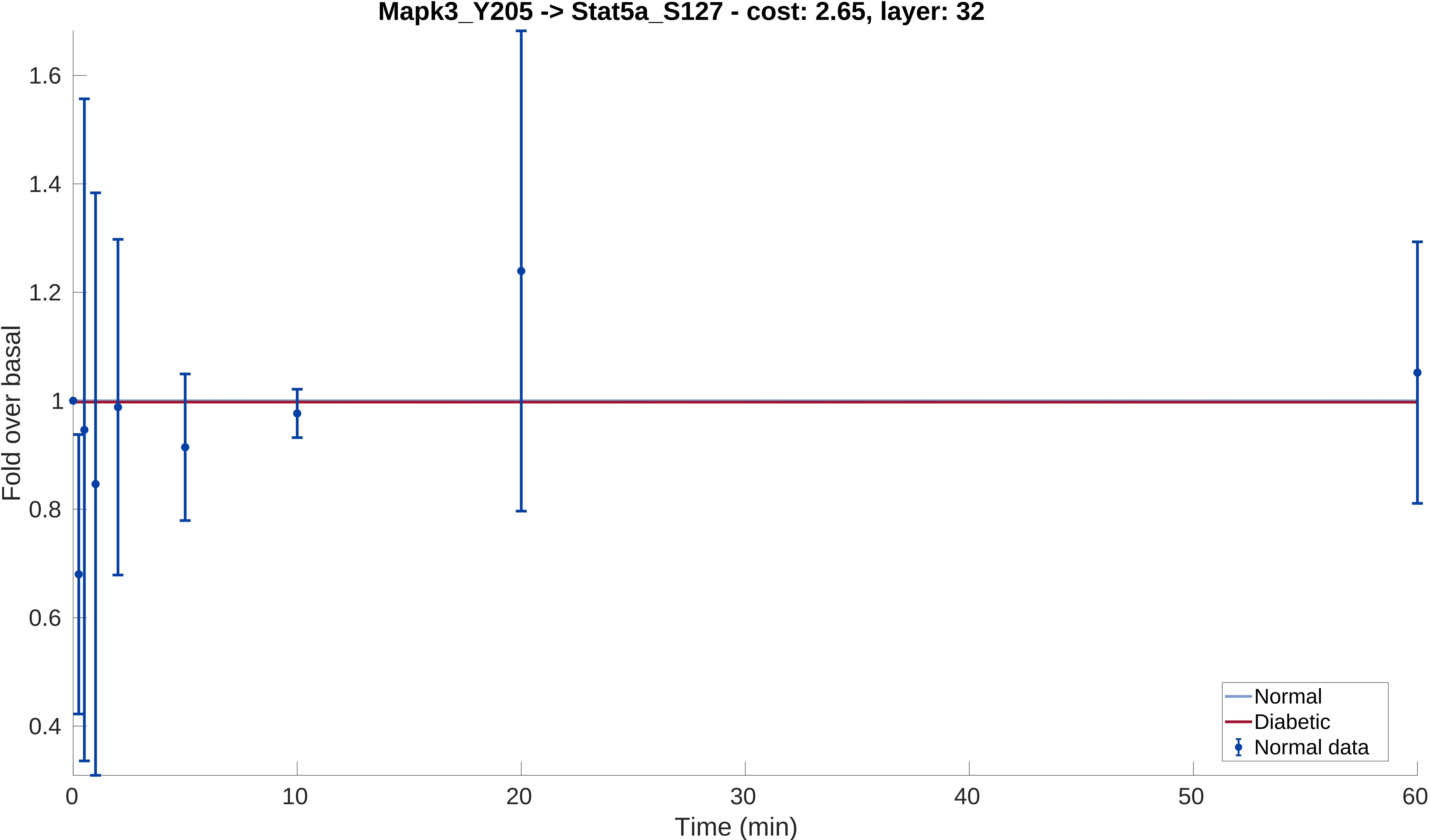
Mapk3_Y205 -> Stat5a_S127 - cost: 2.65, layer: 32
Fold over basal
1.6
1.4
1.2
1
0.8
0.6
0.4
0
10
20
30
40
50
60
Time (min)
Normal
Diabetic
Normal data

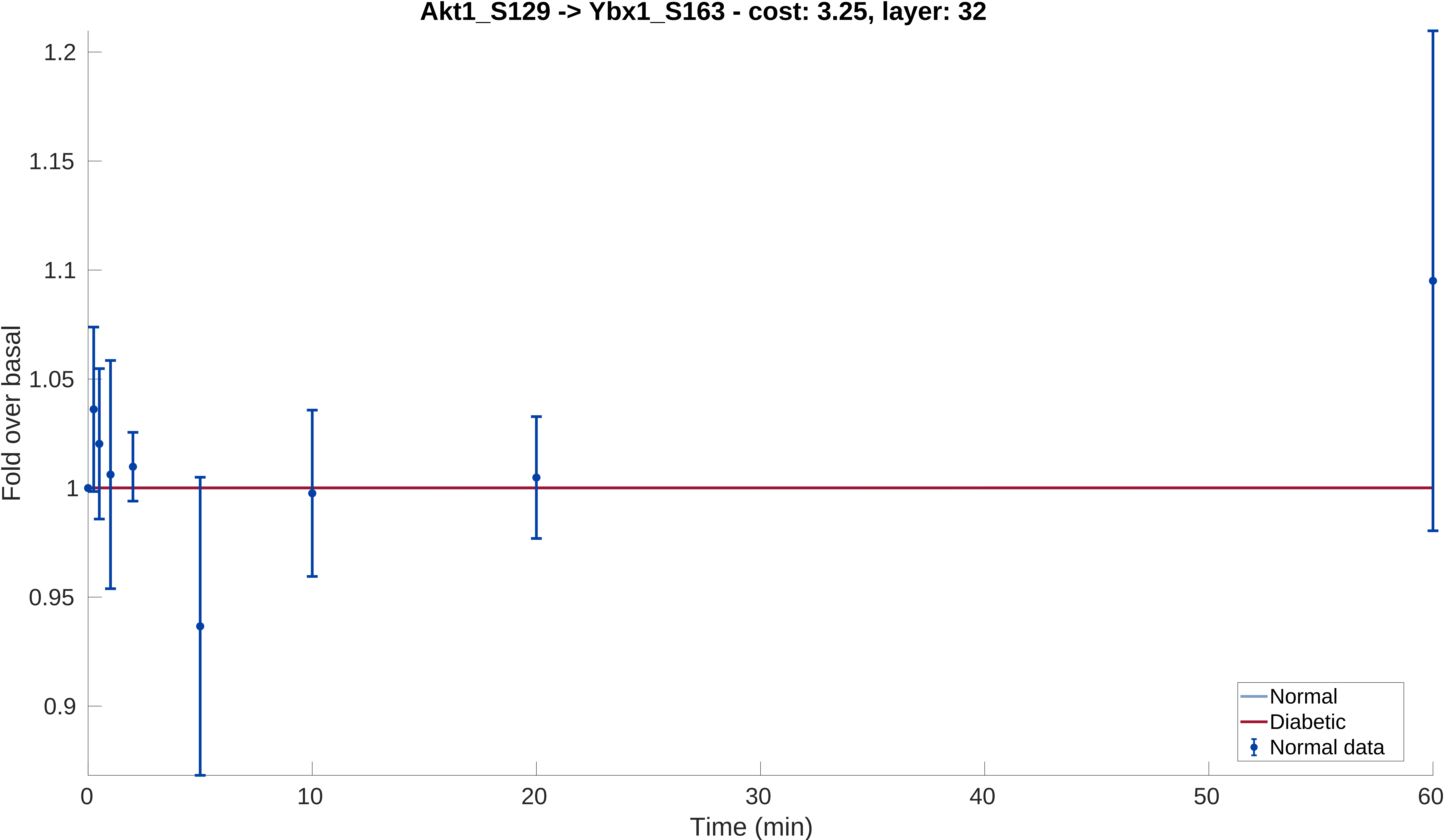

Akt1_S129 -> Ybx1_S163 - cost: 3.25, layer: 32
Fold over basal
1.2
1.15
1.1
1.05
1
0.95
0.9
0
10
20
30
40
50
60
Time (min)
Normal
Diabetic
Normal data

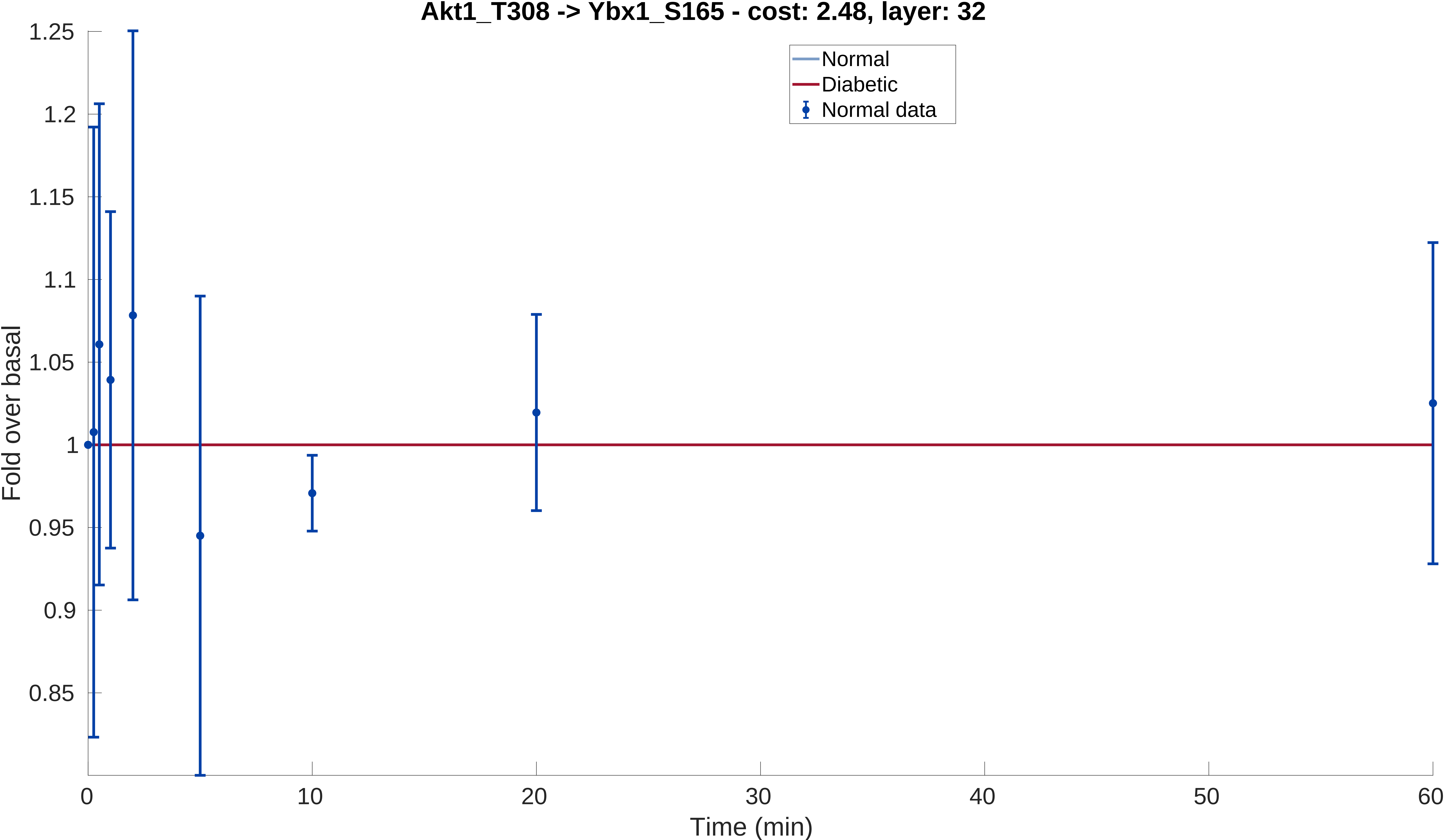

Akt1_T308 -> Ybx1_S165 - cost: 2.48, layer: 32
Normal
Diabetic
Normal data
Fold over basal
1.25
1.2
1.15
1.1
1.05
1
0.95
0.9
0.85
0
10
20
30
40
50
60
Time (min)

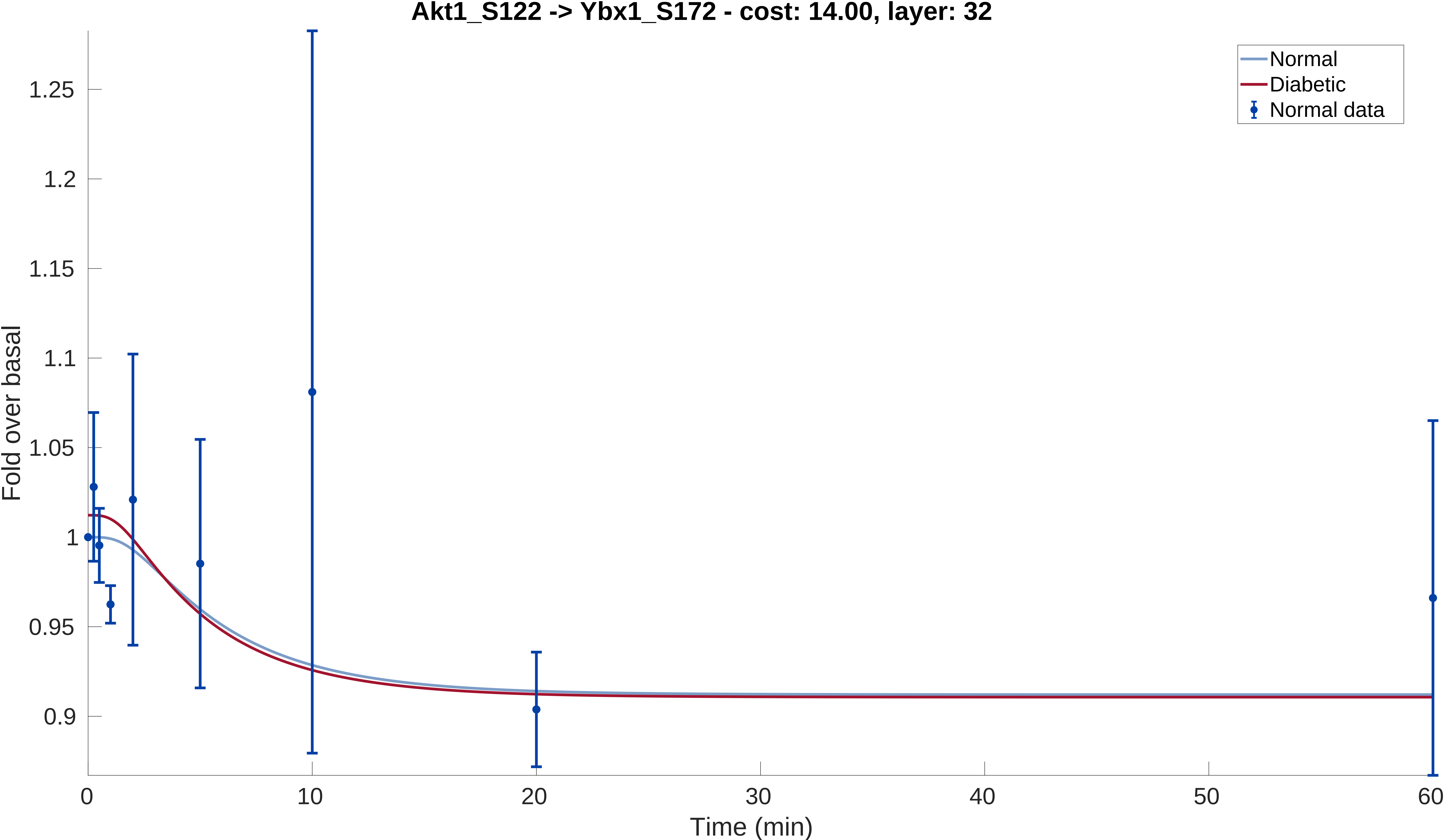

Akt1_S122 -> Ybx1_S172 - cost: 14.00, layer: 32
Normal
Diabetic
Normal data
Fold over basal
1.25
1.2
1.15
1.1
1.05
1
0.95
0.9
0
10
20
30
40
50
60
Time (min)

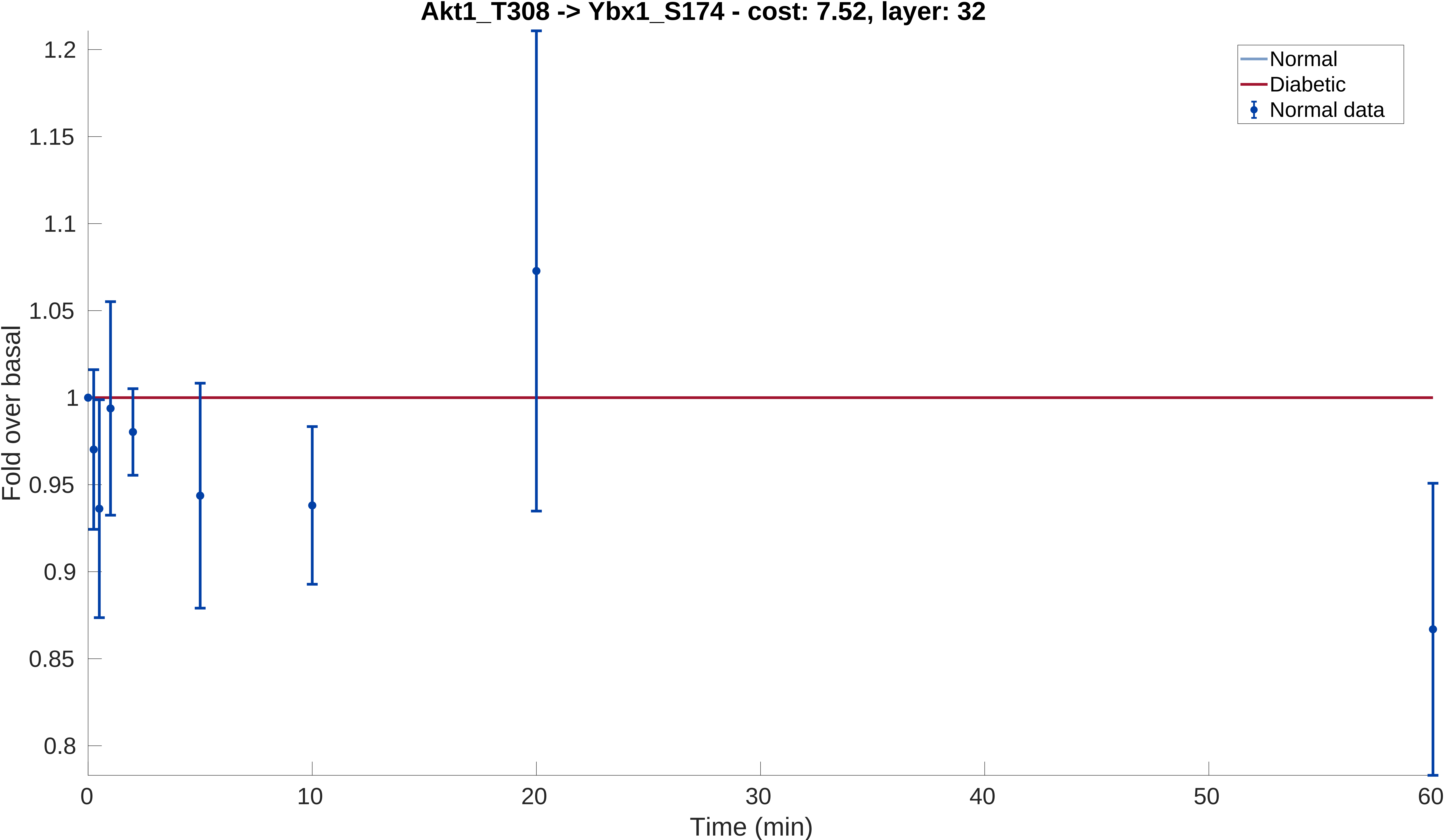

Akt1_T308 -> Ybx1_S174 - cost: 7.52, layer: 32
Normal
Diabetic
Normal data
Fold over basal
1.2
1.15
1.1
1.05
1
0.95
0.9
0.85
0.8
0
10
20
30
40
50
60
Time (min)

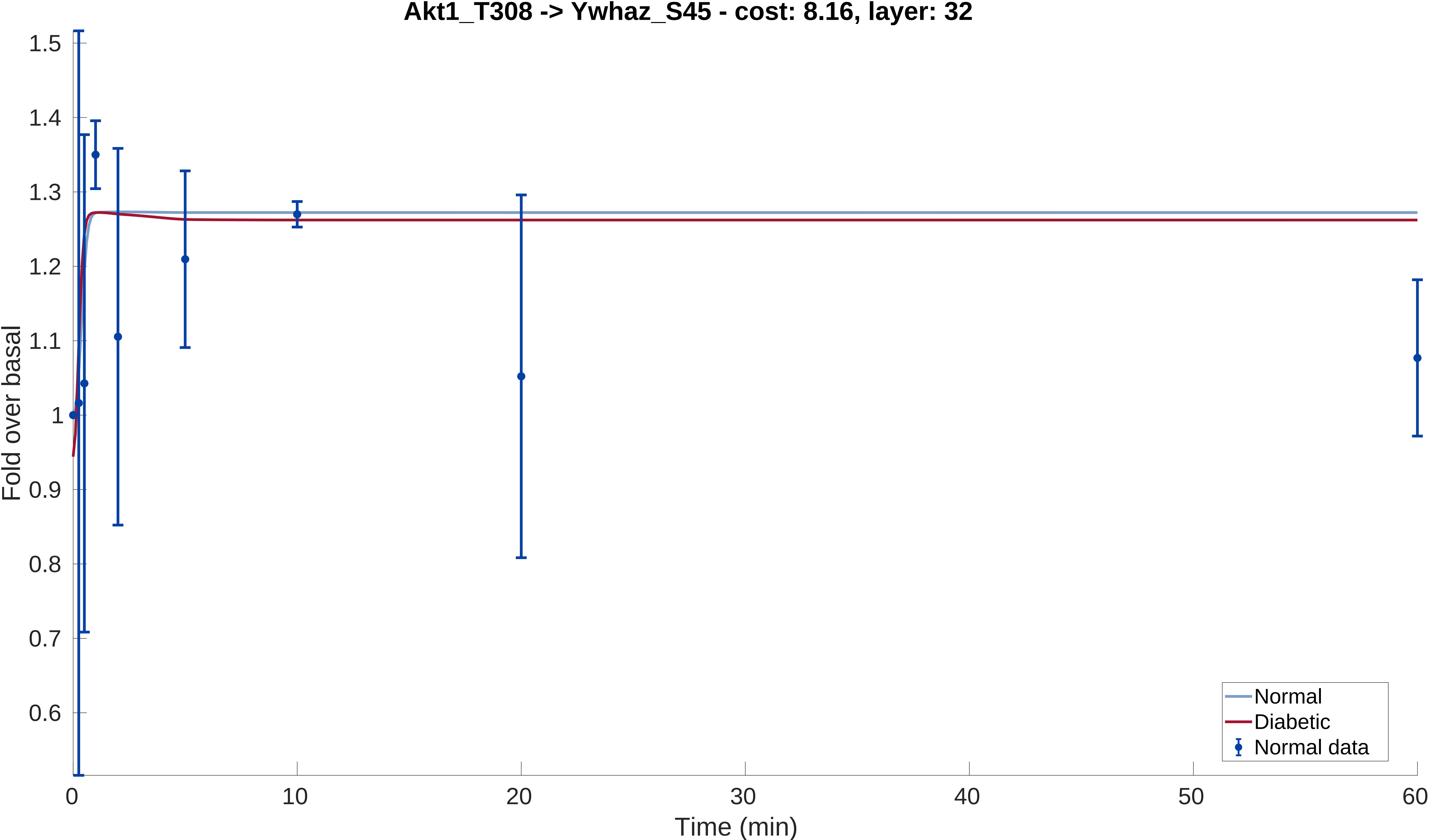

Akt1_T308 -> Ywhaz_S45 - cost: 8.16, layer: 32
Fold over basal
1.5
1.4
1.3
1.2
1.1
1
0.9
0.8
0.7
0.6
0
10
20
30
40
50
60
Time (min)
Normal
Diabetic
Normal data

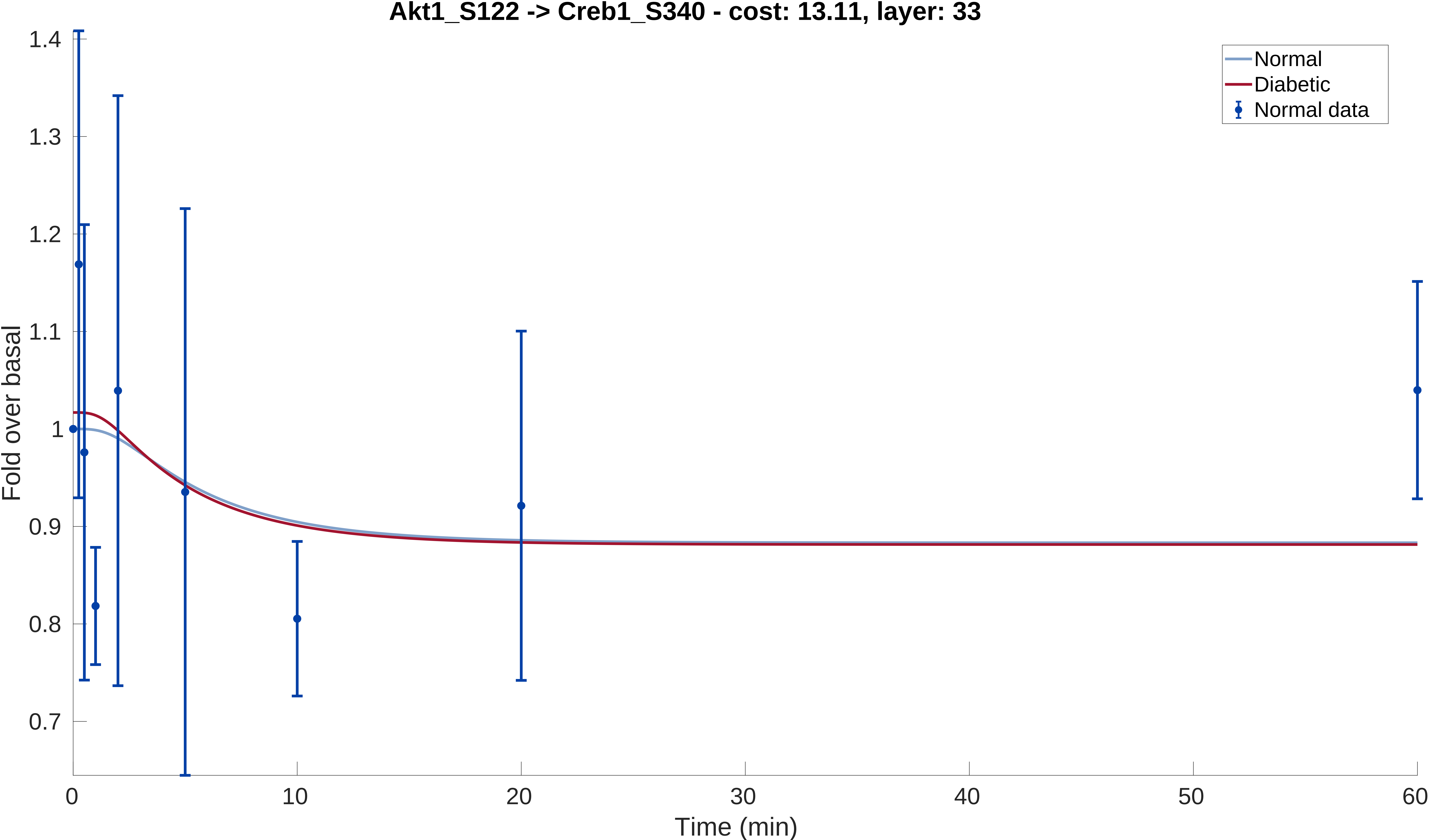

Akt1_S122 -> Creb1_S340 - cost: 13.11, layer: 33
Normal
Diabetic
Normal data
Fold over basal
1.4
1.3
1.2
1.1
1
0.9
0.8
0.7
0
10
20
30
40
50
60
Time (min)

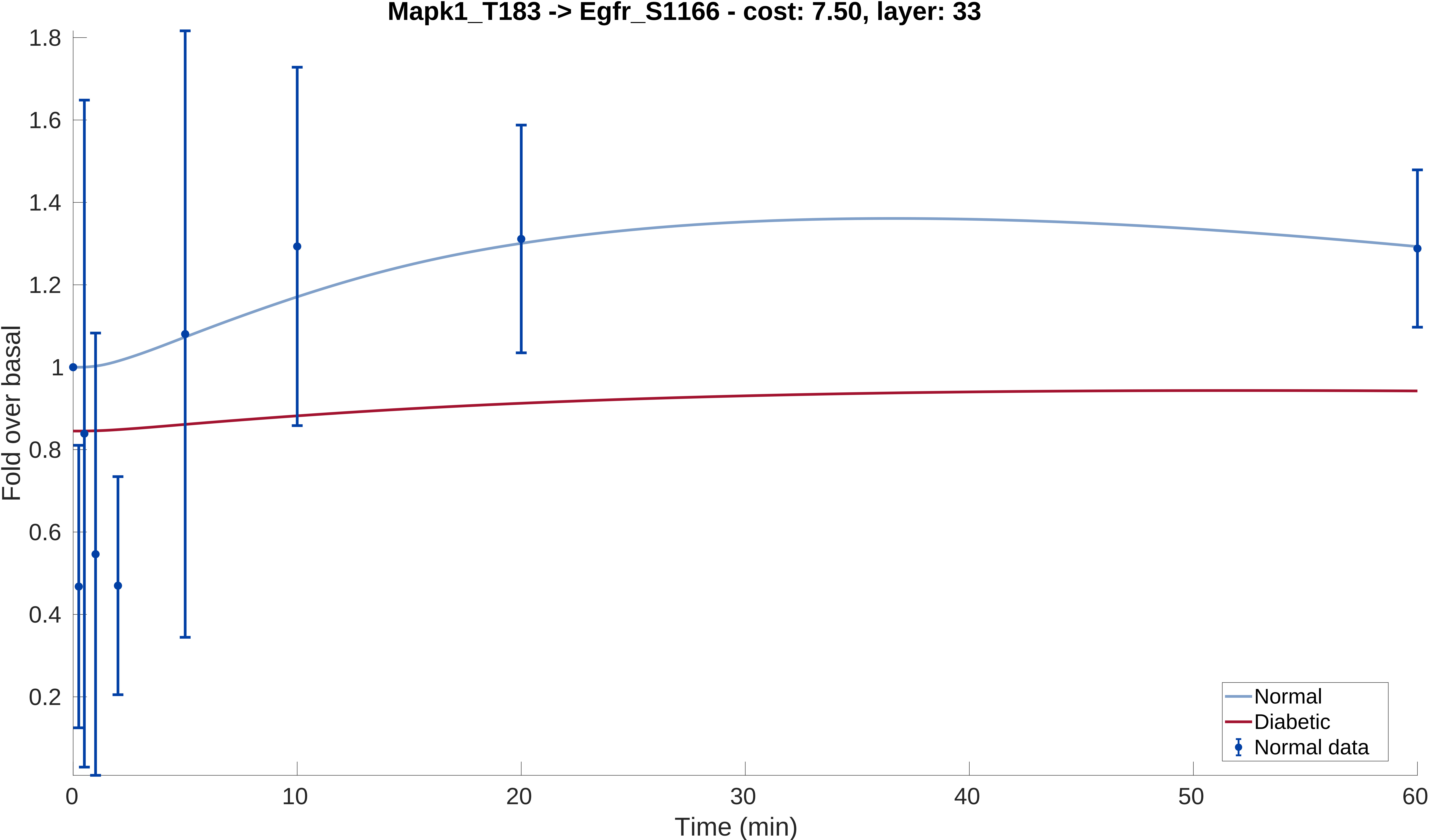

Mapk1_T183 -> Egfr_S1166 - cost: 7.50, layer: 33
Fold over basal
Time (min)
0
10
20
30
40
50
60
0.2
0.4
0.6
0.8
1
1.2
1.4
1.6
1.8
Normal
Diabetic
Normal data

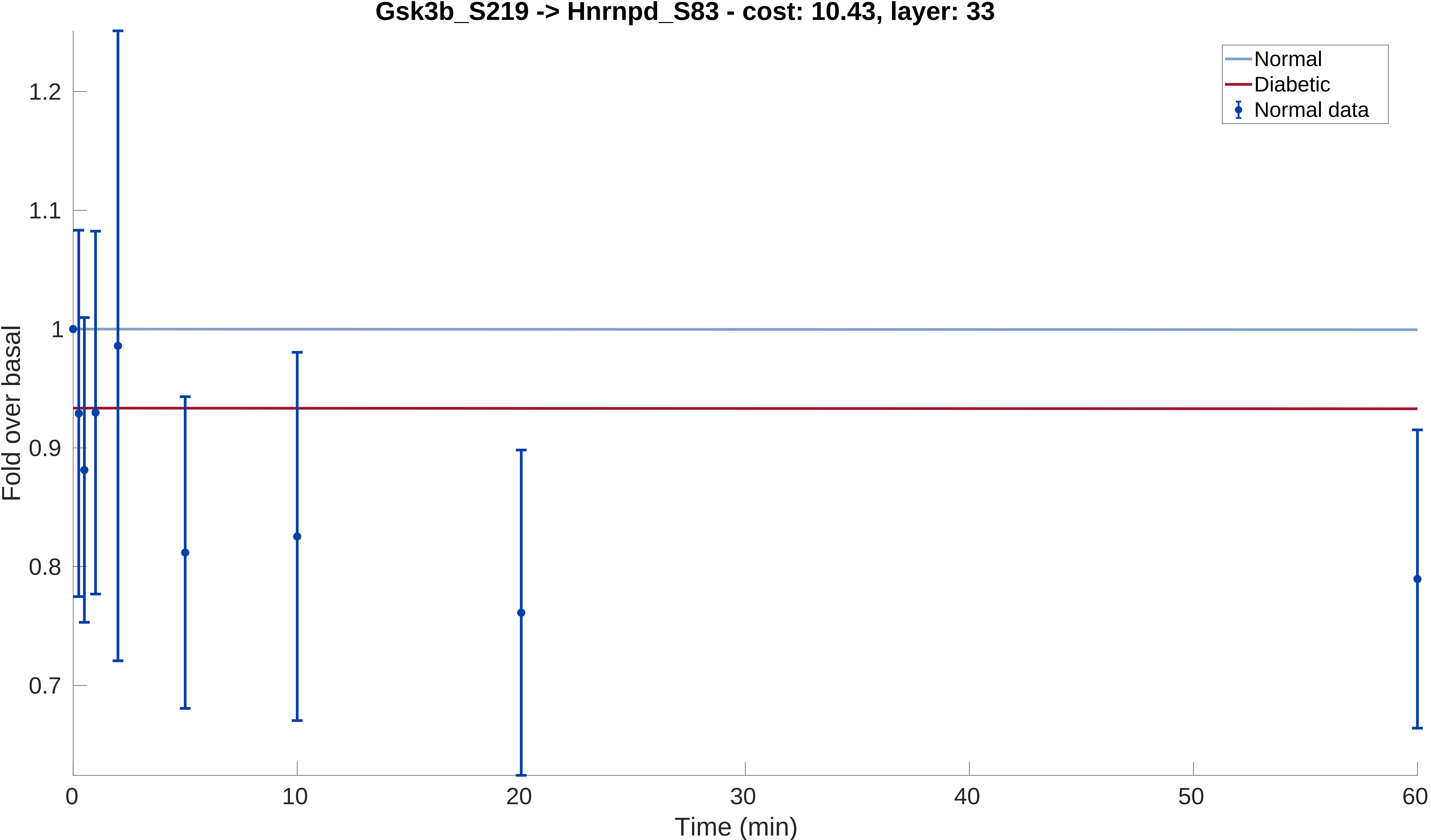
Gsk3b_S219 -> Hnrnpd_S83 - cost: 10.43, layer: 33
Normal
Diabetic
Normal data
Fold over basal
1.2
1.1
1
0.9
0.8
0.7
0
10
20
30
40
50
60
Time (min)

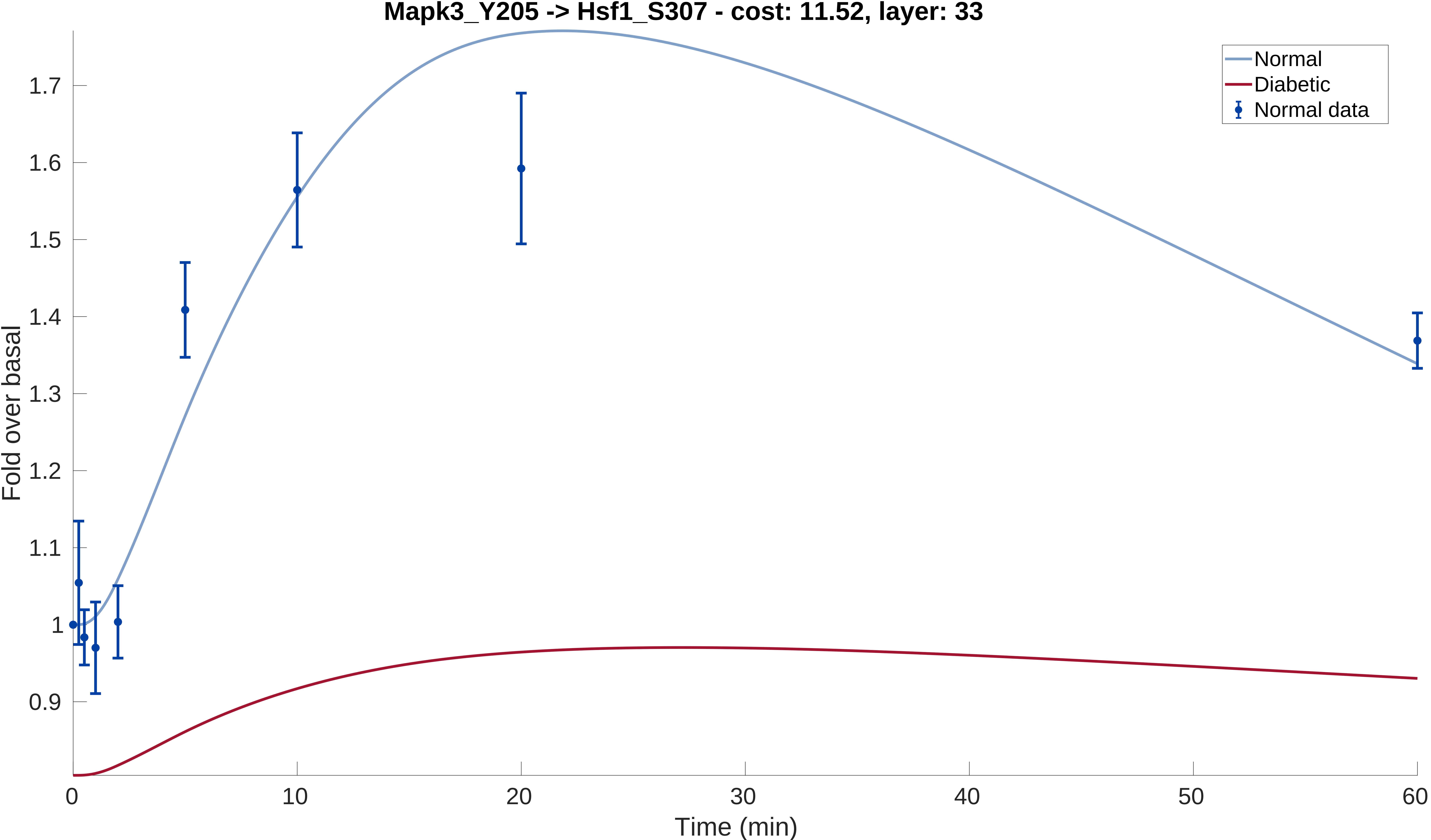

Mapk3_Y205 -> Hsf1_S307 - cost: 11.52, layer: 33
Normal
Diabetic
Normal data
Fold over basal
Time (min)
0.9
1
1.1
1.2
1.3
1.4
1.5
1.6
1.7
0
10
20
30
40
50
60

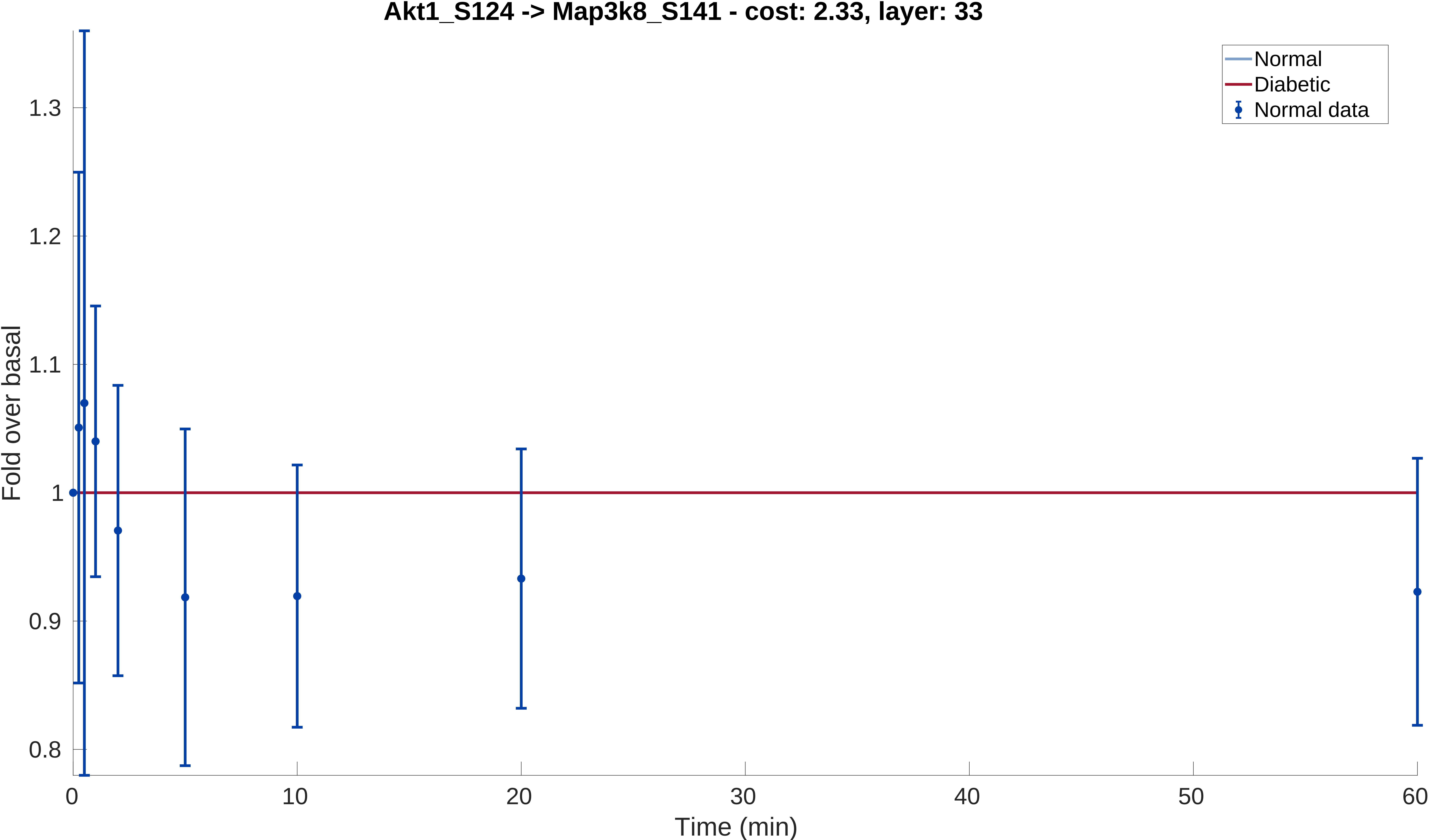

Akt1_S124 -> Map3k8_S141 - cost: 2.33, layer: 33
Normal
Diabetic
Normal data
Fold over basal
1.3
1.2
1.1
1
0.9
0.8
0
10
20
30
40
50
60
Time (min)

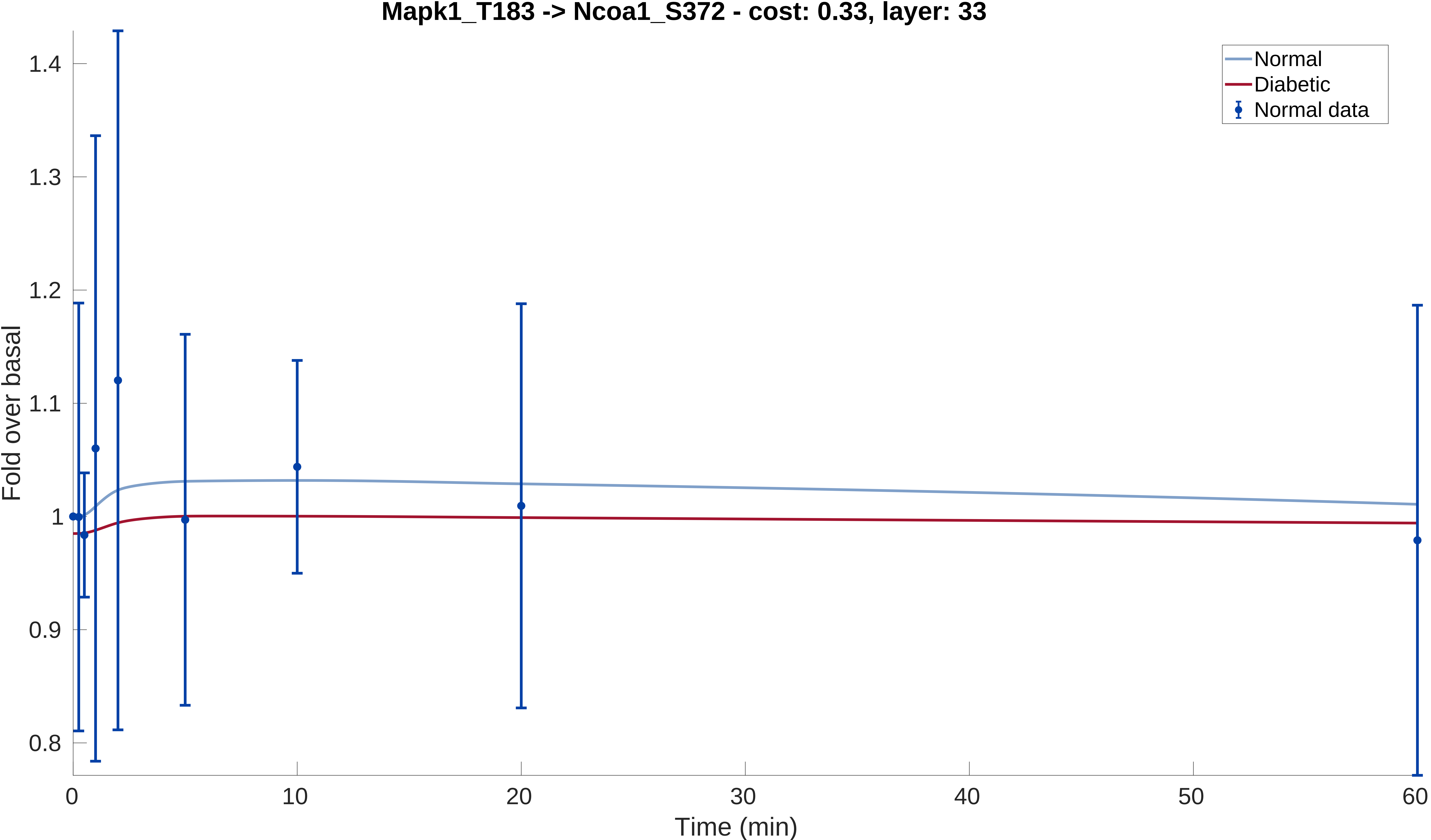
Mapk1_T183 -> Ncoa1_S372 - cost: 0.33, layer: 33
Normal
Diabetic
Normal data
Fold over basal
1.4
1.3
1.2
1.1
1
0.9
0.8
0
10
20
30
40
50
60
Time (min)

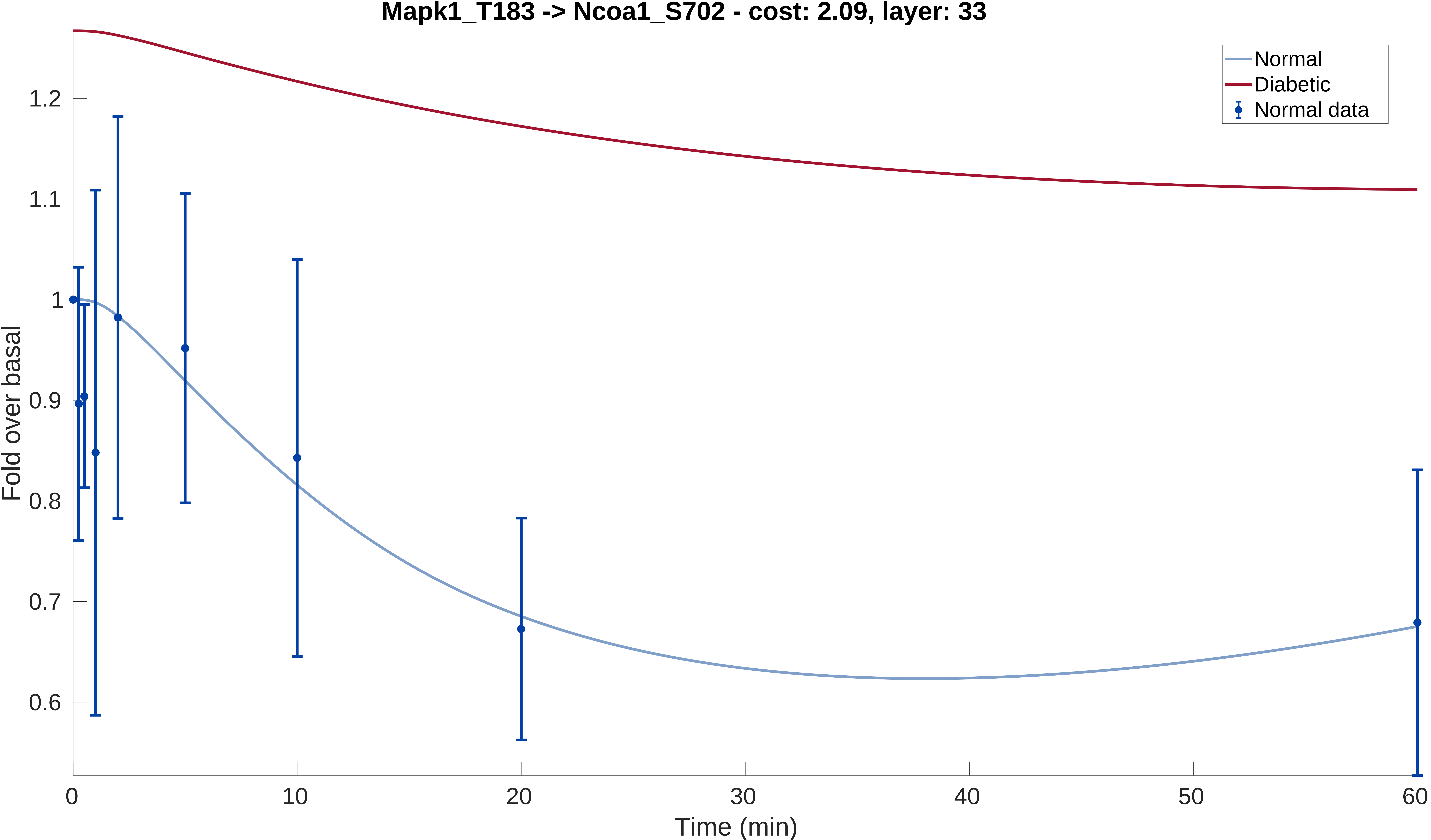

Mapk1_T183 -> Ncoa1_S702 - cost: 2.09, layer: 33
Normal
Diabetic
Normal data
Fold over basal
Time (min)
1.2
1.1
1
0.9
0.8
0.7
0.6
0
10
20
30
40
50
60

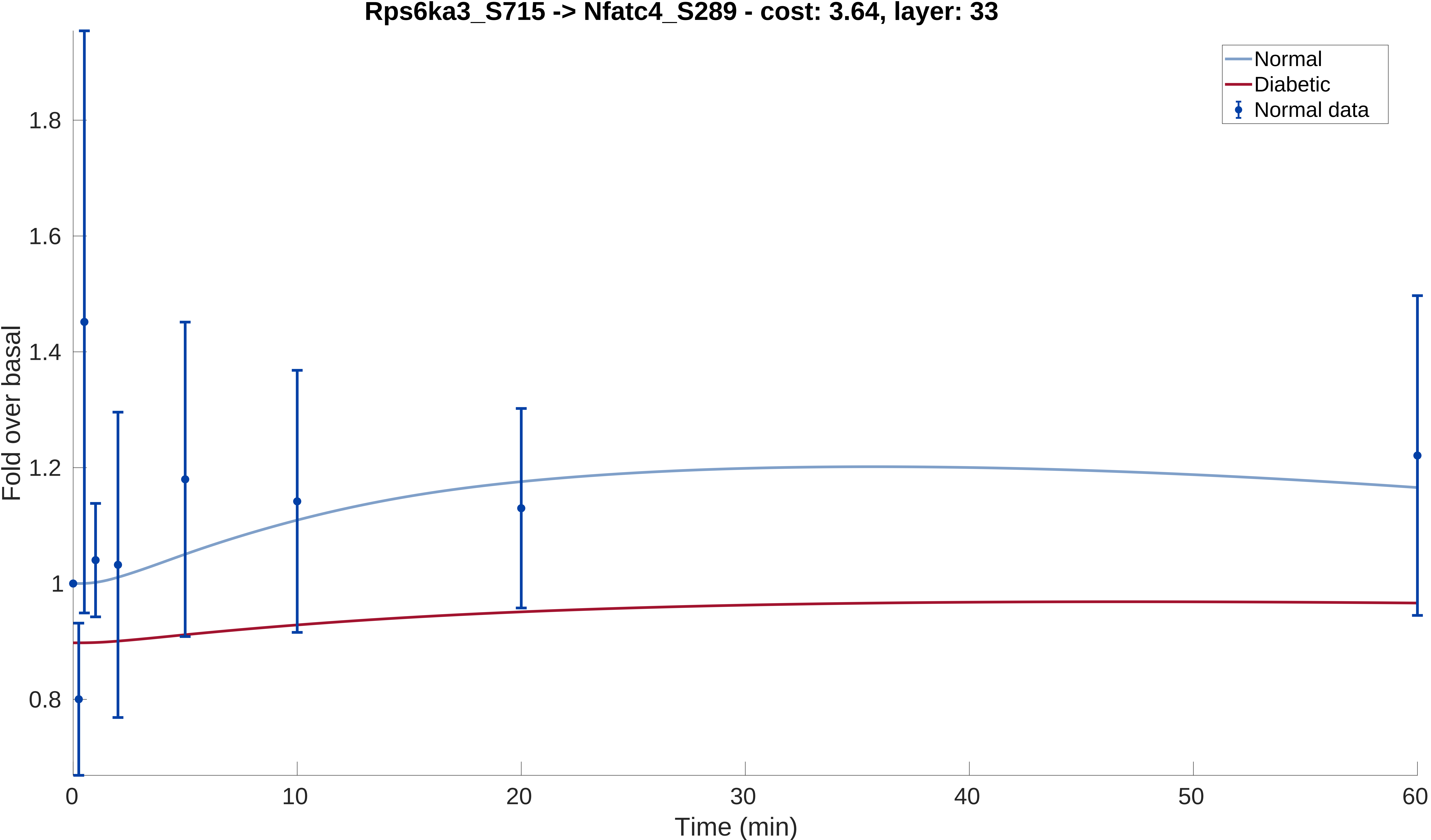

Rps6ka3_S715 -> Nfatc4_S289 - cost: 3.64, layer: 33
Normal
Diabetic
Normal data
Fold over basal
Time (min)
0.8
1
1.2
1.4
1.6
1.8
0
10
20
30
40
50
60

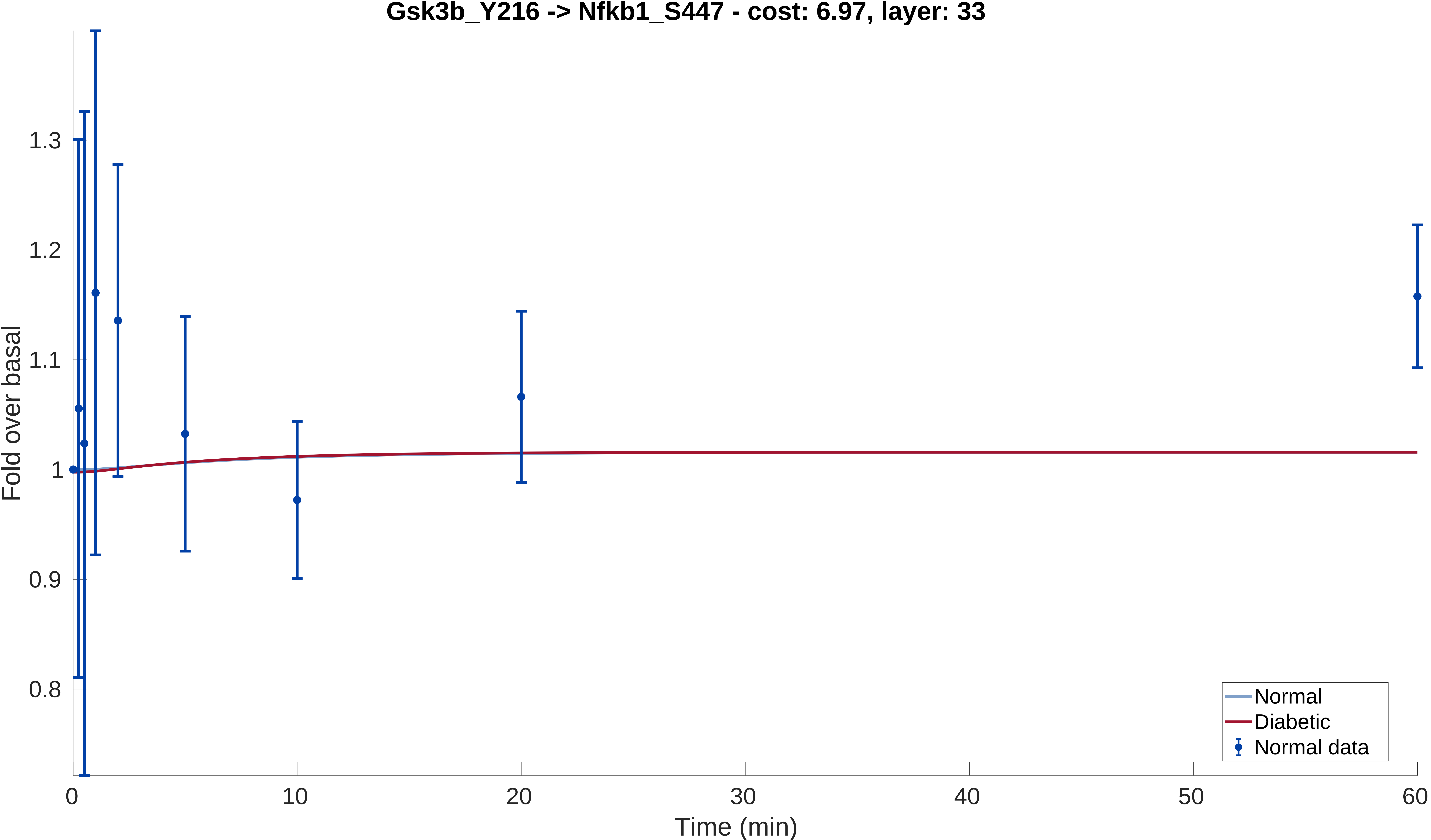

Gsk3b_Y216 -> Nfkb1_S447 - cost: 6.97, layer: 33
Fold over basal
1.3
1.2
1.1
1
0.9
0.8
0
10
20
30
40
50
60
Time (min)
Normal
Diabetic
Normal data

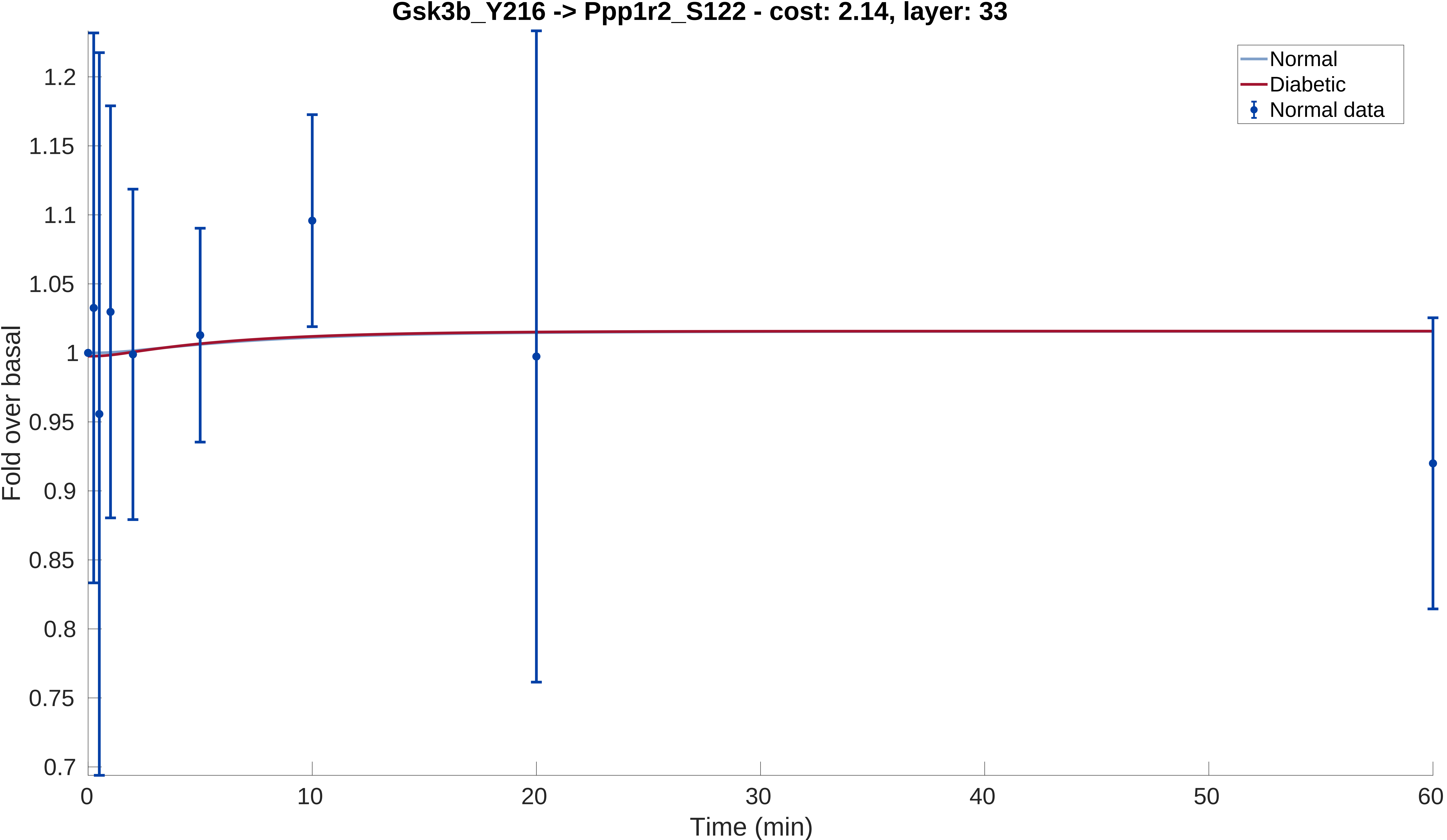

Gsk3b_Y216 -> Ppp1r2_S122 - cost: 2.14, layer: 33
Normal
Diabetic
Normal data
Fold over basal
1.2
1.15
1.1
1.05
1
0.95
0.9
0.85
0.8
0.75
0.7
0
10
20
30
40
50
60
Time (min)

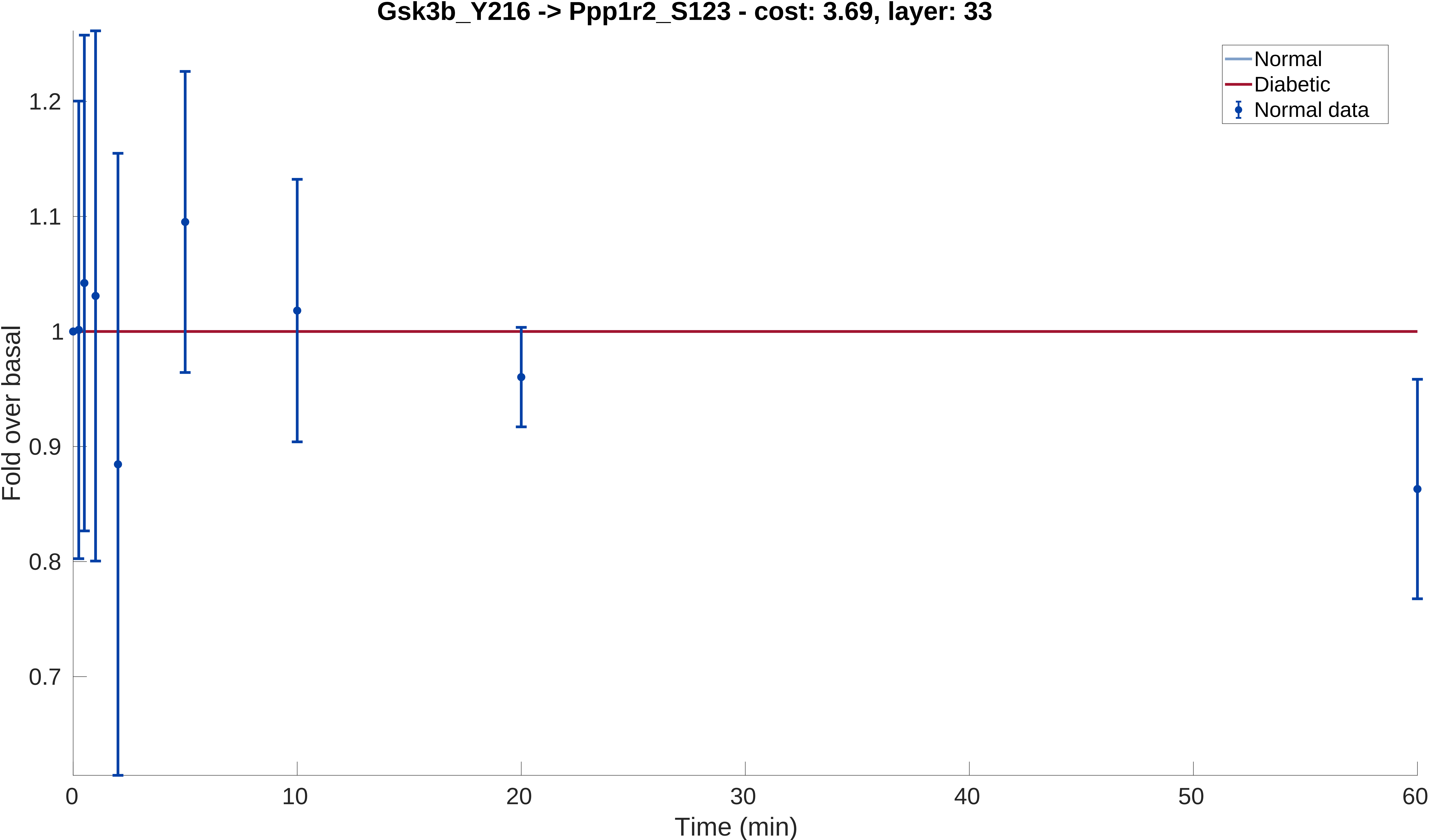

Gsk3b_Y216 -> Ppp1r2_S123 - cost: 3.69, layer: 33
Normal
Diabetic
Normal data
Fold over basal
1.2
1.1
1
0.9
0.8
0.7
0
10
20
30
40
50
60
Time (min)

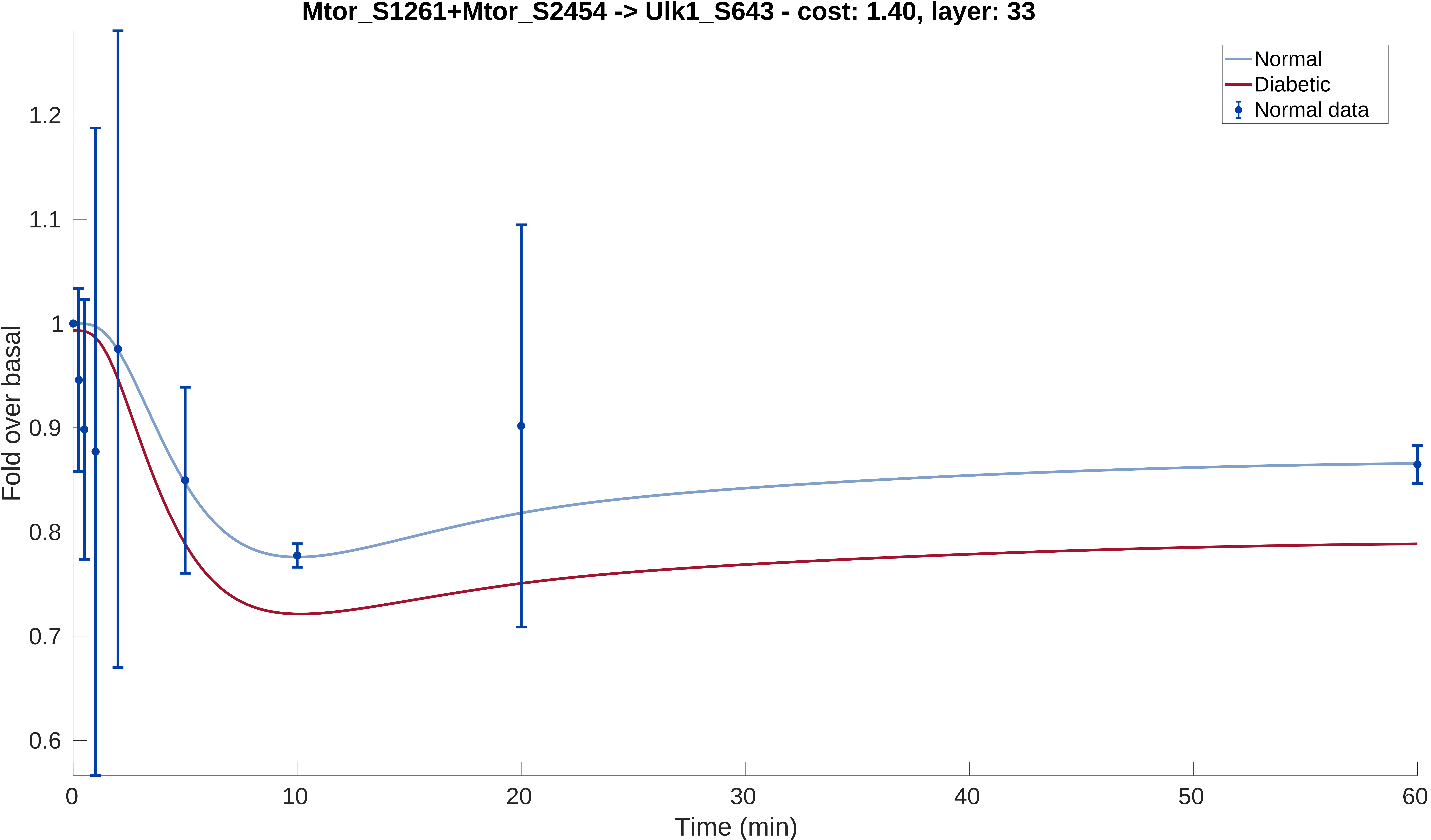

Mtor_S1261+Mtor_S2454 -> Ulk1_S643 - cost: 1.40, layer: 33
Normal
Diabetic
Normal data
Fold over basal
Time (min)
1.2
1.1
1
0.9
0.8
0.7
0.6
0
10
20
30
40
50
60

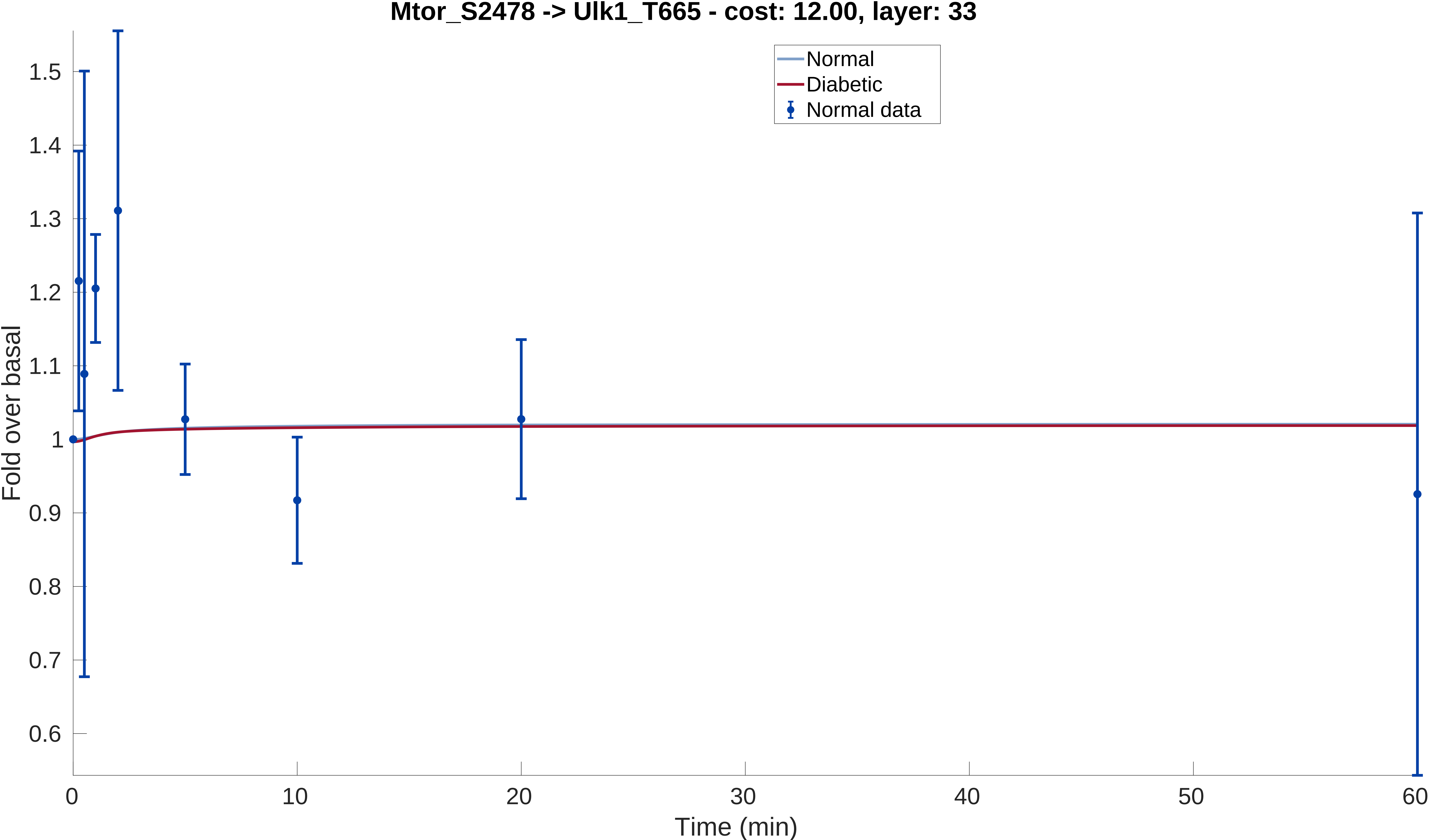

Mtor_S2478 -> Ulk1_T665 - cost: 12.00, layer: 33
Normal
Diabetic
Normal data
1.5
1.4
1.3
1.2
1.1
1
0.9
0.8
0.7
0.6
Fold over basal
0
10
20
30
40
50
60
Time (min)

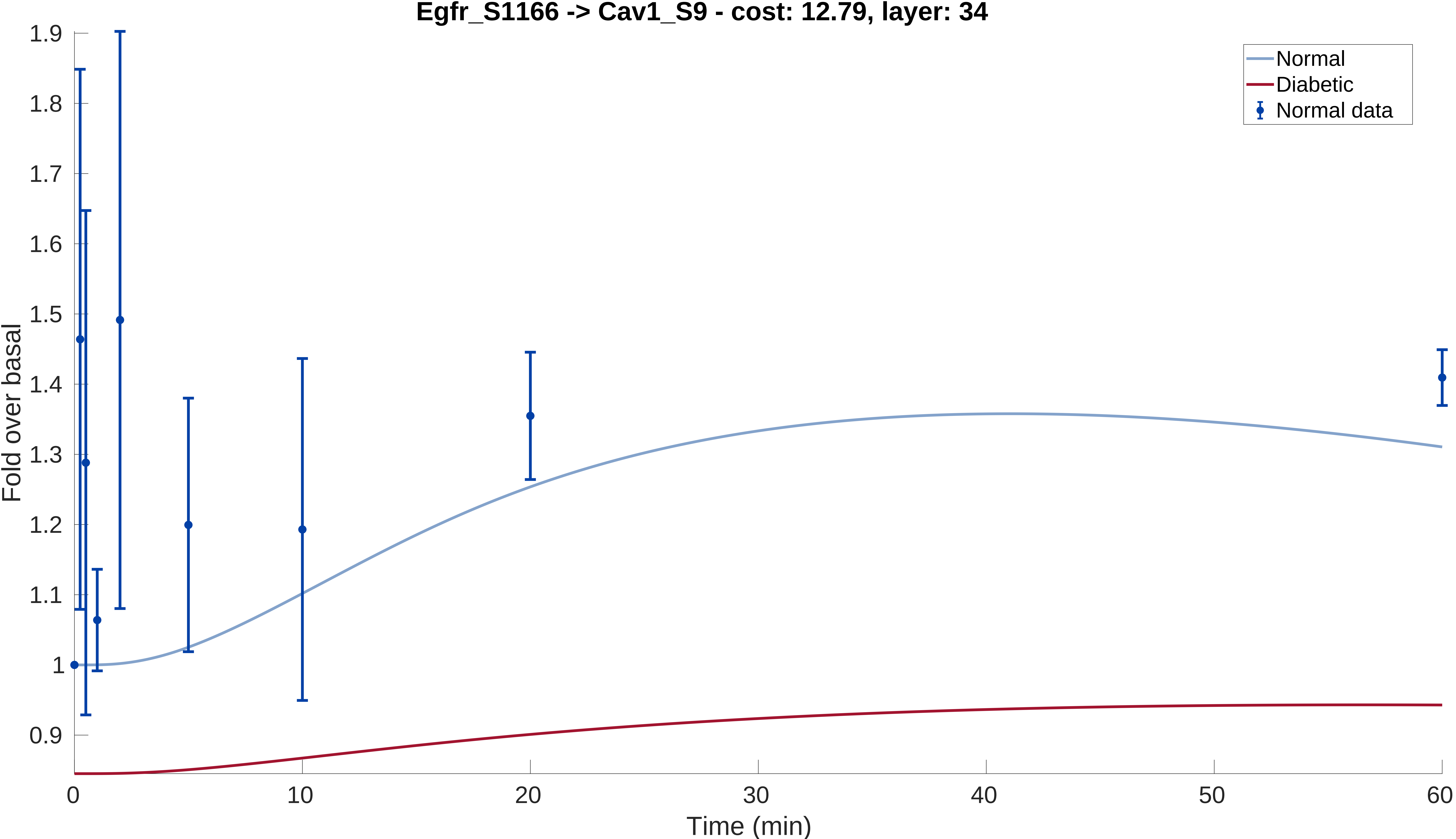
Egfr_S1166 -> Cav1_S9 - cost: 12.79, layer: 34
Fold over basal
Time (min)
Normal
Diabetic
Normal data
1.9
1.8
1.7
1.6
1.5
1.4
1.3
1.2
1.1
1
0.9
0
10
20
30
40
50
60

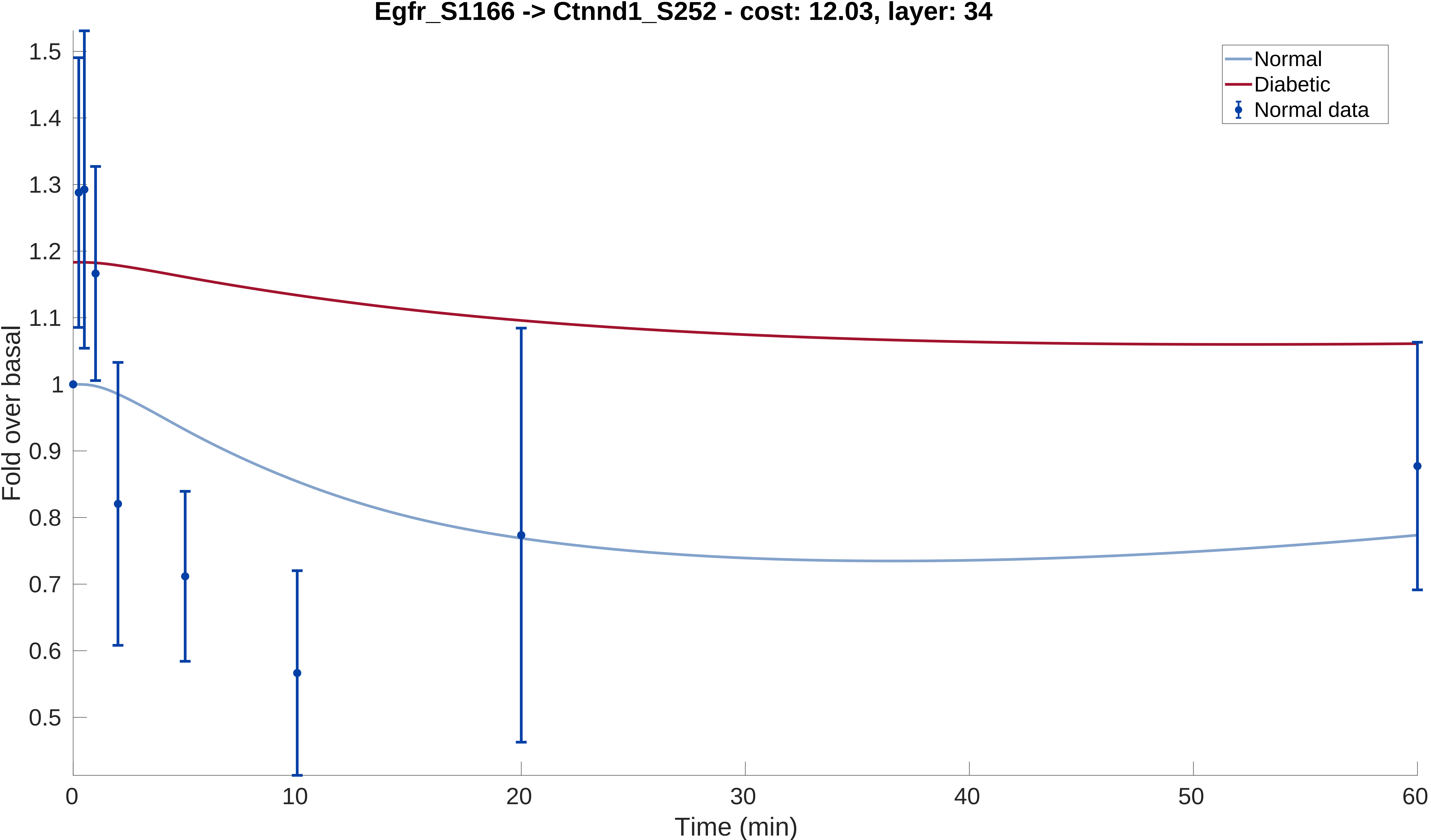
Egfr_S1166 -> Ctnnd1_S252 - cost: 12.03, layer: 34
Normal
Diabetic
Normal data
Fold over basal
Time (min)
0.5
0.6
0.7
0.8
0.9
1
1.1
1.2
1.3
1.4
1.5
0
10
20
30
40
50
60

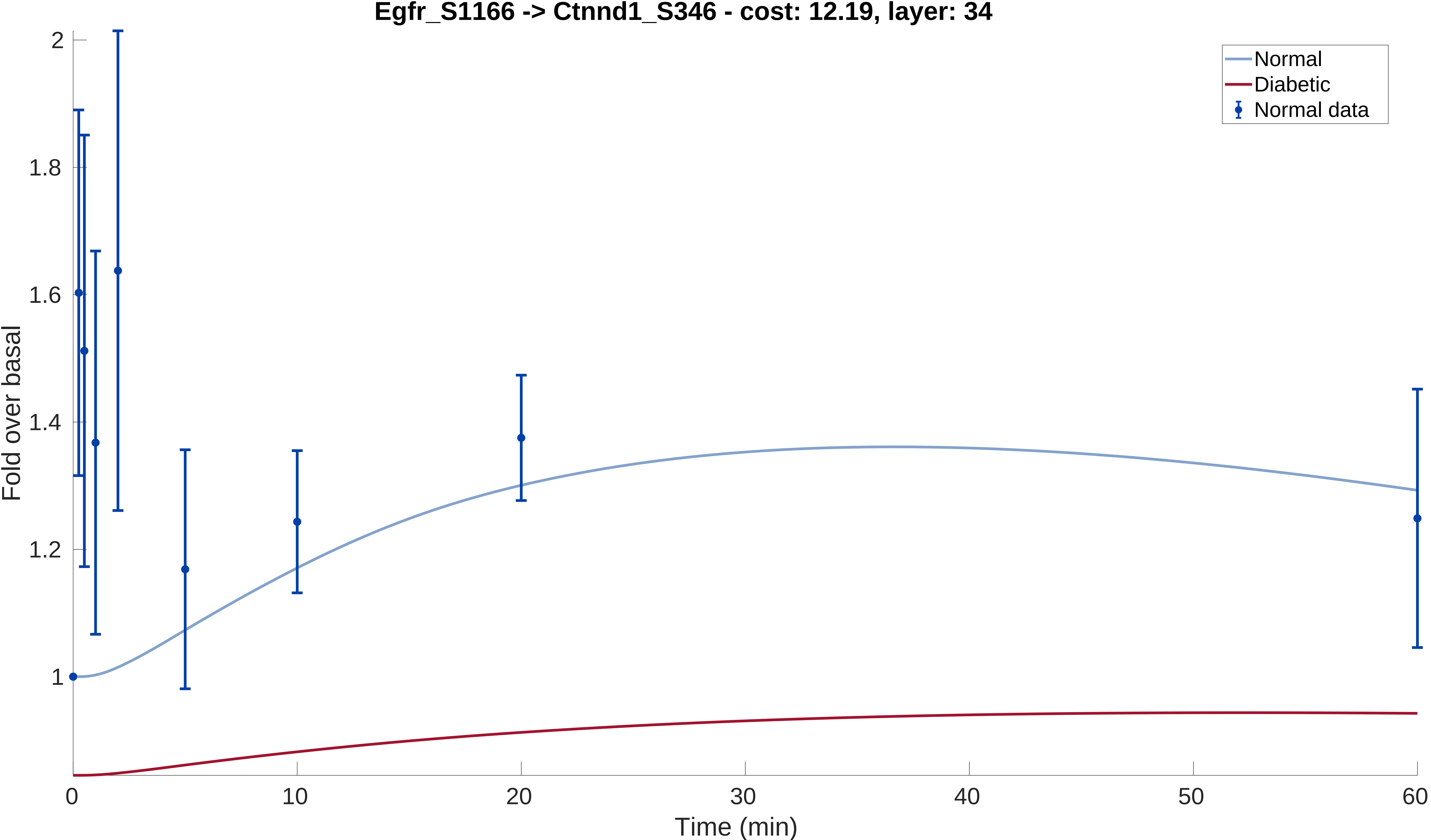
Egfr_S1166 -> Ctnnd1_S346 - cost: 12.19, layer: 34
Normal
Diabetic
Normal data
Fold over basal
2
1.8
1.6
1.4
1.2
1
0
10
20
30
40
50
60
Time (min)

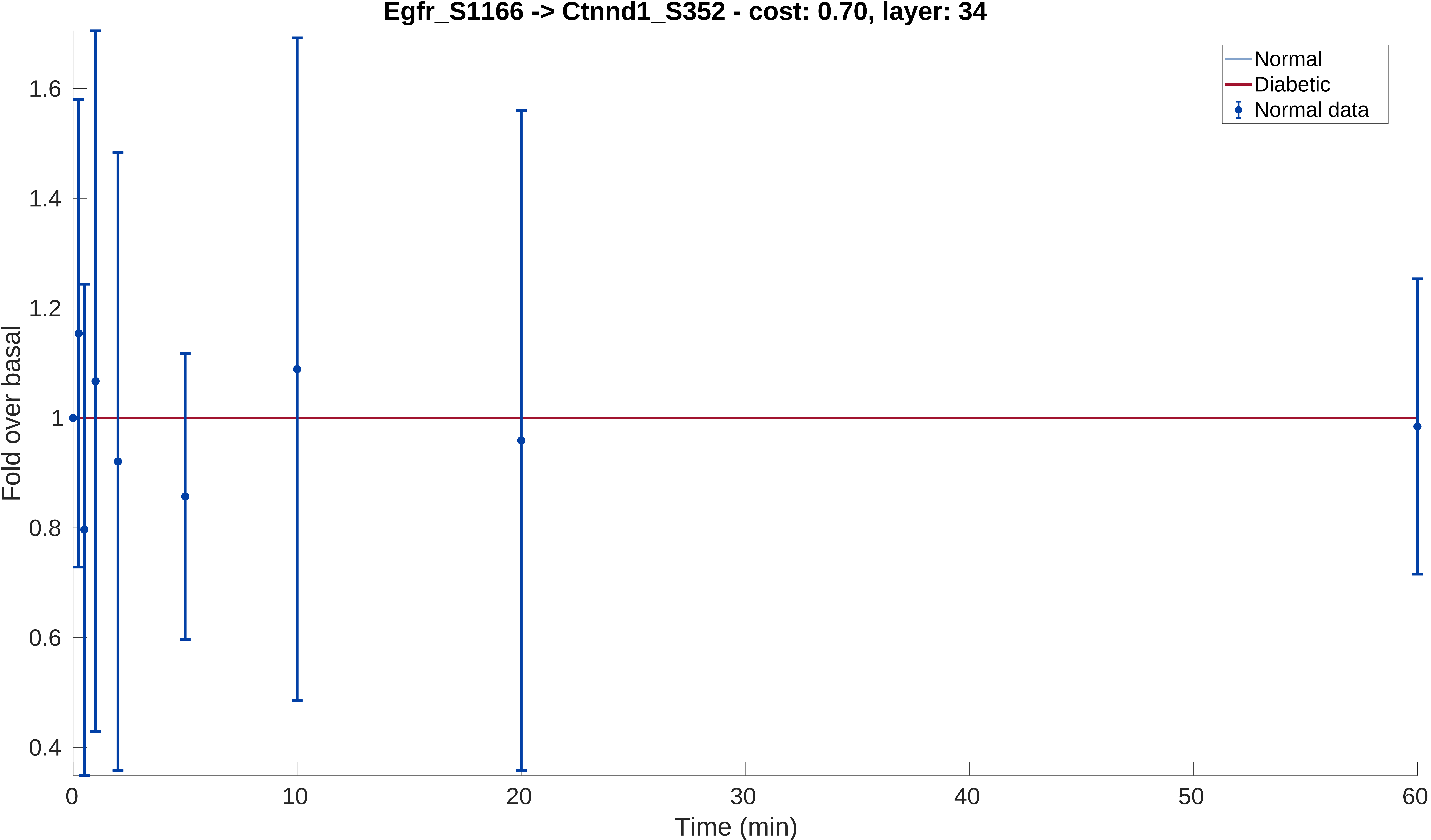

Egfr_S1166 -> Ctnnd1_S352 - cost: 0.70, layer: 34
Normal
Diabetic
Normal data
Fold over basal
1.6
1.4
1.2
1
0.8
0.6
0.4
0
10
20
30
40
50
60
Time (min)

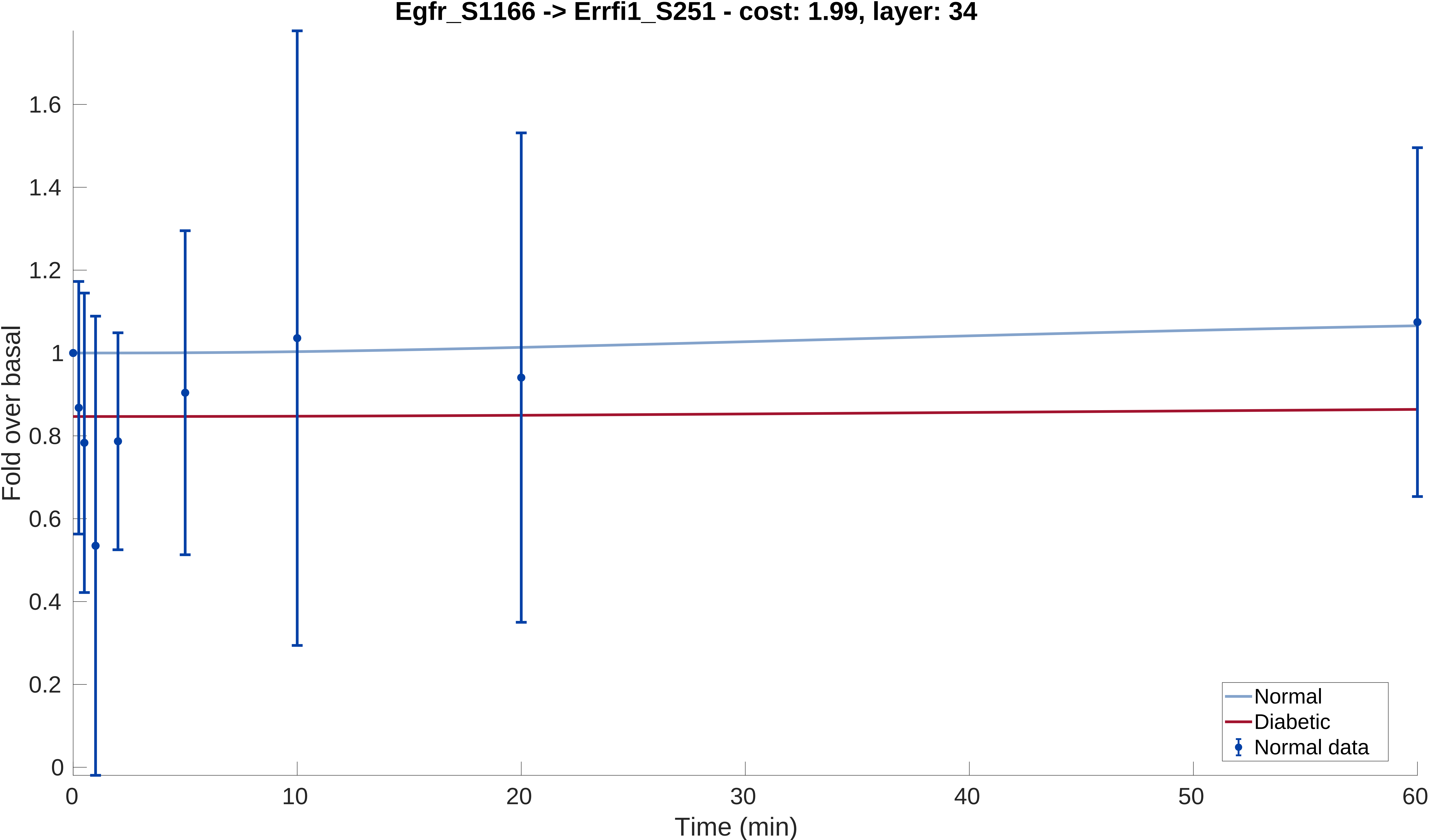

Egfr_S1166 -> Errfi1_S251 - cost: 1.99, layer: 34
Fold over basal
1.6
1.4
1.2
1
0.8
0.6
0.4
0.2
0
0
10
20
30
40
50
60
Time (min)
Normal
Diabetic
Normal data

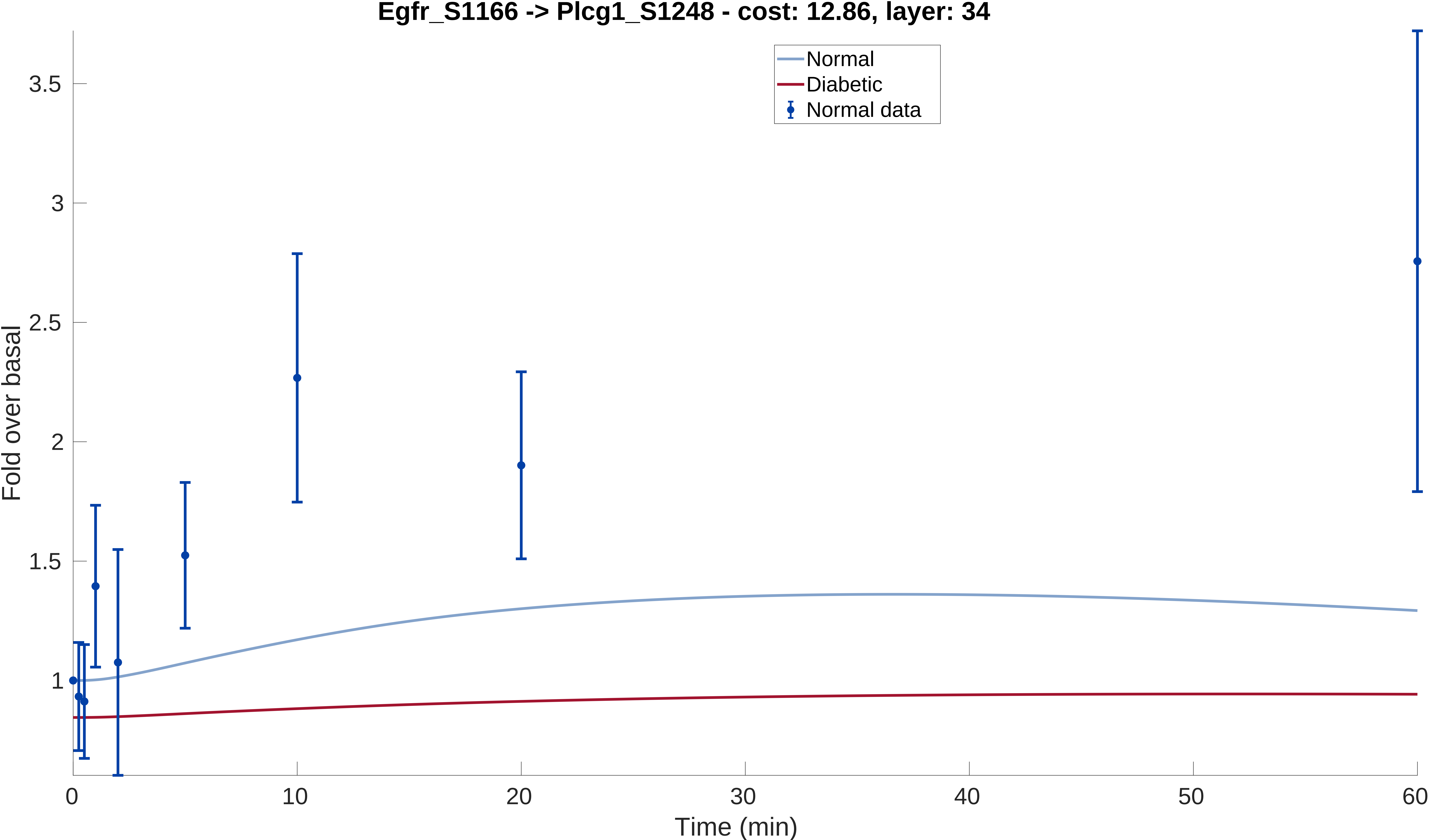

Egfr_S1166 -> Plcg1_S1248 - cost: 12.86, layer: 34
Normal
Diabetic
Normal data
Fold over basal
3.5
3
2.5
2
1.5
1
0
10
20
30
40
50
60
Time (min)

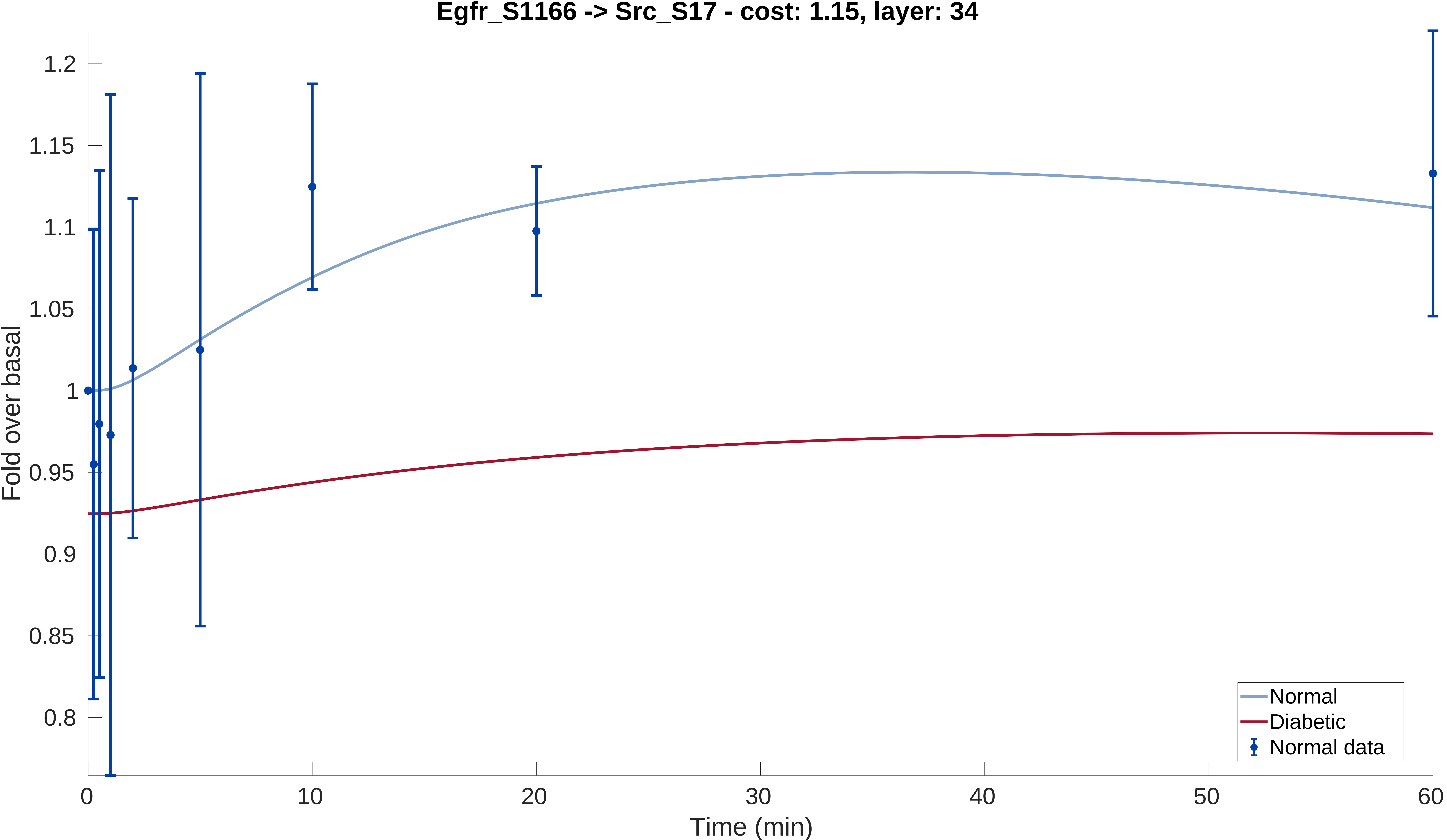

Egfr_S1166 -> Src_S17 - cost: 1.15, layer: 34
Fold over basal
1.2
1.15
1.1
1.05
1
0.95
0.9
0.85
0.8
0
10
20
30
40
50
60
Time (min)
Normal
Diabetic
Normal data

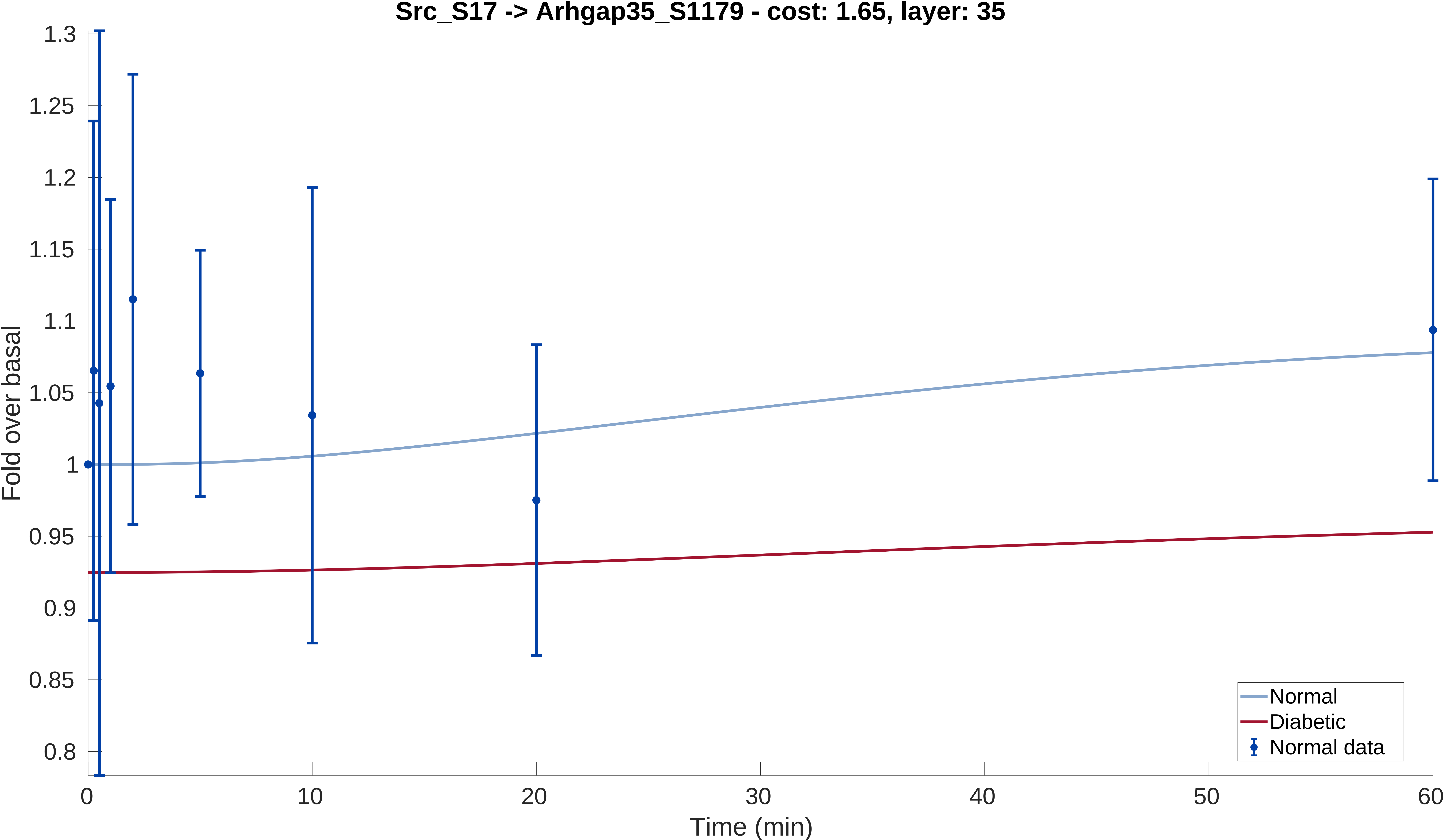

Src_S17 -> Arhgap35_S1179 - cost: 1.65, layer: 35
Fold over basal
Time (min)
1.3
1.25
1.2
1.15
1.1
1.05
1
0.95
0.9
0.85
0.8
0
10
20
30
40
50
60
Normal
Diabetic
Normal data

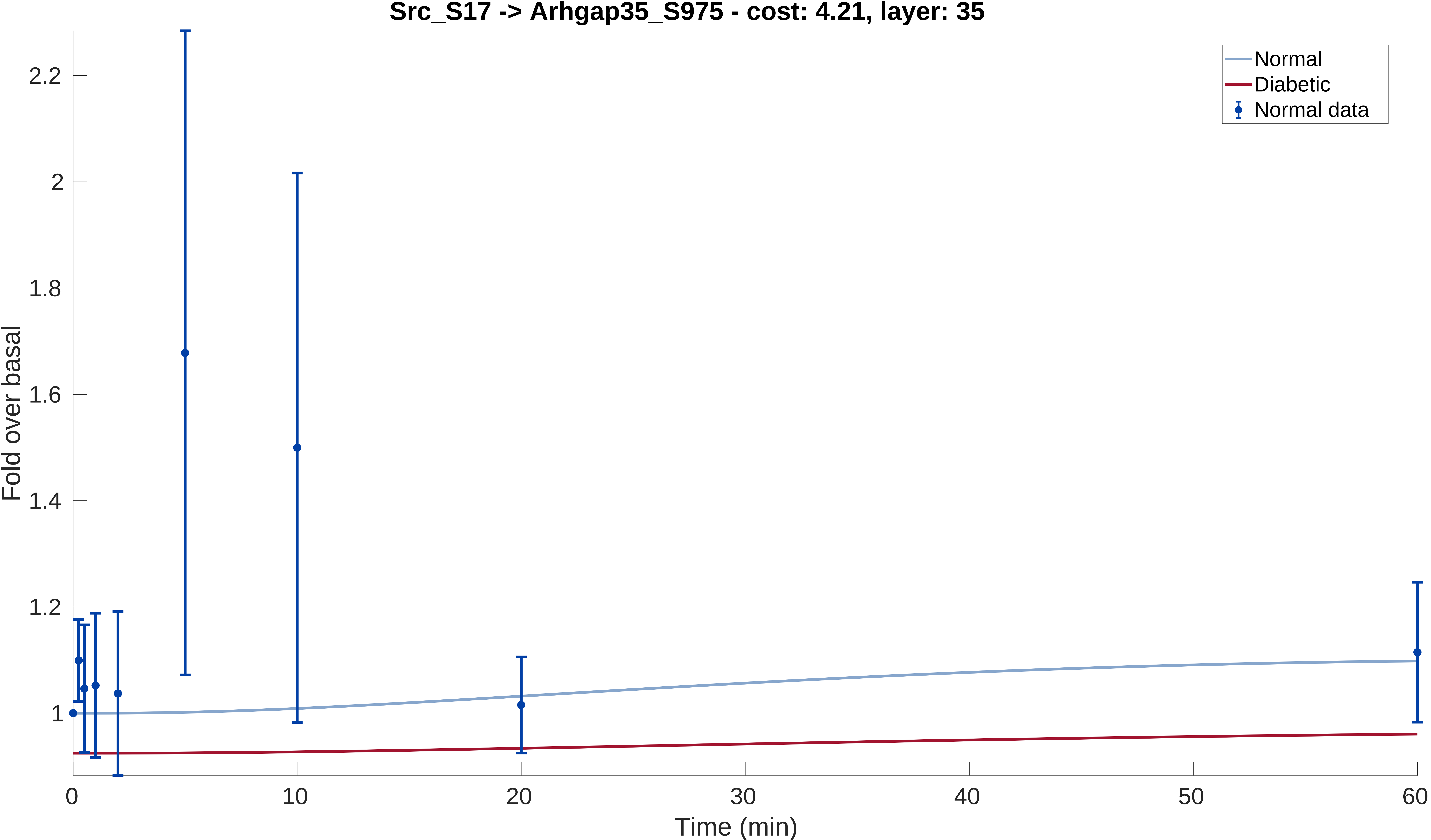

Src_S17 -> Arhgap35_S975 - cost: 4.21, layer: 35
Normal
Diabetic
Normal data
Fold over basal
2.2
2
1.8
1.6
1.4
1.2
1
0
10
20
30
40
50
60
Time (min)

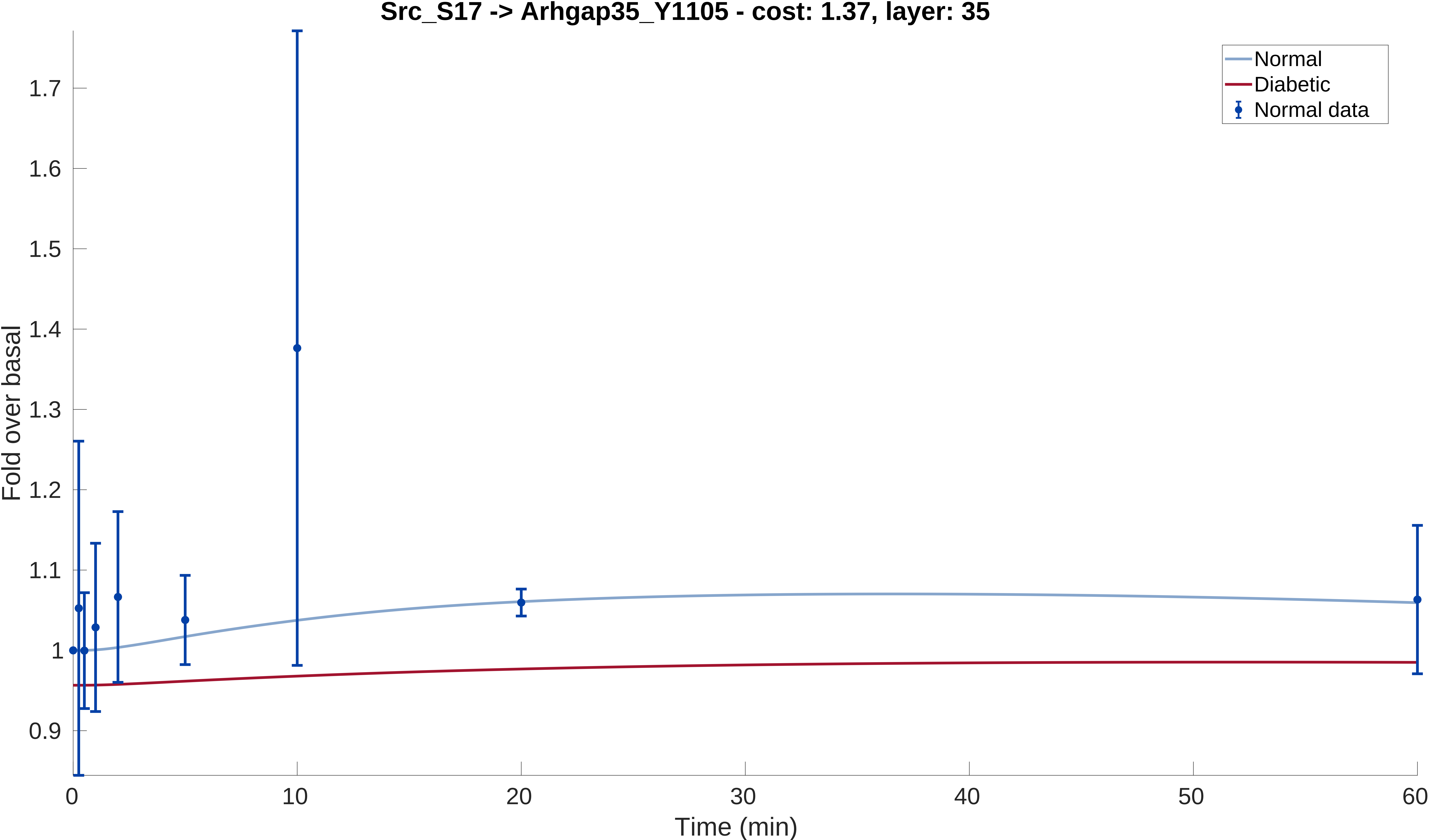

Src_S17 -> Arhgap35_Y1105 - cost: 1.37, layer: 35
Normal
Diabetic
Normal data
Fold over basal
Time (min)
0.9
1
1.1
1.2
1.3
1.4
1.5
1.6
1.7
0
10
20
30
40
50
60

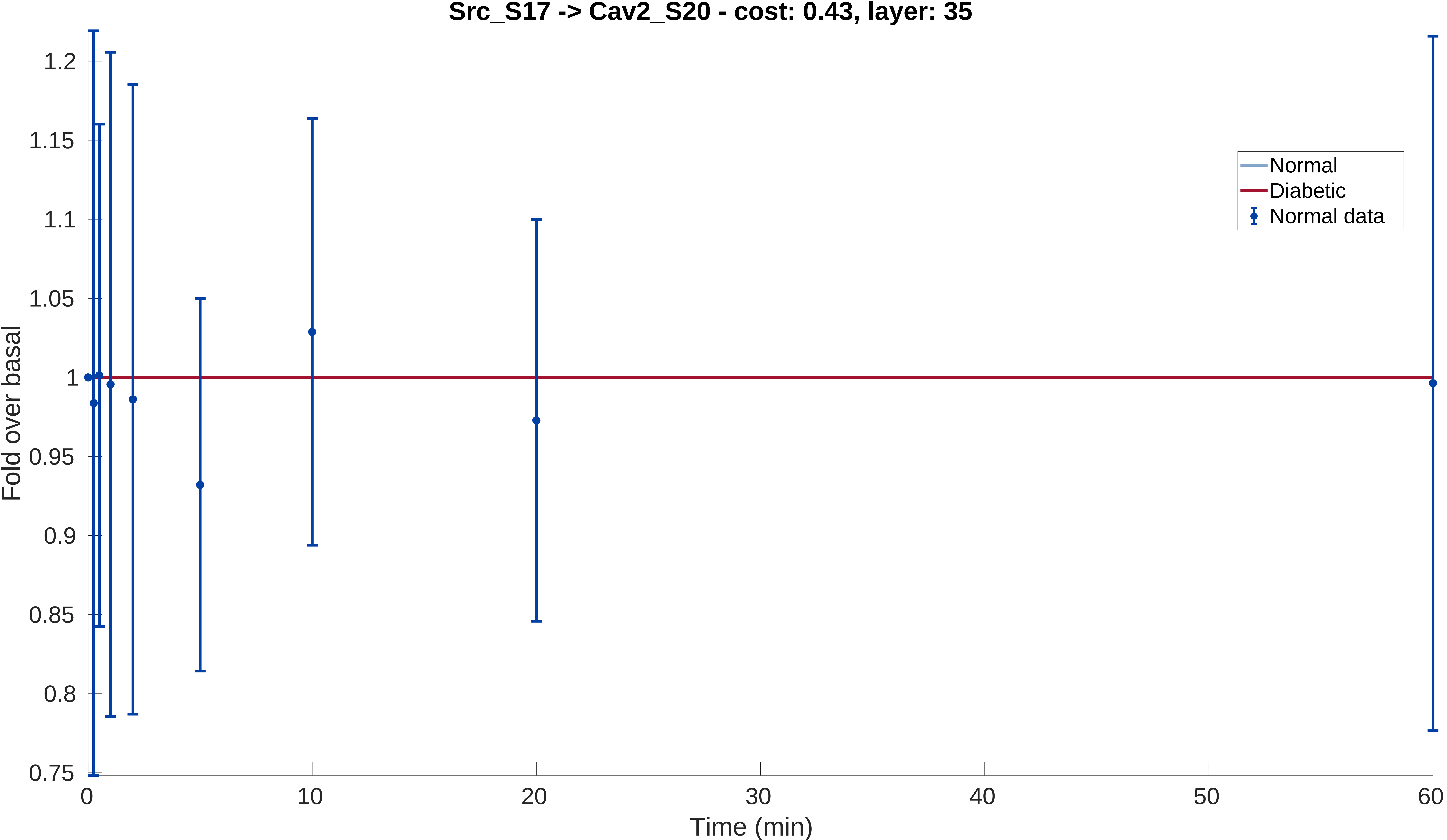

Src_S17 -> Cav2_S20 - cost: 0.43, layer: 35
Normal
Diabetic
Normal data
Fold over basal
Time (min)
1.2
1.15
1.1
1.05
1
0.95
0.9
0.85
0.8
0.75
0
10
20
30
40
50
60

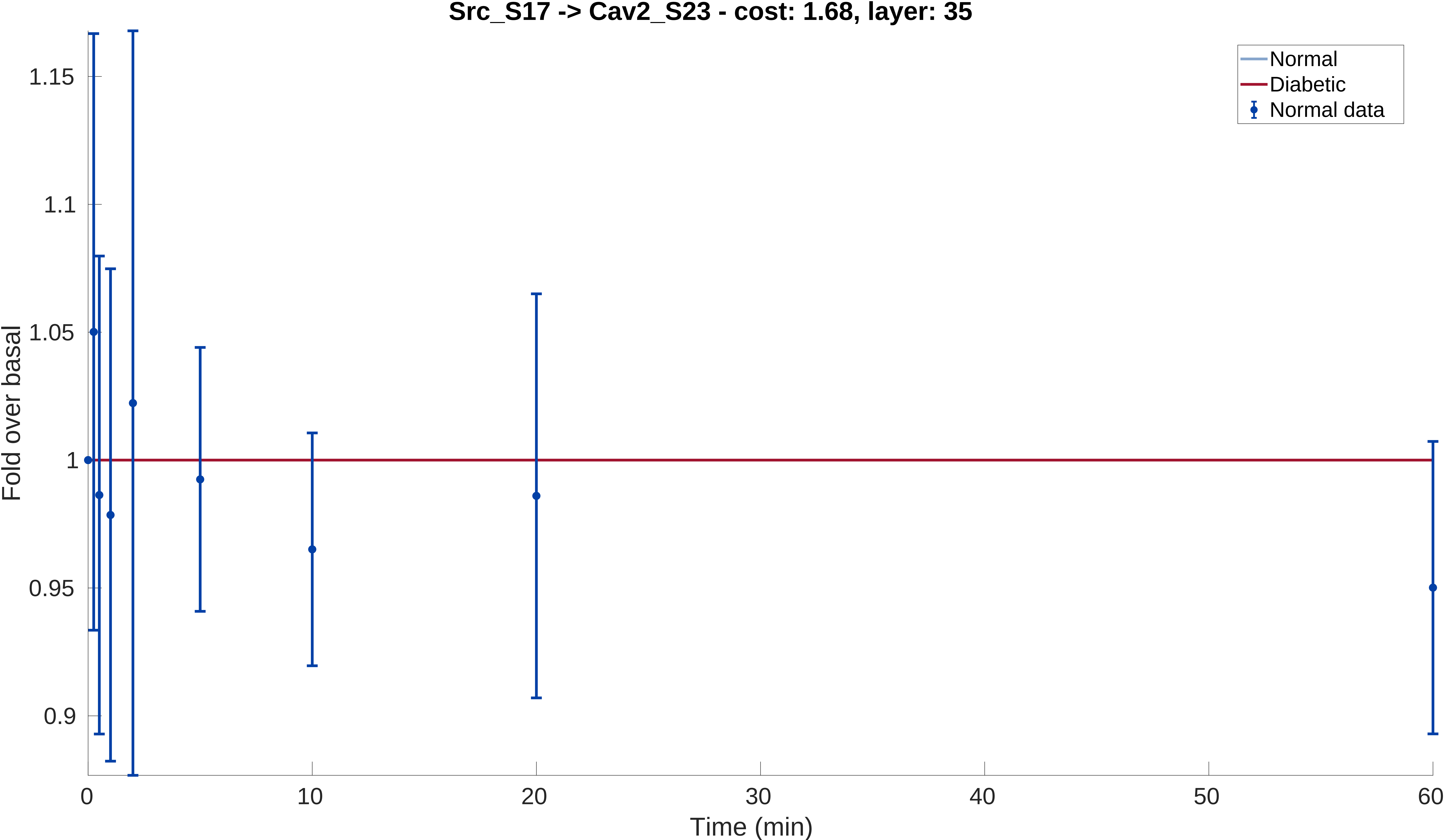
Src_S17 -> Cav2_S23 - cost: 1.68, layer: 35
Normal
Diabetic
Normal data
Fold over basal
1.15
1.1
1.05
1
0.95
0.9
Time (min)
0
10
20
30
40
50
60

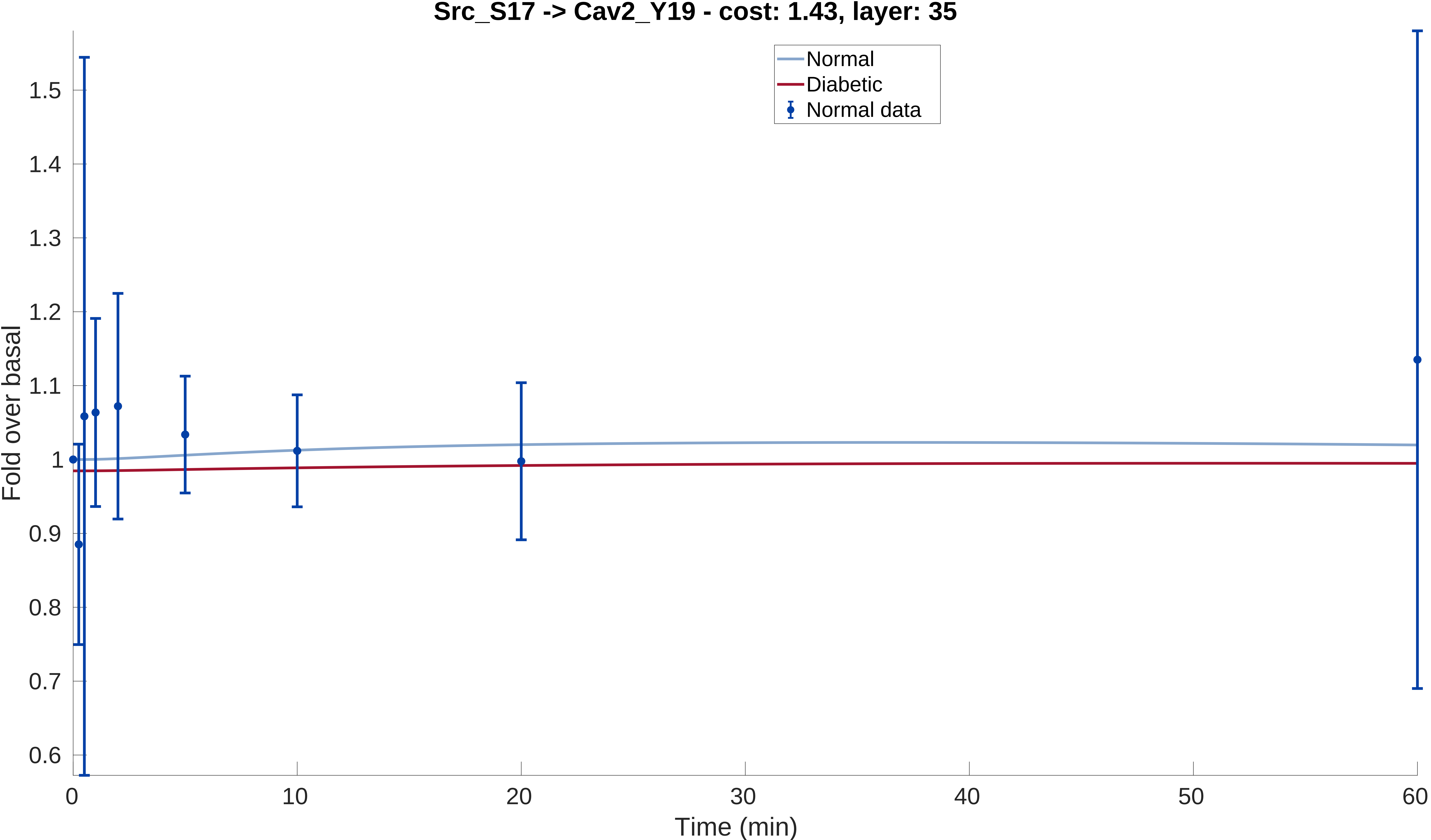

Src_S17 -> Cav2_Y19 - cost: 1.43, layer: 35
Normal
Diabetic
Normal data
Fold over basal
1.5
1.4
1.3
1.2
1.1
1
0.9
0.8
0.7
0.6
0
10
20
30
40
50
60
Time (min)

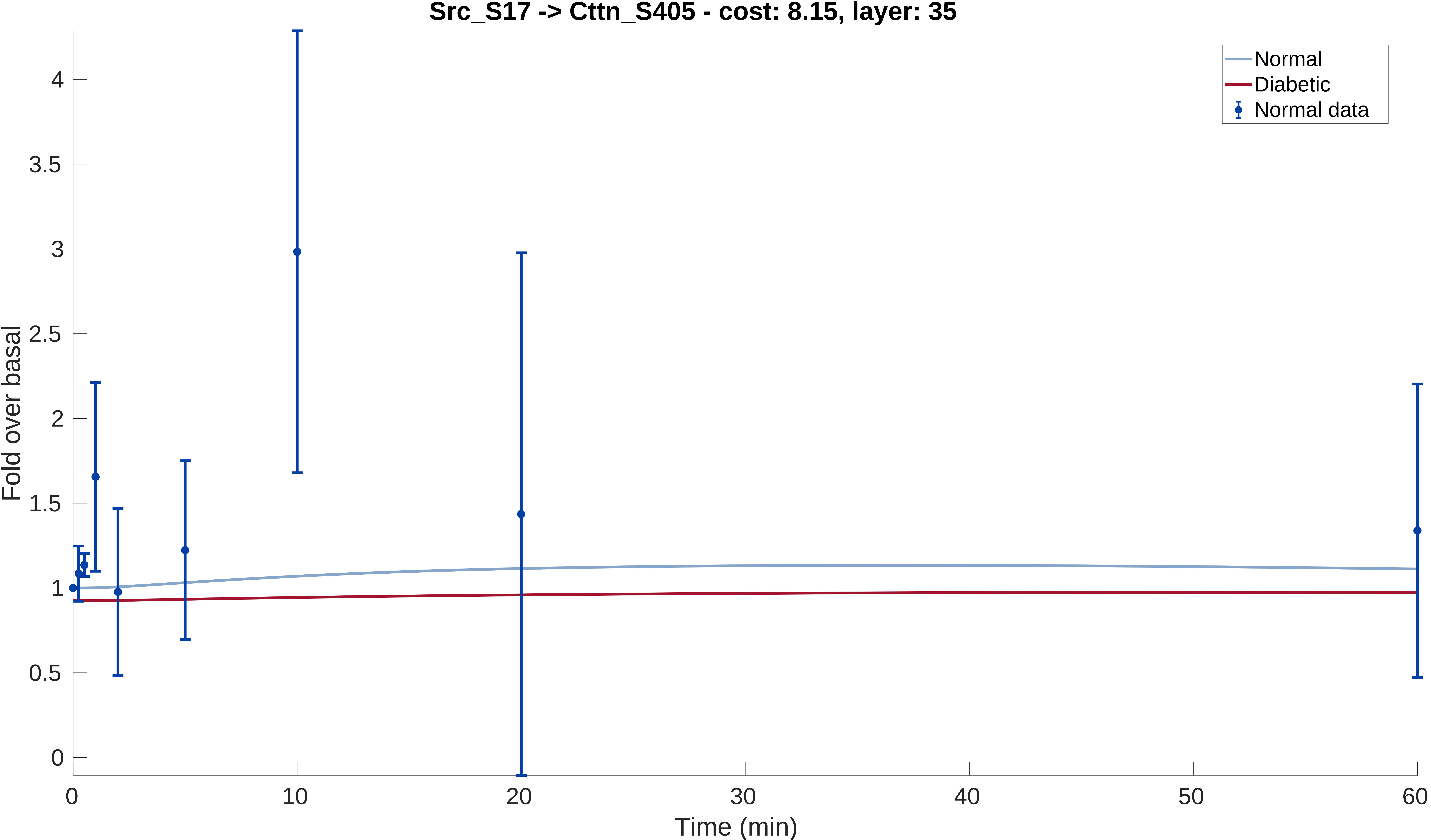

Src_S17 -> Cttn_S405 - cost: 8.15, layer: 35
Normal
Diabetic
Normal data
Fold over basal
Time (min)
0
0.5
1
1.5
2
2.5
3
3.5
4
0
10
20
30
40
50
60

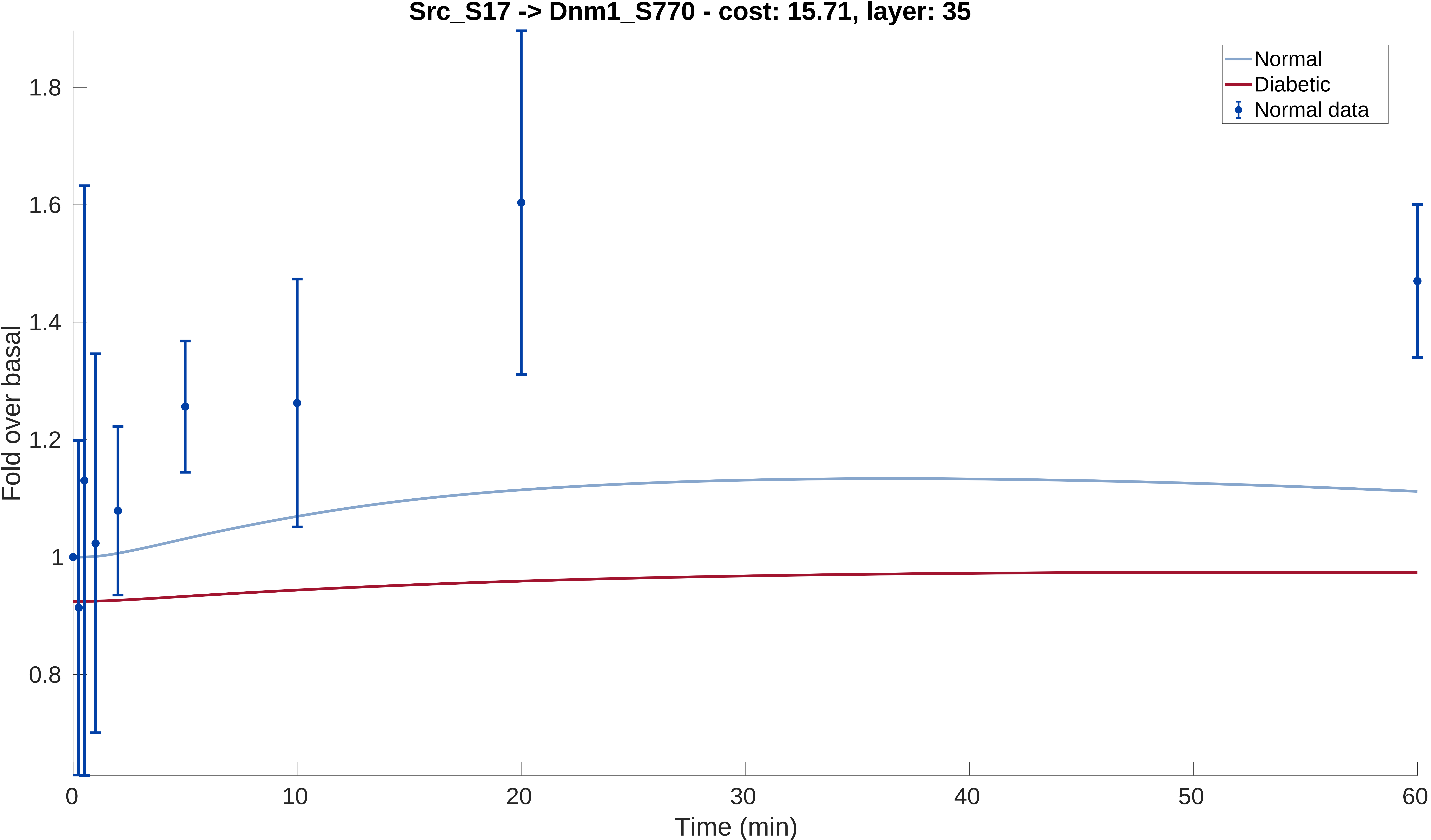

Src_S17 -> Dnm1_S770 - cost: 15.71, layer: 35
Fold over basal
Time (min)
Normal
Diabetic
Normal data
0.8
1
1.2
1.4
1.6
1.8
0
10
20
30
40
50
60

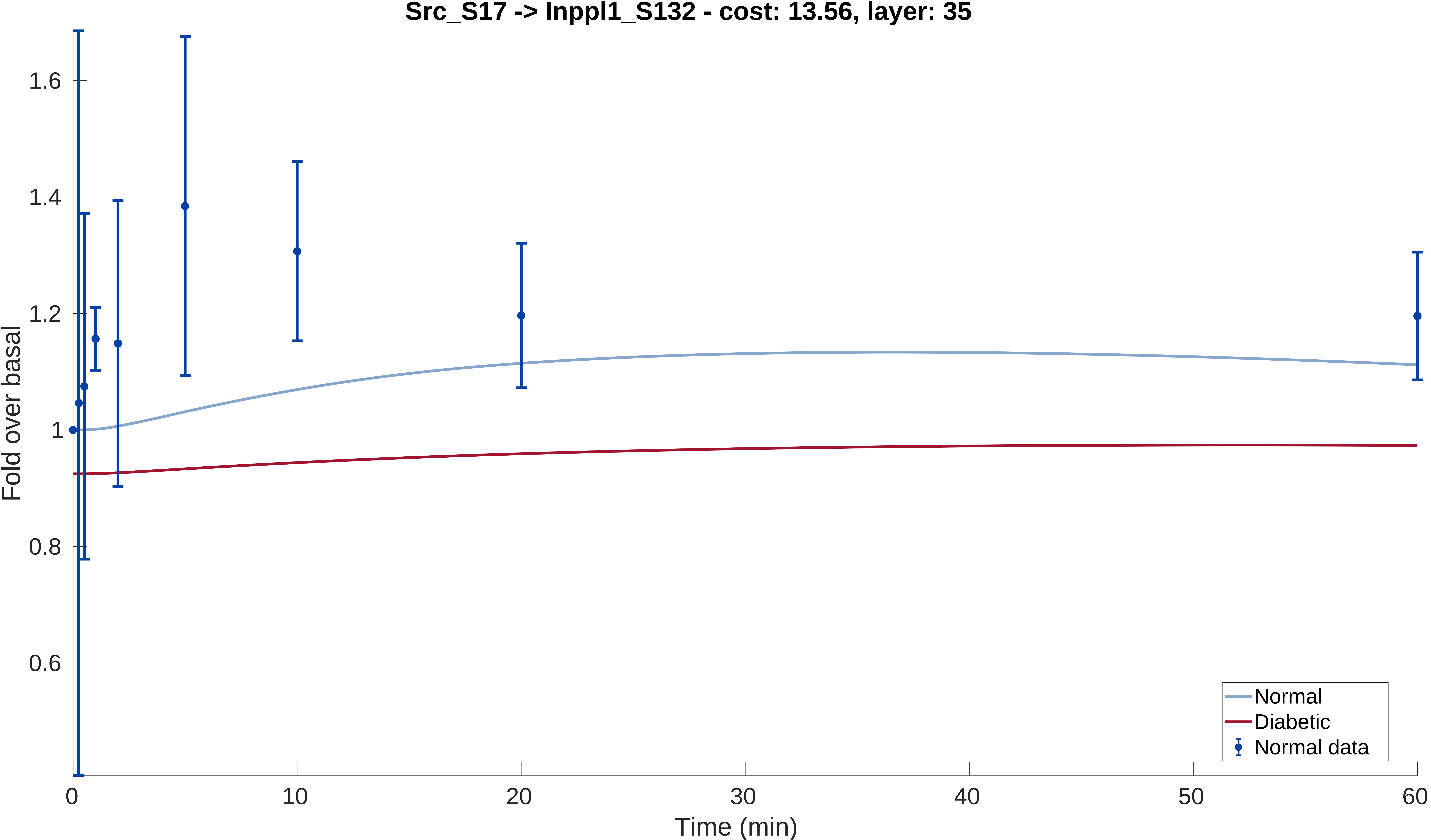

Src_S17 -> Inppl1_S132 - cost: 13.56, layer: 35
1.6
1.4
1.2
1
0.8
0.6
Fold over basal
0
10
20
30
40
50
60
Time (min)
Normal
Diabetic
Normal data

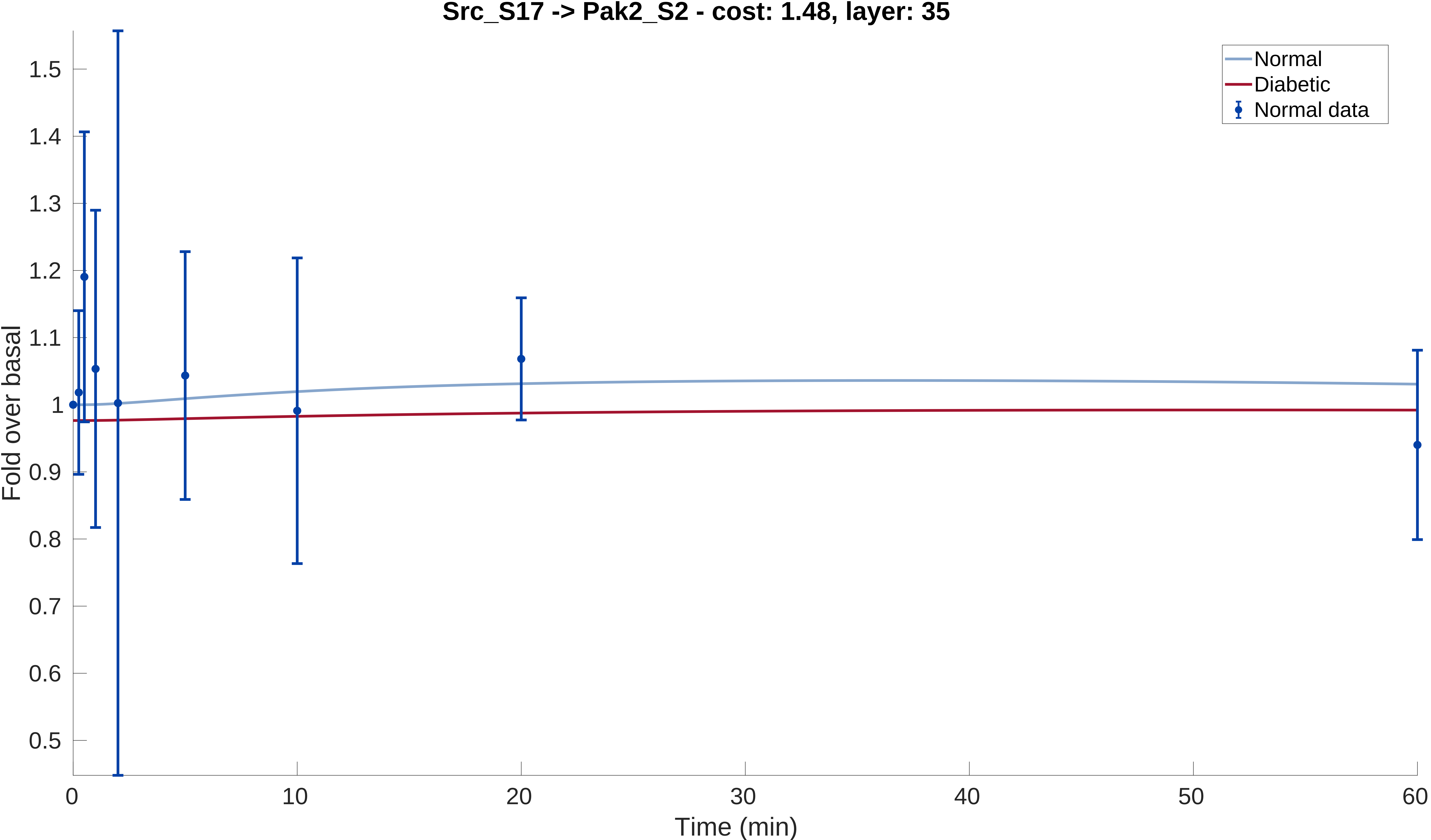

Src_S17 -> Pak2_S2 - cost: 1.48, layer: 35
Normal
Diabetic
Normal data
Fold over basal
1.5
1.4
1.3
1.2
1.1
1
0.9
0.8
0.7
0.6
0.5
0
10
20
30
40
50
60
Time (min)

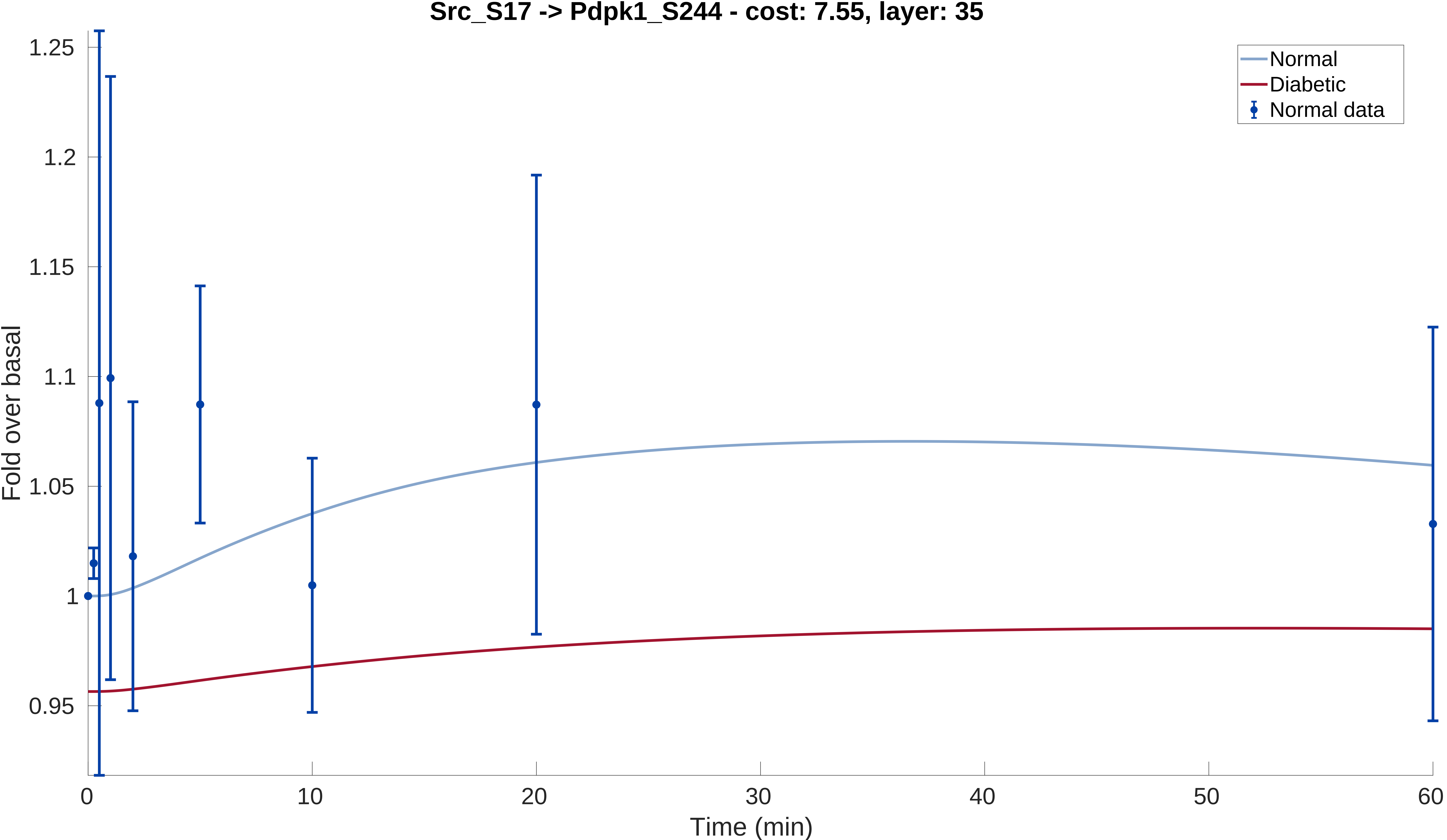

Src_S17 -> Pdpk1_S244 - cost: 7.55, layer: 35
Normal
Diabetic
Normal data
Fold over basal
1.25
1.2
1.15
1.1
1.05
1
0.95
0
10
20
30
40
50
60
Time (min)

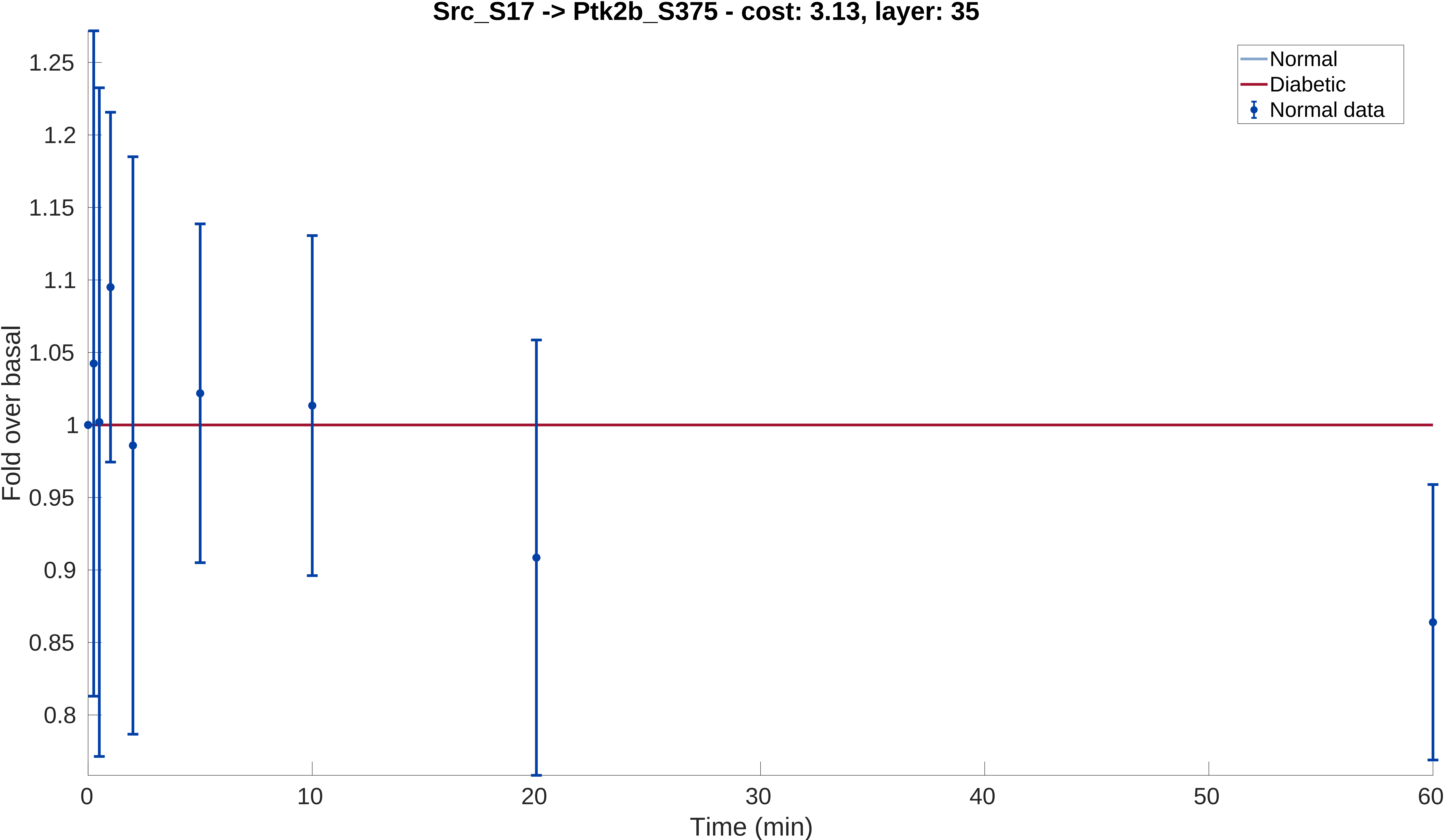

Src_S17 -> Ptk2b_S375 - cost: 3.13, layer: 35
Normal
Diabetic
Normal data
Fold over basal
1.25
1.2
1.15
1.1
1.05
1
0.95
0.9
0.85
0.8
0
10
20
30
40
50
60
Time (min)

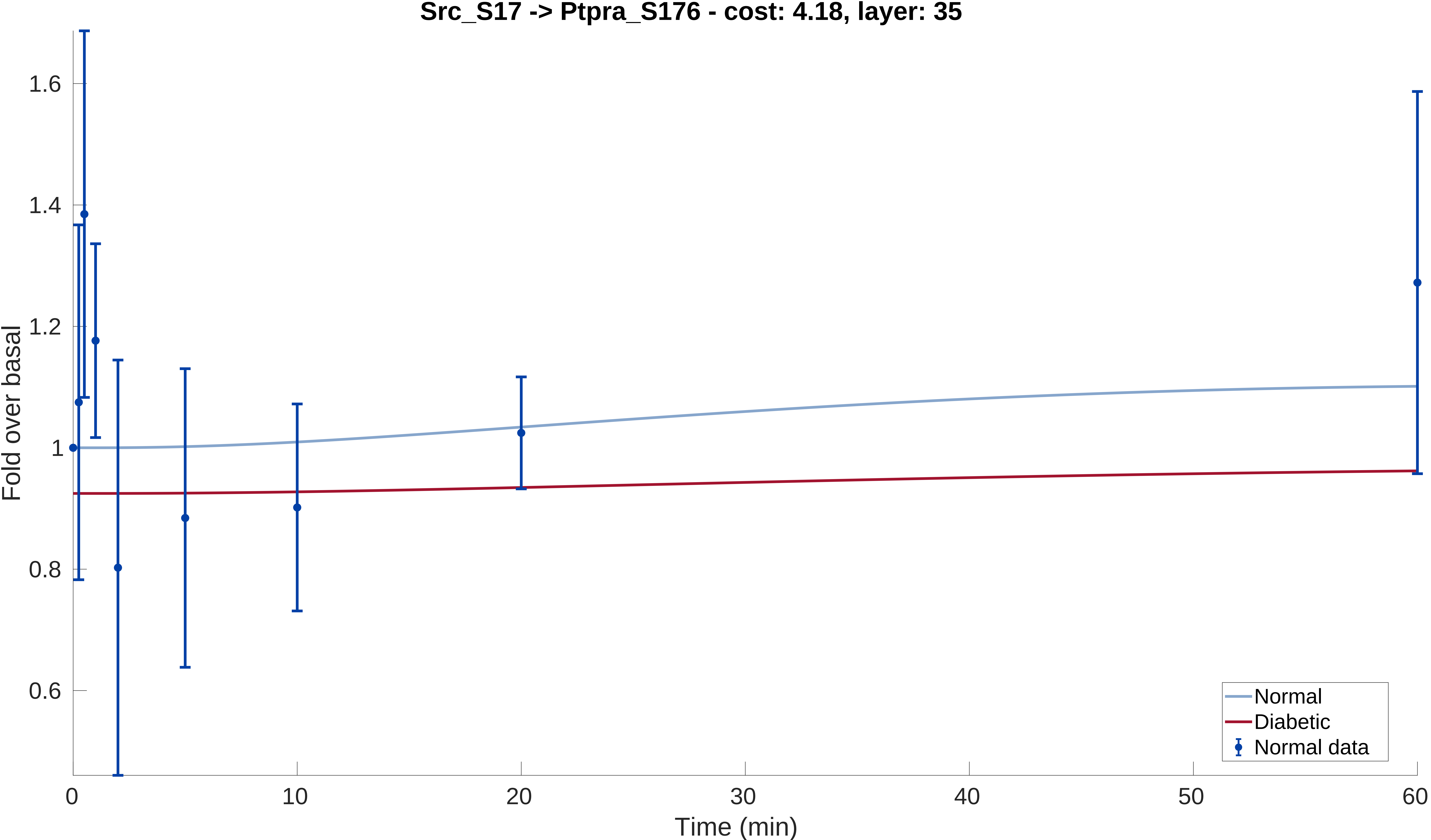

Src_S17 -> Ptpra_S176 - cost: 4.18, layer: 35
Fold over basal
1.6
1.4
1.2
1
0.8
0.6
0
10
20
30
40
50
60
Time (min)
Normal
Diabetic
Normal data

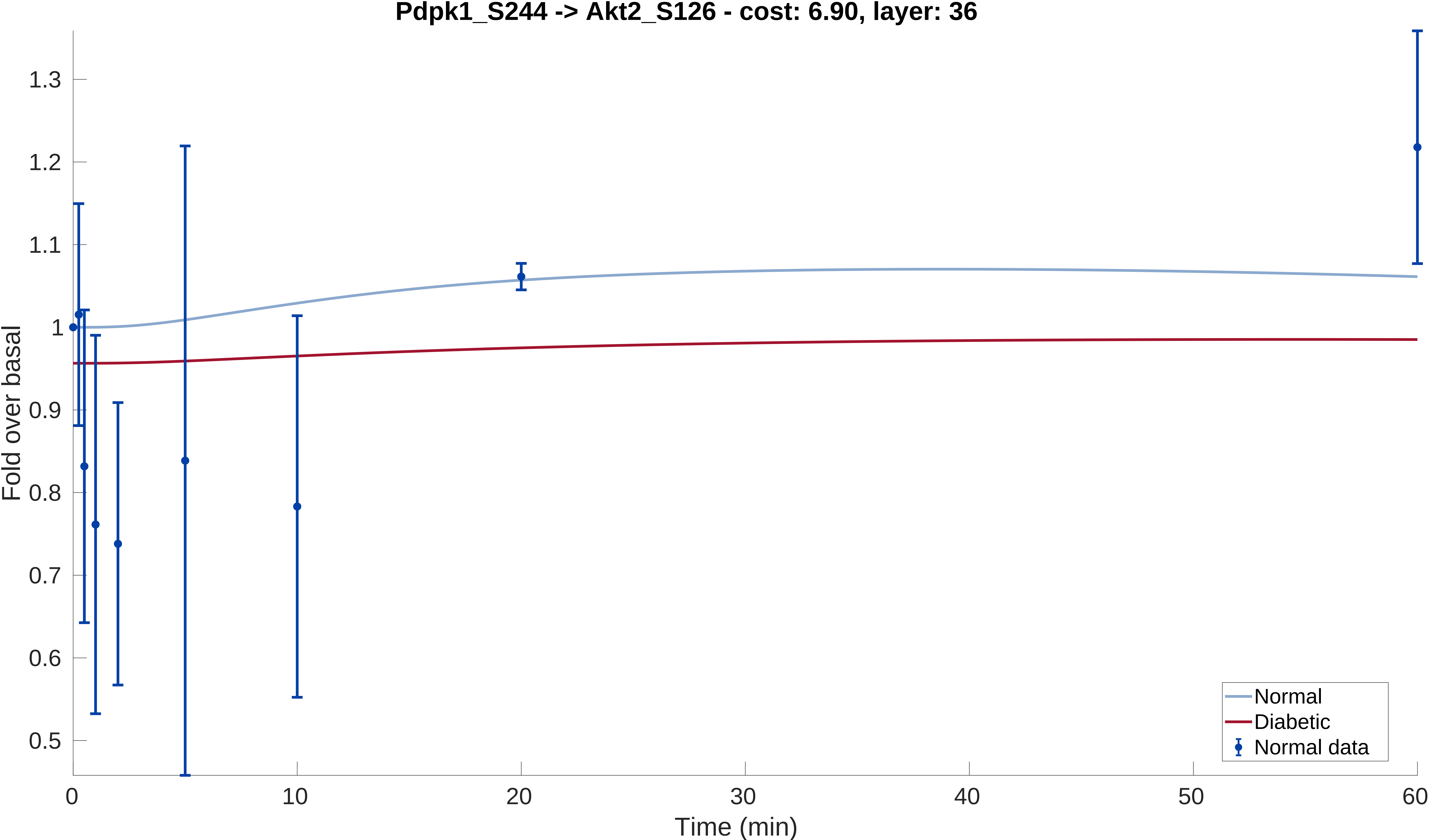
Pdpk1_S244 -> Akt2_S126 - cost: 6.90, layer: 36
Fold over basal
1.3
1.2
1.1
1
0.9
0.8
0.7
0.6
0.5
0
10
20
30
40
50
60
Time (min)
Normal
Diabetic
Normal data

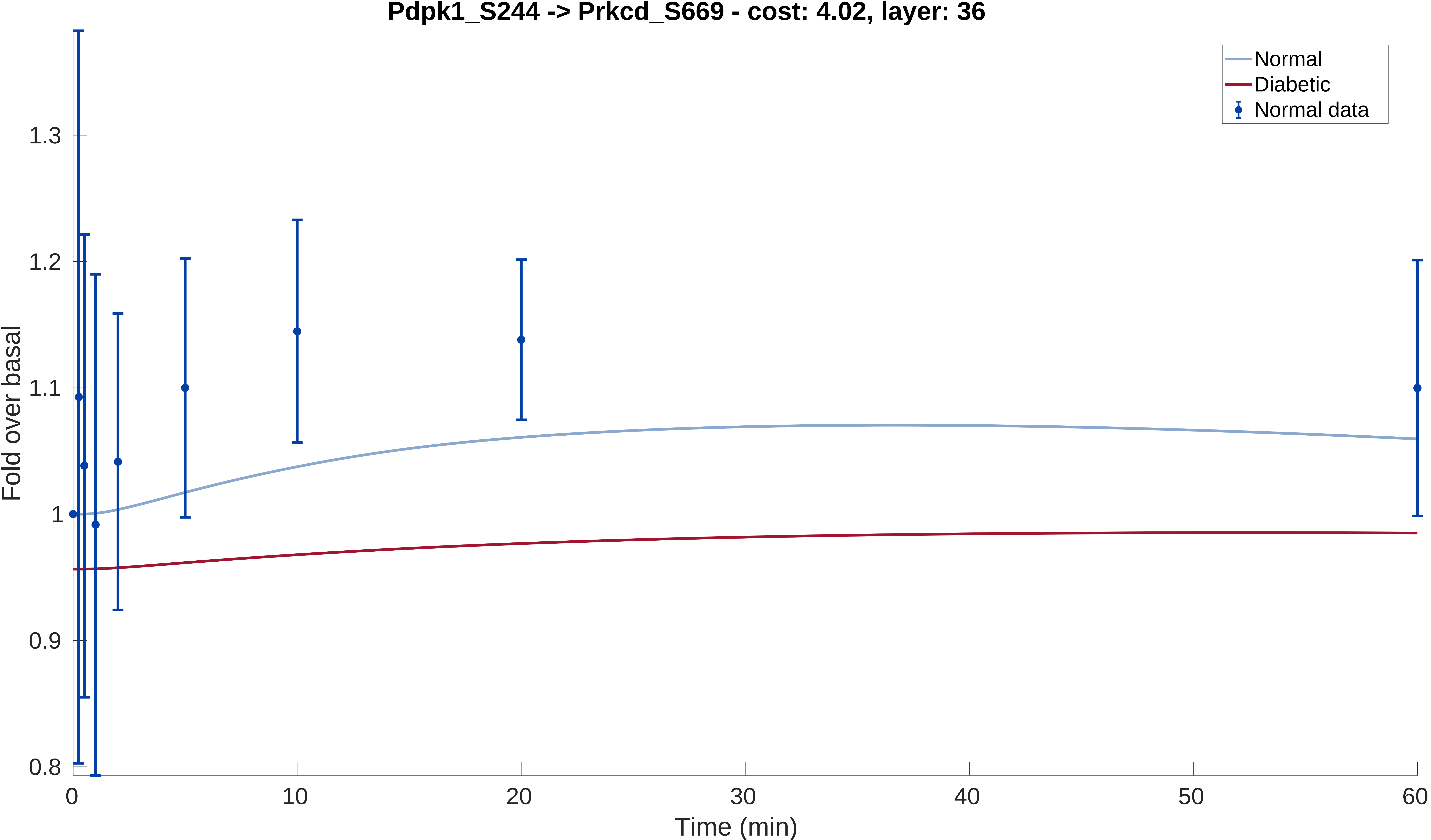

Pdpk1_S244 -> Prkcd_S669 - cost: 4.02, layer: 36
Normal
Diabetic
Normal data
Fold over basal
Time (min)
0.8
0.9
1
1.1
1.2
1.3
0
10
20
30
40
50
60

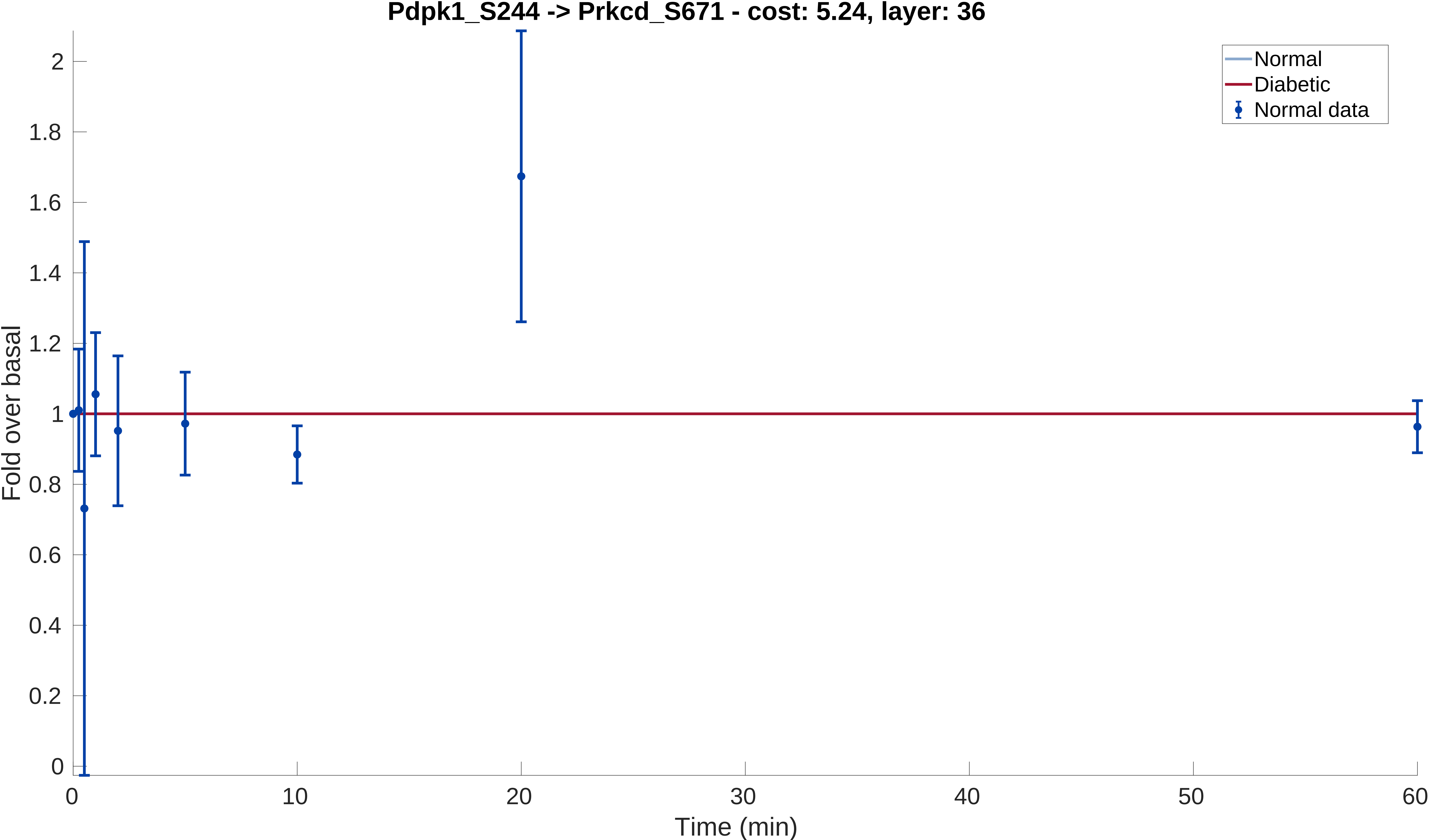

Pdpk1_S244 -> Prkcd_S671 - cost: 5.24, layer: 36
Normal
Diabetic
Normal data
Fold over basal
Time (min)
0
0.2
0.4
0.6
0.8
1
1.2
1.4
1.6
1.8
2
0
10
20
30
40
50
60

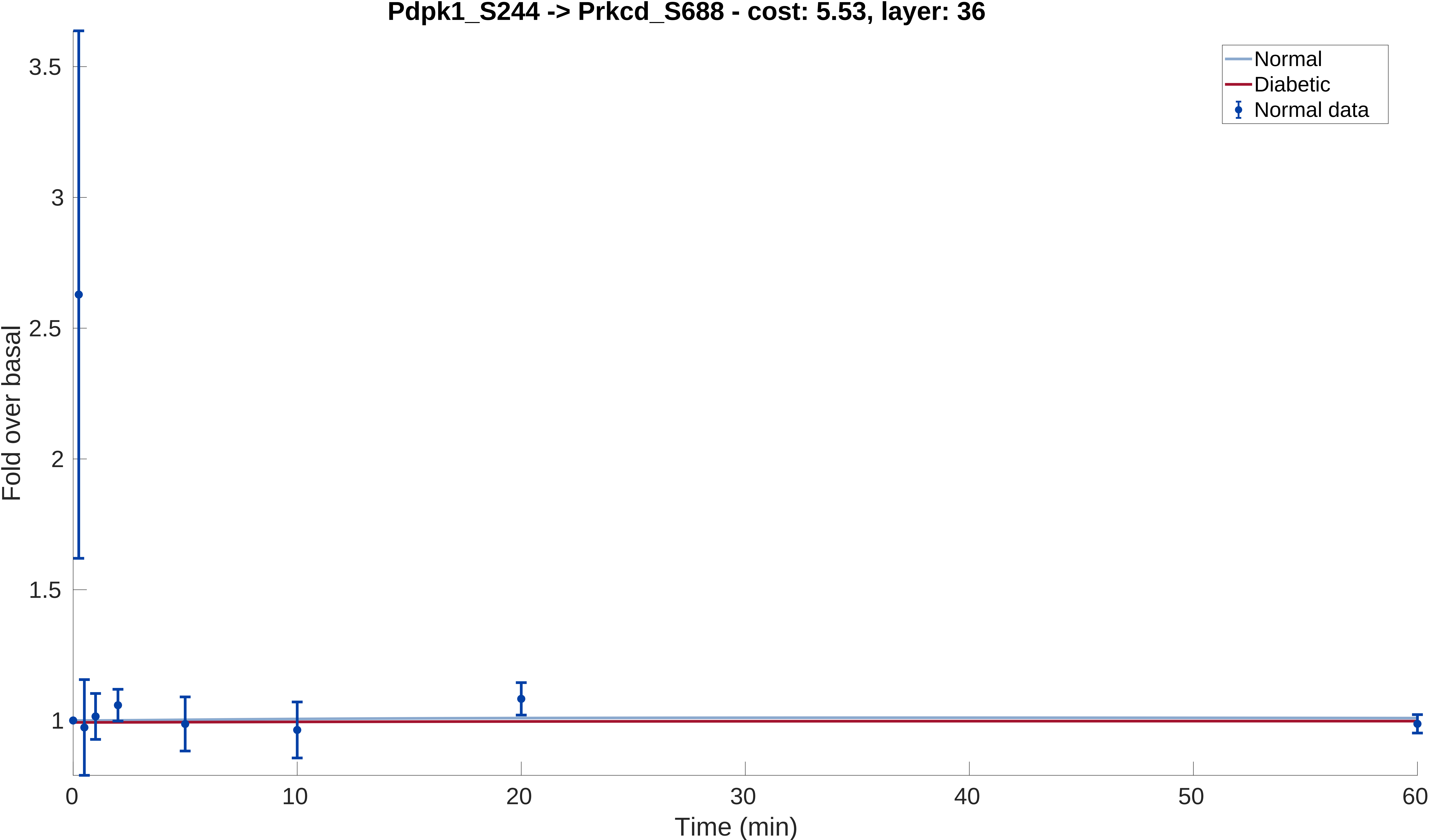

Pdpk1_S244 -> Prkcd_S688 - cost: 5.53, layer: 36
Normal
Diabetic
Normal data
Fold over basal
3.5
3
2.5
2
1.5
1
0
10
20
30
40
50
60
Time (min)

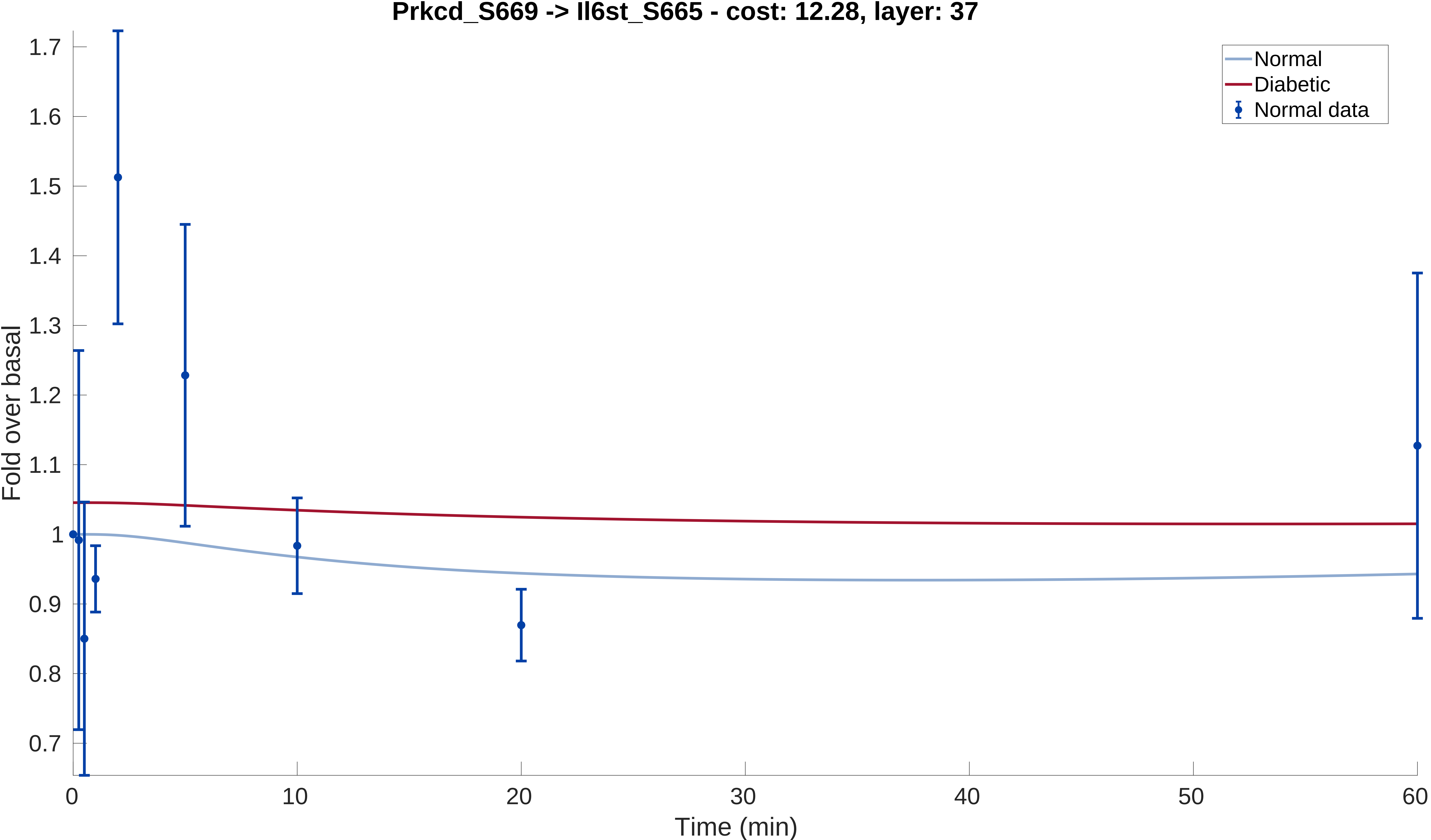

Prkcd_S669 -> Il6st_S665 - cost: 12.28, layer: 37
Normal
Diabetic
Normal data
Fold over basal
1.7
1.6
1.5
1.4
1.3
1.2
1.1
1
0.9
0.8
0.7
0
10
20
30
40
50
60
Time (min)

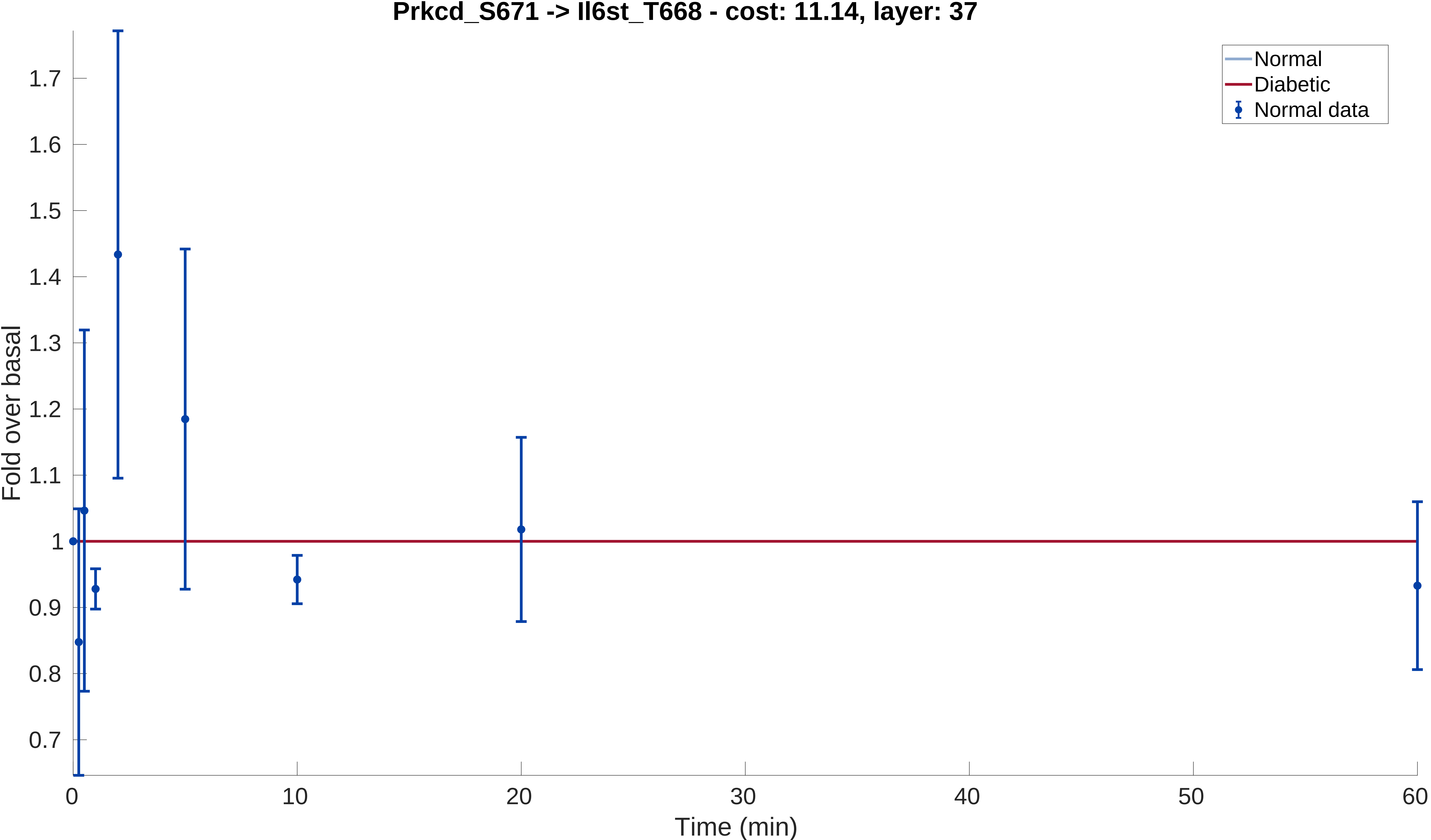

Prkcd_S671 -> Il6st_T668 - cost: 11.14, layer: 37
Normal
Diabetic
Normal data
Fold over basal
1.7
1.6
1.5
1.4
1.3
1.2
1.1
1
0.9
0.8
0.7
0
10
20
30
40
50
60
Time (min)

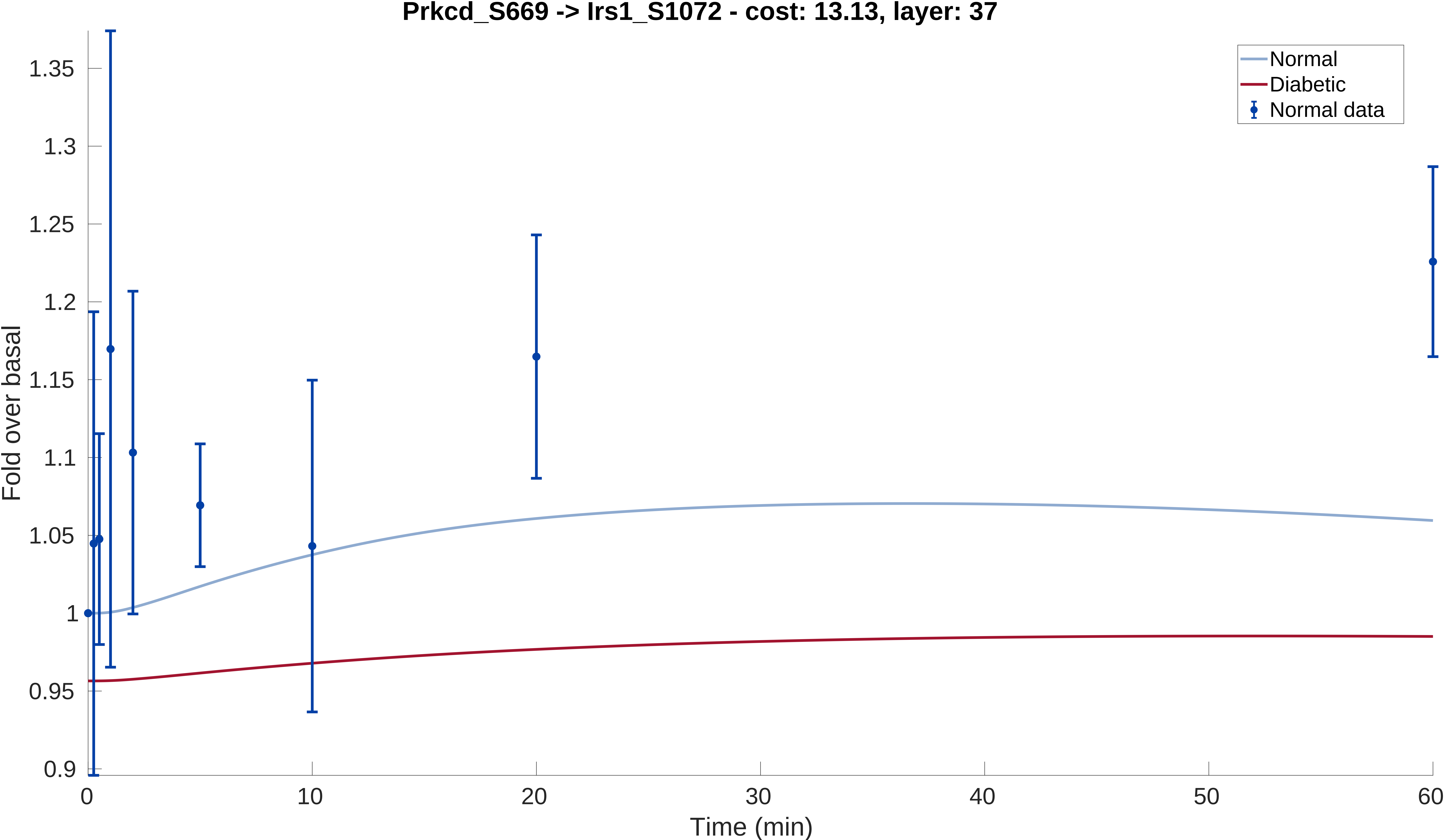

Prkcd_S669 -> Irs1_S1072 - cost: 13.13, layer: 37
Normal
Diabetic
Normal data
Fold over basal
1.35
1.3
1.25
1.2
1.15
1.1
1.05
1
0.95
0.9
0
10
20
30
40
50
60
Time (min)

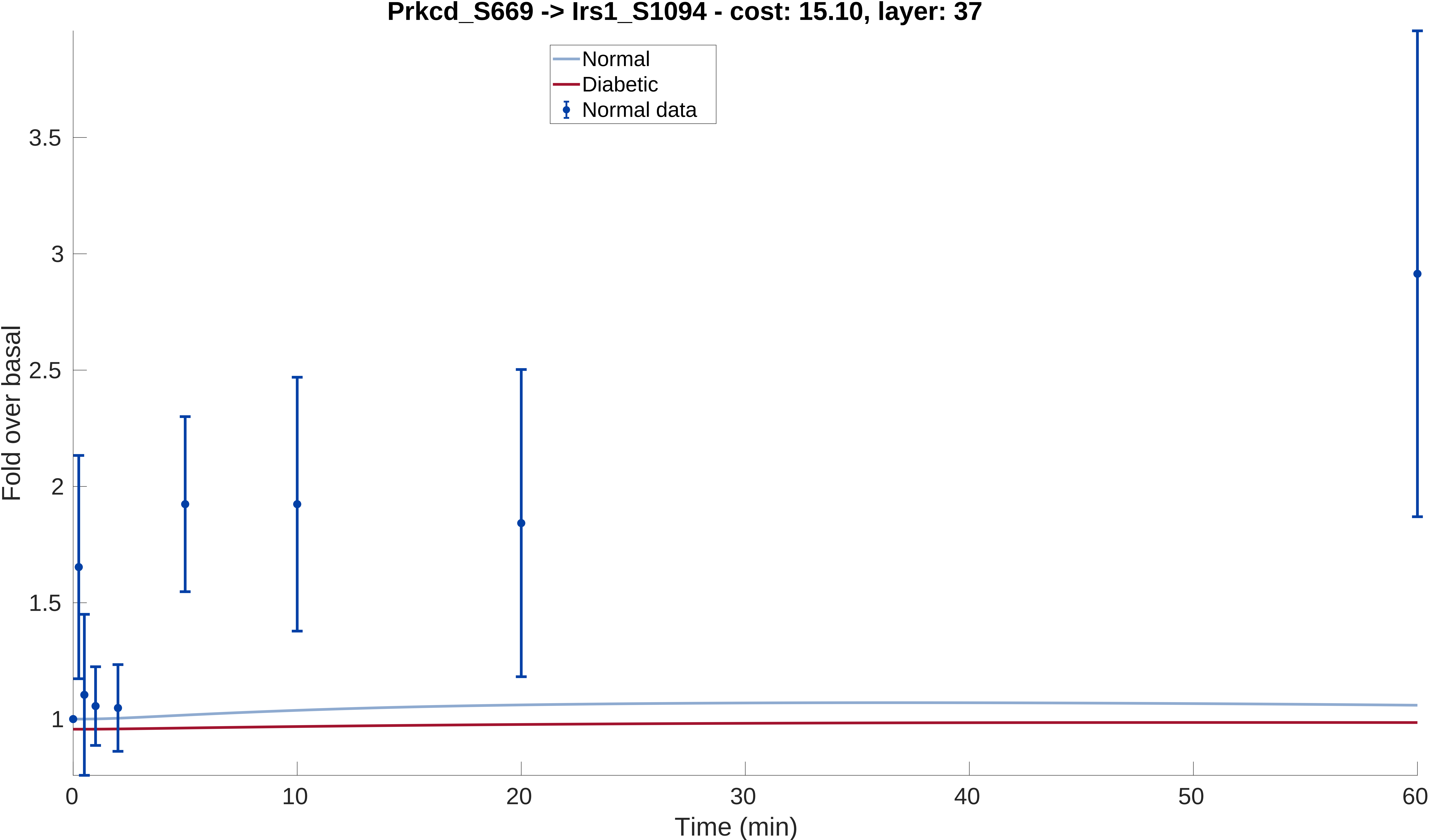

Prkcd_S669 -> Irs1_S1094 - cost: 15.10, layer: 37
Normal
Diabetic
Normal data
Fold over basal
3.5
3
2.5
2
1.5
1
0
10
20
30
40
50
60
Time (min)

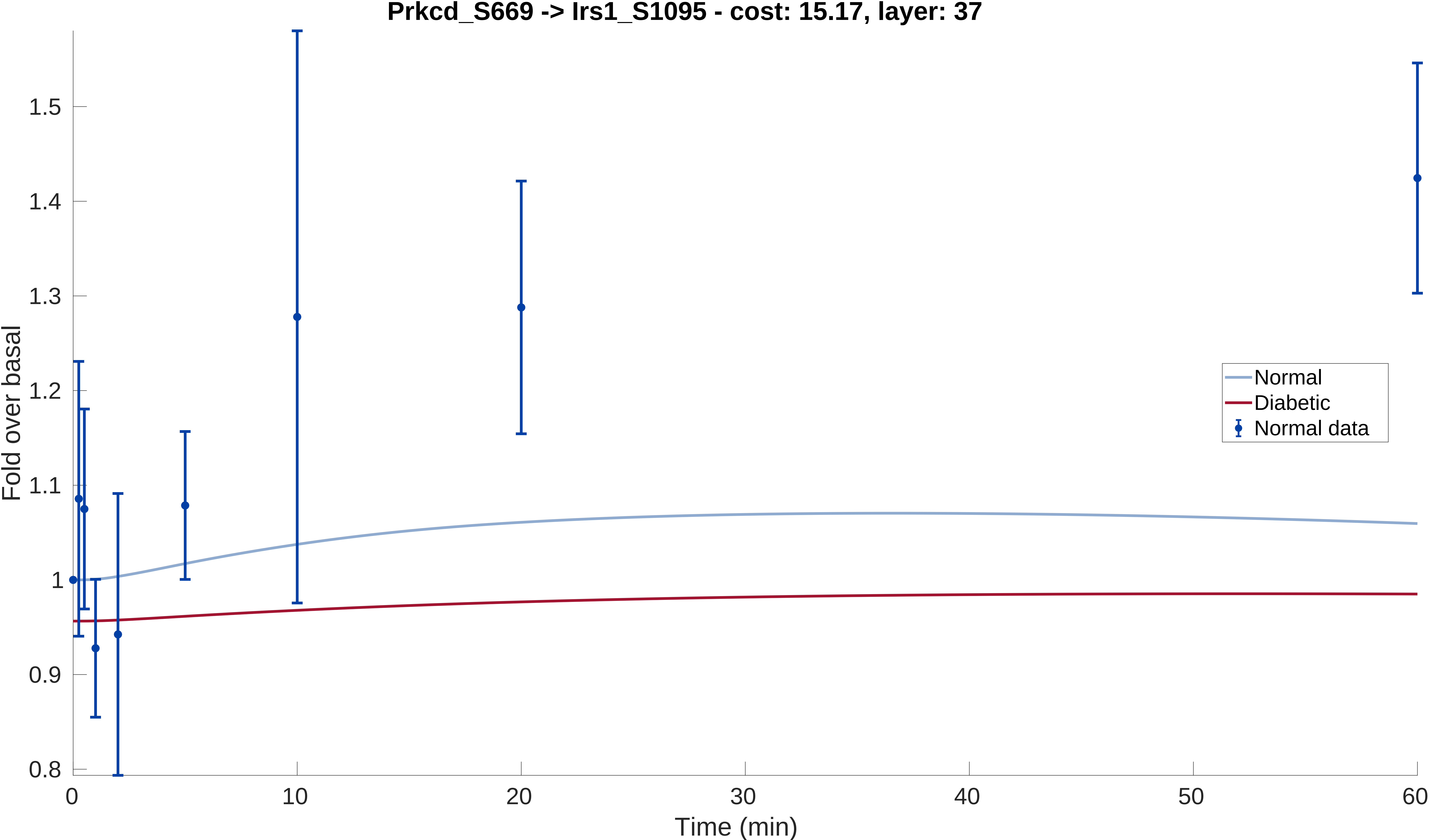
Prkcd_S669 -> Irs1_S1095 - cost: 15.17, layer: 37
Fold over basal
1.5
1.4
1.3
1.2
1.1
1
0.9
0.8
0
10
20
30
40
50
60
Time (min)
Normal
Diabetic
Normal data

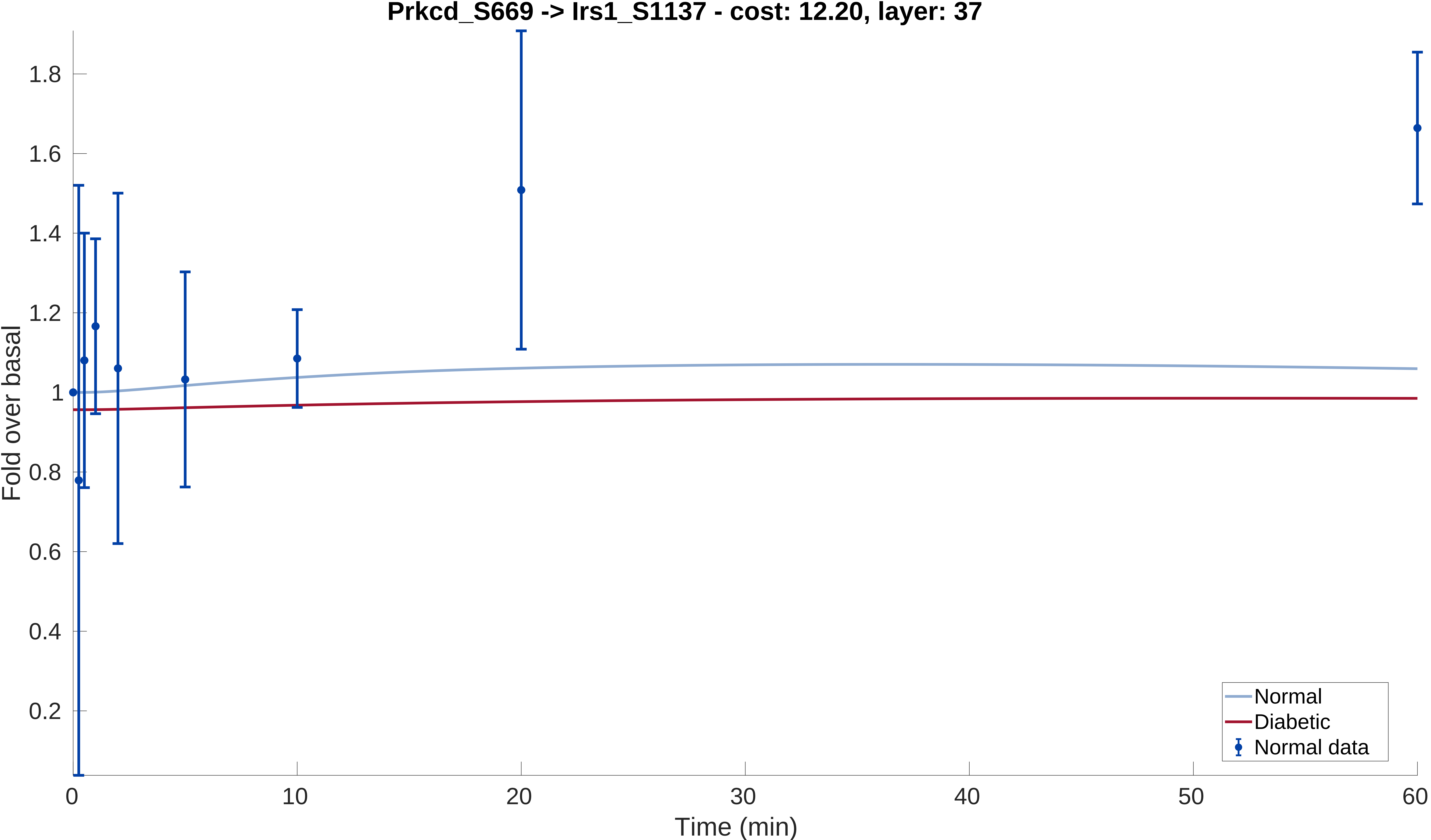

Prkcd_S669 -> Irs1_S1137 - cost: 12.20, layer: 37
Fold over basal
1.8
1.6
1.4
1.2
1
0.8
0.6
0.4
0.2
0
10
20
30
40
50
60
Time (min)
Normal
Diabetic
Normal data

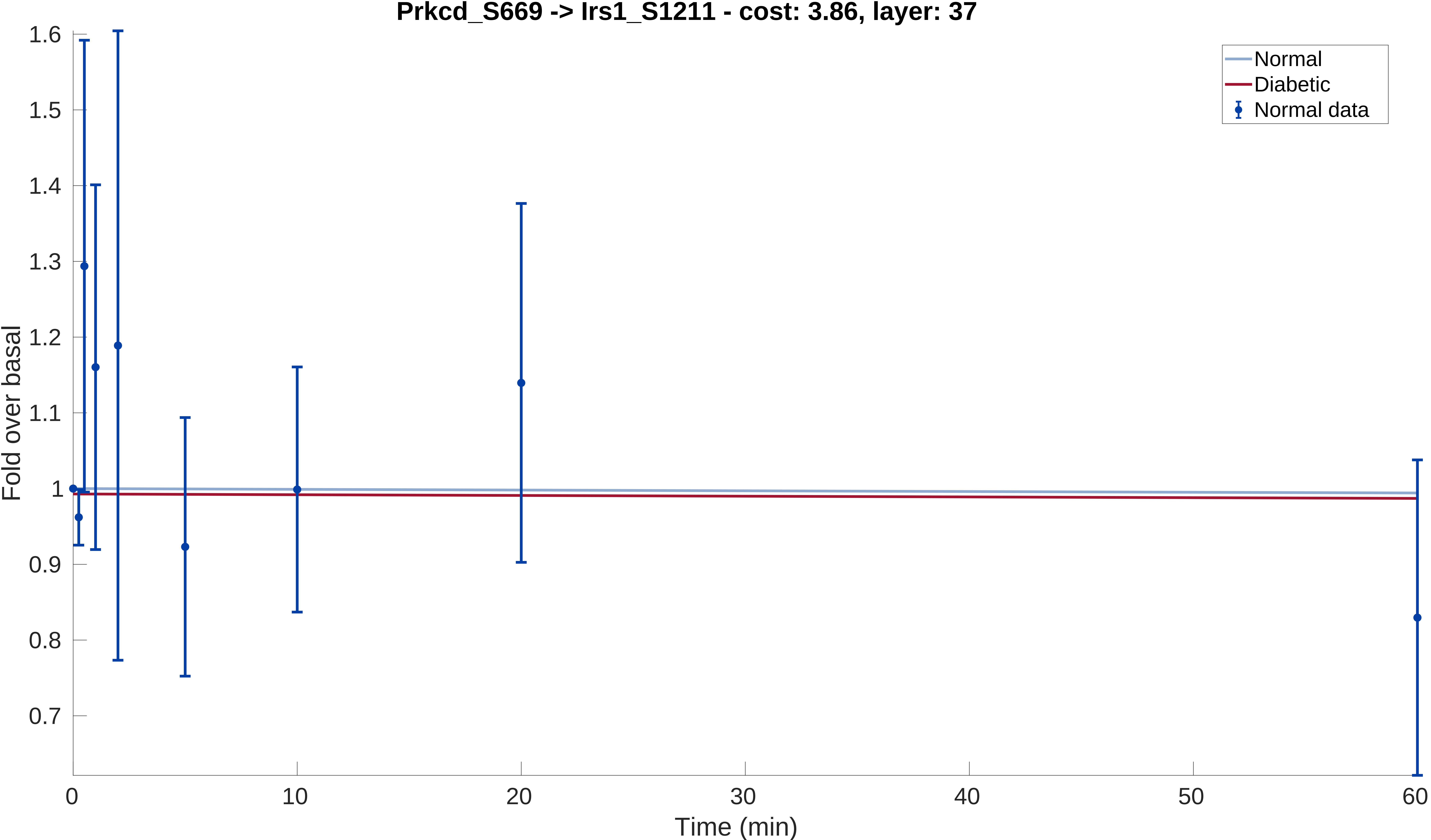

Prkcd_S669 -> Irs1_S1211 - cost: 3.86, layer: 37
Normal
Diabetic
Normal data
Fold over basal
1.6
1.5
1.4
1.3
1.2
1.1
1
0.9
0.8
0.7
0
10
20
30
40
50
60
Time (min)

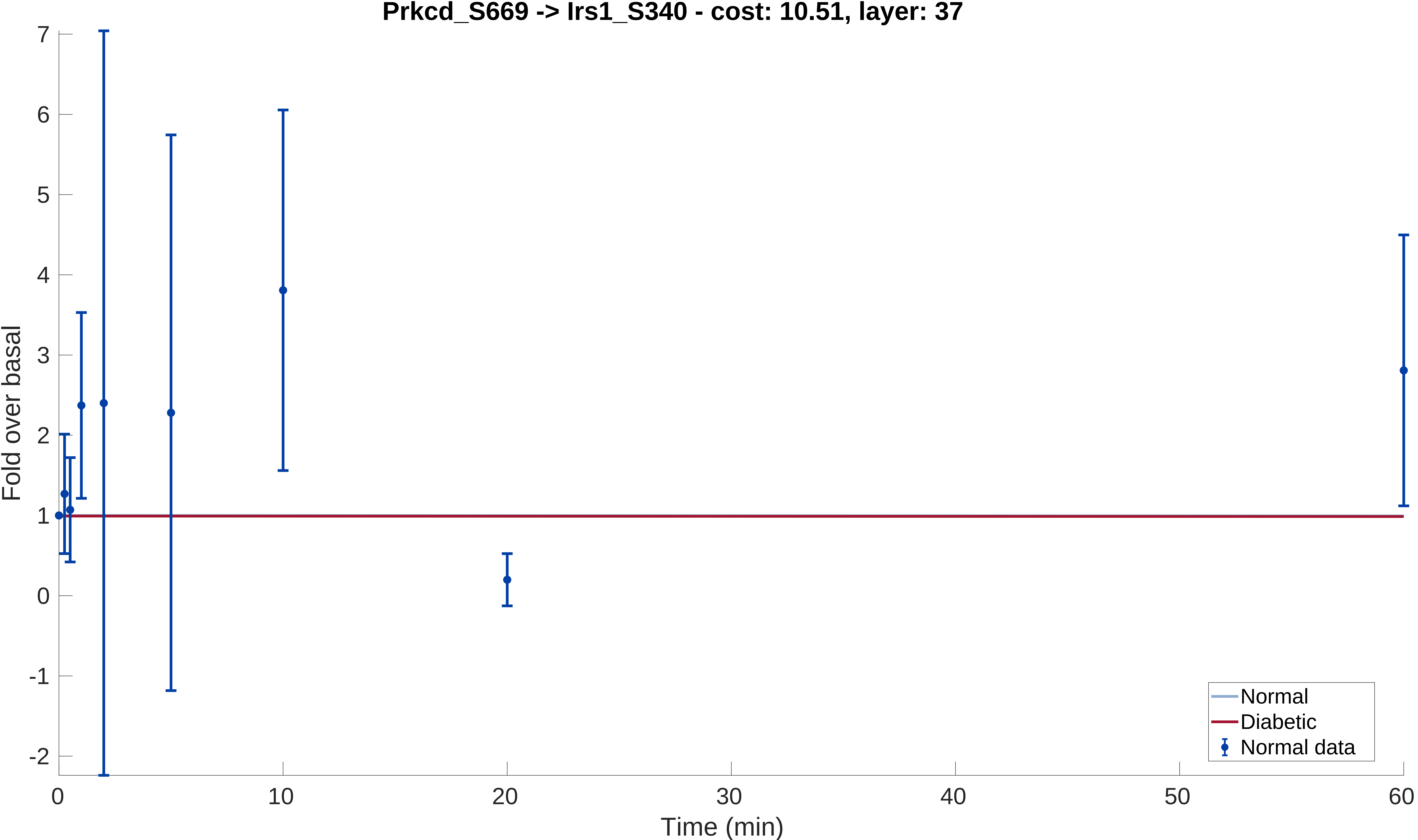

Prkcd_S669 -> Irs1_S340 - cost: 10.51, layer: 37
Fold over basal
Time (min)
7
6
5
4
3
2
1
0
-1
-2
0
10
20
30
40
50
60
Normal
Diabetic
Normal data

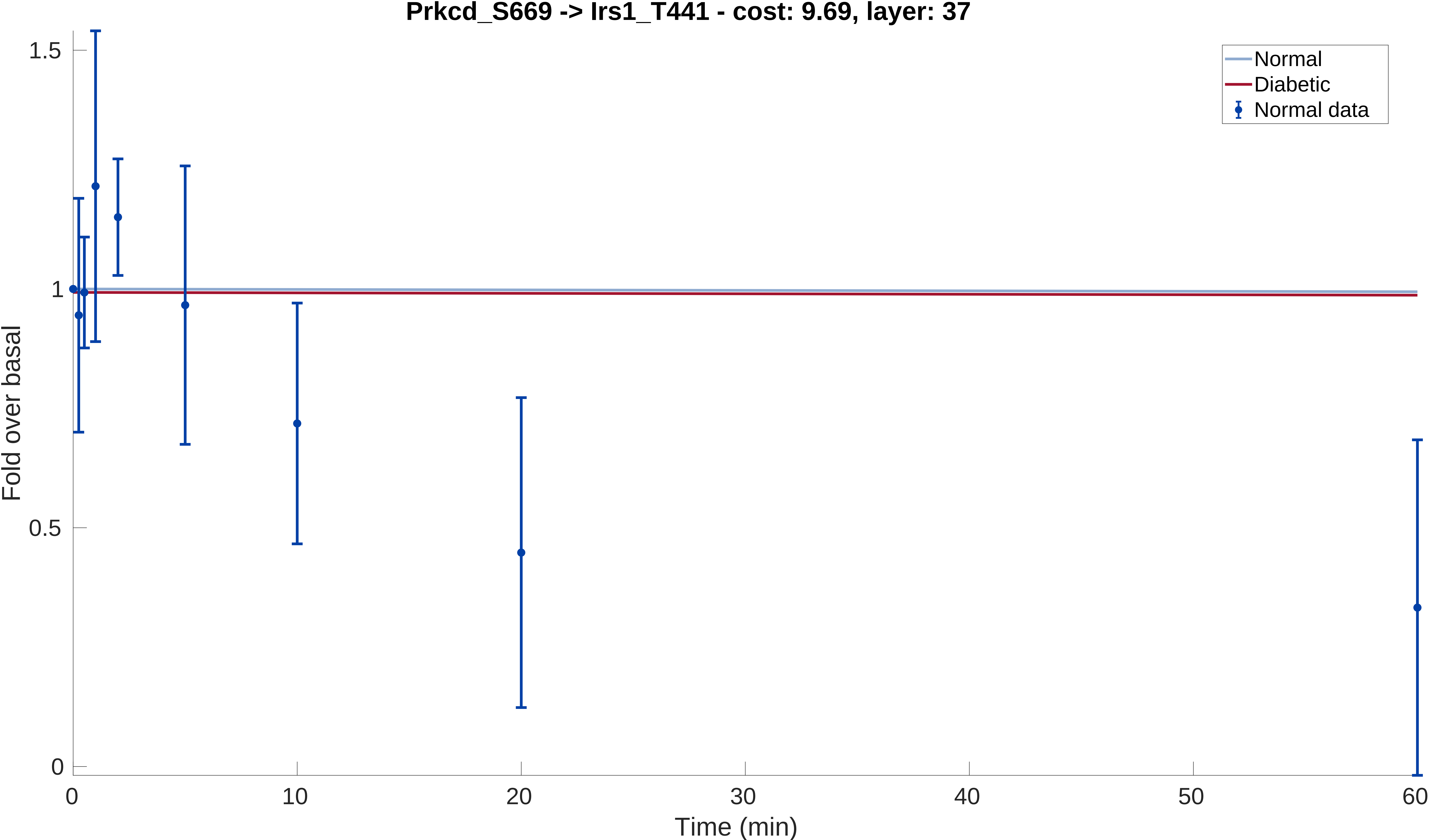

Prkcd_S669 -> Irs1_T441 - cost: 9.69, layer: 37
Normal
Diabetic
Normal data
Fold over basal
Time (min)
1.5
1
0.5
0
0
10
20
30
40
50
60

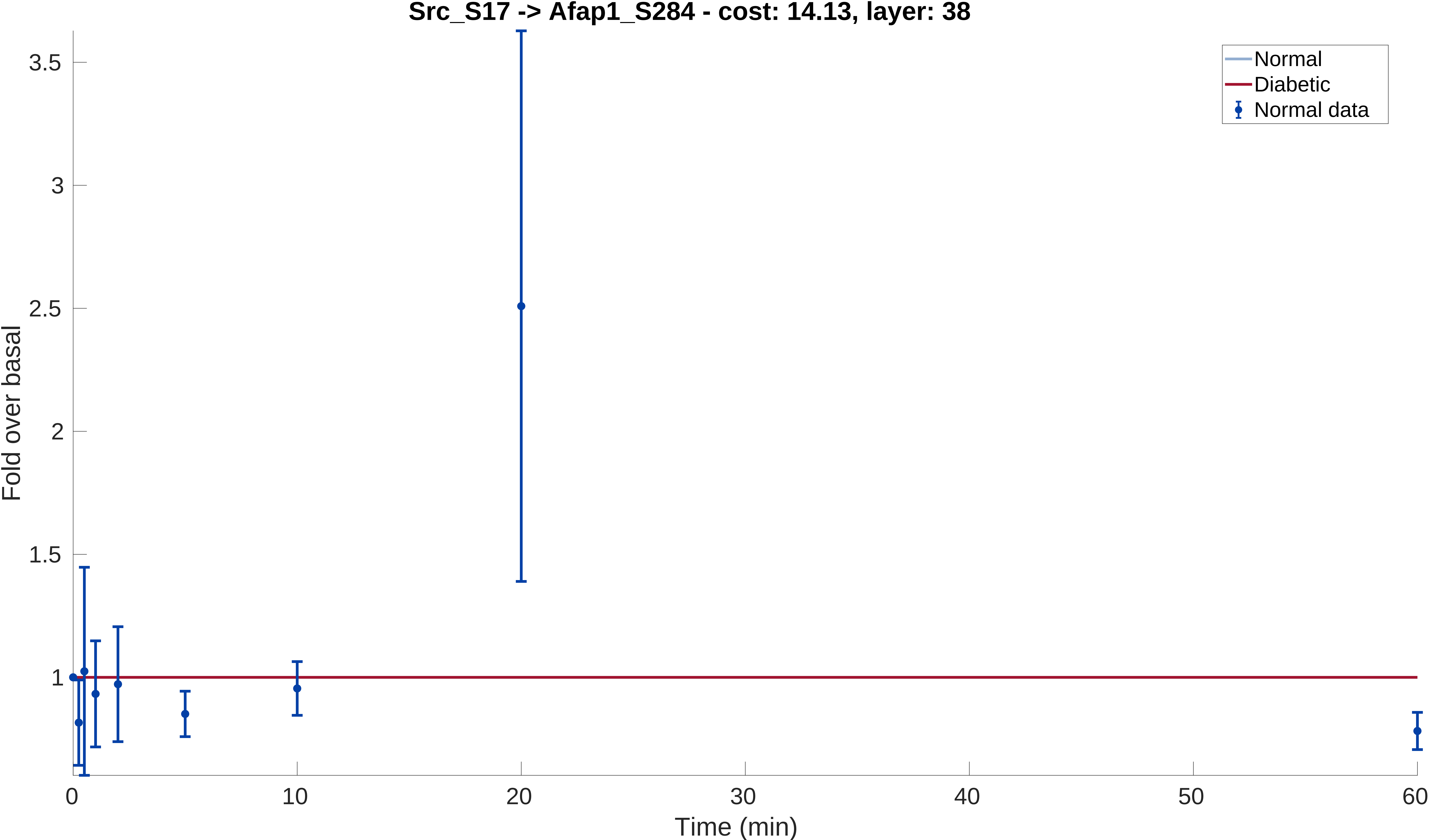

Src_S17 -> Afap1_S284 - cost: 14.13, layer: 38
Normal
Diabetic
Normal data
Fold over basal
3.5
3
2.5
2
1.5
1
0
10
20
30
40
50
60
Time (min)

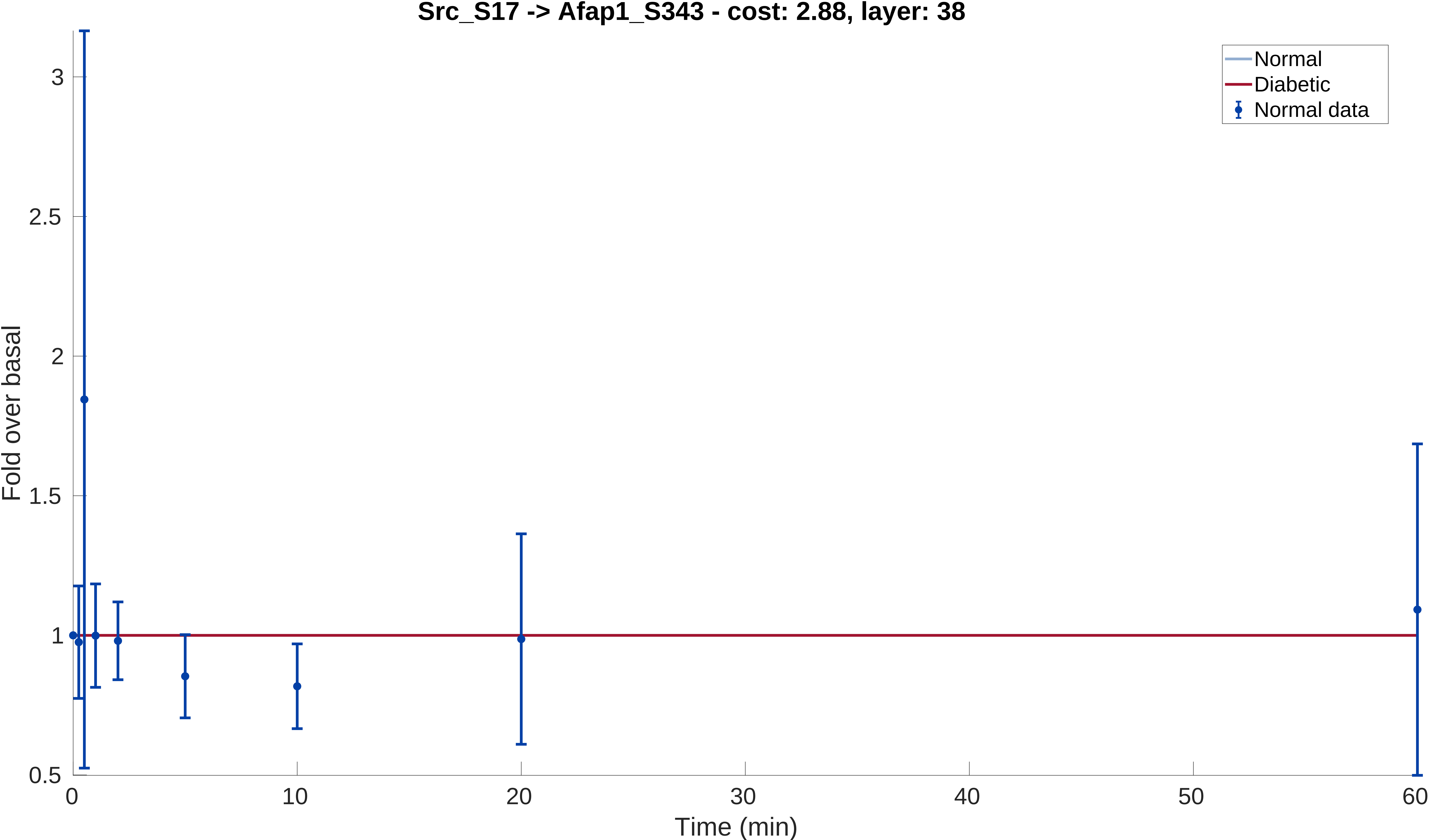

Src_S17 -> Afap1_S343 - cost: 2.88, layer: 38
Normal
Diabetic
Normal data
Fold over basal
0.5
1
1.5
2
2.5
3
0
10
20
30
40
50
60
Time (min)

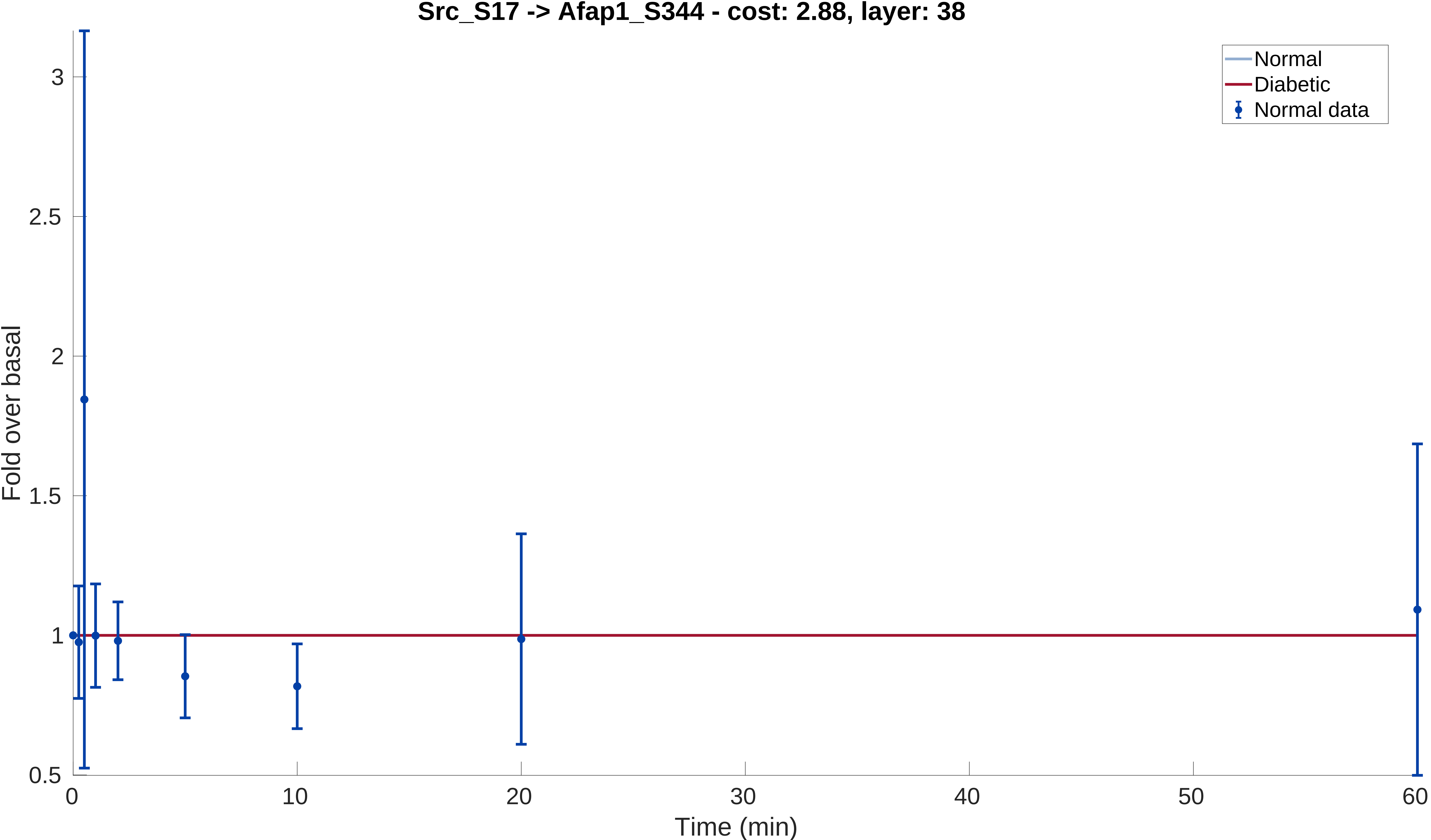
Src_S17 -> Afap1_S344 - cost: 2.88, layer: 38
Normal
Diabetic
Normal data
Fold over basal
3
2.5
2
1.5
1
0.5
0
10
20
30
40
50
60
Time (min)

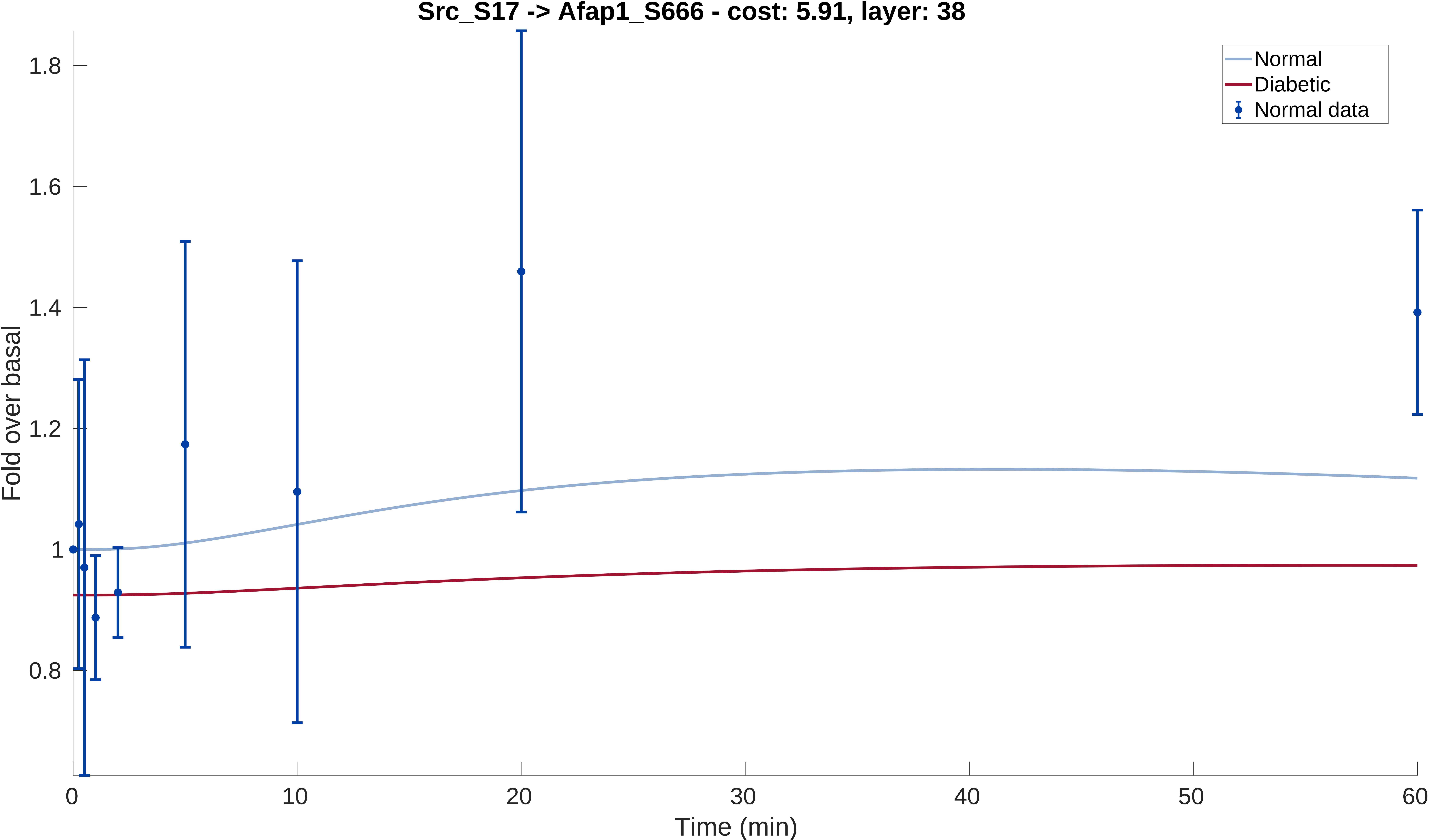

Src_S17 -> Afap1_S666 - cost: 5.91, layer: 38
Normal
Diabetic
Normal data
1.8
1.6
1.4
1.2
1
0.8
Fold over basal
0
10
20
30
40
50
60
Time (min)

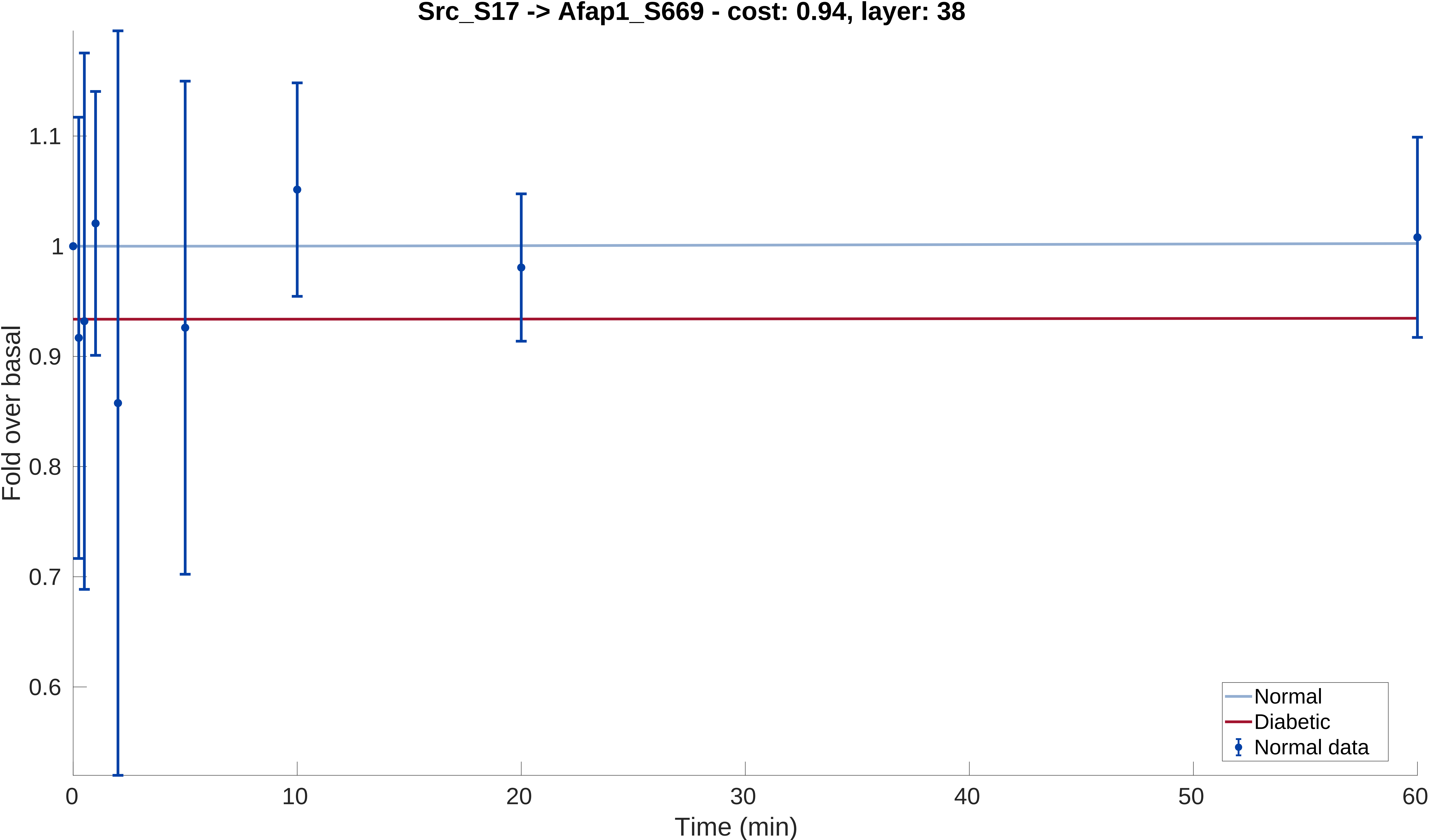

Src_S17 -> Afap1_S669 - cost: 0.94, layer: 38
Fold over basal
1.1
1
0.9
0.8
0.7
0.6
0
10
20
30
40
50
60
Time (min)
Normal
Diabetic
Normal data

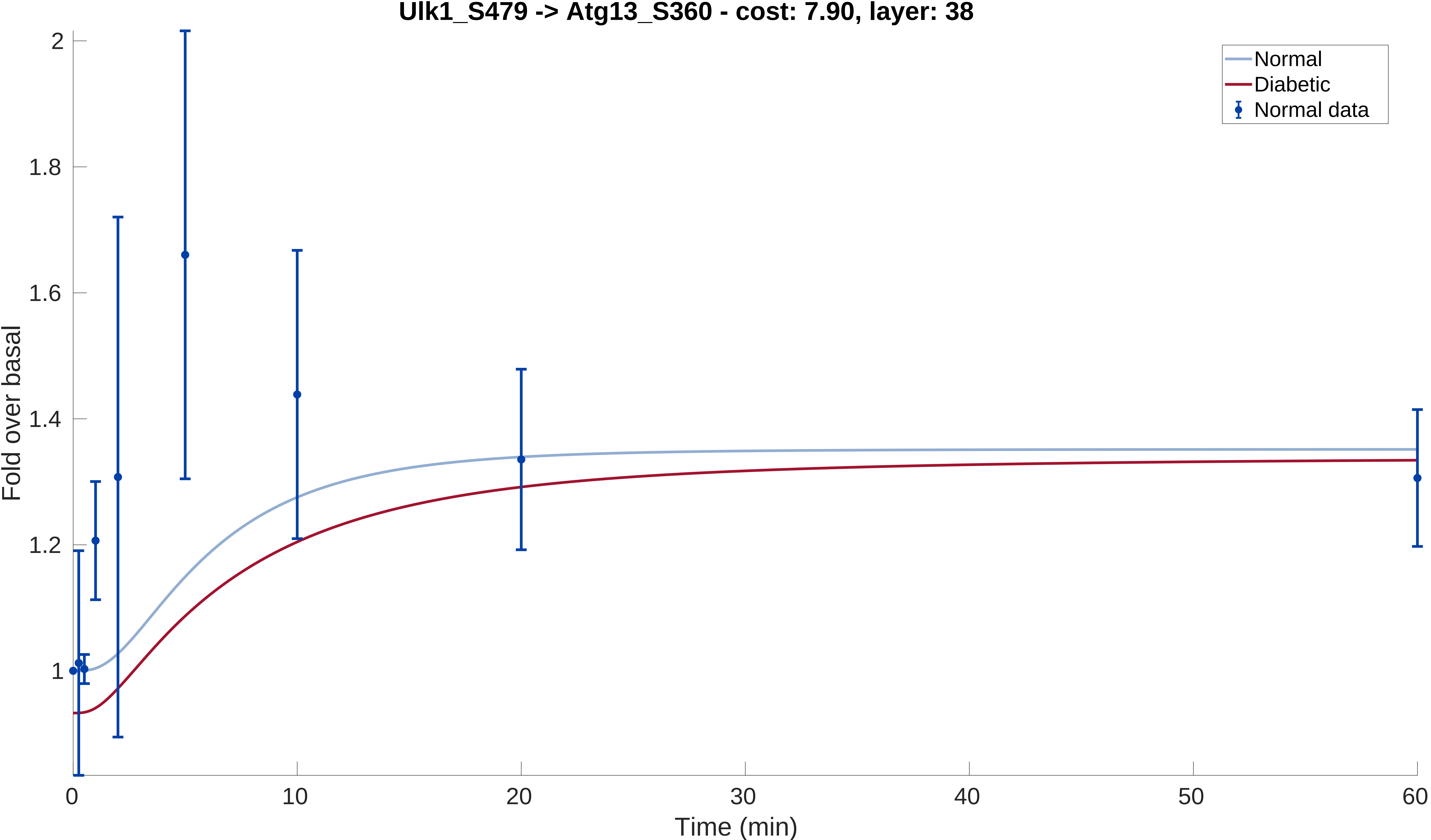

Ulk1_S479 -> Atg13_S360 - cost: 7.90, layer: 38
Fold over basal
2
1.8
1.6
1.4
1.2
1
0
10
20
30
40
50
60
Time (min)
Normal
Diabetic
Normal data

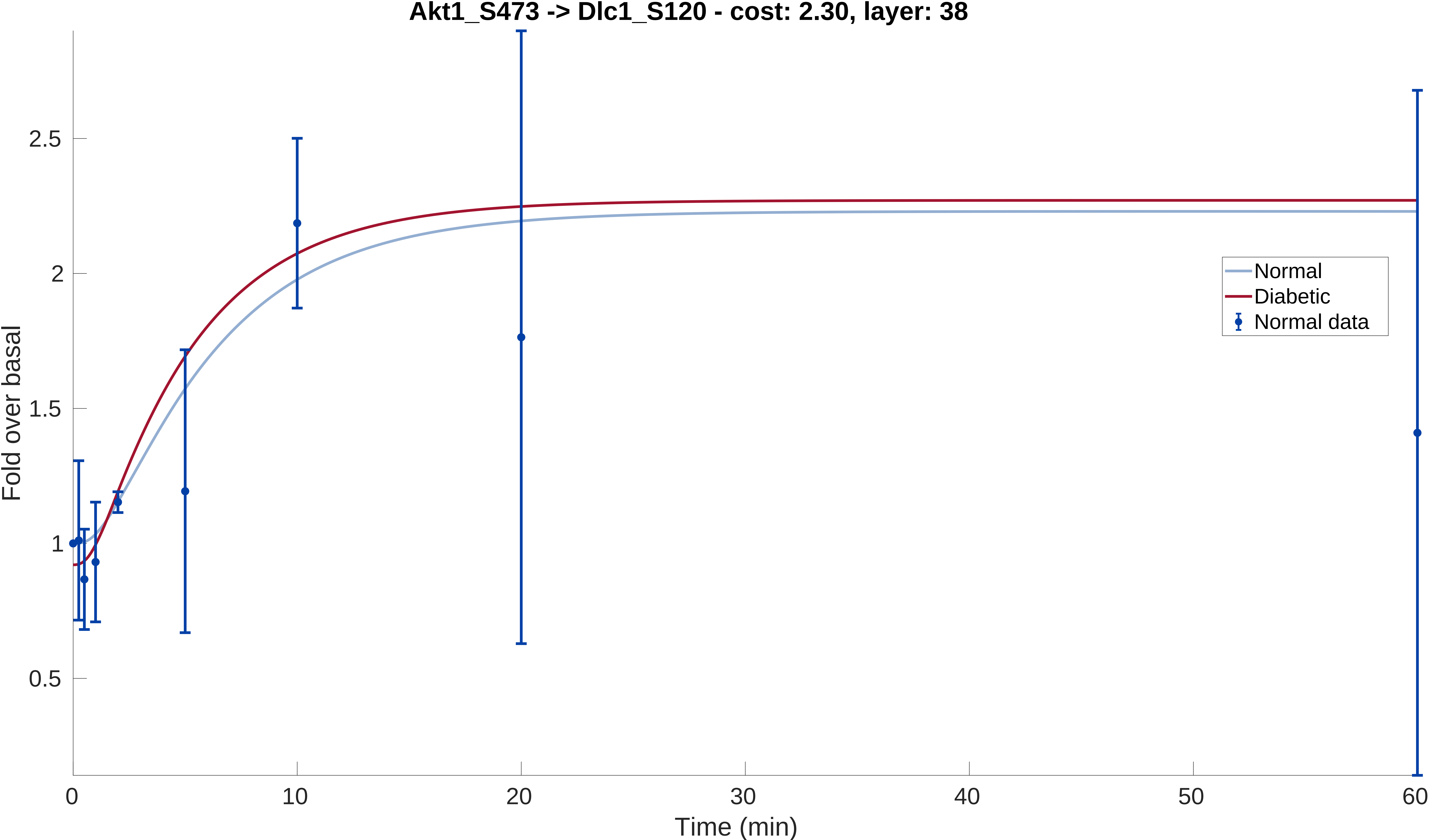

Akt1_S473 -> Dlc1_S120 - cost: 2.30, layer: 38
Fold over basal
2.5
2
1.5
1
0.5
0
10
20
30
40
50
60
Time (min)
Normal
Diabetic
Normal data

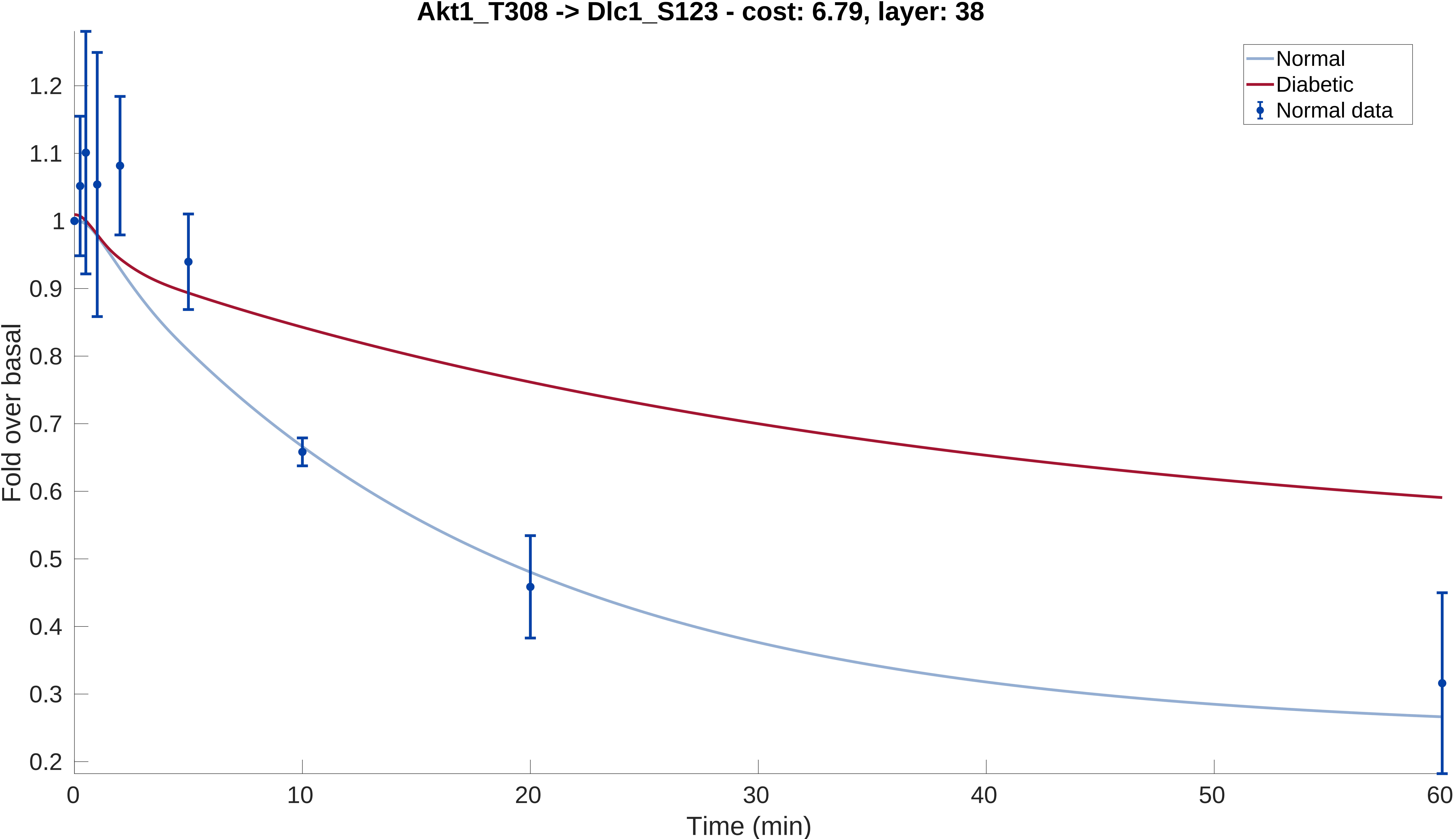
Akt1_T308 -> Dlc1_S123 - cost: 6.79, layer: 38
Normal
Diabetic
Normal data
Fold over basal
Time (min)
0
10
20
30
40
50
60
0.2
0.3
0.4
0.5
0.6
0.7
0.8
0.9
1
1.1
1.2

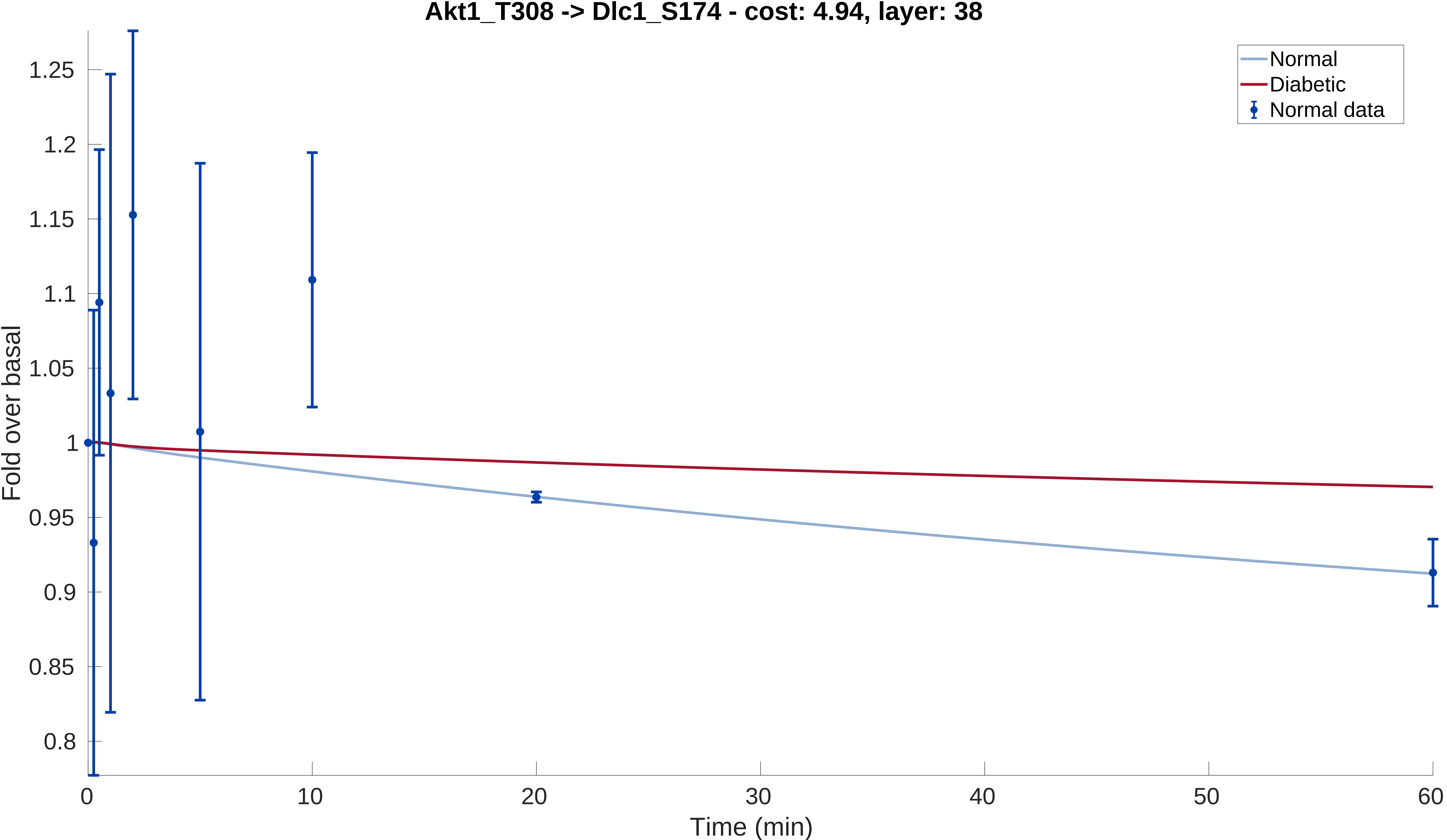

Akt1_T308 -> Dlc1_S174 - cost: 4.94, layer: 38
Normal
Diabetic
Normal data
Fold over basal
Time (min)
1.25
1.2
1.15
1.1
1.05
1
0.95
0.9
0.85
0.8
0
10
20
30
40
50
60

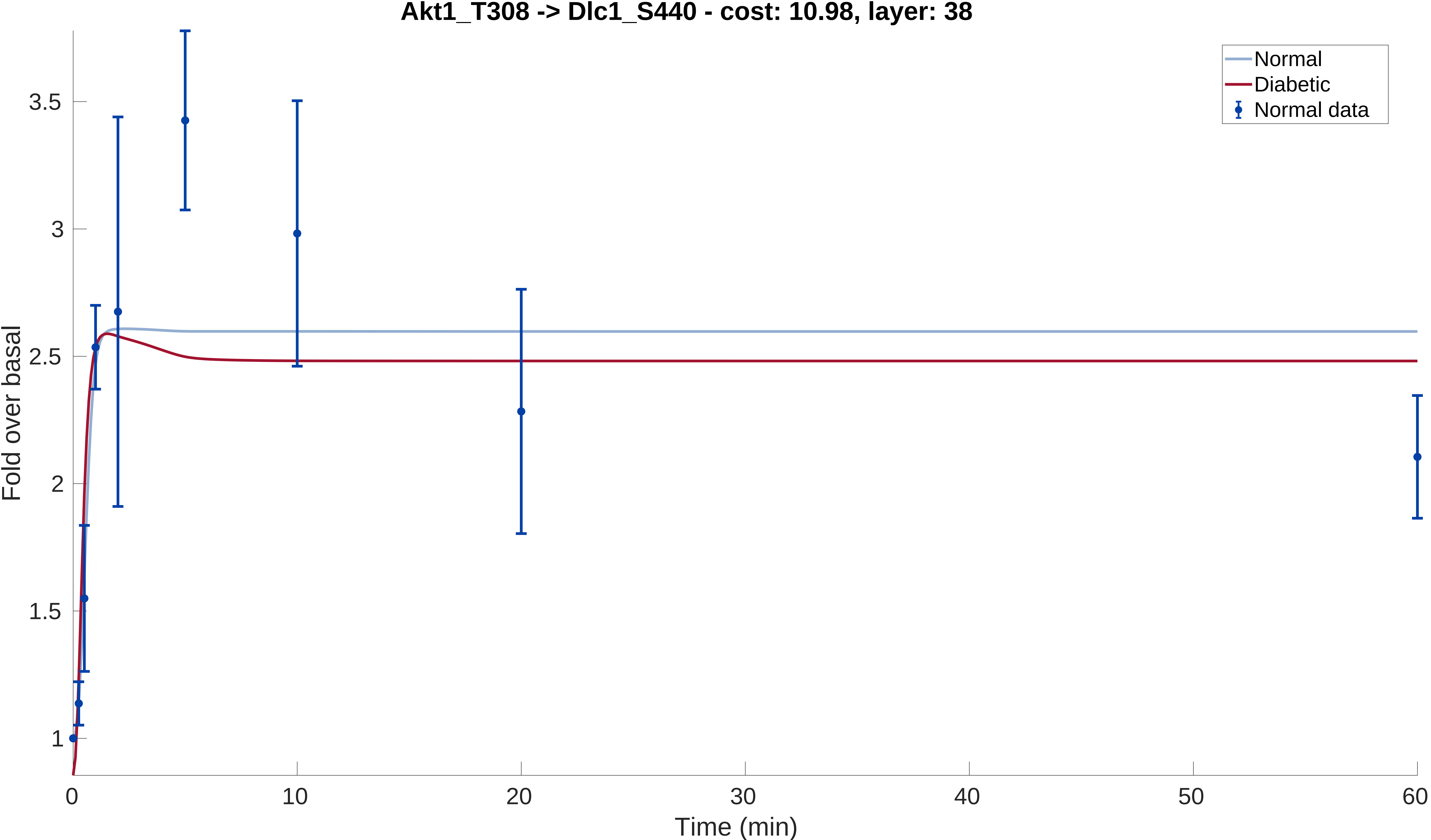

Akt1_T308 -> Dlc1_S440 - cost: 10.98, layer: 38
Normal
Diabetic
Normal data
Fold over basal
3.5
3
2.5
2
1.5
1
0
10
20
30
40
50
60
Time (min)

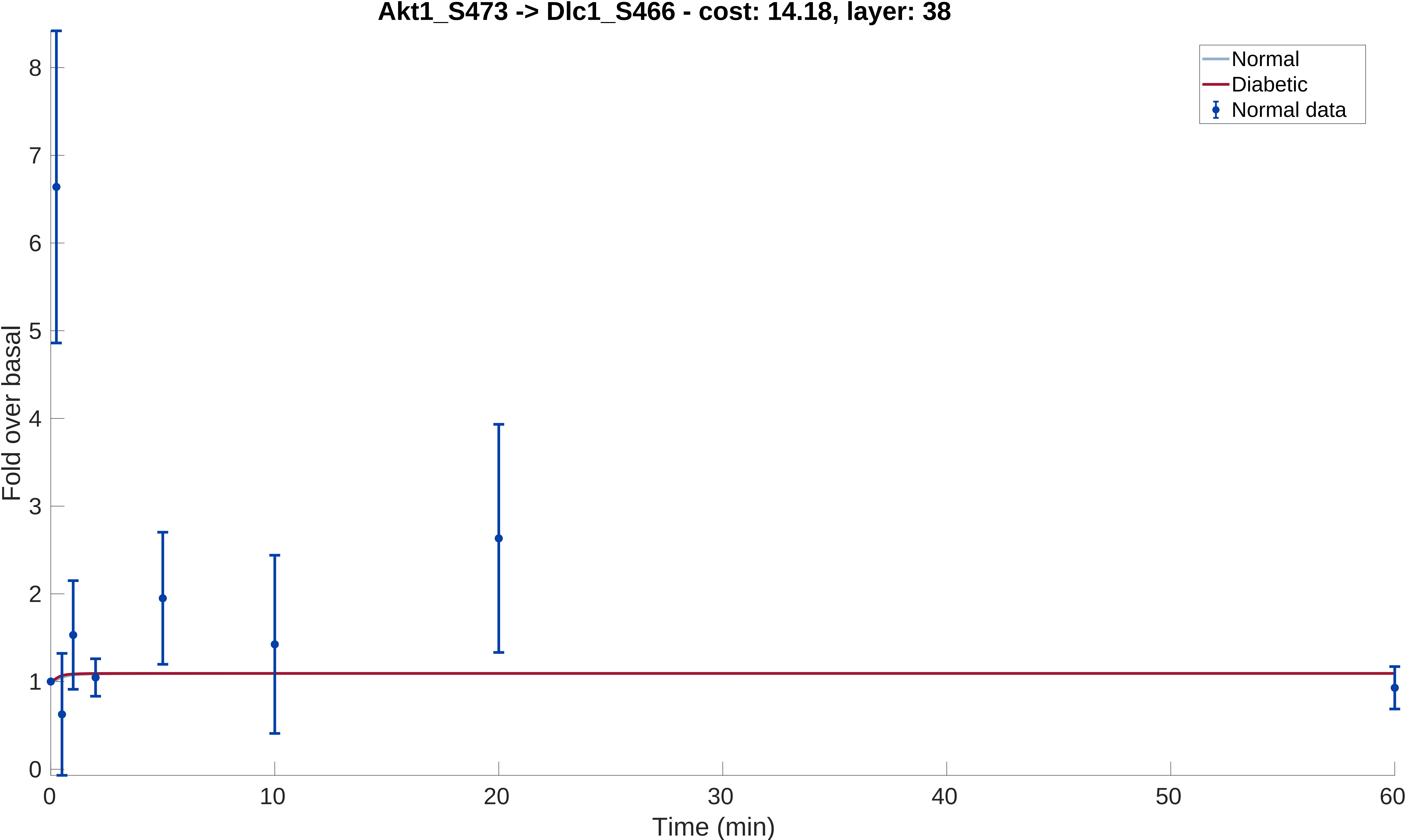

Akt1_S473 -> Dlc1_S466 - cost: 14.18, layer: 38
Normal
Diabetic
Normal data
Fold over basal
Time (min)
0
1
2
3
4
5
6
7
8
0
10
20
30
40
50
60

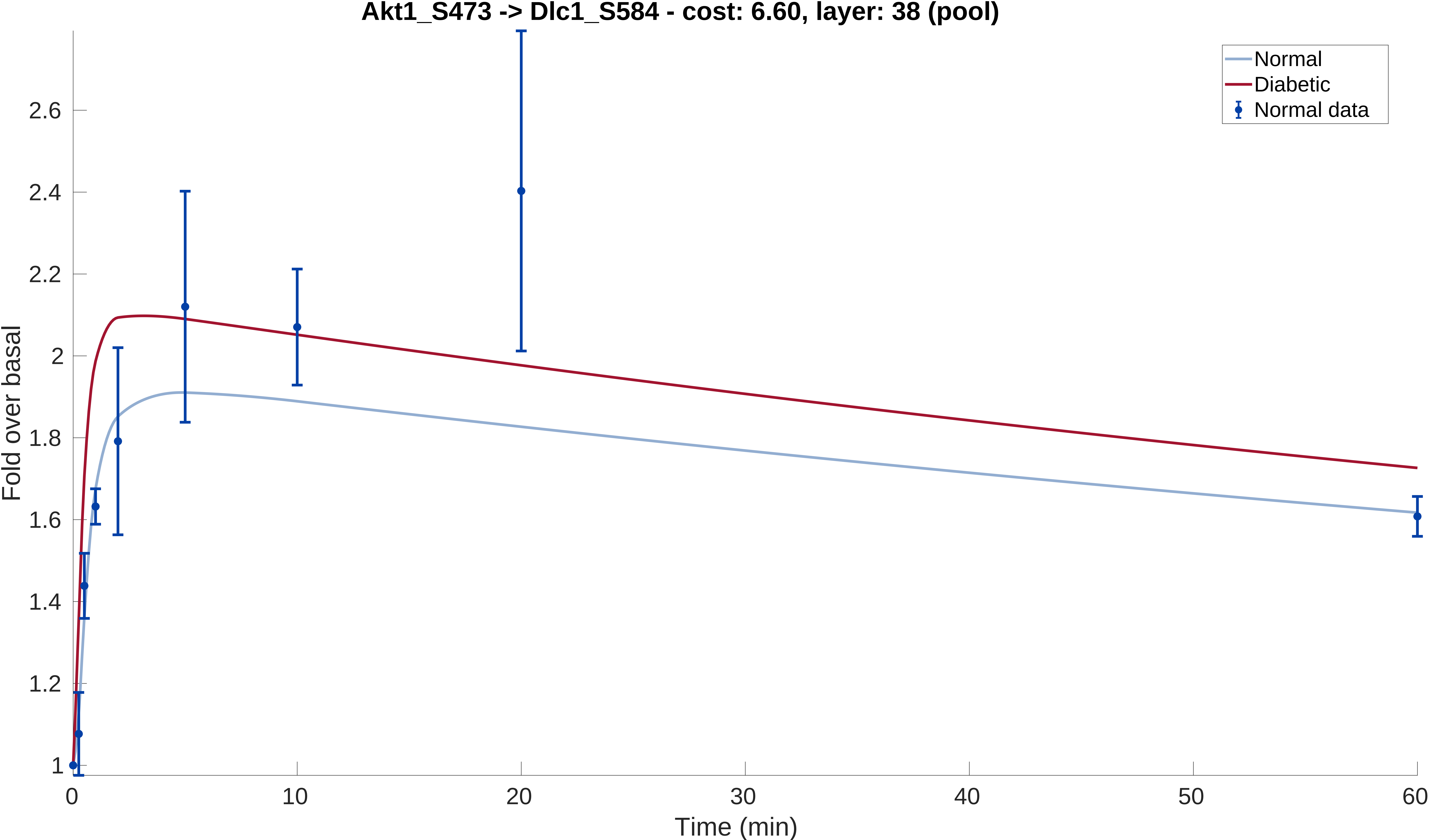

Akt1_S473 -> Dlc1_S584 - cost: 6.60, layer: 38 (pool)
Normal
Diabetic
Normal data
Fold over basal
1
1.2
1.4
1.6
1.8
2
2.2
2.4
2.6
0
10
20
30
40
50
60
Time (min)

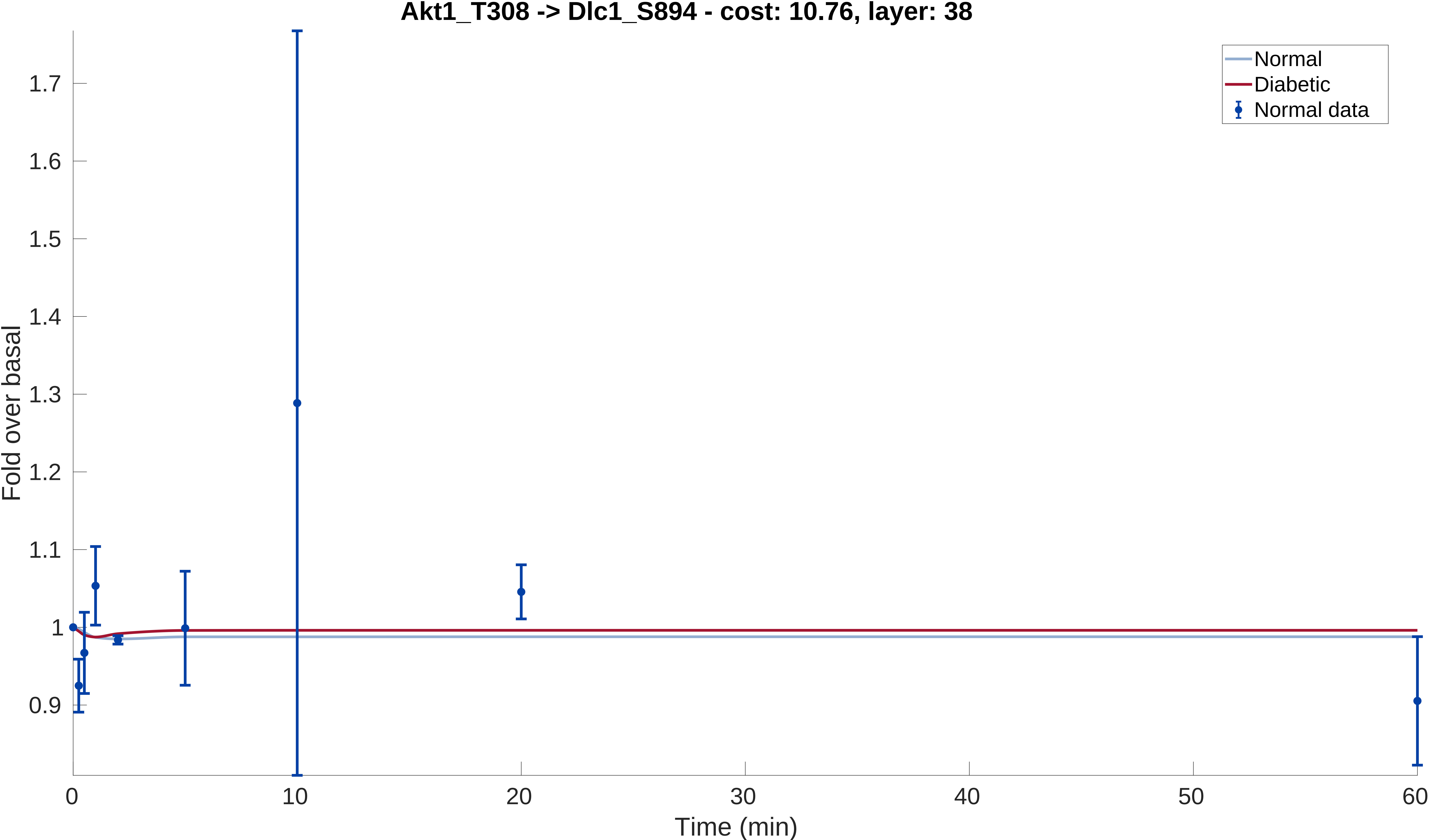

Akt1_T308 -> Dlc1_S894 - cost: 10.76, layer: 38
Normal
Diabetic
Normal data
Fold over basal
0.9
1
1.1
1.2
1.3
1.4
1.5
1.6
1.7
0
10
20
30
40
50
60
Time (min)

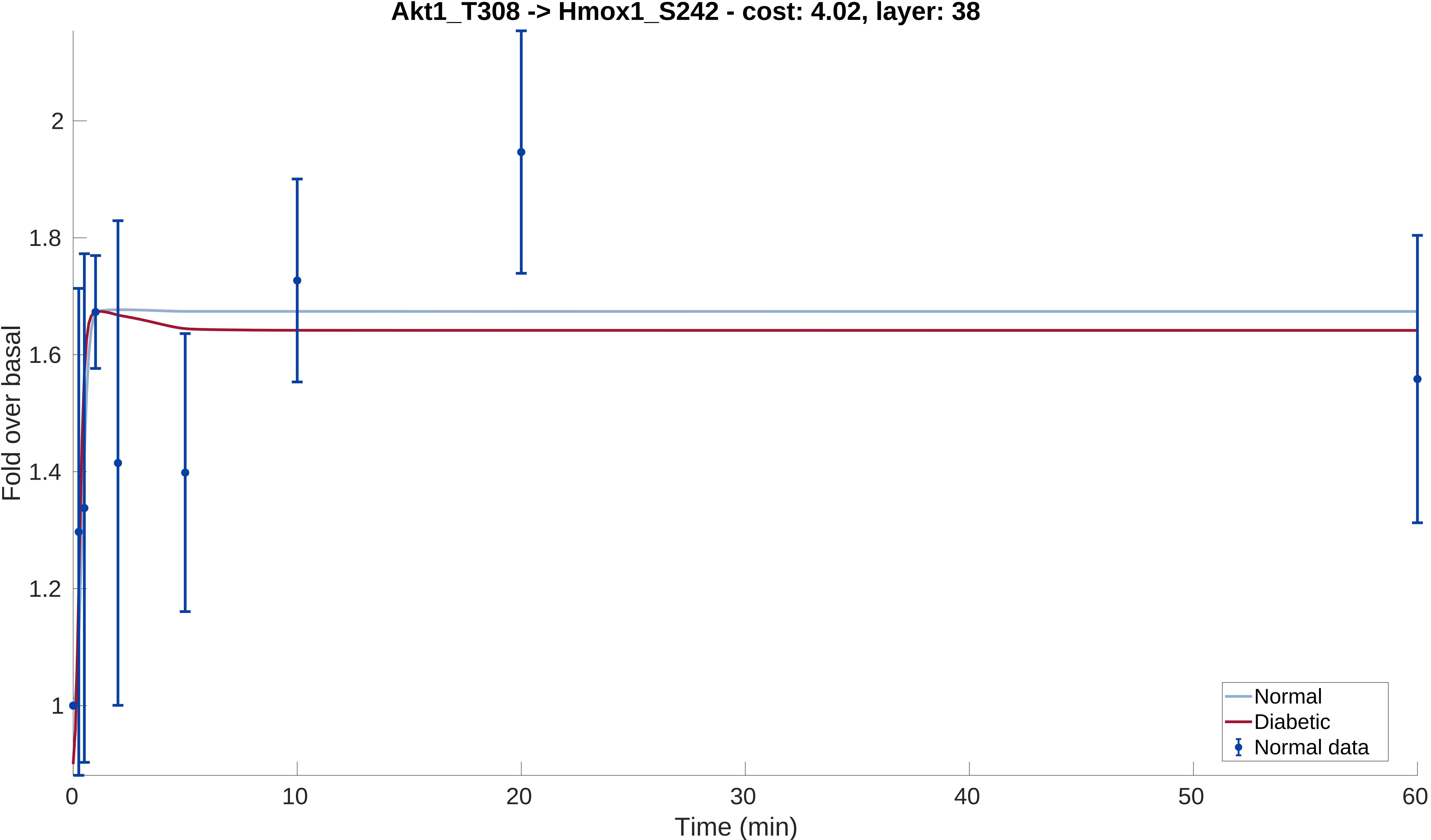

Akt1_T308 -> Hmox1_S242 - cost: 4.02, layer: 38
Fold over basal
2
1.8
1.6
1.4
1.2
1
0
10
20
30
40
50
60
Time (min)
Normal
Diabetic
Normal data

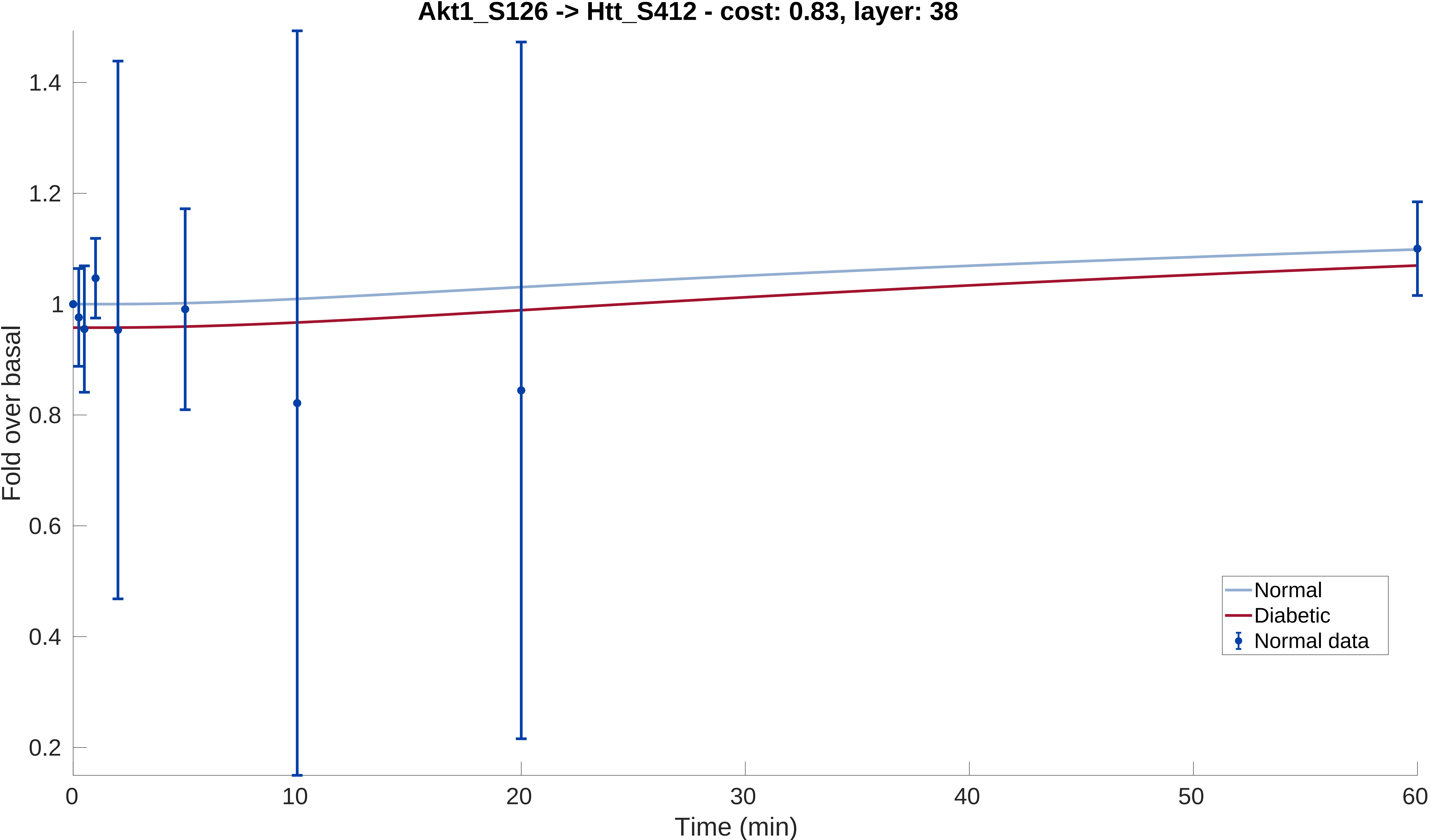

Akt1_S126 -> Htt_S412 - cost: 0.83, layer: 38
Fold over basal
1.4
1.2
1
0.8
0.6
0.4
0.2
0
10
20
30
40
50
60
Time (min)
Normal
Diabetic
Normal data

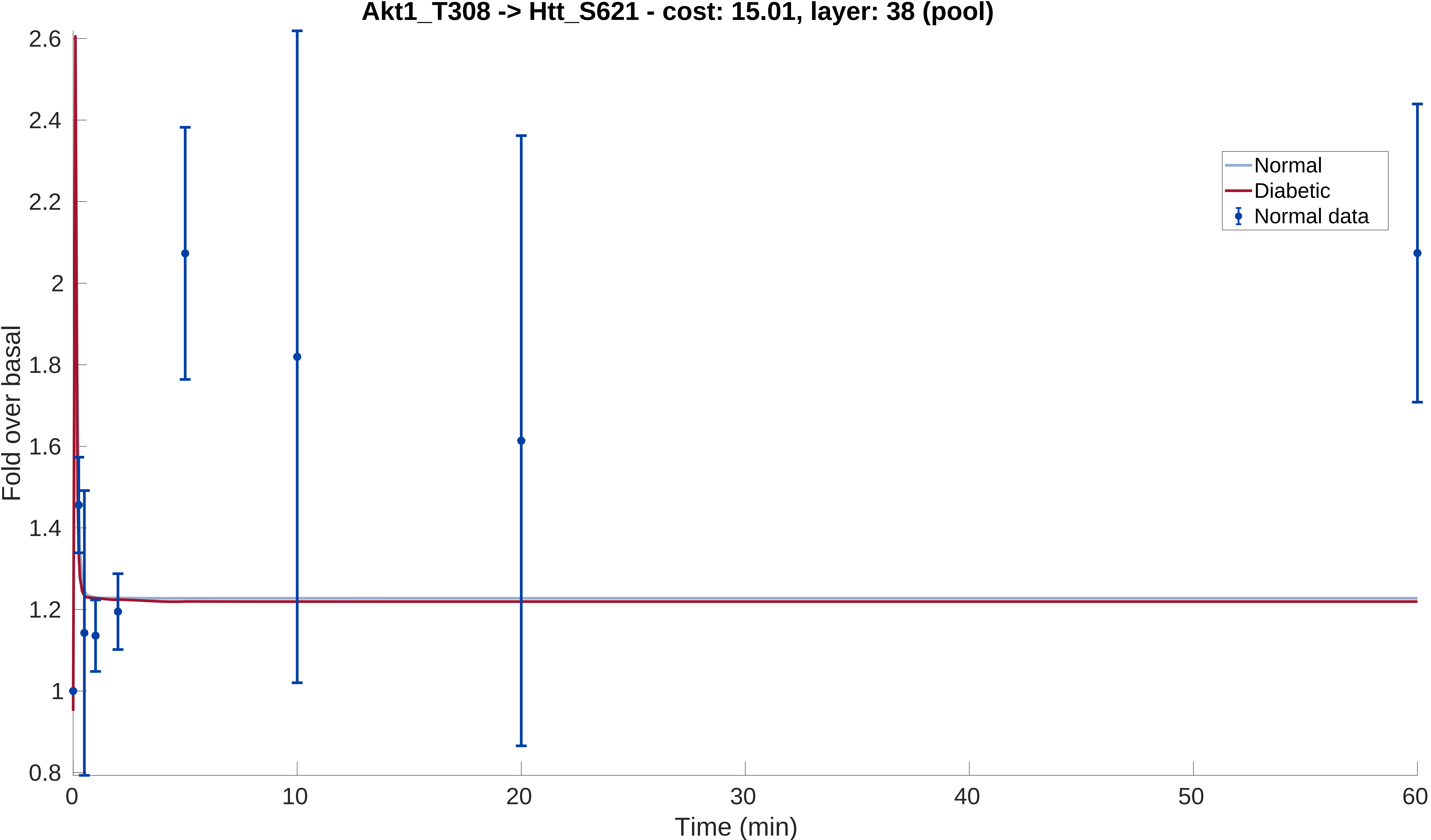
Akt1_T308 -> Htt_S621 - cost: 15.01, layer: 38 (pool)
Fold over basal
2.6
2.4
2.2
2
1.8
1.6
1.4
1.2
1
0.8
0
10
20
30
40
50
60
Time (min)
Normal
Diabetic
Normal data

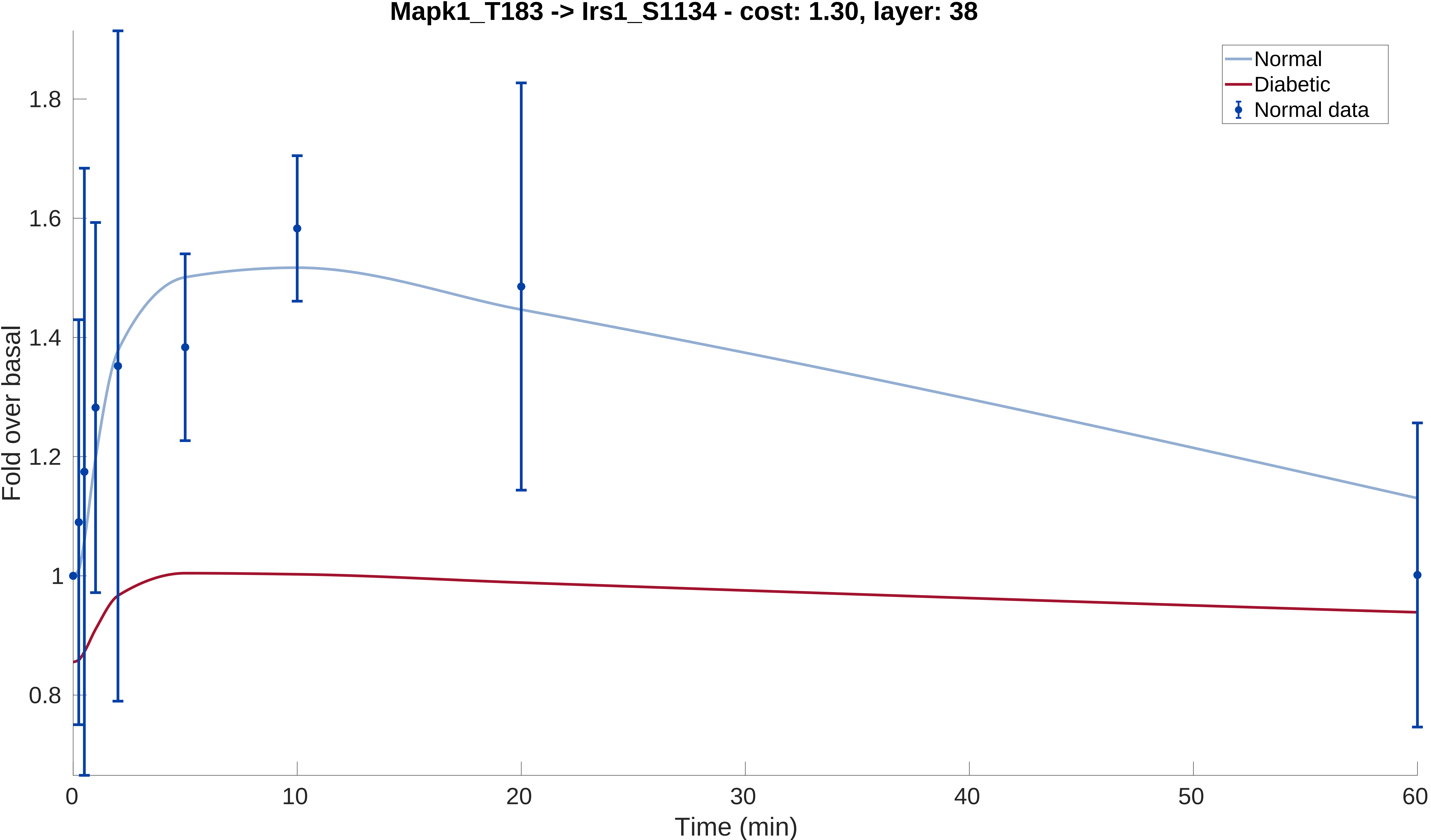

Mapk1_T183 -> Irs1_S1134 - cost: 1.30, layer: 38
Normal
Diabetic
Normal data
Fold over basal
1.8
1.6
1.4
1.2
1
0.8
0
10
20
30
40
50
60
Time (min)

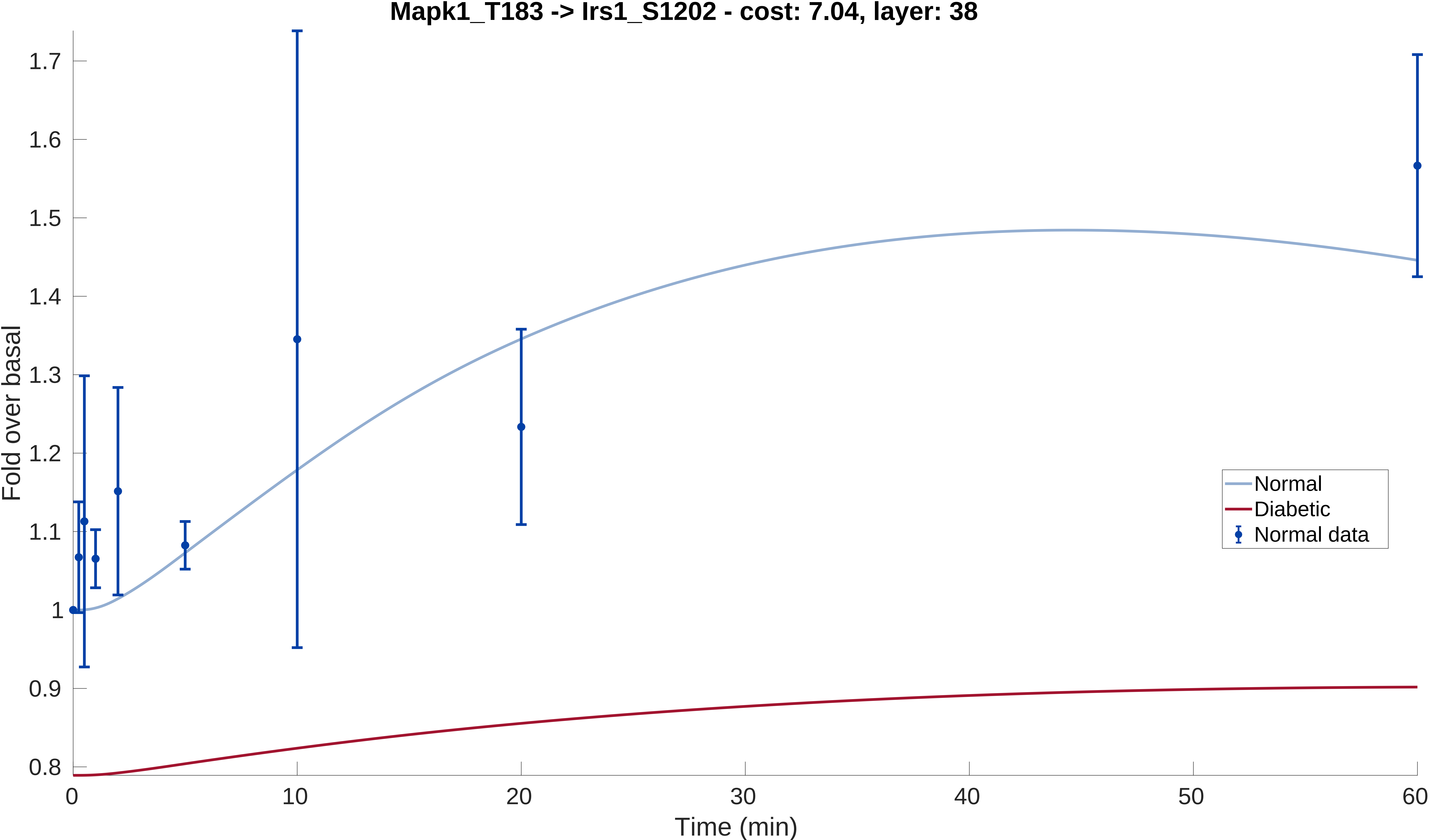
Mapk1_T183 -> Irs1_S1202 - cost: 7.04, layer: 38
Fold over basal
1.7
1.6
1.5
1.4
1.3
1.2
1.1
1
0.9
0.8
0
10
20
30
40
50
60
Time (min)
Normal
Diabetic
Normal data

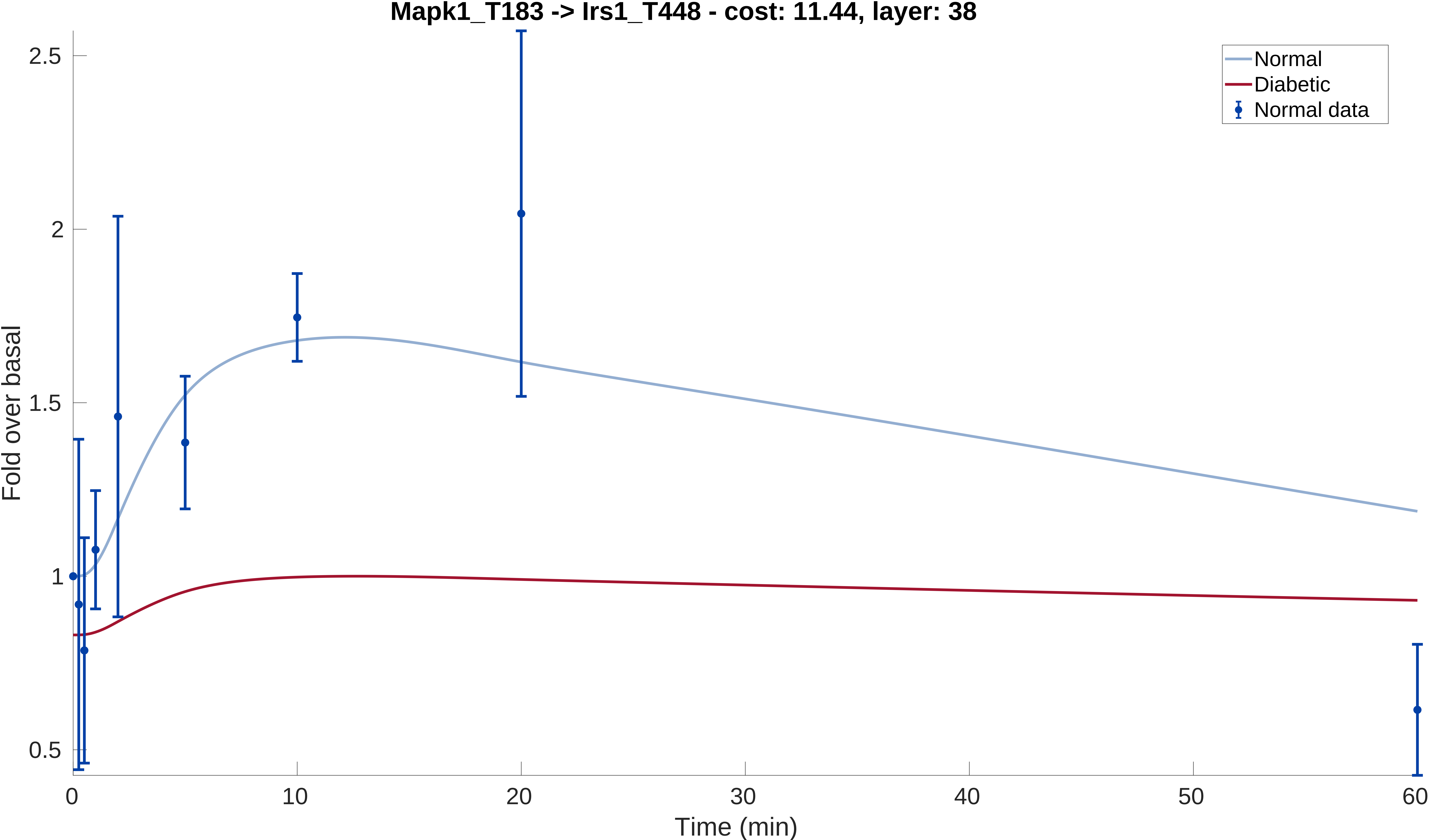

Mapk1_T183 -> Irs1_T448 - cost: 11.44, layer: 38
Normal
Diabetic
Normal data
Fold over basal
2.5
2
1.5
1
0.5
0
10
20
30
40
50
60
Time (min)

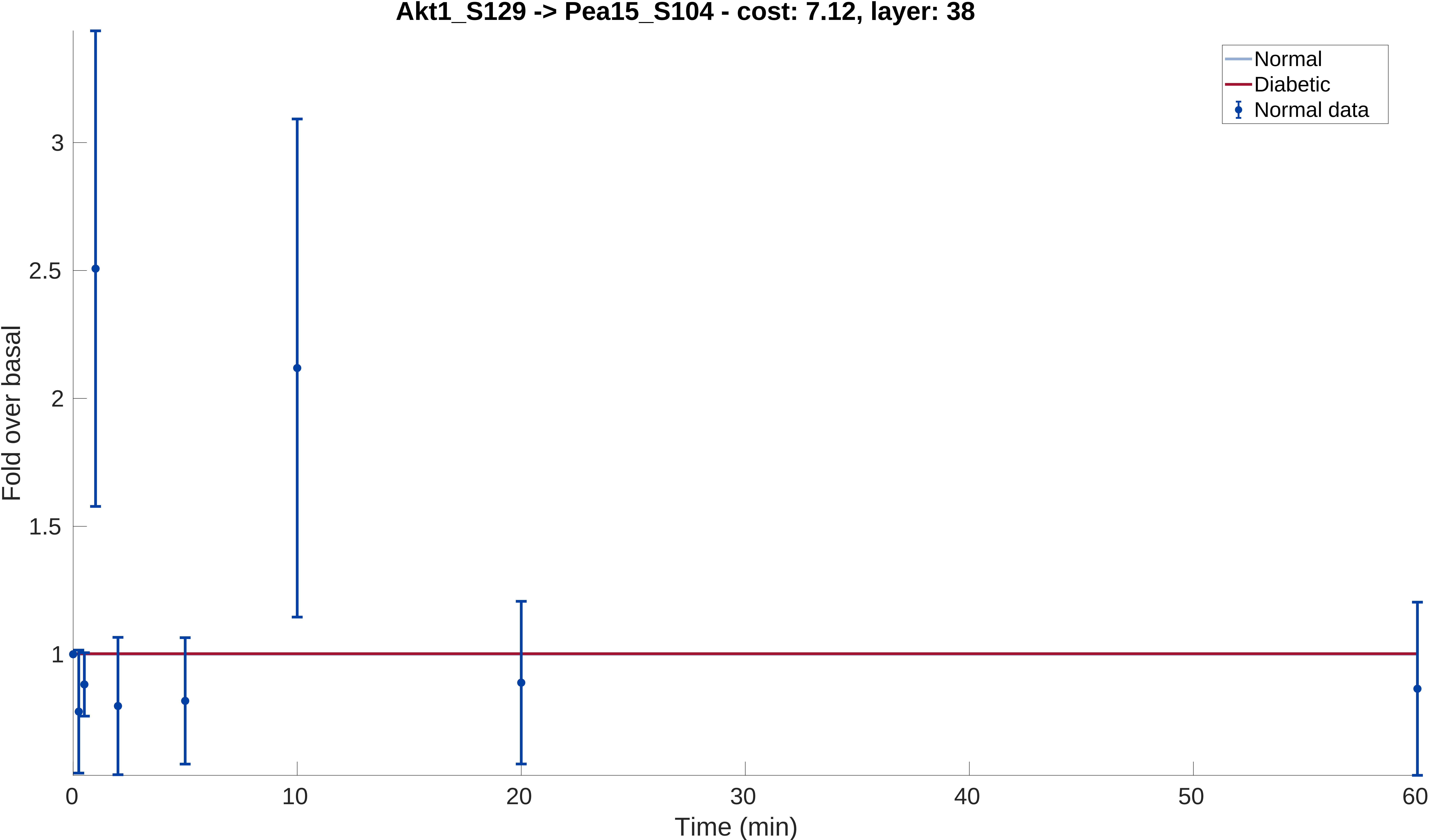
Akt1_S129 -> Pea15_S104 - cost: 7.12, layer: 38
Normal
Diabetic
Normal data
Fold over basal
3
2.5
2
1.5
1
0
10
20
30
40
50
60
Time (min)

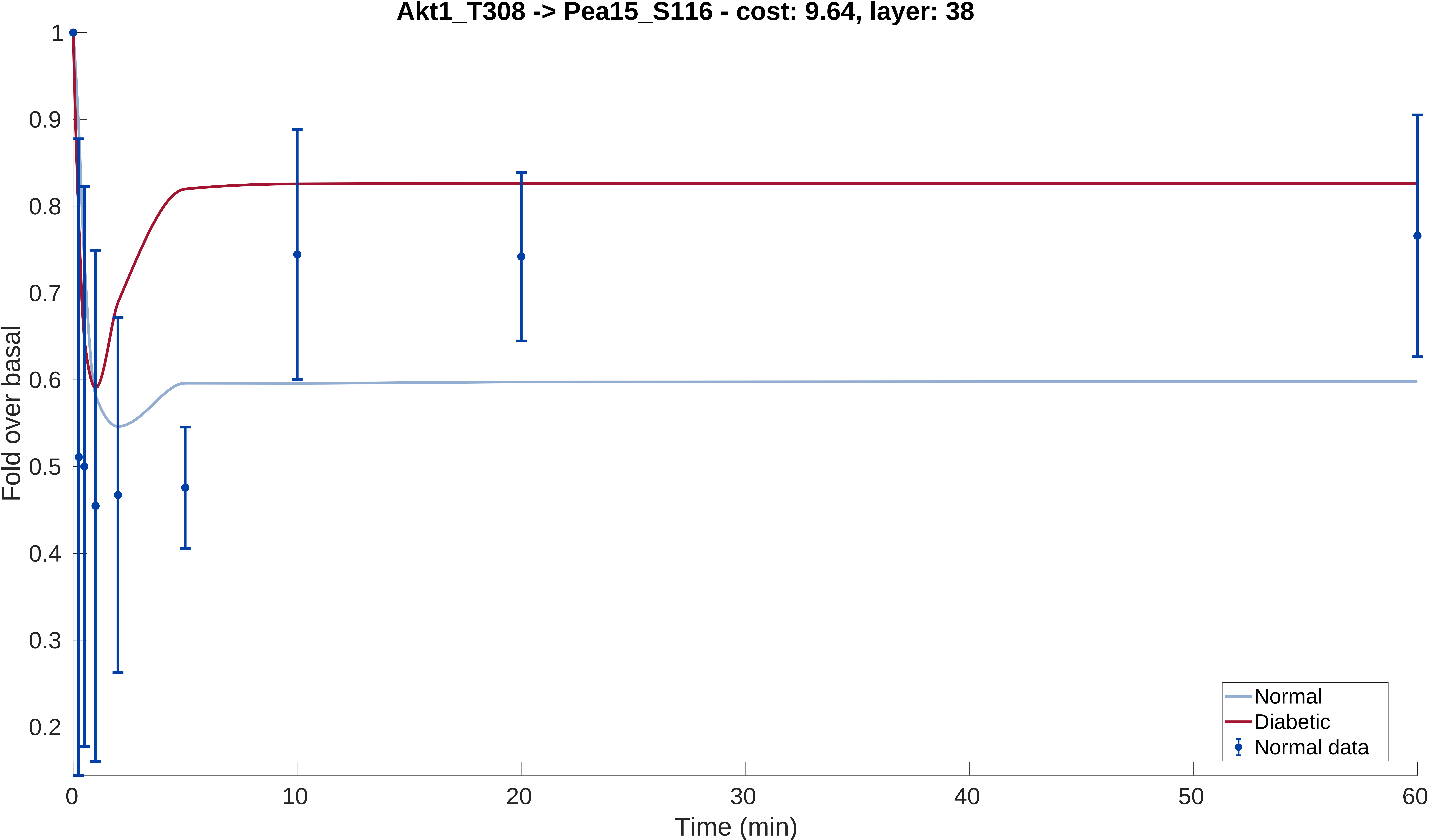

Akt1_T308 -> Pea15_S116 - cost: 9.64, layer: 38
Fold over basal
Time (min)
1
0.9
0.8
0.7
0.6
0.5
0.4
0.3
0.2
0
10
20
30
40
50
60
Normal
Diabetic
Normal data

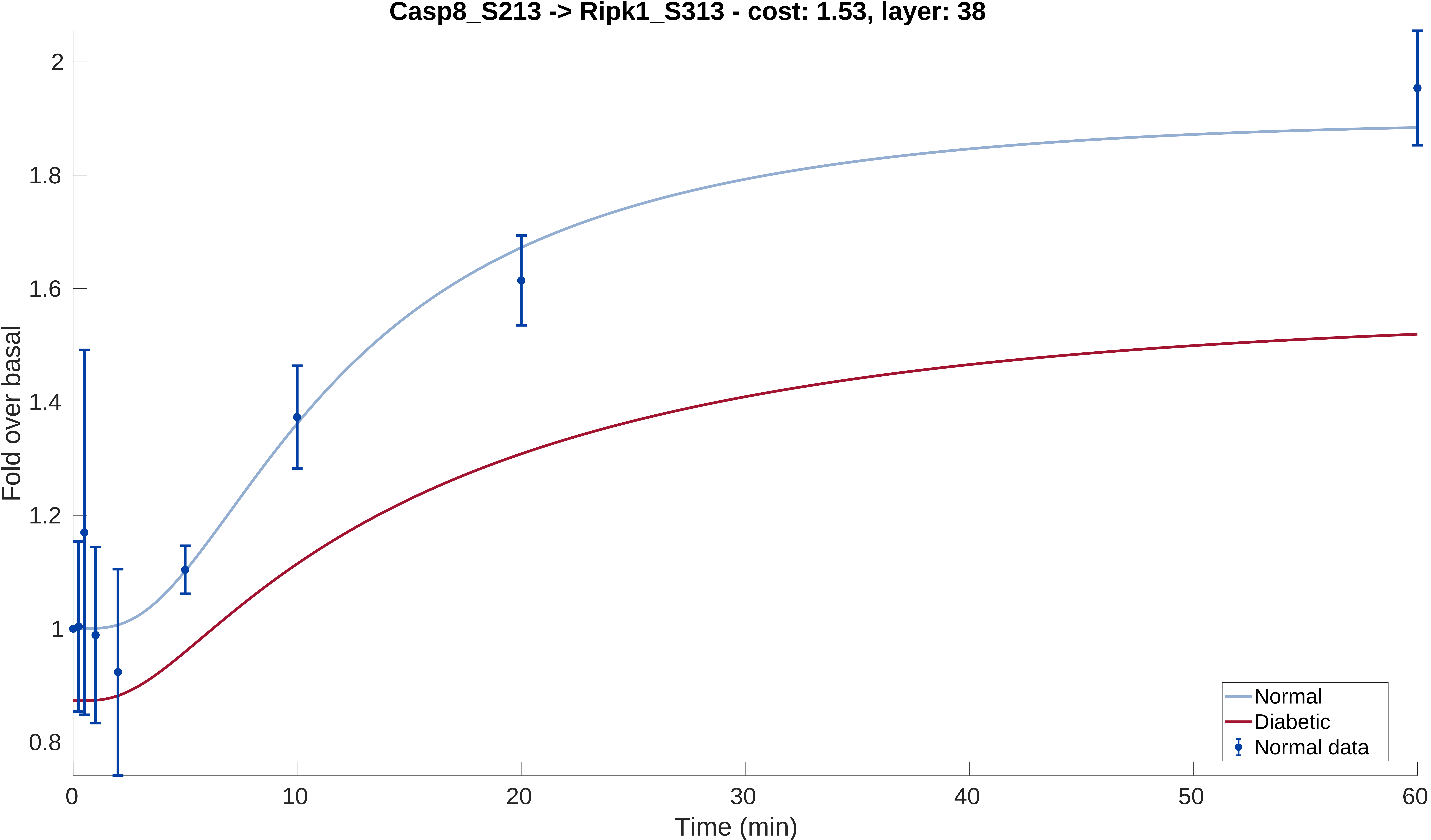

Casp8_S213 -> Ripk1_S313 - cost: 1.53, layer: 38
Fold over basal
Time (min)
0.8
1
1.2
1.4
1.6
1.8
2
0
10
20
30
40
50
60
Normal
Diabetic
Normal data

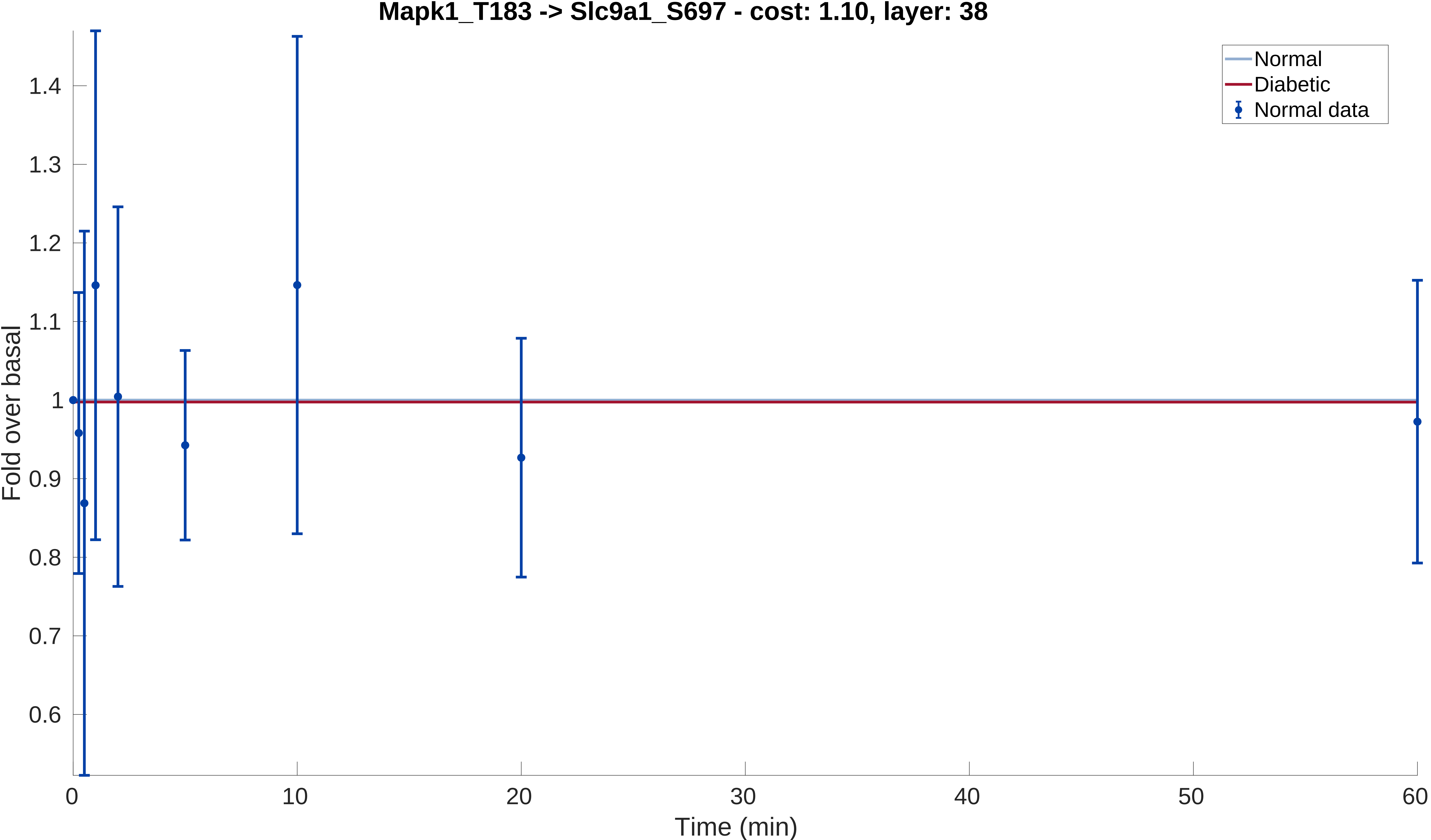

Mapk1_T183 -> Slc9a1_S697 - cost: 1.10, layer: 38
Normal
Diabetic
Normal data
Fold over basal
Time (min)
1.4
1.3
1.2
1.1
1
0.9
0.8
0.7
0.6
0
10
20
30
40
50
60

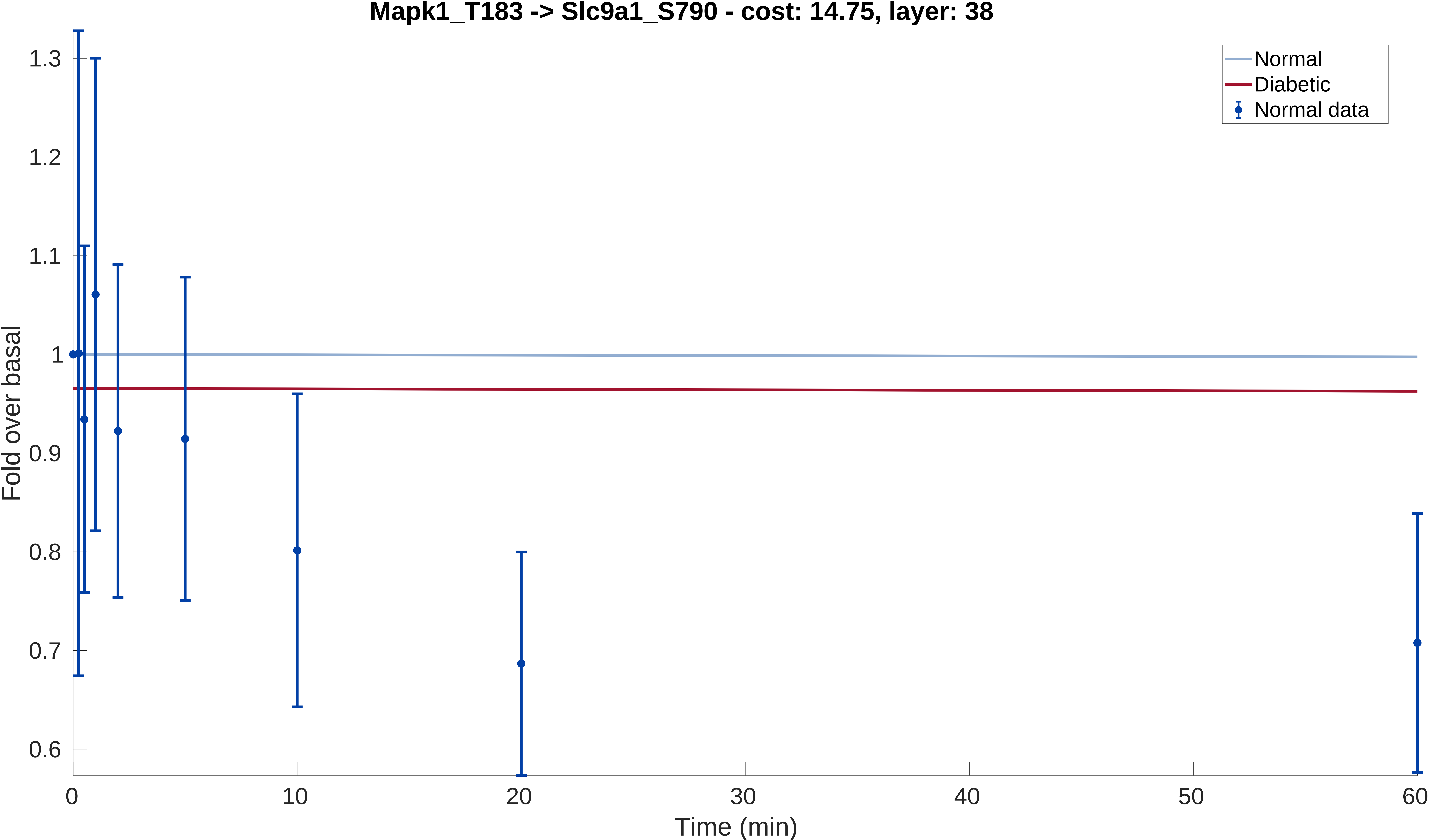

Mapk1_T183 -> Slc9a1_S790 - cost: 14.75, layer: 38
Normal
Diabetic
Normal data
Fold over basal
Time (min)
1.3
1.2
1.1
1
0.9
0.8
0.7
0.6
0
10
20
30
40
50
60

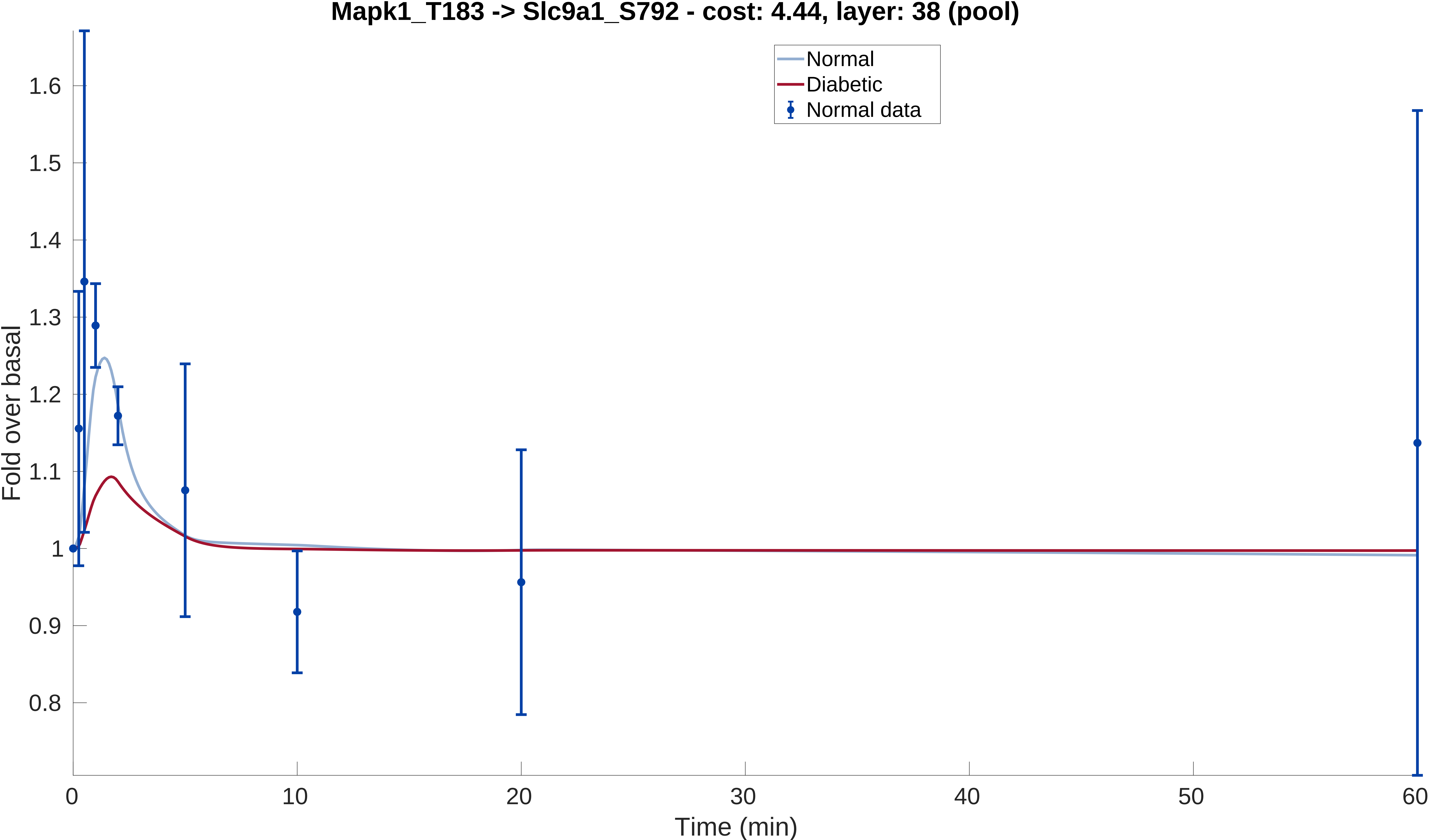

Mapk1_T183 -> Slc9a1_S792 - cost: 4.44, layer: 38 (pool)
Normal
Diabetic
Normal data
Fold over basal
1.6
1.5
1.4
1.3
1.2
1.1
1
0.9
0.8
0
10
20
30
40
50
60
Time (min)

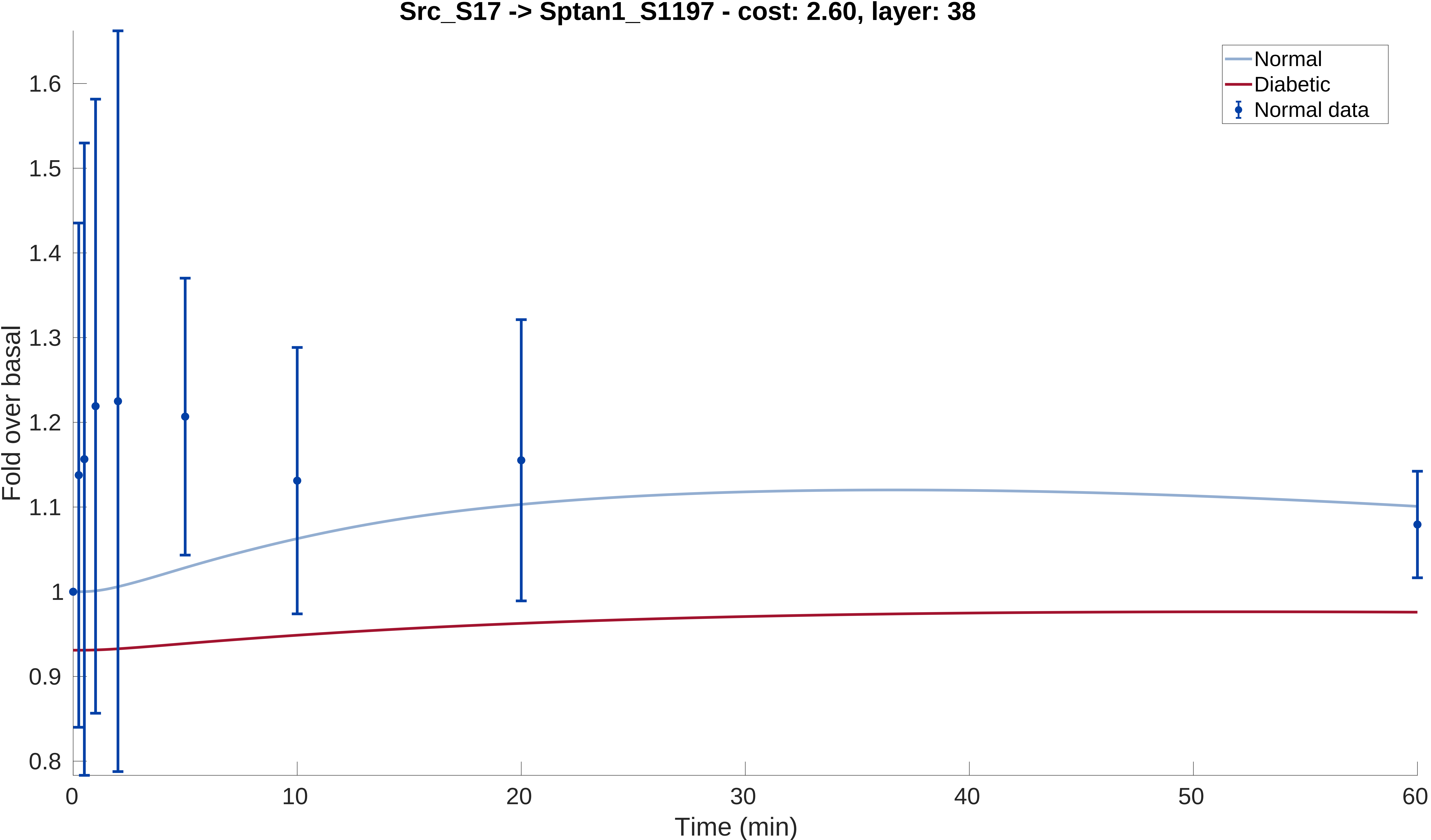
Src_S17 -> Sptan1_S1197 - cost: 2.60, layer: 38
Normal
Diabetic
Normal data
Fold over basal
1.6
1.5
1.4
1.3
1.2
1.1
1
0.9
0.8
0
10
20
30
40
50
60
Time (min)

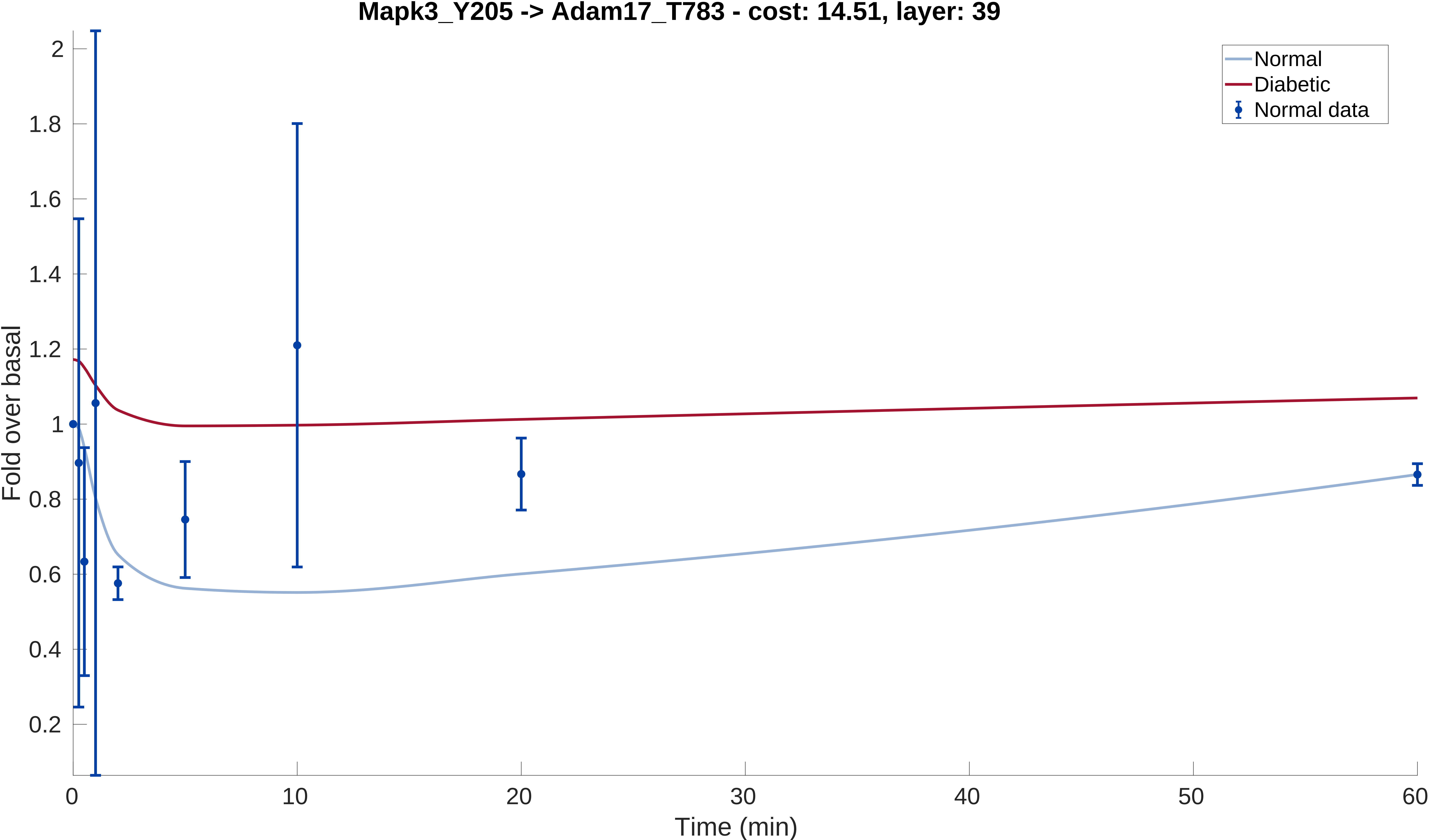

Mapk3_Y205 -> Adam17_T783 - cost: 14.51, layer: 39
Normal
Diabetic
Normal data
Fold over basal
Time (min)
0
10
20
30
40
50
60
0.2
0.4
0.6
0.8
1
1.2
1.4
1.6
1.8
2

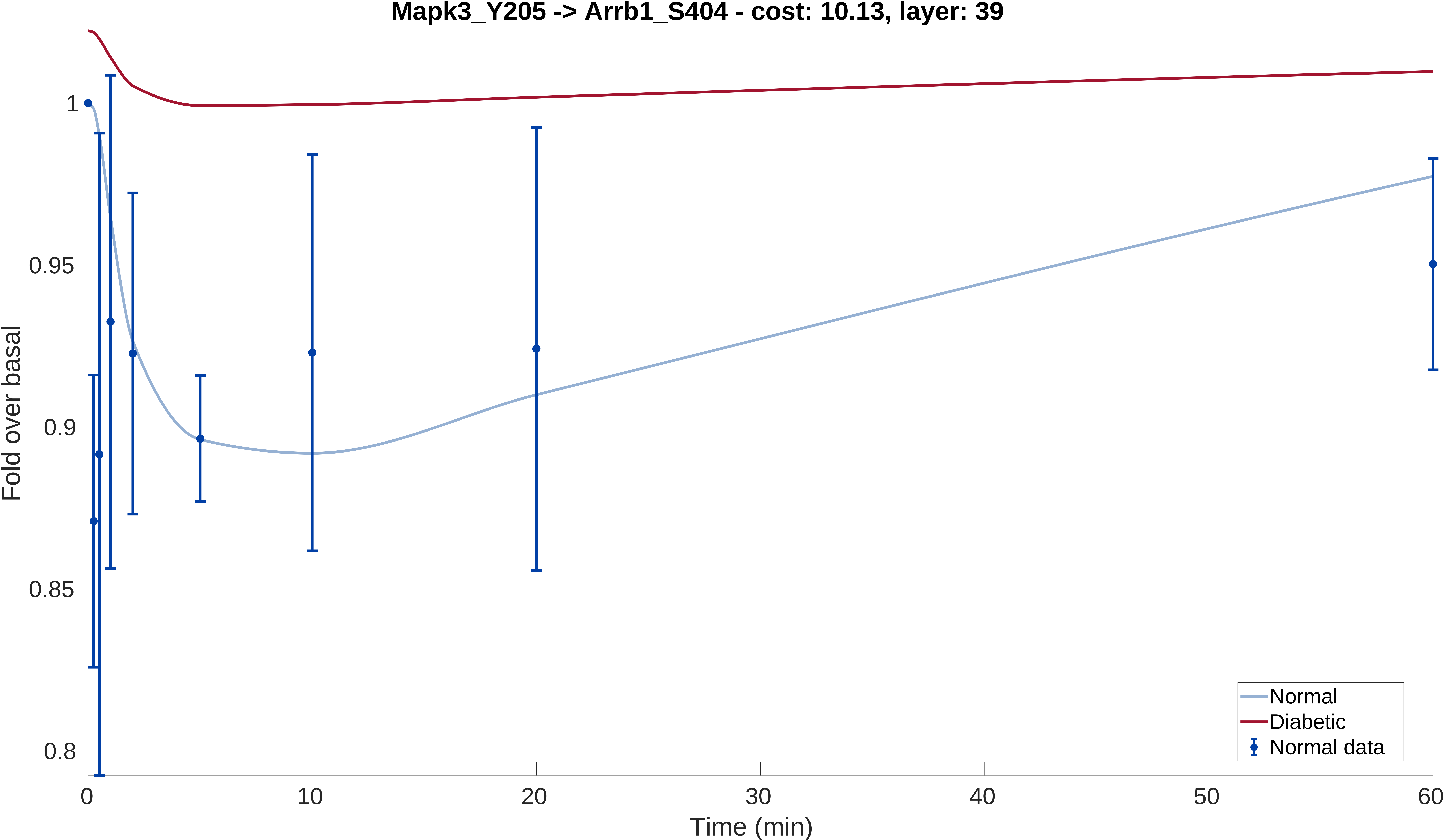
Mapk3_Y205 -> Arrb1_S404 - cost: 10.13, layer: 39
Fold over basal
1
0.95
0.9
0.85
0.8
0
10
20
30
40
50
60
Time (min)
Normal
Diabetic
Normal data

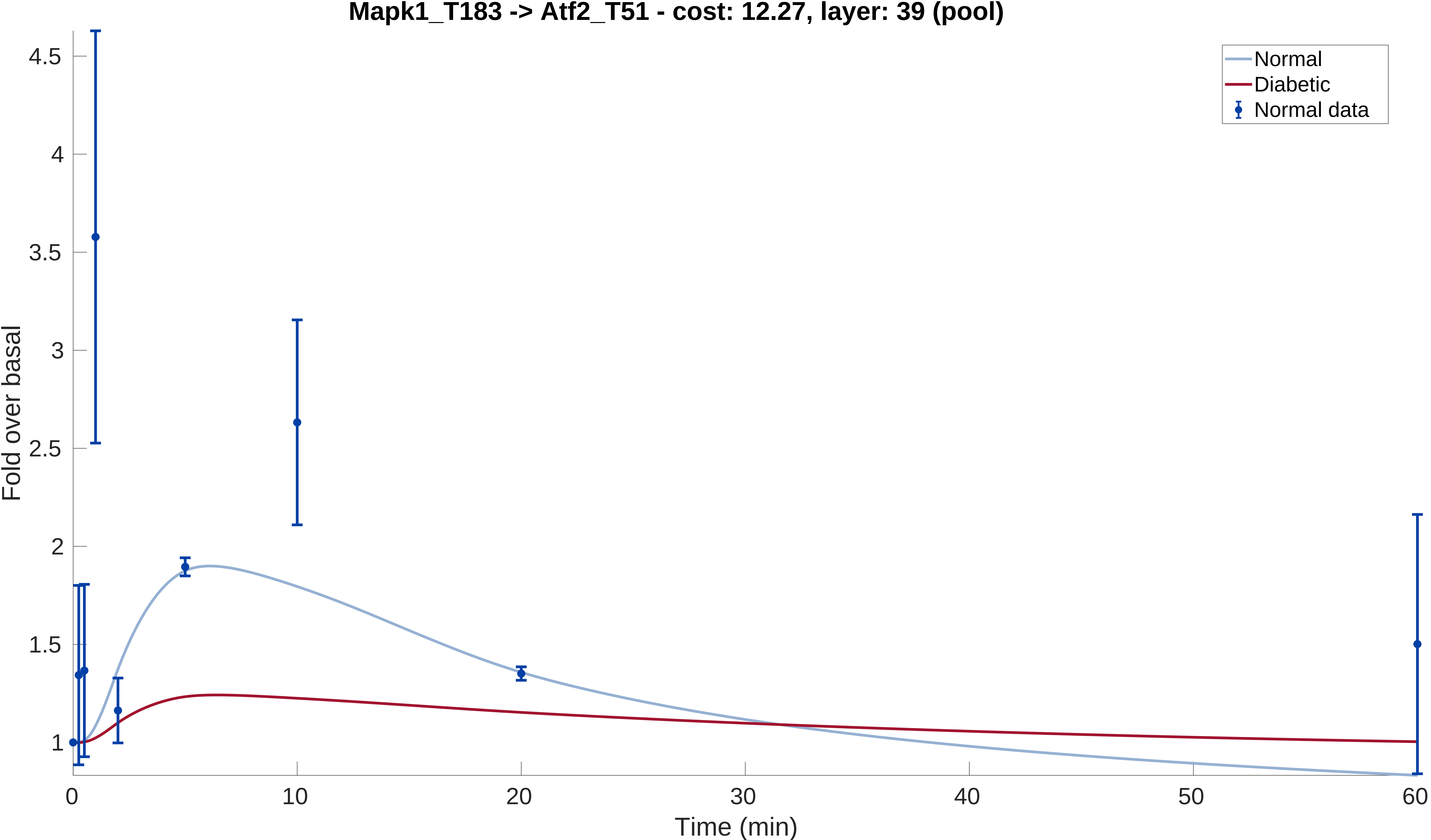

Mapk1_T183 -> Atf2_T51 - cost: 12.27, layer: 39 (pool)
Normal
Diabetic
Normal data
Fold over basal
4.5
4
3.5
3
2.5
2
1.5
1
0
10
20
30
40
50
60
Time (min)

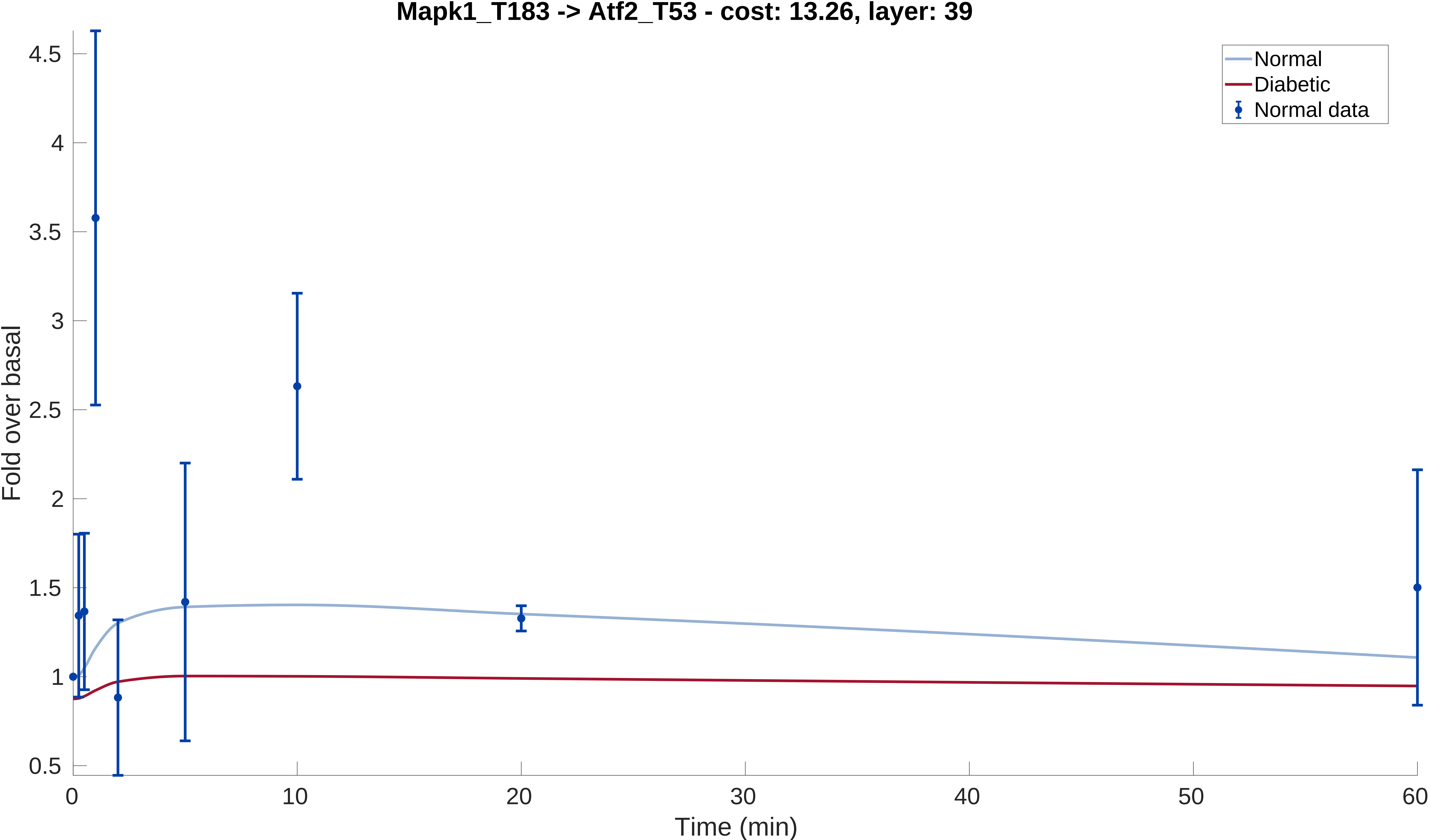
Mapk1_T183 -> Atf2_T53 - cost: 13.26, layer: 39
Normal
Diabetic
Normal data
Fold over basal
4.5
4
3.5
3
2.5
2
1.5
1
0.5
0
10
20
30
40
50
60
Time (min)

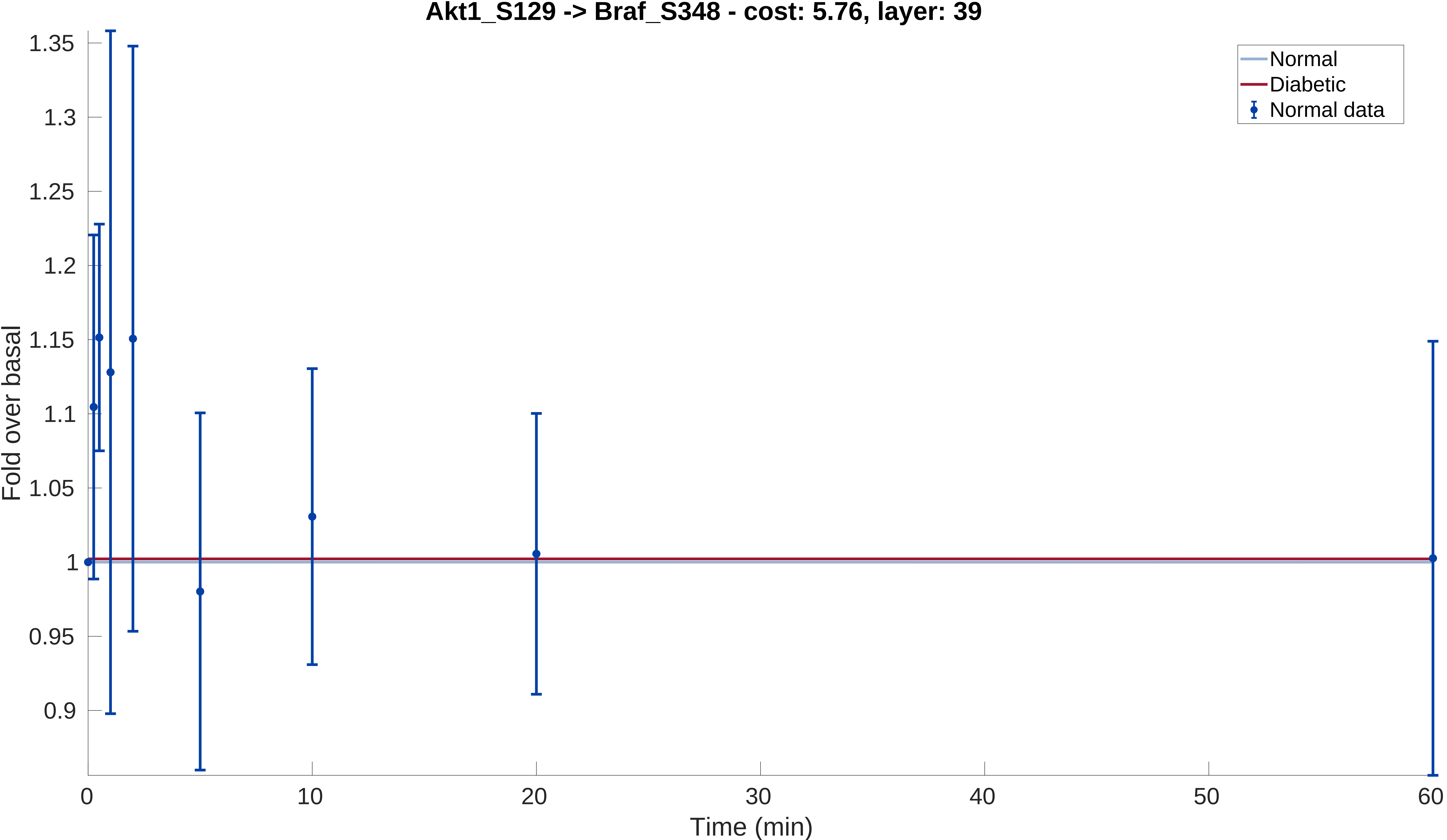

Akt1_S129 -> Braf_S348 - cost: 5.76, layer: 39
Normal
Diabetic
Normal data
Fold over basal
Time (min)
1.35
1.3
1.25
1.2
1.15
1.1
1.05
1
0.95
0.9
0
10
20
30
40
50
60

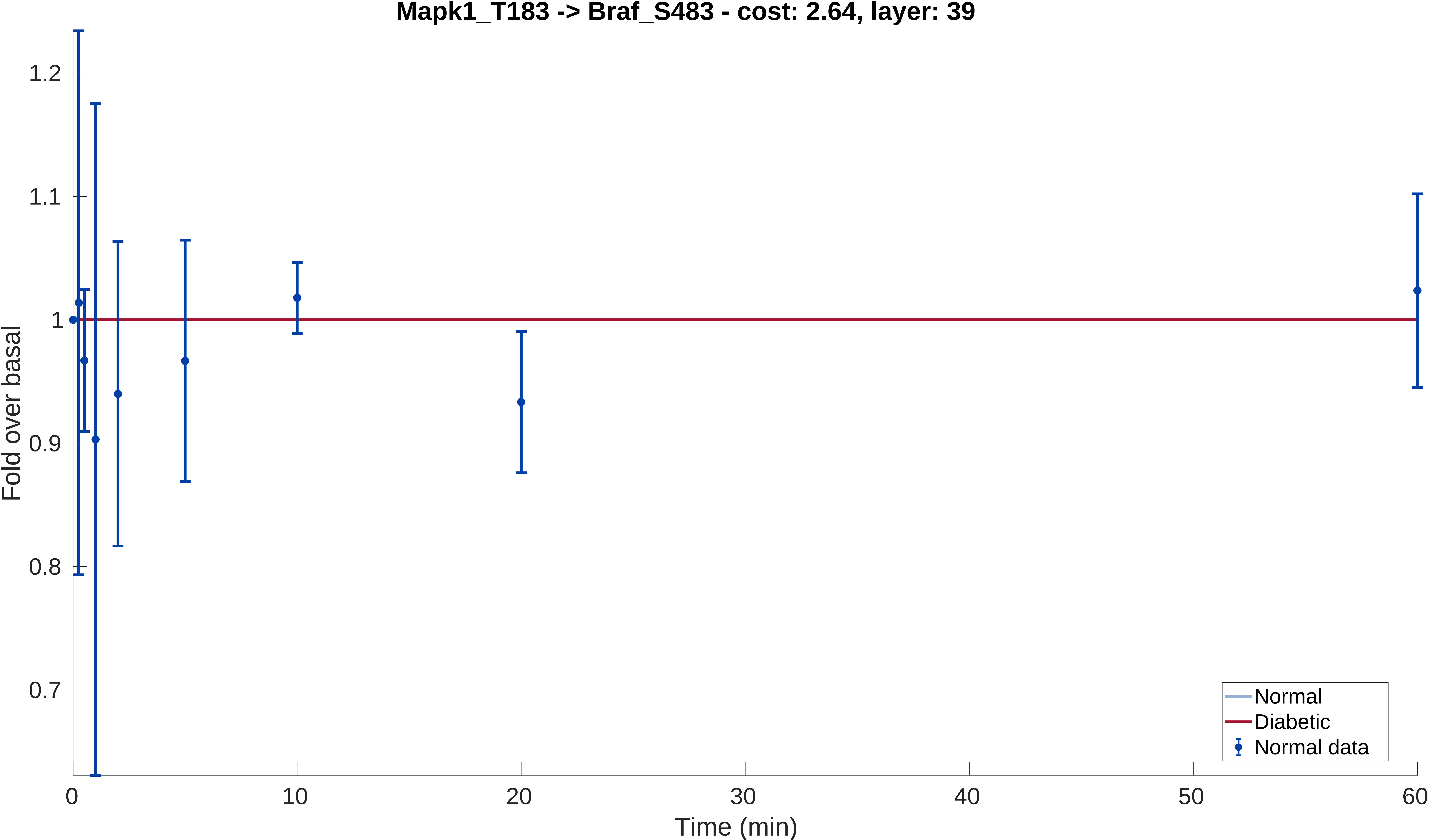

Mapk1_T183 -> Braf_S483 - cost: 2.64, layer: 39
Fold over basal
1.2
1.1
1
0.9
0.8
0.7
Time (min)
0
10
20
30
40
50
60
Normal
Diabetic
Normal data

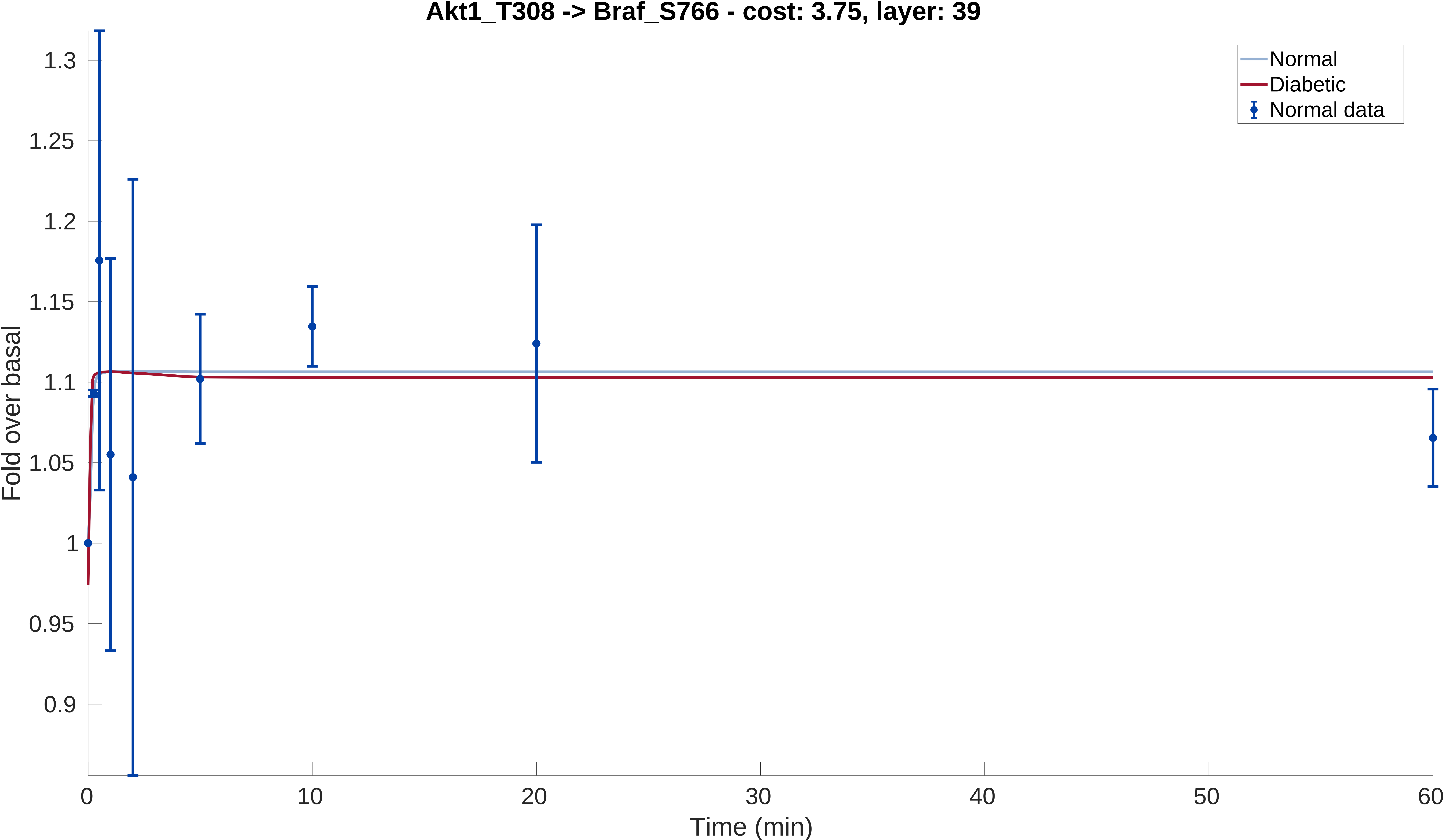

Akt1_T308 -> Braf_S766 - cost: 3.75, layer: 39
Normal
Diabetic
Normal data
Fold over basal
Time (min)
1.3
1.25
1.2
1.15
1.1
1.05
1
0.95
0.9
0
10
20
30
40
50
60

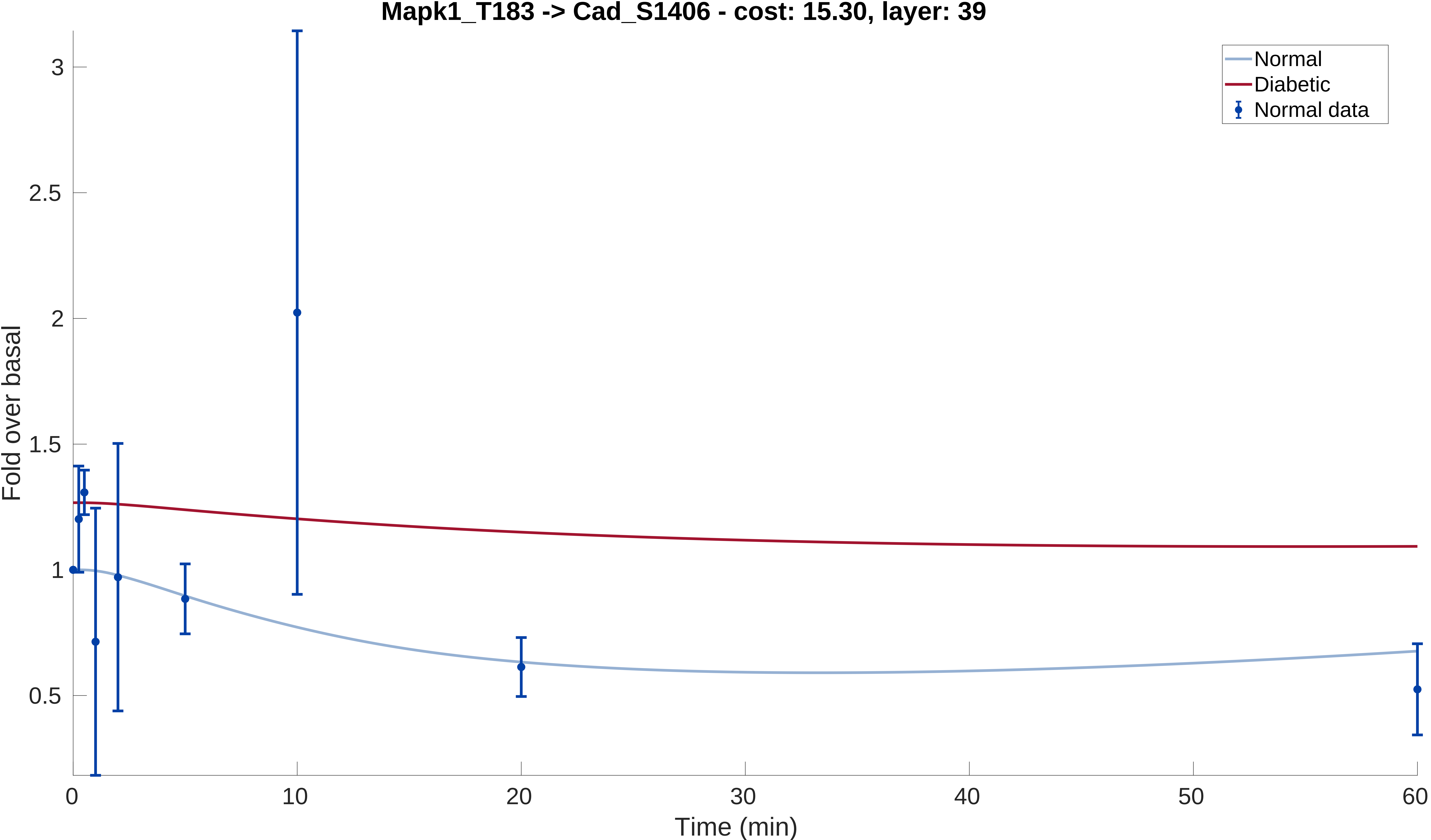

Mapk1_T183 -> Cad_S1406 - cost: 15.30, layer: 39
Normal
Diabetic
Normal data
Fold over basal
0.5
1
1.5
2
2.5
3
0
10
20
30
40
50
60
Time (min)

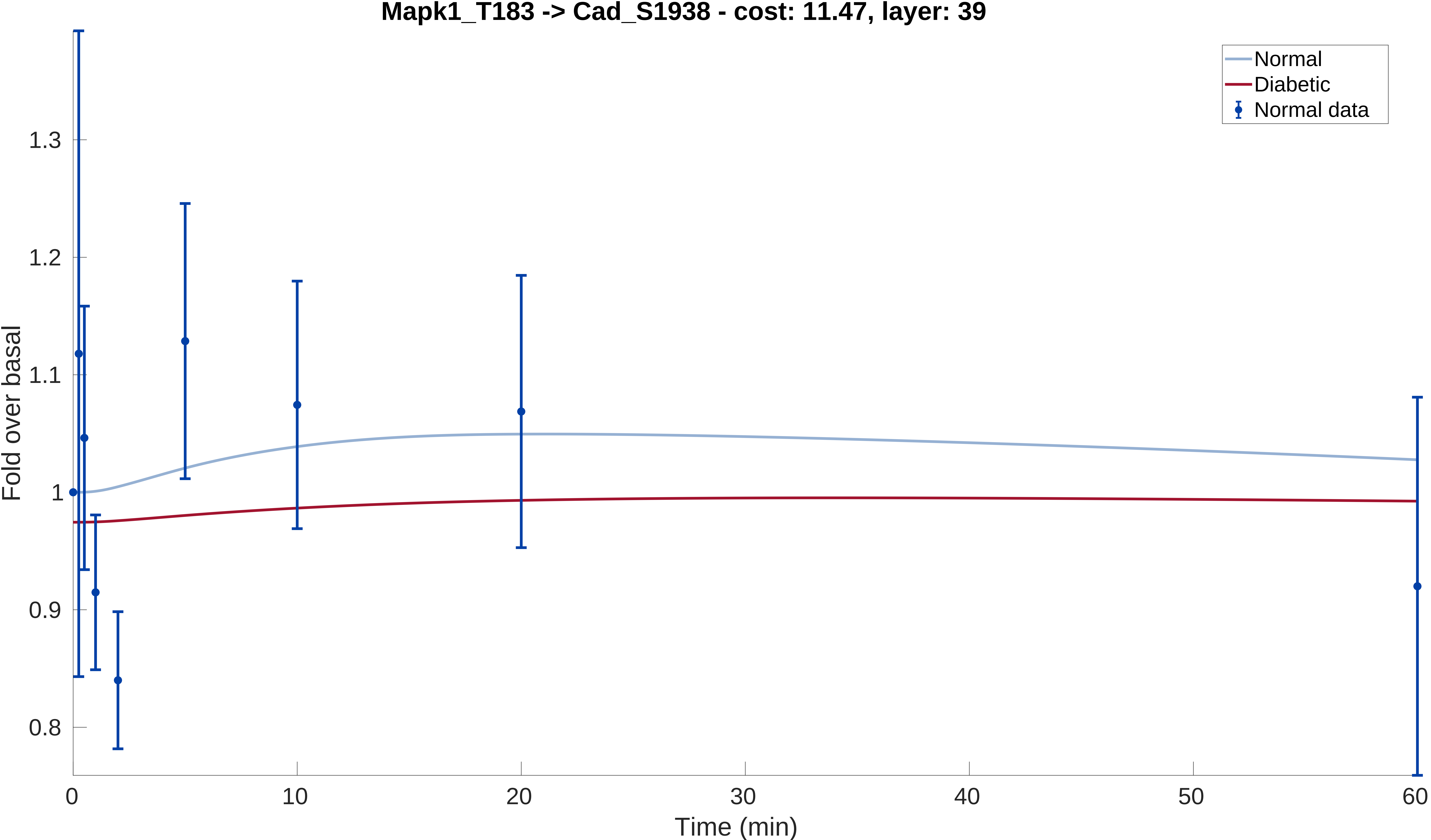

Mapk1_T183 -> Cad_S1938 - cost: 11.47, layer: 39
Normal
Diabetic
Normal data
1.3
1.2
1.1
1
0.9
0.8
Fold over basal
0
10
20
30
40
50
60
Time (min)

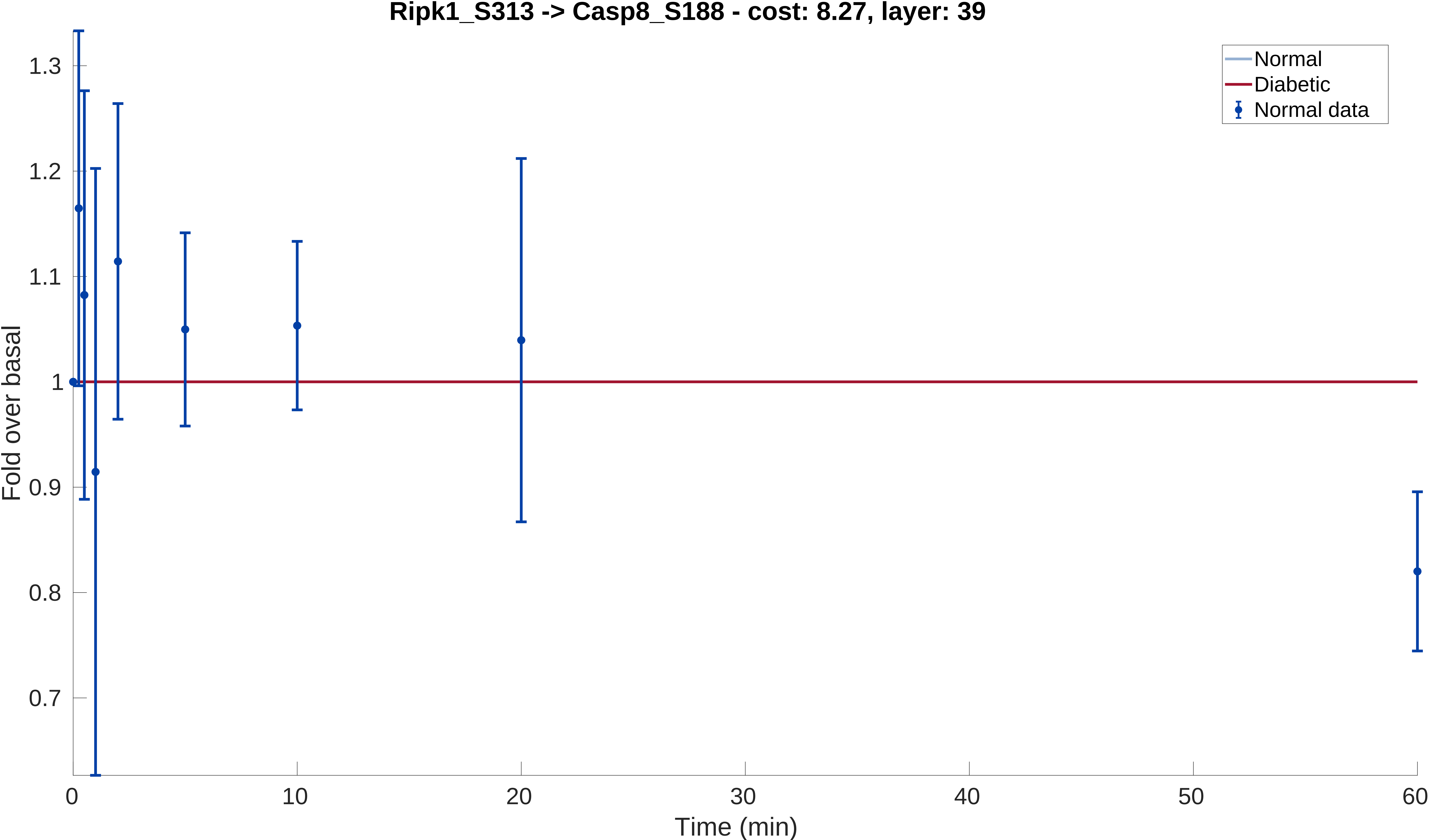

Ripk1_S313 -> Casp8_S188 - cost: 8.27, layer: 39
Normal
Diabetic
Normal data
Fold over basal
Time (min)
1.3
1.2
1.1
1
0.9
0.8
0.7
0
10
20
30
40
50
60

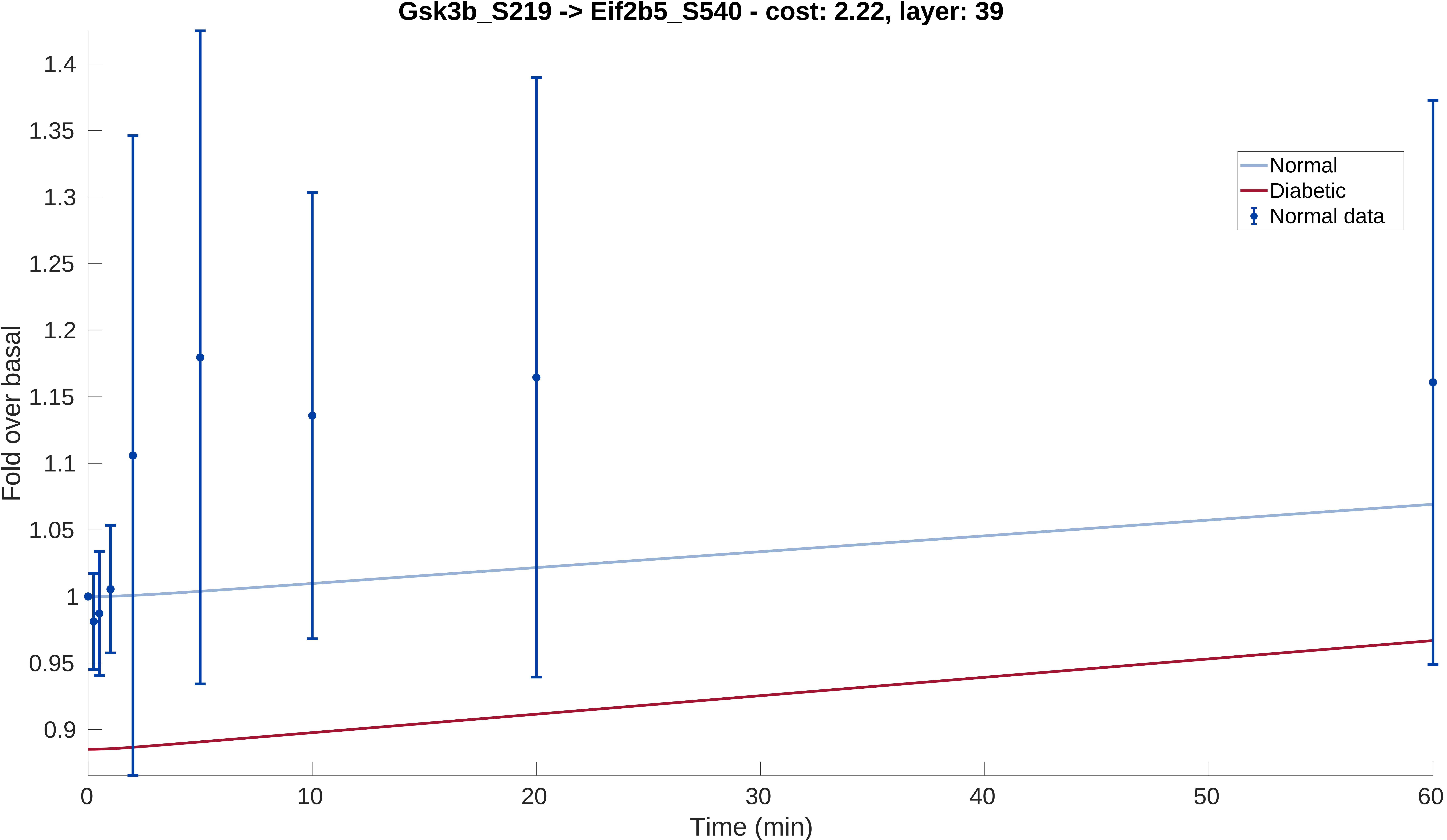
Gsk3b_S219 -> Eif2b5_S540 - cost: 2.22, layer: 39
Fold over basal
1.4
1.35
1.3
1.25
1.2
1.15
1.1
1.05
1
0.95
0.9
0
10
20
30
40
50
60
Time (min)
Normal
Diabetic
Normal data

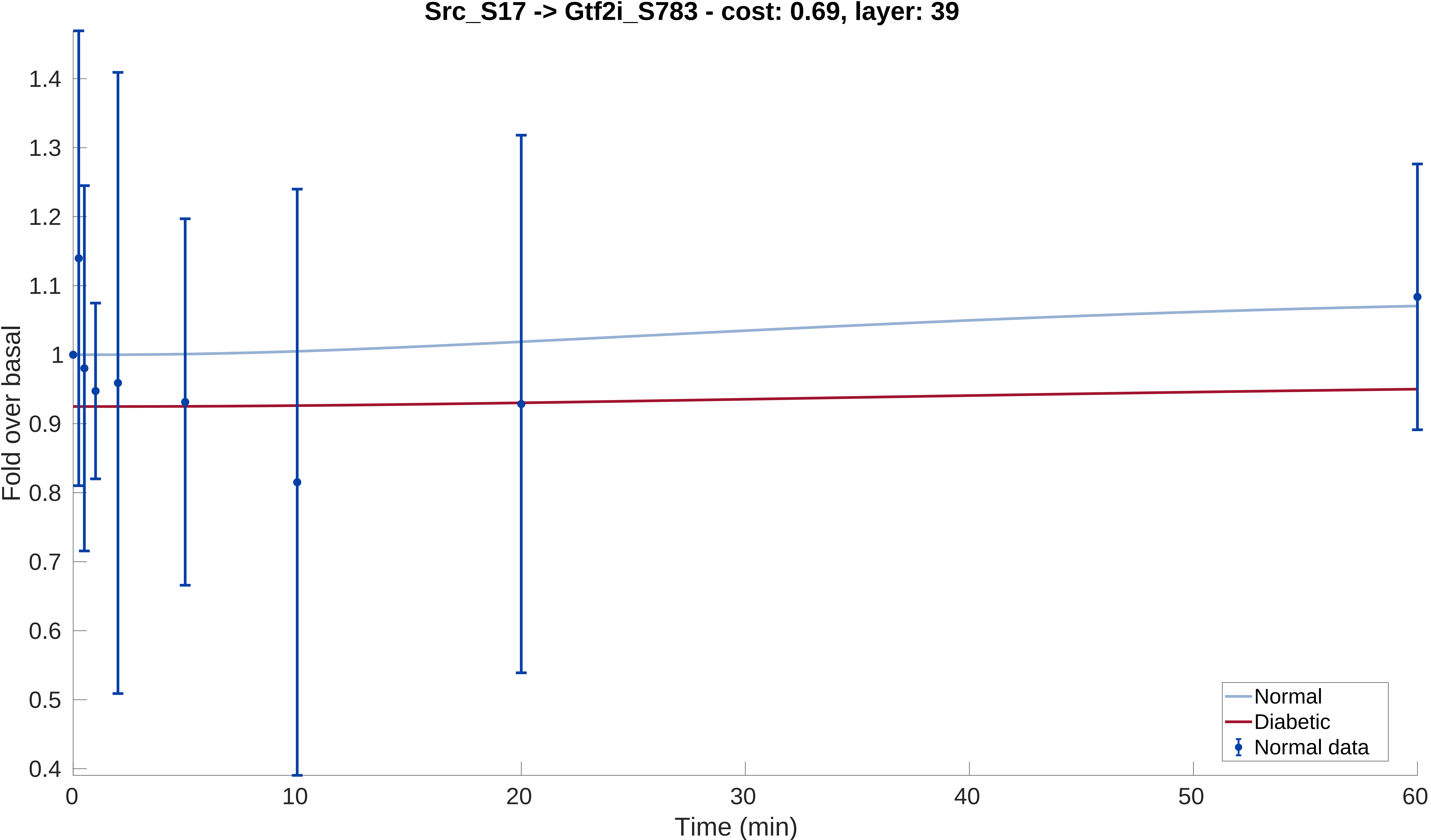

Src_S17 -> Gtf2i_S783 - cost: 0.69, layer: 39
Fold over basal
Time (min)
0
10
20
30
40
50
60
0.4
0.5
0.6
0.7
0.8
0.9
1
1.1
1.2
1.3
1.4
Normal
Diabetic
Normal data

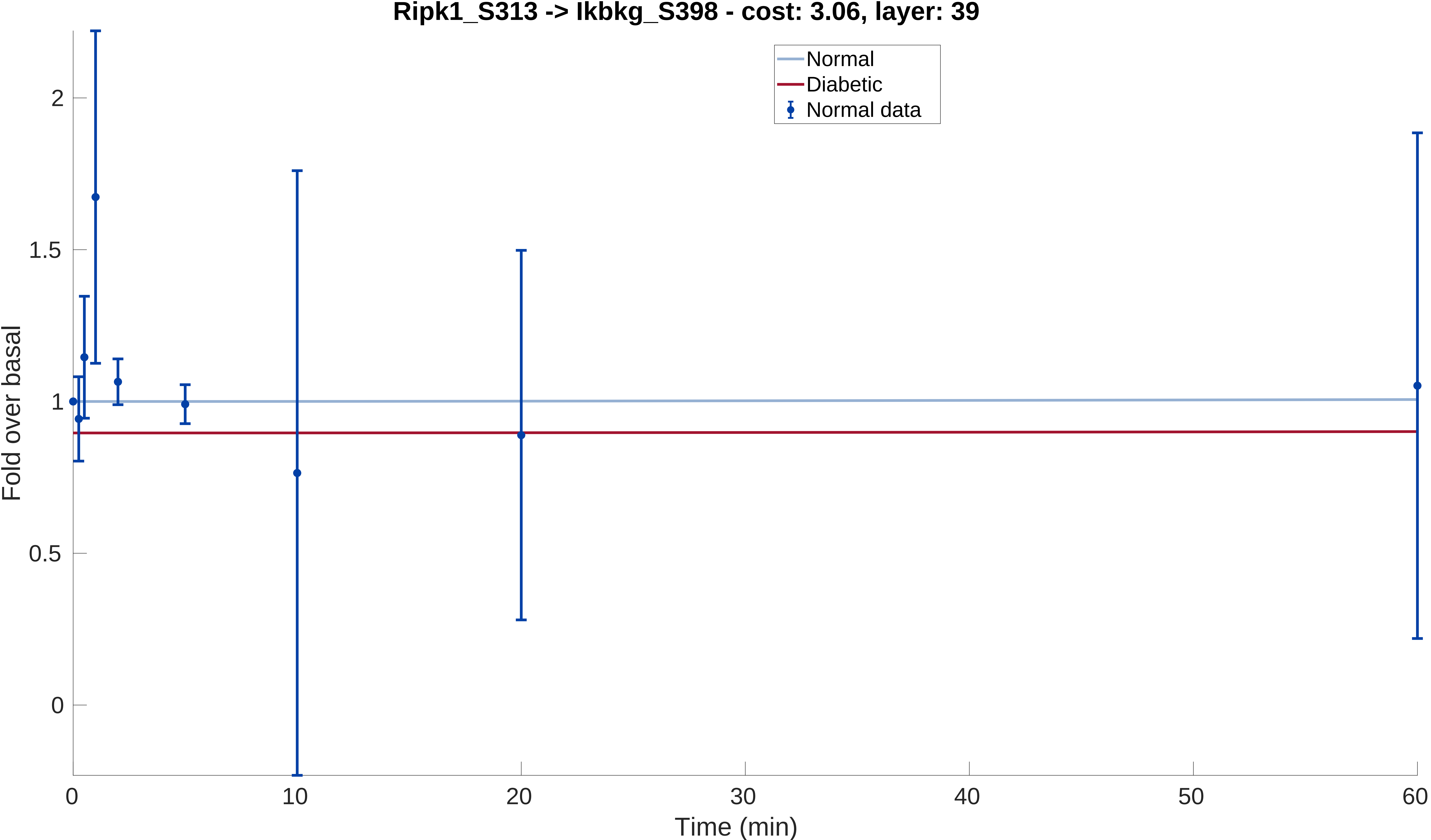

Ripk1_S313 -> Ikbkg_S398 - cost: 3.06, layer: 39
Normal
Diabetic
Normal data
Fold over basal
Time (min)
2
1.5
1
0.5
0
0
10
20
30
40
50
60

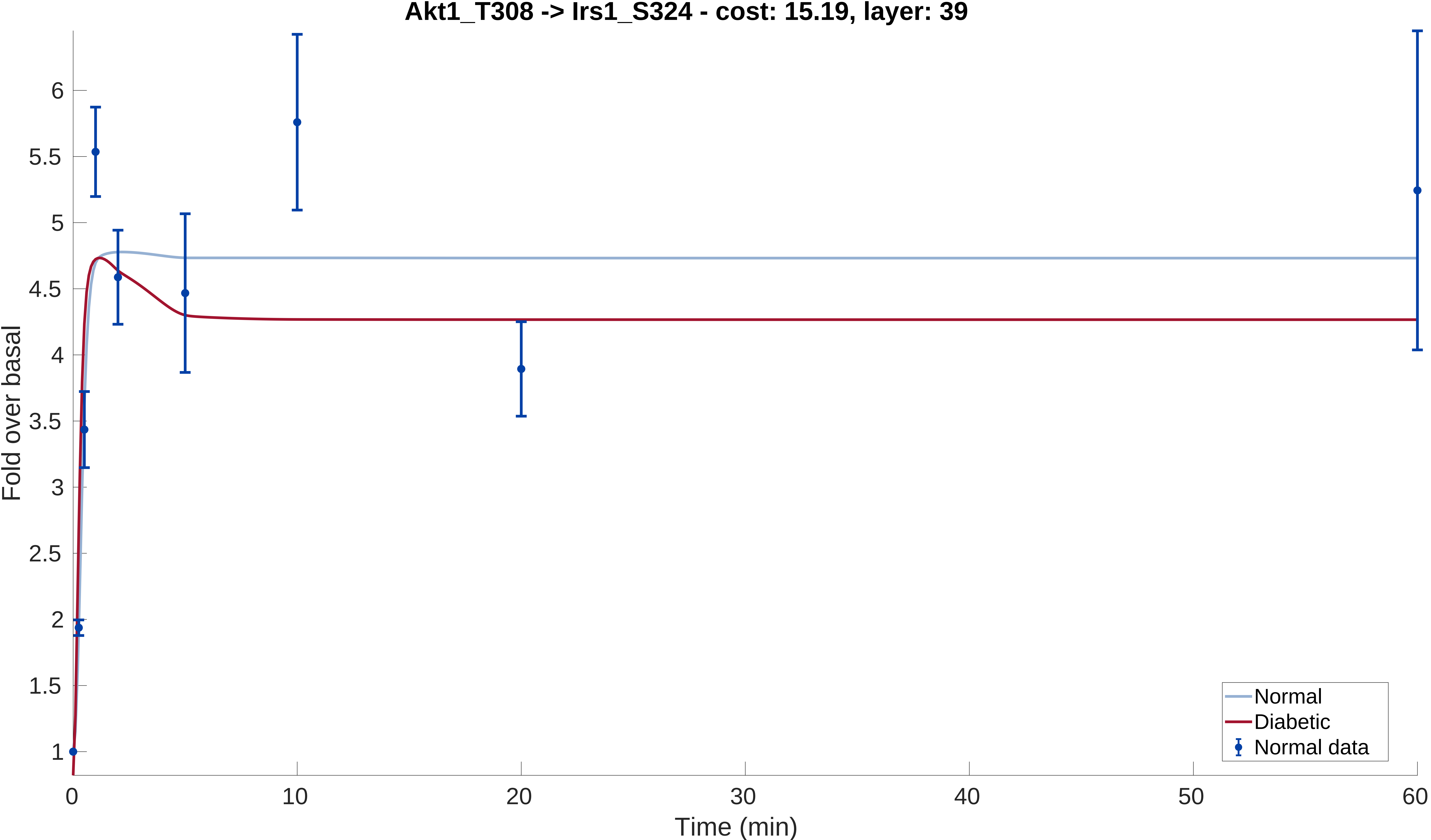
Akt1_T308 -> Irs1_S324 - cost: 15.19, layer: 39
Fold over basal
6
5.5
5
4.5
4
3.5
3
2.5
2
1.5
1
0
10
20
30
40
50
60
Time (min)
Normal
Diabetic
Normal data

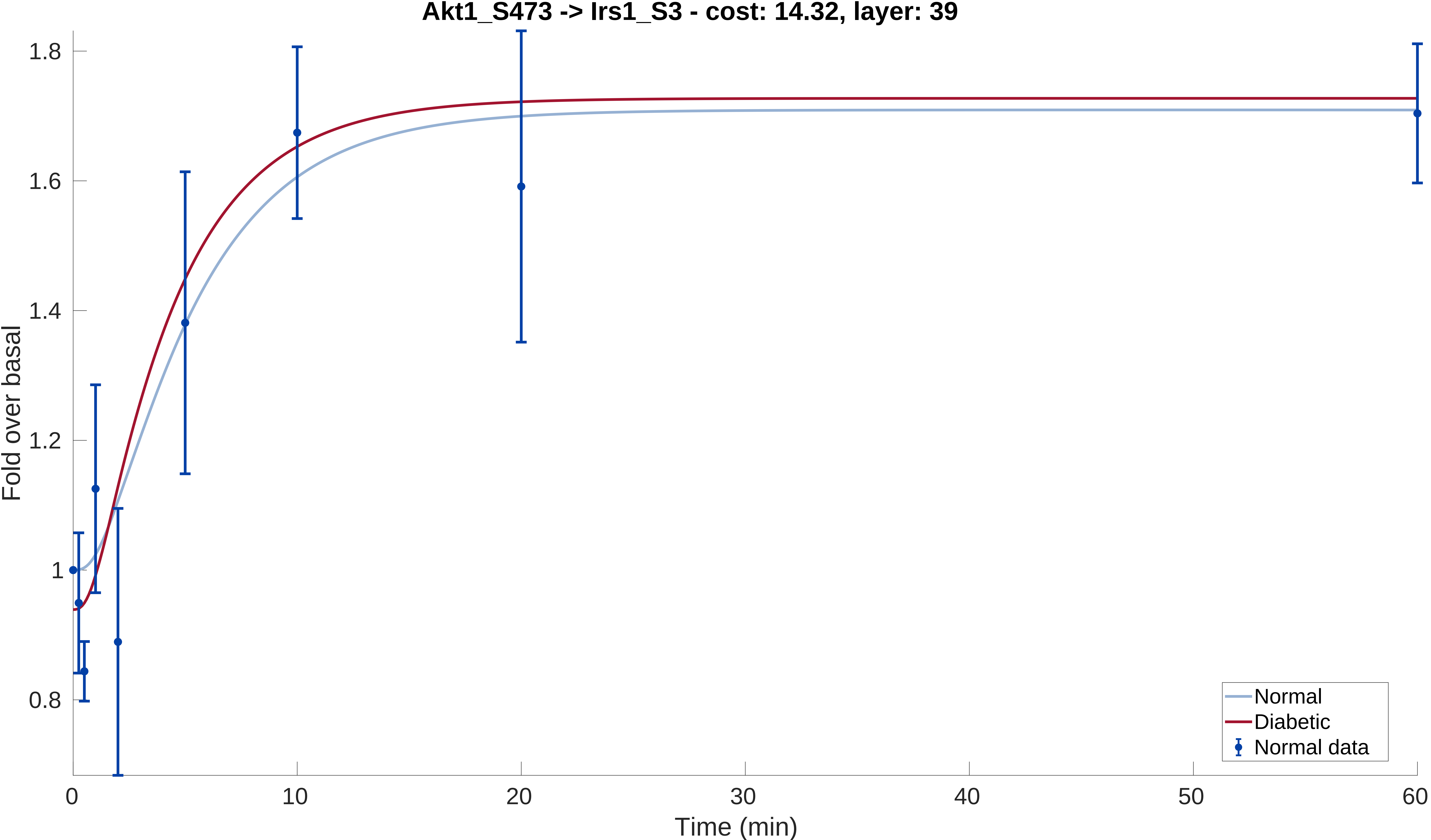

Akt1_S473 -> Irs1_S3 - cost: 14.32, layer: 39
Fold over basal
1.8
1.6
1.4
1.2
1
0.8
0
10
20
30
40
50
60
Time (min)
Normal
Diabetic
Normal data

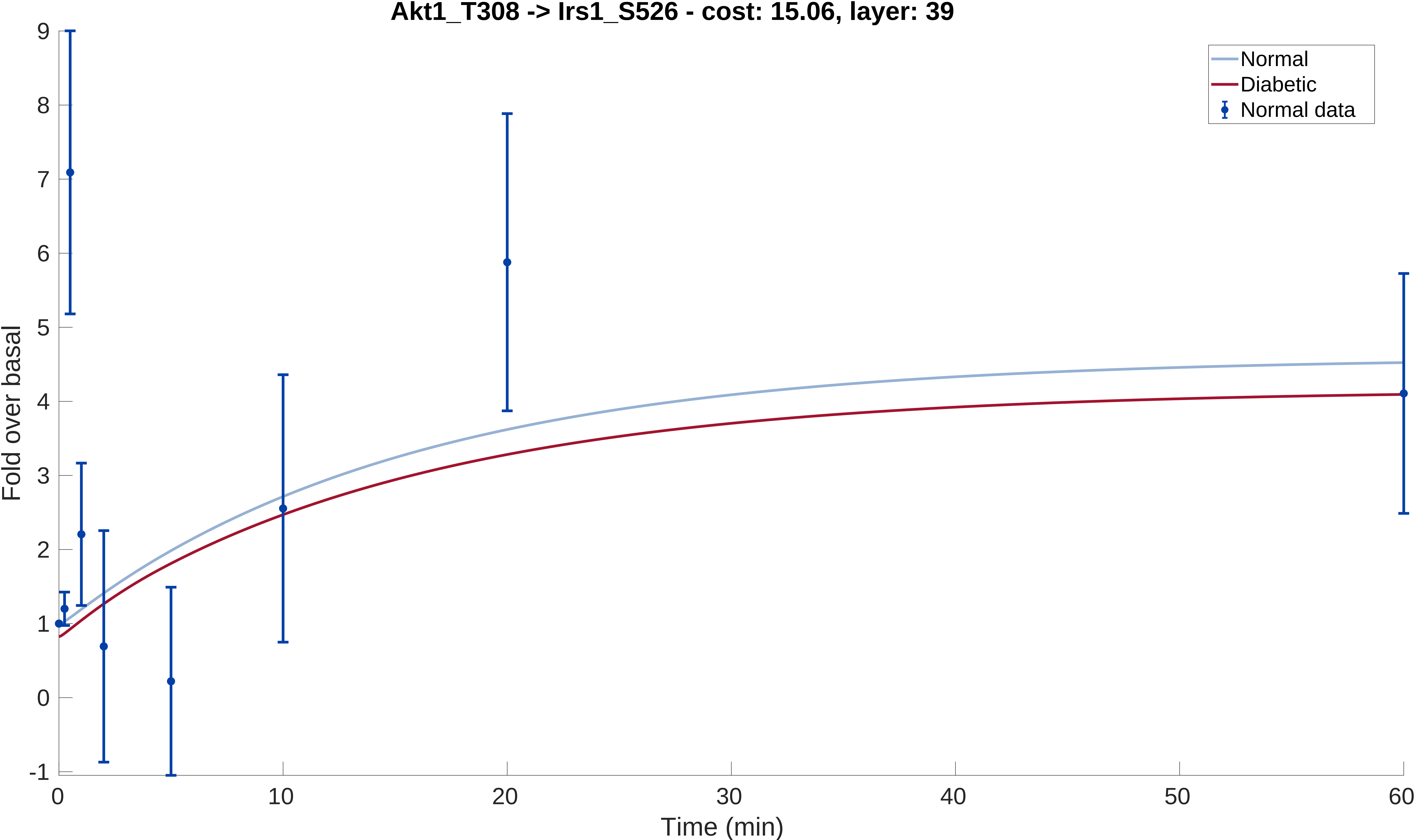

Akt1_T308 -> Irs1_S526 - cost: 15.06, layer: 39
Normal
Diabetic
Normal data
Fold over basal
Time (min)
9
8
7
6
5
4
3
2
1
0
-1
0
10
20
30
40
50
60

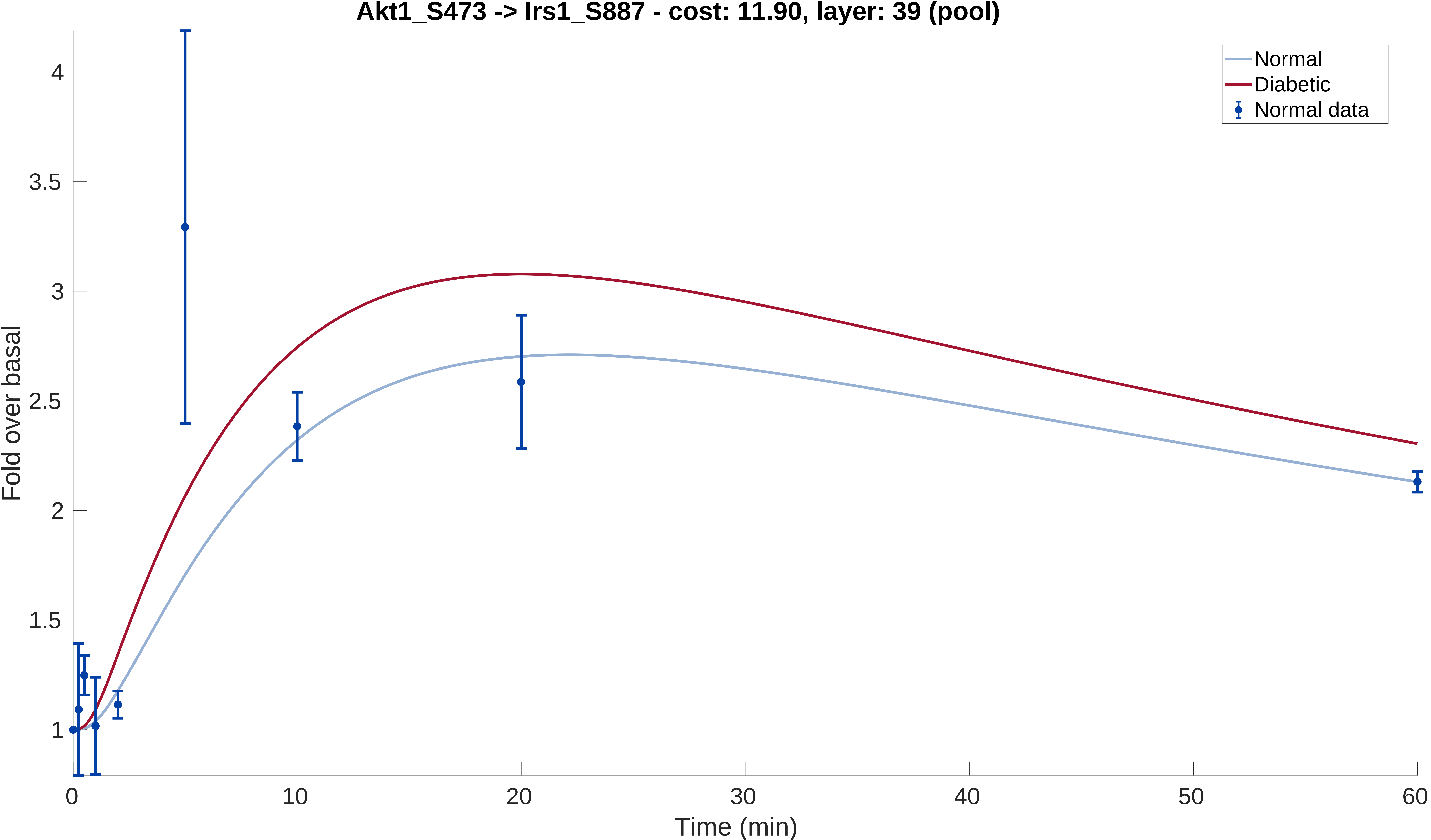

Akt1_S473 -> Irs1_S887 - cost: 11.90, layer: 39 (pool)
Normal
Diabetic
Normal data
Fold over basal
Time (min)
4
3.5
3
2.5
2
1.5
1
0
10
20
30
40
50
60

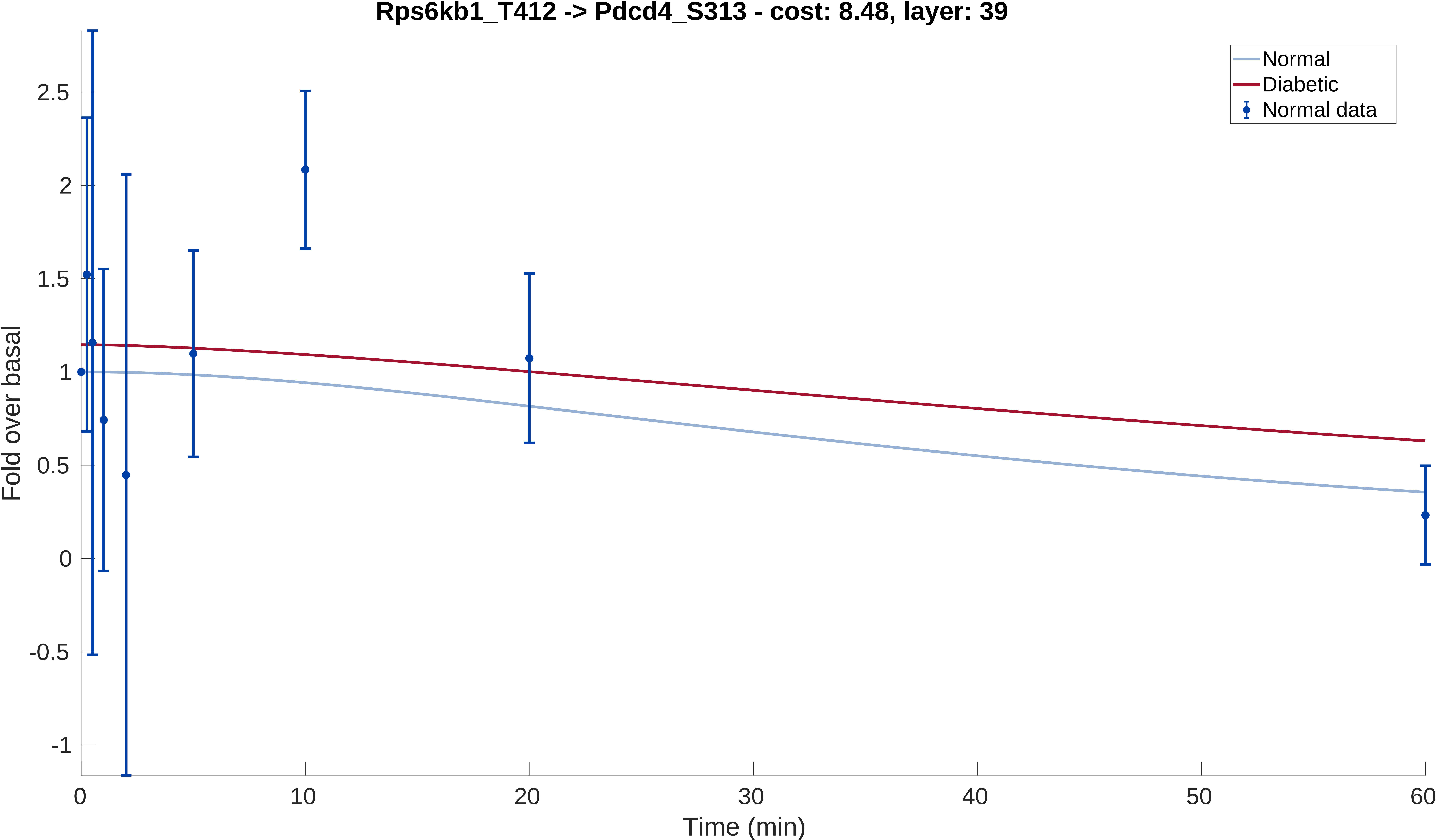

Rps6kb1_T412 -> Pdcd4_S313 - cost: 8.48, layer: 39
Normal
Diabetic
Normal data
Fold over basal
Time (min)
2.5
2
1.5
1
0.5
0
-0.5
-1
0
10
20
30
40
50
60

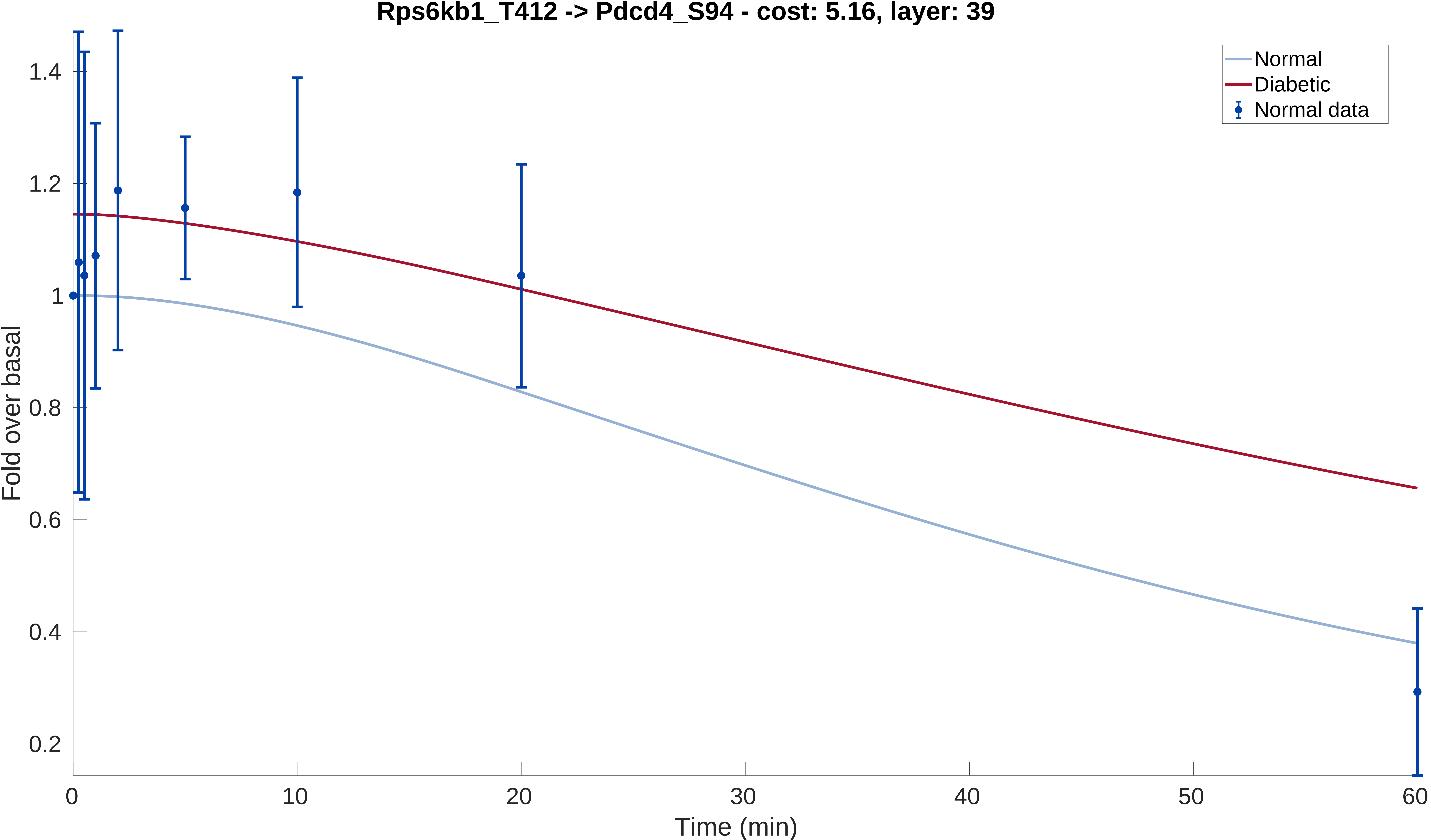

Rps6kb1_T412 -> Pdcd4_S94 - cost: 5.16, layer: 39
Normal
Diabetic
Normal data
Fold over basal
0.2
0.4
0.6
0.8
1
1.2
1.4
0
10
20
30
40
50
60
Time (min)

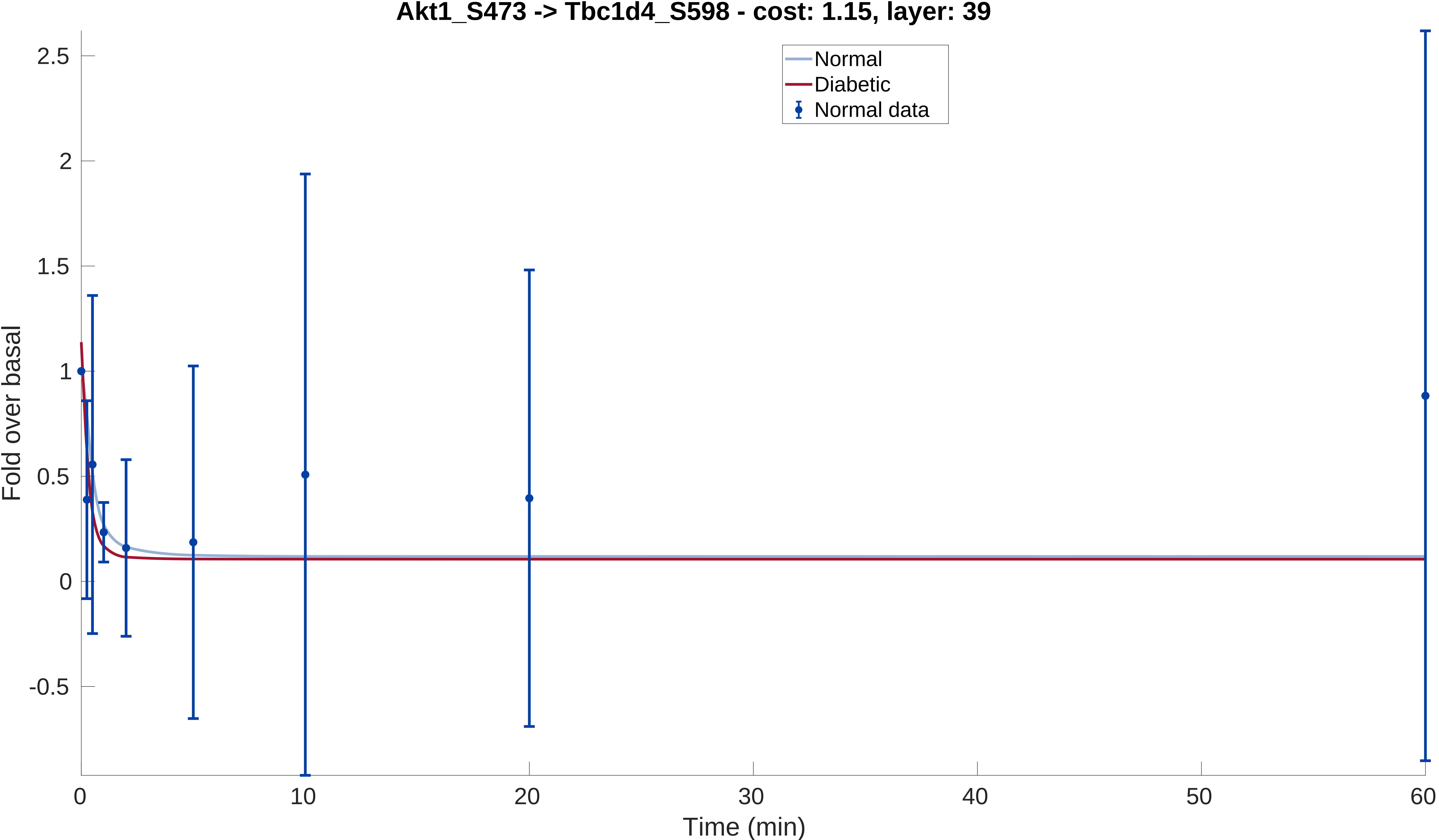

Akt1_S473 -> Tbc1d4_S598 - cost: 1.15, layer: 39
Normal
Diabetic
Normal data
Fold over basal
Time (min)
2.5
2
1.5
1
0.5
0
-0.5
0
10
20
30
40
50
60

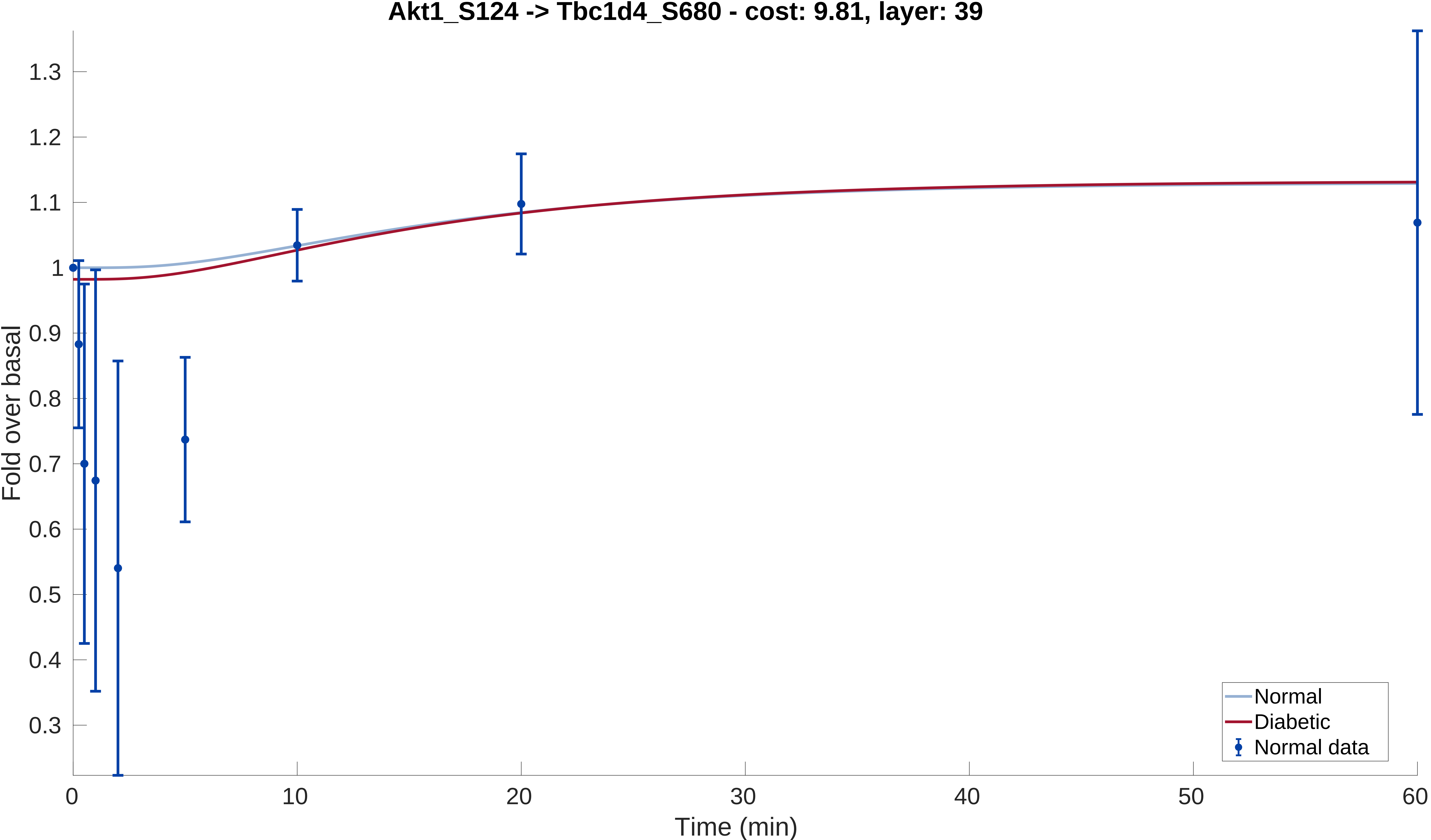
Akt1_S124 -> Tbc1d4_S680 - cost: 9.81, layer: 39
Fold over basal
Time (min)
1.3
1.2
1.1
1
0.9
0.8
0.7
0.6
0.5
0.4
0.3
0
10
20
30
40
50
60
Normal
Diabetic
Normal data

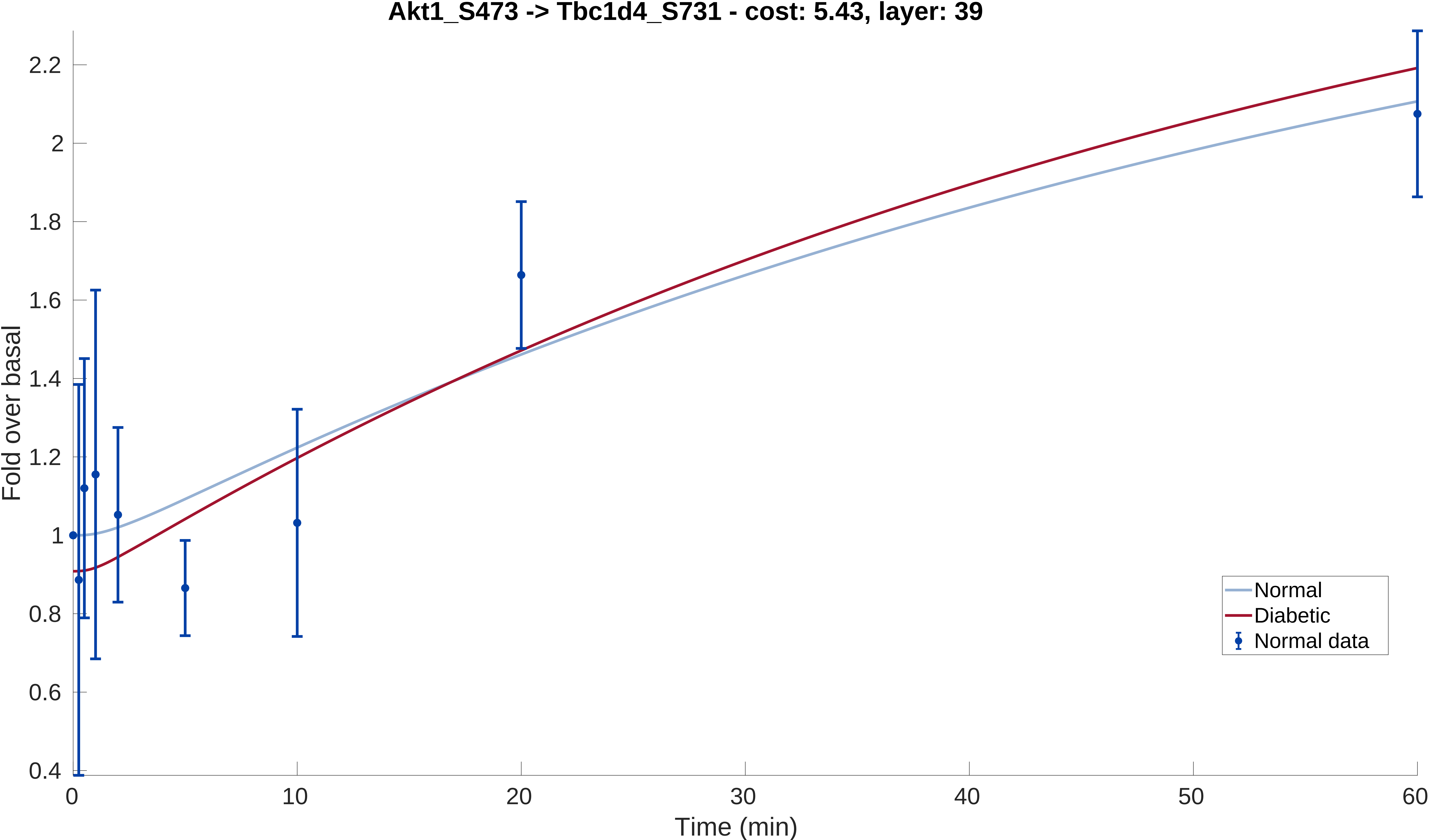

Akt1_S473 -> Tbc1d4_S731 - cost: 5.43, layer: 39
Fold over basal
2.2
2
1.8
1.6
1.4
1.2
1
0.8
0.6
0.4
0
10
20
30
40
50
60
Time (min)
Normal
Diabetic
Normal data

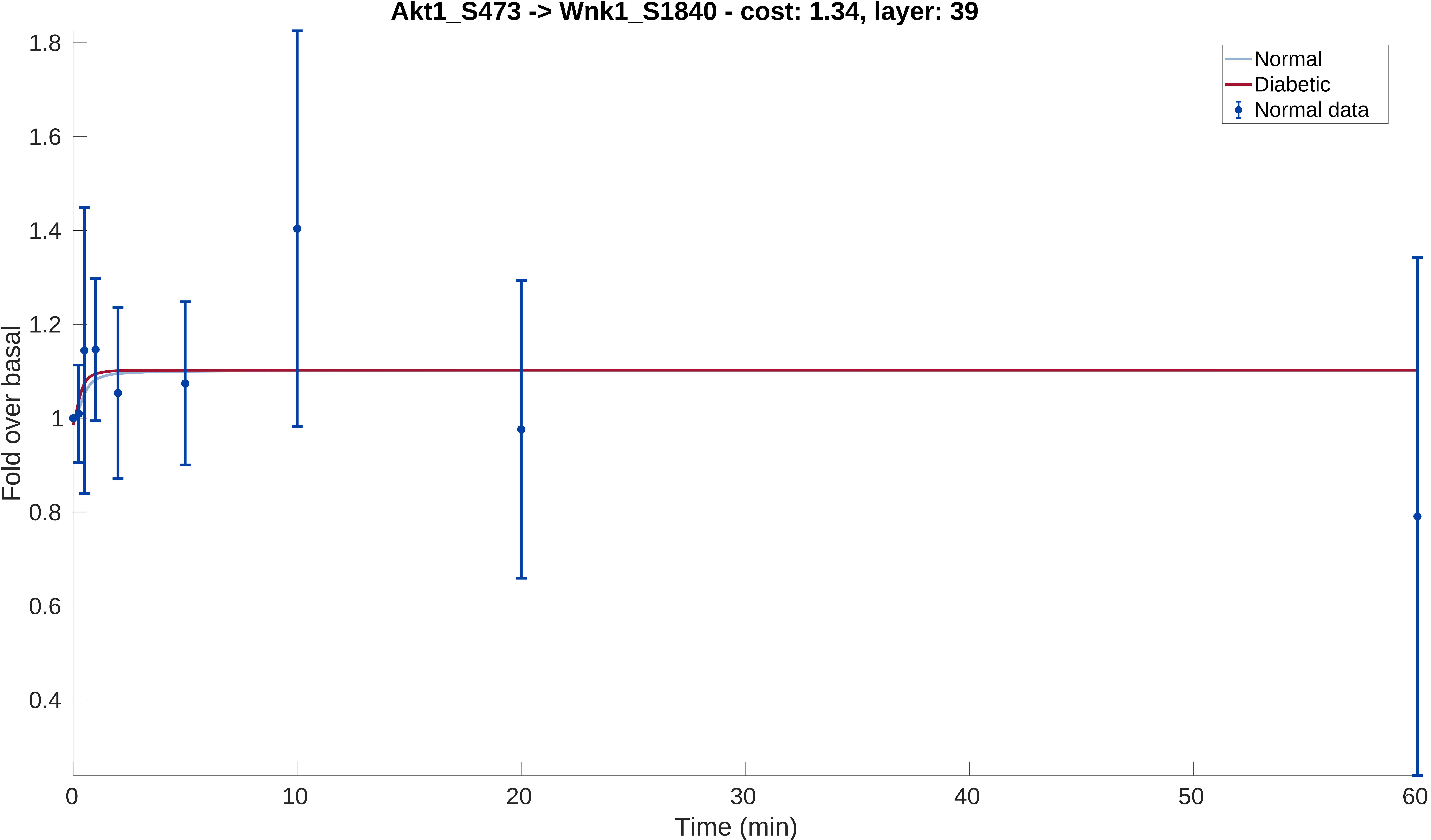
Akt1_S473 -> Wnk1_S1840 - cost: 1.34, layer: 39
Fold over basal
1.8
1.6
1.4
1.2
1
0.8
0.6
0.4
0
10
20
30
40
50
60
Time (min)
Normal
Diabetic
Normal data

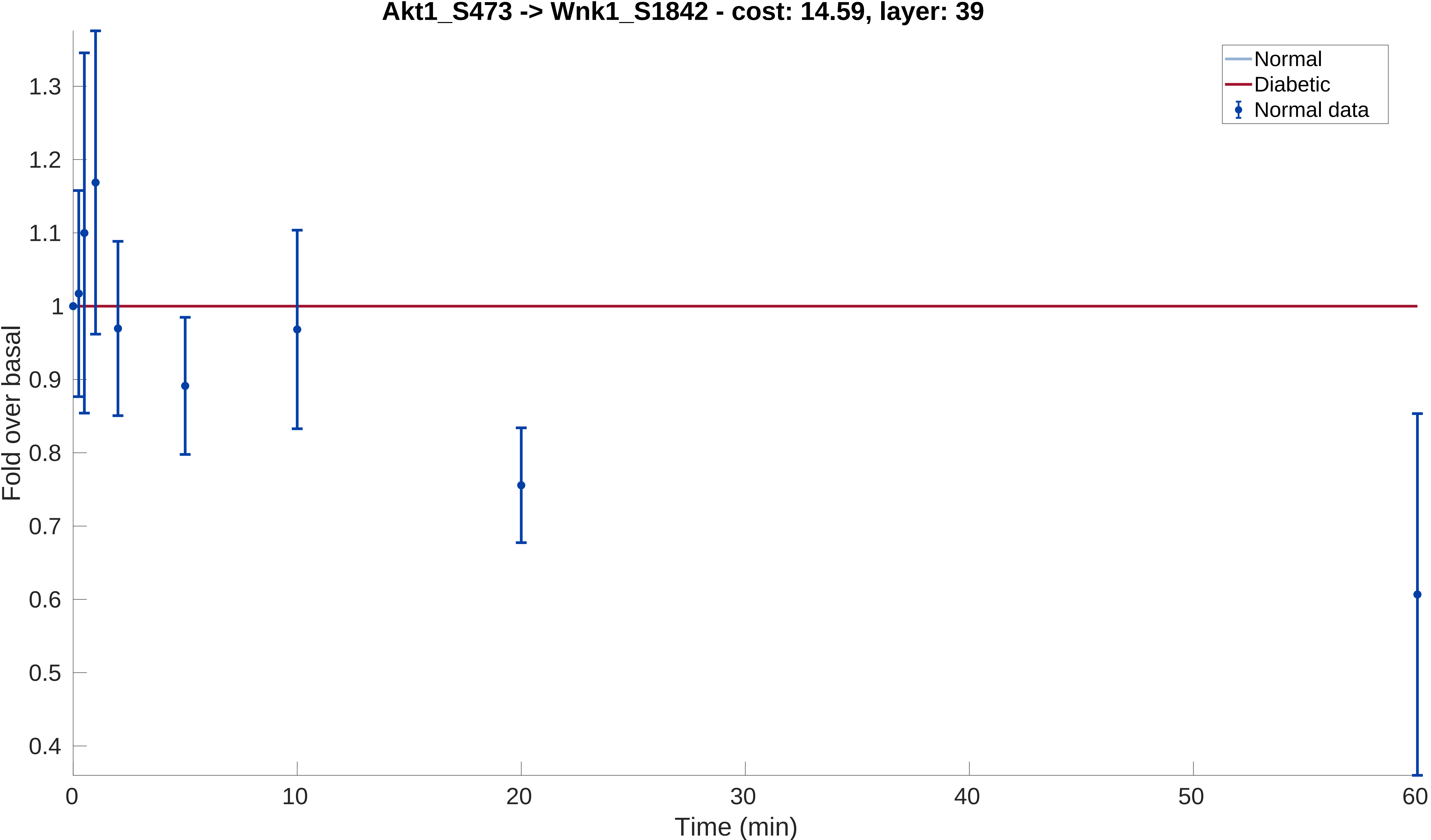
Akt1_S473 -> Wnk1_S1842 - cost: 14.59, layer: 39
Normal
Diabetic
Normal data
Fold over basal
1.3
1.2
1.1
1
0.9
0.8
0.7
0.6
0.5
0.4
0
10
20
30
40
50
60
Time (min)

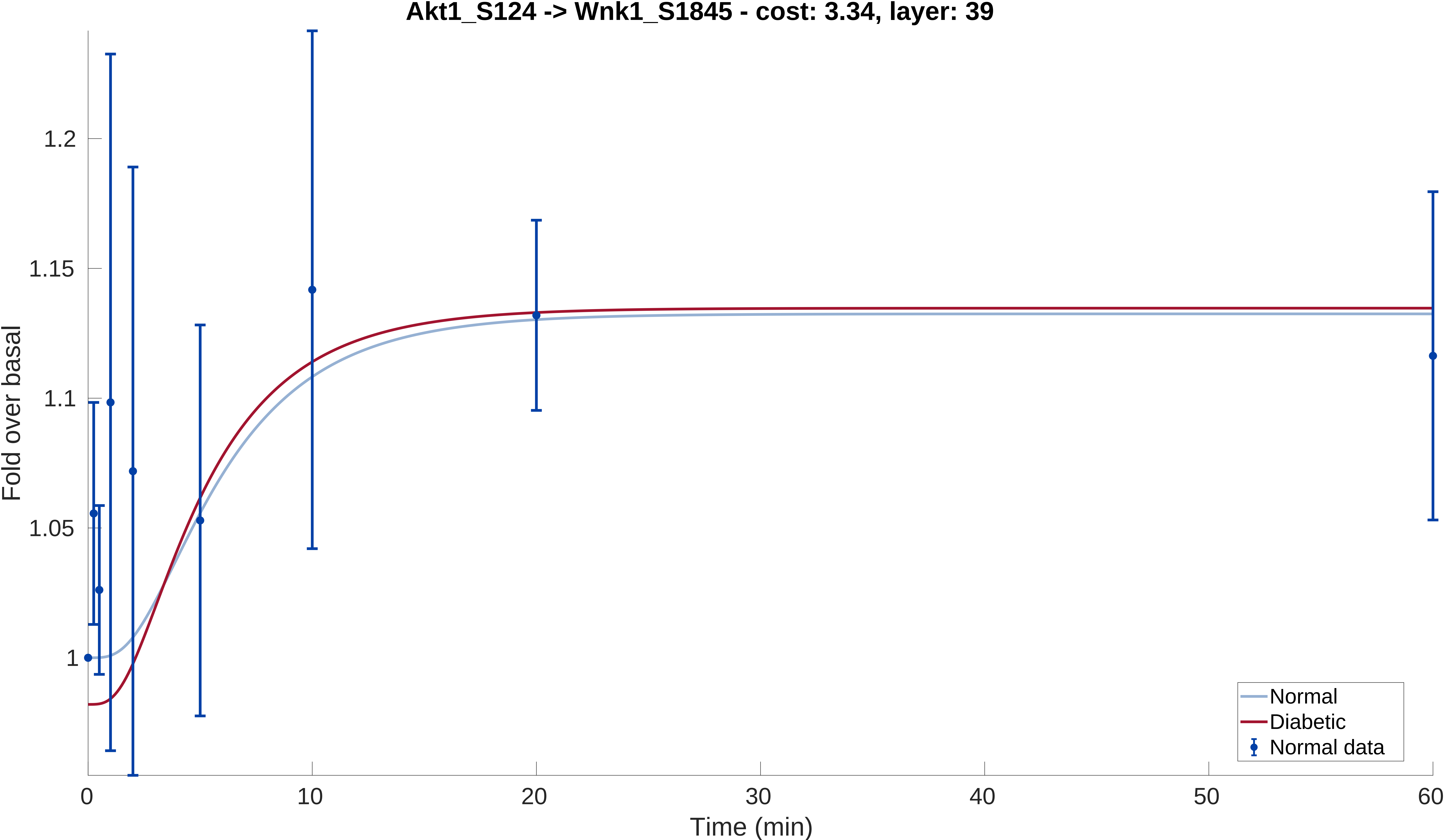
Akt1_S124 -> Wnk1_S1845 - cost: 3.34, layer: 39
Fold over basal
1.2
1.15
1.1
1.05
1
0
10
20
30
40
50
60
Time (min)
Normal
Diabetic
Normal data

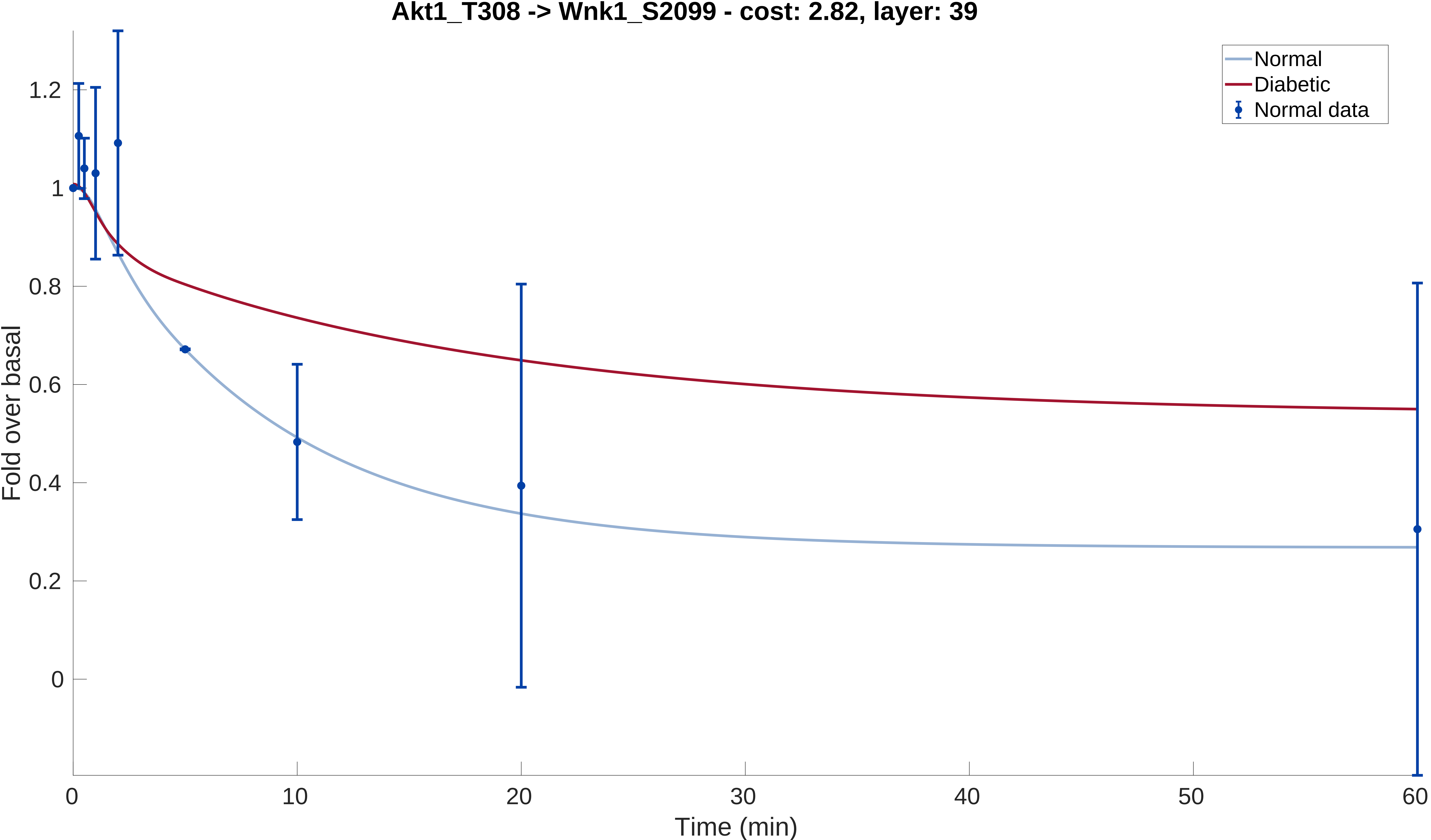

Akt1_T308 -> Wnk1_S2099 - cost: 2.82, layer: 39
Normal
Diabetic
Normal data
Fold over basal
Time (min)
0
0.2
0.4
0.6
0.8
1
1.2
0
10
20
30
40
50
60

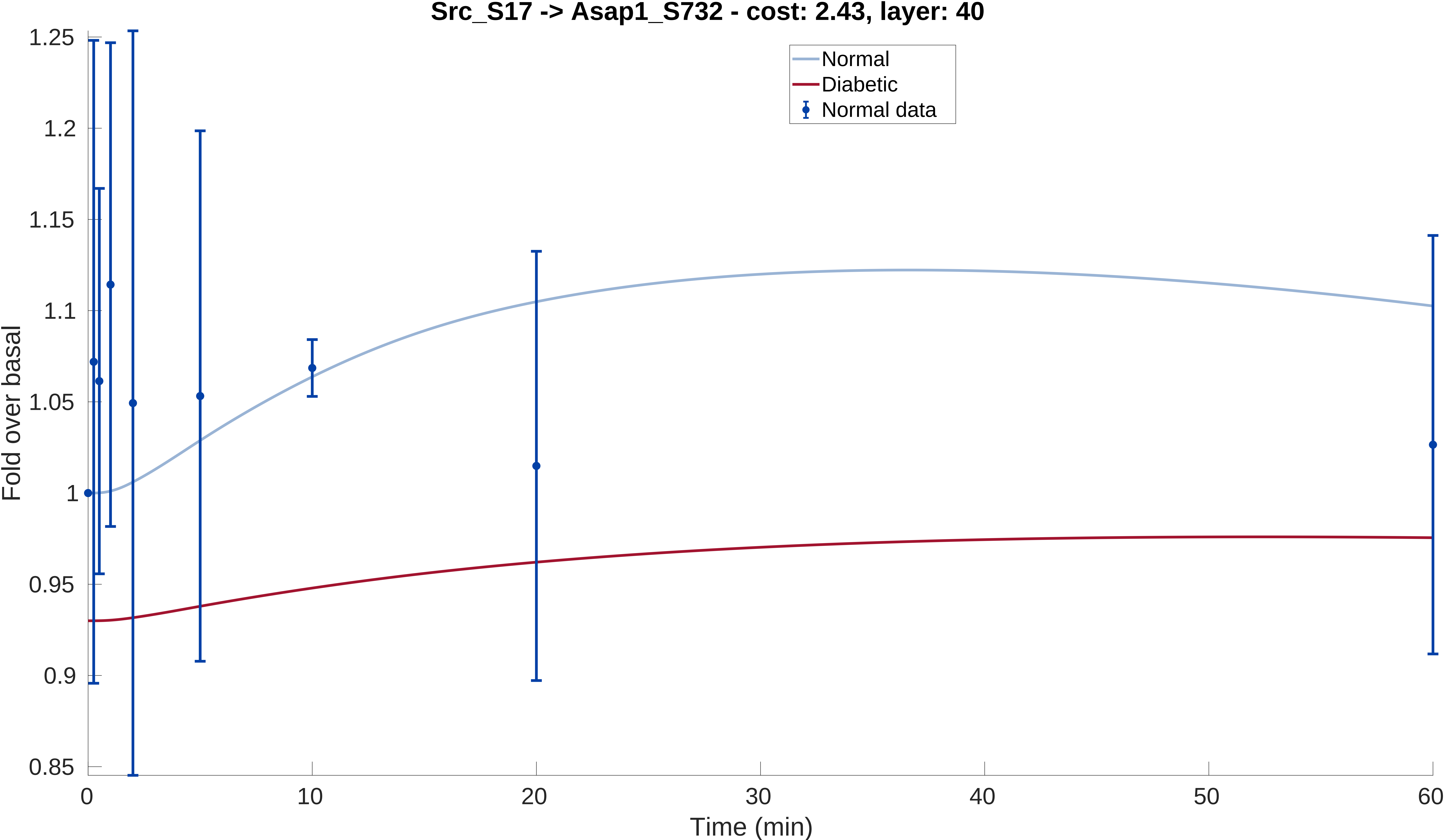
Src_S17 -> Asap1_S732 - cost: 2.43, layer: 40
Normal
Diabetic
Normal data
Fold over basal
Time (min)
1.25
1.2
1.15
1.1
1.05
1
0.95
0.9
0.85
0
10
20
30
40
50
60

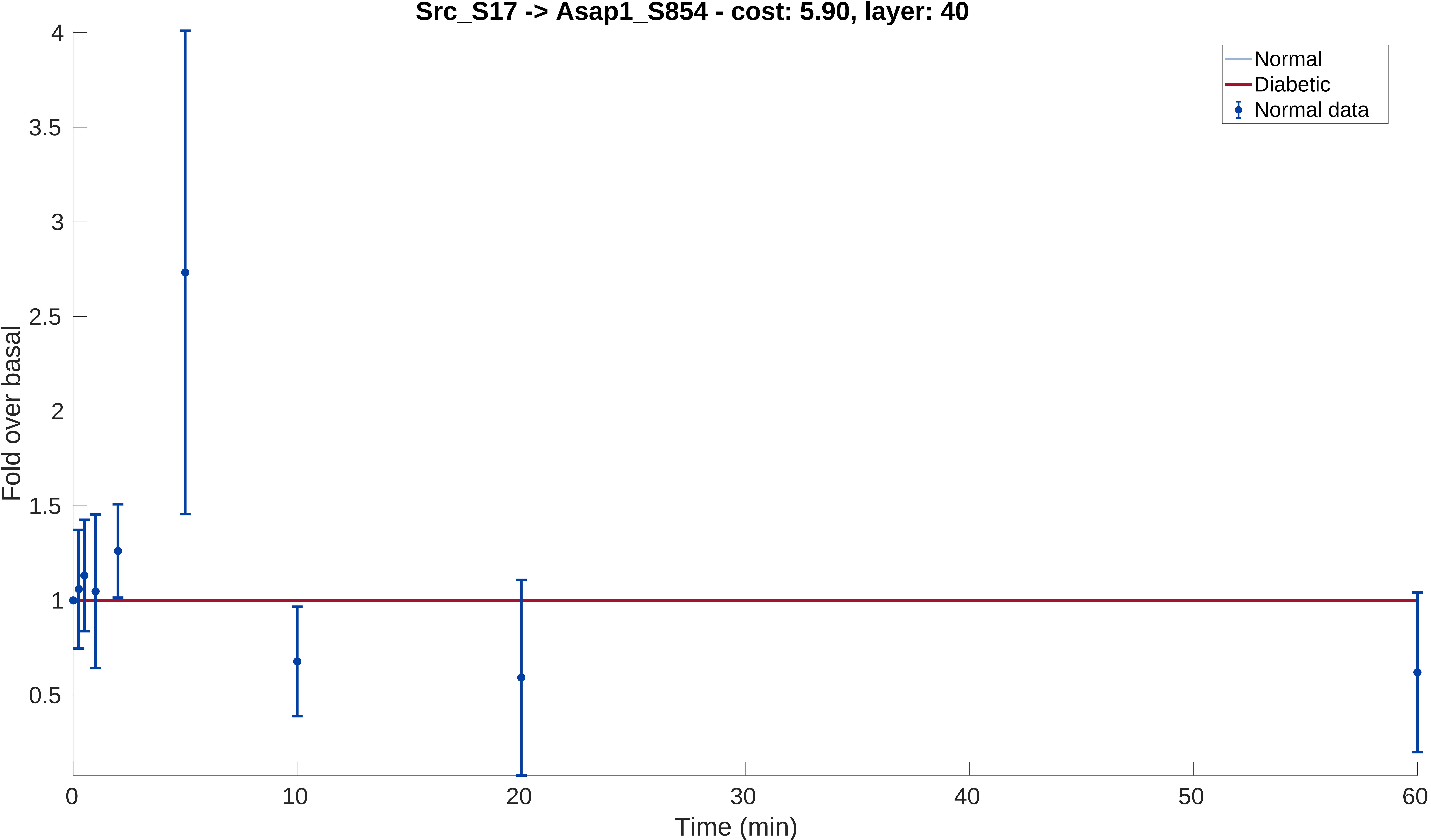

Src_S17 -> Asap1_S854 - cost: 5.90, layer: 40
Normal
Diabetic
Normal data
Fold over basal
Time (min)
0
10
20
30
40
50
60
0.5
1
1.5
2
2.5
3
3.5
4

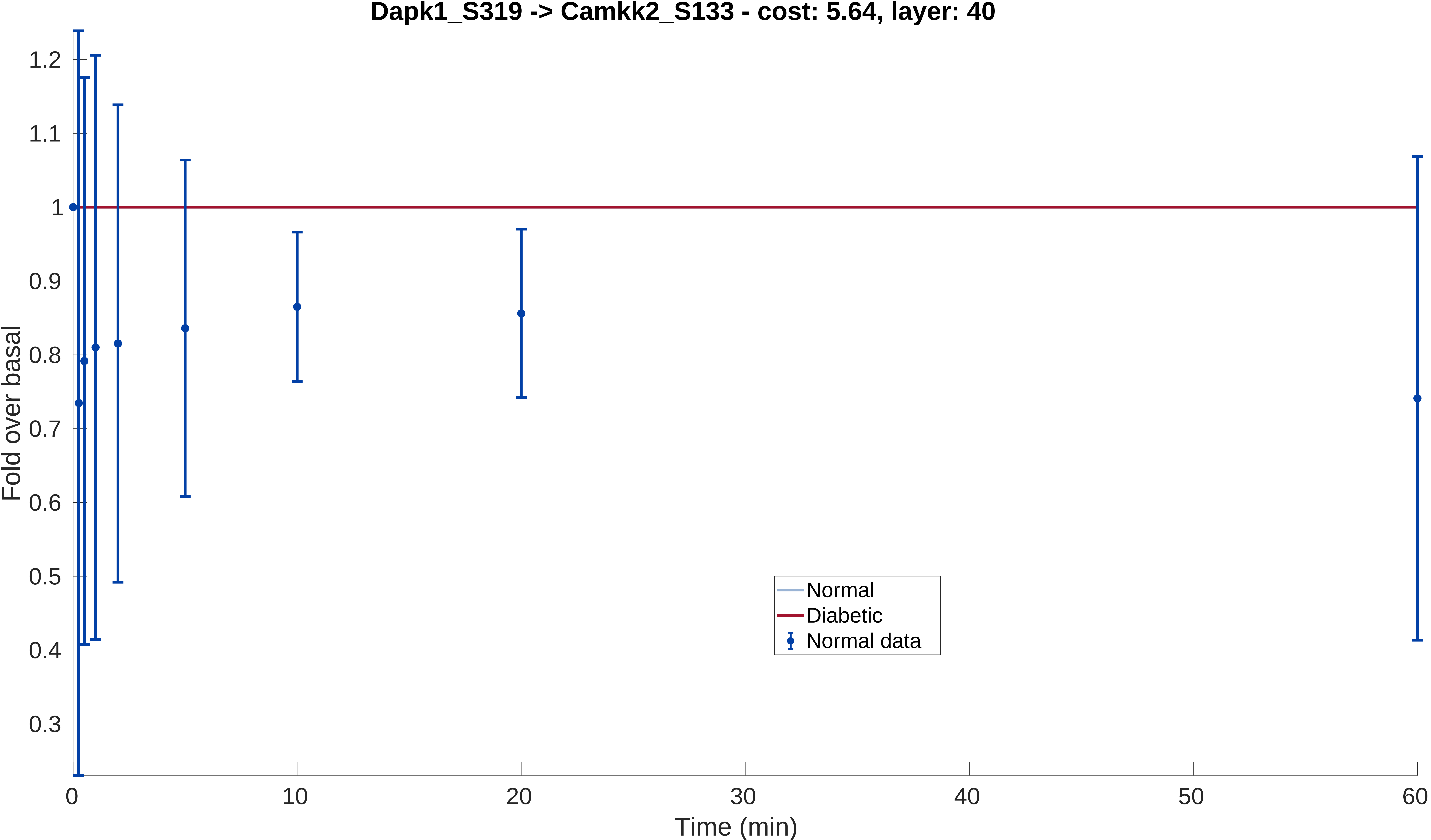

Dapk1_S319 -> Camkk2_S133 - cost: 5.64, layer: 40
Fold over basal
1.2
1.1
1
0.9
0.8
0.7
0.6
0.5
0.4
0.3
0
10
20
30
40
50
60
Time (min)
Normal
Diabetic
Normal data

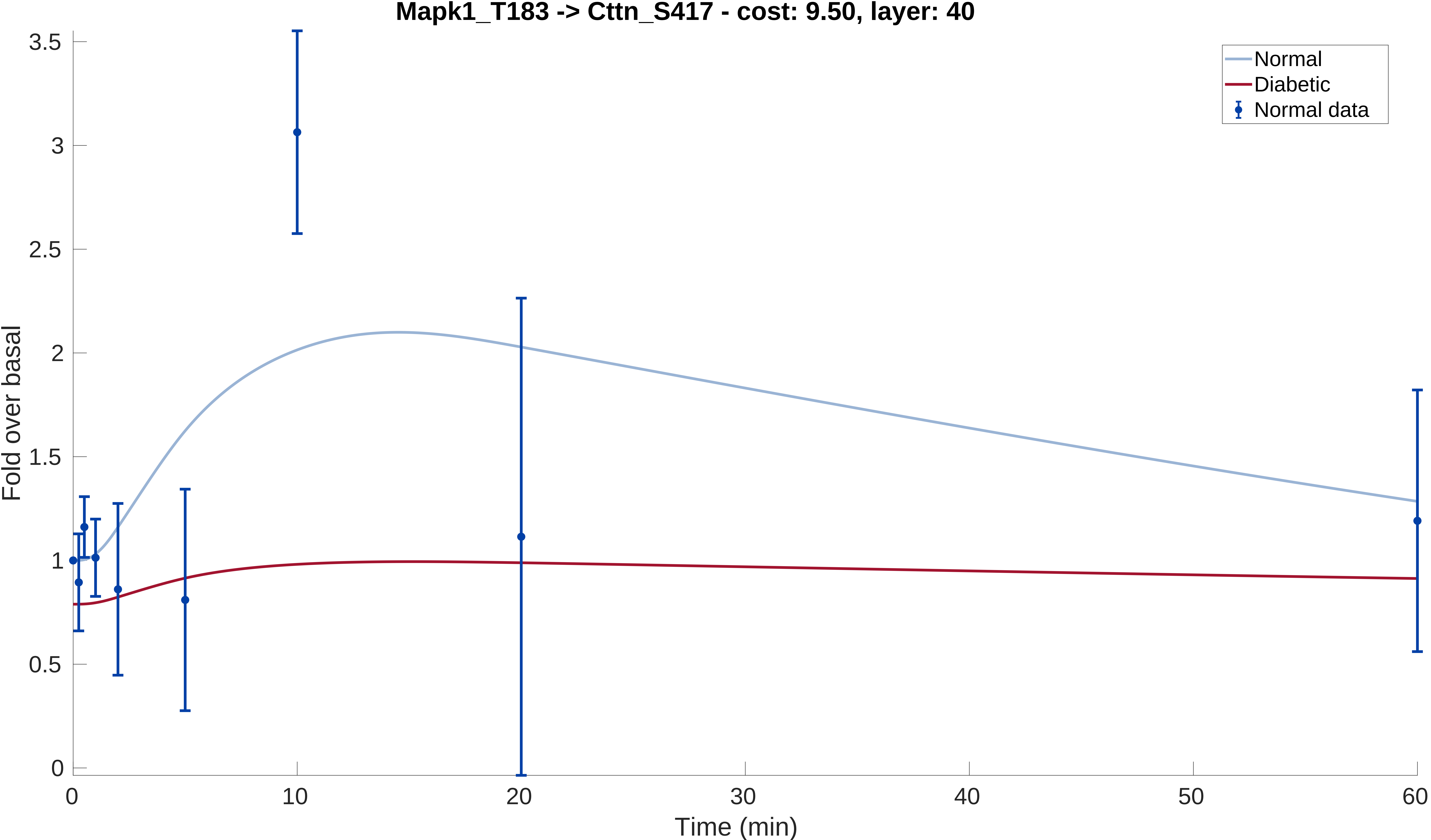
Mapk1_T183 -> Cttn_S417 - cost: 9.50, layer: 40
Normal
Diabetic
Normal data
Fold over basal
Time (min)
0
0.5
1
1.5
2
2.5
3
3.5
0
10
20
30
40
50
60

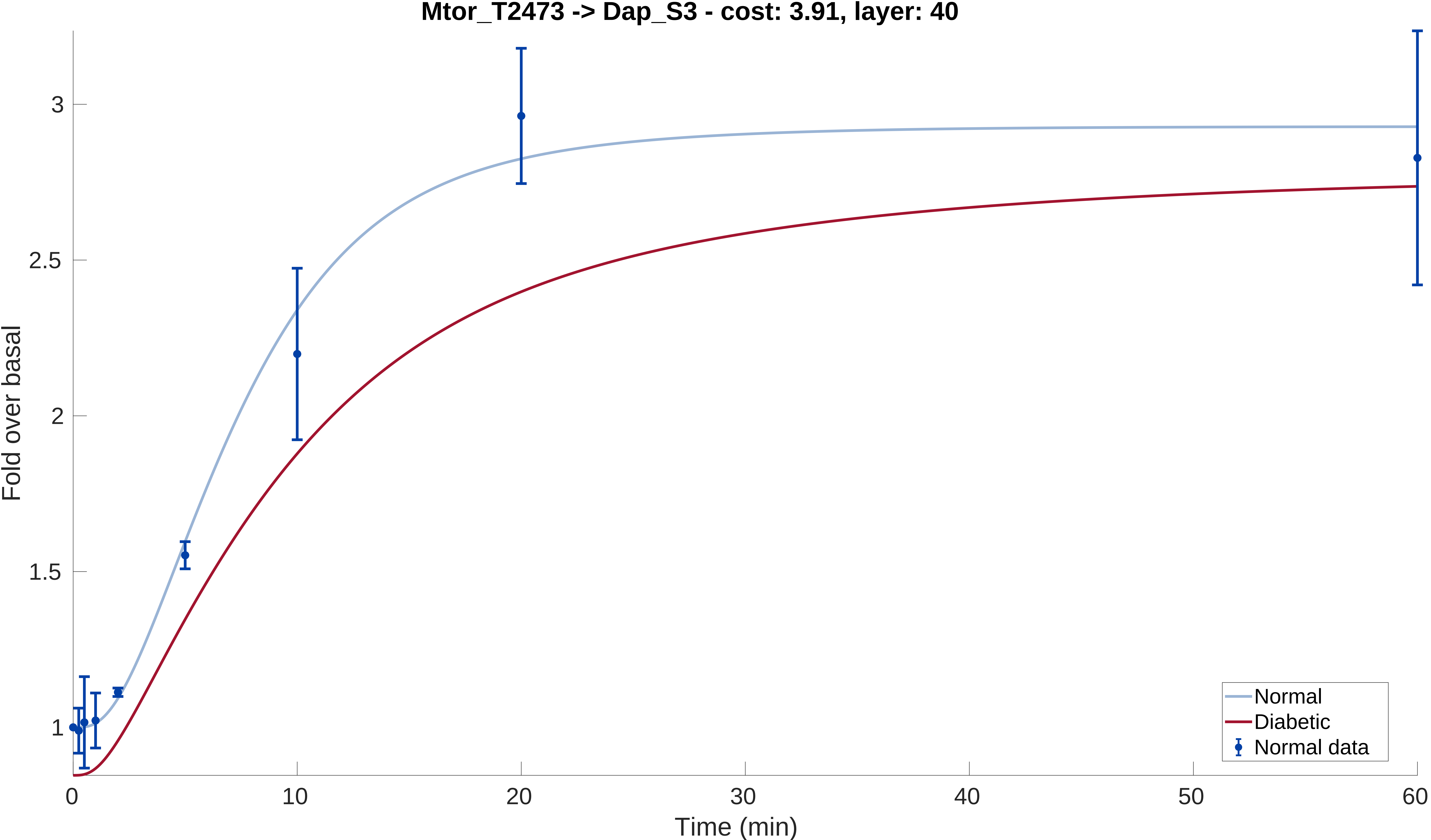

Mtor_T2473 -> Dap_S3 - cost: 3.91, layer: 40
Fold over basal
Time (min)
0
10
20
30
40
50
60
1
1.5
2
2.5
3
Normal
Diabetic
Normal data

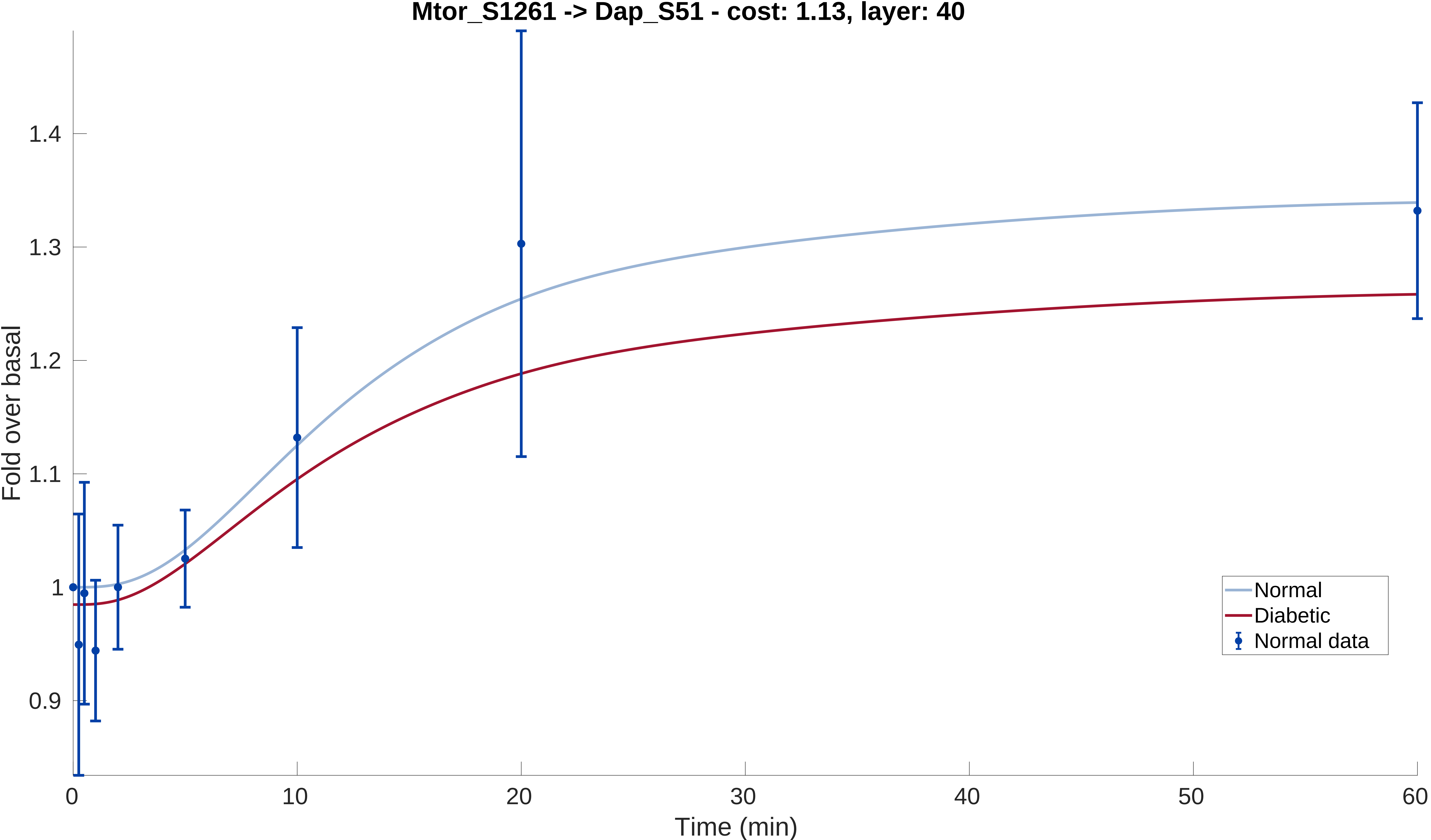
Mtor_S1261 -> Dap_S51 - cost: 1.13, layer: 40
Fold over basal
Time (min)
1.4
1.3
1.2
1.1
1
0.9
0
10
20
30
40
50
60
Normal
Diabetic
Normal data

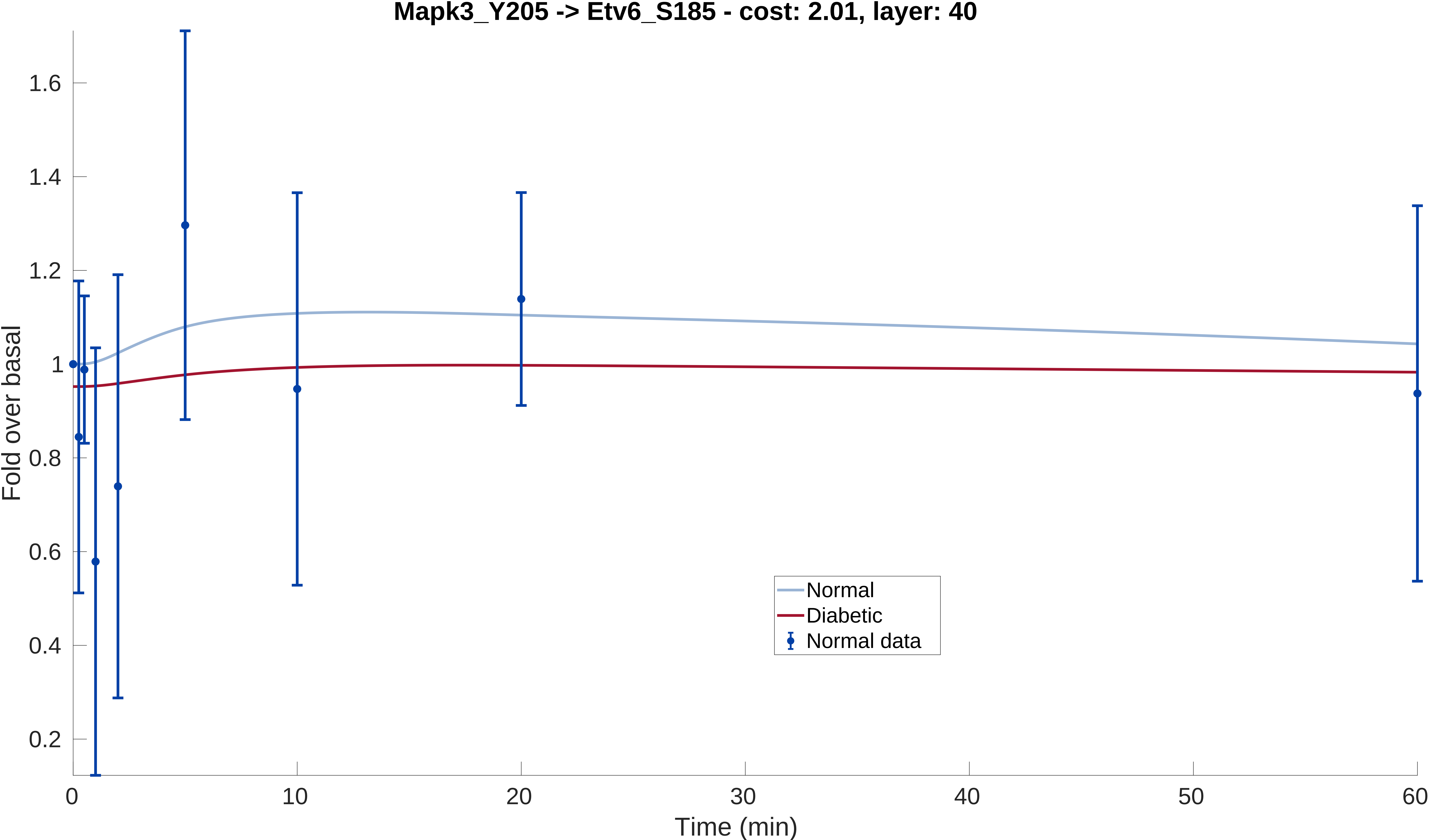

Mapk3_Y205 -> Etv6_S185 - cost: 2.01, layer: 40
Fold over basal
Time (min)
0
10
20
30
40
50
60
0.2
0.4
0.6
0.8
1
1.2
1.4
1.6
Normal
Diabetic
Normal data

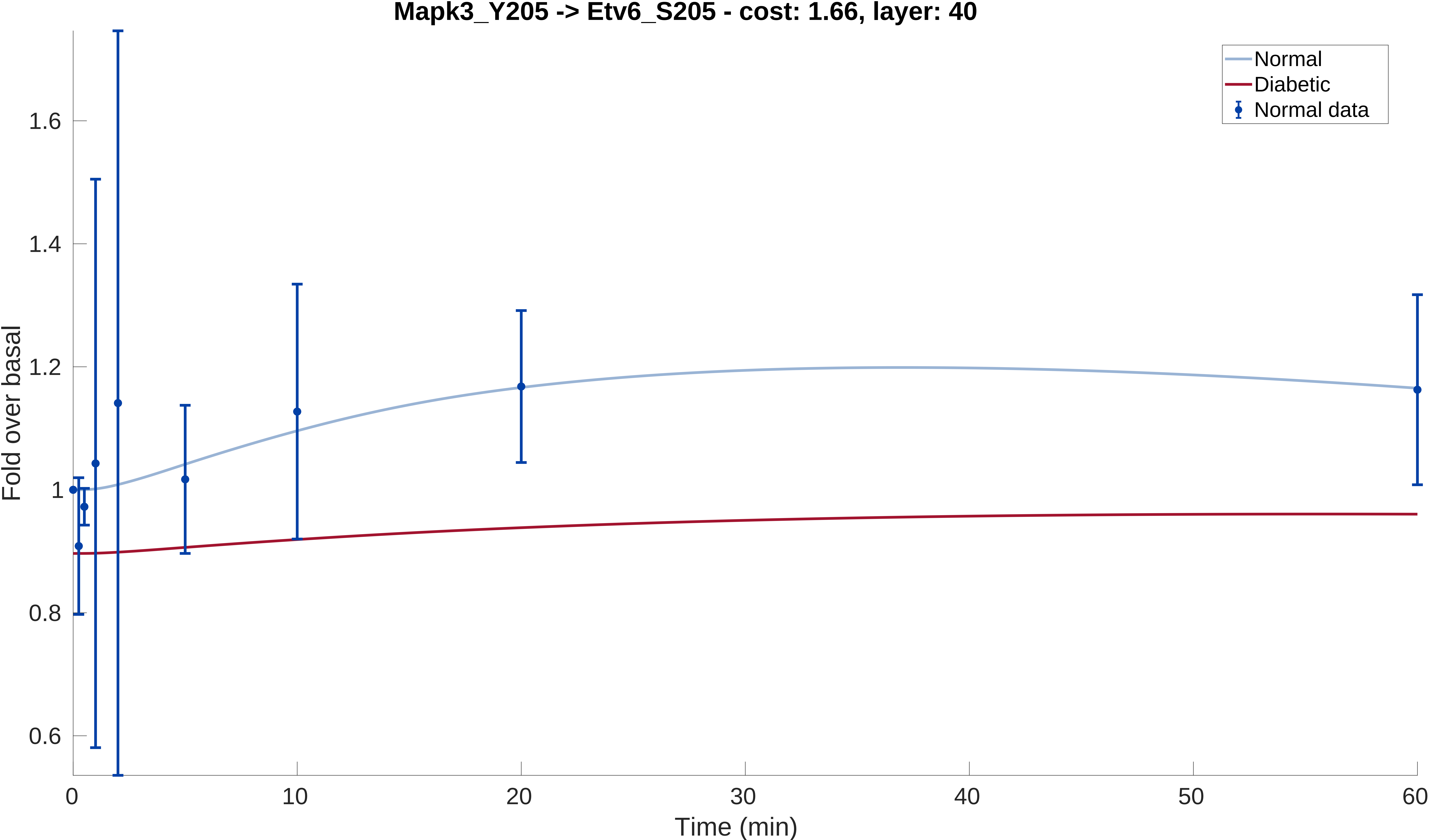

Mapk3_Y205 -> Etv6_S205 - cost: 1.66, layer: 40
Normal
Diabetic
Normal data
Fold over basal
1.6
1.4
1.2
1
0.8
0.6
0
10
20
30
40
50
60
Time (min)

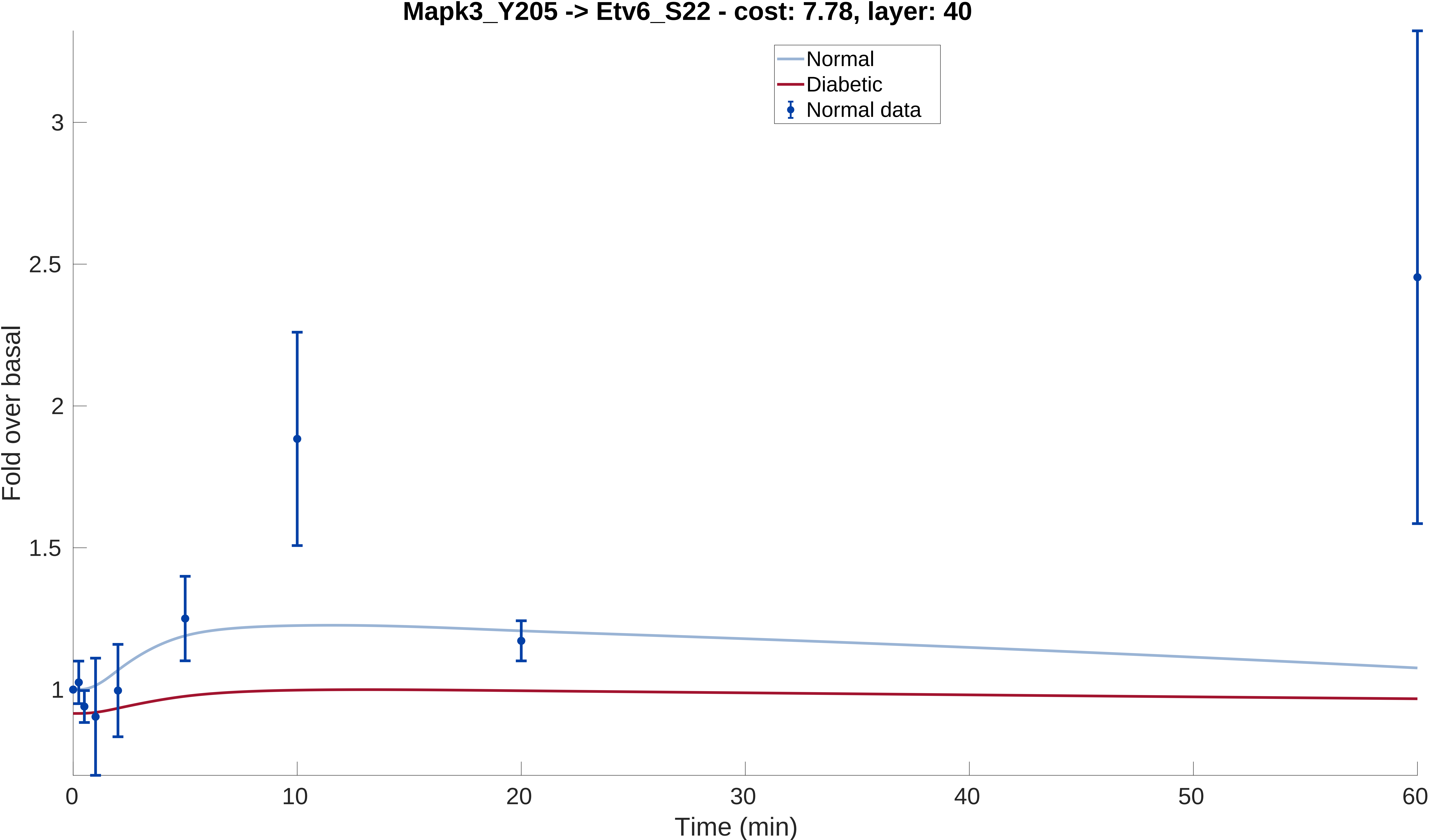

Mapk3_Y205 -> Etv6_S22 - cost: 7.78, layer: 40
Normal
Diabetic
Normal data
Fold over basal
Time (min)
3
2.5
2
1.5
1
0
10
20
30
40
50
60

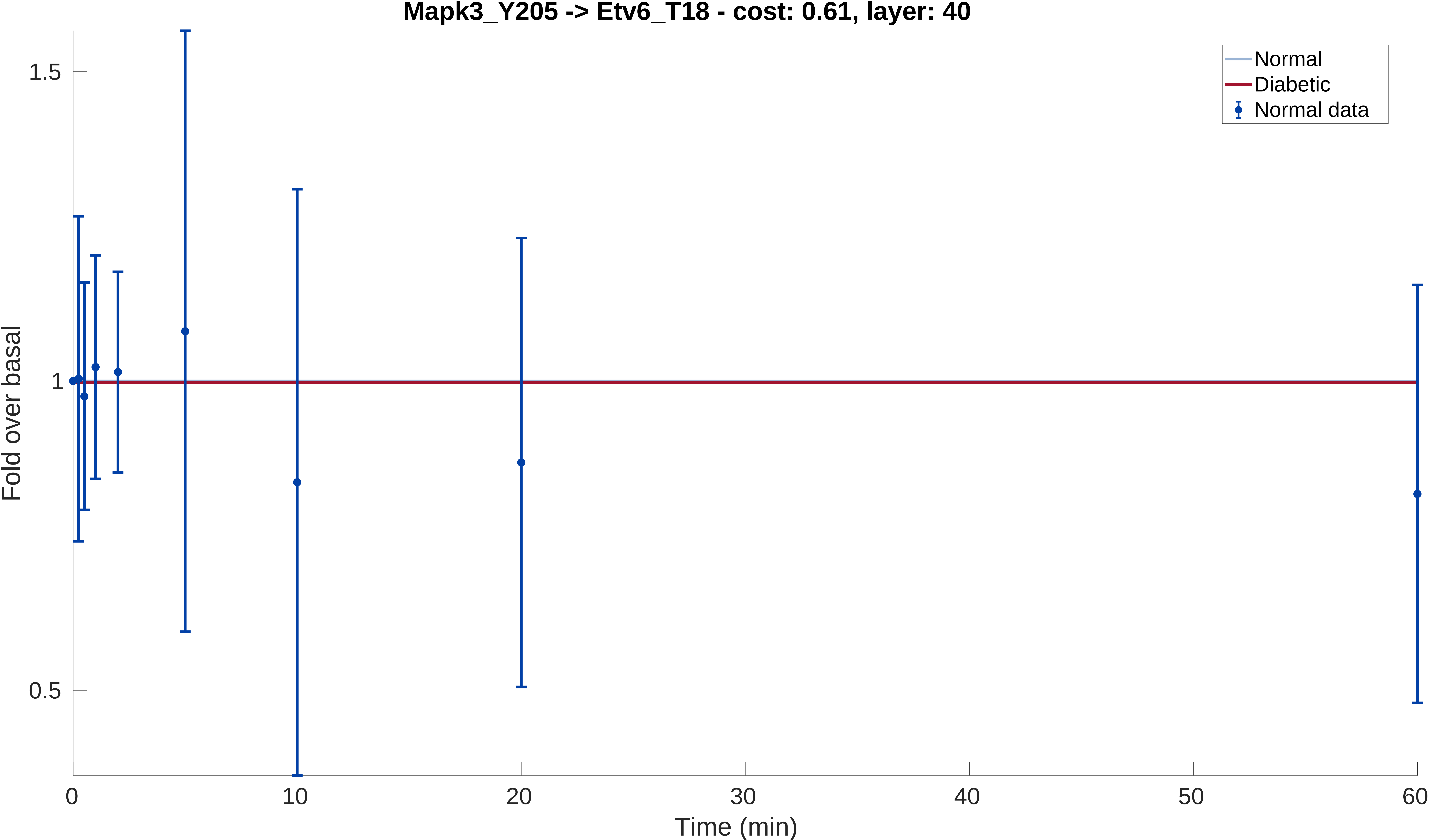
Mapk3_Y205 -> Etv6_T18 - cost: 0.61, layer: 40
Normal
Diabetic
Normal data
Fold over basal
1.5
1
0.5
0
10
20
30
40
50
60
Time (min)

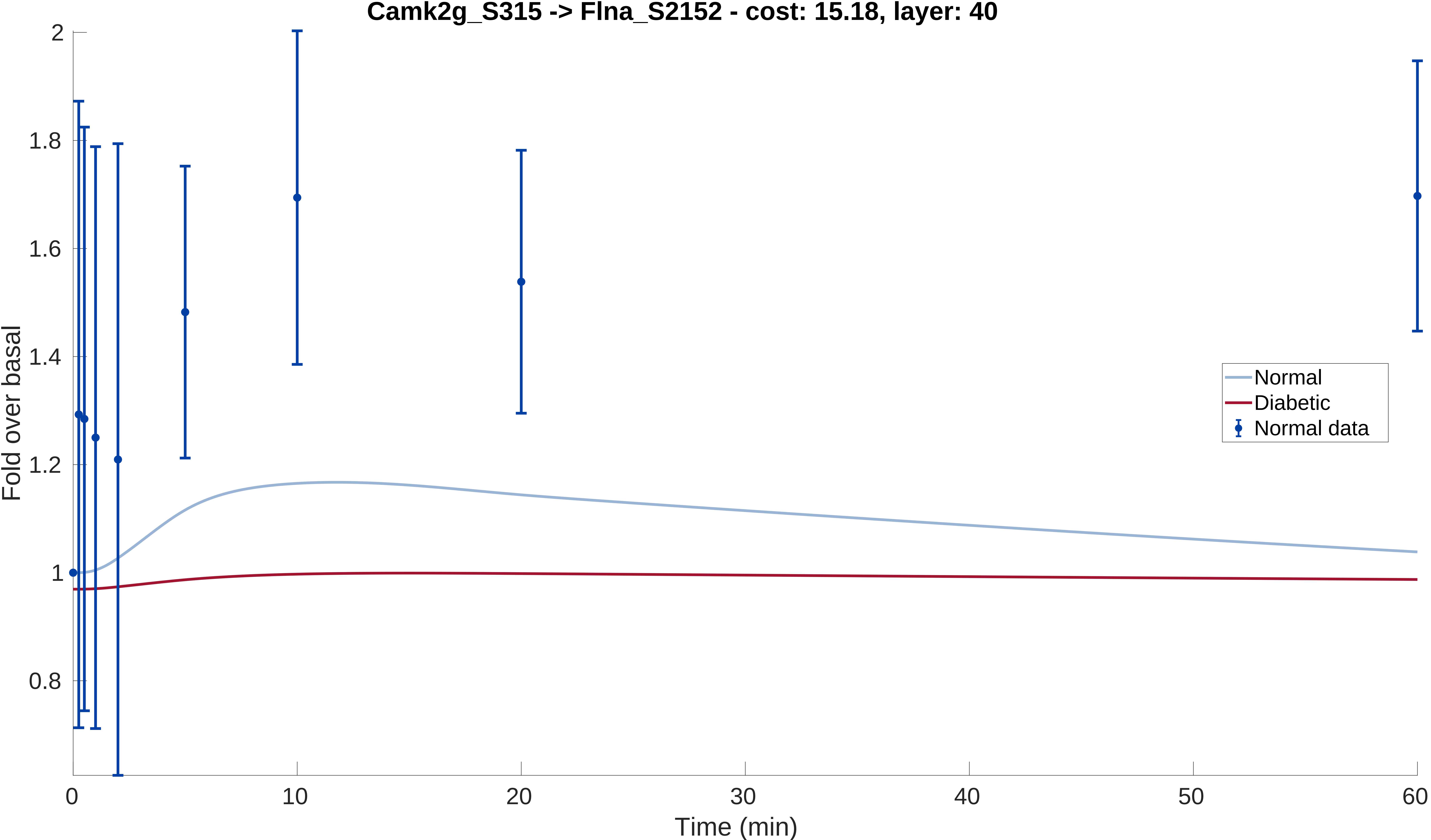

Camk2g_S315 -> Flna_S2152 - cost: 15.18, layer: 40
2
1.8
1.6
1.4
1.2
1
0.8
Fold over basal
0
10
20
30
40
50
60
Time (min)
Normal
Diabetic
Normal data

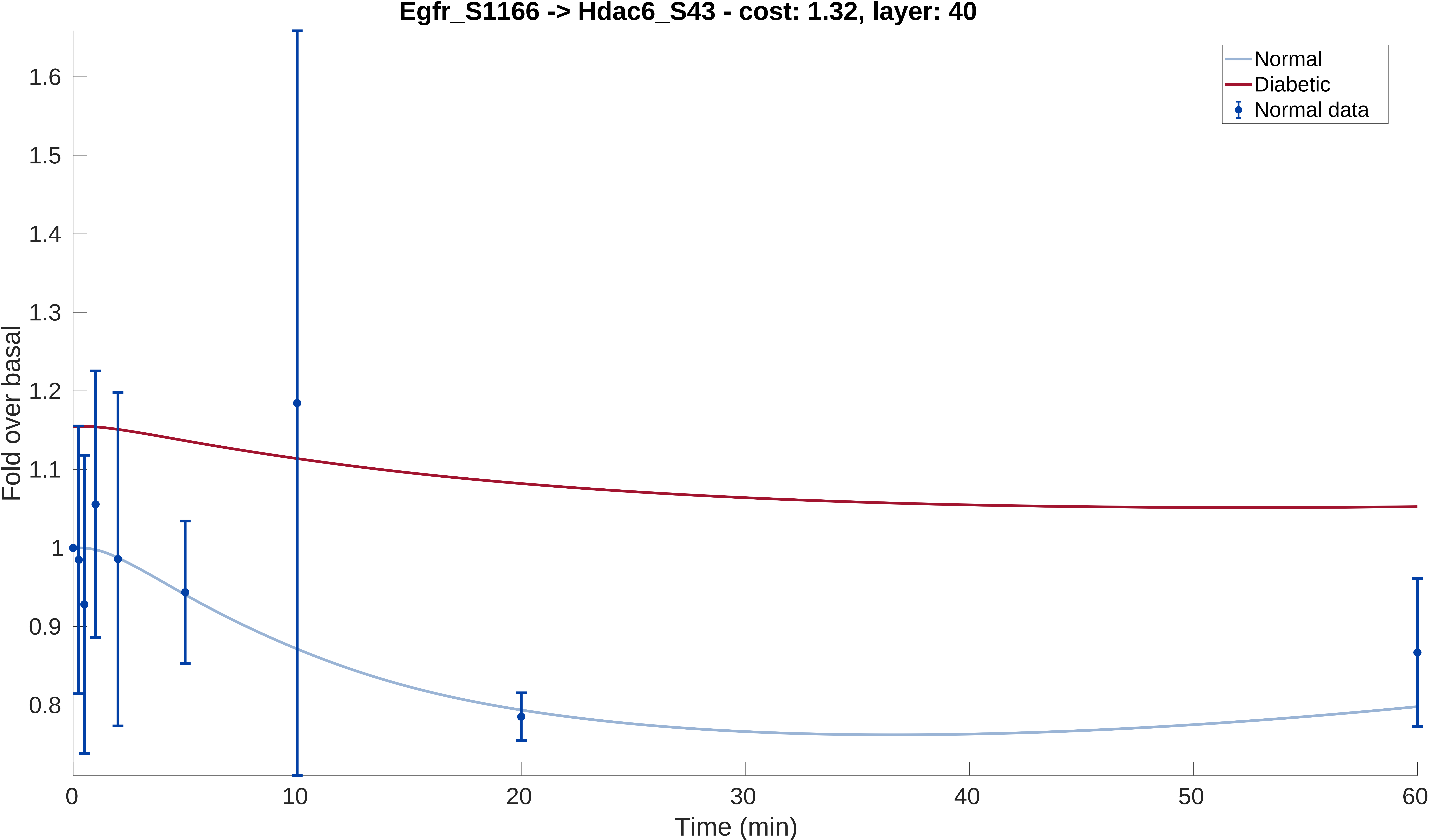

Egfr_S1166 -> Hdac6_S43 - cost: 1.32, layer: 40
Normal
Diabetic
Normal data
Fold over basal
1.6
1.5
1.4
1.3
1.2
1.1
1
0.9
0.8
0
10
20
30
40
50
60
Time (min)

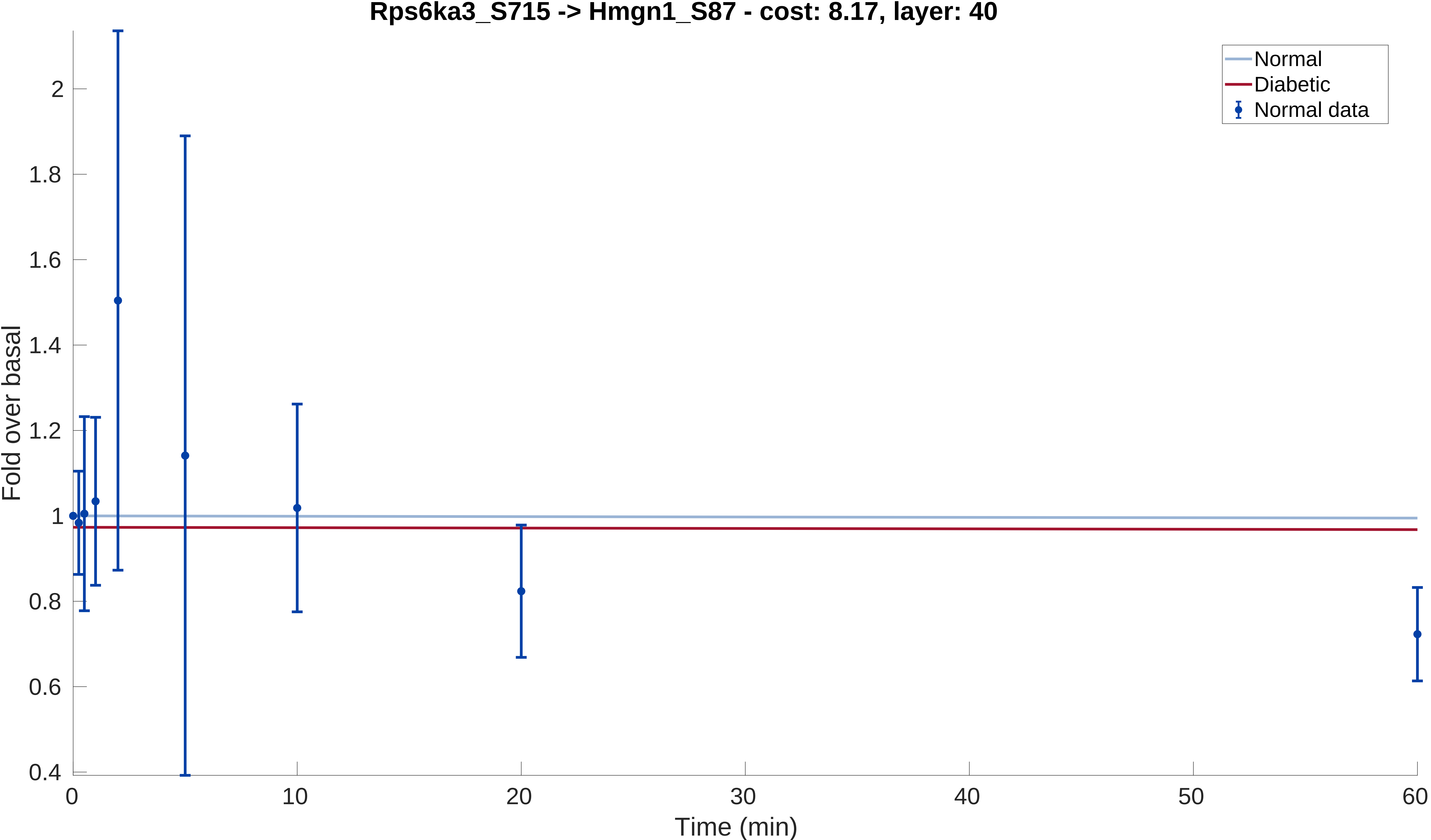

Rps6ka3_S715 -> Hmgn1_S87 - cost: 8.17, layer: 40
Normal
Diabetic
Normal data
Fold over basal
Time (min)
0.4
0.6
0.8
1
1.2
1.4
1.6
1.8
2
0
10
20
30
40
50
60

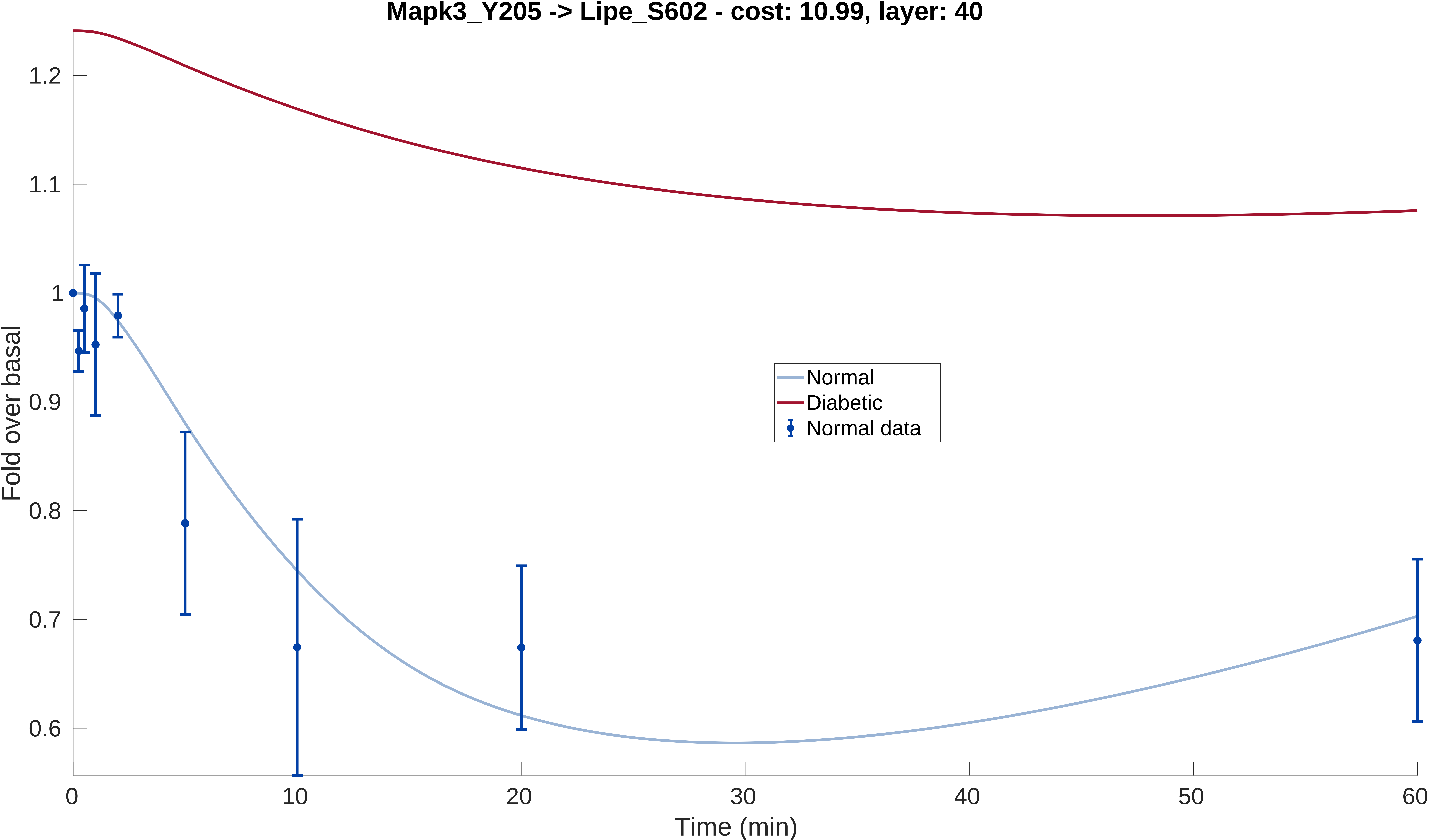

Mapk3_Y205 -> Lipe_S602 - cost: 10.99, layer: 40
Fold over basal
1.2
1.1
1
0.9
0.8
0.7
0.6
0
10
20
30
40
50
60
Time (min)
Normal
Diabetic
Normal data

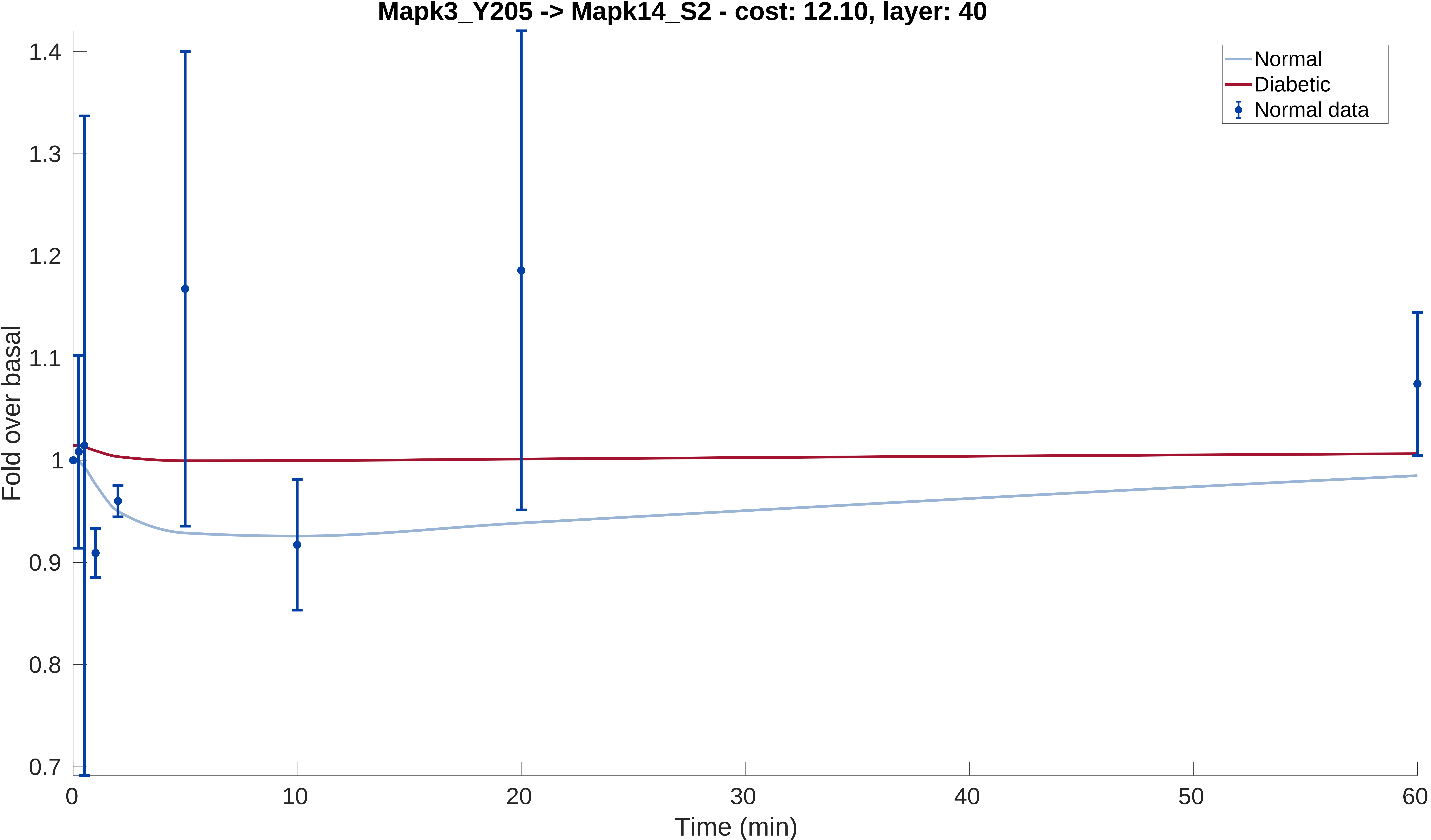

Mapk3_Y205 -> Mapk14_S2 - cost: 12.10, layer: 40
Normal
Diabetic
Normal data
Fold over basal
Time (min)
1.4
1.3
1.2
1.1
1
0.9
0.8
0.7
0
10
20
30
40
50
60

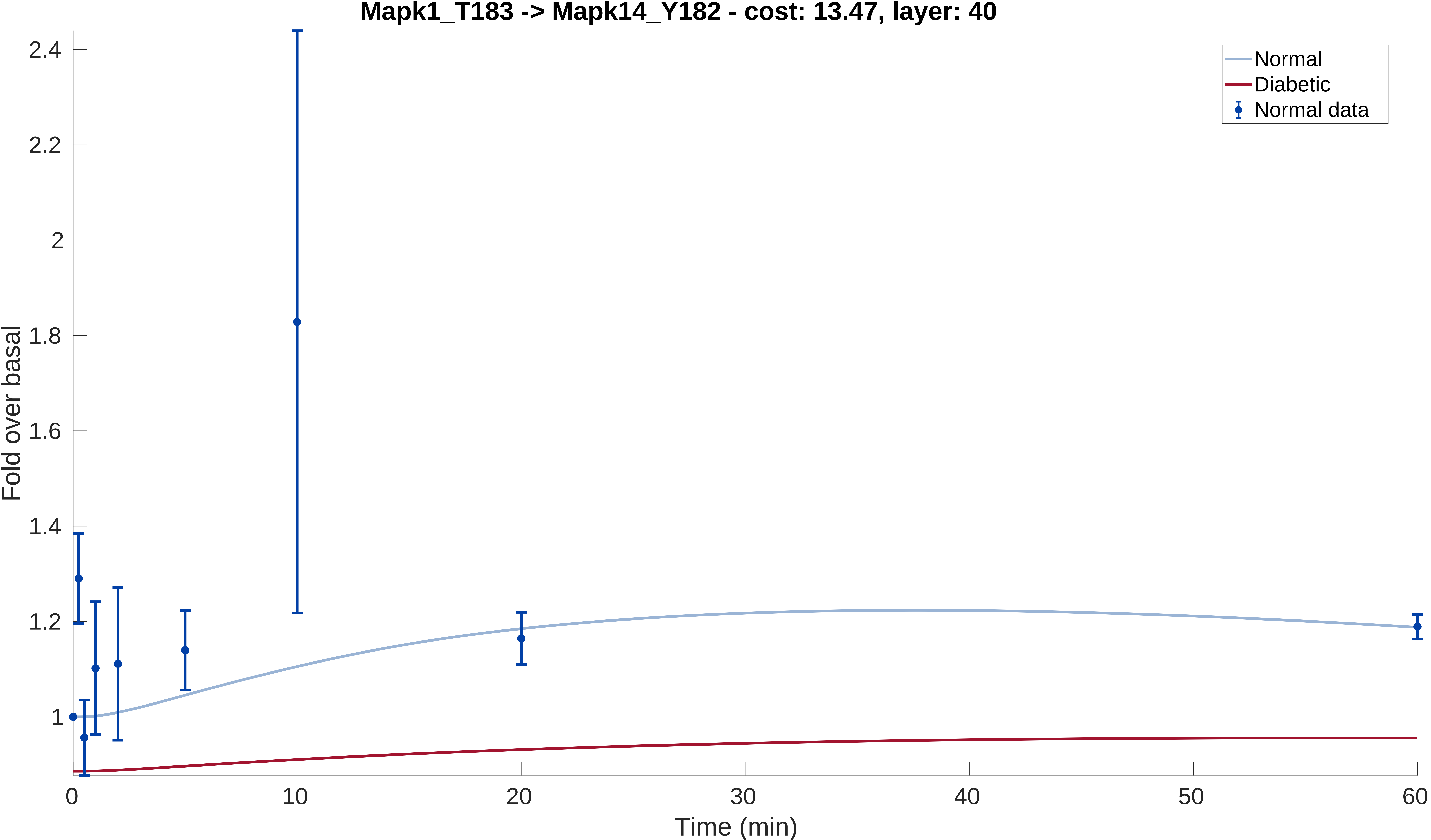
Mapk1_T183 -> Mapk14_Y182 - cost: 13.47, layer: 40
Fold over basal
Time (min)
Normal
Diabetic
Normal data
2.4
2.2
2
1.8
1.6
1.4
1.2
1
0
10
20
30
40
50
60

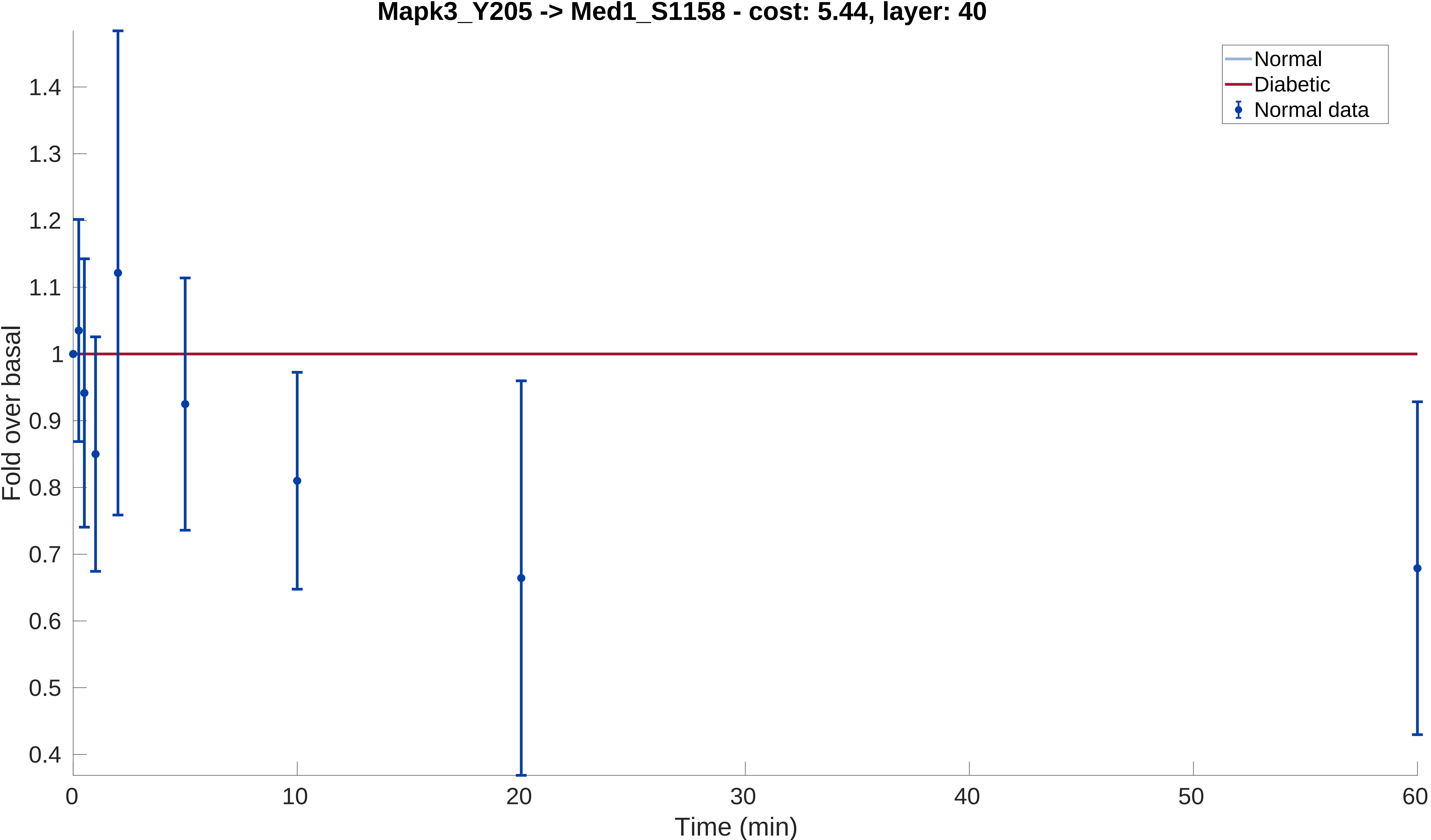

Mapk3_Y205 -> Med1_S1158 - cost: 5.44, layer: 40
Normal
Diabetic
Normal data
Fold over basal
1.4
1.3
1.2
1.1
1
0.9
0.8
0.7
0.6
0.5
0.4
0
10
20
30
40
50
60
Time (min)

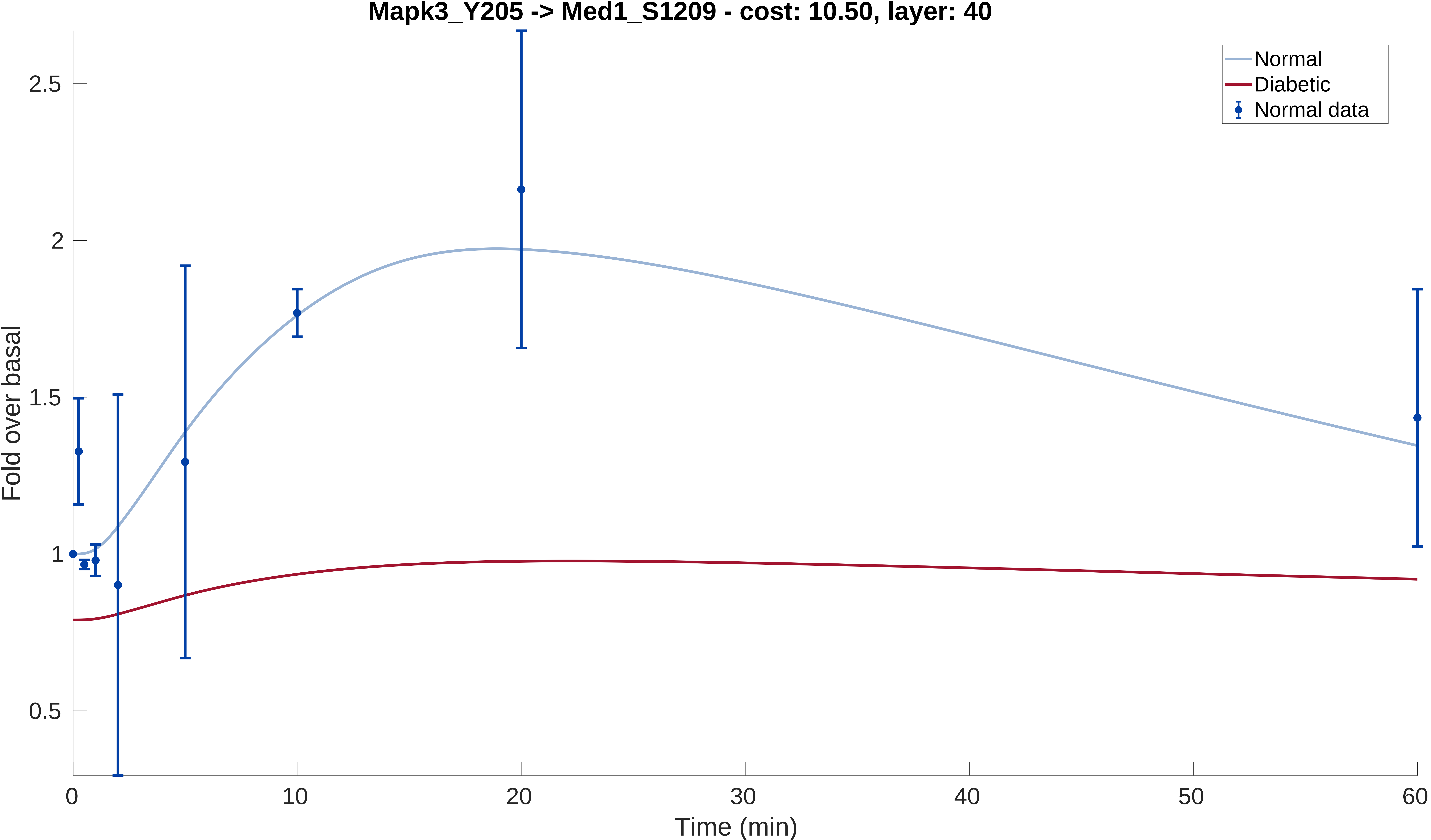

Mapk3_Y205 -> Med1_S1209 - cost: 10.50, layer: 40
Normal
Diabetic
Normal data
Fold over basal
2.5
2
1.5
1
0.5
0
10
20
30
40
50
60
Time (min)

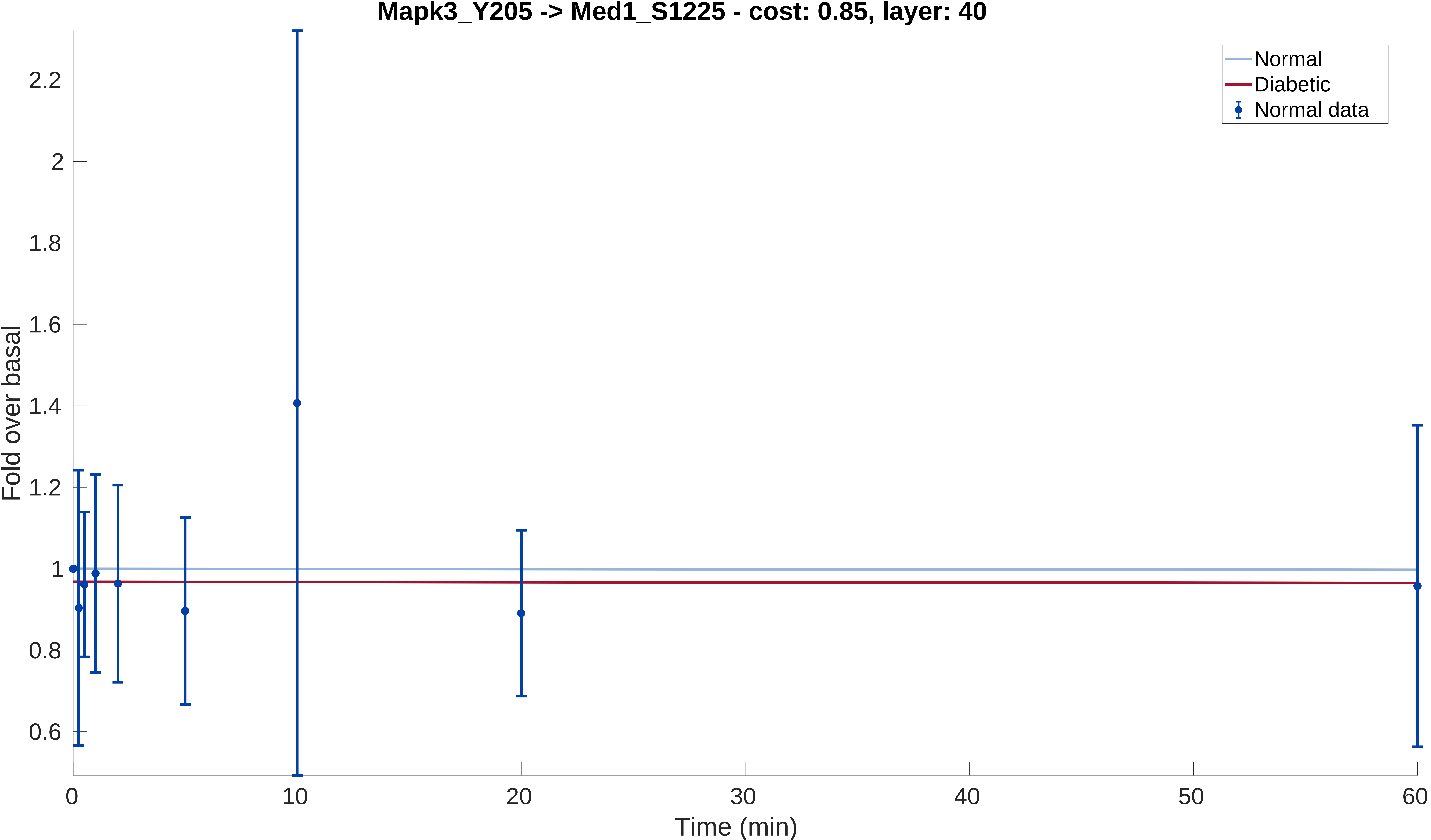

Mapk3_Y205 -> Med1_S1225 - cost: 0.85, layer: 40
Normal
Diabetic
Normal data
Fold over basal
2.2
2
1.8
1.6
1.4
1.2
1
0.8
0.6
0
10
20
30
40
50
60
Time (min)

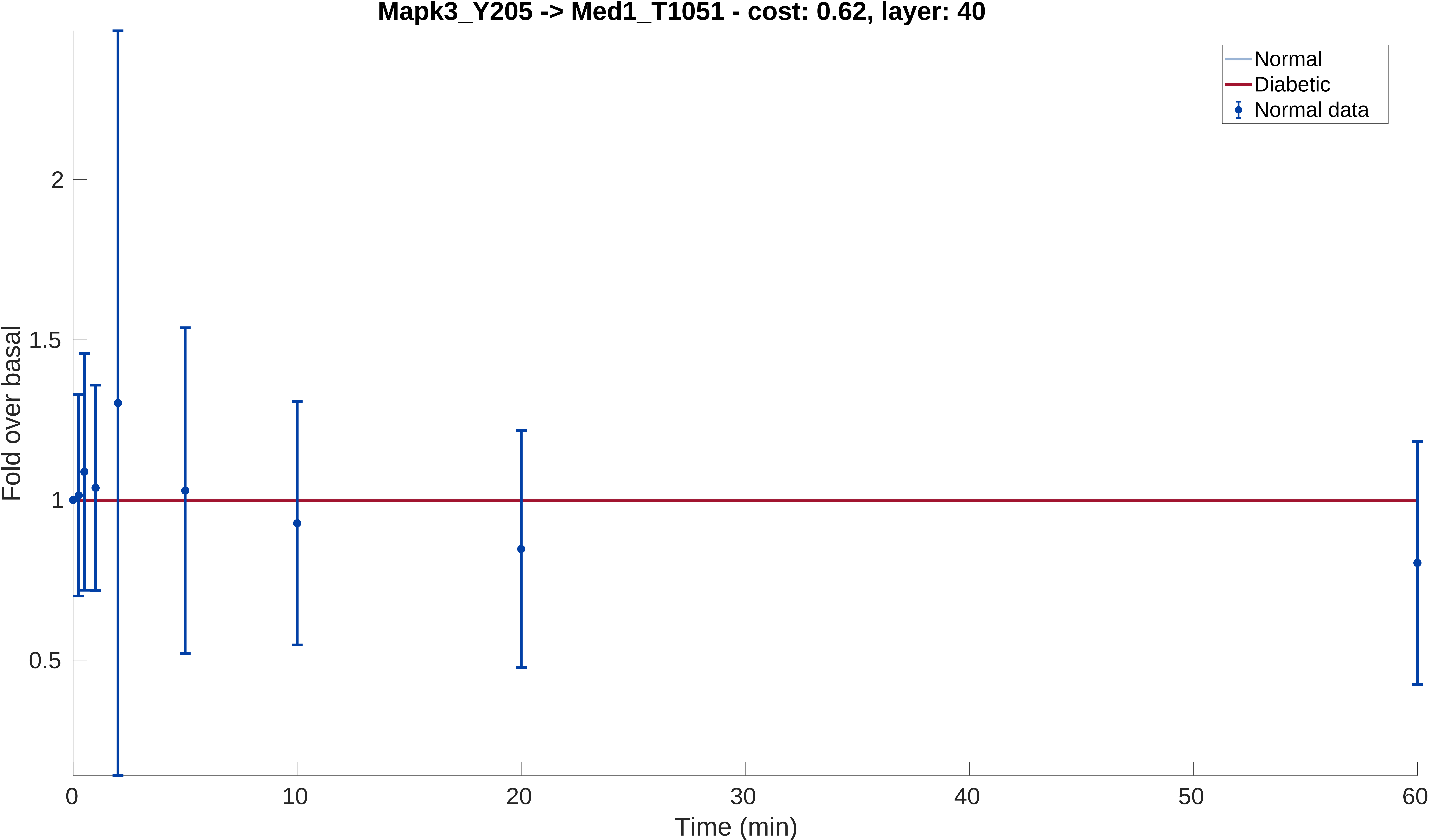

Mapk3_Y205 -> Med1_T1051 - cost: 0.62, layer: 40
Normal
Diabetic
Normal data
Fold over basal
2
1.5
1
0.5
0
10
20
30
40
50
60
Time (min)

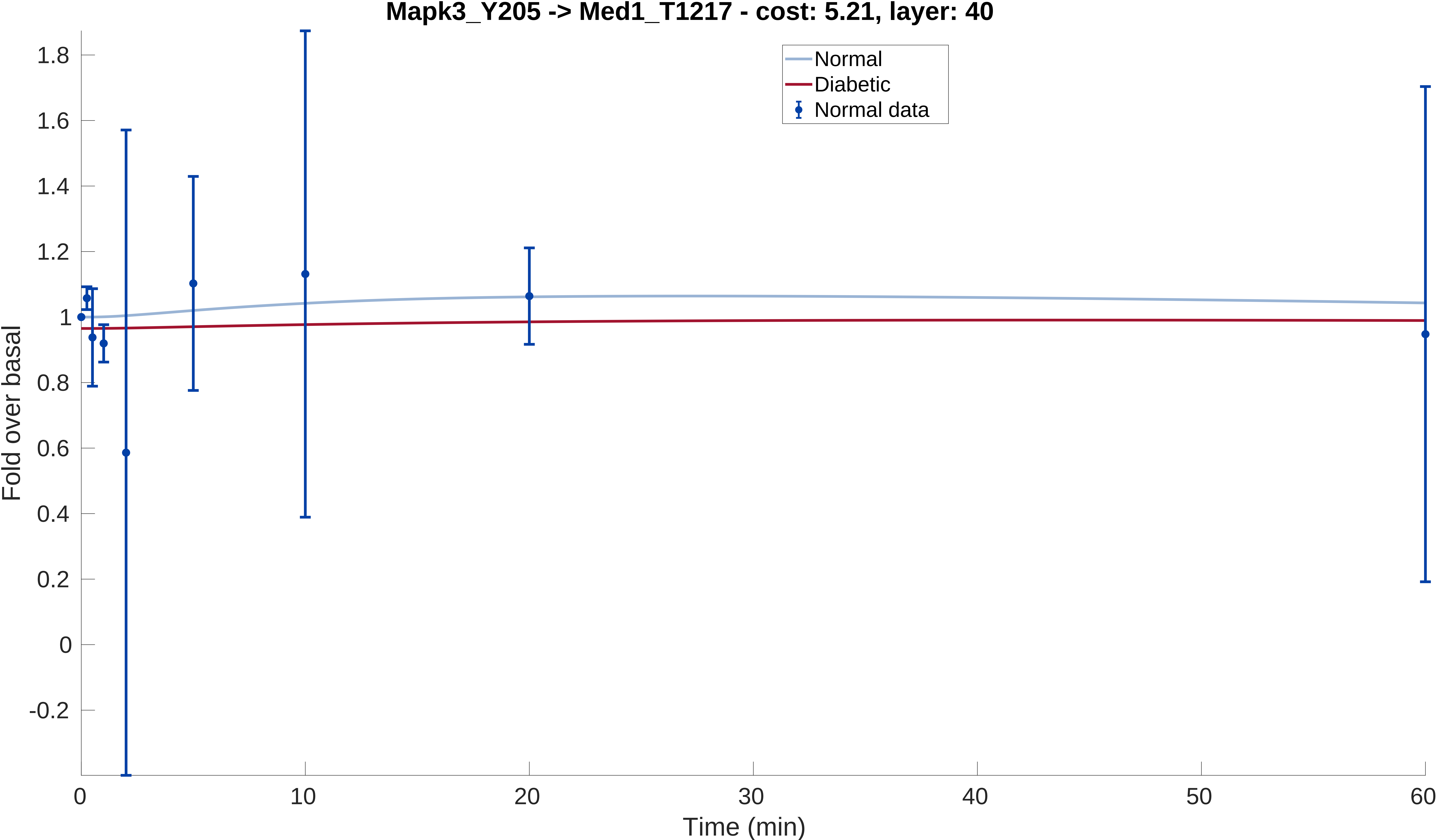

Mapk3_Y205 -> Med1_T1217 - cost: 5.21, layer: 40
Normal
Diabetic
Normal data
Fold over basal
Time (min)
1.8
1.6
1.4
1.2
1
0.8
0.6
0.4
0.2
0
-0.2
0
10
20
30
40
50
60

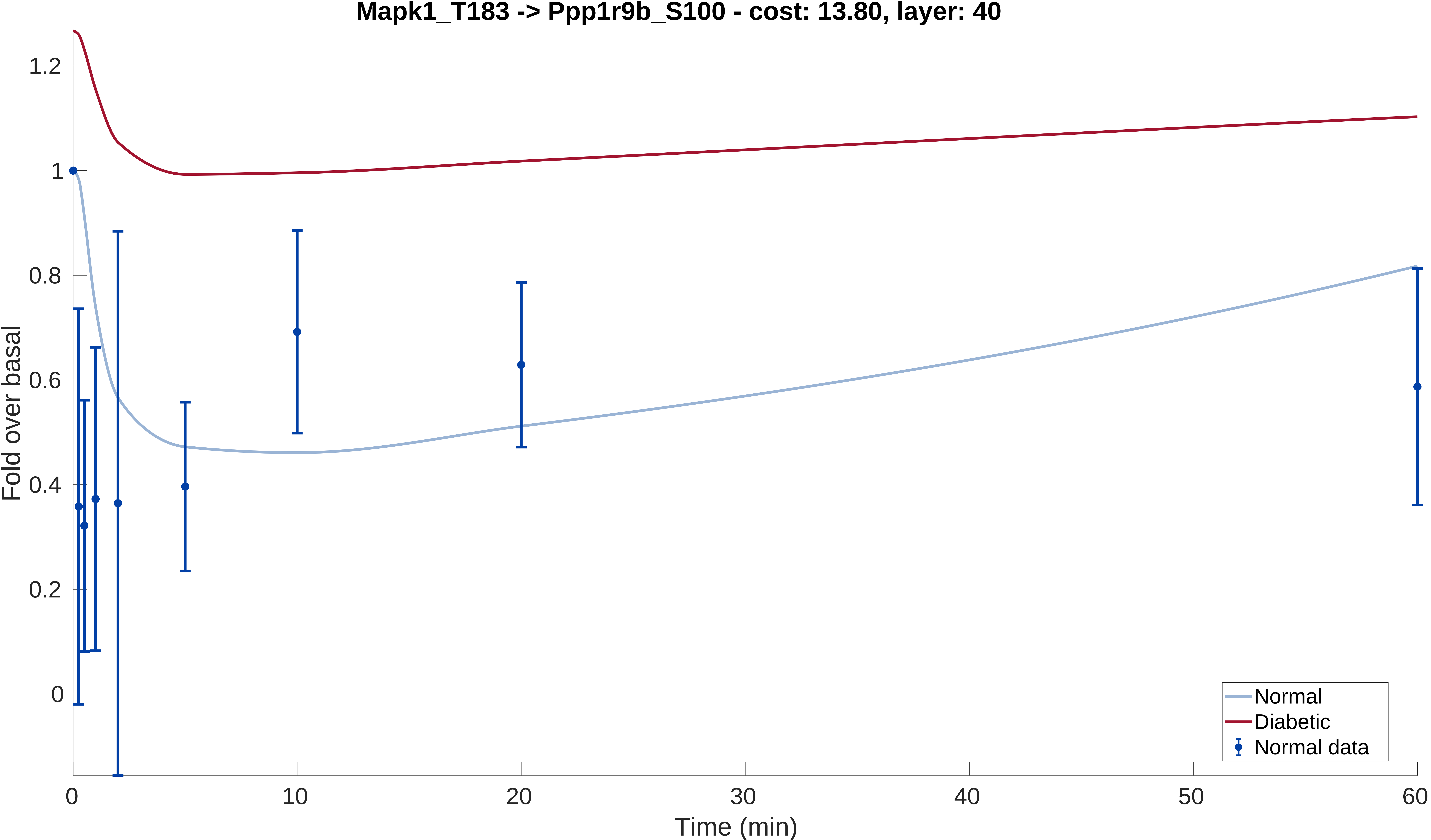
Mapk1_T183 -> Ppp1r9b_S100 - cost: 13.80, layer: 40
1.2
1
0.8
0.6
0.4
0.2
0
Fold over basal
0
10
20
30
40
50
60
Time (min)
Normal
Diabetic
Normal data

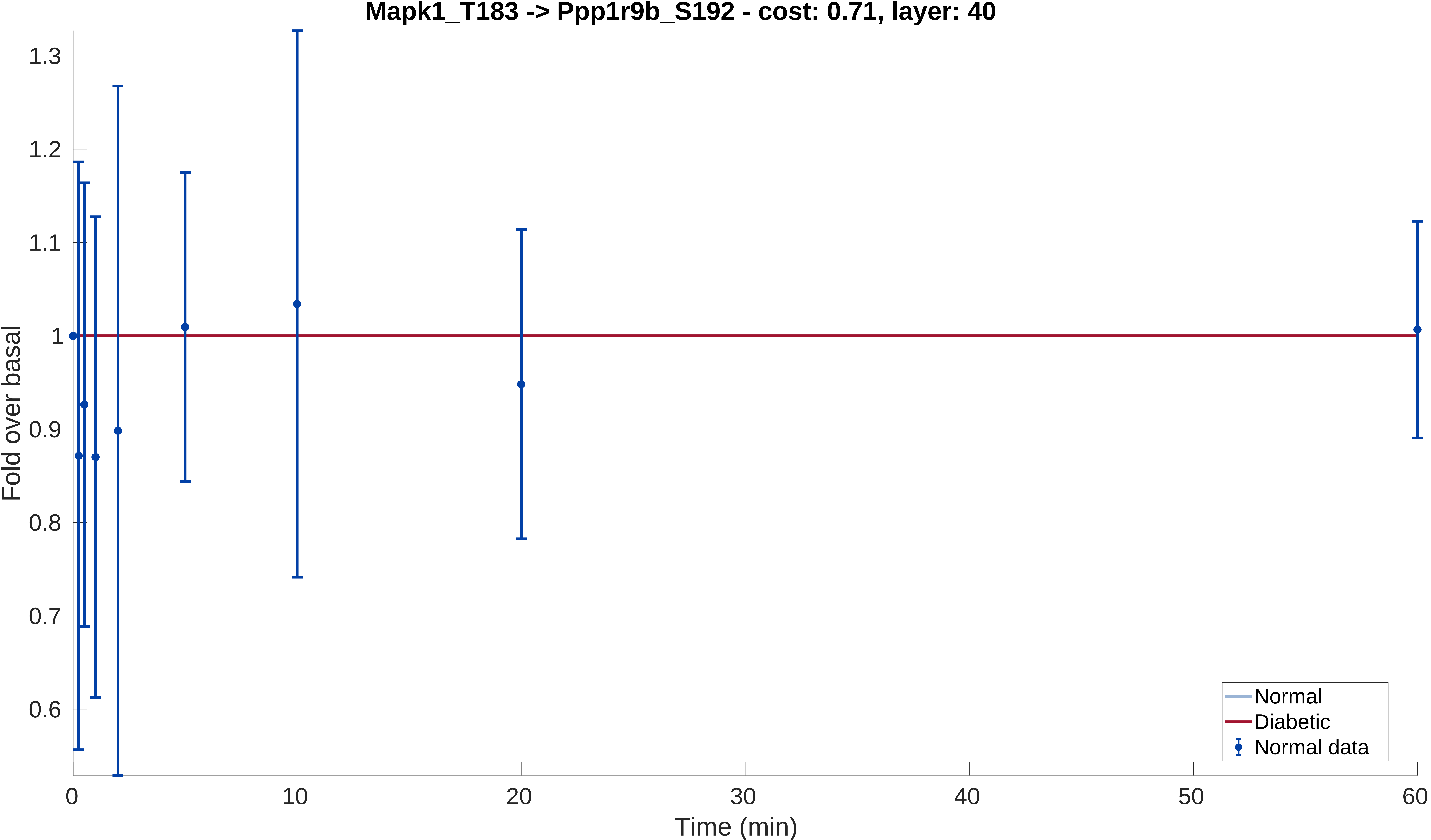

Mapk1_T183 -> Ppp1r9b_S192 - cost: 0.71, layer: 40
Fold over basal
1.3
1.2
1.1
1
0.9
0.8
0.7
0.6
0
10
20
30
40
50
60
Time (min)
Normal
Diabetic
Normal data

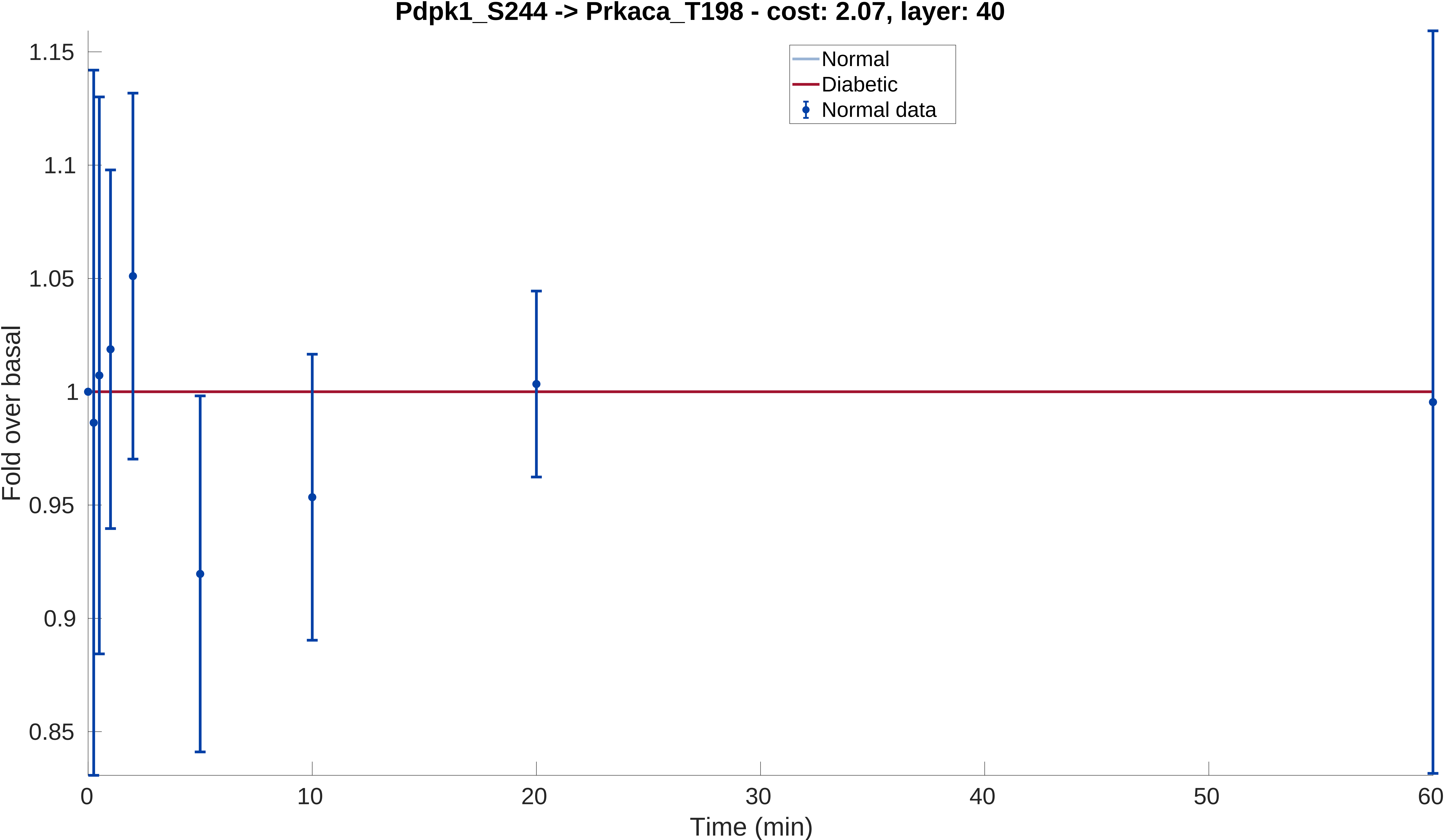

Pdpk1_S244 -> Prkaca_T198 - cost: 2.07, layer: 40
Normal
Diabetic
Normal data
Fold over basal
Time (min)
1.15
1.1
1.05
1
0.95
0.9
0.85
0
10
20
30
40
50
60

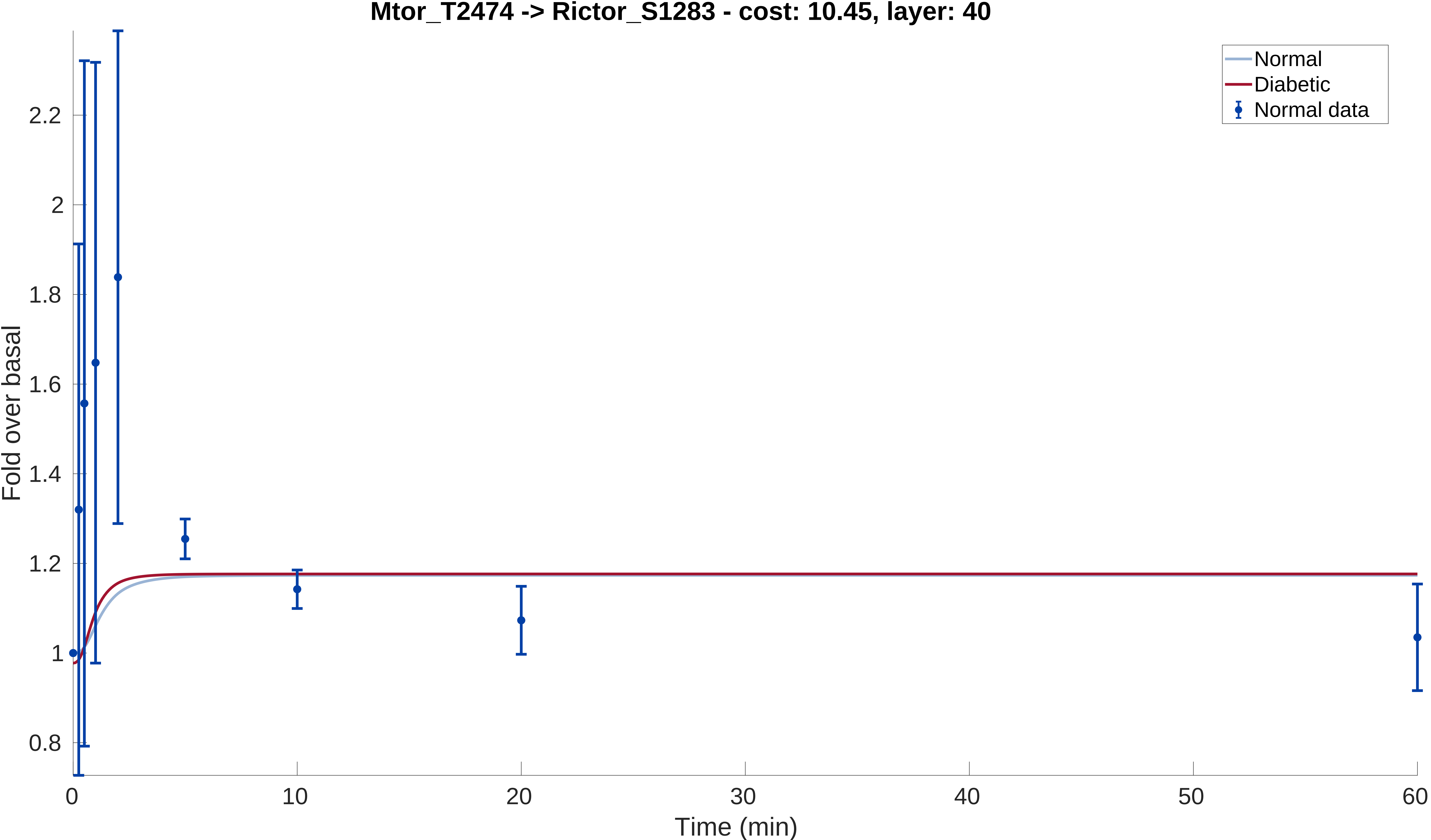

Mtor_T2474 -> Rictor_S1283 - cost: 10.45, layer: 40
Normal
Diabetic
Normal data
Fold over basal
2.2
2
1.8
1.6
1.4
1.2
1
0.8
0
10
20
30
40
50
60
Time (min)

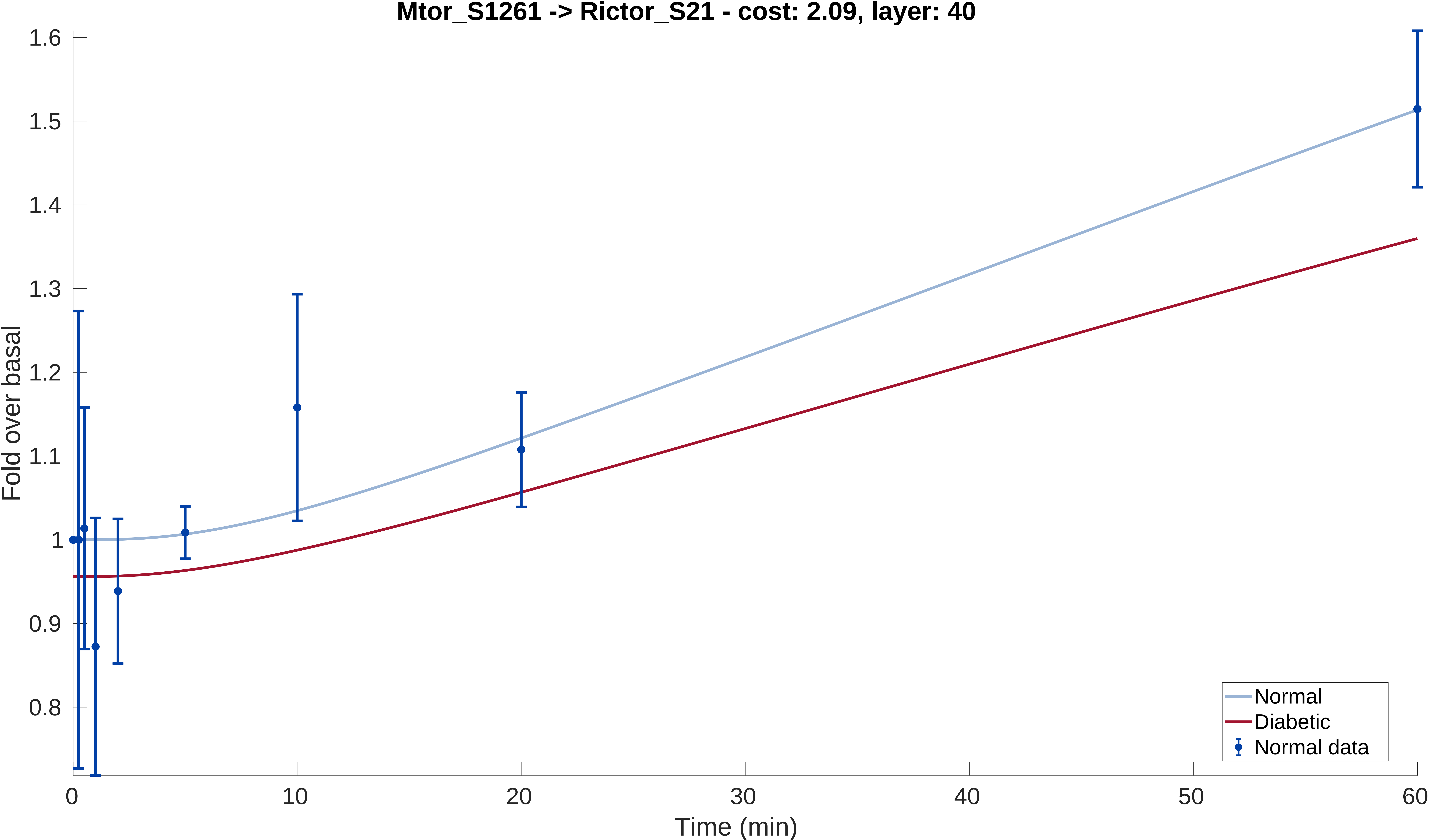
Mtor_S1261 -> Rictor_S21 - cost: 2.09, layer: 40
Fold over basal
1.6
1.5
1.4
1.3
1.2
1.1
1
0.9
0.8
0
10
20
30
40
50
60
Time (min)
Normal
Diabetic
Normal data

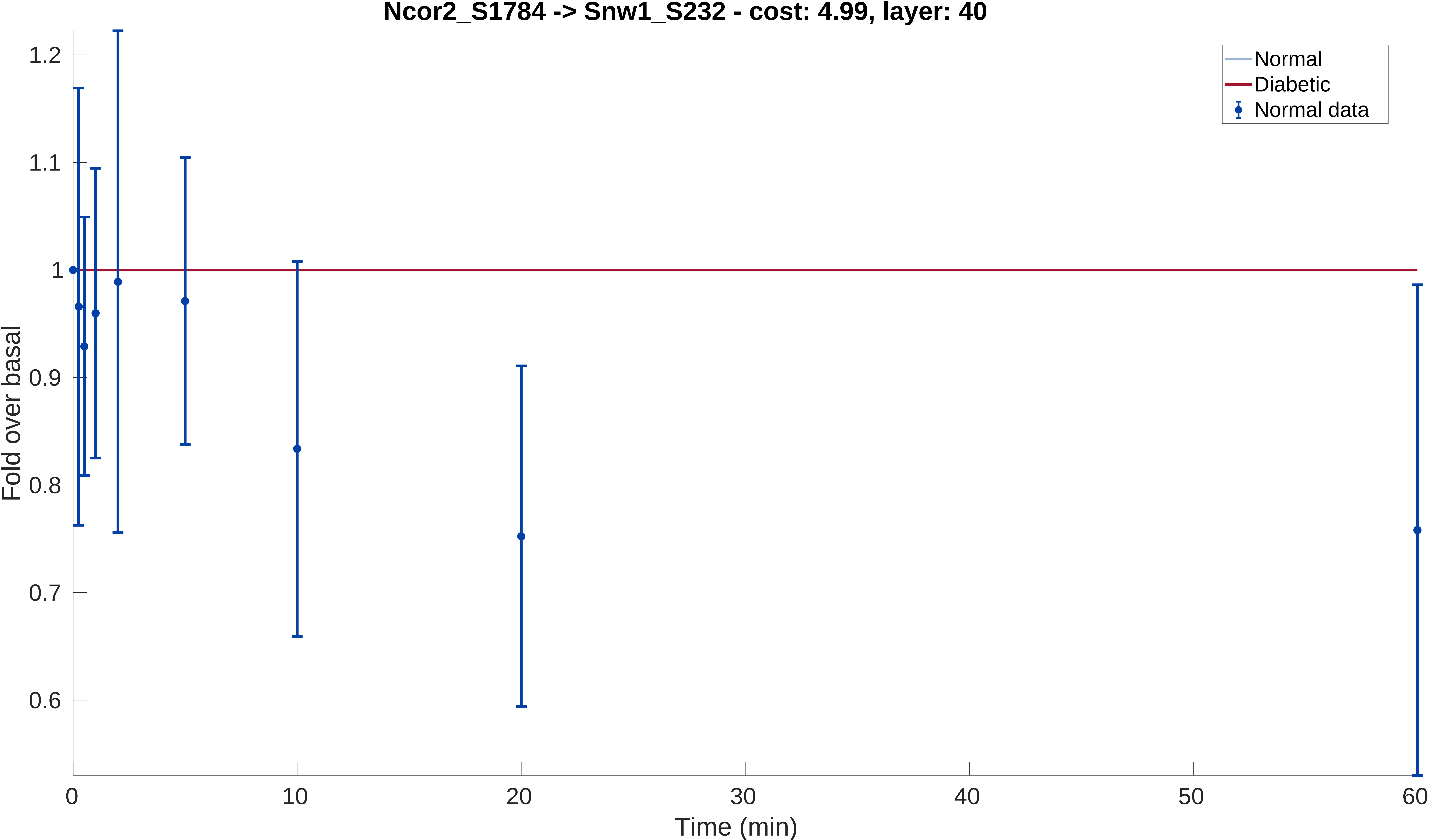

Ncor2_S1784 -> Snw1_S232 - cost: 4.99, layer: 40
Normal
Diabetic
Normal data
Fold over basal
1.2
1.1
1
0.9
0.8
0.7
0.6
0
10
20
30
40
50
60
Time (min)

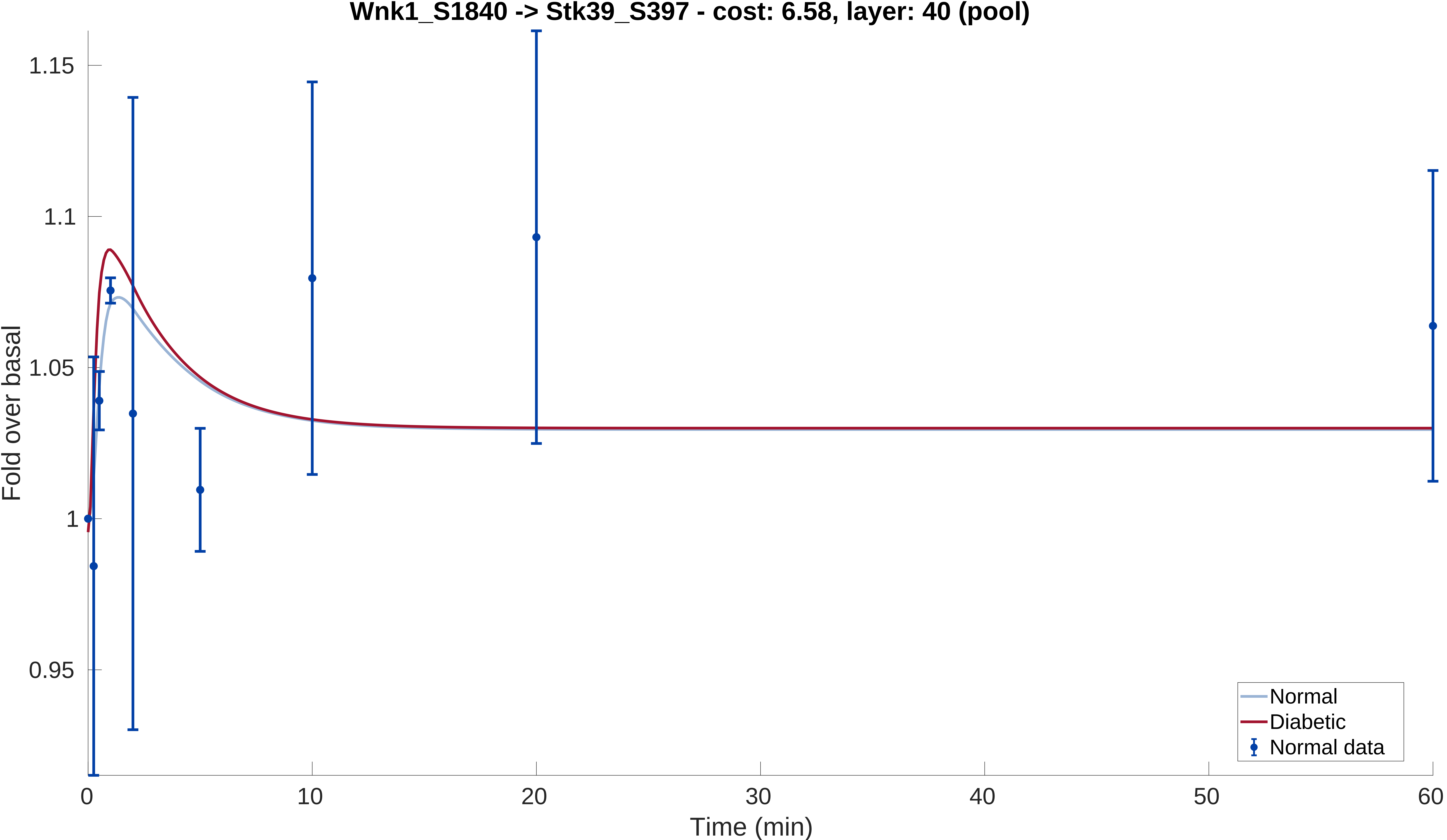

Wnk1_S1840 -> Stk39_S397 - cost: 6.58, layer: 40 (pool)
1.15
1.1
1.05
1
0.95
Fold over basal
0
10
20
30
40
50
60
Time (min)
Normal
Diabetic
Normal data

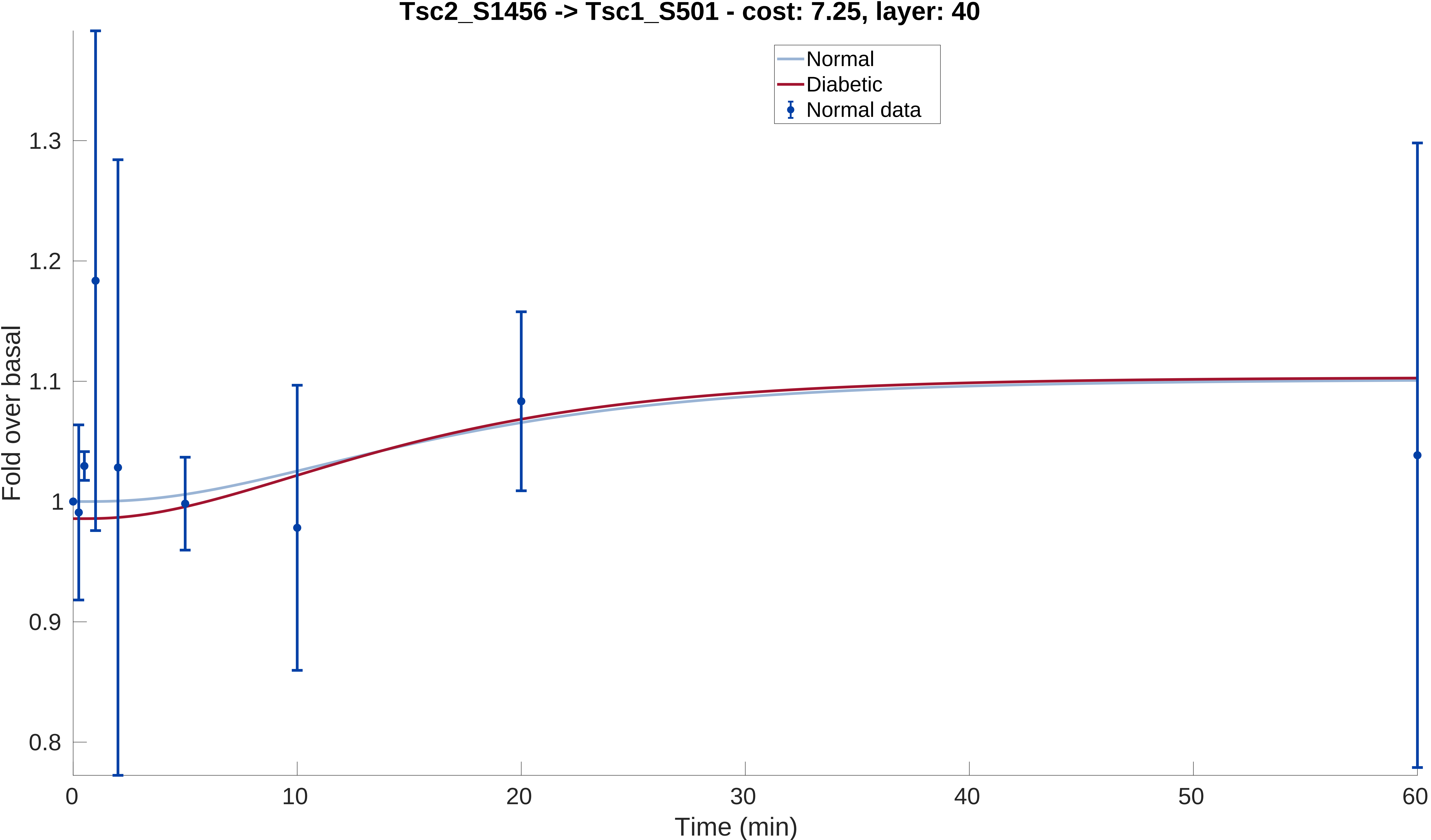

Tsc2_S1456 -> Tsc1_S501 - cost: 7.25, layer: 40
Normal
Diabetic
Normal data
Fold over basal
1.3
1.2
1.1
1
0.9
0.8
0
10
20
30
40
50
60
Time (min)

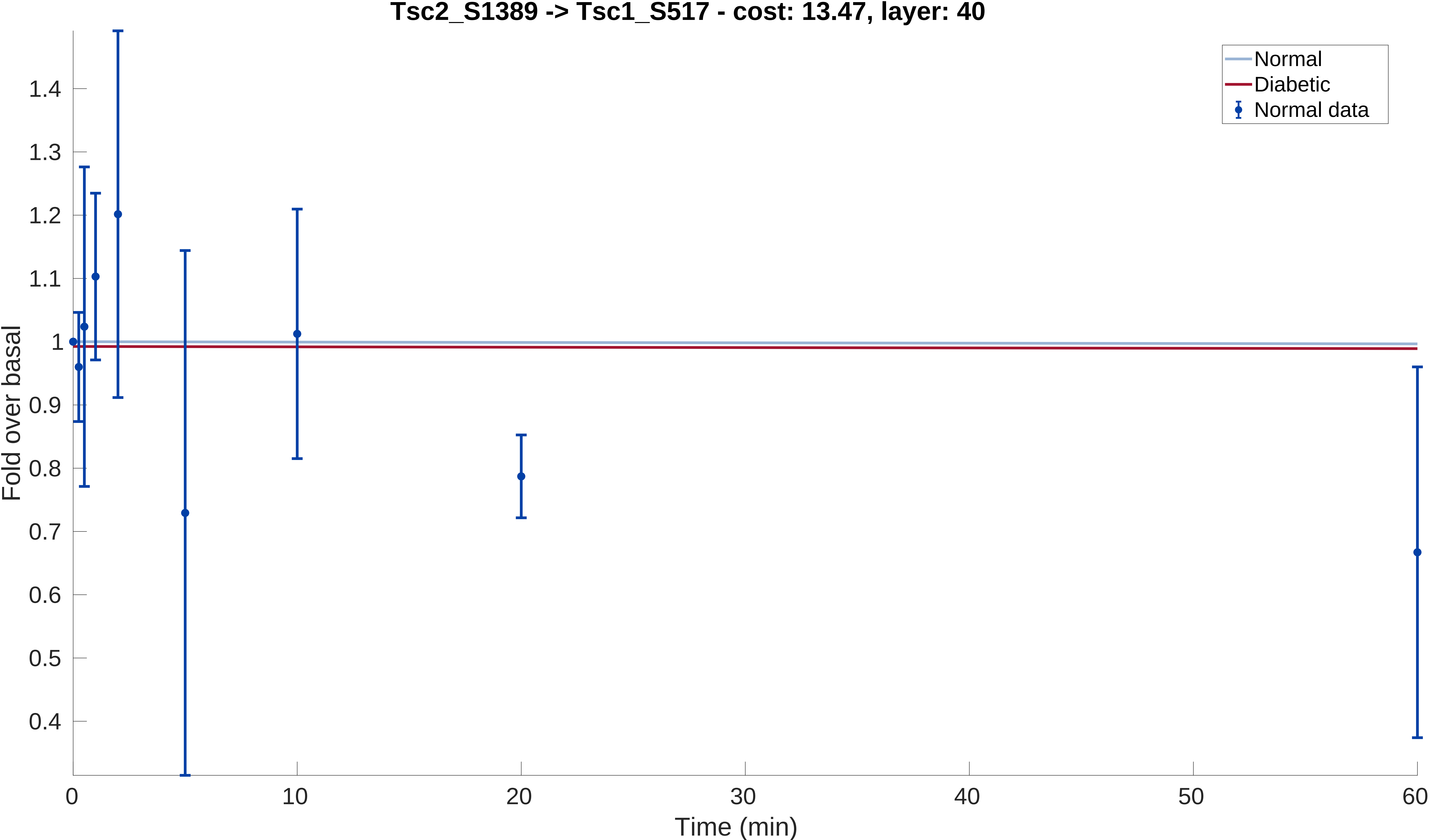
Tsc2_S1389 -> Tsc1_S517 - cost: 13.47, layer: 40
Normal
Diabetic
Normal data
Fold over basal
1.4
1.3
1.2
1.1
1
0.9
0.8
0.7
0.6
0.5
0.4
0
10
20
30
40
50
60
Time (min)

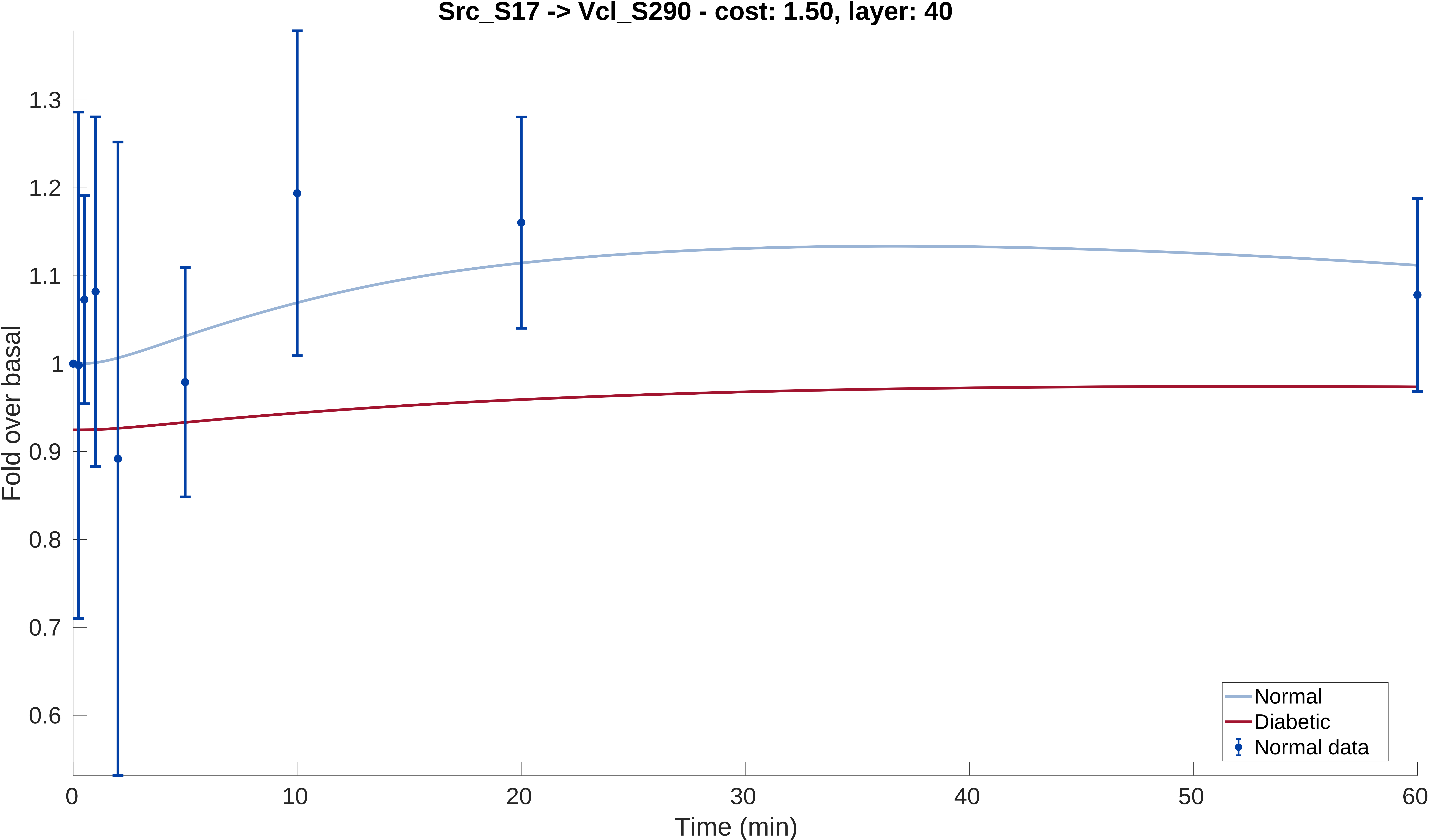
Src_S17 -> Vcl_S290 - cost: 1.50, layer: 40
Fold over basal
1.3
1.2
1.1
1
0.9
0.8
0.7
0.6
0
10
20
30
40
50
60
Time (min)
Normal
Diabetic
Normal data

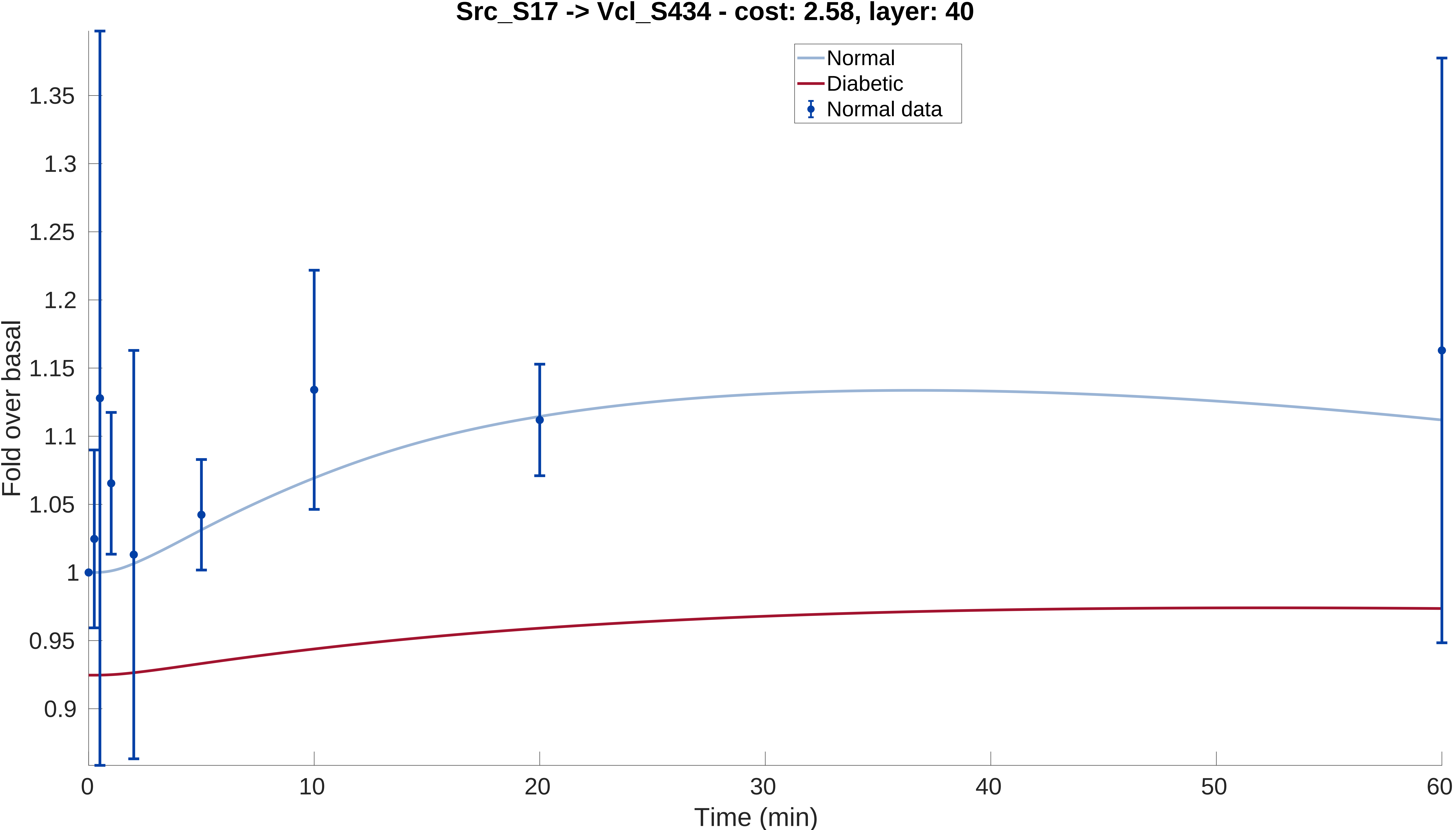

Src_S17 -> Vcl_S434 - cost: 2.58, layer: 40
Normal
Diabetic
Normal data
Fold over basal
Time (min)
1.35
1.3
1.25
1.2
1.15
1.1
1.05
1
0.95
0.9
0
10
20
30
40
50
60

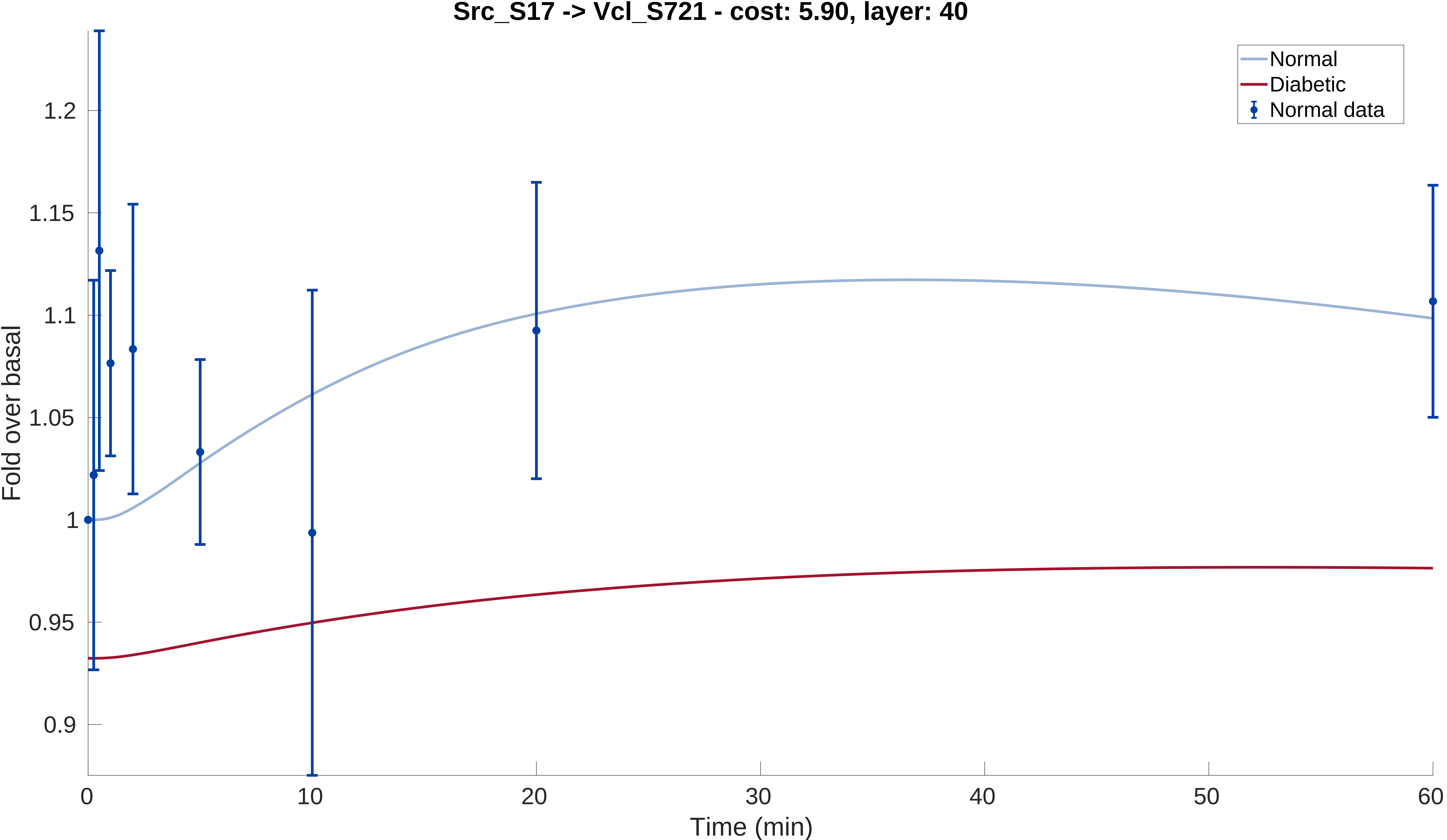

Src_S17 -> Vcl_S721 - cost: 5.90, layer: 40
Normal
Diabetic
Normal data
Fold over basal
Time (min)
1.2
1.15
1.1
1.05
1
0.95
0.9
0
10
20
30
40
50
60

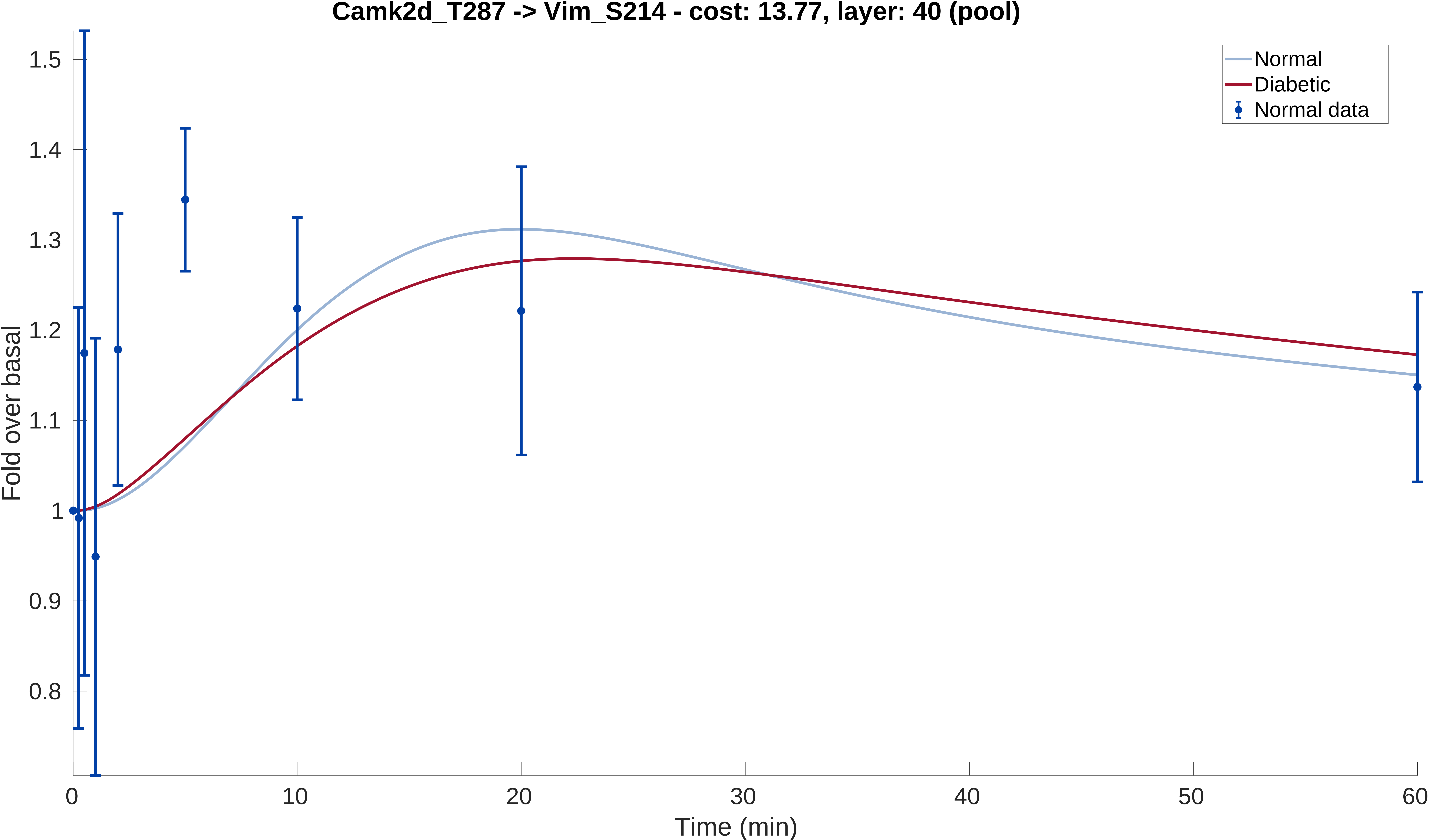

Camk2d_T287 -> Vim_S214 - cost: 13.77, layer: 40 (pool)
Normal
Diabetic
Normal data
Fold over basal
1.5
1.4
1.3
1.2
1.1
1
0.9
0.8
0
10
20
30
40
50
60
Time (min)

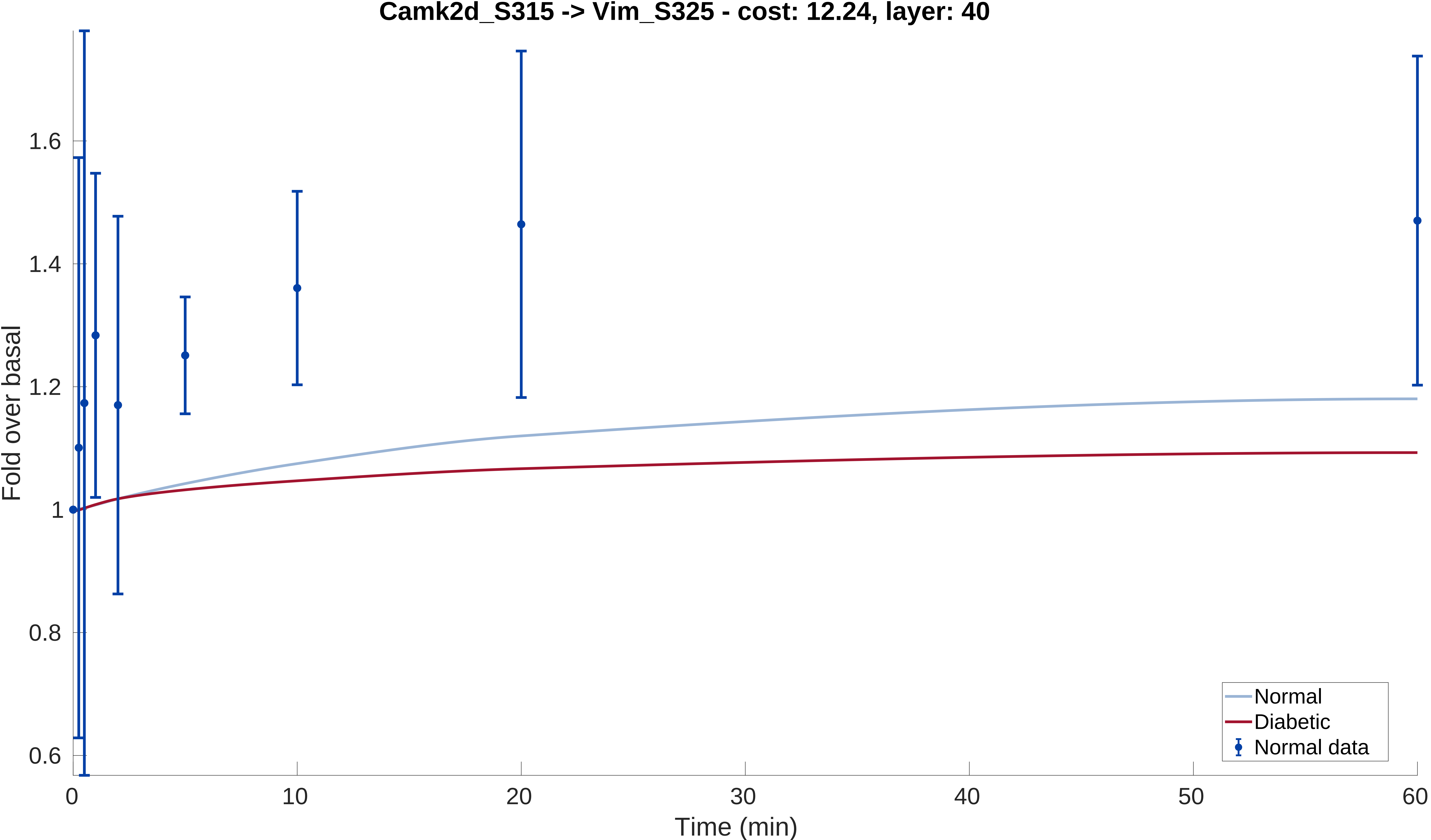

Camk2d_S315 -> Vim_S325 - cost: 12.24, layer: 40
Fold over basal
Time (min)
0.6
0.8
1
1.2
1.4
1.6
0
10
20
30
40
50
60
Normal
Diabetic
Normal data

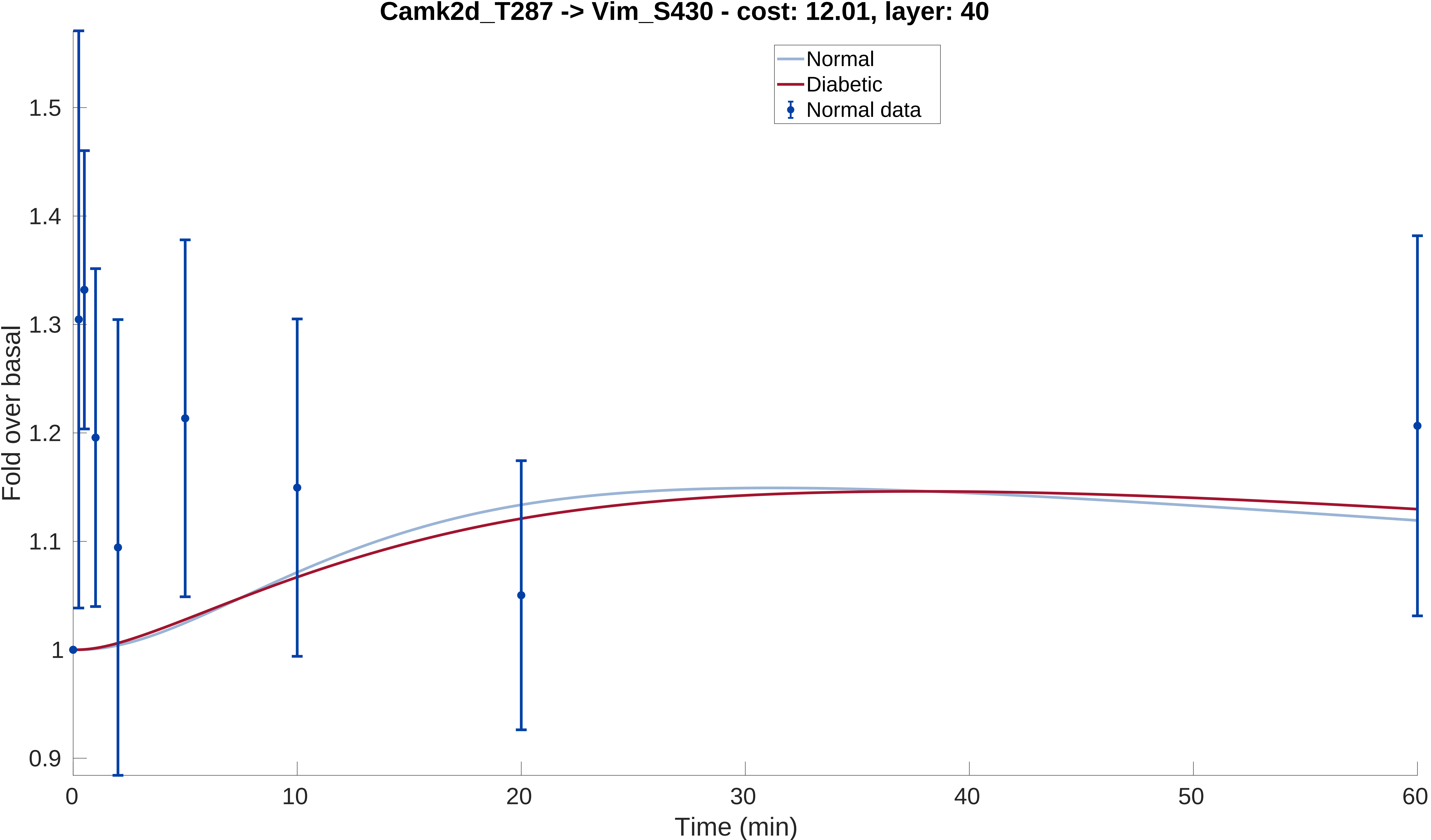
Camk2d_T287 -> Vim_S430 - cost: 12.01, layer: 40
Normal
Diabetic
Normal data
Fold over basal
1.5
1.4
1.3
1.2
1.1
1
0.9
0
10
20
30
40
50
60
Time (min)

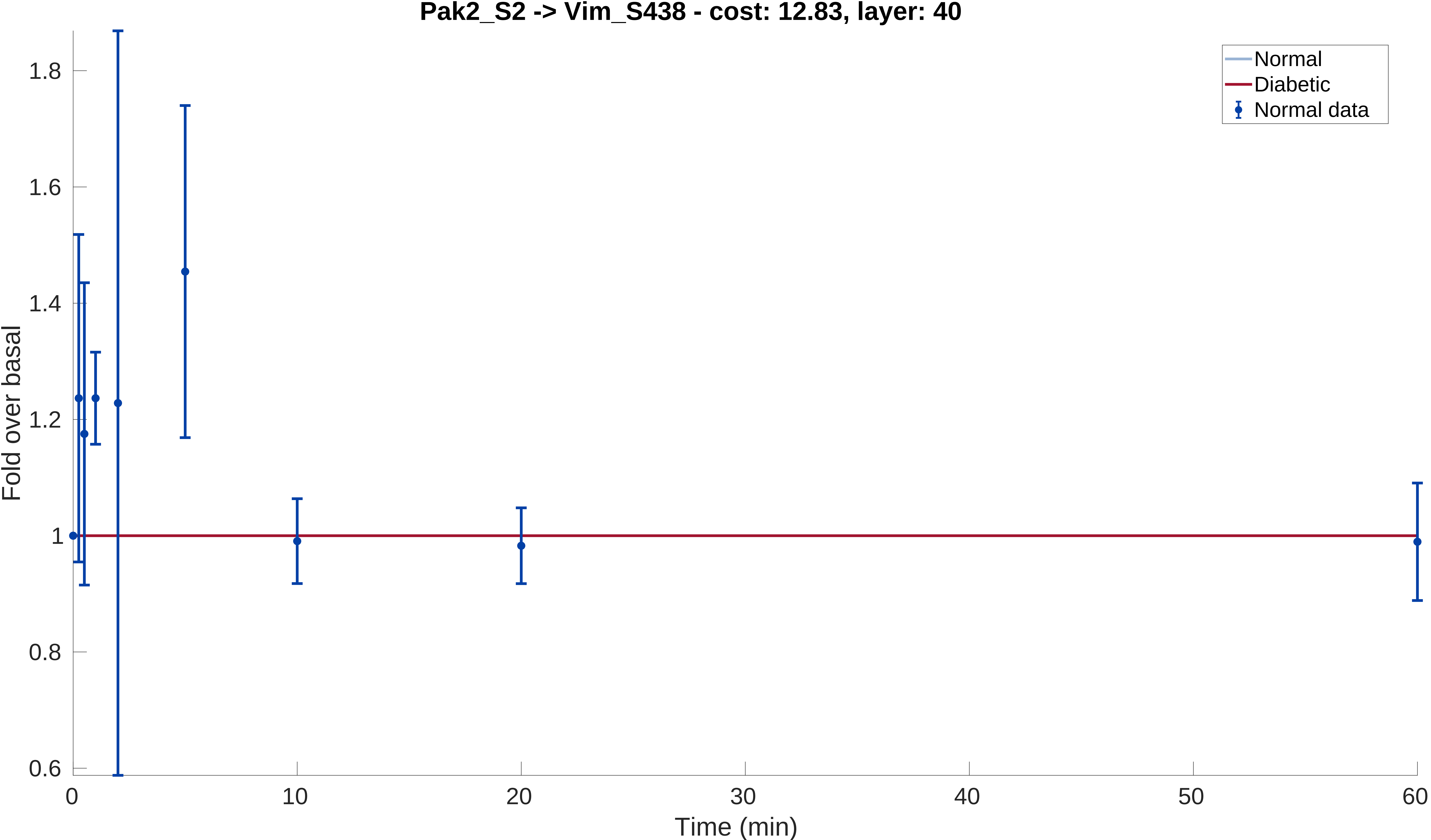
Pak2_S2 -> Vim_S438 - cost: 12.83, layer: 40
Normal
Diabetic
Normal data
Fold over basal
1.8
1.6
1.4
1.2
1
0.8
0.6
0
10
20
30
40
50
60
Time (min)

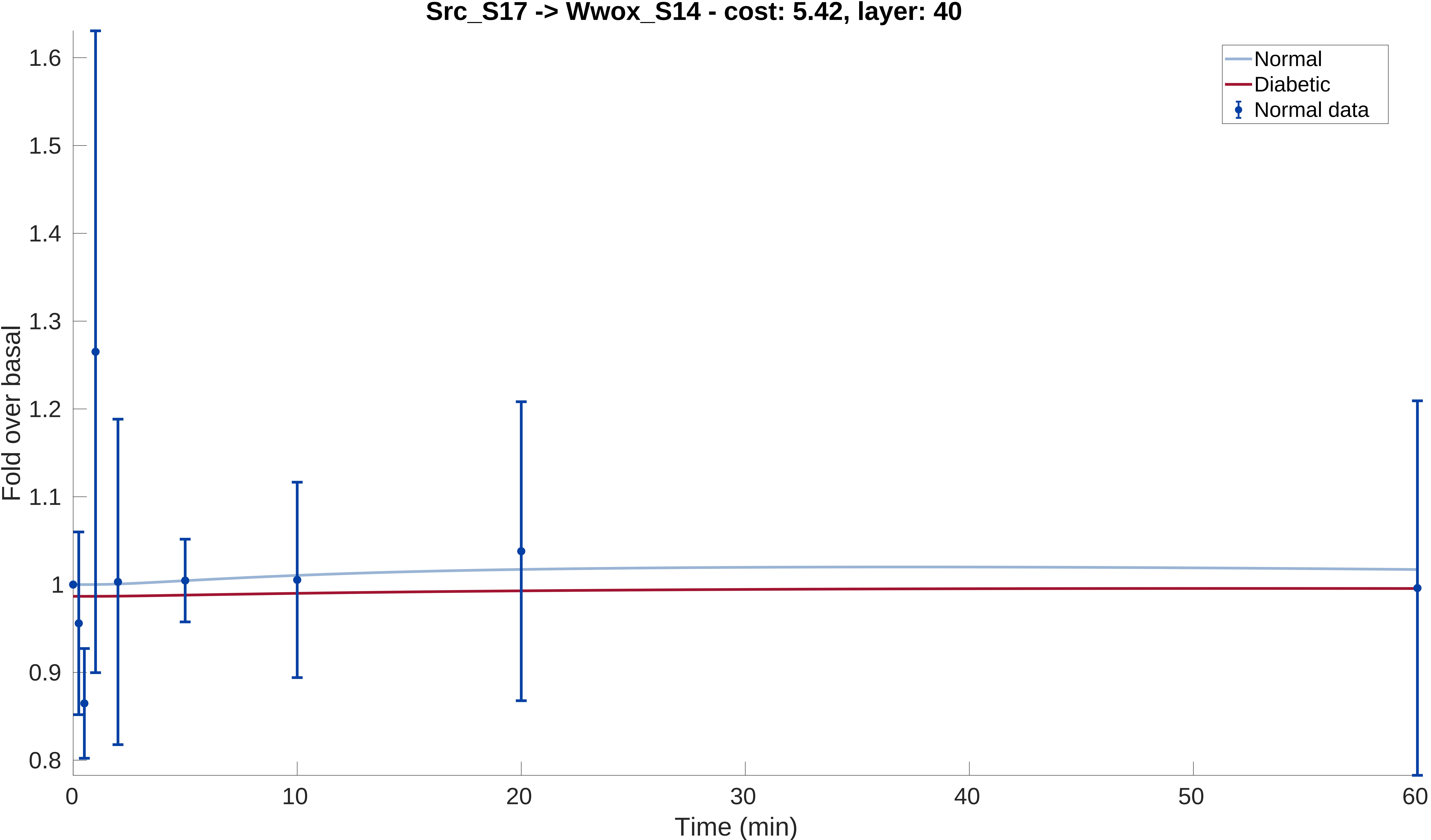

Src_S17 -> Wwox_S14 - cost: 5.42, layer: 40
Normal
Diabetic
Normal data
Fold over basal
1.6
1.5
1.4
1.3
1.2
1.1
1
0.9
0.8
0
10
20
30
40
50
60
Time (min)

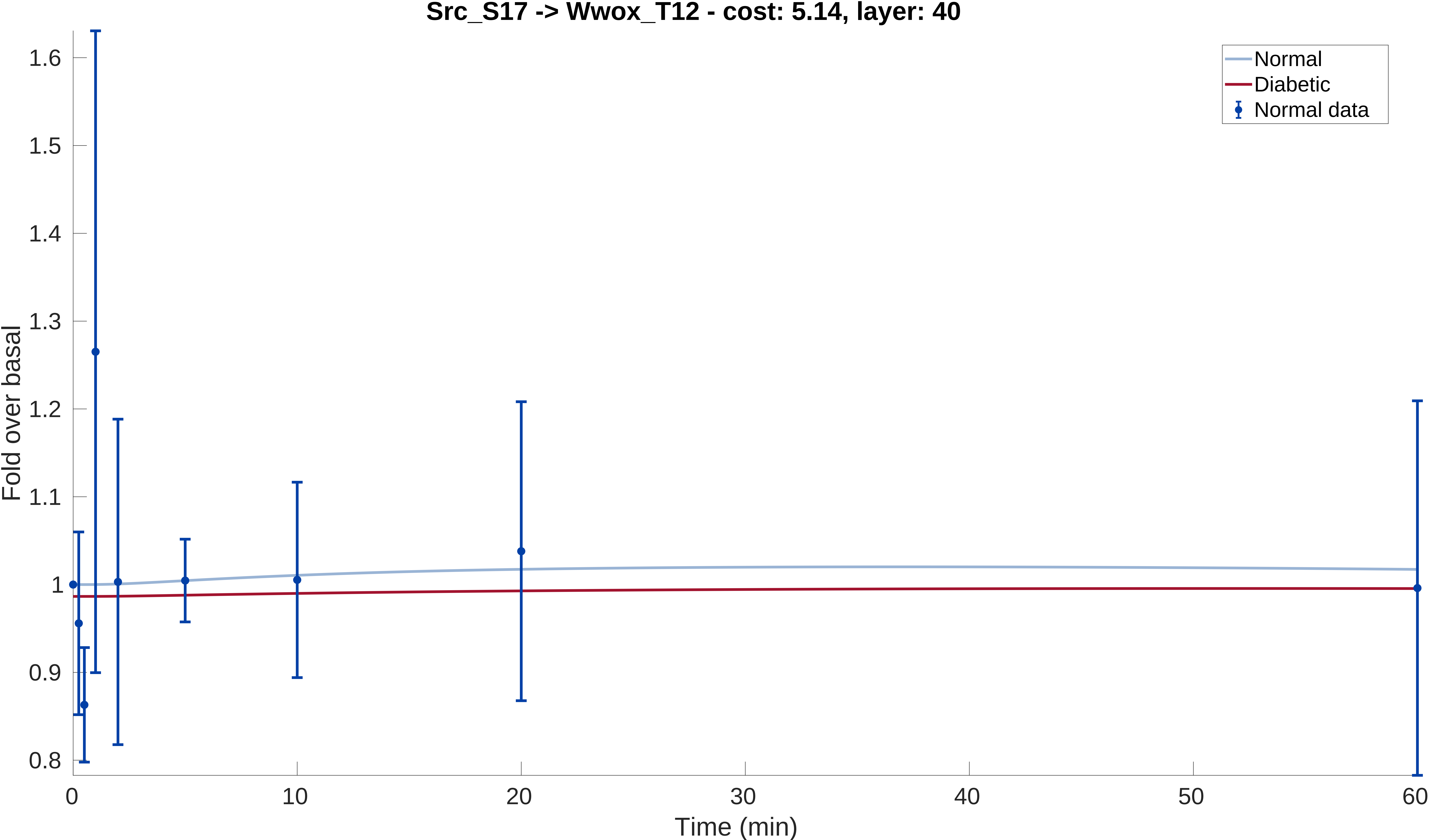
Src_S17 -> Wwox_T12 - cost: 5.14, layer: 40
Normal
Diabetic
Normal data
Fold over basal
1.6
1.5
1.4
1.3
1.2
1.1
1
0.9
0.8
0
10
20
30
40
50
60
Time (min)

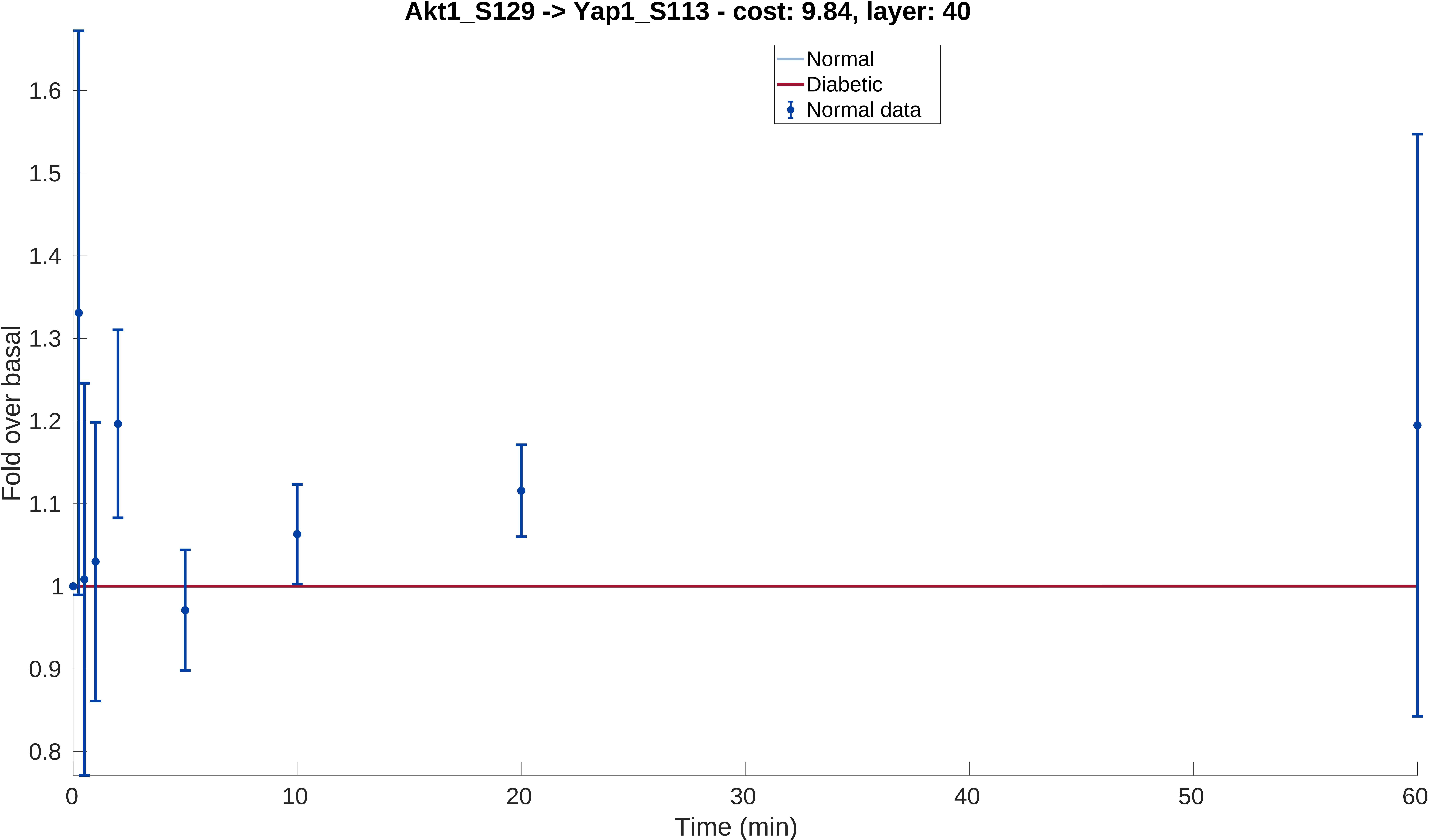
Akt1_S129 -> Yap1_S113 - cost: 9.84, layer: 40
Normal
Diabetic
Normal data
Fold over basal
1.6
1.5
1.4
1.3
1.2
1.1
1
0.9
0.8
0
10
20
30
40
50
60
Time (min)

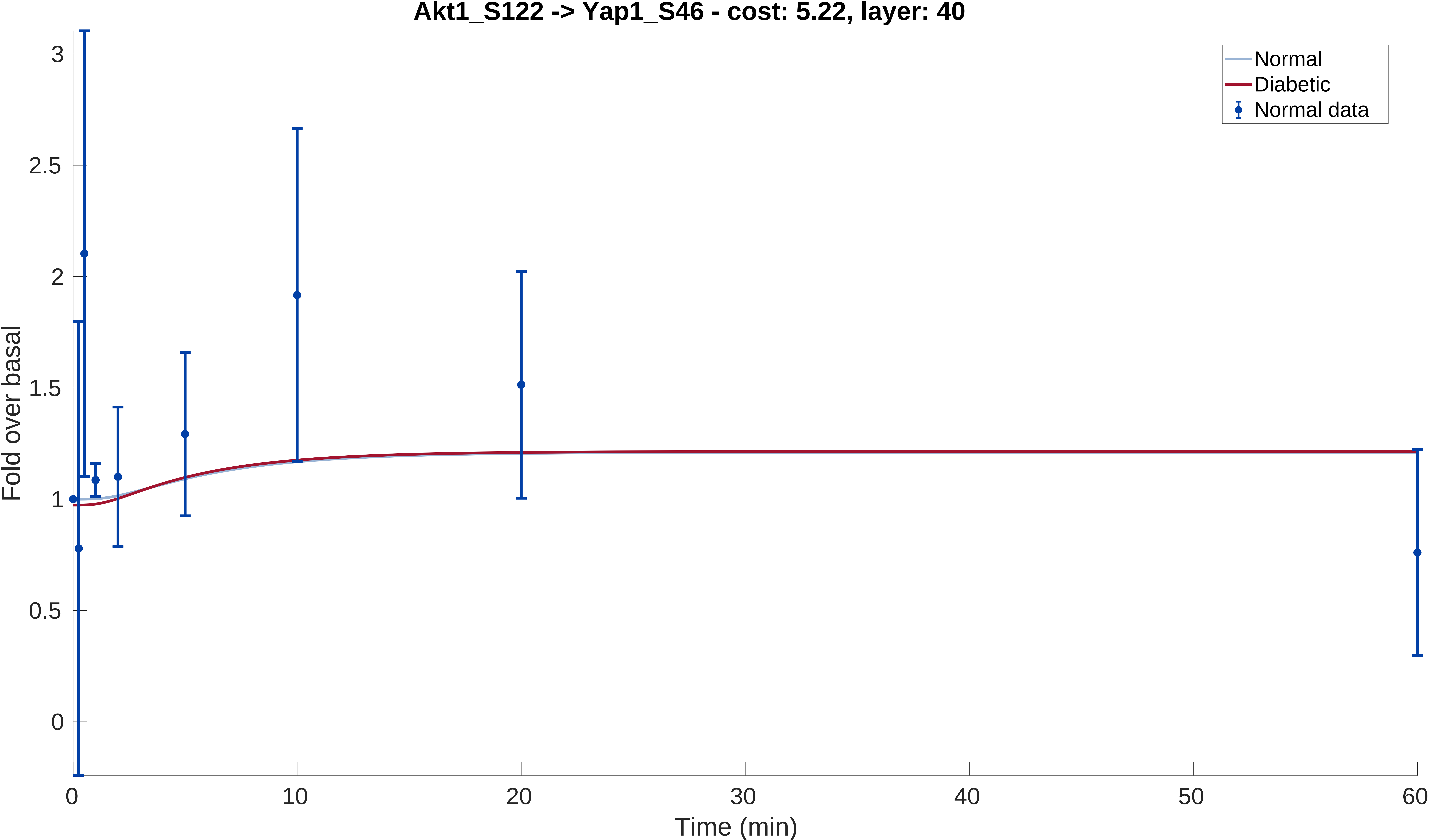

Akt1_S122 -> Yap1_S46 - cost: 5.22, layer: 40
Normal
Diabetic
Normal data
Fold over basal
Time (min)
0
0.5
1
1.5
2
2.5
3
0
10
20
30
40
50
60

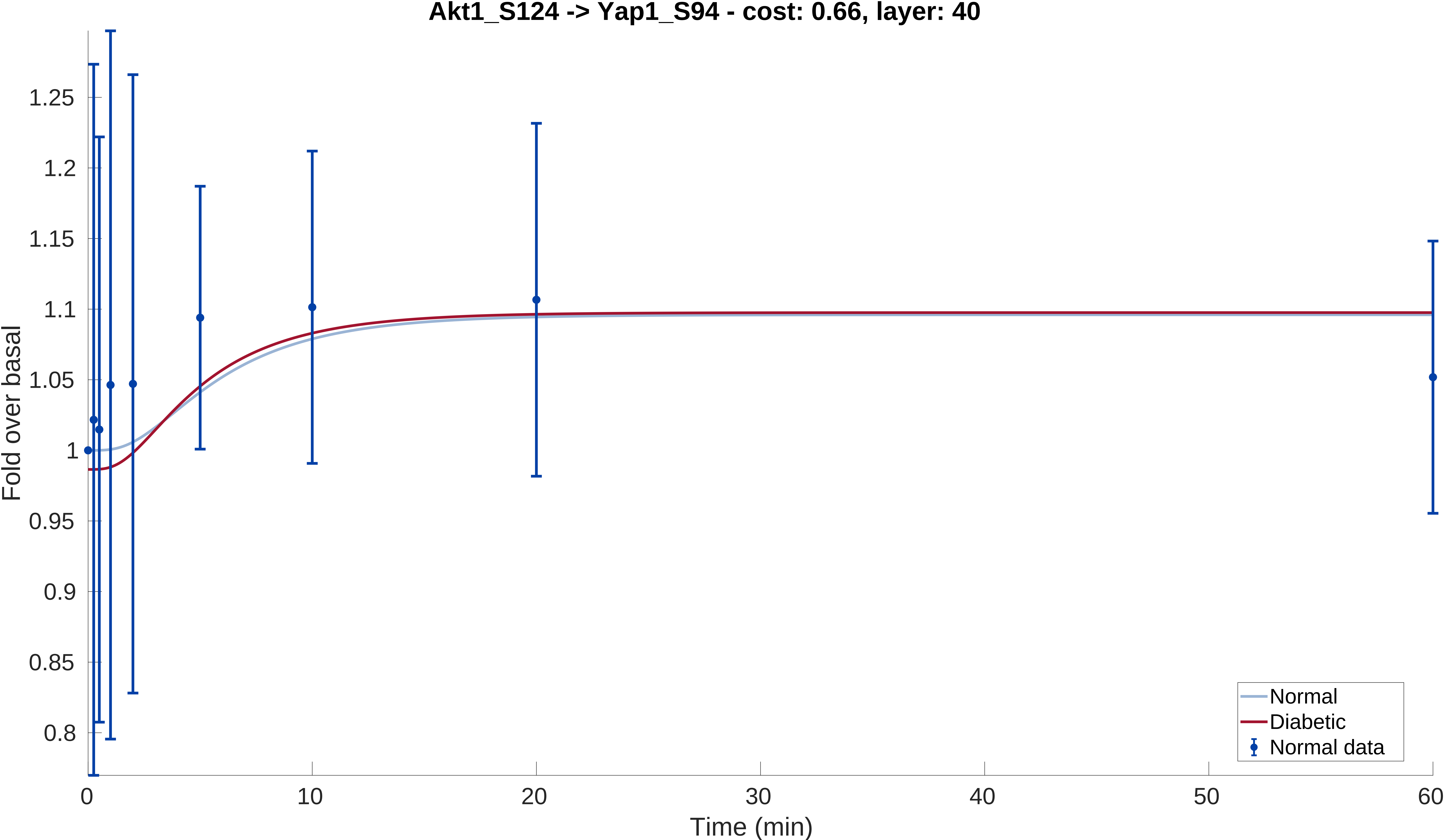

Akt1_S124 -> Yap1_S94 - cost: 0.66, layer: 40
Fold over basal
1.25
1.2
1.15
1.1
1.05
1
0.95
0.9
0.85
0.8
0
10
20
30
40
50
60
Time (min)
Normal
Diabetic
Normal data

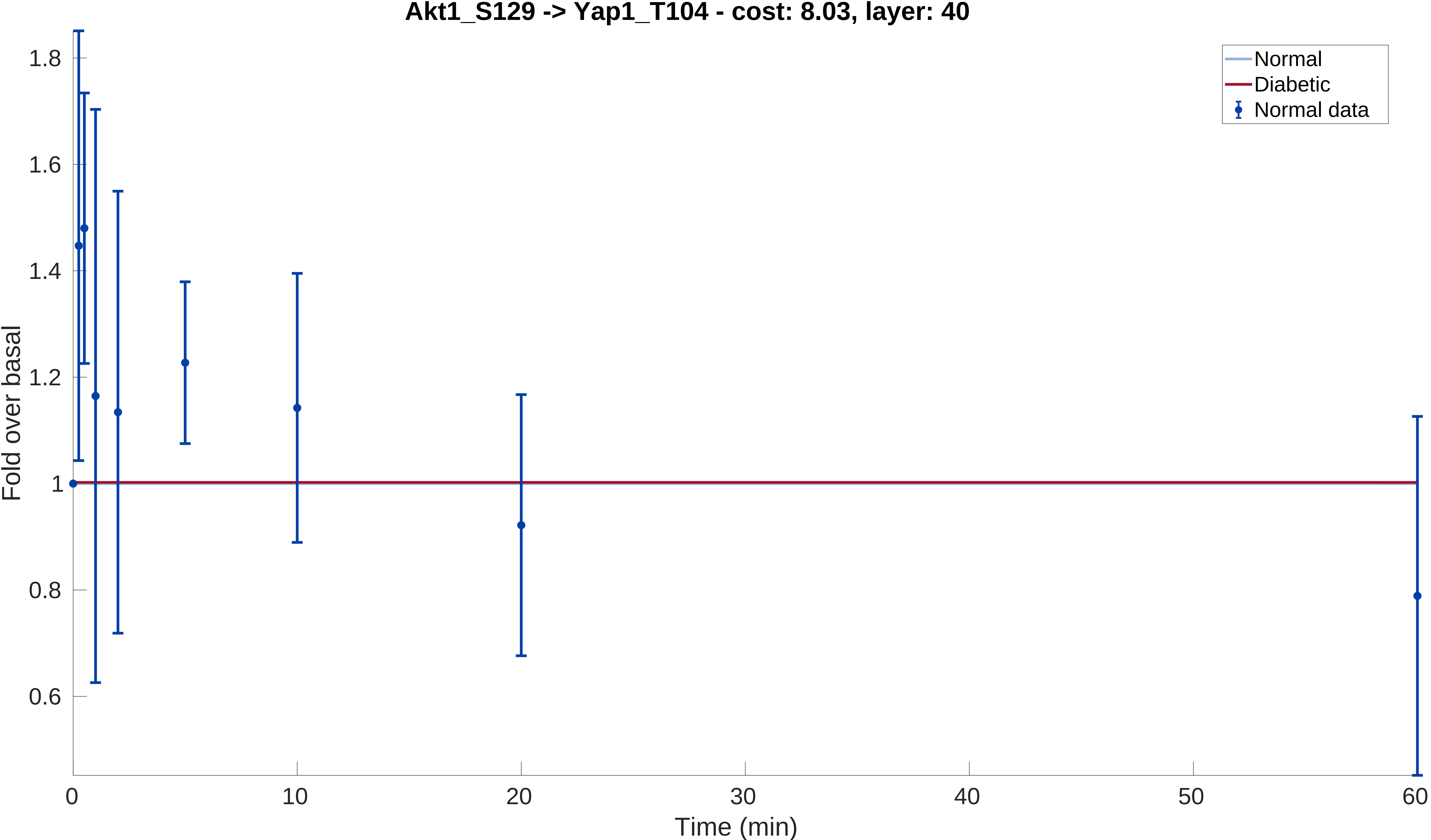
Akt1_S129 -> Yap1_T104 - cost: 8.03, layer: 40
Normal
Diabetic
Normal data
Fold over basal
1.8
1.6
1.4
1.2
1
0.8
0.6
0
10
20
30
40
50
60
Time (min)

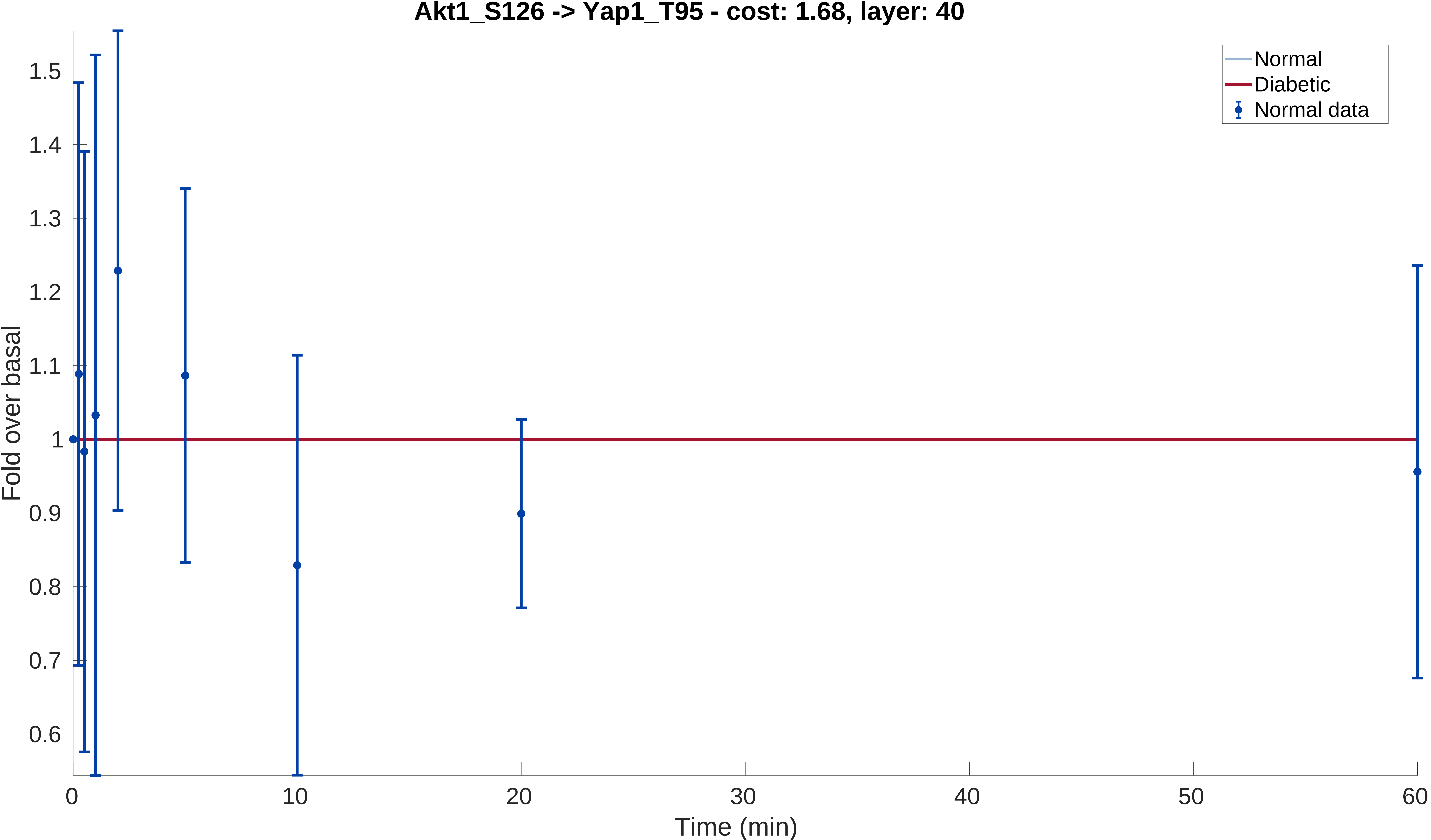

Akt1_S126 -> Yap1_T95 - cost: 1.68, layer: 40
Normal
Diabetic
Normal data
Fold over basal
1.5
1.4
1.3
1.2
1.1
1
0.9
0.8
0.7
0.6
0
10
20
30
40
50
60
Time (min)

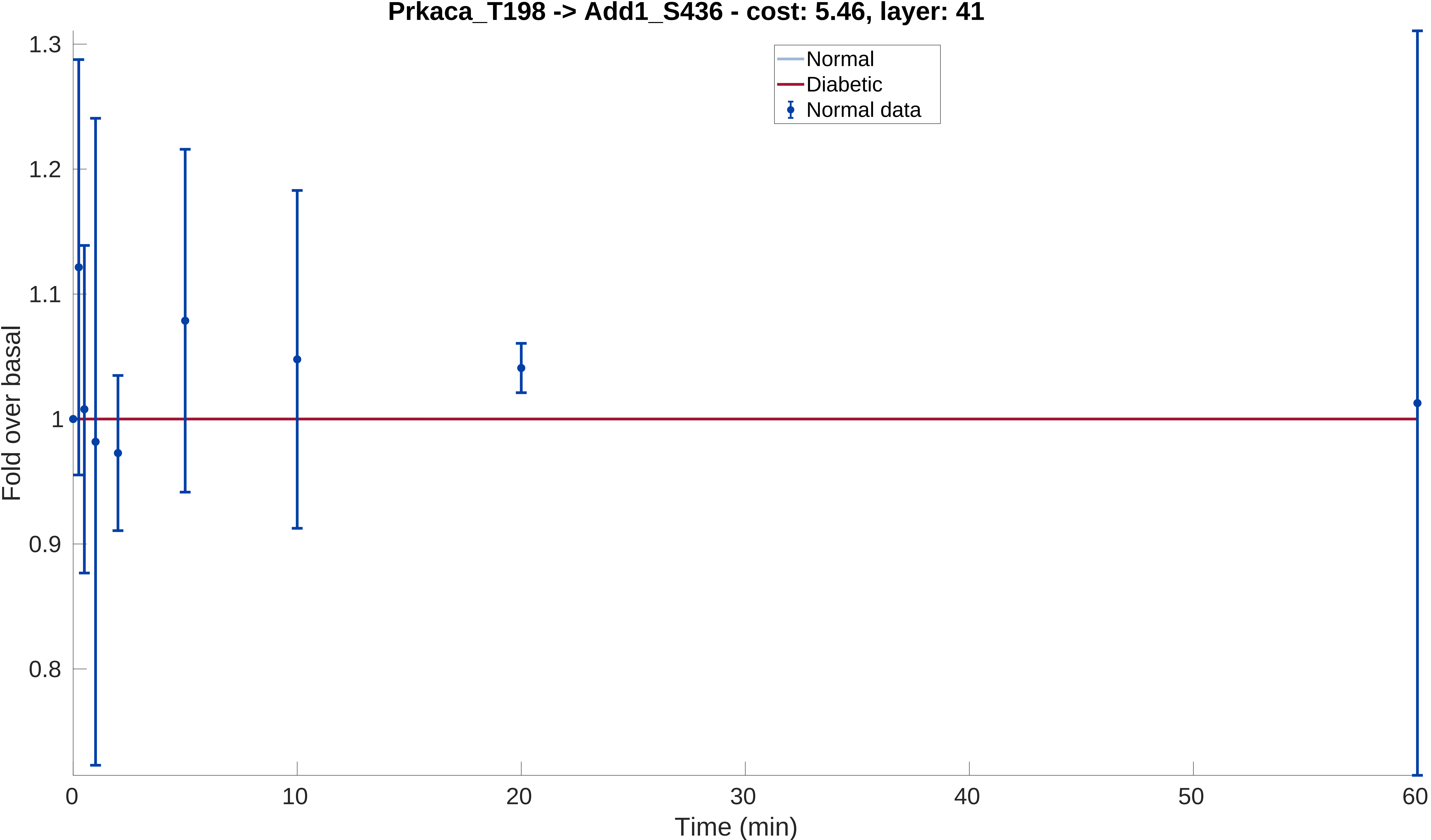

Prkaca_T198 -> Add1_S436 - cost: 5.46, layer: 41
Normal
Diabetic
Normal data
Fold over basal
Time (min)
1.3
1.2
1.1
1
0.9
0.8
0
10
20
30
40
50
60

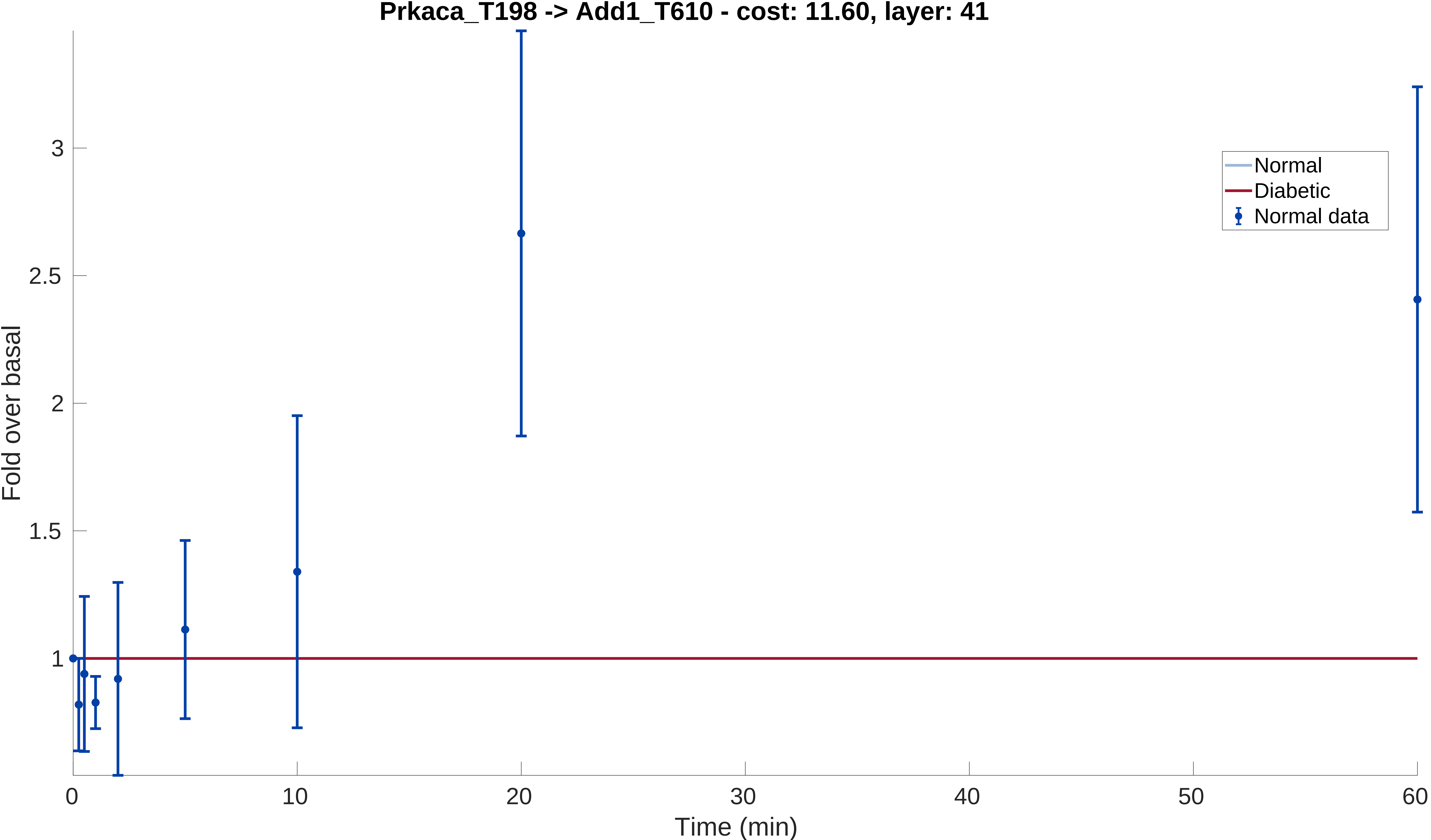

Prkaca_T198 -> Add1_T610 - cost: 11.60, layer: 41
Fold over basal
3
2.5
2
1.5
1
0
10
20
30
40
50
60
Time (min)
Normal
Diabetic
Normal data

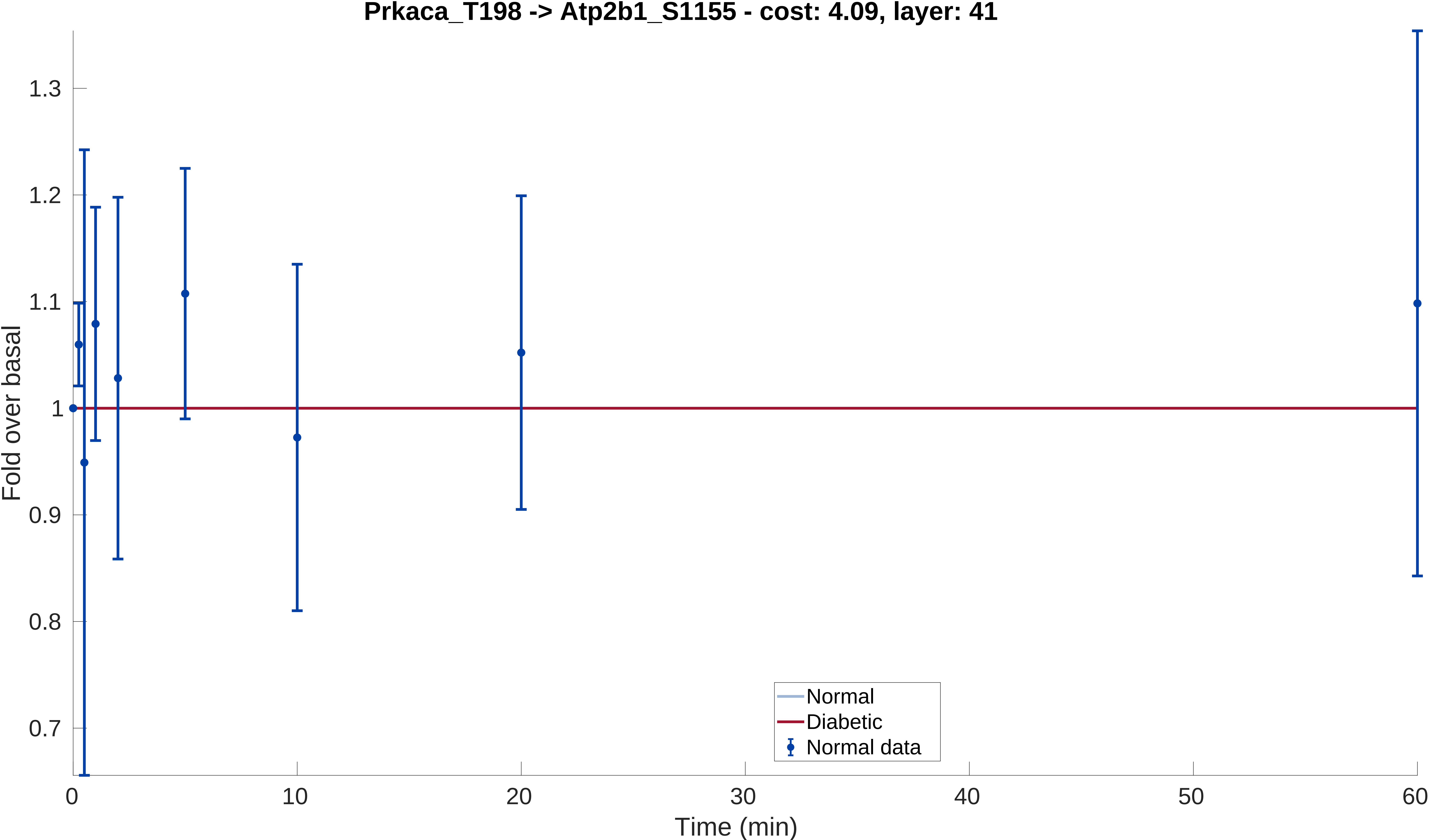

Prkaca_T198 -> Atp2b1_S1155 - cost: 4.09, layer: 41
Fold over basal
1.3
1.2
1.1
1
0.9
0.8
0.7
0
10
20
30
40
50
60
Time (min)
Normal
Diabetic
Normal data

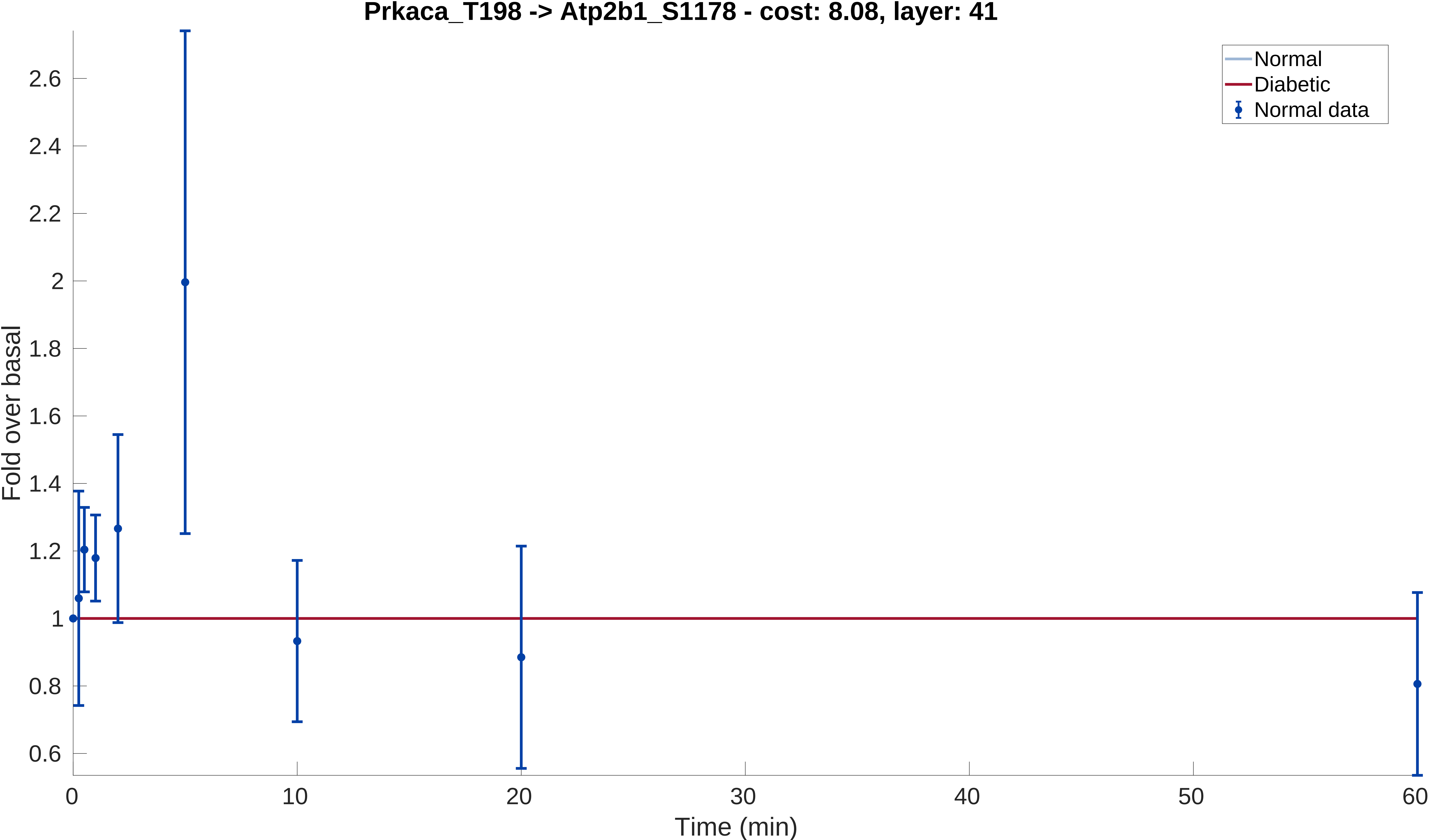
Prkaca_T198 -> Atp2b1_S1178 - cost: 8.08, layer: 41
Normal
Diabetic
Normal data
Fold over basal
2.6
2.4
2.2
2
1.8
1.6
1.4
1.2
1
0.8
0.6
0
10
20
30
40
50
60
Time (min)

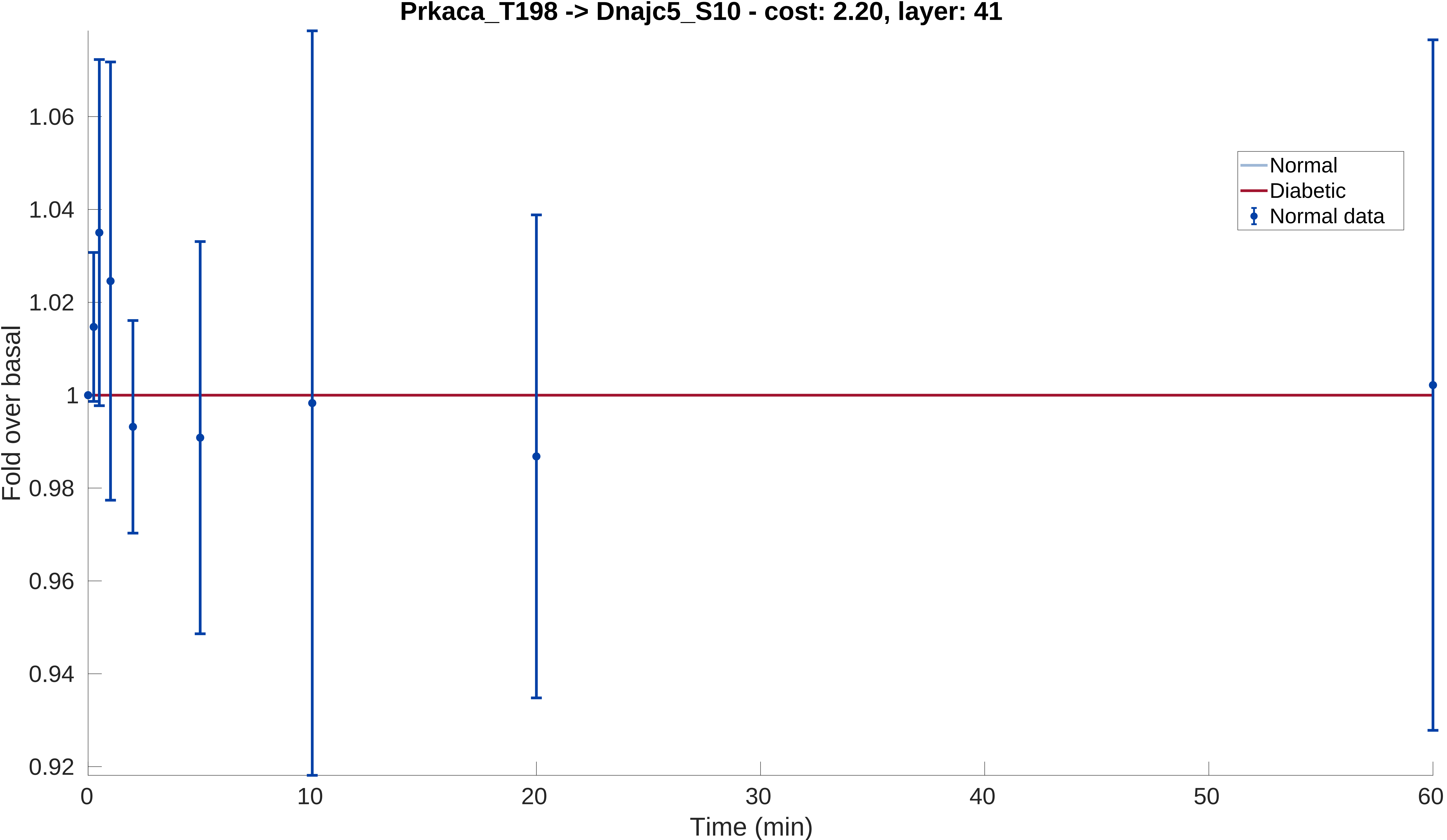

Prkaca_T198 -> Dnajc5_S10 - cost: 2.20, layer: 41
Fold over basal
1.06
1.04
1.02
1
0.98
0.96
0.94
0.92
0
10
20
30
40
50
60
Time (min)
Normal
Diabetic
Normal data

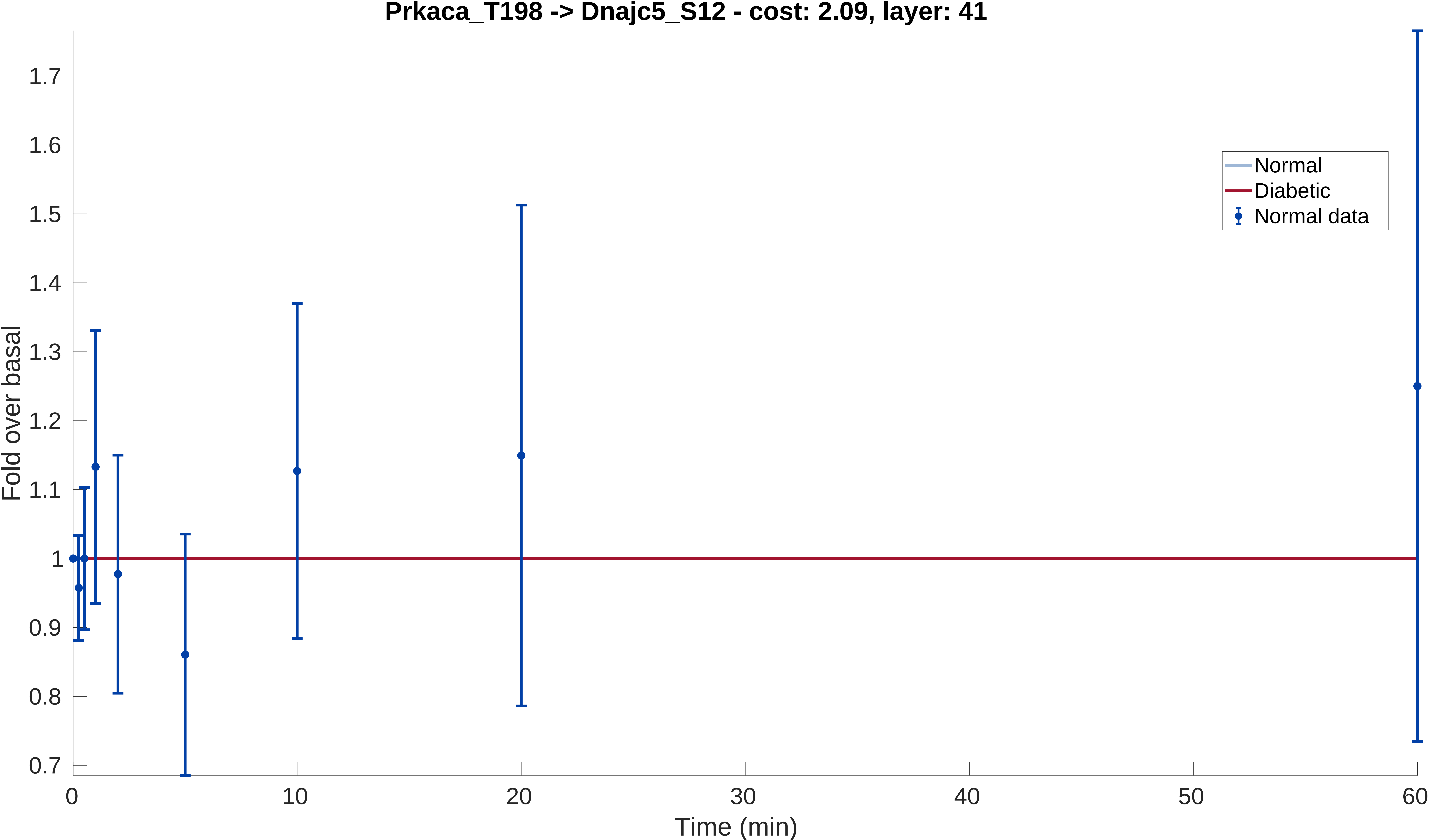

Prrkaca_T198 -> Dnajc5_S12 - cost: 2.09, layer: 41
Fold over basal
Time (min)
Normal
Diabetic
Normal data
0.7
0.8
0.9
1
1.1
1.2
1.3
1.4
1.5
1.6
1.7
0
10
20
30
40
50
60

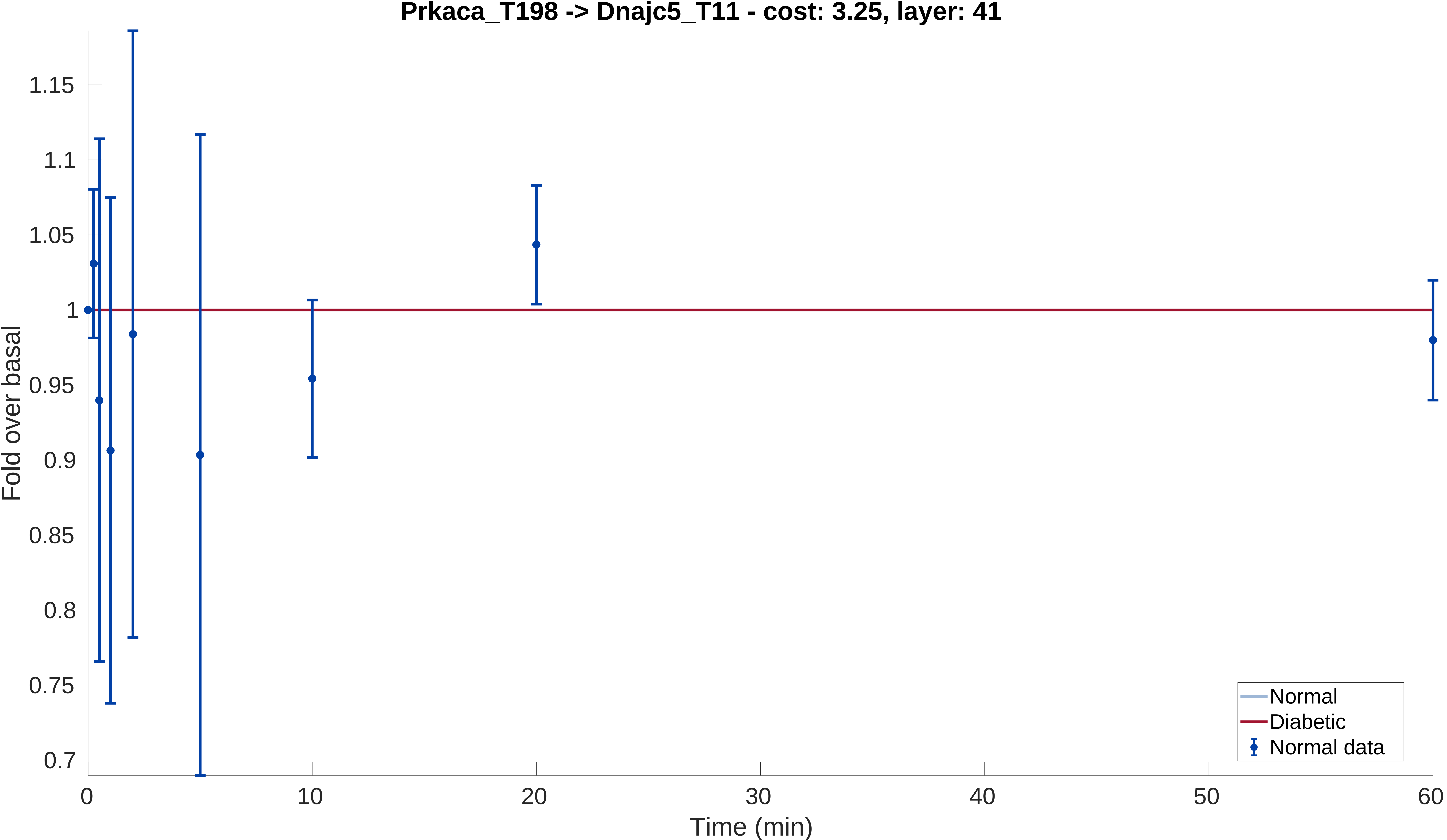

Prkaca_T198 -> Dnajc5_T11 - cost: 3.25, layer: 41
Fold over basal
1.15
1.1
1.05
1
0.95
0.9
0.85
0.8
0.75
0.7
0
10
20
30
40
50
60
Time (min)
Normal
Diabetic
Normal data

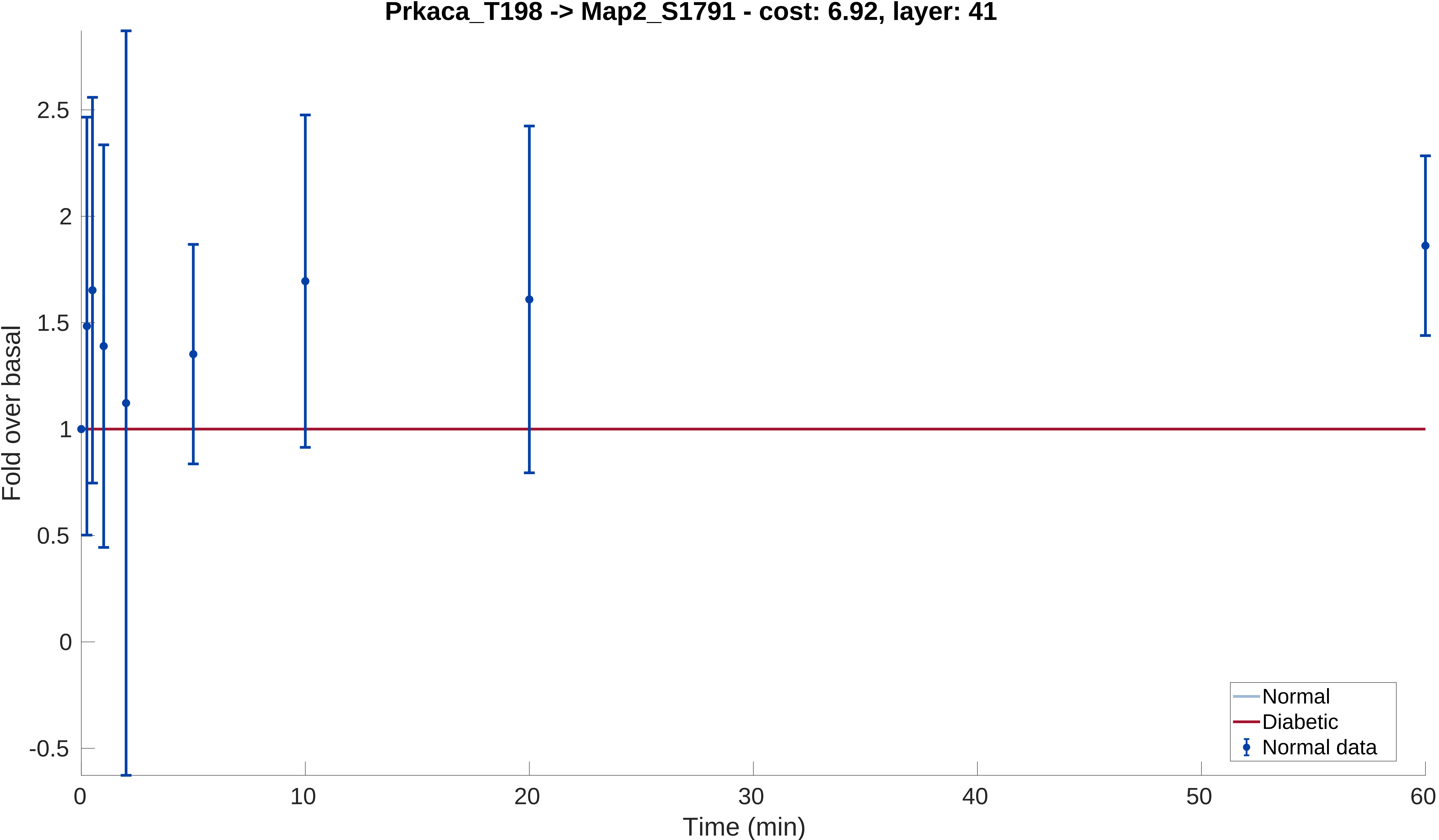

Prkaca_T198 -> Map2_S1791 - cost: 6.92, layer: 41
Fold over basal
2.5
2
1.5
1
0.5
0
-0.5
0
10
20
30
40
50
60
Time (min)
Normal
Diabetic
Normal data

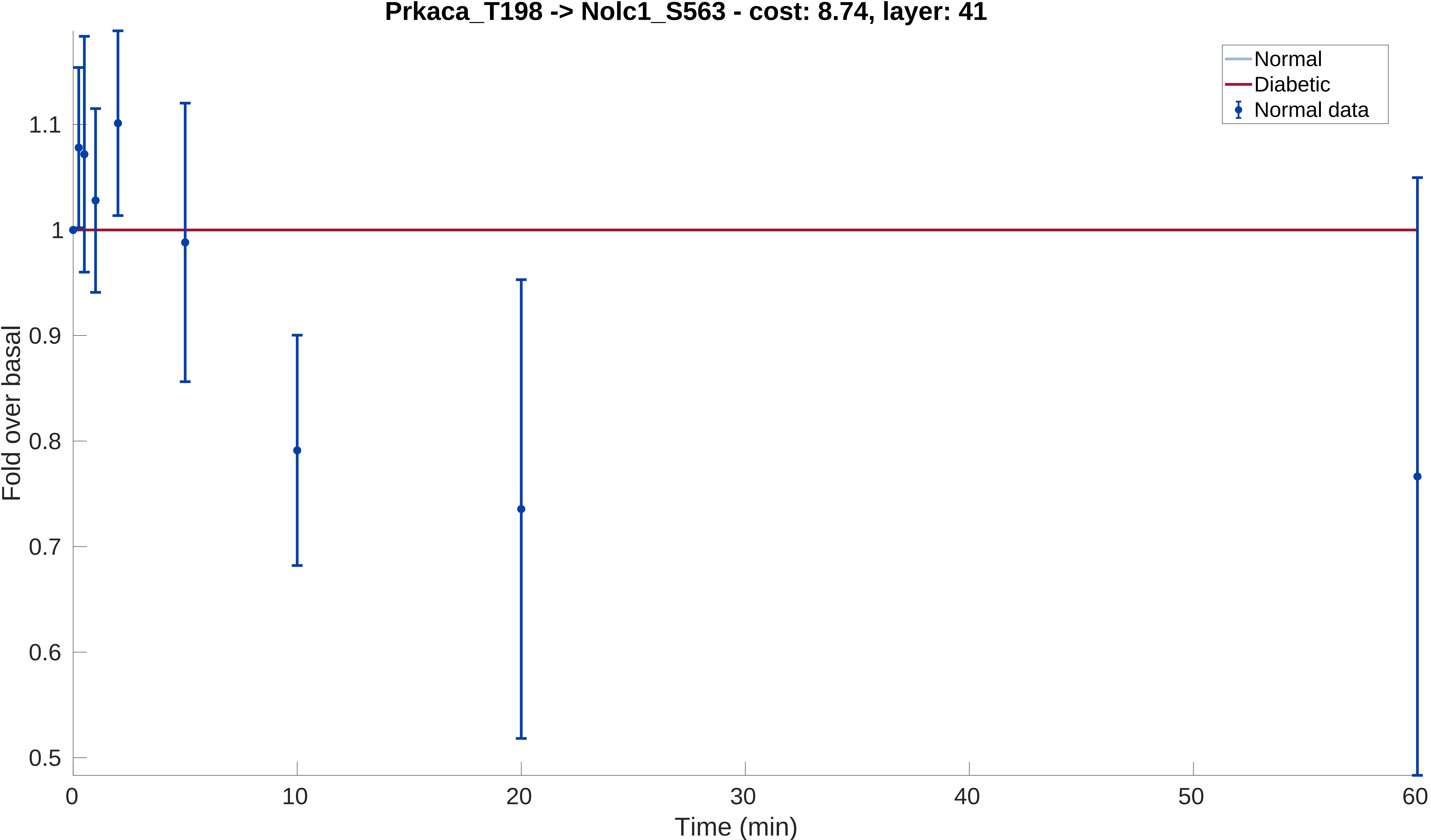

Prkaca_T198 -> Nolc1_S563 - cost: 8.74, layer: 41
Normal
Diabetic
Normal data
Fold over basal
1.1
1
0.9
0.8
0.7
0.6
0.5
0
10
20
30
40
50
60
Time (min)

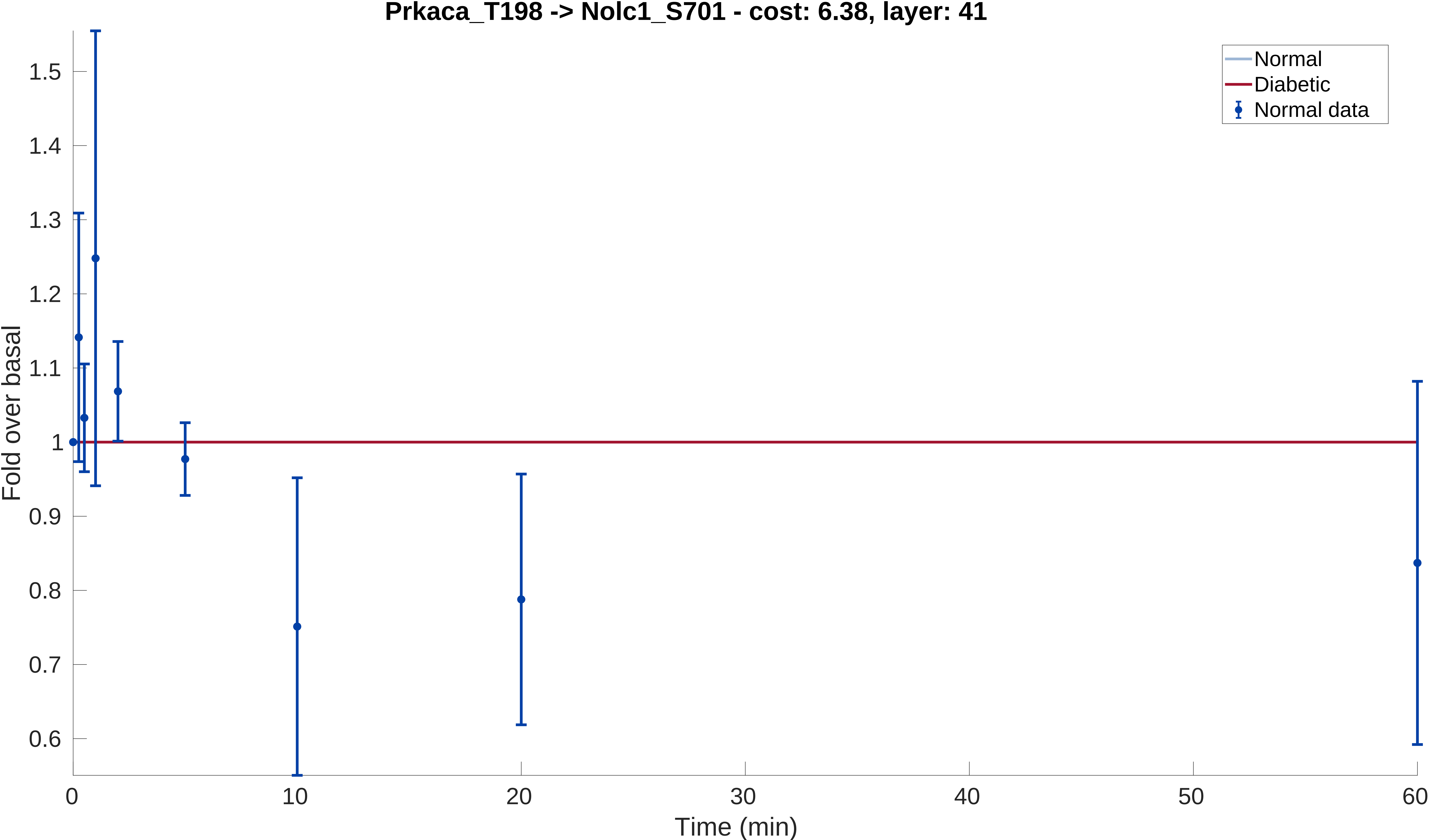

Prkaca_T198 -> Nolc1_S701 - cost: 6.38, layer: 41
Normal
Diabetic
Normal data
Fold over basal
1.5
1.4
1.3
1.2
1.1
1
0.9
0.8
0.7
0.6
0
10
20
30
40
50
60
Time (min)

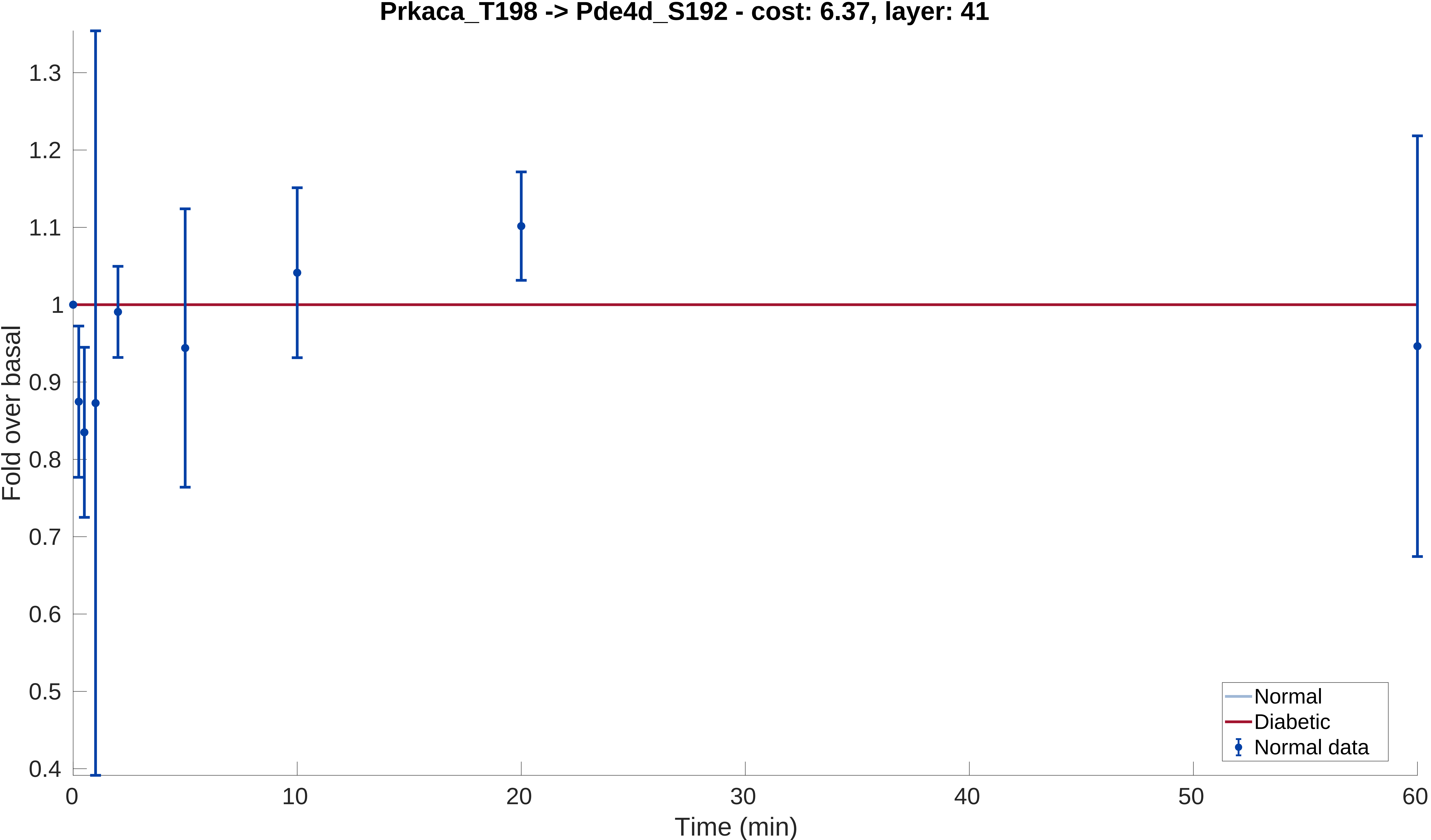

Prkaca_T198 -> Pde4d_S192 - cost: 6.37, layer: 41
Fold over basal
1.3
1.2
1.1
1
0.9
0.8
0.7
0.6
0.5
0.4
0
10
20
30
40
50
60
Time (min)
Normal
Diabetic
Normal data

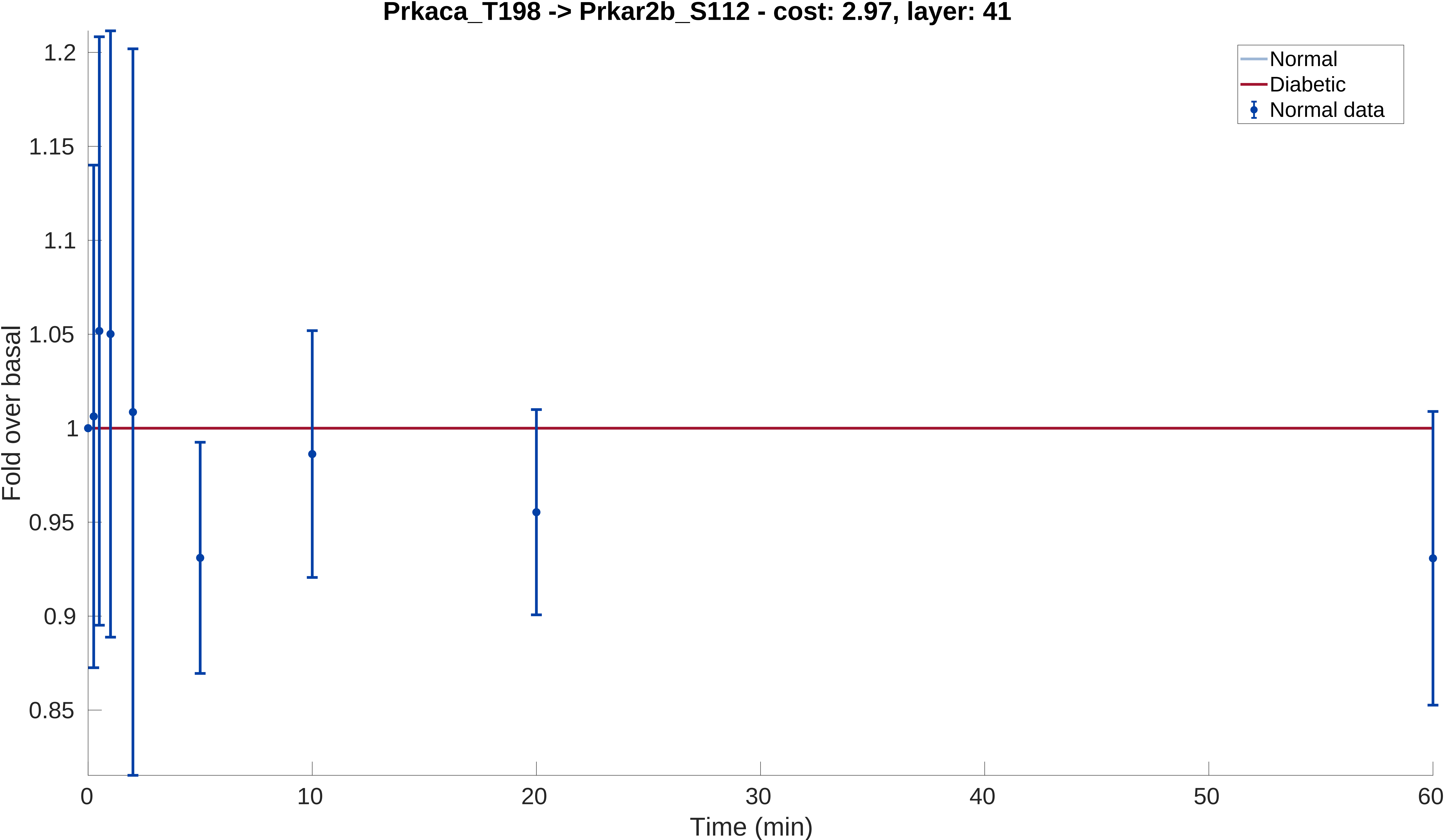

Prkaca_T198 -> Prkar2b_S112 - cost: 2.97, layer: 41
Normal
Diabetic
Normal data
Fold over basal
1.2
1.15
1.1
1.05
1
0.95
0.9
0.85
0
10
20
30
40
50
60
Time (min)

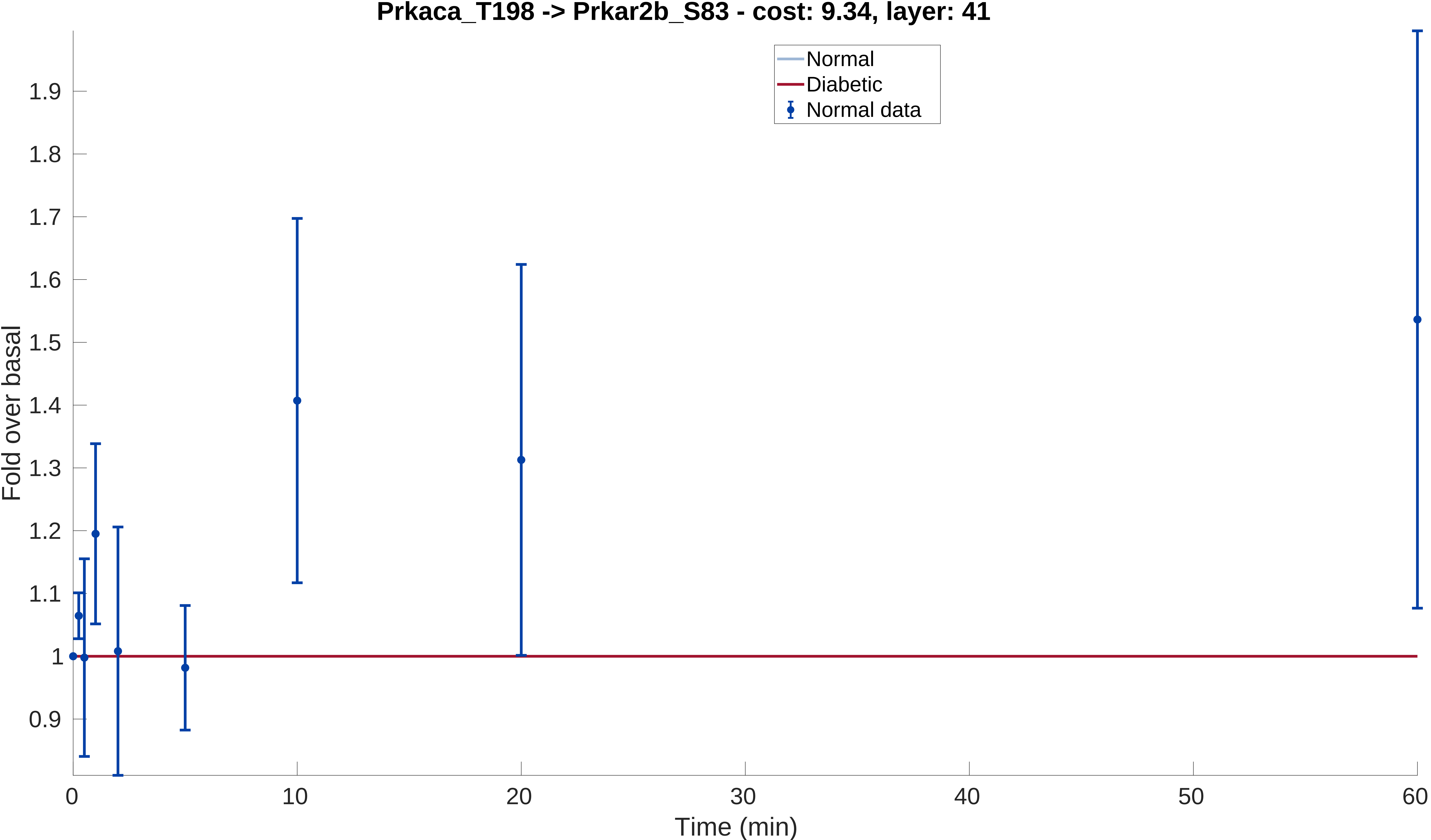

Prrkaca_T198 -> Prkar2b_S83 - cost: 9.34, layer: 41
Normal
Diabetic
Normal data
Fold over basal
1.9
1.8
1.7
1.6
1.5
1.4
1.3
1.2
1.1
1
0.9
0
10
20
30
40
50
60
Time (min)

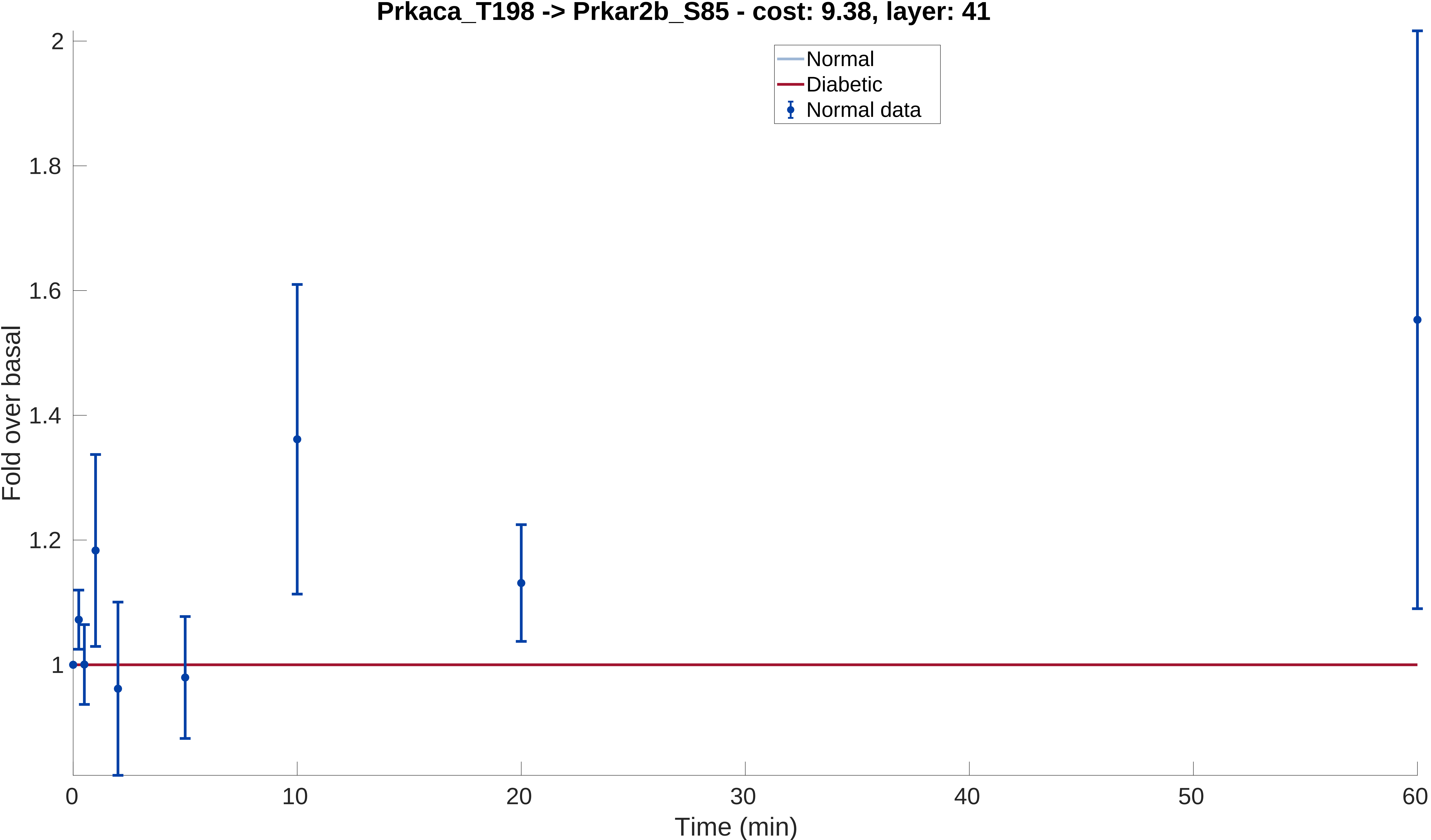
Prkaca_T198 -> Prkar2b_S85 - cost: 9.38, layer: 41
Normal
Diabetic
Normal data
Fold over basal
Time (min)
2
1.8
1.6
1.4
1.2
1
0
10
20
30
40
50
60

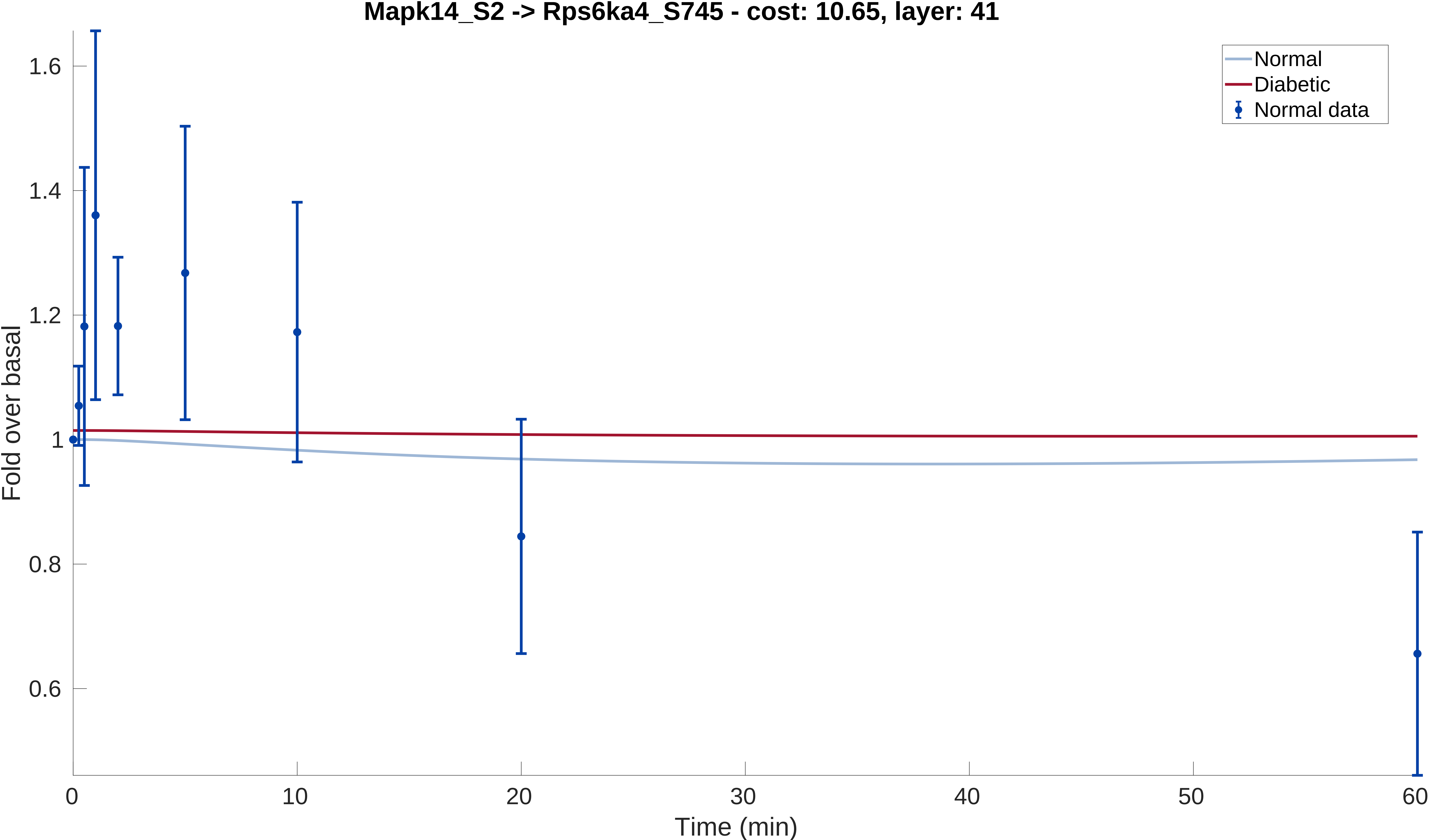
Mapk14_S2 -> Rps6ka4_S745 - cost: 10.65, layer: 41
Normal
Diabetic
Normal data
1.6
1.4
1.2
1
0.8
0.6
Fold over basal
0
10
20
30
40
50
60
Time (min)

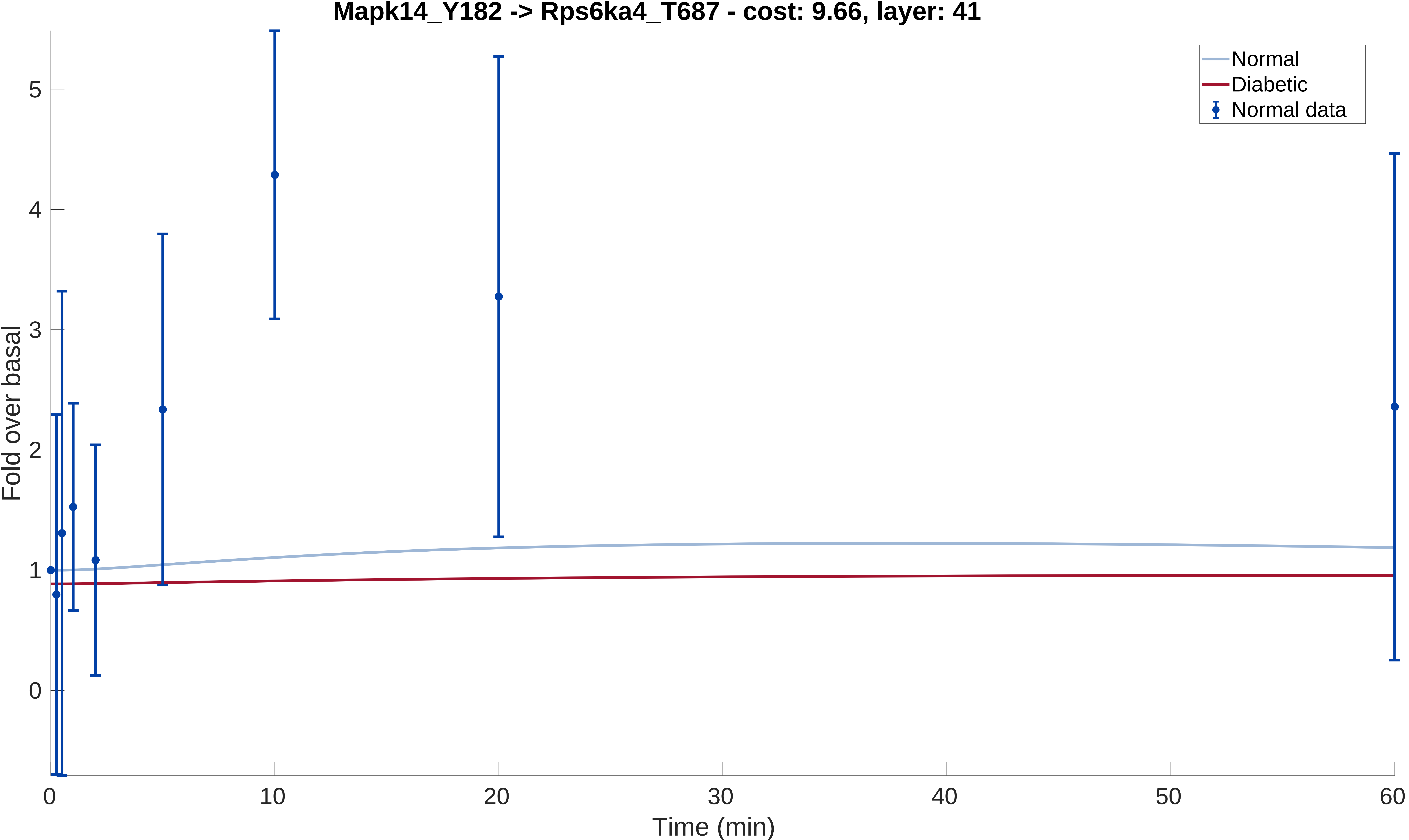

Mapk14_Y182 -> Rps6ka4_T687 - cost: 9.66, layer: 41
Normal
Diabetic
Normal data
Fold over basal
Time (min)
0
1
2
3
4
5
0
10
20
30
40
50
60

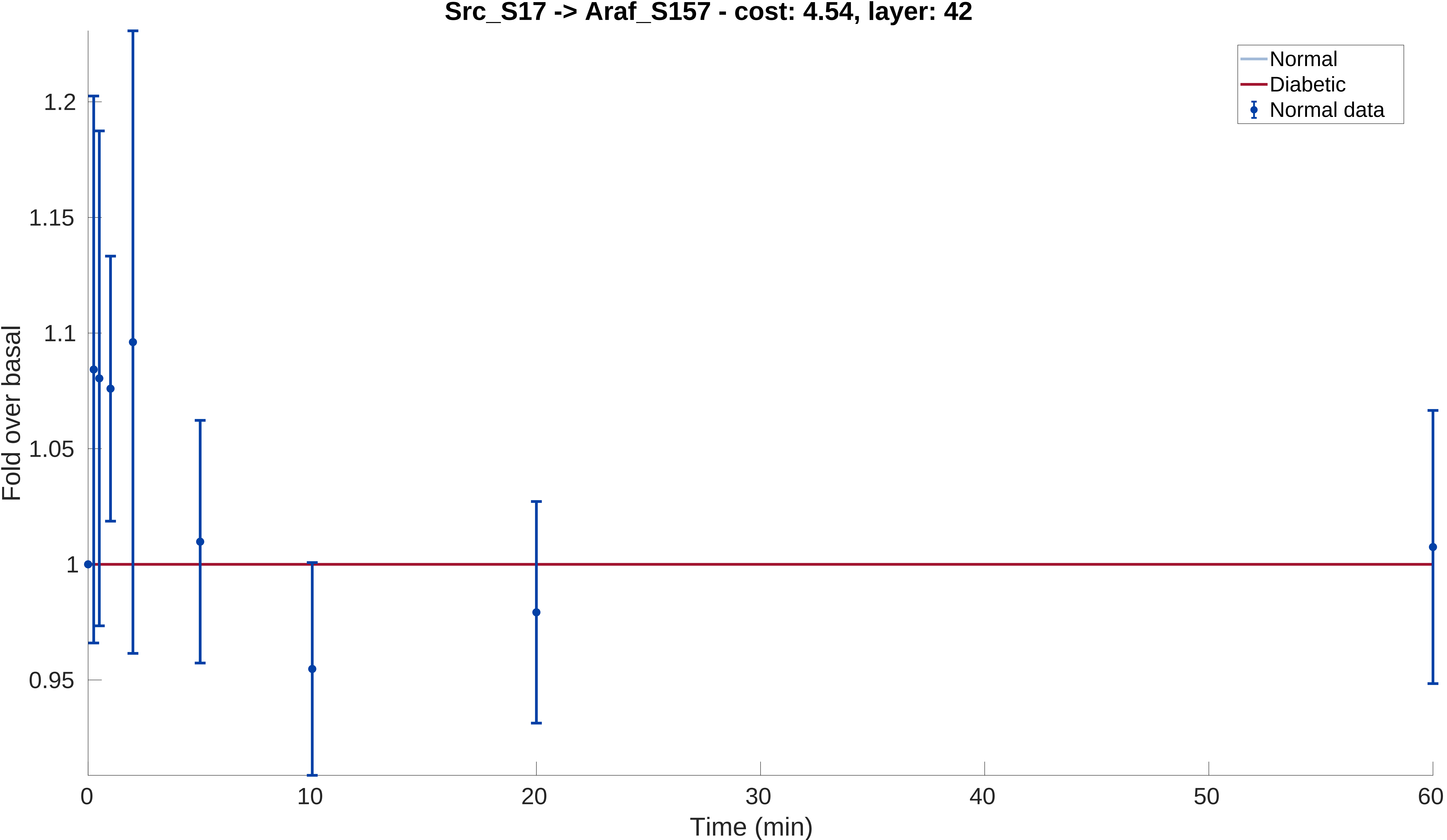

Src_S17 -> Araf_S157 - cost: 4.54, layer: 42
Normal
Diabetic
Normal data
Fold over basal
1.2
1.15
1.1
1.05
1
0.95
0
10
20
30
40
50
60
Time (min)

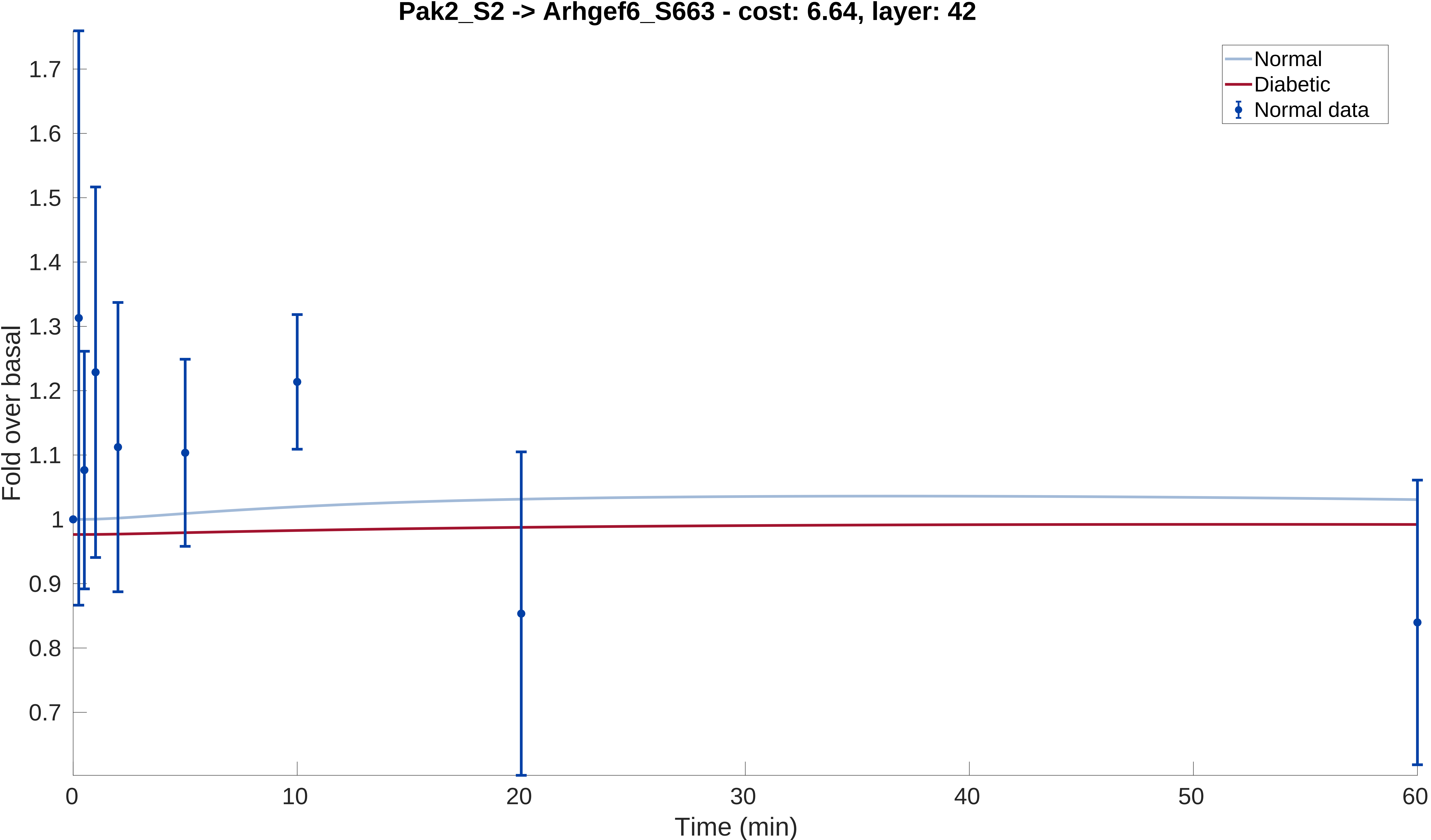

Pak2_S2 -> Arhgef6_S663 - cost: 6.64, layer: 42
Normal
Diabetic
Normal data
Fold over basal
1.7
1.6
1.5
1.4
1.3
1.2
1.1
1
0.9
0.8
0.7
0
10
20
30
40
50
60
Time (min)

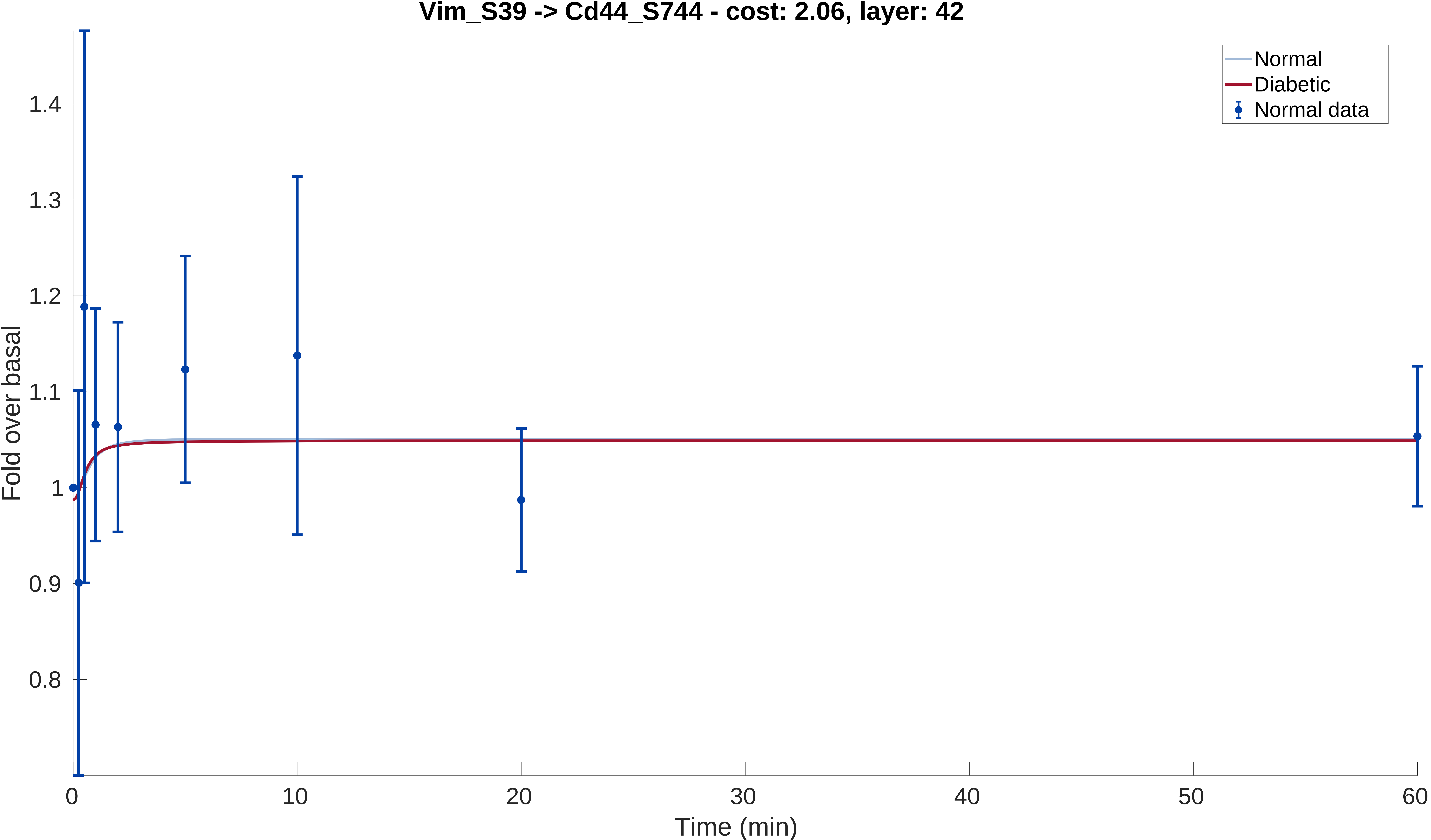

Vim_S39 -> Cd44_S744 - cost: 2.06, layer: 42
Normal
Diabetic
Normal data
Fold over basal
1.4
1.3
1.2
1.1
1
0.9
0.8
0
10
20
30
40
50
60
Time (min)

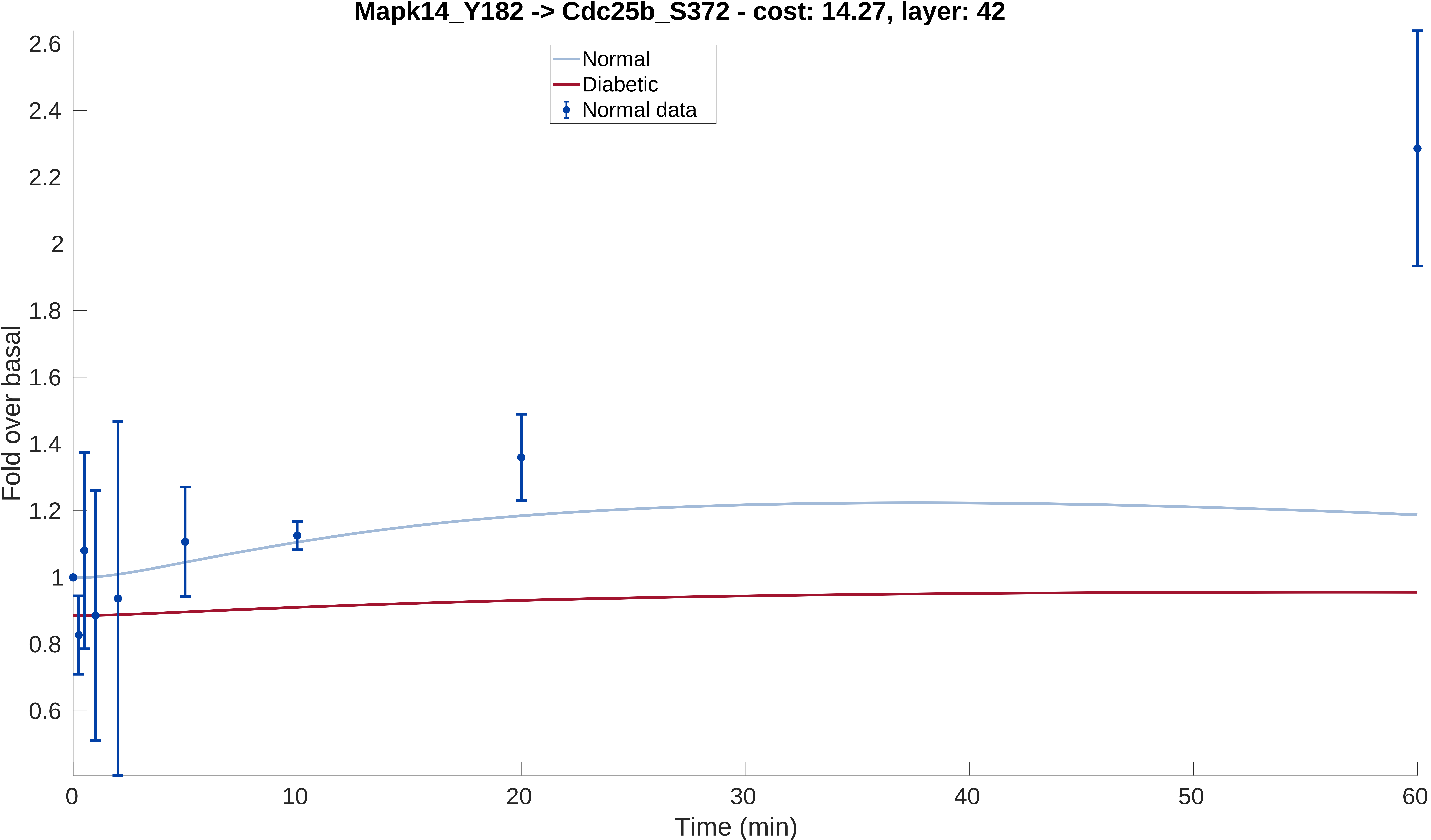
Mapk14_Y182 -> Cdc25b_S372 - cost: 14.27, layer: 42
Normal
Diabetic
Normal data
Fold over basal
Time (min)
2.6
2.4
2.2
2
1.8
1.6
1.4
1.2
1
0.8
0.6
0
10
20
30
40
50
60

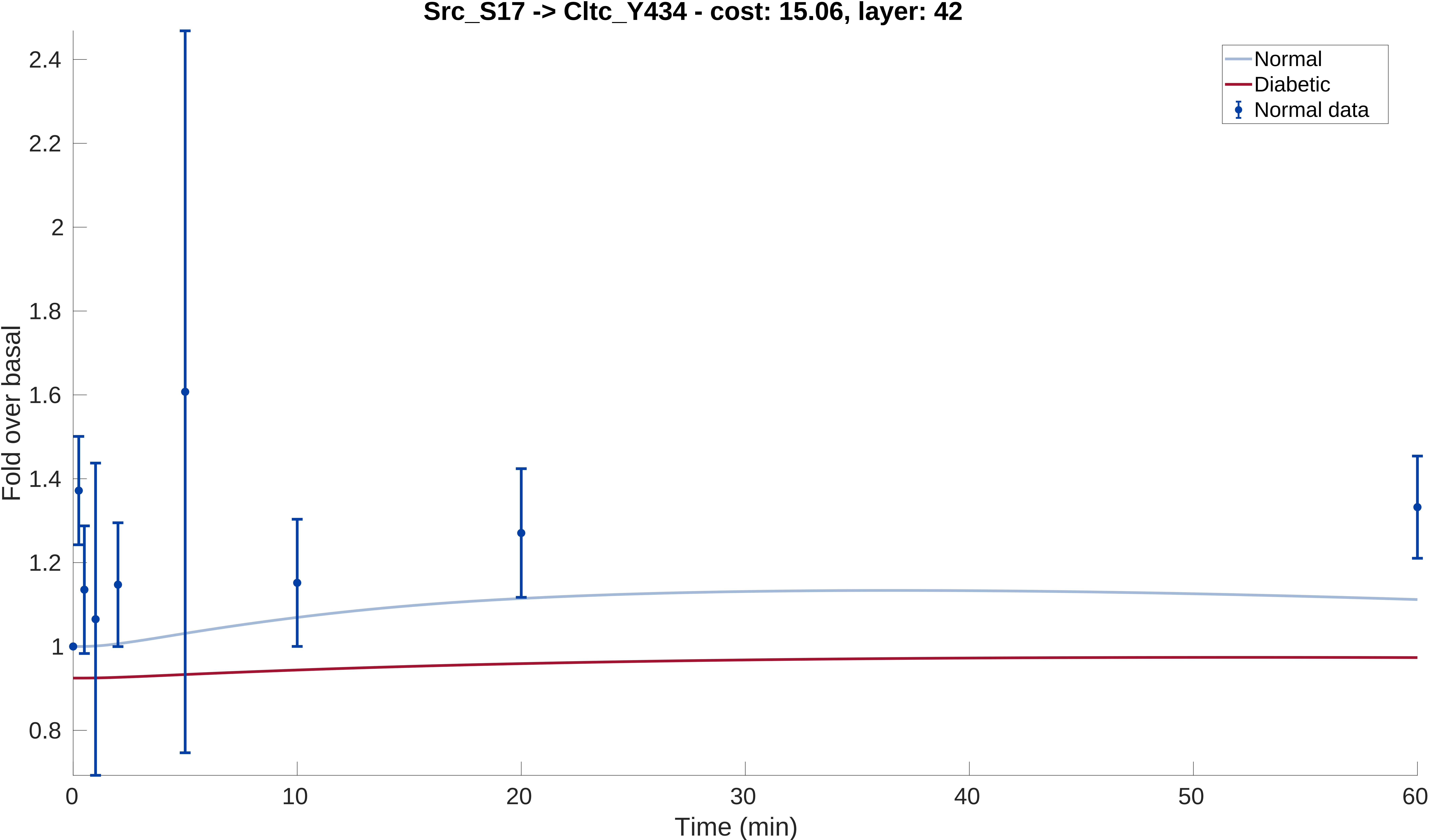
Src_S17 -> Cltc_Y434 - cost: 15.06, layer: 42
Normal
Diabetic
Normal data
Fold over basal
Time (min)
2.4
2.2
2
1.8
1.6
1.4
1.2
1
0.8
0
10
20
30
40
50
60

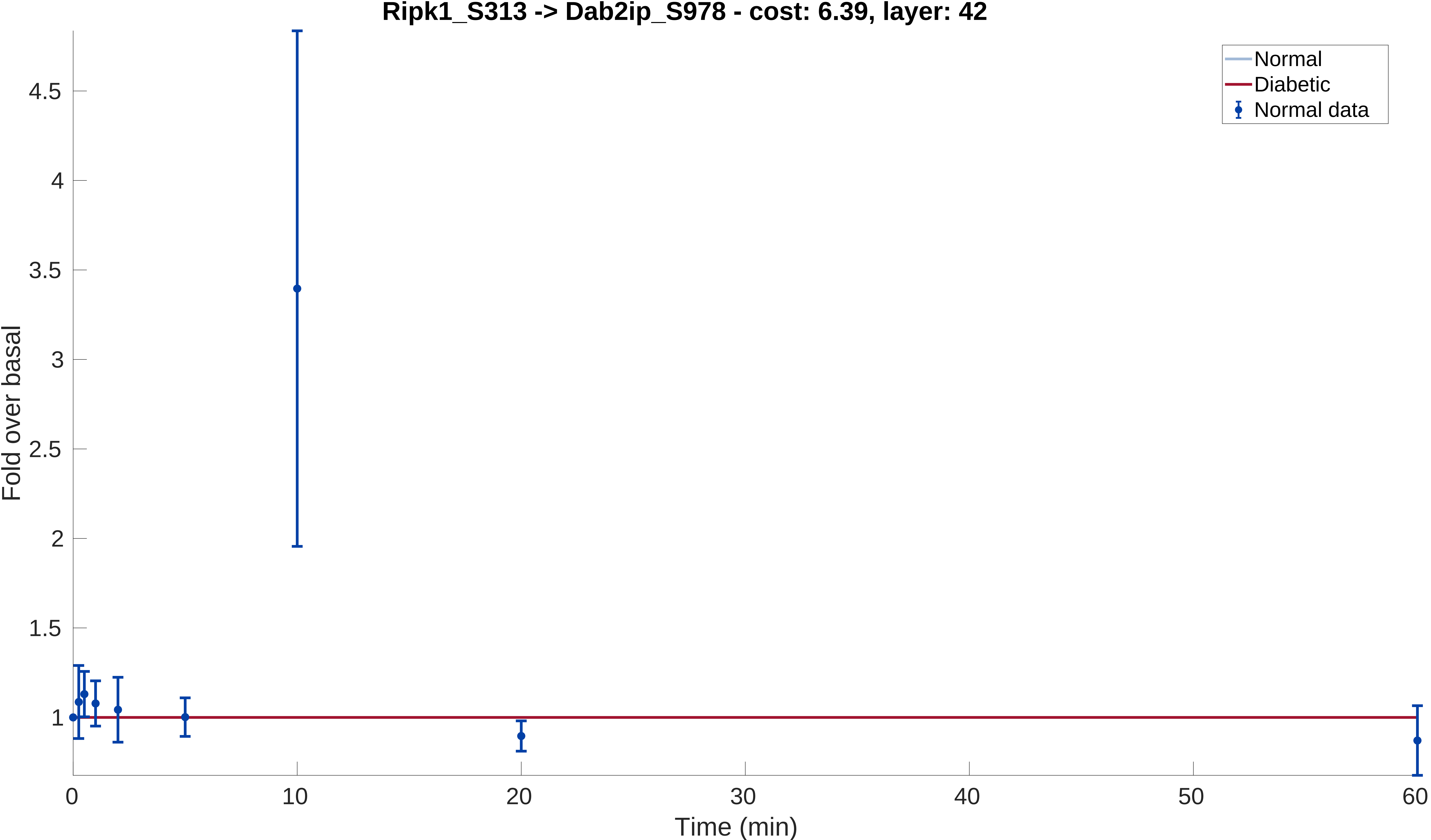

Ripk1_S313 -> Dab2ip_S978 - cost: 6.39, layer: 42
Normal
Diabetic
Normal data
Fold over basal
Time (min)
4.5
4
3.5
3
2.5
2
1.5
1
0
10
20
30
40
50
60

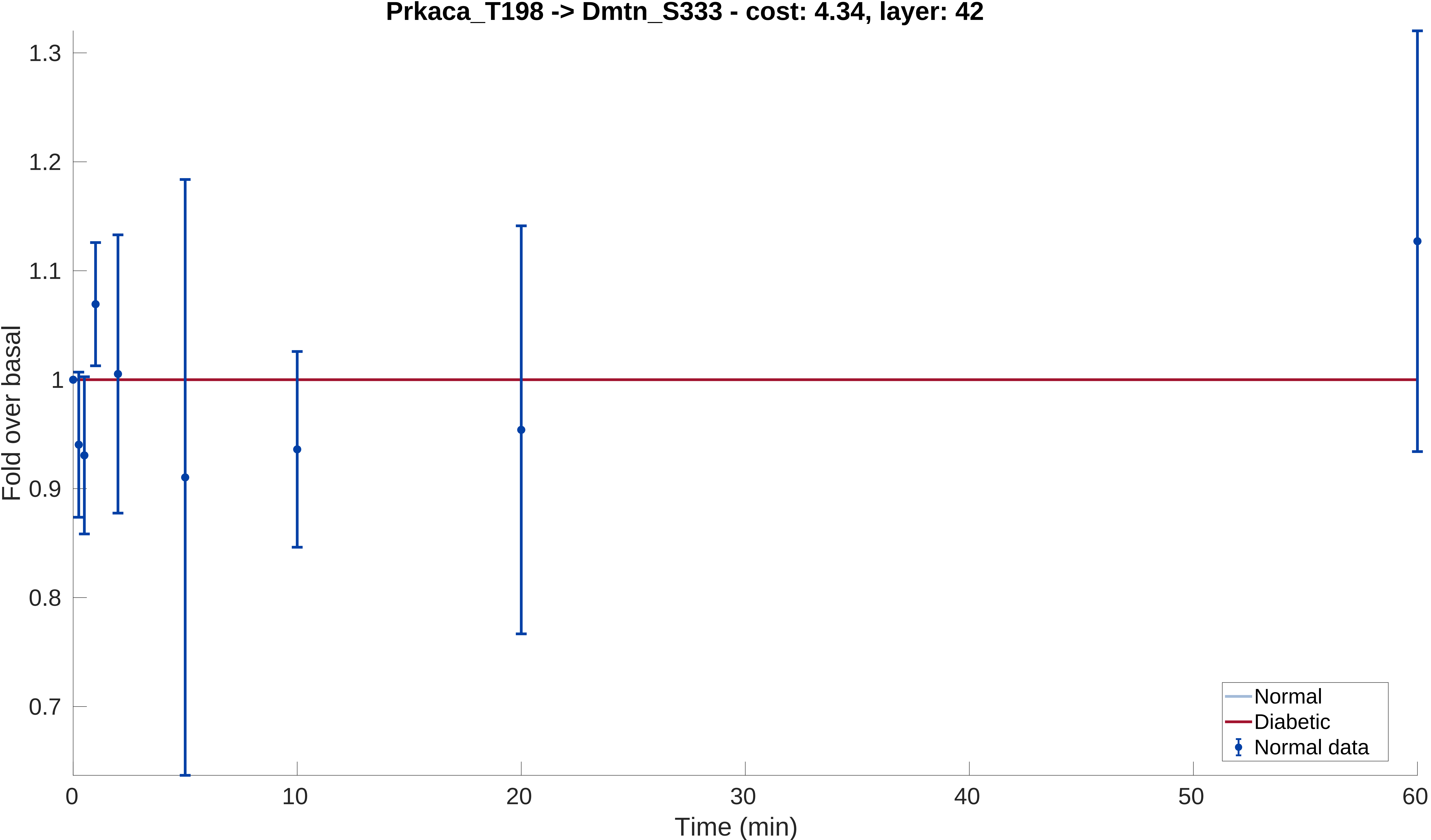

Prkaca_T198 -> Dmtn_S333 - cost: 4.34, layer: 42
Fold over basal
1.3
1.2
1.1
1
0.9
0.8
0.7
0
10
20
30
40
50
60
Time (min)
Normal
Diabetic
Normal data

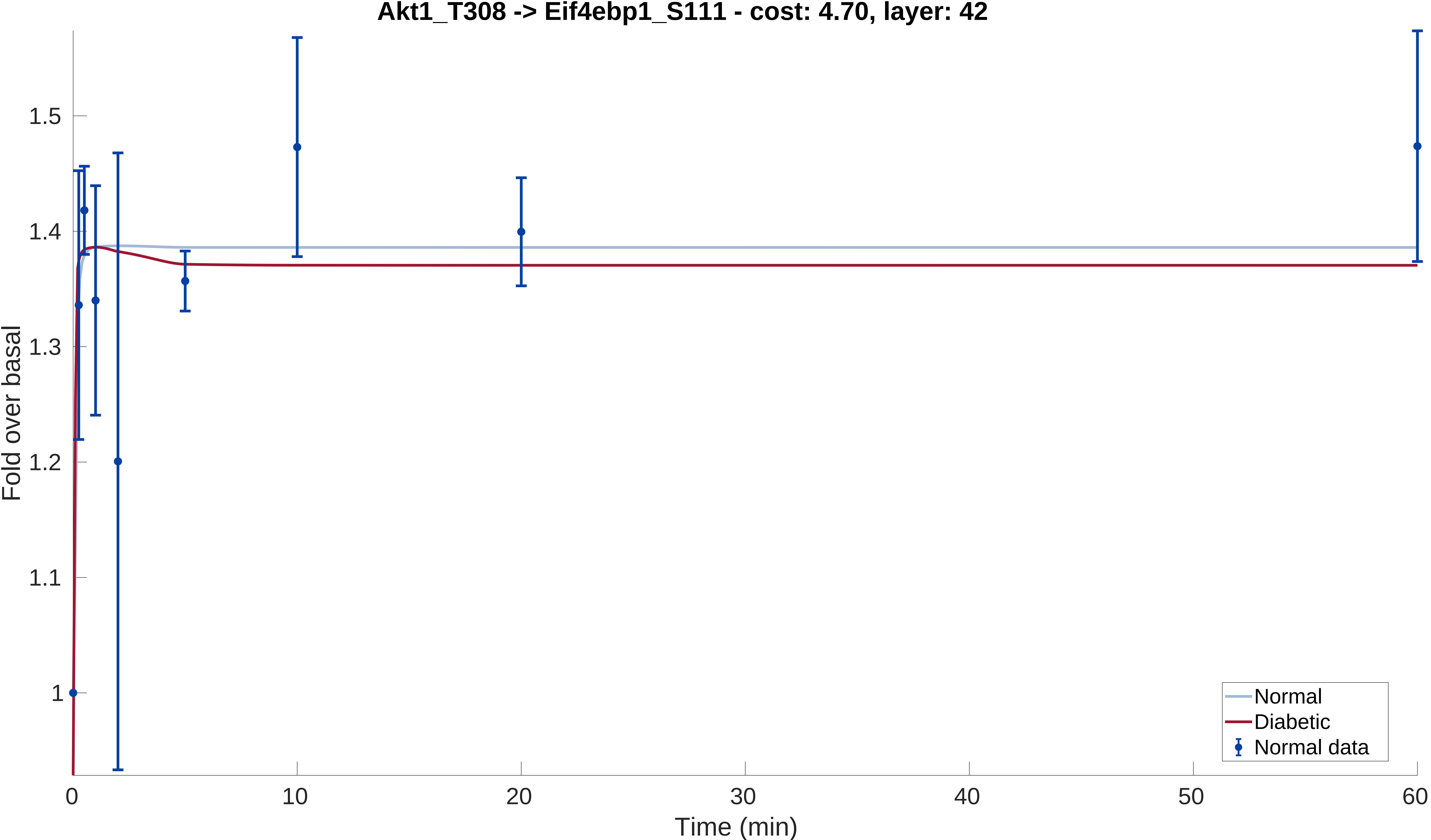

Akt1_T308 -> Eif4ebp1_S111 - cost: 4.70, layer: 42
Fold over basal
1.5
1.4
1.3
1.2
1.1
1
0
10
20
30
40
50
60
Time (min)
Normal
Diabetic
Normal data

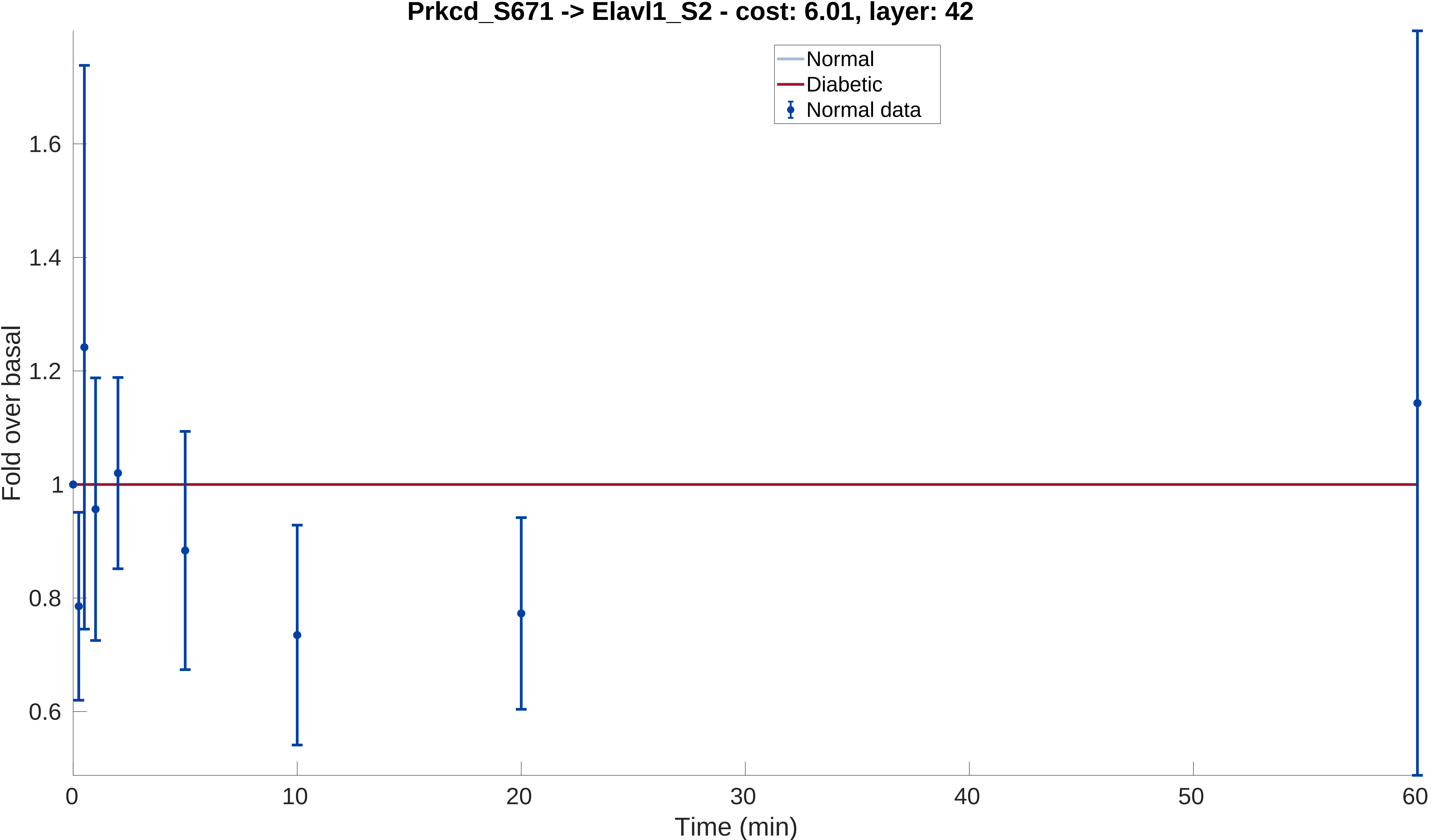
Prkcd_S671 -> Elavl1_S2 - cost: 6.01, layer: 42
Normal
Diabetic
Normal data
Fold over basal
1.6
1.4
1.2
1
0.8
0.6
0
10
20
30
40
50
60
Time (min)

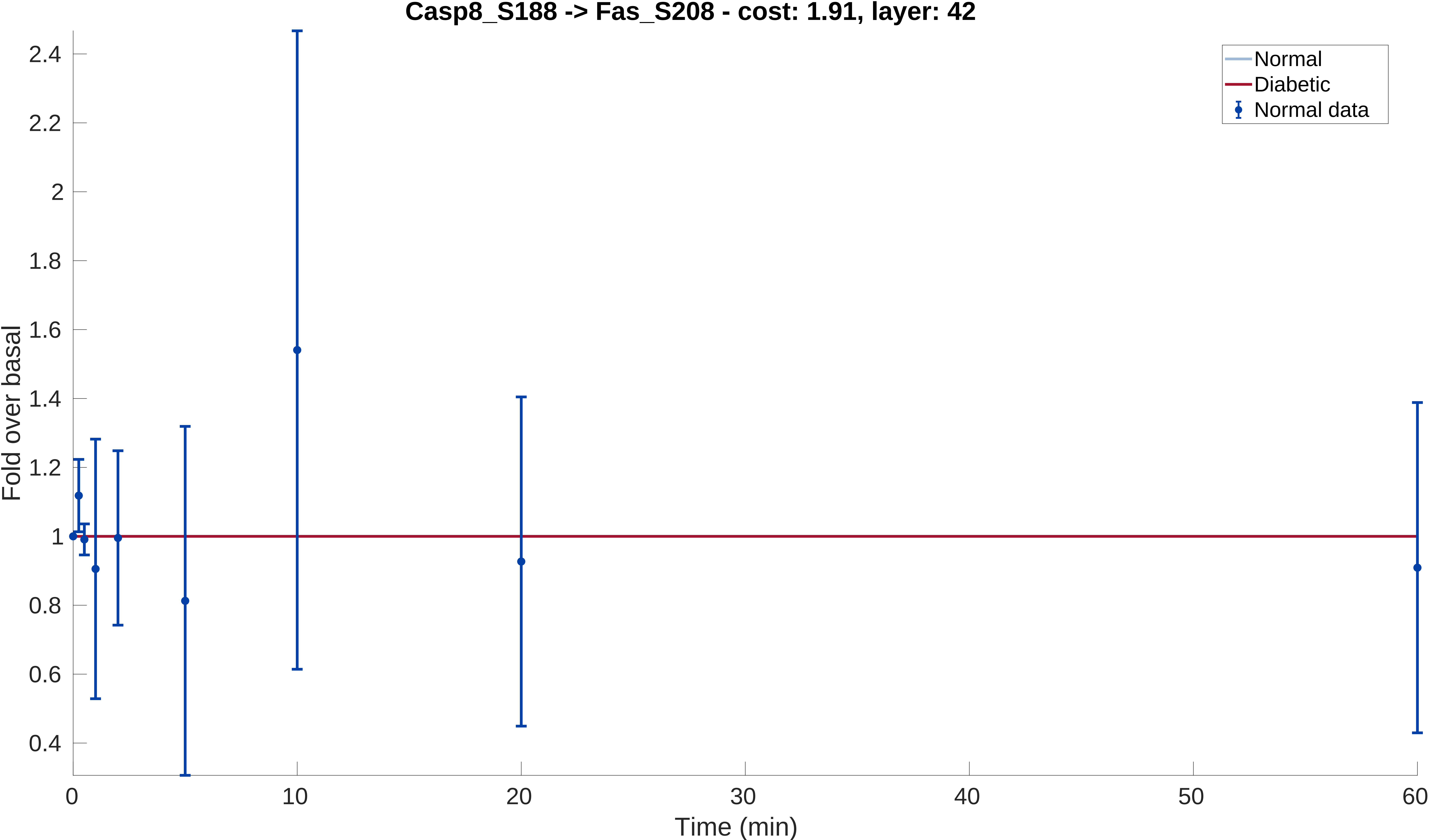

Casp8_S188 -> Fas_S208 - cost: 1.91, layer: 42
Normal
Diabetic
Normal data
Fold over basal
2.4
2.2
2
1.8
1.6
1.4
1.2
1
0.8
0.6
0.4
0
10
20
30
40
50
60
Time (min)

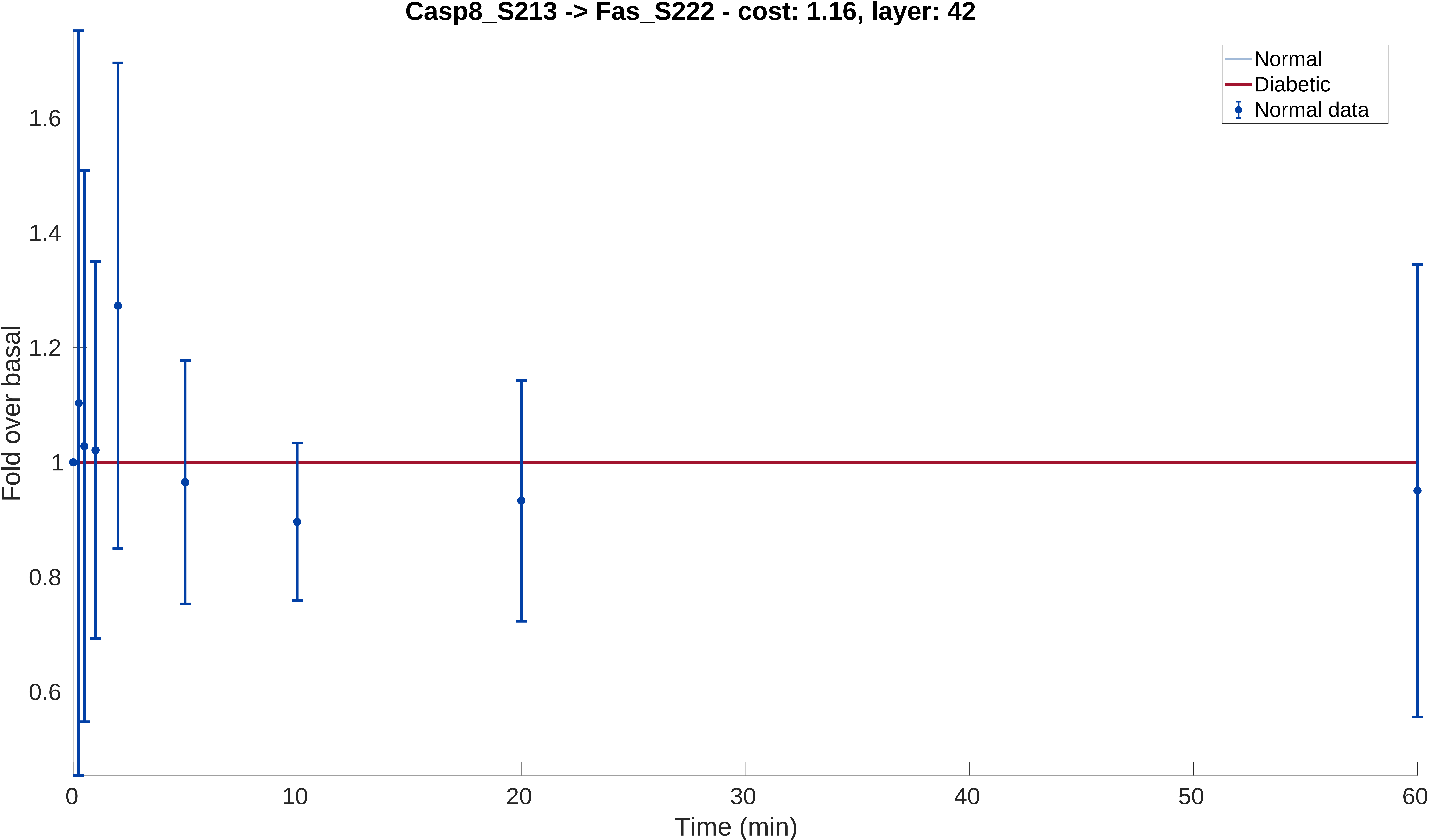

Casp8_S213 -> Fas_S222 - cost: 1.16, layer: 42
Normal
Diabetic
Normal data
Fold over basal
Time (min)
0.6
0.8
1
1.2
1.4
1.6
0
10
20
30
40
50
60

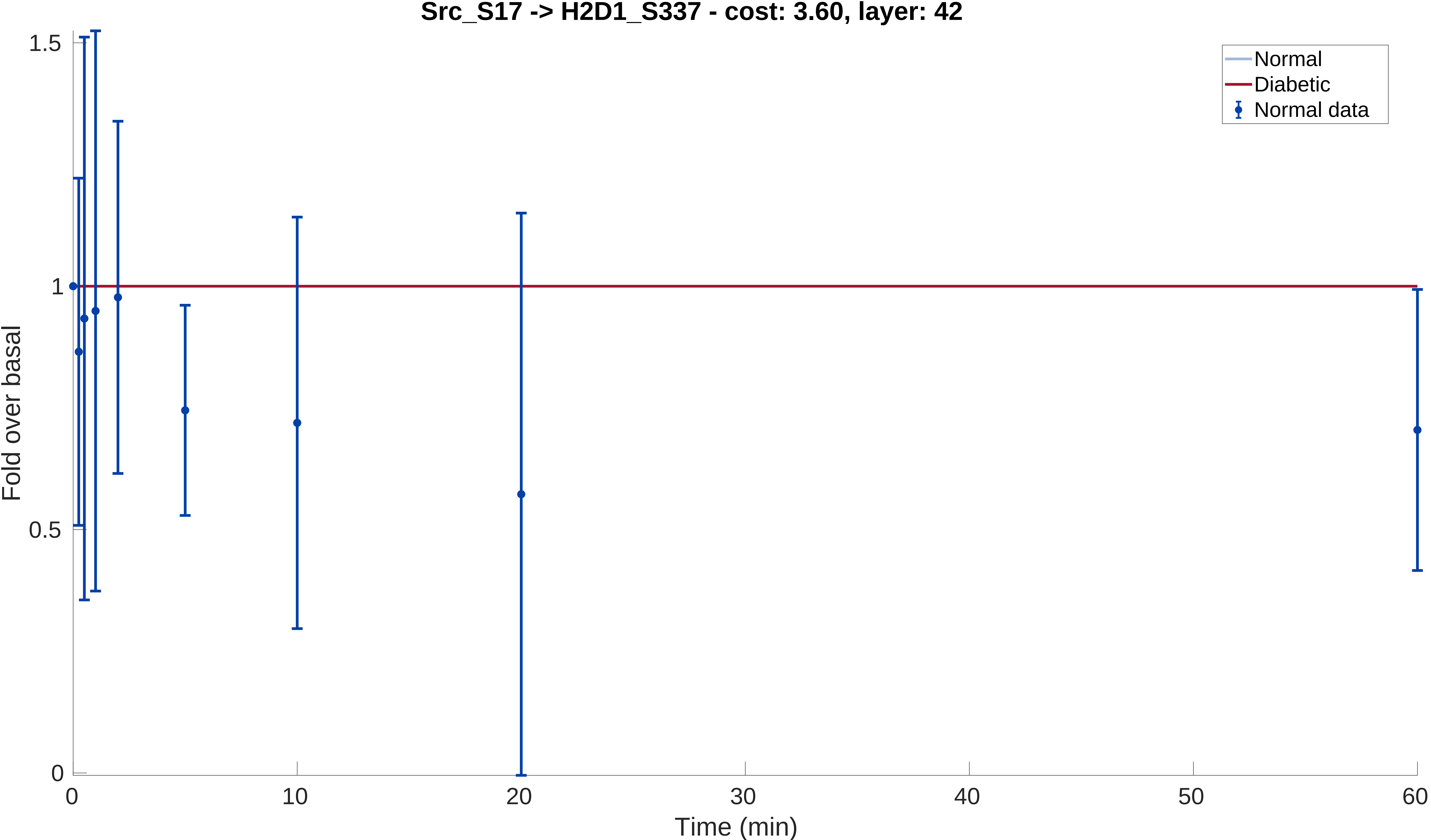
Src_S17 -> H2D1_S337 - cost: 3.60, layer: 42
Normal
Diabetic
Normal data
Fold over basal
1.5
1
0.5
0
0
10
20
30
40
50
60
Time (min)

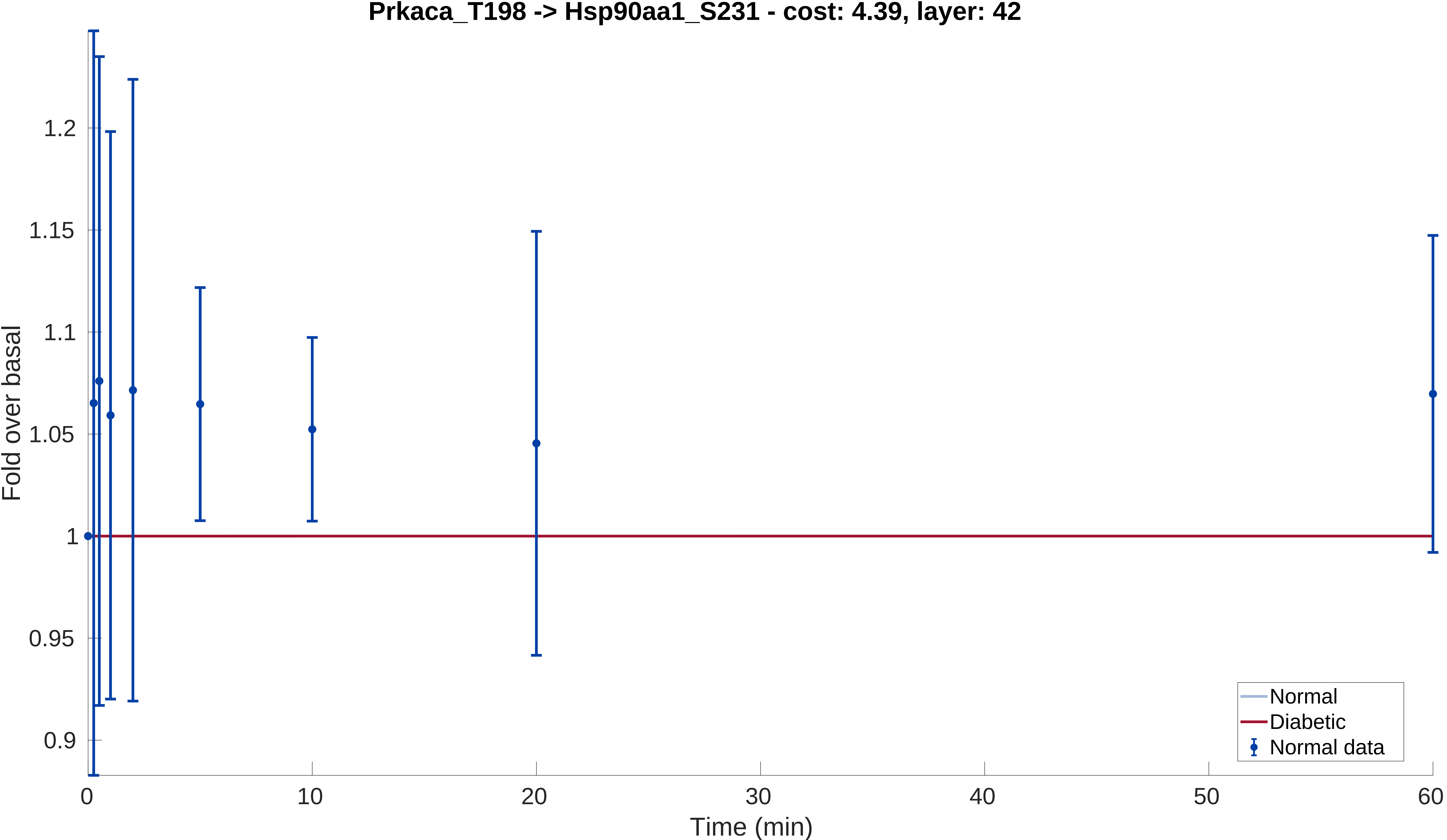

Prkaca_T198 -> Hsp90aa1_S231 - cost: 4.39, layer: 42
Fold over basal
1.2
1.15
1.1
1.05
1
0.95
0.9
0
10
20
30
40
50
60
Time (min)
Normal
Diabetic
Normal data

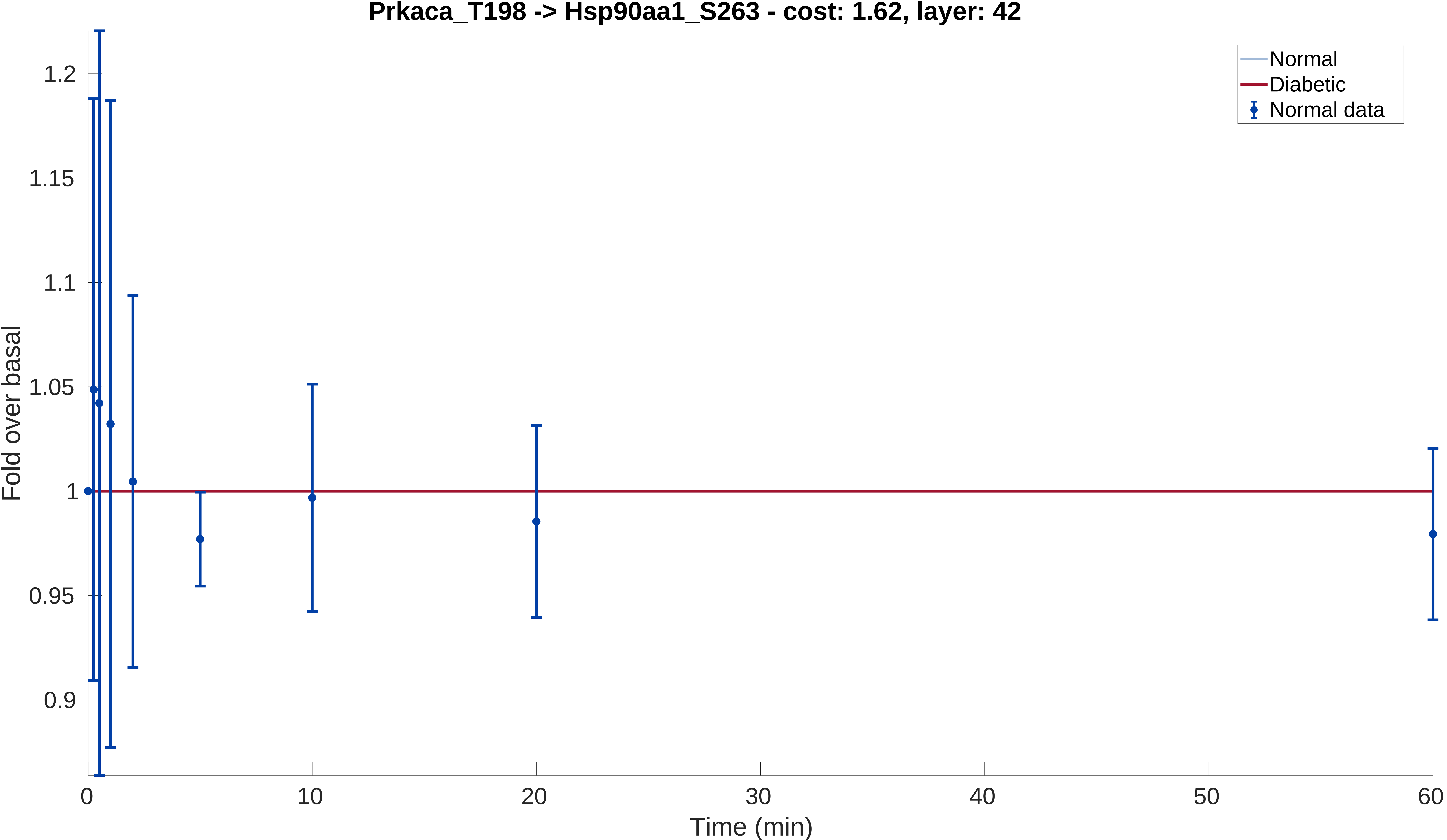

Prkaca_T198 -> Hsp90aa1_S263 - cost: 1.62, layer: 42
Normal
Diabetic
Normal data
Fold over basal
1.2
1.15
1.1
1.05
1
0.95
0.9
0
10
20
30
40
50
60
Time (min)

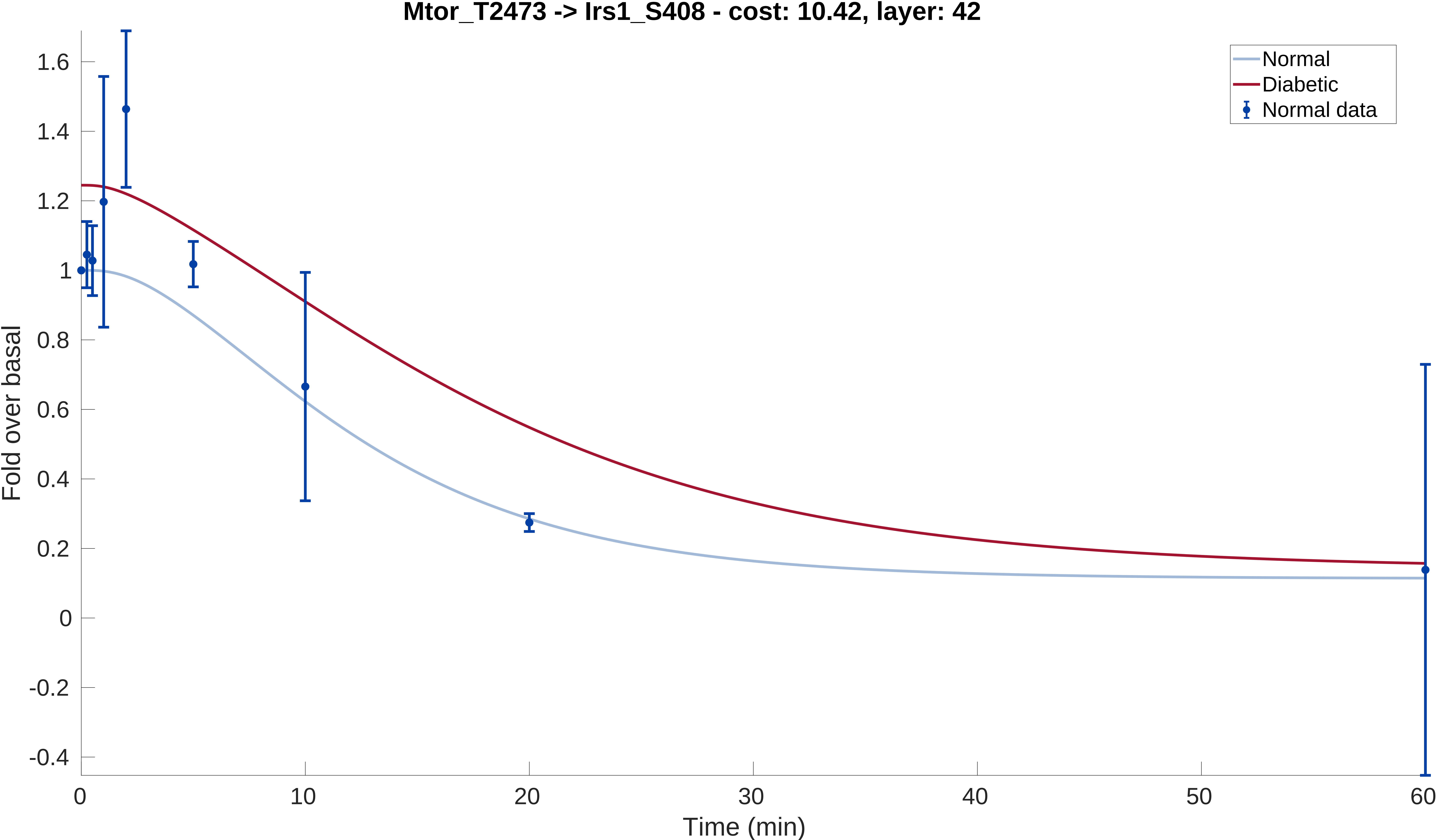
Mtor_T2473 -> Irs1_S408 - cost: 10.42, layer: 42
Normal
Diabetic
Normal data
Fold over basal
Time (min)
1.6
1.4
1.2
1
0.8
0.6
0.4
0.2
0
-0.2
-0.4
0
10
20
30
40
50
60

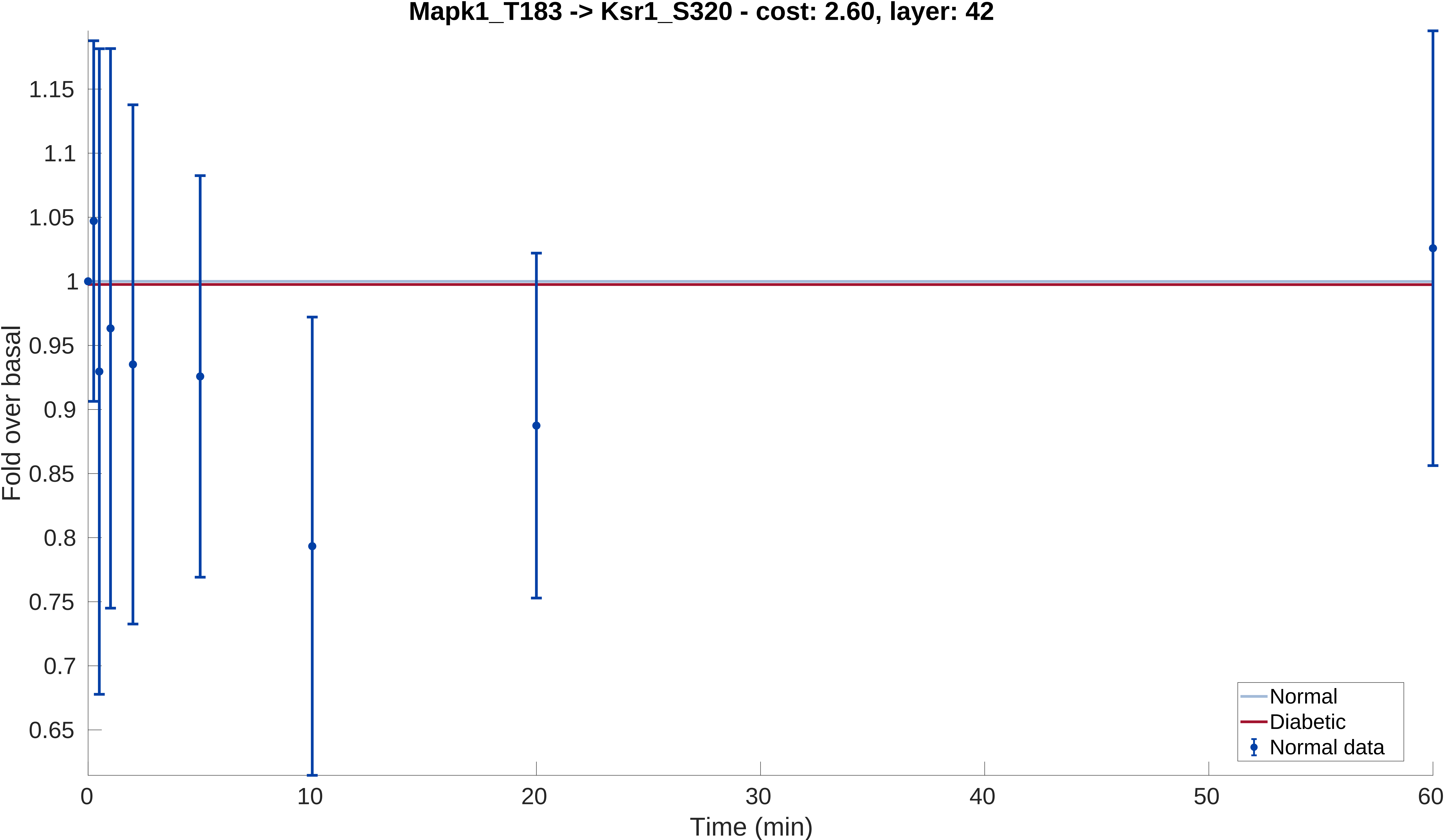
Mapk1_T183 -> Ksr1_S320 - cost: 2.60, layer: 42
Fold over basal
1.15
1.1
1.05
1
0.95
0.9
0.85
0.8
0.75
0.7
0.65
0
10
20
30
40
50
60
Time (min)
Normal
Diabetic
Normal data

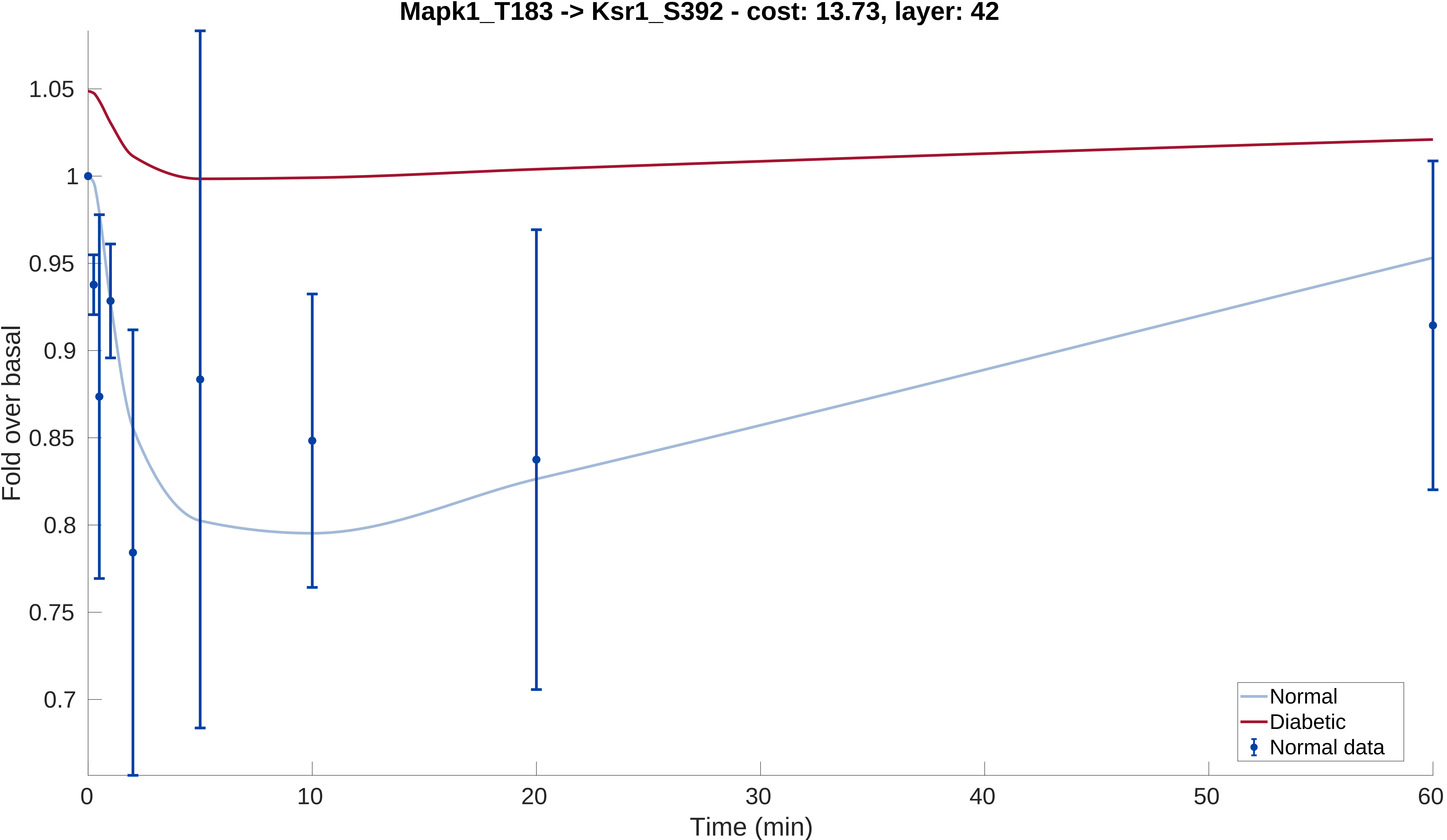

Mapk1_T183 -> Ksr1_S392 - cost: 13.73, layer: 42
1.05
1
0.95
0.9
0.85
0.8
0.75
0.7
Fold over basal
0
10
20
30
40
50
60
Time (min)
Normal
Diabetic
Normal data

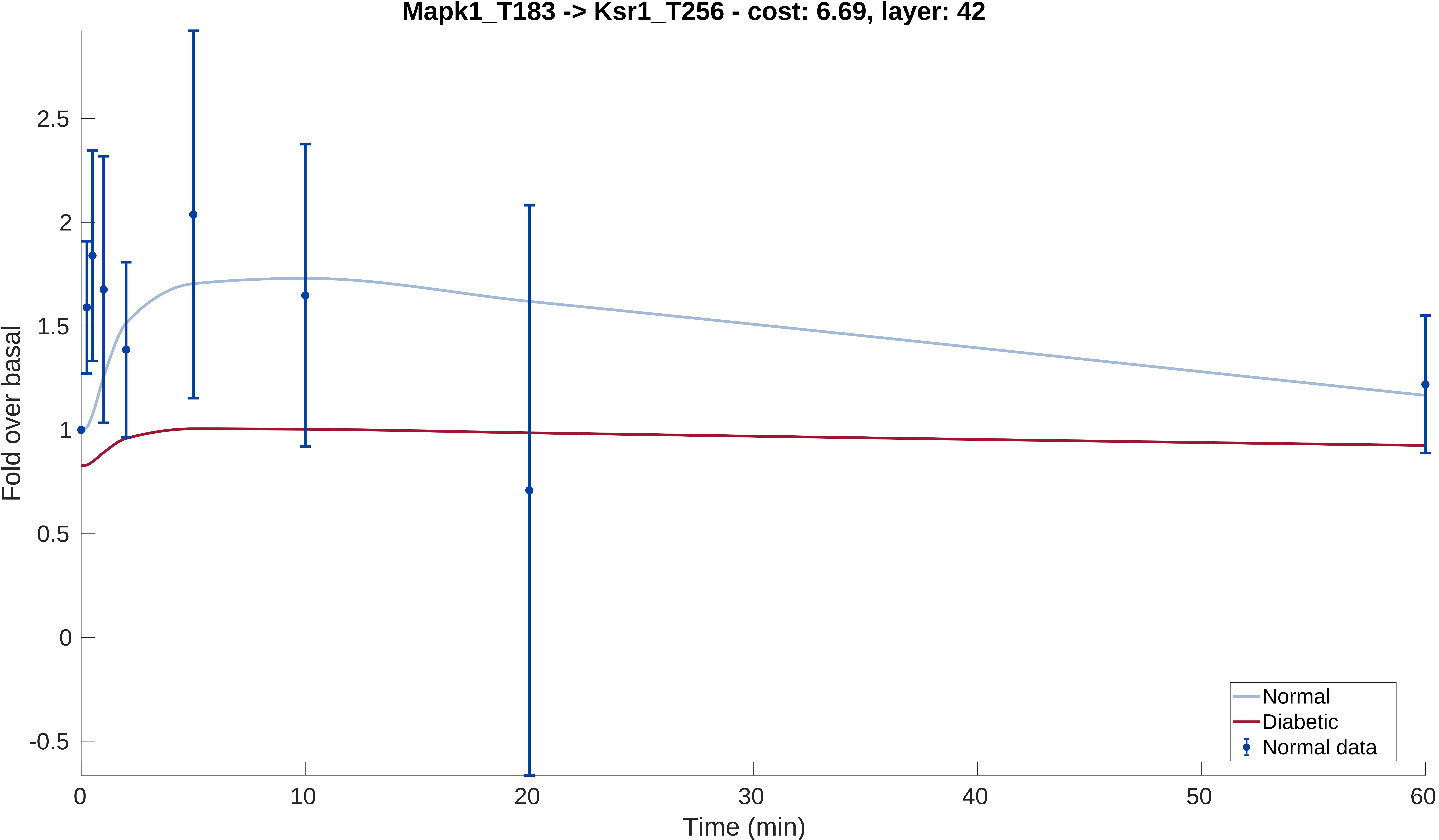

Mapk1_T183 -> Ksr1_T256 - cost: 6.69, layer: 42
Fold over basal
Time (min)
Normal
Diabetic
Normal data
2.5
2
1.5
1
0.5
0
-0.5
0
10
20
30
40
50
60

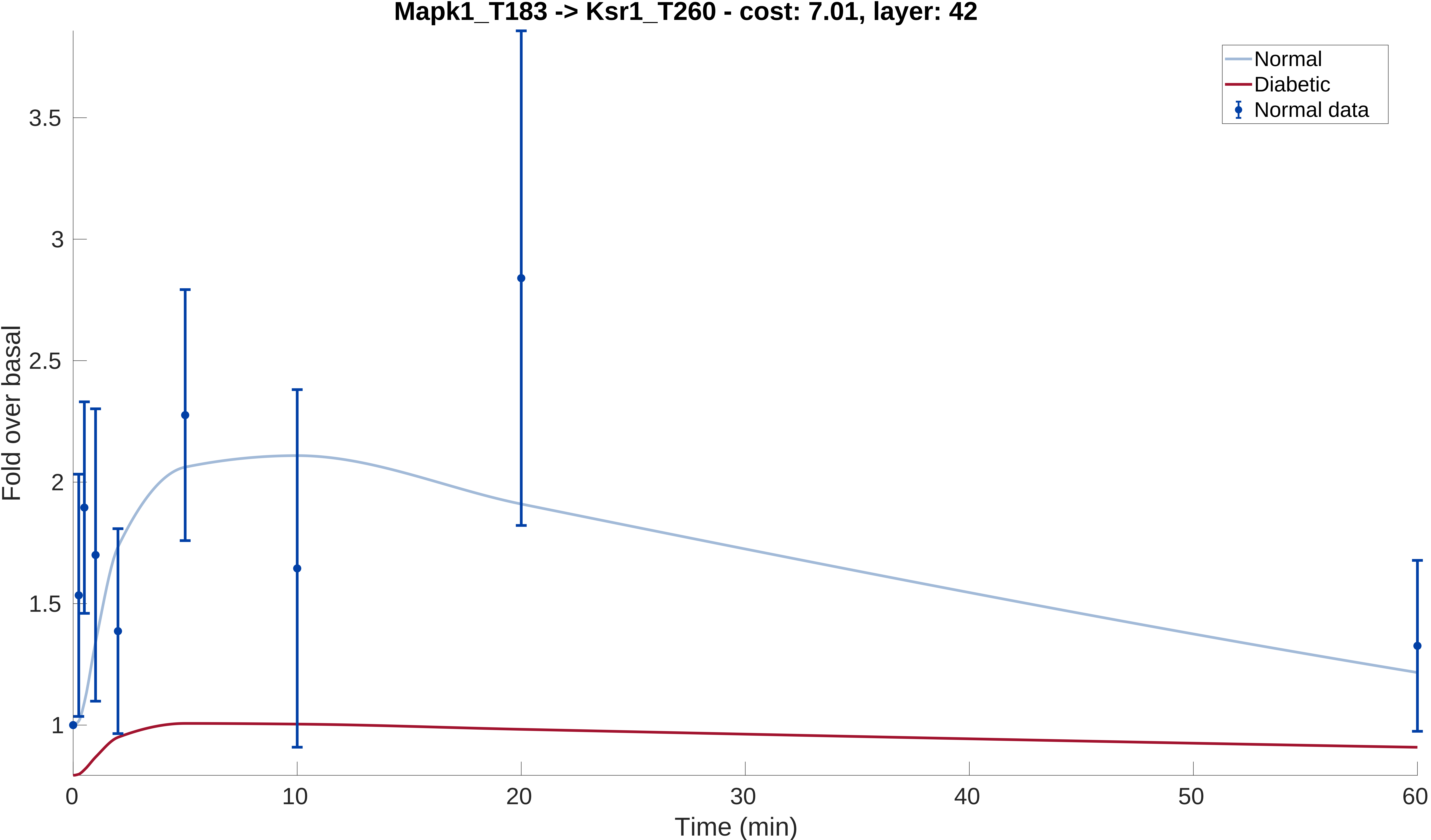

Mapk1_T183 -> Ksr1_T260 - cost: 7.01, layer: 42
Normal
Diabetic
Normal data
Fold over basal
3.5
3
2.5
2
1.5
1
0
10
20
30
40
50
60
Time (min)

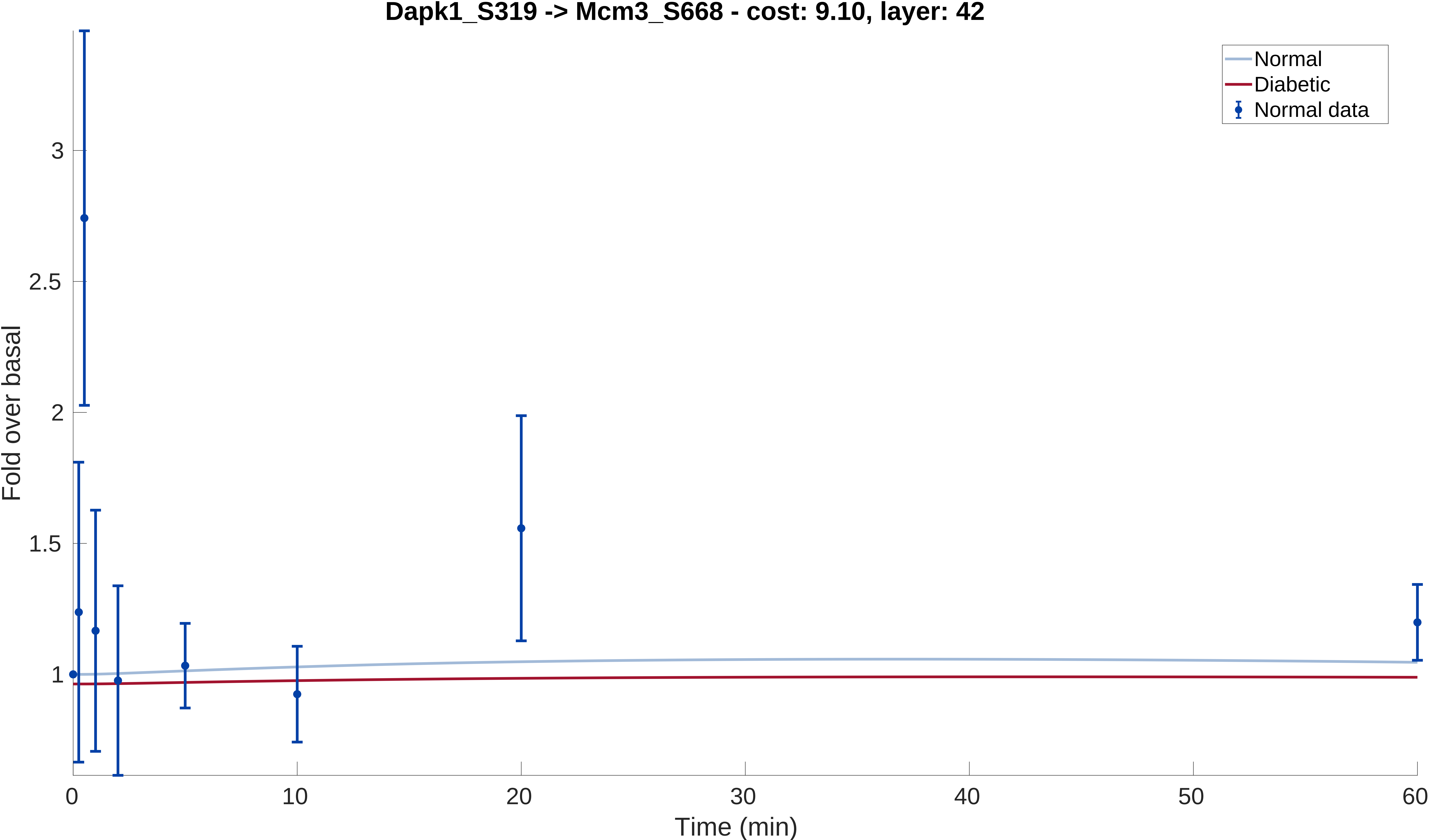

Dapk1_S319 -> Mcm3_S668 - cost: 9.10, layer: 42
Normal
Diabetic
Normal data
Fold over basal
3
2.5
2
1.5
1
0
10
20
30
40
50
60
Time (min)

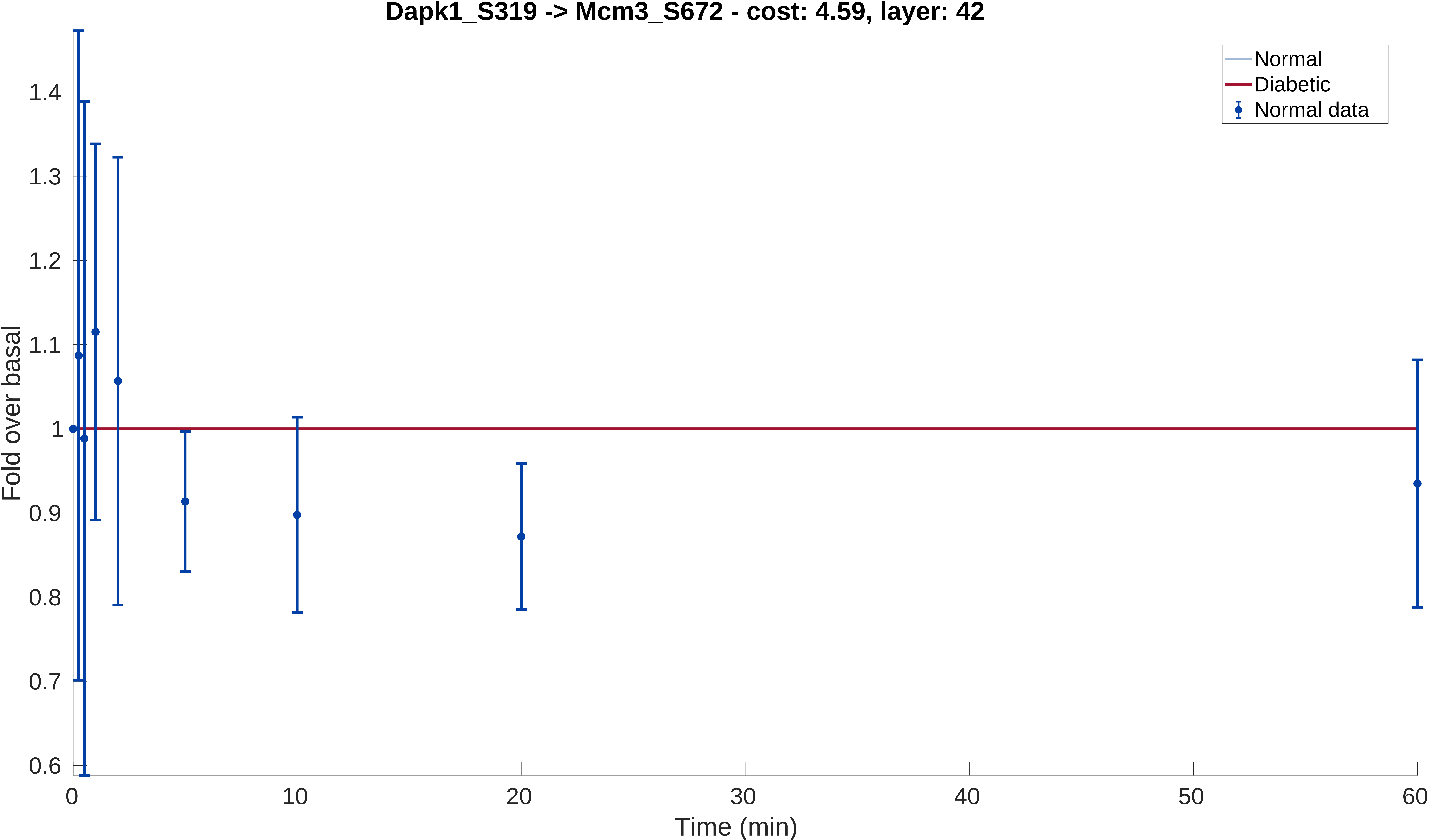
Dapk1_S319 -> Mcm3_S672 - cost: 4.59, layer: 42
Normal
Diabetic
Normal data
Fold over basal
1.4
1.3
1.2
1.1
1
0.9
0.8
0.7
0.6
0
10
20
30
40
50
60
Time (min)

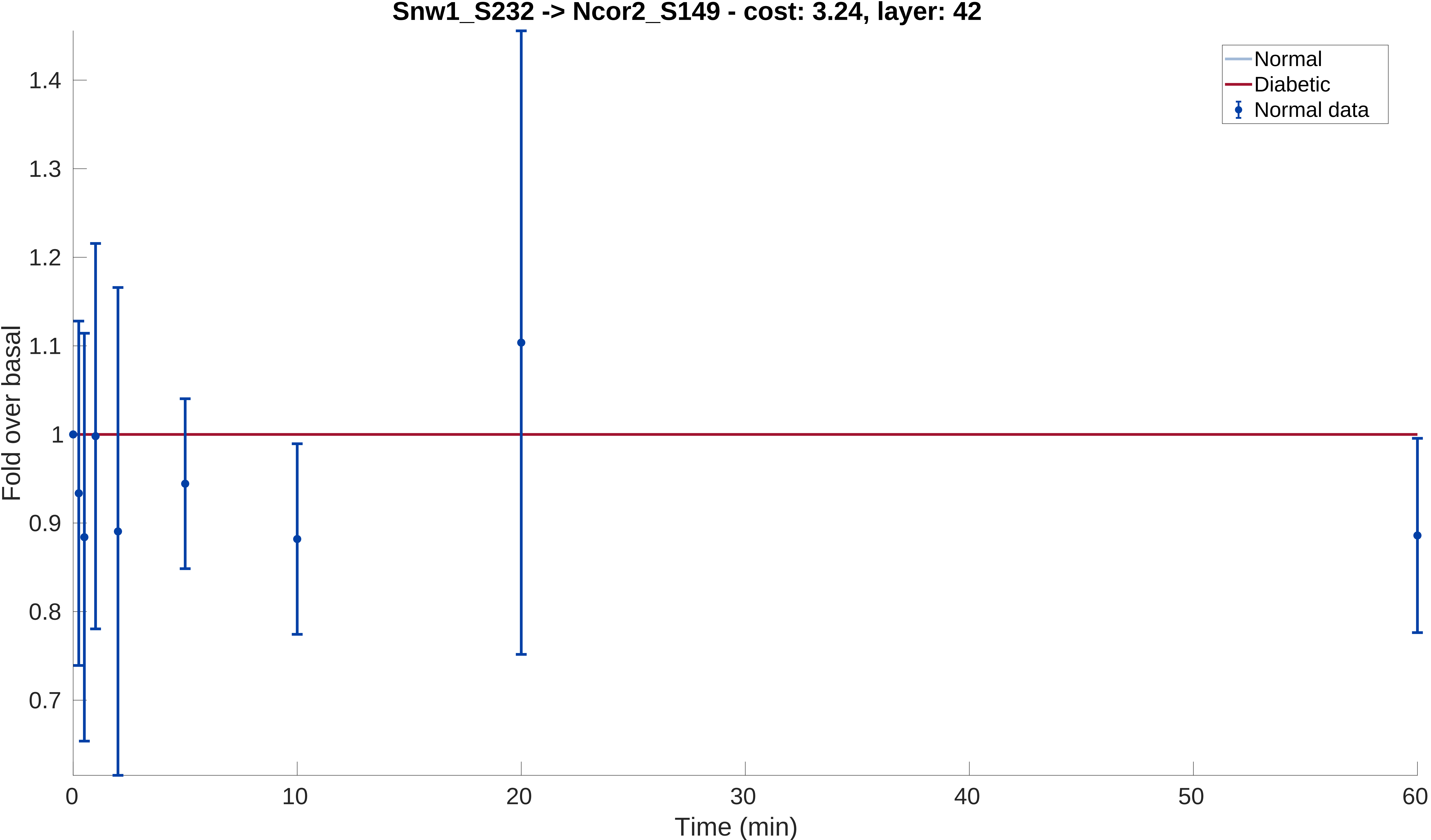

Snw1_S232 -> Ncor2_S149 - cost: 3.24, layer: 42
Normal
Diabetic
Normal data
Fold over basal
1.4
1.3
1.2
1.1
1
0.9
0.8
0.7
0
10
20
30
40
50
60
Time (min)

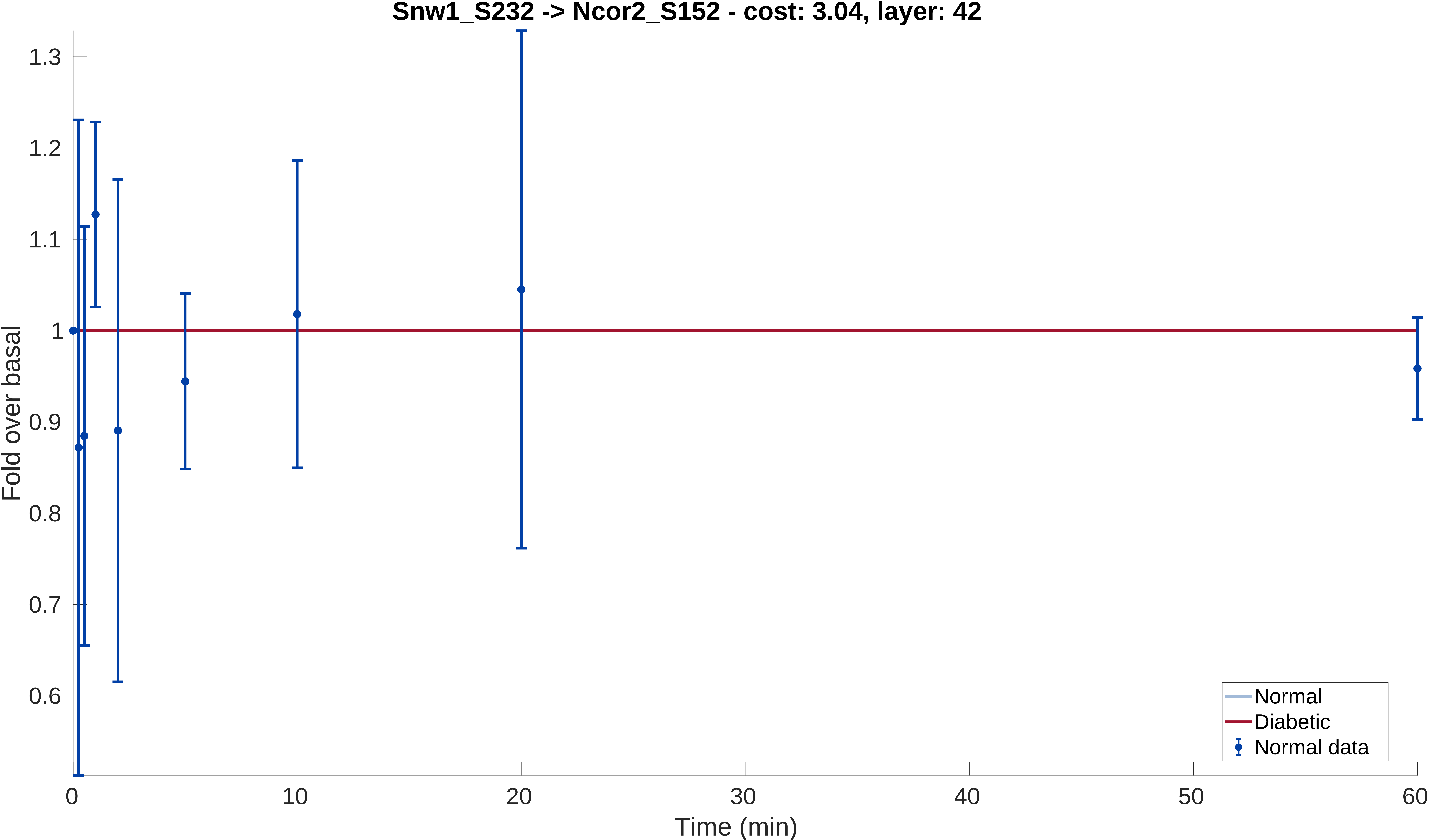
Snw1_S232 -> Ncor2_S152 - cost: 3.04, layer: 42
Fold over basal
1.3
1.2
1.1
1
0.9
0.8
0.7
0.6
0
10
20
30
40
50
60
Time (min)
Normal
Diabetic
Normal data

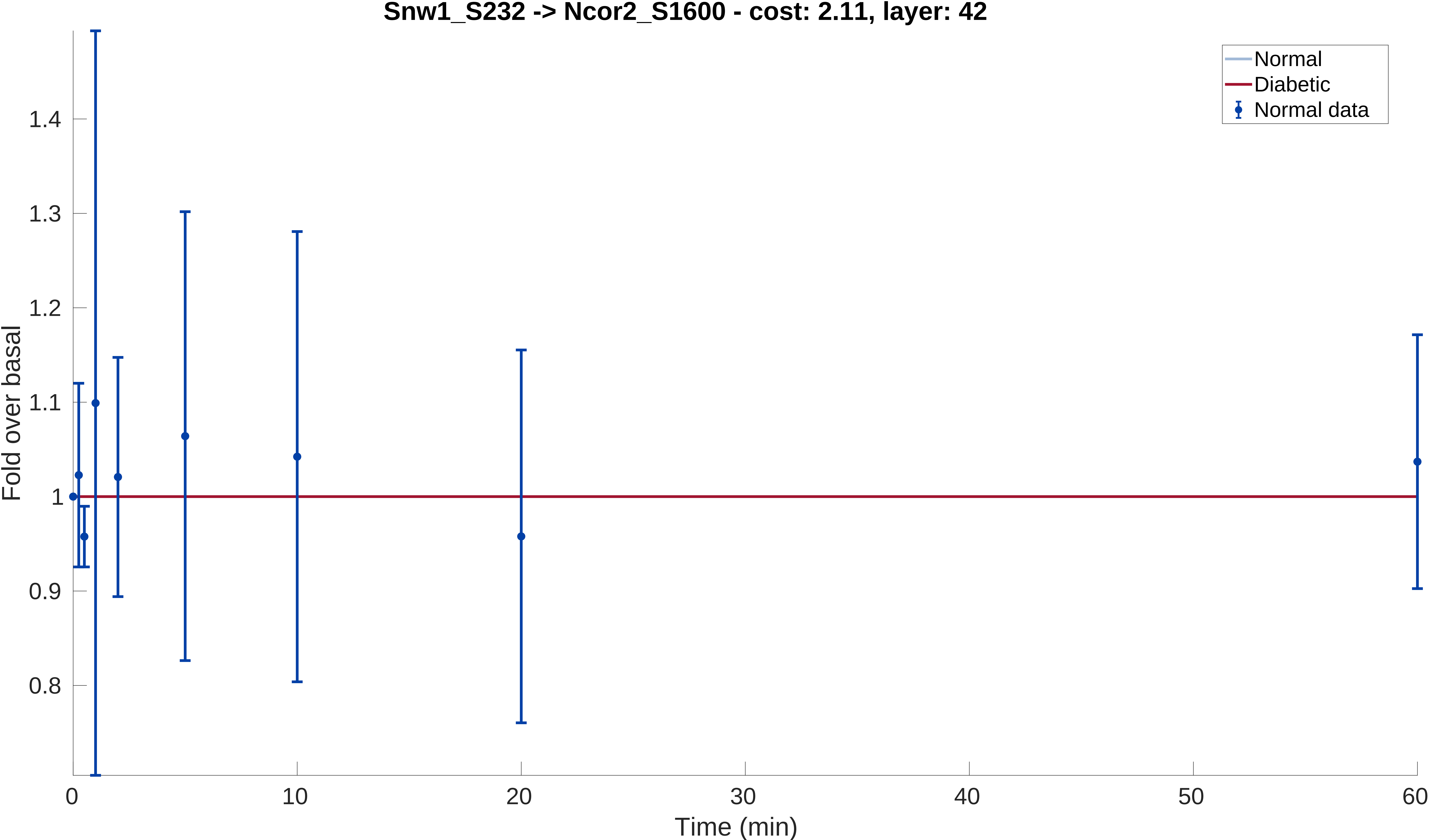
Snw1_S232 -> Ncor2_S1600 - cost: 2.11, layer: 42
Normal
Diabetic
Normal data
Fold over basal
1.4
1.3
1.2
1.1
1
0.9
0.8
0
10
20
30
40
50
60
Time (min)

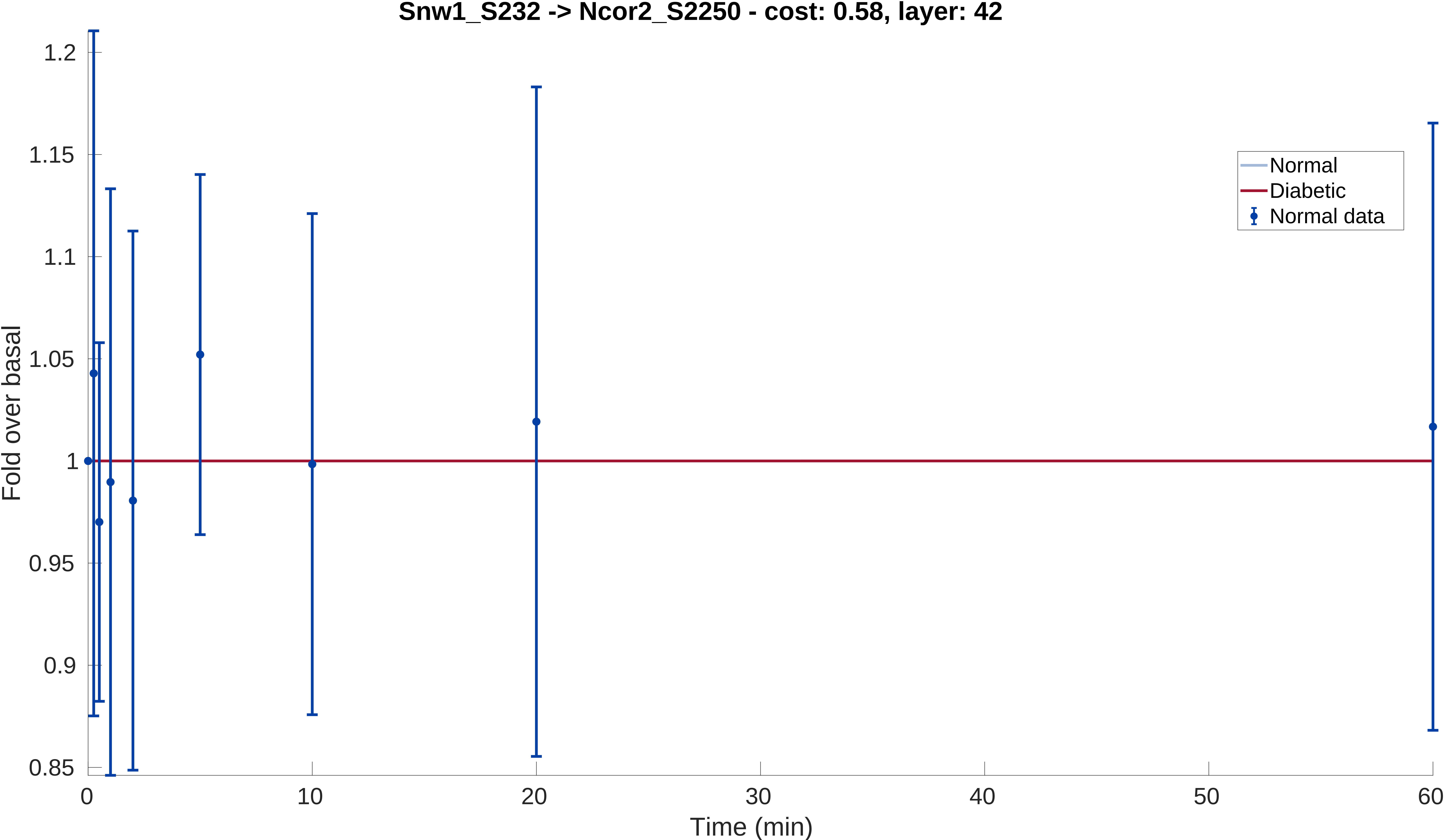

Snw1_S232 -> Ncor2_S2250 - cost: 0.58, layer: 42
Fold over basal
1.2
1.15
1.1
1.05
1
0.95
0.9
0.85
0
10
20
30
40
50
60
Time (min)
Normal
Diabetic
Normal data

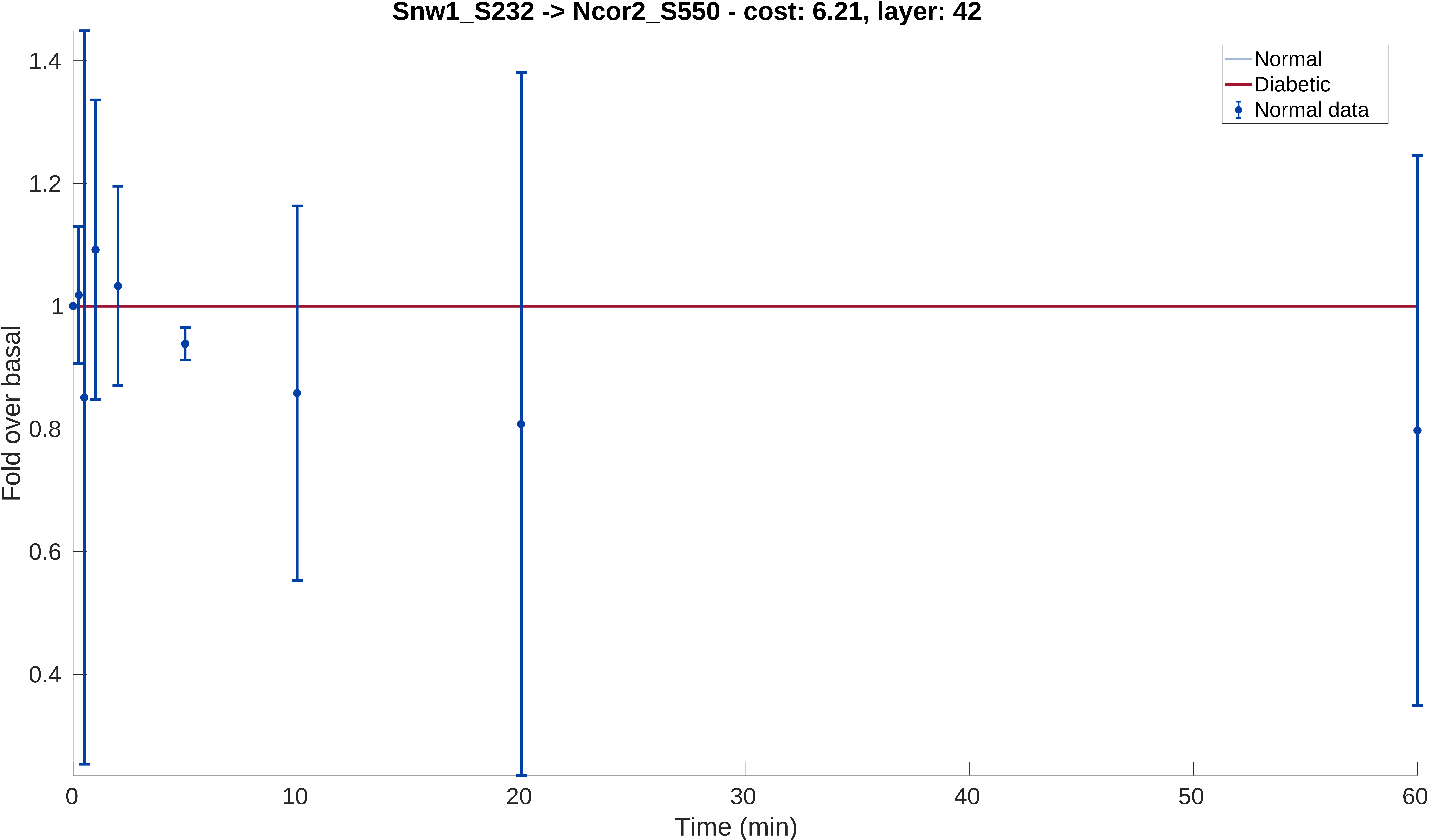

Snw1_S232 -> Ncor2_S550 - cost: 6.21, layer: 42
Normal
Diabetic
Normal data
Fold over basal
1.4
1.2
1
0.8
0.6
0.4
0
10
20
30
40
50
60
Time (min)

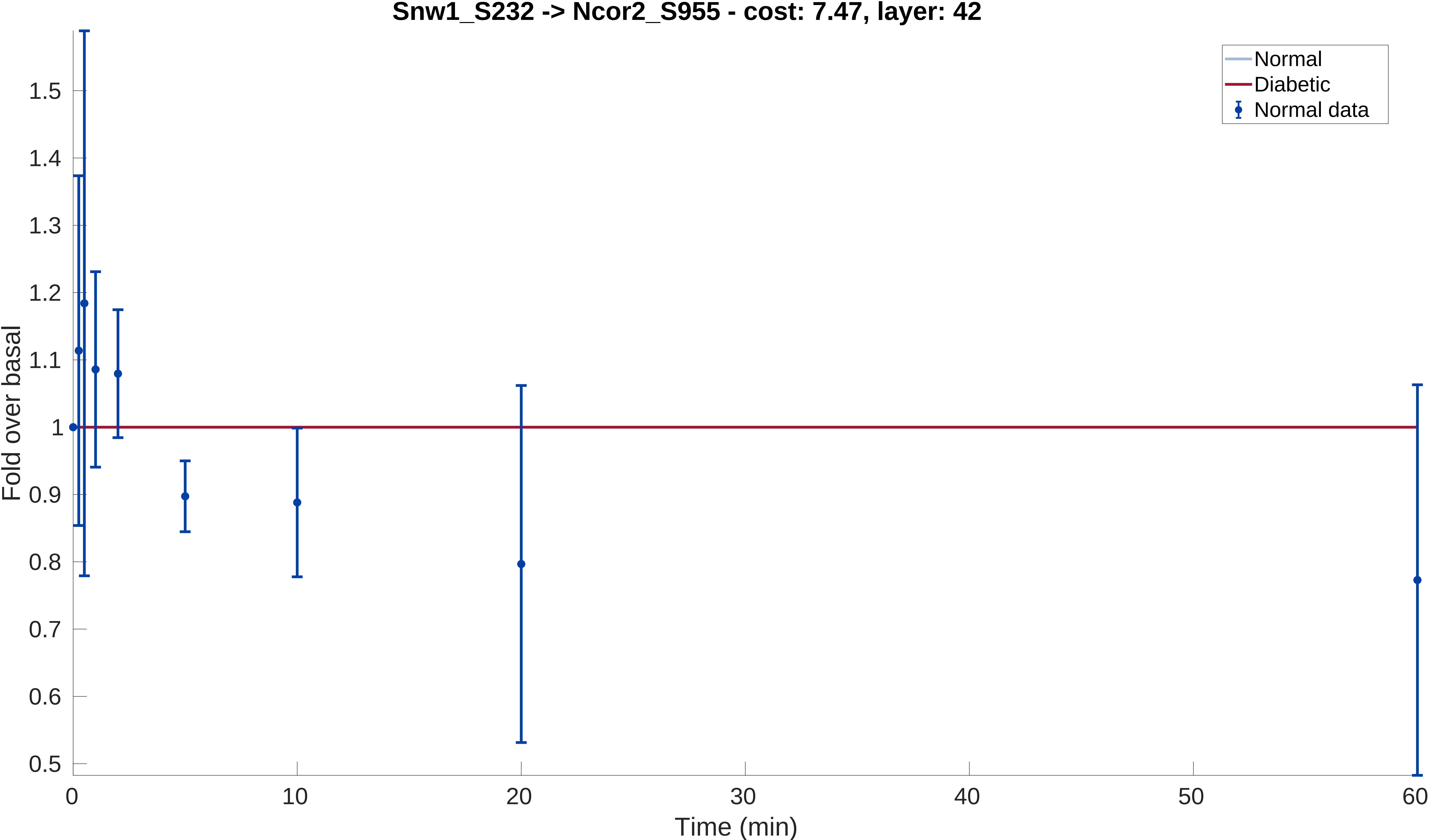
Snw1_S232 -> Ncor2_S955 - cost: 7.47, layer: 42
Normal
Diabetic
Normal data
Fold over basal
1.5
1.4
1.3
1.2
1.1
1
0.9
0.8
0.7
0.6
0.5
0
10
20
30
40
50
60
Time (min)

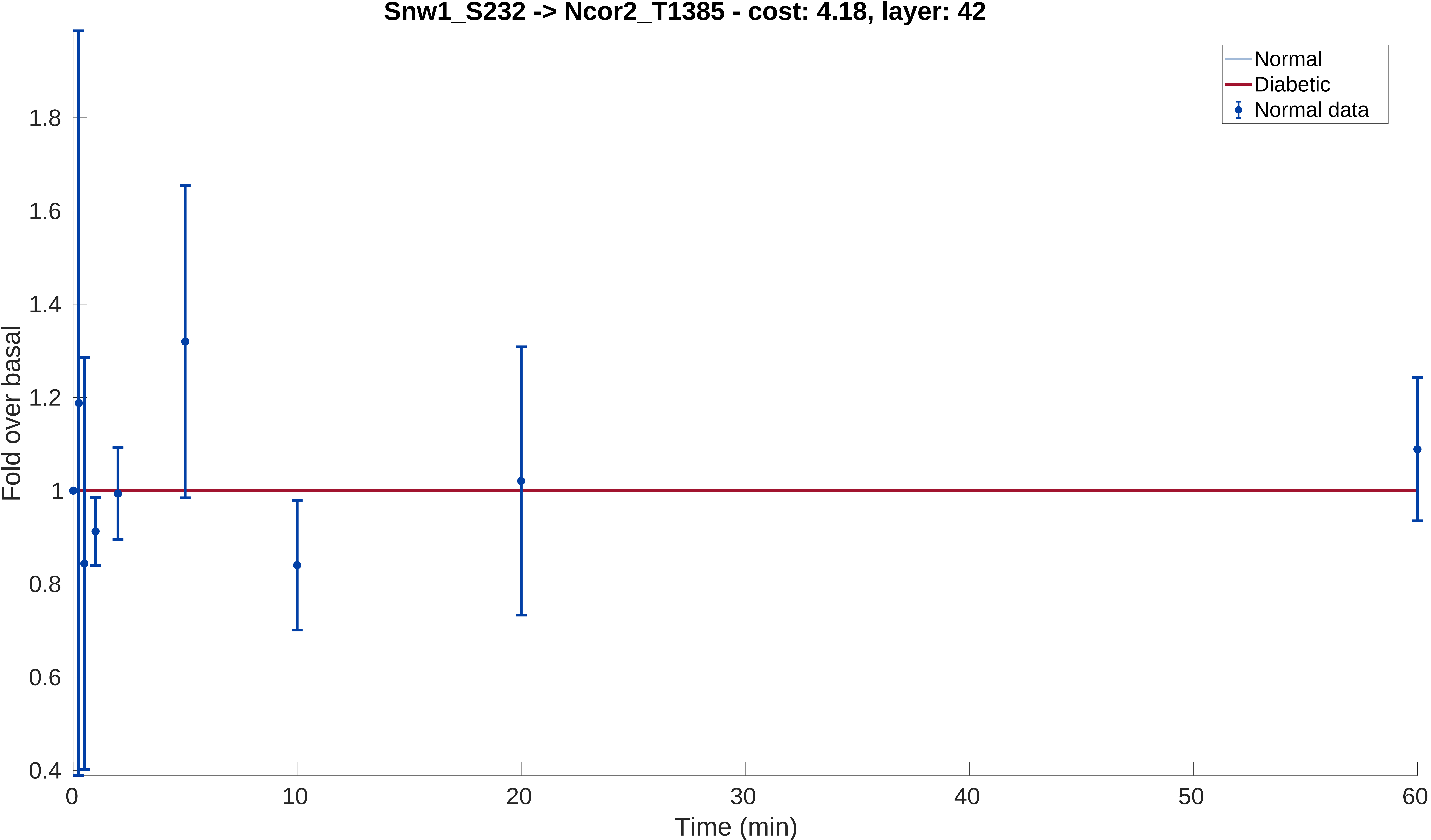
Snw1_S232 -> Ncor2_T1385 - cost: 4.18, layer: 42
Normal
Diabetic
Normal data
Fold over basal
1.8
1.6
1.4
1.2
1
0.8
0.6
0.4
0
10
20
30
40
50
60
Time (min)

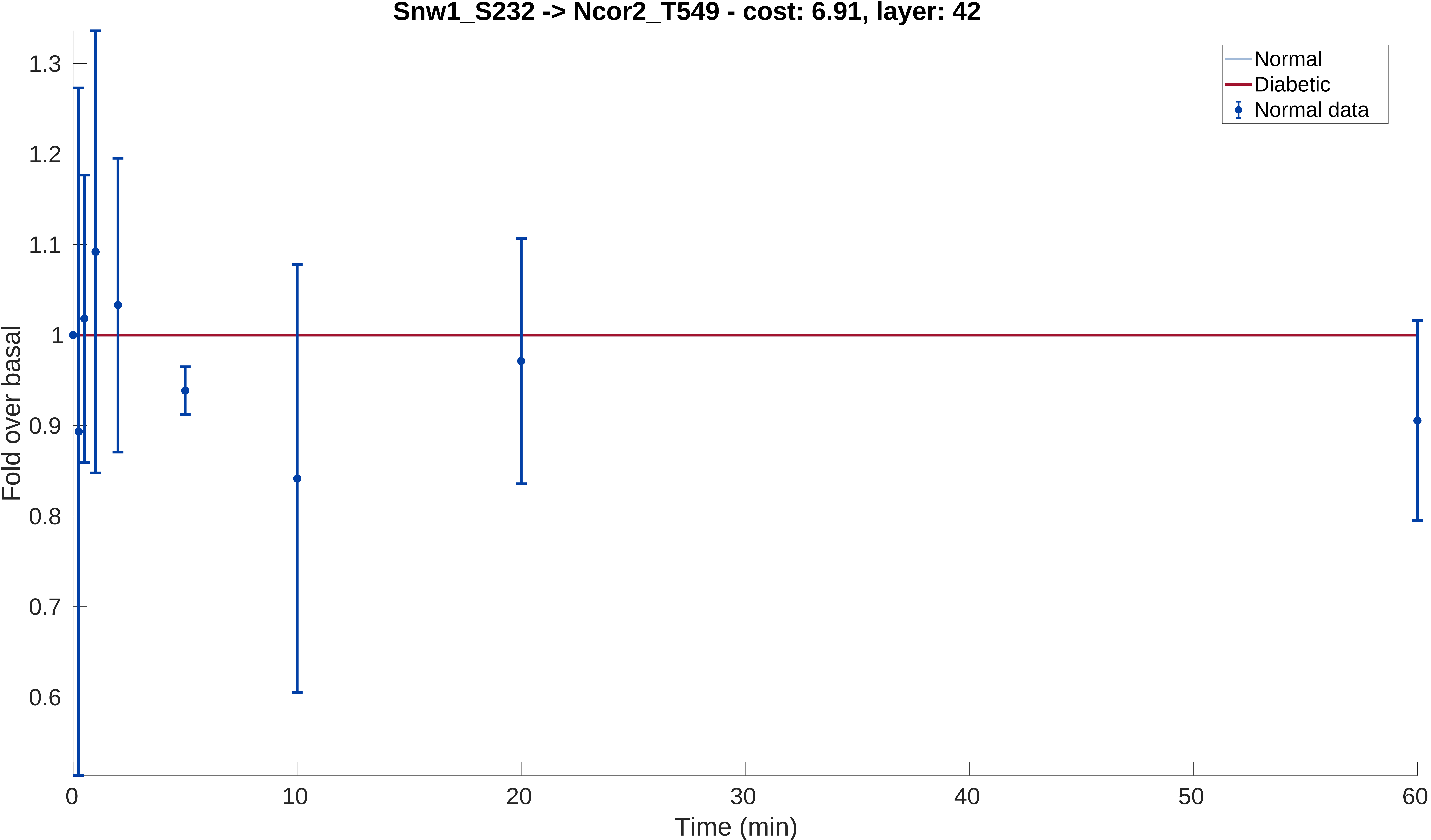

Snw1_S232 -> Ncor2_T549 - cost: 6.91, layer: 42
Normal
Diabetic
Normal data
Fold over basal
Time (min)
1.3
1.2
1.1
1
0.9
0.8
0.7
0.6
0
10
20
30
40
50
60

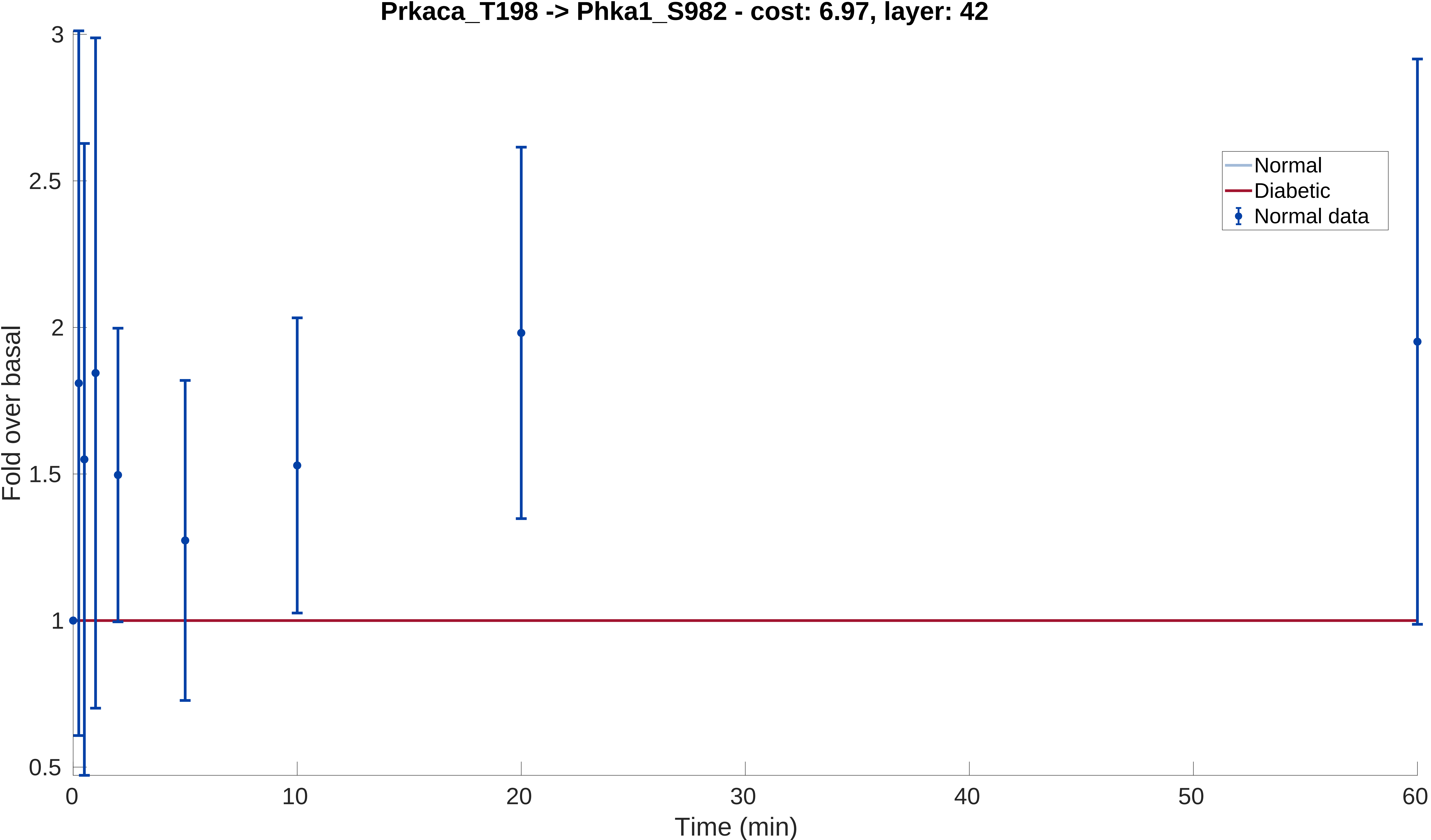

Prkaca_T198 -> Phka1_S982 - cost: 6.97, layer: 42
Fold over basal
Time (min)
Normal
Diabetic
Normal data
0.5
1
1.5
2
2.5
3
0
10
20
30
40
50
60

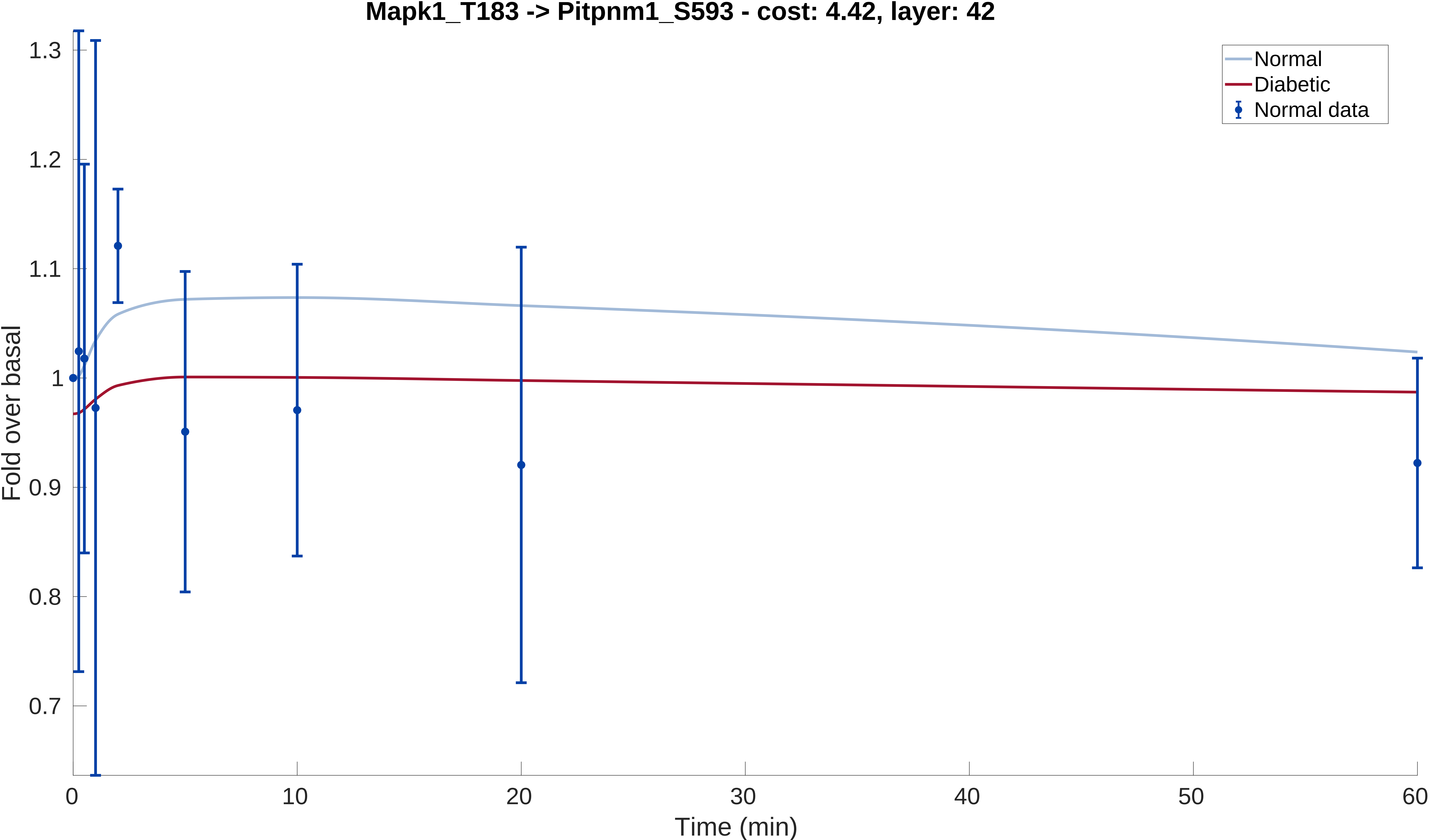

Mapk1_T183 -> Pitpnm1_S593 - cost: 4.42, layer: 42
Normal
Diabetic
Normal data
Fold over basal
1.3
1.2
1.1
1
0.9
0.8
0.7
0
10
20
30
40
50
60
Time (min)

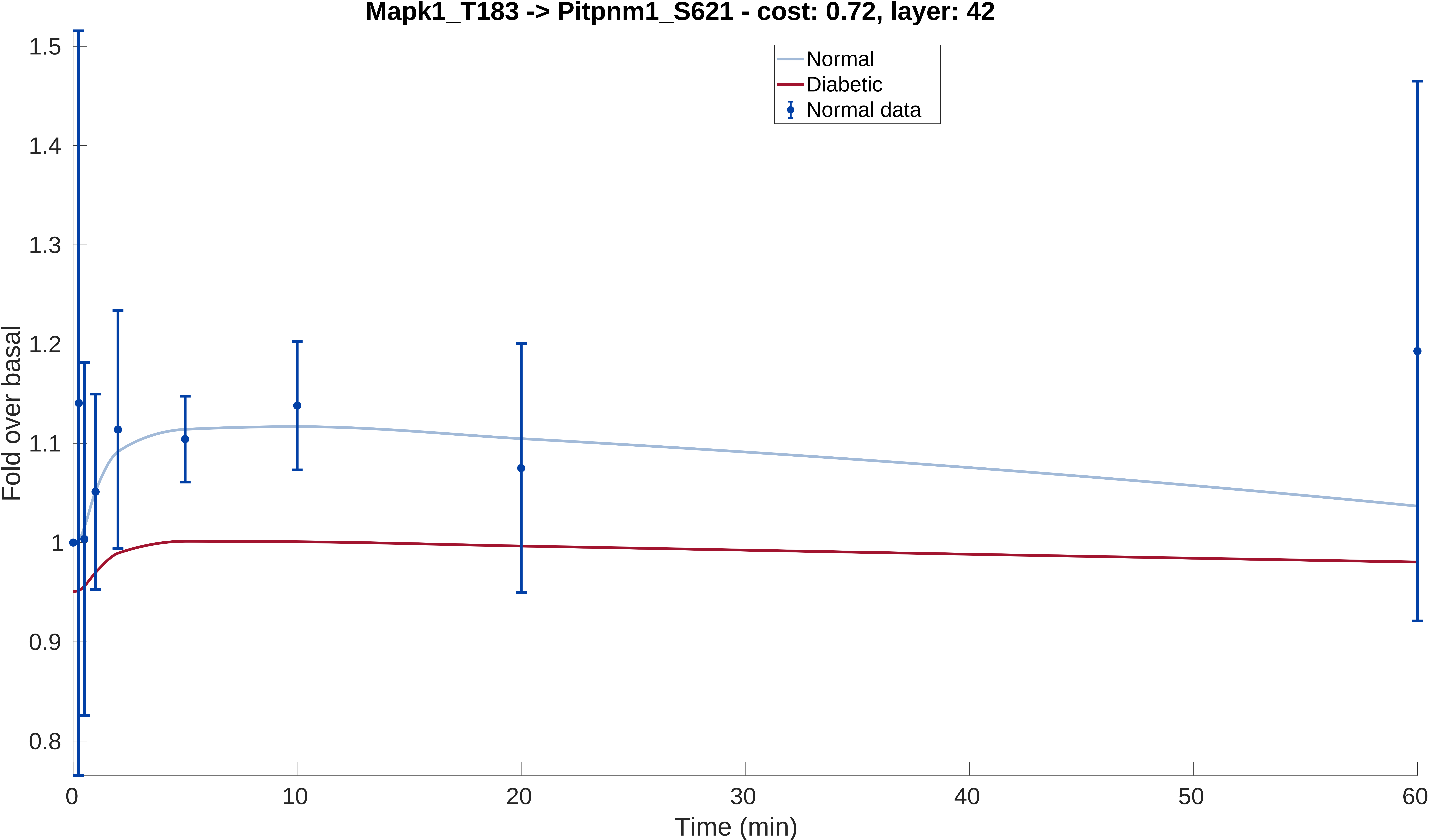

Mapk1_T183 -> Pitpnm1_S621 - cost: 0.72, layer: 42
Normal
Diabetic
Normal data
Fold over basal
1.5
1.4
1.3
1.2
1.1
1
0.9
0.8
0
10
20
30
40
50
60
Time (min)

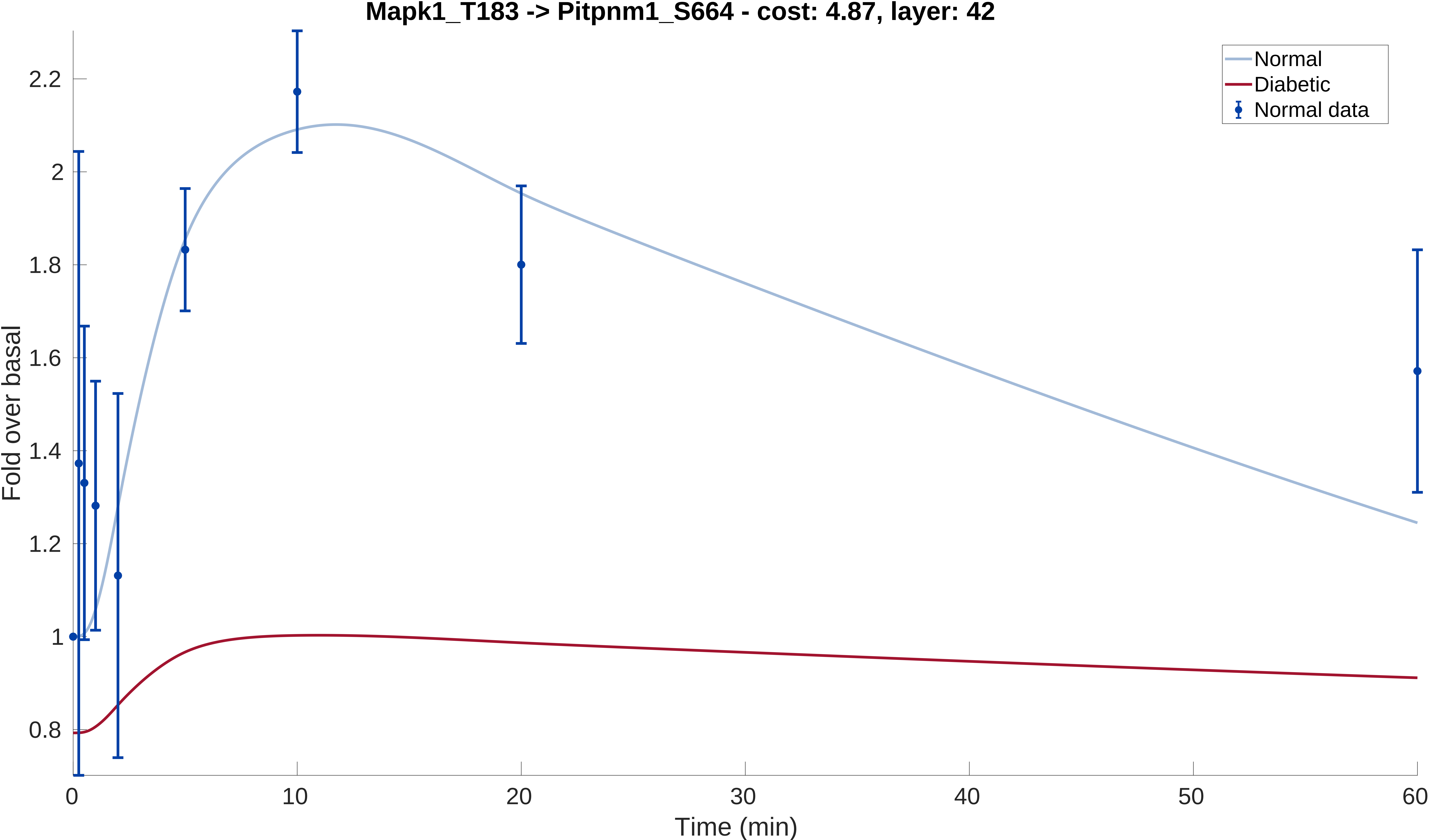

Mapk1_T183 -> Pitpnm1_S664 - cost: 4.87, layer: 42
Normal
Diabetic
Normal data
Fold over basal
0.8
1
1.2
1.4
1.6
1.8
2
2.2
0
10
20
30
40
50
60
Time (min)

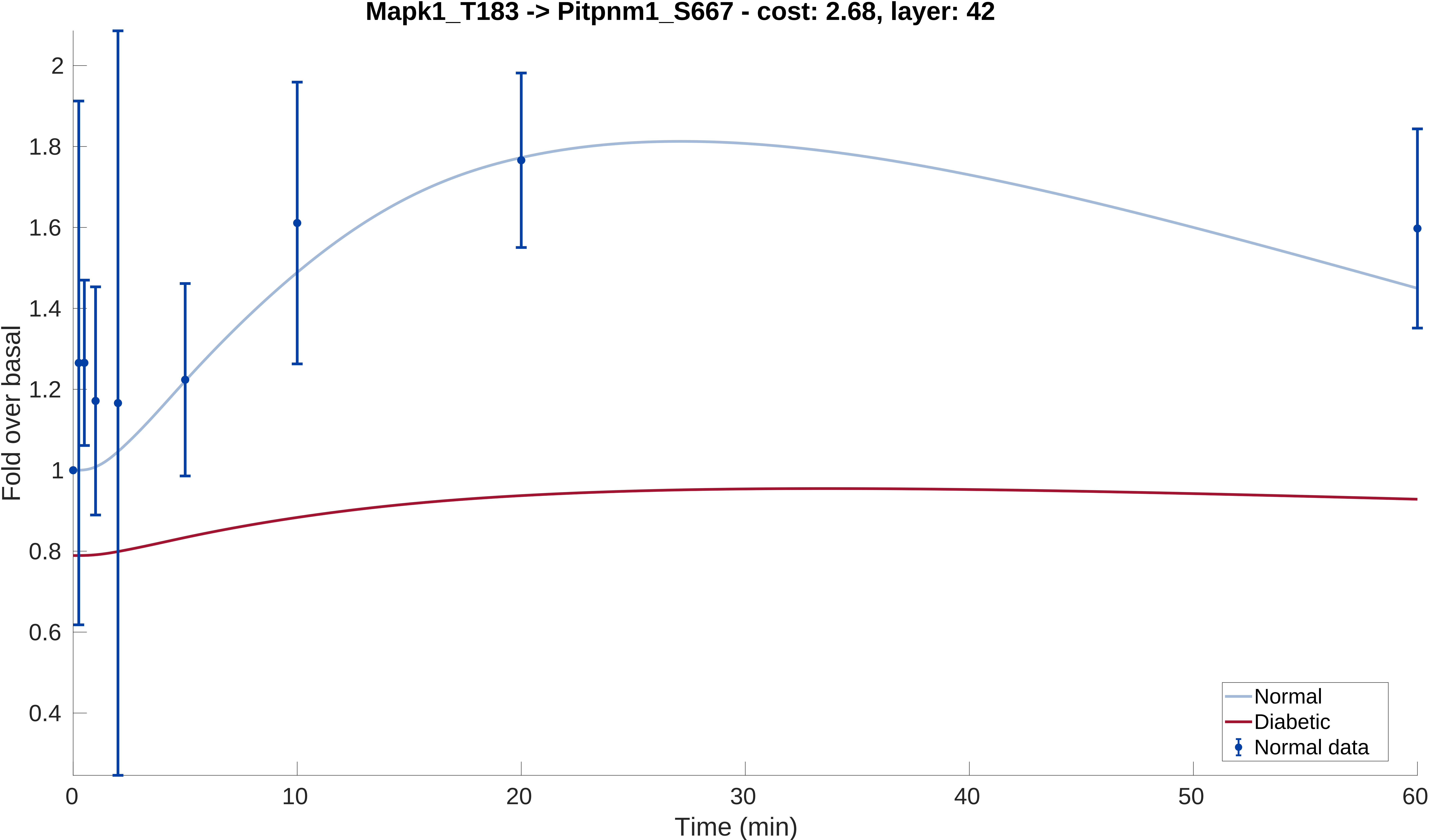
Mapk1_T183 -> Pitpnm1_S667 - cost: 2.68, layer: 42
Fold over basal
Time (min)
0
10
20
30
40
50
60
0.4
0.6
0.8
1
1.2
1.4
1.6
1.8
2
Normal
Diabetic
Normal data

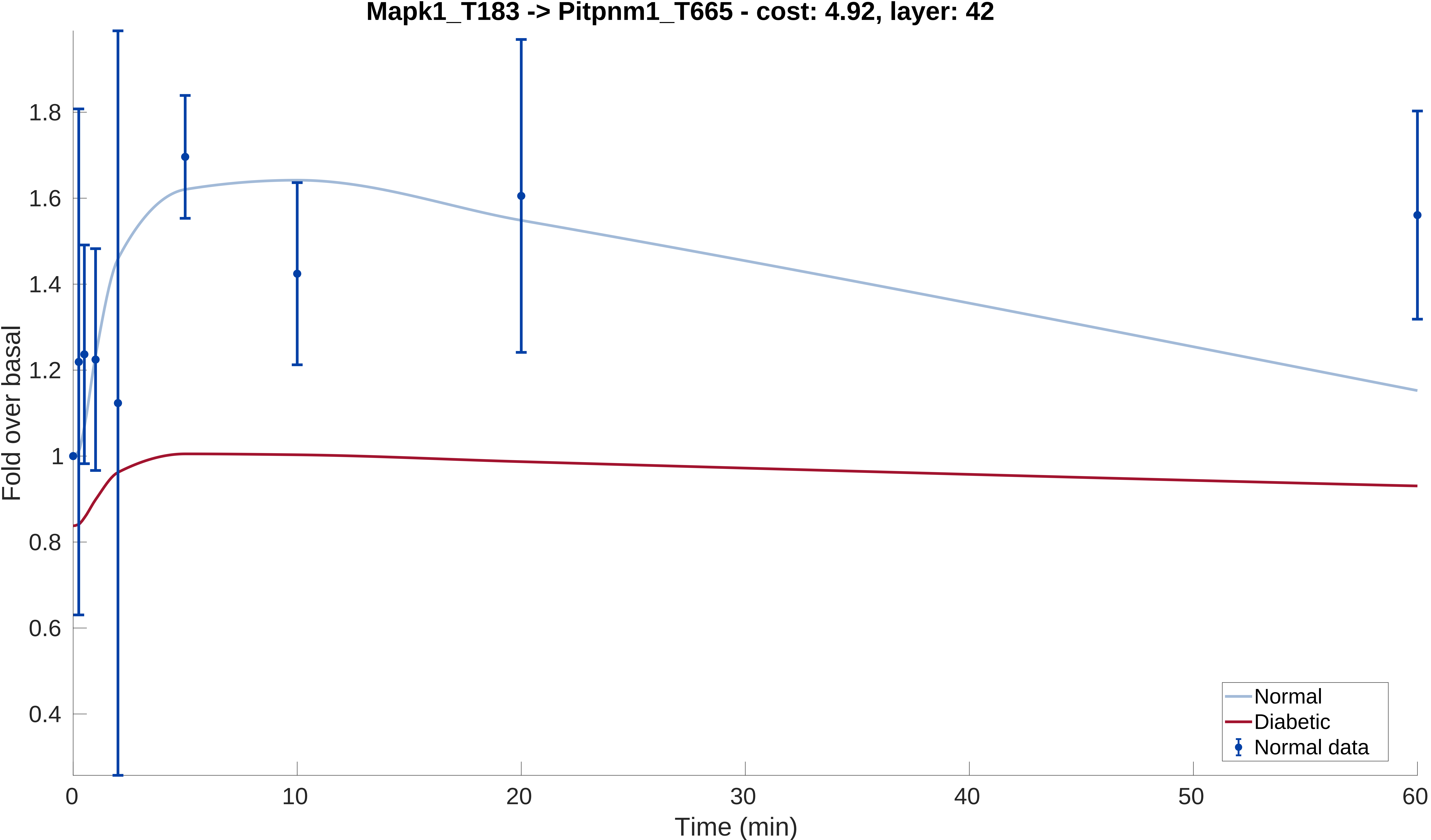

Mapk1_T183 -> Pitpnm1_T665 - cost: 4.92, layer: 42
Fold over basal
1.8
1.6
1.4
1.2
1
0.8
0.6
0.4
0
10
20
30
40
50
60
Time (min)
Normal
Diabetic
Normal data

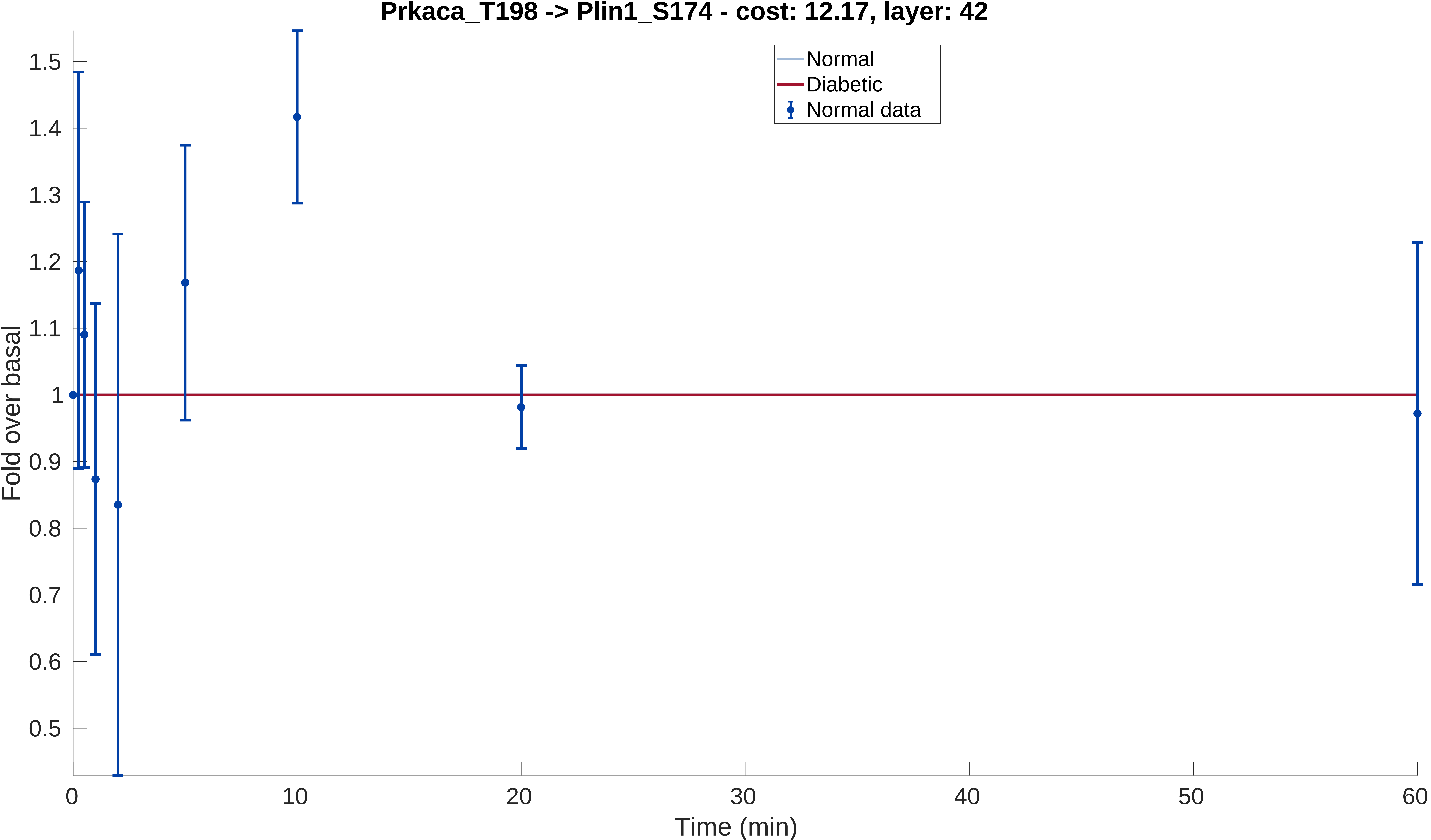

Prkaca_T198 -> Plin1_S174 - cost: 12.17, layer: 42
Normal
Diabetic
Normal data
Fold over basal
1.5
1.4
1.3
1.2
1.1
1
0.9
0.8
0.7
0.6
0.5
0
10
20
30
40
50
60
Time (min)

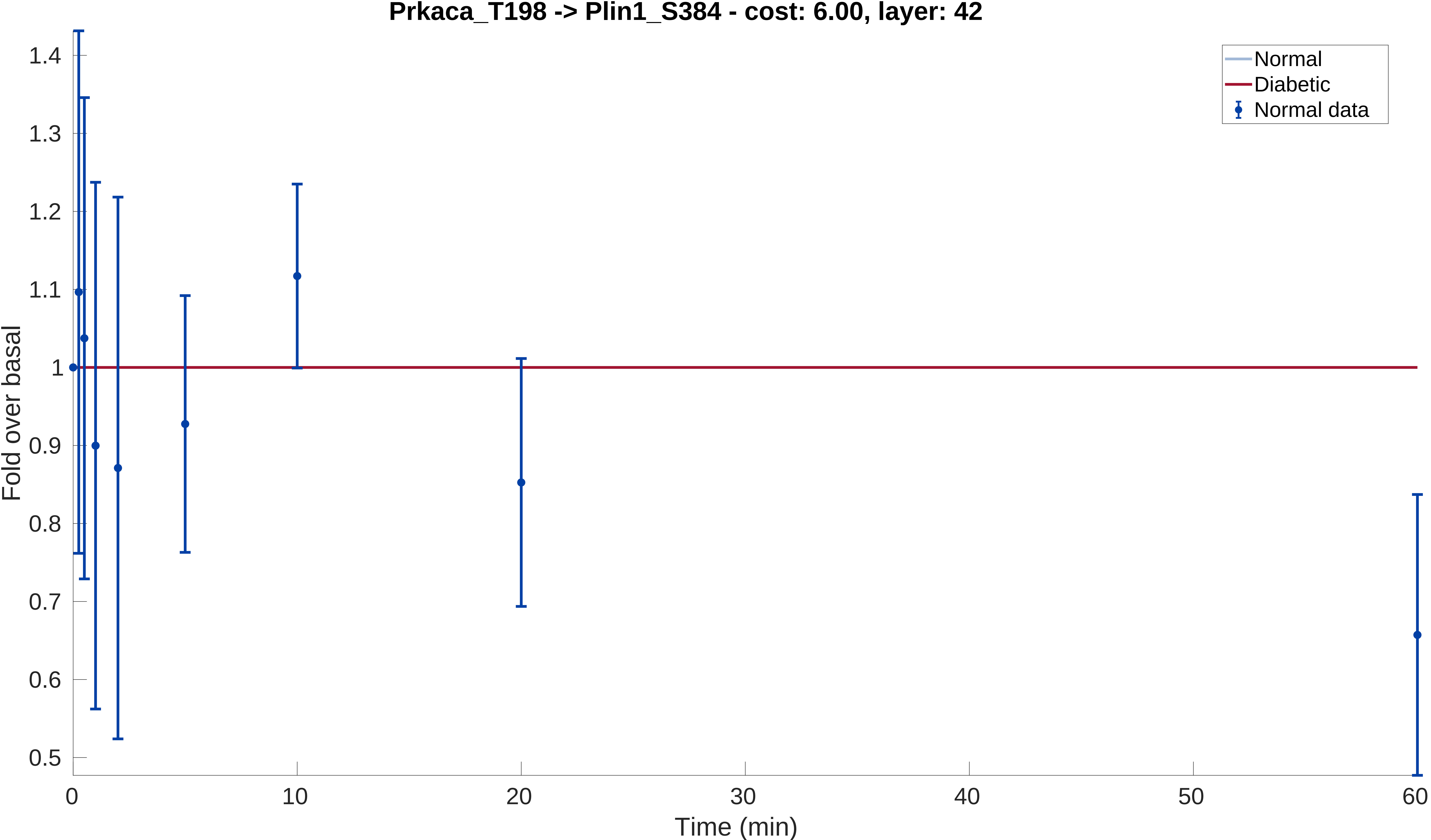

Prkaca_T198 -> Plin1_S384 - cost: 6.00, layer: 42
Normal
Diabetic
Normal data
Fold over basal
1.4
1.3
1.2
1.1
1
0.9
0.8
0.7
0.6
0.5
0
10
20
30
40
50
60
Time (min)

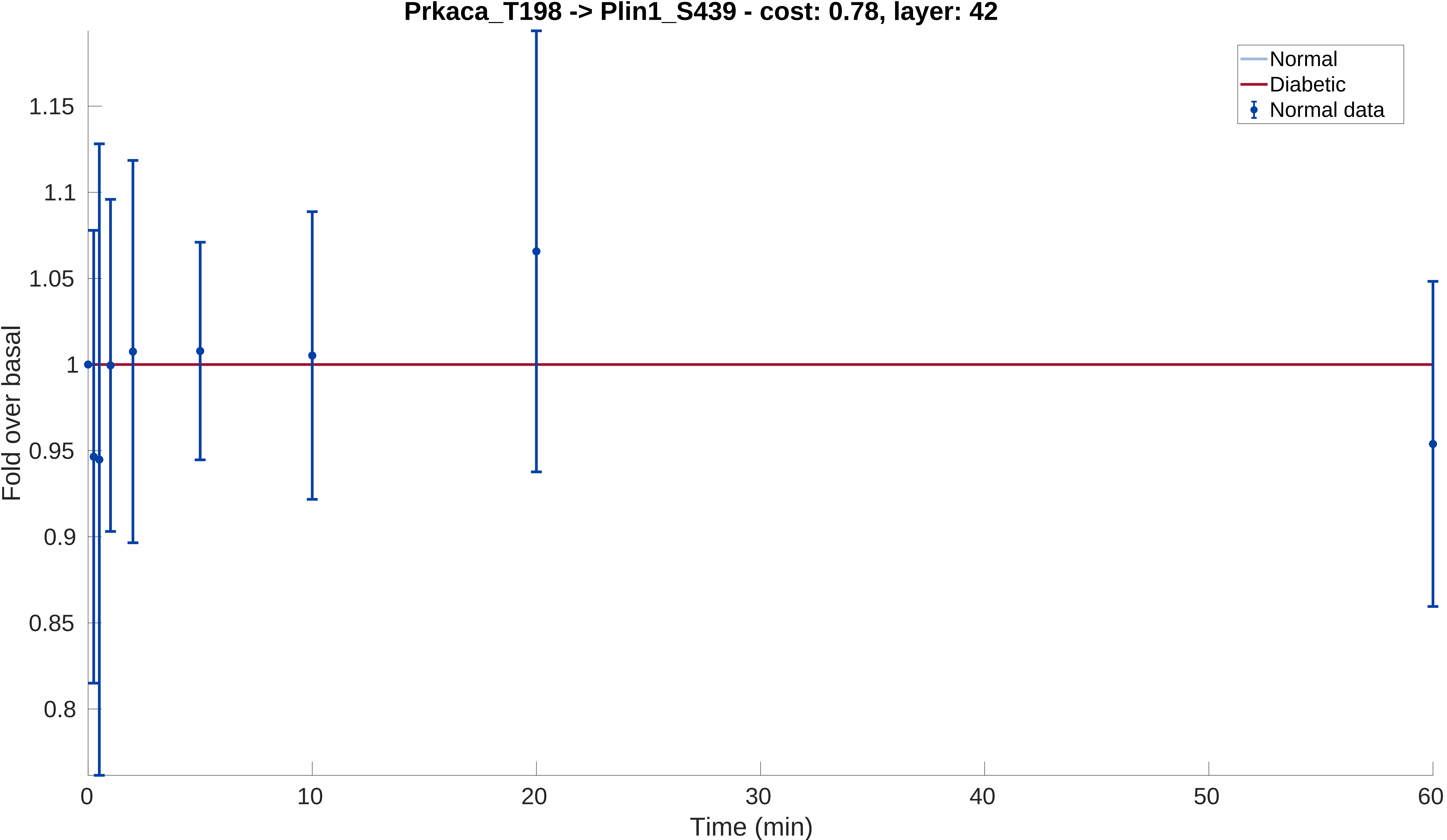

Prkaca_T198 -> Plin1_S439 - cost: 0.78, layer: 42
Normal
Diabetic
Normal data
Fold over basal
1.15
1.1
1.05
1
0.95
0.9
0.85
0.8
0
10
20
30
40
50
60
Time (min)

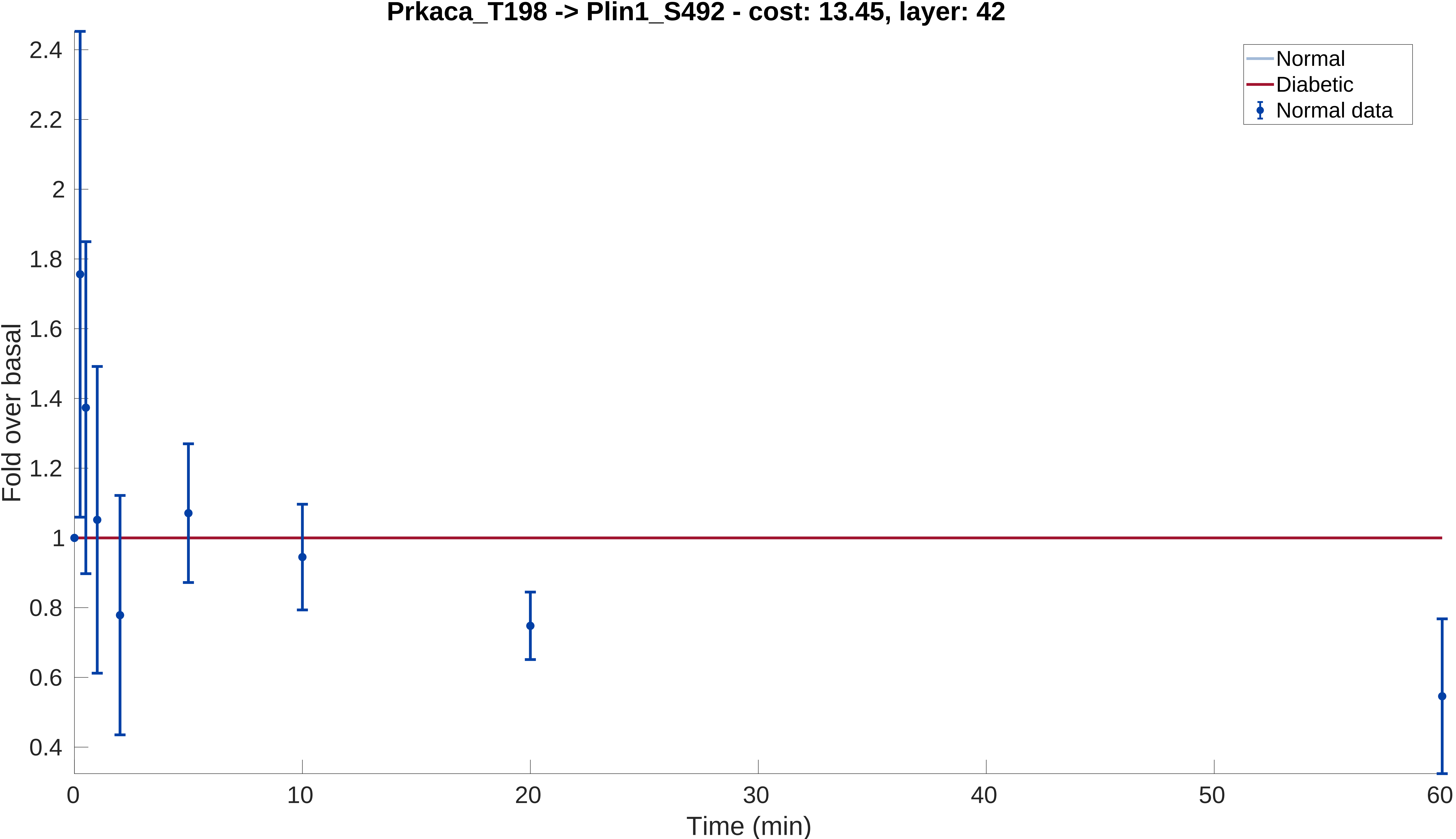

Prkaca_T198 -> Plin1_S492 - cost: 13.45, layer: 42
Normal
Diabetic
Normal data
Fold over basal
2.4
2.2
2
1.8
1.6
1.4
1.2
1
0.8
0.6
0.4
0
10
20
30
40
50
60
Time (min)

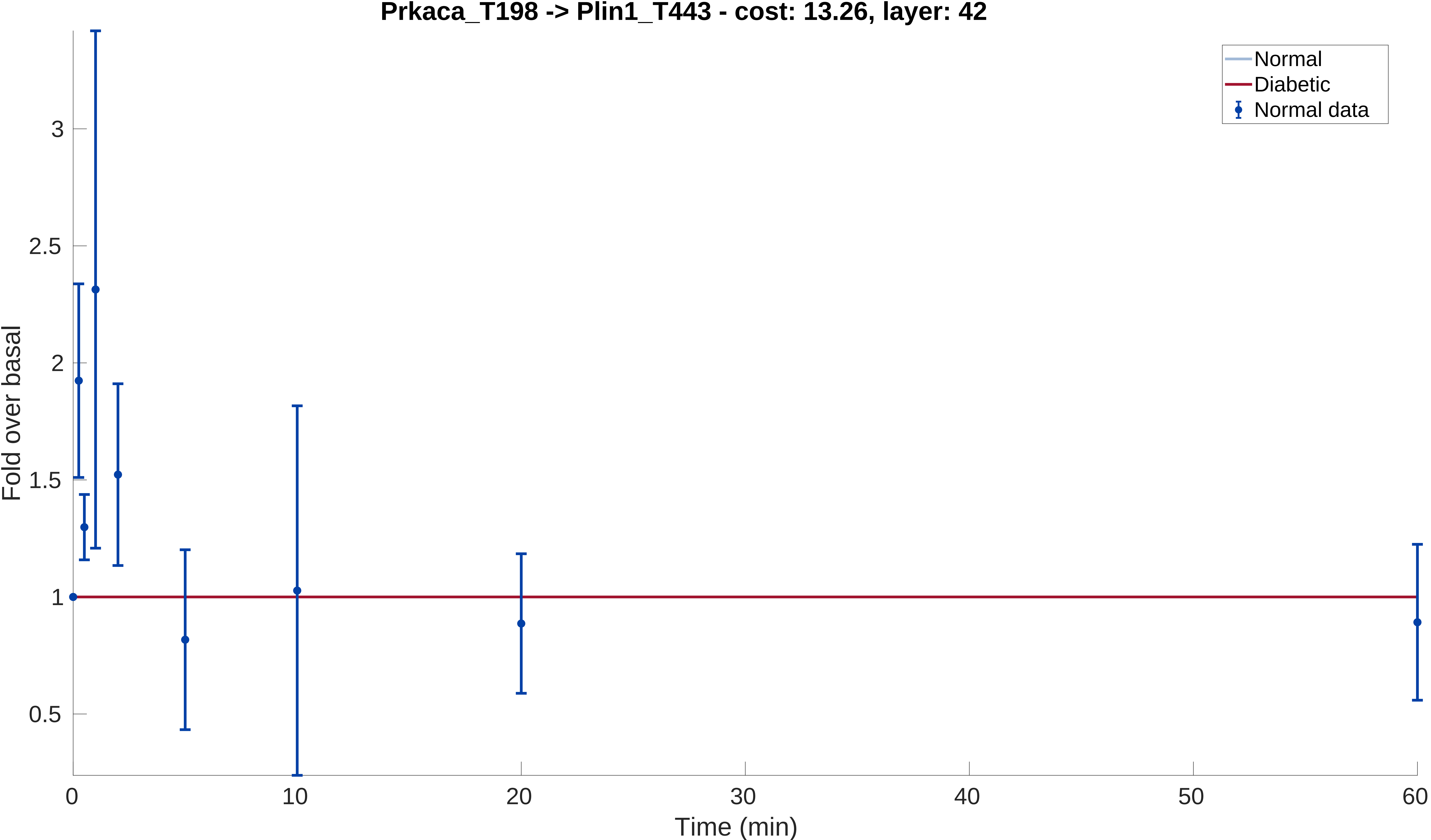
Prkaca_T198 -> Plin1_T443 - cost: 13.26, layer: 42
Normal
Diabetic
Normal data
Fold over basal
0.5
1
1.5
2
2.5
3
0
10
20
30
40
50
60
Time (min)

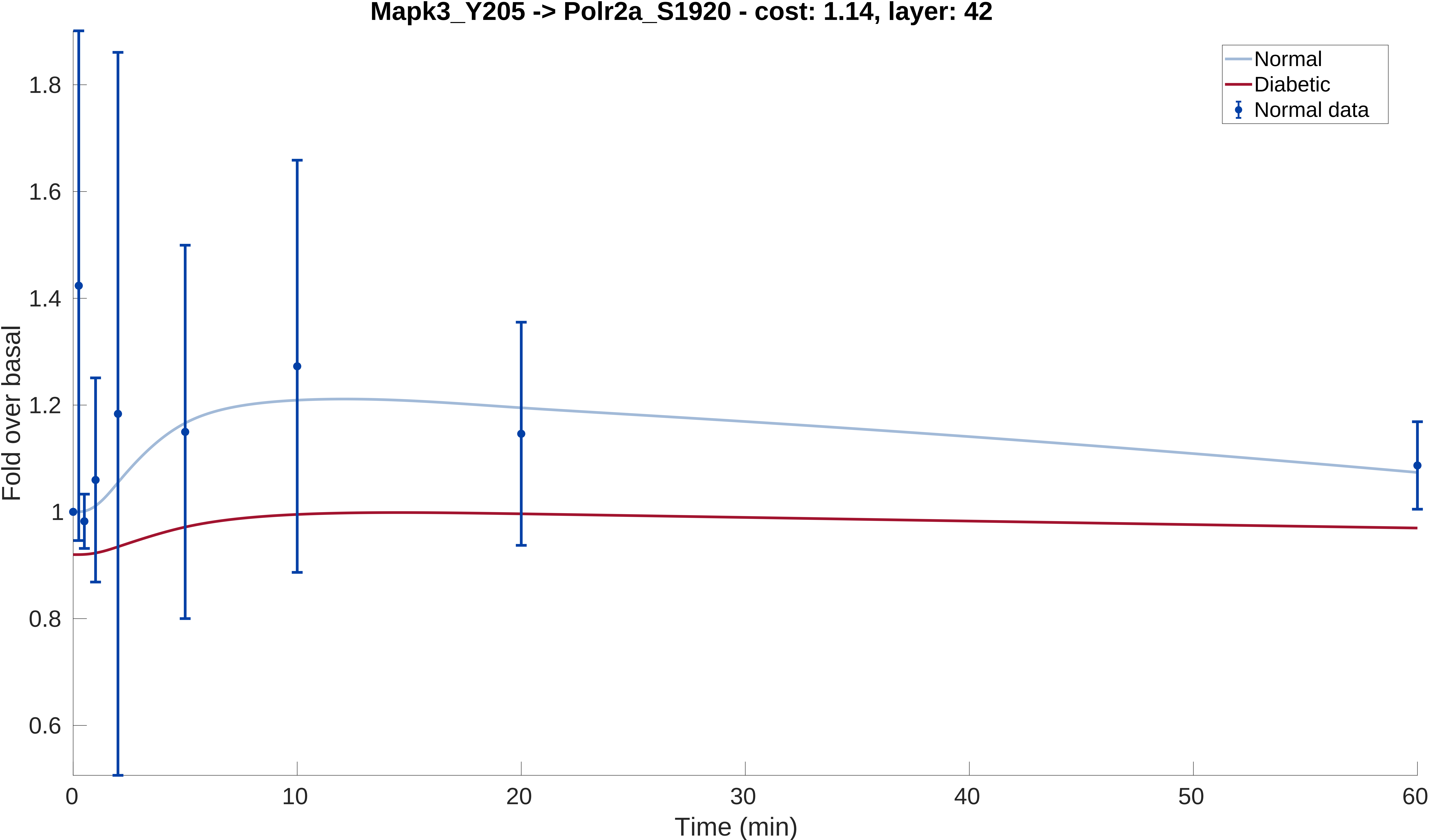

Mapk3_Y205 -> Polr2a_S1920 - cost: 1.14, layer: 42
Normal
Diabetic
Normal data
Fold over basal
1.8
1.6
1.4
1.2
1
0.8
0.6
0
10
20
30
40
50
60
Time (min)

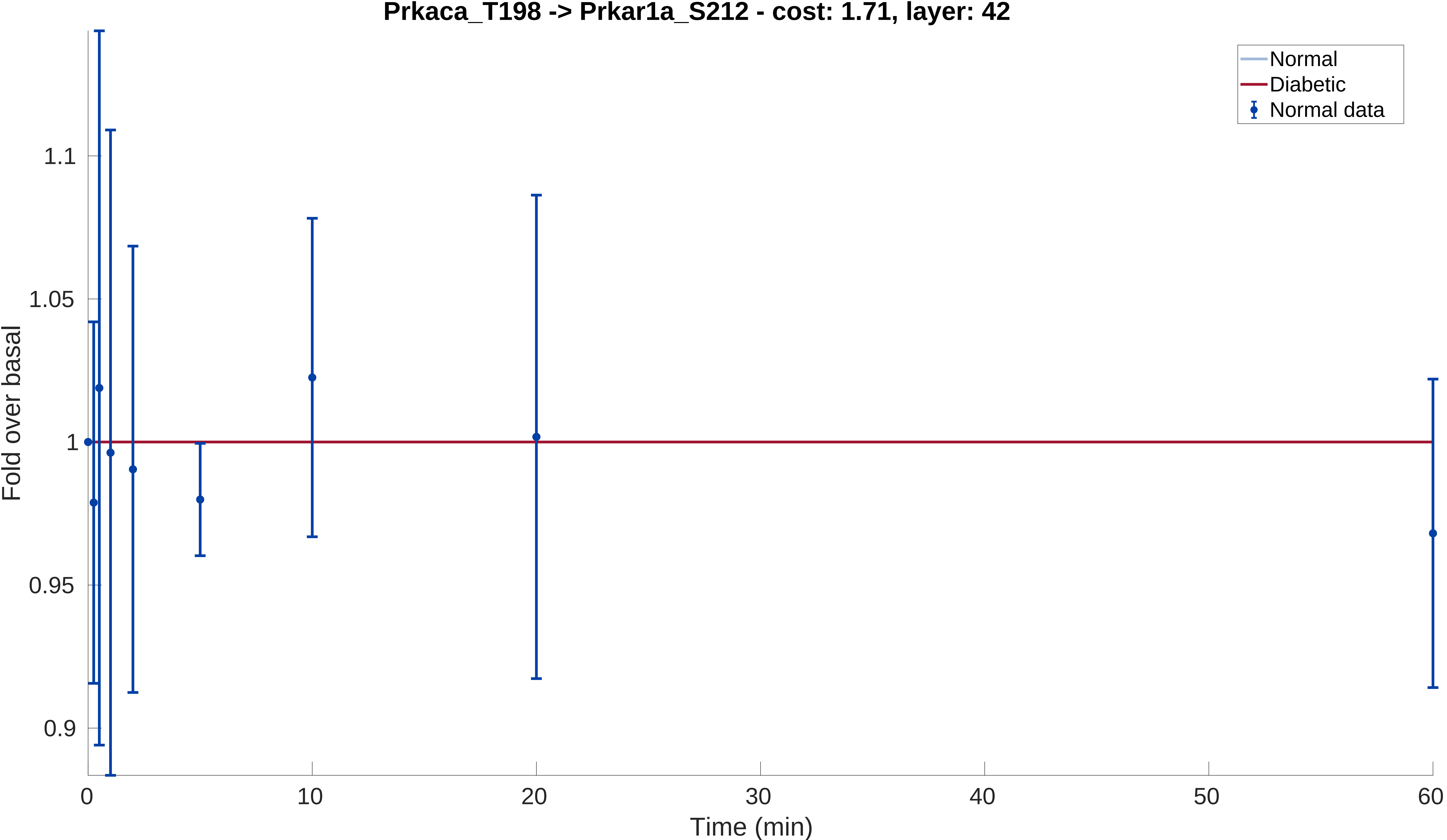

Prkaca_T198 -> Prkar1a_S212 - cost: 1.71, layer: 42
Normal
Diabetic
Normal data
Fold over basal
1.1
1.05
1
0.95
0.9
0
10
20
30
40
50
60
Time (min)

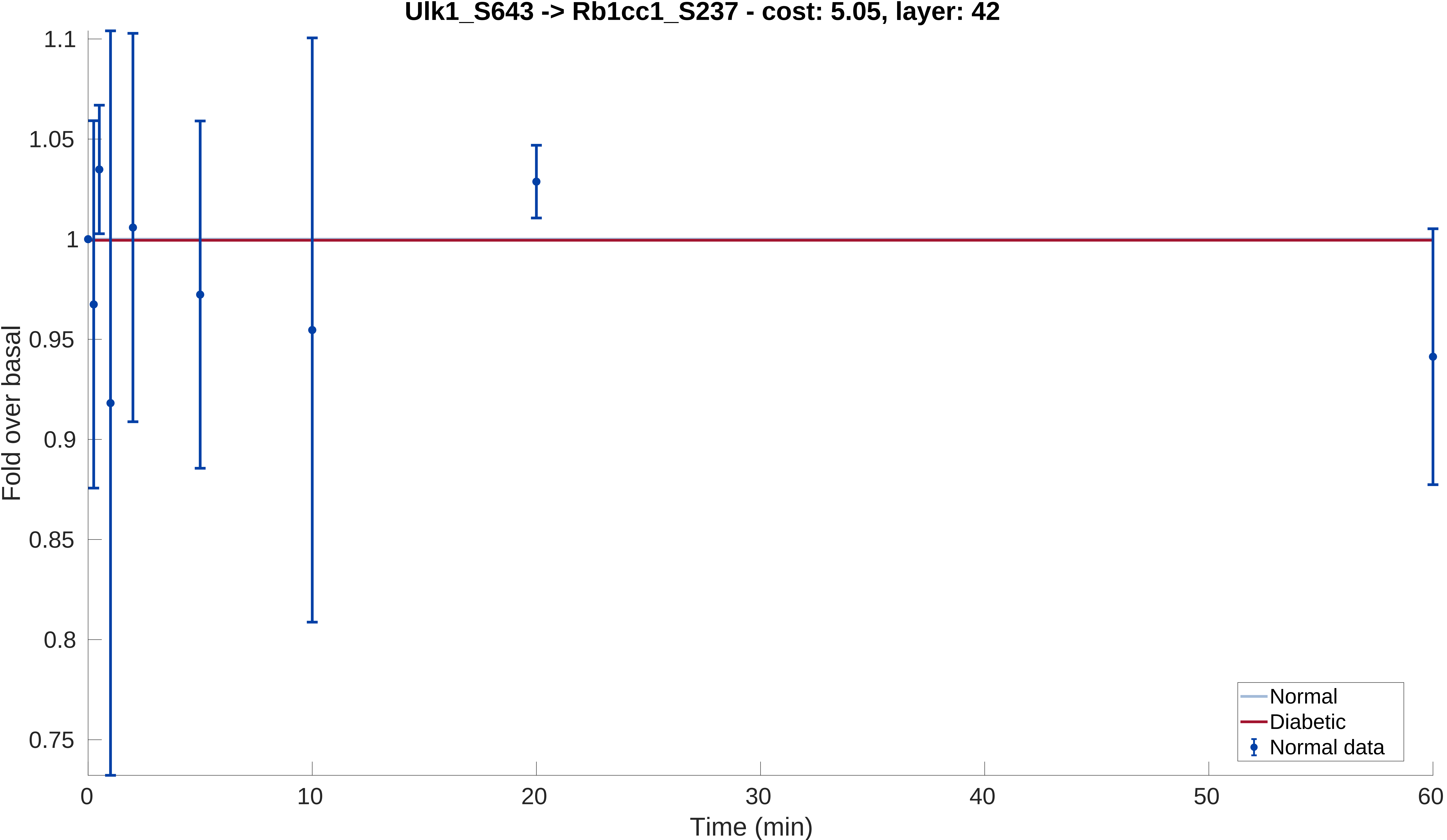

Ulk1_S643 -> Rb1cc1_S237 - cost: 5.05, layer: 42
Fold over basal
Time (min)
1.1
1.05
1
0.95
0.9
0.85
0.8
0.75
0
10
20
30
40
50
60
Normal
Diabetic
Normal data

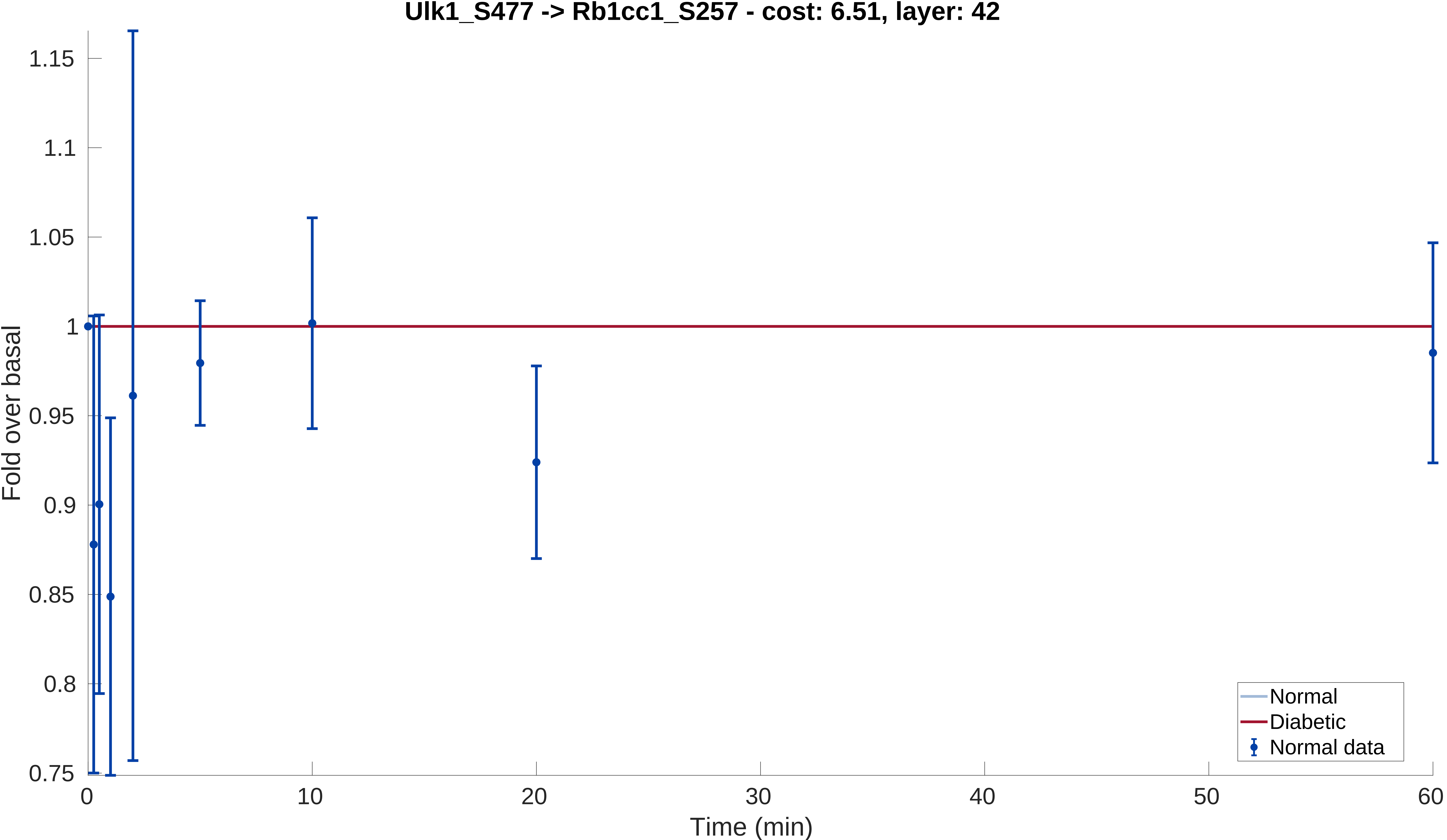

Ulk1_S477 -> Rb1cc1_S257 - cost: 6.51, layer: 42
Fold over basal
1.15
1.1
1.05
1
0.95
0.9
0.85
0.8
0.75
0
10
20
30
40
50
60
Time (min)
Normal
Diabetic
Normal data

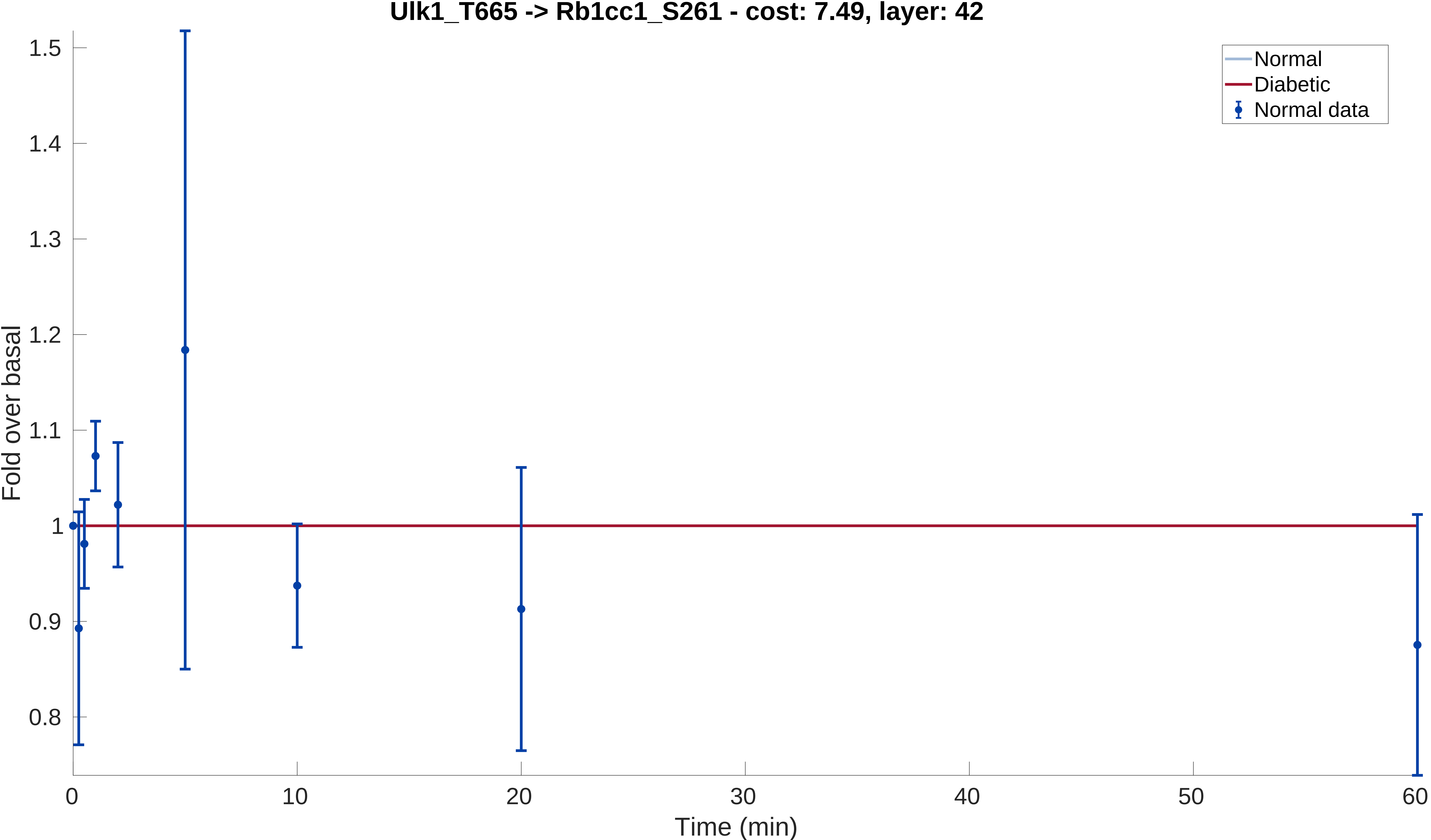

Ulk1_T665 -> Rb1cc1_S261 - cost: 7.49, layer: 42
Normal
Diabetic
Normal data
Fold over basal
1.5
1.4
1.3
1.2
1.1
1
0.9
0.8
0
10
20
30
40
50
60
Time (min)

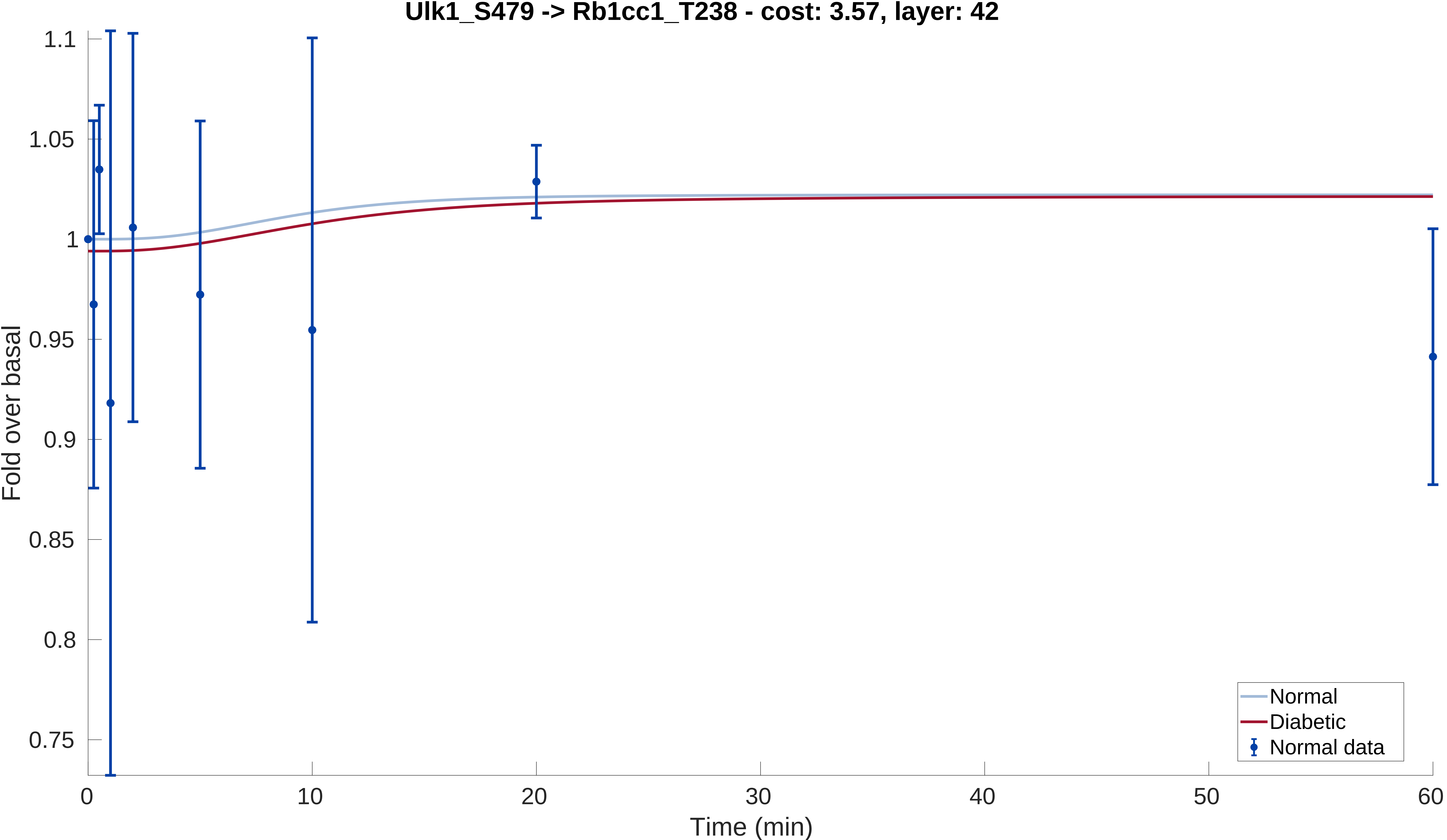

Ulk1_S479 -> Rb1cc1_T238 - cost: 3.57, layer: 42
Fold over basal
1.1
1.05
1
0.95
0.9
0.85
0.8
0.75
0
10
20
30
40
50
60
Time (min)
Normal
Diabetic
Normal data

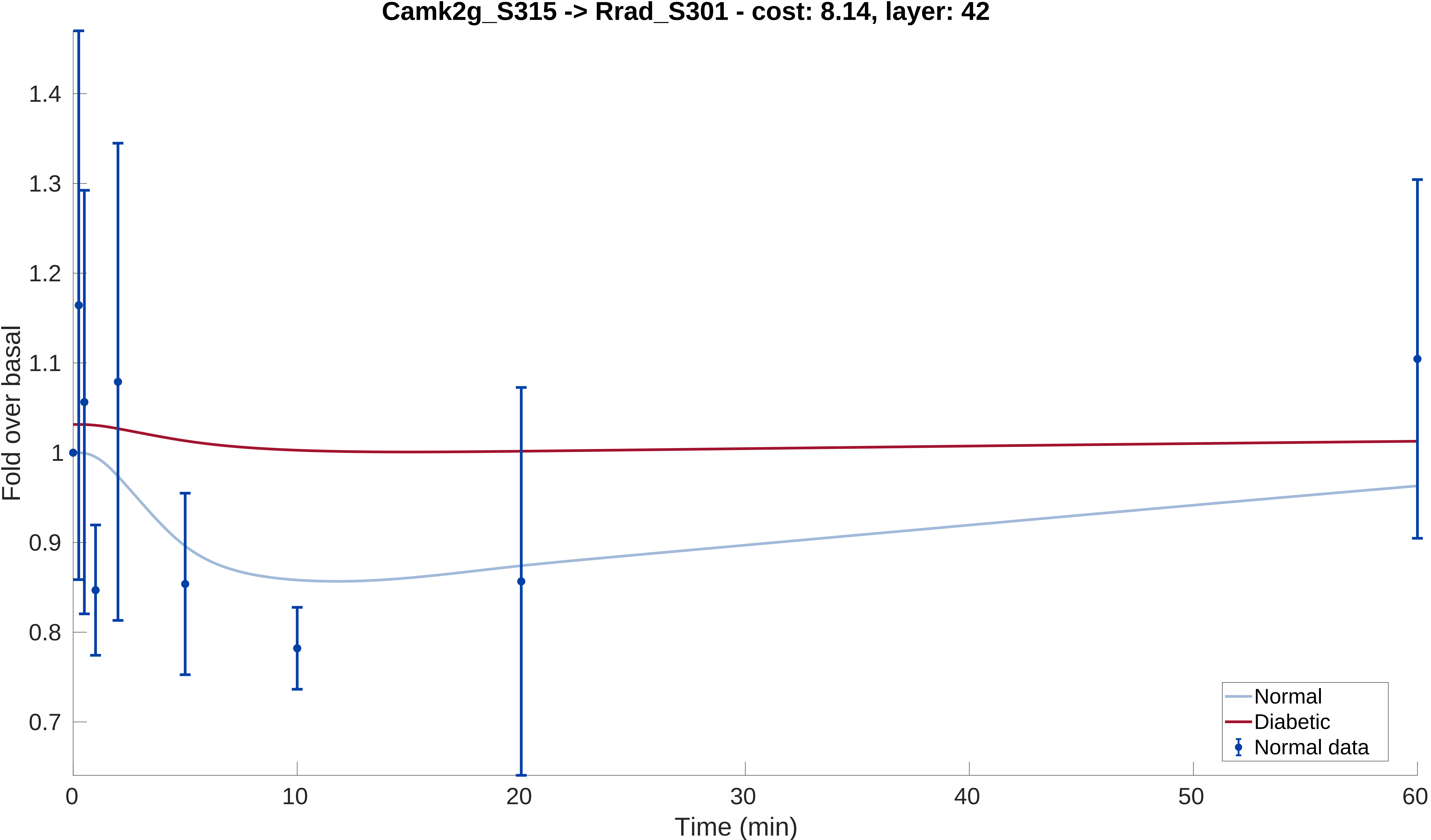

Camk2g_S315 -> Rrad_S301 - cost: 8.14, layer: 42
1.4
1.3
1.2
1.1
1
0.9
0.8
0.7
Fold over basal
0
10
20
30
40
50
60
Time (min)
Normal
Diabetic
Normal data

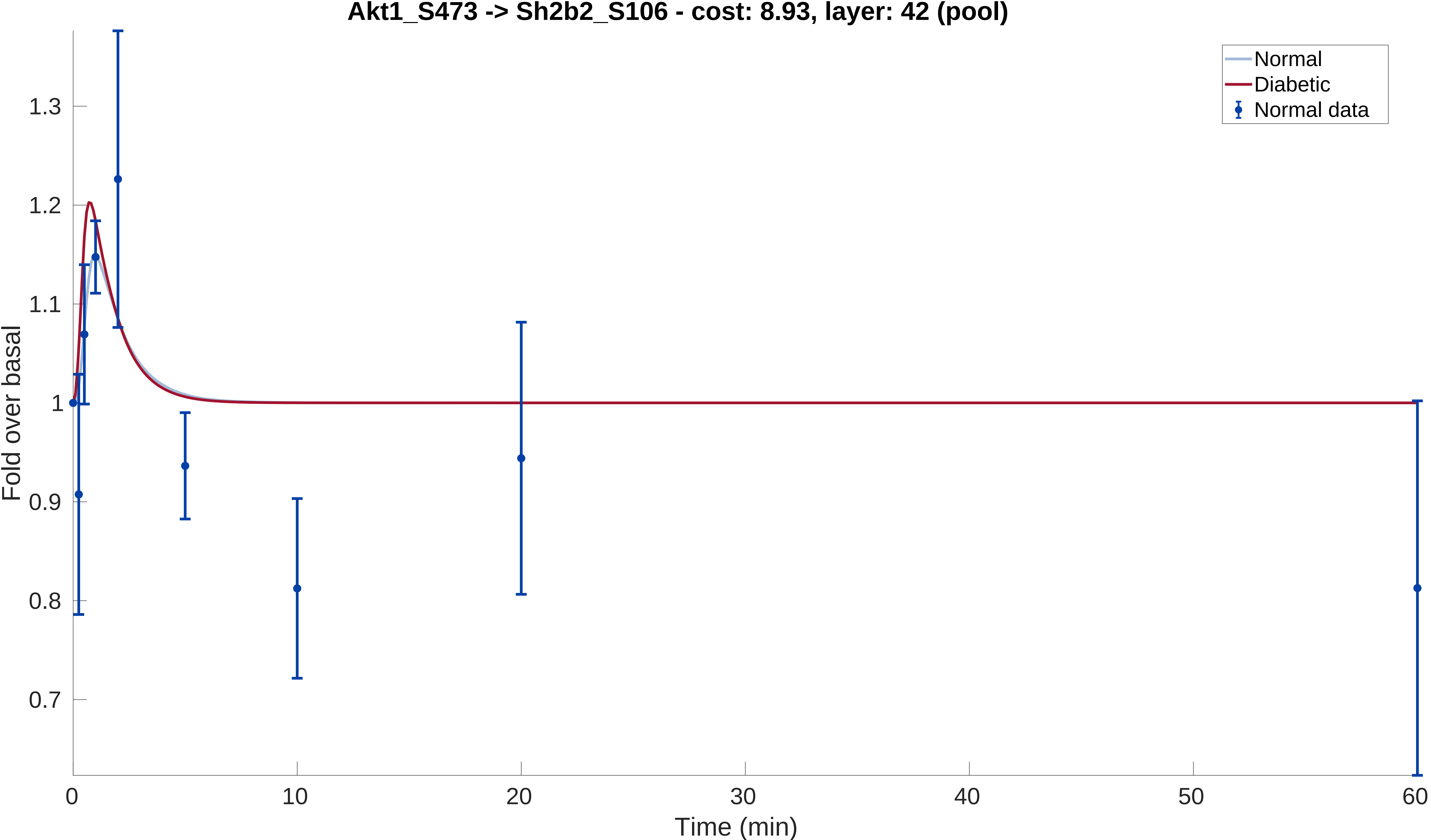

Akt1_S473 -> Sh2b2_S106 - cost: 8.93, layer: 42 (pool)
Normal
Diabetic
Normal data
Fold over basal
Time (min)
1.3
1.2
1.1
1
0.9
0.8
0.7
0
10
20
30
40
50
60

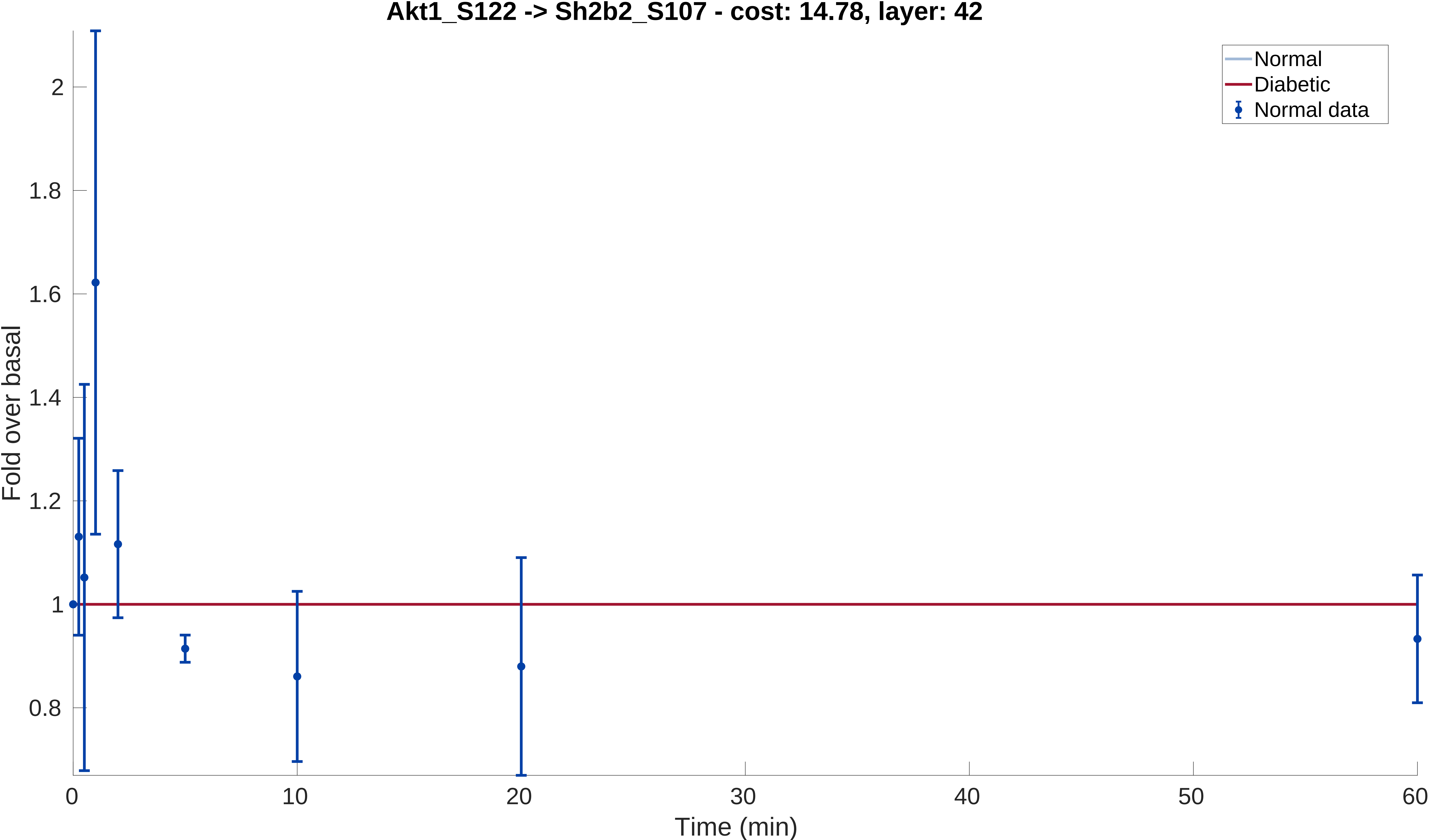

Akt1_S122 -> Sh2b2_S107 - cost: 14.78, layer: 42
Normal
Diabetic
Normal data
Fold over basal
2
1.8
1.6
1.4
1.2
1
0.8
0
10
20
30
40
50
60
Time (min)

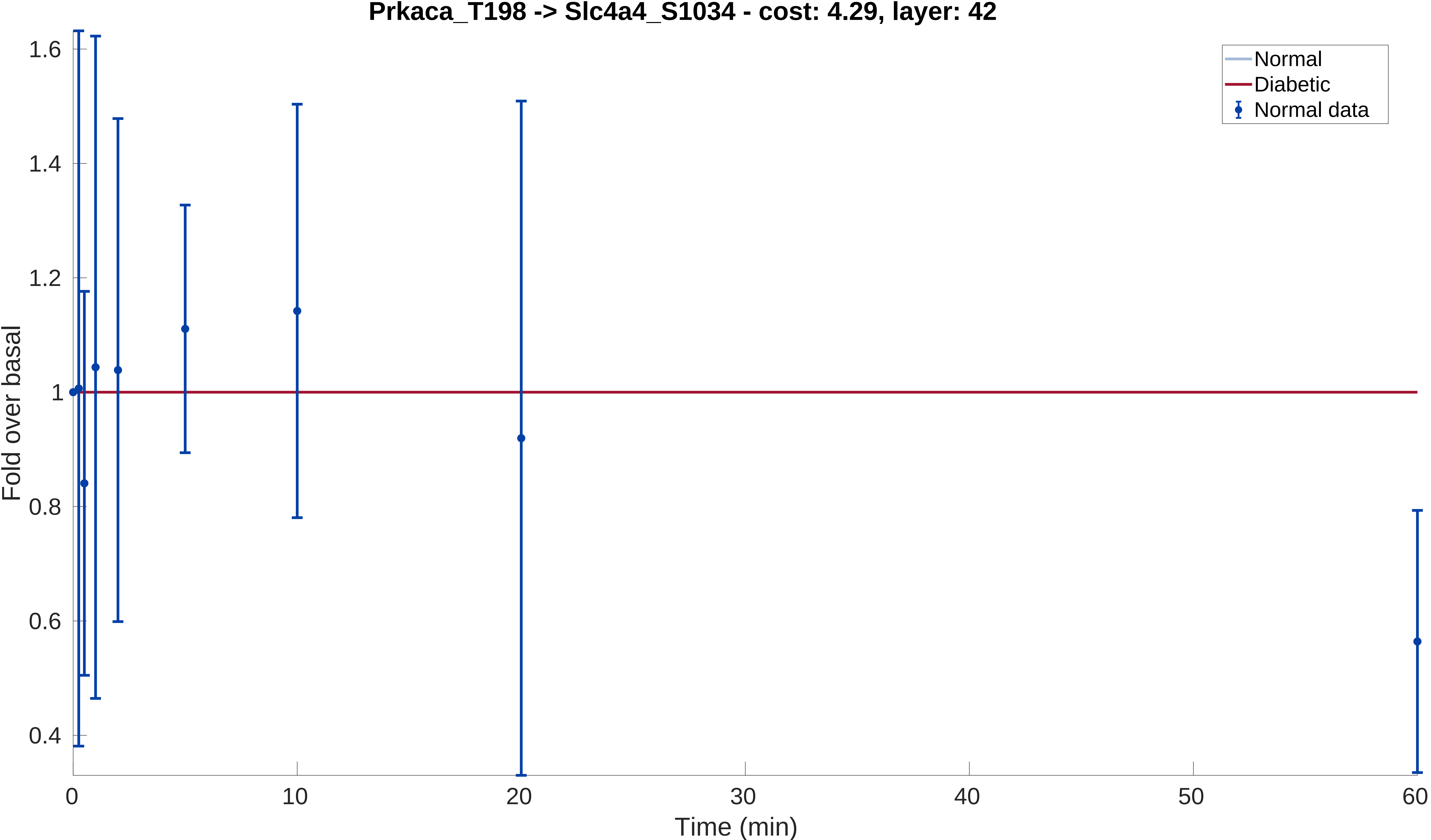

Prkaca_T198 -> Slc4a4_S1034 - cost: 4.29, layer: 42
Normal
Diabetic
Normal data
Fold over basal
Time (min)
1.6
1.4
1.2
1
0.8
0.6
0.4
0
10
20
30
40
50
60

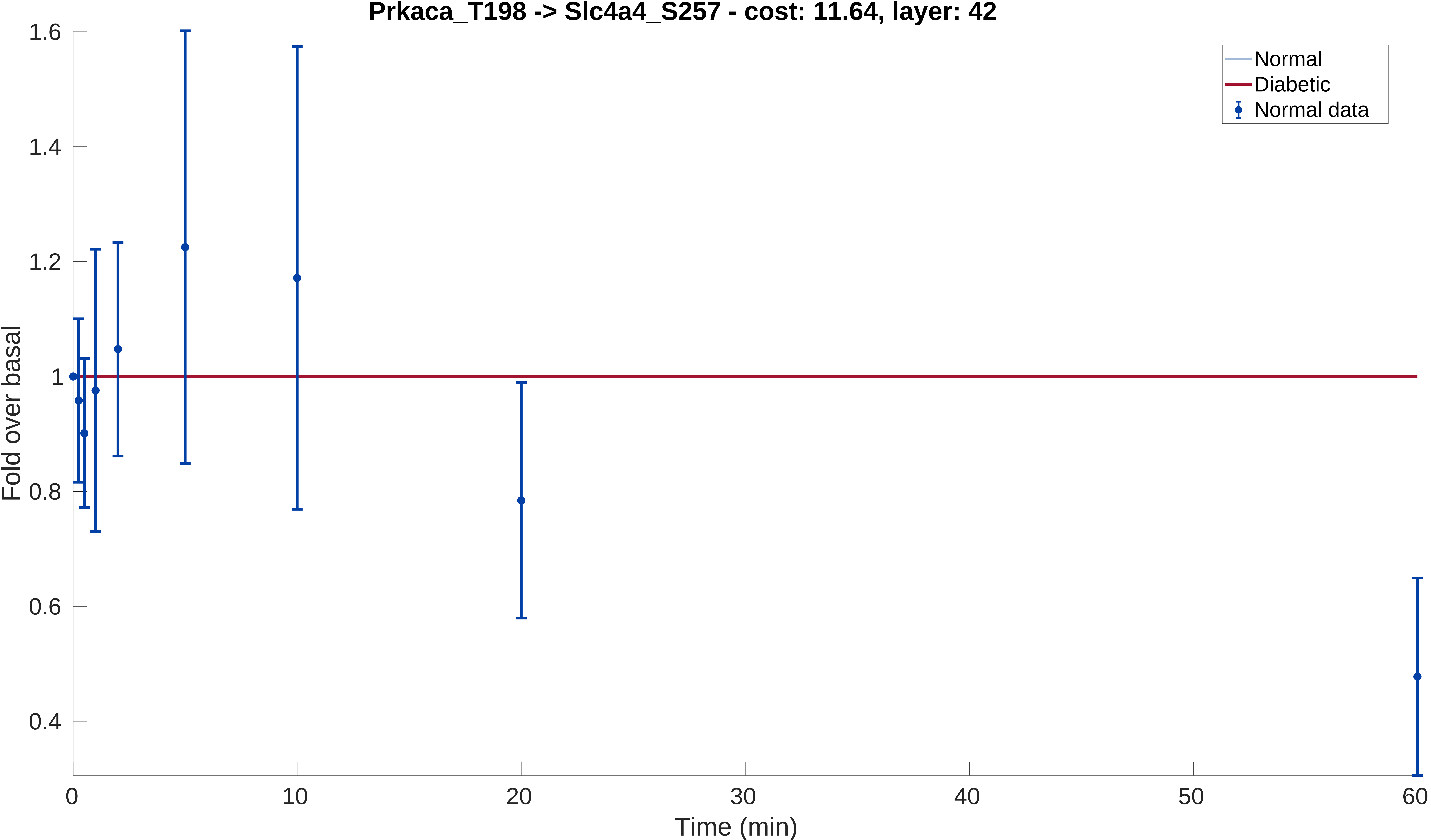
Prkaca_T198 -> Slc4a4_S257 - cost: 11.64, layer: 42
Normal
Diabetic
Normal data
Fold over basal
Time (min)
0.4
0.6
0.8
1
1.2
1.4
1.6
0
10
20
30
40
50
60

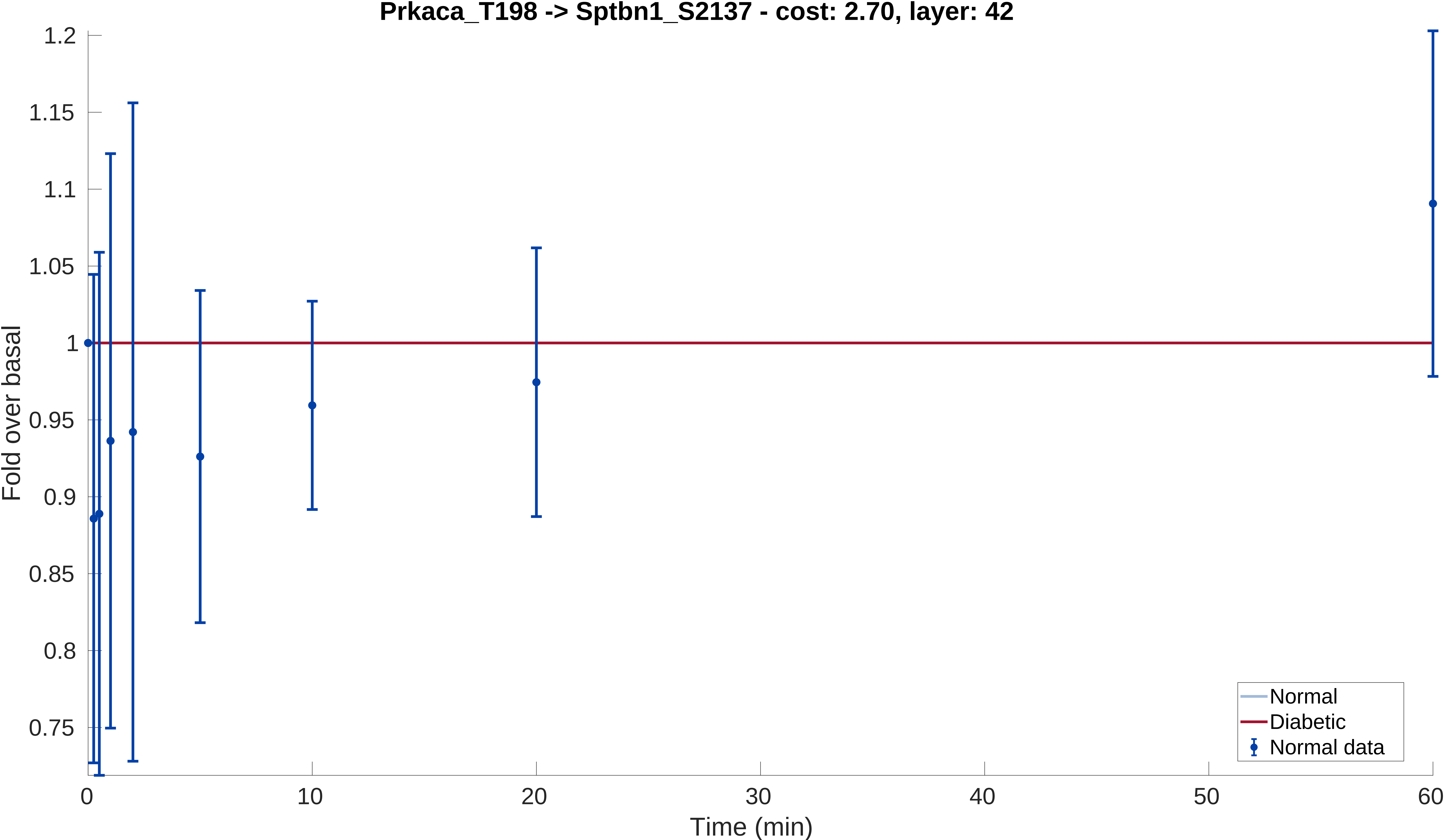

Prkaca_T198 -> Sptbn1_S2137 - cost: 2.70, layer: 42
Fold over basal
1.2
1.15
1.1
1.05
1
0.95
0.9
0.85
0.8
0.75
0
10
20
30
40
50
60
Time (min)
Normal
Diabetic
Normal data

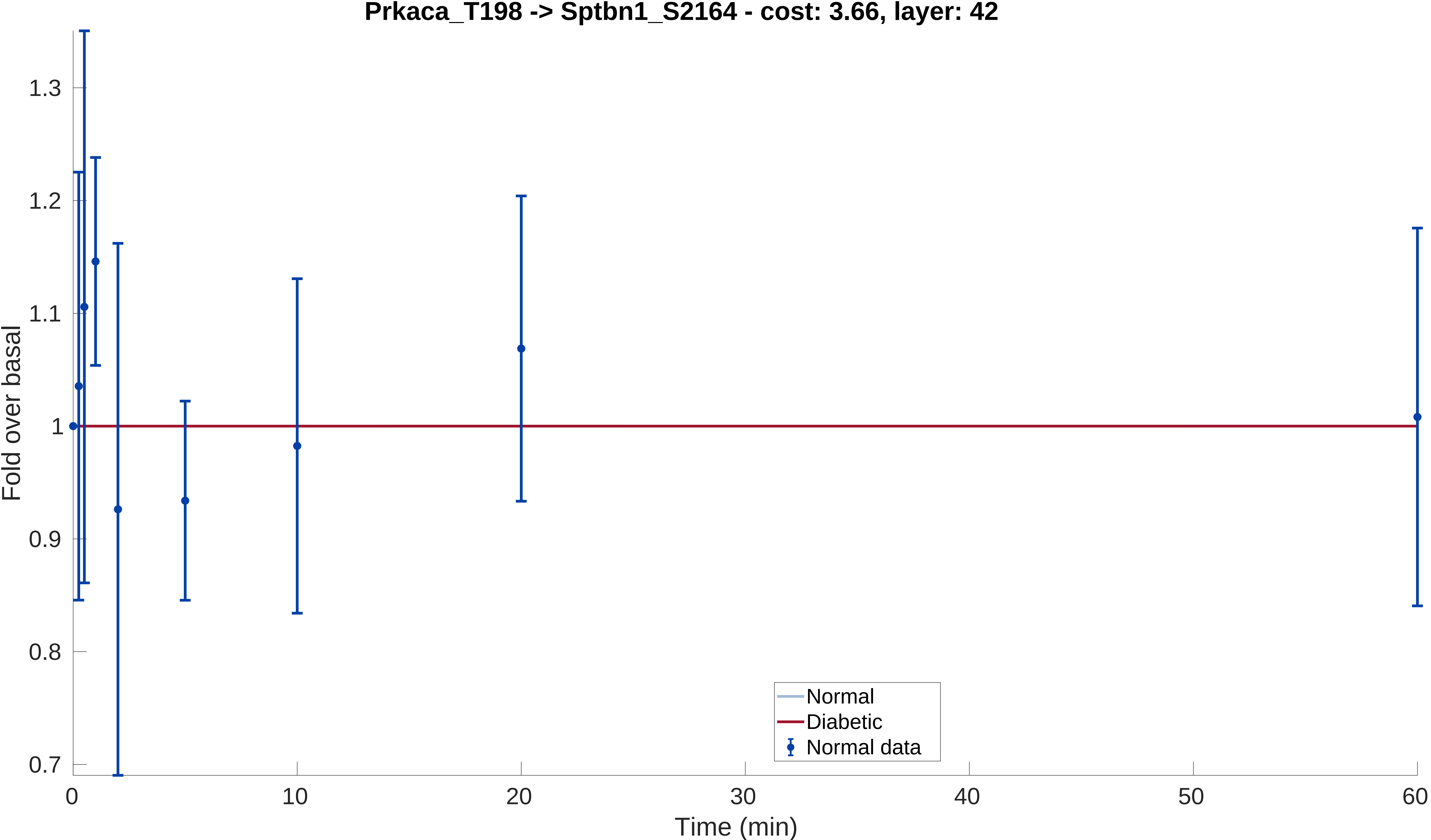

Prkaca_T198 -> Sptbn1_S2164 - cost: 3.66, layer: 42
Fold over basal
1.3
1.2
1.1
1
0.9
0.8
0.7
0
10
20
30
40
50
60
Time (min)
Normal
Diabetic
Normal data

### Prkaca_T198 -> Sptbn1_S2168 - cost: 1.66, layer: 42

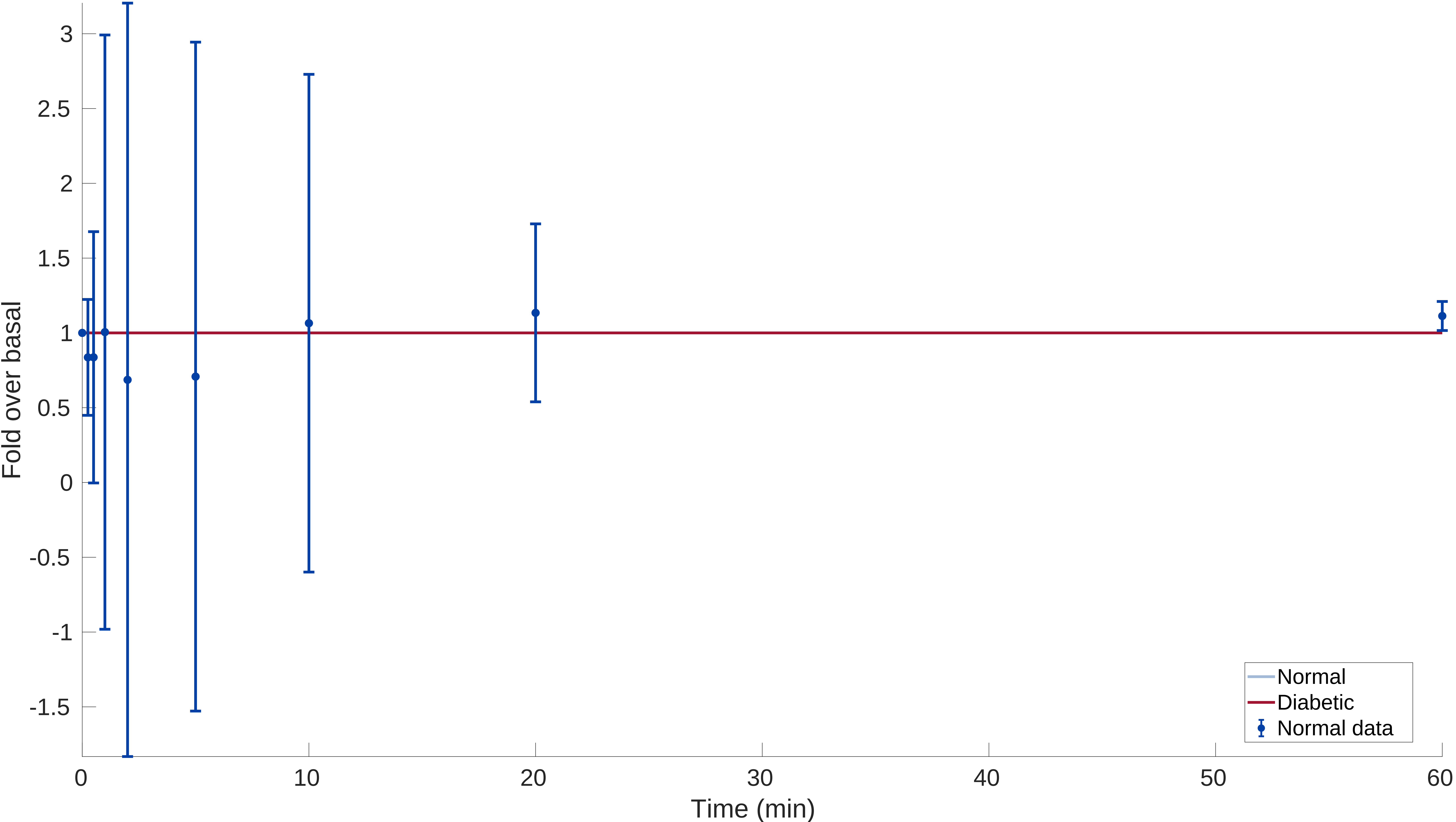

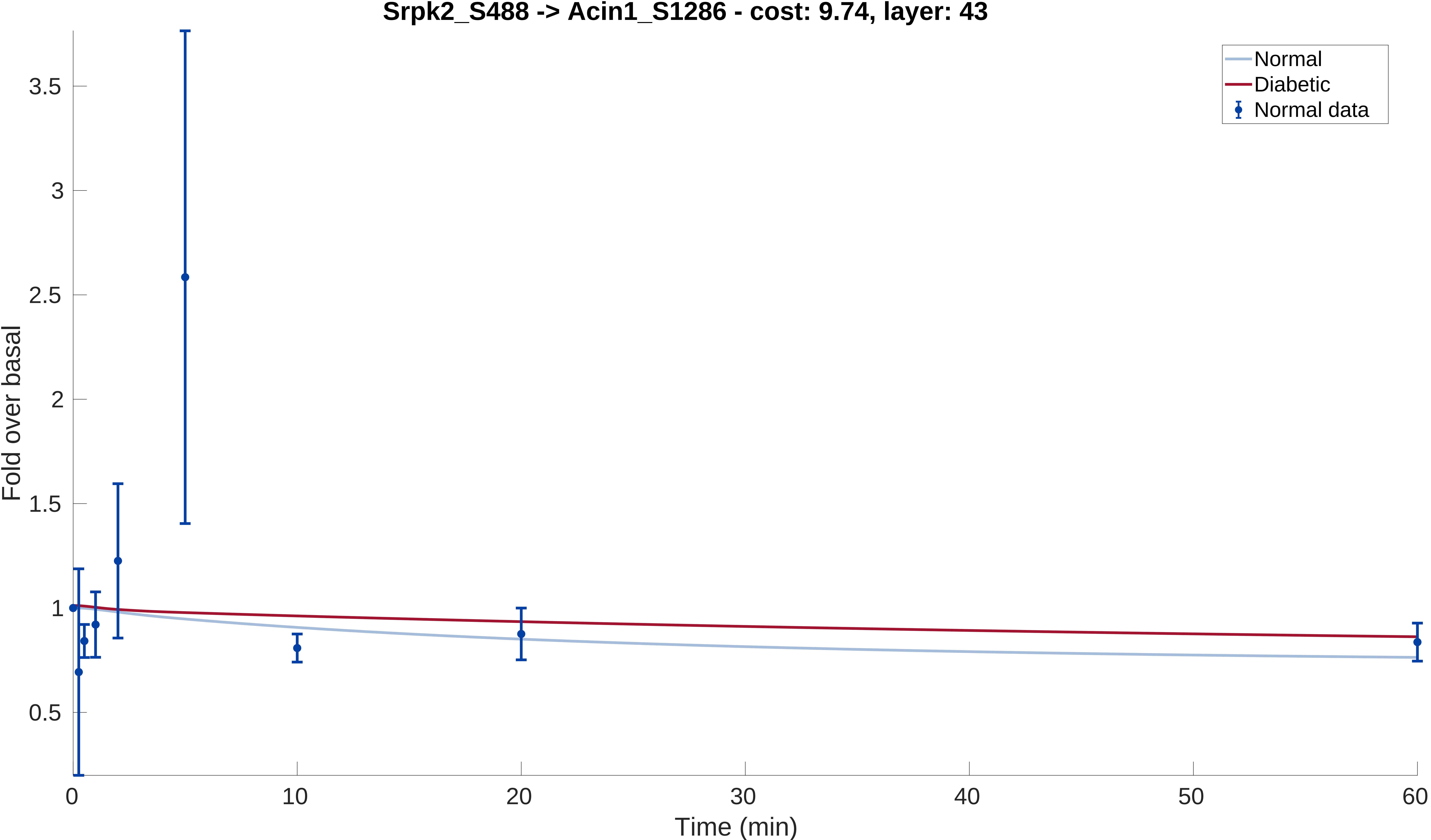

Srpk2_S488 -> Acin1_S1286 - cost: 9.74, layer: 43
Normal
Diabetic
Normal data
Fold over basal
3.5
3
2.5
2
1.5
1
0.5
0
10
20
30
40
50
60
Time (min)

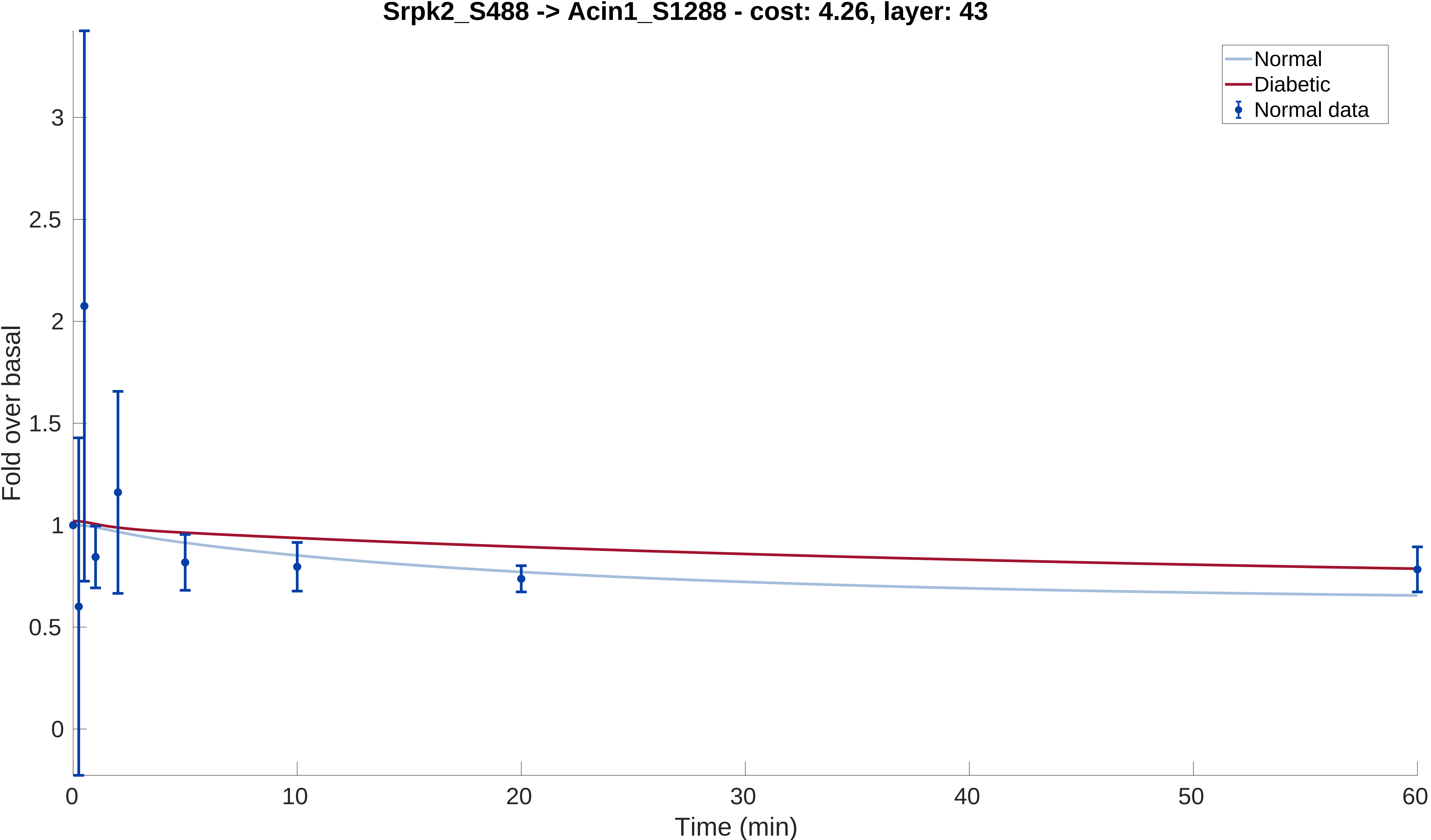

Srpk2_S488 -> Acin1_S1288 - cost: 4.26, layer: 43
Normal
Diabetic
Normal data
Fold over basal
0
0.5
1
1.5
2
2.5
3
0
10
20
30
40
50
60
Time (min)

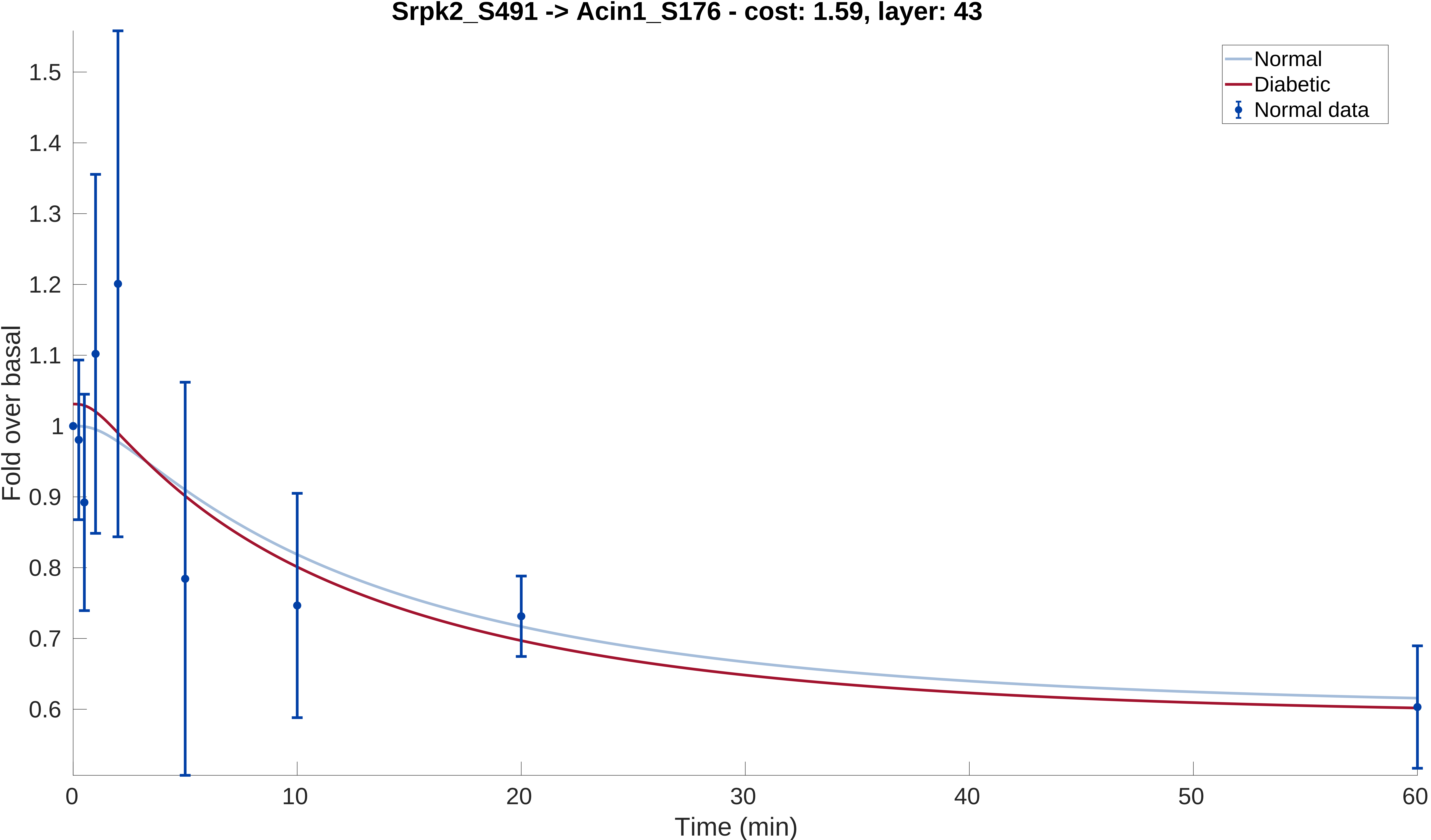
Srpk2_S491 -> Acin1_S176 - cost: 1.59, layer: 43
Normal
Diabetic
Normal data
Fold over basal
1.5
1.4
1.3
1.2
1.1
1
0.9
0.8
0.7
0.6
0
10
20
30
40
50
60
Time (min)

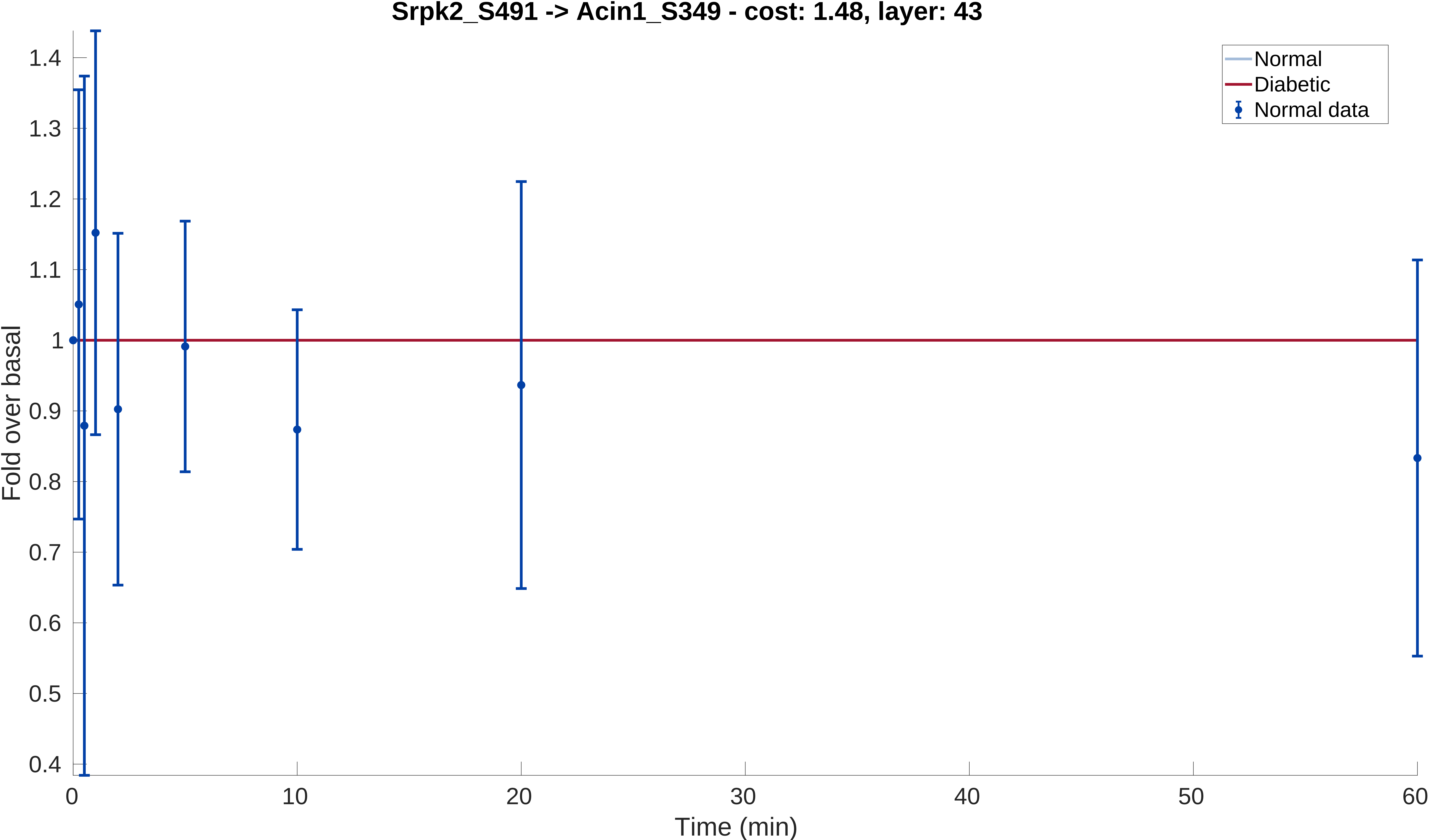

Srpk2_S491 -> Acin1_S349 - cost: 1.48, layer: 43
Normal
Diabetic
Normal data
Fold over basal
1.4
1.3
1.2
1.1
1
0.9
0.8
0.7
0.6
0.5
0.4
0
10
20
30
40
50
60
Time (min)

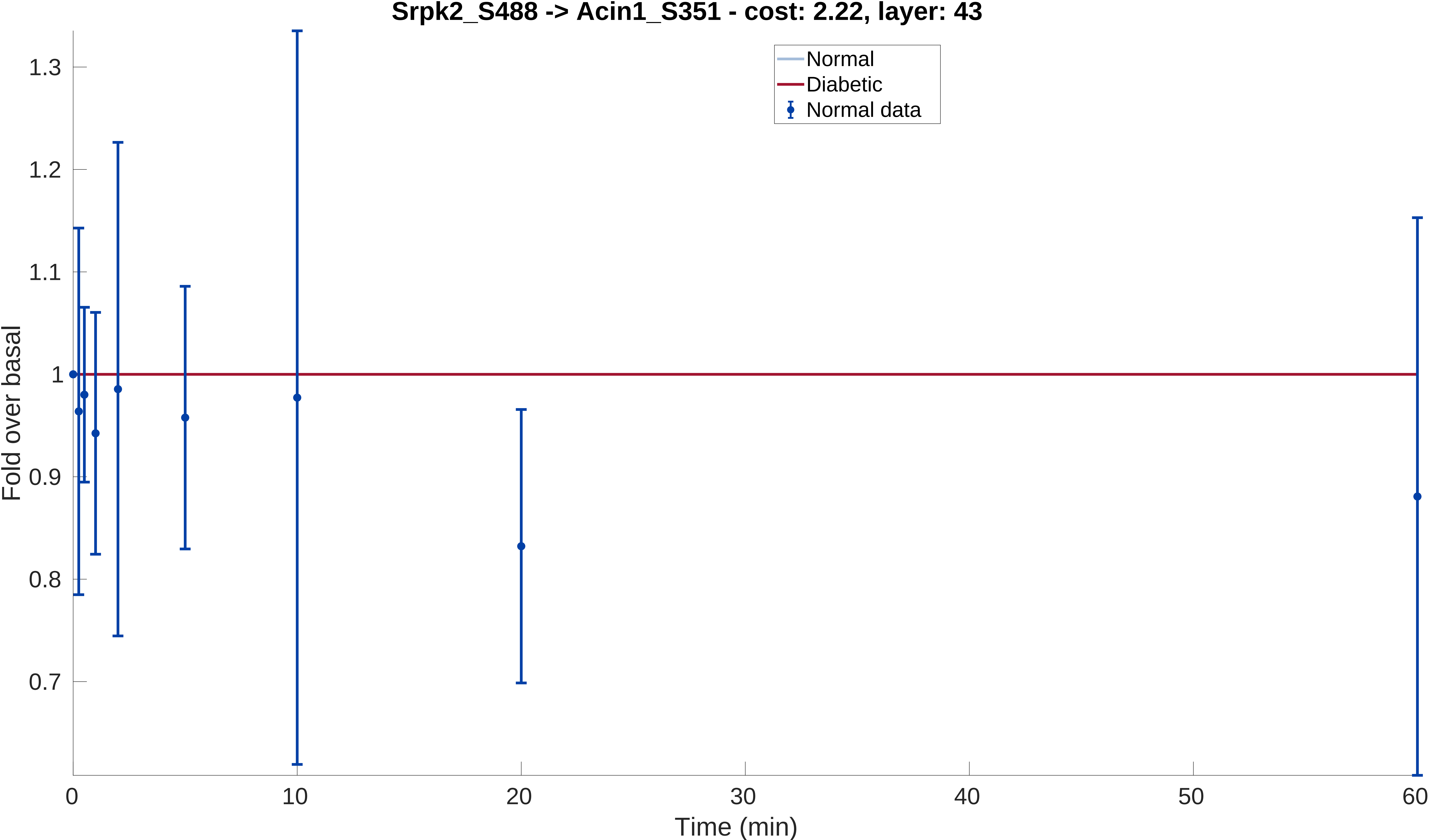

Srpk2_S488 -> Acin1_S351 - cost: 2.22, layer: 43
Normal
Diabetic
Normal data
Fold over basal
Time (min)
1.3
1.2
1.1
1
0.9
0.8
0.7
0
10
20
30
40
50
60

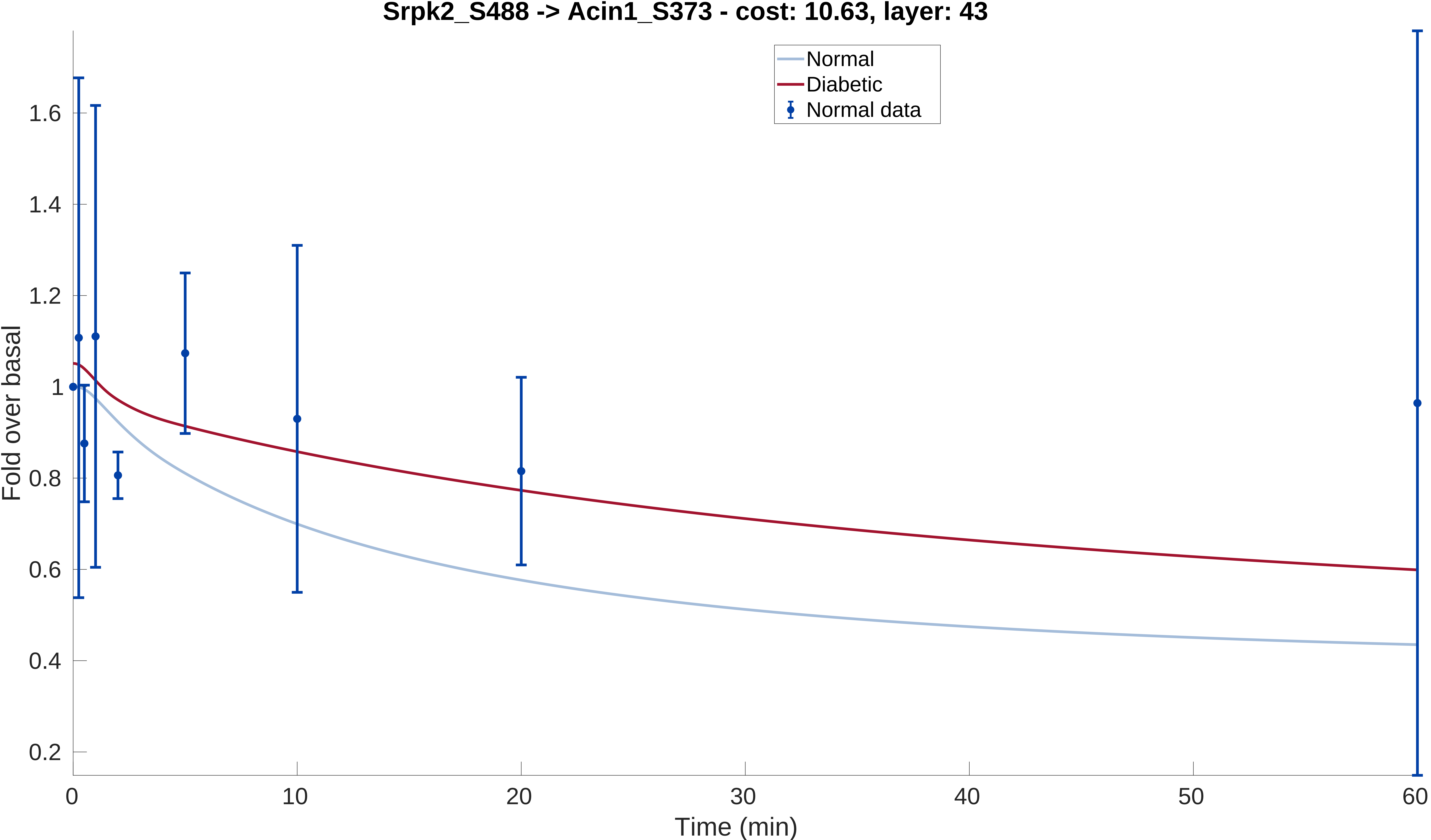

Srpk2_S488 -> Acin1_S373 - cost: 10.63, layer: 43
Normal
Diabetic
Normal data
Fold over basal
1.6
1.4
1.2
1
0.8
0.6
0.4
0.2
0
10
20
30
40
50
60
Time (min)

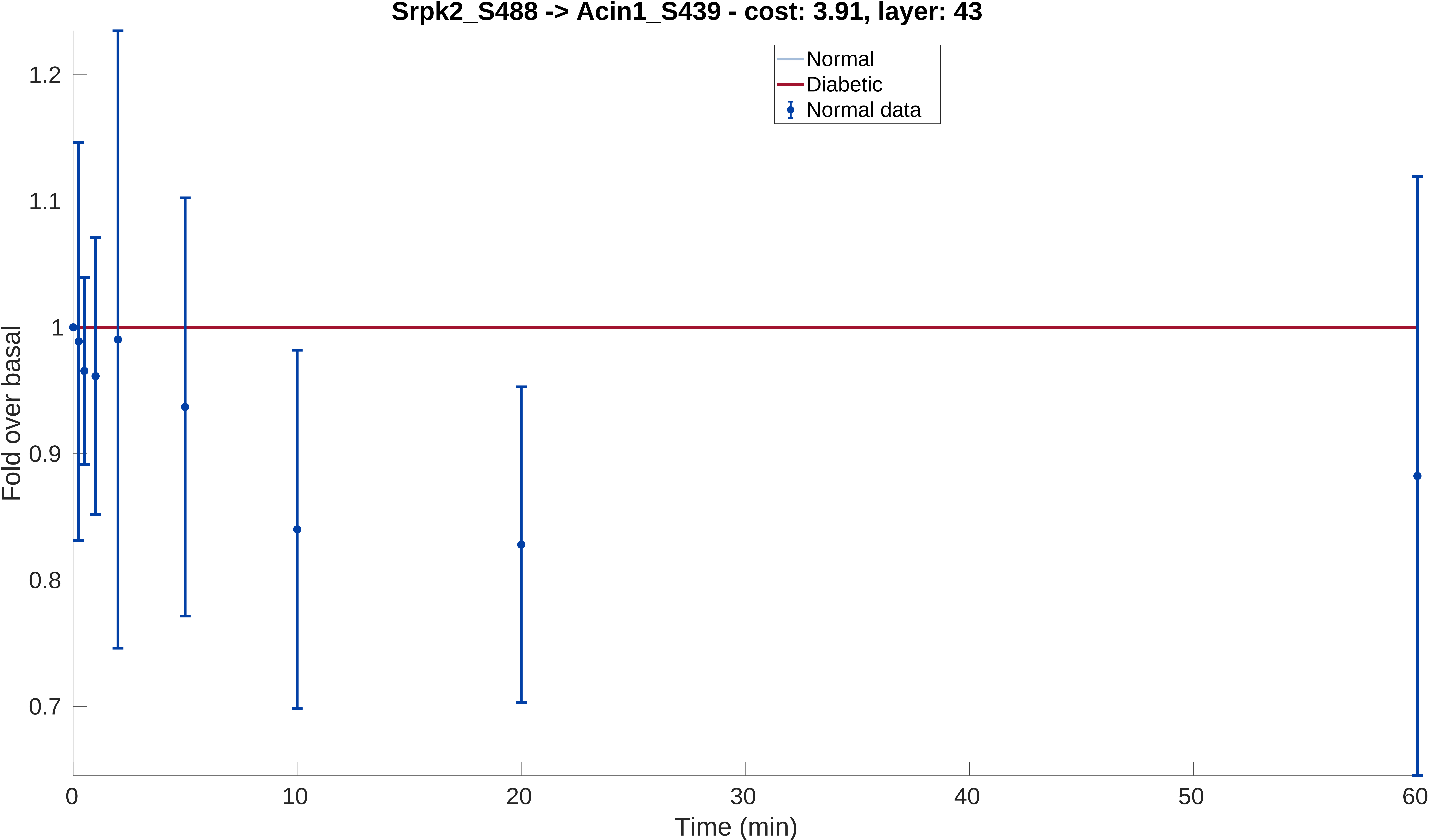
Srpk2_S488 -> Acin1_S439 - cost: 3.91, layer: 43
Normal
Diabetic
Normal data
Fold over basal
Time (min)
0.7
0.8
0.9
1
1.1
1.2
0
10
20
30
40
50
60

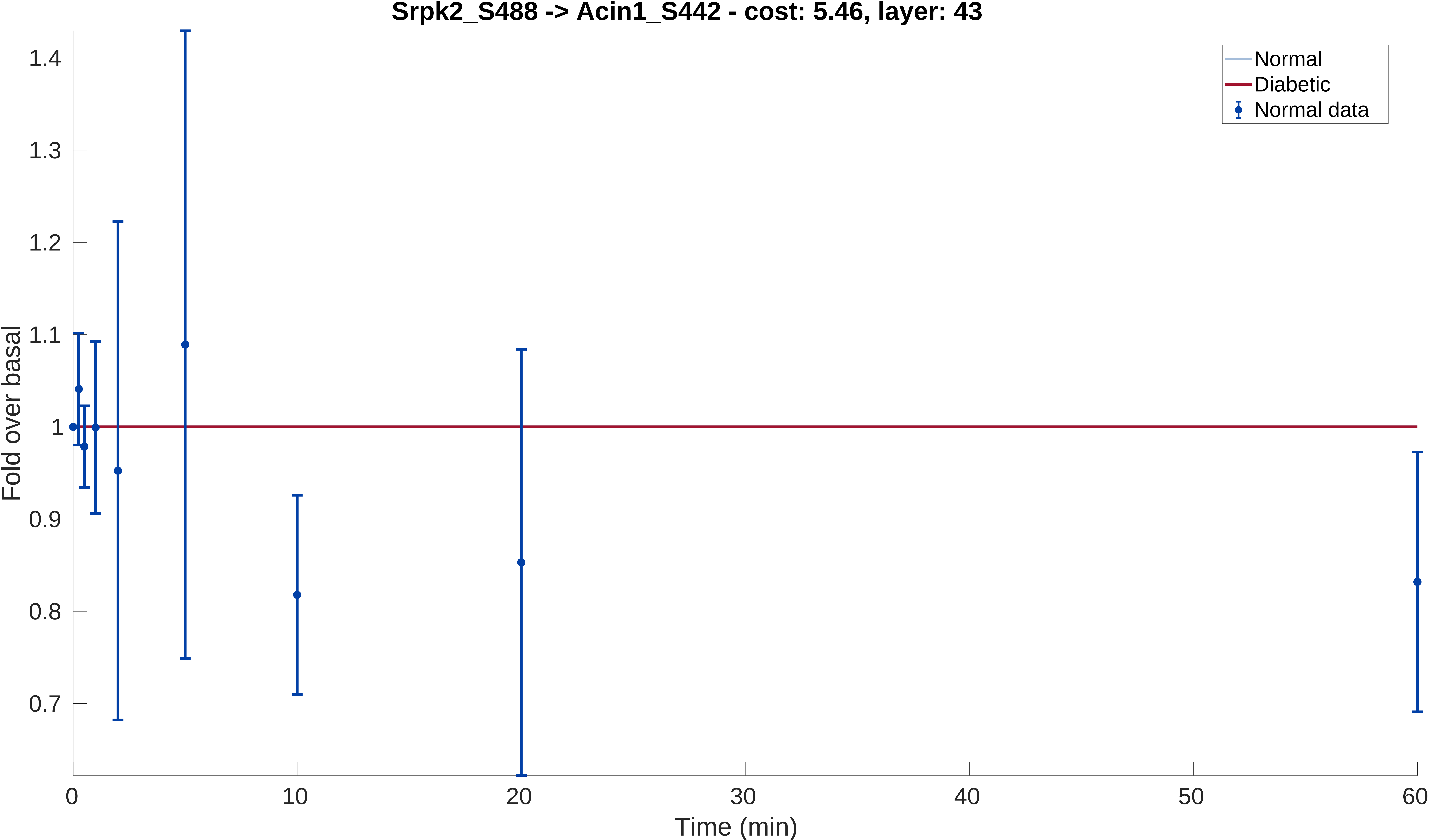

Srpk2_S488 -> Acin1_S442 - cost: 5.46, layer: 43
Normal
Diabetic
Normal data
Fold over basal
1.4
1.3
1.2
1.1
1
0.9
0.8
0.7
0
10
20
30
40
50
60
Time (min)

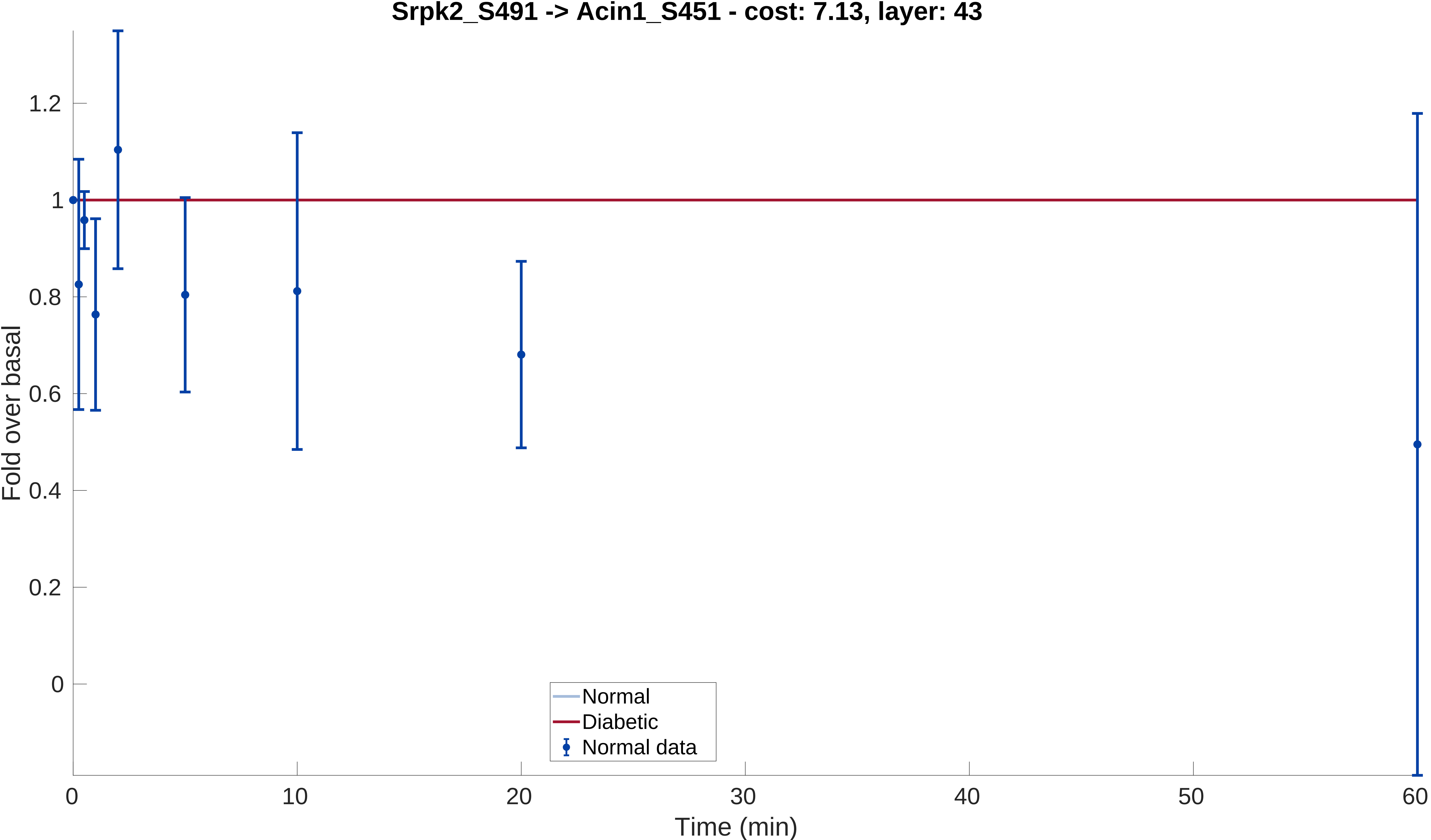

Srpk2_S491 -> Acin1_S451 - cost: 7.13, layer: 43
1.2
1
0.8
0.6
0.4
0.2
0
Fold over basal
0
10
20
30
40
50
60
Time (min)
Normal
Diabetic
Normal data

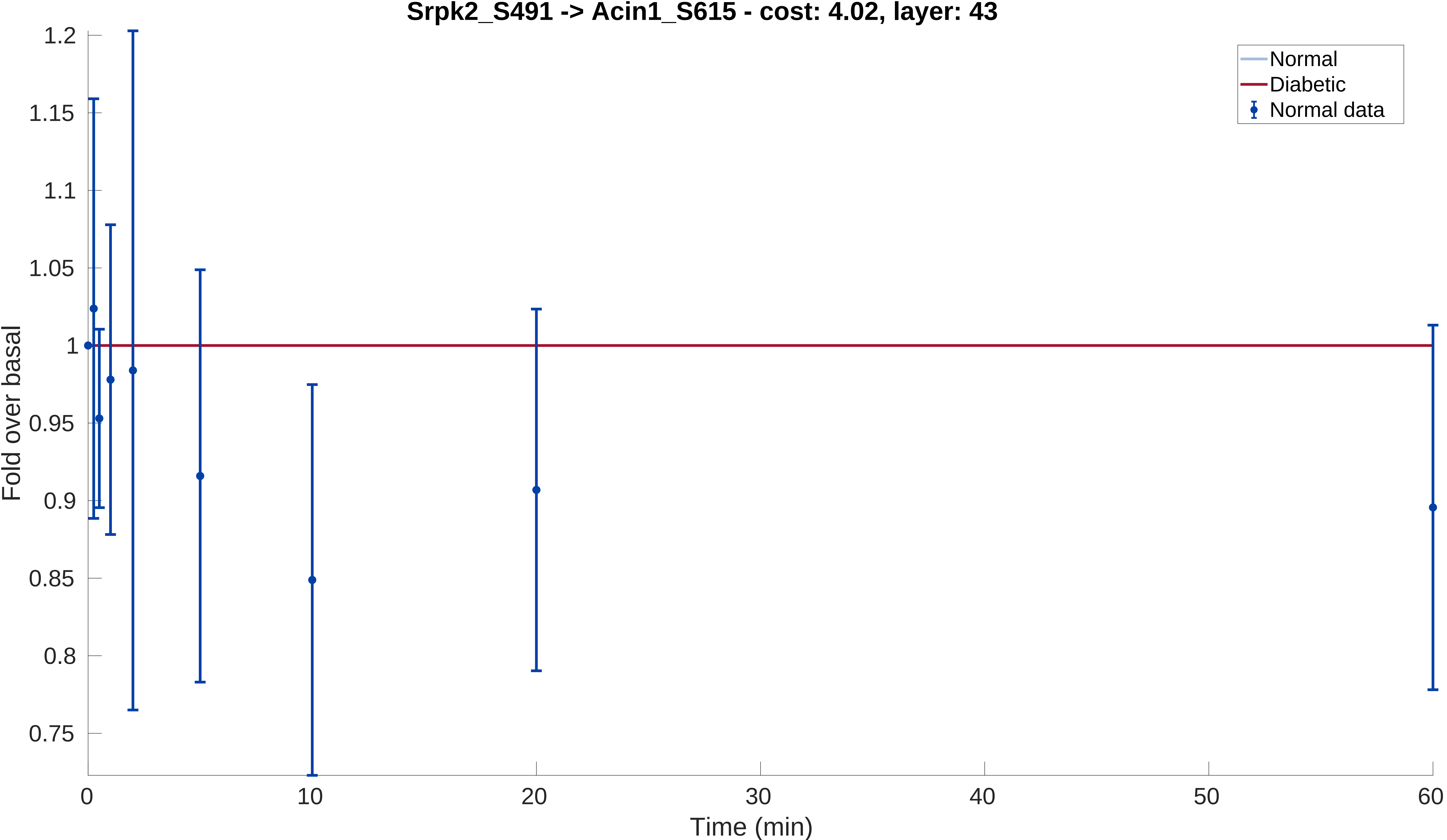

Srpk2_S491 -> Acin1_S615 - cost: 4.02, layer: 43
Normal
Diabetic
Normal data
Fold over basal
Time (min)
1.2
1.15
1.1
1.05
1
0.95
0.9
0.85
0.8
0.75
0
10
20
30
40
50
60

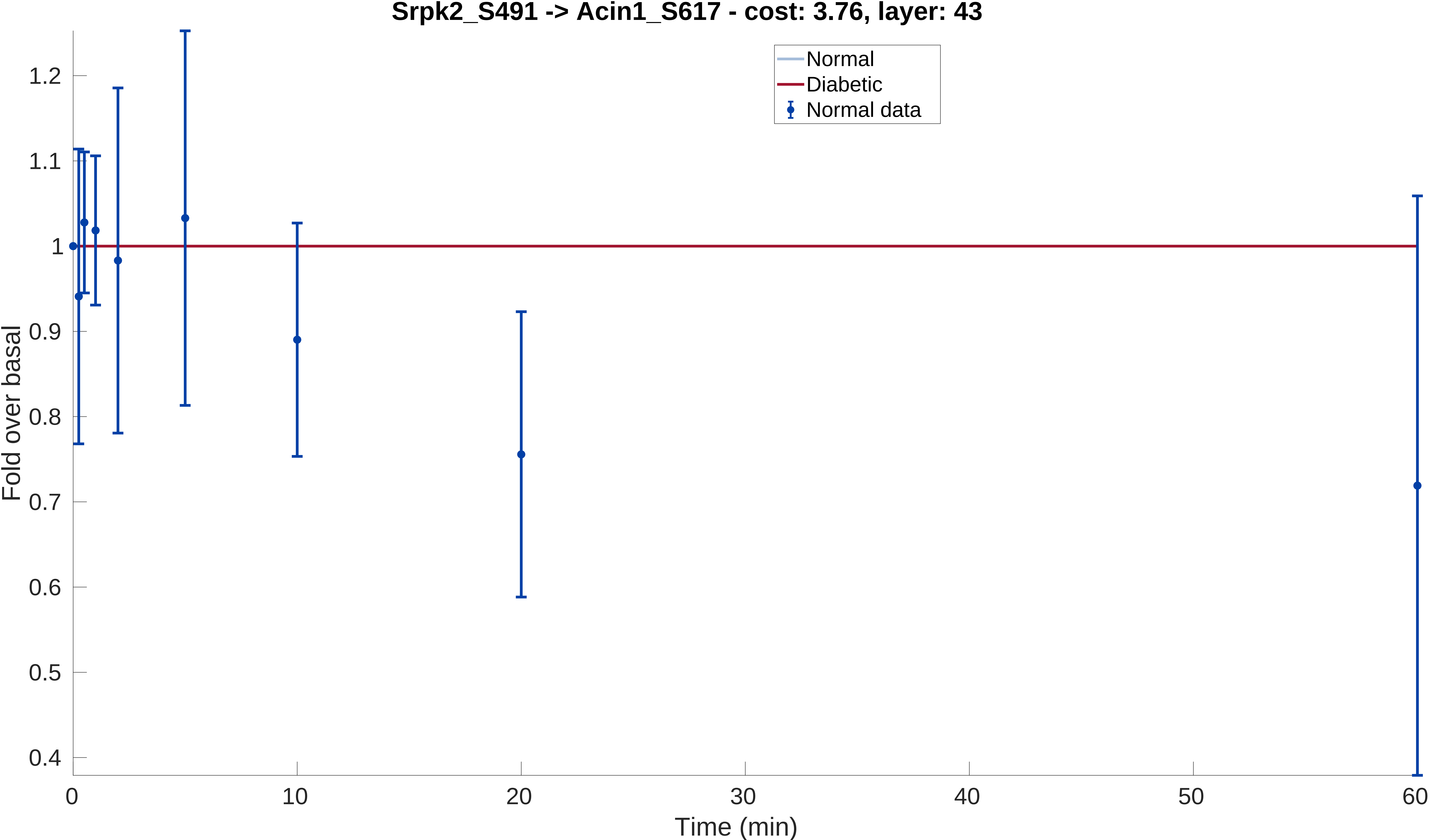

Srpk2_S491 -> Acin1_S617 - cost: 3.76, layer: 43
Normal
Diabetic
Normal data
Fold over basal
Time (min)
0.4
0.5
0.6
0.7
0.8
0.9
1
1.1
1.2
0
10
20
30
40
50
60

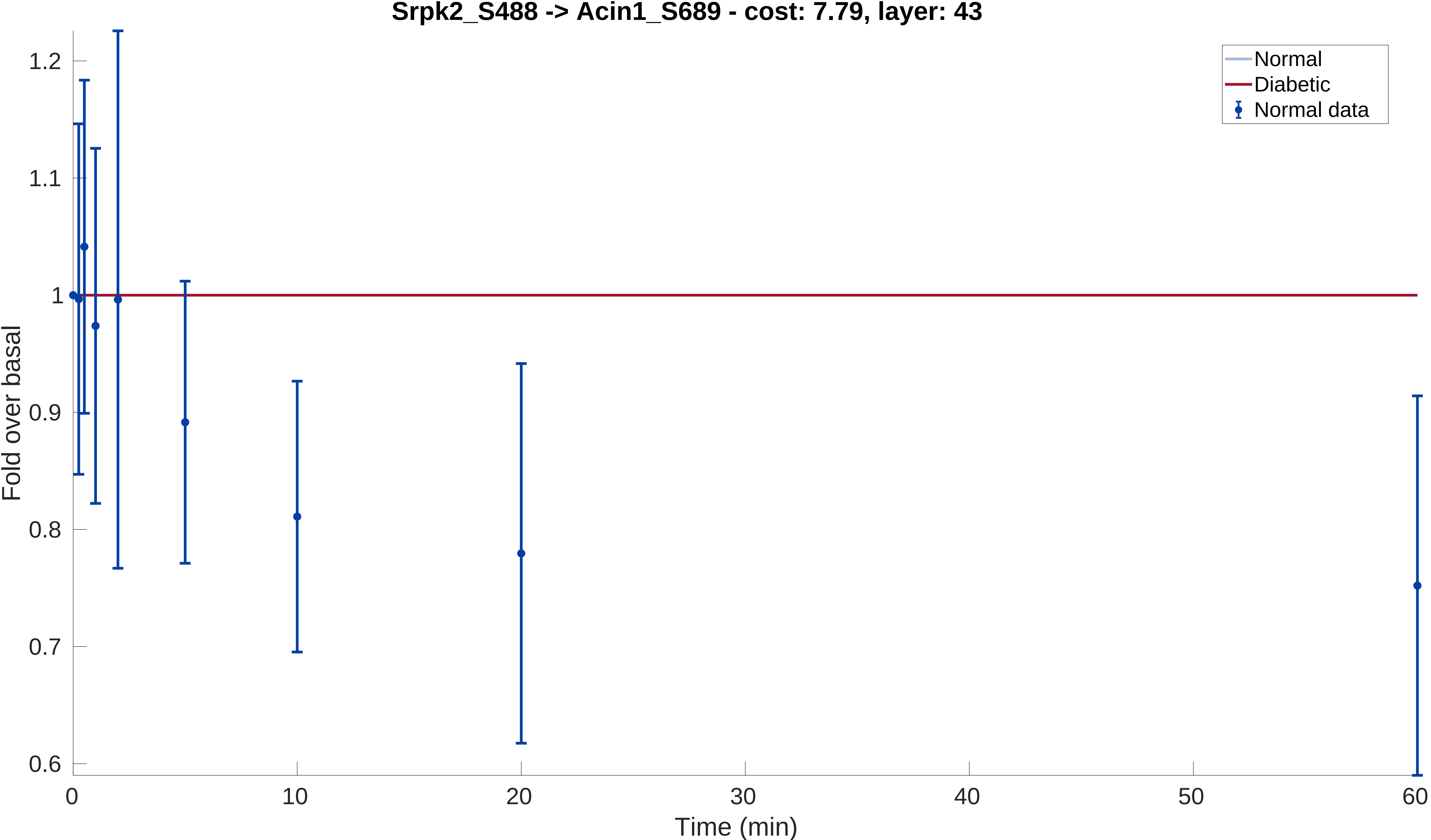

Srpk2_S488 -> Acin1_S689 - cost: 7.79, layer: 43
Normal
Diabetic
Normal data
Fold over basal
1.2
1.1
1
0.9
0.8
0.7
0.6
0
10
20
30
40
50
60
Time (min)

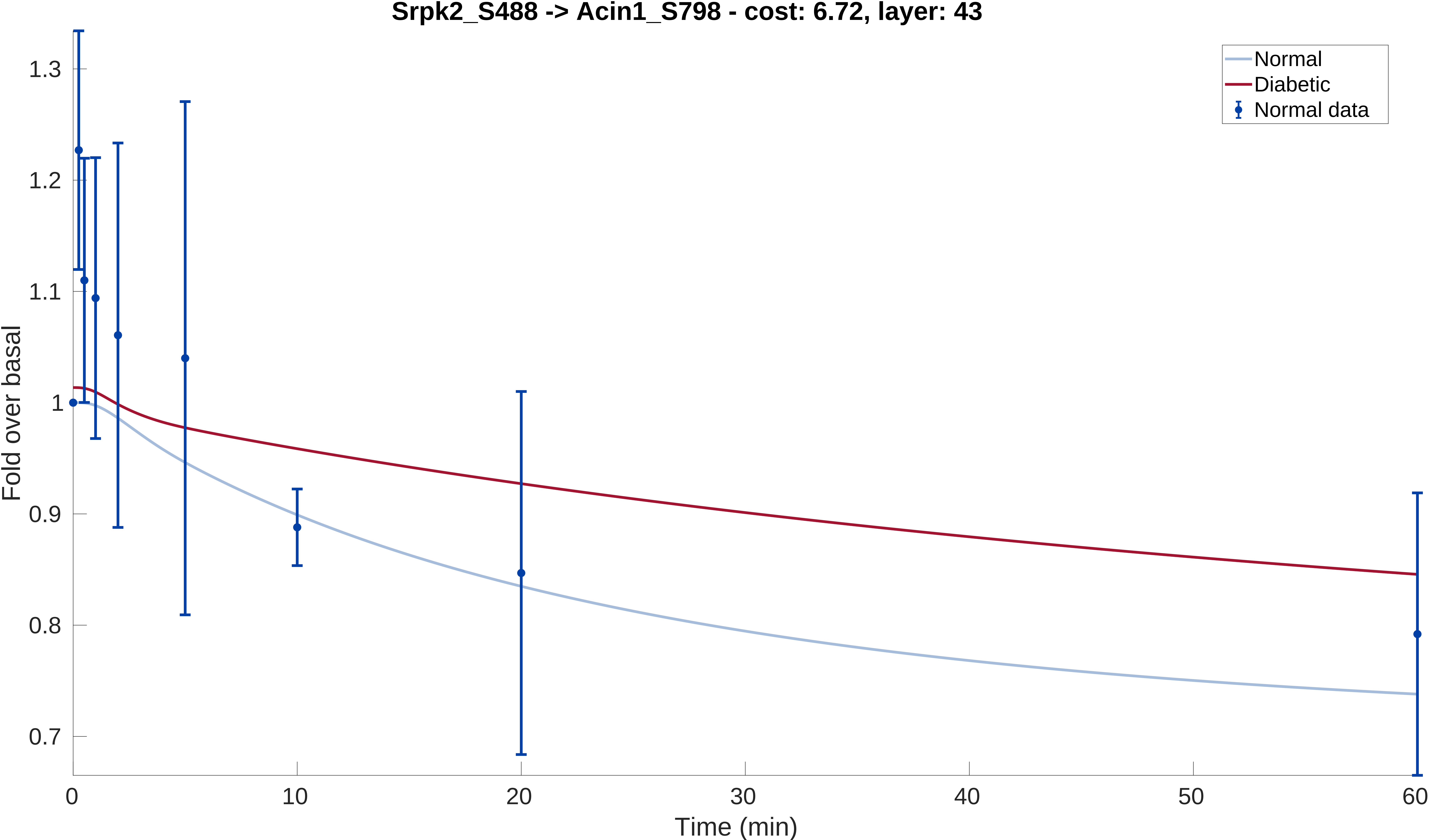

Srpk2_S488 -> Acin1_S798 - cost: 6.72, layer: 43
Normal
Diabetic
Normal data
Fold over basal
1.3
1.2
1.1
1
0.9
0.8
0.7
0
10
20
30
40
50
60
Time (min)

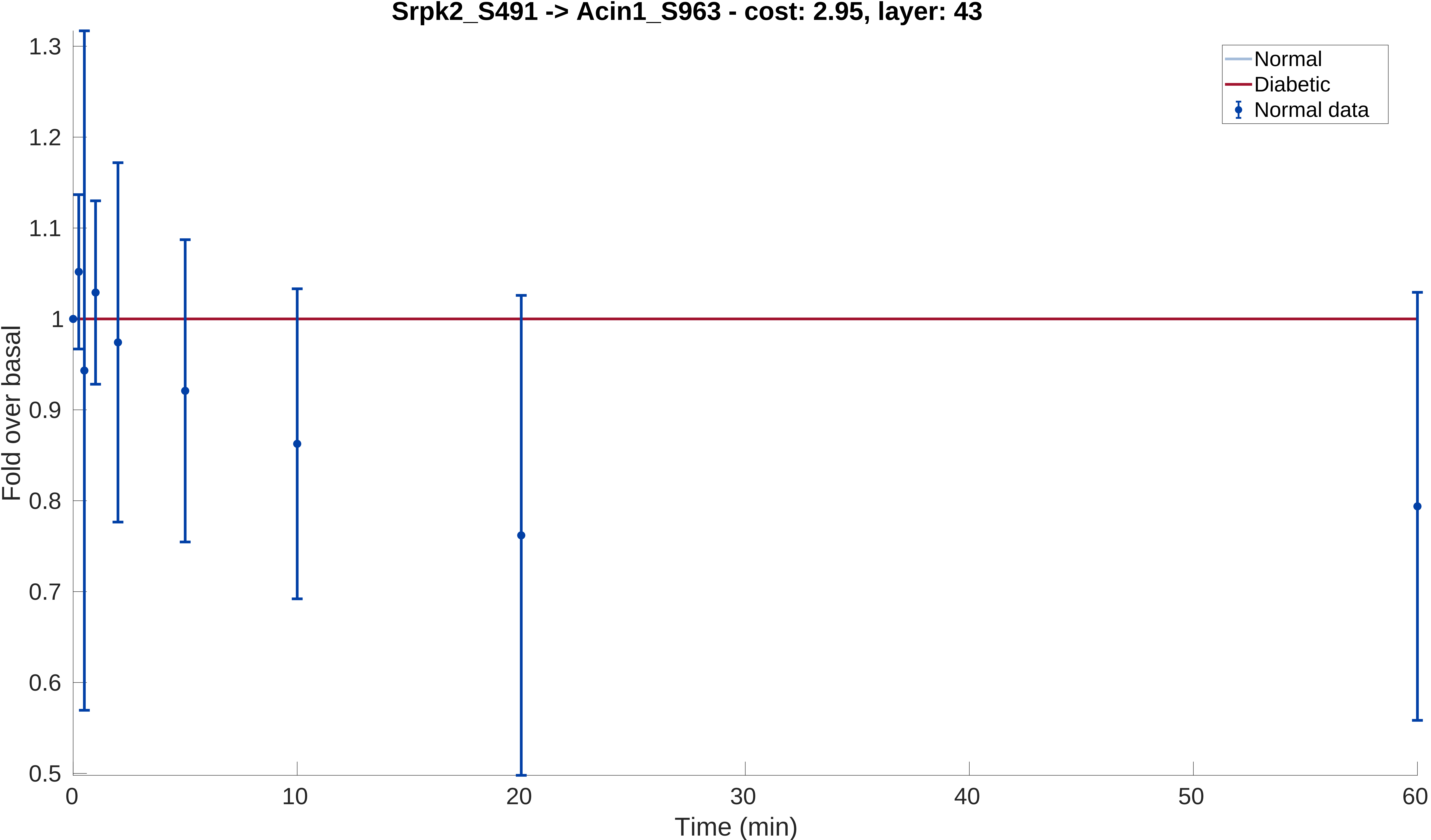
Srpk2_S491 -> Acin1_S963 - cost: 2.95, layer: 43
Normal
Diabetic
Normal data
Fold over basal
1.3
1.2
1.1
1
0.9
0.8
0.7
0.6
0.5
0
10
20
30
40
50
60
Time (min)

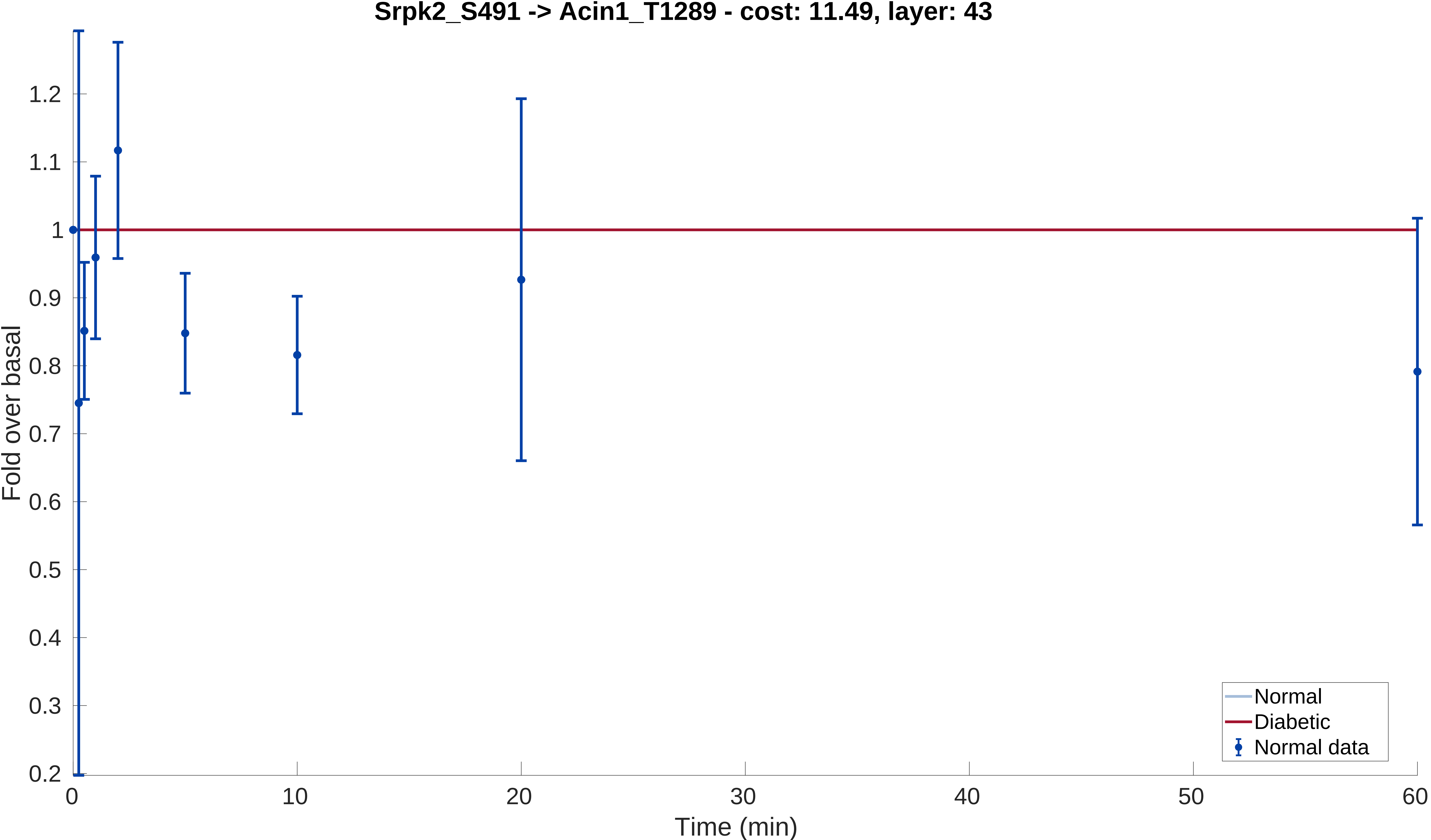
Srpk2_S491 -> Acin1_T1289 - cost: 11.49, layer: 43
Fold over basal
Time (min)
0.2
0.3
0.4
0.5
0.6
0.7
0.8
0.9
1
1.1
1.2
0
10
20
30
40
50
60
Normal
Diabetic
Normal data

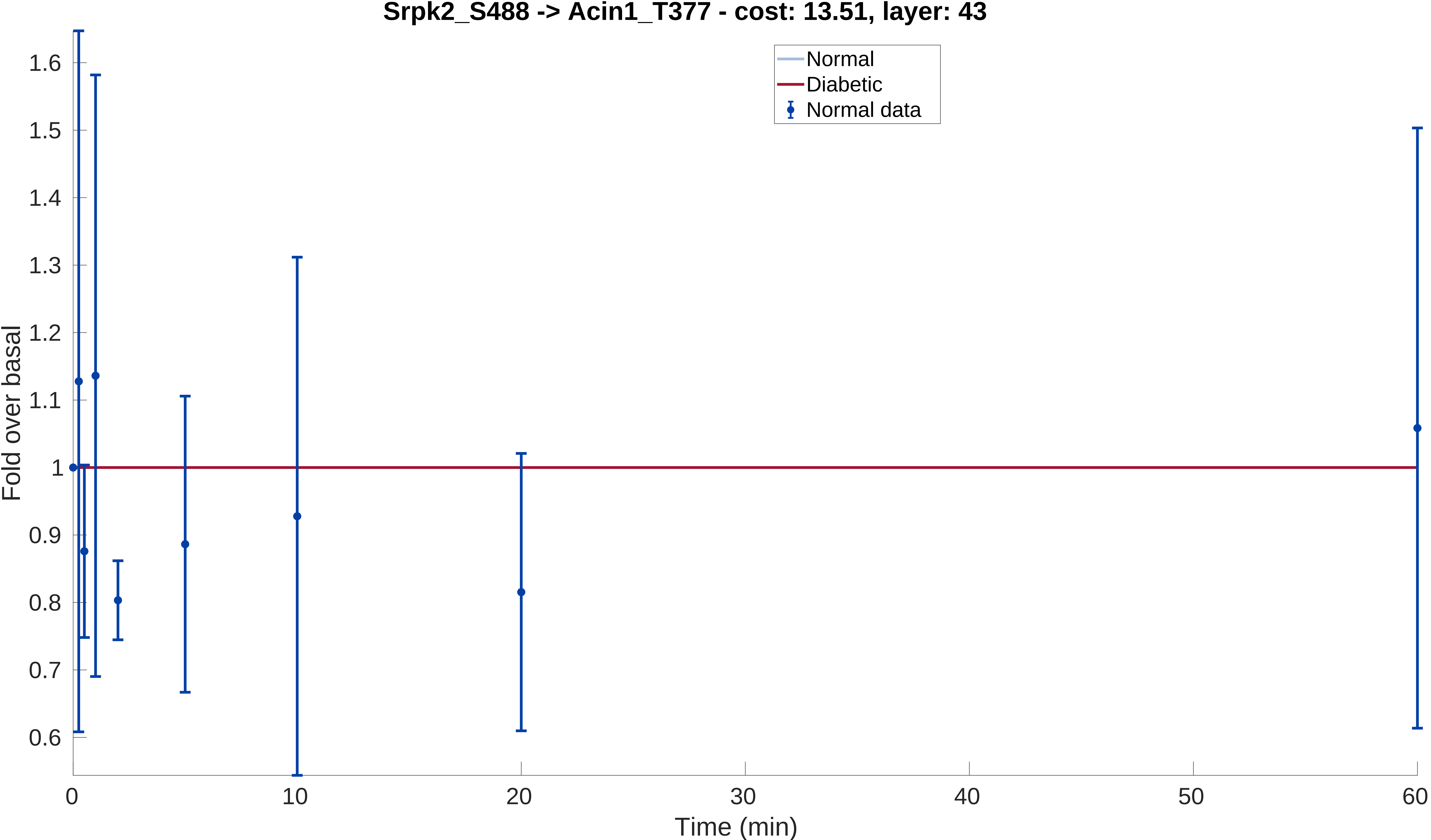

Srpk2_S488 -> Acin1_T377 - cost: 13.51, layer: 43
Normal
Diabetic
Normal data
Fold over basal
1.6
1.5
1.4
1.3
1.2
1.1
1
0.9
0.8
0.7
0.6
0
10
20
30
40
50
60
Time (min)

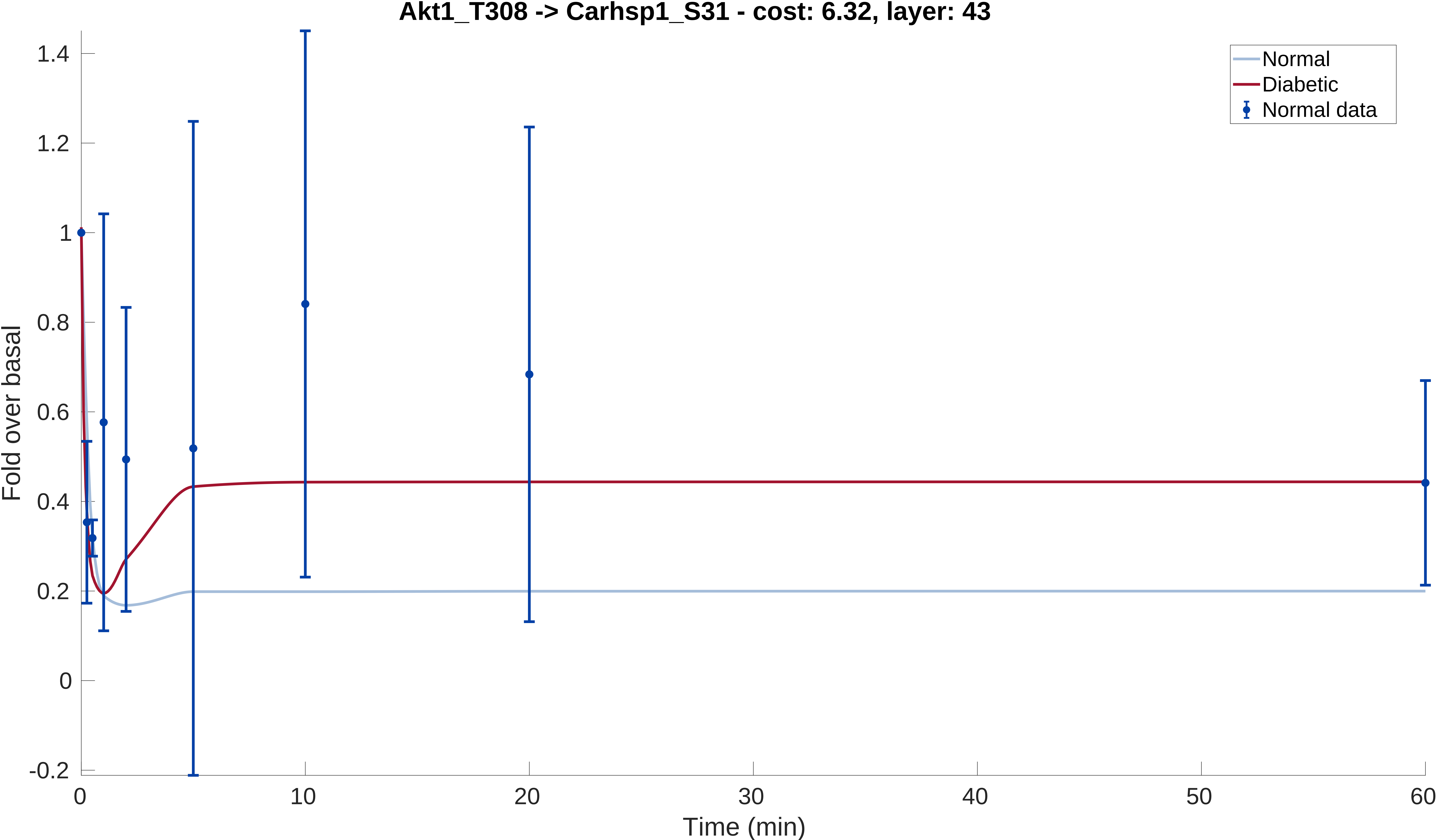

Akt1_T308 -> Carhsp1_S31 - cost: 6.32, layer: 43
Normal
Diabetic
Normal data
Fold over basal
Time (min)
1.4
1.2
1
0.8
0.6
0.4
0.2
0
-0.2
0
10
20
30
40
50
60

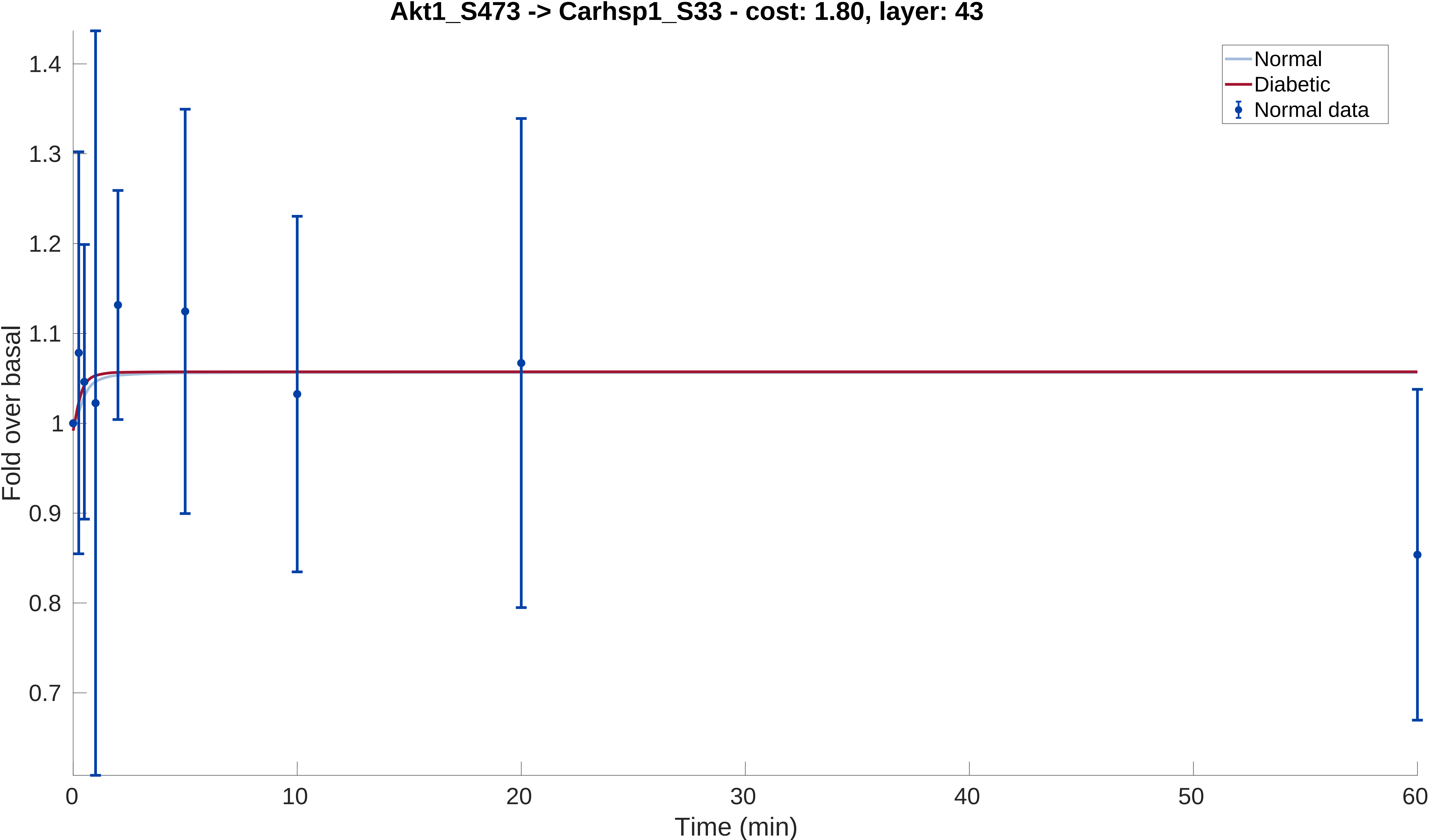

Akt1_S473 -> Carhsp1_S33 - cost: 1.80, layer: 43
Normal
Diabetic
Normal data
Fold over basal
1.4
1.3
1.2
1.1
1
0.9
0.8
0.7
0
10
20
30
40
50
60
Time (min)

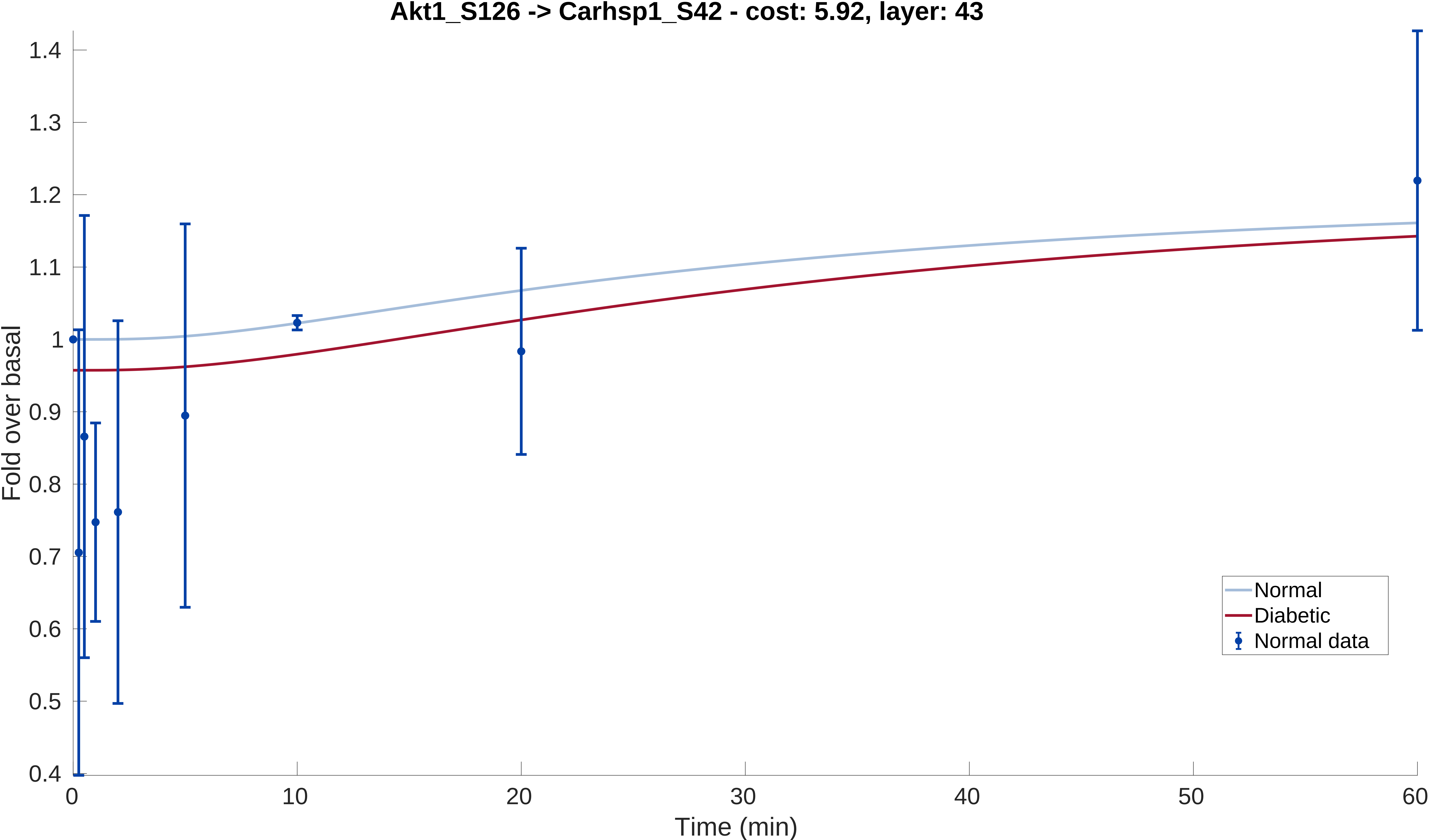

Akt1_S126 -> Carhsp1_S42 - cost: 5.92, layer: 43
Fold over basal
1.4
1.3
1.2
1.1
1
0.9
0.8
0.7
0.6
0.5
0.4
0
10
20
30
40
50
60
Time (min)
Normal
Diabetic
Normal data

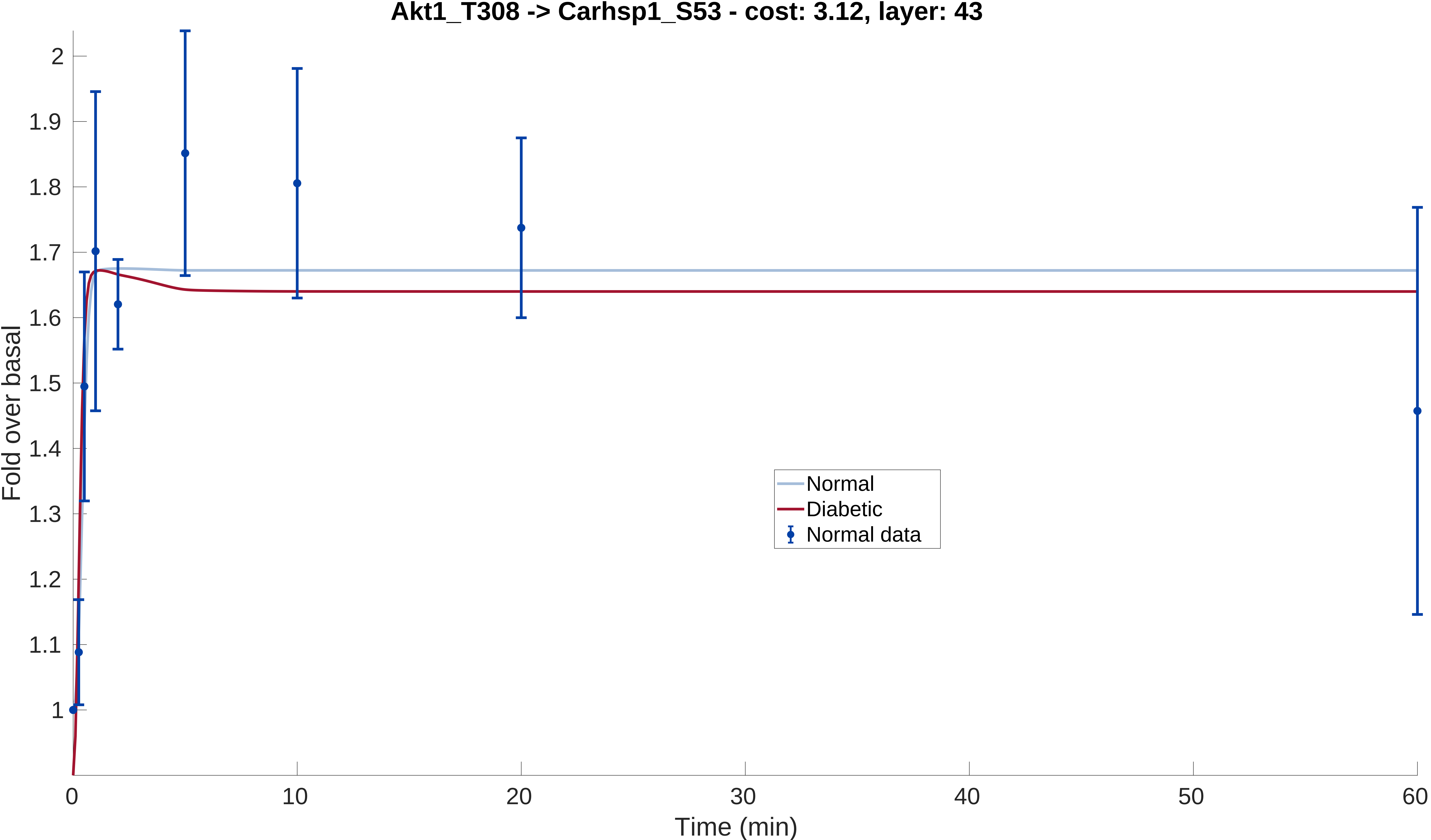
Akt1_T308 -> Carhsp1_S53 - cost: 3.12, layer: 43
Fold over basal
Time (min)
2
1.9
1.8
1.7
1.6
1.5
1.4
1.3
1.2
1.1
1
0
10
20
30
40
50
60
Normal
Diabetic
Normal data

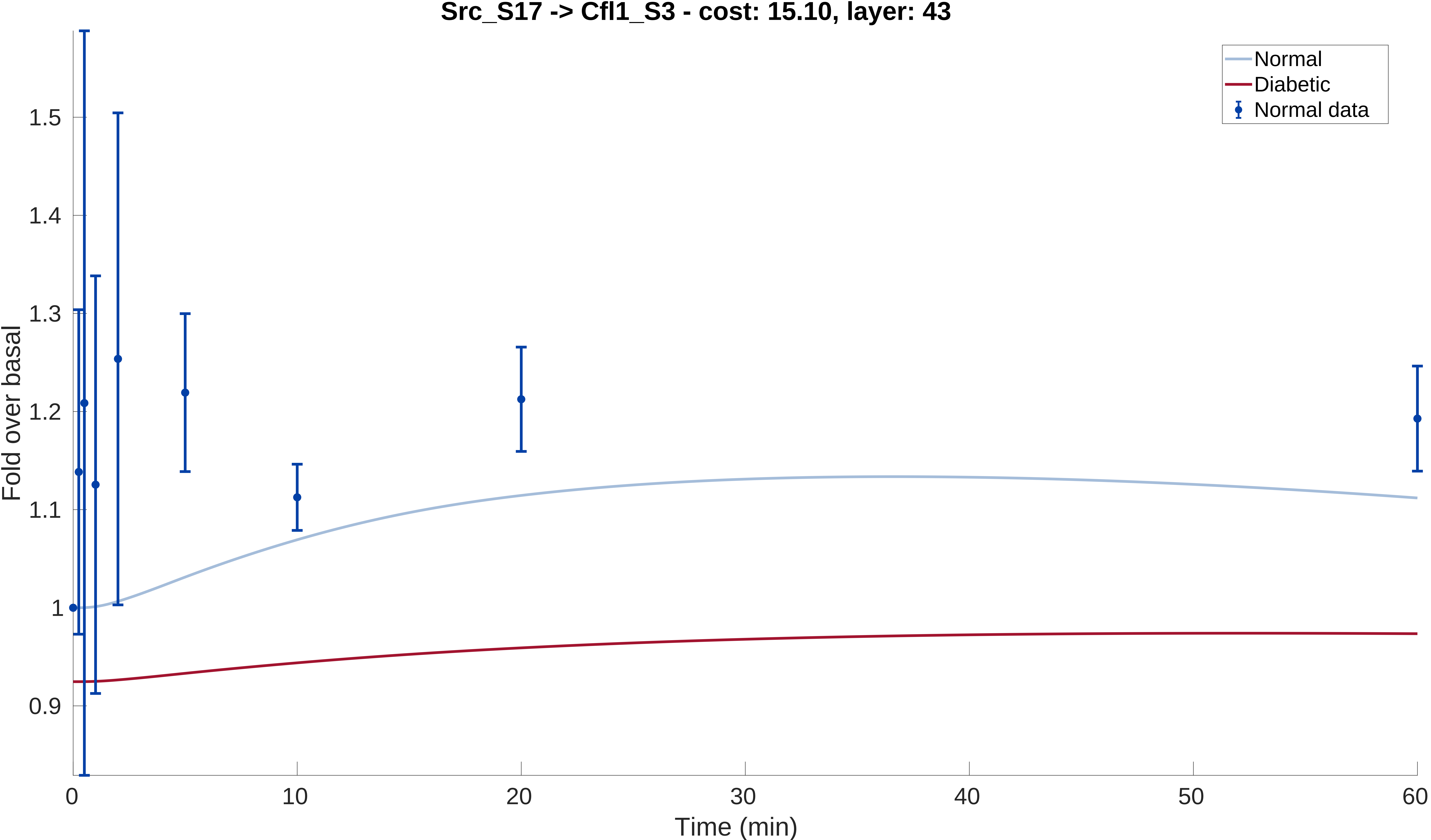
Src_S17 -> Cfl1_S3 - cost: 15.10, layer: 43
Normal
Diabetic
Normal data
Fold over basal
1.5
1.4
1.3
1.2
1.1
1
0.9
0
10
20
30
40
50
60
Time (min)

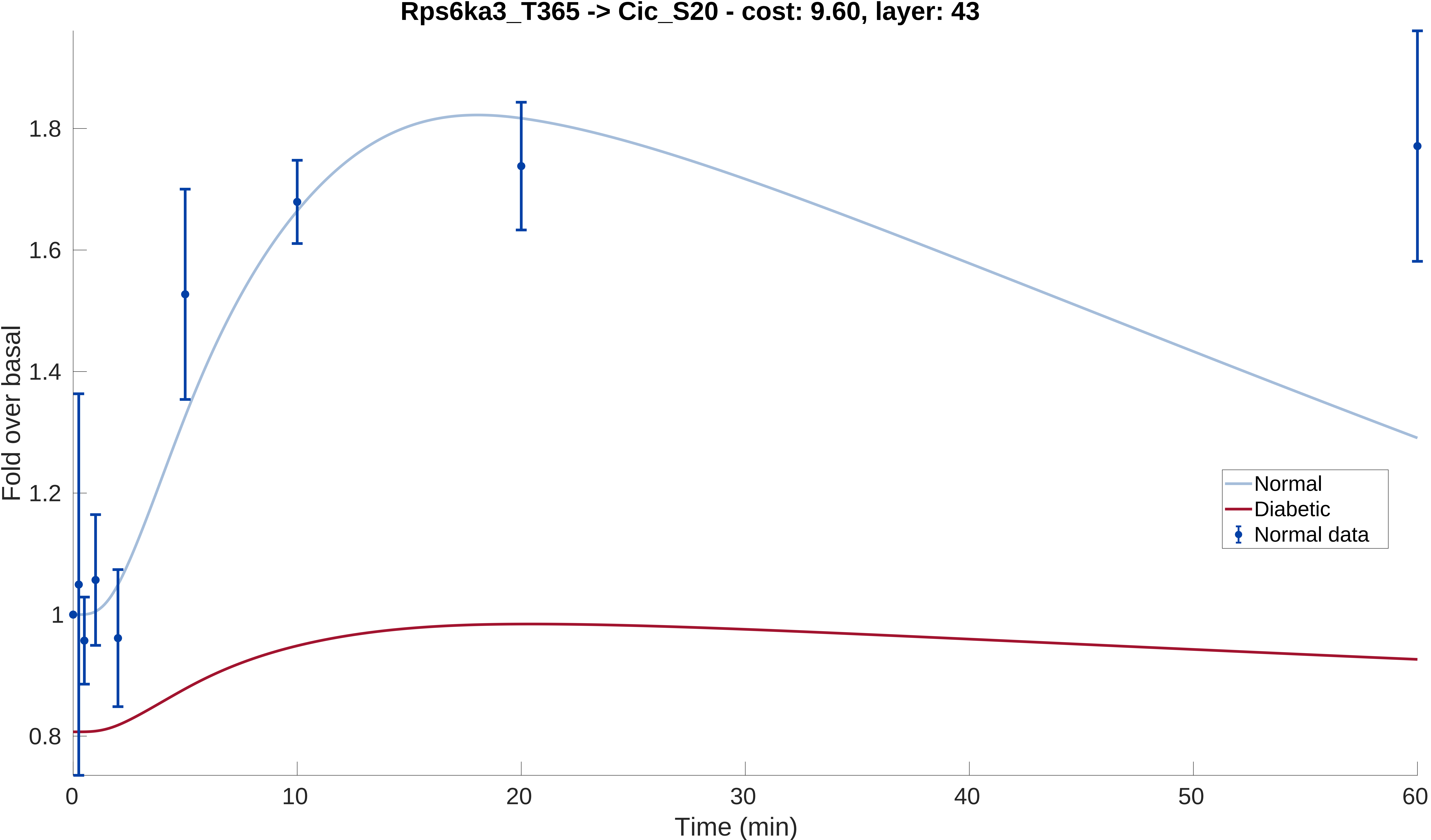
Rps6ka3_T365 -> Cic_S20 - cost: 9.60, layer: 43
Fold over basal
1.8
1.6
1.4
1.2
1
0.8
0
10
20
30
40
50
60
Time (min)
Normal
Diabetic
Normal data

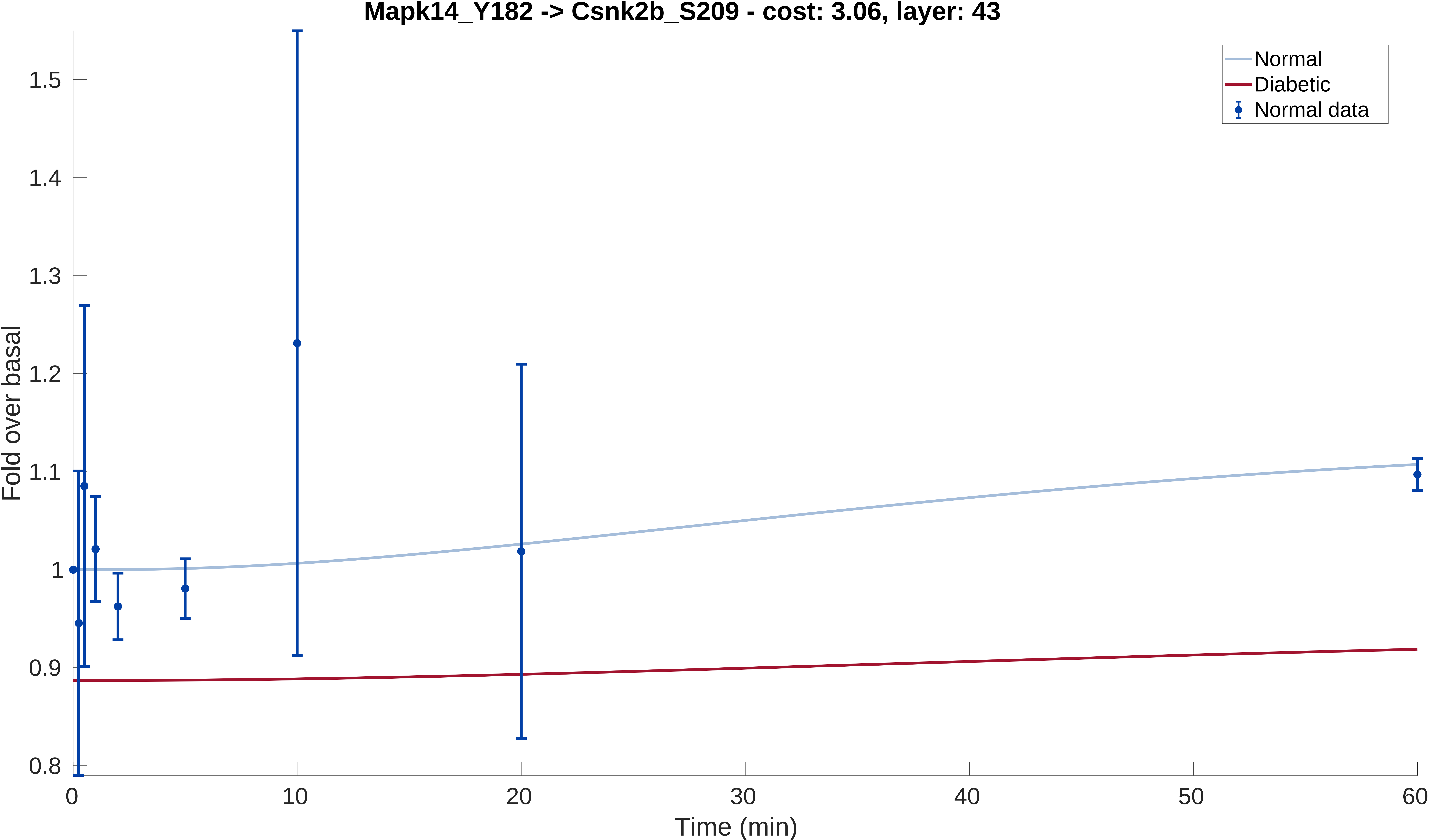

Mapk14_Y182 -> Csnk2b_S209 - cost: 3.06, layer: 43
Normal
Diabetic
Normal data
Fold over basal
Time (min)
0.8
0.9
1
1.1
1.2
1.3
1.4
1.5
0
10
20
30
40
50
60

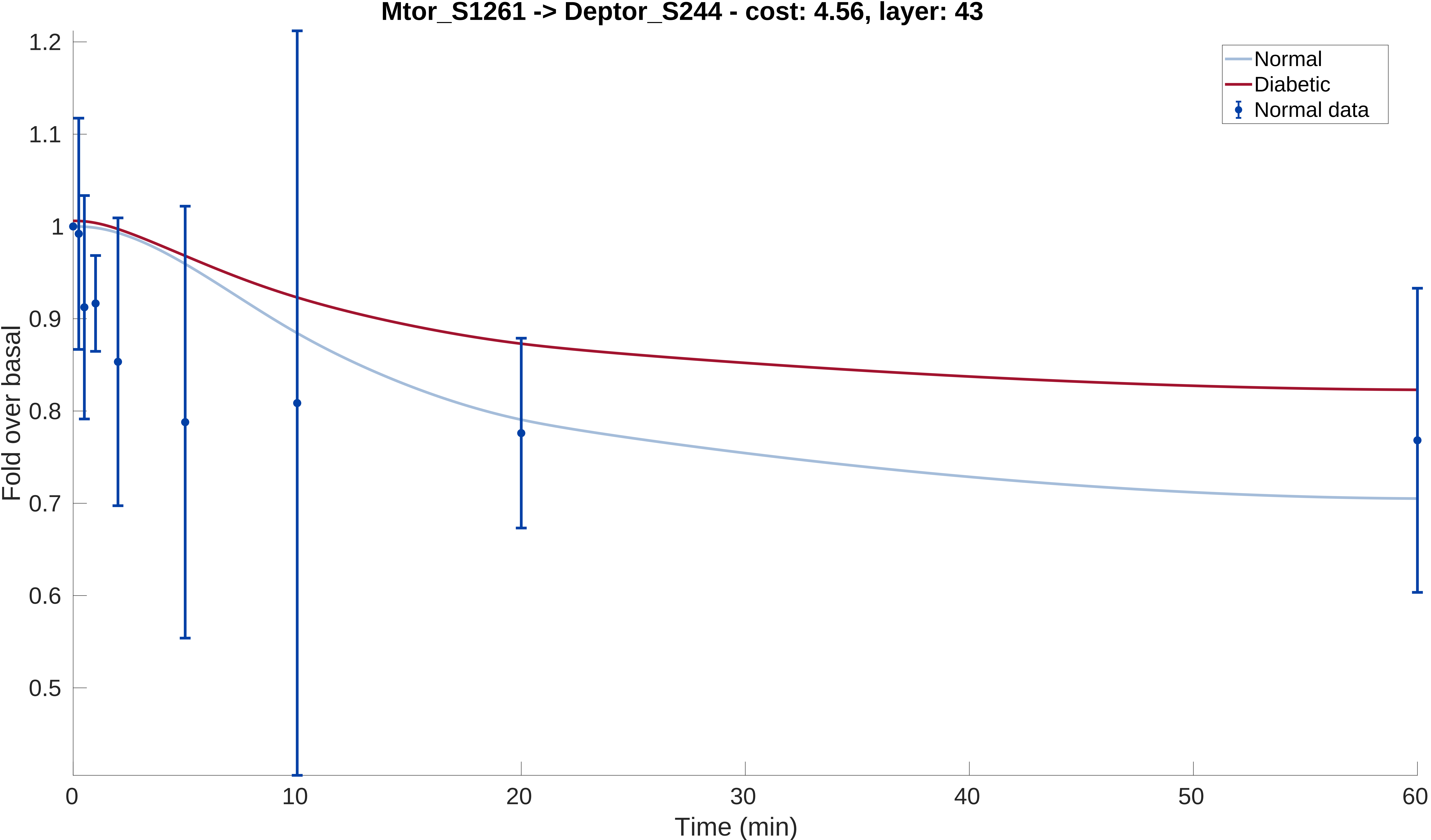

Mtor_S1261 -> Deptor_S244 - cost: 4.56, layer: 43
Normal
Diabetic
Normal data
Fold over basal
1.2
1.1
1
0.9
0.8
0.7
0.6
0.5
0
10
20
30
40
50
60
Time (min)

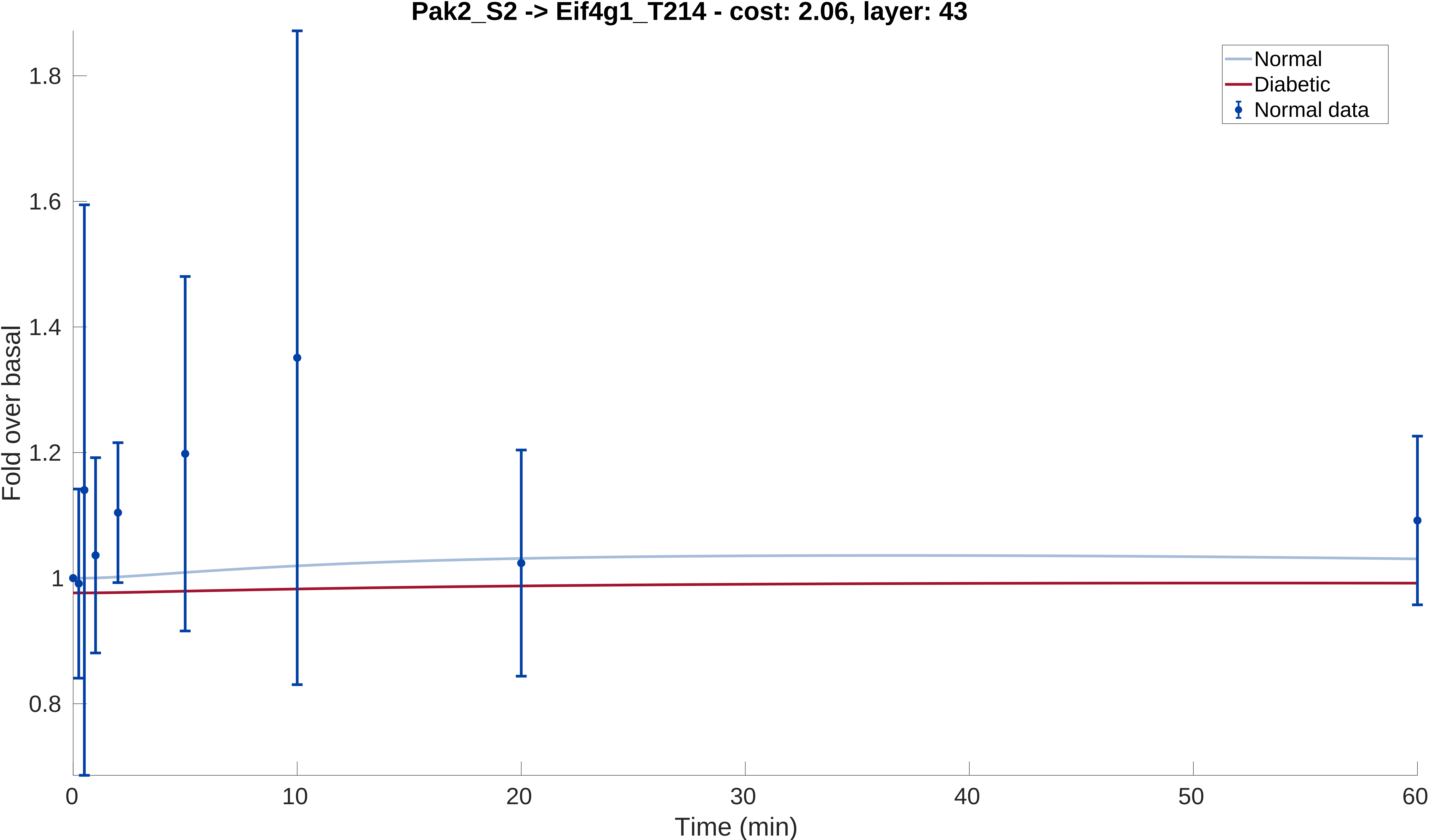

Pak2_S2 -> Eif4g1_T214 - cost: 2.06, layer: 43
Normal
Diabetic
Normal data
Fold over basal
1.8
1.6
1.4
1.2
1
0.8
0
10
20
30
40
50
60
Time (min)

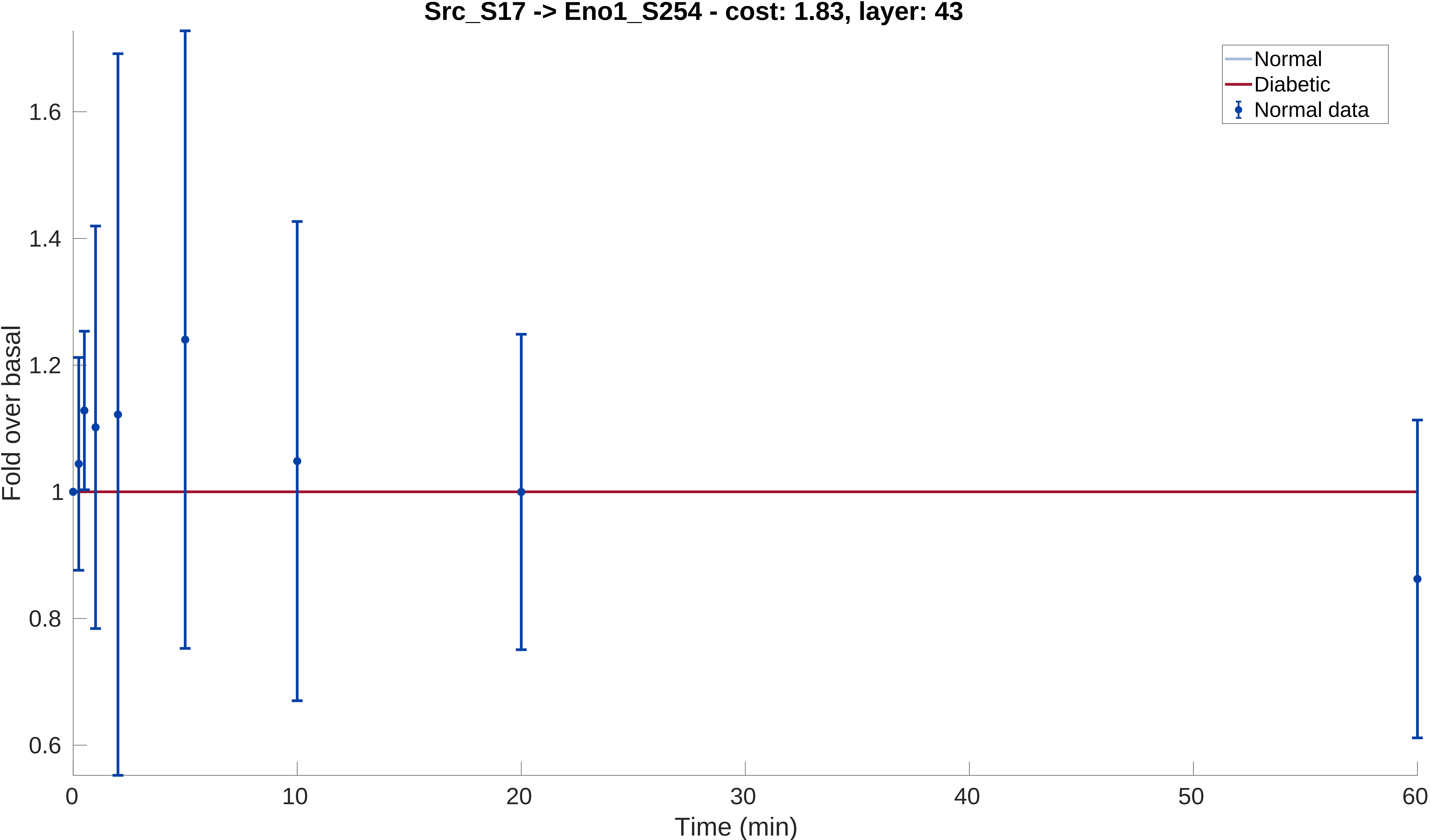

Src_S17 -> Eno1_S254 - cost: 1.83, layer: 43
Normal
Diabetic
Normal data
Fold over basal
Time (min)
0.6
0.8
1
1.2
1.4
1.6
0
10
20
30
40
50
60

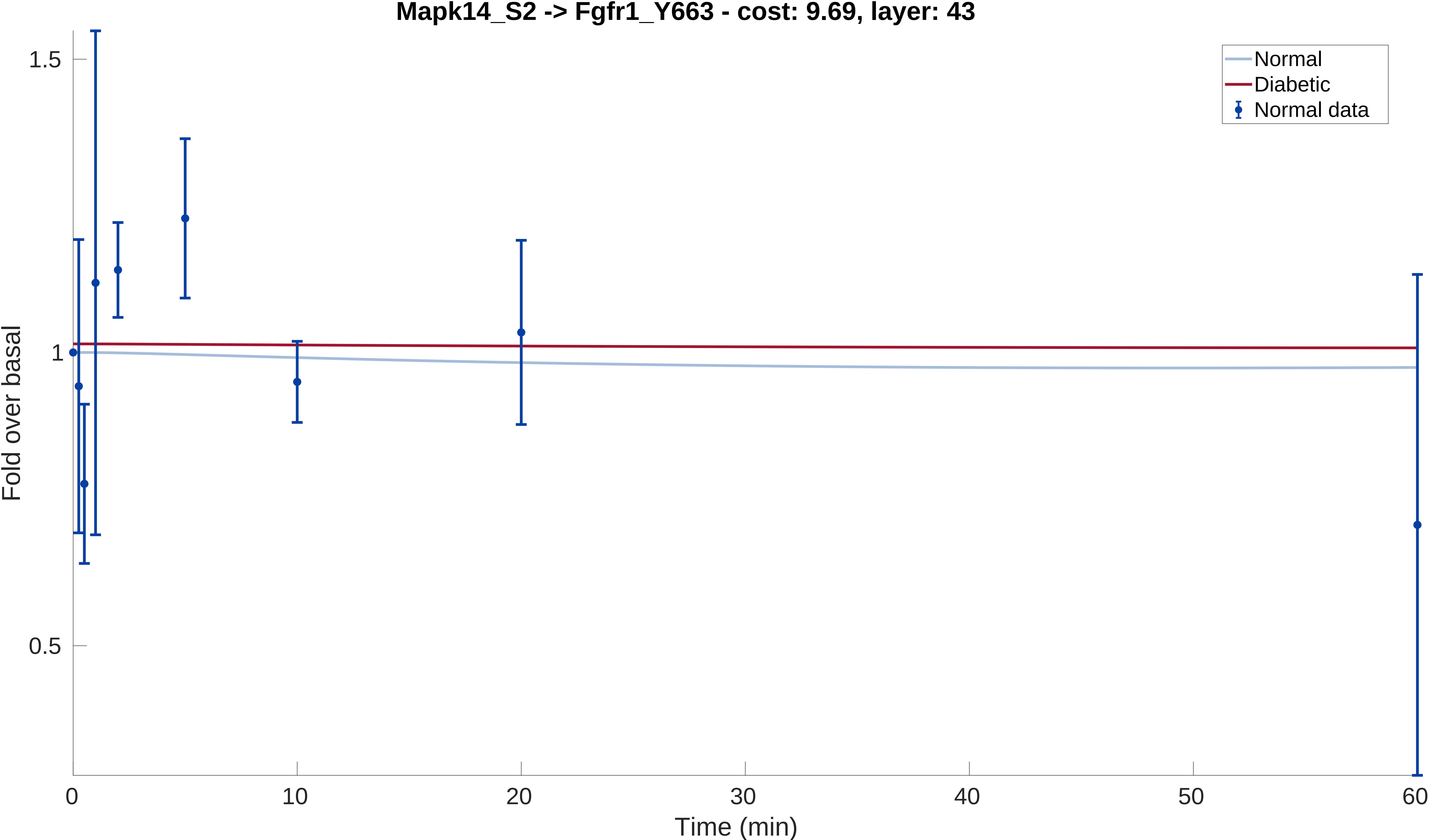
Mapk14_S2 -> Fgfr1_Y663 - cost: 9.69, layer: 43
Normal
Diabetic
Normal data
Fold over basal
1.5
1
0.5
0
10
20
30
40
50
60
Time (min)

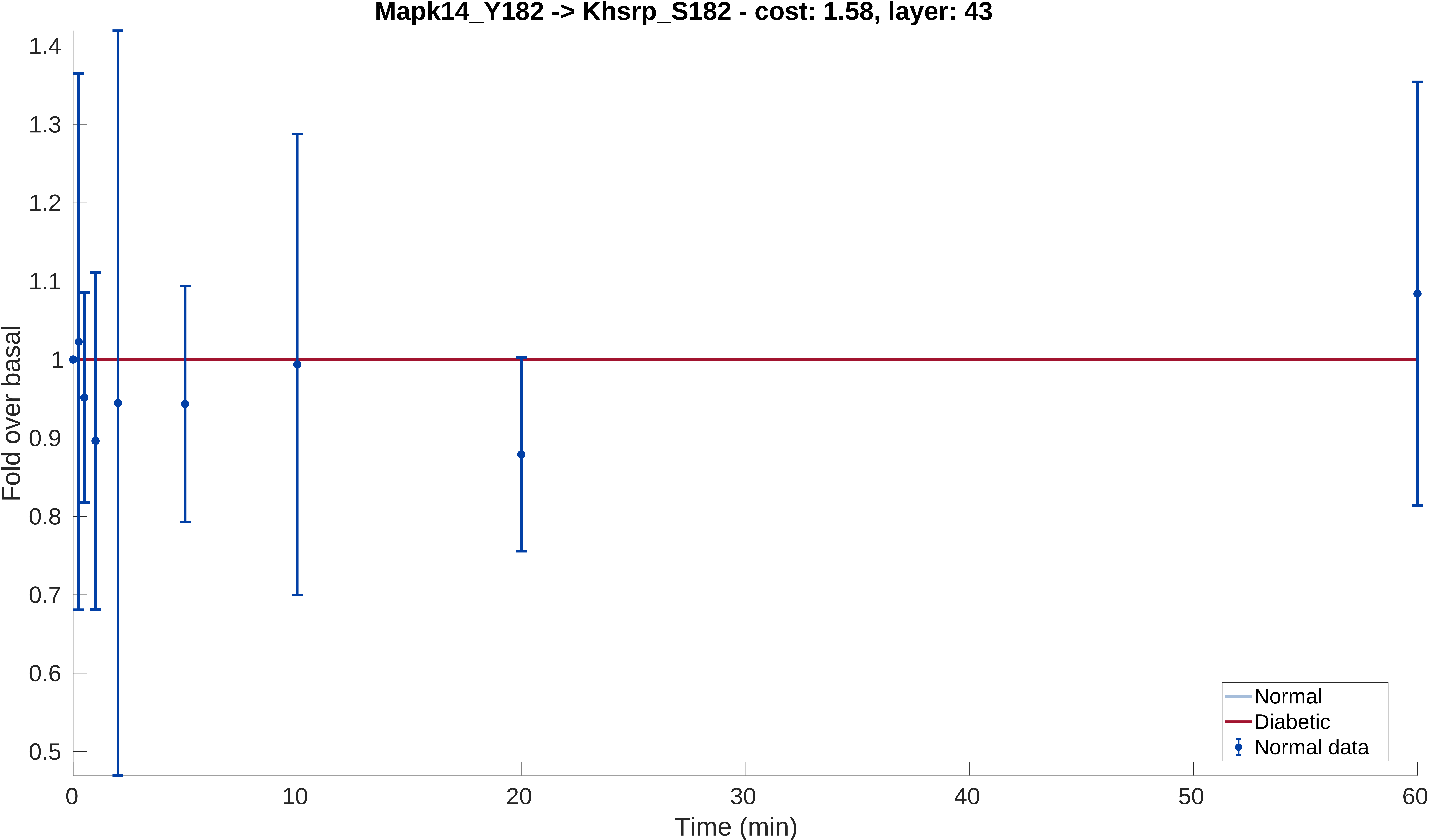

Mapk14_Y182 -> Khsrp_S182 - cost: 1.58, layer: 43
Fold over basal
1.4
1.3
1.2
1.1
1
0.9
0.8
0.7
0.6
0.5
0
10
20
30
40
50
60
Time (min)
Normal
Diabetic
Normal data

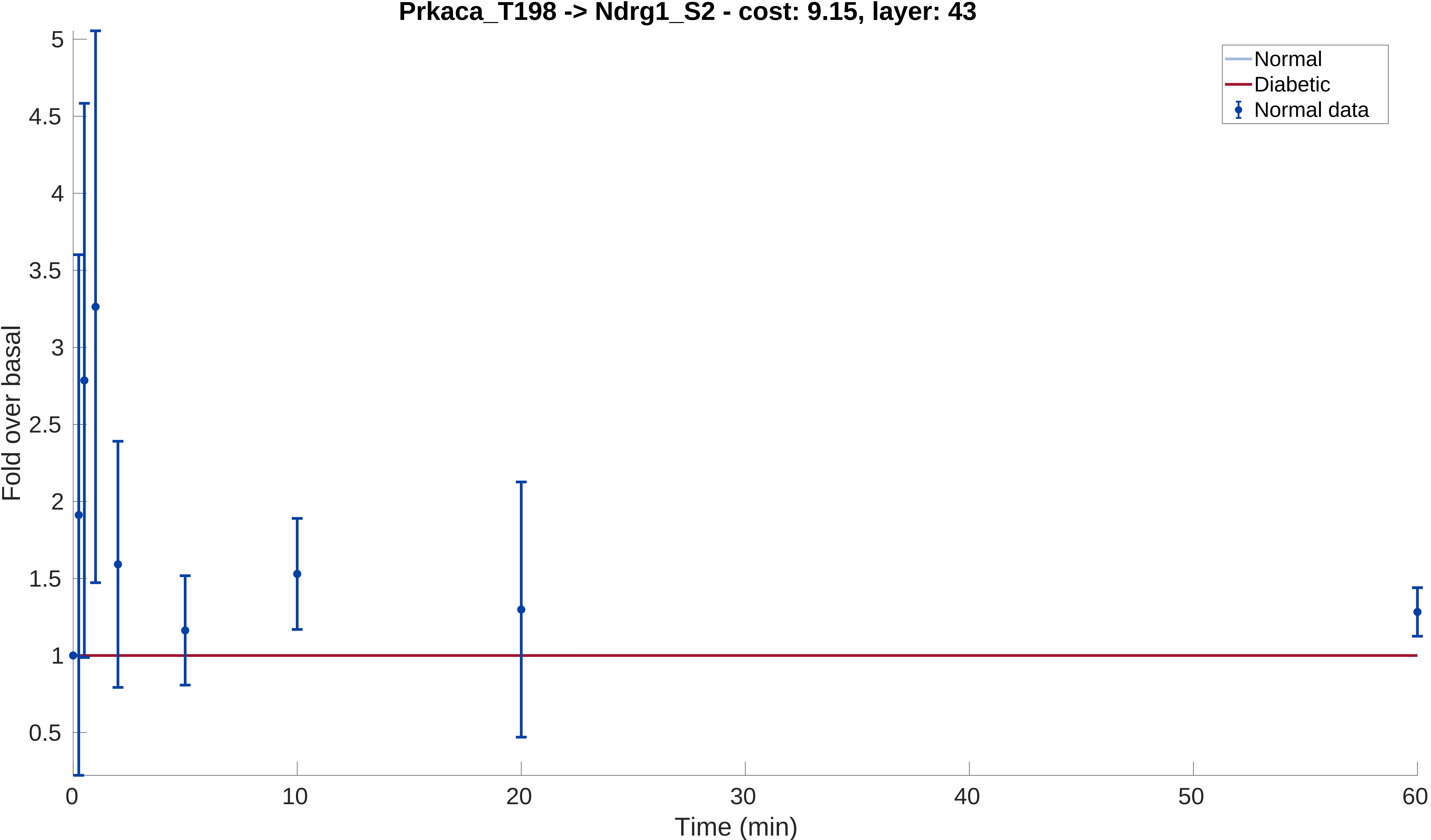

Prkaca_T198 -> Ndrg1_S2 - cost: 9.15, layer: 43
Normal
Diabetic
Normal data
Fold over basal
Time (min)
0
10
20
30
40
50
60
0.5
1
1.5
2
2.5
3
3.5
4
4.5
5

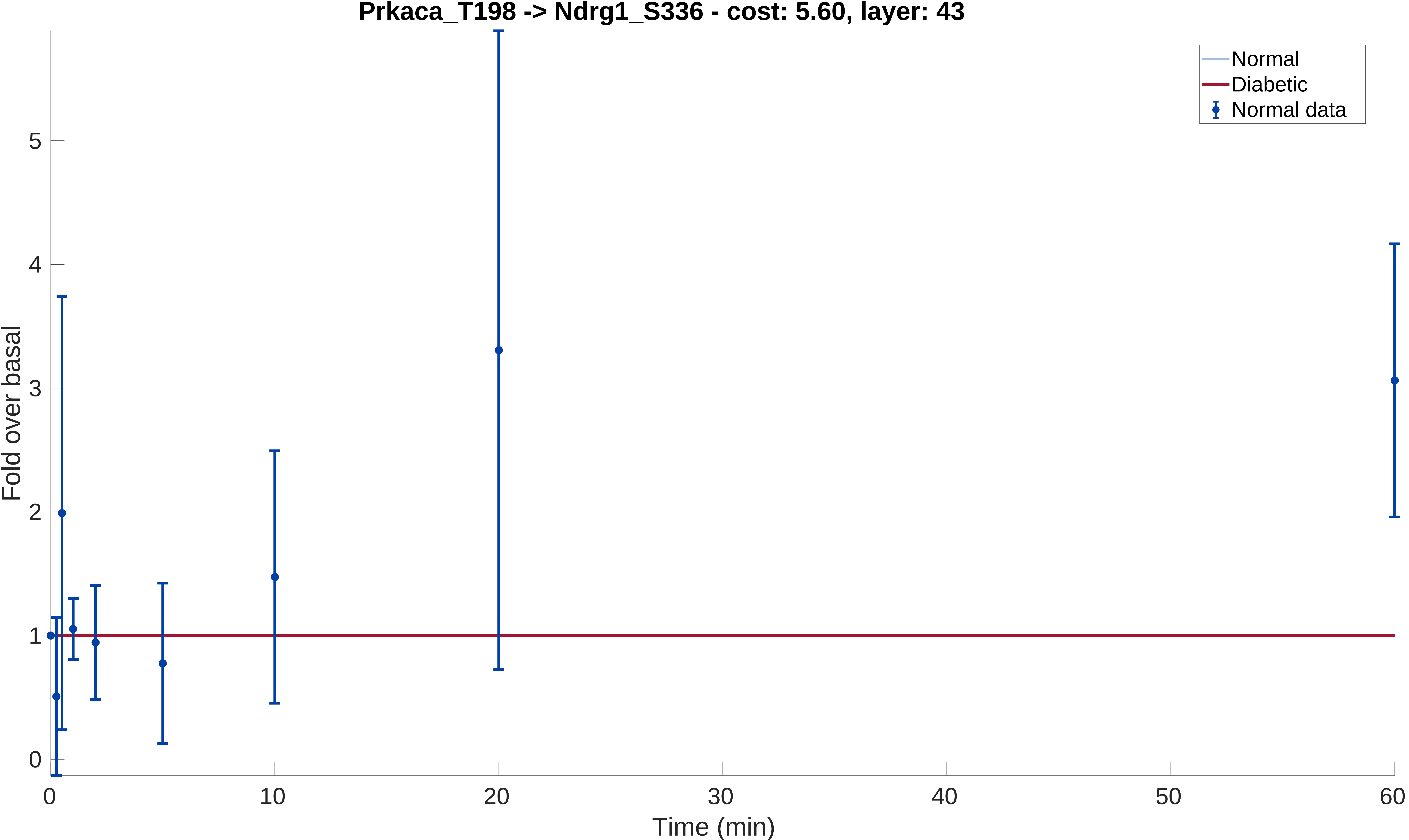

Prkaca_T198 -> Ndrg1_S336 - cost: 5.60, layer: 43
Normal
Diabetic
Normal data
Fold over basal
0
1
2
3
4
5
0
10
20
30
40
50
60
Time (min)

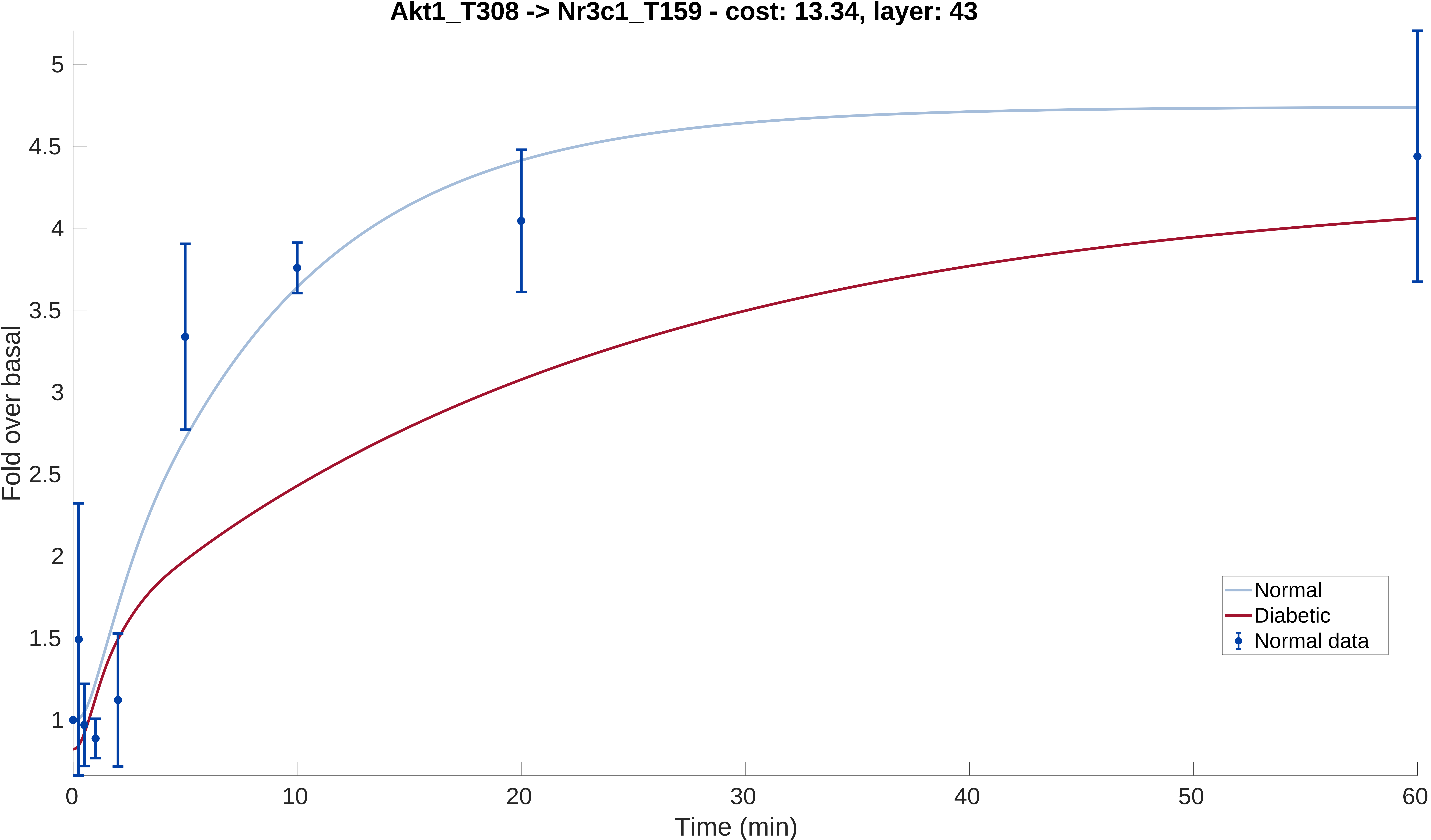

Akt1_T308 -> Nr3c1_T159 - cost: 13.34, layer: 43
Fold over basal
Time (min)
5
4.5
4
3.5
3
2.5
2
1.5
1
0
10
20
30
40
50
60
Normal
Diabetic
Normal data

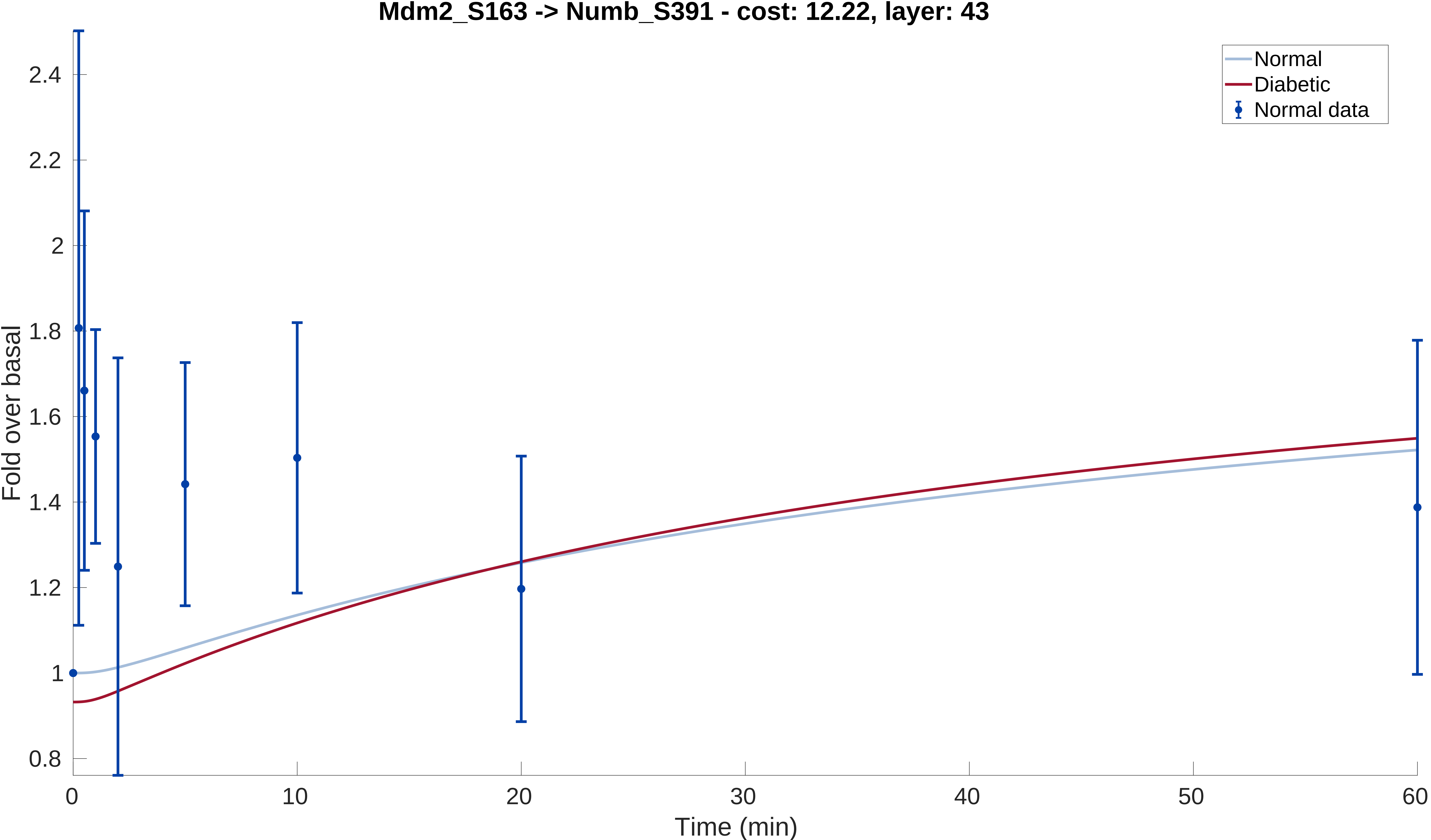

Mdm2_S163 -> Numb_S391 - cost: 12.22, layer: 43
Normal
Diabetic
Normal data
Fold over basal
2.4
2.2
2
1.8
1.6
1.4
1.2
1
0.8
0
10
20
30
40
50
60
Time (min)

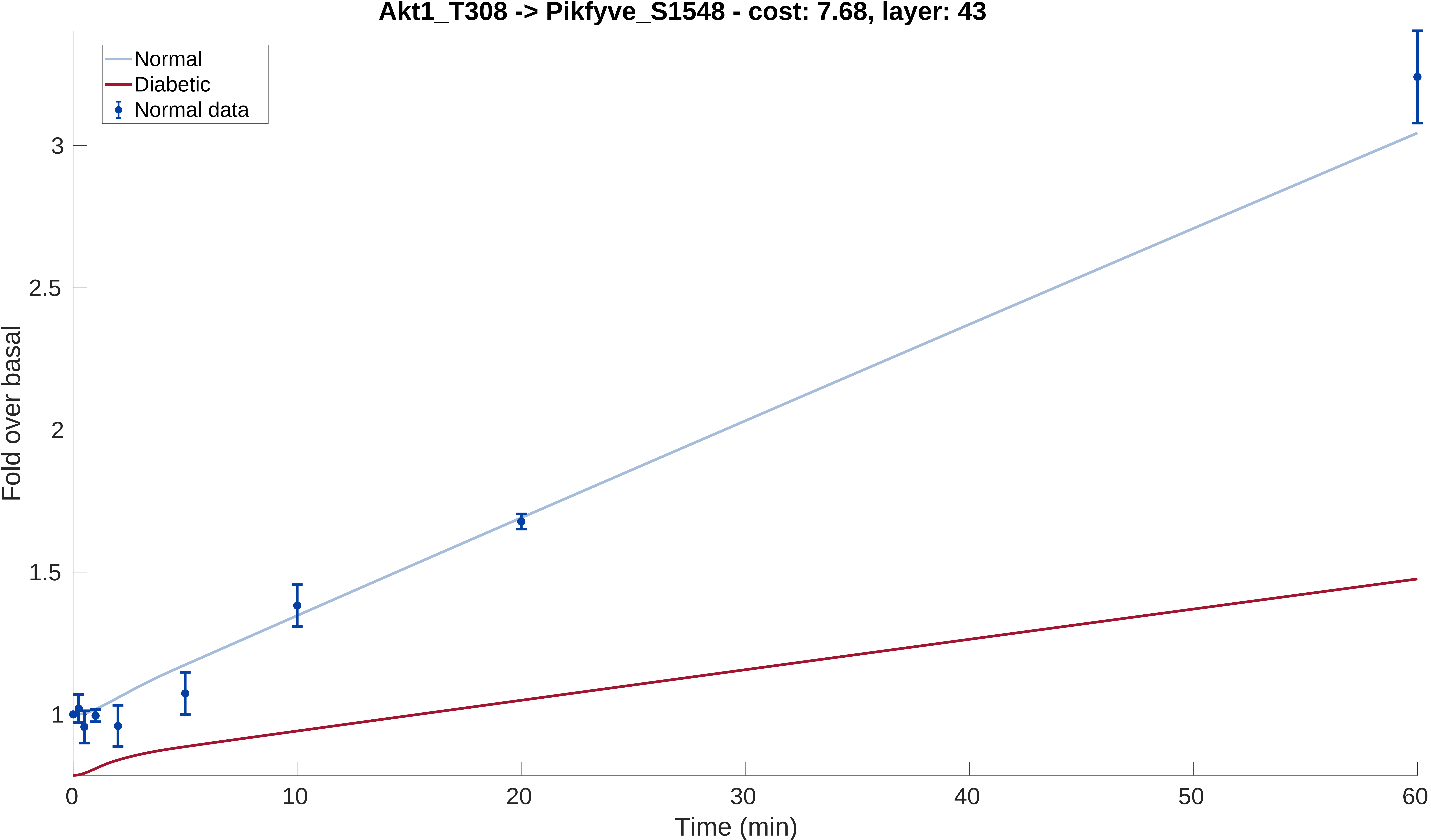

Akt1_T308 -> Pikfyve_S1548 - cost: 7.68, layer: 43
Normal
Diabetic
Normal data
3
2.5
2
1.5
1
0
10
20
30
40
50
60
Time (min)
Fold over basal

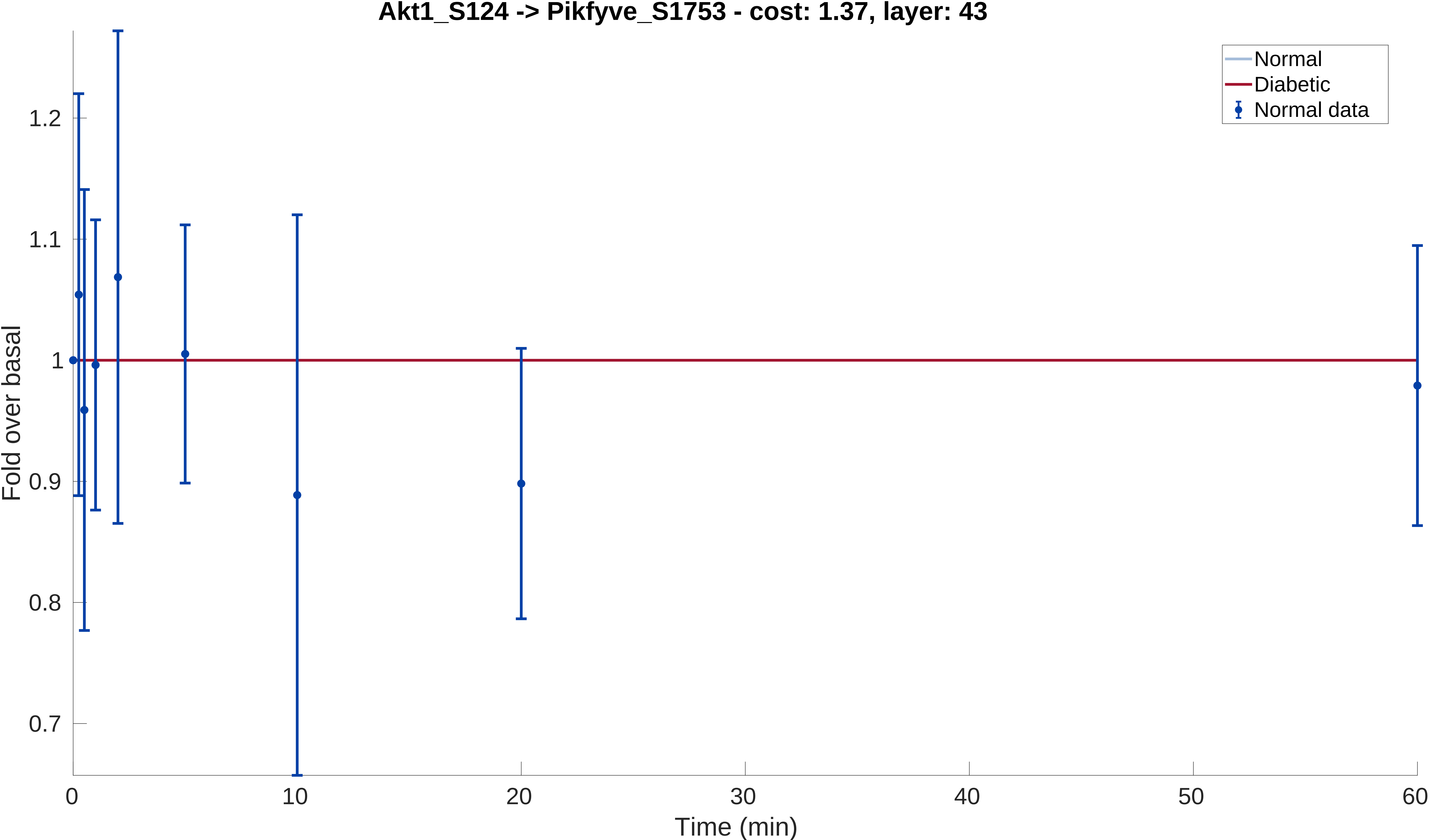

Akt1_S124 -> Pikfyve_S1753 - cost: 1.37, layer: 43
Normal
Diabetic
Normal data
Fold over basal
1.2
1.1
1
0.9
0.8
0.7
0
10
20
30
40
50
60
Time (min)

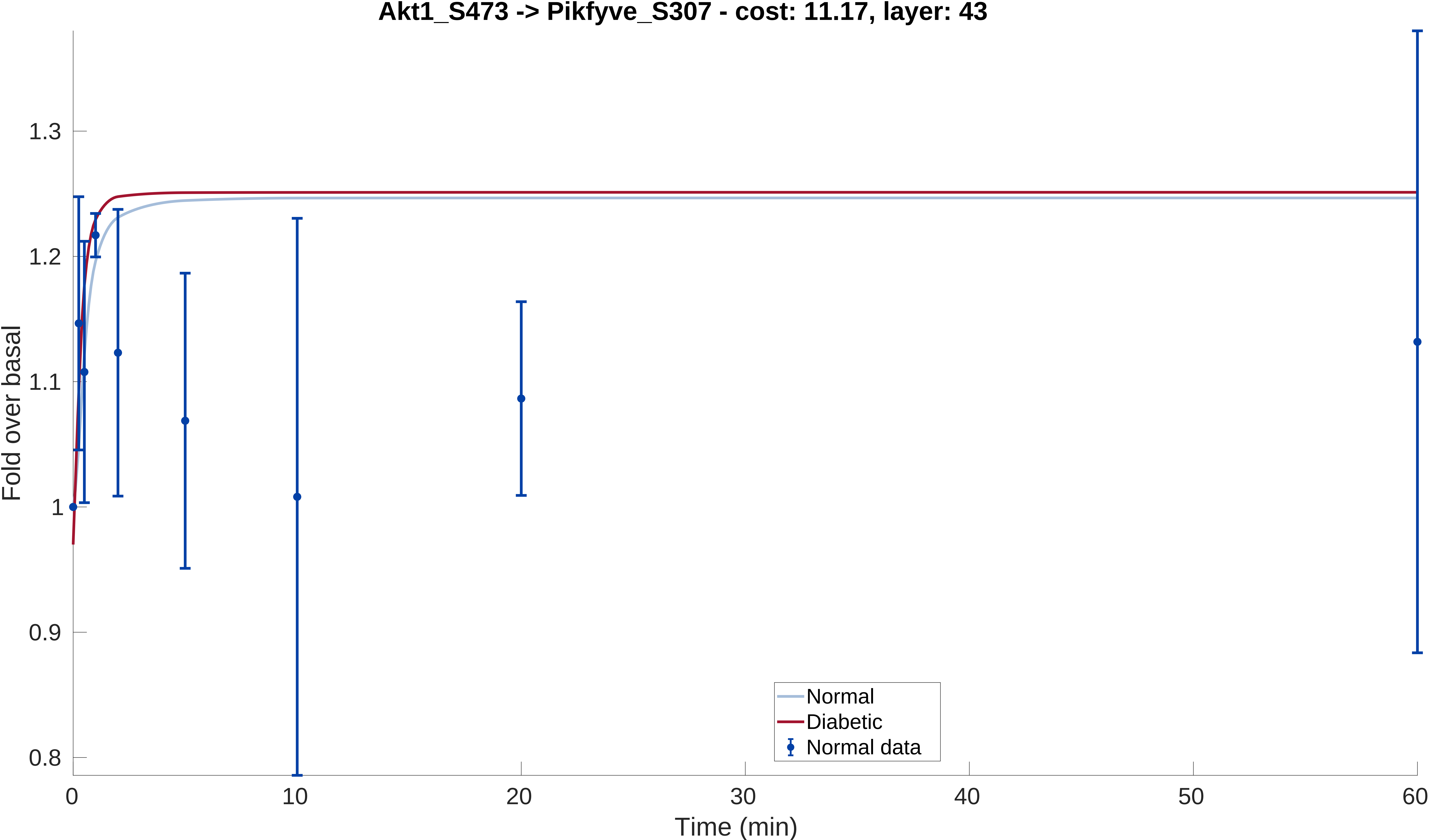

Akt1_S473 -> Pikfyve_S307 - cost: 11.17, layer: 43
Fold over basal
1.3
1.2
1.1
1
0.9
0.8
0
10
20
30
40
50
60
Time (min)
Normal
Diabetic
Normal data

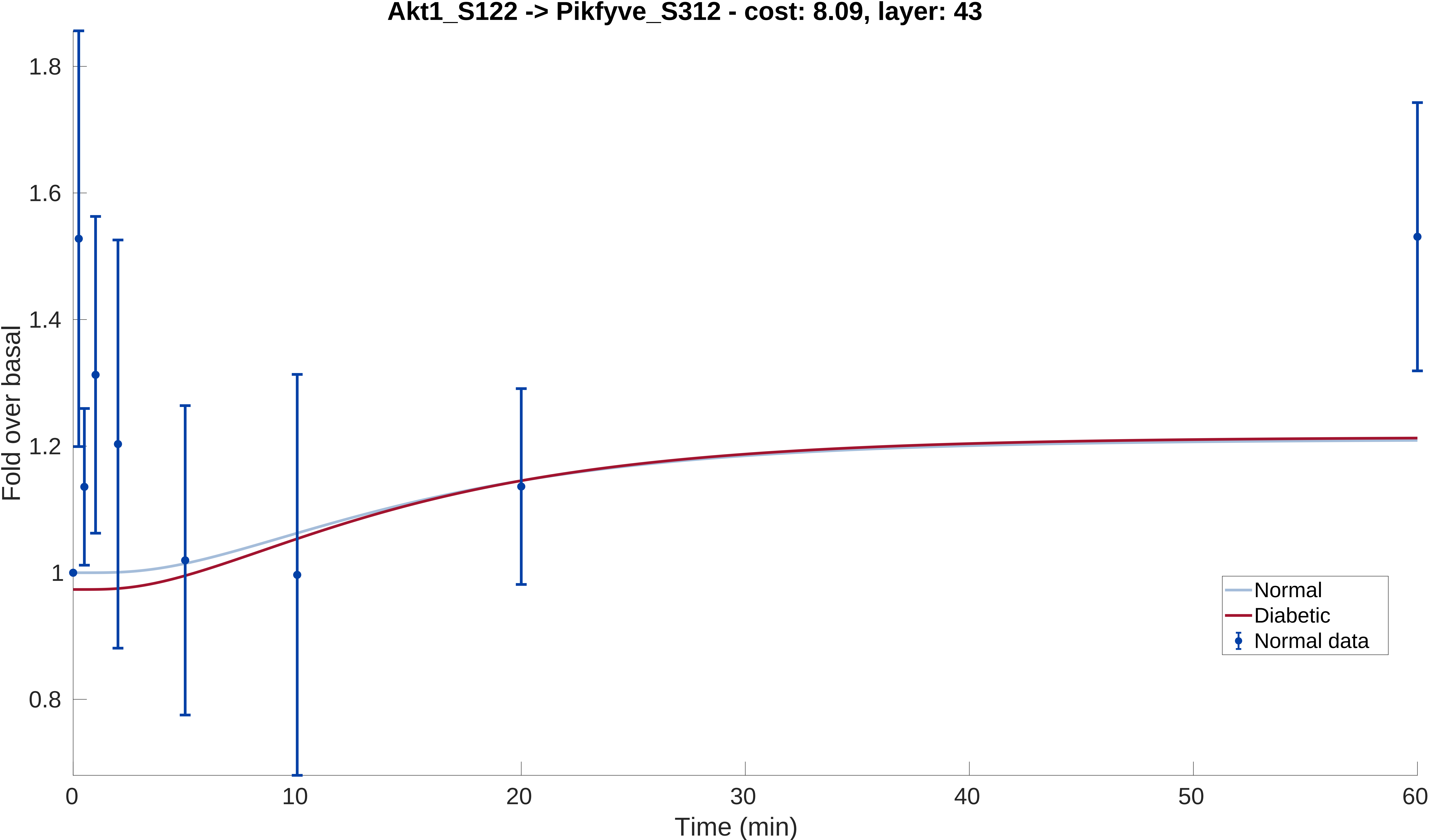

Akt1_S122 -> Pikfyve_S312 - cost: 8.09, layer: 43
Fold over basal
1.8
1.6
1.4
1.2
1
0.8
0
10
20
30
40
50
60
Time (min)
Normal
Diabetic
Normal data

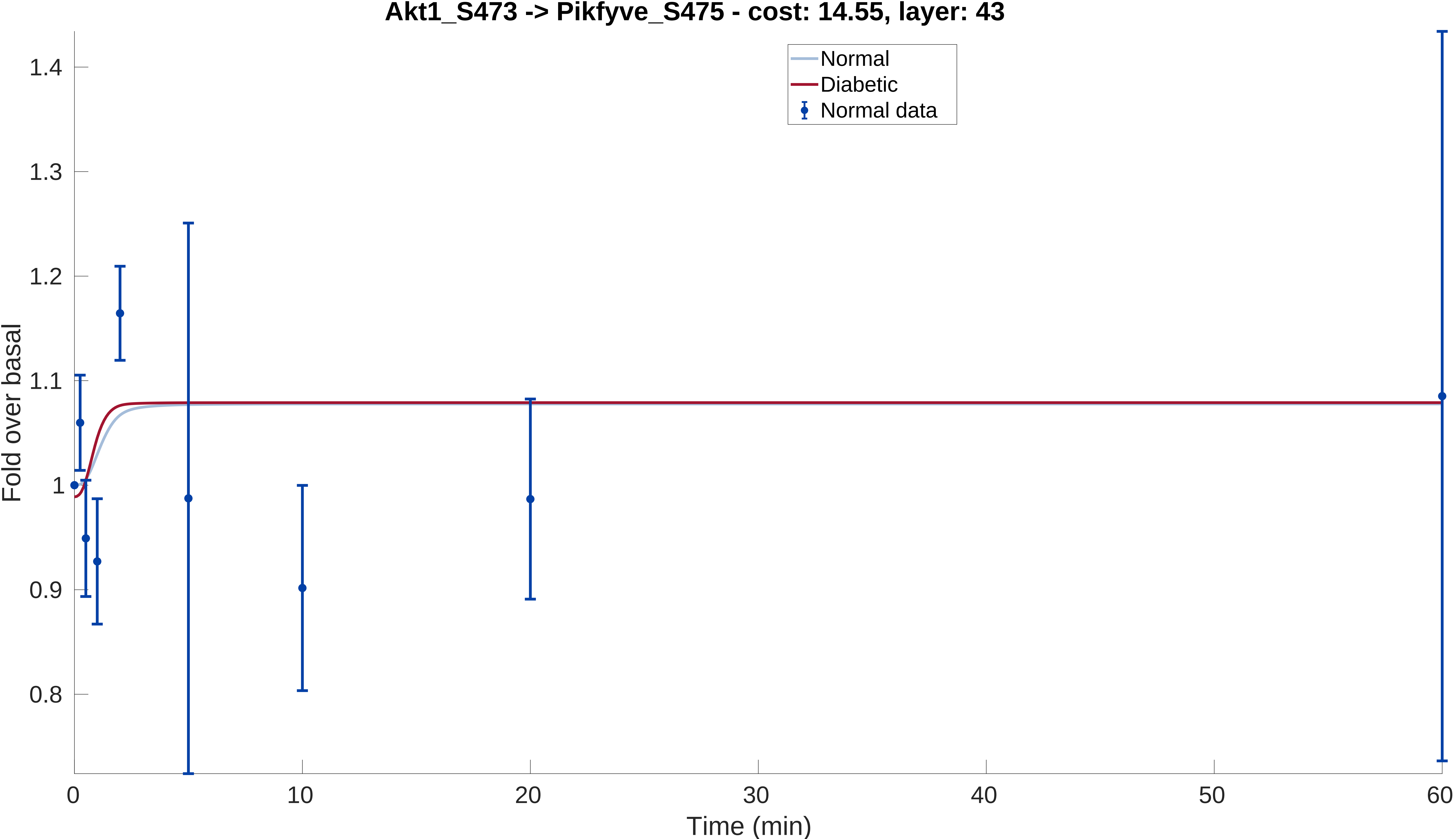
Akt1_S473 -> Pikfyve_S475 - cost: 14.55, layer: 43
Normal
Diabetic
Normal data
Fold over basal
1.4
1.3
1.2
1.1
1
0.9
0.8
0
10
20
30
40
50
60
Time (min)

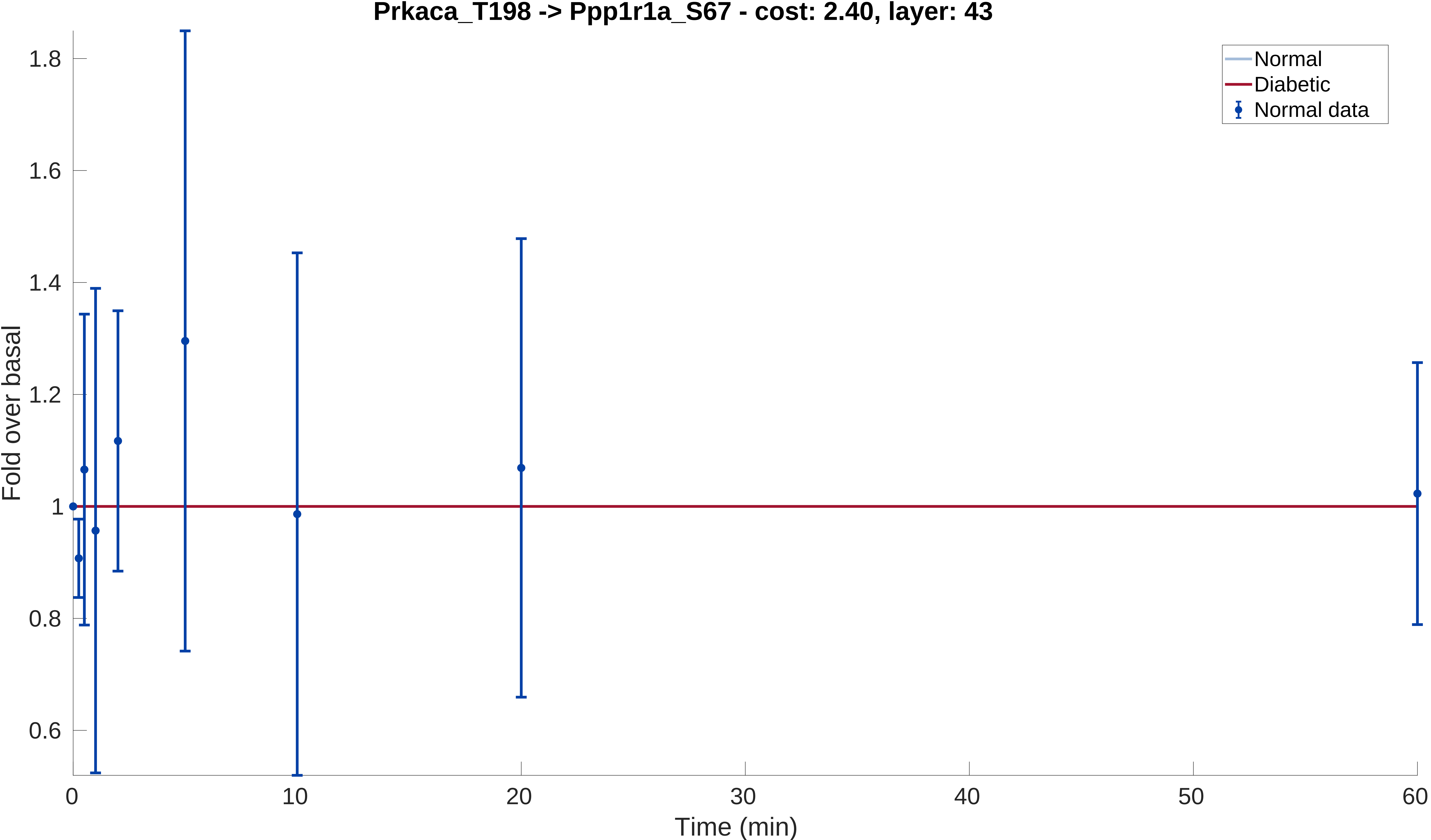

Prkaca_T198 -> Ppp1r1a_S67 - cost: 2.40, layer: 43
Normal
Diabetic
Normal data
Fold over basal
1.8
1.6
1.4
1.2
1
0.8
0.6
0
10
20
30
40
50
60
Time (min)

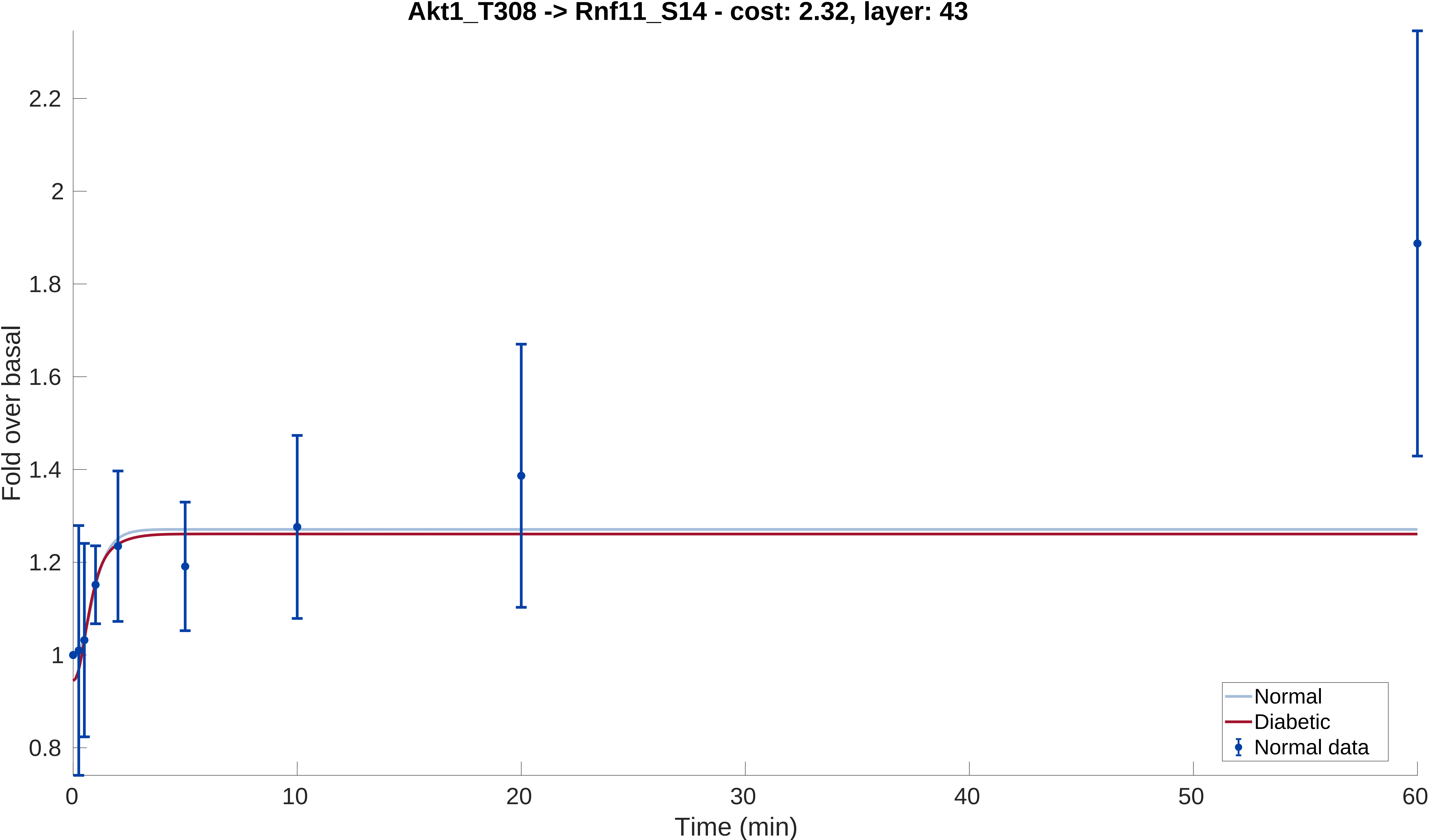

Akt1_T308 -> Rnf11_S14 - cost: 2.32, layer: 43
Fold over basal
Time (min)
0.8
1
1.2
1.4
1.6
1.8
2
2.2
0
10
20
30
40
50
60
Normal
Diabetic
Normal data

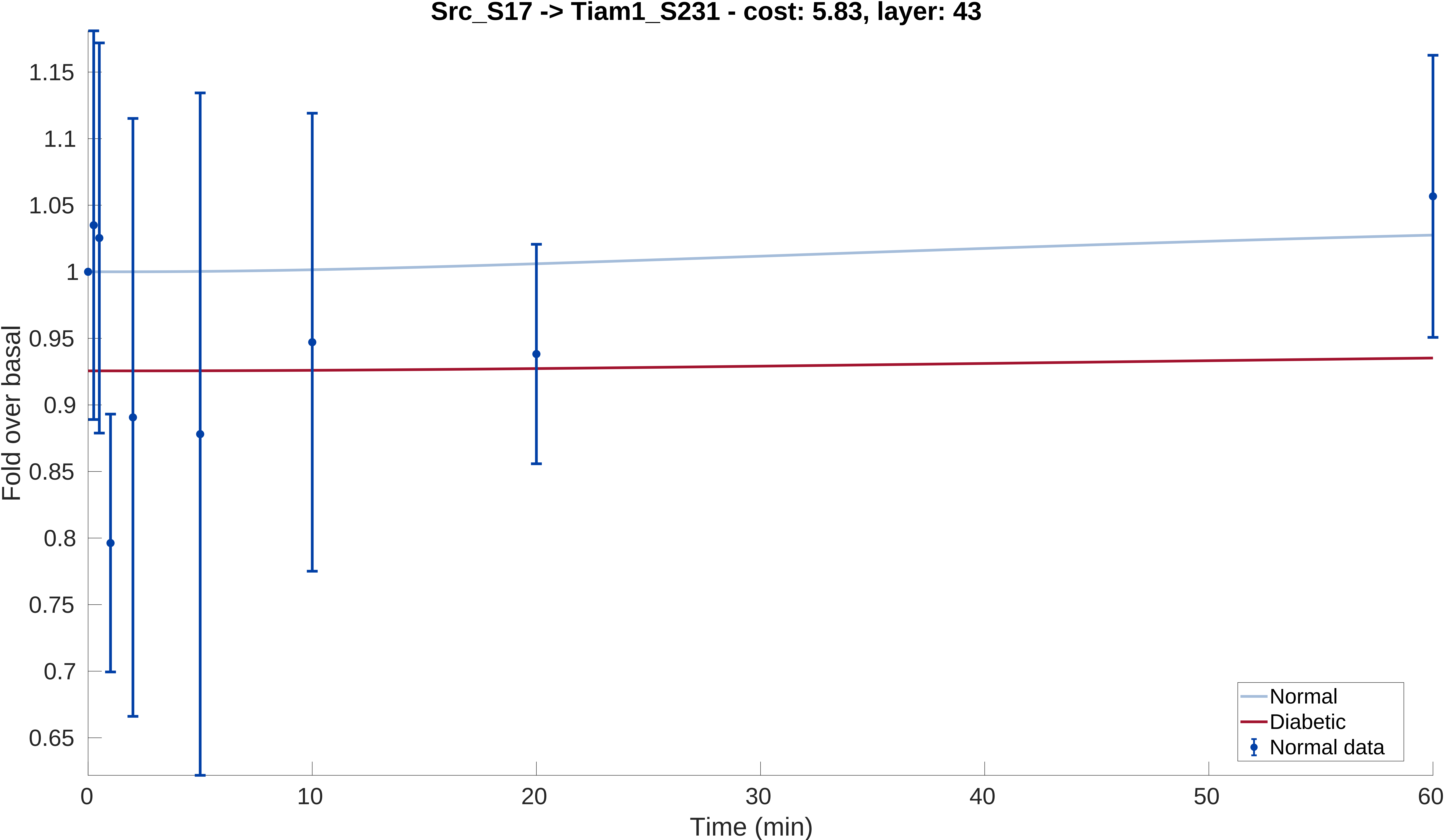
Src_S17 -> Tiam1_S231 - cost: 5.83, layer: 43
Fold over basal
Time (min)
1.15
1.1
1.05
1
0.95
0.9
0.85
0.8
0.75
0.7
0.65
0
10
20
30
40
50
60
Normal
Diabetic
Normal data

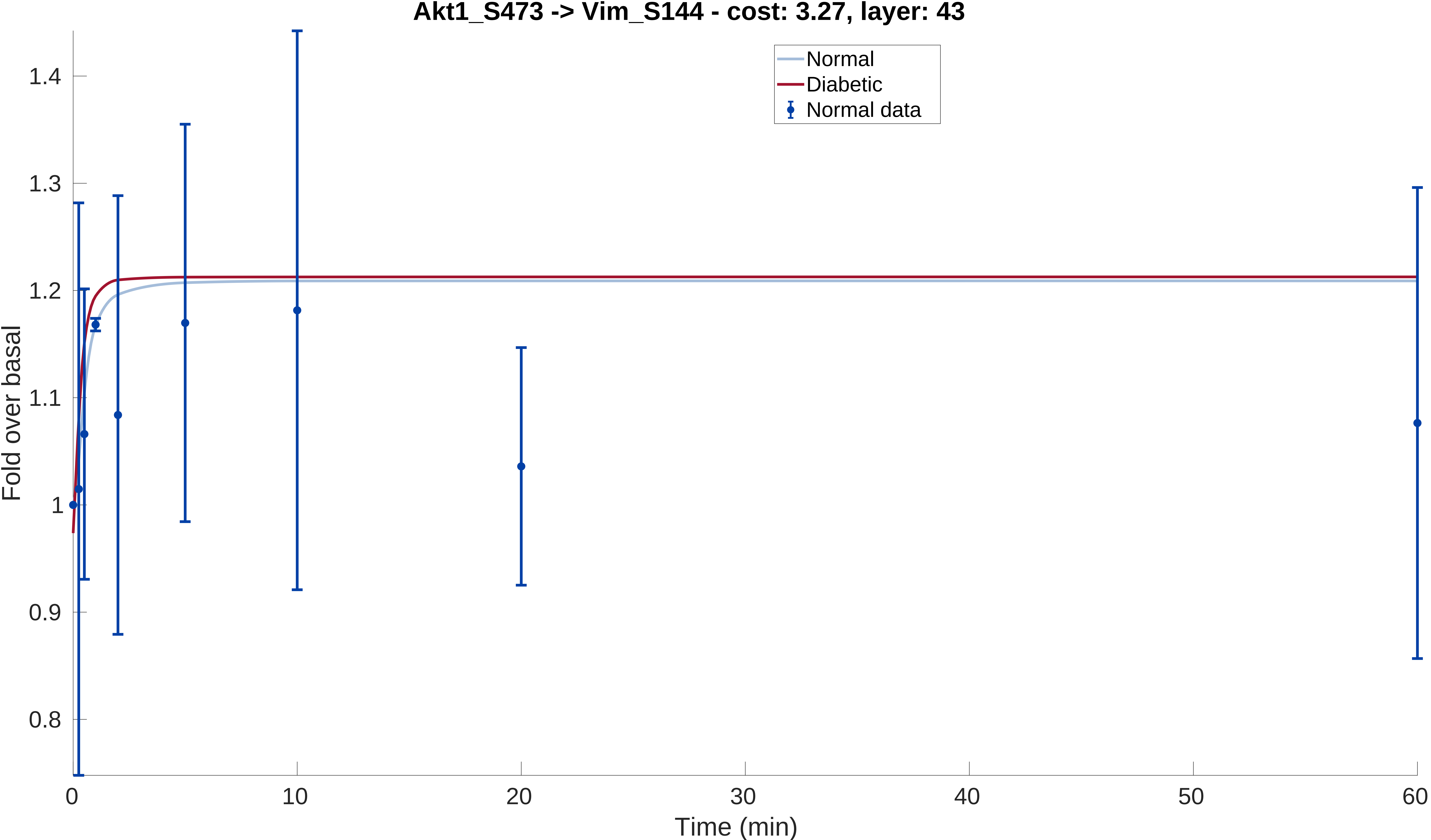

Akt1_S473 -> Vim_S144 - cost: 3.27, layer: 43
Normal
Diabetic
Normal data
Fold over basal
Time (min)
1.4
1.3
1.2
1.1
1
0.9
0.8
0
10
20
30
40
50
60

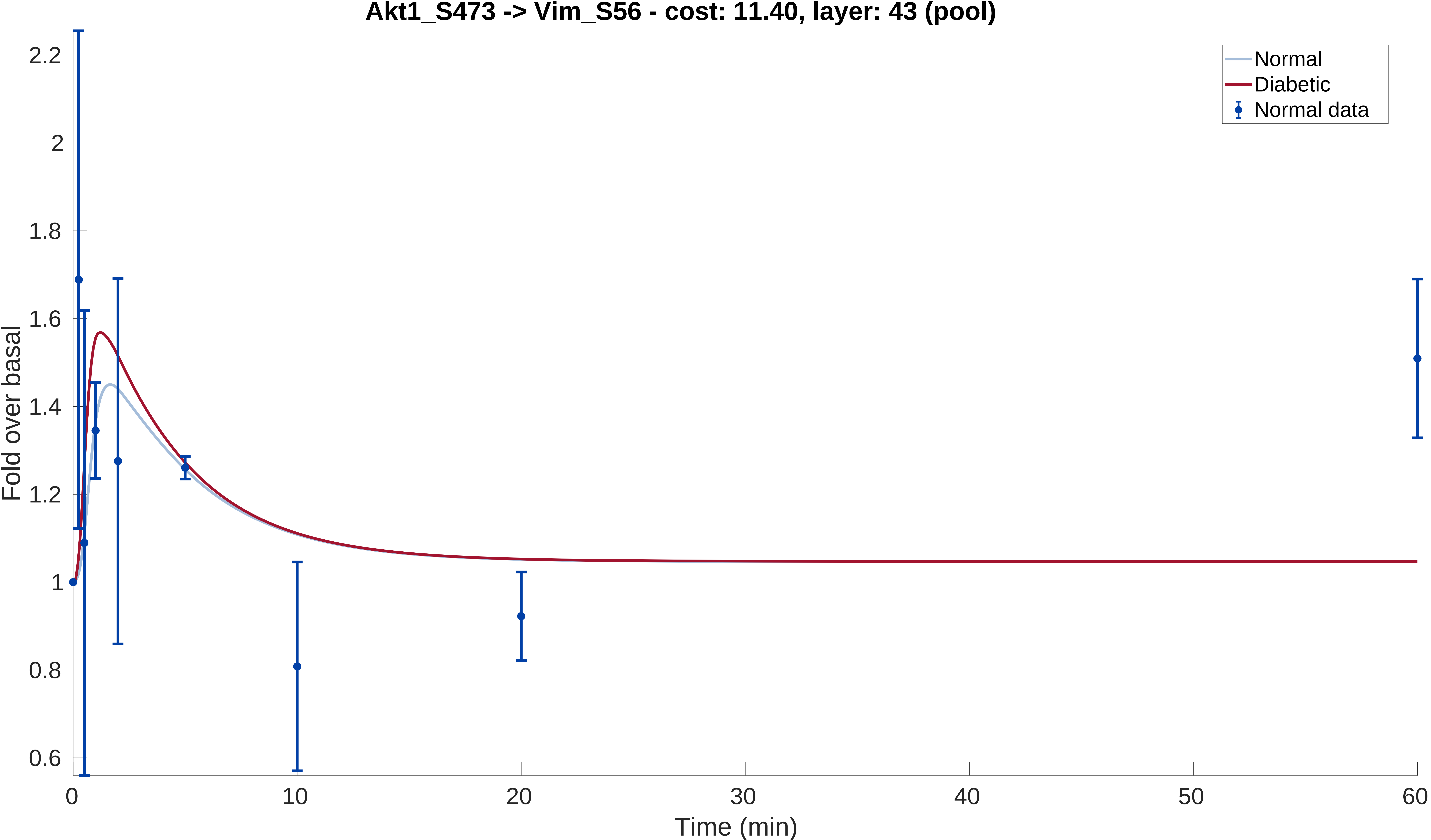
Akt1_S473 -> Vim_S56 - cost: 11.40, layer: 43 (pool)
Normal
Diabetic
Normal data
Fold over basal
Time (min)
0.6
0.8
1
1.2
1.4
1.6
1.8
2
2.2
0
10
20
30
40
50
60

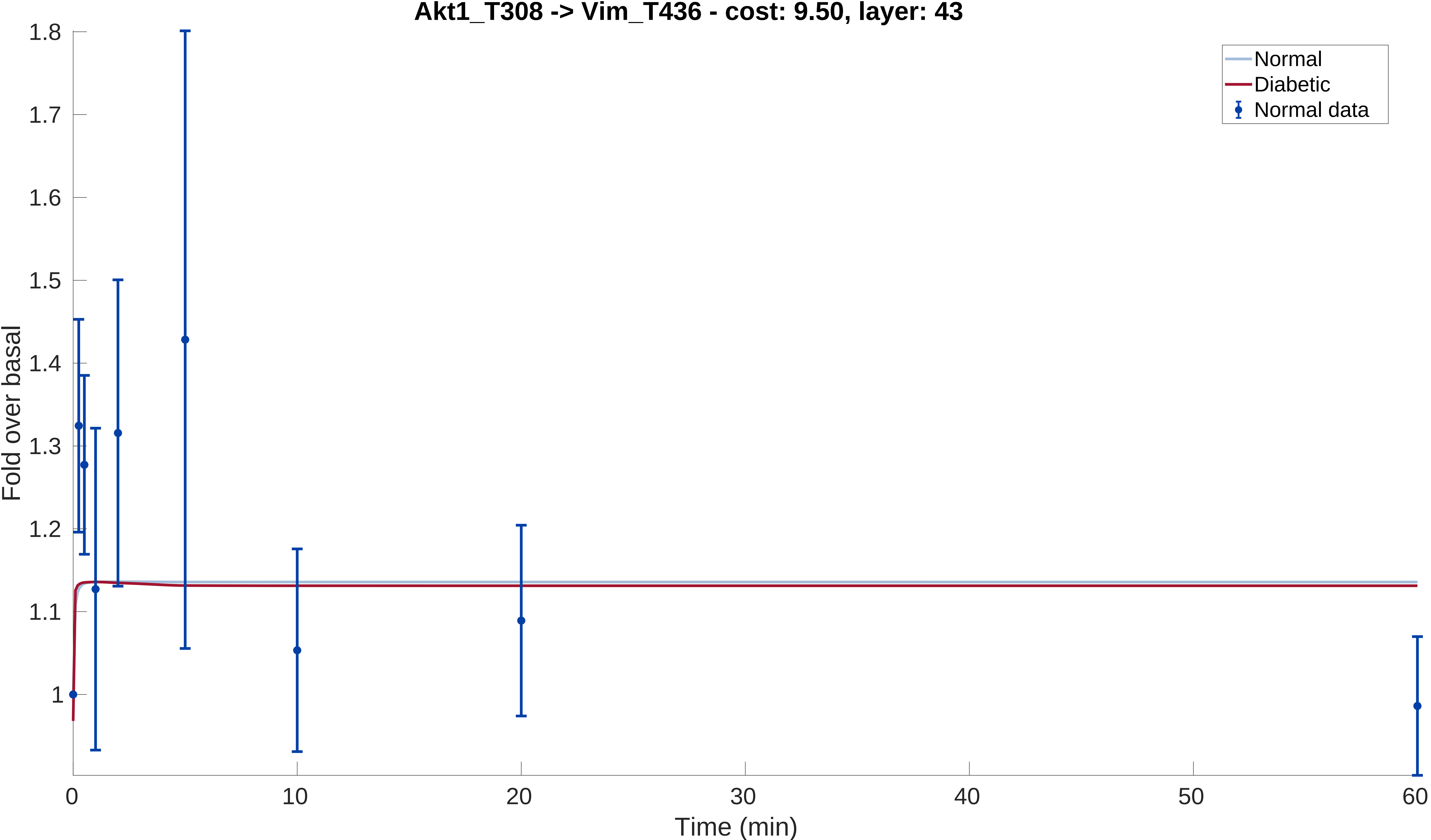

Akt1_T308 -> Vim_T436 - cost: 9.50, layer: 43
Normal
Diabetic
Normal data
Fold over basal
Time (min)
1.8
1.7
1.6
1.5
1.4
1.3
1.2
1.1
1
0
10
20
30
40
50
60

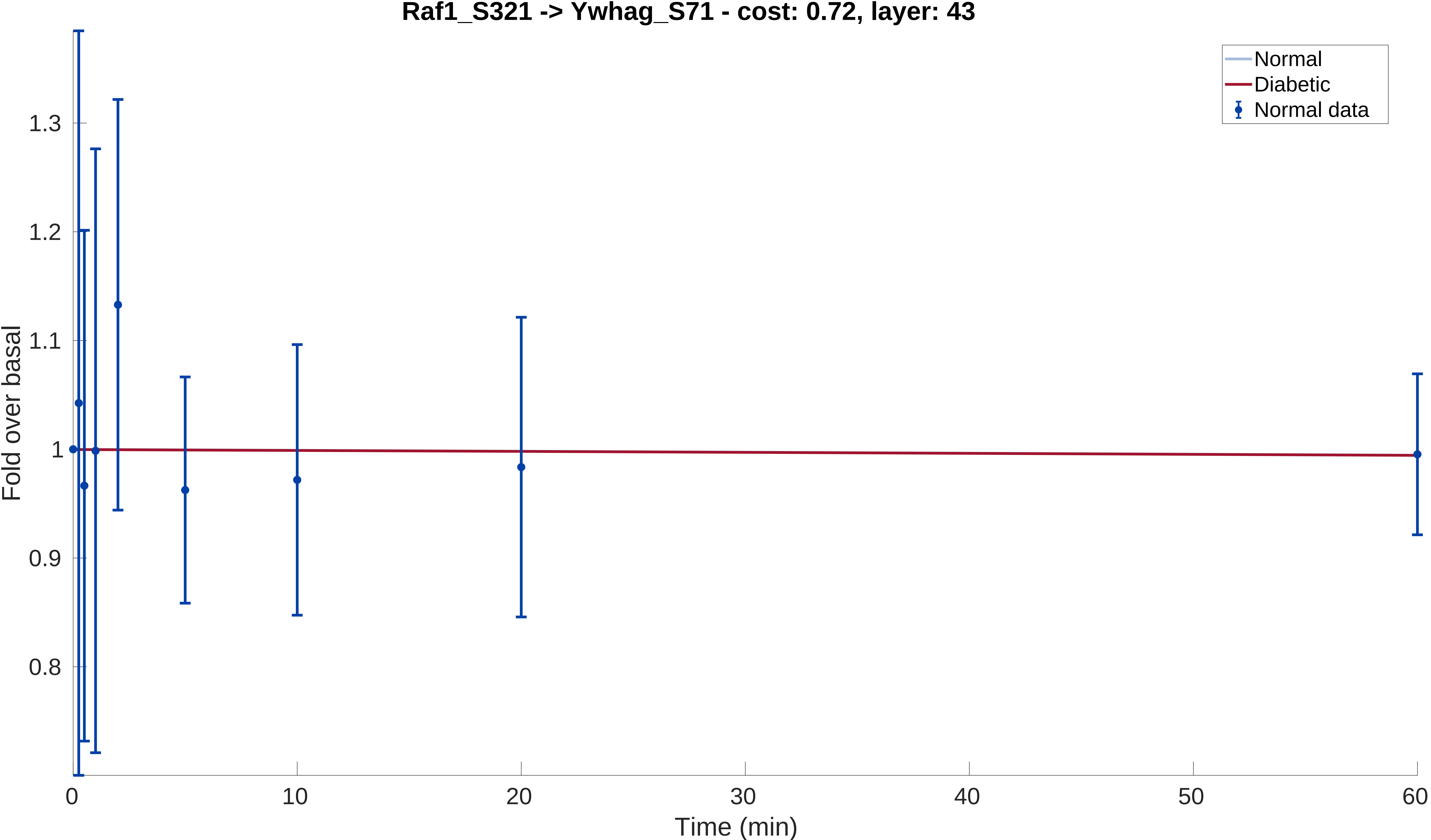
Raf1_S321 -> Ywhag_S71 - cost: 0.72, layer: 43
Normal
Diabetic
Normal data
Fold over basal
1.3
1.2
1.1
1
0.9
0.8
0
10
20
30
40
50
60
Time (min)

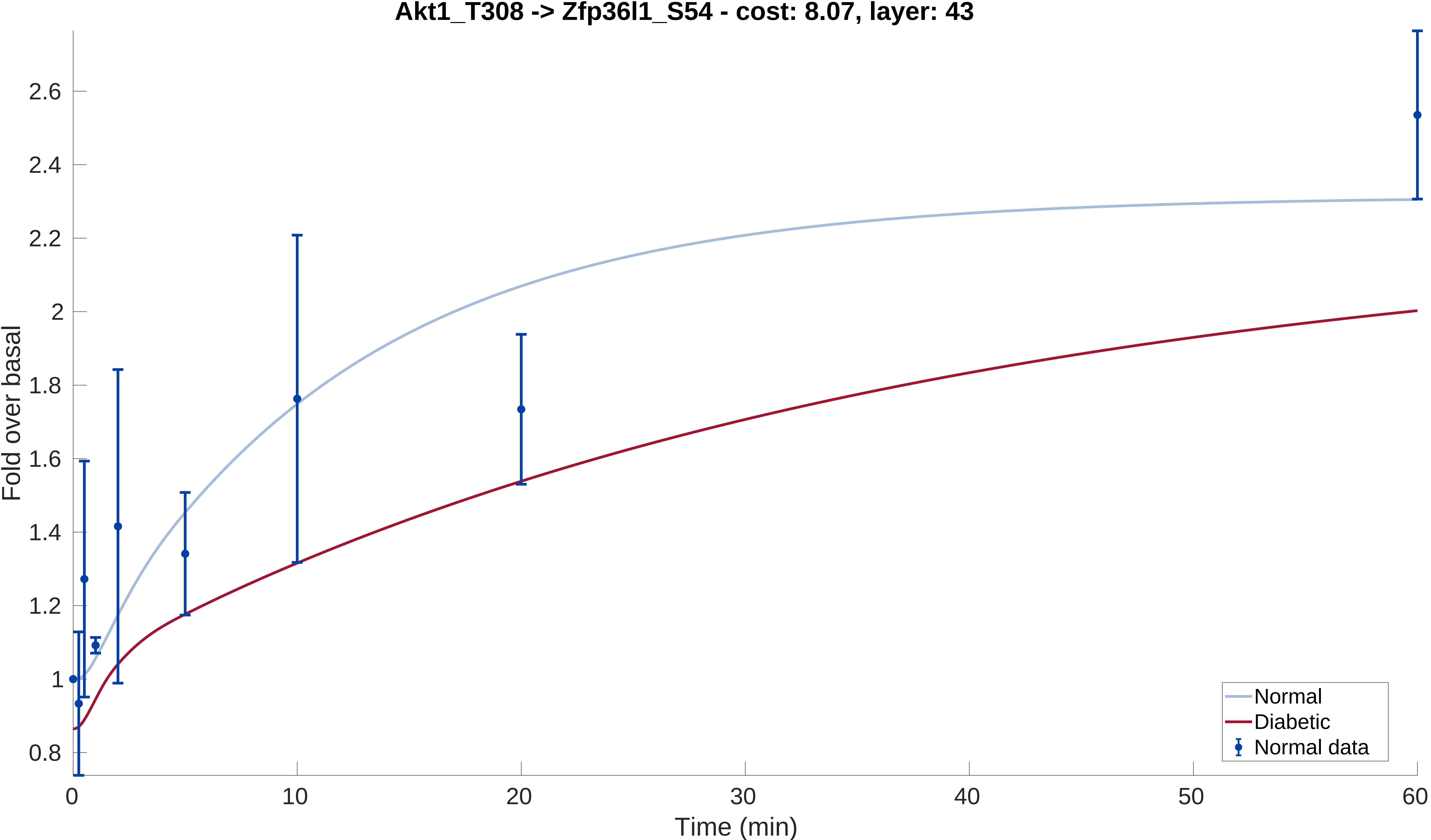

Akt1_T308 -> Zfp36l1_S54 - cost: 8.07, layer: 43
Fold over basal
0.8
1
1.2
1.4
1.6
1.8
2
2.2
2.4
2.6
0
10
20
30
40
50
60
Time (min)
Normal
Diabetic
Normal data

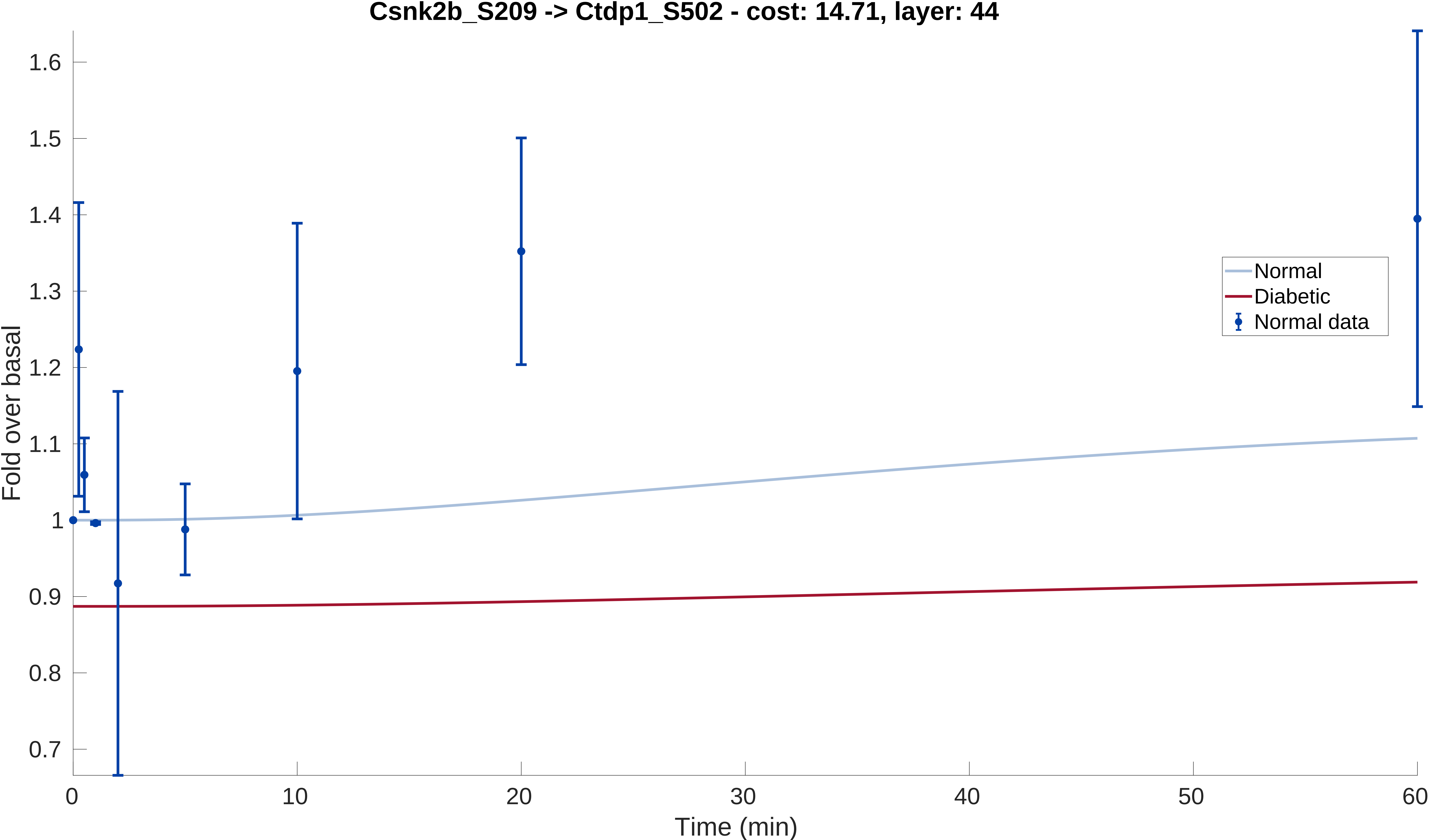
Csnk2b_S209 -> Ctdp1_S502 - cost: 14.71, layer: 44
Fold over basal
Time (min)
Normal
Diabetic
Normal data
1.6
1.5
1.4
1.3
1.2
1.1
1
0.9
0.8
0.7
0
10
20
30
40
50
60

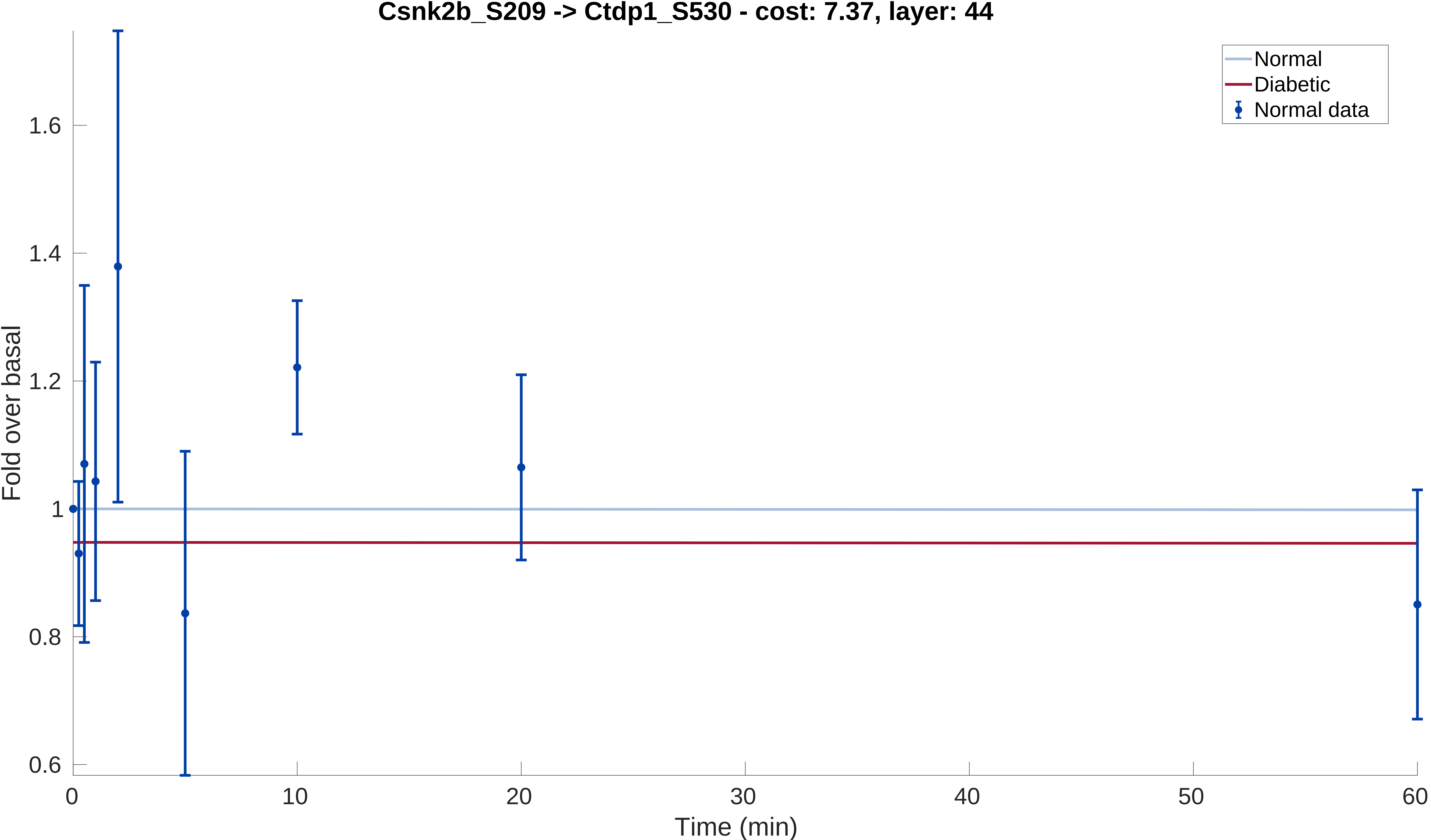

Csnk2b_S209 -> Ctdp1_S530 - cost: 7.37, layer: 44
Normal
Diabetic
Normal data
Fold over basal
1.6
1.4
1.2
1
0.8
0.6
0
10
20
30
40
50
60
Time (min)

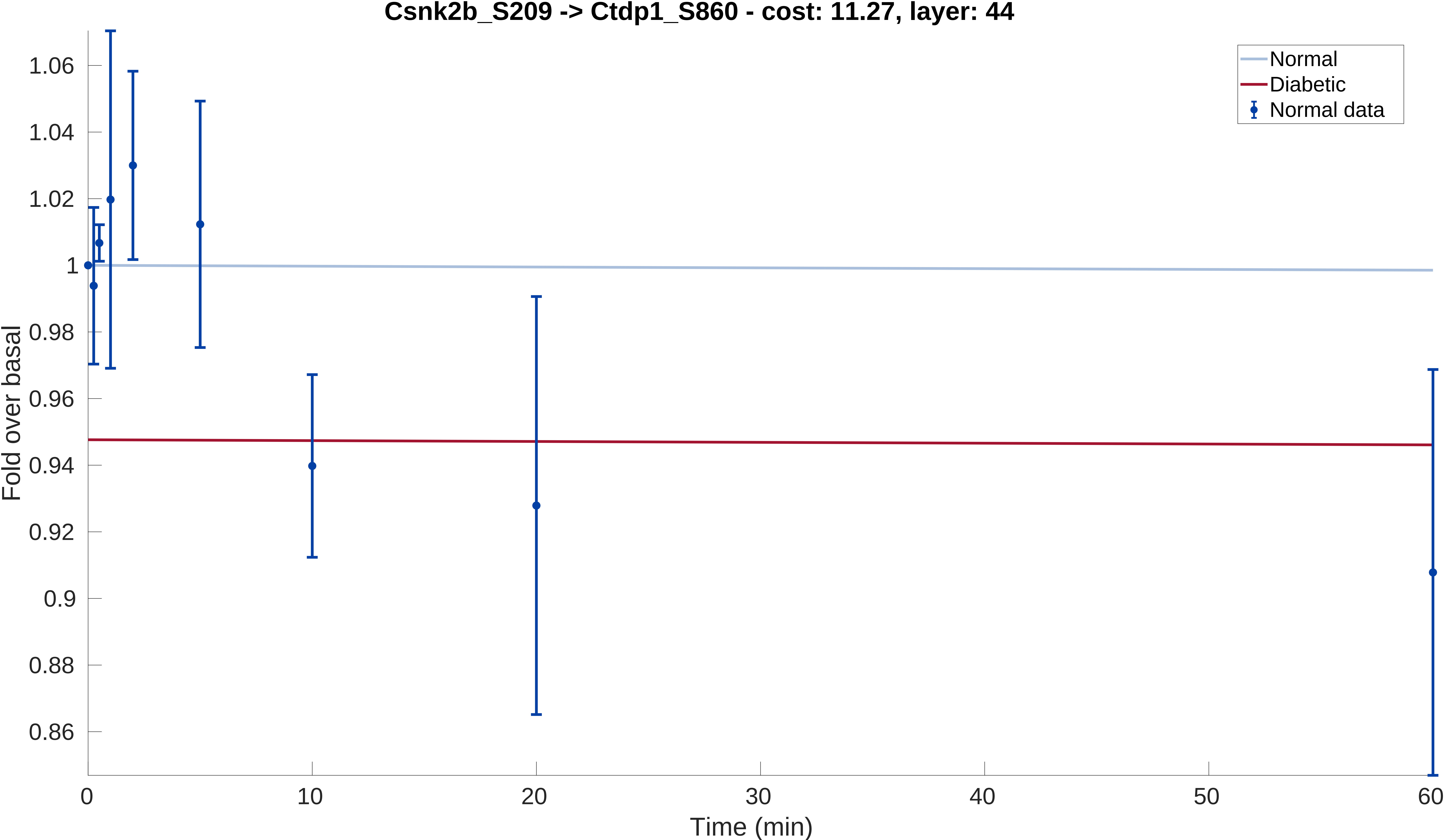

Csnk2b_S209 -> Ctdp1_S860 - cost: 11.27, layer: 44
Normal
Diabetic
Normal data
Fold over basal
1.06
1.04
1.02
1
0.98
0.96
0.94
0.92
0.9
0.88
0.86
0
10
20
30
40
50
60
Time (min)

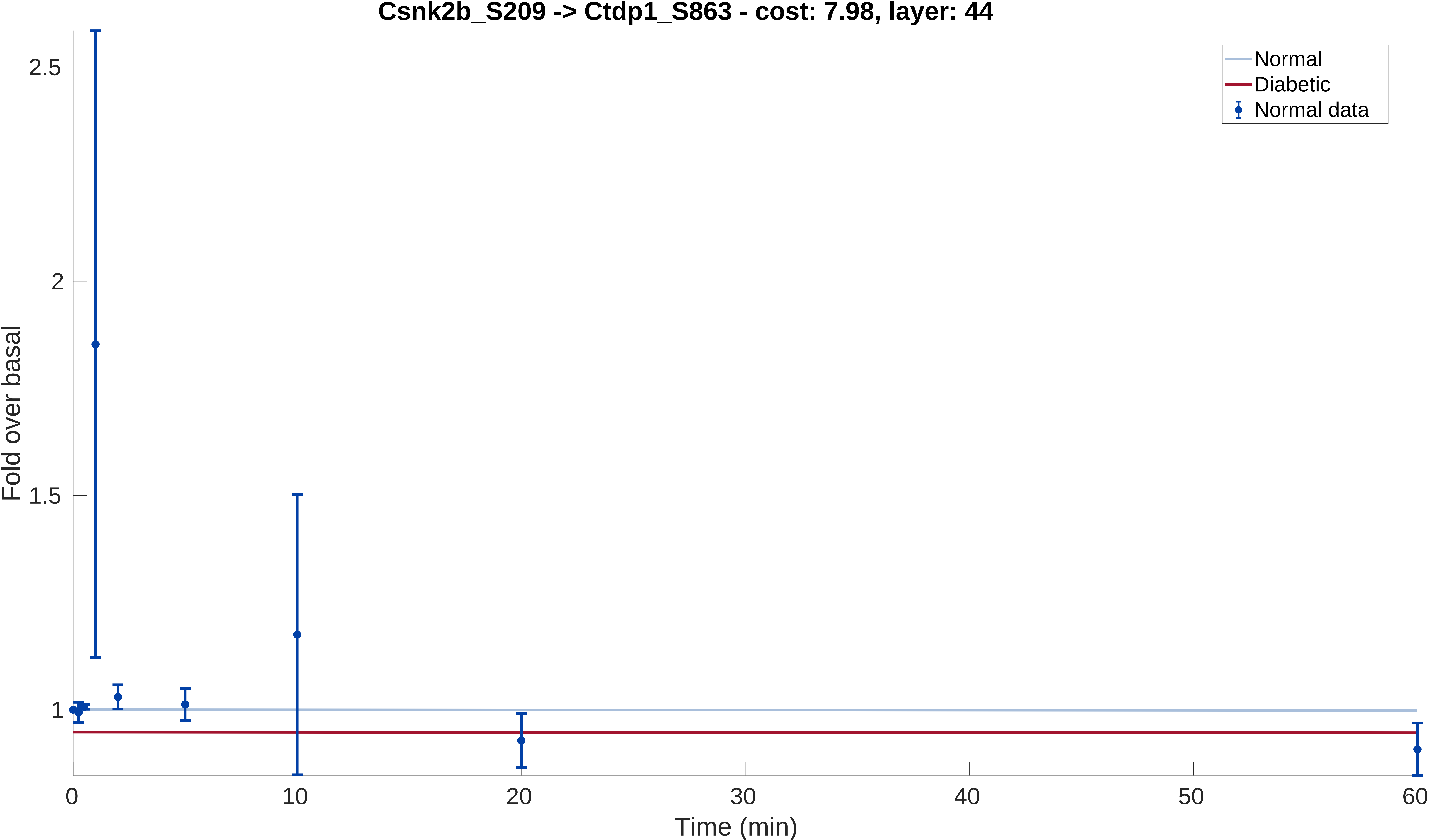
Csnk2b_S209 -> Ctdp1_S863 - cost: 7.98, layer: 44
Normal
Diabetic
Normal data
Fold over basal
2.5
2
1.5
1
0
10
20
30
40
50
60
Time (min)

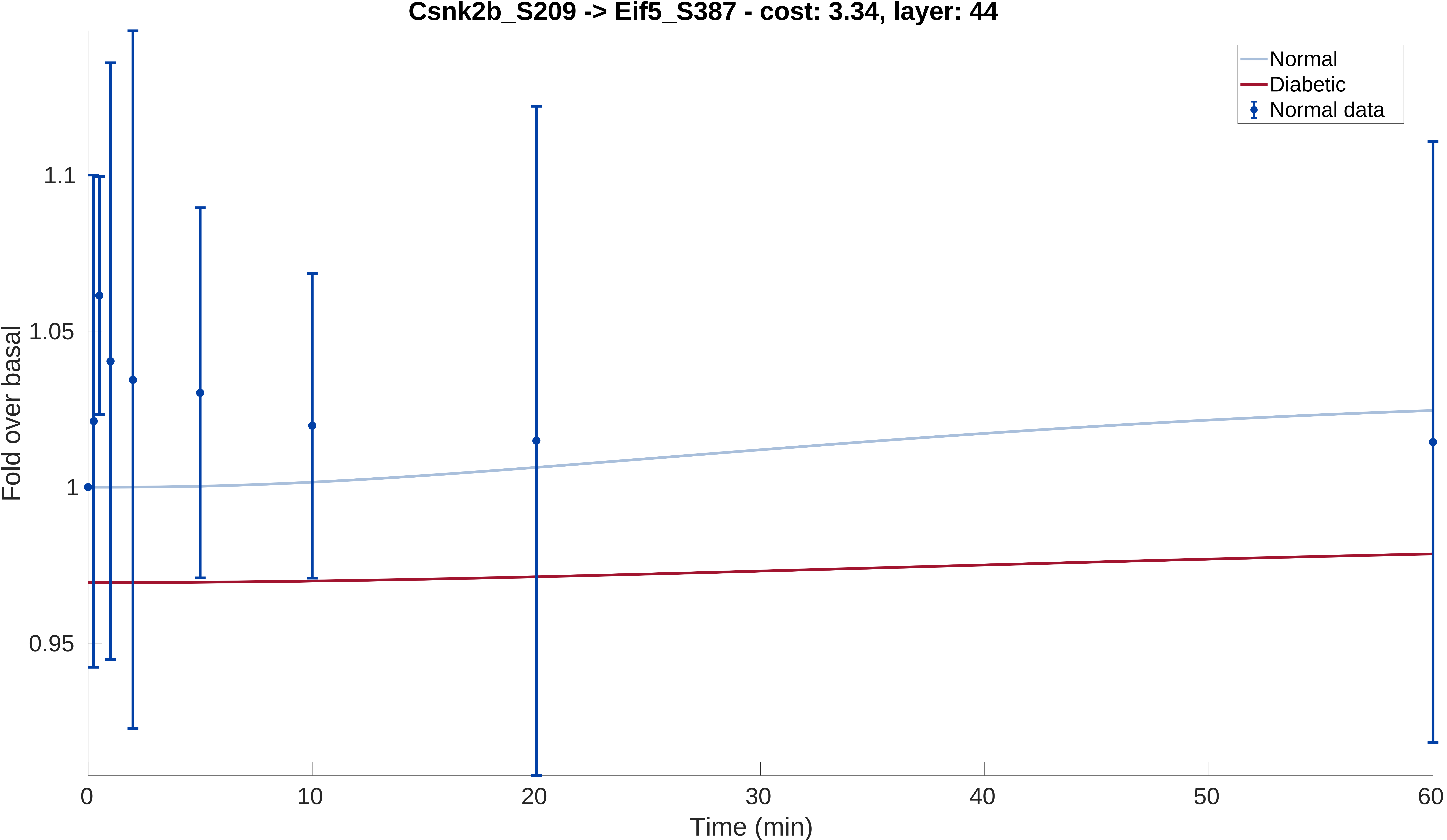
Csnk2b_S209 -> Eif5_S387 - cost: 3.34, layer: 44
Normal
Diabetic
Normal data
Fold over basal
1.1
1.05
1
0.95
0
10
20
30
40
50
60
Time (min)

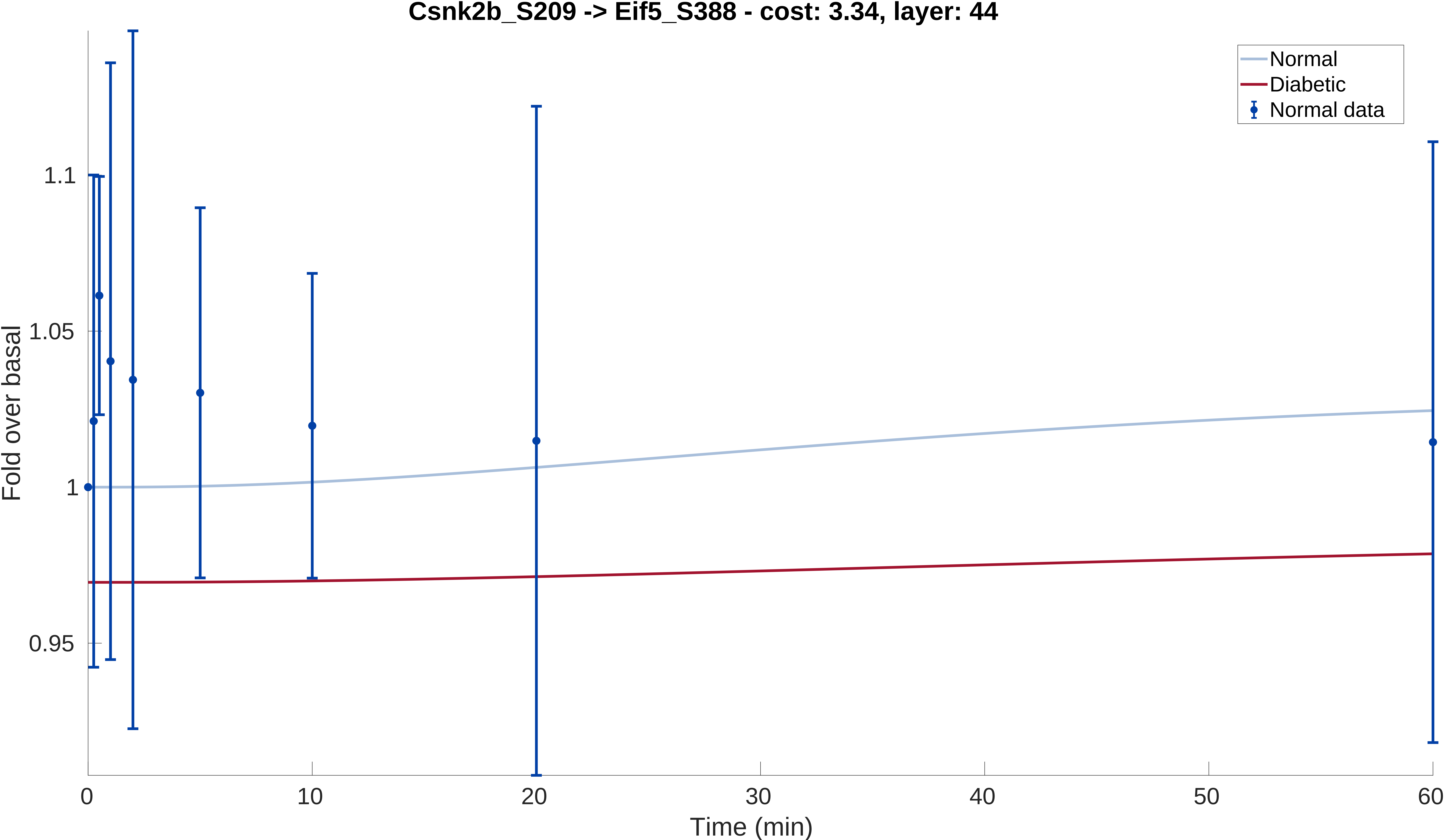
Csnk2b_S209 -> Eif5_S388 - cost: 3.34, layer: 44
Normal
Diabetic
Normal data
Fold over basal
1.1
1.05
1
0.95
0
10
20
30
40
50
60
Time (min)

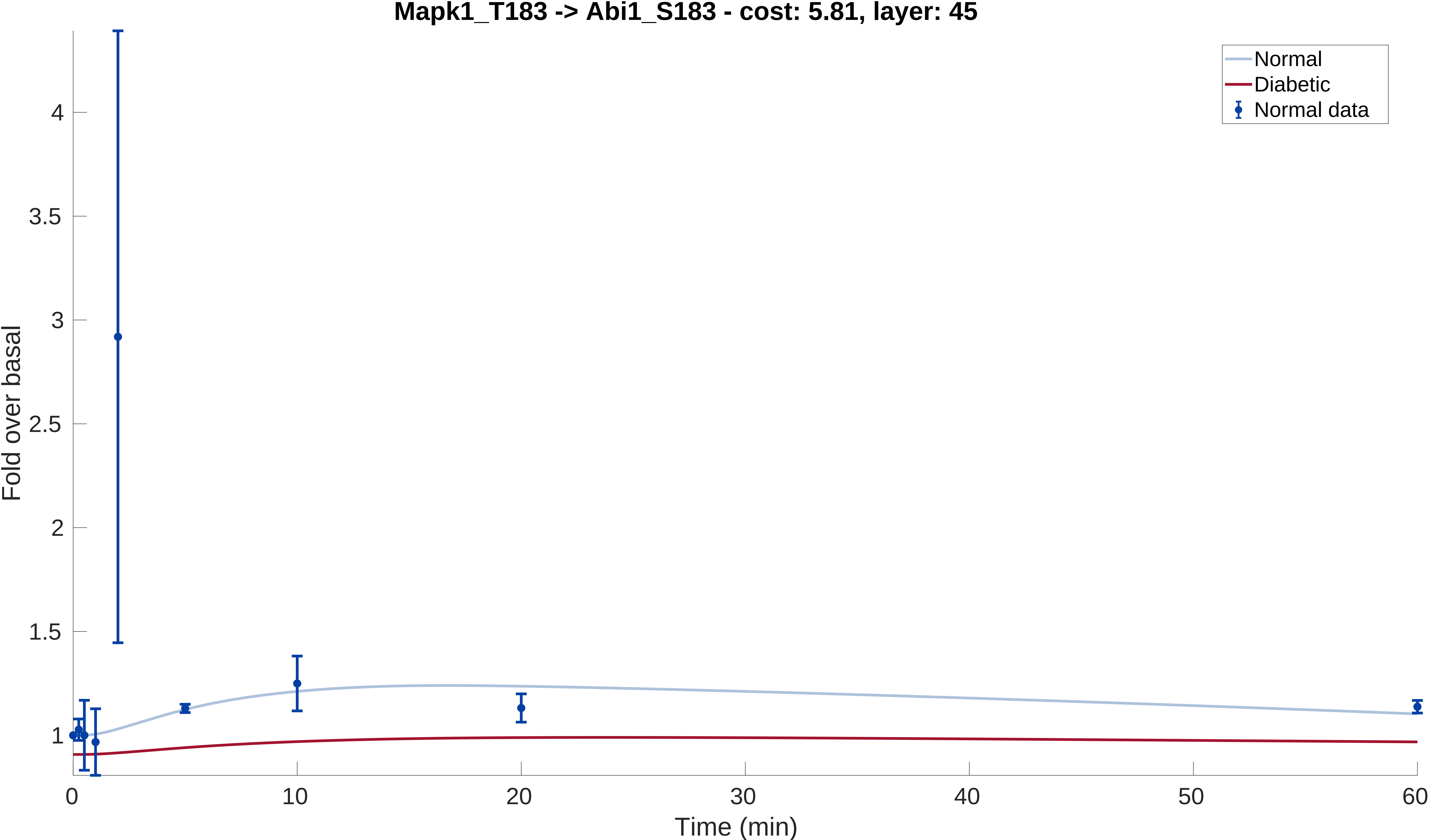
Mapk1_T183 -> Abi1_S183 - cost: 5.81, layer: 45
Normal
Diabetic
Normal data
Fold over basal
4
3.5
3
2.5
2
1.5
1
0
10
20
30
40
50
60
Time (min)

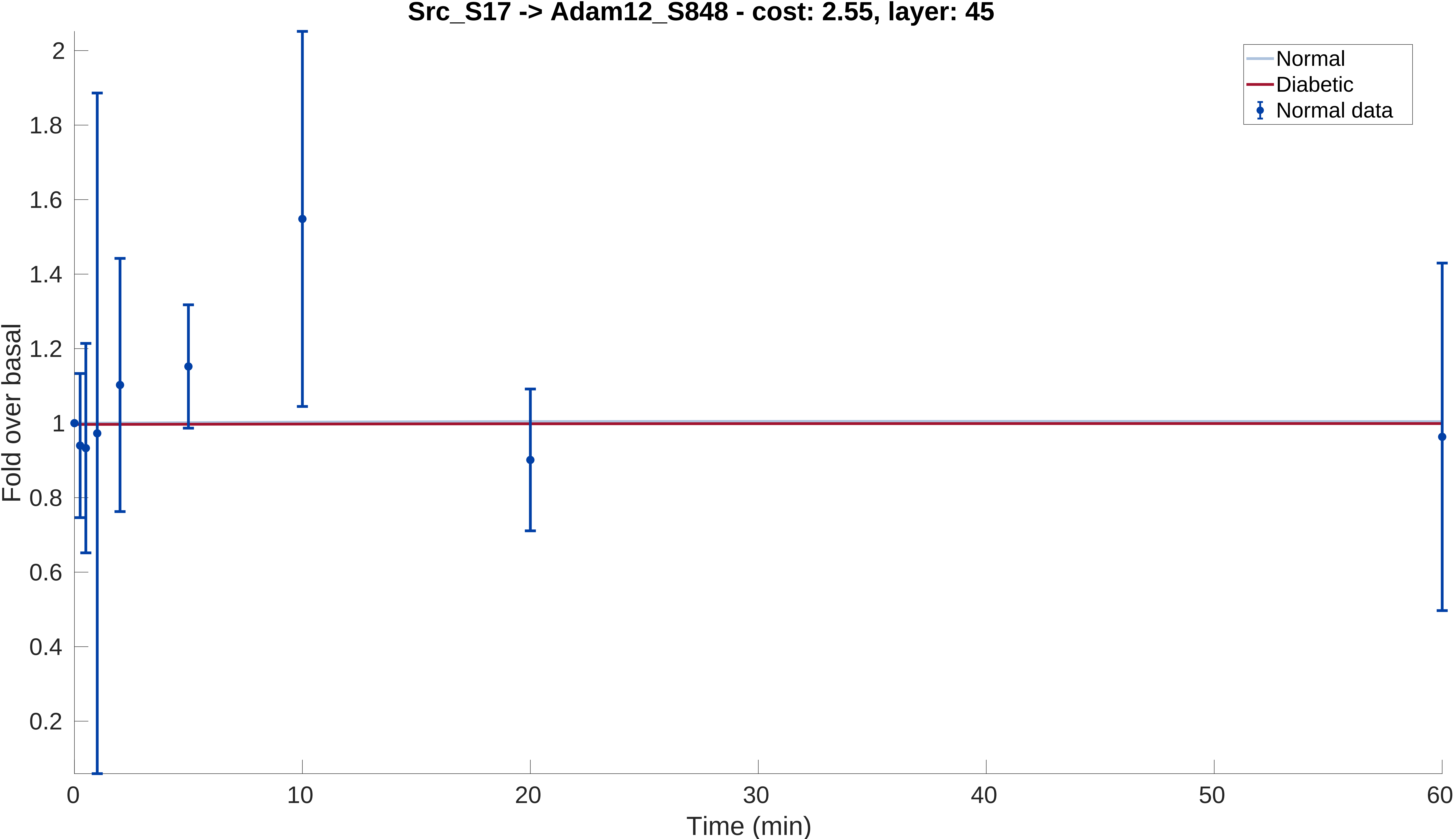

Src_S17 -> Adam12_S848 - cost: 2.55, layer: 45
Normal
Diabetic
Normal data
Fold over basal
2
1.8
1.6
1.4
1.2
1
0.8
0.6
0.4
0.2
0
10
20
30
40
50
60
Time (min)

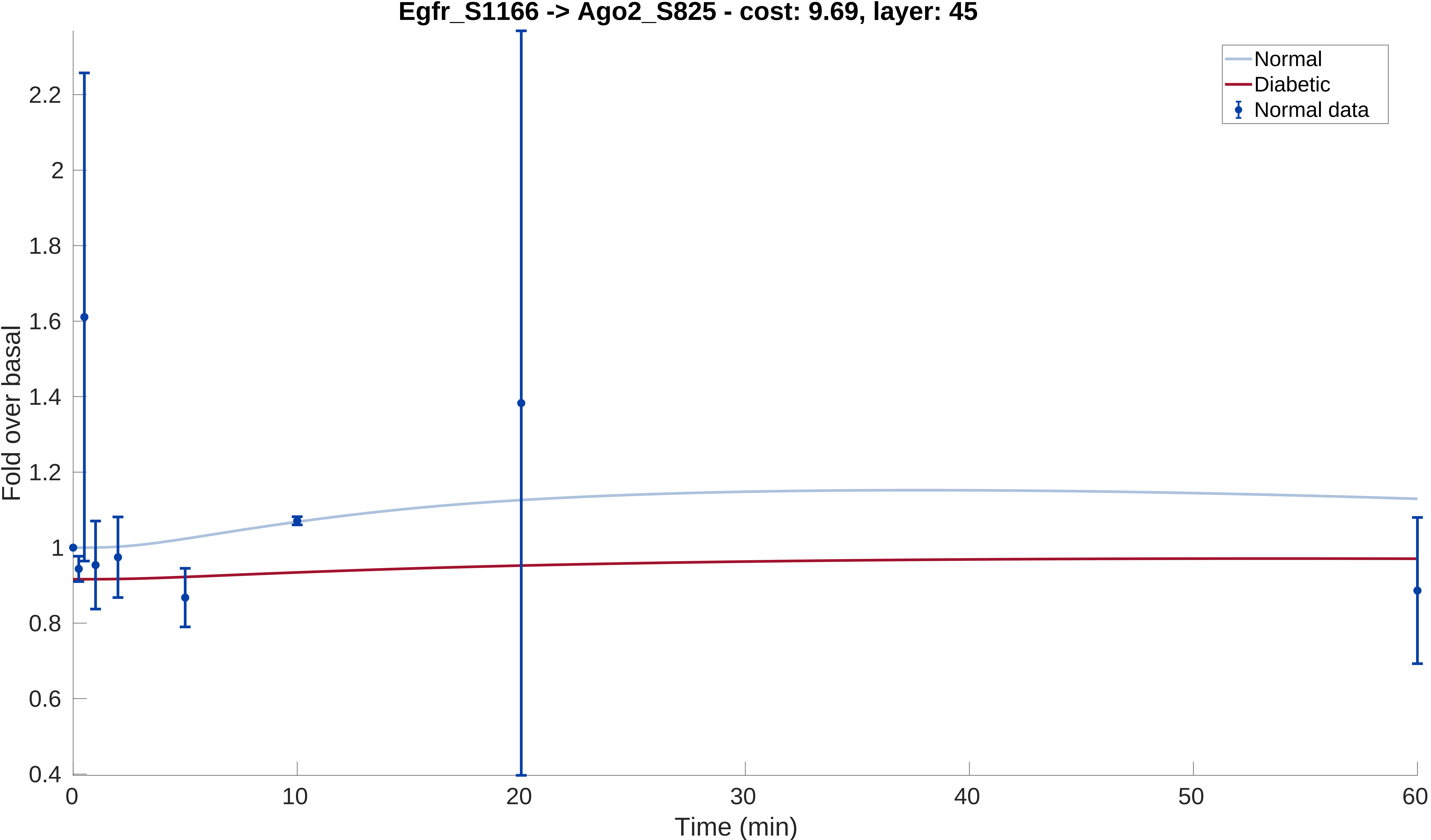

Egfr_S1166 -> Ago2_S825 - cost: 9.69, layer: 45
Normal
Diabetic
Normal data
Fold over basal
Time (min)
0.4
0.6
0.8
1
1.2
1.4
1.6
1.8
2
2.2
0
10
20
30
40
50
60

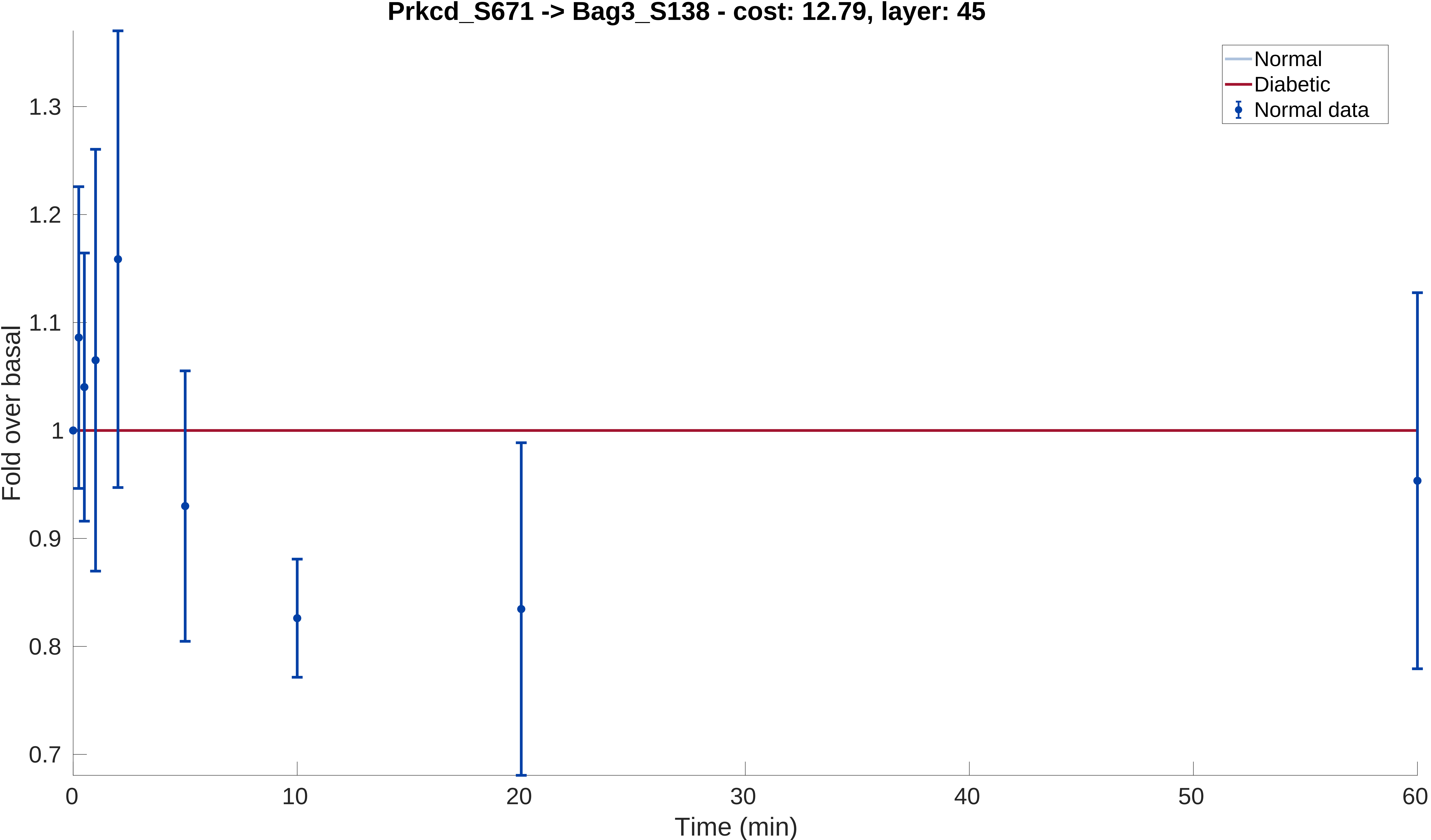
Prkcd_S671 -> Bag3_S138 - cost: 12.79, layer: 45
Normal
Diabetic
Normal data
Fold over basal
1.3
1.2
1.1
1
0.9
0.8
0.7
0
10
20
30
40
50
60
Time (min)

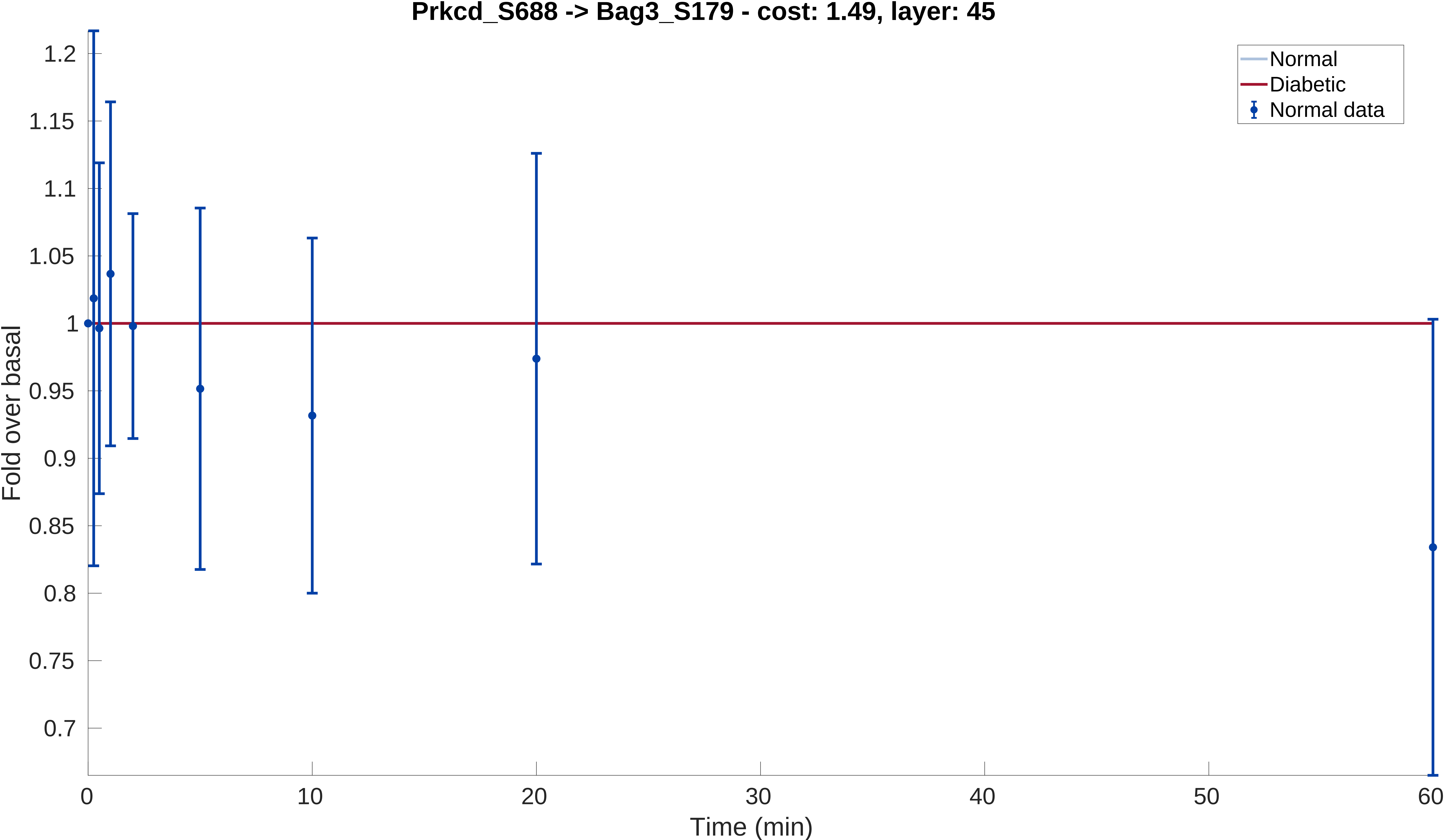

Prkcd_S688 -> Bag3_S179 - cost: 1.49, layer: 45
Normal
Diabetic
Normal data
Fold over basal
1.2
1.15
1.1
1.05
1
0.95
0.9
0.85
0.8
0.75
0.7
0
10
20
30
40
50
60
Time (min)

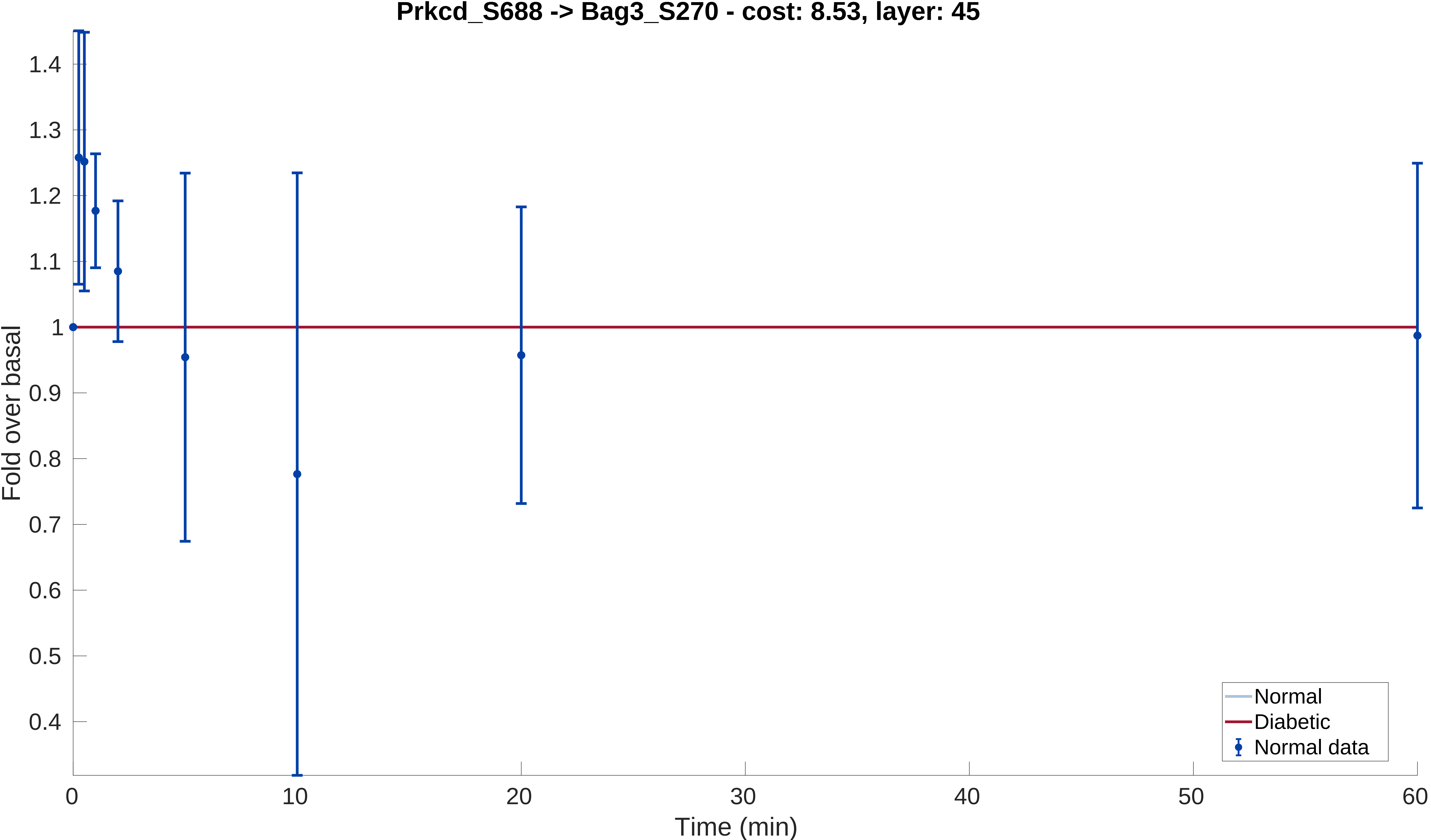

Prkcd_S688 -> Bag3_S270 - cost: 8.53, layer: 45
Fold over basal
1.4
1.3
1.2
1.1
1
0.9
0.8
0.7
0.6
0.5
0.4
Normal
Diabetic
Normal data
0
10
20
30
40
50
60
Time (min)

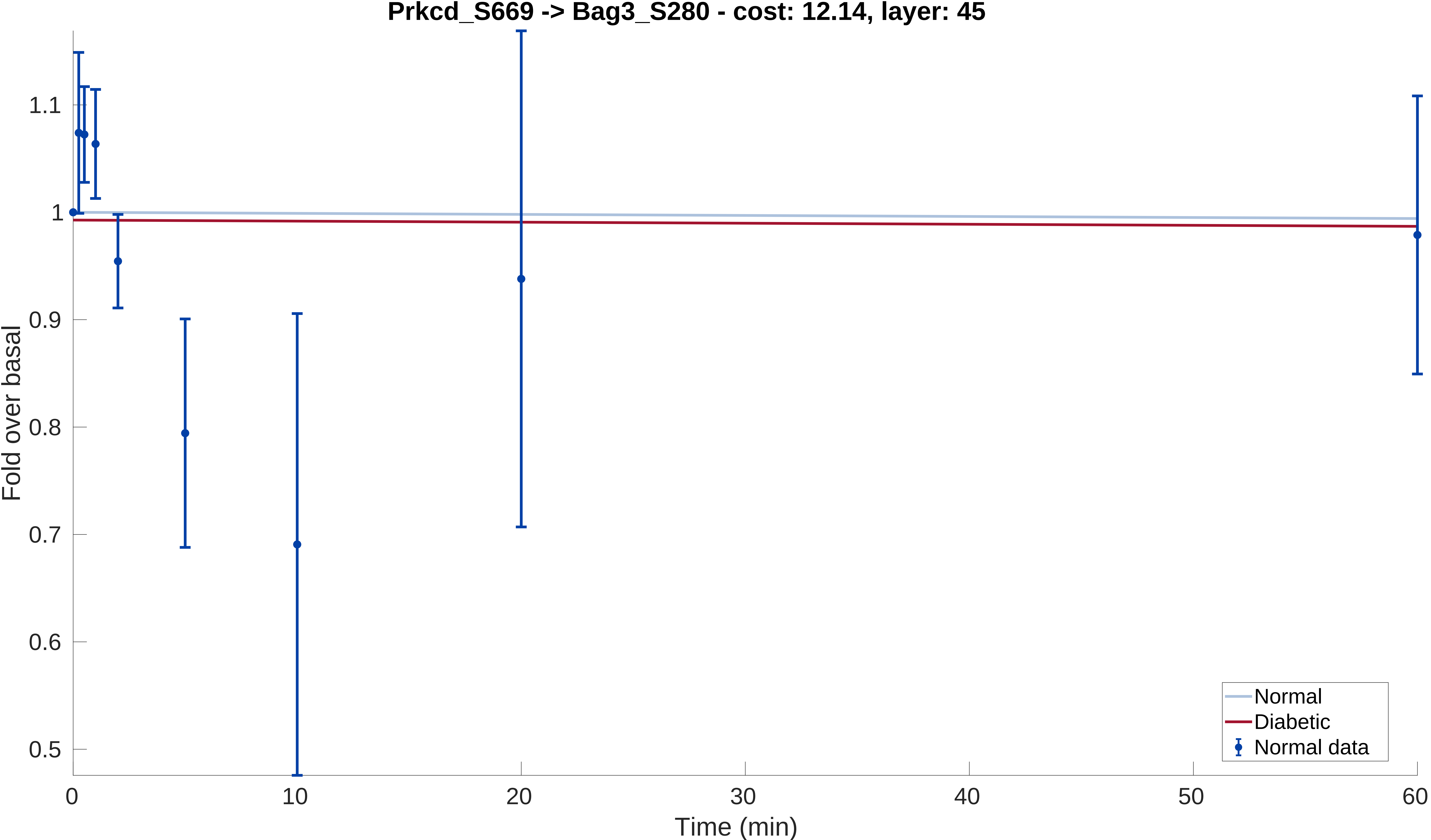

Prkcd_S669 -> Bag3_S280 - cost: 12.14, layer: 45
Fold over basal
1.1
1
0.9
0.8
0.7
0.6
0.5
0
10
20
30
40
50
60
Time (min)
Normal
Diabetic
Normal data

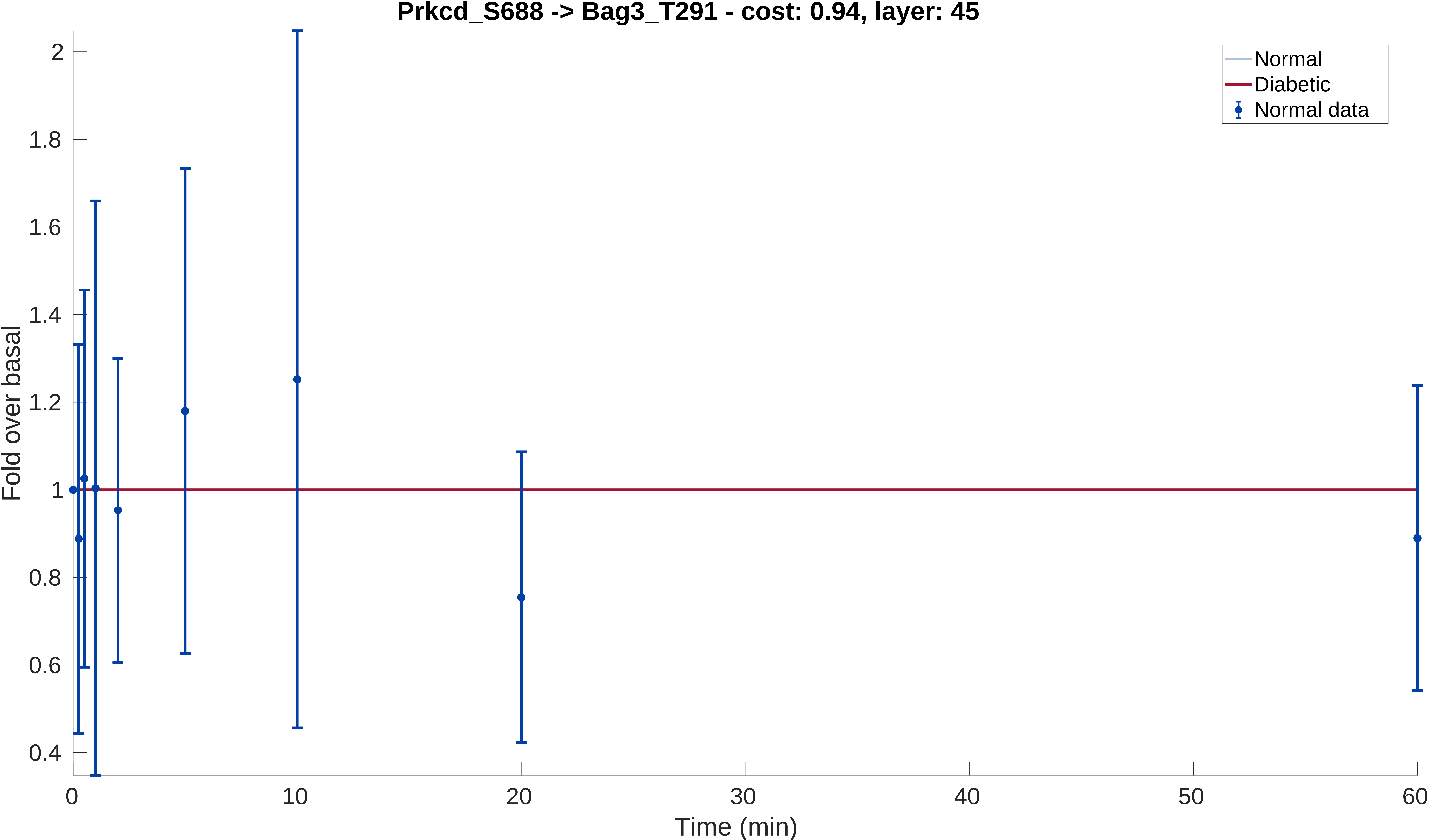

Prkcd_S688 -> Bag3_T291 - cost: 0.94, layer: 45
Normal
Diabetic
Normal data
Fold over basal
2
1.8
1.6
1.4
1.2
1
0.8
0.6
0.4
0
10
20
30
40
50
60
Time (min)

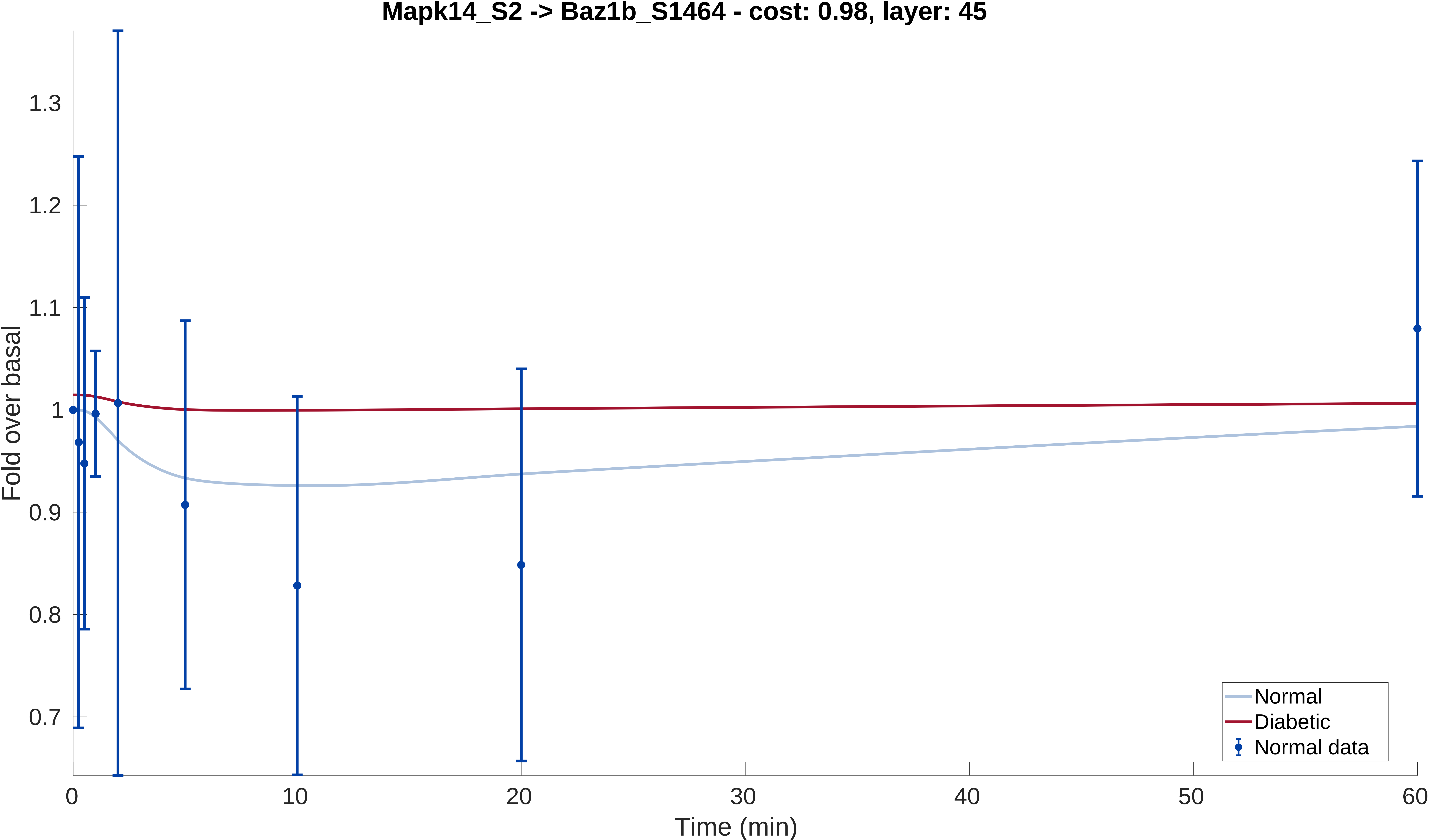
Mapk14_S2 -> Baz1b_S1464 - cost: 0.98, layer: 45
Fold over basal
1.3
1.2
1.1
1
0.9
0.8
0.7
0
10
20
30
40
50
60
Time (min)
Normal
Diabetic
Normal data

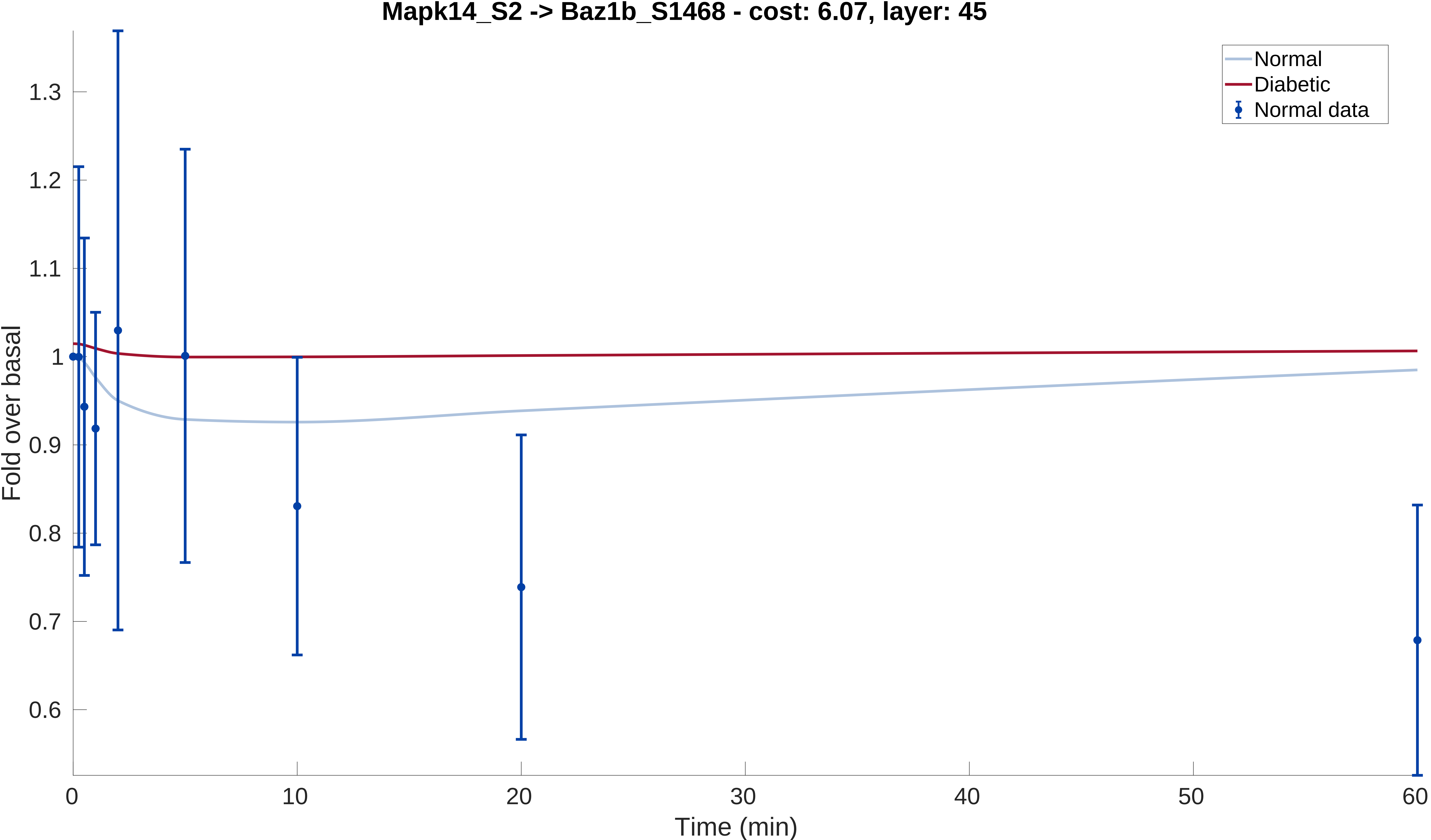
Mapk14_S2 -> Baz1b_S1468 - cost: 6.07, layer: 45
Normal
Diabetic
Normal data
Fold over basal
Time (min)
1.3
1.2
1.1
1
0.9
0.8
0.7
0.6
0
10
20
30
40
50
60

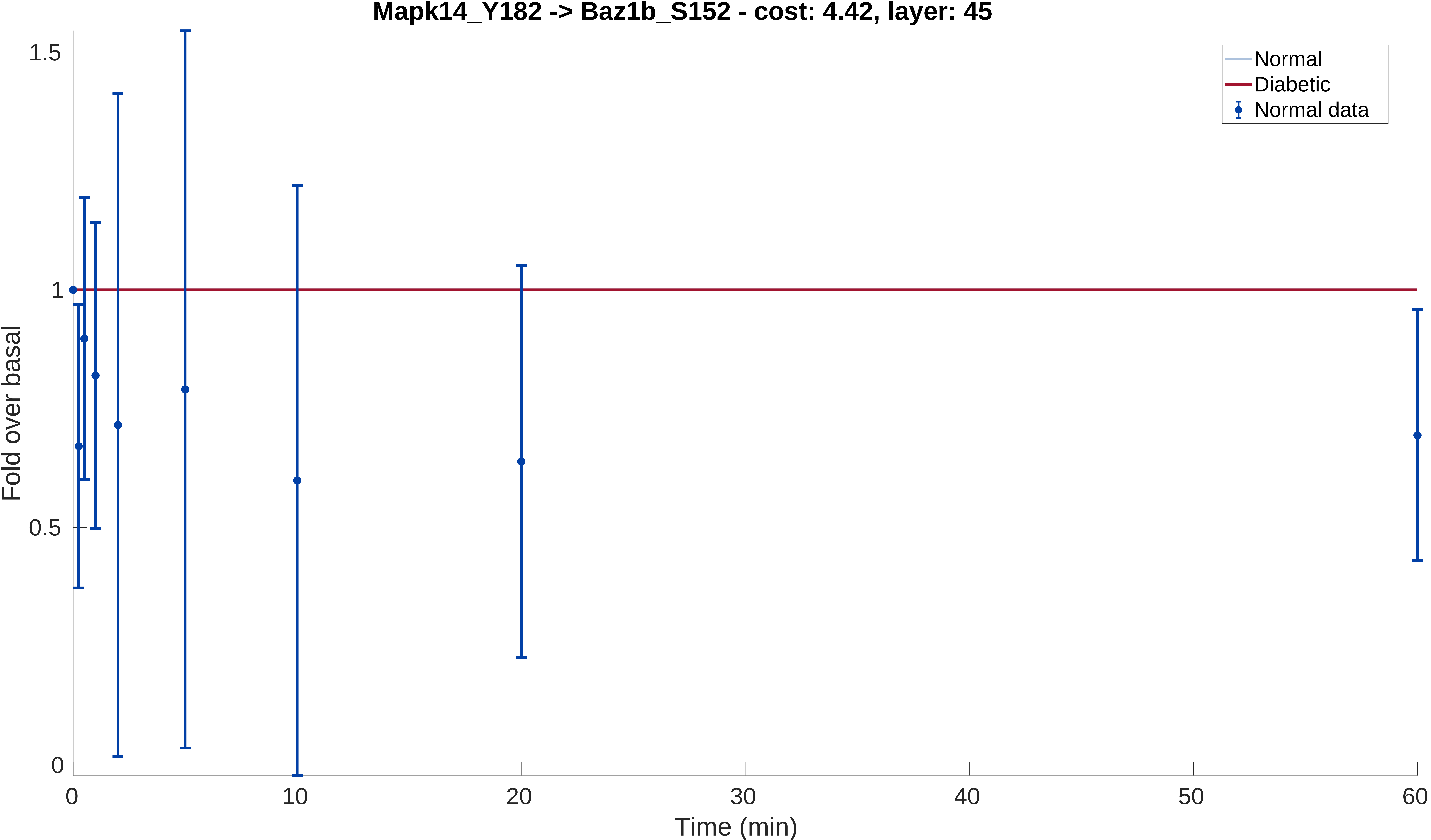
Mapk14_Y182 -> Baz1b_S152 - cost: 4.42, layer: 45
Normal
Diabetic
Normal data
Fold over basal
Time (min)
0
0.5
1
1.5
0
10
20
30
40
50
60

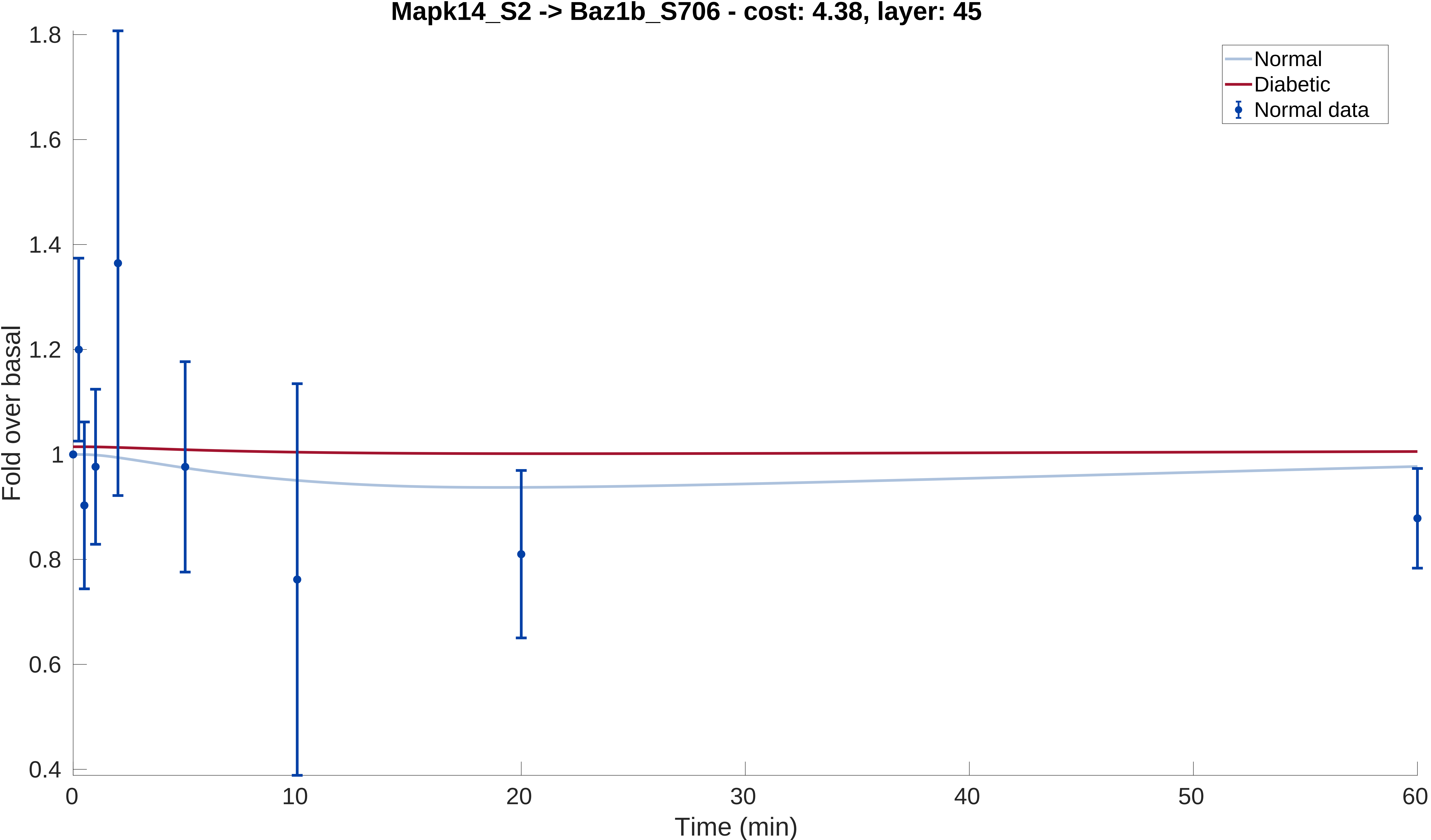

Mapk14_S2 -> Baz1b_S706 - cost: 4.38, layer: 45
Normal
Diabetic
Normal data
Fold over basal
1.8
1.6
1.4
1.2
1
0.8
0.6
0.4
0
10
20
30
40
50
60
Time (min)

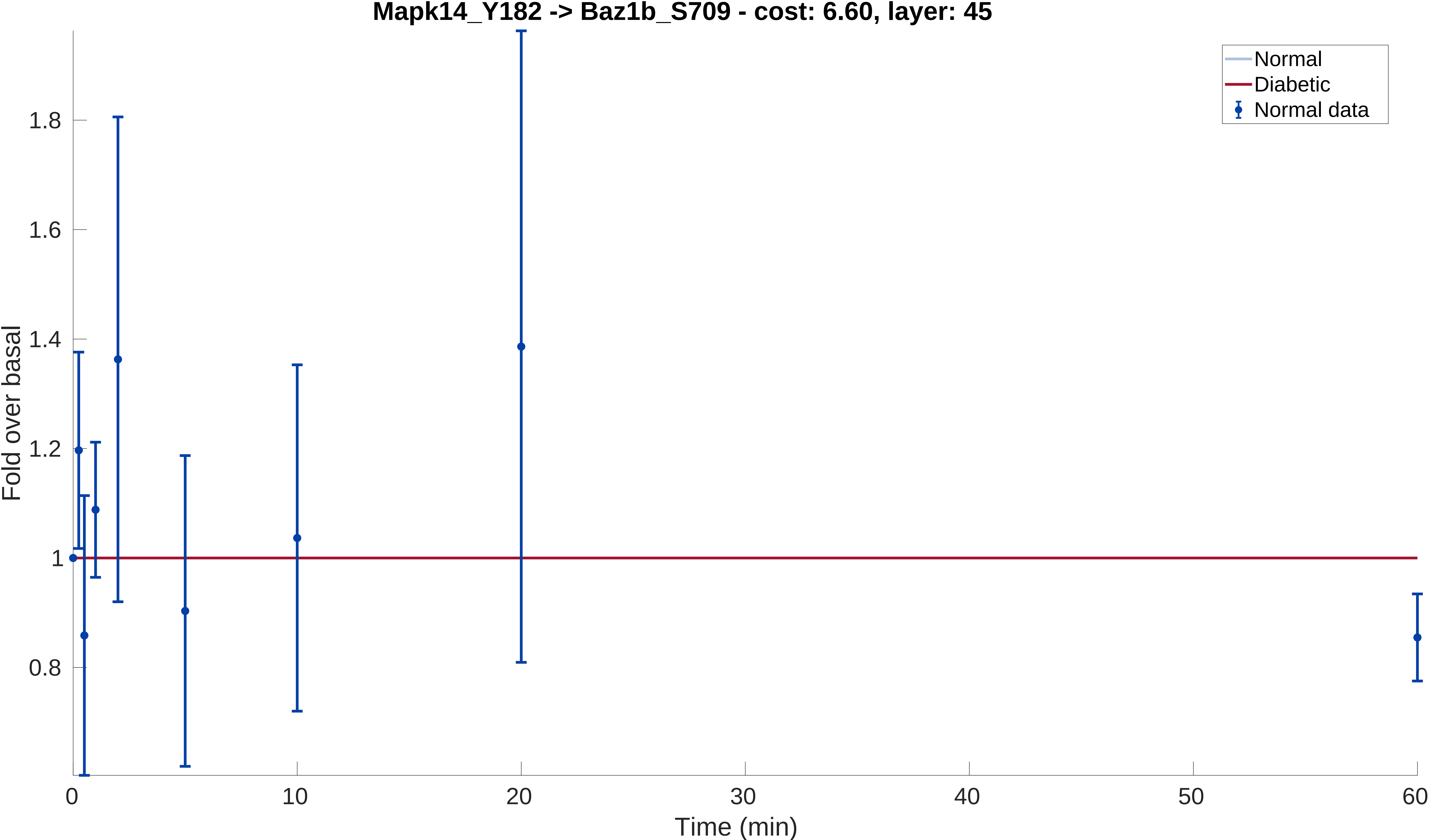
Mapk14_Y182 -> Baz1b_S709 - cost: 6.60, layer: 45
Normal
Diabetic
Normal data
Fold over basal
1.8
1.6
1.4
1.2
1
0.8
0
10
20
30
40
50
60
Time (min)

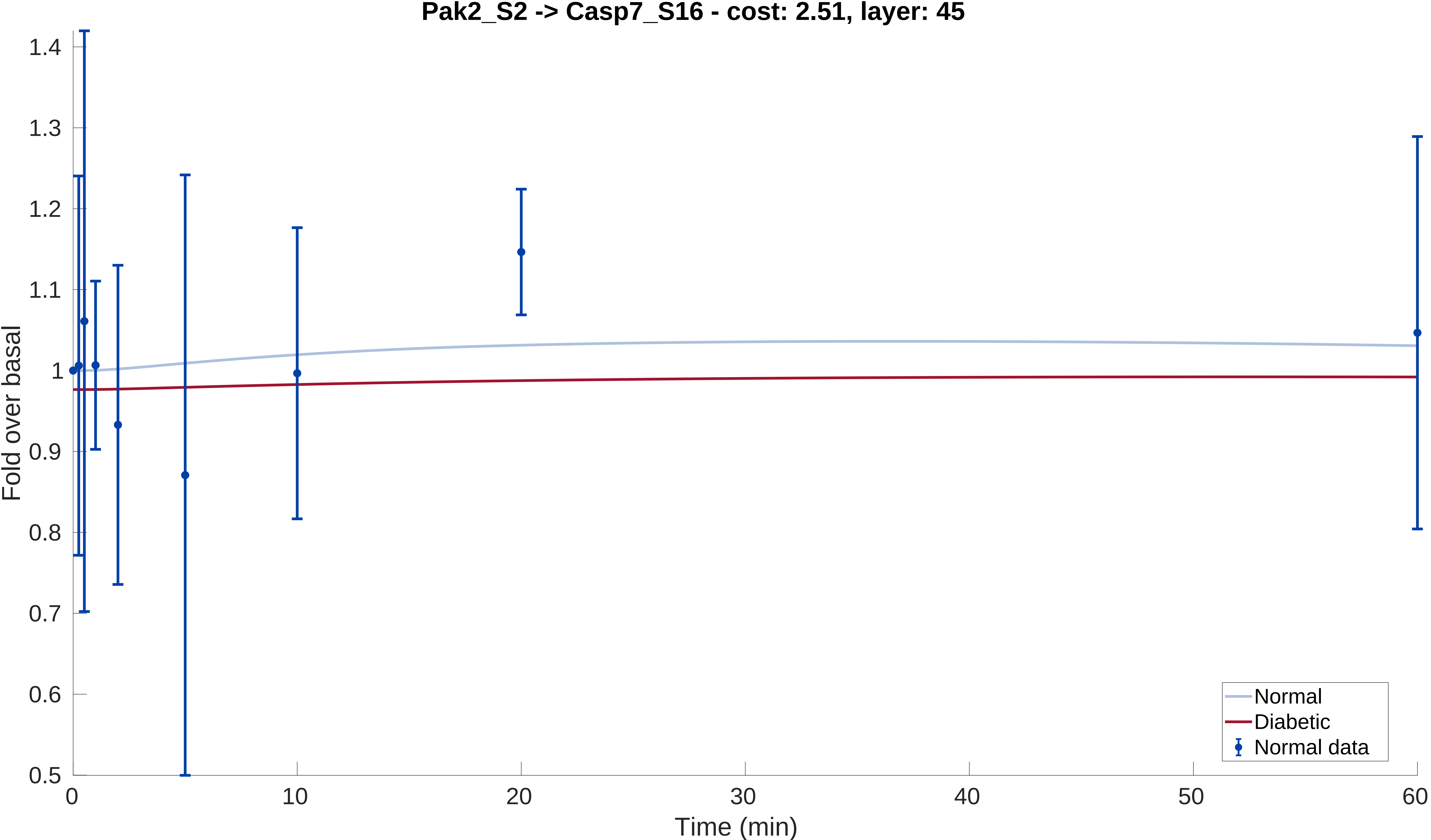

Pak2_S2 -> Casp7_S16 - cost: 2.51, layer: 45
Fold over basal
1.4
1.3
1.2
1.1
1
0.9
0.8
0.7
0.6
0.5
0
10
20
30
40
50
60
Time (min)
Normal
Diabetic
Normal data

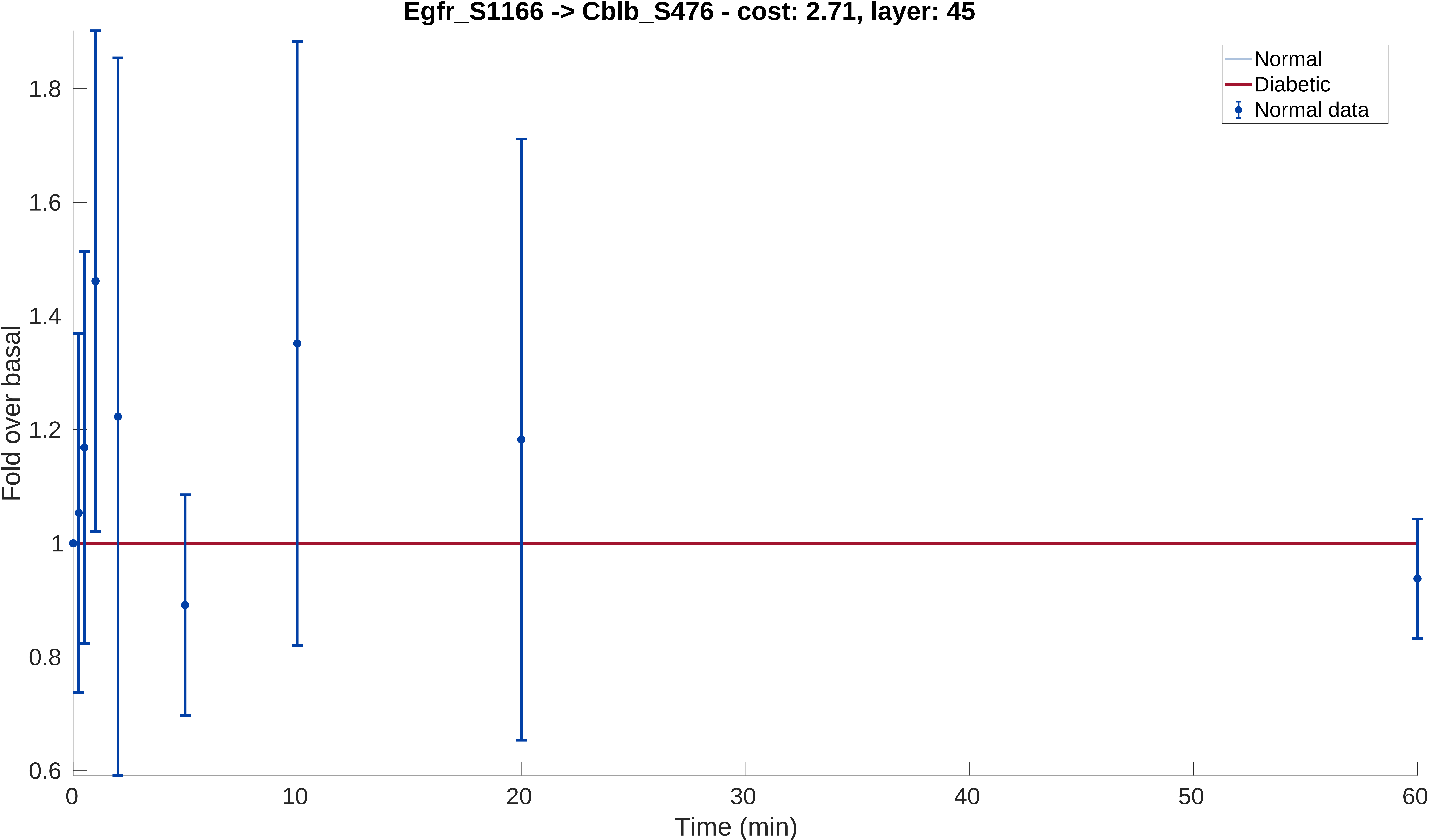
Egfr_S1166 -> Cblb_S476 - cost: 2.71, layer: 45
Normal
Diabetic
Normal data
Fold over basal
1.8
1.6
1.4
1.2
1
0.8
0.6
0
10
20
30
40
50
60
Time (min)

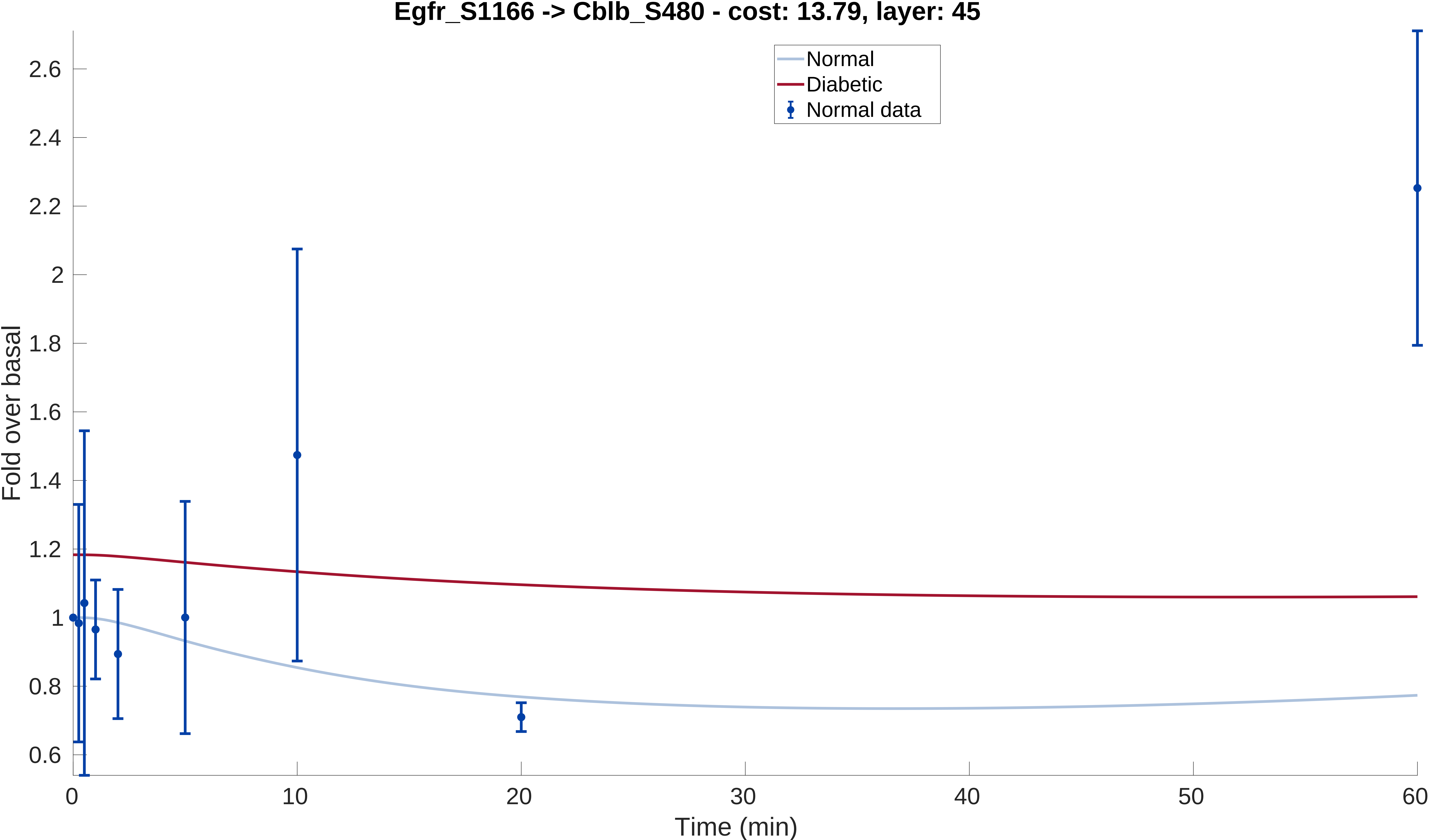

Egfr_S1166 -> Cblb_S480 - cost: 13.79, layer: 45
Normal
Diabetic
Normal data
Fold over basal
2.6
2.4
2.2
2
1.8
1.6
1.4
1.2
1
0.8
0.6
0
10
20
30
40
50
60
Time (min)

#### Egfr_S1166 -> Cblb_S521 - cost: 14.19, layer: 45

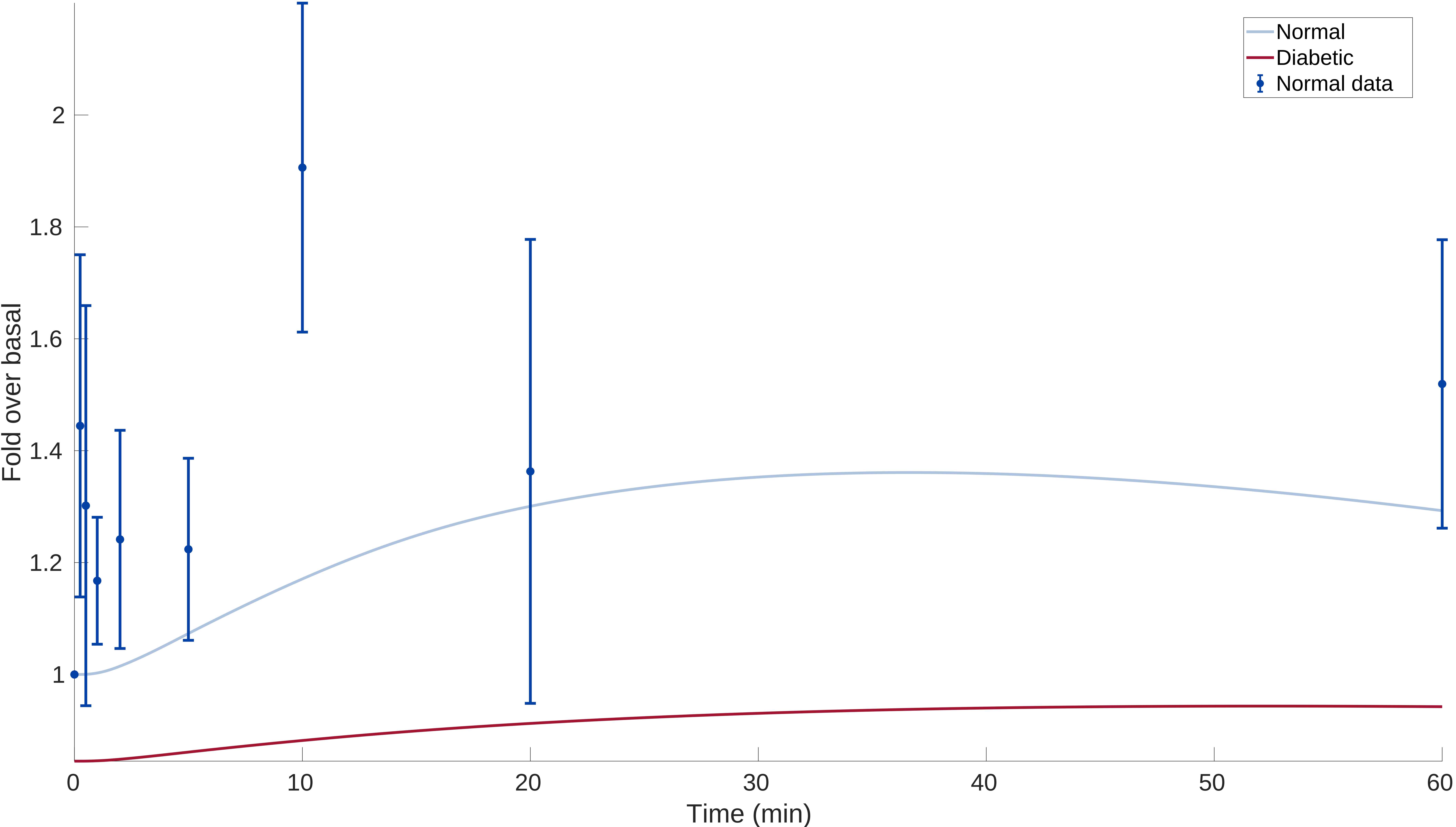

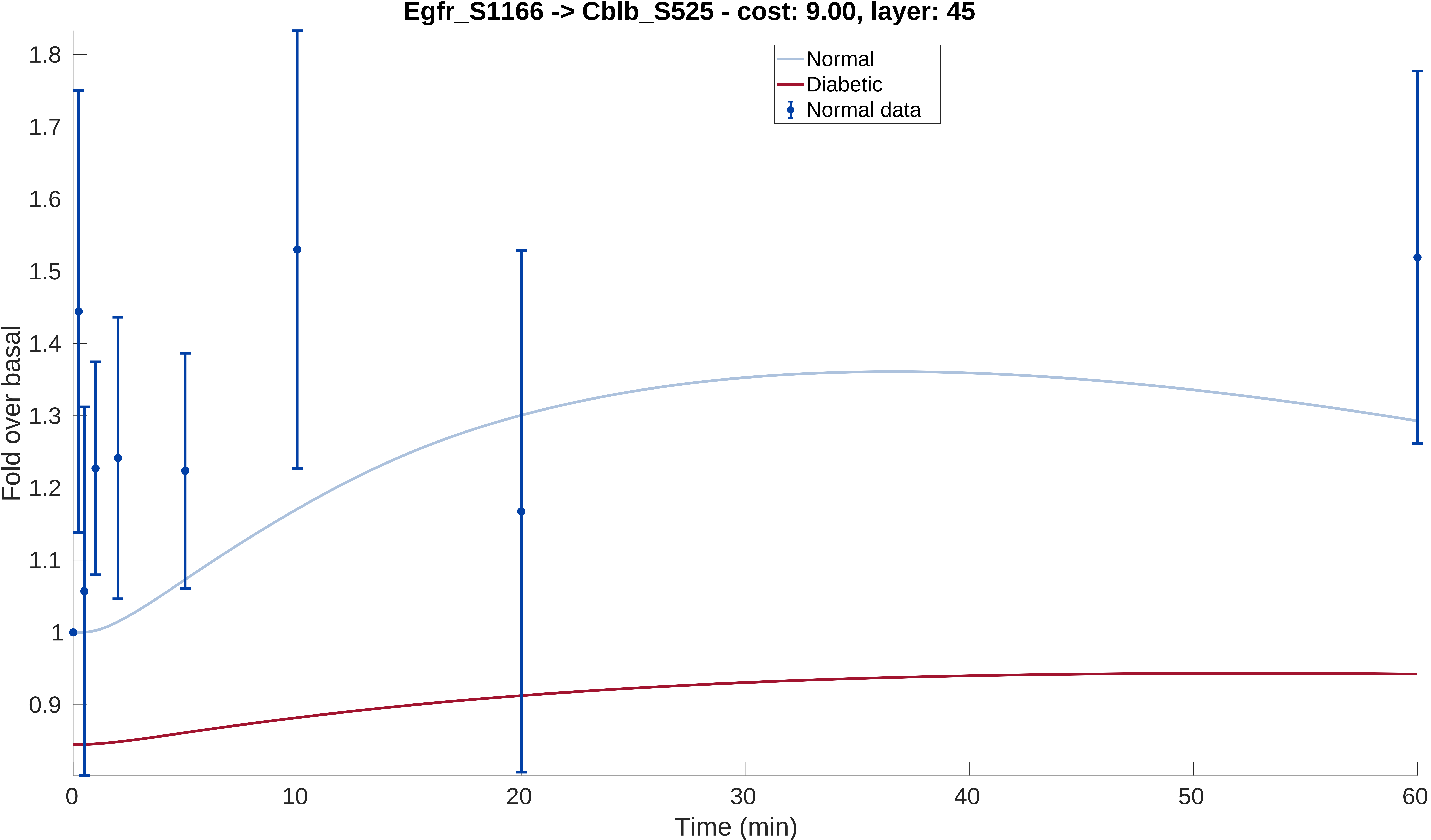

Egfr_S1166 -> Cblb_S525 - cost: 9.00, layer: 45
Normal
Diabetic
Normal data
Fold over basal
Time (min)
0
10
20
30
40
50
60
0.9
1
1.1
1.2
1.3
1.4
1.5
1.6
1.7
1.8

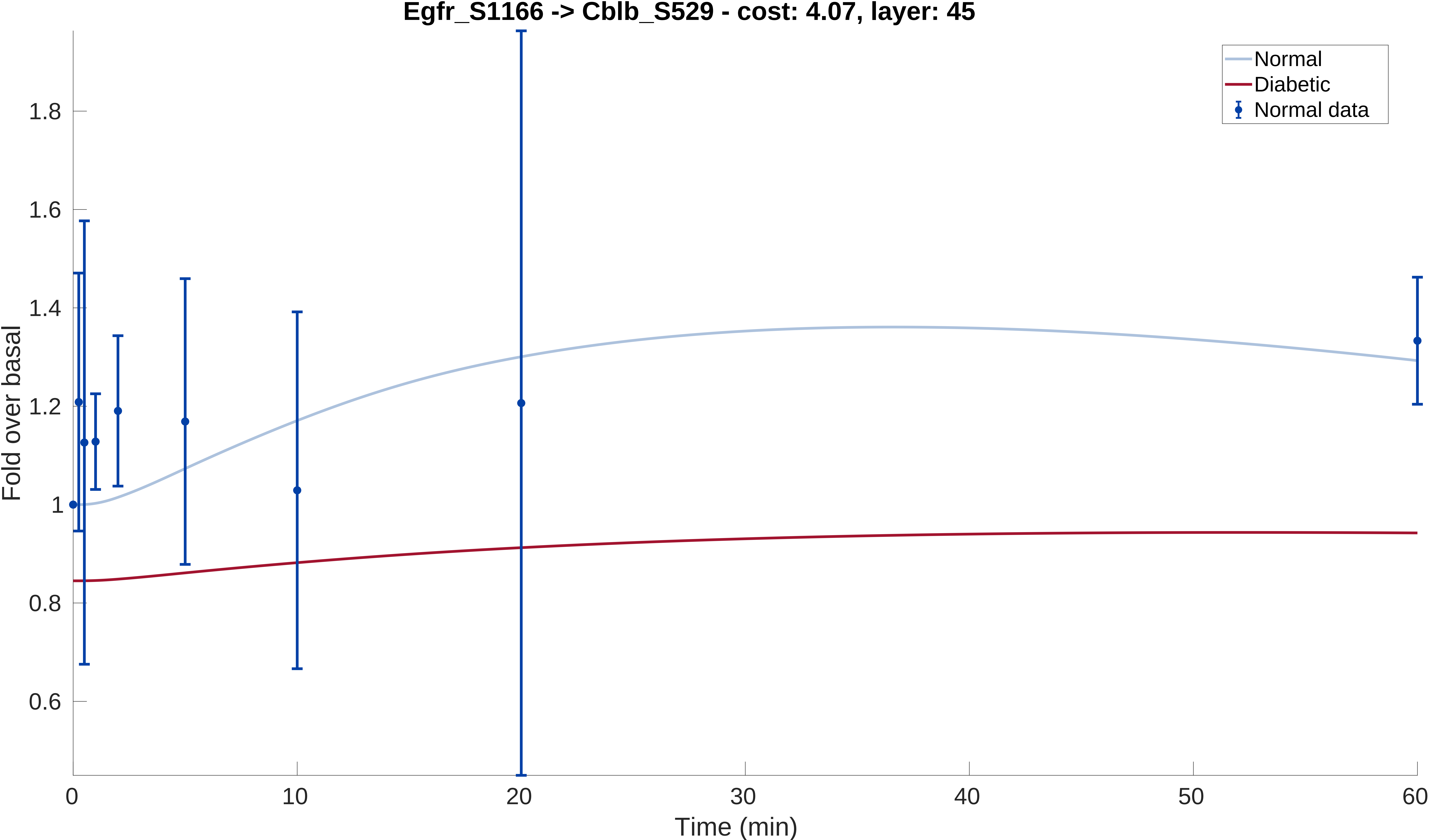

Egfr_S1166 -> Cblb_S529 - cost: 4.07, layer: 45
Normal
Diabetic
Normal data
Fold over basal
1.8
1.6
1.4
1.2
1
0.8
0.6
0
10
20
30
40
50
60
Time (min)

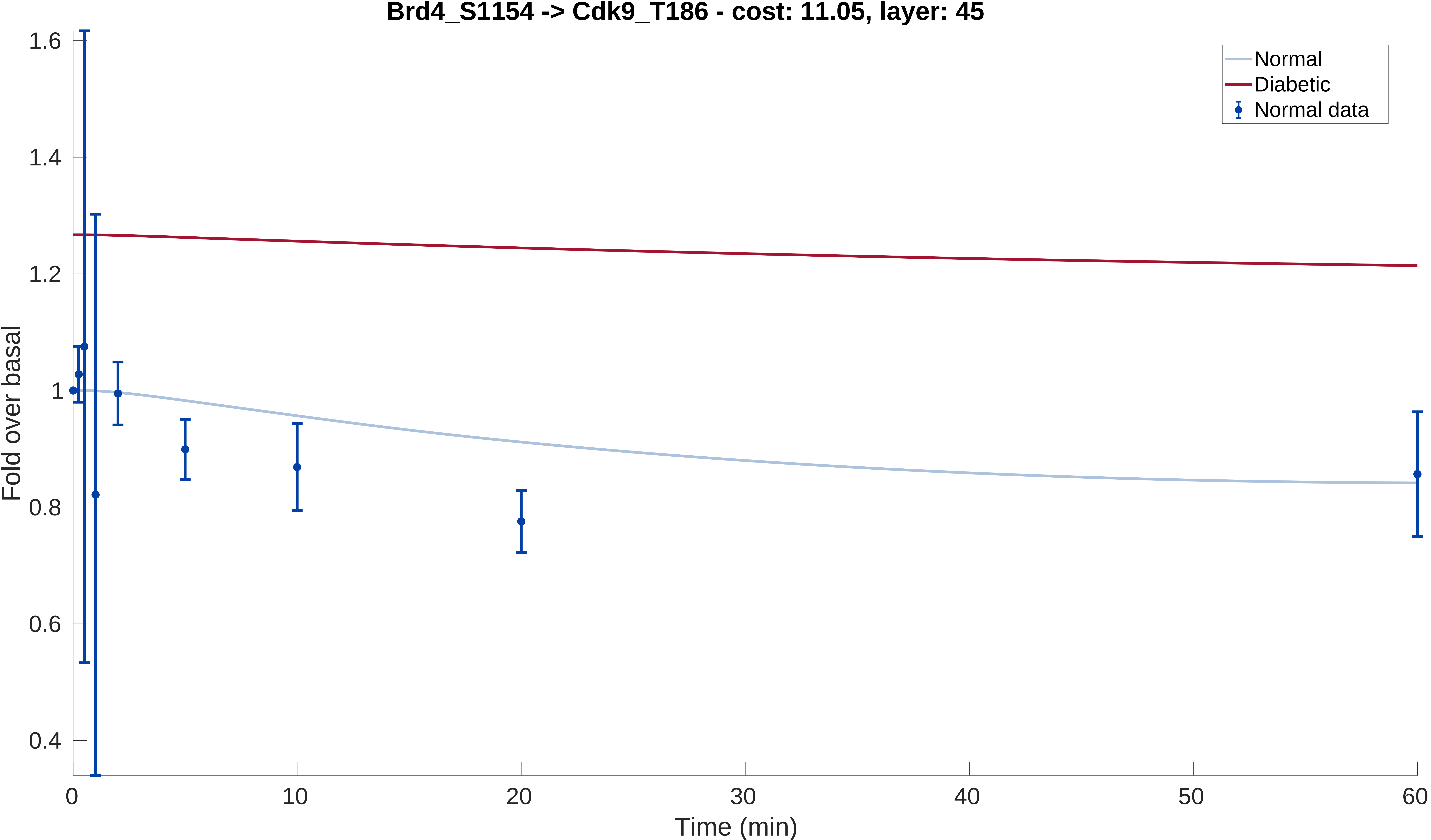

Brd4_S1154 -> Cdk9_T186 - cost: 11.05, layer: 45
Normal
Diabetic
Normal data
Fold over basal
1.6
1.4
1.2
1
0.8
0.6
0.4
0
10
20
30
40
50
60
Time (min)

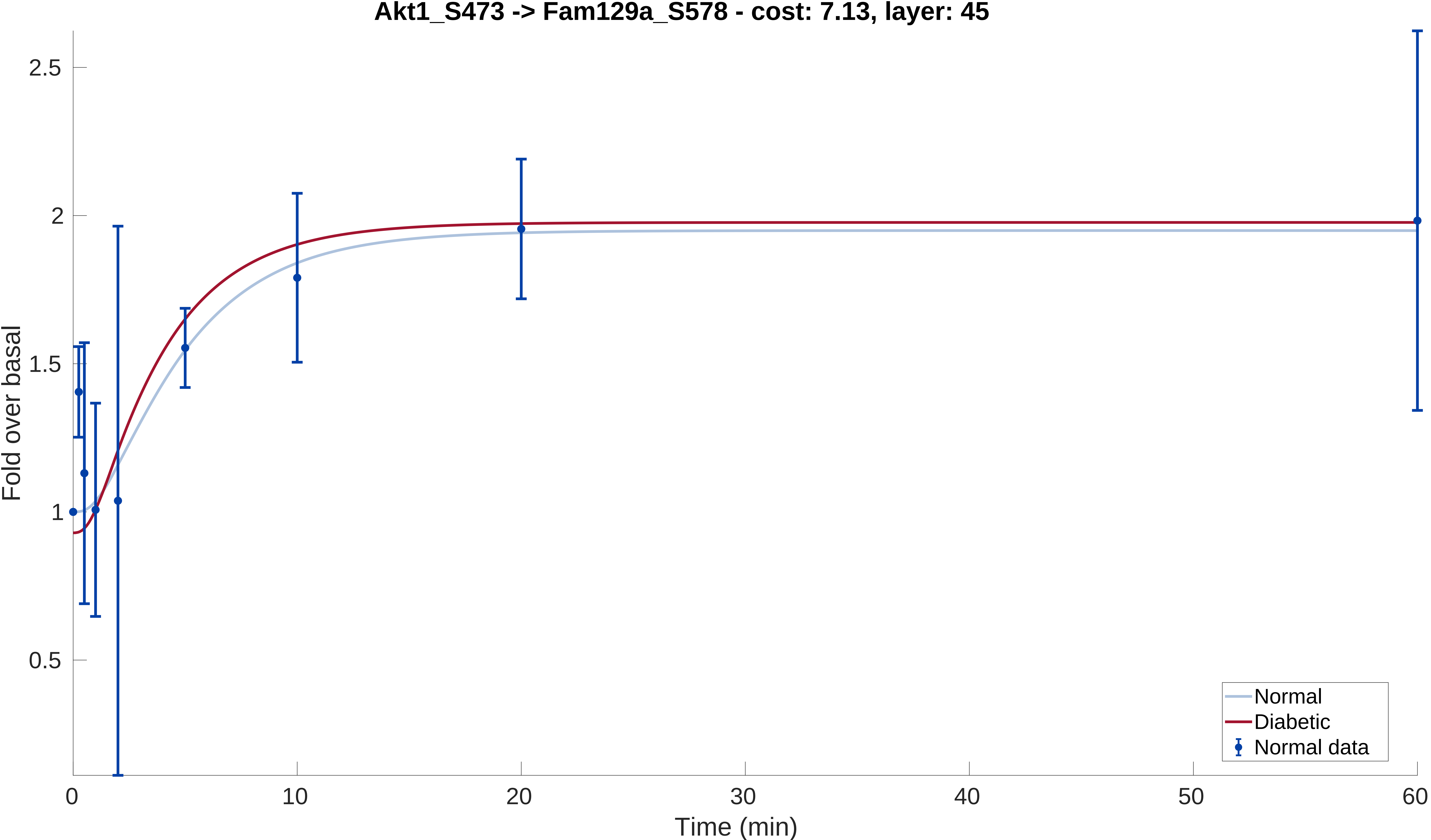

Akt1_S473 -> Fam129a_S578 - cost: 7.13, layer: 45
Fold over basal
2.5
2
1.5
1
0.5
0
10
20
30
40
50
60
Time (min)
Normal
Diabetic
Normal data

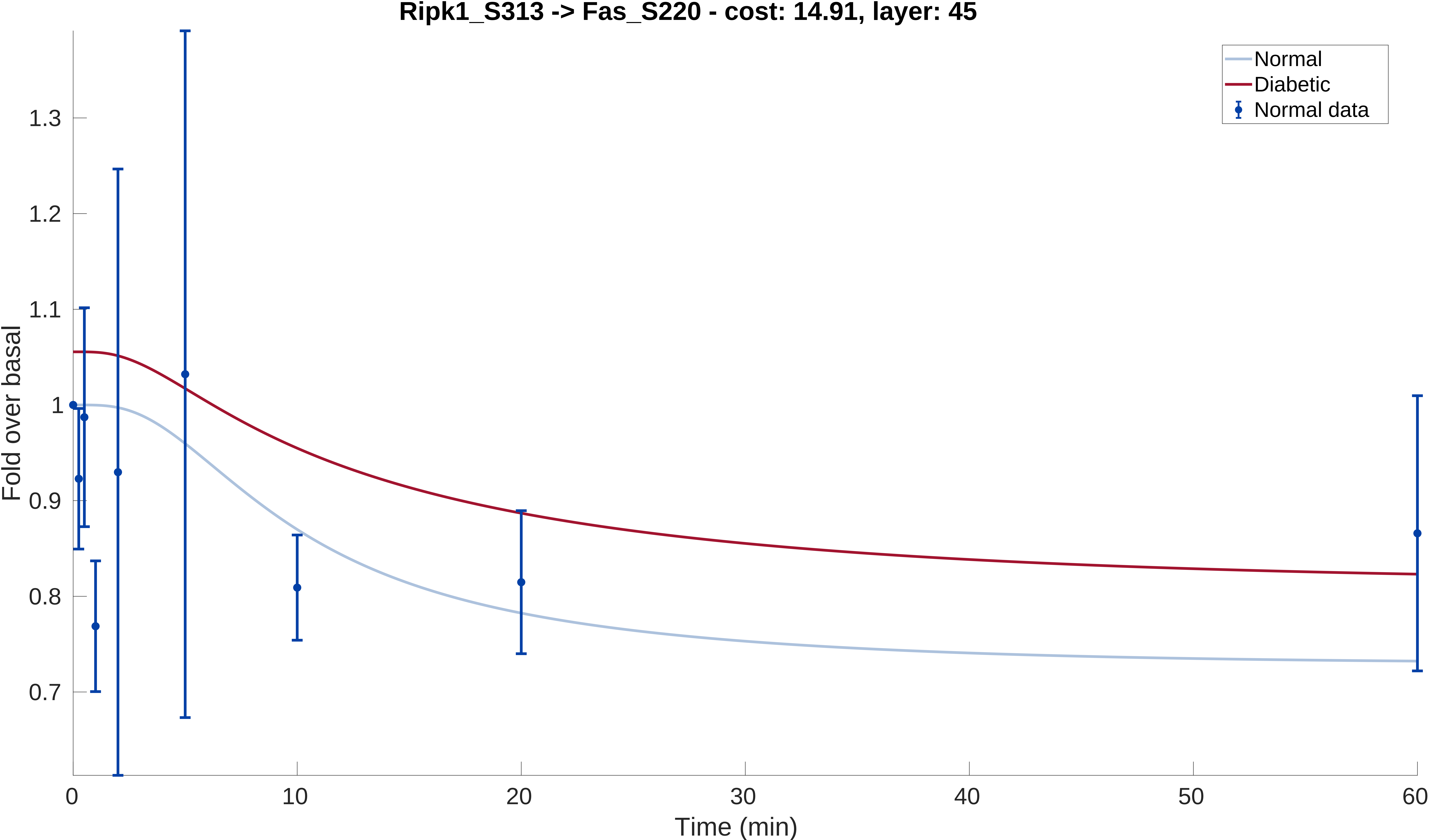

Ripk1_S313 -> Fas_S220 - cost: 14.91, layer: 45
Normal
Diabetic
Normal data
1.3
1.2
1.1
1
0.9
0.8
0.7
Fold over basal
0
10
20
30
40
50
60
Time (min)

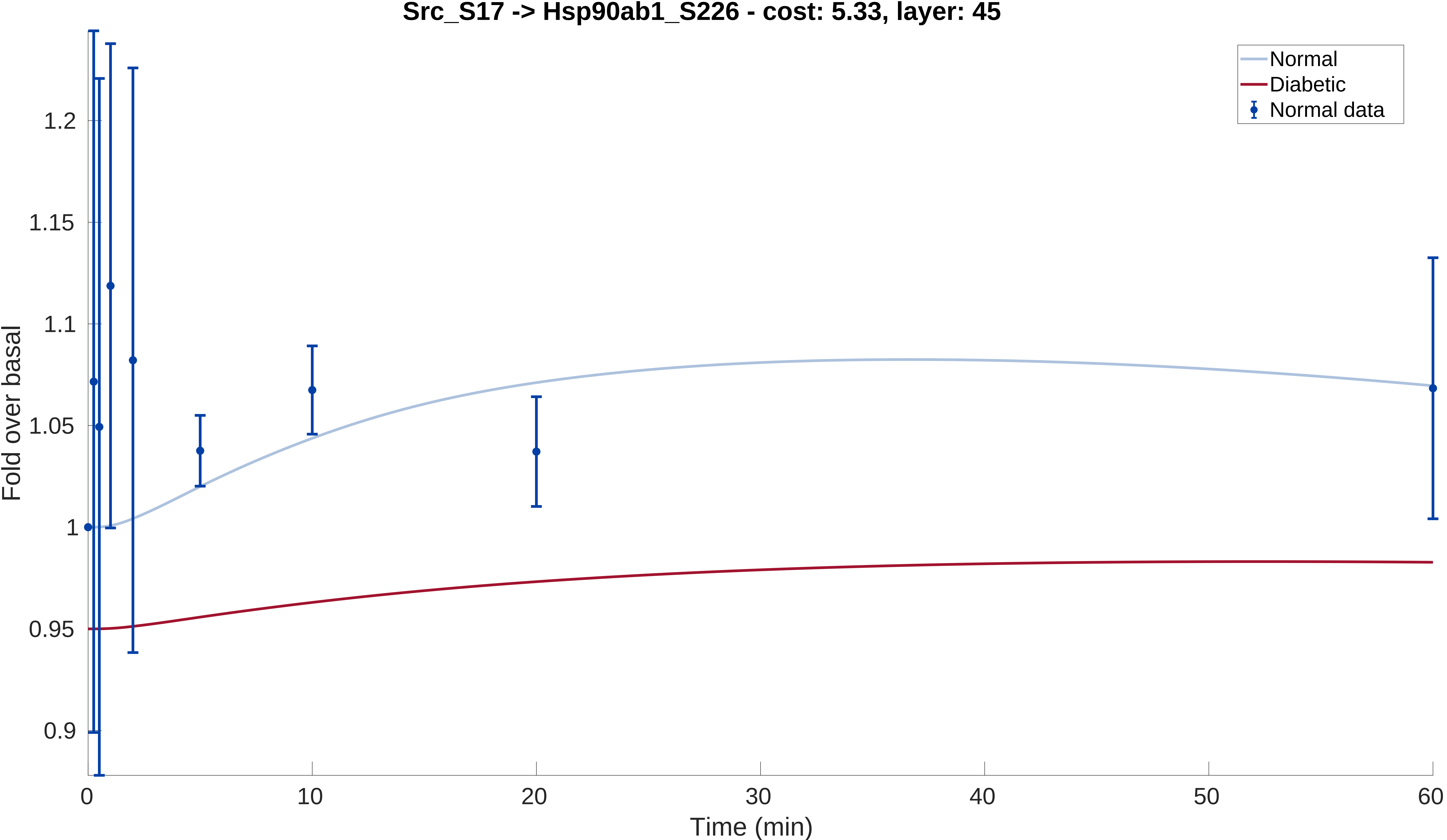
Src_S17 -> Hsp90ab1_S226 - cost: 5.33, layer: 45
Normal
Diabetic
Normal data
Fold over basal
1.2
1.15
1.1
1.05
1
0.95
0.9
0
10
20
30
40
50
60
Time (min)

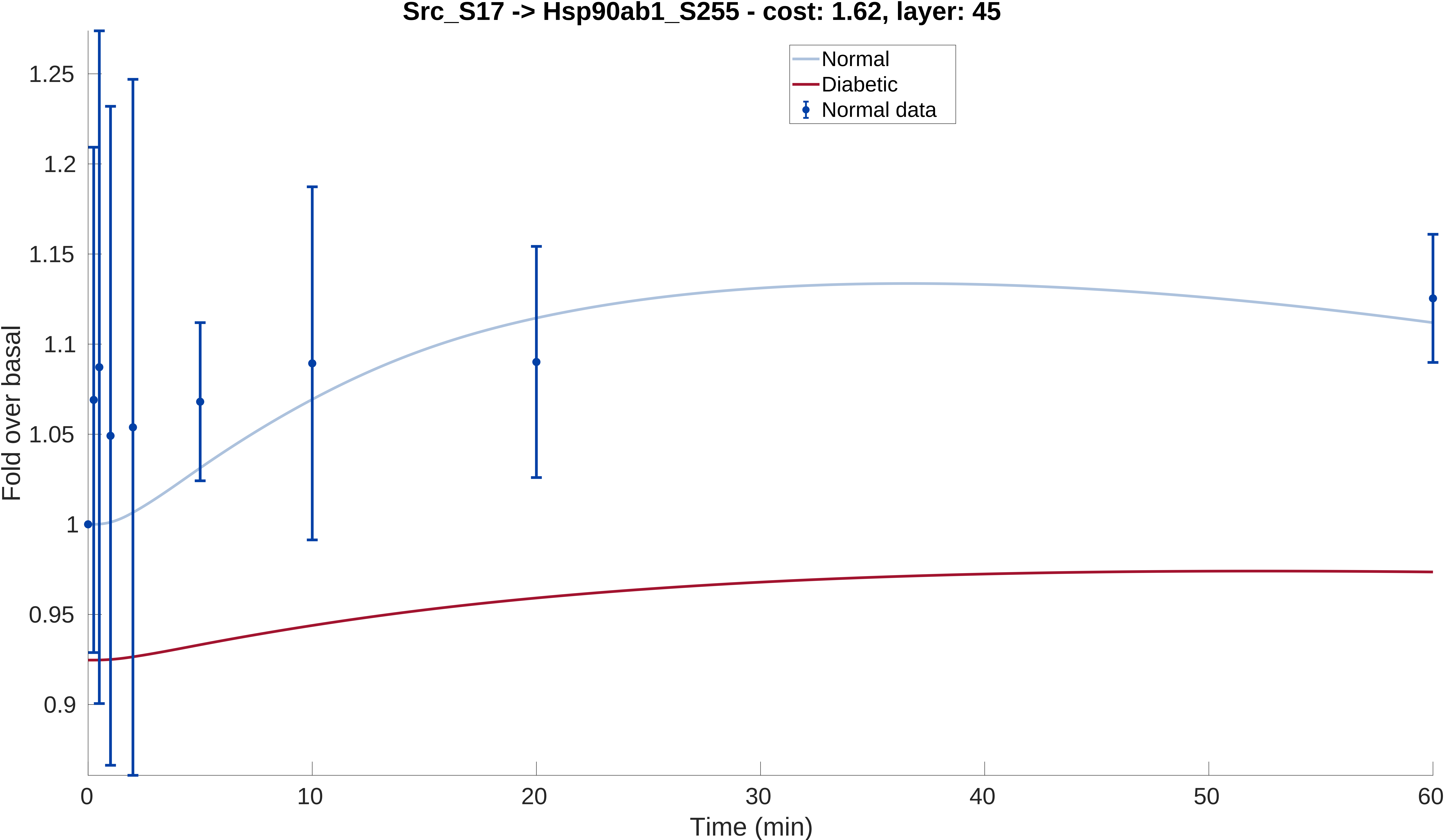
Src_S17 -> Hsp90ab1_S255 - cost: 1.62, layer: 45
Normal
Diabetic
Normal data
Fold over basal
1.25
1.2
1.15
1.1
1.05
1
0.95
0.9
0
10
20
30
40
50
60
Time (min)

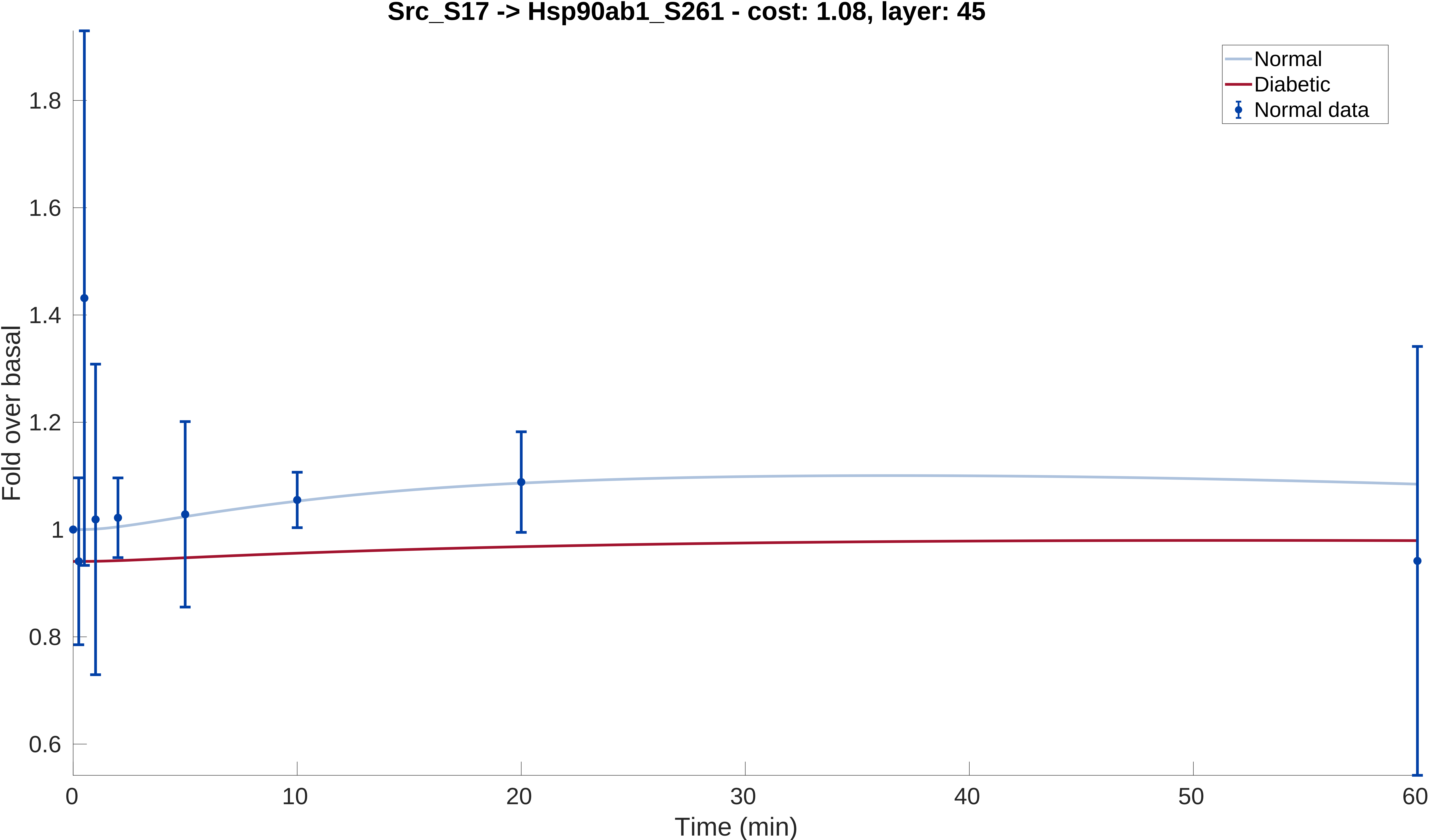
Src_S17 -> Hsp90ab1_S261 - cost: 1.08, layer: 45
Normal
Diabetic
Normal data
Fold over basal
1.8
1.6
1.4
1.2
1
0.8
0.6
0
10
20
30
40
50
60
Time (min)

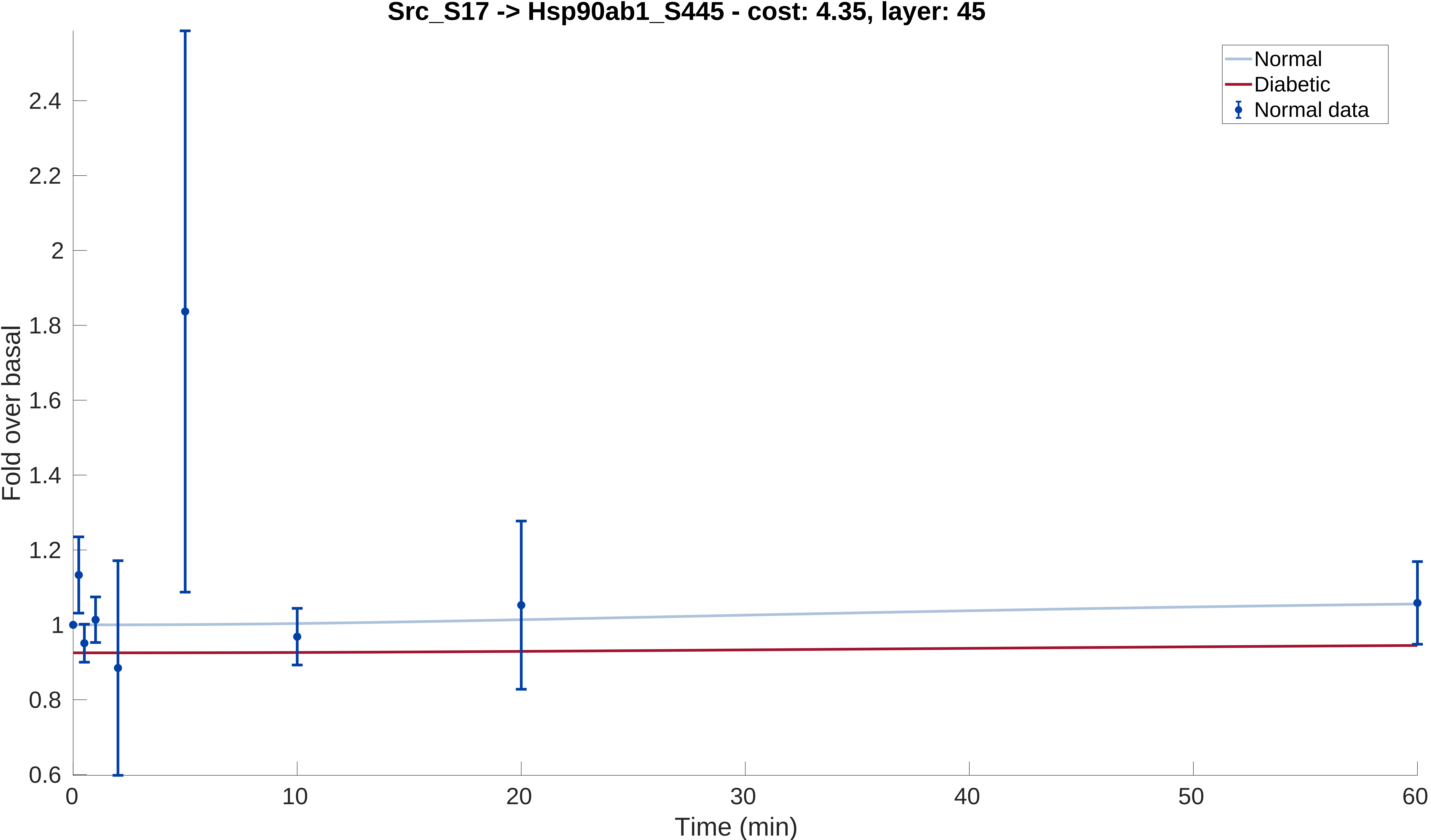

Src_S17 -> Hsp90ab1_S445 - cost: 4.35, layer: 45
Normal
Diabetic
Normal data
Fold over basal
2.4
2.2
2
1.8
1.6
1.4
1.2
1
0.8
0.6
0
10
20
30
40
50
60
Time (min)

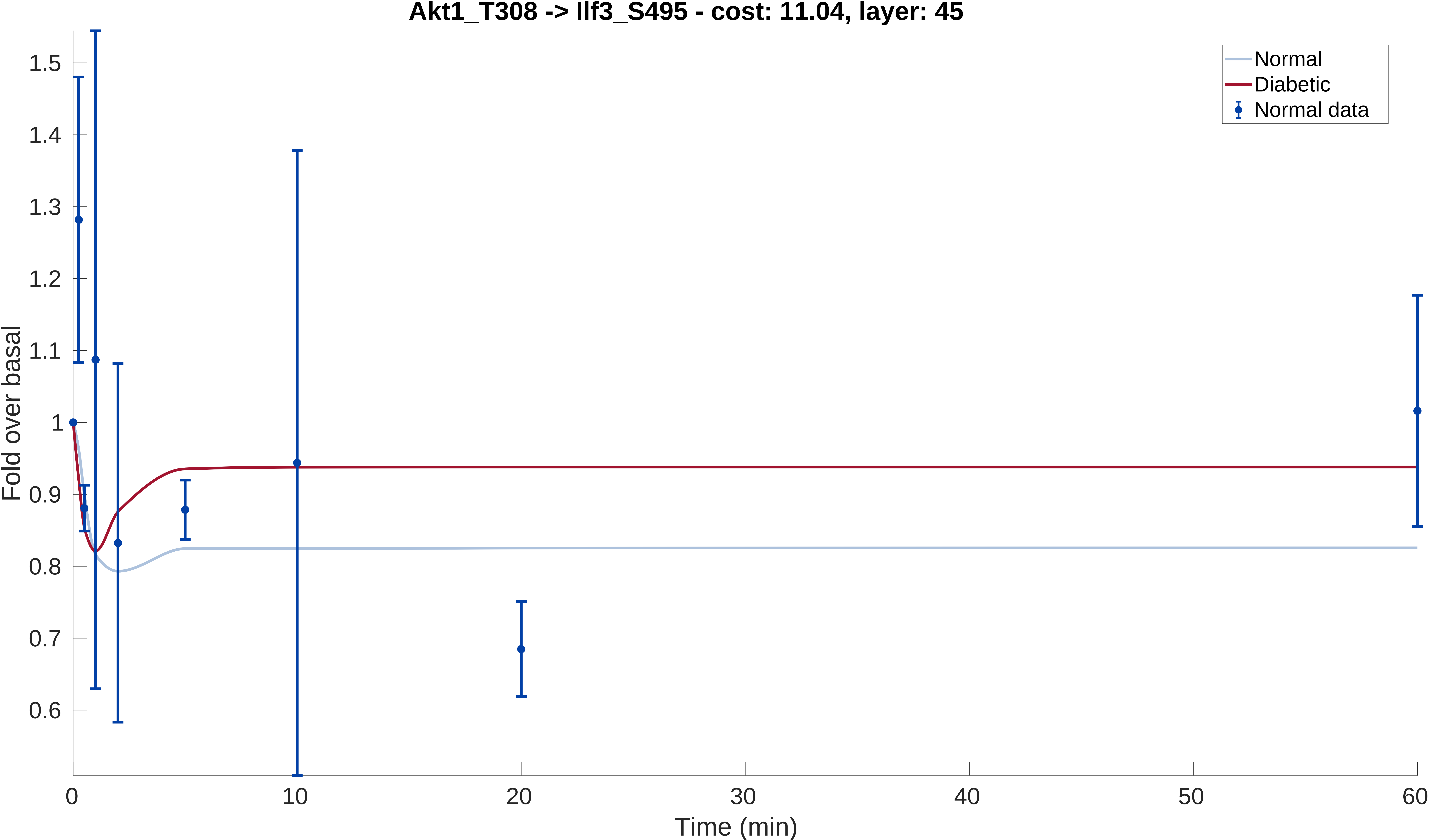
Akt1_T308 -> Ilf3_S495 - cost: 11.04, layer: 45
Normal
Diabetic
Normal data
Fold over basal
1.5
1.4
1.3
1.2
1.1
1
0.9
0.8
0.7
0.6
0
10
20
30
40
50
60
Time (min)

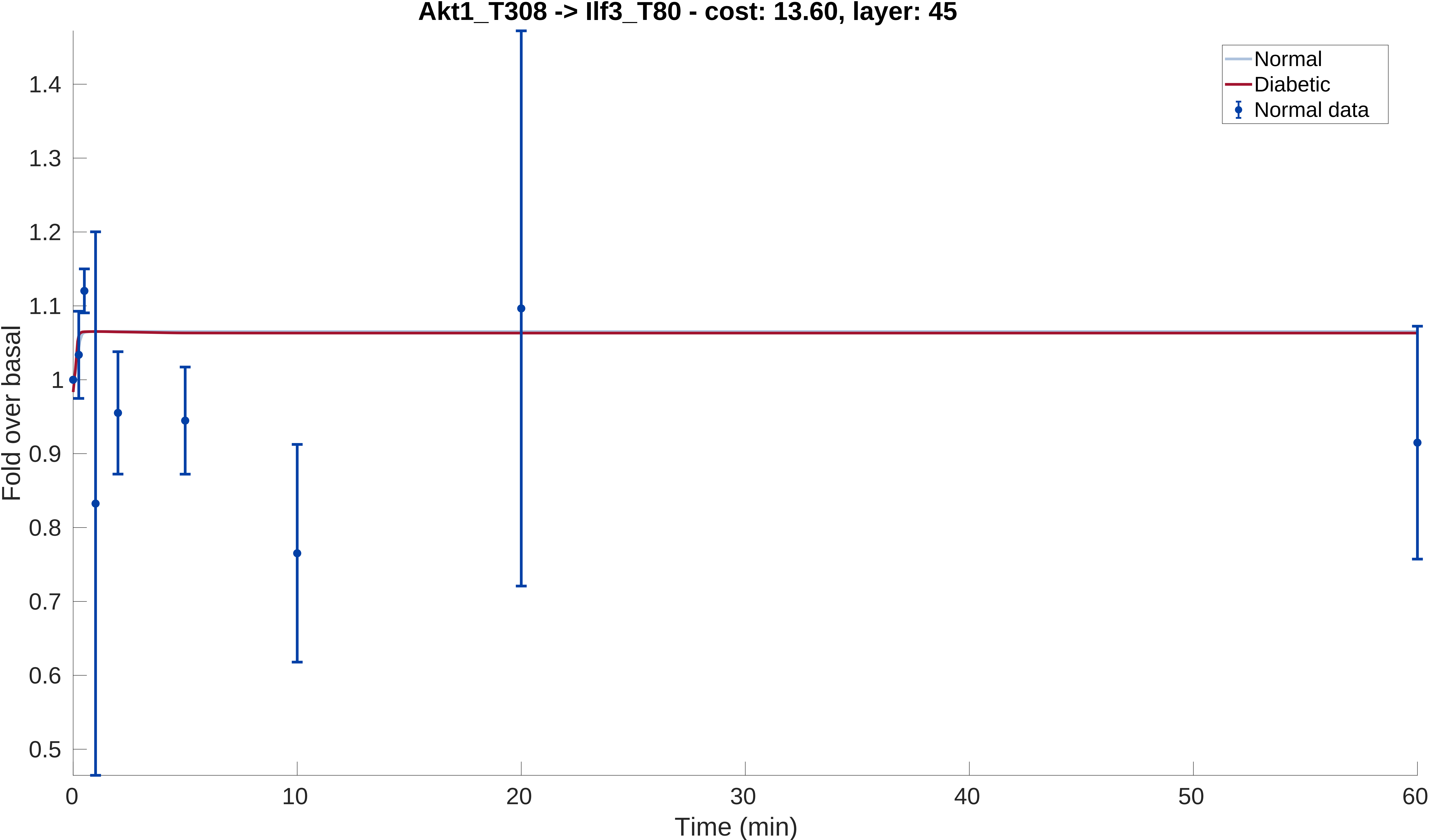

Akt1_T308 -> Ilf3_T80 - cost: 13.60, layer: 45
Normal
Diabetic
Normal data
Fold over basal
1.4
1.3
1.2
1.1
1
0.9
0.8
0.7
0.6
0.5
Time (min)
0
10
20
30
40
50
60

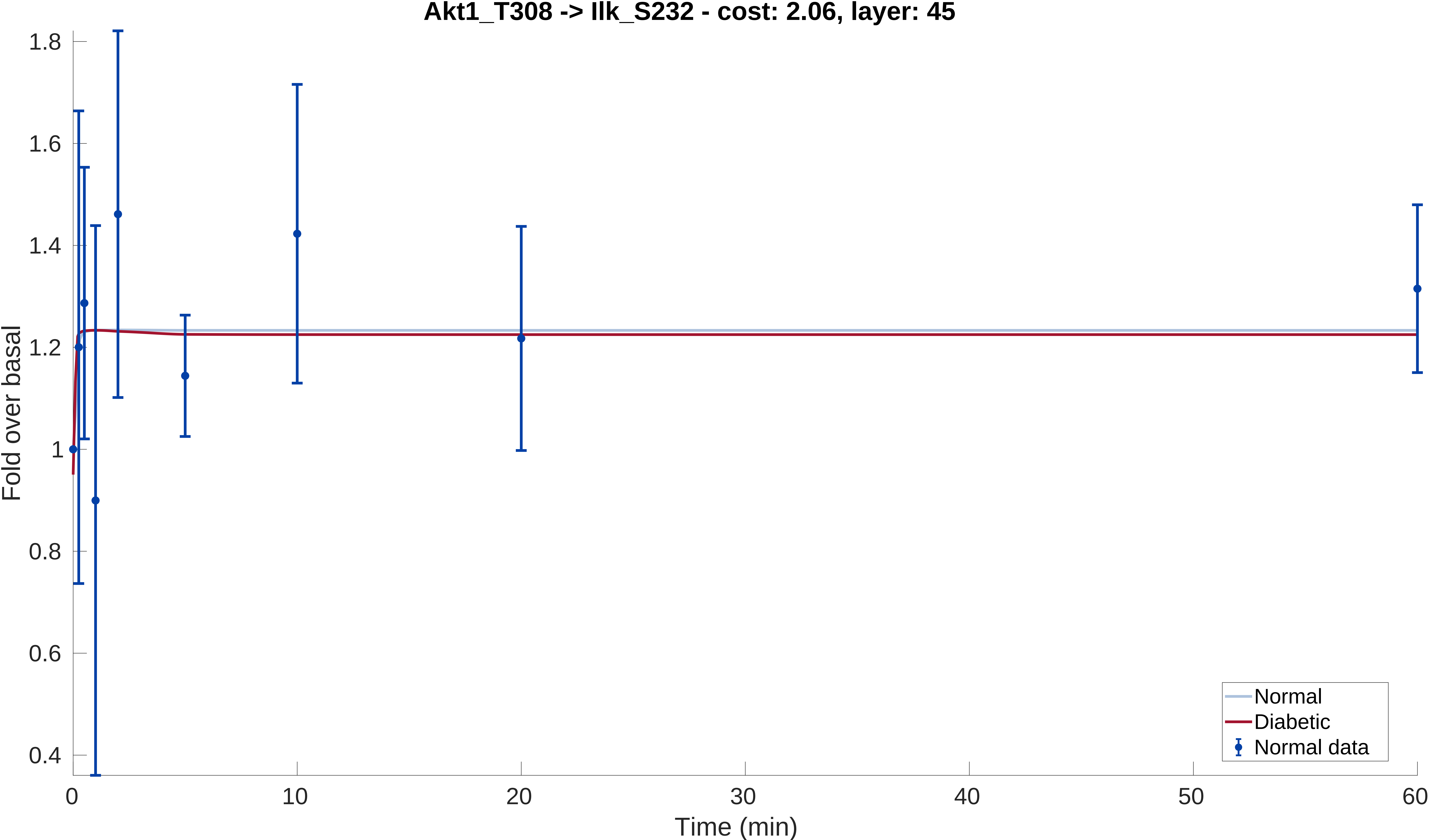
Akt1_T308 -> Ilk_S232 - cost: 2.06, layer: 45
1.8
1.6
1.4
1.2
1
0.8
0.6
0.4
Fold over basal
0
10
20
30
40
50
60
Time (min)
Normal
Diabetic
Normal data

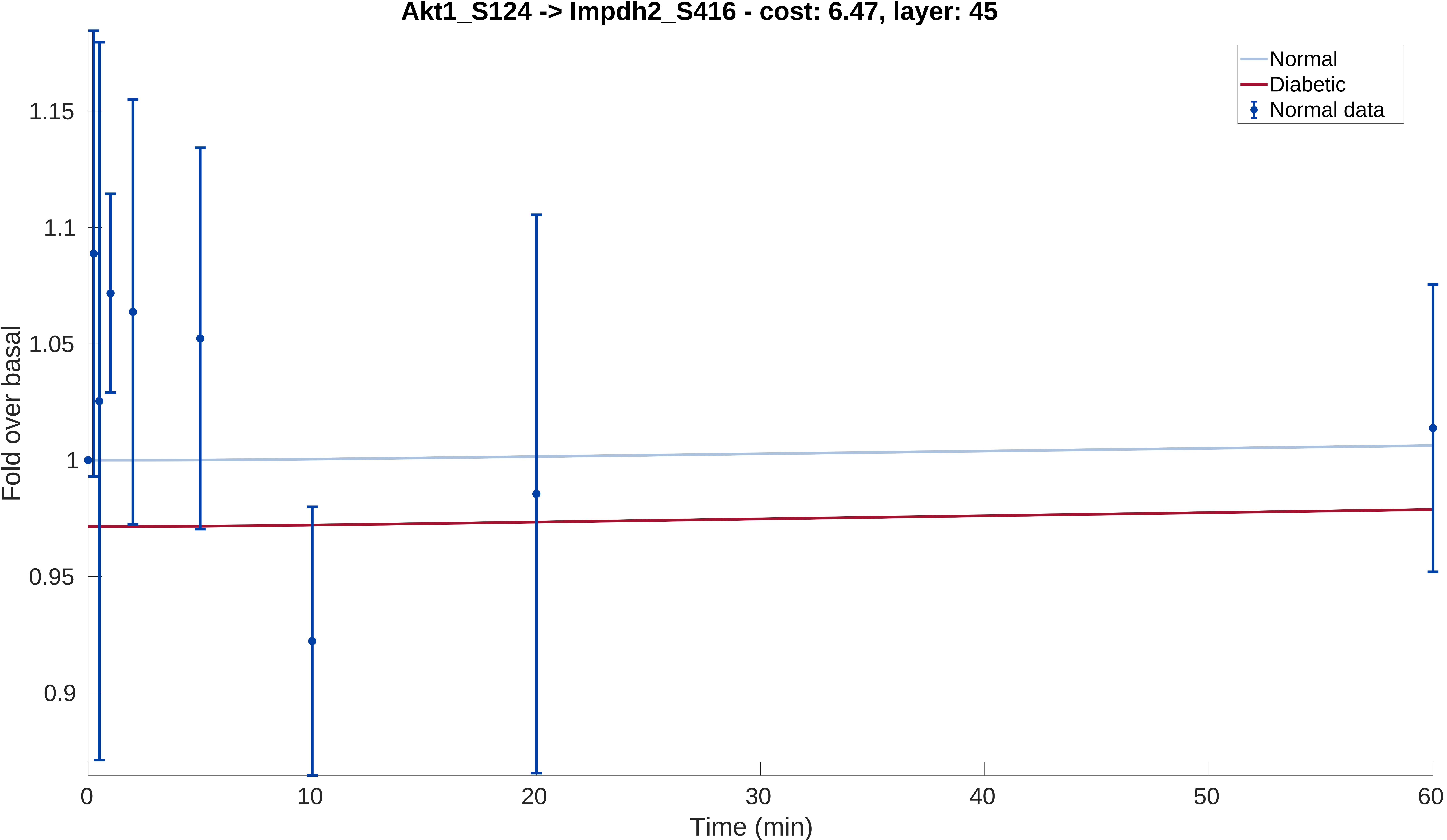
Akt1_S124 -> Impdh2_S416 - cost: 6.47, layer: 45
Normal
Diabetic
Normal data
Fold over basal
1.15
1.1
1.05
1
0.95
0.9
0
10
20
30
40
50
60
Time (min)

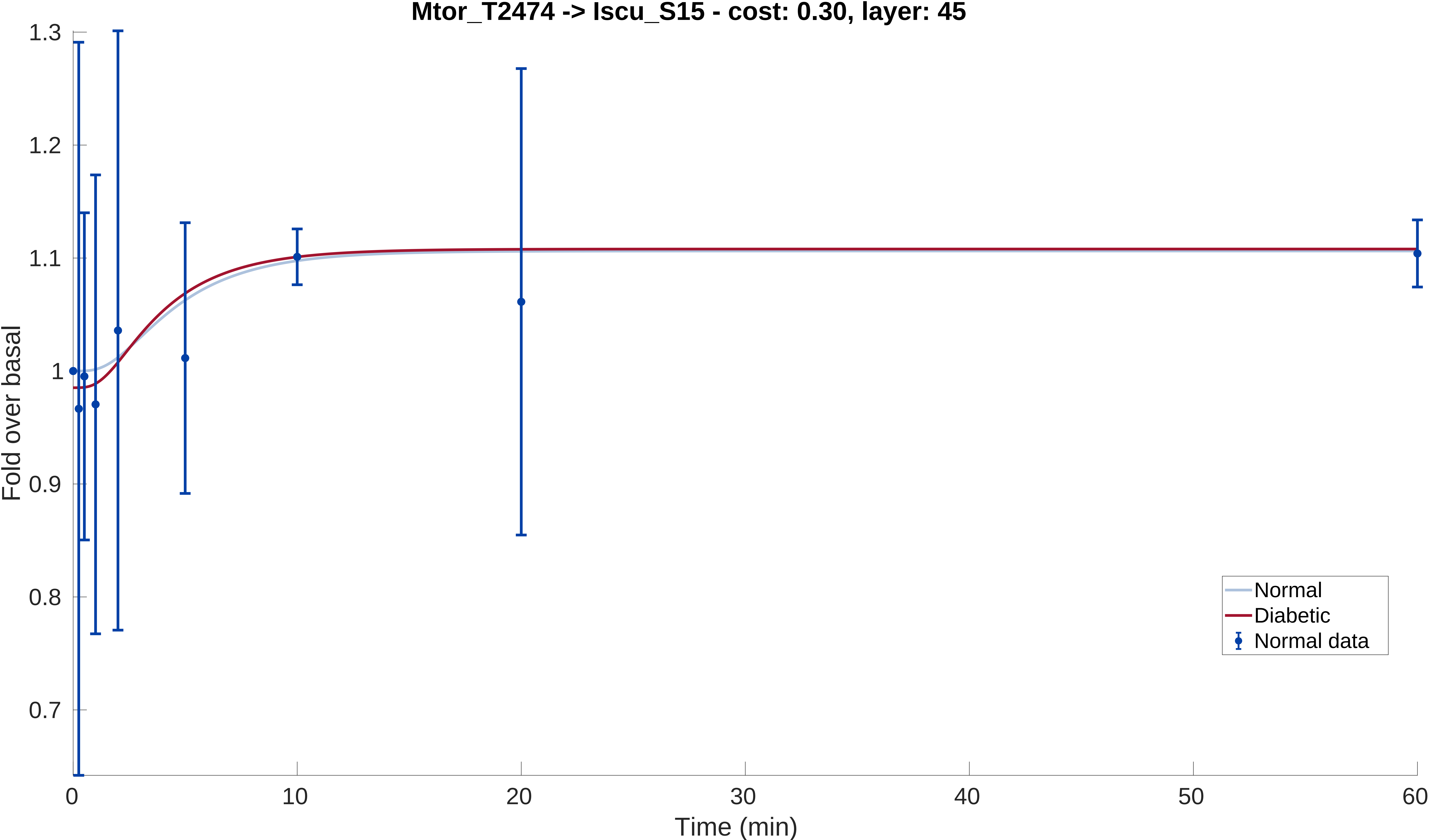

Mtor_T2474 -> Iscu_S15 - cost: 0.30, layer: 45
1.3
1.2
1.1
1
0.9
0.8
0.7
Fold over basal
0
10
20
30
40
50
60
Time (min)
Normal
Diabetic
Normal data

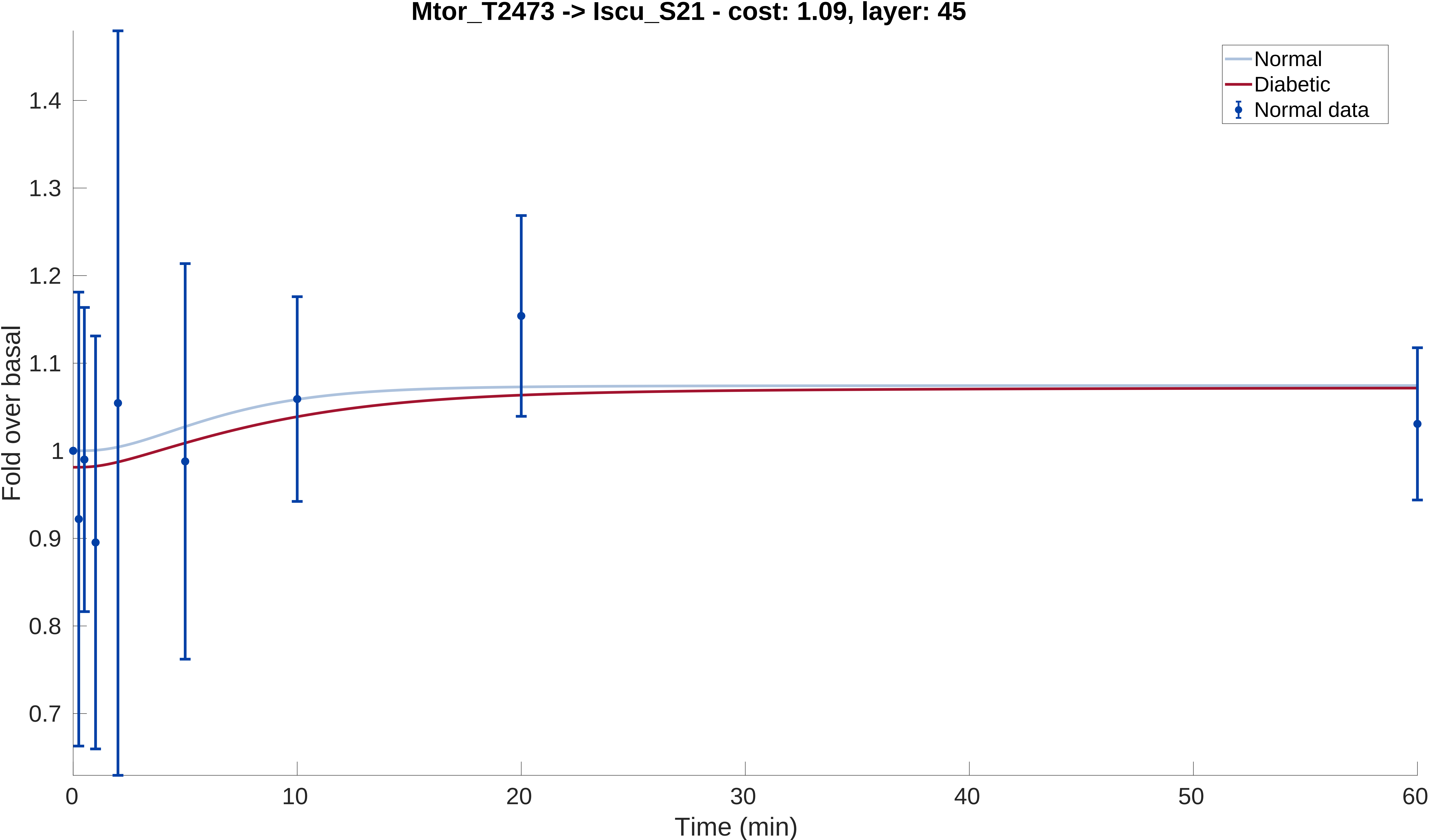

Mtor_T2473 -> Iscu_S21 - cost: 1.09, layer: 45
Normal
Diabetic
Normal data
Fold over basal
1.4
1.3
1.2
1.1
1
0.9
0.8
0.7
0
10
20
30
40
50
60
Time (min)

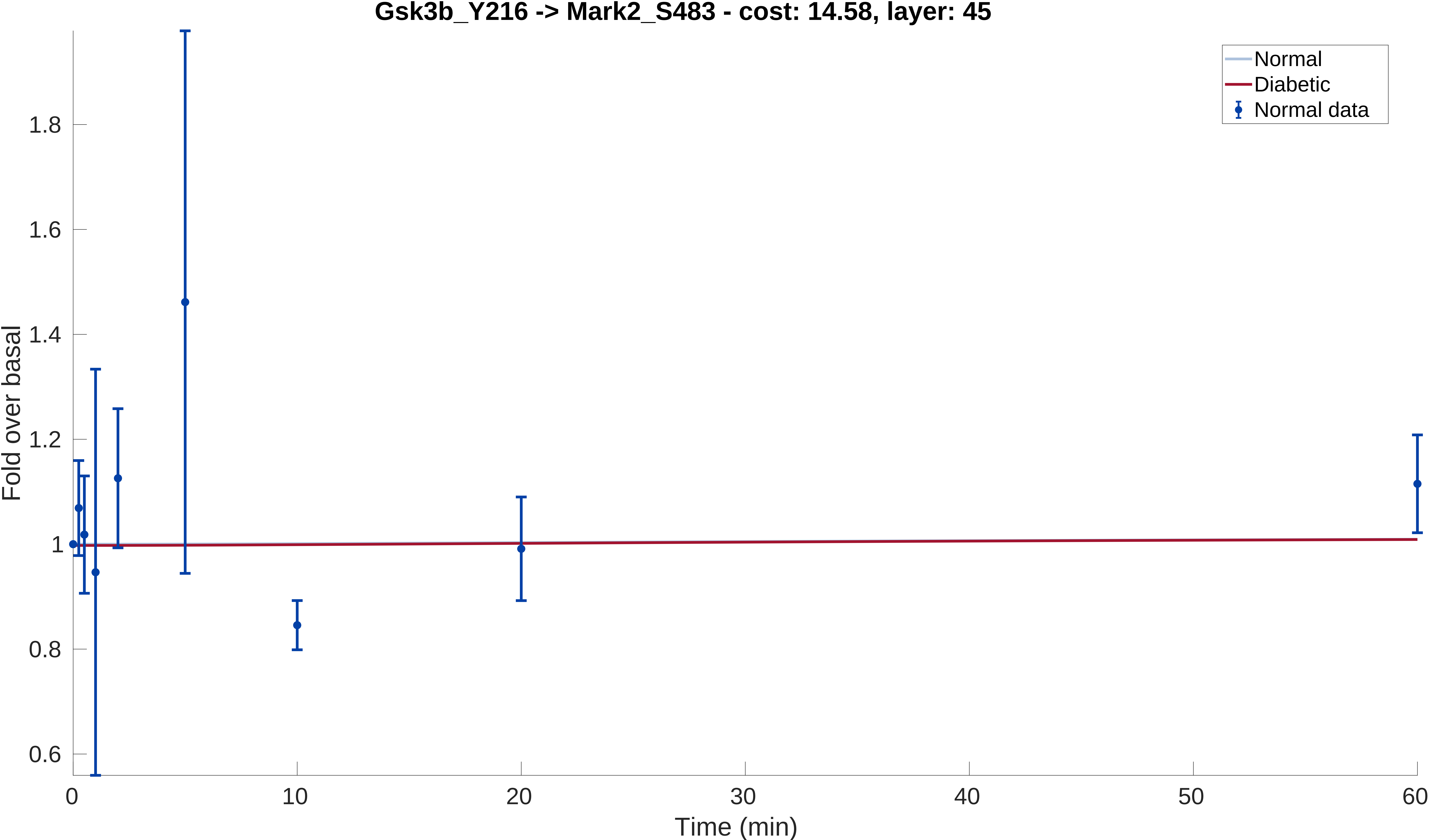

Gsk3b_Y216 -> Mark2_S483 - cost: 14.58, layer: 45
Normal
Diabetic
Normal data
Fold over basal
Time (min)
0.6
0.8
1
1.2
1.4
1.6
1.8
0
10
20
30
40
50
60

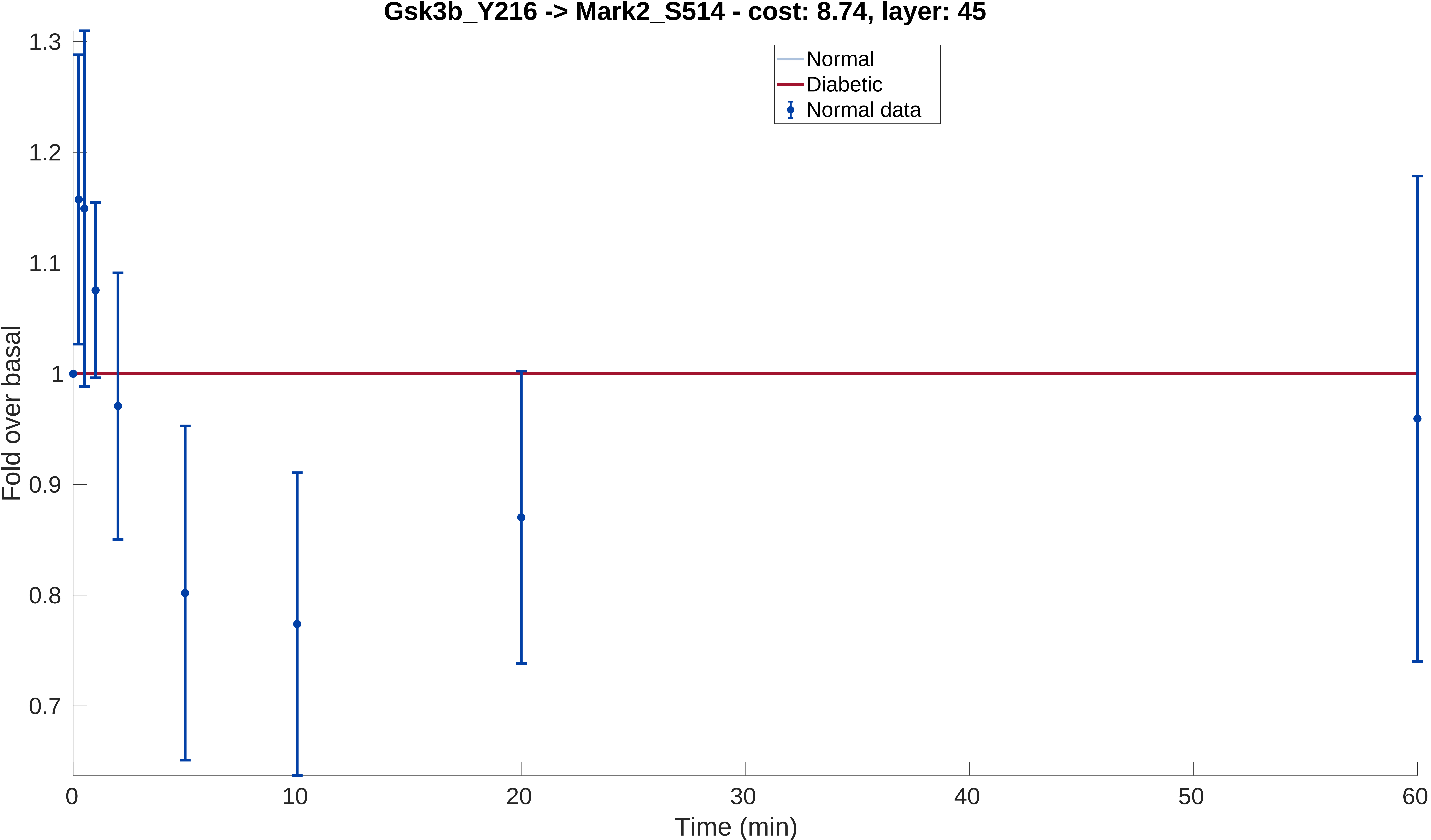

Gsk3b_Y216 -> Mark2_S514 - cost: 8.74, layer: 45
Normal
Diabetic
Normal data
Fold over basal
1.3
1.2
1.1
1
0.9
0.8
0.7
0
10
20
30
40
50
60
Time (min)

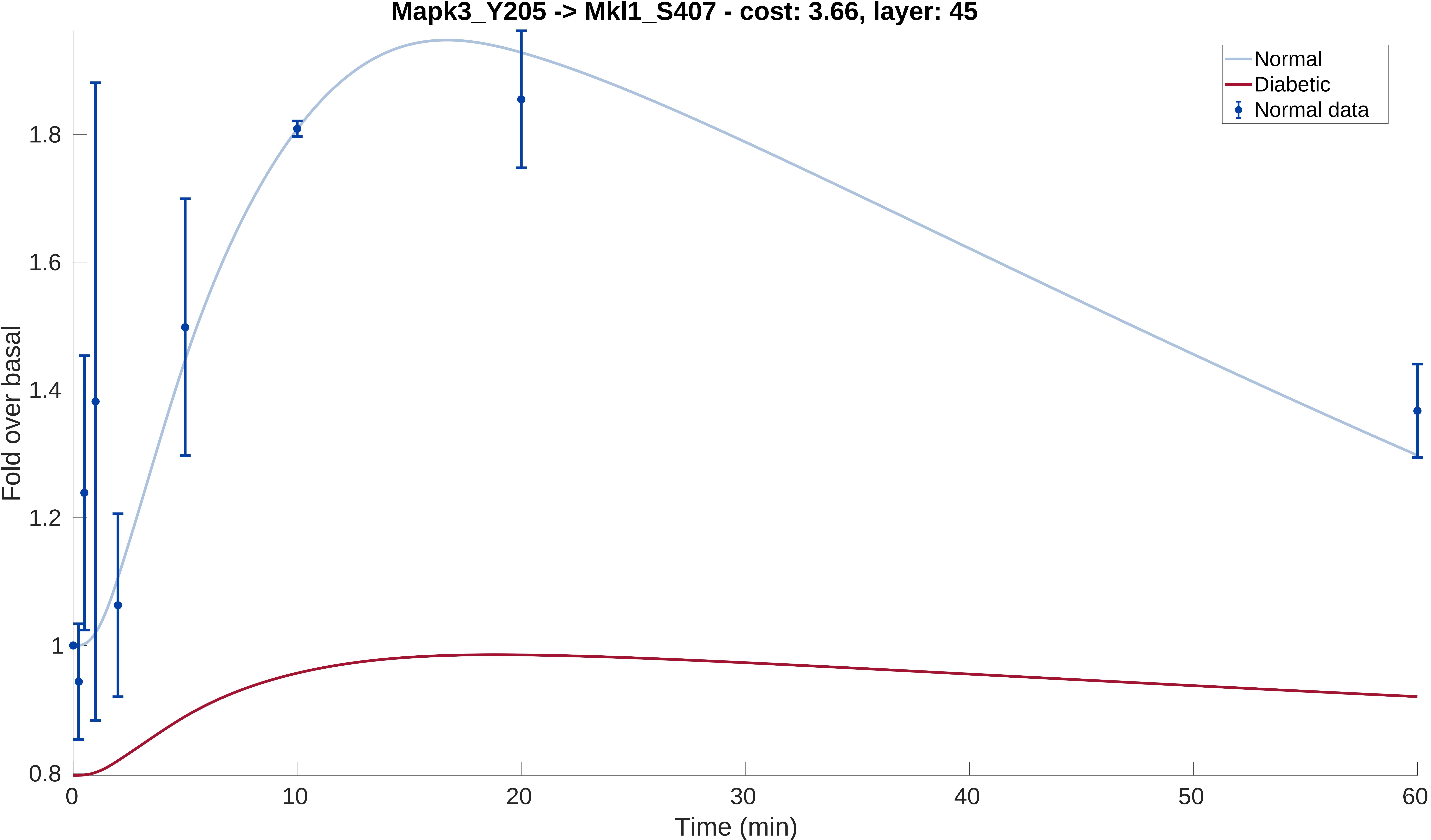

Mapk3_Y205 -> Mkl1_S407 - cost: 3.66, layer: 45
Normal
Diabetic
Normal data
Fold over basal
0.8
1
1.2
1.4
1.6
1.8
0
10
20
30
40
50
60
Time (min)

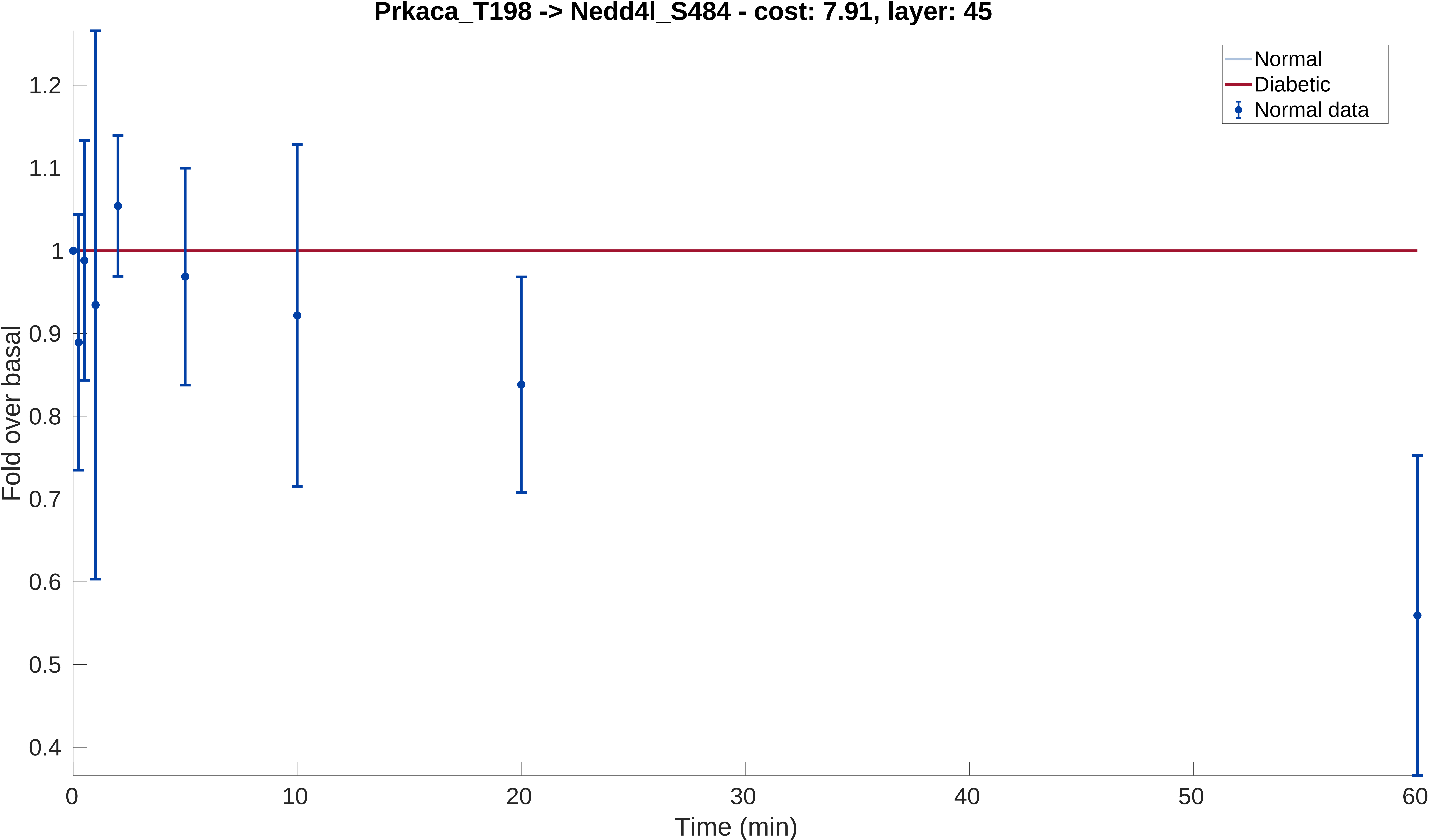

Prkaca_T198 -> Nedd4l_S484 - cost: 7.91, layer: 45
Normal
Diabetic
Normal data
Fold over basal
1.2
1.1
1
0.9
0.8
0.7
0.6
0.5
0.4
0
10
20
30
40
50
60
Time (min)

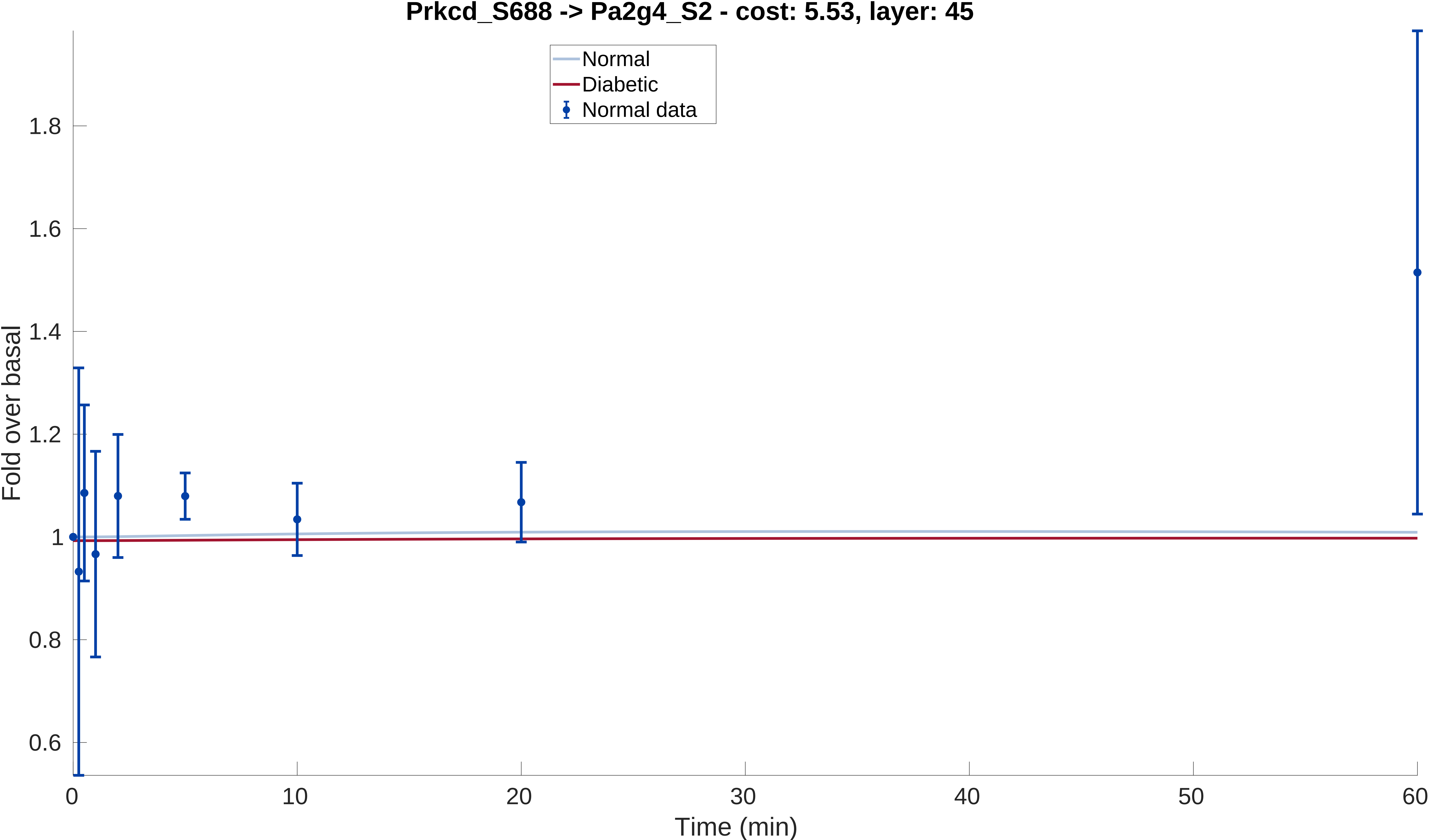

Prkcd_S688 -> Pa2g4_S2 - cost: 5.53, layer: 45
Normal
Diabetic
Normal data
Fold over basal
1.8
1.6
1.4
1.2
1
0.8
0.6
0
10
20
30
40
50
60
Time (min)

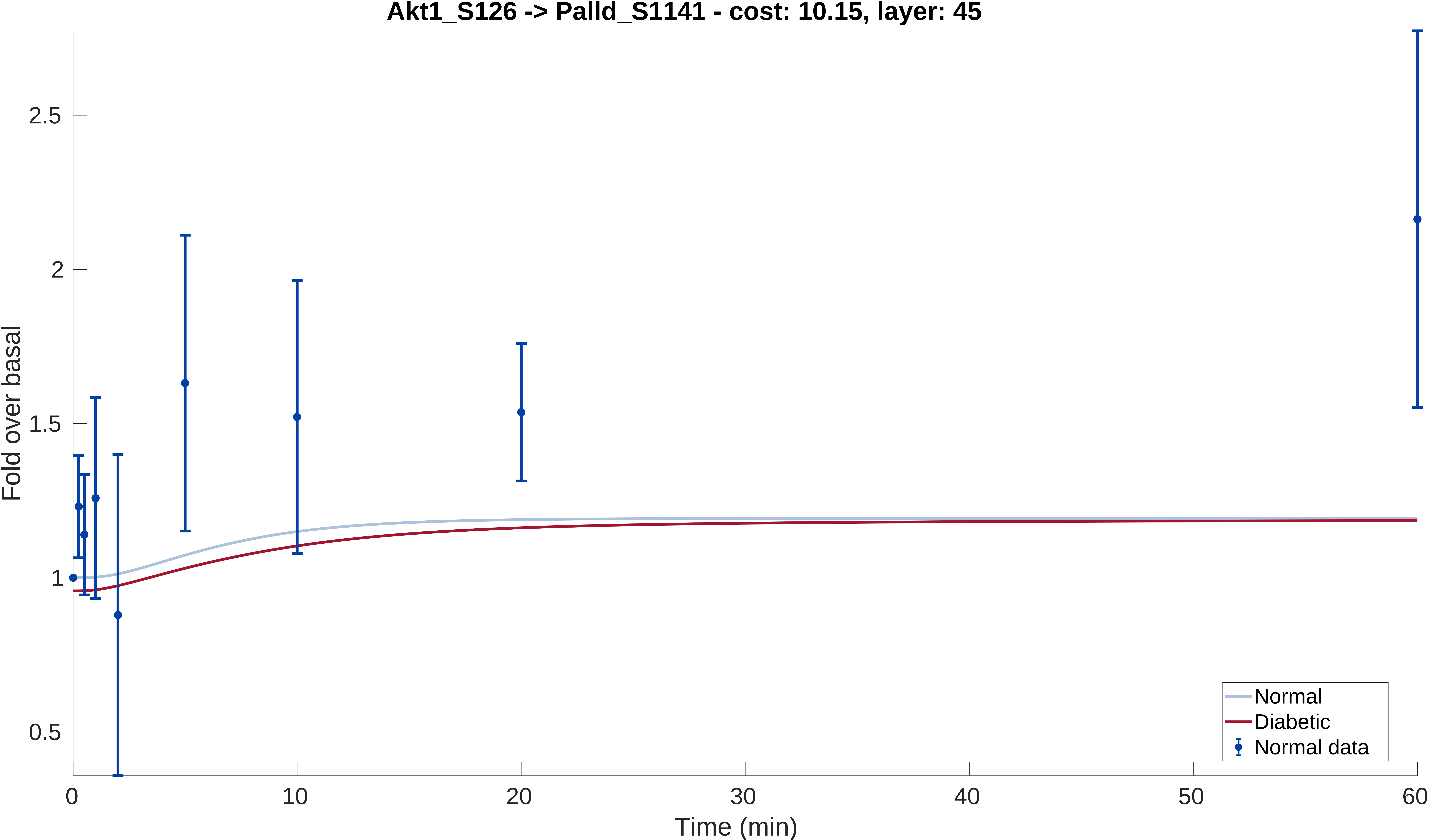
Akt1_S126 -> Palld_S1141 - cost: 10.15, layer: 45
Fold over basal
2.5
2
1.5
1
0.5
0
10
20
30
40
50
60
Time (min)
Normal
Diabetic
Normal data

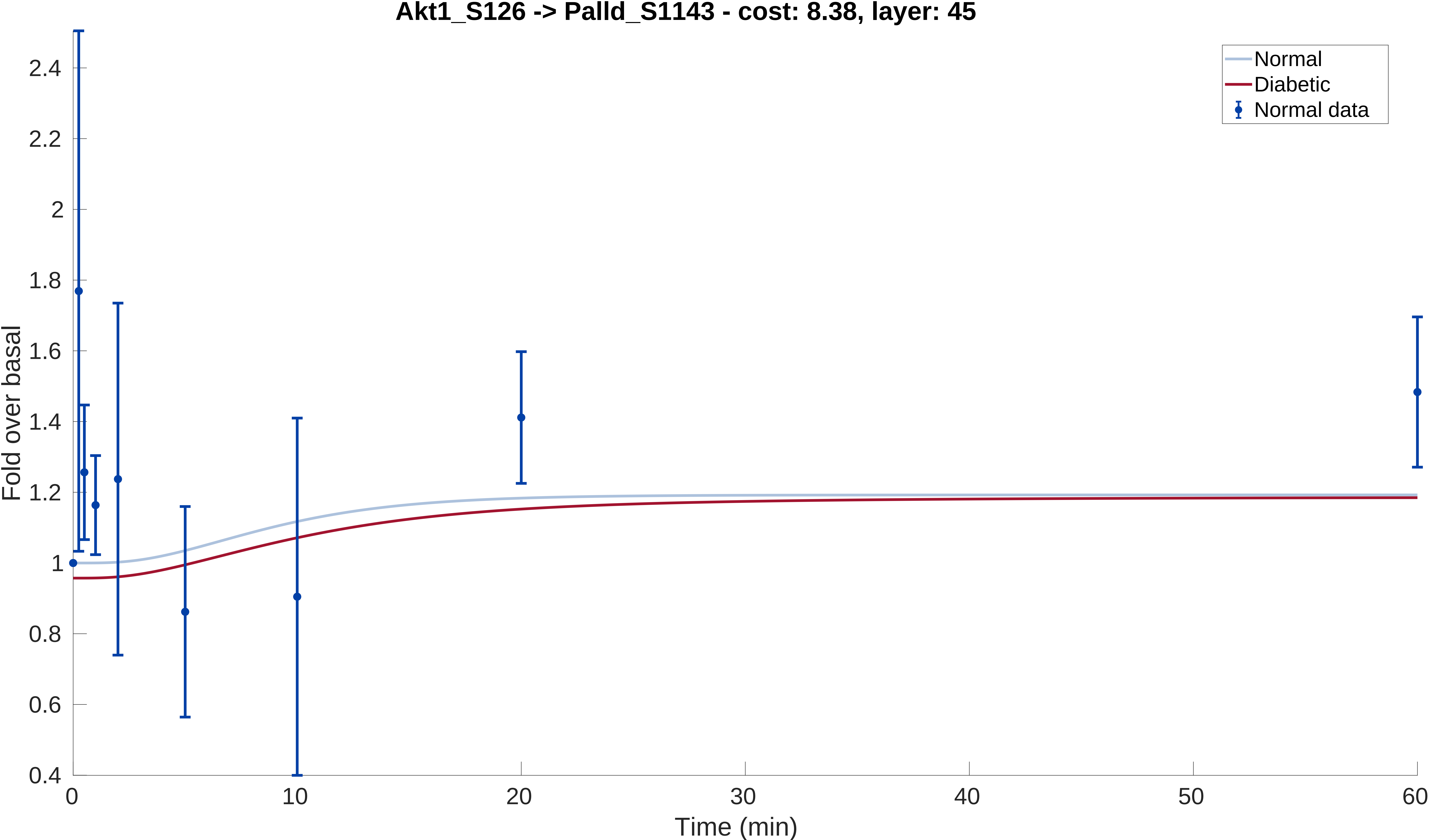

Akt1_S126 -> Palld_S1143 - cost: 8.38, layer: 45
Normal
Diabetic
Normal data
Fold over basal
2.4
2.2
2
1.8
1.6
1.4
1.2
1
0.8
0.6
0.4
0
10
20
30
40
50
60
Time (min)

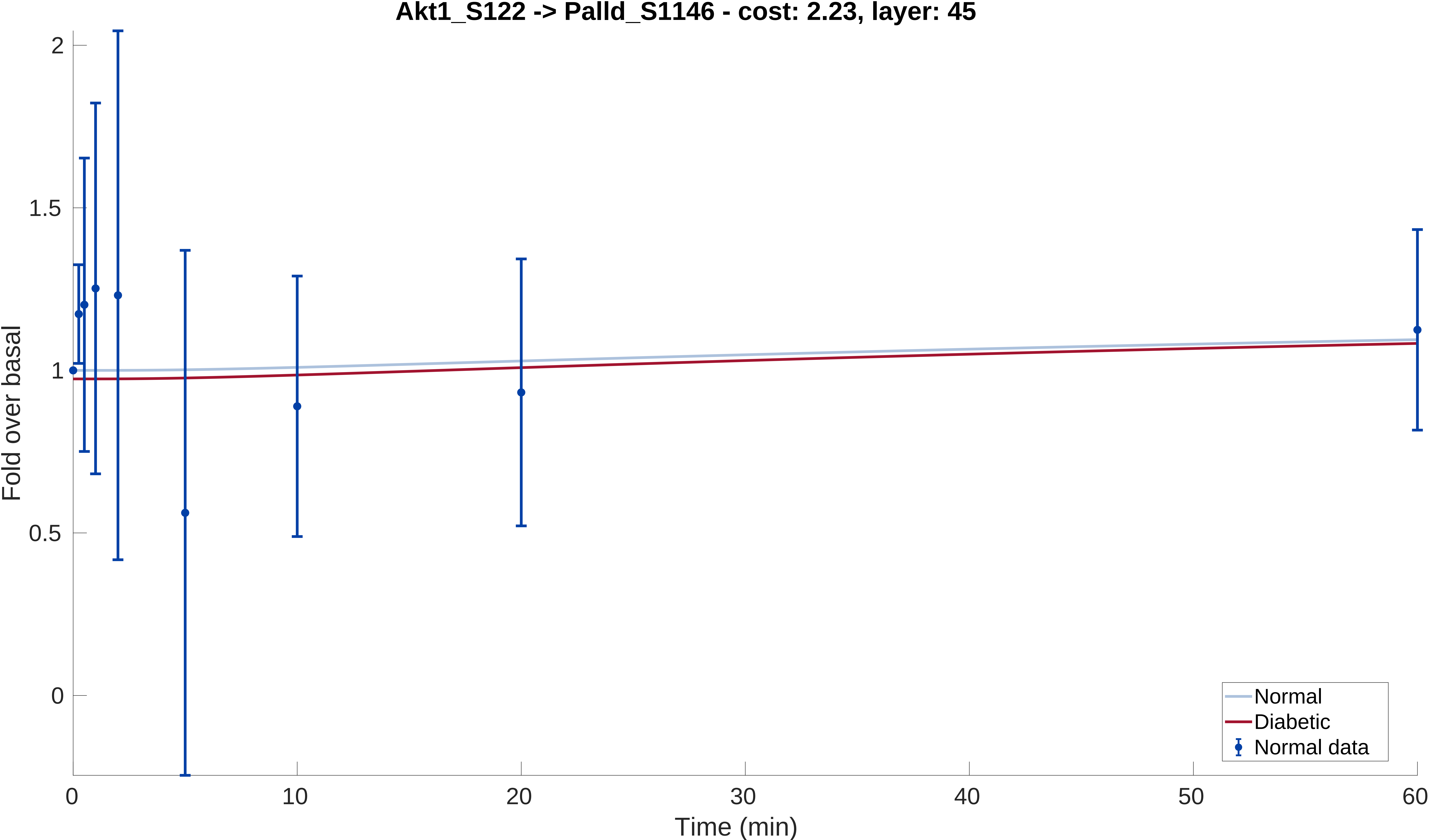

Akt1_S122 -> Palld_S1146 - cost: 2.23, layer: 45
Fold over basal
Time (min)
0
0.5
1
1.5
2
0
10
20
30
40
50
60
Normal
Diabetic
Normal data

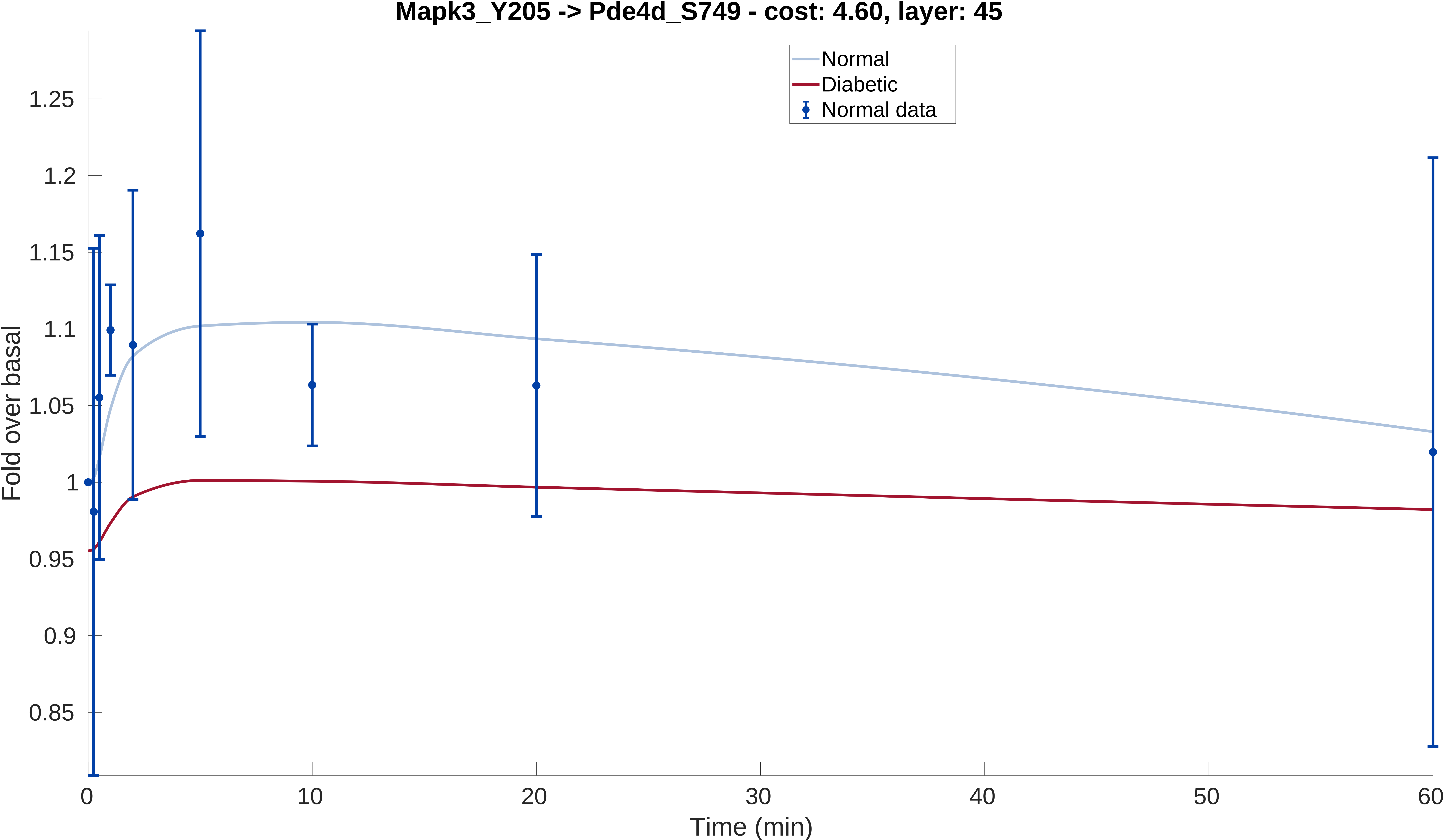

Mapk3_Y205 -> Pde4d_S749 - cost: 4.60, layer: 45
Normal
Diabetic
Normal data
Fold over basal
Time (min)
1.25
1.2
1.15
1.1
1.05
1
0.95
0.9
0.85
0
10
20
30
40
50
60

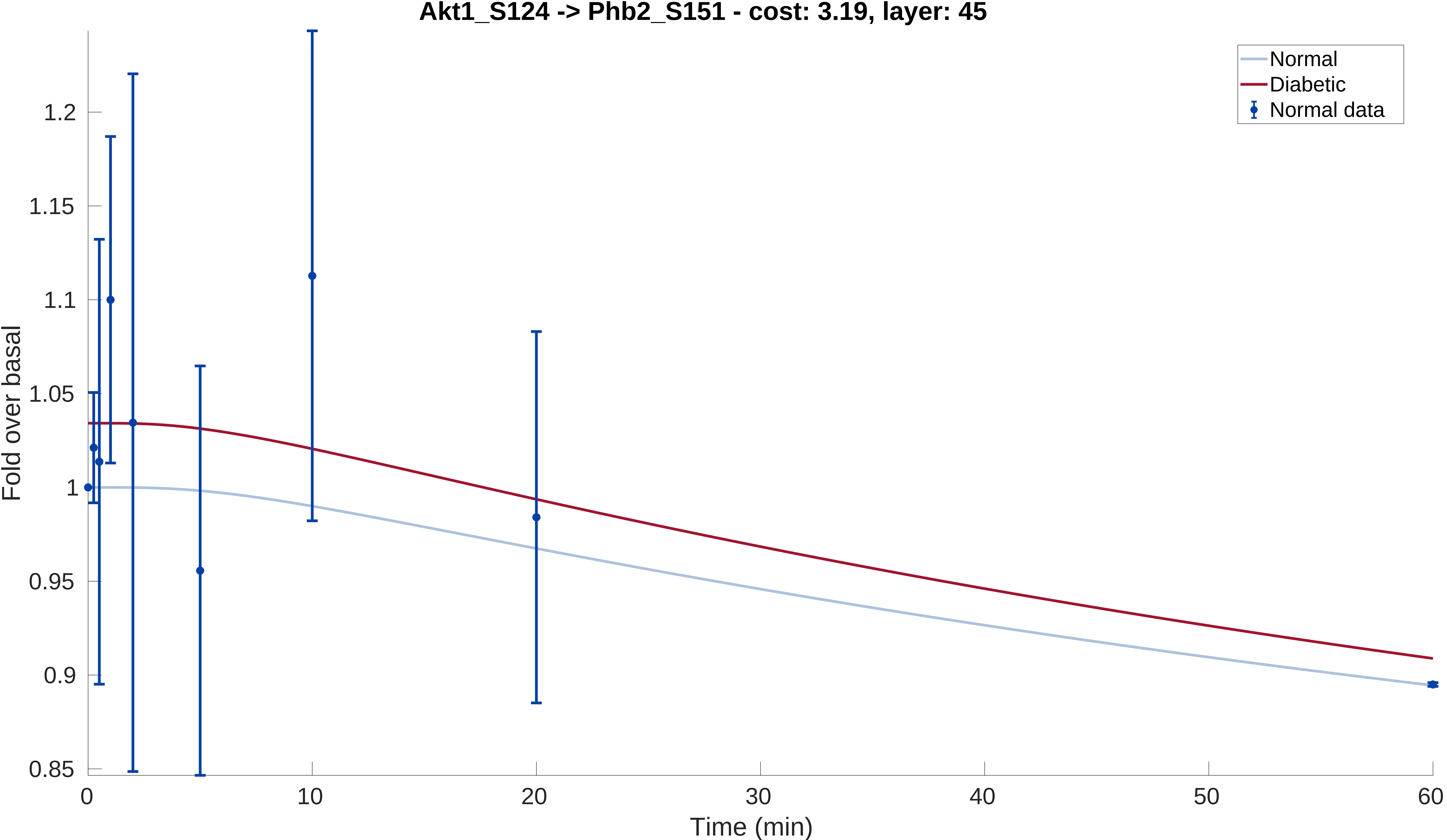
Akt1_S124 -> Phb2_S151 - cost: 3.19, layer: 45
Normal
Diabetic
Normal data
Fold over basal
1.2
1.15
1.1
1.05
1
0.95
0.9
0.85
0
10
20
30
40
50
60
Time (min)

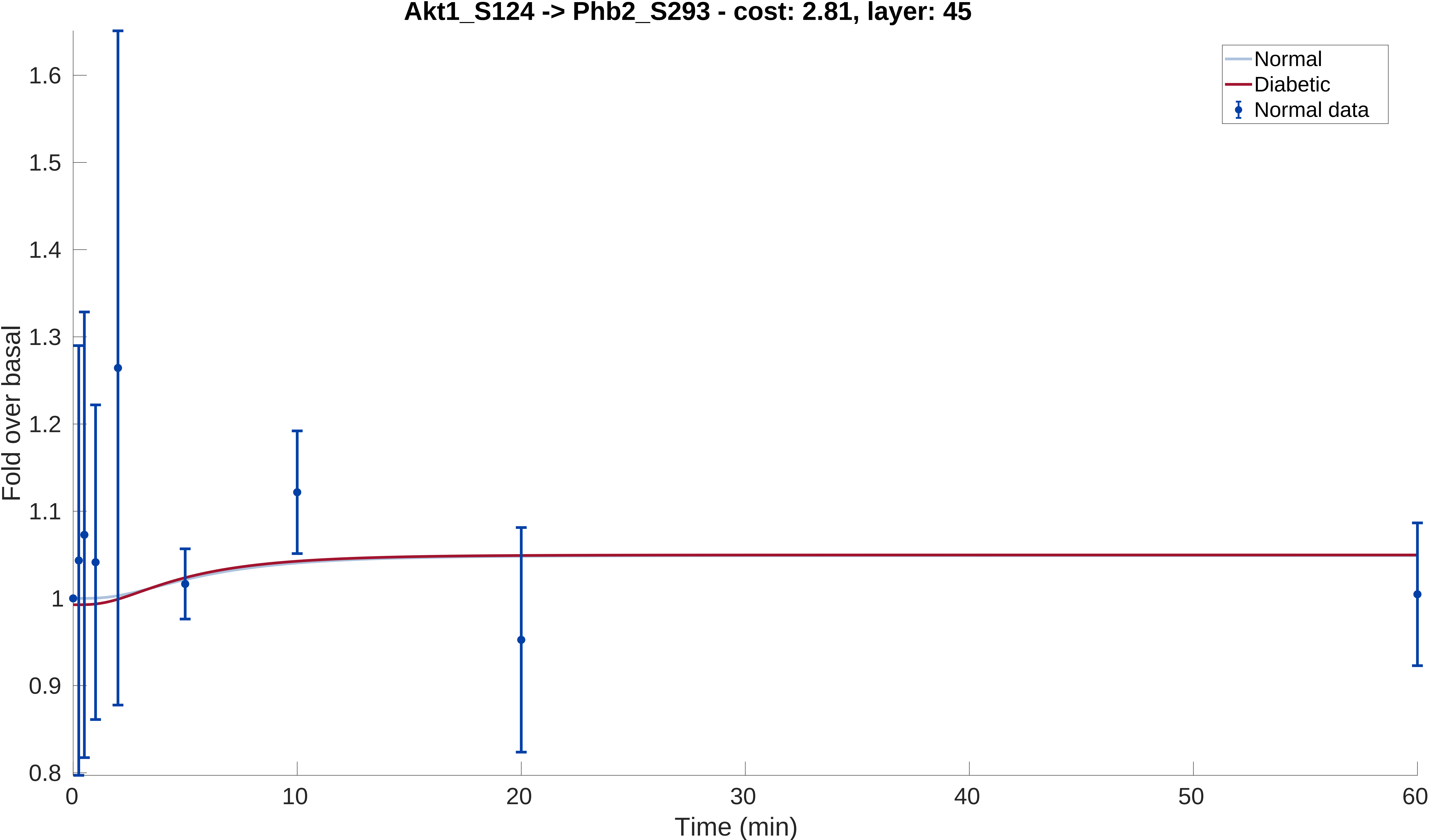

Akt1_S124 -> Phb2_S293 - cost: 2.81, layer: 45
Normal
Diabetic
Normal data
Fold over basal
1.6
1.5
1.4
1.3
1.2
1.1
1
0.9
0.8
0
10
20
30
40
50
60
Time (min)

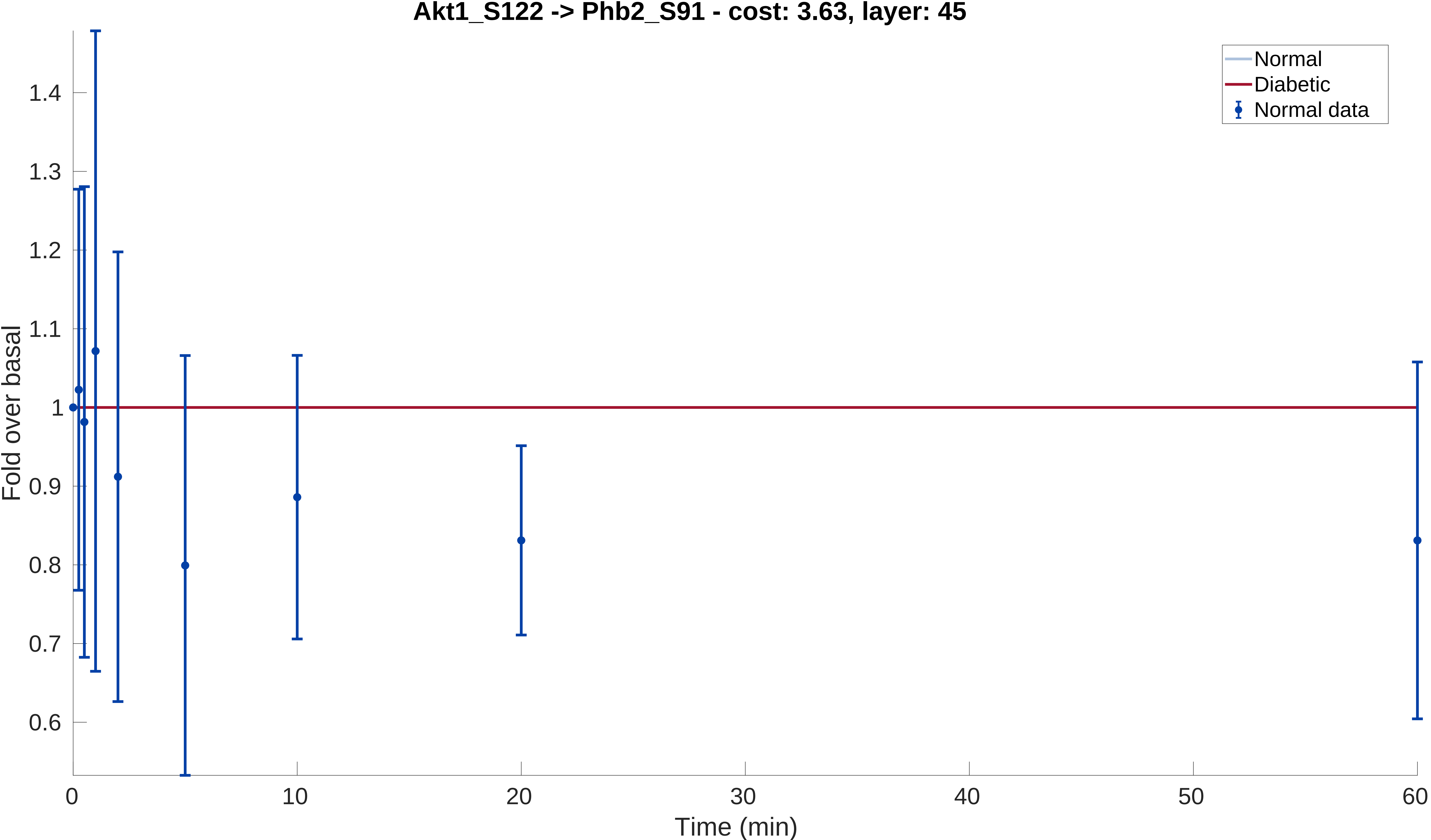
Akt1_S122 -> Phb2_S91 - cost: 3.63, layer: 45
Normal
Diabetic
Normal data
Fold over basal
Time (min)
1.4
1.3
1.2
1.1
1
0.9
0.8
0.7
0.6
0
10
20
30
40
50
60

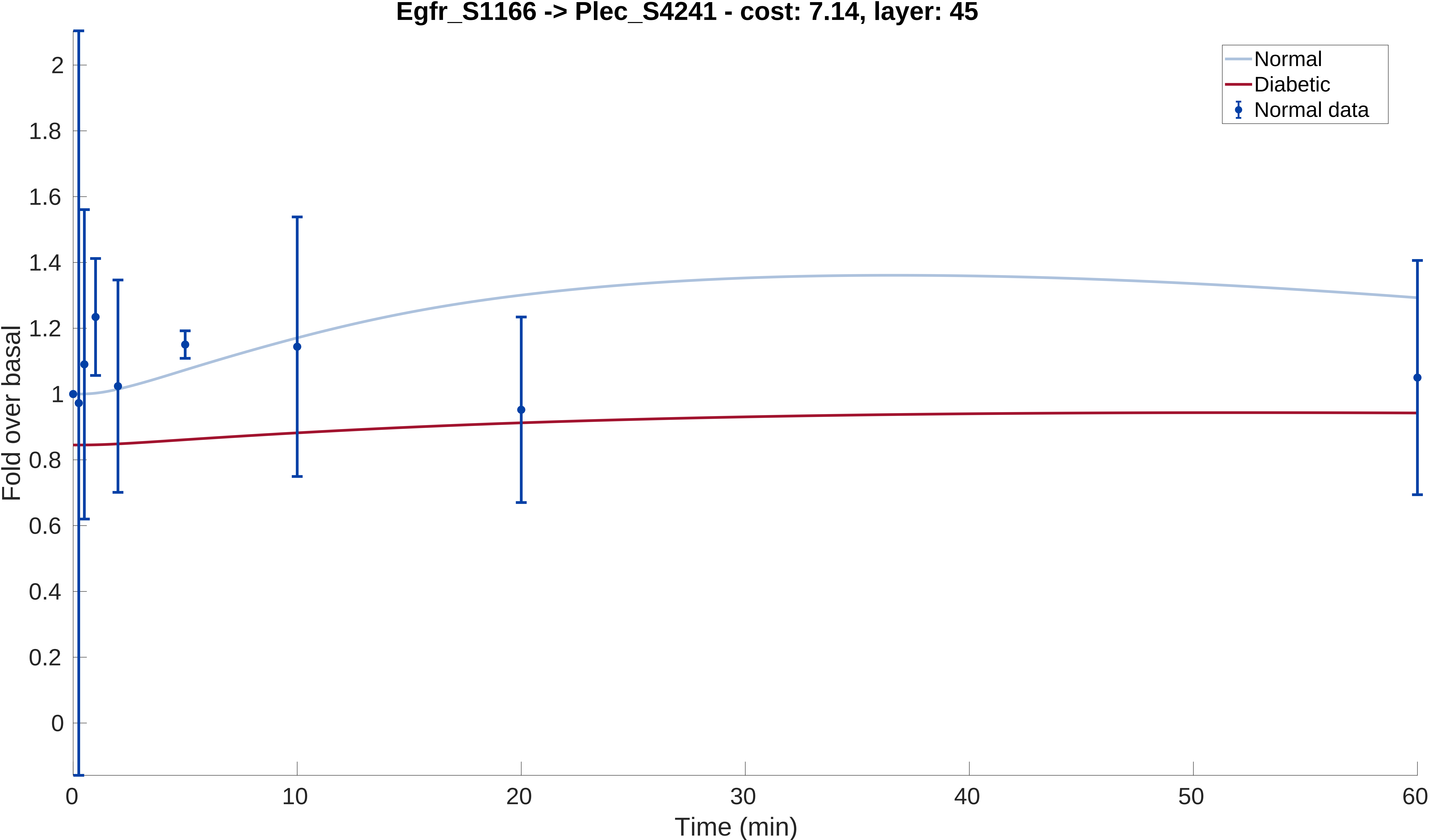

Egfr_S1166 -> Plec_S4241 - cost: 7.14, layer: 45
Normal
Diabetic
Normal data
Fold over basal
Time (min)
0
0.2
0.4
0.6
0.8
1
1.2
1.4
1.6
1.8
2
0
10
20
30
40
50
60

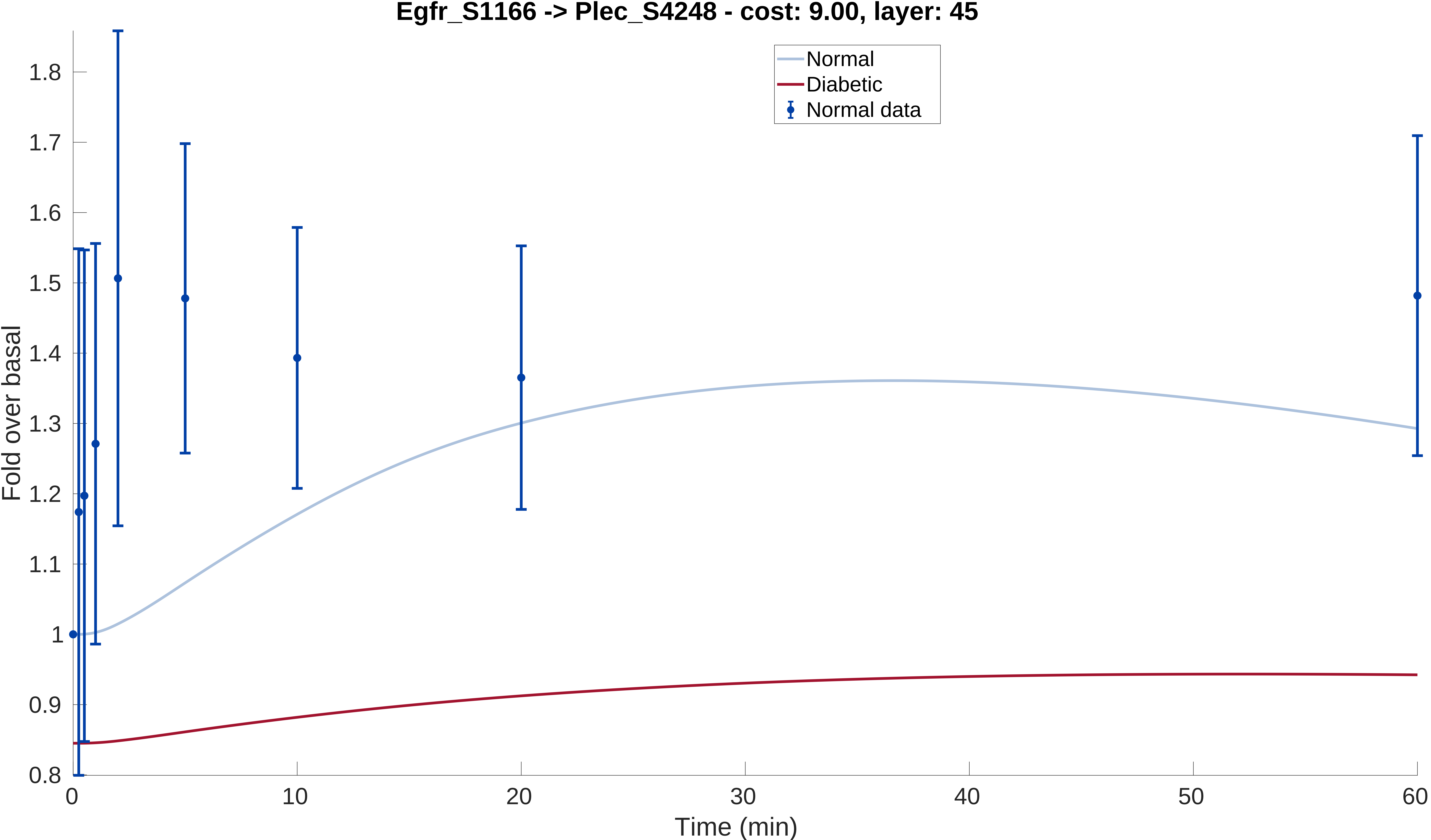

Egfr_S1166 -> Plec_S4248 - cost: 9.00, layer: 45
Normal
Diabetic
Normal data
Fold over basal
1.8
1.7
1.6
1.5
1.4
1.3
1.2
1.1
1
0.9
0.8
0
10
20
30
40
50
60
Time (min)

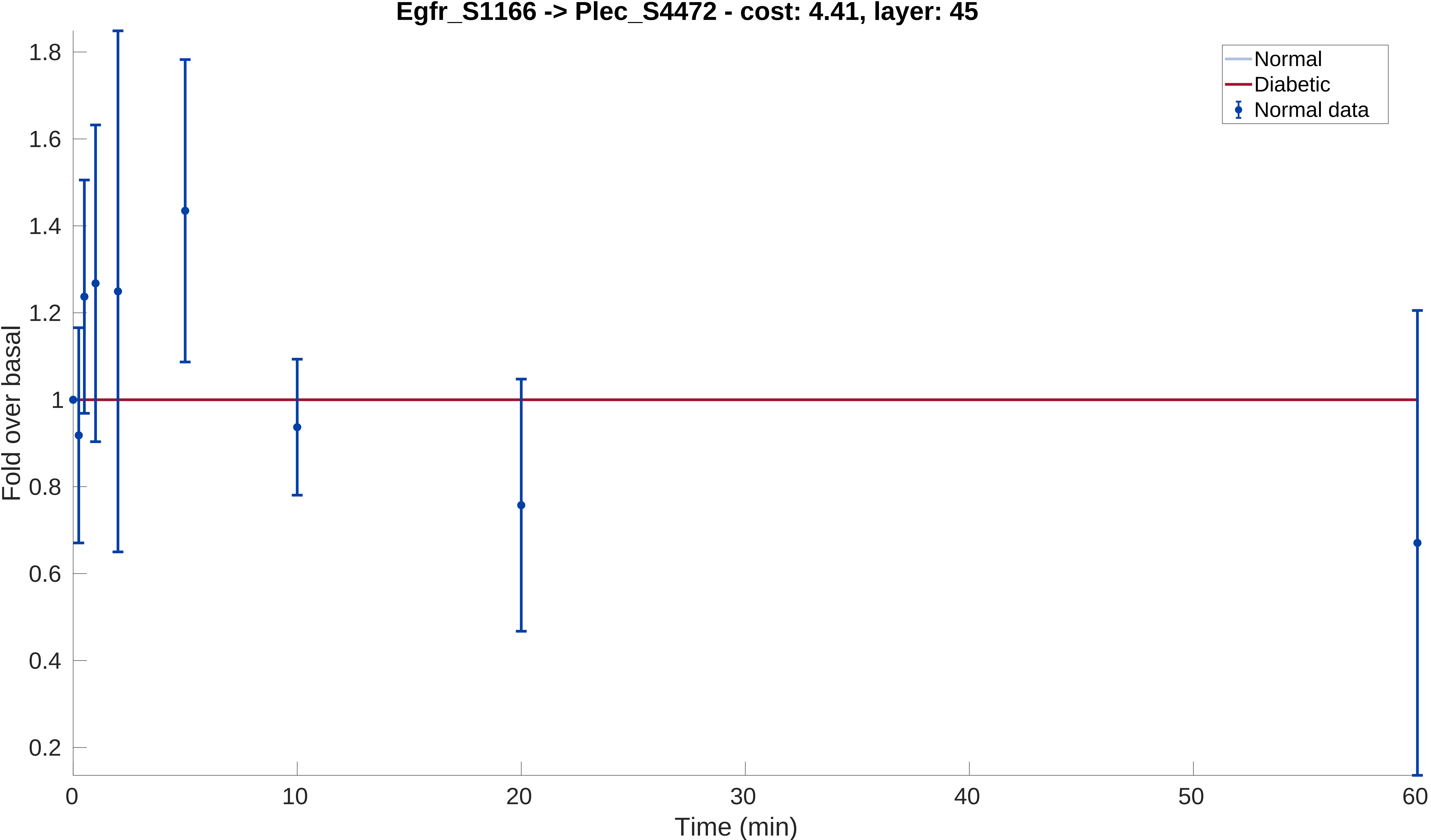
Egfr_S1166 -> Plec_S4472 - cost: 4.41, layer: 45
Normal
Diabetic
Normal data
Fold over basal
1.8
1.6
1.4
1.2
1
0.8
0.6
0.4
0.2
0
10
20
30
40
50
60
Time (min)

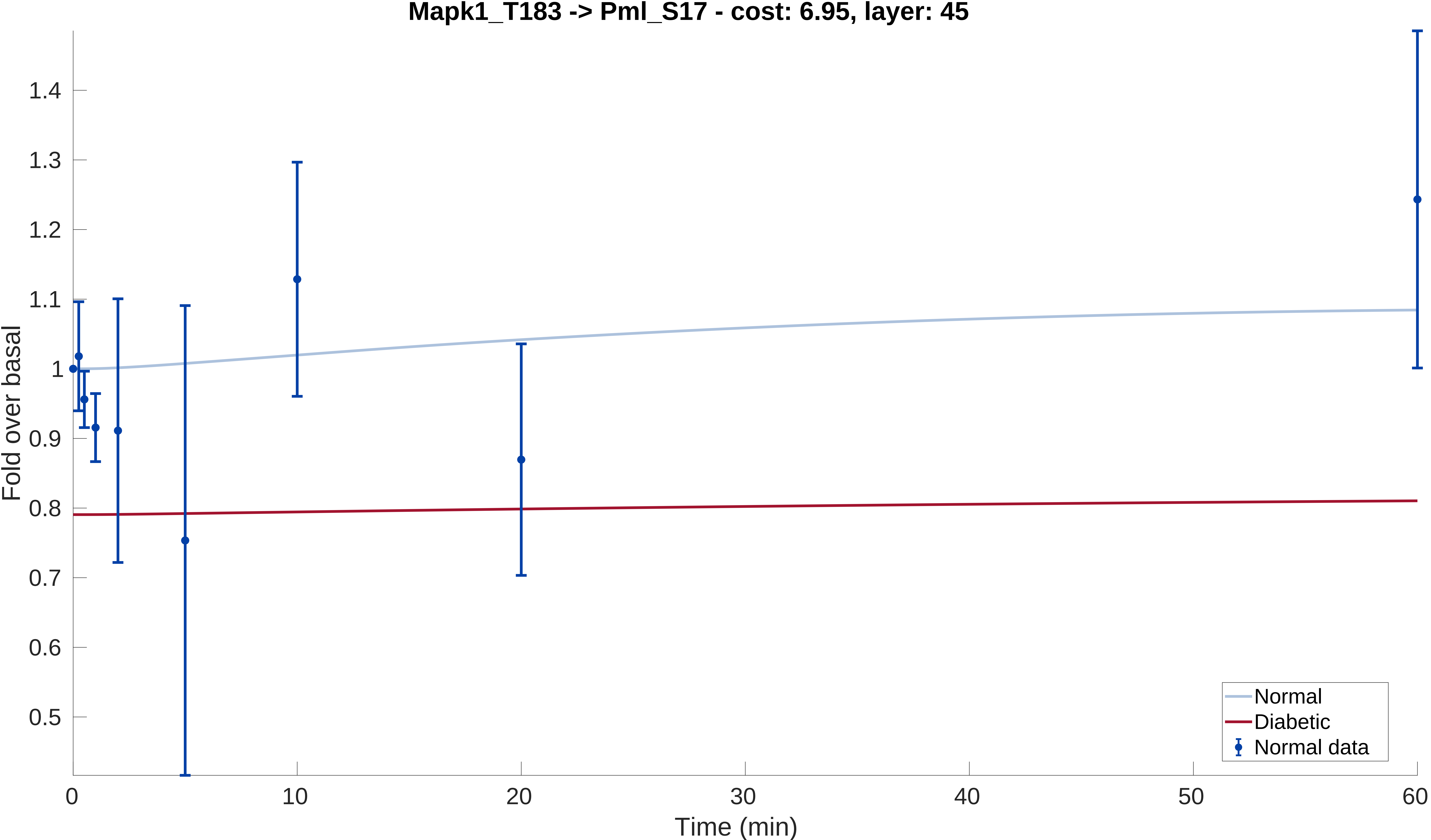
Mapk1_T183 -> Pml_S17 - cost: 6.95, layer: 45
Fold over basal
1.4
1.3
1.2
1.1
1
0.9
0.8
0.7
0.6
0.5
0
10
20
30
40
50
60
Time (min)
Normal
Diabetic
Normal data

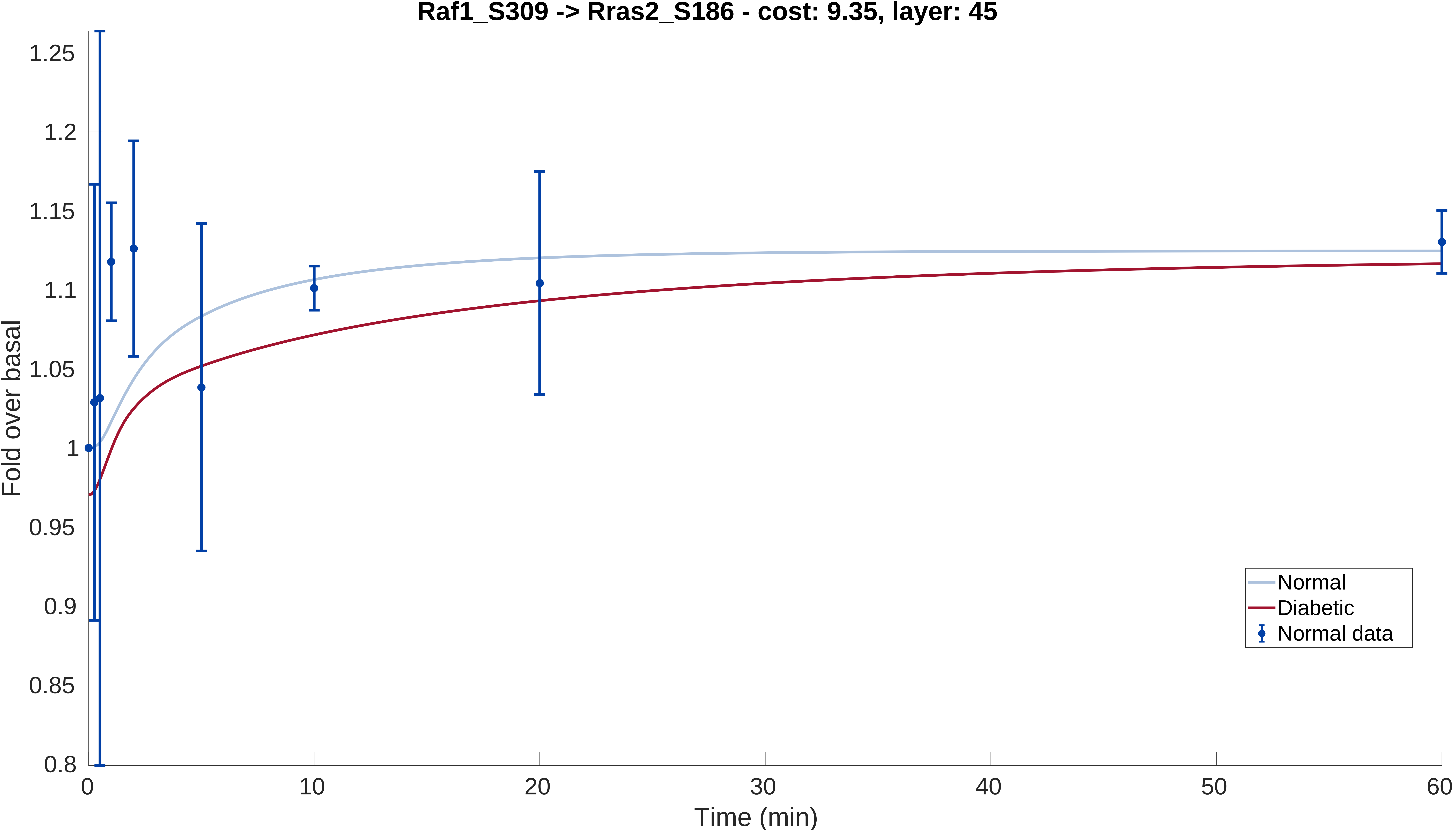

Raf1_S309 -> Rras2_S186 - cost: 9.35, layer: 45
Fold over basal
Time (min)
1.25
1.2
1.15
1.1
1.05
1
0.95
0.9
0.85
0.8
0
10
20
30
40
50
60
Normal
Diabetic
Normal data

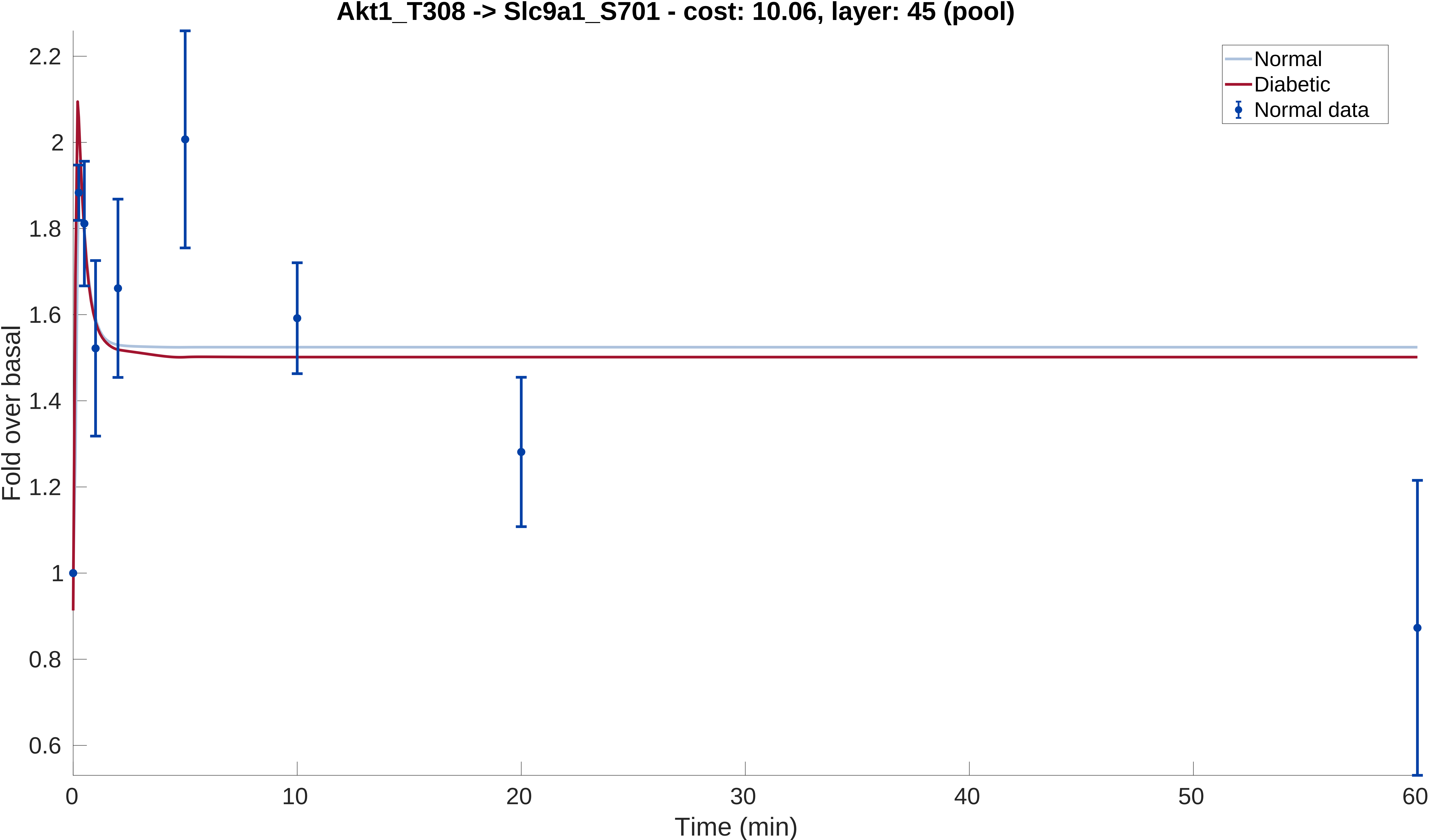

Akt1_T308 -> Slc9a1_S701 - cost: 10.06, layer: 45 (pool)
Normal
Diabetic
Normal data
Fold over basal
2.2
2
1.8
1.6
1.4
1.2
1
0.8
0.6
0
10
20
30
40
50
60
Time (min)

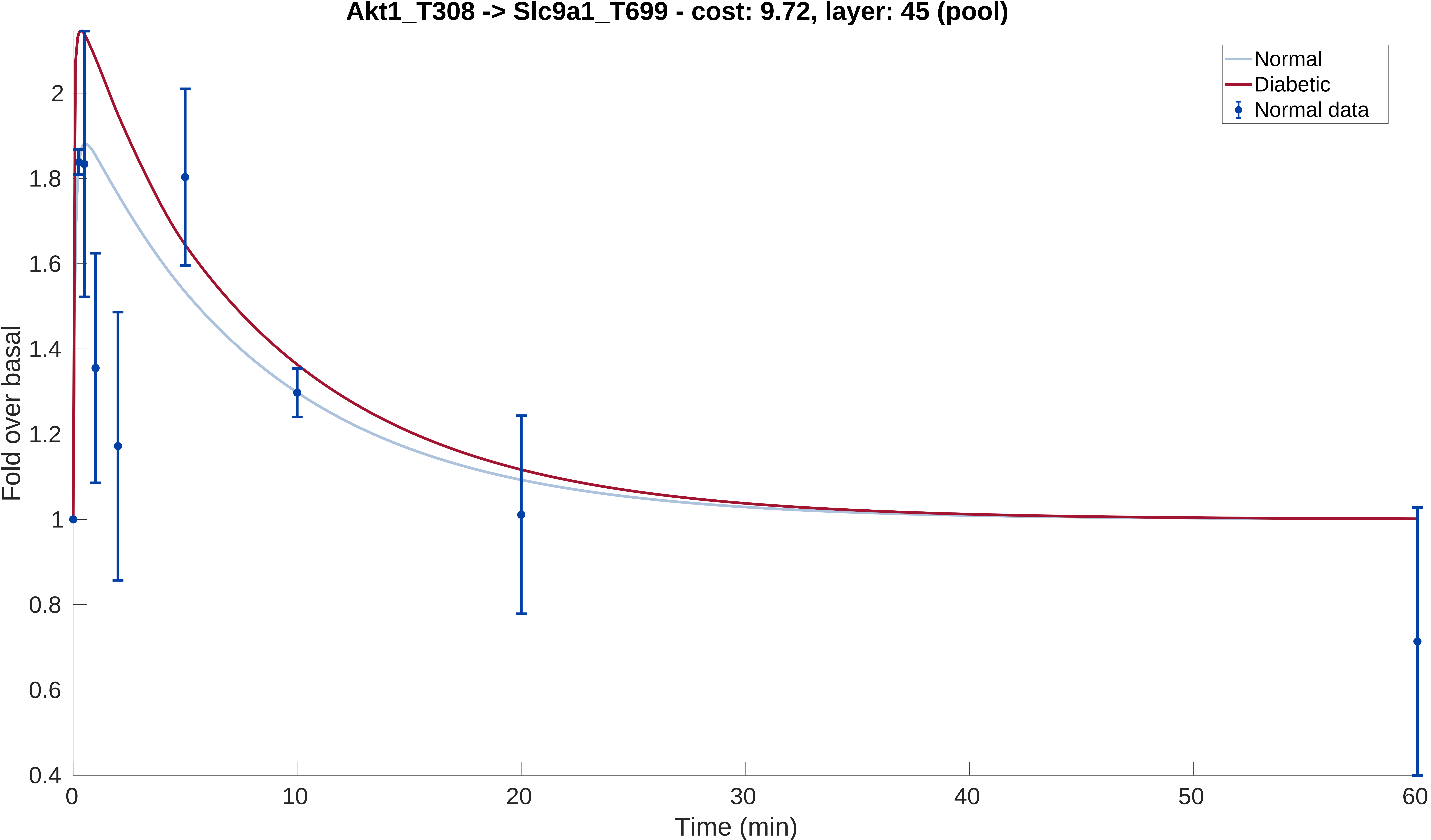

Akt1_T308 -> Slc9a1_T699 - cost: 9.72, layer: 45 (pool)
Normal
Diabetic
Normal data
Fold over basal
Time (min)
0.4
0.6
0.8
1
1.2
1.4
1.6
1.8
2
0
10
20
30
40
50
60

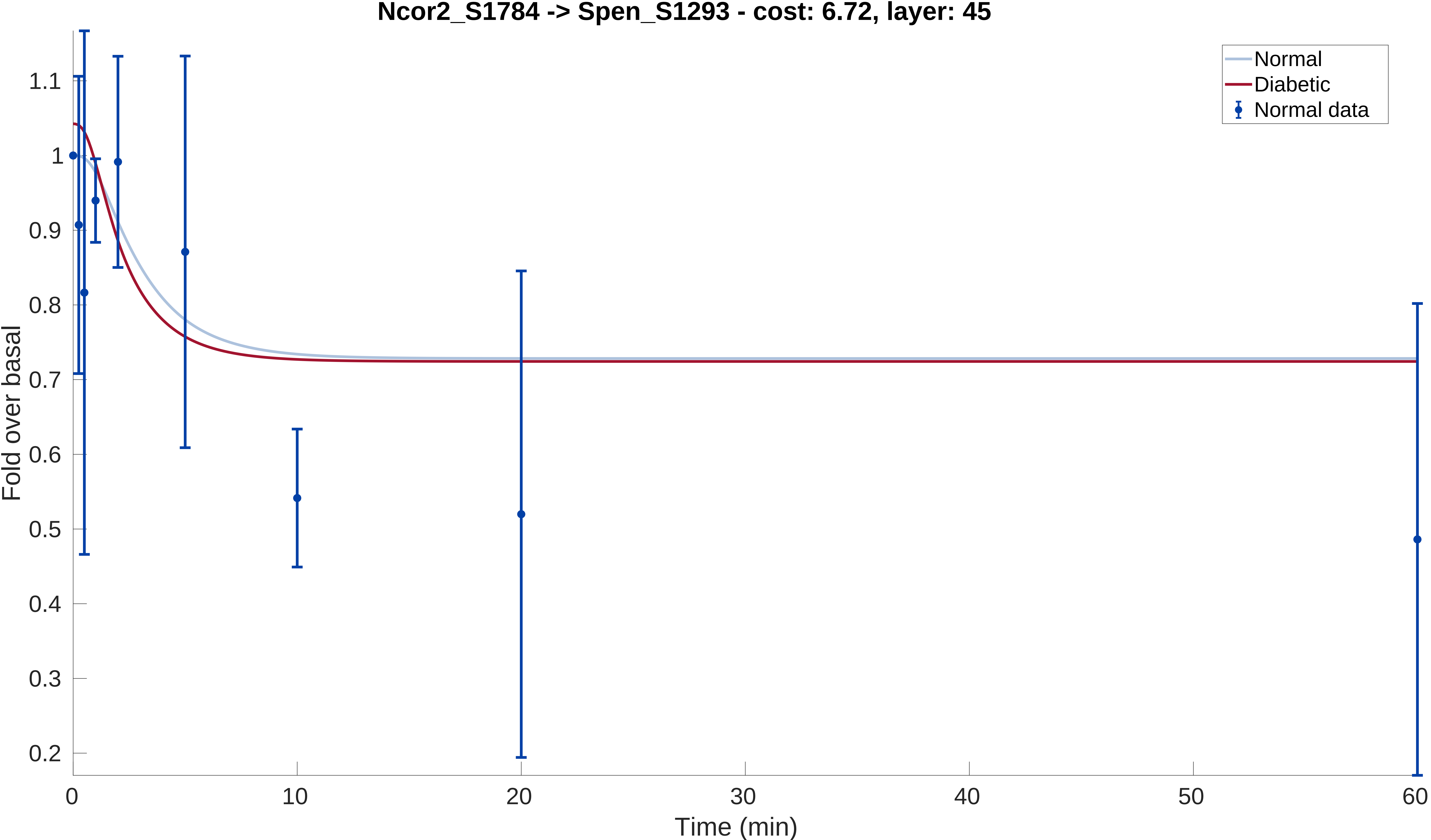

Ncor2_S1784 -> Spen_S1293 - cost: 6.72, layer: 45
Normal
Diabetic
Normal data
Fold over basal
Time (min)
0.2
0.3
0.4
0.5
0.6
0.7
0.8
0.9
1
1.1
0
10
20
30
40
50
60

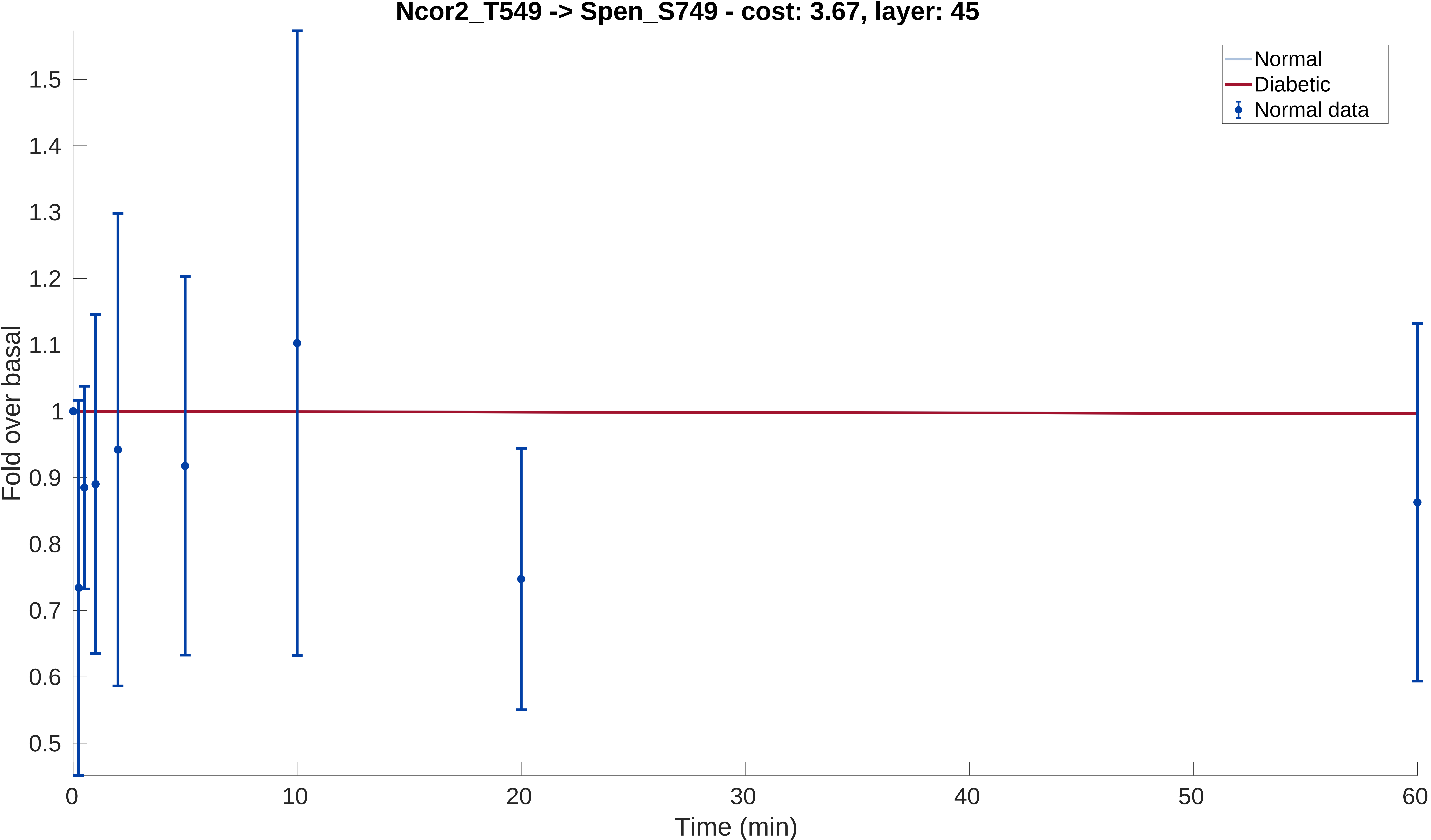

Ncor2_T549 -> Spen_S749 - cost: 3.67, layer: 45
Normal
Diabetic
Normal data
Fold over basal
1.5
1.4
1.3
1.2
1.1
1
0.9
0.8
0.7
0.6
0.5
0
10
20
30
40
50
60
Time (min)

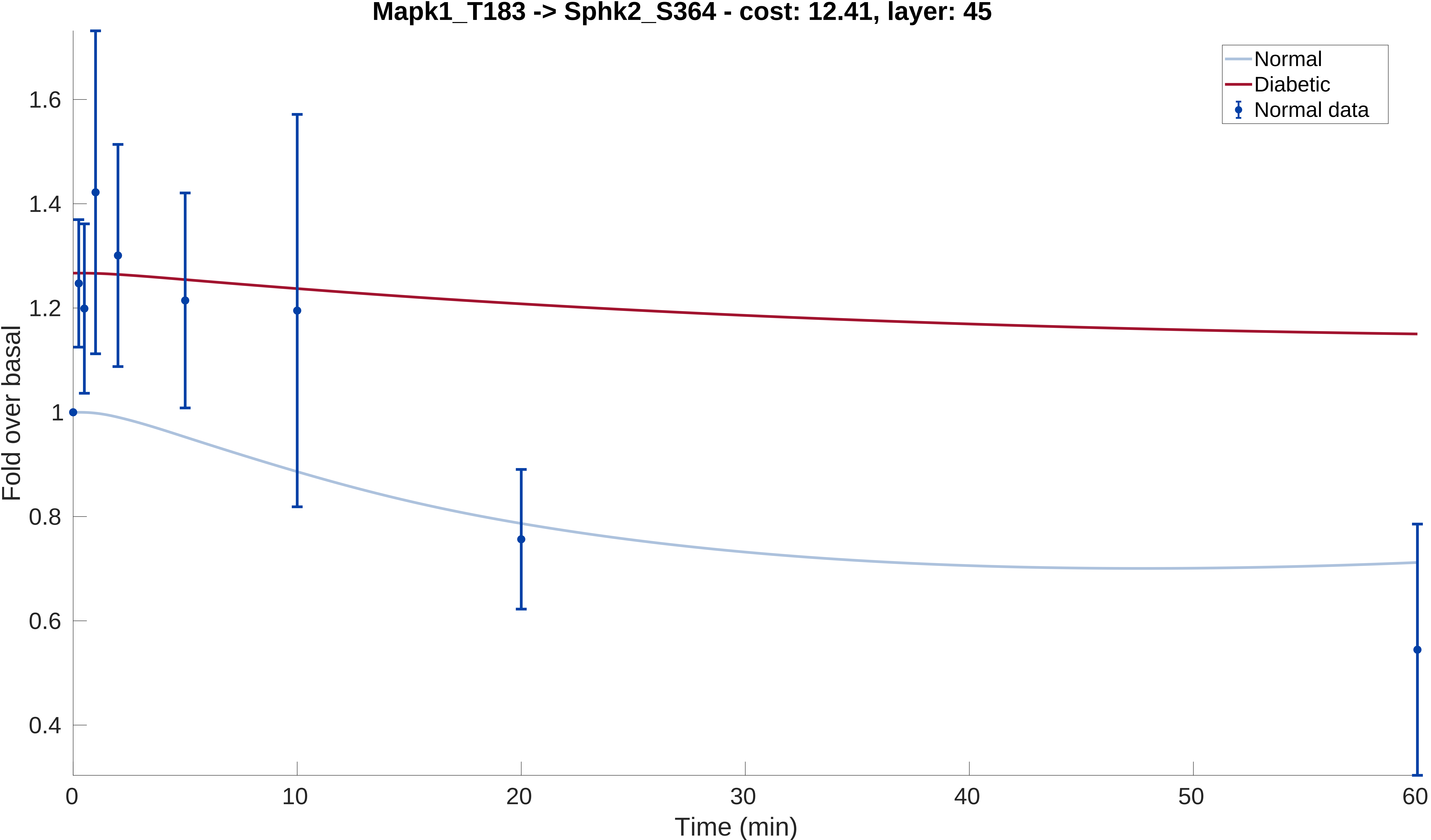

Mapk1_T183 -> Sphk2_S364 - cost: 12.41, layer: 45
Normal
Diabetic
Normal data
1.6
1.4
1.2
1
0.8
0.6
0.4
Fold over basal
0
10
20
30
40
50
60
Time (min)

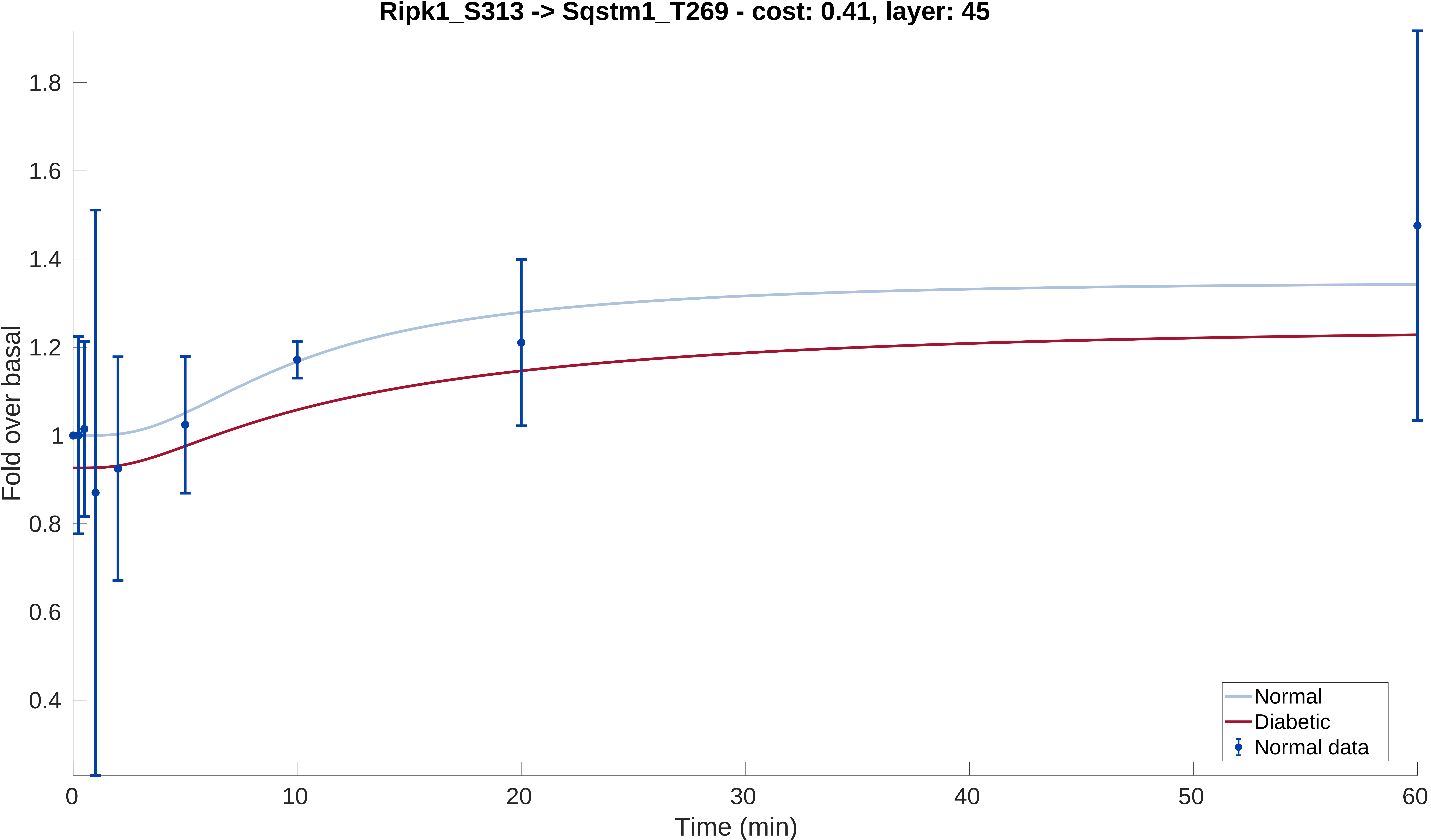

Ripk1_S313 -> Sqstm1_T269 - cost: 0.41, layer: 45
Fold over basal
1.8
1.6
1.4
1.2
1
0.8
0.6
0.4
0
10
20
30
40
50
60
Time (min)
Normal
Diabetic
Normal data

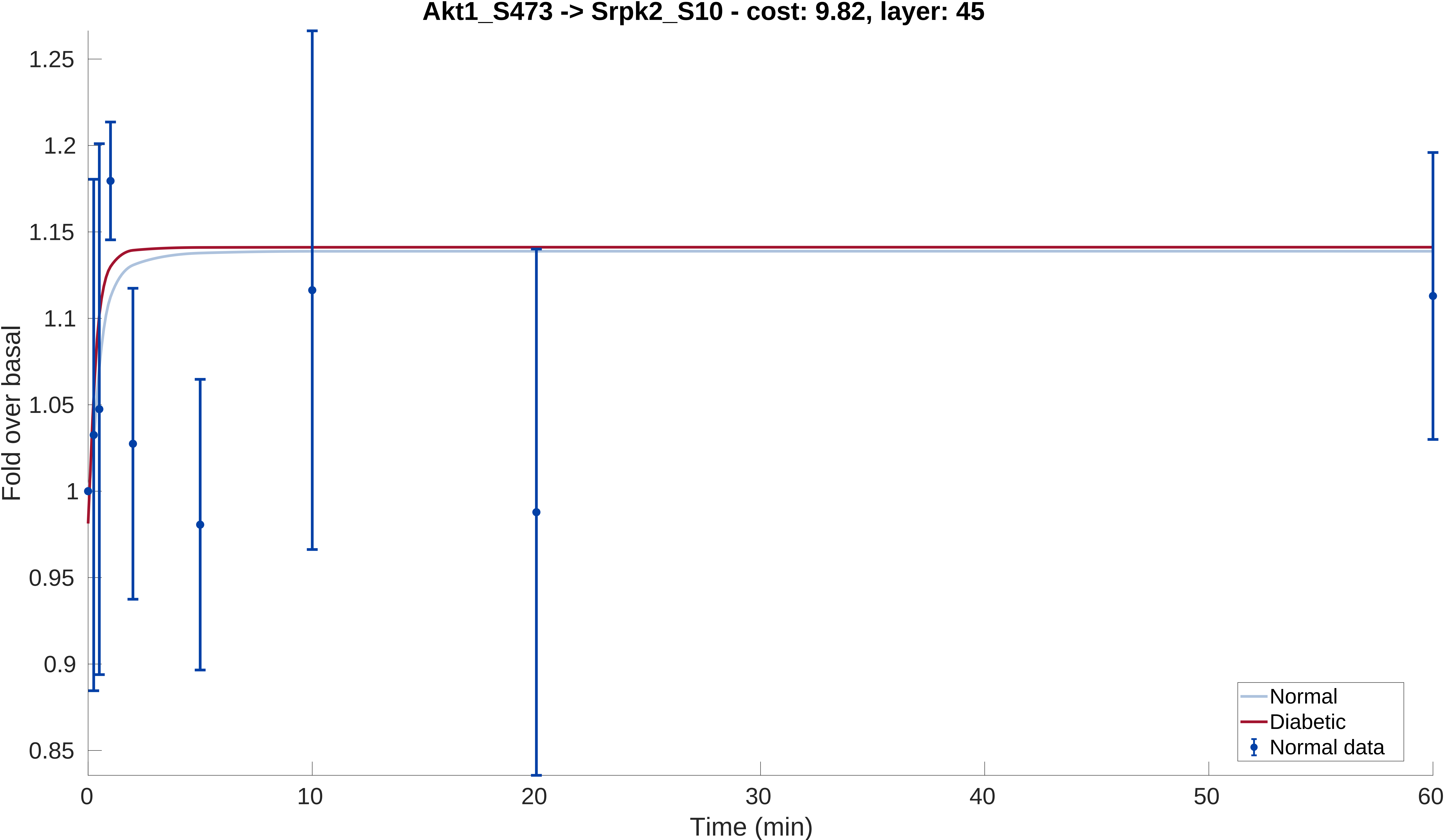

Akt1_S473 -> Srpk2_S10 - cost: 9.82, layer: 45
Fold over basal
1.25
1.2
1.15
1.1
1.05
1
0.95
0.9
0.85
0
10
20
30
40
50
60
Time (min)
Normal
Diabetic
Normal data

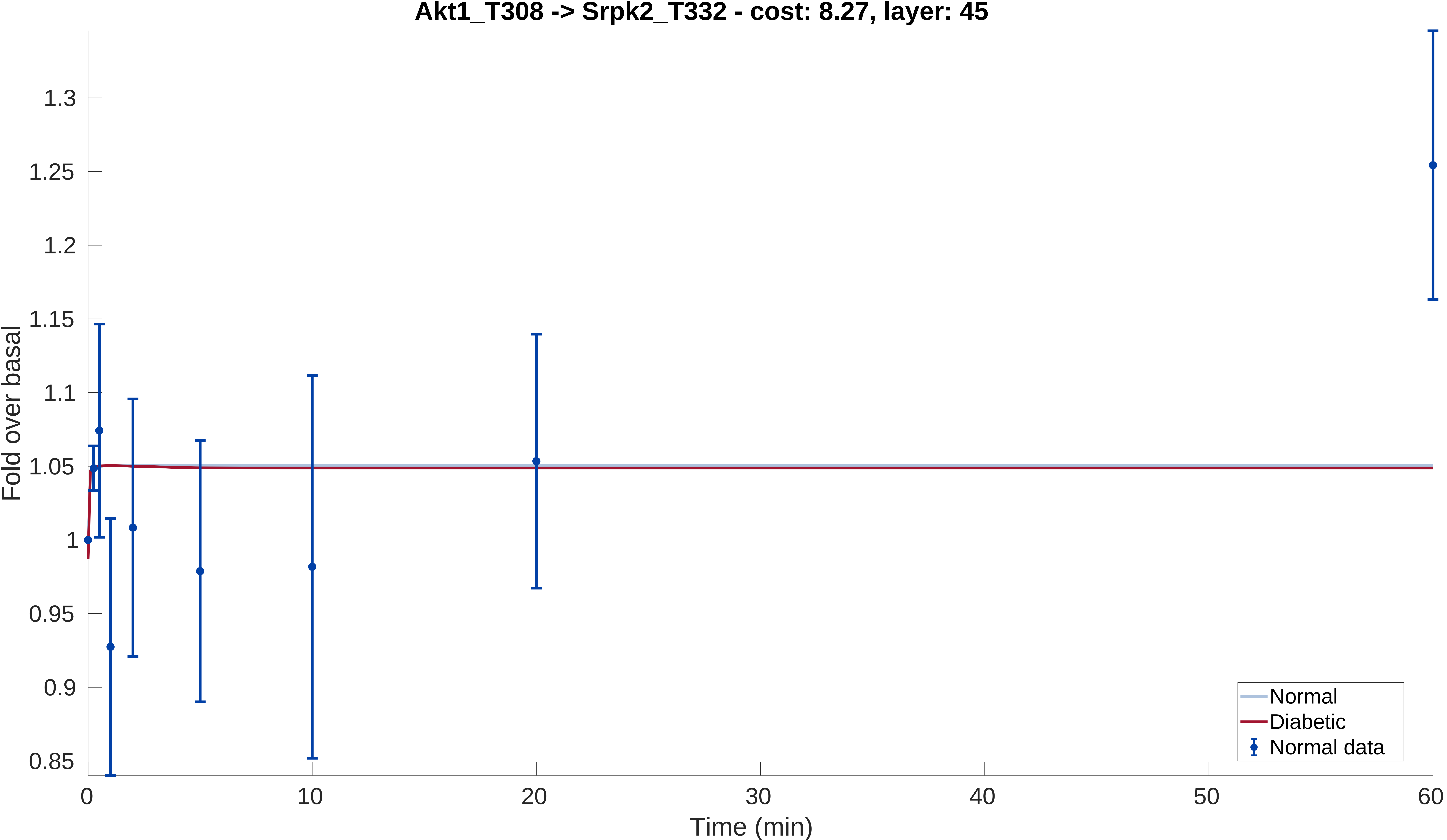

Akt1_T308 -> Srpk2_T332 - cost: 8.27, layer: 45
Fold over basal
1.3
1.25
1.2
1.15
1.1
1.05
1
0.95
0.9
0.85
0
10
20
30
40
50
60
Time (min)
Normal
Diabetic
Normal data

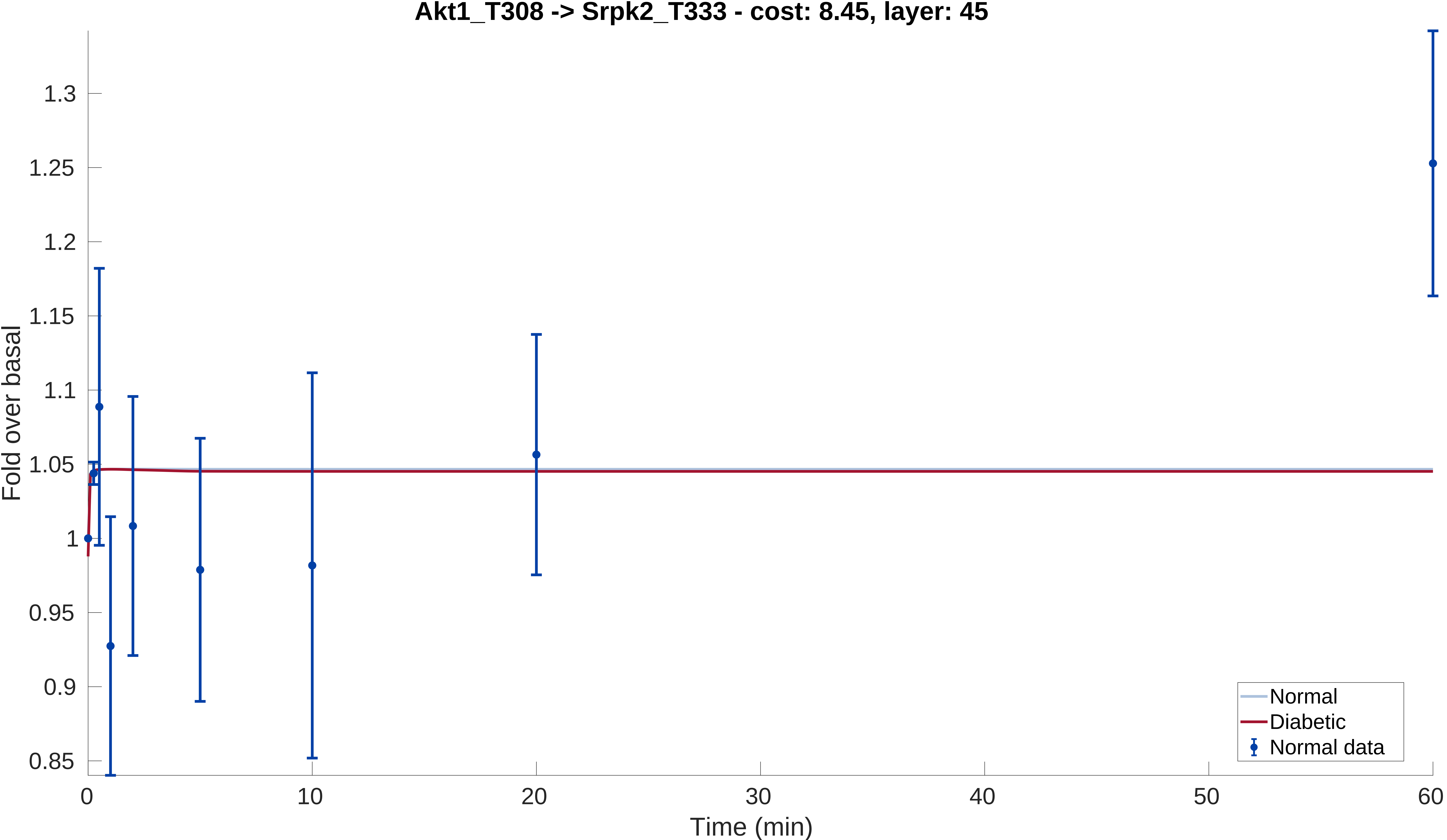

Akt1_T308 -> Srpk2_T333 - cost: 8.45, layer: 45
Fold over basal
1.3
1.25
1.2
1.15
1.1
1.05
1
0.95
0.9
0.85
0
10
20
30
40
50
60
Time (min)
Normal
Diabetic
Normal data

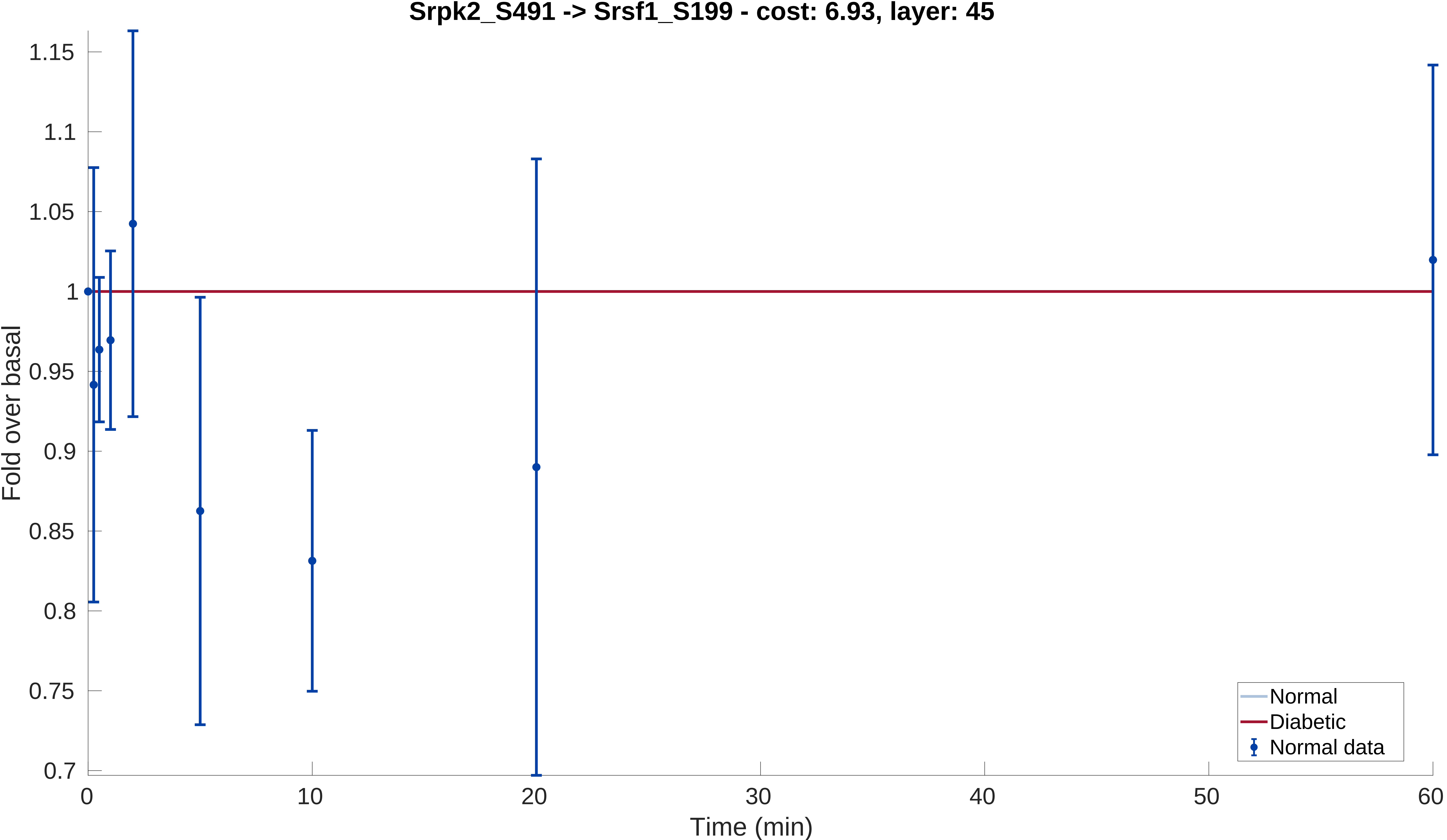

Srpk2_S491 -> Srsf1_S199 - cost: 6.93, layer: 45
Fold over basal
1.15
1.1
1.05
1
0.95
0.9
0.85
0.8
0.75
0.7
0
10
20
30
40
50
60
Time (min)
Normal
Diabetic
Normal data

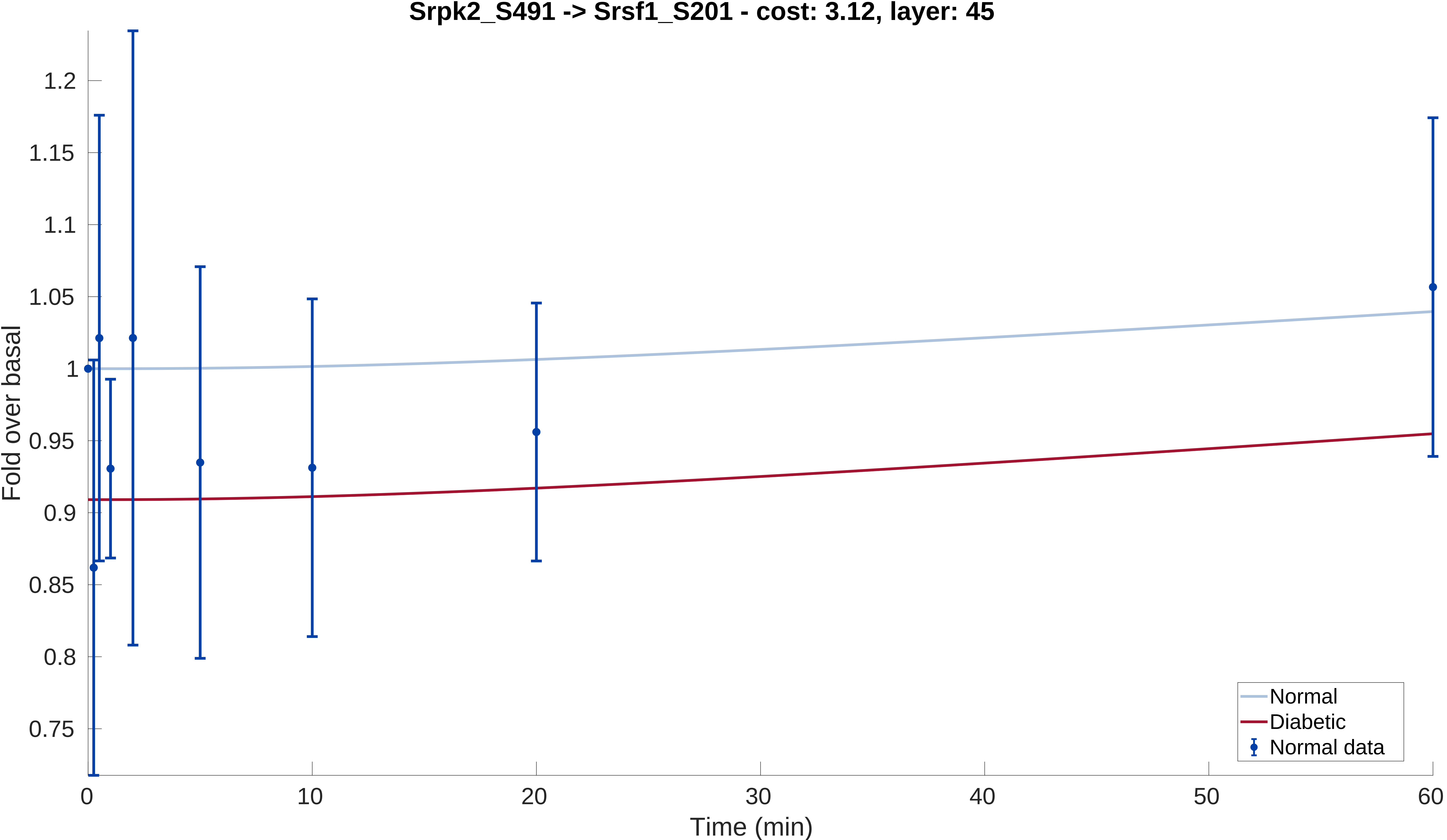
Srpk2_S491 -> Srsf1_S201 - cost: 3.12, layer: 45
Fold over basal
1.2
1.15
1.1
1.05
1
0.95
0.9
0.85
0.8
0.75
0
10
20
30
40
50
60
Time (min)
Normal
Diabetic
Normal data

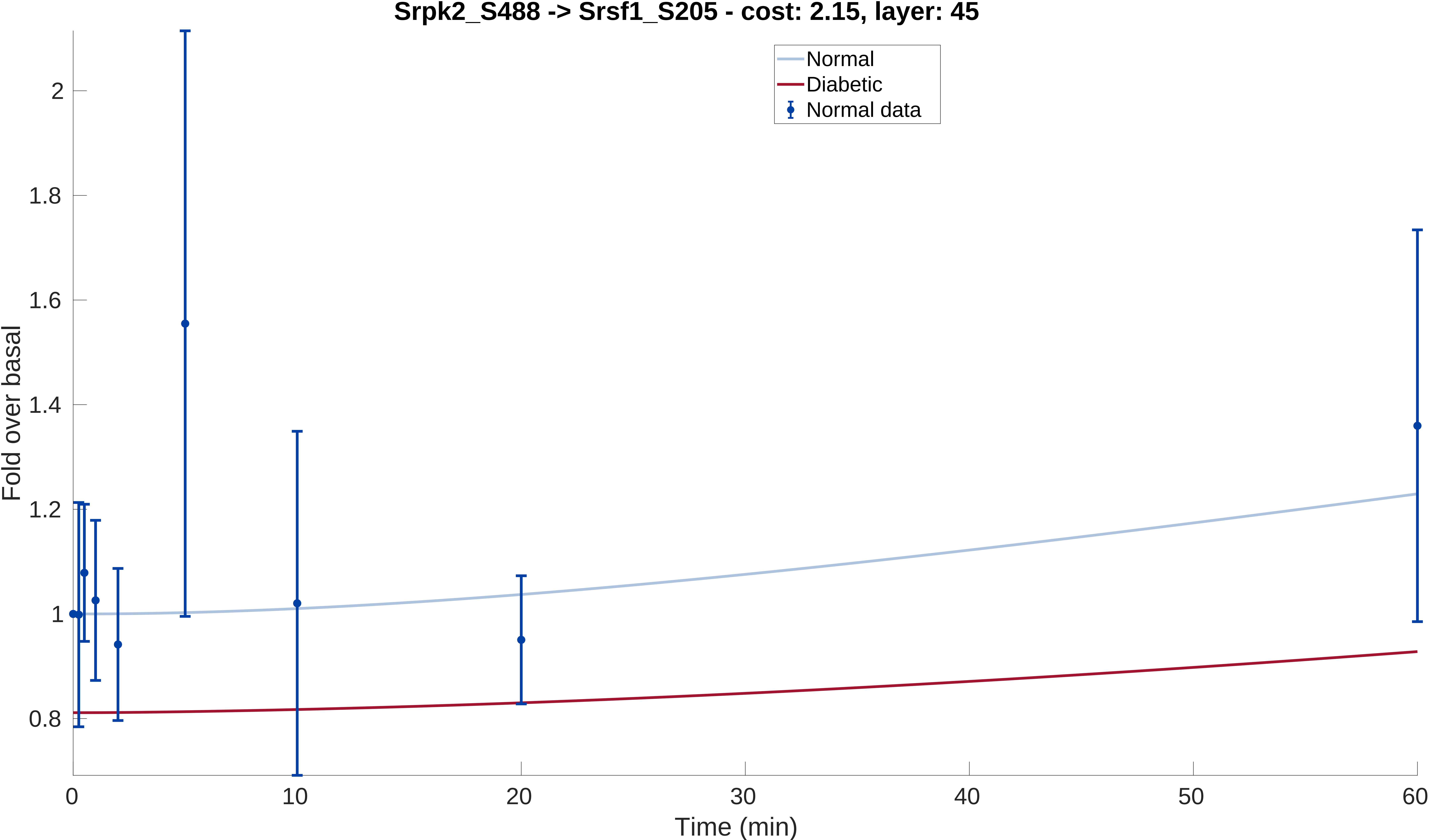
Srpk2_S488 -> Srsf1_S205 - cost: 2.15, layer: 45
Normal
Diabetic
Normal data
Fold over basal
2
1.8
1.6
1.4
1.2
1
0.8
0
10
20
30
40
50
60
Time (min)

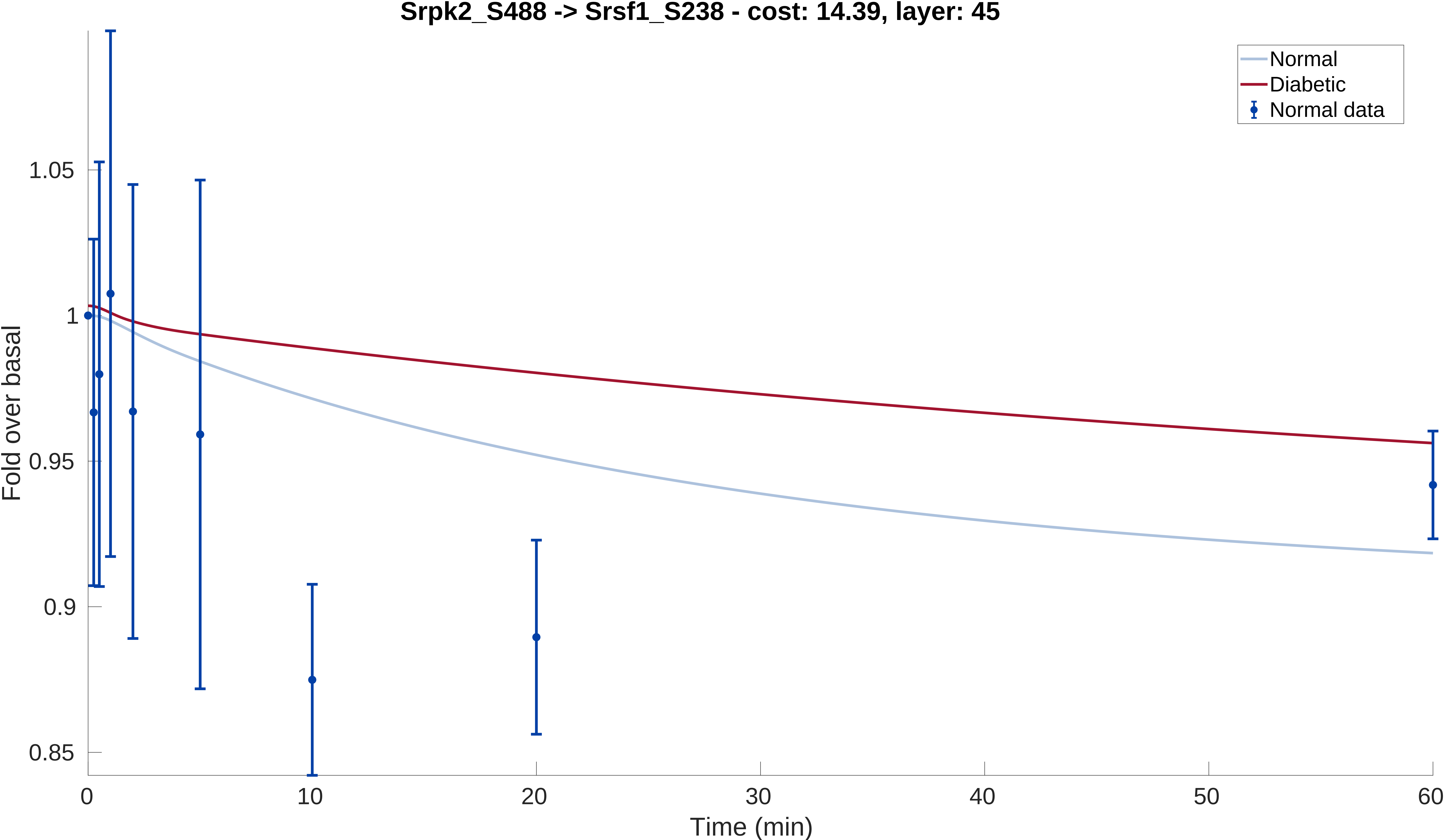
Srpk2_S488 -> Srsf1_S238 - cost: 14.39, layer: 45
Normal
Diabetic
Normal data
Fold over basal
1.05
1
0.95
0.9
0.85
0
10
20
30
40
50
60
Time (min)

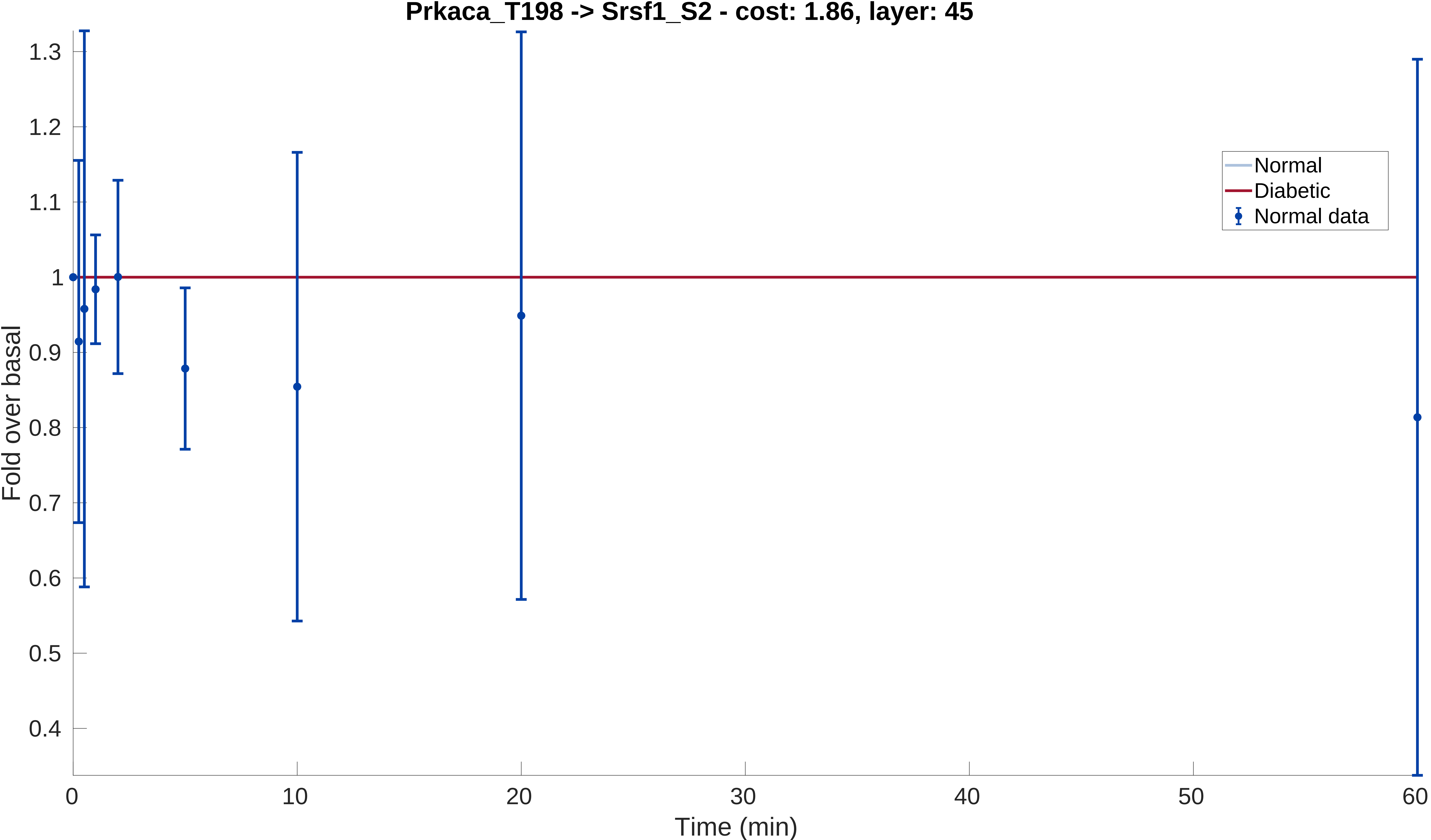

Prkaca_T198 -> Srsf1_S2 - cost: 1.86, layer: 45
Normal
Diabetic
Normal data
Fold over basal
Time (min)
1.3
1.2
1.1
1
0.9
0.8
0.7
0.6
0.5
0.4
0
10
20
30
40
50
60

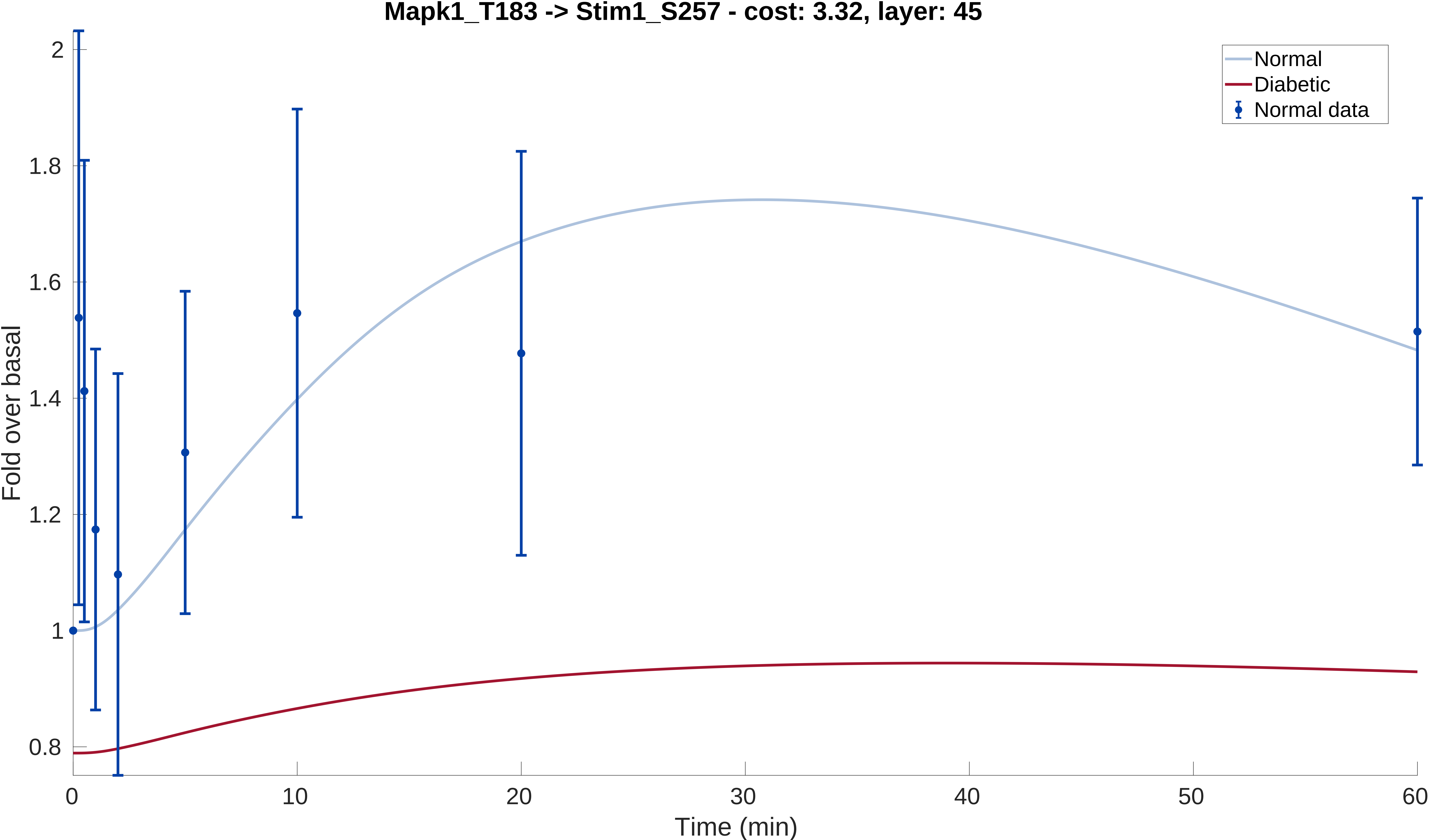

Mapk1_T183 -> Stim1_S257 - cost: 3.32, layer: 45
Normal
Diabetic
Normal data
Fold over basal
2
1.8
1.6
1.4
1.2
1
0.8
0
10
20
30
40
50
60
Time (min)

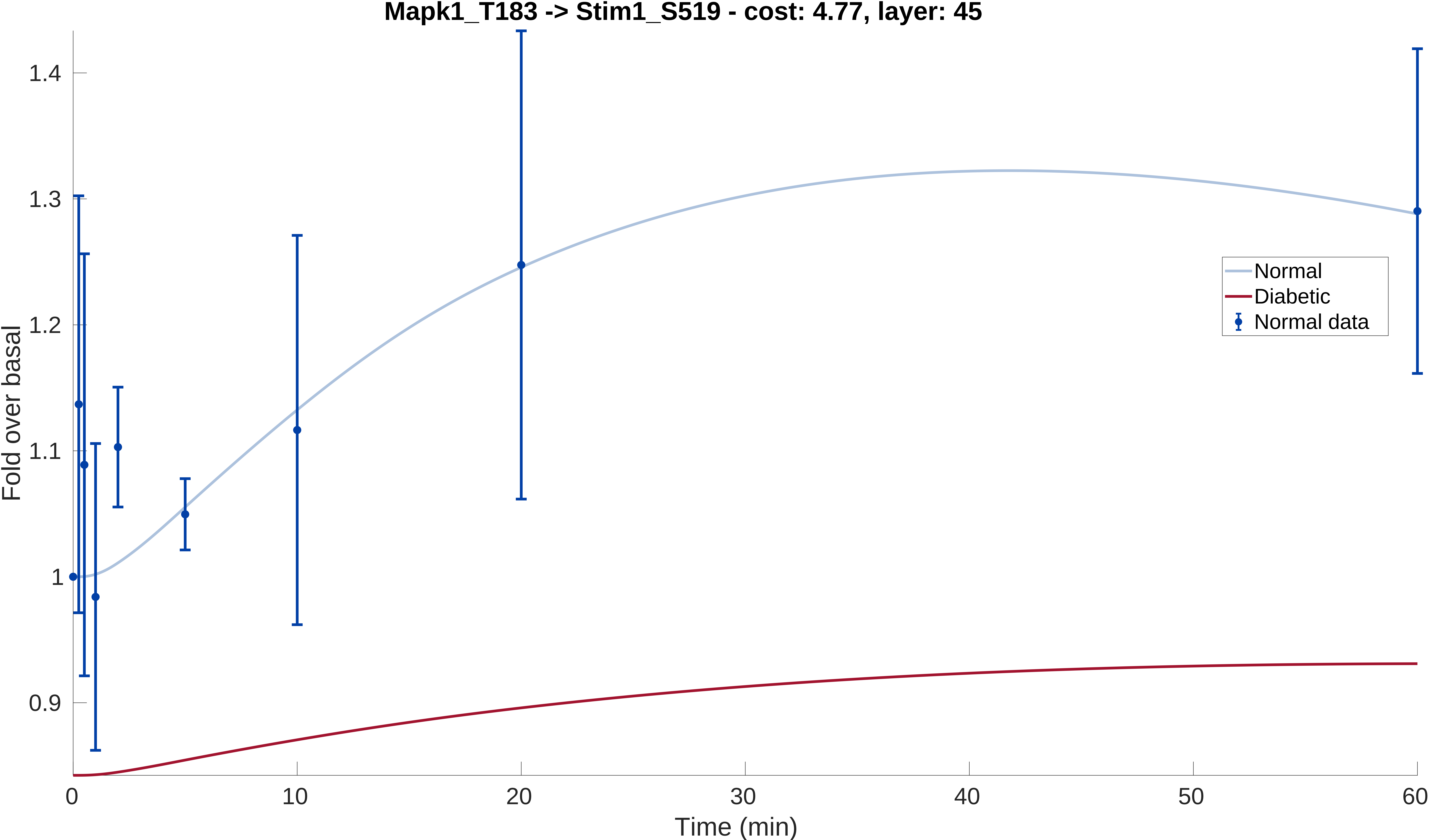

Mapk1_T183 -> Stim1_S519 - cost: 4.77, layer: 45
Fold over basal
Time (min)
1.4
1.3
1.2
1.1
1
0.9
0
10
20
30
40
50
60
Normal
Diabetic
Normal data

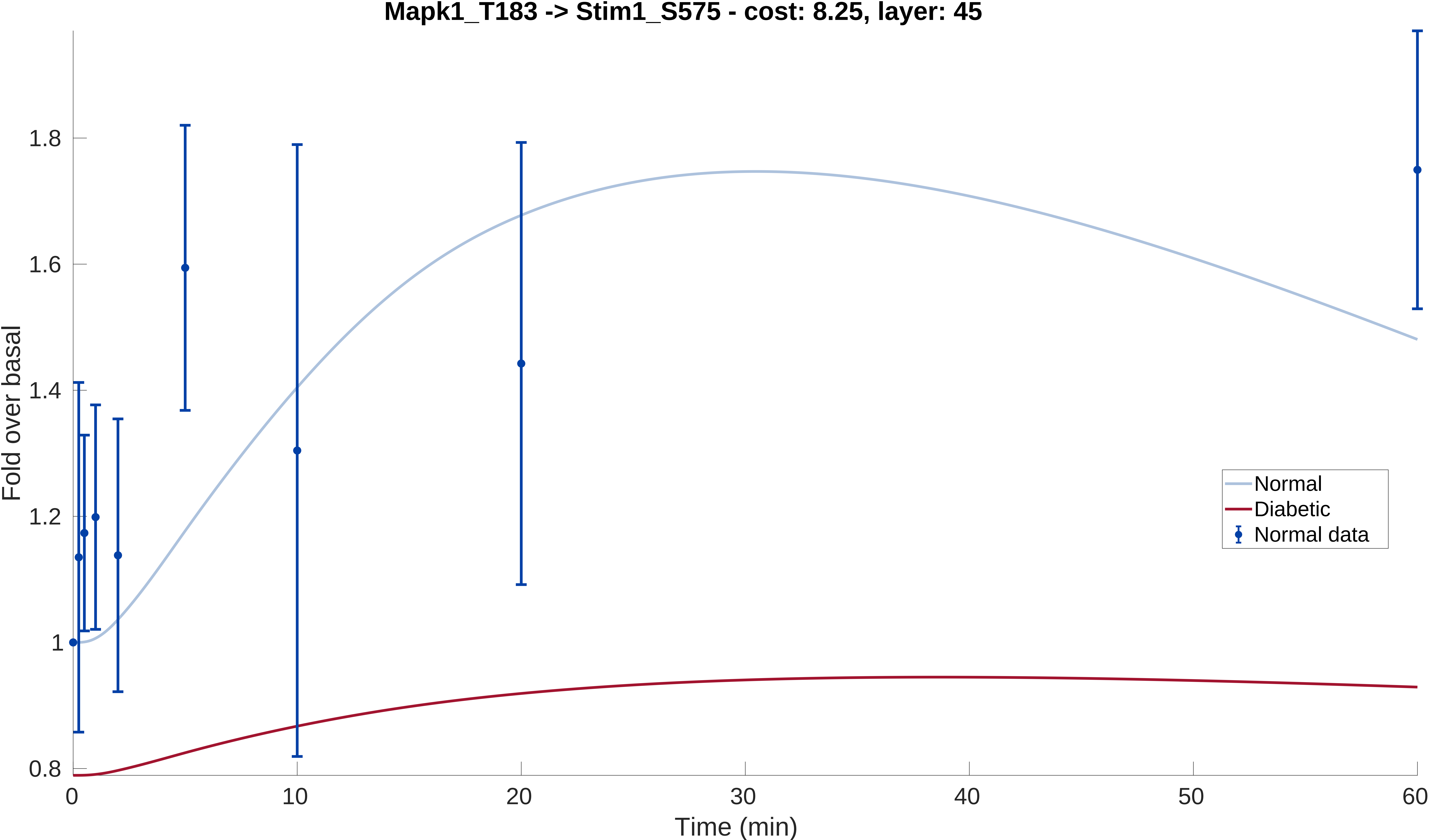

Mapk1_T183 -> Stim1_S575 - cost: 8.25, layer: 45
Fold over basal
Time (min)
0.8
1
1.2
1.4
1.6
1.8
0
10
20
30
40
50
60
Normal
Diabetic
Normal data

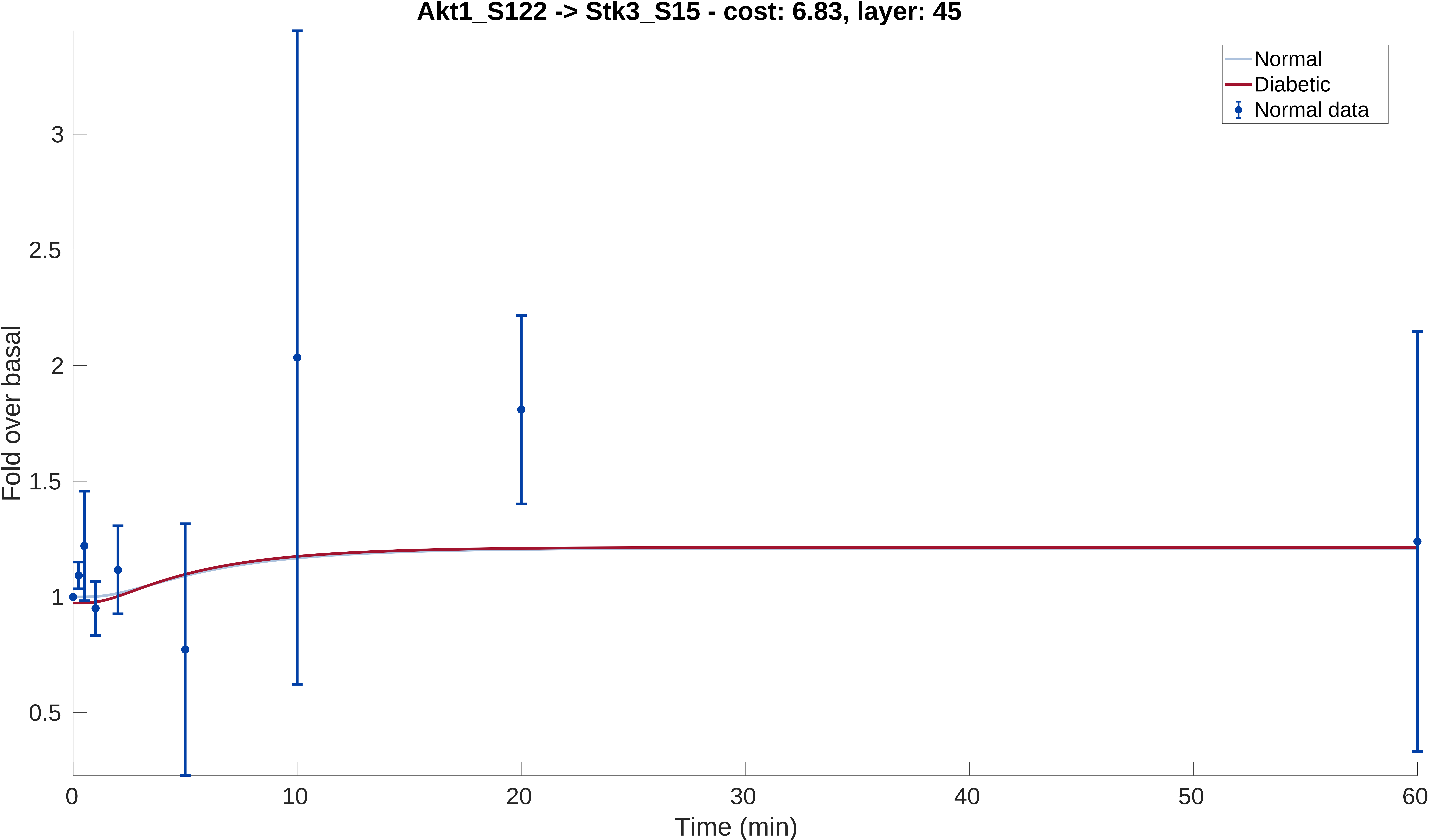

Akt1_S122 -> Stk3_S15 - cost: 6.83, layer: 45
Normal
Diabetic
Normal data
Fold over basal
0.5
1
1.5
2
2.5
3
0
10
20
30
40
50
60
Time (min)

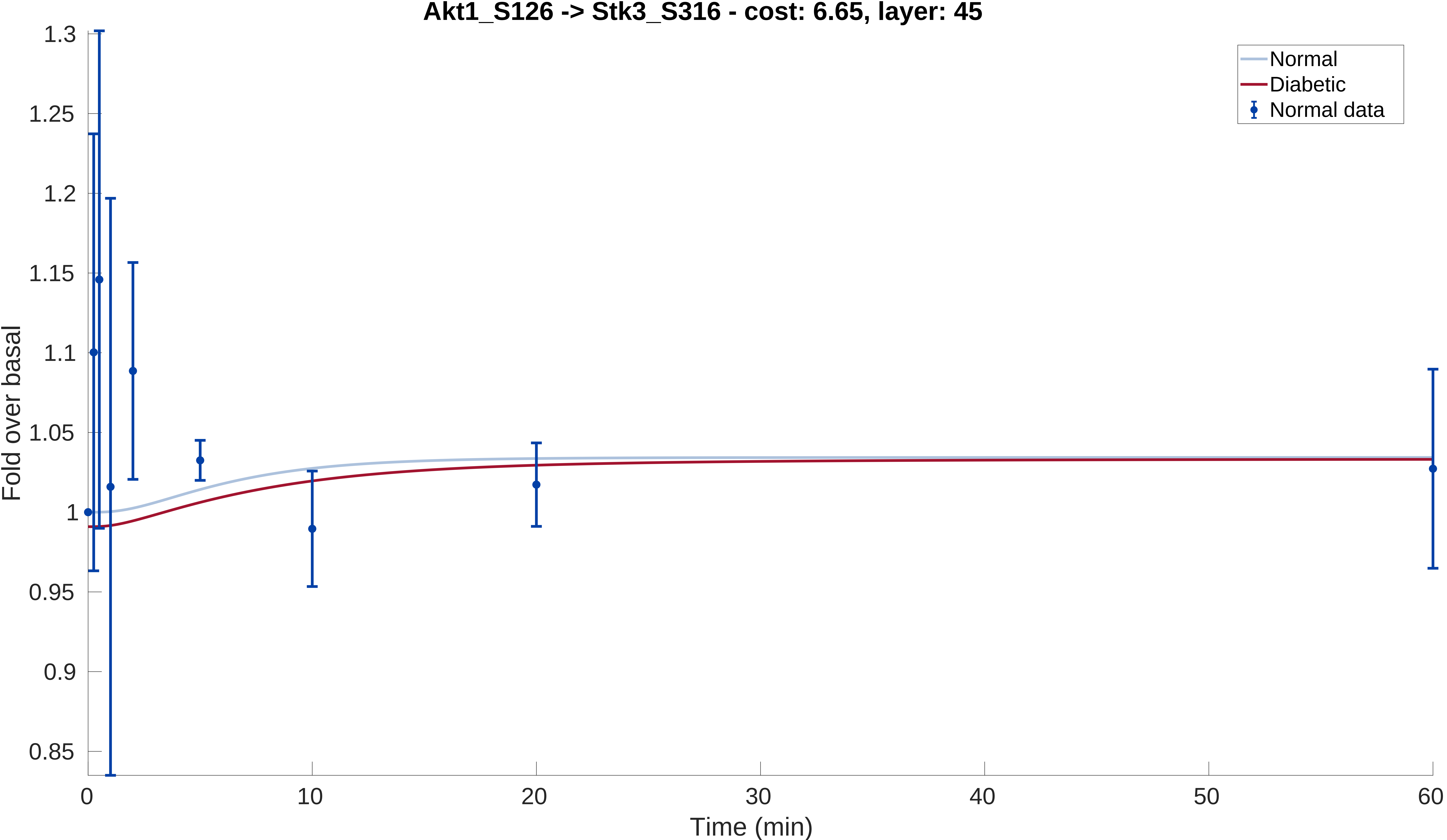
Akt1_S126 -> Stk3_S316 - cost: 6.65, layer: 45
Normal
Diabetic
Normal data
Fold over basal
1.3
1.25
1.2
1.15
1.1
1.05
1
0.95
0.9
0.85
0
10
20
30
40
50
60
Time (min)

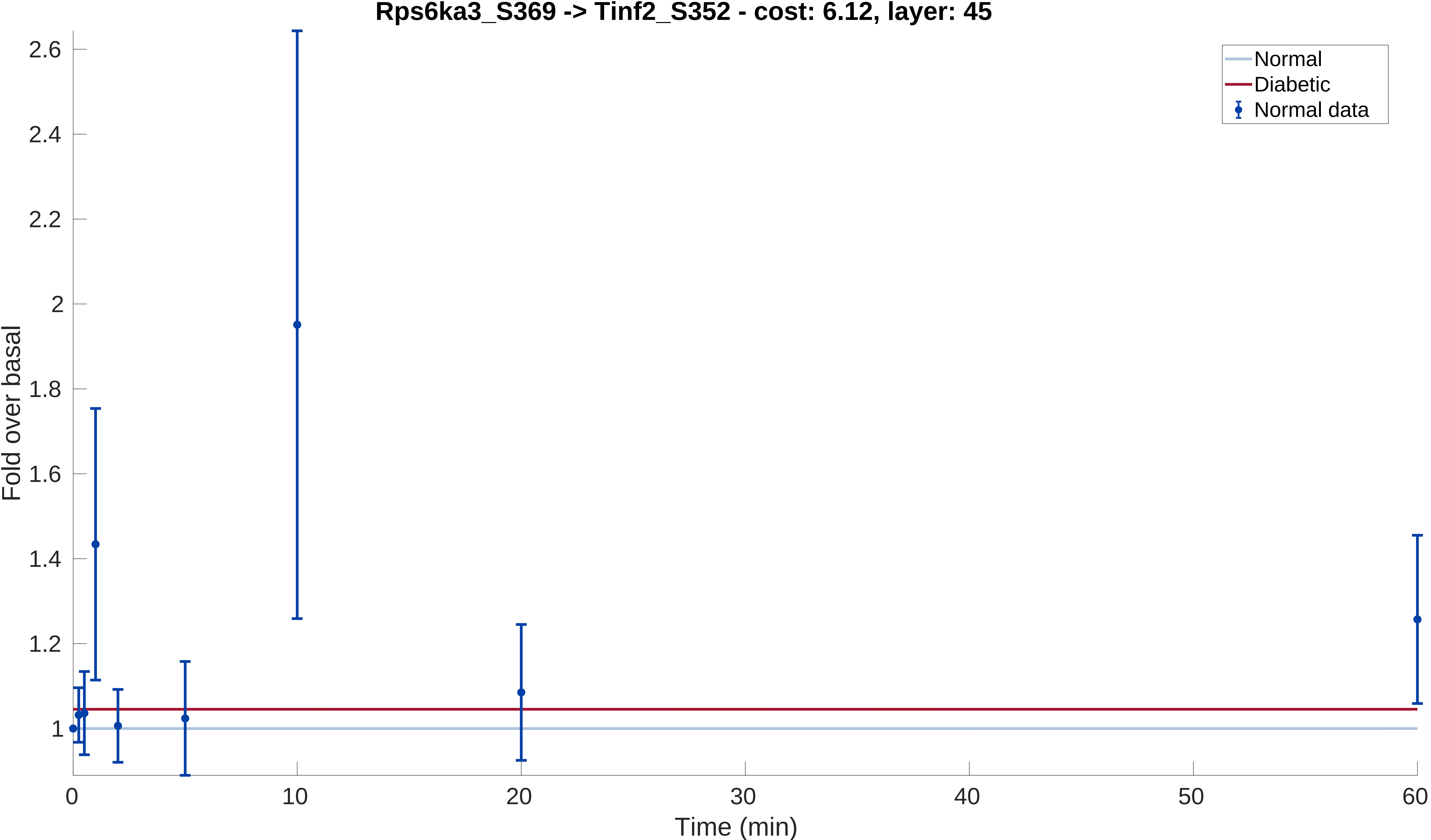

Rps6ka3_S369 -> Tinf2_S352 - cost: 6.12, layer: 45
Normal
Diabetic
Normal data
Fold over basal
2.6
2.4
2.2
2
1.8
1.6
1.4
1.2
1
0
10
20
30
40
50
60
Time (min)

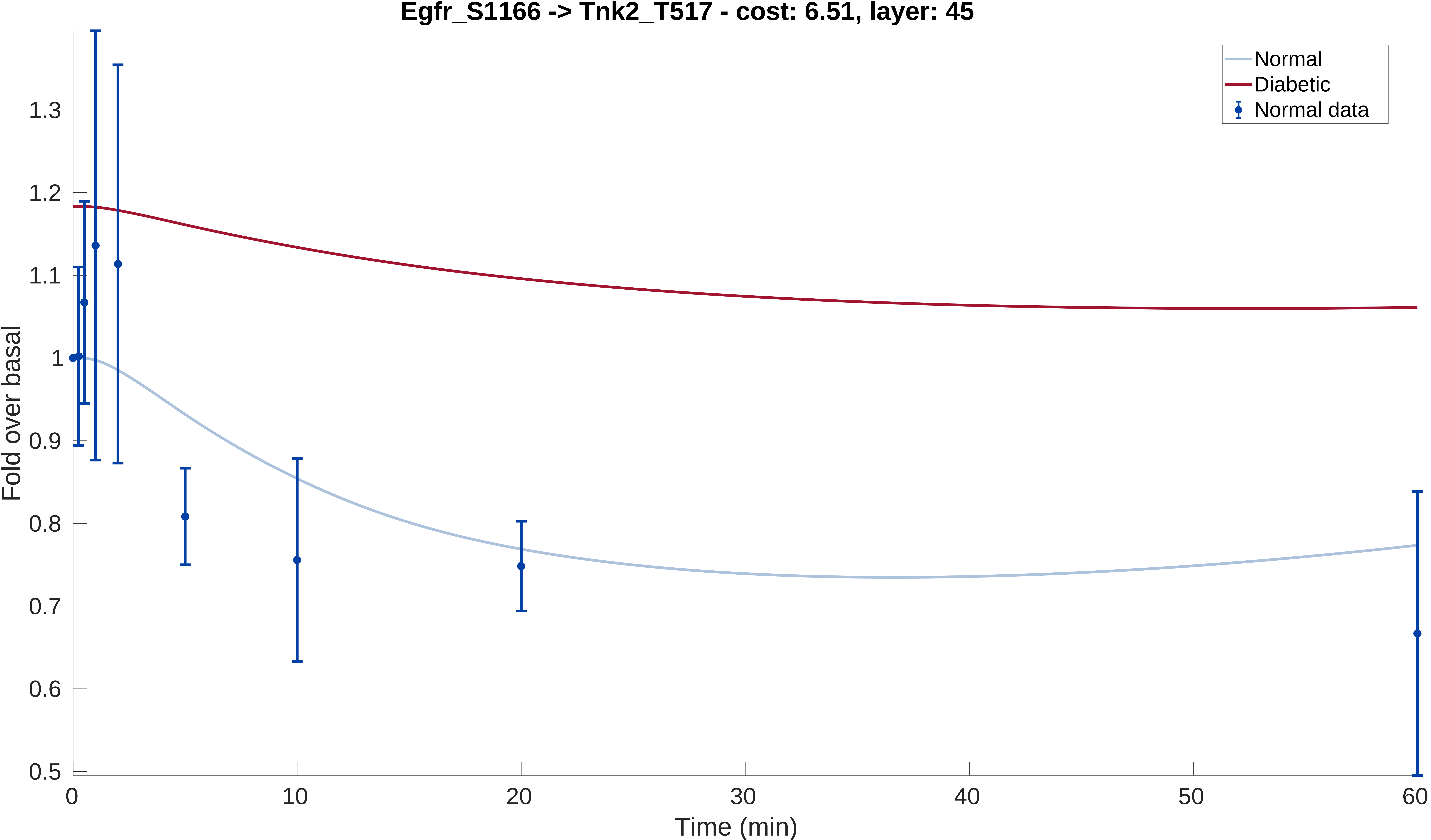
Egfr_S1166 -> Tnk2_T517 - cost: 6.51, layer: 45
Normal
Diabetic
Normal data
Fold over basal
Time (min)
0.5
0.6
0.7
0.8
0.9
1
1.1
1.2
1.3
0
10
20
30
40
50
60

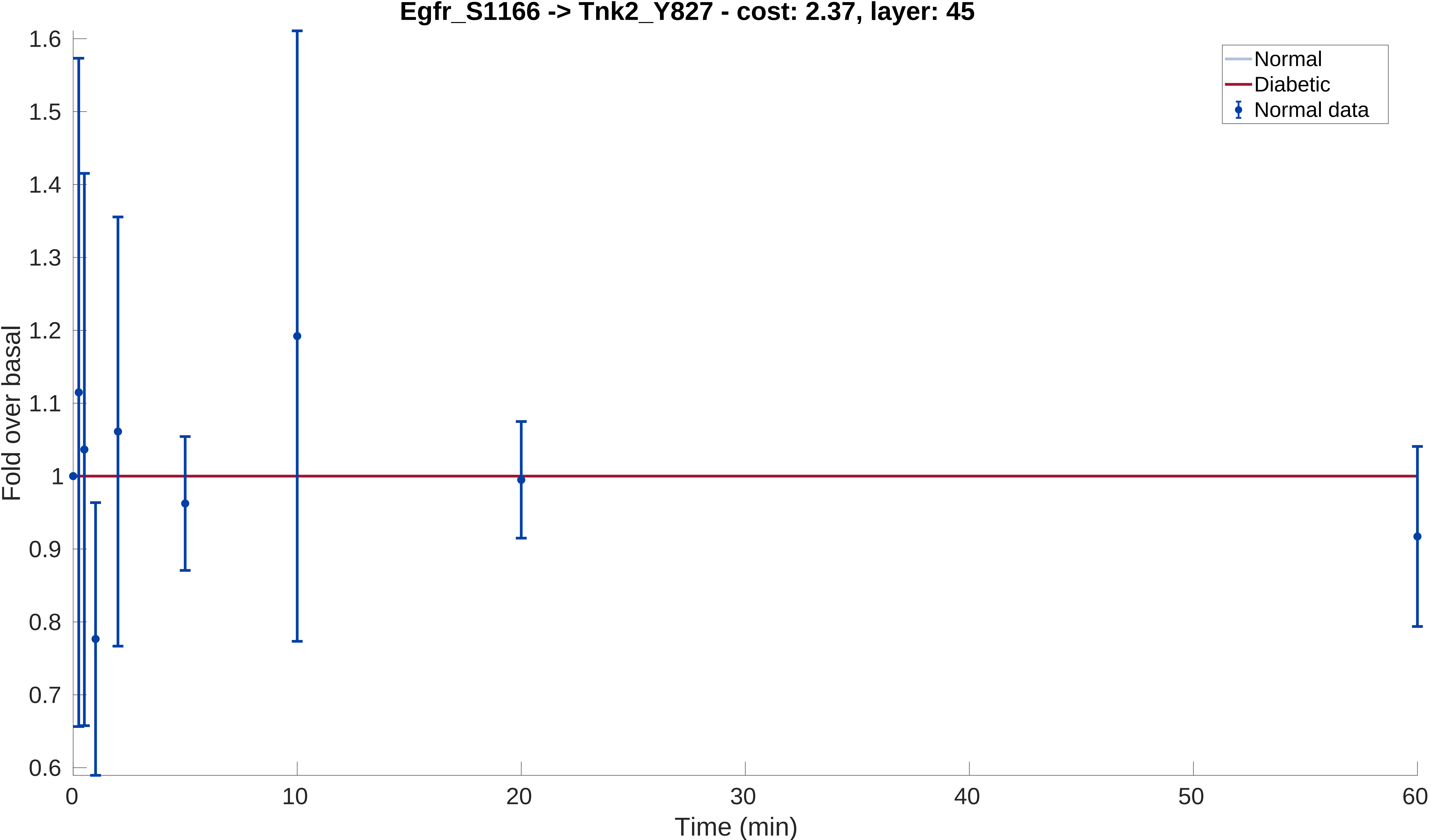

Egfr_S1166 -> Tnk2_Y827 - cost: 2.37, layer: 45
Normal
Diabetic
Normal data
Fold over basal
Time (min)
1.6
1.5
1.4
1.3
1.2
1.1
1
0.9
0.8
0.7
0.6
0
10
20
30
40
50
60

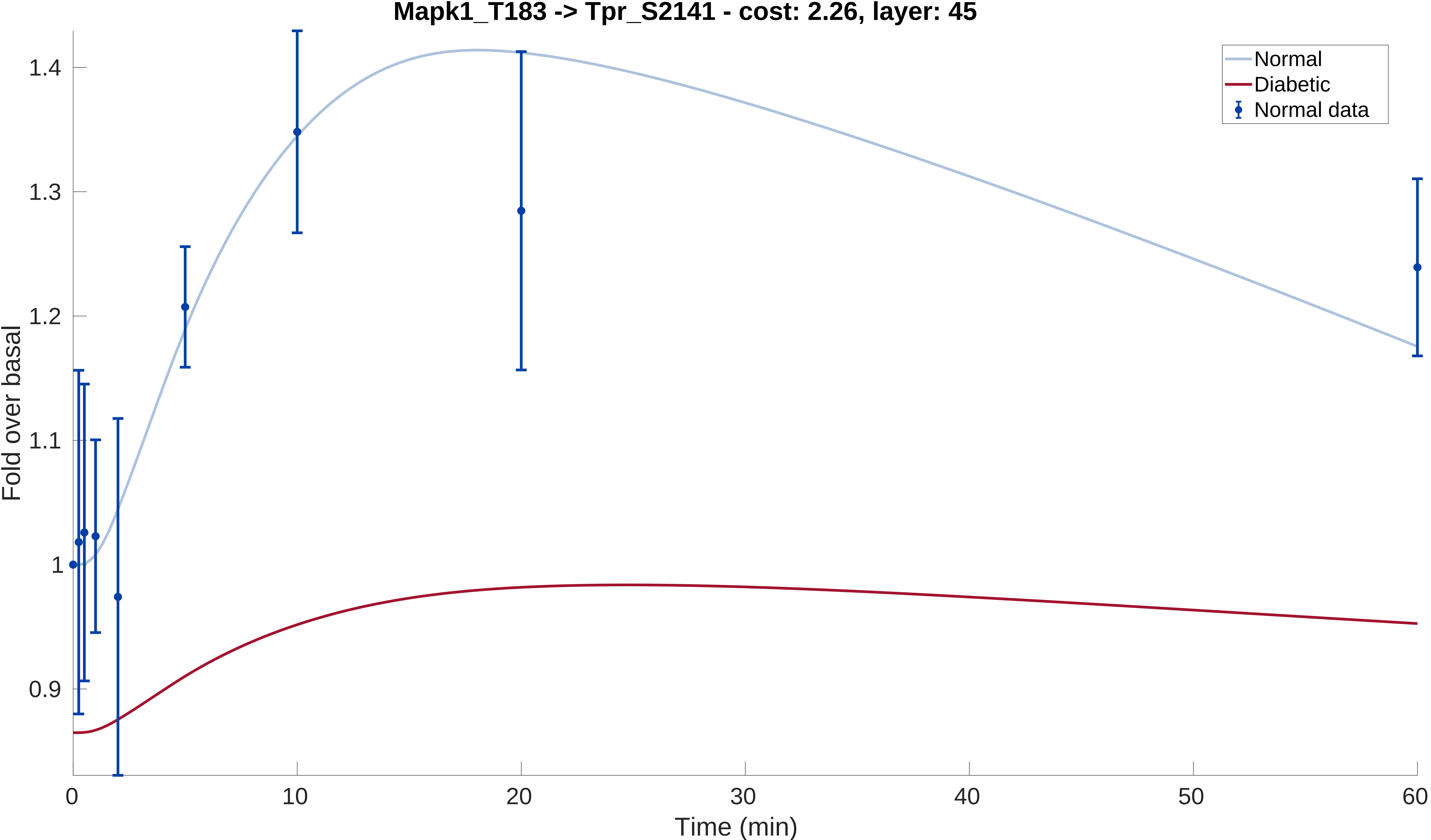

Mapk1_T183 -> Tpr_S2141 - cost: 2.26, layer: 45
Normal
Diabetic
Normal data
Fold over basal
Time (min)
0.9
1
1.1
1.2
1.3
1.4
0
10
20
30
40
50
60

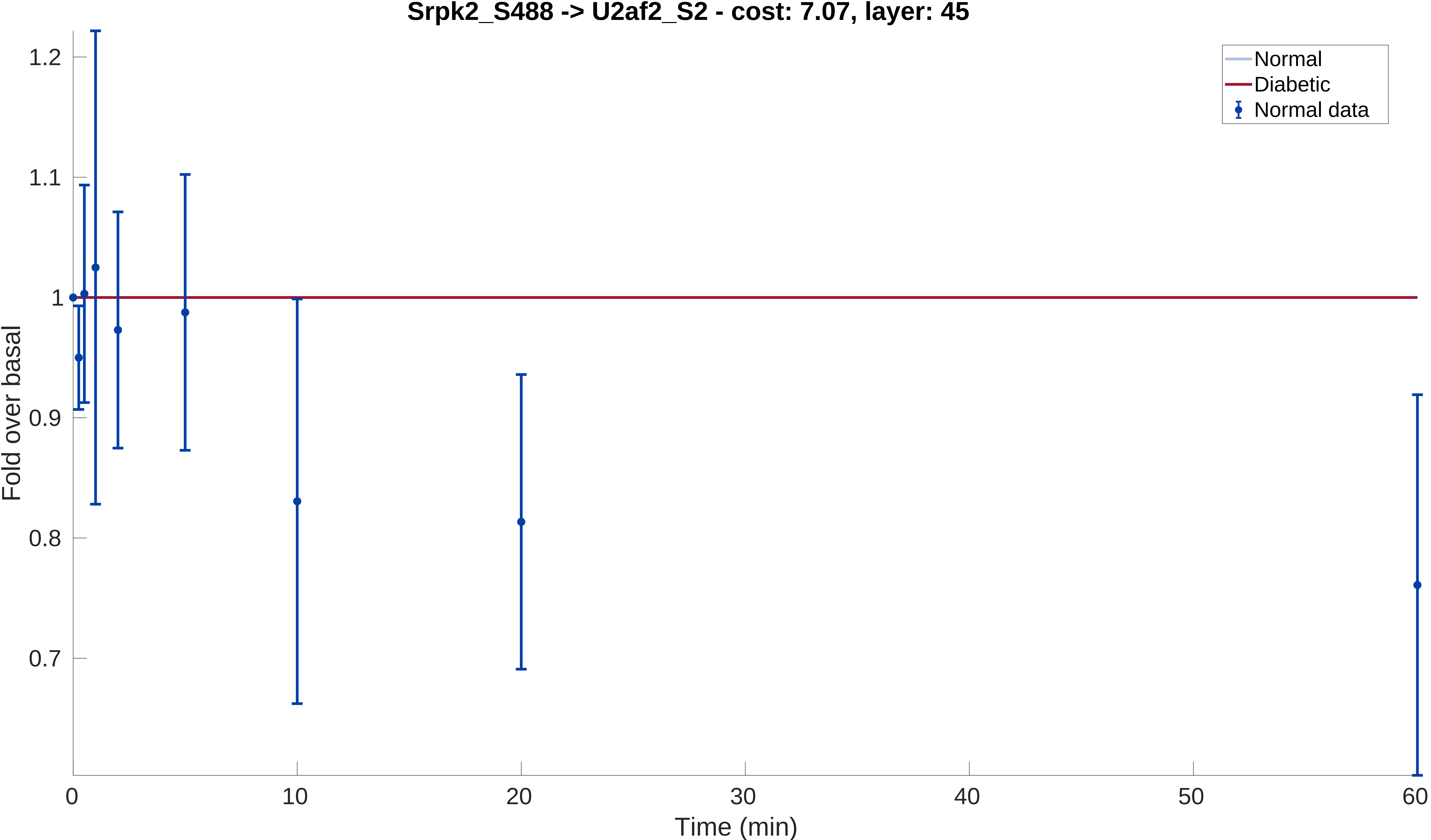

Srpk2_S488 -> U2af2_S2 - cost: 7.07, layer: 45
Normal
Diabetic
Normal data
Fold over basal
1.2
1.1
1
0.9
0.8
0.7
0
10
20
30
40
50
60
Time (min)

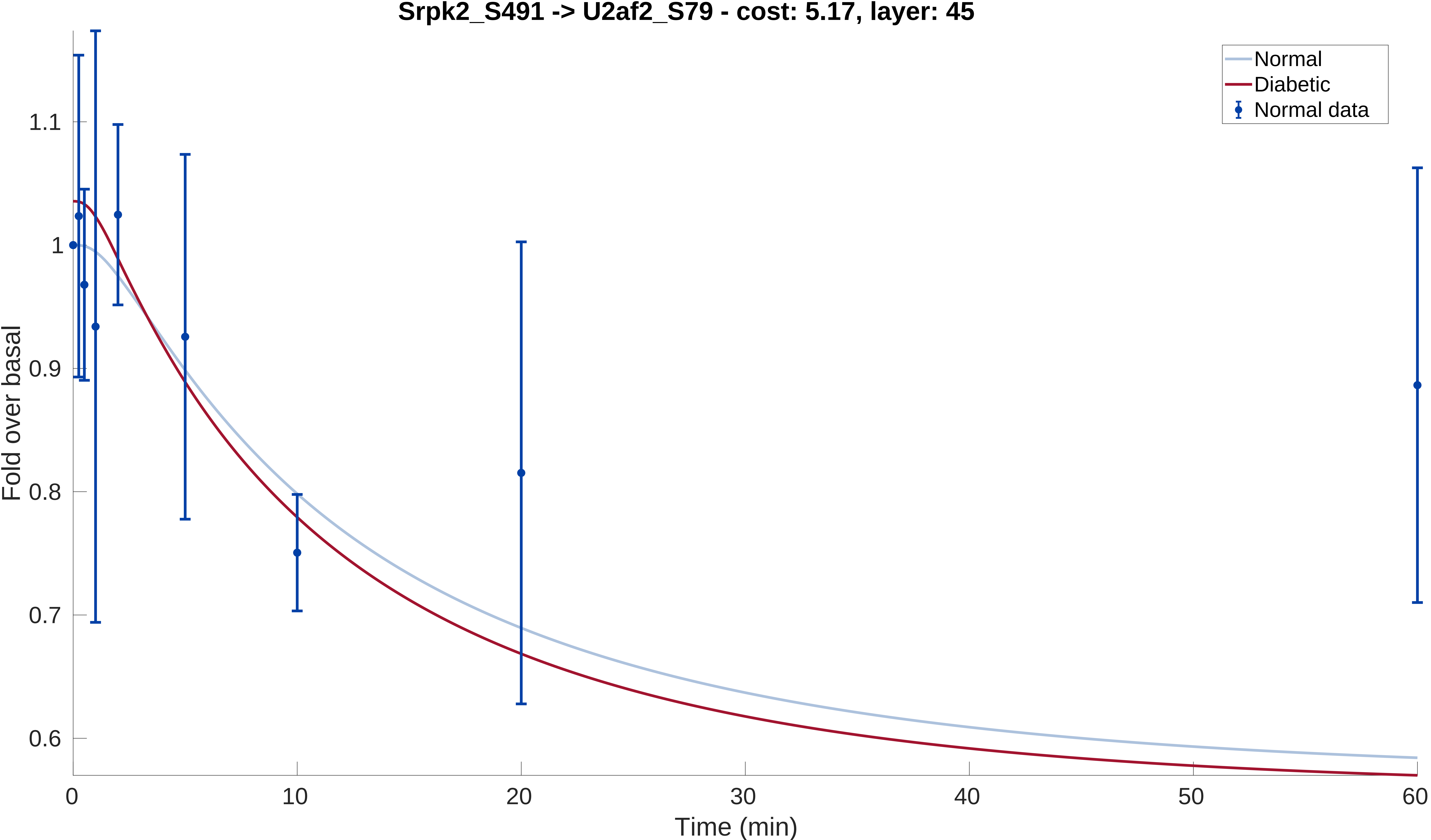

Srpk2_S491 -> U2af2_S79 - cost: 5.17, layer: 45
Normal
Diabetic
Normal data
Fold over basal
Time (min)
1.1
1
0.9
0.8
0.7
0.6
0
10
20
30
40
50
60

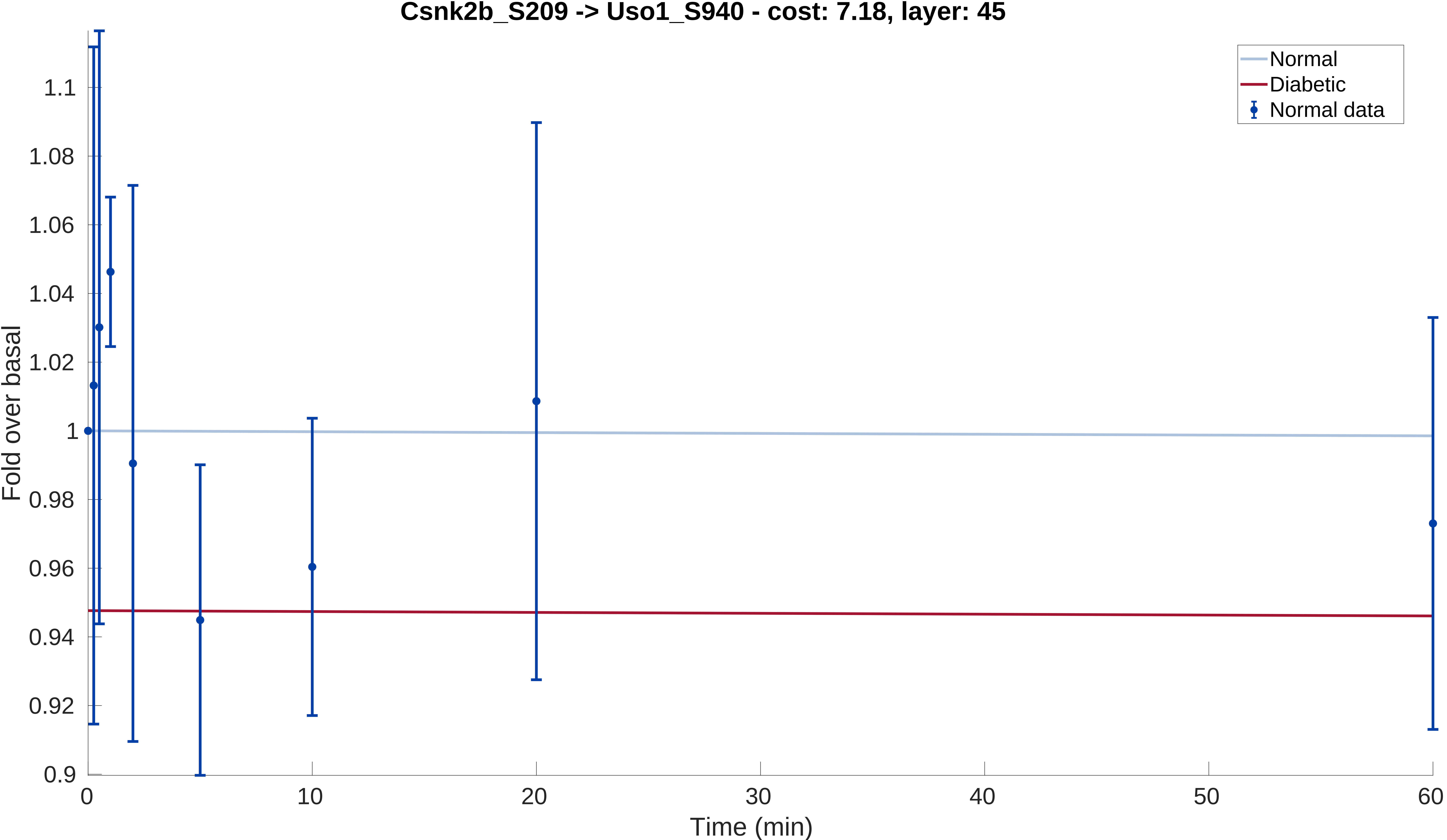

Csnk2b_S209 -> Uso1_S940 - cost: 7.18, layer: 45
Normal
Diabetic
Normal data
Fold over basal
1.1
1.08
1.06
1.04
1.02
1
0.98
0.96
0.94
0.92
0.9
0
10
20
30
40
50
60
Time (min)

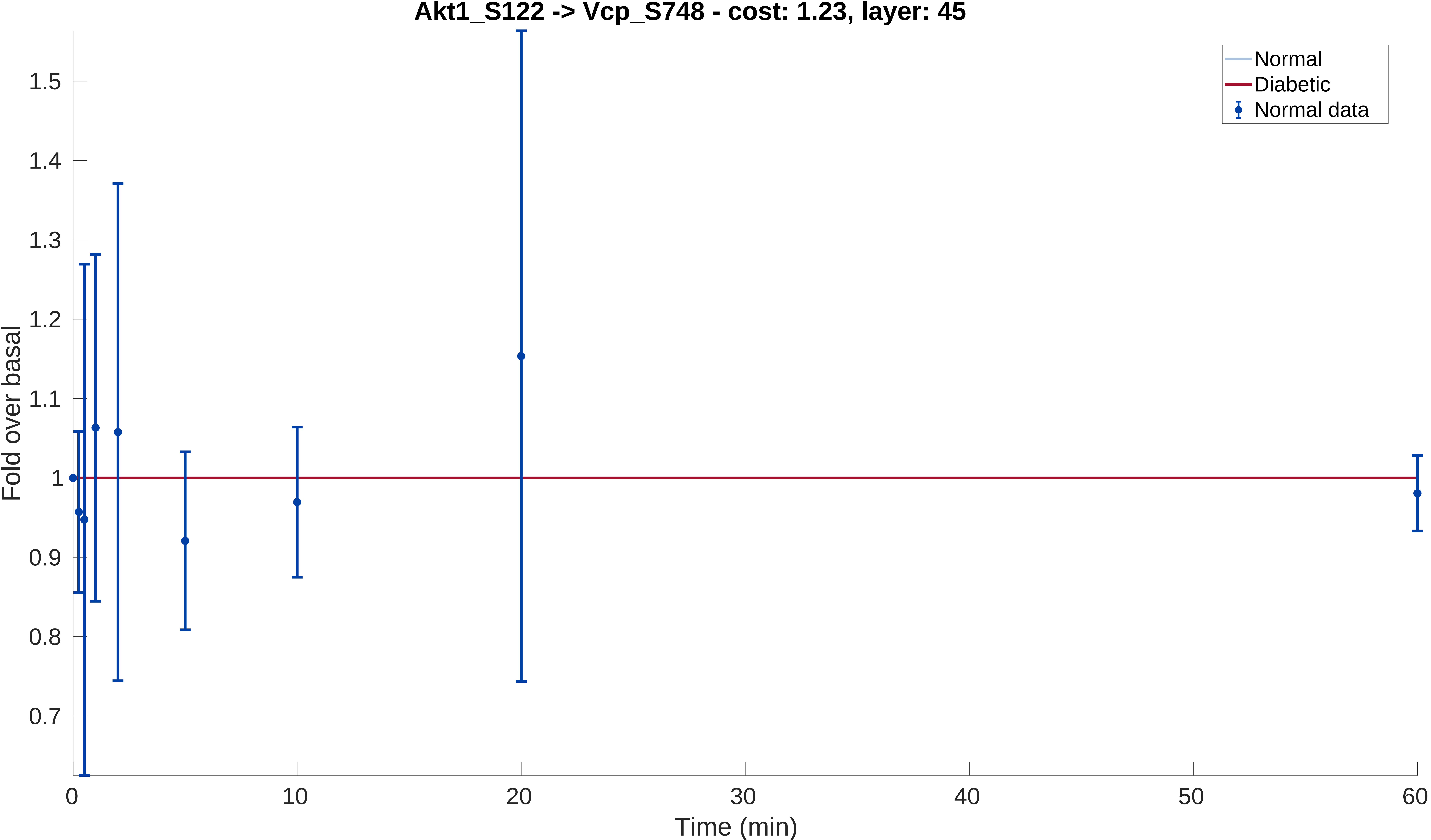
Akt1_S122 -> Vcp_S748 - cost: 1.23, layer: 45
Normal
Diabetic
Normal data
Fold over basal
1.5
1.4
1.3
1.2
1.1
1
0.9
0.8
0.7
0
10
20
30
40
50
60
Time (min)

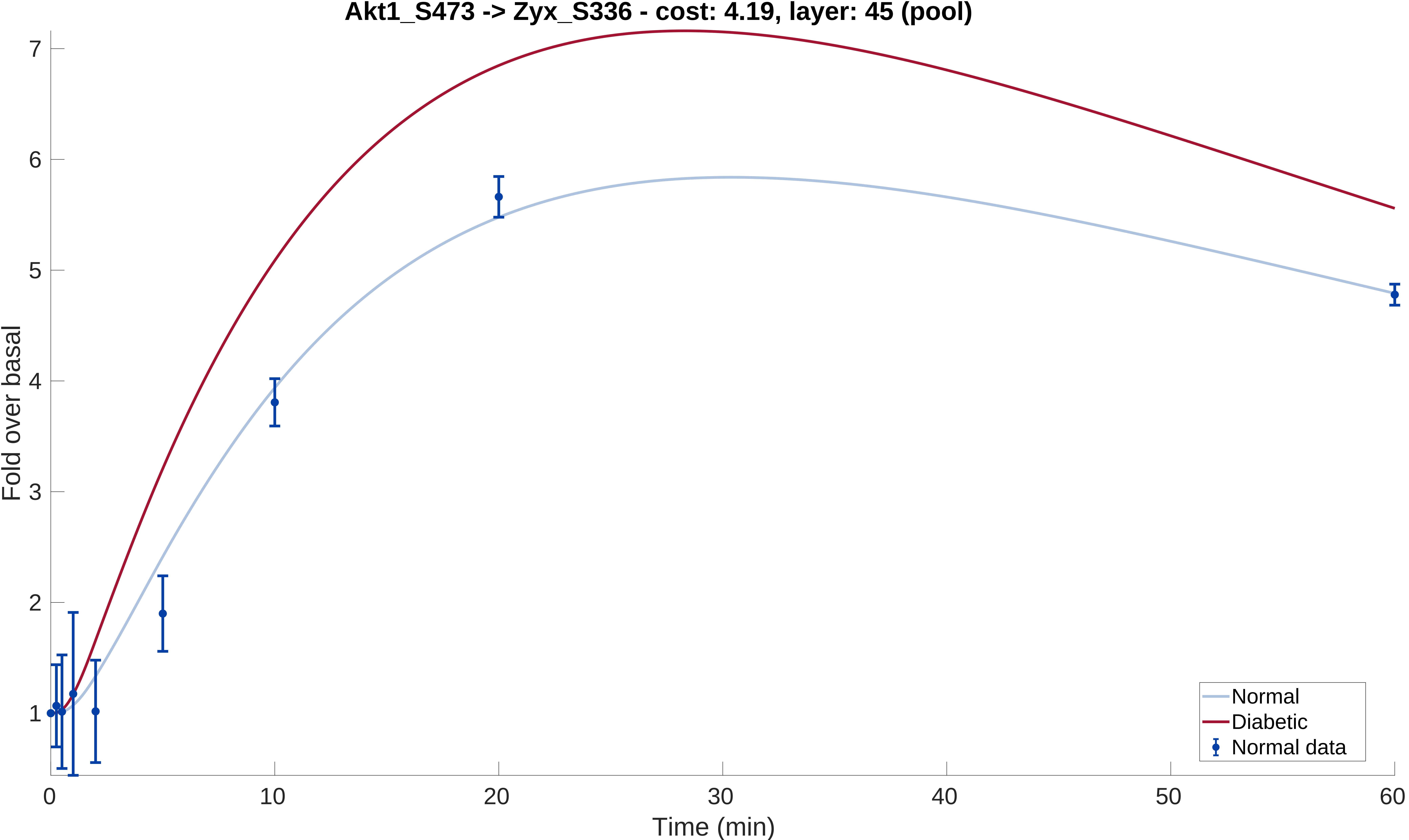
Akt1_S473 -> Zyx_S336 - cost: 4.19, layer: 45 (pool)
Fold over basal
Time (min)
0
10
20
30
40
50
60
1
2
3
4
5
6
7
Normal
Diabetic
Normal data

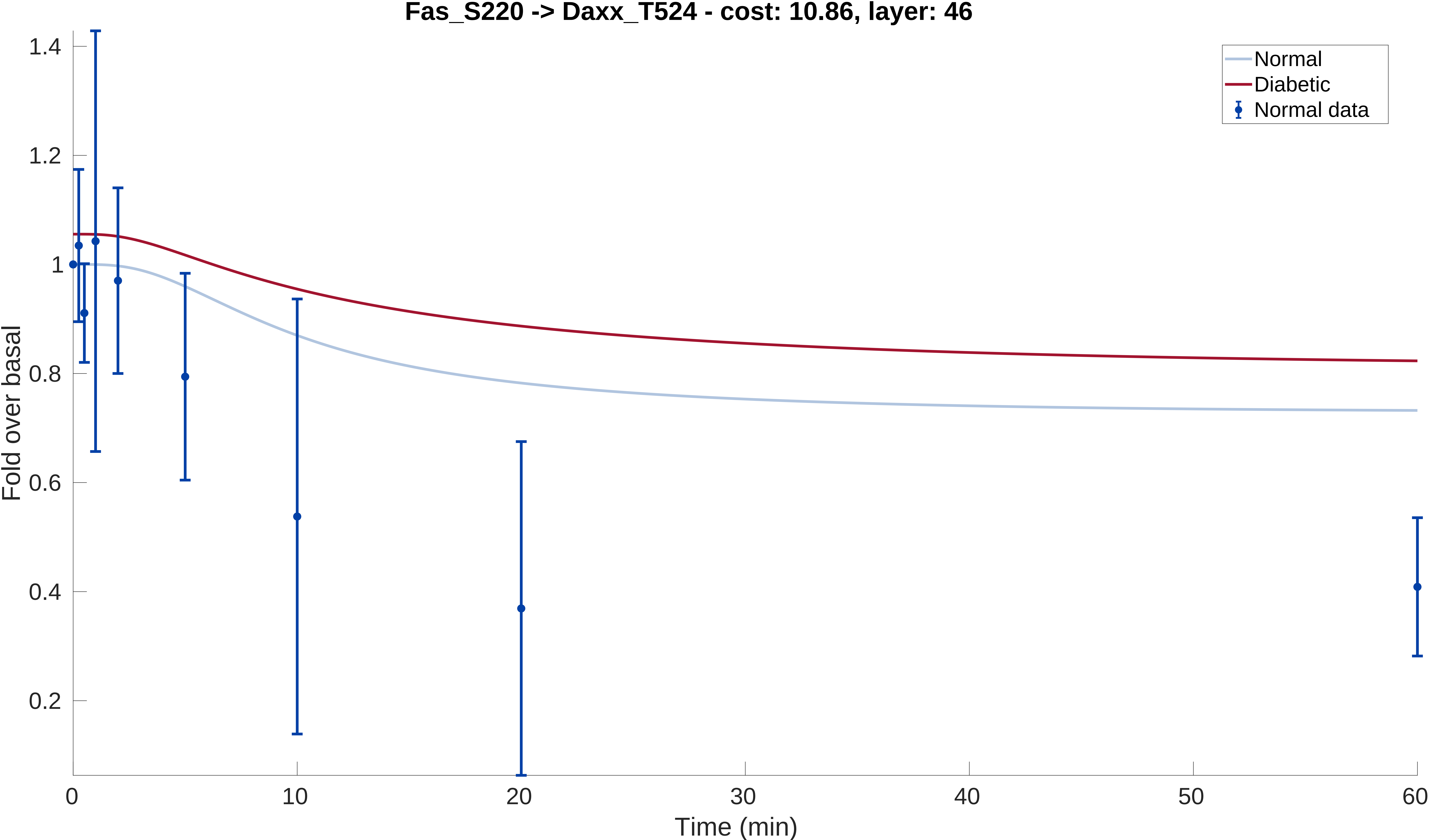
Fas_S220 -> Daxx_T524 - cost: 10.86, layer: 46
Normal
Diabetic
Normal data
Fold over basal
Time (min)
0.2
0.4
0.6
0.8
1
1.2
1.4
0
10
20
30
40
50
60

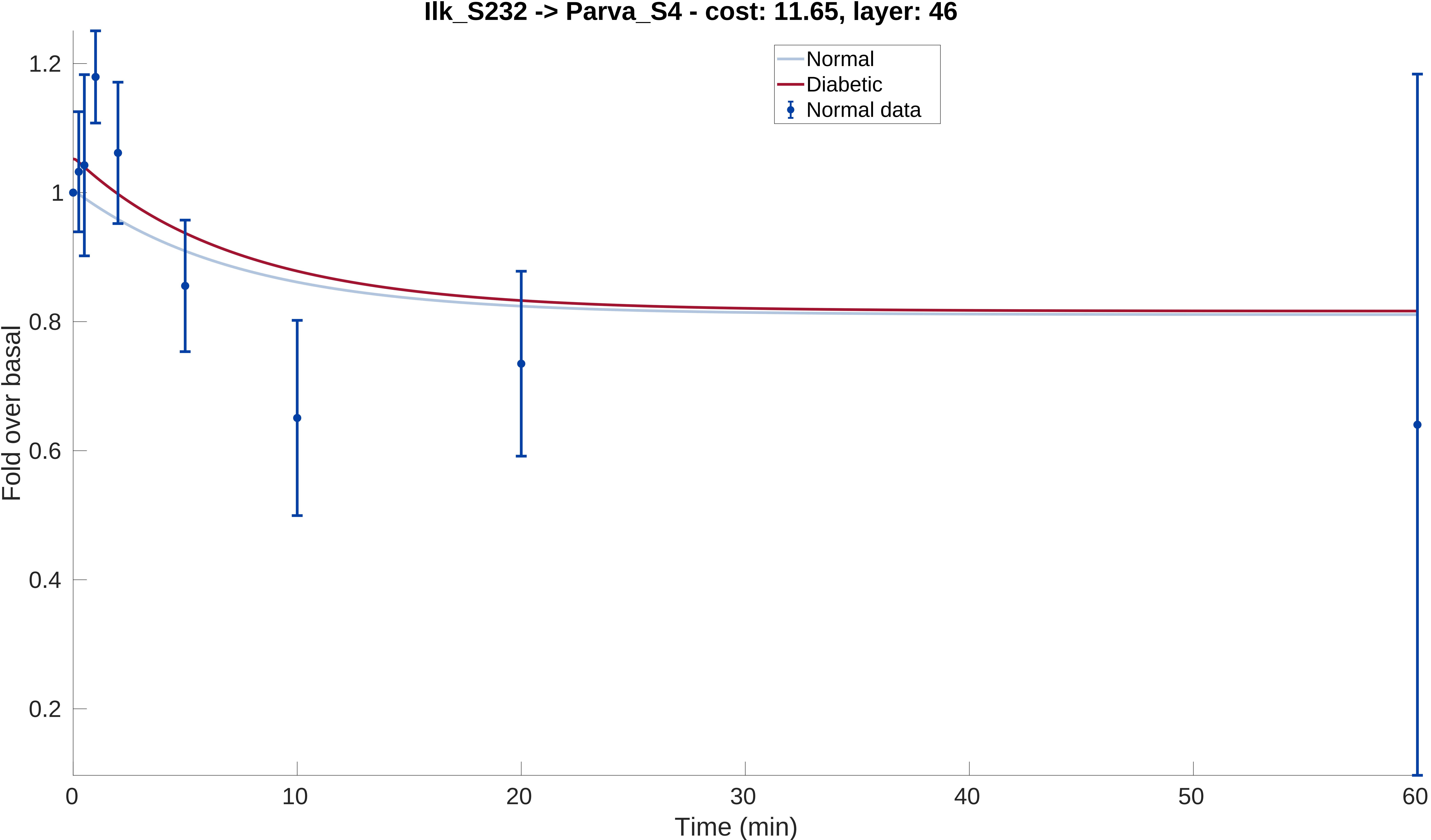

Ilk_S232 -> Parva_S4 - cost: 11.65, layer: 46
Normal
Diabetic
Normal data
Fold over basal
Time (min)
1.2
1
0.8
0.6
0.4
0.2
0
10
20
30
40
50
60

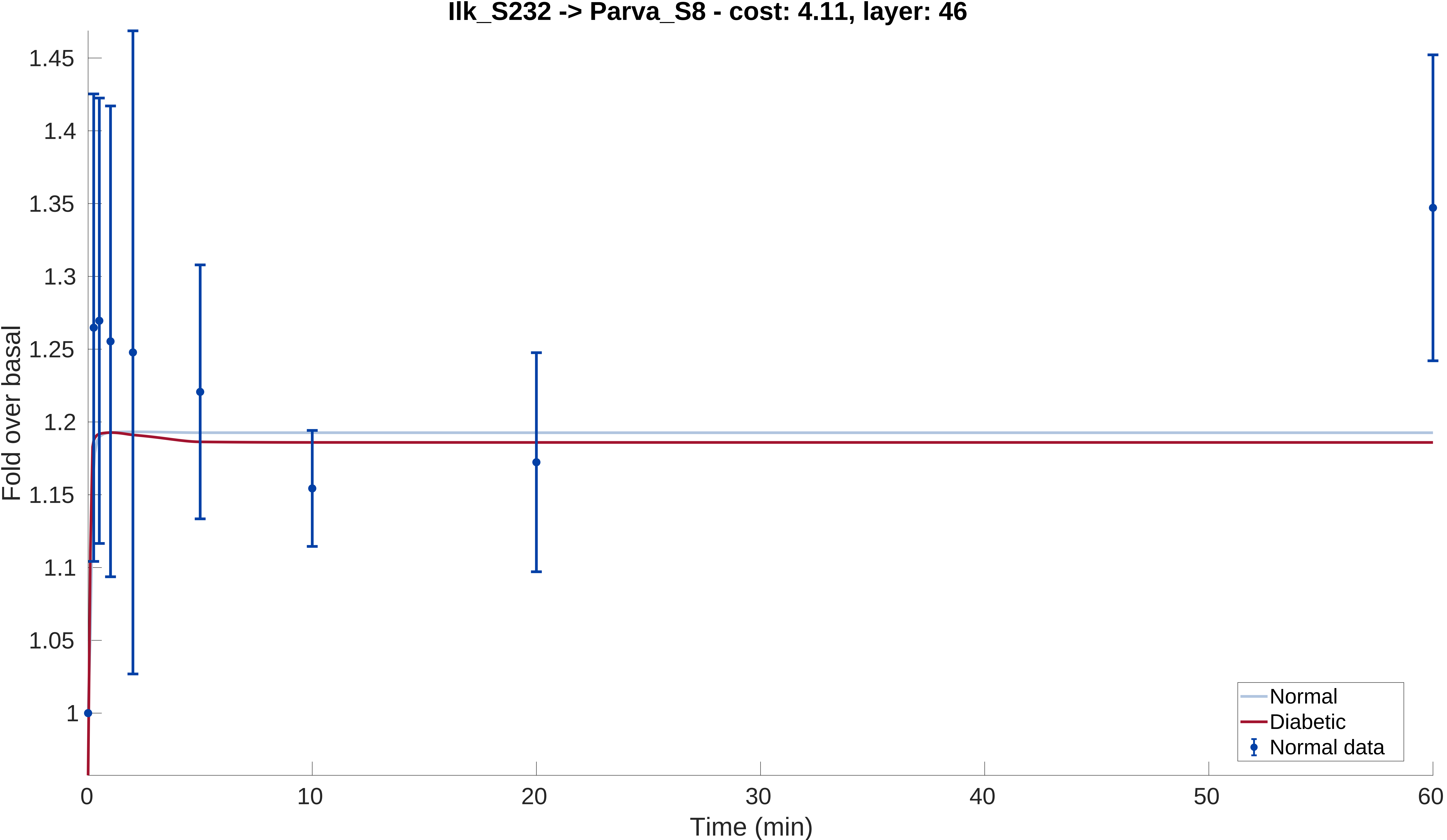

Ilk_S232 -> Parva_S8 - cost: 4.11, layer: 46
Fold over basal
1.45
1.4
1.35
1.3
1.25
1.2
1.15
1.1
1.05
1
0
10
20
30
40
50
60
Time (min)
Normal
Diabetic
Normal data

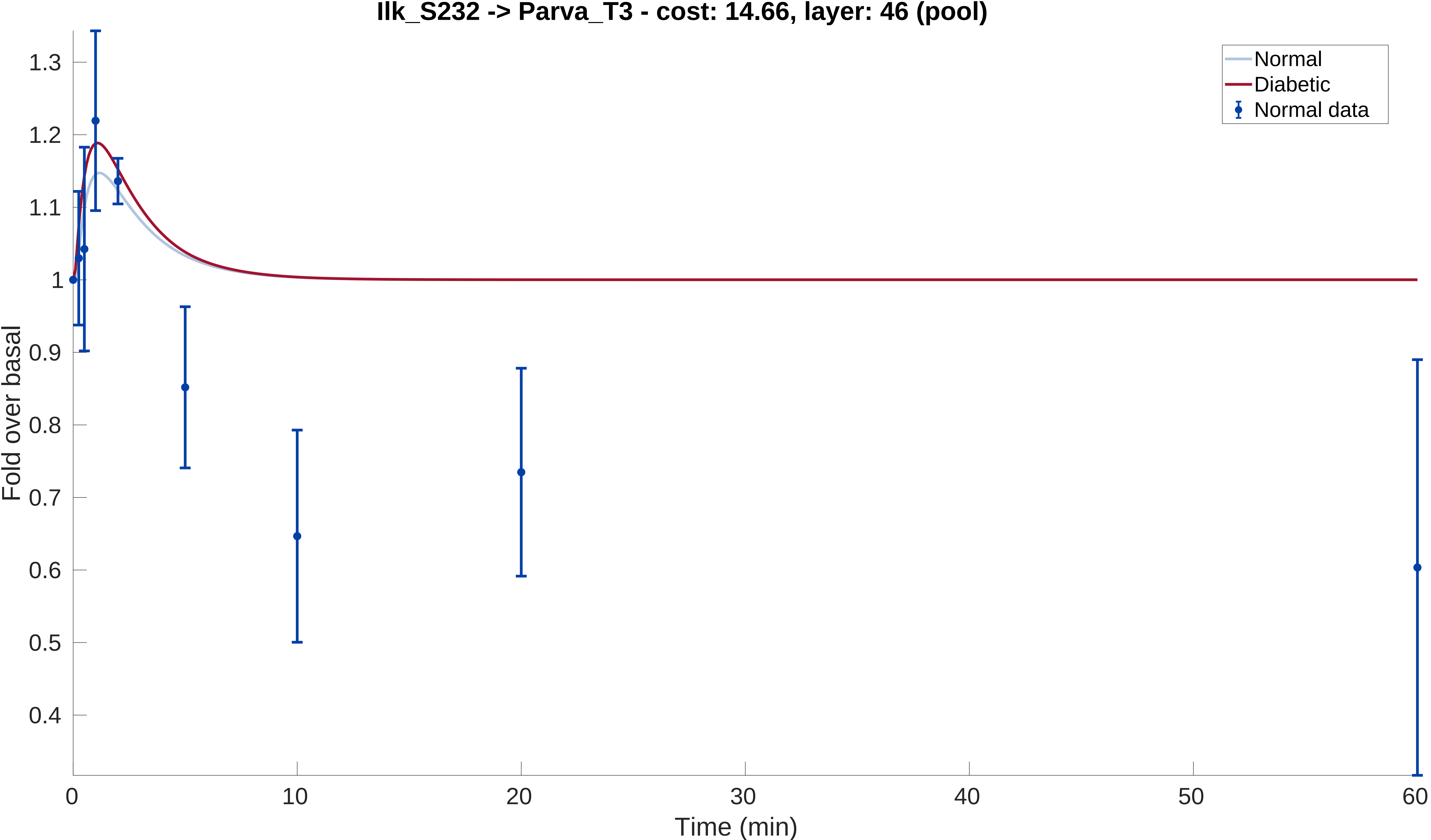

Ilk_S232 -> Parva_T3 - cost: 14.66, layer: 46 (pool)
Normal
Diabetic
Normal data
Fold over basal
Time (min)
1.3
1.2
1.1
1
0.9
0.8
0.7
0.6
0.5
0.4
0
10
20
30
40
50
60

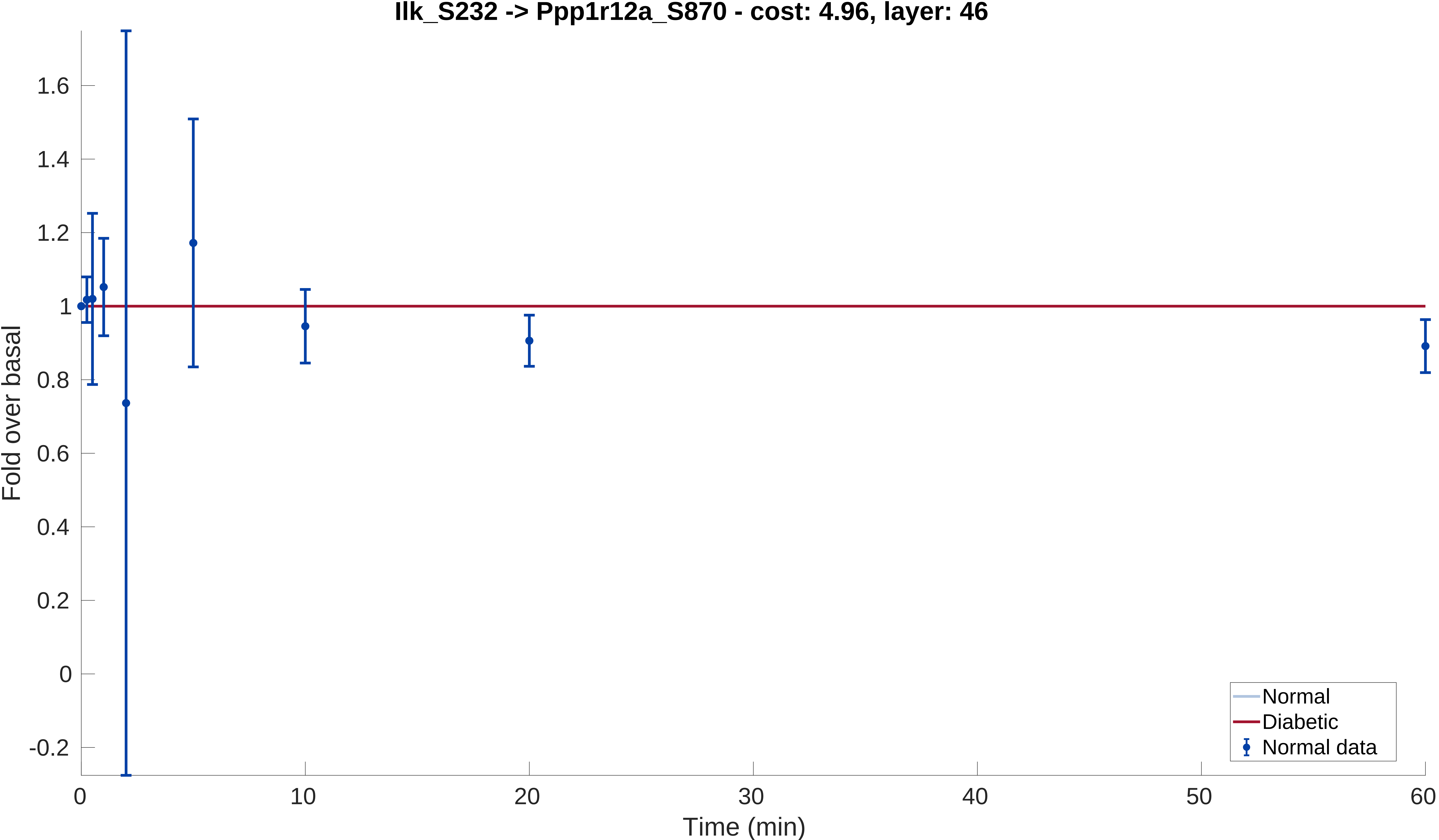

Ilk_S232 -> Ppp1r12a_S870 - cost: 4.96, layer: 46
Fold over basal
1.6
1.4
1.2
1
0.8
0.6
0.4
0.2
0
-0.2
0
10
20
30
40
50
60
Time (min)
Normal
Diabetic
Normal data

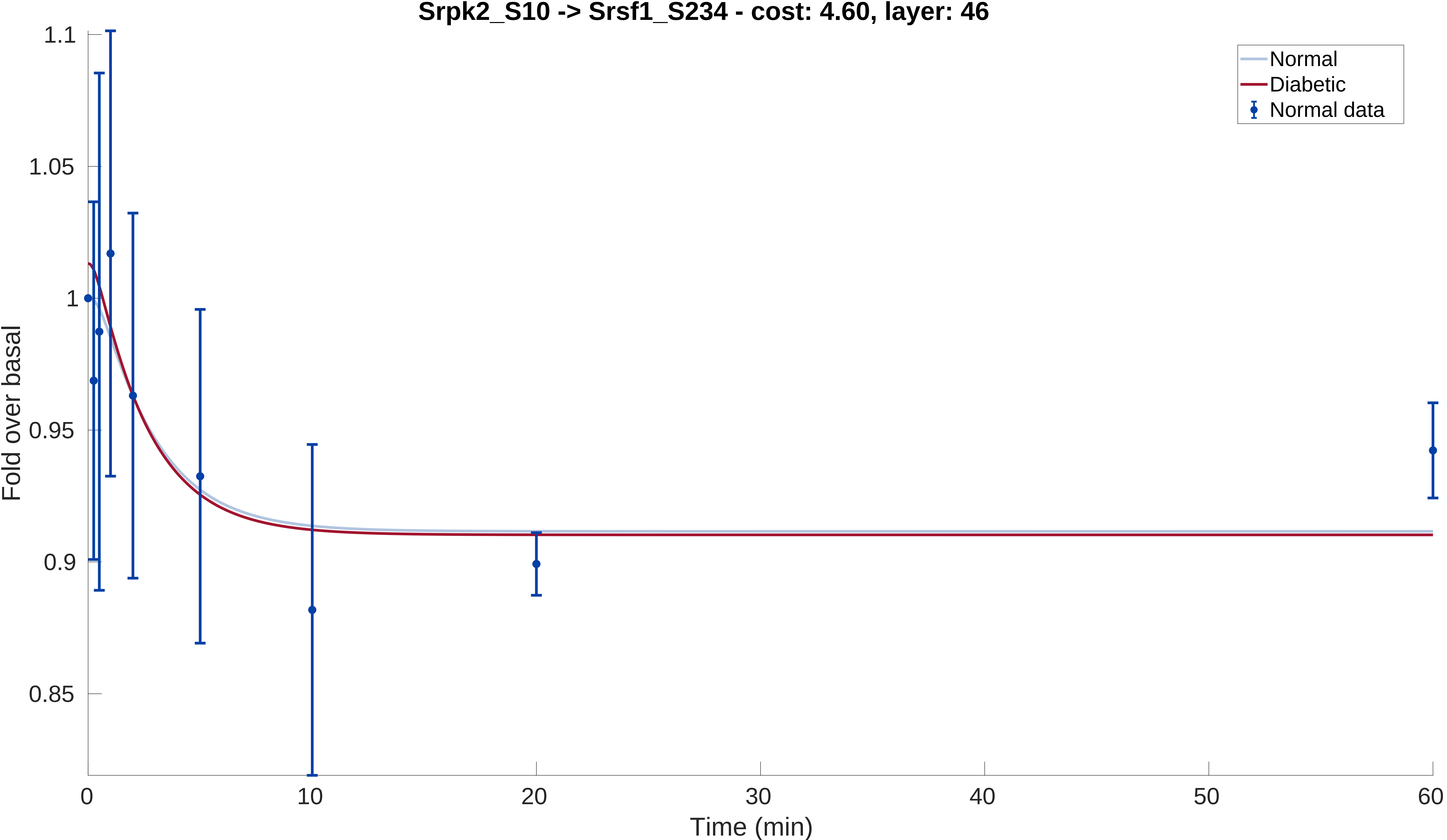
Srpk2_S10 -> Srsf1_S234 - cost: 4.60, layer: 46
Normal
Diabetic
Normal data
Fold over basal
Time (min)
1.1
1.05
1
0.95
0.9
0.85
0
10
20
30
40
50
60

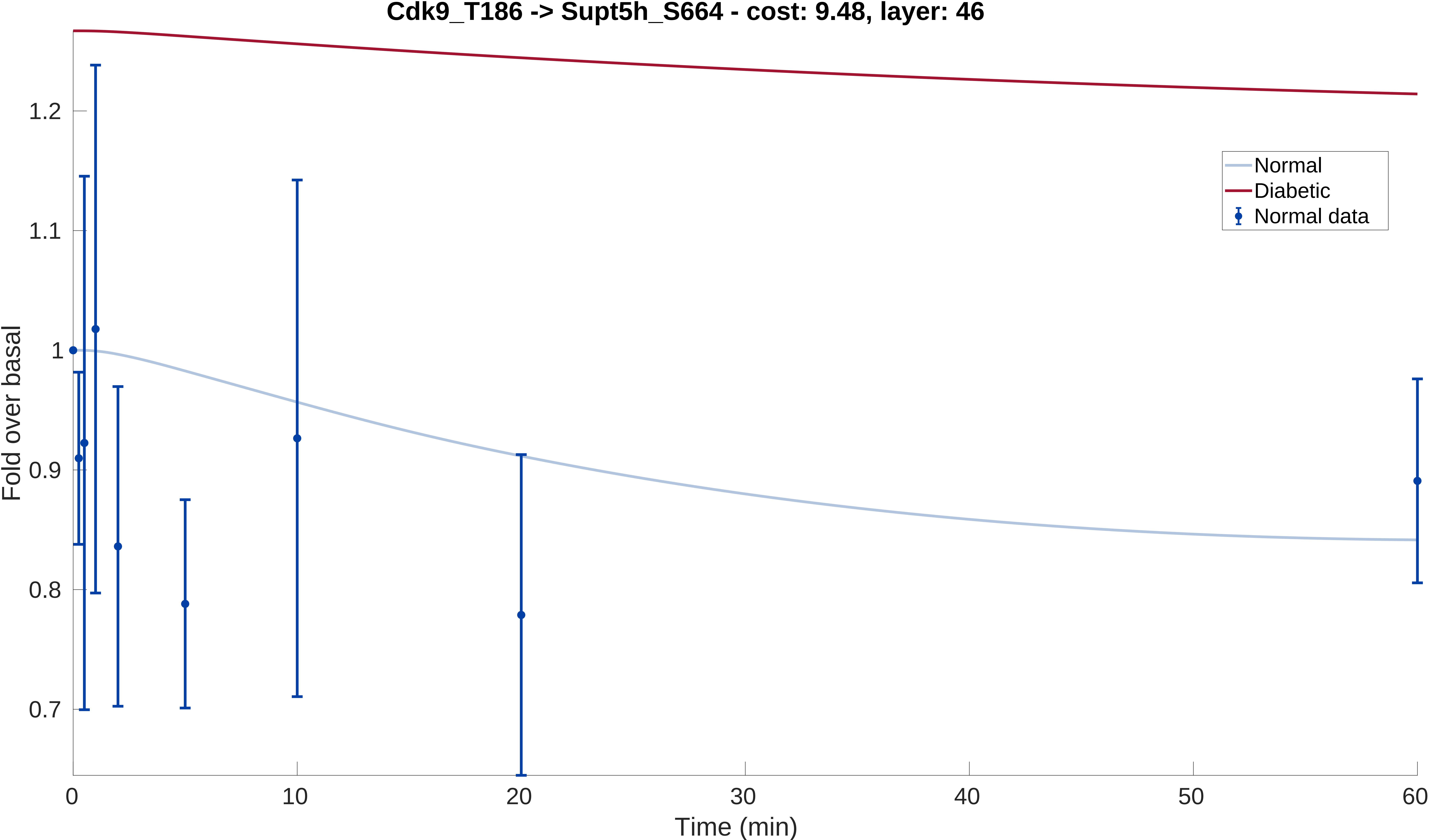

Cdk9_T186 -> Supt5h_S664 - cost: 9.48, layer: 46
Normal
Diabetic
Normal data
Fold over basal
Time (min)
1.2
1.1
1
0.9
0.8
0.7
0
10
20
30
40
50
60

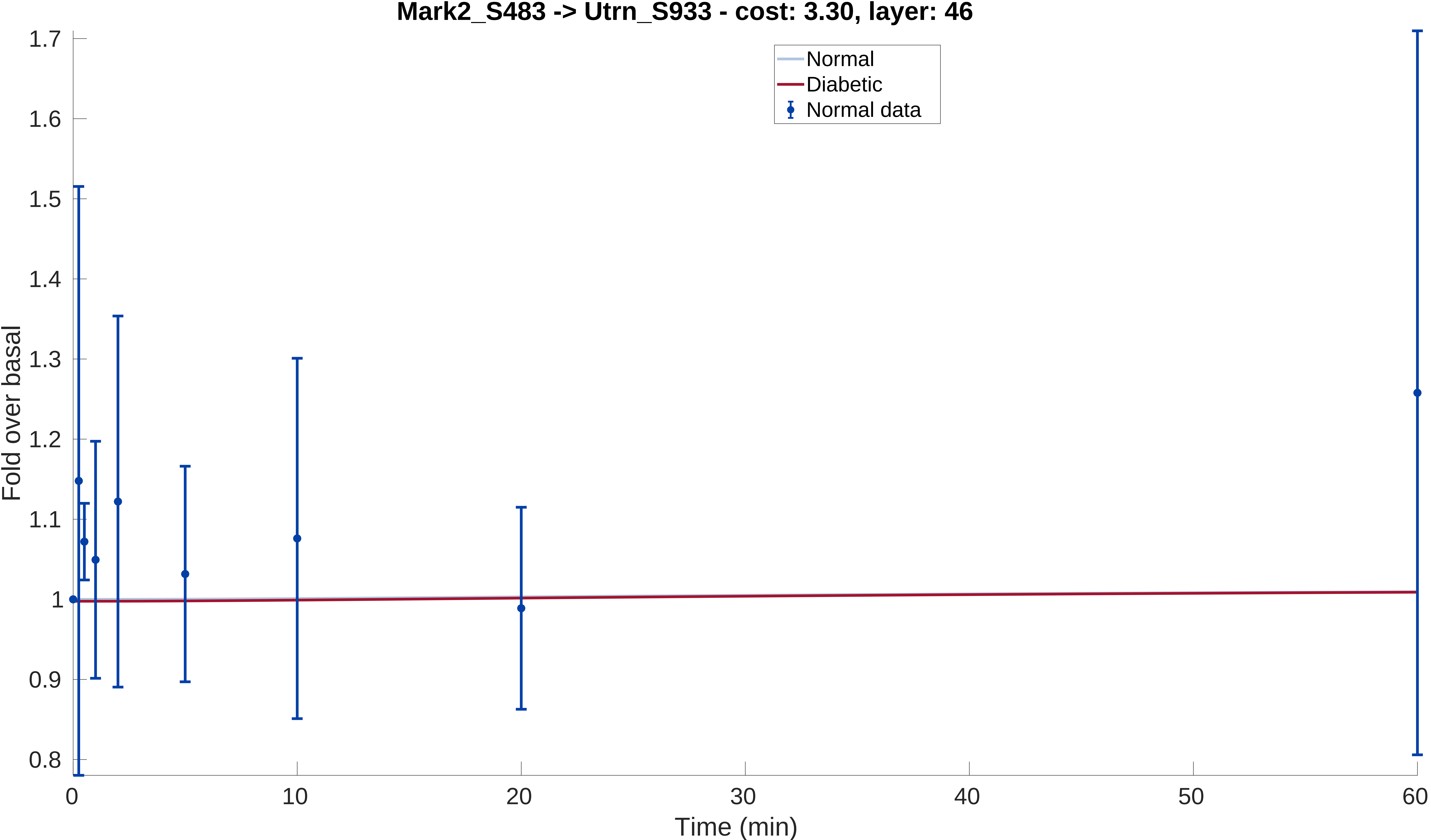

Mark2_S483 -> Utrn_S933 - cost: 3.30, layer: 46
Normal
Diabetic
Normal data
Fold over basal
Time (min)
1.7
1.6
1.5
1.4
1.3
1.2
1.1
1
0.9
0.8
0
10
20
30
40
50
60

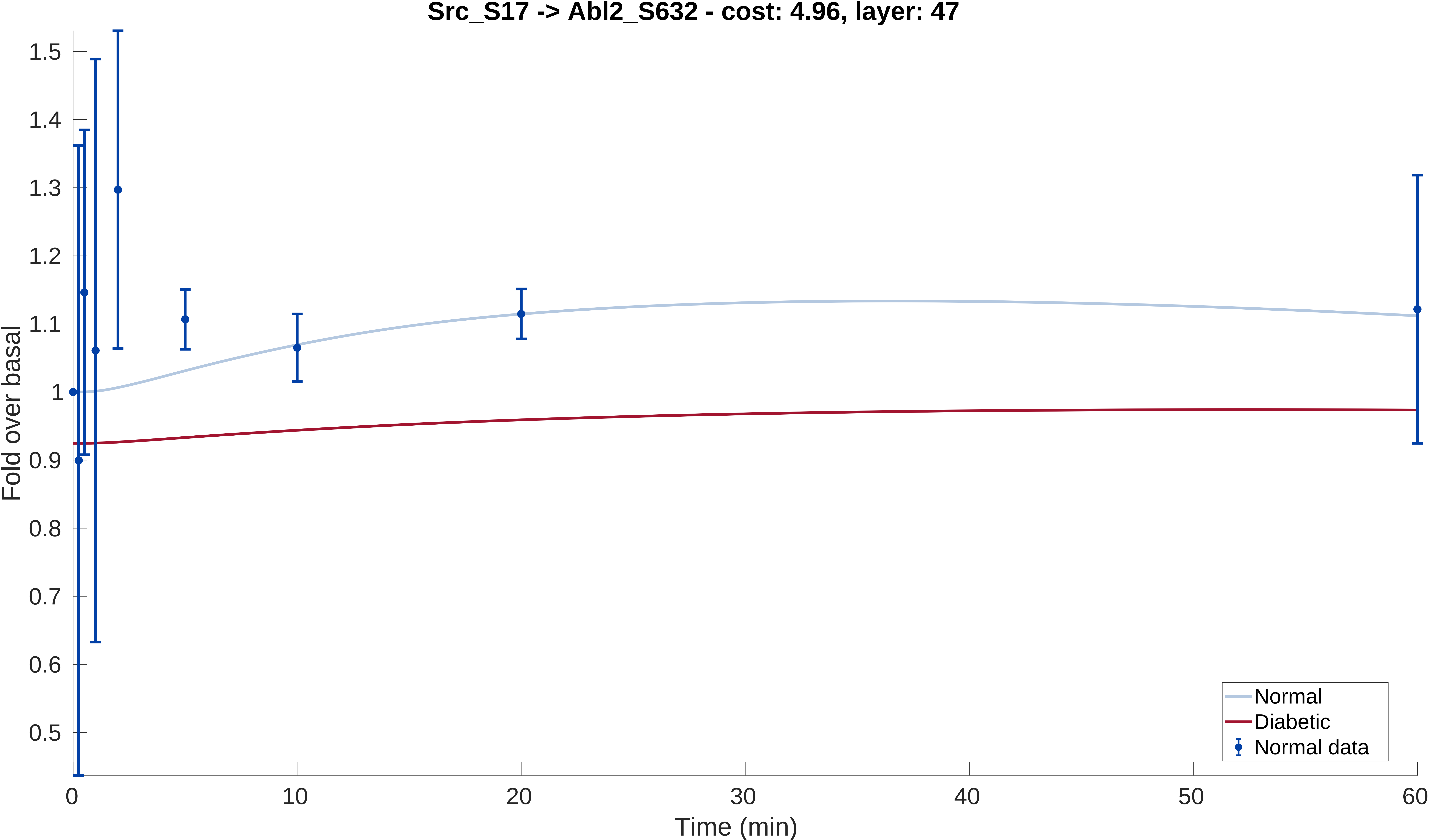

Src_S17 -> Abl2_S632 - cost: 4.96, layer: 47
Fold over basal
1.5
1.4
1.3
1.2
1.1
1
0.9
0.8
0.7
0.6
0.5
0
10
20
30
40
50
60
Time (min)
Normal
Diabetic
Normal data

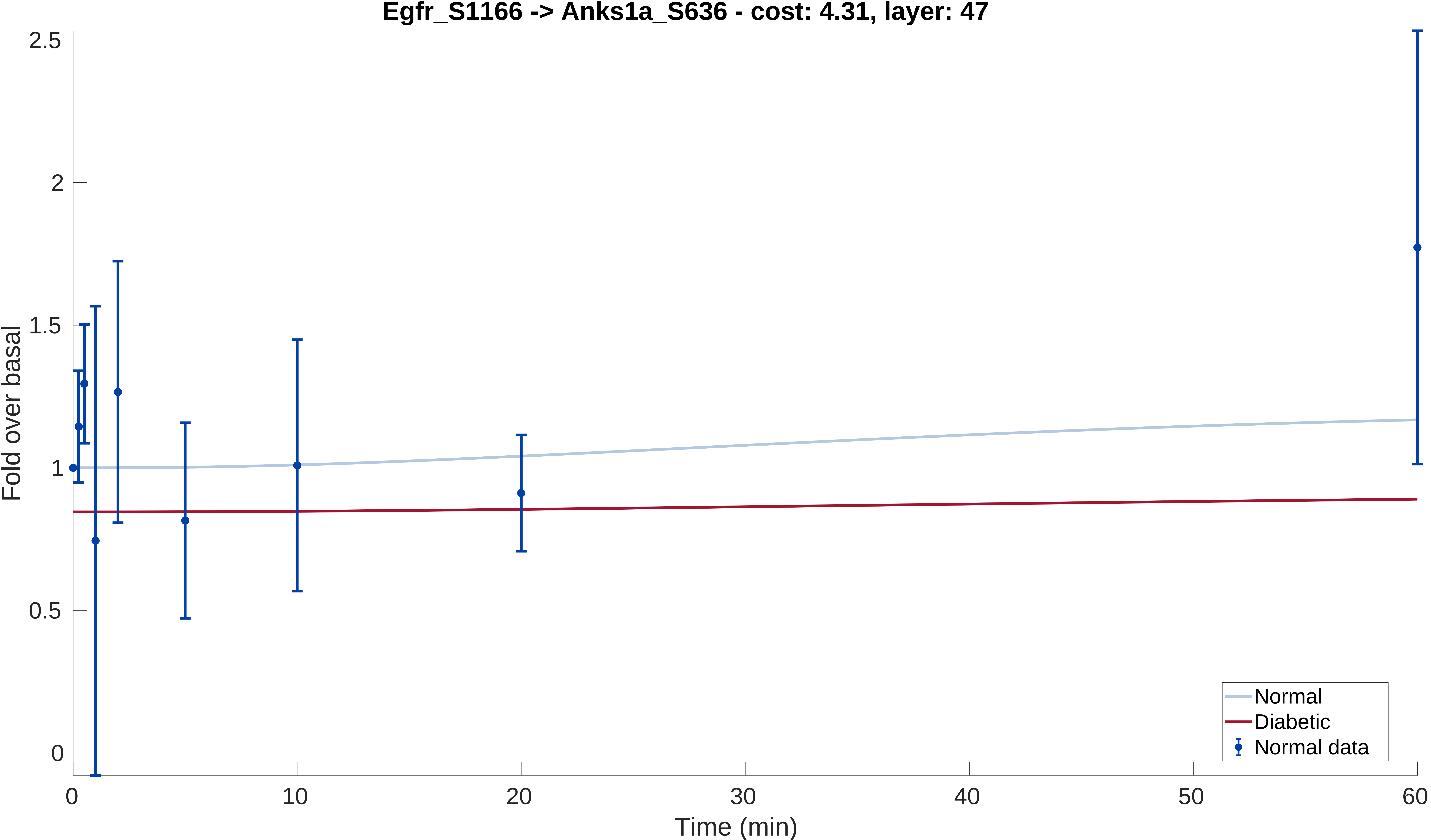

Egfr_S1166 -> Anks1a_S636 - cost: 4.31, layer: 47
Fold over basal
Time (min)
0
0.5
1
1.5
2
2.5
0
10
20
30
40
50
60
Normal
Diabetic
Normal data

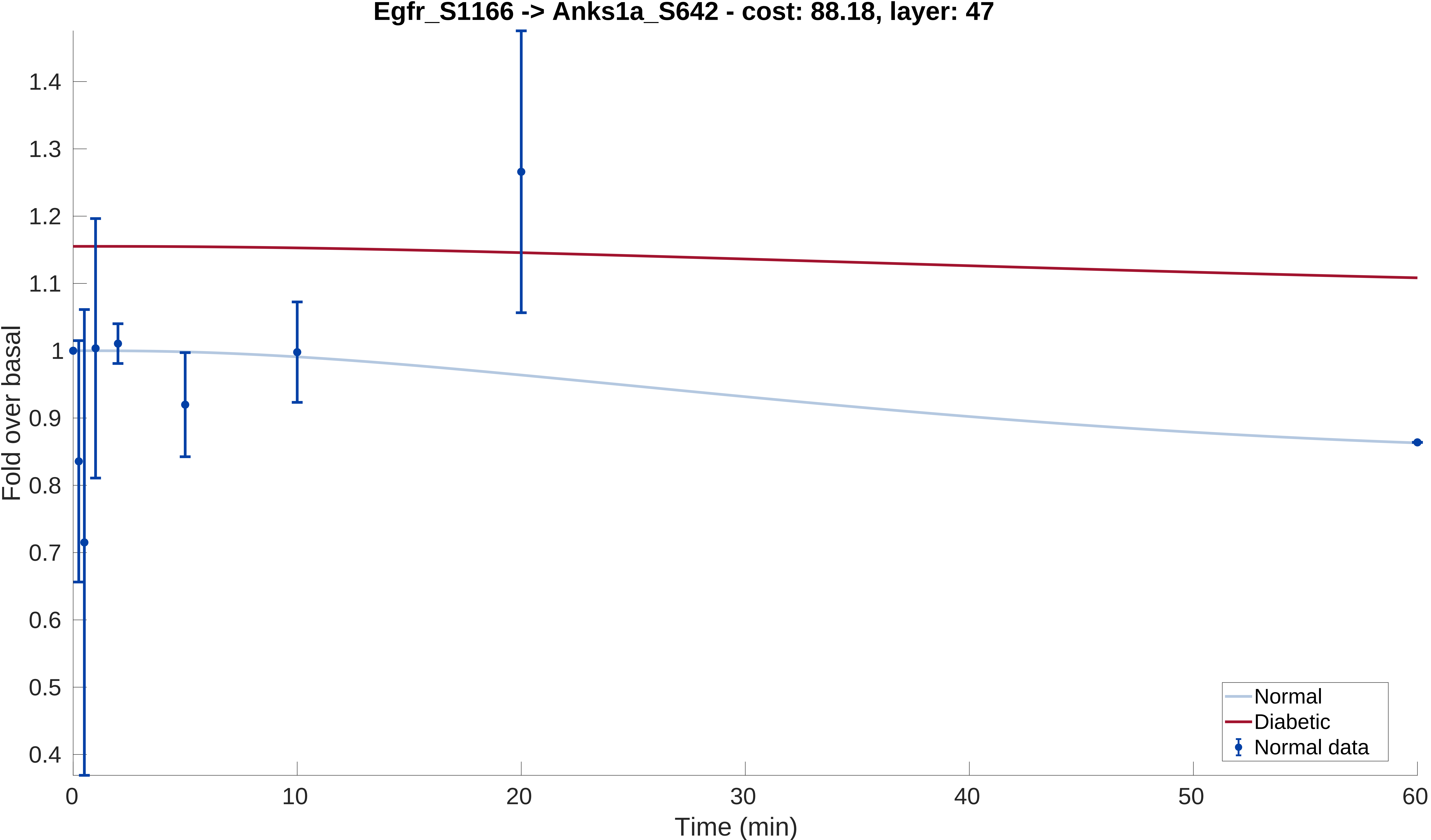

Egfr_S1166 -> Anks1a_S642 - cost: 88.18, layer: 47
Fold over basal
1.4
1.3
1.2
1.1
1
0.9
0.8
0.7
0.6
0.5
0.4
0
10
20
30
40
50
60
Time (min)
Normal
Diabetic
Normal data

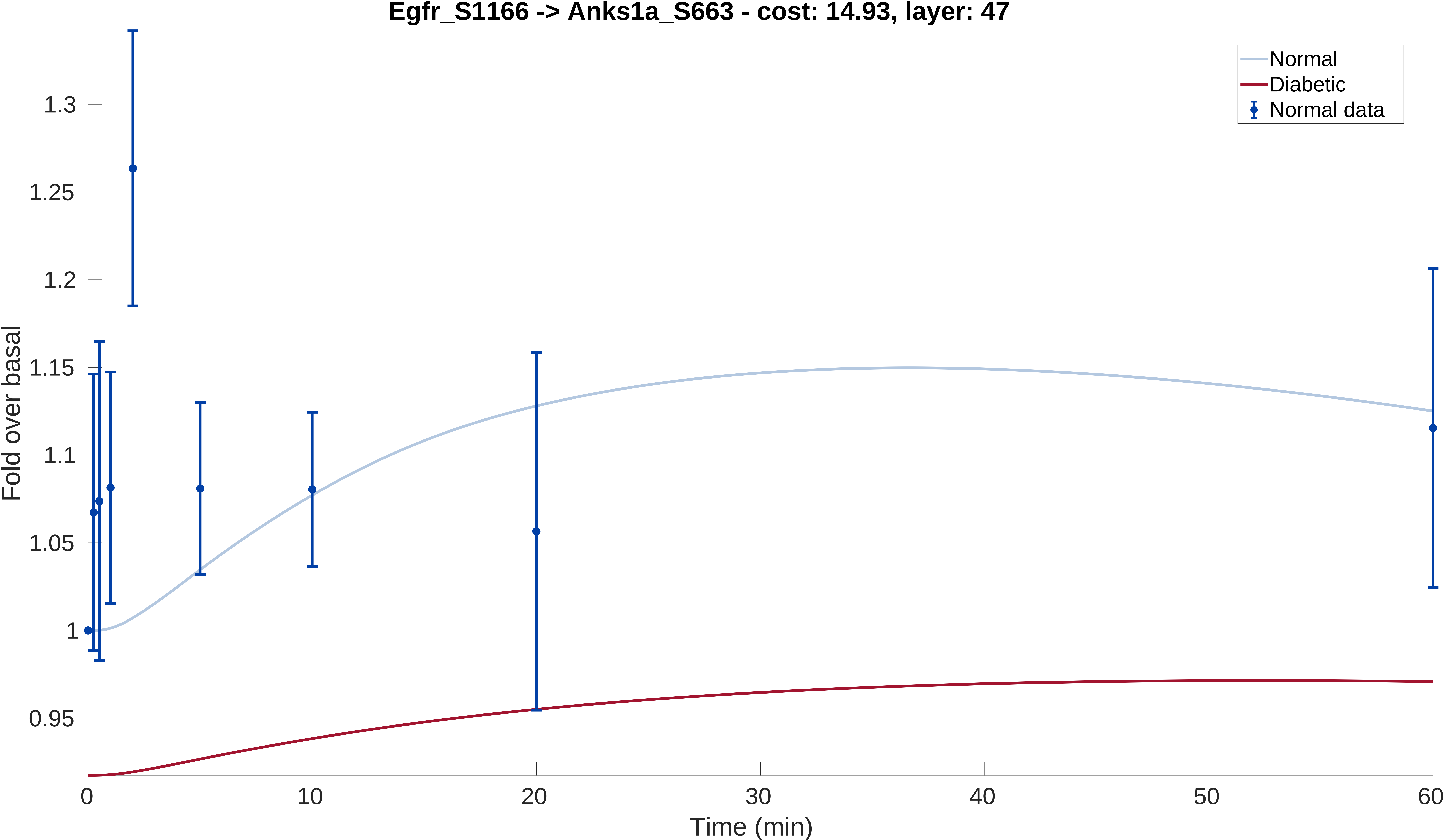

Egfr_S1166 -> Anks1a_S663 - cost: 14.93, layer: 47
Normal
Diabetic
Normal data
Fold over basal
1.3
1.25
1.2
1.15
1.1
1.05
1
0.95
0
10
20
30
40
50
60
Time (min)

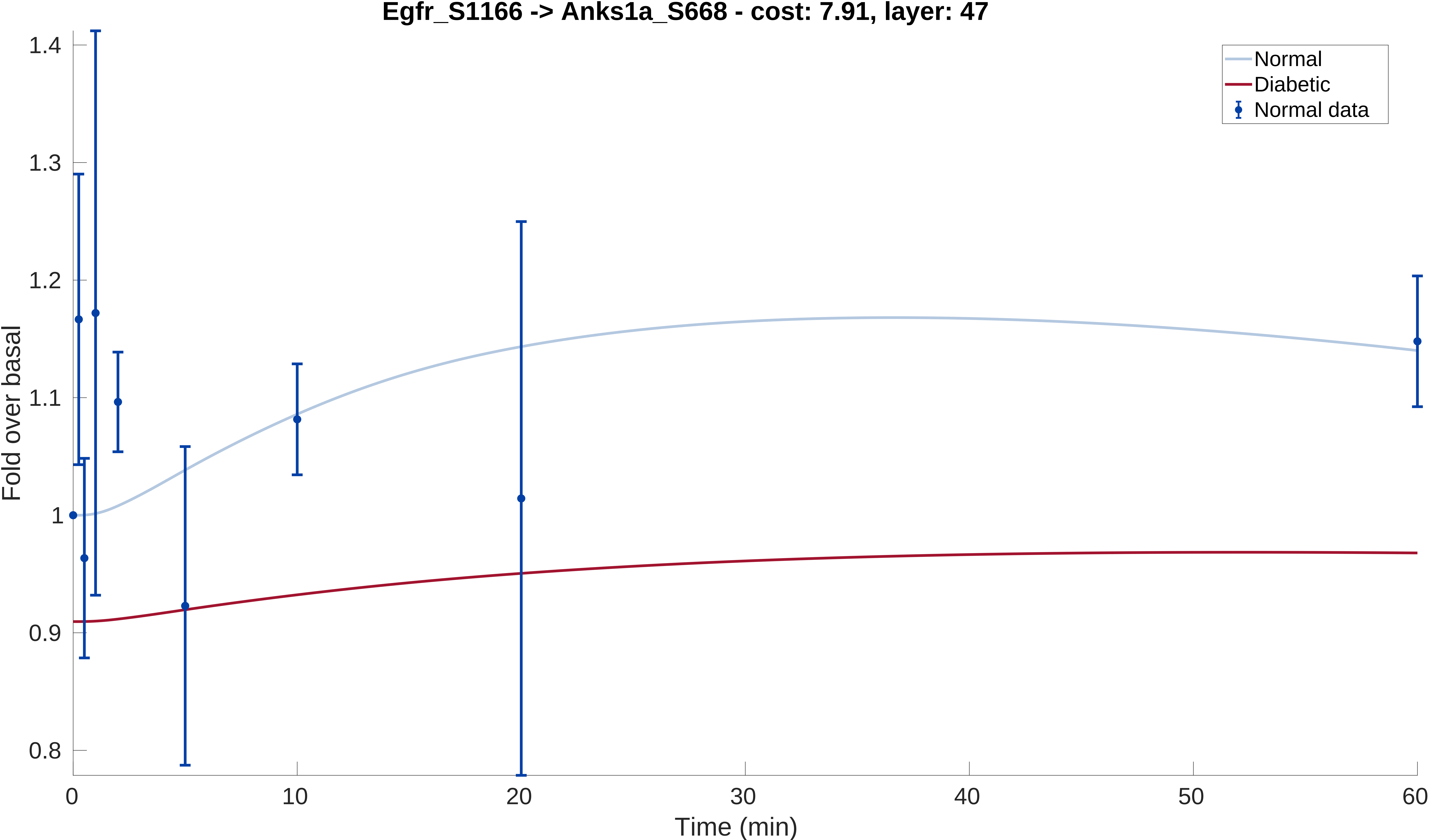

Egfr_S1166 -> Anks1a_S668 - cost: 7.91, layer: 47
Normal
Diabetic
Normal data
Fold over basal
1.4
1.3
1.2
1.1
1
0.9
0.8
0
10
20
30
40
50
60
Time (min)

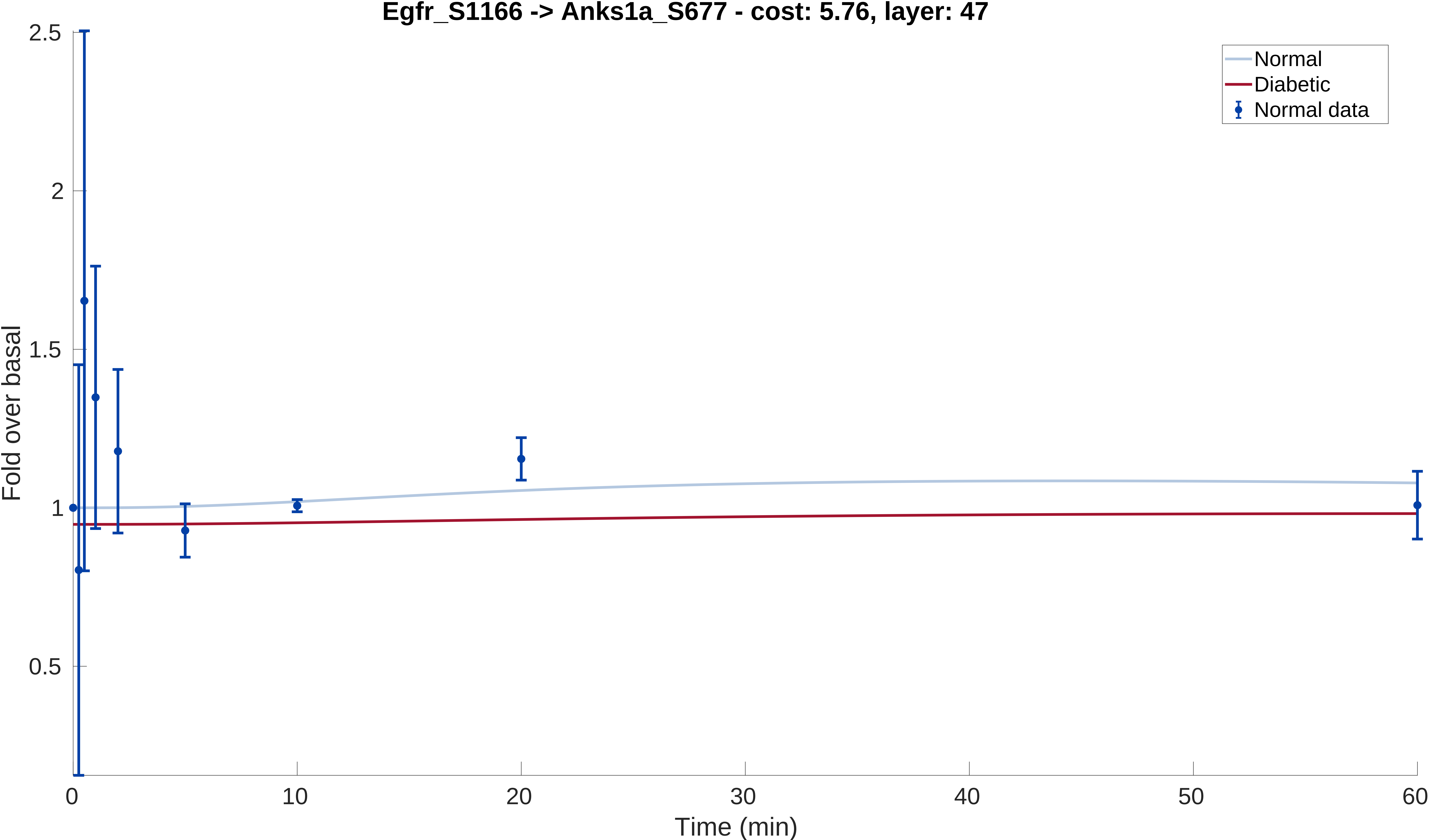
Egfr_S1166 -> Anks1a_S677 - cost: 5.76, layer: 47
Fold over basal
Time (min)
Normal
Diabetic
Normal data
0.5
1
1.5
2
2.5
0
10
20
30
40
50
60

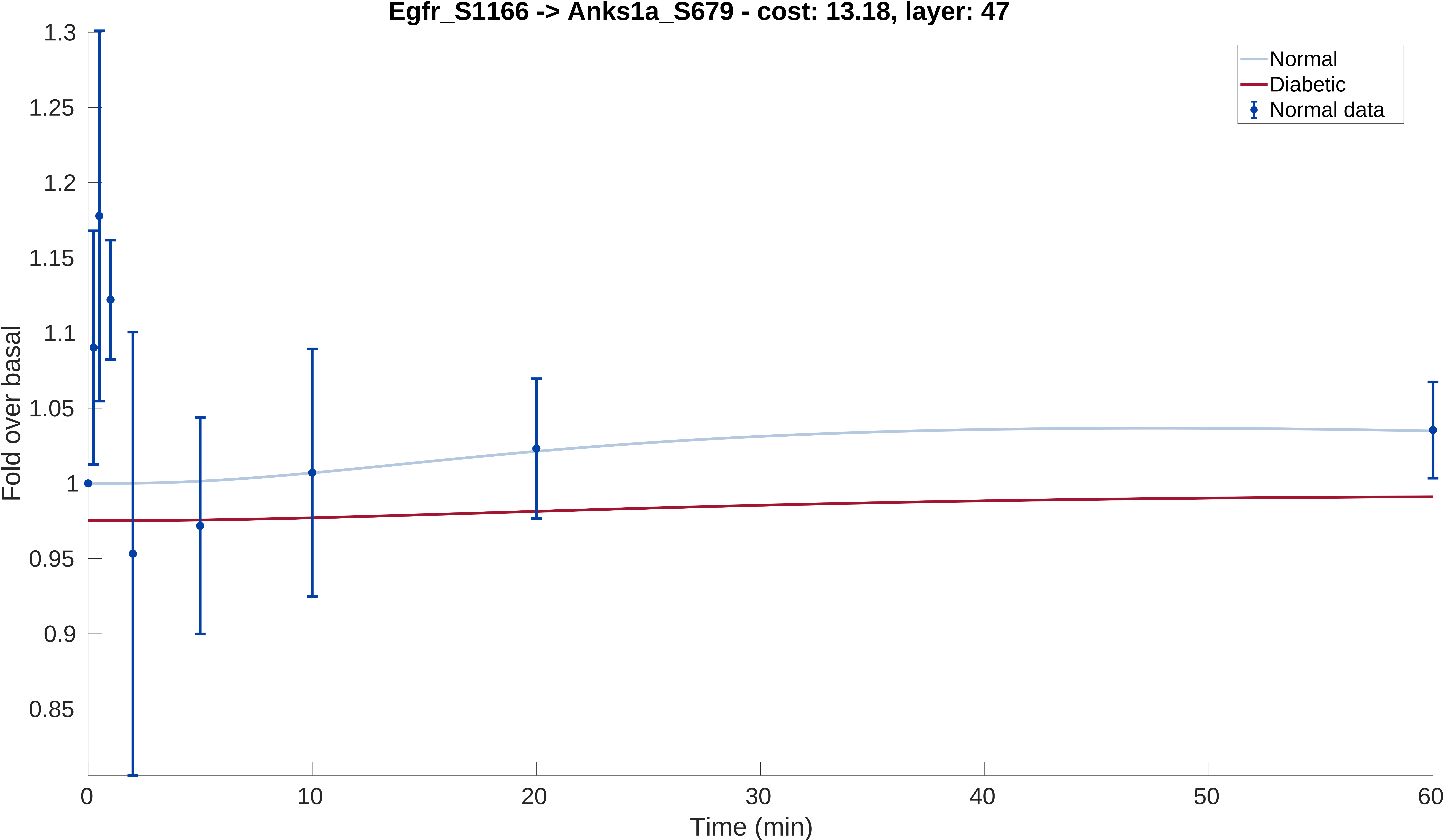

Egfr_S1166 -> Anks1a_S679 - cost: 13.18, layer: 47
Normal
Diabetic
Normal data
Fold over basal
1.3
1.25
1.2
1.15
1.1
1.05
1
0.95
0.9
0.85
0
10
20
30
40
50
60
Time (min)

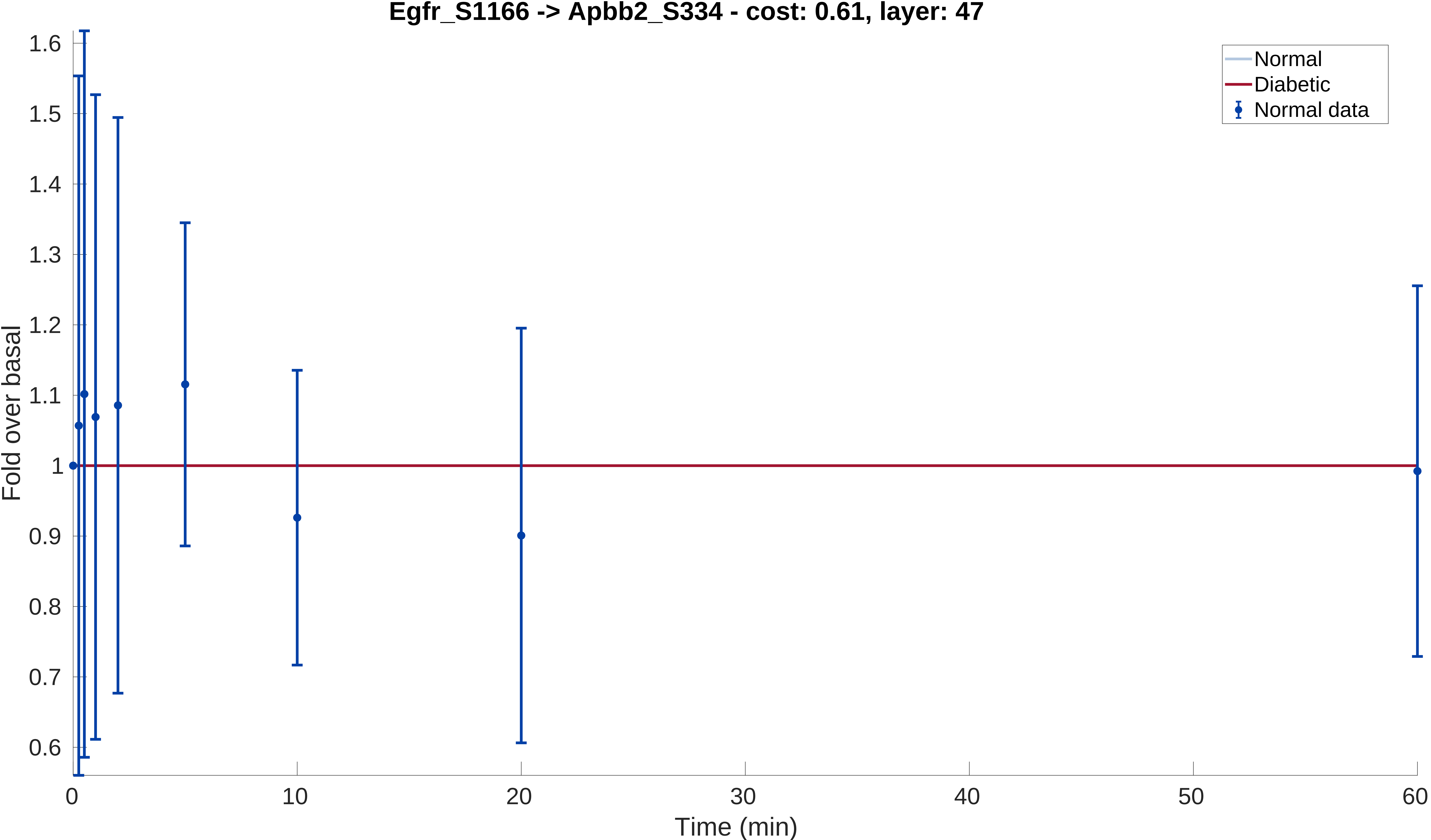

Egfr_S1166 -> Apbb2_S334 - cost: 0.61, layer: 47
Normal
Diabetic
Normal data
Fold over basal
Time (min)
1.6
1.5
1.4
1.3
1.2
1.1
1
0.9
0.8
0.7
0.6
0
10
20
30
40
50
60

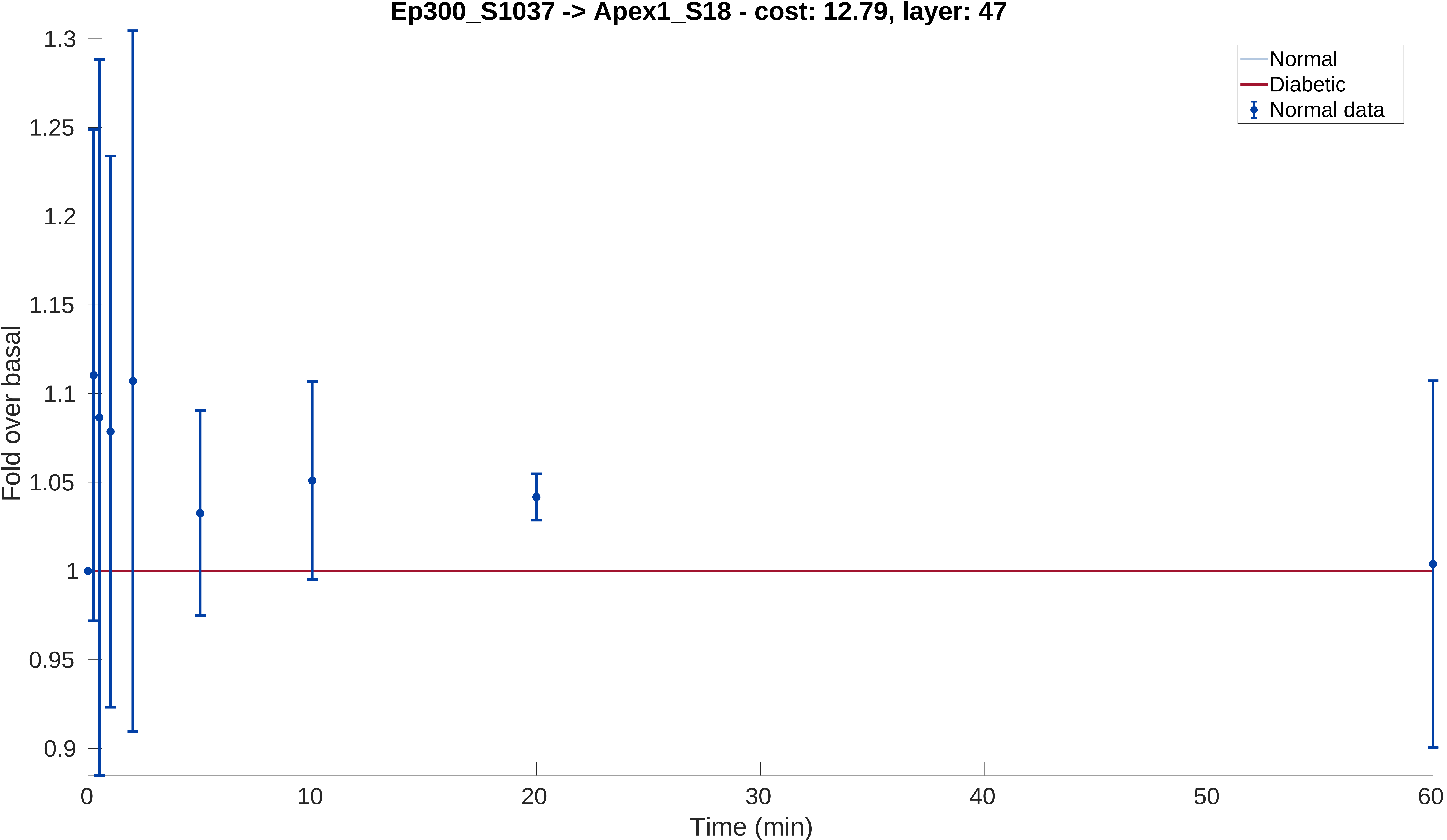

Ep300_S1037 -> Apex1_S18 - cost: 12.79, layer: 47
Normal
Diabetic
Normal data
Fold over basal
Time (min)
1.3
1.25
1.2
1.15
1.1
1.05
1
0.95
0.9
0
10
20
30
40
50
60

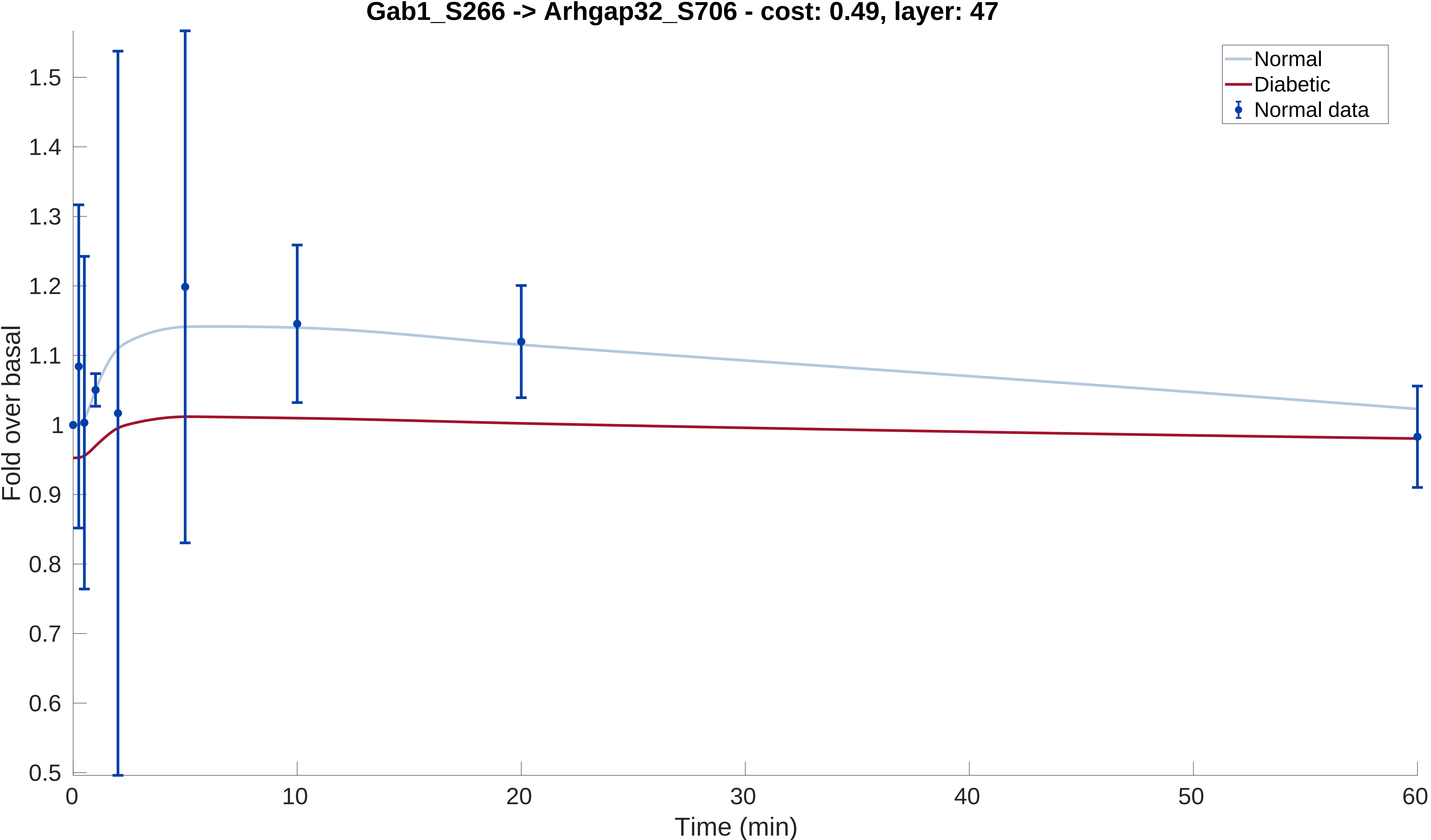

Gab1_S266 -> Arhgap32_S706 - cost: 0.49, layer: 47
Normal
Diabetic
Normal data
Fold over basal
1.5
1.4
1.3
1.2
1.1
1
0.9
0.8
0.7
0.6
0.5
0
10
20
30
40
50
60
Time (min)

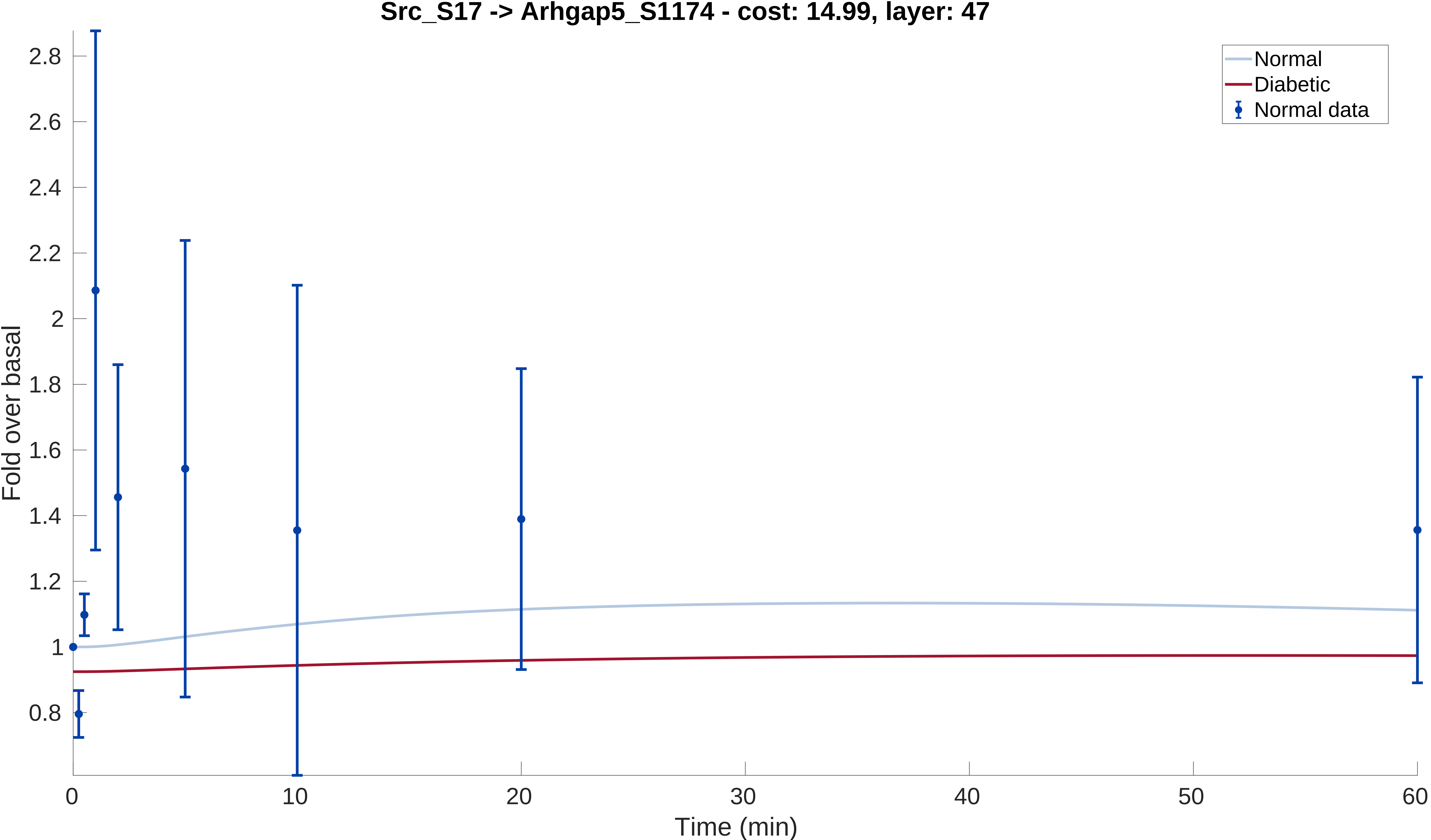

Src_S17 -> Arhgap5_S1174 - cost: 14.99, layer: 47
Fold over basal
Time (min)
2.8
2.6
2.4
2.2
2
1.8
1.6
1.4
1.2
1
0.8
0
10
20
30
40
50
60
Normal
Diabetic
Normal data

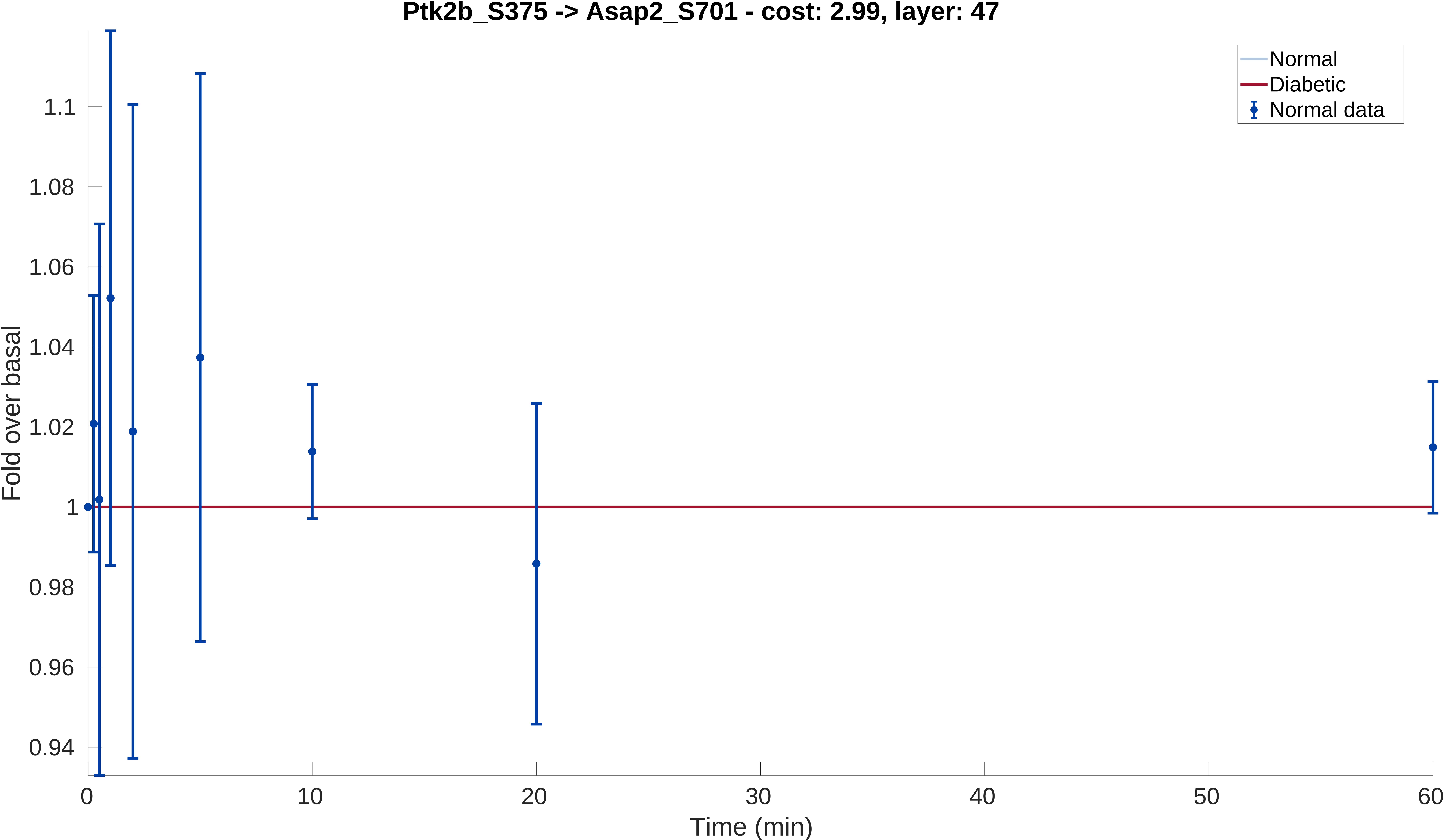

Ptk2b_S375 -> Asap2_S701 - cost: 2.99, layer: 47
Normal
Diabetic
Normal data
Fold over basal
1.1
1.08
1.06
1.04
1.02
1
0.98
0.96
0.94
0
10
20
30
40
50
60
Time (min)

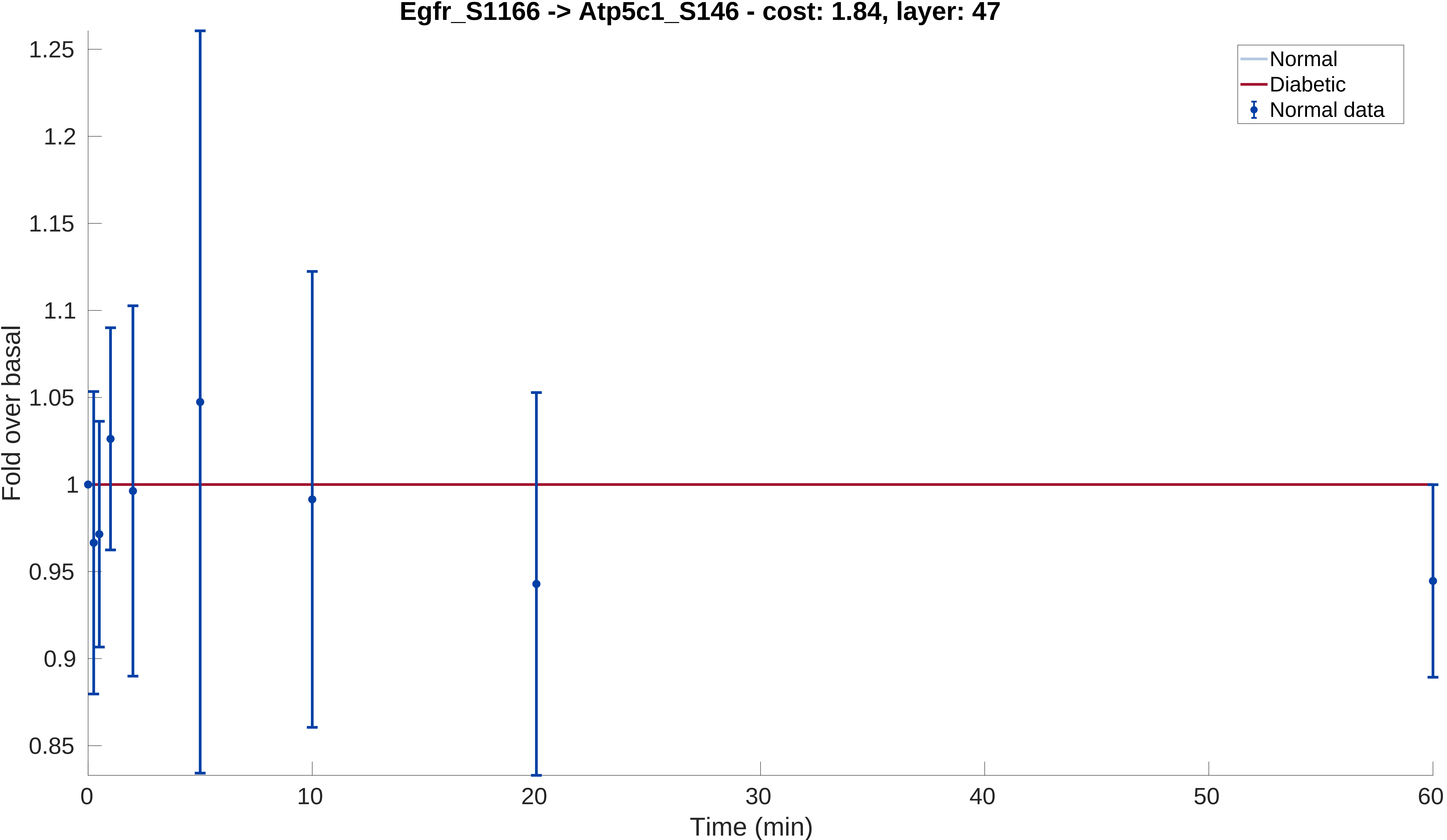

Egfr_S1166 -> Atp5c1_S146 - cost: 1.84, layer: 47
Normal
Diabetic
Normal data
Fold over basal
1.25
1.2
1.15
1.1
1.05
1
0.95
0.9
0.85
0
10
20
30
40
50
60
Time (min)

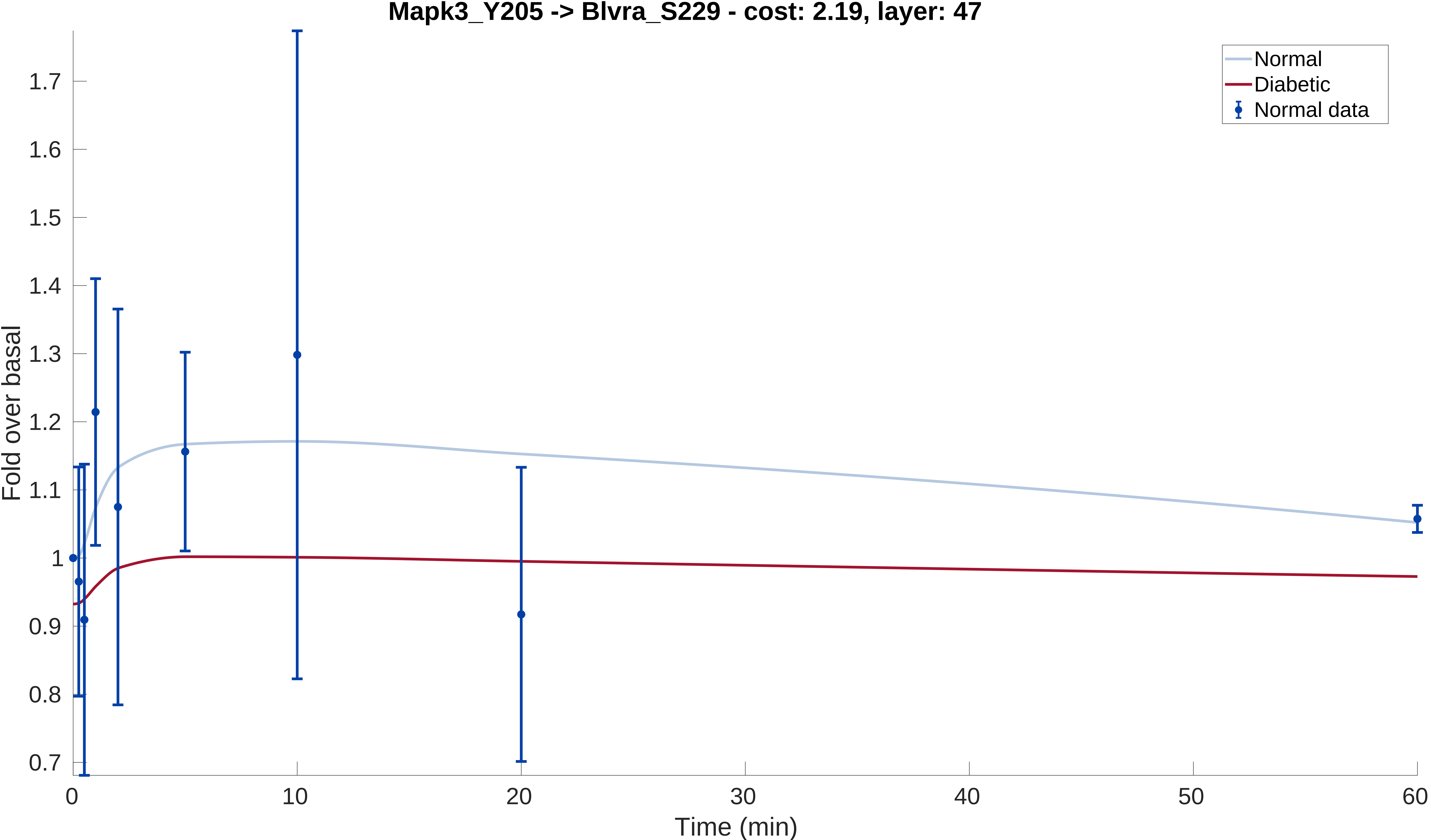
Mapk3_Y205 -> Blvra_S229 - cost: 2.19, layer: 47
Normal
Diabetic
Normal data
Fold over basal
0.7
0.8
0.9
1
1.1
1.2
1.3
1.4
1.5
1.6
1.7
0
10
20
30
40
50
60
Time (min)

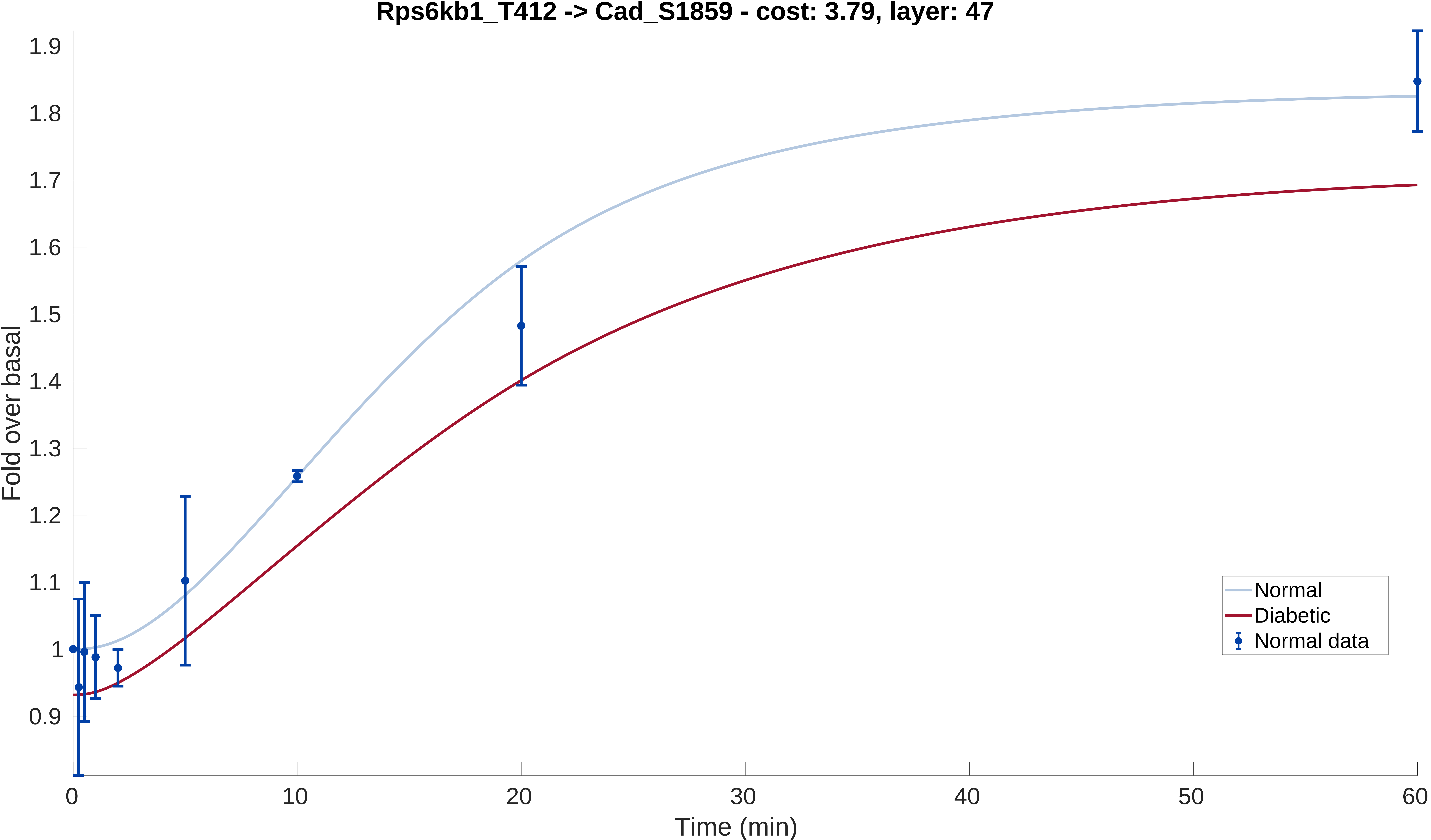
Rps6kb1_T412 -> Cad_S1859 - cost: 3.79, layer: 47
Fold over basal
1.9
1.8
1.7
1.6
1.5
1.4
1.3
1.2
1.1
1
0.9
0
10
20
30
40
50
60
Time (min)
Normal
Diabetic
Normal data

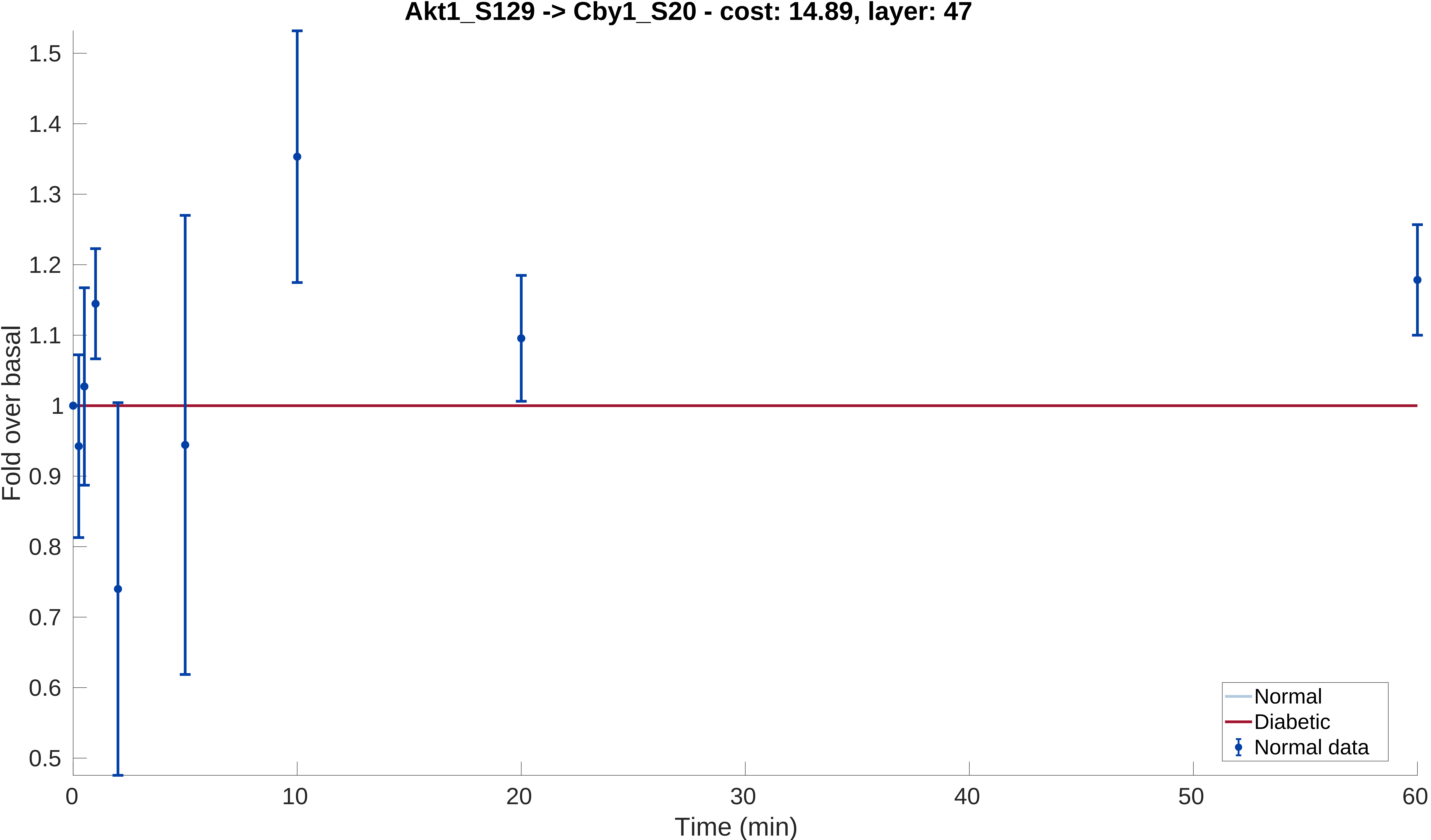

Akt1_S129 -> Cby1_S20 - cost: 14.89, layer: 47
Fold over basal
1.5
1.4
1.3
1.2
1.1
1
0.9
0.8
0.7
0.6
0.5
0
10
20
30
40
50
60
Time (min)
Normal
Diabetic
Normal data

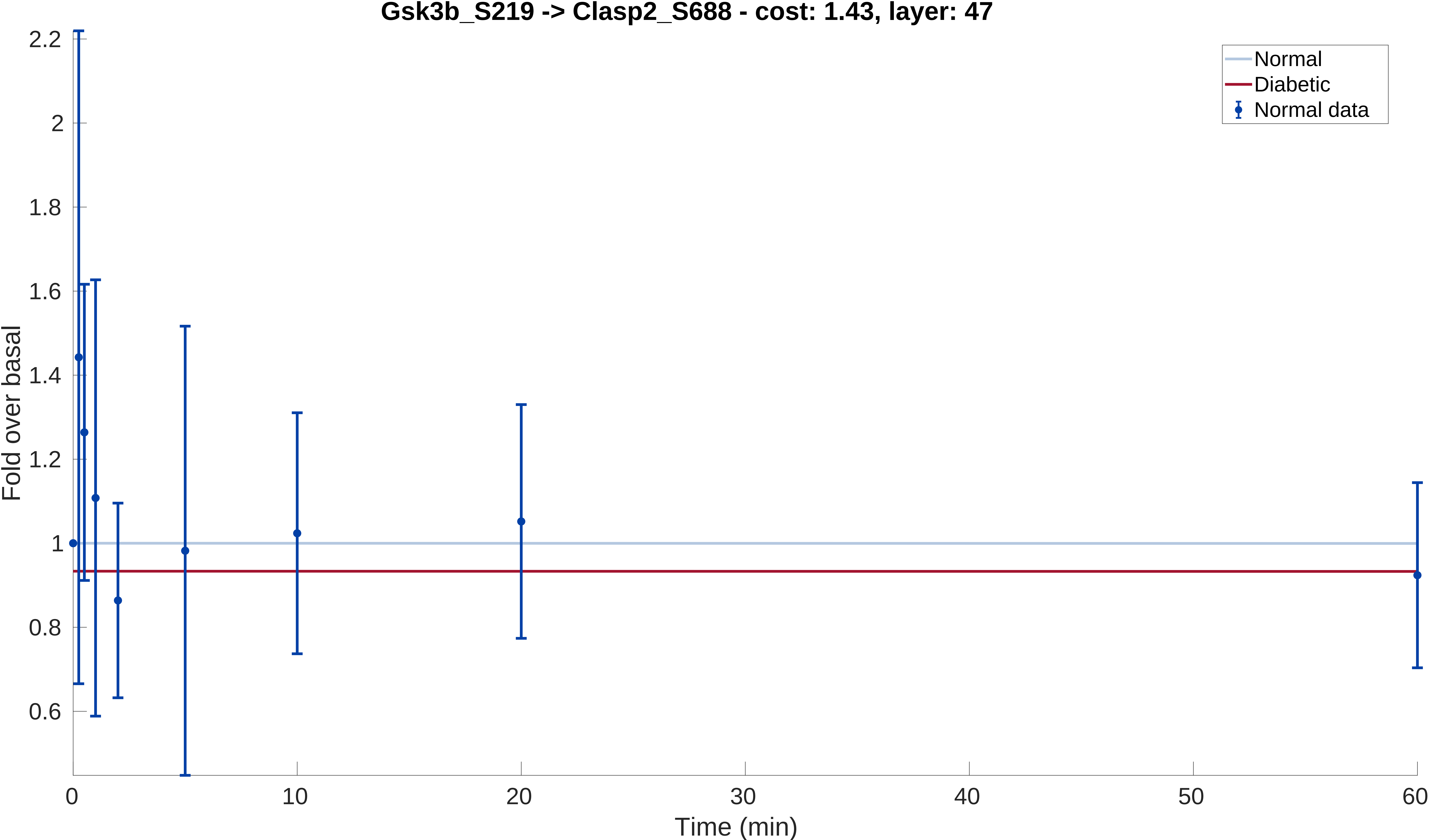

Gsk3b_S219 -> Clasp2_S688 - cost: 1.43, layer: 47
Normal
Diabetic
Normal data
Fold over basal
2.2
2
1.8
1.6
1.4
1.2
1
0.8
0.6
0
10
20
30
40
50
60
Time (min)

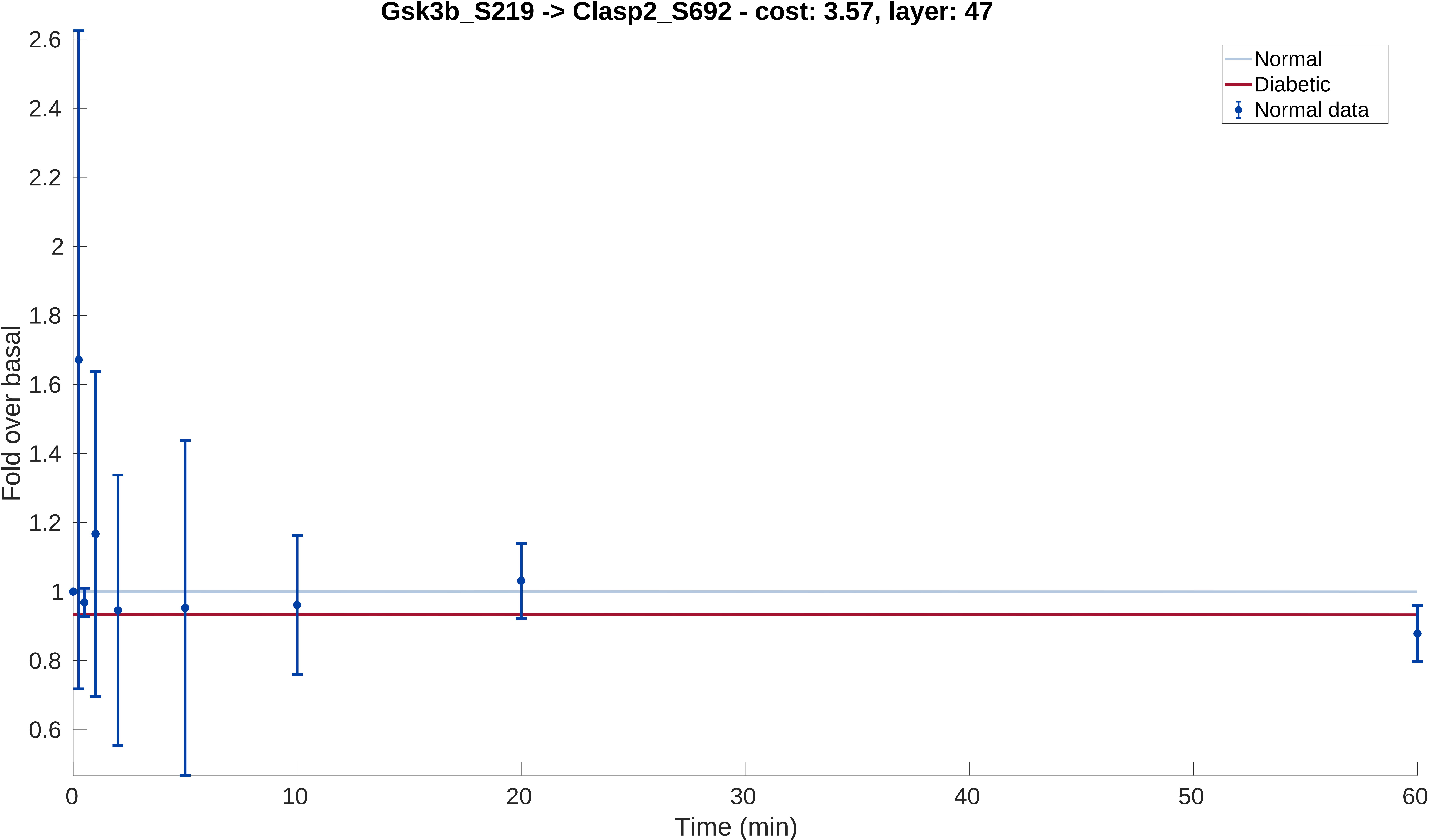

Gsk3b_S219 -> Clasp2_S692 - cost: 3.57, layer: 47
Normal
Diabetic
Normal data
Fold over basal
2.6
2.4
2.2
2
1.8
1.6
1.4
1.2
1
0.8
0.6
0
10
20
30
40
50
60
Time (min)

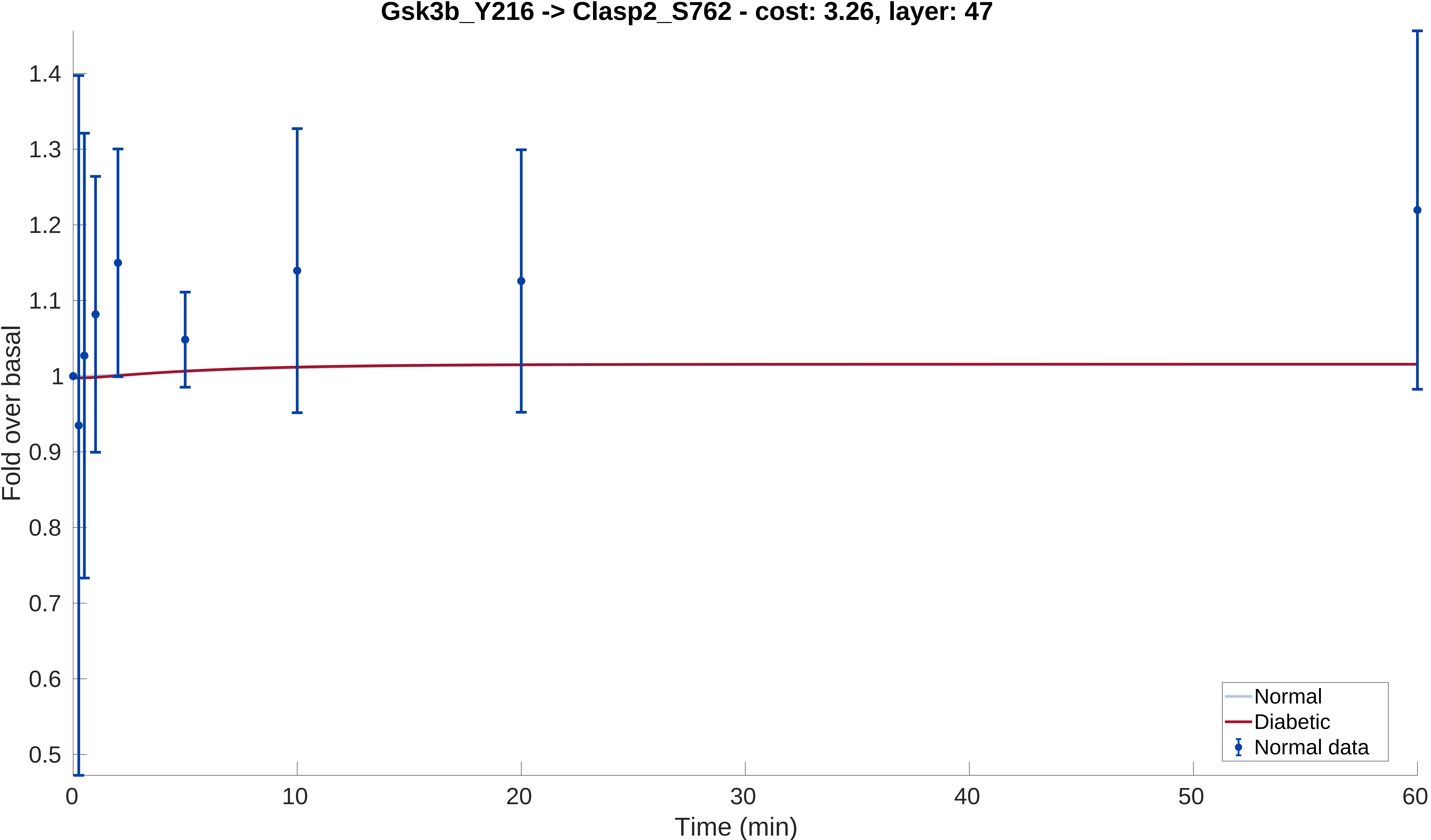
Gsk3b_Y216 -> Clasp2_S762 - cost: 3.26, layer: 47
Fold over basal
Time (min)
1.4
1.3
1.2
1.1
1
0.9
0.8
0.7
0.6
0.5
0
10
20
30
40
50
60
Normal
Diabetic
Normal data

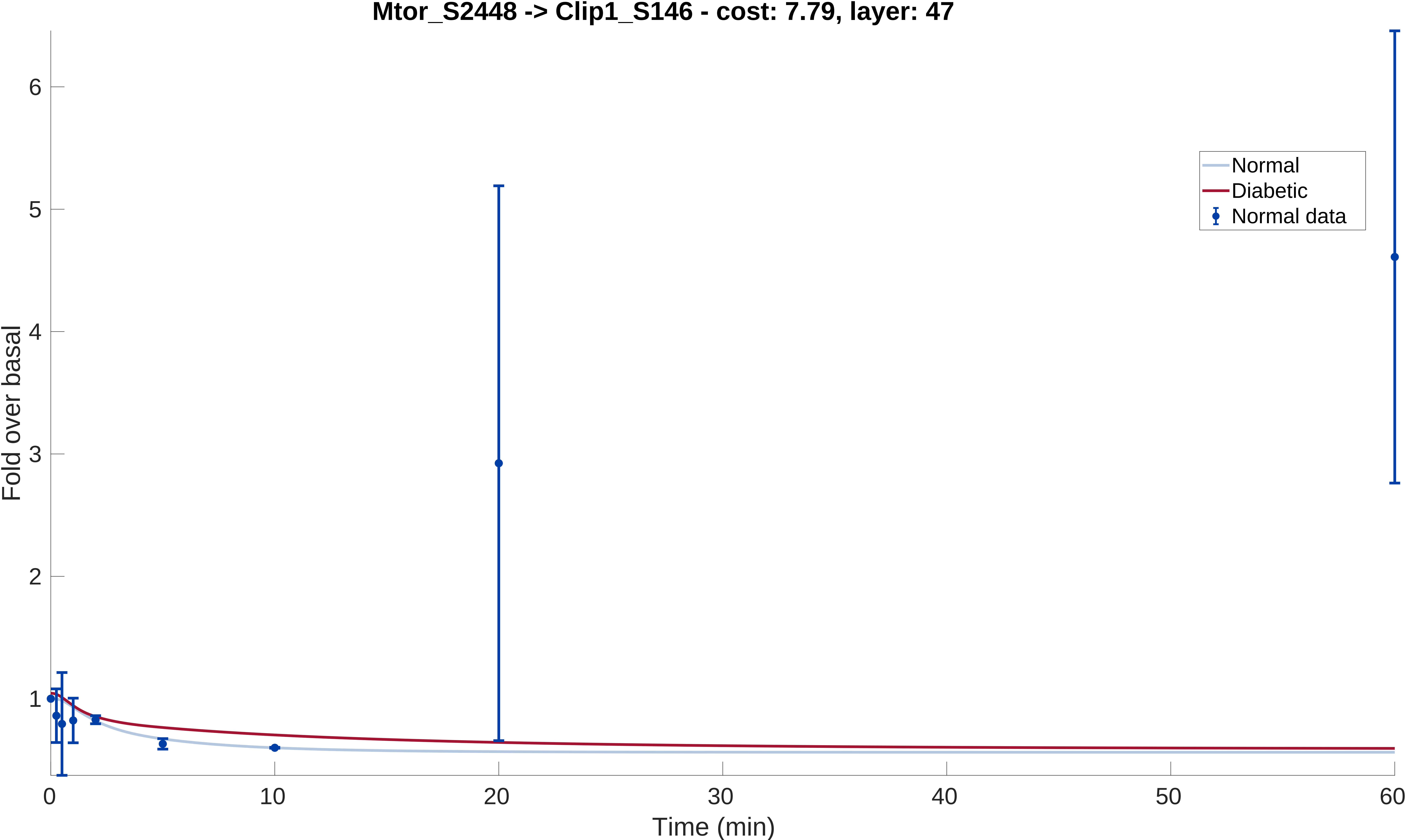
Mtor_S2448 -> Clip1_S146 - cost: 7.79, layer: 47
Fold over basal
Time (min)
Normal
Diabetic
Normal data
0
10
20
30
40
50
60
1
2
3
4
5
6

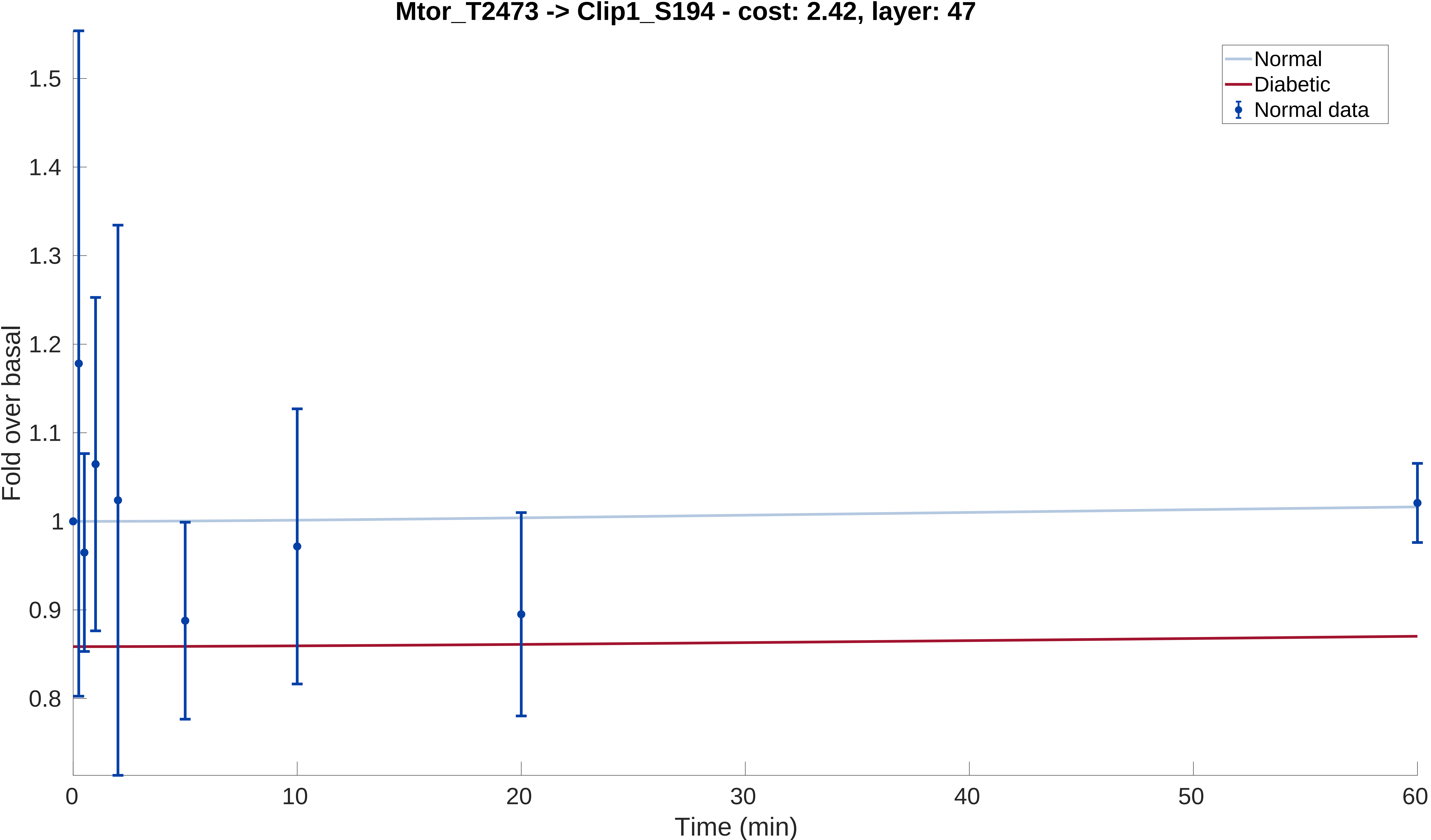

Mtor_T2473 -> Clip1_S194 - cost: 2.42, layer: 47
Normal
Diabetic
Normal data
Fold over basal
1.5
1.4
1.3
1.2
1.1
1
0.9
0.8
0
10
20
30
40
50
60
Time (min)

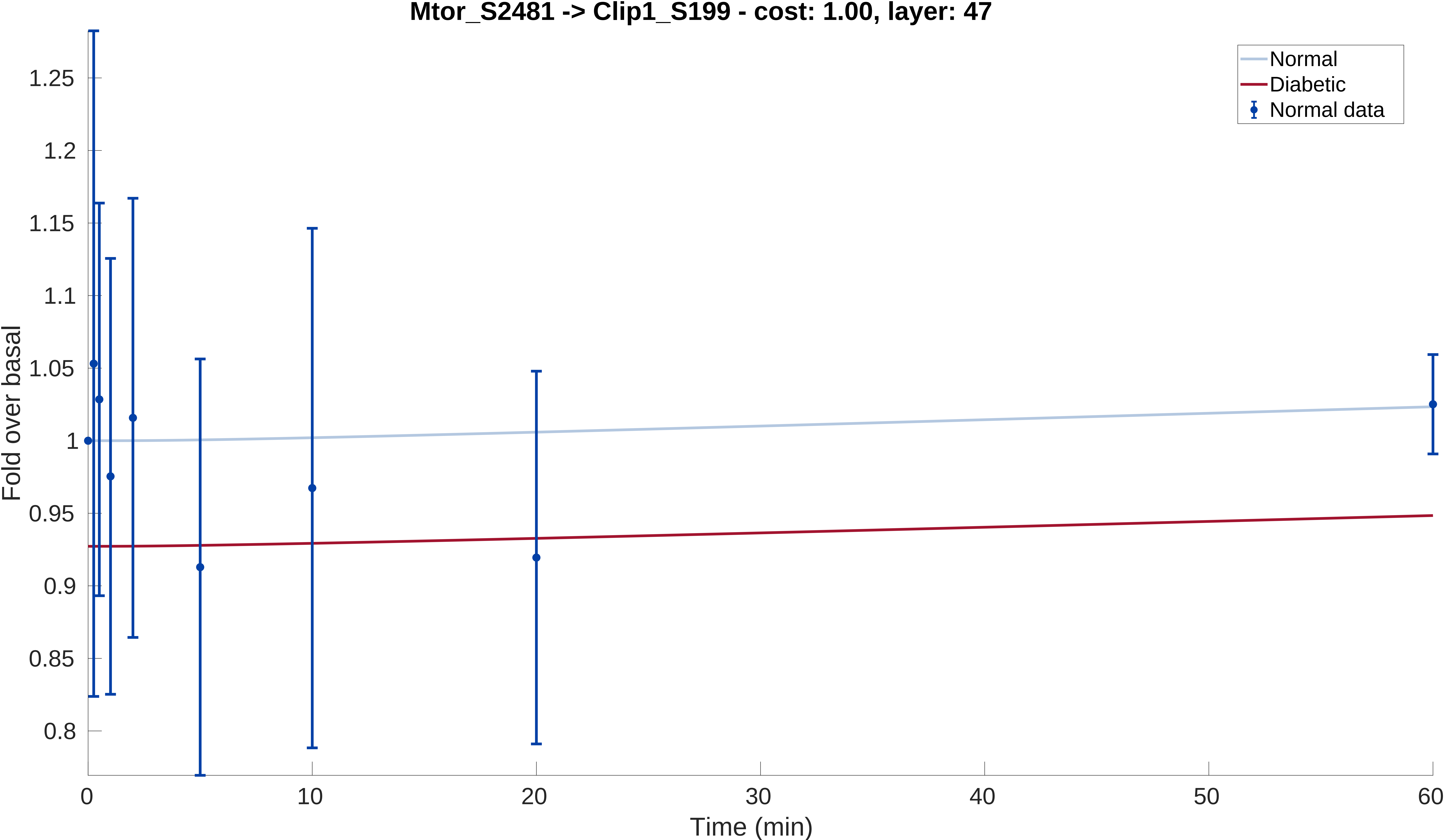

Mtor_S2481 -> Clip1_S199 - cost: 1.00, layer: 47
Normal
Diabetic
Normal data
Fold over basal
1.25
1.2
1.15
1.1
1.05
1
0.95
0.9
0.85
0.8
0
10
20
30
40
50
60
Time (min)

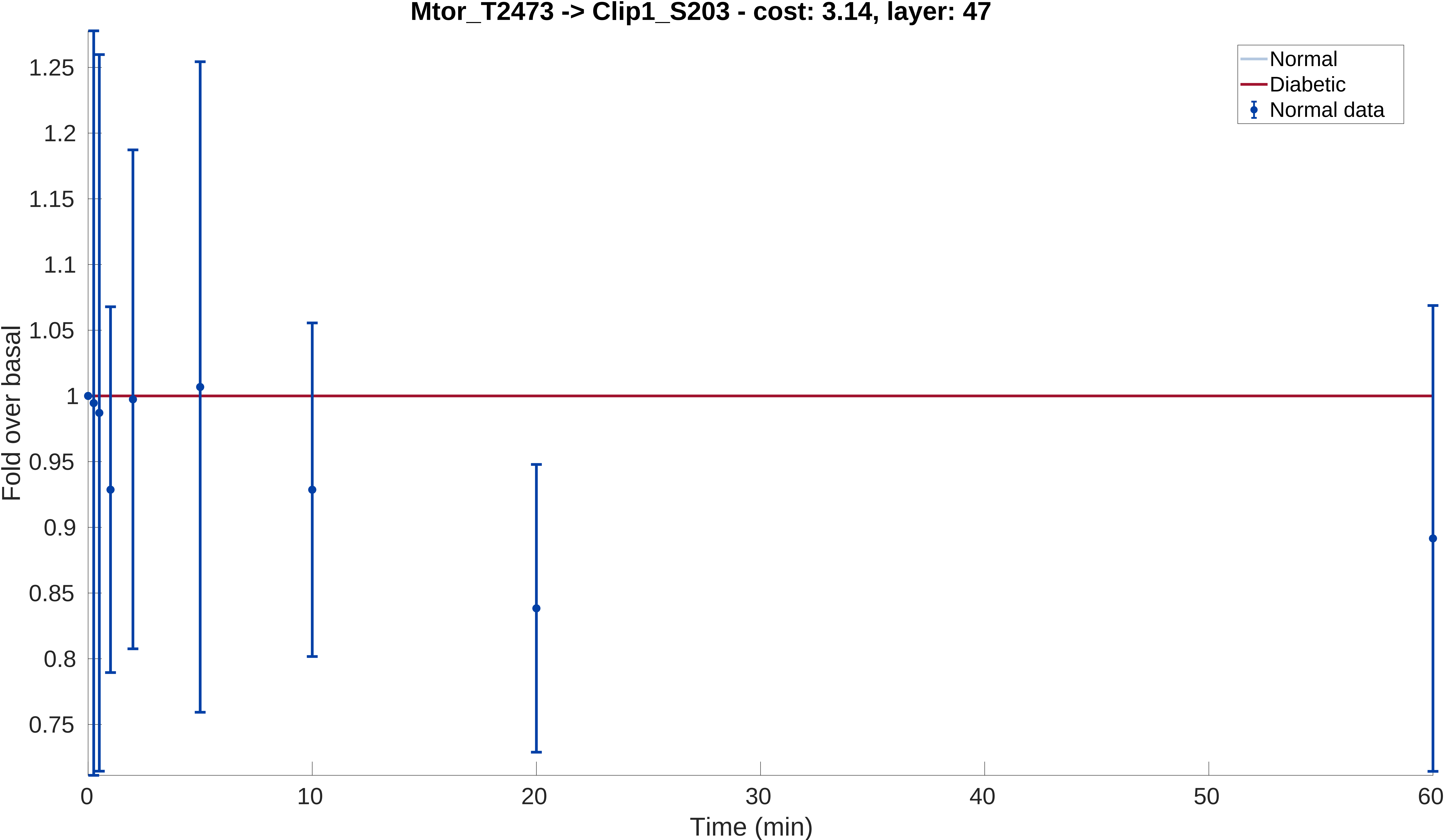

Mtor_T2473 -> Clip1_S203 - cost: 3.14, layer: 47
Normal
Diabetic
Normal data
Fold over basal
Time (min)
1.25
1.2
1.15
1.1
1.05
1
0.95
0.9
0.85
0.8
0.75
0
10
20
30
40
50
60

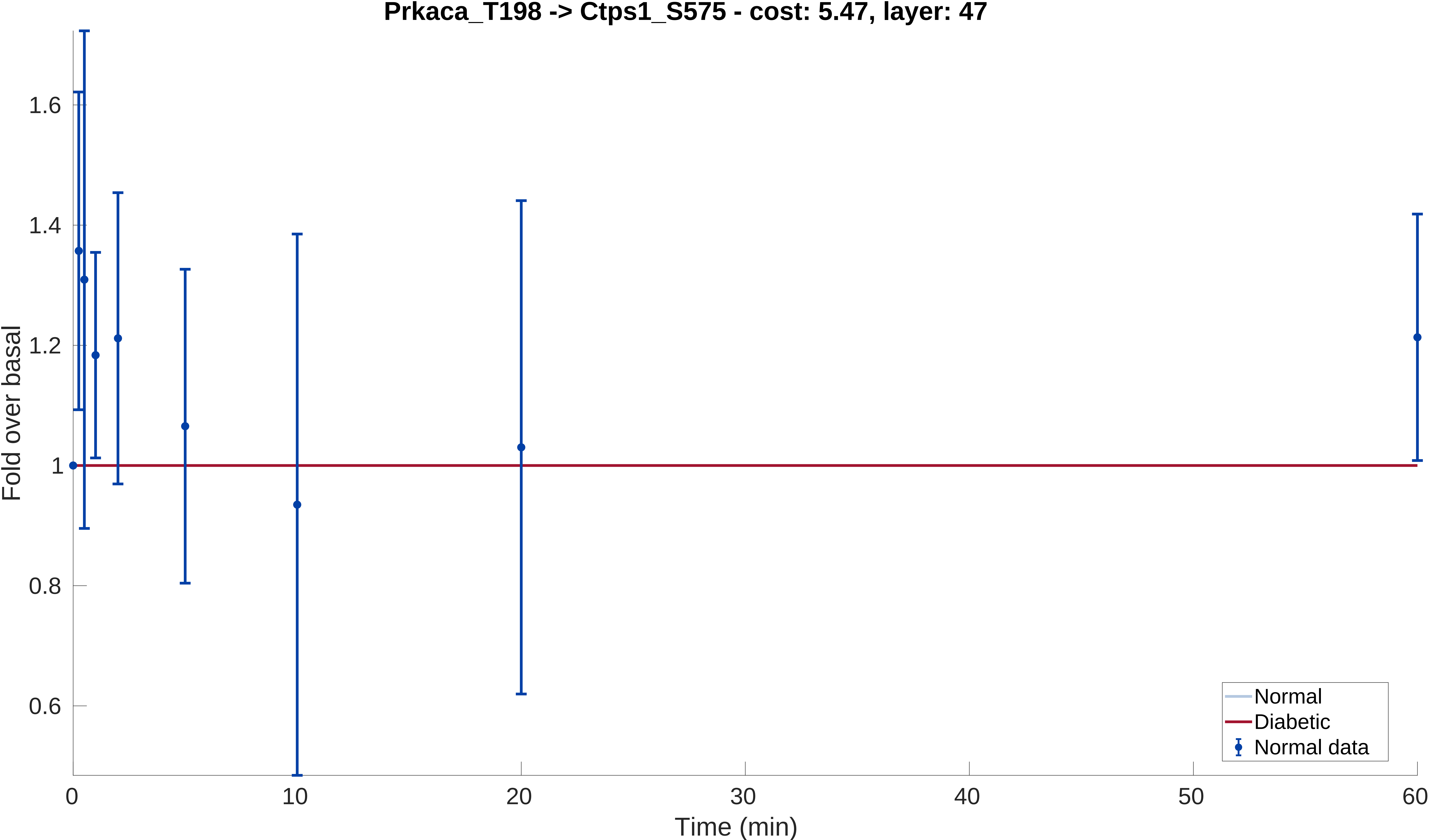

Prrkaca_T198 -> Ctps1_S575 - cost: 5.47, layer: 47
Fold over basal
1.6
1.4
1.2
1
0.8
0.6
0
10
20
30
40
50
60
Time (min)
Normal
Diabetic
Normal data

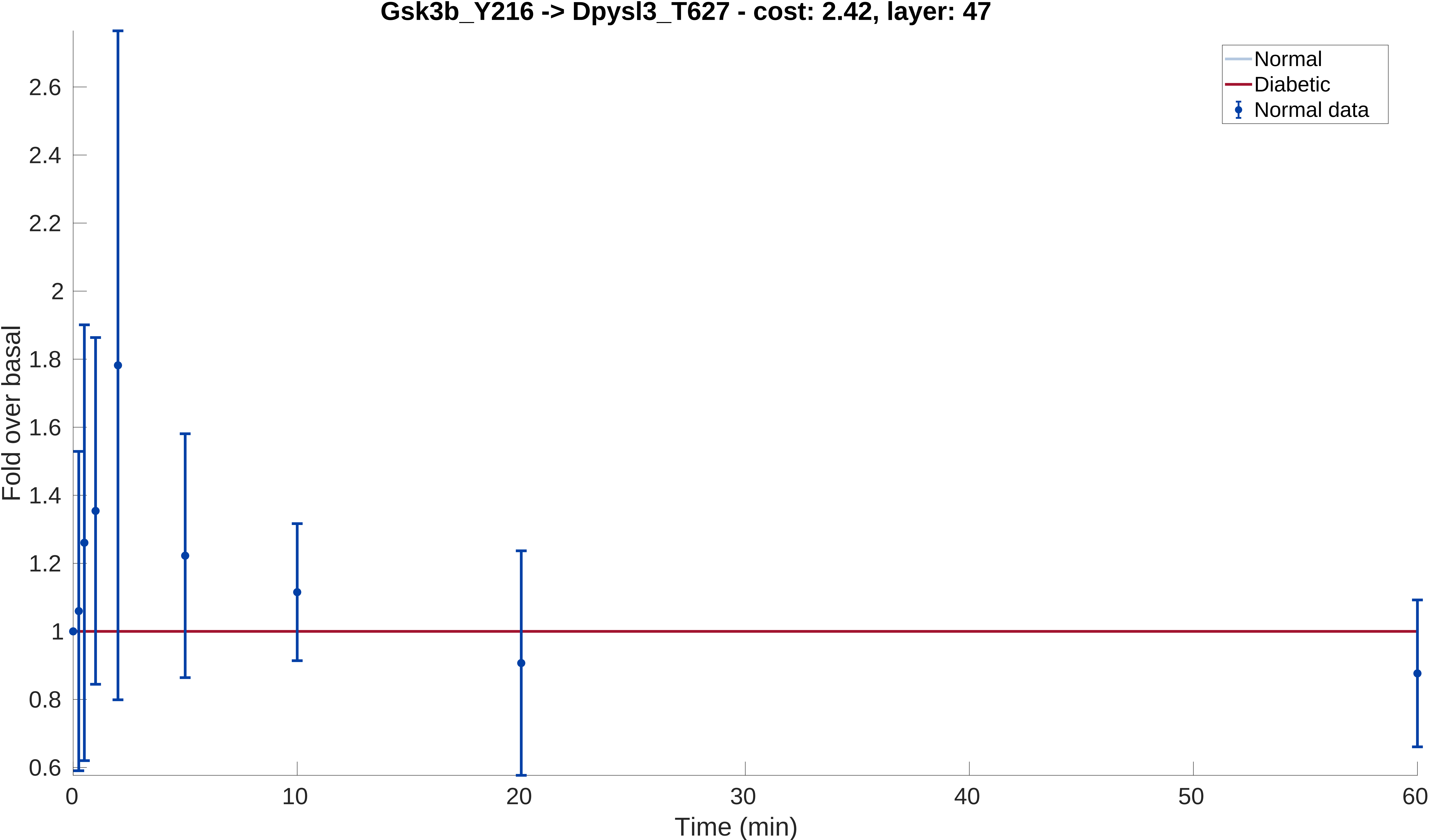

Gsk3b_Y216 -> Dpysl3_T627 - cost: 2.42, layer: 47
Normal
Diabetic
Normal data
Fold over basal
Time (min)
0
10
20
30
40
50
60
0.6
0.8
1
1.2
1.4
1.6
1.8
2
2.2
2.4
2.6

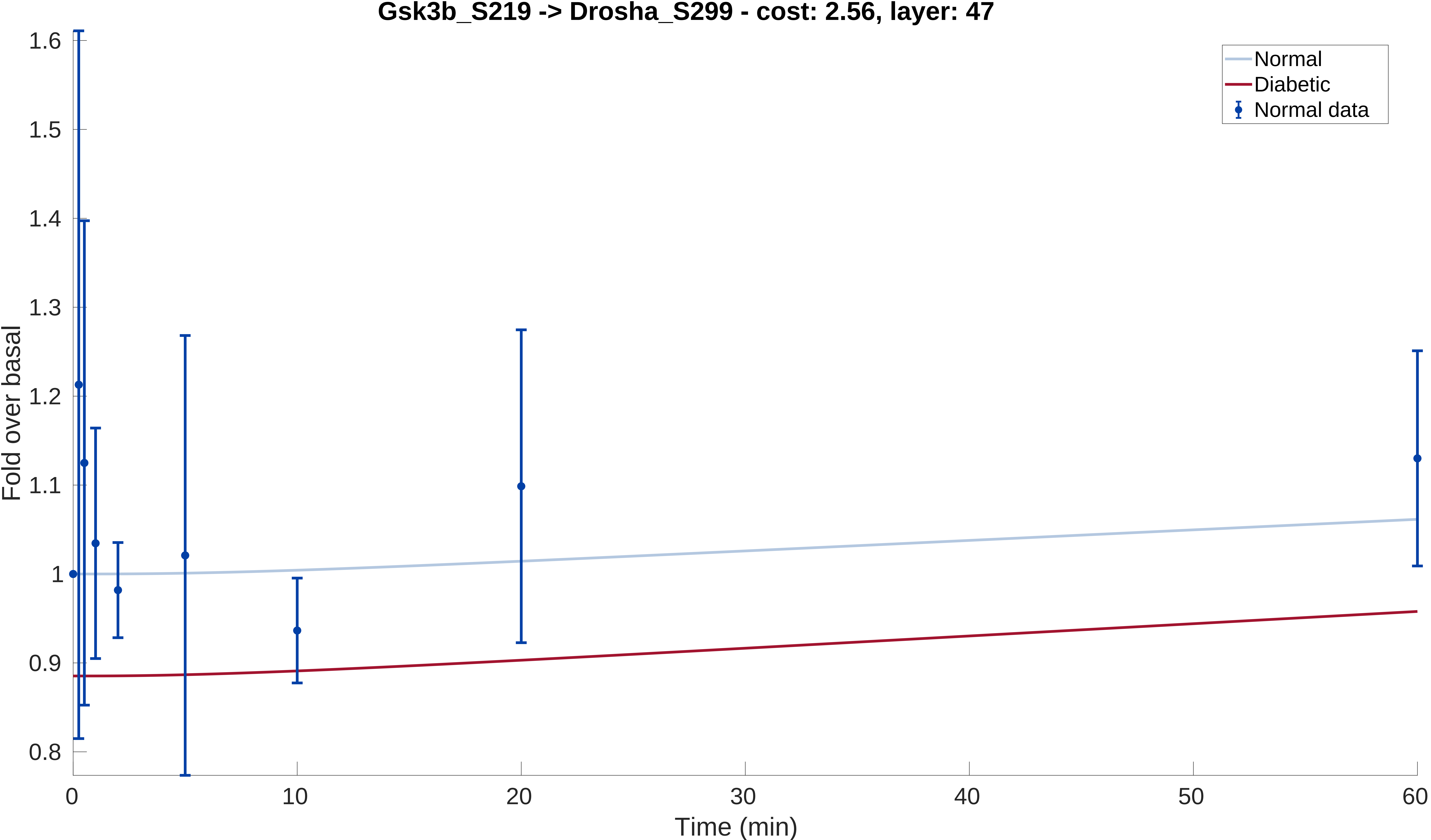

Gsk3b_S219 -> Drosha_S299 - cost: 2.56, layer: 47
Normal
Diabetic
Normal data
Fold over basal
1.6
1.5
1.4
1.3
1.2
1.1
1
0.9
0.8
0
10
20
30
40
50
60
Time (min)

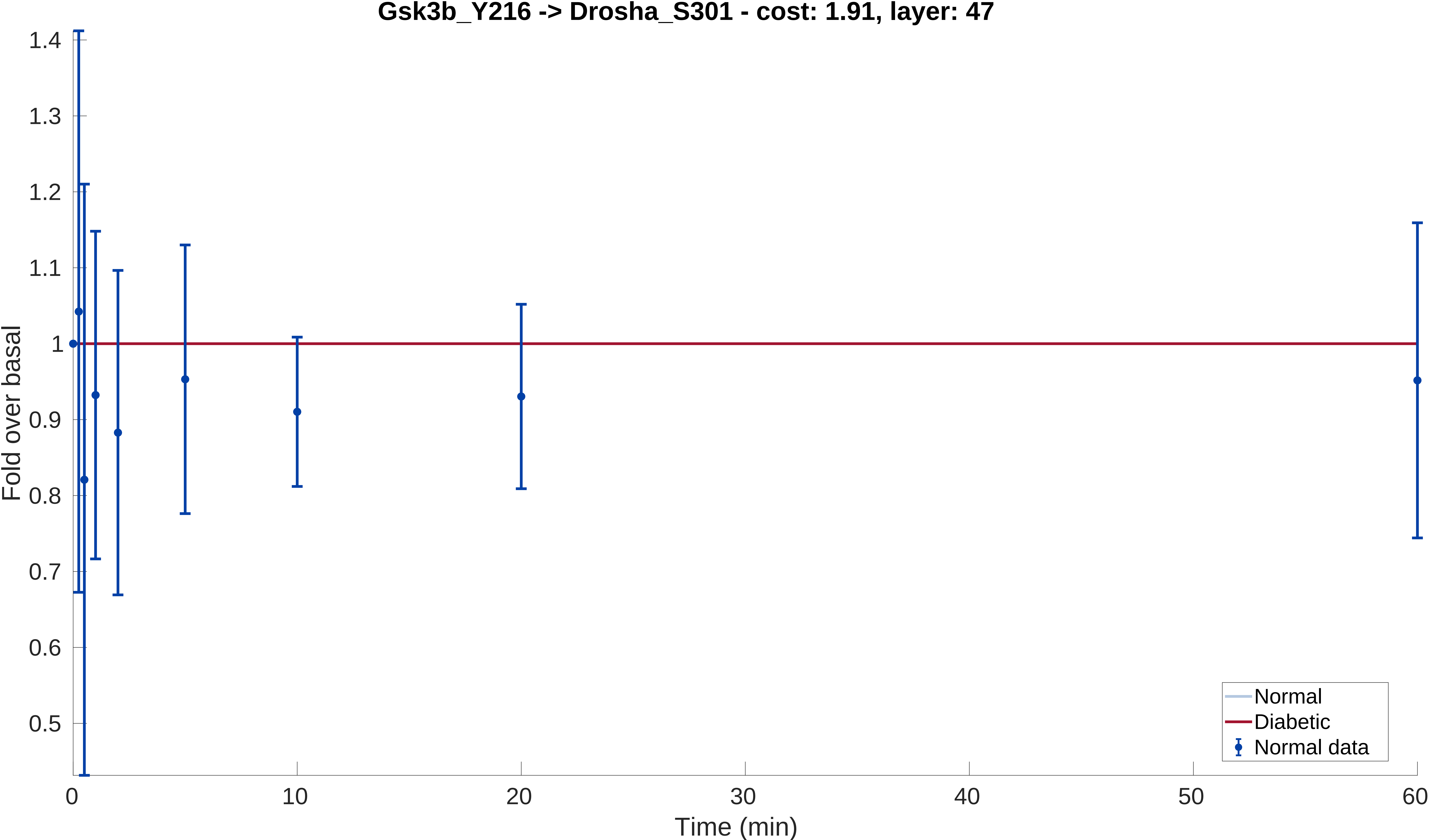

Gsk3b_Y216 -> Drosha_S301 - cost: 1.91, layer: 47
Fold over basal
1.4
1.3
1.2
1.1
1
0.9
0.8
0.7
0.6
0.5
0
10
20
30
40
50
60
Time (min)
Normal
Diabetic
Normal data

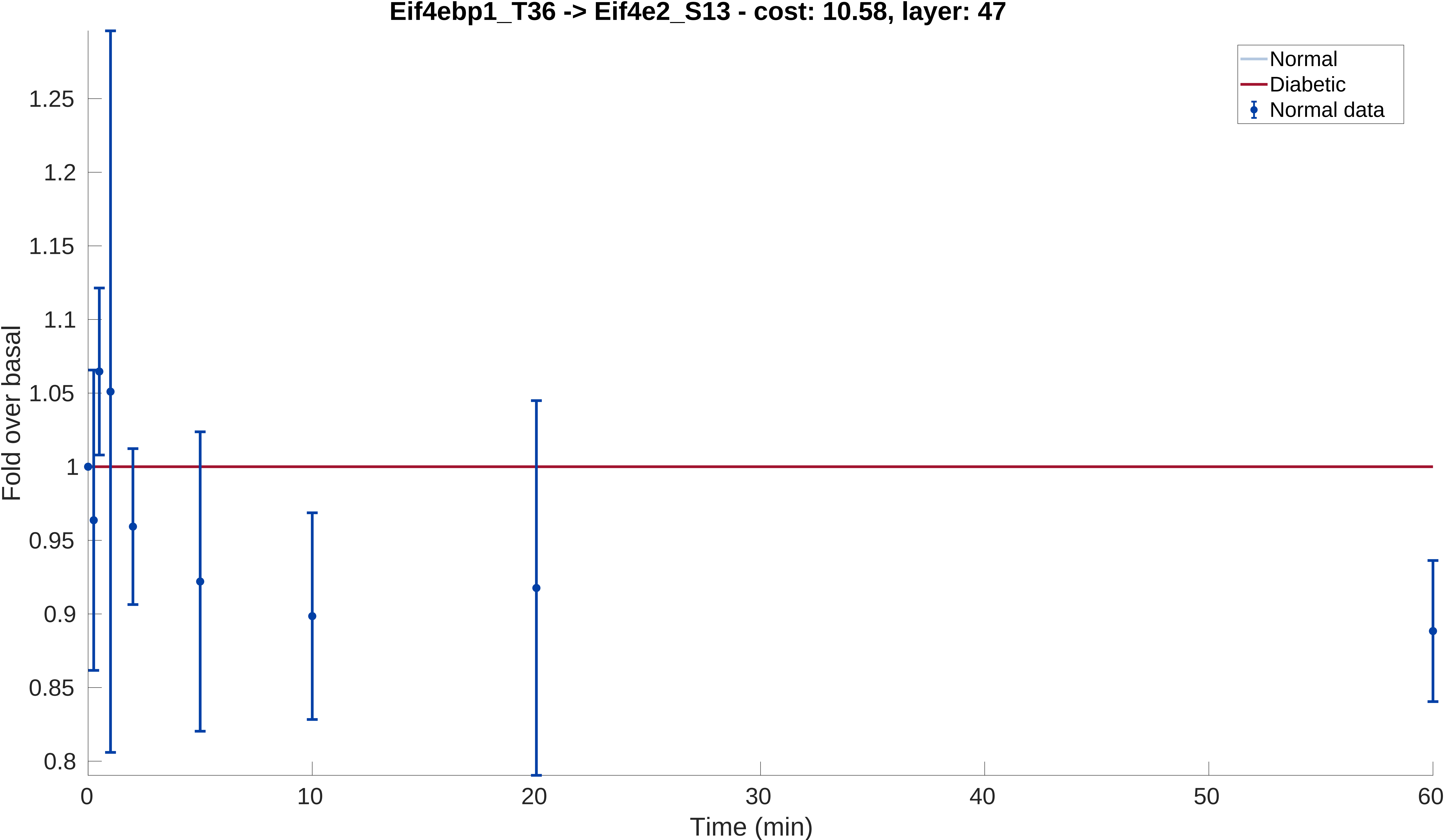
Eif4ebp1_T36 -> Eif4e2_S13 - cost: 10.58, layer: 47
Normal
Diabetic
Normal data
Fold over basal
1.25
1.2
1.15
1.1
1.05
1
0.95
0.9
0.85
0.8
0
10
20
30
40
50
60
Time (min)

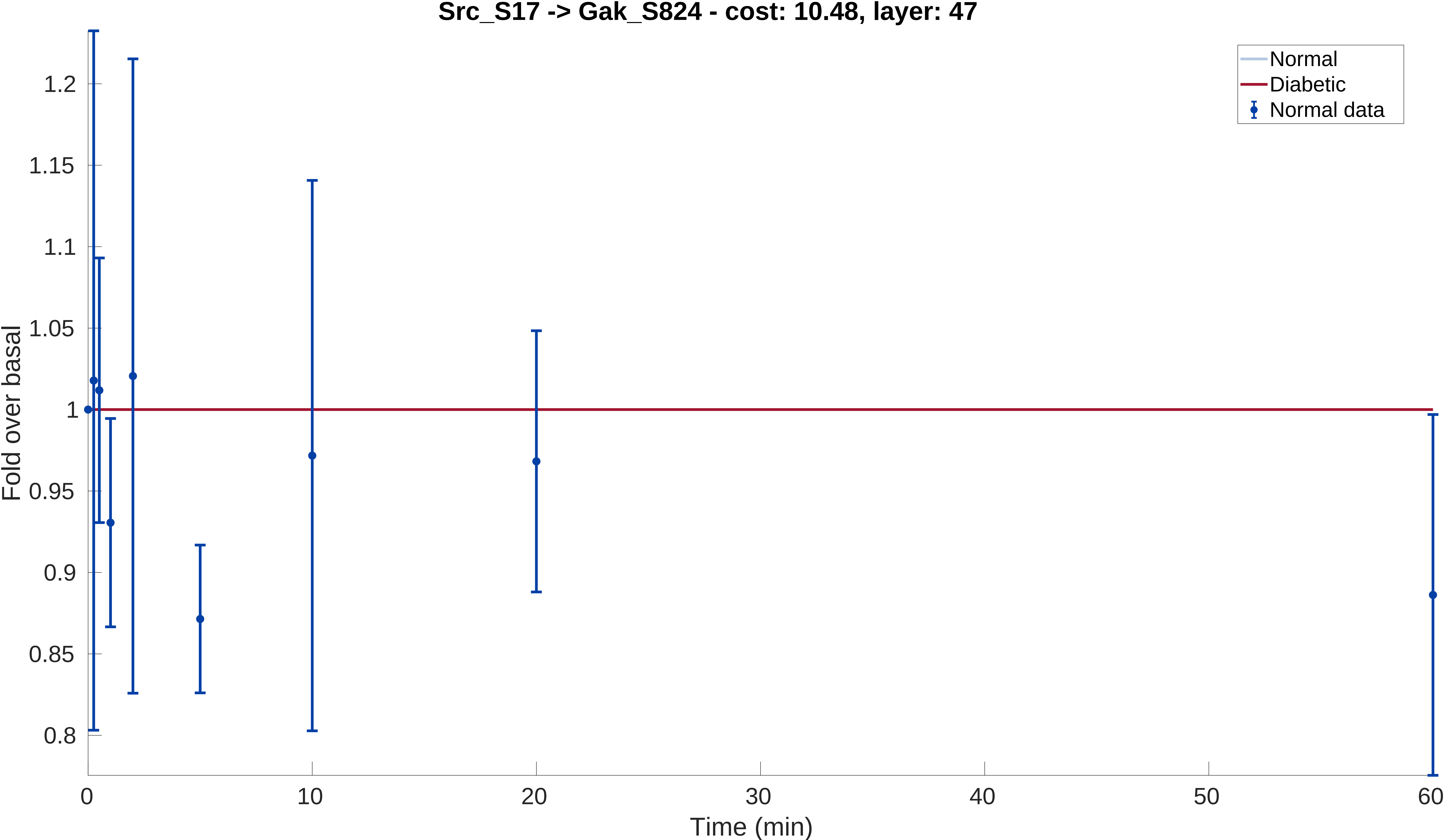
Src_S17 -> Gak_S824 - cost: 10.48, layer: 47
Normal
Diabetic
Normal data
Fold over basal
1.2
1.15
1.1
1.05
1
0.95
0.9
0.85
0.8
0
10
20
30
40
50
60
Time (min)

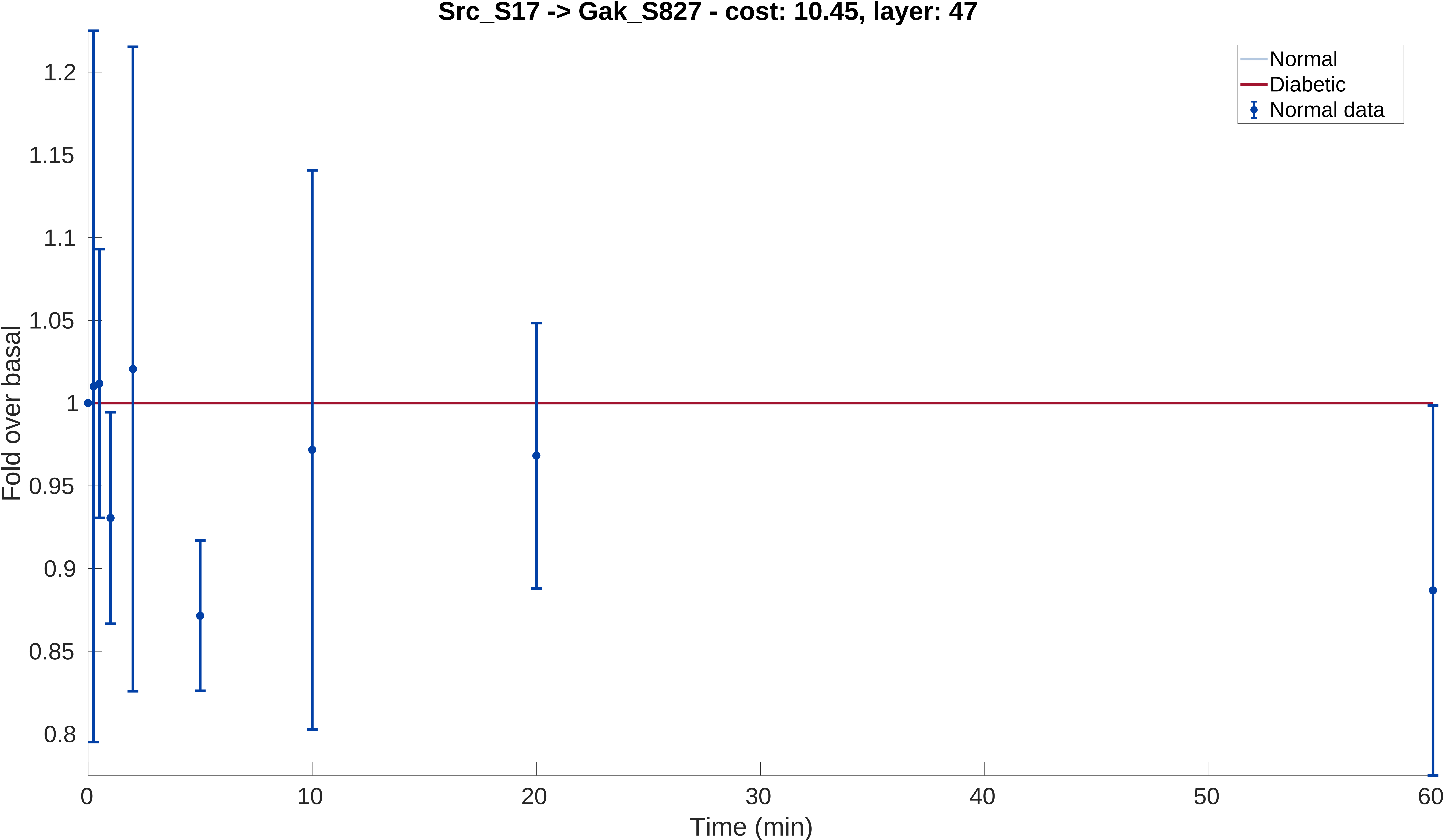

Src_S17 -> Gak_S827 - cost: 10.45, layer: 47
Normal
Diabetic
Normal data
Fold over basal
1.2
1.15
1.1
1.05
1
0.95
0.9
0.85
0.8
0
10
20
30
40
50
60
Time (min)

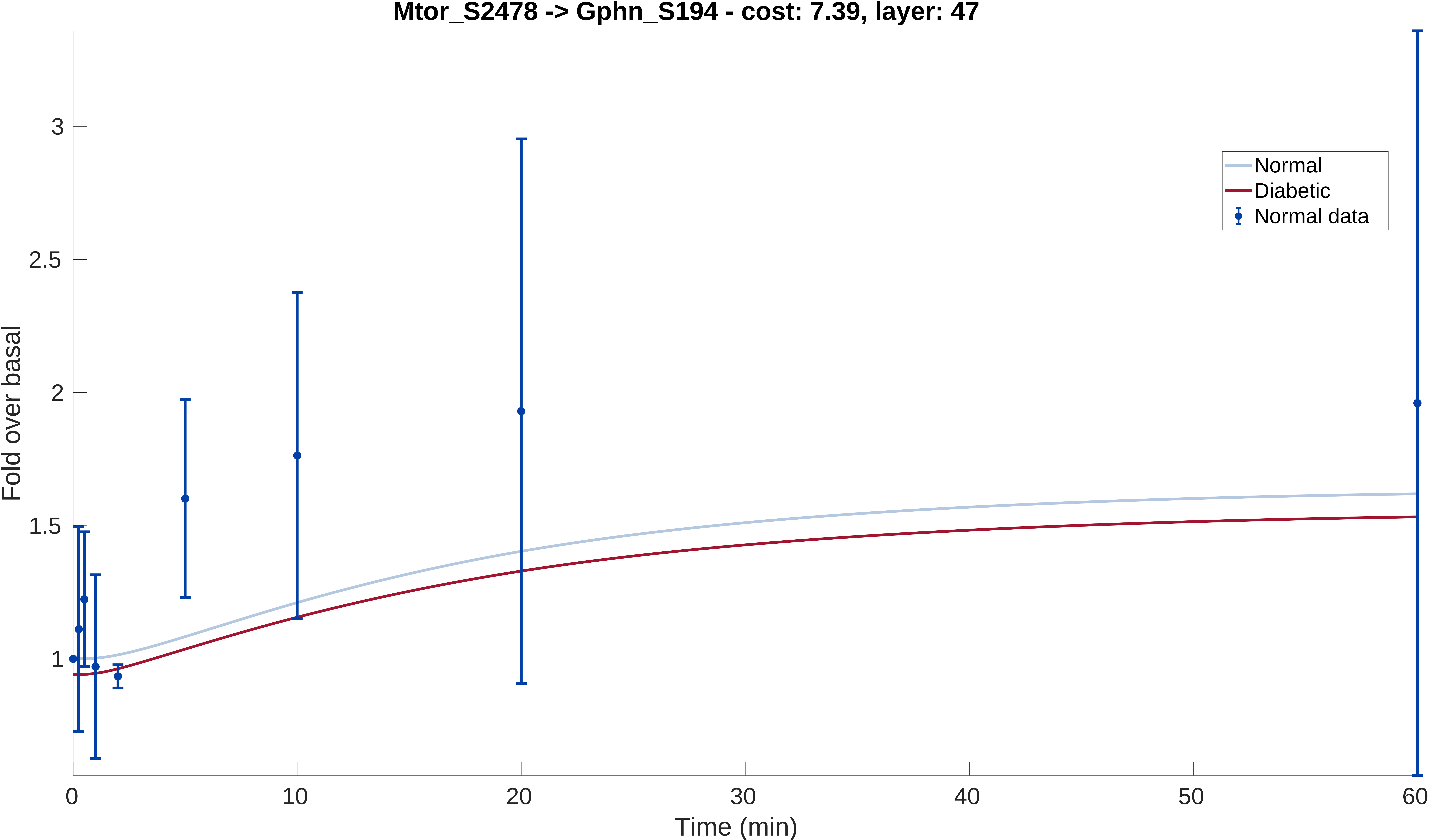

Mtor_S2478 -> Gphn_S194 - cost: 7.39, layer: 47
Normal
Diabetic
Normal data
Fold over basal
3
2.5
2
1.5
1
0
10
20
30
40
50
60
Time (min)

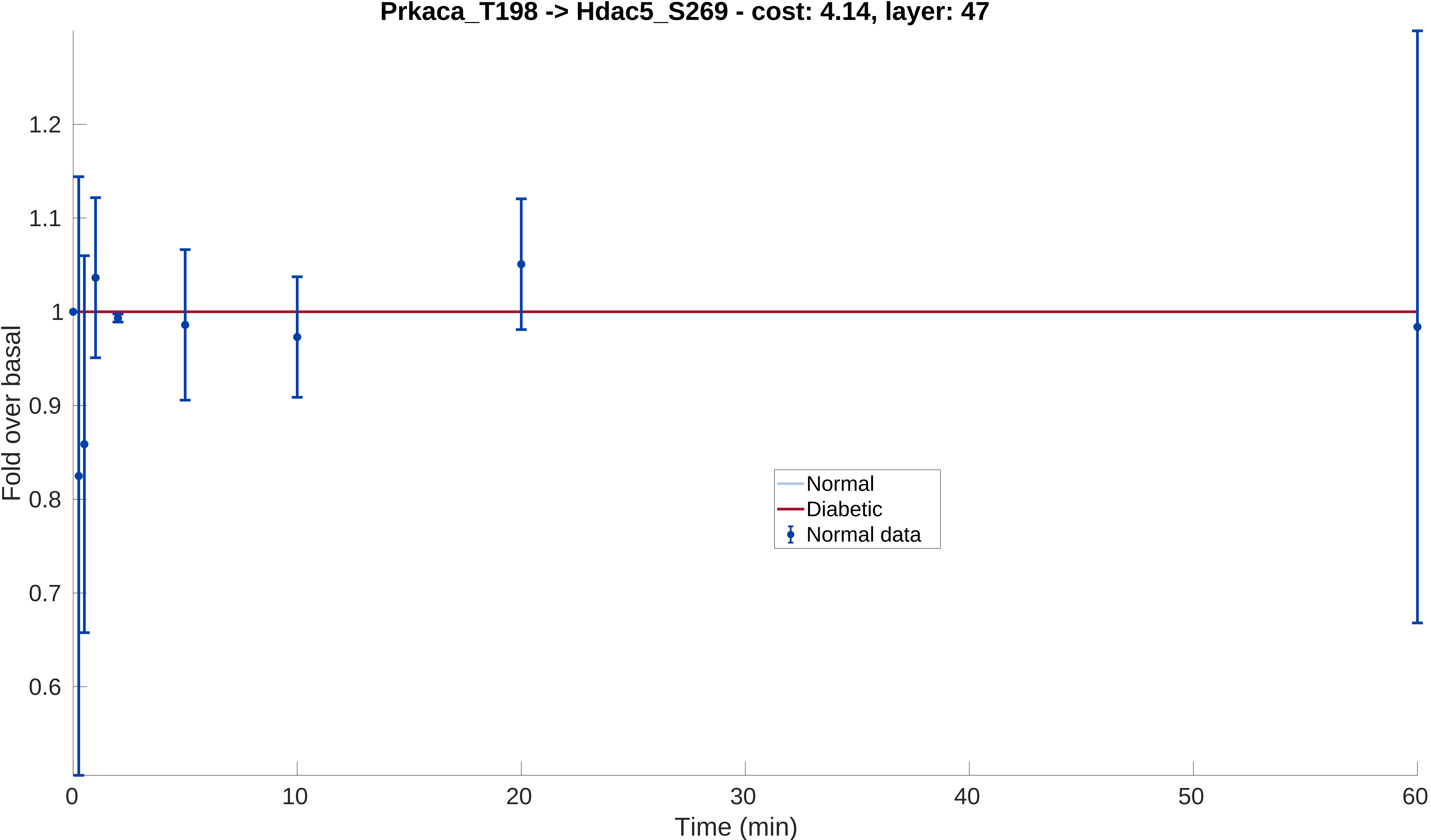
Prkaca_T198 -> Hdac5_S269 - cost: 4.14, layer: 47
Fold over basal
1.2
1.1
1
0.9
0.8
0.7
0.6
0
10
20
30
40
50
60
Time (min)
Normal
Diabetic
Normal data

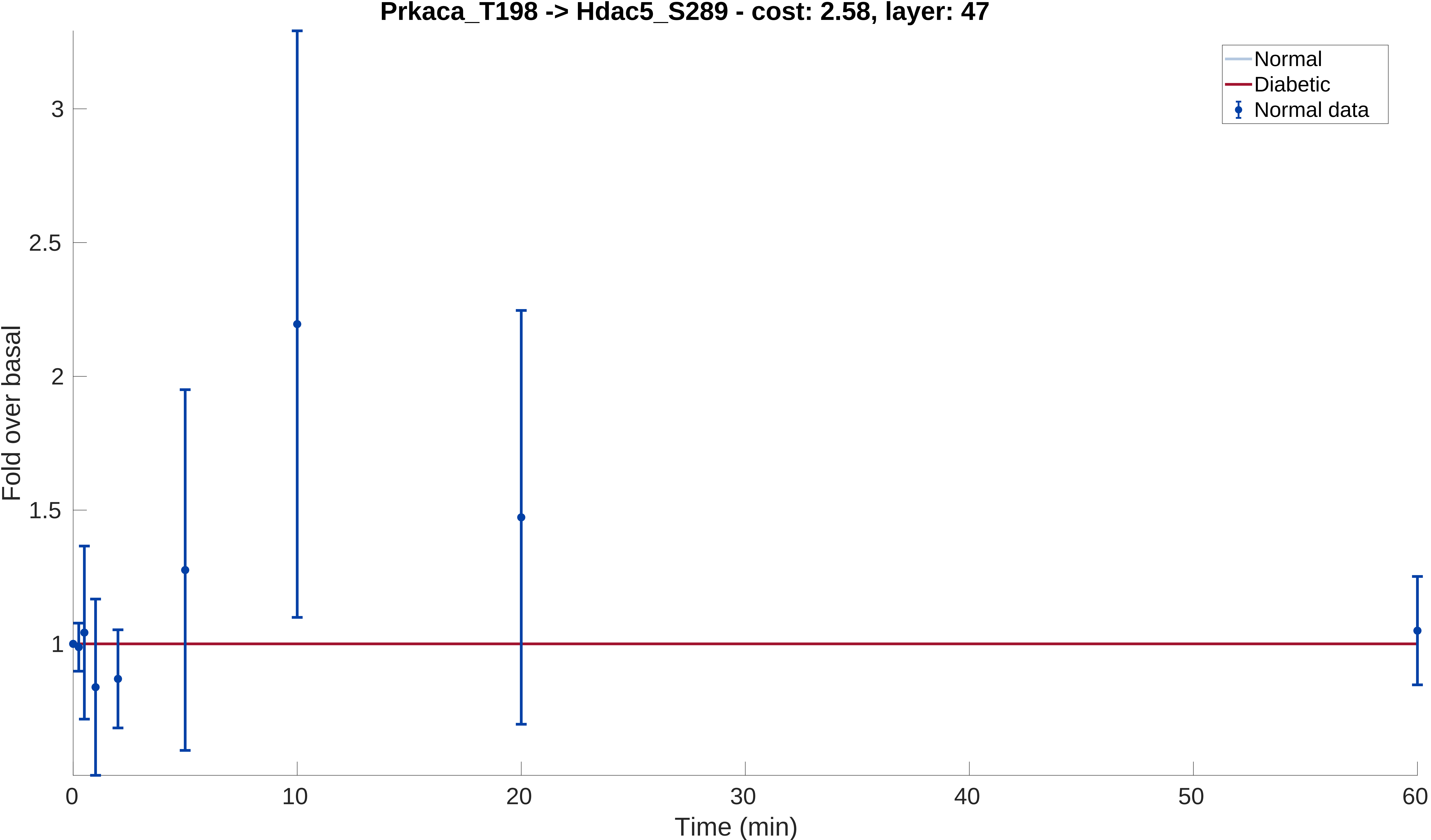

Prkaca_T198 -> Hdac5_S289 - cost: 2.58, layer: 47
Normal
Diabetic
Normal data
Fold over basal
3
2.5
2
1.5
1
0
10
20
30
40
50
60
Time (min)

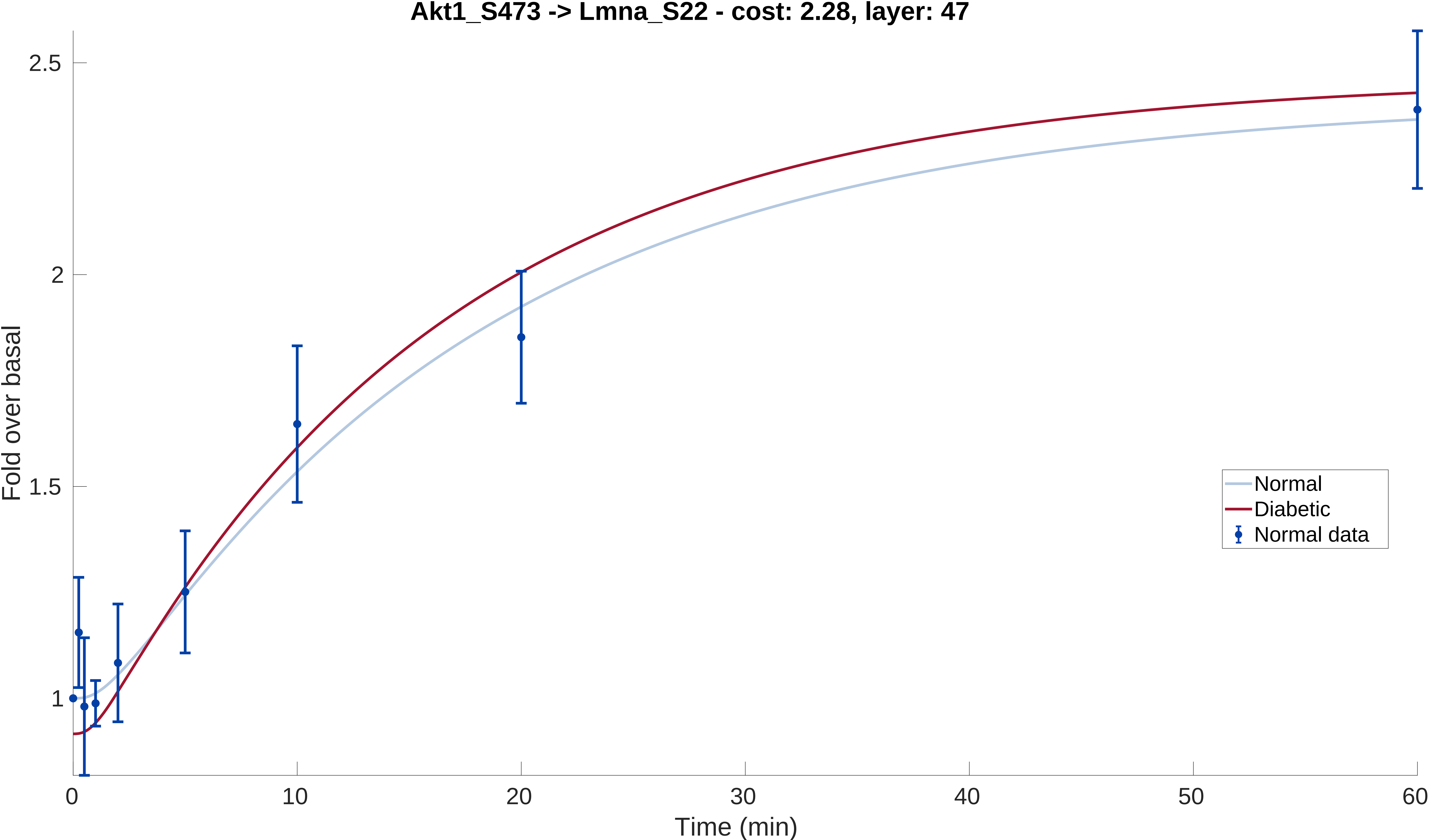

Akt1_S473 -> Lmna_S22 - cost: 2.28, layer: 47
2.5
2
1.5
1
Fold over basal
0
10
20
30
40
50
60
Time (min)
Normal
Diabetic
Normal data

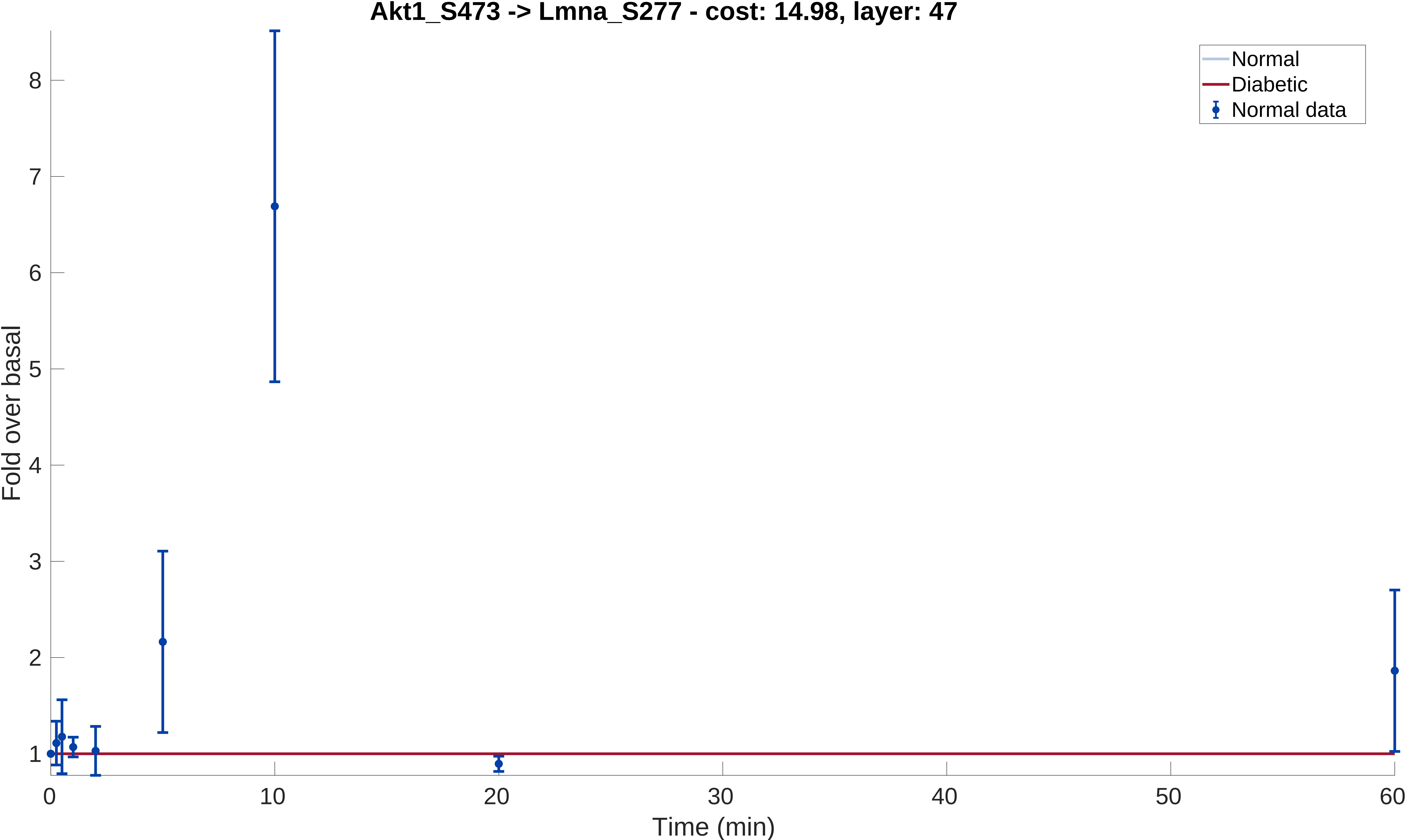

Akt1_S473 -> Lmna_S277 - cost: 14.98, layer: 47
Normal
Diabetic
Normal data
Fold over basal
1
2
3
4
5
6
7
8
0
10
20
30
40
50
60
Time (min)

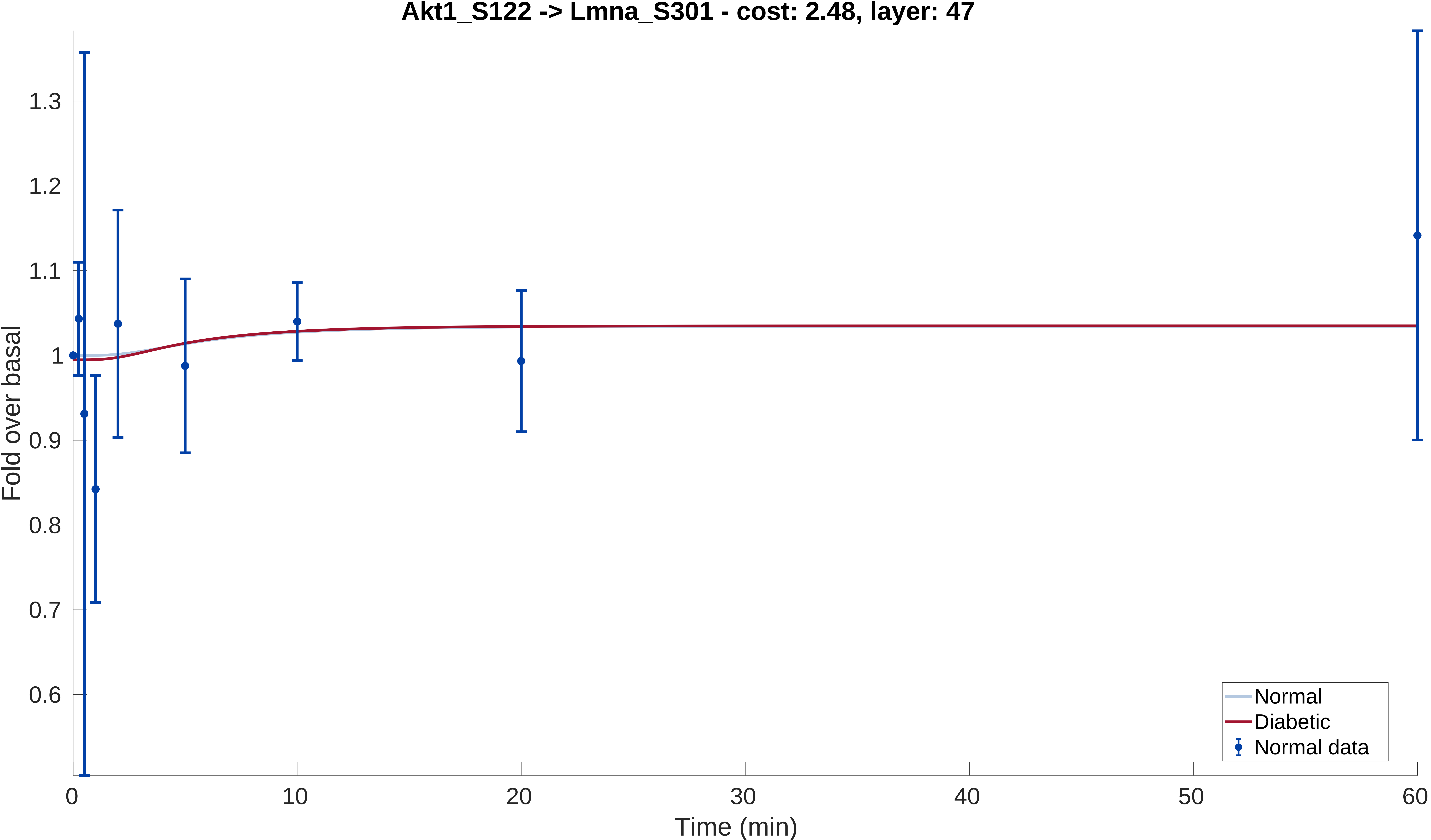

Akt1_S122 -> Lmna_S301 - cost: 2.48, layer: 47
Fold over basal
1.3
1.2
1.1
1
0.9
0.8
0.7
0.6
0
10
20
30
40
50
60
Time (min)
Normal
Diabetic
Normal data

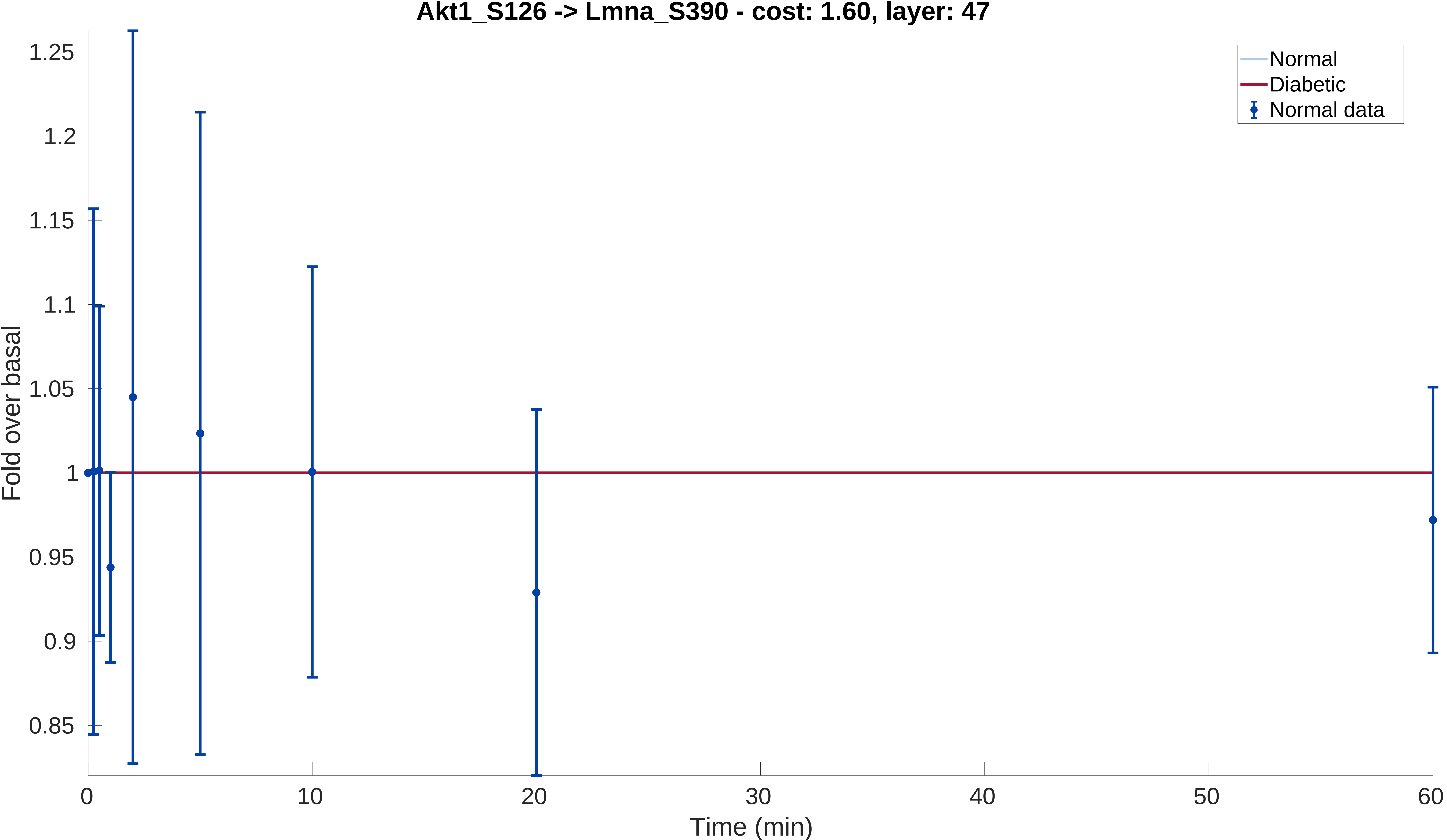

Akt1_S126 -> Lmna_S390 - cost: 1.60, layer: 47
Normal
Diabetic
Normal data
Fold over basal
1.25
1.2
1.15
1.1
1.05
1
0.95
0.9
0.85
0
10
20
30
40
50
60
Time (min)

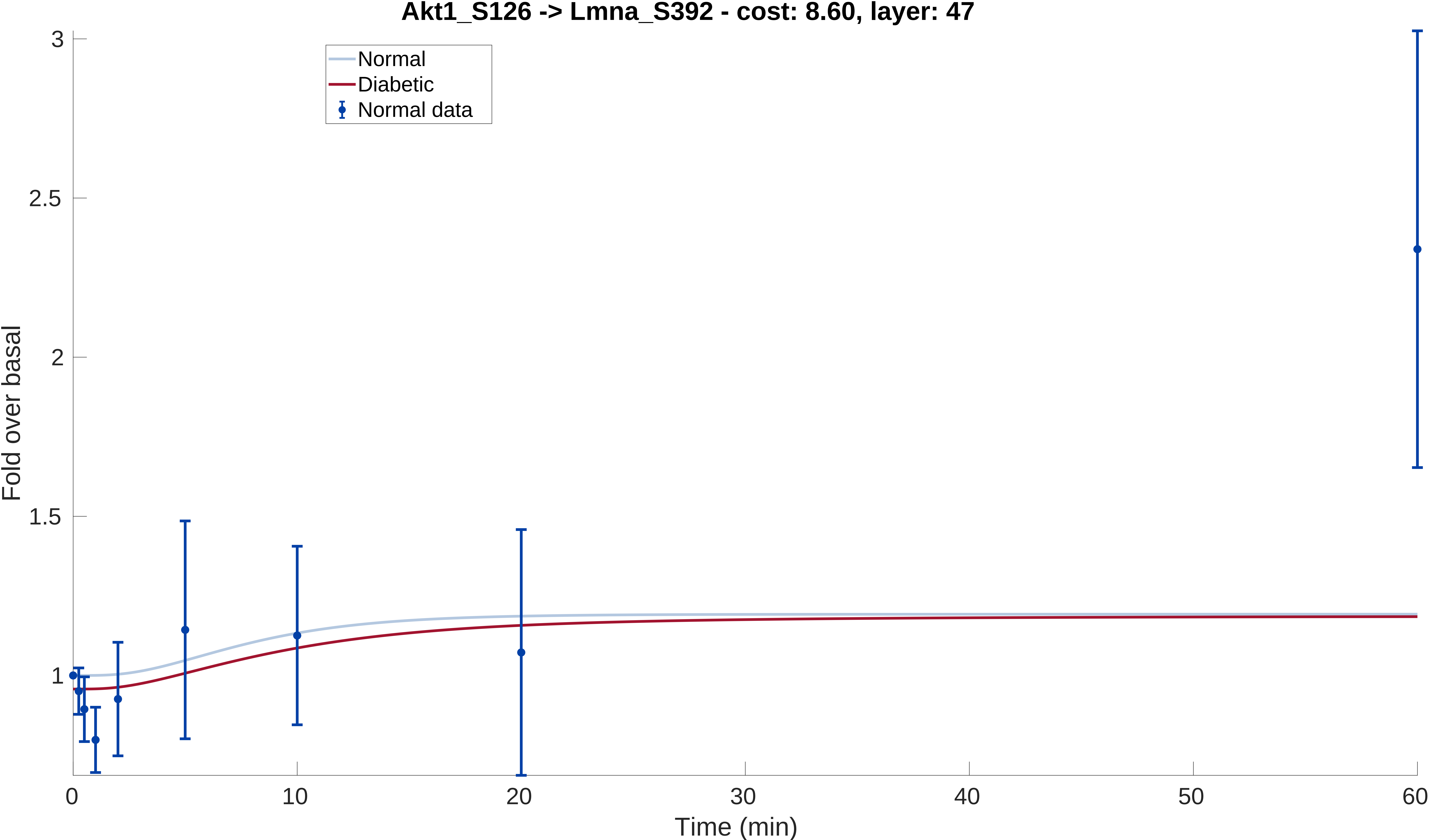

Akt1_S126 -> Lmna_S392 - cost: 8.60, layer: 47
Normal
Diabetic
Normal data
Fold over basal
Time (min)
3
2.5
2
1.5
1
0
10
20
30
40
50
60

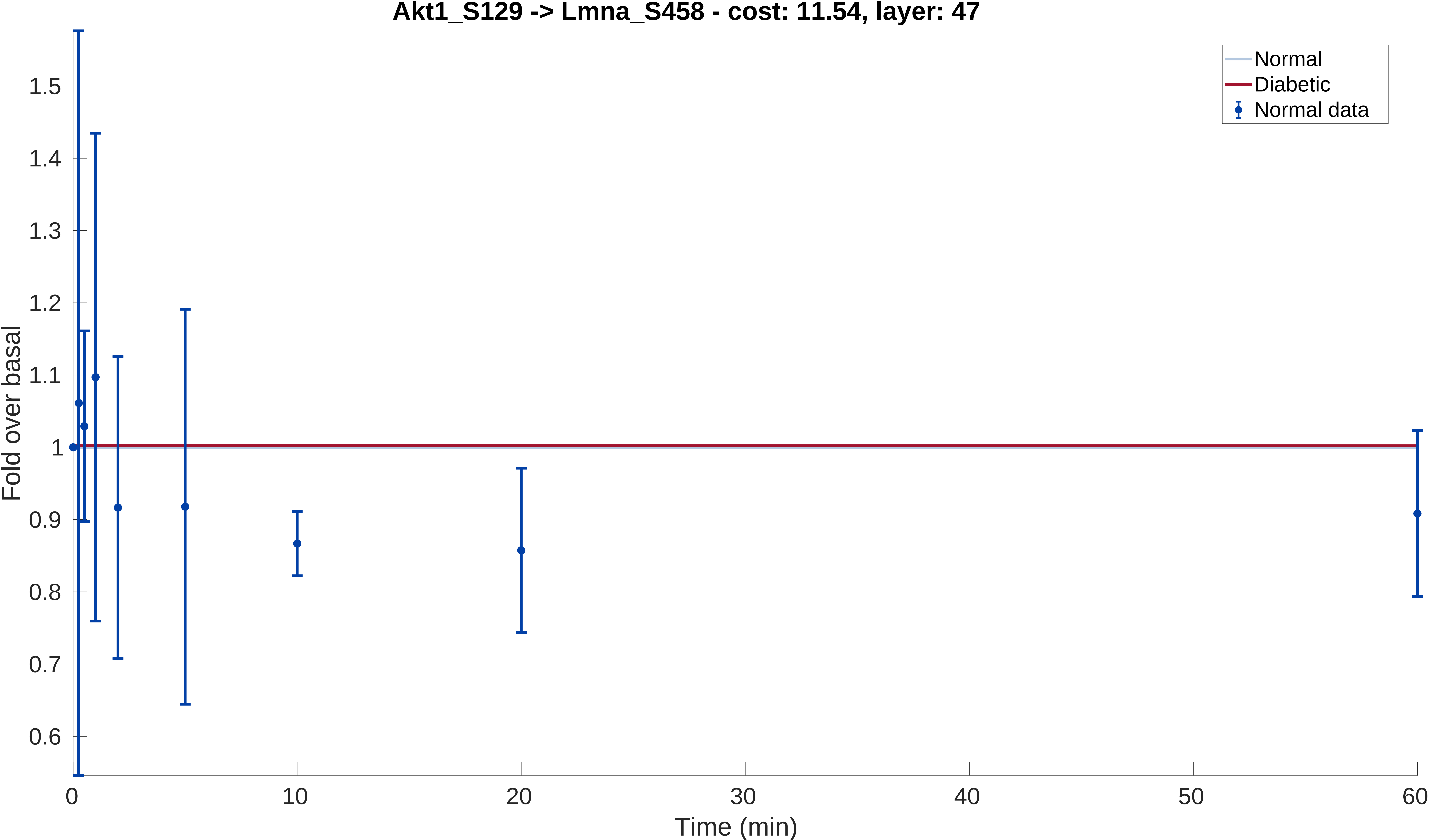
Akt1_S129 -> Lmna_S458 - cost: 11.54, layer: 47
Normal
Diabetic
Normal data
Fold over basal
Time (min)
0.6
0.7
0.8
0.9
1
1.1
1.2
1.3
1.4
1.5
0
10
20
30
40
50
60

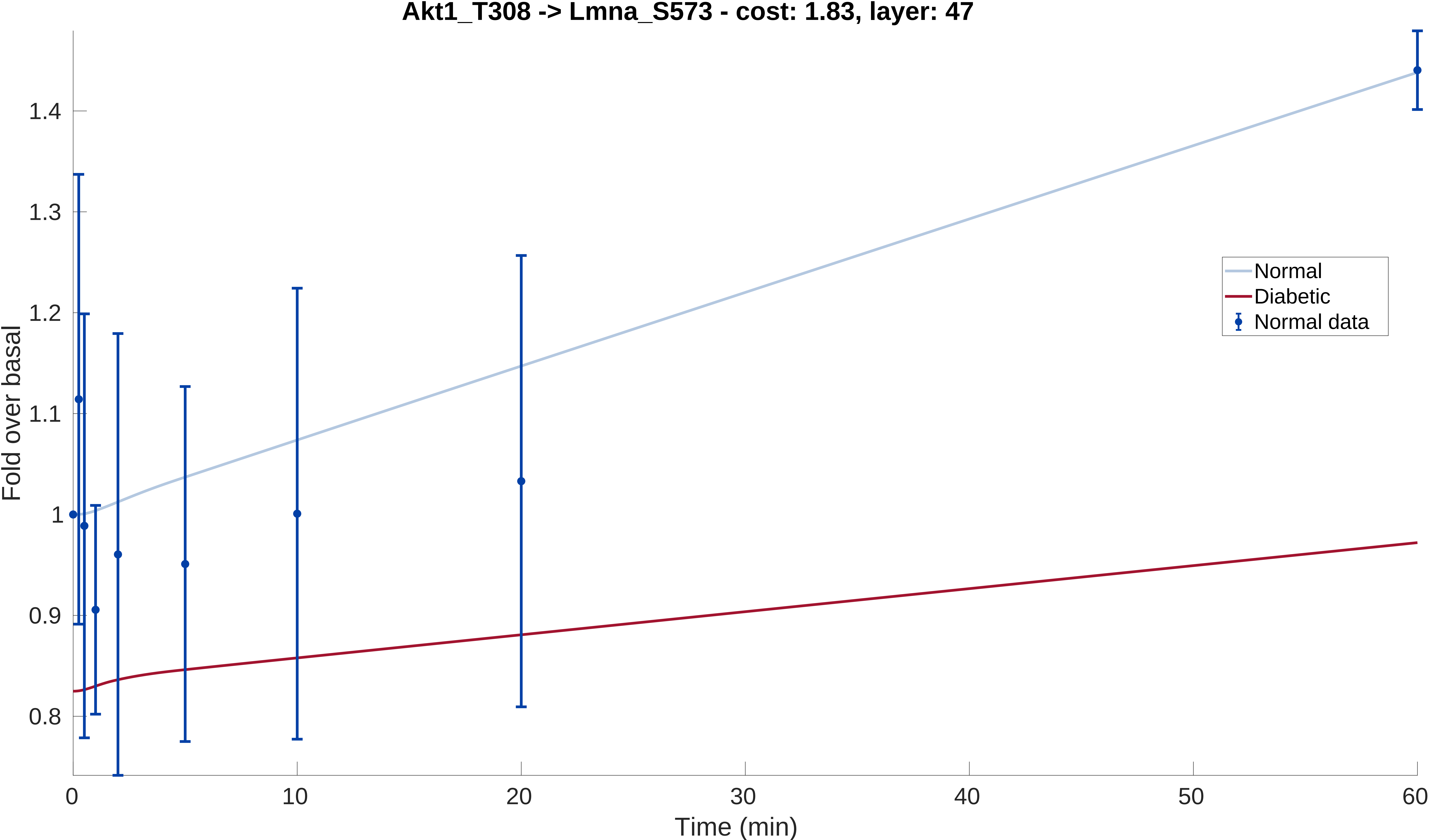

Akt1_T308 -> Lmna_S573 - cost: 1.83, layer: 47
Fold over basal
Time (min)
Normal
Diabetic
Normal data
1.4
1.3
1.2
1.1
1
0.9
0.8
0
10
20
30
40
50
60

#### Akt1_T308 -> Lmna_S575 - cost: 3.51, layer: 47

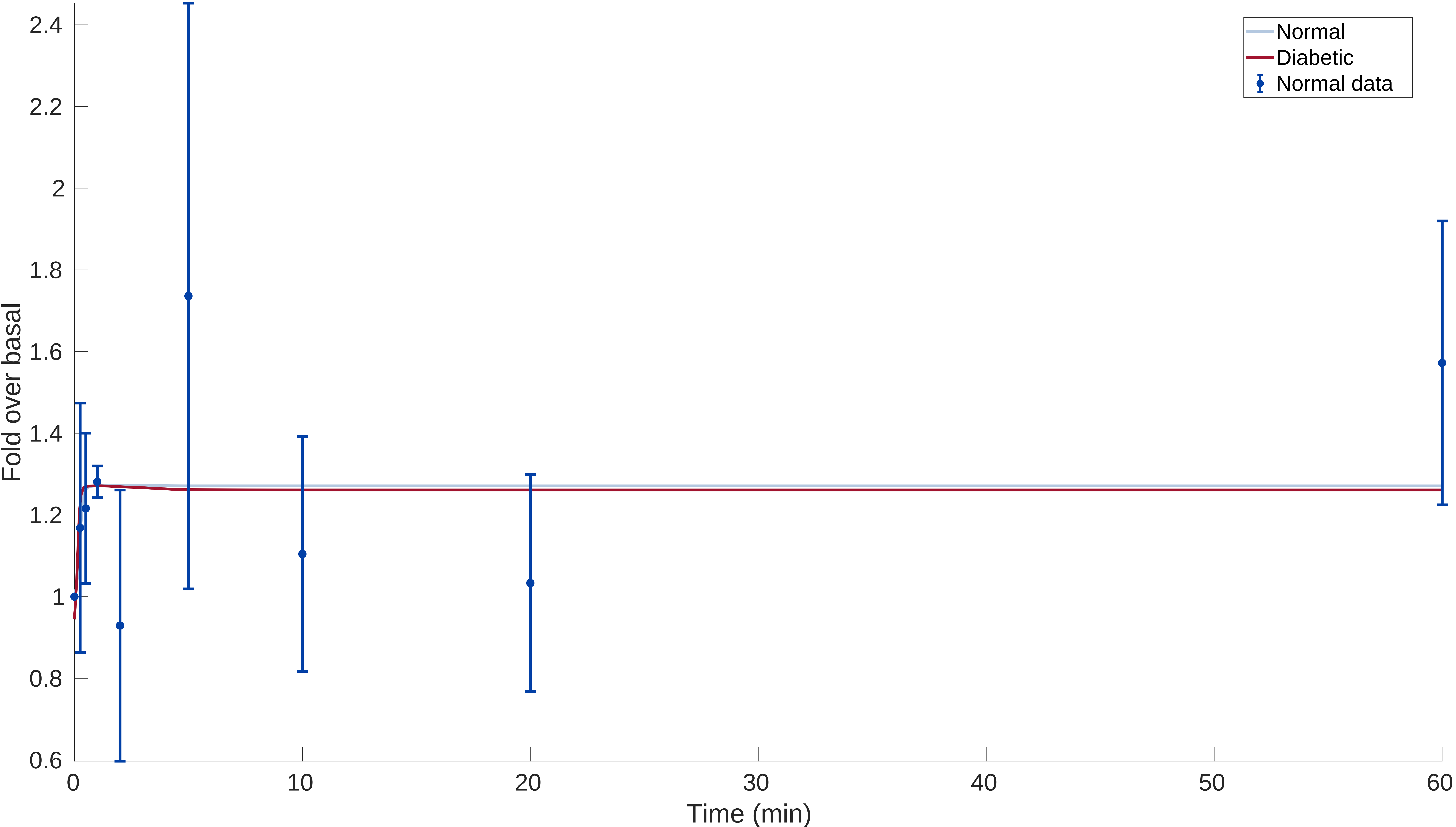

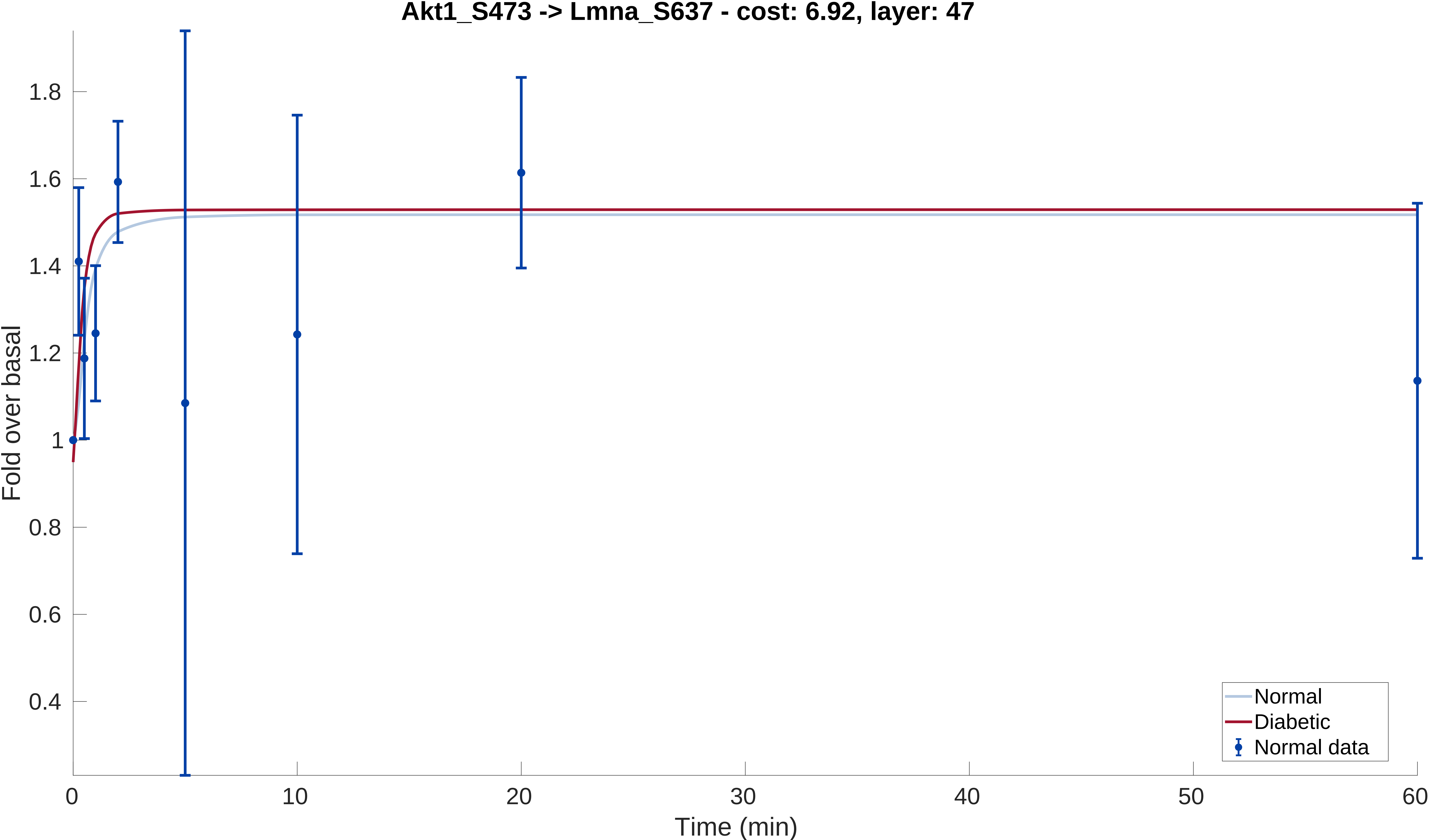
Akt1_S473 -> Lmna_S637 - cost: 6.92, layer: 47
Fold over basal
Time (min)
0.4
0.6
0.8
1
1.2
1.4
1.6
1.8
0
10
20
30
40
50
60
Normal
Diabetic
Normal data

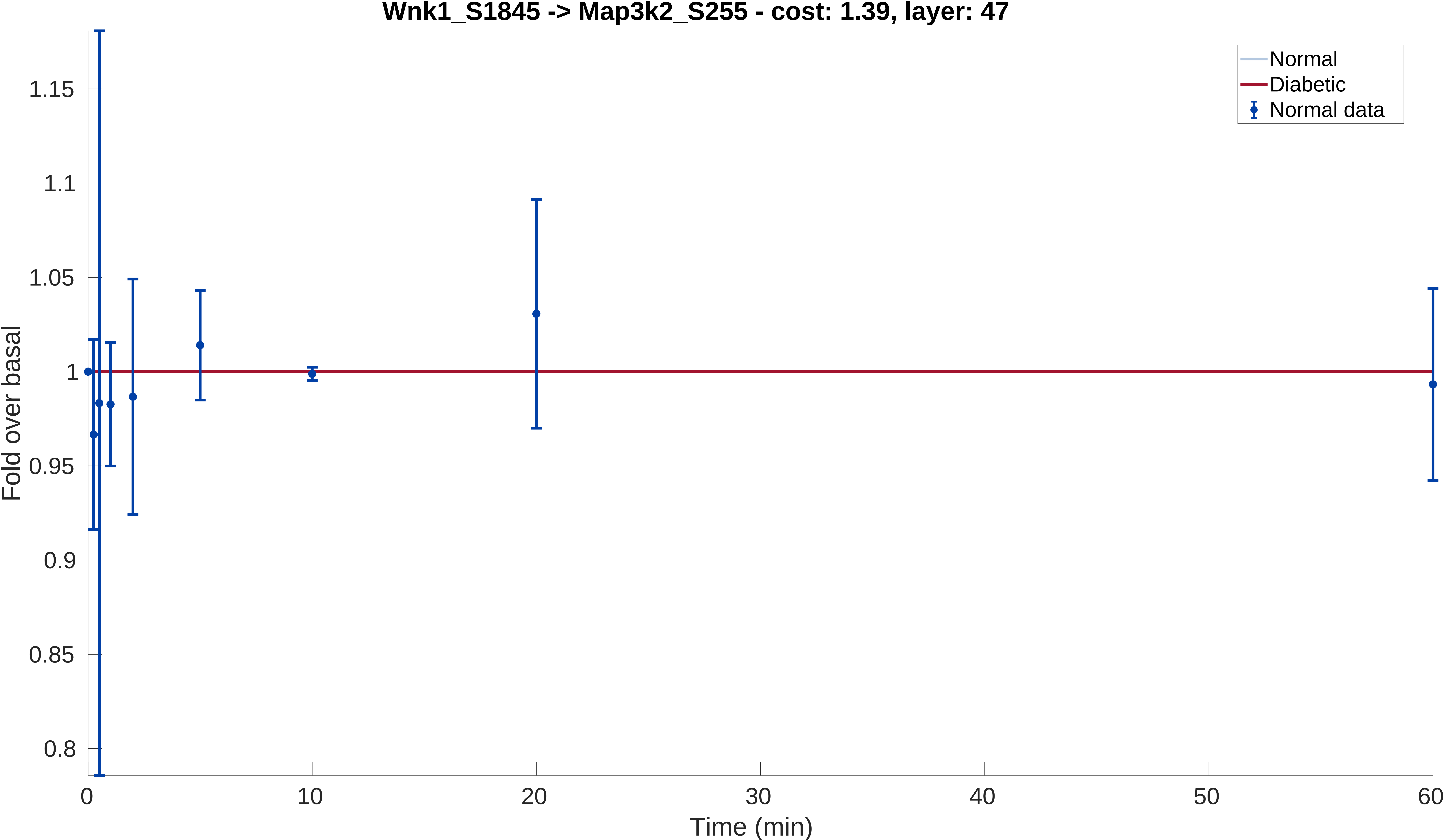

Wnk1_S1845 -> Map3k2_S255 - cost: 1.39, layer: 47
Normal
Diabetic
Normal data
1.15
1.1
1.05
1
0.95
0.9
0.85
0.8
Fold over basal
0
10
20
30
40
50
60
Time (min)

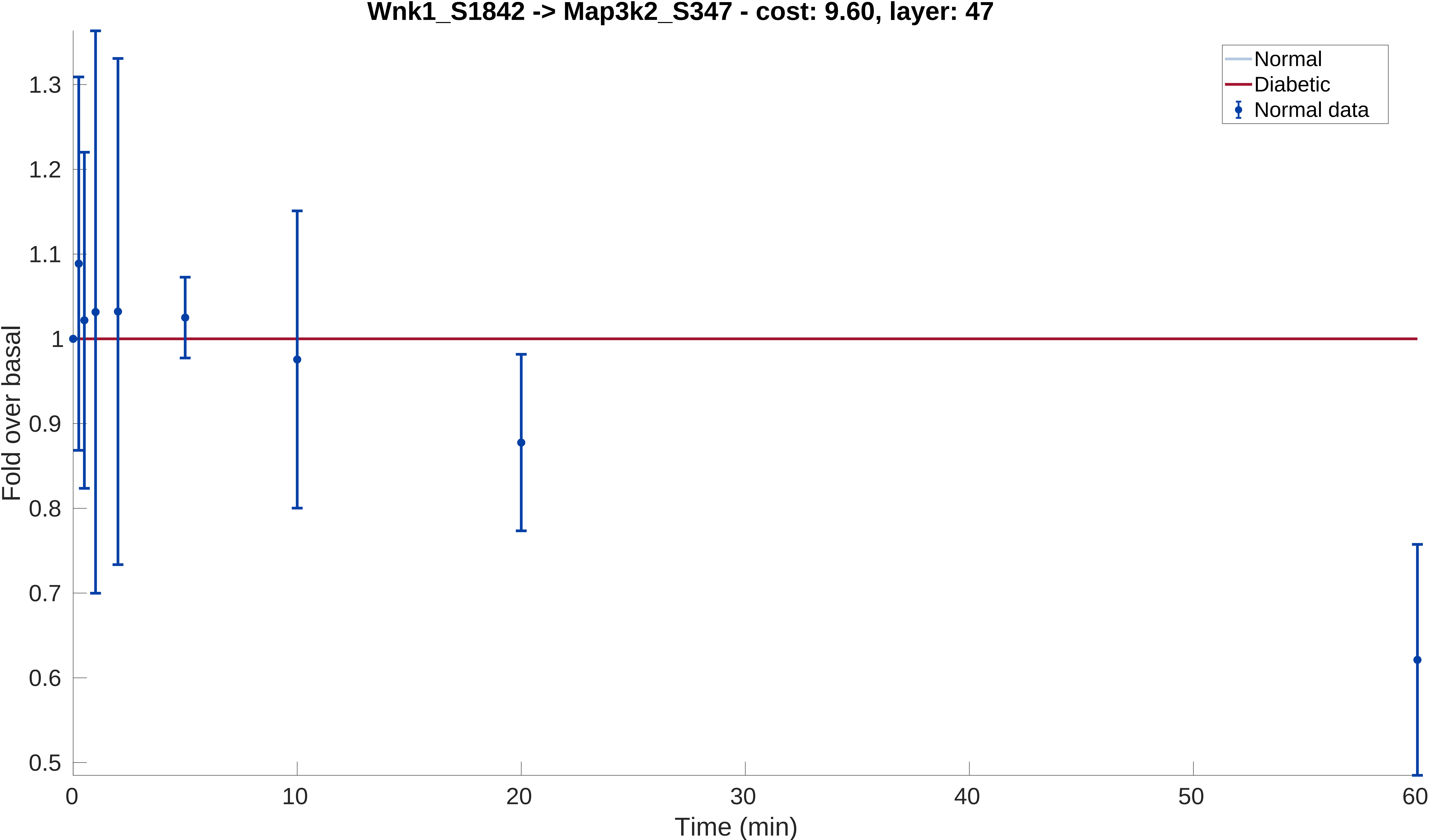

Wnk1_S1842 -> Map3k2_S347 - cost: 9.60, layer: 47
Normal
Diabetic
Normal data
Fold over basal
1.3
1.2
1.1
1
0.9
0.8
0.7
0.6
0.5
0
10
20
30
40
50
60
Time (min)

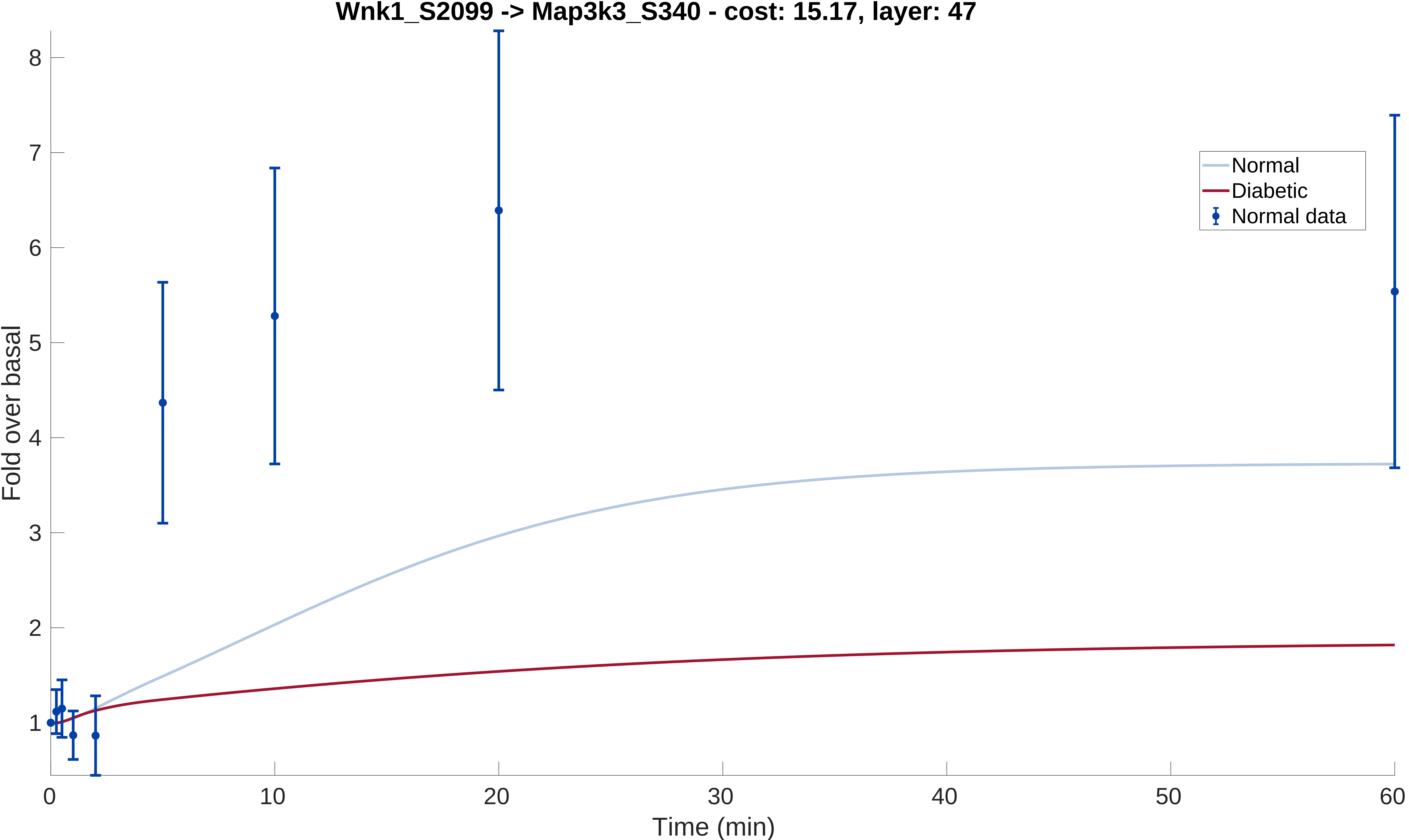
Wnk1_S2099 -> Map3k3_S340 - cost: 15.17, layer: 47
Fold over basal
Time (min)
0
10
20
30
40
50
60
1
2
3
4
5
6
7
8
Normal
Diabetic
Normal data

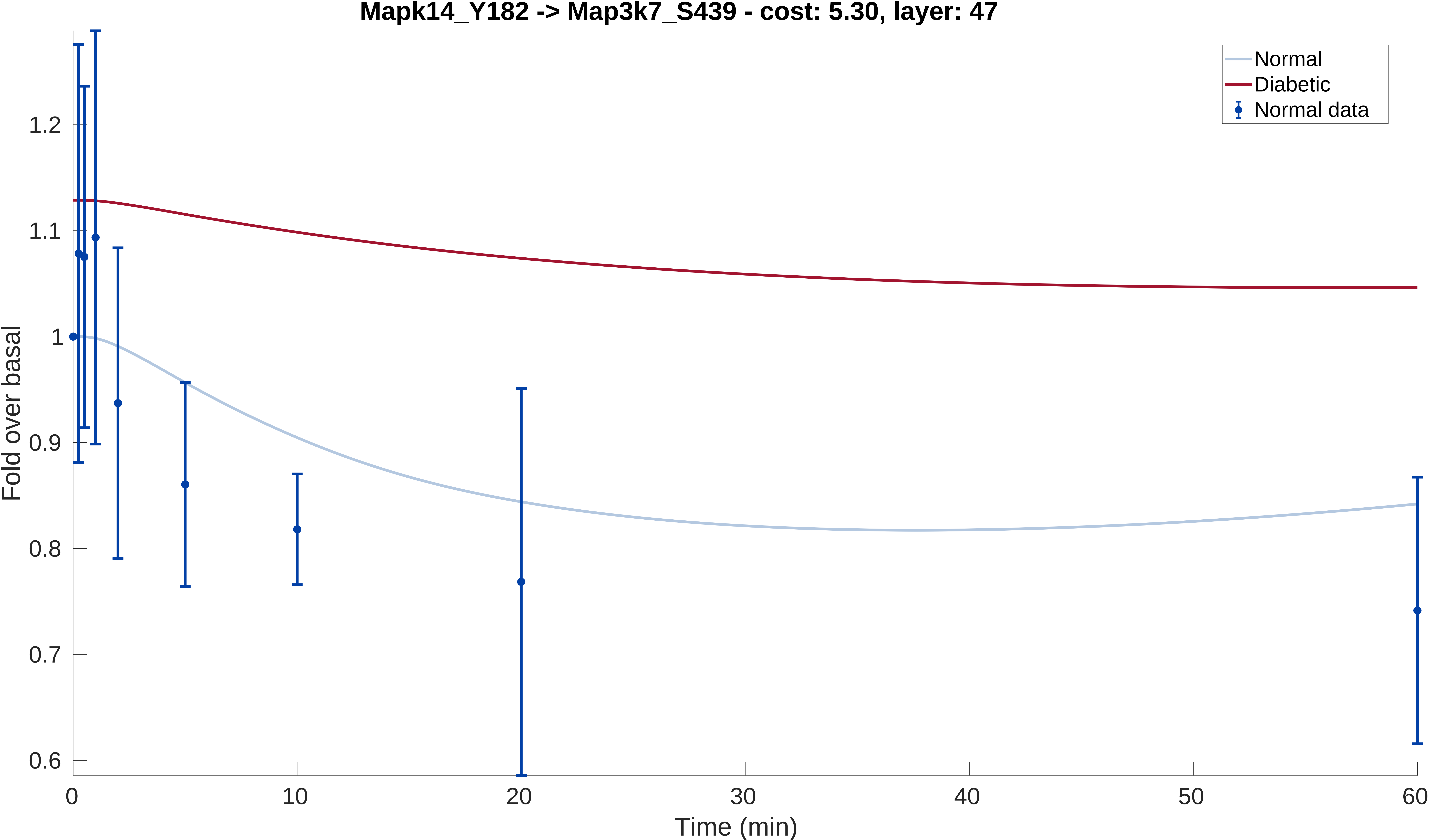
Mapk14_Y182 -> Map3k7_S439 - cost: 5.30, layer: 47
Normal
Diabetic
Normal data
Fold over basal
Time (min)
0.6
0.7
0.8
0.9
1
1.1
1.2
0
10
20
30
40
50
60

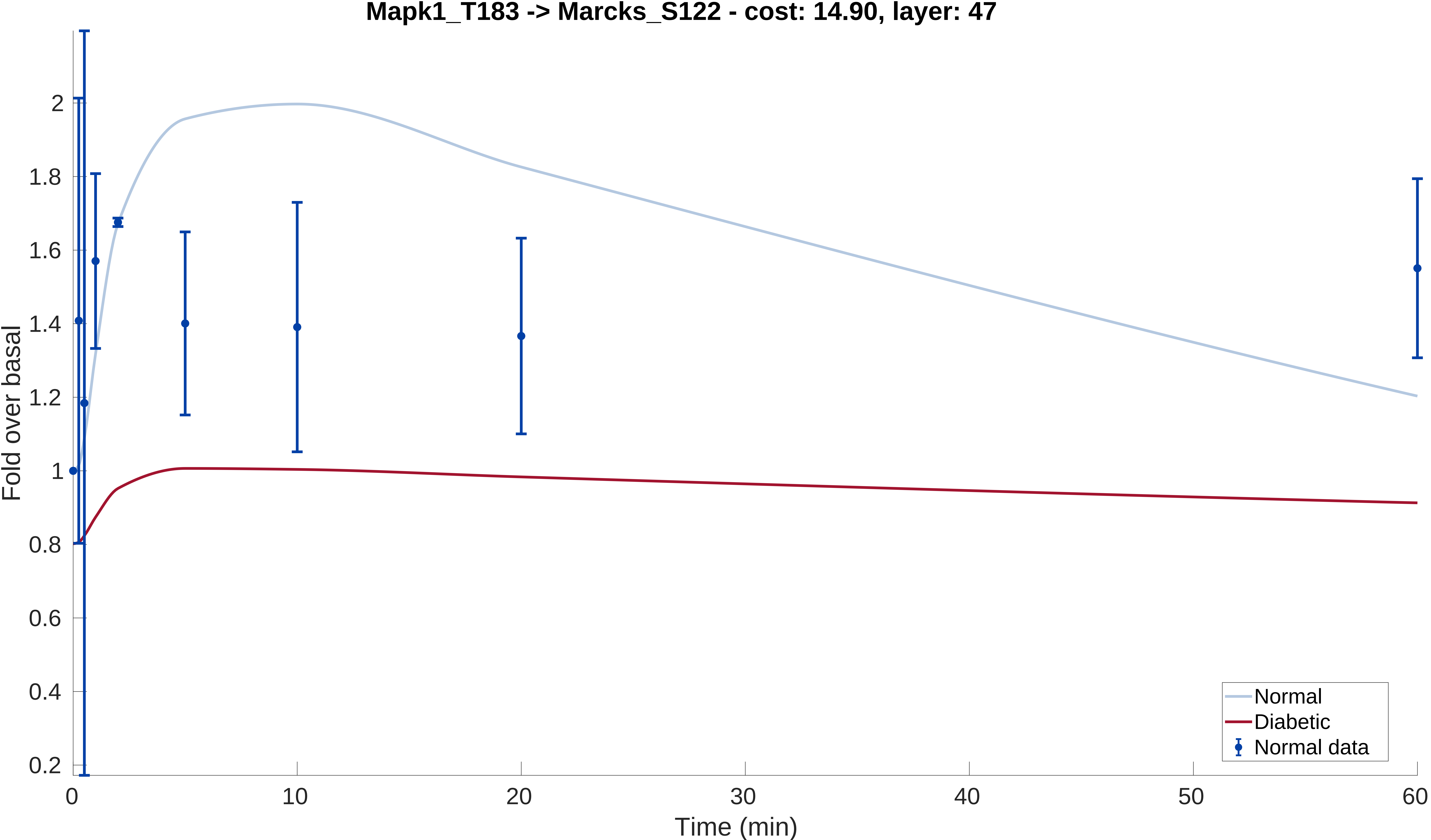

Mapk1_T183 -> Marcks_S122 - cost: 14.90, layer: 47
Fold over basal
2
1.8
1.6
1.4
1.2
1
0.8
0.6
0.4
0.2
0
10
20
30
40
50
60
Time (min)
Normal
Diabetic
Normal data

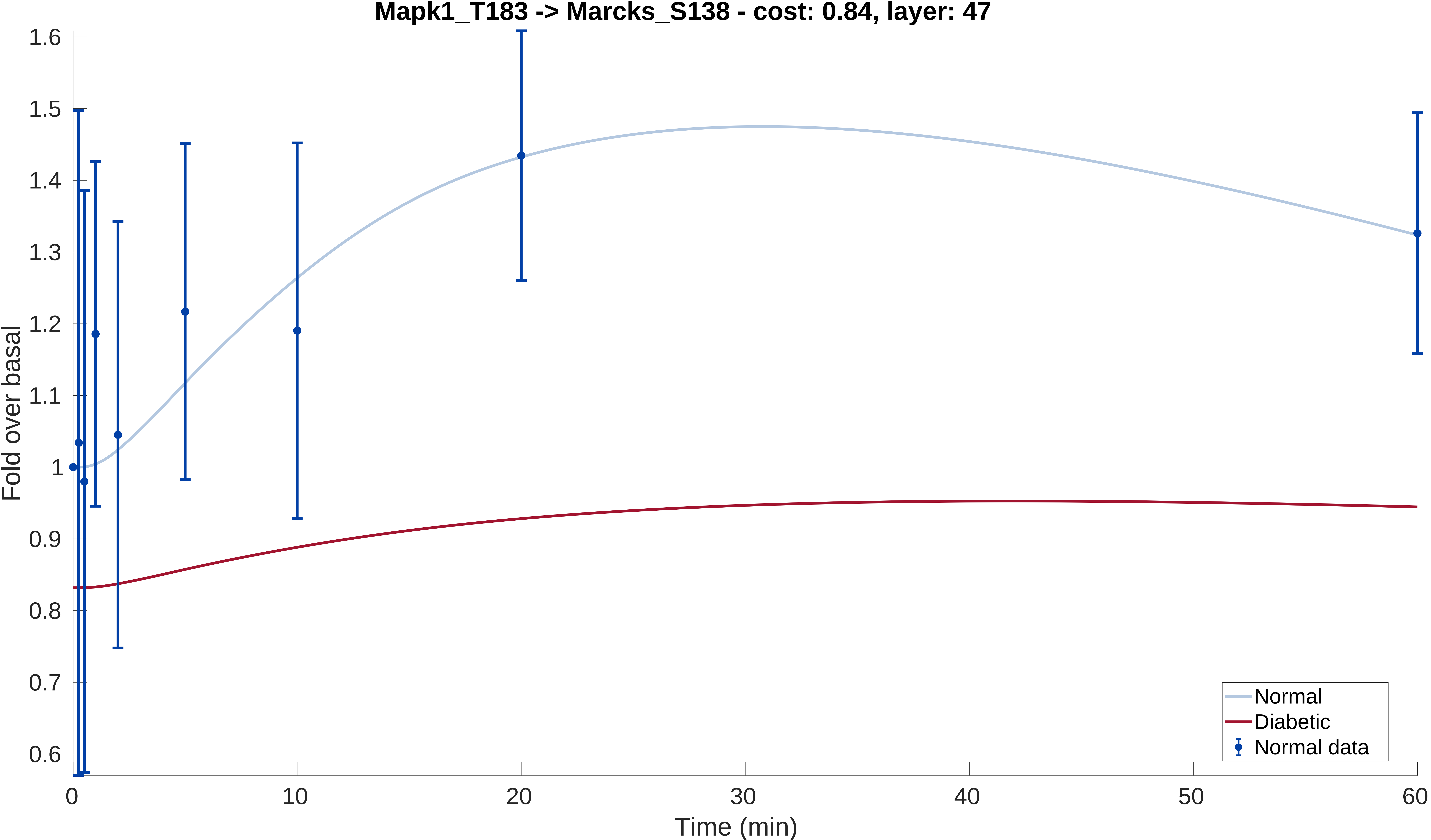

Mapk1_T183 -> Marcks_S138 - cost: 0.84, layer: 47
Fold over basal
Time (min)
1.6
1.5
1.4
1.3
1.2
1.1
1
0.9
0.8
0.7
0.6
0
10
20
30
40
50
60
Normal
Diabetic
Normal data

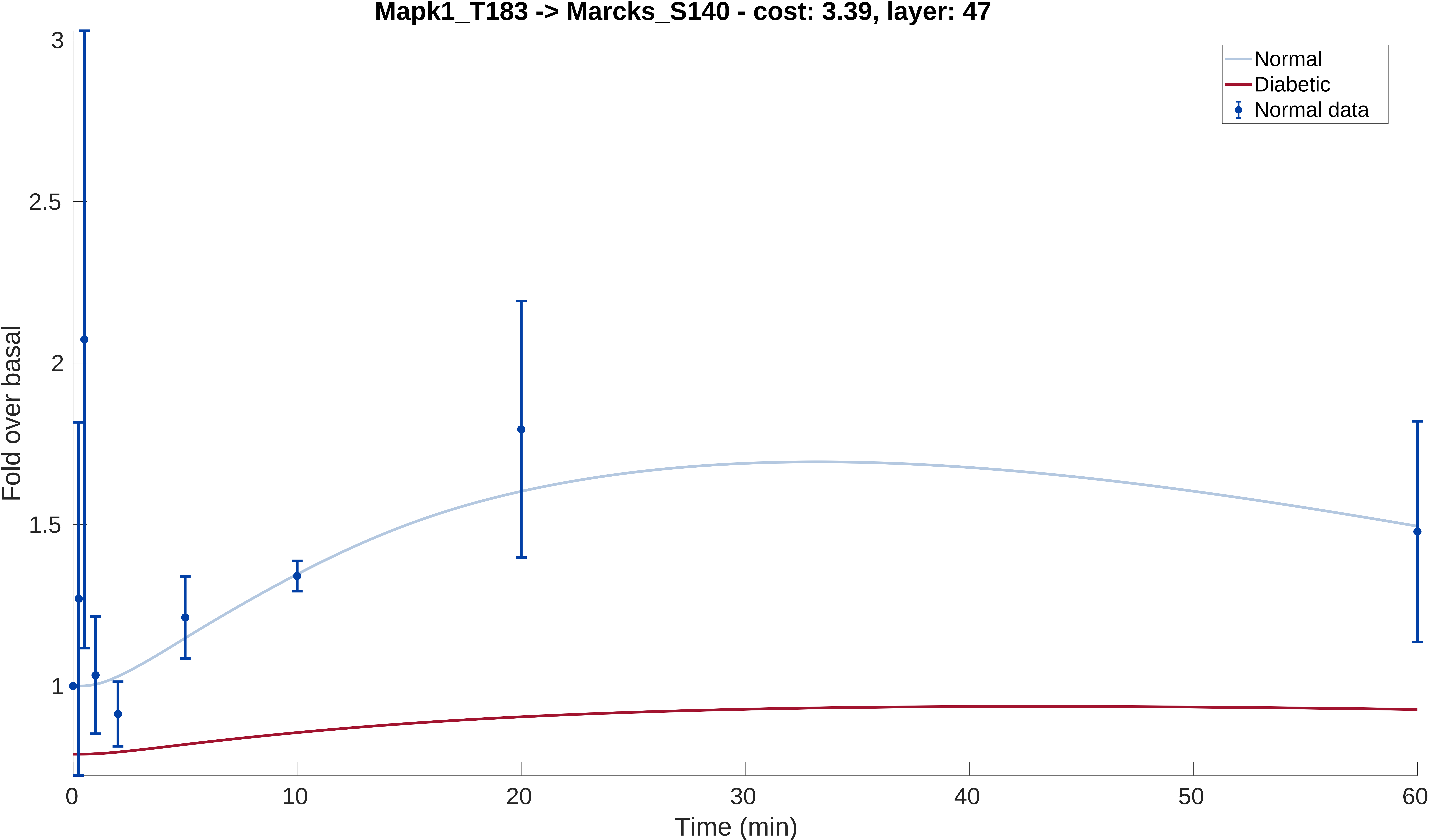

Mapk1_T183 -> Marcks_S140 - cost: 3.39, layer: 47
Normal
Diabetic
Normal data
Fold over basal
3
2.5
2
1.5
1
0
10
20
30
40
50
60
Time (min)

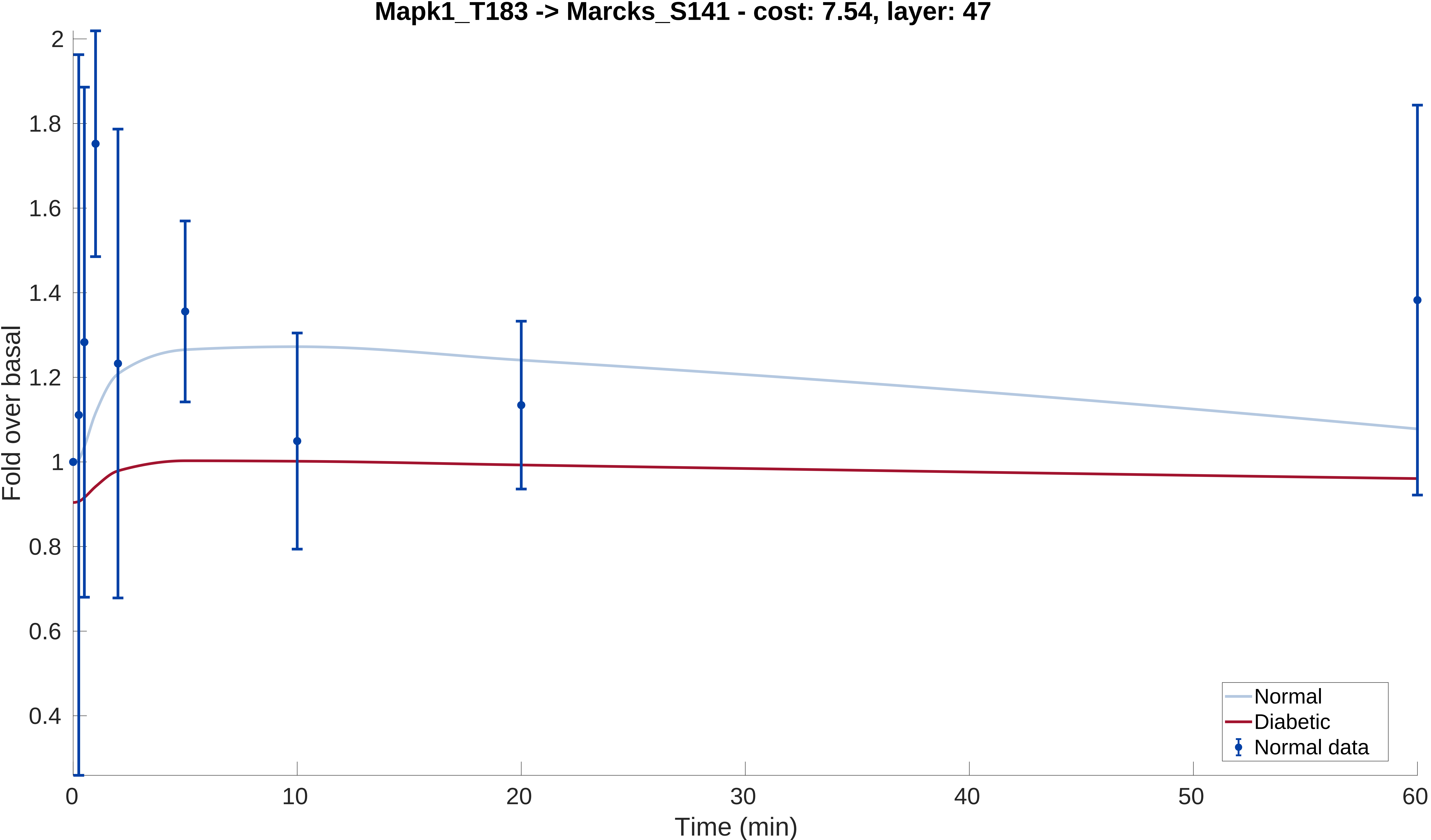

Mapk1_T183 -> Marcks_S141 - cost: 7.54, layer: 47
Fold over basal
2
1.8
1.6
1.4
1.2
1
0.8
0.6
0.4
0
10
20
30
40
50
60
Time (min)
Normal
Diabetic
Normal data

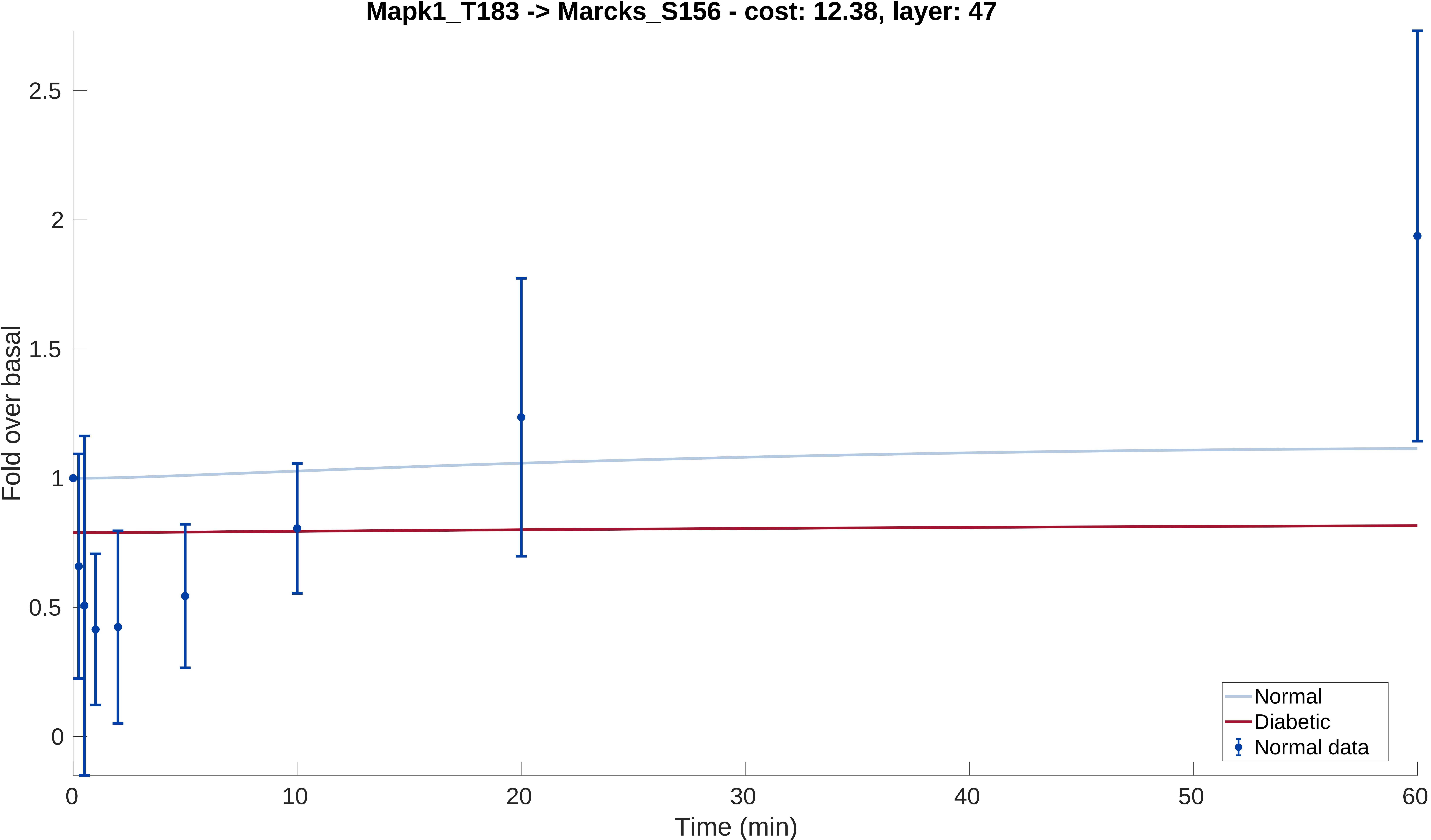

Mapk1_T183 -> Marcks_S156 - cost: 12.38, layer: 47
Fold over basal
2.5
2
1.5
1
0.5
0
0
10
20
30
40
50
60
Time (min)
Normal
Diabetic
Normal data

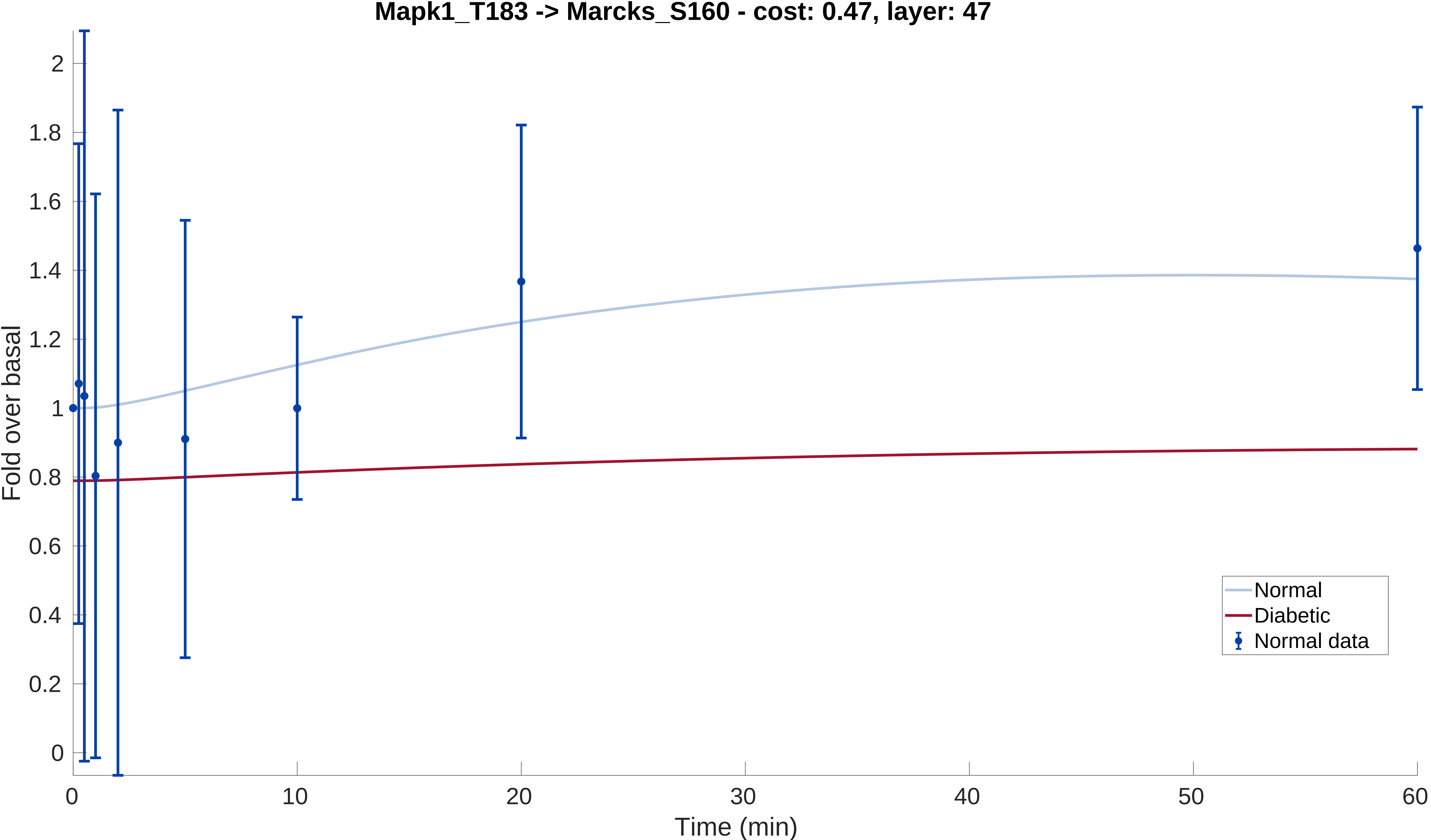
Mapk1_T183 -> Marcks_S160 - cost: 0.47, layer: 47
Fold over basal
Time (min)
0
0.2
0.4
0.6
0.8
1
1.2
1.4
1.6
1.8
2
0
10
20
30
40
50
60
Normal
Diabetic
Normal data

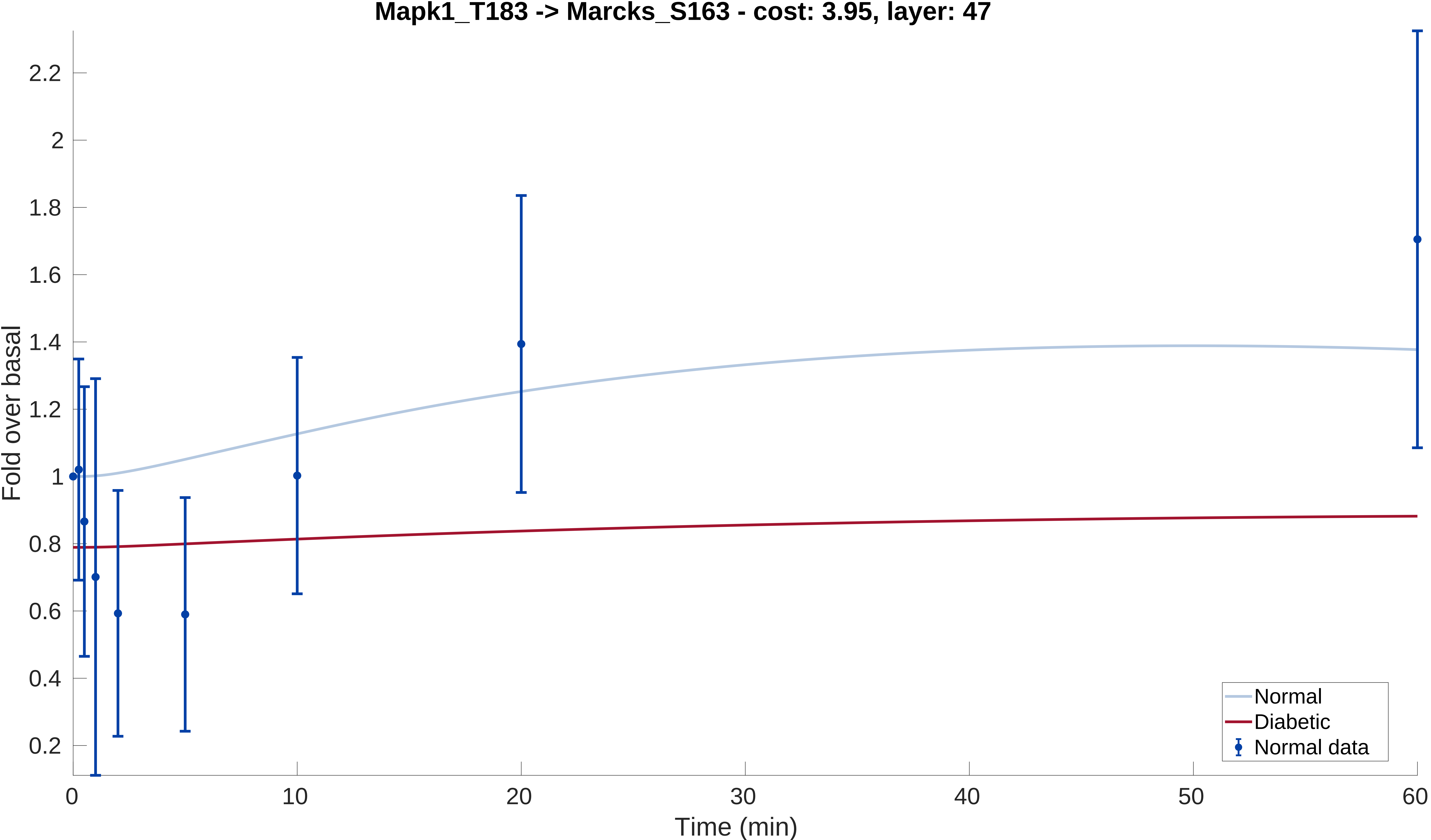

Mapk1_T183 -> Marcks_S163 - cost: 3.95, layer: 47
Fold over basal
Time (min)
2.2
2
1.8
1.6
1.4
1.2
1
0.8
0.6
0.4
0.2
0
10
20
30
40
50
60
Normal
Diabetic
Normal data

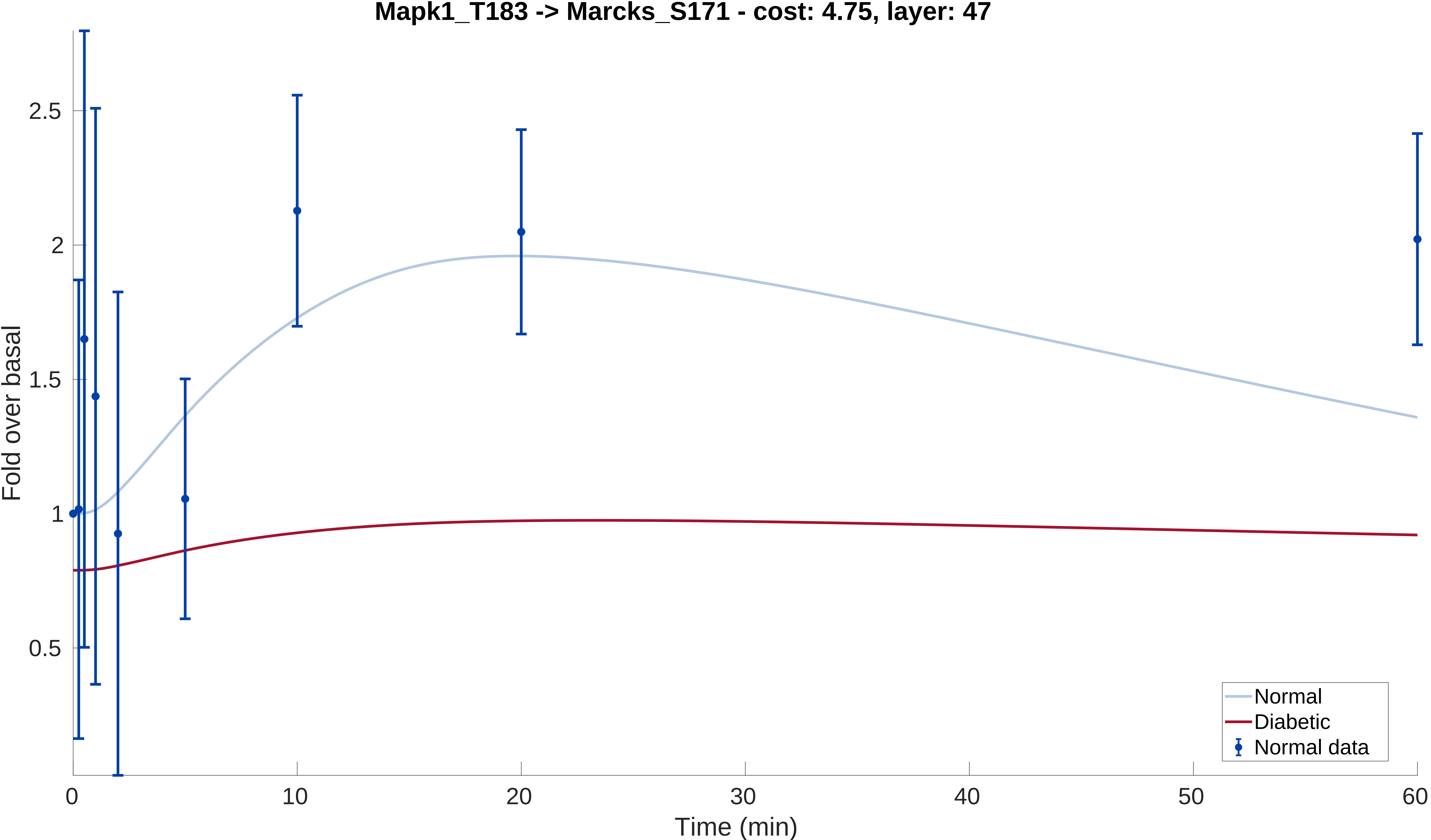

Mapk1_T183 -> Marcks_S171 - cost: 4.75, layer: 47
Fold over basal
Time (min)
Normal
Diabetic
Normal data
0.5
1
1.5
2
2.5
0
10
20
30
40
50
60

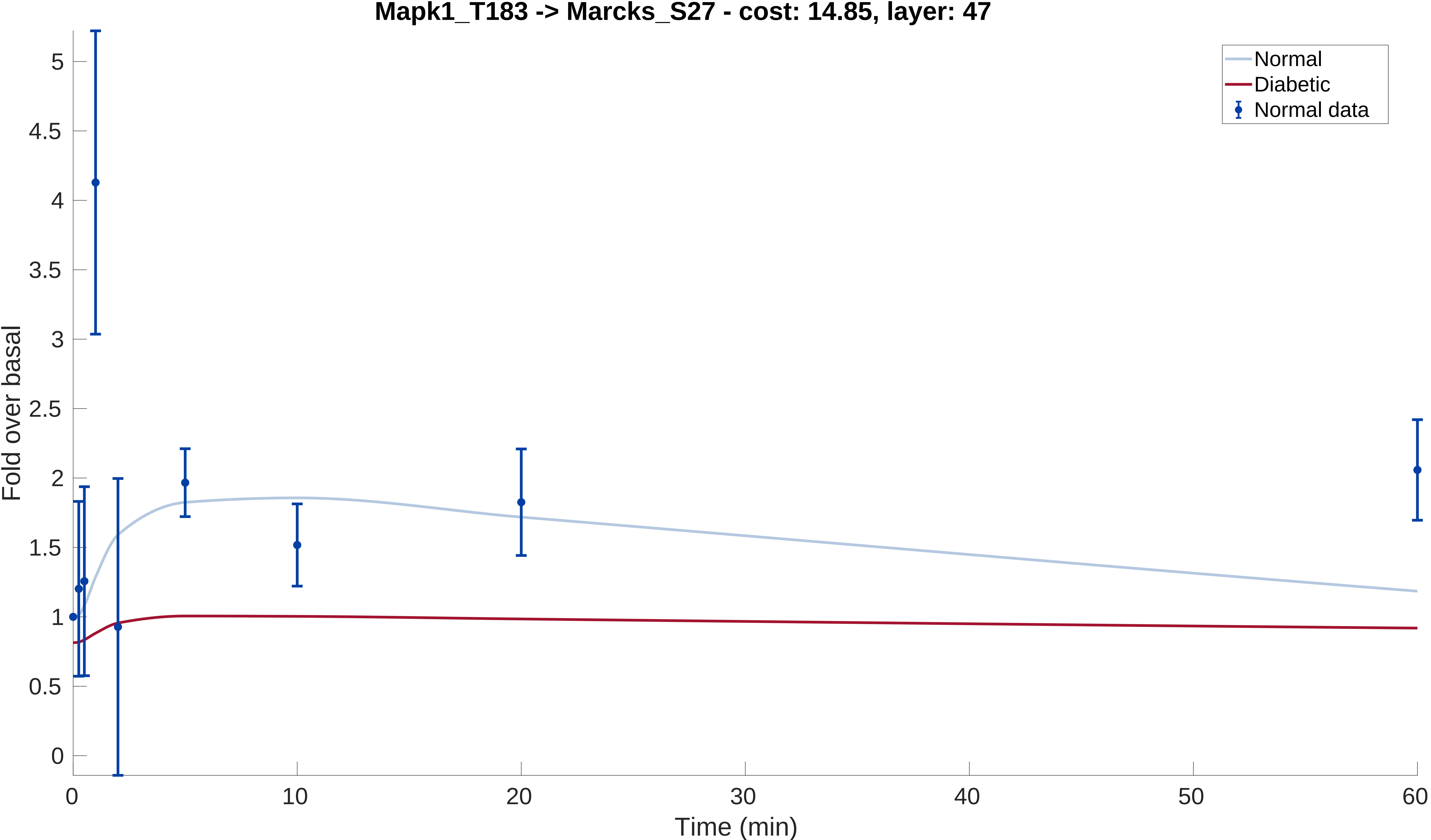

Mapk1_T183 -> Marcks_S27 - cost: 14.85, layer: 47
Normal
Diabetic
Normal data
Fold over basal
Time (min)
0
0.5
1
1.5
2
2.5
3
3.5
4
4.5
5
0
10
20
30
40
50
60

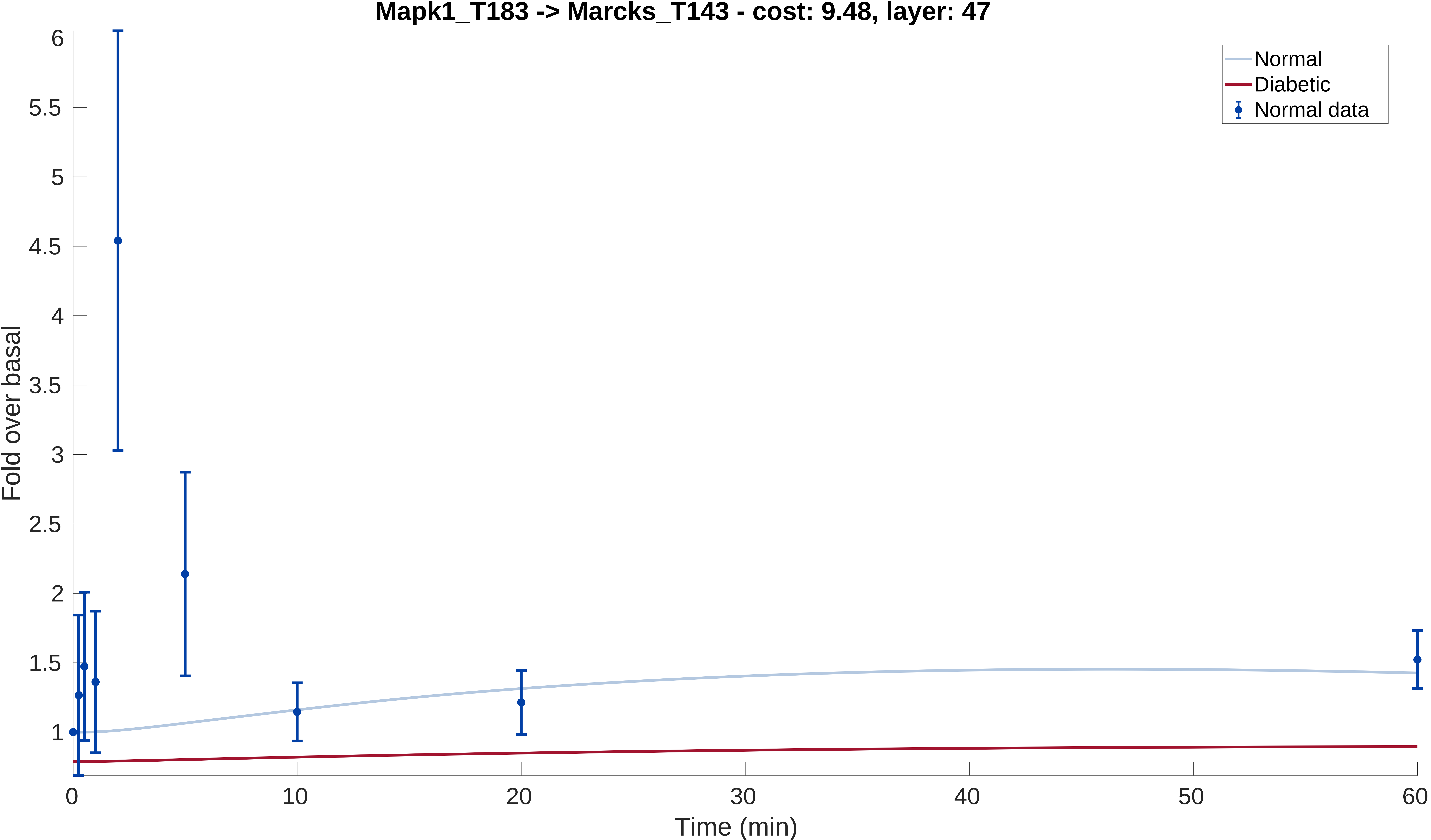

Mapk1_T183 -> Marcks_T143 - cost: 9.48, layer: 47
Fold over basal
Time (min)
Normal
Diabetic
Normal data
0
10
20
30
40
50
60
1
1.5
2
2.5
3
3.5
4
4.5
5
5.5
6

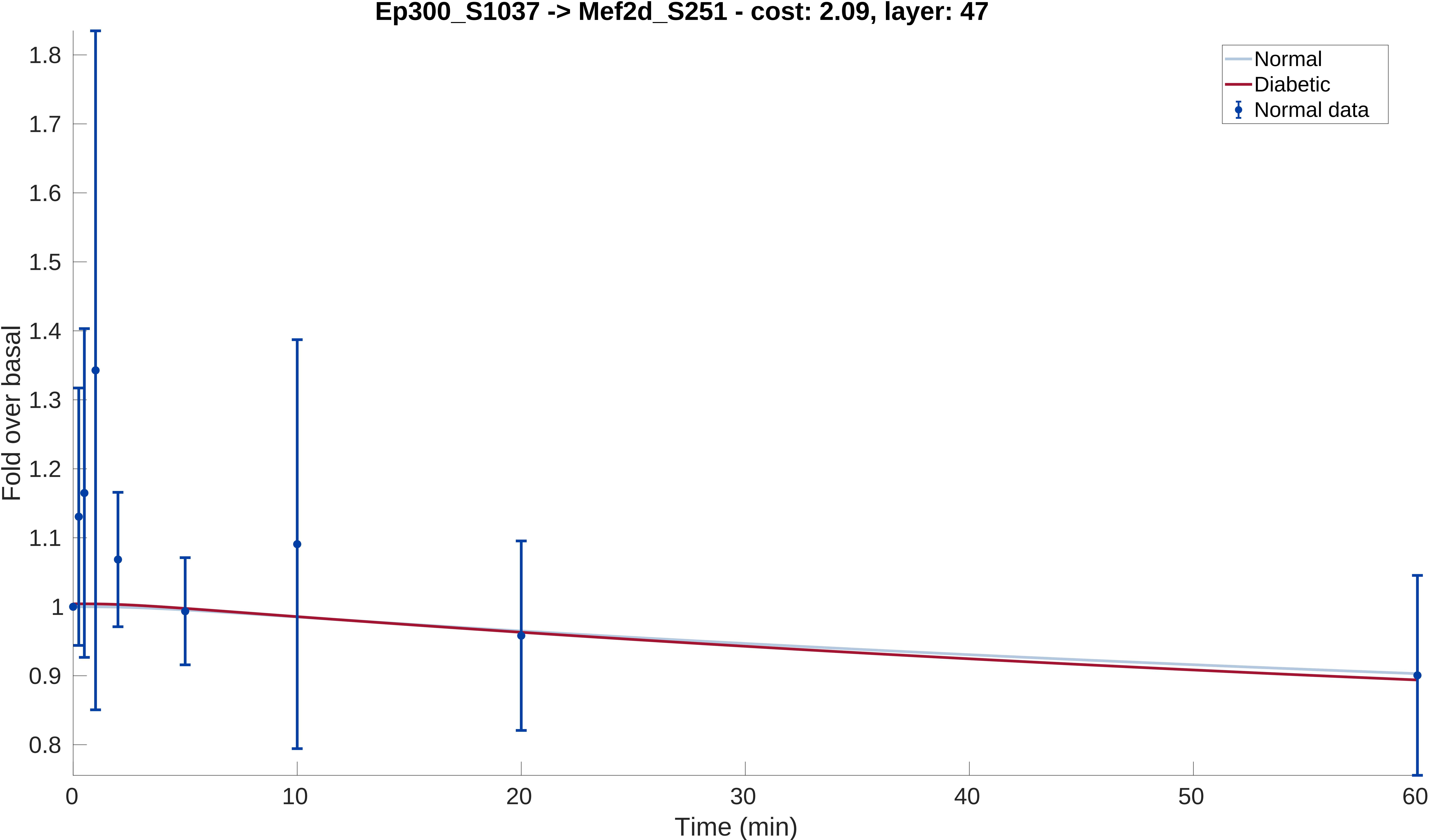
Ep300_S1037 -> Mef2d_S251 - cost: 2.09, layer: 47
Fold over basal
1.8
1.7
1.6
1.5
1.4
1.3
1.2
1.1
1
0.9
0.8
0
10
20
30
40
50
60
Time (min)
Normal
Diabetic
Normal data

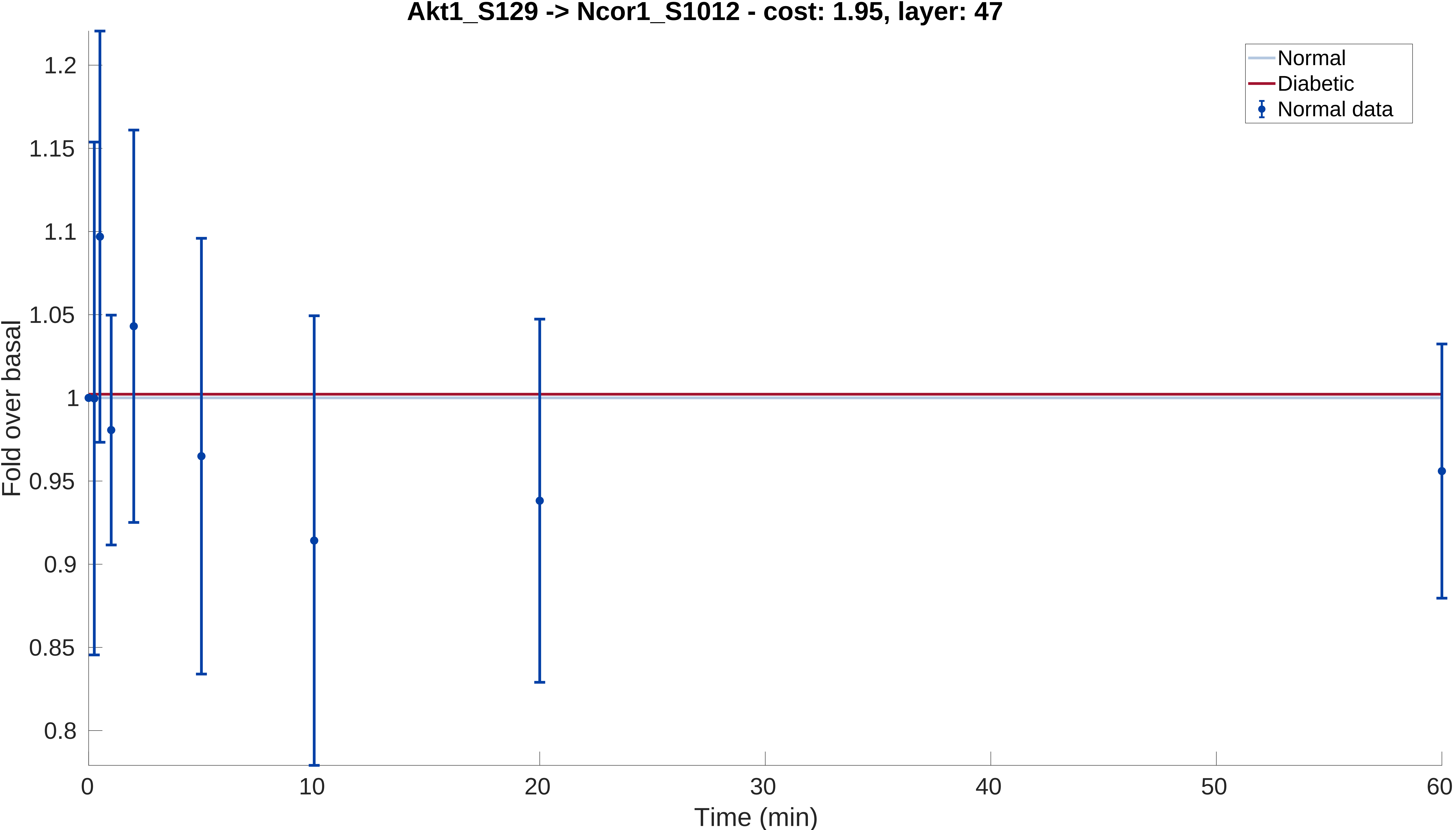

Akt1_S129 -> Ncor1_S1012 - cost: 1.95, layer: 47
Normal
Diabetic
Normal data
Fold over basal
Time (min)
1.2
1.15
1.1
1.05
1
0.95
0.9
0.85
0.8
0
10
20
30
40
50
60

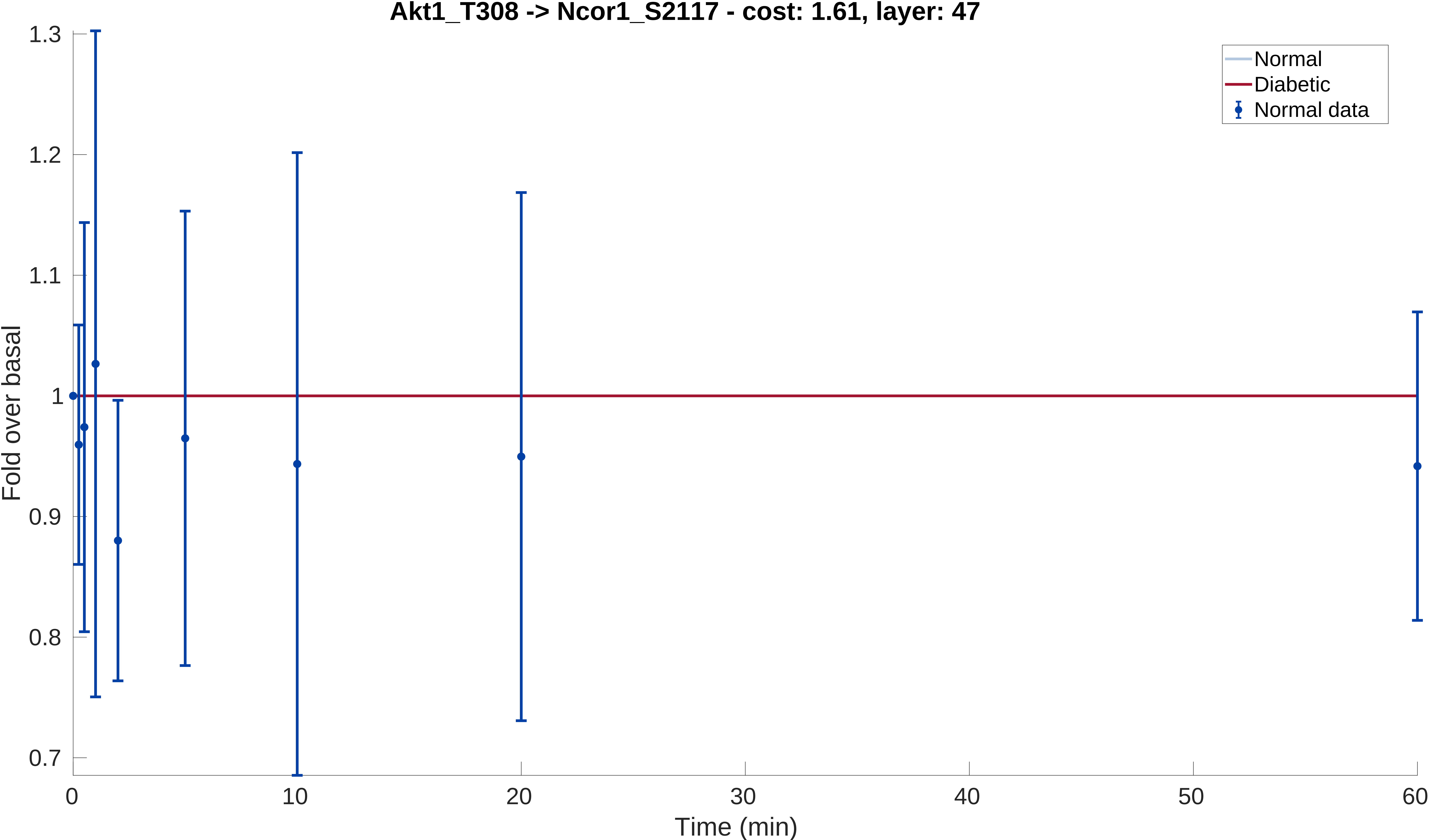

Akt1_T308 -> Ncor1_S2117 - cost: 1.61, layer: 47
Normal
Diabetic
Normal data
Fold over basal
1.3
1.2
1.1
1
0.9
0.8
0.7
0
10
20
30
40
50
60
Time (min)

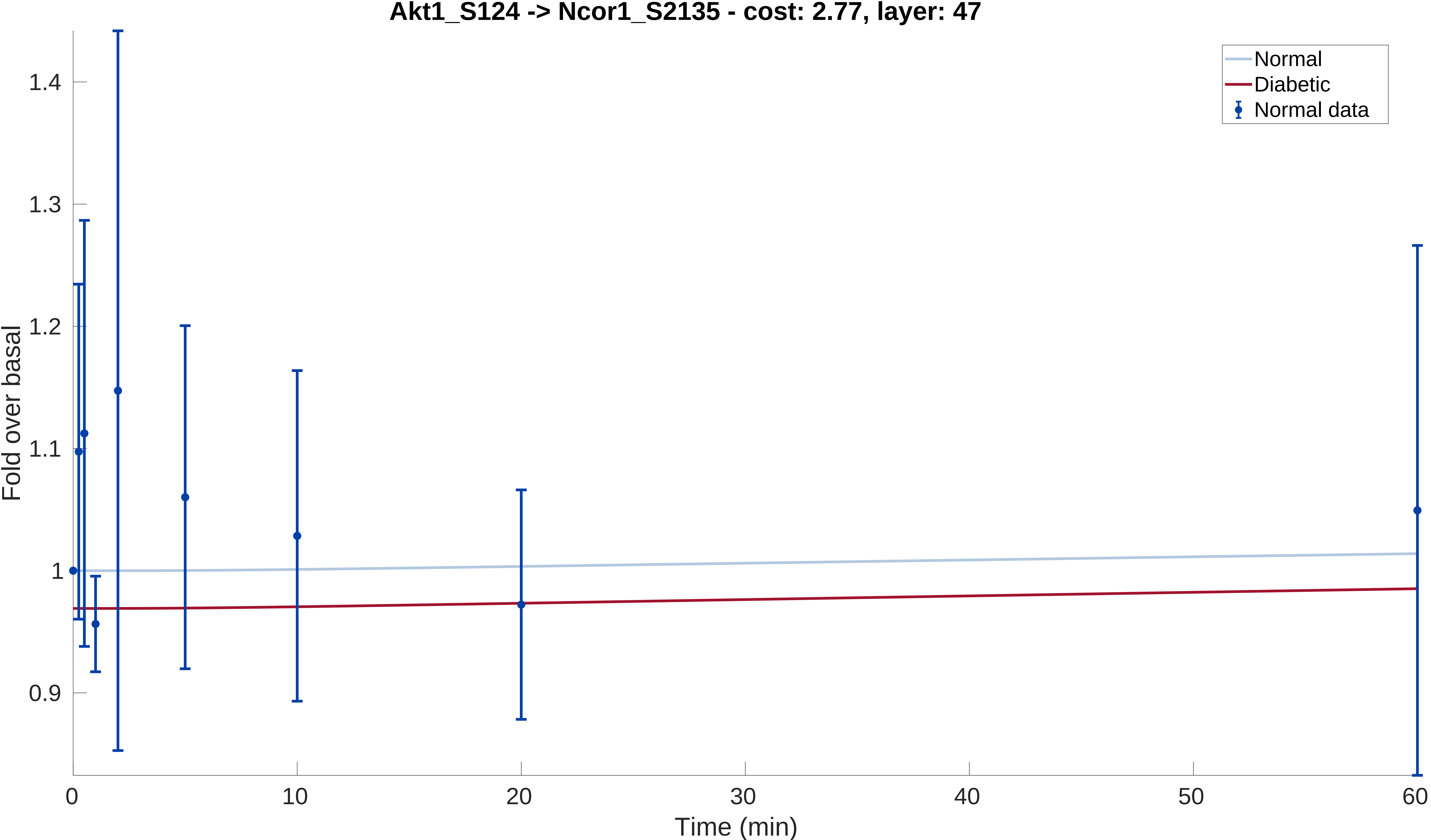

Akt1_S124 -> Ncor1_S2135 - cost: 2.77, layer: 47
Normal
Diabetic
Normal data
Fold over basal
1.4
1.3
1.2
1.1
1
0.9
0
10
20
30
40
50
60
Time (min)

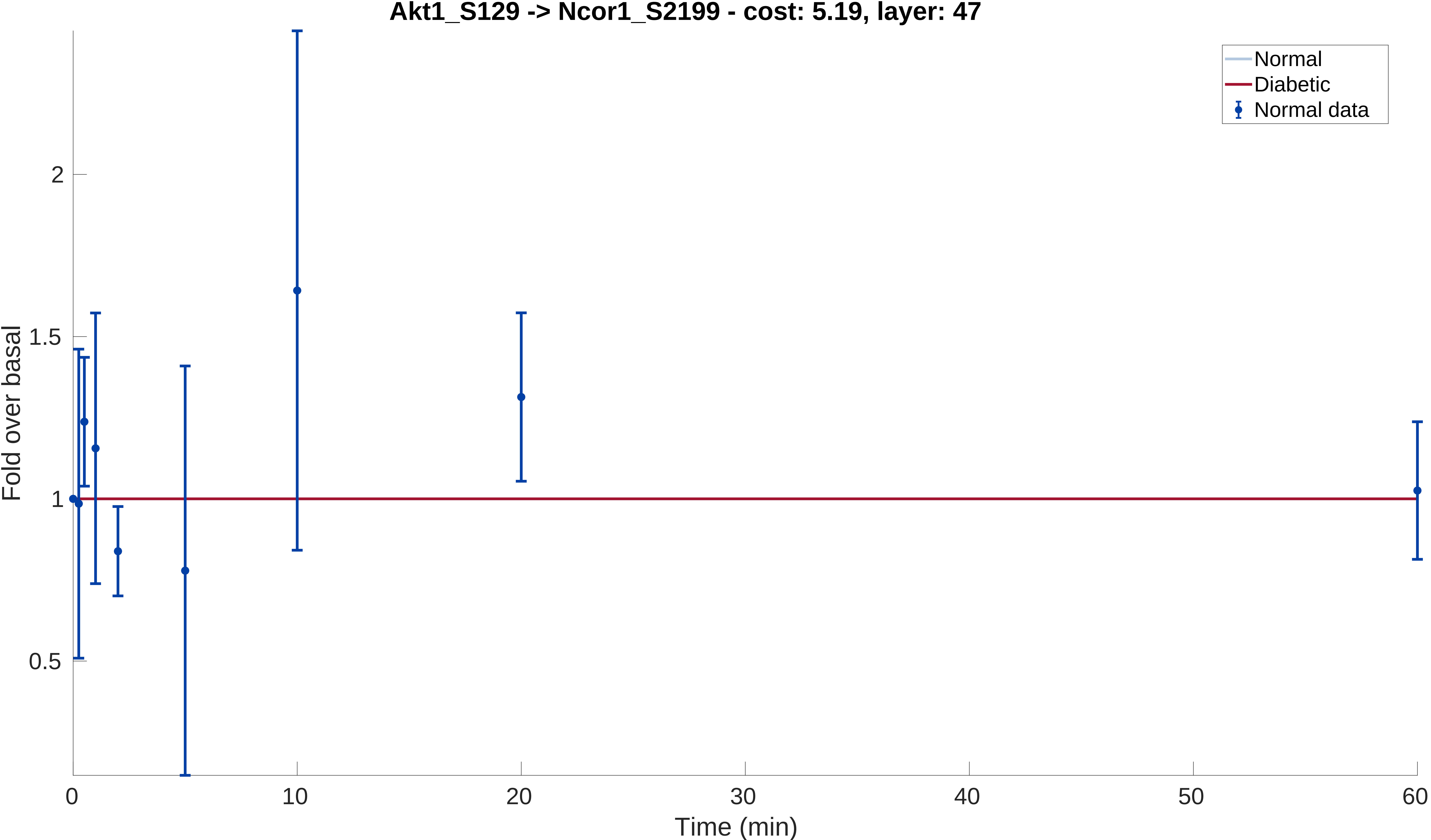

Akt1_S129 -> Ncor1_S2199 - cost: 5.19, layer: 47
Normal
Diabetic
Normal data
Fold over basal
0.5
1
1.5
2
0
10
20
30
40
50
60
Time (min)

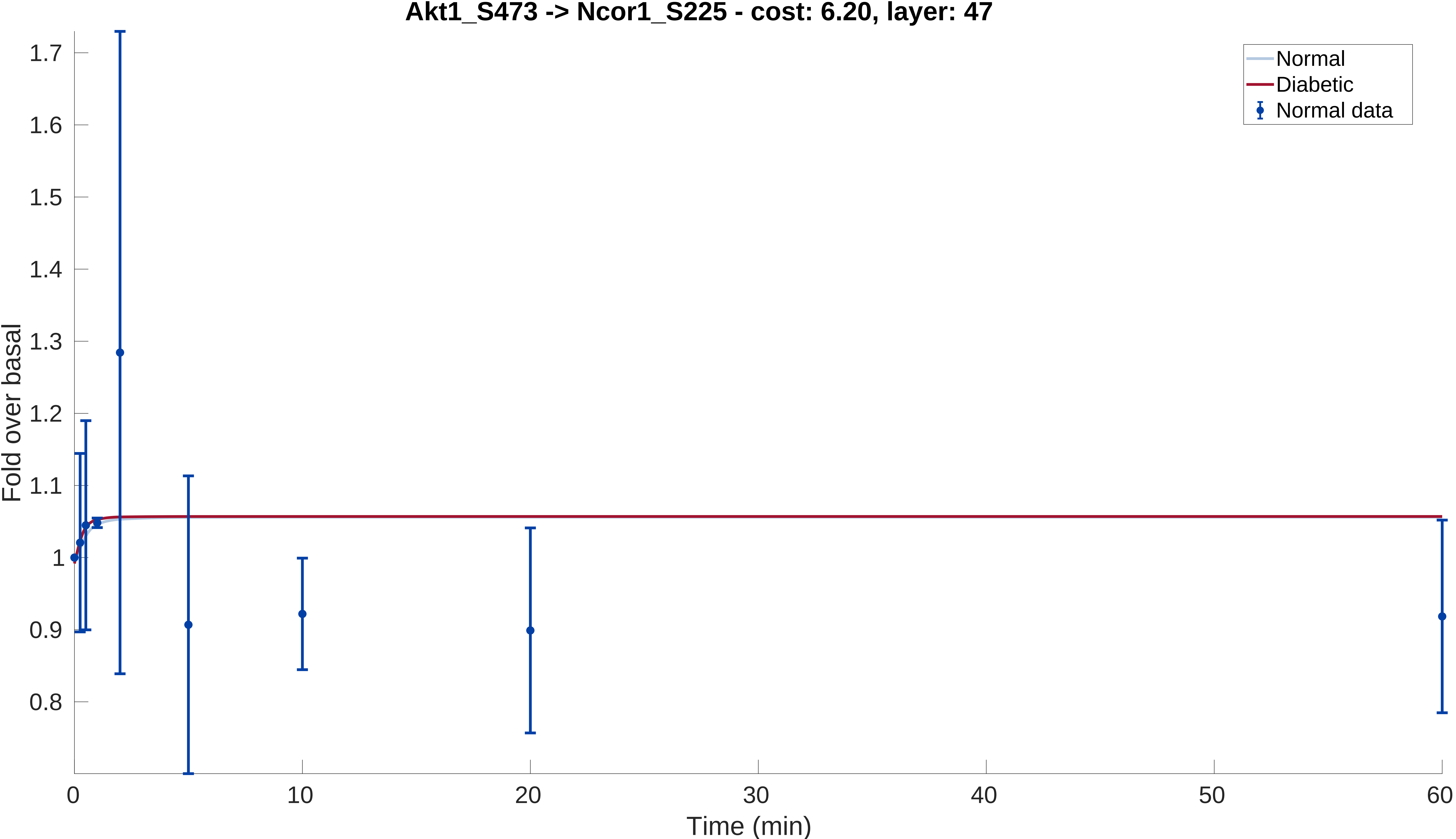

Akt1_S473 -> Ncor1_S225 - cost: 6.20, layer: 47
Normal
Diabetic
Normal data
Fold over basal
Time (min)
1.7
1.6
1.5
1.4
1.3
1.2
1.1
1
0.9
0.8
0
10
20
30
40
50
60

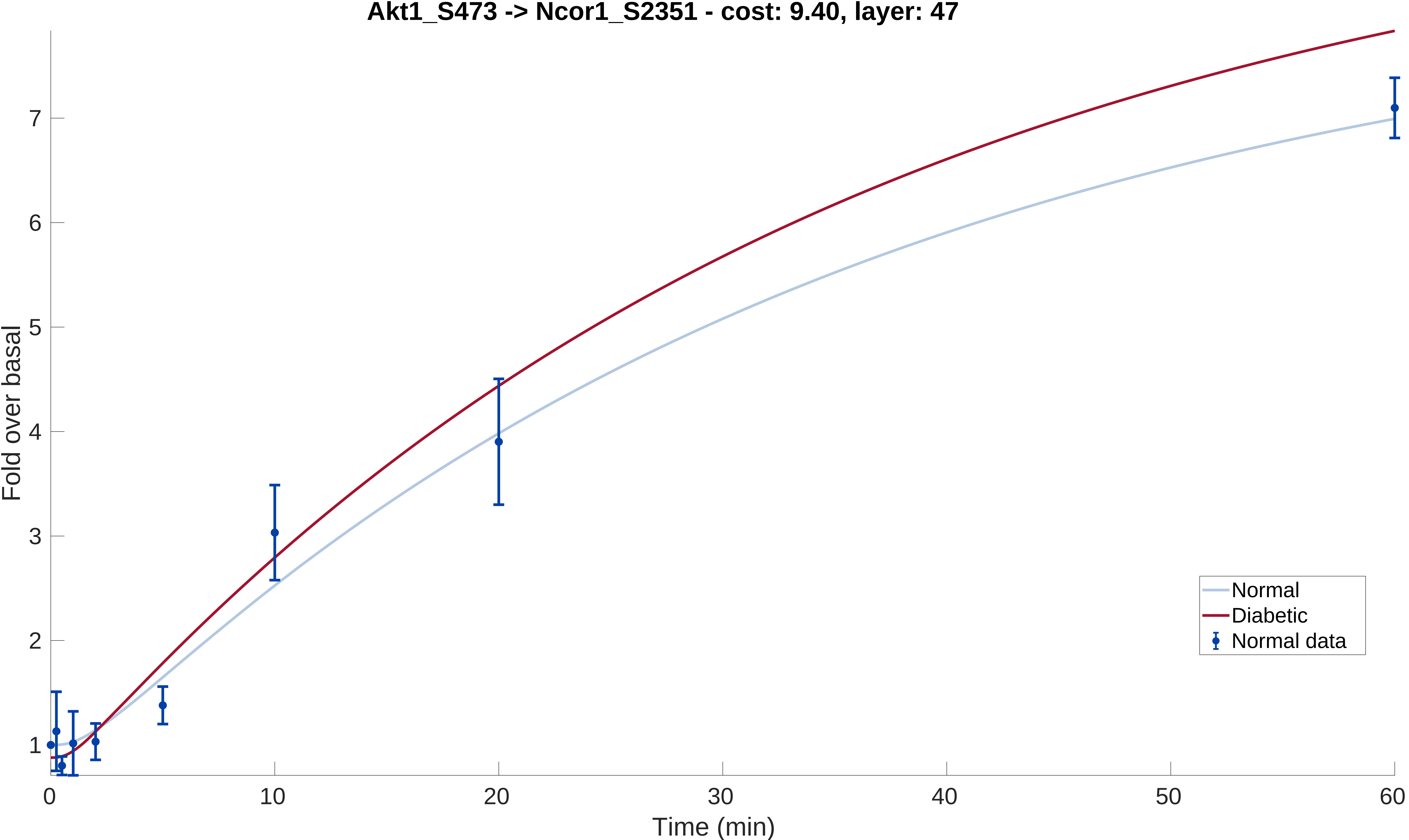

Akt1_S473 -> Ncor1_S2351 - cost: 9.40, layer: 47
Fold over basal
Time (min)
1
2
3
4
5
6
7
0
10
20
30
40
50
60
Normal
Diabetic
Normal data

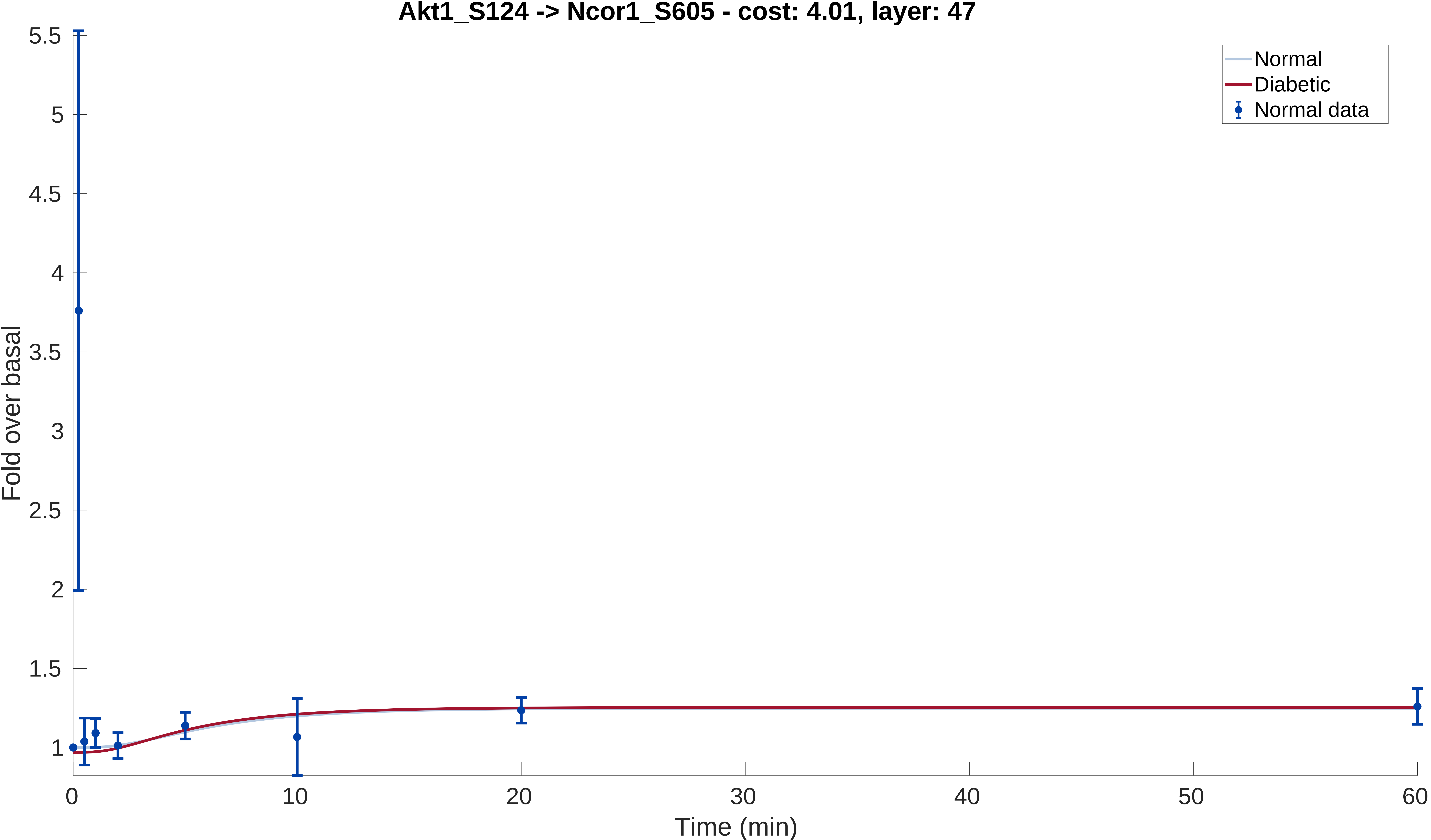

Akt1_S124 -> Ncor1_S605 - cost: 4.01, layer: 47
Normal
Diabetic
Normal data
Fold over basal
5.5
5
4.5
4
3.5
3
2.5
2
1.5
1
0
10
20
30
40
50
60
Time (min)

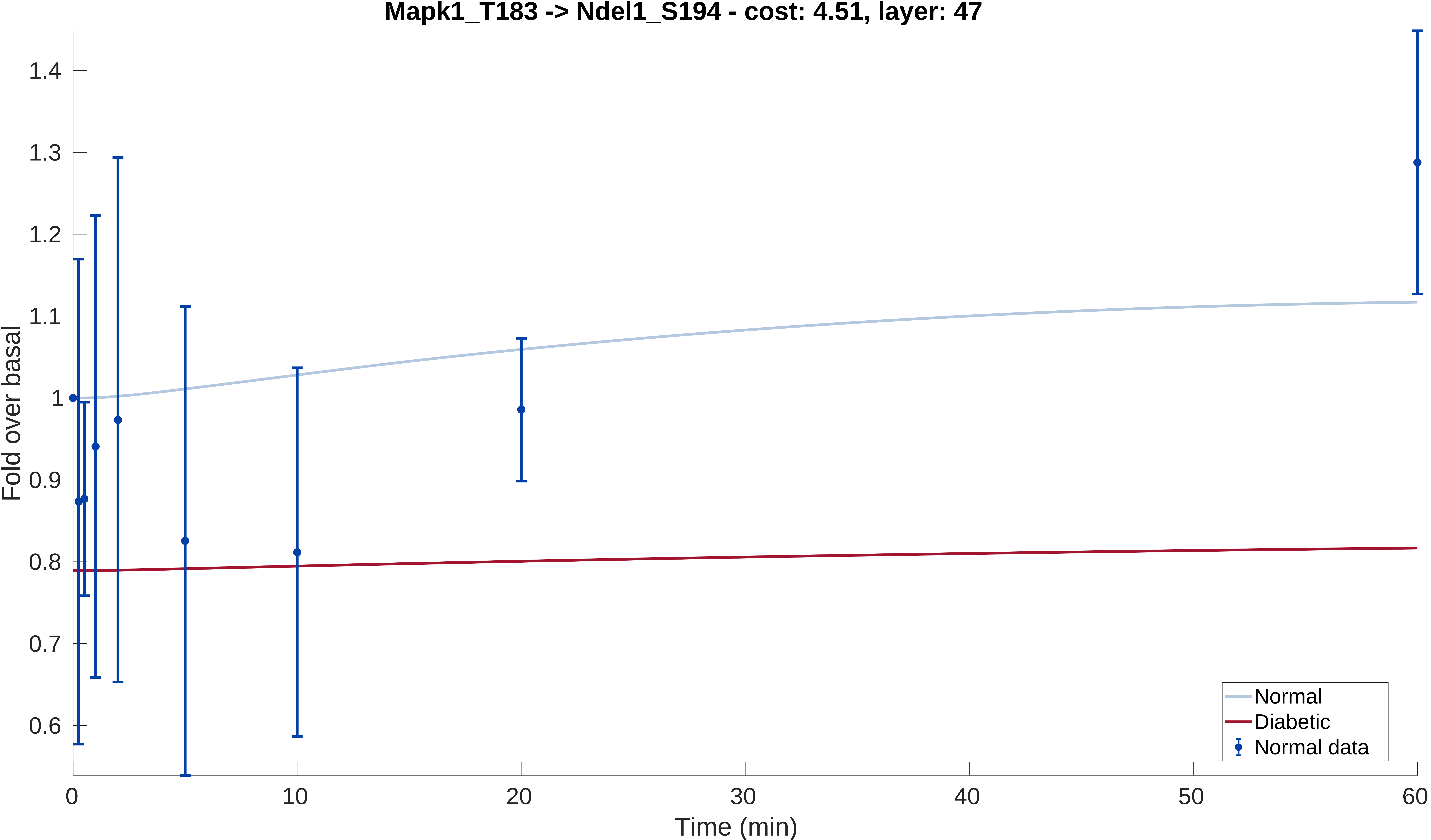
Mapk1_T183 -> Ndel1_S194 - cost: 4.51, layer: 47
Fold over basal
Time (min)
1.4
1.3
1.2
1.1
1
0.9
0.8
0.7
0.6
0
10
20
30
40
50
60
Normal
Diabetic
Normal data

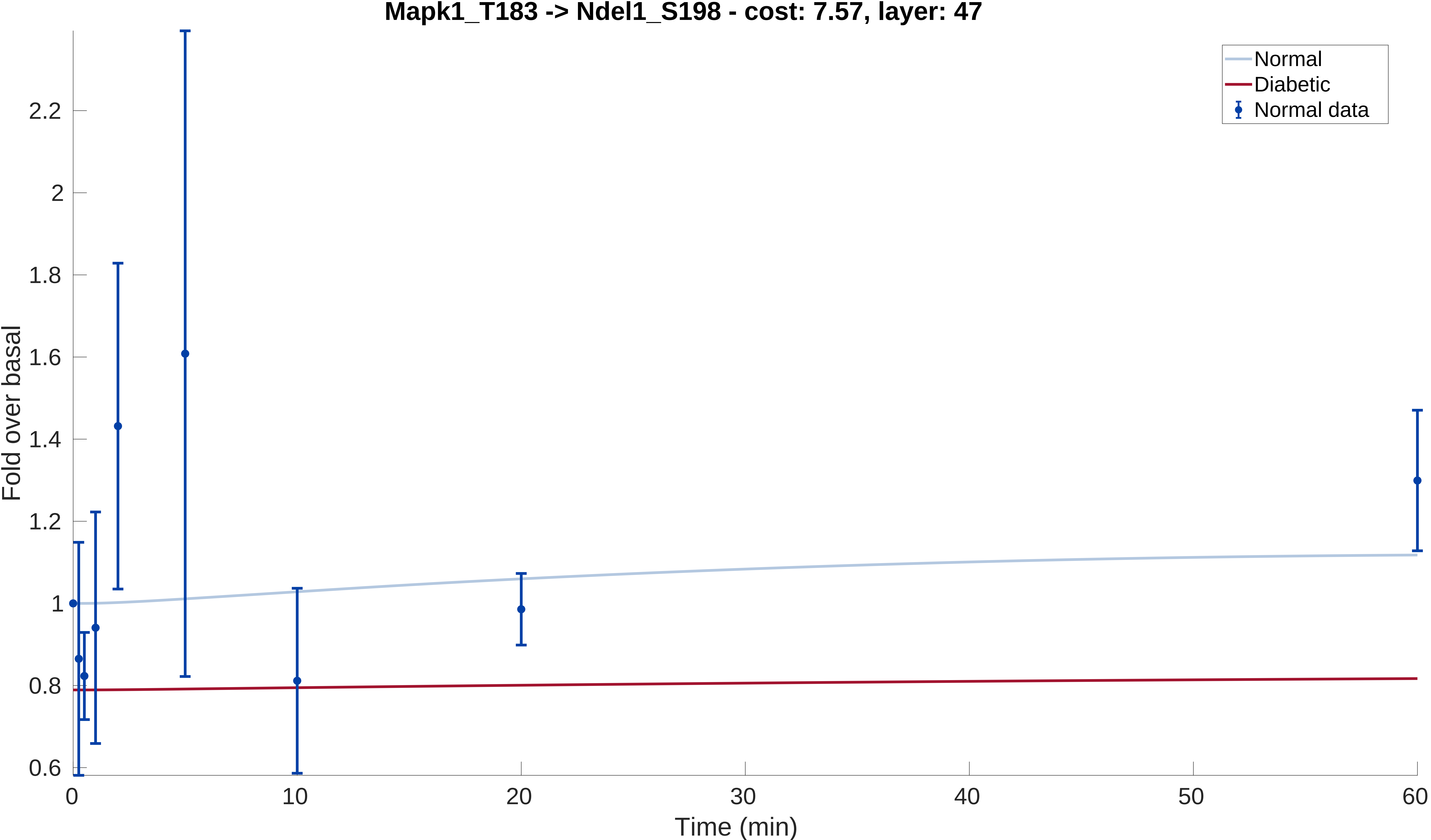

Mapk1_T183 -> Ndel1_S198 - cost: 7.57, layer: 47
Normal
Diabetic
Normal data
Fold over basal
Time (min)
0.6
0.8
1
1.2
1.4
1.6
1.8
2
2.2
0
10
20
30
40
50
60

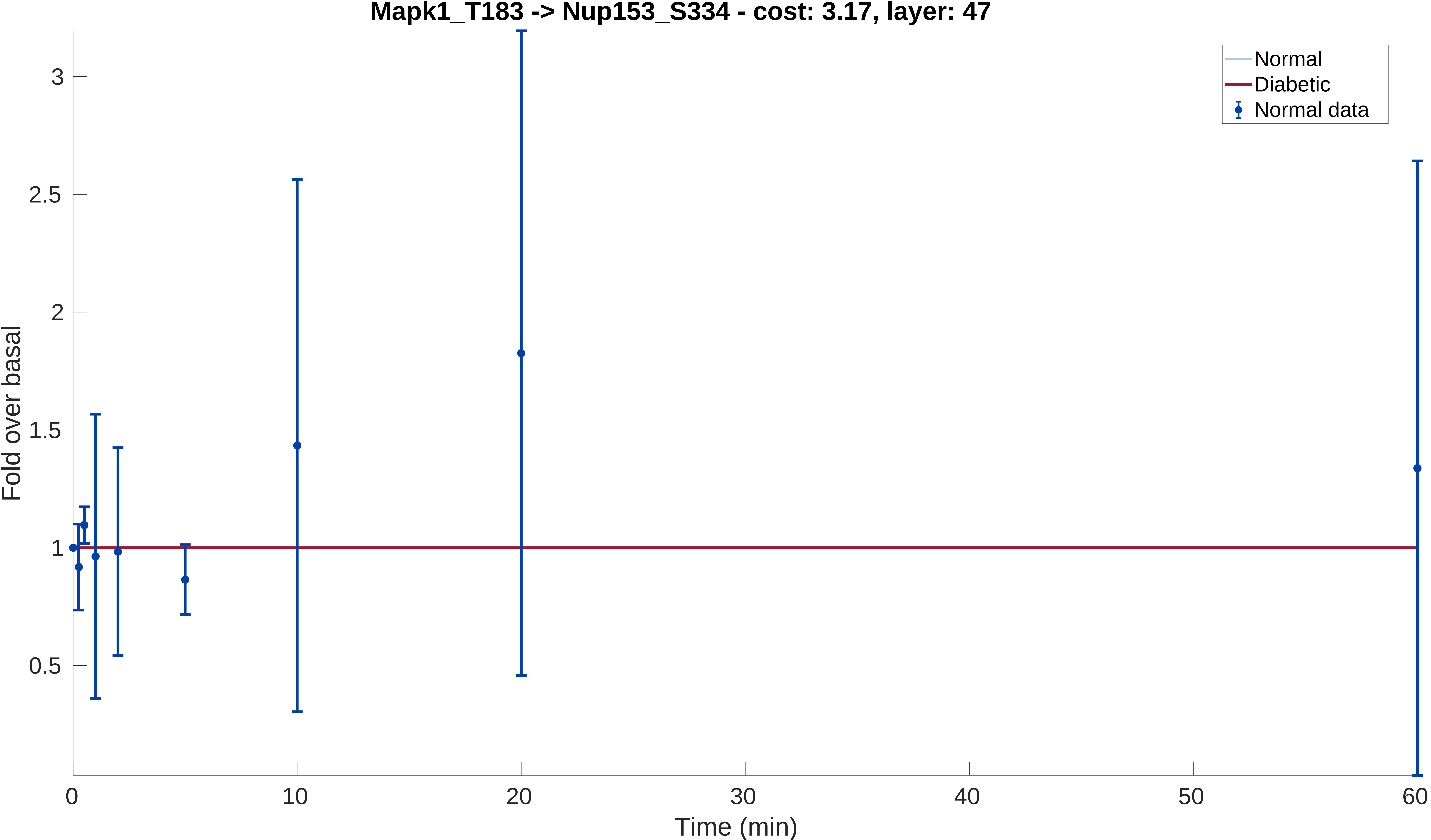

Mapk1_T183 -> Nup153_S334 - cost: 3.17, layer: 47
Normal
Diabetic
Normal data
3
2.5
2
1.5
1
0.5
Fold over basal
0
10
20
30
40
50
60
Time (min)

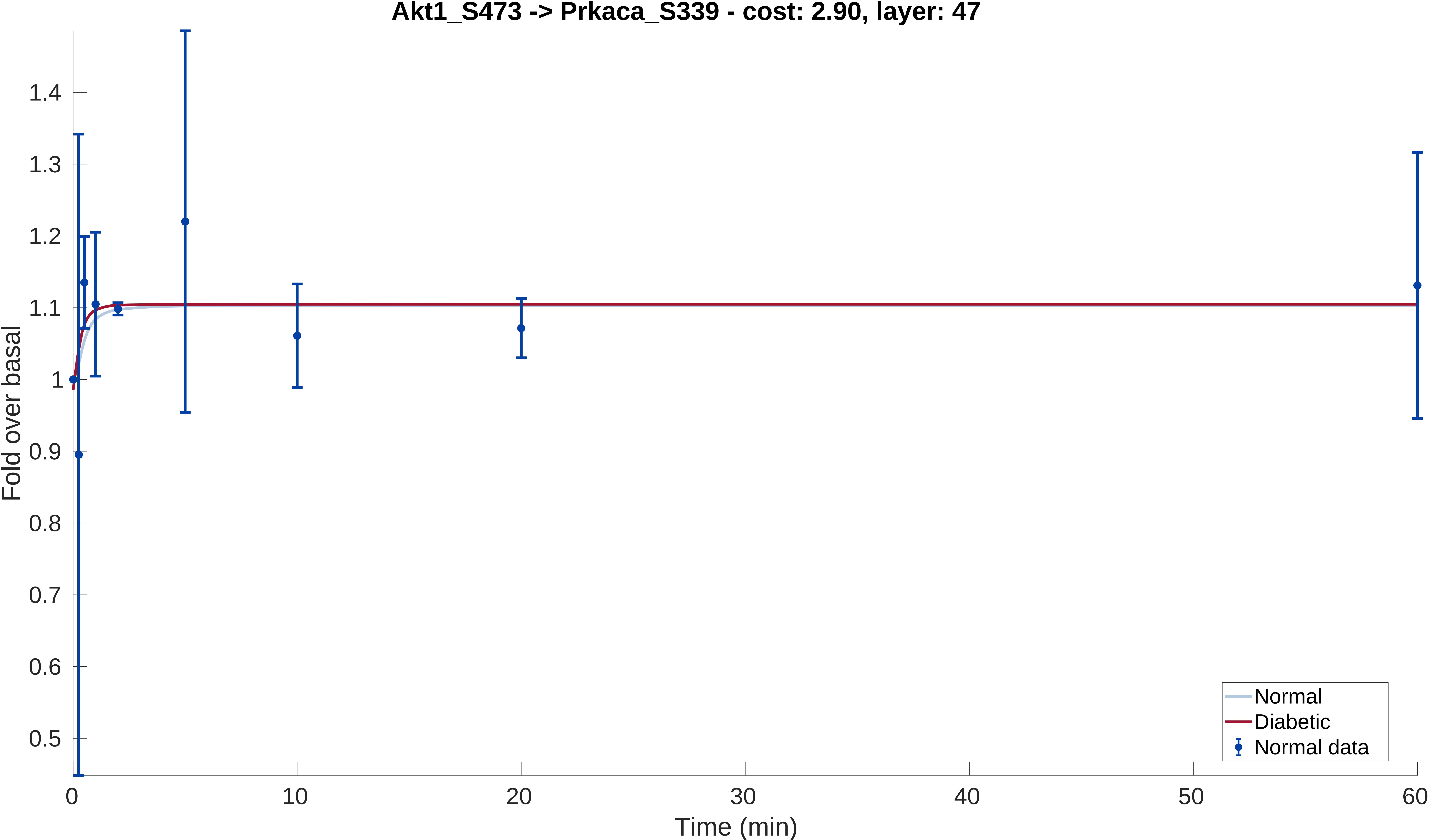

Akt1_S473 -> Prkaca_S339 - cost: 2.90, layer: 47
Fold over basal
1.4
1.3
1.2
1.1
1
0.9
0.8
0.7
0.6
0.5
0
10
20
30
40
50
60
Time (min)
Normal
Diabetic
Normal data

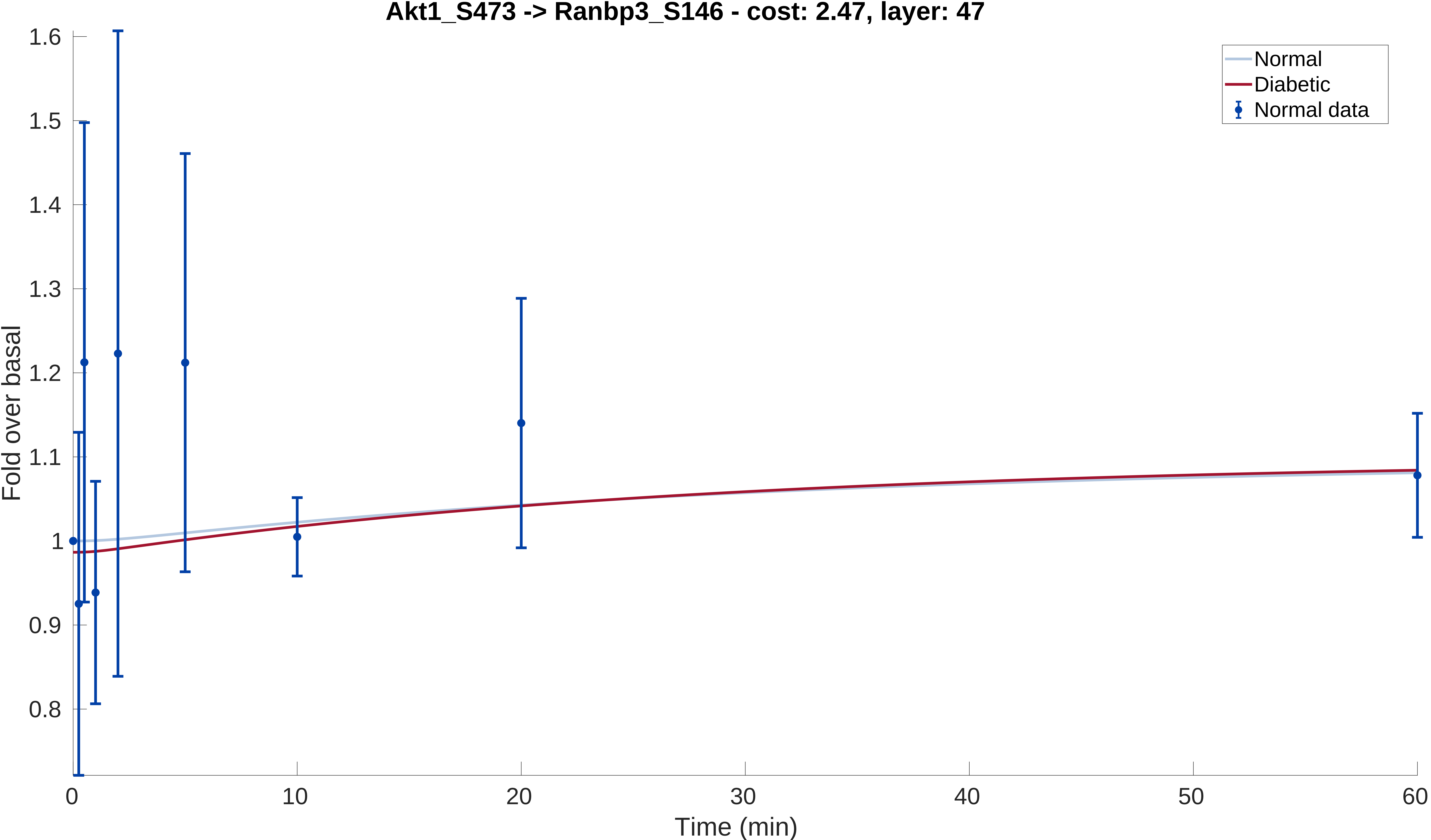

Akt1_S473 -> Ranbp3_S146 - cost: 2.47, layer: 47
Normal
Diabetic
Normal data
Fold over basal
1.6
1.5
1.4
1.3
1.2
1.1
1
0.9
0.8
0
10
20
30
40
50
60
Time (min)

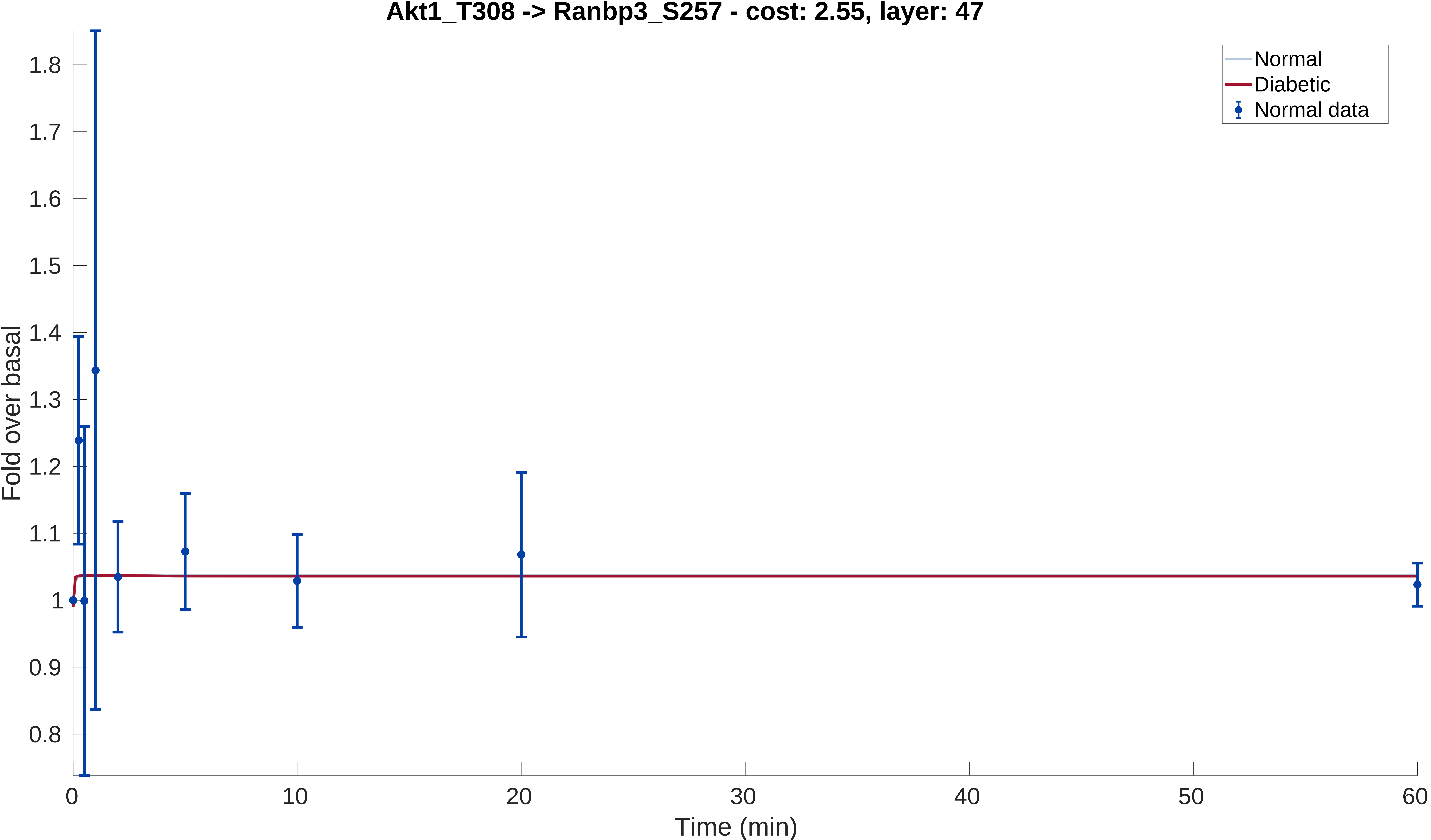

Akt1_T308 -> Ranbp3_S257 - cost: 2.55, layer: 47
Normal
Diabetic
Normal data
Fold over basal
1.8
1.7
1.6
1.5
1.4
1.3
1.2
1.1
1
0.9
0.8
0
10
20
30
40
50
60
Time (min)

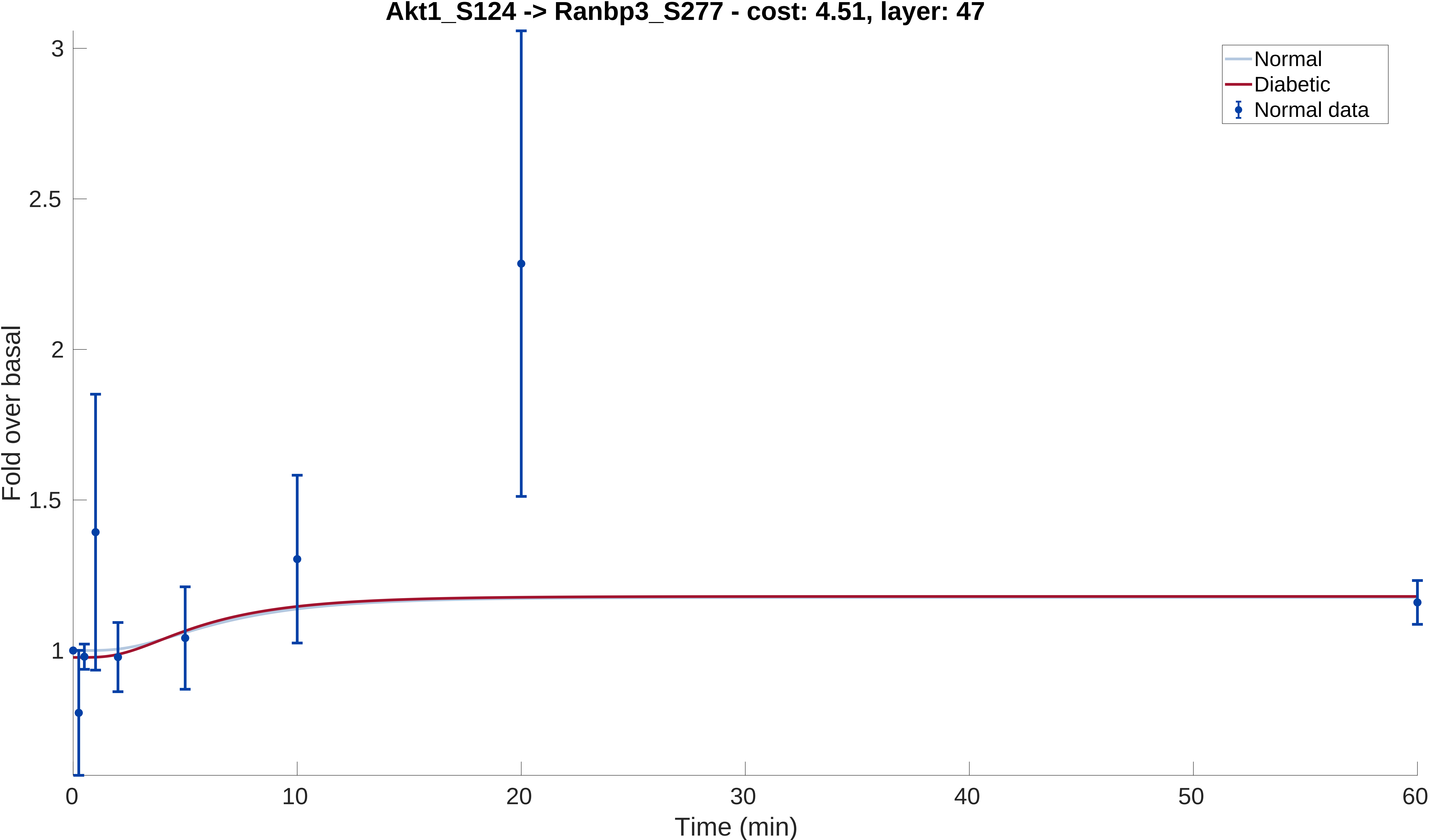

Akt1_S124 -> Ranbp3_S277 - cost: 4.51, layer: 47
Normal
Diabetic
Normal data
Fold over basal
3
2.5
2
1.5
1
0
10
20
30
40
50
60
Time (min)

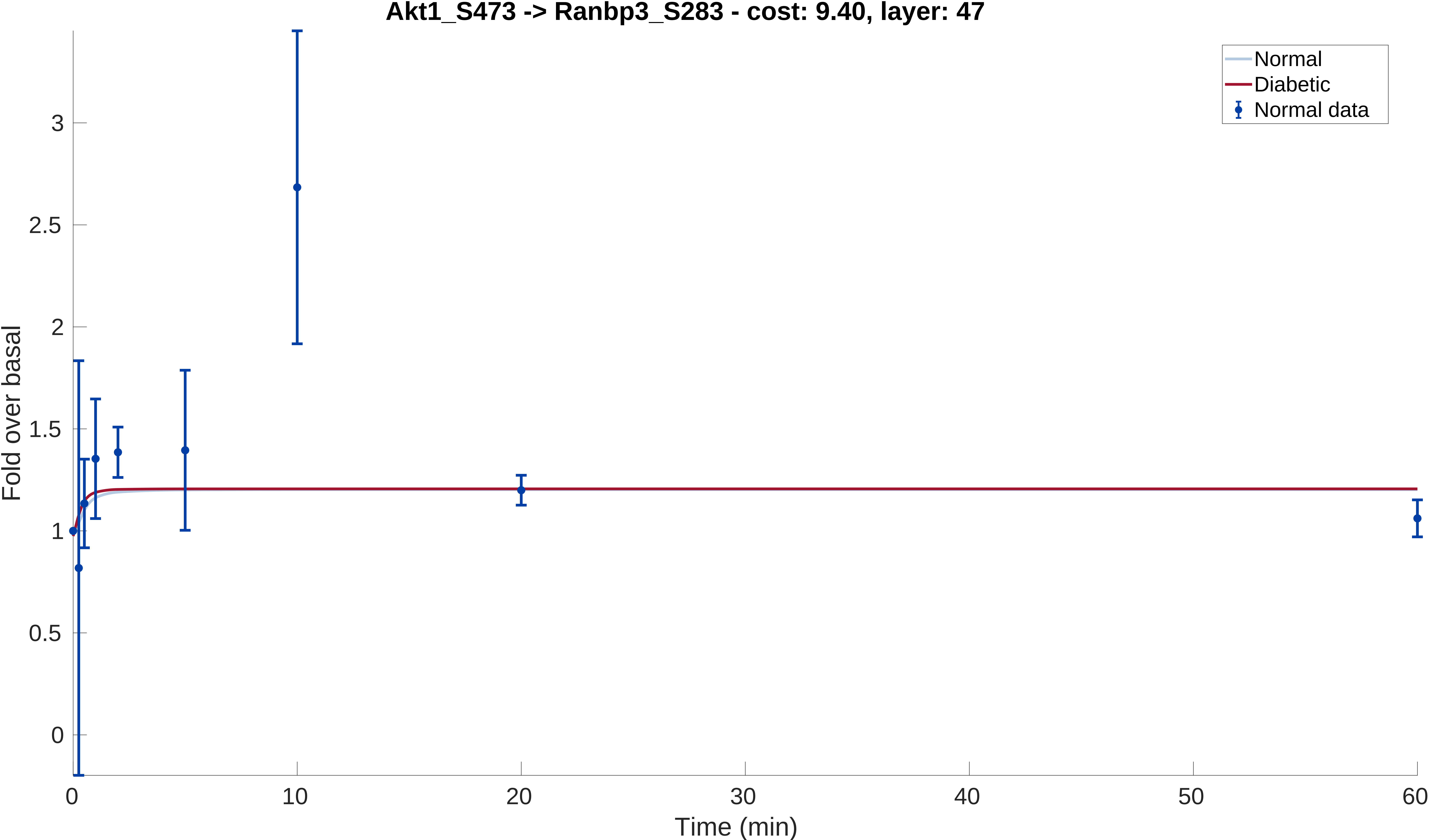
Akt1_S473 -> Ranbp3_S283 - cost: 9.40, layer: 47
Normal
Diabetic
Normal data
Fold over basal
0
0.5
1
1.5
2
2.5
3
0
10
20
30
40
50
60
Time (min)

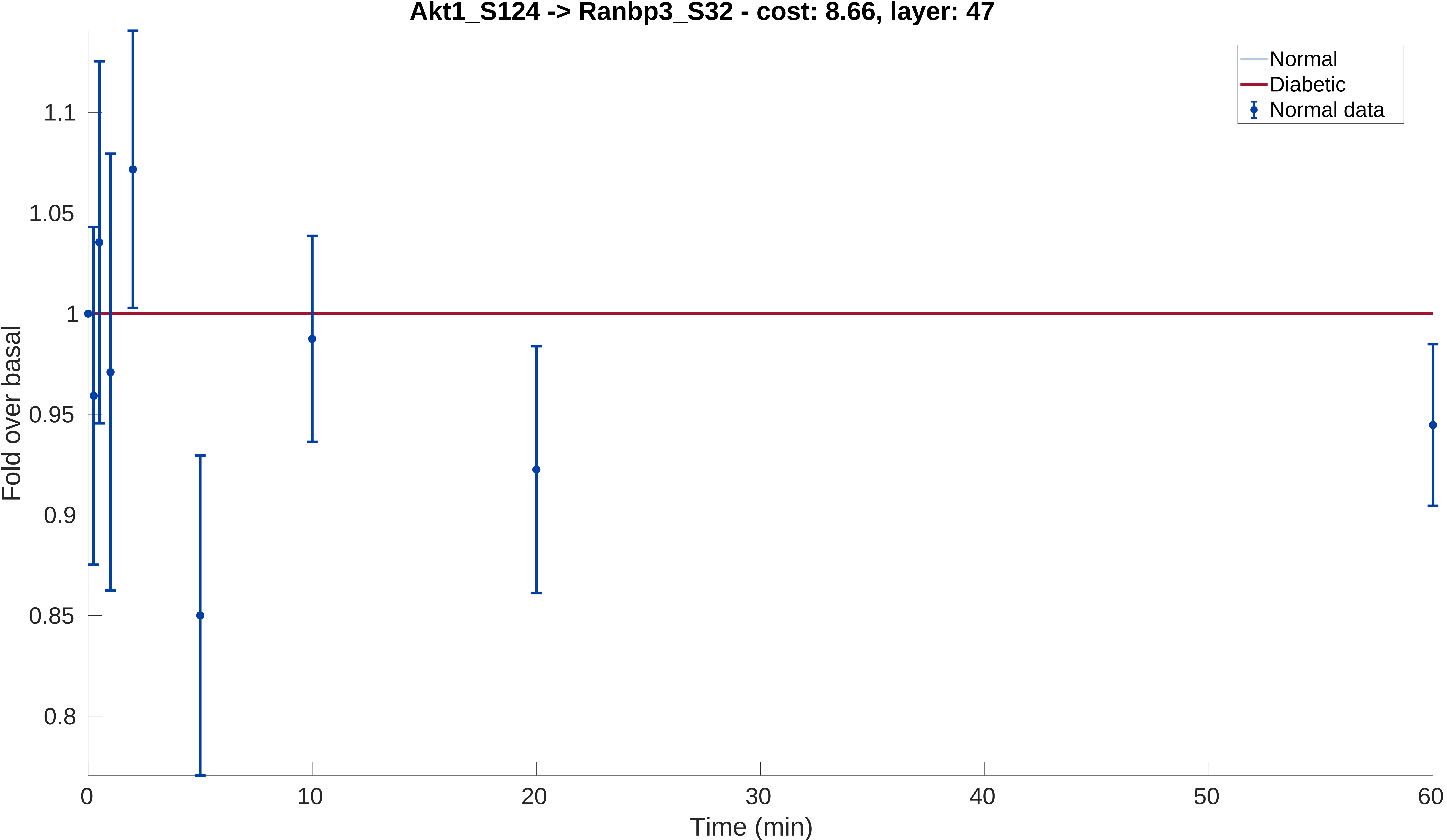
Akt1_S124 -> Ranbp3_S32 - cost: 8.66, layer: 47
Normal
Diabetic
Normal data
Fold over basal
1.1
1.05
1
0.95
0.9
0.85
0.8
0
10
20
30
40
50
60
Time (min)

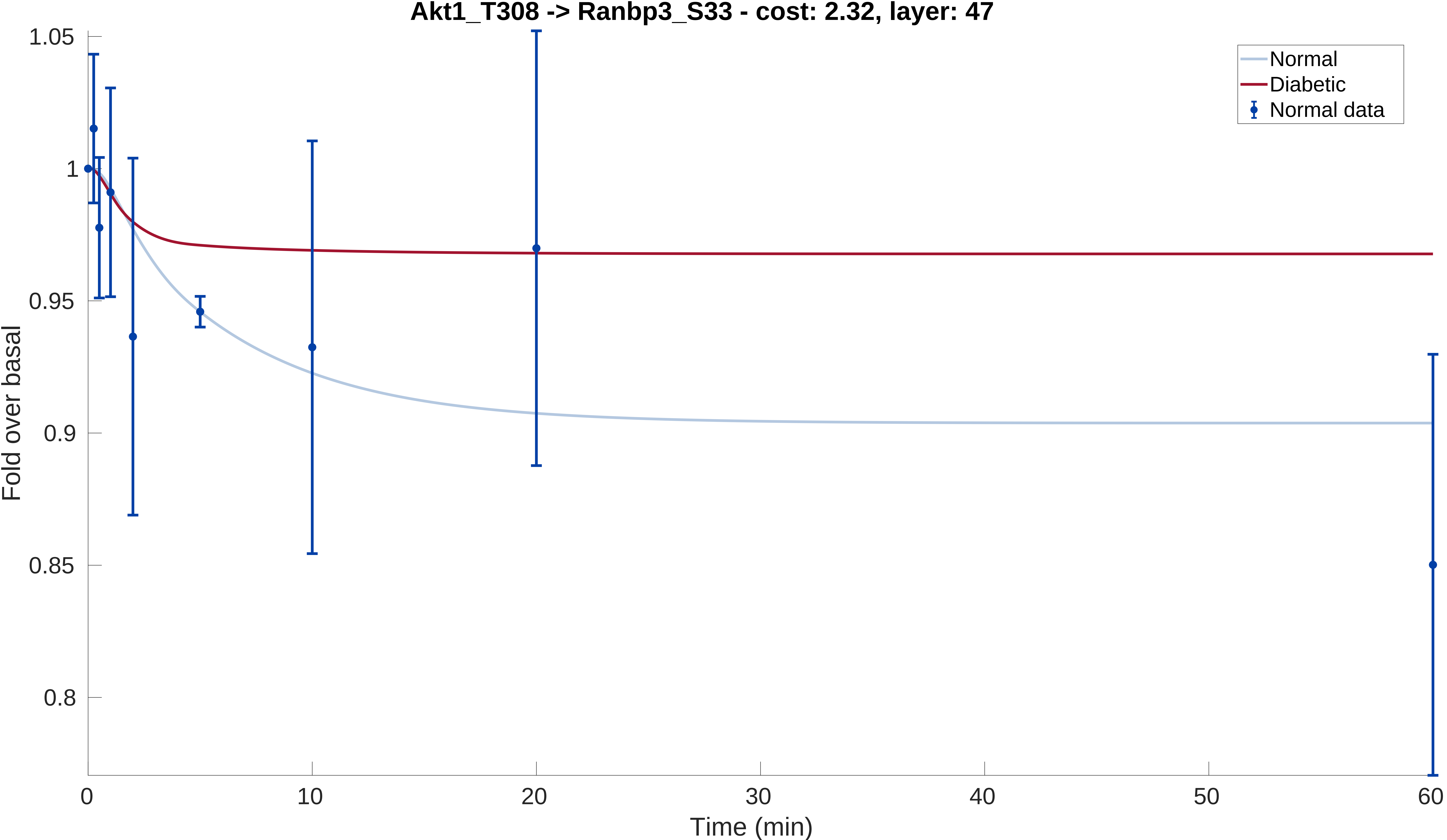

Akt1_T308 -> Ranbp3_S33 - cost: 2.32, layer: 47
Normal
Diabetic
Normal data
Fold over basal
1.05
1
0.95
0.9
0.85
0.8
0
10
20
30
40
50
60
Time (min)

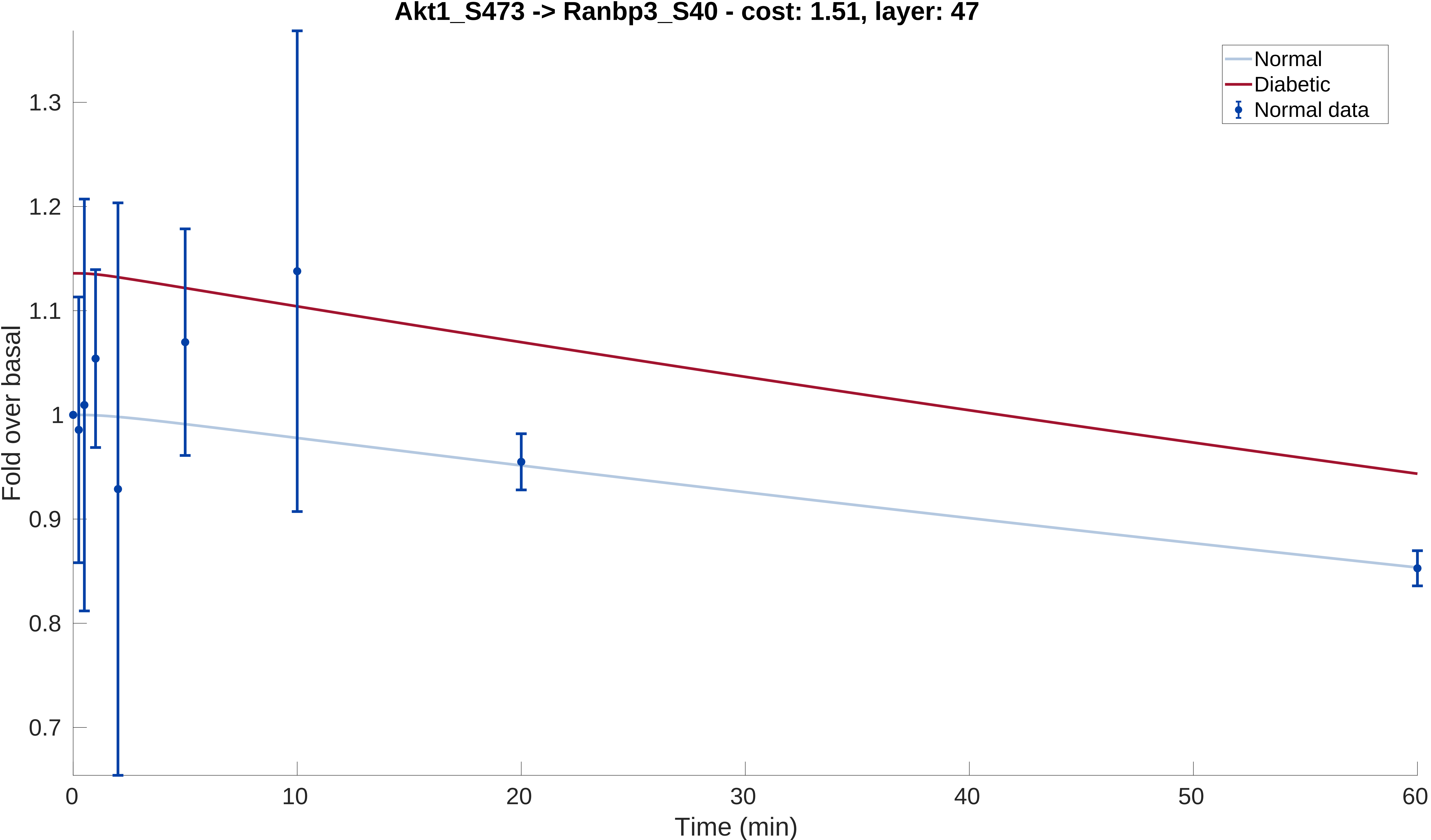

Akt1_S473 -> Ranbp3_S40 - cost: 1.51, layer: 47
Normal
Diabetic
Normal data
1.3
1.2
1.1
1
0.9
0.8
0.7
Fold over basal
0
10
20
30
40
50
60
Time (min)

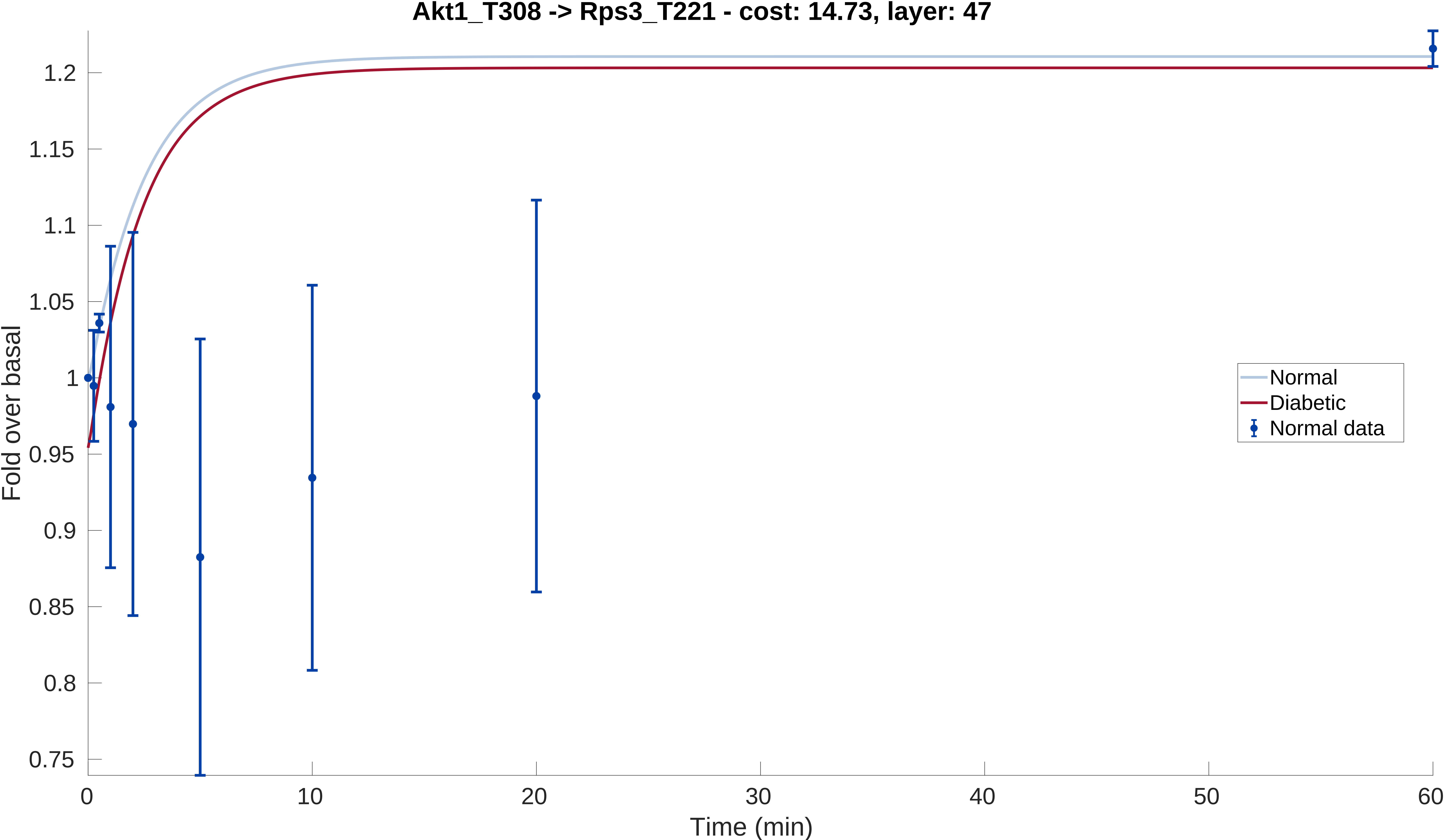

Akt1_T308 -> Rps3_T221 - cost: 14.73, layer: 47
Fold over basal
1.2
1.15
1.1
1.05
1
0.95
0.9
0.85
0.8
0.75
0
10
20
30
40
50
60
Time (min)
Normal
Diabetic
Normal data

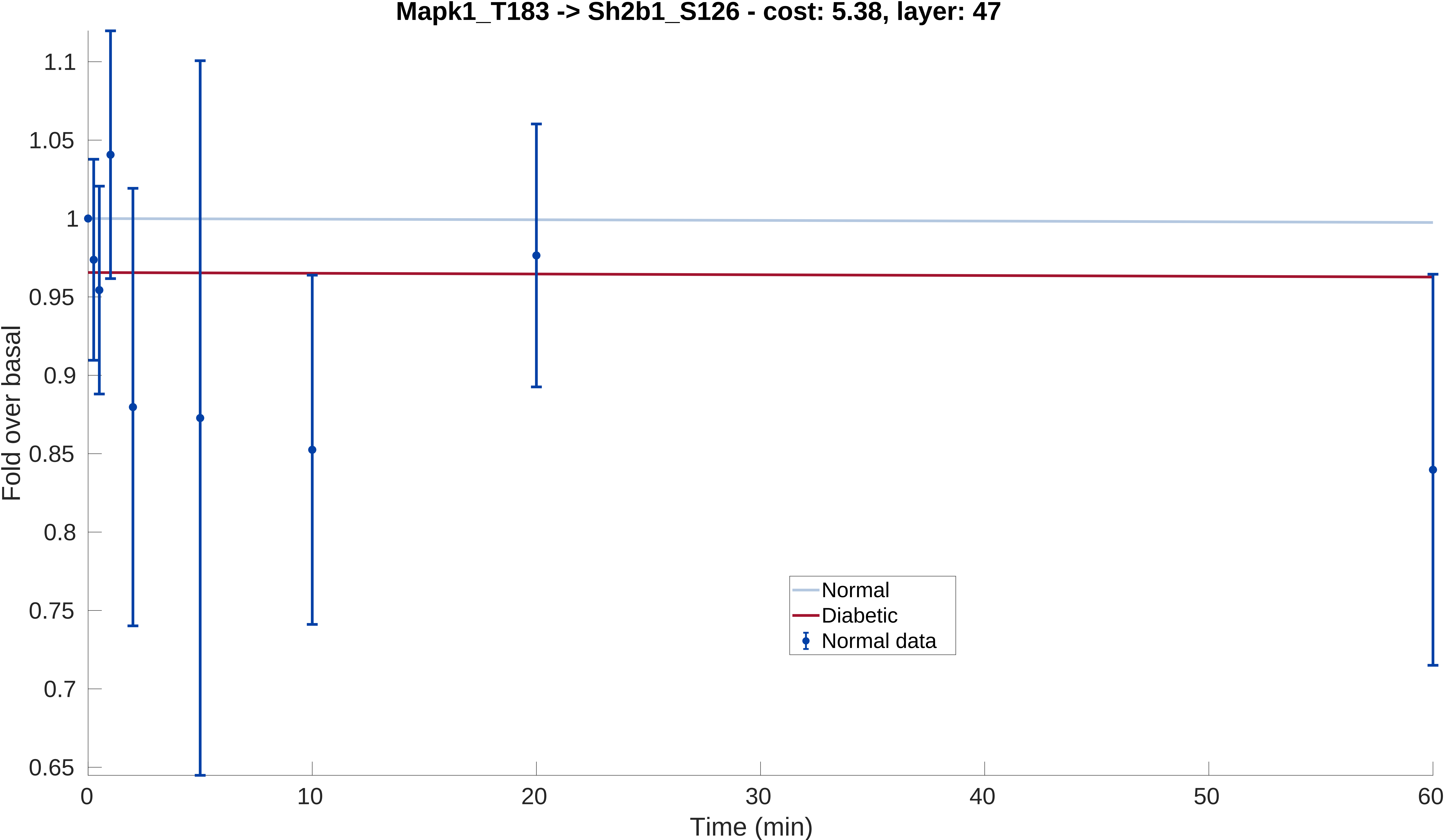

Mapk1_T183 -> Sh2b1_S126 - cost: 5.38, layer: 47
Fold over basal
1.1
1.05
1
0.95
0.9
0.85
0.8
0.75
0.7
0.65
0
10
20
30
40
50
60
Time (min)
Normal
Diabetic
Normal data

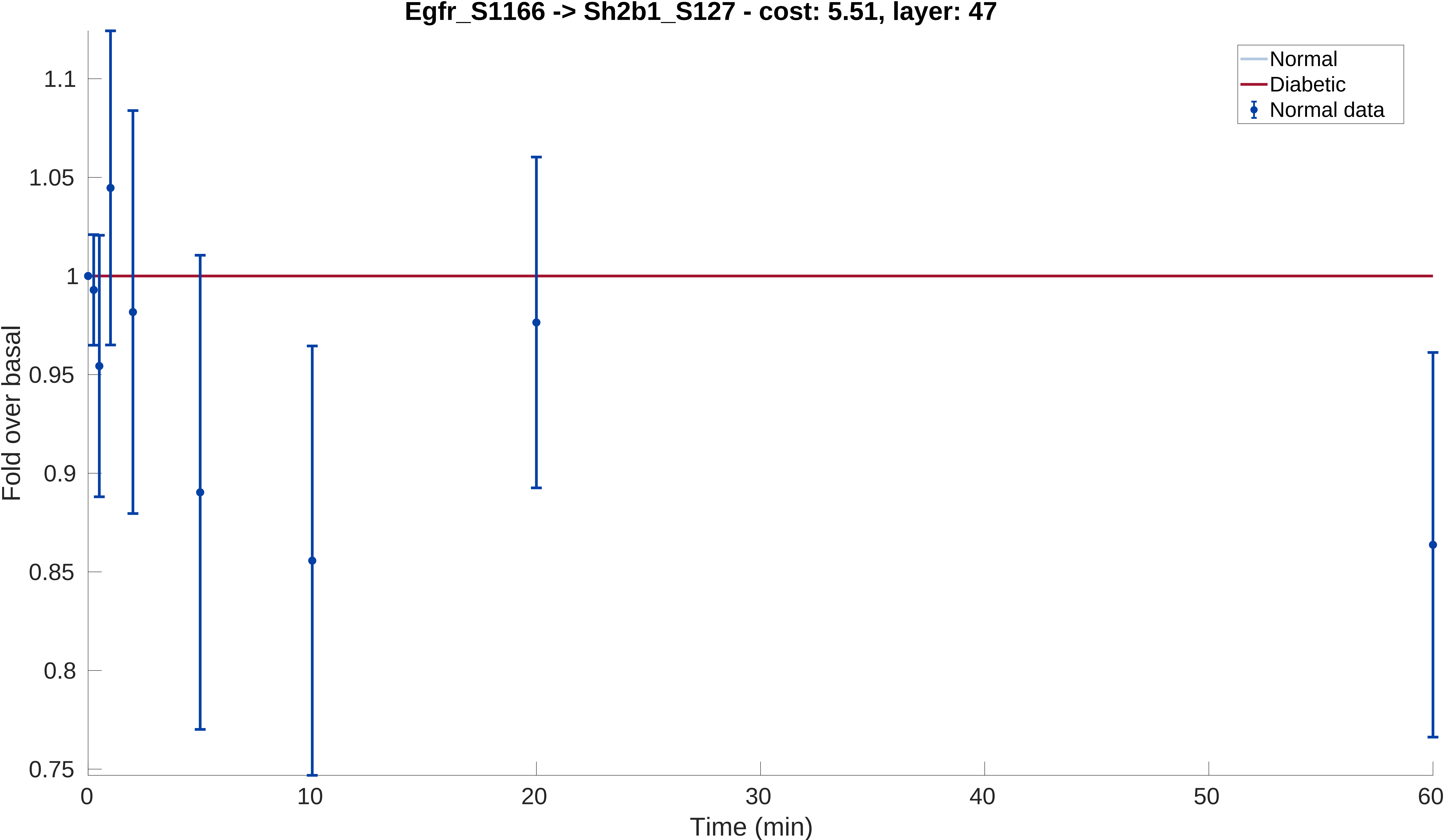

Egfr_S1166 -> Sh2b1_S127 - cost: 5.51, layer: 47
Normal
Diabetic
Normal data
Fold over basal
1.1
1.05
1
0.95
0.9
0.85
0.8
0.75
0
10
20
30
40
50
60
Time (min)

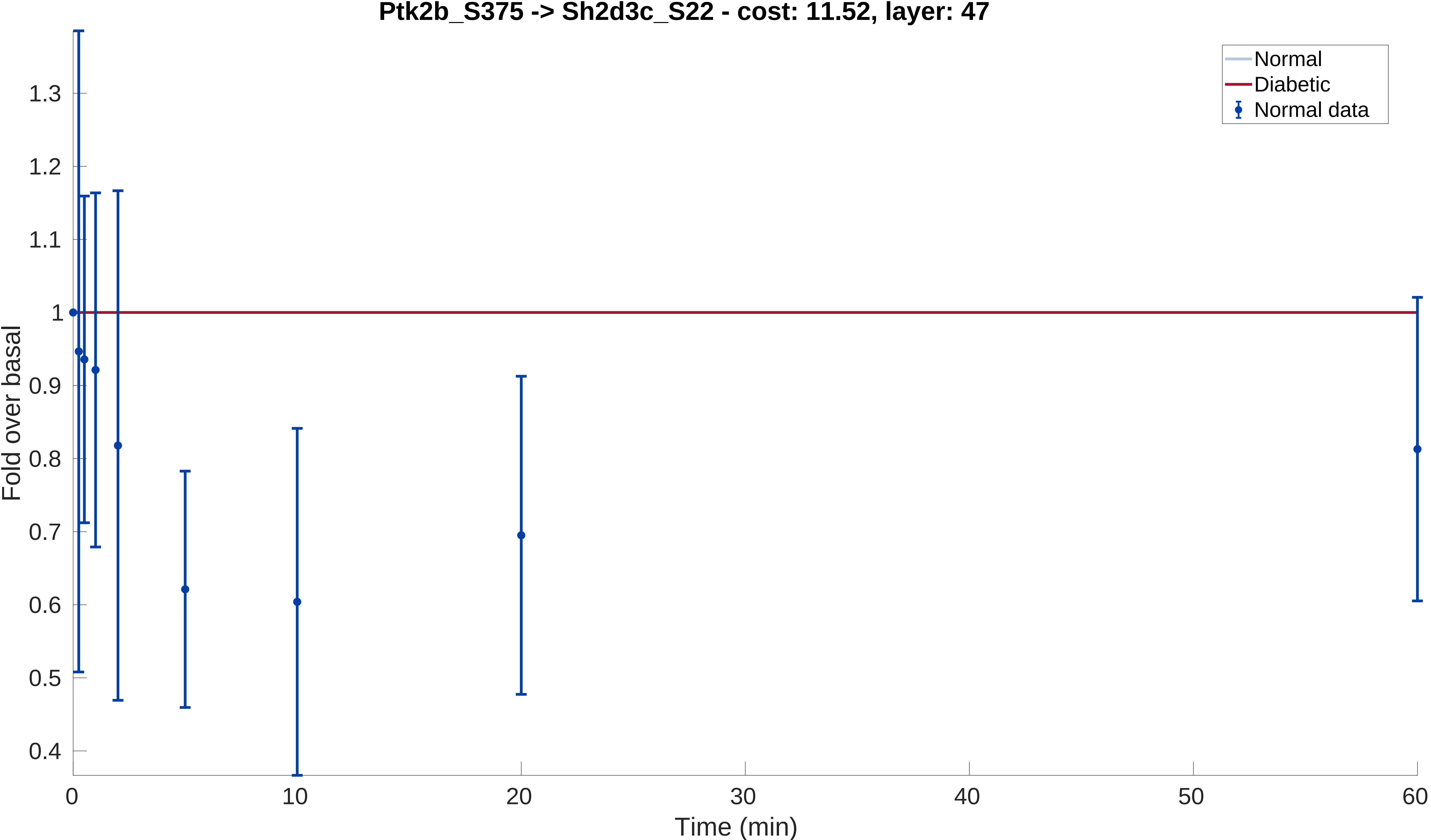

Ptk2b_S375 -> Sh2d3c_S22 - cost: 11.52, layer: 47
Normal
Diabetic
Normal data
Fold over basal
1.3
1.2
1.1
1
0.9
0.8
0.7
0.6
0.5
0.4
0
10
20
30
40
50
60
Time (min)

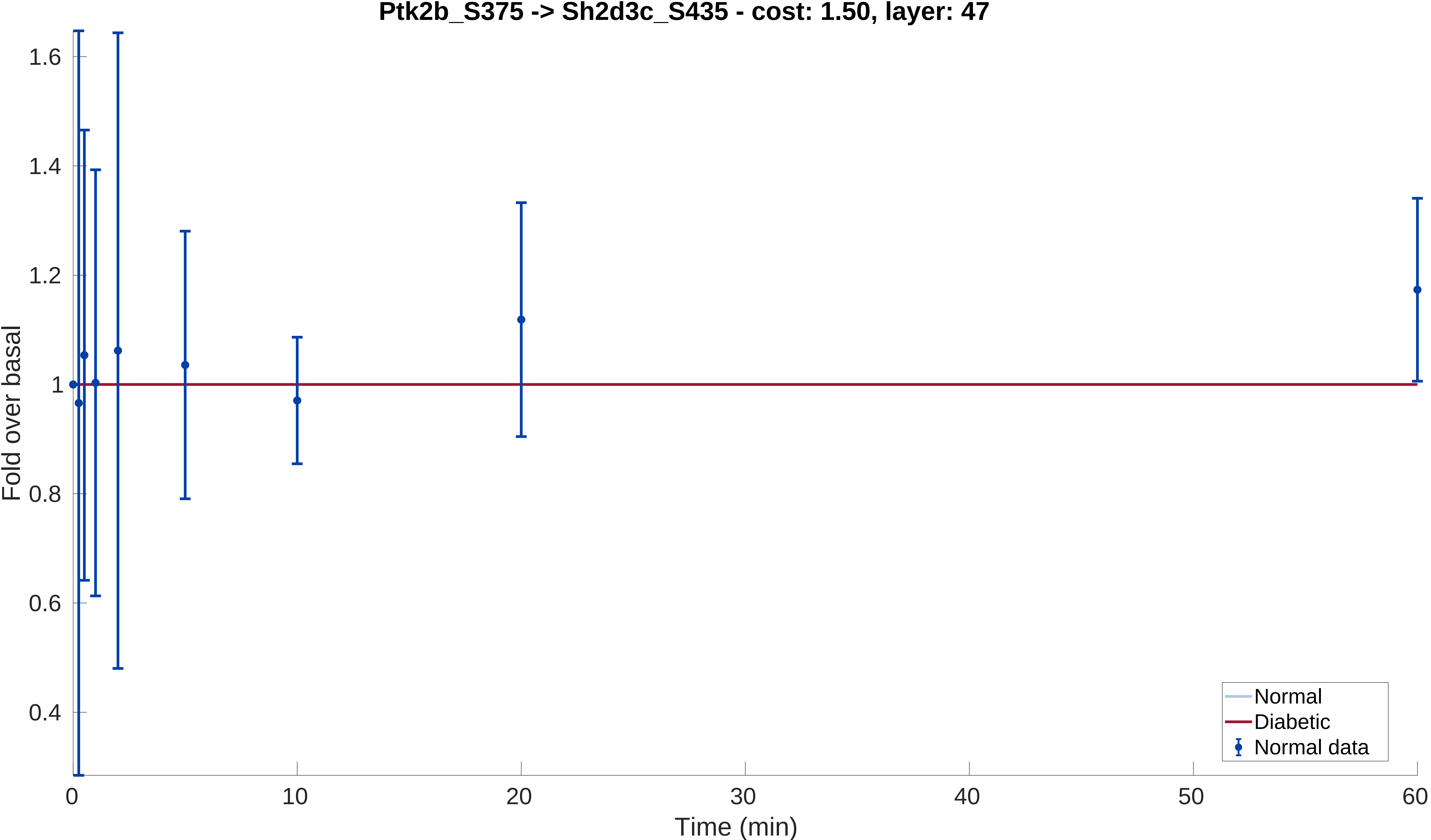

Ptk2b_S375 -> Sh2d3c_S435 - cost: 1.50, layer: 47
1.6
1.4
1.2
1
0.8
0.6
0.4
Fold over basal
0
10
20
30
40
50
60
Time (min)
Normal
Diabetic
Normal data

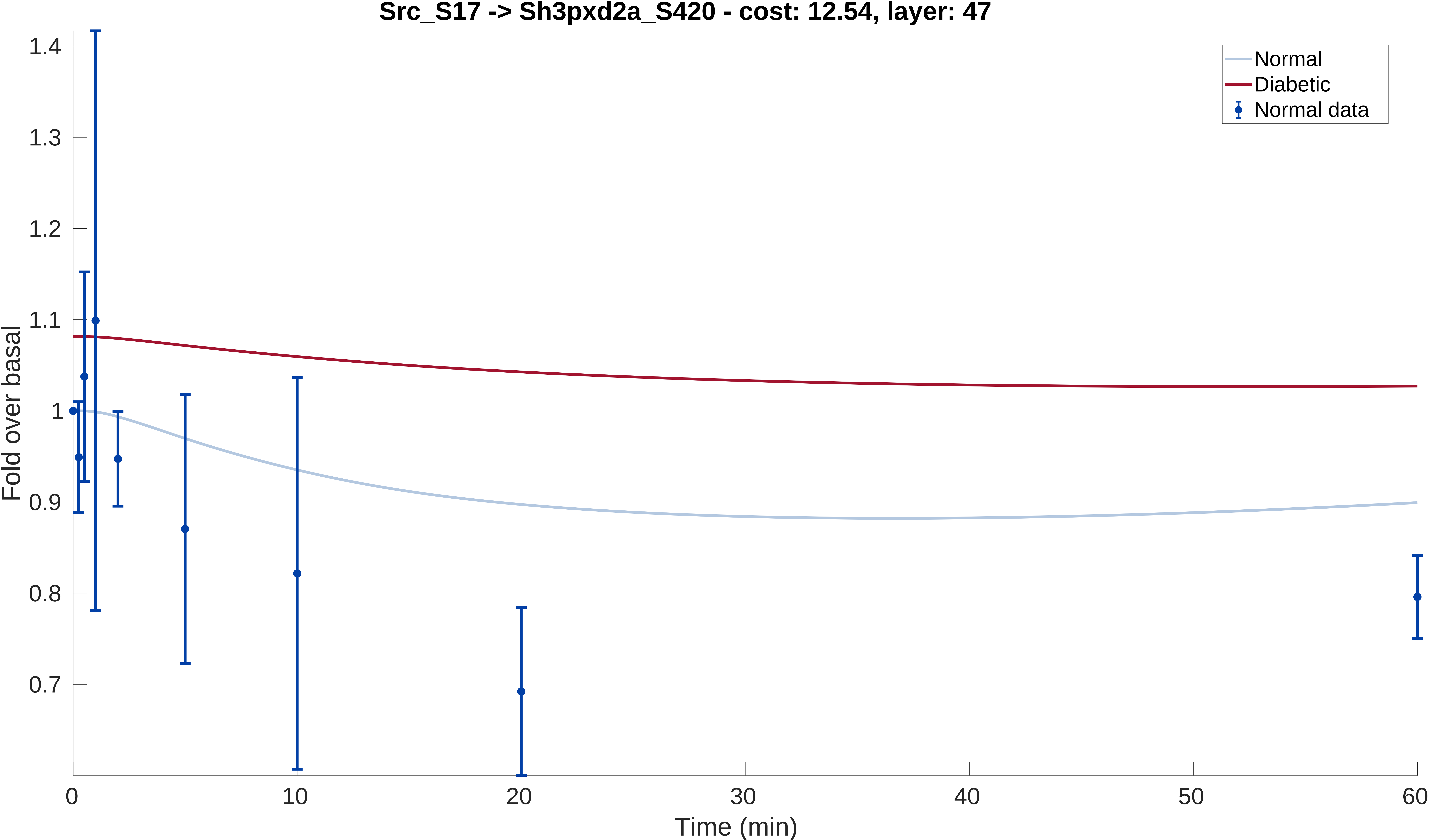

Src_S17 -> Sh3pxd2a_S420 - cost: 12.54, layer: 47
Normal
Diabetic
Normal data
Fold over basal
Time (min)
1.4
1.3
1.2
1.1
1
0.9
0.8
0.7
0
10
20
30
40
50
60

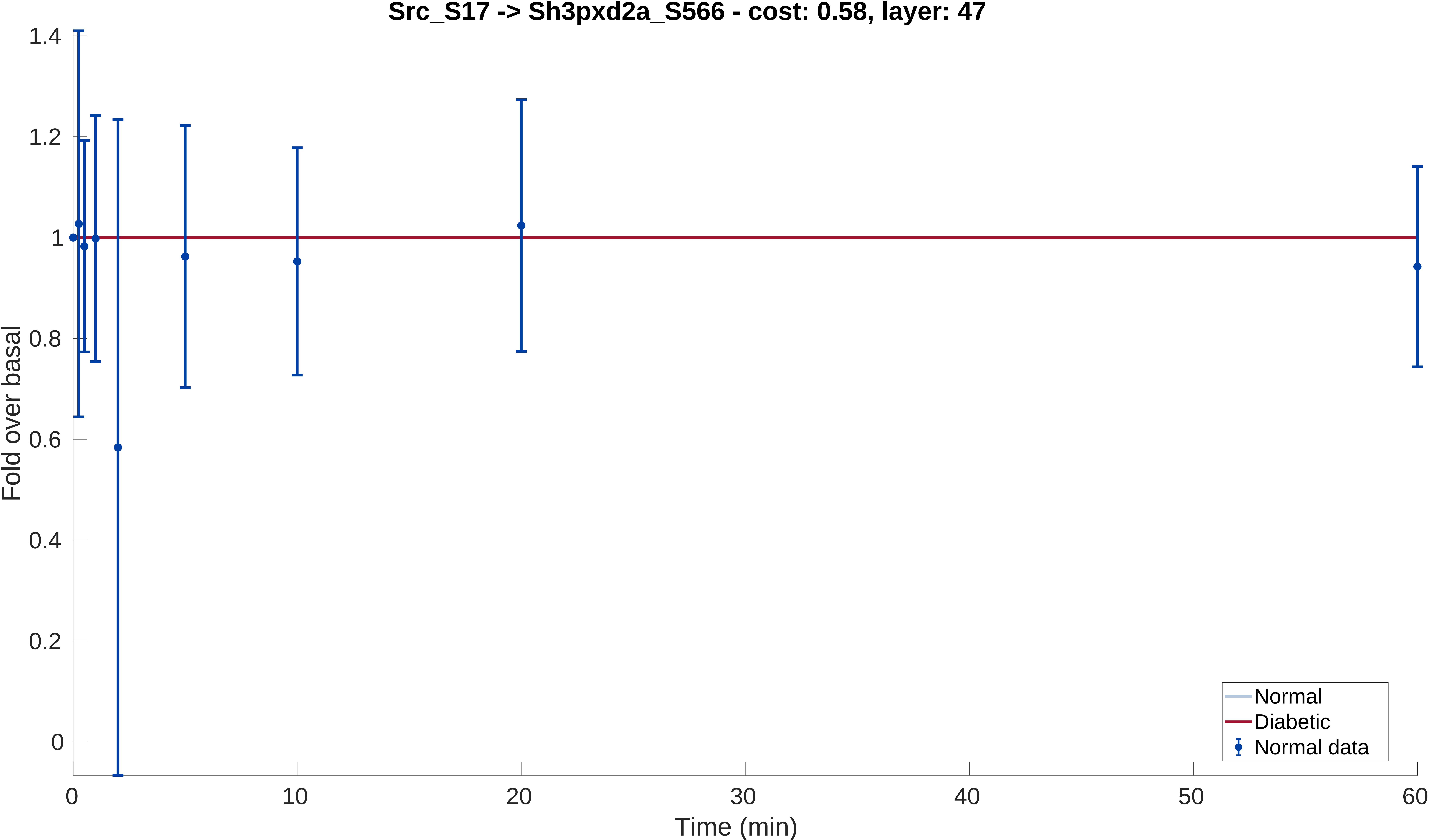

Src_S17 -> Sh3pxd2a_S566 - cost: 0.58, layer: 47
Fold over basal
Time (min)
0
0.2
0.4
0.6
0.8
1
1.2
1.4
0
10
20
30
40
50
60
Normal
Diabetic
Normal data

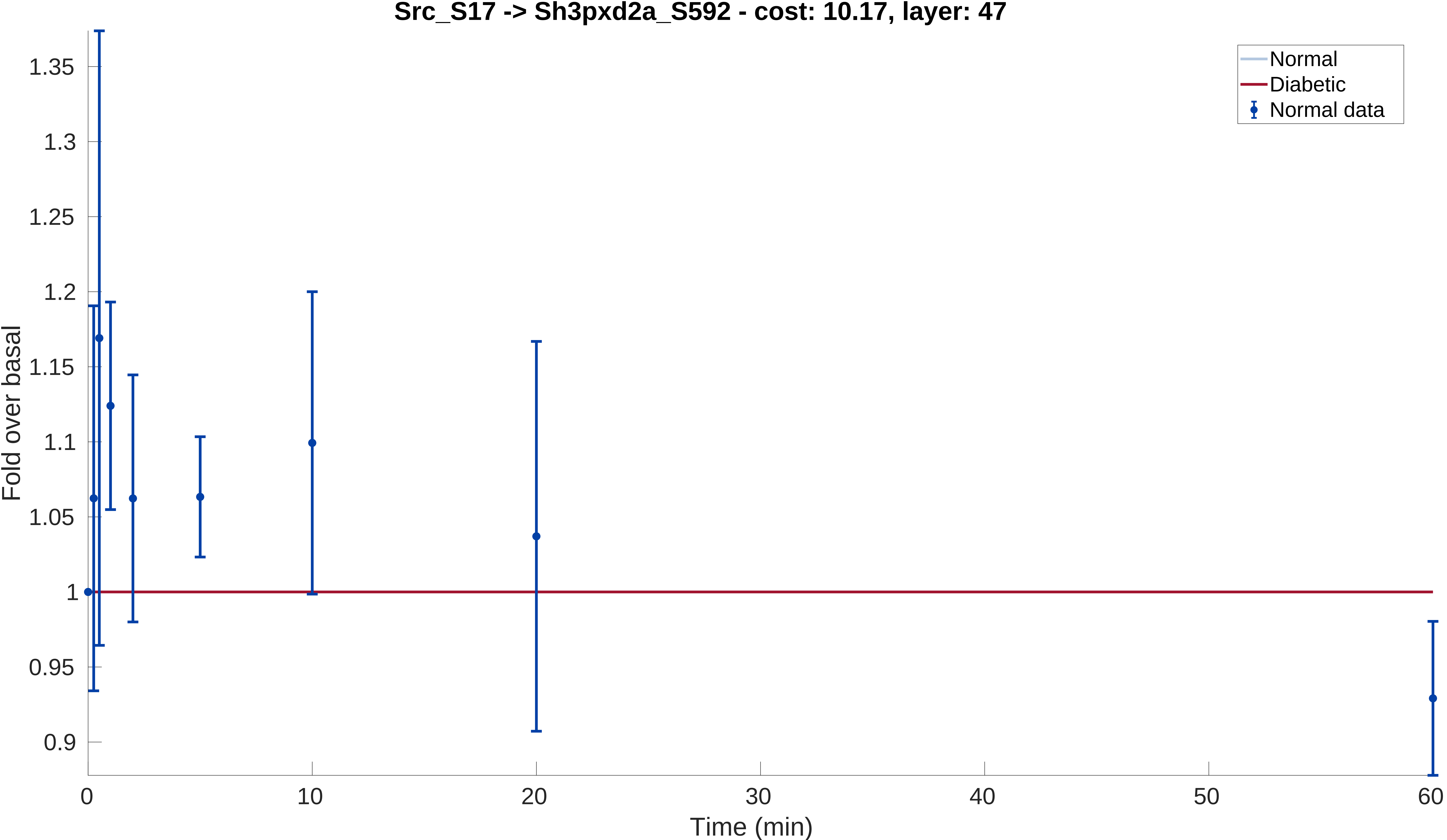

Src_S17 -> Sh3pxd2a_S592 - cost: 10.17, layer: 47
Normal
Diabetic
Normal data
Fold over basal
1.35
1.3
1.25
1.2
1.15
1.1
1.05
1
0.95
0.9
0
10
20
30
40
50
60
Time (min)

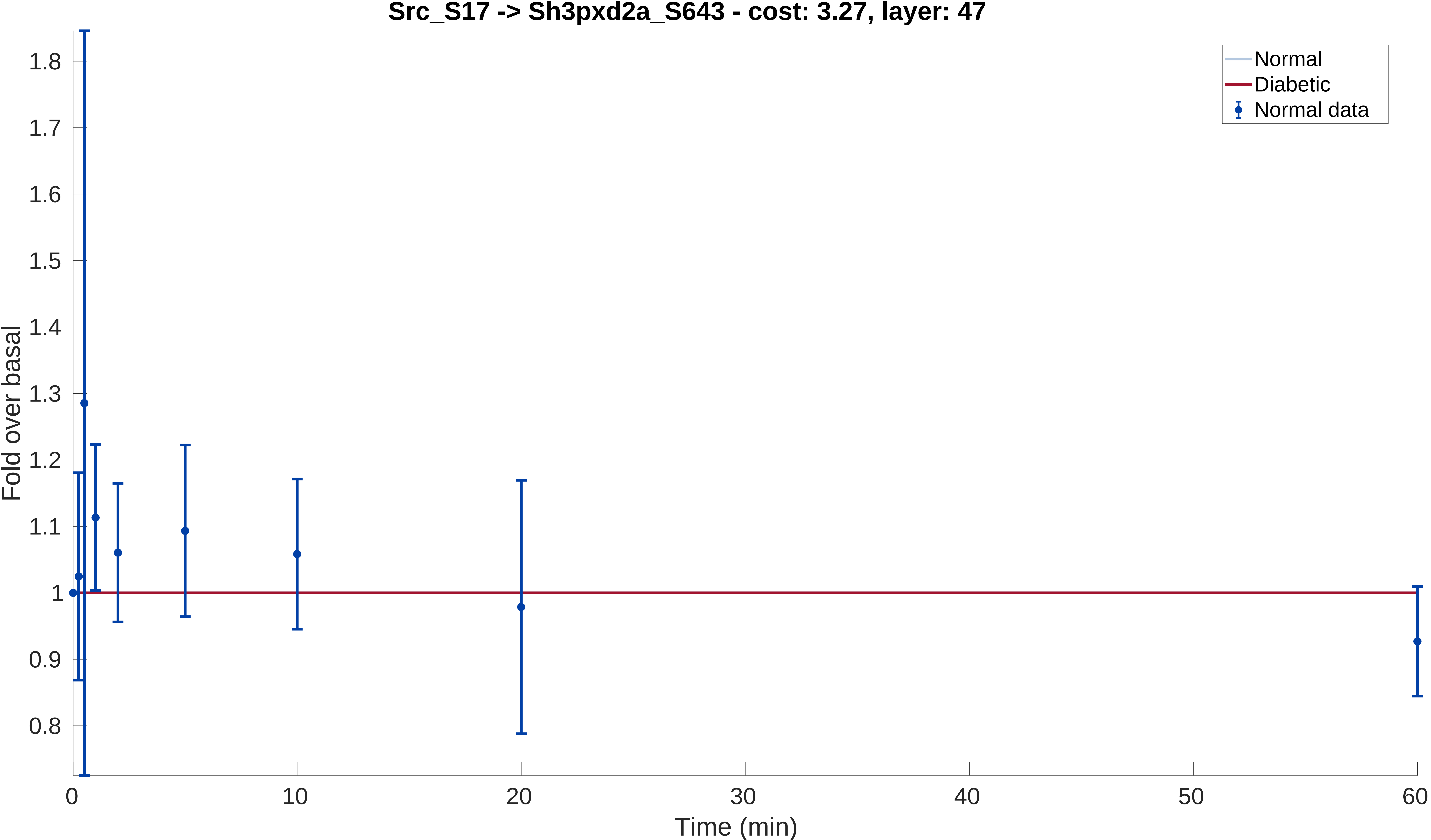

Src_S17 -> Sh3pxd2a_S643 - cost: 3.27, layer: 47
Normal
Diabetic
Normal data
Fold over basal
1.8
1.7
1.6
1.5
1.4
1.3
1.2
1.1
1
0.9
0.8
0
10
20
30
40
50
60
Time (min)

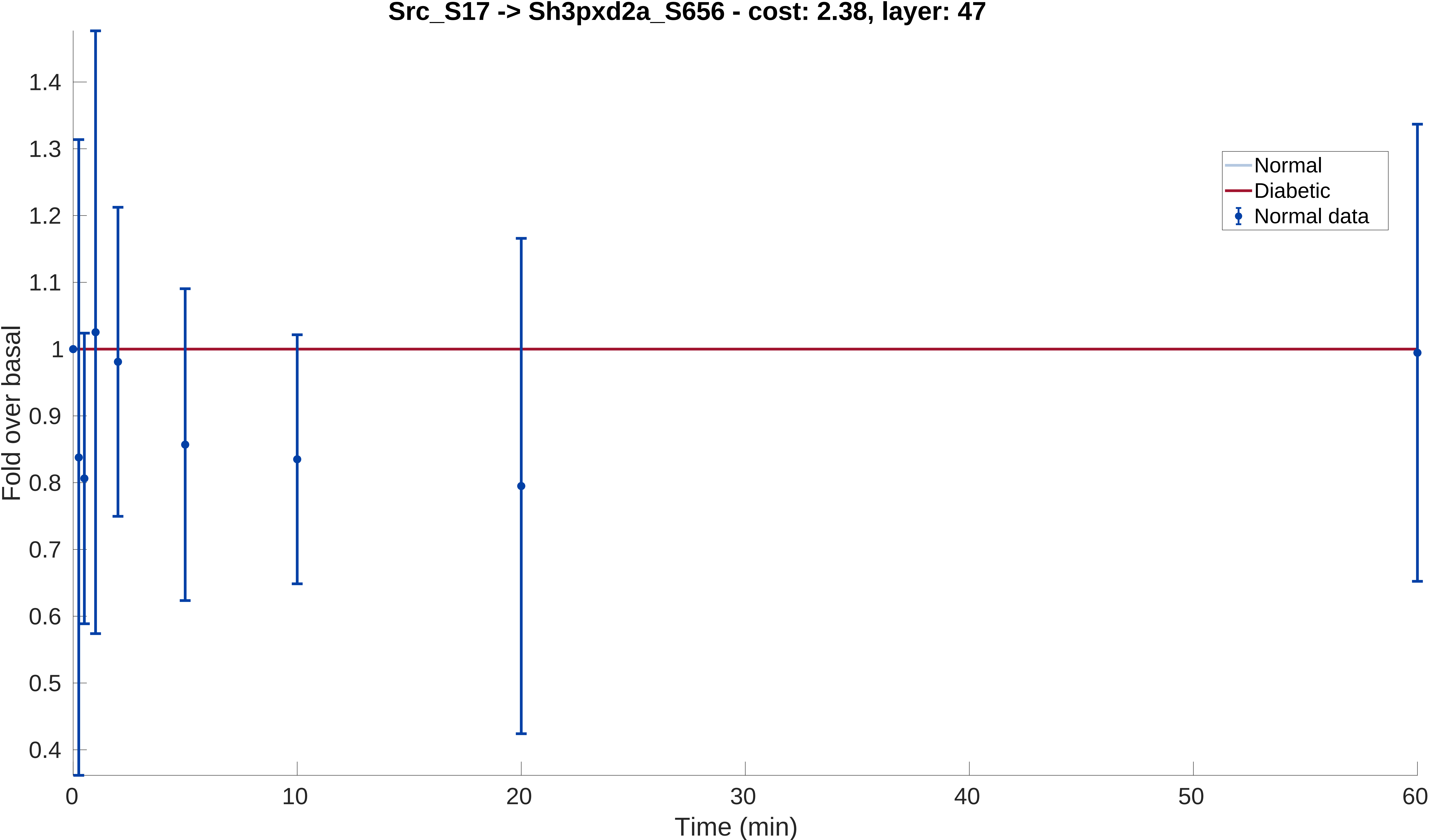

Src_S17 -> Sh3pxd2a_S656 - cost: 2.38, layer: 47
Fold over basal
1.4
1.3
1.2
1.1
1
0.9
0.8
0.7
0.6
0.5
0.4
0
10
20
30
40
50
60
Time (min)
Normal
Diabetic
Normal data

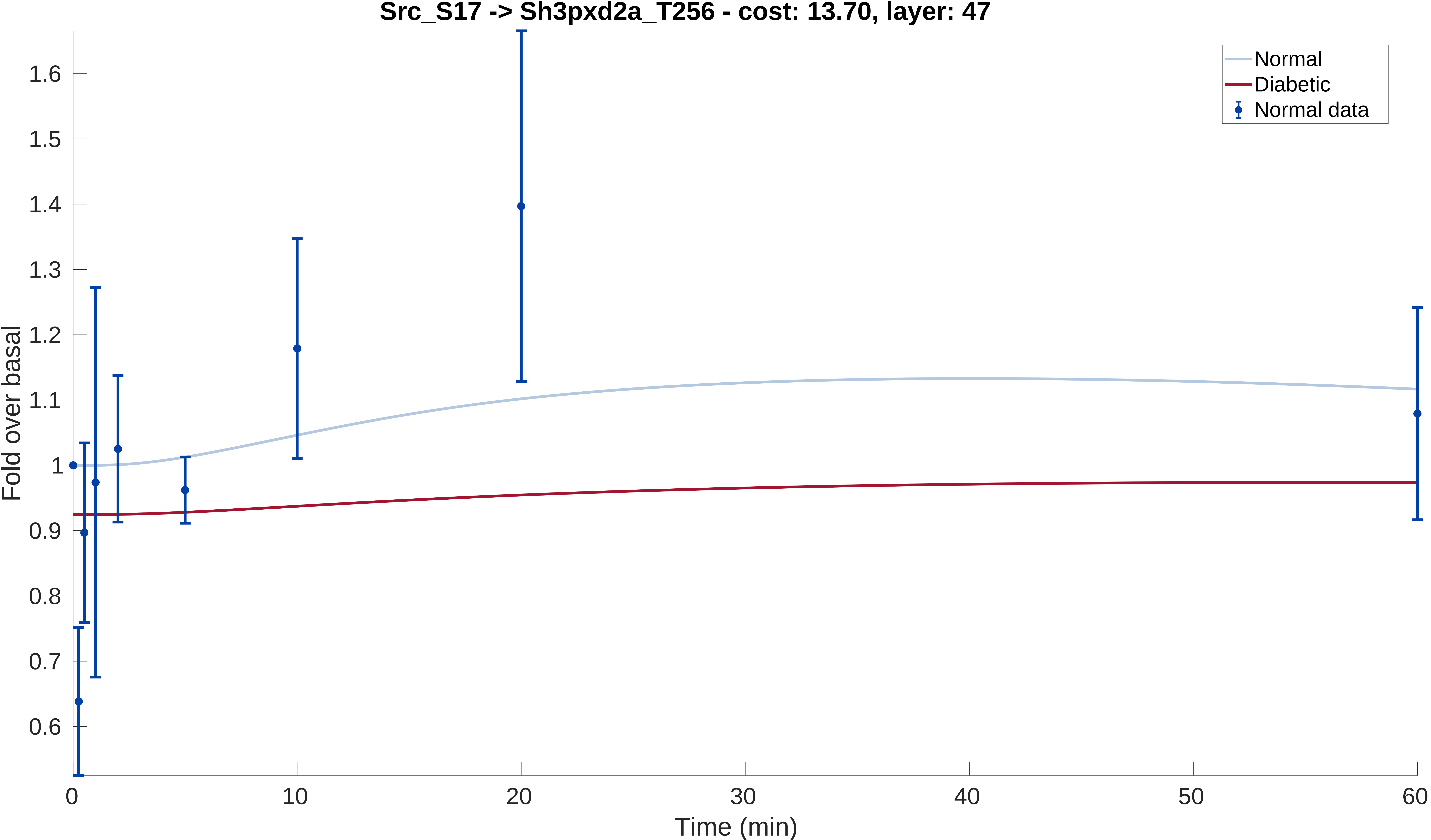

Src_S17 -> Sh3pxd2a_T256 - cost: 13.70, layer: 47
Normal
Diabetic
Normal data
Fold over basal
1.6
1.5
1.4
1.3
1.2
1.1
1
0.9
0.8
0.7
0.6
0
10
20
30
40
50
60
Time (min)

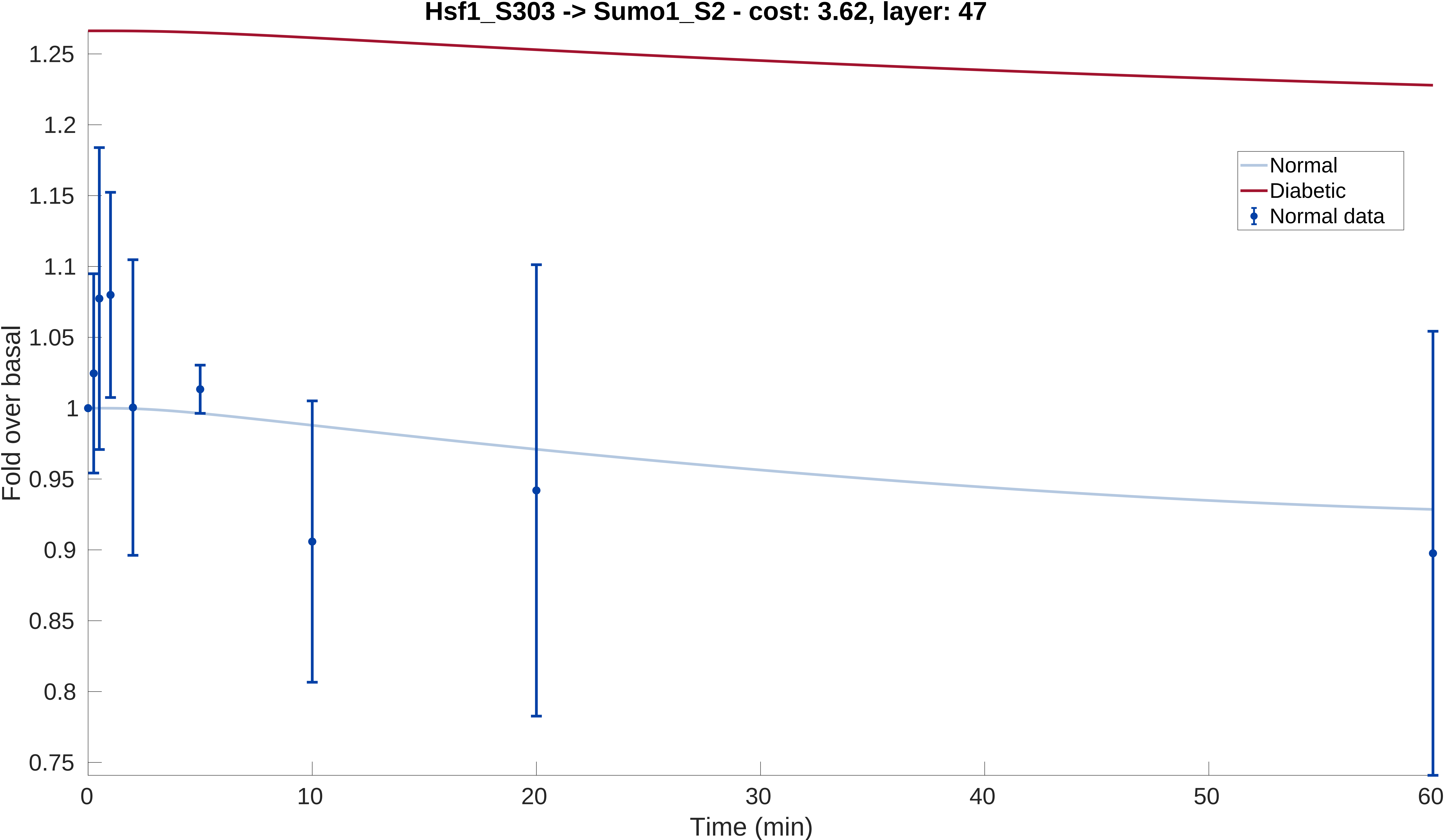

Hsf1_S303 -> Sumo1_S2 - cost: 3.62, layer: 47
Normal
Diabetic
Normal data
Fold over basal
Time (min)
1.25
1.2
1.15
1.1
1.05
1
0.95
0.9
0.85
0.8
0.75
0
10
20
30
40
50
60

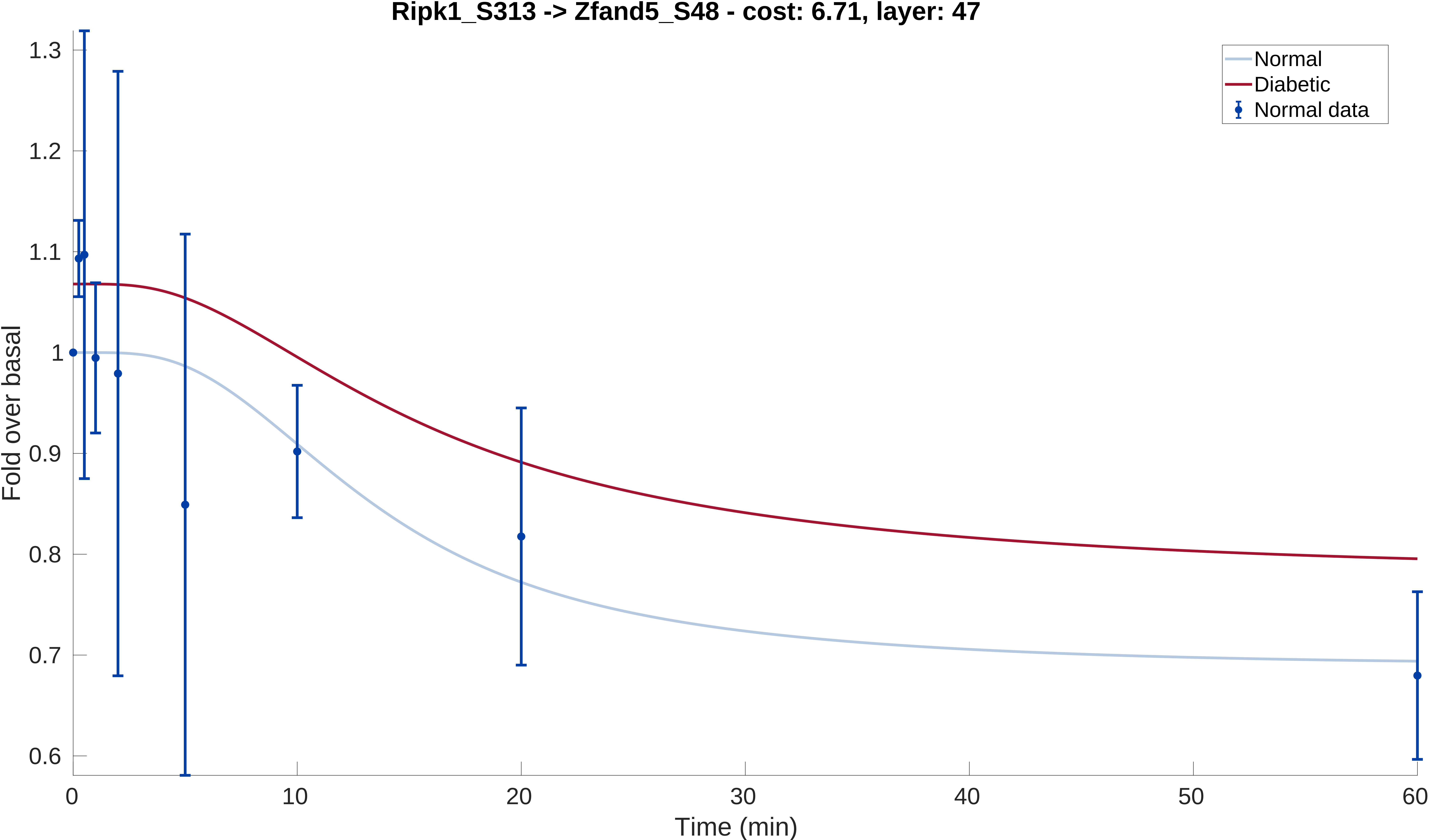

Ripk1_S313 -> Zfand5_S48 - cost: 6.71, layer: 47
Normal
Diabetic
Normal data
Fold over basal
Time (min)
1.3
1.2
1.1
1
0.9
0.8
0.7
0.6
0
10
20
30
40
50
60

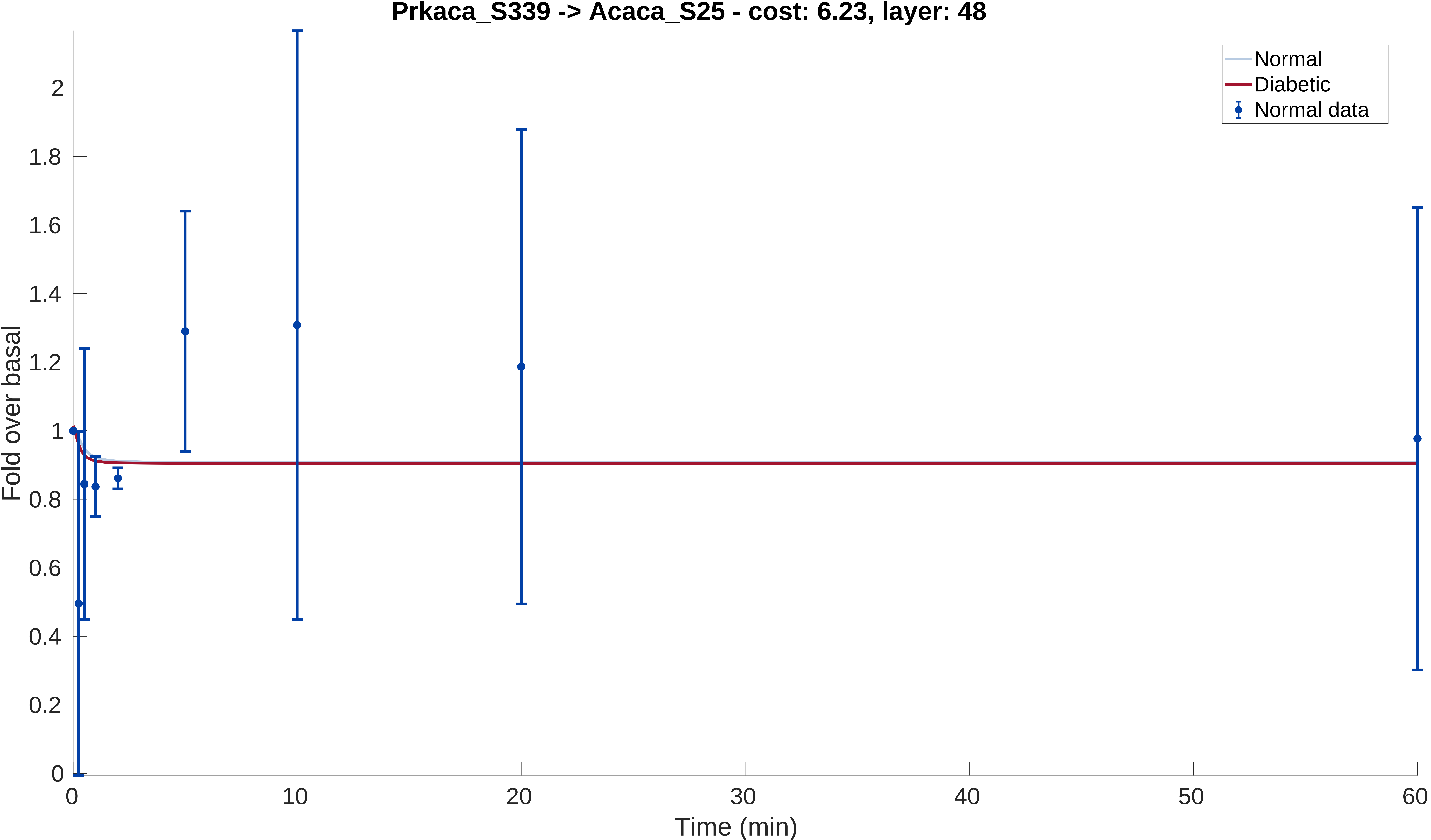

Prkaca_S339 -> Acaca_S25 - cost: 6.23, layer: 48
Normal
Diabetic
Normal data
Fold over basal
Time (min)
0
0.2
0.4
0.6
0.8
1
1.2
1.4
1.6
1.8
2
0
10
20
30
40
50
60

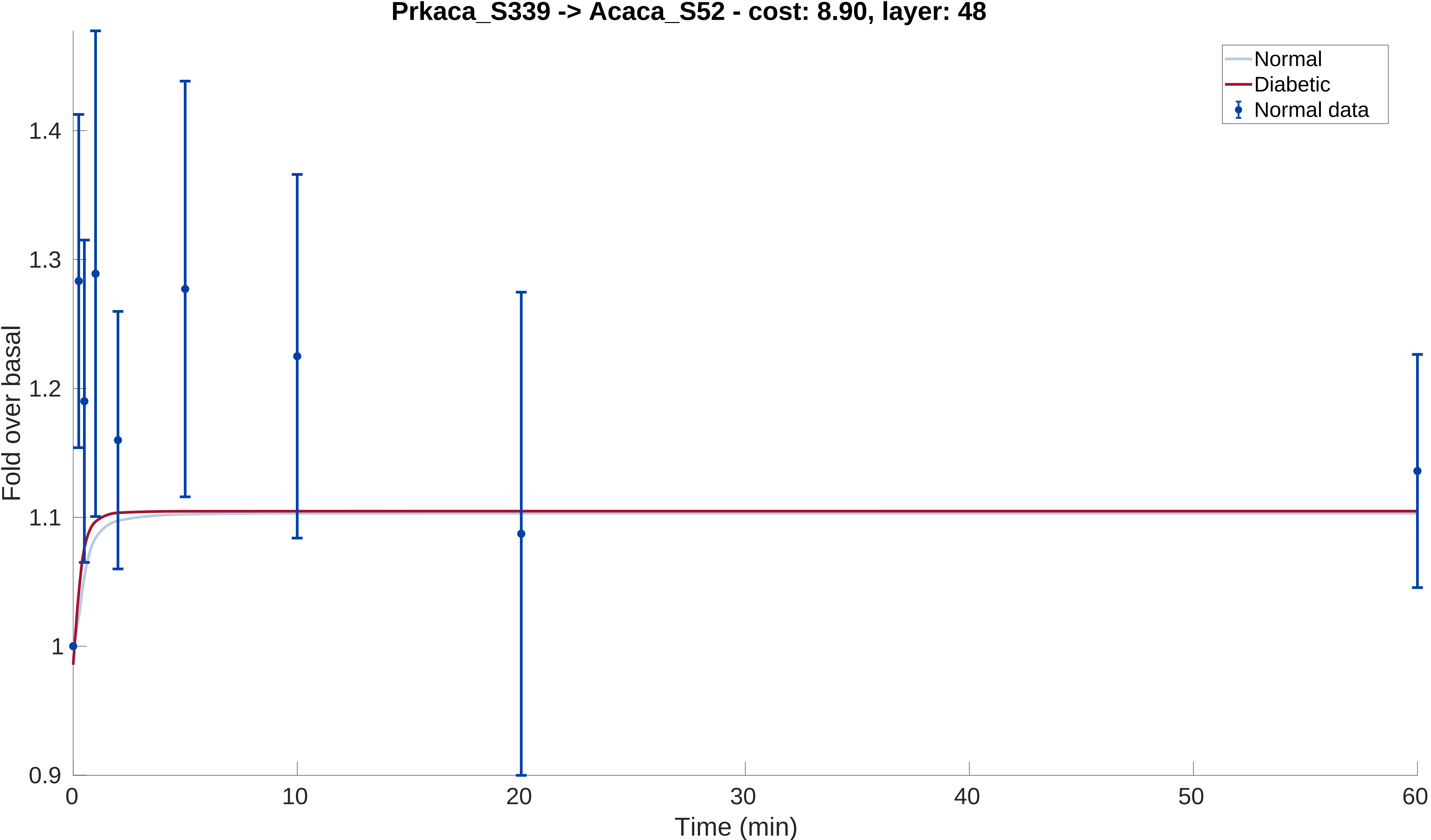
Prkaca_S339 -> Acaca_S52 - cost: 8.90, layer: 48
Normal
Diabetic
Normal data
Fold over basal
1.4
1.3
1.2
1.1
1
0.9
0
10
20
30
40
50
60
Time (min)

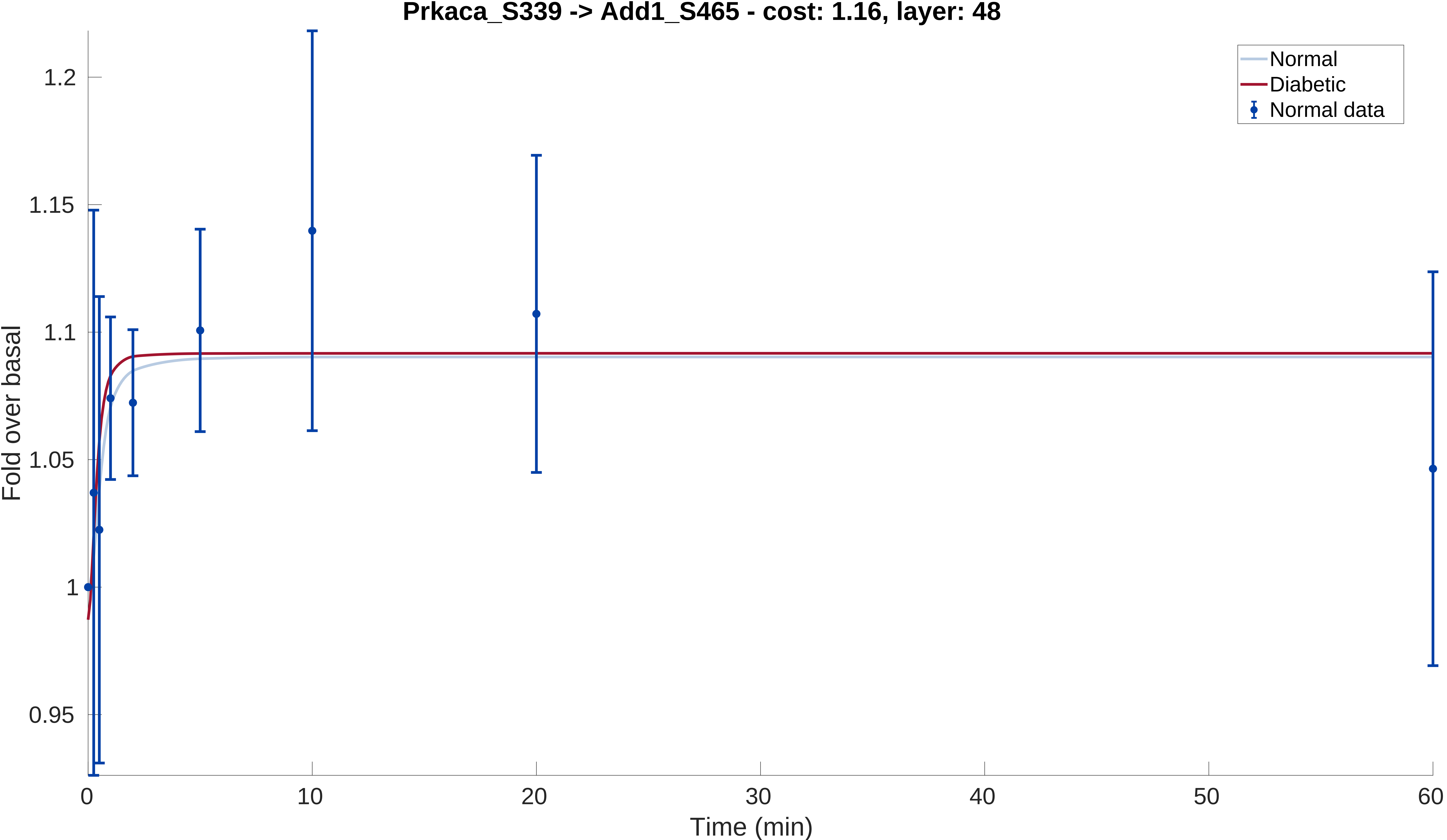

Prkaca_S339 -> Add1_S465 - cost: 1.16, layer: 48
Normal
Diabetic
Normal data
Fold over basal
Time (min)
1.2
1.15
1.1
1.05
1
0.95
0
10
20
30
40
50
60

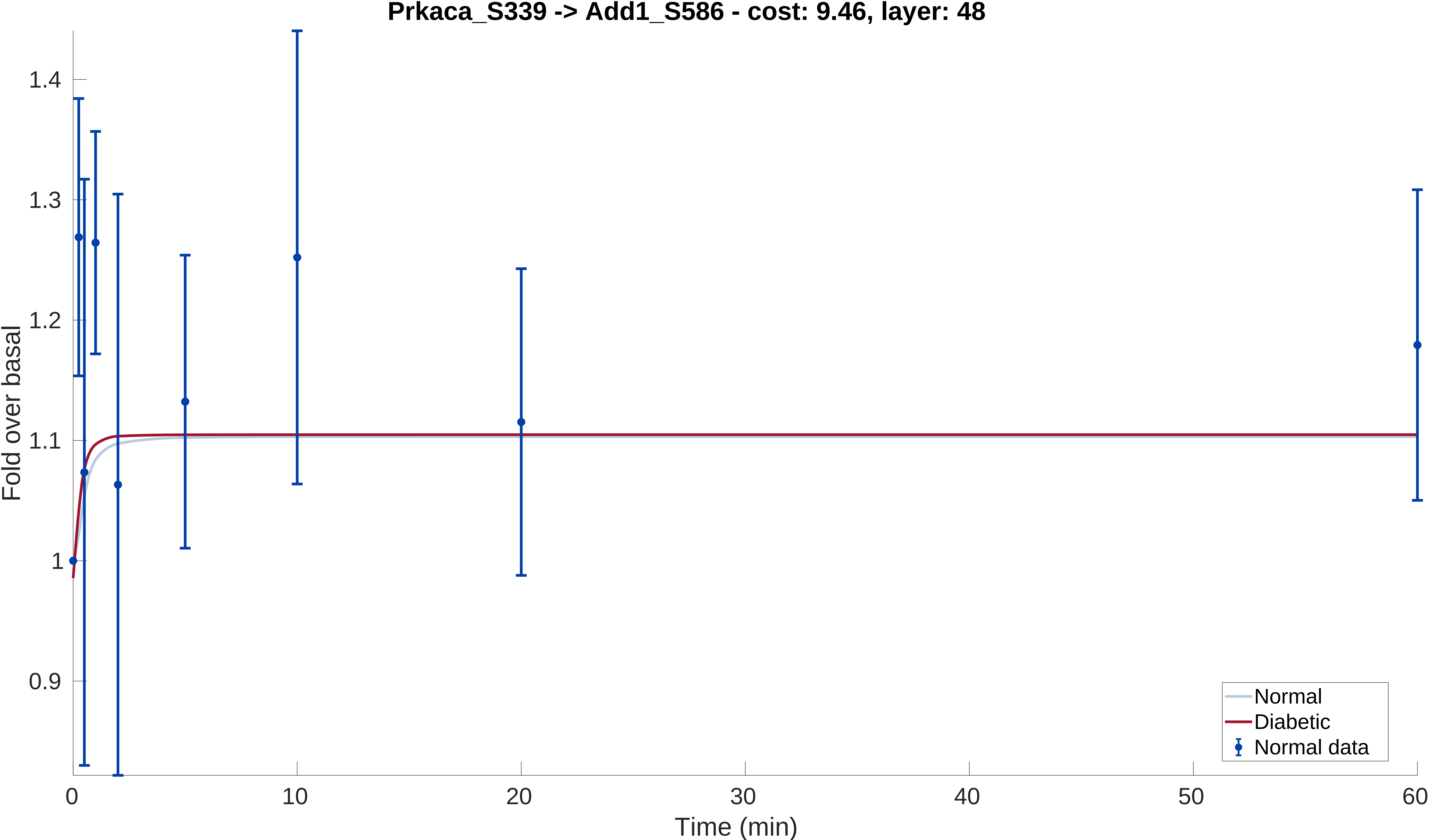

Prkaca_S339 -> Add1_S586 - cost: 9.46, layer: 48
Fold over basal
1.4
1.3
1.2
1.1
1
0.9
0
10
20
30
40
50
60
Time (min)
Normal
Diabetic
Normal data

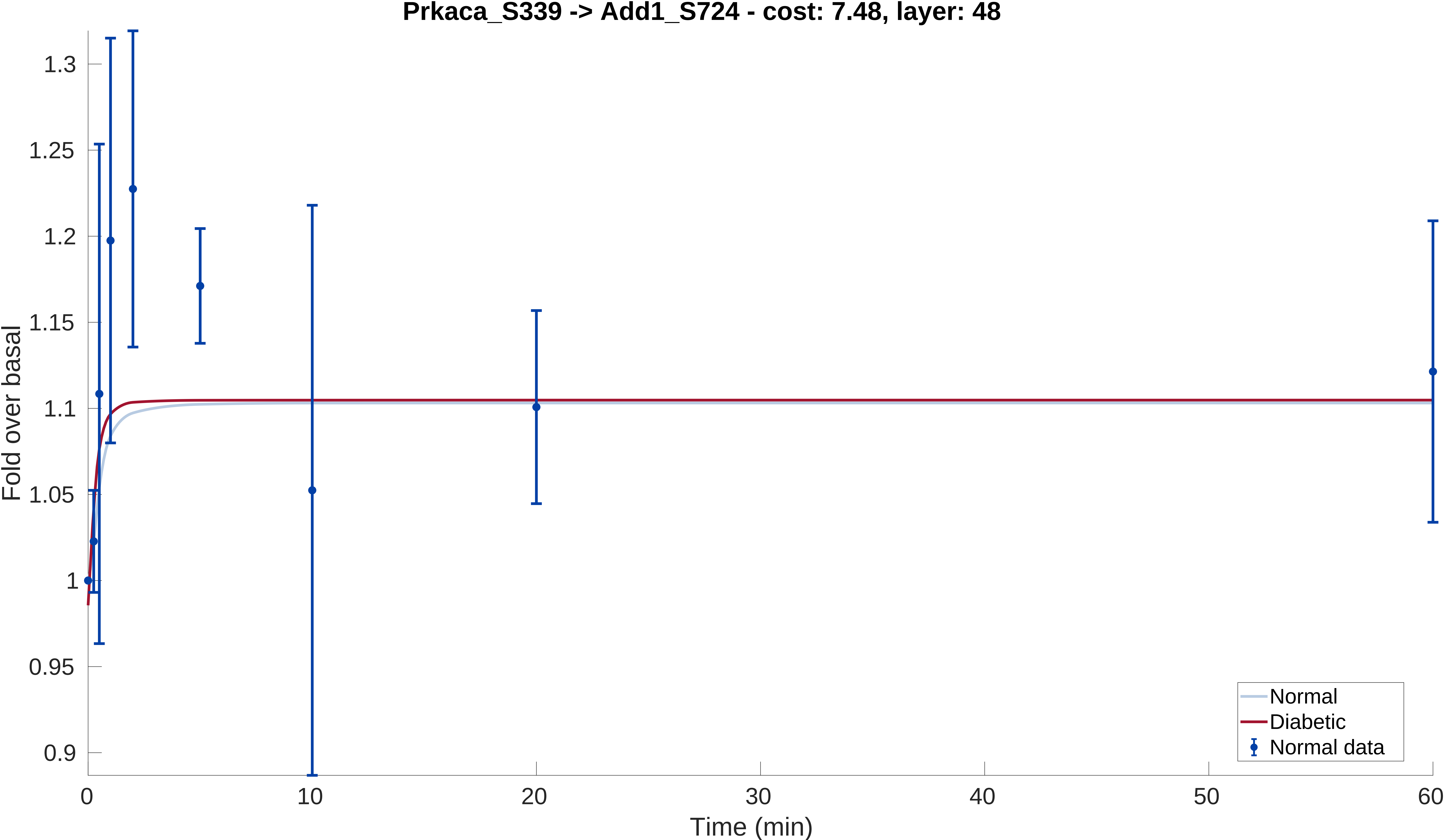
Prkaca_S339 -> Add1_S724 - cost: 7.48, layer: 48
Fold over basal
1.3
1.25
1.2
1.15
1.1
1.05
1
0.95
0.9
0
10
20
30
40
50
60
Time (min)
Normal
Diabetic
Normal data

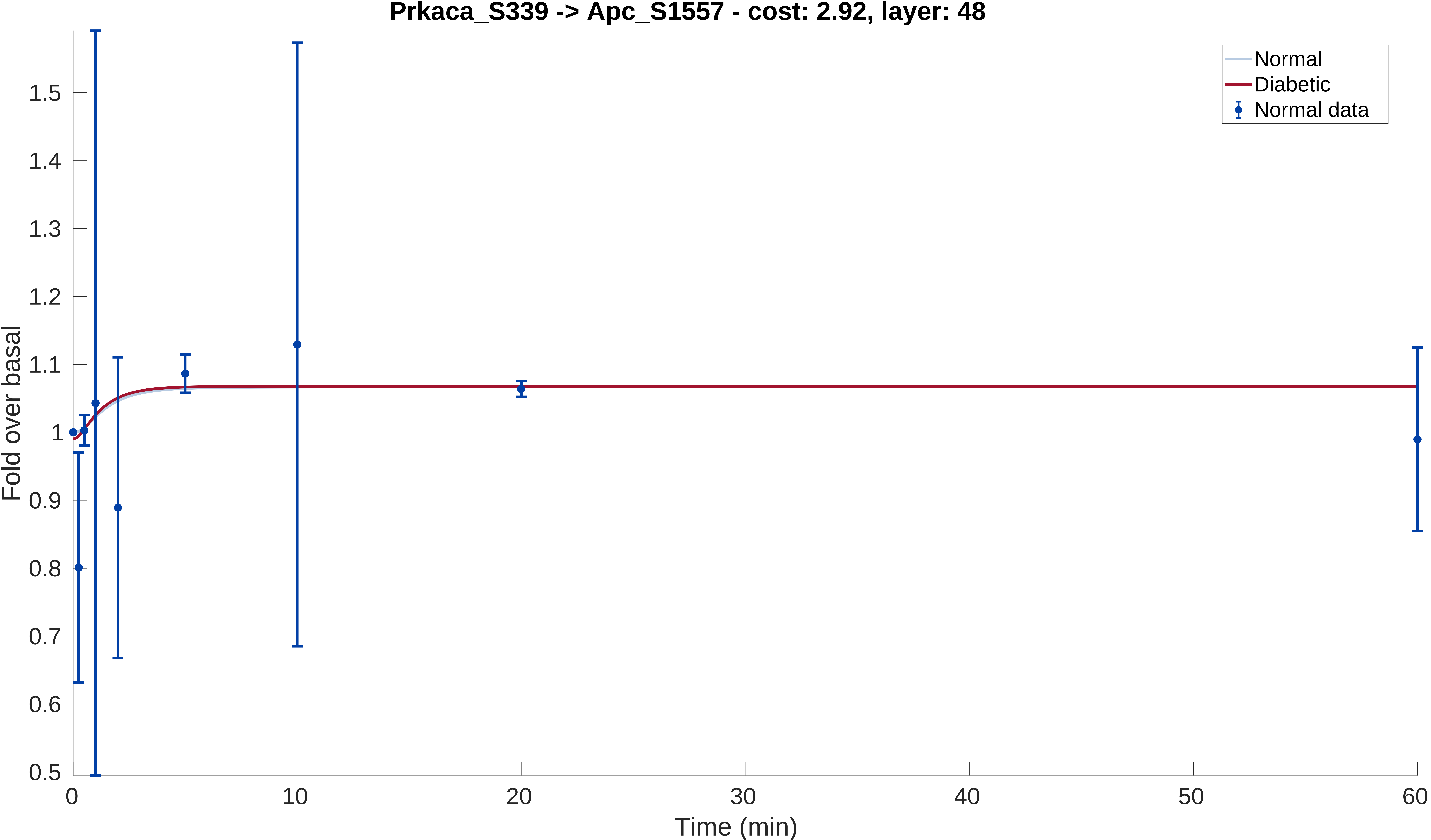

Prkaca_S339 -> Apc_S1557 - cost: 2.92, layer: 48
Normal
Diabetic
Normal data
Fold over basal
1.5
1.4
1.3
1.2
1.1
1
0.9
0.8
0.7
0.6
0.5
0
10
20
30
40
50
60
Time (min)

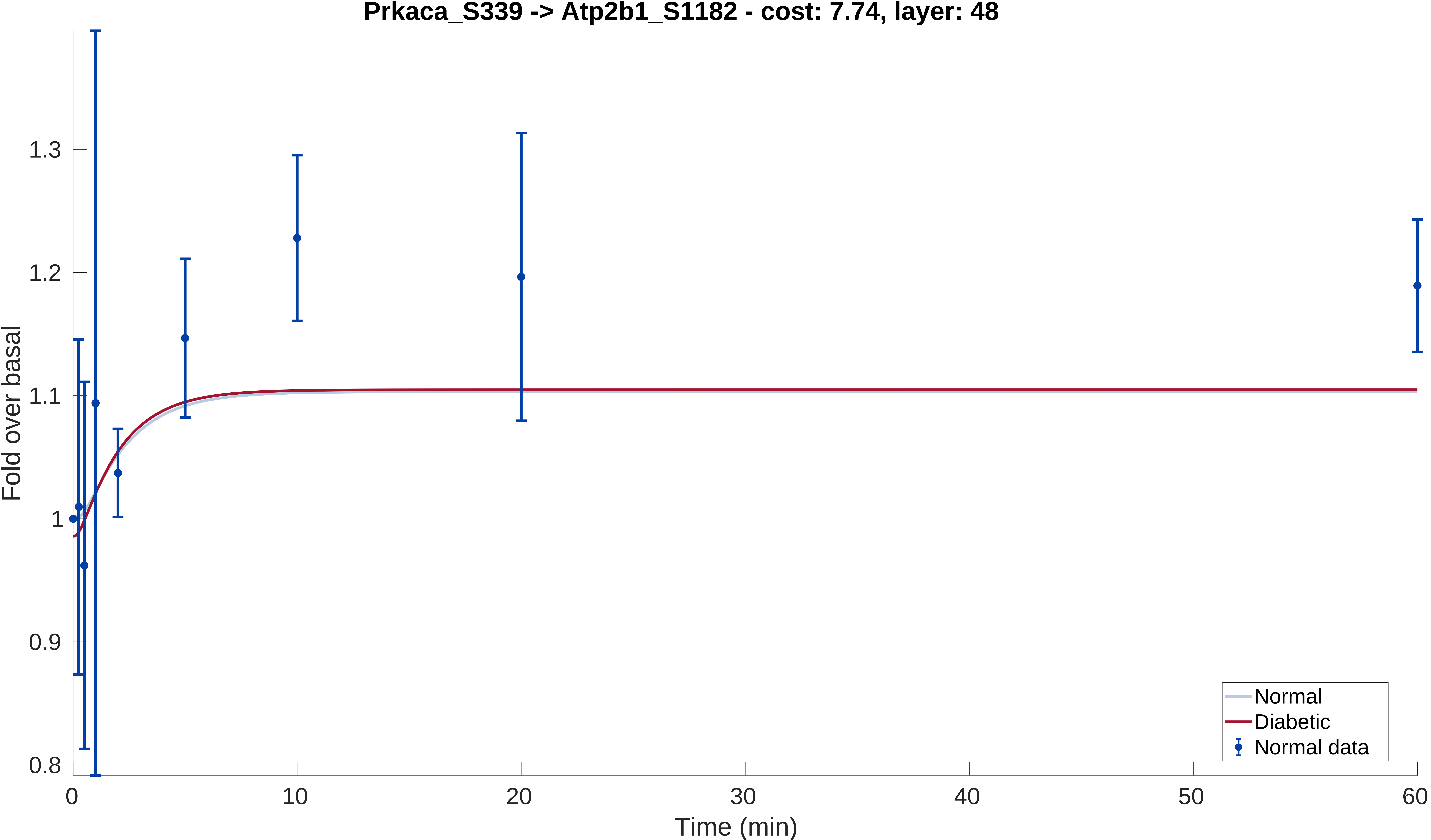

Prkaca_S339 -> Atp2b1_S1182 - cost: 7.74, layer: 48
Fold over basal
Time (min)
0.8
0.9
1
1.1
1.2
1.3
0
10
20
30
40
50
60
Normal
Diabetic
Normal data

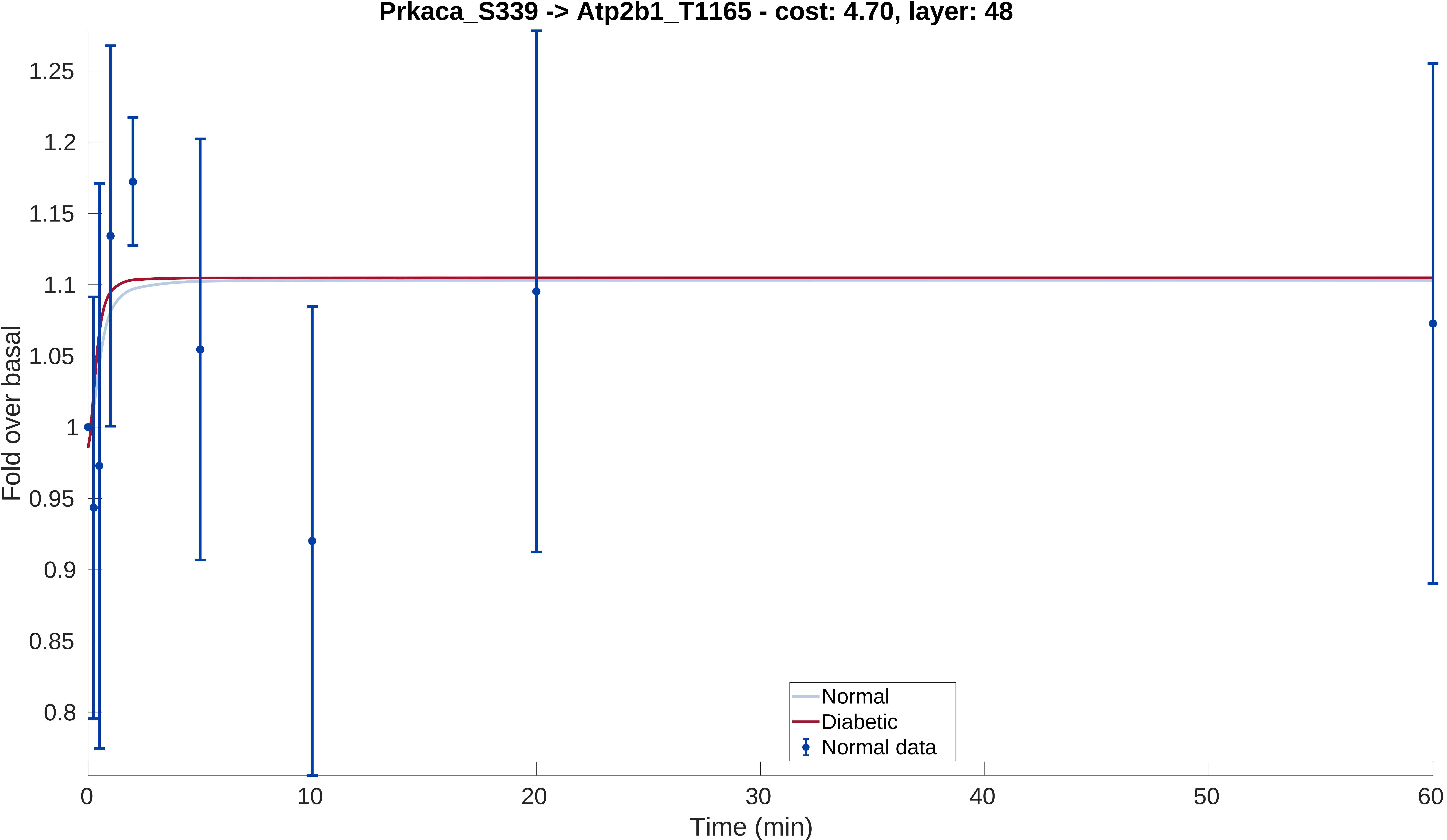

Prkaca_S339 -> Atp2b1_T1165 - cost: 4.70, layer: 48
Fold over basal
1.25
1.2
1.15
1.1
1.05
1
0.95
0.9
0.85
0.8
0
10
20
30
40
50
60
Time (min)
Normal
Diabetic
Normal data

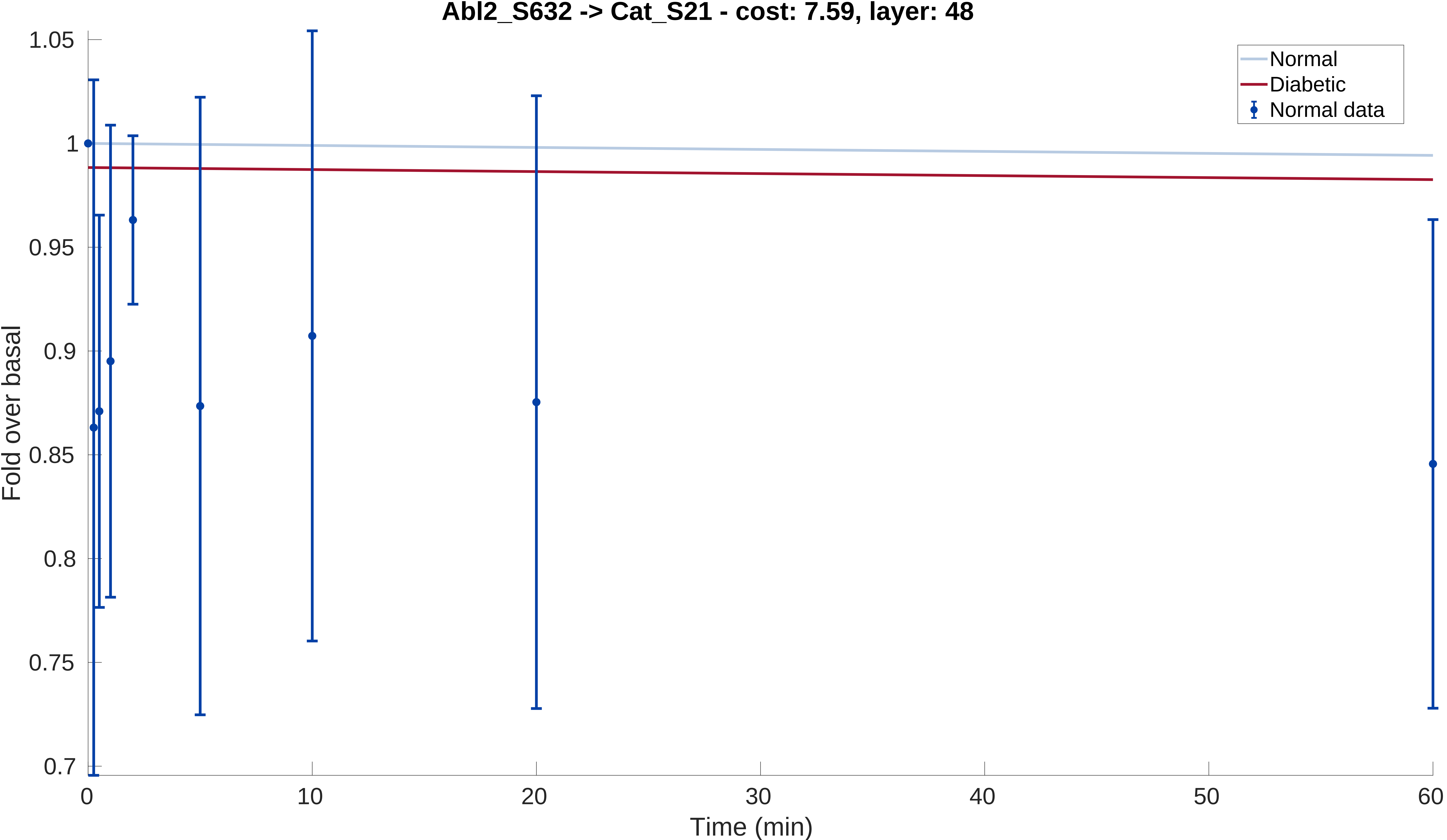

Abl2_S632 -> Cat_S21 - cost: 7.59, layer: 48
Normal
Diabetic
Normal data
Fold over basal
Time (min)
1.05
1
0.95
0.9
0.85
0.8
0.75
0.7
0
10
20
30
40
50
60

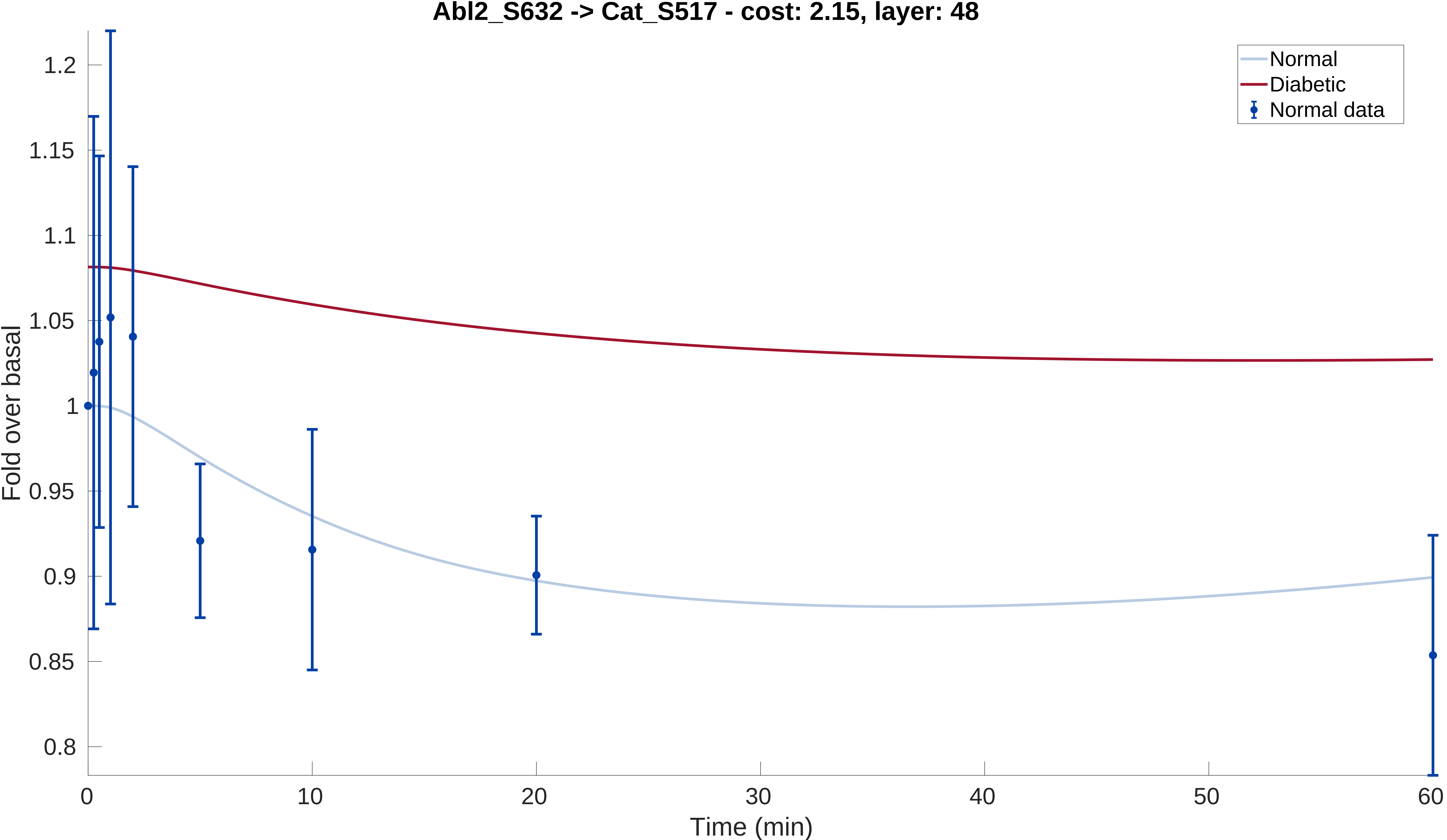

Abl2_S632 -> Cat_S517 - cost: 2.15, layer: 48
Normal
Diabetic
Normal data
Fold over basal
Time (min)
1.2
1.15
1.1
1.05
1
0.95
0.9
0.85
0.8
0
10
20
30
40
50
60

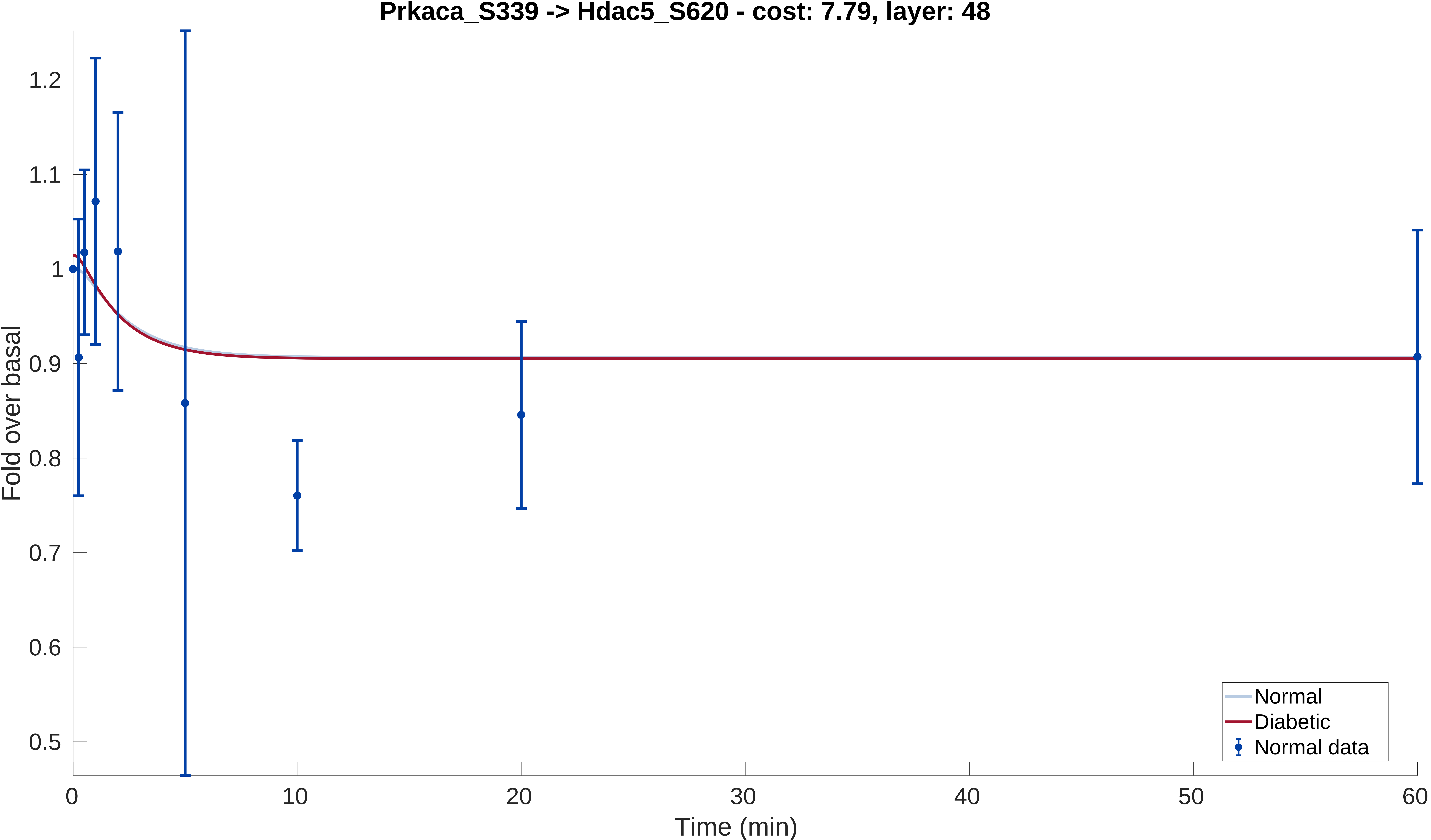

Prkaca_S339 -> Hdac5_S620 - cost: 7.79, layer: 48
Fold over basal
1.2
1.1
1
0.9
0.8
0.7
0.6
0.5
0
10
20
30
40
50
60
Time (min)
Normal
Diabetic
Normal data

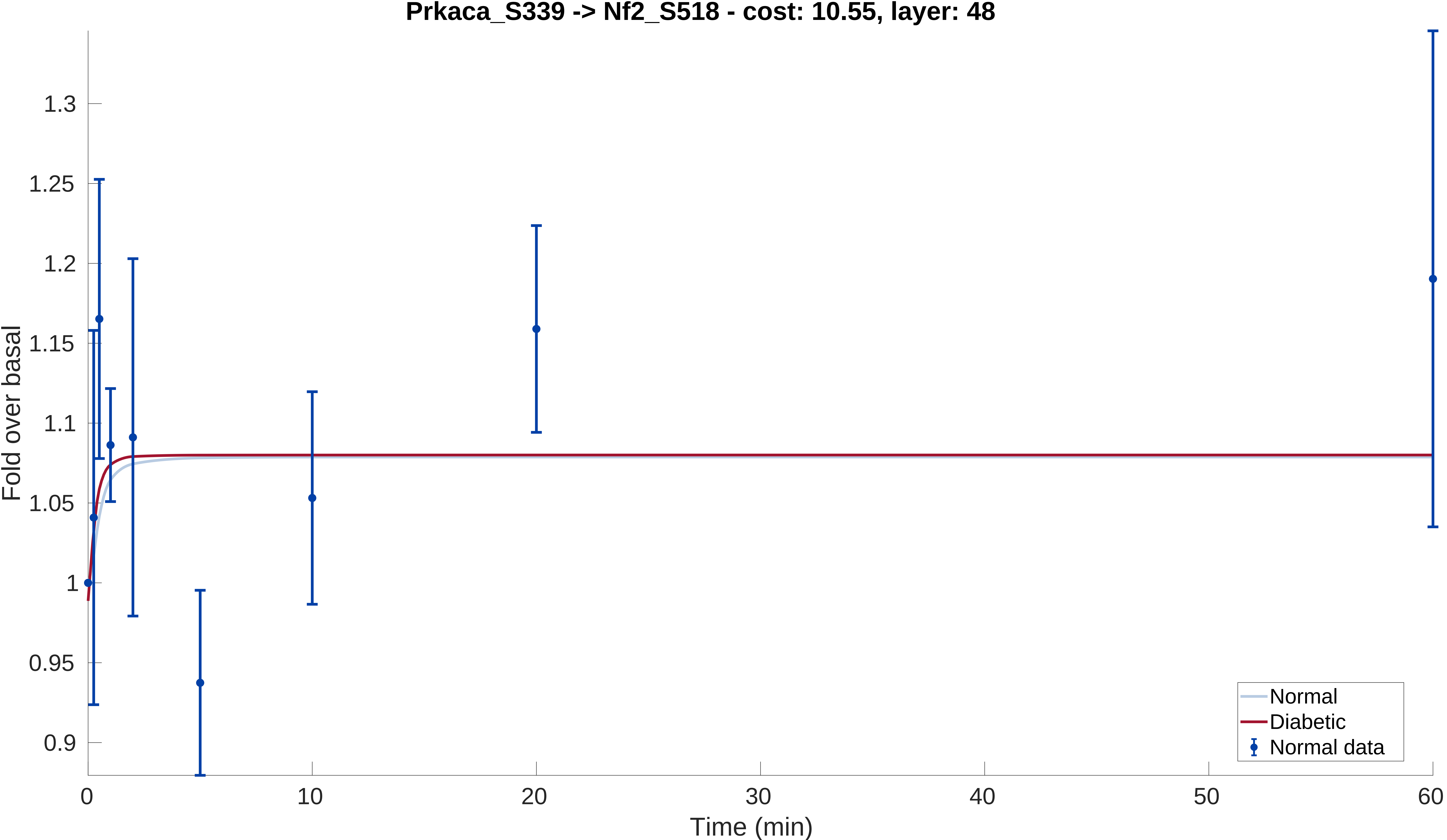

Prkaca_S339 -> Nf2_S518 - cost: 10.55, layer: 48
Fold over basal
1.3
1.25
1.2
1.15
1.1
1.05
1
0.95
0.9
0
10
20
30
40
50
60
Time (min)
Normal
Diabetic
Normal data

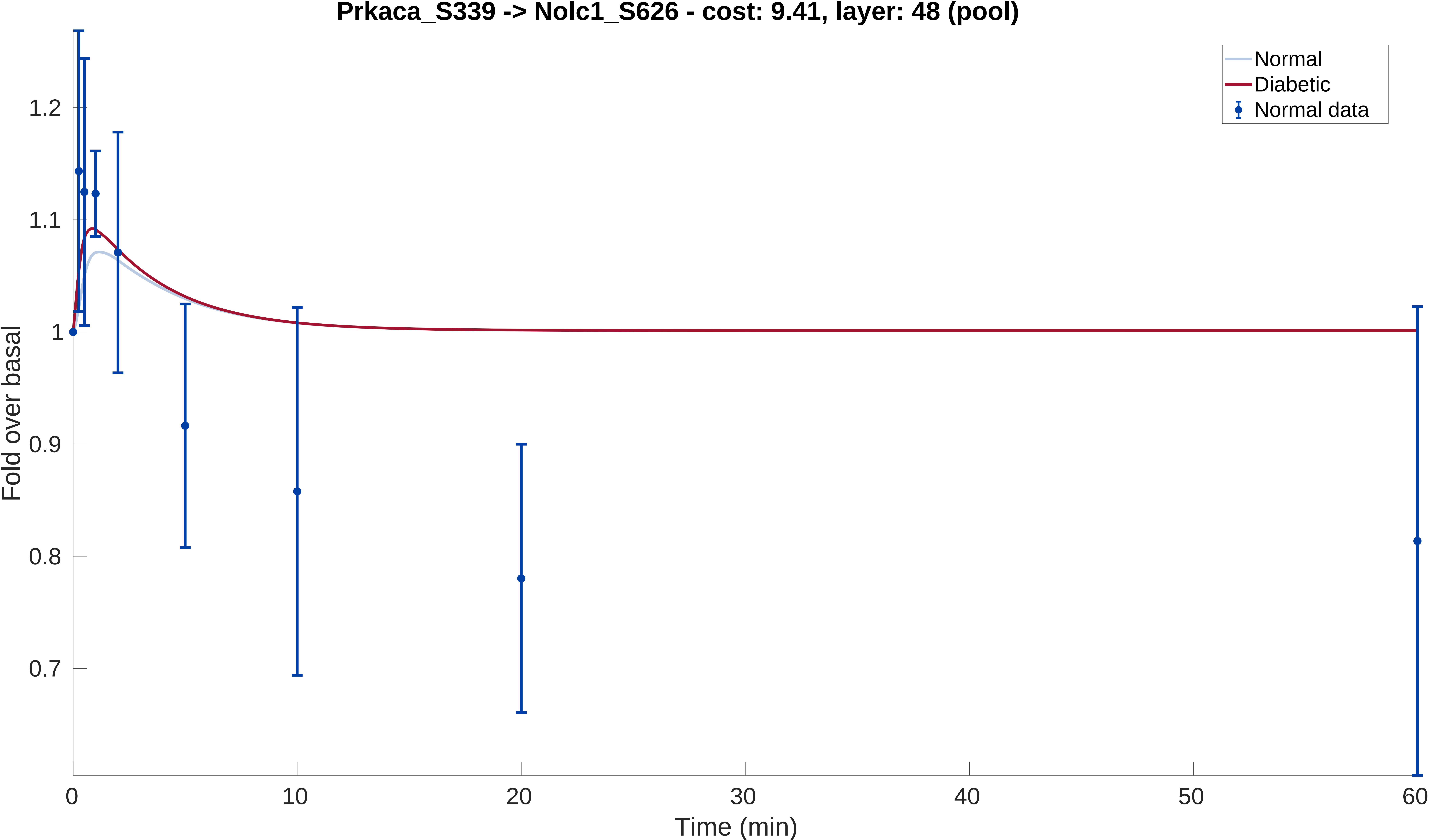

Prkaca_S339 -> Nolc1_S626 - cost: 9.41, layer: 48 (pool)
Normal
Diabetic
Normal data
Fold over basal
1.2
1.1
1
0.9
0.8
0.7
0
10
20
30
40
50
60
Time (min)

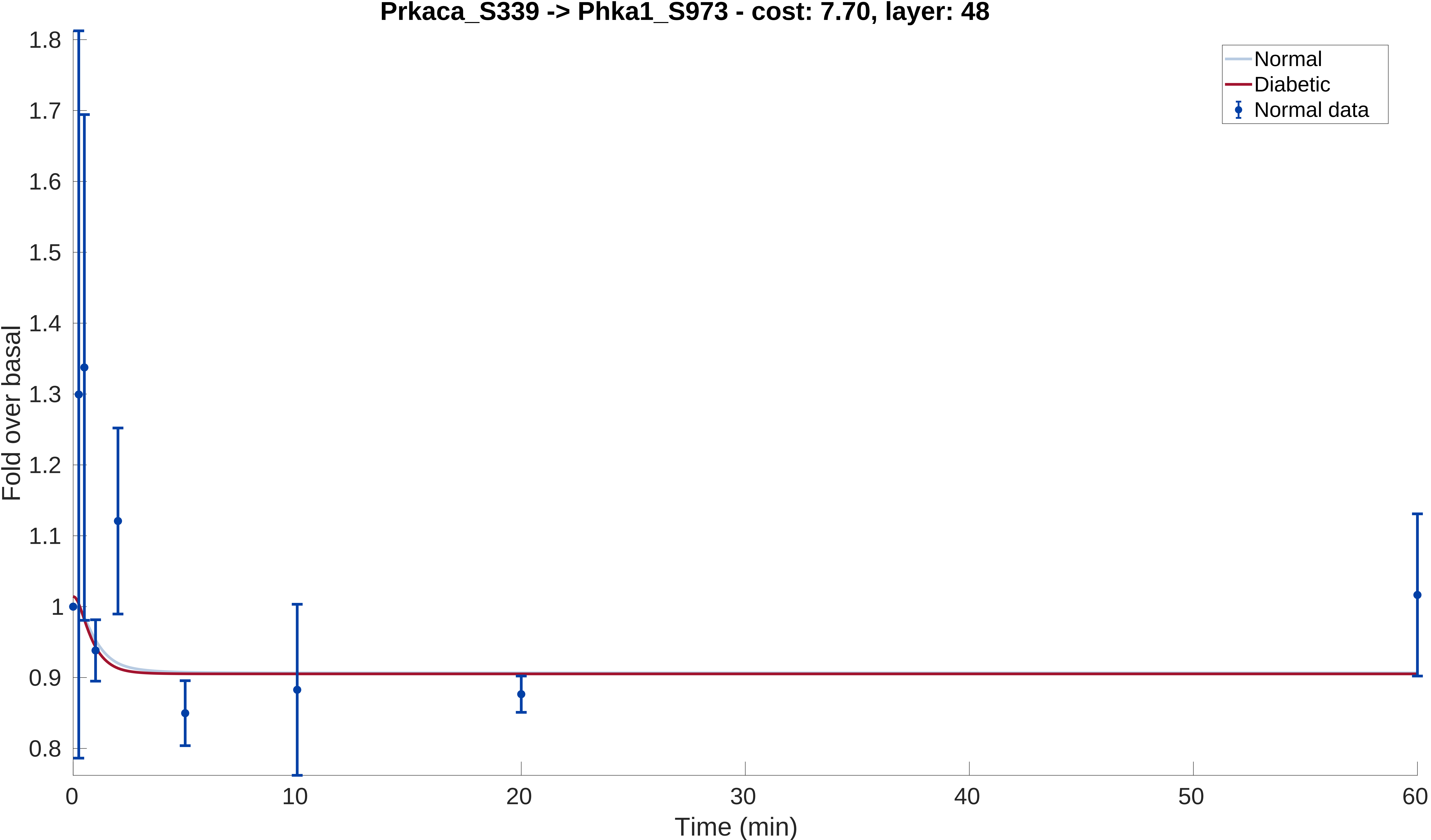
Prkaca_S339 -> Phka1_S973 - cost: 7.70, layer: 48
Normal
Diabetic
Normal data
Fold over basal
1.8
1.7
1.6
1.5
1.4
1.3
1.2
1.1
1
0.9
0.8
0
10
20
30
40
50
60
Time (min)

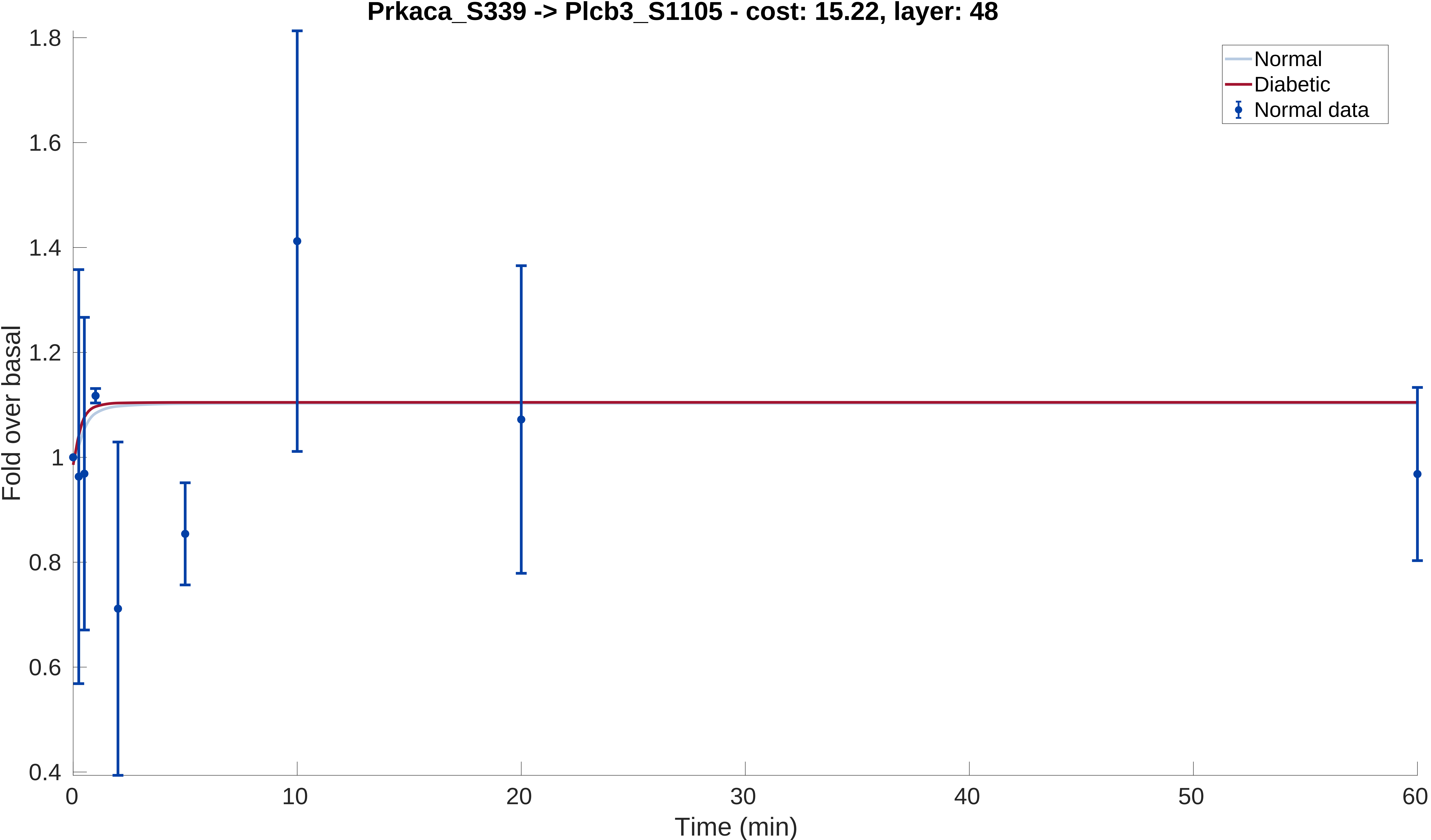

Prkaca_S339 -> Plcb3_S1105 - cost: 15.22, layer: 48
Normal
Diabetic
Normal data
Fold over basal
Time (min)
1.8
1.6
1.4
1.2
1
0.8
0.6
0.4
0
10
20
30
40
50
60

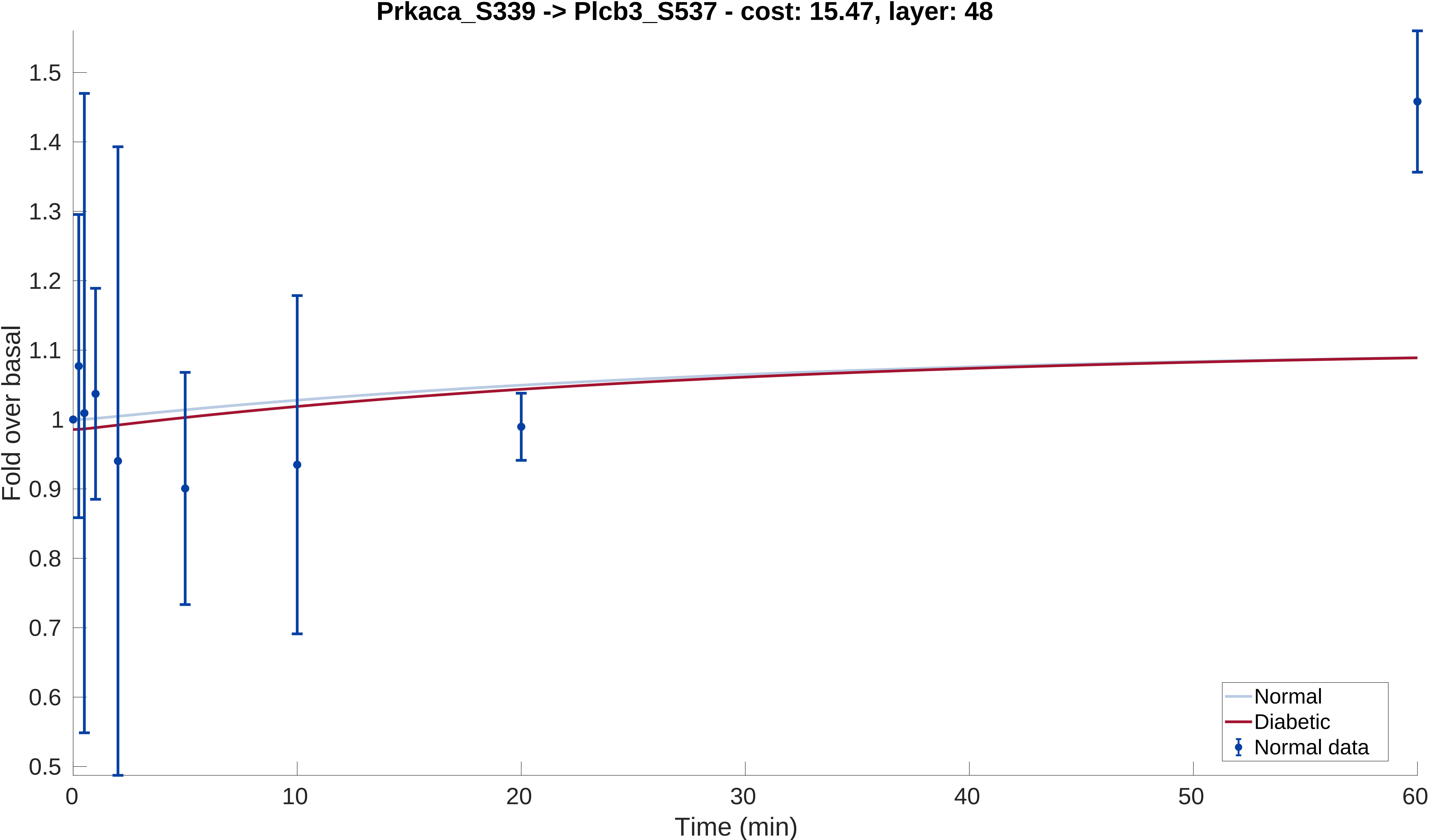
Prkaca_S339 -> Plcb3_S537 - cost: 15.47, layer: 48
Fold over basal
1.5
1.4
1.3
1.2
1.1
1
0.9
0.8
0.7
0.6
0.5
0
10
20
30
40
50
60
Time (min)
Normal
Diabetic
Normal data

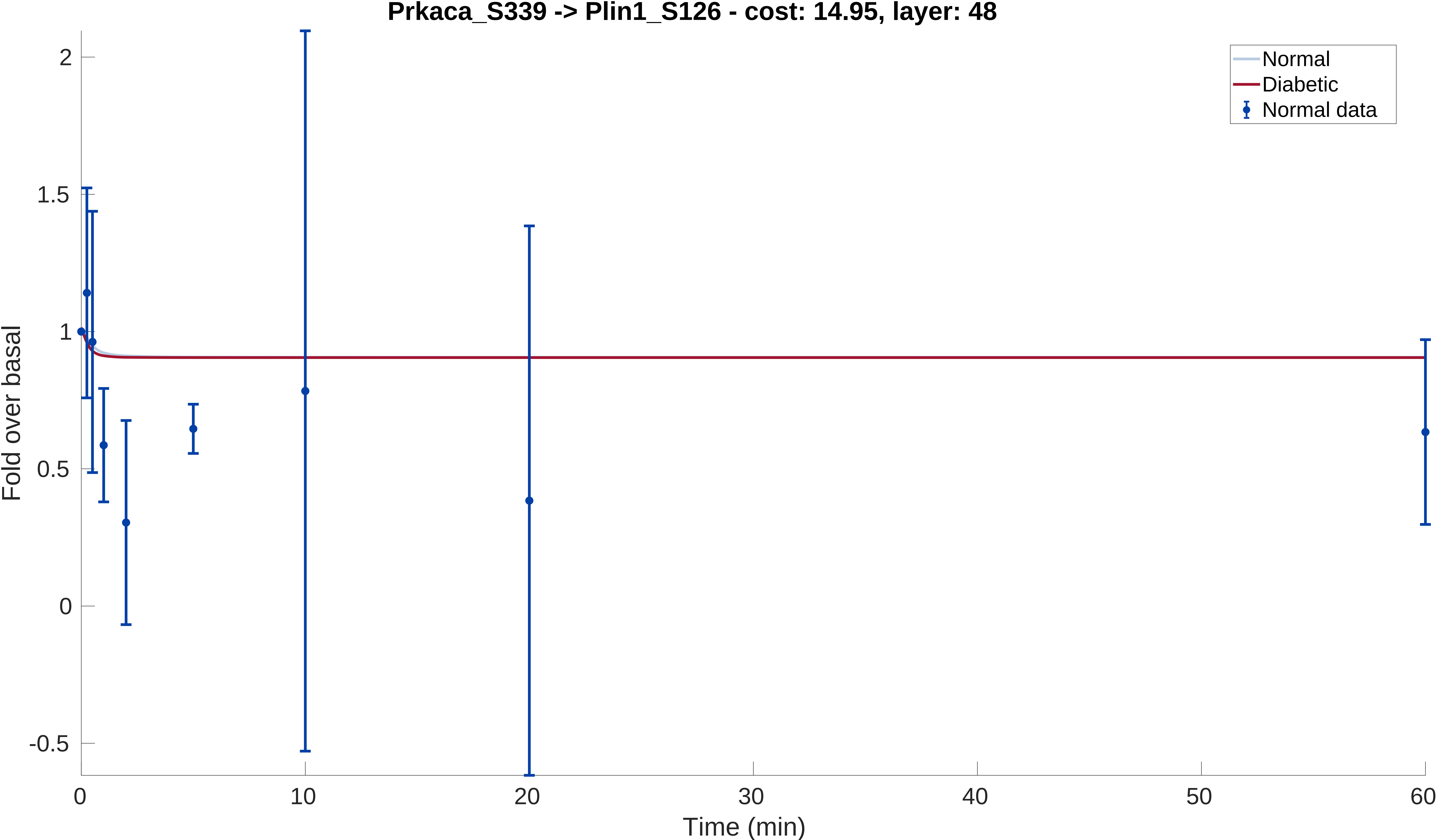
Prkaca_S339 -> Plin1_S126 - cost: 14.95, layer: 48
Normal
Diabetic
Normal data
Fold over basal
Time (min)
2
1.5
1
0.5
0
-0.5
0
10
20
30
40
50
60

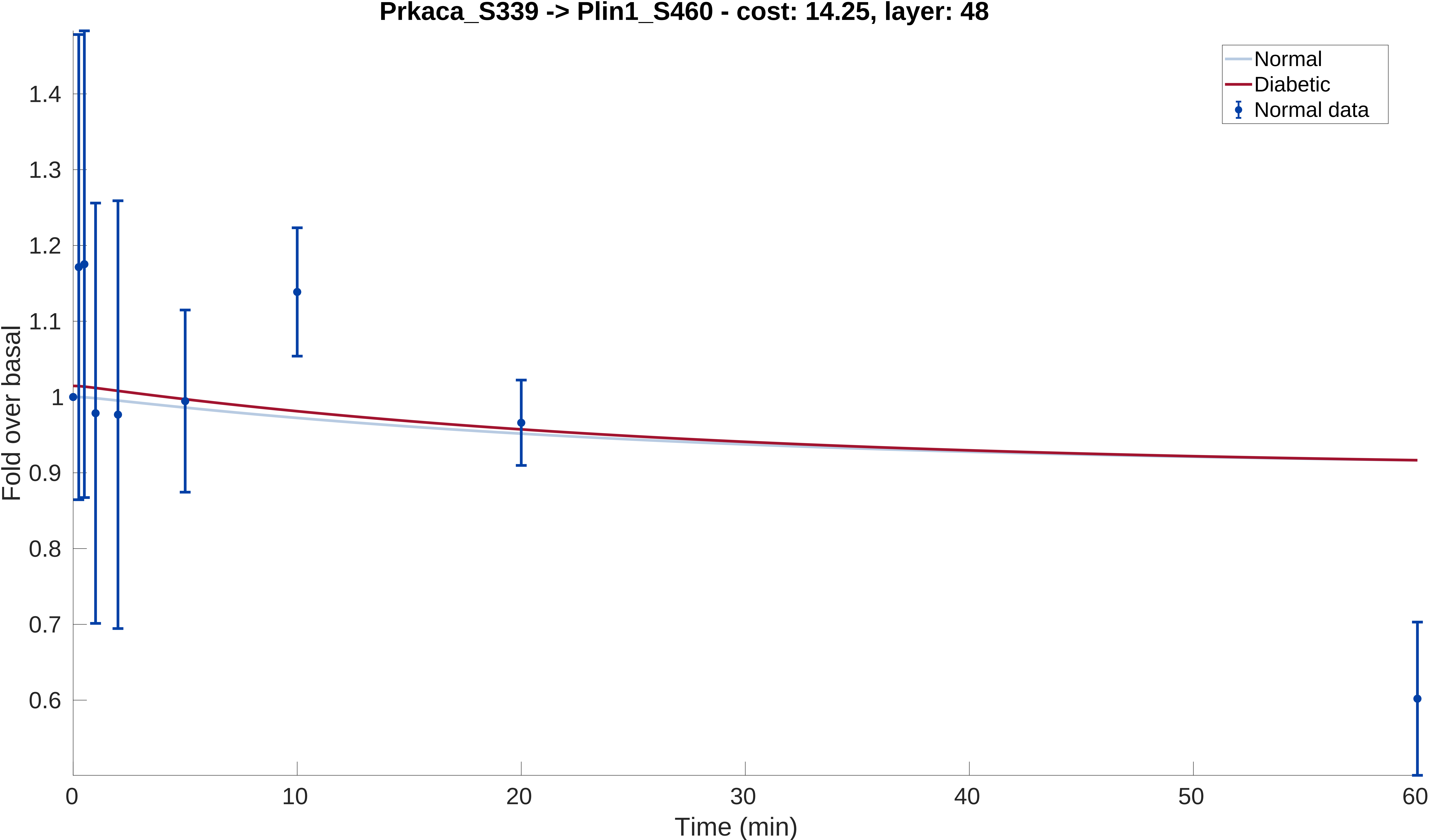

Prrkaca_S339 -> Plin1_S460 - cost: 14.25, layer: 48
Normal
Diabetic
Normal data
1.4
1.3
1.2
1.1
1
0.9
0.8
0.7
0.6
Fold over basal
0
10
20
30
40
50
60
Time (min)

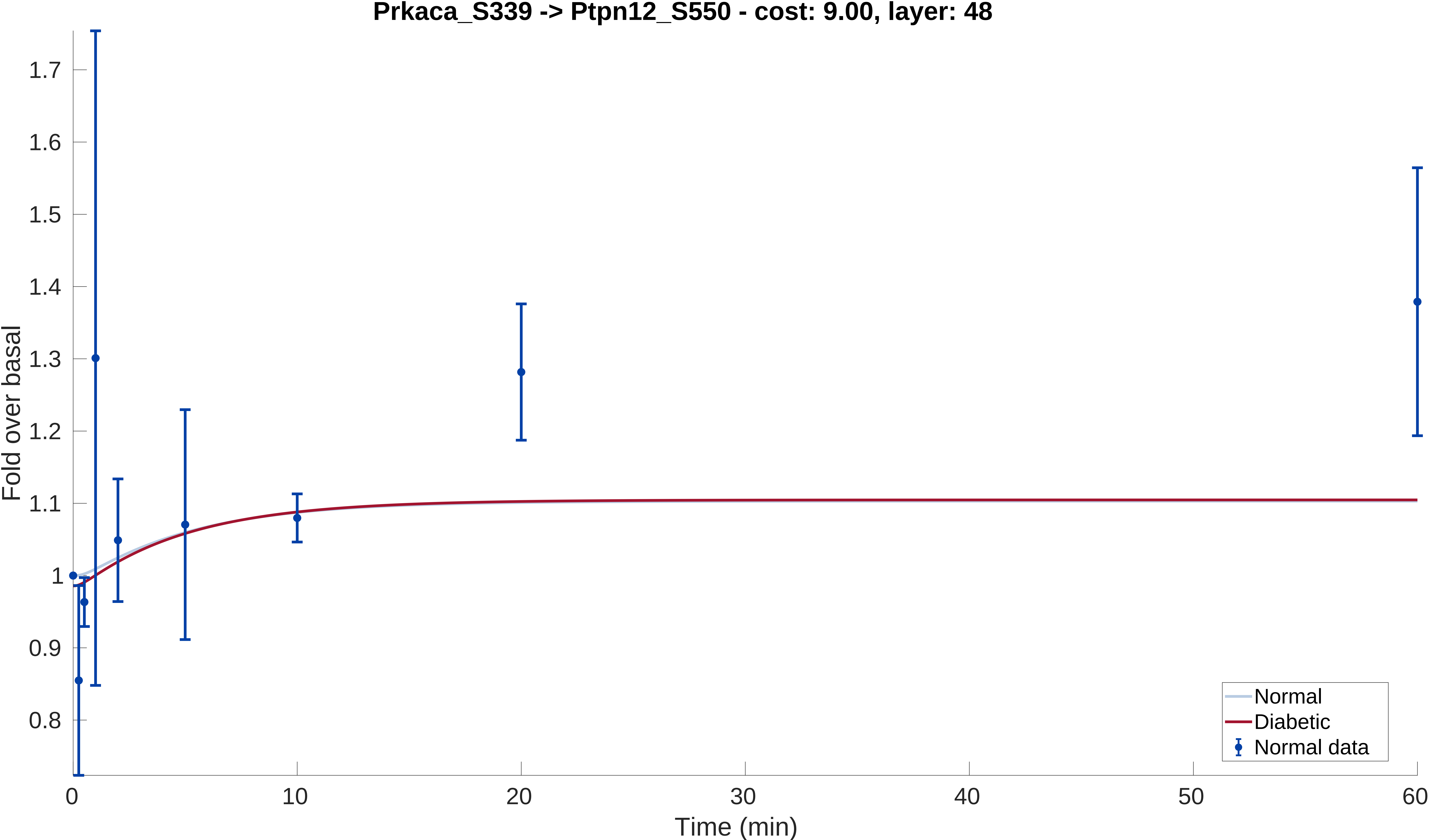

Prkaca_S339 -> Ptpn12_S550 - cost: 9.00, layer: 48
Fold over basal
1.7
1.6
1.5
1.4
1.3
1.2
1.1
1
0.9
0.8
0
10
20
30
40
50
60
Time (min)
Normal
Diabetic
Normal data

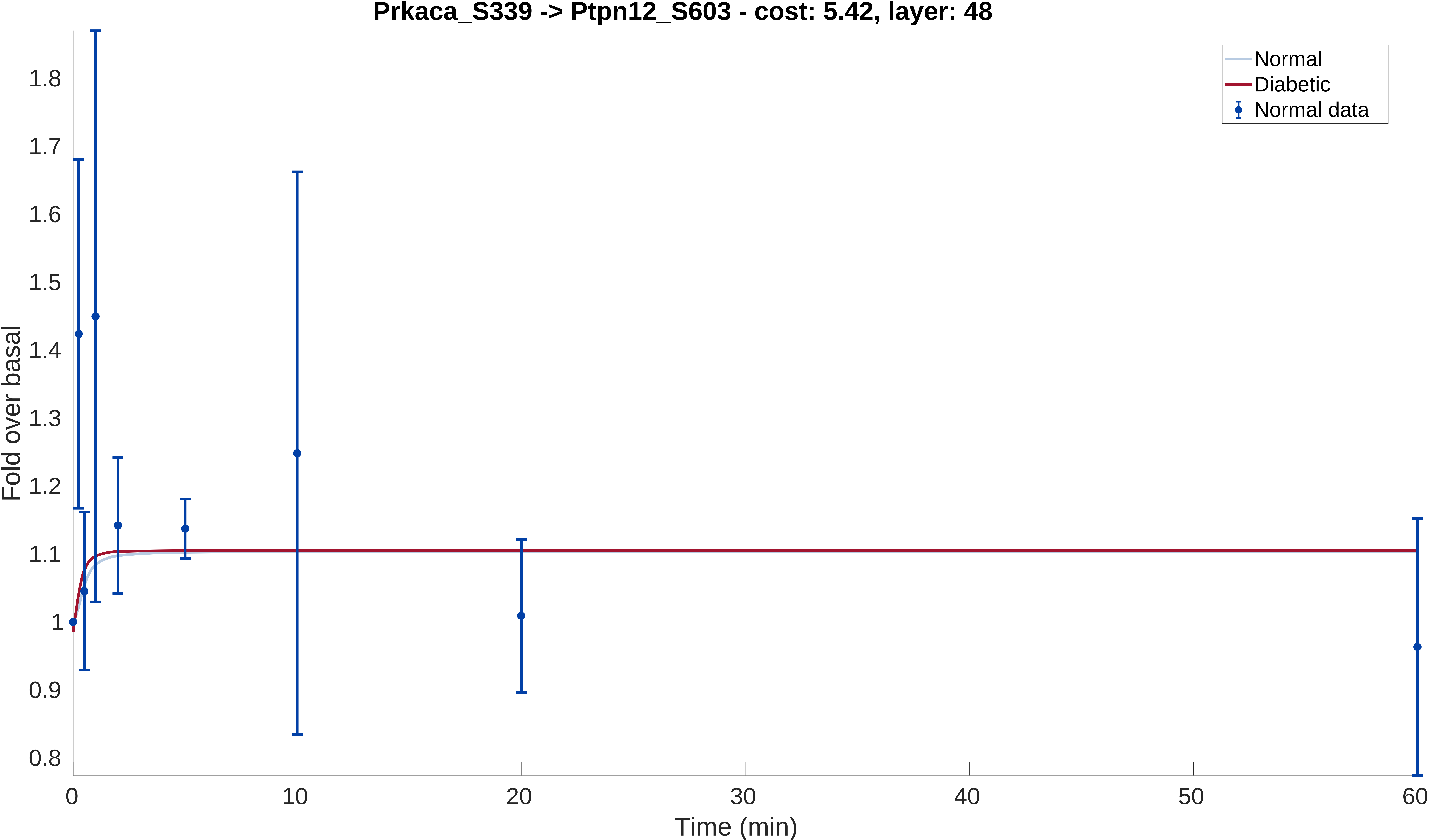

Prrkaca_S339 -> Ptpn12_S603 - cost: 5.42, layer: 48
Normal
Diabetic
Normal data
Fold over basal
Time (min)
0.8
0.9
1
1.1
1.2
1.3
1.4
1.5
1.6
1.7
1.8
0
10
20
30
40
50
60

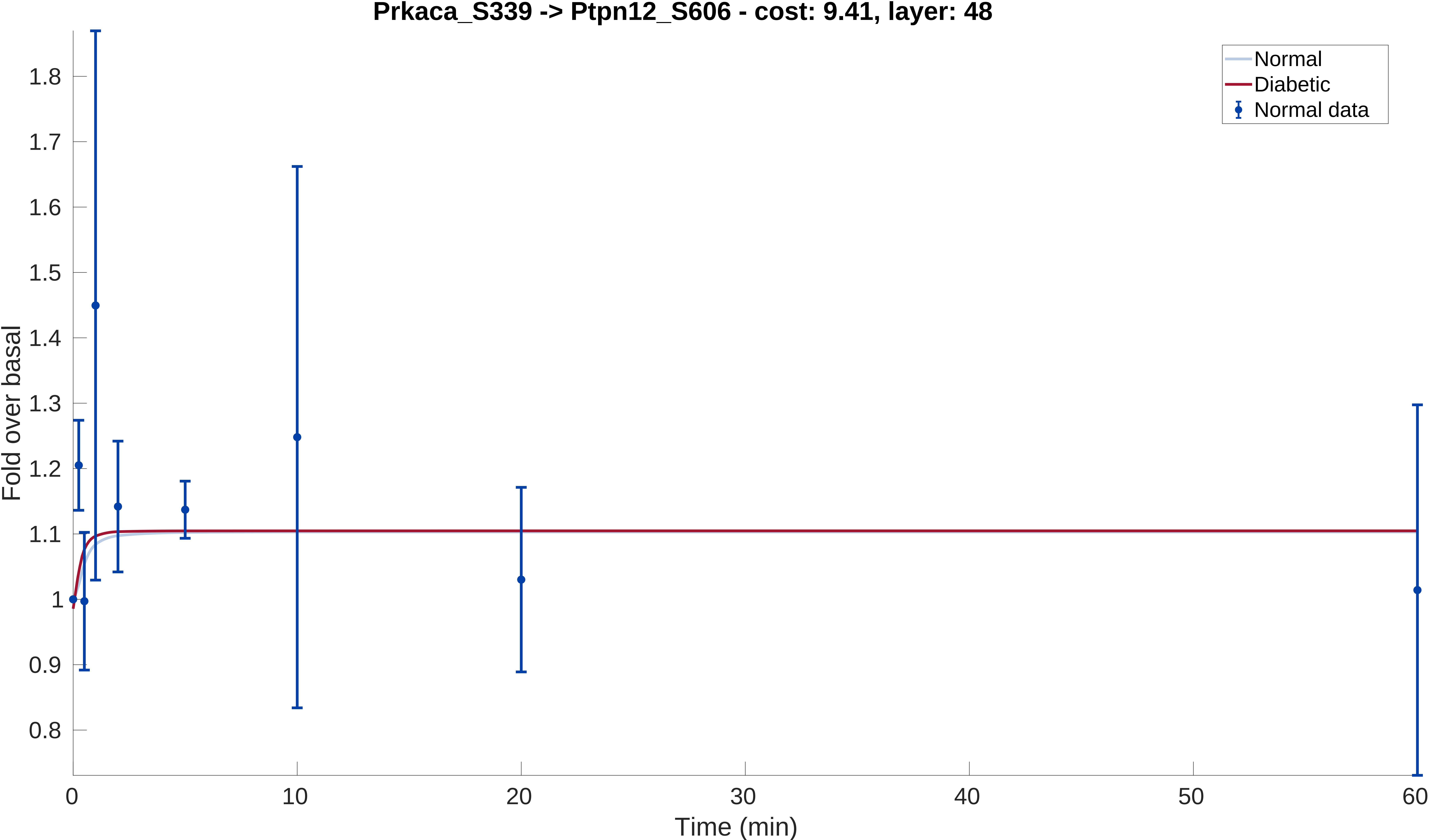

Prkaca_S339 -> Ptpn12_S606 - cost: 9.41, layer: 48
Normal
Diabetic
Normal data
Fold over basal
Time (min)
0.8
0.9
1
1.1
1.2
1.3
1.4
1.5
1.6
1.7
1.8
0
10
20
30
40
50
60

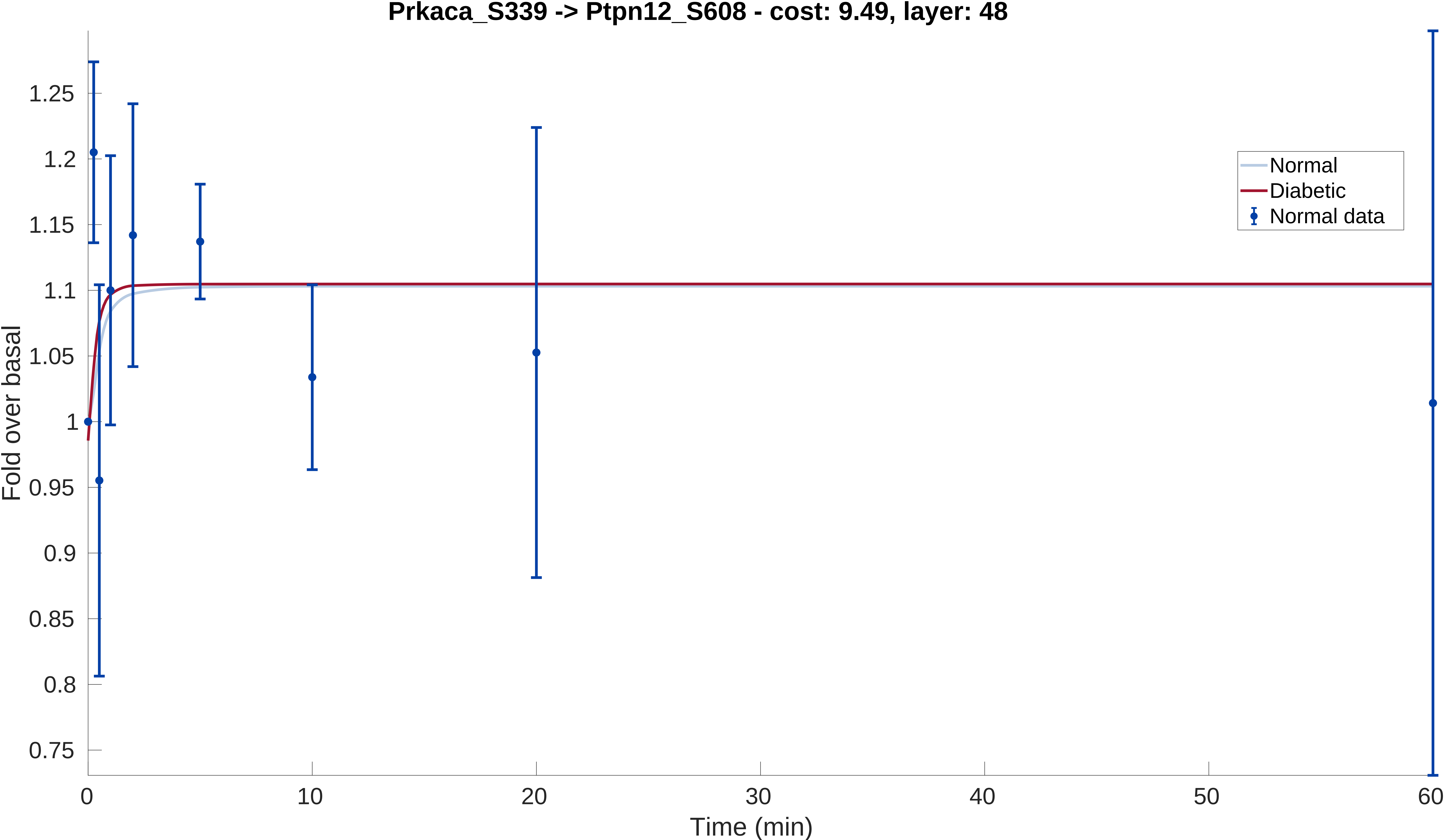

Prkaca_S339 -> Ptpn12_S608 - cost: 9.49, layer: 48
Normal
Diabetic
Normal data
Fold over basal
1.25
1.2
1.15
1.1
1.05
1
0.95
0.9
0.85
0.8
0.75
0
10
20
30
40
50
60
Time (min)

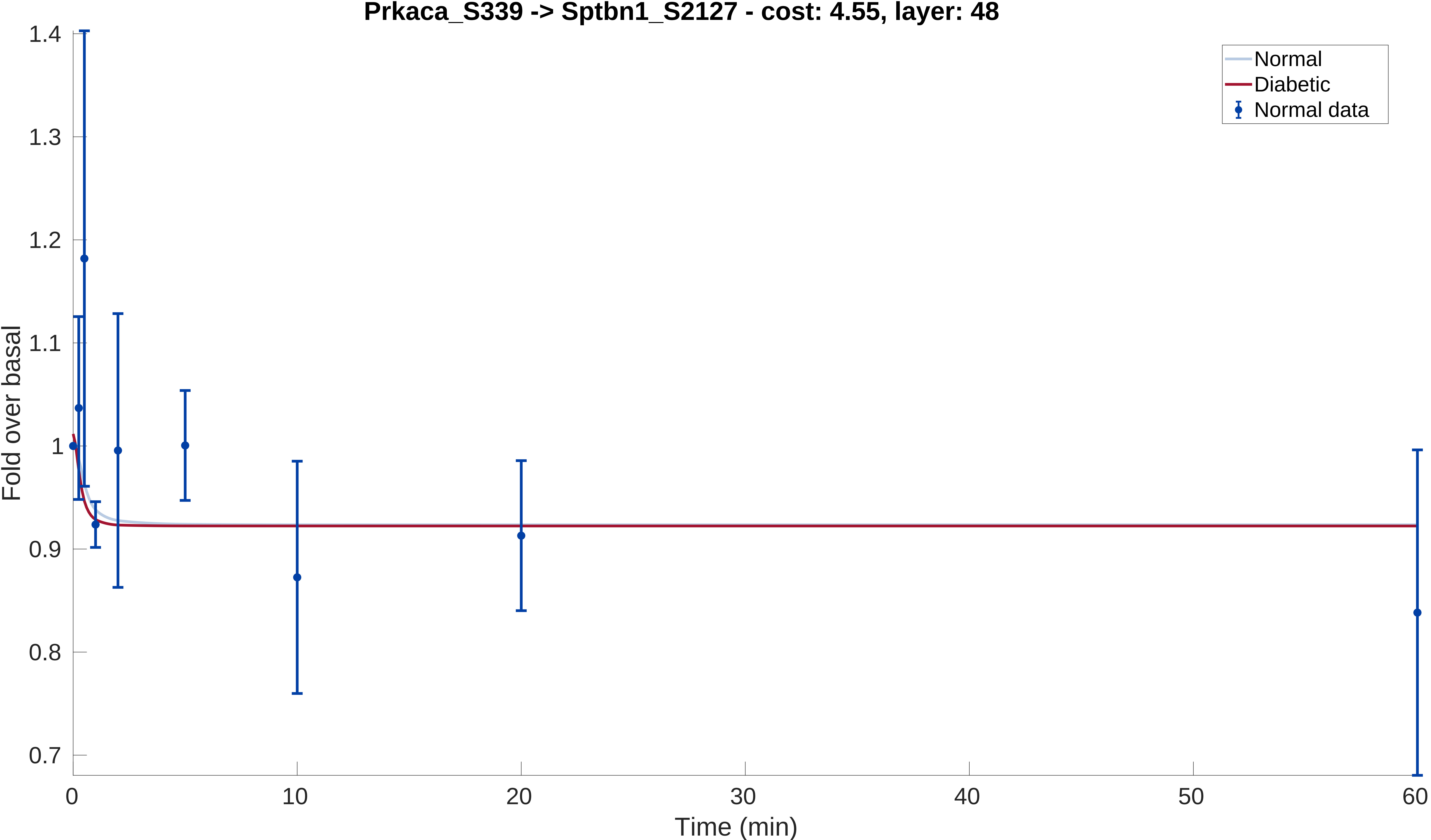
Prkaca_S339 -> Sptbn1_S2127 - cost: 4.55, layer: 48
Normal
Diabetic
Normal data
Fold over basal
1.4
1.3
1.2
1.1
1
0.9
0.8
0.7
0
10
20
30
40
50
60
Time (min)

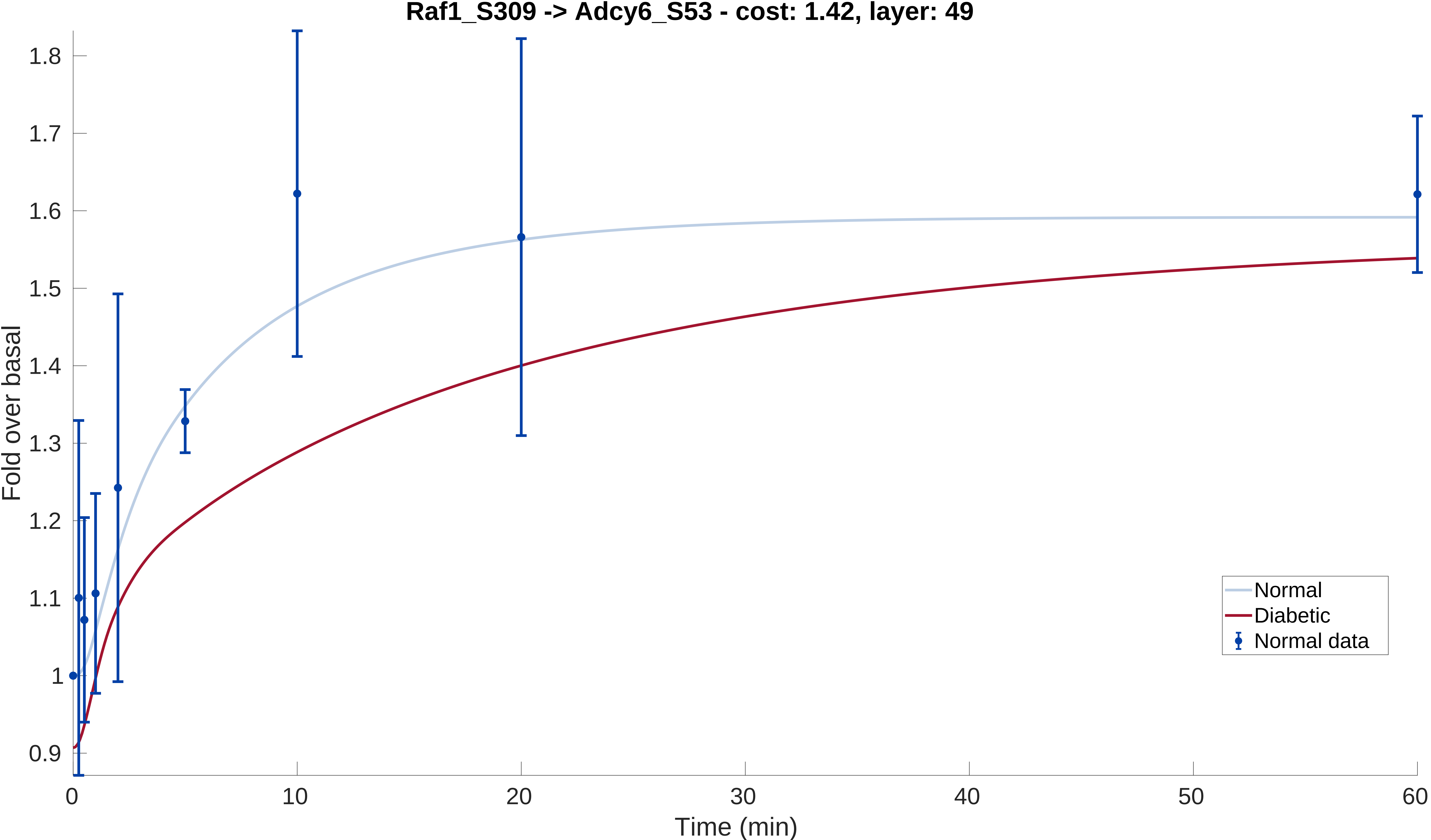
Raf1_S309 -> Adcy6_S53 - cost: 1.42, layer: 49
Fold over basal
0.9
1
1.1
1.2
1.3
1.4
1.5
1.6
1.7
1.8
0
10
20
30
40
50
60
Time (min)
Normal
Diabetic
Normal data

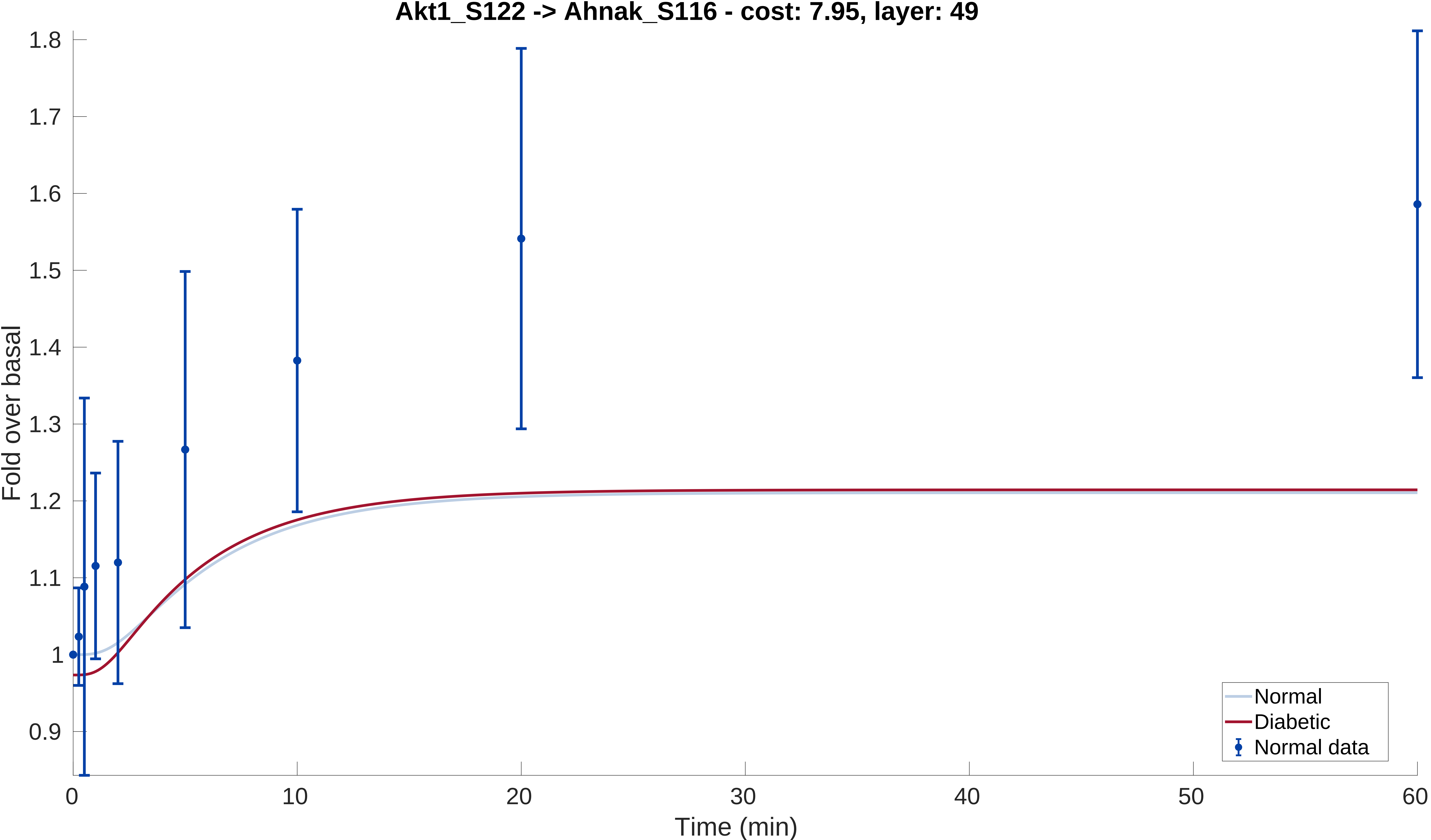
Akt1_S122 -> Ahnak_S116 - cost: 7.95, layer: 49
Fold over basal
1.8
1.7
1.6
1.5
1.4
1.3
1.2
1.1
1
0.9
0
10
20
30
40
50
60
Time (min)
Normal
Diabetic
Normal data

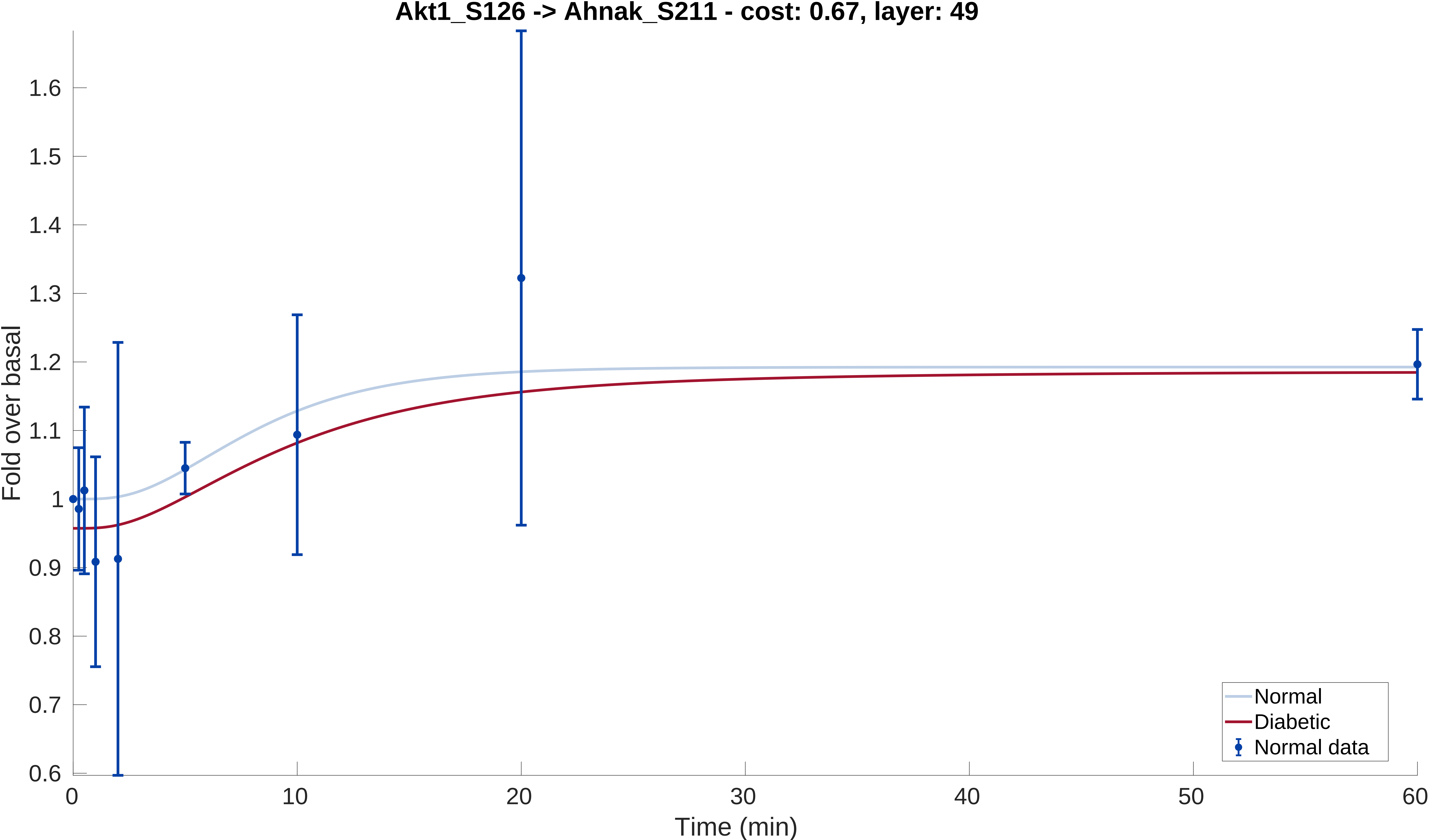

Akt1_S126 -> Ahnak_S211 - cost: 0.67, layer: 49
Fold over basal
1.6
1.5
1.4
1.3
1.2
1.1
1
0.9
0.8
0.7
0.6
0
10
20
30
40
50
60
Time (min)
Normal
Diabetic
Normal data

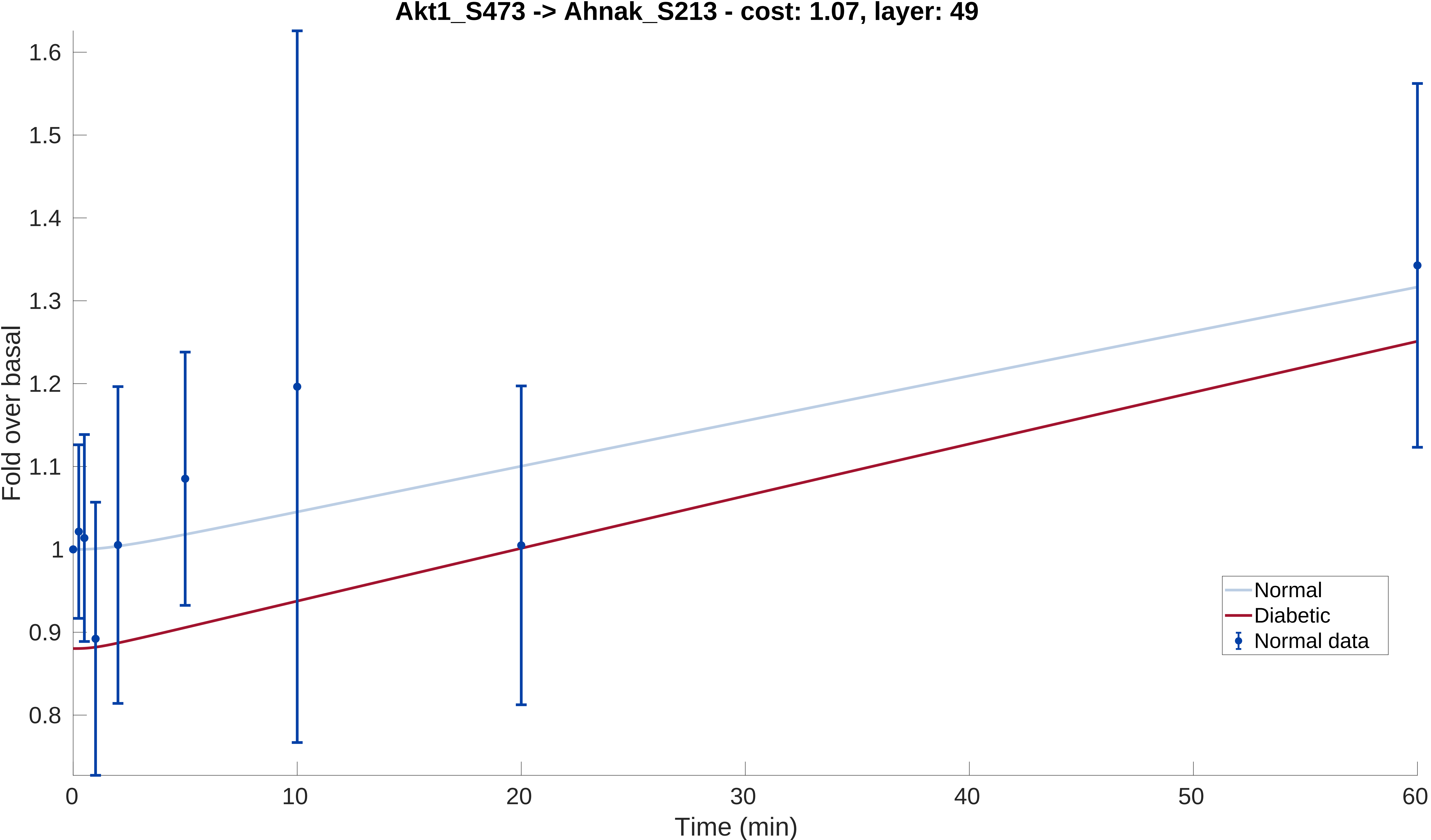
Akt1_S473 -> Ahnak_S213 - cost: 1.07, layer: 49
Fold over basal
Time (min)
0
10
20
30
40
50
60
0.8
0.9
1
1.1
1.2
1.3
1.4
1.5
1.6
Normal
Diabetic
Normal data

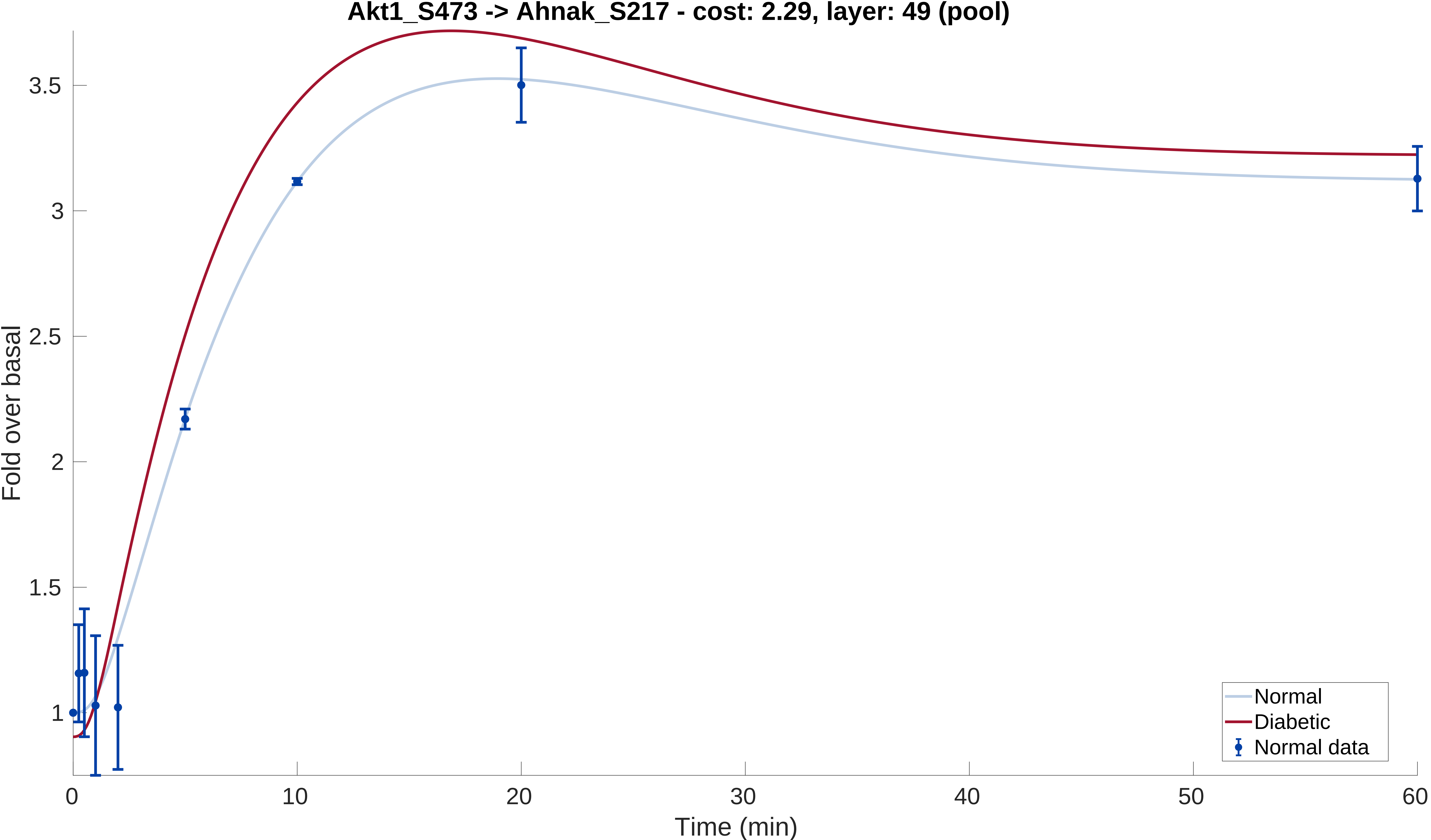

Akt1_S473 -> Ahnak_S217 - cost: 2.29, layer: 49 (pool)
Fold over basal
1
1.5
2
2.5
3
3.5
0
10
20
30
40
50
60
Time (min)
Normal
Diabetic
Normal data

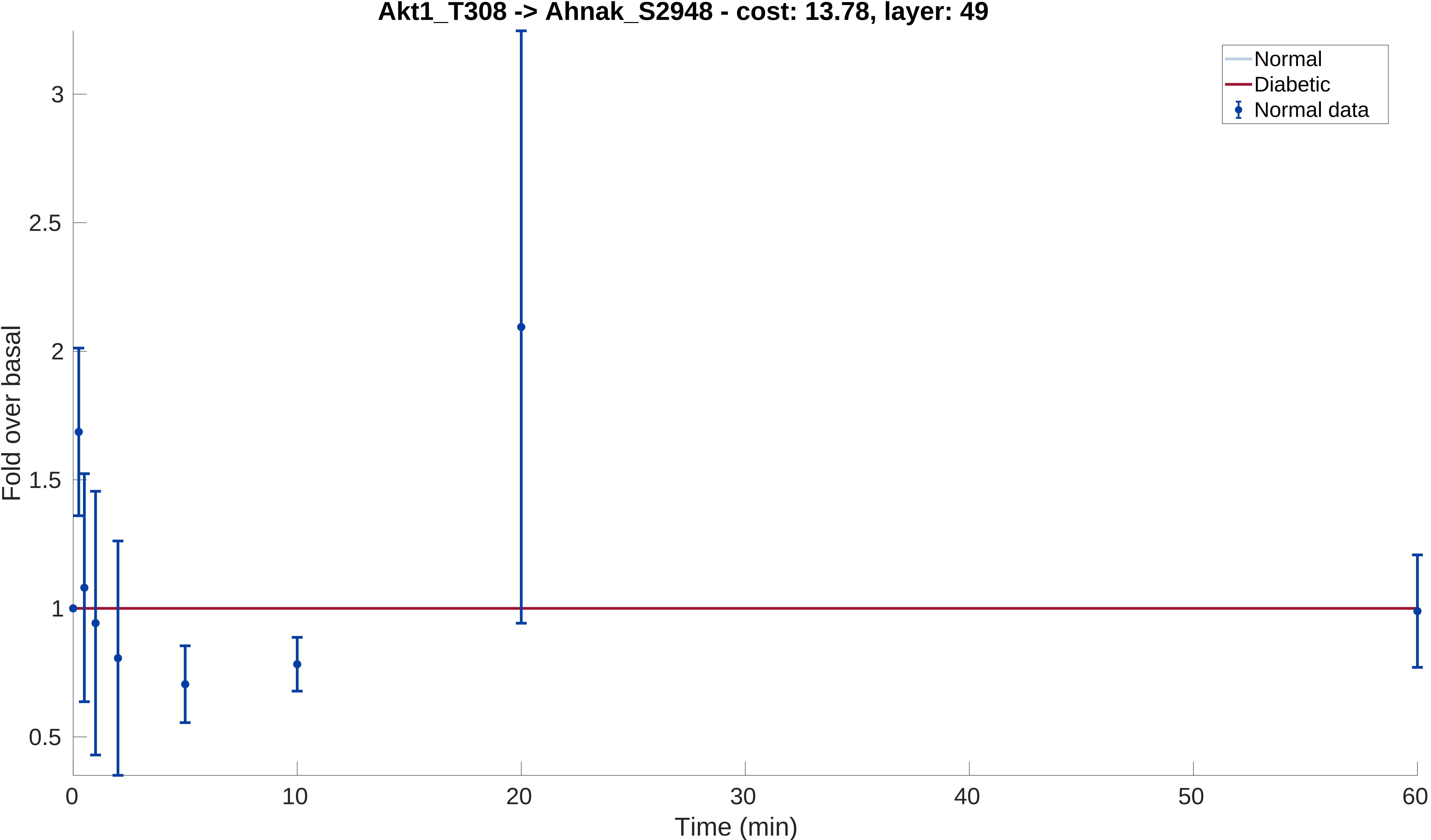

Akt1_T308 -> Ahnak_S2948 - cost: 13.78, layer: 49
Normal
Diabetic
Normal data
Fold over basal
3
2.5
2
1.5
1
0.5
0
10
20
30
40
50
60
Time (min)

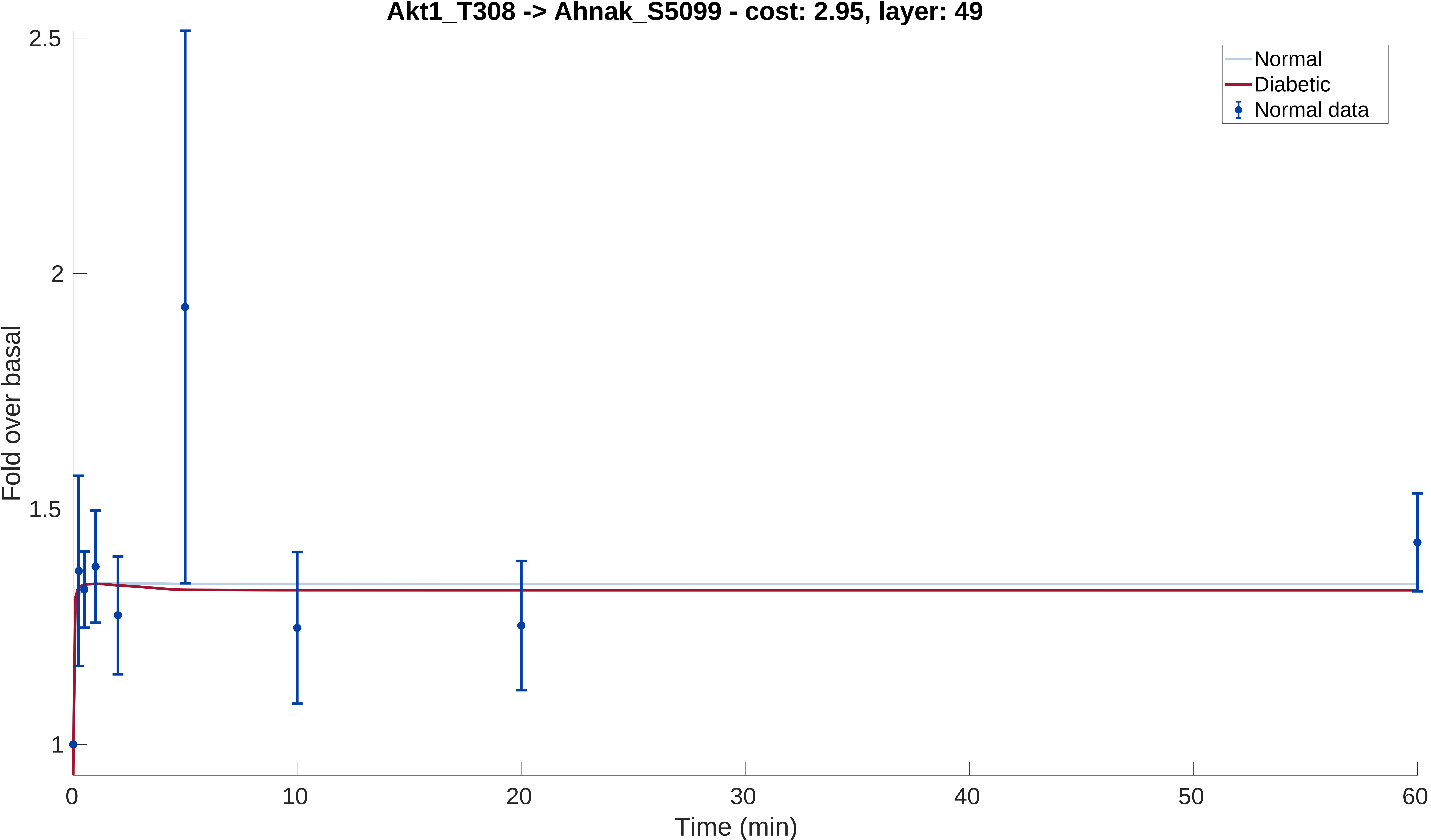

Akt1_T308 -> Ahnak_S5099 - cost: 2.95, layer: 49
Normal
Diabetic
Normal data
Fold over basal
Time (min)
2.5
2
1.5
1
0
10
20
30
40
50
60

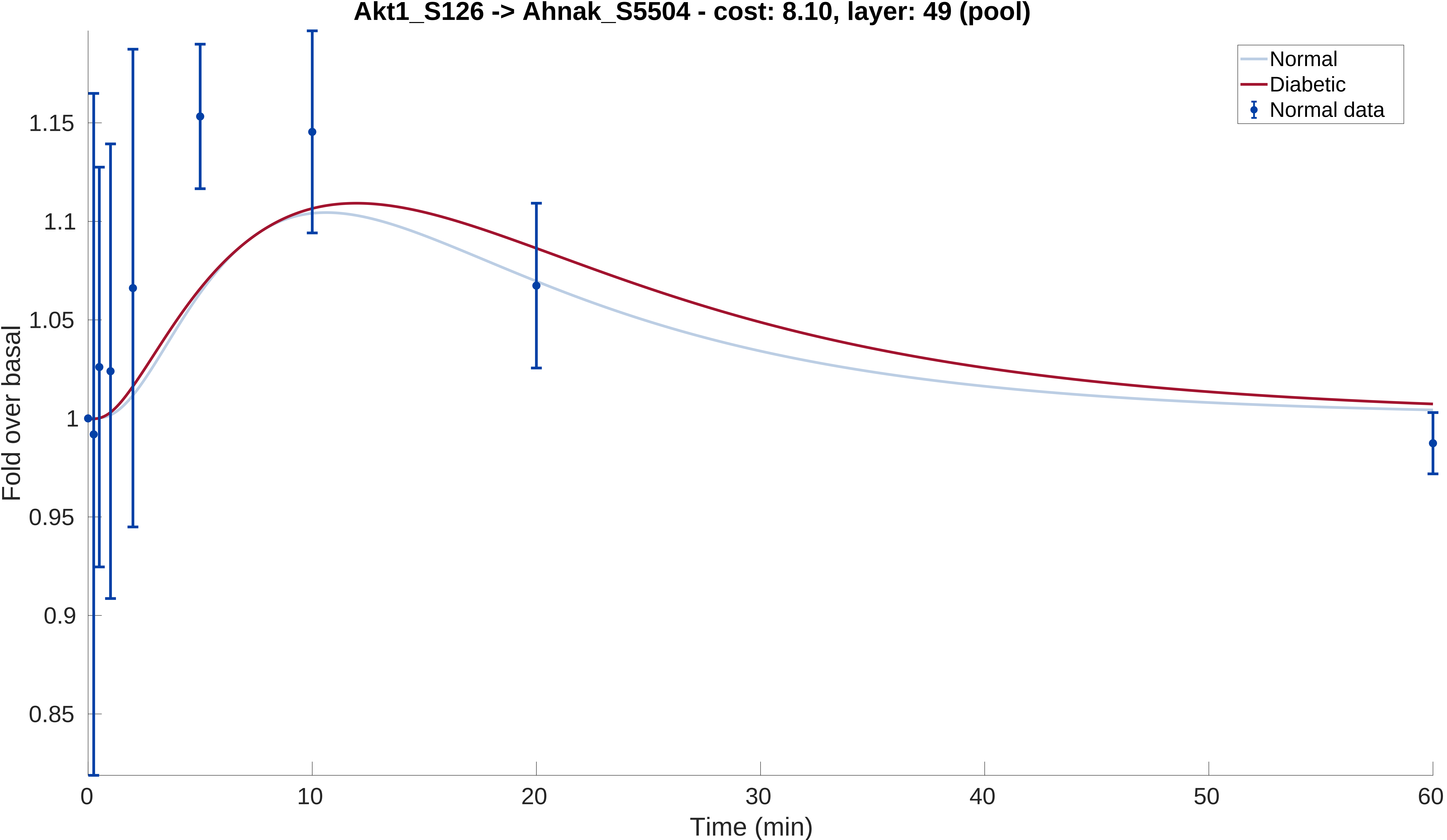

Akt1_S126 -> Ahnak_S5504 - cost: 8.10, layer: 49 (pool)
Normal
Diabetic
Normal data
Fold over basal
1.15
1.1
1.05
1
0.95
0.9
0.85
0
10
20
30
40
50
60
Time (min)

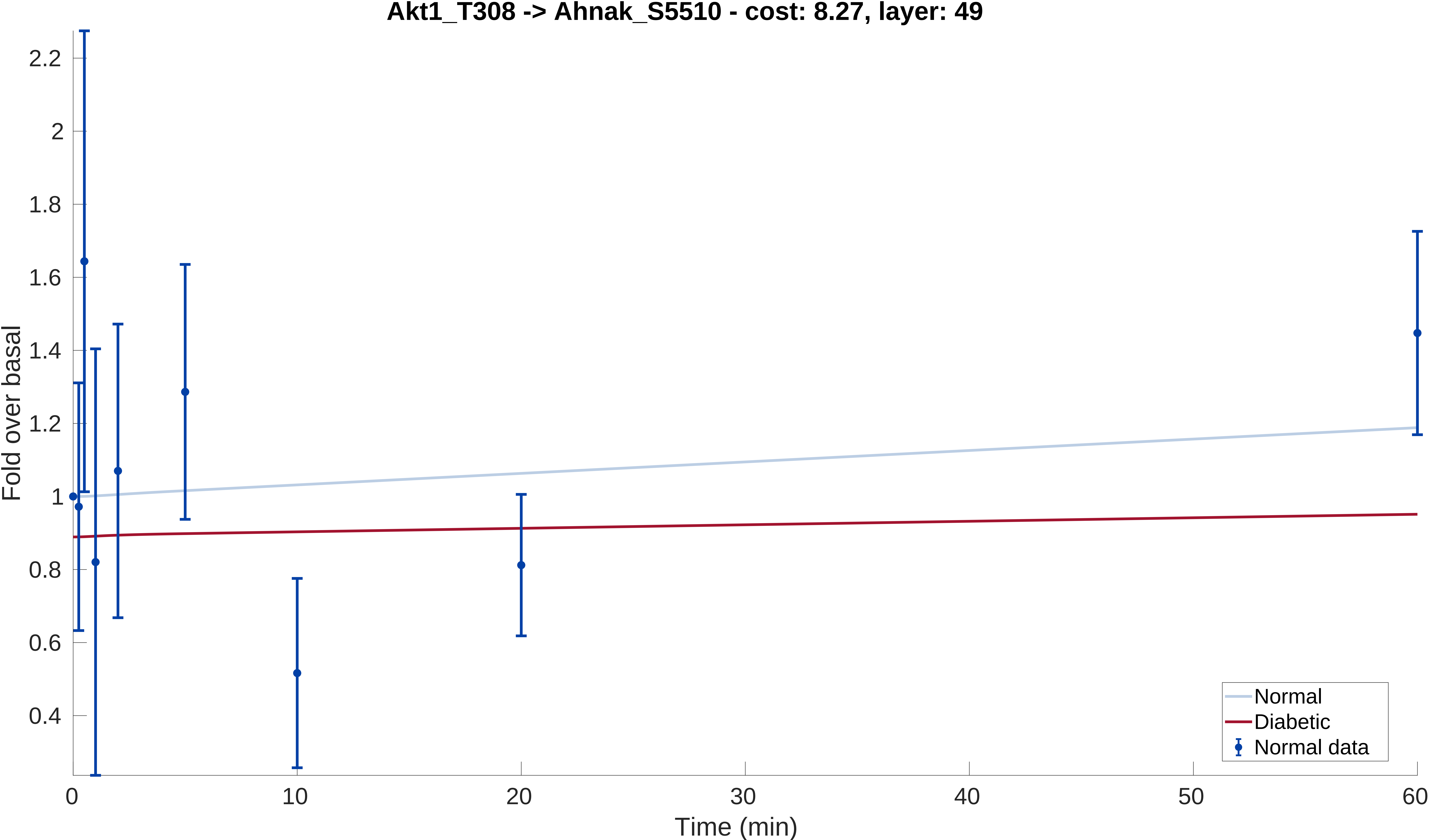

Akt1_T308 -> Ahnak_S5510 - cost: 8.27, layer: 49
Fold over basal
2.2
2
1.8
1.6
1.4
1.2
1
0.8
0.6
0.4
0
10
20
30
40
50
60
Time (min)
Normal
Diabetic
Normal data

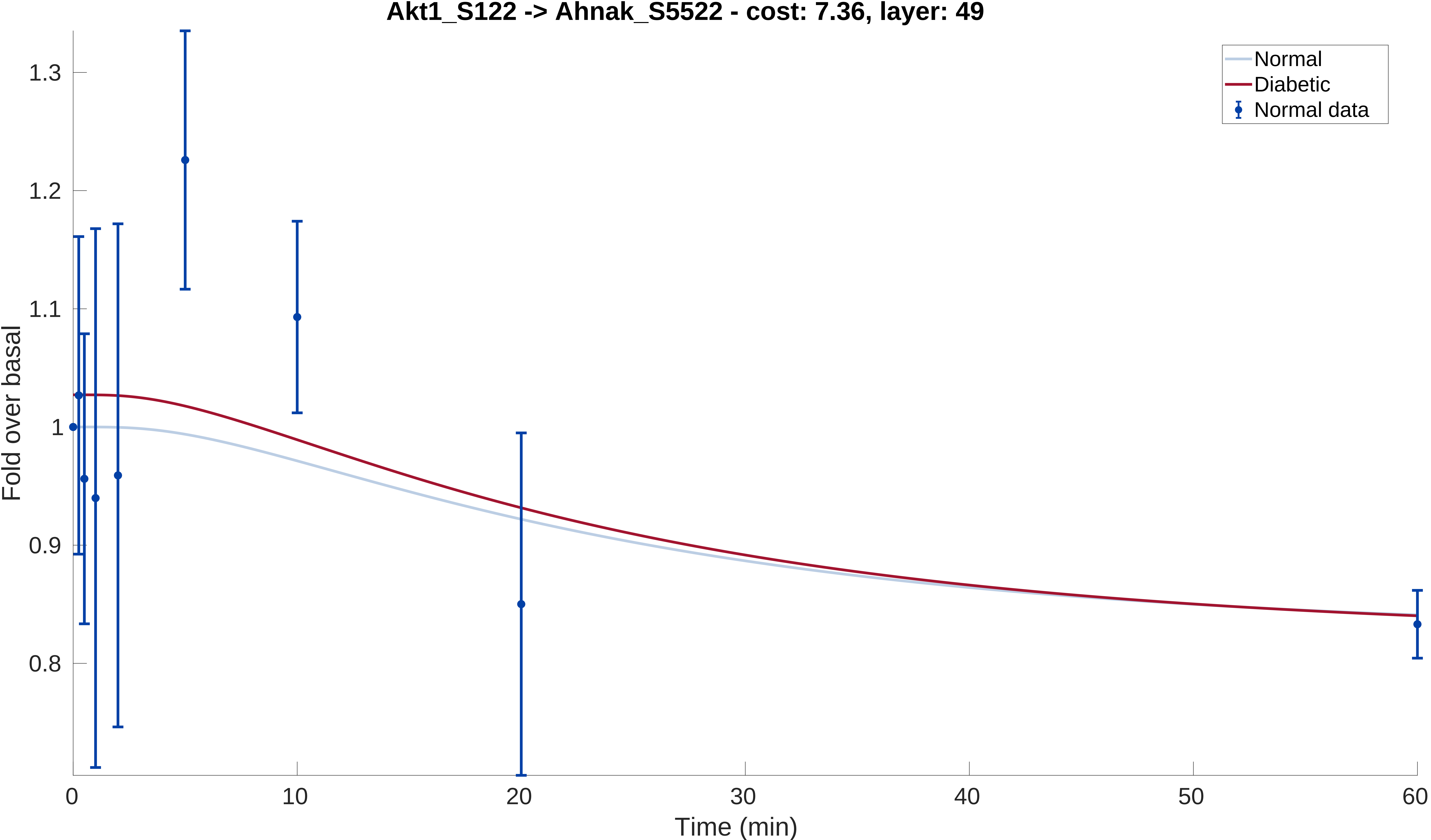

Akt1_S122 -> Ahnak_S5522 - cost: 7.36, layer: 49
Normal
Diabetic
Normal data
Fold over basal
1.3
1.2
1.1
1
0.9
0.8
0
10
20
30
40
50
60
Time (min)

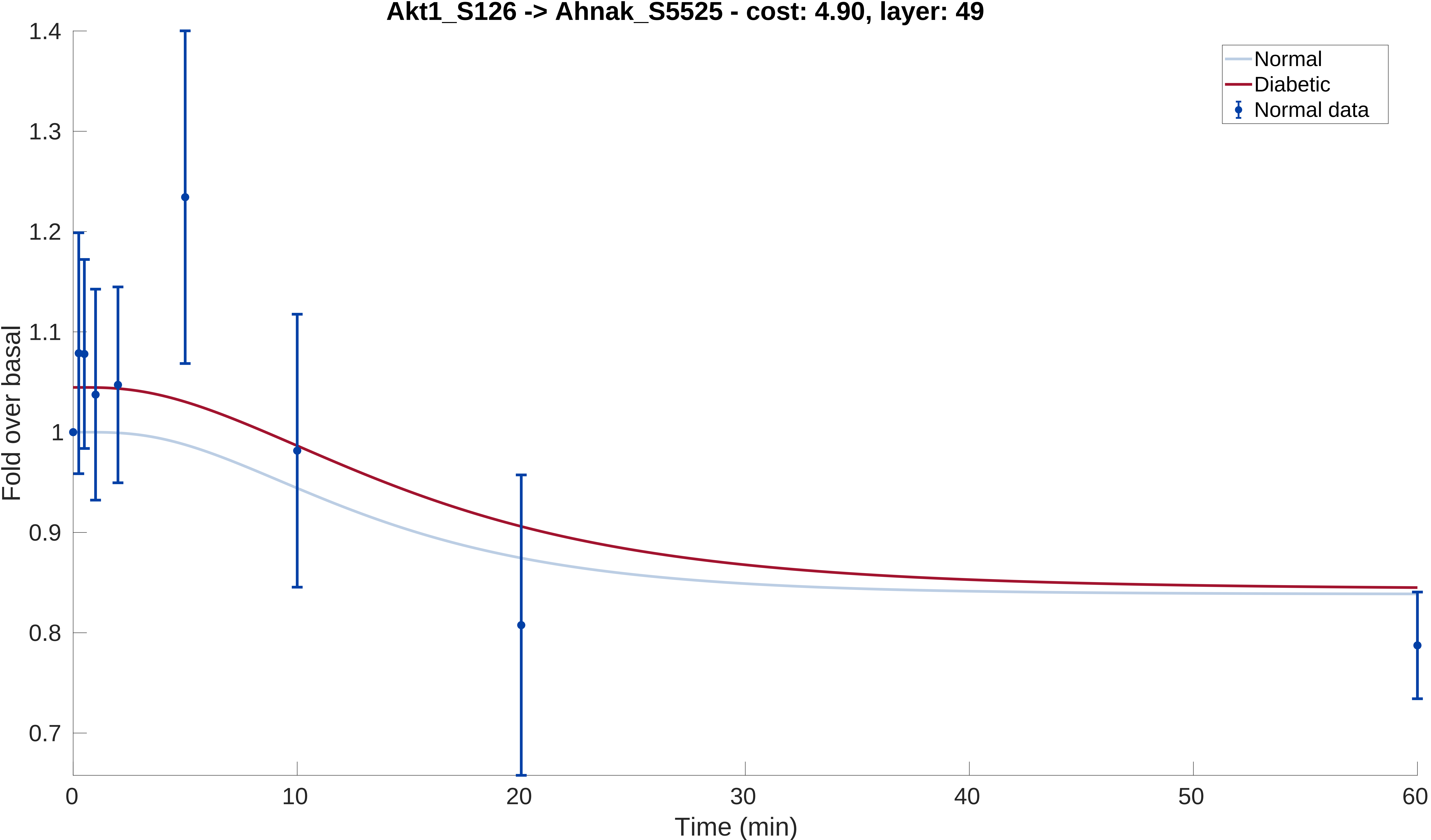

Akt1_S126 -> Ahnak_S5525 - cost: 4.90, layer: 49
Normal
Diabetic
Normal data
Fold over basal
1.4
1.3
1.2
1.1
1
0.9
0.8
0.7
0
10
20
30
40
50
60
Time (min)

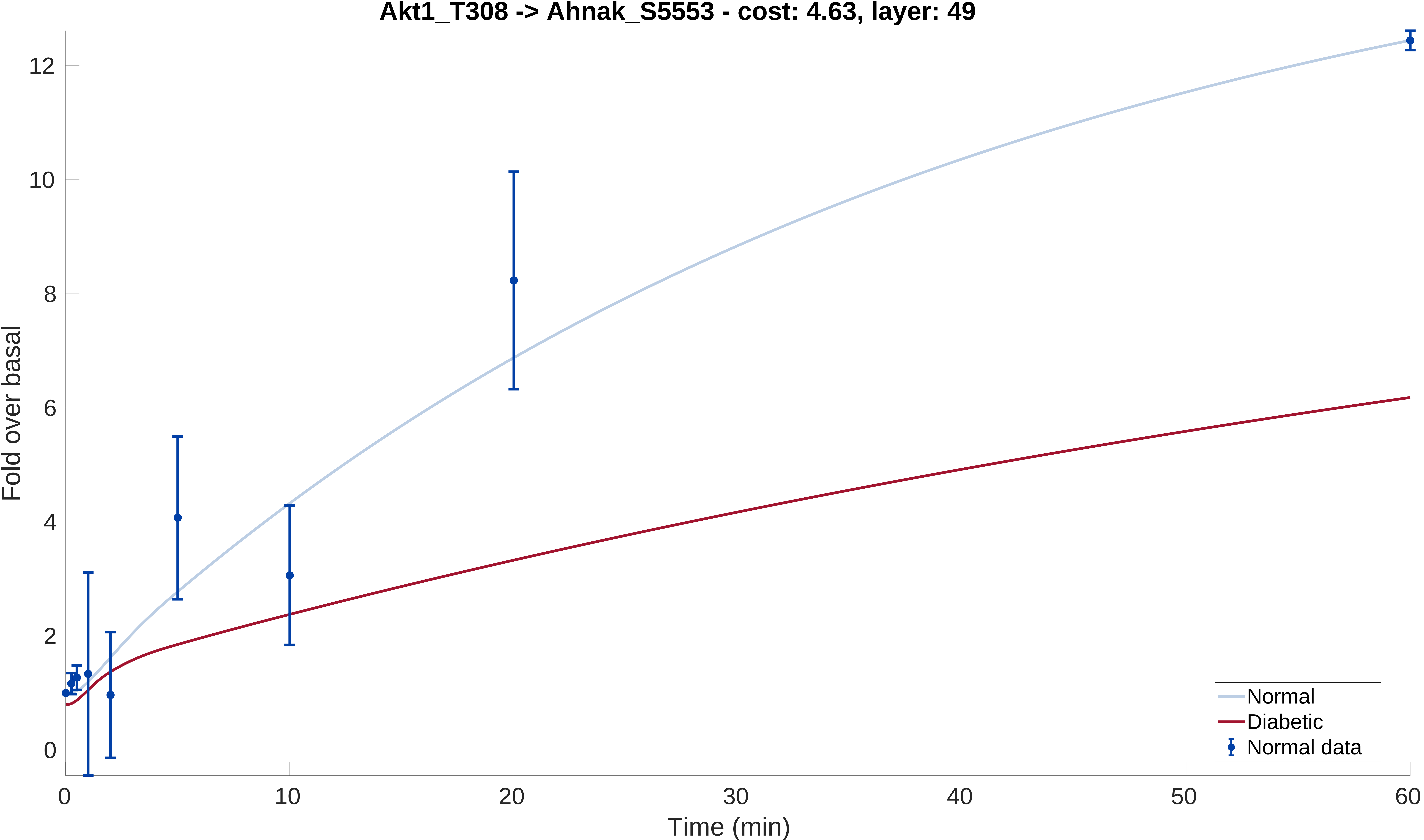

Akt1_T308 -> Ahnak_S5553 - cost: 4.63, layer: 49
Fold over basal
Time (min)
0
2
4
6
8
10
12
0
10
20
30
40
50
60
Normal
Diabetic
Normal data

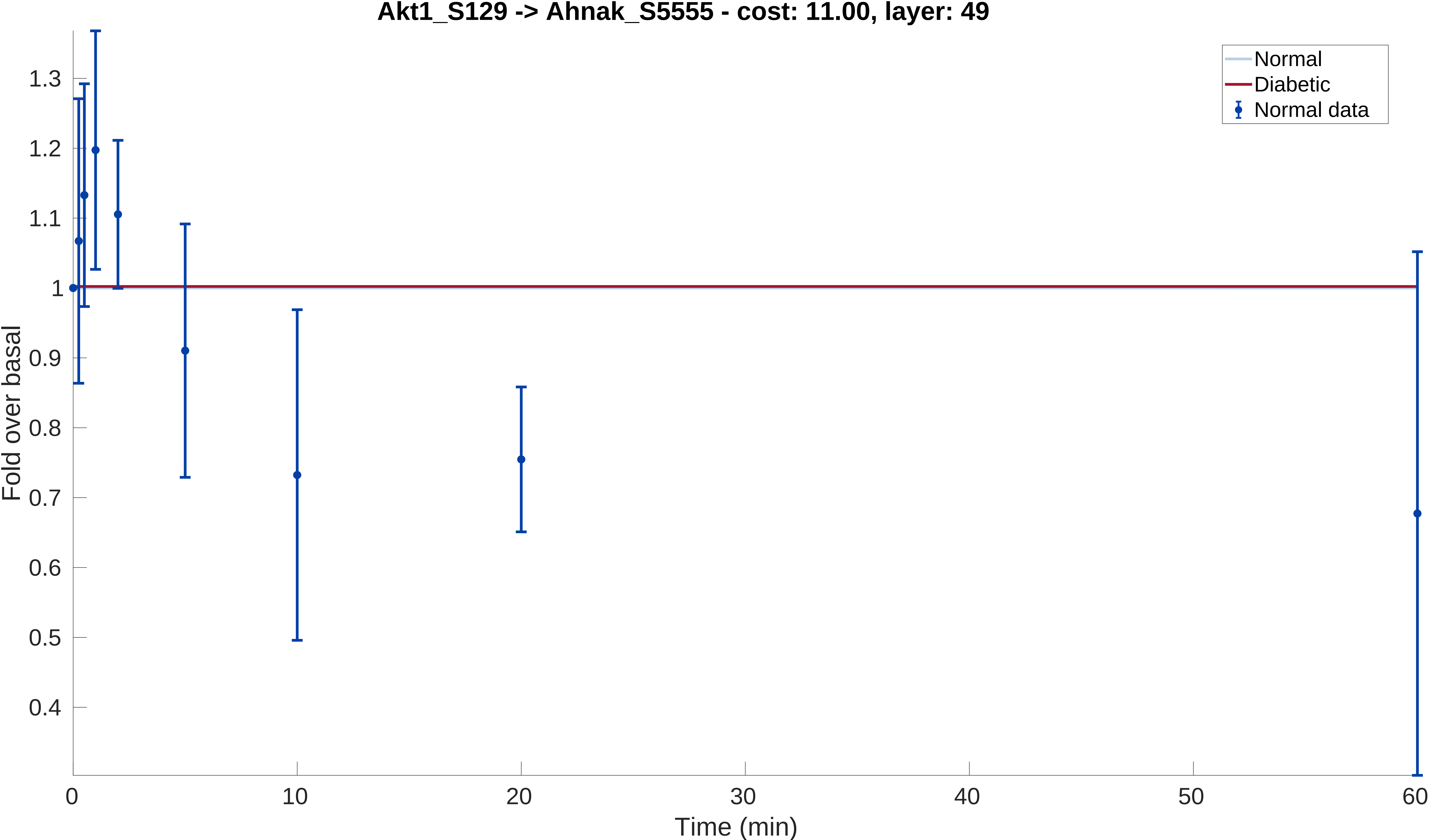

Akt1_S129 -> Ahnak_S5555 - cost: 11.00, layer: 49
Normal
Diabetic
Normal data
Fold over basal
1.3
1.2
1.1
1
0.9
0.8
0.7
0.6
0.5
0.4
0
10
20
30
40
50
60
Time (min)

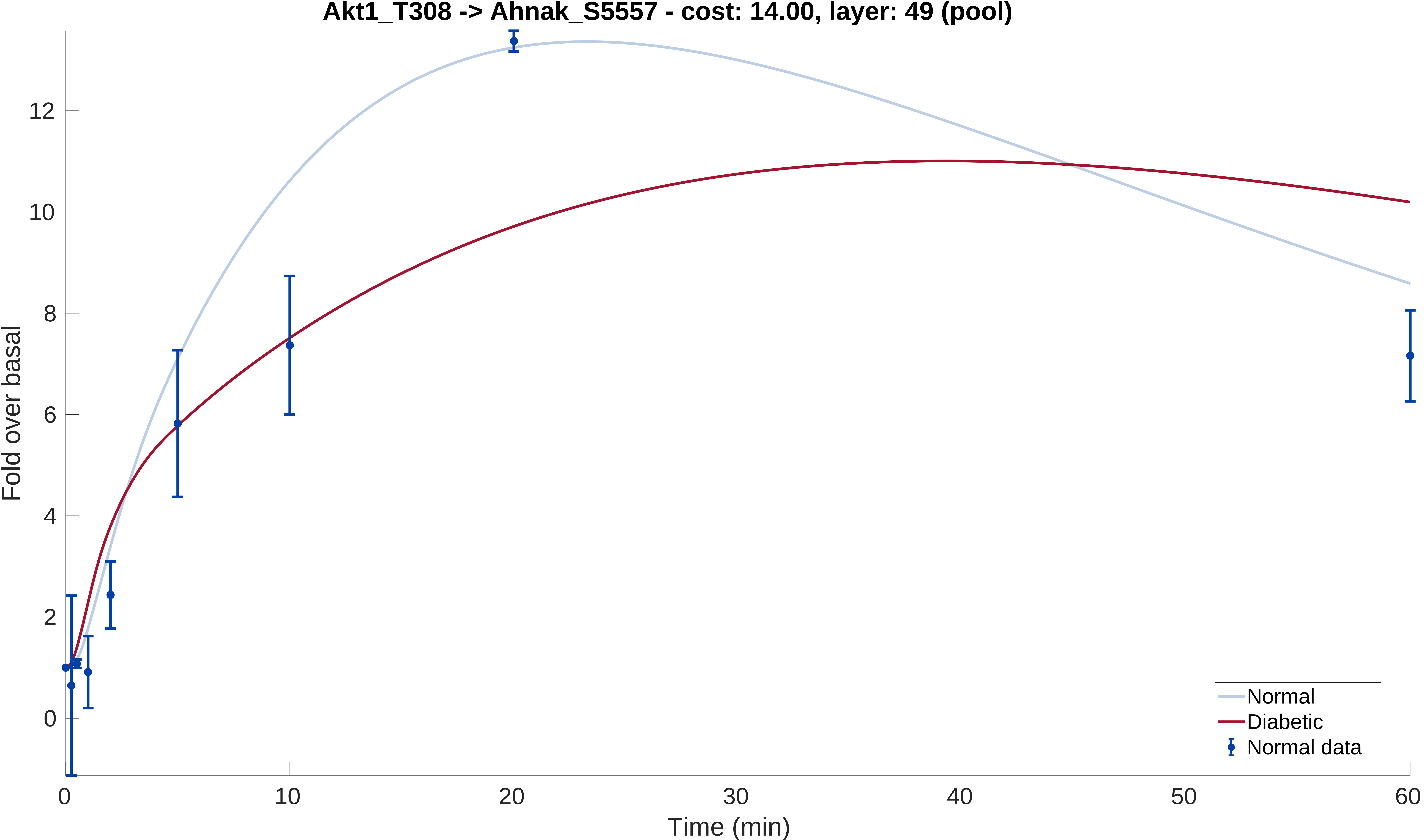
Akt1_T308 -> Ahnak_S5557 - cost: 14.00, layer: 49 (pool)
Fold over basal
Time (min)
0
2
4
6
8
10
12
0
10
20
30
40
50
60
Normal
Diabetic
Normal data

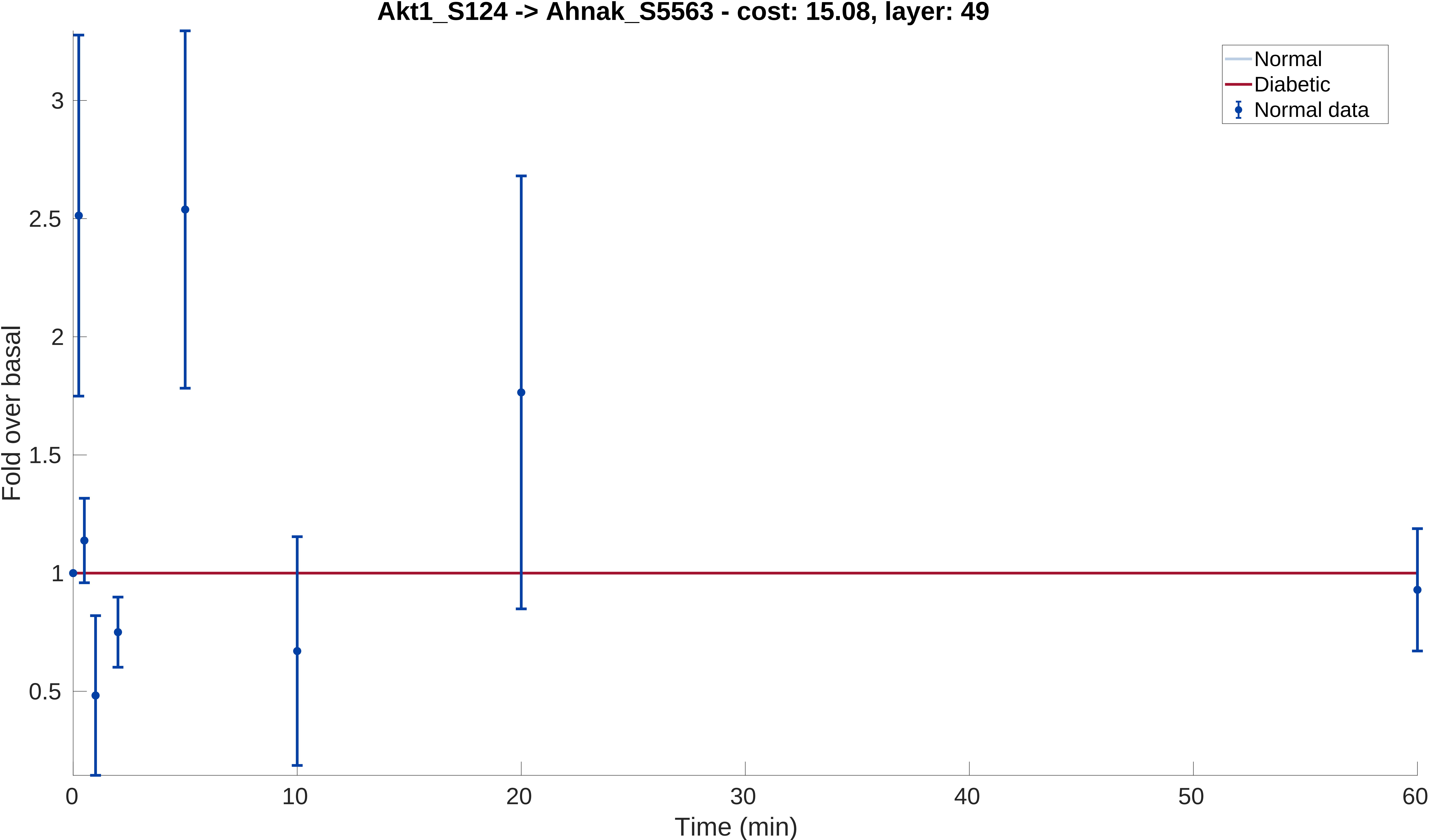

Akt1_S124 -> Ahnak_S5563 - cost: 15.08, layer: 49
Normal
Diabetic
Normal data
Fold over basal
3
2.5
2
1.5
1
0.5
0
10
20
30
40
50
60
Time (min)

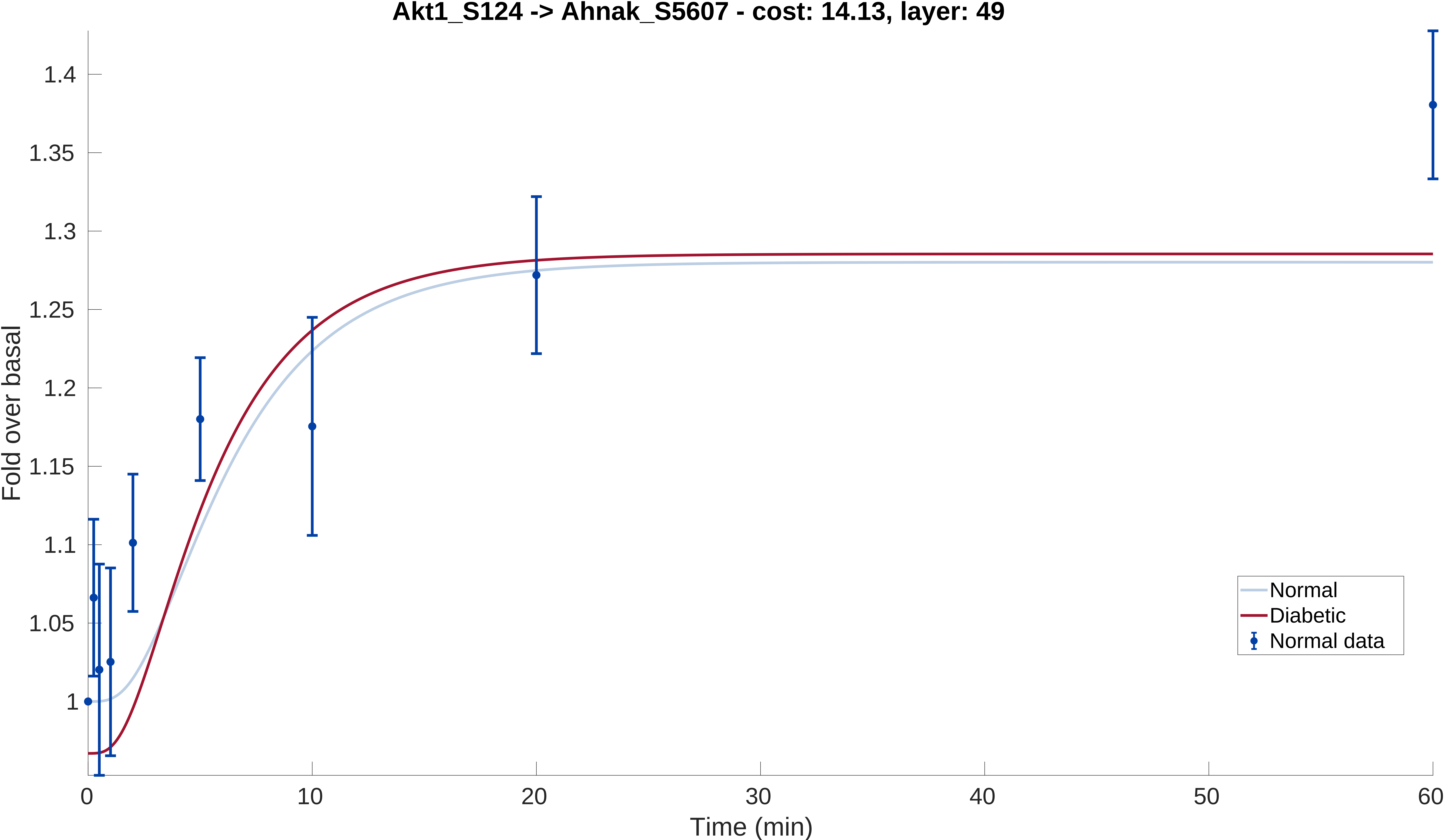
Akt1_S124 -> Ahnak_S5607 - cost: 14.13, layer: 49
Fold over basal
1.4
1.35
1.3
1.25
1.2
1.15
1.1
1.05
1
0
10
20
30
40
50
60
Time (min)
Normal
Diabetic
Normal data

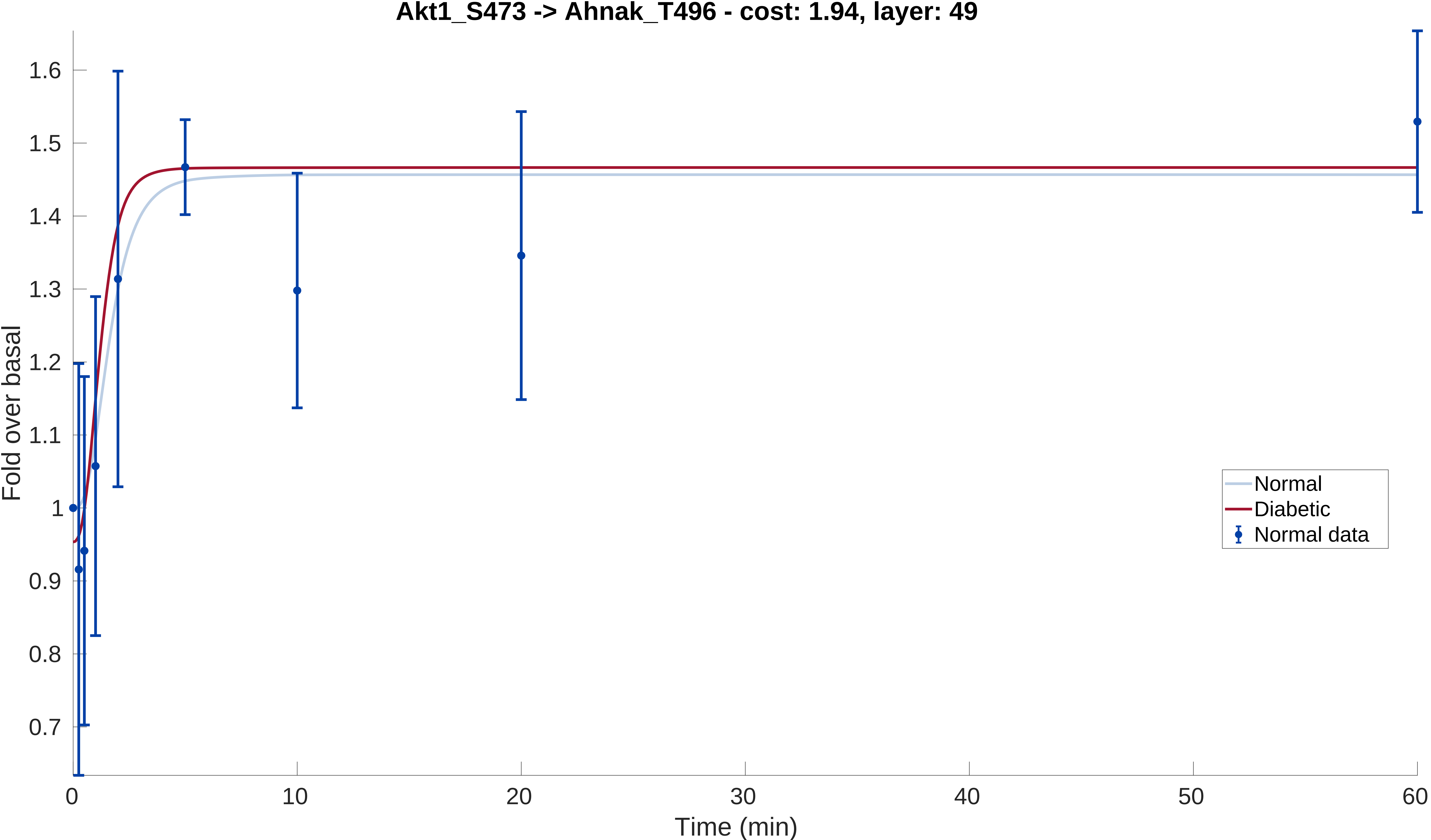

Akt1_S473 -> Ahnak_T496 - cost: 1.94, layer: 49
Fold over basal
Time (min)
0.7
0.8
0.9
1
1.1
1.2
1.3
1.4
1.5
1.6
0
10
20
30
40
50
60
Normal
Diabetic
Normal data

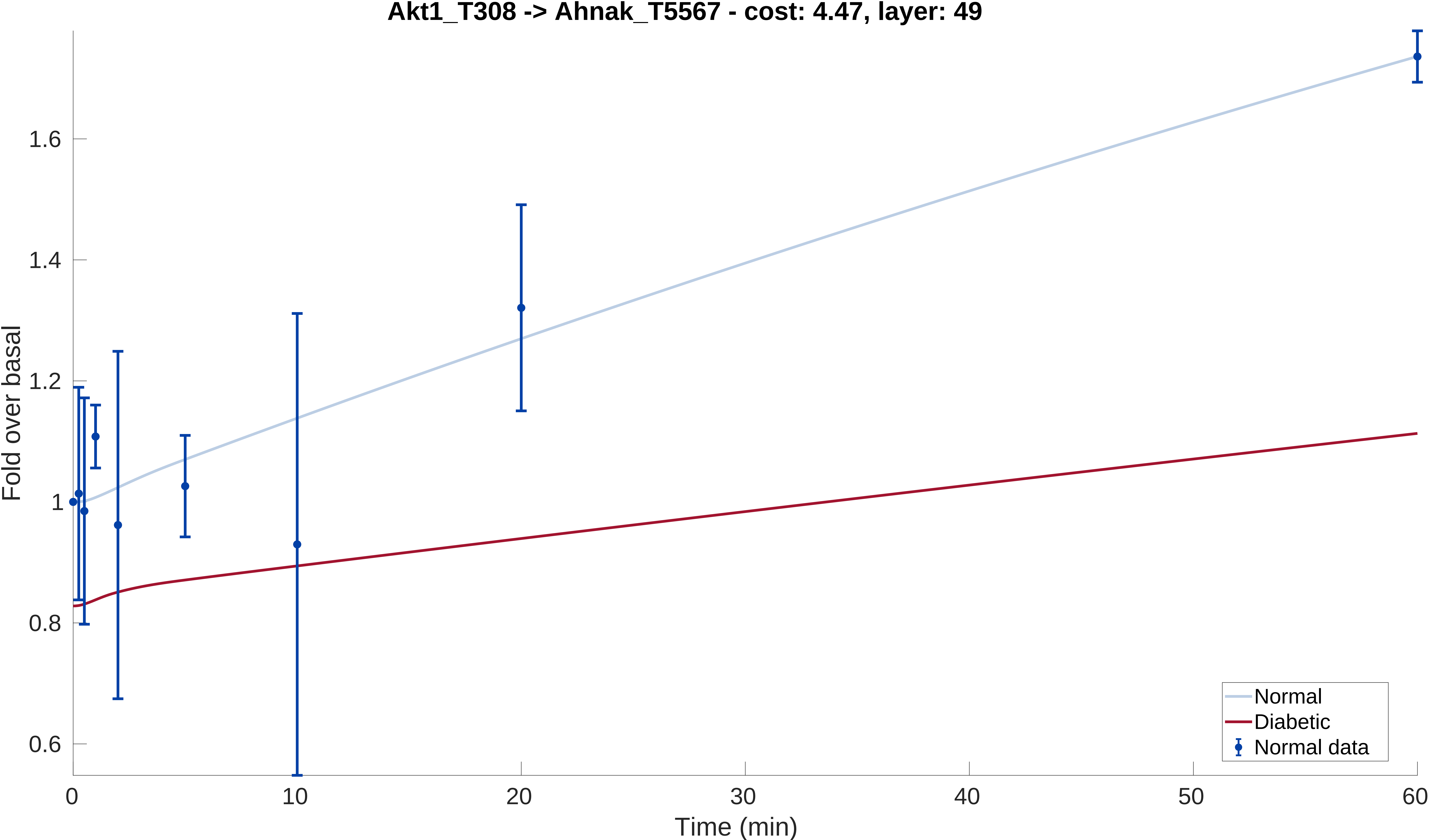

Akt1_T308 -> Ahnak_T5567 - cost: 4.47, layer: 49
Fold over basal
1.6
1.4
1.2
1
0.8
0.6
0
10
20
30
40
50
60
Time (min)
Normal
Diabetic
Normal data

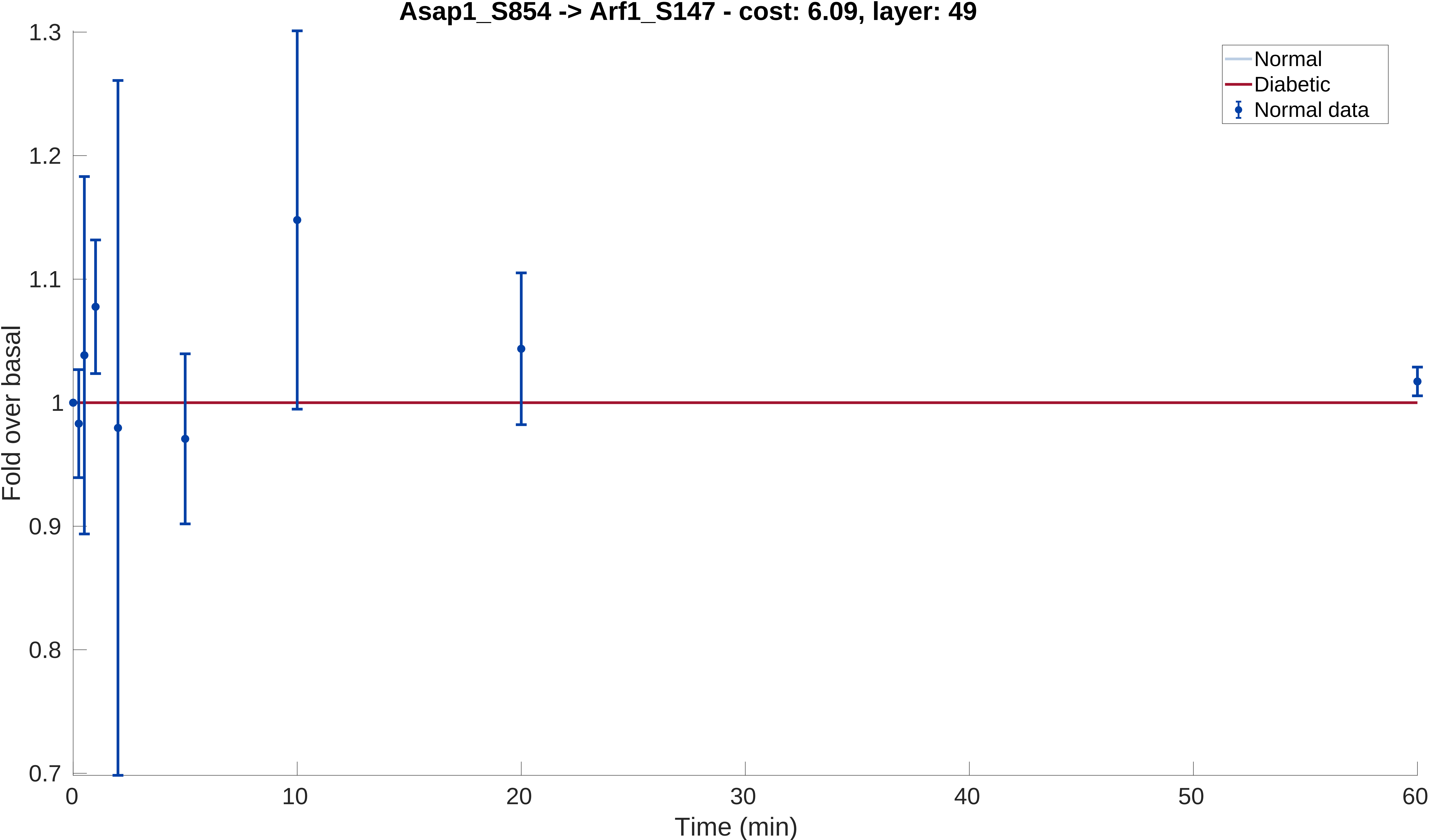

Asap1_S854 -> Arf1_S147 - cost: 6.09, layer: 49
Normal
Diabetic
Normal data
Fold over basal
Time (min)
1.3
1.2
1.1
1
0.9
0.8
0.7
0
10
20
30
40
50
60

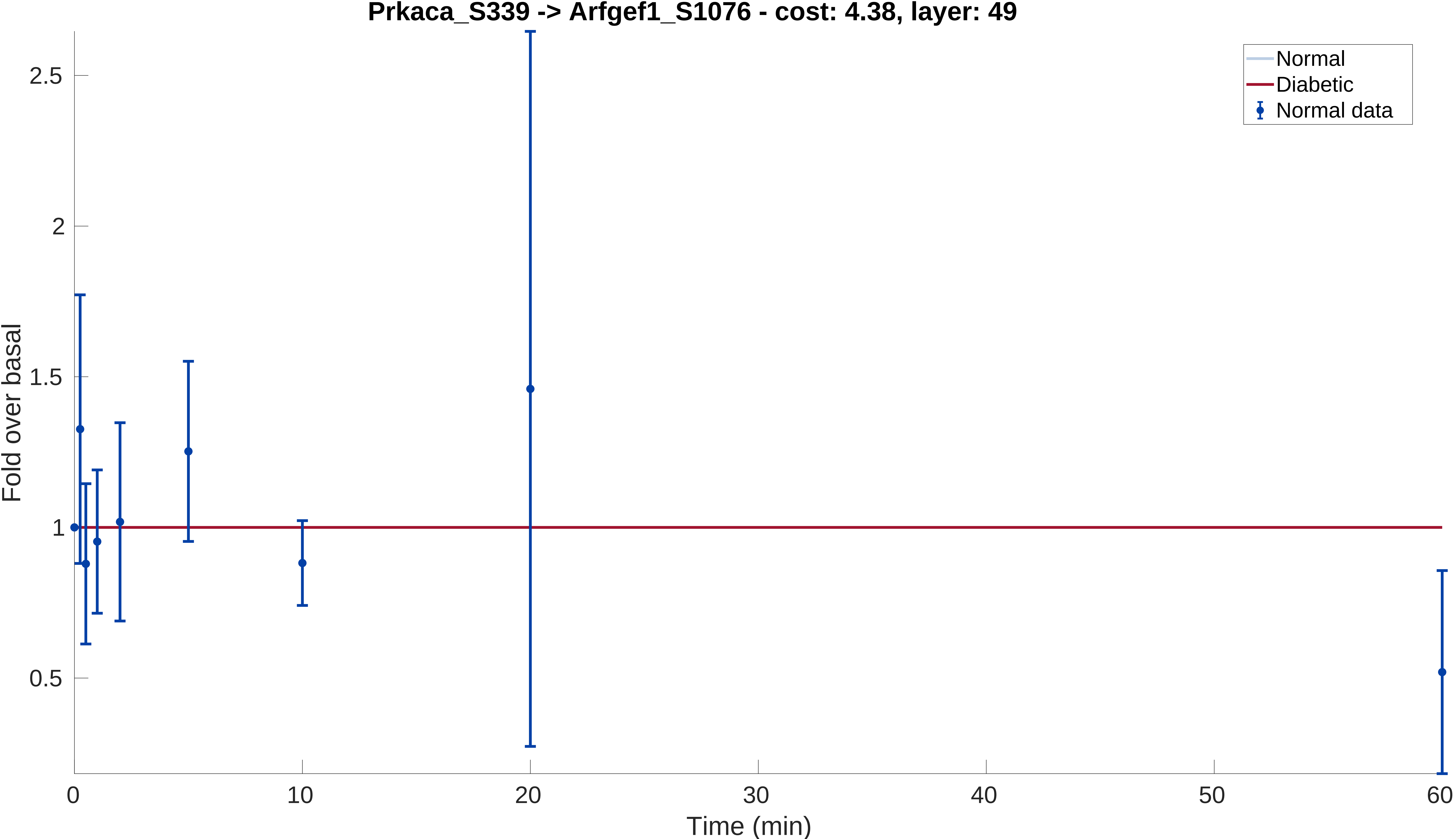
Prkaca_S339 -> Arfgef1_S1076 - cost: 4.38, layer: 49
Normal
Diabetic
Normal data
Fold over basal
2.5
2
1.5
1
0.5
0
10
20
30
40
50
60
Time (min)

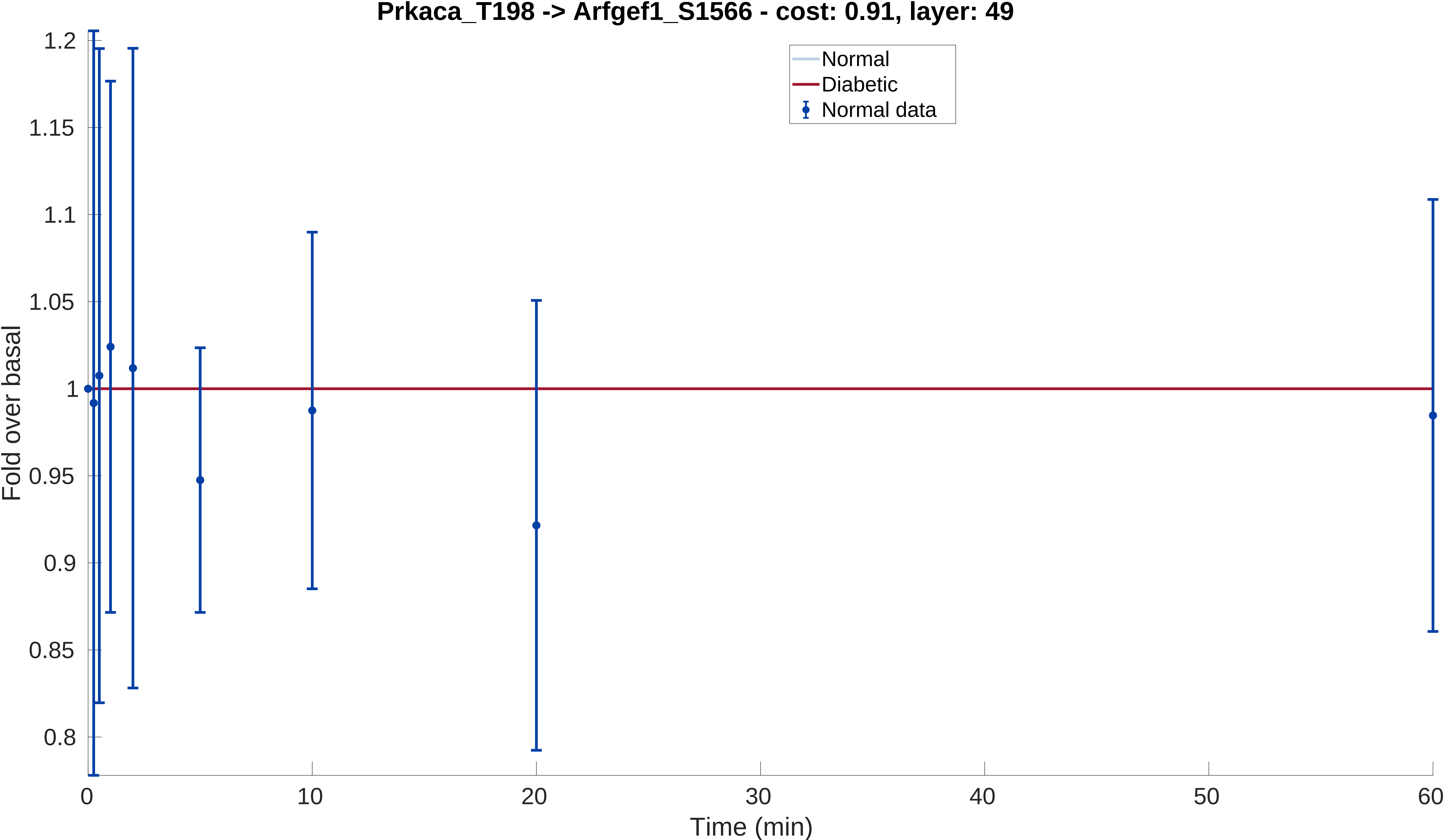

Prkaca_T198 -> Arfgef1_S1566 - cost: 0.91, layer: 49
Normal
Diabetic
Normal data
Fold over basal
Time (min)
1.2
1.15
1.1
1.05
1
0.95
0.9
0.85
0.8
0
10
20
30
40
50
60

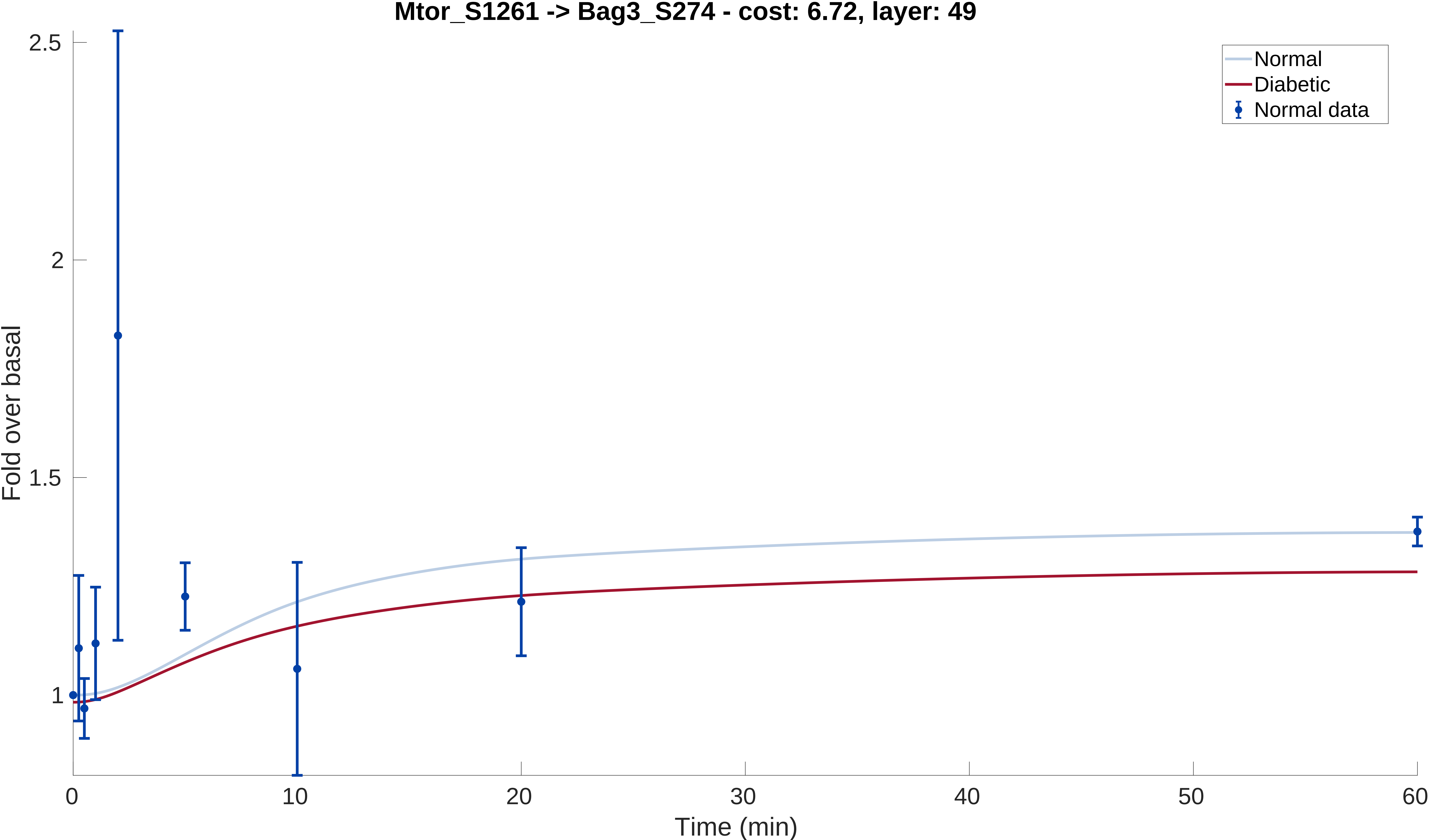

Mtor_S1261 -> Bag3_S274 - cost: 6.72, layer: 49
Normal
Diabetic
Normal data
Fold over basal
2.5
2
1.5
1
0
10
20
30
40
50
60
Time (min)

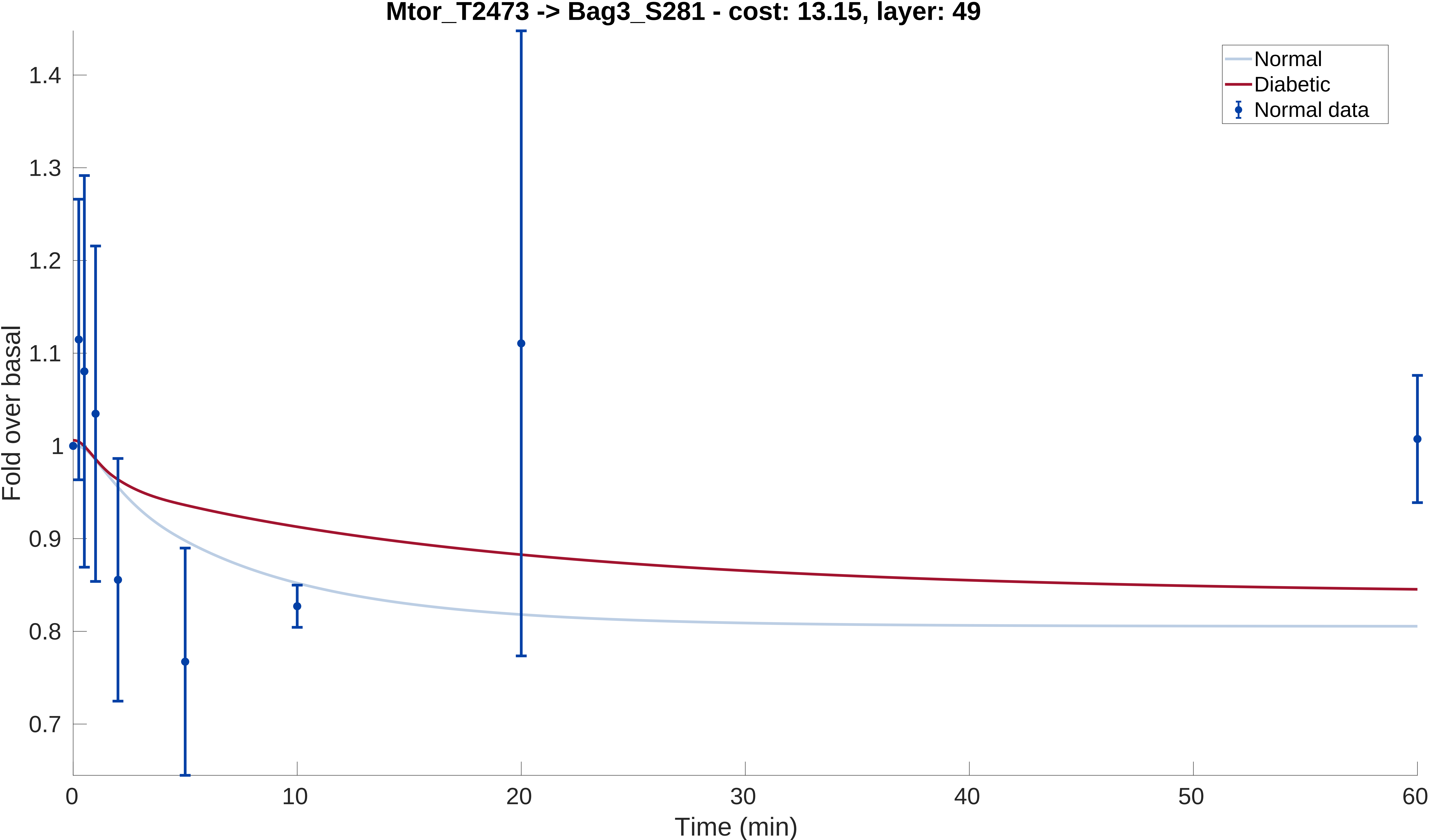
Mtor_T2473 -> Bag3_S281 - cost: 13.15, layer: 49
Normal
Diabetic
Normal data
Fold over basal
1.4
1.3
1.2
1.1
1
0.9
0.8
0.7
0
10
20
30
40
50
60
Time (min)

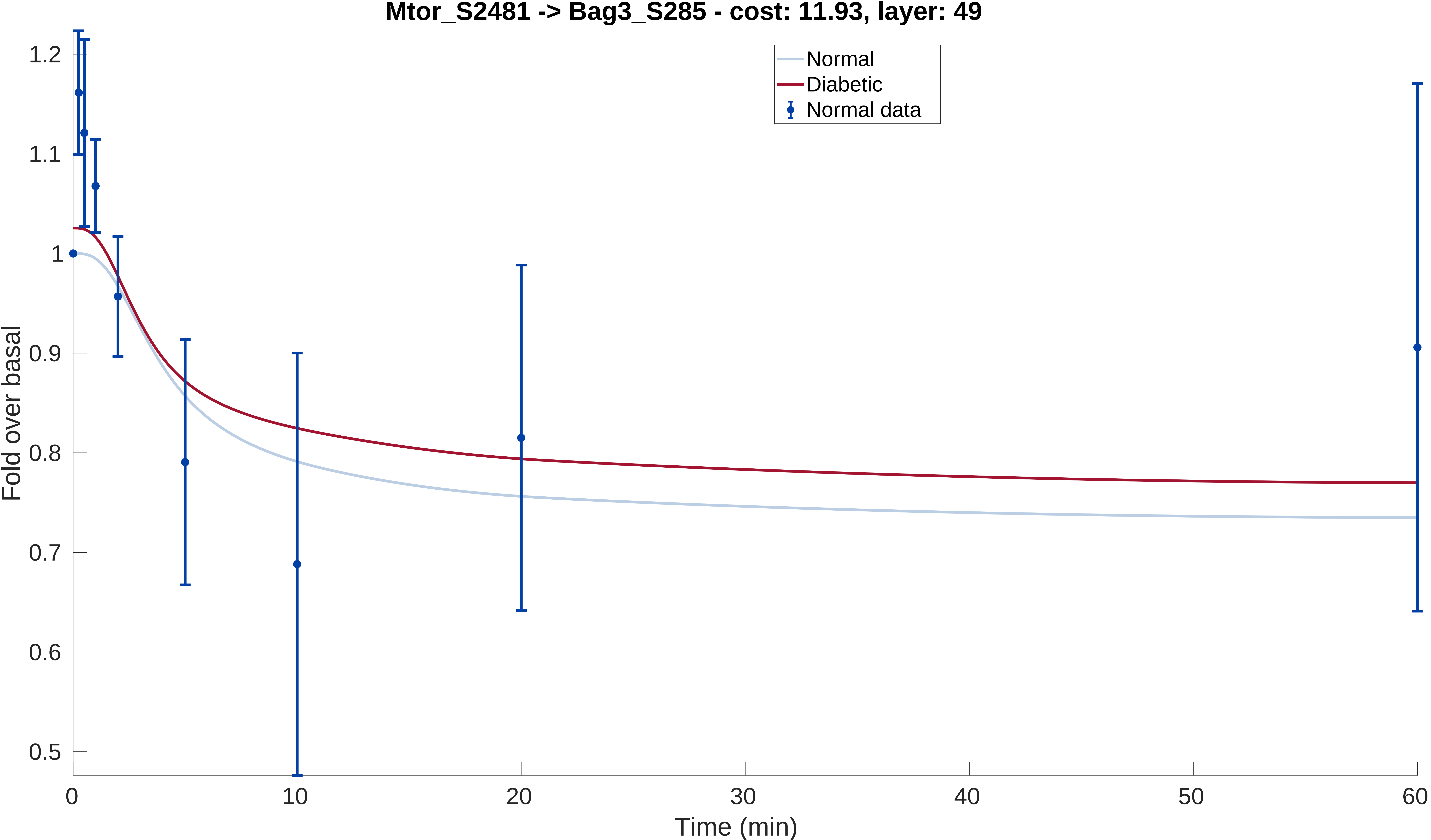
Mtor_S2481 -> Bag3_S285 - cost: 11.93, layer: 49
Normal
Diabetic
Normal data
Fold over basal
Time (min)
1.2
1.1
1
0.9
0.8
0.7
0.6
0.5
0
10
20
30
40
50
60

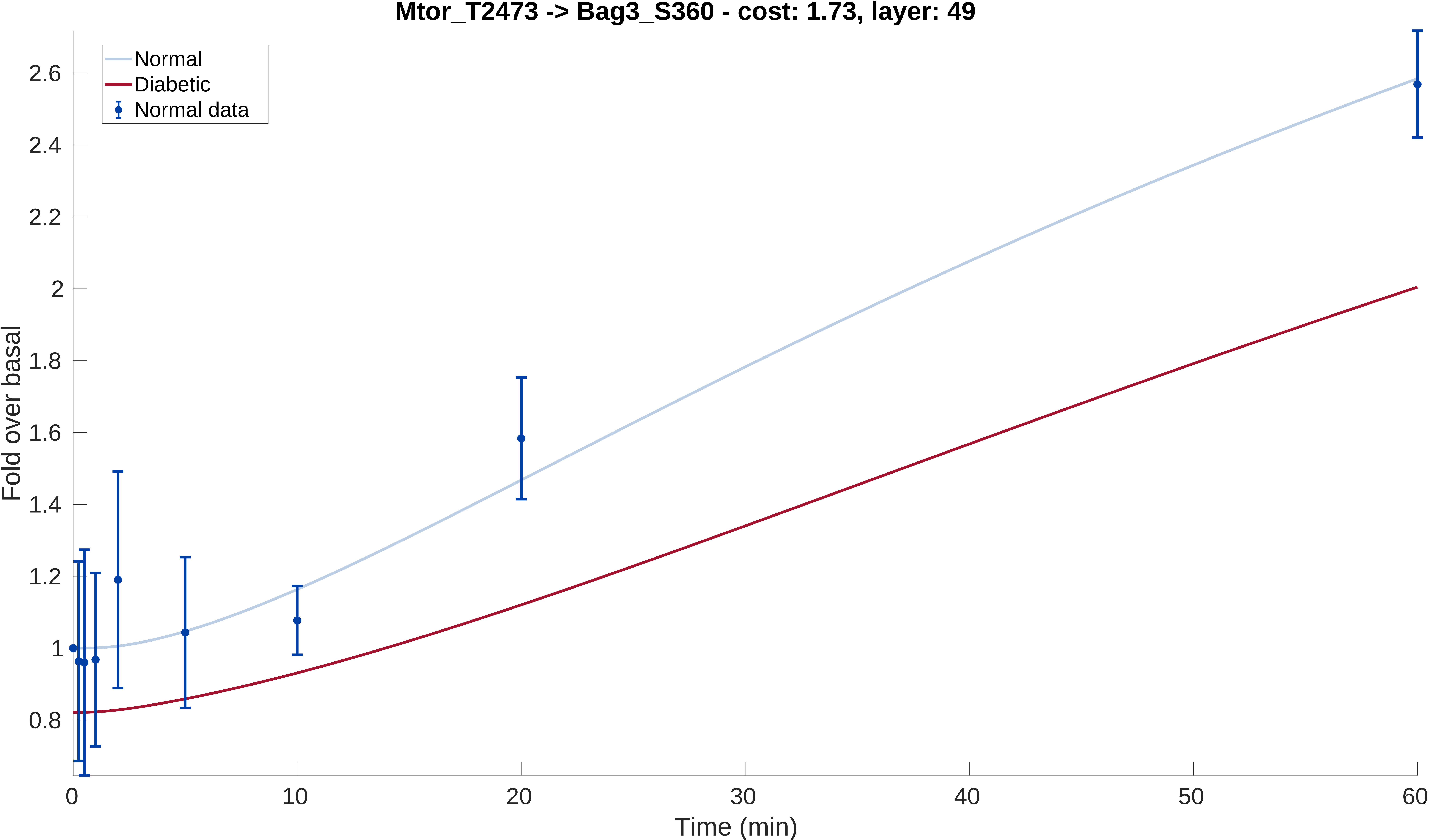

Mtor_T2473 -> Bag3_S360 - cost: 1.73, layer: 49
Normal
Diabetic
Normal data
Fold over basal
2.6
2.4
2.2
2
1.8
1.6
1.4
1.2
1
0.8
0
10
20
30
40
50
60
Time (min)

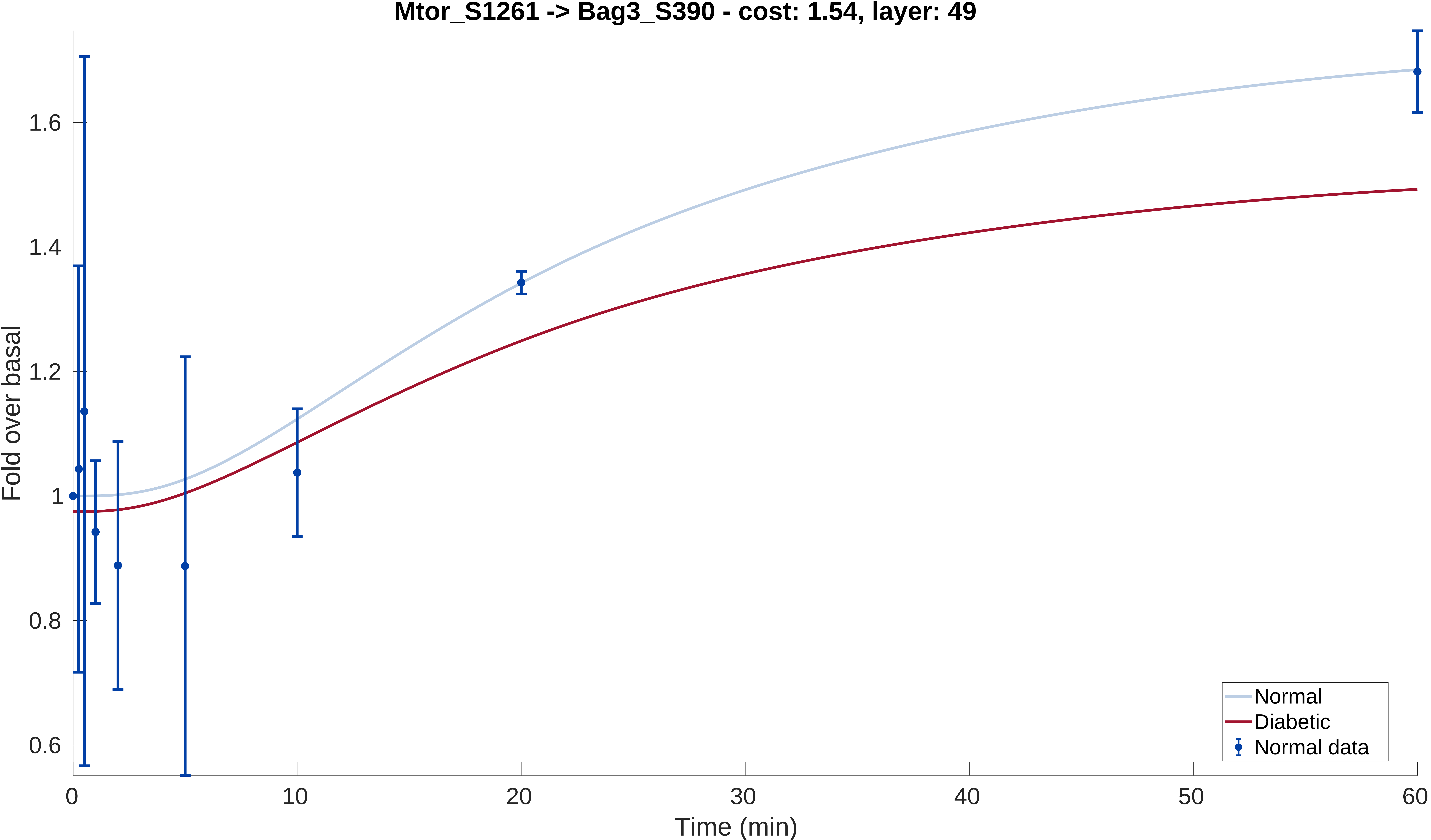
Mtor_S1261 -> Bag3_S390 - cost: 1.54, layer: 49
Fold over basal
0.6
0.8
1
1.2
1.4
1.6
0
10
20
30
40
50
60
Time (min)
Normal
Diabetic
Normal data

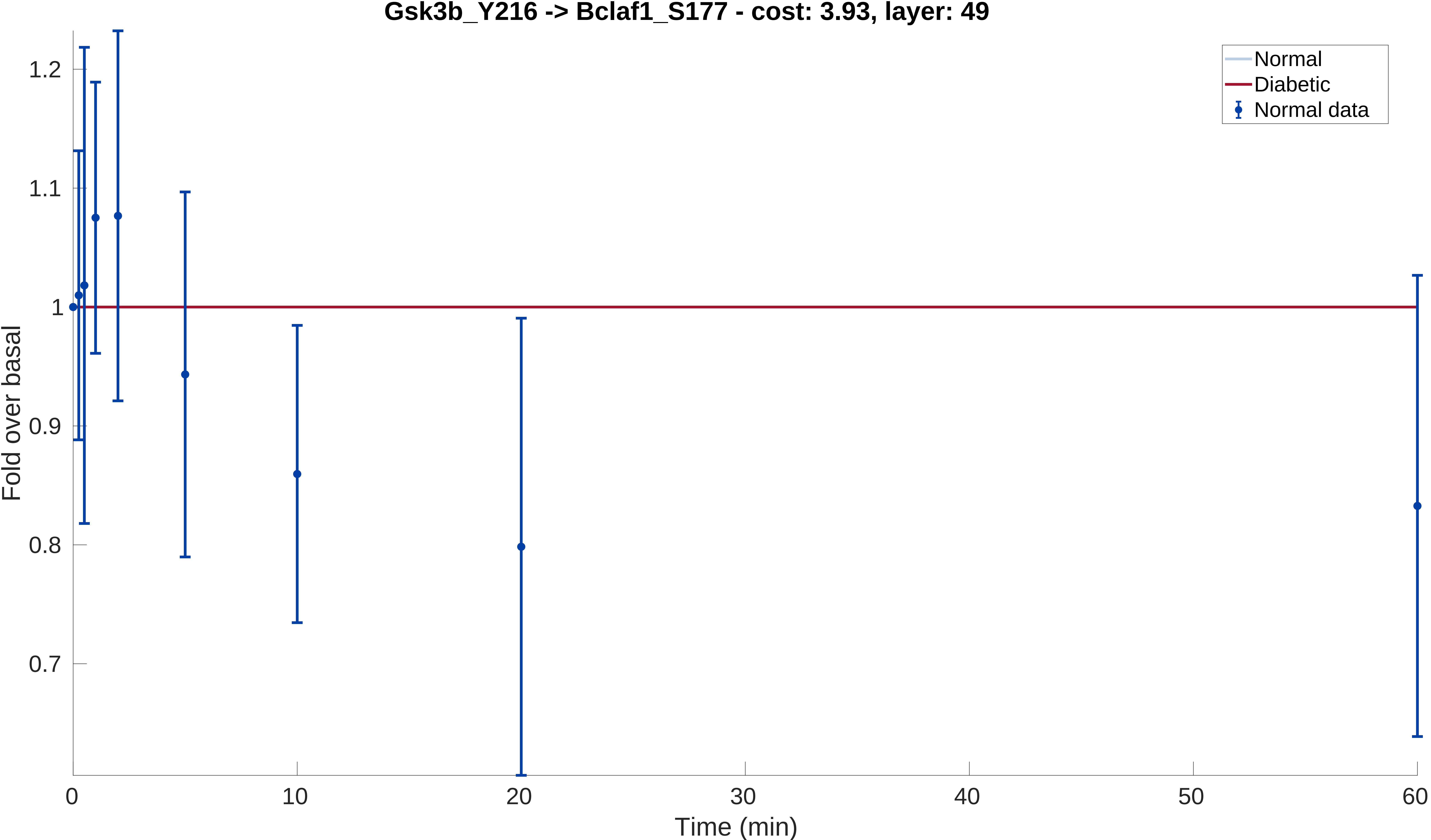

Gsk3b_Y216 -> Bclaf1_S177 - cost: 3.93, layer: 49
Normal
Diabetic
Normal data
Fold over basal
Time (min)
1.2
1.1
1
0.9
0.8
0.7
0
10
20
30
40
50
60

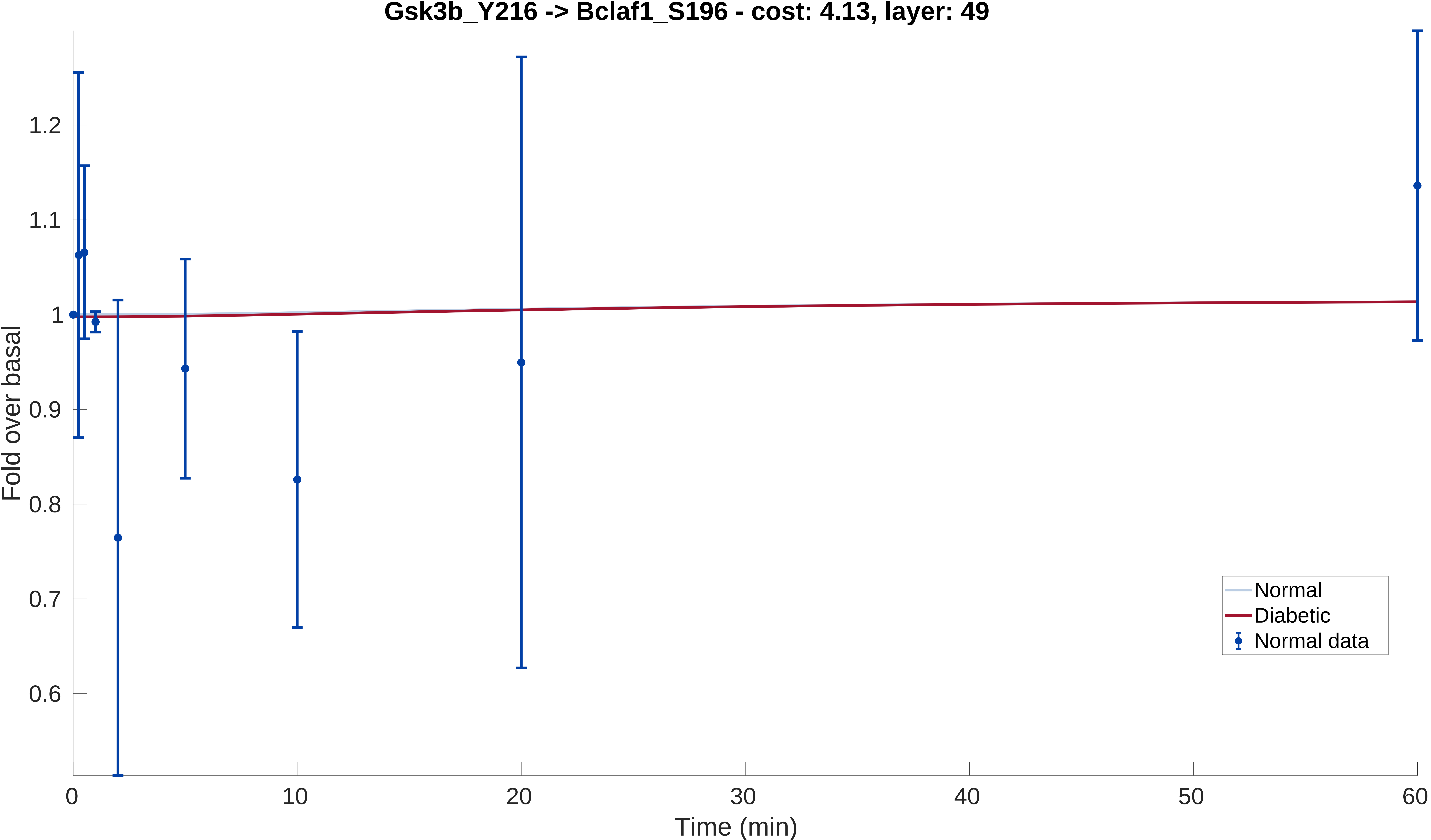

Gsk3b_Y216 -> Bclaf1_S196 - cost: 4.13, layer: 49
Fold over basal
1.2
1.1
1
0.9
0.8
0.7
0.6
0
10
20
30
40
50
60
Time (min)
Normal
Diabetic
Normal data

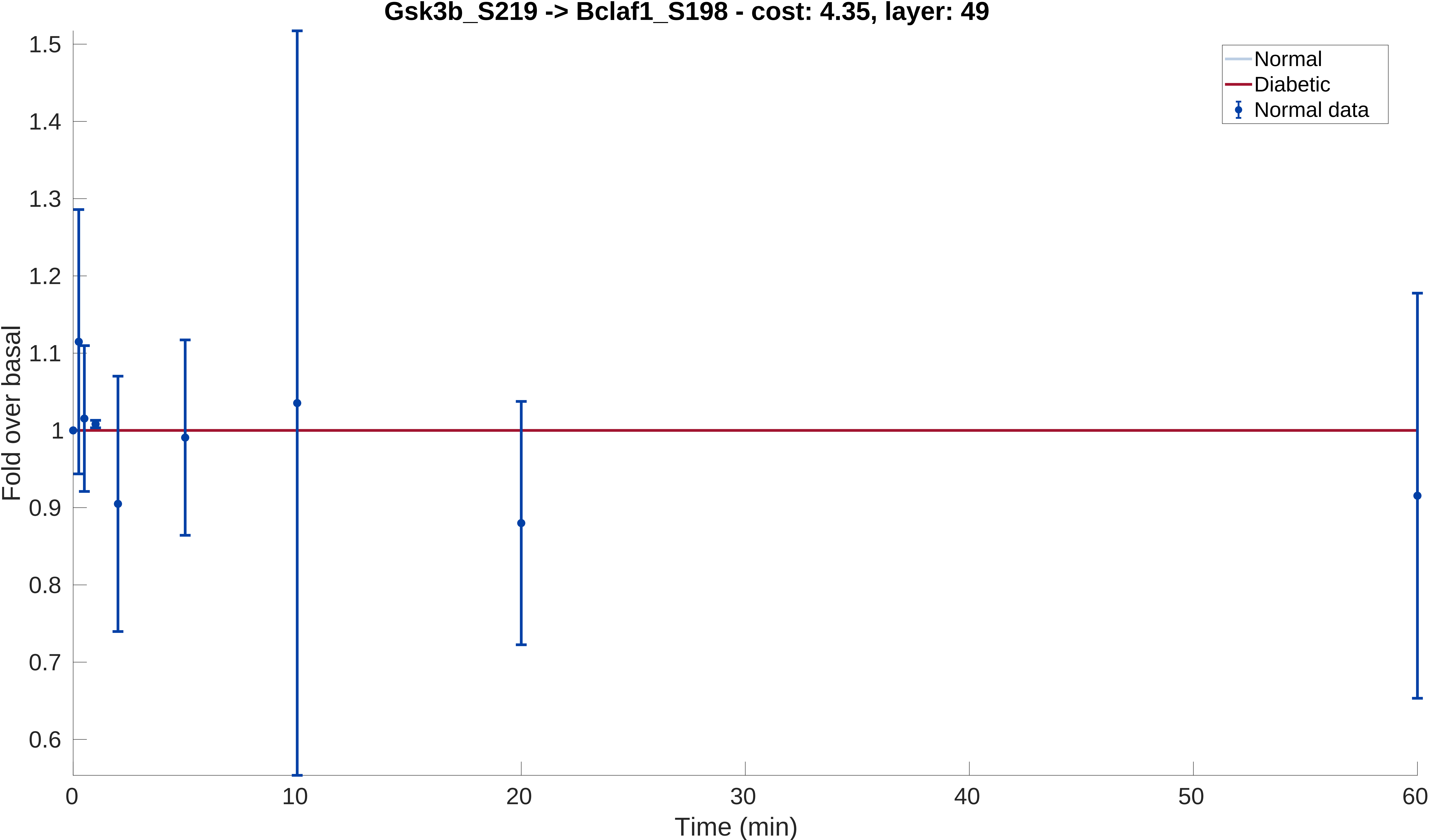

Gsk3b_S219 -> Bclaf1_S198 - cost: 4.35, layer: 49
Normal
Diabetic
Normal data
Fold over basal
Time (min)
1.5
1.4
1.3
1.2
1.1
1
0.9
0.8
0.7
0.6
0
10
20
30
40
50
60

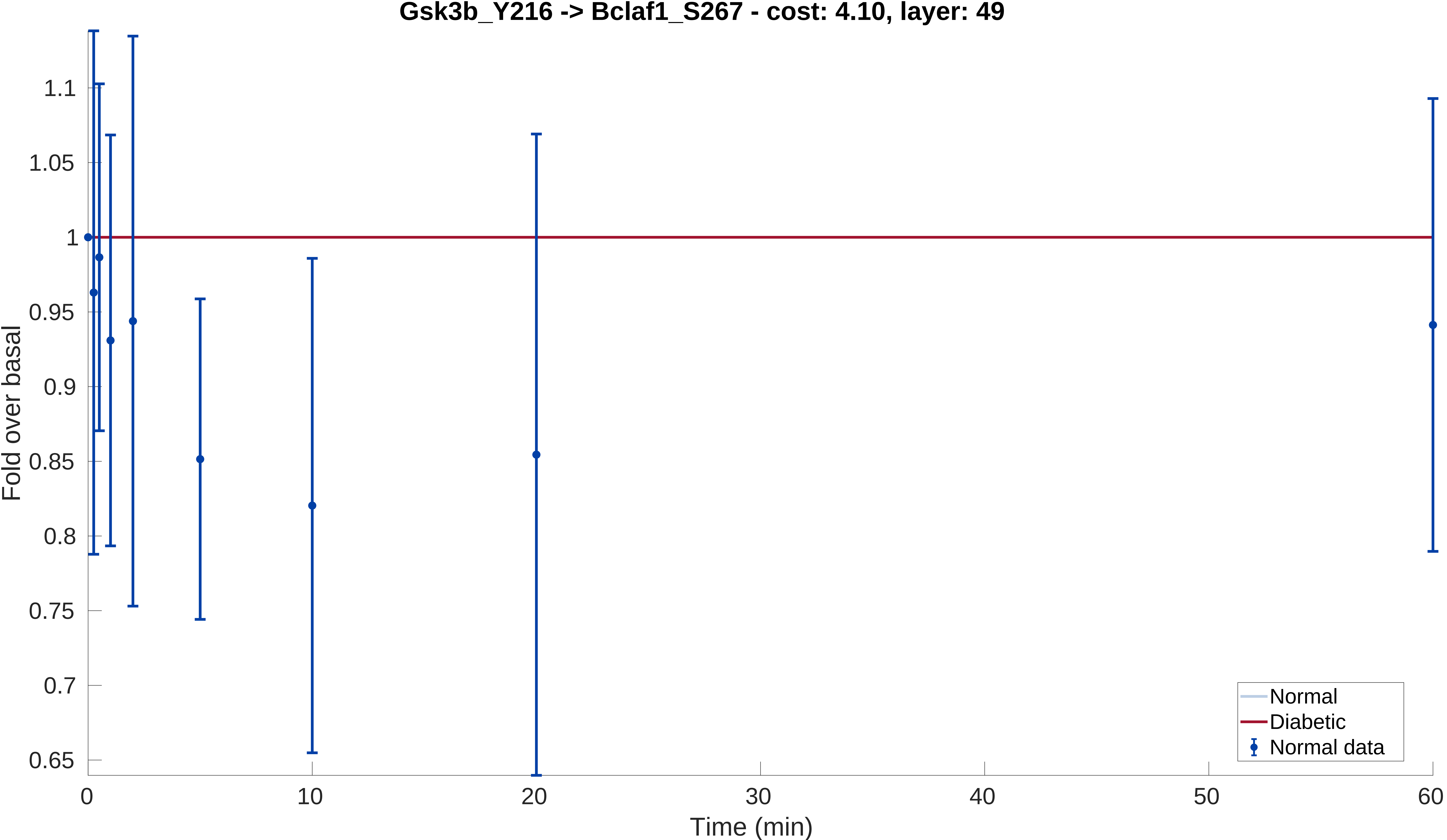

Gsk3b_Y216 -> Bclaf1_S267 - cost: 4.10, layer: 49
Fold over basal
1.1
1.05
1
0.95
0.9
0.85
0.8
0.75
0.7
0.65
0
10
20
30
40
50
60
Time (min)
Normal
Diabetic
Normal data

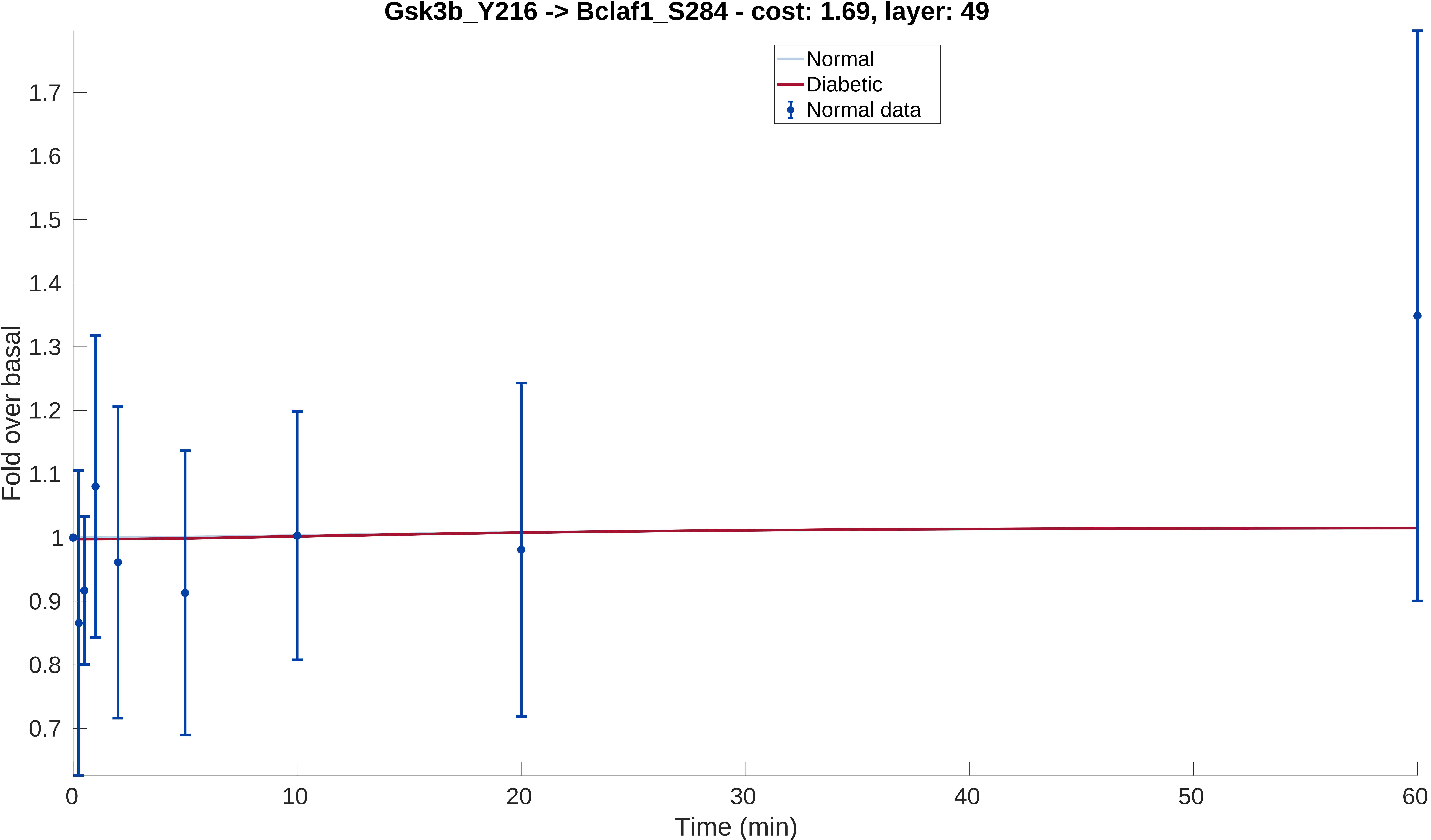
Gsk3b_Y216 -> Bclaf1_S284 - cost: 1.69, layer: 49
Normal
Diabetic
Normal data
Fold over basal
1.7
1.6
1.5
1.4
1.3
1.2
1.1
1
0.9
0.8
0.7
0
10
20
30
40
50
60
Time (min)

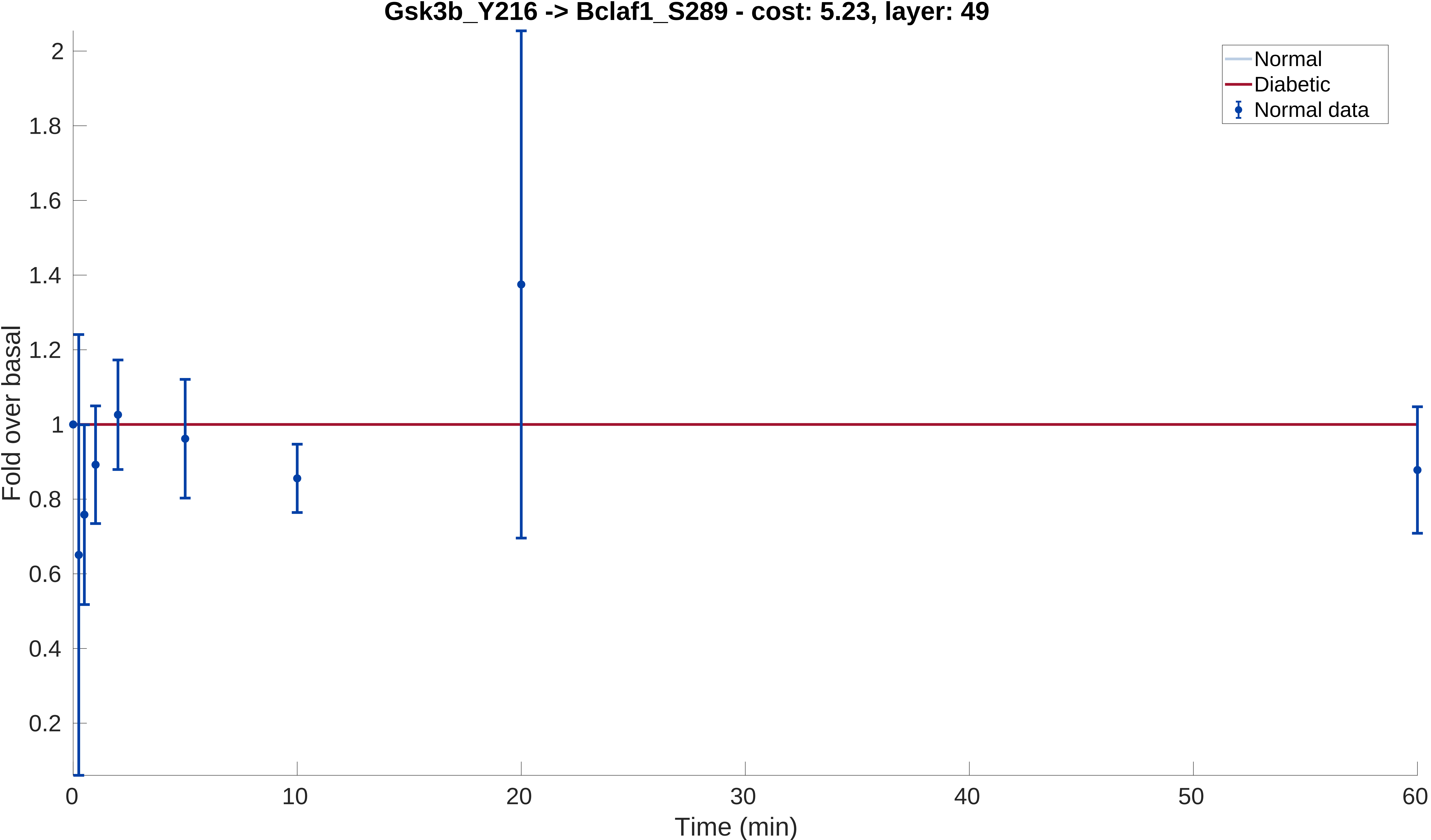

Gsk3b_Y216 -> Bclaf1_S289 - cost: 5.23, layer: 49
Normal
Diabetic
Normal data
Fold over basal
Time (min)
2
1.8
1.6
1.4
1.2
1
0.8
0.6
0.4
0.2
0
10
20
30
40
50
60

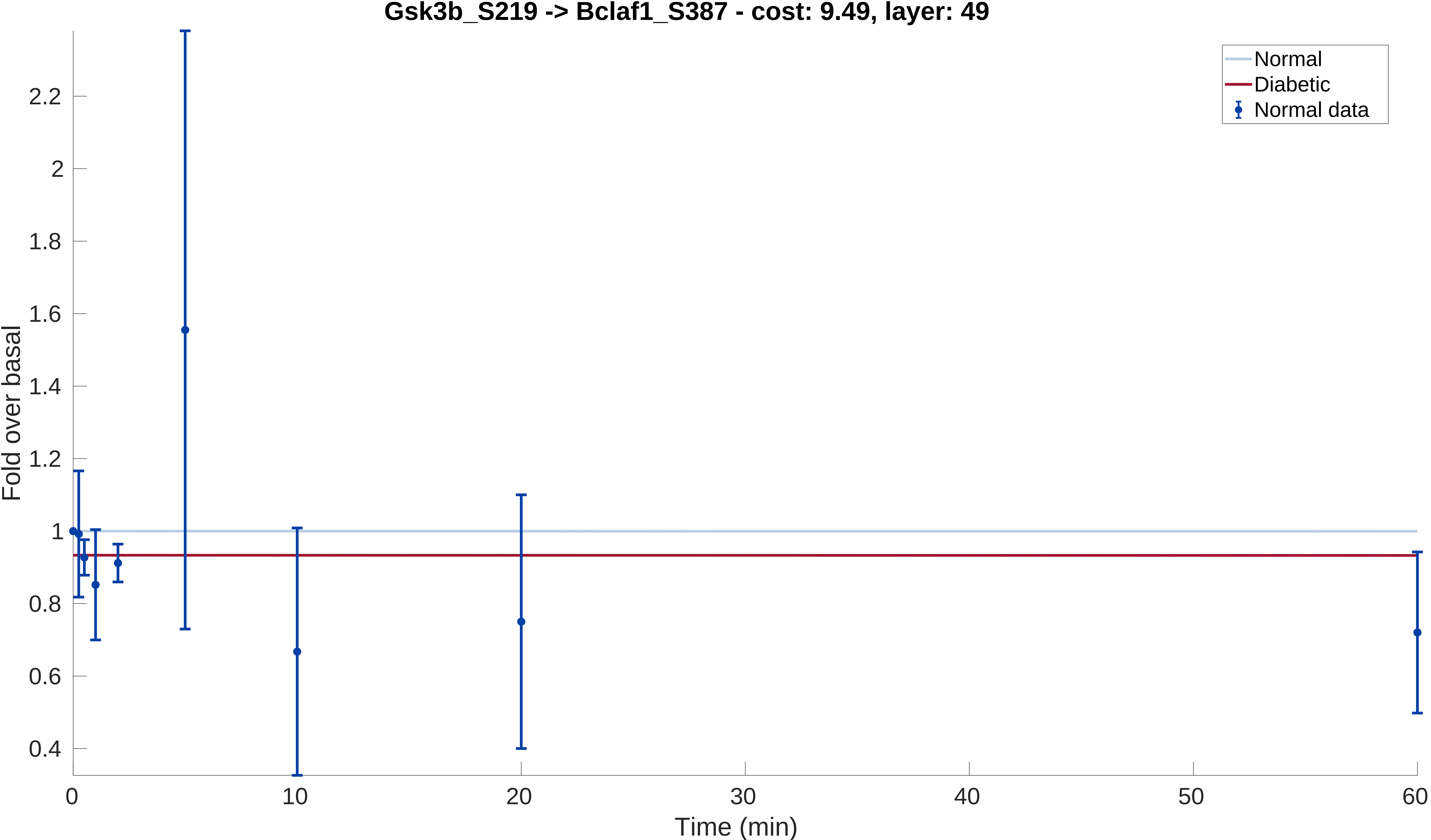

Gsk3b_S219 -> Bclaf1_S387 - cost: 9.49, layer: 49
Normal
Diabetic
Normal data
Fold over basal
2.2
2
1.8
1.6
1.4
1.2
1
0.8
0.6
0.4
0
10
20
30
40
50
60
Time (min)

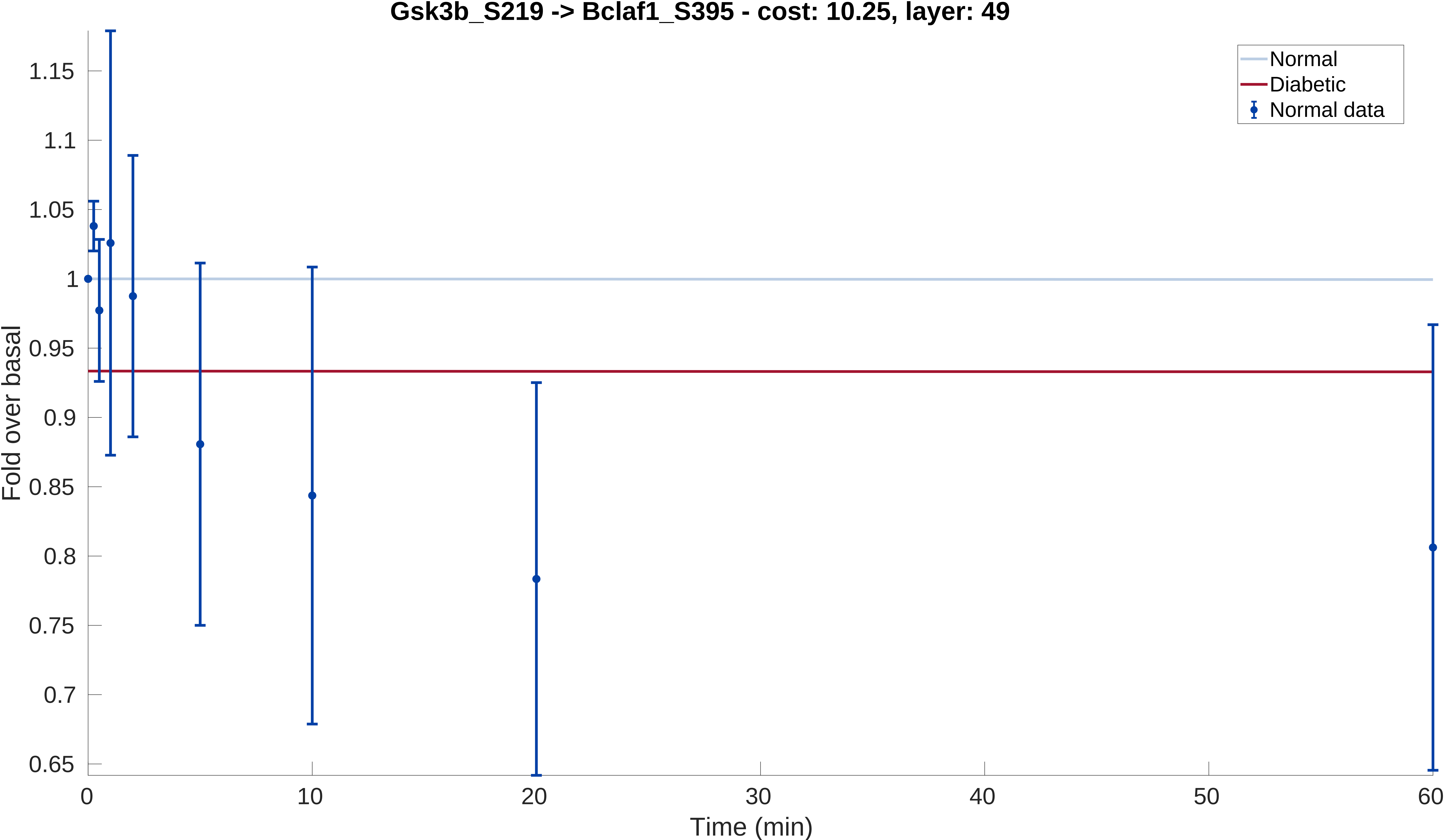
Gsk3b_S219 -> Bclaf1_S395 - cost: 10.25, layer: 49
Normal
Diabetic
Normal data
Fold over basal
Time (min)
1.15
1.1
1.05
1
0.95
0.9
0.85
0.8
0.75
0.7
0.65
0
10
20
30
40
50
60

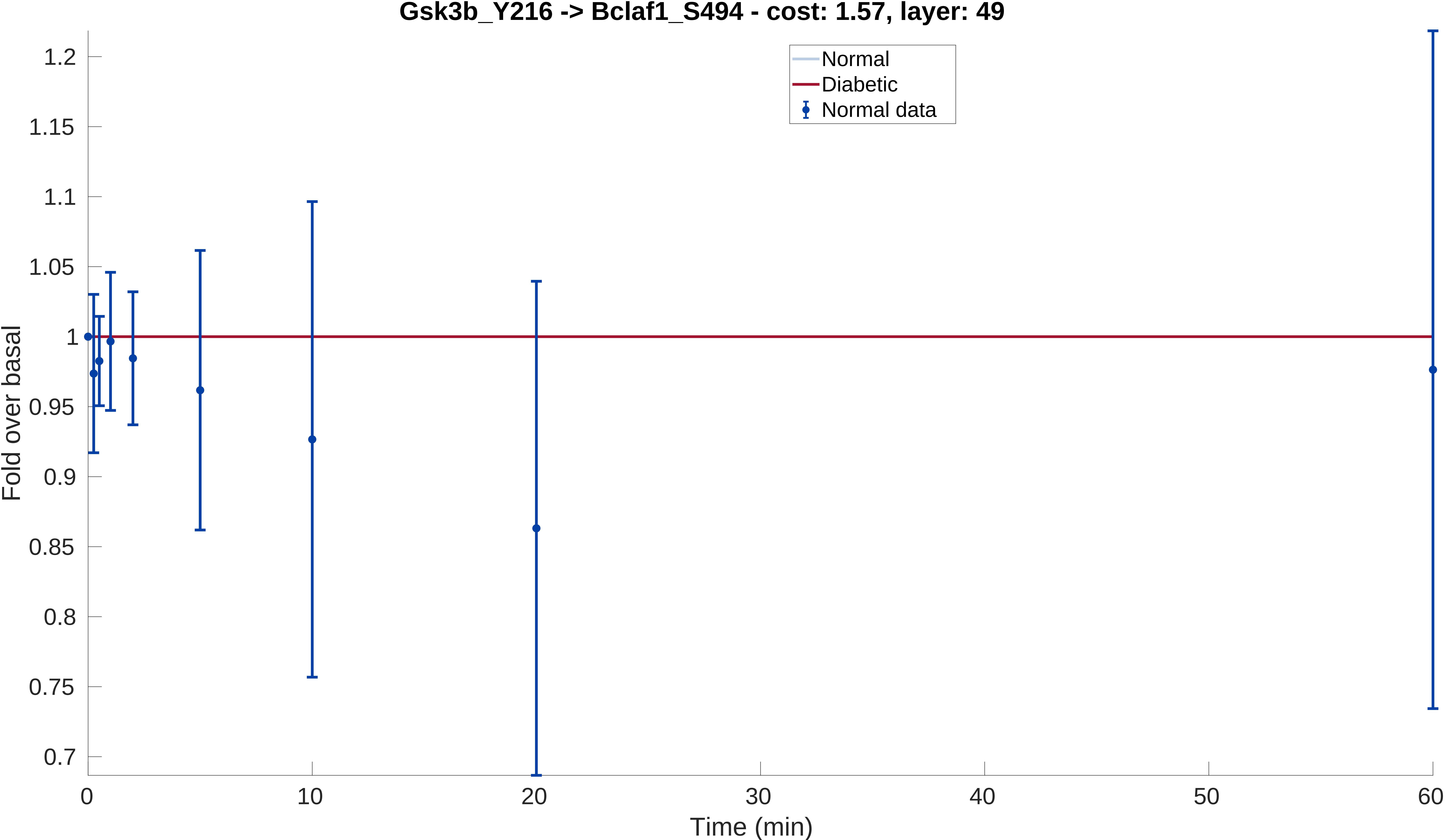

Gsk3b_Y216 -> Bclaf1_S494 - cost: 1.57, layer: 49
Normal
Diabetic
Normal data
Fold over basal
1.2
1.15
1.1
1.05
1
0.95
0.9
0.85
0.8
0.75
0.7
0
10
20
30
40
50
60
Time (min)

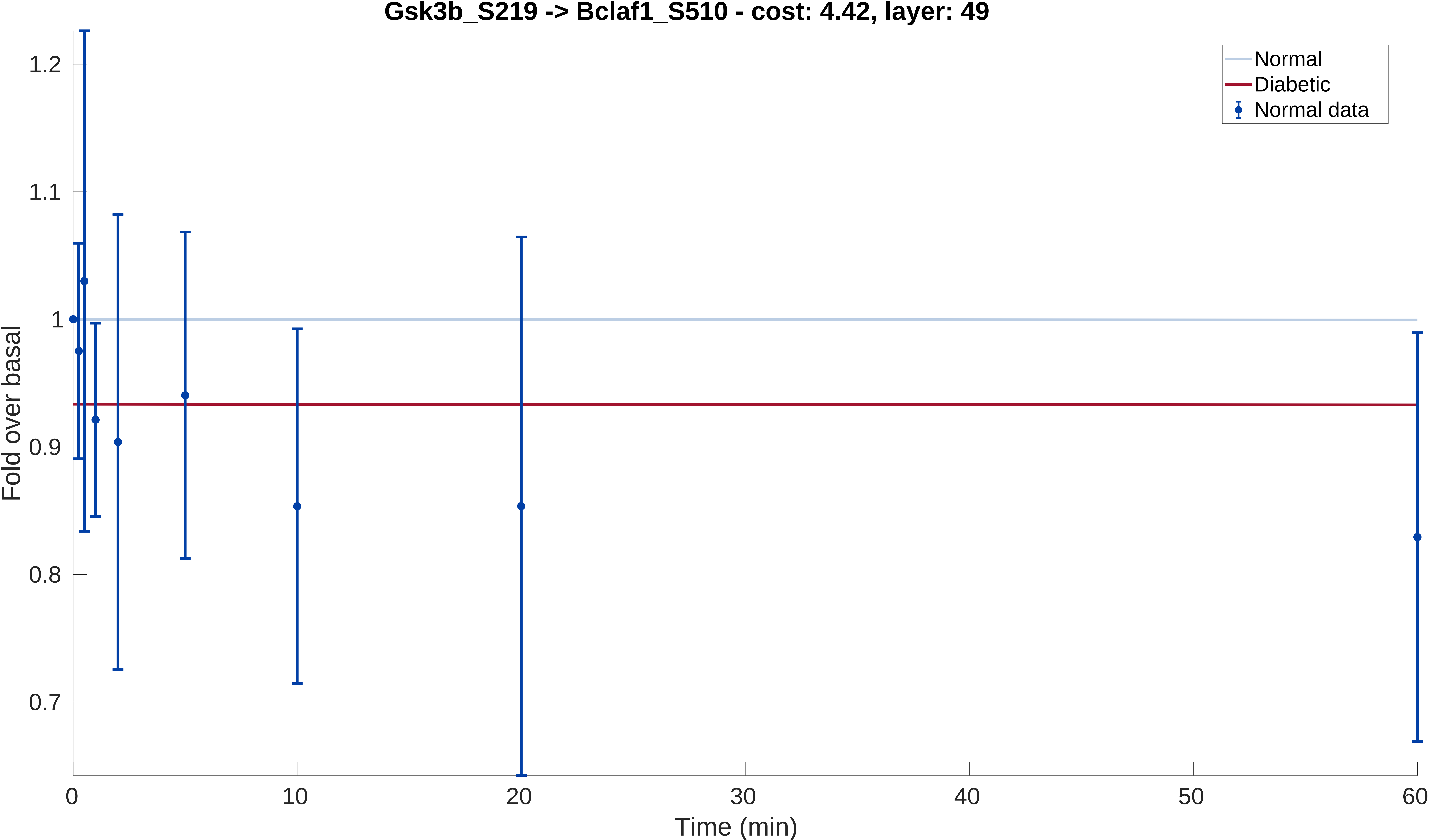

Gsk3b_S219 -> Bclaf1_S510 - cost: 4.42, layer: 49
Normal
Diabetic
Normal data
Fold over basal
Time (min)
1.2
1.1
1
0.9
0.8
0.7
0
10
20
30
40
50
60

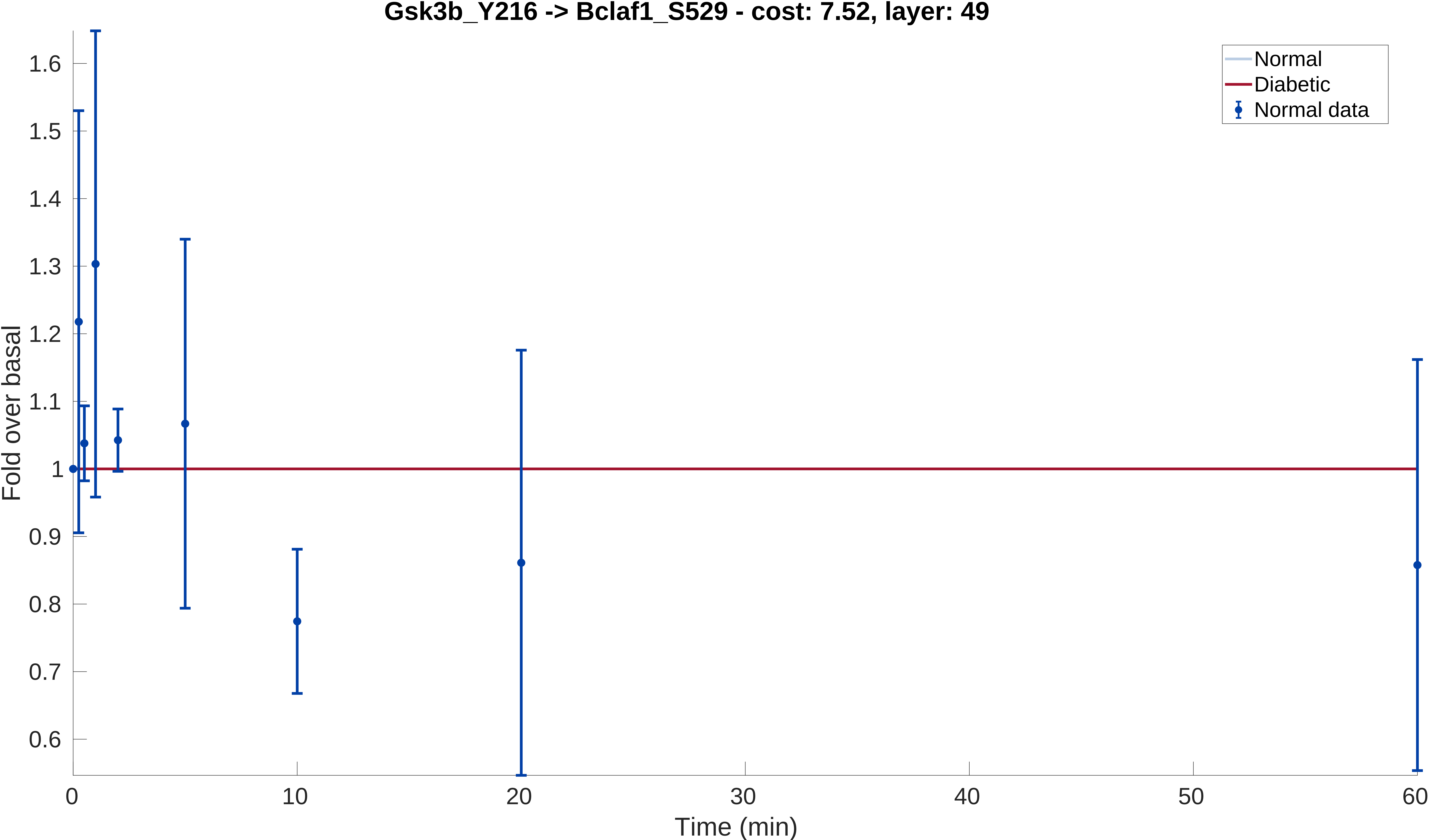

Gsk3b_Y216 -> Bclaf1_S529 - cost: 7.52, layer: 49
Normal
Diabetic
Normal data
Fold over basal
1.6
1.5
1.4
1.3
1.2
1.1
1
0.9
0.8
0.7
0.6
0
10
20
30
40
50
60
Time (min)

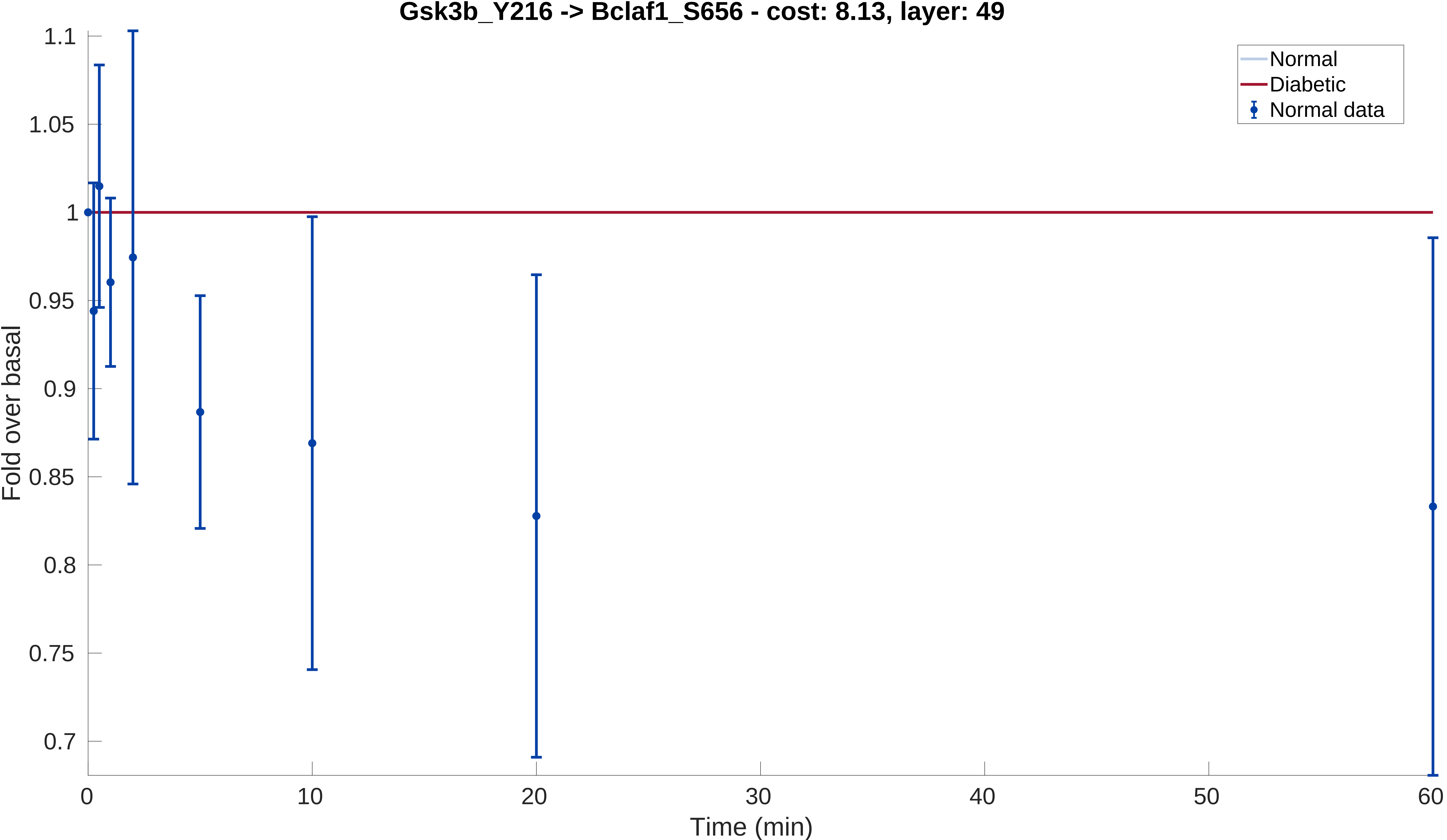

Gsk3b_Y216 -> Bclaf1_S656 - cost: 8.13, layer: 49
Normal
Diabetic
Normal data
Fold over basal
Time (min)
1.1
1.05
1
0.95
0.9
0.85
0.8
0.75
0.7
0
10
20
30
40
50
60

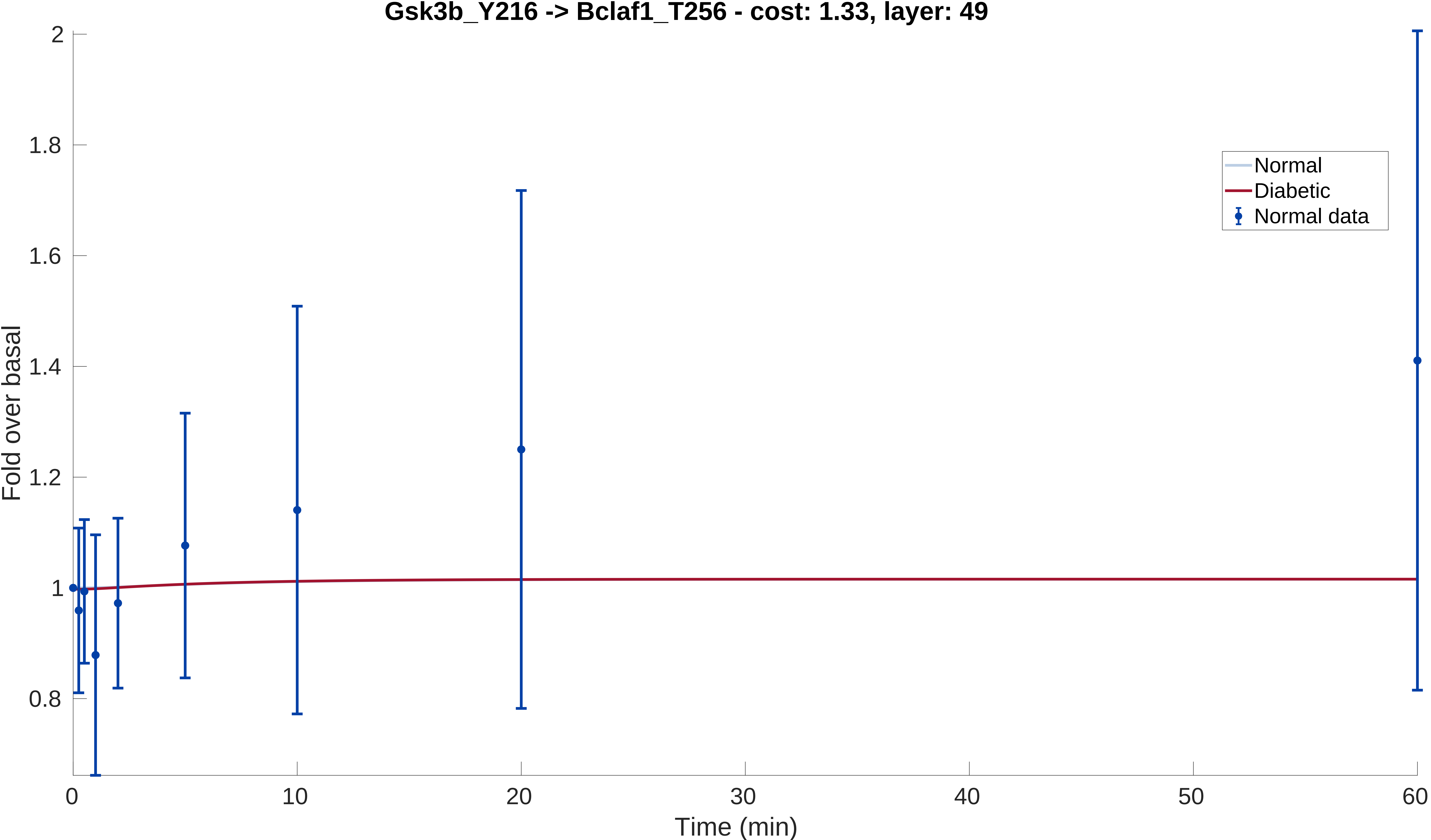

Gsk3b_Y216 -> Bclaf1_T256 - cost: 1.33, layer: 49
Fold over basal
Time (min)
0
10
20
30
40
50
60
0.8
1
1.2
1.4
1.6
1.8
2
Normal
Diabetic
Normal data

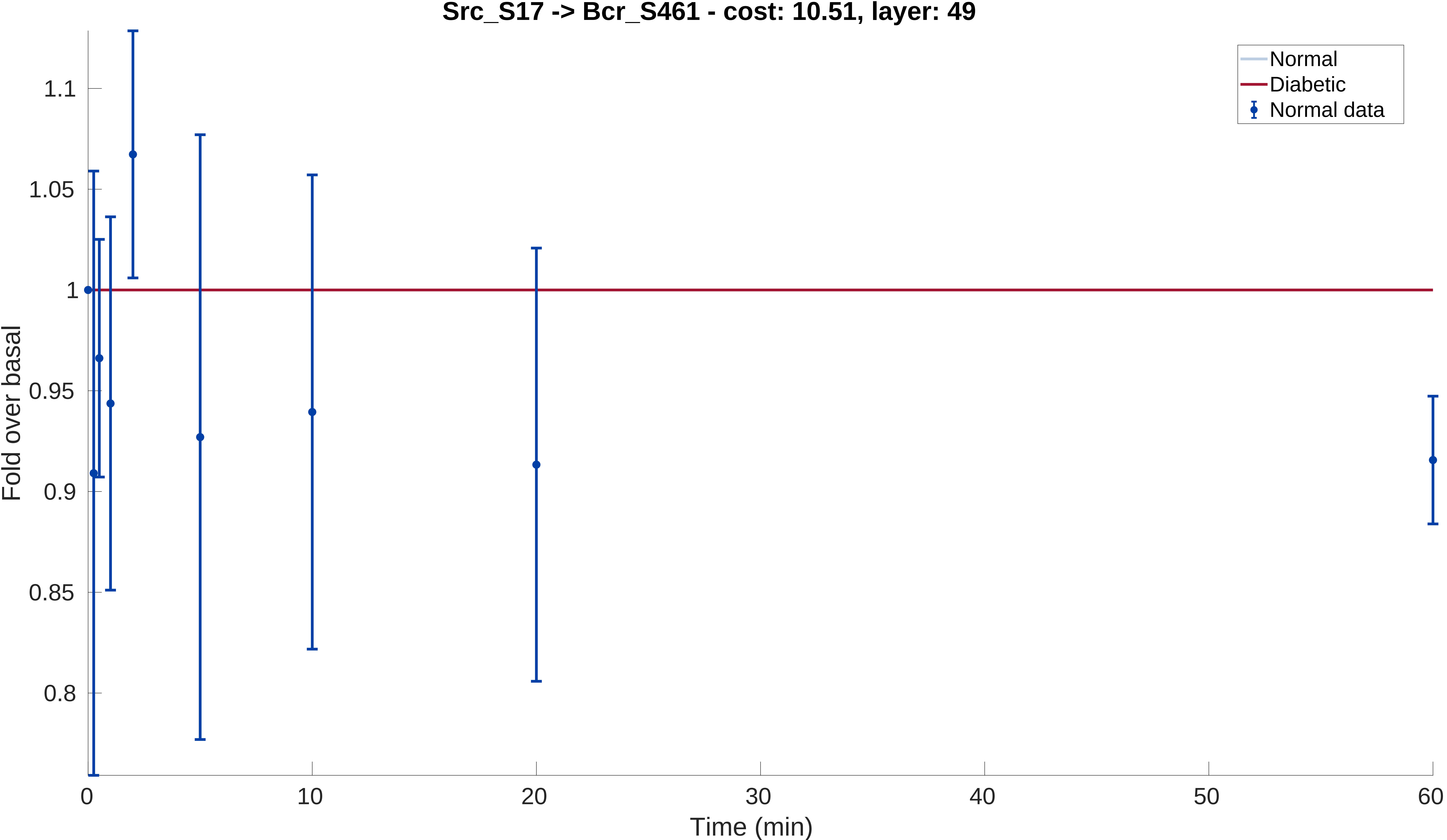

Src_S17 -> Bcr_S461 - cost: 10.51, layer: 49
Normal
Diabetic
Normal data
1.1
1.05
1
0.95
0.9
0.85
0.8
Fold over basal
0
10
20
30
40
50
60
Time (min)

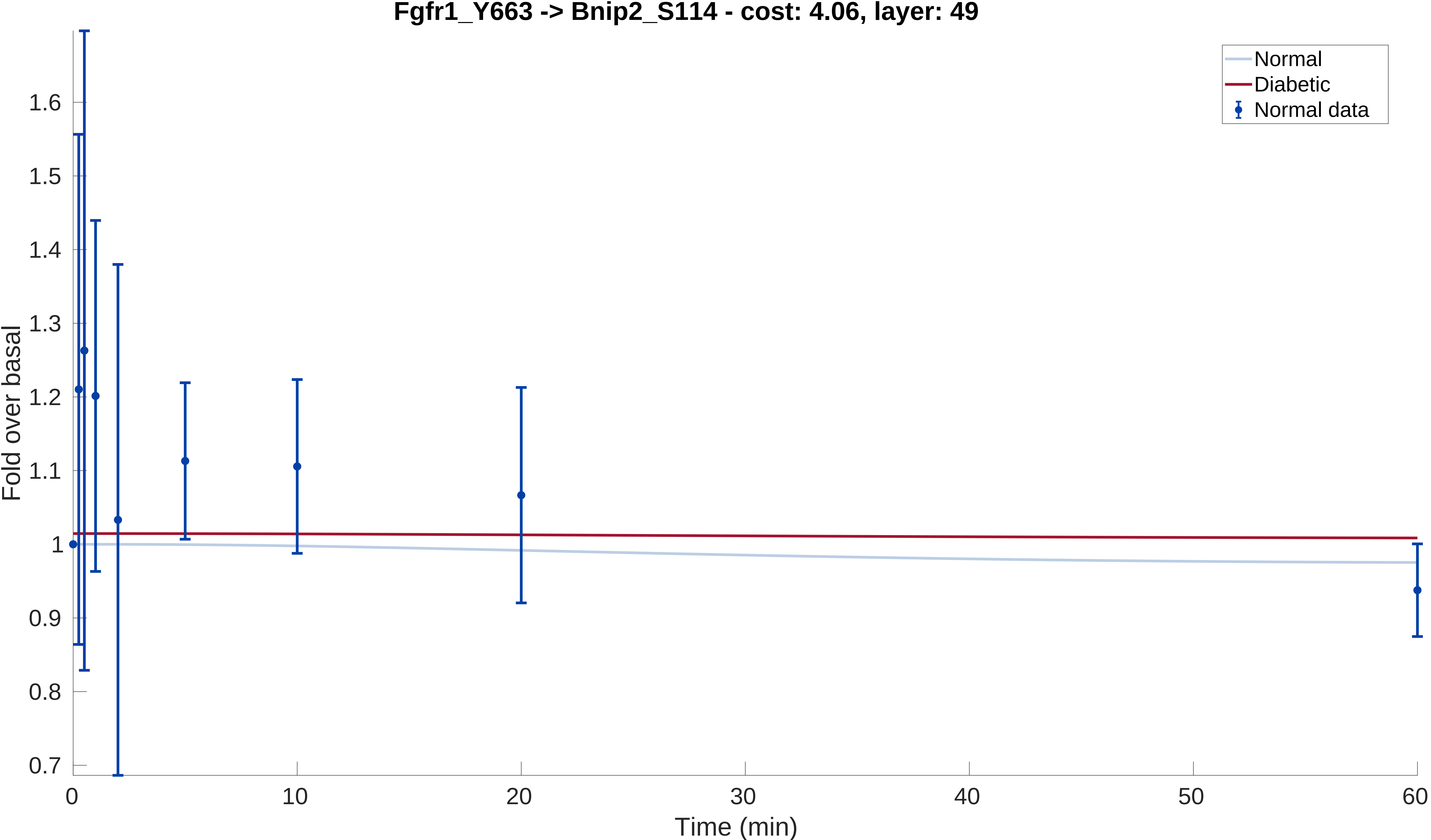

Fgfr1_Y663 -> Bnip2_S114 - cost: 4.06, layer: 49
Normal
Diabetic
Normal data
Fold over basal
1.6
1.5
1.4
1.3
1.2
1.1
1
0.9
0.8
0.7
0
10
20
30
40
50
60
Time (min)

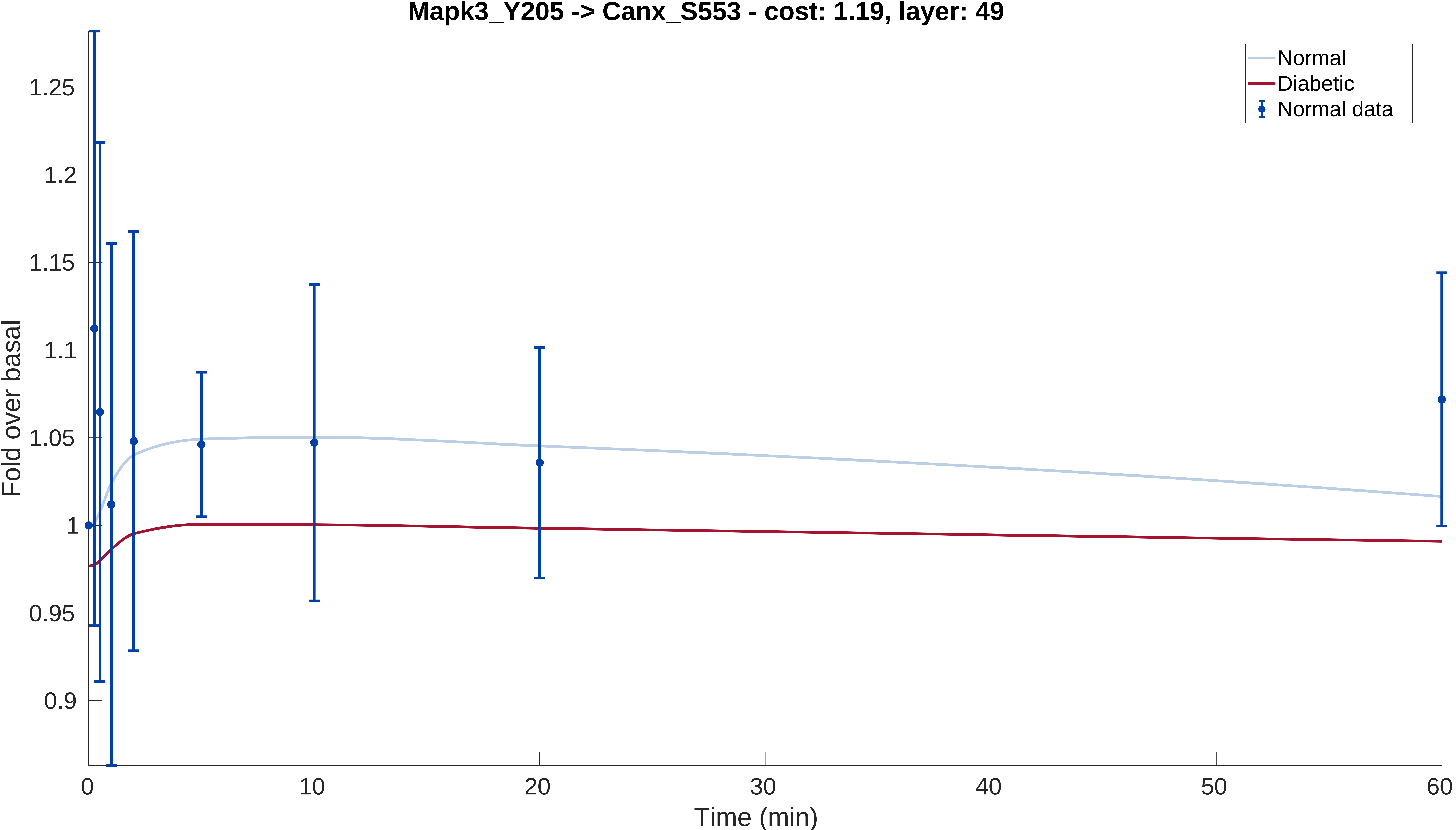

Mapk3_Y205 -> Canx_S553 - cost: 1.19, layer: 49
Normal
Diabetic
Normal data
Fold over basal
1.25
1.2
1.15
1.1
1.05
1
0.95
0.9
0
10
20
30
40
50
60
Time (min)

Mapk3_Y205 -> Canx_S563 - cost: 10.55, layer: 49

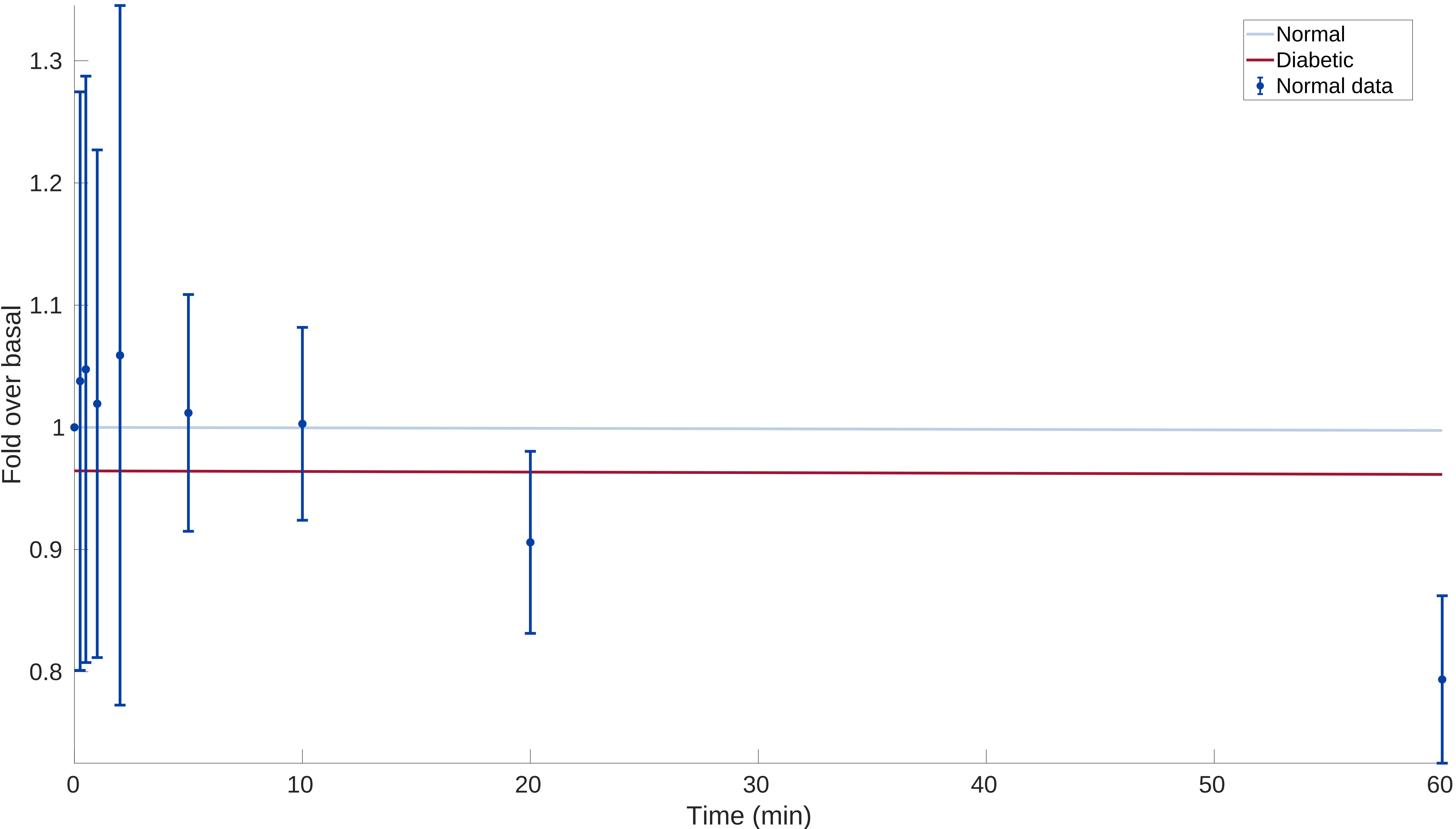

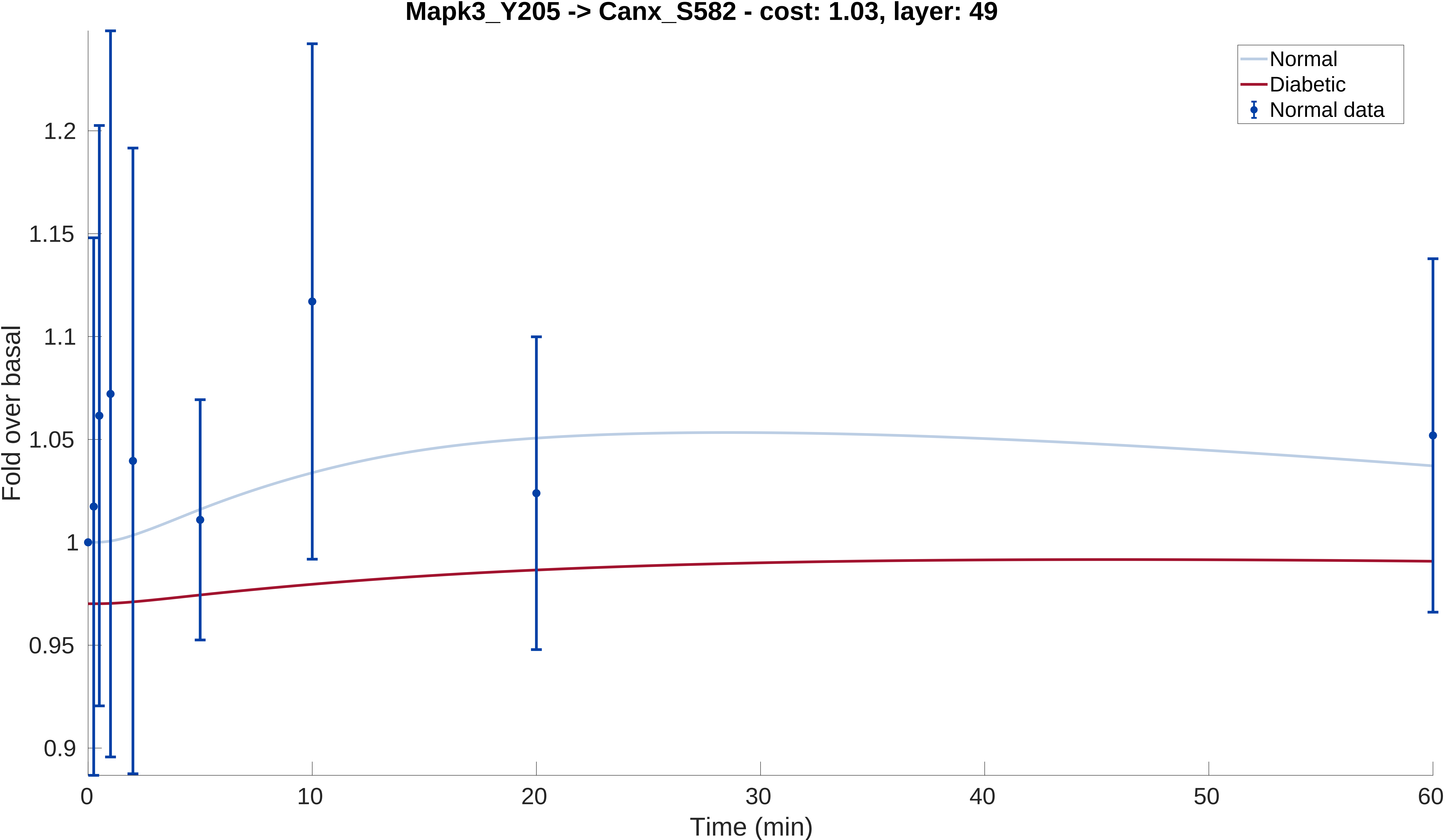

Mapk3_Y205 -> Canx_S582 - cost: 1.03, layer: 49
Normal
Diabetic
Normal data
Fold over basal
1.2
1.15
1.1
1.05
1
0.95
0.9
0
10
20
30
40
50
60
Time (min)

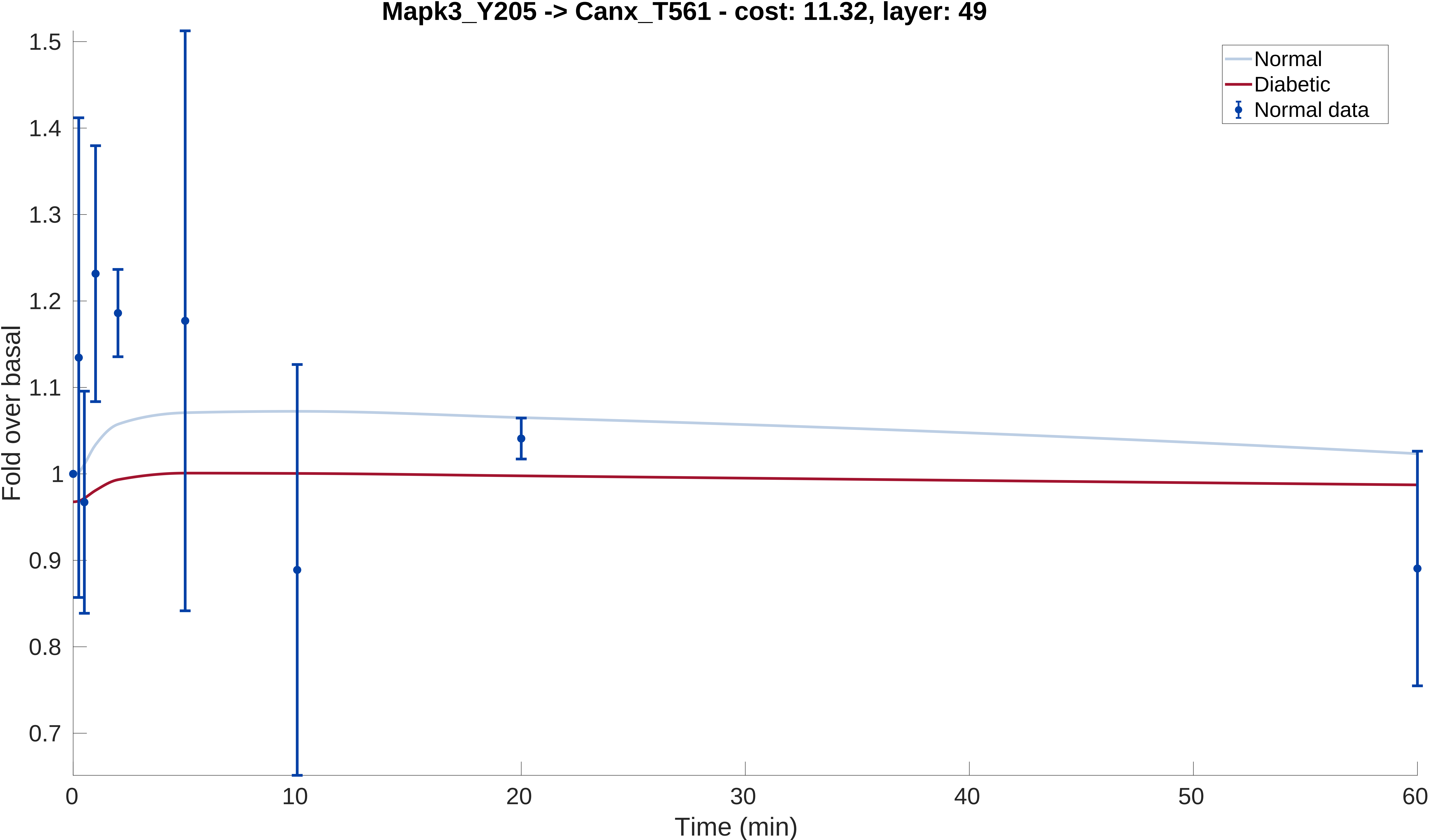

Mapk3_Y205 -> Canx_T561 - cost: 11.32, layer: 49
Normal
Diabetic
Normal data
Fold over basal
1.5
1.4
1.3
1.2
1.1
1
0.9
0.8
0.7
0
10
20
30
40
50
60
Time (min)

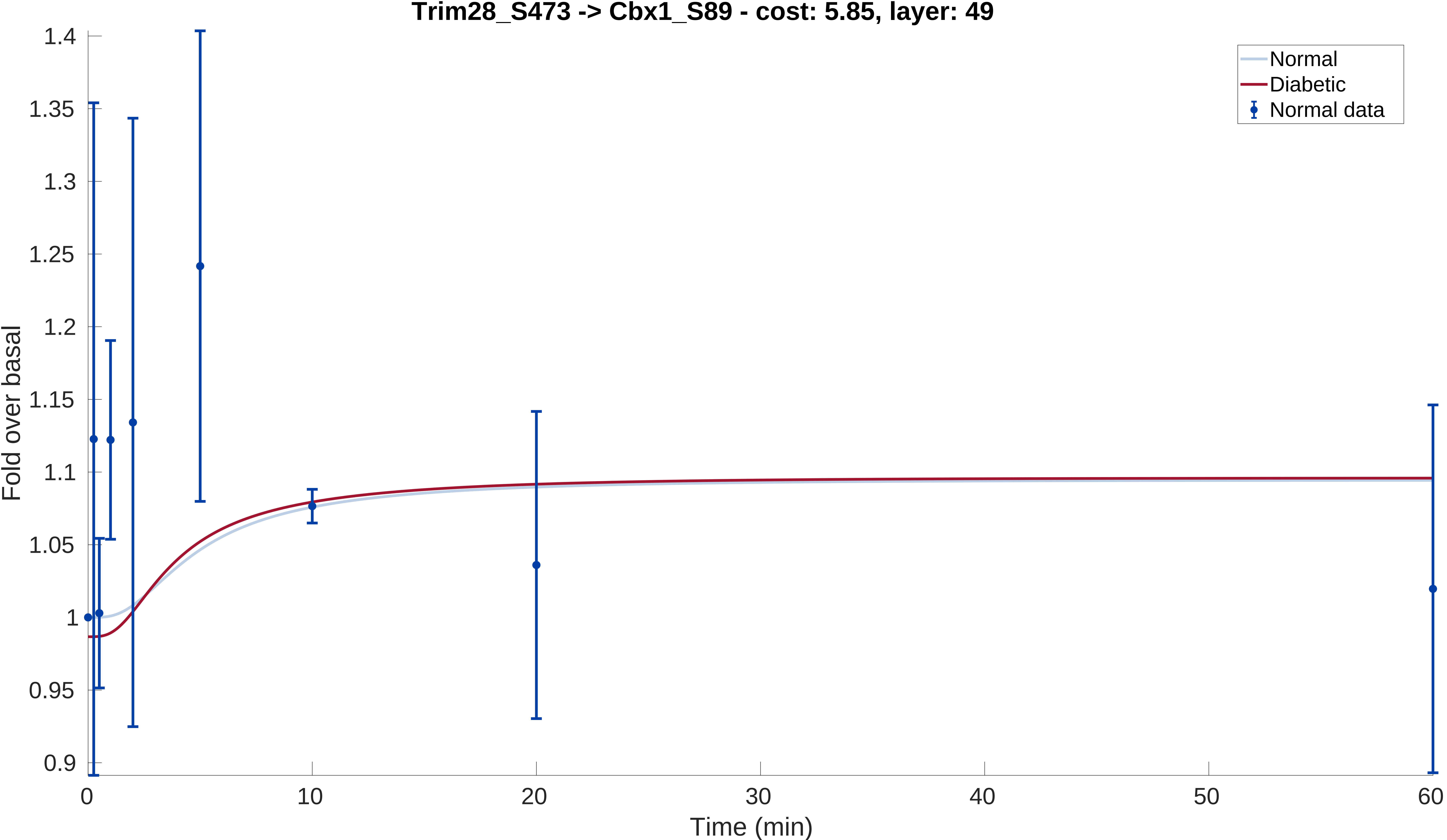

Trim28_S473 -> Cbx1_S89 - cost: 5.85, layer: 49
Normal
Diabetic
Normal data
Fold over basal
1.4
1.35
1.3
1.25
1.2
1.15
1.1
1.05
1
0.95
0.9
0
10
20
30
40
50
60
Time (min)

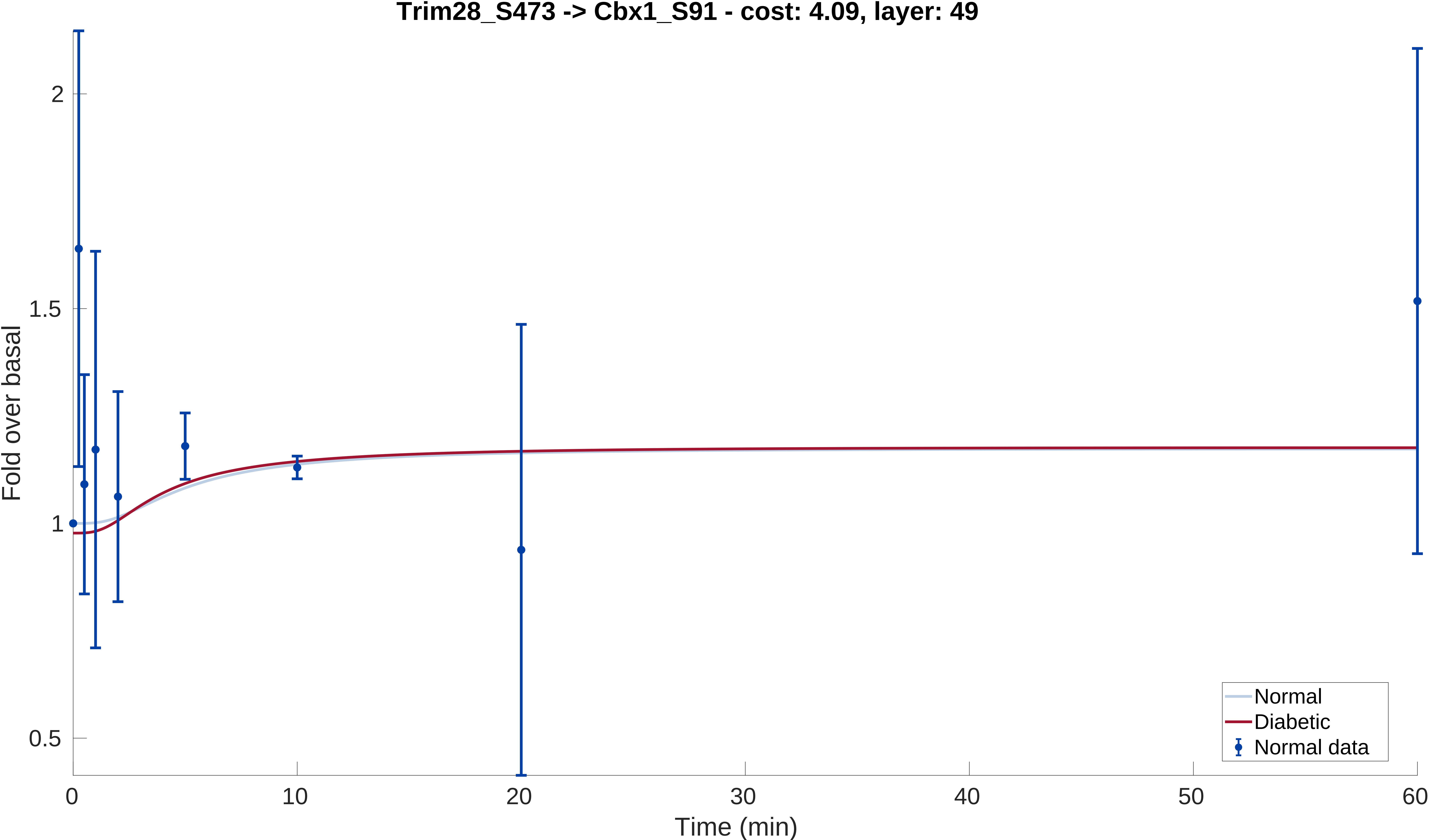

Trim28_S473 -> Cbx1_S91 - cost: 4.09, layer: 49
Fold over basal
2
1.5
1
0.5
0
10
20
30
40
50
60
Time (min)
Normal
Diabetic
Normal data

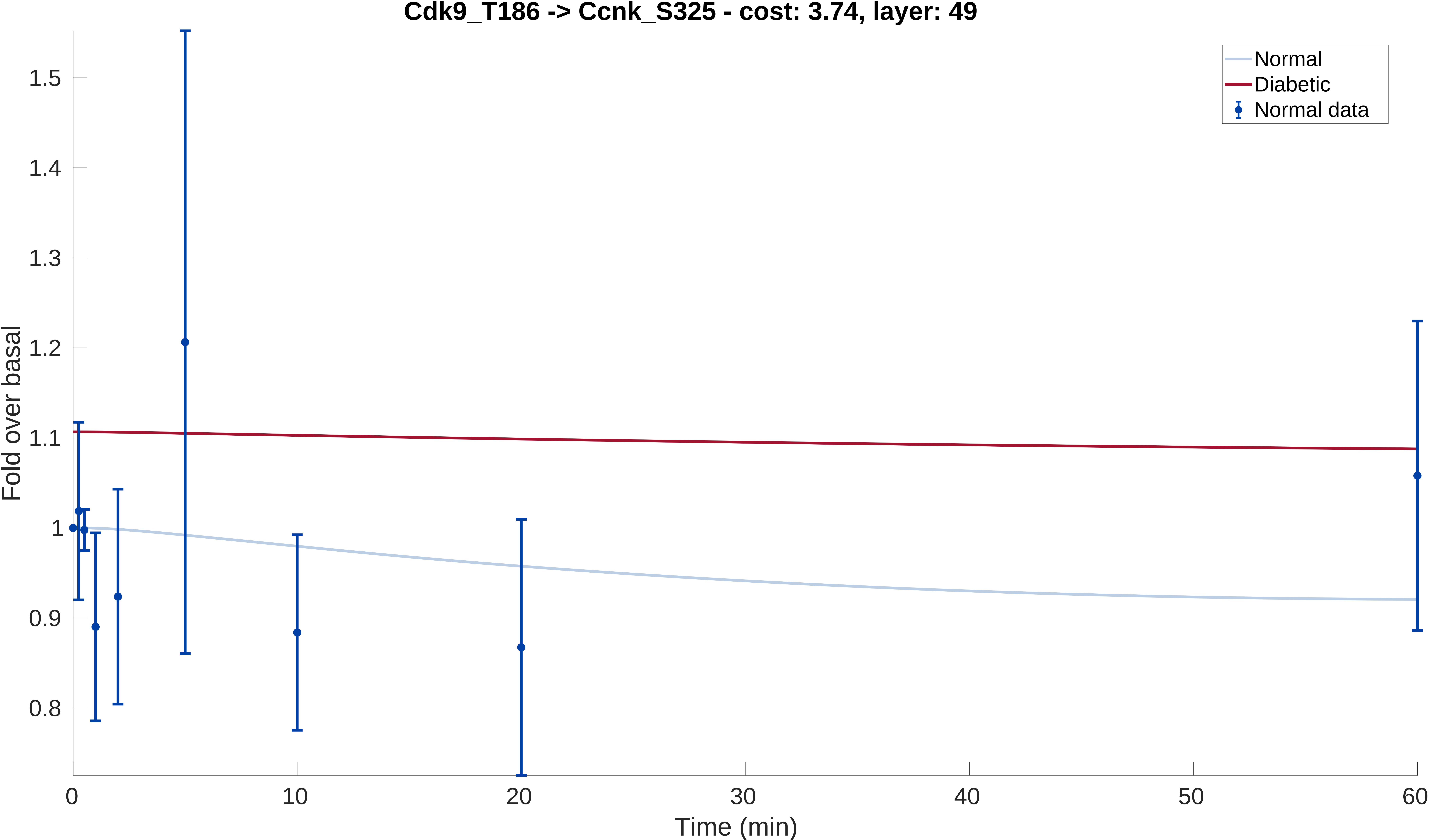

Cdk9_T186 -> Ccnk_S325 - cost: 3.74, layer: 49
Normal
Diabetic
Normal data
Fold over basal
1.5
1.4
1.3
1.2
1.1
1
0.9
0.8
0
10
20
30
40
50
60
Time (min)

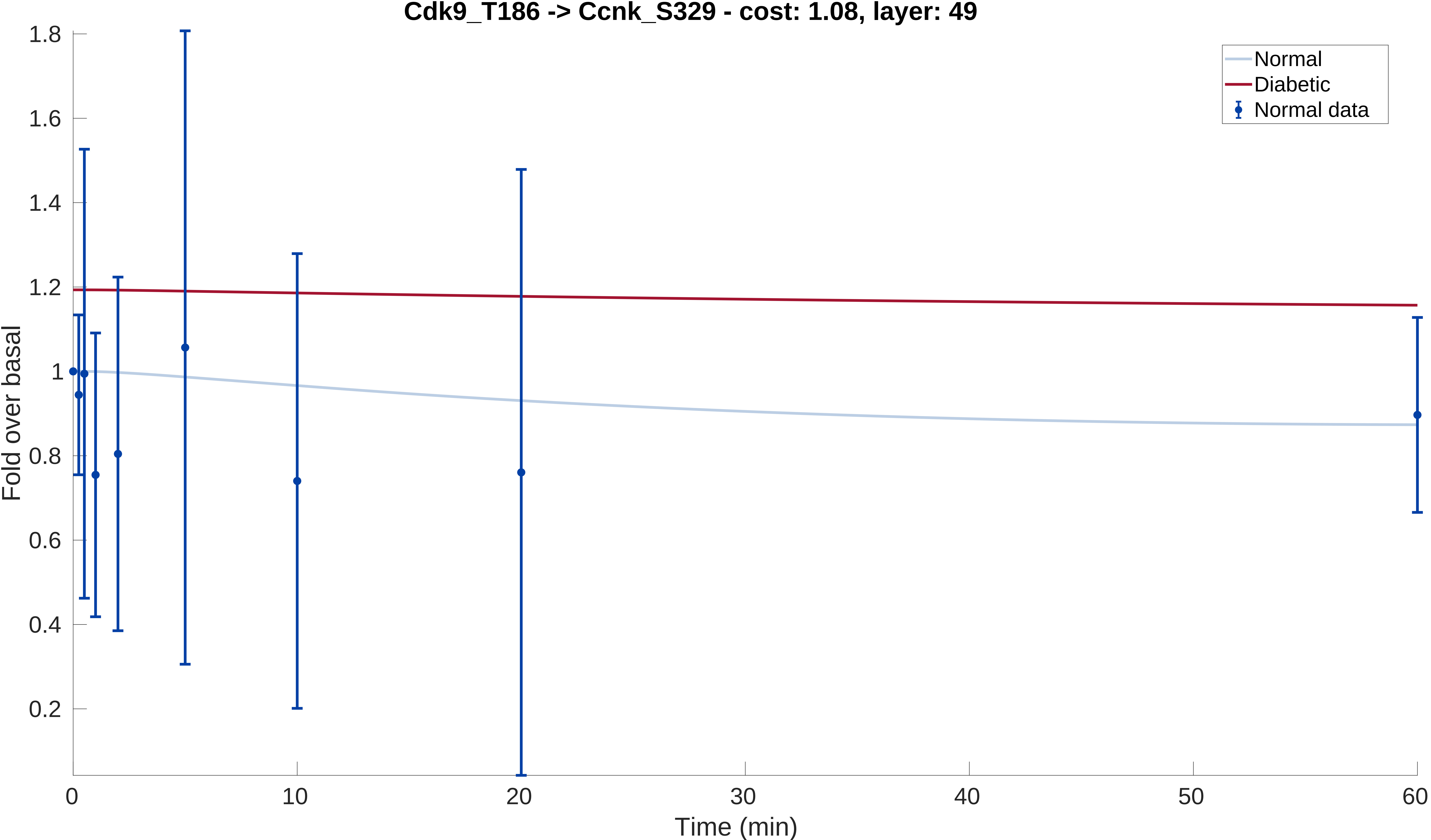

Cdk9_T186 -> Ccnk_S329 - cost: 1.08, layer: 49
Normal
Diabetic
Normal data
Fold over basal
1.8
1.6
1.4
1.2
1
0.8
0.6
0.4
0.2
0
10
20
30
40
50
60
Time (min)

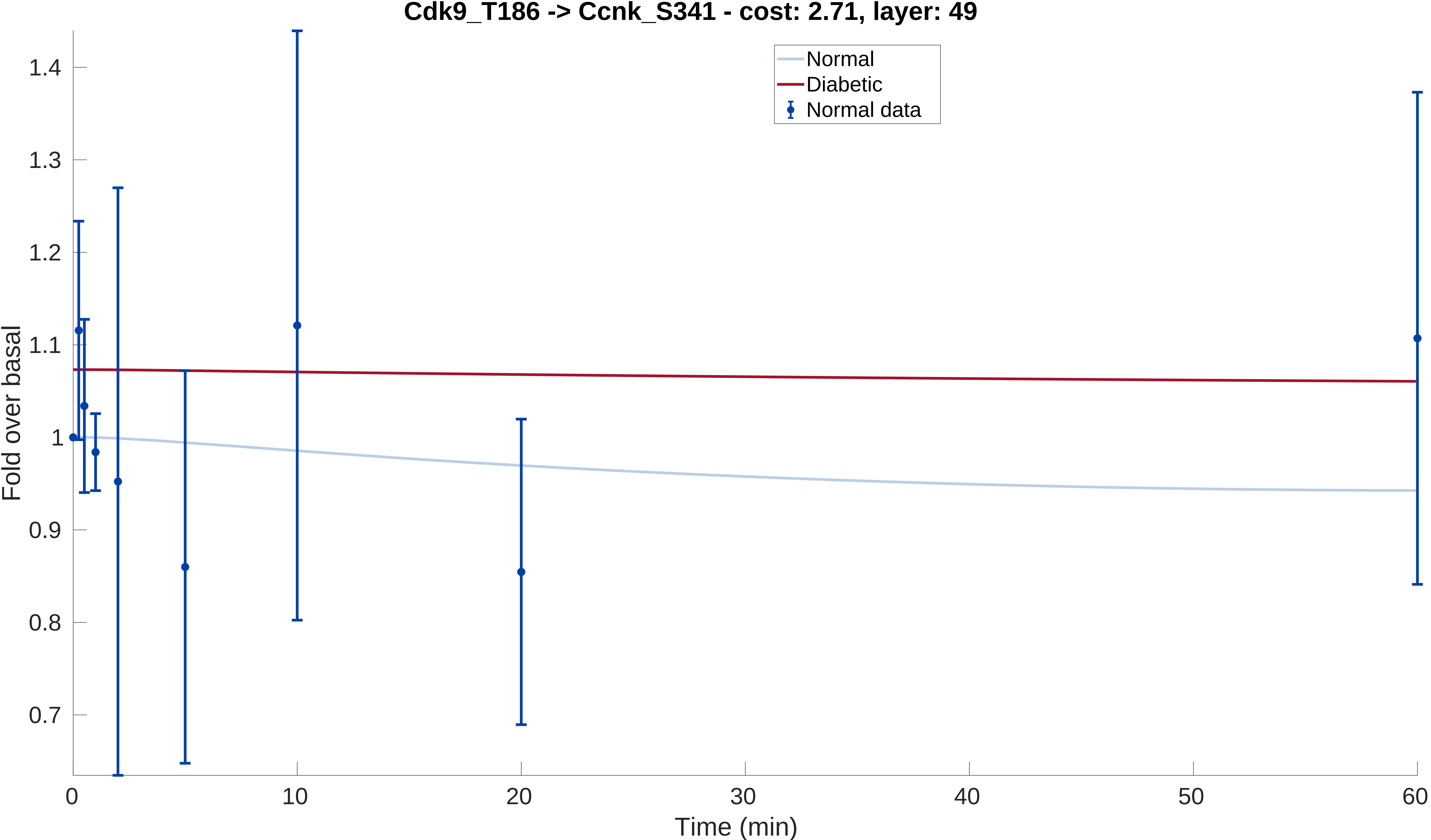

Cdk9_T186 -> Ccnk_S341 - cost: 2.71, layer: 49
Normal
Diabetic
Normal data
Fold over basal
1.4
1.3
1.2
1.1
1
0.9
0.8
0.7
0
10
20
30
40
50
60
Time (min)

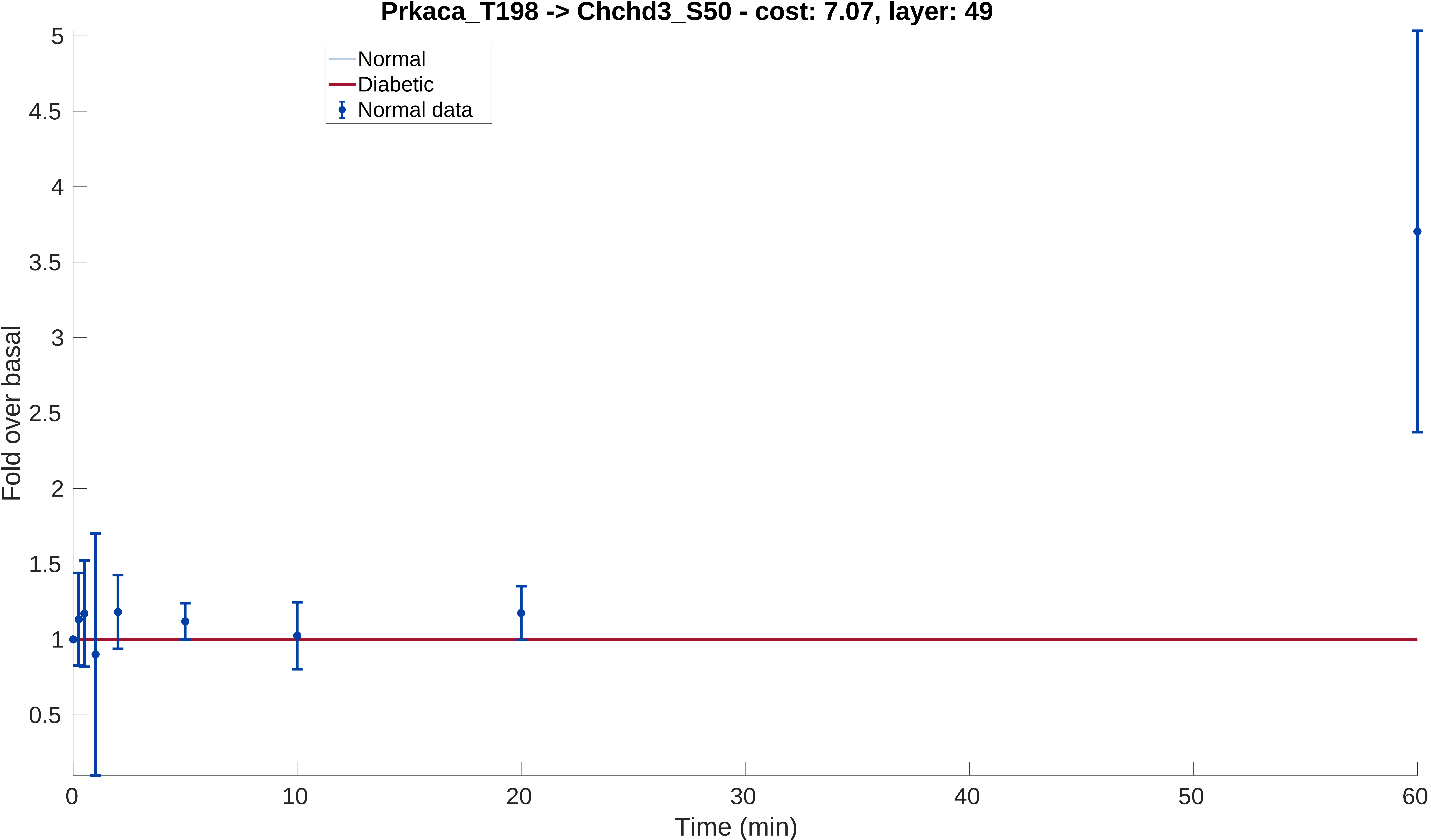

Prkaca_T198 -> Chchd3_S50 - cost: 7.07, layer: 49
Normal
Diabetic
Normal data
Fold over basal
Time (min)
0.5
1
1.5
2
2.5
3
3.5
4
4.5
5
0
10
20
30
40
50
60

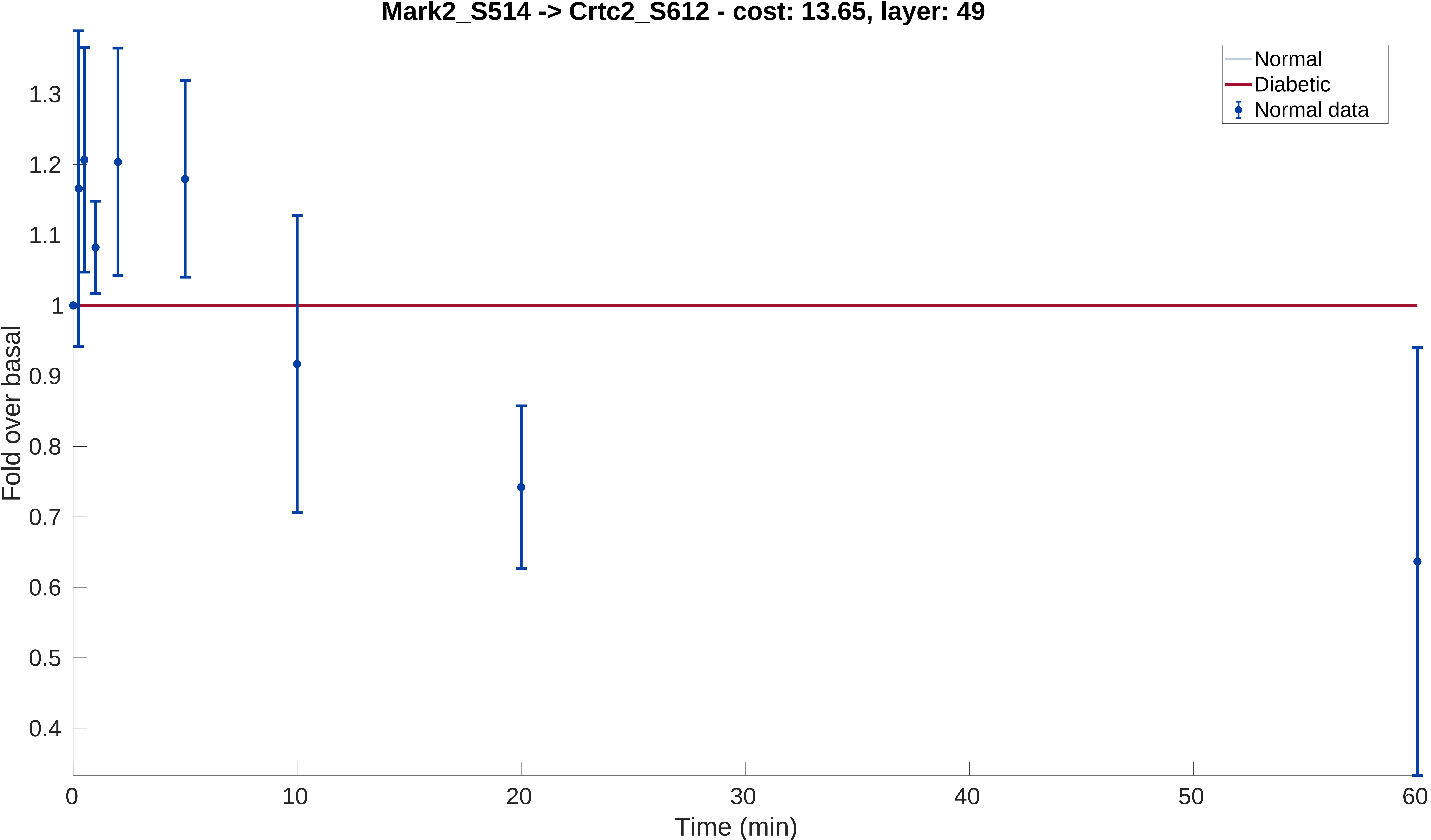
Mark2_S514 -> Crtc2_S612 - cost: 13.65, layer: 49
Normal
Diabetic
Normal data
Fold over basal
Time (min)
0.4
0.5
0.6
0.7
0.8
0.9
1
1.1
1.2
1.3
0
10
20
30
40
50
60

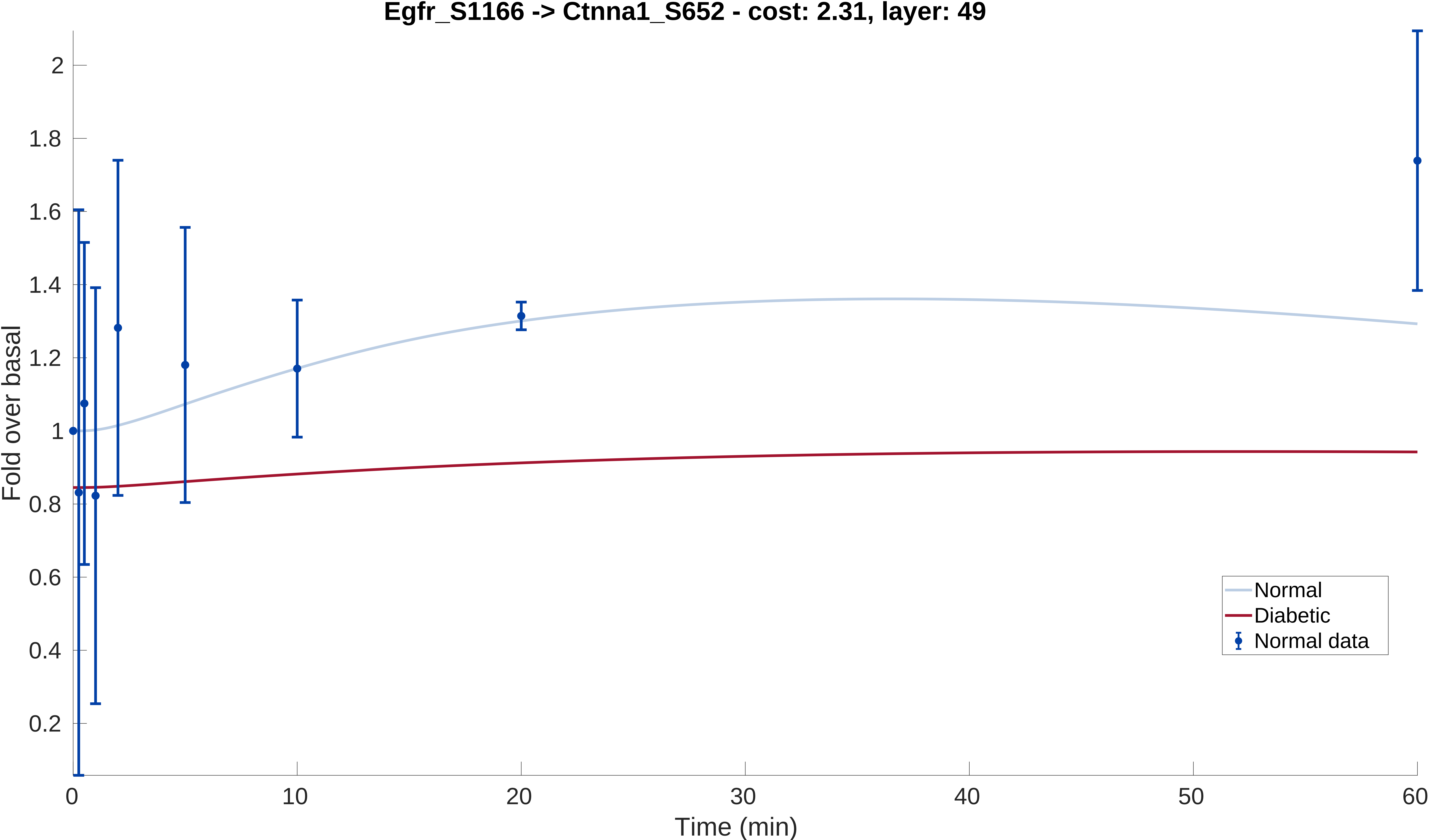

Egfr_S1166 -> Ctnna1_S652 - cost: 2.31, layer: 49
Fold over basal
Time (min)
2
1.8
1.6
1.4
1.2
1
0.8
0.6
0.4
0.2
0
10
20
30
40
50
60
Normal
Diabetic
Normal data

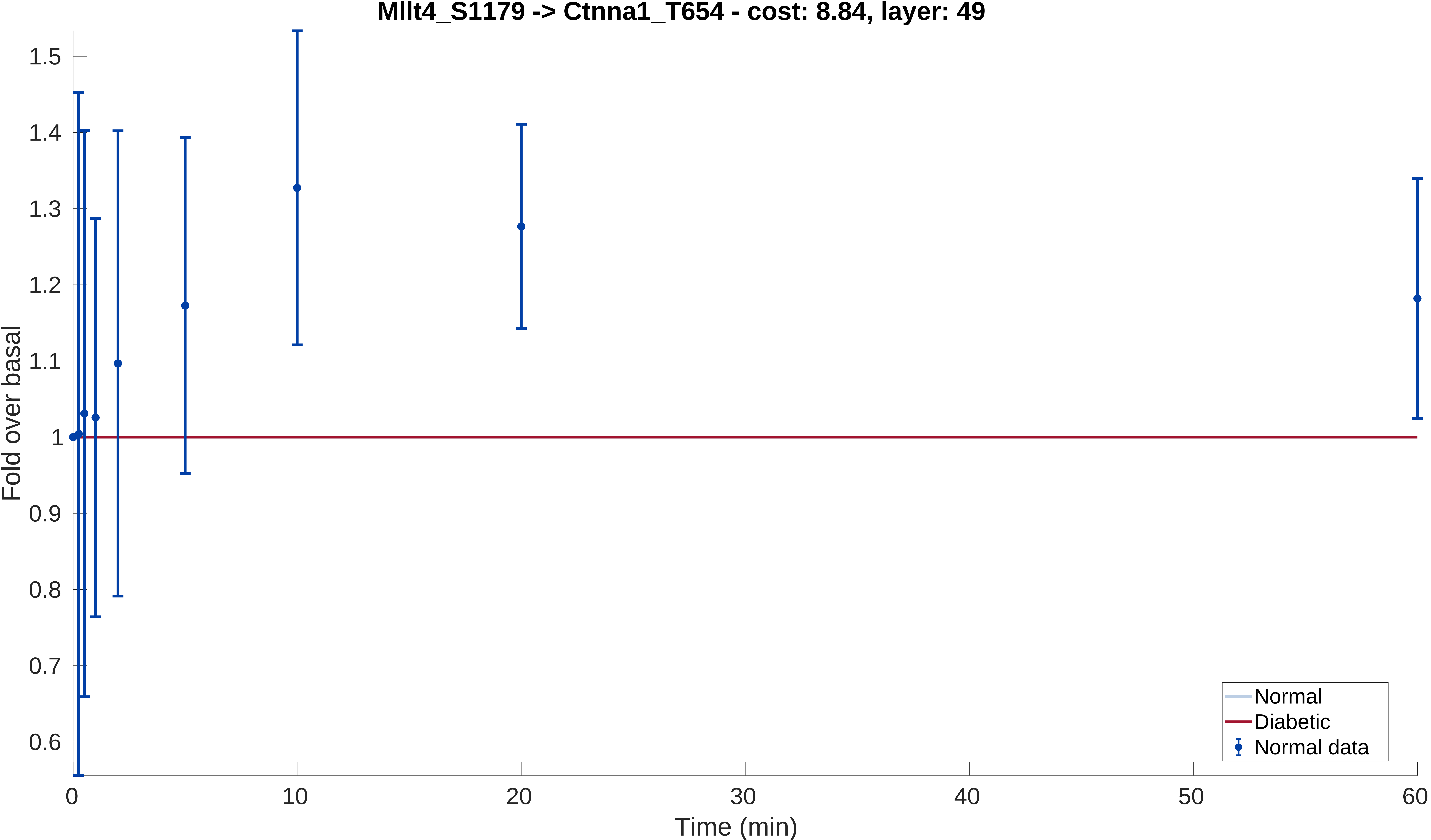
Mllt4_S1179 -> Ctnna1_T654 - cost: 8.84, layer: 49
Fold over basal
1.5
1.4
1.3
1.2
1.1
1
0.9
0.8
0.7
0.6
0
10
20
30
40
50
60
Time (min)
Normal
Diabetic
Normal data

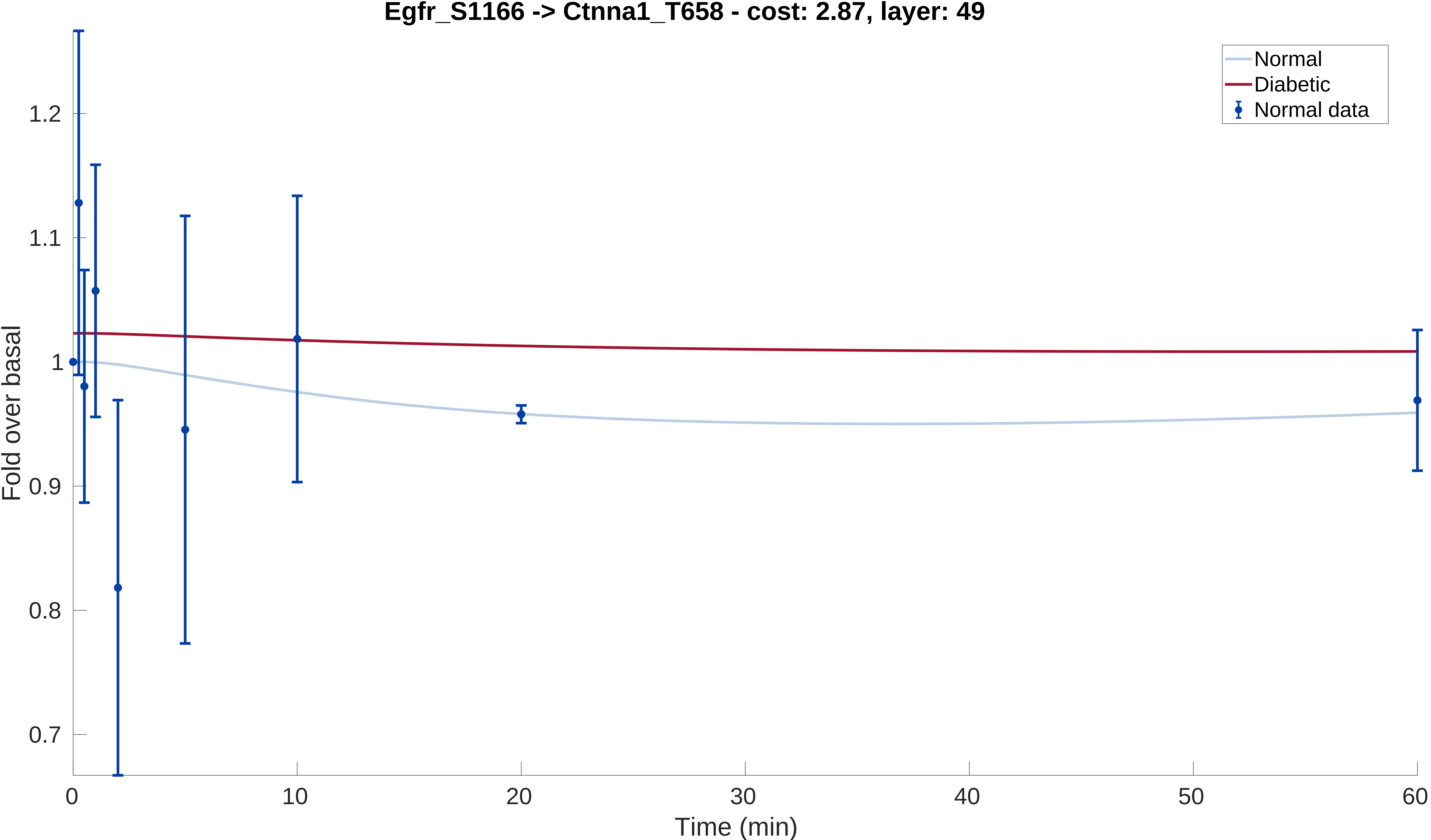

Egfr_S1166 -> Ctnna1_T658 - cost: 2.87, layer: 49
Normal
Diabetic
Normal data
1.2
1.1
1
0.9
0.8
0.7
Fold over basal
0
10
20
30
40
50
60
Time (min)

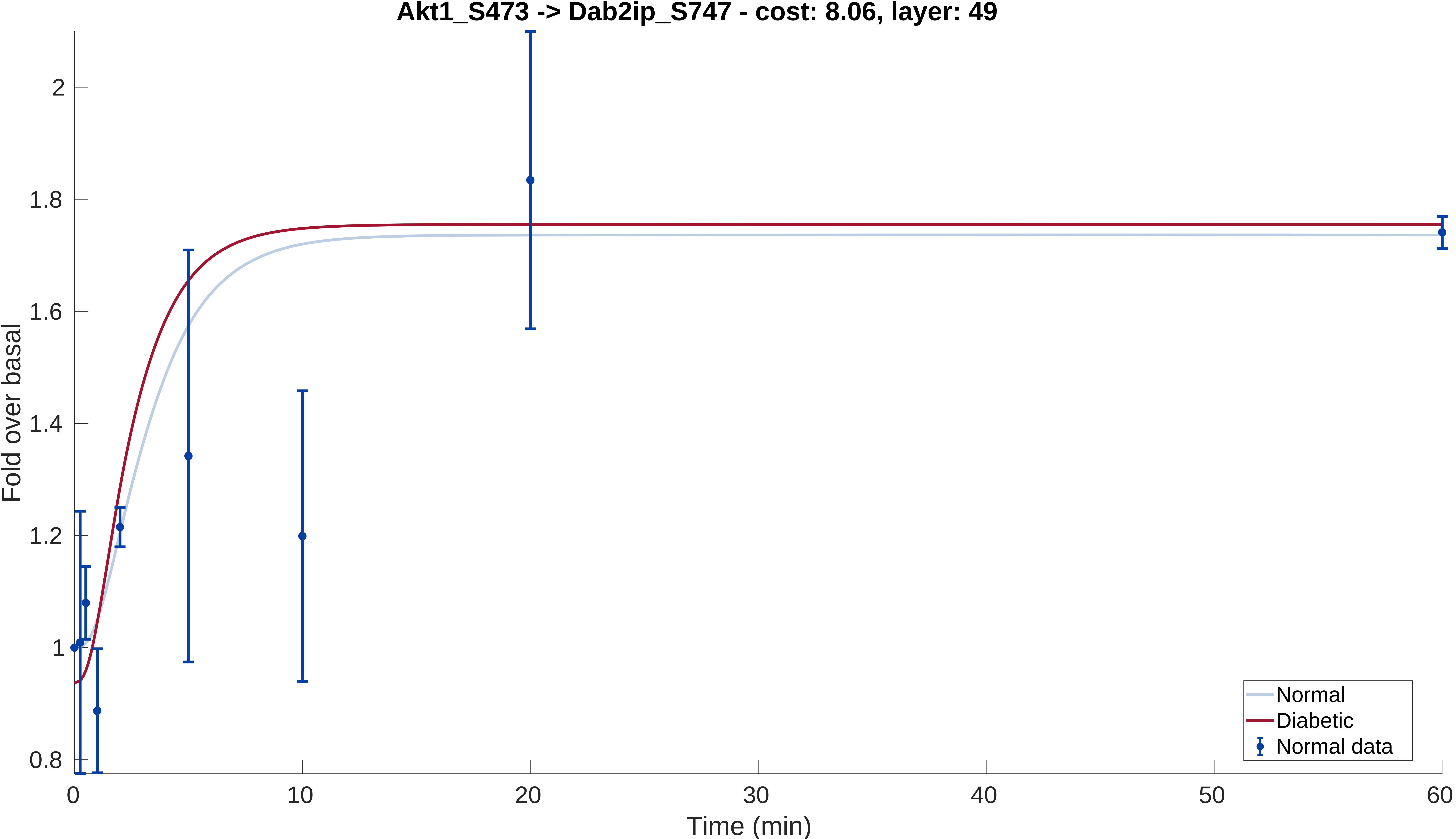
Akt1_S473 -> Dab2ip_S747 - cost: 8.06, layer: 49
Fold over basal
2
1.8
1.6
1.4
1.2
1
0.8
0
10
20
30
40
50
60
Time (min)
Normal
Diabetic
Normal data

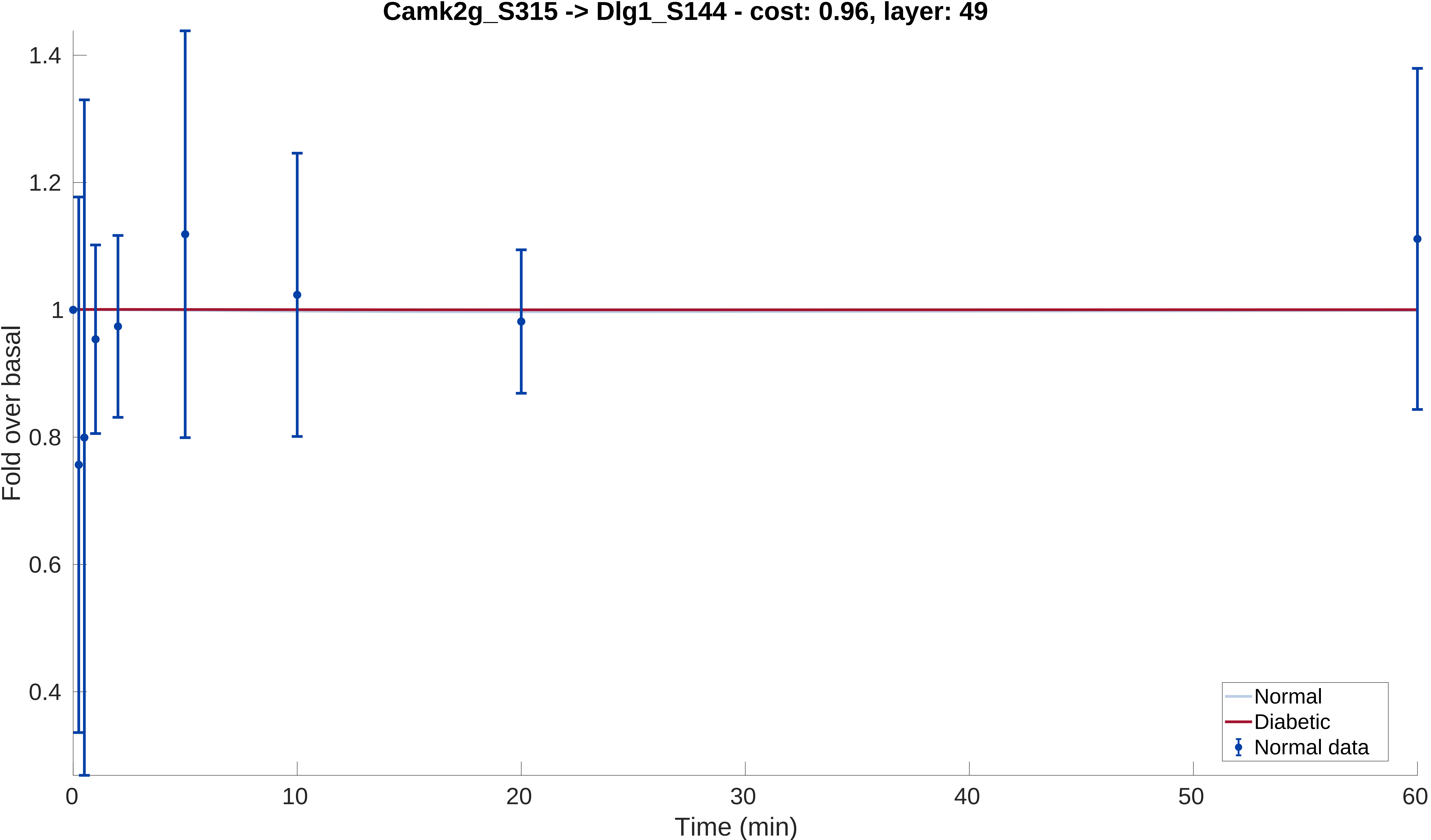

Camk2g_S315 -> Dlg1_S144 - cost: 0.96, layer: 49
1.4
1.2
1
0.8
0.6
0.4
Fold over basal
0
10
20
30
40
50
60
Time (min)
Normal
Diabetic
Normal data

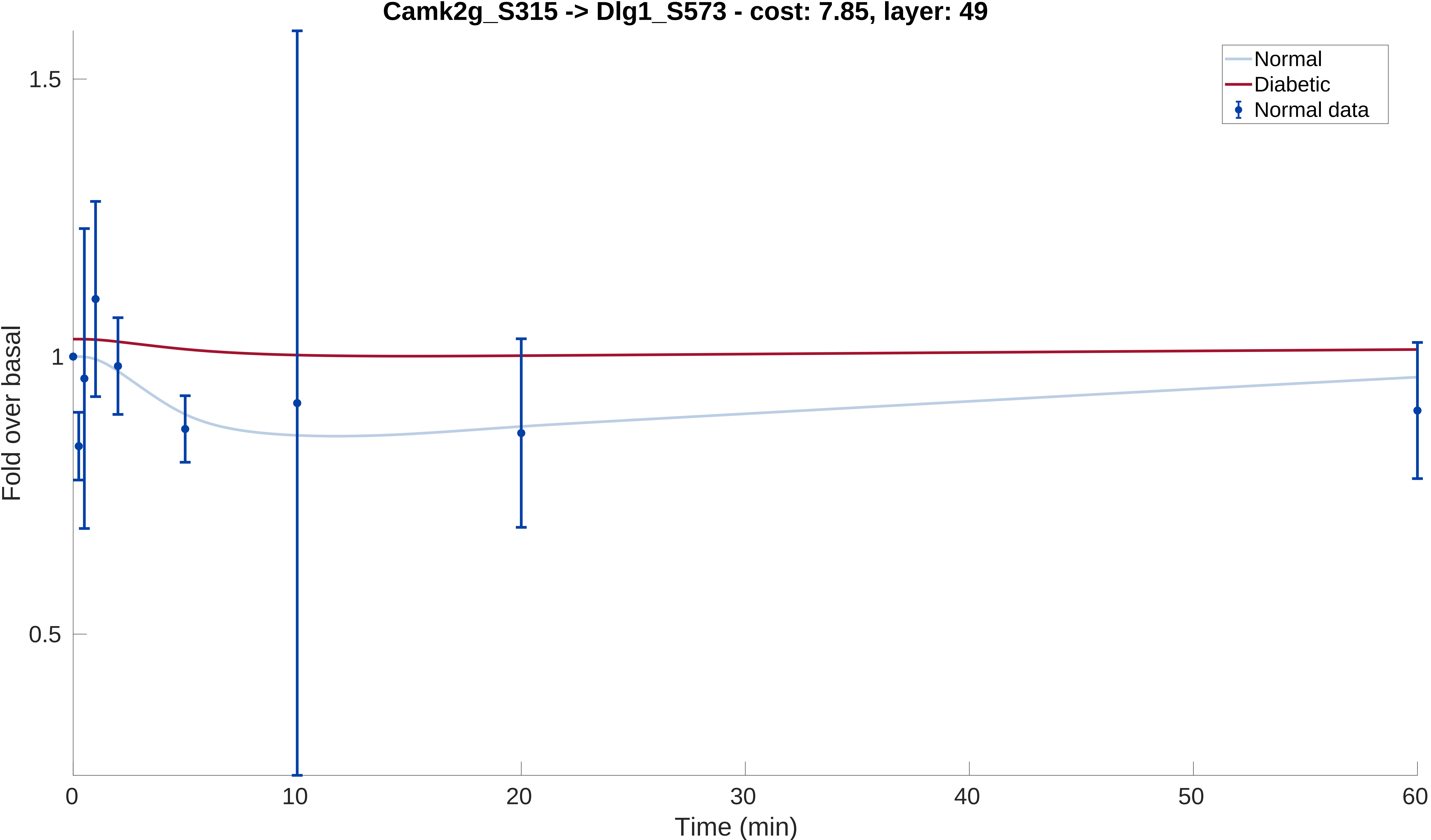

Camk2g_S315 -> Dlg1_S573 - cost: 7.85, layer: 49
Normal
Diabetic
Normal data
Fold over basal
1.5
1
0.5
0
10
20
30
40
50
60
Time (min)

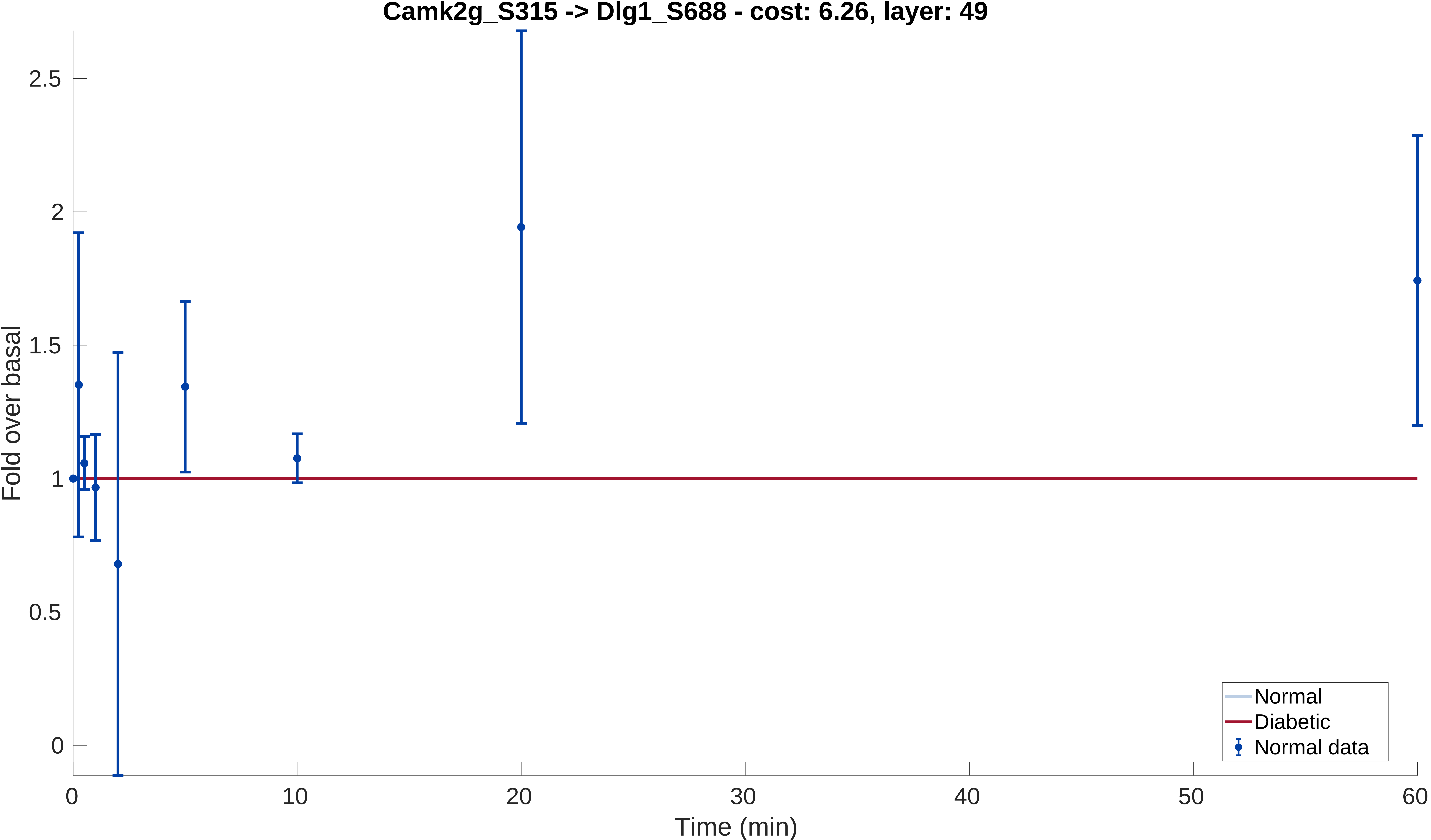
Camk2g_S315 -> Dlg1_S688 - cost: 6.26, layer: 49
Fold over basal
2.5
2
1.5
1
0.5
0
Time (min)
0
10
20
30
40
50
60
Normal
Diabetic
Normal data

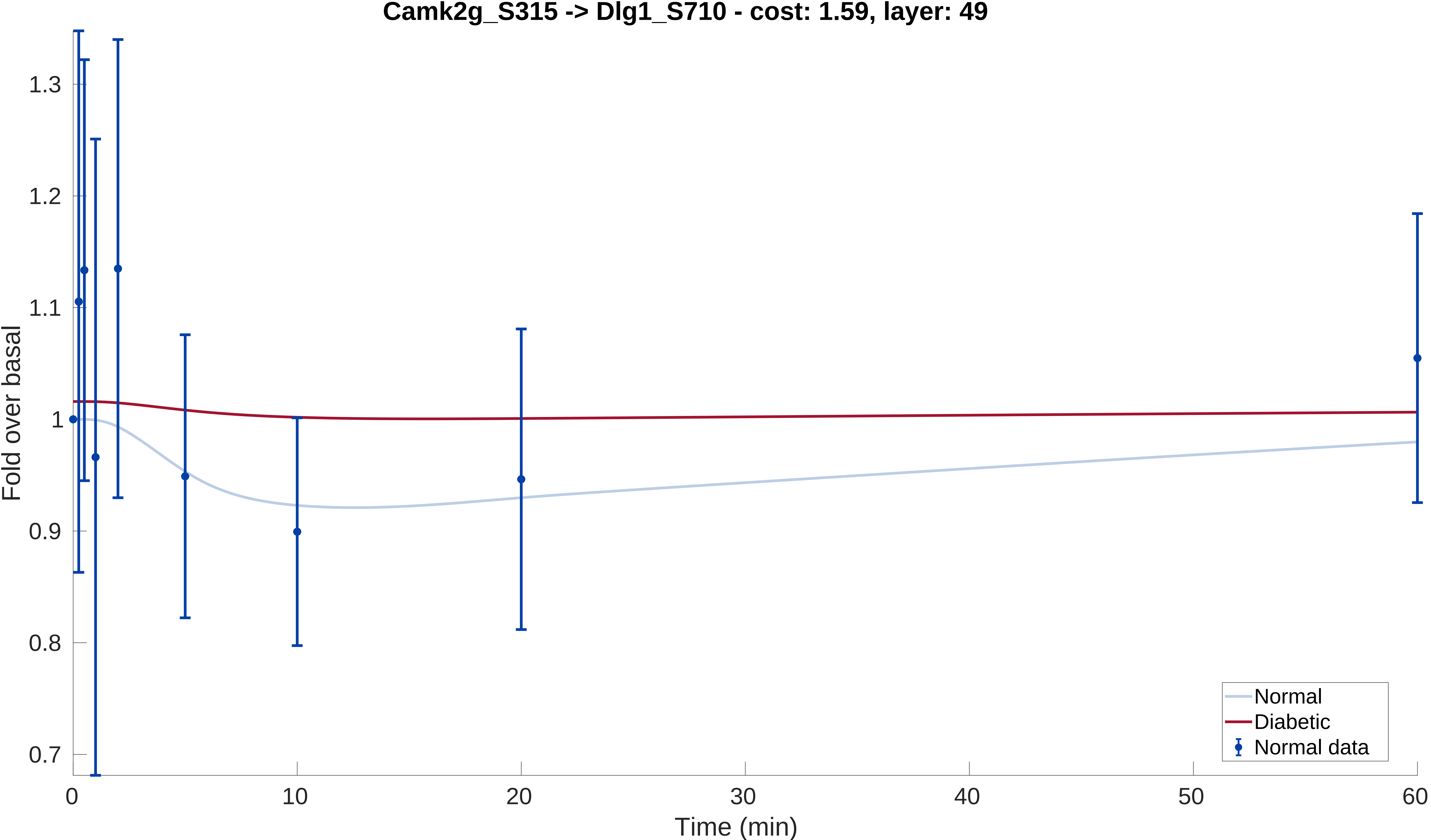

Camk2g_S315 -> Dlg1_S710 - cost: 1.59, layer: 49
Fold over basal
1.3
1.2
1.1
1
0.9
0.8
0.7
0
10
20
30
40
50
60
Time (min)
Normal
Diabetic
Normal data

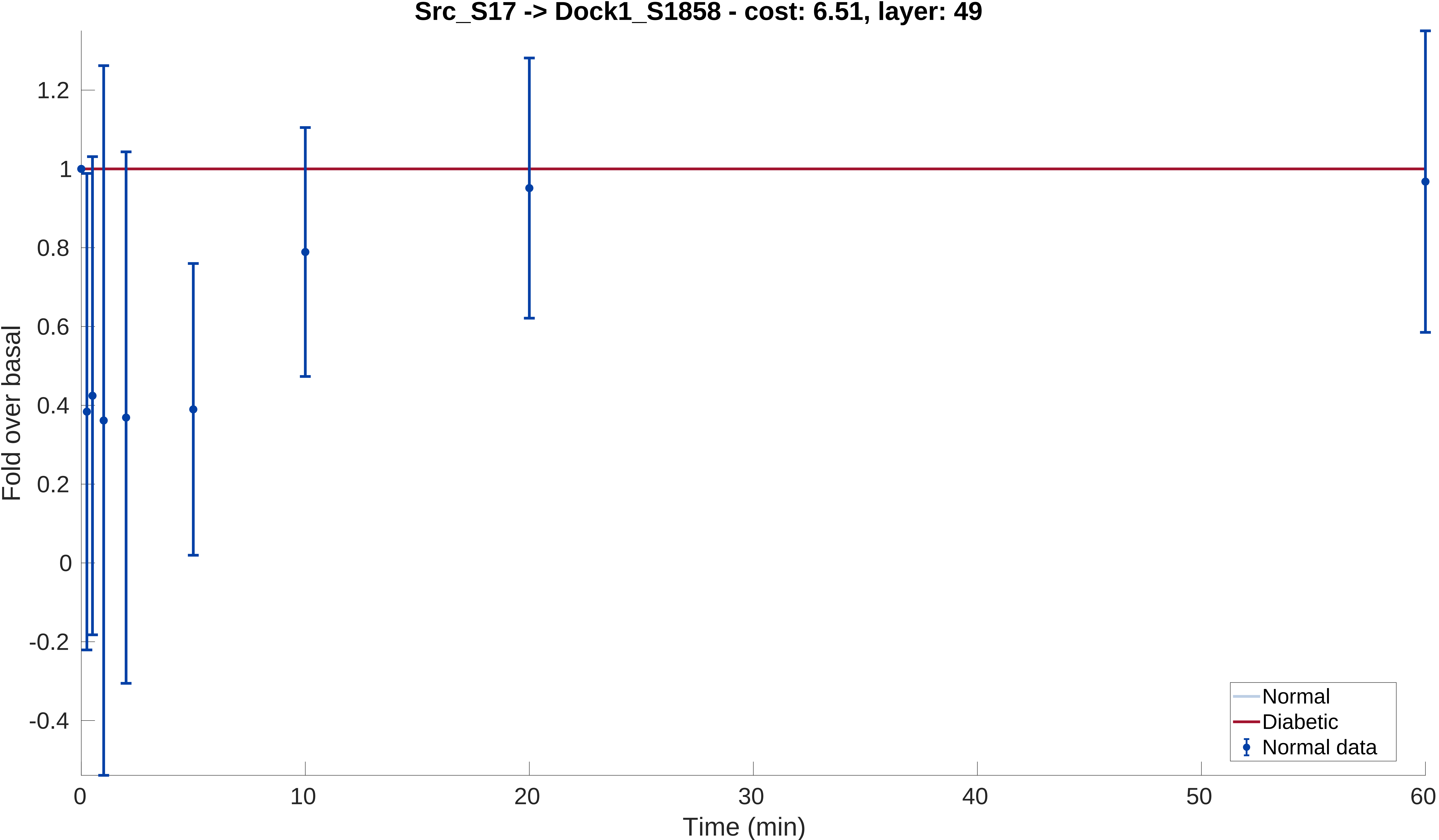

Src_S17 -> Dock1_S1858 - cost: 6.51, layer: 49
Fold over basal
1.2
1
0.8
0.6
0.4
0.2
0
-0.2
-0.4
0
10
20
30
40
50
60
Time (min)
Normal
Diabetic
Normal data

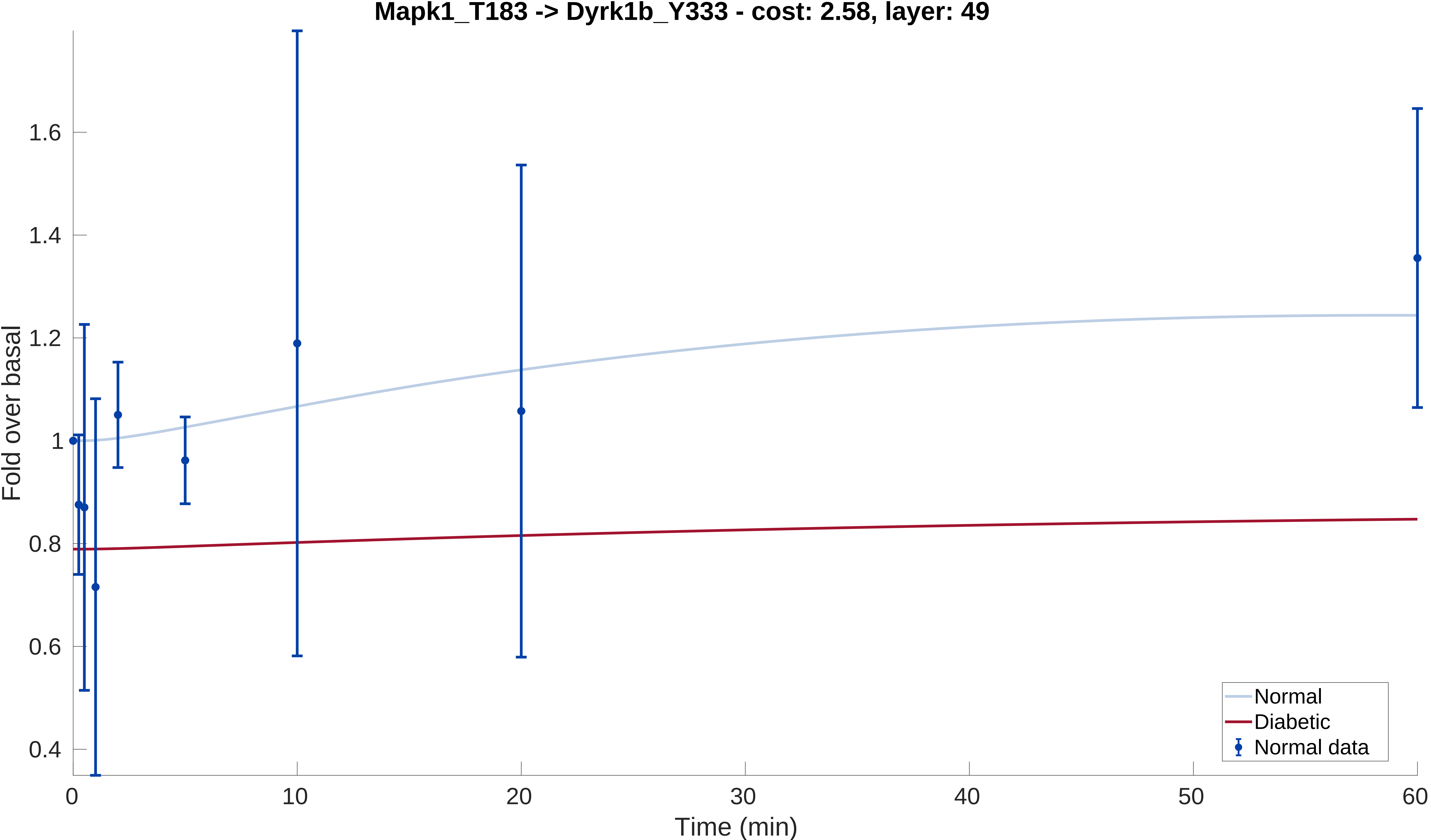
Mapk1_T183 -> Dyrk1b_Y333 - cost: 2.58, layer: 49
Fold over basal
1.6
1.4
1.2
1
0.8
0.6
0.4
0
10
20
30
40
50
60
Time (min)
Normal
Diabetic
Normal data

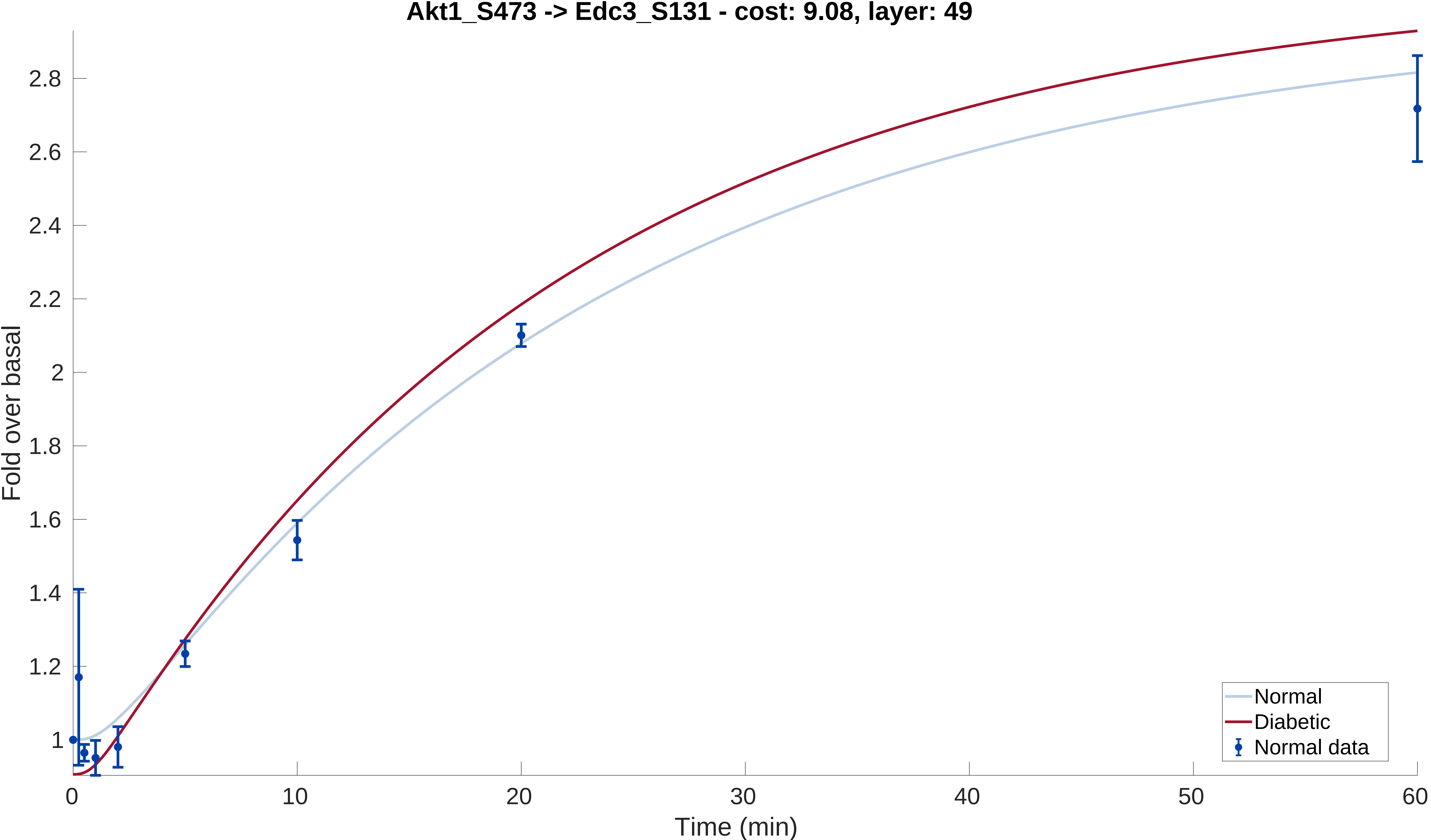

Akt1_S473 -> Edc3_S131 - cost: 9.08, layer: 49
Fold over basal
2.8
2.6
2.4
2.2
2
1.8
1.6
1.4
1.2
1
0
10
20
30
40
50
60
Time (min)
Normal
Diabetic
Normal data

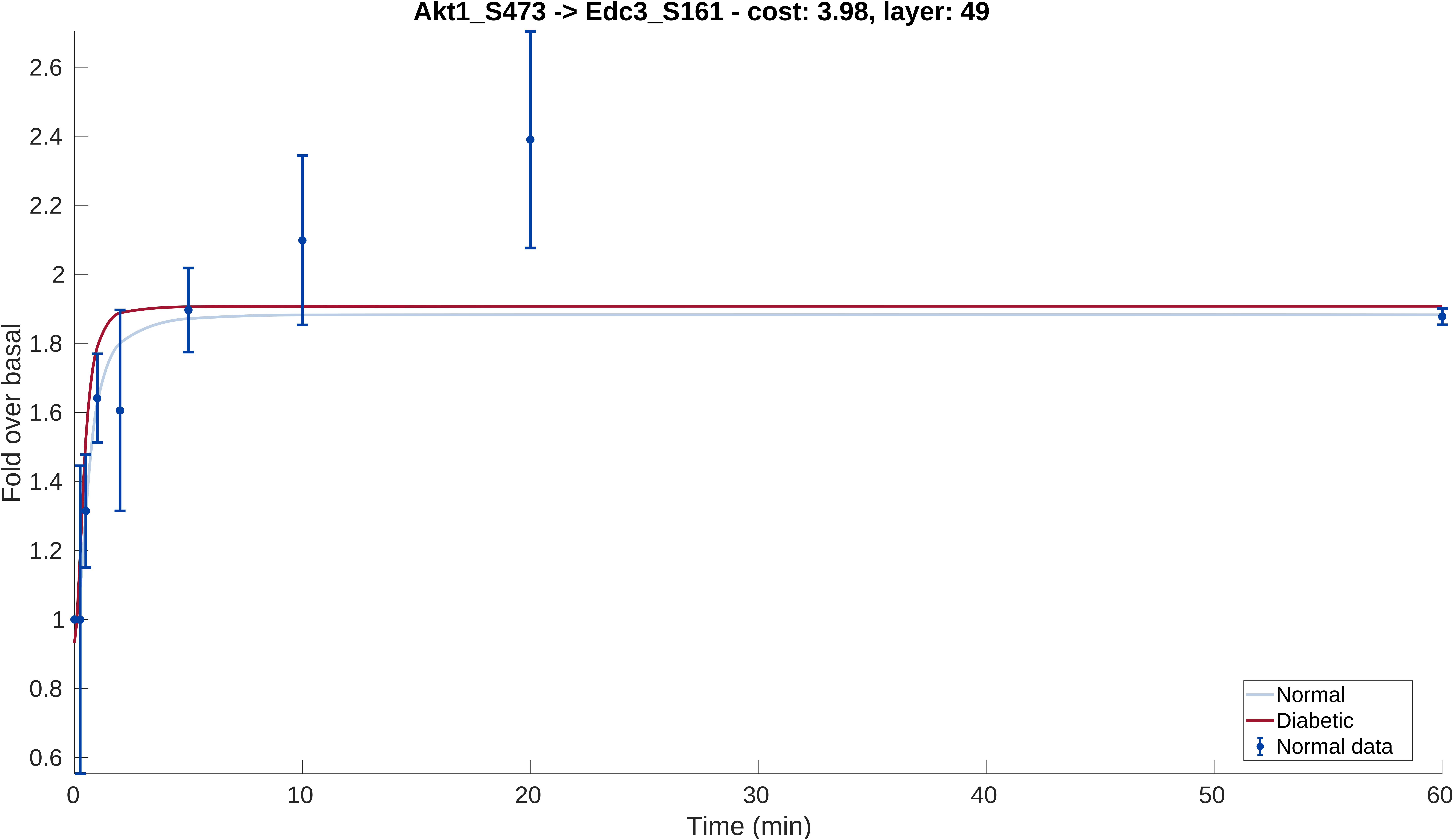

Akt1_S473 -> Edc3_S161 - cost: 3.98, layer: 49
Fold over basal
2.6
2.4
2.2
2
1.8
1.6
1.4
1.2
1
0.8
0.6
0
10
20
30
40
50
60
Time (min)
Normal
Diabetic
Normal data

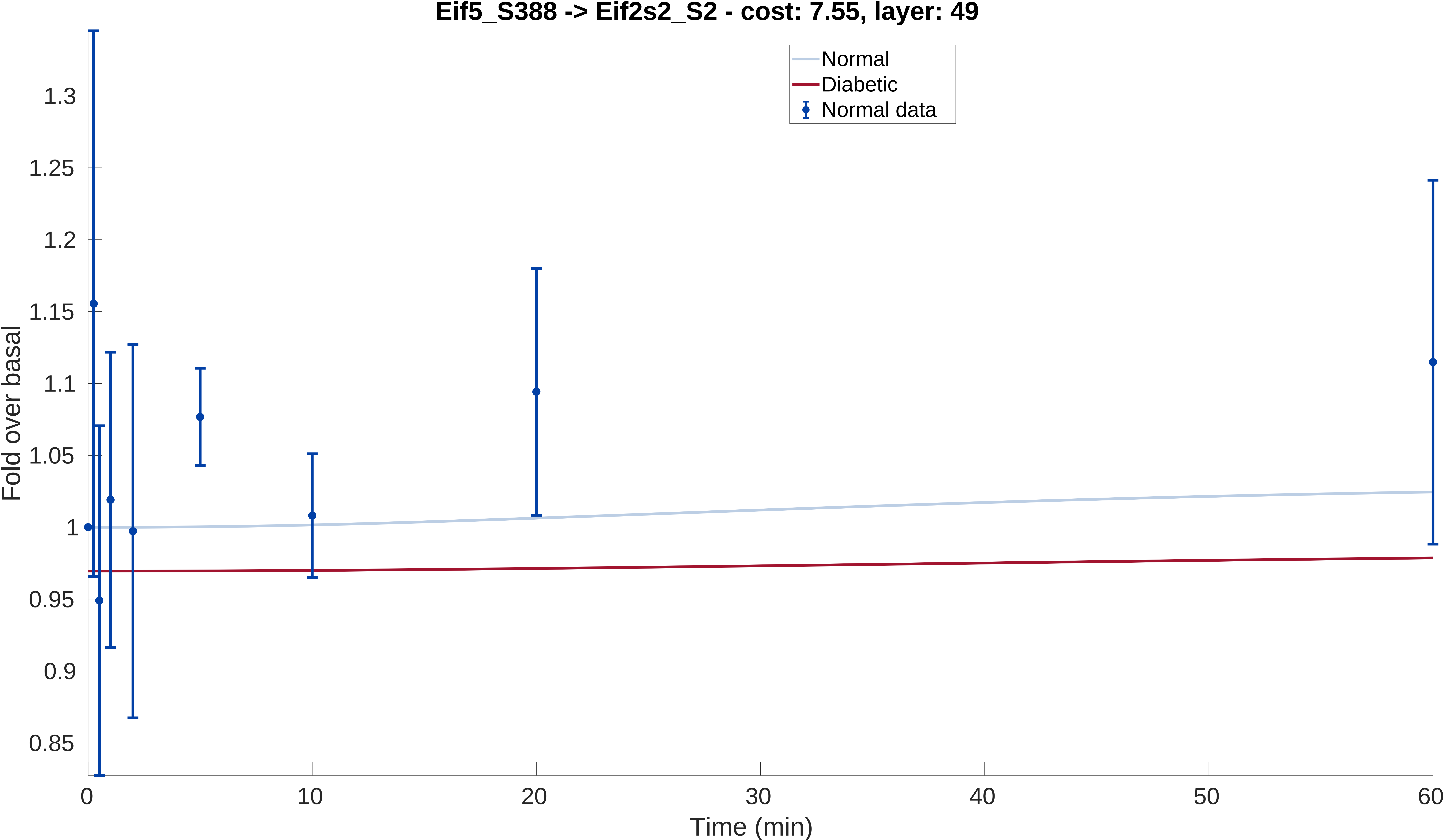

Eif5_S388 -> Eif2s2_S2 - cost: 7.55, layer: 49
Normal
Diabetic
Normal data
Fold over basal
1.3
1.25
1.2
1.15
1.1
1.05
1
0.95
0.9
0.85
0
10
20
30
40
50
60
Time (min)

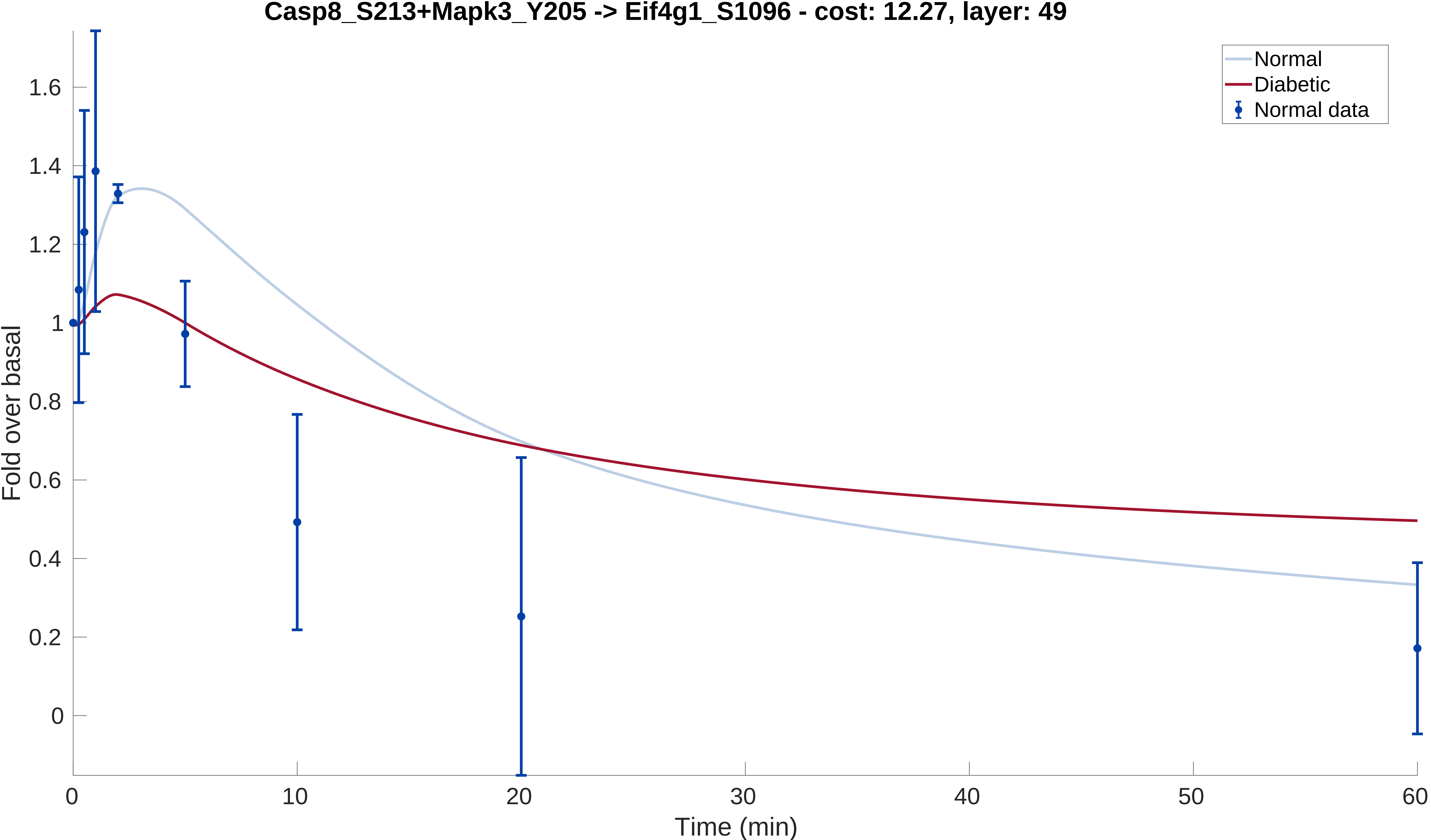

Casp8_S213+Mapk3_Y205 -> Eif4g1_S1096 - cost: 12.27, layer: 49
Normal
Diabetic
Normal data
Fold over basal
Time (min)
0
10
20
30
40
50
60
0
0.2
0.4
0.6
0.8
1
1.2
1.4
1.6

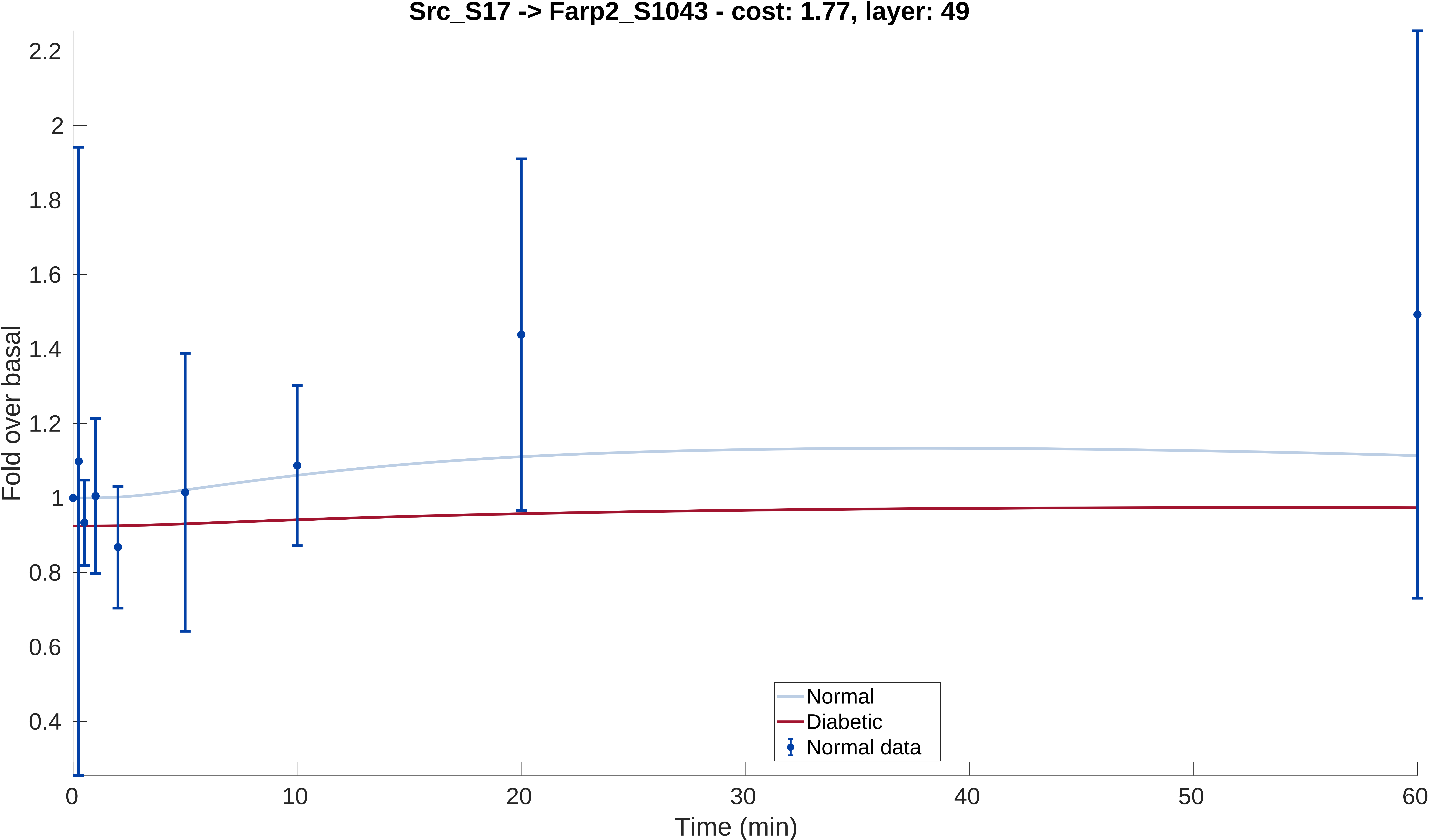

Src_S17 -> Farp2_S1043 - cost: 1.77, layer: 49
2.2
2
1.8
1.6
1.4
1.2
1
0.8
0.6
0.4
Fold over basal
0
10
20
30
40
50
60
Time (min)
Normal
Diabetic
Normal data

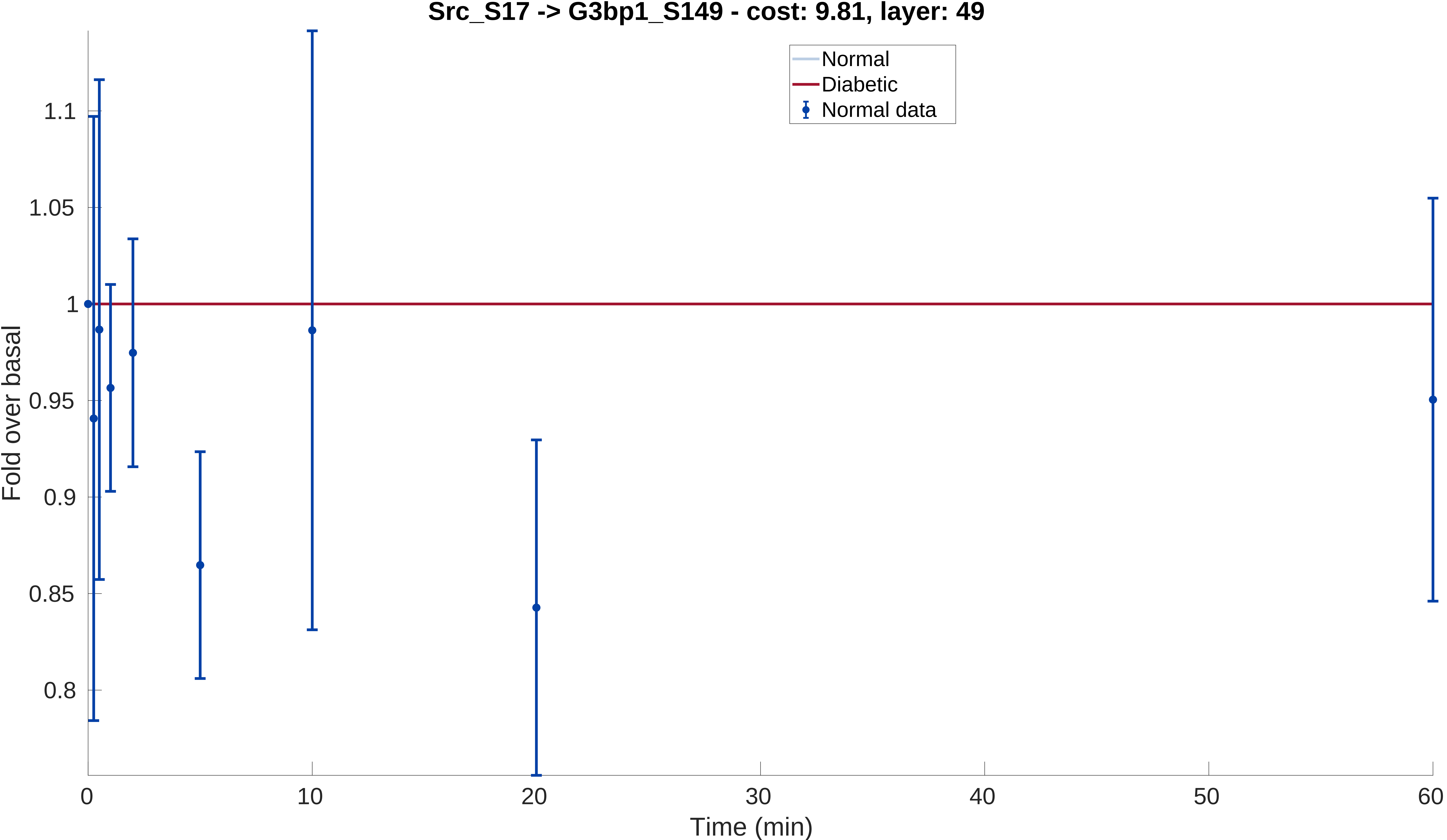

Src_S17 -> G3bp1_S149 - cost: 9.81, layer: 49
Normal
Diabetic
Normal data
Fold over basal
1.1
1.05
1
0.95
0.9
0.85
0.8
0
10
20
30
40
50
60
Time (min)

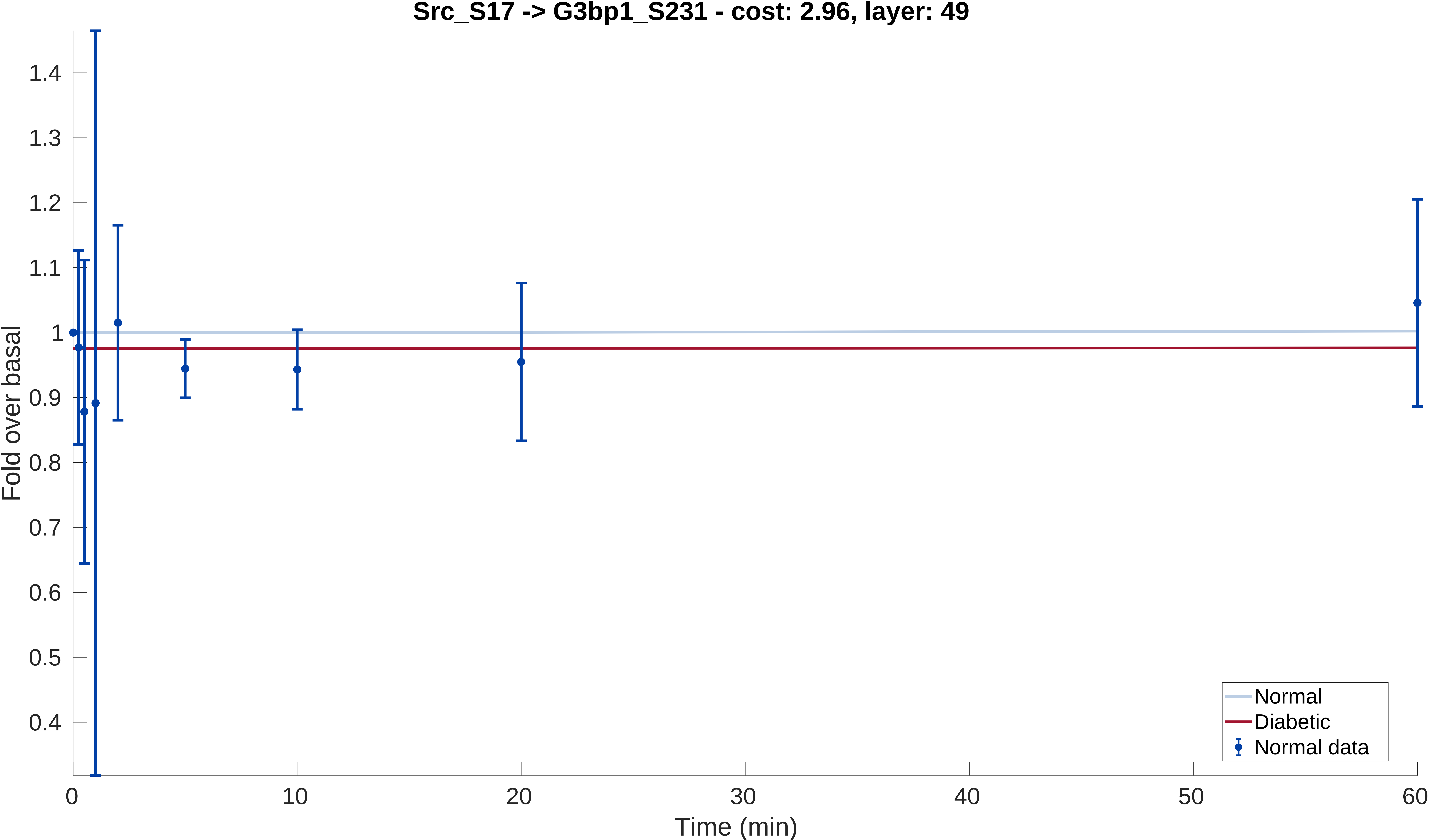

Src_S17 -> G3bp1_S231 - cost: 2.96, layer: 49
Fold over basal
1.4
1.3
1.2
1.1
1
0.9
0.8
0.7
0.6
0.5
0.4
0
10
20
30
40
50
60
Time (min)
Normal
Diabetic
Normal data

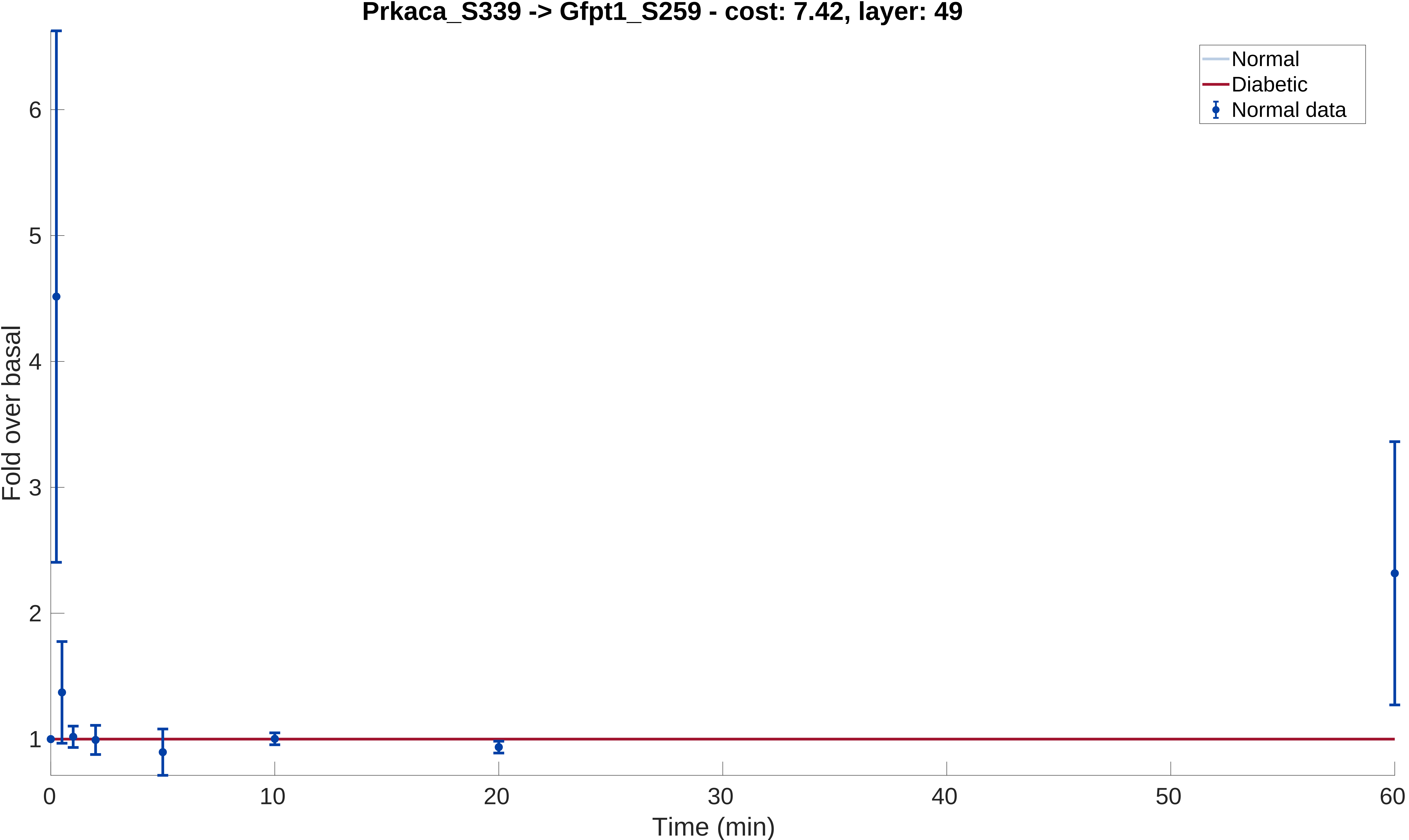

Prkaca_S339 -> Gfpt1_S259 - cost: 7.42, layer: 49
Normal
Diabetic
Normal data
Fold over basal
Time (min)
1
2
3
4
5
6
0
10
20
30
40
50
60

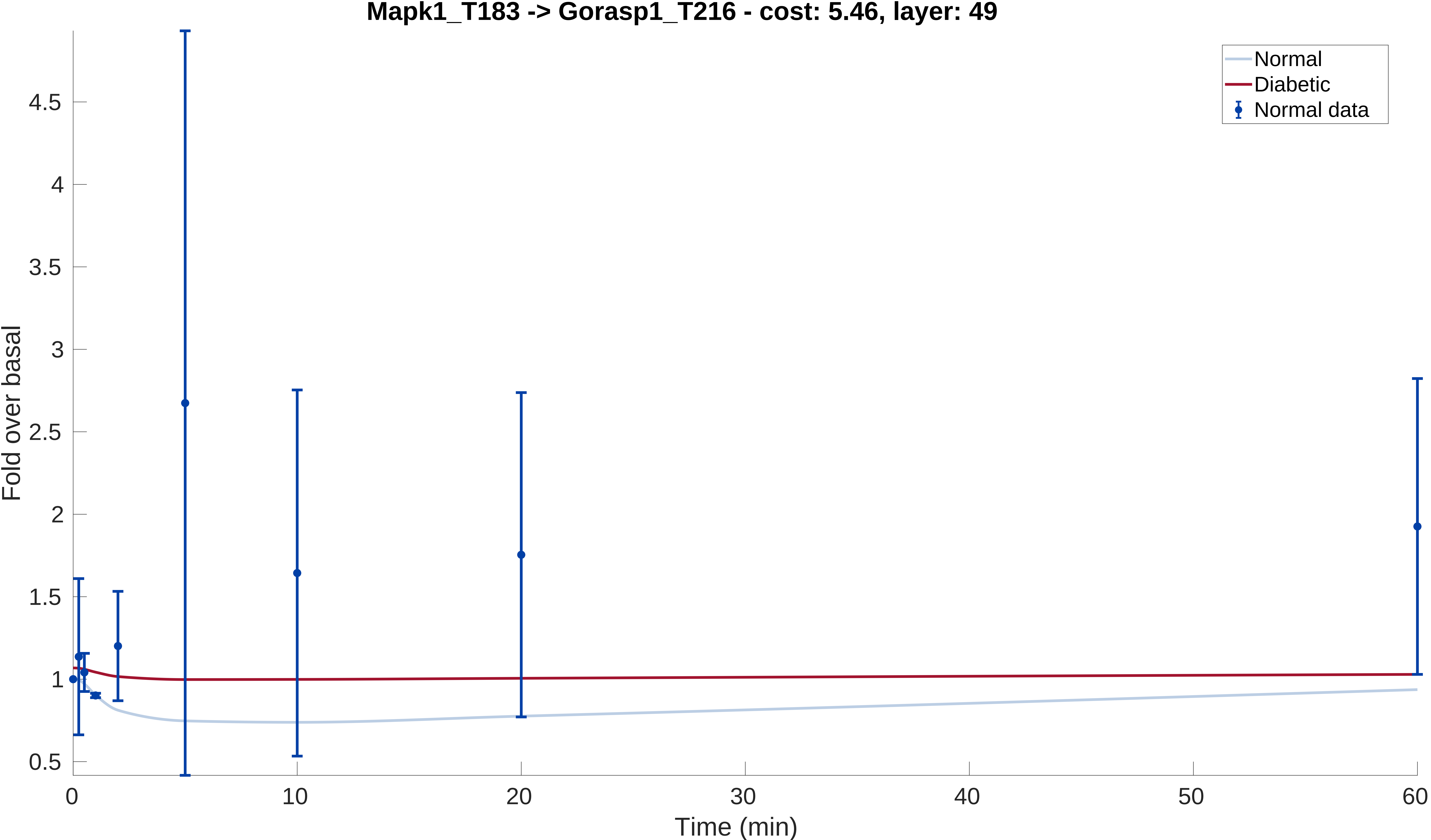

Mapk1_T183 -> Gorasp1_T216 - cost: 5.46, layer: 49
Normal
Diabetic
Normal data
Fold over basal
Time (min)
0.5
1
1.5
2
2.5
3
3.5
4
4.5
0
10
20
30
40
50
60

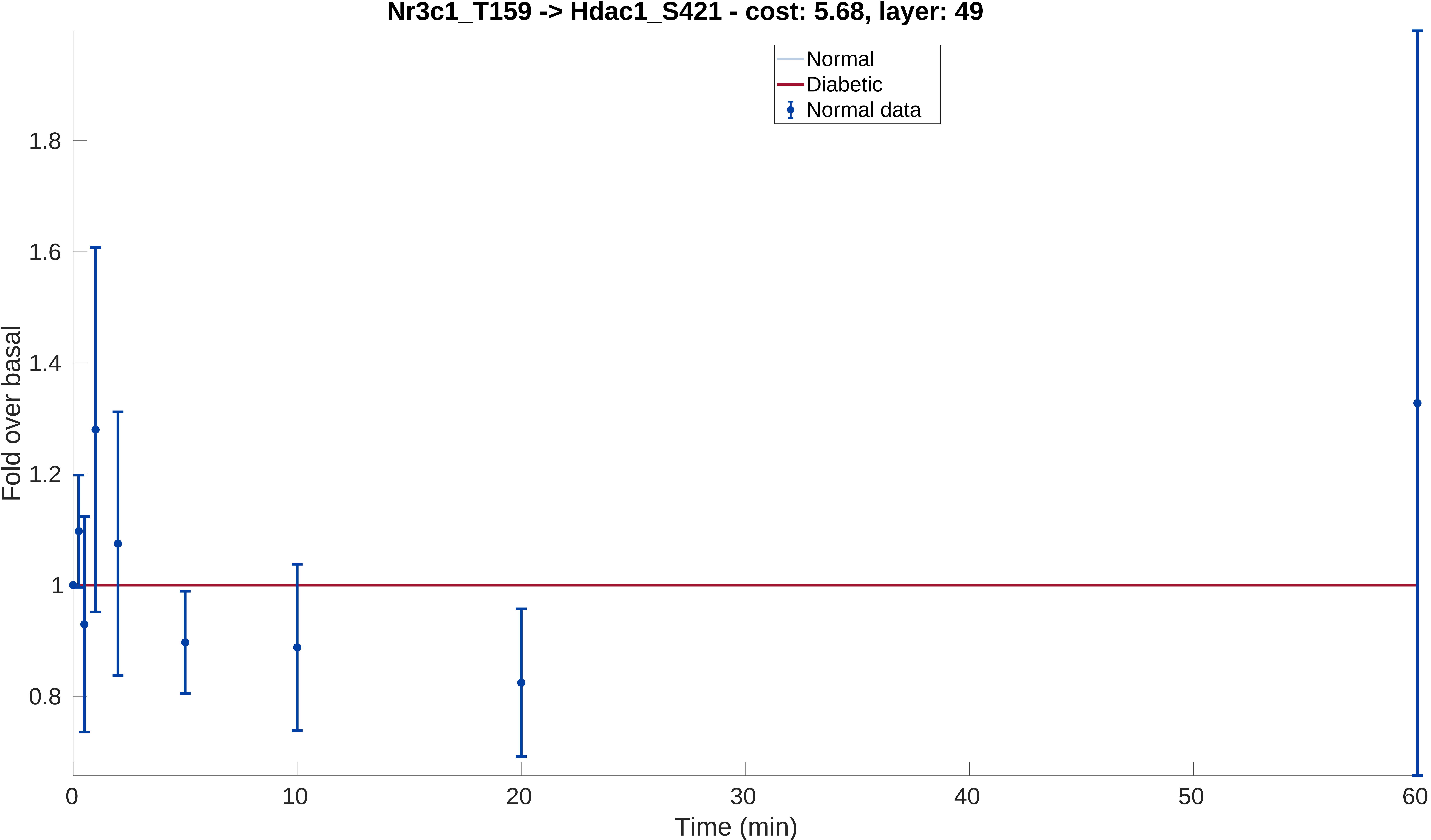

Nr3c1_T159 -> Hdac1_S421 - cost: 5.68, layer: 49
Normal
Diabetic
Normal data
Fold over basal
1.8
1.6
1.4
1.2
1
0.8
0
10
20
30
40
50
60
Time (min)

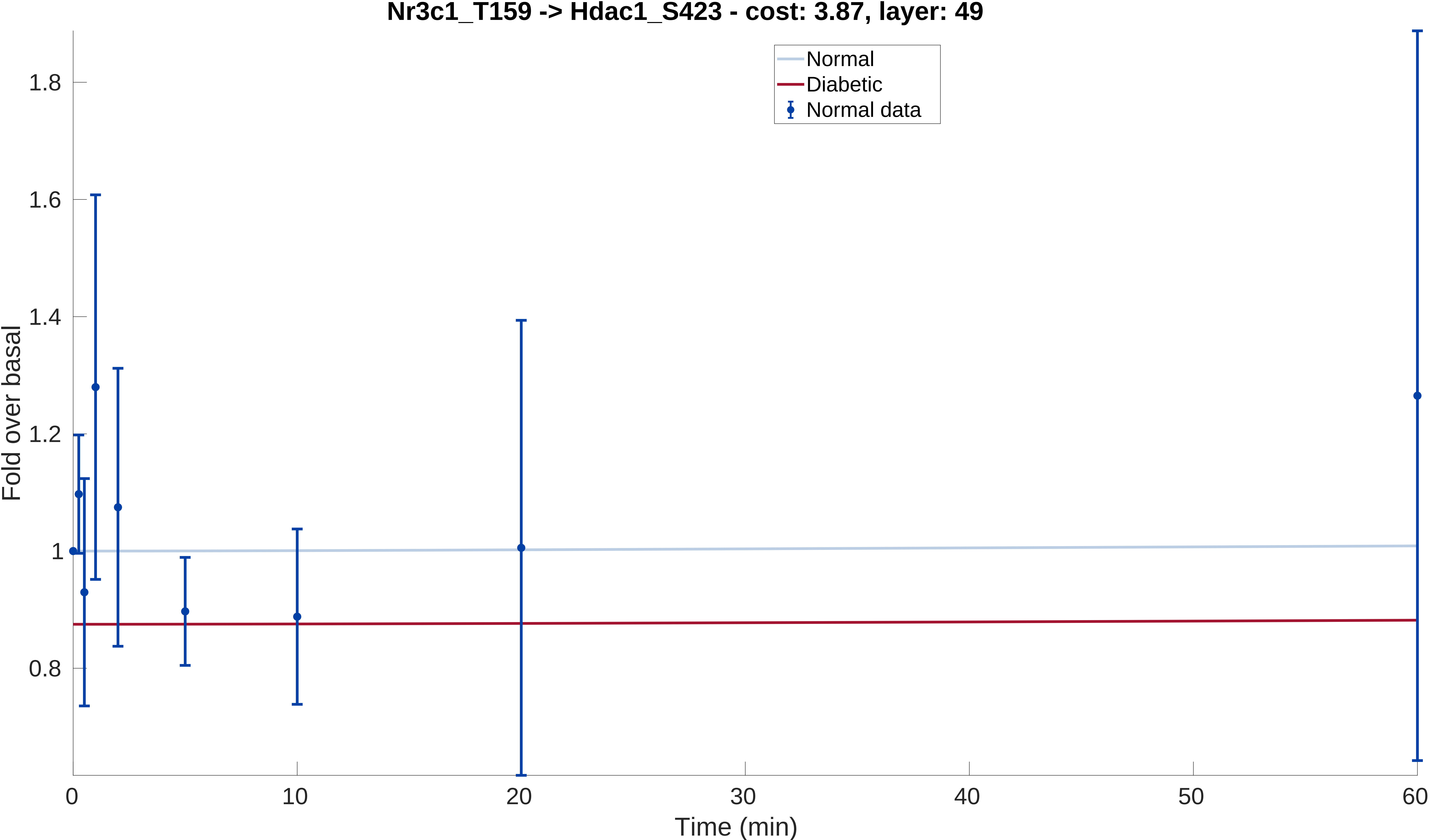

Nr3c1_T159 -> Hdac1_S423 - cost: 3.87, layer: 49
Normal
Diabetic
Normal data
Fold over basal
1.8
1.6
1.4
1.2
1
0.8
0
10
20
30
40
50
60
Time (min)

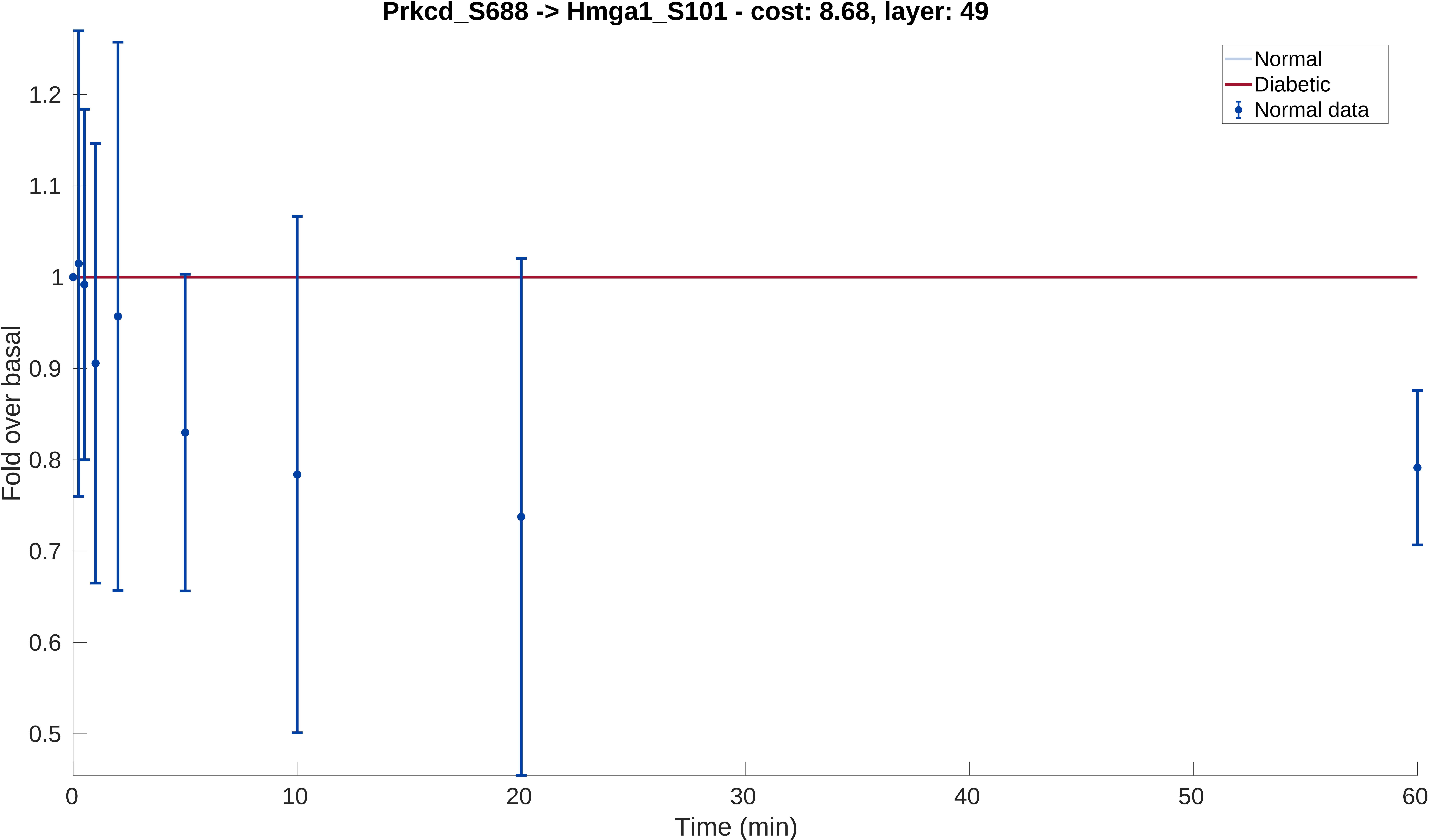
Prkcd_S688 -> Hmga1_S101 - cost: 8.68, layer: 49
Normal
Diabetic
Normal data
Fold over basal
1.2
1.1
1
0.9
0.8
0.7
0.6
0.5
0
10
20
30
40
50
60
Time (min)

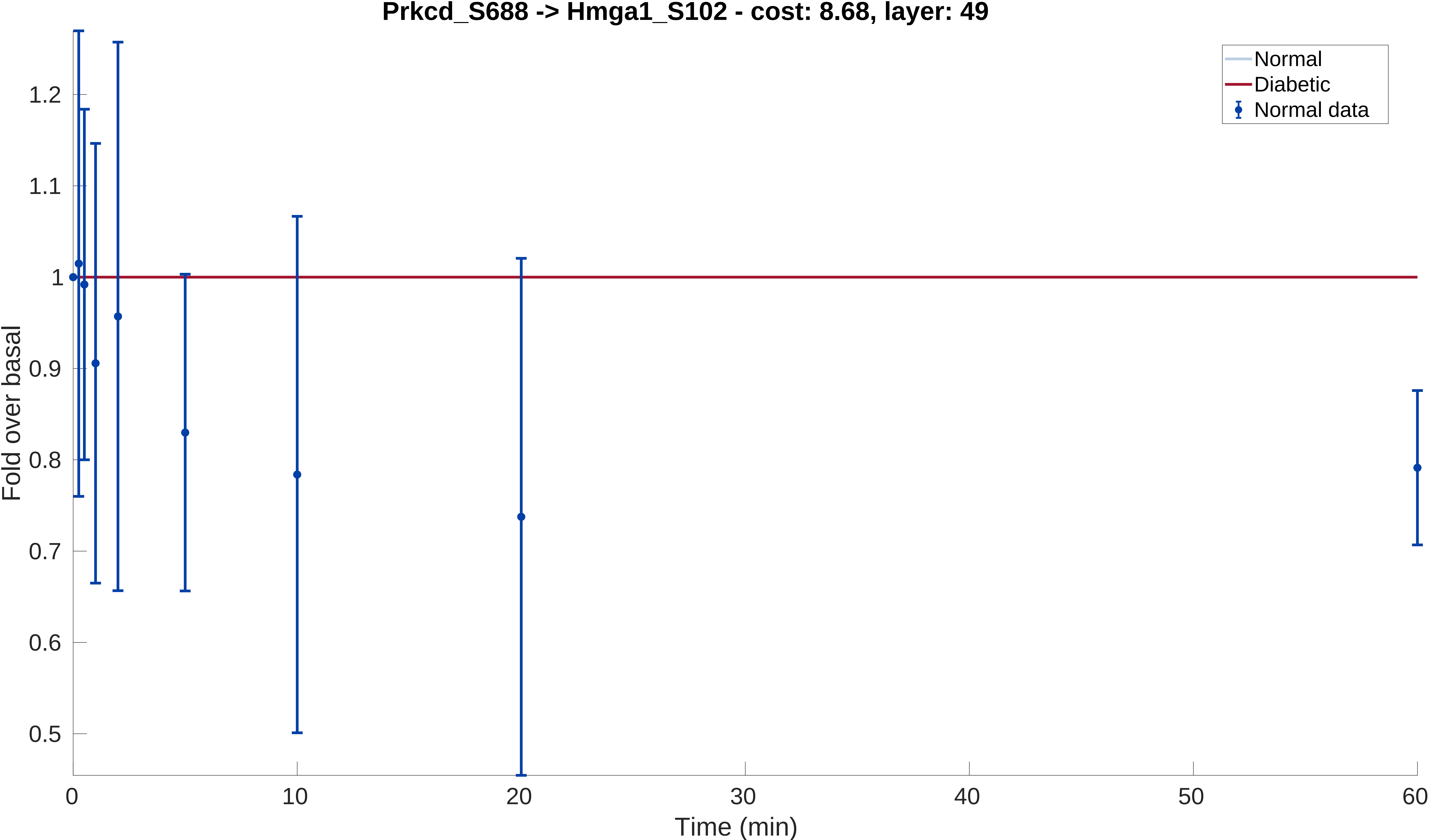
Prkcd_S688 -> Hmga1_S102 - cost: 8.68, layer: 49
Normal
Diabetic
Normal data
Fold over basal
1.2
1.1
1
0.9
0.8
0.7
0.6
0.5
0
10
20
30
40
50
60
Time (min)

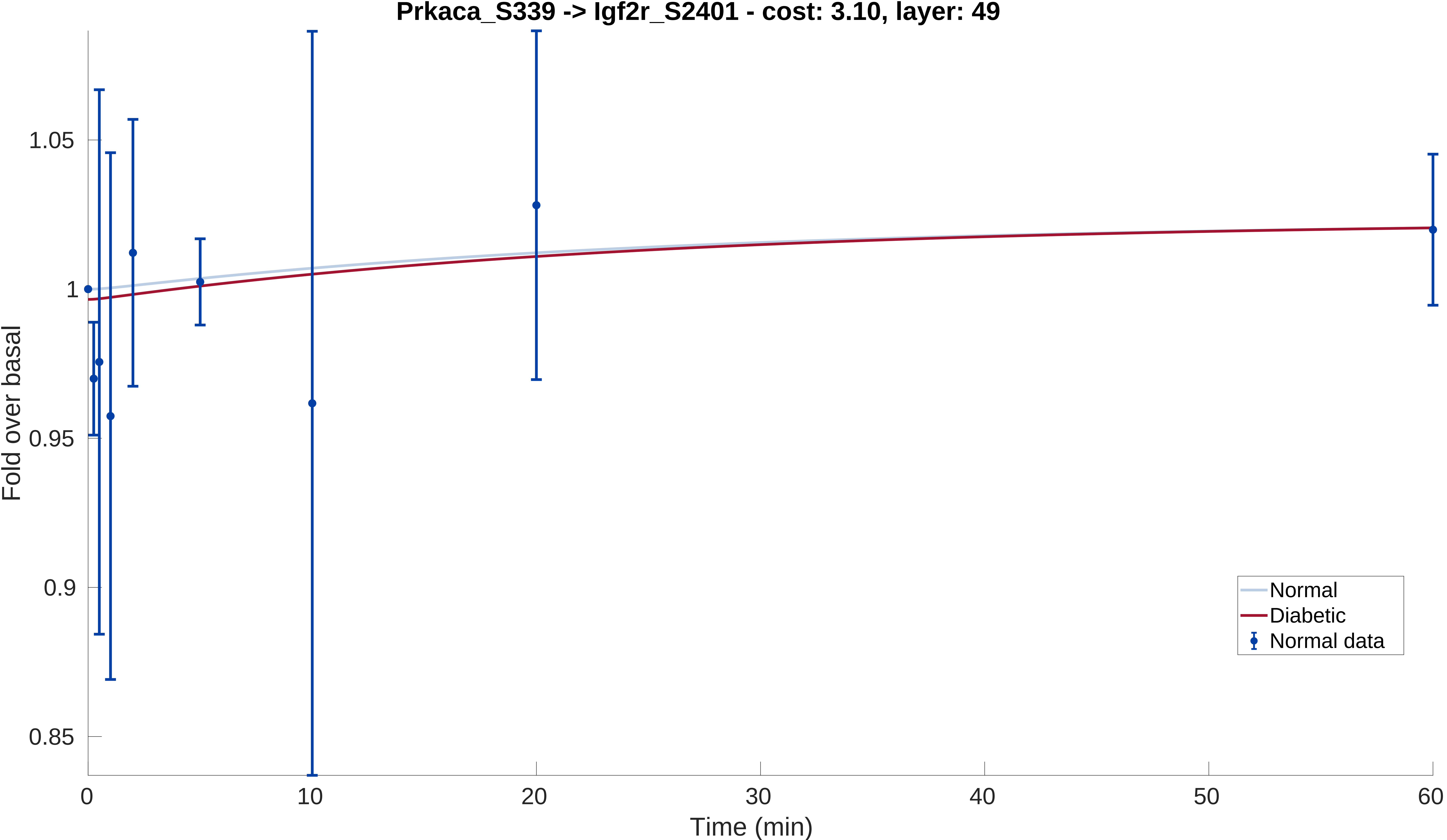

Prkaca_S339 -> Igf2r_S2401 - cost: 3.10, layer: 49
Fold over basal
1.05
1
0.95
0.9
0.85
0
10
20
30
40
50
60
Time (min)
Normal
Diabetic
Normal data

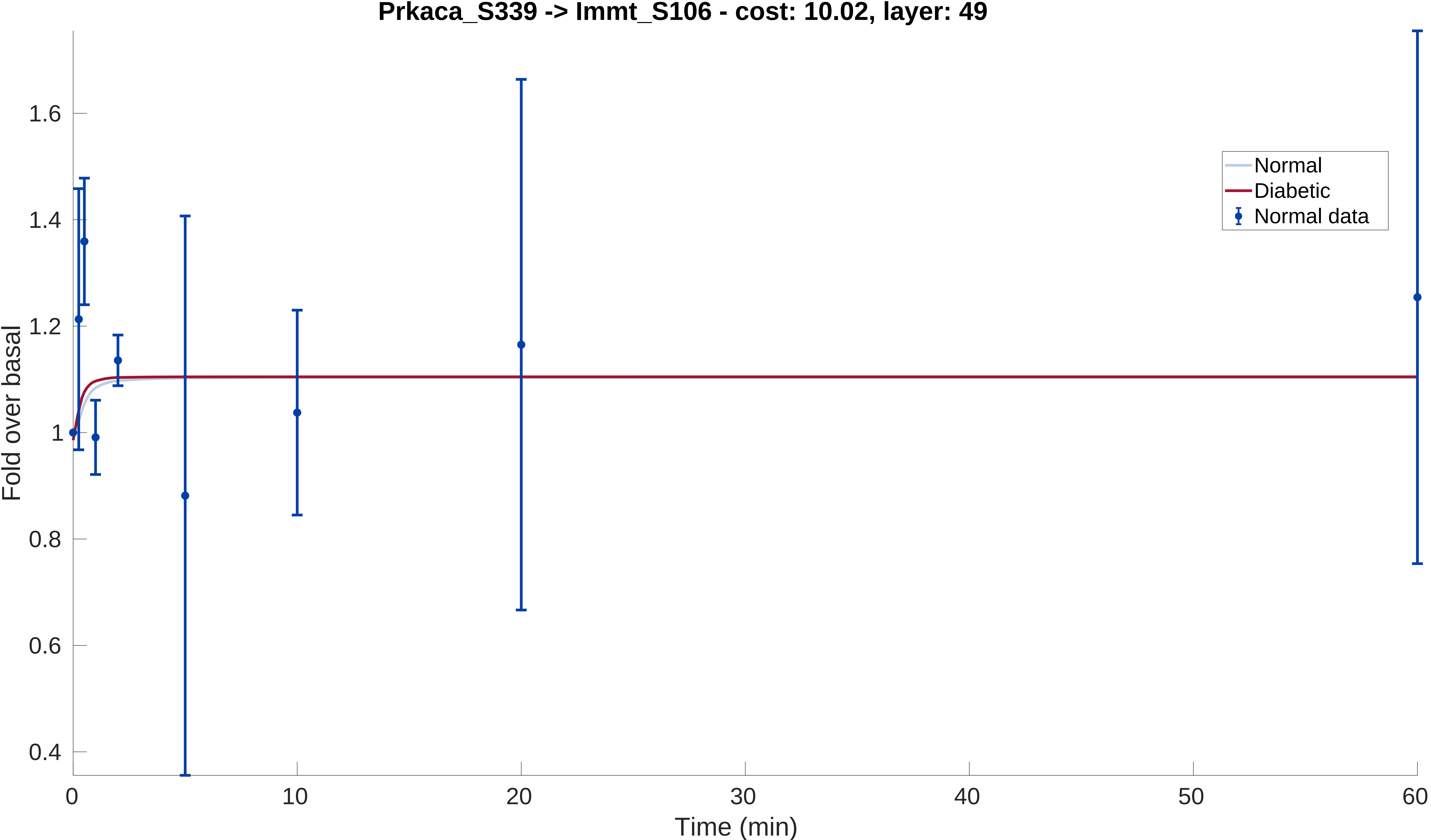
Prkaca_S339 -> Immt_S106 - cost: 10.02, layer: 49
Fold over basal
1.6
1.4
1.2
1
0.8
0.6
0.4
0
10
20
30
40
50
60
Time (min)
Normal
Diabetic
Normal data

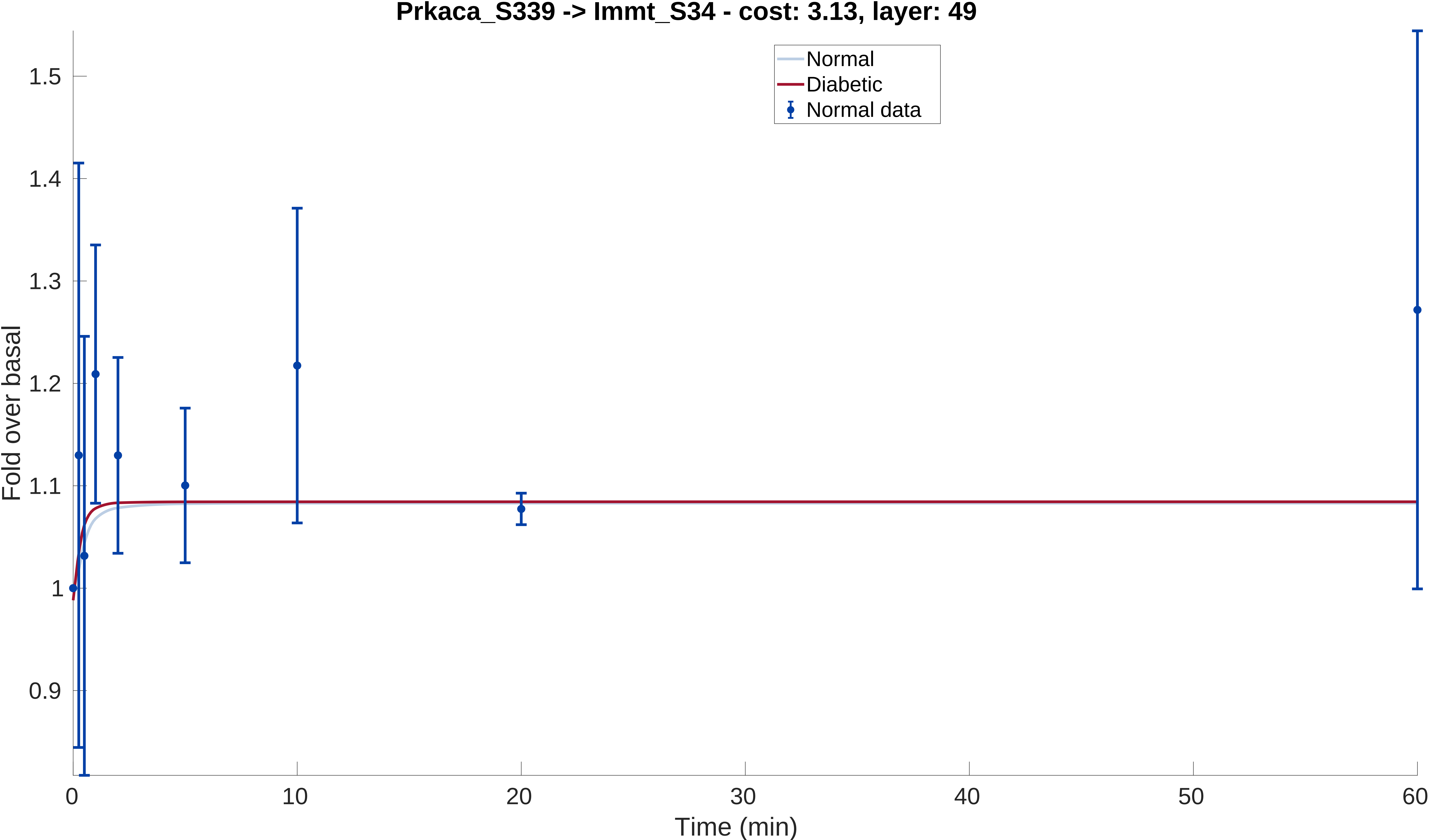

Prkaca_S339 -> Immt_S34 - cost: 3.13, layer: 49
Normal
Diabetic
Normal data
Fold over basal
1.5
1.4
1.3
1.2
1.1
1
0.9
0
10
20
30
40
50
60
Time (min)

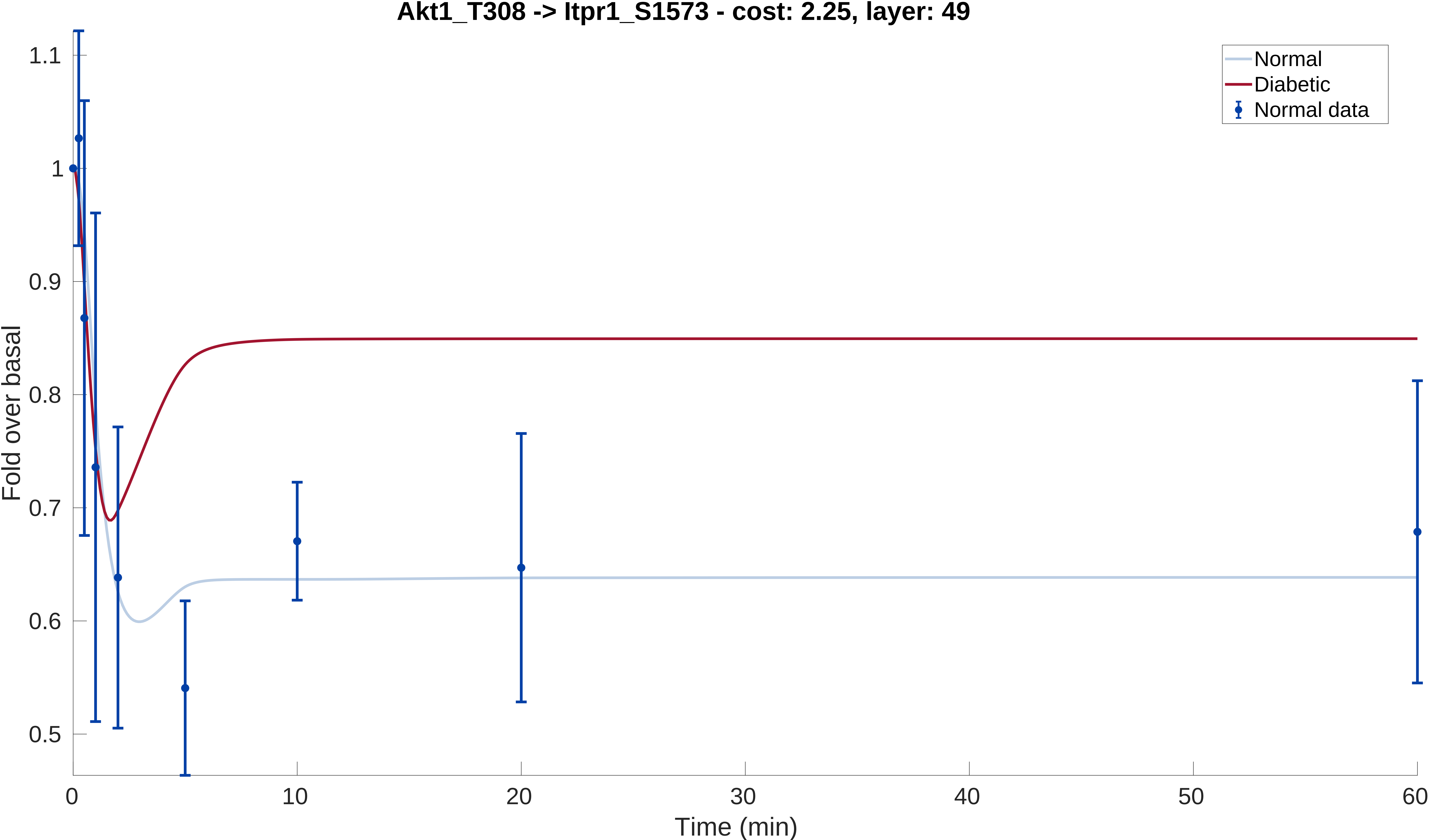

Akt1_T308 -> Itpr1_S1573 - cost: 2.25, layer: 49
Normal
Diabetic
Normal data
Fold over basal
1.1
1
0.9
0.8
0.7
0.6
0.5
0
10
20
30
40
50
60
Time (min)

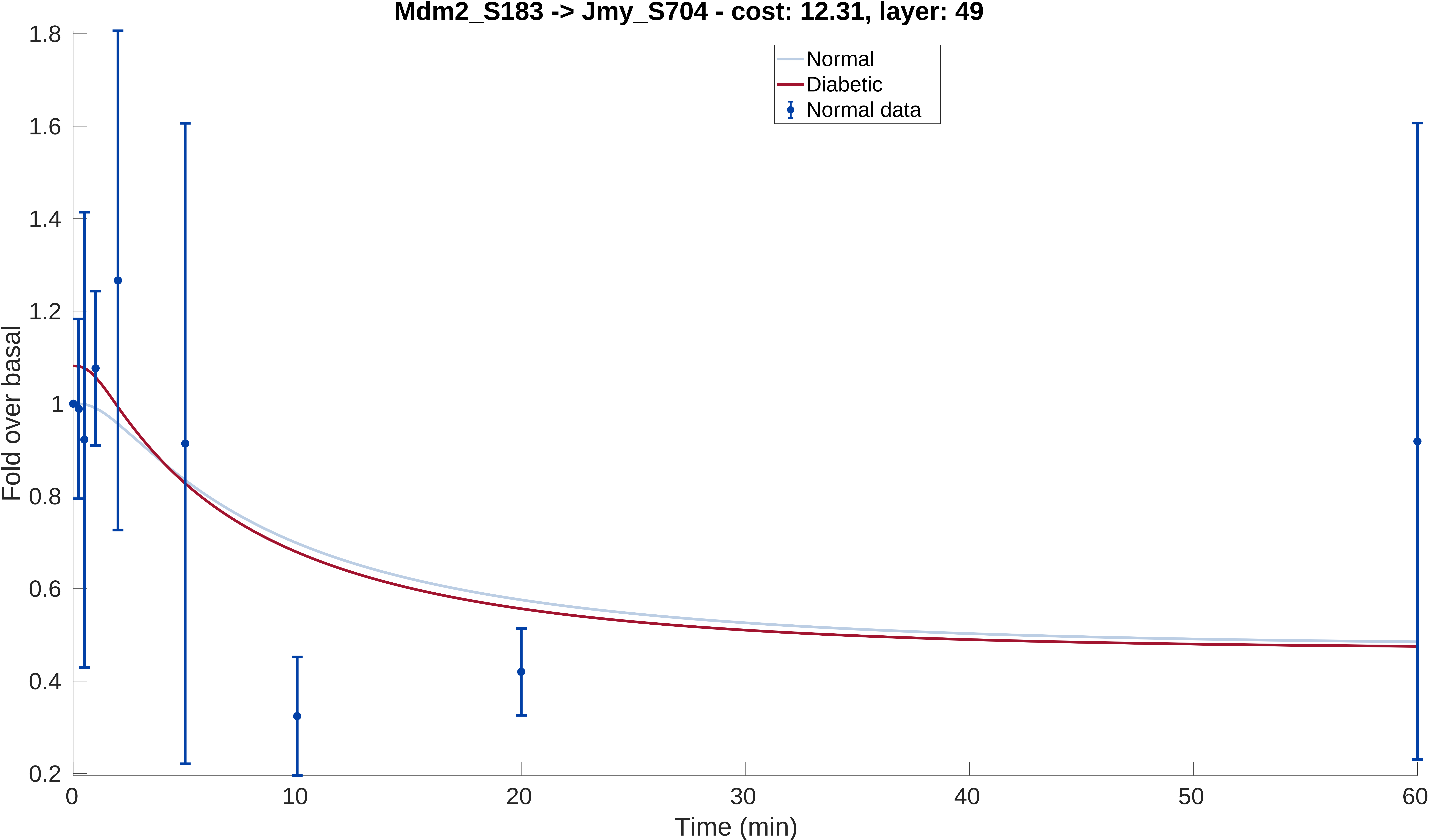

Mdm2_S183 -> Jmy_S704 - cost: 12.31, layer: 49
Normal
Diabetic
Normal data
Fold over basal
Time (min)
1.8
1.6
1.4
1.2
1
0.8
0.6
0.4
0.2
0
10
20
30
40
50
60

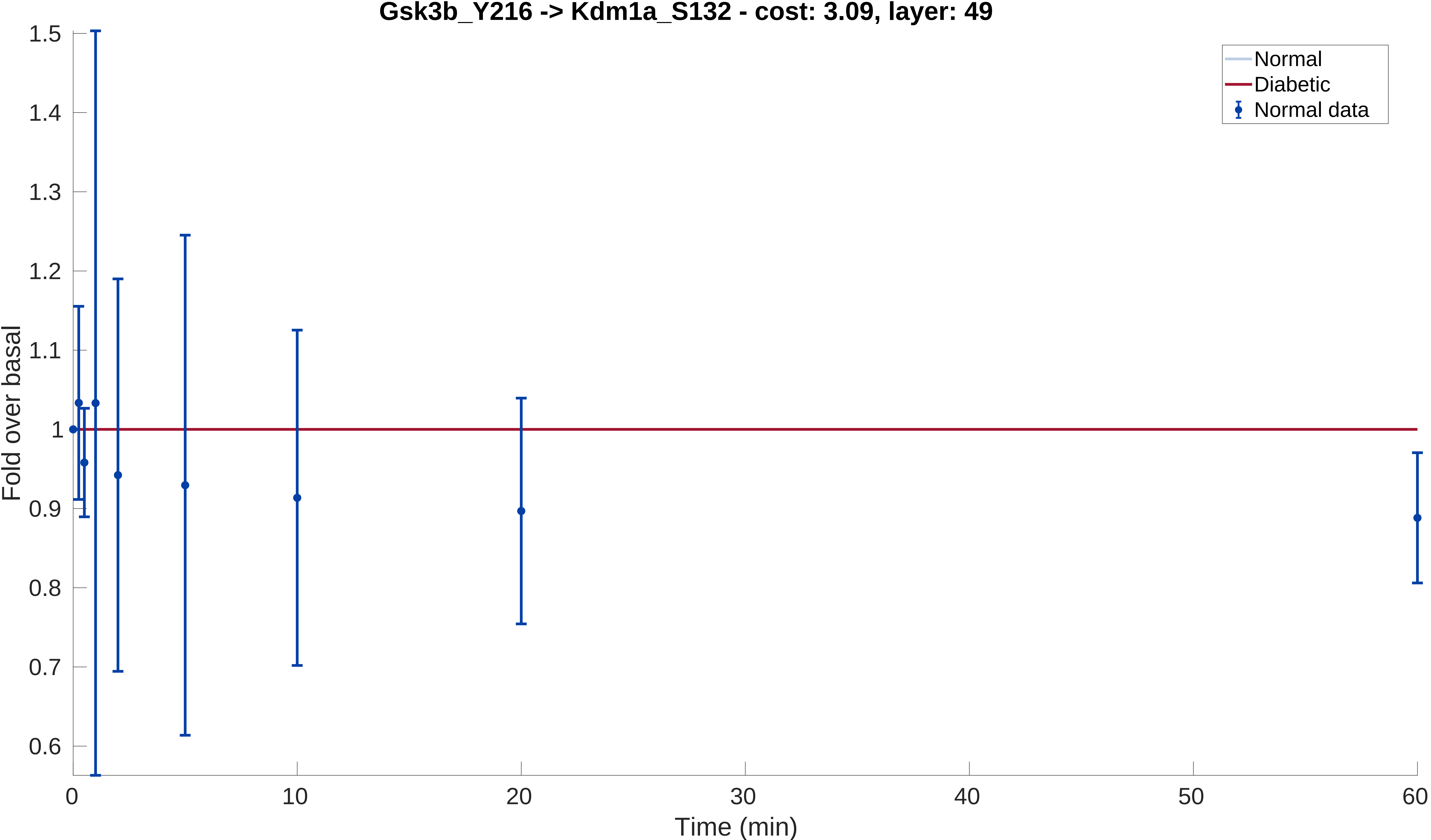

Gsk3b_Y216 -> Kdm1a_S132 - cost: 3.09, layer: 49
Normal
Diabetic
Normal data
Fold over basal
Time (min)
1.5
1.4
1.3
1.2
1.1
1
0.9
0.8
0.7
0.6
0
10
20
30
40
50
60

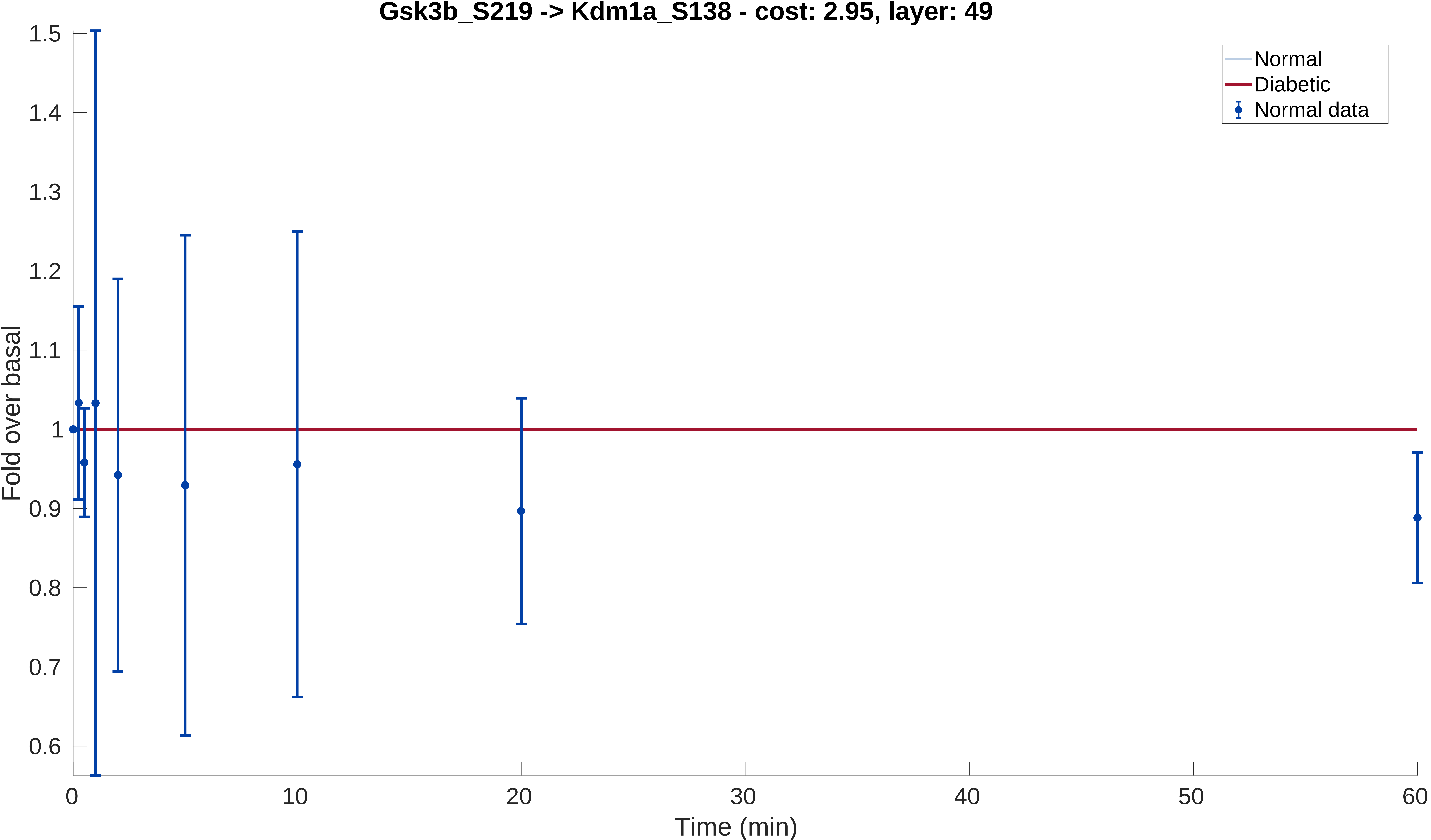

Gsk3b_S219 -> Kdm1a_S138 - cost: 2.95, layer: 49
Fold over basal
Time (min)
Normal
Diabetic
Normal data
1.5
1.4
1.3
1.2
1.1
1
0.9
0.8
0.7
0.6
0
10
20
30
40
50
60

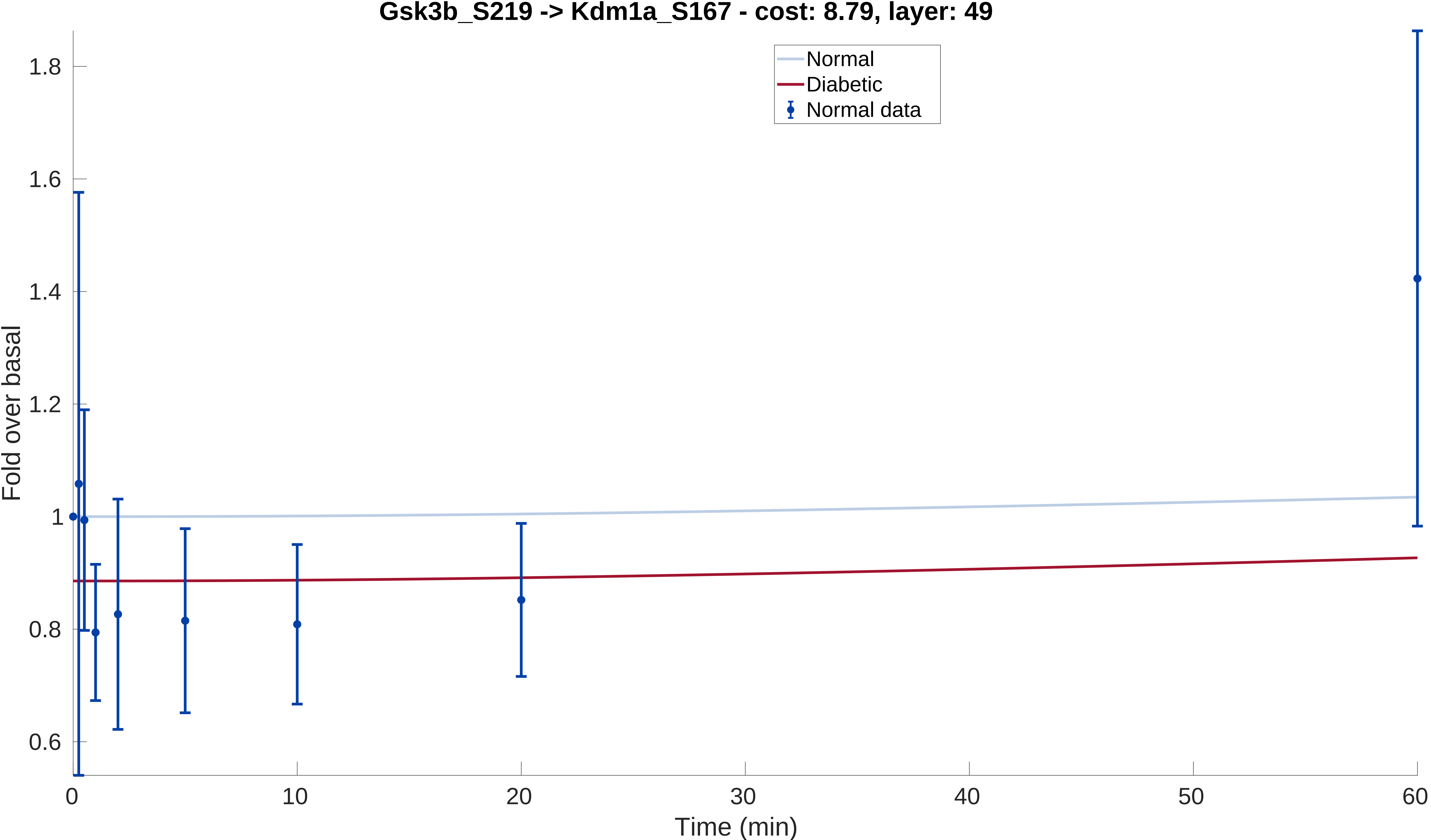

Gsk3b_S219 -> Kdm1a_S167 - cost: 8.79, layer: 49
Normal
Diabetic
Normal data
Fold over basal
Time (min)
0.6
0.8
1
1.2
1.4
1.6
1.8
0
10
20
30
40
50
60

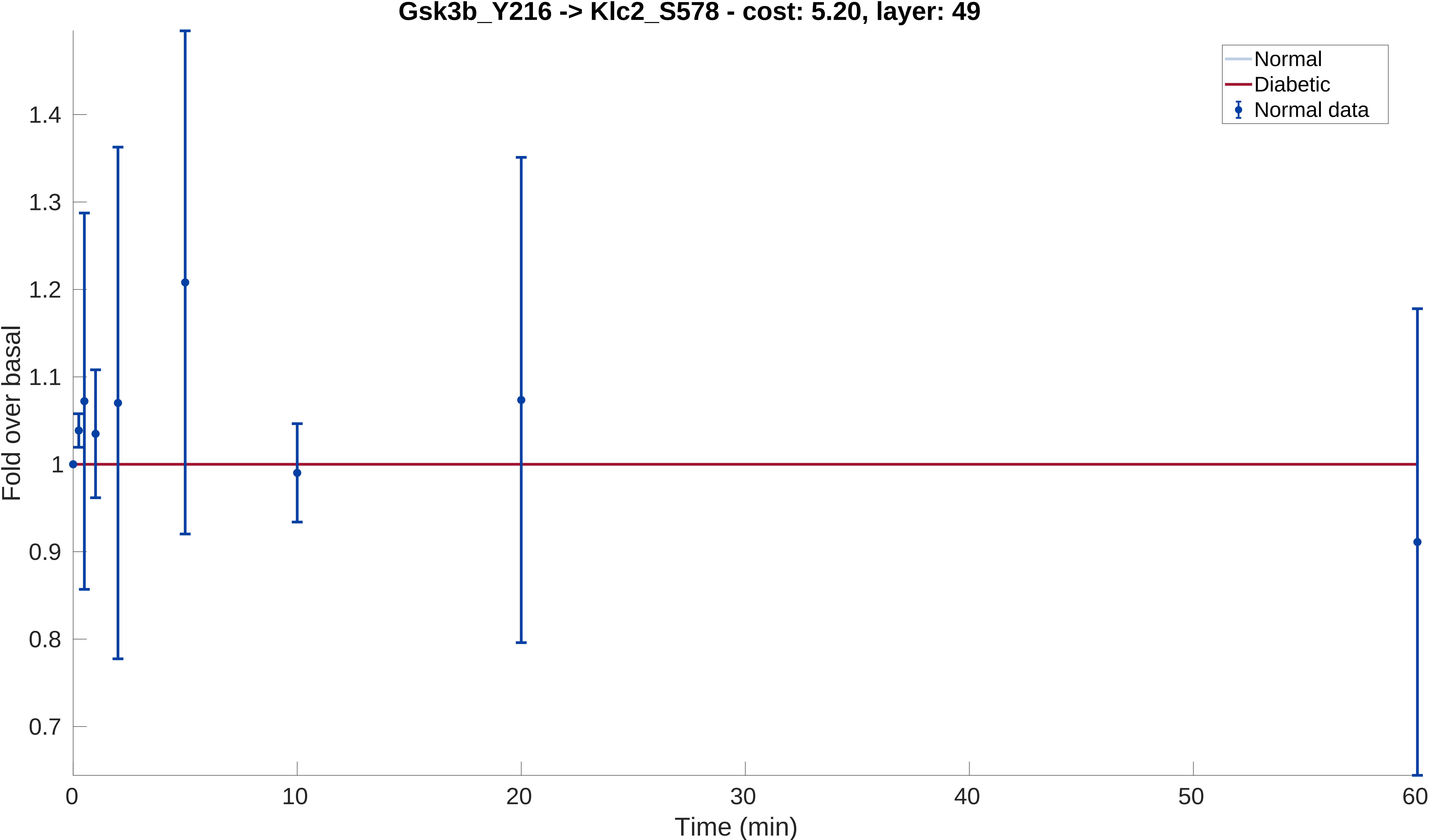
Gsk3b_Y216 -> Klc2_S578 - cost: 5.20, layer: 49
Normal
Diabetic
Normal data
Fold over basal
Time (min)
1.4
1.3
1.2
1.1
1
0.9
0.8
0.7
0
10
20
30
40
50
60

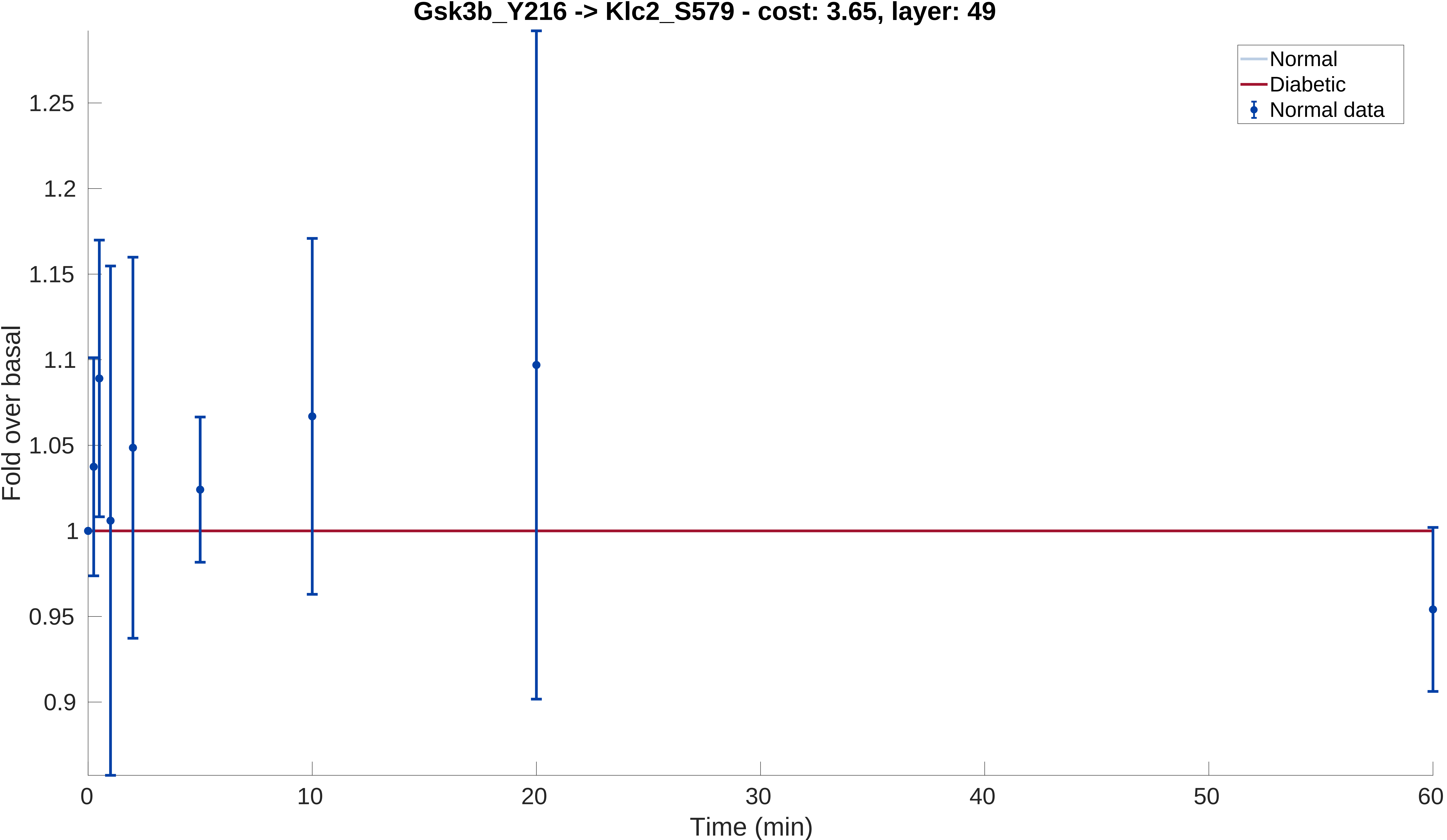

Gsk3b_Y216 -> Klc2_S579 - cost: 3.65, layer: 49
Normal
Diabetic
Normal data
Fold over basal
1.25
1.2
1.15
1.1
1.05
1
0.95
0.9
0
10
20
30
40
50
60
Time (min)

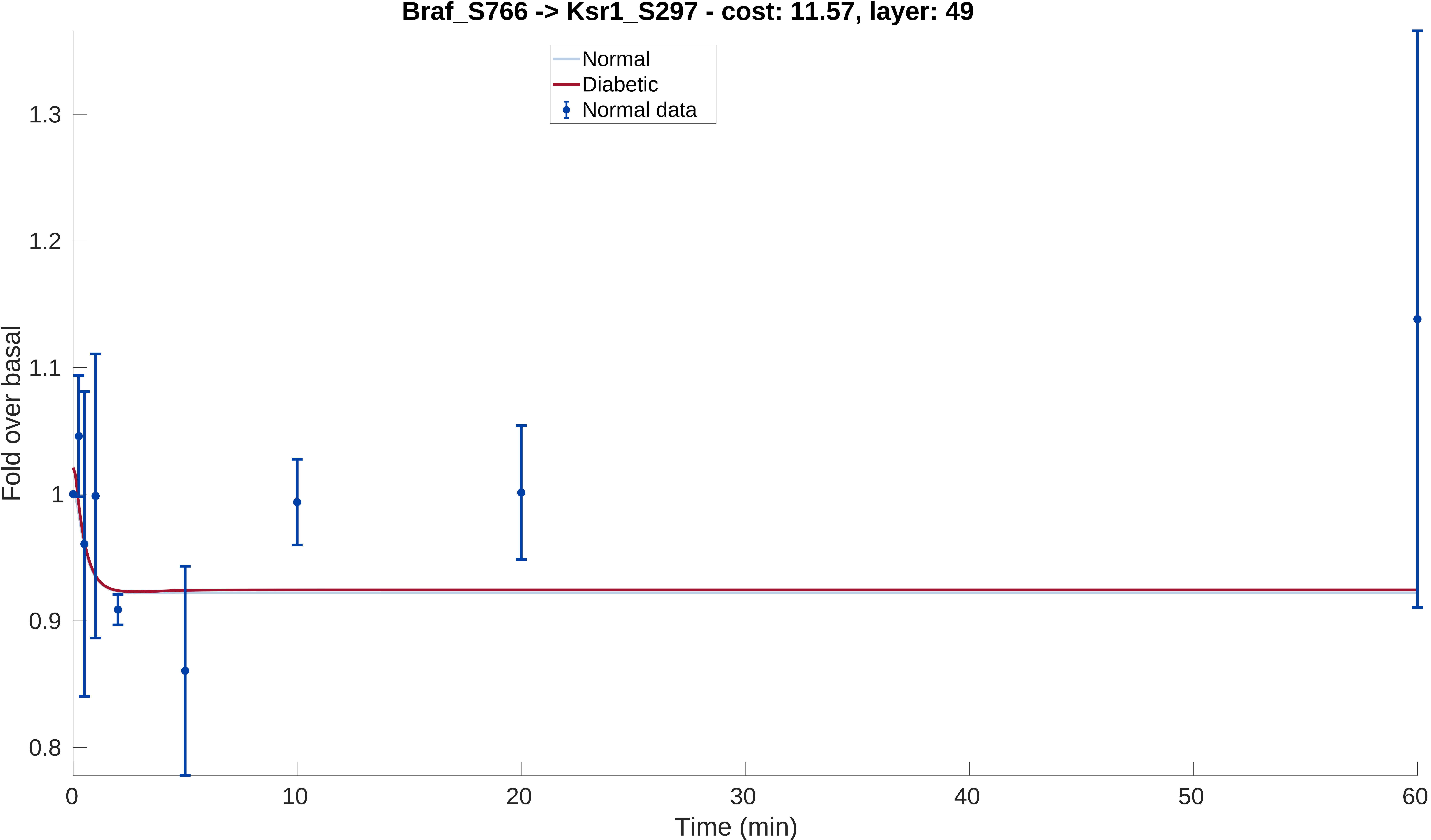

Braf_S766 -> Ksr1_S297 - cost: 11.57, layer: 49
Normal
Diabetic
Normal data
Fold over basal
1.3
1.2
1.1
1
0.9
0.8
0
10
20
30
40
50
60
Time (min)

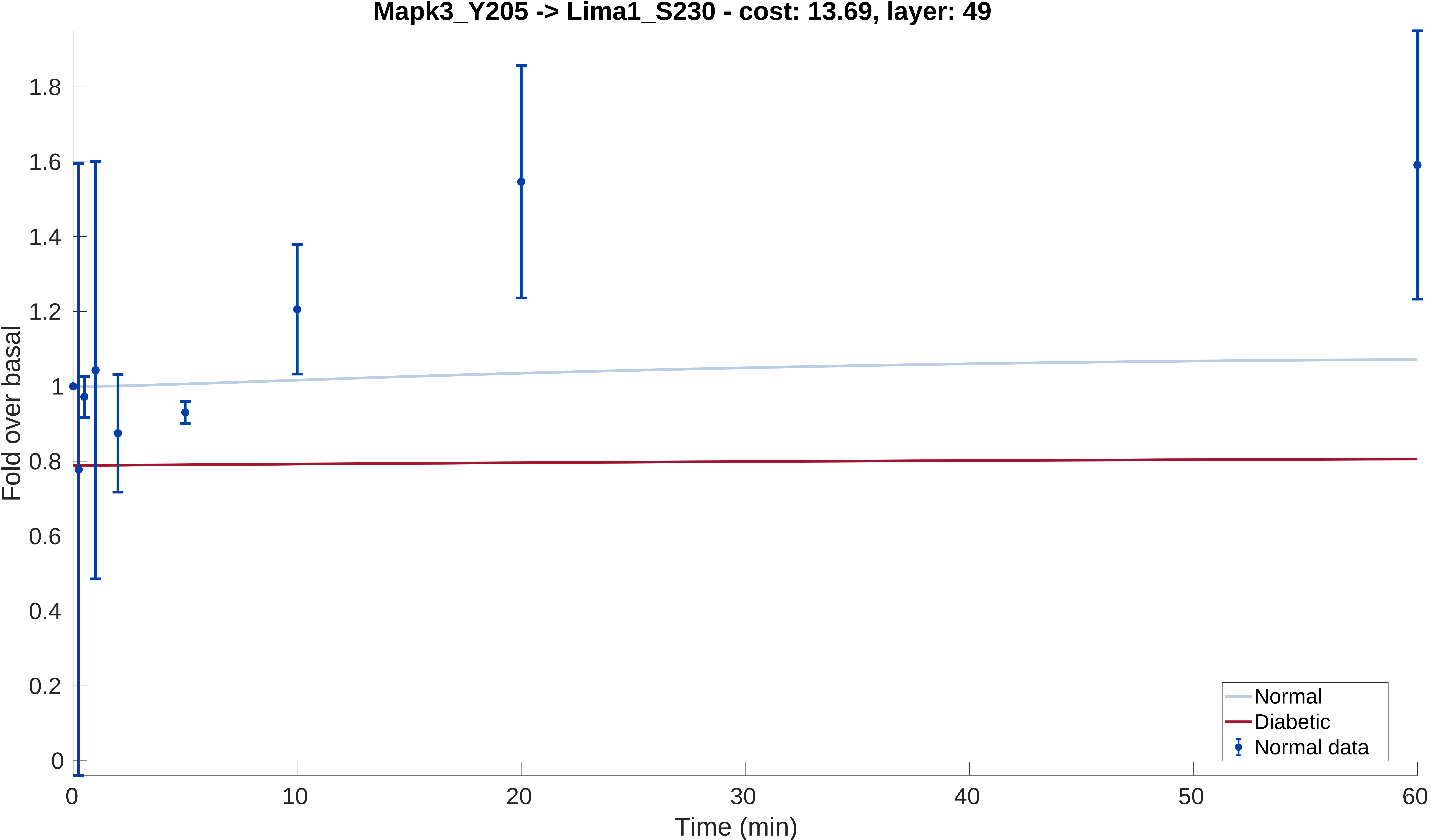

Mapk3_Y205 -> Lima1_S230 - cost: 13.69, layer: 49
Fold over basal
Time (min)
0
0.2
0.4
0.6
0.8
1
1.2
1.4
1.6
1.8
0
10
20
30
40
50
60
Normal
Diabetic
Normal data

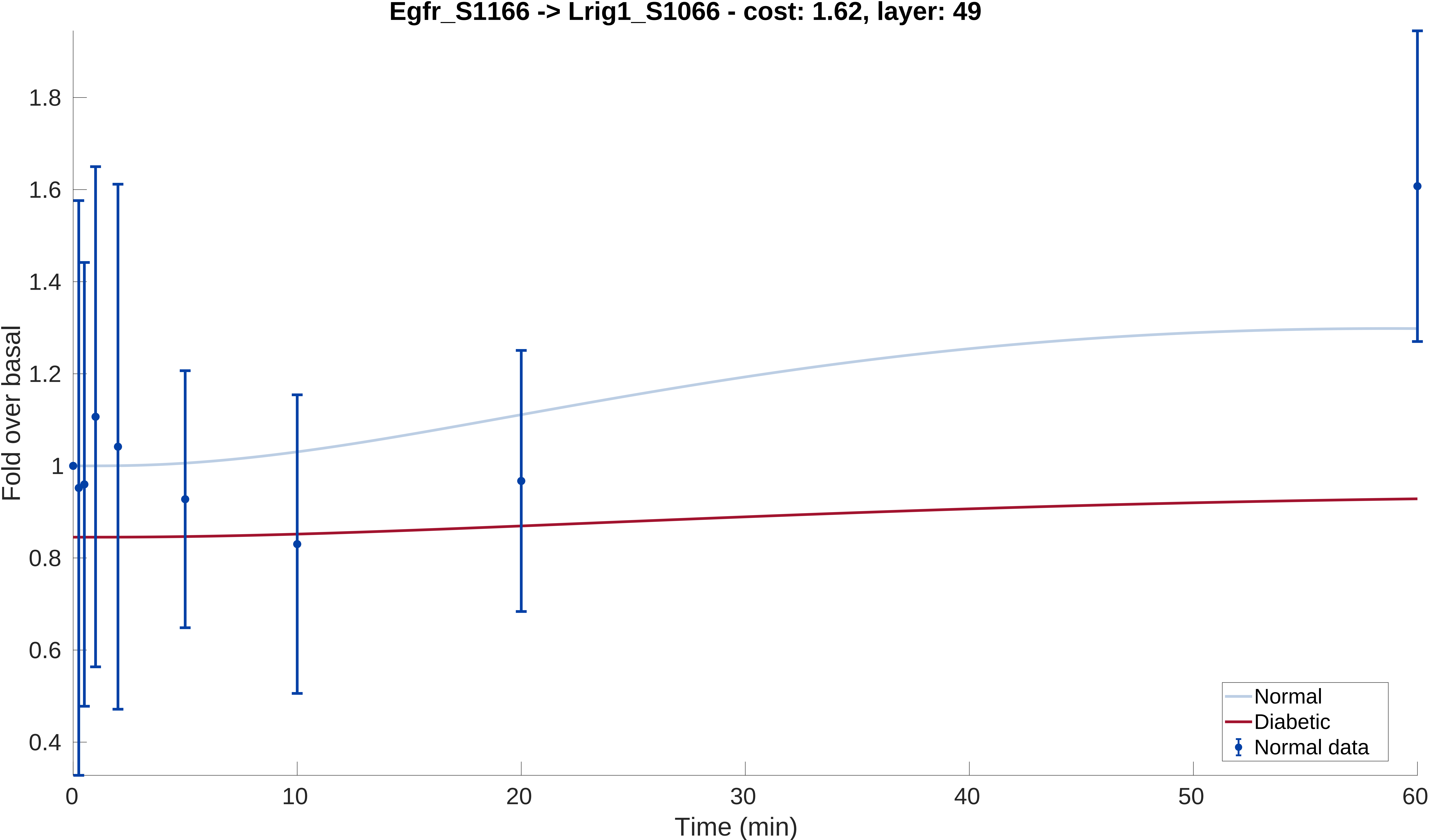
Egfr_S1166 -> Lrig1_S1066 - cost: 1.62, layer: 49
Fold over basal
1.8
1.6
1.4
1.2
1
0.8
0.6
0.4
0
10
20
30
40
50
60
Time (min)
Normal
Diabetic
Normal data

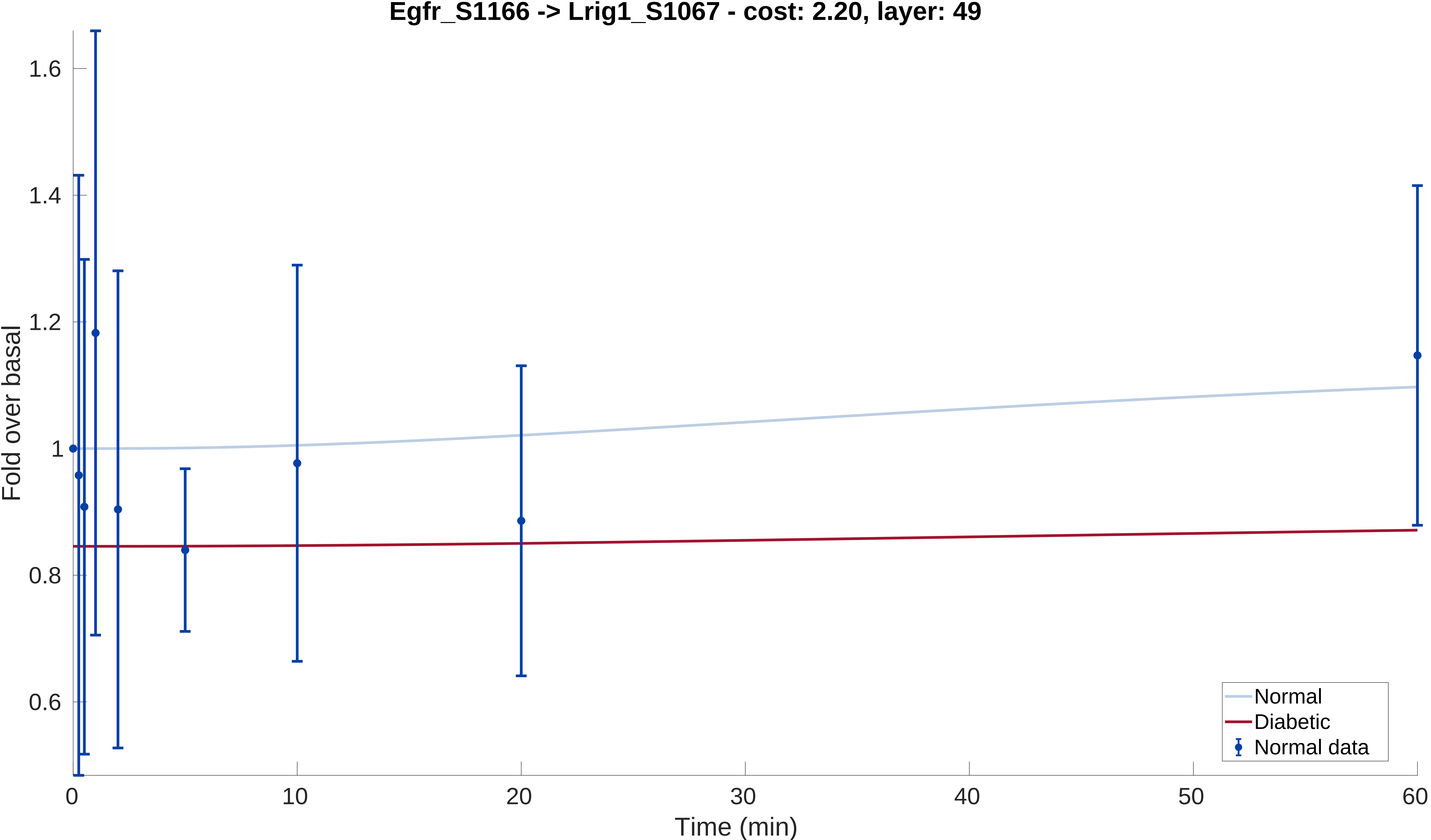

Egfr_S1166 -> Lrig1_S1067 - cost: 2.20, layer: 49
Fold over basal
1.6
1.4
1.2
1
0.8
0.6
0
10
20
30
40
50
60
Time (min)
Normal
Diabetic
Normal data

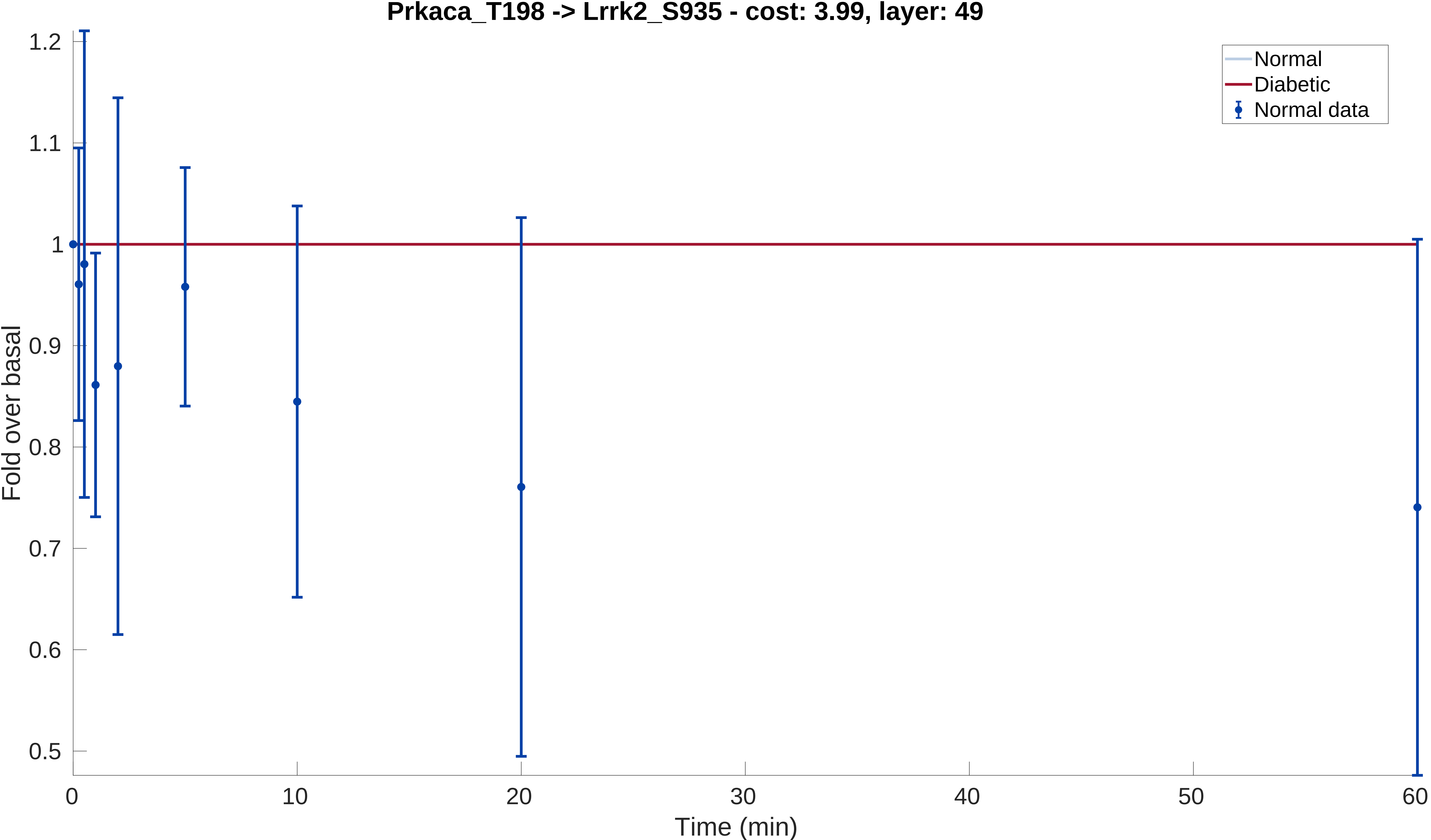
Prkaca_T198 -> Lrrk2_S935 - cost: 3.99, layer: 49
Normal
Diabetic
Normal data
Fold over basal
Time (min)
1.2
1.1
1
0.9
0.8
0.7
0.6
0.5
0
10
20
30
40
50
60

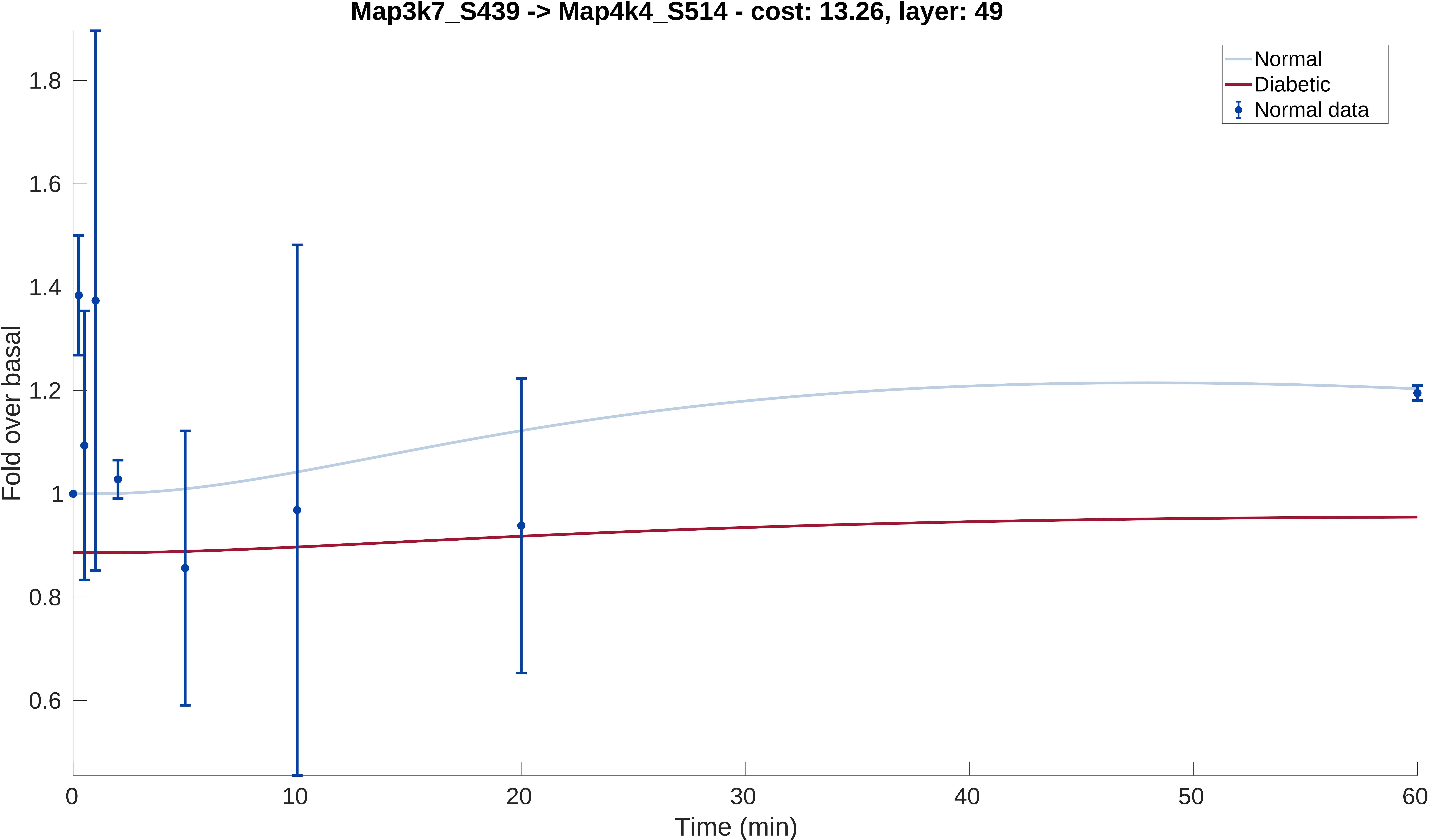
Map3k7_S439 -> Map4k4_S514 - cost: 13.26, layer: 49
Normal
Diabetic
Normal data
Fold over basal
1.8
1.6
1.4
1.2
1
0.8
0.6
0
10
20
30
40
50
60
Time (min)

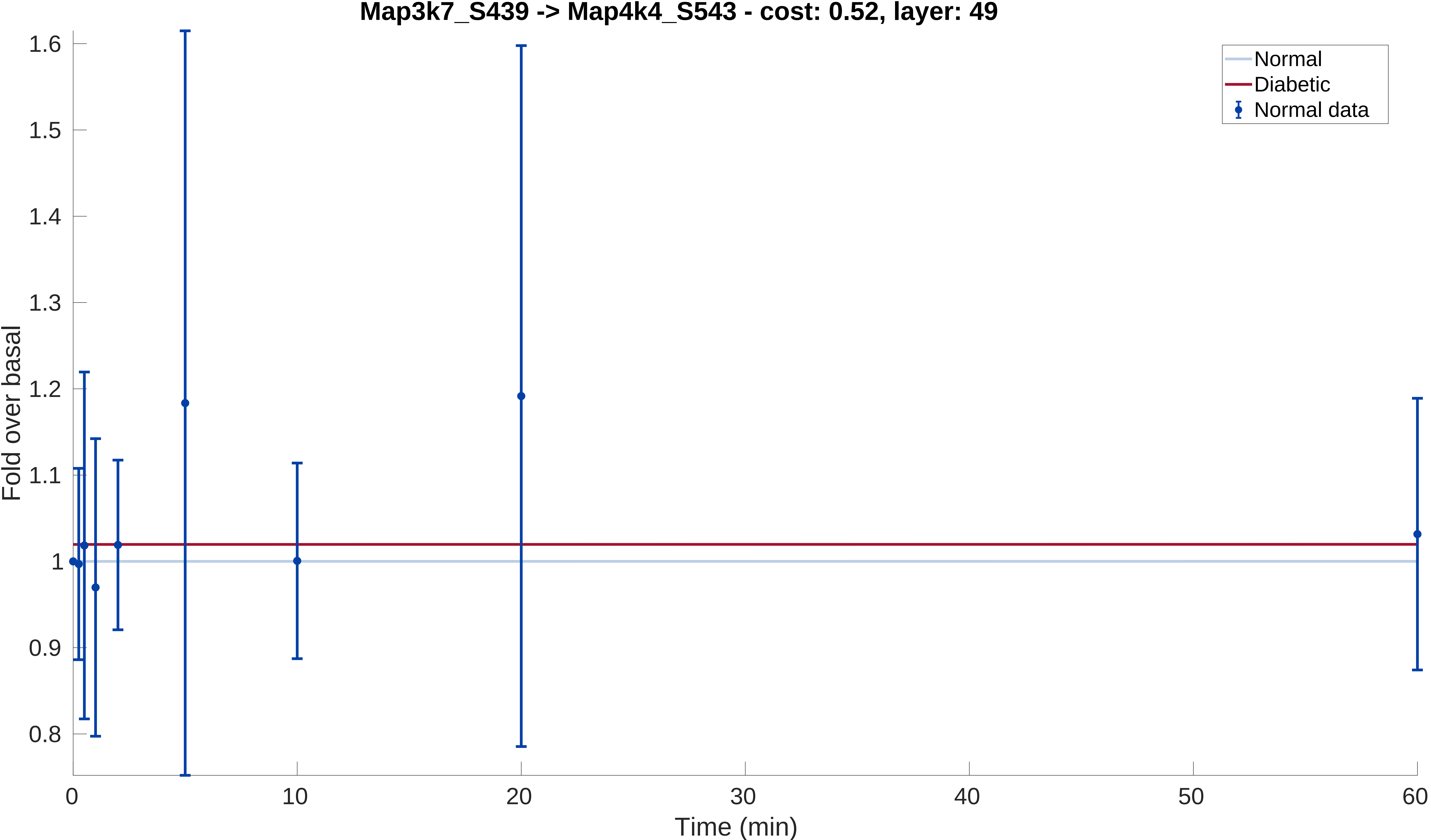

Map3k7_S439 -> Map4k4_S543 - cost: 0.52, layer: 49
Normal
Diabetic
Normal data
Fold over basal
1.6
1.5
1.4
1.3
1.2
1.1
1
0.9
0.8
0
10
20
30
40
50
60
Time (min)

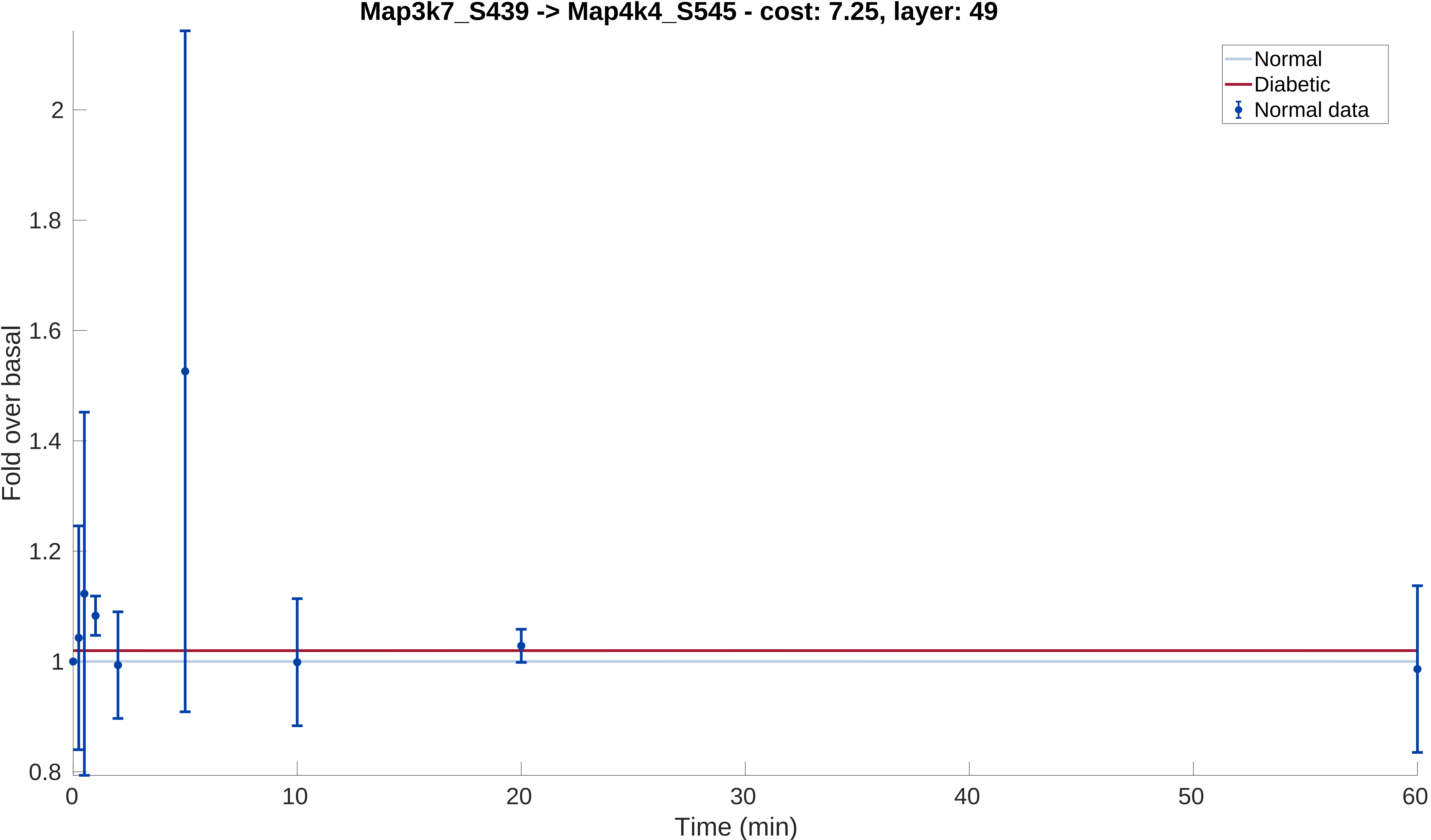

Map3k7_S439 -> Map4k4_S545 - cost: 7.25, layer: 49
Normal
Diabetic
Normal data
Fold over basal
0.8
1
1.2
1.4
1.6
1.8
2
0
10
20
30
40
50
60
Time (min)

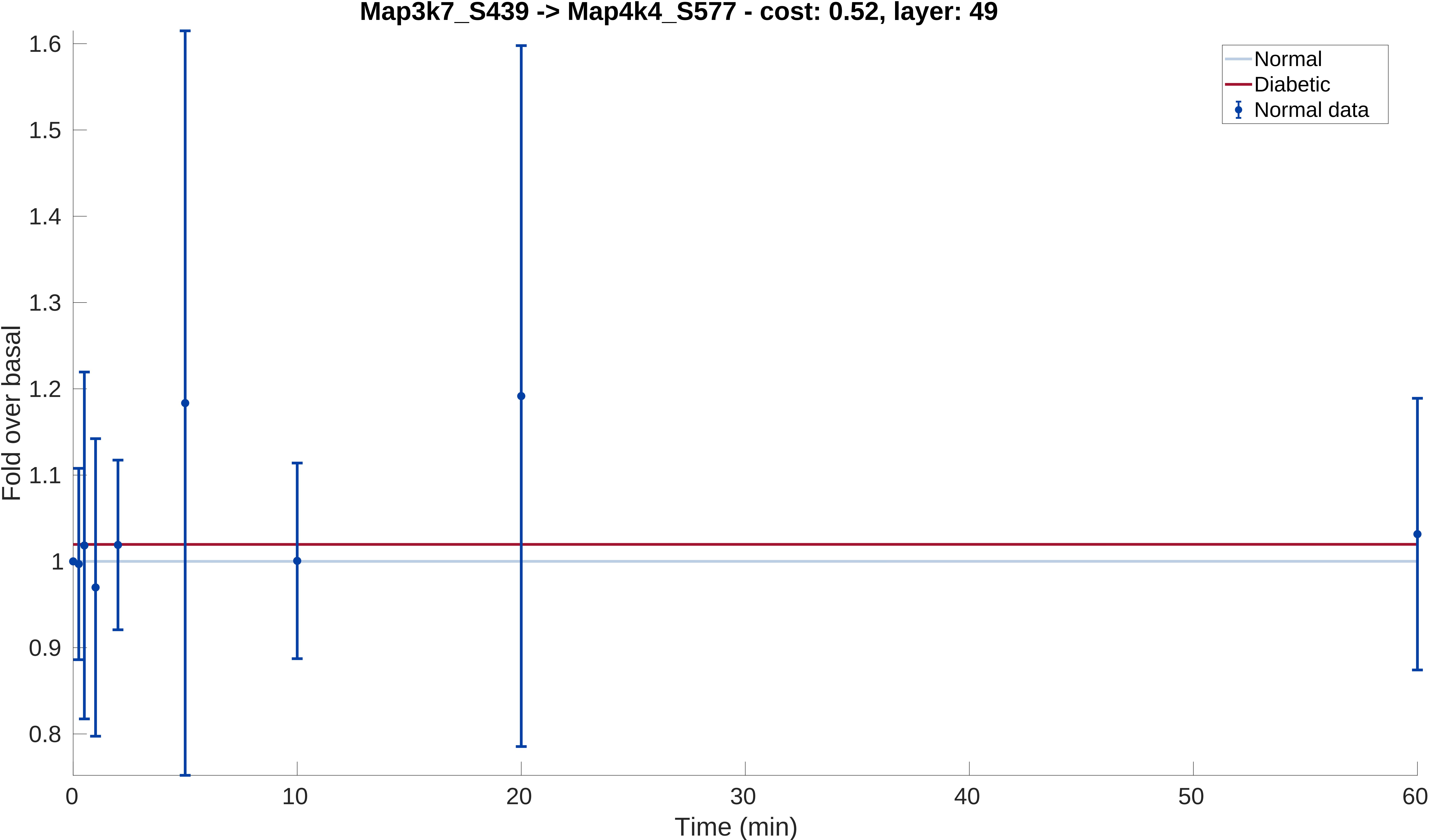

Map3k7_S439 -> Map4k4_S577 - cost: 0.52, layer: 49
Normal
Diabetic
Normal data
Fold over basal
1.6
1.5
1.4
1.3
1.2
1.1
1
0.9
0.8
0
10
20
30
40
50
60
Time (min)

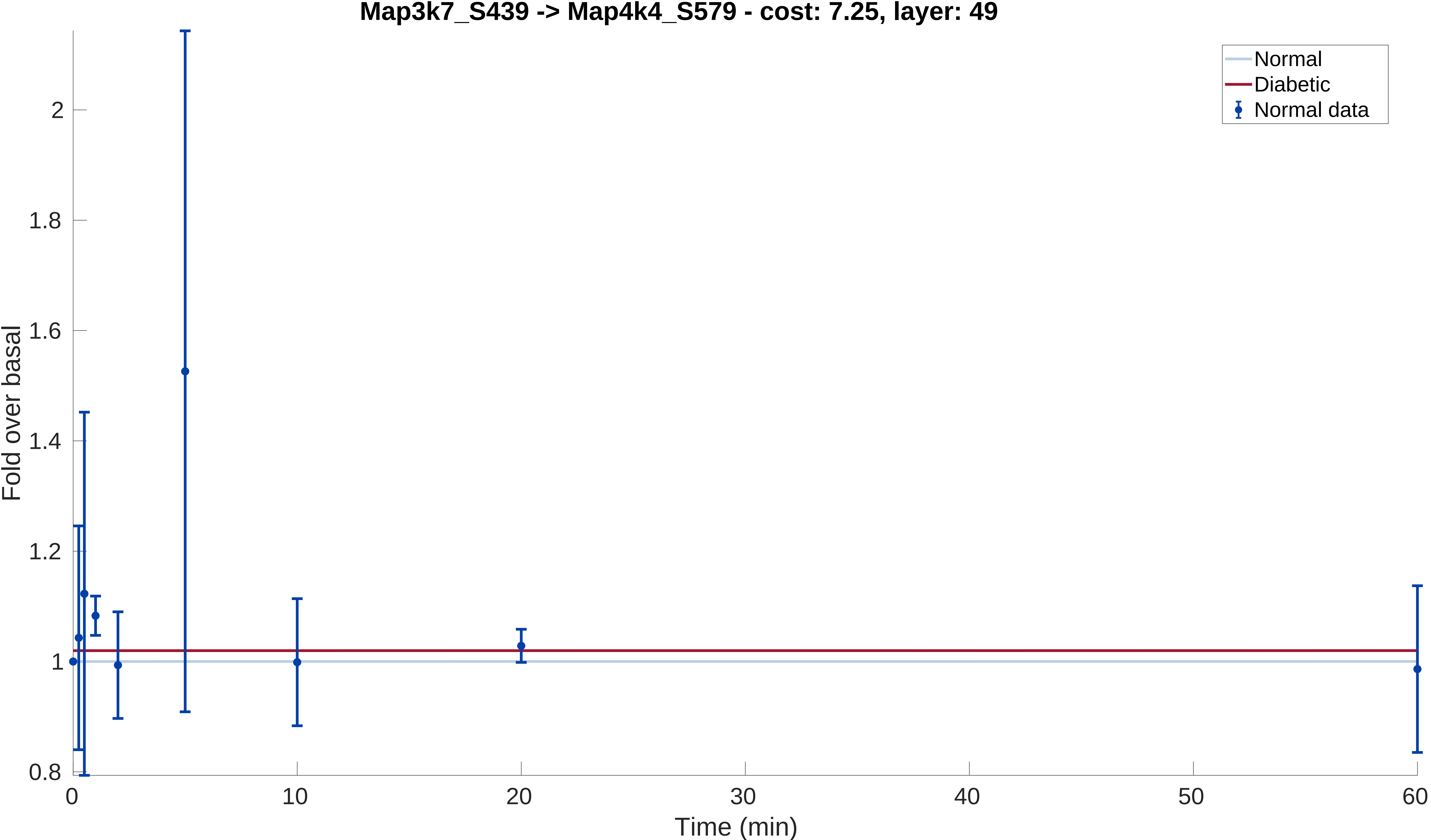

Map3k7_S439 -> Map4k4_S579 - cost: 7.25, layer: 49
Normal
Diabetic
Normal data
Fold over basal
0.8
1
1.2
1.4
1.6
1.8
2
0
10
20
30
40
50
60
Time (min)

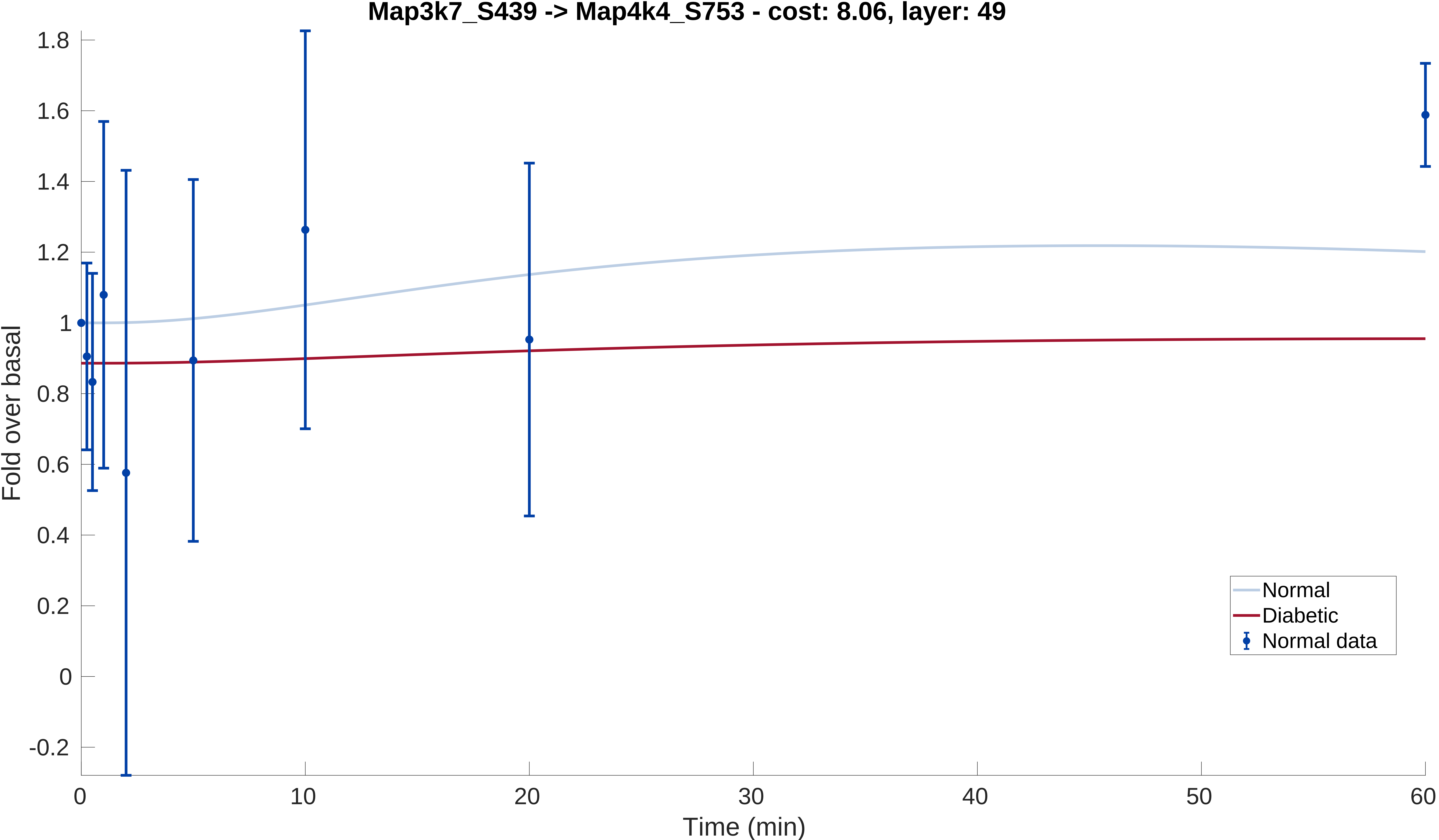

Map3k7_S439 -> Map4k4_S753 - cost: 8.06, layer: 49
Fold over basal
1.8
1.6
1.4
1.2
1
0.8
0.6
0.4
0.2
0
-0.2
0
10
20
30
40
50
60
Time (min)
Normal
Diabetic
Normal data

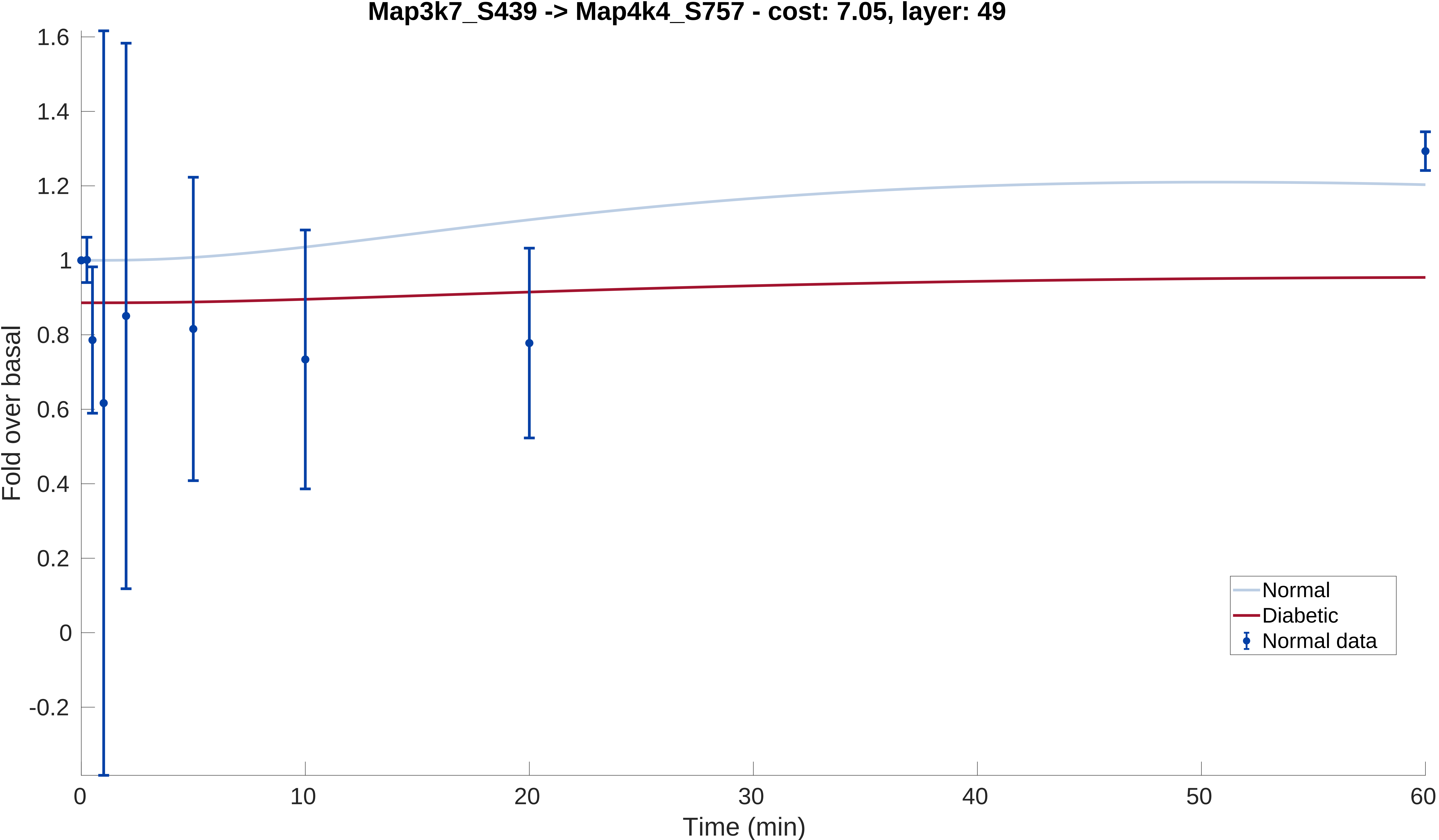

Map3k7_S439 -> Map4k4_S757 - cost: 7.05, layer: 49
Fold over basal
1.6
1.4
1.2
1
0.8
0.6
0.4
0.2
0
-0.2
0
10
20
30
40
50
60
Time (min)
Normal
Diabetic
Normal data

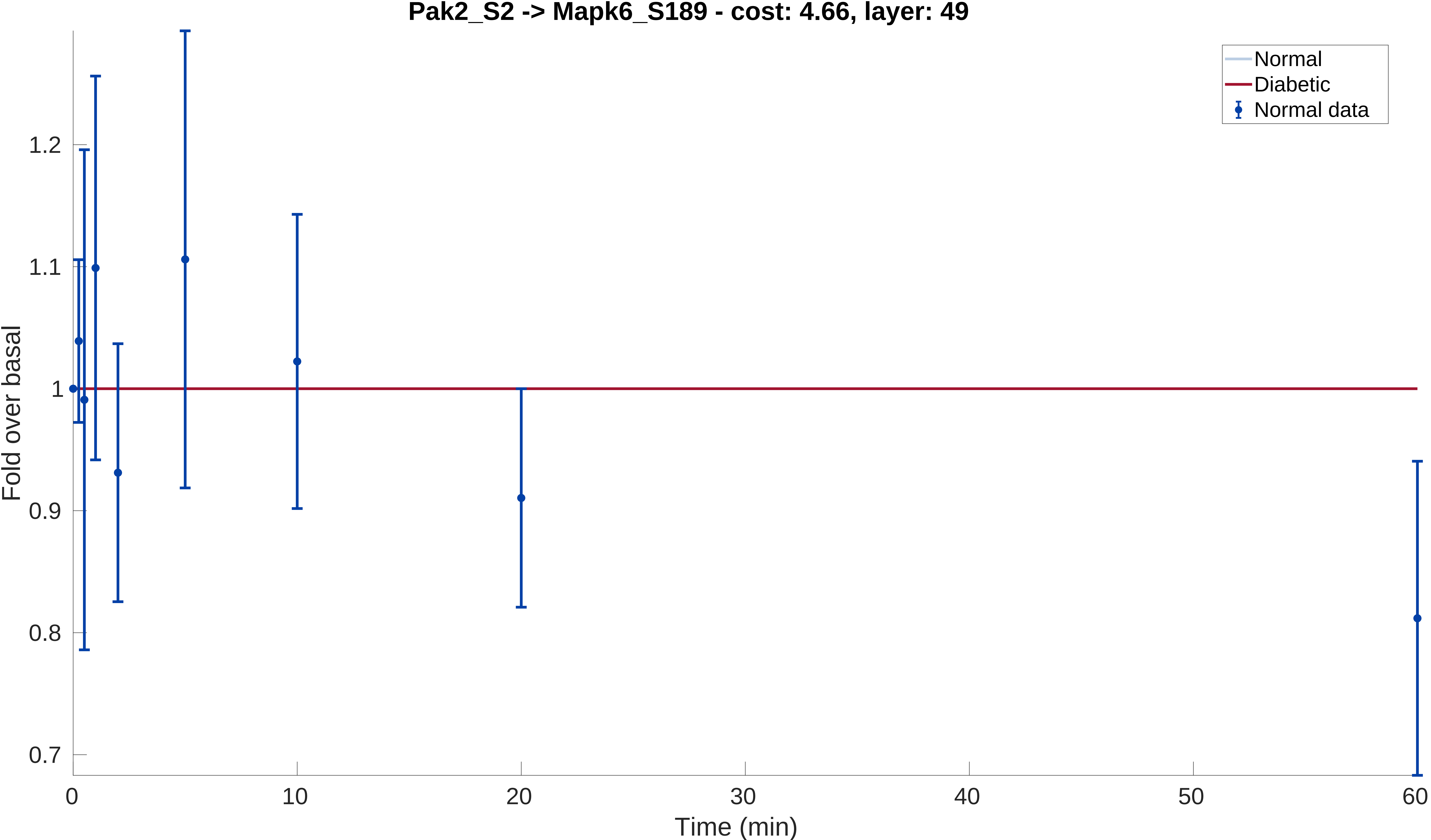

Pak2_S2 -> Mapk6_S189 - cost: 4.66, layer: 49
Normal
Diabetic
Normal data
Fold over basal
1.2
1.1
1
0.9
0.8
0.7
0
10
20
30
40
50
60
Time (min)

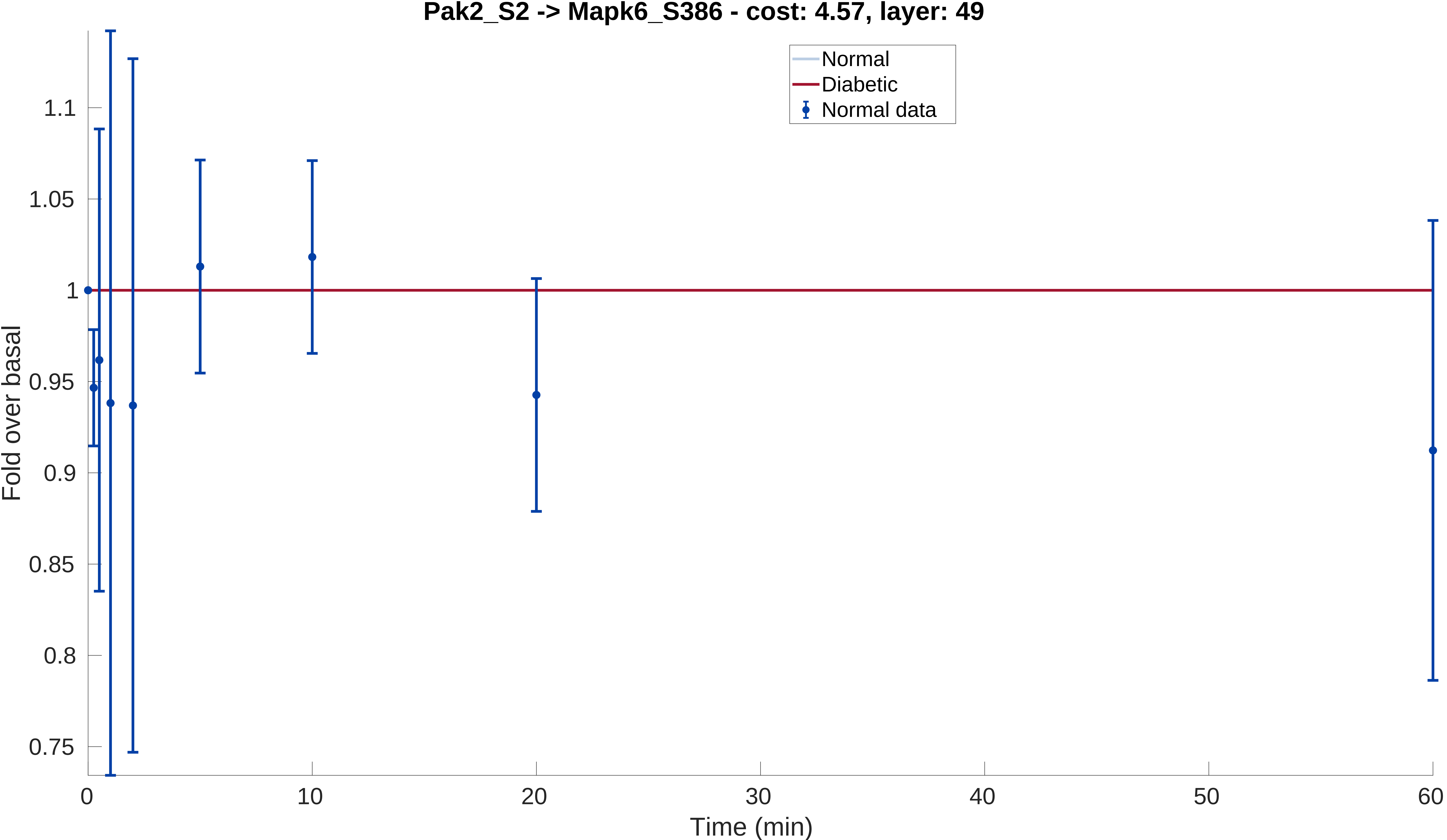

Pak2_S2 -> Mapk6_S386 - cost: 4.57, layer: 49
Normal
Diabetic
Normal data
Fold over basal
1.1
1.05
1
0.95
0.9
0.85
0.8
0.75
0
10
20
30
40
50
60
Time (min)

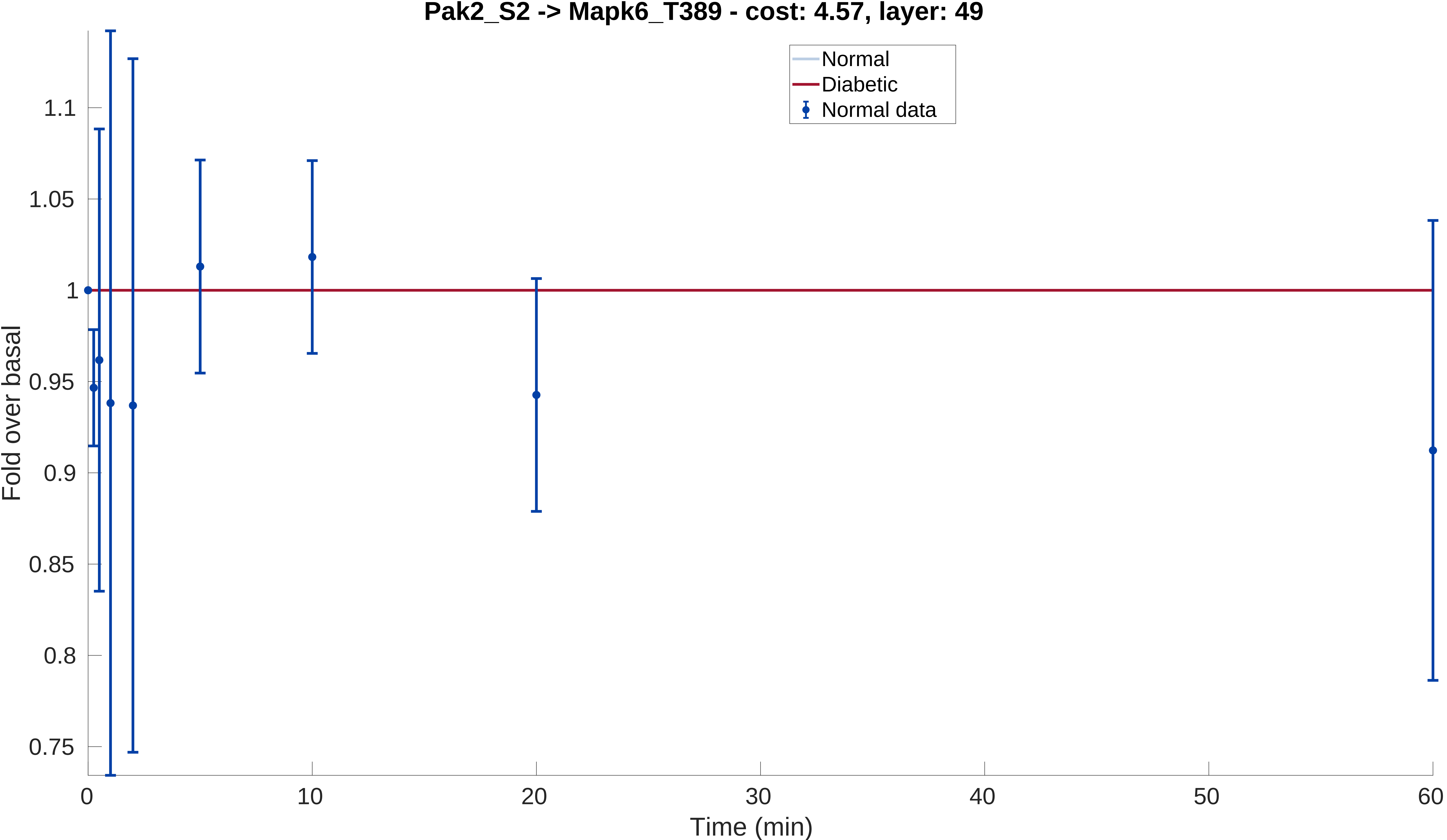

Pak2_S2 -> Mapk6_T389 - cost: 4.57, layer: 49
Normal
Diabetic
Normal data
Fold over basal
1.1
1.05
1
0.95
0.9
0.85
0.8
0.75
0
10
20
30
40
50
60
Time (min)

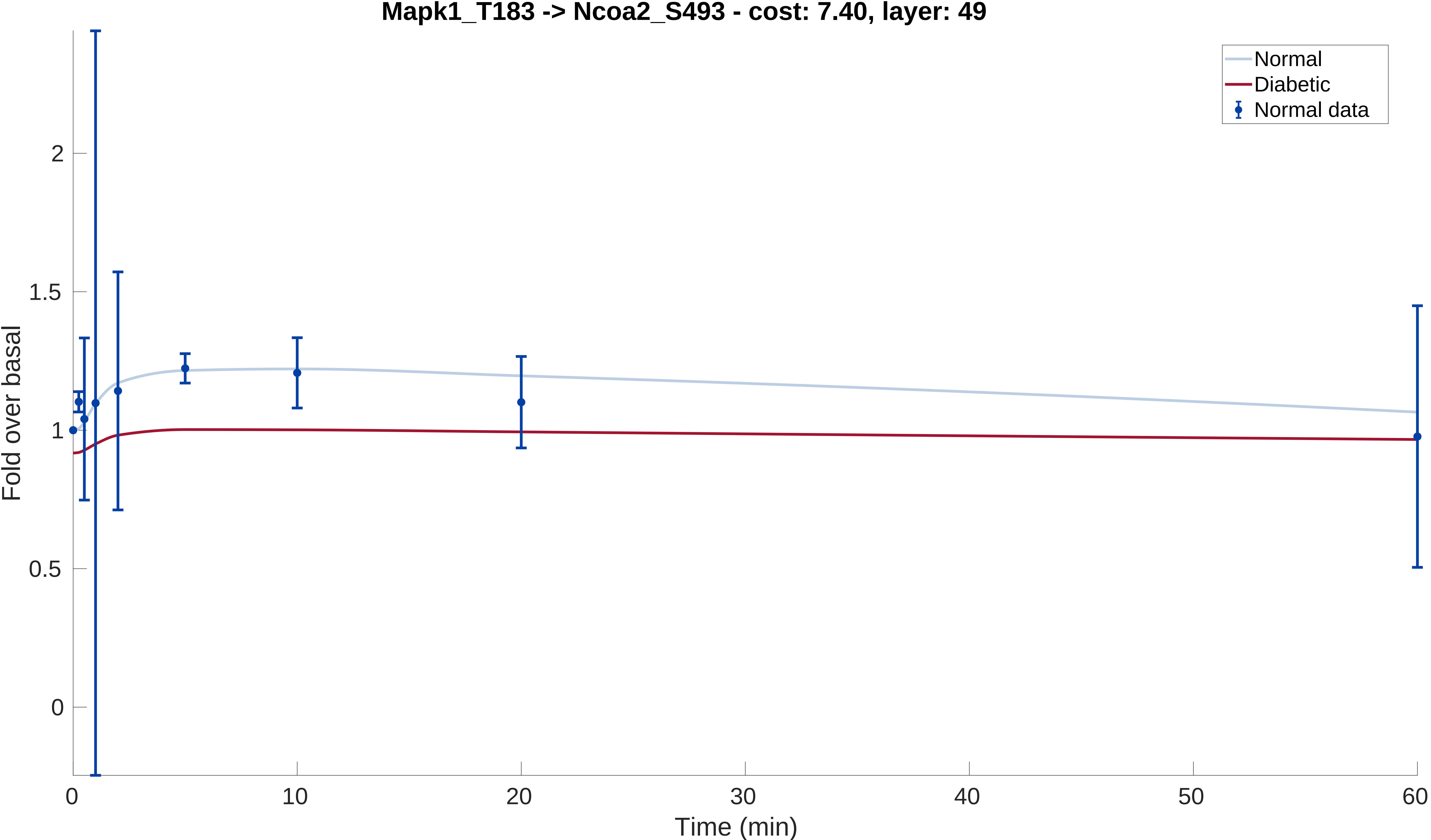

Mapk1_T183 -> Ncoa2_S493 - cost: 7.40, layer: 49
Normal
Diabetic
Normal data
Fold over basal
2
1.5
1
0.5
0
0
10
20
30
40
50
60
Time (min)

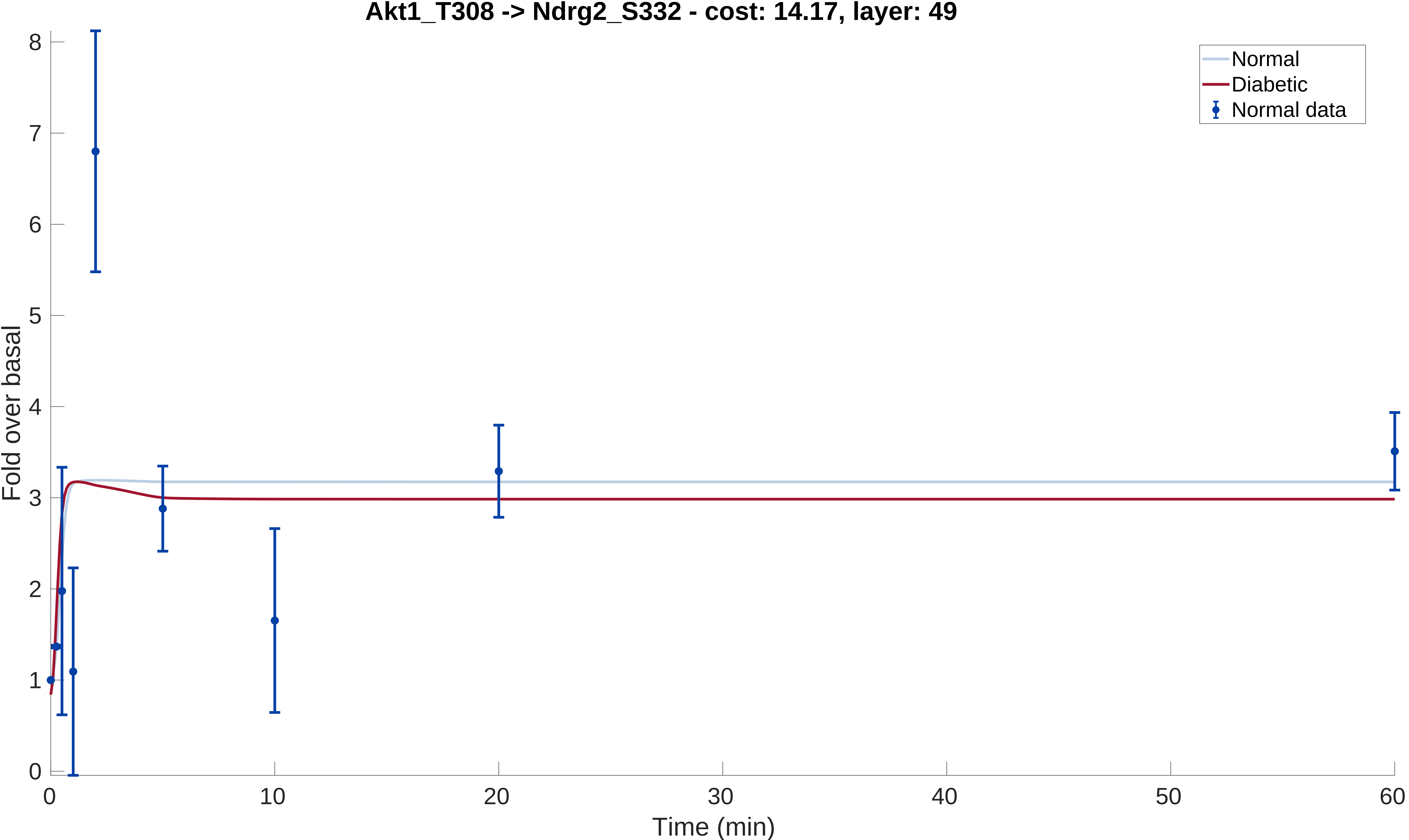

Akt1_T308 -> Ndrg2_S332 - cost: 14.17, layer: 49
Normal
Diabetic
Normal data
Fold over basal
0
1
2
3
4
5
6
7
8
0
10
20
30
40
50
60
Time (min)

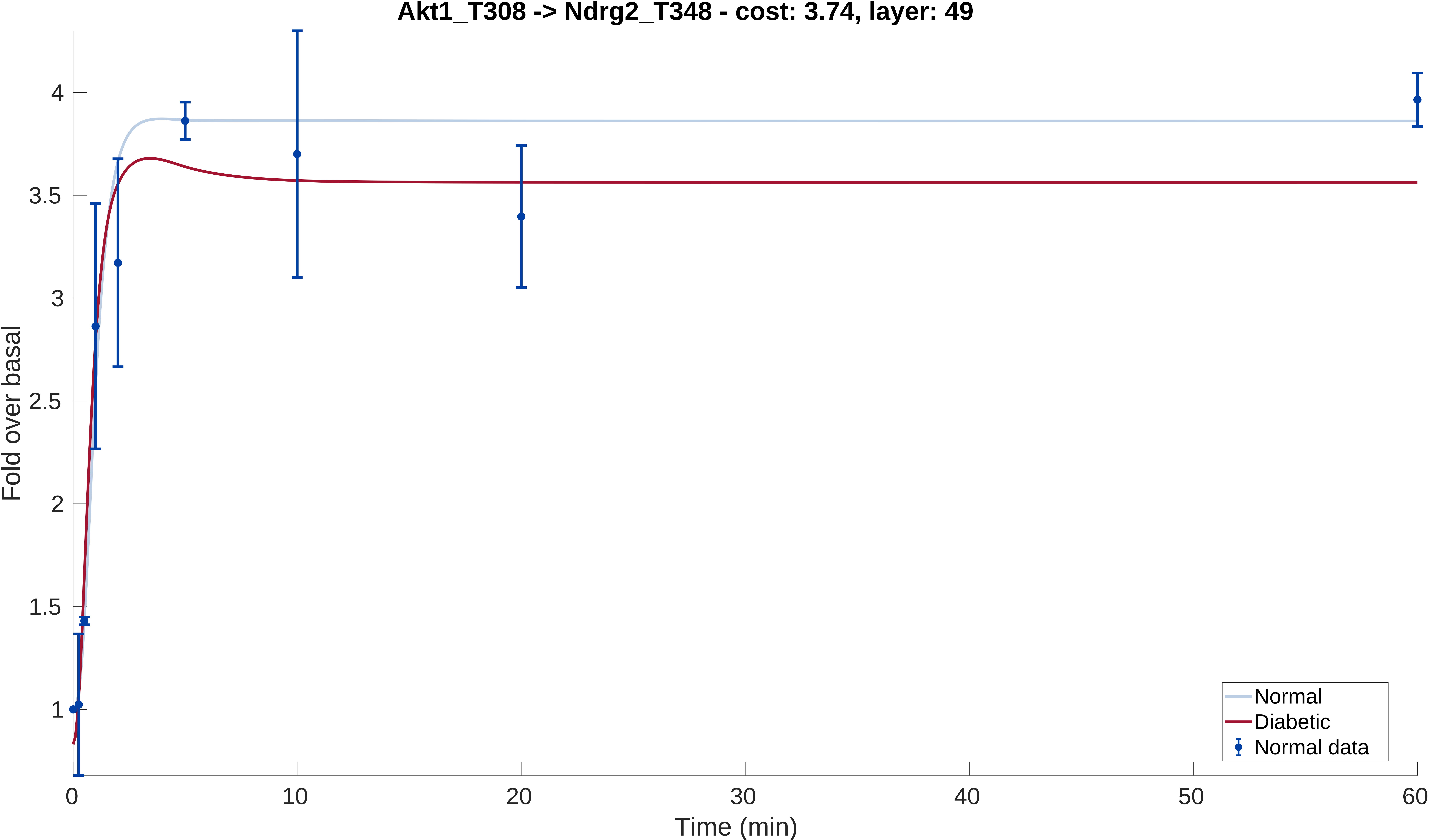
Akt1_T308 -> Ndrg2_T348 - cost: 3.74, layer: 49
Fold over basal
Time (min)
0
10
20
30
40
50
60
1
1.5
2
2.5
3
3.5
4
Normal
Diabetic
Normal data

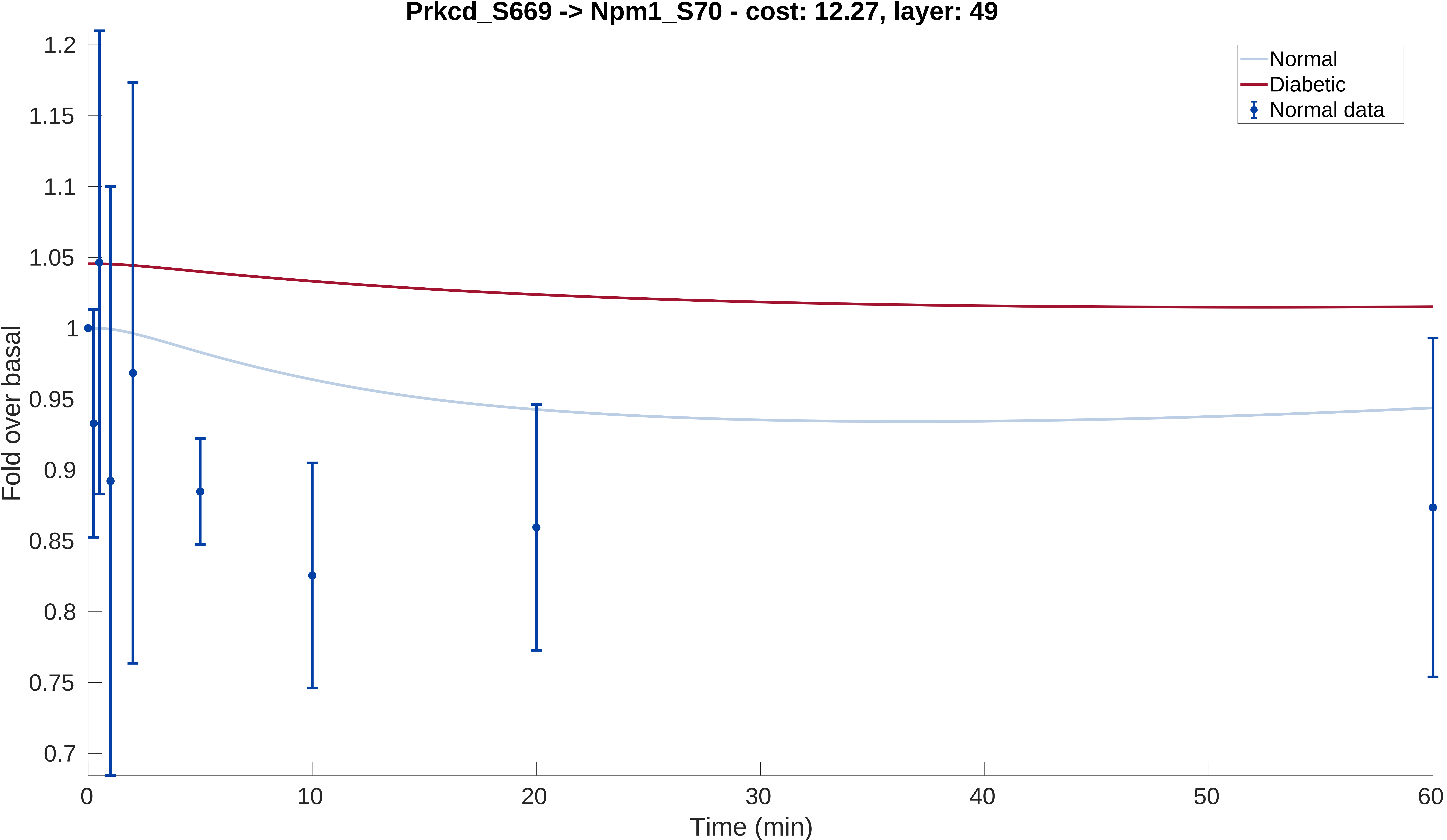
Prkcd_S669 -> Npm1_S70 - cost: 12.27, layer: 49
Normal
Diabetic
Normal data
Fold over basal
Time (min)
1.2
1.15
1.1
1.05
1
0.95
0.9
0.85
0.8
0.75
0.7
0
10
20
30
40
50
60

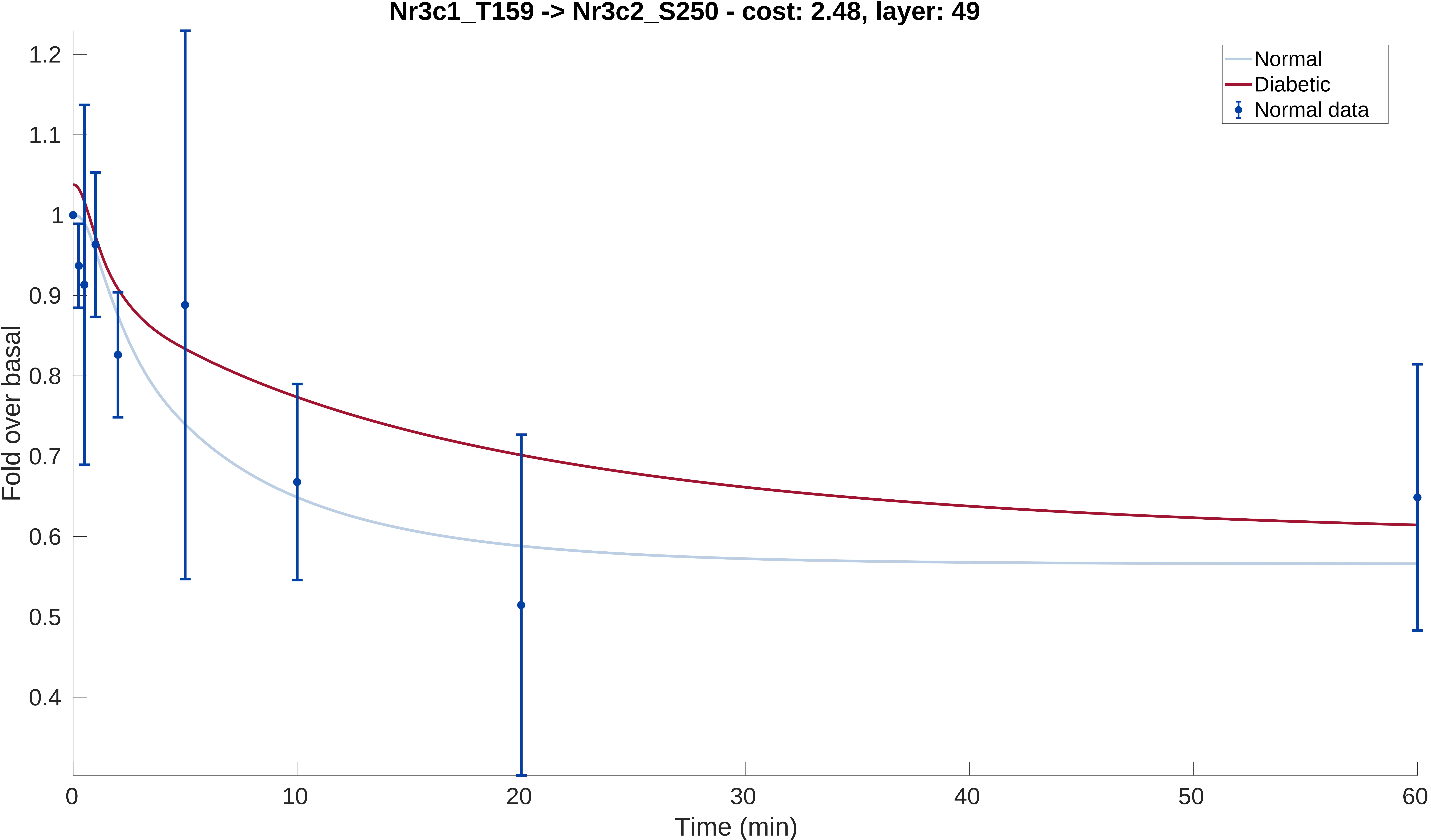

Nr3c1_T159 -> Nr3c2_S250 - cost: 2.48, layer: 49
Normal
Diabetic
Normal data
Fold over basal
1.2
1.1
1
0.9
0.8
0.7
0.6
0.5
0.4
0
10
20
30
40
50
60
Time (min)

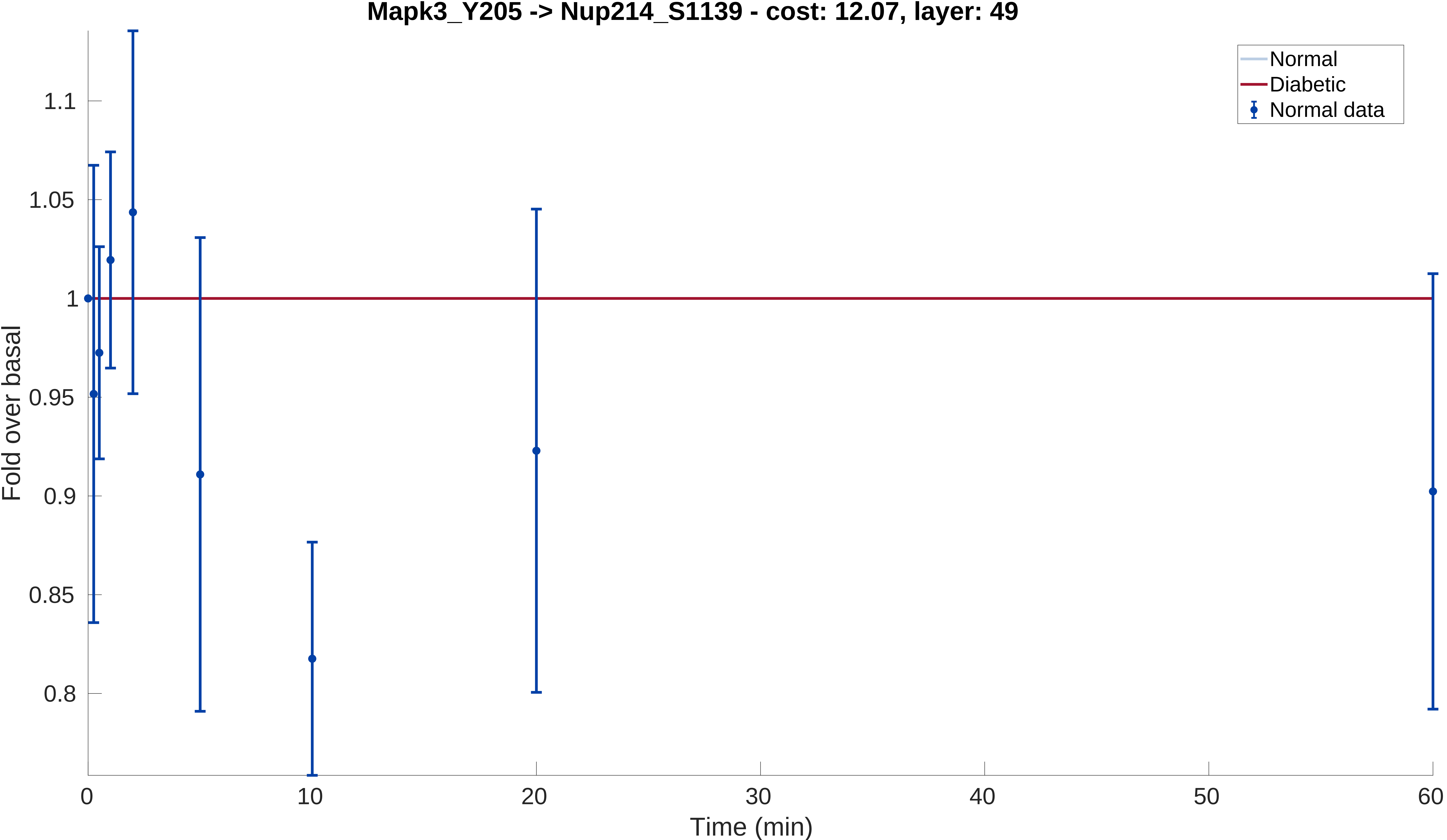
Mapk3_Y205 -> Nup214_S1139 - cost: 12.07, layer: 49
Normal
Diabetic
Normal data
Fold over basal
1.1
1.05
1
0.95
0.9
0.85
0.8
0
10
20
30
40
50
60
Time (min)

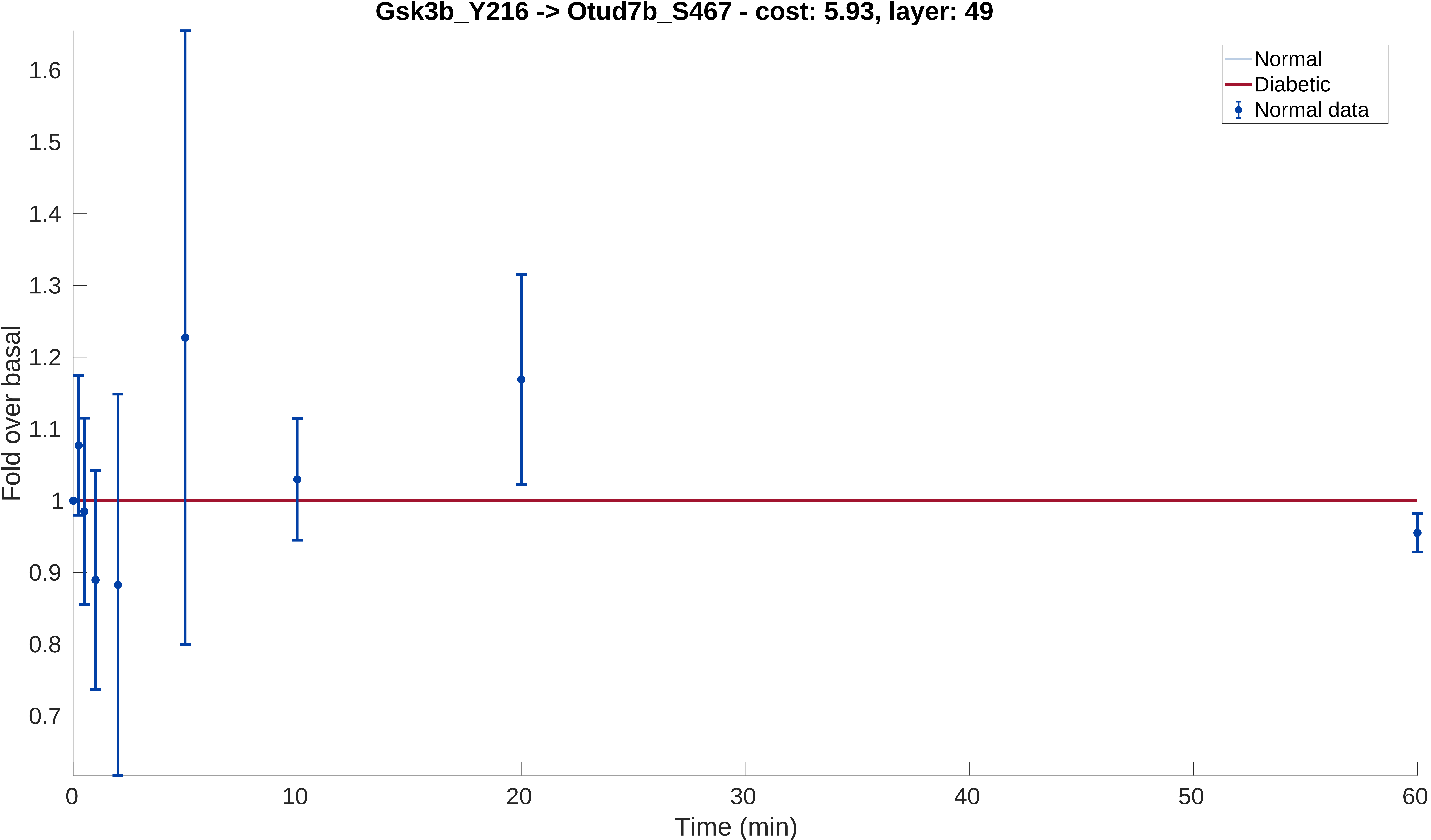
Gsk3b_Y216 -> Otud7b_S467 - cost: 5.93, layer: 49
Normal
Diabetic
Normal data
Fold over basal
1.6
1.5
1.4
1.3
1.2
1.1
1
0.9
0.8
0.7
0
10
20
30
40
50
60
Time (min)

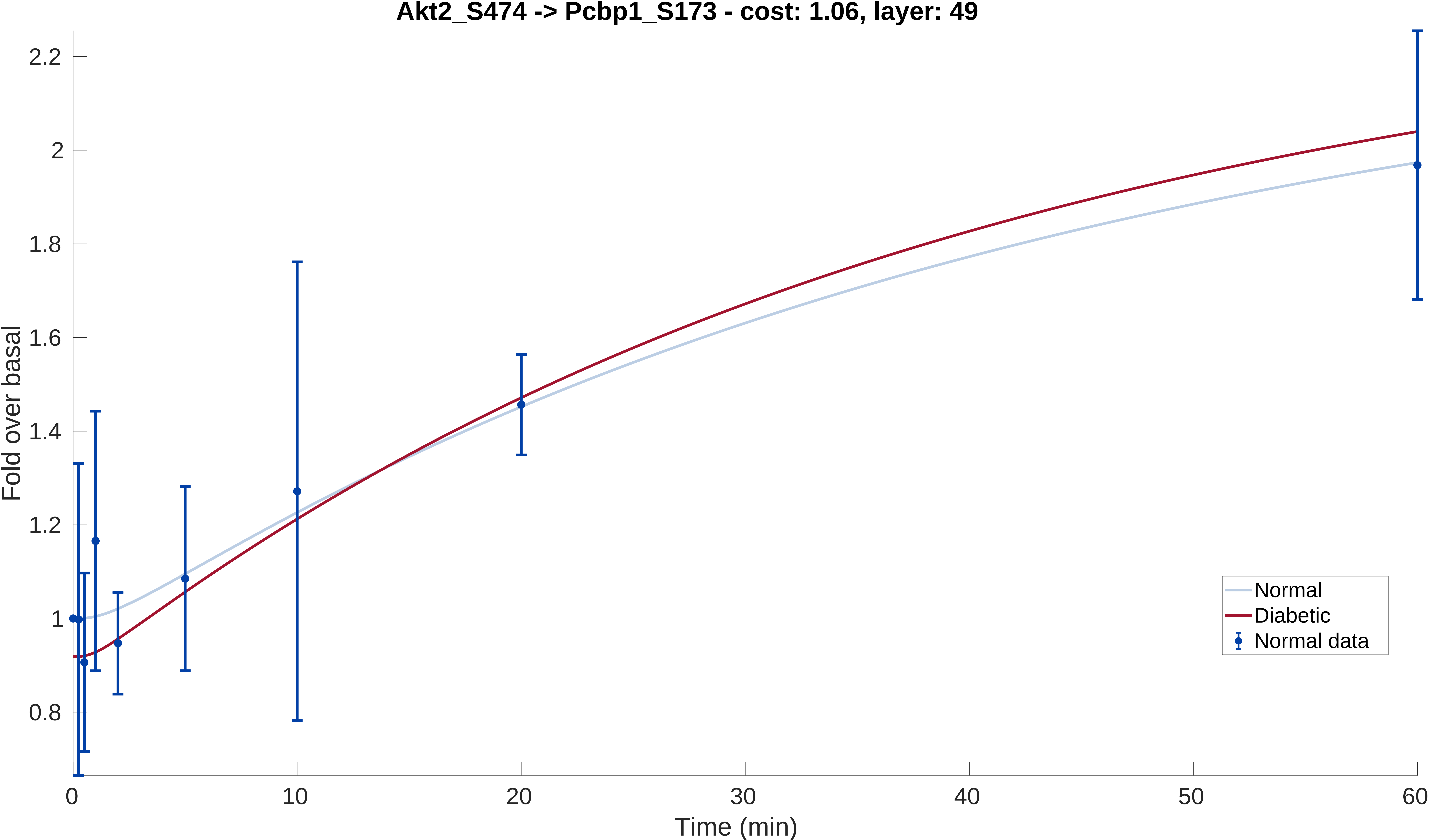

Akt2_S474 -> Pcbp1_S173 - cost: 1.06, layer: 49
Fold over basal
2.2
2
1.8
1.6
1.4
1.2
1
0.8
0
10
20
30
40
50
60
Time (min)
Normal
Diabetic
Normal data

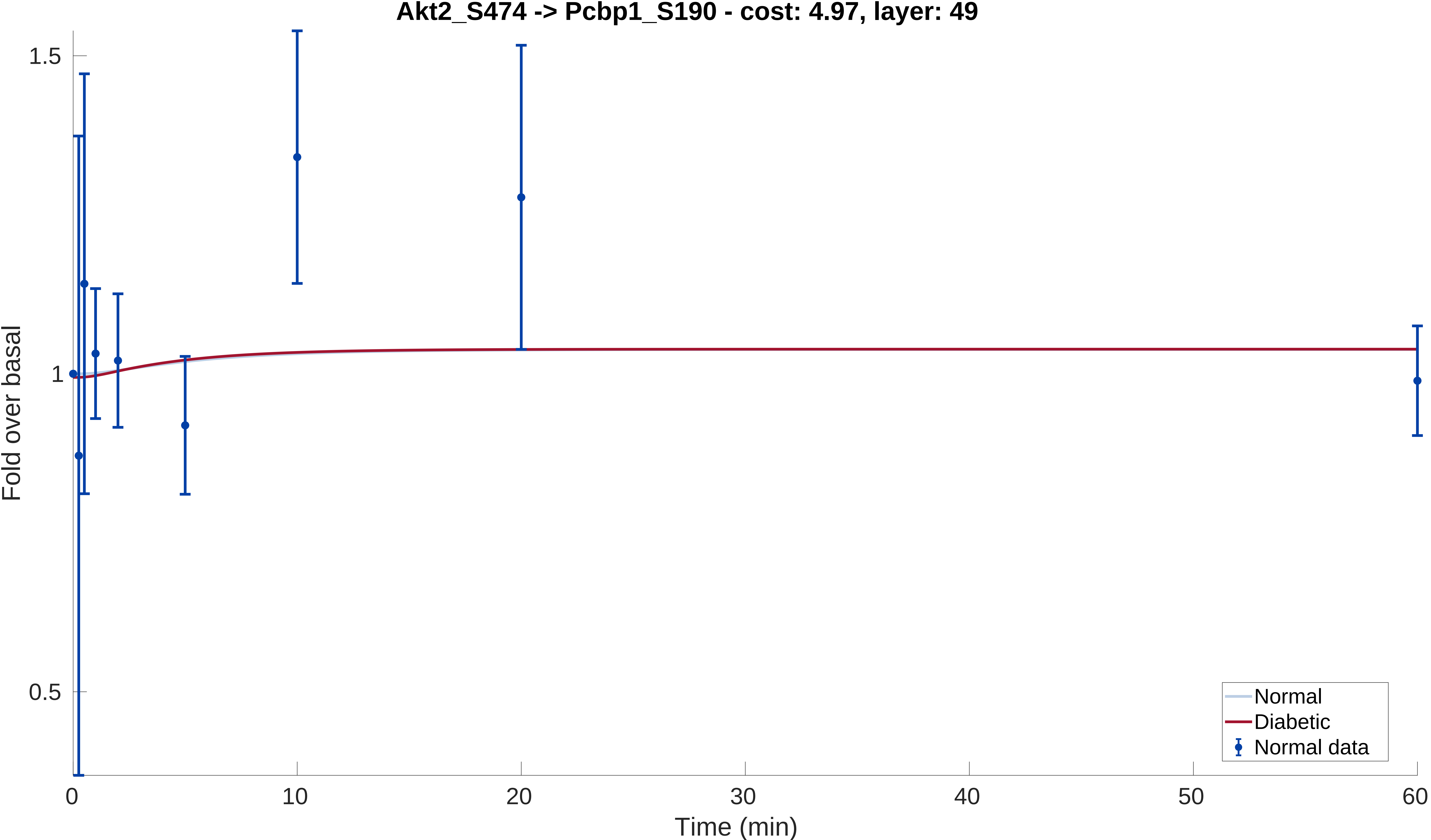

Akt2_S474 -> Pcbp1_S190 - cost: 4.97, layer: 49
1.5
1
0.5
Fold over basal
0
10
20
30
40
50
60
Time (min)
Normal
Diabetic
Normal data

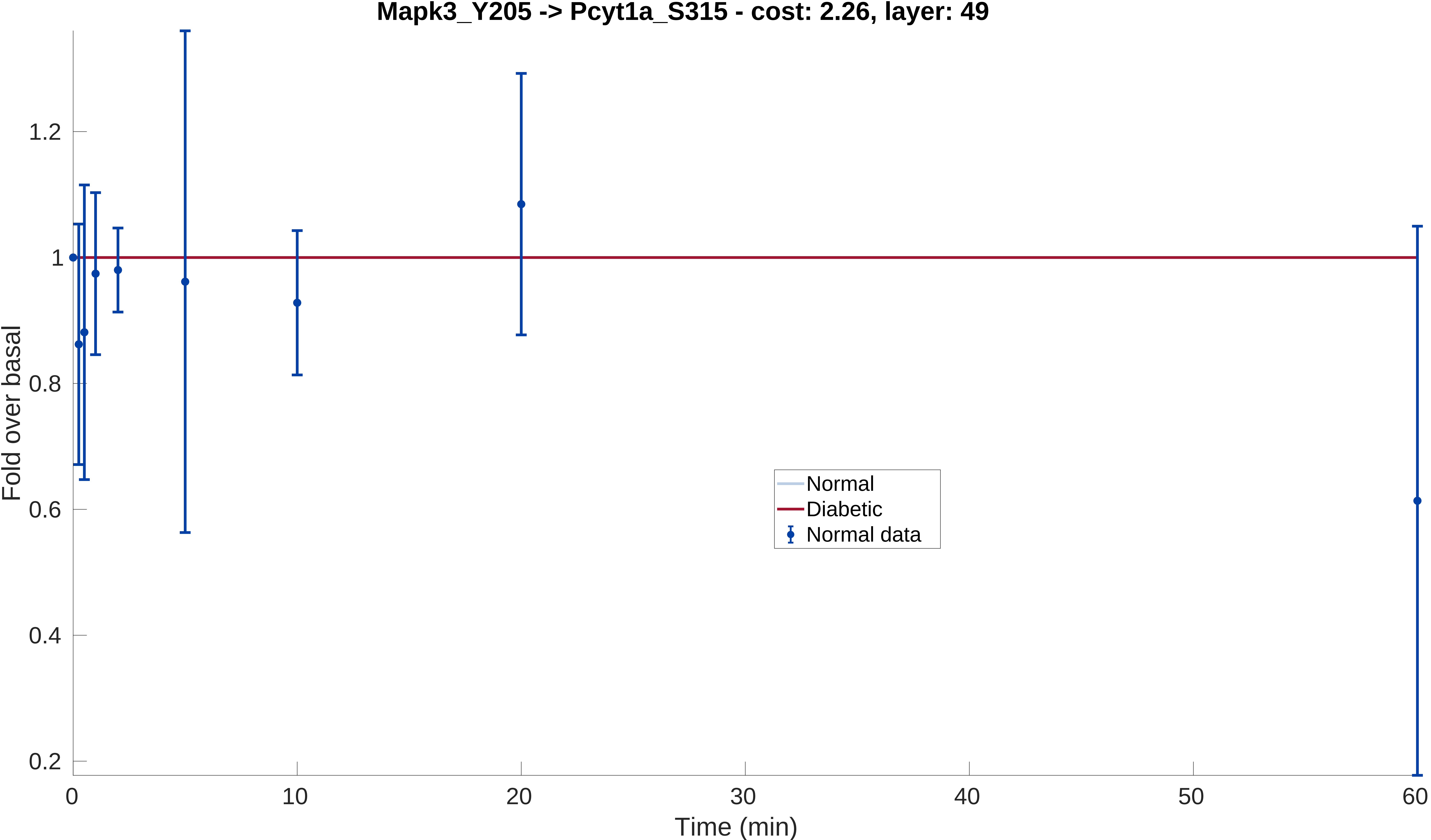

Mapk3_Y205 -> Pcyt1a_S315 - cost: 2.26, layer: 49
Fold over basal
1.2
1
0.8
0.6
0.4
0.2
0
10
20
30
40
50
60
Time (min)
Normal
Diabetic
Normal data

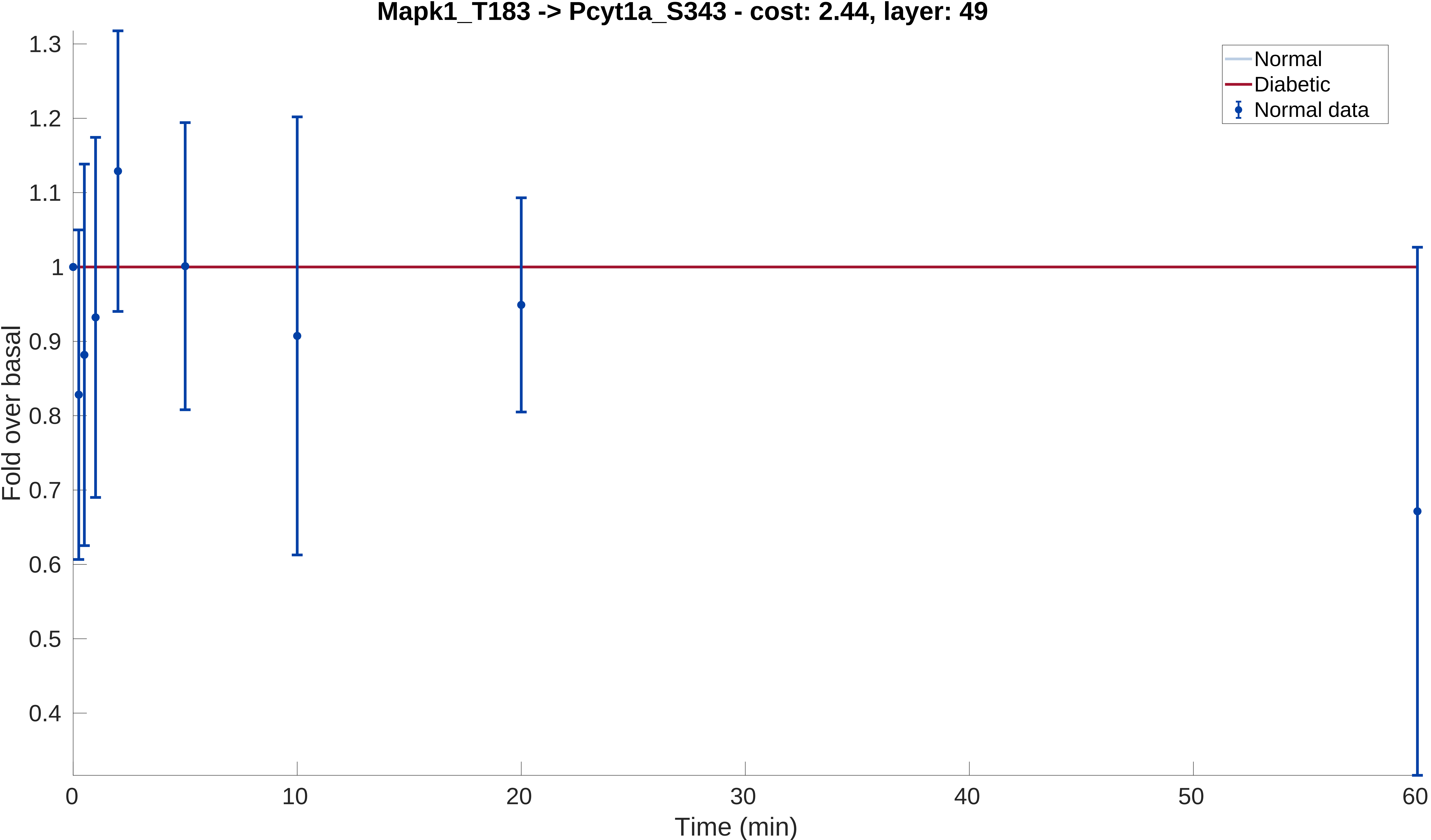

Mapk1_T183 -> Pcyt1a_S343 - cost: 2.44, layer: 49
Normal
Diabetic
Normal data
Fold over basal
Time (min)
0
10
20
30
40
50
60
0.4
0.5
0.6
0.7
0.8
0.9
1
1.1
1.2
1.3

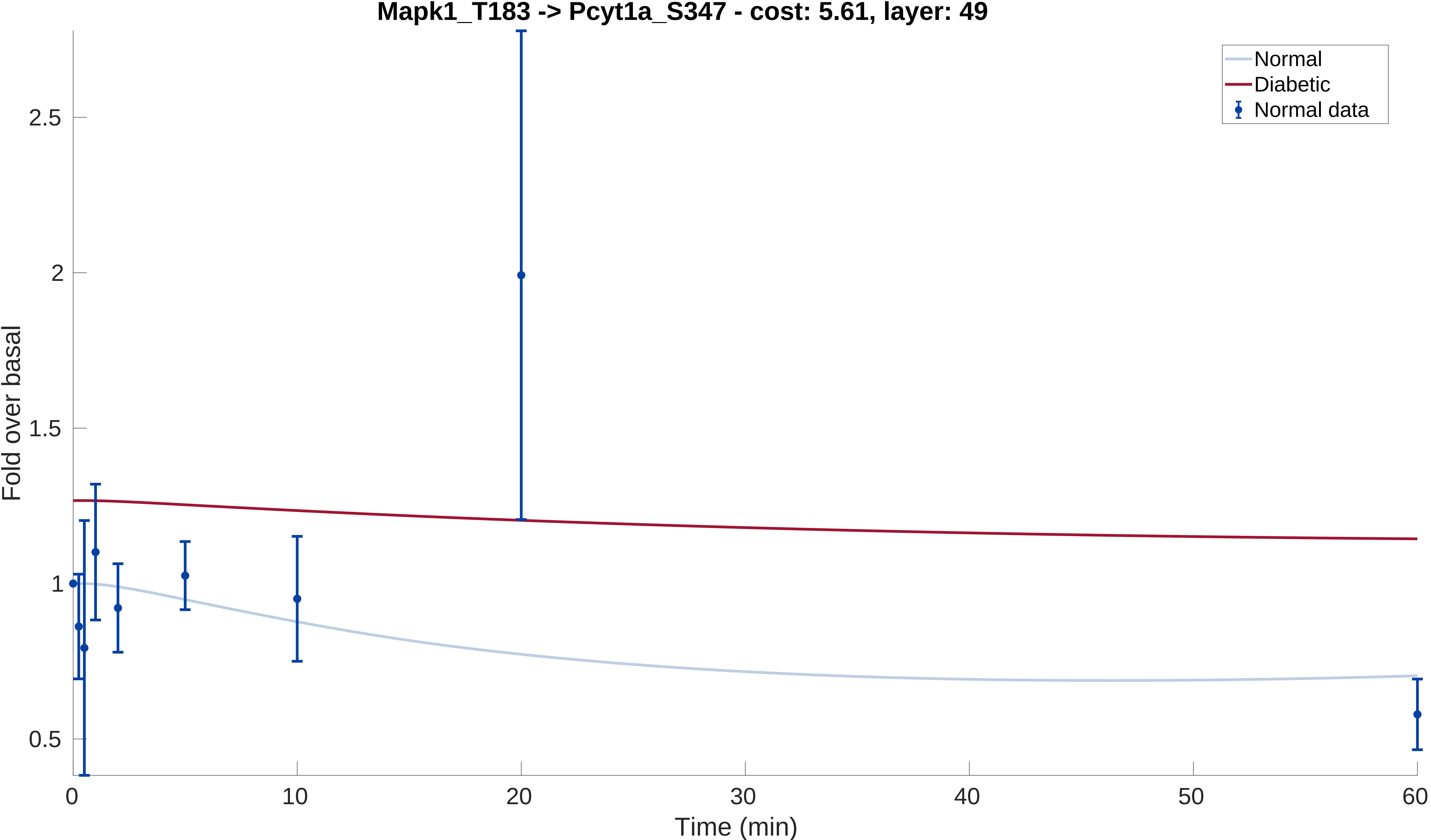
Mapk1_T183 -> Pcyt1a_S347 - cost: 5.61, layer: 49
Normal
Diabetic
Normal data
Fold over basal
0.5
1
1.5
2
2.5
0
10
20
30
40
50
60
Time (min)

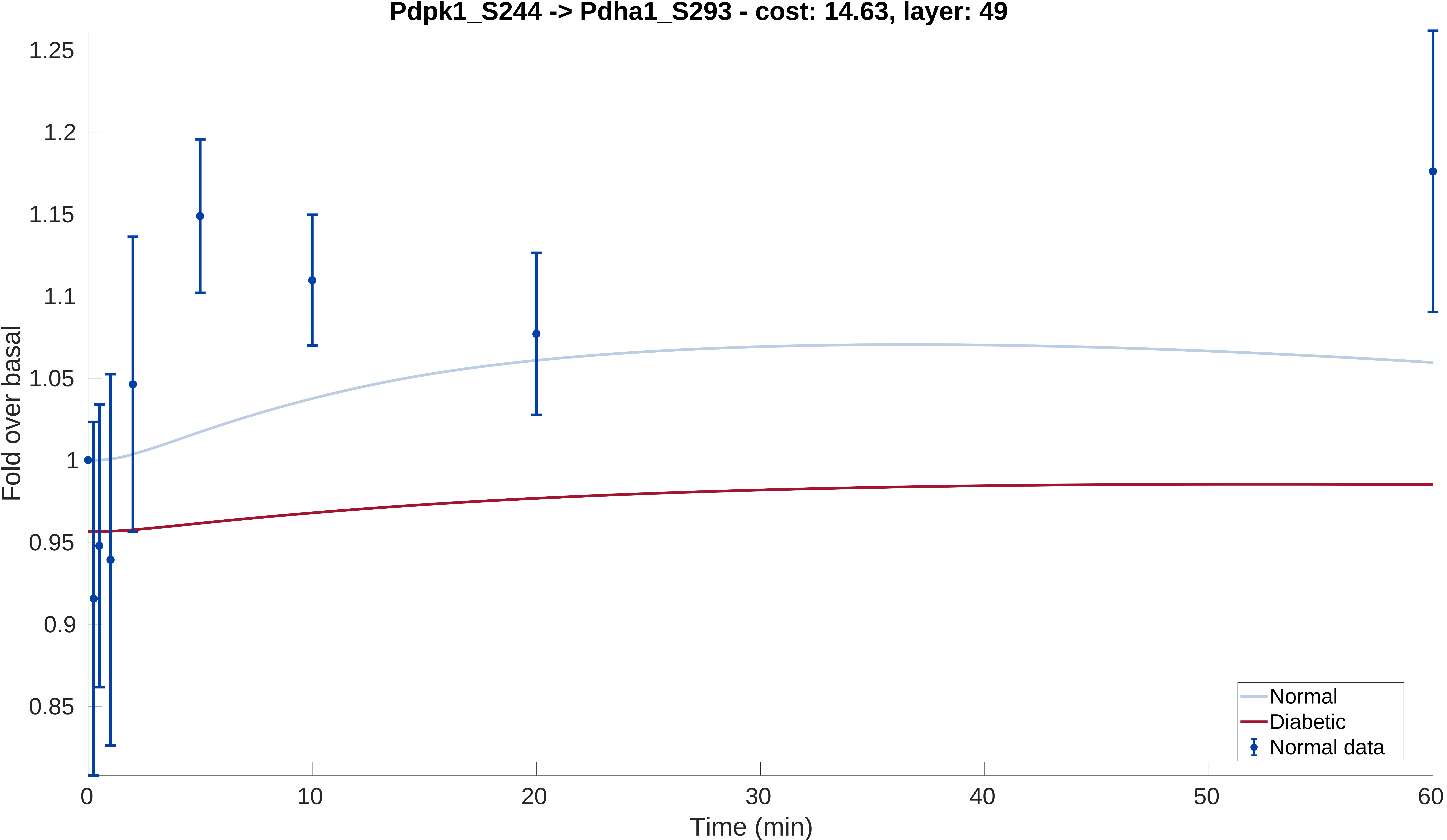

Pdpk1_S244 -> Pdha1_S293 - cost: 14.63, layer: 49
Fold over basal
1.25
1.2
1.15
1.1
1.05
1
0.95
0.9
0.85
0
10
20
30
40
50
60
Time (min)
Normal
Diabetic
Normal data

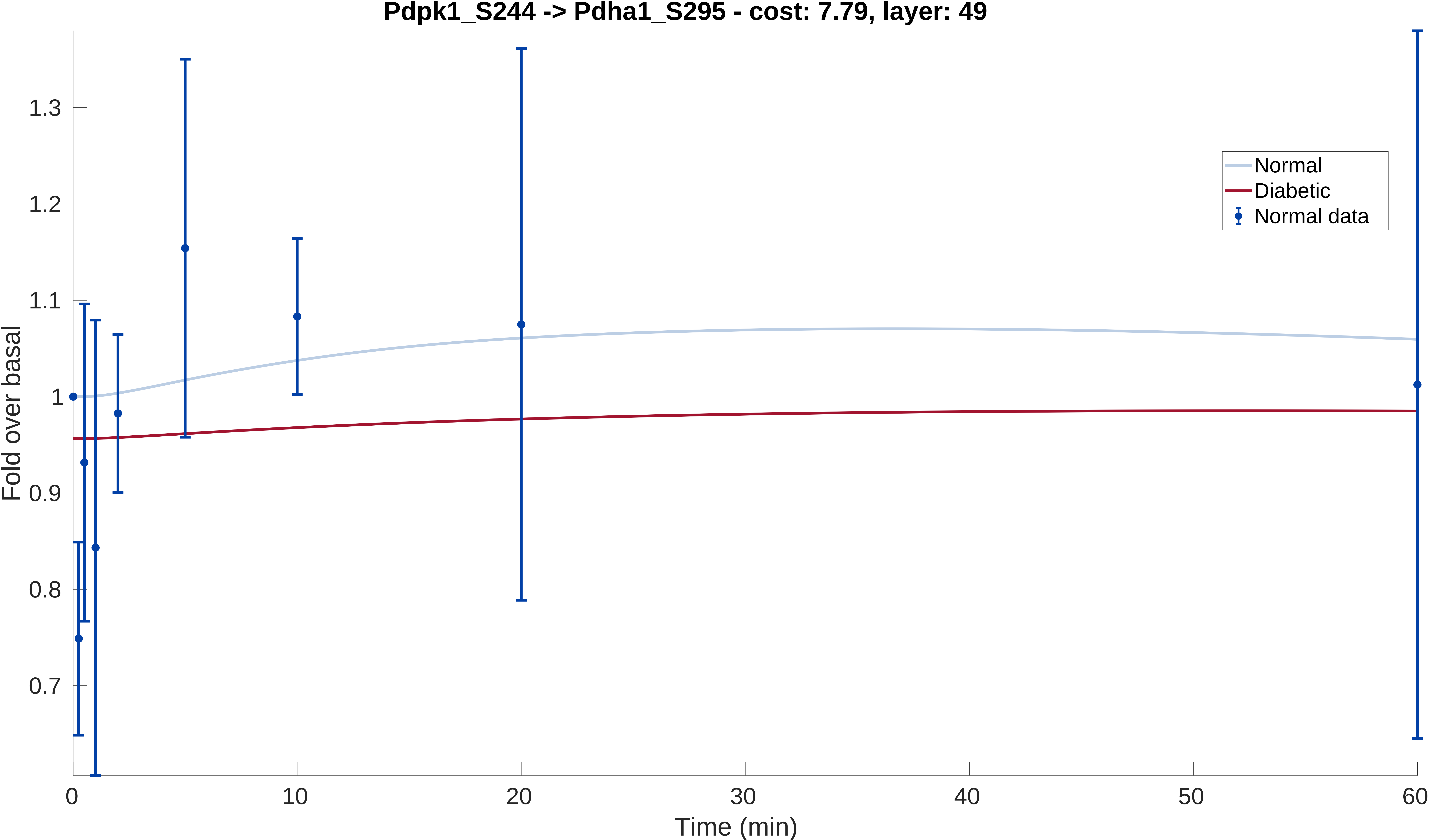

Pdpk1_S244 -> Pdha1_S295 - cost: 7.79, layer: 49
1.3
1.2
1.1
1
0.9
0.8
0.7
Fold over basal
0
10
20
30
40
50
60
Time (min)
Normal
Diabetic
Normal data

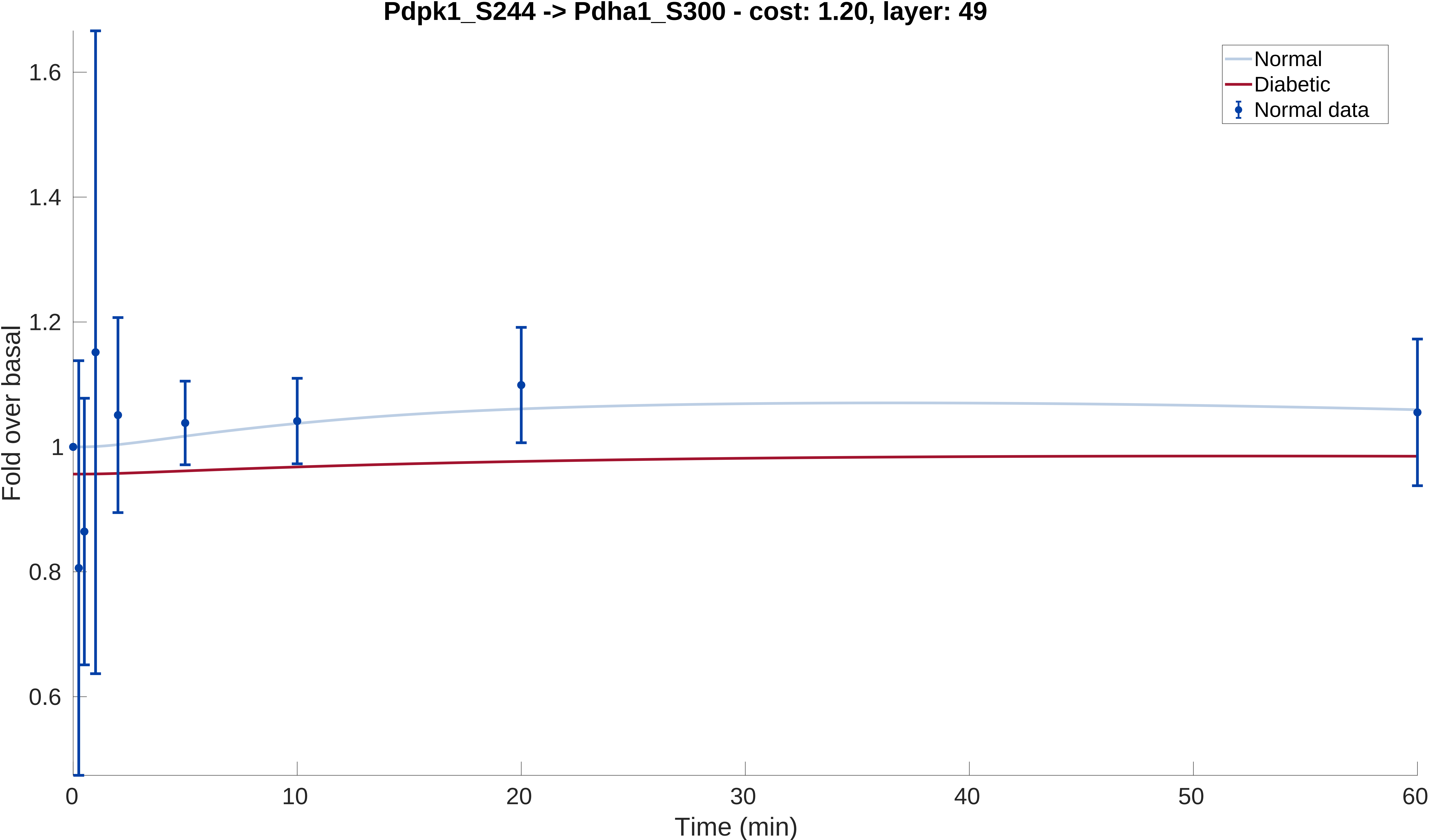

Pdpk1_S244 -> Pdha1_S300 - cost: 1.20, layer: 49
Normal
Diabetic
Normal data
Fold over basal
1.6
1.4
1.2
1
0.8
0.6
0
10
20
30
40
50
60
Time (min)

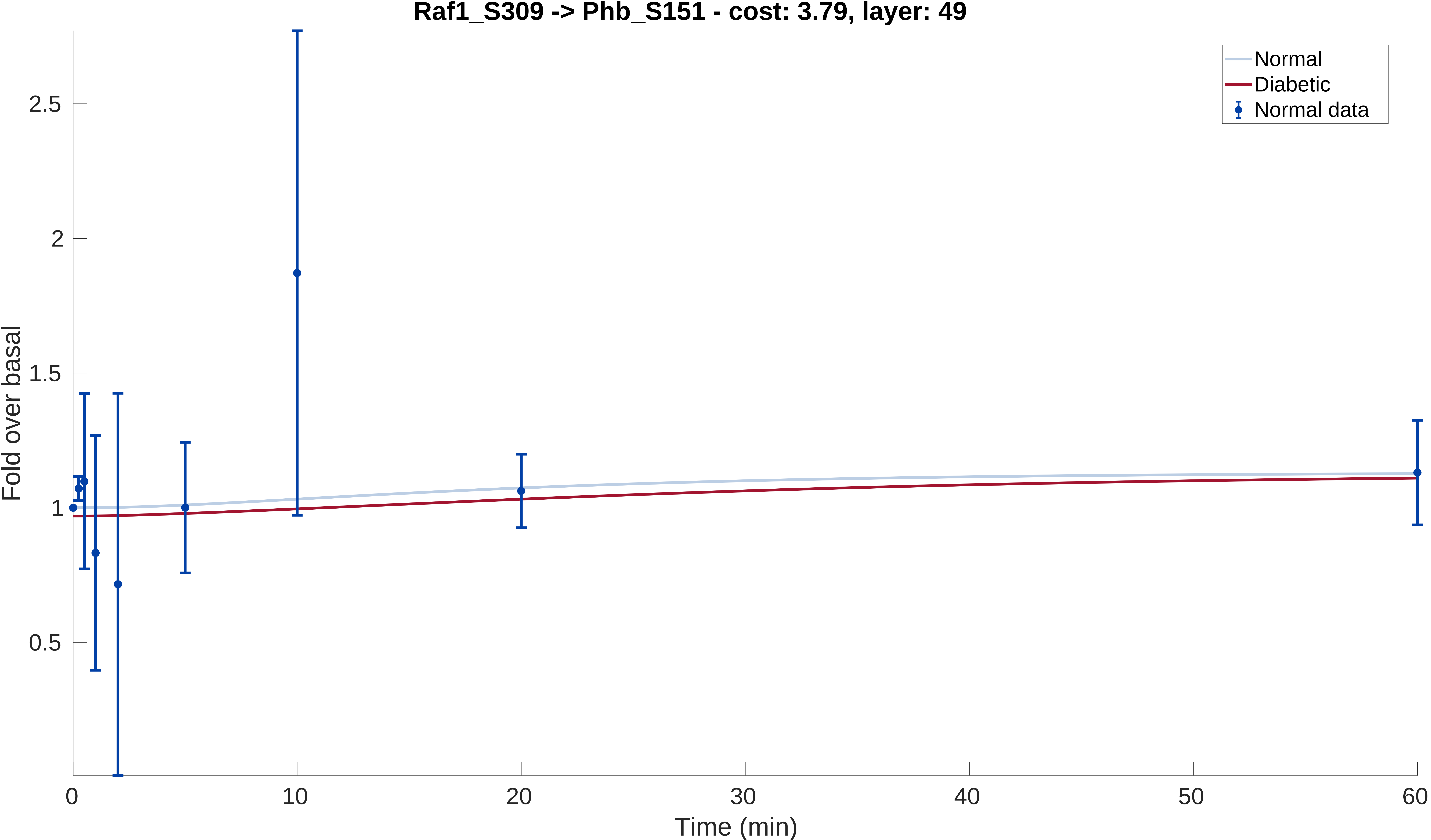

Raf1_S309 -> Phb_S151 - cost: 3.79, layer: 49
Normal
Diabetic
Normal data
Fold over basal
2.5
2
1.5
1
0.5
0
10
20
30
40
50
60
Time (min)

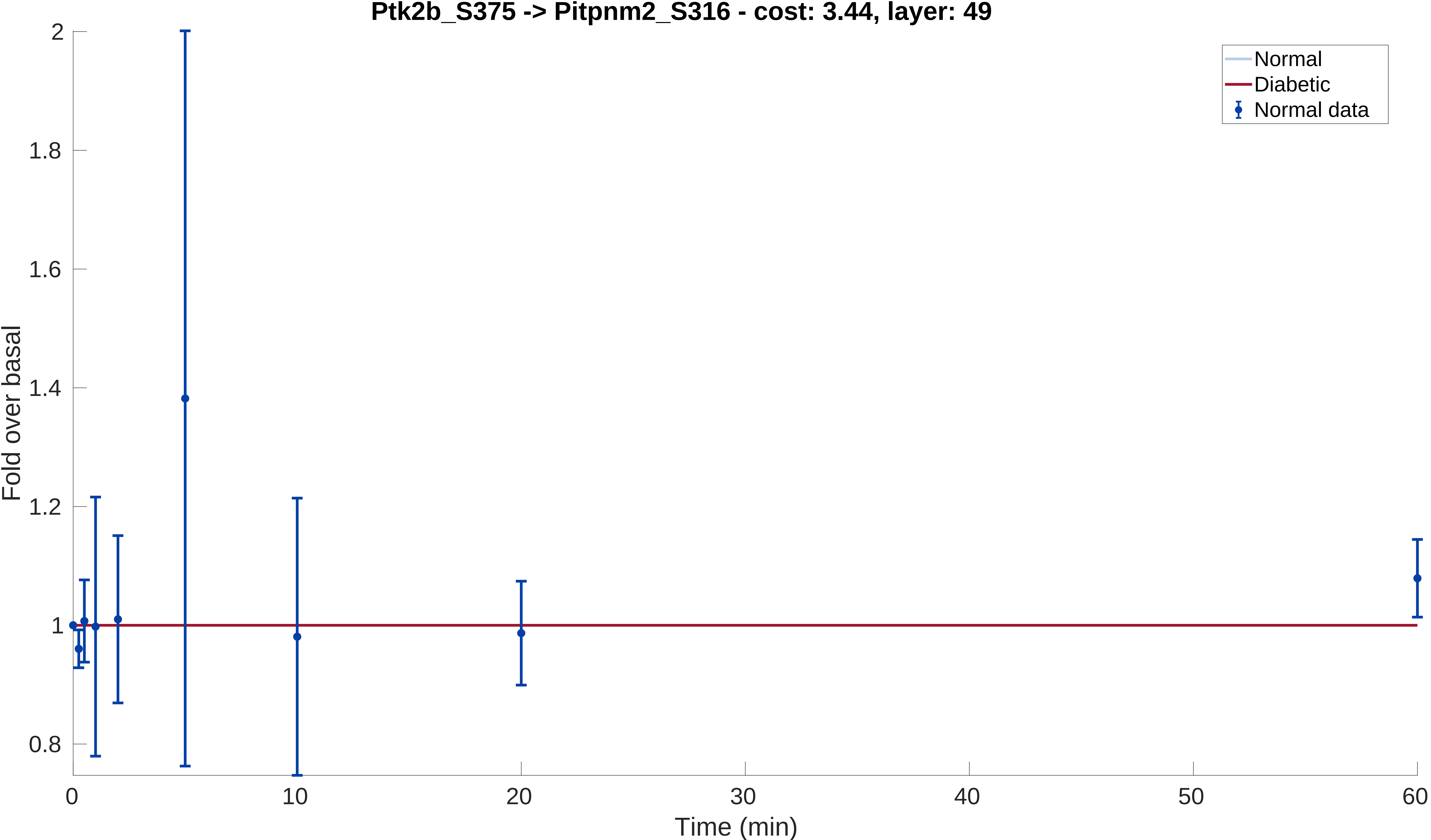

Ptk2b_S375 -> Pitpnm2_S316 - cost: 3.44, layer: 49
Normal
Diabetic
Normal data
Fold over basal
2
1.8
1.6
1.4
1.2
1
0.8
0
10
20
30
40
50
60
Time (min)

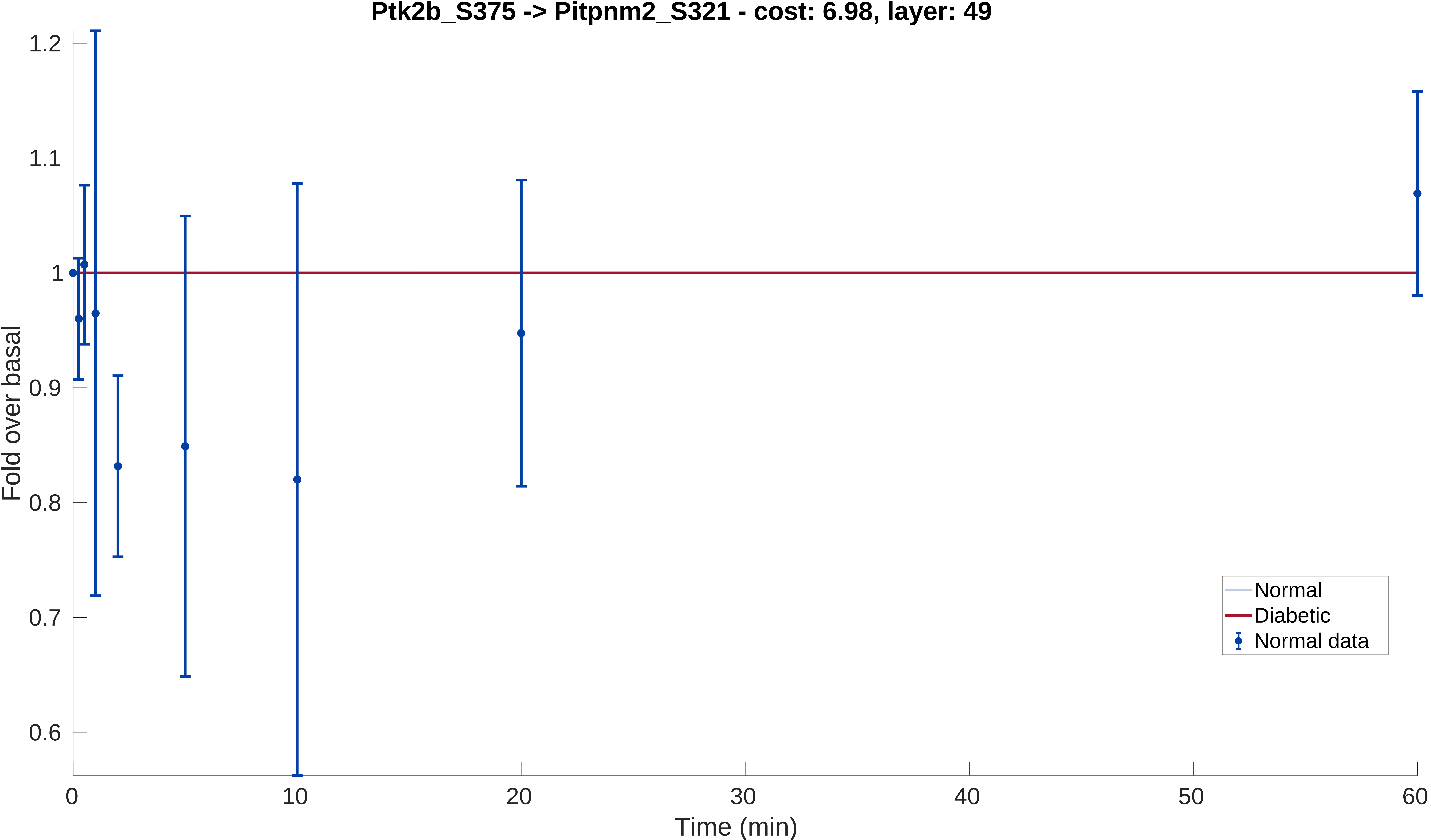

Ptk2b_S375 -> Pitpnm2_S321 - cost: 6.98, layer: 49
1.2
1.1
1
0.9
0.8
0.7
0.6
Fold over basal
0
10
20
30
40
50
60
Time (min)
Normal
Diabetic
Normal data

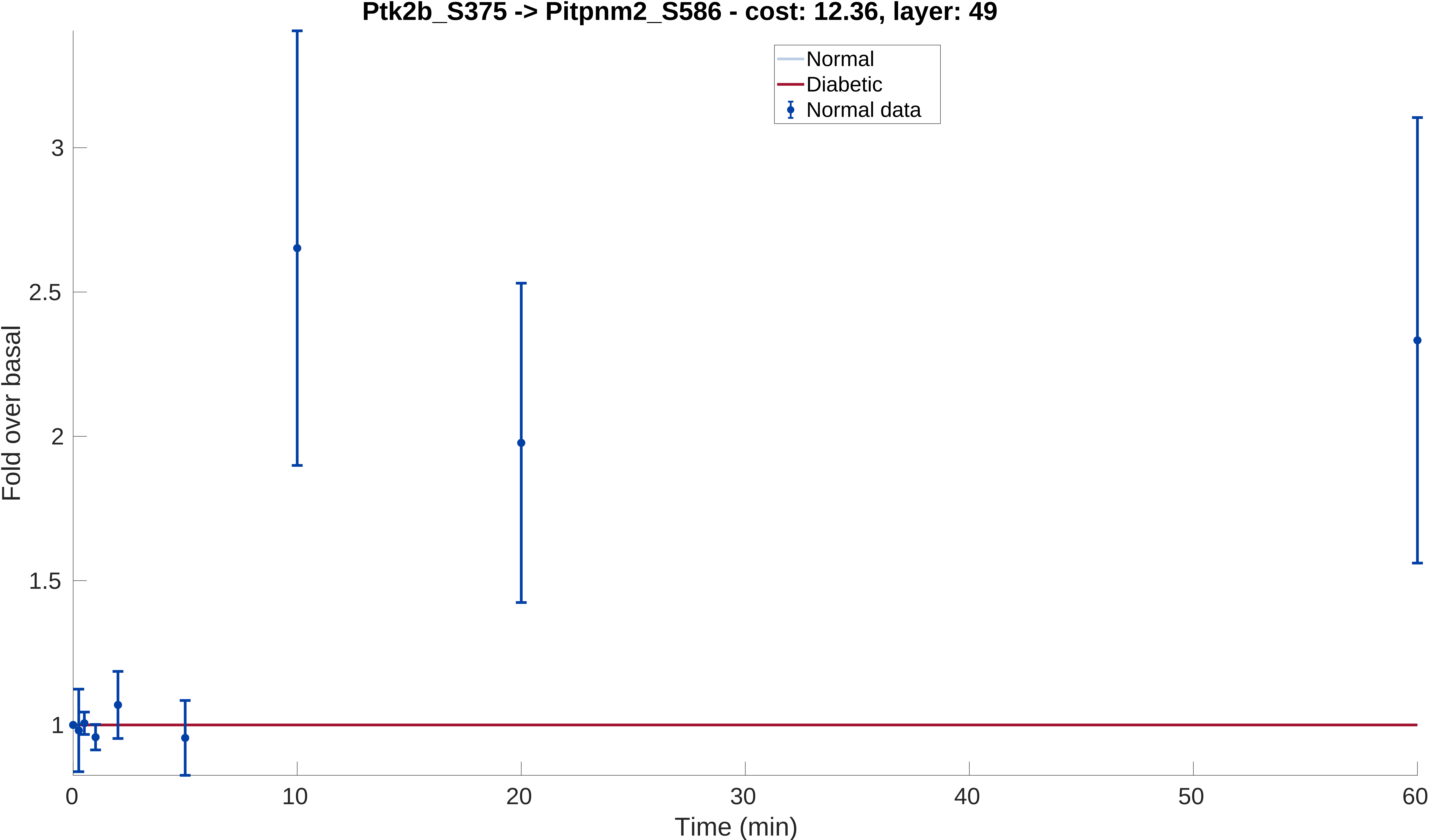
Ptk2b_S375 -> Pitpnm2_S586 - cost: 12.36, layer: 49
Normal
Diabetic
Normal data
Fold over basal
3
2.5
2
1.5
1
0
10
20
30
40
50
60
Time (min)

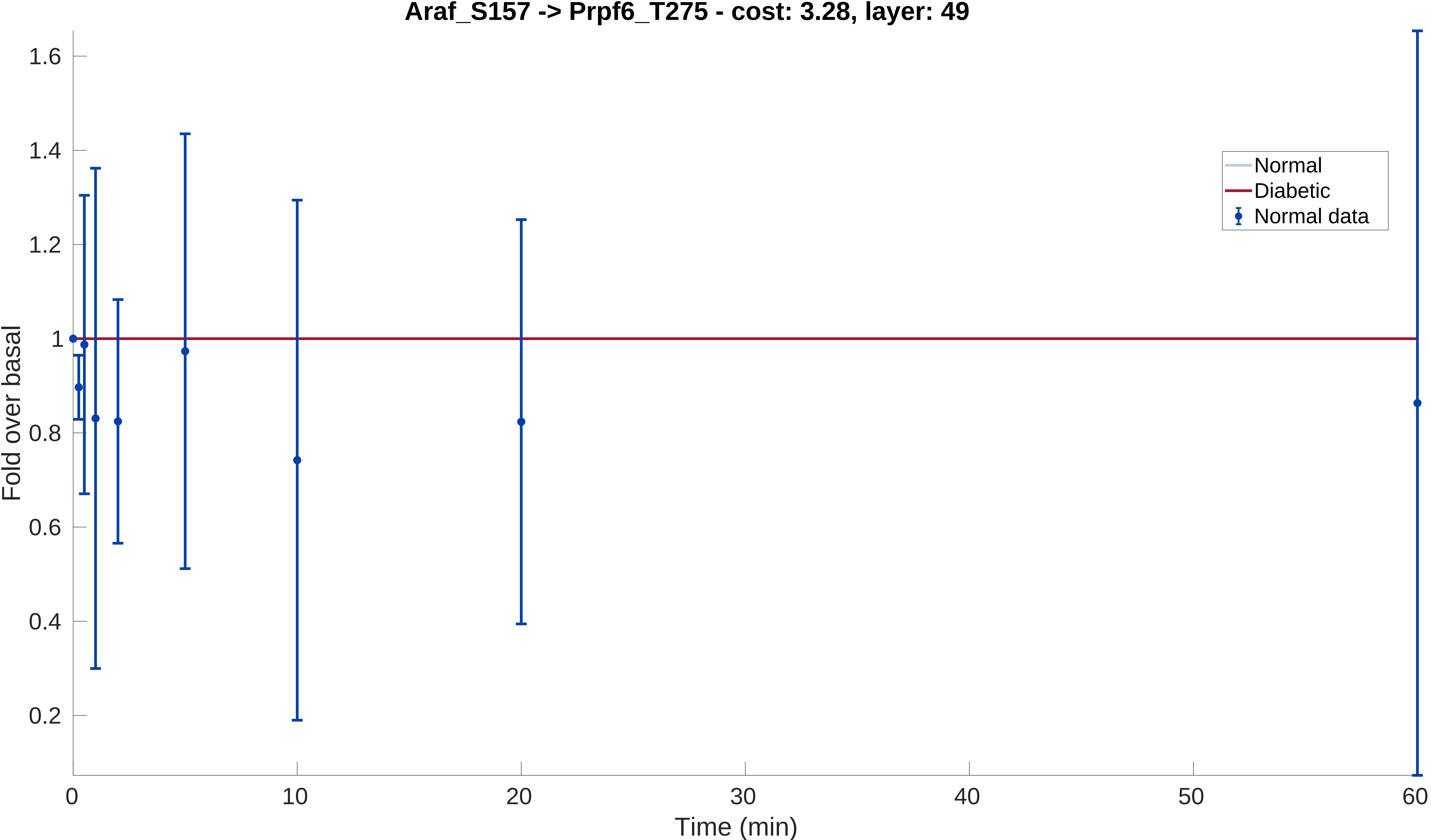

Araf_S157 -> Prpf6_T275 - cost: 3.28, layer: 49
Fold over basal
Time (min)
0
10
20
30
40
50
60
0.2
0.4
0.6
0.8
1
1.2
1.4
1.6
Normal
Diabetic
Normal data

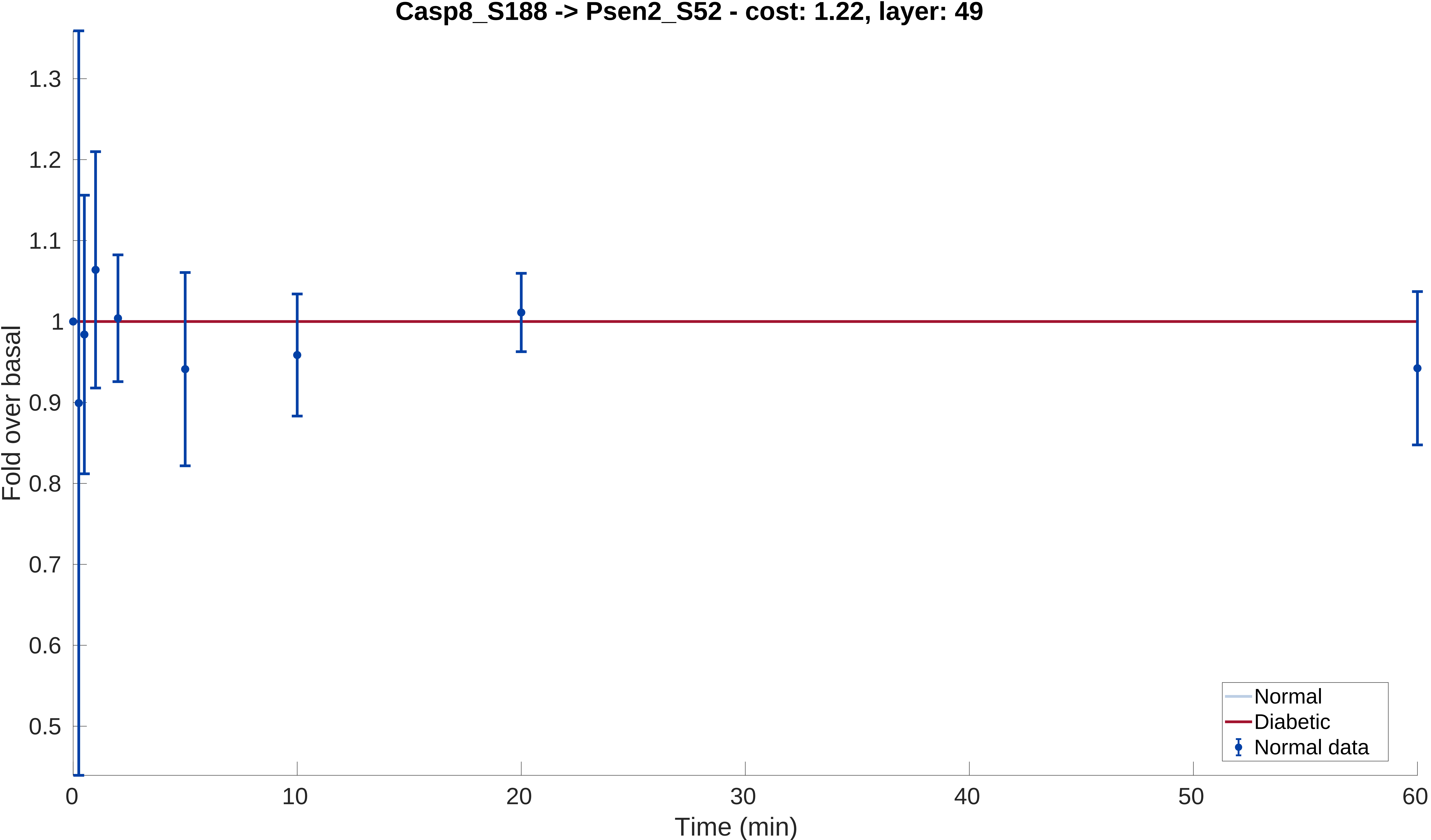

Casp8_S188 -> Psen2_S52 - cost: 1.22, layer: 49
Fold over basal
1.3
1.2
1.1
1
0.9
0.8
0.7
0.6
0.5
0
10
20
30
40
50
60
Time (min)
Normal
Diabetic
Normal data

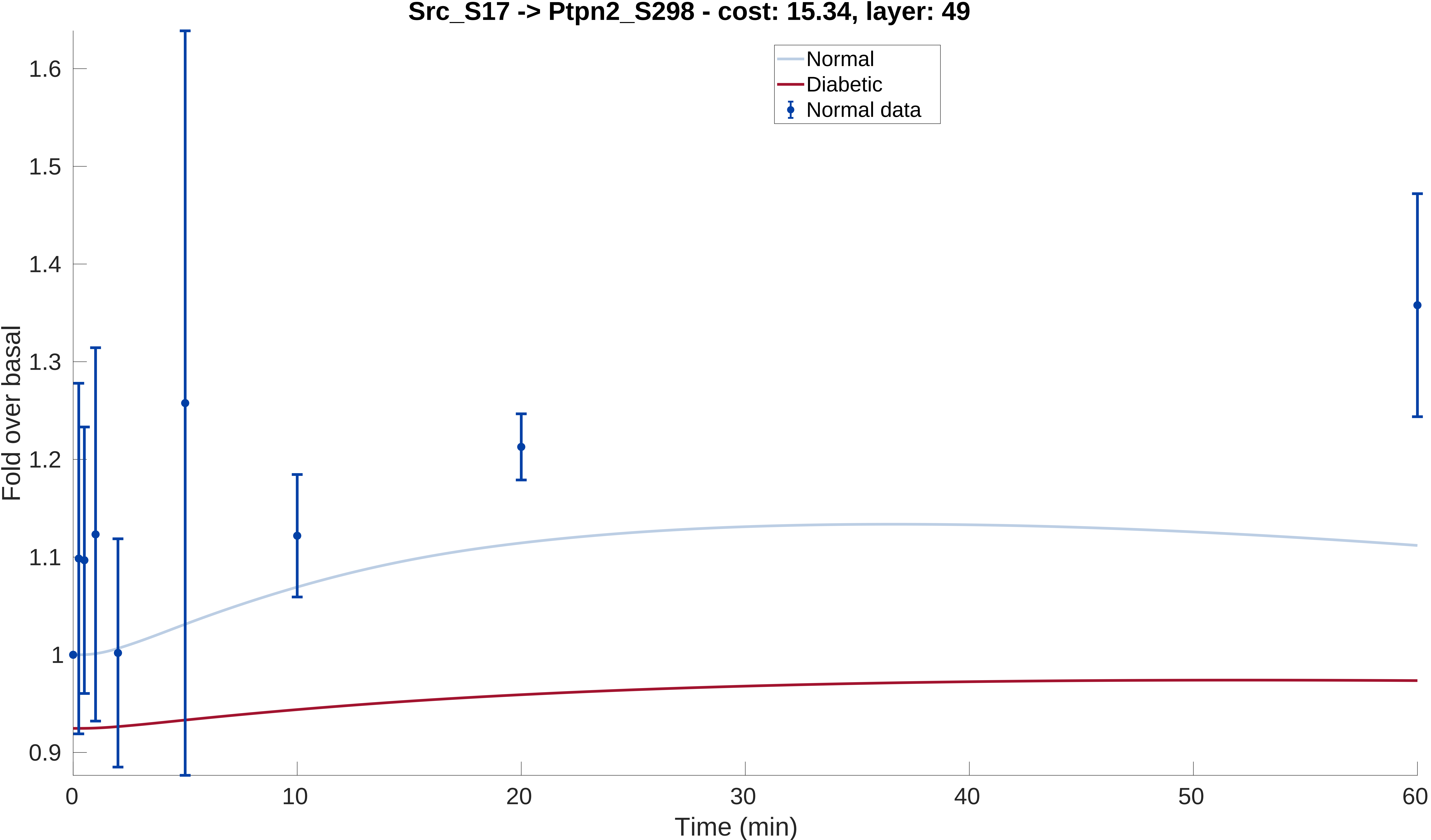
Src_S17 -> Ptpn2_S298 - cost: 15.34, layer: 49
Normal
Diabetic
Normal data
Fold over basal
Time (min)
0.9
1
1.1
1.2
1.3
1.4
1.5
1.6
0
10
20
30
40
50
60

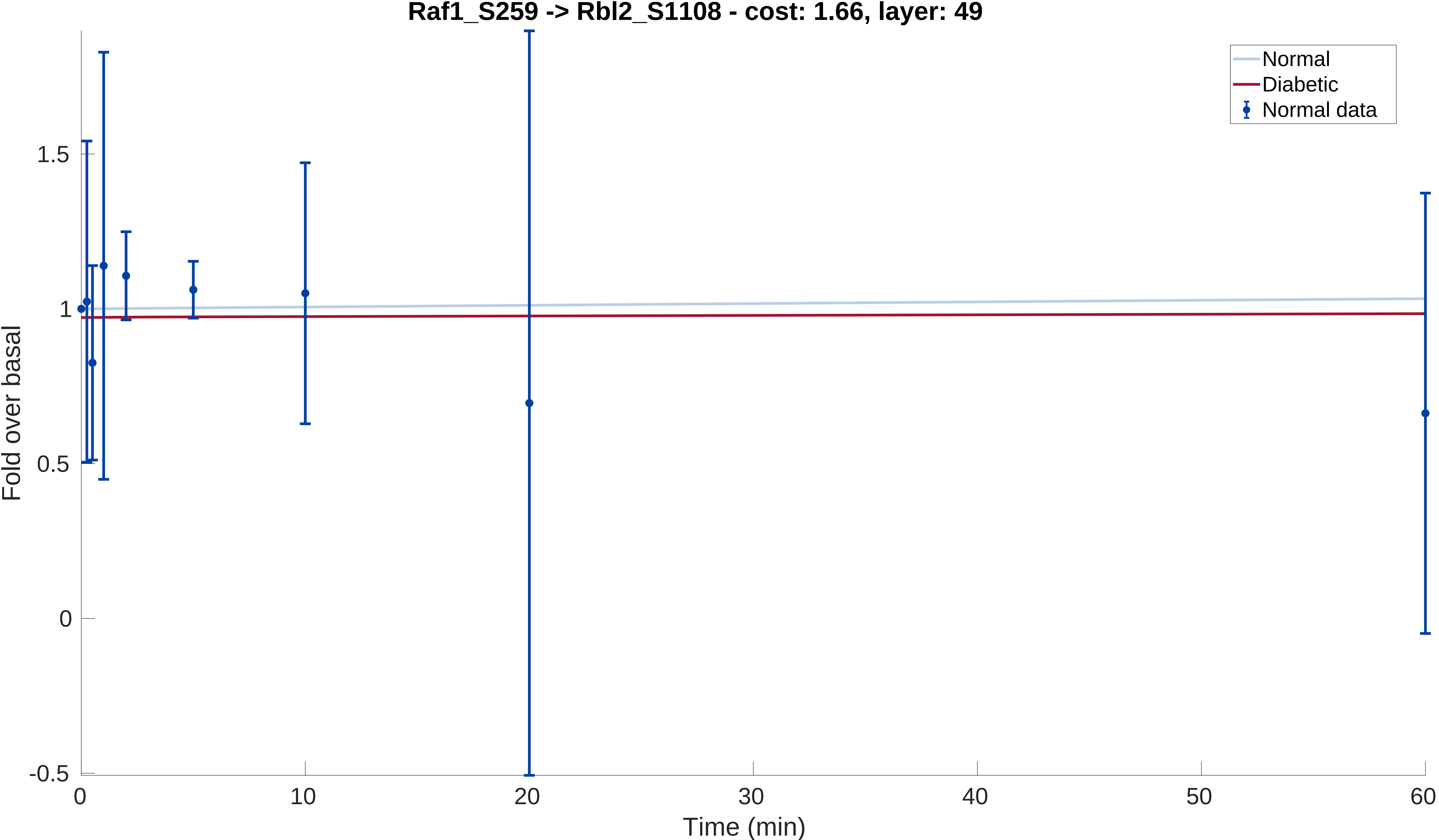

Raf1_S259 -> Rbl2_S1108 - cost: 1.66, layer: 49
Normal
Diabetic
Normal data
Fold over basal
1.5
1
0.5
0
-0.5
0
10
20
30
40
50
60
Time (min)

#### Raf1_S321 -> Rbl2_S37 - cost: 6.68, layer: 49

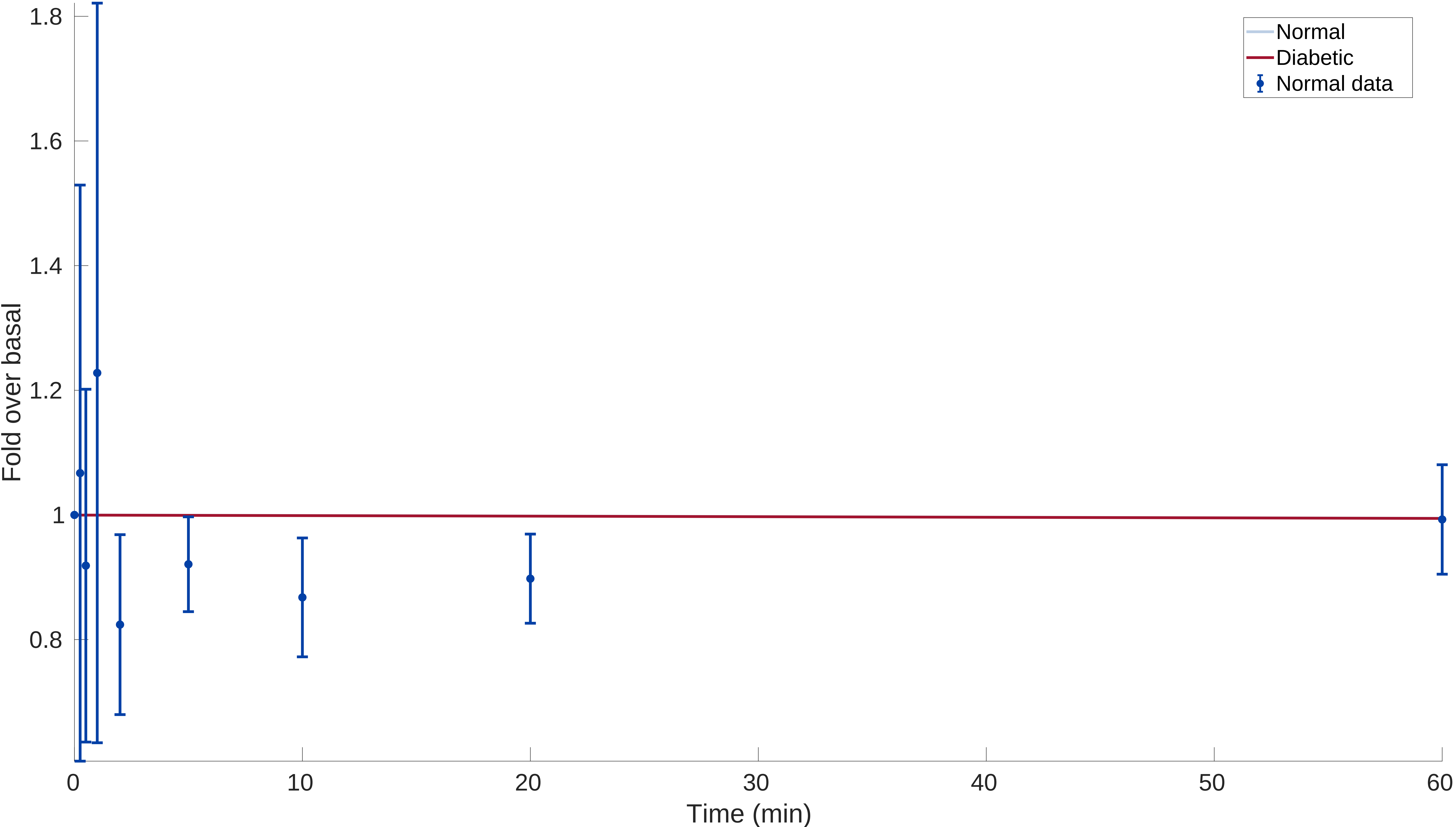

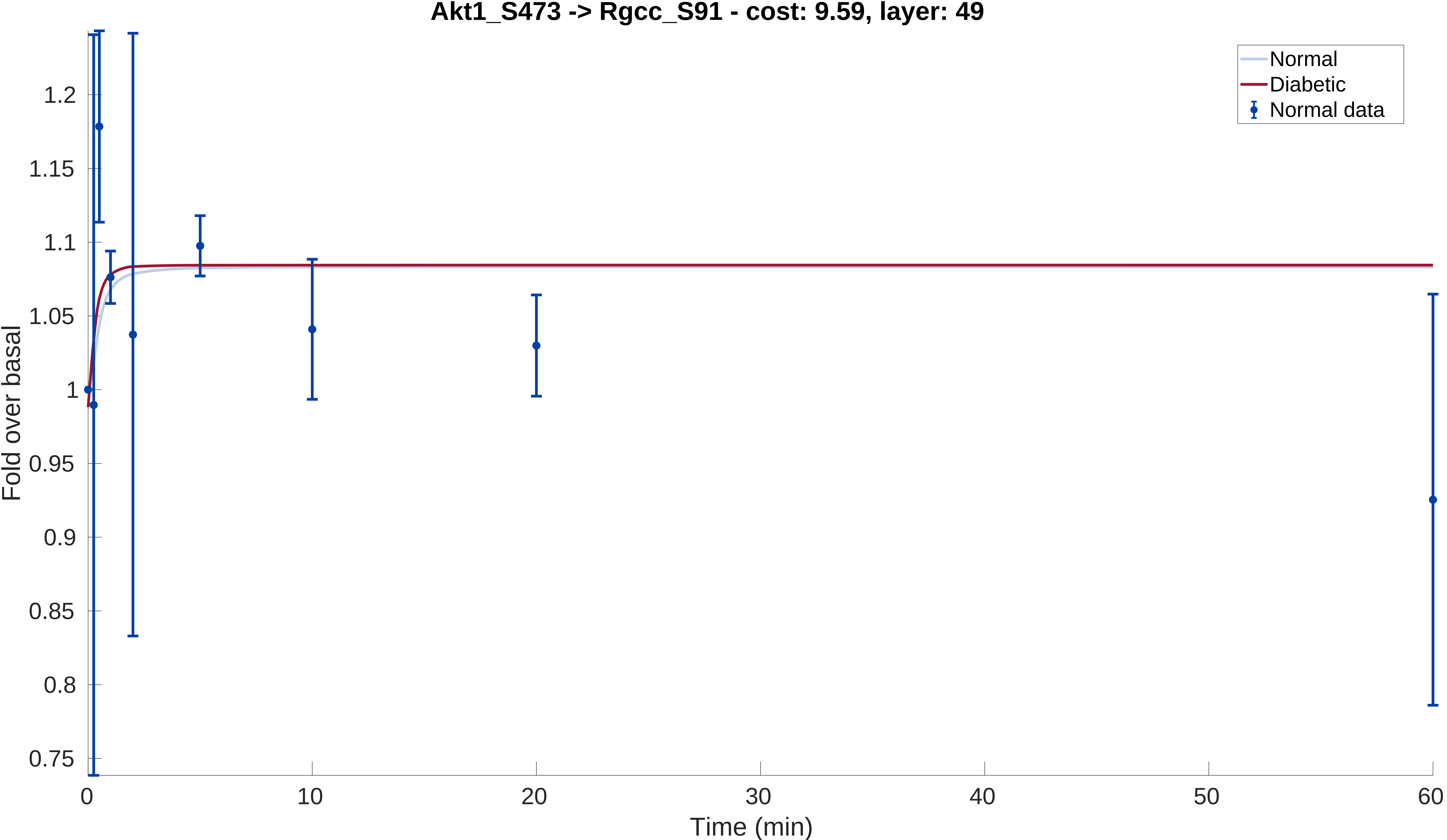

Akt1_S473 -> Rgcc_S91 - cost: 9.59, layer: 49
Normal
Diabetic
Normal data
Fold over basal
Time (min)
0
10
20
30
40
50
60
0.75
0.8
0.85
0.9
0.95
1
1.05
1.1
1.15
1.2

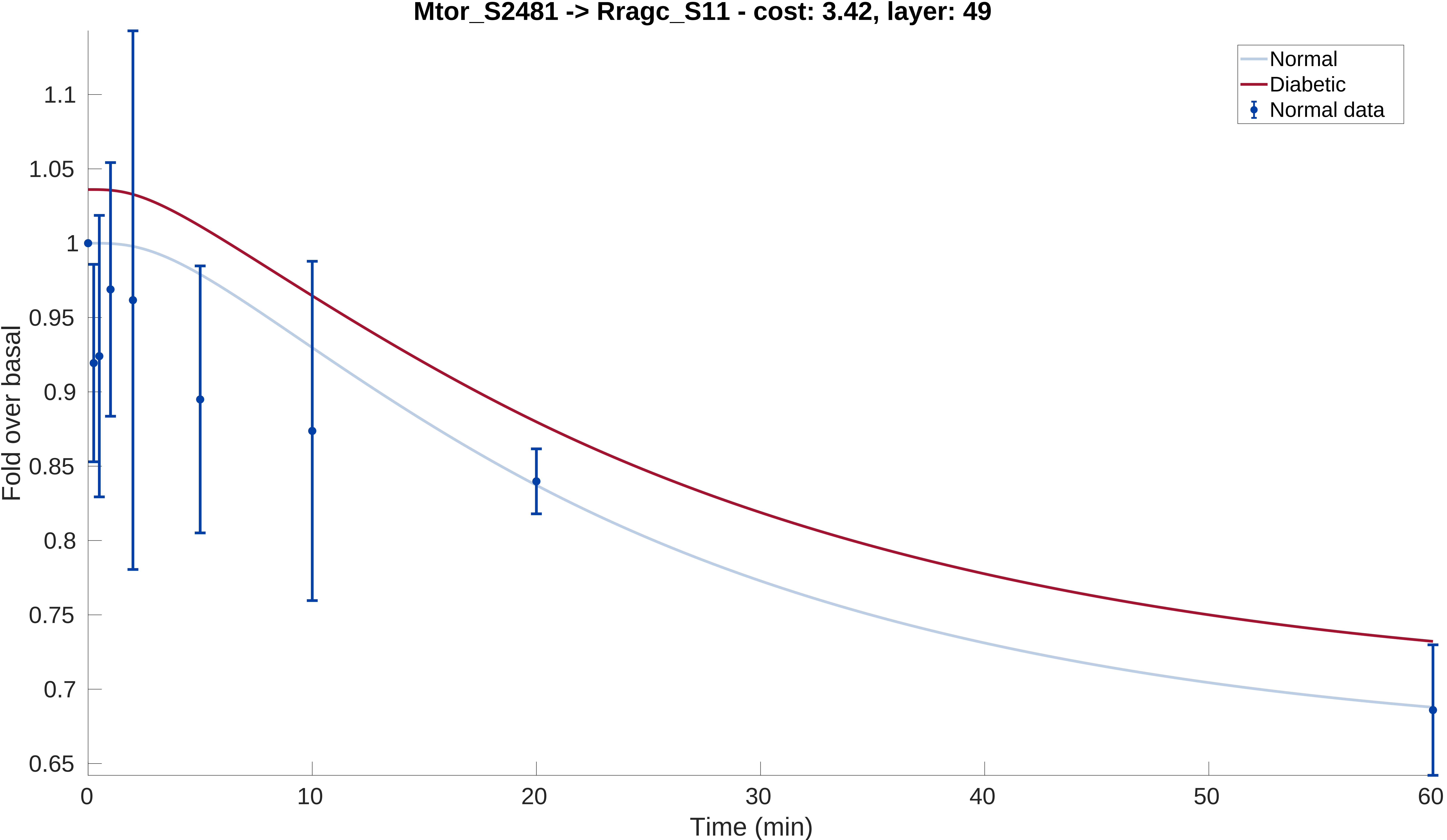
Mtor_S2481 -> Rragc_S11 - cost: 3.42, layer: 49
Normal
Diabetic
Normal data
Fold over basal
Time (min)
1.1
1.05
1
0.95
0.9
0.85
0.8
0.75
0.7
0.65
0
10
20
30
40
50
60

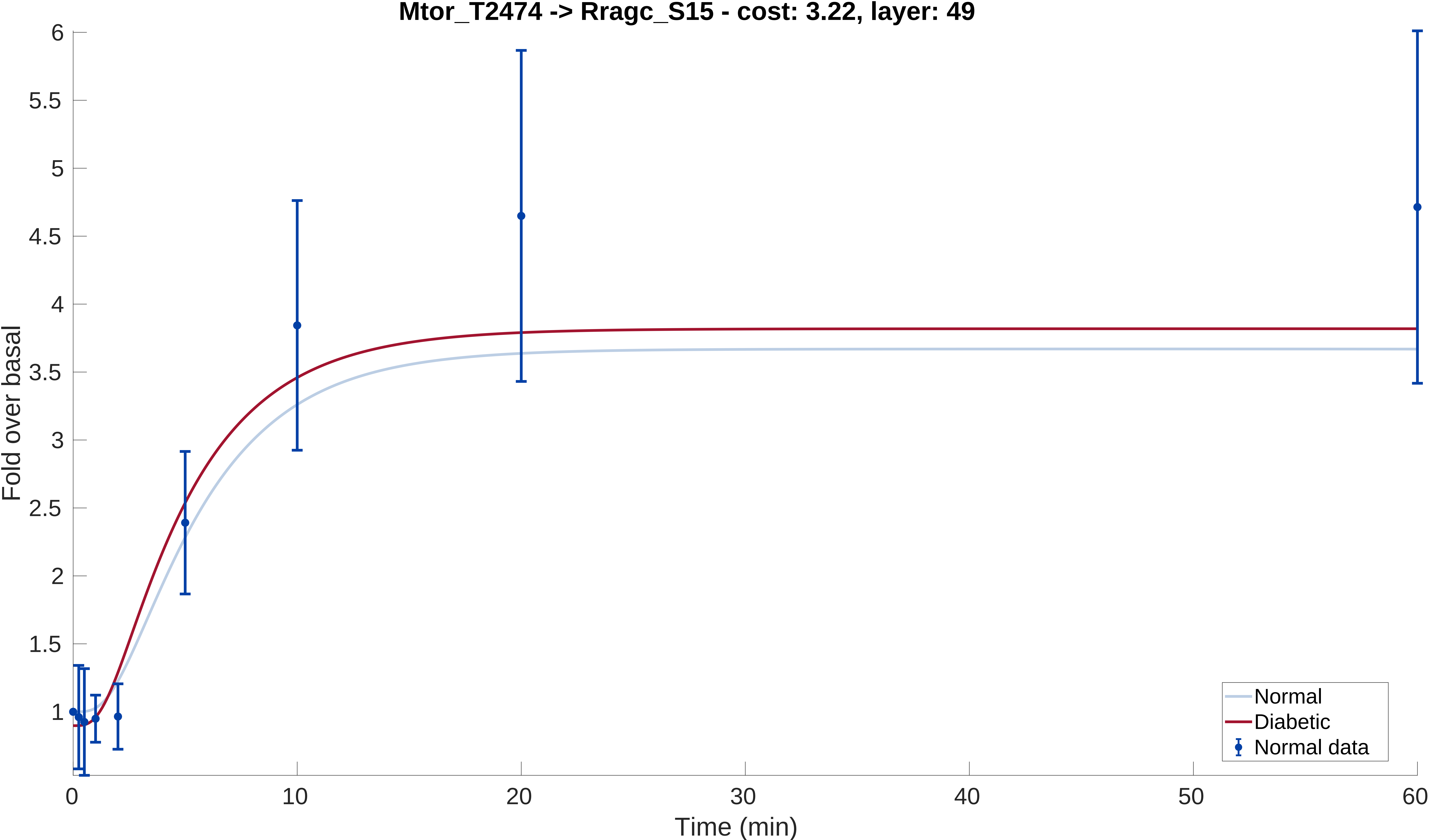

Mtor_T2474 -> Rragc_S15 - cost: 3.22, layer: 49
Fold over basal
Time (min)
0
10
20
30
40
50
60
1
1.5
2
2.5
3
3.5
4
4.5
5
5.5
6
Normal
Diabetic
Normal data

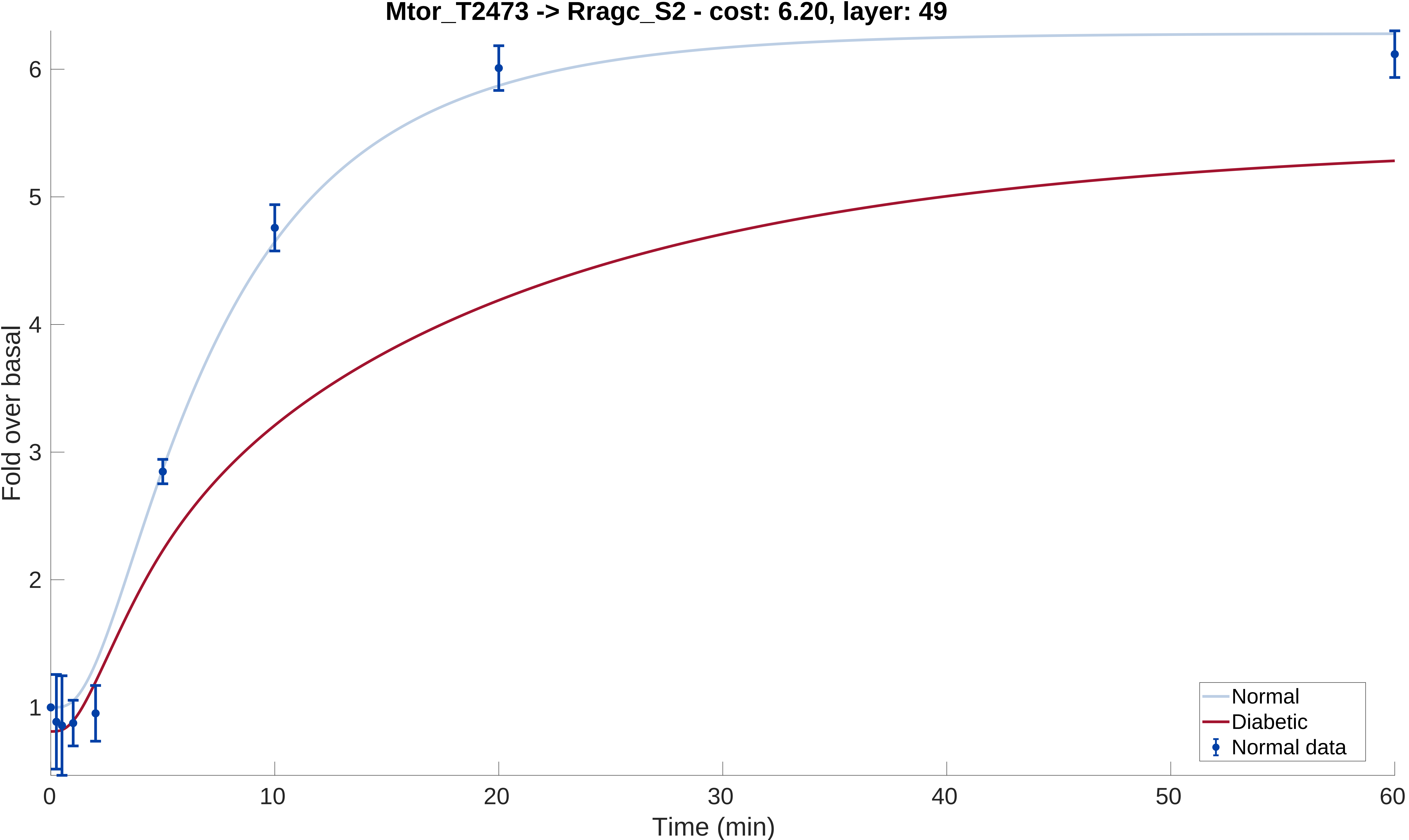

Mtor_T2473 -> Rragc_S2 - cost: 6.20, layer: 49
Fold over basal
Time (min)
0
10
20
30
40
50
60
1
2
3
4
5
6
Normal
Diabetic
Normal data

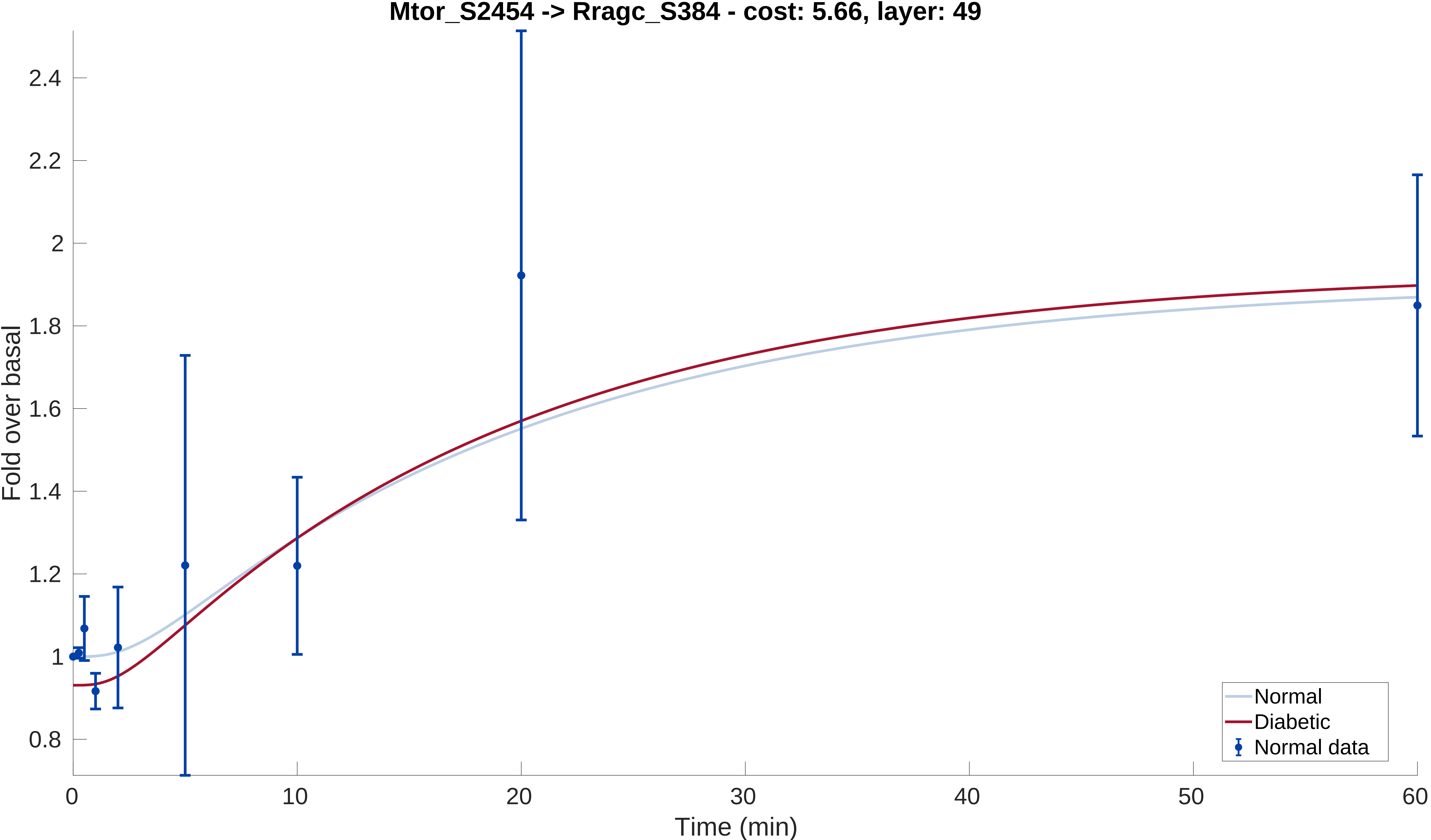

Mtor_S2454 -> Rragc_S384 - cost: 5.66, layer: 49
Fold over basal
2.4
2.2
2
1.8
1.6
1.4
1.2
1
0.8
0
10
20
30
40
50
60
Time (min)
Normal
Diabetic
Normal data

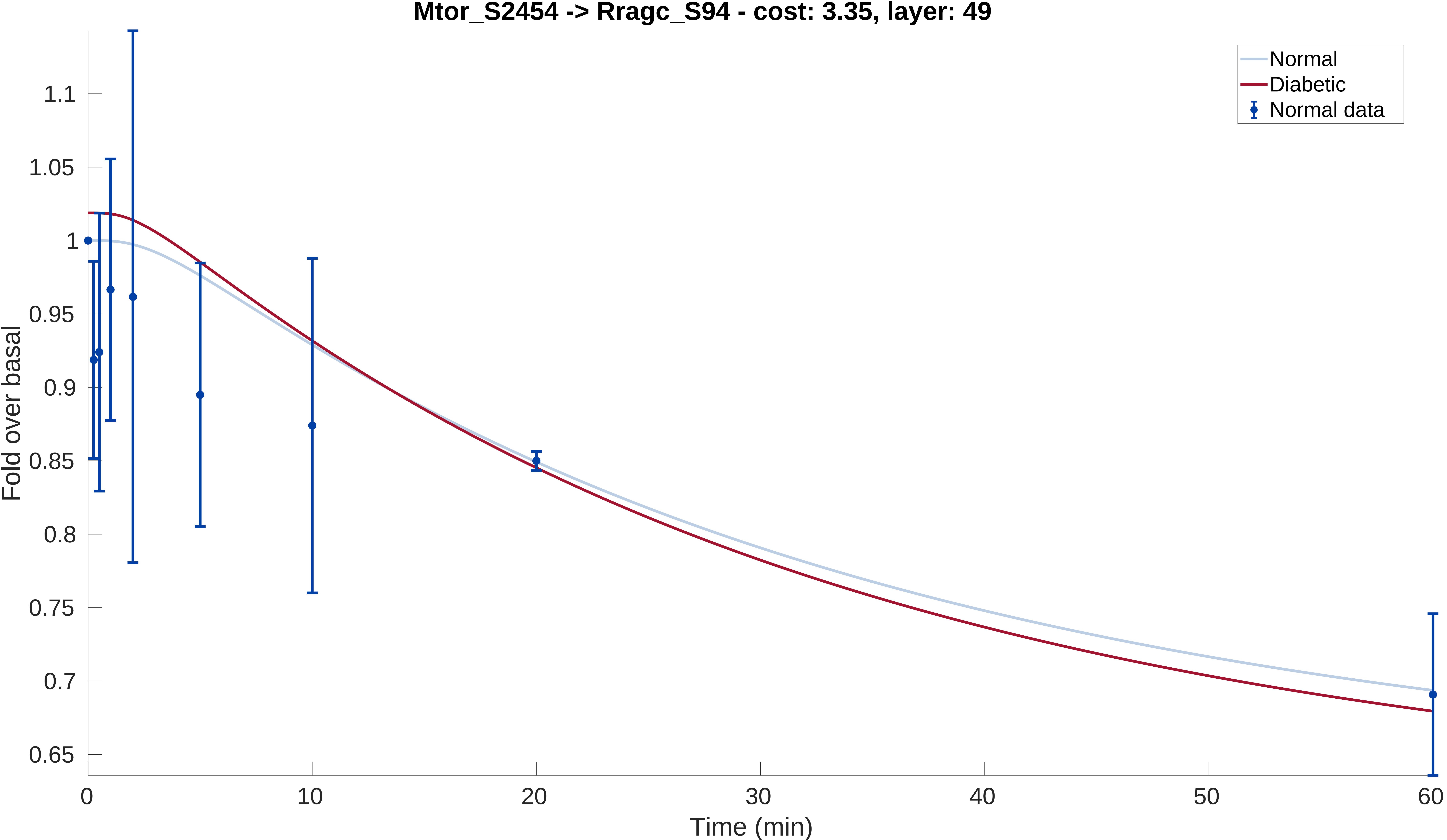

Mtor_S2454 -> Rragc_S94 - cost: 3.35, layer: 49
Normal
Diabetic
Normal data
Fold over basal
1.1
1.05
1
0.95
0.9
0.85
0.8
0.75
0.7
0.65
0
10
20
30
40
50
60
Time (min)

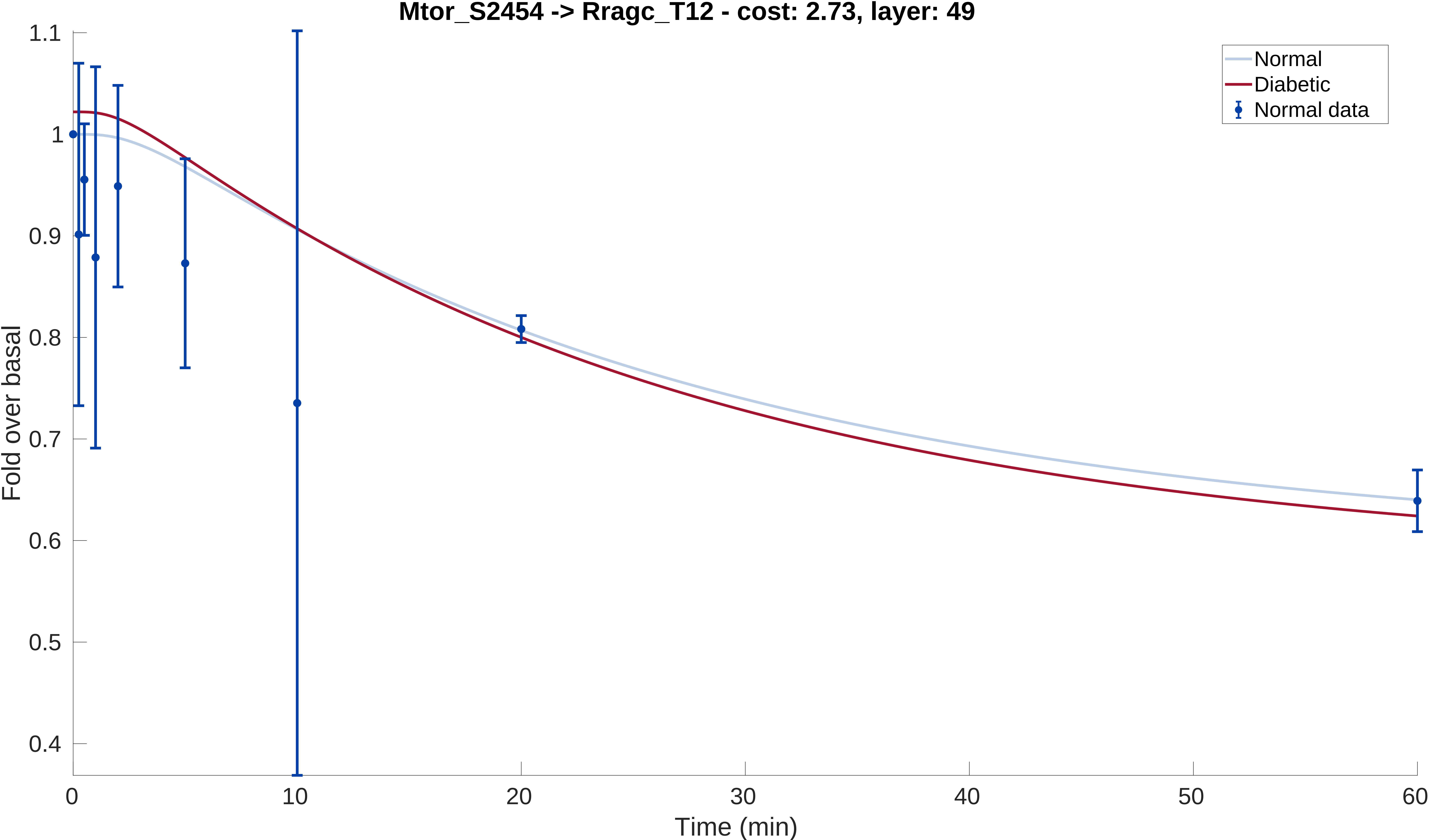

Mtor_S2454 -> Rragc_T12 - cost: 2.73, layer: 49
Fold over basal
Time (min)
Normal
Diabetic
Normal data
1.1
1
0.9
0.8
0.7
0.6
0.5
0.4
0
10
20
30
40
50
60

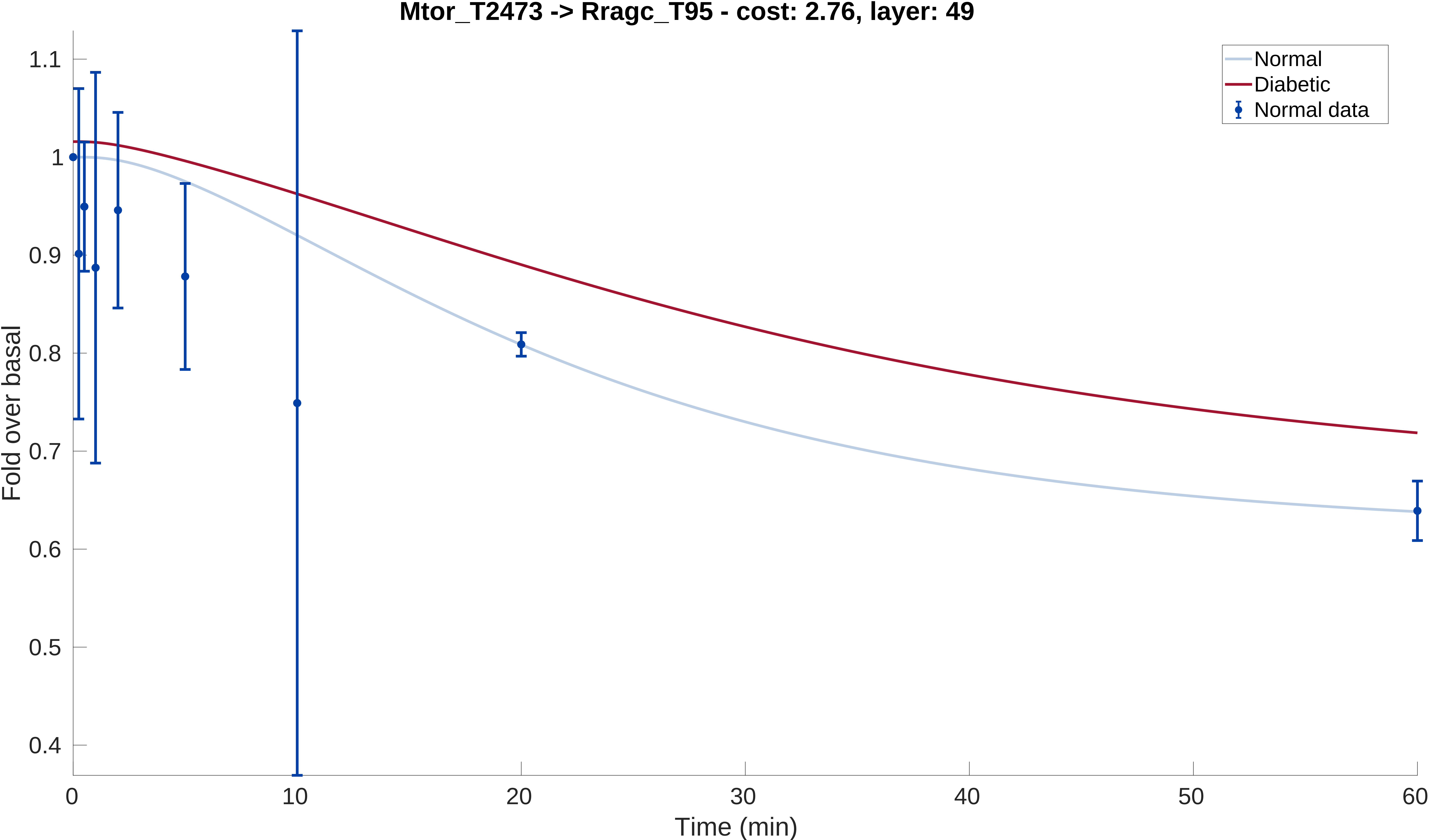

Mtor_T2473 -> Rragc_T95 - cost: 2.76, layer: 49
Normal
Diabetic
Normal data
Fold over basal
1.1
1
0.9
0.8
0.7
0.6
0.5
0.4
0
10
20
30
40
50
60
Time (min)

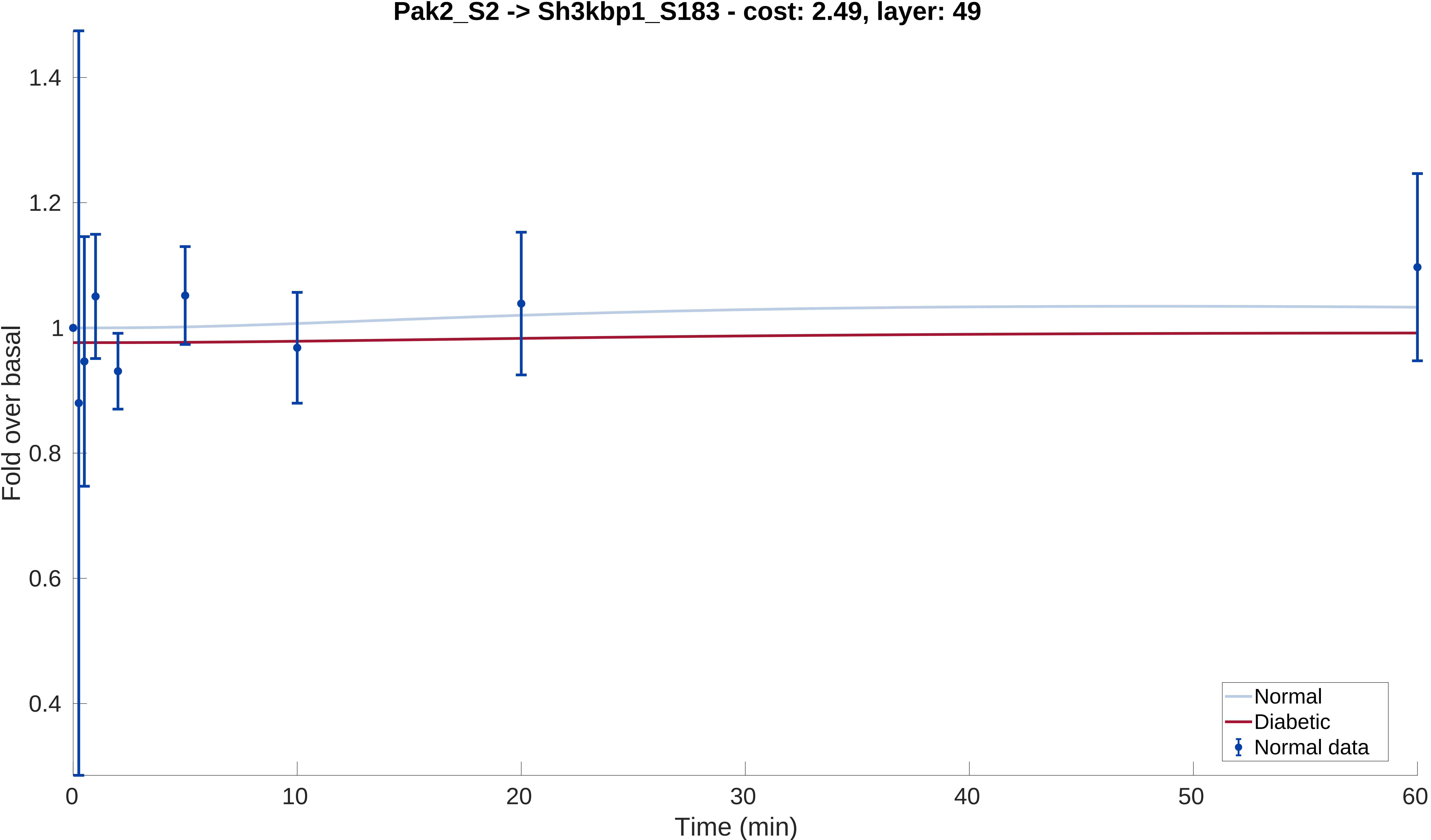

Pak2_S2 -> Sh3kbp1_S183 - cost: 2.49, layer: 49
Fold over basal
1.4
1.2
1
0.8
0.6
0.4
0
10
20
30
40
50
60
Time (min)
Normal
Diabetic
Normal data

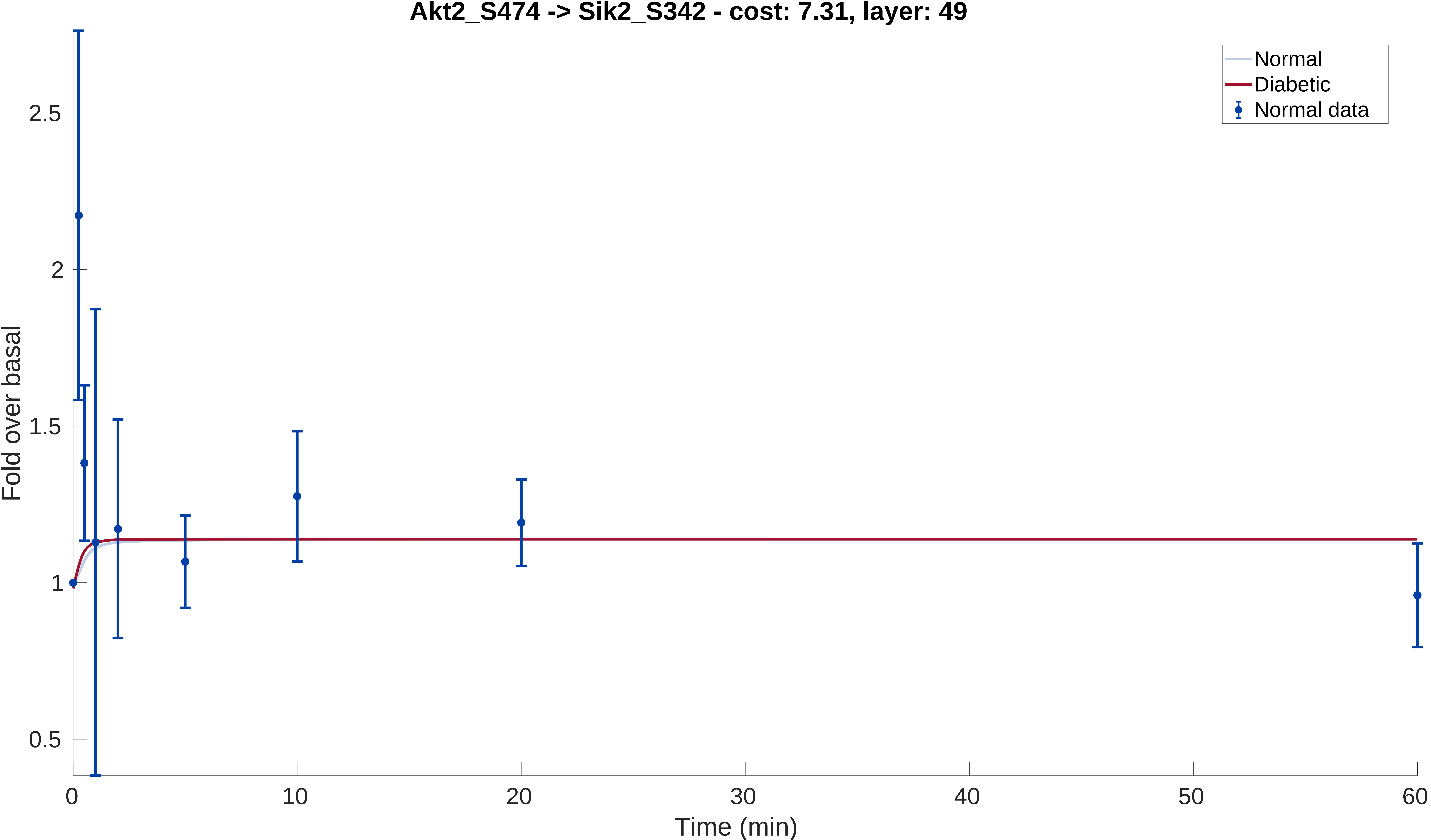

Akt2_S474 -> Sik2_S342 - cost: 7.31, layer: 49
Normal
Diabetic
Normal data
Fold over basal
2.5
2
1.5
1
0.5
0
10
20
30
40
50
60
Time (min)

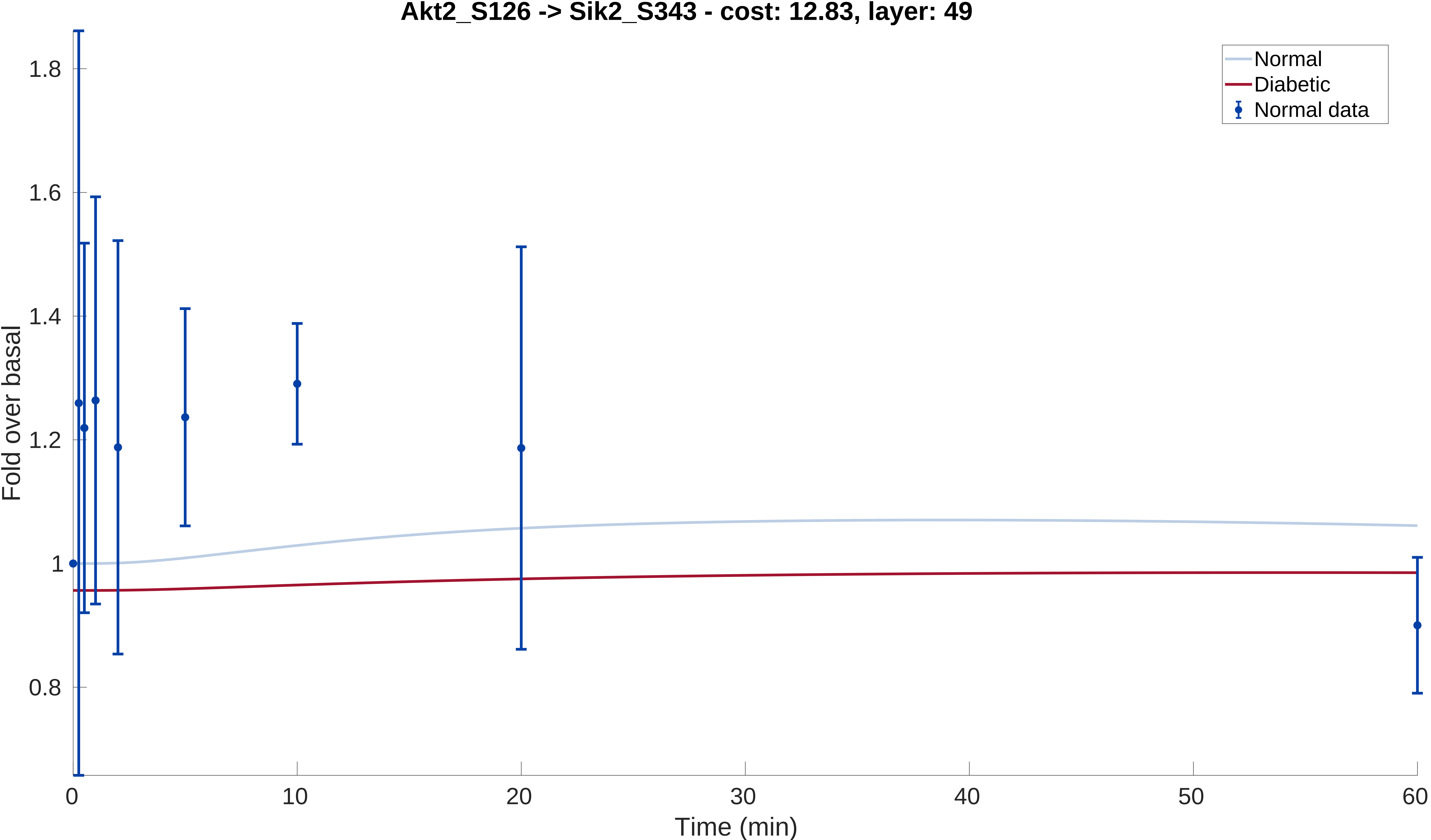

Akt2_S126 -> Sik2_S343 - cost: 12.83, layer: 49
Normal
Diabetic
Normal data
1.8
1.6
1.4
1.2
1
0.8
Fold over basal
0
10
20
30
40
50
60
Time (min)

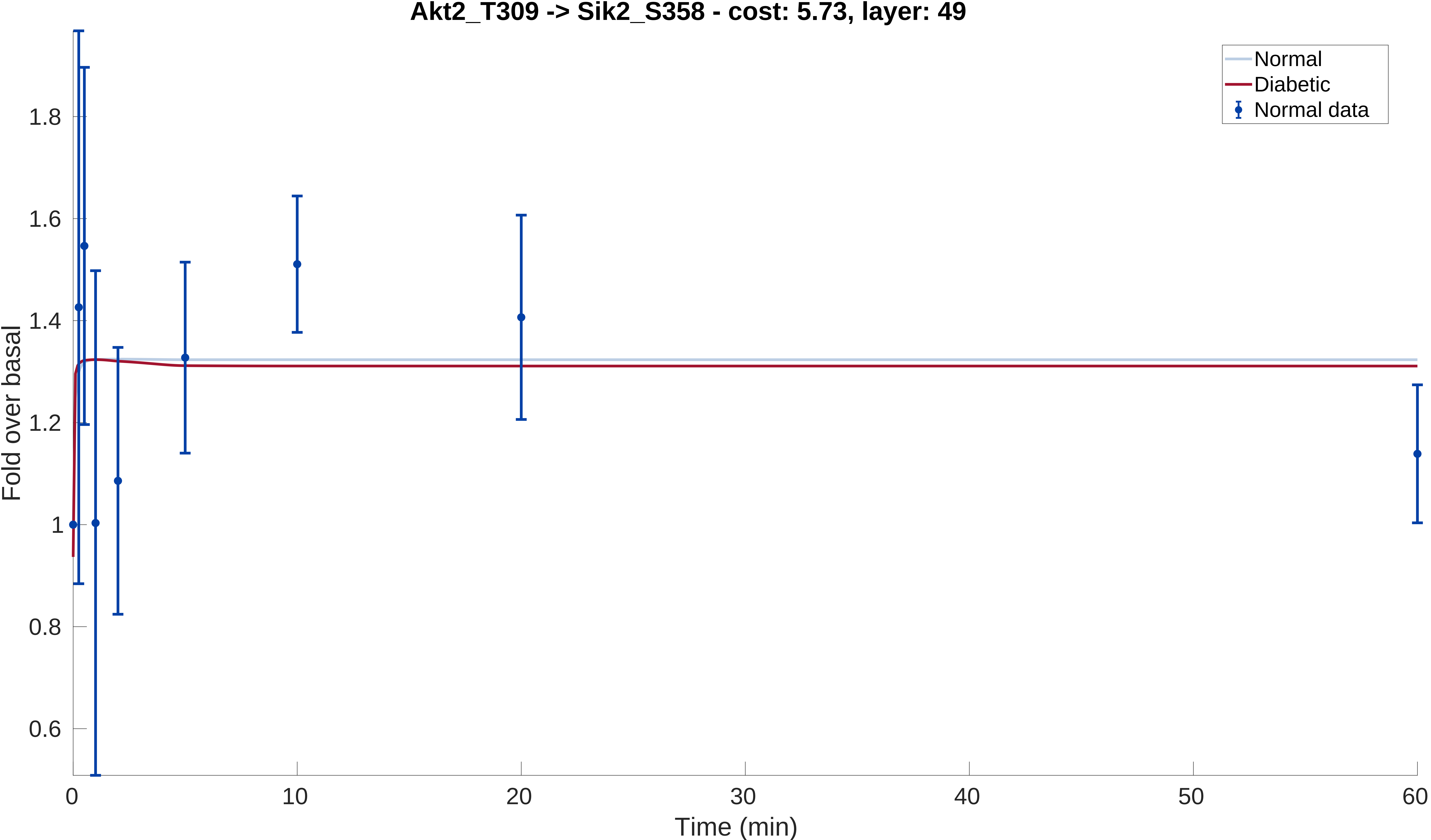
Akt2_T309 -> Sik2_S358 - cost: 5.73, layer: 49
Normal
Diabetic
Normal data
Fold over basal
1.8
1.6
1.4
1.2
1
0.8
0.6
0
10
20
30
40
50
60
Time (min)

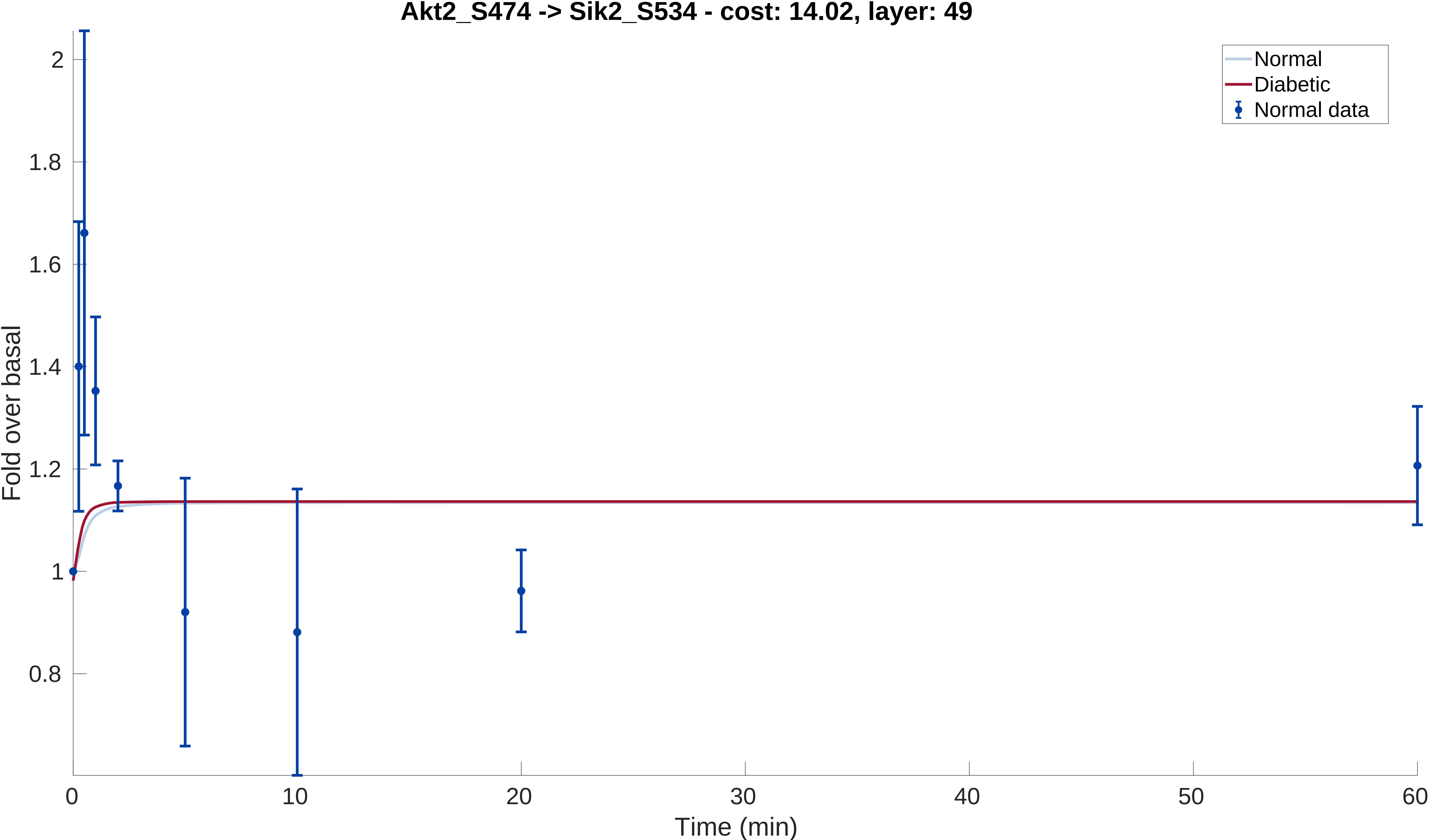

Akt2_S474 -> Sik2_S534 - cost: 14.02, layer: 49
Normal
Diabetic
Normal data
Fold over basal
2
1.8
1.6
1.4
1.2
1
0.8
0
10
20
30
40
50
60
Time (min)

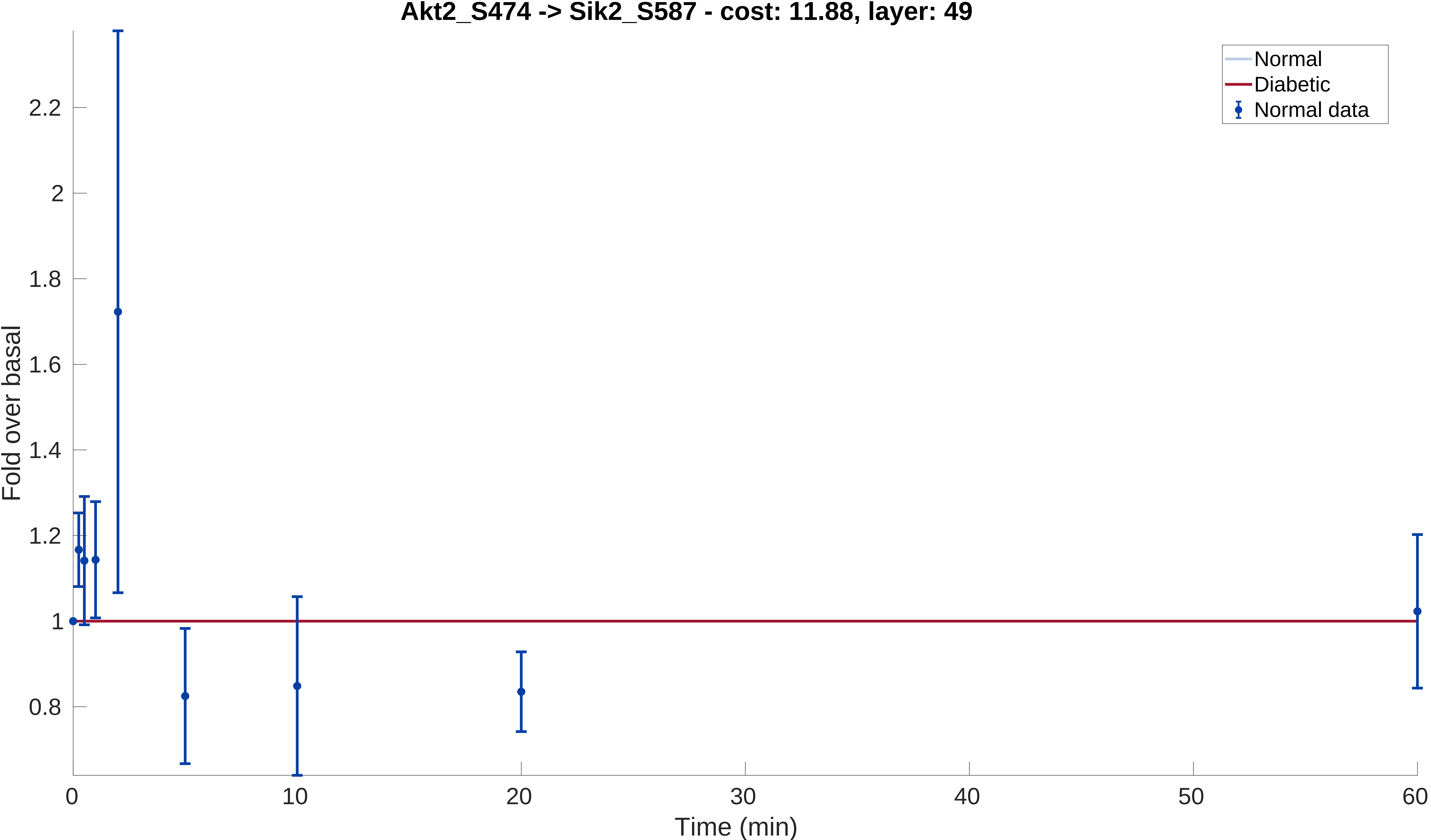

Akt2_S474 -> Sik2_S587 - cost: 11.88, layer: 49
Normal
Diabetic
Normal data
Fold over basal
2.2
2
1.8
1.6
1.4
1.2
1
0.8
0
10
20
30
40
50
60
Time (min)

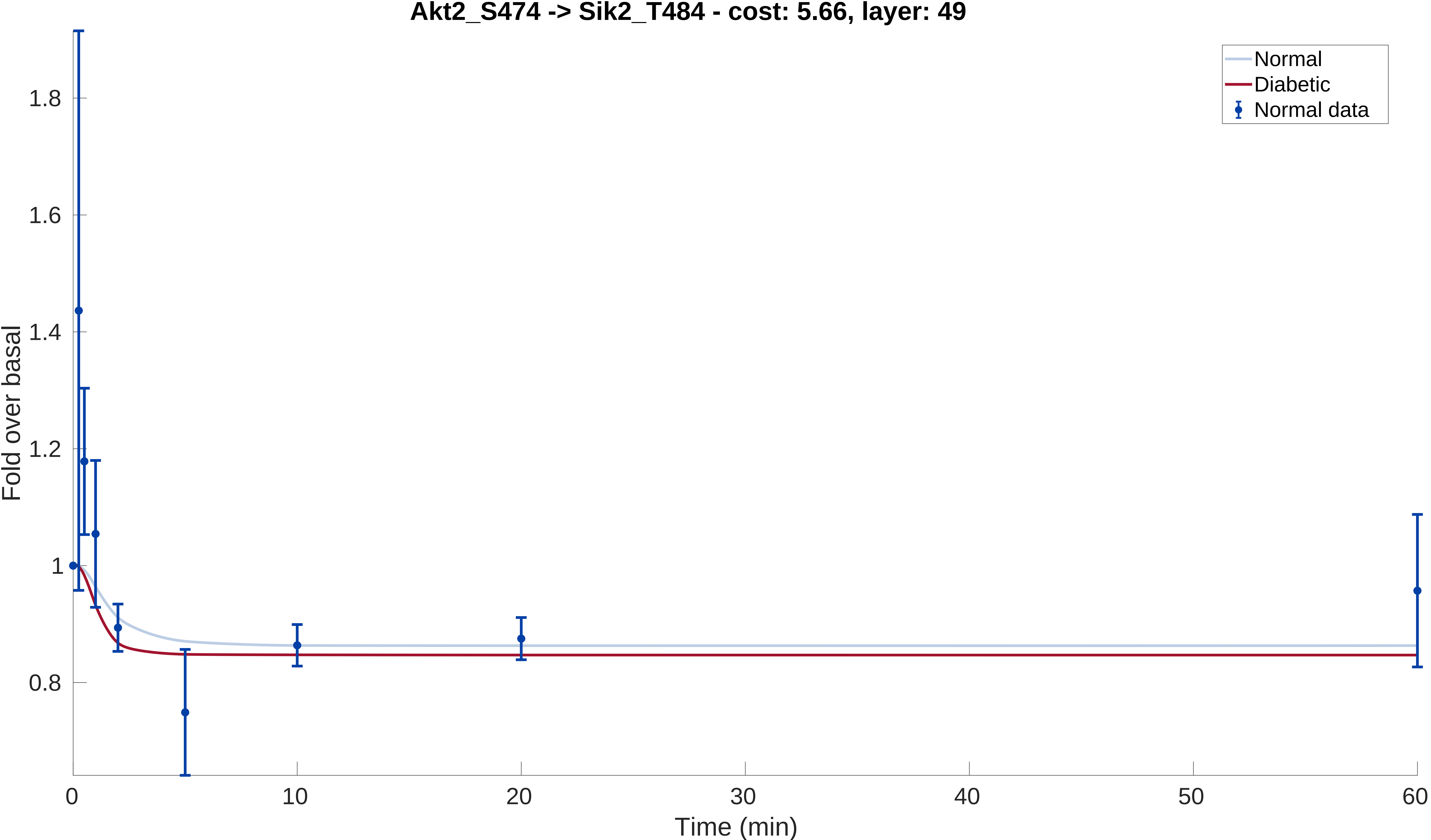
Akt2_S474 -> Sik2_T484 - cost: 5.66, layer: 49
Normal
Diabetic
Normal data
Fold over basal
1.8
1.6
1.4
1.2
1
0.8
0
10
20
30
40
50
60
Time (min)

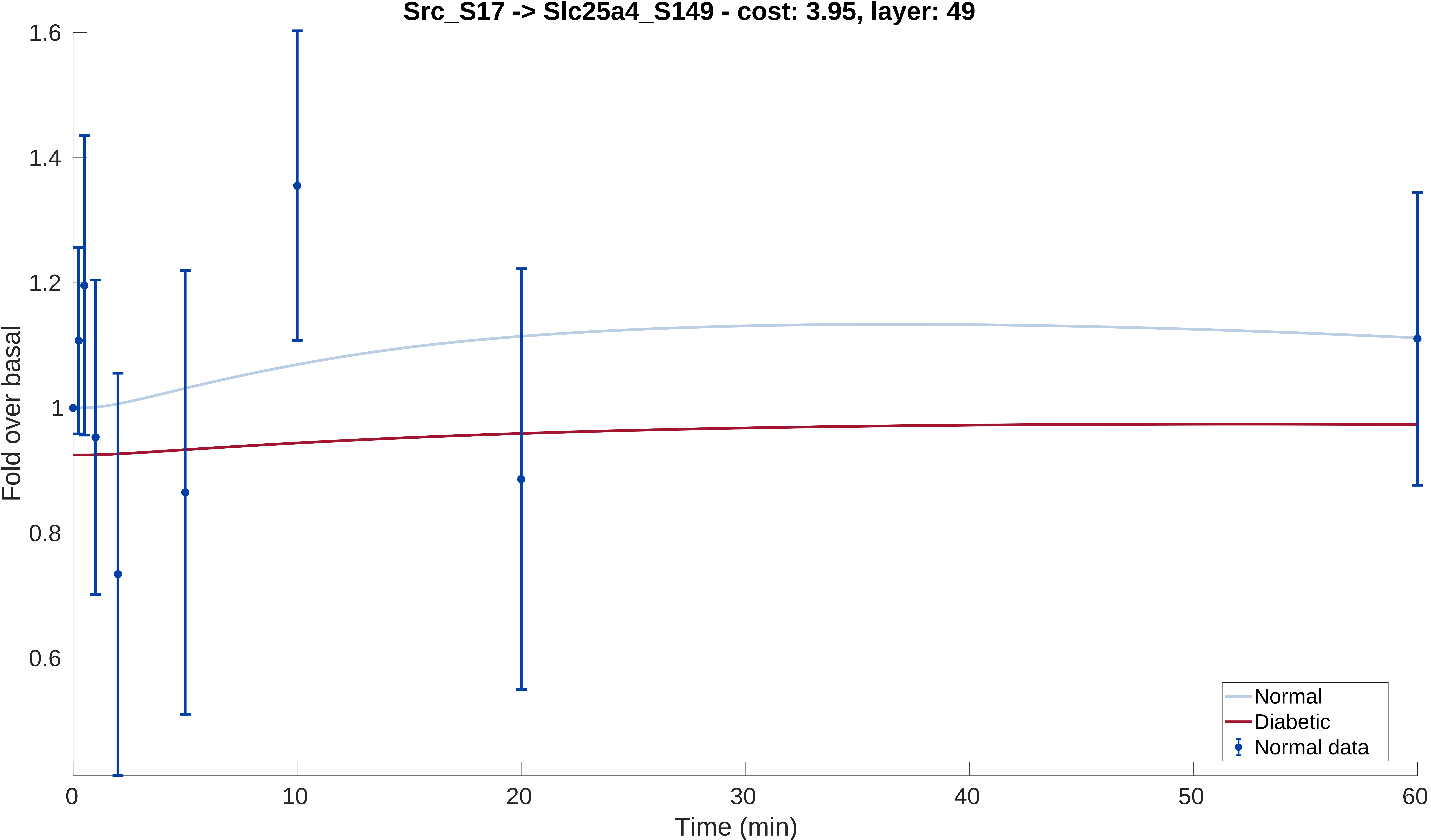

Src_S17 -> Slc25a4_S149 - cost: 3.95, layer: 49
1.6
1.4
1.2
1
0.8
0.6
Fold over basal
0
10
20
30
40
50
60
Time (min)
Normal
Diabetic
Normal data

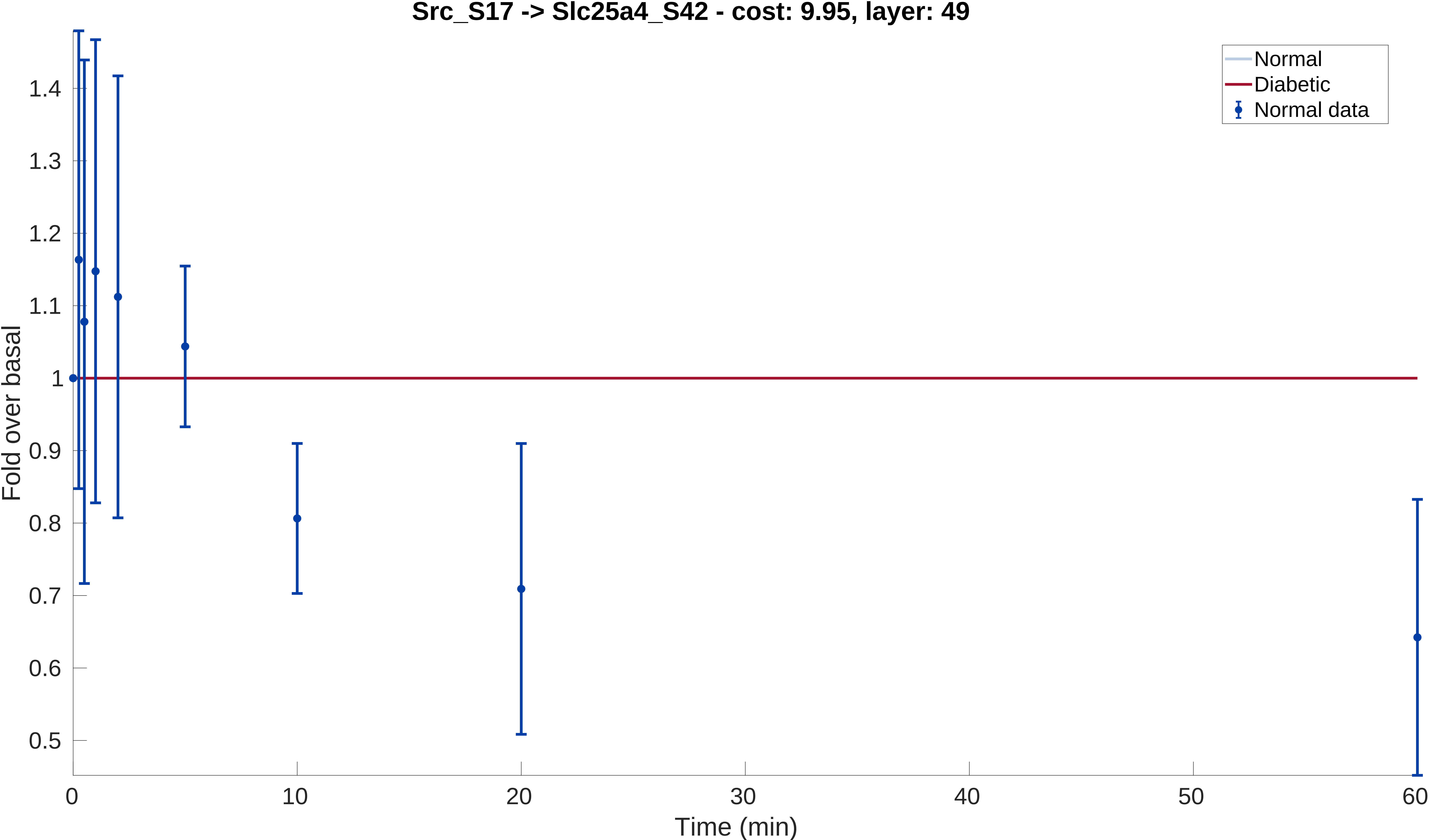

Src_S17 -> Slc25a4_S42 - cost: 9.95, layer: 49
Normal
Diabetic
Normal data
Fold over basal
1.4
1.3
1.2
1.1
1
0.9
0.8
0.7
0.6
0.5
0
10
20
30
40
50
60
Time (min)

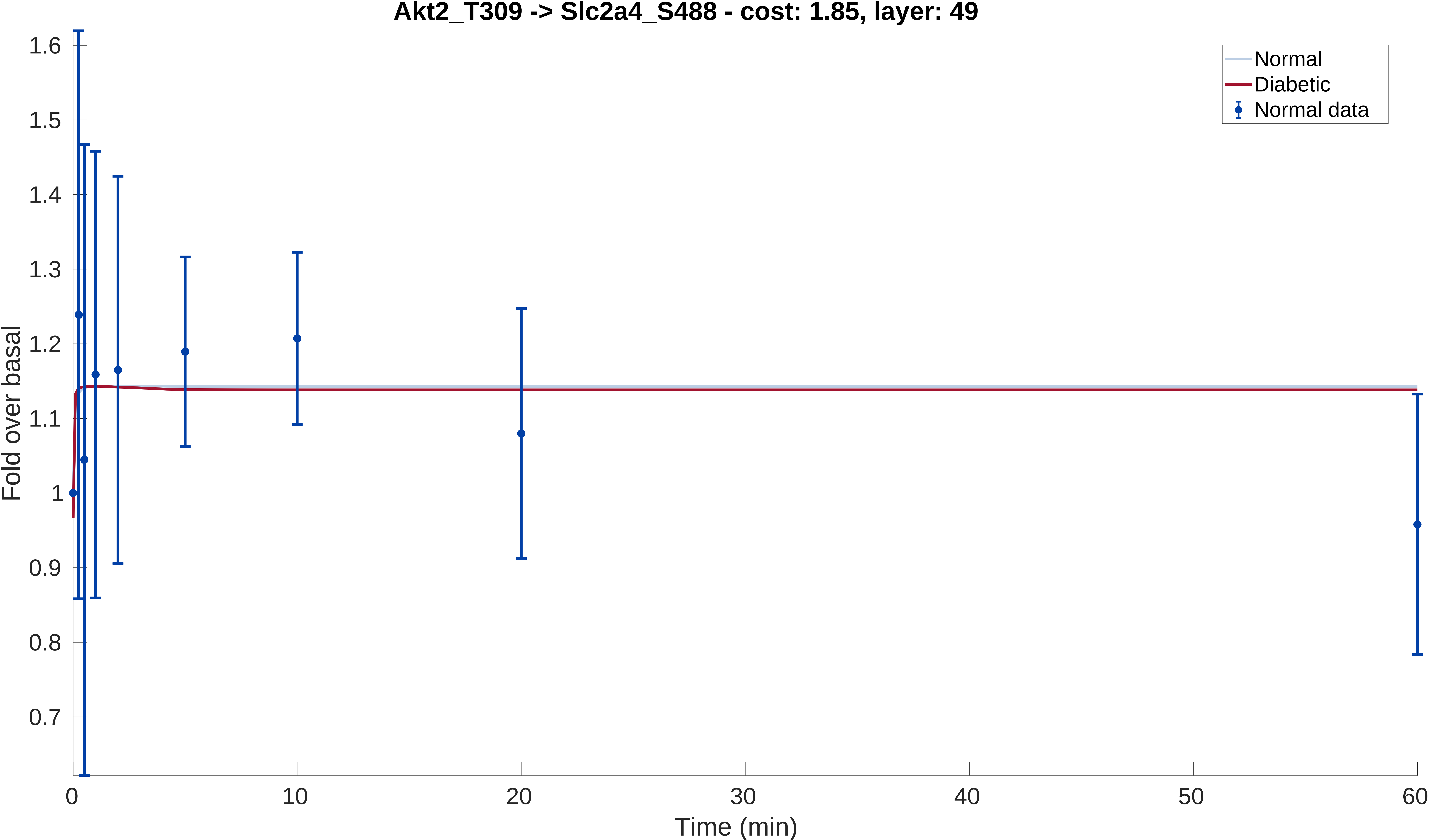

Akt2_T309 -> Slc2a4_S488 - cost: 1.85, layer: 49
Normal
Diabetic
Normal data
Fold over basal
1.6
1.5
1.4
1.3
1.2
1.1
1
0.9
0.8
0.7
0
10
20
30
40
50
60
Time (min)

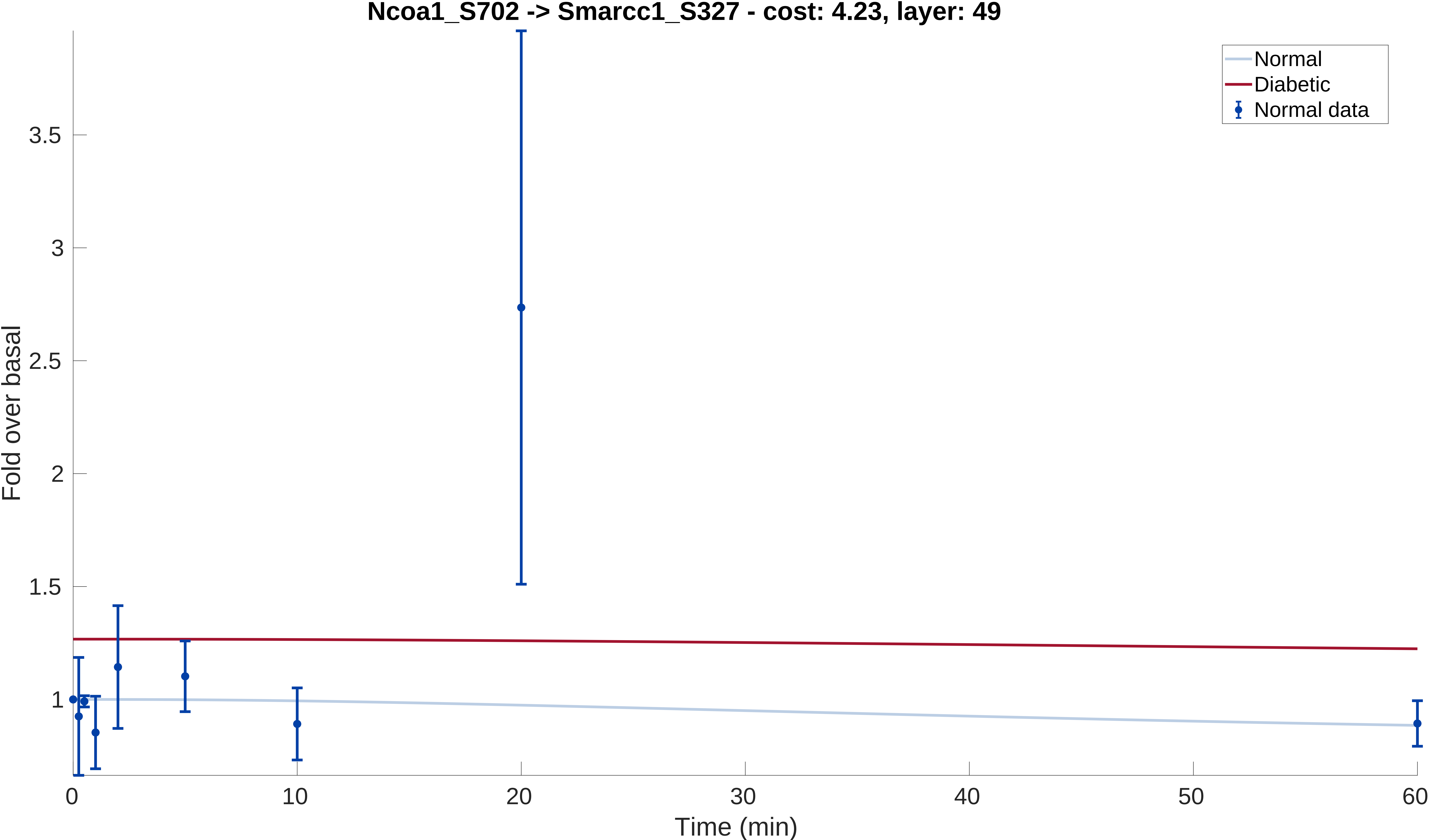

Ncoa1_S702 -> Smarcc1_S327 - cost: 4.23, layer: 49
Normal
Diabetic
Normal data
Fold over basal
3.5
3
2.5
2
1.5
1
0
10
20
30
40
50
60
Time (min)

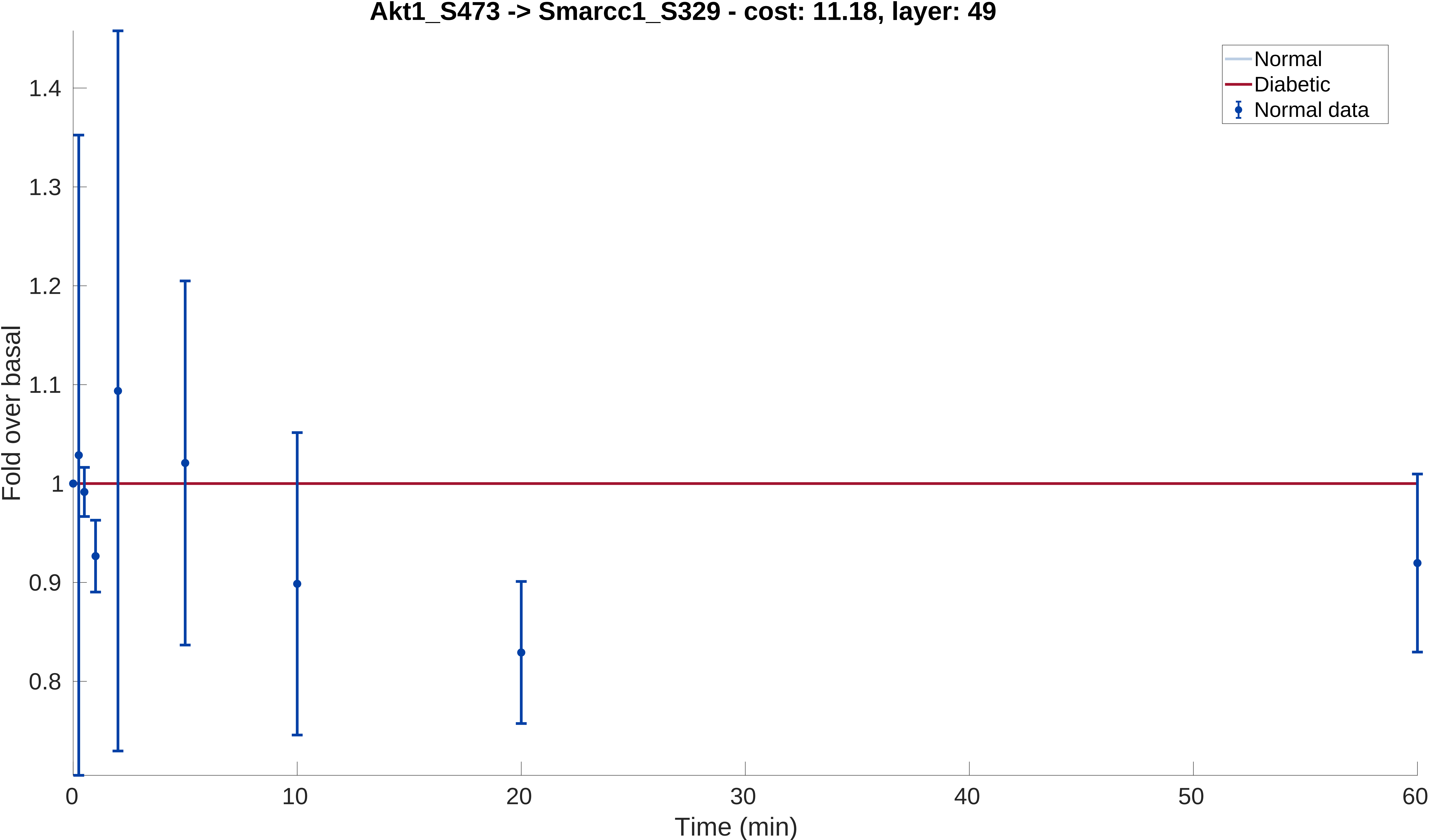

Akt1_S473 -> Smarcc1_S329 - cost: 11.18, layer: 49
Normal
Diabetic
Normal data
Fold over basal
1.4
1.3
1.2
1.1
1
0.9
0.8
0
10
20
30
40
50
60
Time (min)

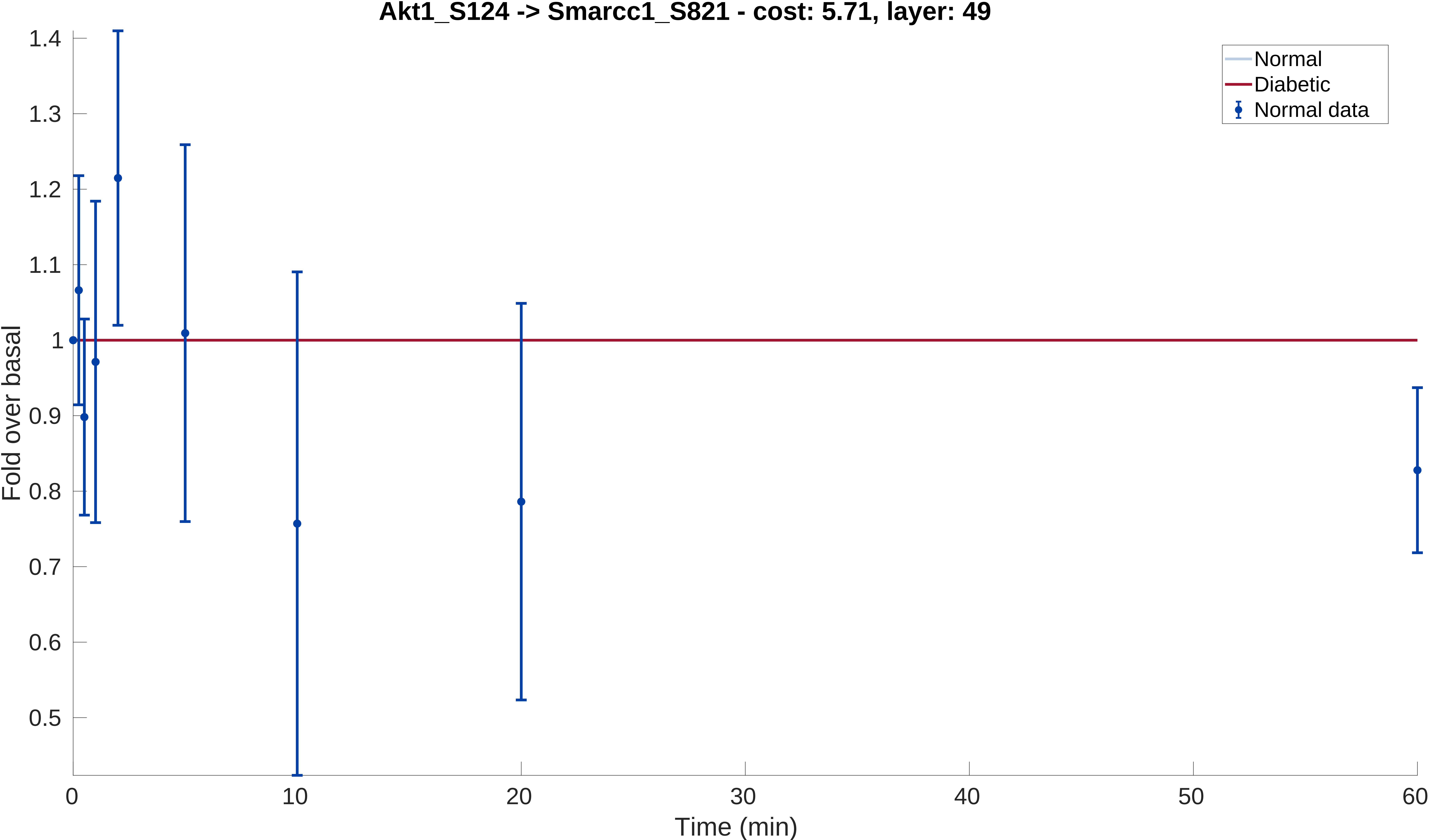
Akt1_S124 -> Smarcc1_S821 - cost: 5.71, layer: 49
Normal
Diabetic
Normal data
Fold over basal
Time (min)
1.4
1.3
1.2
1.1
1
0.9
0.8
0.7
0.6
0.5
0
10
20
30
40
50
60

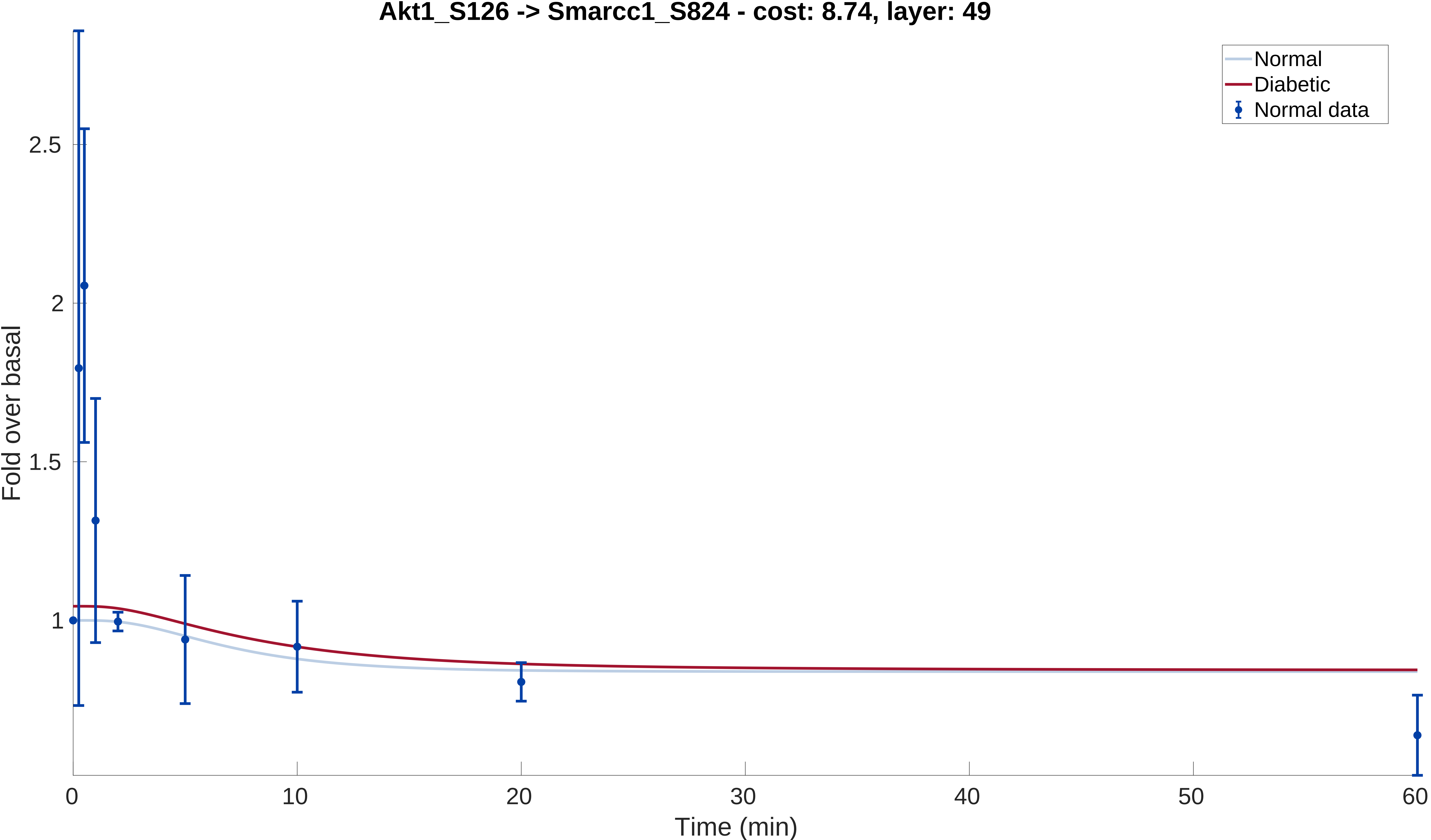

Akt1_S126 -> Smarcc1_S824 - cost: 8.74, layer: 49
Normal
Diabetic
Normal data
2.5
2
1.5
1
Fold over basal
0
10
20
30
40
50
60
Time (min)

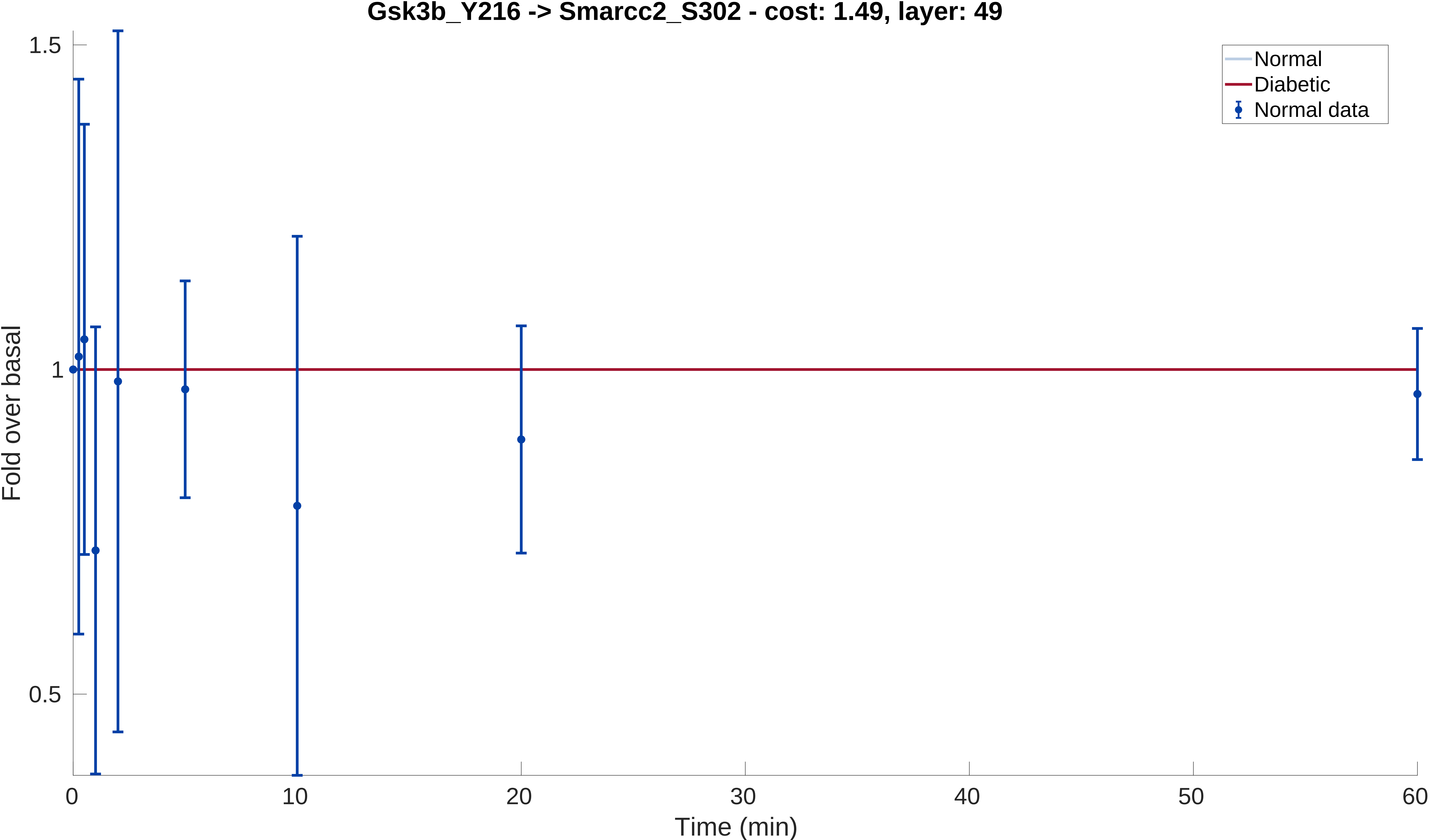
Gsk3b_Y216 -> Smarcc2_S302 - cost: 1.49, layer: 49
Normal
Diabetic
Normal data
1.5
1
0.5
Fold over basal
0
10
20
30
40
50
60
Time (min)

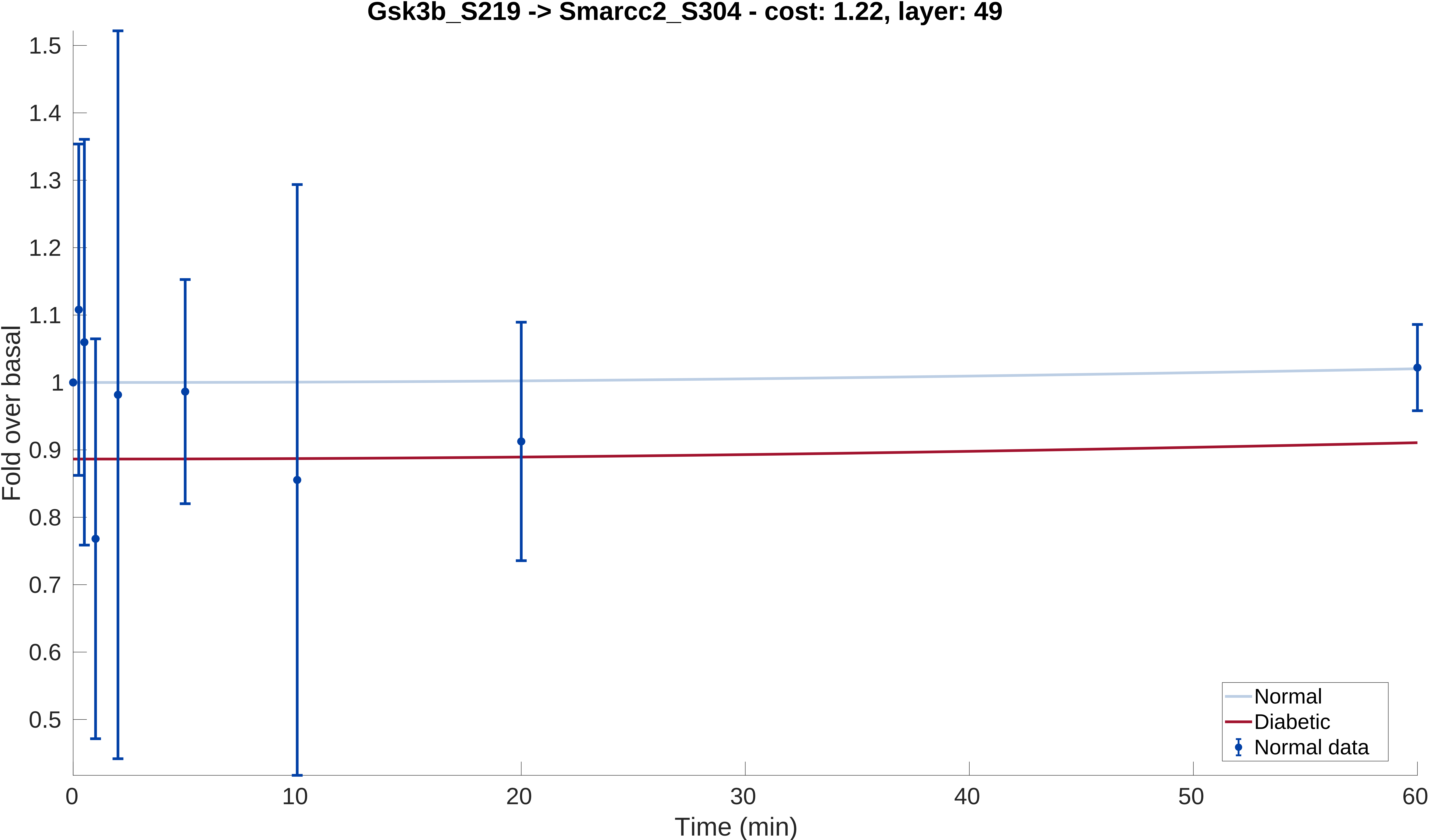

Gsk3b_S219 -> Smarcc2_S304 - cost: 1.22, layer: 49
Fold over basal
Time (min)
1.5
1.4
1.3
1.2
1.1
1
0.9
0.8
0.7
0.6
0.5
0
10
20
30
40
50
60
Normal
Diabetic
Normal data

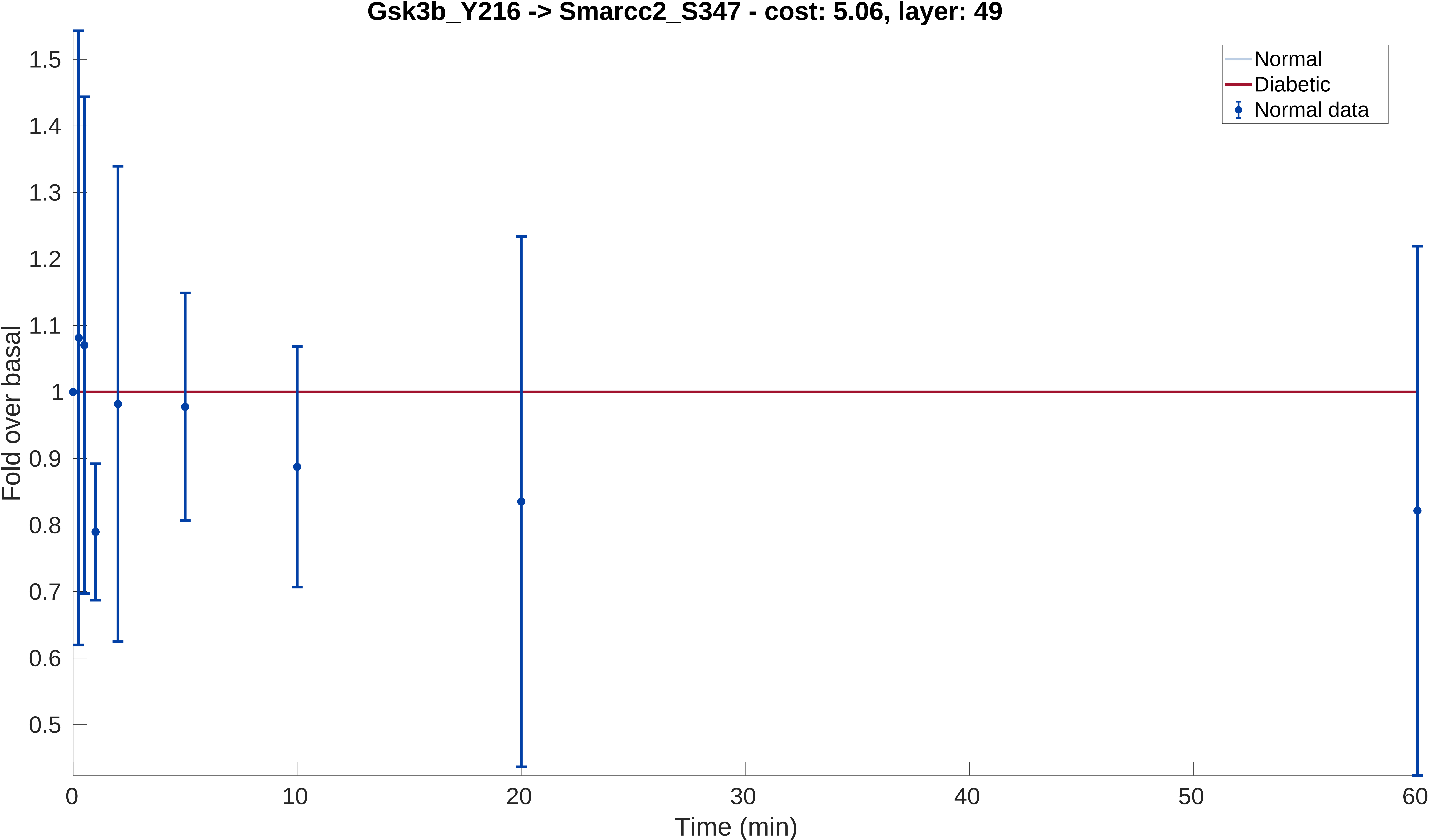

Gsk3b_Y216 -> Smarcc2_S347 - cost: 5.06, layer: 49
Normal
Diabetic
Normal data
Fold over basal
Time (min)
1.5
1.4
1.3
1.2
1.1
1
0.9
0.8
0.7
0.6
0.5
0
10
20
30
40
50
60

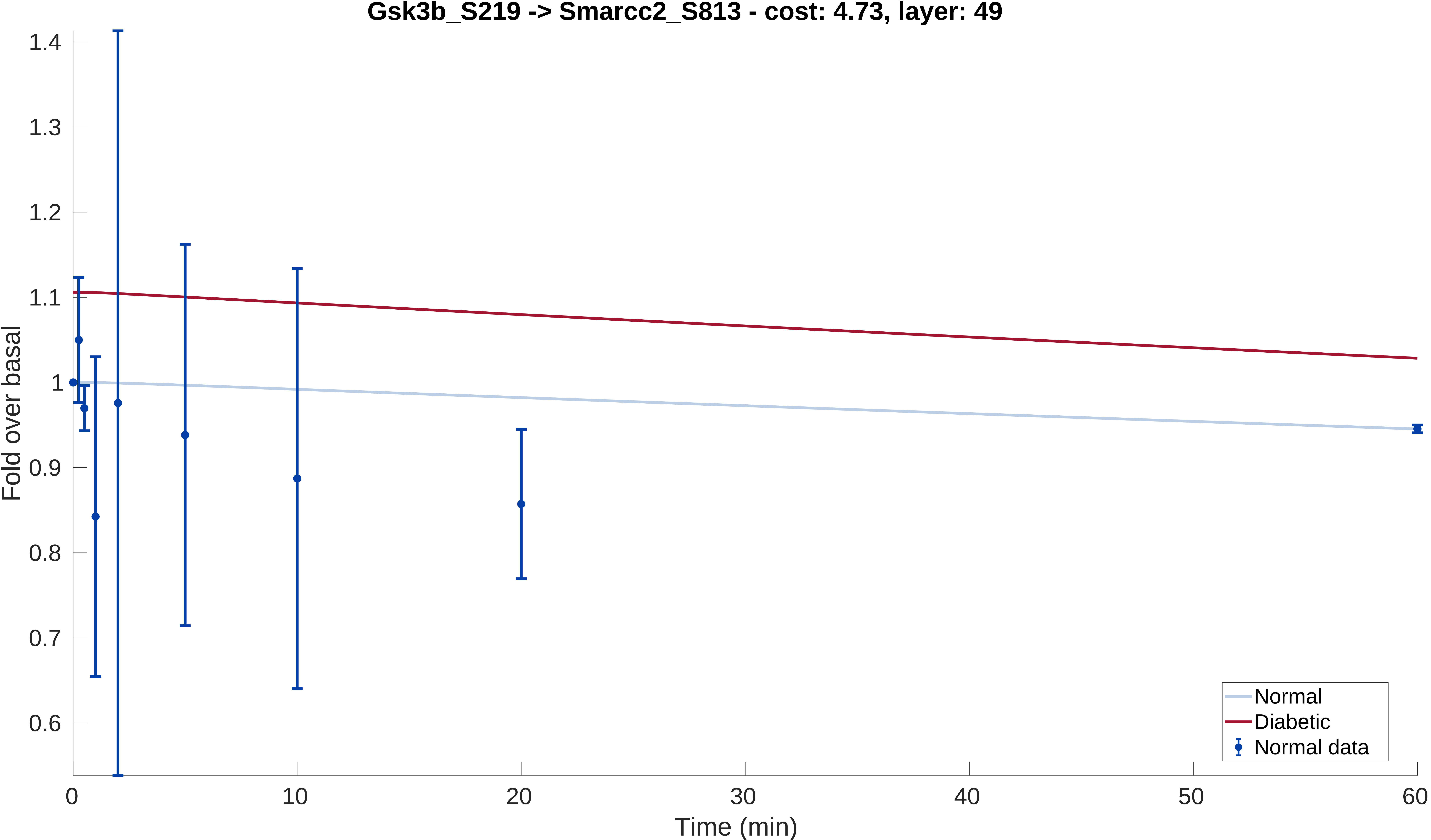

Gsk3b_S219 -> Smarcc2_S813 - cost: 4.73, layer: 49
1.4
1.3
1.2
1.1
1
0.9
0.8
0.7
0.6
Fold over basal
0
10
20
30
40
50
60
Time (min)
Normal
Diabetic
Normal data

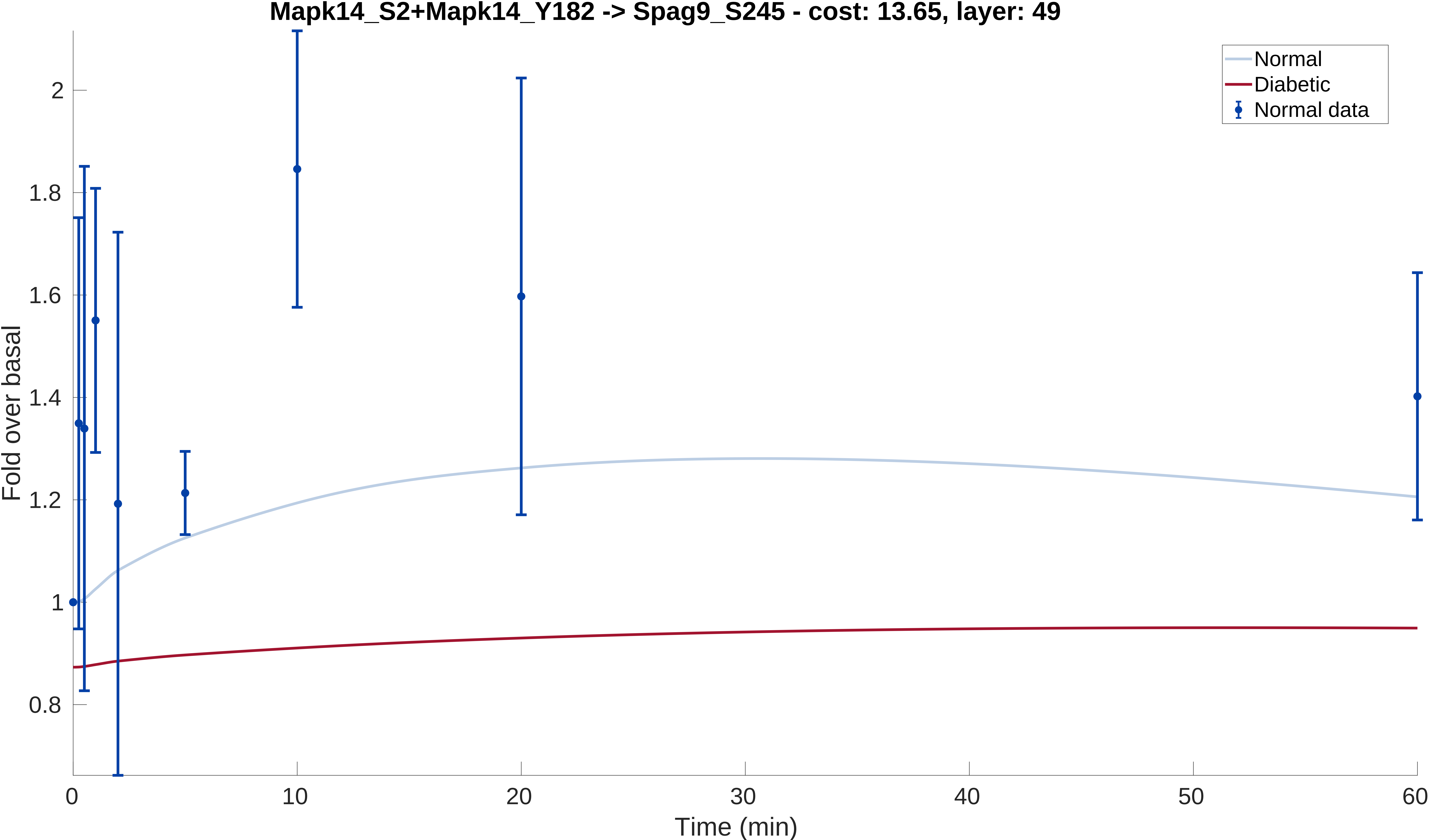

Mapk14_S2+Mapk14_Y182 -> Spag9_S245 - cost: 13.65, layer: 49
Normal
Diabetic
Normal data
Fold over basal
2
1.8
1.6
1.4
1.2
1
0.8
0
10
20
30
40
50
60
Time (min)

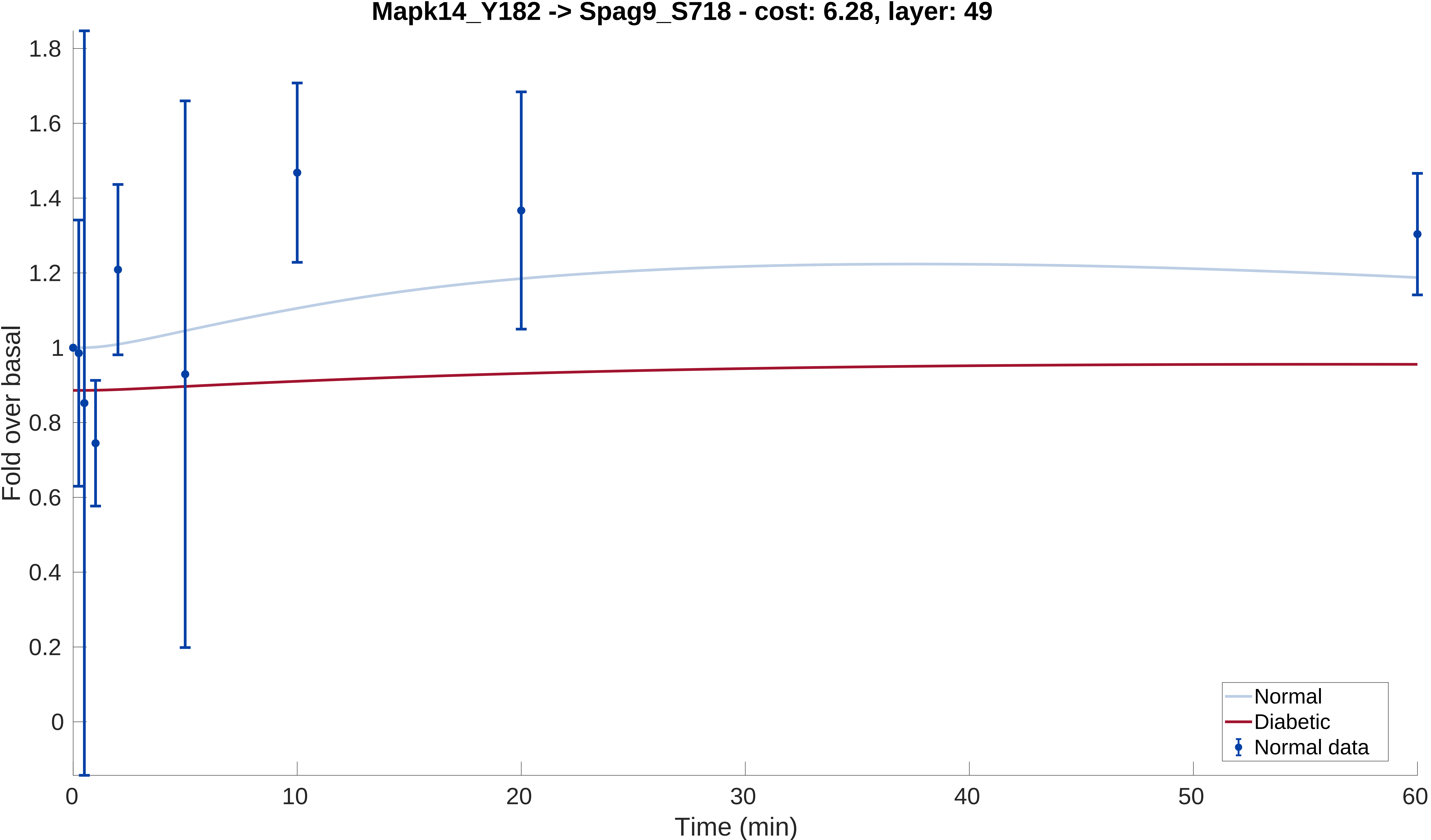

Mapk14_Y182 -> Spag9_S718 - cost: 6.28, layer: 49
Fold over basal
1.8
1.6
1.4
1.2
1
0.8
0.6
0.4
0.2
0
0
10
20
30
40
50
60
Time (min)
Normal
Diabetic
Normal data

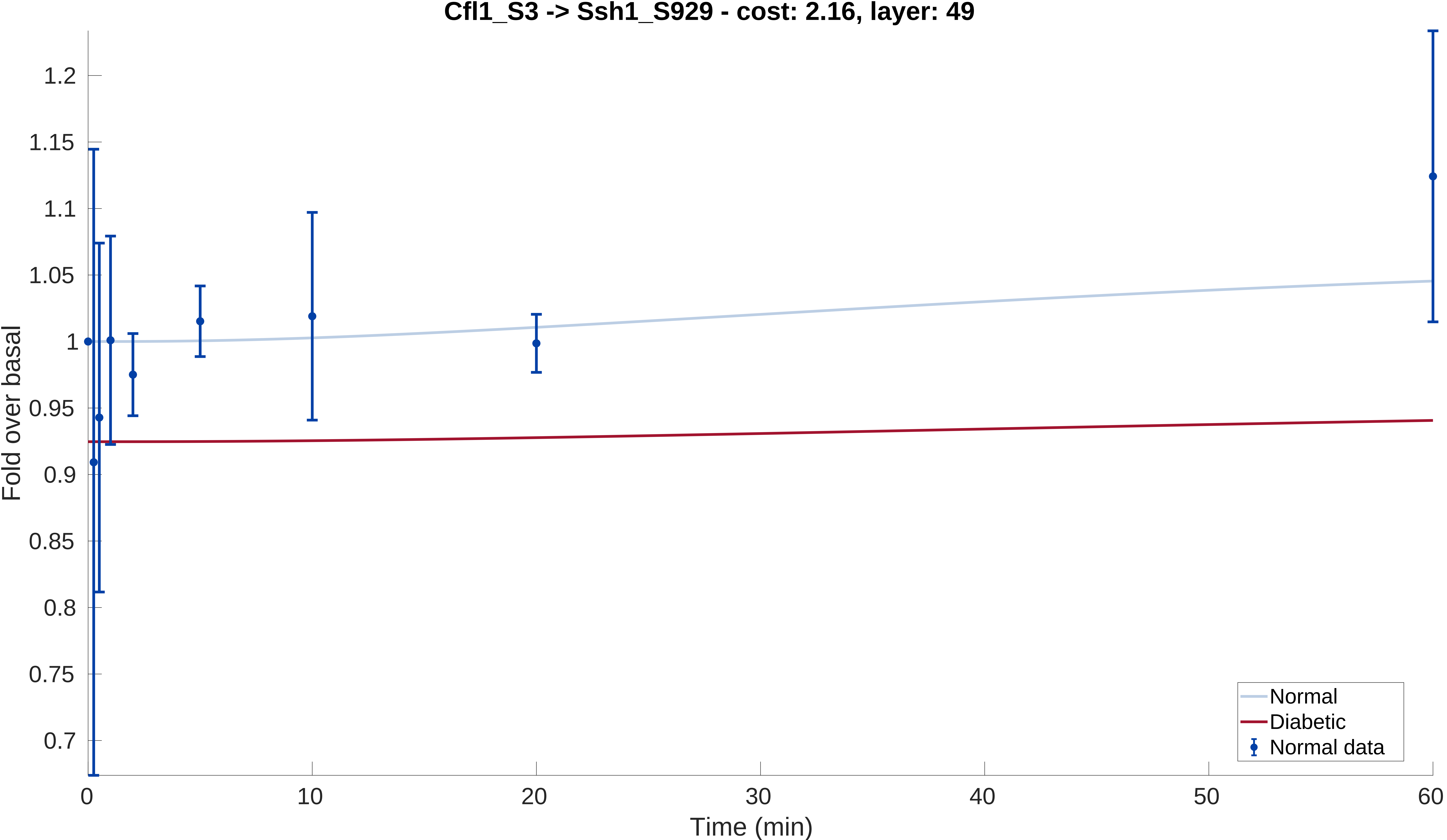

Cfl1_S3 -> Ssh1_S929 - cost: 2.16, layer: 49
1.2
1.15
1.1
1.05
1
0.95
0.9
0.85
0.8
0.75
0.7
Fold over basal
0
10
20
30
40
50
60
Time (min)
Normal
Diabetic
Normal data

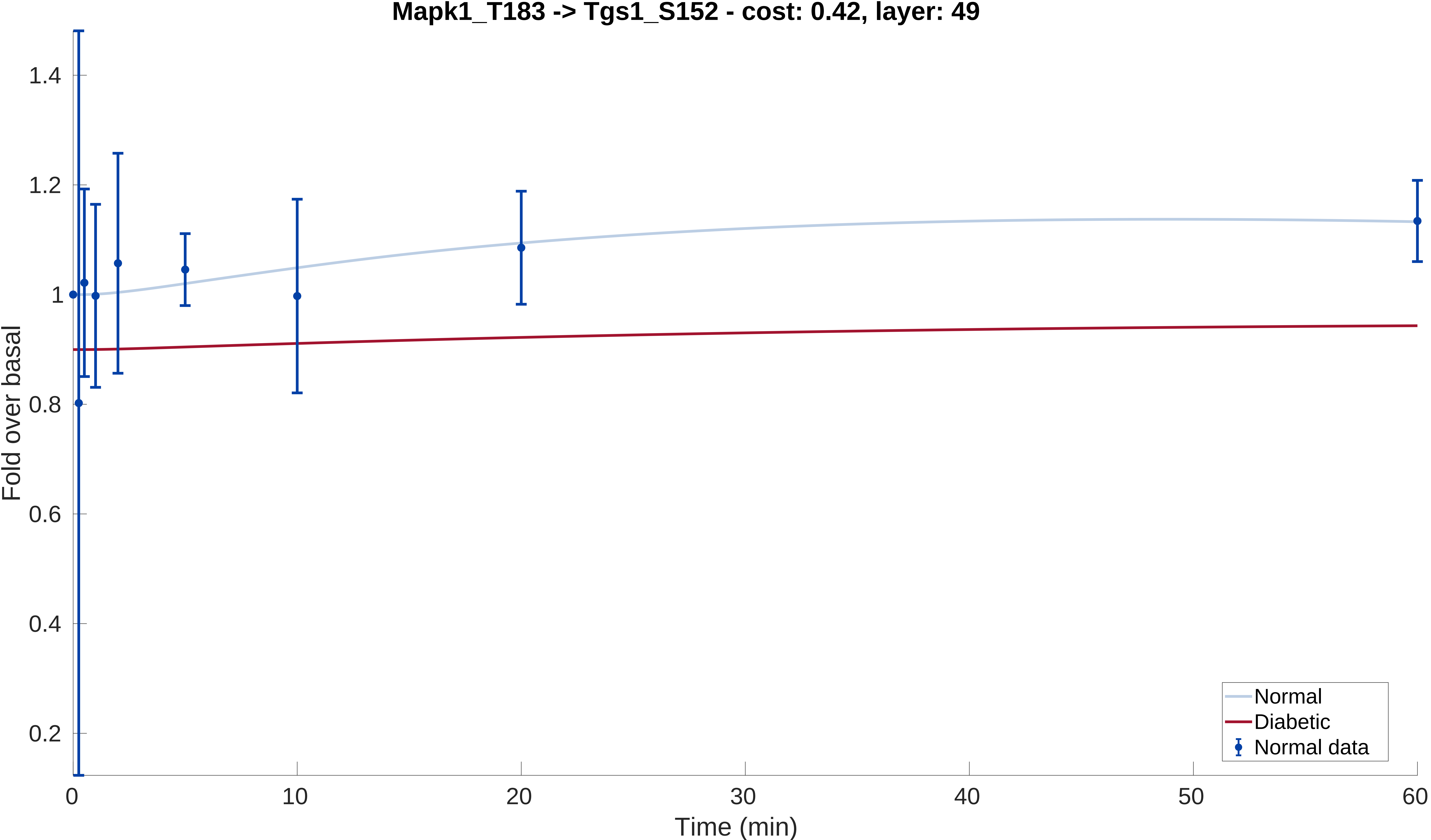
Mapk1_T183 -> Tgs1_S152 - cost: 0.42, layer: 49
Fold over basal
1.4
1.2
1
0.8
0.6
0.4
0.2
0
10
20
30
40
50
60
Time (min)
Normal
Diabetic
Normal data

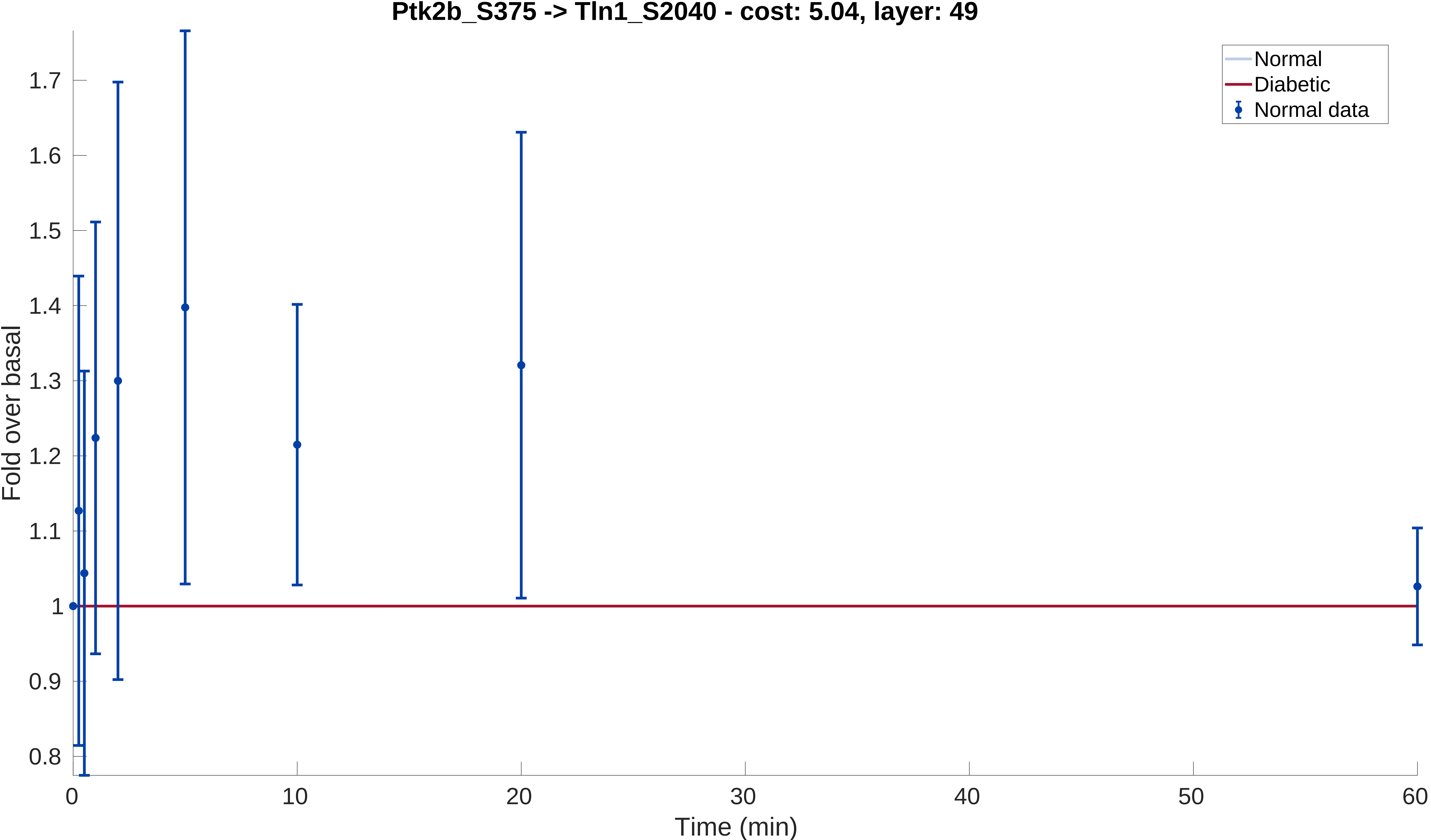

Ptk2b_S375 -> Tln1_S2040 - cost: 5.04, layer: 49
Normal
Diabetic
Normal data
Fold over basal
1.7
1.6
1.5
1.4
1.3
1.2
1.1
1
0.9
0.8
0
10
20
30
40
50
60
Time (min)

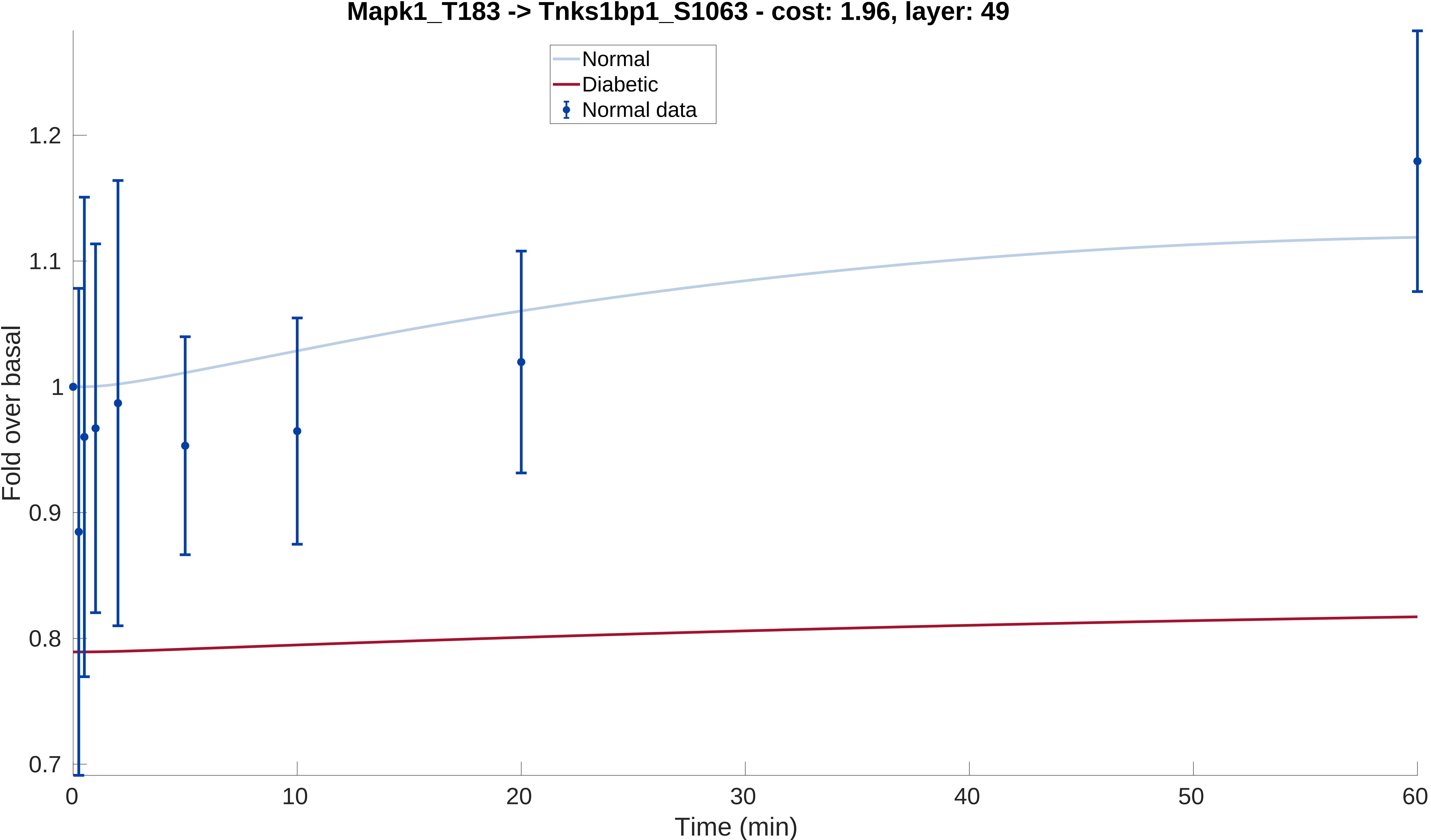

Mapk1_T183 -> Tnks1bp1_S1063 - cost: 1.96, layer: 49
Normal
Diabetic
Normal data
Fold over basal
1.2
1.1
1
0.9
0.8
0.7
0
10
20
30
40
50
60
Time (min)

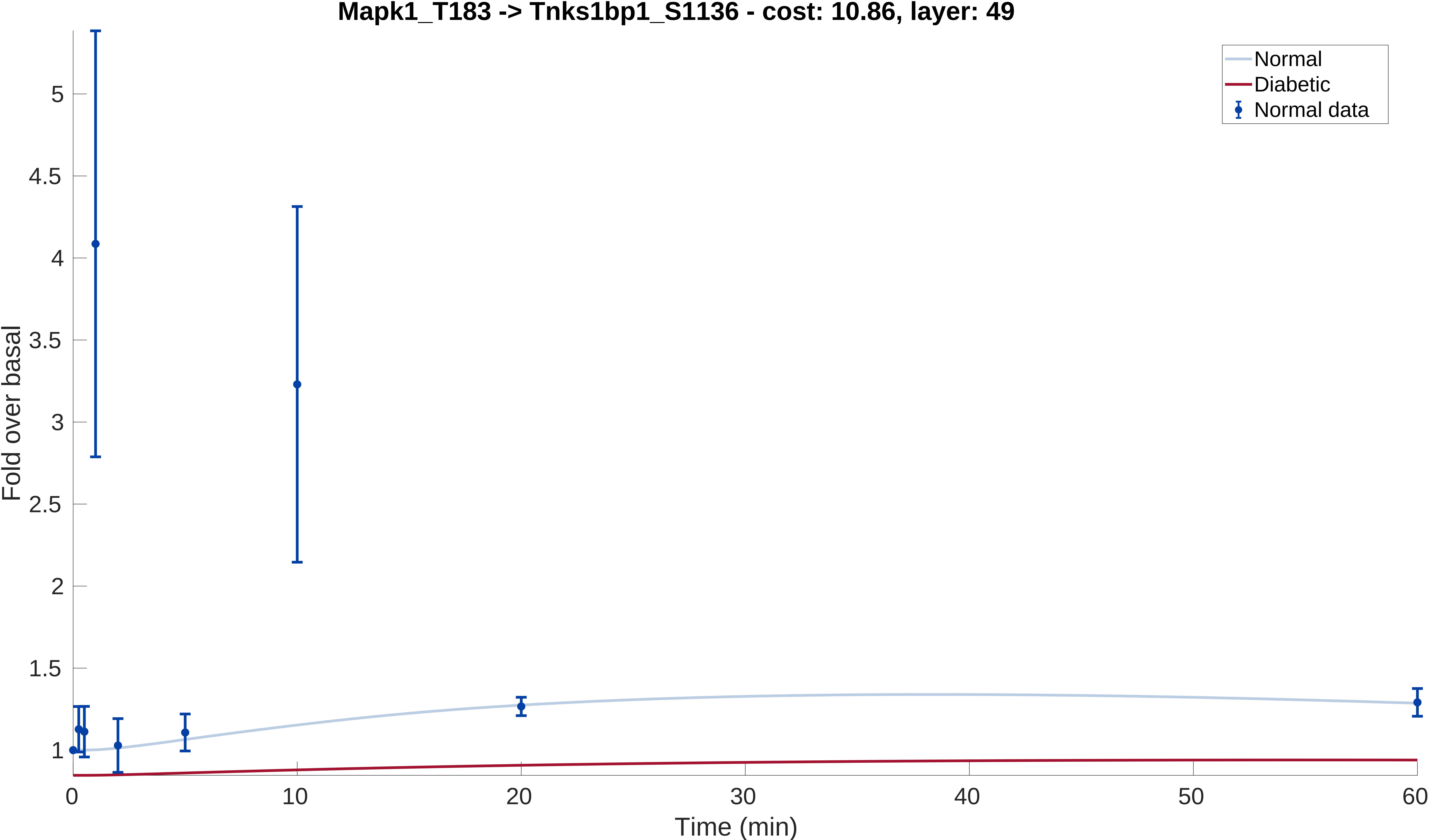
Mapk1_T183 -> Tnks1bp1_S1136 - cost: 10.86, layer: 49
Fold over basal
Time (min)
Normal
Diabetic
Normal data
5
4.5
4
3.5
3
2.5
2
1.5
1
0
10
20
30
40
50
60

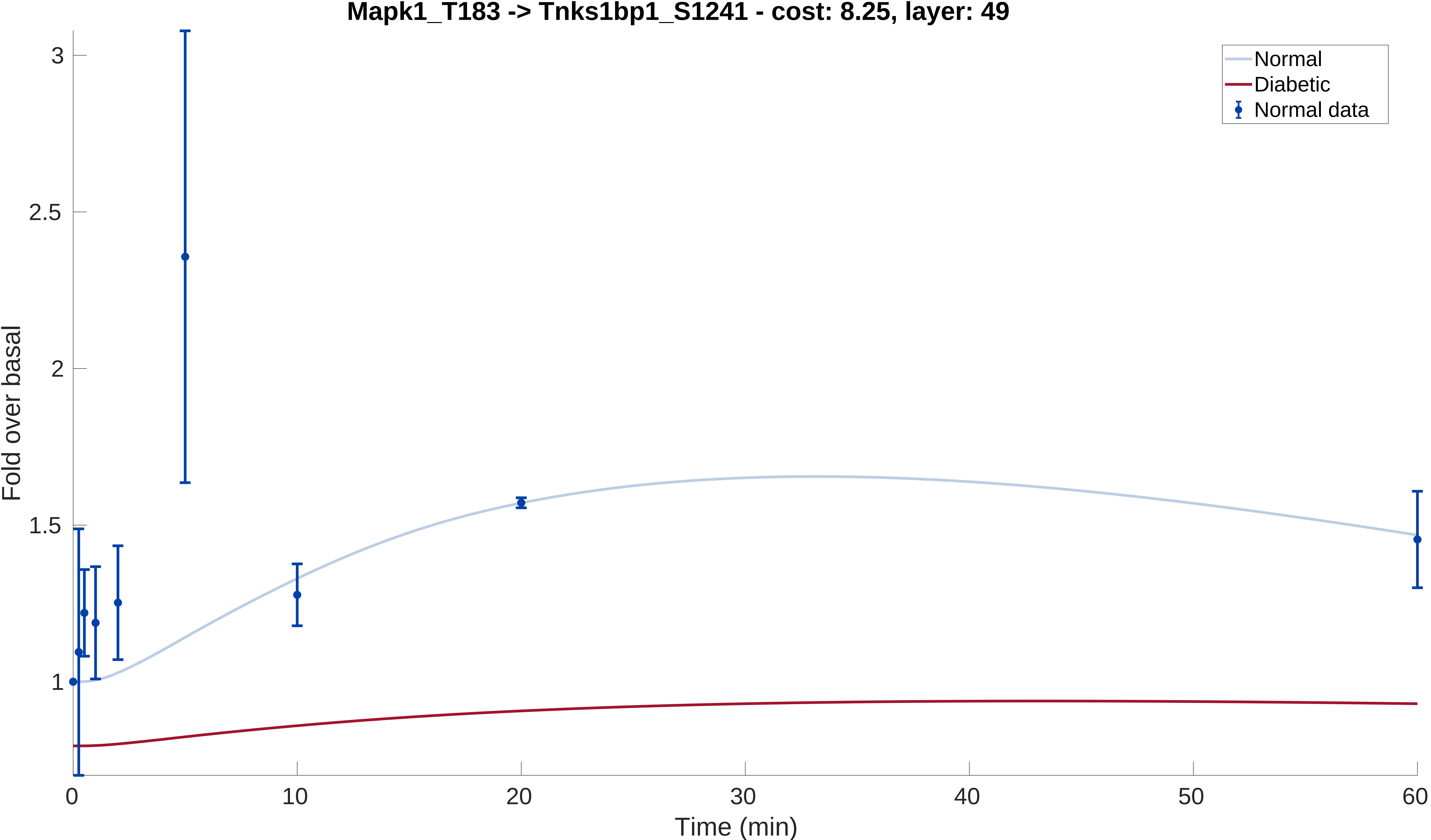

Mapk1_T183 -> Tnks1bp1_S1241 - cost: 8.25, layer: 49
Normal
Diabetic
Normal data
Fold over basal
Time (min)
3
2.5
2
1.5
1
0
10
20
30
40
50
60

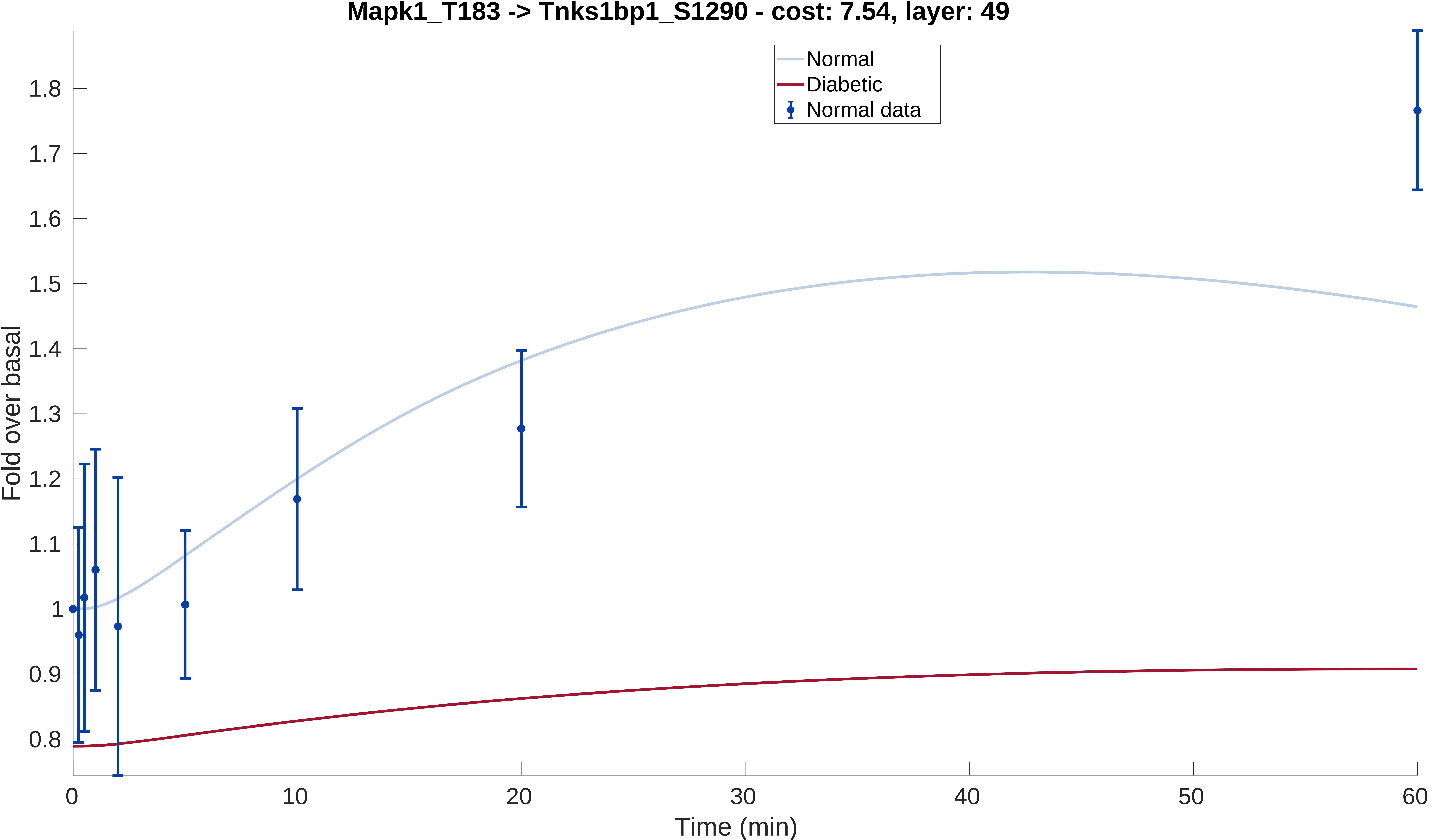

Mapk1_T183 -> Tnks1bp1_S1290 - cost: 7.54, layer: 49
Normal
Diabetic
Normal data
Fold over basal
Time (min)
0.8
0.9
1
1.1
1.2
1.3
1.4
1.5
1.6
1.7
1.8
0
10
20
30
40
50
60

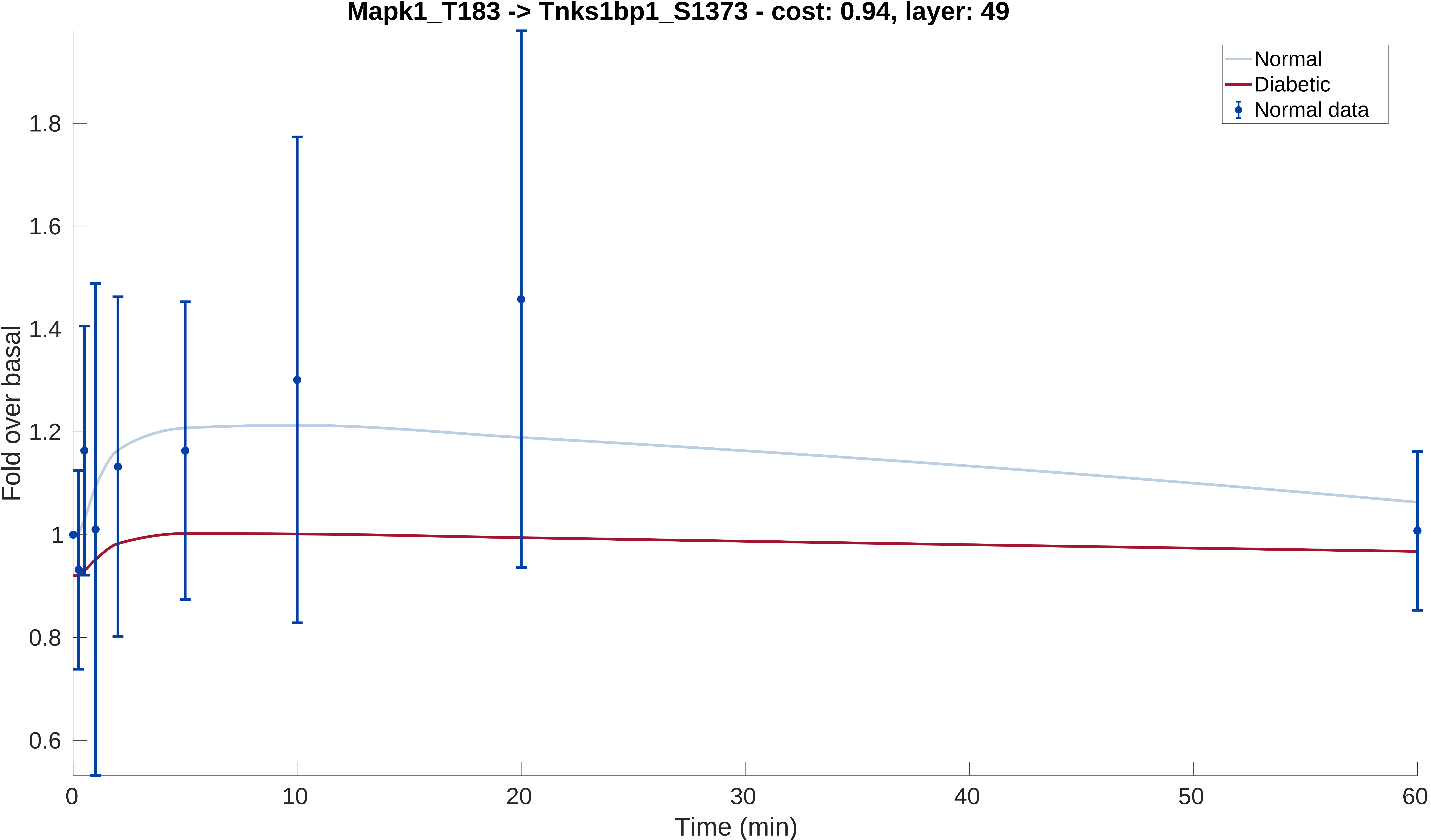
Mapk1_T183 -> Tnks1bp1_S1373 - cost: 0.94, layer: 49
Normal
Diabetic
Normal data
Fold over basal
Time (min)
0.6
0.8
1
1.2
1.4
1.6
1.8
0
10
20
30
40
50
60

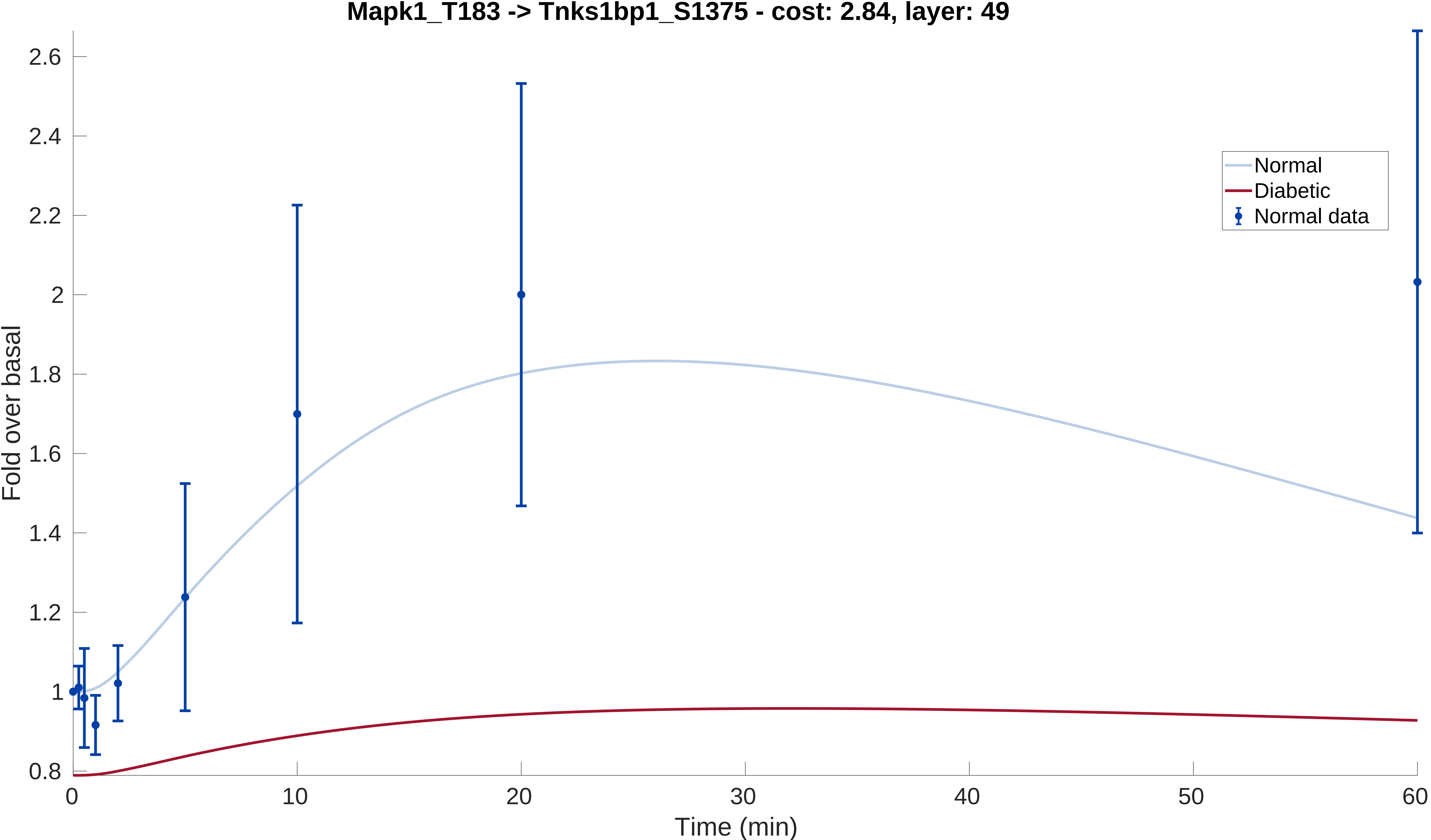

Mapk1_T183 -> Tnks1bp1_S1375 - cost: 2.84, layer: 49
Fold over basal
Time (min)
Normal
Diabetic
Normal data
2.6
2.4
2.2
2
1.8
1.6
1.4
1.2
1
0.8
0
10
20
30
40
50
60

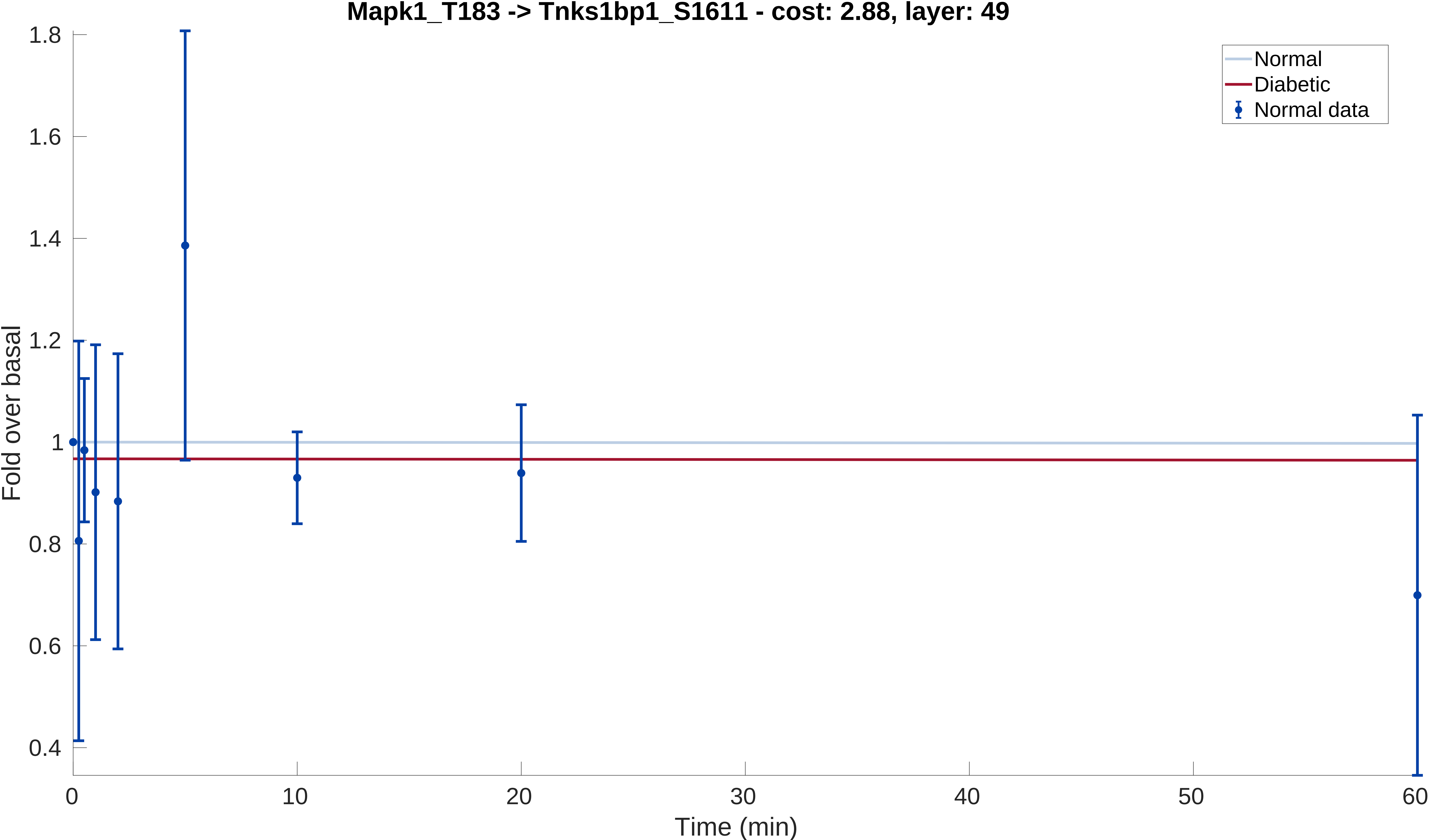
Mapk1_T183 -> Tnks1bp1_S1611 - cost: 2.88, layer: 49
Fold over basal
Time (min)
Normal
Diabetic
Normal data
0.4
0.6
0.8
1
1.2
1.4
1.6
1.8
0
10
20
30
40
50
60

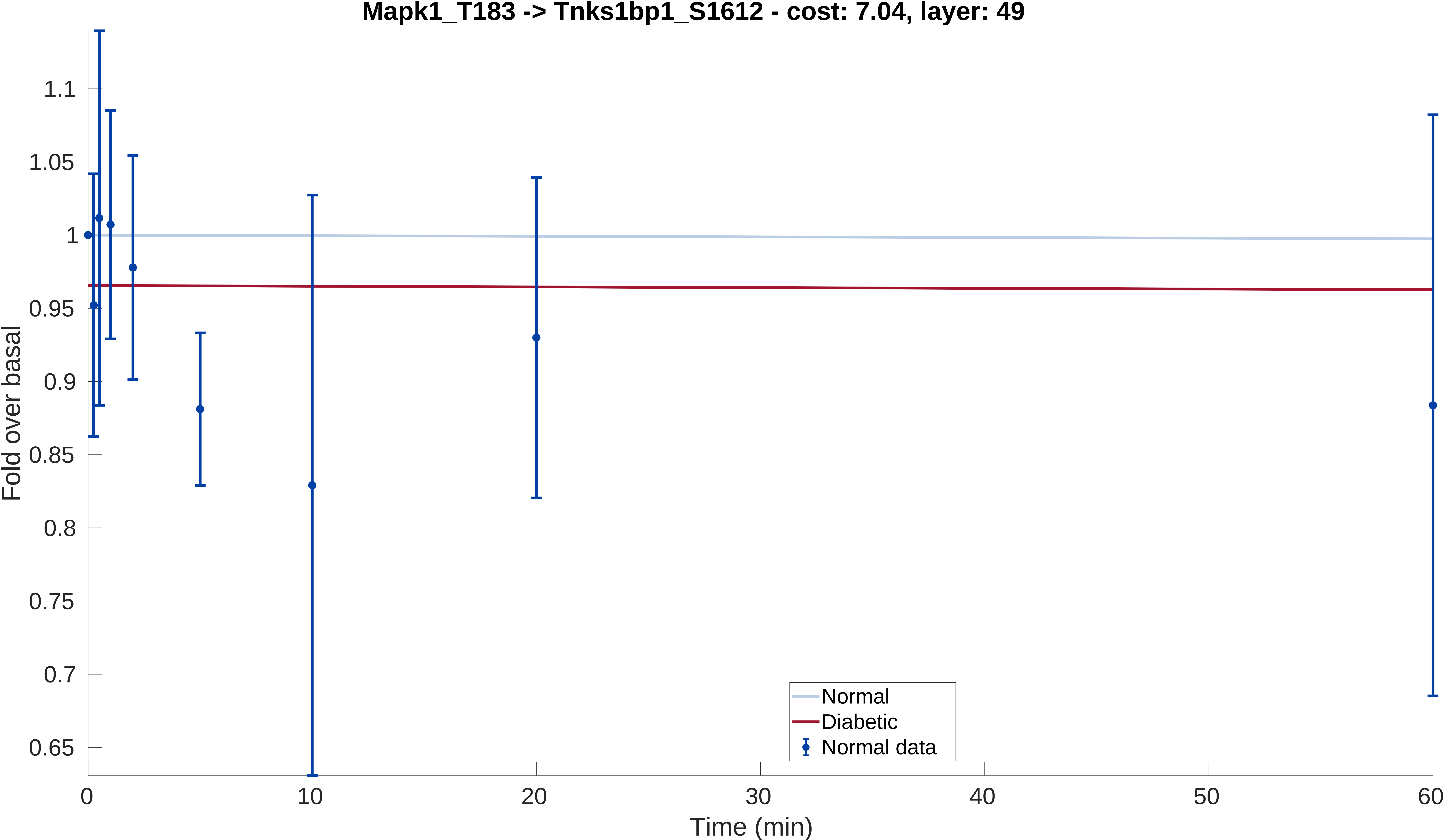

Mapk1_T183 -> Tnks1bp1_S1612 - cost: 7.04, layer: 49
Fold over basal
1.1
1.05
1
0.95
0.9
0.85
0.8
0.75
0.7
0.65
0
10
20
30
40
50
60
Time (min)
Normal
Diabetic
Normal data

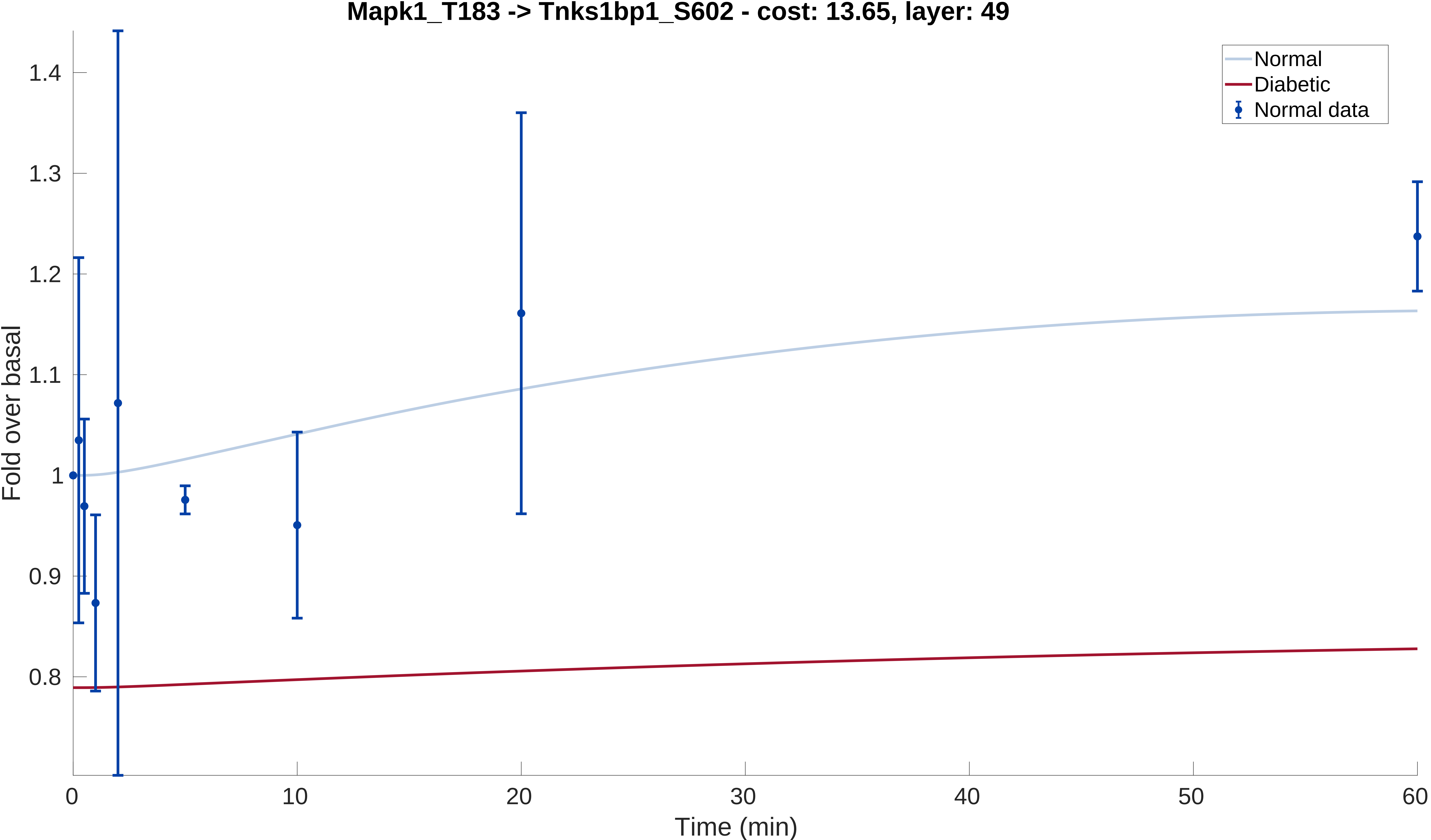
Mapk1_T183 -> Tnks1bp1_S602 - cost: 13.65, layer: 49
Normal
Diabetic
Normal data
Fold over basal
Time (min)
1.4
1.3
1.2
1.1
1
0.9
0.8
0
10
20
30
40
50
60

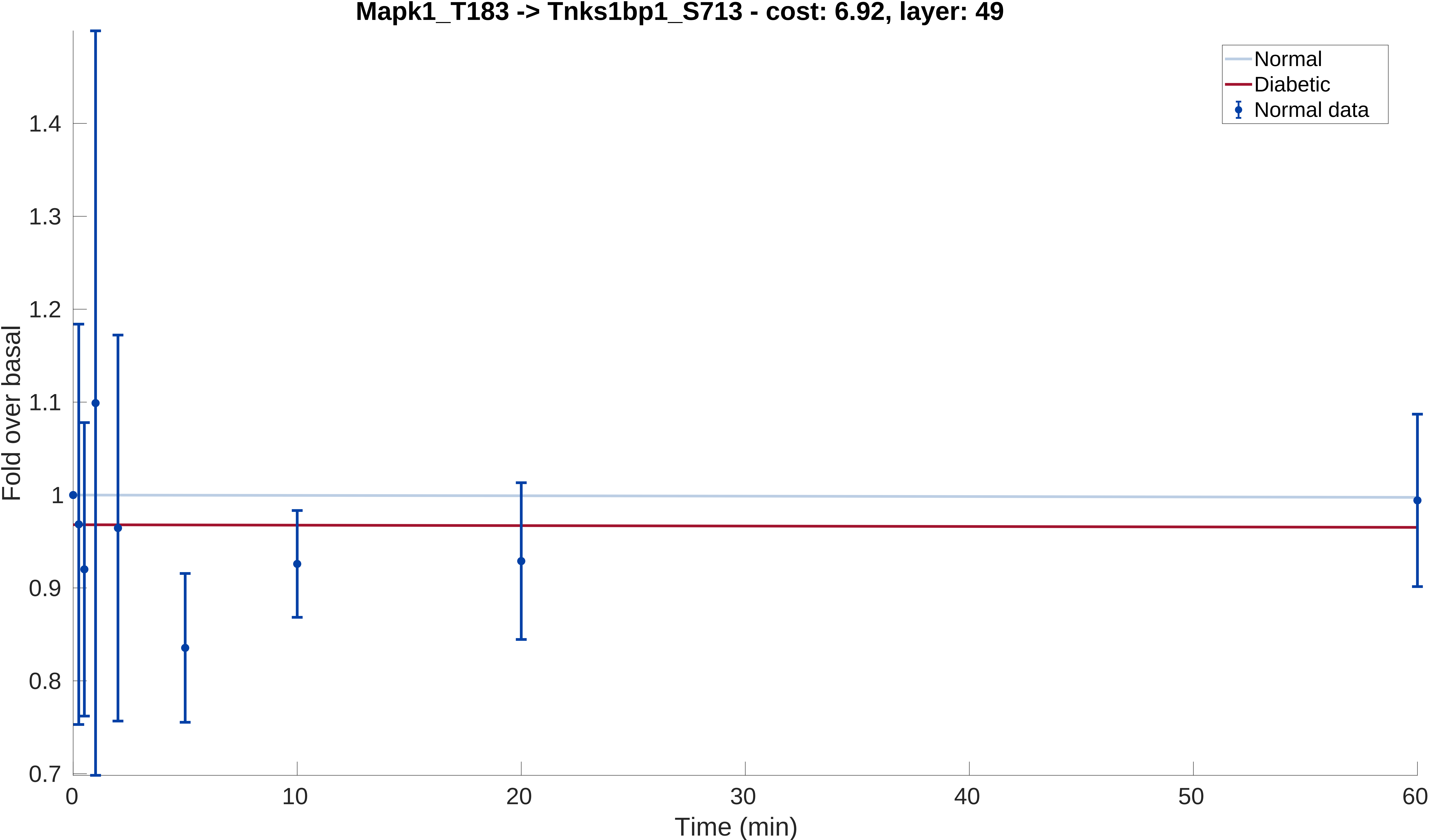

Mapk1_T183 -> Tnks1bp1_S713 - cost: 6.92, layer: 49
Normal
Diabetic
Normal data
Fold over basal
1.4
1.3
1.2
1.1
1
0.9
0.8
0.7
0
10
20
30
40
50
60
Time (min)

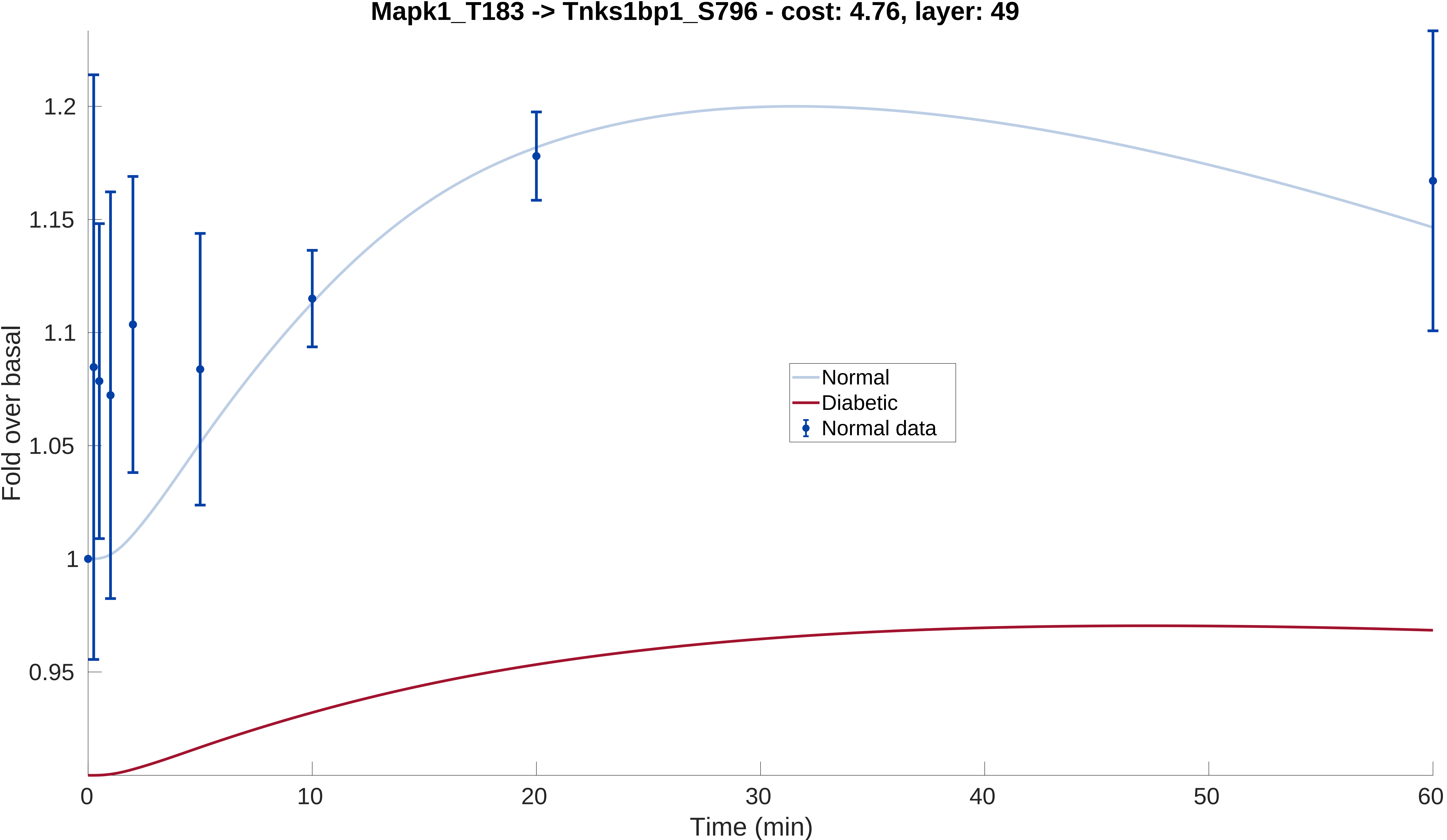

Mapk1_T183 -> Tnks1bp1_S796 - cost: 4.76, layer: 49
Fold over basal
1.2
1.15
1.1
1.05
1
0.95
0
10
20
30
40
50
60
Time (min)
Normal
Diabetic
Normal data

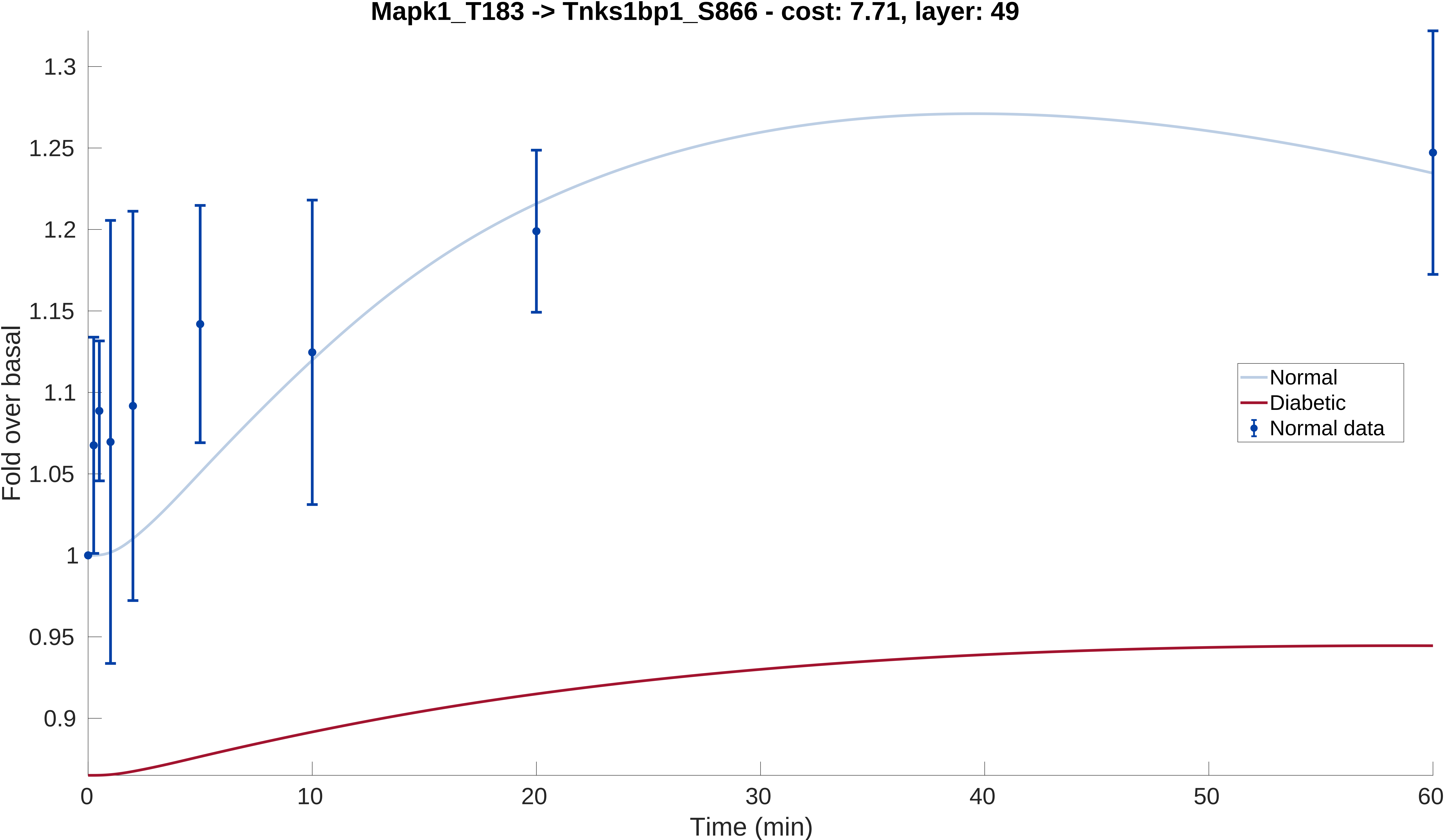
Mapk1_T183 -> Tnks1bp1_S866 - cost: 7.71, layer: 49
Fold over basal
Time (min)
1.3
1.25
1.2
1.15
1.1
1.05
1
0.95
0.9
0
10
20
30
40
50
60
Normal
Diabetic
Normal data

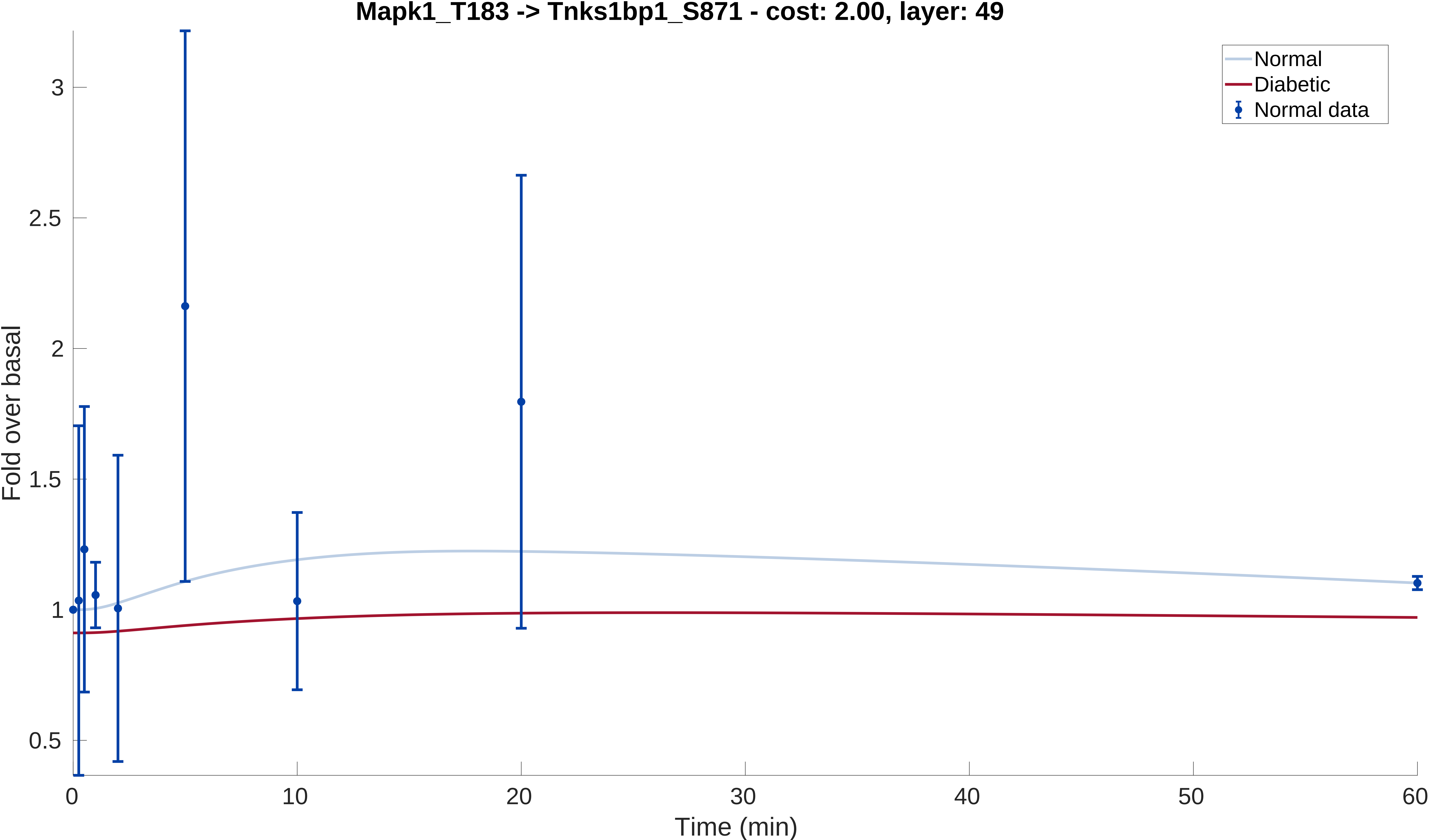

Mapk1_T183 -> Tnks1bp1_S871 - cost: 2.00, layer: 49
Normal
Diabetic
Normal data
Fold over basal
Time (min)
0
10
20
30
40
50
60
0.5
1
1.5
2
2.5
3

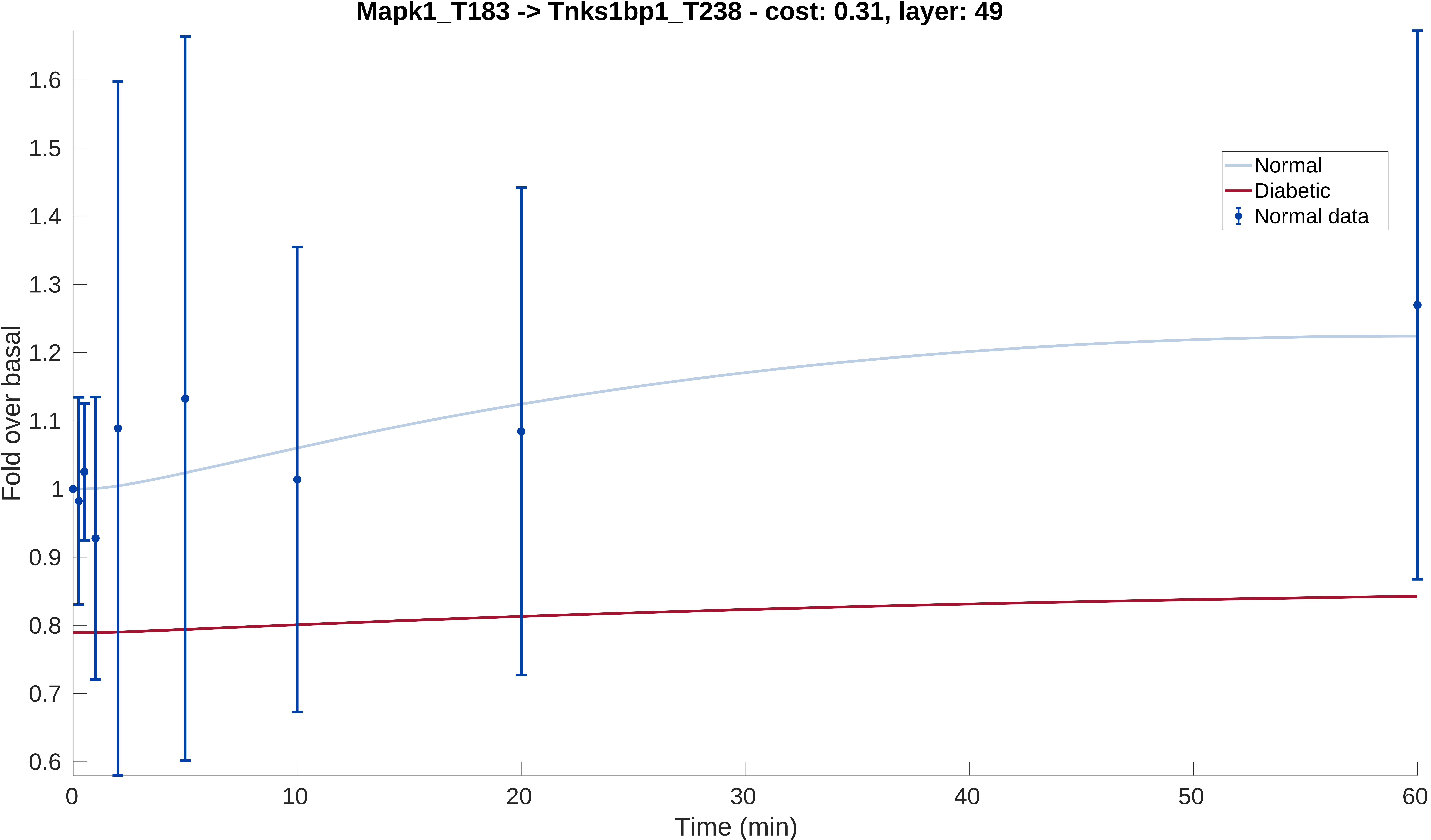
Mapk1_T183 -> Tnks1bp1_T238 - cost: 0.31, layer: 49
Normal
Diabetic
Normal data
Fold over basal
Time (min)
0.6
0.7
0.8
0.9
1
1.1
1.2
1.3
1.4
1.5
1.6
0
10
20
30
40
50
60

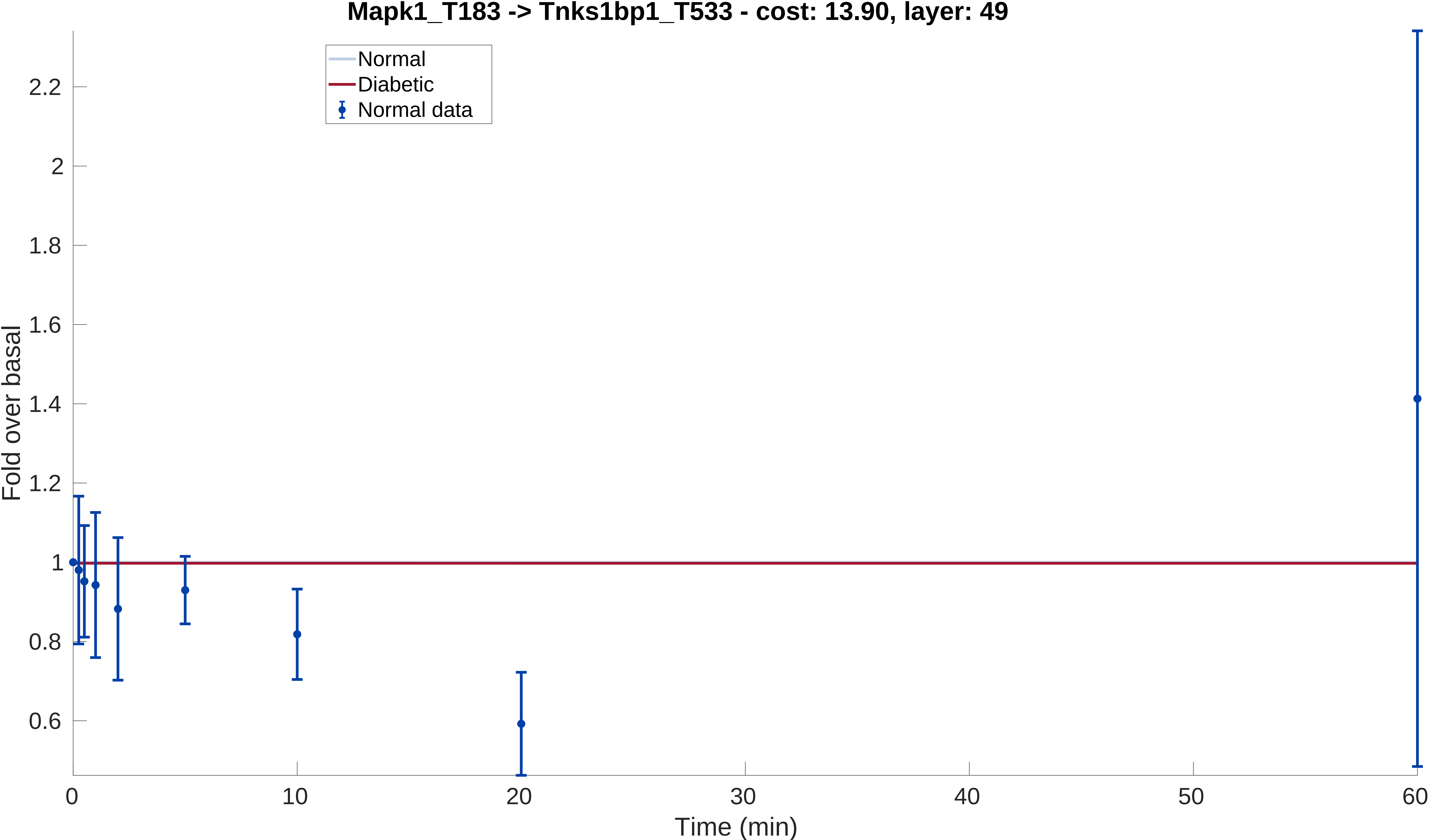

Mapk1_T183 -> Tnks1bp1_T533 - cost: 13.90, layer: 49
Normal
Diabetic
Normal data
Fold over basal
Time (min)
2.2
2
1.8
1.6
1.4
1.2
1
0.8
0.6
0
10
20
30
40
50
60

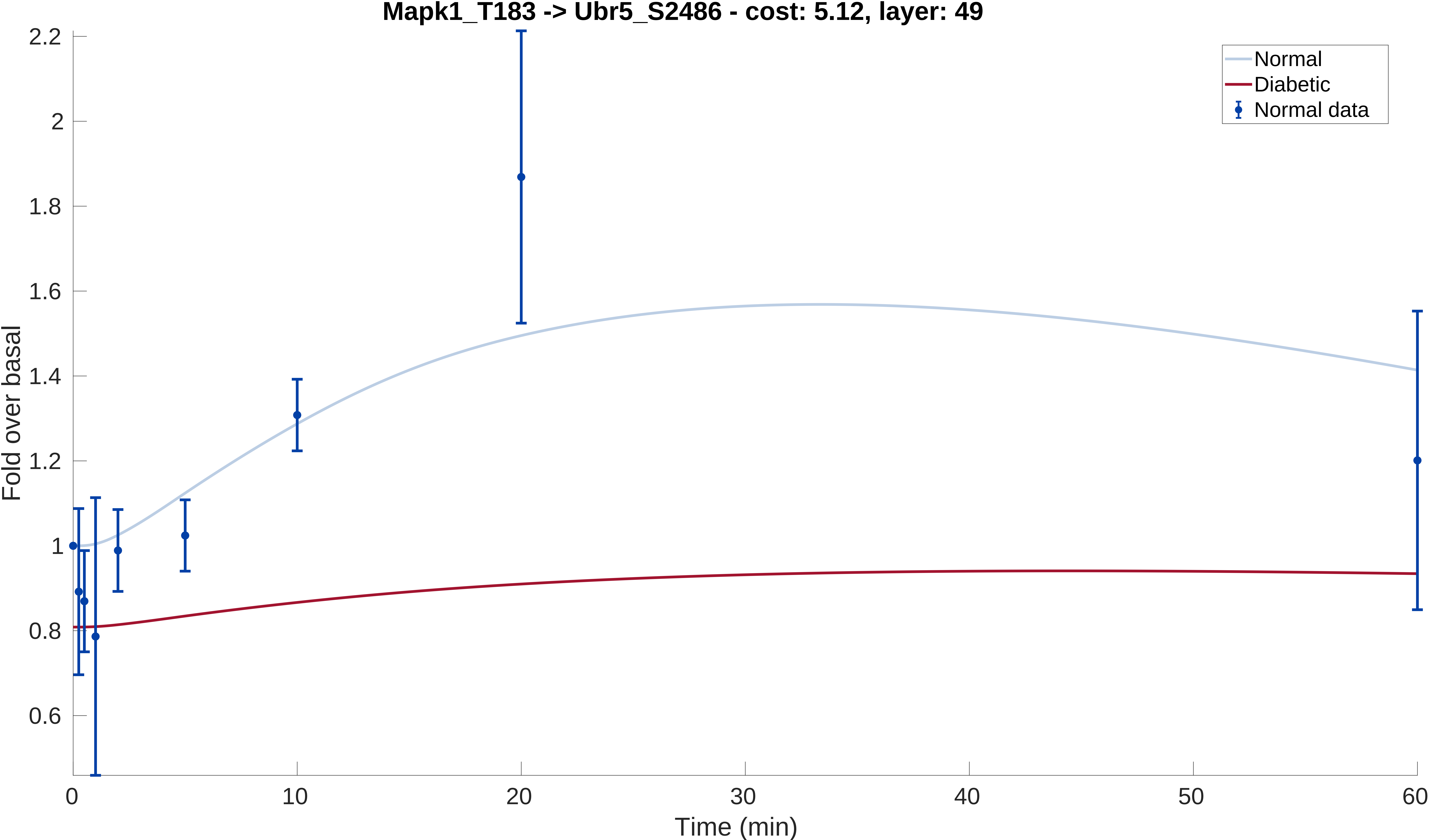

Mapk1_T183 -> Ubr5_S2486 - cost: 5.12, layer: 49
Normal
Diabetic
Normal data
Fold over basal
Time (min)
2.2
2
1.8
1.6
1.4
1.2
1
0.8
0.6
0
10
20
30
40
50
60

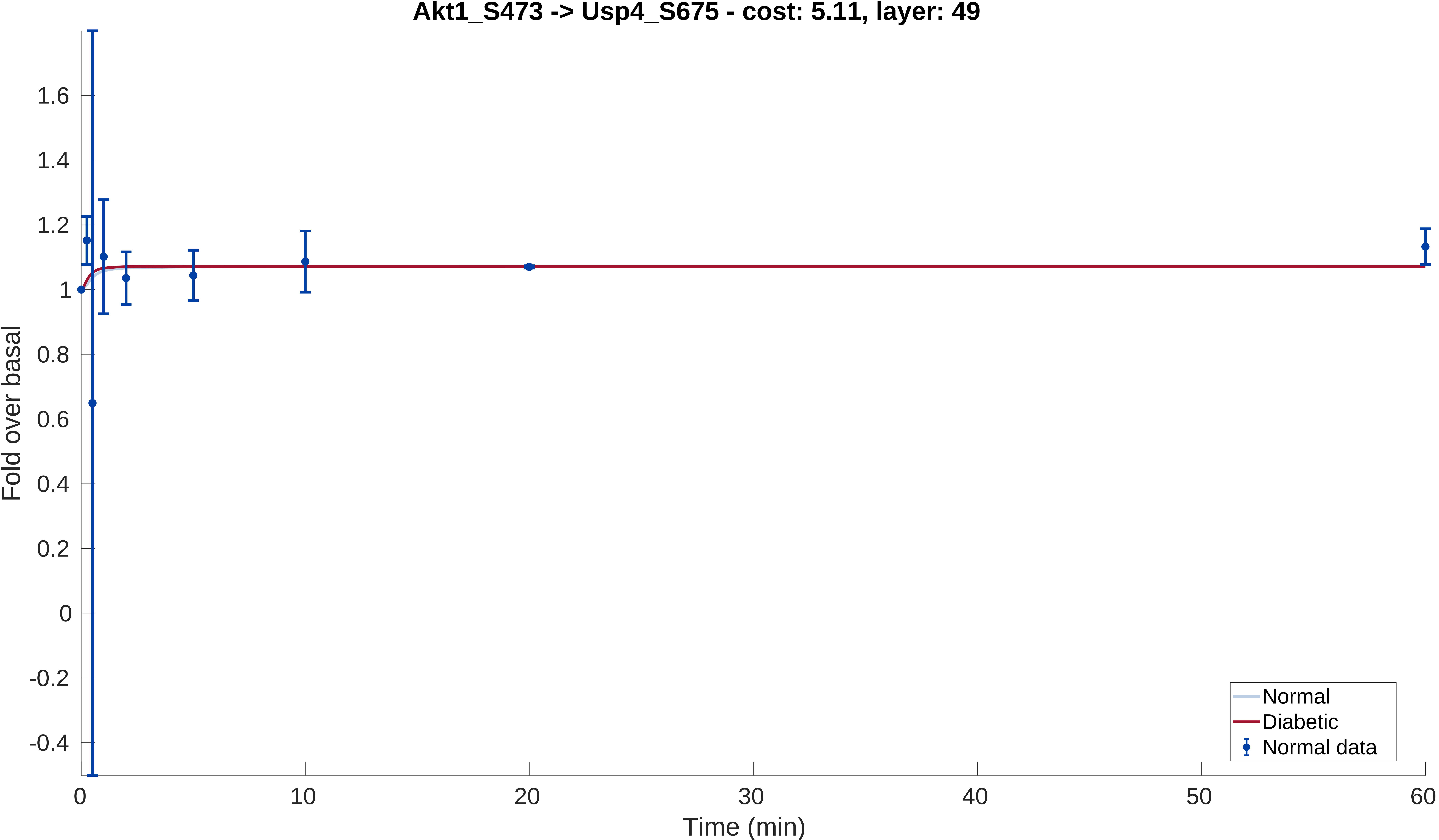
Akt1_S473 -> Usp4_S675 - cost: 5.11, layer: 49
Fold over basal
1.6
1.4
1.2
1
0.8
0.6
0.4
0.2
0
-0.2
-0.4
0
10
20
30
40
50
60
Time (min)
Normal
Diabetic
Normal data

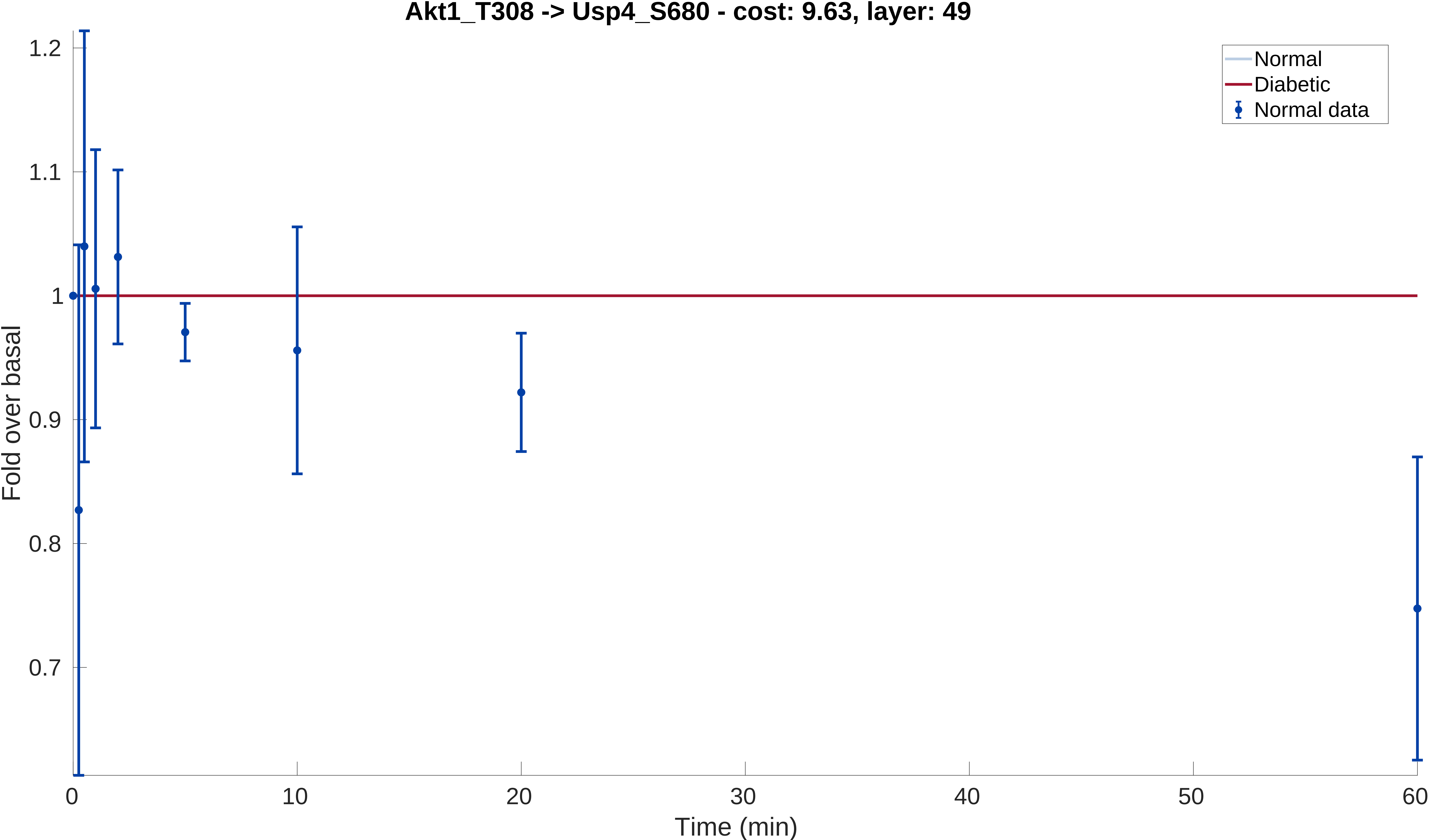

Akt1_T308 -> Usp4_S680 - cost: 9.63, layer: 49
Normal
Diabetic
Normal data
Fold over basal
Time (min)
1.2
1.1
1
0.9
0.8
0.7
0
10
20
30
40
50
60

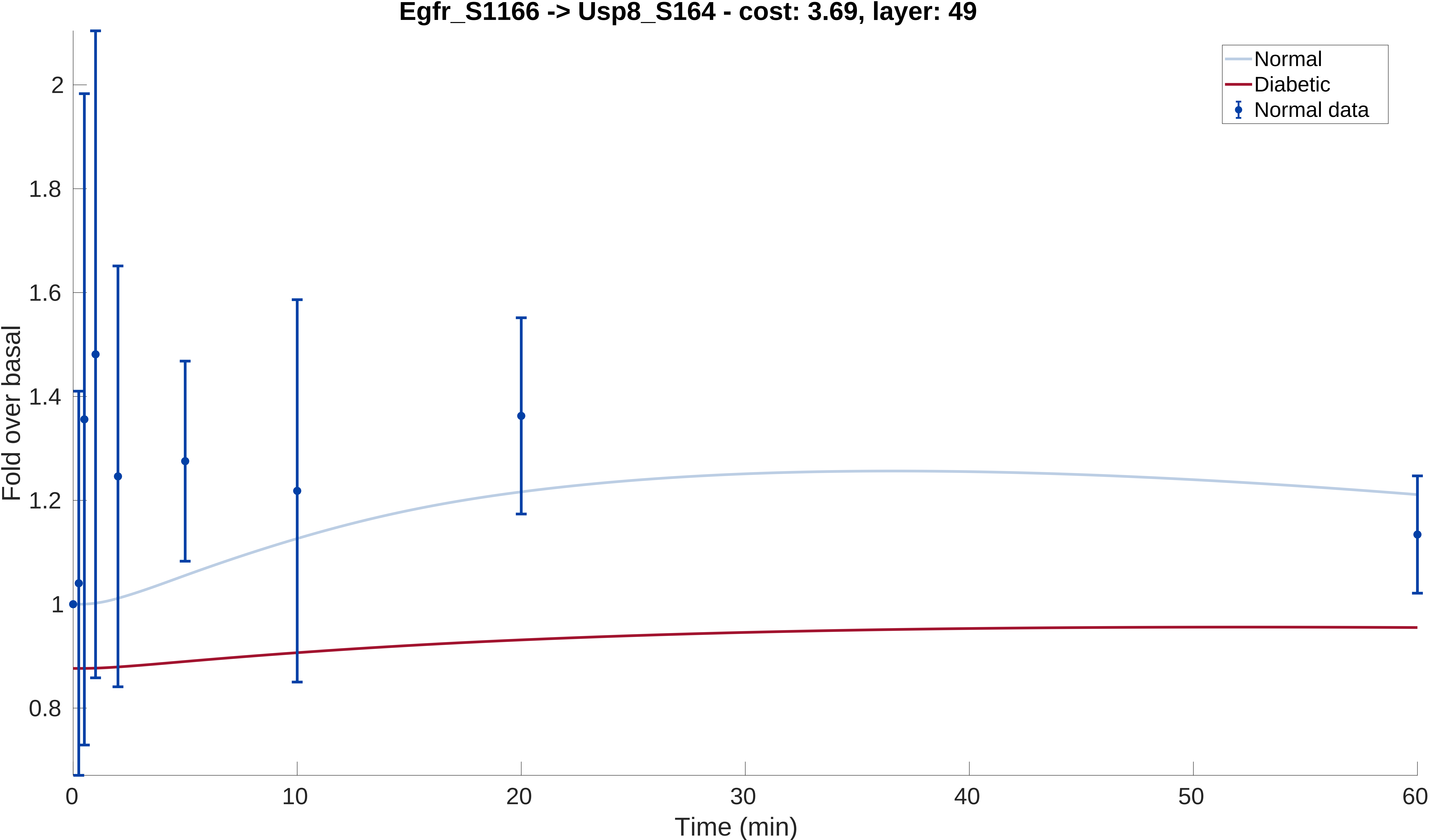

Egfr_S1166 -> Usp8_S164 - cost: 3.69, layer: 49
Normal
Diabetic
Normal data
Fold over basal
Time (min)
0.8
1
1.2
1.4
1.6
1.8
2
0
10
20
30
40
50
60

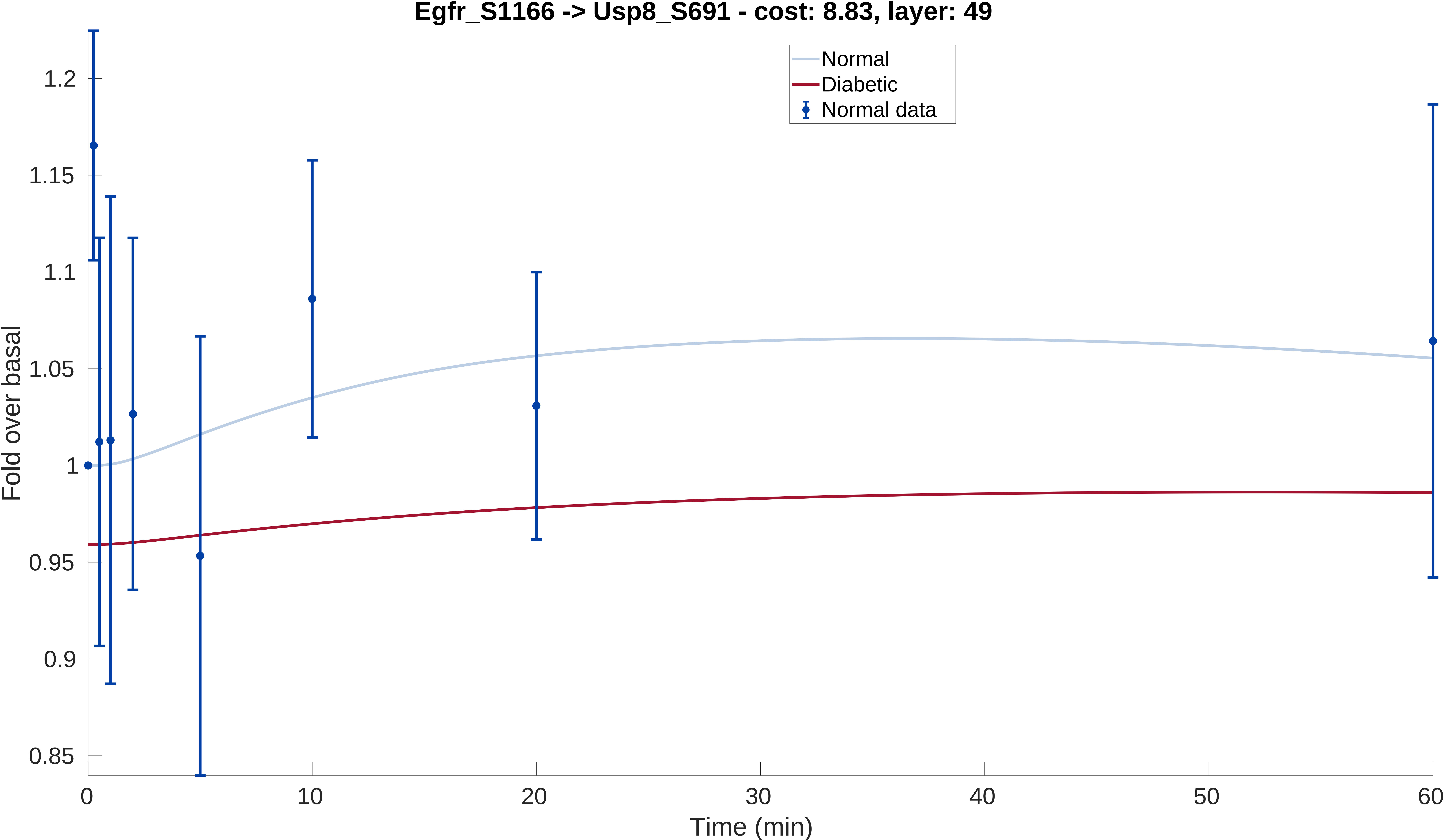

Egfr_S1166 -> Usp8_S691 - cost: 8.83, layer: 49
Normal
Diabetic
Normal data
Fold over basal
1.2
1.15
1.1
1.05
1
0.95
0.9
0.85
0
10
20
30
40
50
60
Time (min)

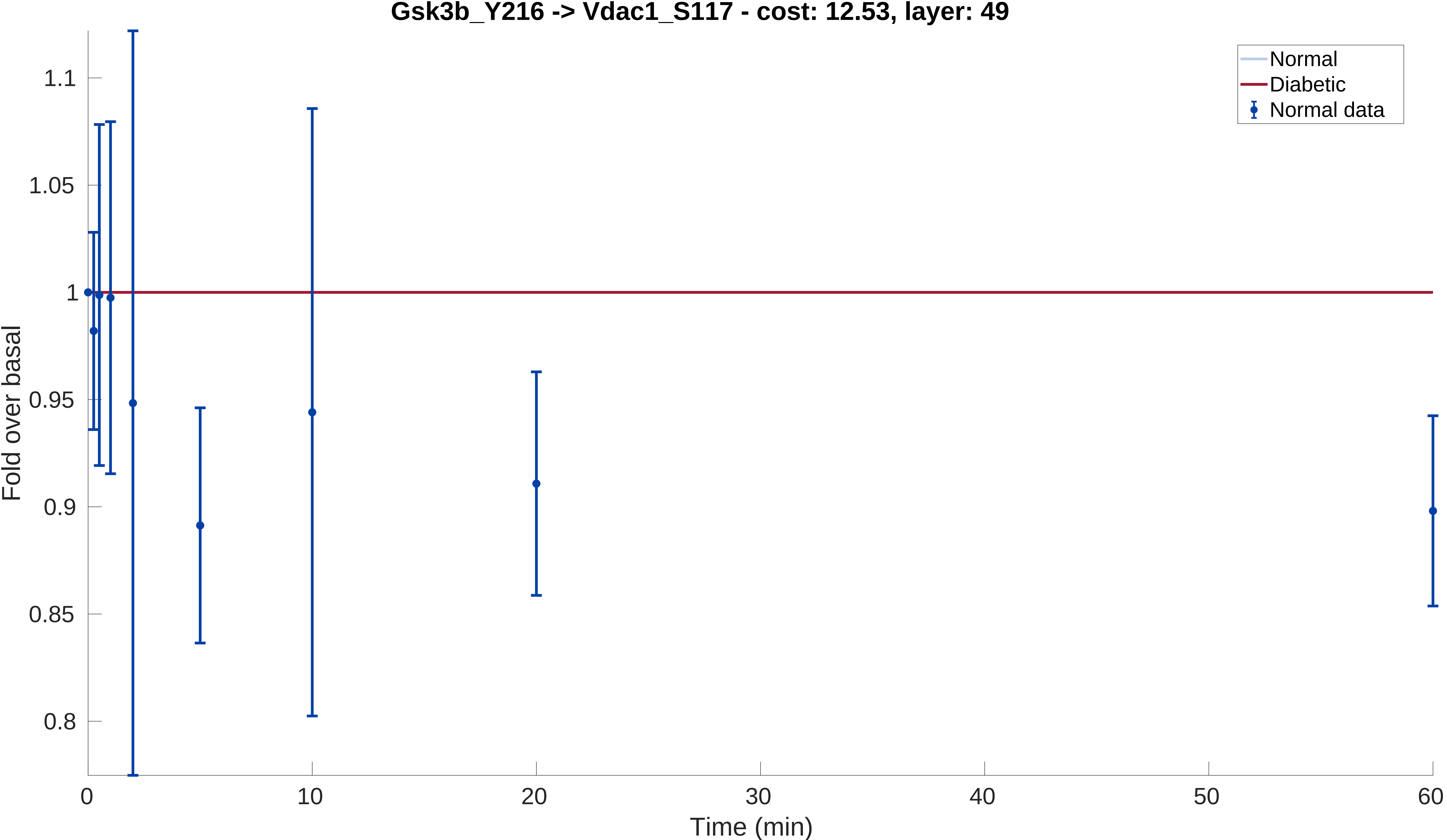
Gsk3b_Y216 -> Vdac1_S117 - cost: 12.53, layer: 49
Normal
Diabetic
Normal data
Fold over basal
1.1
1.05
1
0.95
0.9
0.85
0.8
0
10
20
30
40
50
60
Time (min)

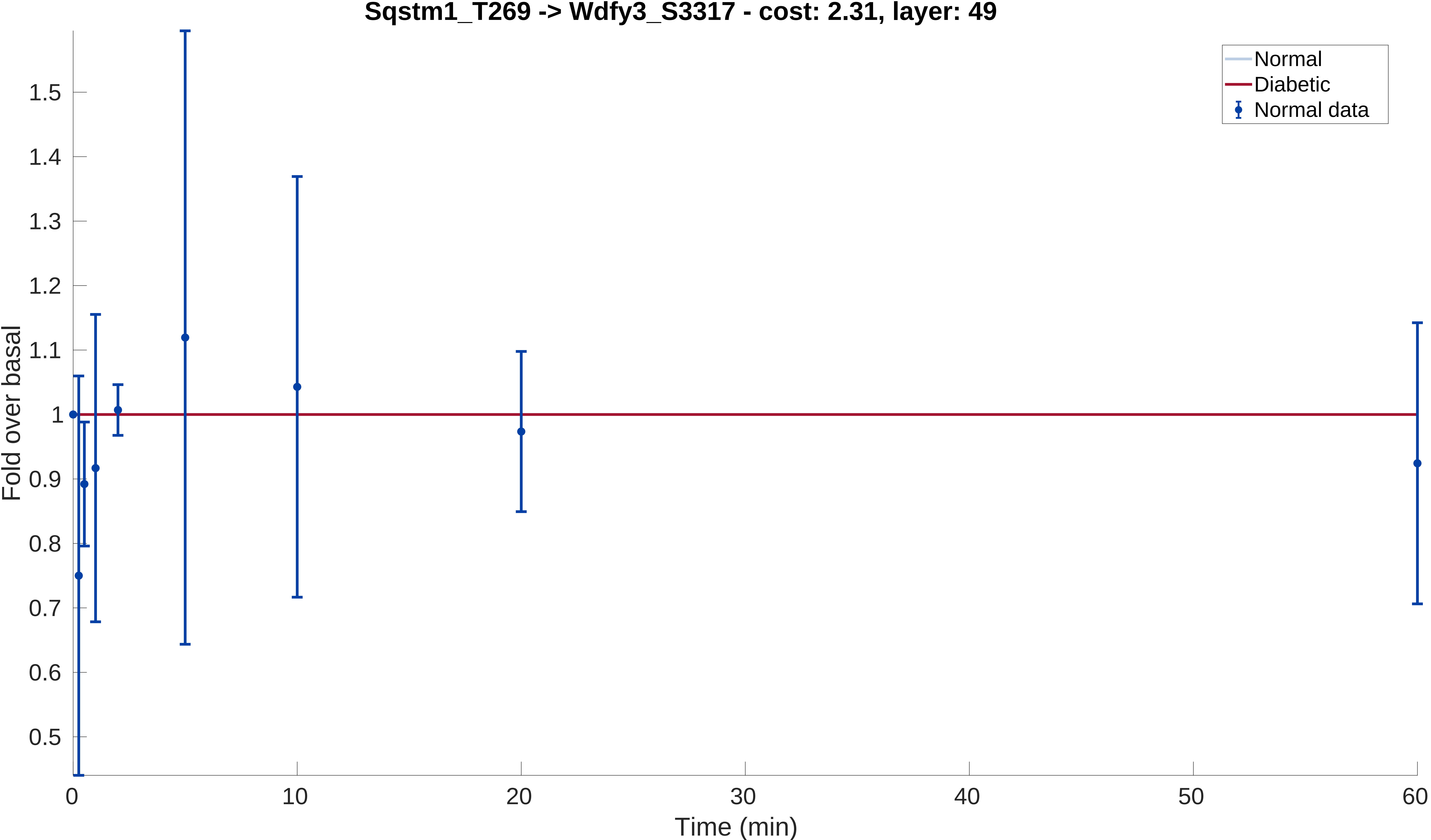
Sqstm1_T269 -> Wdfy3_S3317 - cost: 2.31, layer: 49
Normal
Diabetic
Normal data
Fold over basal
Time (min)
1.5
1.4
1.3
1.2
1.1
1
0.9
0.8
0.7
0.6
0.5
0
10
20
30
40
50
60

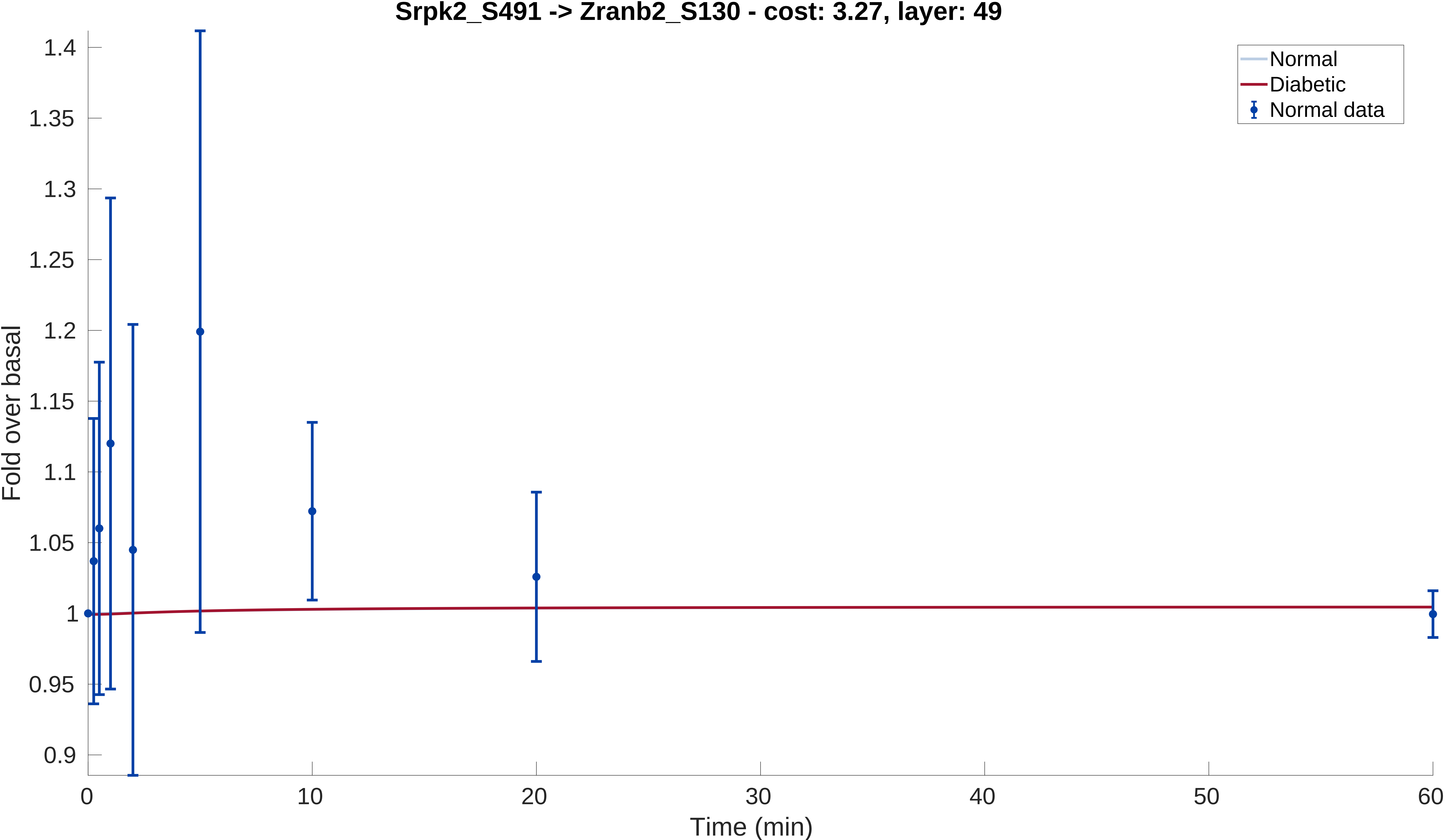

Srpk2_S491 -> Zranb2_S130 - cost: 3.27, layer: 49
Normal
Diabetic
Normal data
Fold over basal
Time (min)
1.4
1.35
1.3
1.25
1.2
1.15
1.1
1.05
1
0.95
0.9
0
10
20
30
40
50
60

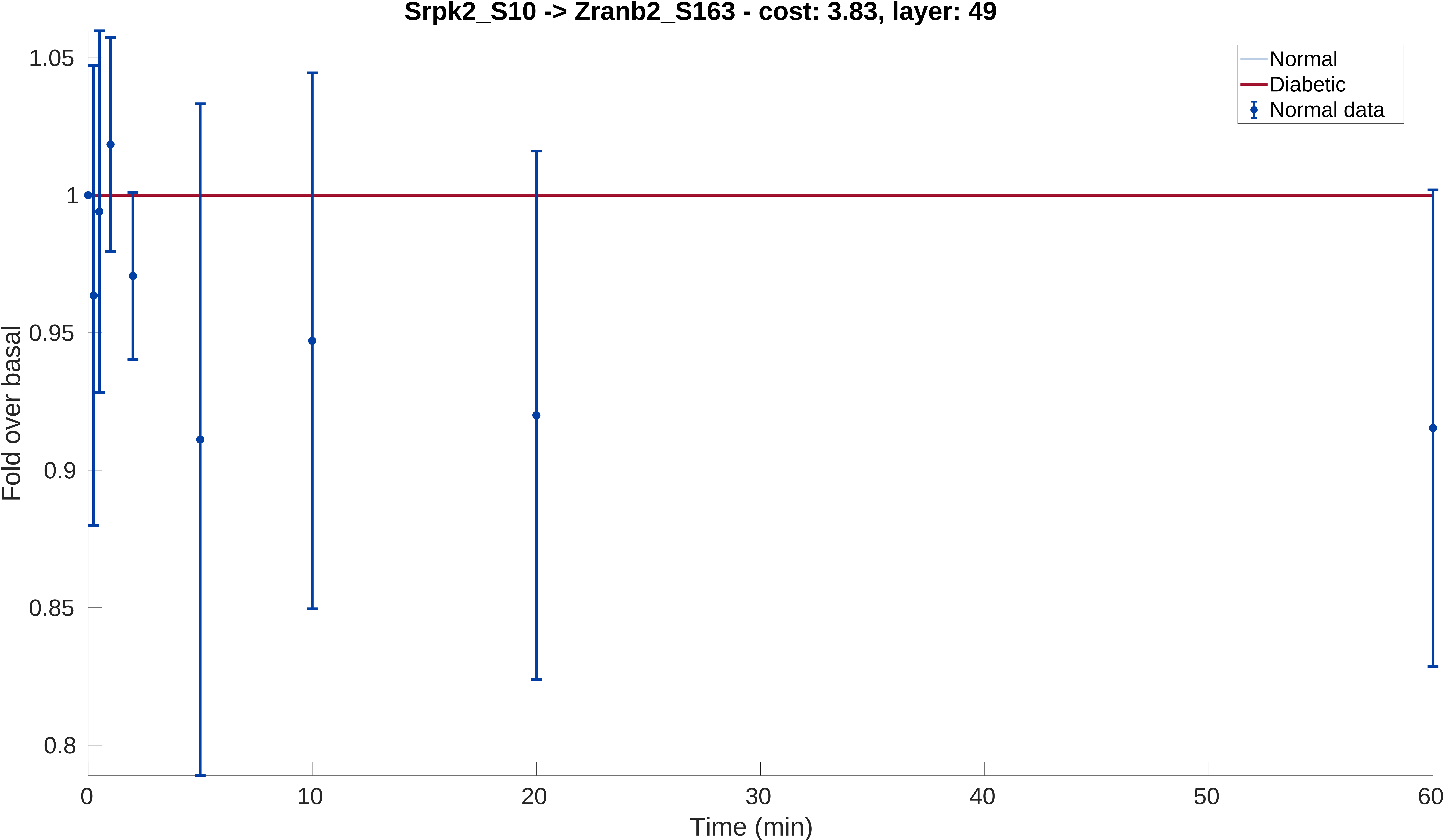

Srpk2_S10 -> Zranb2_S163 - cost: 3.83, layer: 49
Normal
Diabetic
Normal data
Fold over basal
Time (min)
1.05
1
0.95
0.9
0.85
0.8
0
10
20
30
40
50
60

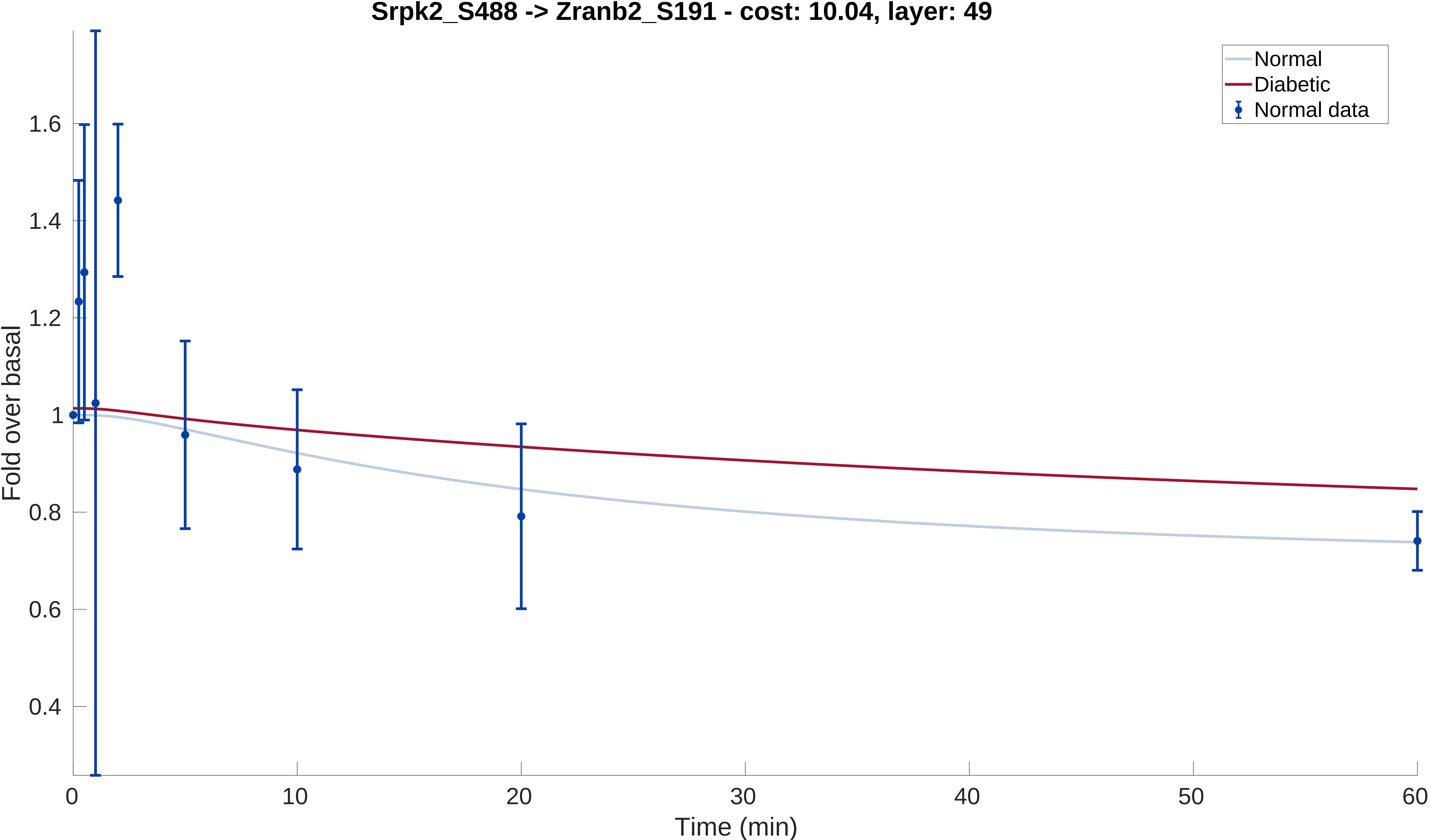

Srpk2_S488 -> Zranb2_S191 - cost: 10.04, layer: 49
Normal
Diabetic
Normal data
1.6
1.4
1.2
1
0.8
0.6
0.4
Fold over basal
0
10
20
30
40
50
60
Time (min)

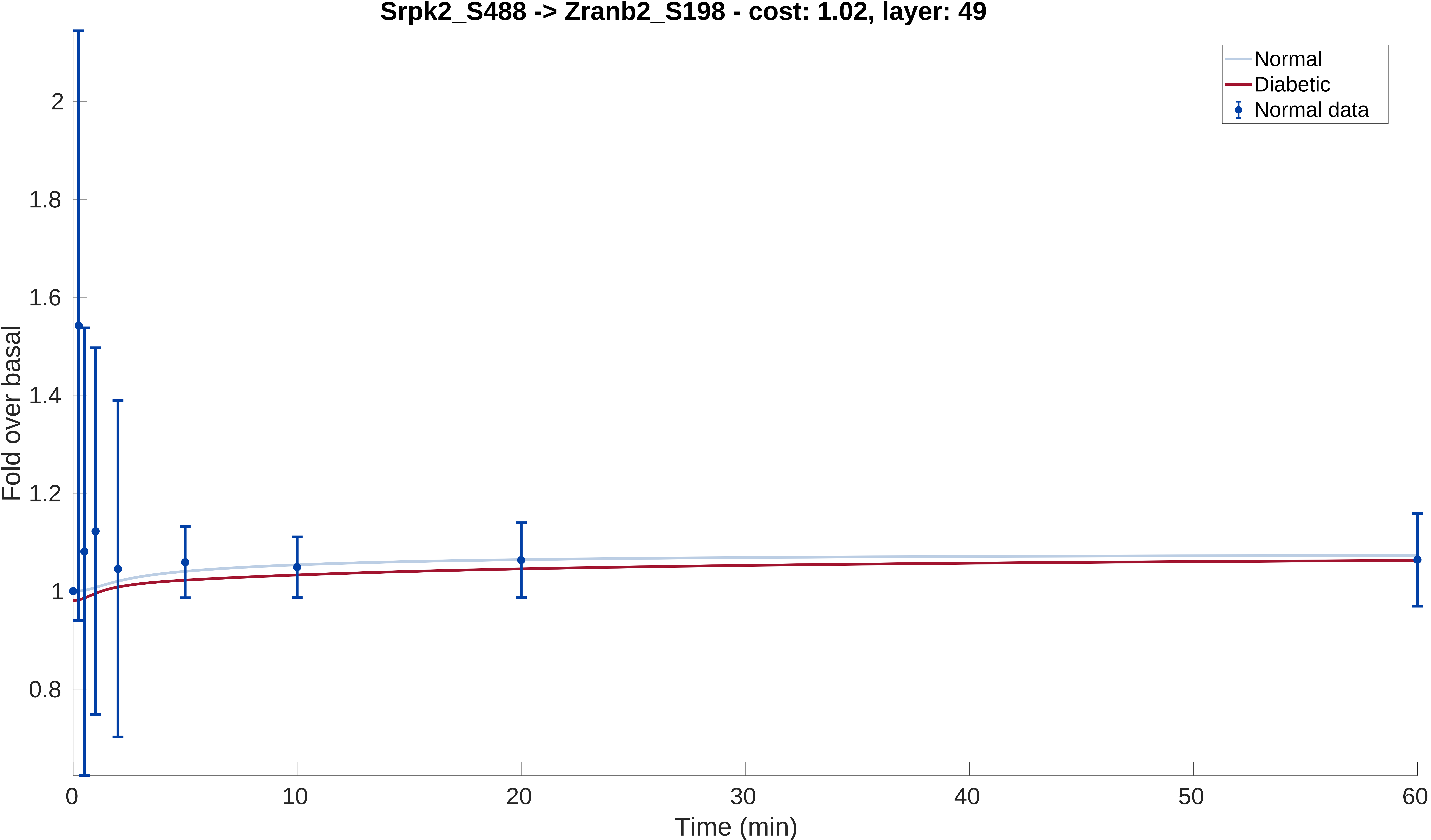

Srpk2_S488 -> Zranb2_S198 - cost: 1.02, layer: 49
Normal
Diabetic
Normal data
Fold over basal
2
1.8
1.6
1.4
1.2
1
0.8
0
10
20
30
40
50
60
Time (min)

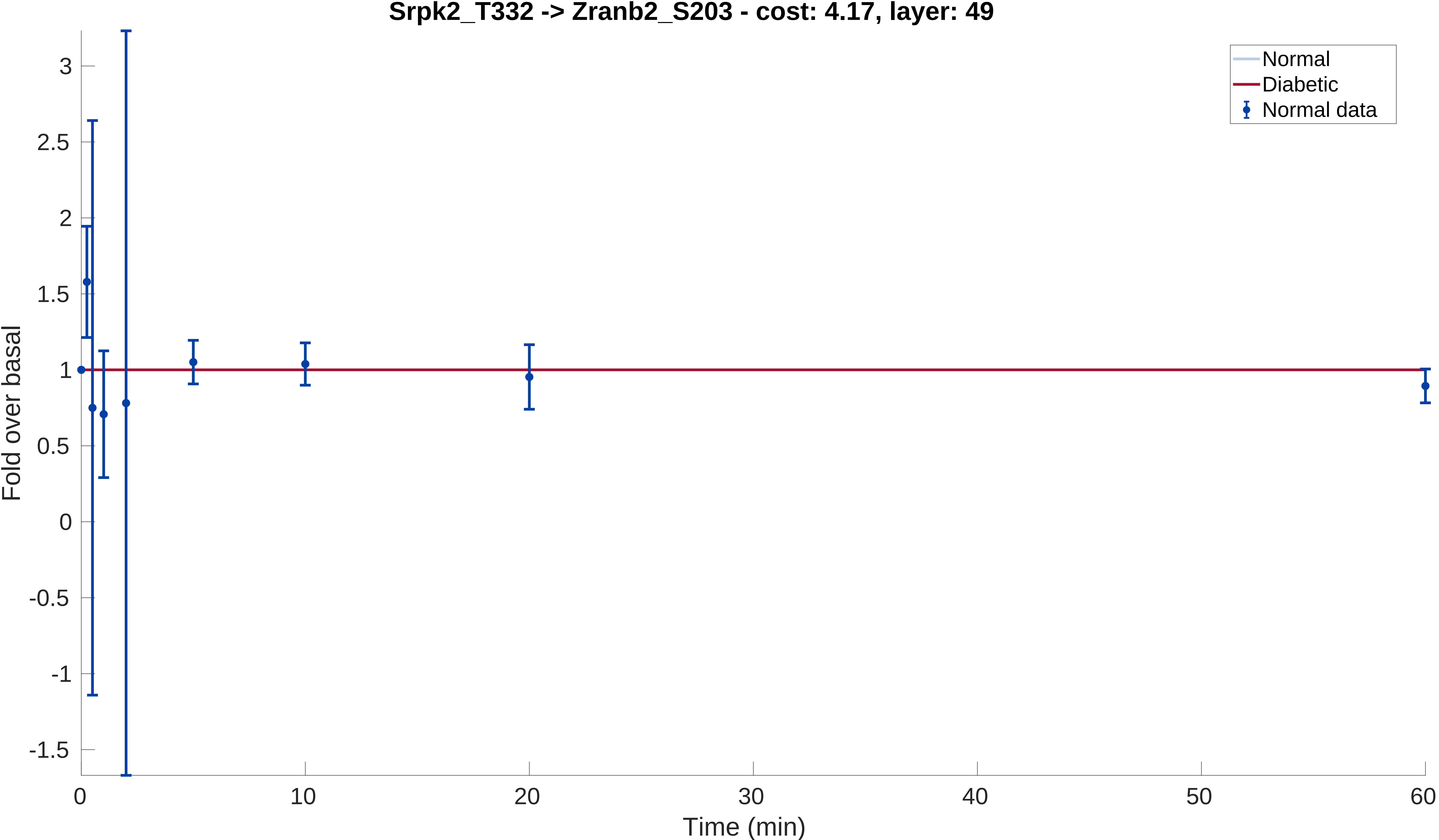

Srpk2_T332 -> Zranb2_S203 - cost: 4.17, layer: 49
Normal
Diabetic
Normal data
Fold over basal
3
2.5
2
1.5
1
0.5
0
-0.5
-1
-1.5
0
10
20
30
40
50
60
Time (min)

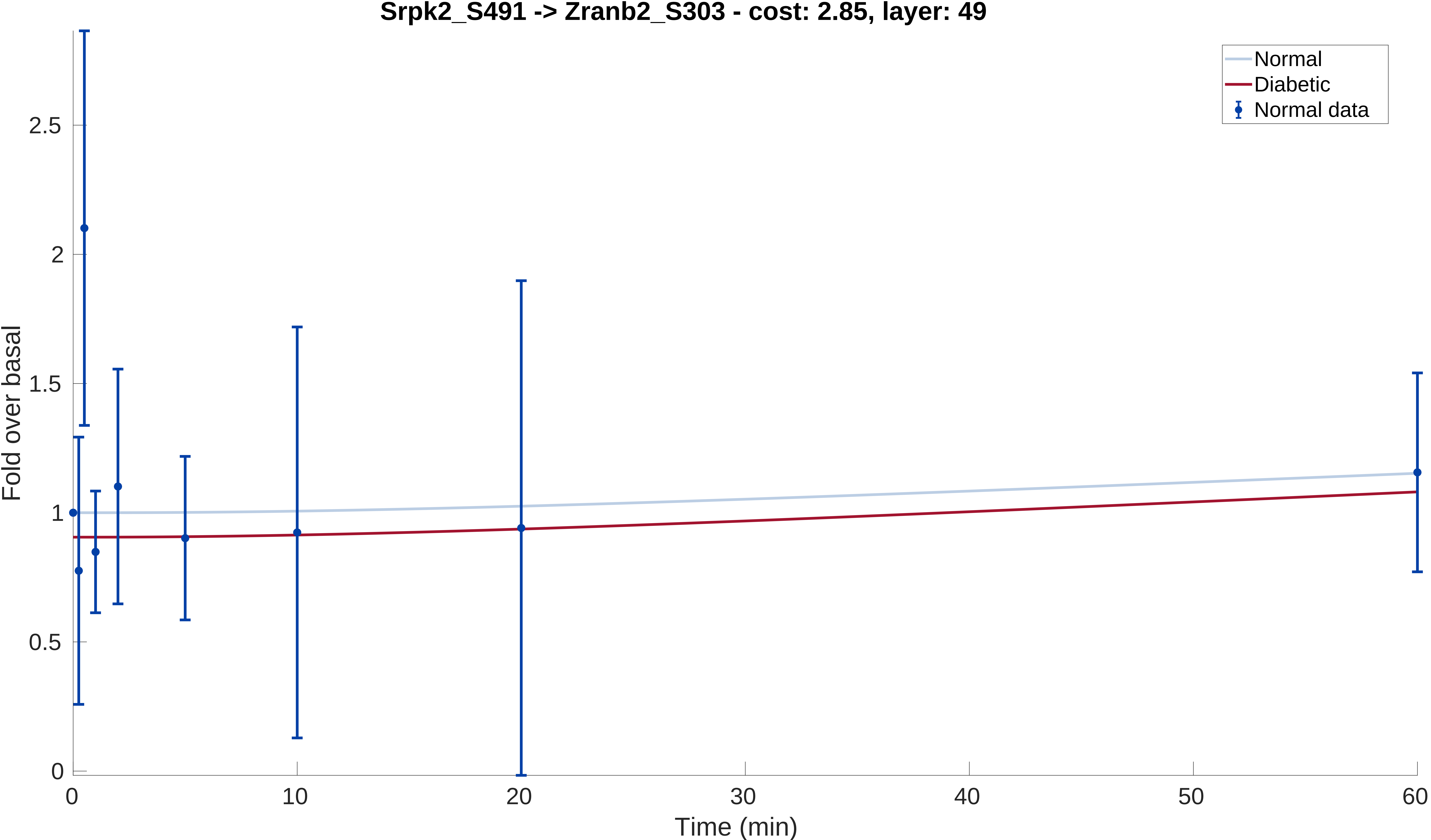
Srpk2_S491 -> Zranb2_S303 - cost: 2.85, layer: 49
Normal
Diabetic
Normal data
Fold over basal
0
0.5
1
1.5
2
2.5
0
10
20
30
40
50
60
Time (min)

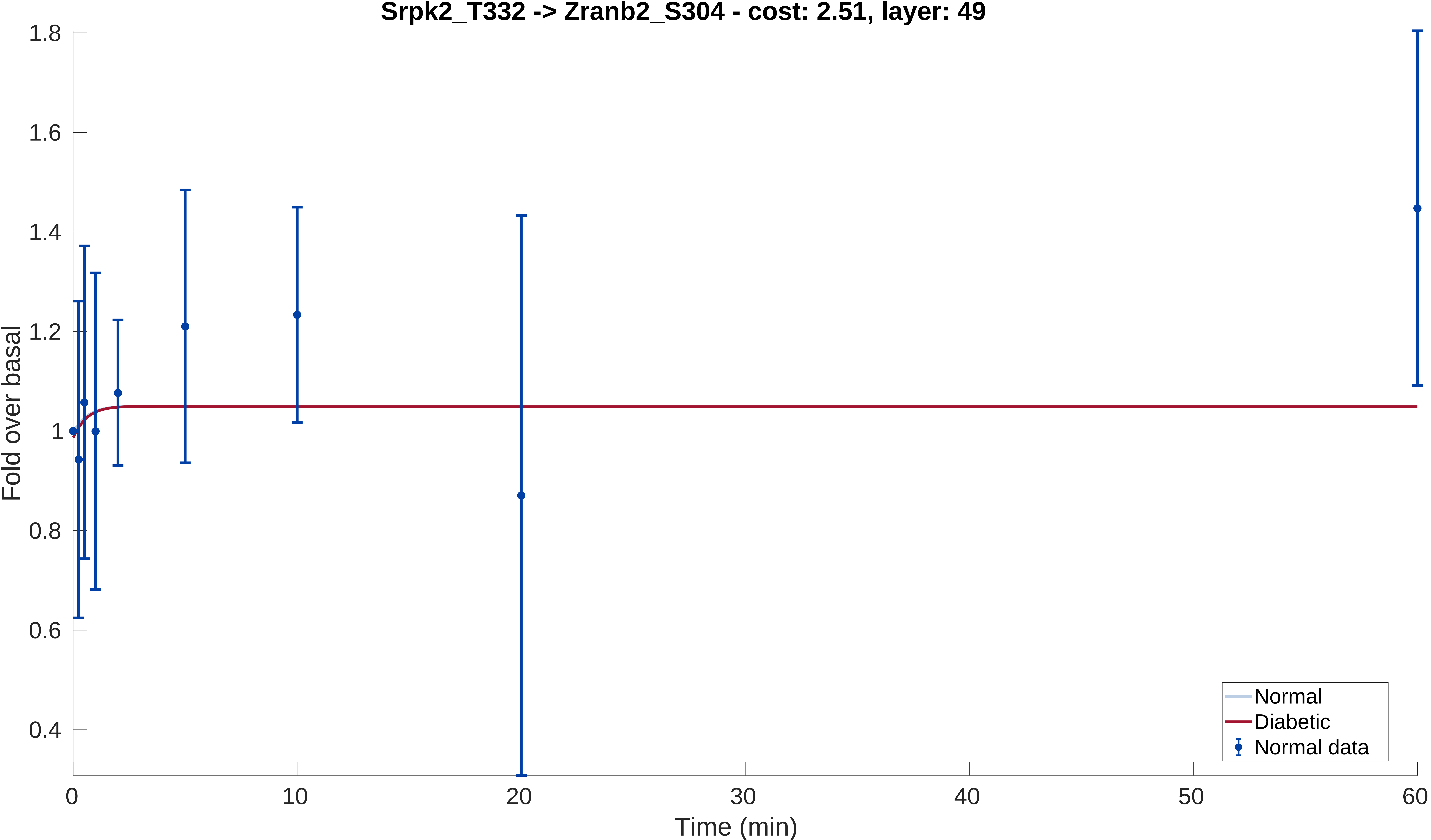

Srpk2_T332 -> Zranb2_S304 - cost: 2.51, layer: 49
Fold over basal
Time (min)
1.8
1.6
1.4
1.2
1
0.8
0.6
0.4
0
10
20
30
40
50
60
Normal
Diabetic
Normal data

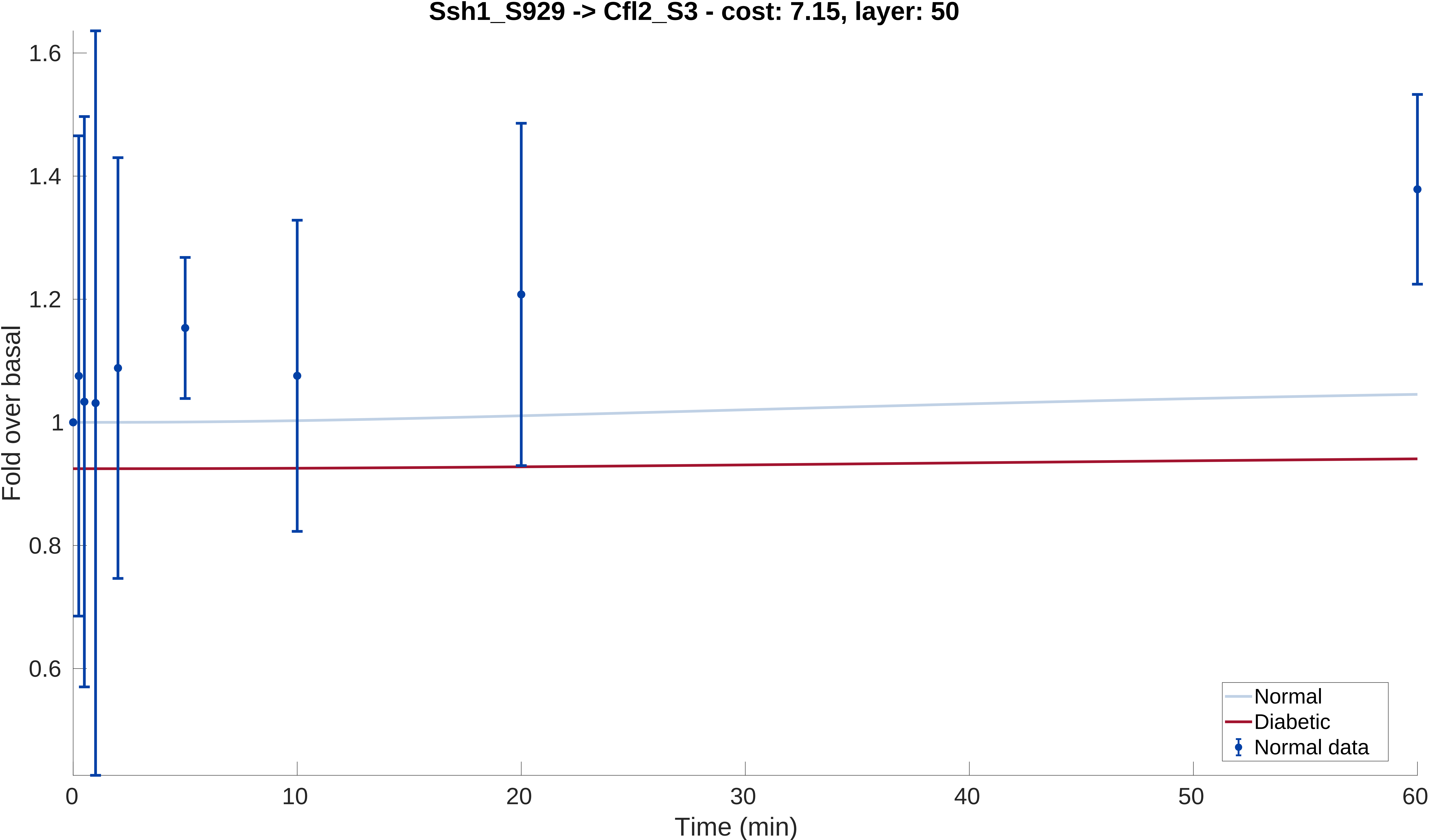

Ssh1_S929 -> Cfl2_S3 - cost: 7.15, layer: 50
Fold over basal
1.6
1.4
1.2
1
0.8
0.6
0
10
20
30
40
50
60
Time (min)
Normal
Diabetic
Normal data

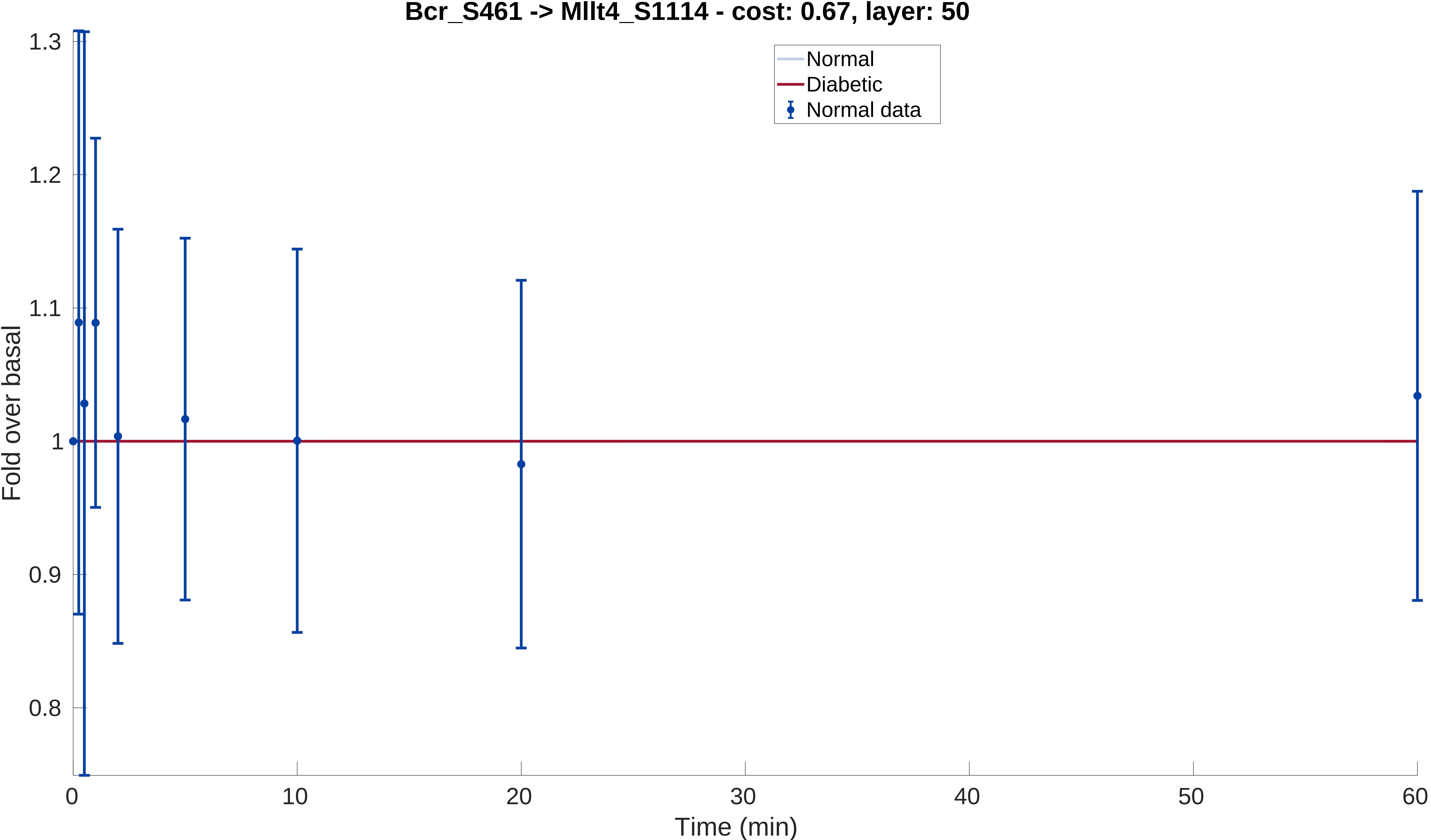
Bcr_S461 -> Mllt4_S1114 - cost: 0.67, layer: 50
Normal
Diabetic
Normal data
Fold over basal
Time (min)
1.3
1.2
1.1
1
0.9
0.8
0
10
20
30
40
50
60

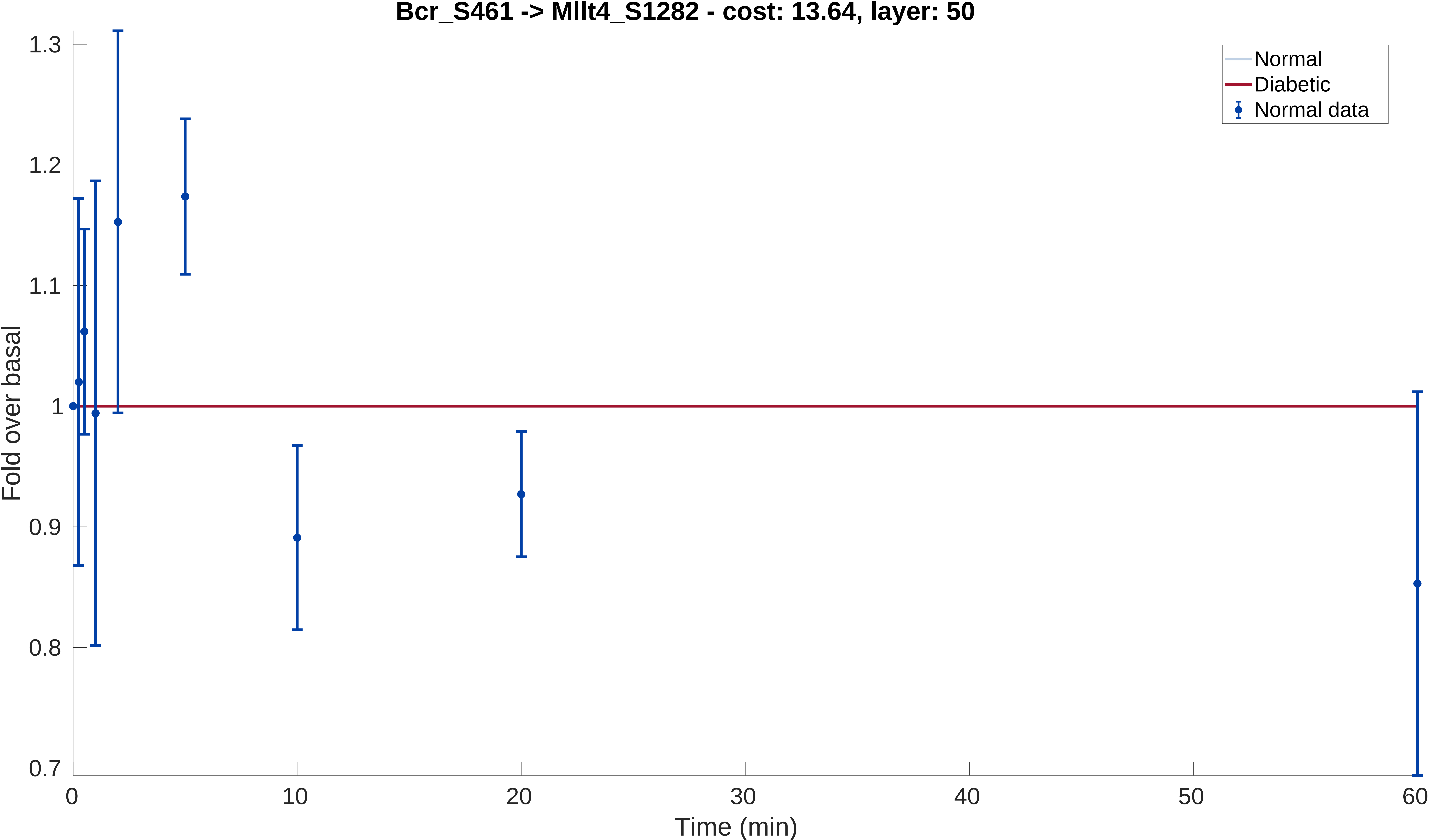

Bcr_S461 -> Mllt4_S1282 - cost: 13.64, layer: 50
Normal
Diabetic
Normal data
Fold over basal
1.3
1.2
1.1
1
0.9
0.8
0.7
0
10
20
30
40
50
60
Time (min)

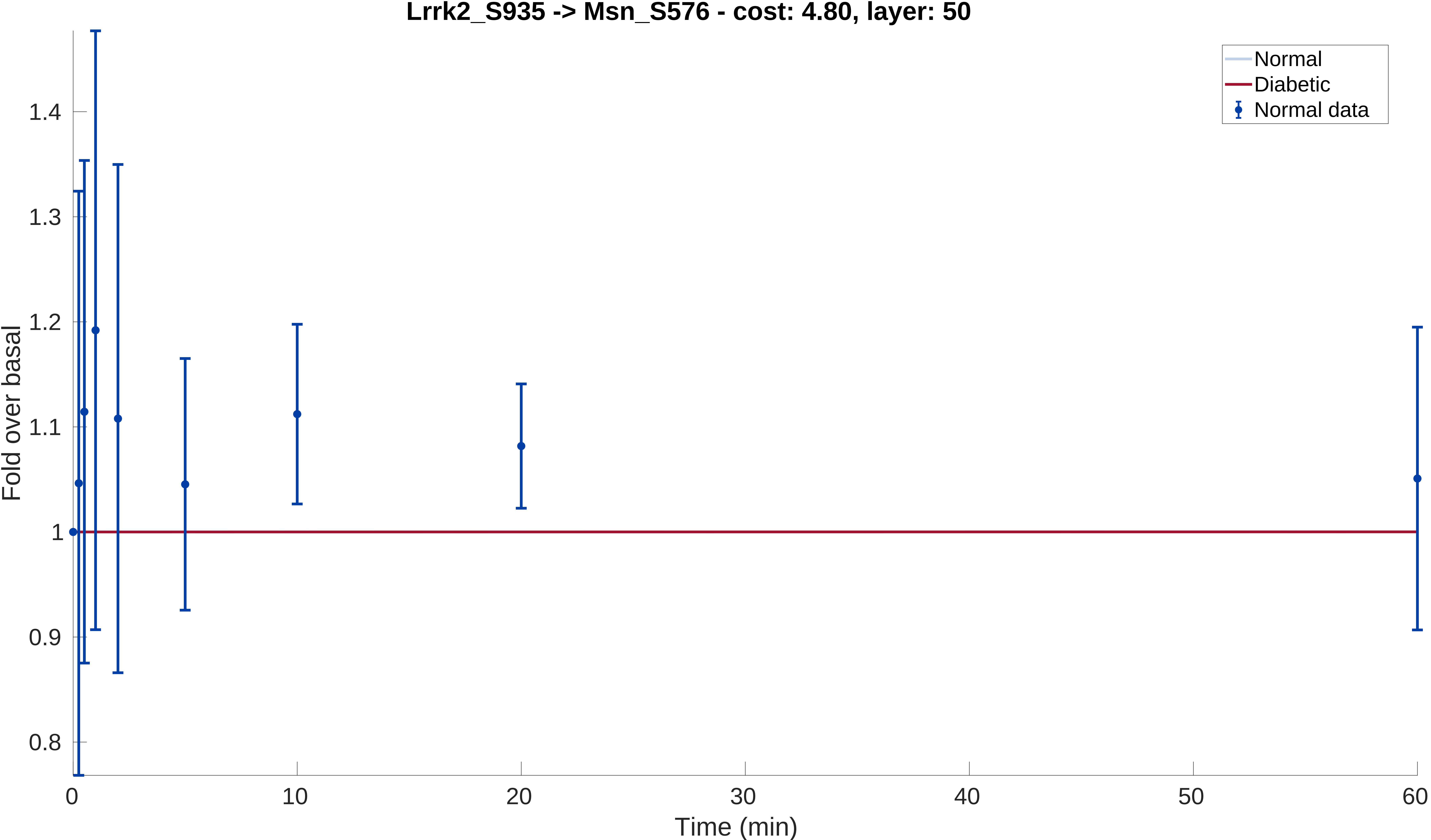

Lrrk2_S935 -> Msn_S576 - cost: 4.80, layer: 50
Normal
Diabetic
Normal data
Fold over basal
1.4
1.3
1.2
1.1
1
0.9
0.8
0
10
20
30
40
50
60
Time (min)

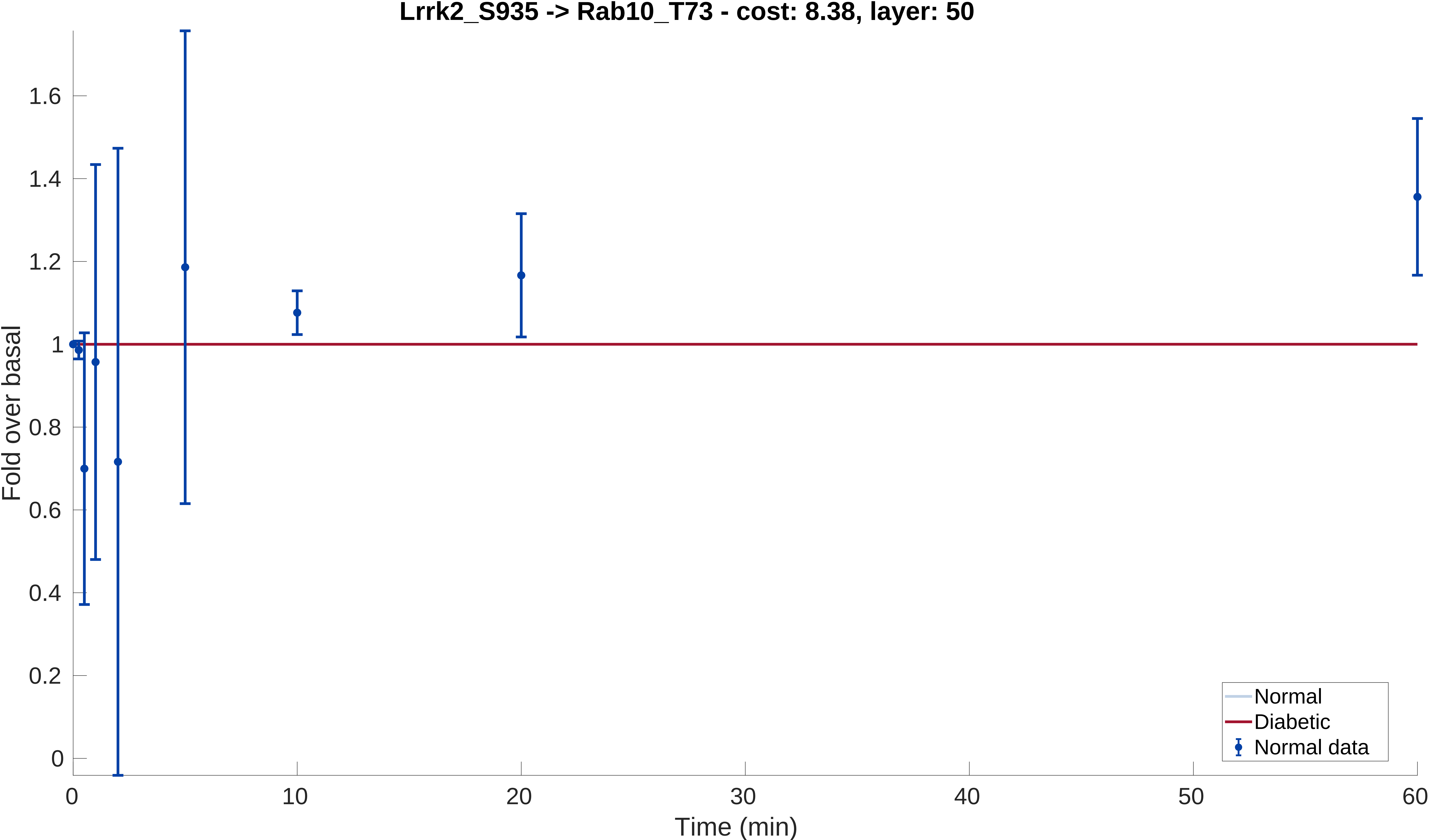

Lrrk2_S935 -> Rab10_T73 - cost: 8.38, layer: 50
Fold over basal
1.6
1.4
1.2
1
0.8
0.6
0.4
0.2
0
0
10
20
30
40
50
60
Time (min)
Normal
Diabetic
Normal data

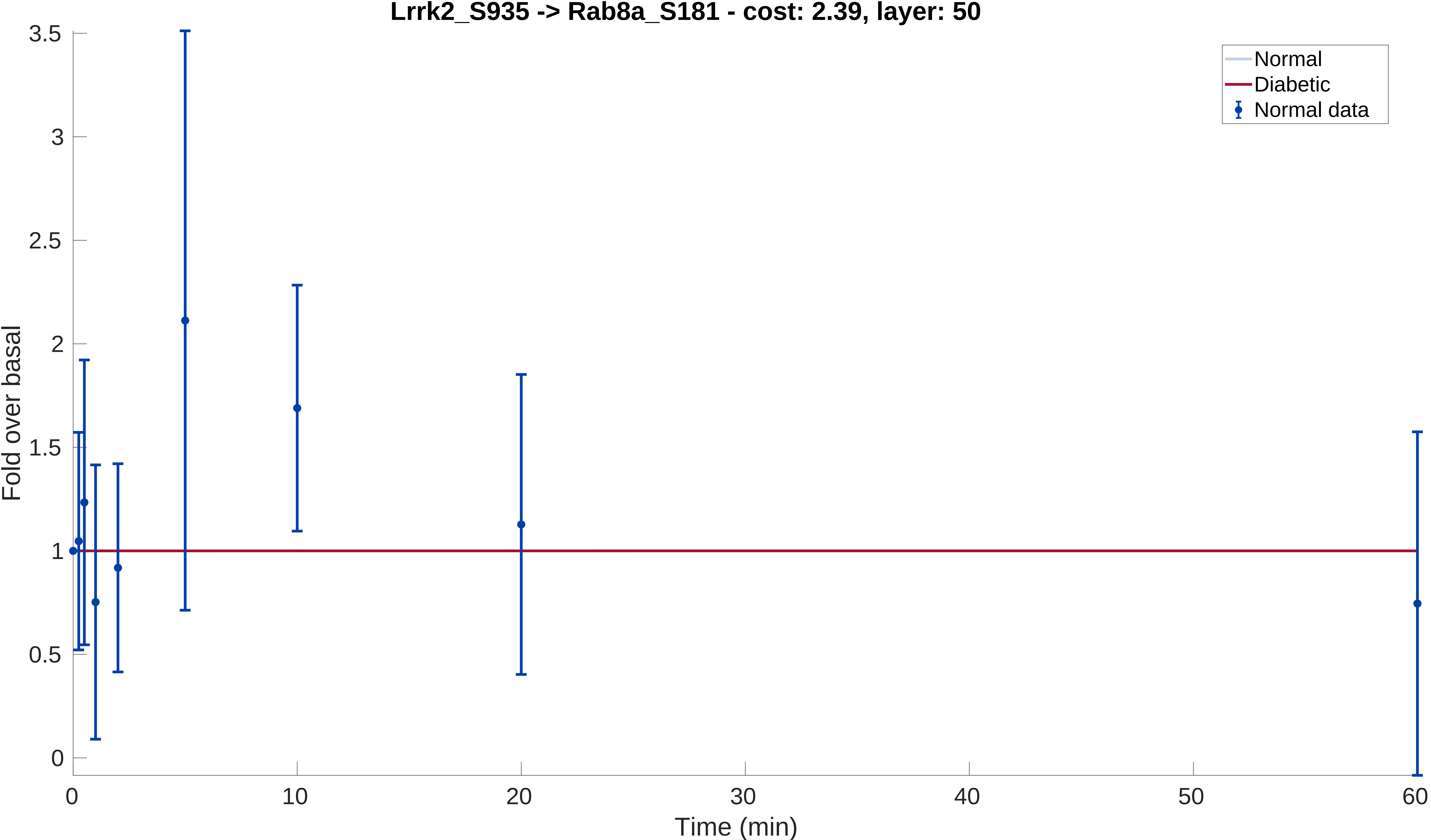

Lrrk2_S935 -> Rab8a_S181 - cost: 2.39, layer: 50
Normal
Diabetic
Normal data
Fold over basal
Time (min)
0
0.5
1
1.5
2
2.5
3
3.5
0
10
20
30
40
50
60

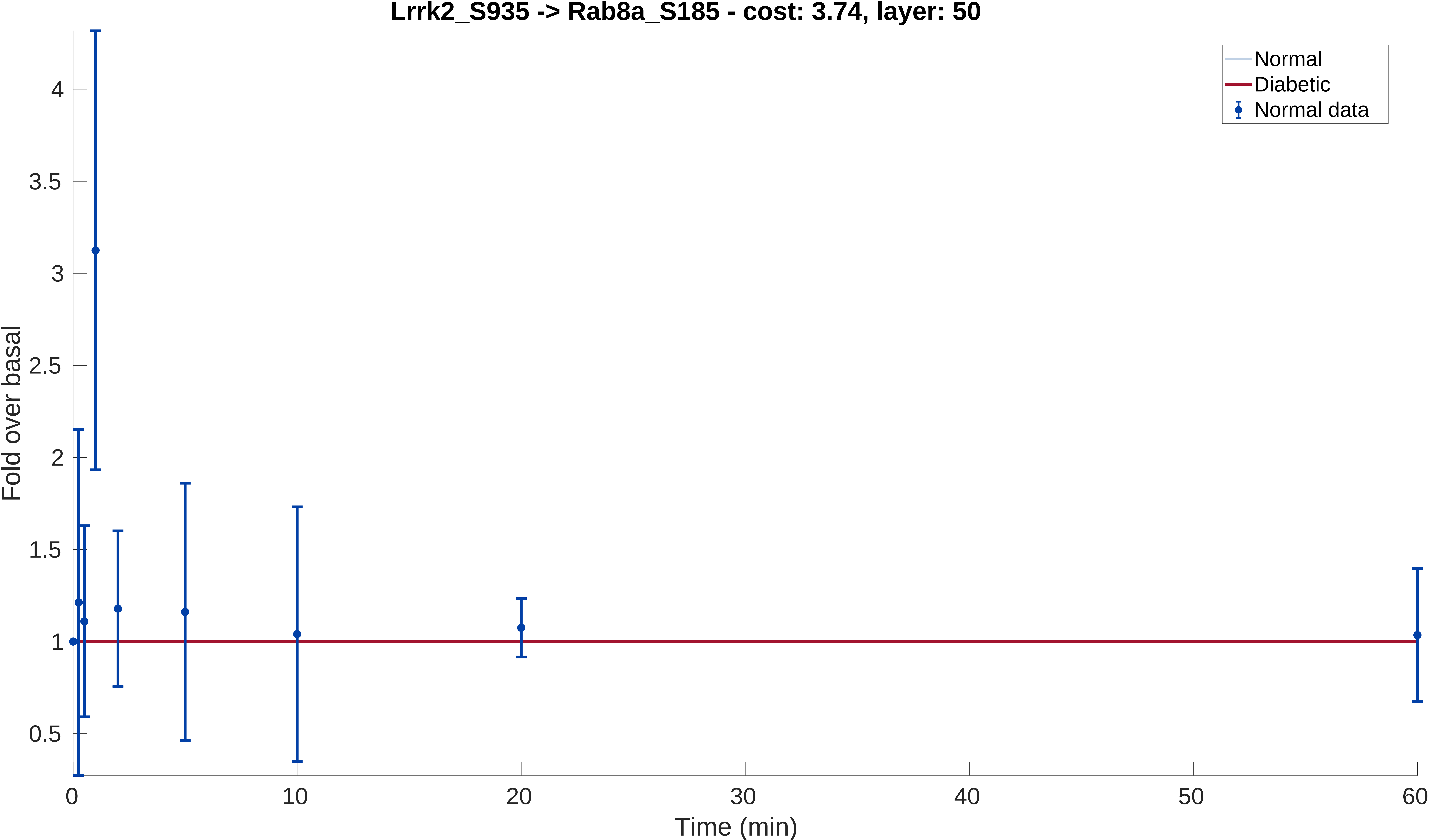
Lrrk2_S935 -> Rab8a_S185 - cost: 3.74, layer: 50
Normal
Diabetic
Normal data
Fold over basal
0.5
1
1.5
2
2.5
3
3.5
4
0
10
20
30
40
50
60
Time (min)

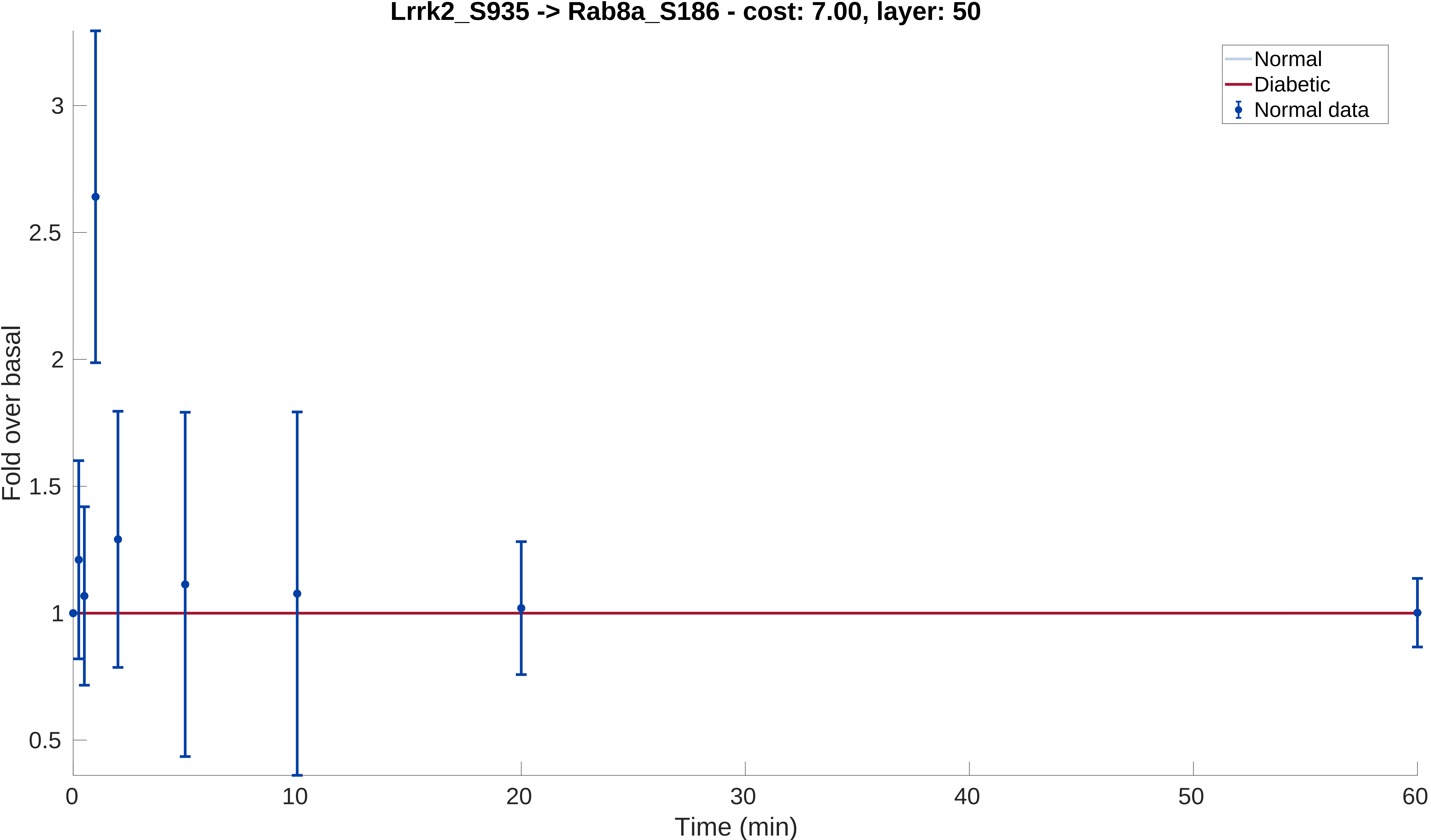
Lrrk2_S935 -> Rab8a_S186 - cost: 7.00, layer: 50
Normal
Diabetic
Normal data
Fold over basal
Time (min)
0.5
1
1.5
2
2.5
3
0
10
20
30
40
50
60

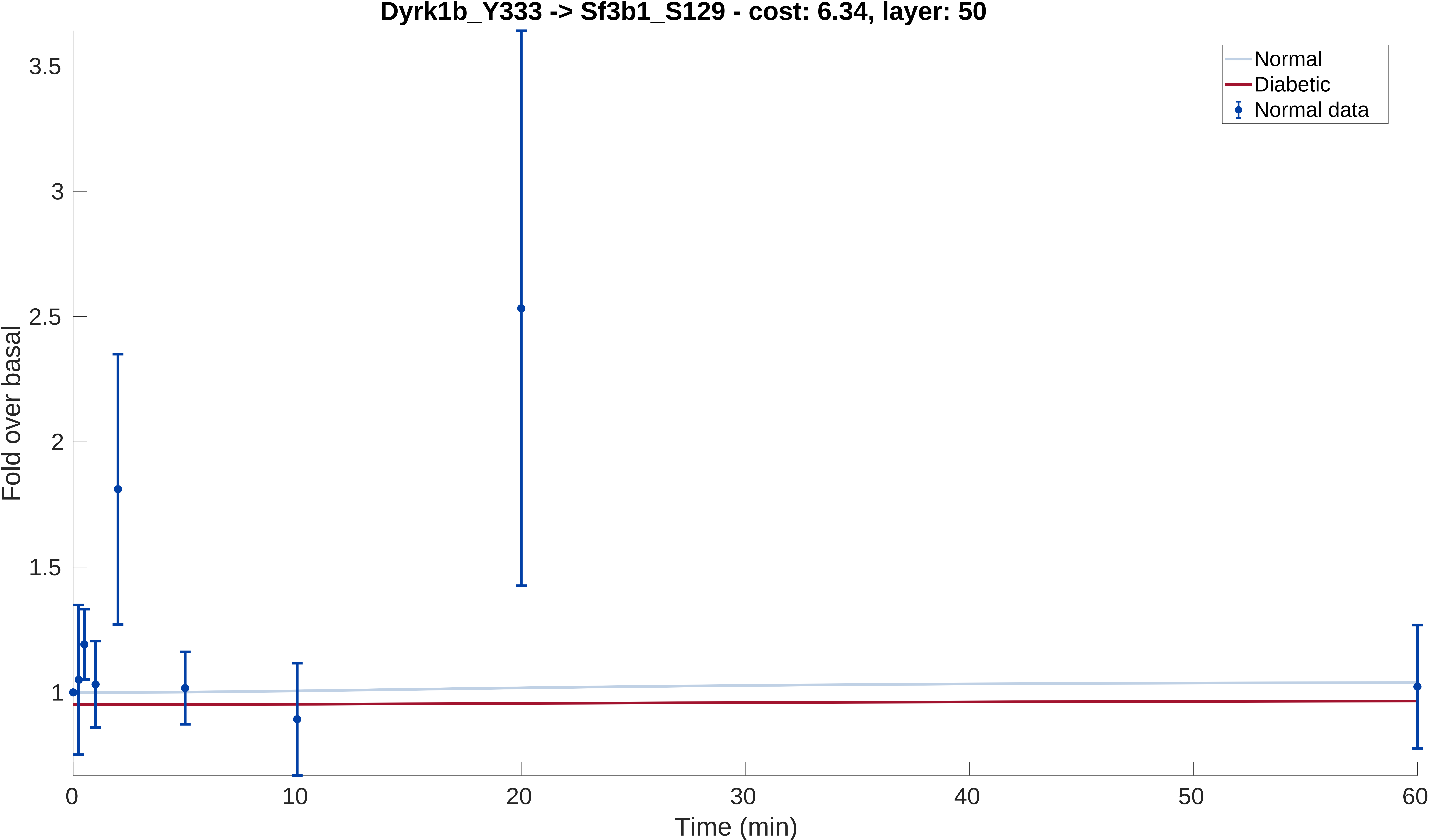
Dyrk1b_Y333 -> Sf3b1_S129 - cost: 6.34, layer: 50
Normal
Diabetic
Normal data
Fold over basal
3.5
3
2.5
2
1.5
1
0
10
20
30
40
50
60
Time (min)

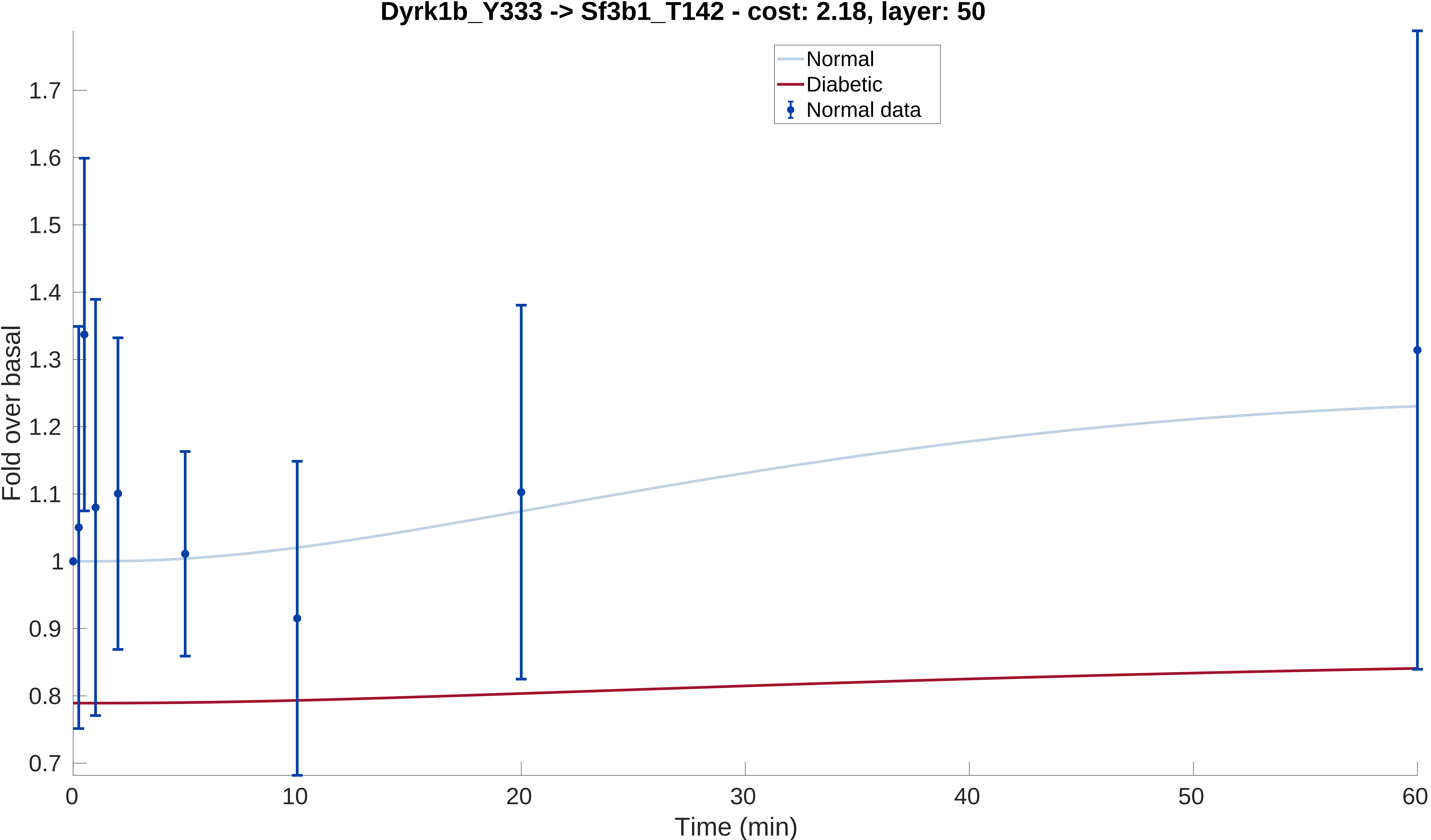
Dyrk1b_Y333 -> Sf3b1_T142 - cost: 2.18, layer: 50
Normal
Diabetic
Normal data
Fold over basal
1.7
1.6
1.5
1.4
1.3
1.2
1.1
1
0.9
0.8
0.7
0
10
20
30
40
50
60
Time (min)

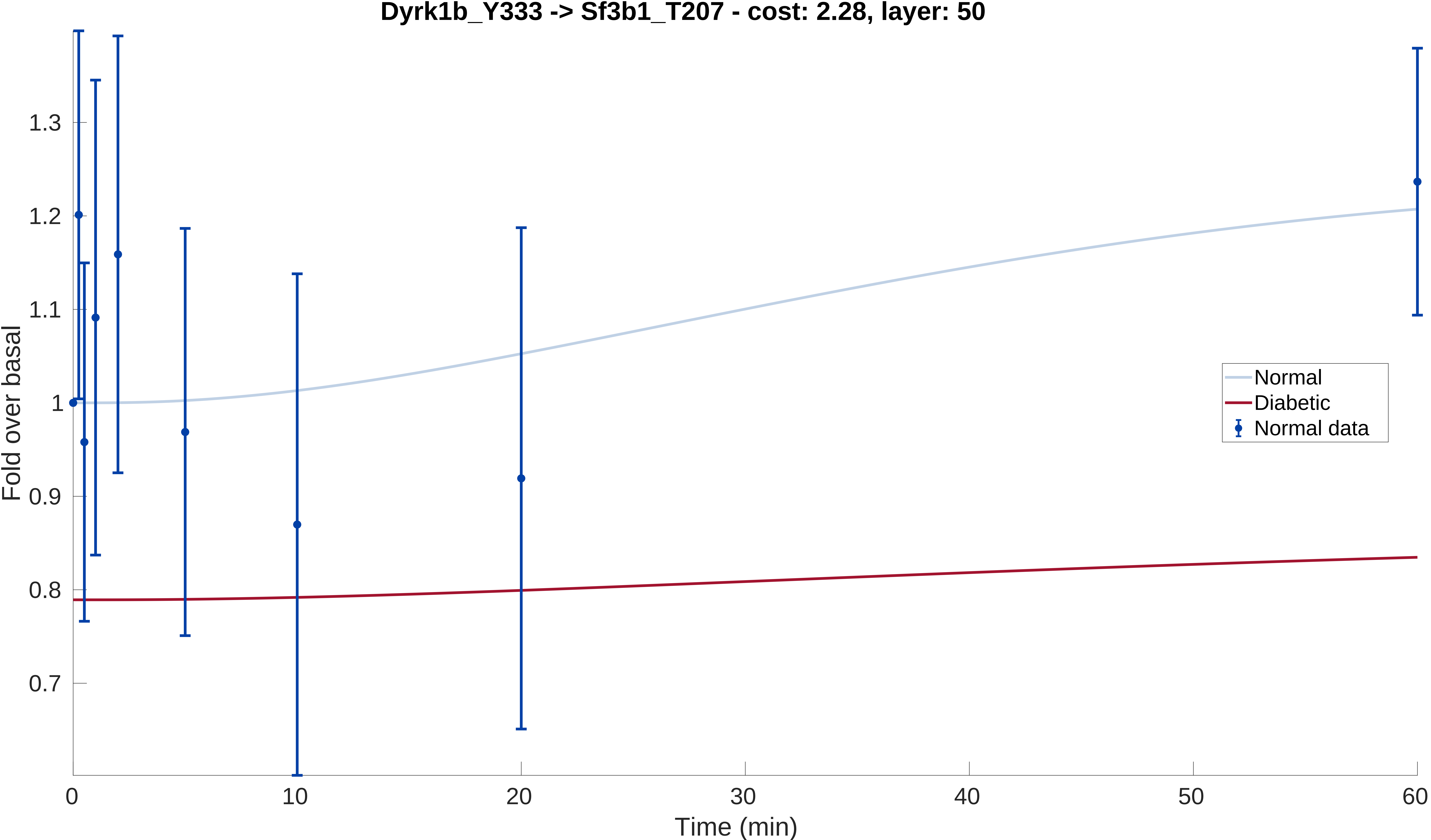

Dyrk1b_Y333 -> Sf3b1_T207 - cost: 2.28, layer: 50
Fold over basal
1.3
1.2
1.1
1
0.9
0.8
0.7
0
10
20
30
40
50
60
Time (min)
Normal
Diabetic
Normal data

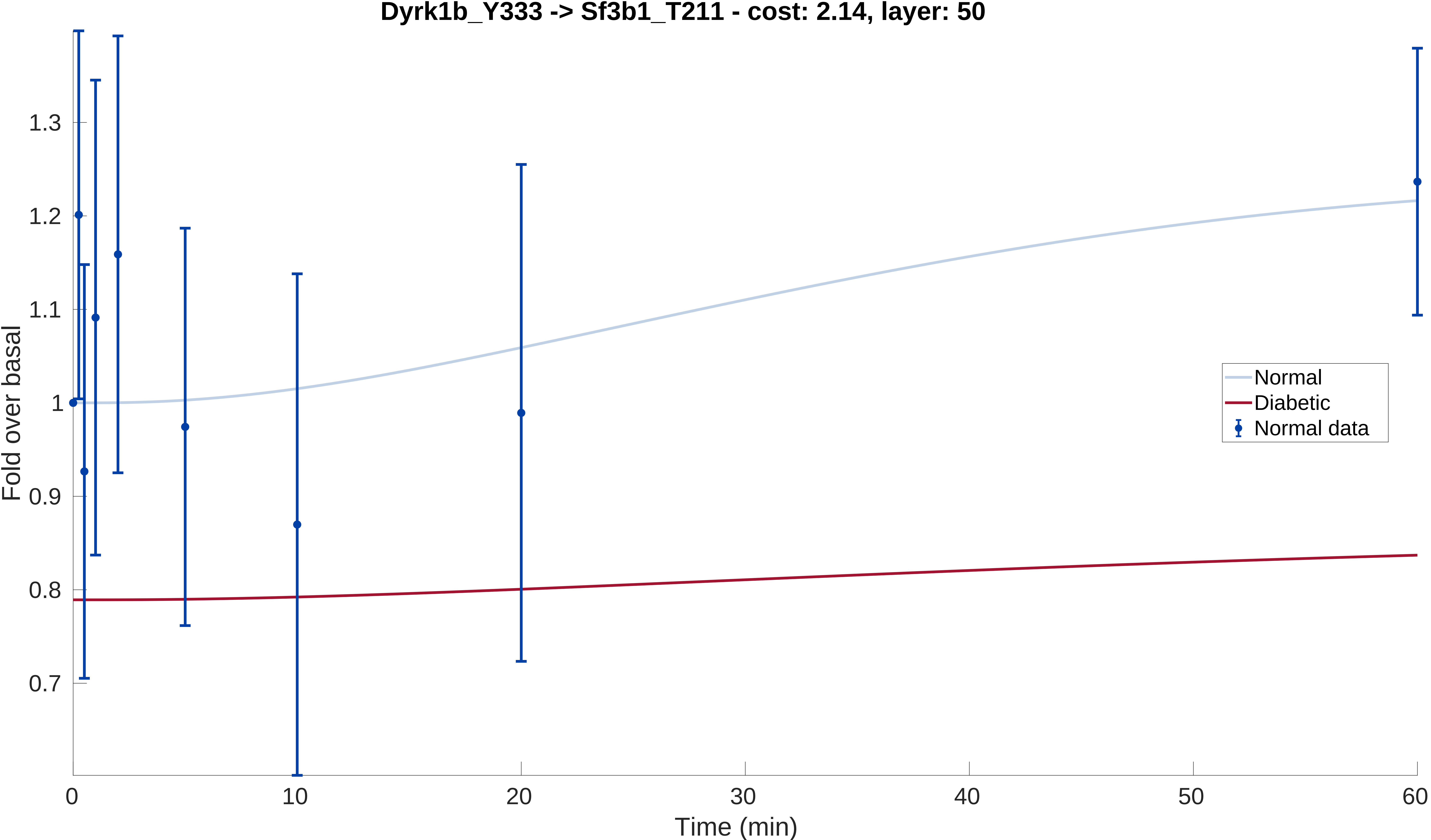

Dyrk1b_Y333 -> Sf3b1_T211 - cost: 2.14, layer: 50
Fold over basal
Time (min)
Normal
Diabetic
Normal data
1.3
1.2
1.1
1
0.9
0.8
0.7
0
10
20
30
40
50
60

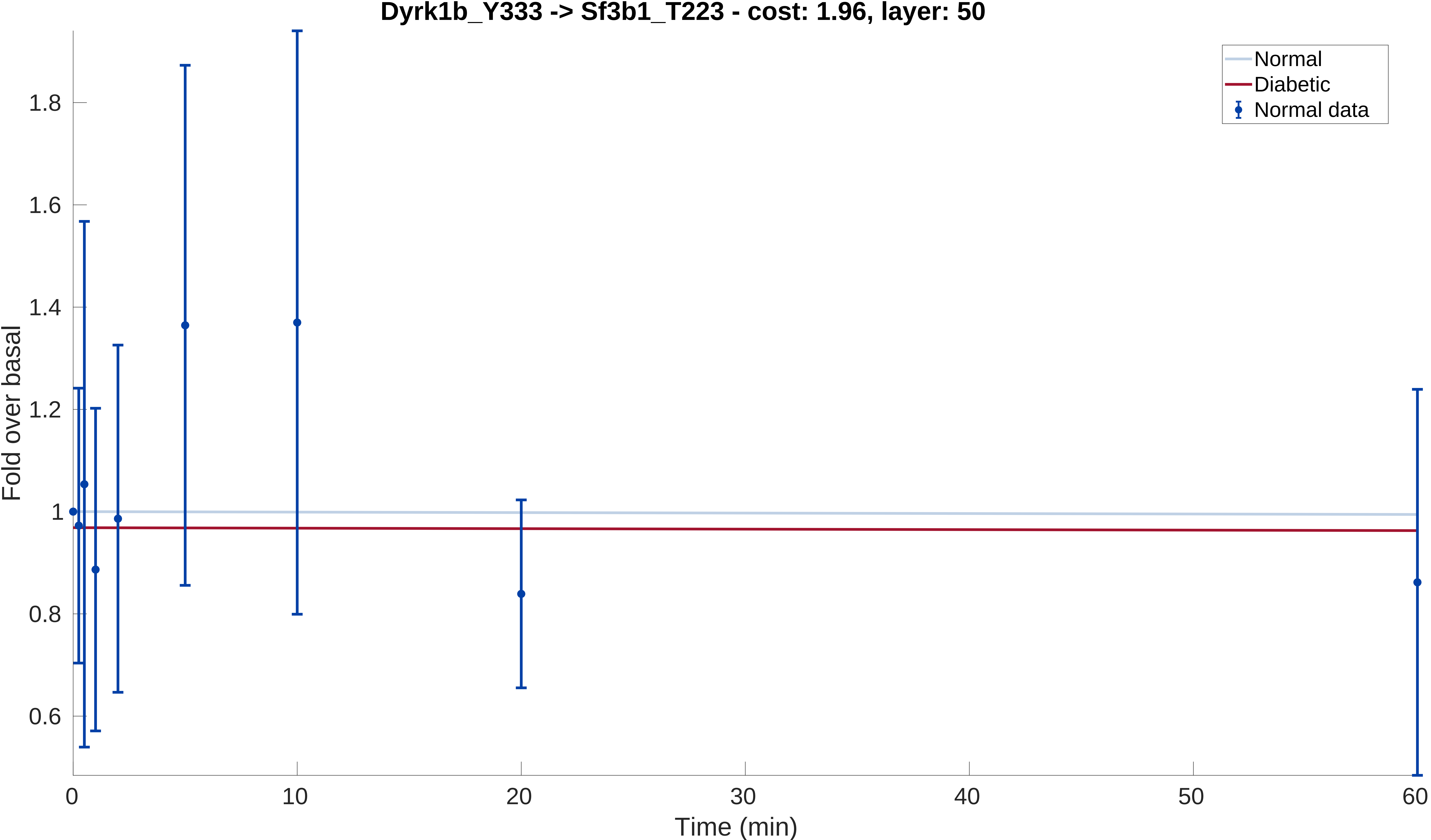

Dyrk1b_Y333 -> Sf3b1_T223 - cost: 1.96, layer: 50
Normal
Diabetic
Normal data
Fold over basal
Time (min)
0
10
20
30
40
50
60
0.6
0.8
1
1.2
1.4
1.6
1.8

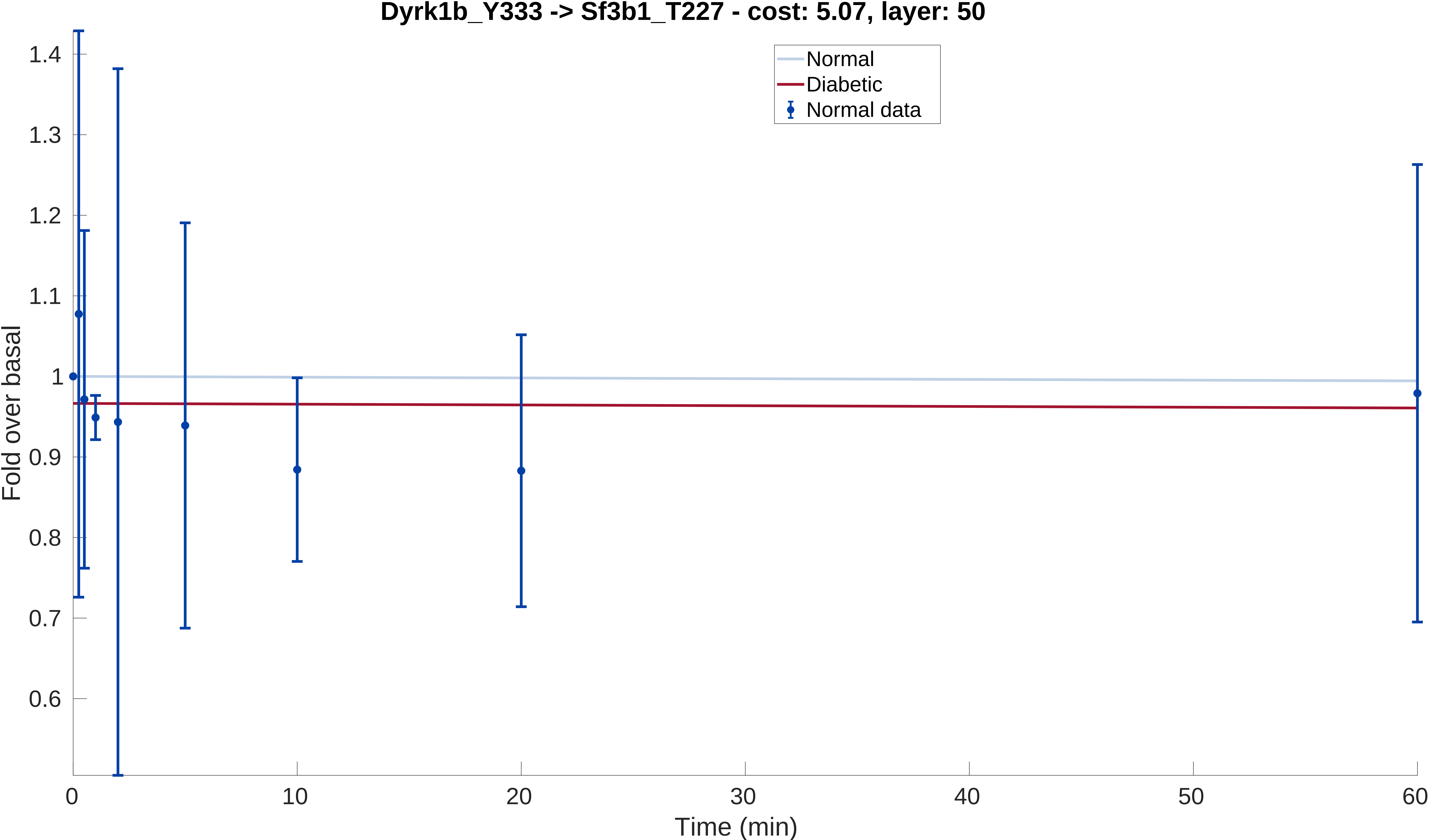

Dyrk1b_Y333 -> Sf3b1_T227 - cost: 5.07, layer: 50
Normal
Diabetic
Normal data
Fold over basal
1.4
1.3
1.2
1.1
1
0.9
0.8
0.7
0.6
0
10
20
30
40
50
60
Time (min)

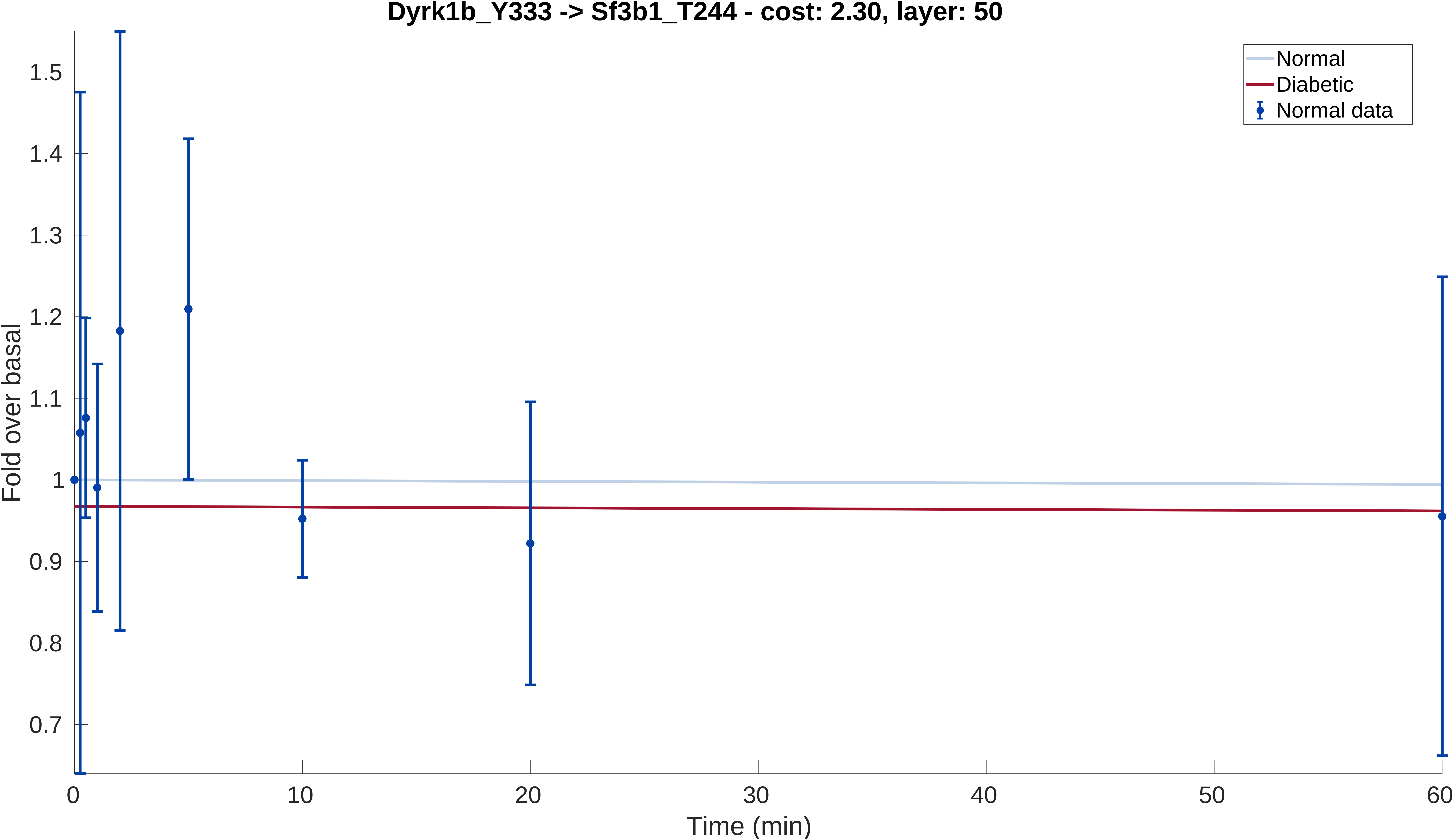

Dyrk1b_Y333 -> Sf3b1_T244 - cost: 2.30, layer: 50
Normal
Diabetic
Normal data
Fold over basal
Time (min)
1.5
1.4
1.3
1.2
1.1
1
0.9
0.8
0.7
0
10
20
30
40
50
60

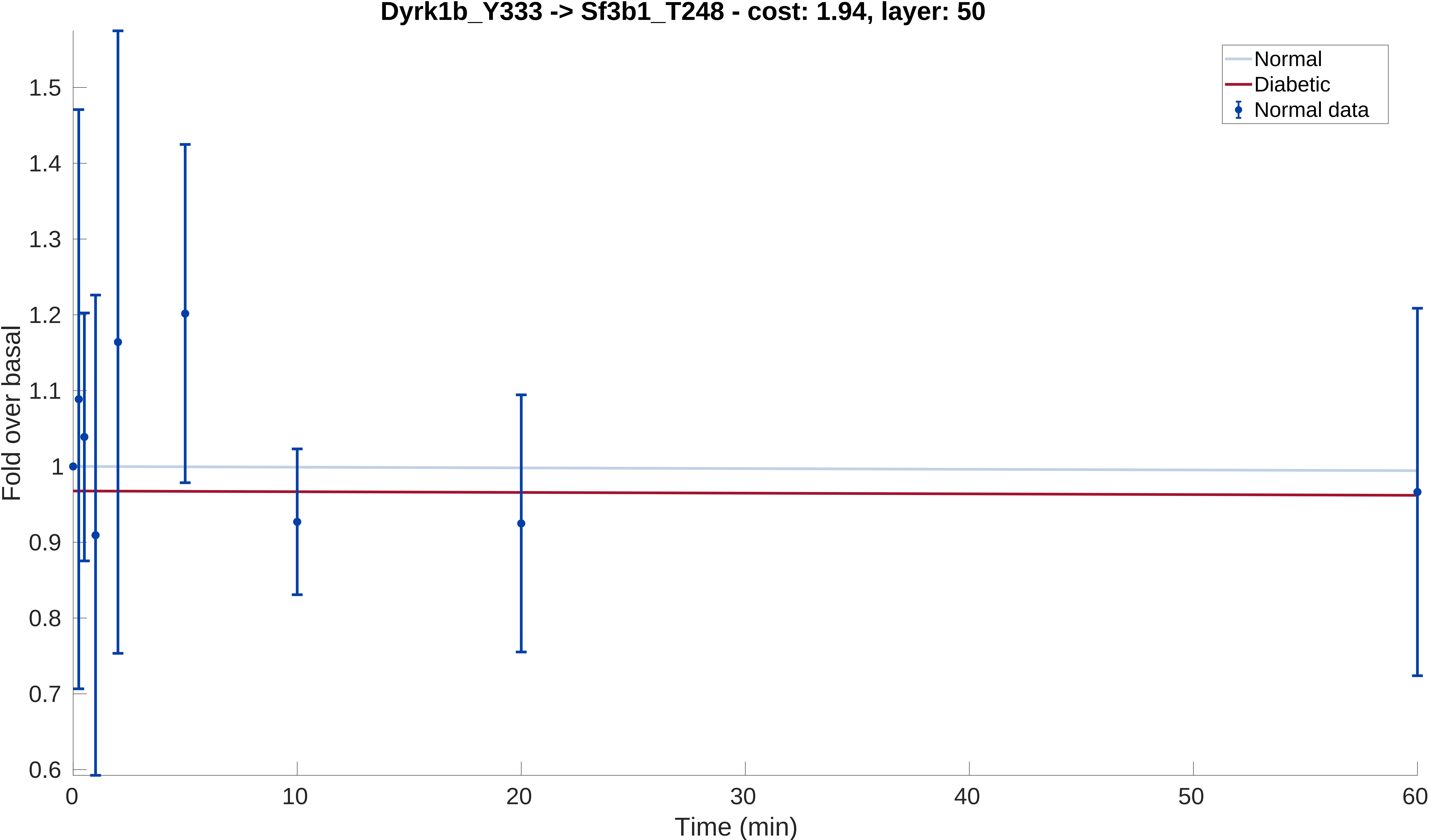
Dyrk1b_Y333 -> Sf3b1_T248 - cost: 1.94, layer: 50
Normal
Diabetic
Normal data
Fold over basal
Time (min)
1.5
1.4
1.3
1.2
1.1
1
0.9
0.8
0.7
0.6
0
10
20
30
40
50
60

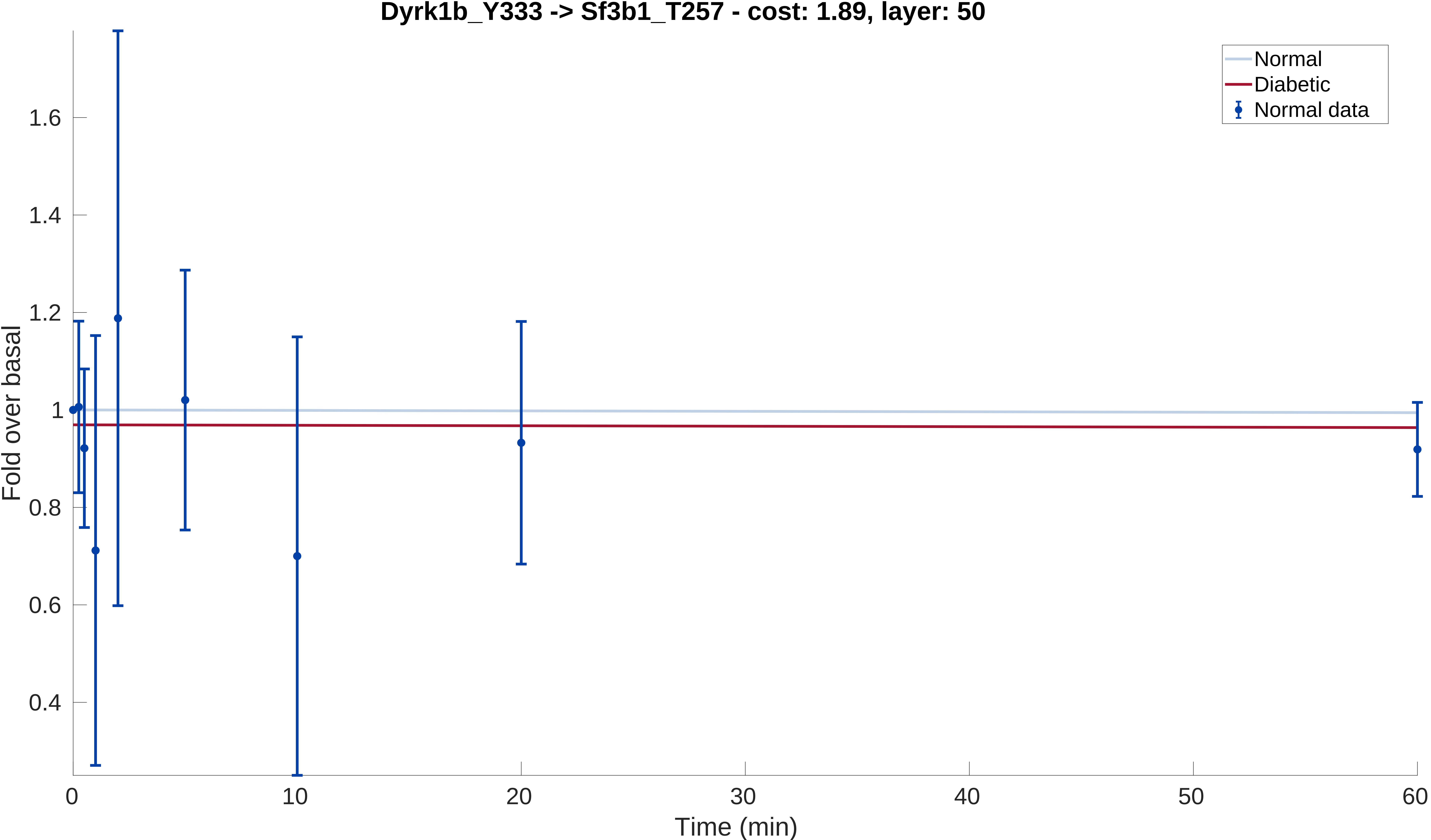

Dyrk1b_Y333 -> Sf3b1_T257 - cost: 1.89, layer: 50
Normal
Diabetic
Normal data
Fold over basal
1.6
1.4
1.2
1
0.8
0.6
0.4
0
10
20
30
40
50
60
Time (min)

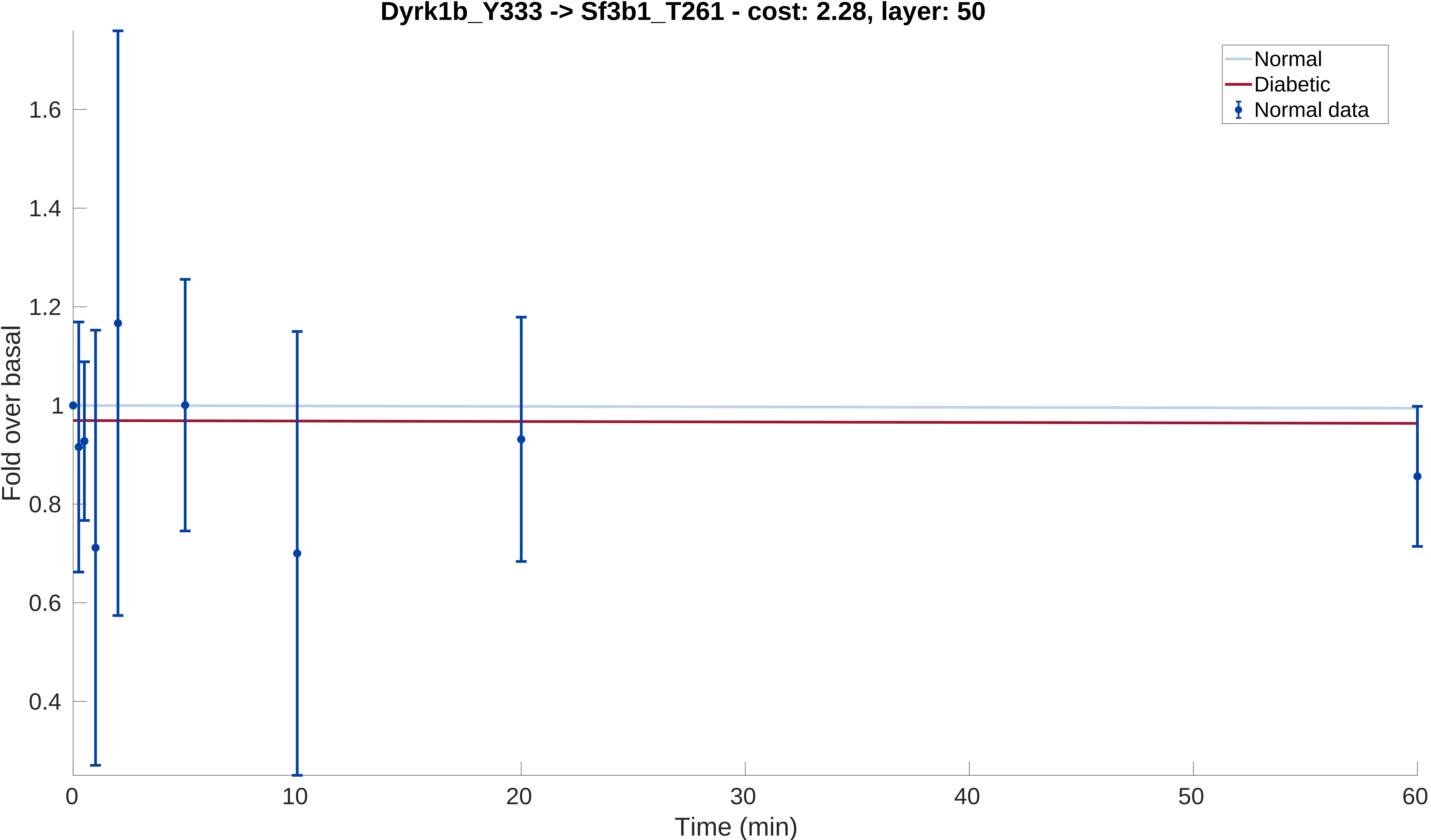

Dyrk1b_Y333 -> Sf3b1_T261 - cost: 2.28, layer: 50
Normal
Diabetic
Normal data
Fold over basal
Time (min)
0.4
0.6
0.8
1
1.2
1.4
1.6
0
10
20
30
40
50
60

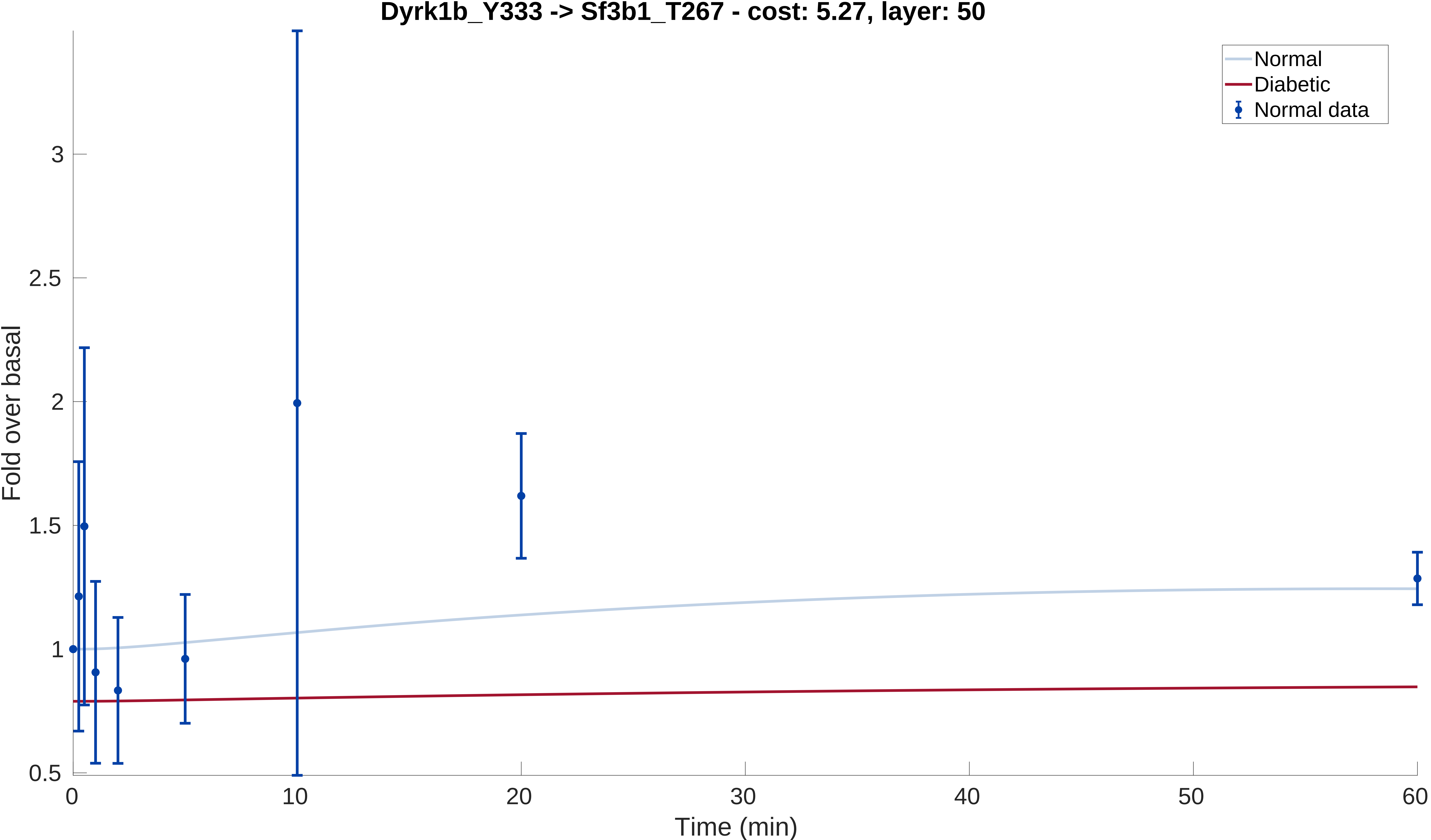

Dyrk1b_Y333 -> Sf3b1_T267 - cost: 5.27, layer: 50
Normal
Diabetic
Normal data
Fold over basal
0.5
1
1.5
2
2.5
3
0
10
20
30
40
50
60
Time (min)

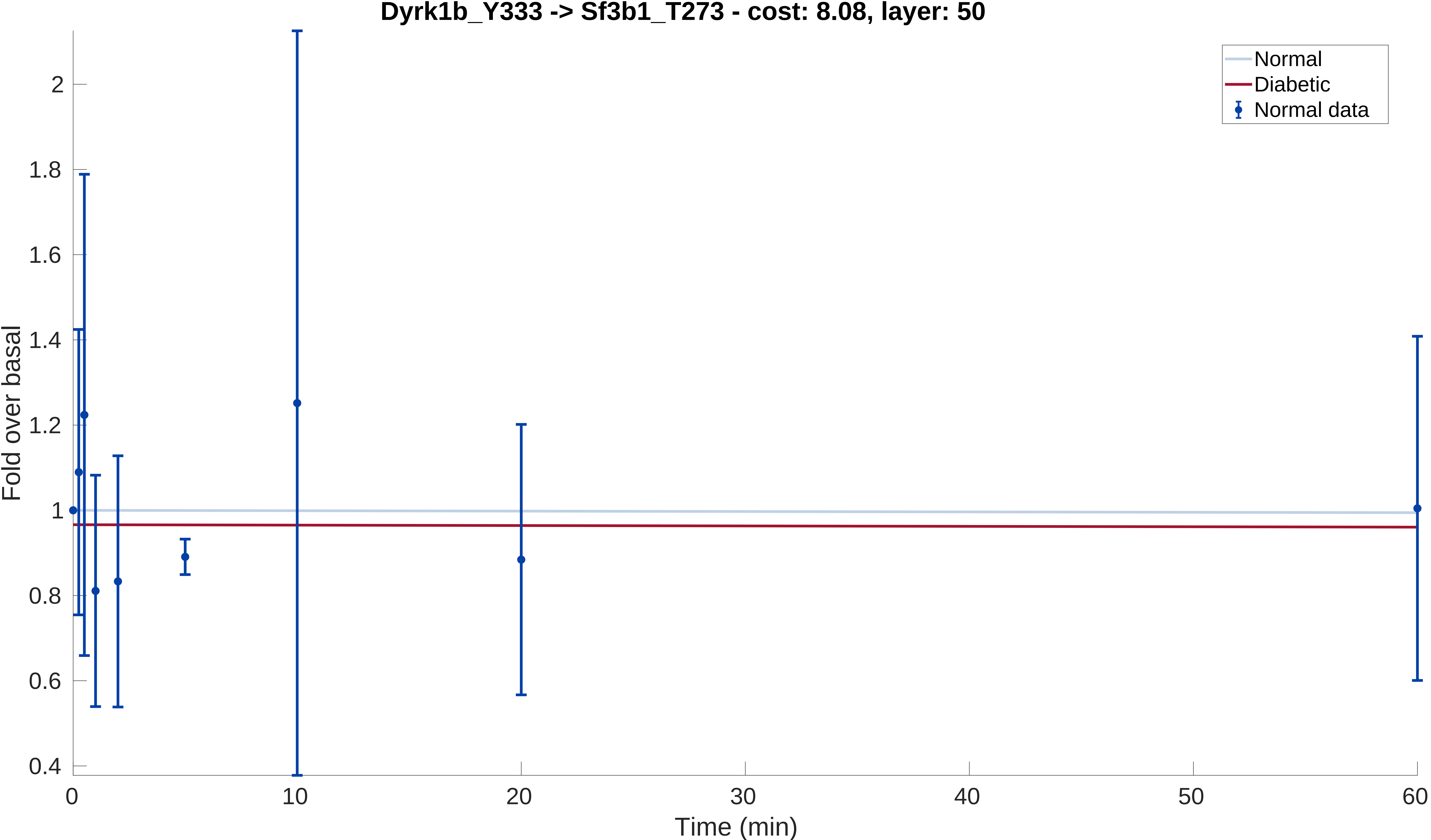

Dyrk1b_Y333 -> Sf3b1_T273 - cost: 8.08, layer: 50
Normal
Diabetic
Normal data
Fold over basal
Time (min)
0
10
20
30
40
50
60
0.4
0.6
0.8
1
1.2
1.4
1.6
1.8
2

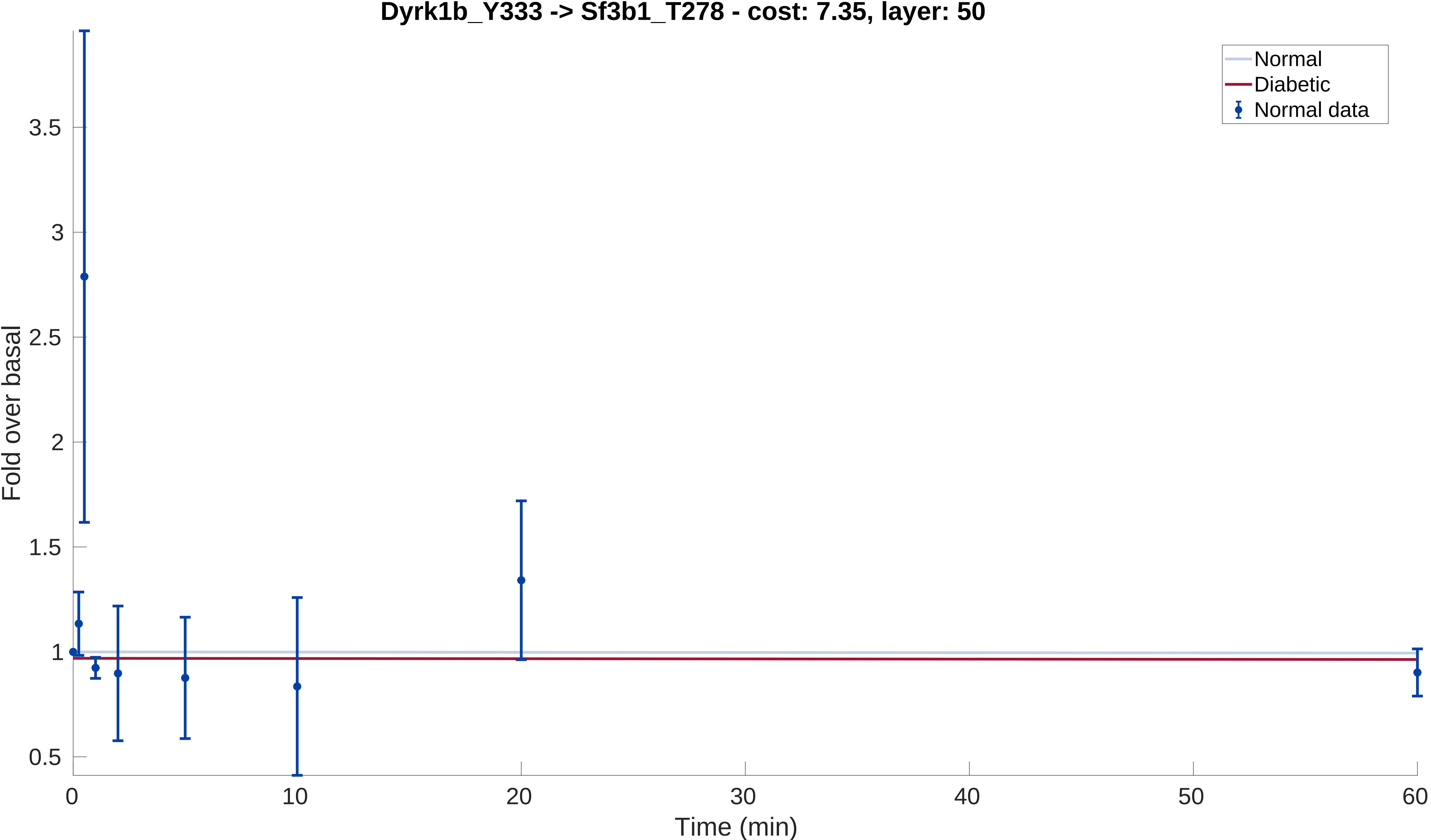
Dyrk1b_Y333 -> Sf3b1_T278 - cost: 7.35, layer: 50
Normal
Diabetic
Normal data
Fold over basal
3.5
3
2.5
2
1.5
1
0.5
0
10
20
30
40
50
60
Time (min)

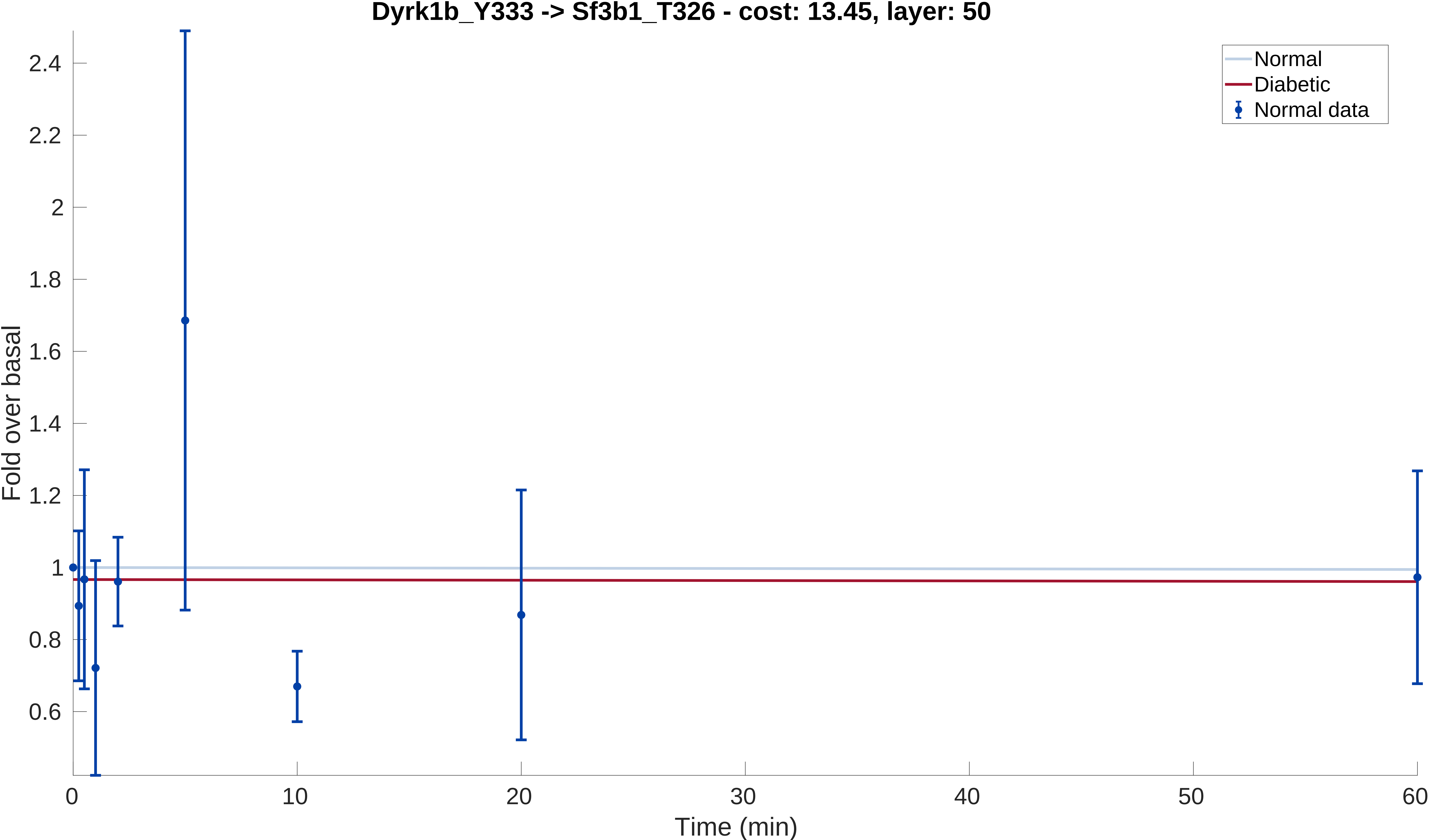
Dyrk1b_Y333 -> Sf3b1_T326 - cost: 13.45, layer: 50
Normal
Diabetic
Normal data
Fold over basal
2.4
2.2
2
1.8
1.6
1.4
1.2
1
0.8
0.6
0
10
20
30
40
50
60
Time (min)

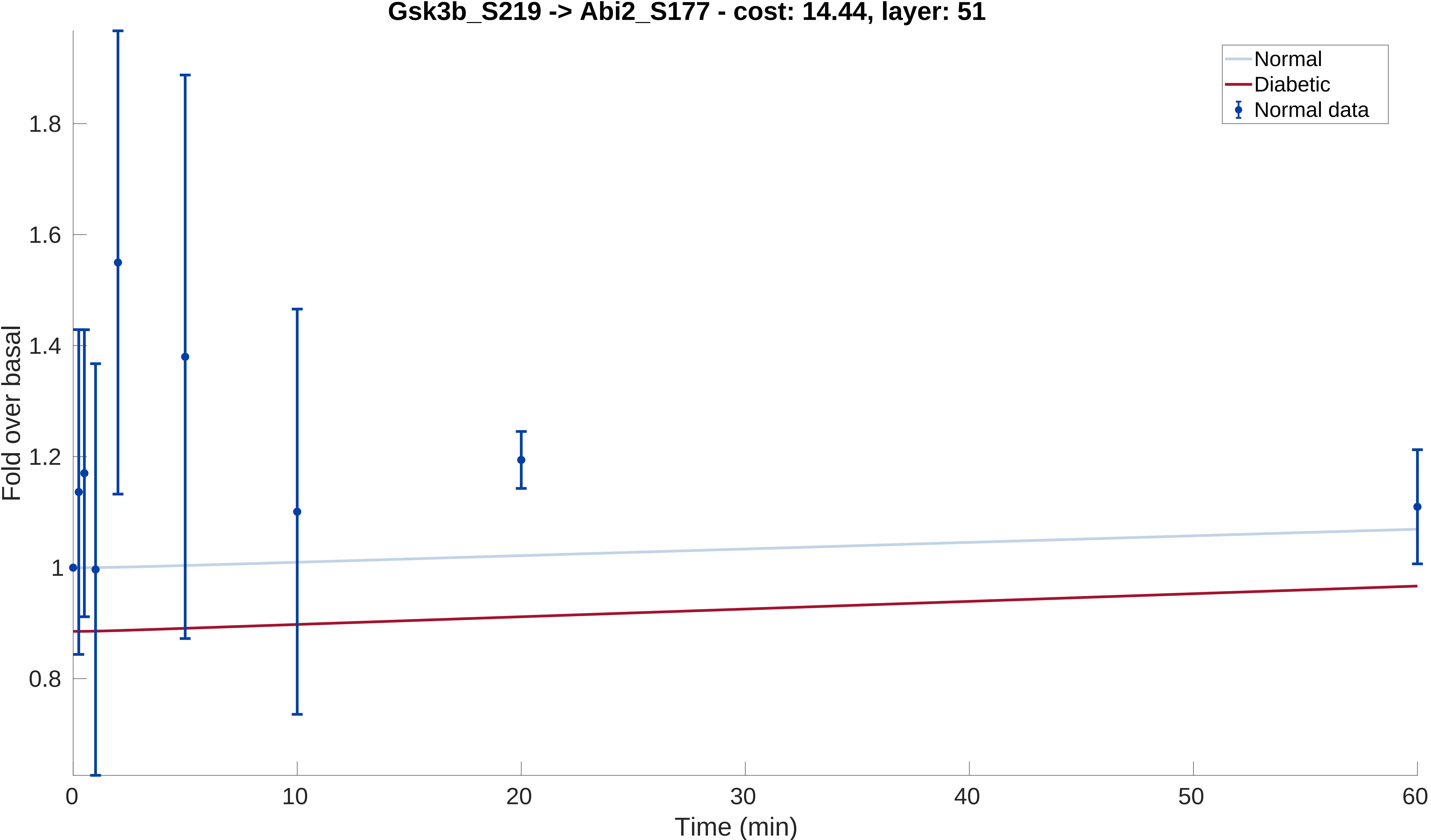

Gsk3b_S219 -> Abi2_S177 - cost: 14.44, layer: 51
Normal
Diabetic
Normal data
Fold over basal
1.8
1.6
1.4
1.2
1
0.8
0
10
20
30
40
50
60
Time (min)

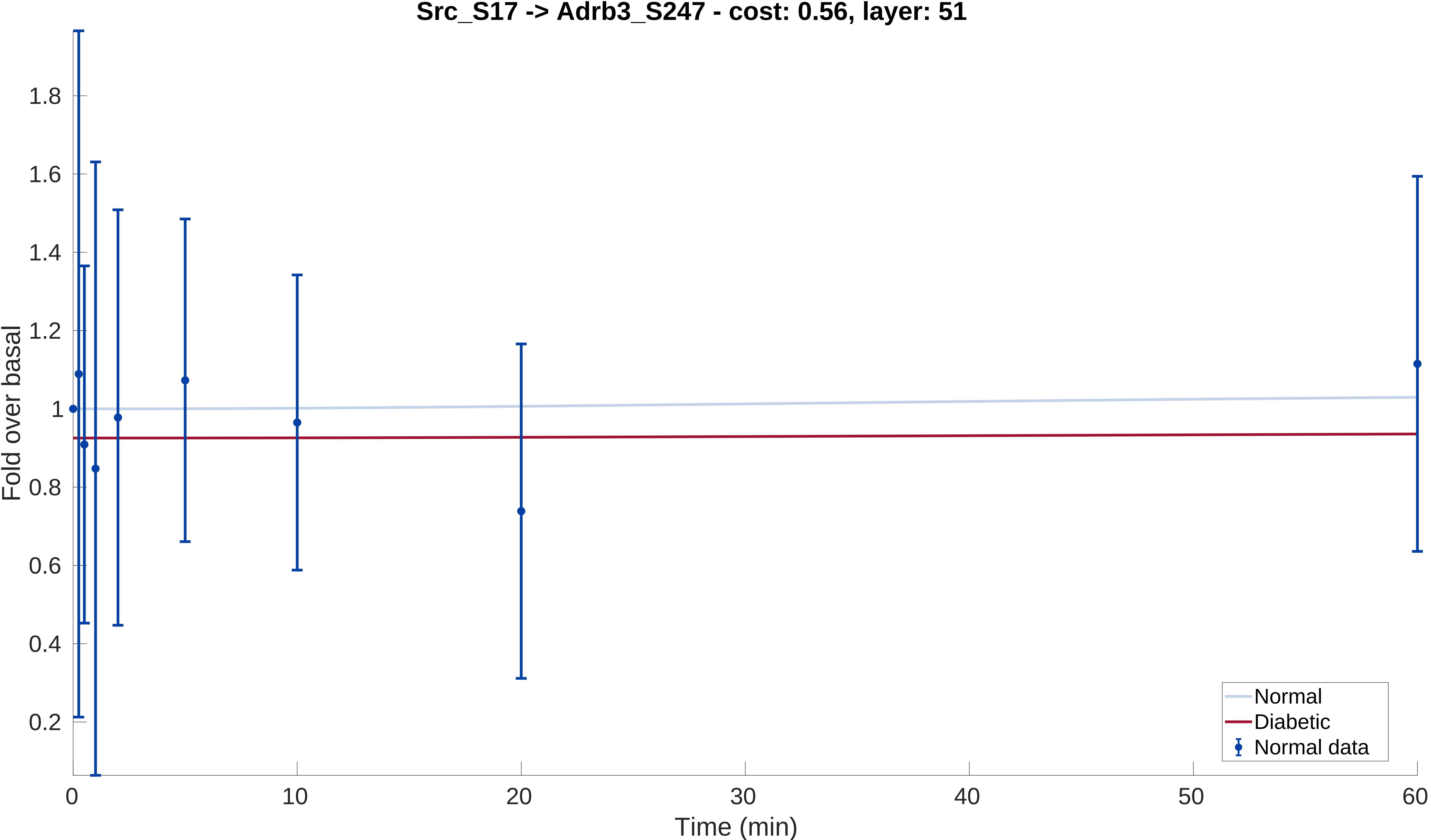

Src_S17 -> Adrb3_S247 - cost: 0.56, layer: 51
Fold over basal
1.8
1.6
1.4
1.2
1
0.8
0.6
0.4
0.2
0
10
20
30
40
50
60
Time (min)
Normal
Diabetic
Normal data

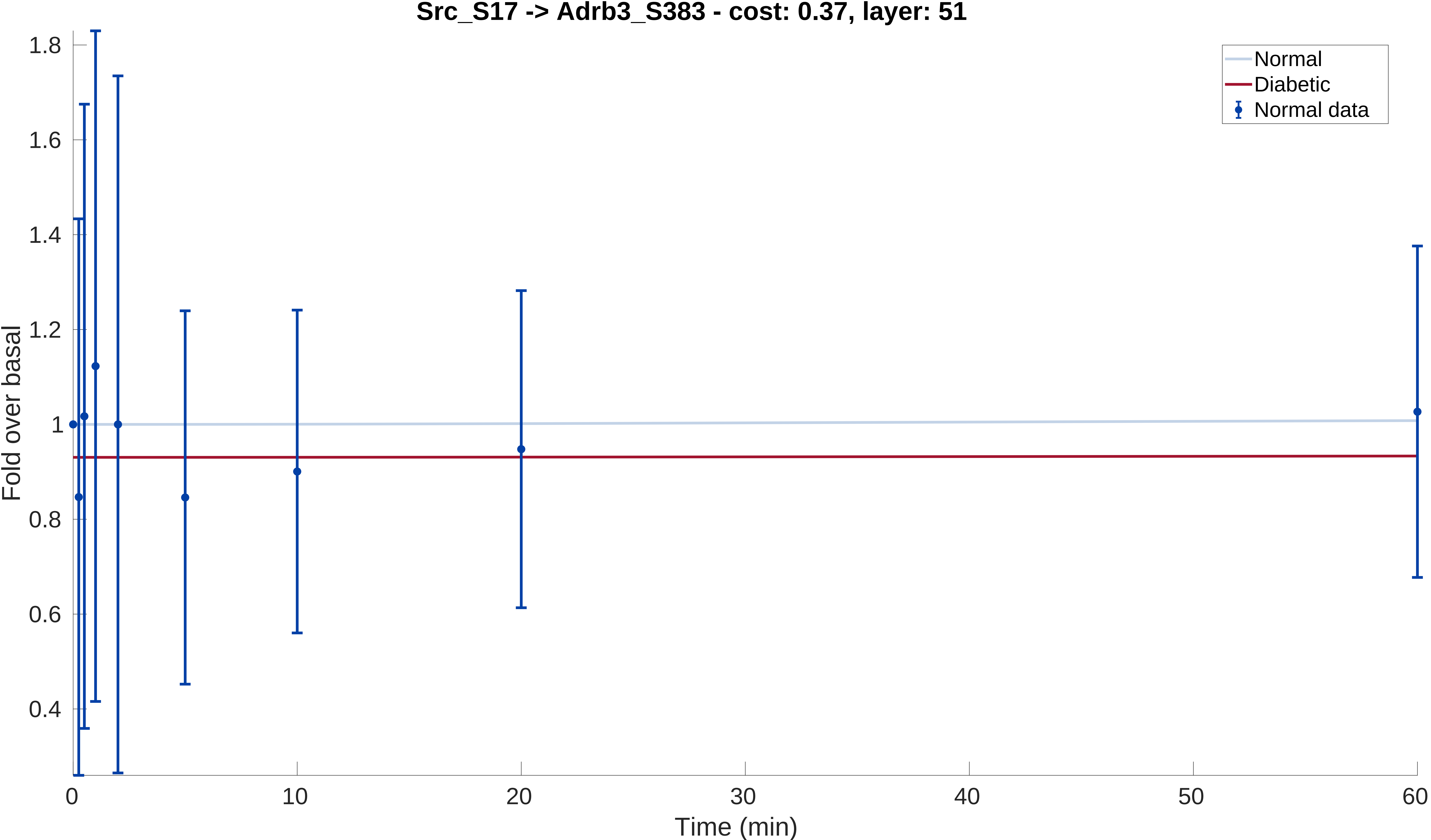

Src_S17 -> Adrb3_S383 - cost: 0.37, layer: 51
Normal
Diabetic
Normal data
Fold over basal
1.8
1.6
1.4
1.2
1
0.8
0.6
0.4
0
10
20
30
40
50
60
Time (min)

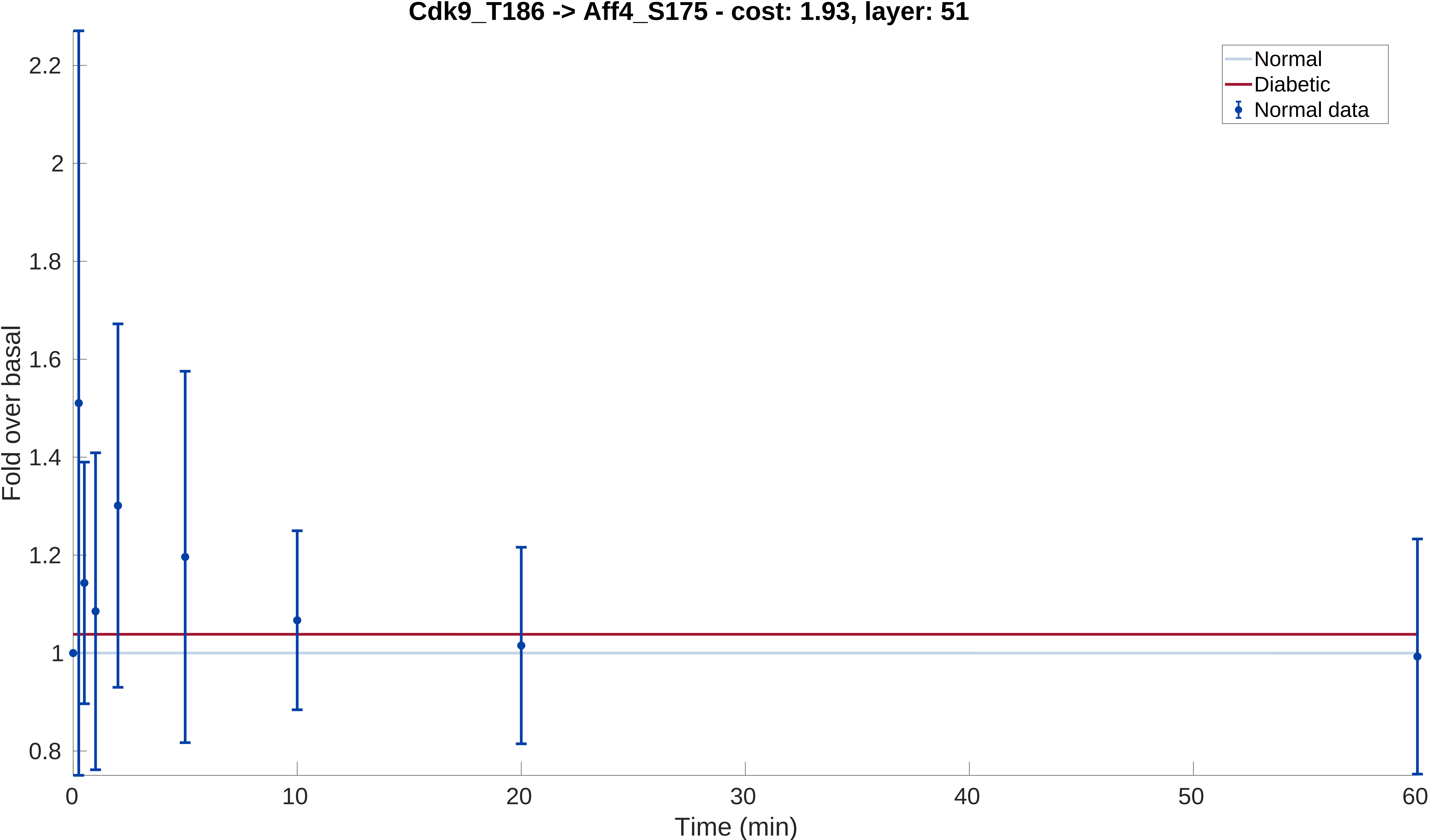

Cdk9_T186 -> Aff4_S175 - cost: 1.93, layer: 51
Normal
Diabetic
Normal data
Fold over basal
2.2
2
1.8
1.6
1.4
1.2
1
0.8
0
10
20
30
40
50
60
Time (min)

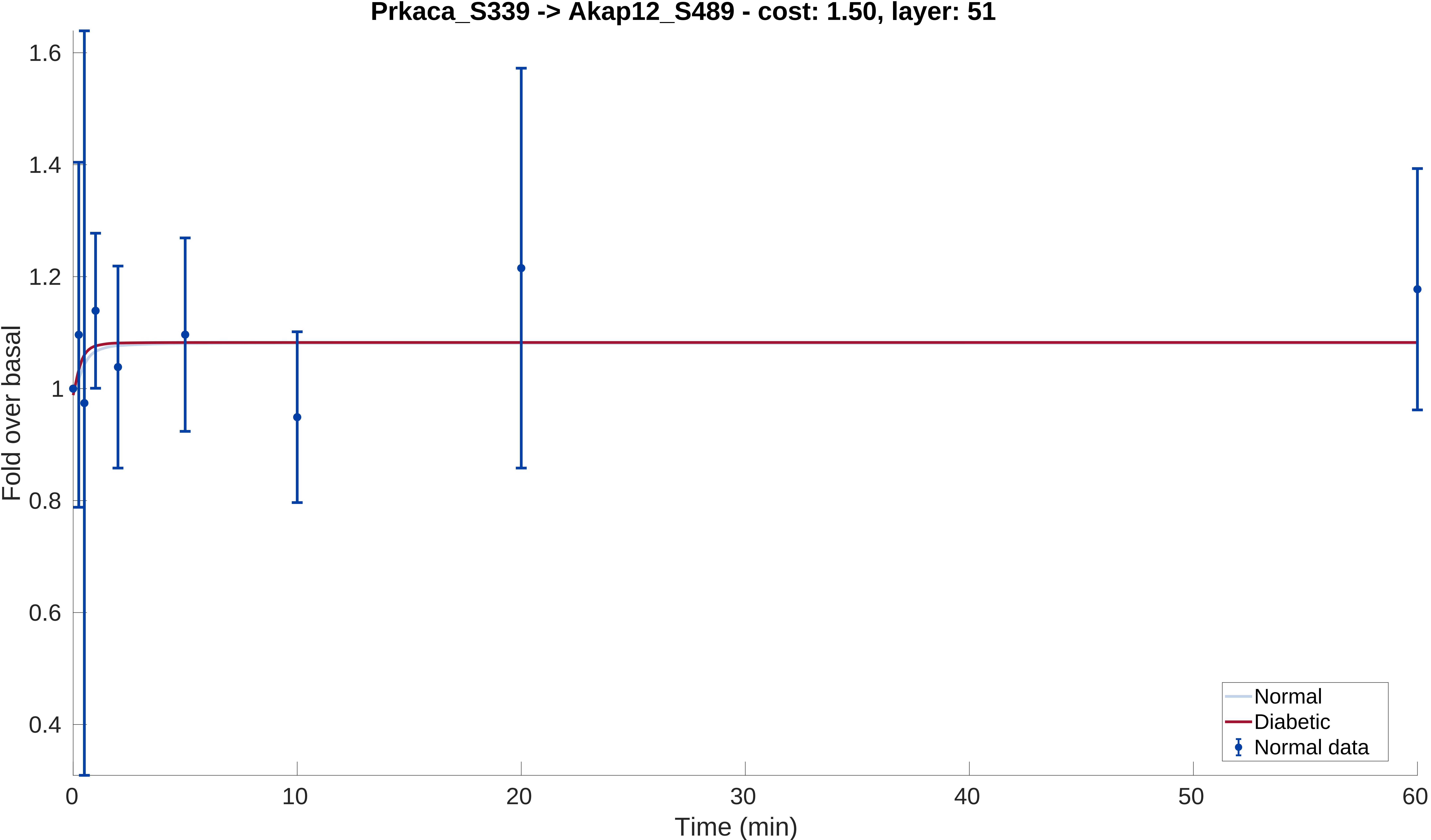
Prkaca_S339 -> Akap12_S489 - cost: 1.50, layer: 51
Fold over basal
1.6
1.4
1.2
1
0.8
0.6
0.4
0
10
20
30
40
50
60
Time (min)
Normal
Diabetic
Normal data

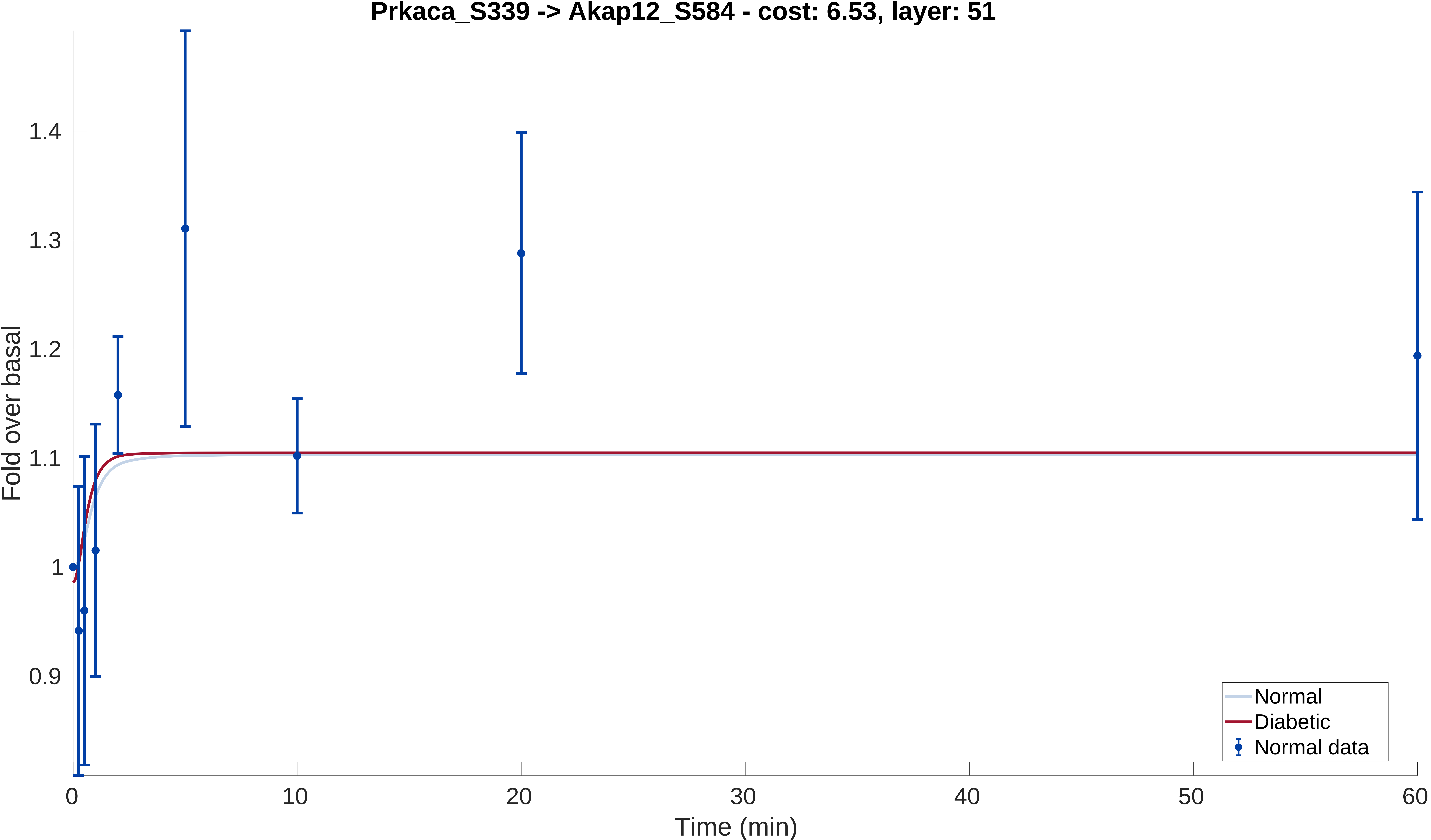
Prkaca_S339 -> Akap12_S584 - cost: 6.53, layer: 51
Fold over basal
1.4
1.3
1.2
1.1
1
0.9
0
10
20
30
40
50
60
Time (min)
Normal
Diabetic
Normal data

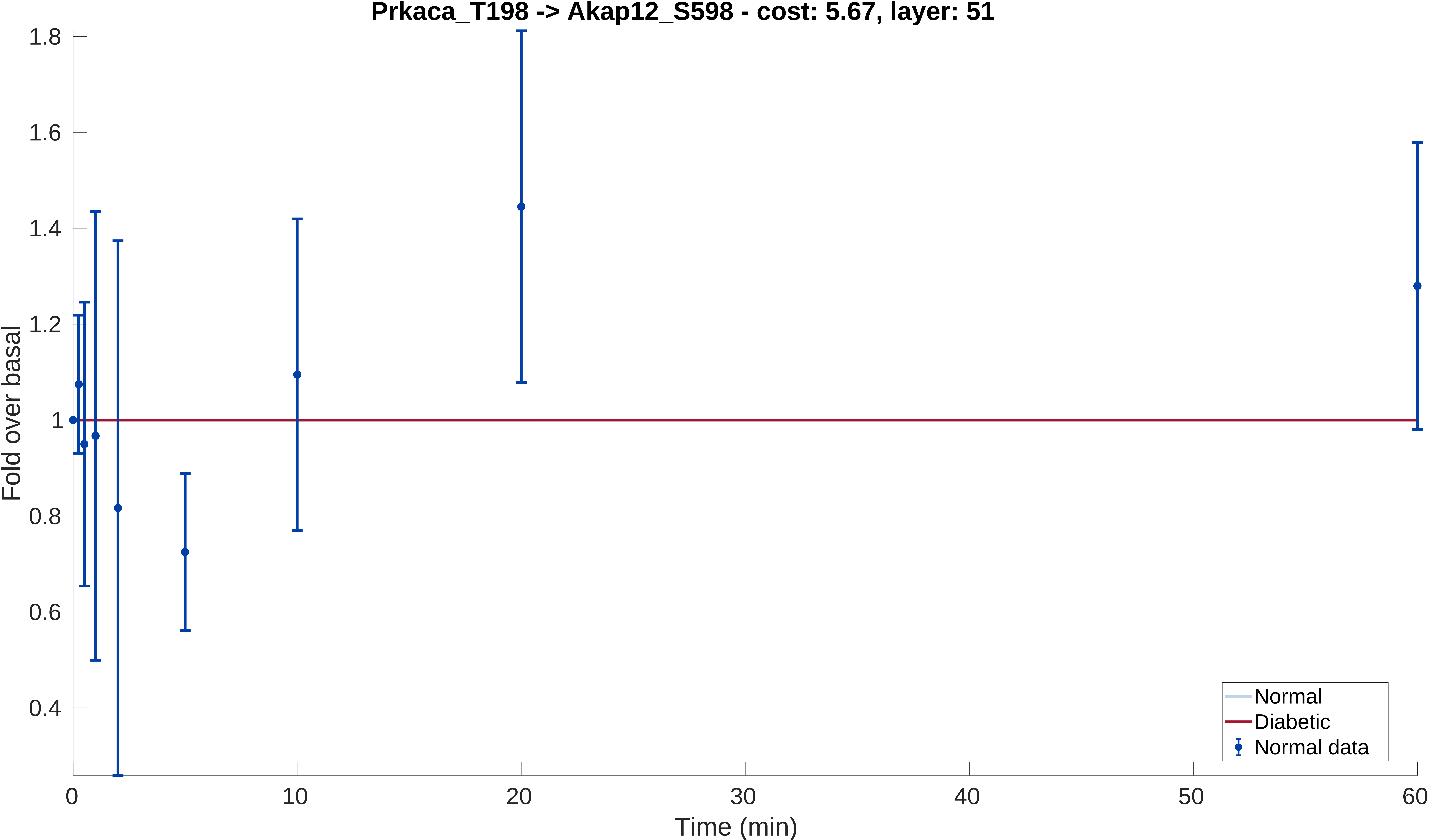

Prkaca_T198 -> Akap12_S598 - cost: 5.67, layer: 51
Fold over basal
Time (min)
Normal
Diabetic
Normal data

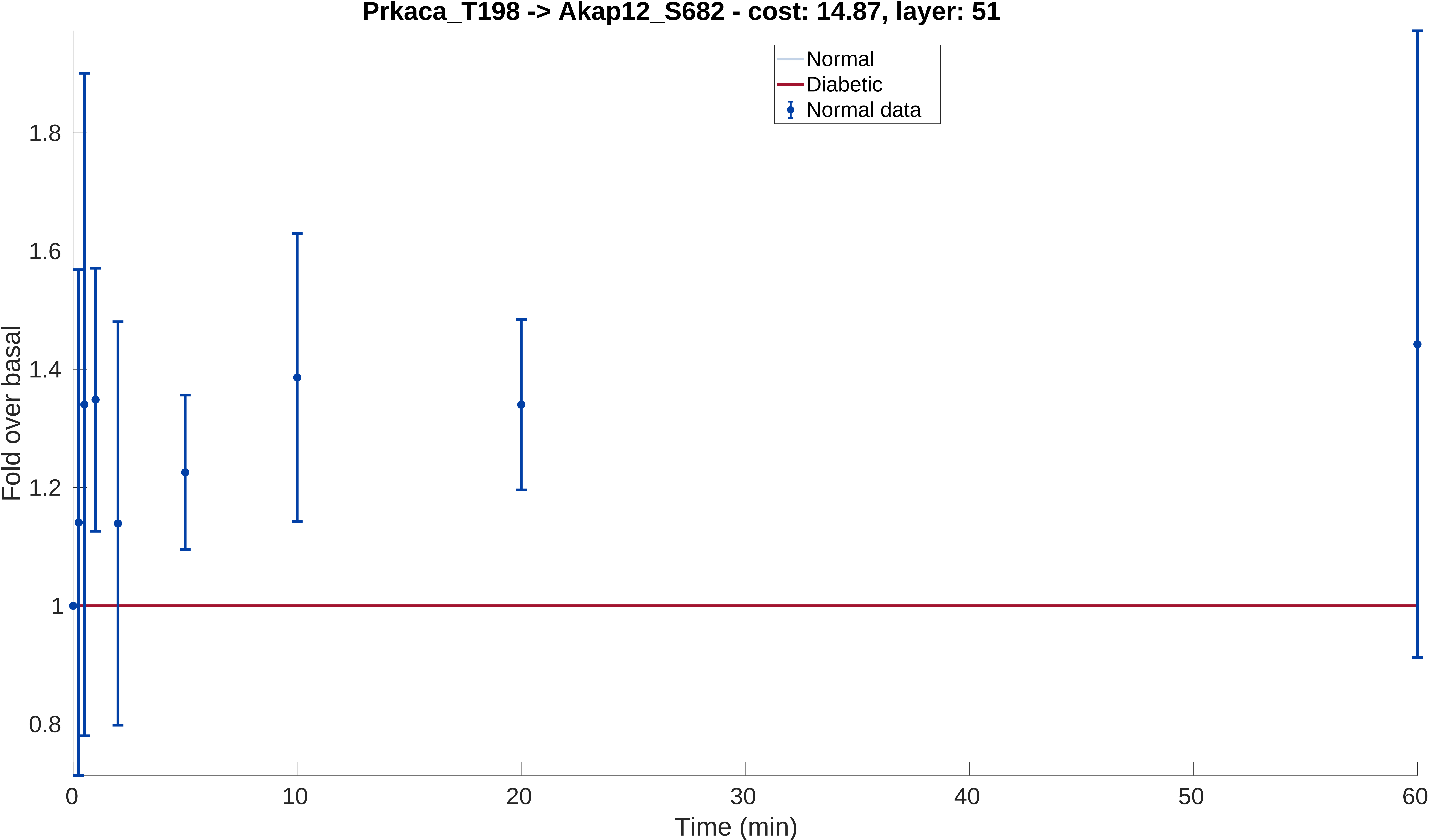

Prkaca_T198 -> Akap12_S682 - cost: 14.87, layer: 51
Normal
Diabetic
Normal data
Fold over basal
1.8
1.6
1.4
1.2
1
0.8
0
10
20
30
40
50
60
Time (min)

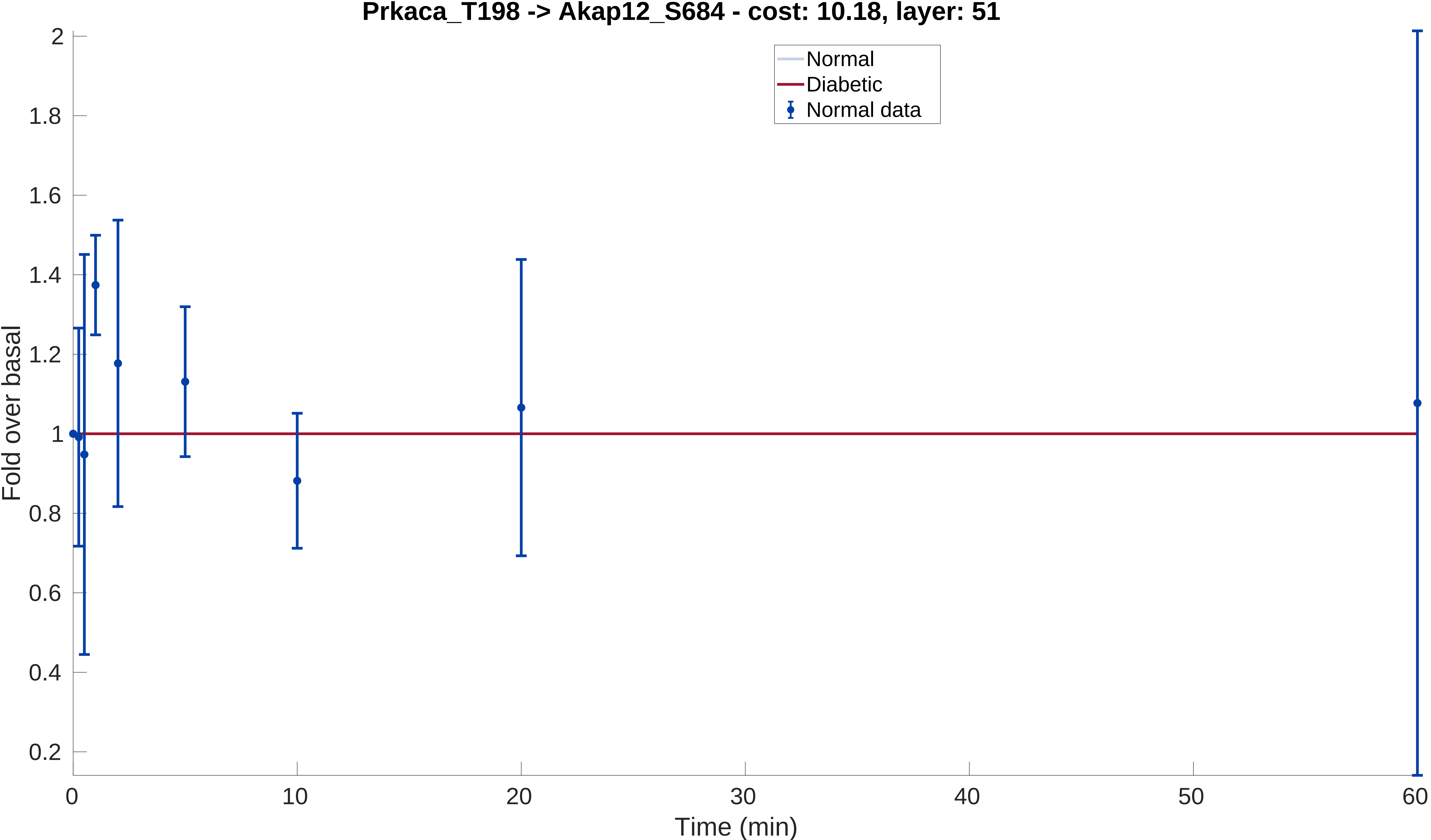

Prrkaca_T198 -> Akap12_S684 - cost: 10.18, layer: 51
Normal
Diabetic
Normal data
Fold over basal
Time (min)
2
1.8
1.6
1.4
1.2
1
0.8
0.6
0.4
0.2
0
10
20
30
40
50
60

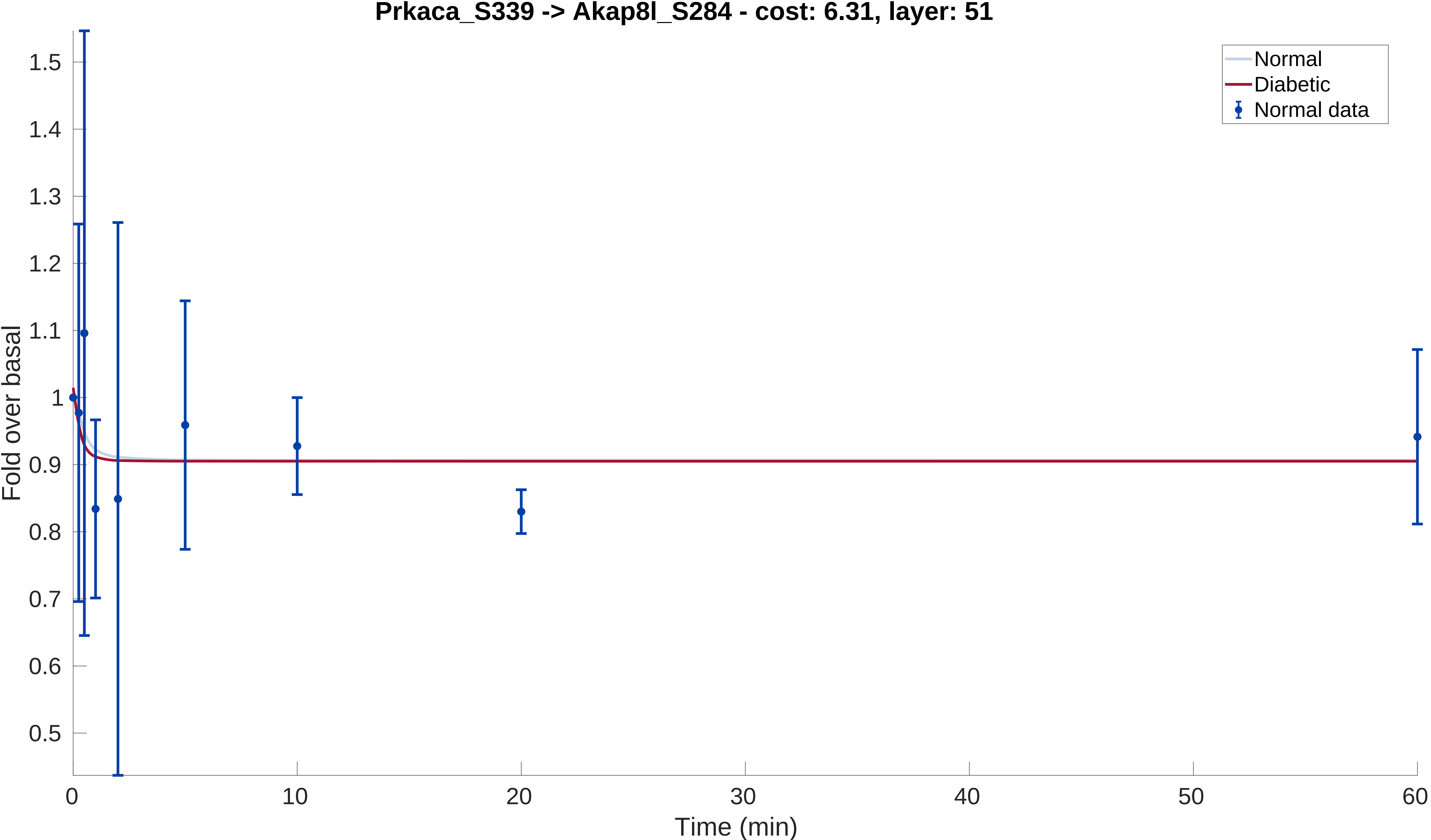
Prkaca_S339 -> Akap8l_S284 - cost: 6.31, layer: 51
Normal
Diabetic
Normal data
Fold over basal
1.5
1.4
1.3
1.2
1.1
1
0.9
0.8
0.7
0.6
0.5
0
10
20
30
40
50
60
Time (min)

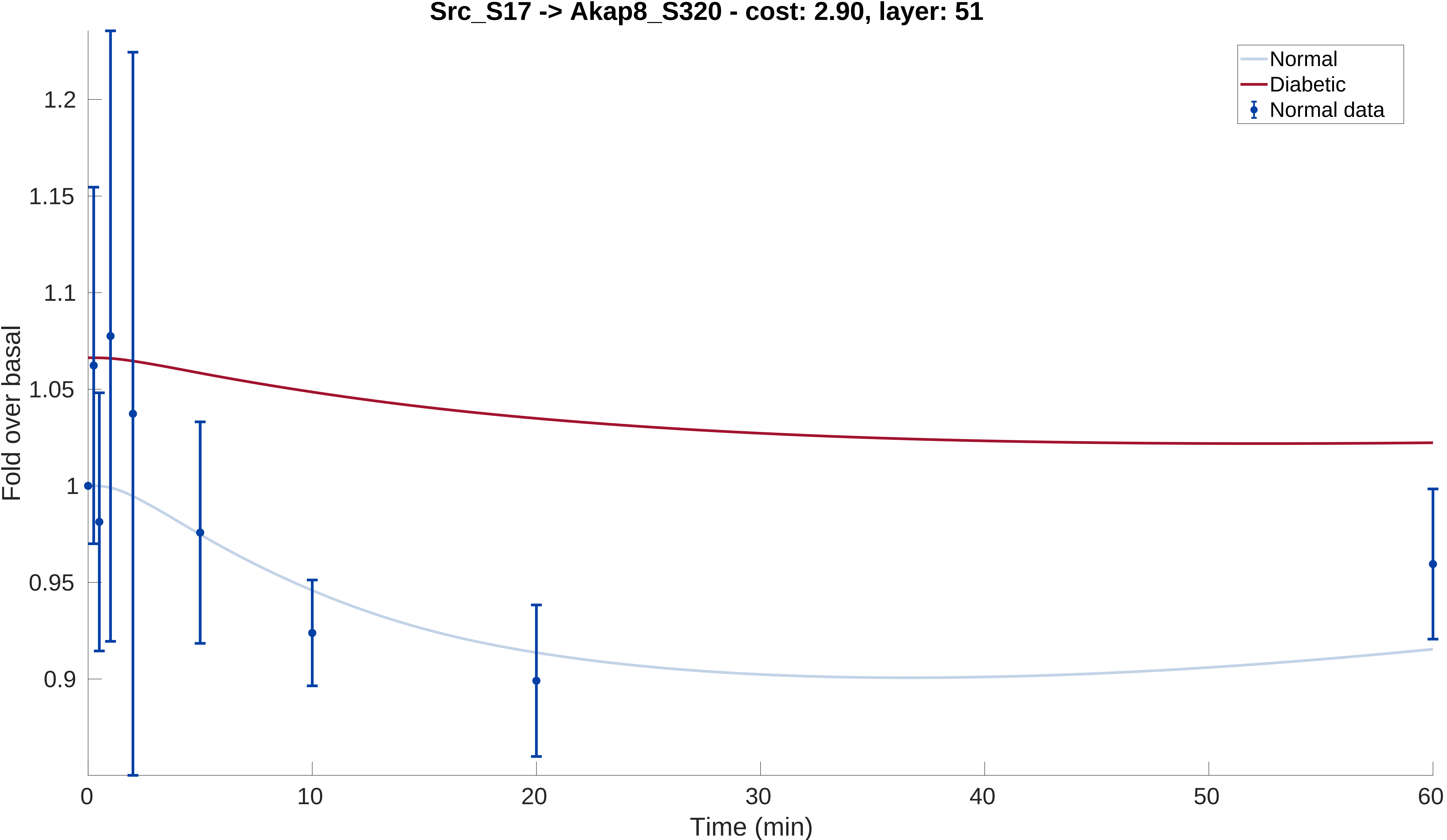

Src_S17 -> Akap8_S320 - cost: 2.90, layer: 51
Normal
Diabetic
Normal data
Fold over basal
1.2
1.15
1.1
1.05
1
0.95
0.9
0
10
20
30
40
50
60
Time (min)

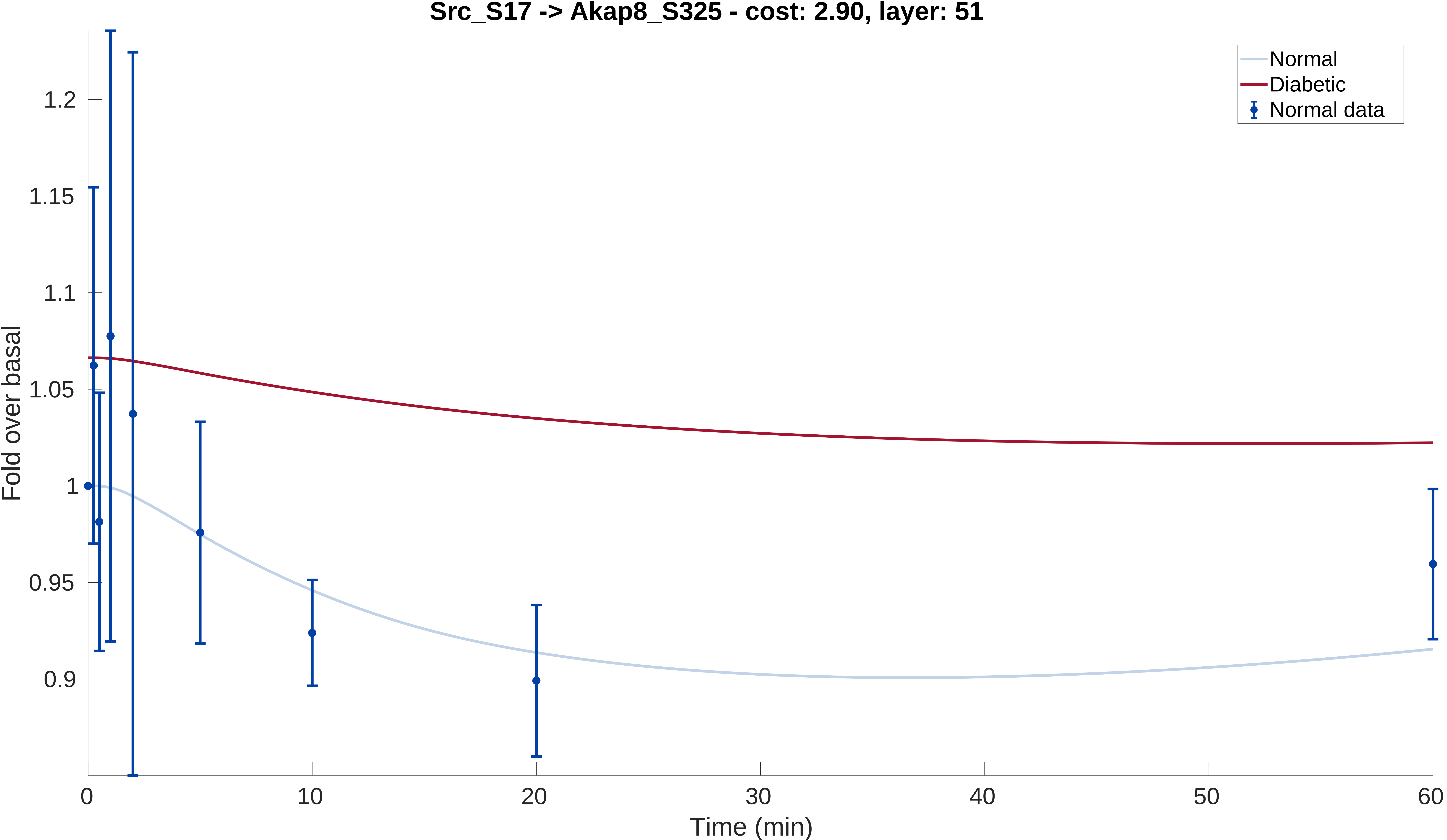

Src_S17 -> Akap8_S325 - cost: 2.90, layer: 51
Normal
Diabetic
Normal data
Fold over basal
1.2
1.15
1.1
1.05
1
0.95
0.9
0
10
20
30
40
50
60
Time (min)

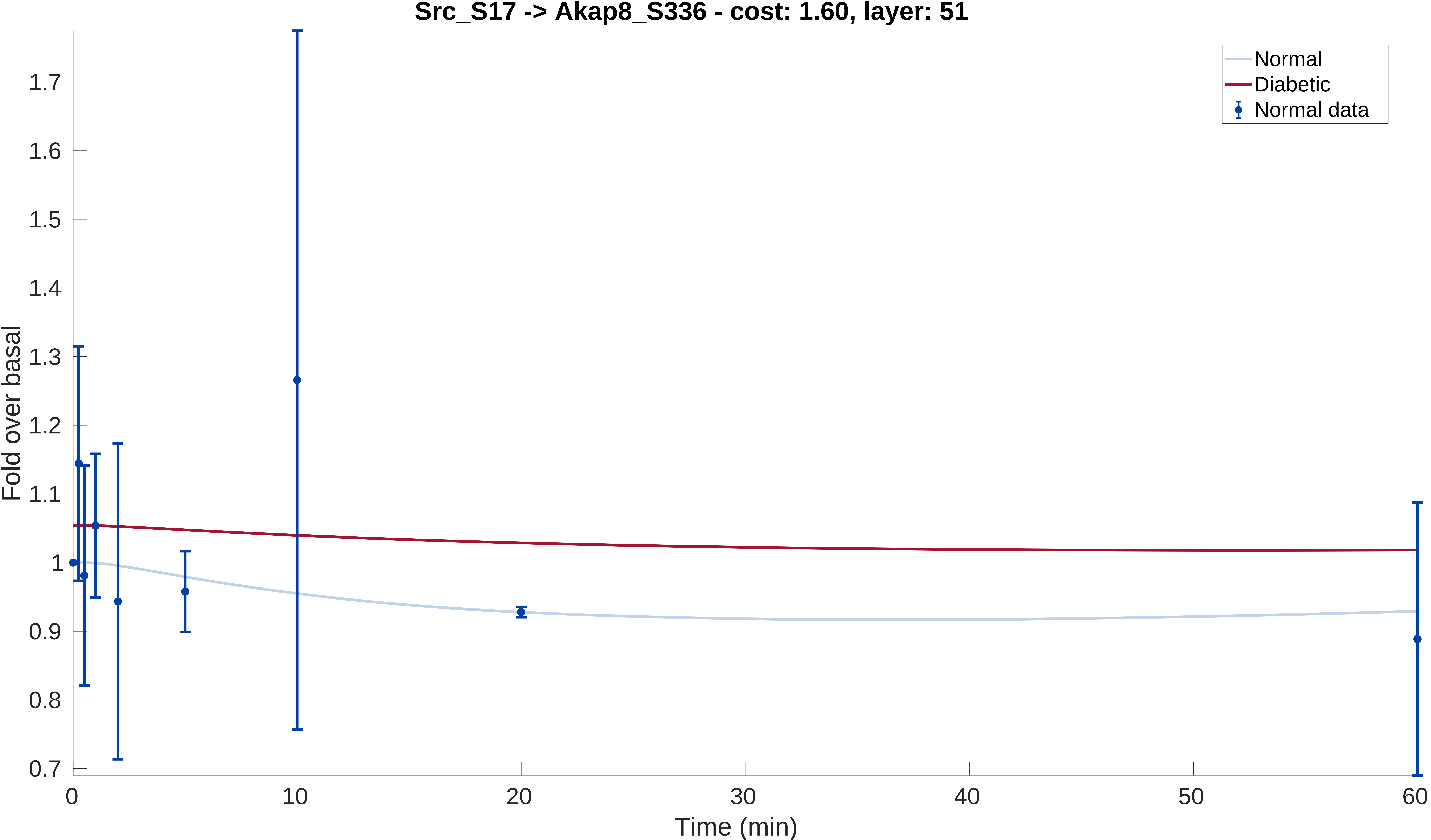

Src_S17 -> Akap8_S336 - cost: 1.60, layer: 51
Normal
Diabetic
Normal data
Fold over basal
Time (min)
1.7
1.6
1.5
1.4
1.3
1.2
1.1
1
0.9
0.8
0.7
0
10
20
30
40
50
60

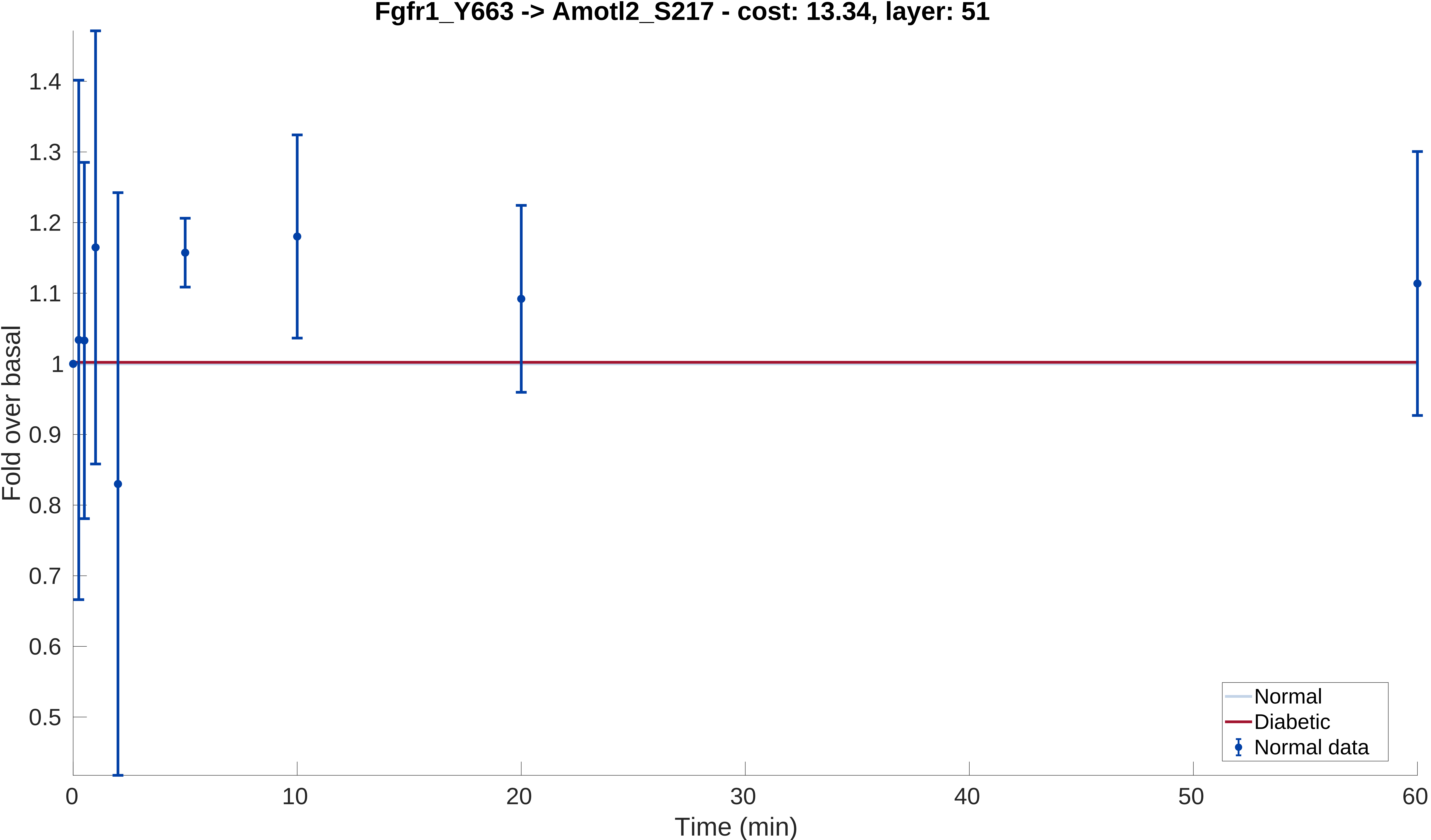

Fgfr1_Y663 -> Amotl2_S217 - cost: 13.34, layer: 51
Fold over basal
1.4
1.3
1.2
1.1
1
0.9
0.8
0.7
0.6
0.5
0
10
20
30
40
50
60
Time (min)
Normal
Diabetic
Normal data

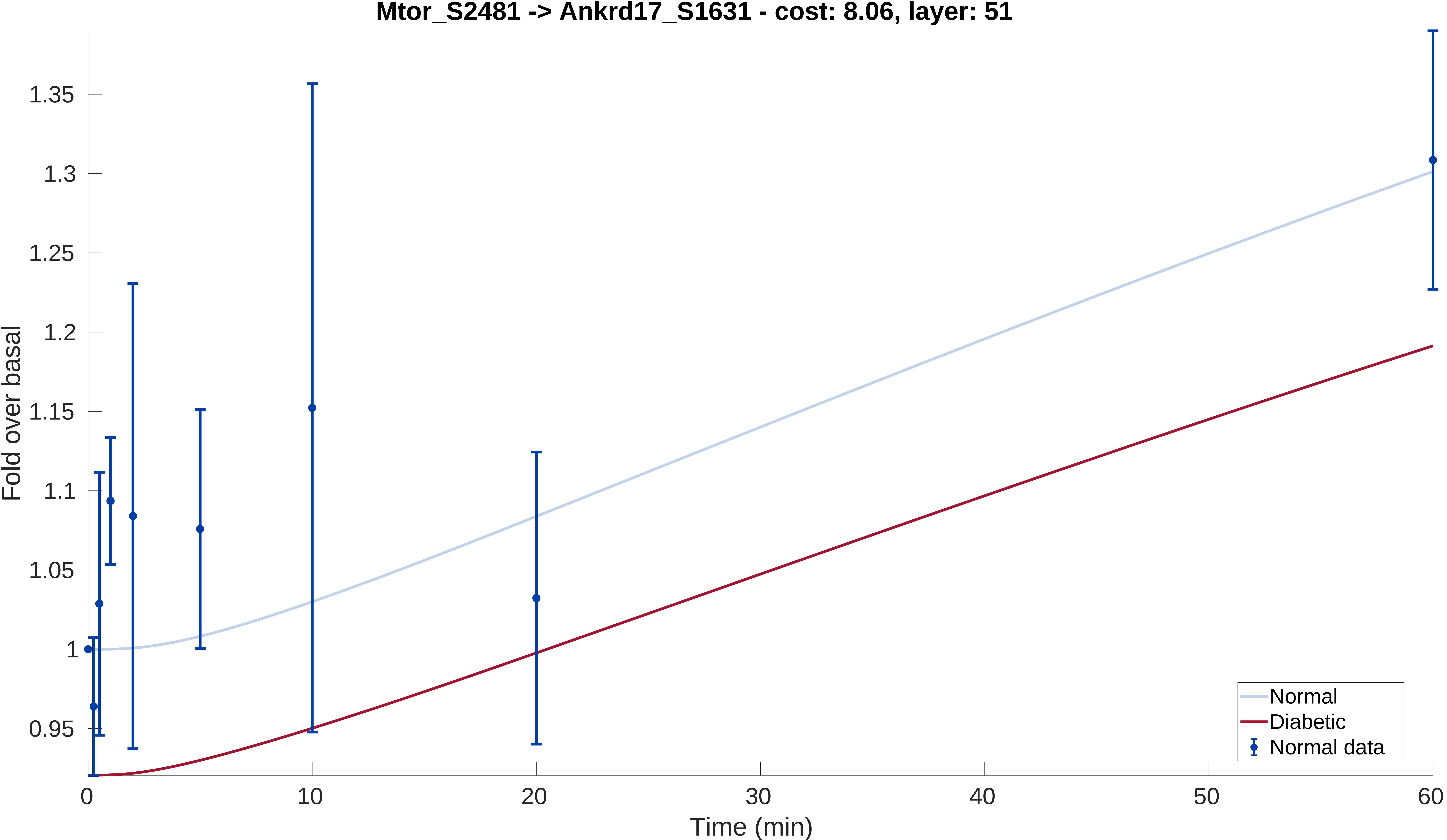

Mtor_S2481 -> Ankrd17_S1631 - cost: 8.06, layer: 51
Fold over basal
1.35
1.3
1.25
1.2
1.15
1.1
1.05
1
0.95
0
10
20
30
40
50
60
Time (min)
Normal
Diabetic
Normal data

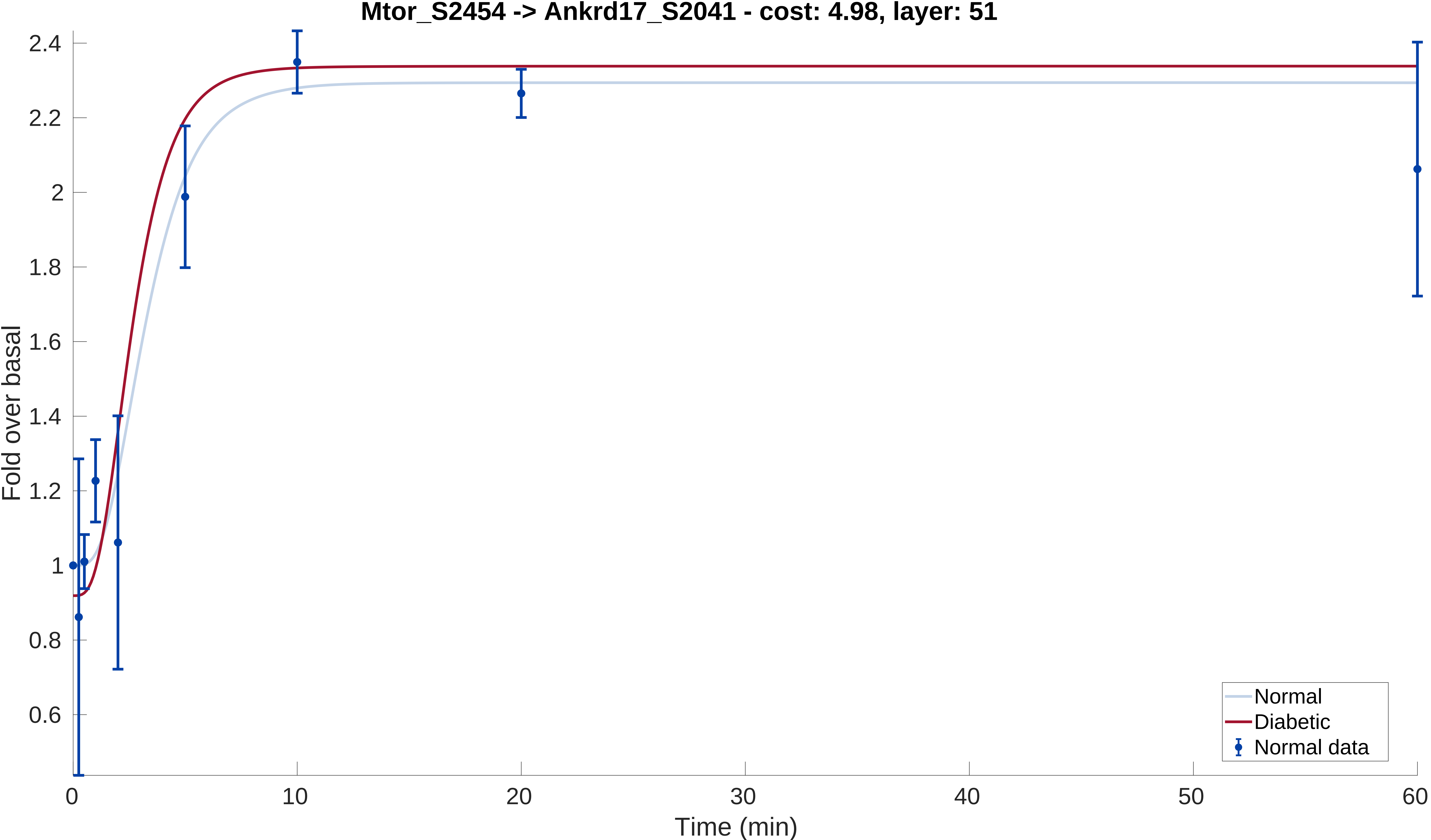
Mtor_S2454 -> Ankrd17_S2041 - cost: 4.98, layer: 51
Fold over basal
Time (min)
Normal
Diabetic
Normal data
2.4
2.2
2
1.8
1.6
1.4
1.2
1
0.8
0.6
0
10
20
30
40
50
60

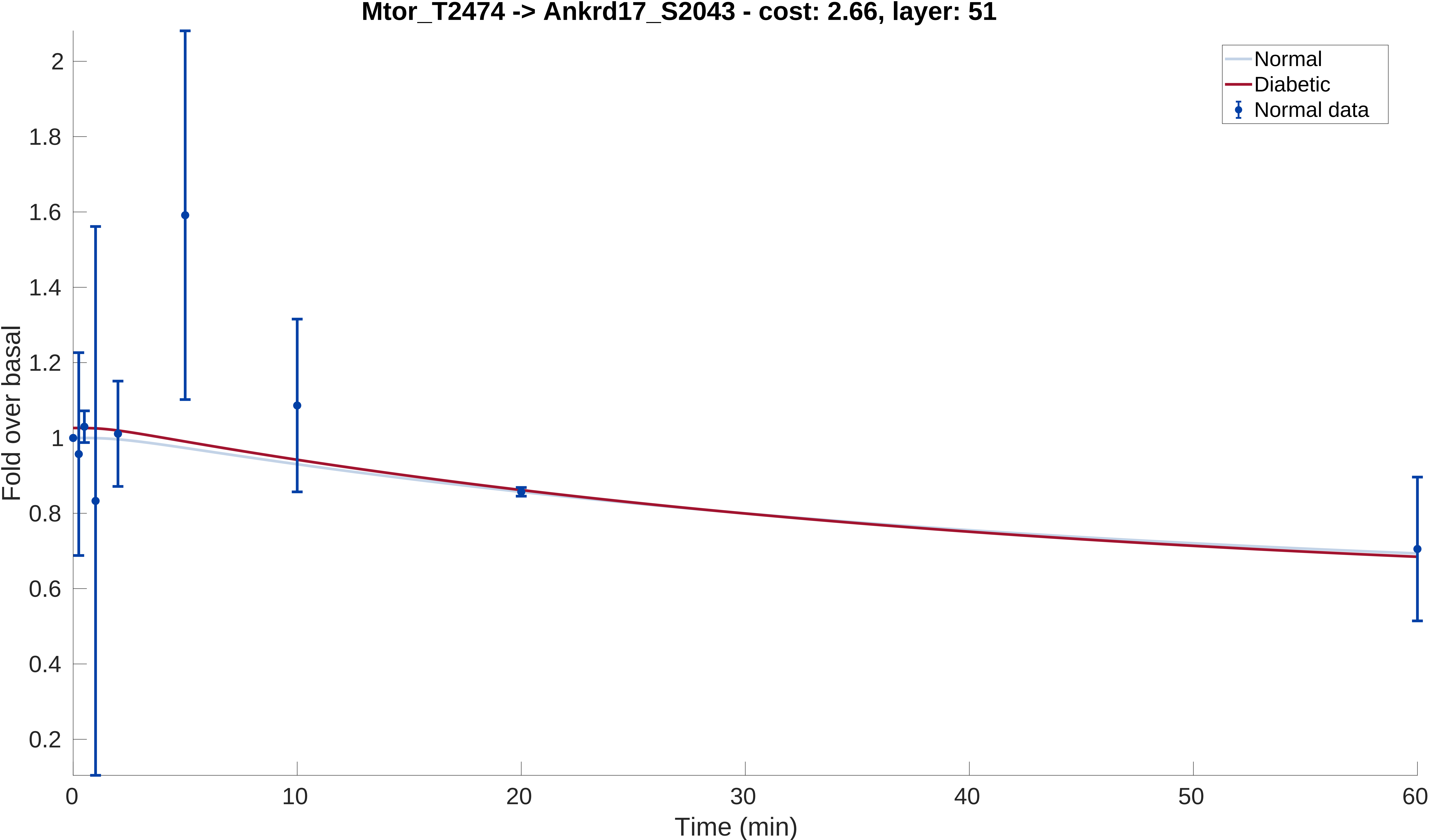
Mtor_T2474 -> Ankrd17_S2043 - cost: 2.66, layer: 51
Normal
Diabetic
Normal data
Fold over basal
Time (min)
0.2
0.4
0.6
0.8
1
1.2
1.4
1.6
1.8
2
0
10
20
30
40
50
60

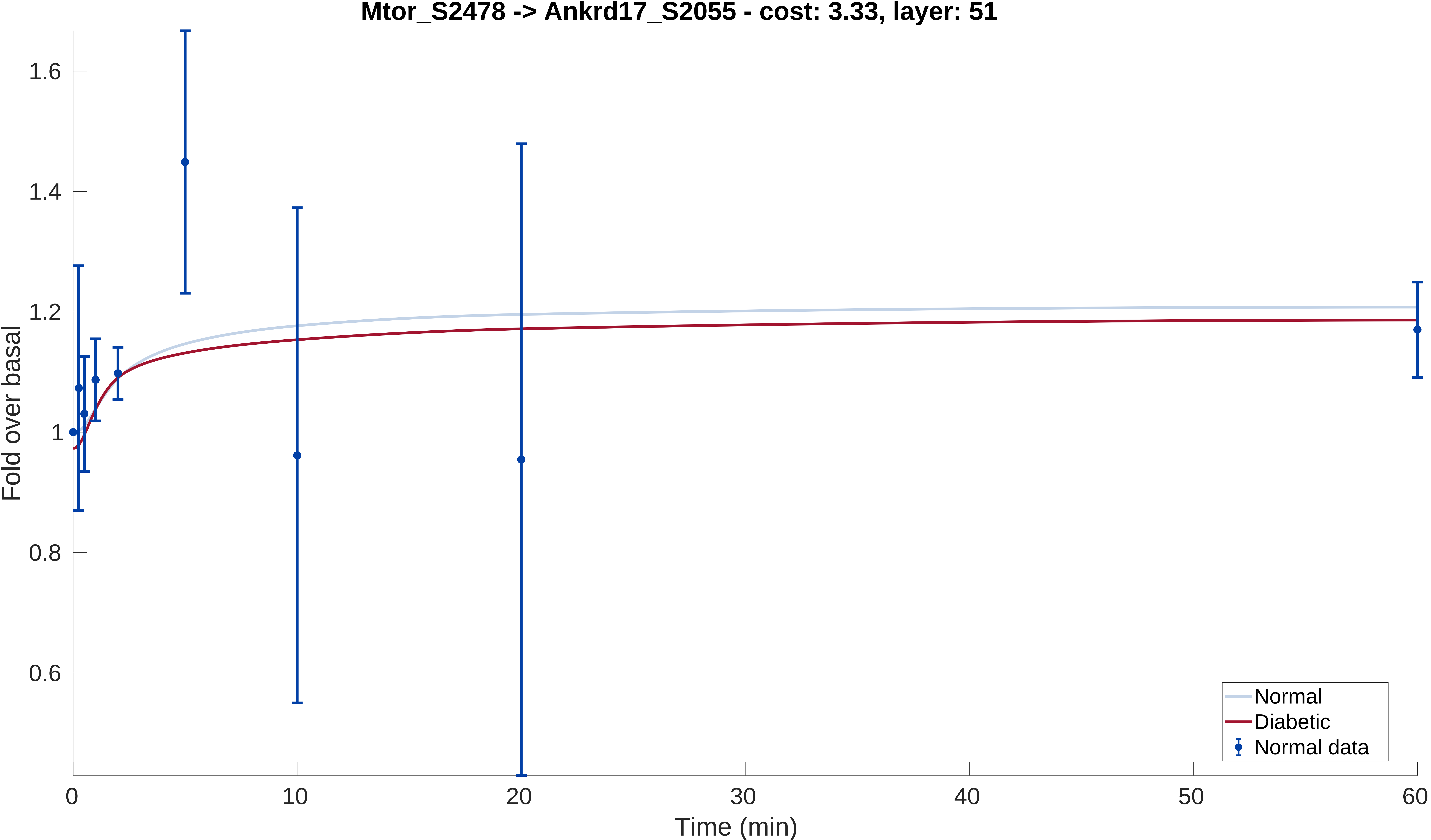

Mtor_S2478 -> Ankrd17_S2055 - cost: 3.33, layer: 51
Fold over basal
1.6
1.4
1.2
1
0.8
0.6
0
10
20
30
40
50
60
Time (min)
Normal
Diabetic
Normal data

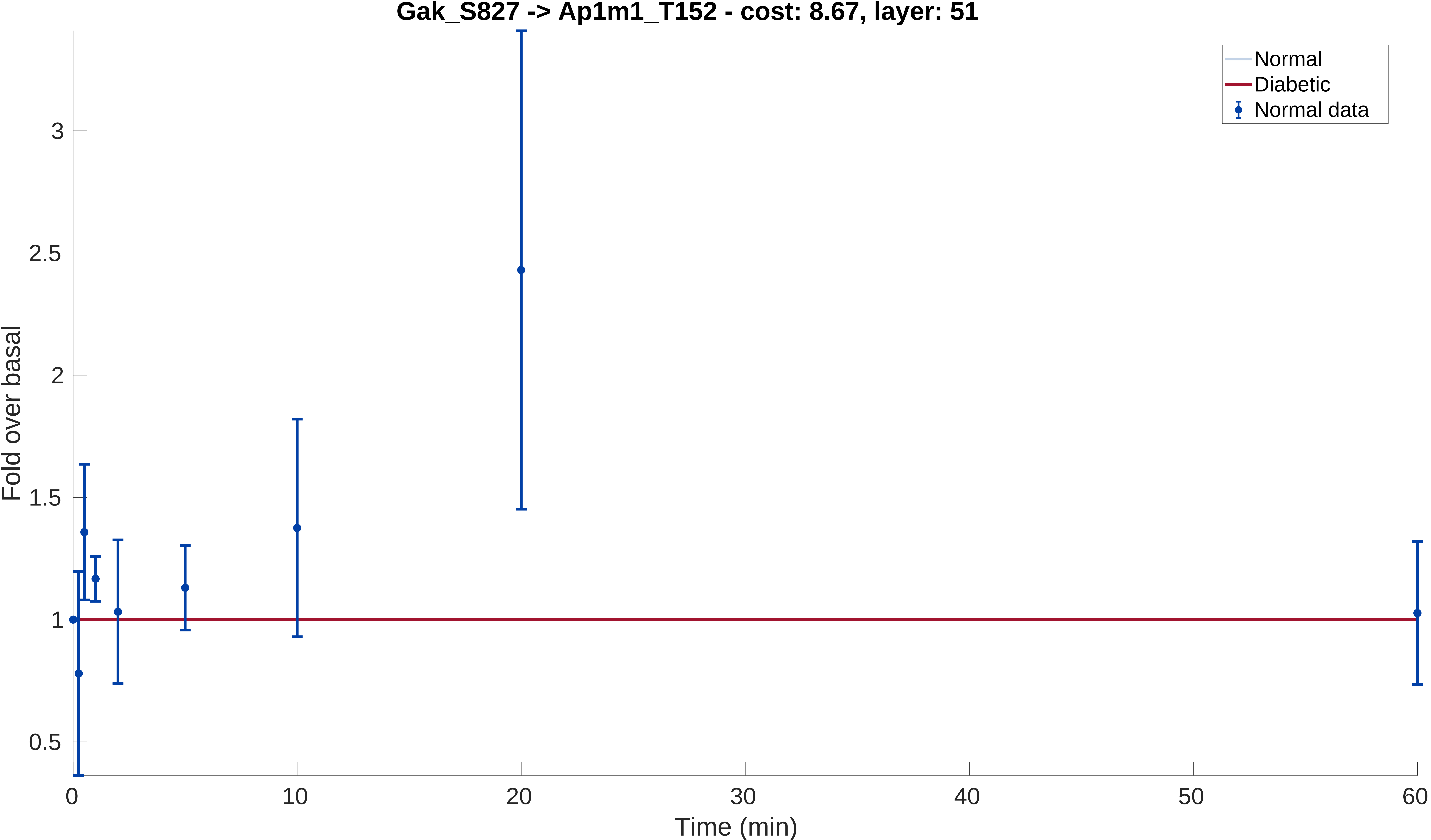

Gak_S827 -> Ap1m1_T152 - cost: 8.67, layer: 51
Normal
Diabetic
Normal data
Fold over basal
3
2.5
2
1.5
1
0.5
0
10
20
30
40
50
60
Time (min)

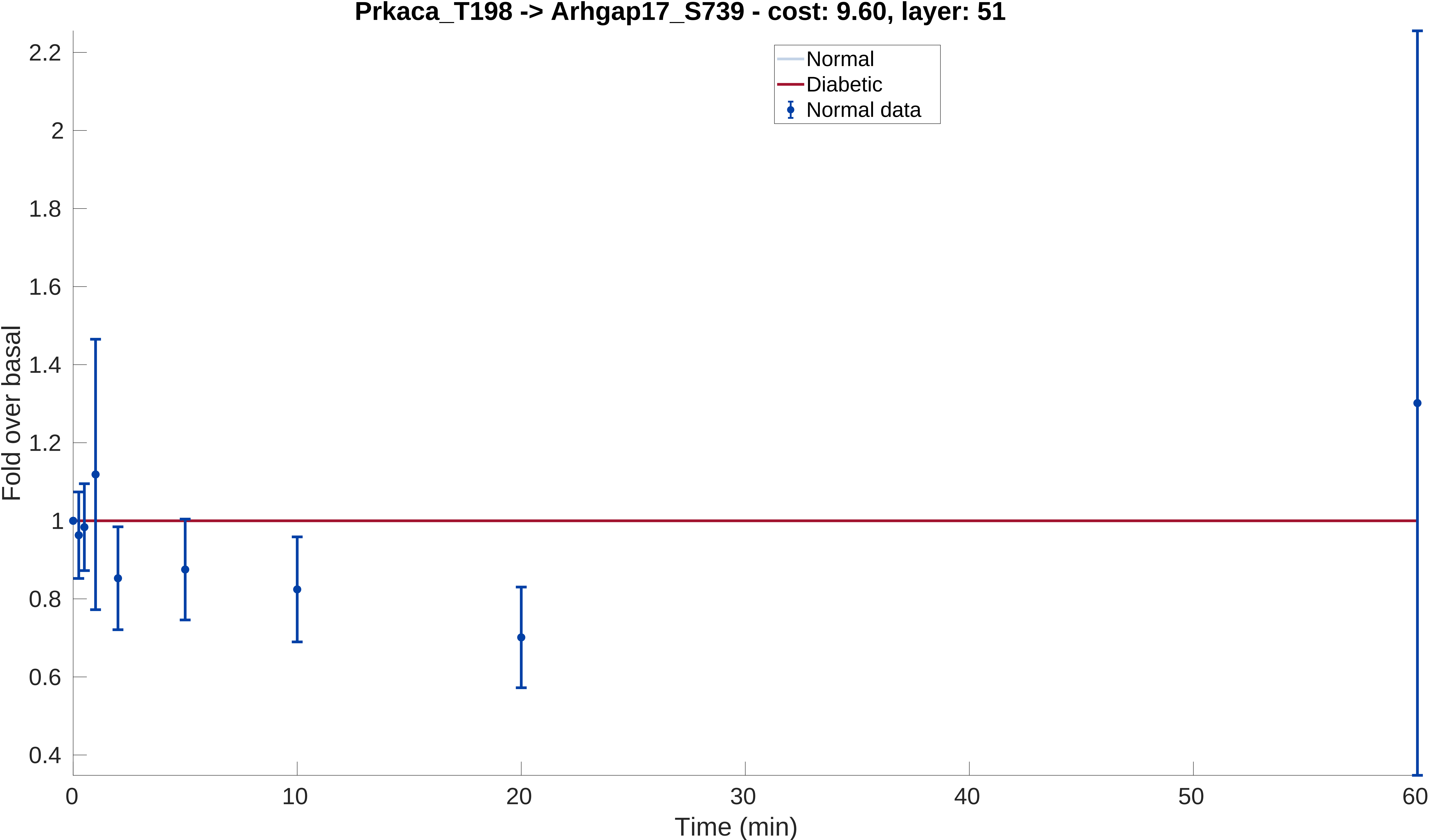

Prkaca_T198 -> Arhgap17_S739 - cost: 9.60, layer: 51
Normal
Diabetic
Normal data
Fold over basal
2.2
2
1.8
1.6
1.4
1.2
1
0.8
0.6
0.4
0
10
20
30
40
50
60
Time (min)

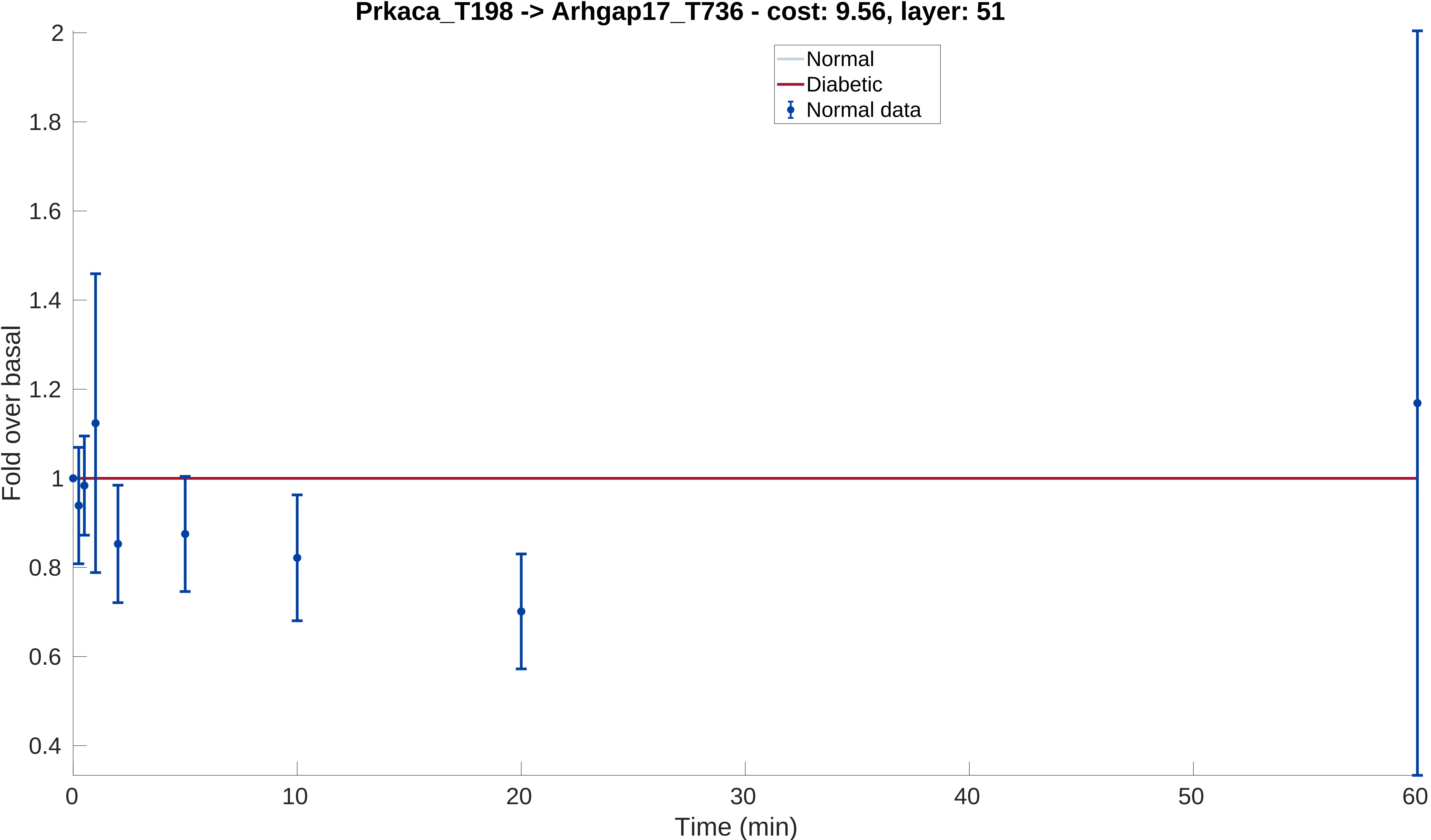

Prkaca_T198 -> Arhgap17_T736 - cost: 9.56, layer: 51
Normal
Diabetic
Normal data
Fold over basal
2
1.8
1.6
1.4
1.2
1
0.8
0.6
0.4
0
10
20
30
40
50
60
Time (min)

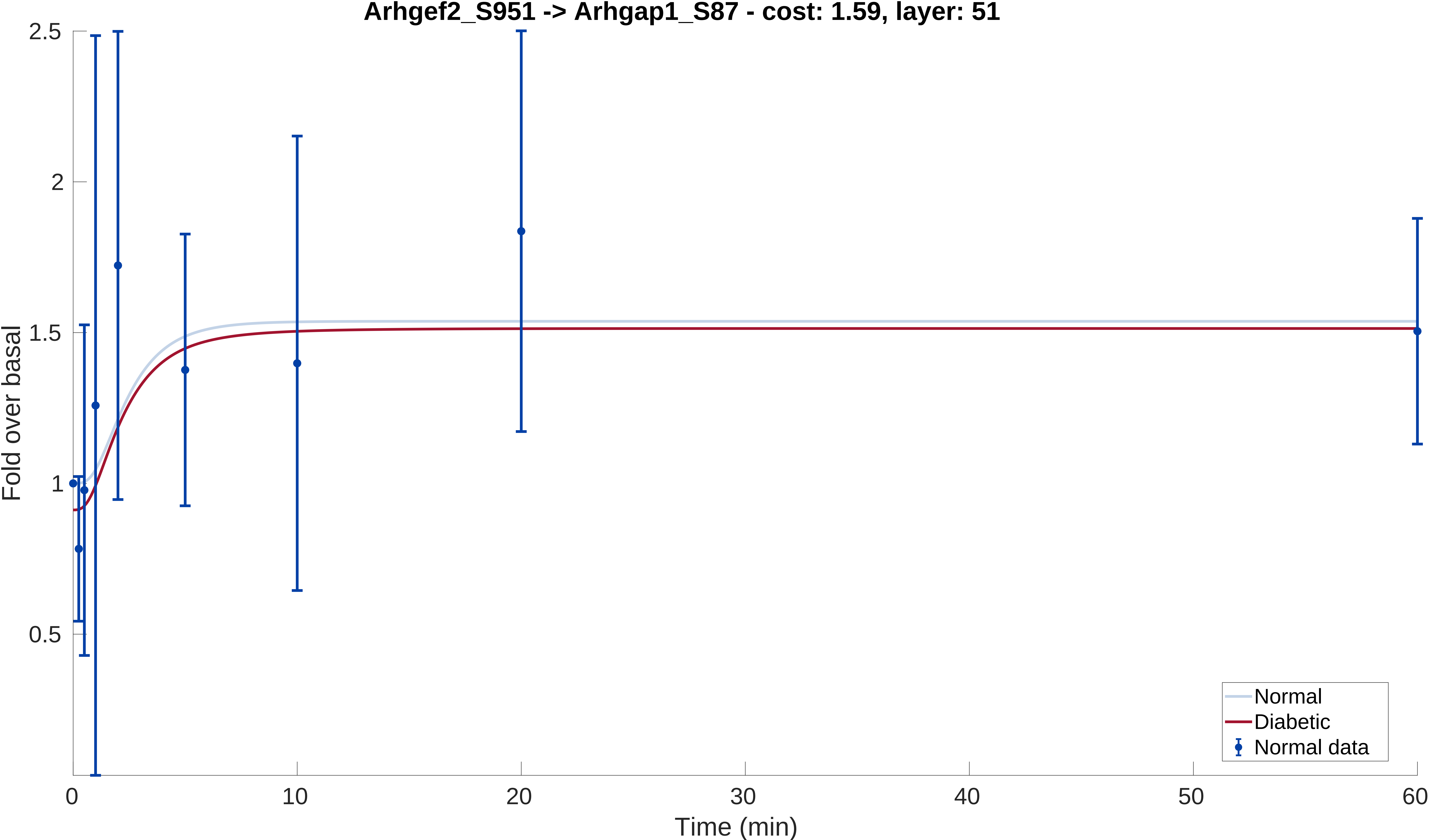

Arhgef2_S951 -> Arhgap1_S87 - cost: 1.59, layer: 51
Fold over basal
Time (min)
0.5
1
1.5
2
2.5
0
10
20
30
40
50
60
Normal
Diabetic
Normal data

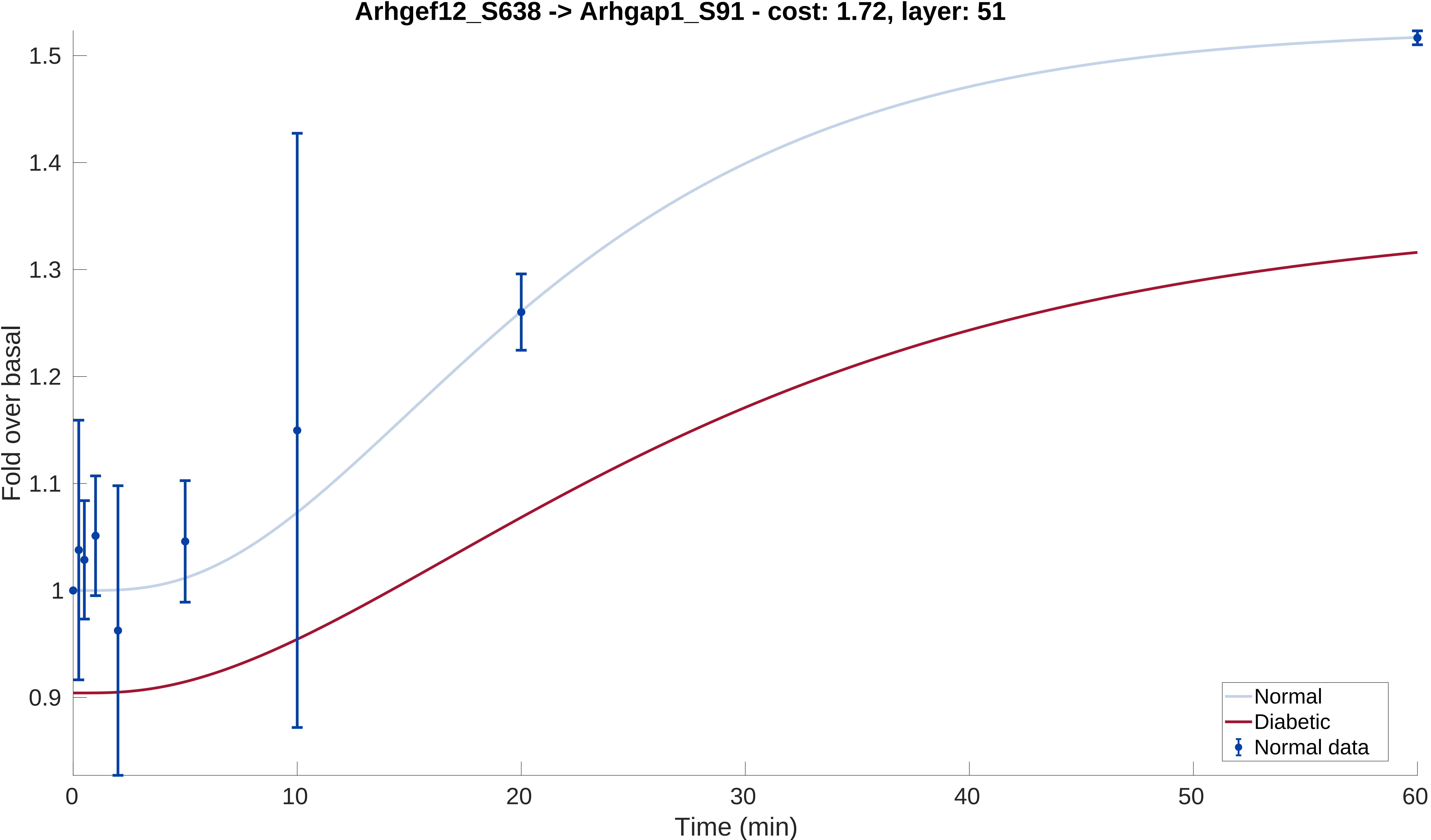

Arhgef12_S638 -> Arhgap1_S91 - cost: 1.72, layer: 51
Fold over basal
Time (min)
1.5
1.4
1.3
1.2
1.1
1
0.9
0
10
20
30
40
50
60
Normal
Diabetic
Normal data

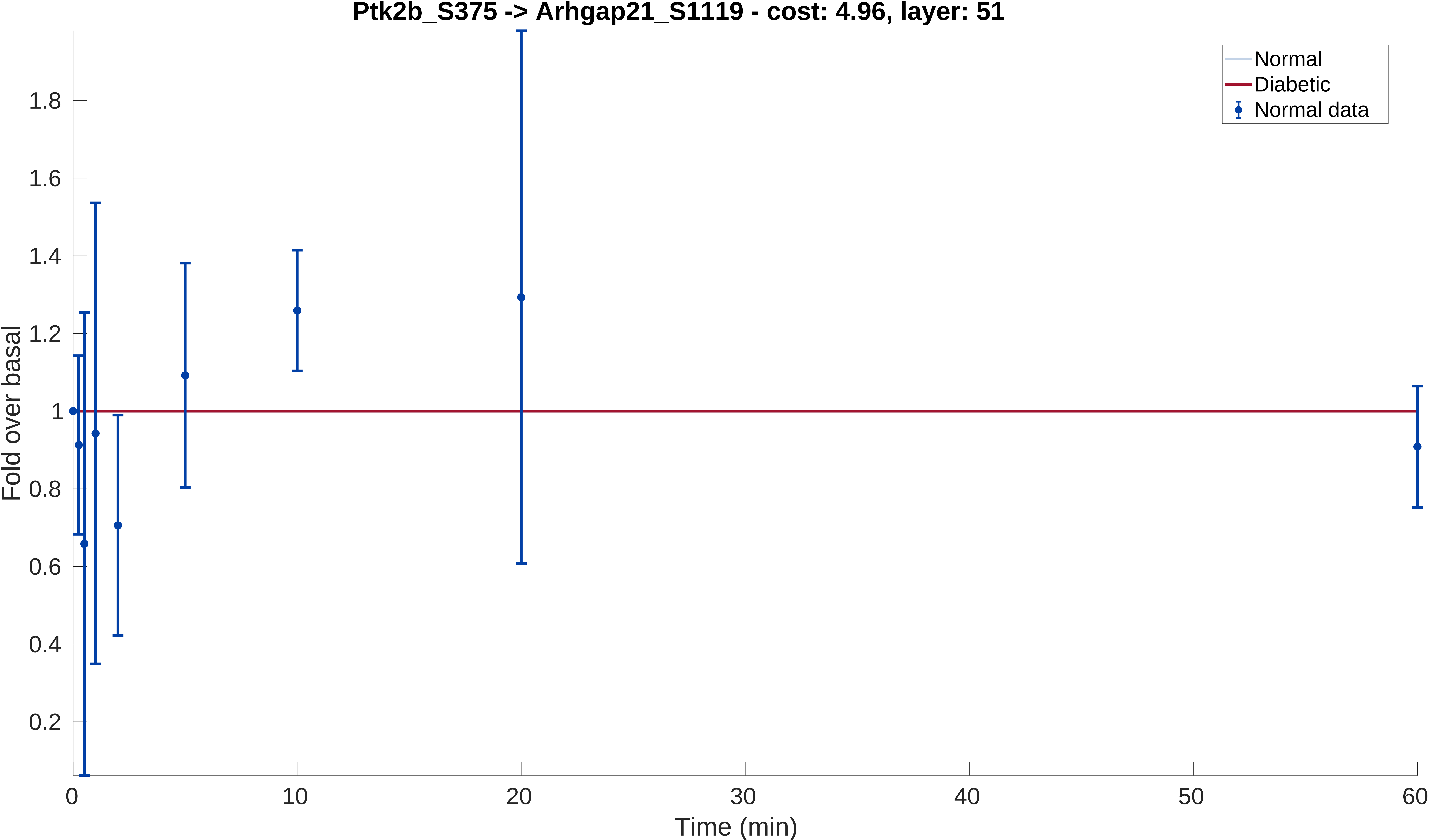
Ptk2b_S375 -> Arhgap21_S1119 - cost: 4.96, layer: 51
Normal
Diabetic
Normal data
Fold over basal
1.8
1.6
1.4
1.2
1
0.8
0.6
0.4
0.2
0
10
20
30
40
50
60
Time (min)

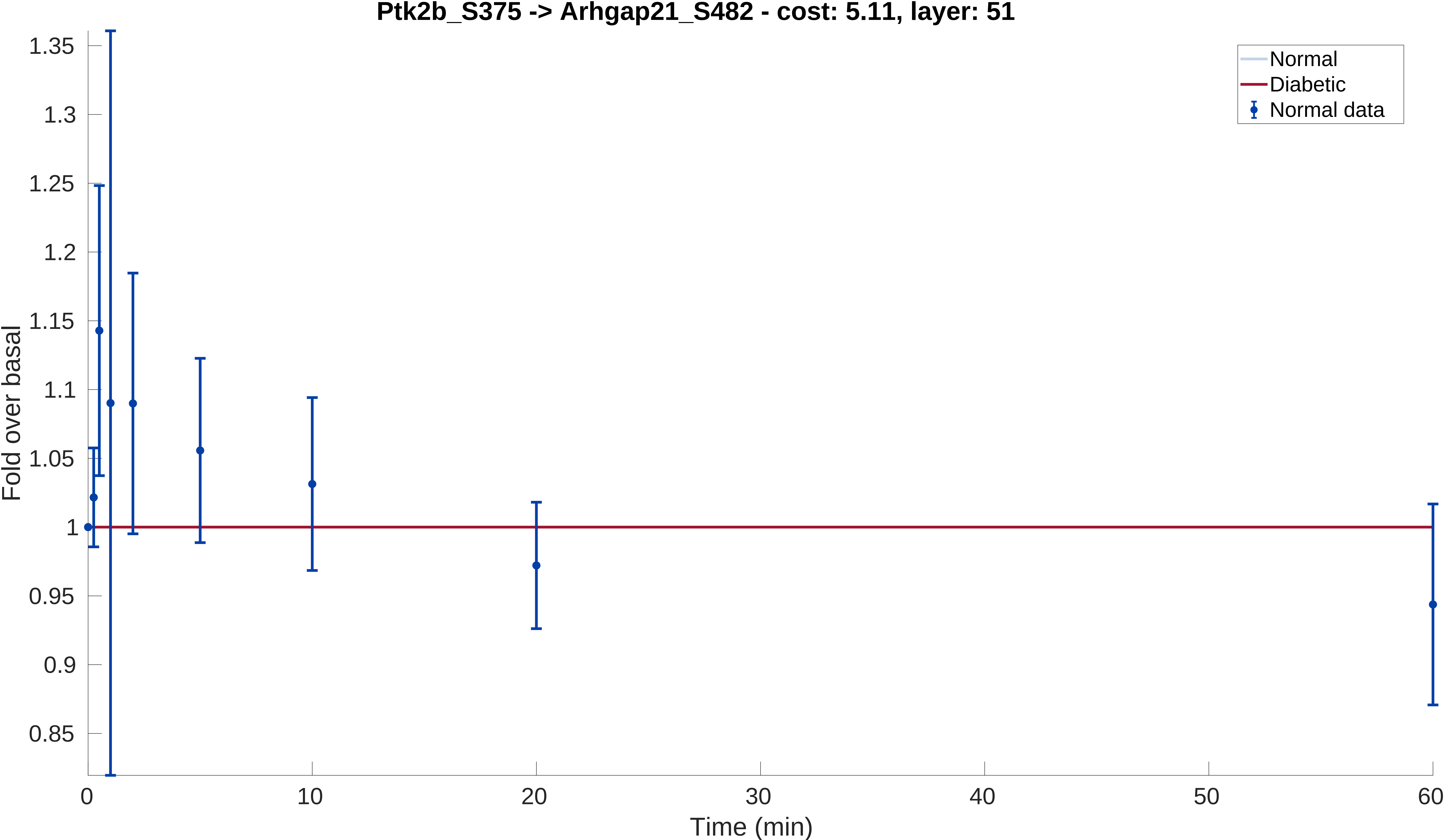

Ptk2b_S375 -> Arhgap21_S482 - cost: 5.11, layer: 51
Normal
Diabetic
Normal data
Fold over basal
1.35
1.3
1.25
1.2
1.15
1.1
1.05
1
0.95
0.9
0.85
0
10
20
30
40
50
60
Time (min)

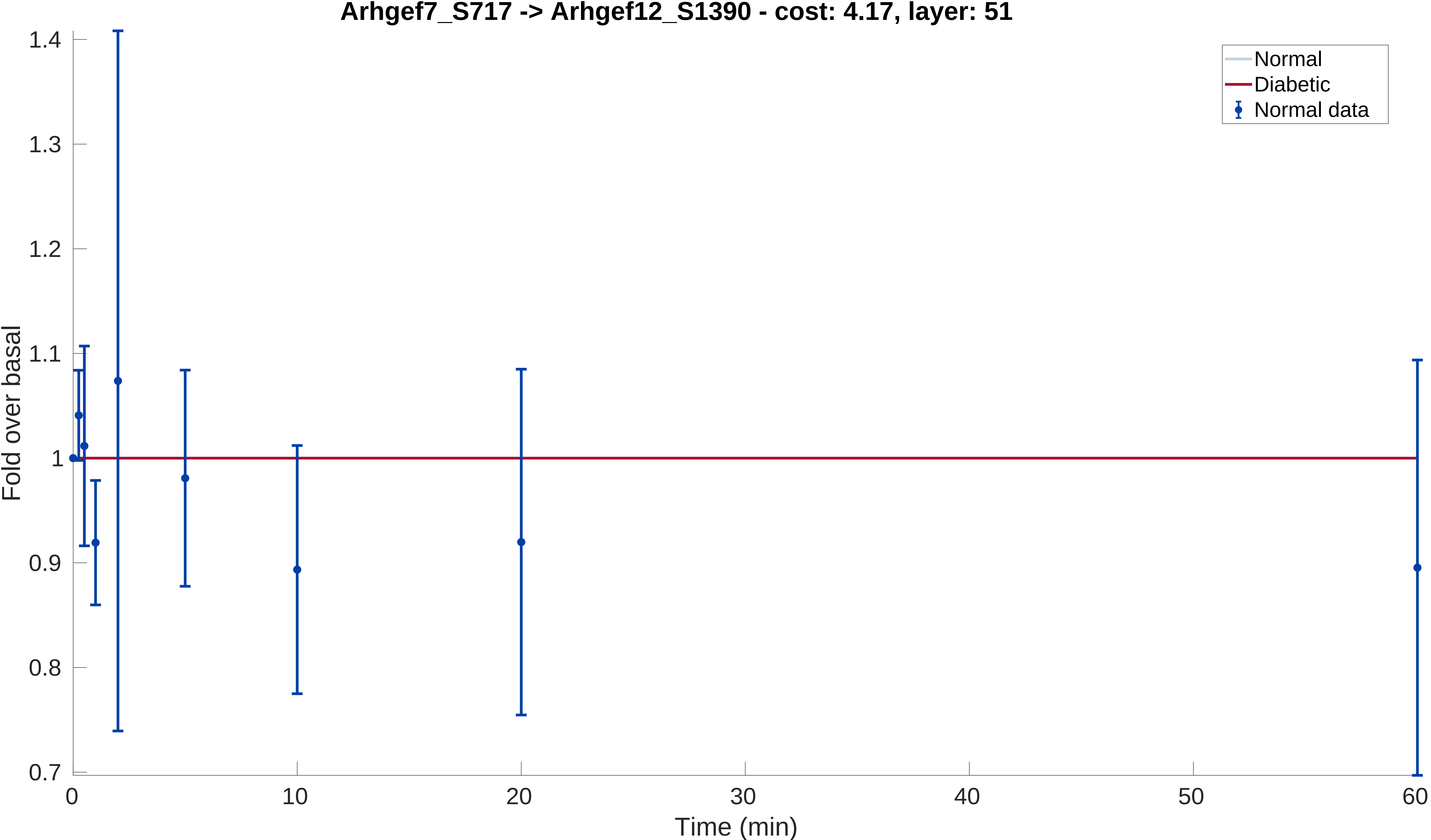

Arhgef7_S717 -> Arhgef12_S1390 - cost: 4.17, layer: 51
Fold over basal
Time (min)
Normal
Diabetic
Normal data
1.4
1.3
1.2
1.1
1
0.9
0.8
0.7
0
10
20
30
40
50
60

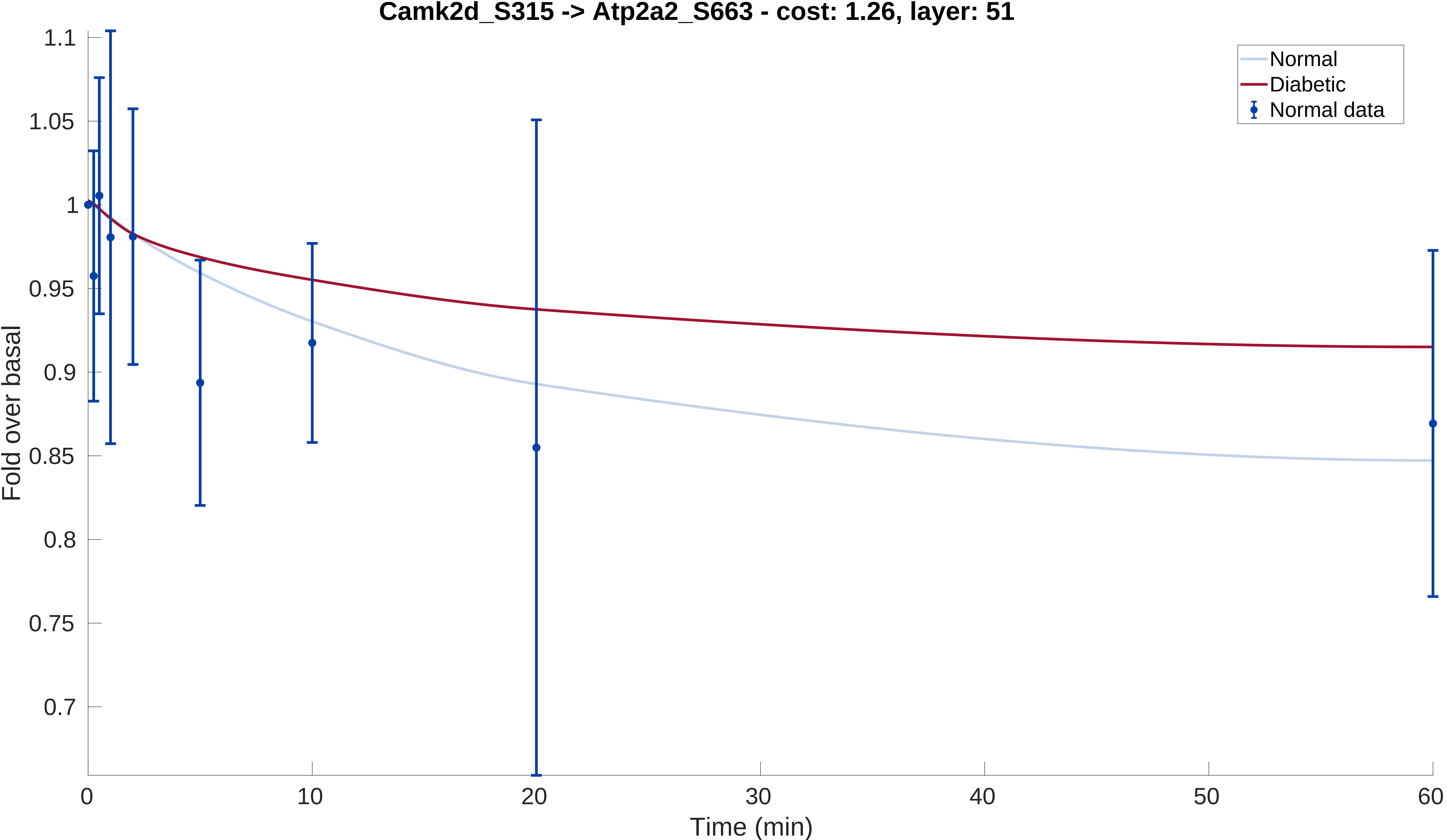
Camk2d_S315 -> Atp2a2_S663 - cost: 1.26, layer: 51
Normal
Diabetic
Normal data
Fold over basal
Time (min)
1.1
1.05
1
0.95
0.9
0.85
0.8
0.75
0.7
0
10
20
30
40
50
60

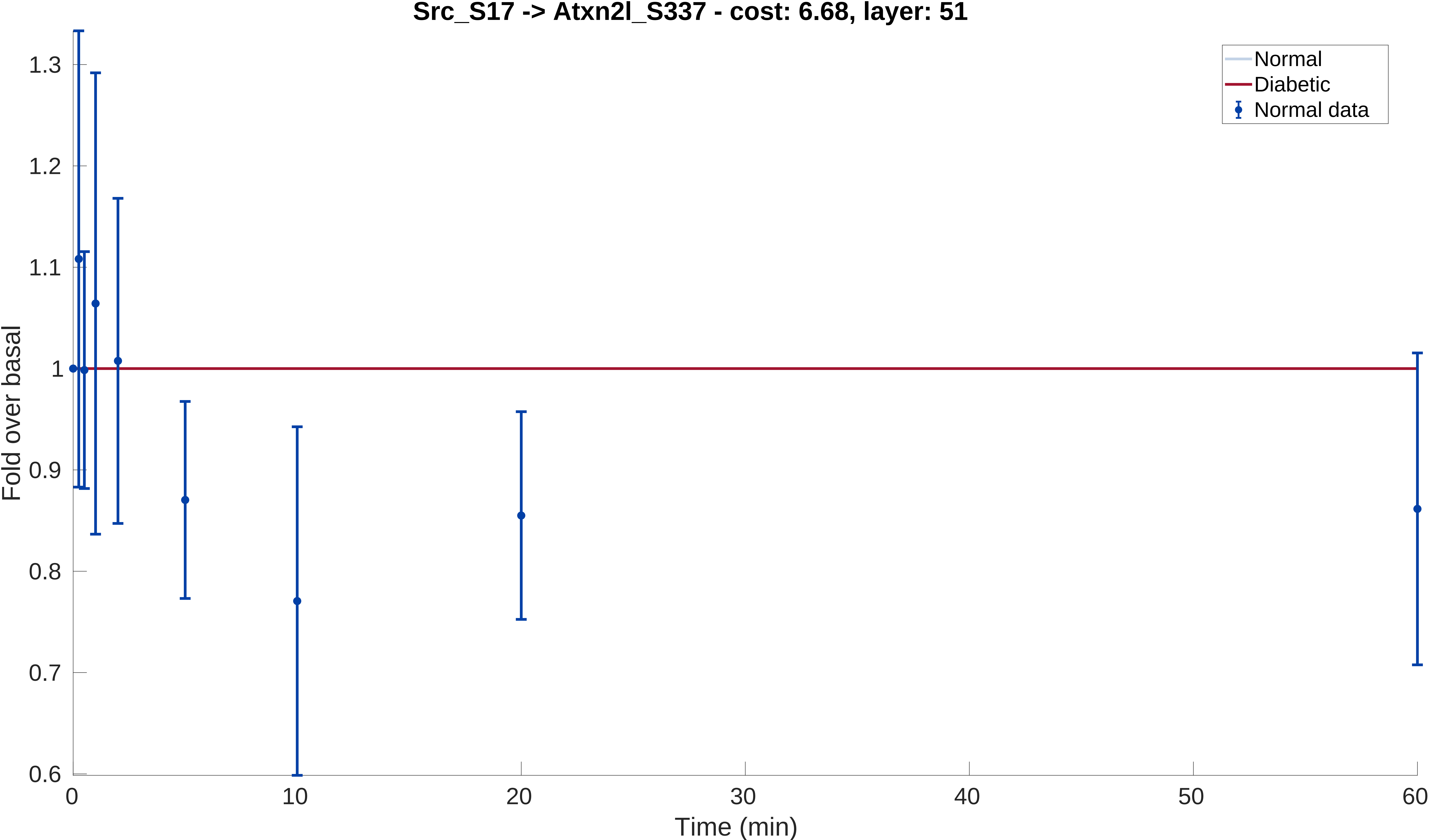
Src_S17 -> Atxn2l_S337 - cost: 6.68, layer: 51
Normal
Diabetic
Normal data
1.3
1.2
1.1
1
0.9
0.8
0.7
0.6
Fold over basal
0
10
20
30
40
50
60
Time (min)

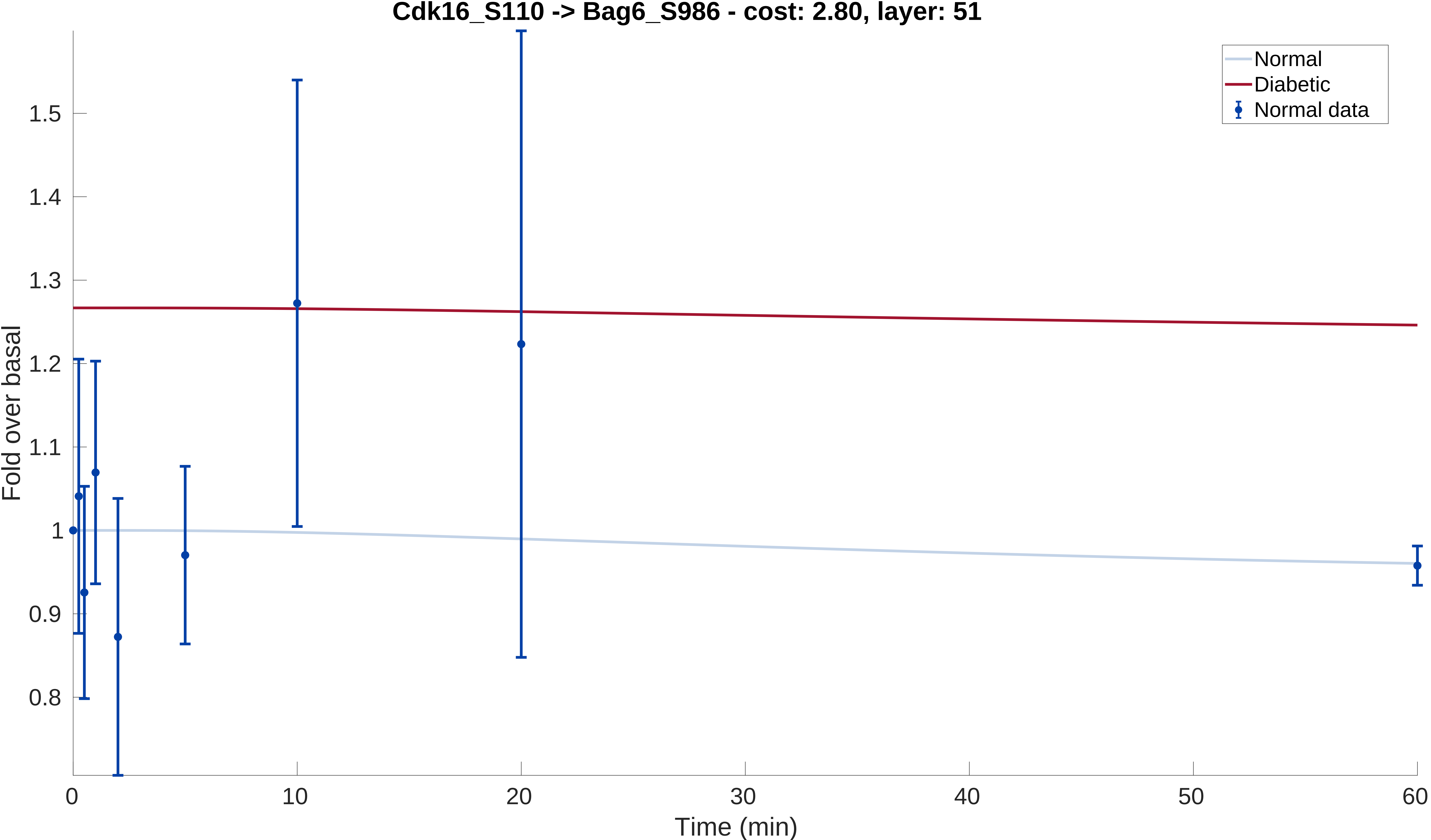

Cdk16_S110 -> Bag6_S986 - cost: 2.80, layer: 51
Normal
Diabetic
Normal data
Fold over basal
1.5
1.4
1.3
1.2
1.1
1
0.9
0.8
0
10
20
30
40
50
60
Time (min)

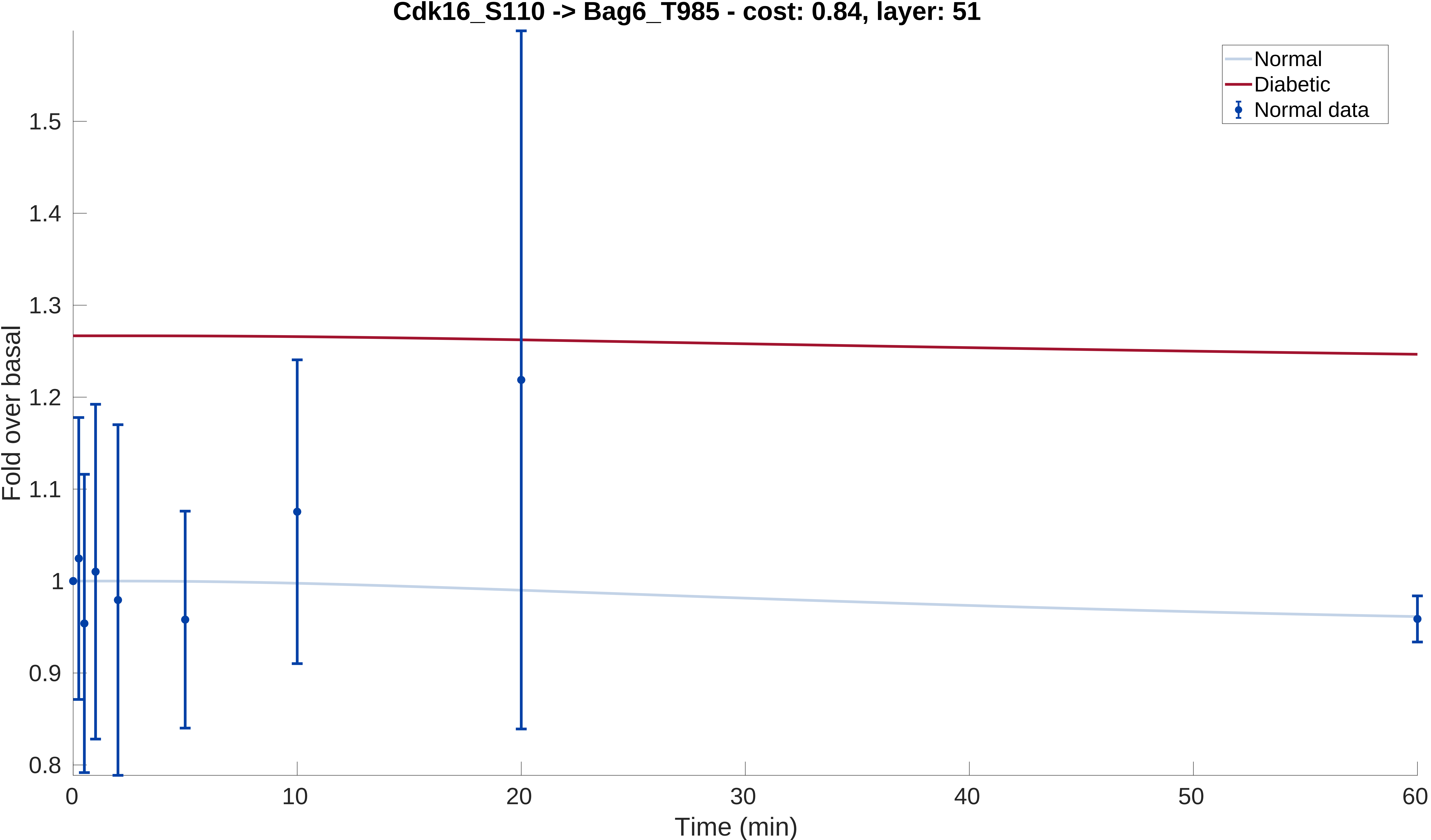

Cdk16_S110 -> Bag6_T985 - cost: 0.84, layer: 51
Normal
Diabetic
Normal data
Fold over basal
Time (min)
0.8
0.9
1
1.1
1.2
1.3
1.4
1.5
0
10
20
30
40
50
60

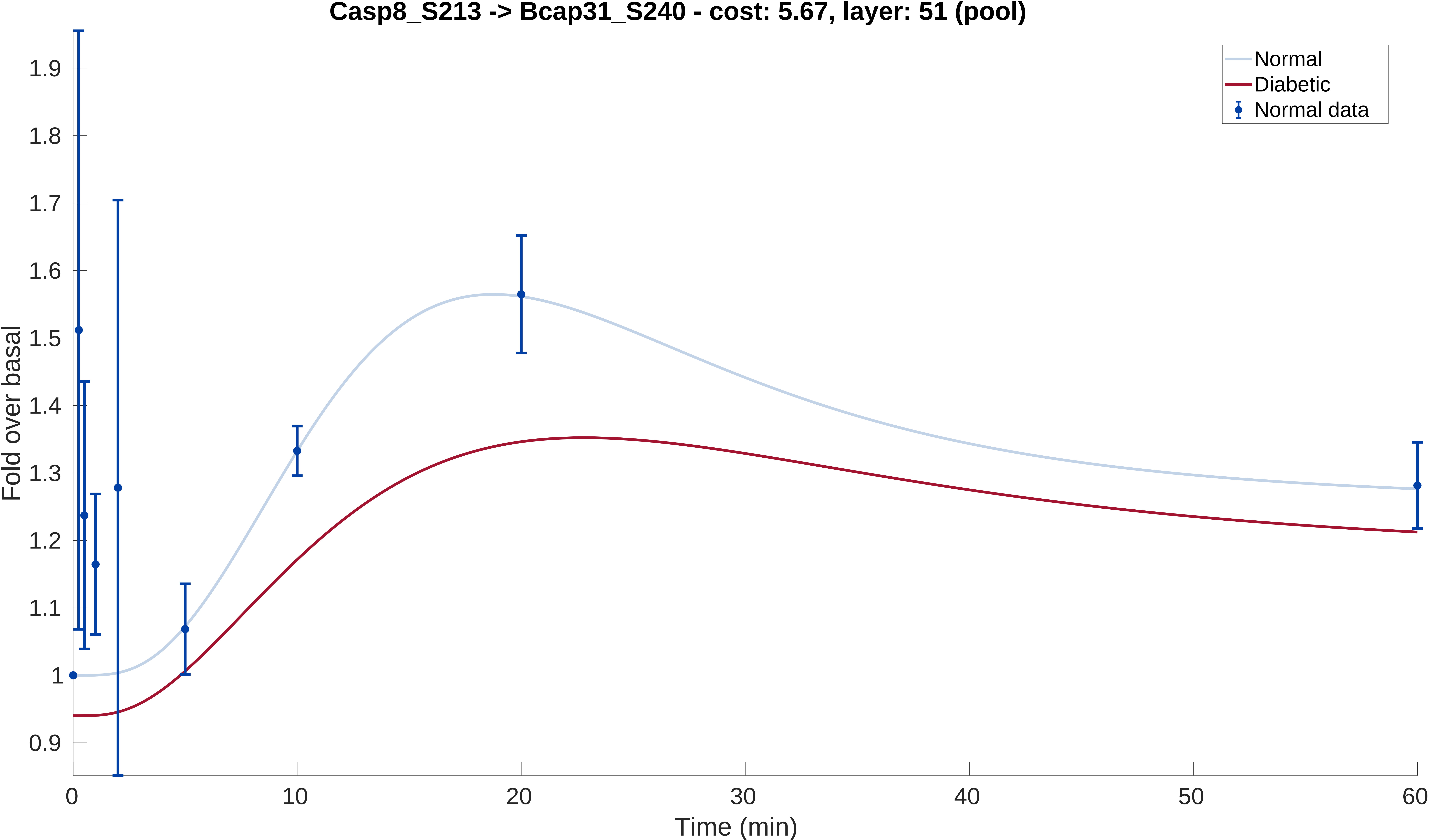

Casp8_S213 -> Bcap31_S240 - cost: 5.67, layer: 51 (pool)
Normal
Diabetic
Normal data
Fold over basal
Time (min)
0.9
1
1.1
1.2
1.3
1.4
1.5
1.6
1.7
1.8
1.9
0
10
20
30
40
50
60

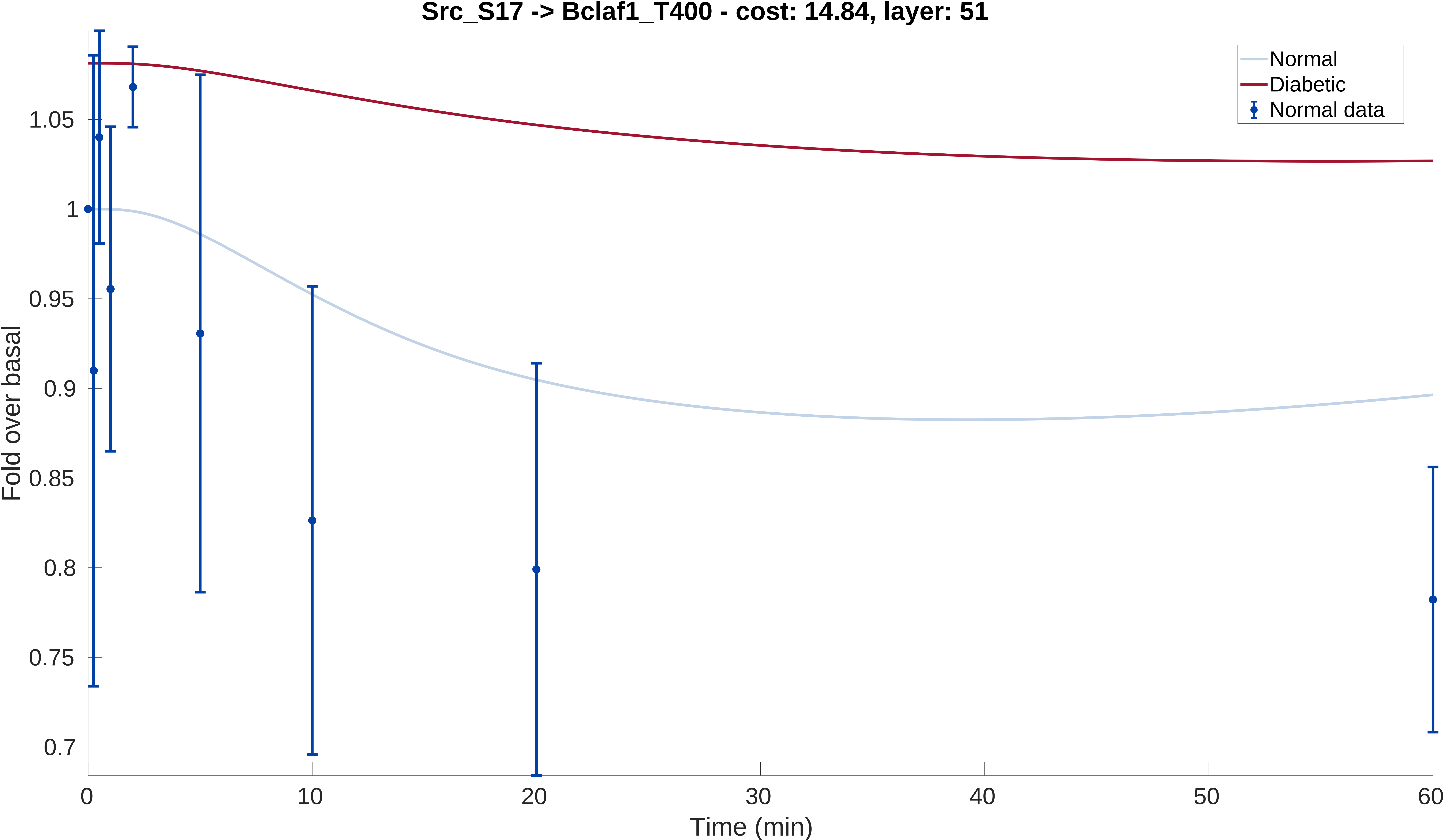
Src_S17 -> Bclaf1_T400 - cost: 14.84, layer: 51
Normal
Diabetic
Normal data
Fold over basal
Time (min)
1.05
1
0.95
0.9
0.85
0.8
0.75
0.7
0
10
20
30
40
50
60

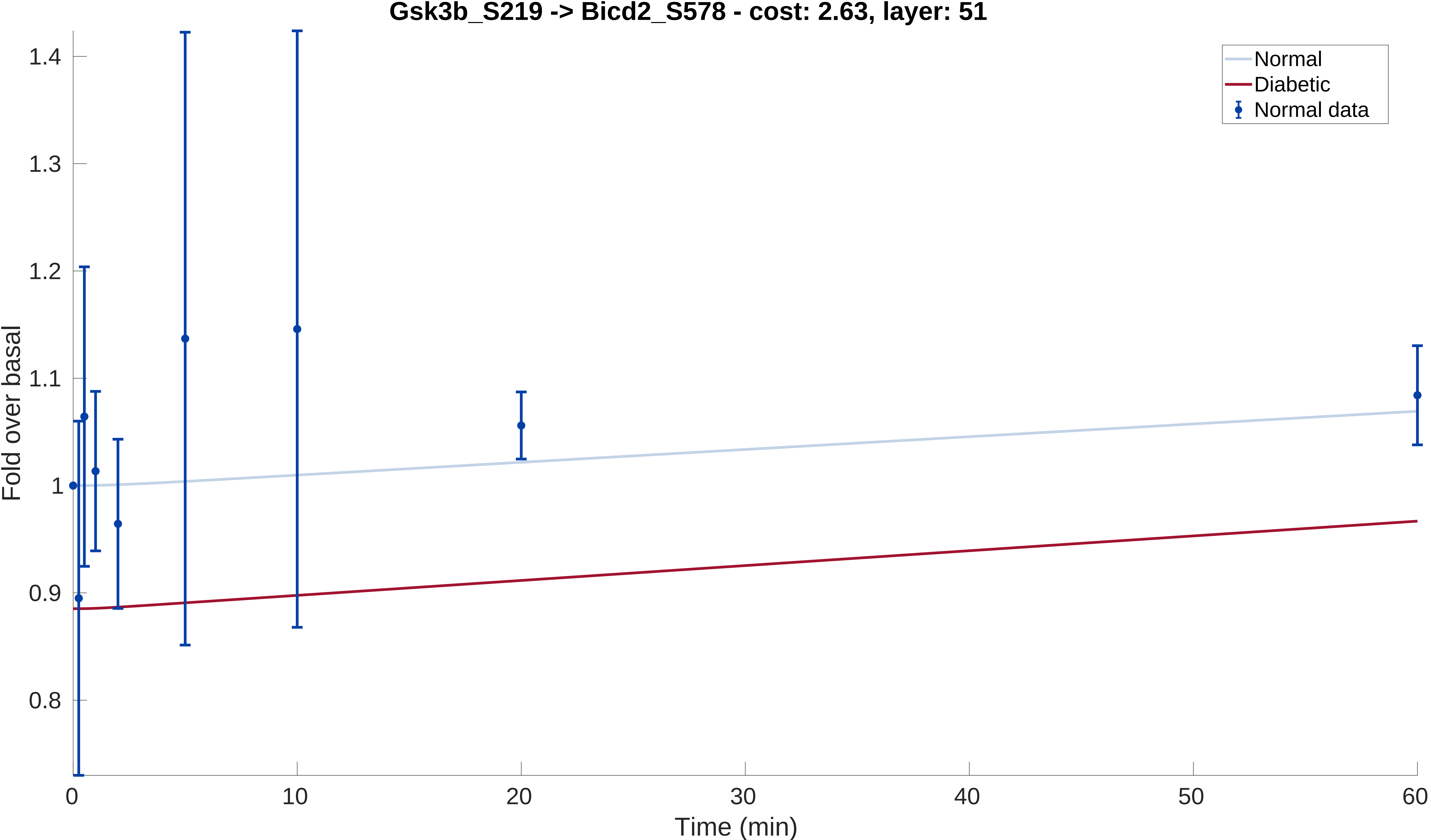

Gsk3b_S219 -> Bicd2_S578 - cost: 2.63, layer: 51
Normal
Diabetic
Normal data
Fold over basal
Time (min)
1.4
1.3
1.2
1.1
1
0.9
0.8
0
10
20
30
40
50
60

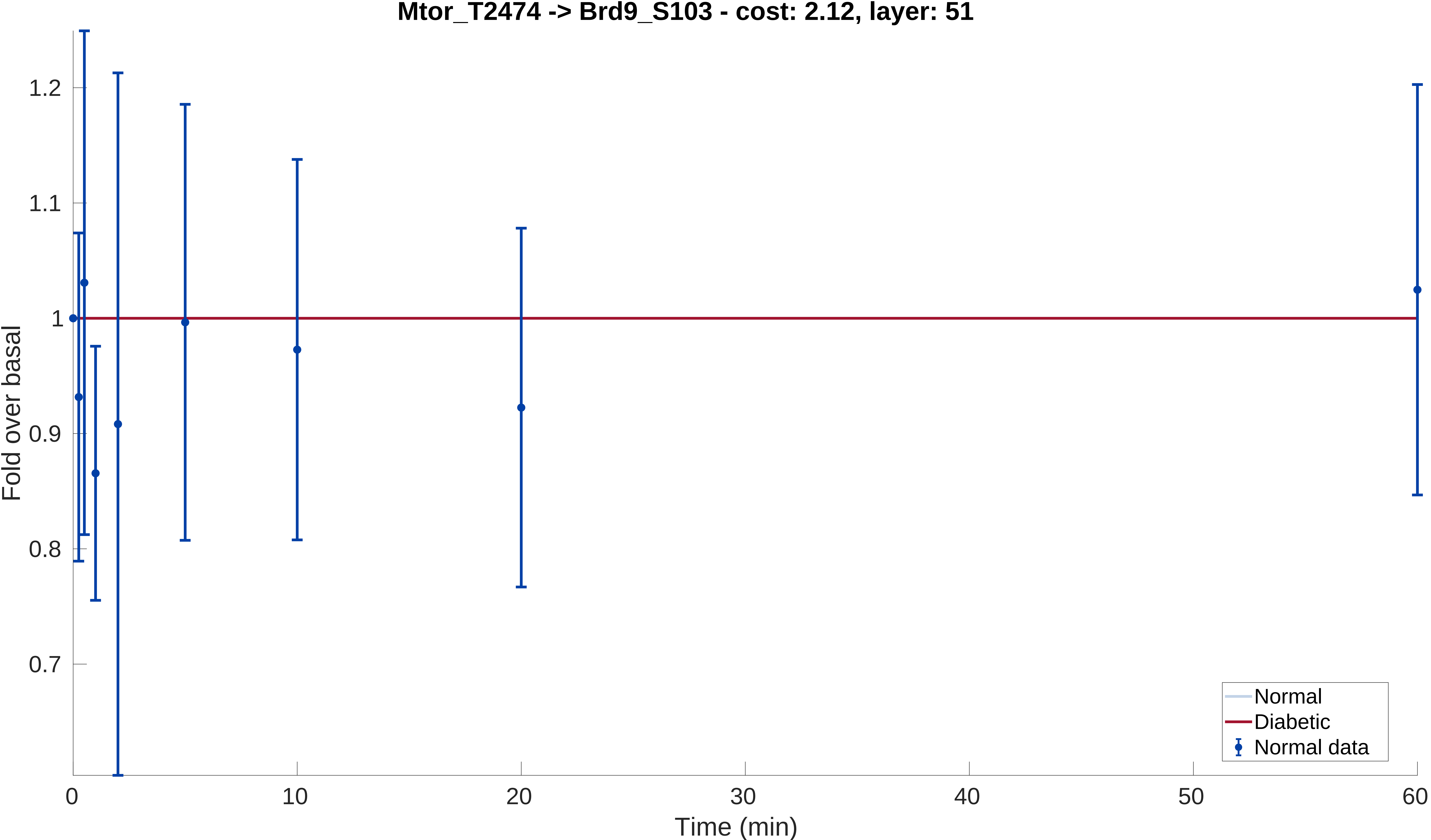
Mtor_T2474 -> Brd9_S103 - cost: 2.12, layer: 51
Fold over basal
1.2
1.1
1
0.9
0.8
0.7
0
10
20
30
40
50
60
Time (min)
Normal
Diabetic
Normal data

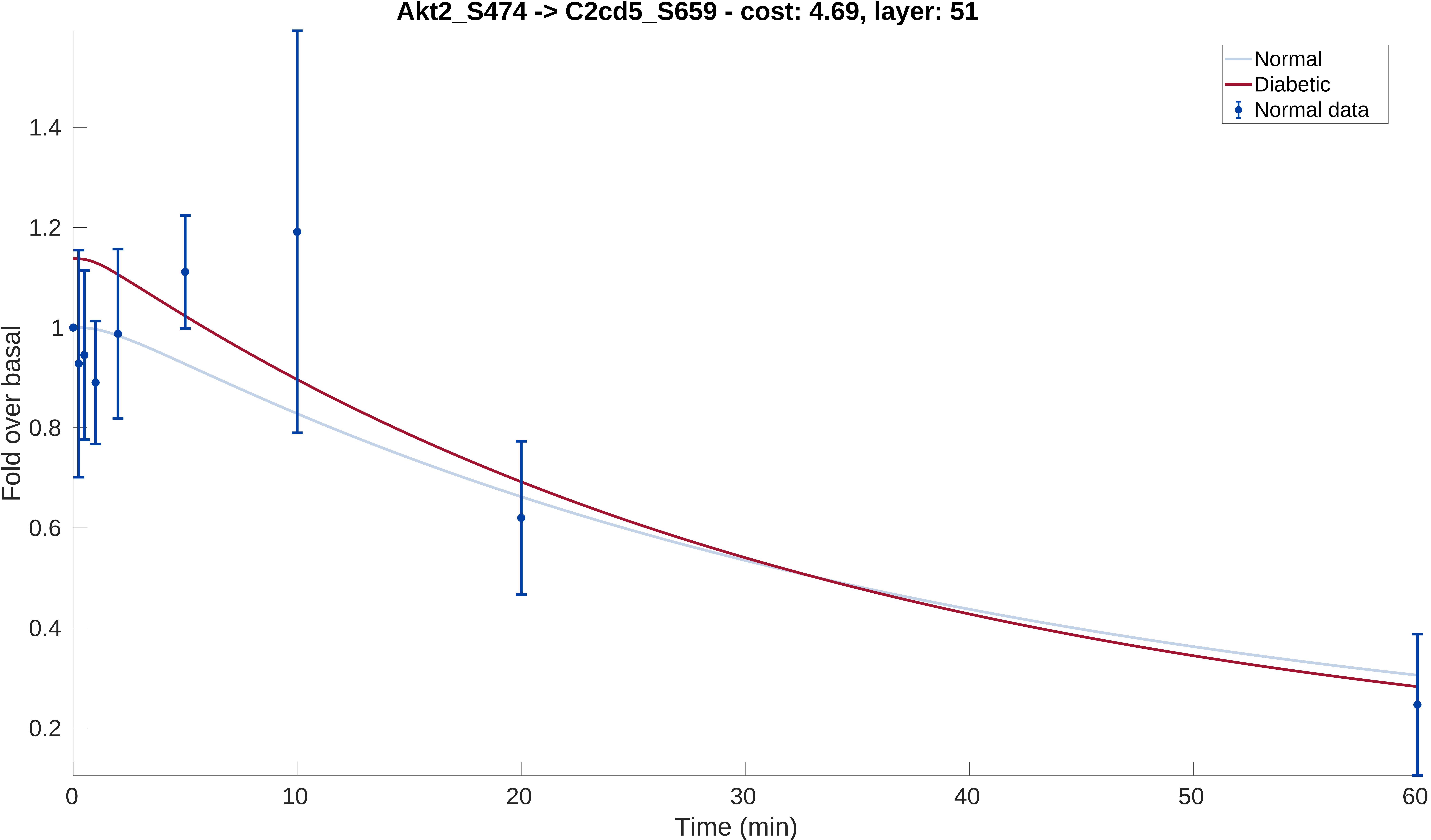
Akt2_S474 -> C2cd5_S659 - cost: 4.69, layer: 51
Normal
Diabetic
Normal data
Fold over basal
1.4
1.2
1
0.8
0.6
0.4
0.2
0
10
20
30
40
50
60
Time (min)

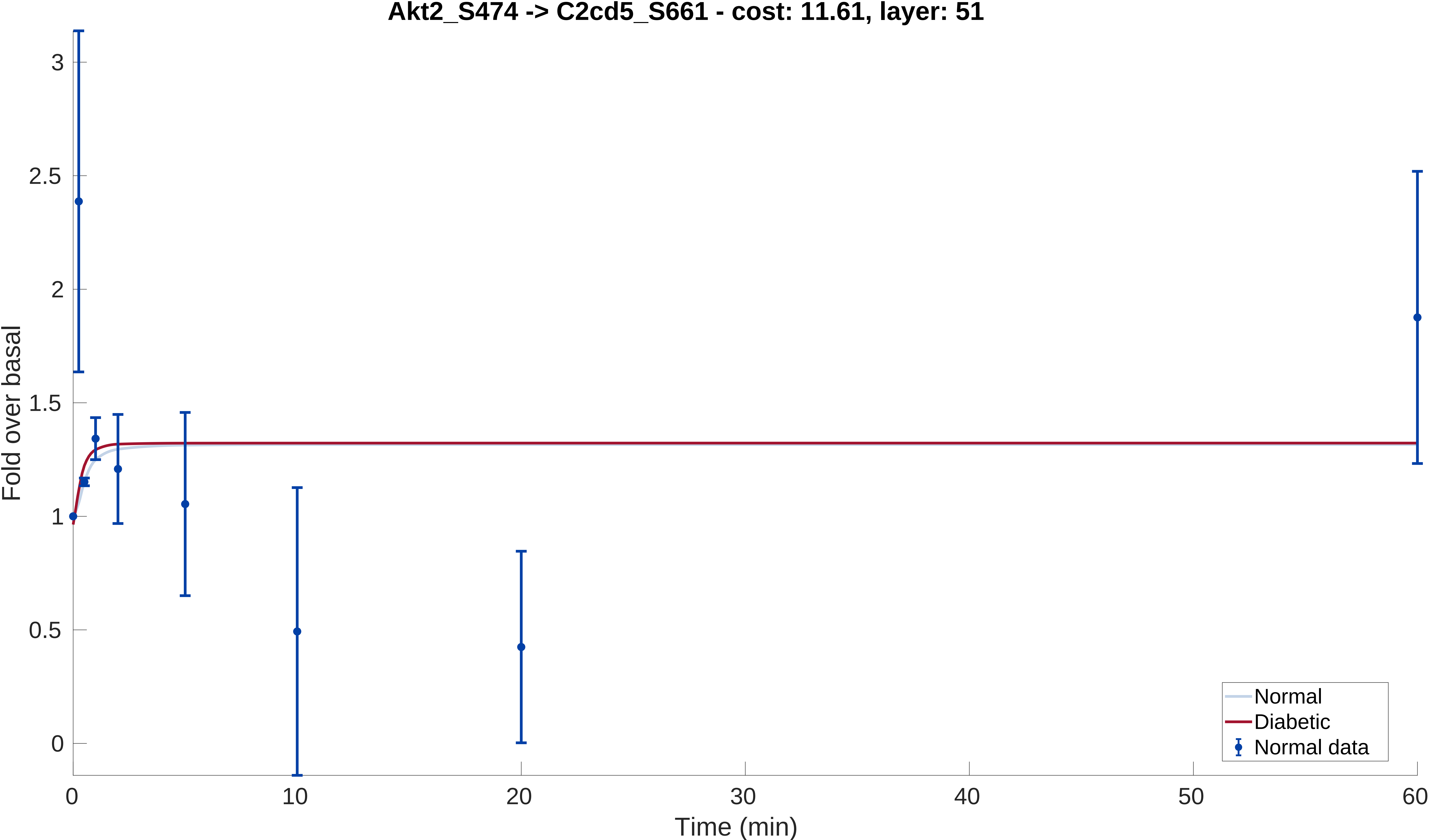

Akt2_S474 -> C2cd5_S661 - cost: 11.61, layer: 51
Fold over basal
Time (min)
0
0.5
1
1.5
2
2.5
3
0
10
20
30
40
50
60
Normal
Diabetic
Normal data

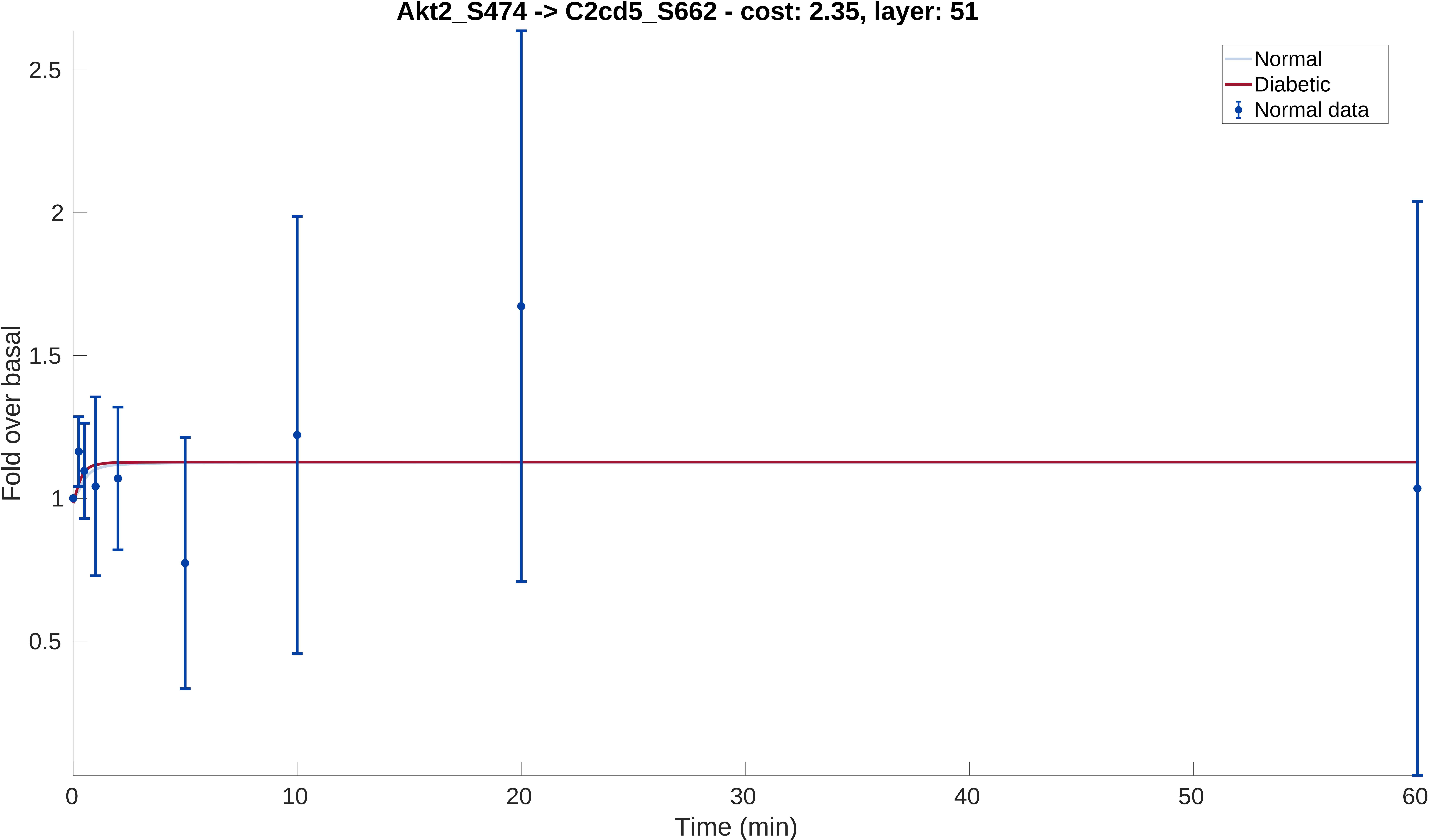

Akt2_S474 -> C2cd5_S662 - cost: 2.35, layer: 51
Normal
Diabetic
Normal data
Fold over basal
2.5
2
1.5
1
0.5
0
10
20
30
40
50
60
Time (min)

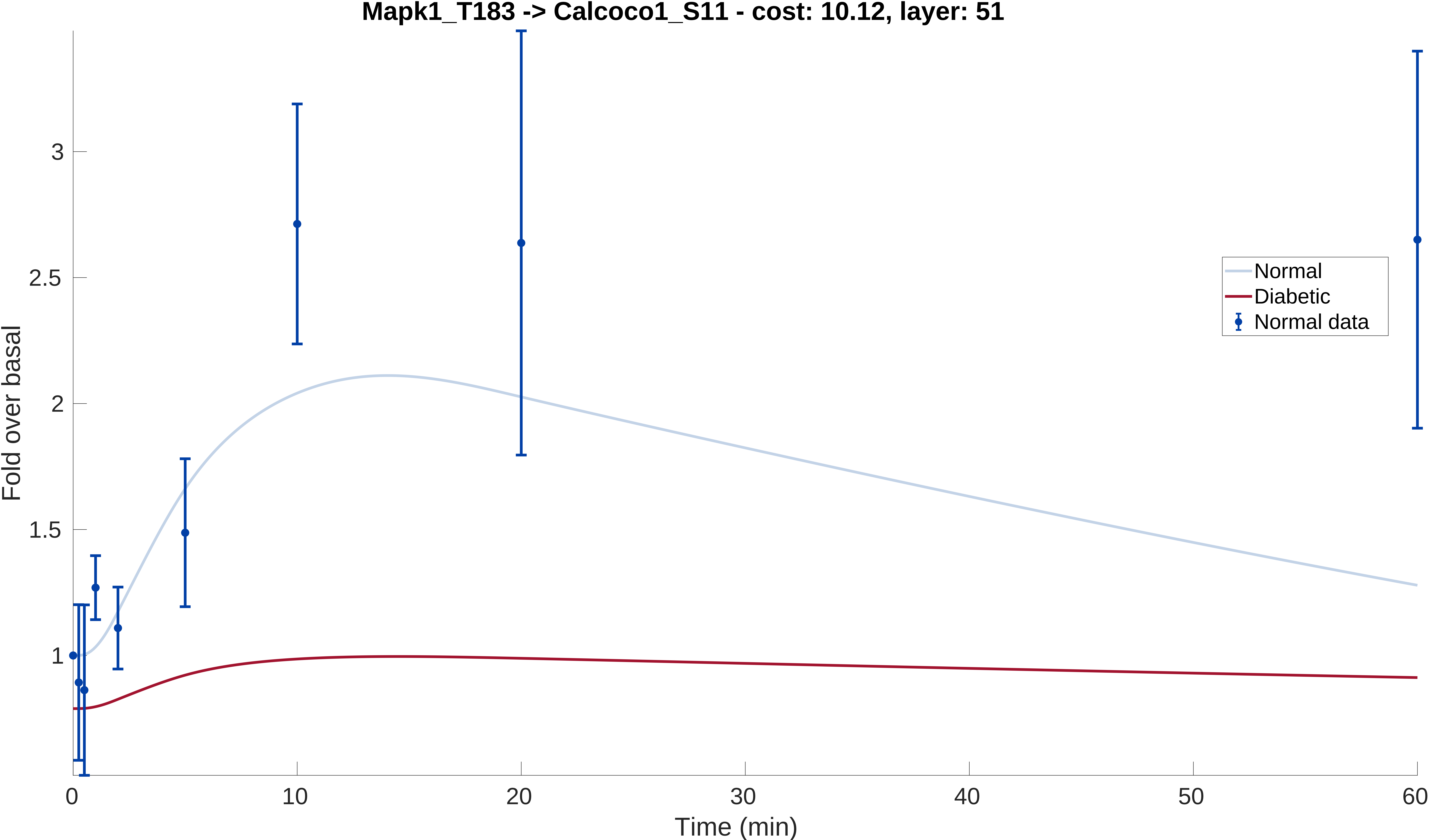
Mapk1_T183 -> Calcoco1_S11 - cost: 10.12, layer: 51
Fold over basal
3
2.5
2
1.5
1
0
10
20
30
40
50
60
Time (min)
Normal
Diabetic
Normal data

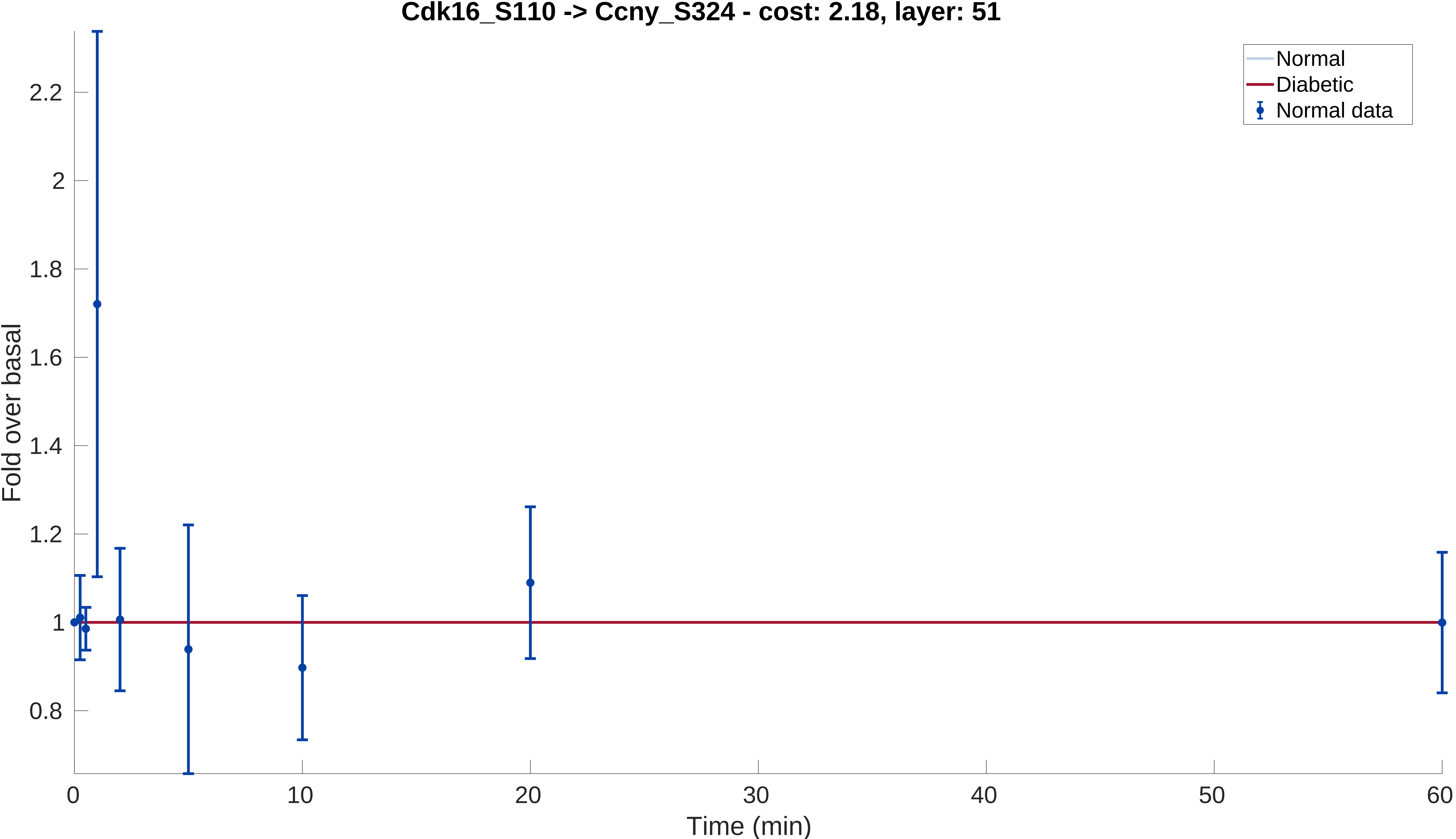
Cdk16_S110 -> Ccny_S324 - cost: 2.18, layer: 51
Normal
Diabetic
Normal data
Fold over basal
0.8
1
1.2
1.4
1.6
1.8
2
2.2
0
10
20
30
40
50
60
Time (min)

Cdk16_S110 -> Ccny_S326 - cost: 1.09, layer: 51

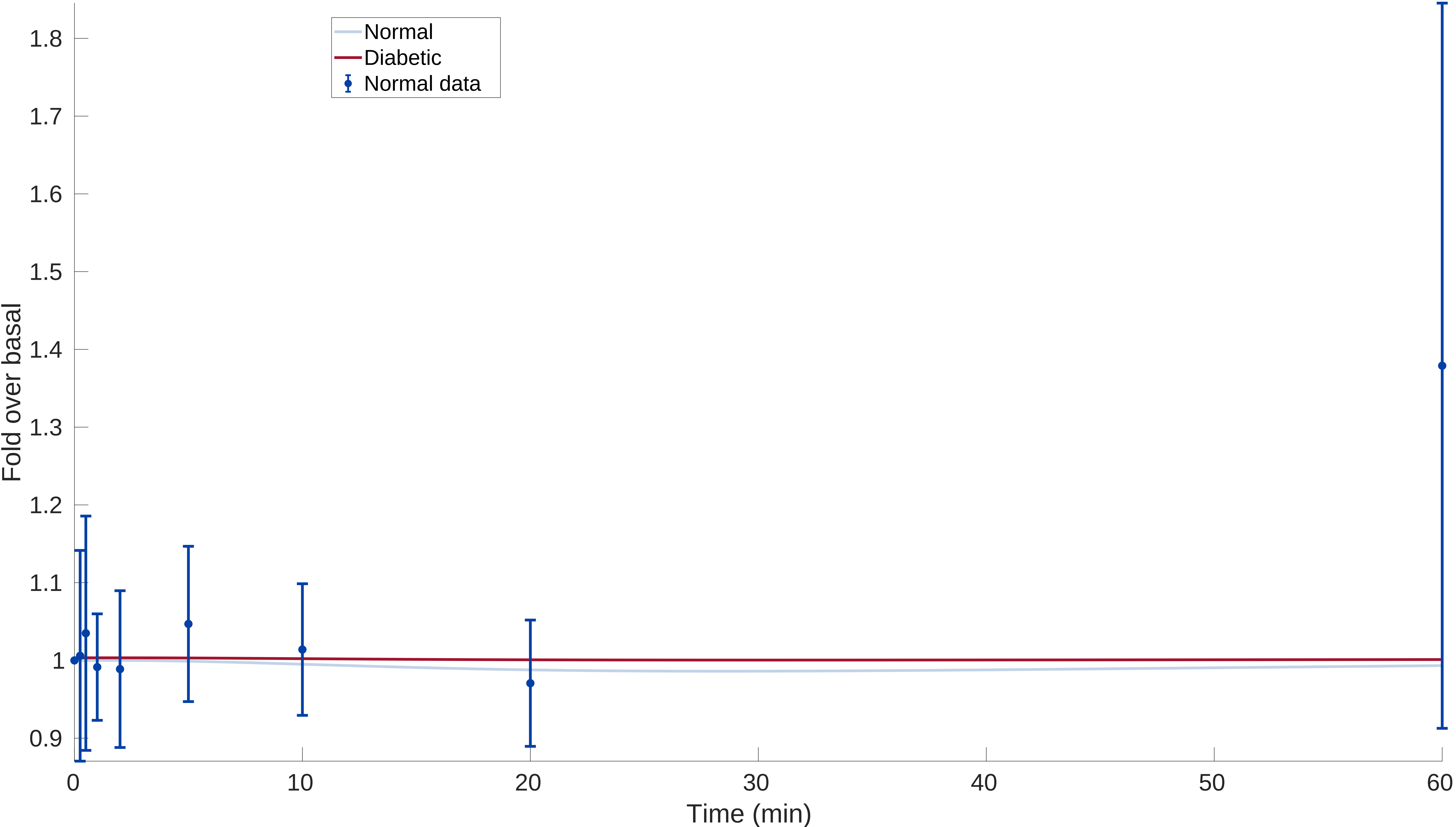

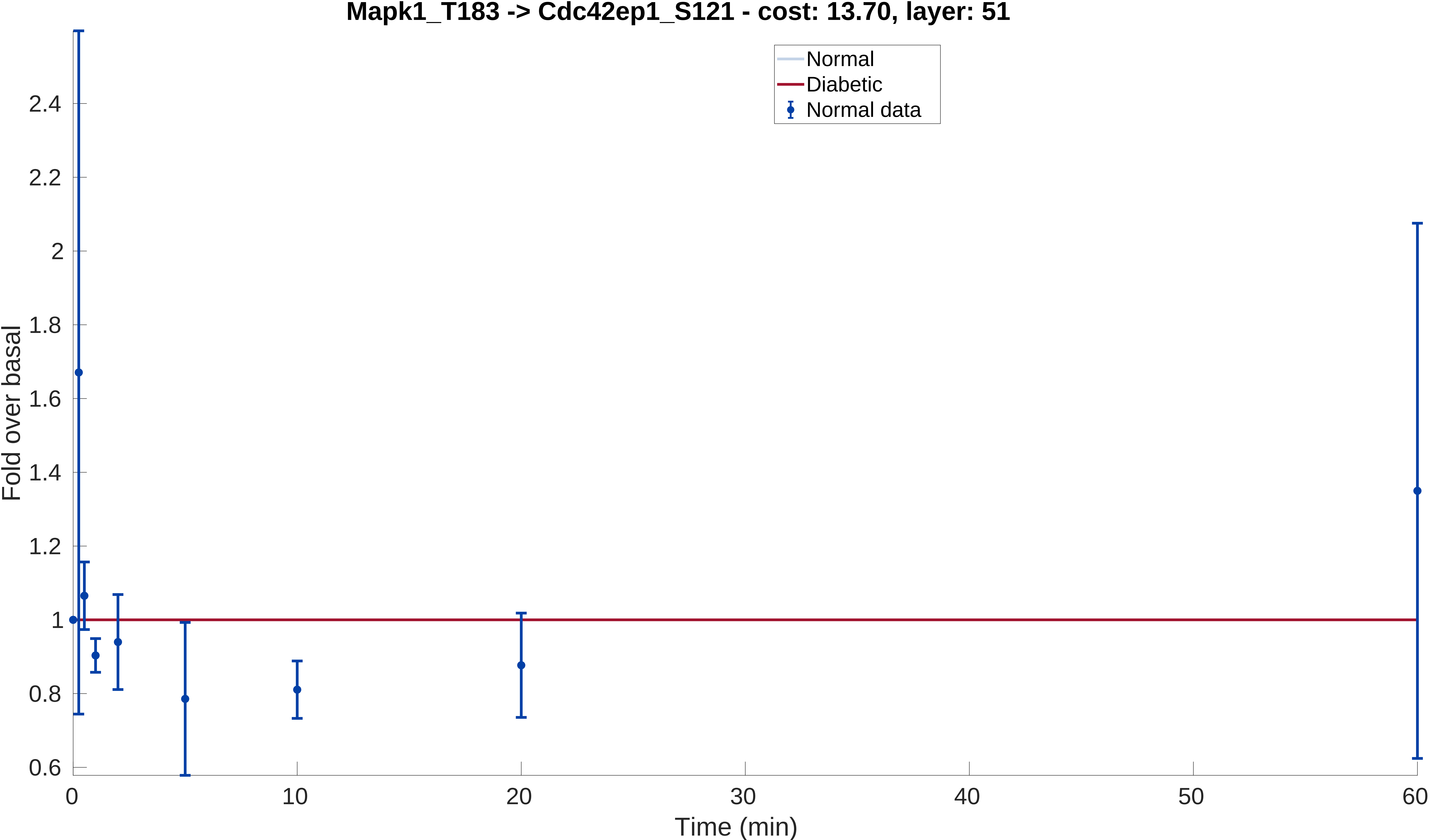

Mapk1_T183 -> Cdc42ep1_S121 - cost: 13.70, layer: 51
Normal
Diabetic
Normal data
Fold over basal
Time (min)
0.6
0.8
1
1.2
1.4
1.6
1.8
2
2.2
2.4
0
10
20
30
40
50
60

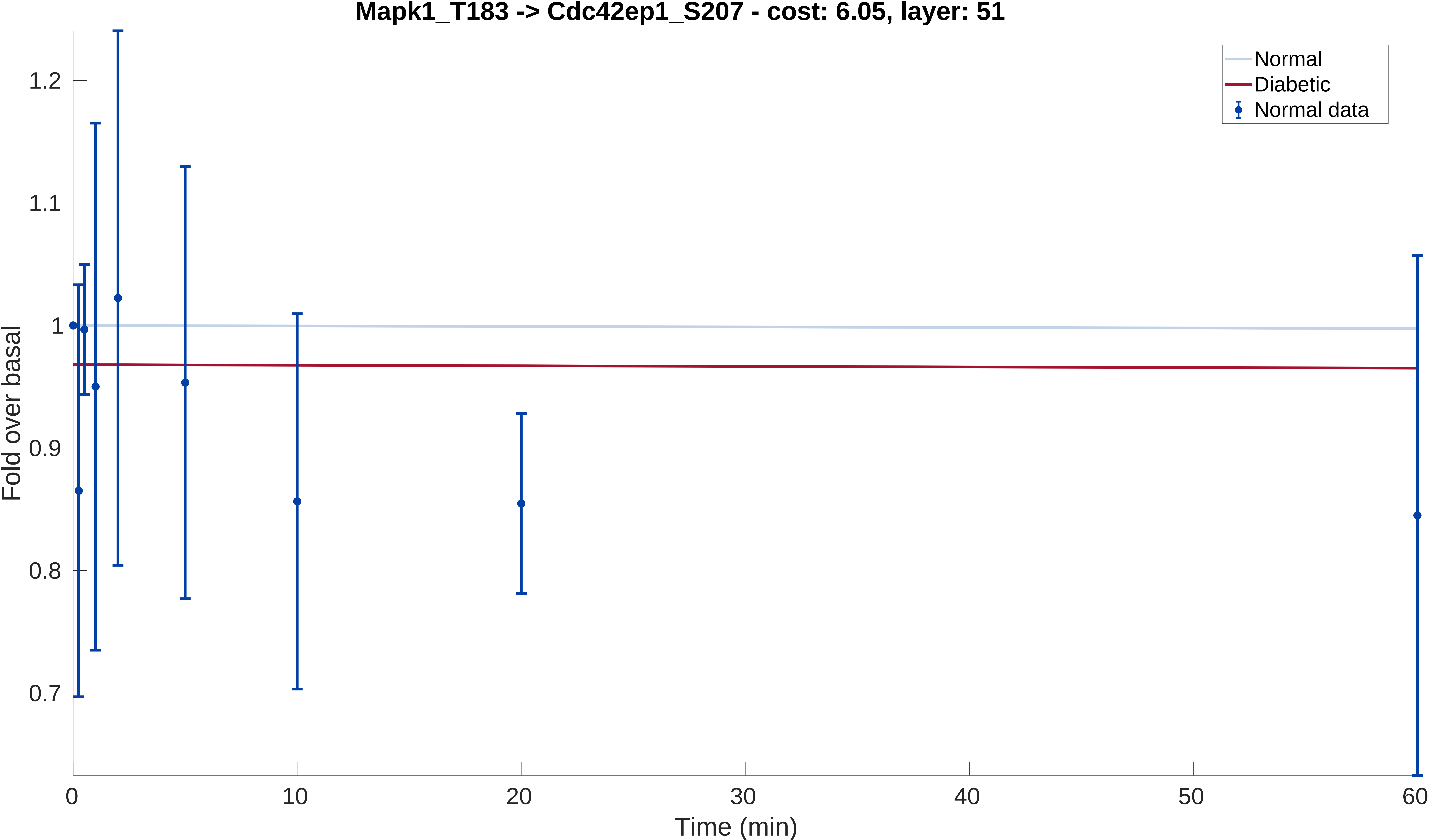

Mapk1_T183 -> Cdc42ep1_S207 - cost: 6.05, layer: 51
Normal
Diabetic
Normal data
1.2
1.1
1
0.9
0.8
0.7
Fold over basal
0
10
20
30
40
50
60
Time (min)

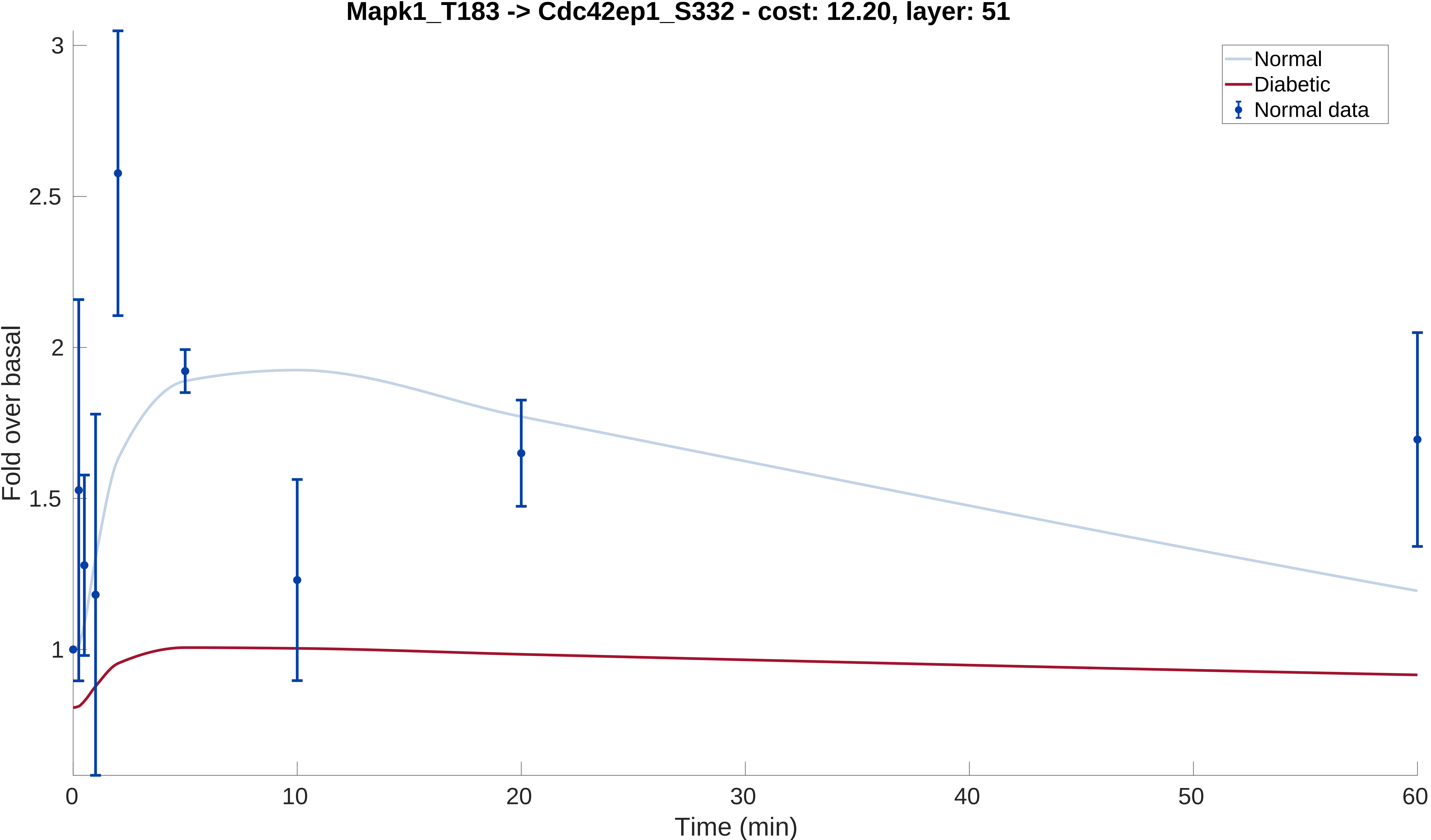

Mapk1_T183 -> Cdc42ep1_S332 - cost: 12.20, layer: 51
Normal
Diabetic
Normal data
Fold over basal
Time (min)
3
2.5
2
1.5
1
0
10
20
30
40
50
60

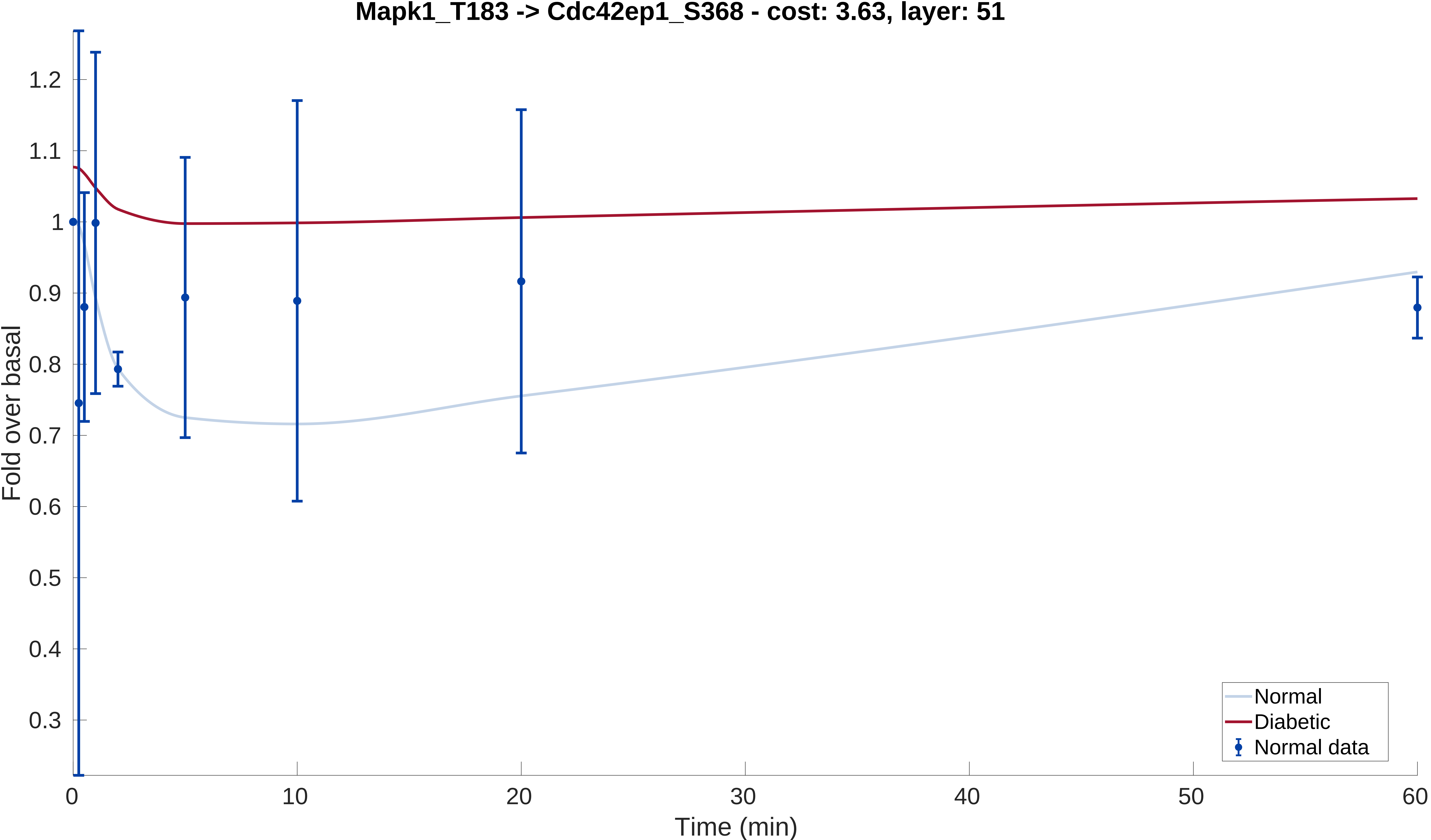

Mapk1_T183 -> Cdc42ep1_S368 - cost: 3.63, layer: 51
Fold over basal
1.2
1.1
1
0.9
0.8
0.7
0.6
0.5
0.4
0.3
0
10
20
30
40
50
60
Time (min)
Normal
Diabetic
Normal data

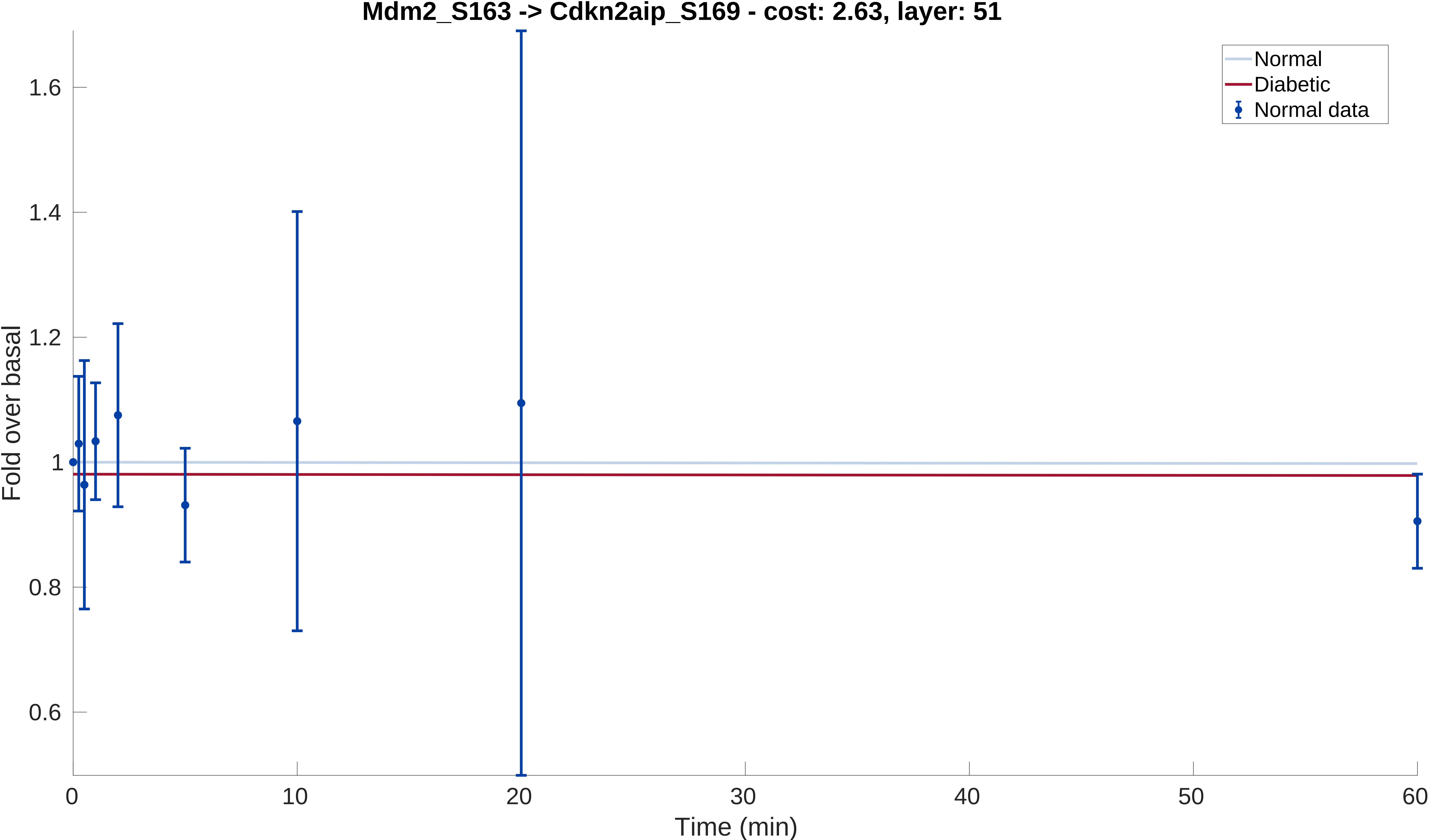

Mdm2_S163 -> Cdkn2aip_S169 - cost: 2.63, layer: 51
Normal
Diabetic
Normal data
Fold over basal
Time (min)
0.6
0.8
1
1.2
1.4
1.6
0
10
20
30
40
50
60

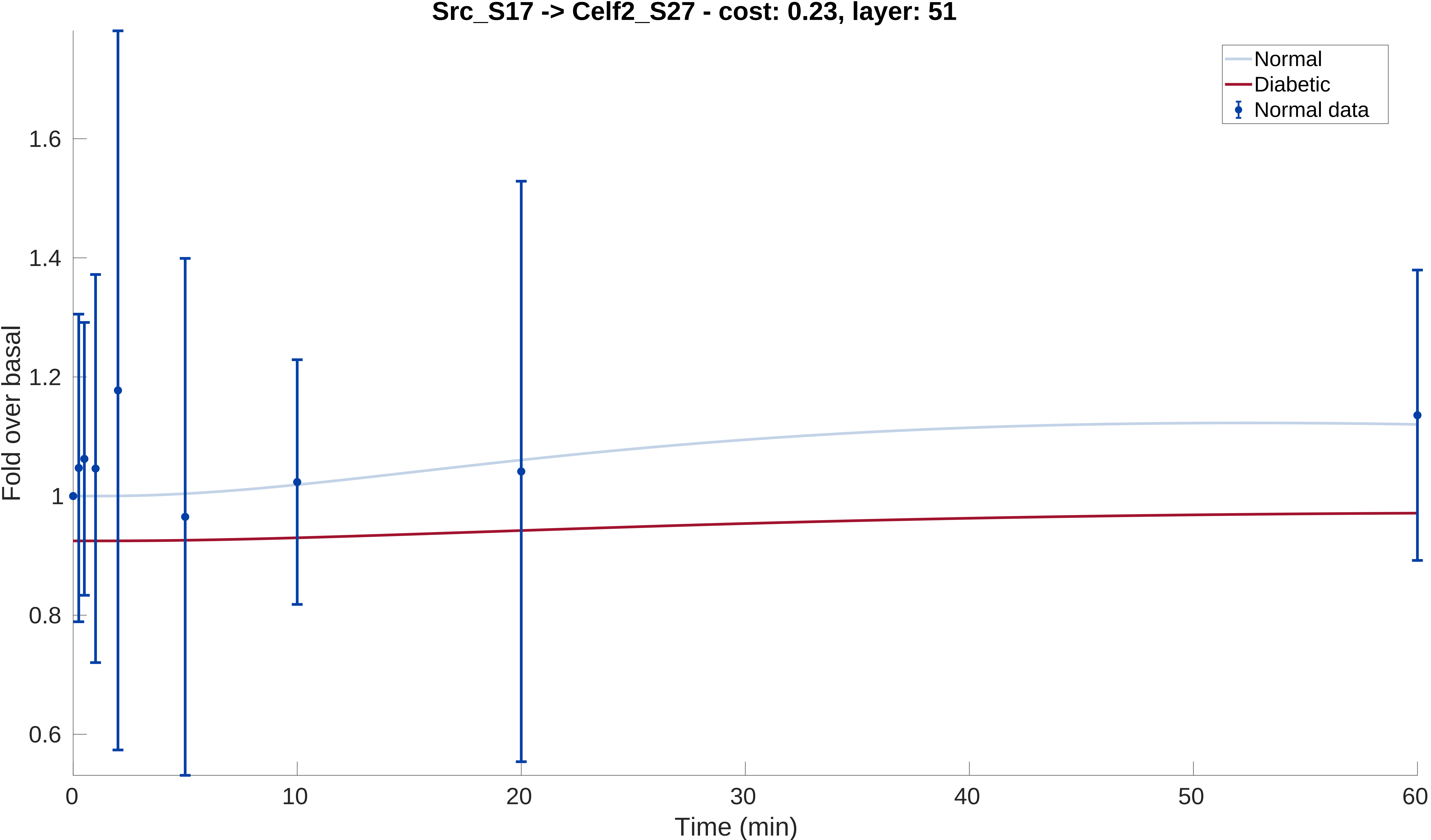
Src_S17 -> Celf2_S27 - cost: 0.23, layer: 51
Normal
Diabetic
Normal data
1.6
1.4
1.2
1
0.8
0.6
Fold over basal
0
10
20
30
40
50
60
Time (min)

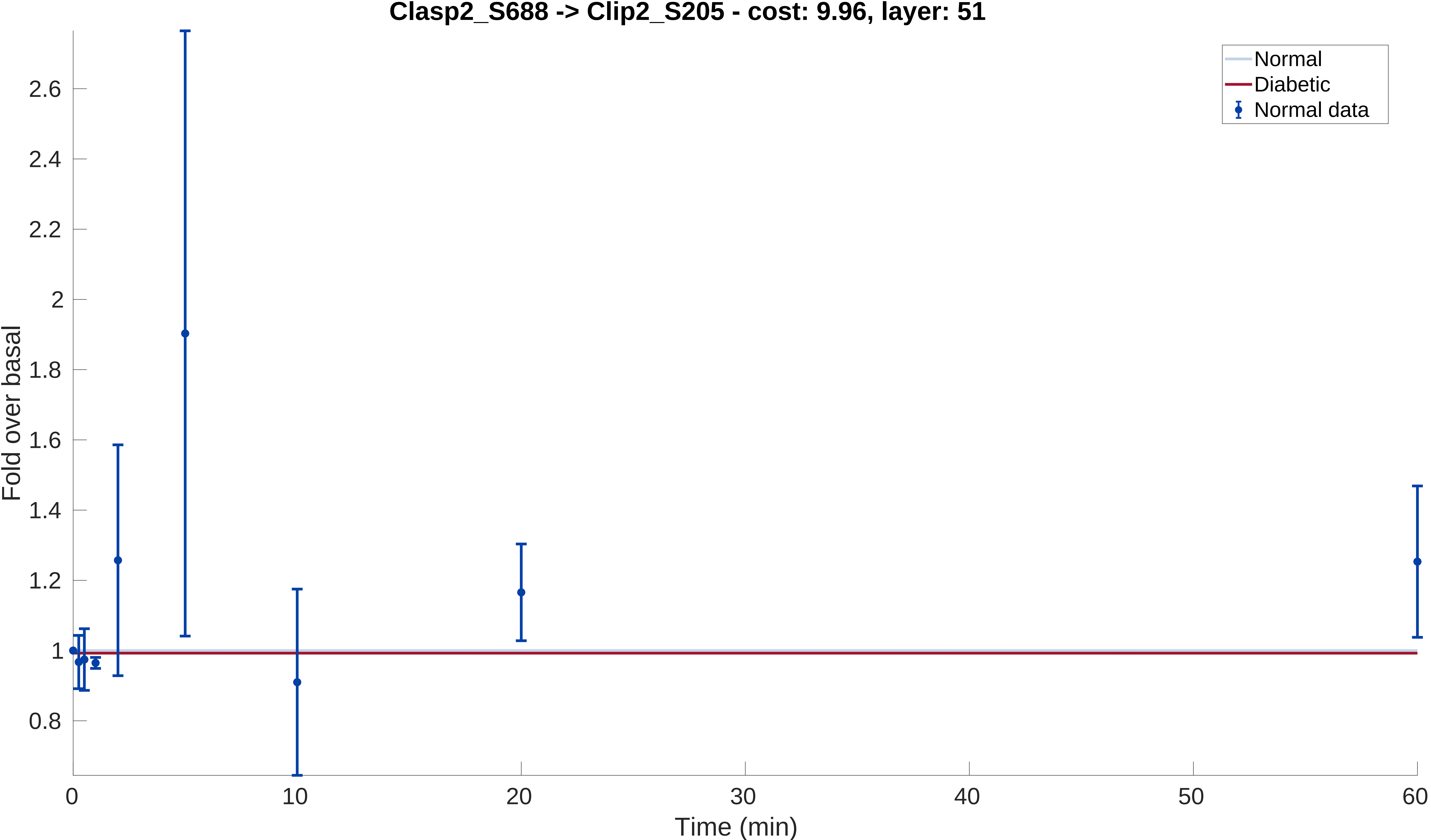
Clasp2_S688 -> Clip2_S205 - cost: 9.96, layer: 51
Normal
Diabetic
Normal data
Fold over basal
2.6
2.4
2.2
2
1.8
1.6
1.4
1.2
1
0.8
0
10
20
30
40
50
60
Time (min)

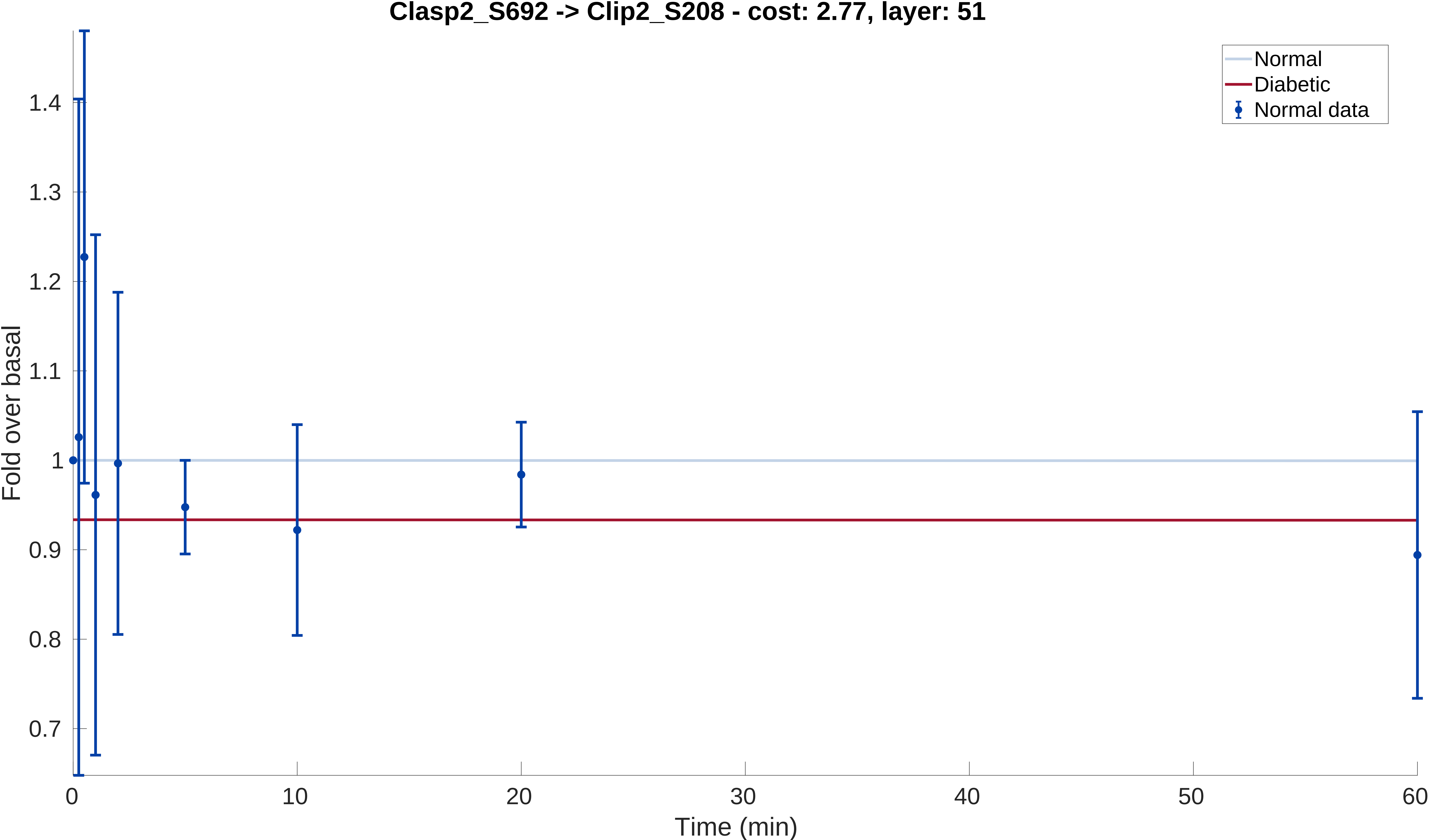

Clasp2_S692 -> Clip2_S208 - cost: 2.77, layer: 51
Normal
Diabetic
Normal data
Fold over basal
1.4
1.3
1.2
1.1
1
0.9
0.8
0.7
0
10
20
30
40
50
60
Time (min)

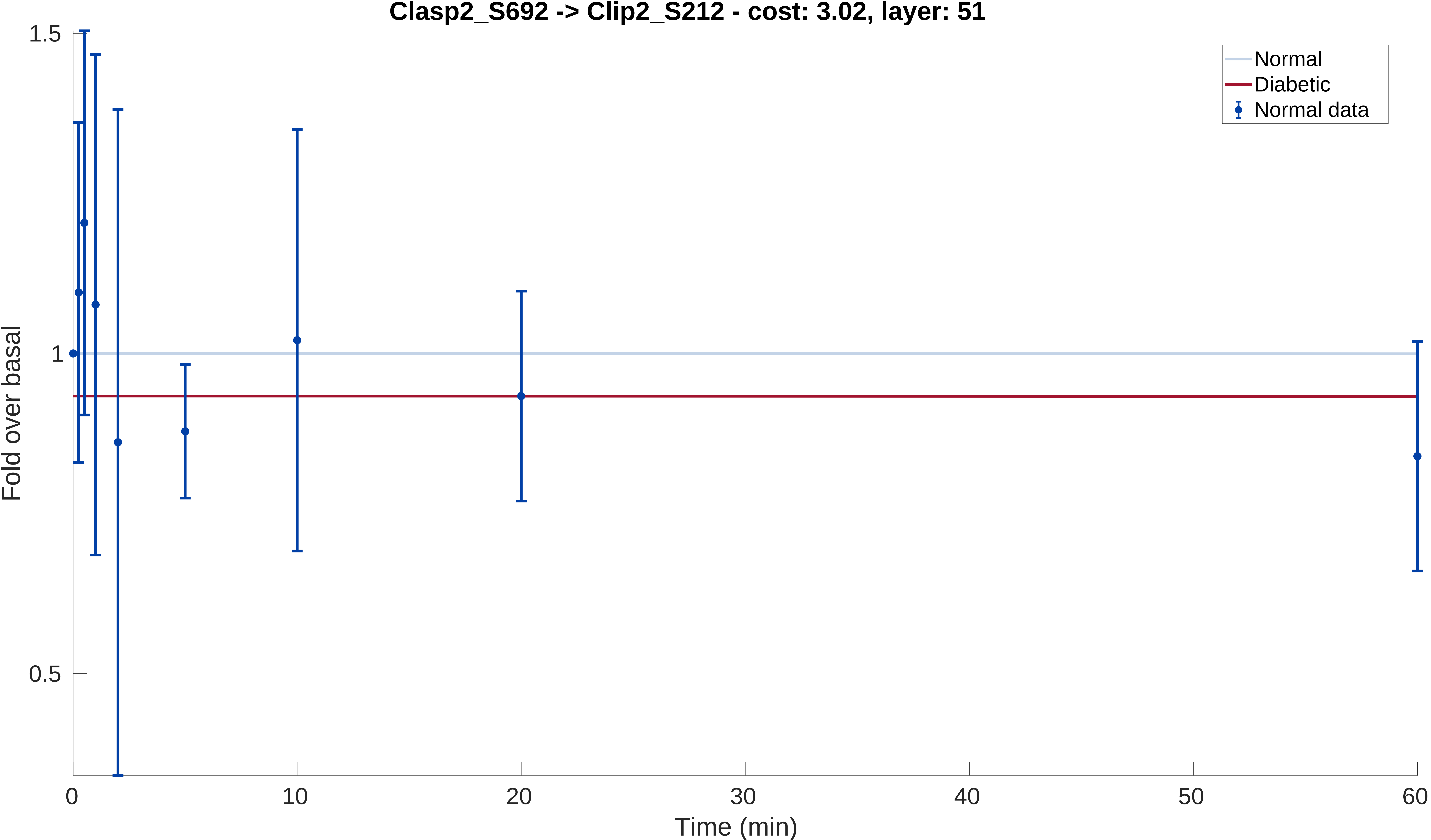

Clasp2_S692 -> Clip2_S212 - cost: 3.02, layer: 51
Normal
Diabetic
Normal data
Fold over basal
1.5
1
0.5
0
10
20
30
40
50
60
Time (min)

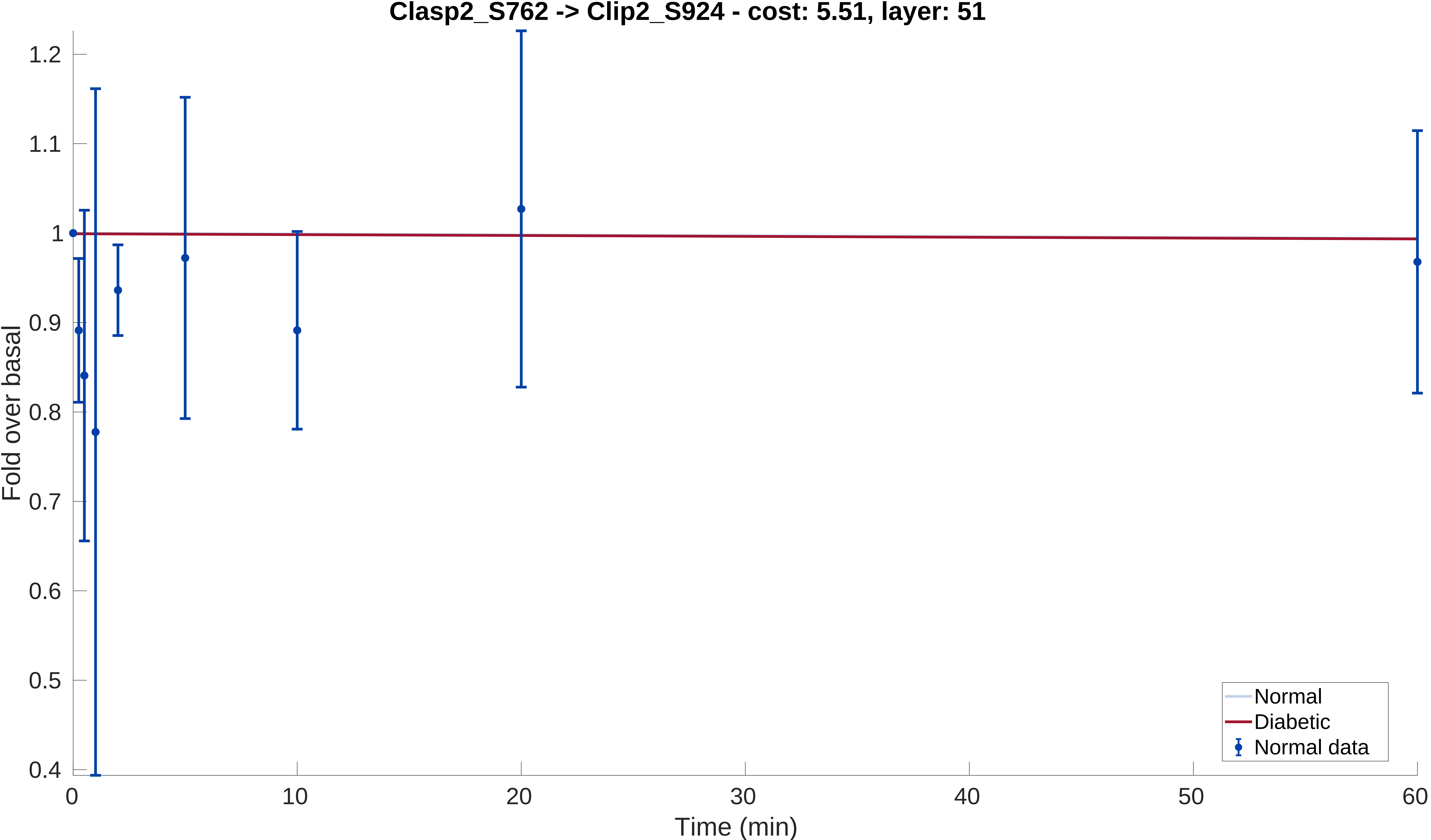

Clasp2_S762 -> Clip2_S924 - cost: 5.51, layer: 51
Fold over basal
1.2
1.1
1
0.9
0.8
0.7
0.6
0.5
0.4
0
10
20
30
40
50
60
Time (min)
Normal
Diabetic
Normal data

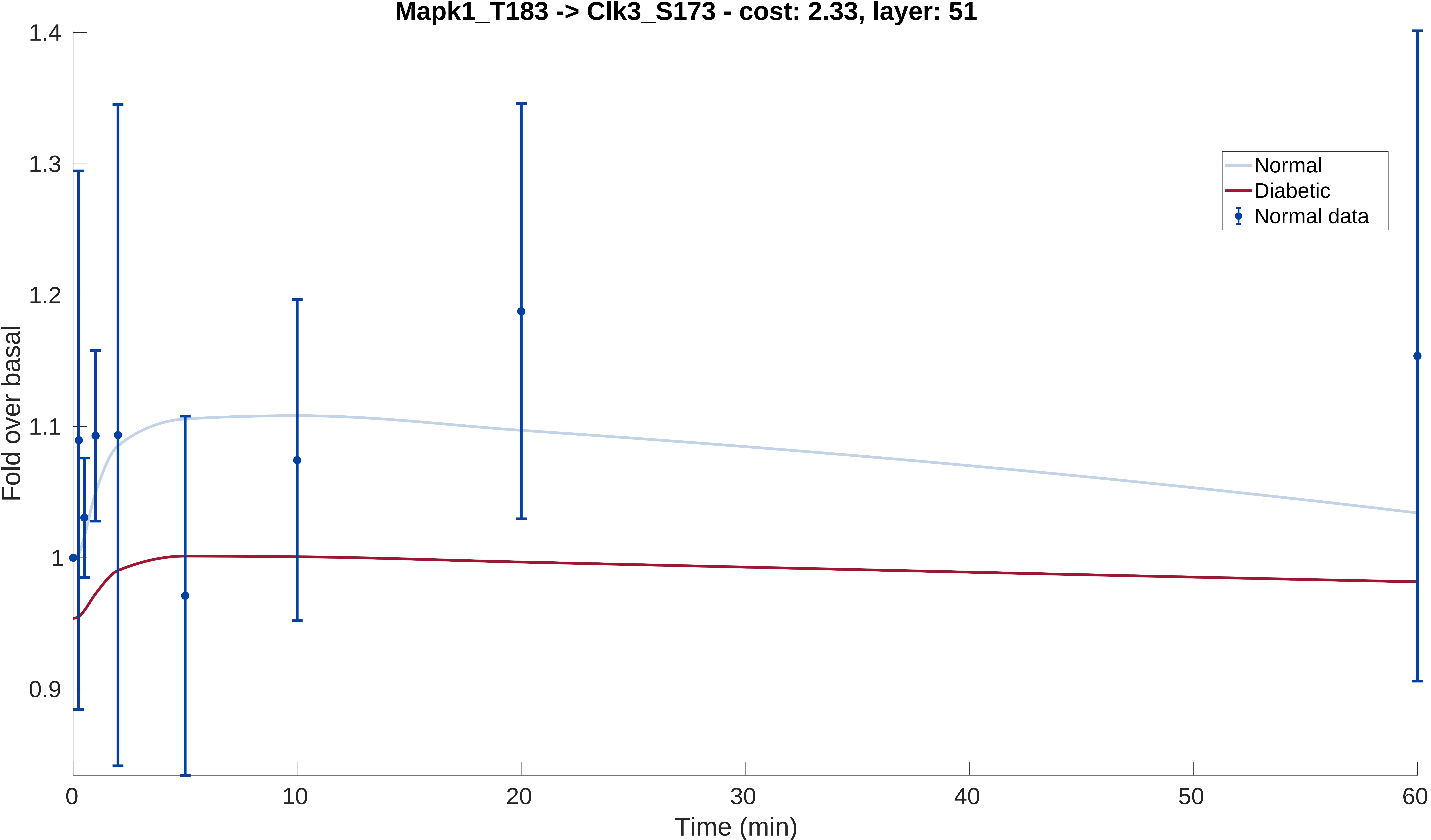
Mapk1_T183 -> Clk3_S173 - cost: 2.33, layer: 51
Fold over basal
Time (min)
Normal
Diabetic
Normal data
0.9
1
1.1
1.2
1.3
1.4
0
10
20
30
40
50
60

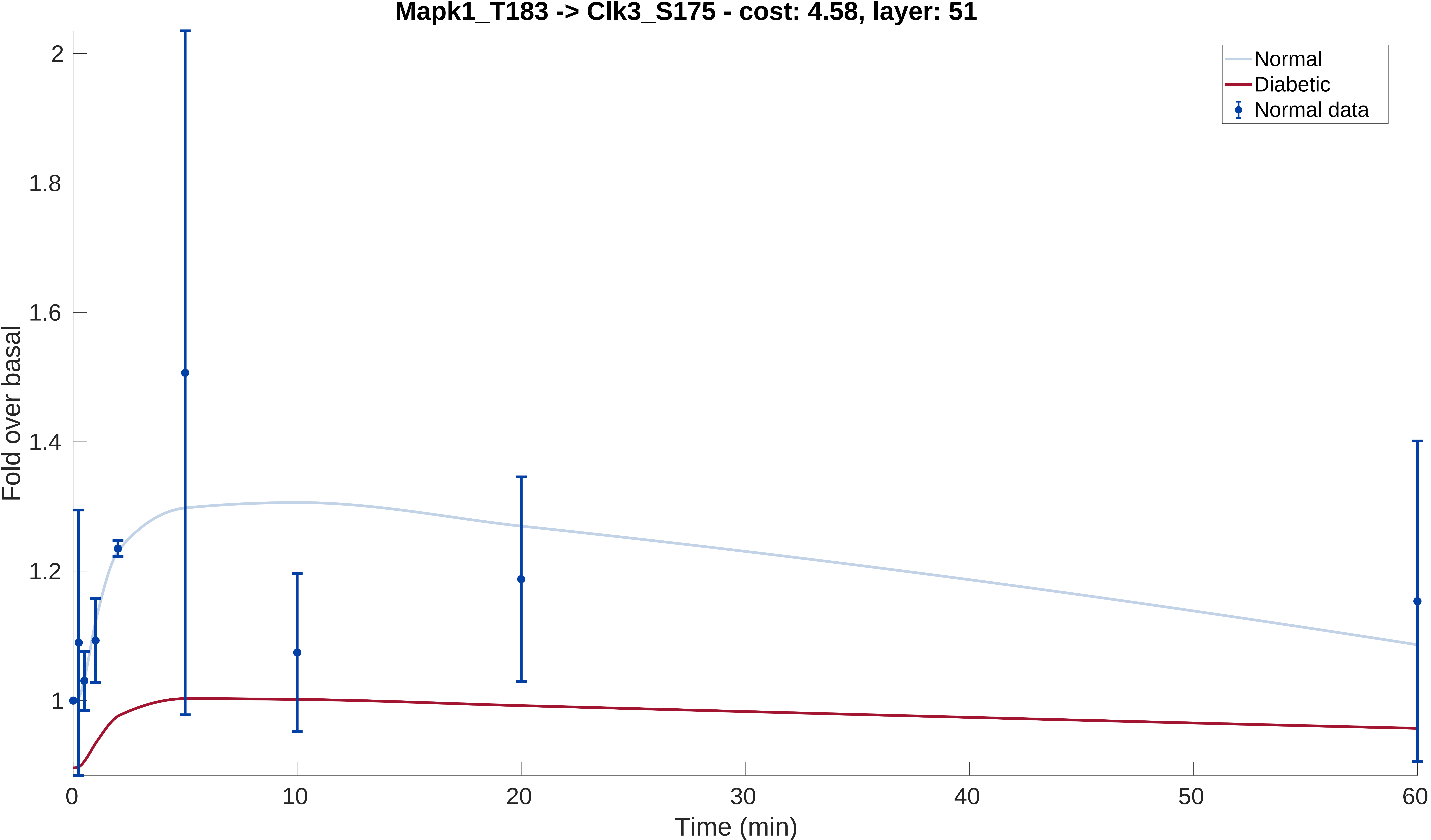

Mapk1_T183 -> Clk3_S175 - cost: 4.58, layer: 51
Normal
Diabetic
Normal data
Fold over basal
Time (min)
2
1.8
1.6
1.4
1.2
1
0
10
20
30
40
50
60

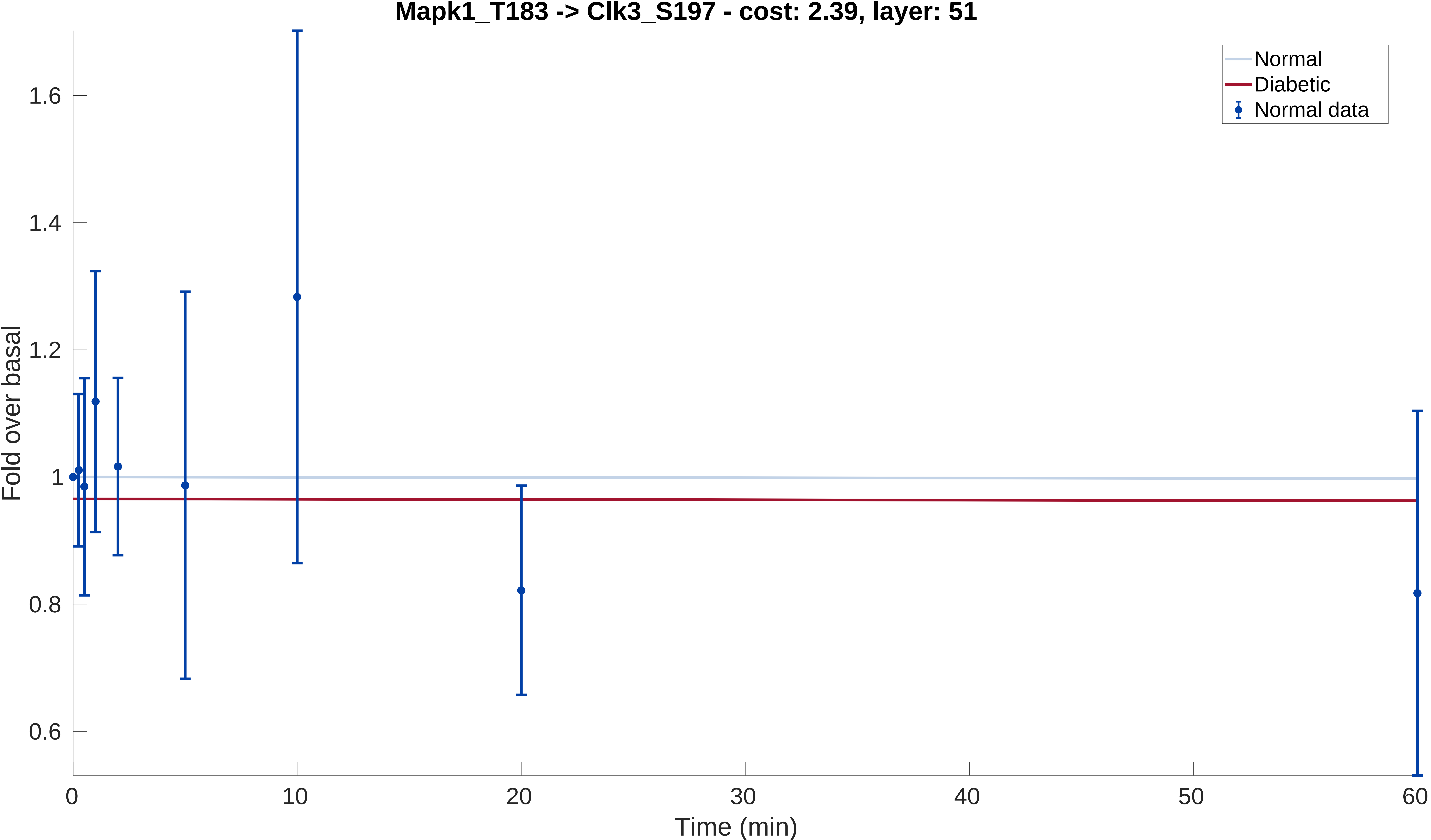
Mapk1_T183 -> Clk3_S197 - cost: 2.39, layer: 51
Normal
Diabetic
Normal data
1.6
1.4
1.2
1
0.8
0.6
Fold over basal
0
10
20
30
40
50
60
Time (min)

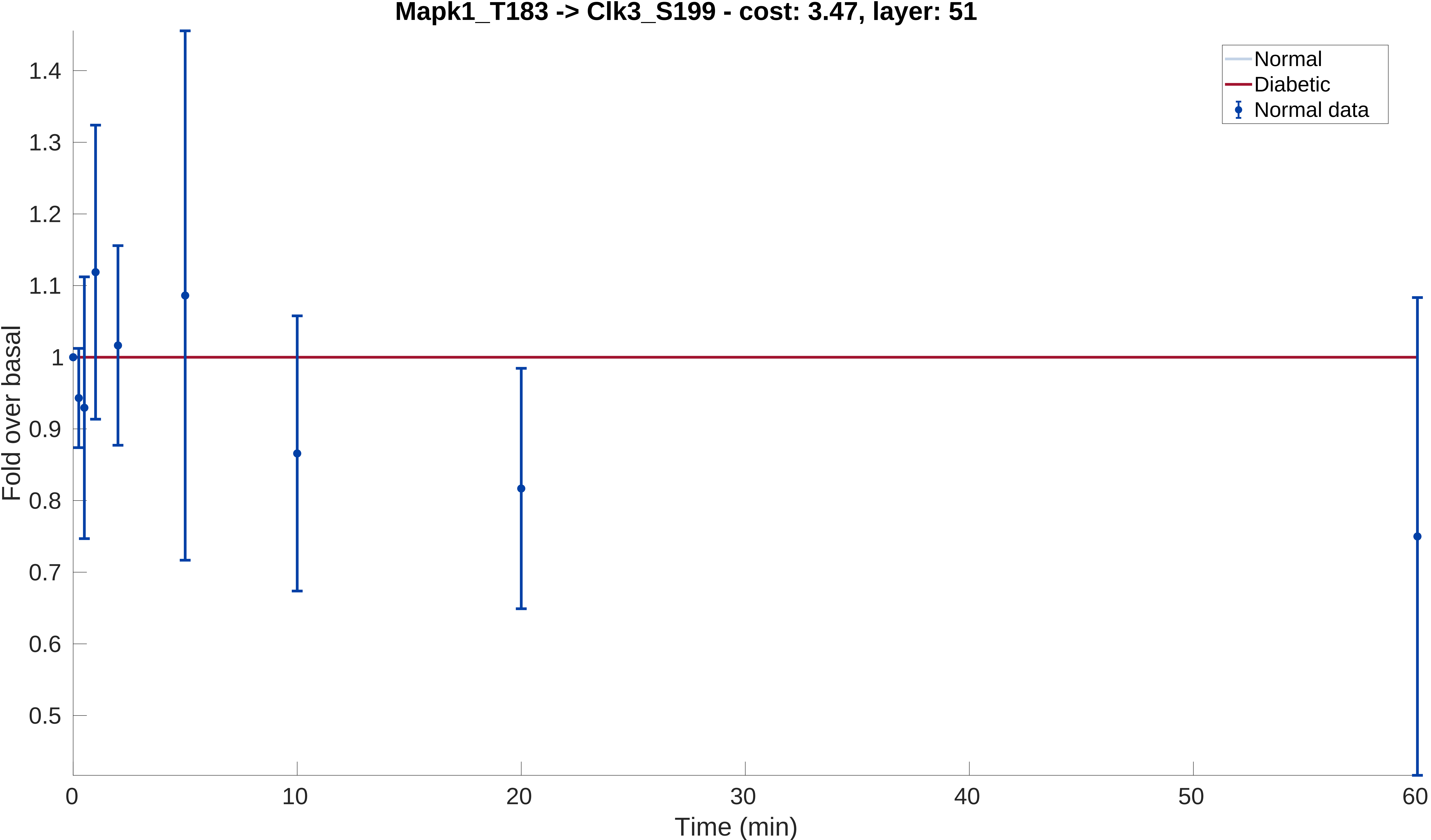

Mapk1_T183 -> Clk3_S199 - cost: 3.47, layer: 51
Normal
Diabetic
Normal data
Fold over basal
Time (min)
1.4
1.3
1.2
1.1
1
0.9
0.8
0.7
0.6
0.5
0
10
20
30
40
50
60

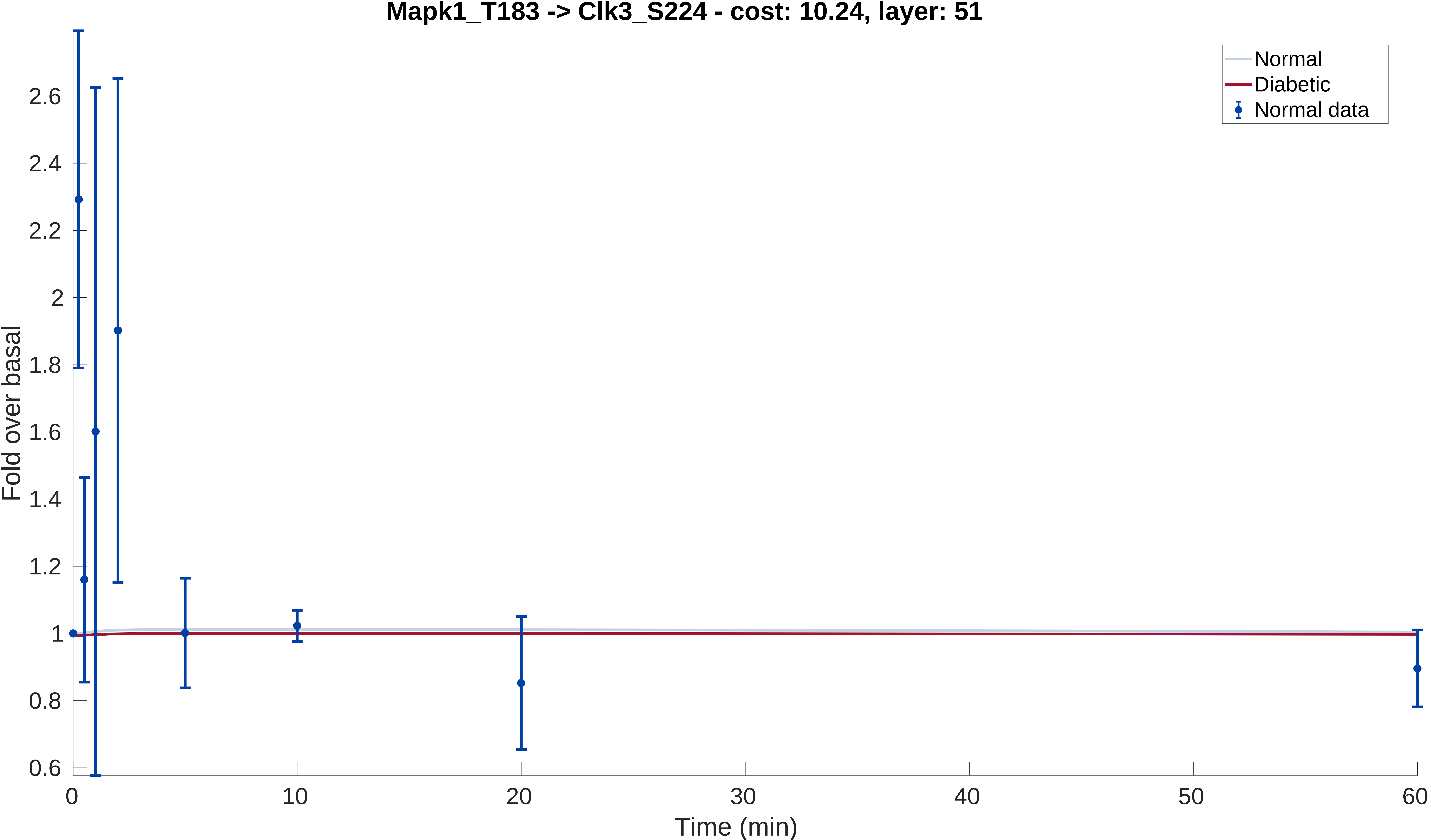
Mapk1_T183 -> Clk3_S224 - cost: 10.24, layer: 51
Normal
Diabetic
Normal data
Fold over basal
2.6
2.4
2.2
2
1.8
1.6
1.4
1.2
1
0.8
0.6
0
10
20
30
40
50
60
Time (min)

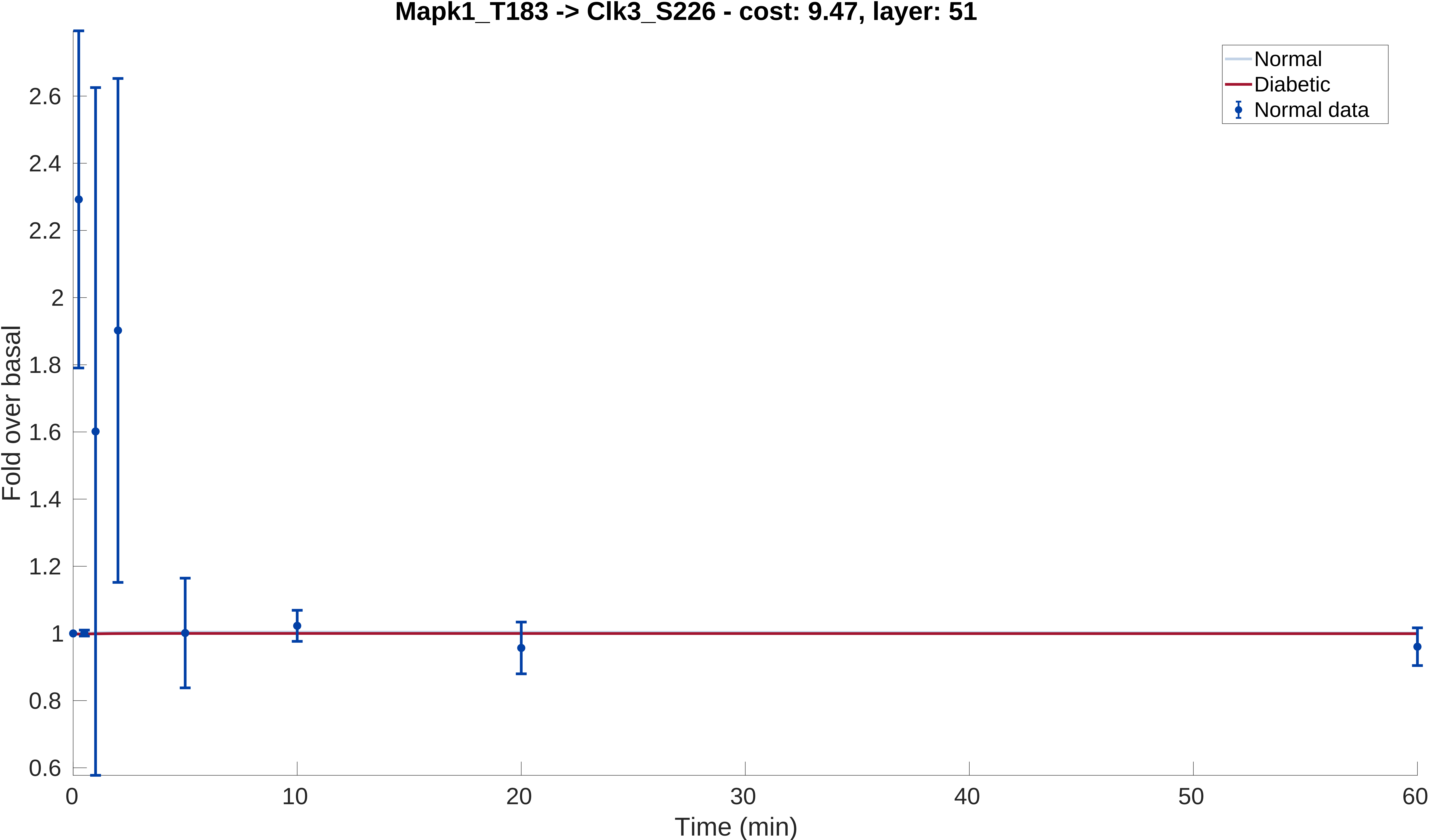

Mapk1_T183 -> Clk3_S226 - cost: 9.47, layer: 51
Normal
Diabetic
Normal data
Fold over basal
2.6
2.4
2.2
2
1.8
1.6
1.4
1.2
1
0.8
0.6
0
10
20
30
40
50
60
Time (min)

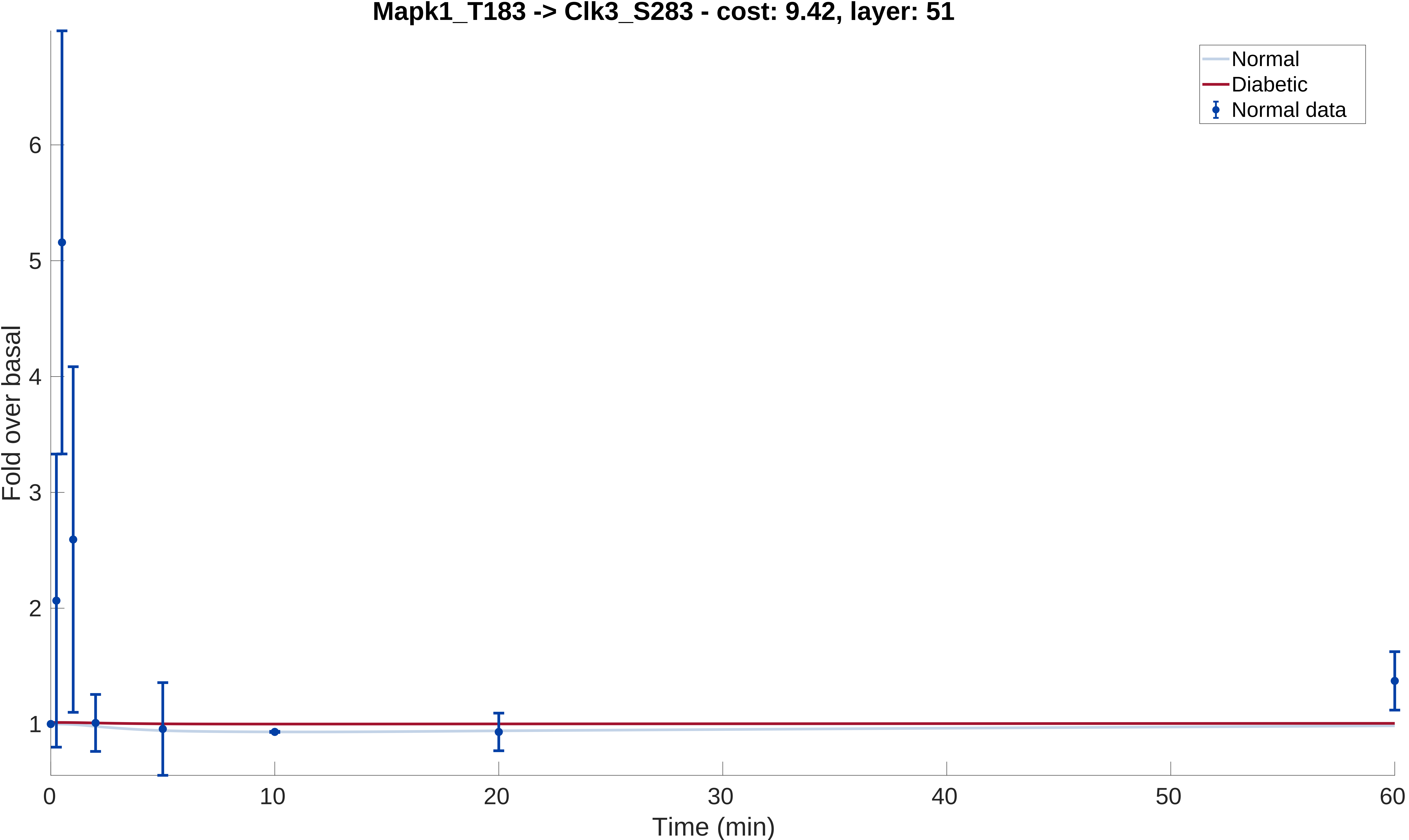

Mapk1_T183 -> Clk3_S283 - cost: 9.42, layer: 51
Normal
Diabetic
Normal data
Fold over basal
1
2
3
4
5
6
0
10
20
30
40
50
60
Time (min)

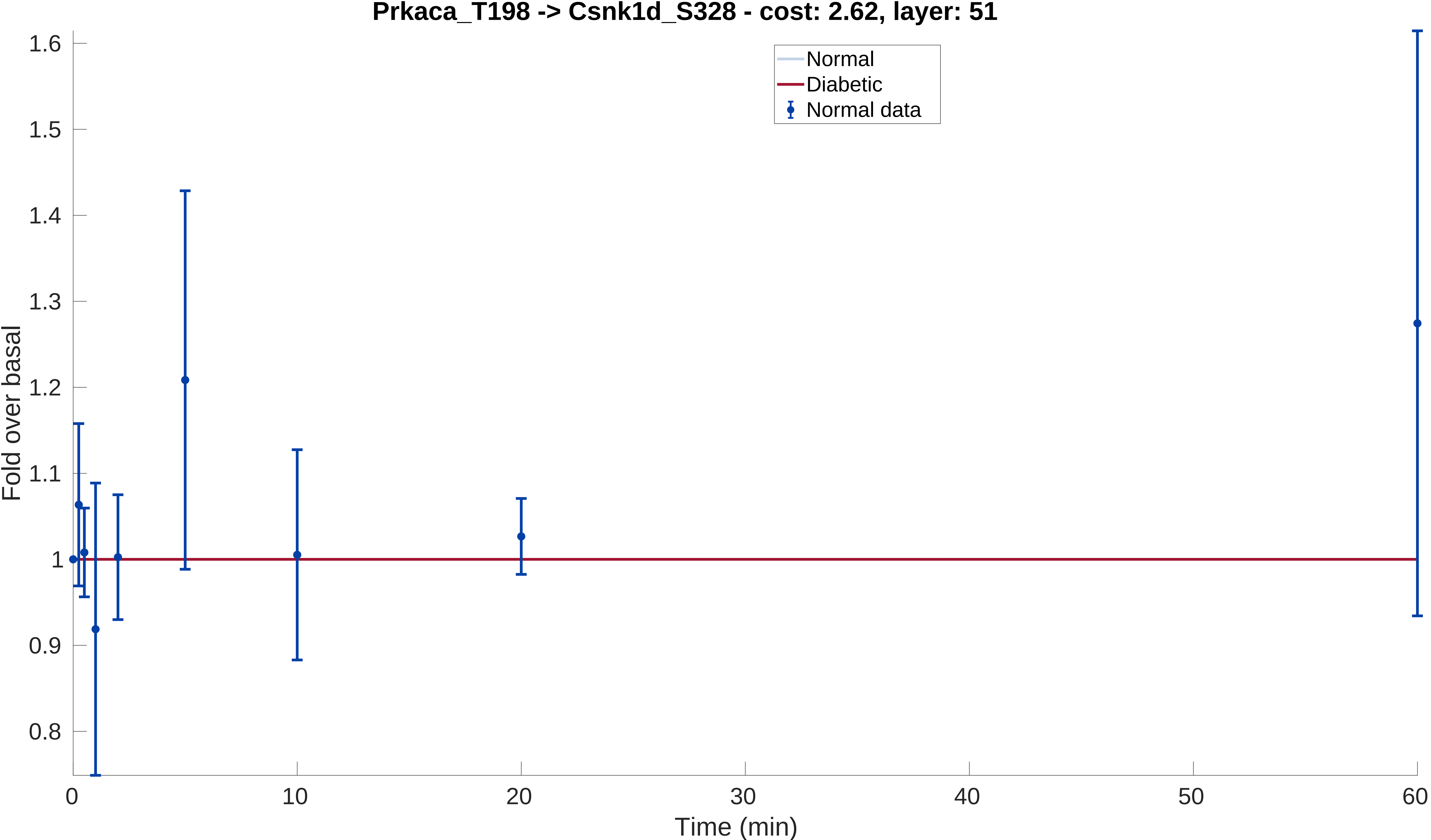

Prkaca_T198 -> Csnk1d_S328 - cost: 2.62, layer: 51
Normal
Diabetic
Normal data
Fold over basal
Time (min)
1.6
1.5
1.4
1.3
1.2
1.1
1
0.9
0.8
0
10
20
30
40
50
60

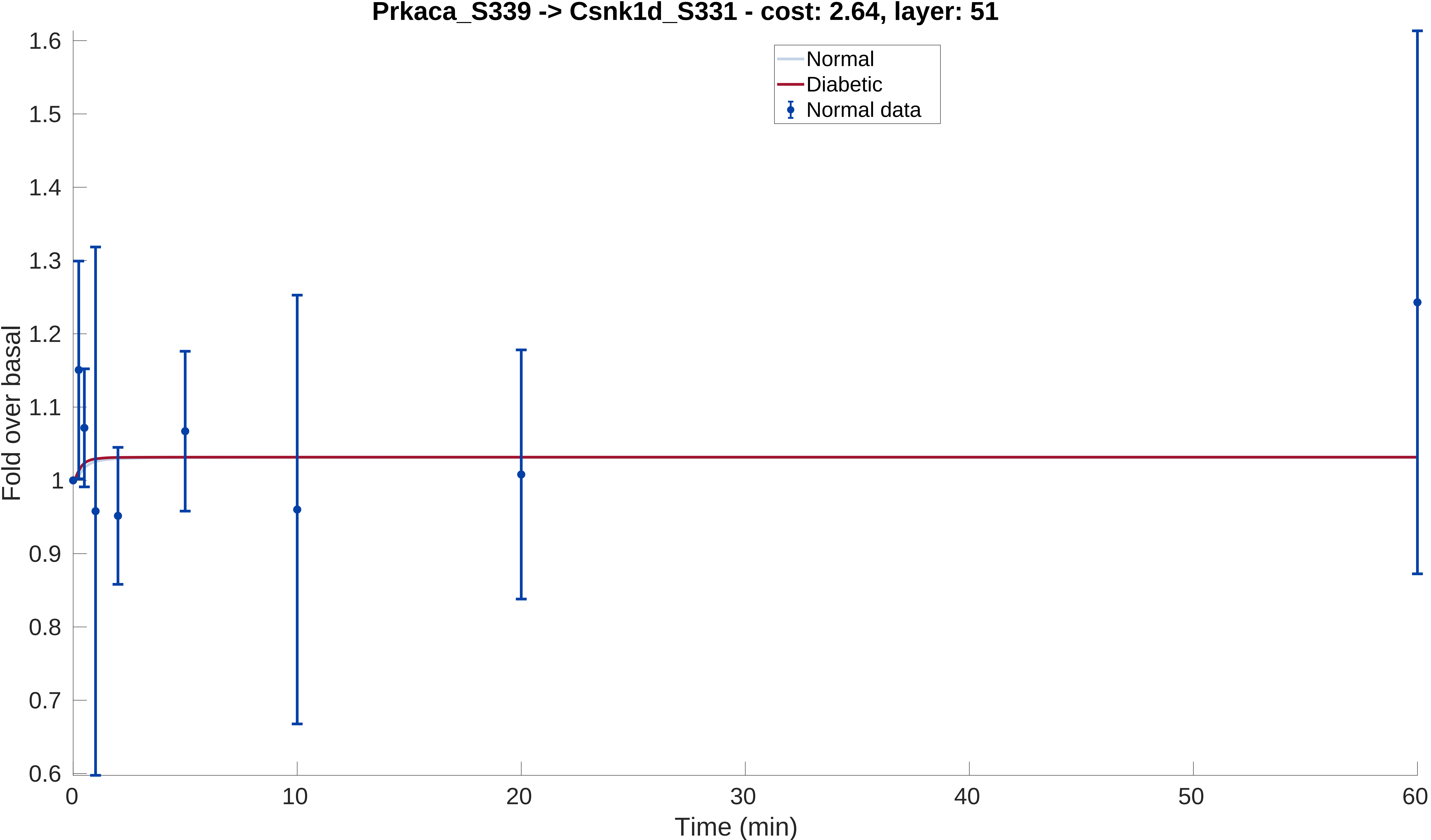

Prkaca_S339 -> Csnk1d_S331 - cost: 2.64, layer: 51
Normal
Diabetic
Normal data
Fold over basal
Time (min)
1.6
1.5
1.4
1.3
1.2
1.1
1
0.9
0.8
0.7
0.6
0
10
20
30
40
50
60

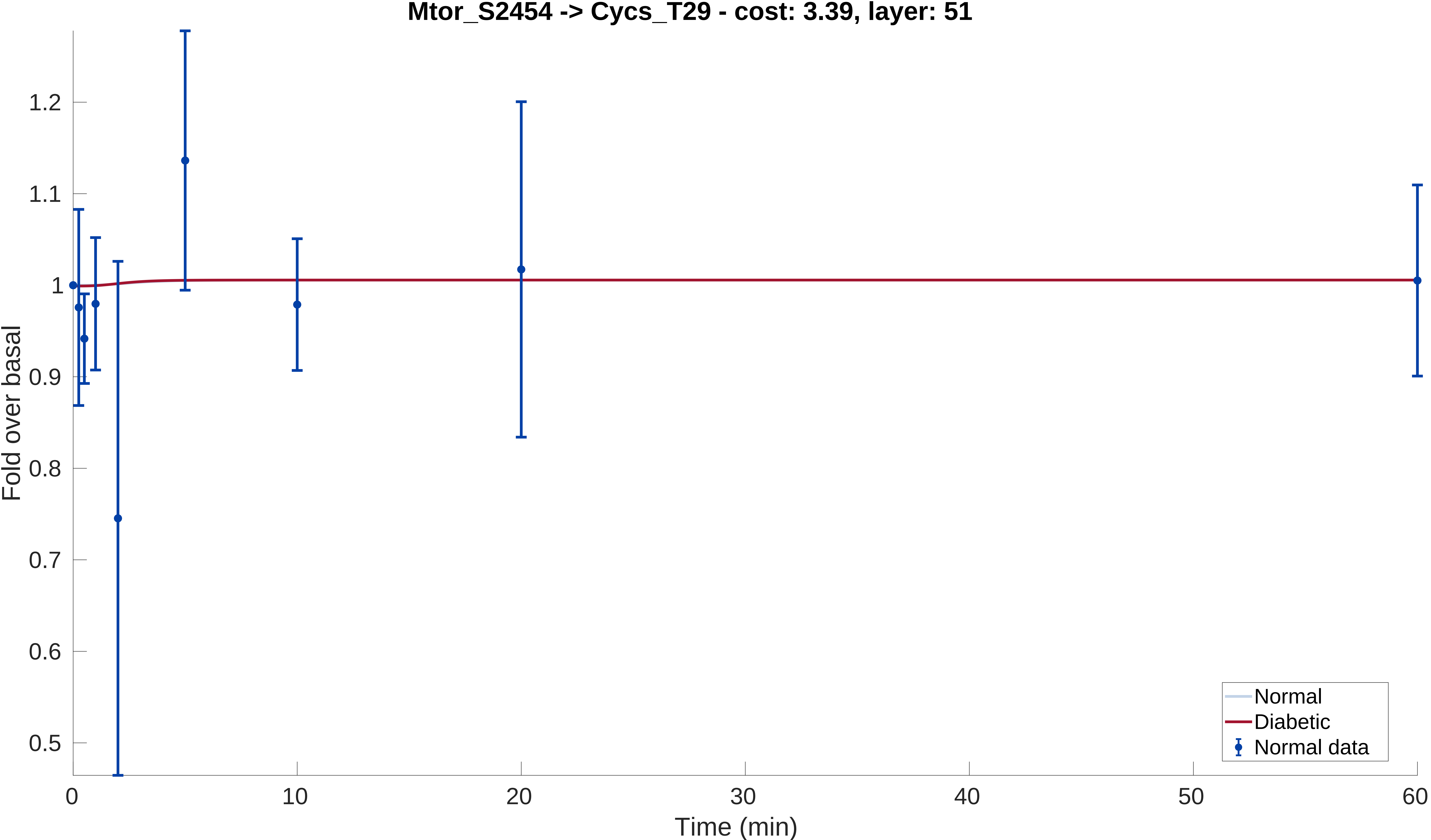

Mtor_S2454 -> Cycs_T29 - cost: 3.39, layer: 51
Fold over basal
Time (min)
1.2
1.1
1
0.9
0.8
0.7
0.6
0.5
0
10
20
30
40
50
60
Normal
Diabetic
Normal data

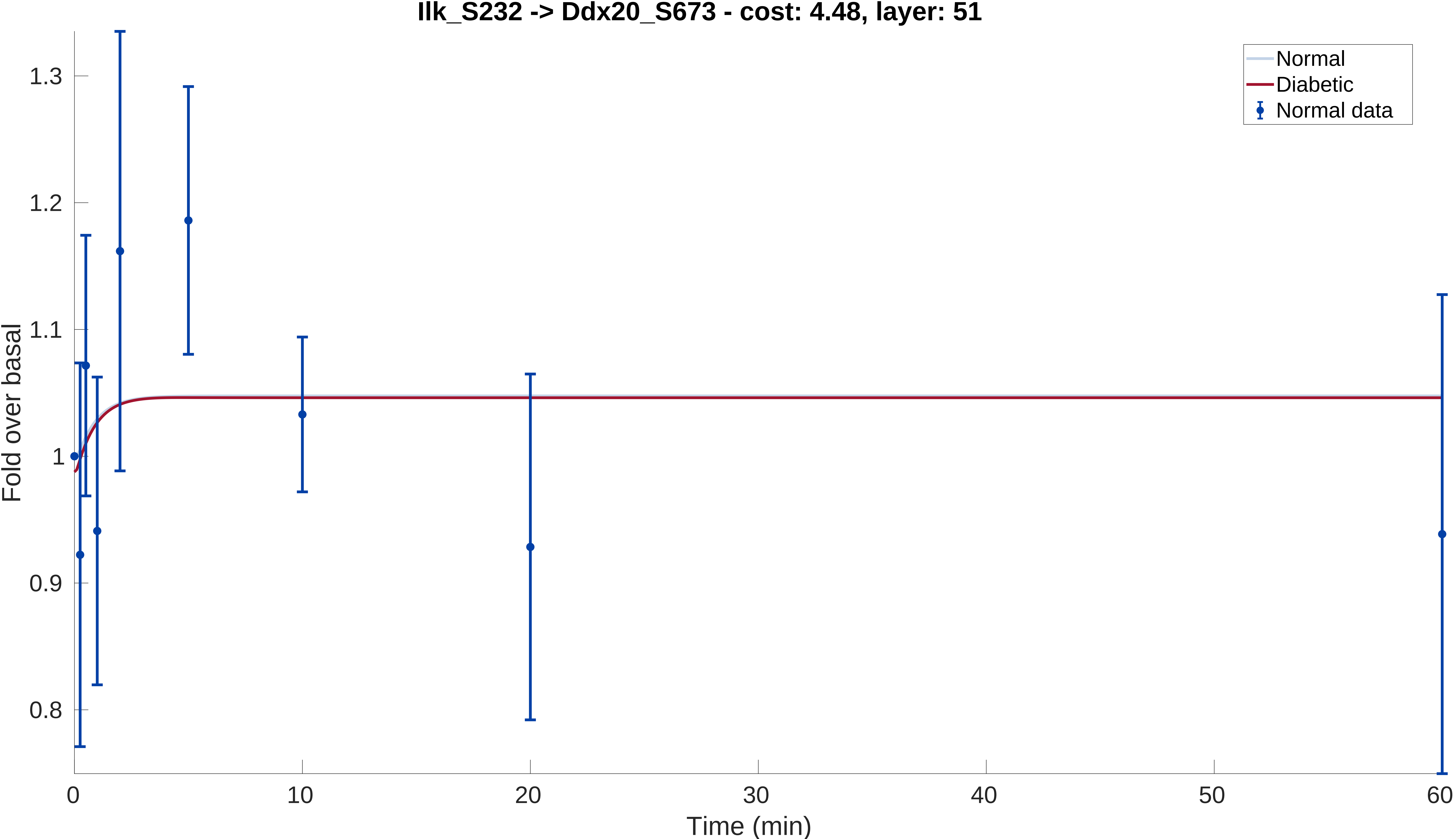

Ilk_S232 -> Ddx20_S673 - cost: 4.48, layer: 51
Normal
Diabetic
Normal data
Fold over basal
1.3
1.2
1.1
1
0.9
0.8
0
10
20
30
40
50
60
Time (min)

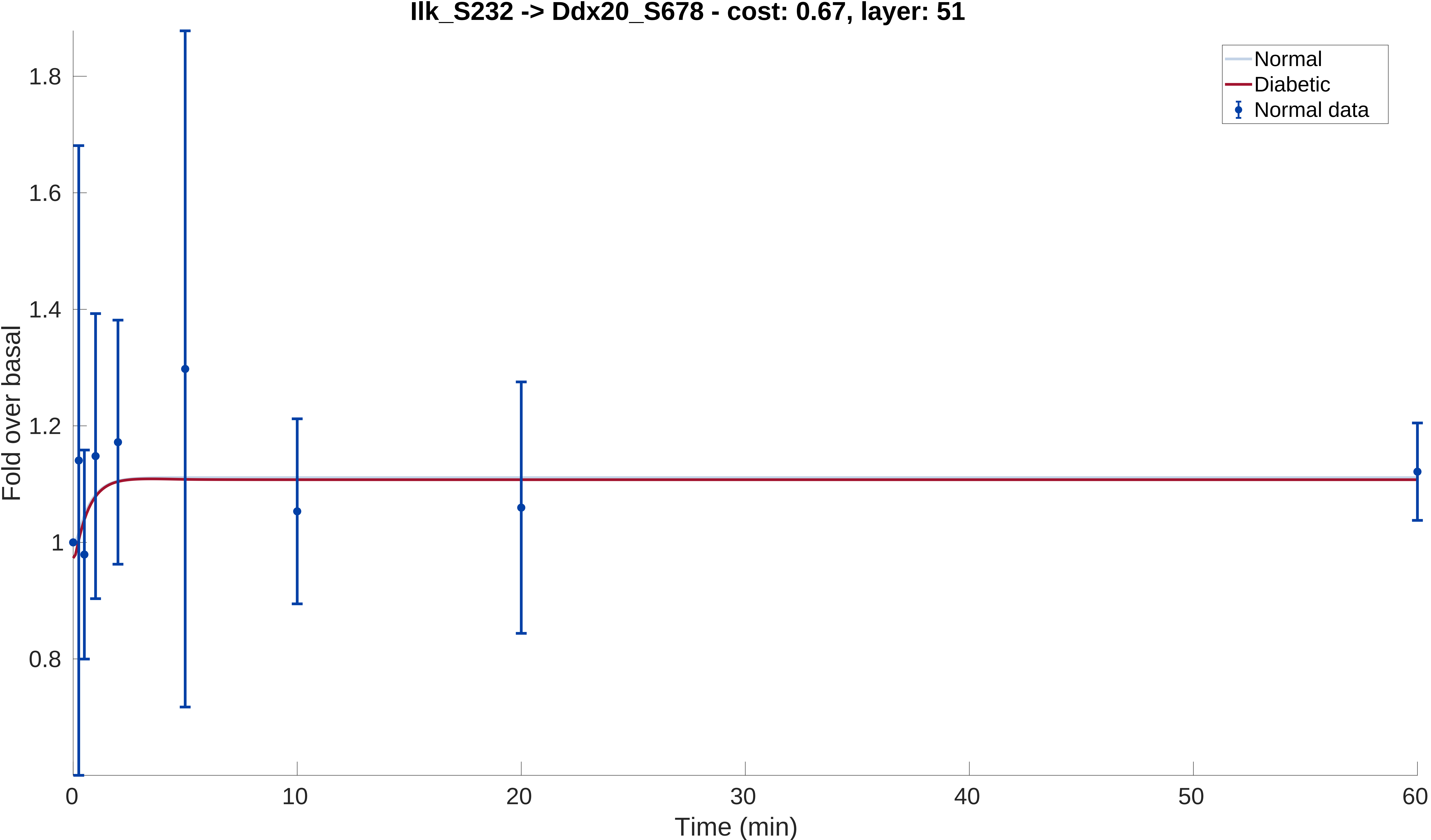

Ilk_S232 -> Ddx20_S678 - cost: 0.67, layer: 51
Normal
Diabetic
Normal data
Fold over basal
1.8
1.6
1.4
1.2
1
0.8
0
10
20
30
40
50
60
Time (min)

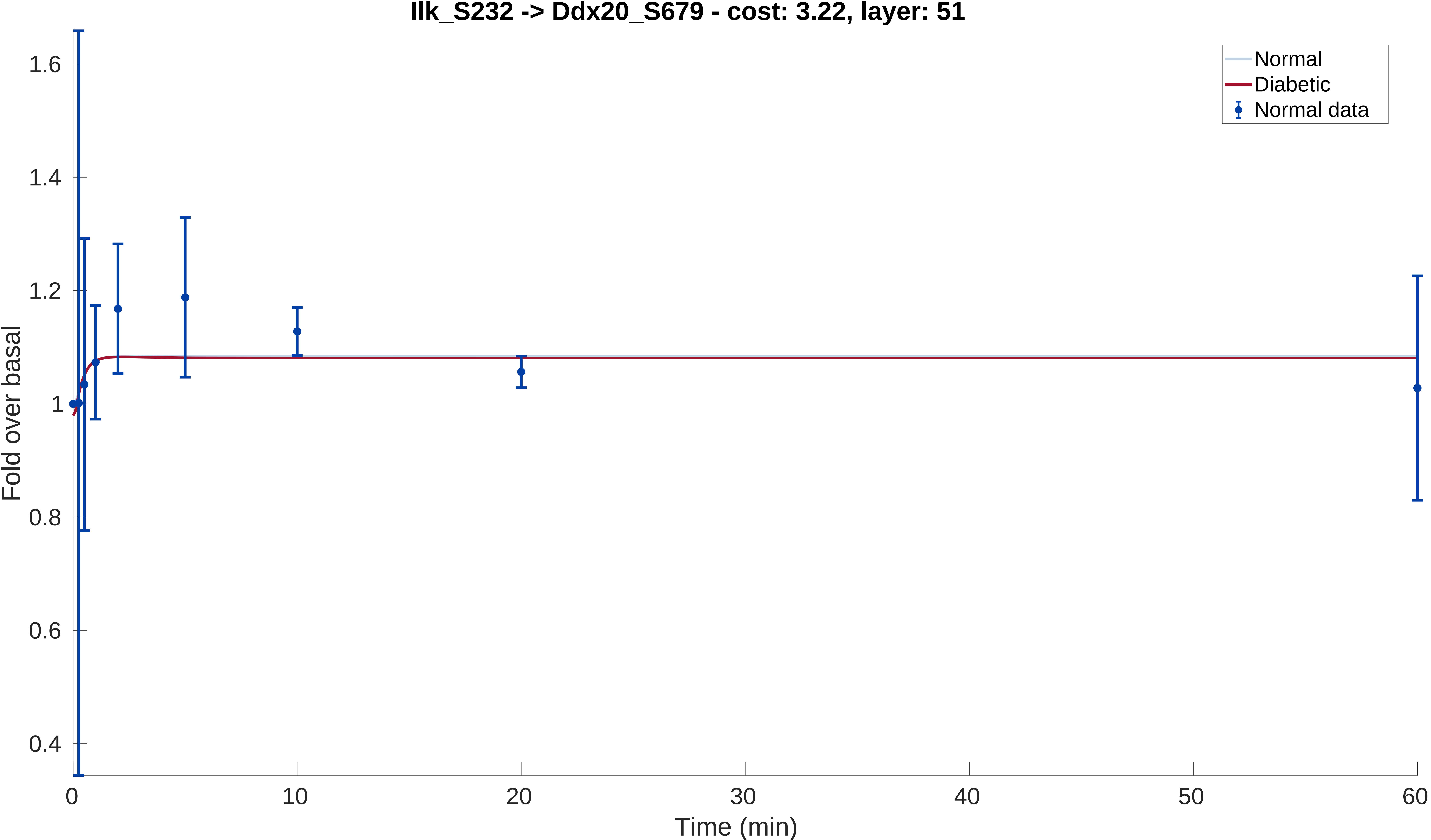

Ilk_S232 -> Ddx20_S679 - cost: 3.22, layer: 51
Normal
Diabetic
Normal data
Fold over basal
1.6
1.4
1.2
1
0.8
0.6
0.4
0
10
20
30
40
50
60
Time (min)

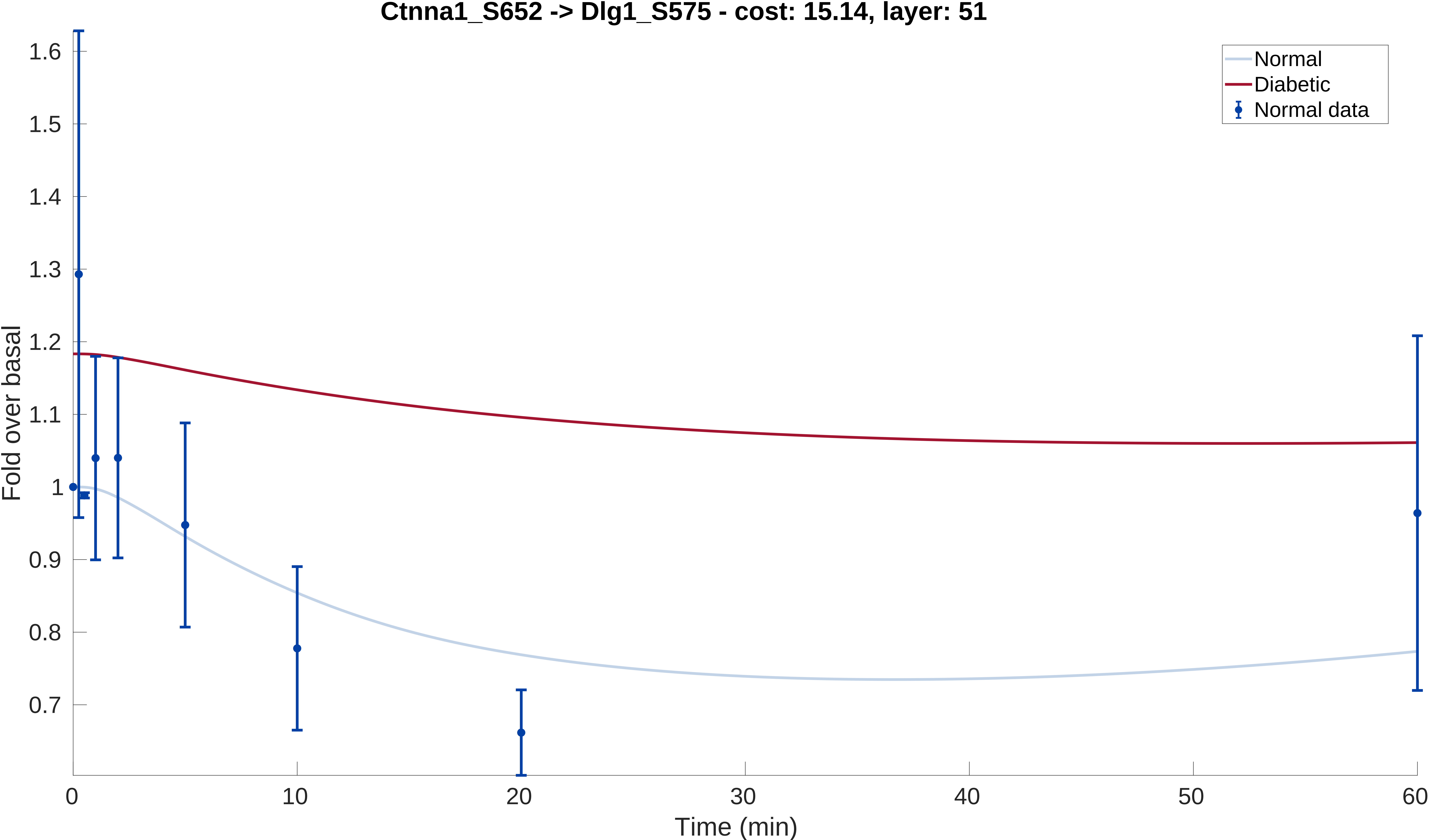
Ctnna1_S652 -> Dlg1_S575 - cost: 15.14, layer: 51
Normal
Diabetic
Normal data
Fold over basal
Time (min)
1.6
1.5
1.4
1.3
1.2
1.1
1
0.9
0.8
0.7
0
10
20
30
40
50
60

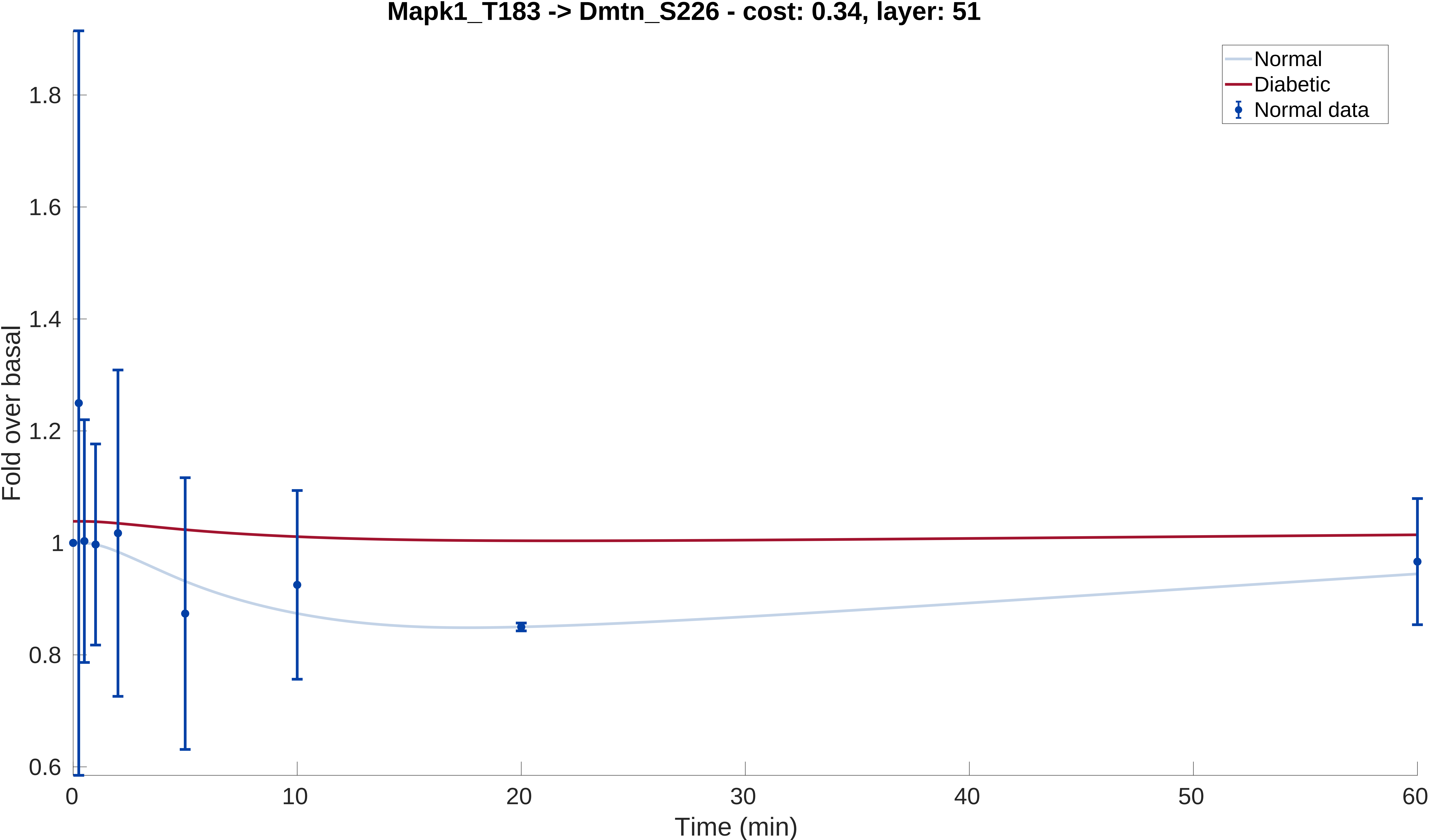

Mapk1_T183 -> Dmtn_S226 - cost: 0.34, layer: 51
Normal
Diabetic
Normal data
Fold over basal
1.8
1.6
1.4
1.2
1
0.8
0.6
0
10
20
30
40
50
60
Time (min)

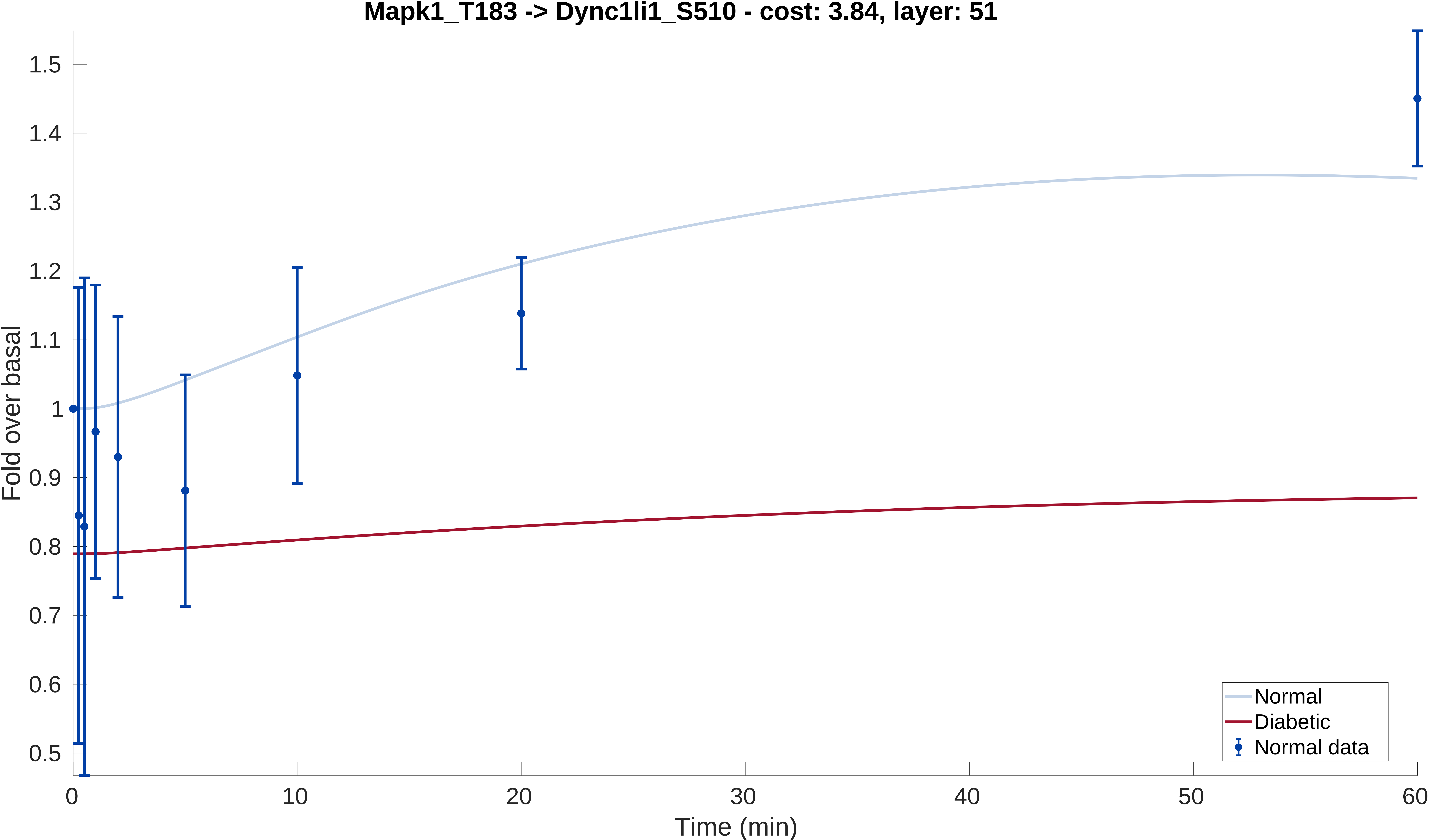
Mapk1_T183 -> Dync1li1_S510 - cost: 3.84, layer: 51
Fold over basal
Time (min)
1.5
1.4
1.3
1.2
1.1
1
0.9
0.8
0.7
0.6
0.5
0
10
20
30
40
50
60
Normal
Diabetic
Normal data

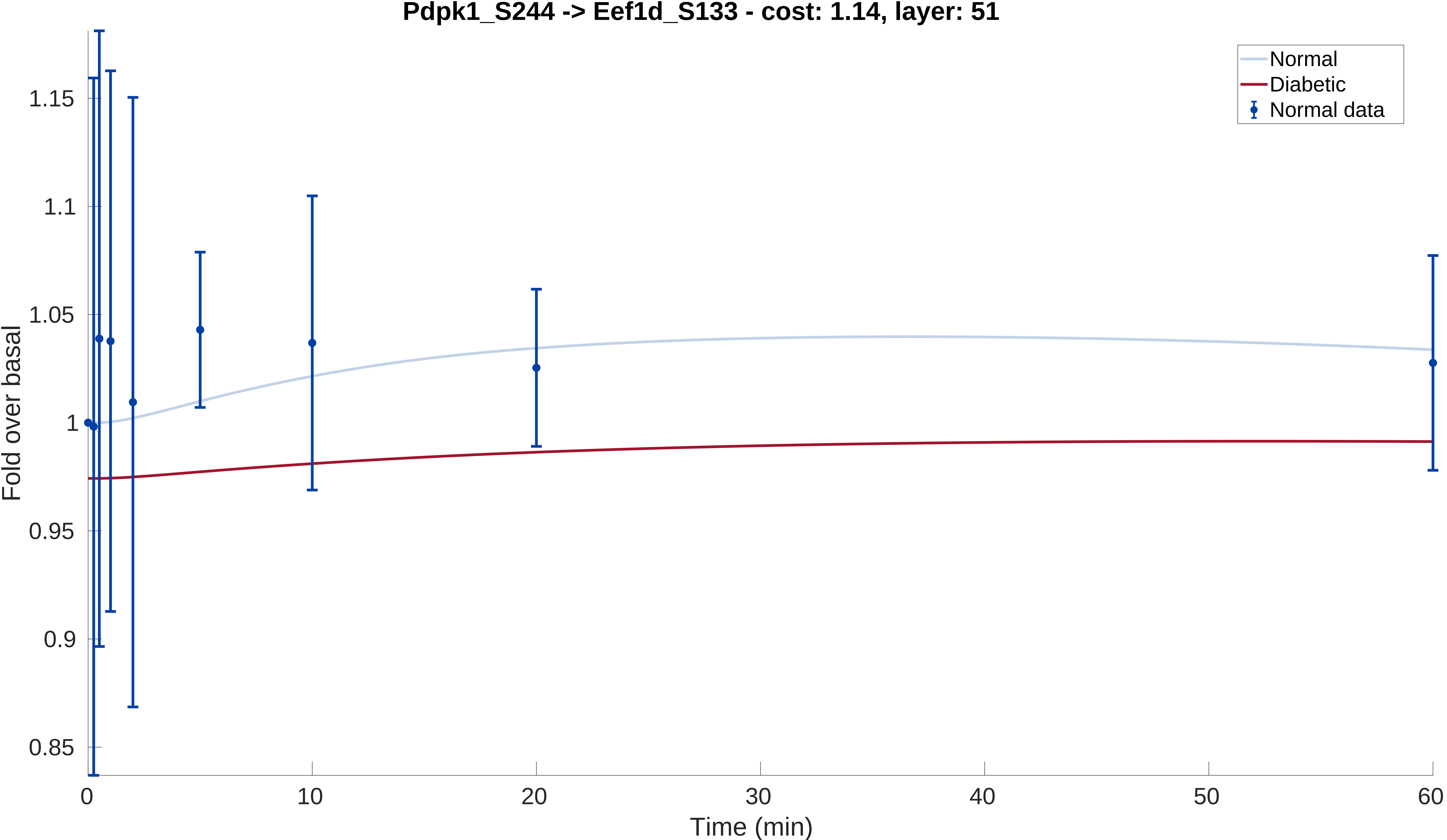

Pdpk1_S244 -> Eef1d_S133 - cost: 1.14, layer: 51
Normal
Diabetic
Normal data
1.15
1.1
1.05
1
0.95
0.9
0.85
Fold over basal
0
10
20
30
40
50
60
Time (min)

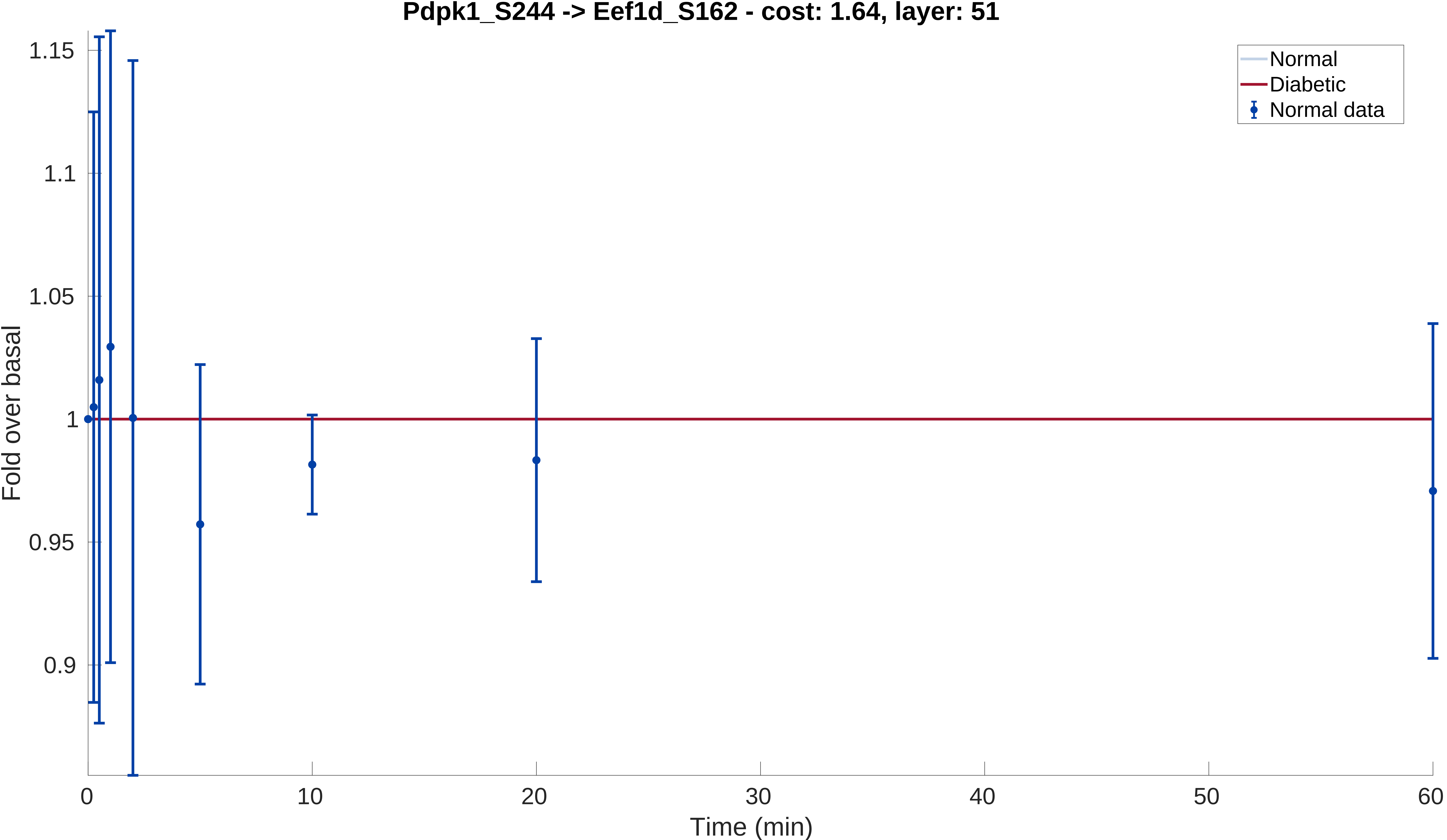

Pdpk1_S244 -> Eef1d_S162 - cost: 1.64, layer: 51
Normal
Diabetic
Normal data
Fold over basal
Time (min)
1.15
1.1
1.05
1
0.95
0.9
0
10
20
30
40
50
60

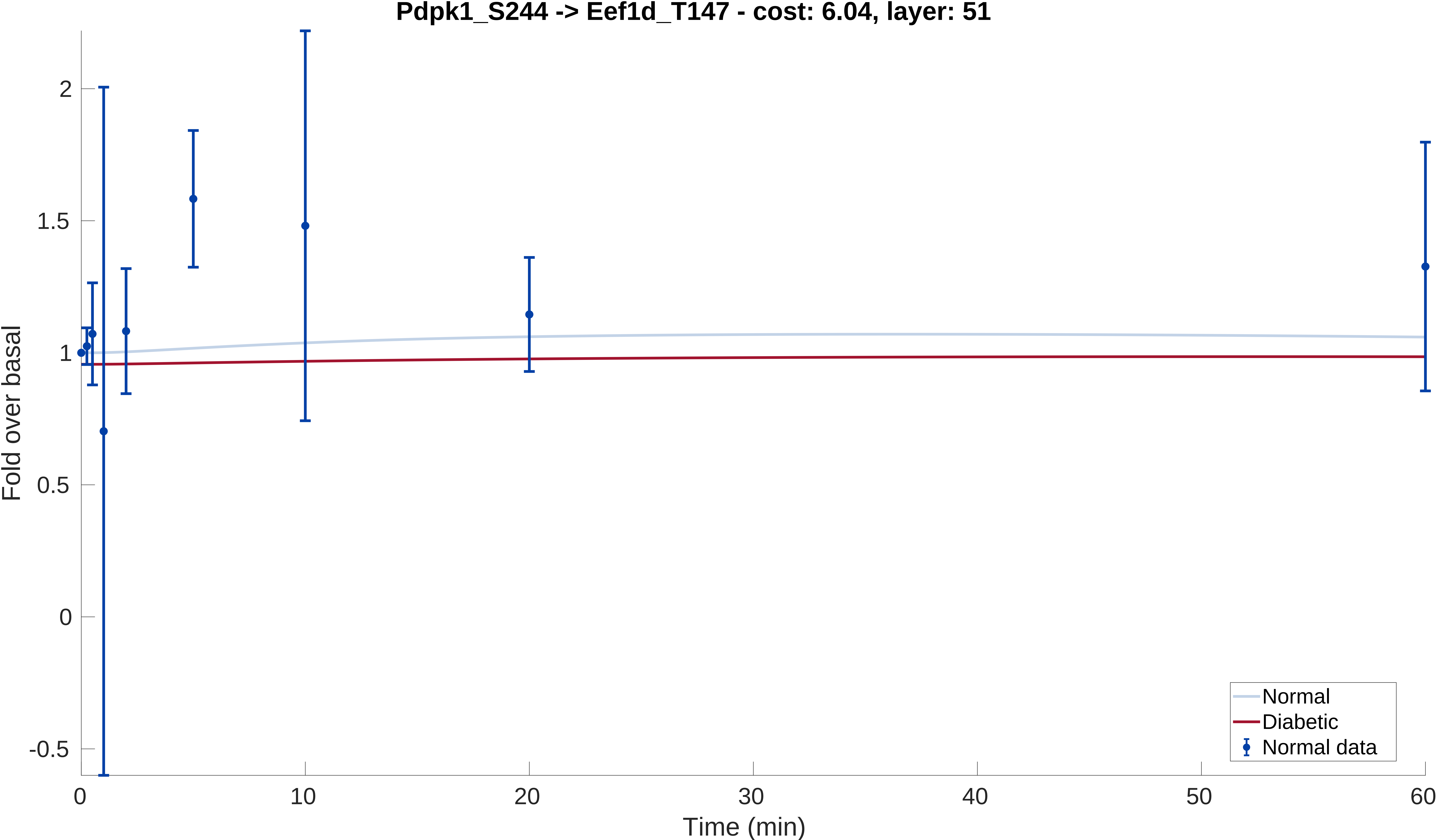

Pdpk1_S244 -> Eef1d_T147 - cost: 6.04, layer: 51
Fold over basal
Time (min)
2
1.5
1
0.5
0
-0.5
0
10
20
30
40
50
60
Normal
Diabetic
Normal data

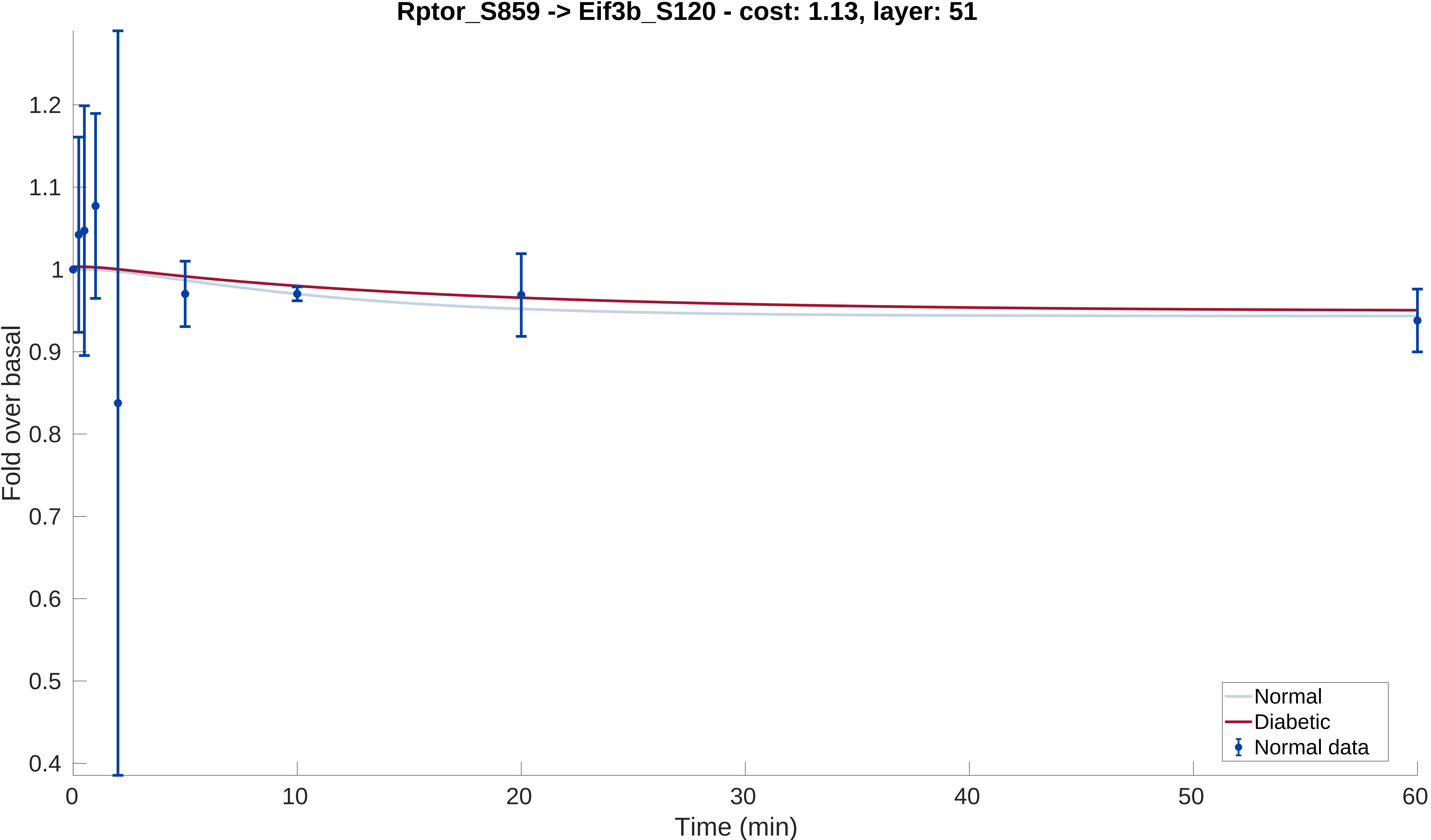

Rptor_S859 -> Eif3b_S120 - cost: 1.13, layer: 51
Fold over basal
1.2
1.1
1
0.9
0.8
0.7
0.6
0.5
0.4
0
10
20
30
40
50
60
Time (min)
Normal
Diabetic
Normal data

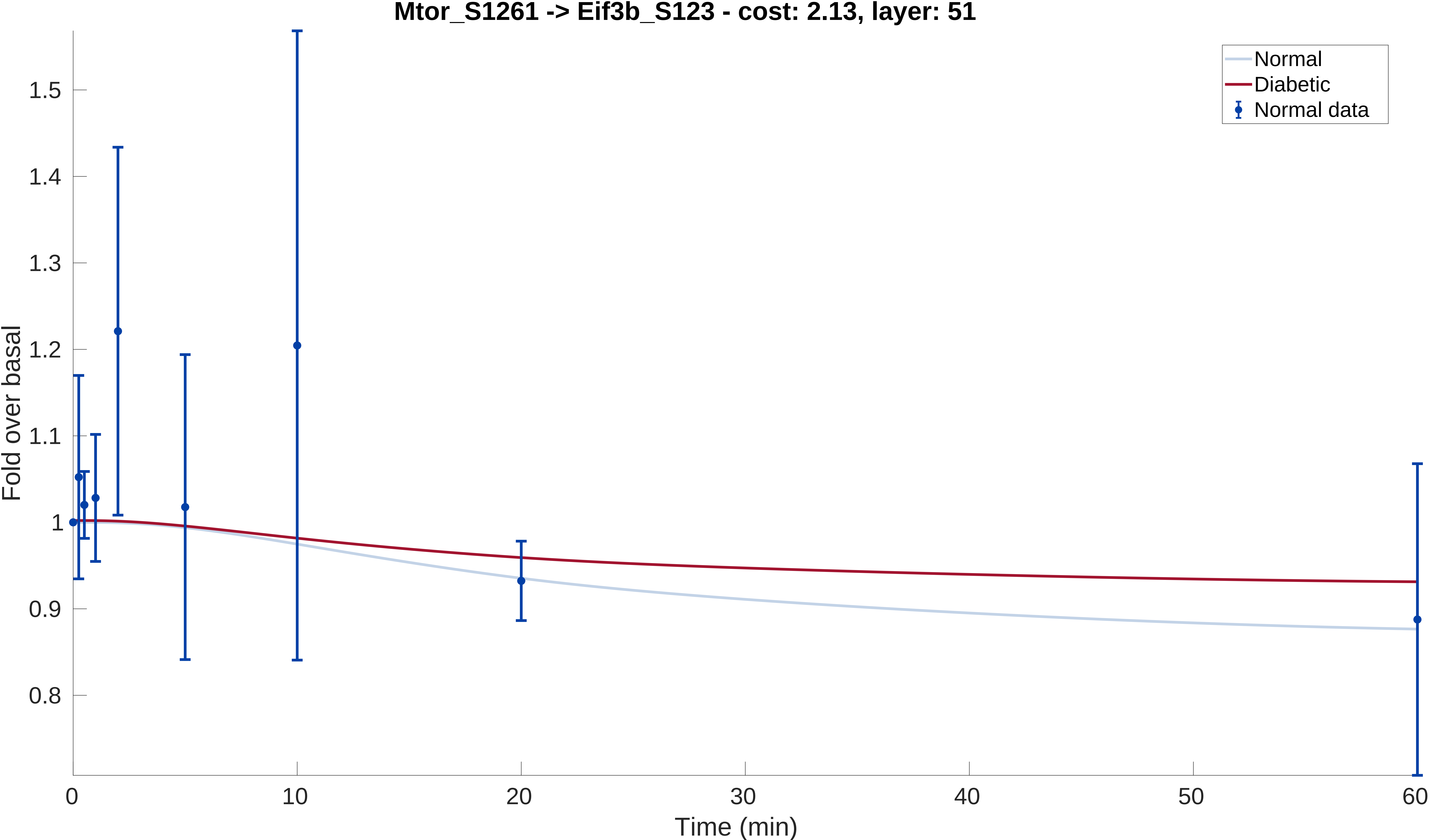

Mtor_S1261 -> Eif3b_S123 - cost: 2.13, layer: 51
Normal
Diabetic
Normal data
Fold over basal
1.5
1.4
1.3
1.2
1.1
1
0.9
0.8
0
10
20
30
40
50
60
Time (min)

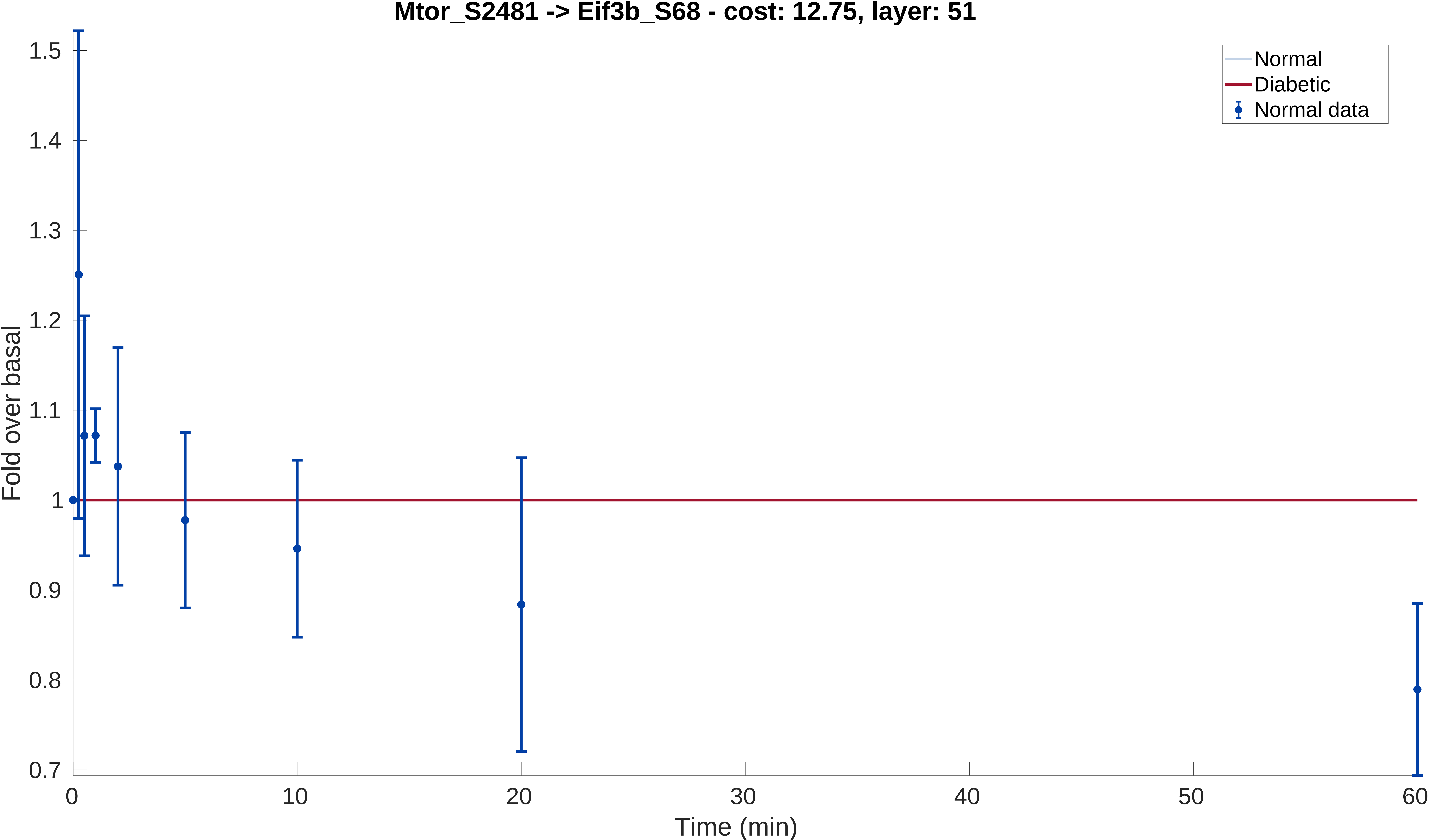

Mtor_S2481 -> Eif3b_S68 - cost: 12.75, layer: 51
Normal
Diabetic
Normal data
Fold over basal
1.5
1.4
1.3
1.2
1.1
1
0.9
0.8
0.7
0
10
20
30
40
50
60
Time (min)

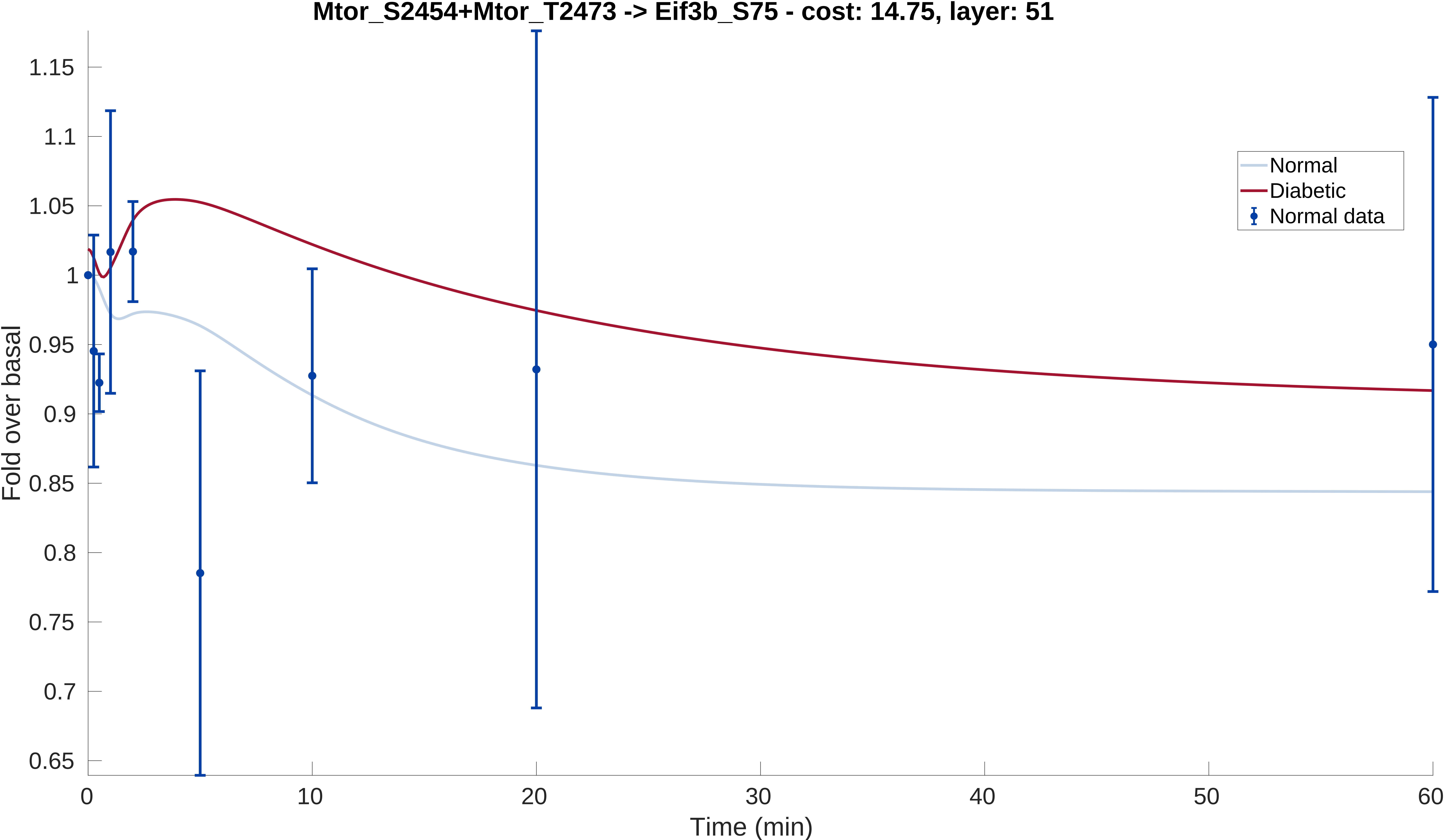

Mtor_S2454+Mtor_T2473 -> Eif3b_S75 - cost: 14.75, layer: 51
Fold over basal
Time (min)
Normal
Diabetic
Normal data
1.15
1.1
1.05
1
0.95
0.9
0.85
0.8
0.75
0.7
0.65
0
10
20
30
40
50
60

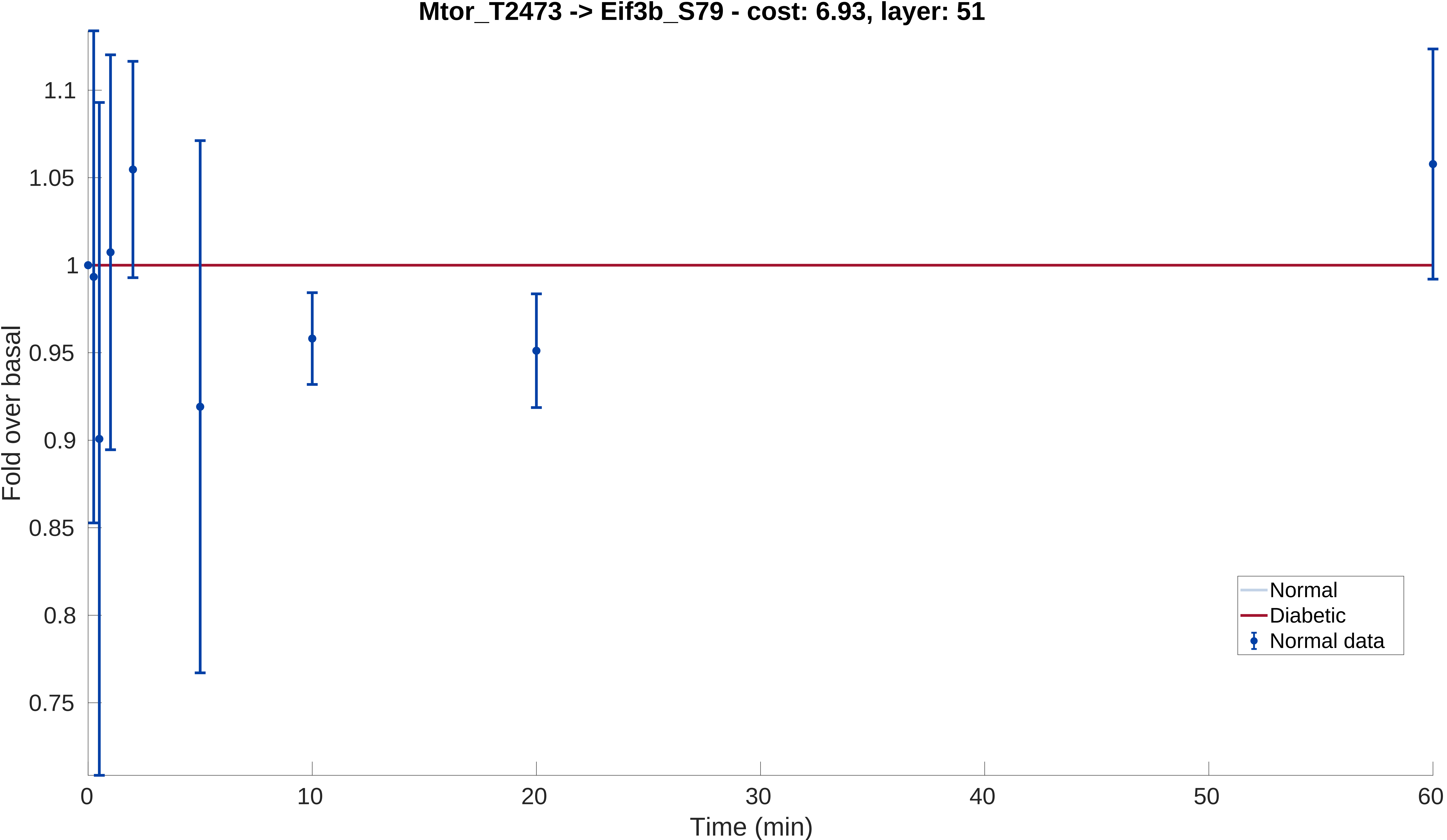

Mtor_T2473 -> Eif3b_S79 - cost: 6.93, layer: 51
1.1
1.05
1
0.95
0.9
0.85
0.8
0.75
Fold over basal
0
10
20
30
40
50
60
Time (min)
Normal
Diabetic
Normal data

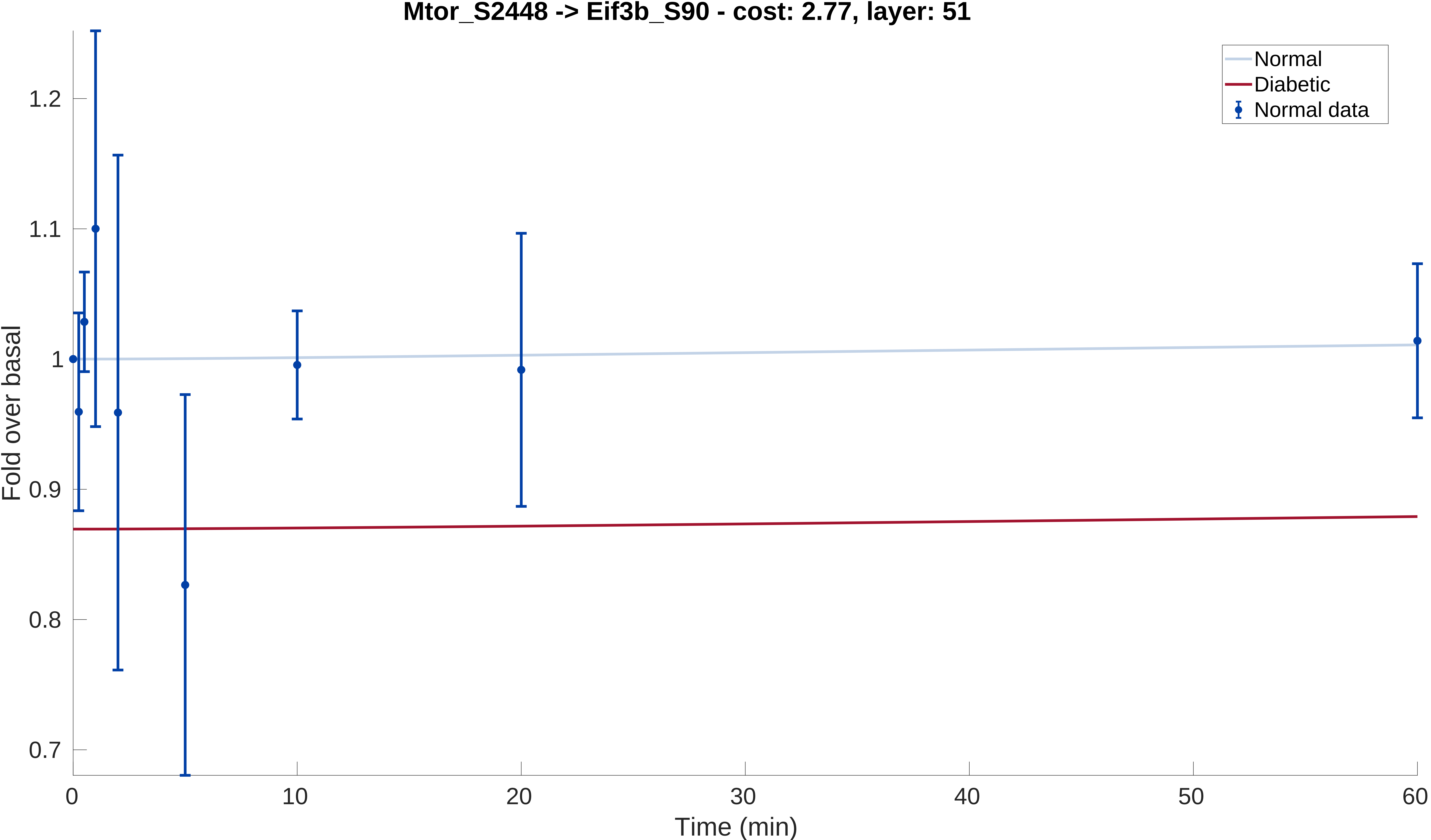

Mtor_S2448 -> Eif3b_S90 - cost: 2.77, layer: 51
Normal
Diabetic
Normal data
Fold over basal
1.2
1.1
1
0.9
0.8
0.7
0
10
20
30
40
50
60
Time (min)

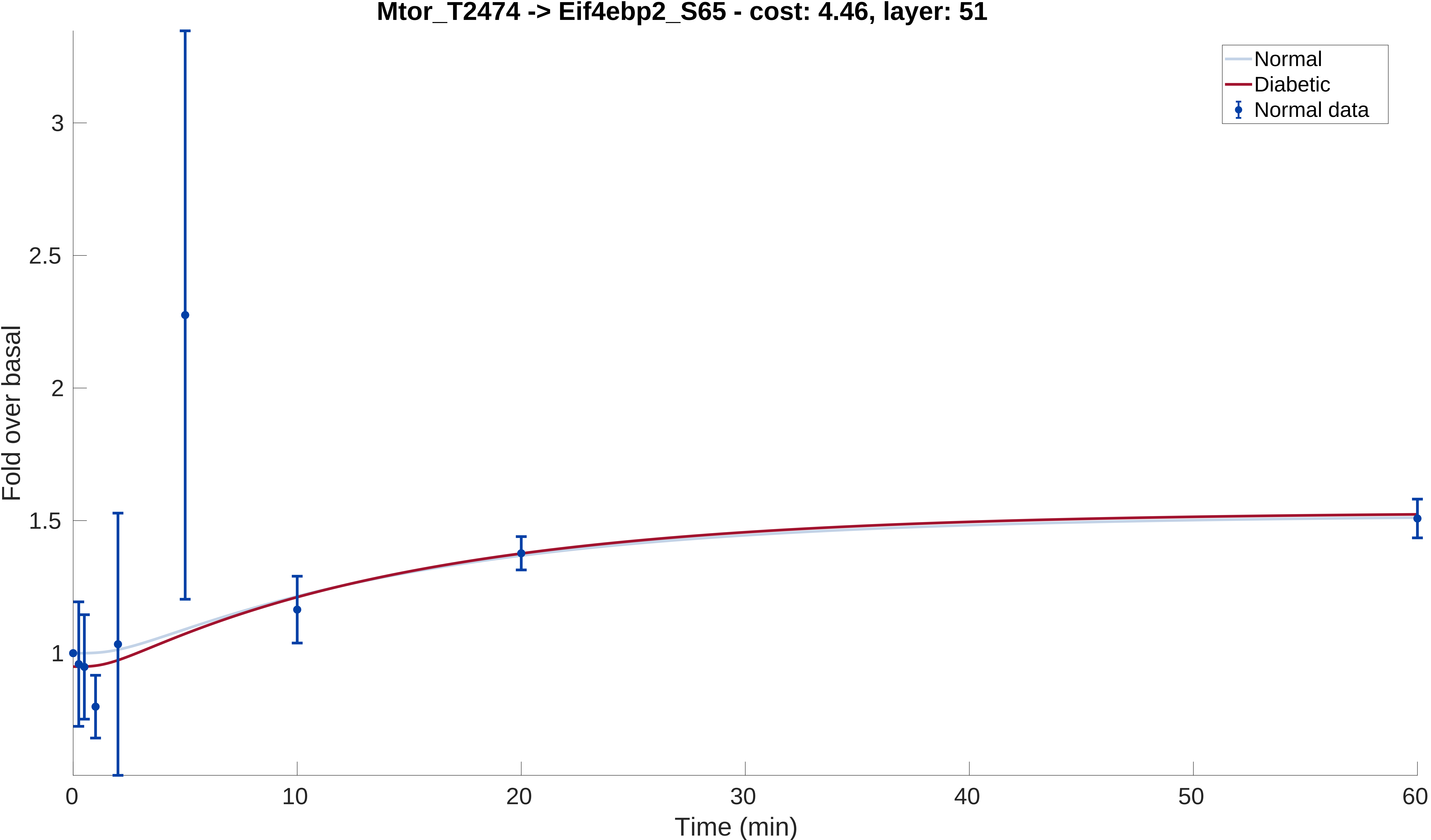
Mtor_T2474 -> Eif4ebp2_S65 - cost: 4.46, layer: 51
Normal
Diabetic
Normal data
Fold over basal
3
2.5
2
1.5
1
0
10
20
30
40
50
60
Time (min)

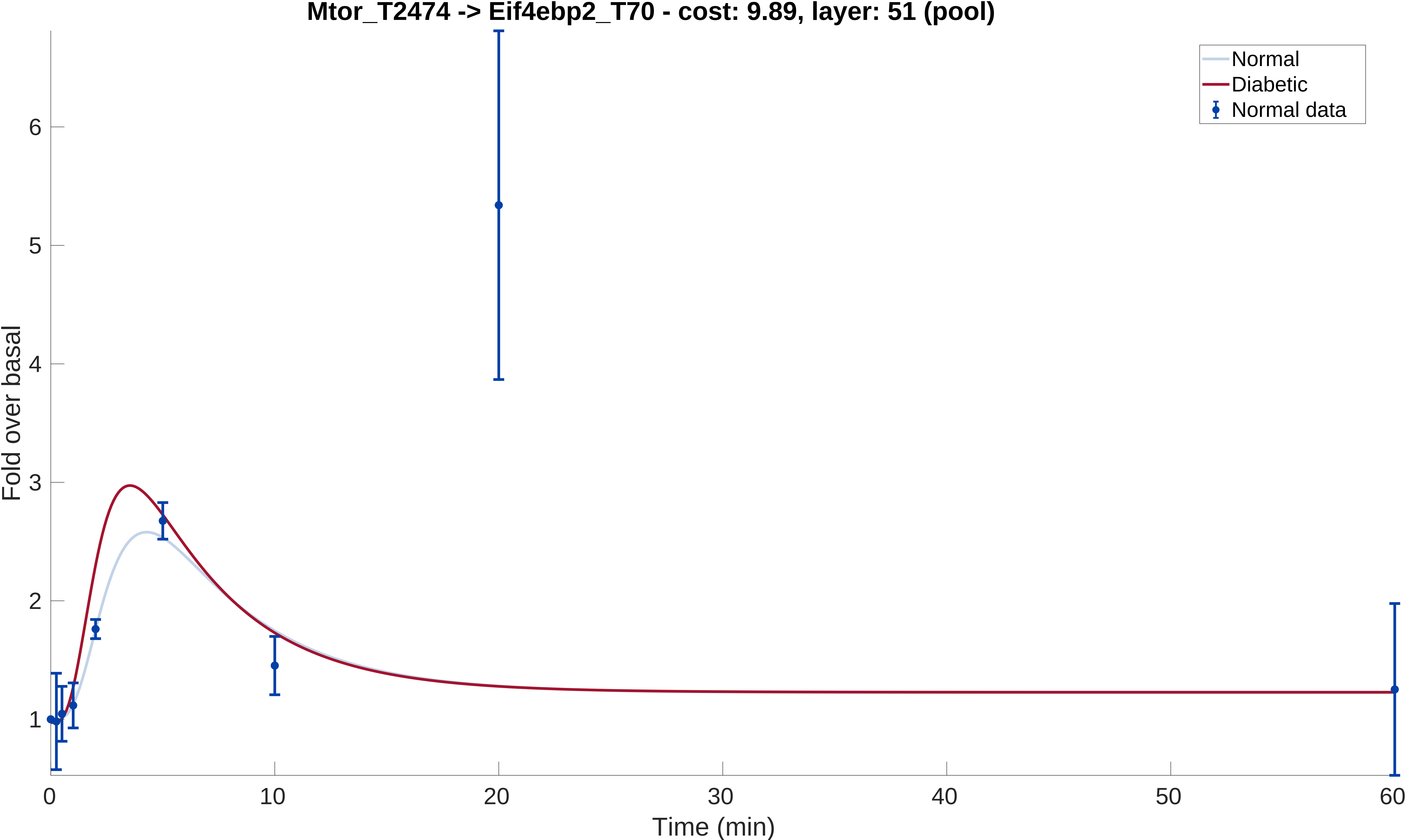
Mtor_T2474 -> Eif4ebp2_T70 - cost: 9.89, layer: 51 (pool)
Normal
Diabetic
Normal data
Fold over basal
Time (min)
1
2
3
4
5
6
0
10
20
30
40
50
60

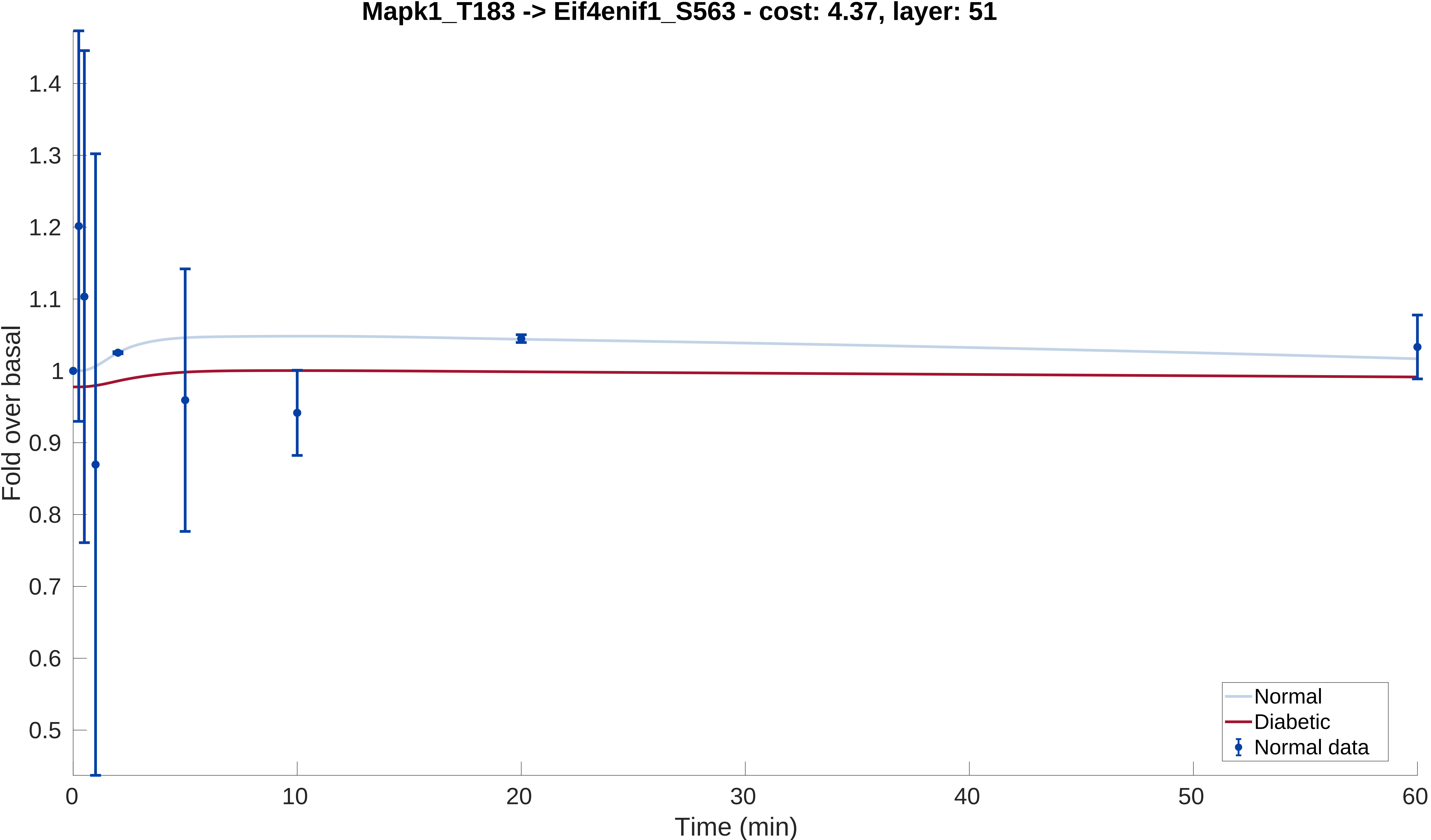
Mapk1_T183 -> Eif4enif1_S563 - cost: 4.37, layer: 51
Fold over basal
1.4
1.3
1.2
1.1
1
0.9
0.8
0.7
0.6
0.5
0
10
20
30
40
50
60
Time (min)
Normal
Diabetic
Normal data

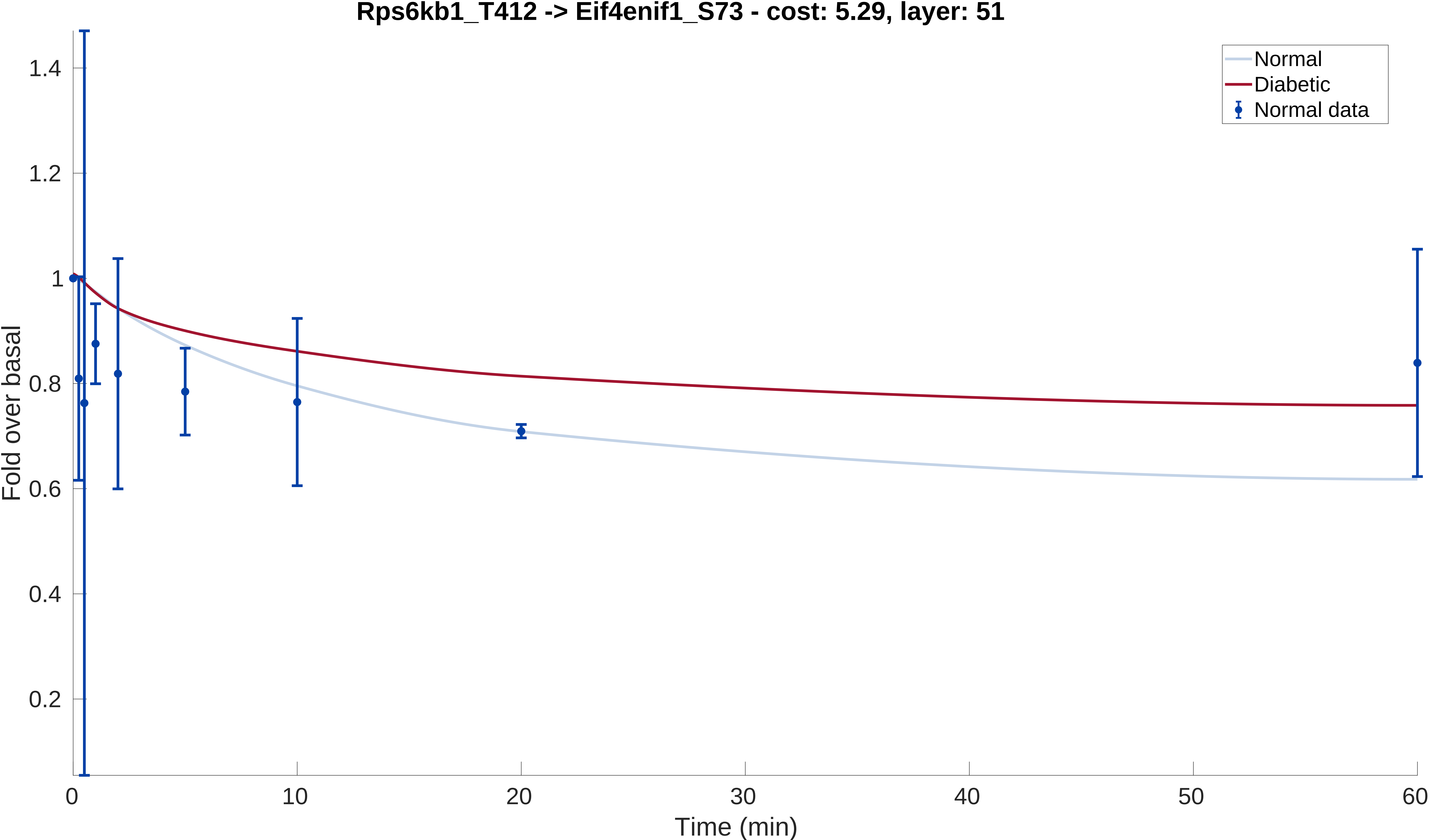

Rps6kb1_T412 -> Eif4enif1_S73 - cost: 5.29, layer: 51
Normal
Diabetic
Normal data
Fold over basal
Time (min)
1.4
1.2
1
0.8
0.6
0.4
0.2
0
10
20
30
40
50
60

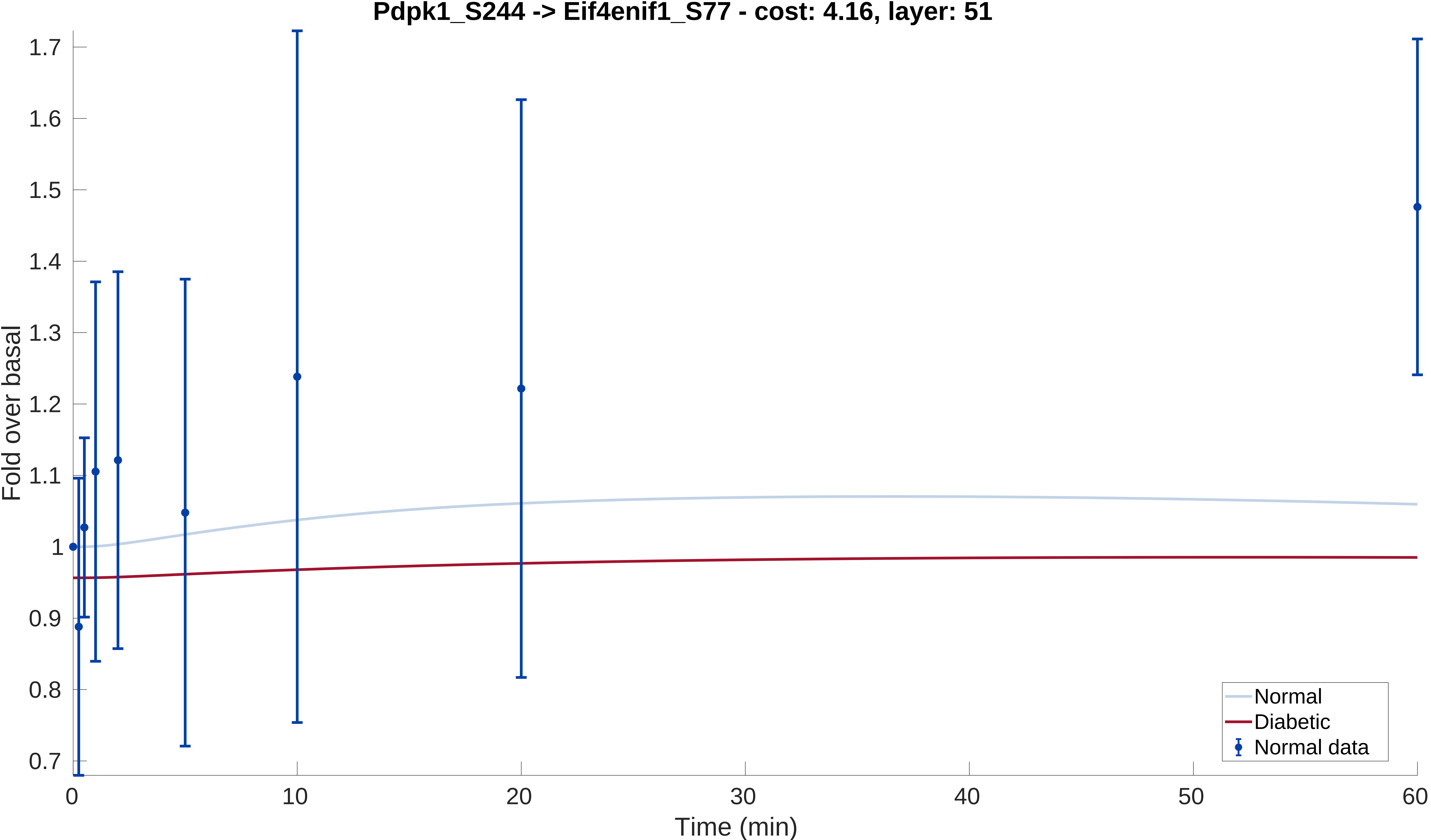
Pdpk1_S244 -> Eif4enif1_S77 - cost: 4.16, layer: 51
Fold over basal
1.7
1.6
1.5
1.4
1.3
1.2
1.1
1
0.9
0.8
0.7
0
10
20
30
40
50
60
Time (min)
Normal
Diabetic
Normal data

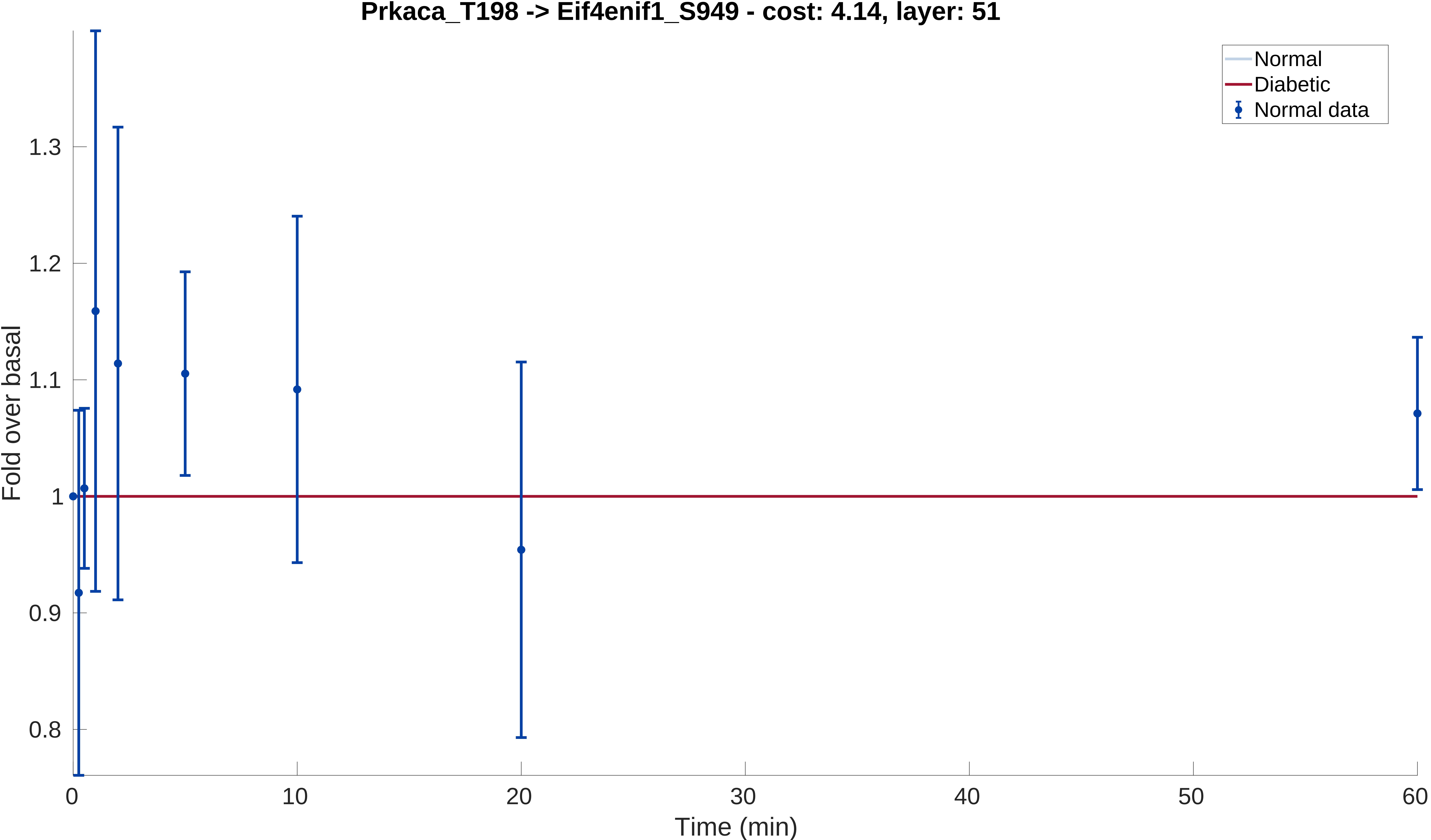

Prkaca_T198 -> Eif4enif1_S949 - cost: 4.14, layer: 51
Normal
Diabetic
Normal data
Fold over basal
1.3
1.2
1.1
1
0.9
0.8
0
10
20
30
40
50
60
Time (min)

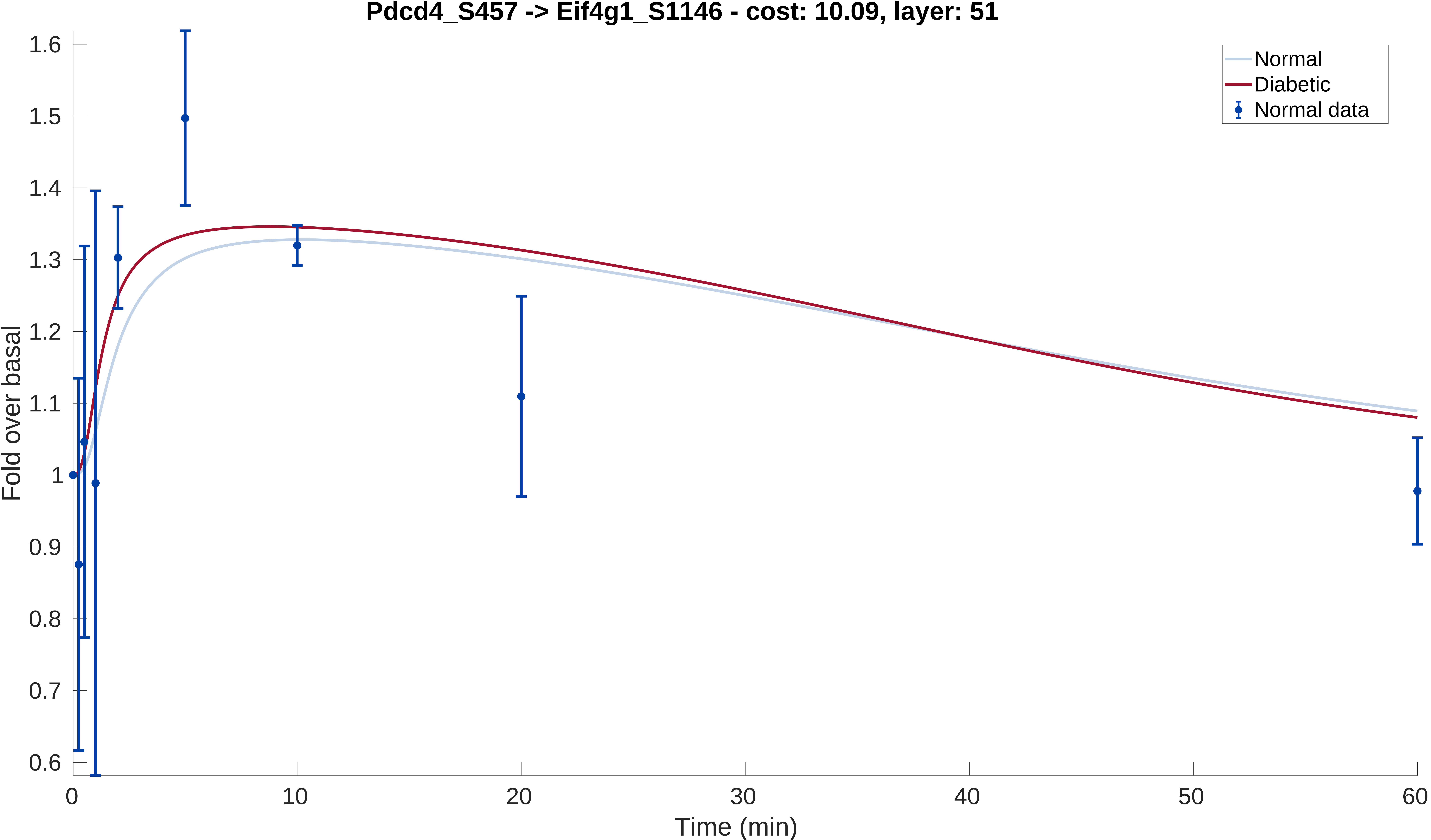

Pdcd4_S457 -> Eif4g1_S1146 - cost: 10.09, layer: 51
Normal
Diabetic
Normal data
Fold over basal
Time (min)
1.6
1.5
1.4
1.3
1.2
1.1
1
0.9
0.8
0.7
0.6
0
10
20
30
40
50
60

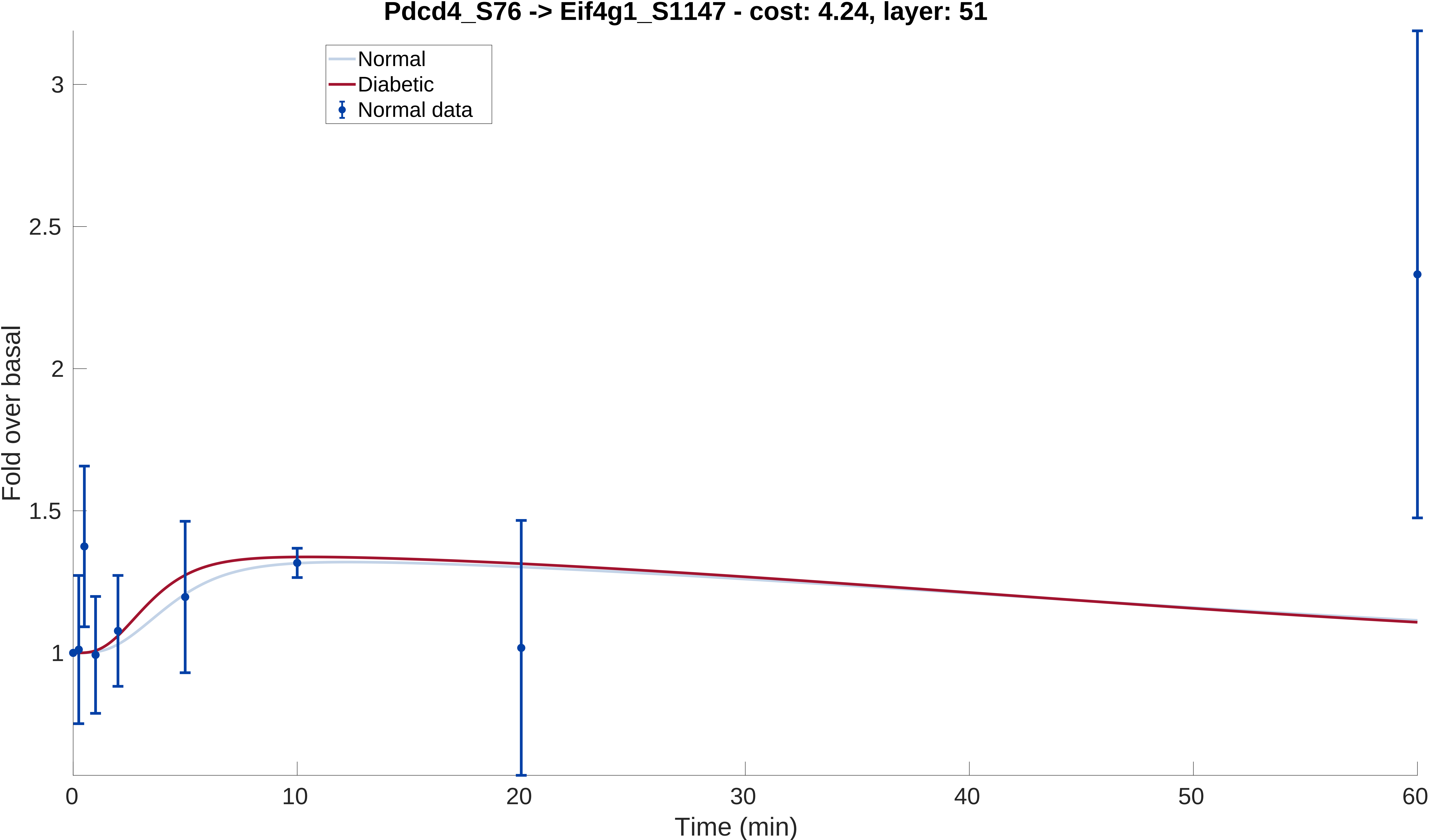
Pdcd4_S76 -> Eif4g1_S1147 - cost: 4.24, layer: 51
Normal
Diabetic
Normal data
Fold over basal
3
2.5
2
1.5
1
0
10
20
30
40
50
60
Time (min)

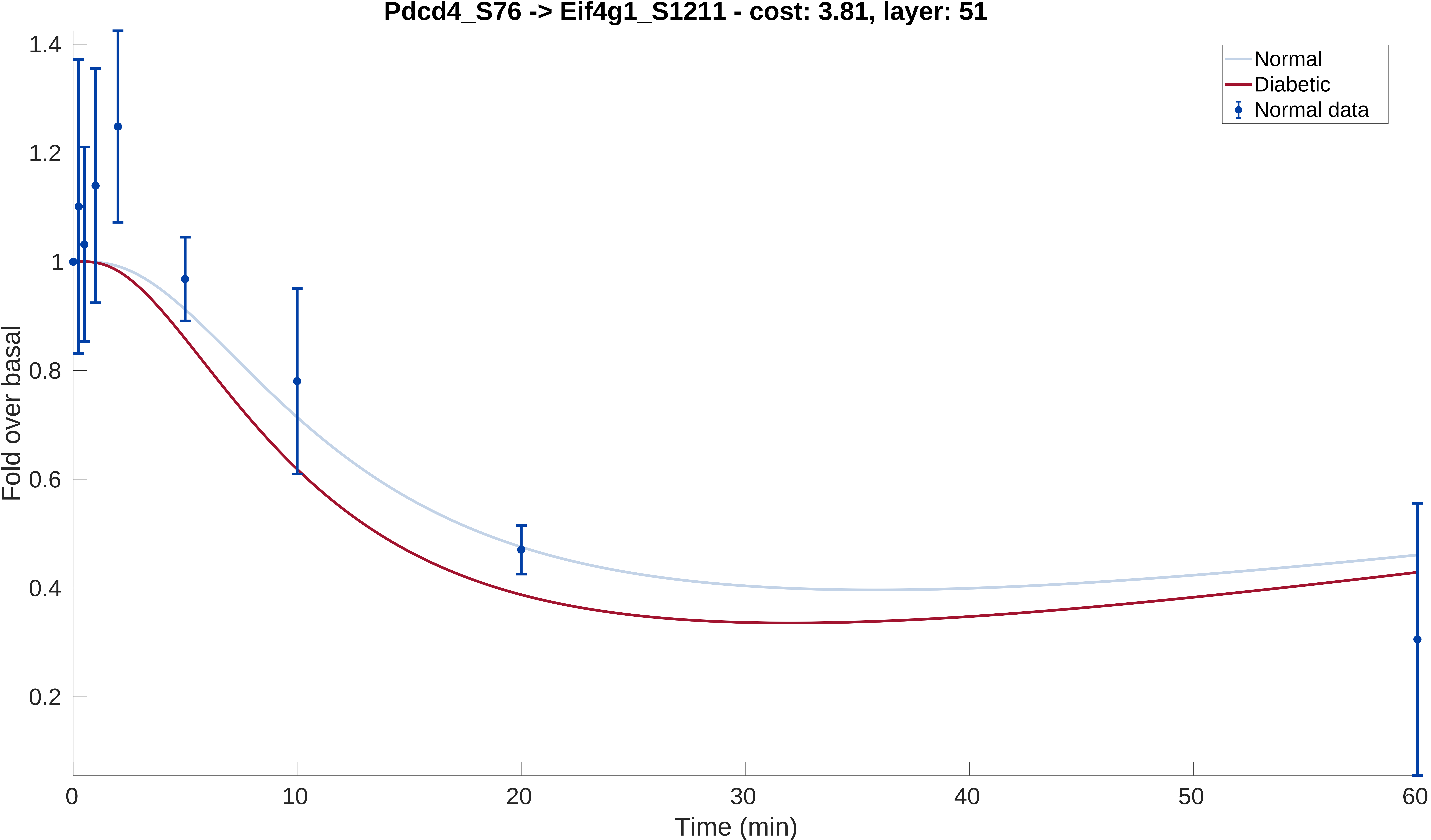
Pdcd4_S76 -> Eif4g1_S1211 - cost: 3.81, layer: 51
Fold over basal
1.4
1.2
1
0.8
0.6
0.4
0.2
0
10
20
30
40
50
60
Time (min)
Normal
Diabetic
Normal data

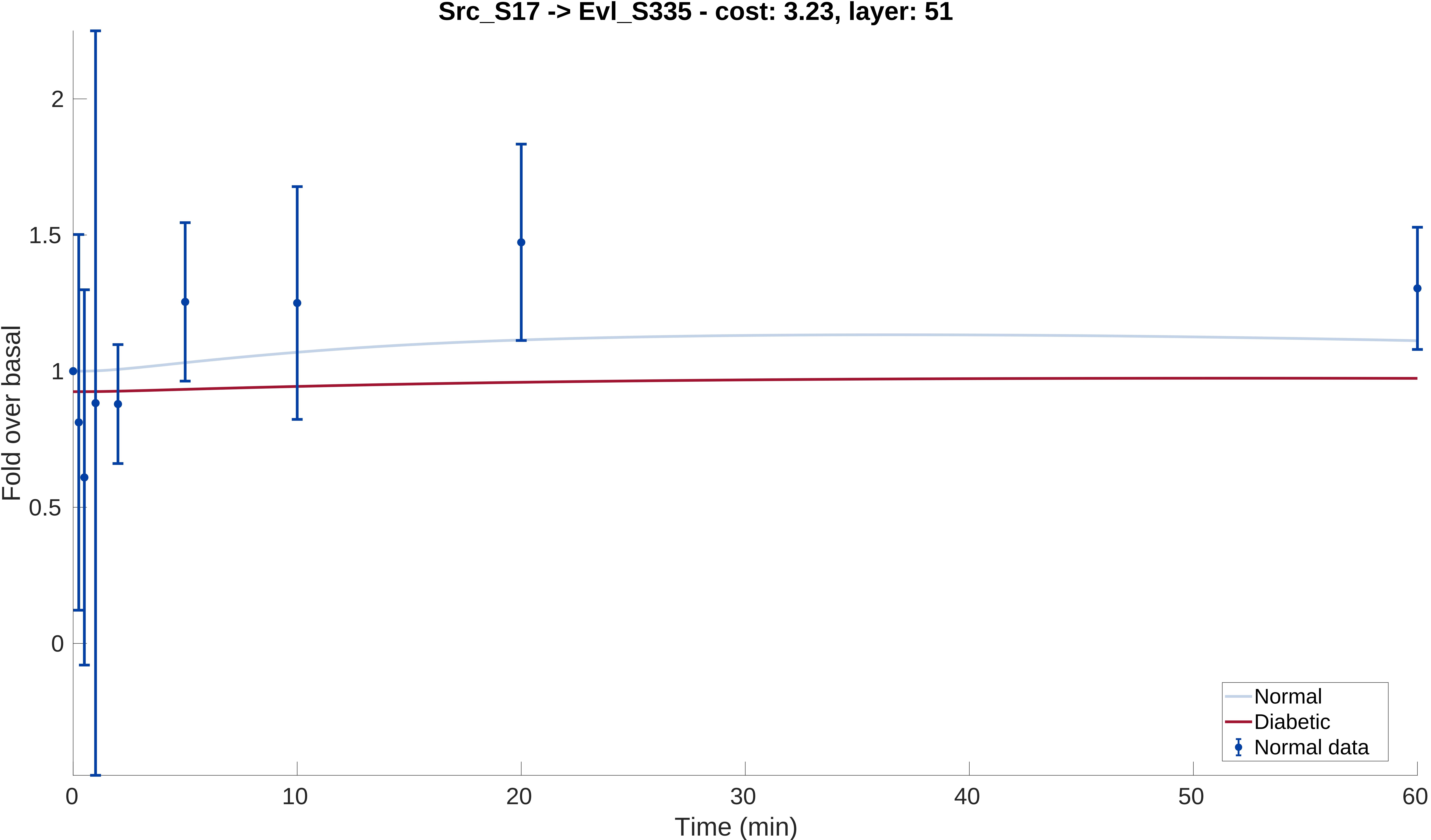

Src_S17 -> Evl_S335 - cost: 3.23, layer: 51
Fold over basal
2
1.5
1
0.5
0
0
10
20
30
40
50
60
Time (min)
Normal
Diabetic
Normal data

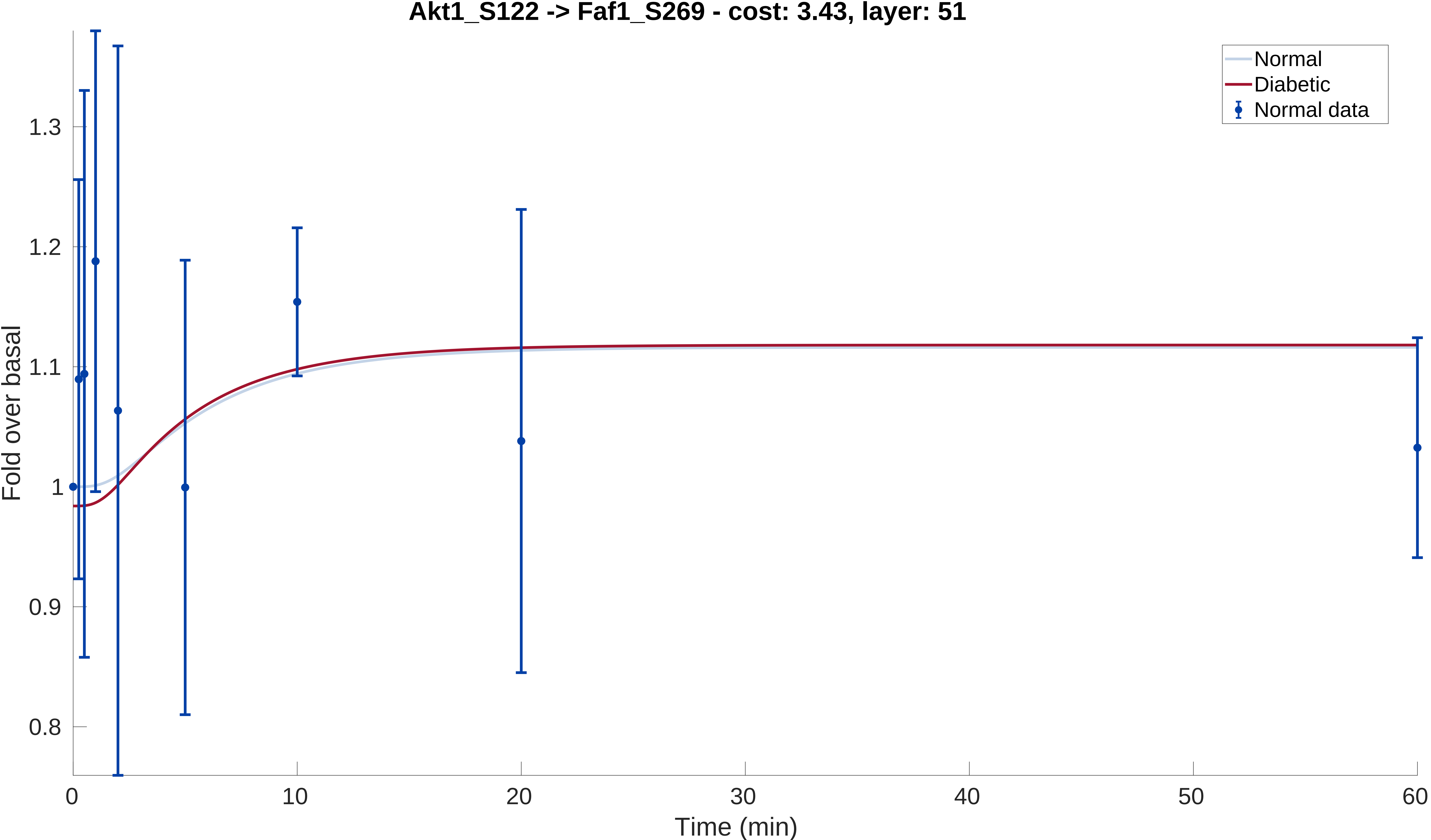

Akt1_S122 -> Faf1_S269 - cost: 3.43, layer: 51
Normal
Diabetic
Normal data
1.3
1.2
1.1
1
0.9
0.8
Fold over basal
0
10
20
30
40
50
60
Time (min)

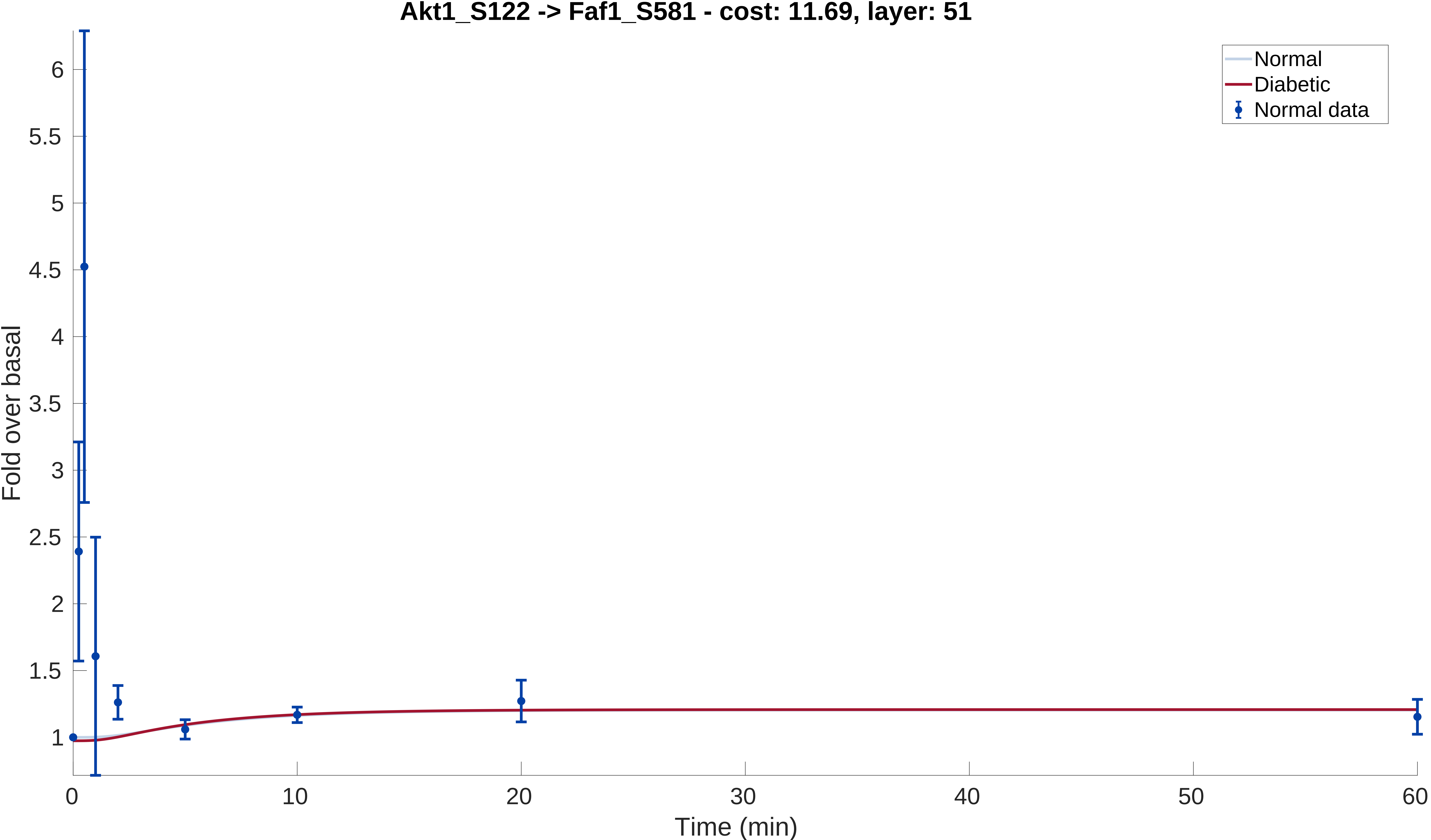

Akt1_S122 -> Faf1_S581 - cost: 11.69, layer: 51
Normal
Diabetic
Normal data
Fold over basal
Time (min)
6
5.5
5
4.5
4
3.5
3
2.5
2
1.5
1
0
10
20
30
40
50
60

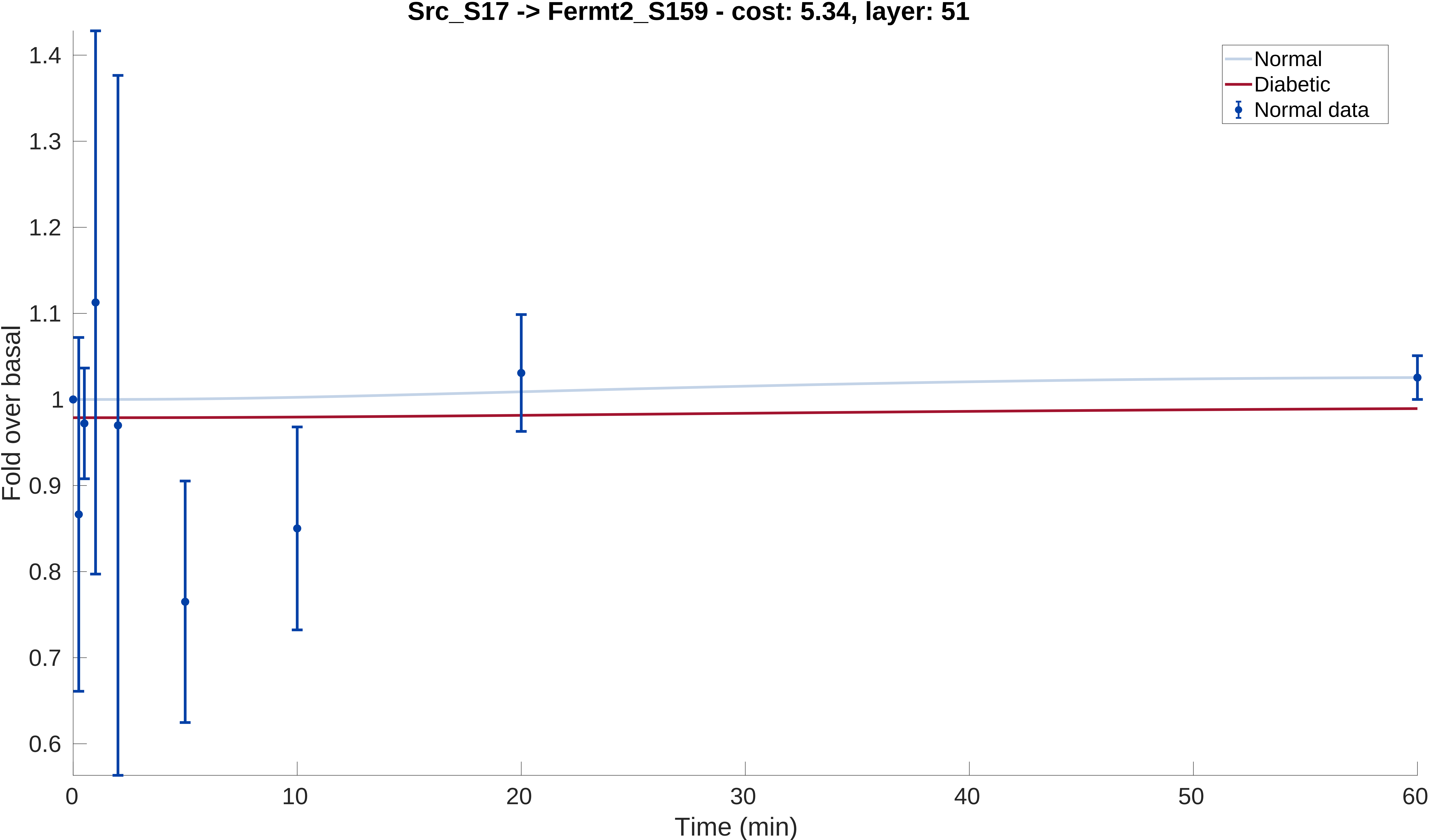

Src_S17 -> Fermt2_S159 - cost: 5.34, layer: 51
Normal
Diabetic
Normal data
Fold over basal
Time (min)
1.4
1.3
1.2
1.1
1
0.9
0.8
0.7
0.6
0
10
20
30
40
50
60

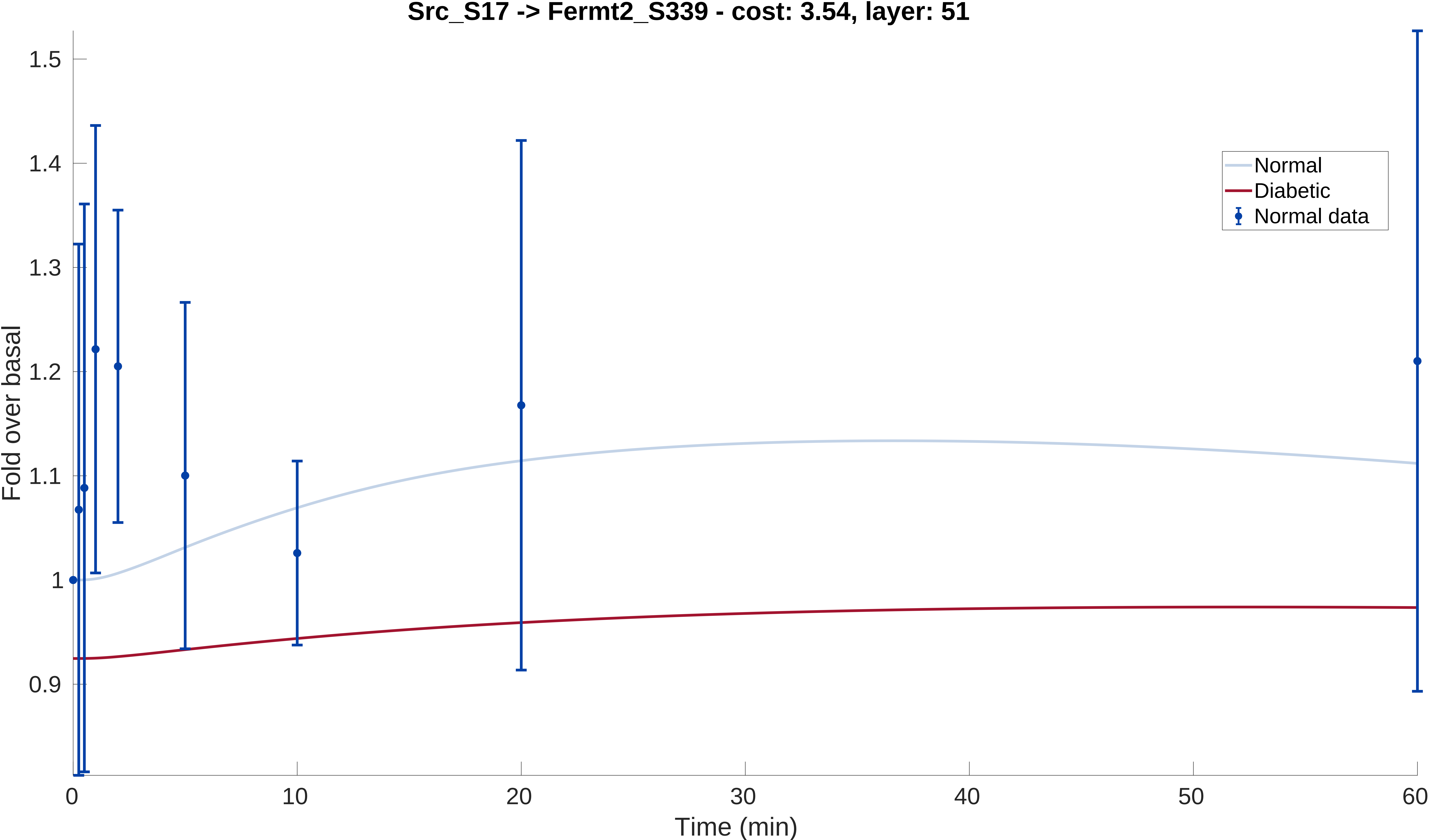

Src_S17 -> Fermt2_S339 - cost: 3.54, layer: 51
Fold over basal
Time (min)
Normal
Diabetic
Normal data
1.5
1.4
1.3
1.2
1.1
1
0.9
0
10
20
30
40
50
60

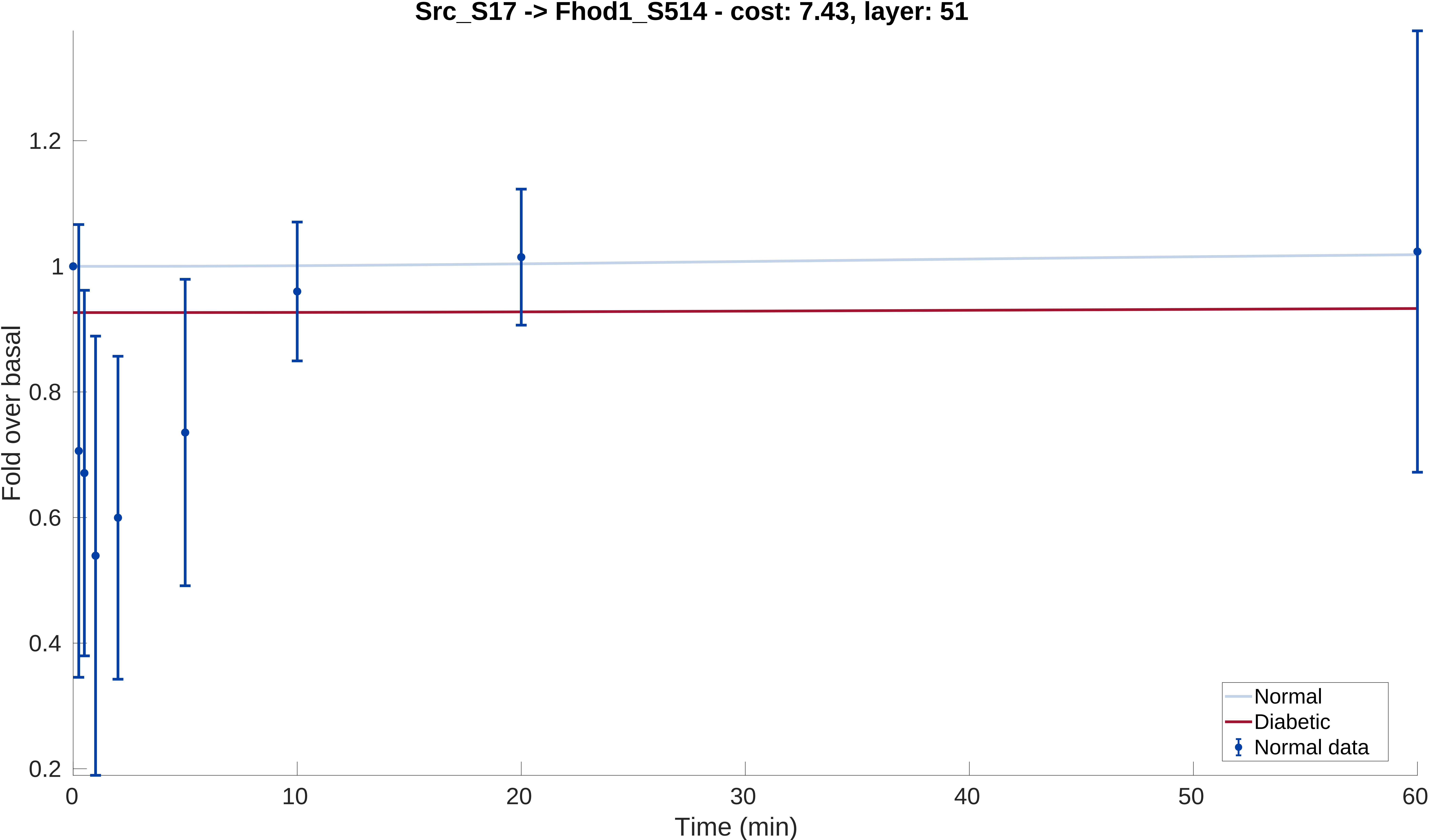

Src_S17 -> Fhod1_S514 - cost: 7.43, layer: 51
Fold over basal
1.2
1
0.8
0.6
0.4
0.2
0
10
20
30
40
50
60
Time (min)
Normal
Diabetic
Normal data

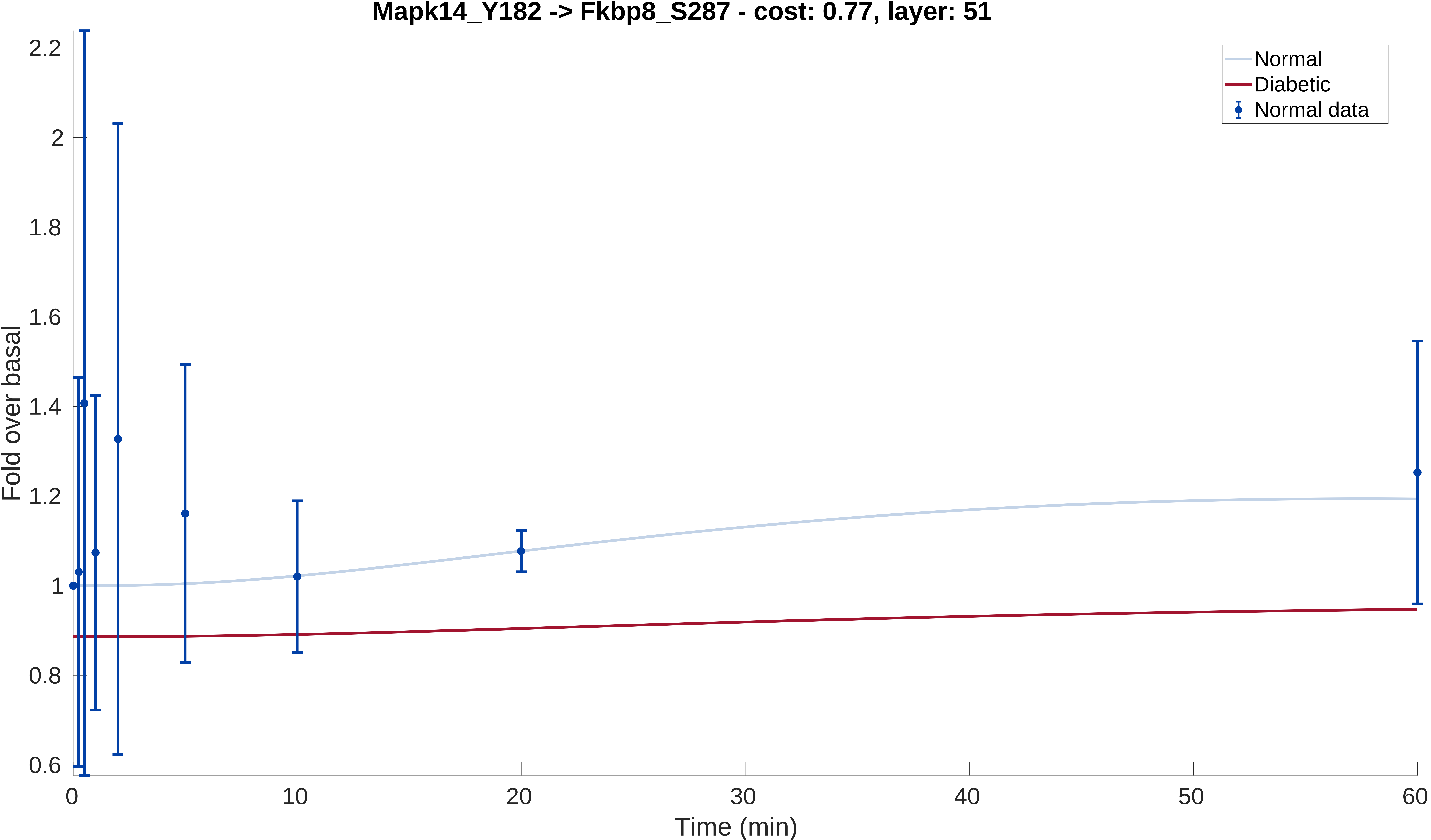

Mapk14_Y182 -> Fkbp8_S287 - cost: 0.77, layer: 51
Normal
Diabetic
Normal data
Fold over basal
0.6
0.8
1
1.2
1.4
1.6
1.8
2
2.2
0
10
20
30
40
50
60
Time (min)

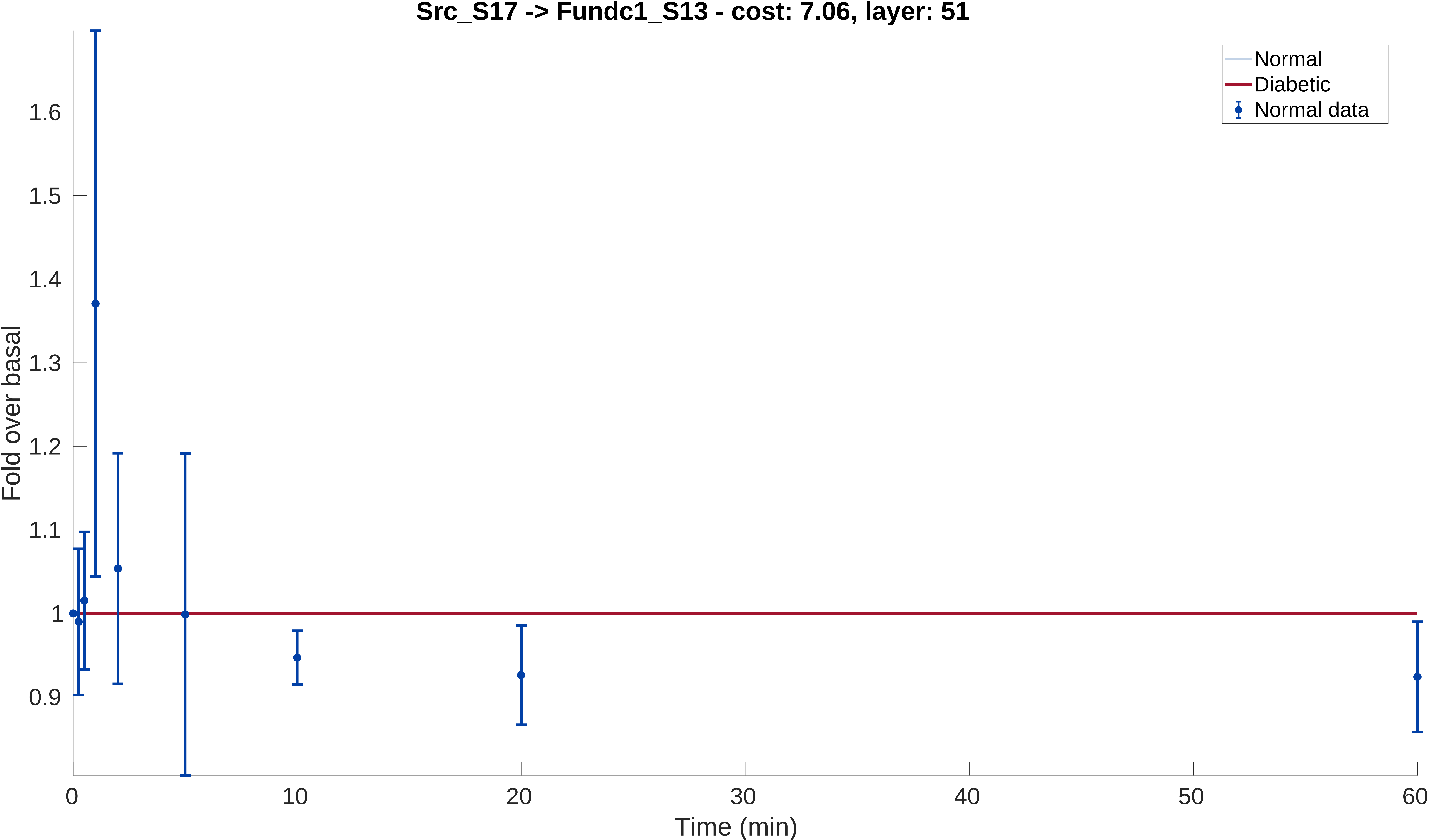

Src_S17 -> Fundc1_S13 - cost: 7.06, layer: 51
Normal
Diabetic
Normal data
Fold over basal
1.6
1.5
1.4
1.3
1.2
1.1
1
0.9
0
10
20
30
40
50
60
Time (min)

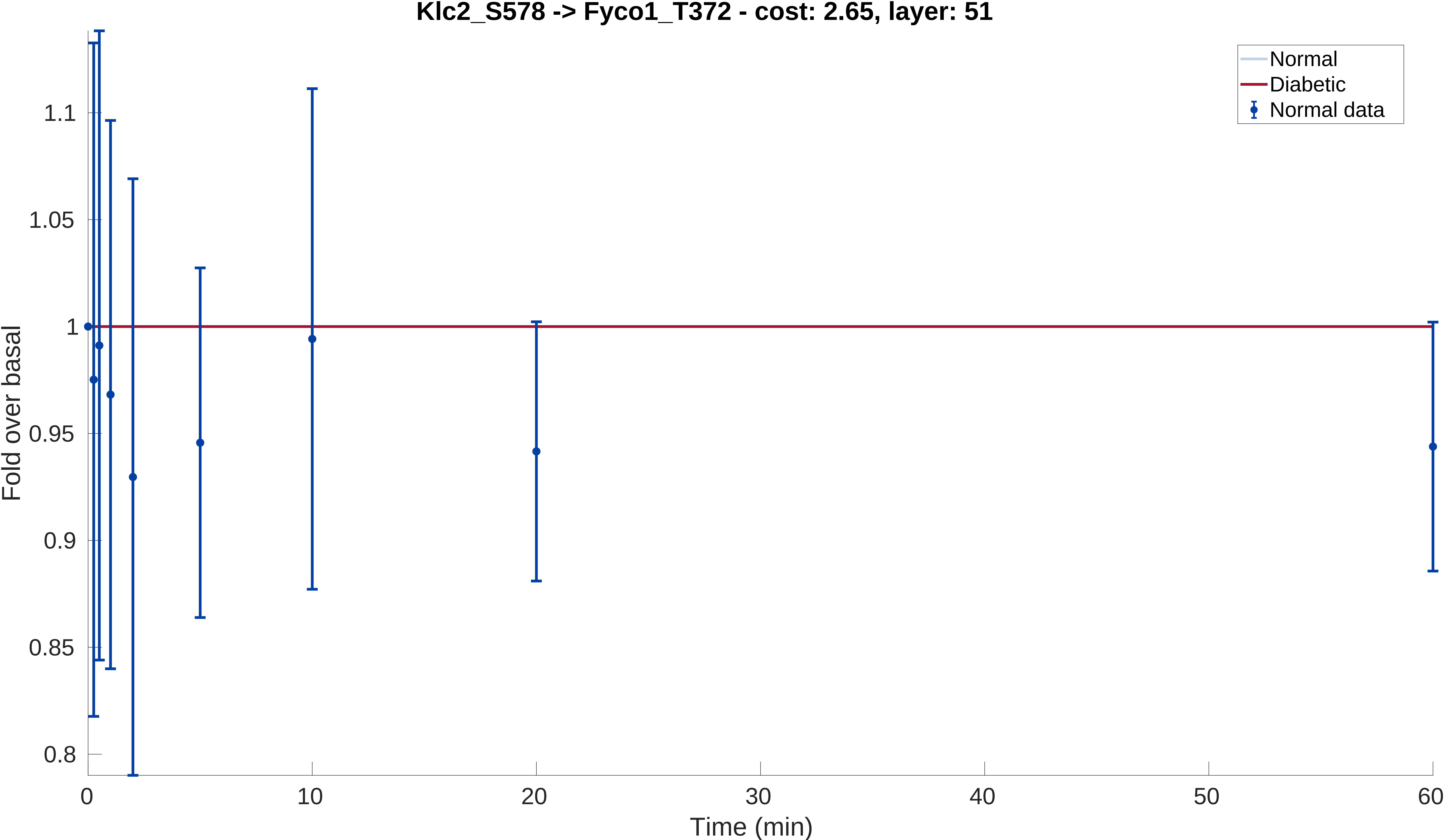

Klc2_S578 -> Fyco1_T372 - cost: 2.65, layer: 51
Normal
Diabetic
Normal data
Fold over basal
Time (min)
1.1
1.05
1
0.95
0.9
0.85
0.8
0
10
20
30
40
50
60

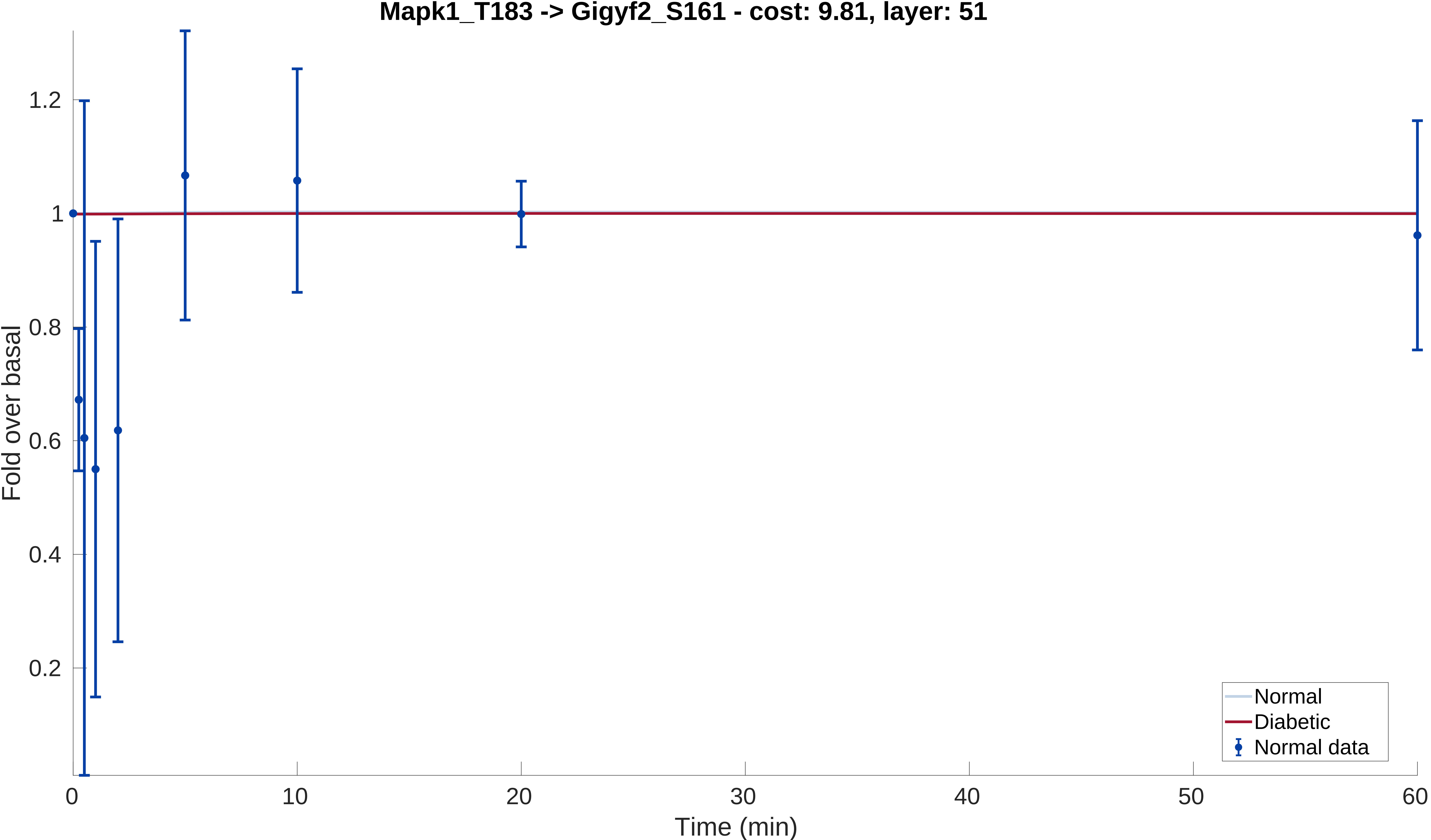

Mapk1_T183 -> Gigyf2_S161 - cost: 9.81, layer: 51
Fold over basal
1.2
1
0.8
0.6
0.4
0.2
0
10
20
30
40
50
60
Time (min)
Normal
Diabetic
Normal data

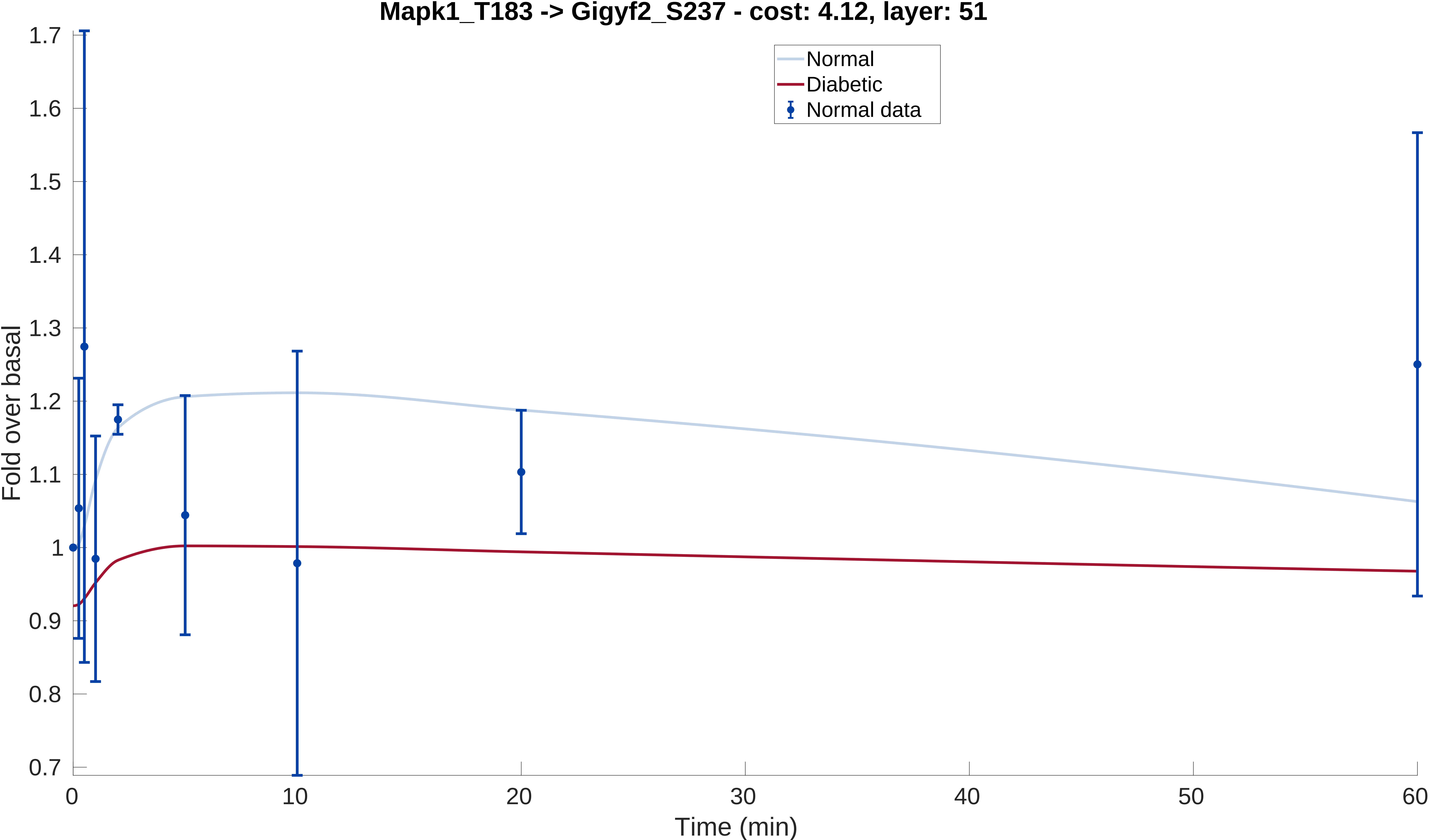

Mapk1_T183 -> Gigyf2_S237 - cost: 4.12, layer: 51
Normal
Diabetic
Normal data
Fold over basal
1.7
1.6
1.5
1.4
1.3
1.2
1.1
1
0.9
0.8
0.7
0
10
20
30
40
50
60
Time (min)

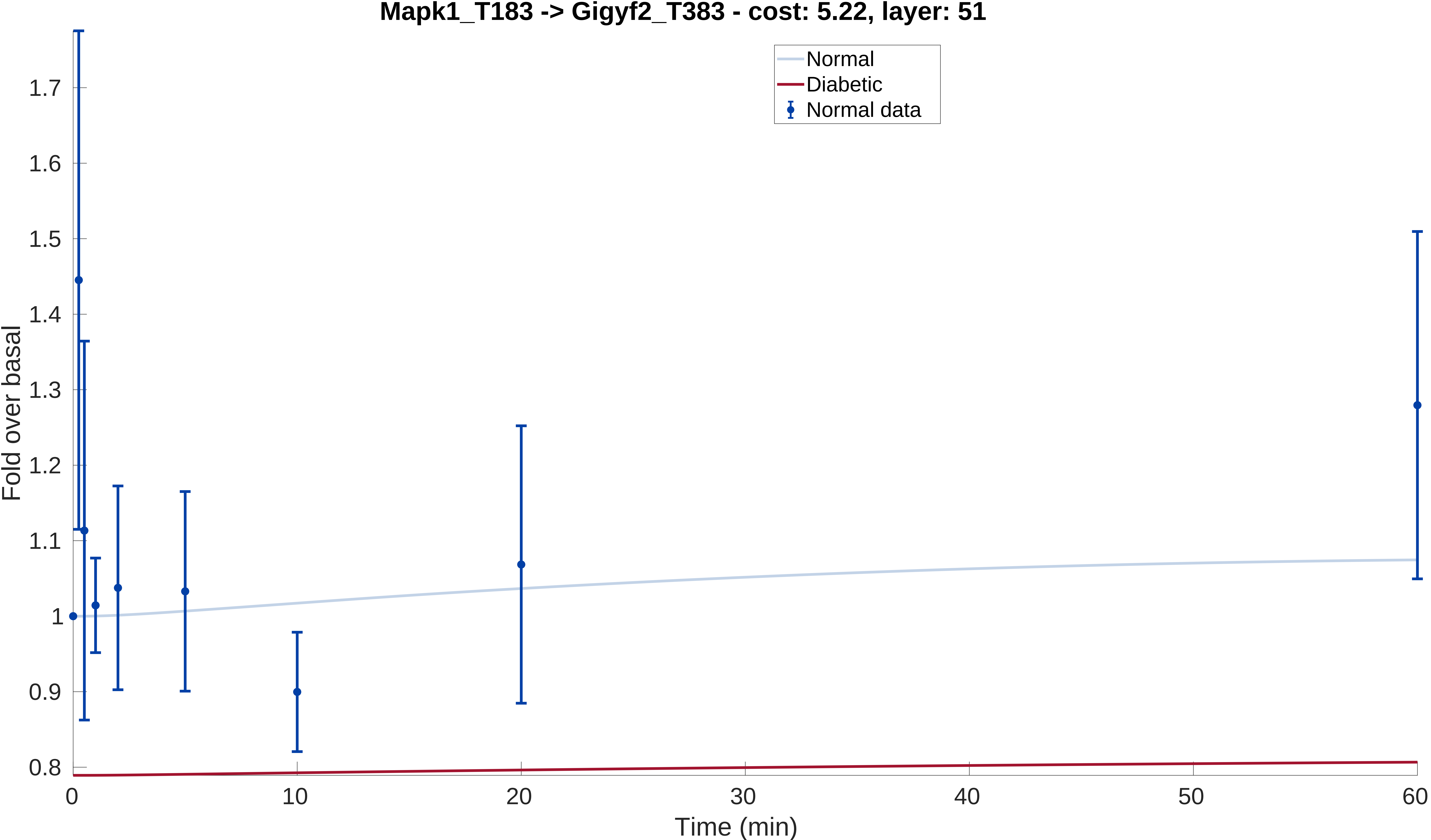

Mapk1_T183 -> Gigyf2_T383 - cost: 5.22, layer: 51
Normal
Diabetic
Normal data
Fold over basal
1.7
1.6
1.5
1.4
1.3
1.2
1.1
1
0.9
0.8
0
10
20
30
40
50
60
Time (min)

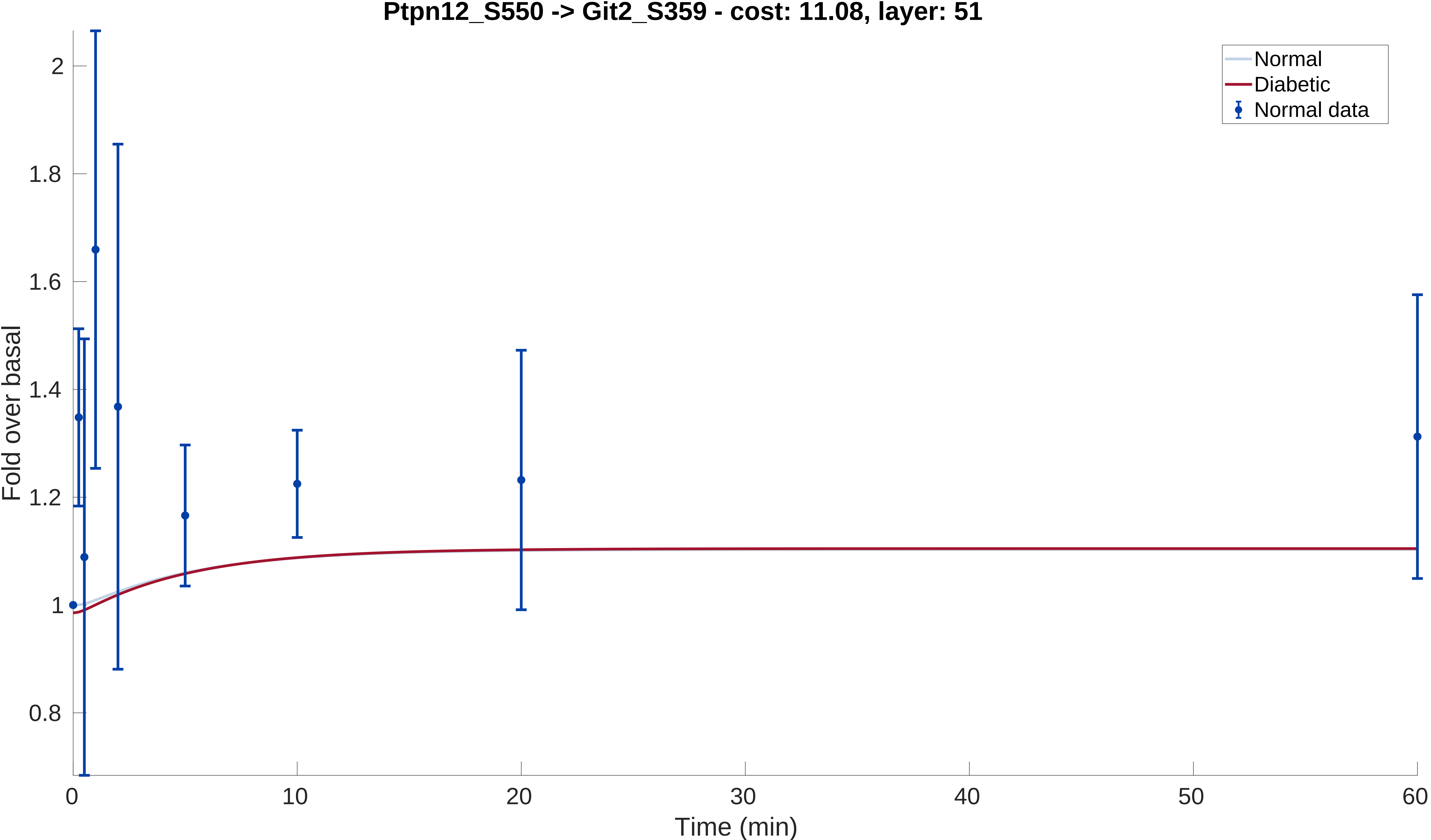

Ptpn12_S550 -> Git2_S359 - cost: 11.08, layer: 51
Normal
Diabetic
Normal data
Fold over basal
Time (min)
0.8
1
1.2
1.4
1.6
1.8
2
0
10
20
30
40
50
60

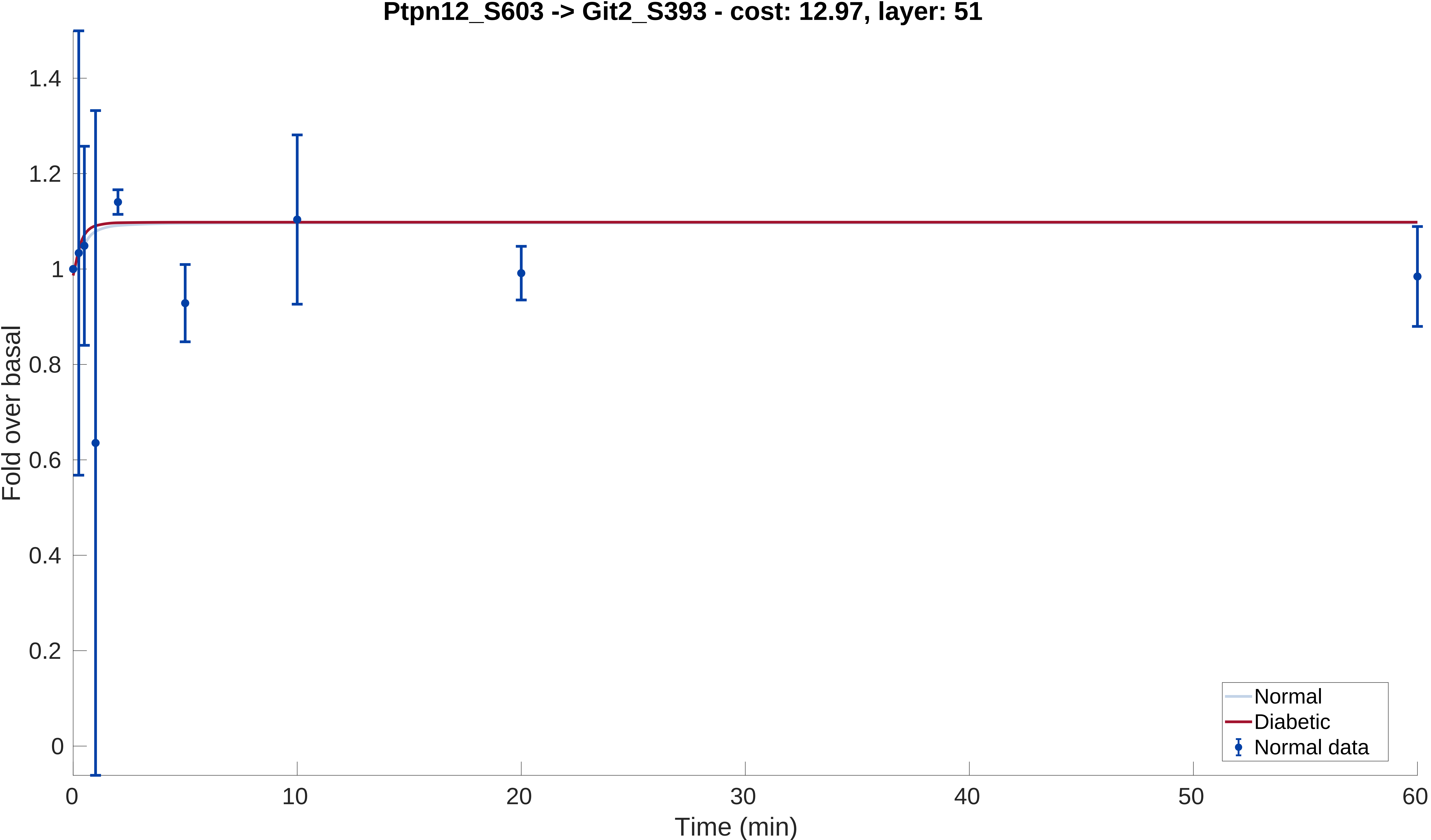

Ptpn12_S603 -> Git2_S393 - cost: 12.97, layer: 51
Fold over basal
1.4
1.2
1
0.8
0.6
0.4
0.2
0
0
10
20
30
40
50
60
Time (min)
Normal
Diabetic
Normal data

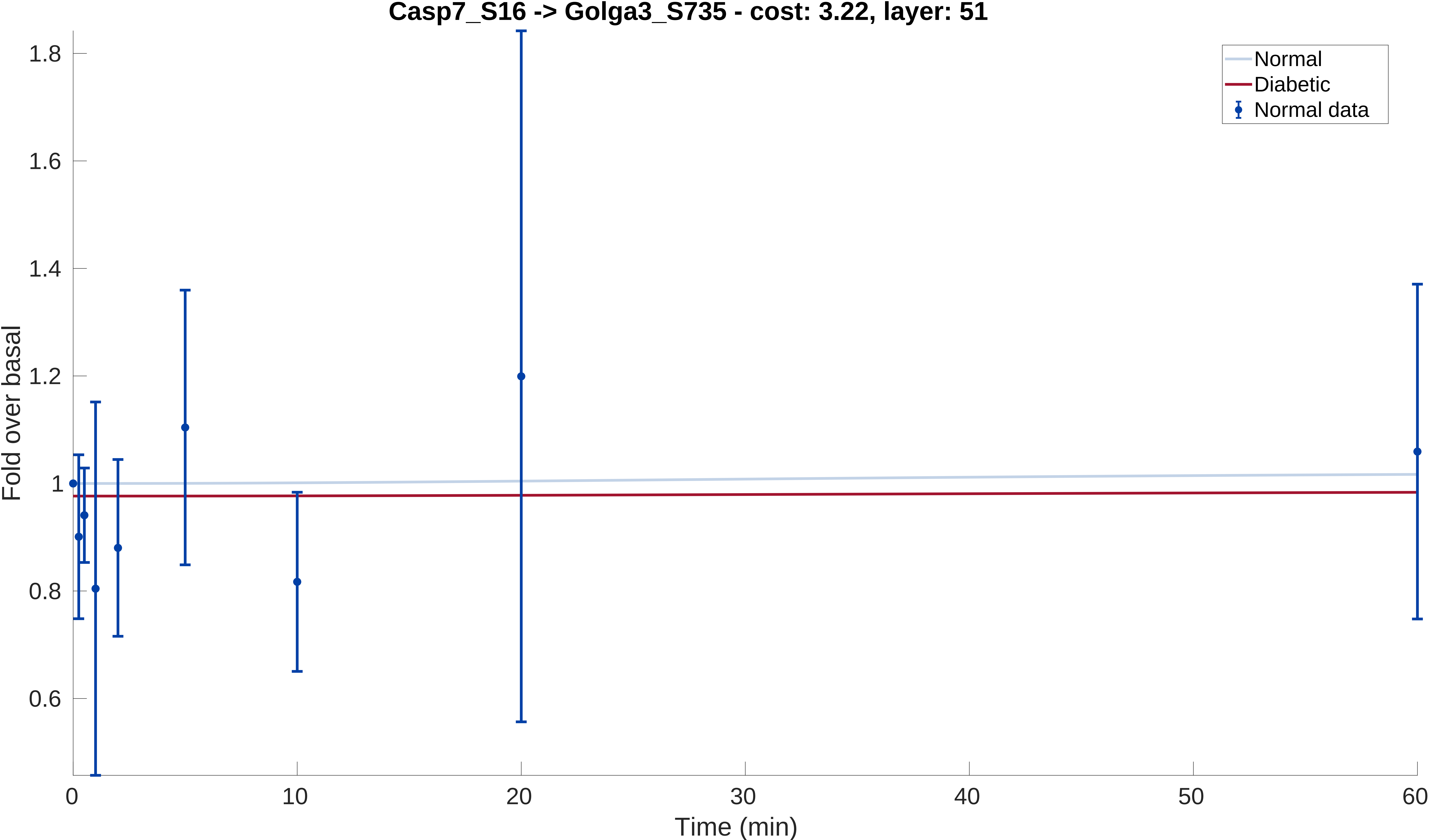

Casp7_S16 -> Golga3_S735 - cost: 3.22, layer: 51
Normal
Diabetic
Normal data
Fold over basal
1.8
1.6
1.4
1.2
1
0.8
0.6
0
10
20
30
40
50
60
Time (min)

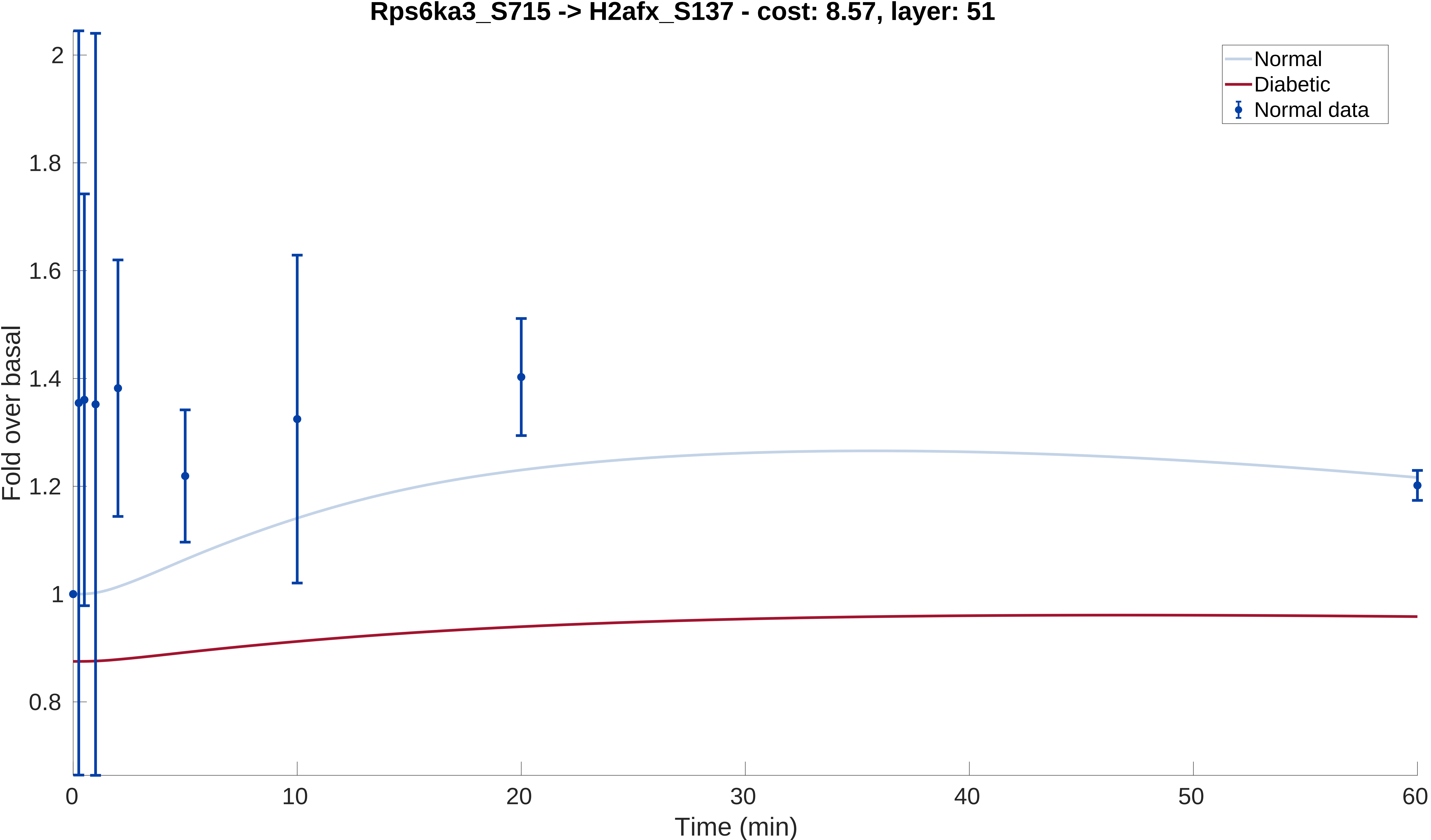
Rps6ka3_S715 -> H2afx_S137 - cost: 8.57, layer: 51
Normal
Diabetic
Normal data
Fold over basal
Time (min)
0.8
1
1.2
1.4
1.6
1.8
2
0
10
20
30
40
50
60

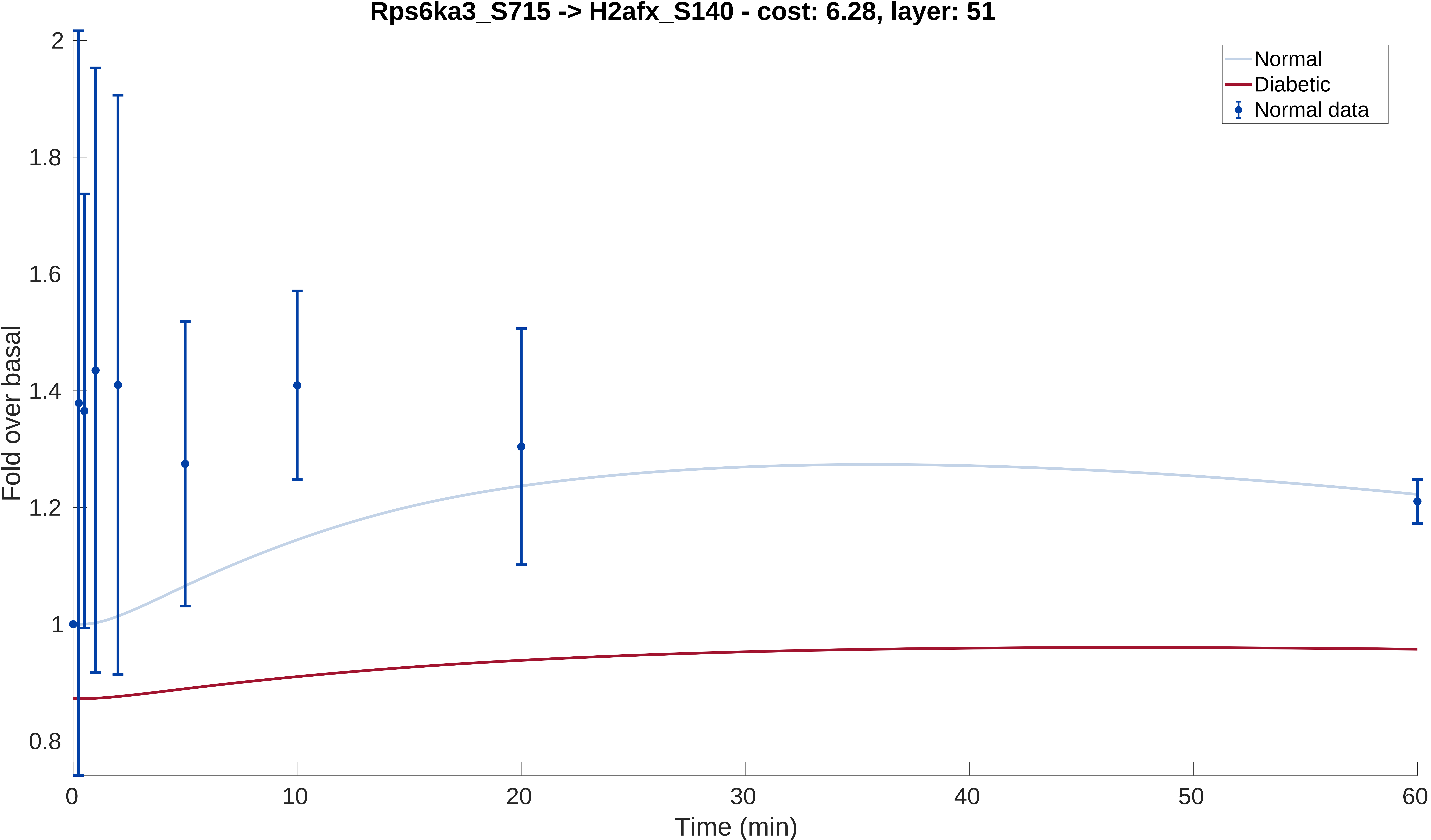
Rps6ka3_S715 -> H2afx_S140 - cost: 6.28, layer: 51
Normal
Diabetic
Normal data
Fold over basal
2
1.8
1.6
1.4
1.2
1
0.8
0
10
20
30
40
50
60
Time (min)

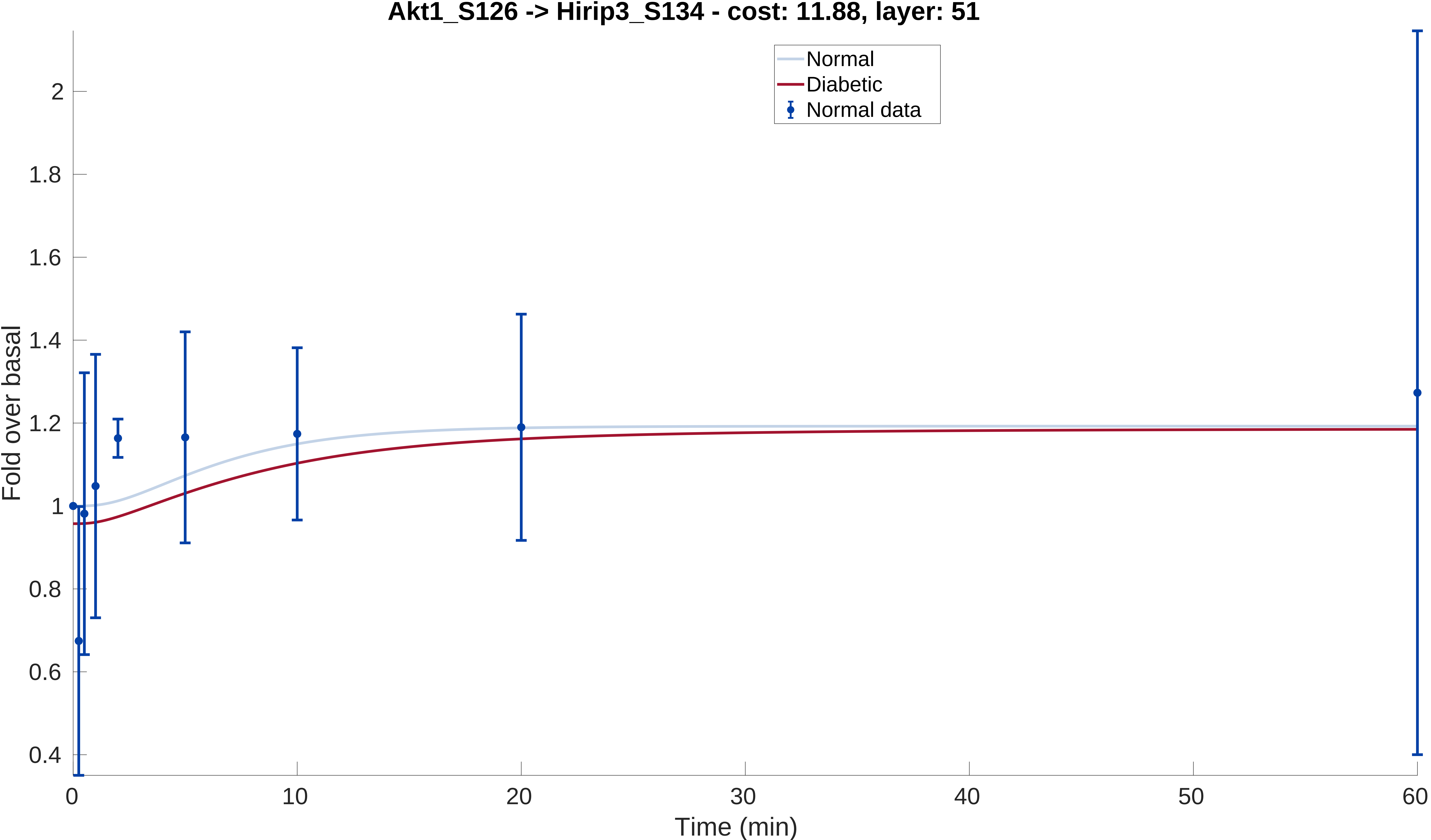

Akt1_S126 -> Hirip3_S134 - cost: 11.88, layer: 51
Normal
Diabetic
Normal data
Fold over basal
2
1.8
1.6
1.4
1.2
1
0.8
0.6
0.4
0
10
20
30
40
50
60
Time (min)

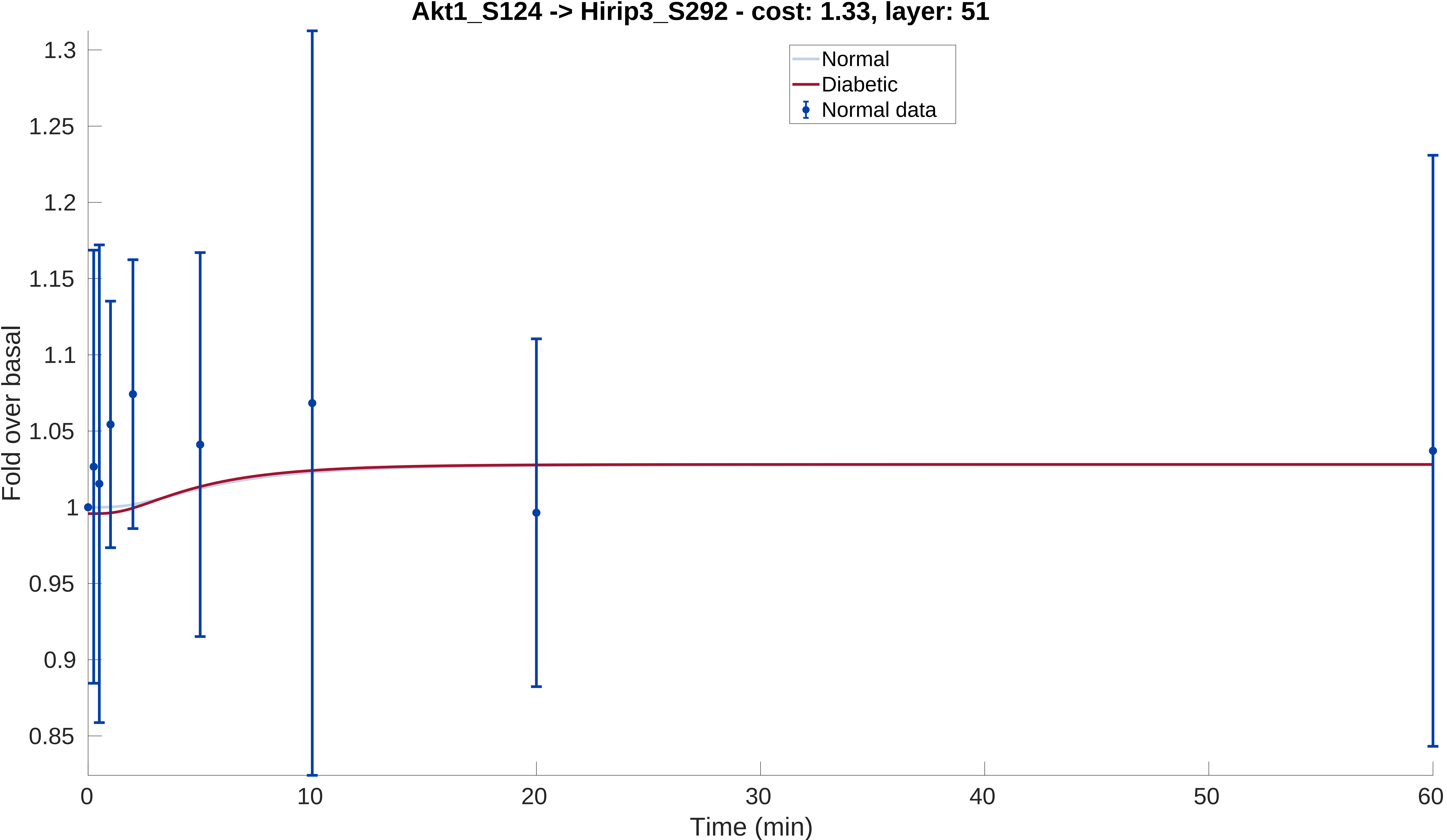
Akt1_S124 -> Hirip3_S292 - cost: 1.33, layer: 51
Normal
Diabetic
Normal data
Fold over basal
1.3
1.25
1.2
1.15
1.1
1.05
1
0.95
0.9
0.85
0
10
20
30
40
50
60
Time (min)

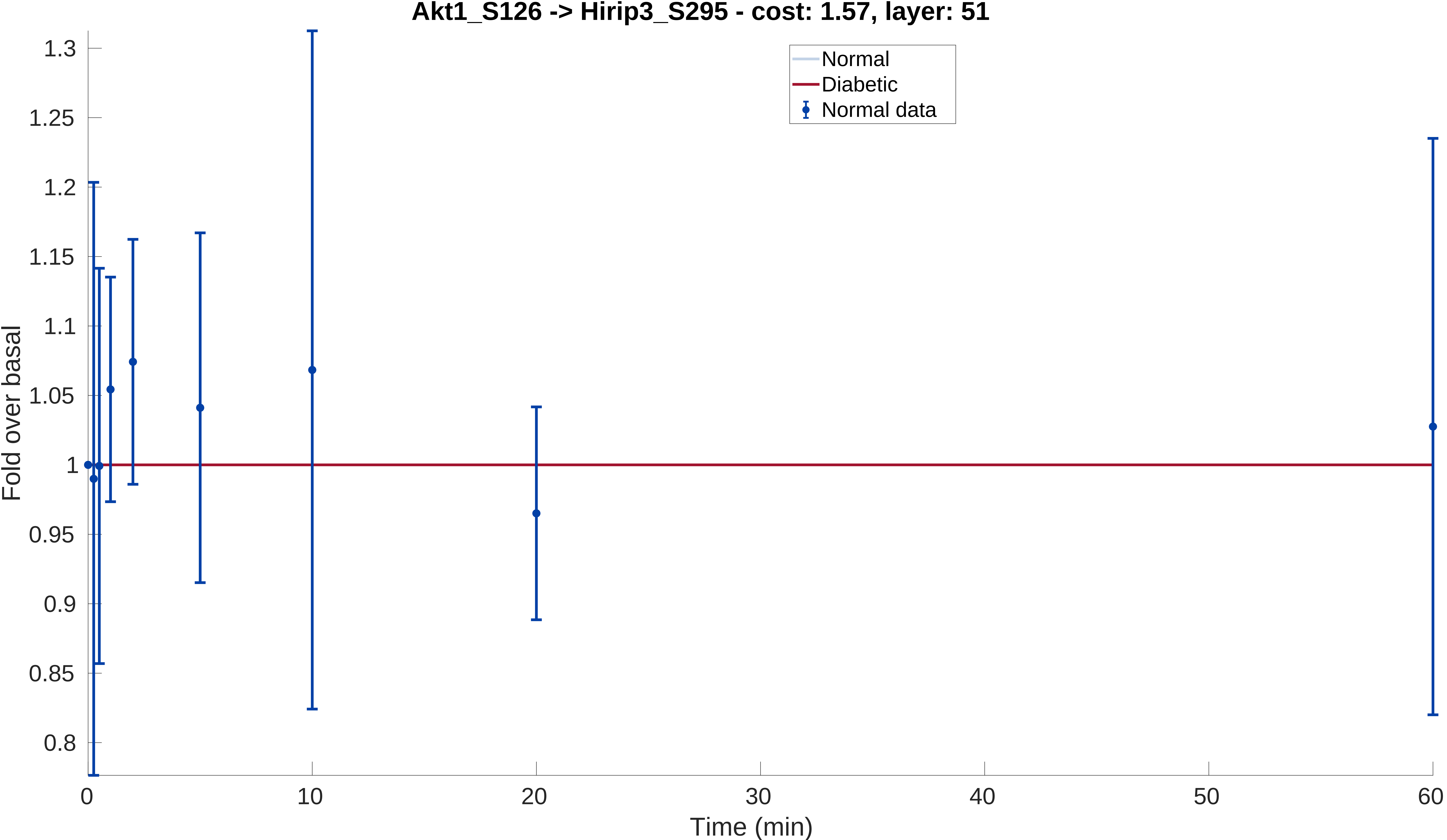

Akt1_S126 -> Hirip3_S295 - cost: 1.57, layer: 51
Normal
Diabetic
Normal data
Fold over basal
1.3
1.25
1.2
1.15
1.1
1.05
1
0.95
0.9
0.85
0.8
0
10
20
30
40
50
60
Time (min)

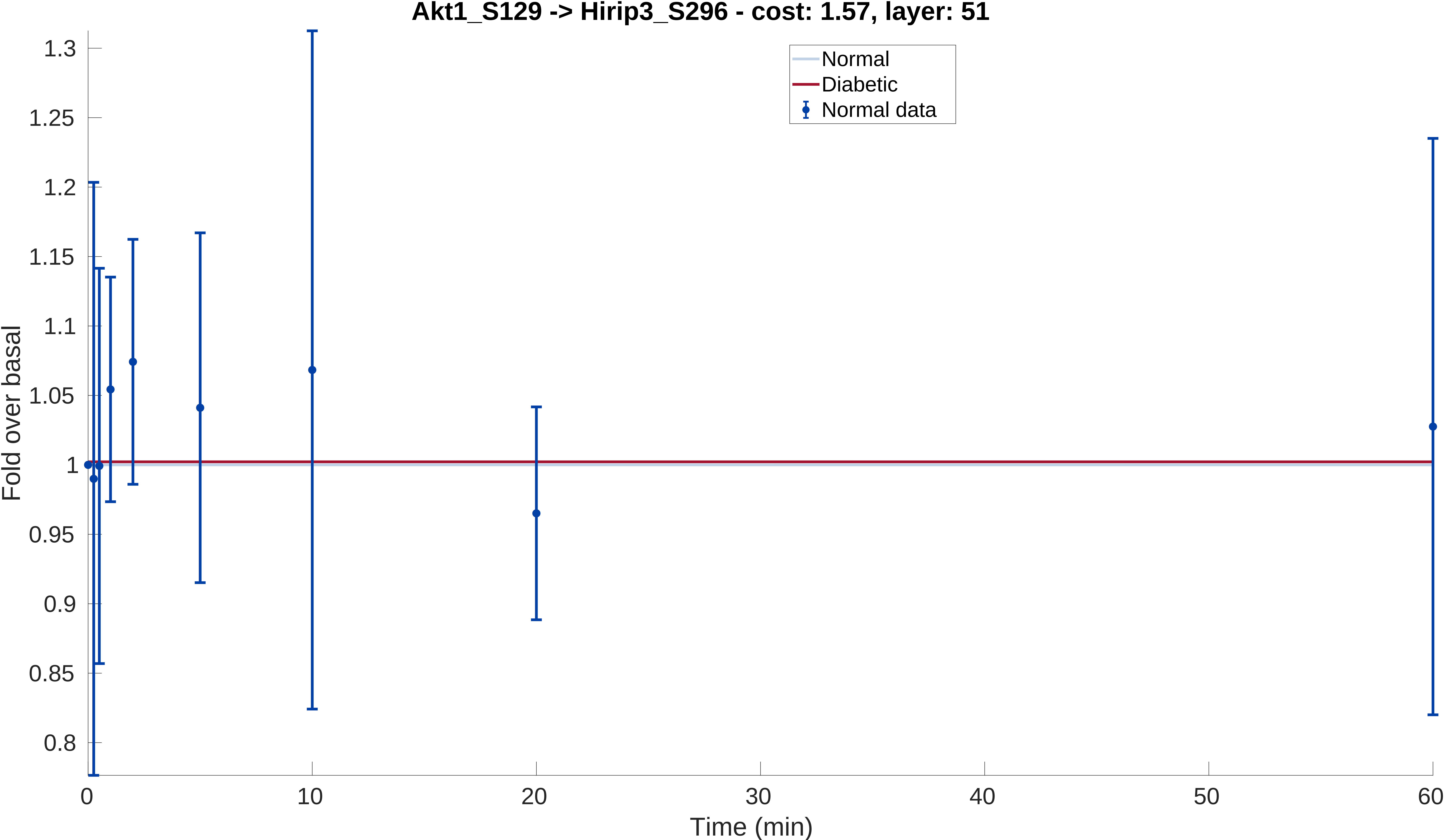

Akt1_S129 -> Hirip3_S296 - cost: 1.57, layer: 51
Normal
Diabetic
Normal data
Fold over basal
Time (min)
1.3
1.25
1.2
1.15
1.1
1.05
1
0.95
0.9
0.85
0.8
0
10
20
30
40
50
60

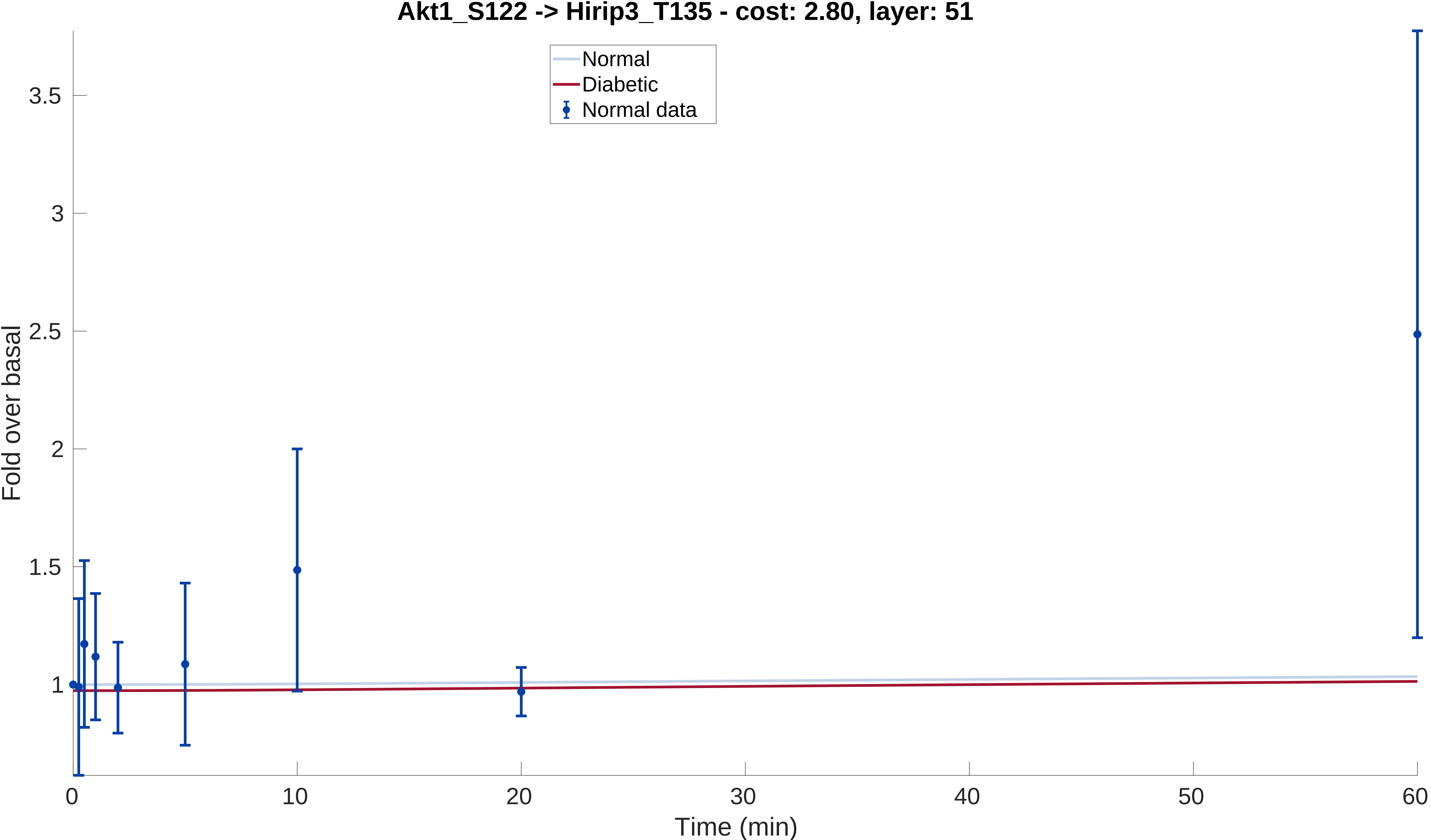

Akt1_S122 -> Hirip3_T135 - cost: 2.80, layer: 51
Normal
Diabetic
Normal data
Fold over basal
3.5
3
2.5
2
1.5
1
0
10
20
30
40
50
60
Time (min)

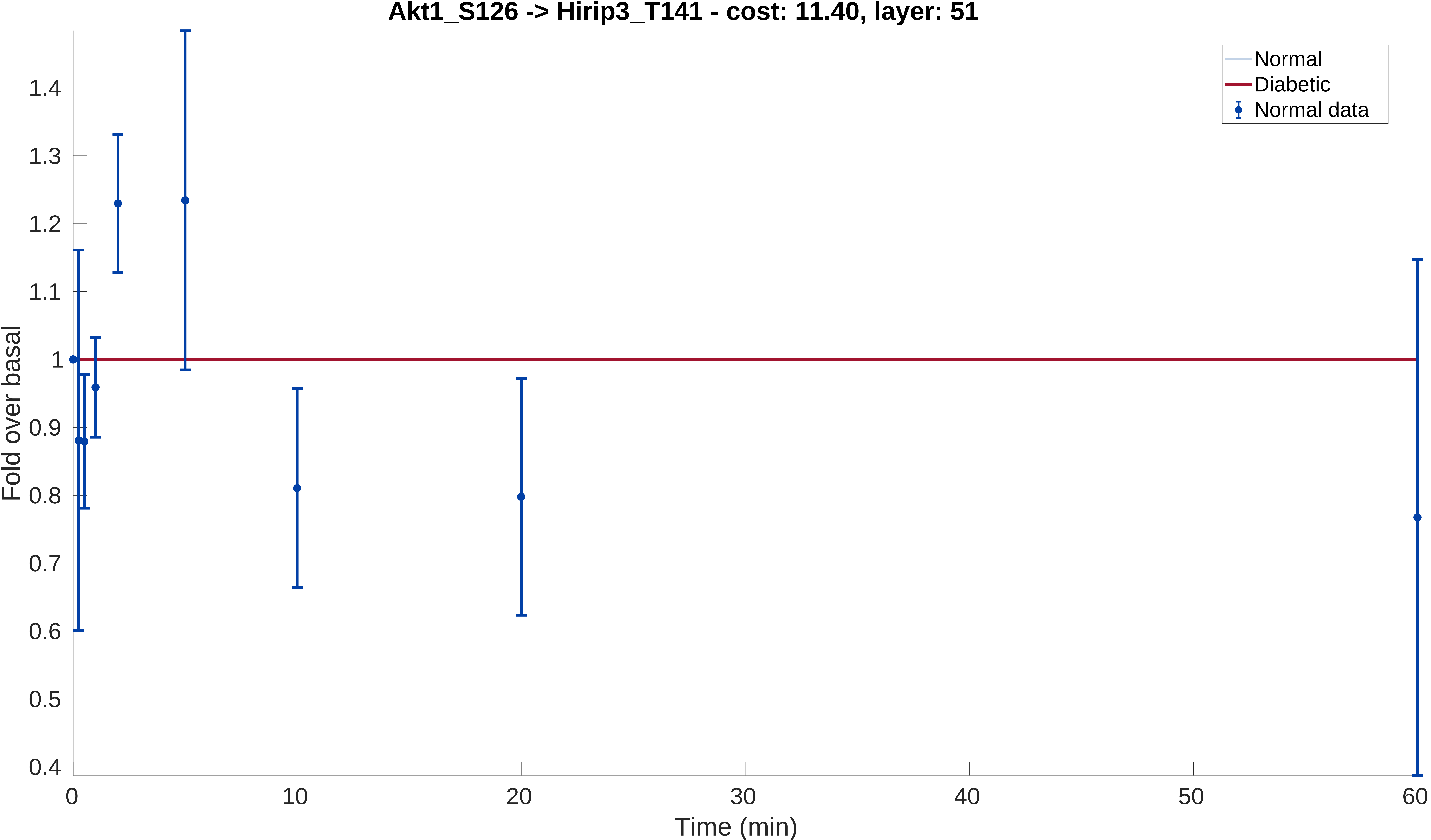

Akt1_S126 -> Hirip3_T141 - cost: 11.40, layer: 51
Normal
Diabetic
Normal data
Fold over basal
1.4
1.3
1.2
1.1
1
0.9
0.8
0.7
0.6
0.5
0.4
0
10
20
30
40
50
60
Time (min)

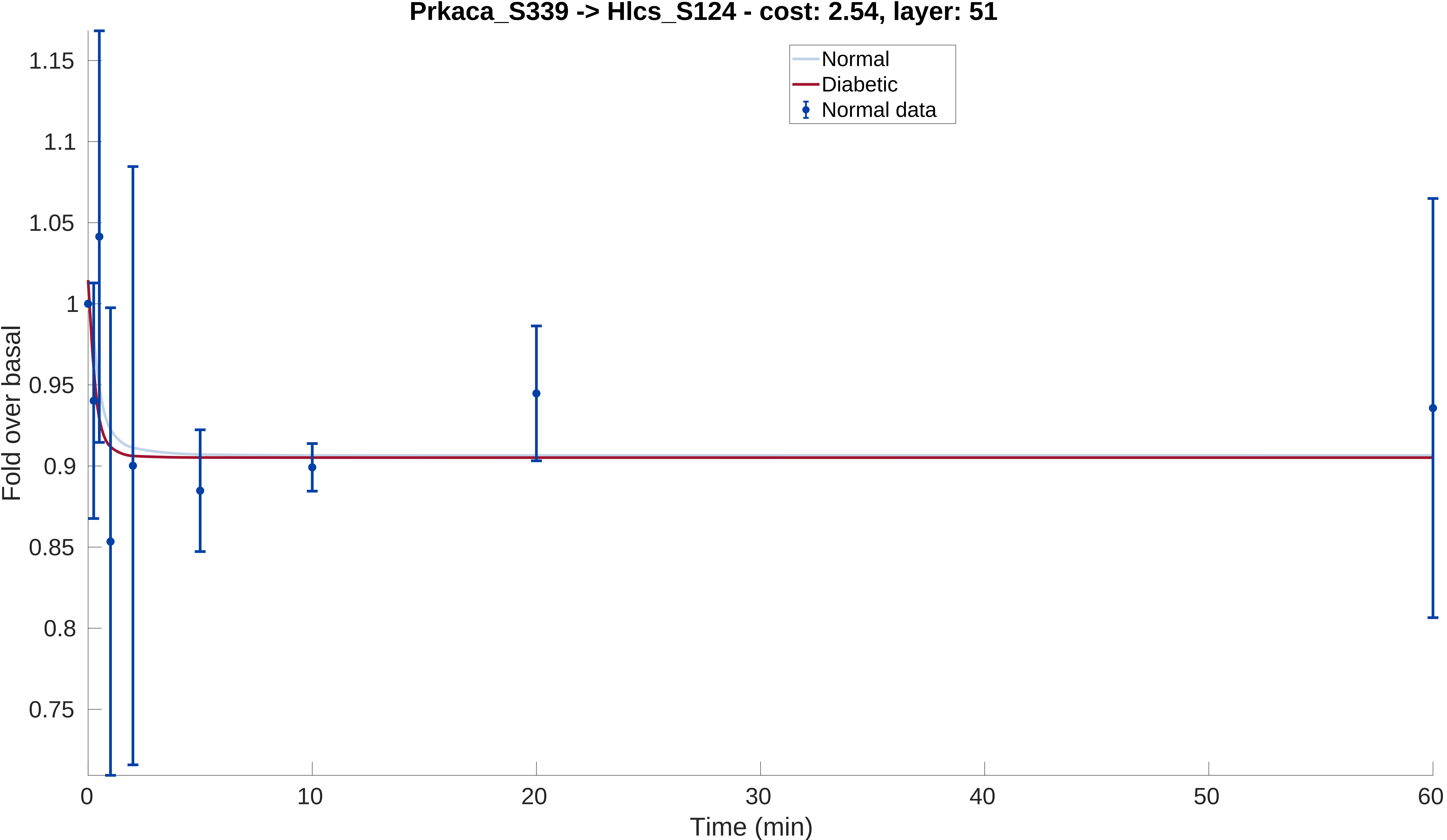
Prkaca_S339 -> Hlcs_S124 - cost: 2.54, layer: 51
Normal
Diabetic
Normal data
Fold over basal
1.15
1.1
1.05
1
0.95
0.9
0.85
0.8
0.75
0
10
20
30
40
50
60
Time (min)

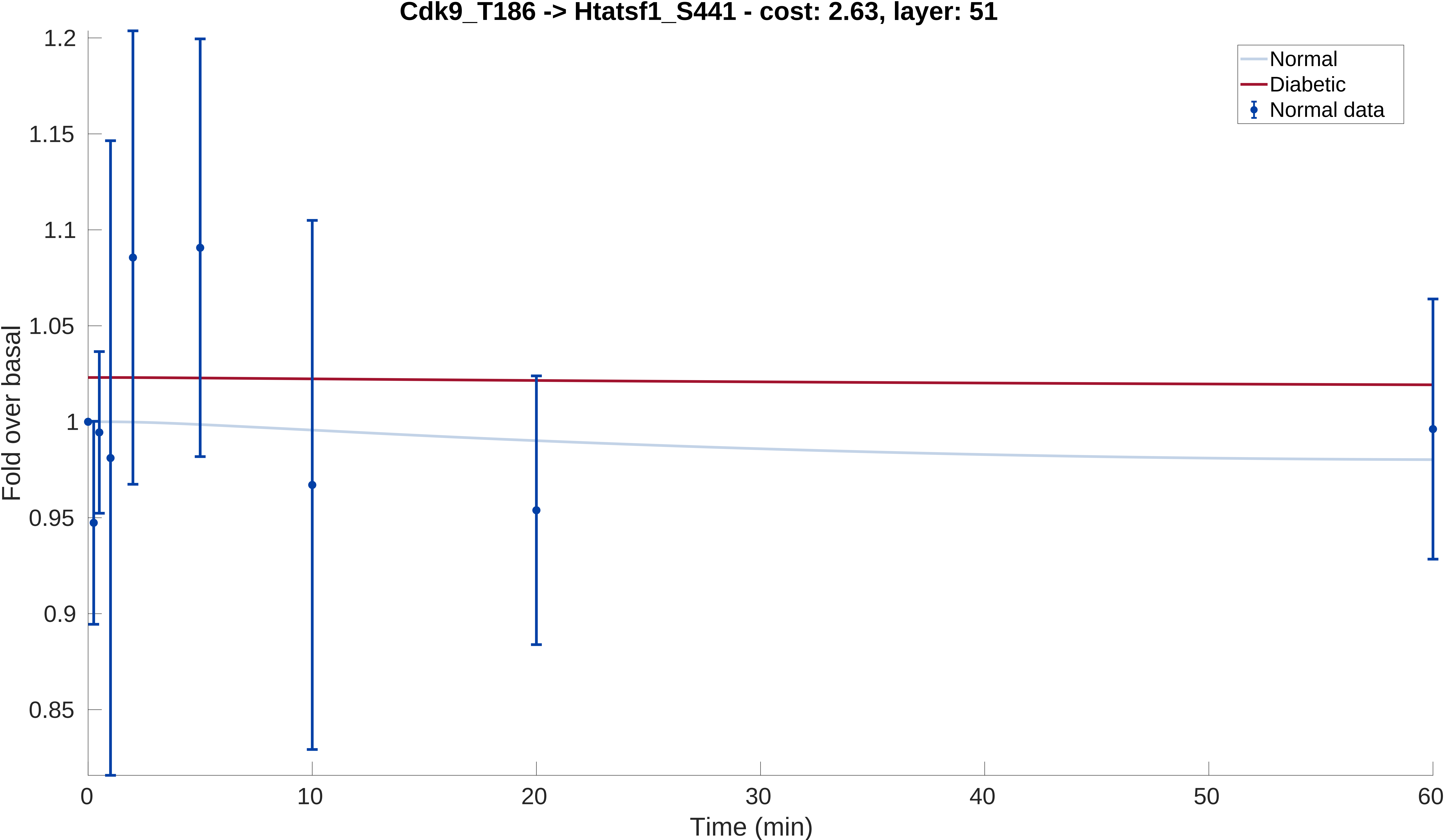

Cdk9_T186 -> Htatsf1_S441 - cost: 2.63, layer: 51
Normal
Diabetic
Normal data
Fold over basal
1.2
1.15
1.1
1.05
1
0.95
0.9
0.85
0
10
20
30
40
50
60
Time (min)

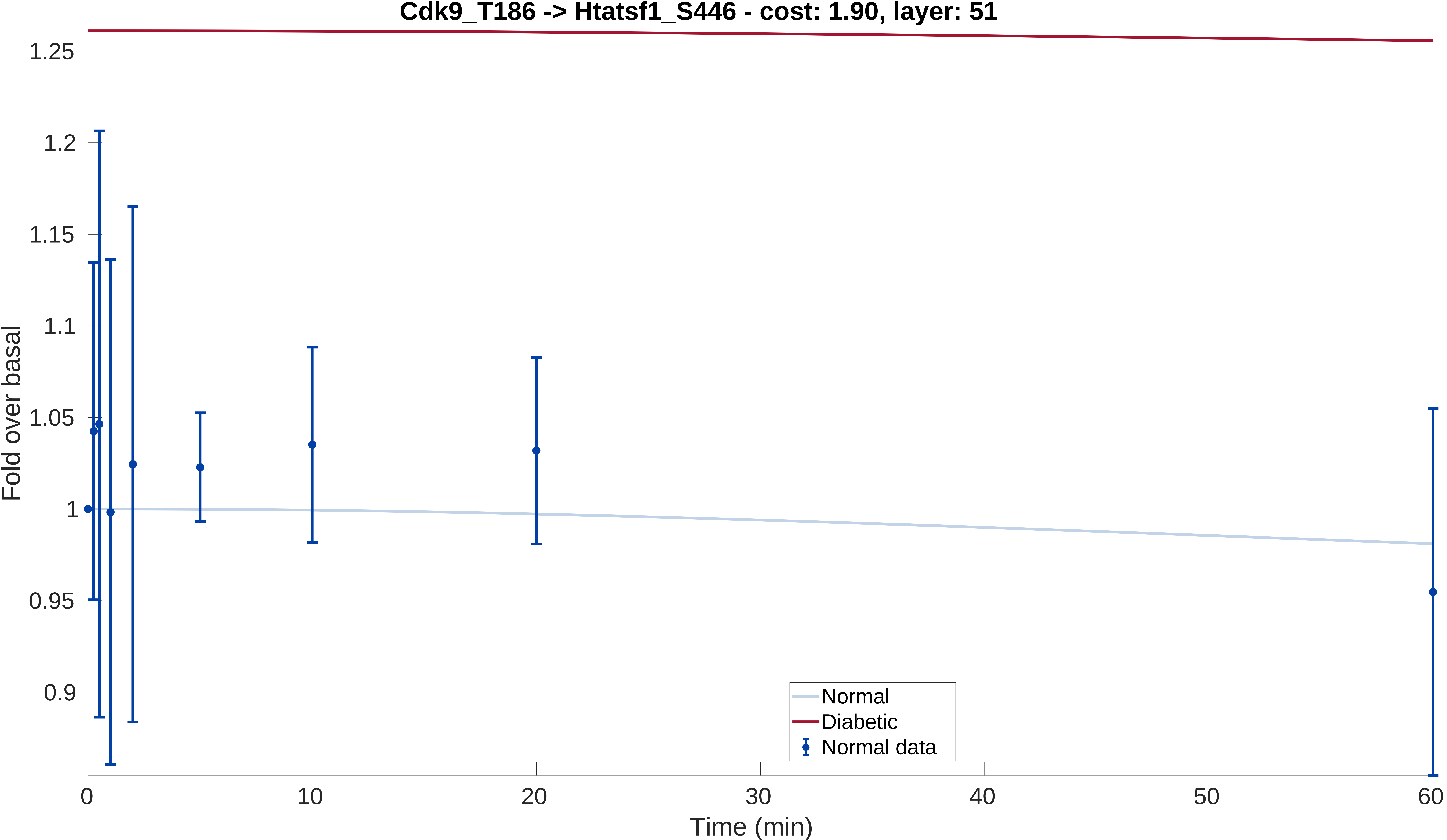

Cdk9_T186 -> Htatsf1_S446 - cost: 1.90, layer: 51
Fold over basal
1.25
1.2
1.15
1.1
1.05
1
0.95
0.9
0
10
20
30
40
50
60
Time (min)
Normal
Diabetic
Normal data

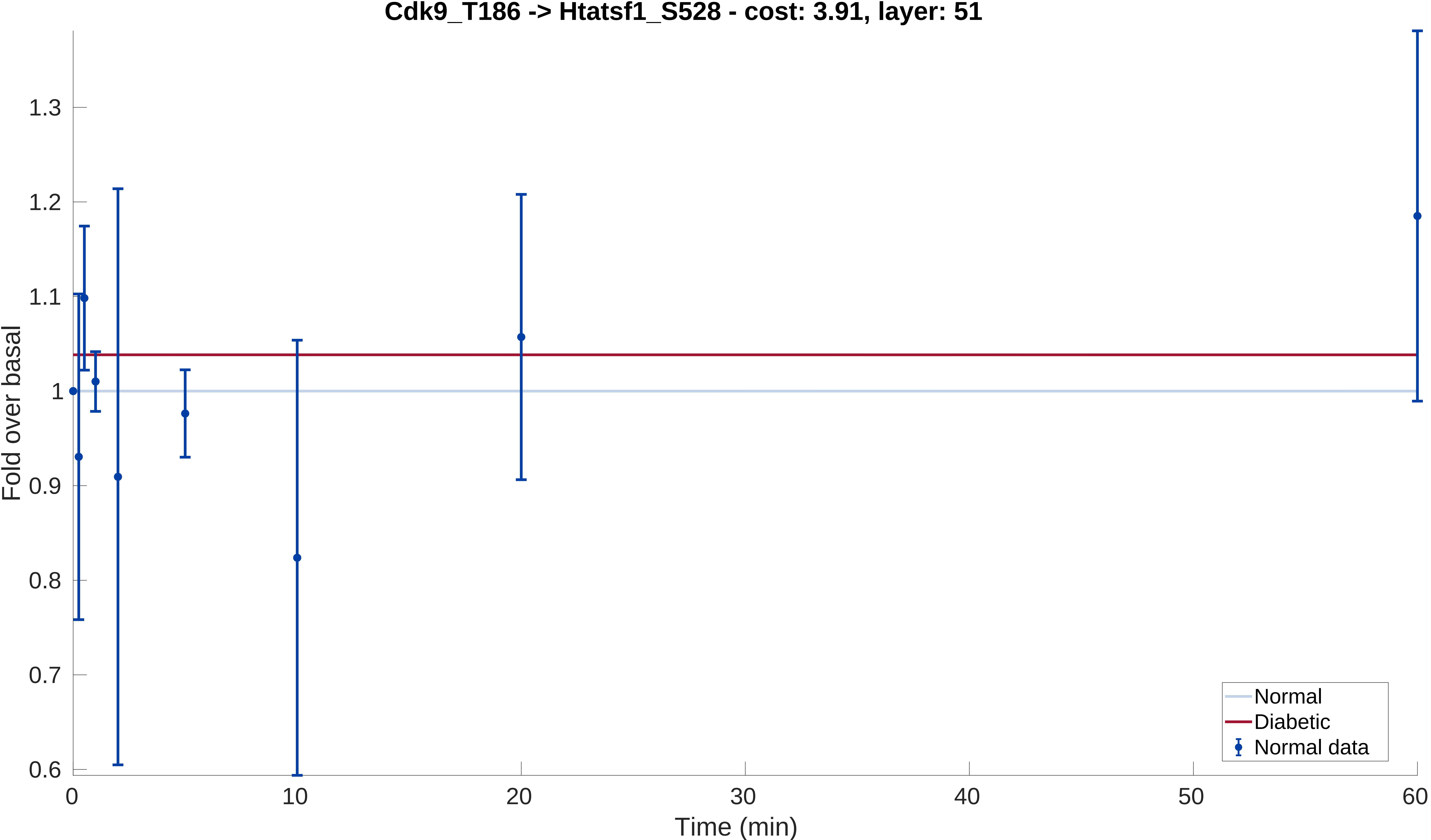

Cdk9_T186 -> Htatsf1_S528 - cost: 3.91, layer: 51
Fold over basal
Time (min)
0.6
0.7
0.8
0.9
1
1.1
1.2
1.3
0
10
20
30
40
50
60
Normal
Diabetic
Normal data

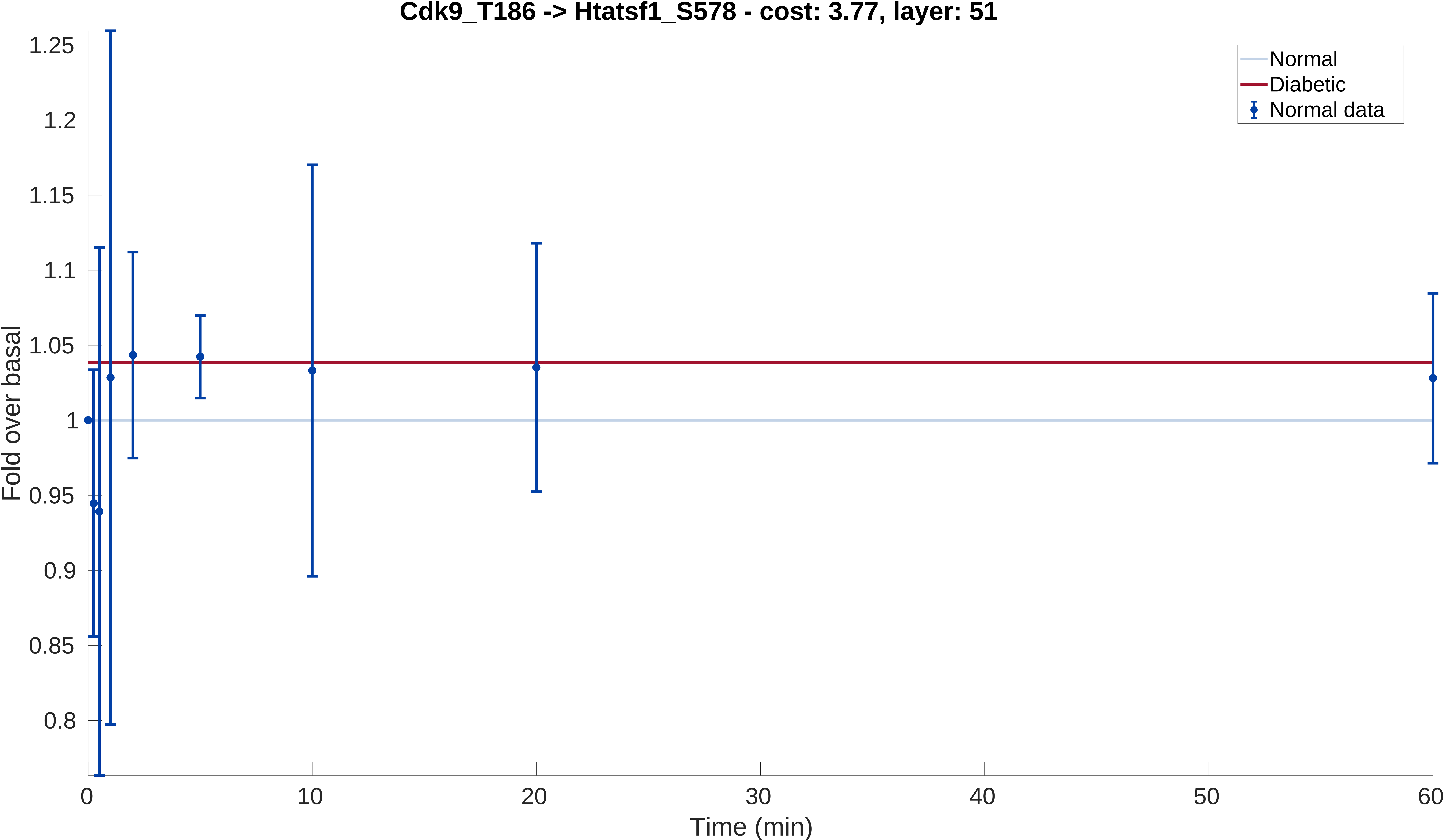

Cdk9_T186 -> Htatsf1_S578 - cost: 3.77, layer: 51
Normal
Diabetic
Normal data
Fold over basal
1.25
1.2
1.15
1.1
1.05
1
0.95
0.9
0.85
0.8
0
10
20
30
40
50
60
Time (min)

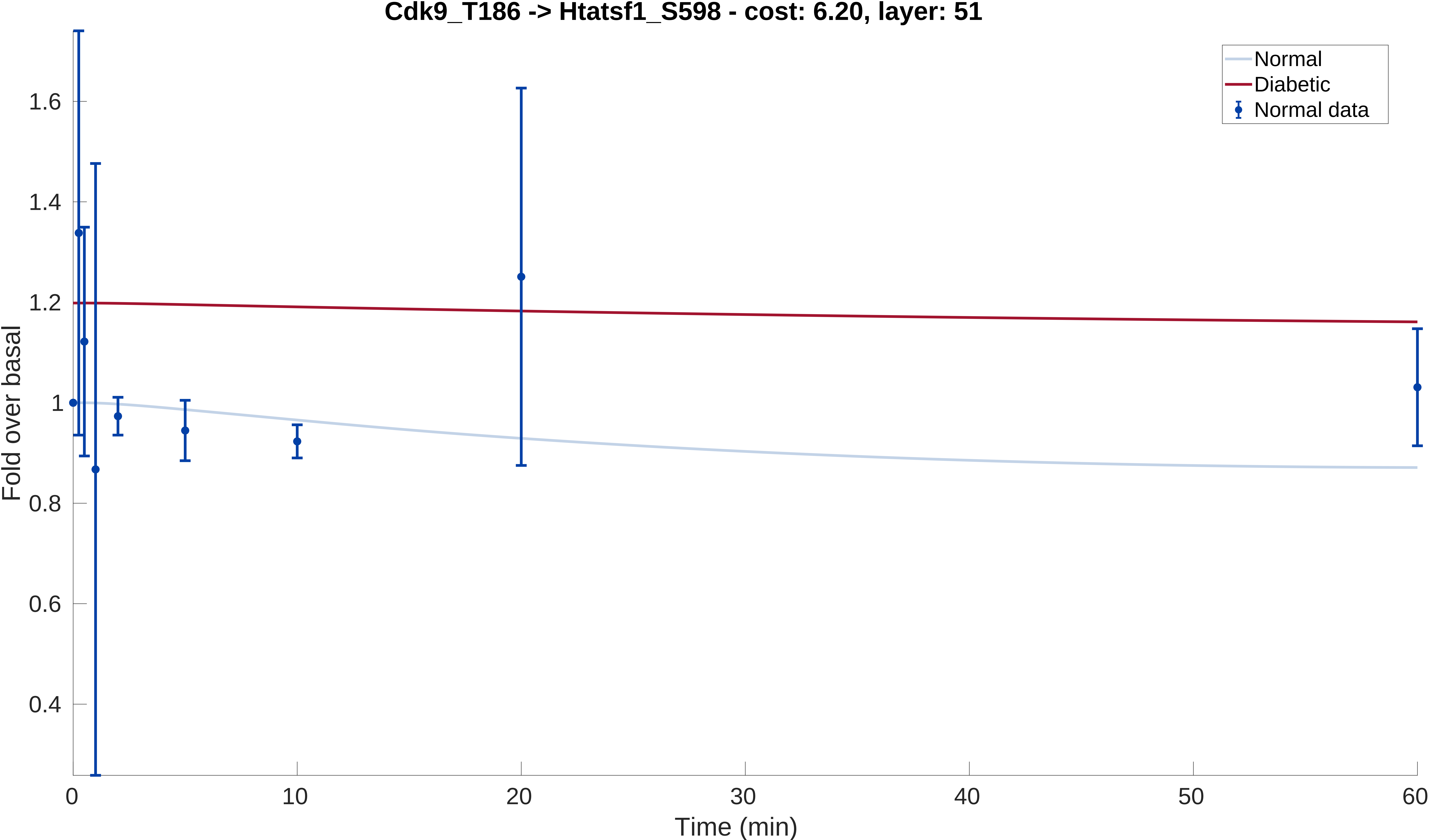

Cdk9_T186 -> Htatsf1_S598 - cost: 6.20, layer: 51
Normal
Diabetic
Normal data
Fold over basal
1.6
1.4
1.2
1
0.8
0.6
0.4
0
10
20
30
40
50
60
Time (min)

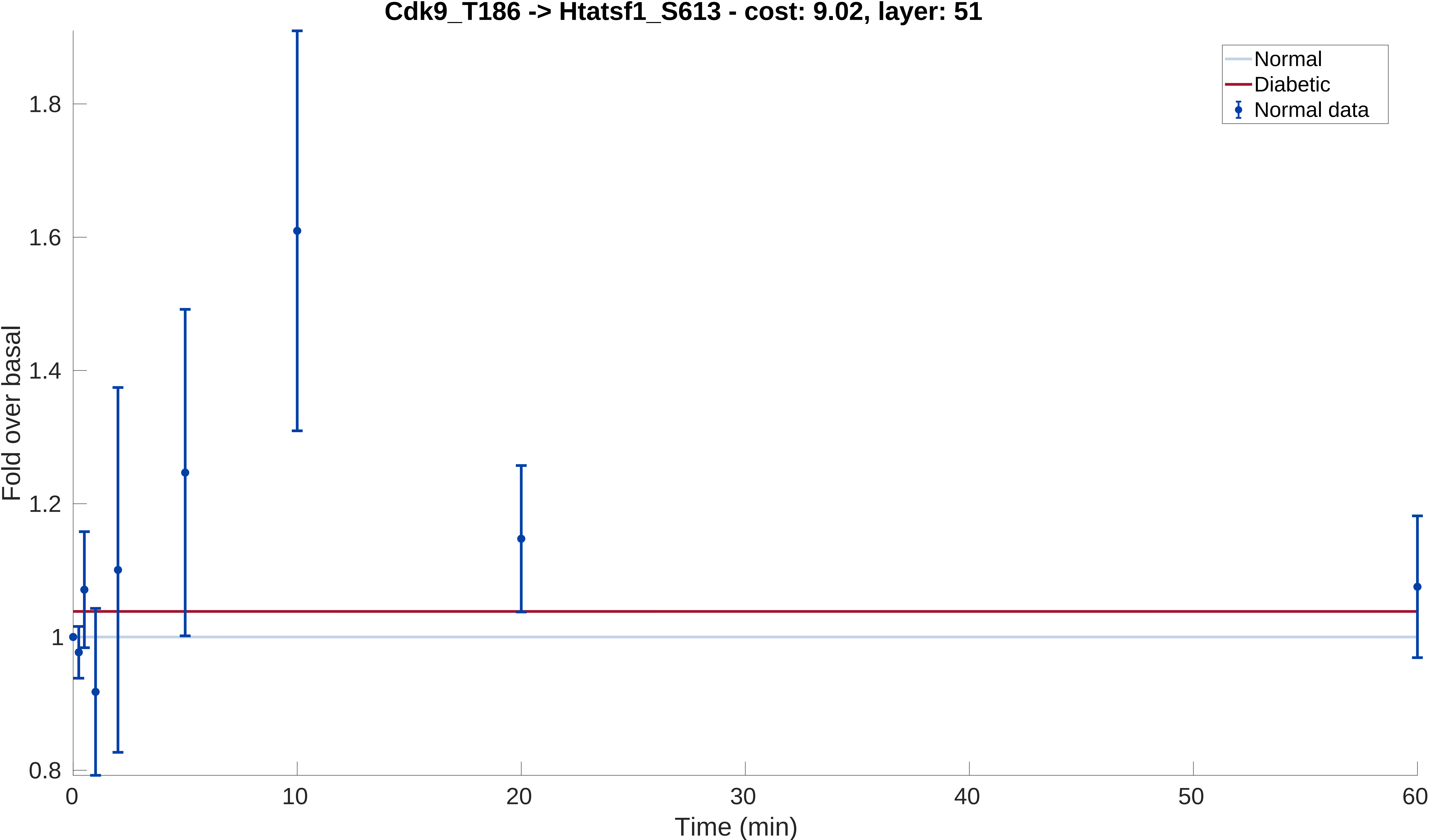
Cdk9_T186 -> Htatsf1_S613 - cost: 9.02, layer: 51
Normal
Diabetic
Normal data
Fold over basal
1.8
1.6
1.4
1.2
1
0.8
0
10
20
30
40
50
60
Time (min)

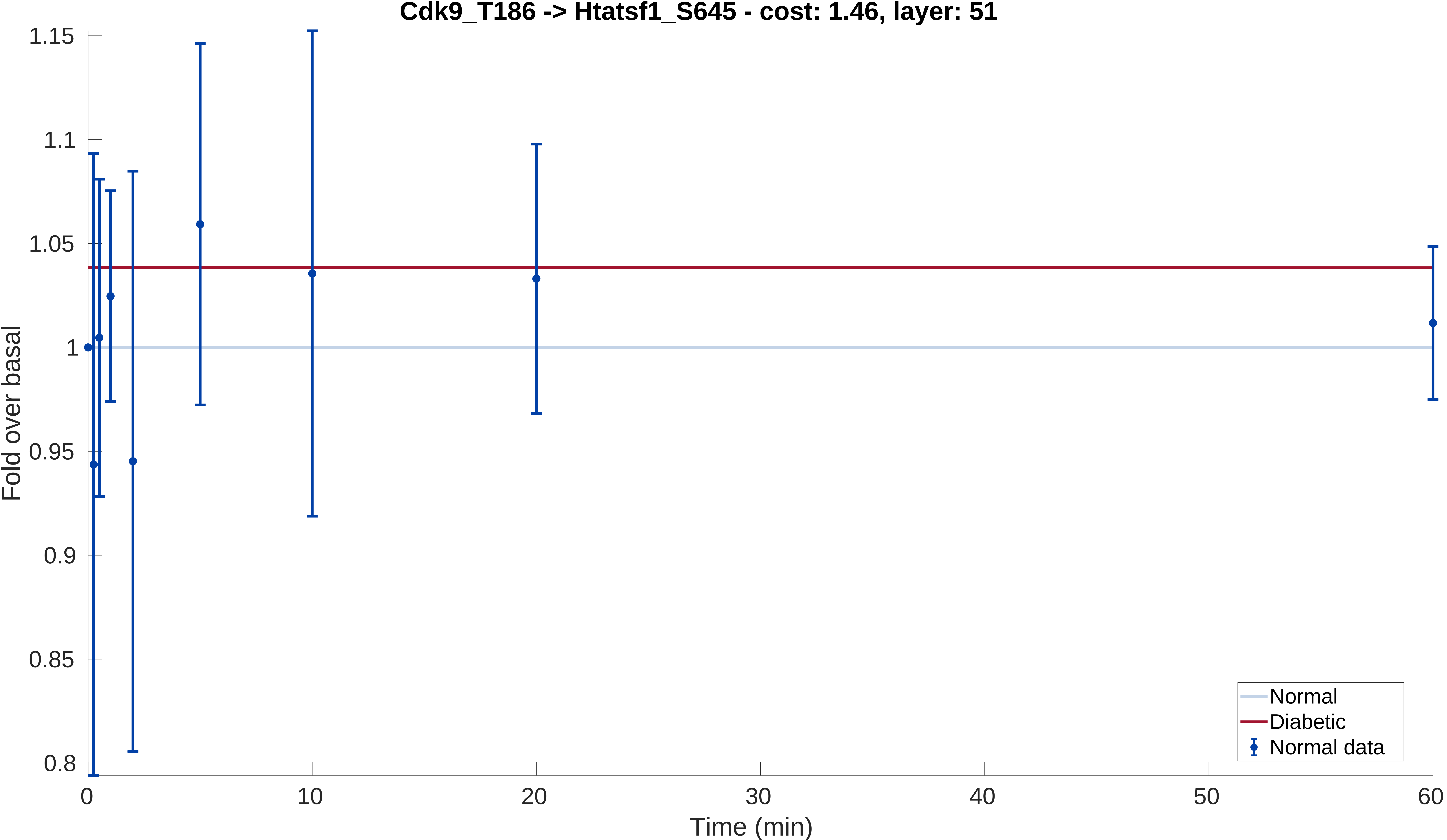

Cdk9_T186 -> Htatsf1_S645 - cost: 1.46, layer: 51
Fold over basal
Time (min)
1.15
1.1
1.05
1
0.95
0.9
0.85
0.8
0
10
20
30
40
50
60
Normal
Diabetic
Normal data

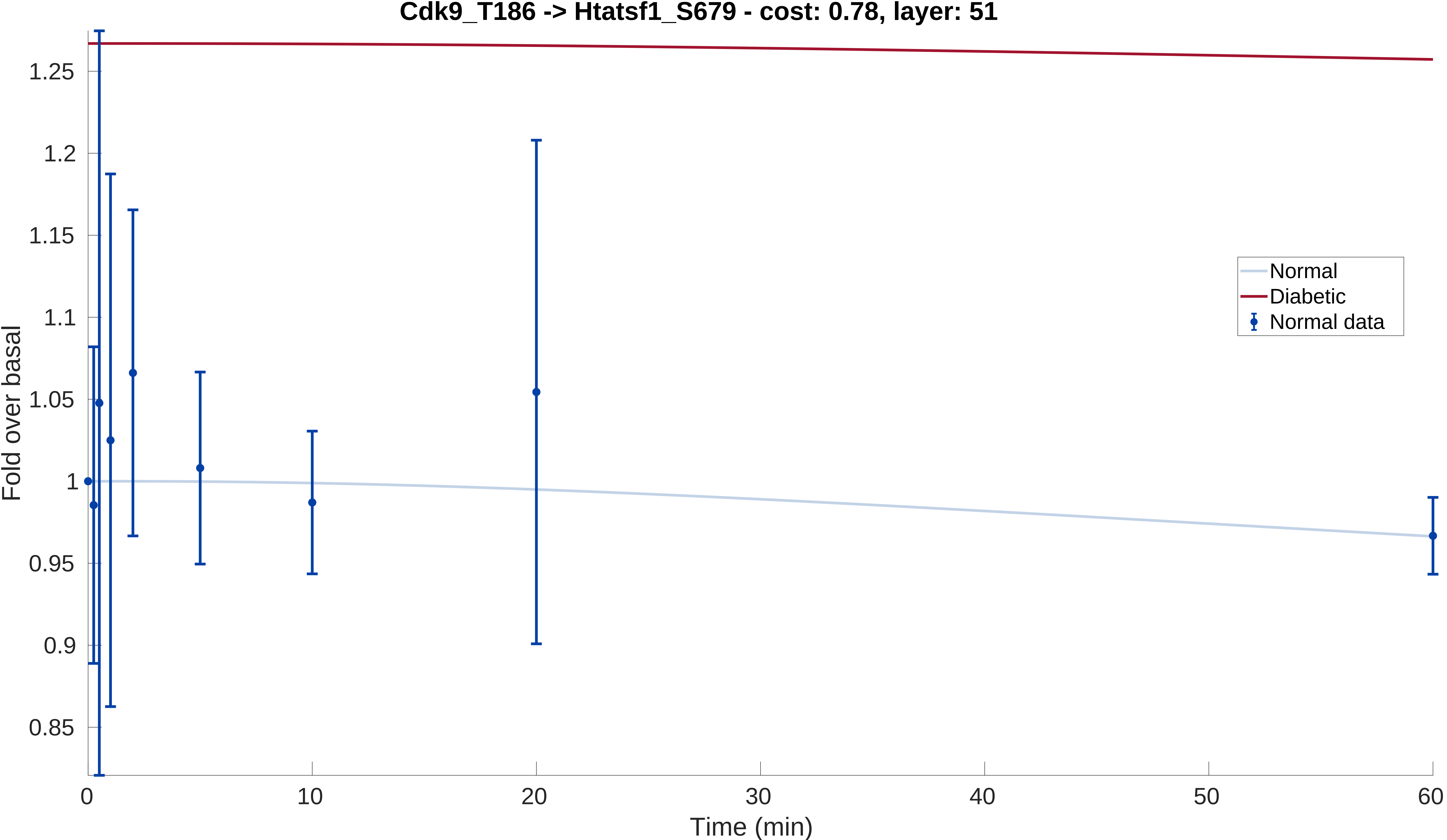

Cdk9_T186 -> Htatsf1_S679 - cost: 0.78, layer: 51
Fold over basal
1.25
1.2
1.15
1.1
1.05
1
0.95
0.9
0.85
0
10
20
30
40
50
60
Time (min)
Normal
Diabetic
Normal data

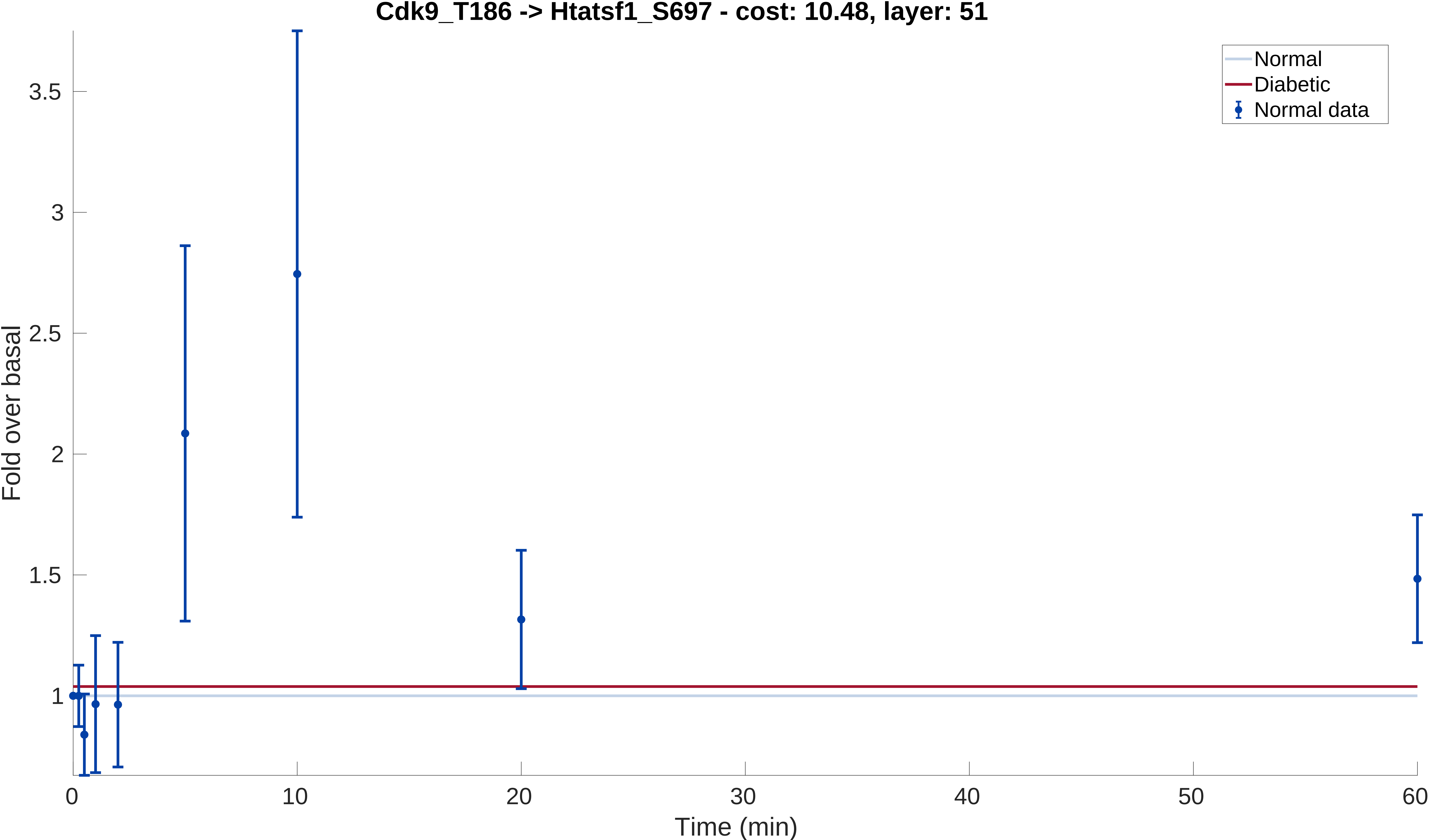

Cdk9_T186 -> Htatsf1_S697 - cost: 10.48, layer: 51
Normal
Diabetic
Normal data
Fold over basal
3.5
3
2.5
2
1.5
1
0
10
20
30
40
50
60
Time (min)

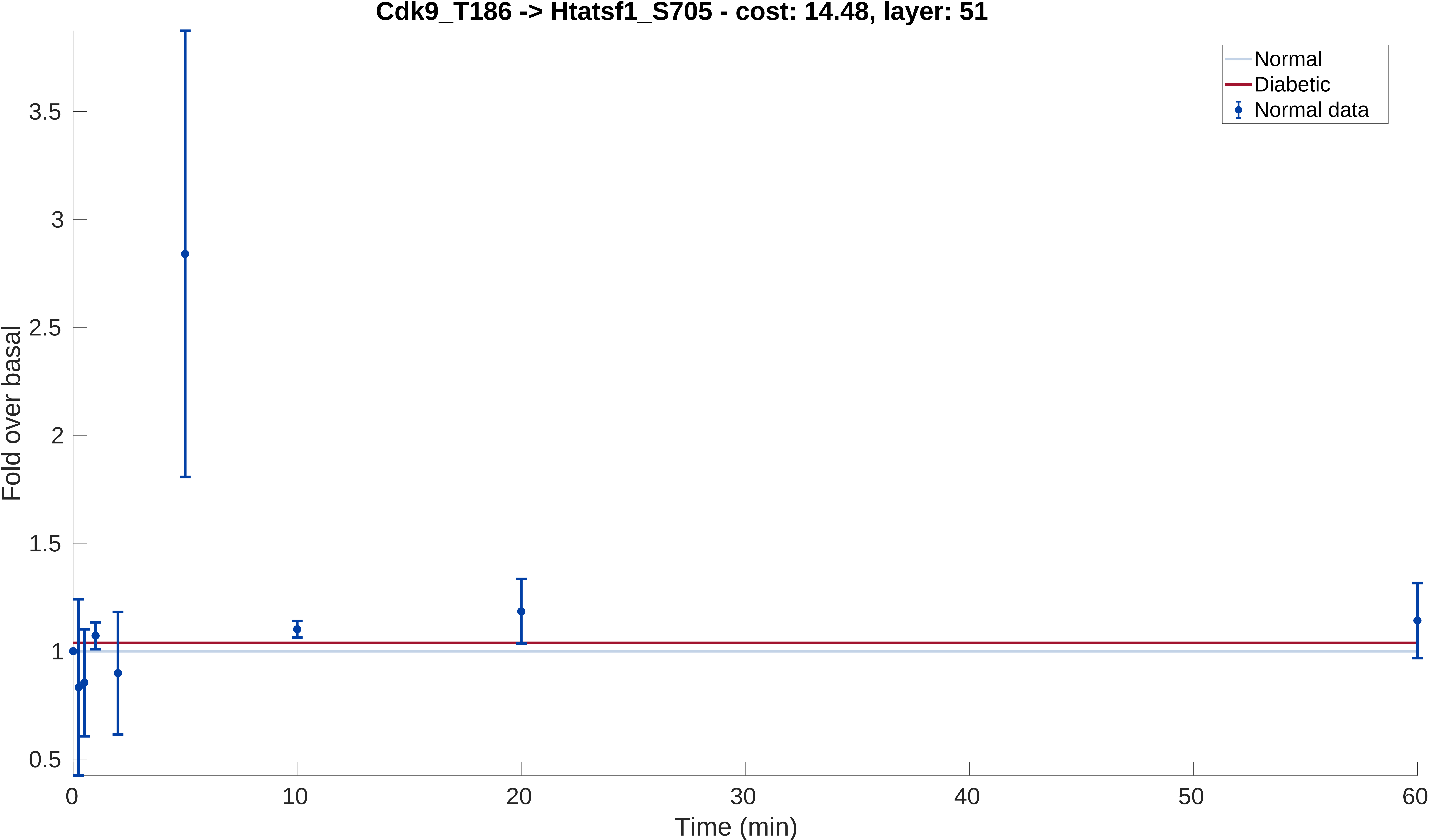

Cdk9_T186 -> Htatsf1_S705 - cost: 14.48, layer: 51
Normal
Diabetic
Normal data
Fold over basal
3.5
3
2.5
2
1.5
1
0.5
0
10
20
30
40
50
60
Time (min)

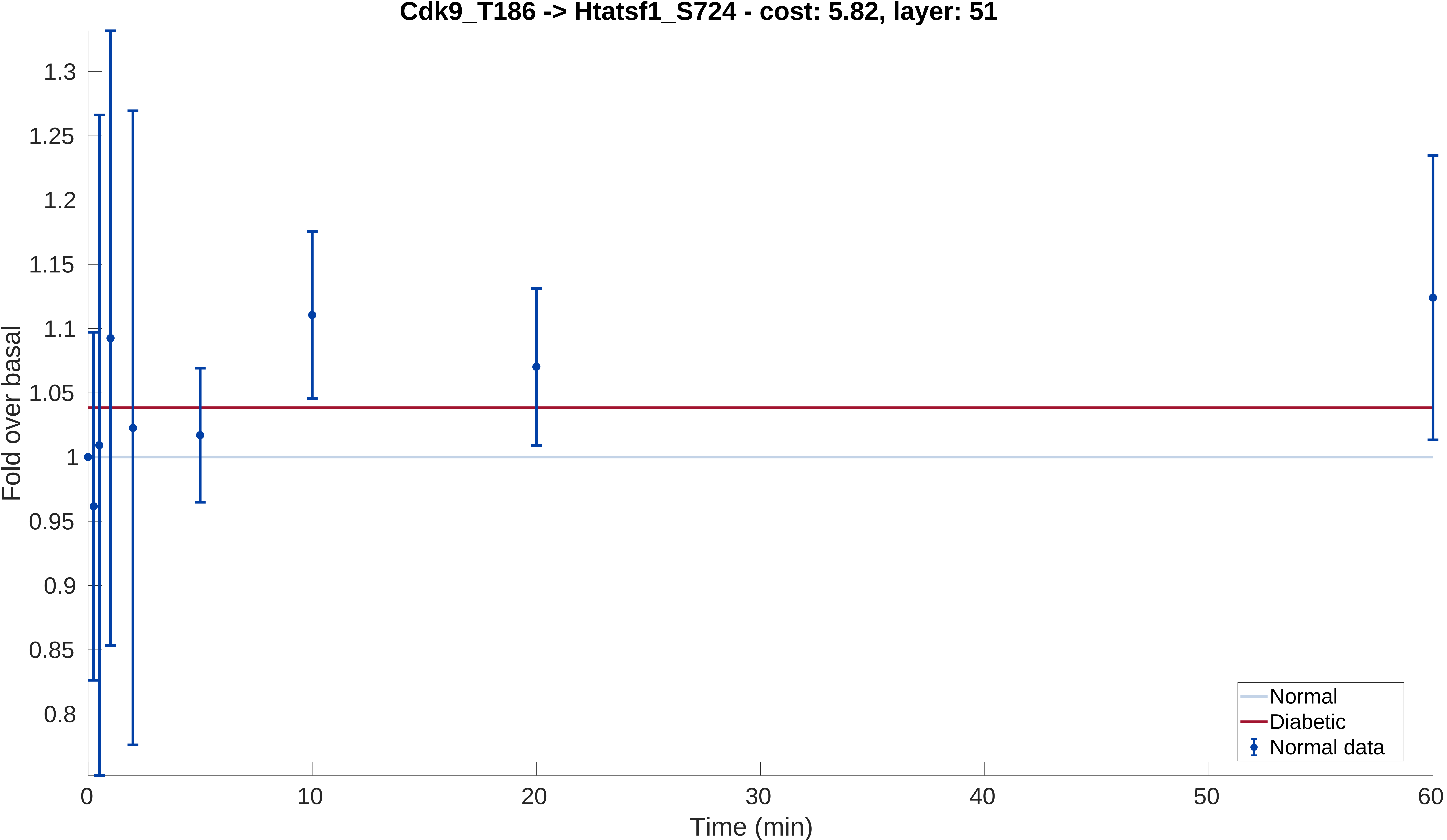
Cdk9_T186 -> Htatsf1_S724 - cost: 5.82, layer: 51
Fold over basal
1.3
1.25
1.2
1.15
1.1
1.05
1
0.95
0.9
0.85
0.8
0
10
20
30
40
50
60
Time (min)
Normal
Diabetic
Normal data

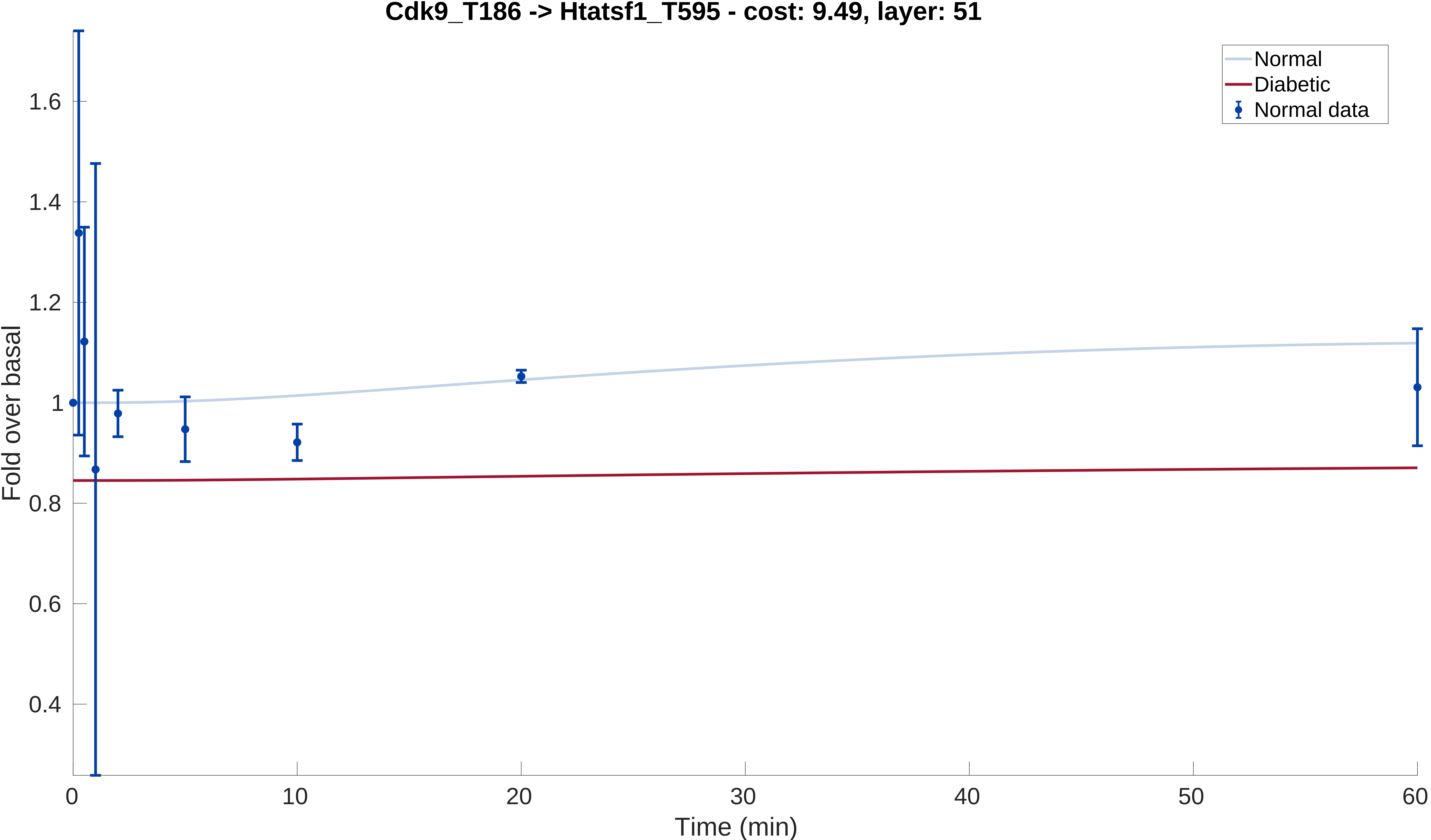

Cdk9_T186 -> Htatsf1_T595 - cost: 9.49, layer: 51
Normal
Diabetic
Normal data
Fold over basal
1.6
1.4
1.2
1
0.8
0.6
0.4
0
10
20
30
40
50
60
Time (min)

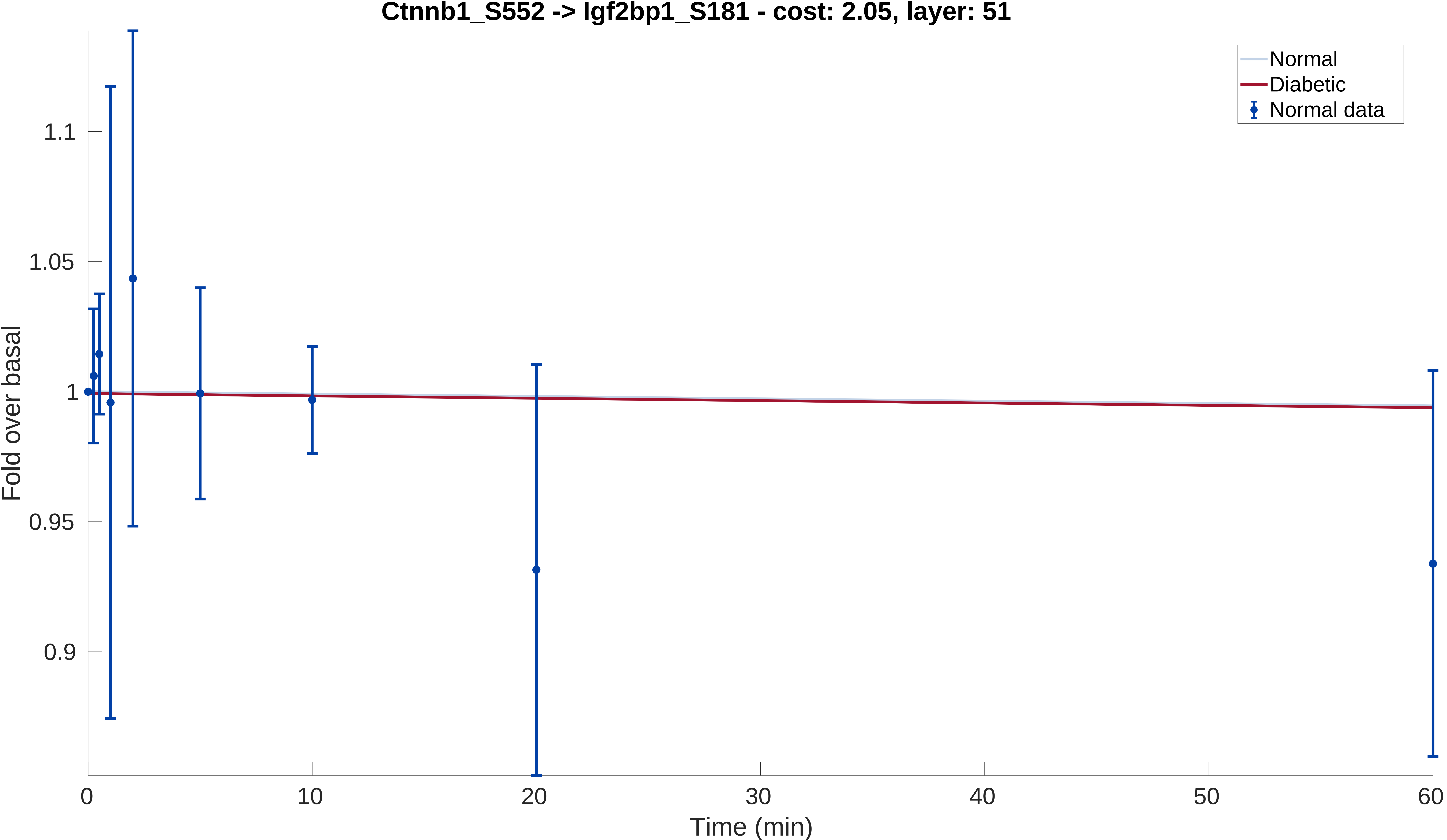

Ctnnb1_S552 -> Igf2bp1_S181 - cost: 2.05, layer: 51
Normal
Diabetic
Normal data
Fold over basal
1.1
1.05
1
0.95
0.9
0
10
20
30
40
50
60
Time (min)

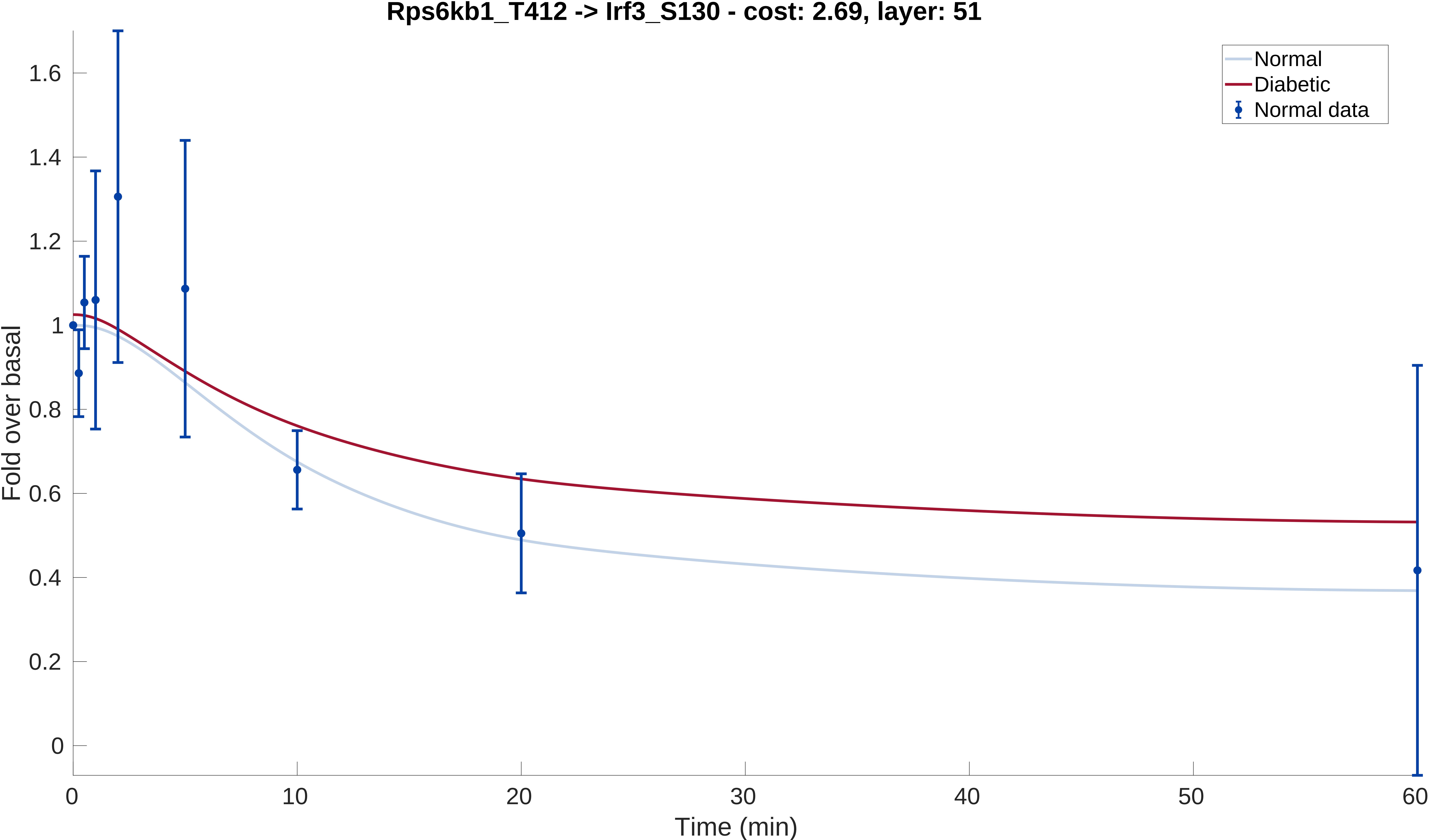

Rps6kb1_T412 -> Irf3_S130 - cost: 2.69, layer: 51
Normal
Diabetic
Normal data
Fold over basal
1.6
1.4
1.2
1
0.8
0.6
0.4
0.2
0
0
10
20
30
40
50
60
Time (min)

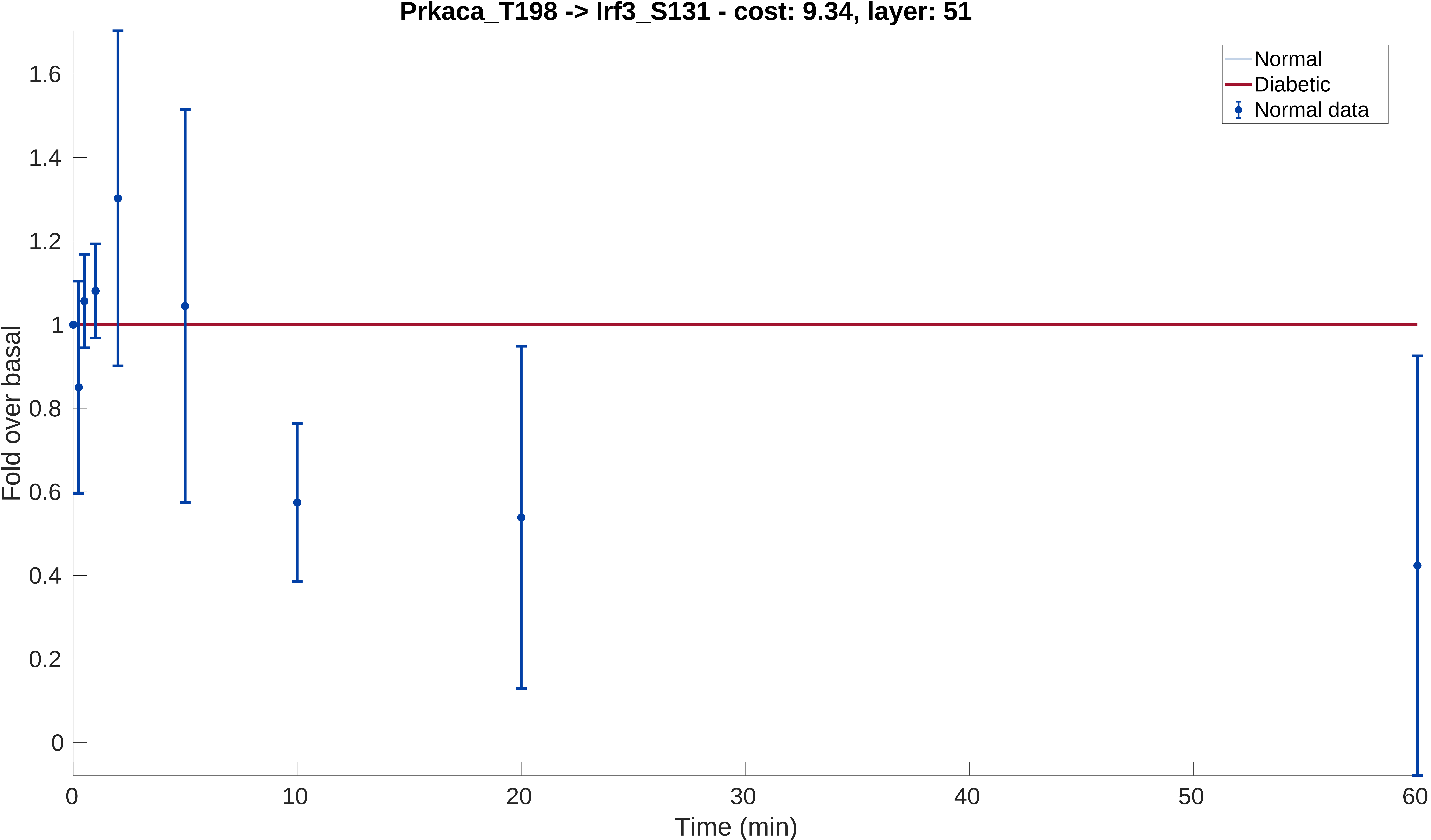

Prkaca_T198 -> Irf3_S131 - cost: 9.34, layer: 51
Normal
Diabetic
Normal data
Fold over basal
1.6
1.4
1.2
1
0.8
0.6
0.4
0.2
0
0
10
20
30
40
50
60
Time (min)

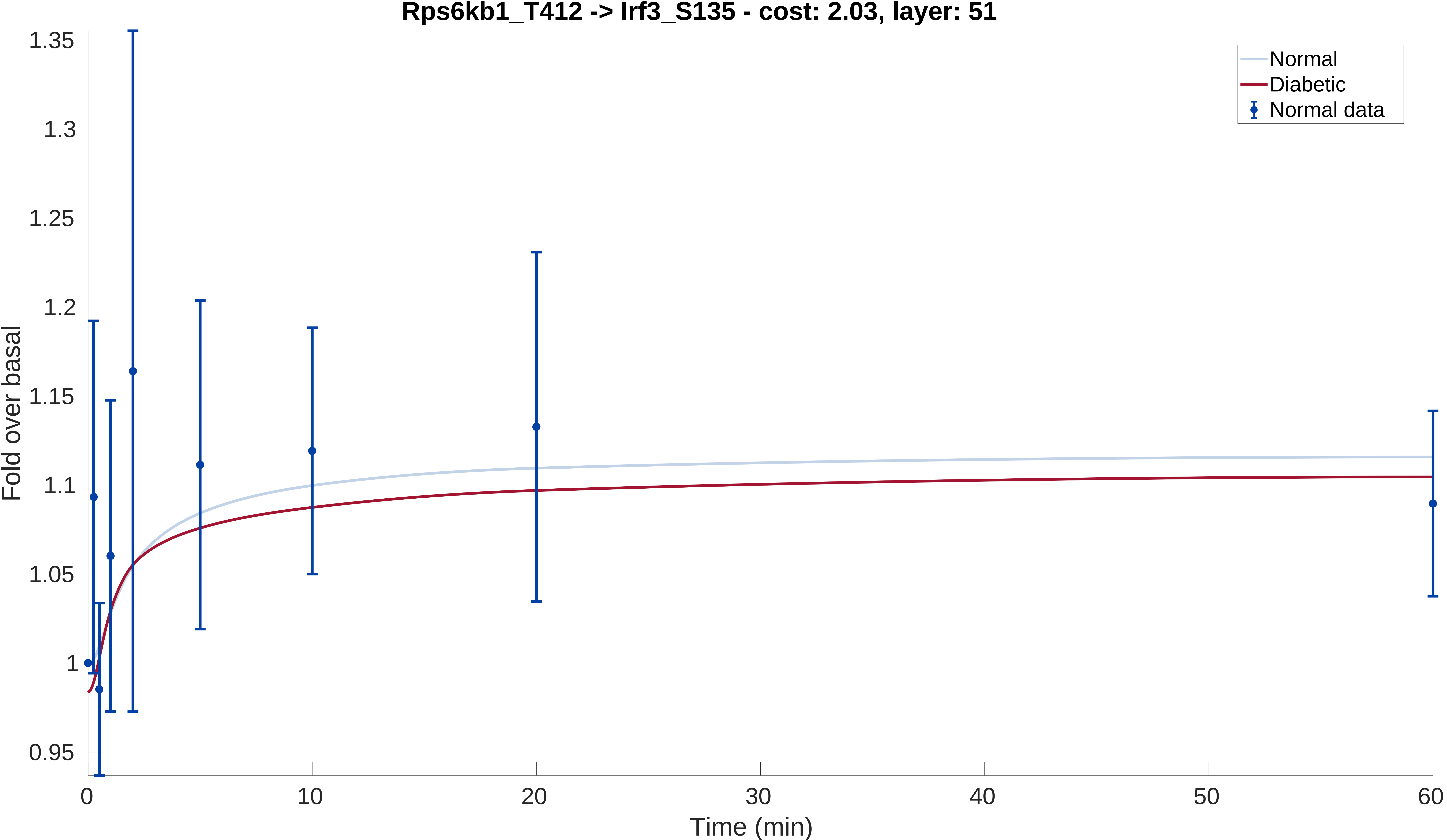
Rps6kb1_T412 -> Irf3_S135 - cost: 2.03, layer: 51
Fold over basal
Time (min)
Normal
Diabetic
Normal data
1.35
1.3
1.25
1.2
1.15
1.1
1.05
1
0.95
0
10
20
30
40
50
60

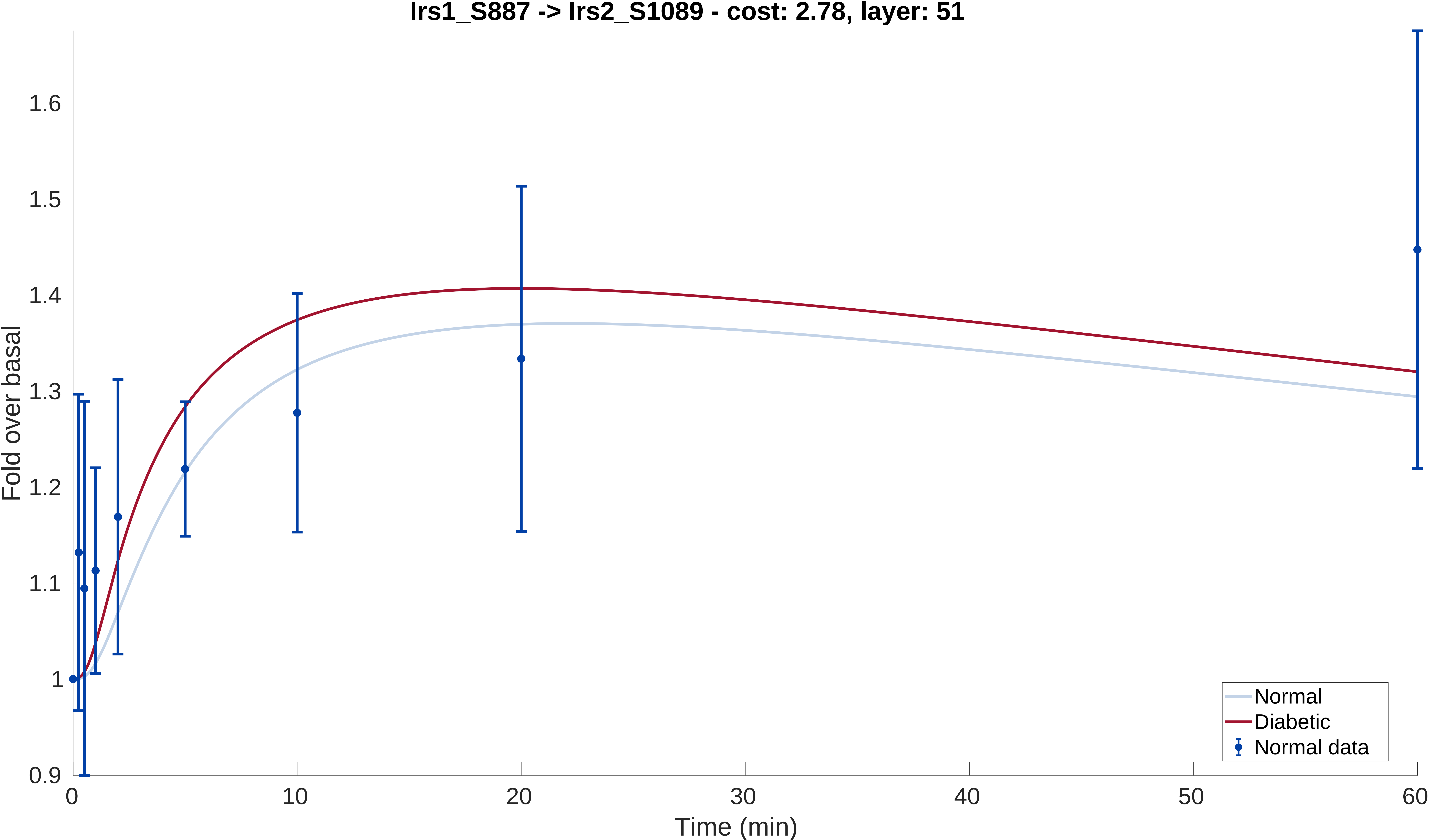
Irs1_S887 -> Irs2_S1089 - cost: 2.78, layer: 51
Fold over basal
1.6
1.5
1.4
1.3
1.2
1.1
1
0.9
0
10
20
30
40
50
60
Time (min)
Normal
Diabetic
Normal data

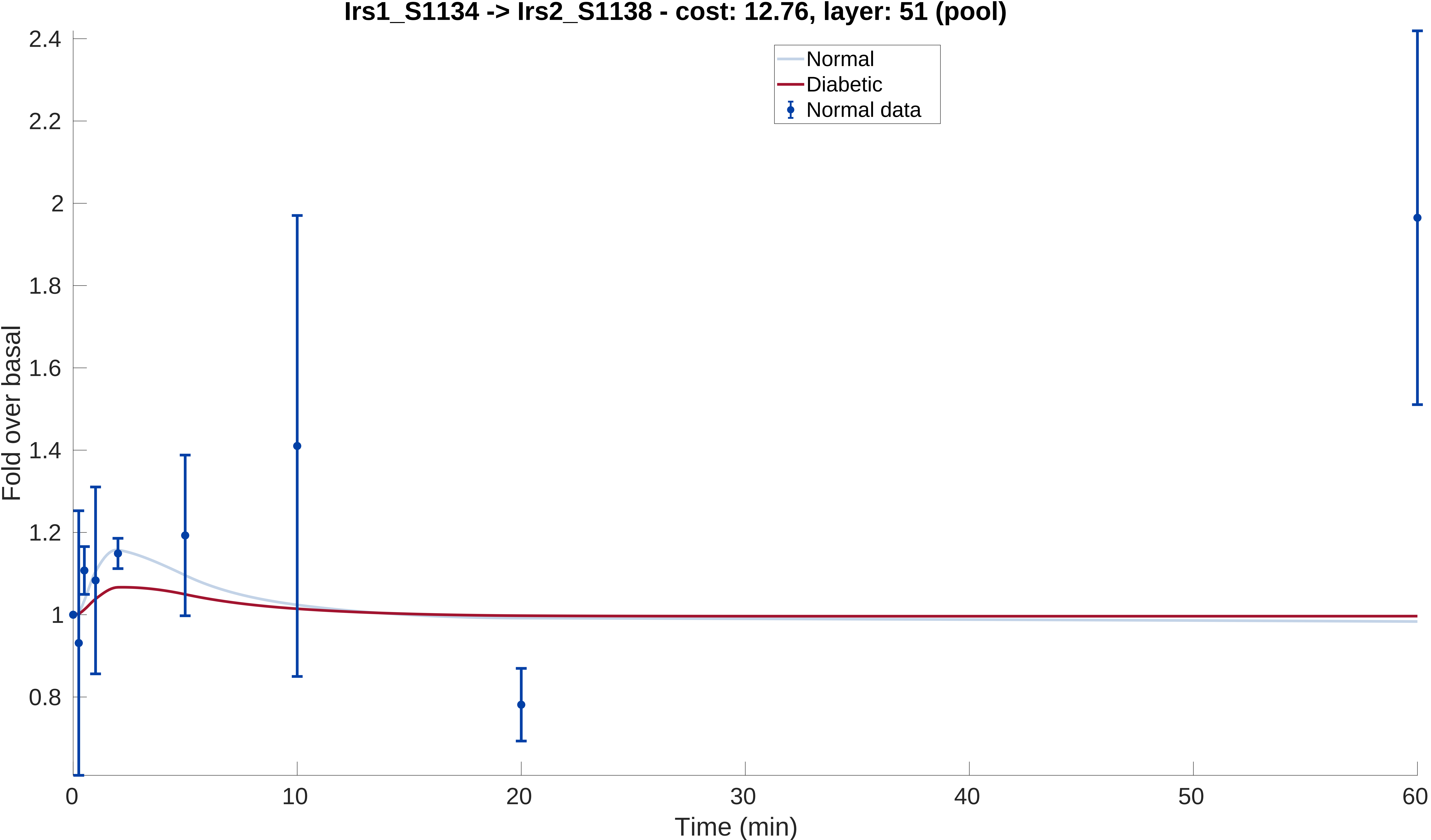

Irs1_S1134 -> Irs2_S1138 - cost: 12.76, layer: 51 (pool)
Normal
Diabetic
Normal data
Fold over basal
Time (min)
2.4
2.2
2
1.8
1.6
1.4
1.2
1
0.8
0
10
20
30
40
50
60

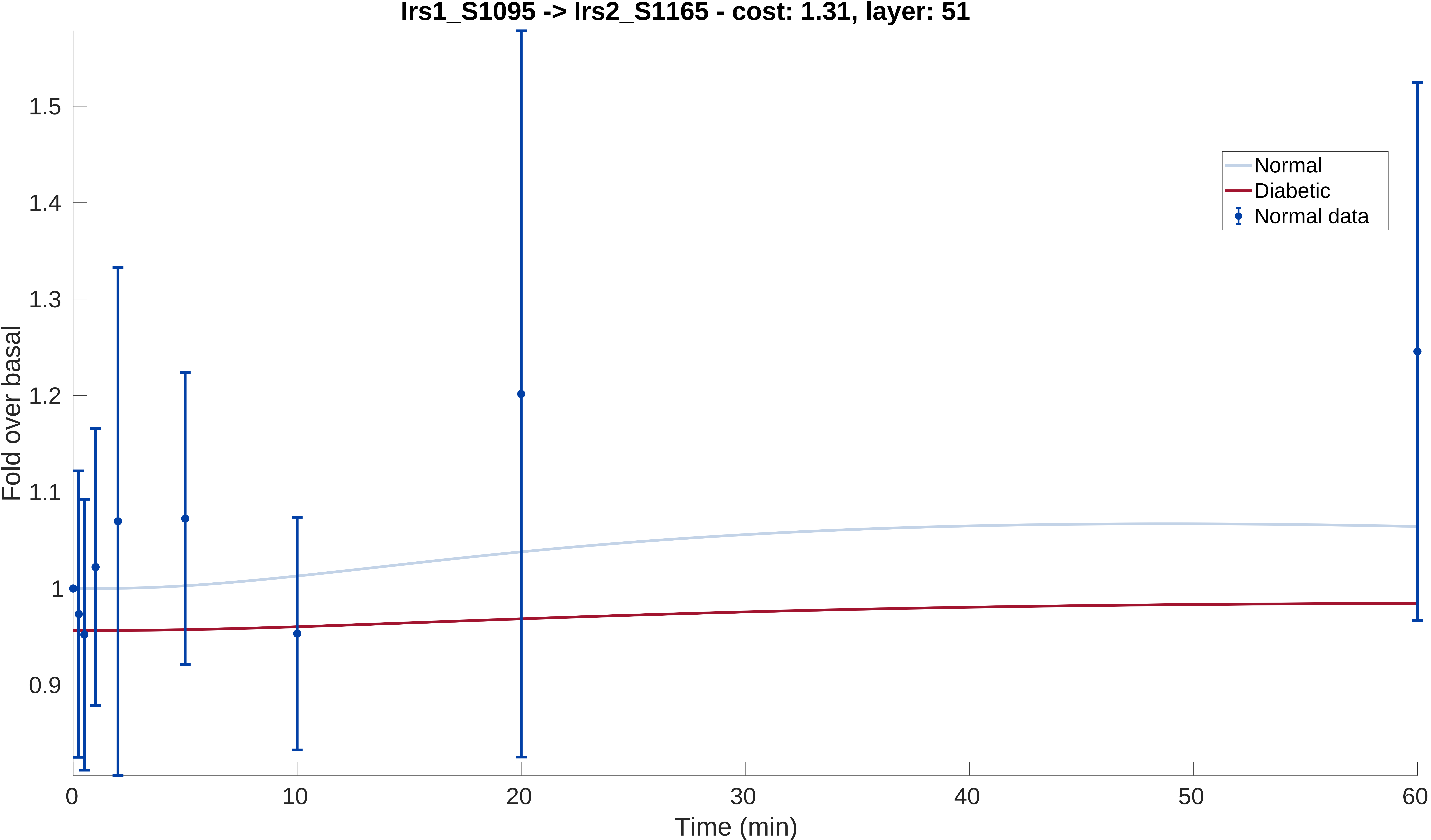
Irs1_S1095 -> Irs2_S1165 - cost: 1.31, layer: 51
Fold over basal
Time (min)
Normal
Diabetic
Normal data
0.9
1
1.1
1.2
1.3
1.4
1.5
0
10
20
30
40
50
60

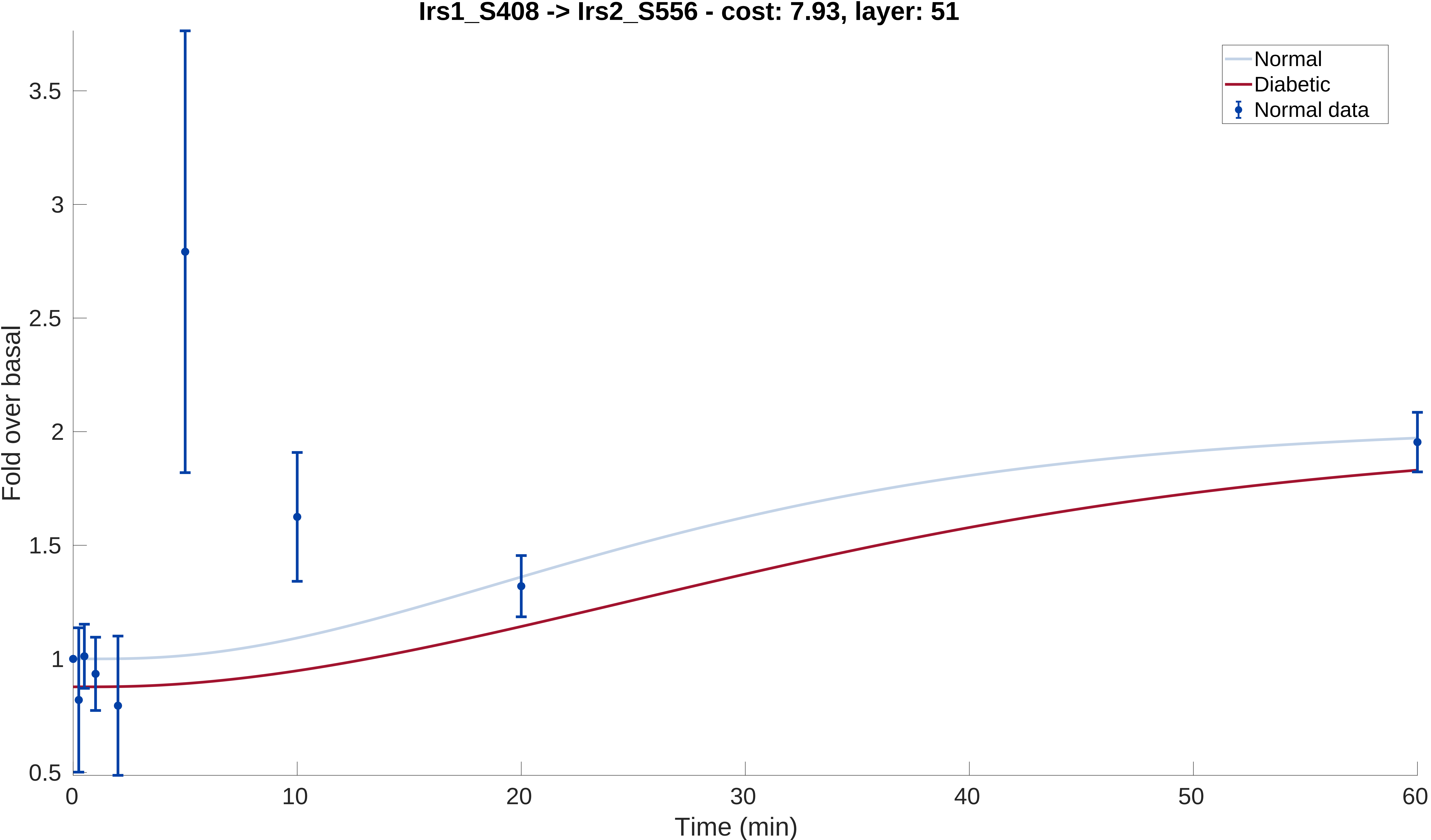

Irs1_S408 -> Irs2_S556 - cost: 7.93, layer: 51
Normal
Diabetic
Normal data
Fold over basal
3.5
3
2.5
2
1.5
1
0.5
0
10
20
30
40
50
60
Time (min)

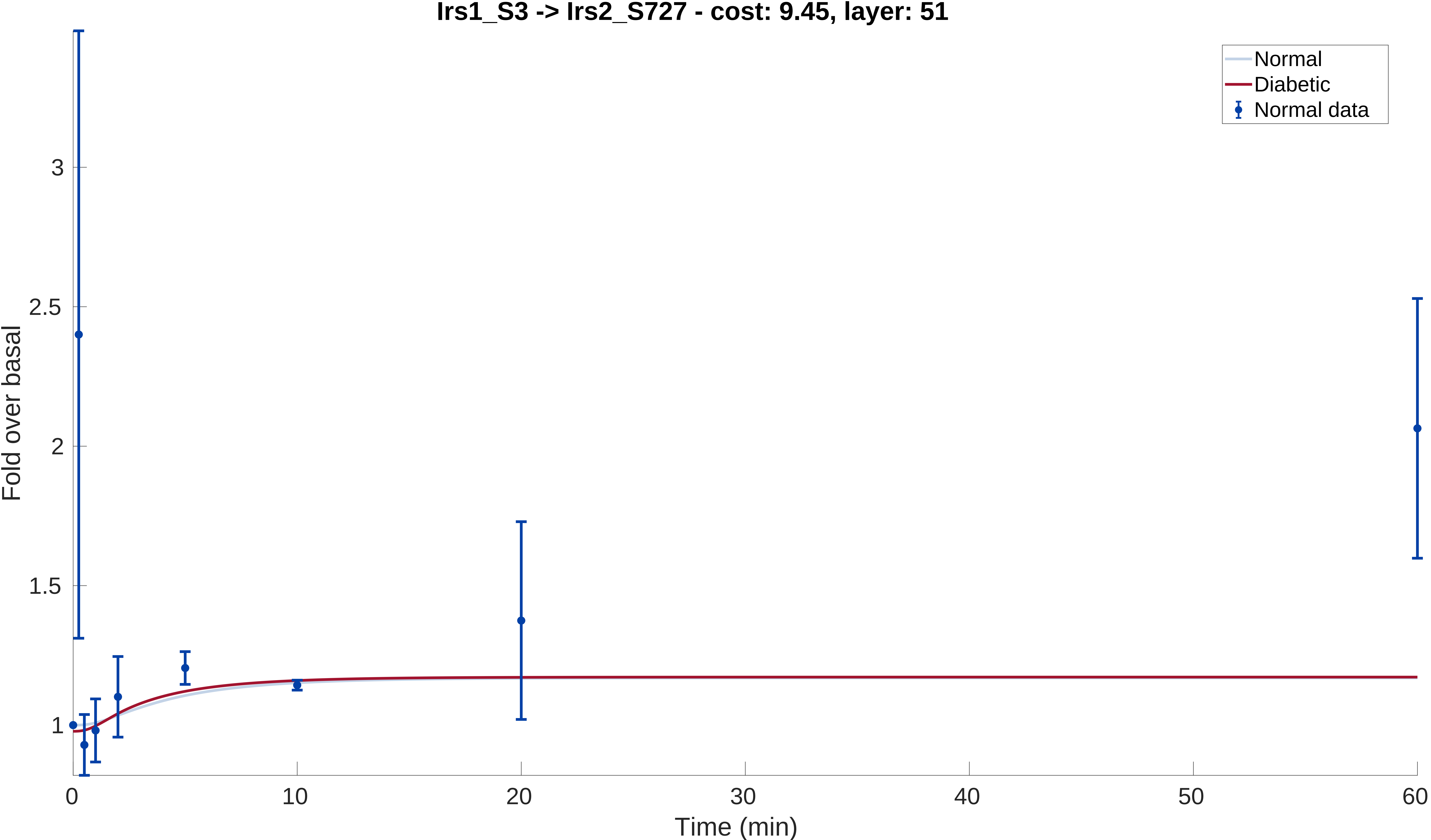

Irs1_S3 -> Irs2_S727 - cost: 9.45, layer: 51
Normal
Diabetic
Normal data
3
2.5
2
1.5
1
Fold over basal
0
10
20
30
40
50
60
Time (min)

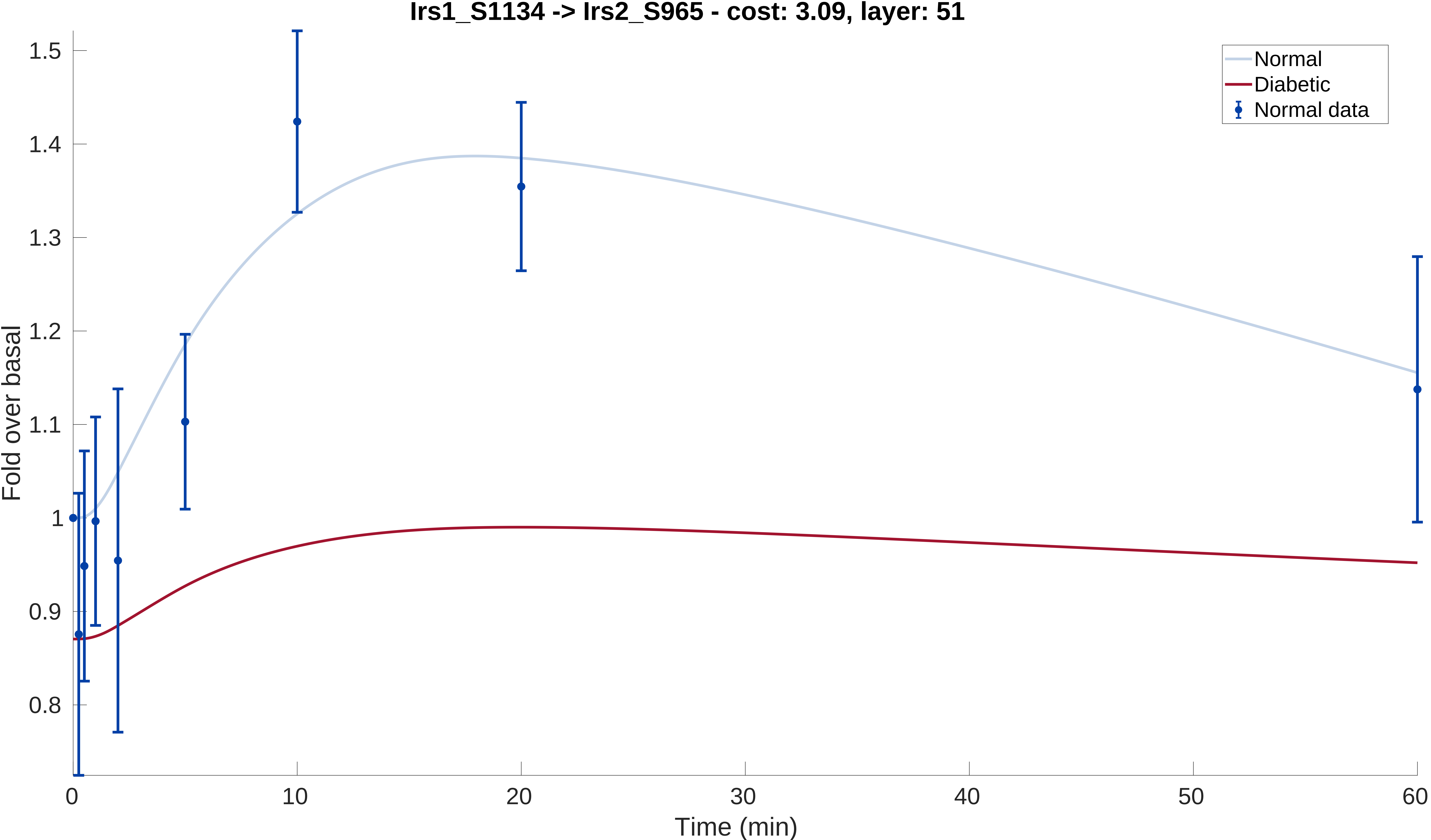

Irs1_S1134 -> Irs2_S965 - cost: 3.09, layer: 51
Normal
Diabetic
Normal data
Fold over basal
1.5
1.4
1.3
1.2
1.1
1
0.9
0.8
0
10
20
30
40
50
60
Time (min)

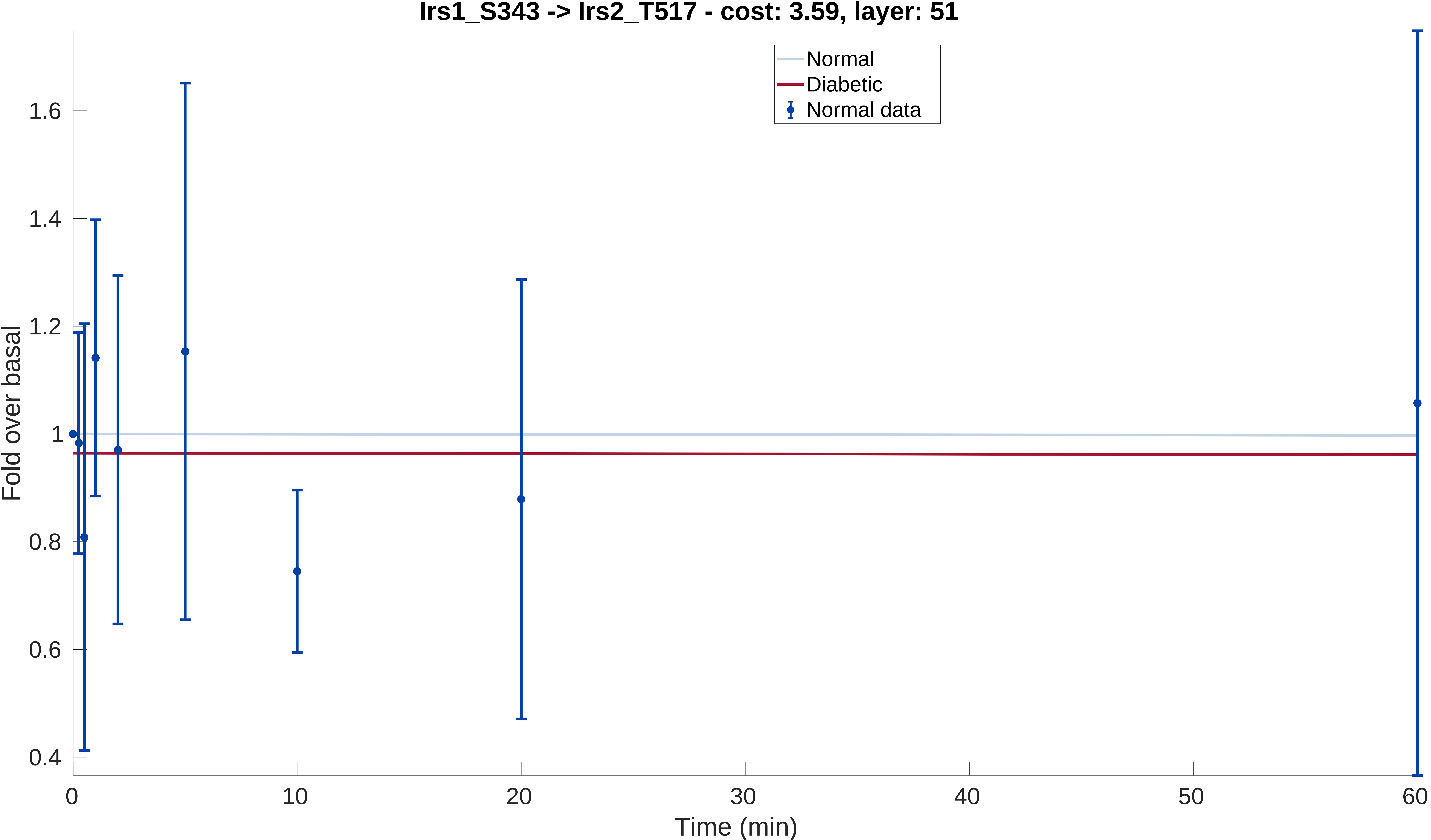
Irs1_S343 -> Irs2_T517 - cost: 3.59, layer: 51
Normal
Diabetic
Normal data
Fold over basal
0.4
0.6
0.8
1
1.2
1.4
1.6
0
10
20
30
40
50
60
Time (min)

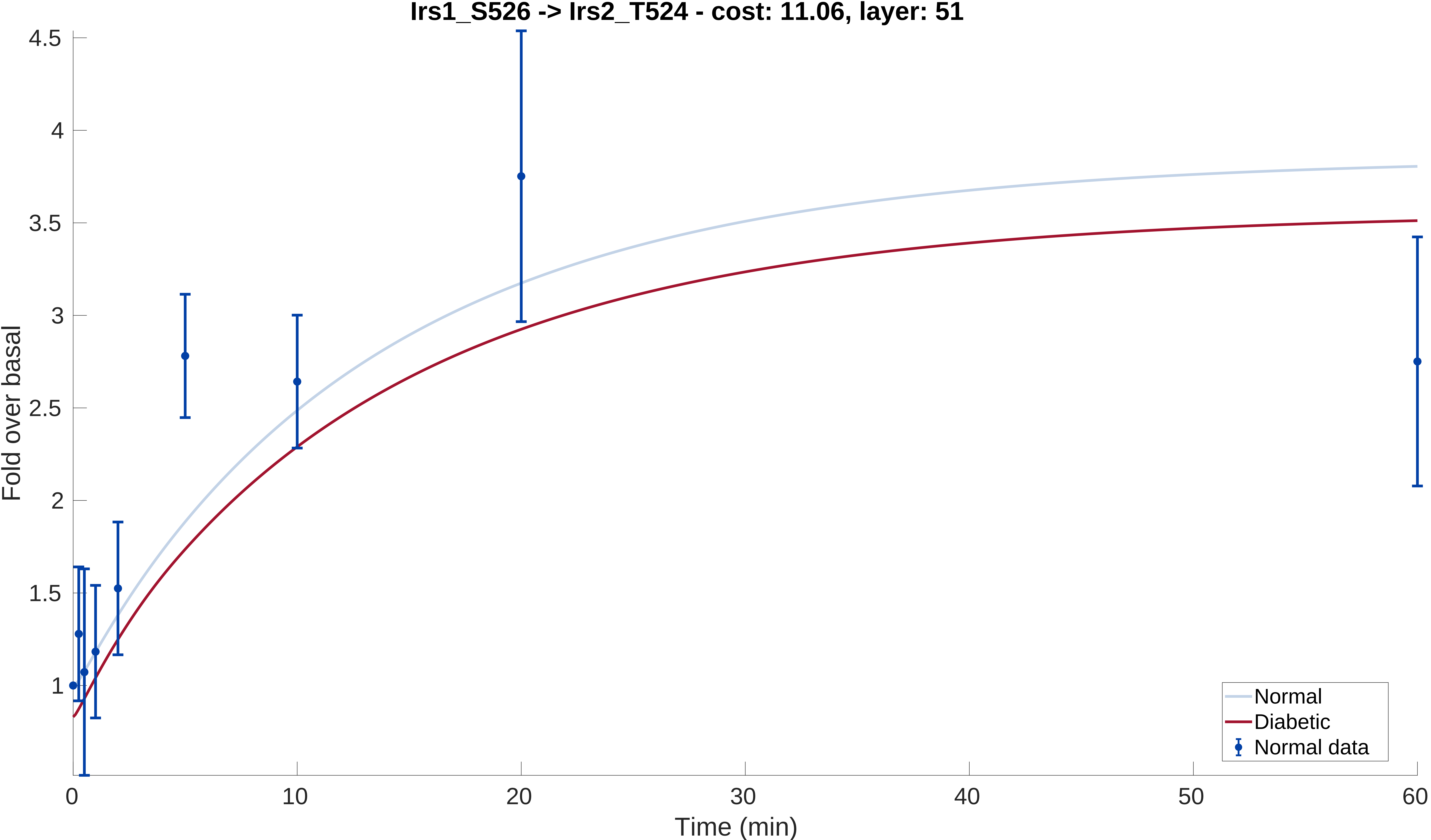

Irs1_S526 -> Irs2_T524 - cost: 11.06, layer: 51
Fold over basal
Time (min)
4.5
4
3.5
3
2.5
2
1.5
1
0
10
20
30
40
50
60
Normal
Diabetic
Normal data

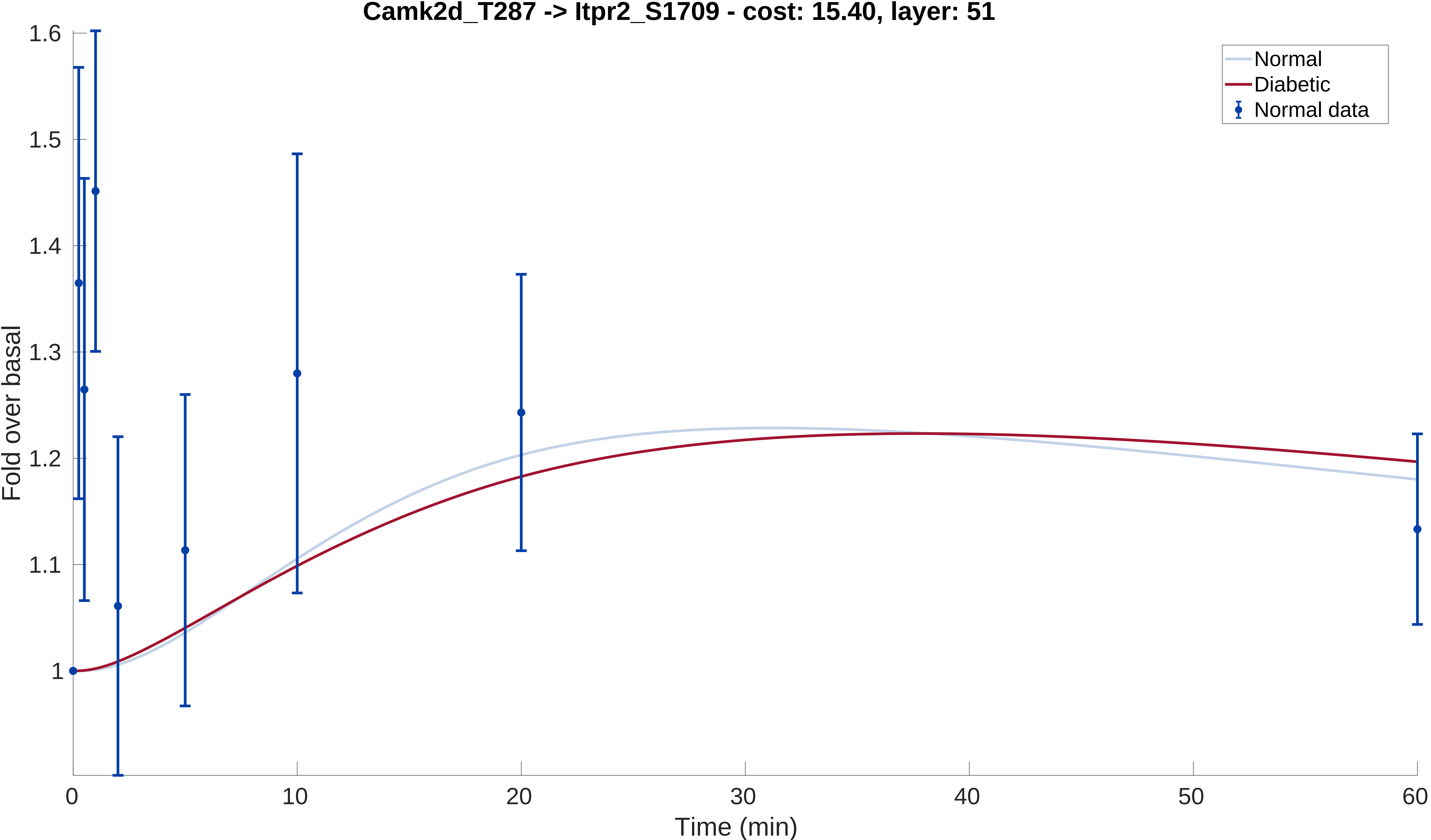
Camk2d_T287 -> Itpr2_S1709 - cost: 15.40, layer: 51
Normal
Diabetic
Normal data
Fold over basal
1.6
1.5
1.4
1.3
1.2
1.1
1
0
10
20
30
40
50
60
Time (min)

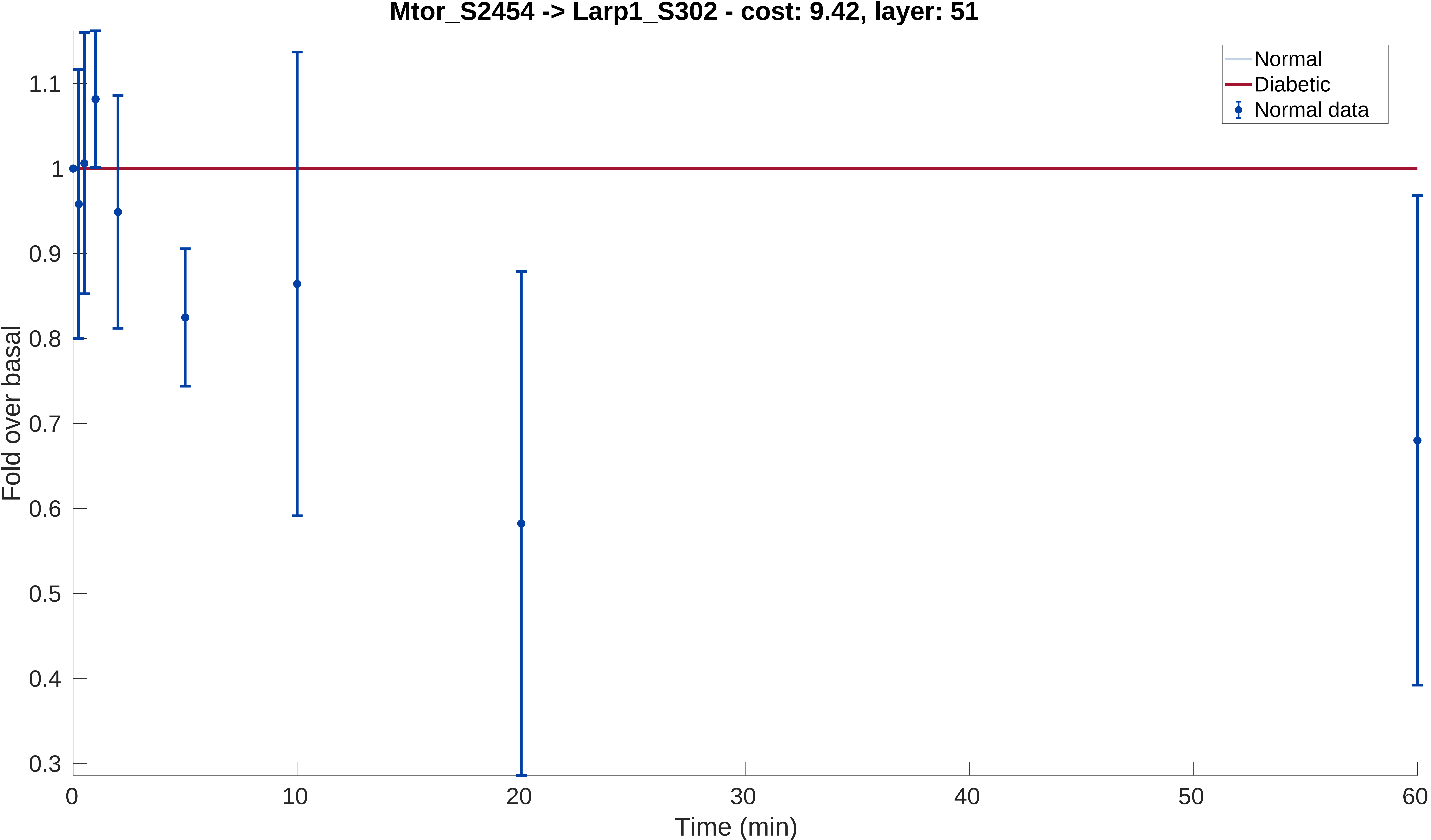

Mtor_S2454 -> Larp1_S302 - cost: 9.42, layer: 51
Normal
Diabetic
Normal data
Fold over basal
Time (min)
1.1
1
0.9
0.8
0.7
0.6
0.5
0.4
0.3
0
10
20
30
40
50
60

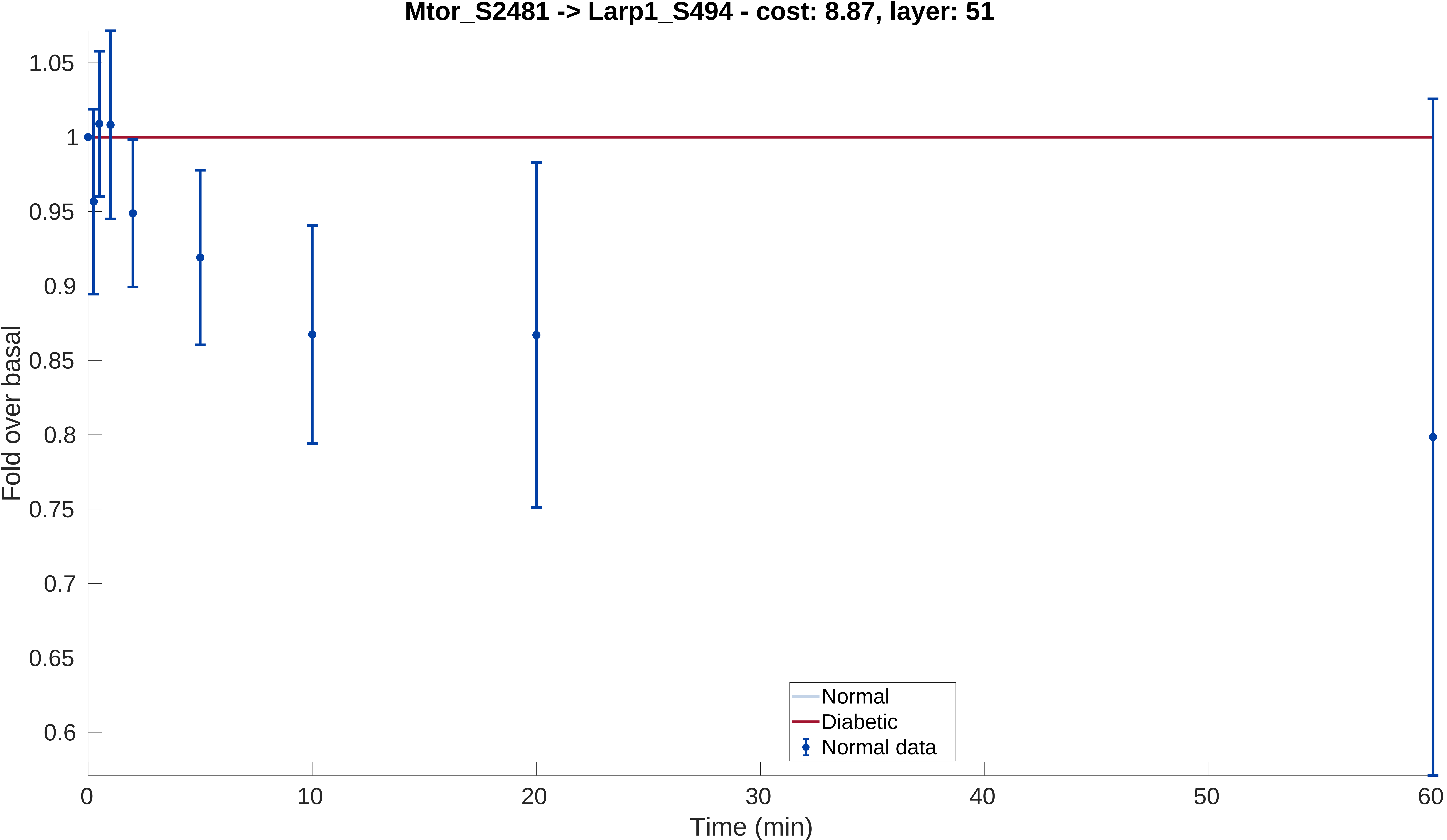
Mtor_S2481 -> Larp1_S494 - cost: 8.87, layer: 51
Fold over basal
1.05
1
0.95
0.9
0.85
0.8
0.75
0.7
0.65
0.6
0
10
20
30
40
50
60
Time (min)
Normal
Diabetic
Normal data

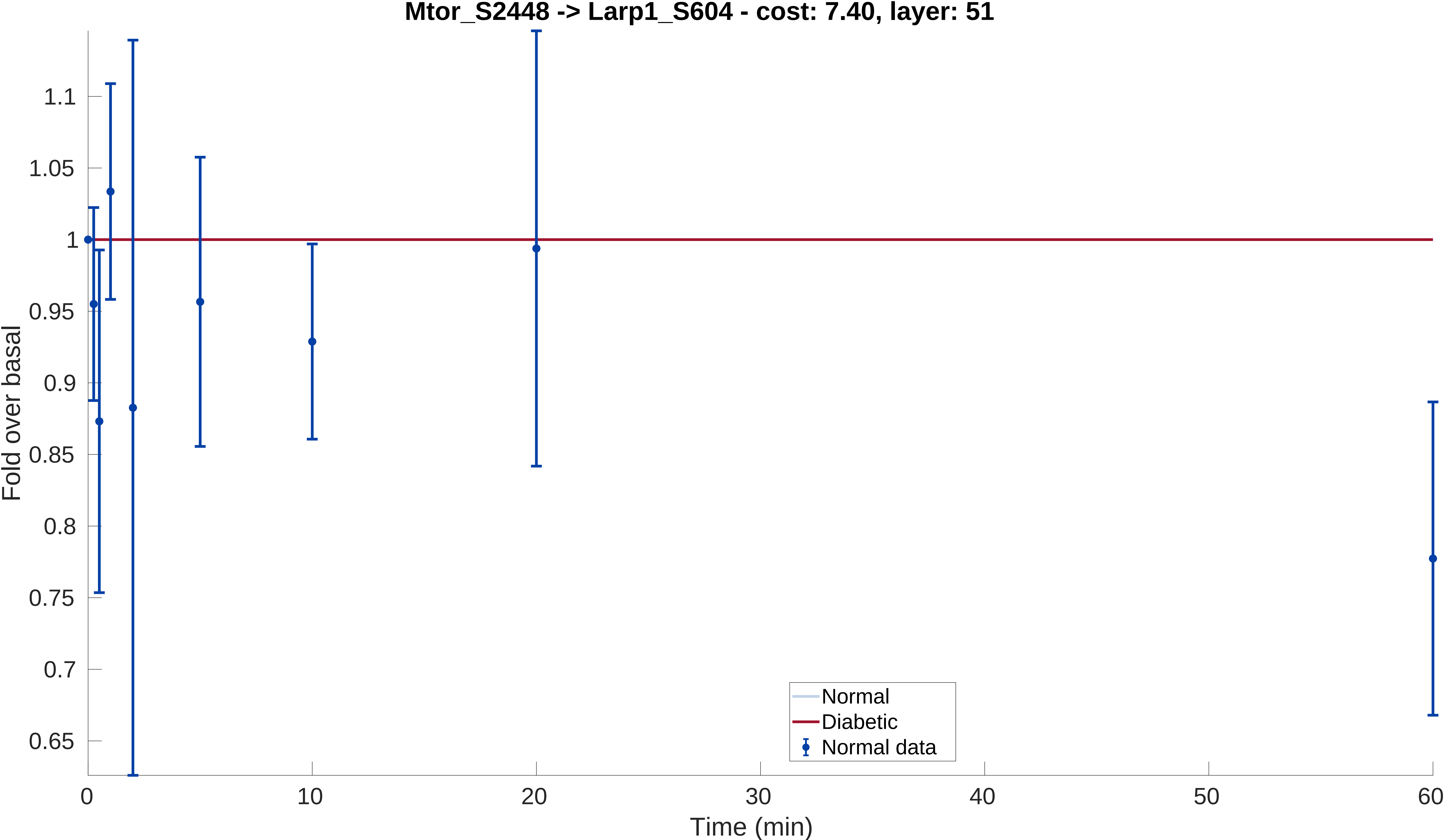

Mtor_S2448 -> Larp1_S604 - cost: 7.40, layer: 51
Fold over basal
Time (min)
Normal
Diabetic
Normal data
1.1
1.05
1
0.95
0.9
0.85
0.8
0.75
0.7
0.65
0
10
20
30
40
50
60

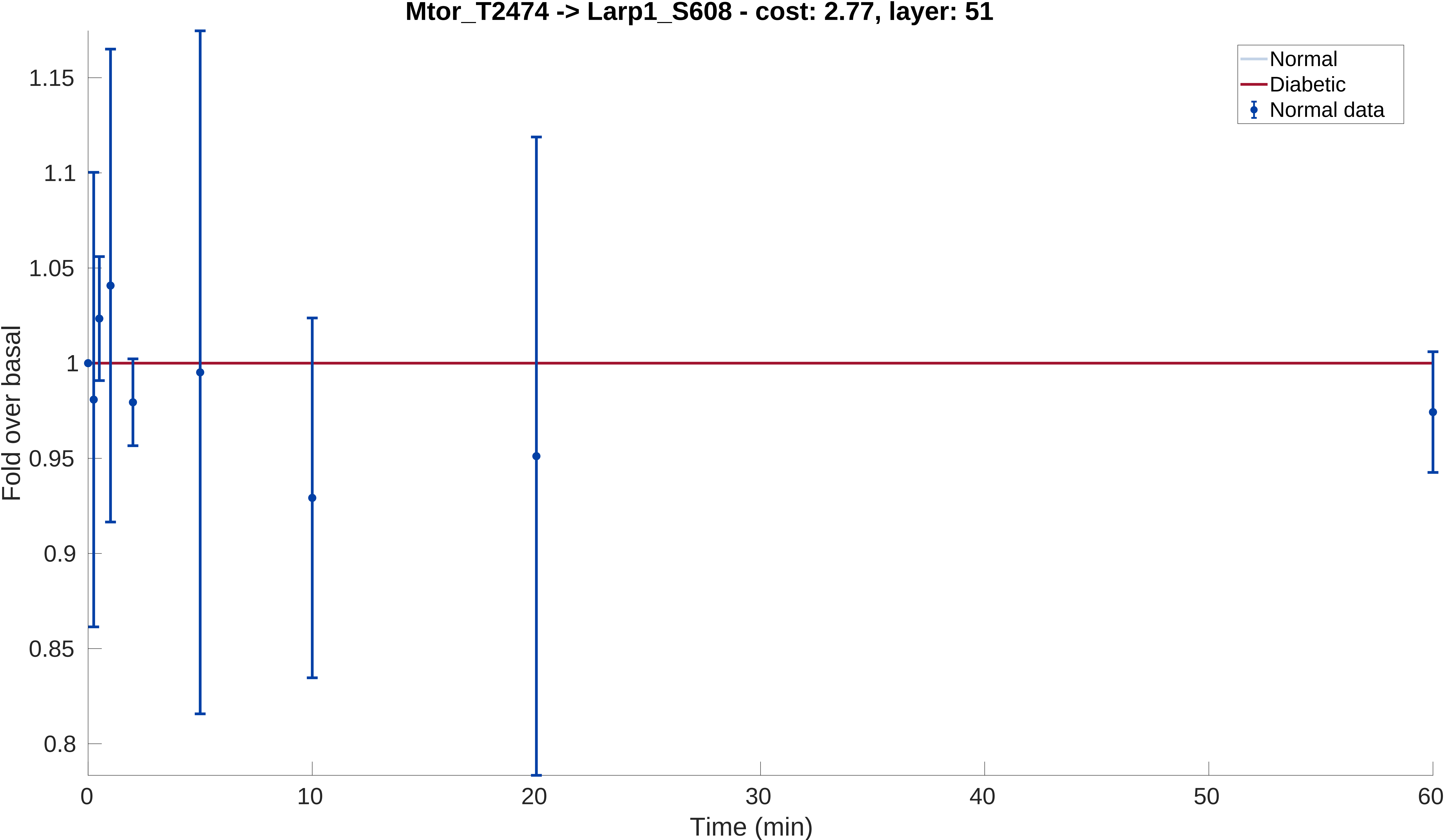

Mtor_T2474 -> Larp1_S608 - cost: 2.77, layer: 51
Normal
Diabetic
Normal data
Fold over basal
1.15
1.1
1.05
1
0.95
0.9
0.85
0.8
0
10
20
30
40
50
60
Time (min)

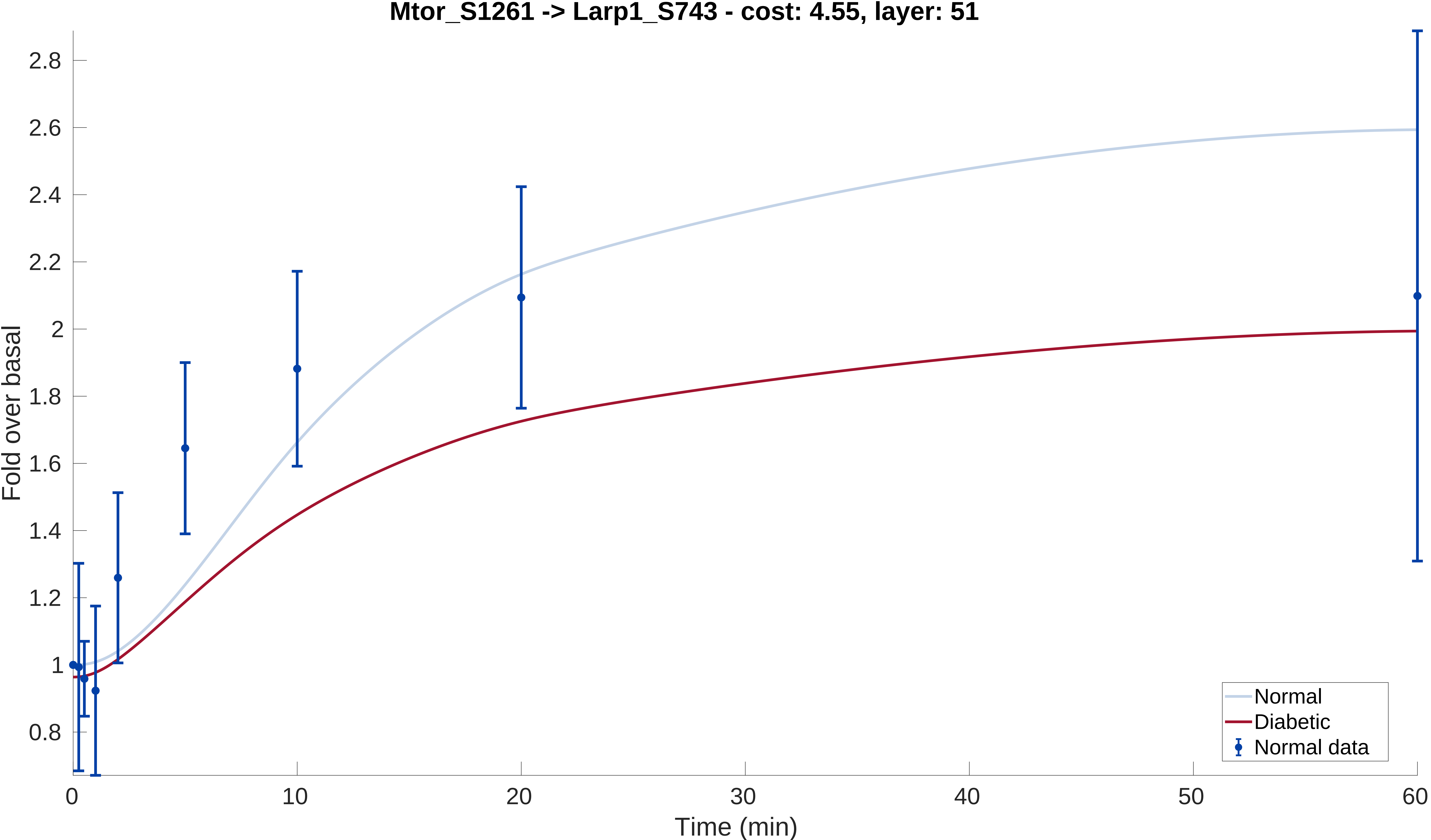

Mtor_S1261 -> Larp1_S743 - cost: 4.55, layer: 51
Fold over basal
2.8
2.6
2.4
2.2
2
1.8
1.6
1.4
1.2
1
0.8
0
10
20
30
40
50
60
Time (min)
Normal
Diabetic
Normal data

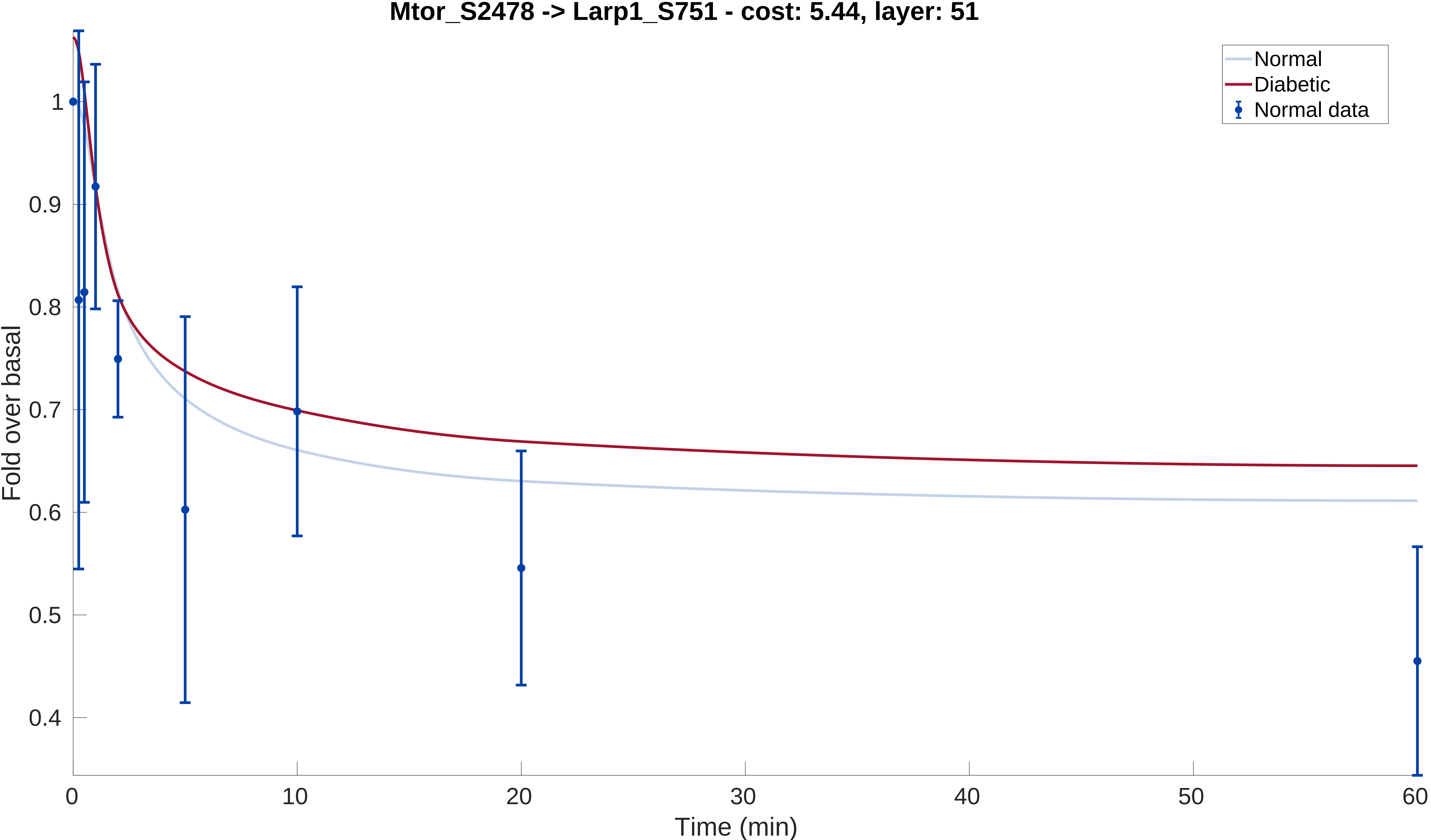

Mtor_S2478 -> Larp1_S751 - cost: 5.44, layer: 51
Normal
Diabetic
Normal data
Fold over basal
1
0.9
0.8
0.7
0.6
0.5
0.4
0
10
20
30
40
50
60
Time (min)

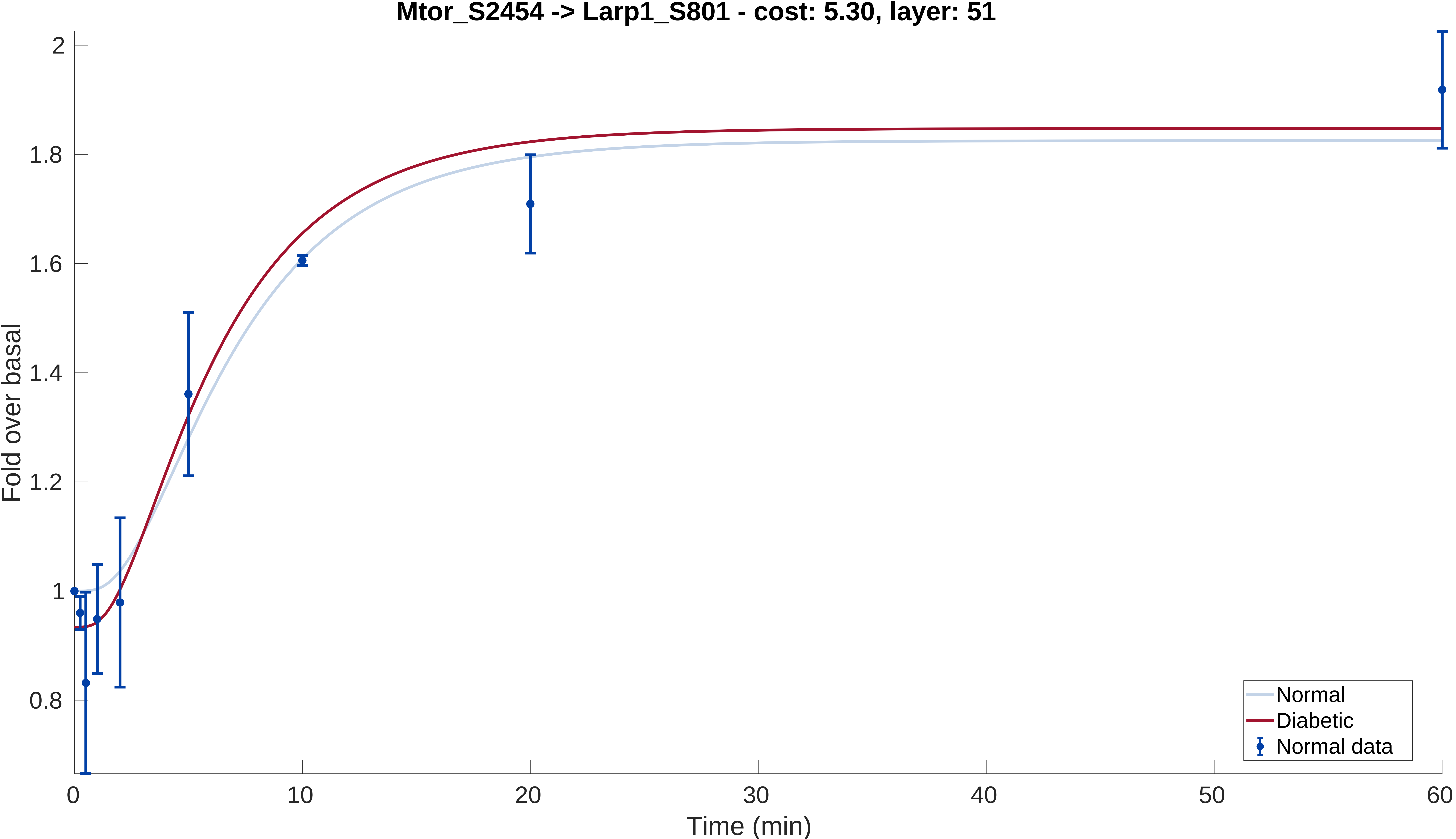

Mtor_S2454 -> Larp1_S801 - cost: 5.30, layer: 51
Fold over basal
Time (min)
0.8
1
1.2
1.4
1.6
1.8
2
0
10
20
30
40
50
60
Normal
Diabetic
Normal data

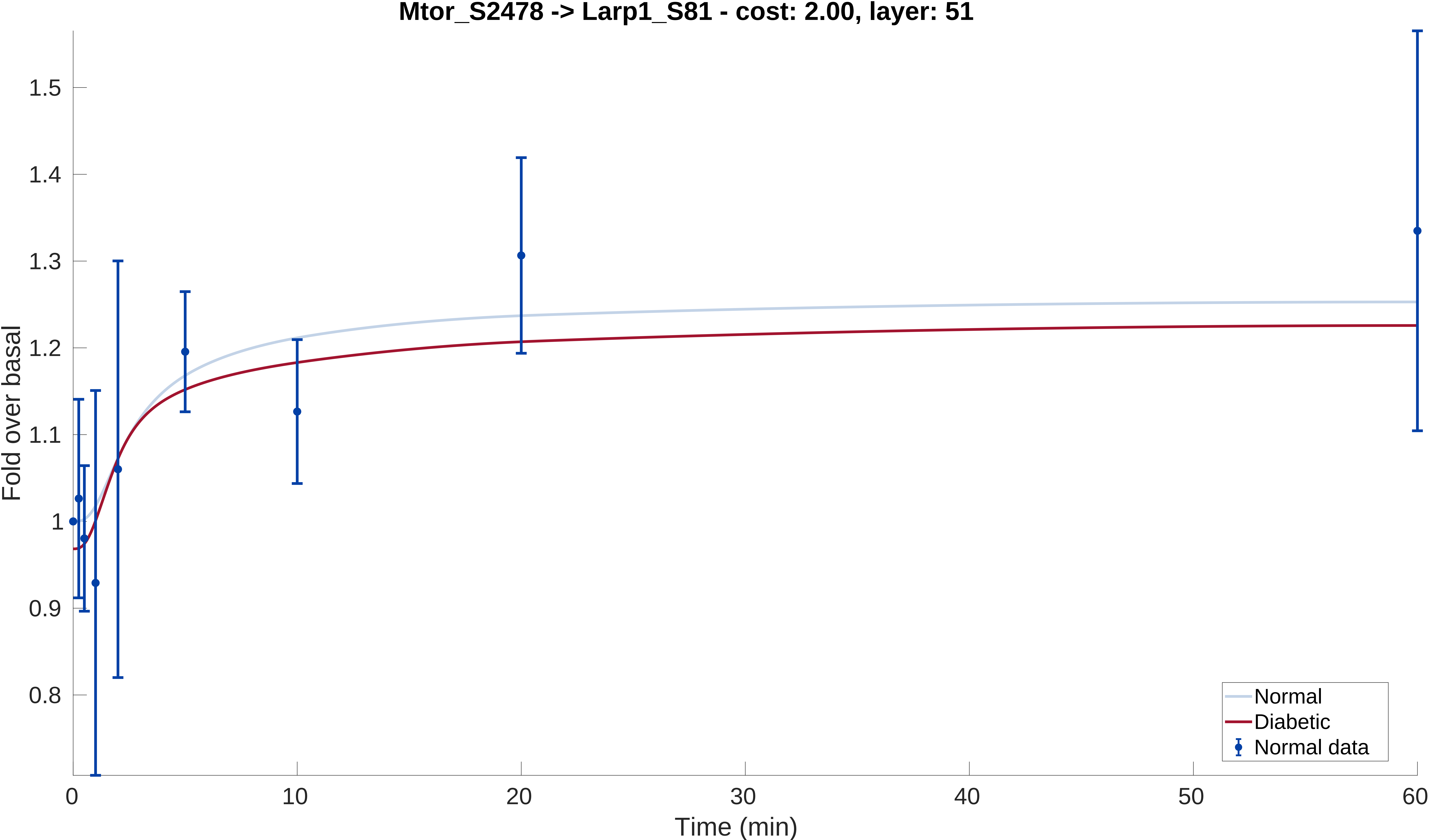

Mtor_S2478 -> Larp1_S81 - cost: 2.00, layer: 51
Fold over basal
1.5
1.4
1.3
1.2
1.1
1
0.9
0.8
0
10
20
30
40
50
60
Time (min)
Normal
Diabetic
Normal data

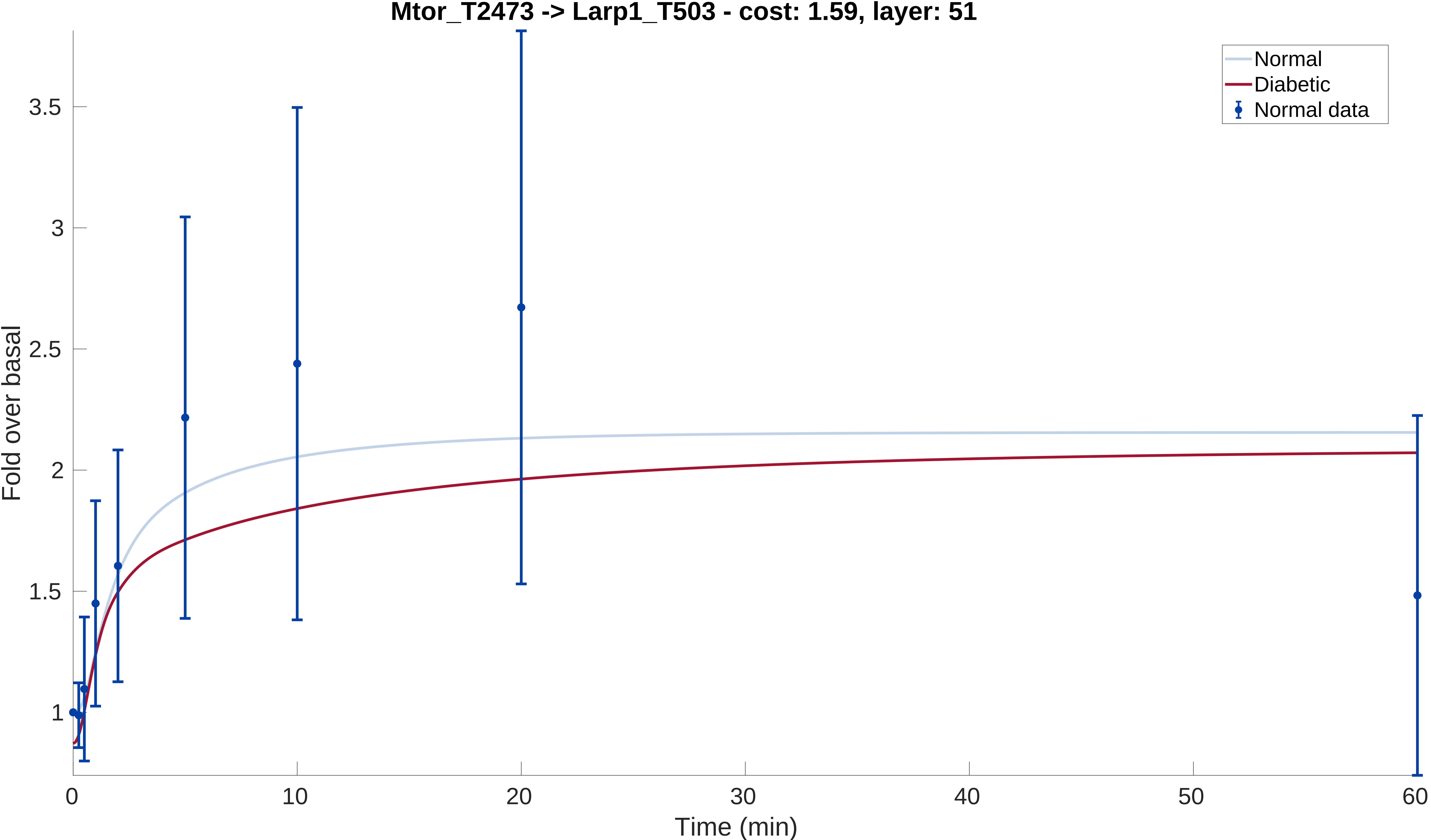

Mtor_T2473 -> Larp1_T503 - cost: 1.59, layer: 51
Normal
Diabetic
Normal data
3.5
3
2.5
2
1.5
1
Fold over basal
0
10
20
30
40
50
60
Time (min)

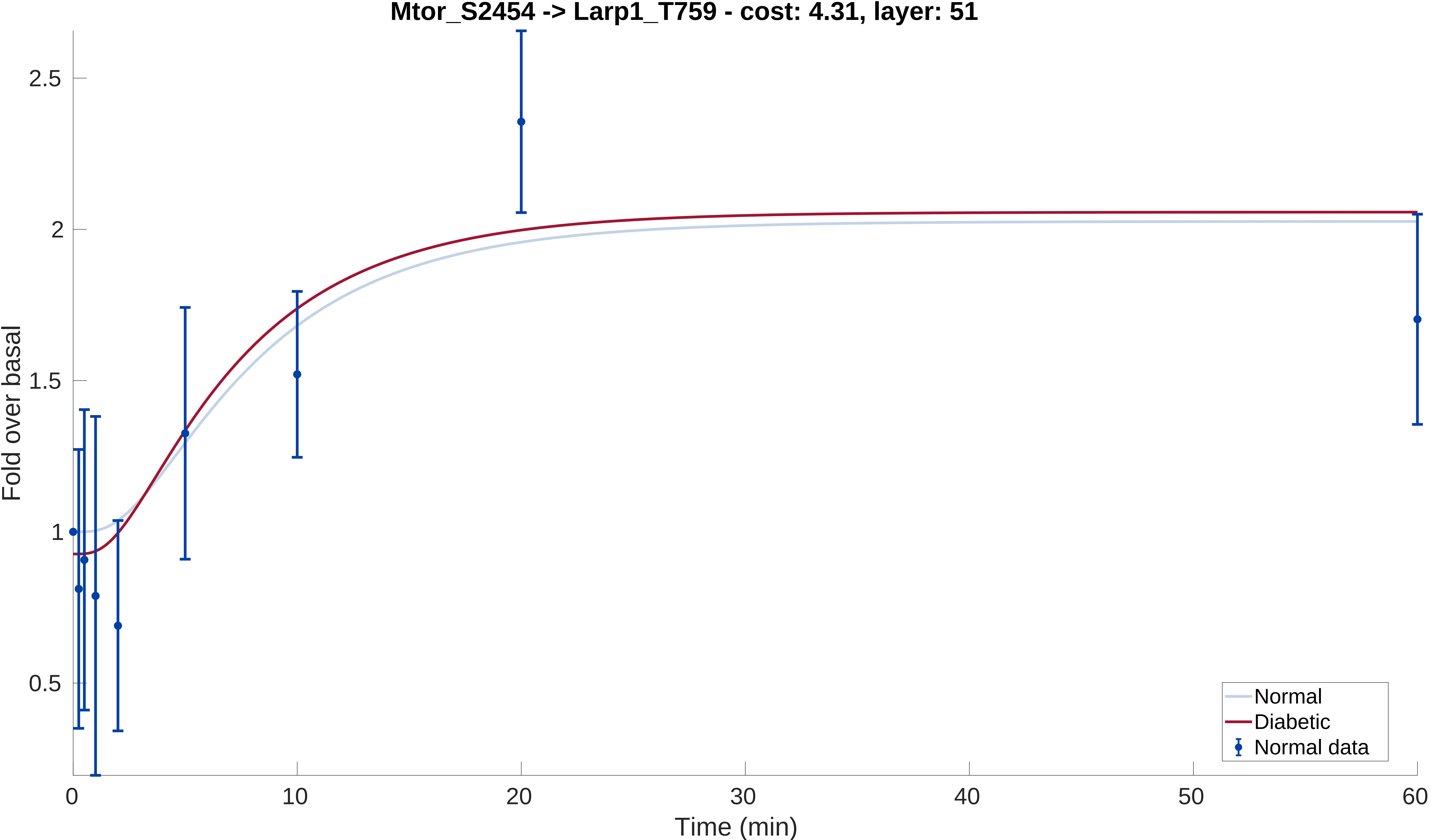
Mtor_S2454 -> Larp1_T759 - cost: 4.31, layer: 51
Fold over basal
2.5
2
1.5
1
0.5
0
10
20
30
40
50
60
Time (min)
Normal
Diabetic
Normal data

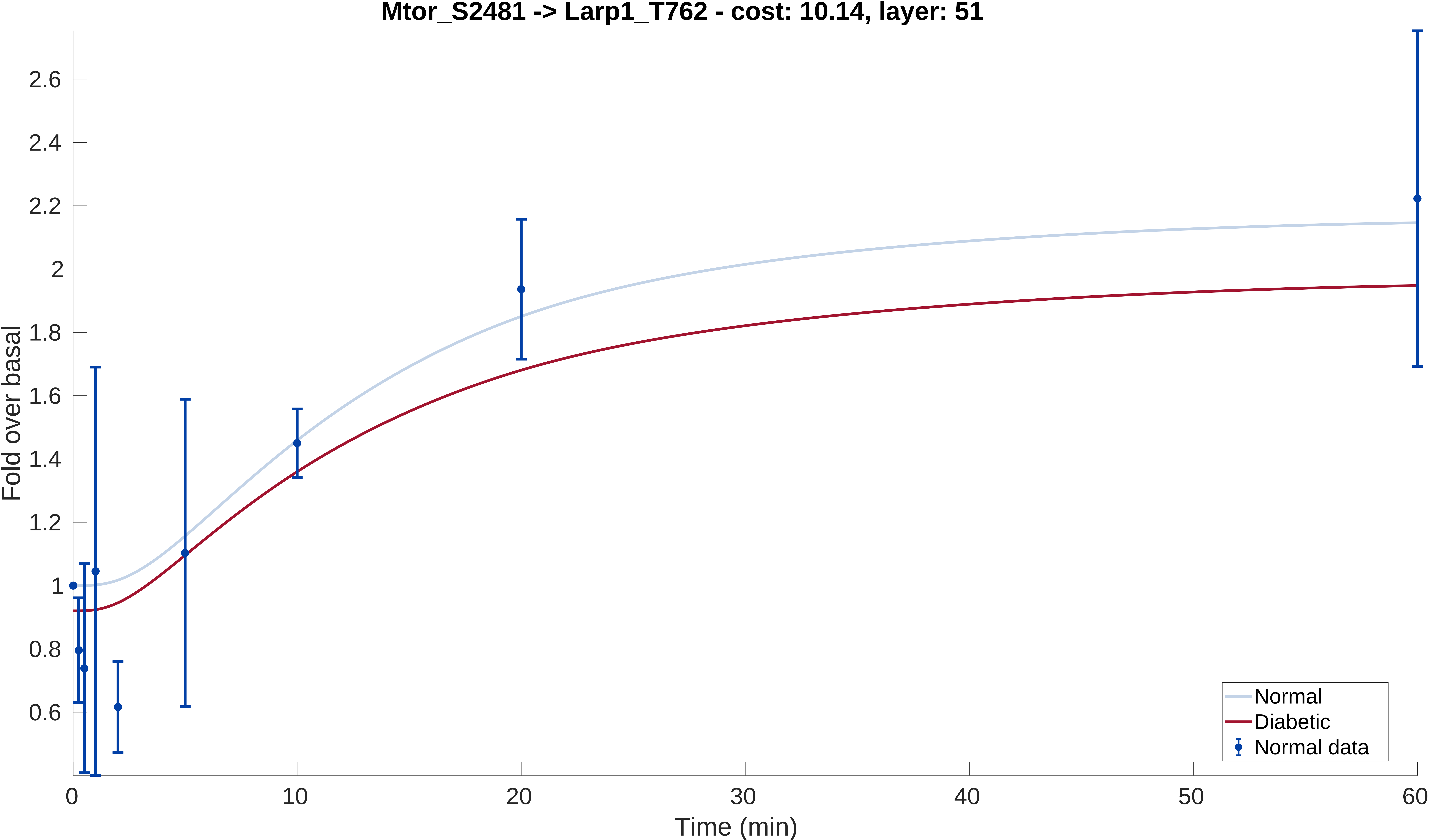
Mtor_S2481 -> Larp1_T762 - cost: 10.14, layer: 51
Fold over basal
2.6
2.4
2.2
2
1.8
1.6
1.4
1.2
1
0.8
0.6
0
10
20
30
40
50
60
Time (min)
Normal
Diabetic
Normal data

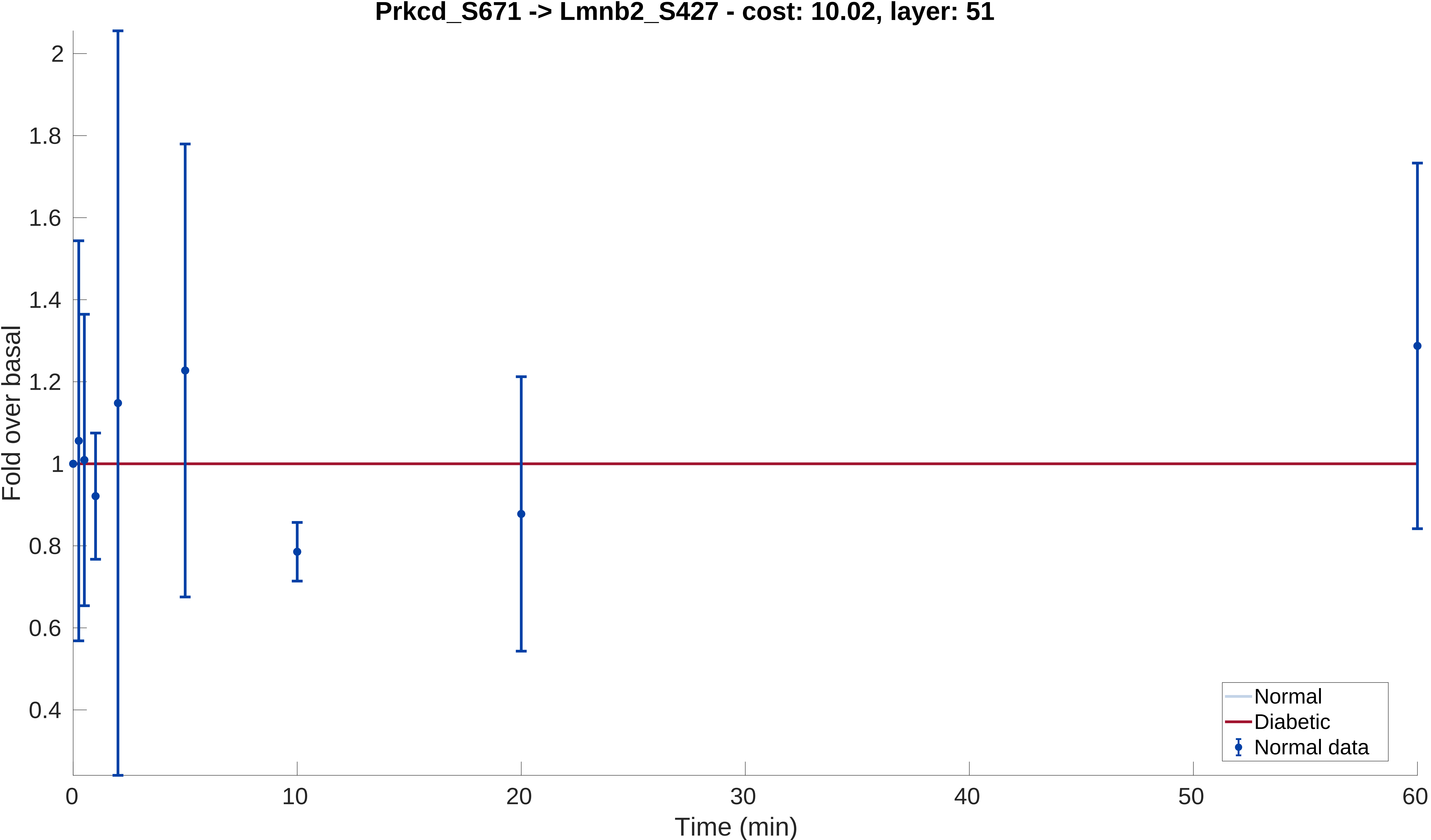
Prkcd_S671 -> Lmnb2_S427 - cost: 10.02, layer: 51
Fold over basal
2
1.8
1.6
1.4
1.2
1
0.8
0.6
0.4
0
10
20
30
40
50
60
Time (min)
Normal
Diabetic
Normal data

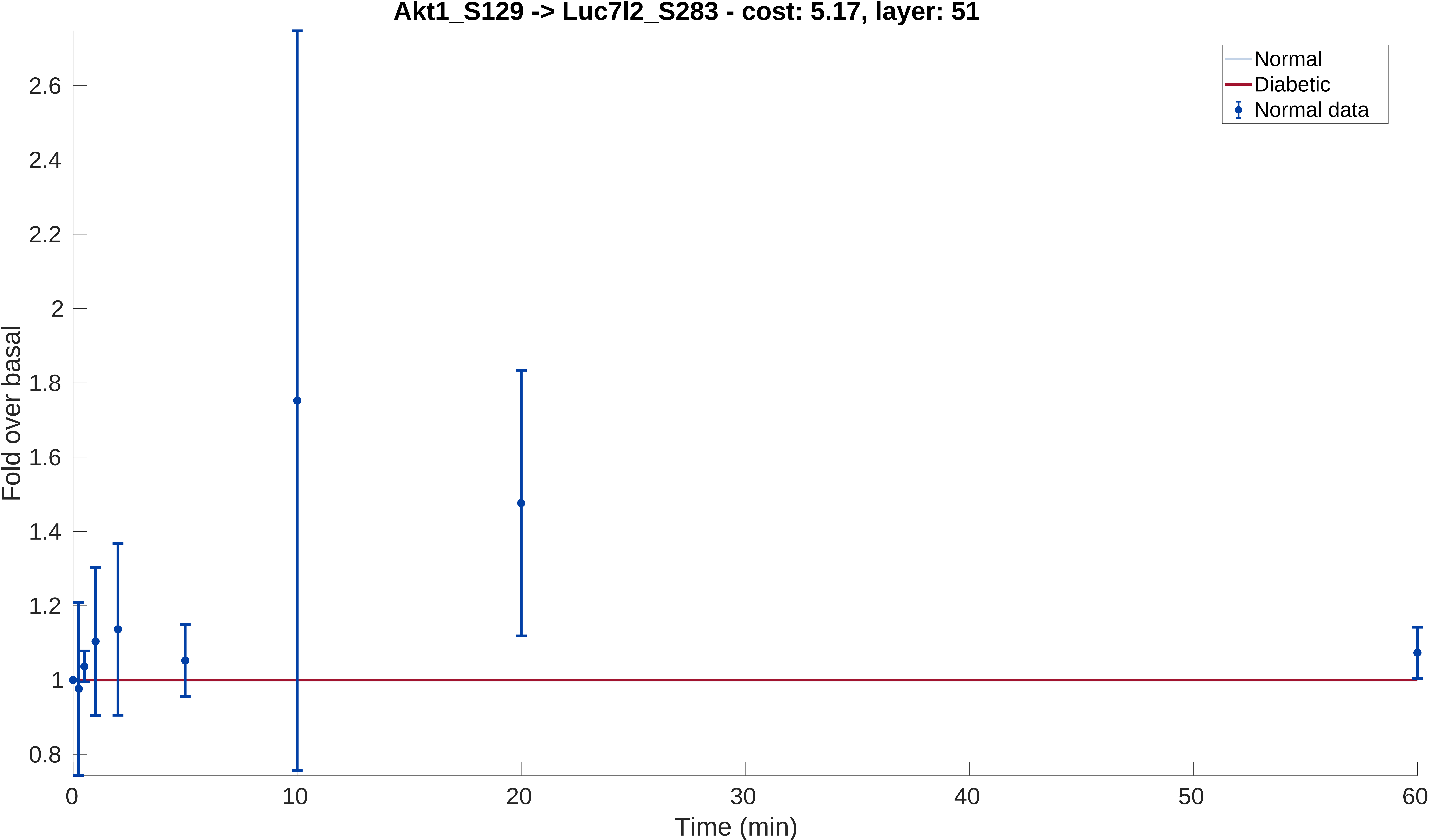
Akt1_S129 -> Luc7l2_S283 - cost: 5.17, layer: 51
Normal
Diabetic
Normal data
Fold over basal
0.8
1
1.2
1.4
1.6
1.8
2
2.2
2.4
2.6
0
10
20
30
40
50
60
Time (min)

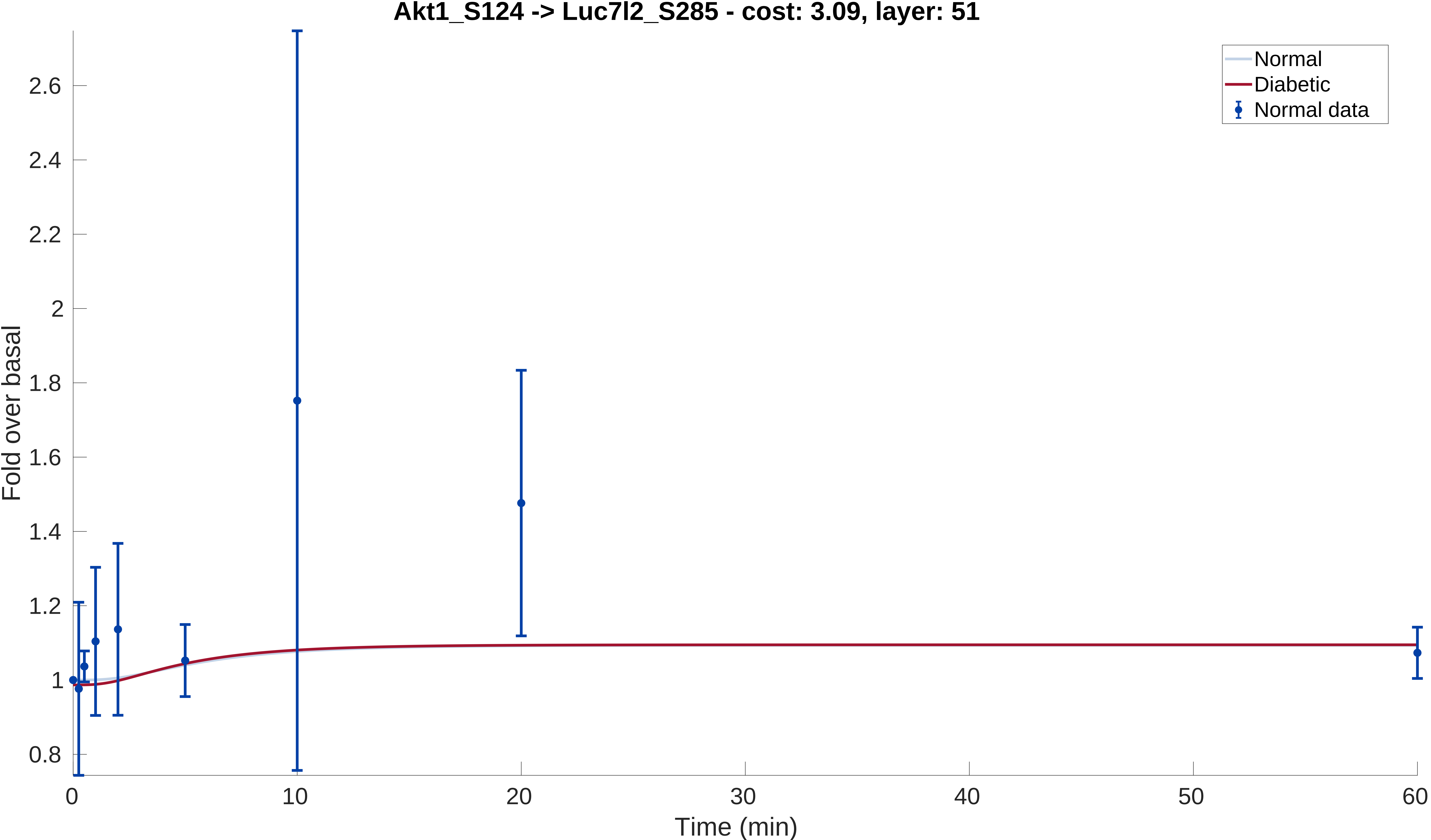
Akt1_S124 -> Luc7l2_S285 - cost: 3.09, layer: 51
Normal
Diabetic
Normal data
Fold over basal
0.8
1
1.2
1.4
1.6
1.8
2
2.2
2.4
2.6
Time (min)
0
10
20
30
40
50
60

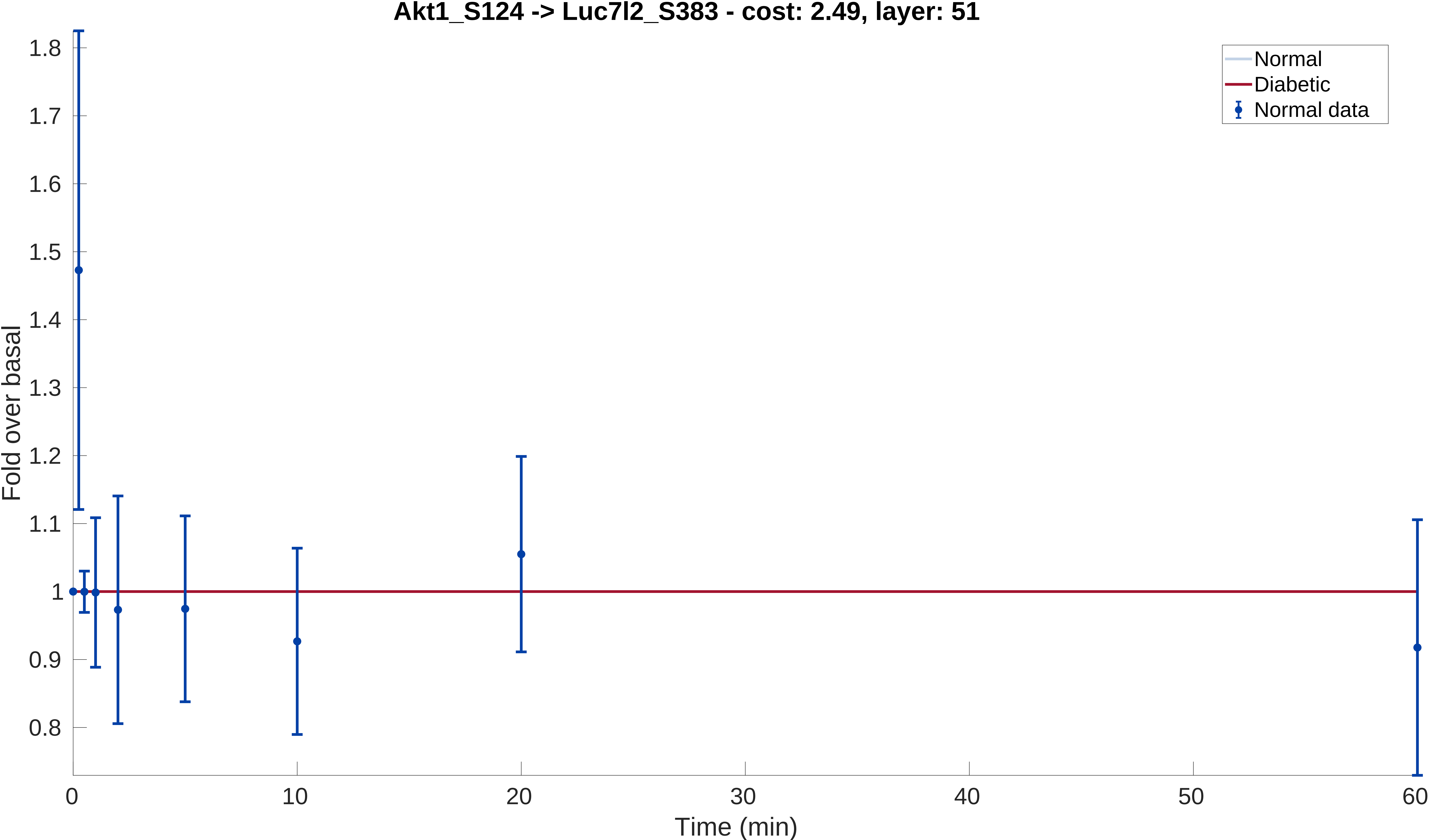

Akt1_S124 -> Luc7l2_S383 - cost: 2.49, layer: 51
Normal
Diabetic
Normal data
Fold over basal
1.8
1.7
1.6
1.5
1.4
1.3
1.2
1.1
1
0.9
0.8
0
10
20
30
40
50
60
Time (min)

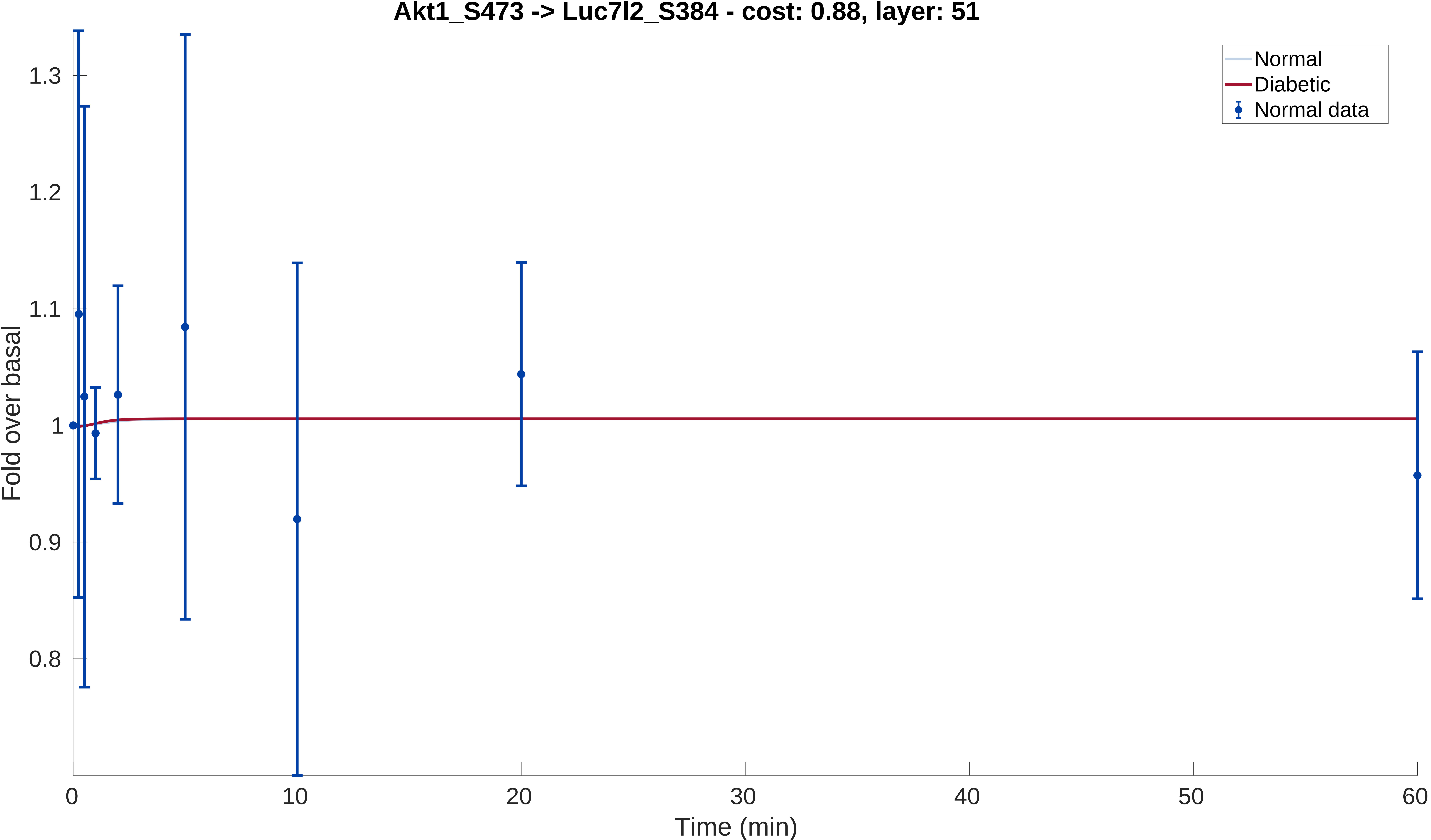

Akt1_S473 -> Luc7l2_S384 - cost: 0.88, layer: 51
Normal
Diabetic
Normal data
Fold over basal
1.3
1.2
1.1
1
0.9
0.8
0
10
20
30
40
50
60
Time (min)

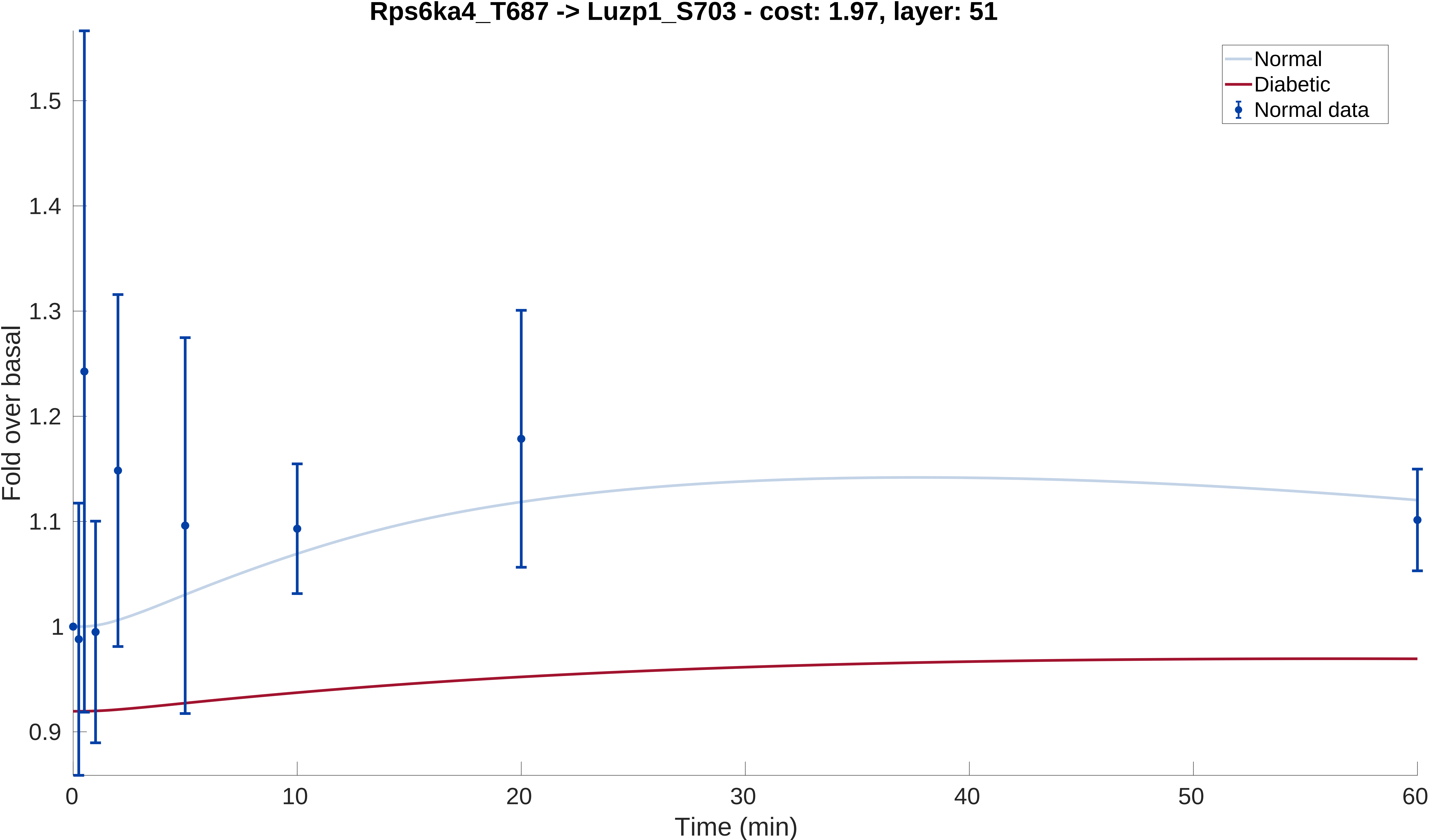

Rps6ka4_T687 -> Luzp1_S703 - cost: 1.97, layer: 51
Normal
Diabetic
Normal data
Fold over basal
Time (min)
1.5
1.4
1.3
1.2
1.1
1
0.9
0
10
20
30
40
50
60

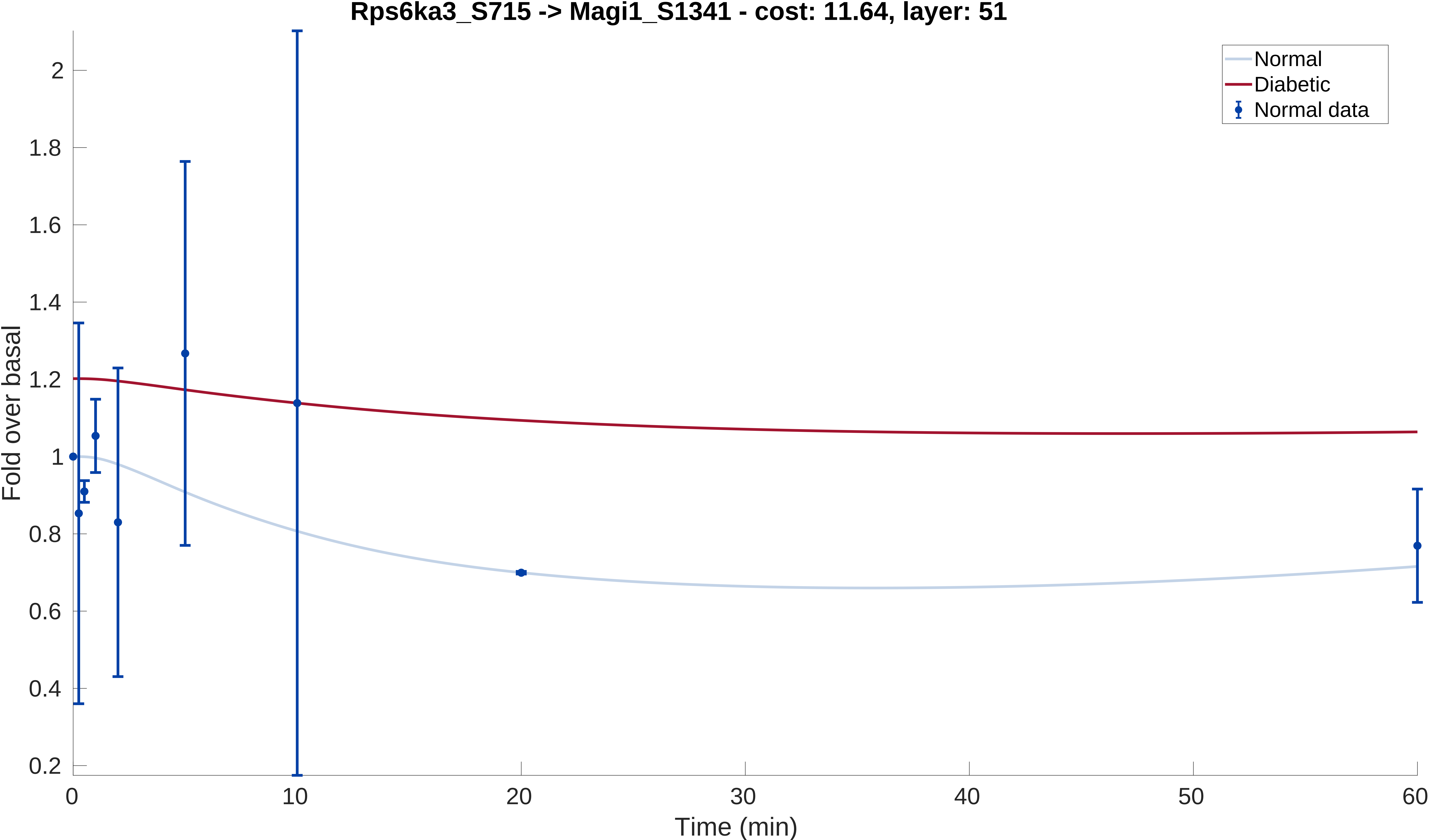

Rps6ka3_S715 -> Magi1_S1341 - cost: 11.64, layer: 51
Normal
Diabetic
Normal data
Fold over basal
2
1.8
1.6
1.4
1.2
1
0.8
0.6
0.4
0.2
0
10
20
30
40
50
60
Time (min)

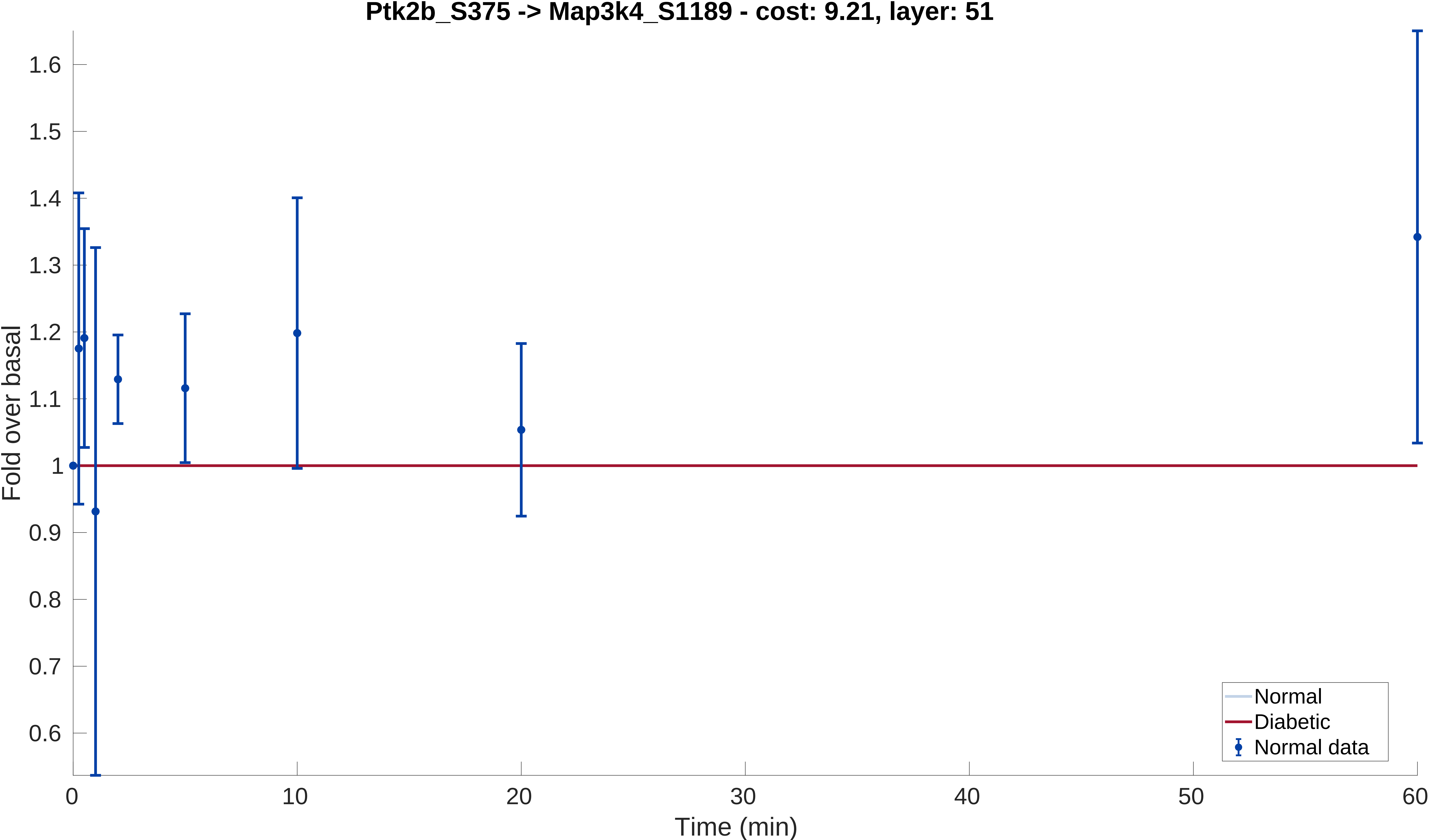

Ptk2b_S375 -> Map3k4_S1189 - cost: 9.21, layer: 51
Fold over basal
1.6
1.5
1.4
1.3
1.2
1.1
1
0.9
0.8
0.7
0.6
0
10
20
30
40
50
60
Time (min)
Normal
Diabetic
Normal data

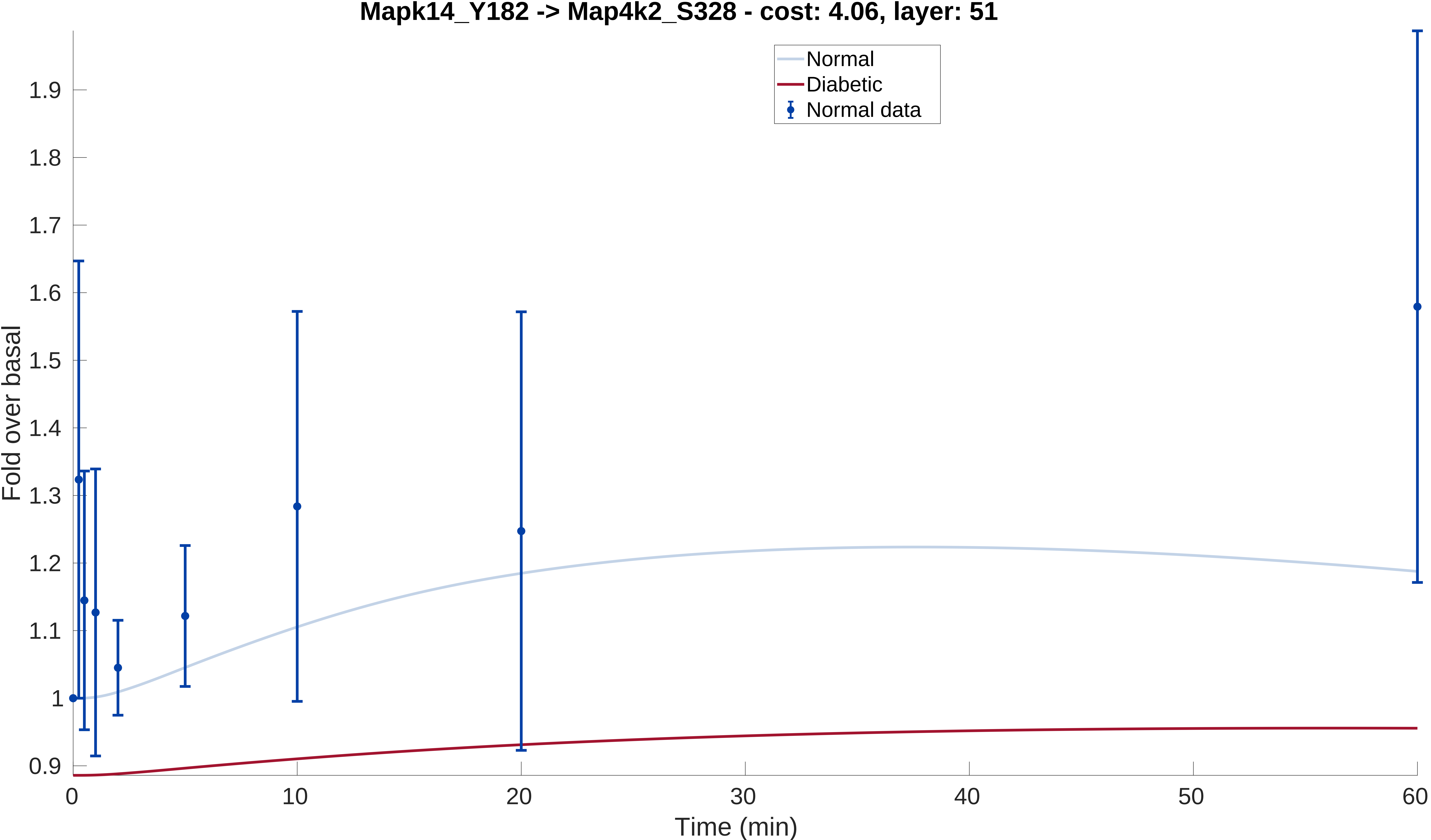

Mapk14_Y182 -> Map4k2_S328 - cost: 4.06, layer: 51
Normal
Diabetic
Normal data
Fold over basal
1.9
1.8
1.7
1.6
1.5
1.4
1.3
1.2
1.1
1
0.9
0
10
20
30
40
50
60
Time (min)

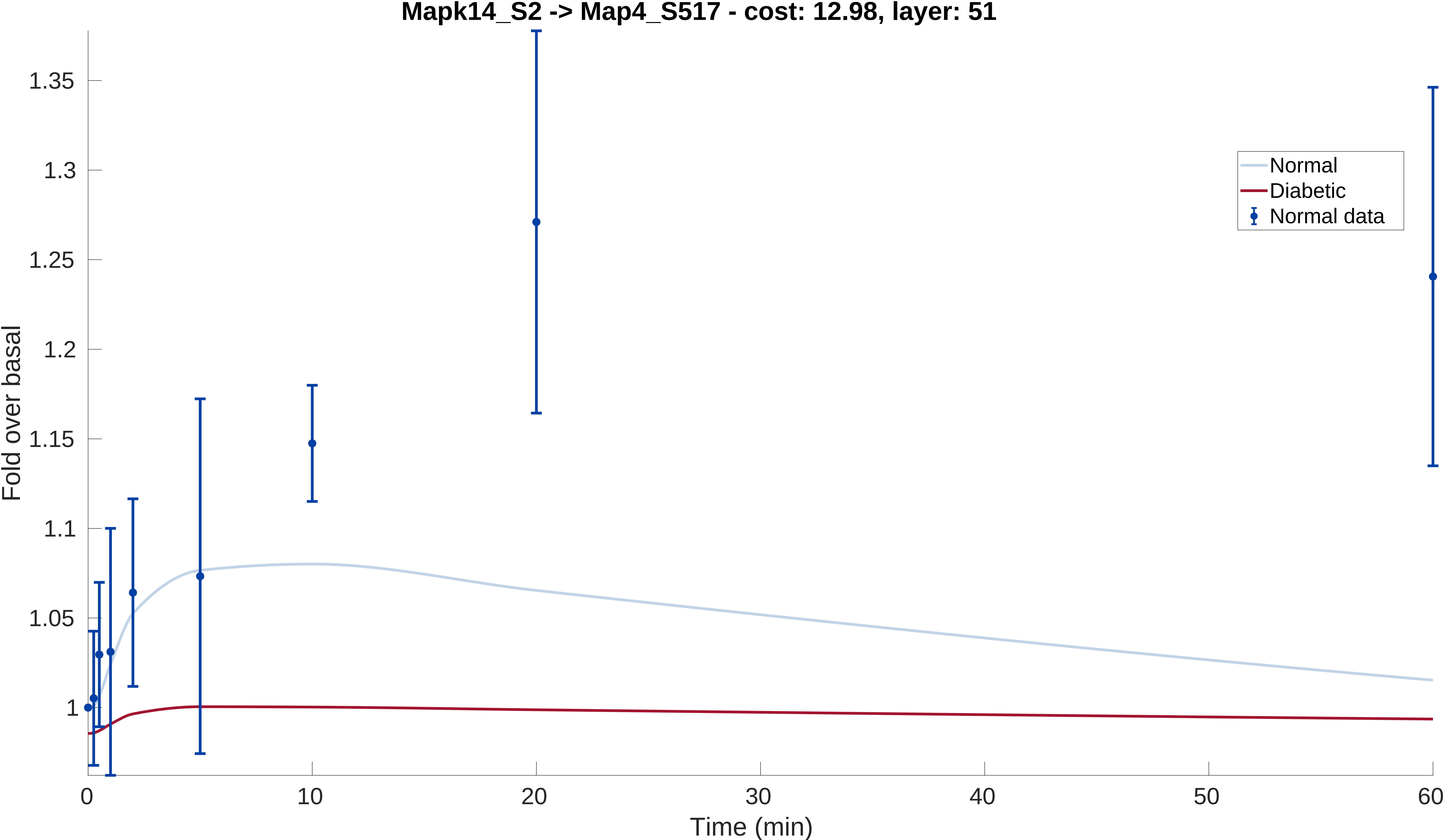

Mapk14_S2 -> Map4_S517 - cost: 12.98, layer: 51
Fold over basal
1.35
1.3
1.25
1.2
1.15
1.1
1.05
1
0
10
20
30
40
50
60
Time (min)
Normal
Diabetic
Normal data

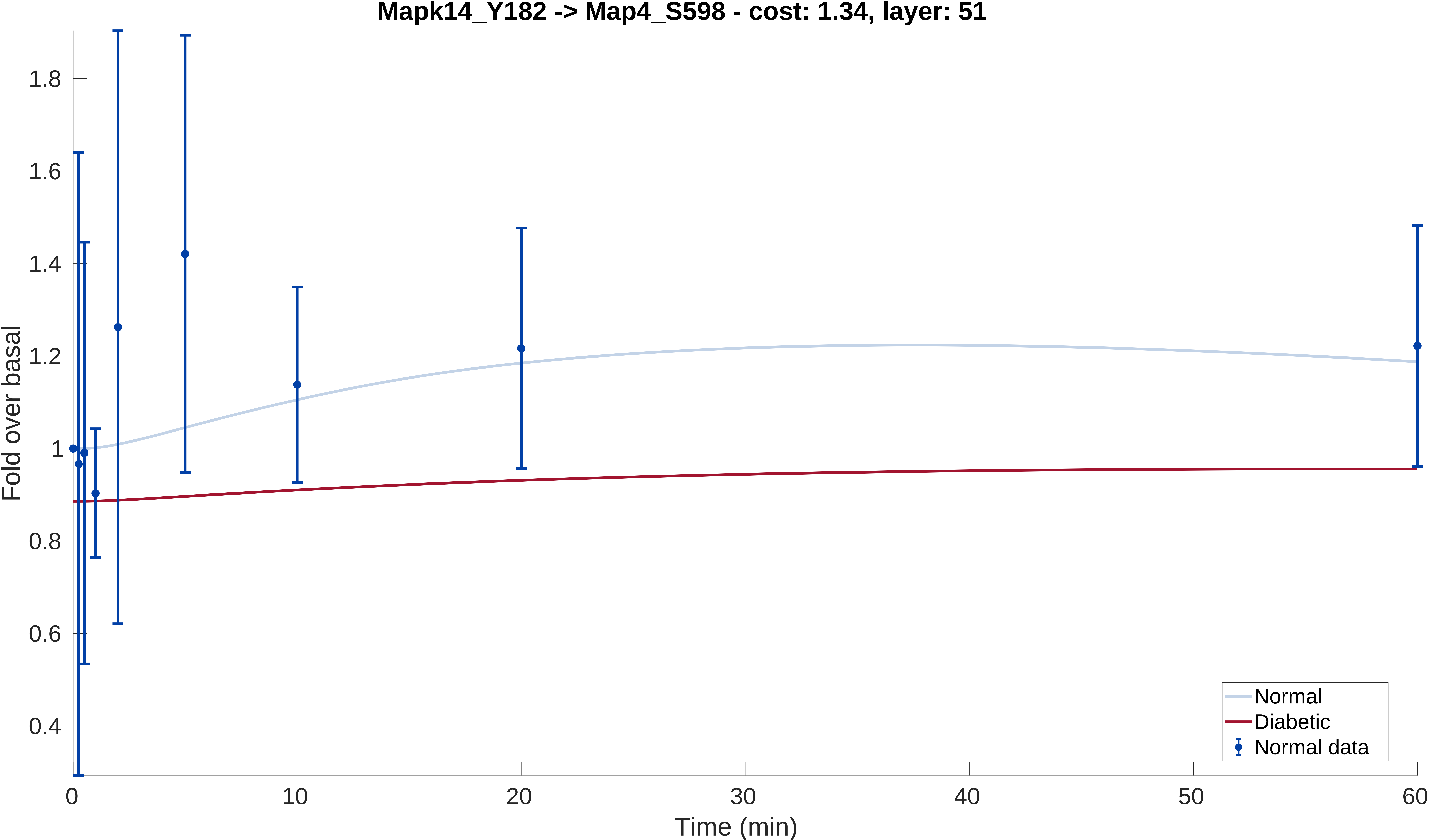

Mapk14_Y182 -> Map4_S598 - cost: 1.34, layer: 51
Fold over basal
1.8
1.6
1.4
1.2
1
0.8
0.6
0.4
0
10
20
30
40
50
60
Time (min)
Normal
Diabetic
Normal data

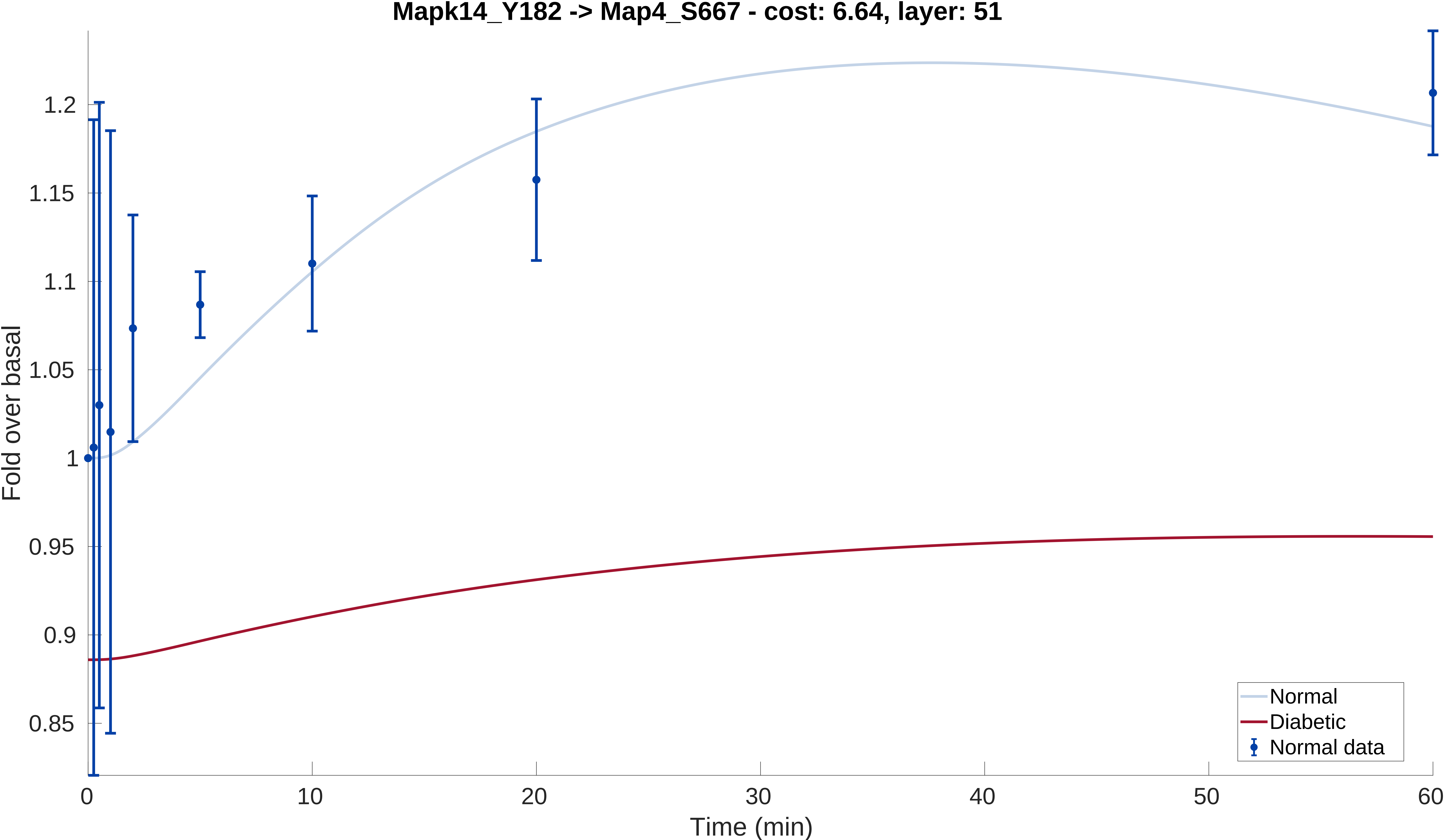
Mapk14_Y182 -> Map4_S667 - cost: 6.64, layer: 51
Fold over basal
1.2
1.15
1.1
1.05
1
0.95
0.9
0.85
0
10
20
30
40
50
60
Time (min)
Normal
Diabetic
Normal data

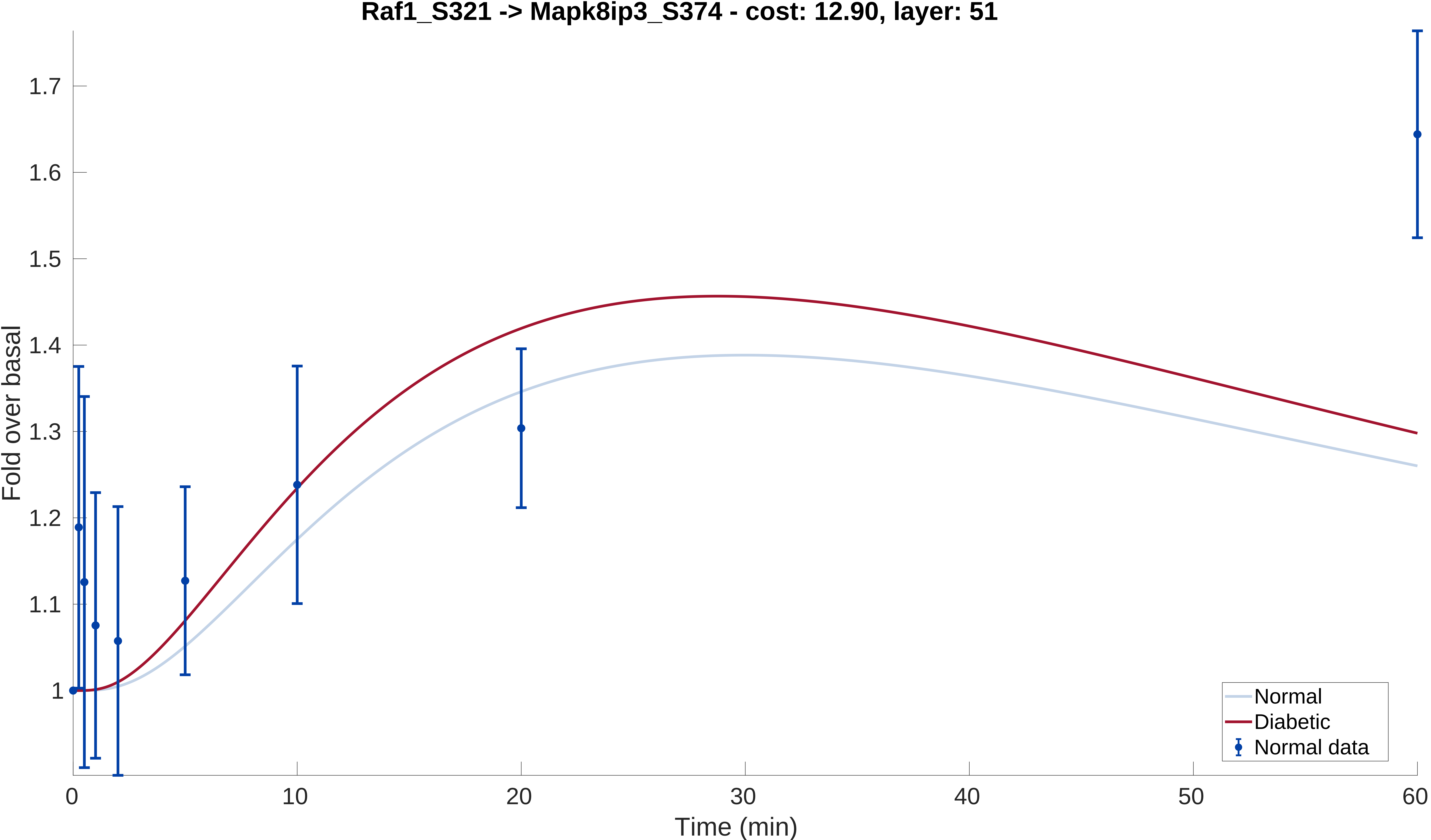

Raf1_S321 -> Mapk8ip3_S374 - cost: 12.90, layer: 51
Fold over basal
1.7
1.6
1.5
1.4
1.3
1.2
1.1
1
0
10
20
30
40
50
60
Time (min)
Normal
Diabetic
Normal data

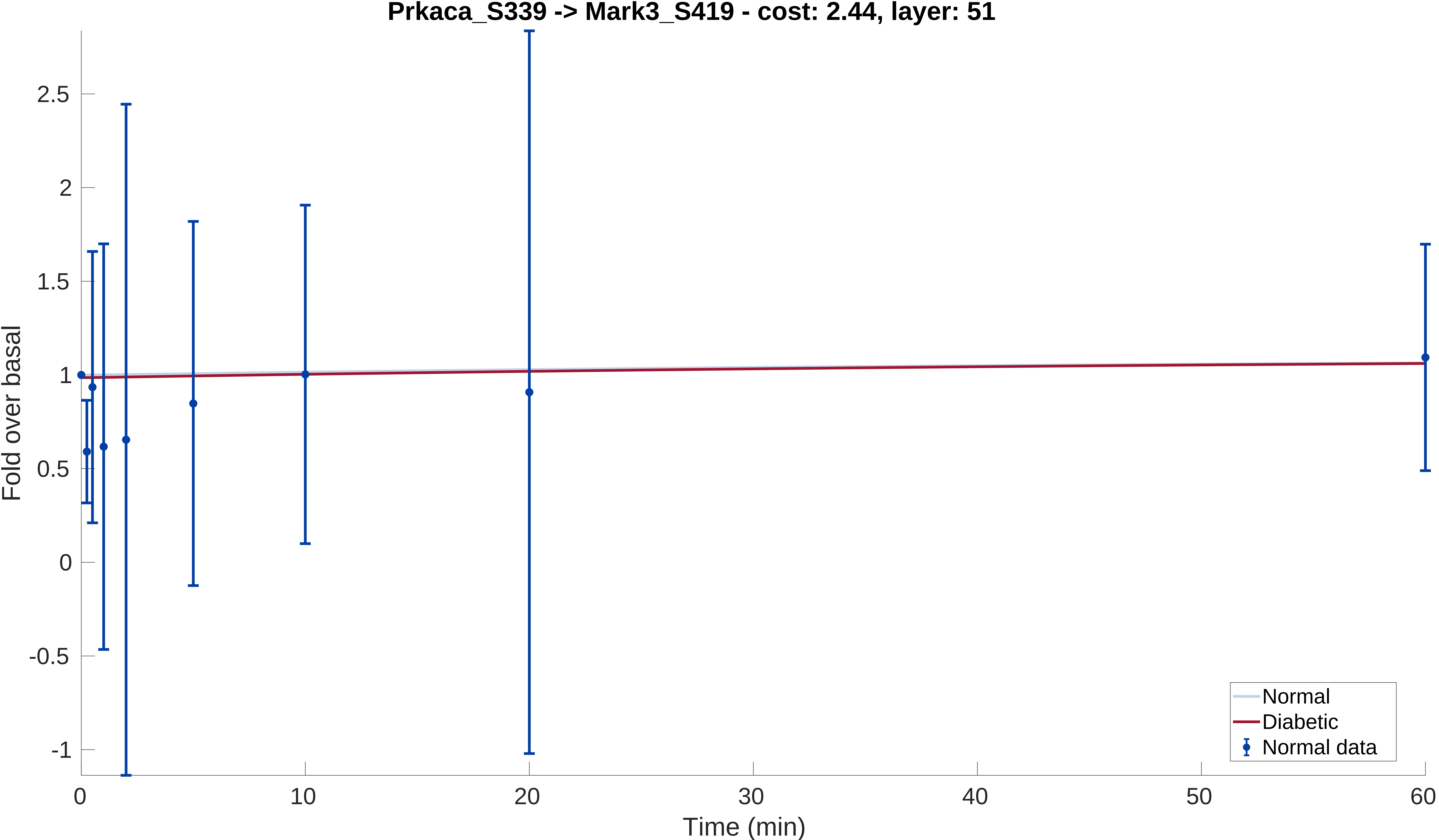
Prkaca_S339 -> Mark3_S419 - cost: 2.44, layer: 51
Fold over basal
Time (min)
Normal
Diabetic
Normal data
2.5
2
1.5
1
0.5
0
-0.5
-1
0
10
20
30
40
50
60

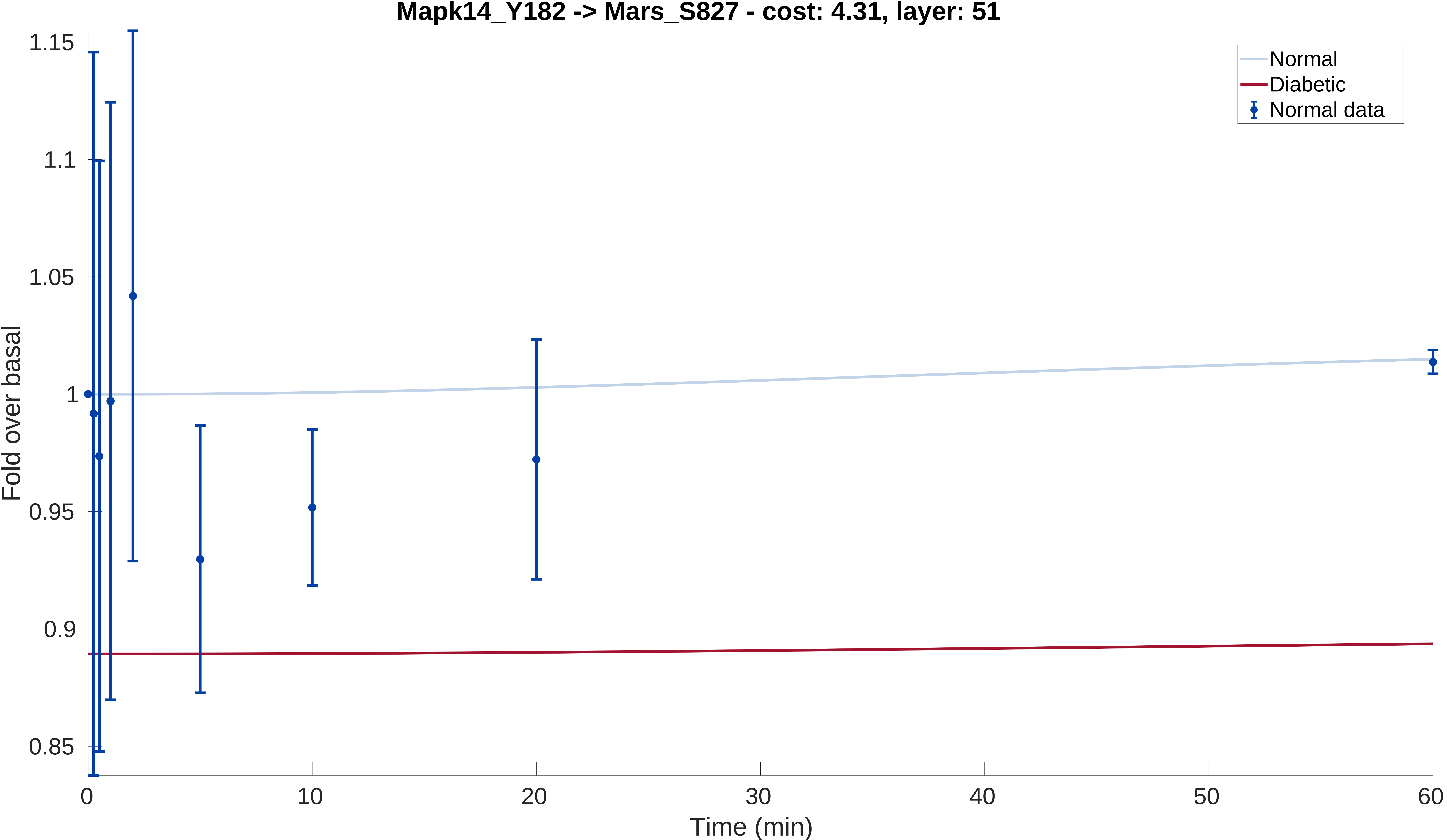

Mapk14_Y182 -> Mars_S827 - cost: 4.31, layer: 51
Normal
Diabetic
Normal data
Fold over basal
1.15
1.1
1.05
1
0.95
0.9
0.85
0
10
20
30
40
50
60
Time (min)

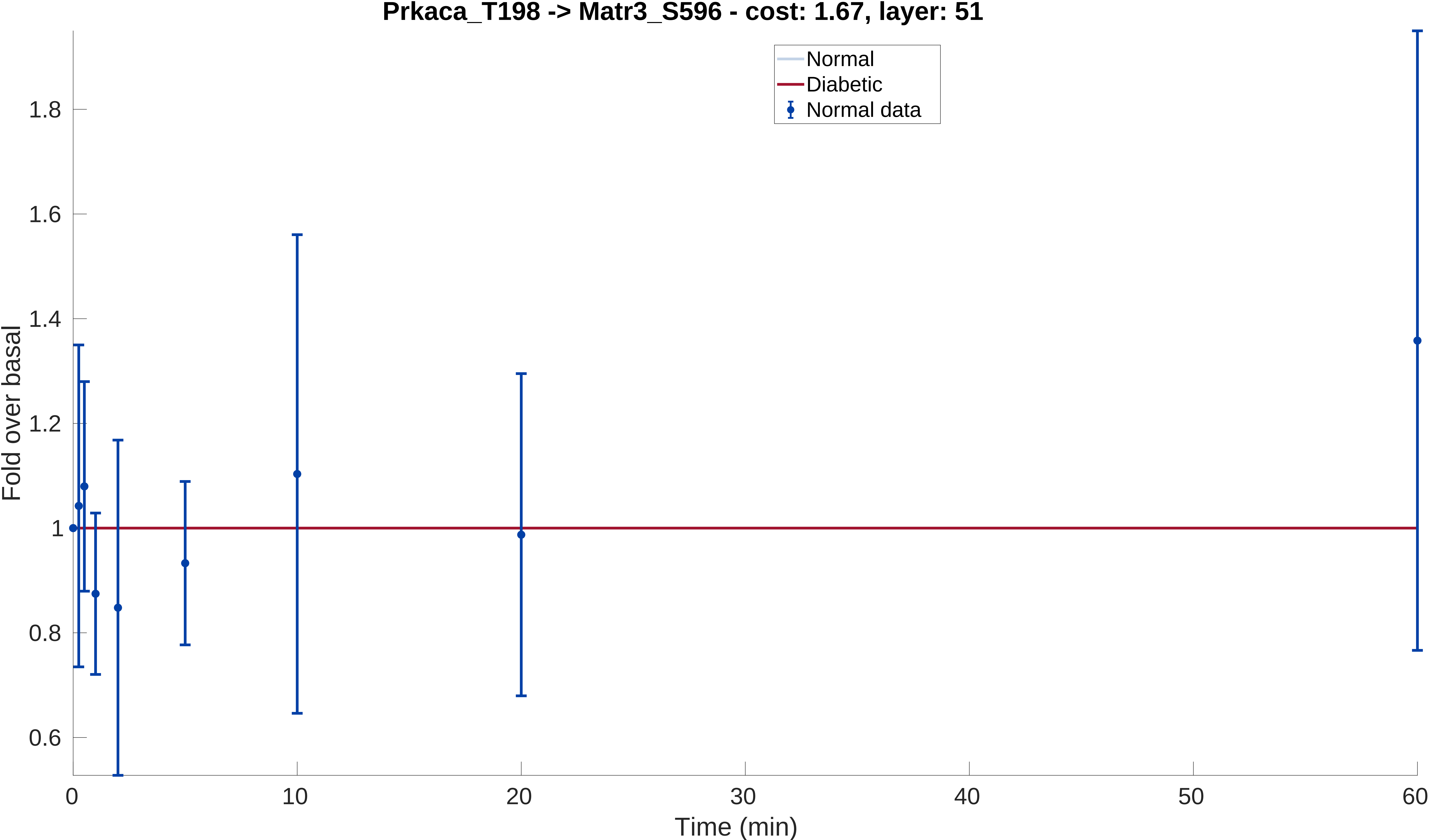

Prkaca_T198 -> Matr3_S596 - cost: 1.67, layer: 51
Normal
Diabetic
Normal data
Fold over basal
1.8
1.6
1.4
1.2
1
0.8
0.6
0
10
20
30
40
50
60
Time (min)

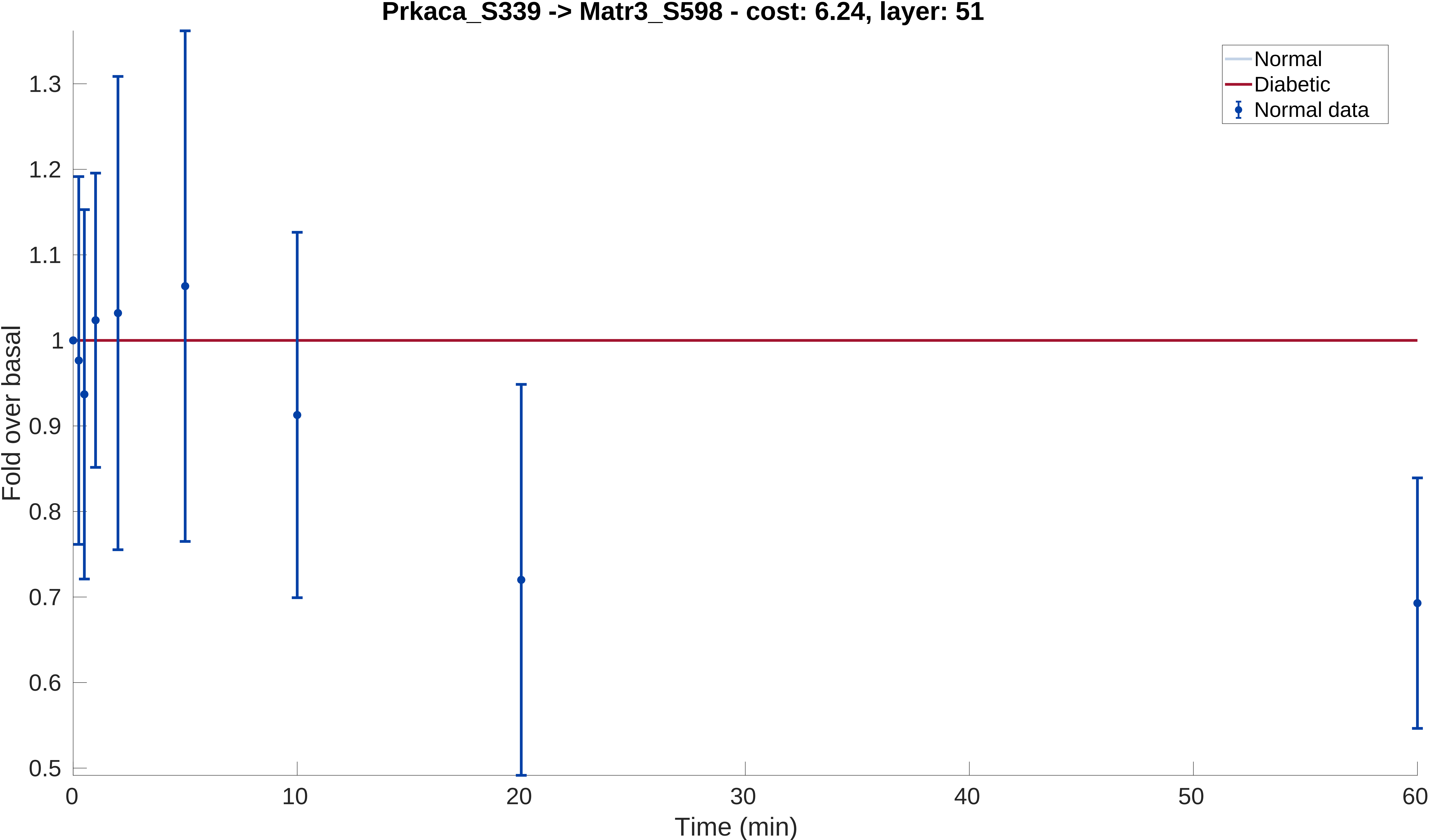
Prkaca_S339 -> Matr3_S598 - cost: 6.24, layer: 51
Normal
Diabetic
Normal data
Fold over basal
1.3
1.2
1.1
1
0.9
0.8
0.7
0.6
0.5
0
10
20
30
40
50
60
Time (min)

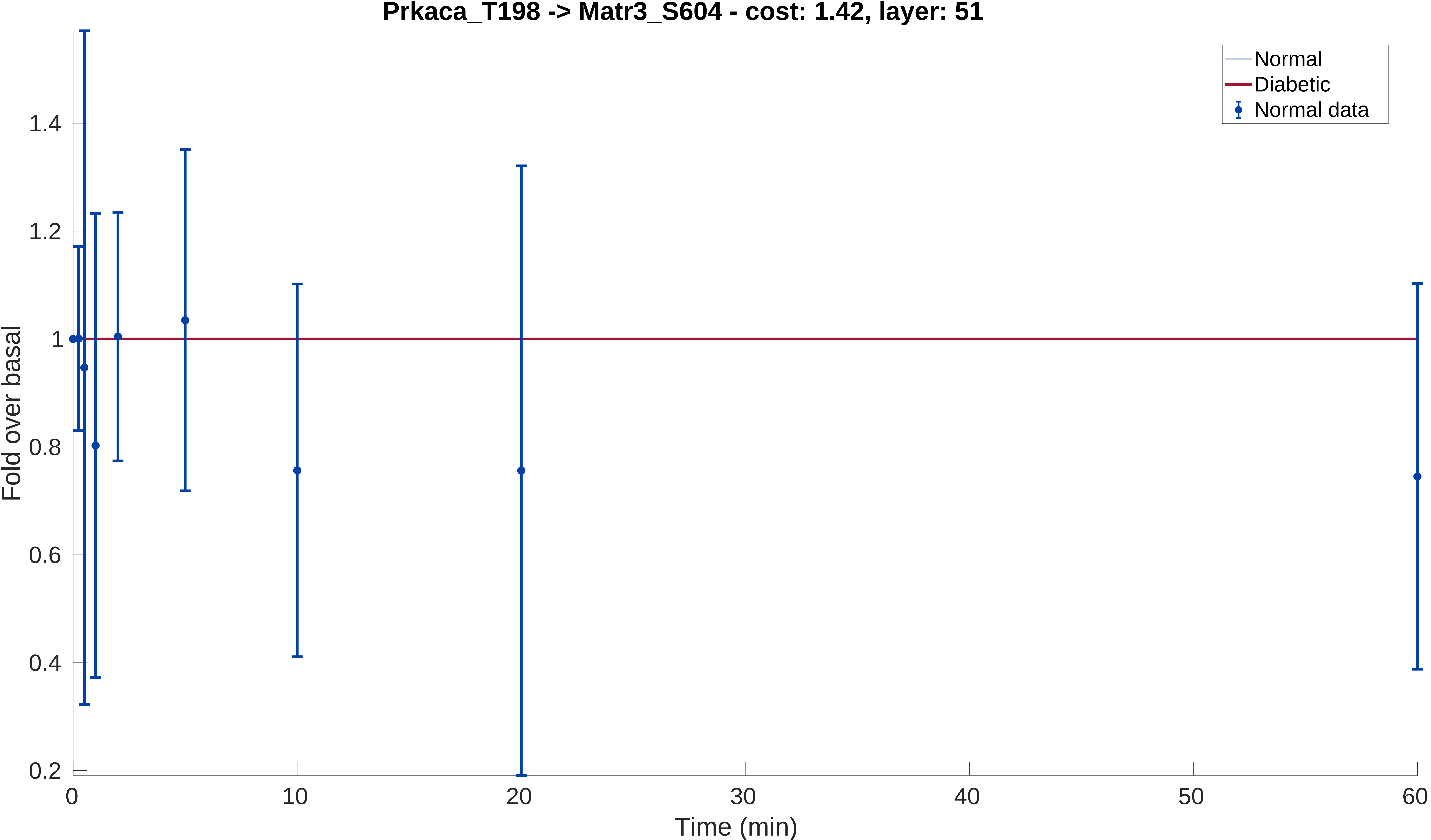

Prkaca_T198 -> Matr3_S604 - cost: 1.42, layer: 51
Normal
Diabetic
Normal data
Fold over basal
1.4
1.2
1
0.8
0.6
0.4
0.2
0
10
20
30
40
50
60
Time (min)

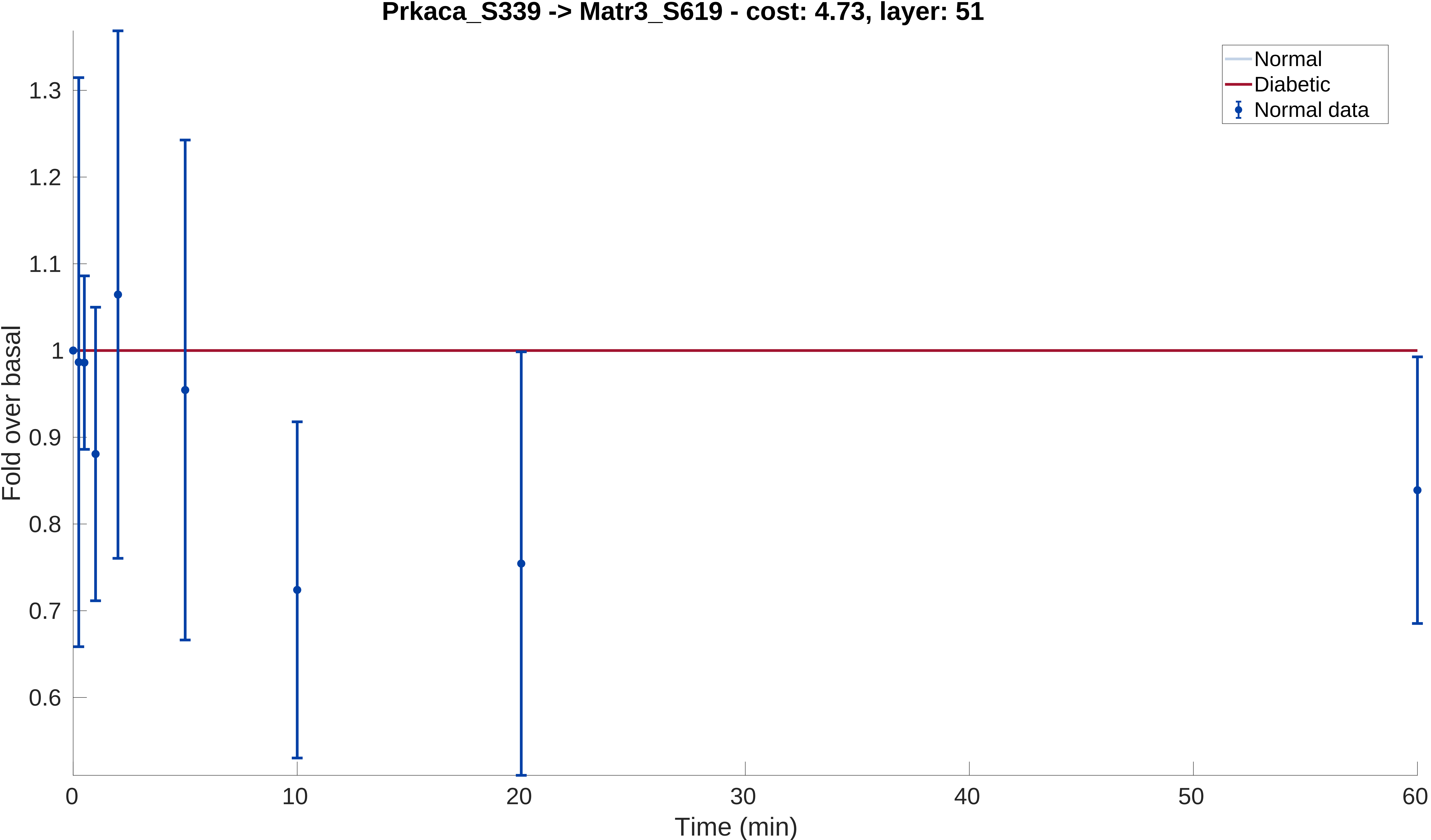
Prkaca_S339 -> Matr3_S619 - cost: 4.73, layer: 51
Normal
Diabetic
Normal data
Fold over basal
1.3
1.2
1.1
1
0.9
0.8
0.7
0.6
0
10
20
30
40
50
60
Time (min)

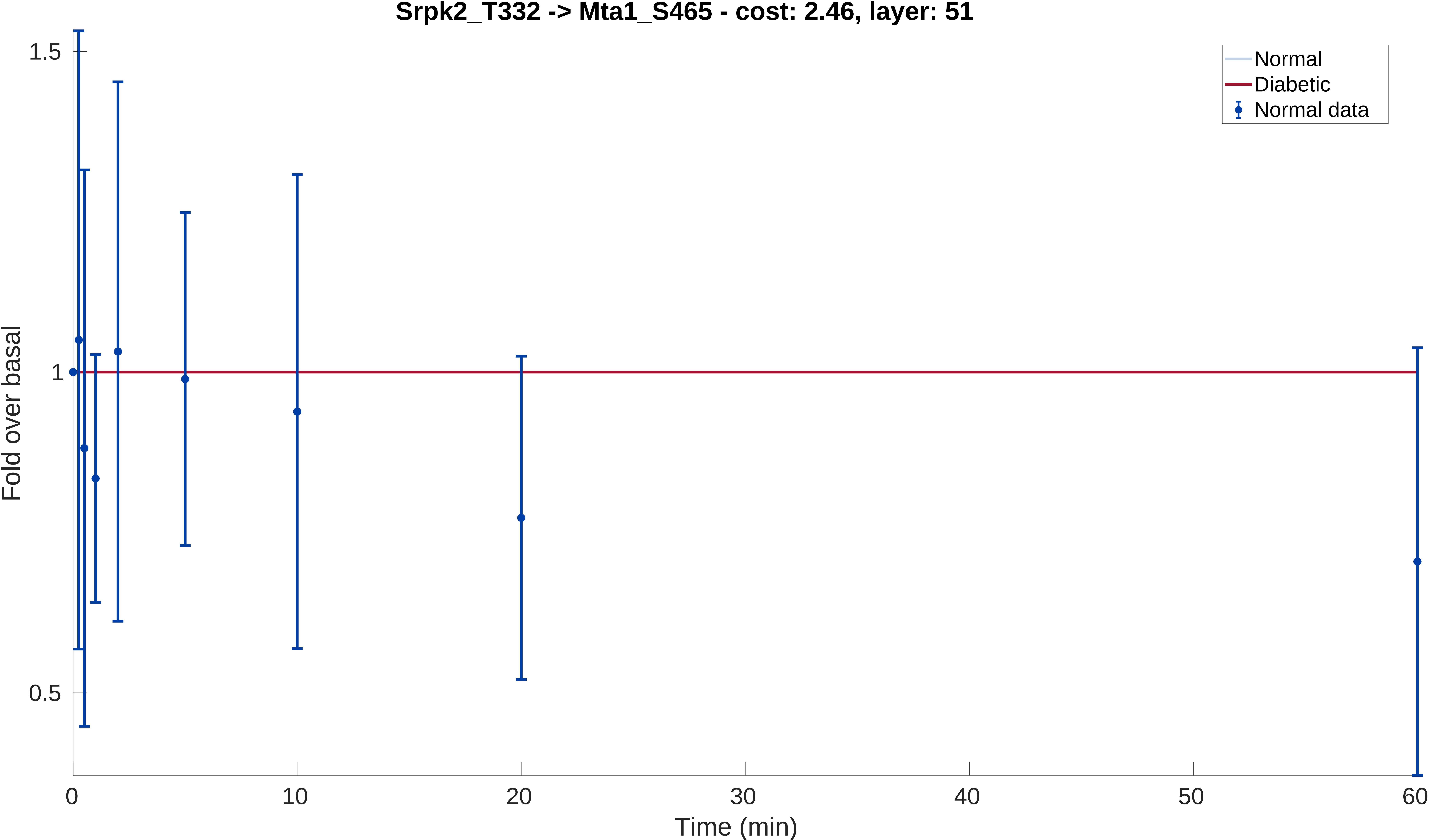

Srpk2_T332 -> Mta1_S465 - cost: 2.46, layer: 51
Normal
Diabetic
Normal data
1.5
1
0.5
Fold over basal
0
10
20
30
40
50
60
Time (min)

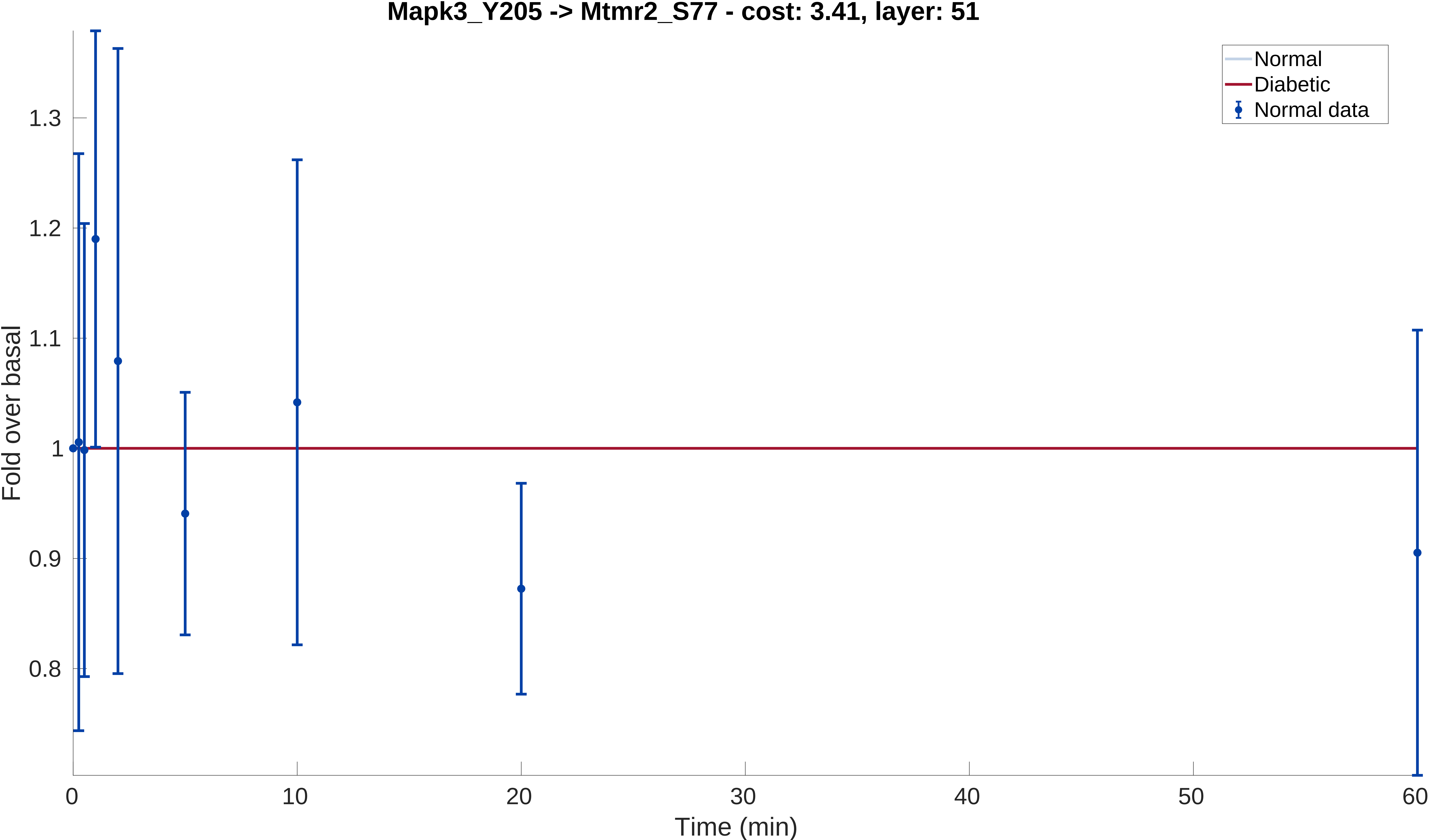
Mapk3_Y205 -> Mtmr2_S77 - cost: 3.41, layer: 51
Normal
Diabetic
Normal data
Fold over basal
1.3
1.2
1.1
1
0.9
0.8
0
10
20
30
40
50
60
Time (min)

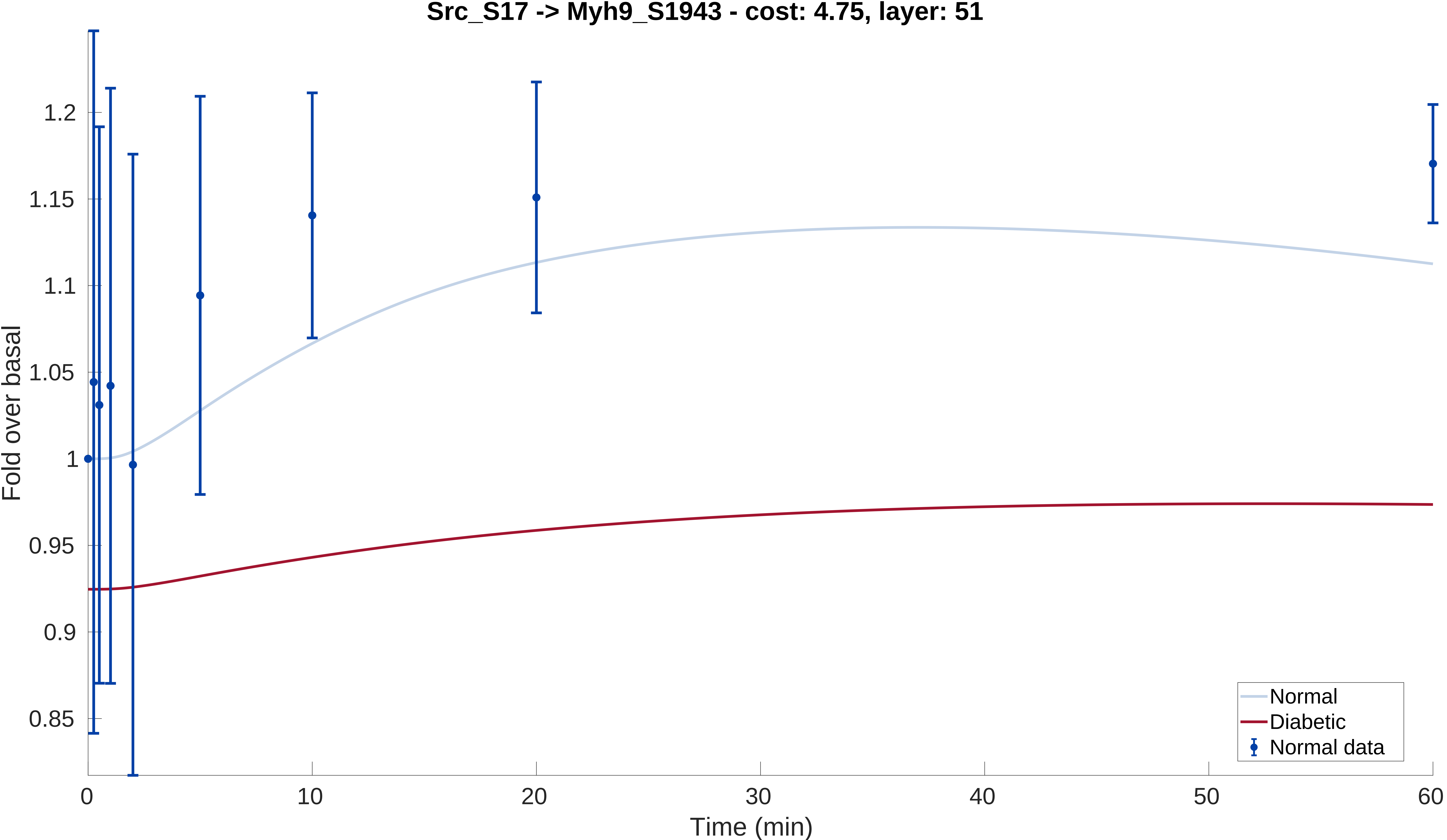

Src_S17 -> Myh9_S1943 - cost: 4.75, layer: 51
Fold over basal
1.2
1.15
1.1
1.05
1
0.95
0.9
0.85
0
10
20
30
40
50
60
Time (min)
Normal
Diabetic
Normal data

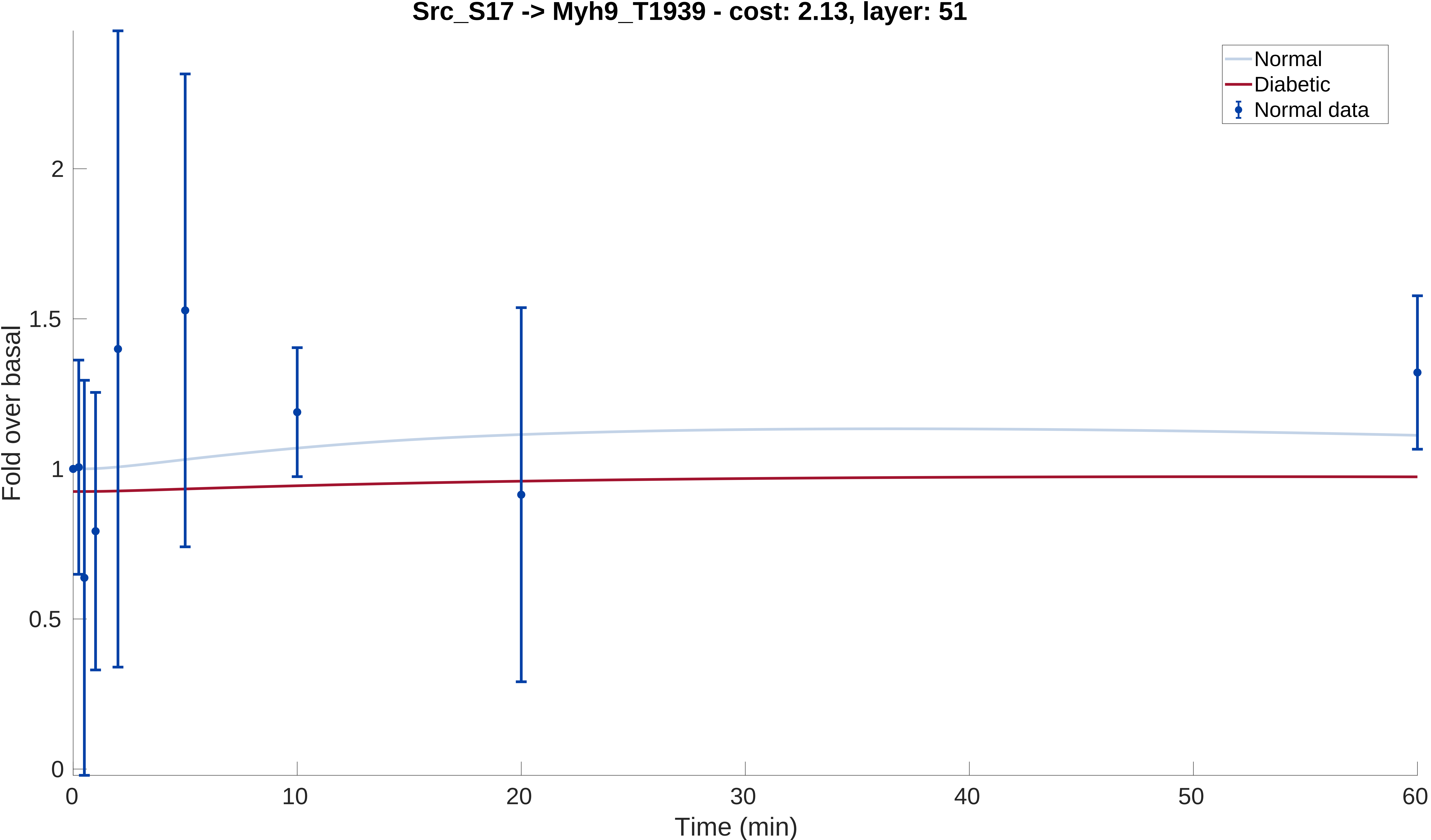
Src_S17 -> Myh9_T1939 - cost: 2.13, layer: 51
Normal
Diabetic
Normal data
Fold over basal
0
0.5
1
1.5
2
0
10
20
30
40
50
60
Time (min)

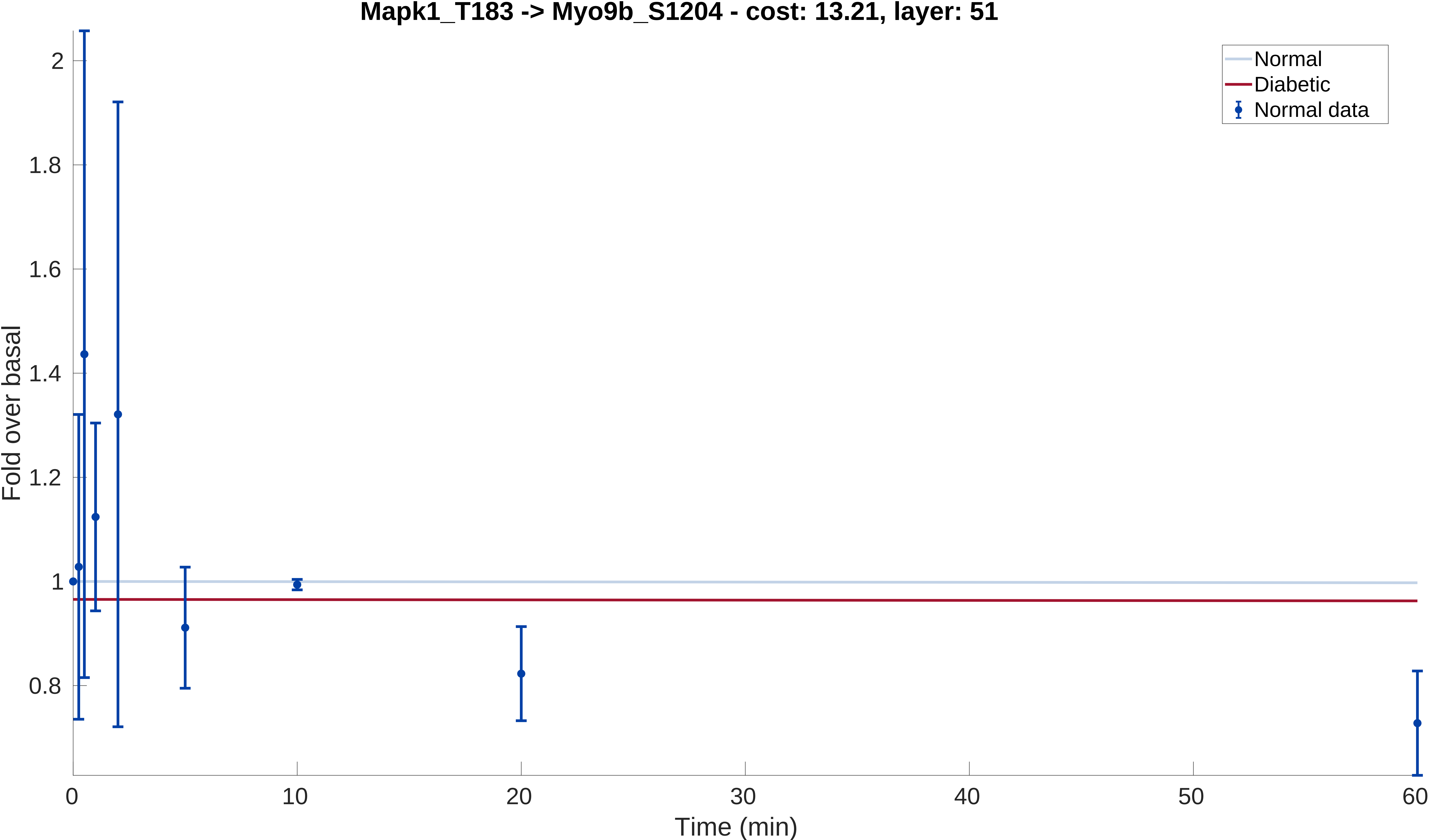

Mapk1_T183 -> Myo9b_S1204 - cost: 13.21, layer: 51
Fold over basal
2
1.8
1.6
1.4
1.2
1
0.8
0
10
20
30
40
50
60
Time (min)
Normal
Diabetic
Normal data

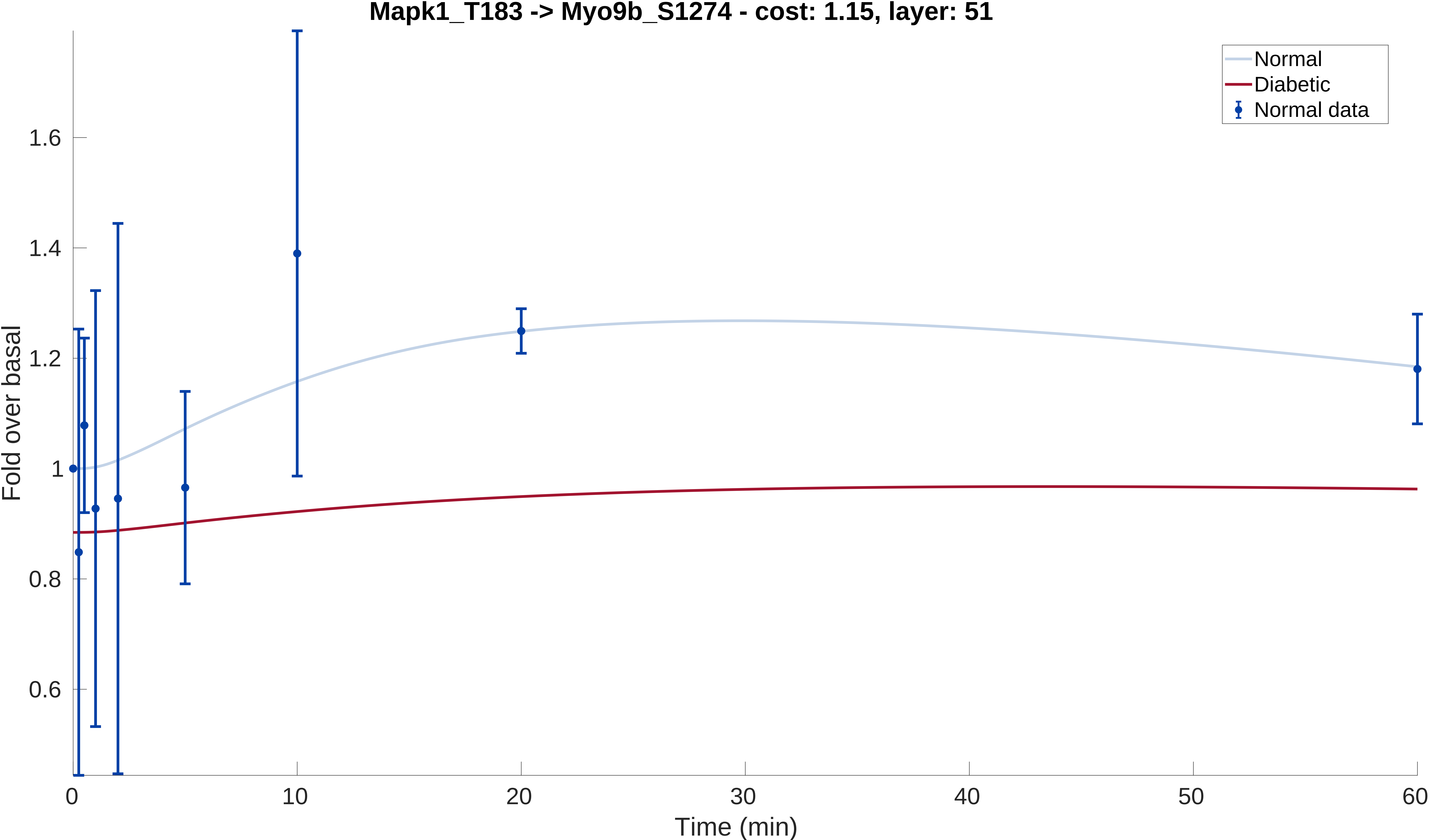

Mapk1_T183 -> Myo9b_S1274 - cost: 1.15, layer: 51
Normal
Diabetic
Normal data
Fold over basal
1.6
1.4
1.2
1
0.8
0.6
0
10
20
30
40
50
60
Time (min)

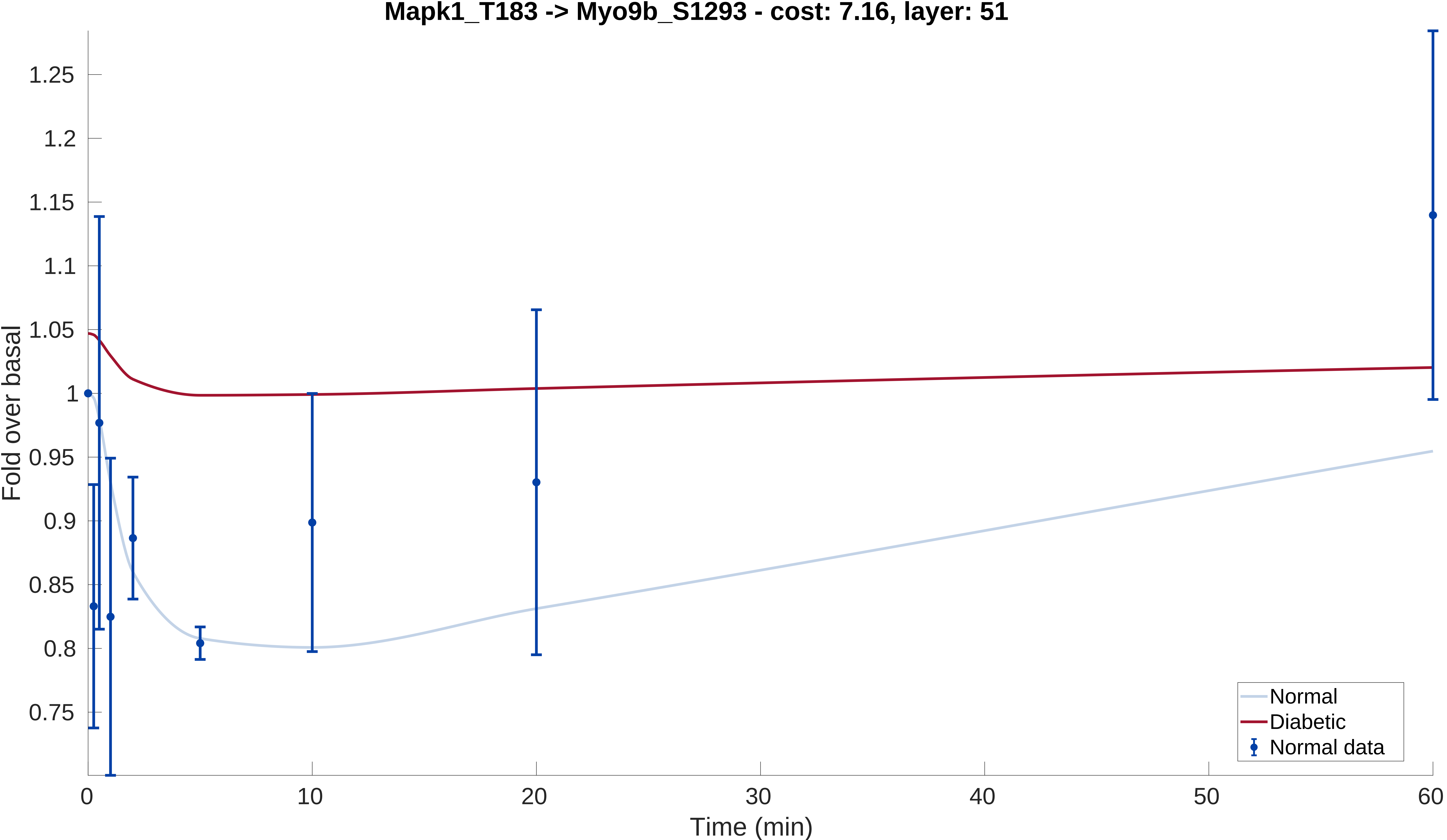

Mapk1_T183 -> Myo9b_S1293 - cost: 7.16, layer: 51
Fold over basal
1.25
1.2
1.15
1.1
1.05
1
0.95
0.9
0.85
0.8
0.75
0
10
20
30
40
50
60
Time (min)
Normal
Diabetic
Normal data

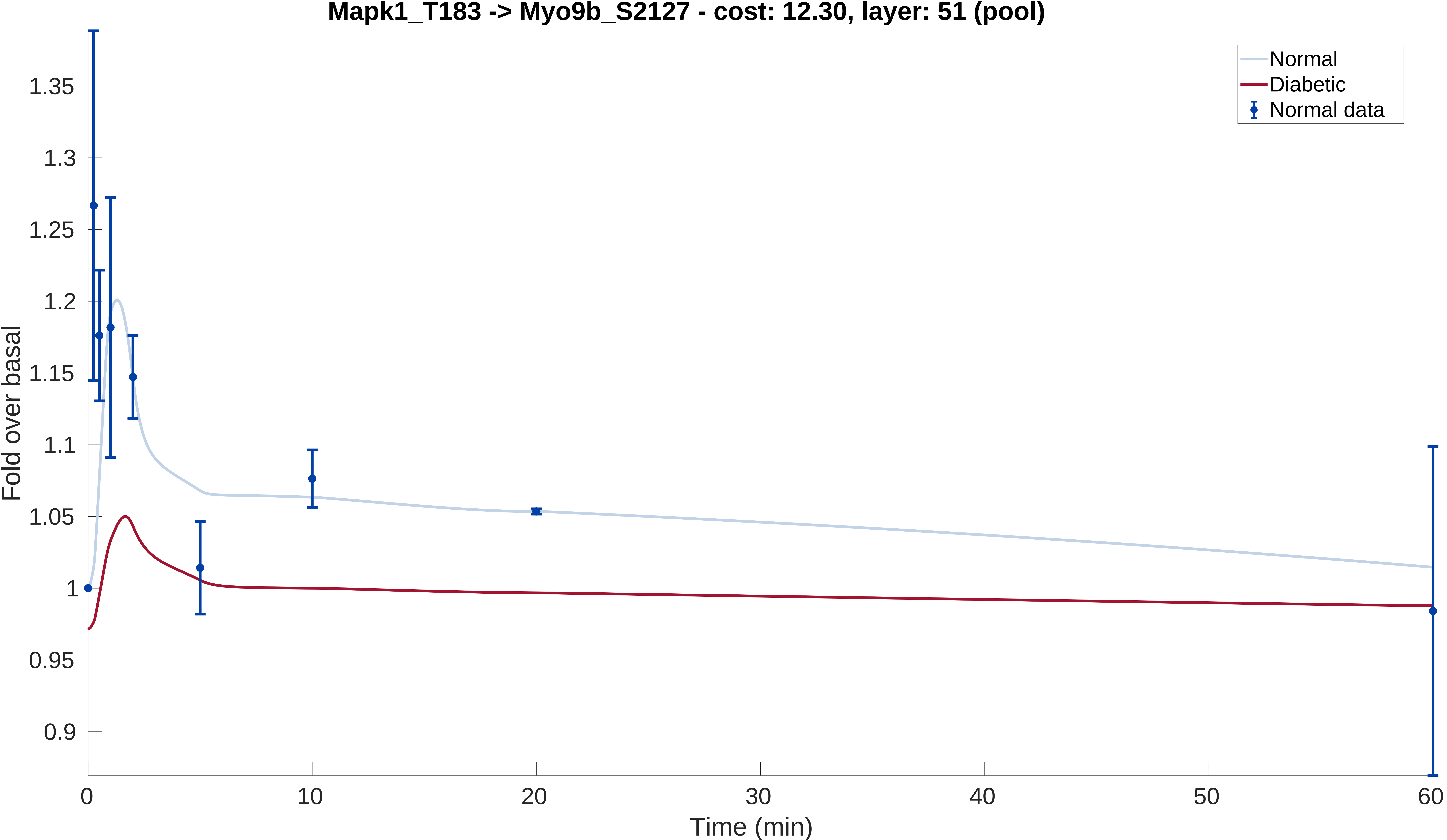

Mapk1_T183 -> Myo9b_S2127 - cost: 12.30, layer: 51 (pool)
Normal
Diabetic
Normal data
Fold over basal
Time (min)
1.35
1.3
1.25
1.2
1.15
1.1
1.05
1
0.95
0.9
0
10
20
30
40
50
60

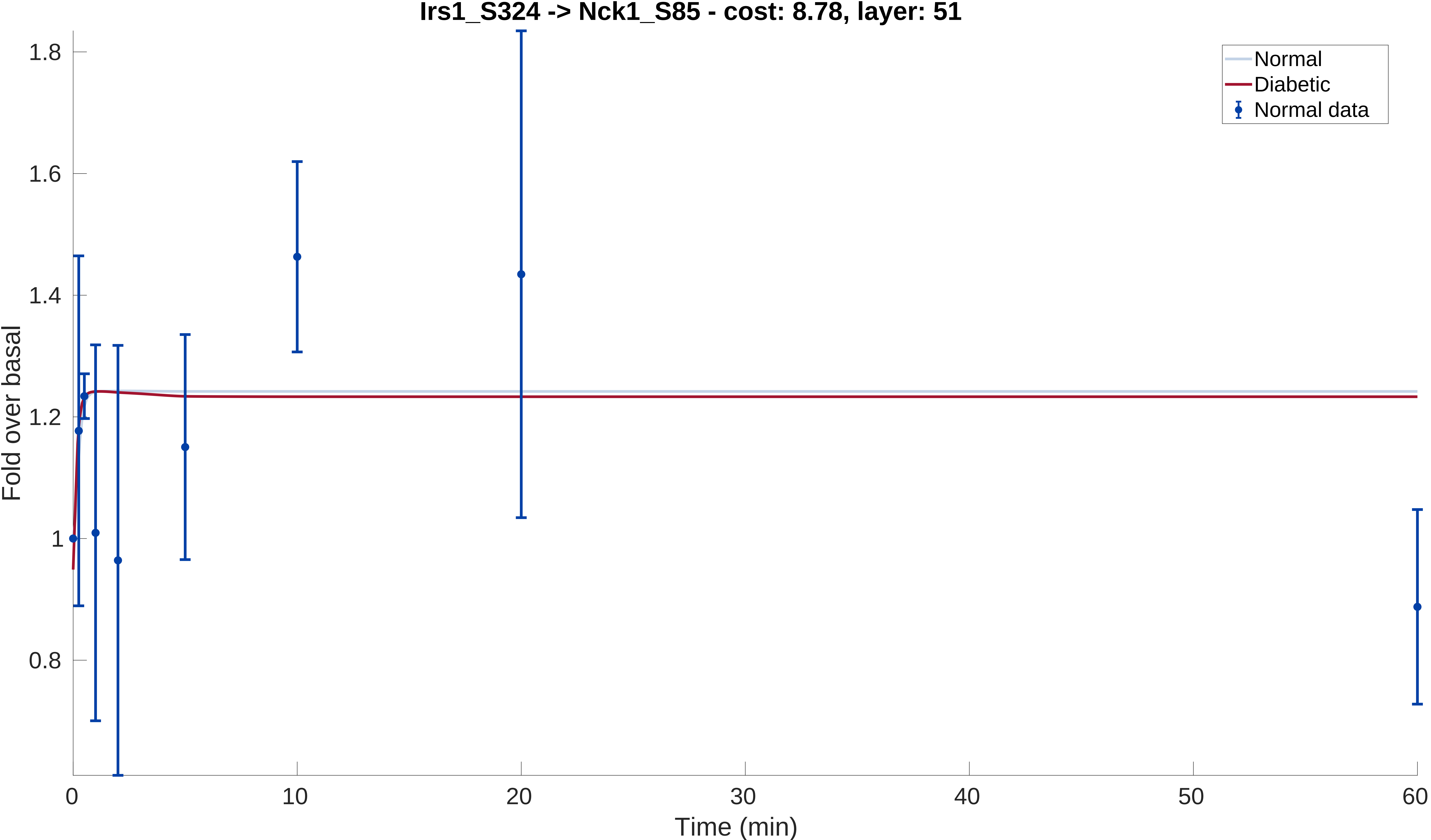

Irs1_S324 -> Nck1_S85 - cost: 8.78, layer: 51
Normal
Diabetic
Normal data
1.8
1.6
1.4
1.2
1
0.8
Fold over basal
0
10
20
30
40
50
60
Time (min)

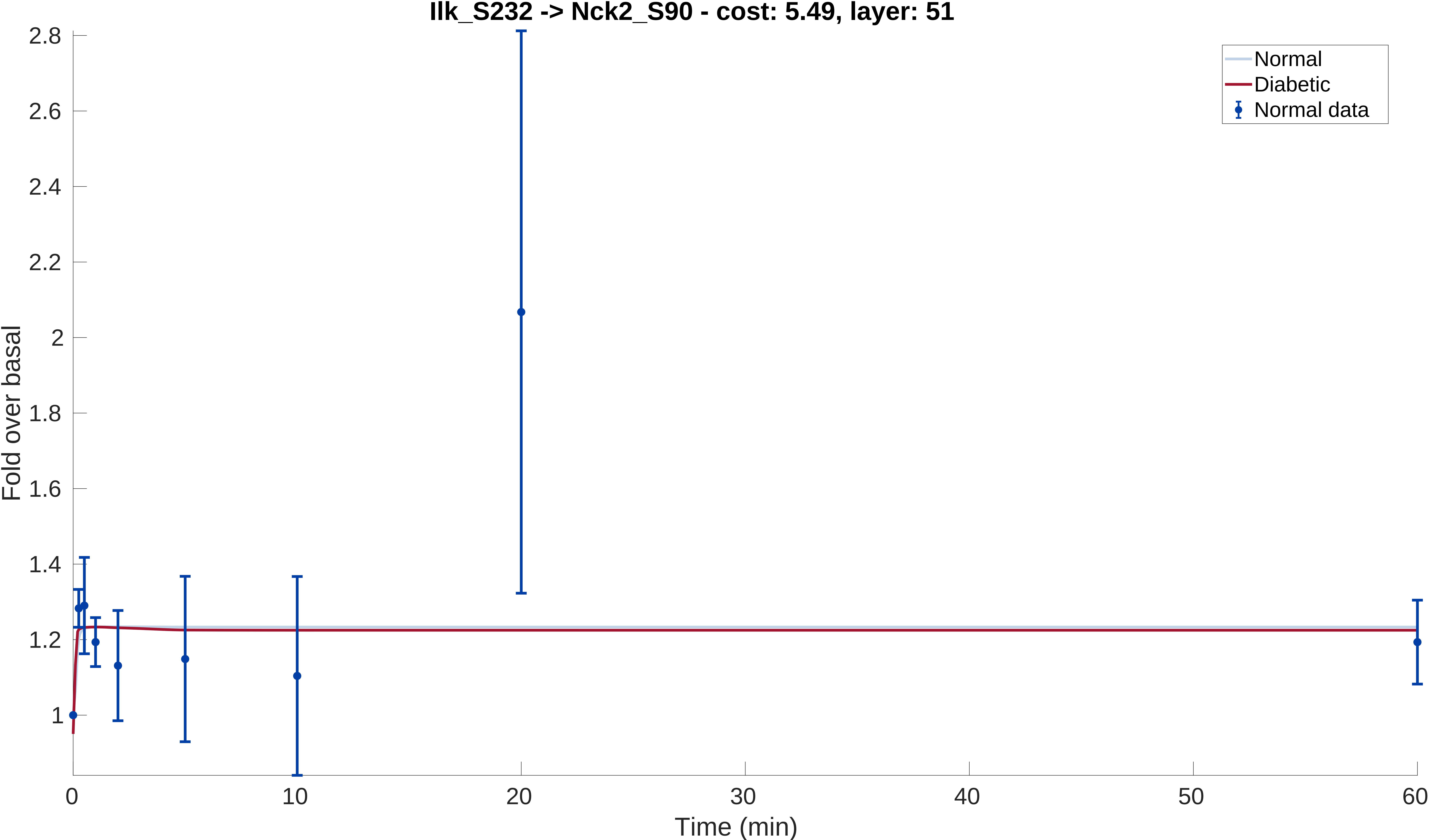

Ilk_S232 -> Nck2_S90 - cost: 5.49, layer: 51
Normal
Diabetic
Normal data
Fold over basal
2.8
2.6
2.4
2.2
2
1.8
1.6
1.4
1.2
1
0
10
20
30
40
50
60
Time (min)

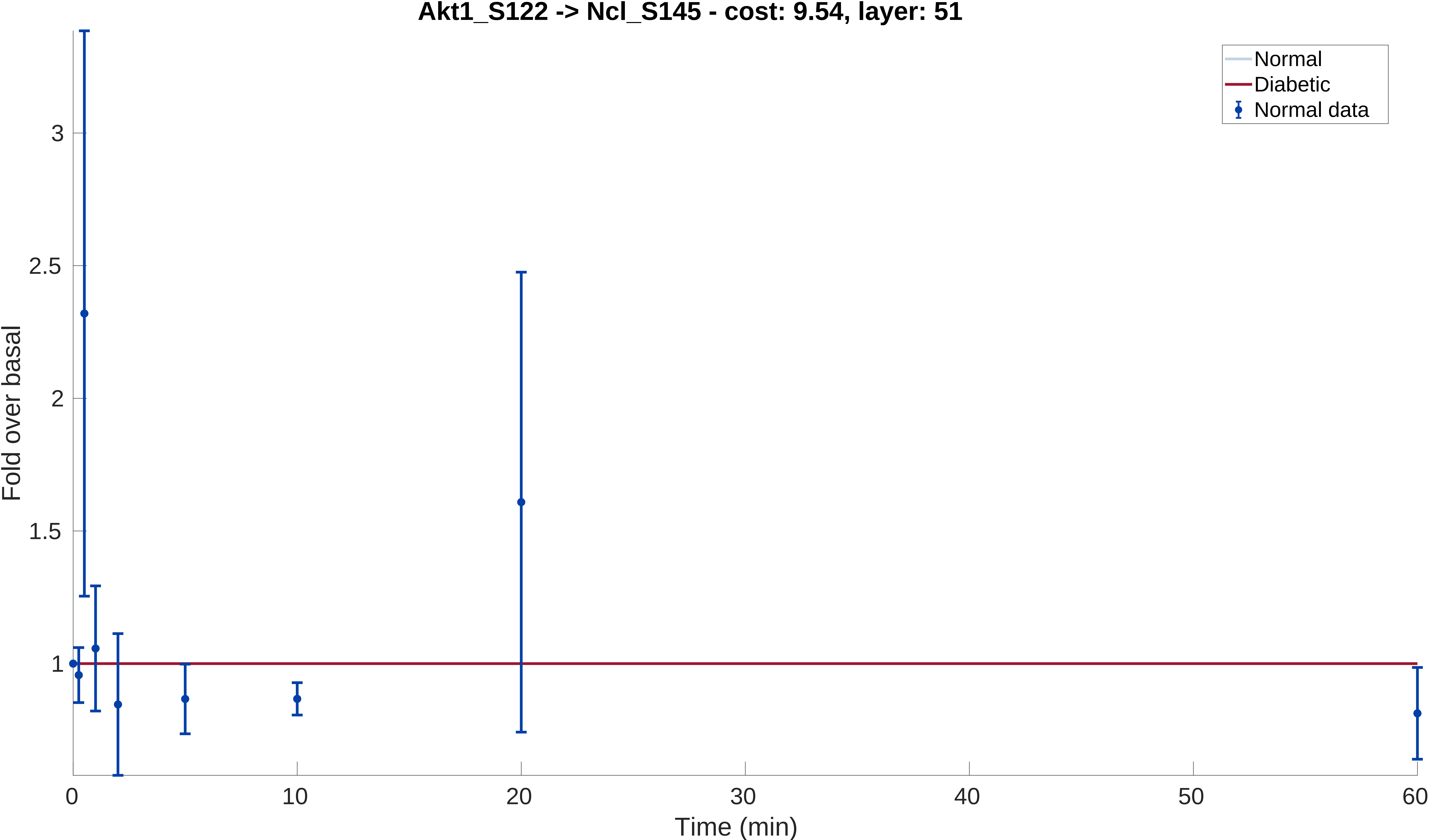

Akt1_S122 -> Ncl_S145 - cost: 9.54, layer: 51
Normal
Diabetic
Normal data
Fold over basal
3
2.5
2
1.5
1
0
10
20
30
40
50
60
Time (min)

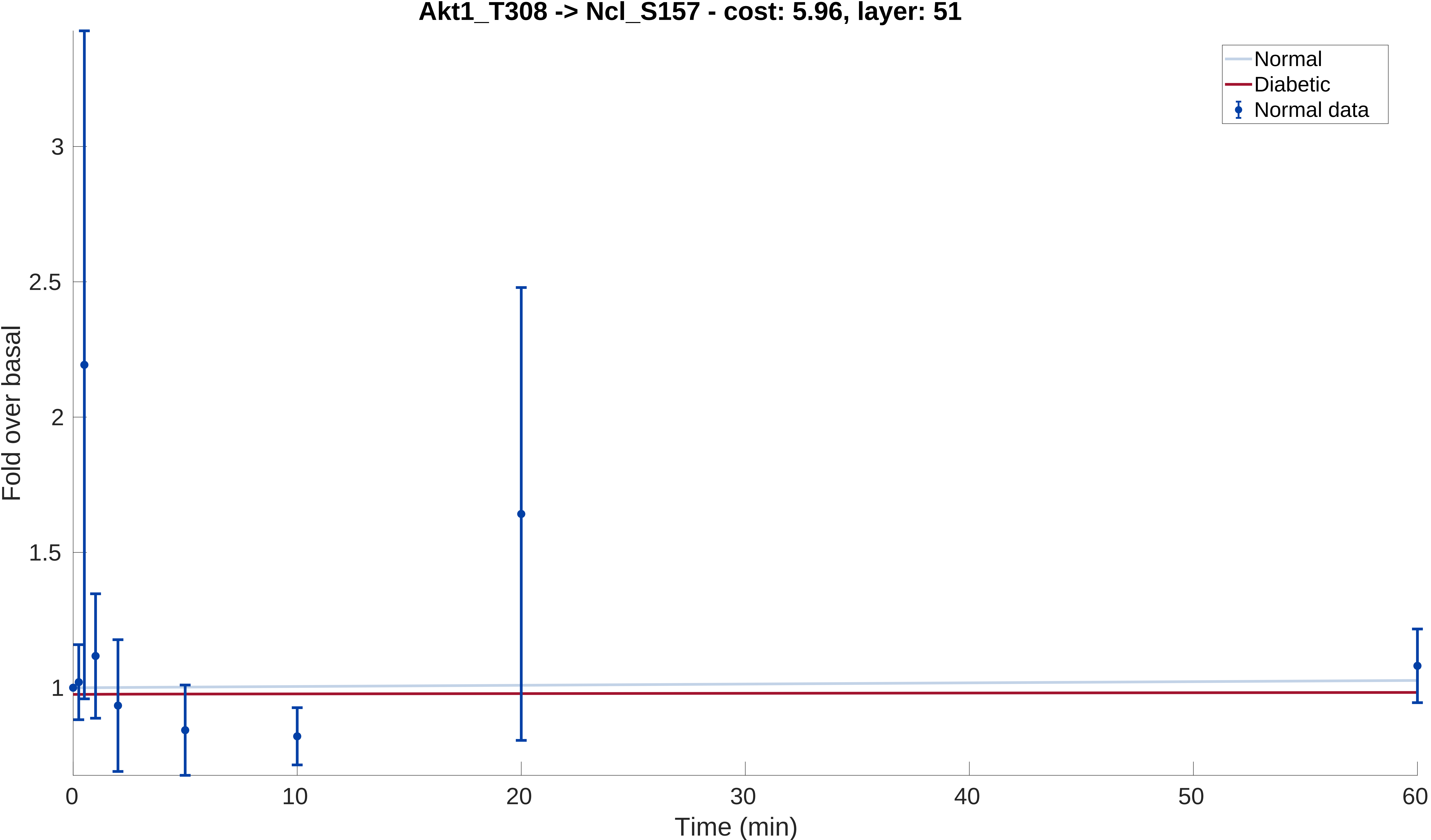
Akt1_T308 -> Ncl_S157 - cost: 5.96, layer: 51
Normal
Diabetic
Normal data
Fold over basal
3
2.5
2
1.5
1
0
10
20
30
40
50
60
Time (min)

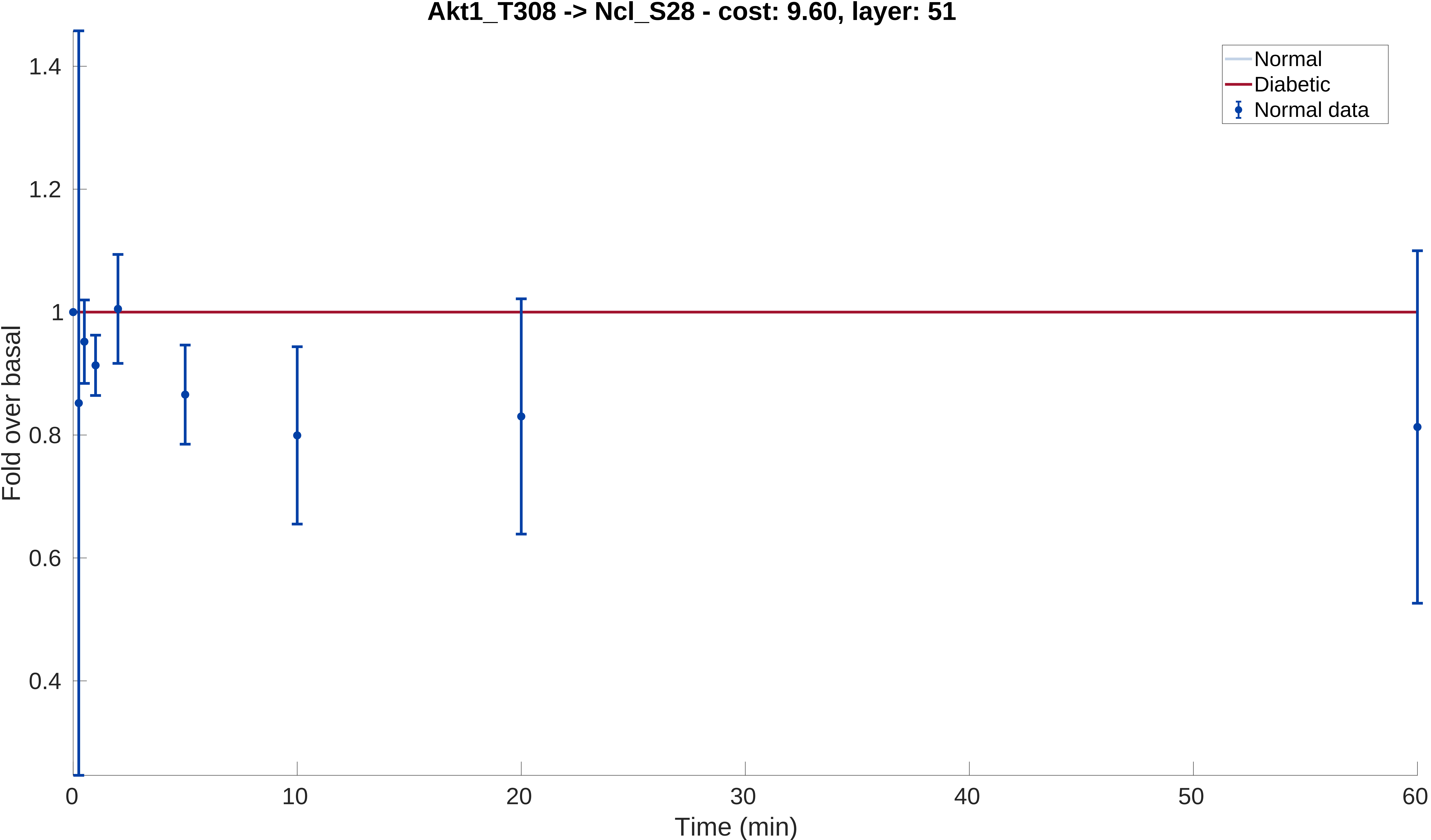
Akt1_T308 -> Ncl_S28 - cost: 9.60, layer: 51
Normal
Diabetic
Normal data
Fold over basal
1.4
1.2
1
0.8
0.6
0.4
0
10
20
30
40
50
60
Time (min)

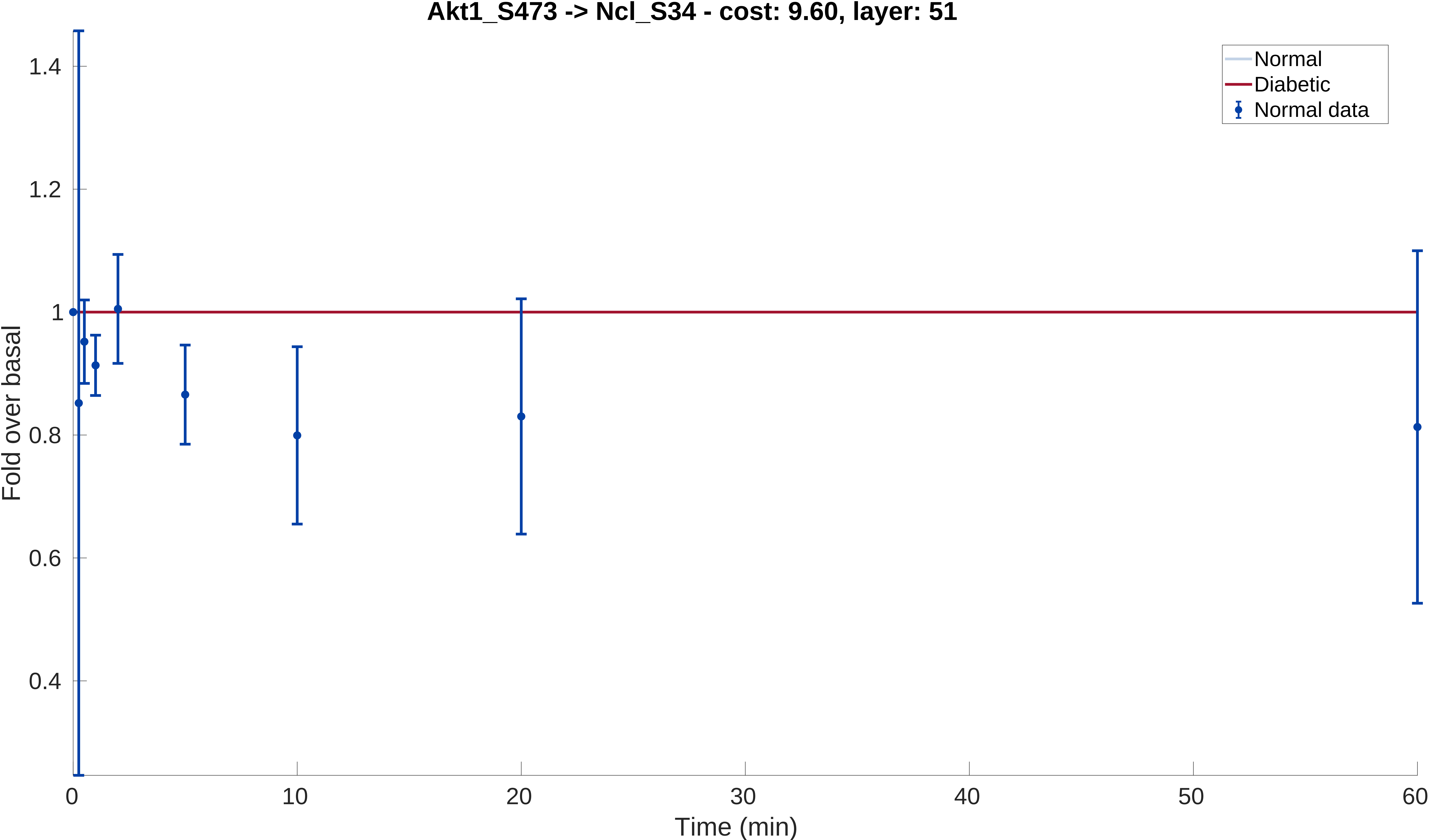

Akt1_S473 -> Ncl_S34 - cost: 9.60, layer: 51
Normal
Diabetic
Normal data
Fold over basal
1.4
1.2
1
0.8
0.6
0.4
0
10
20
30
40
50
60
Time (min)

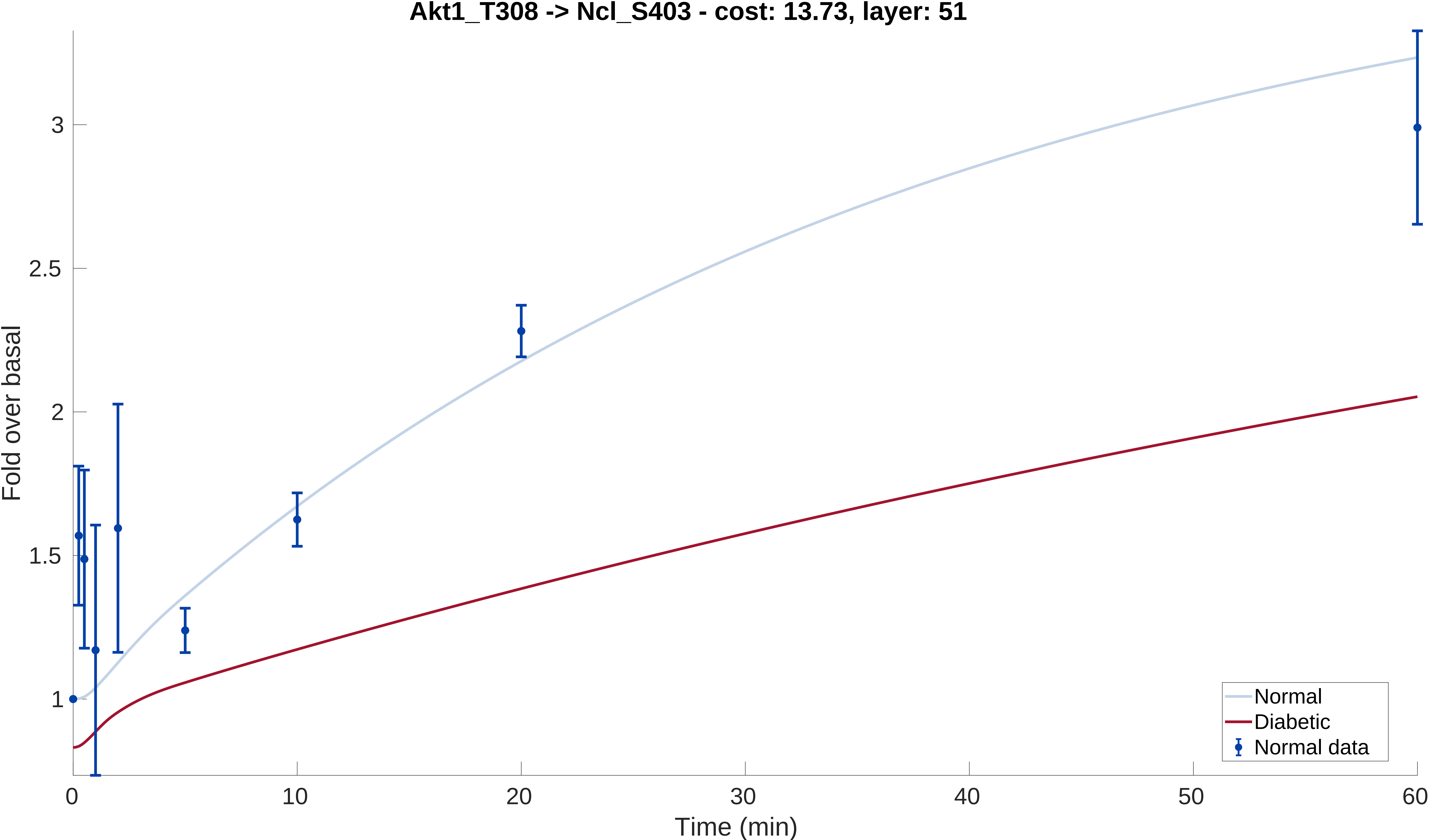
Akt1_T308 -> Ncl_S403 - cost: 13.73, layer: 51
Fold over basal
Time (min)
0
10
20
30
40
50
60
1
1.5
2
2.5
3
Normal
Diabetic
Normal data

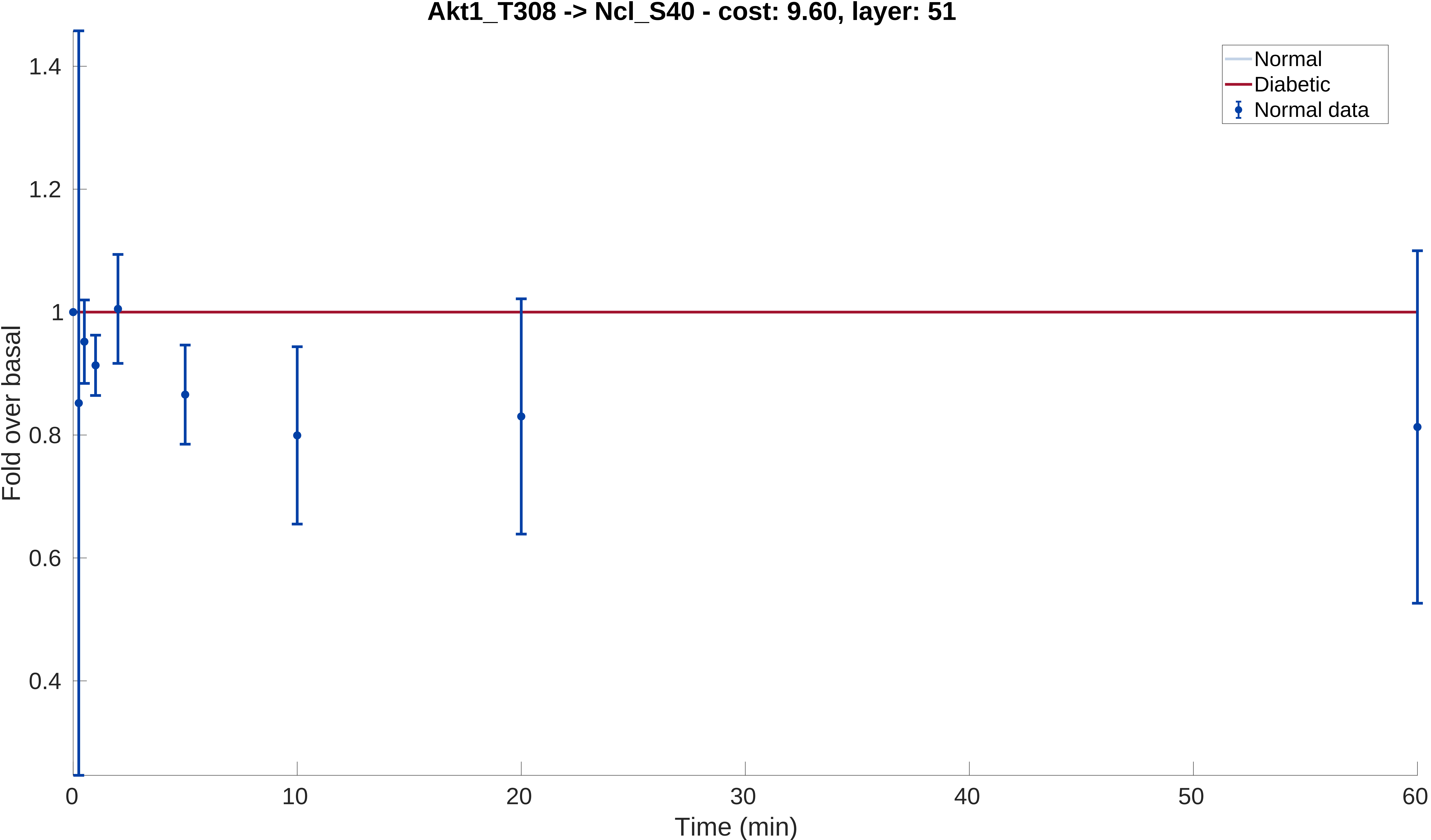
Akt1_T308 -> Ncl_S40 - cost: 9.60, layer: 51
Normal
Diabetic
Normal data
Fold over basal
1.4
1.2
1
0.8
0.6
0.4
0
10
20
30
40
50
60
Time (min)

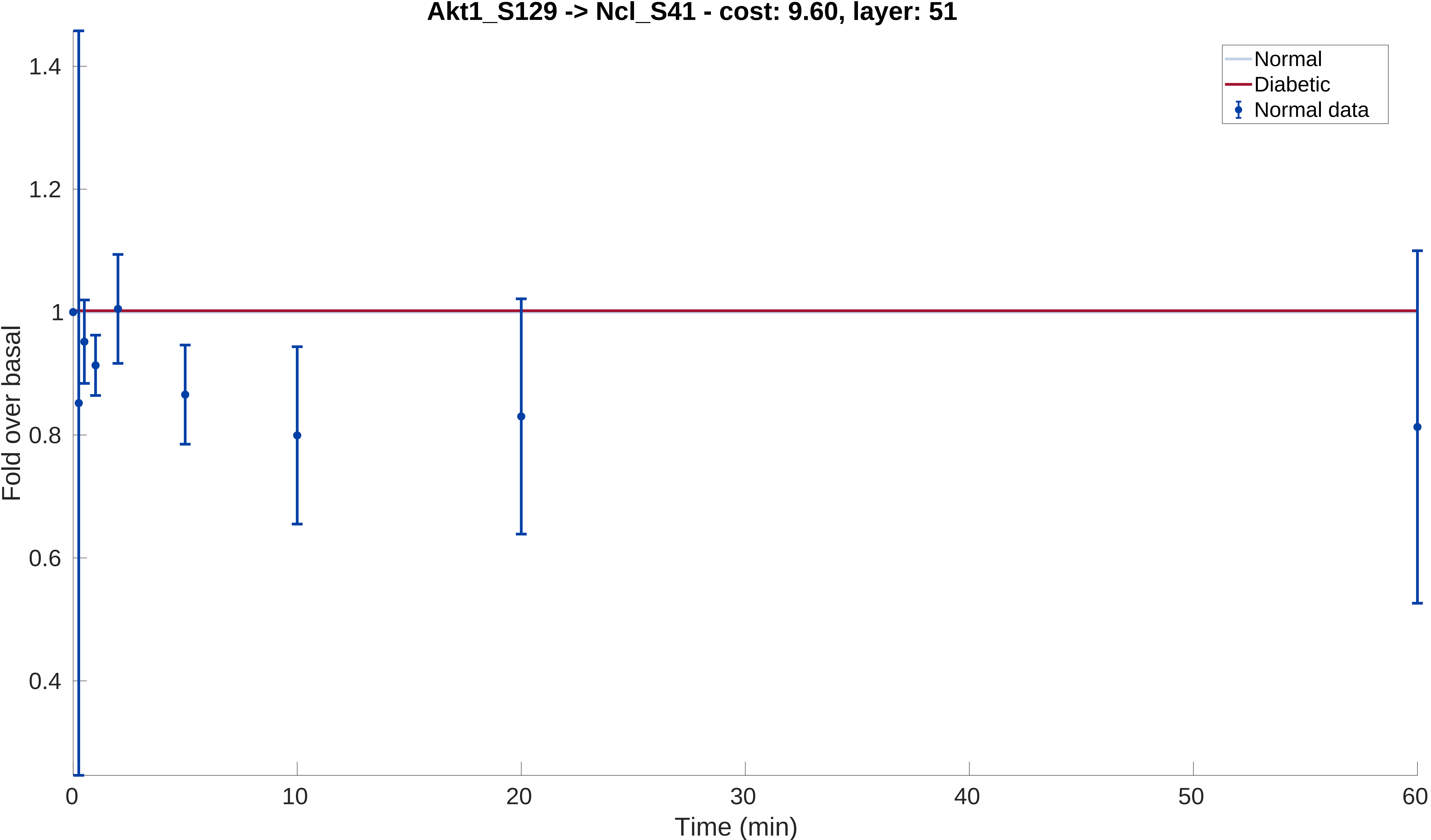

Akt1_S129 -> Ncl_S41 - cost: 9.60, layer: 51
Normal
Diabetic
Normal data
Fold over basal
1.4
1.2
1
0.8
0.6
0.4
0
10
20
30
40
50
60
Time (min)

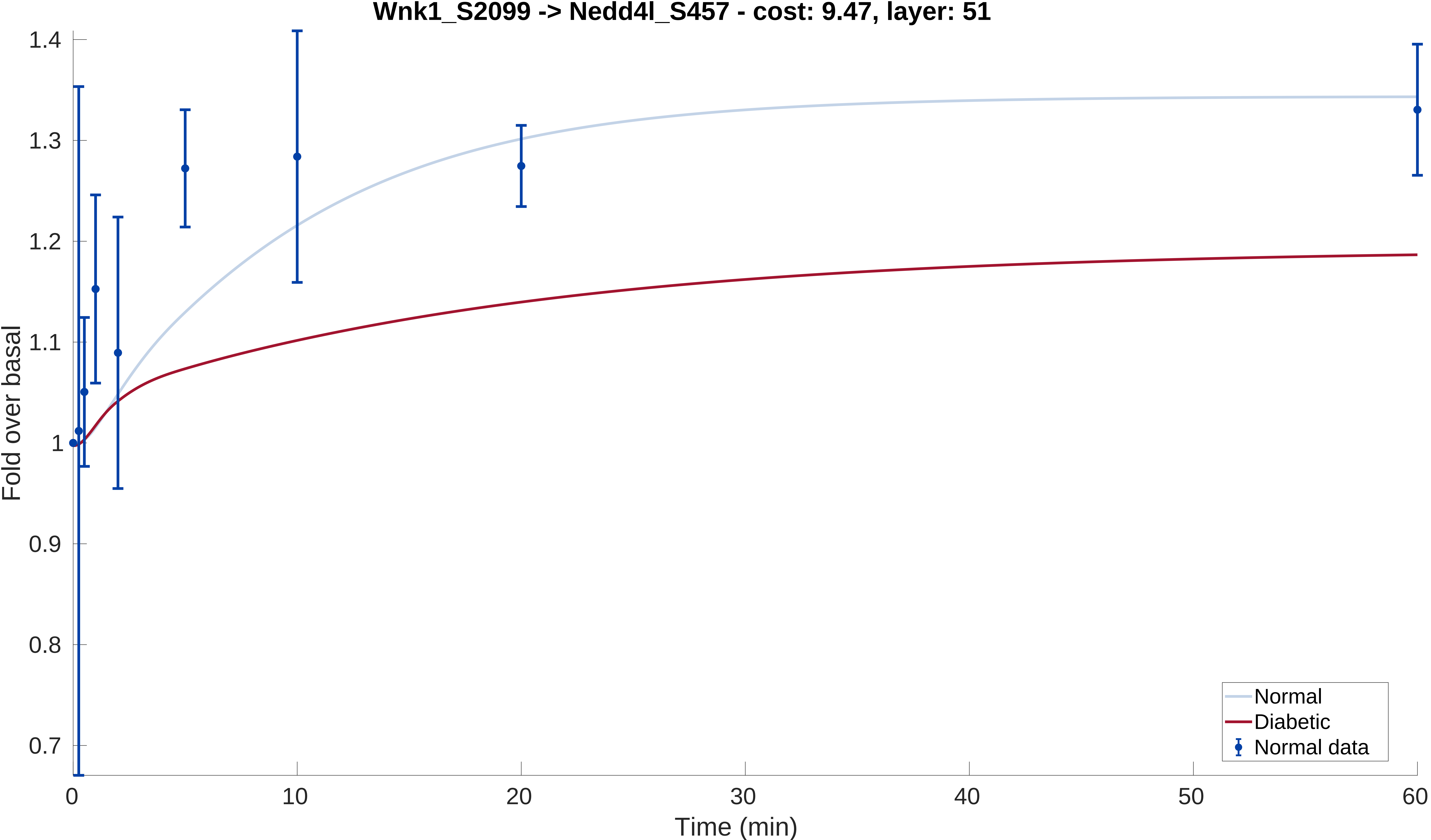
Wnk1_S2099 -> Nedd4l_S457 - cost: 9.47, layer: 51
Fold over basal
1.4
1.3
1.2
1.1
1
0.9
0.8
0.7
0
10
20
30
40
50
60
Time (min)
Normal
Diabetic
Normal data

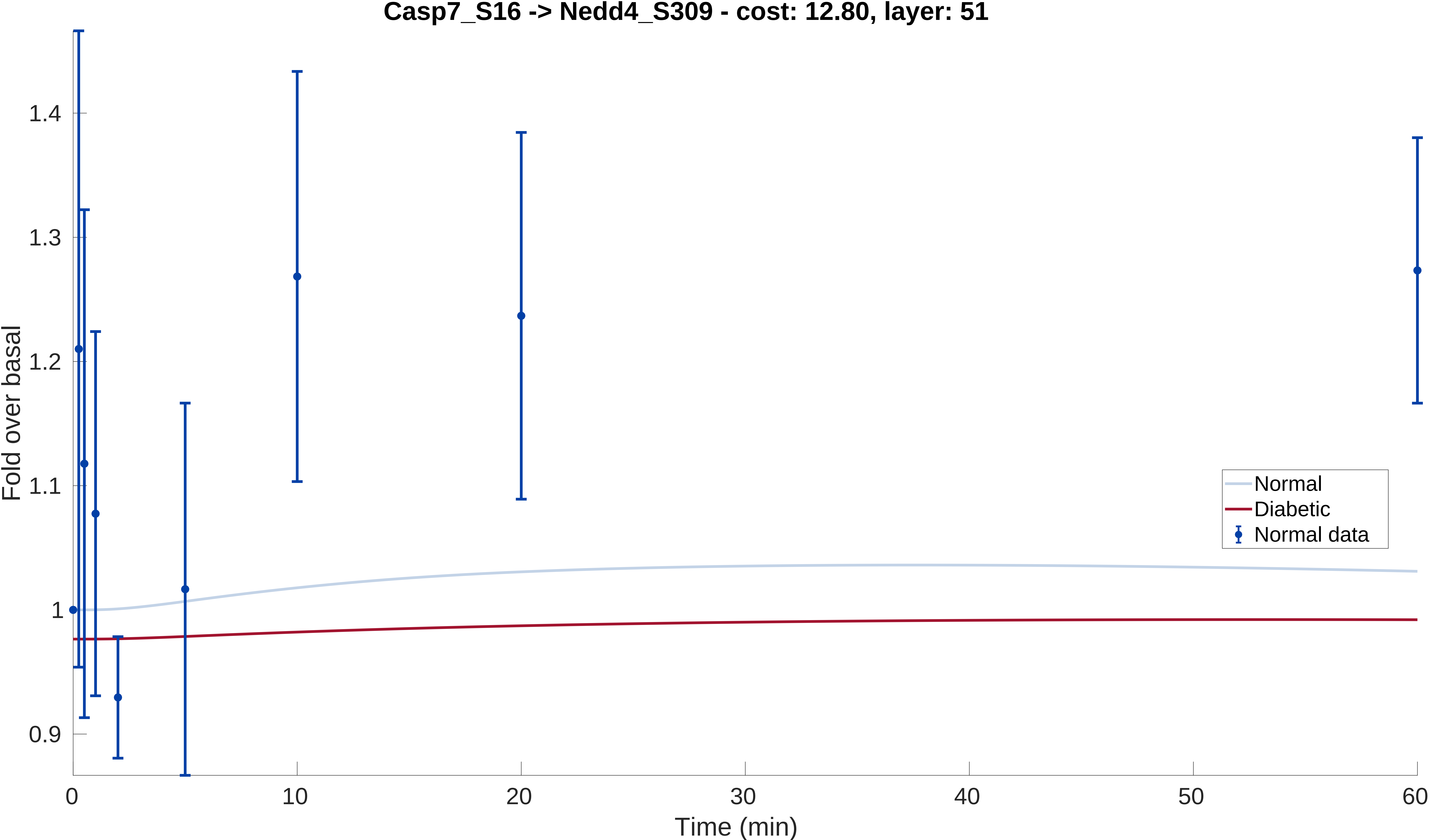
Casp7_S16 -> Nedd4_S309 - cost: 12.80, layer: 51
Fold over basal
1.4
1.3
1.2
1.1
1
0.9
0
10
20
30
40
50
60
Time (min)
Normal
Diabetic
Normal data

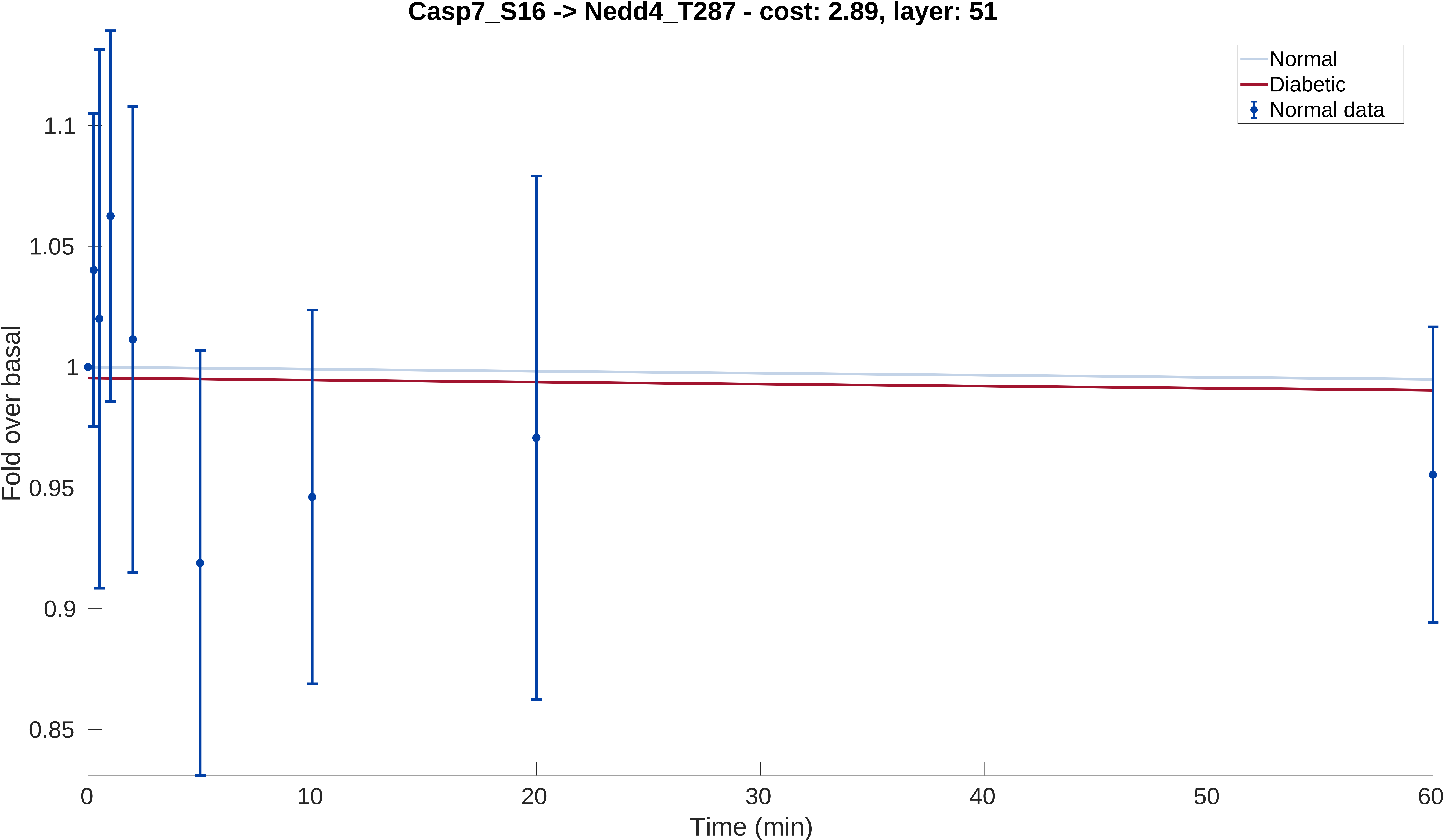

Casp7_S16 -> Nedd4_T287 - cost: 2.89, layer: 51
Normal
Diabetic
Normal data
Fold over basal
1.1
1.05
1
0.95
0.9
0.85
0
10
20
30
40
50
60
Time (min)

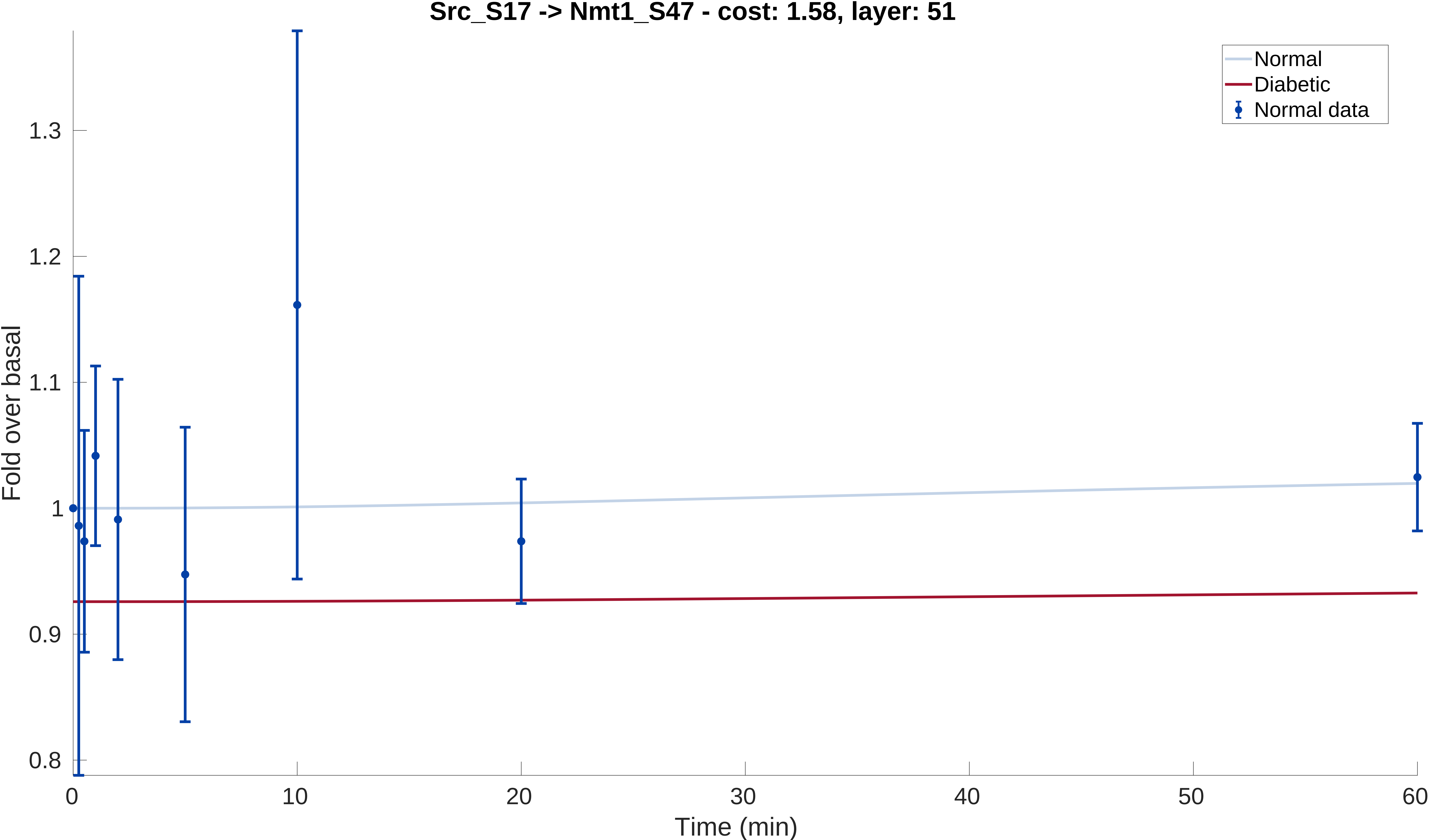

Src_S17 -> Nmt1_S47 - cost: 1.58, layer: 51
Normal
Diabetic
Normal data
Fold over basal
1.3
1.2
1.1
1
0.9
0.8
0
10
20
30
40
50
60
Time (min)

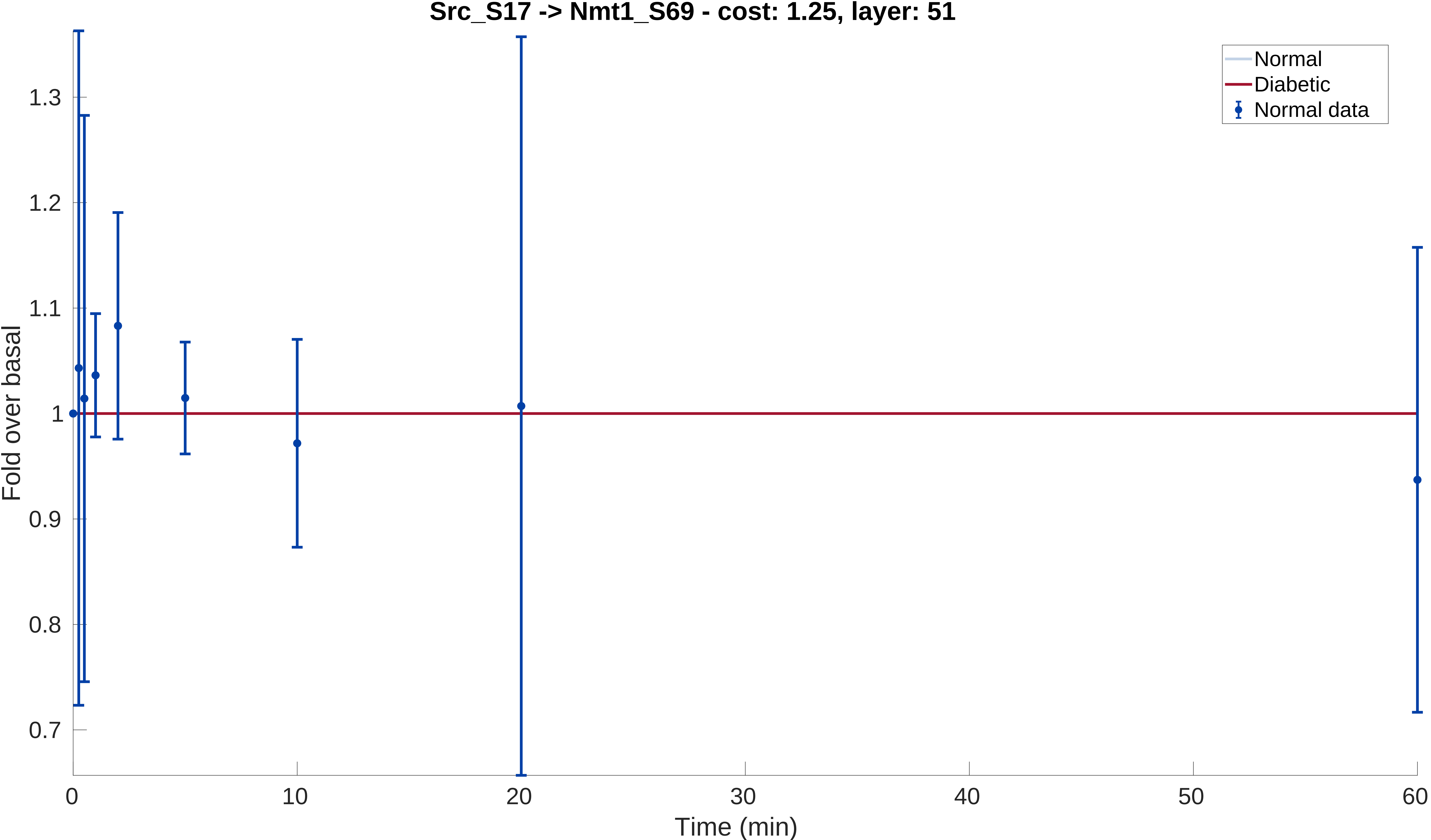

Src_S17 -> Nmt1_S69 - cost: 1.25, layer: 51
Normal
Diabetic
Normal data
Fold over basal
1.3
1.2
1.1
1
0.9
0.8
0.7
0
10
20
30
40
50
60
Time (min)

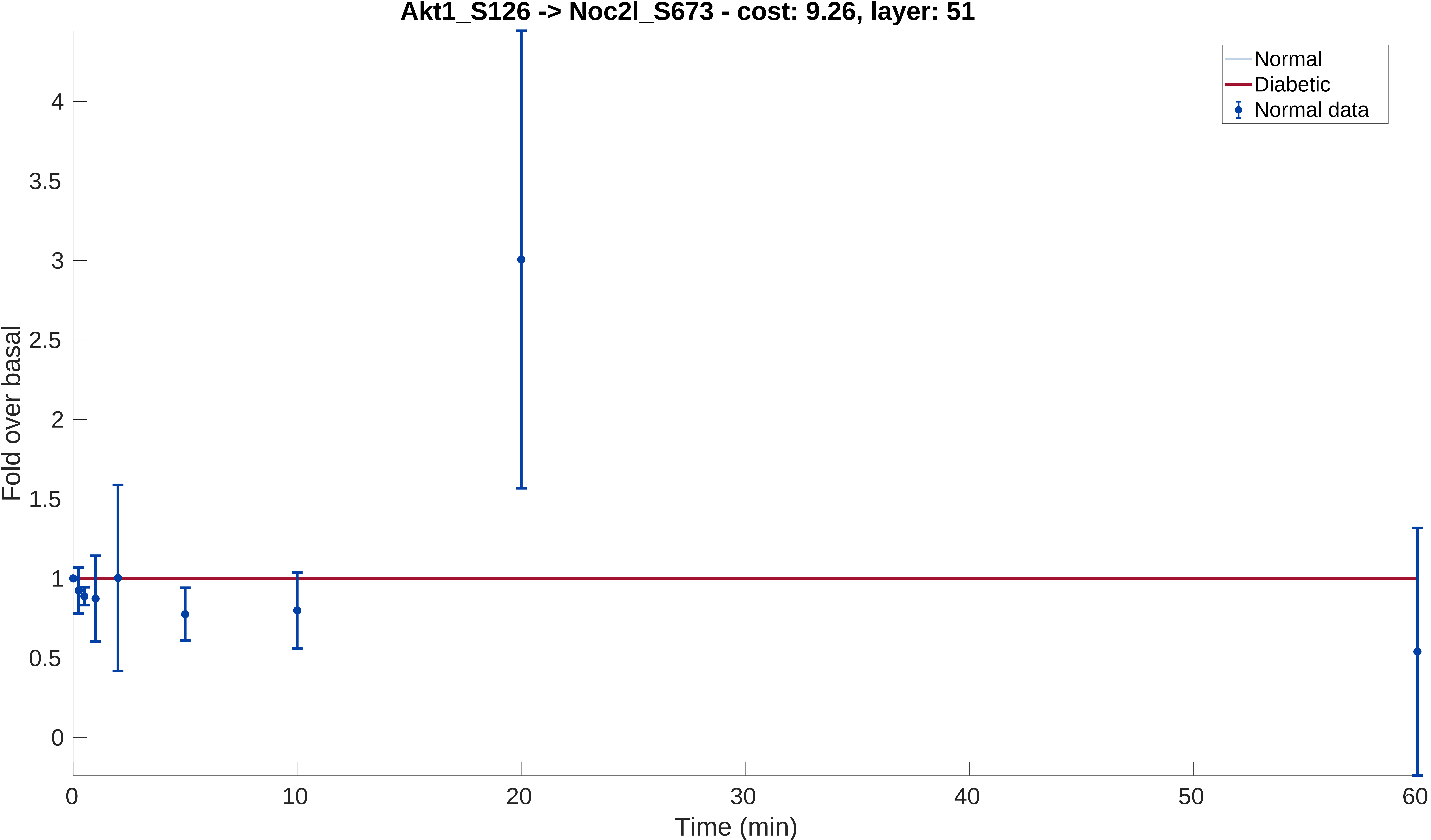

Akt1_S126 -> Noc2l_S673 - cost: 9.26, layer: 51
Normal
Diabetic
Normal data
Fold over basal
4
3.5
3
2.5
2
1.5
1
0.5
0
0
10
20
30
40
50
60
Time (min)

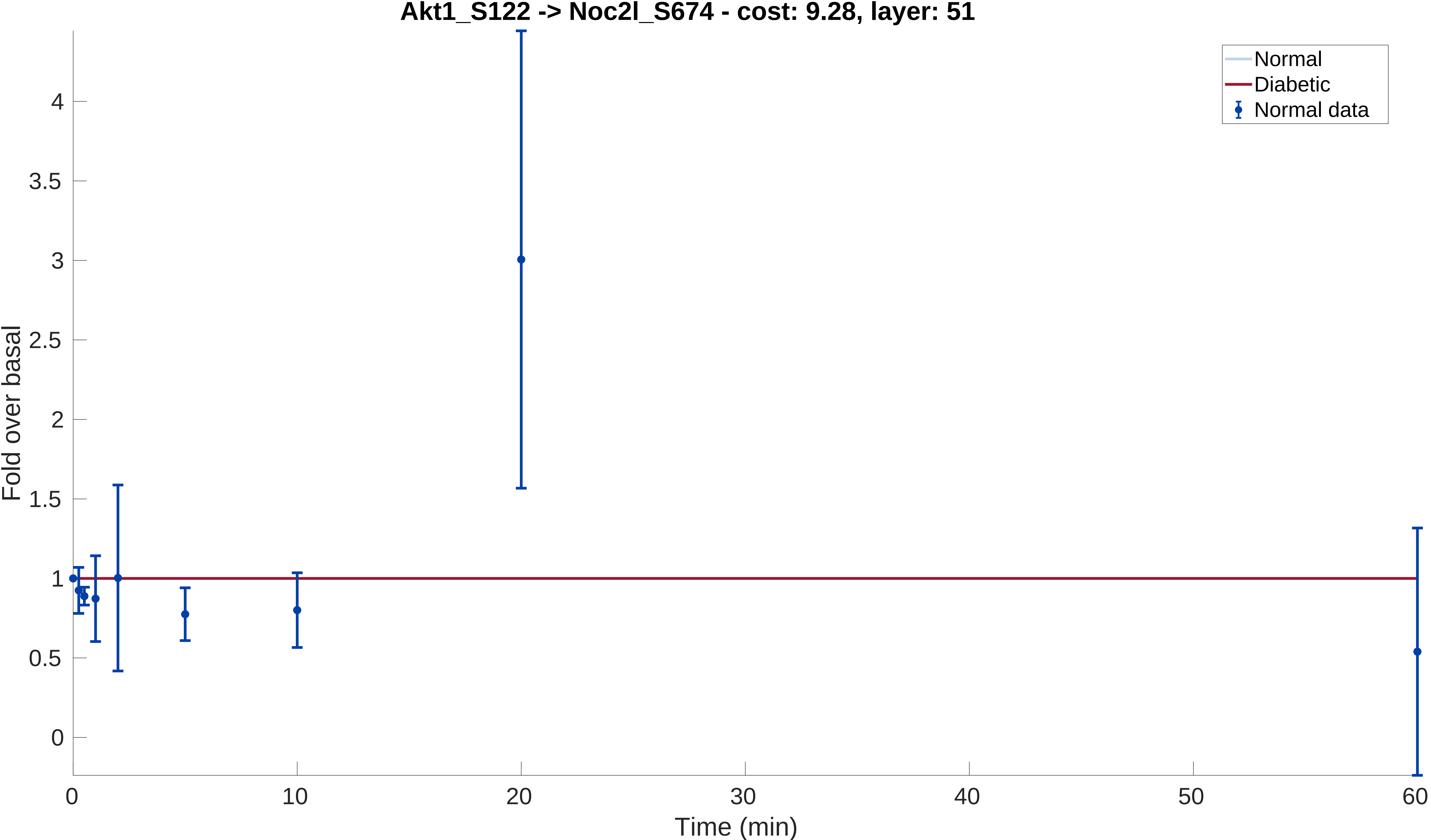

Akt1_S122 -> Noc2l_S674 - cost: 9.28, layer: 51
Normal
Diabetic
Normal data
Fold over basal
0
0.5
1
1.5
2
2.5
3
3.5
4
0
10
20
30
40
50
60
Time (min)

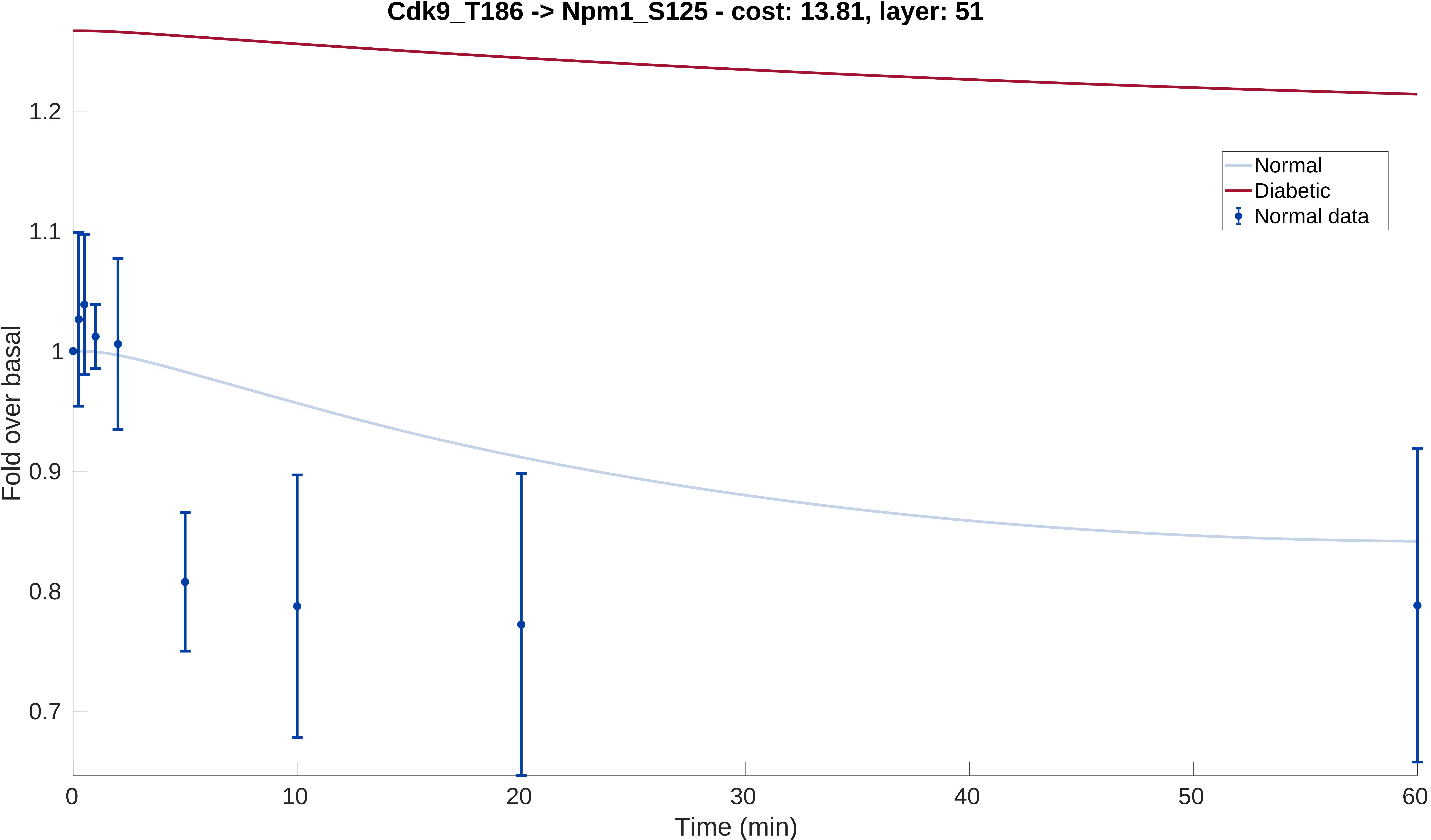
Cdk9_T186 -> Npm1_S125 - cost: 13.81, layer: 51
Fold over basal
Time (min)
Normal
Diabetic
Normal data
1.2
1.1
1
0.9
0.8
0.7
0
10
20
30
40
50
60

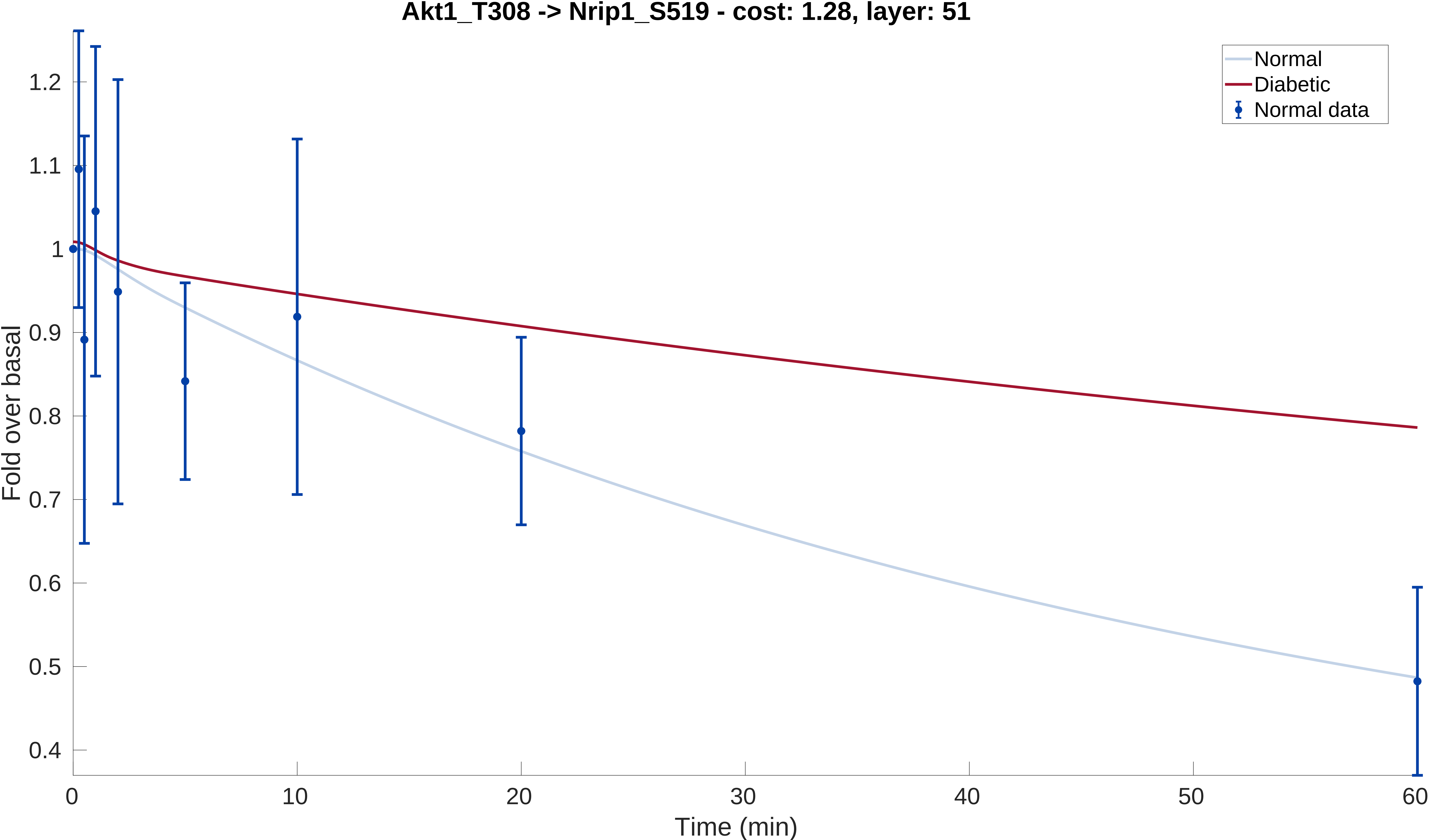

Akt1_T308 -> Nrip1_S519 - cost: 1.28, layer: 51
Normal
Diabetic
Normal data
1.2
1.1
1
0.9
0.8
0.7
0.6
0.5
0.4
Fold over basal
0
10
20
30
40
50
60
Time (min)

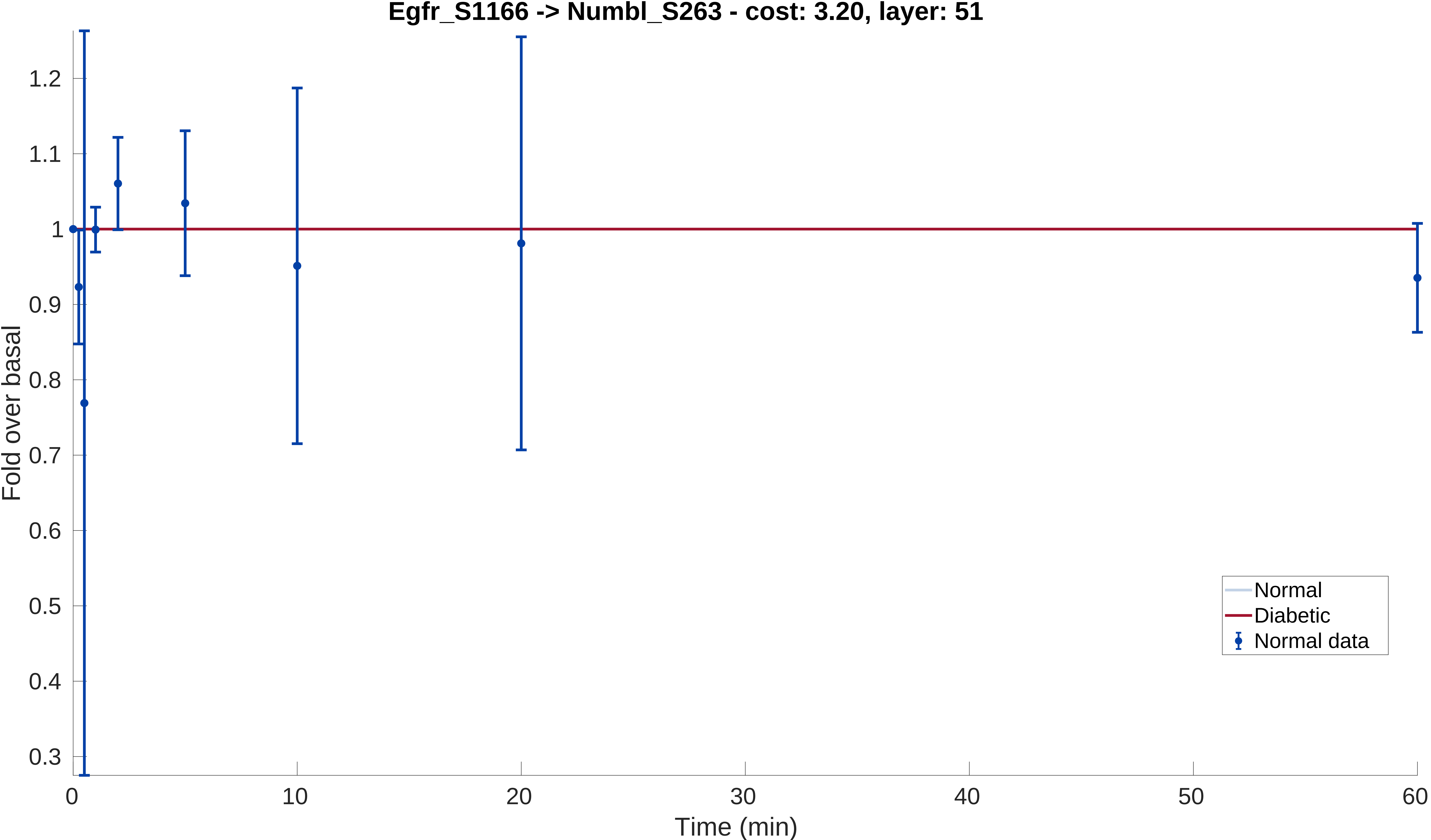

Egfr_S1166 -> Numbl_S263 - cost: 3.20, layer: 51
Fold over basal
1.2
1.1
1
0.9
0.8
0.7
0.6
0.5
0.4
0.3
0
10
20
30
40
50
60
Time (min)
Normal
Diabetic
Normal data

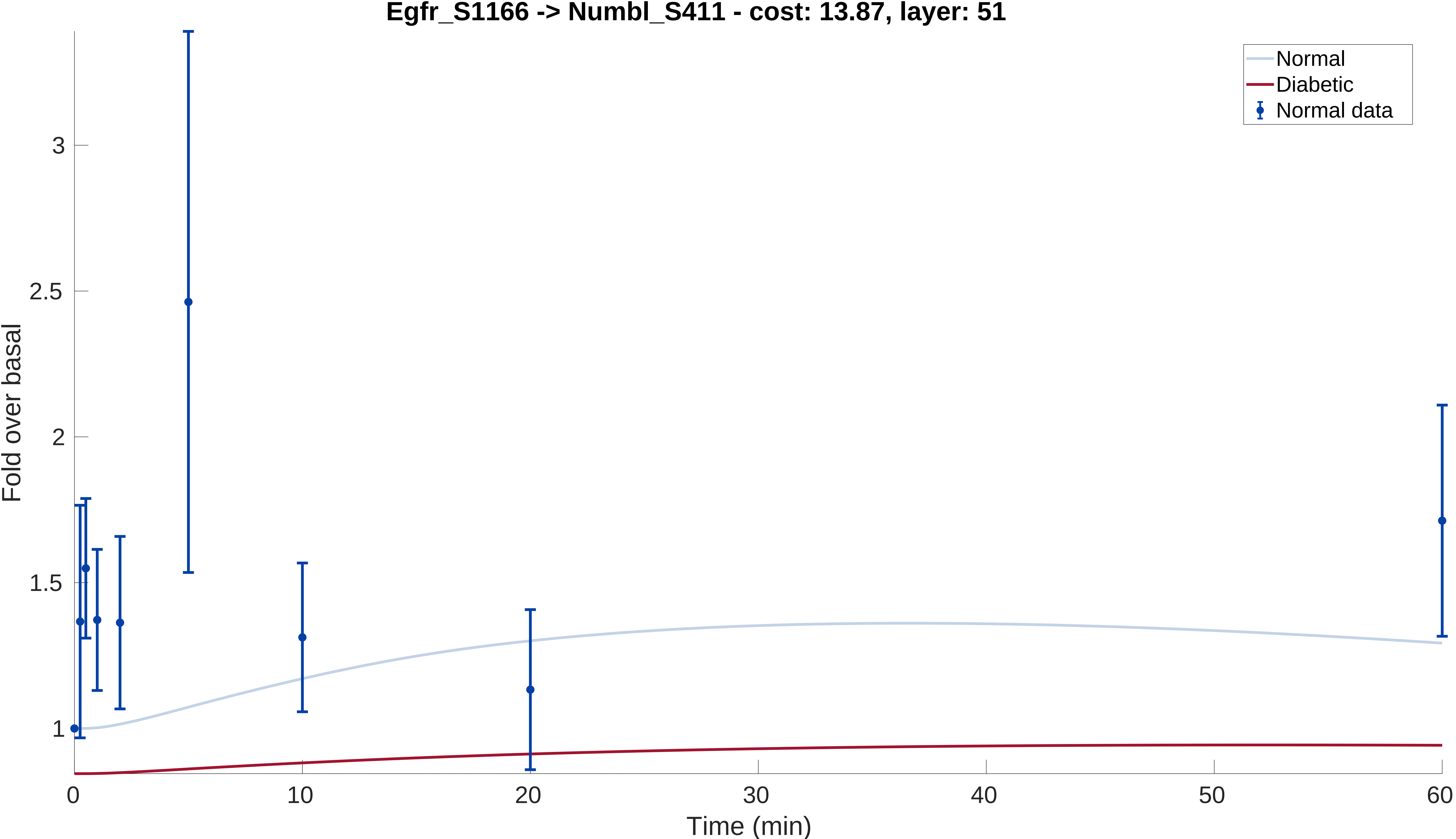

Egfr_S1166 -> Numbl_S411 - cost: 13.87, layer: 51
Normal
Diabetic
Normal data
Fold over basal
1
1.5
2
2.5
3
0
10
20
30
40
50
60
Time (min)

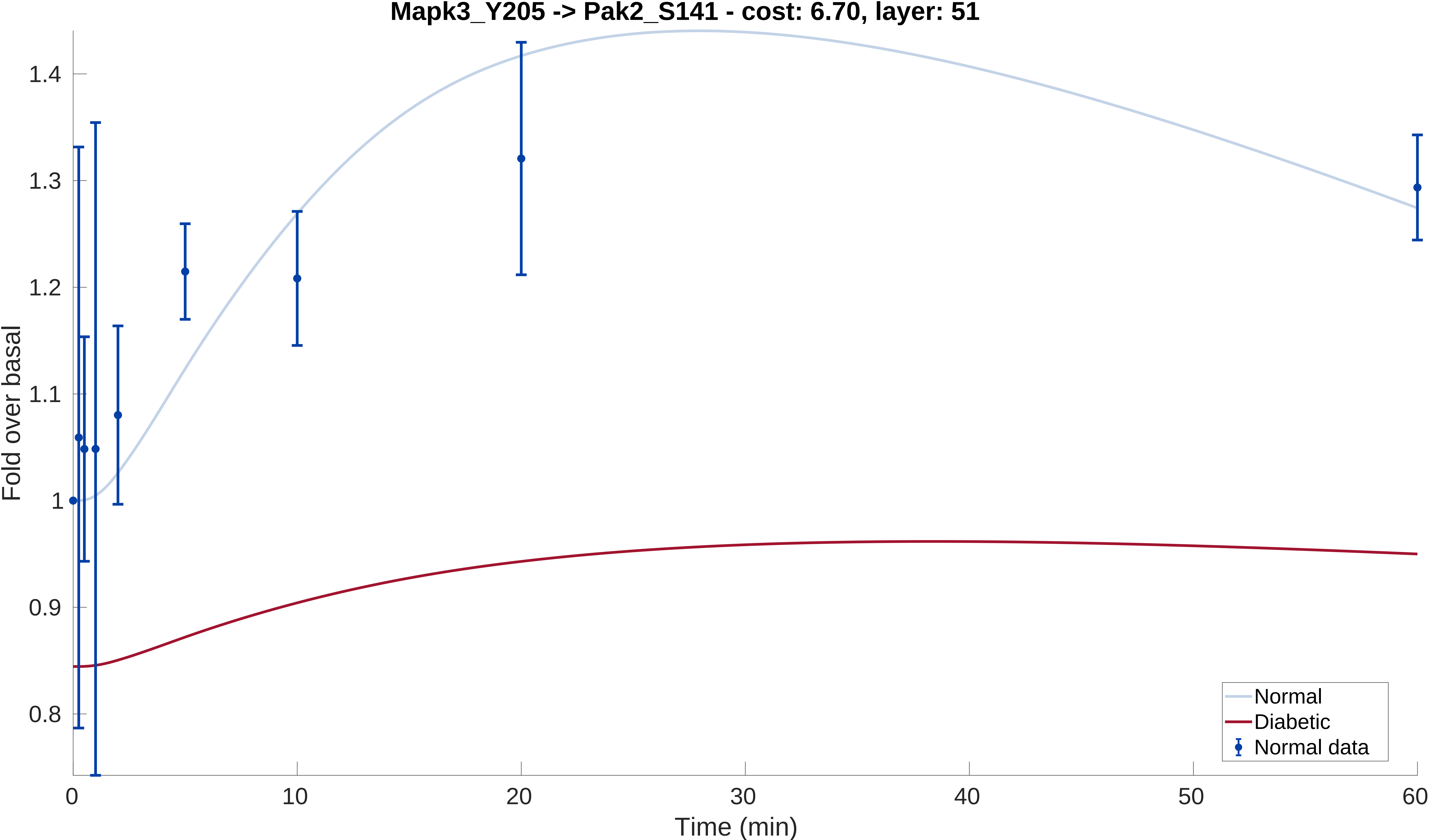
Mapk3_Y205 -> Pak2_S141 - cost: 6.70, layer: 51
Fold over basal
1.4
1.3
1.2
1.1
1
0.9
0.8
0
10
20
30
40
50
60
Time (min)
Normal
Diabetic
Normal data

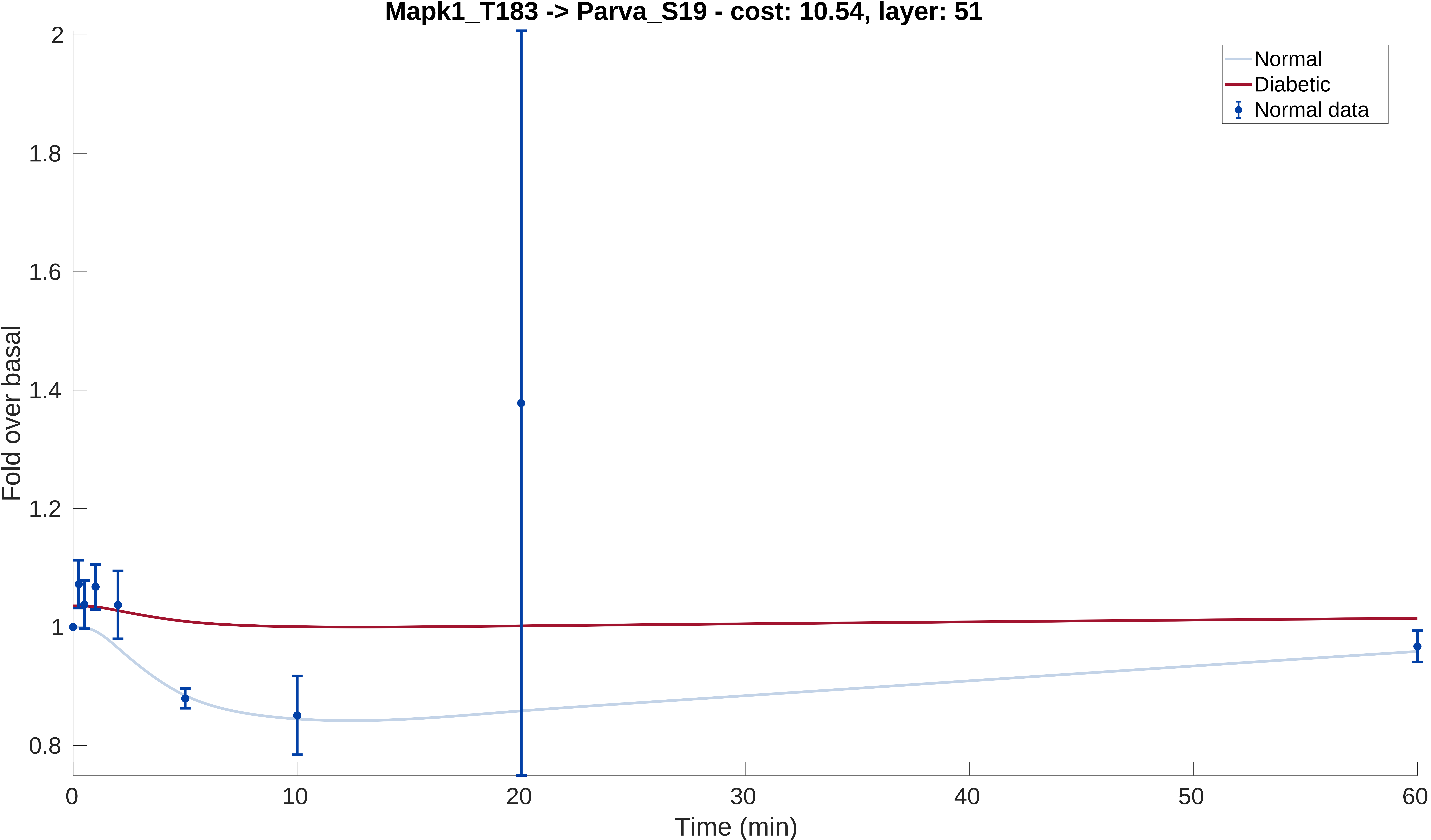

Mapk1_T183 -> Parva_S19 - cost: 10.54, layer: 51
Fold over basal
Time (min)
Normal
Diabetic
Normal data
0.8
1
1.2
1.4
1.6
1.8
2
0
10
20
30
40
50
60

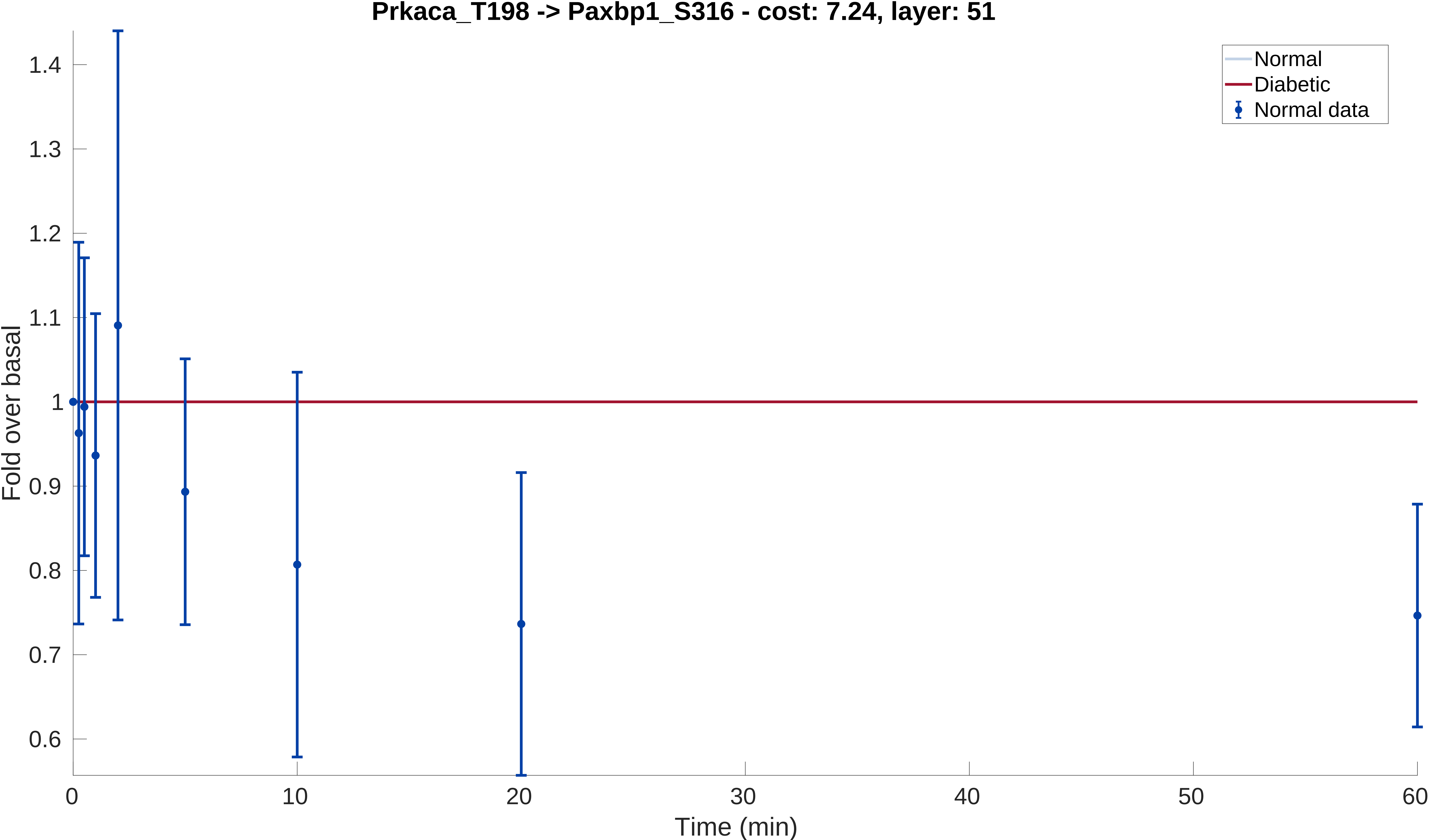

Prkaca_T198 -> Paxbp1_S316 - cost: 7.24, layer: 51
Normal
Diabetic
Normal data
Fold over basal
1.4
1.3
1.2
1.1
1
0.9
0.8
0.7
0.6
0
10
20
30
40
50
60
Time (min)

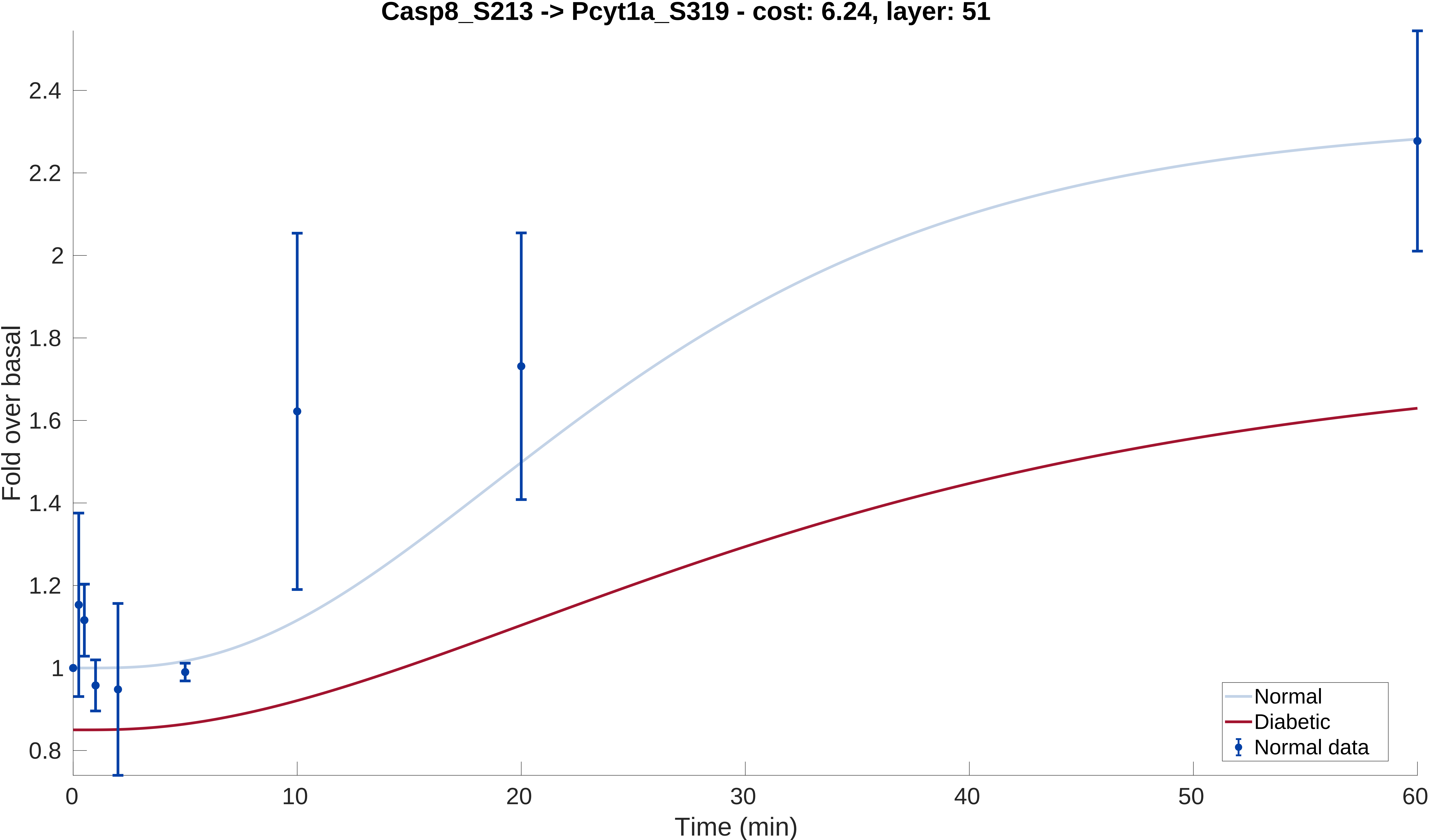

Casp8_S213 -> Pcyt1a_S319 - cost: 6.24, layer: 51
Fold over basal
Time (min)
0.8
1
1.2
1.4
1.6
1.8
2
2.2
2.4
0
10
20
30
40
50
60
Normal
Diabetic
Normal data

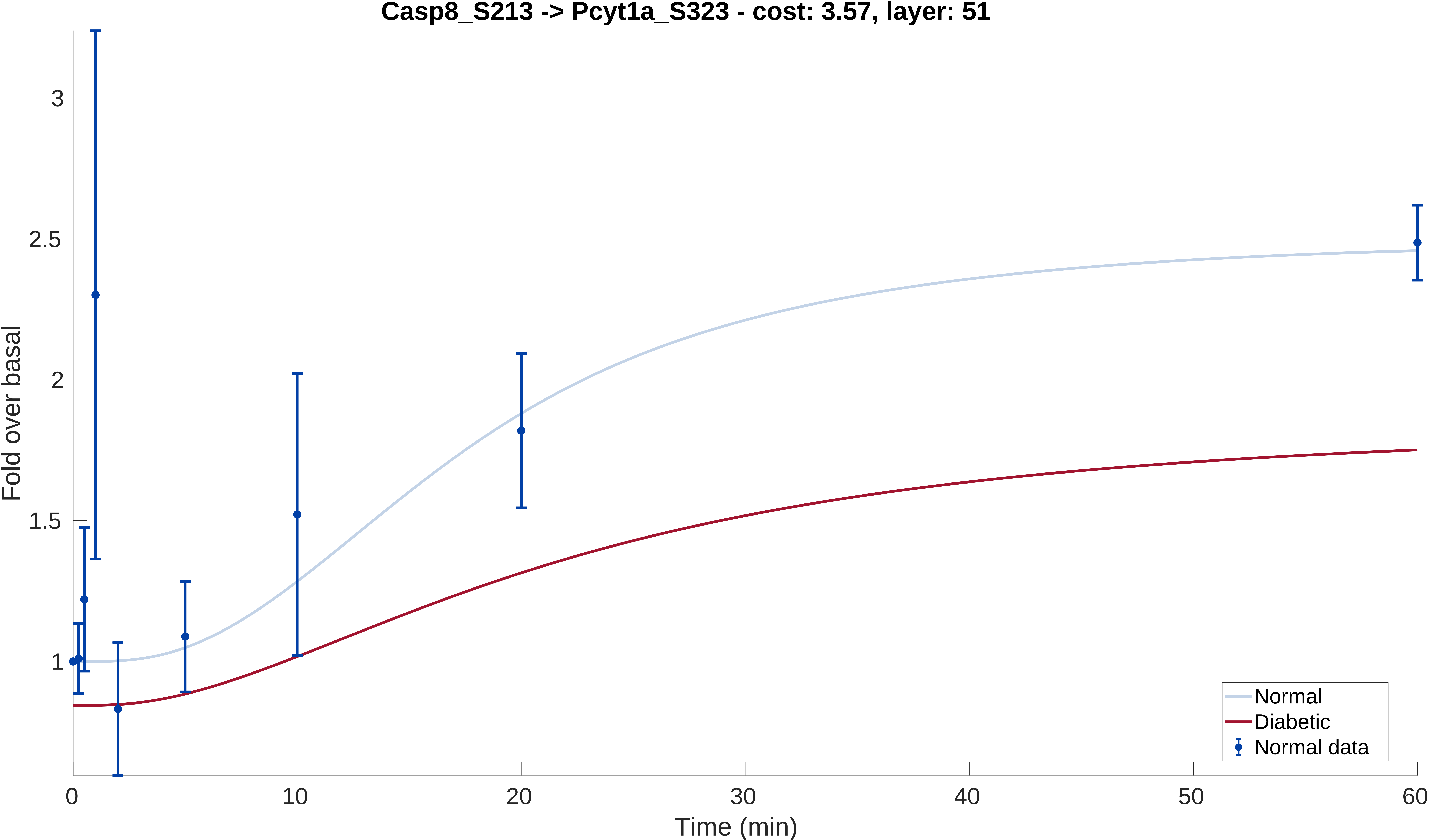

Casp8_S213 -> Pcyt1a_S323 - cost: 3.57, layer: 51
Fold over basal
Time (min)
0
10
20
30
40
50
60
1
1.5
2
2.5
3
Normal
Diabetic
Normal data

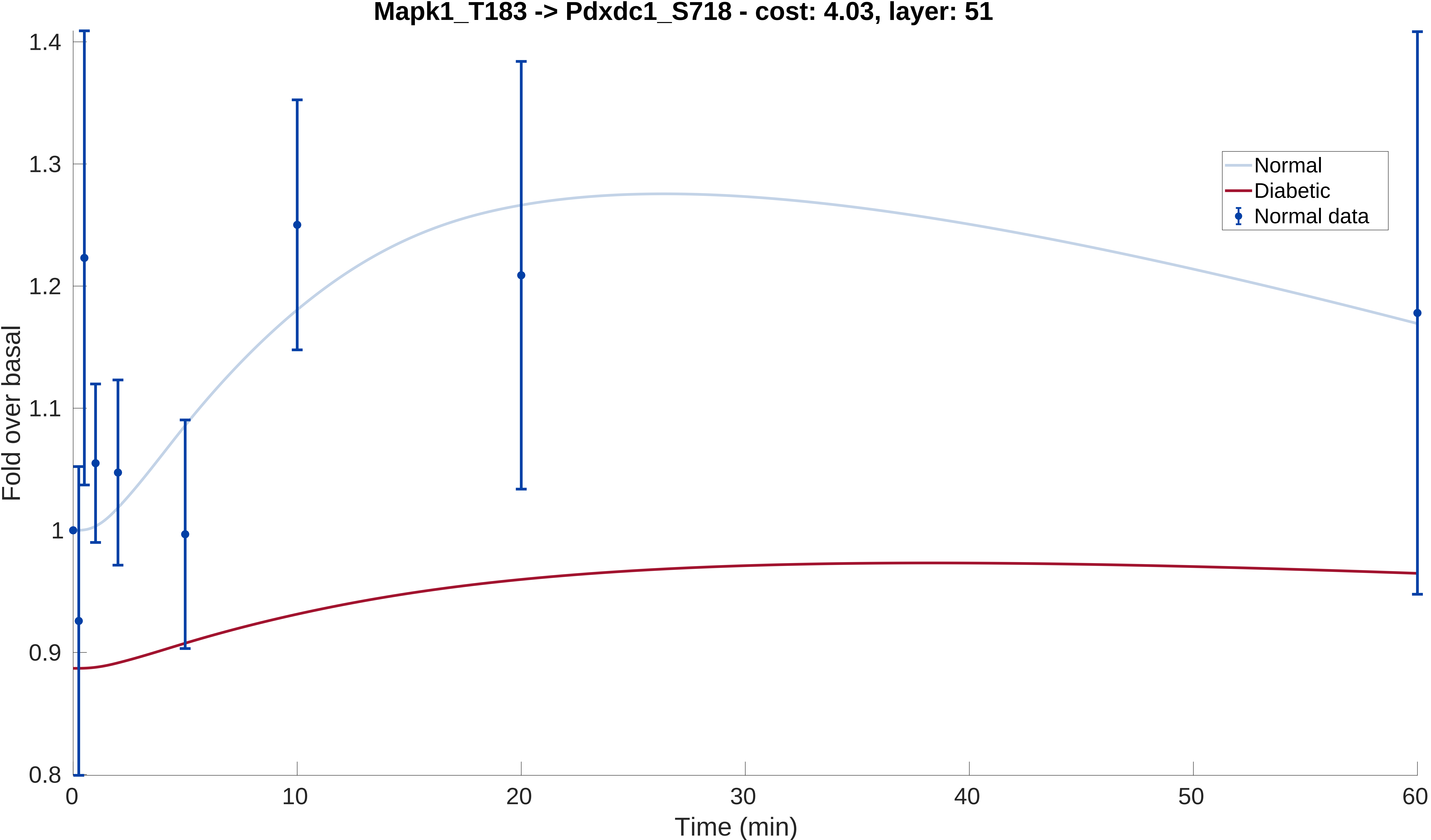

Mapk1_T183 -> Pdxdc1_S718 - cost: 4.03, layer: 51
Fold over basal
1.4
1.3
1.2
1.1
1
0.9
0.8
0
10
20
30
40
50
60
Time (min)
Normal
Diabetic
Normal data

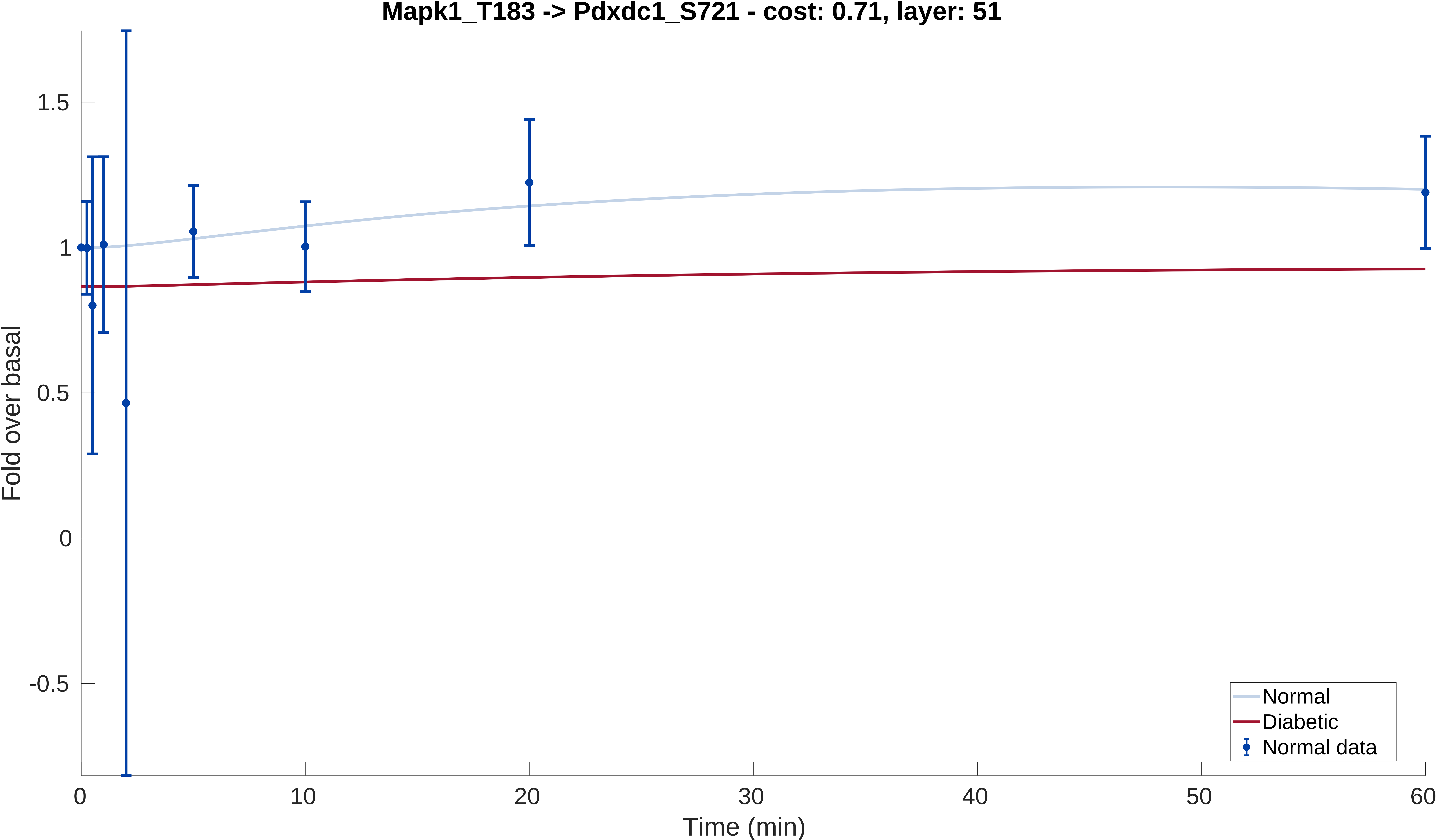

Mapk1_T183 -> Pdxdc1_S721 - cost: 0.71, layer: 51
1.5
1
0.5
0
-0.5
Fold over basal
0
10
20
30
40
50
60
Time (min)
Normal
Diabetic
Normal data

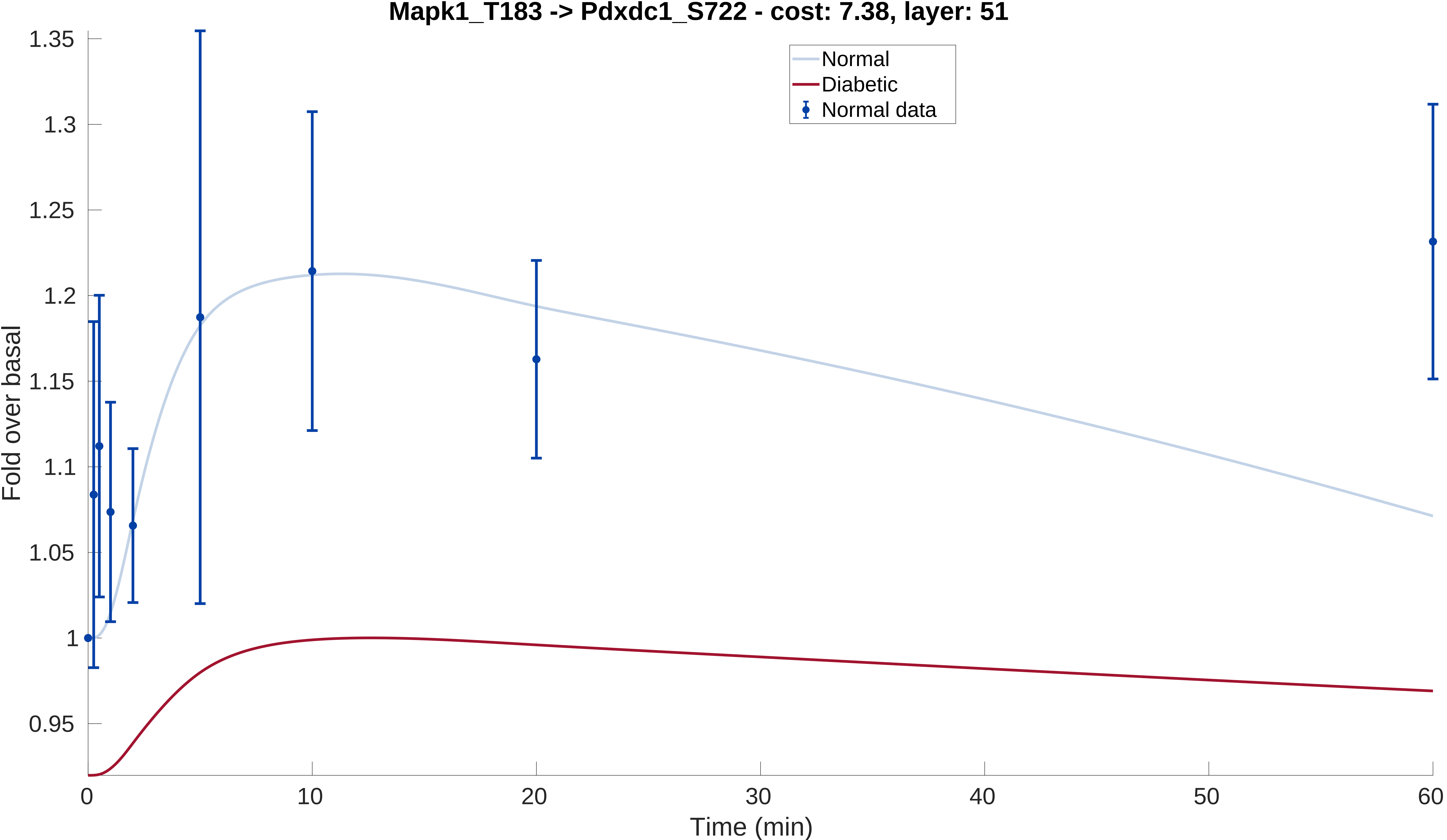

Mapk1_T183 -> Pdxdc1_S722 - cost: 7.38, layer: 51
Normal
Diabetic
Normal data
Fold over basal
Time (min)
1.35
1.3
1.25
1.2
1.15
1.1
1.05
1
0.95
0
10
20
30
40
50
60

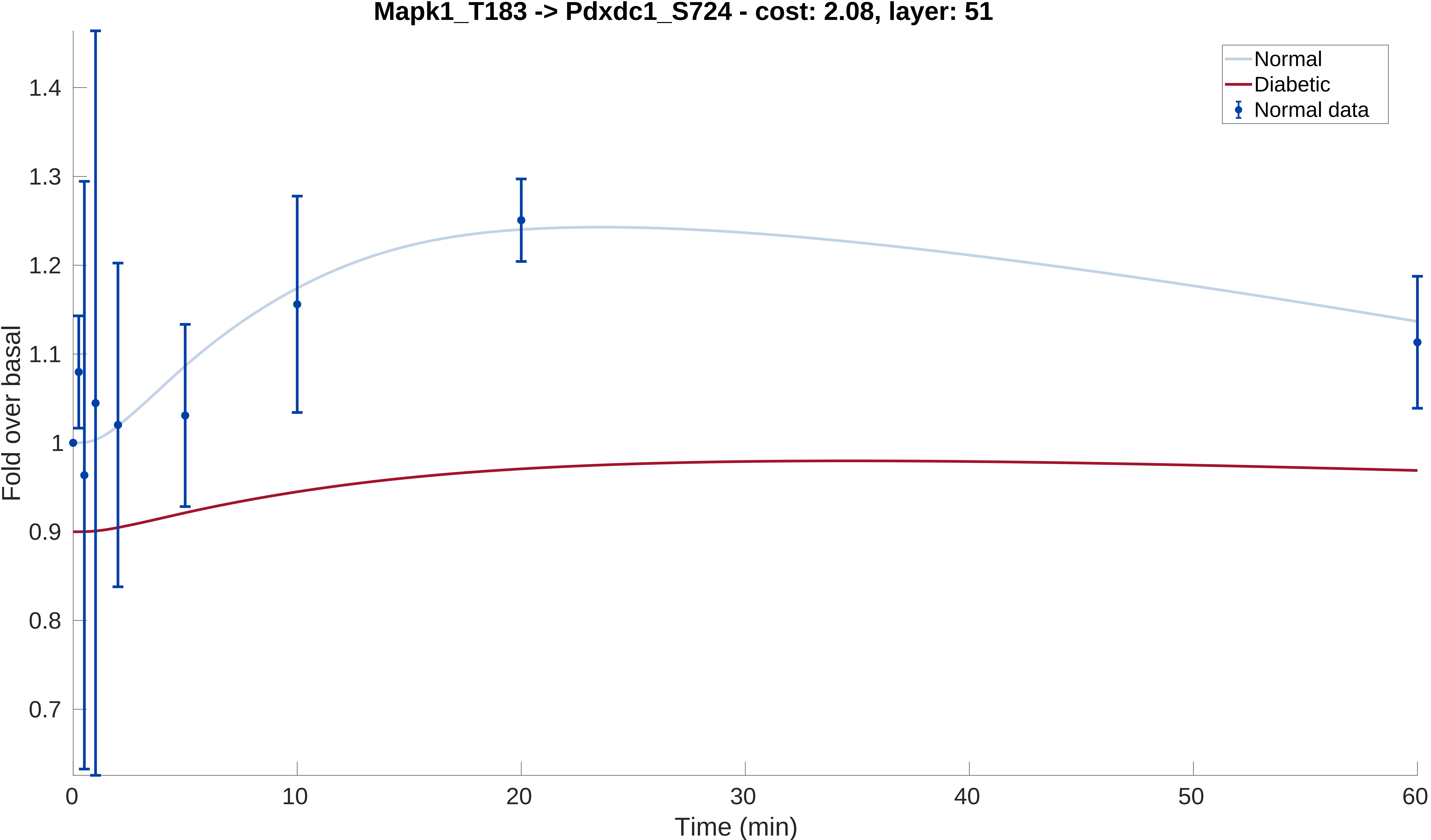

Mapk1_T183 -> Pdxdc1_S724 - cost: 2.08, layer: 51
Normal
Diabetic
Normal data
Fold over basal
Time (min)
1.4
1.3
1.2
1.1
1
0.9
0.8
0.7
0
10
20
30
40
50
60

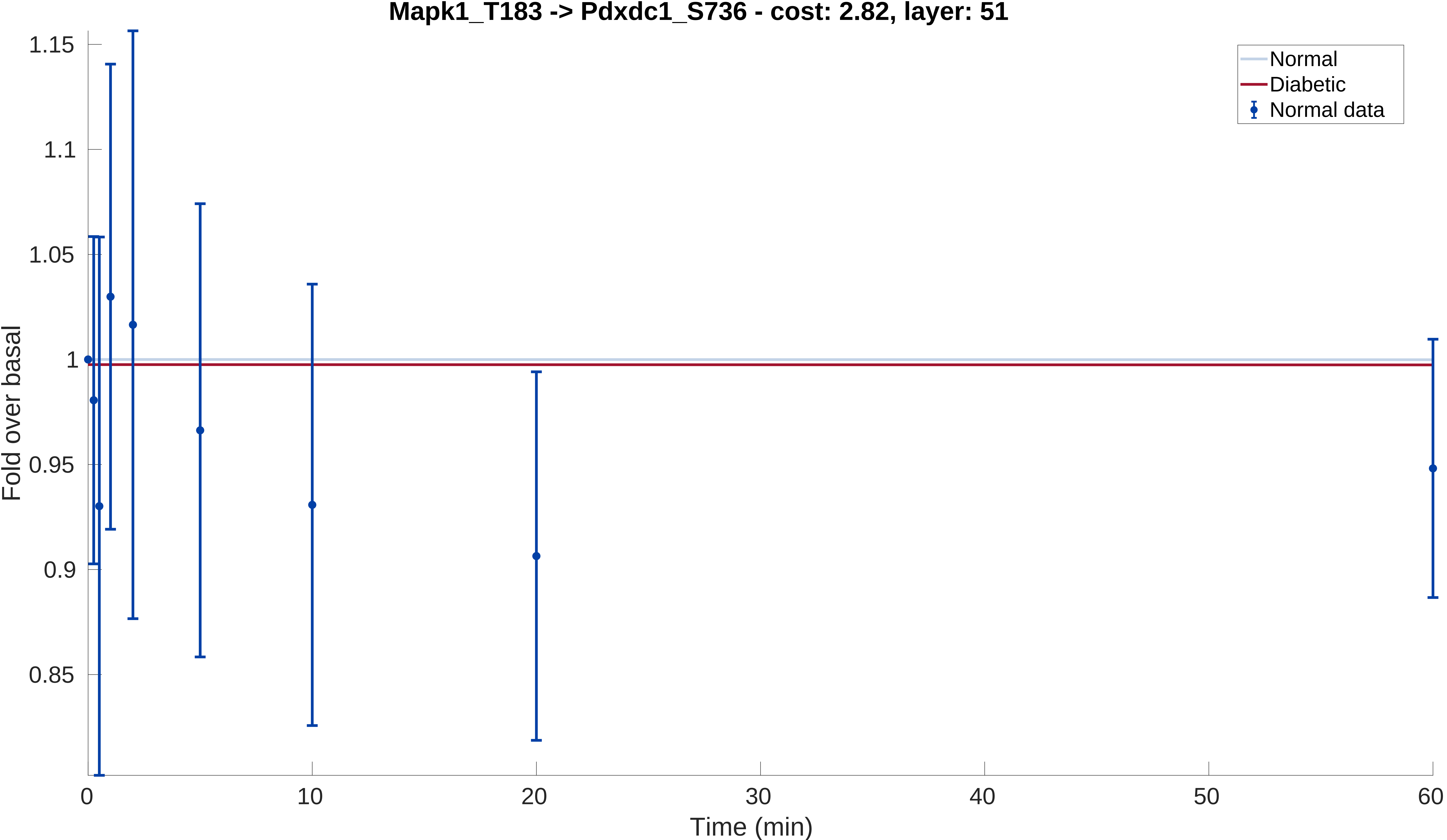

Mapk1_T183 -> Pdxdc1_S736 - cost: 2.82, layer: 51
Normal
Diabetic
Normal data
Fold over basal
1.15
1.1
1.05
1
0.95
0.9
0.85
0
10
20
30
40
50
60
Time (min)

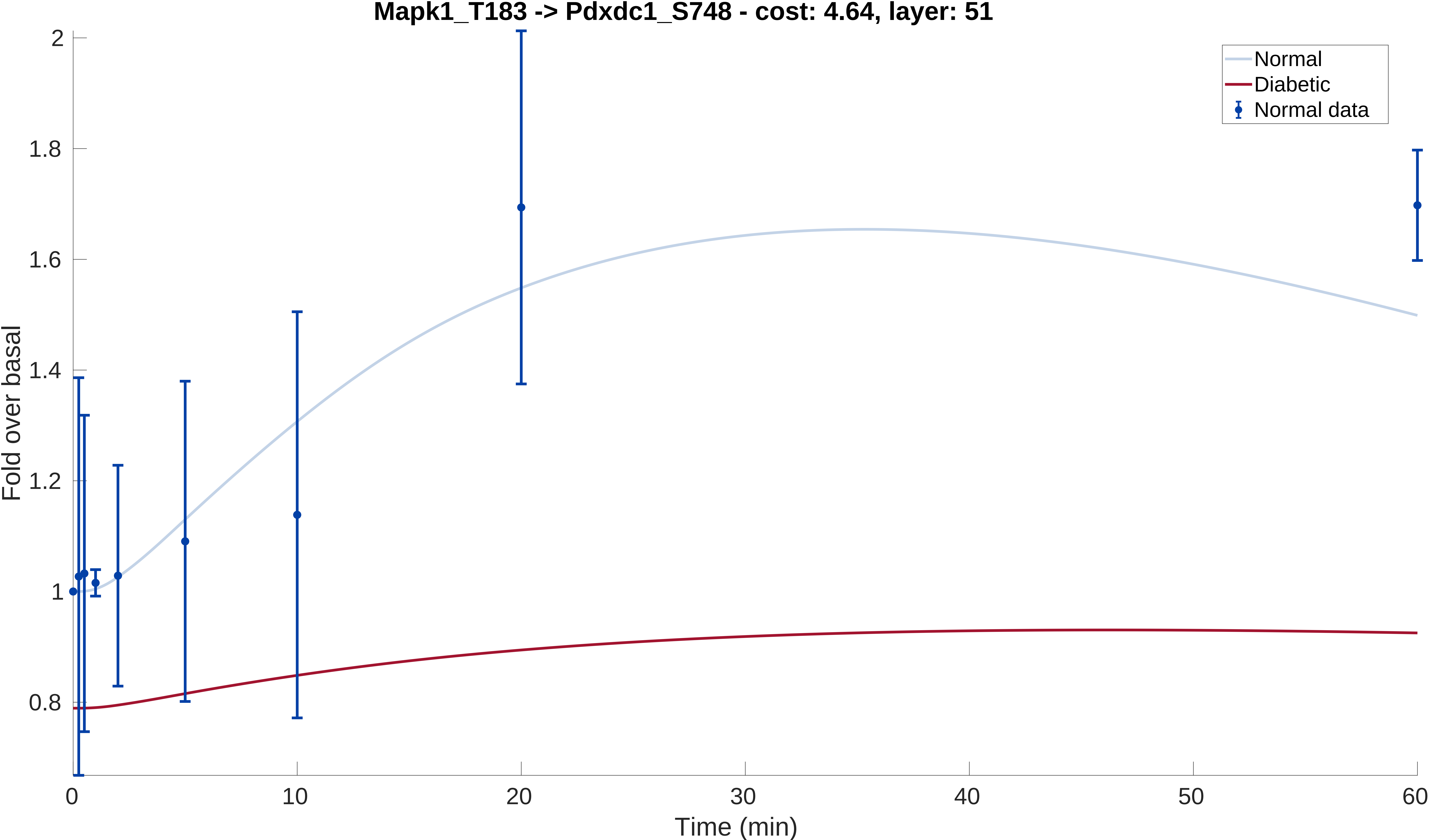

Mapk1_T183 -> Pdxdc1_S748 - cost: 4.64, layer: 51
Normal
Diabetic
Normal data
Fold over basal
Time (min)
2
1.8
1.6
1.4
1.2
1
0.8
0
10
20
30
40
50
60

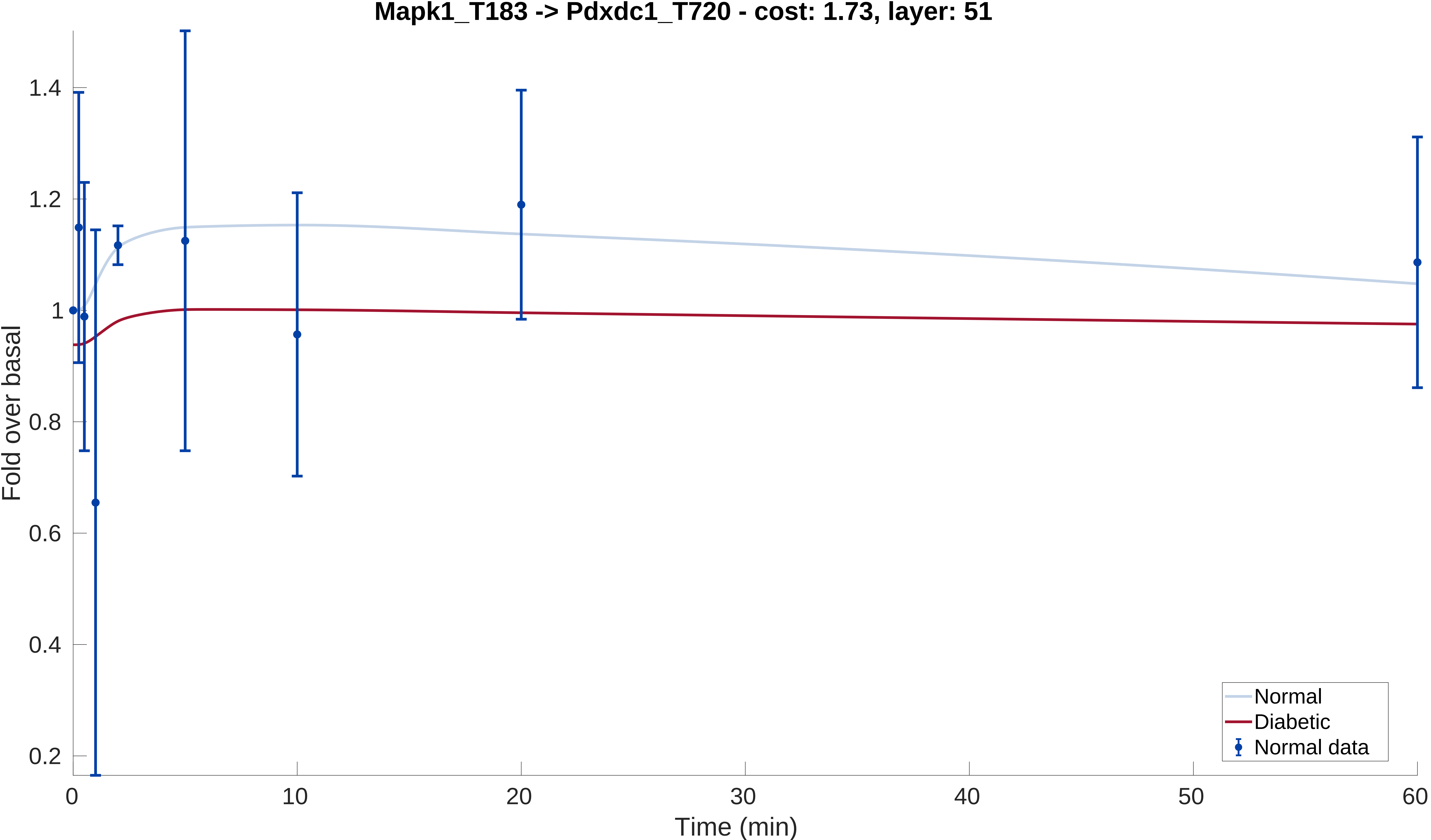

Mapk1_T183 -> Pdxdc1_T720 - cost: 1.73, layer: 51
Fold over basal
1.4
1.2
1
0.8
0.6
0.4
0.2
0
10
20
30
40
50
60
Time (min)
Normal
Diabetic
Normal data

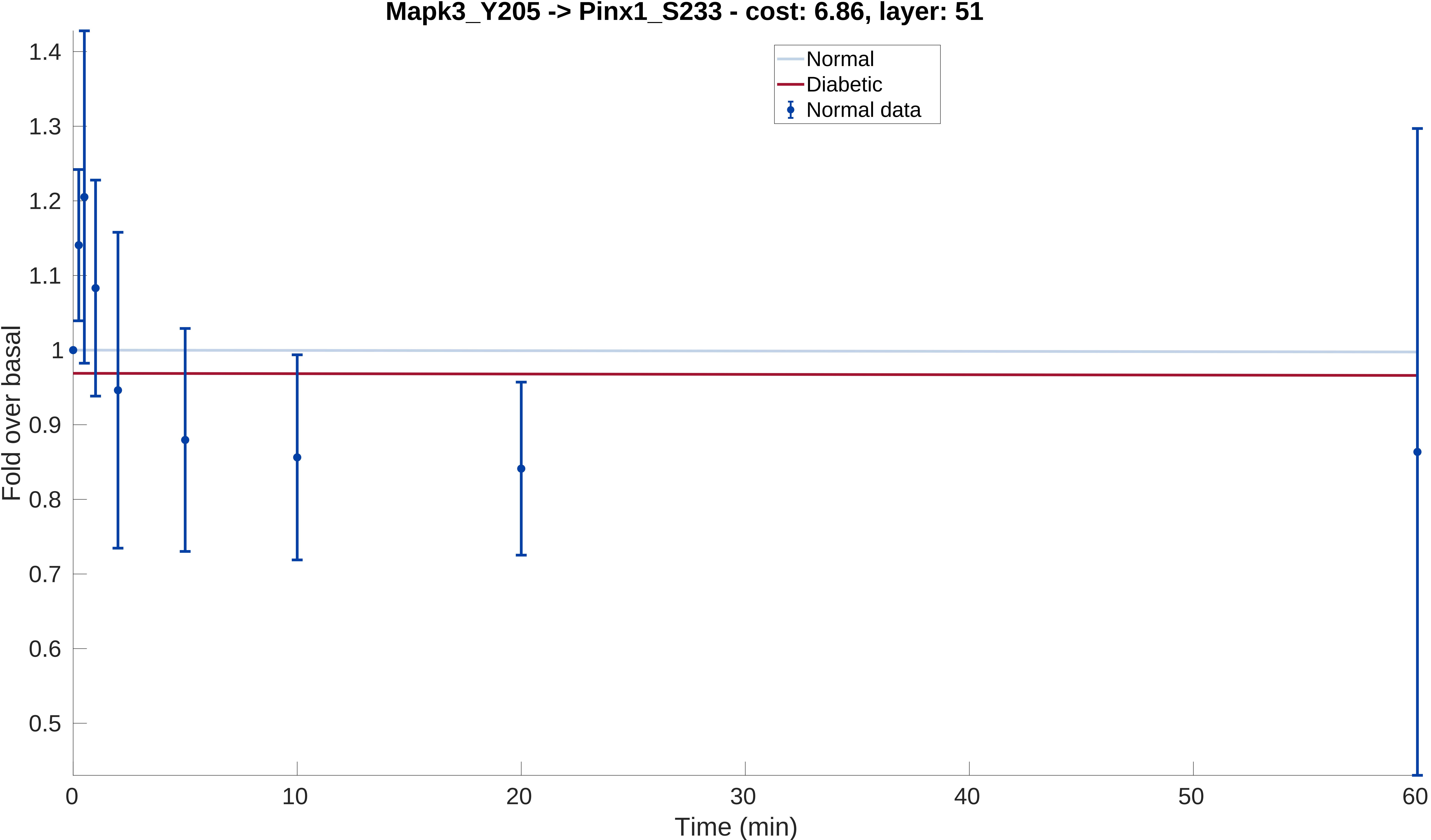

Mapk3_Y205 -> Pinx1_S233 - cost: 6.86, layer: 51
Normal
Diabetic
Normal data
Fold over basal
Time (min)
1.4
1.3
1.2
1.1
1
0.9
0.8
0.7
0.6
0.5
0
10
20
30
40
50
60

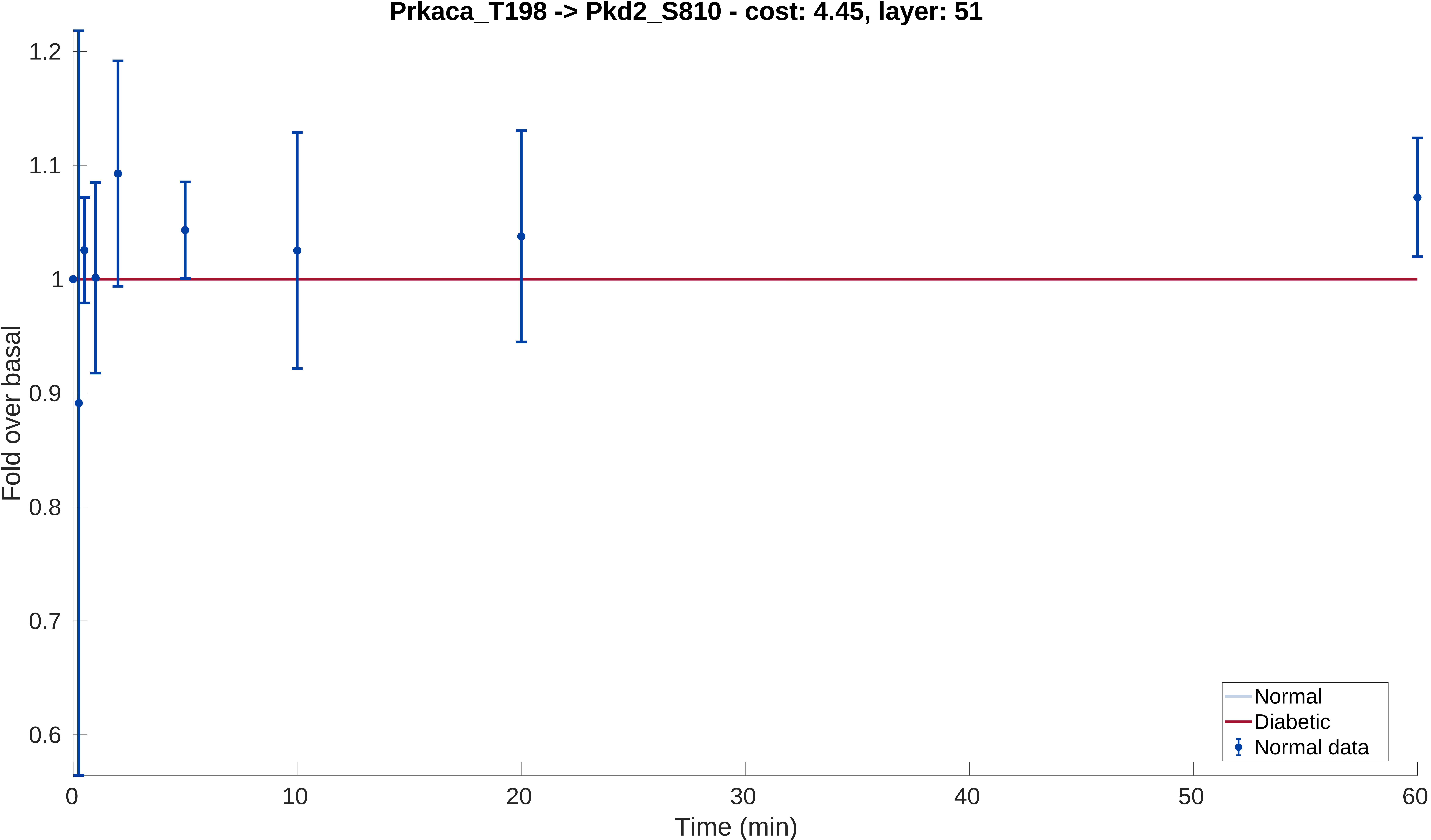
Prkaca_T198 -> Pkd2_S810 - cost: 4.45, layer: 51
Fold over basal
1.2
1.1
1
0.9
0.8
0.7
0.6
0
10
20
30
40
50
60
Time (min)
Normal
Diabetic
Normal data

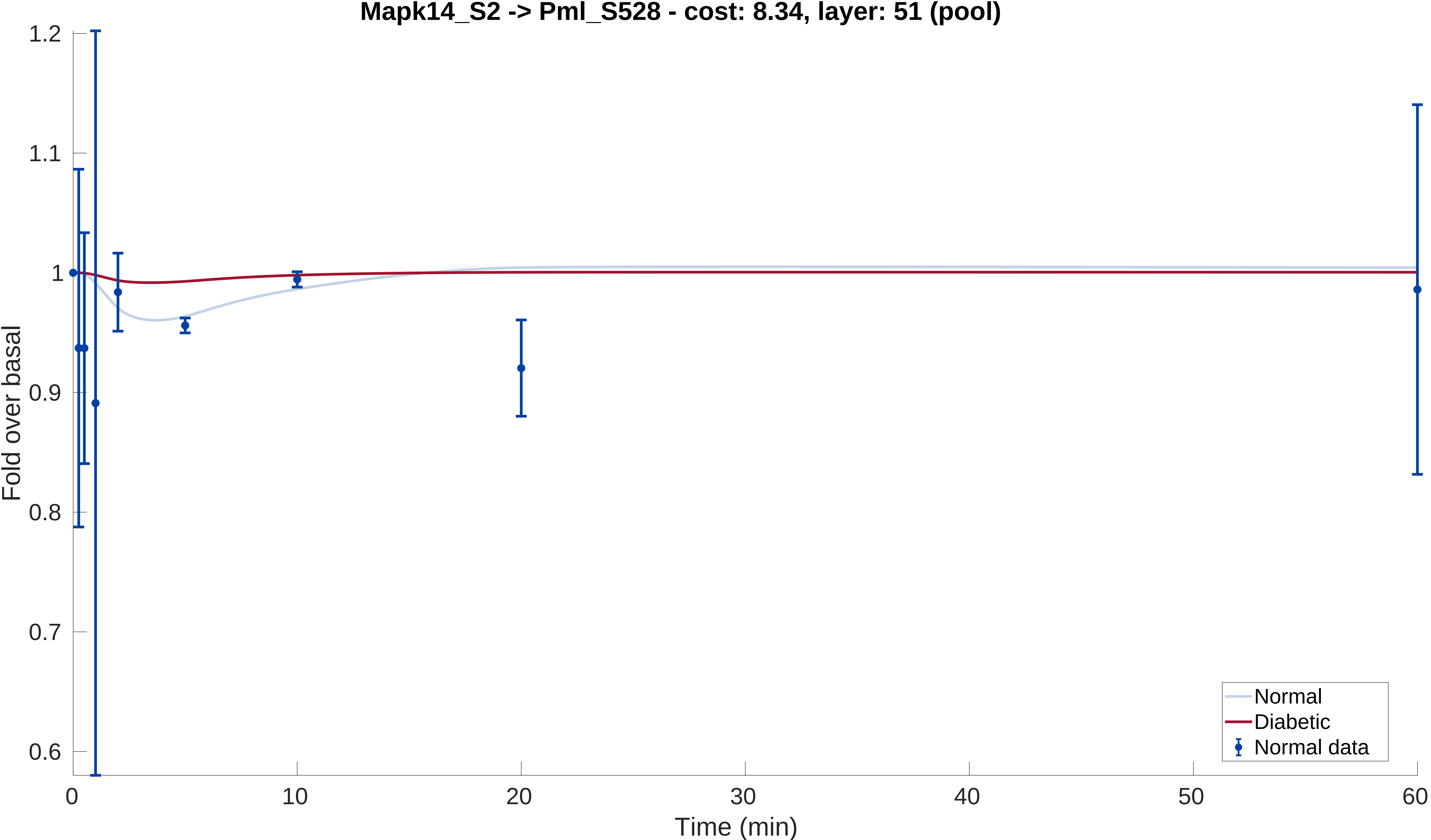

Mapk14_S2 -> Pml_S528 - cost: 8.34, layer: 51 (pool)
1.2
1.1
1
0.9
0.8
0.7
0.6
Fold over basal
0
10
20
30
40
50
60
Time (min)
Normal
Diabetic
Normal data

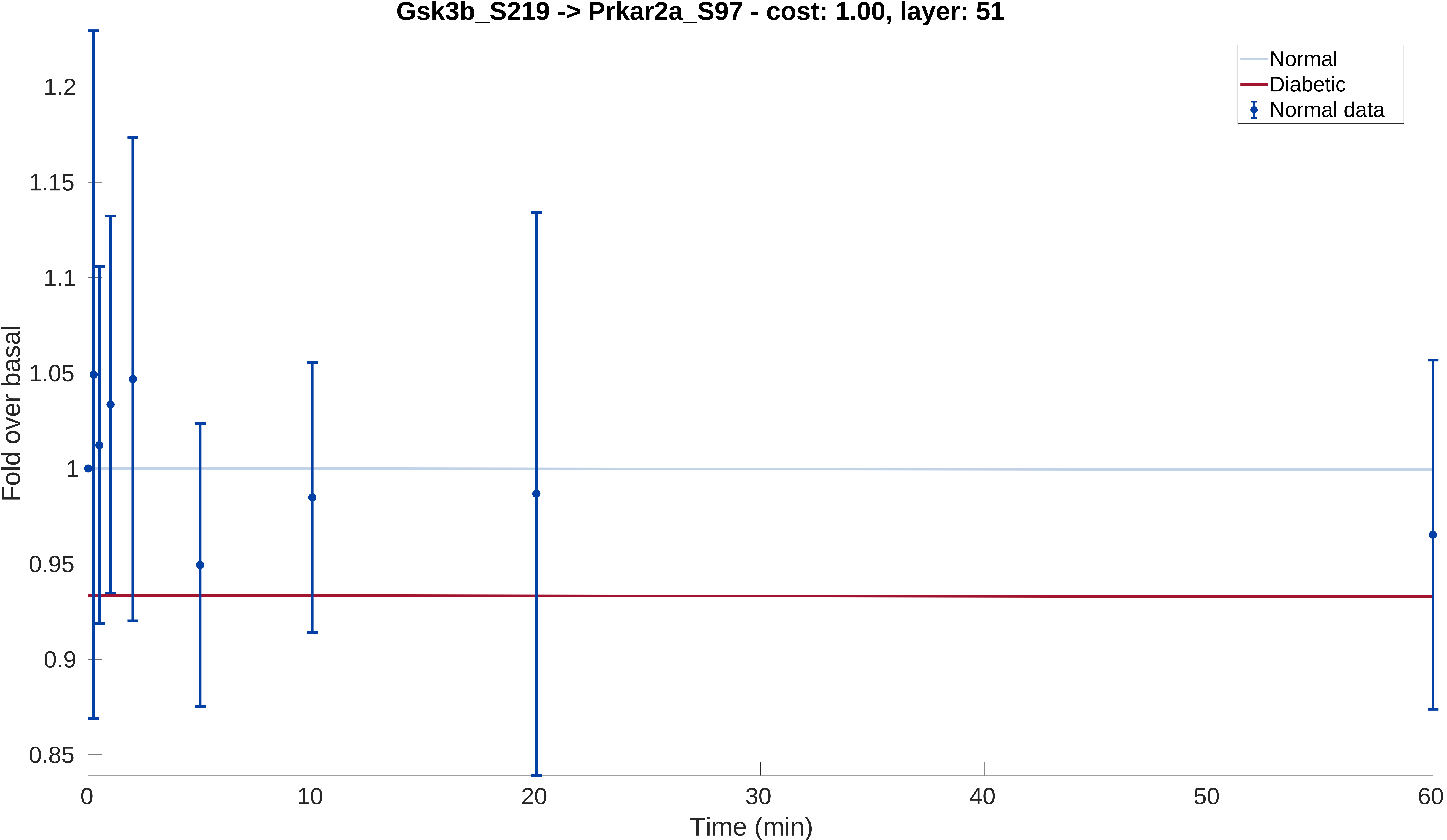

Gsk3b_S219 -> Prkar2a_S97 - cost: 1.00, layer: 51
Normal
Diabetic
Normal data
1.2
1.15
1.1
1.05
1
0.95
0.9
0.85
Fold over basal
0
10
20
30
40
50
60
Time (min)

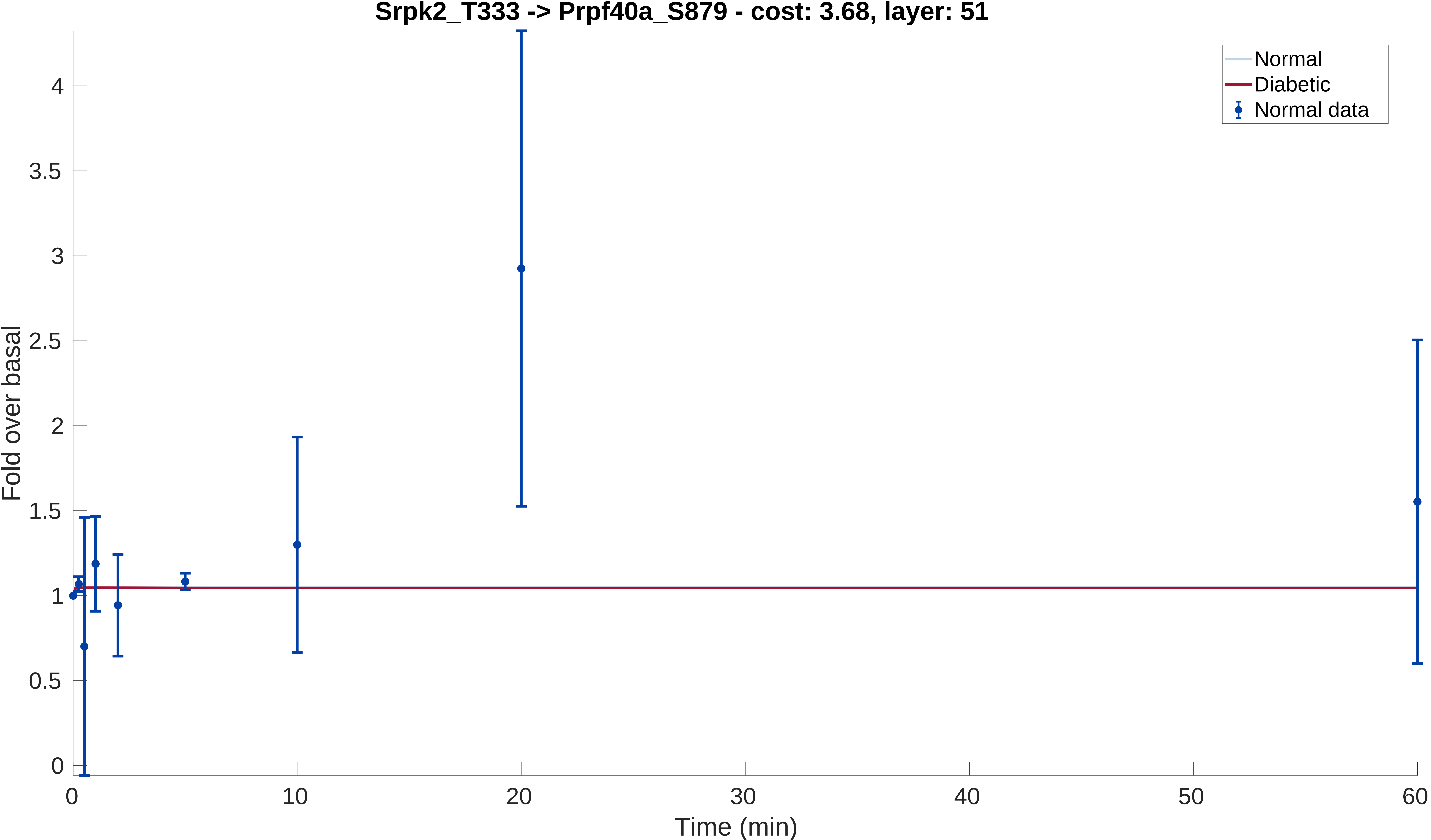

Srpk2_T333 -> Prpf40a_S879 - cost: 3.68, layer: 51
Normal
Diabetic
Normal data
Fold over basal
0
0.5
1
1.5
2
2.5
3
3.5
4
0
10
20
30
40
50
60
Time (min)

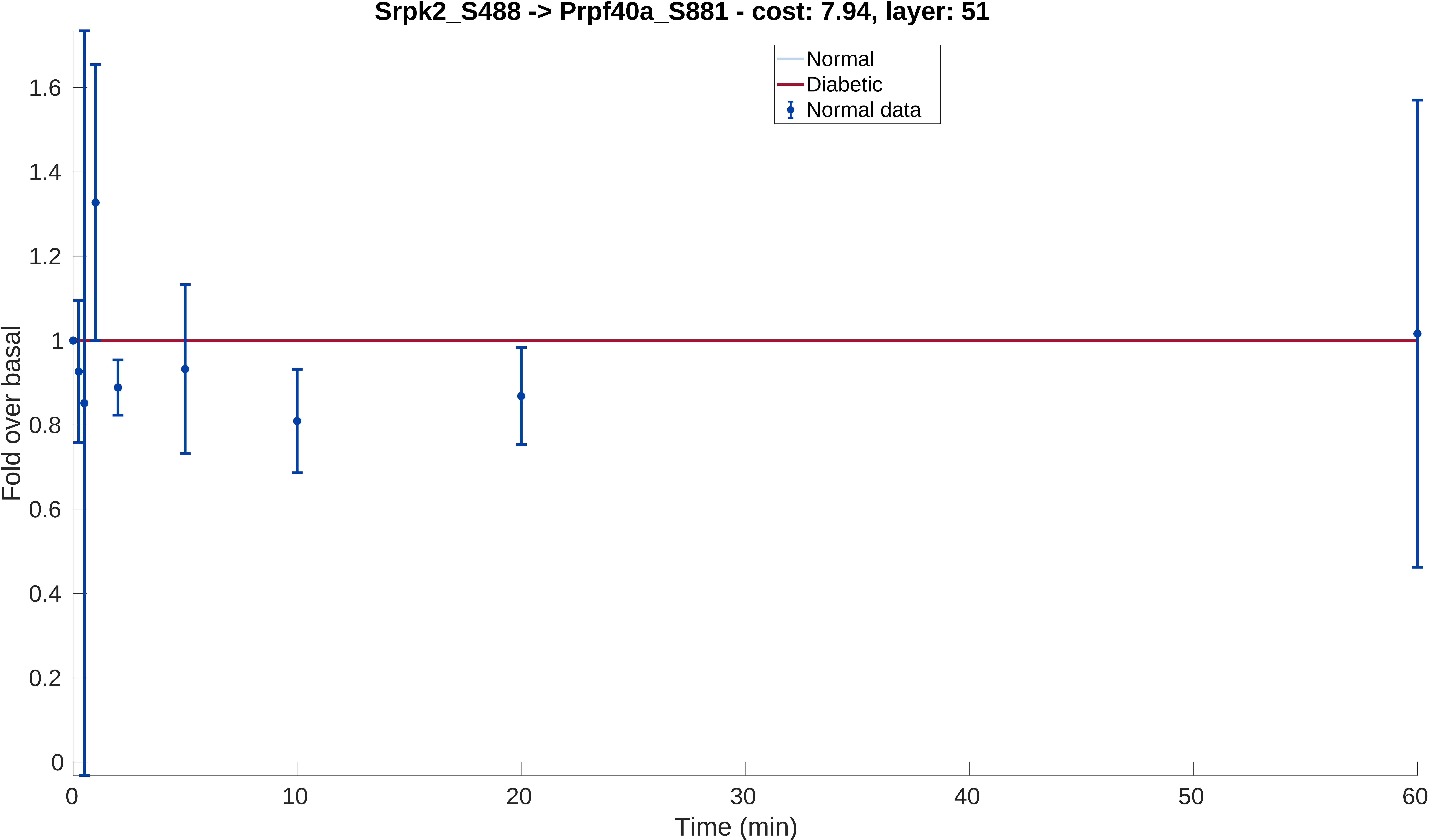

Srpk2_S488 -> Prpf40a_S881 - cost: 7.94, layer: 51
Normal
Diabetic
Normal data
Fold over basal
0
0.2
0.4
0.6
0.8
1
1.2
1.4
1.6
0
10
20
30
40
50
60
Time (min)

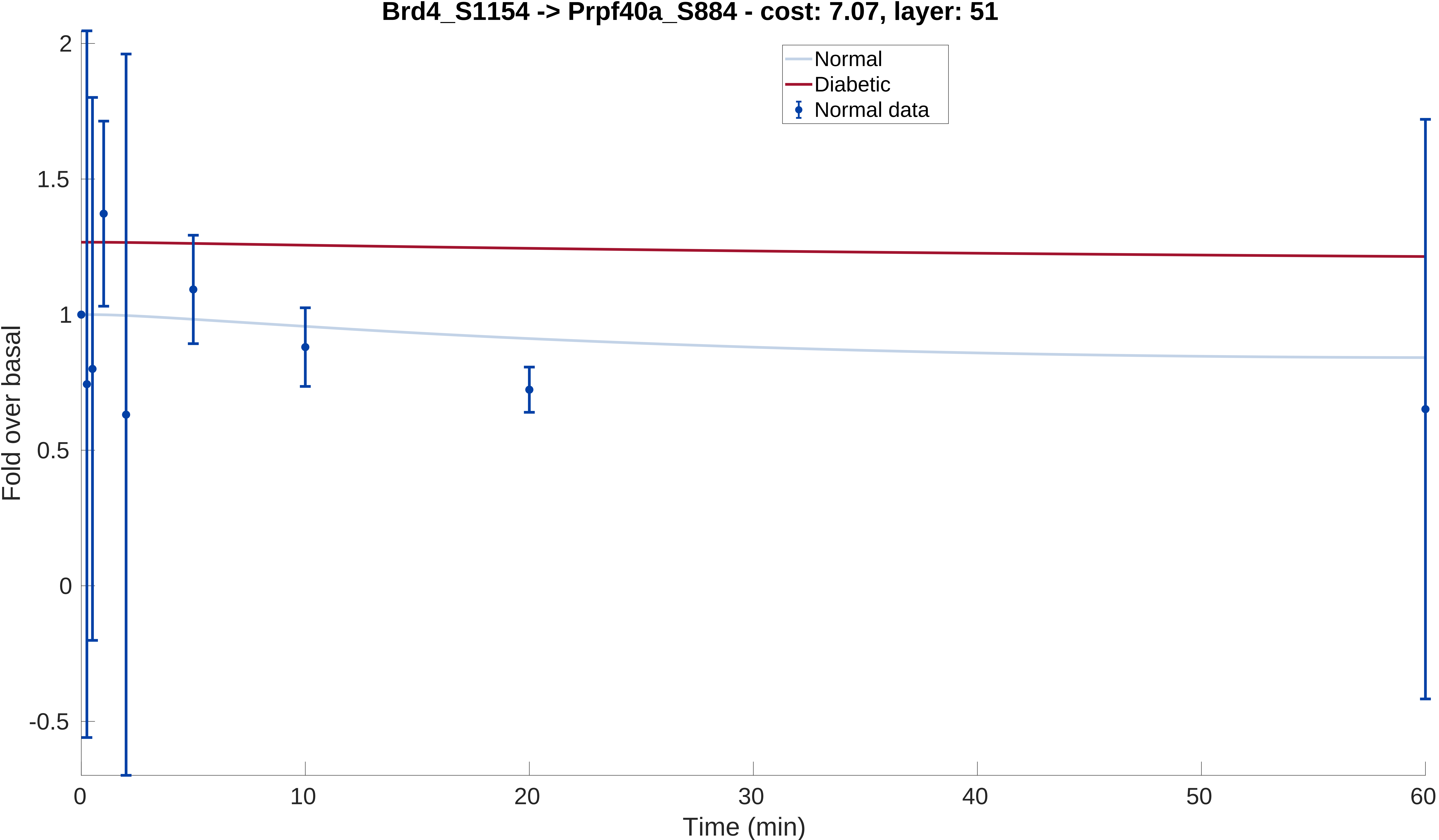
Brd4_S1154 -> Prpf40a_S884 - cost: 7.07, layer: 51
Normal
Diabetic
Normal data
Fold over basal
2
1.5
1
0.5
0
-0.5
0
10
20
30
40
50
60
Time (min)

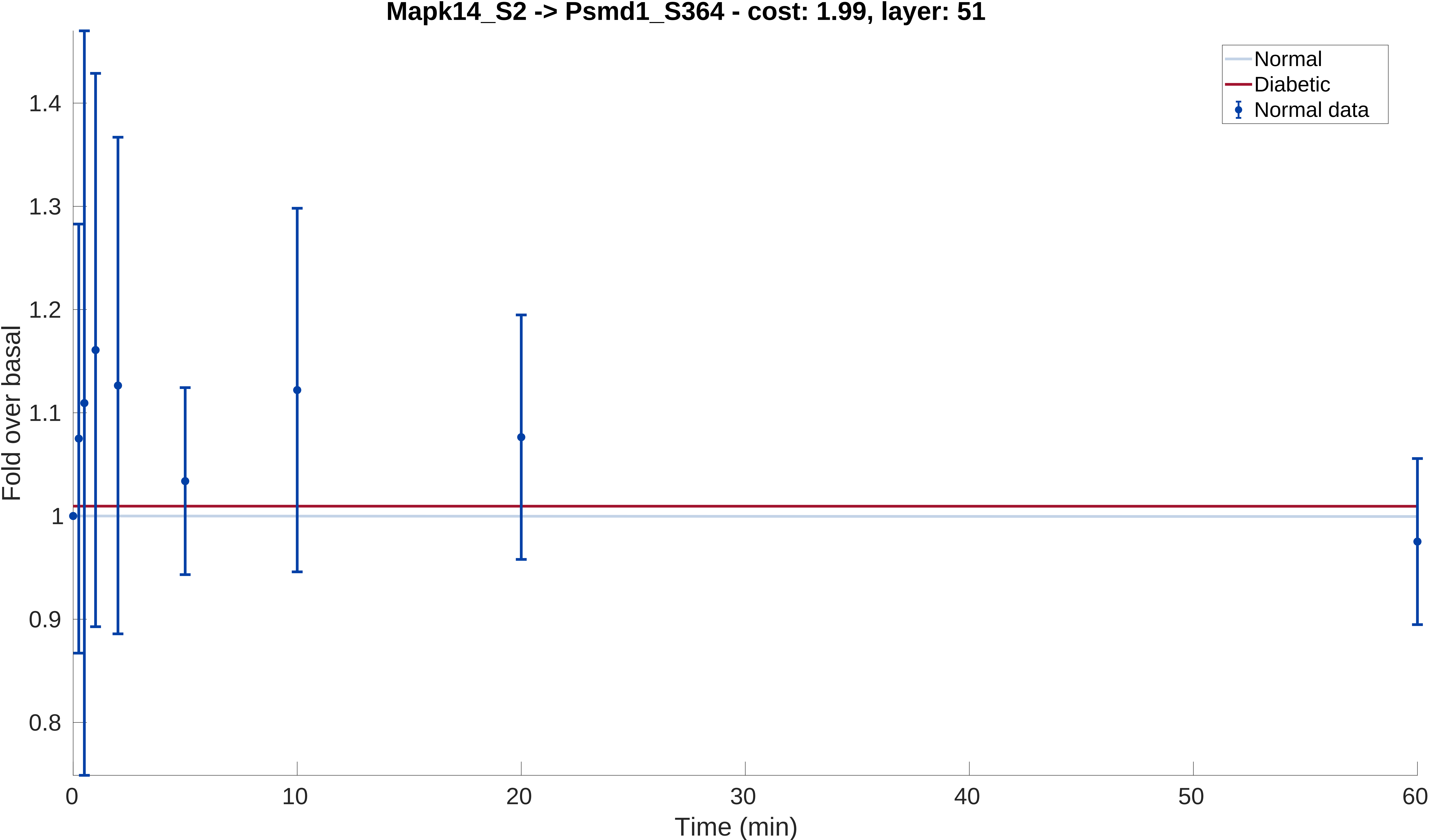

Mapk14_S2 -> Psmd1_S364 - cost: 1.99, layer: 51
Normal
Diabetic
Normal data
Fold over basal
1.4
1.3
1.2
1.1
1
0.9
0.8
0
10
20
30
40
50
60
Time (min)

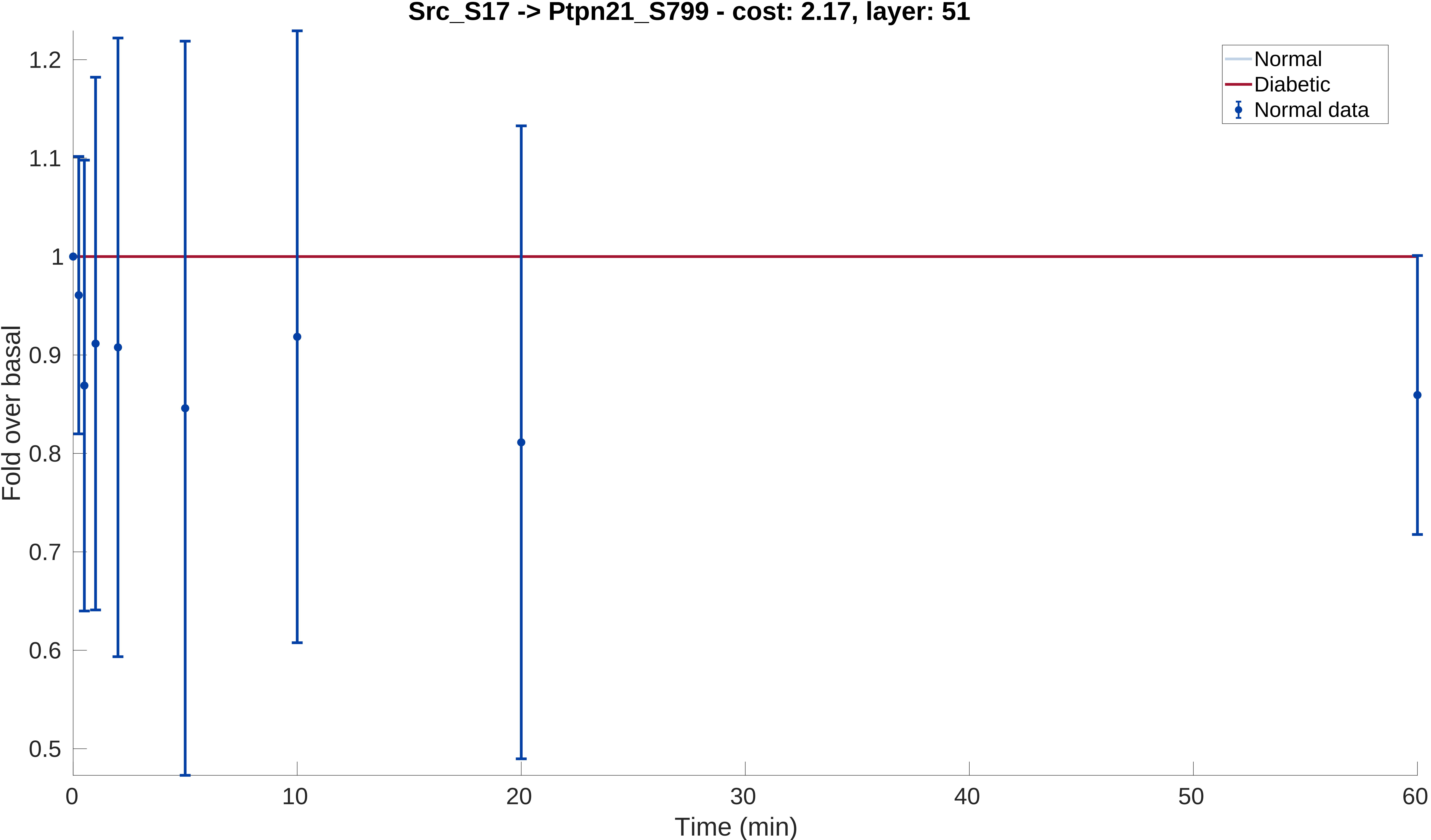
Src_S17 -> Ptpn21_S799 - cost: 2.17, layer: 51
Fold over basal
Time (min)
Normal
Diabetic
Normal data
1.2
1.1
1
0.9
0.8
0.7
0.6
0.5
0
10
20
30
40
50
60

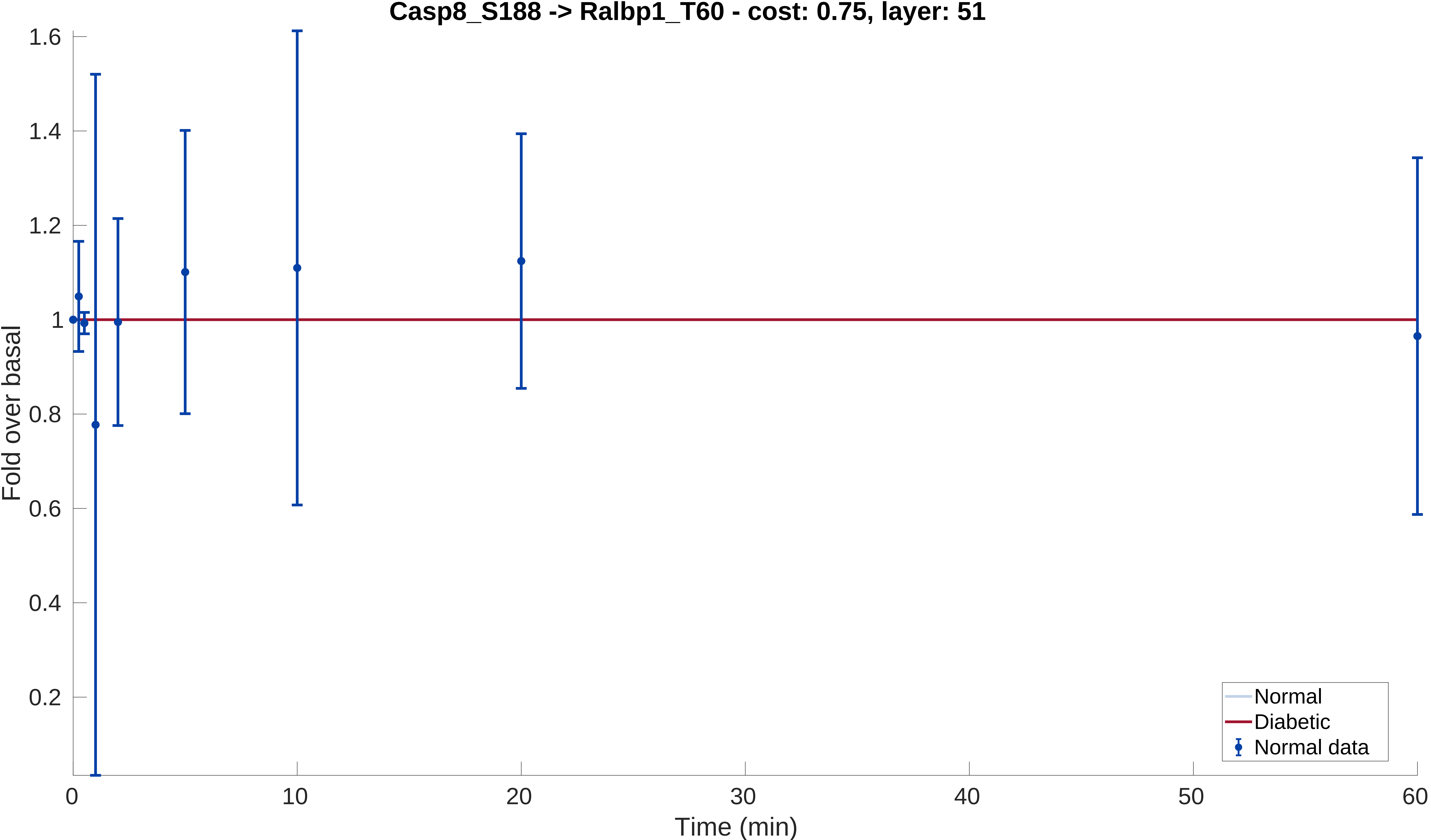

Casp8_S188 -> Ralbp1_T60 - cost: 0.75, layer: 51
Fold over basal
Time (min)
1.6
1.4
1.2
1
0.8
0.6
0.4
0.2
0
10
20
30
40
50
60
Normal
Diabetic
Normal data

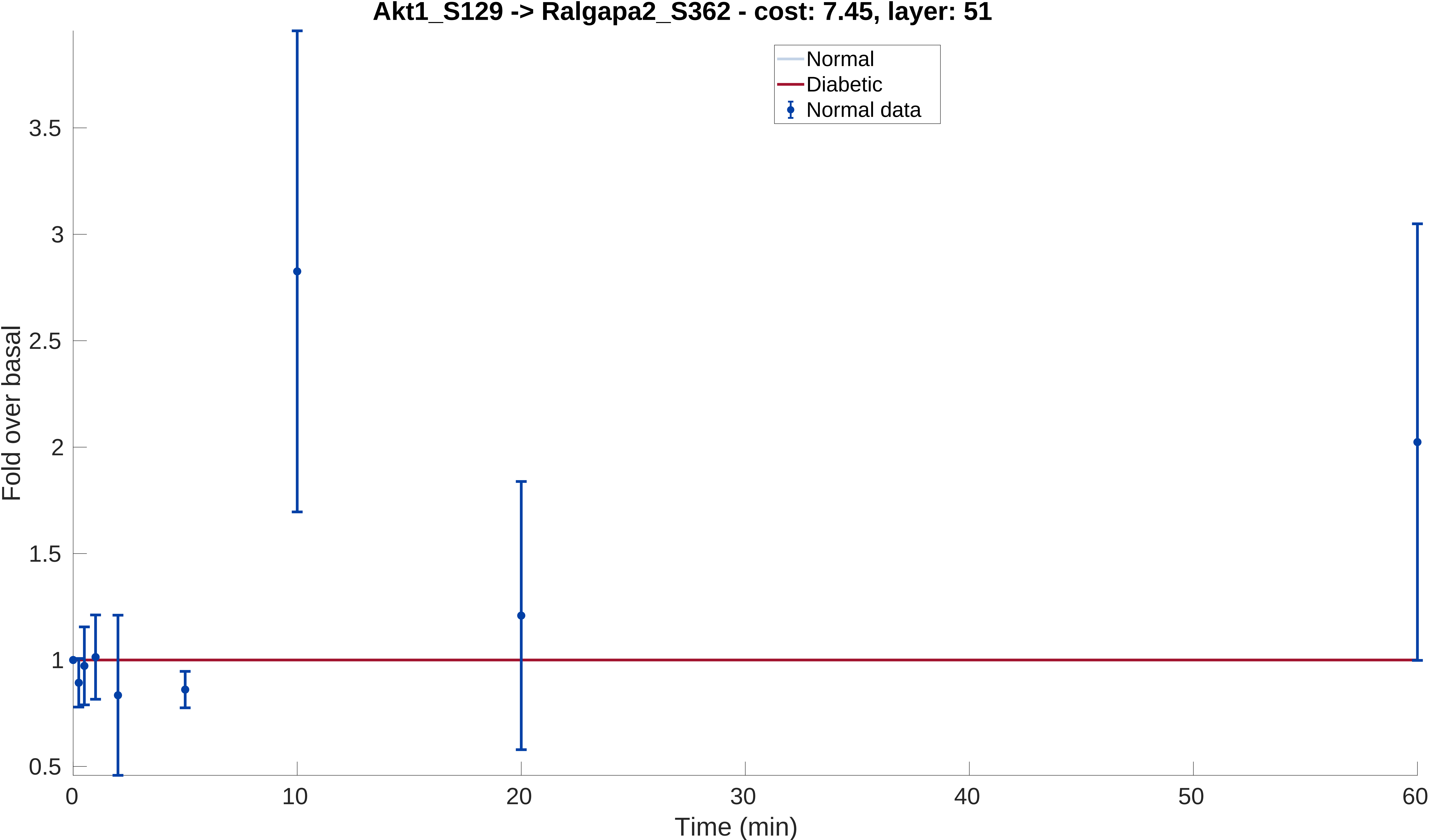
Akt1_S129 -> Ralgapa2_S362 - cost: 7.45, layer: 51
Normal
Diabetic
Normal data
3.5
3
2.5
2
1.5
1
0.5
Fold over basal
0
10
20
30
40
50
60
Time (min)

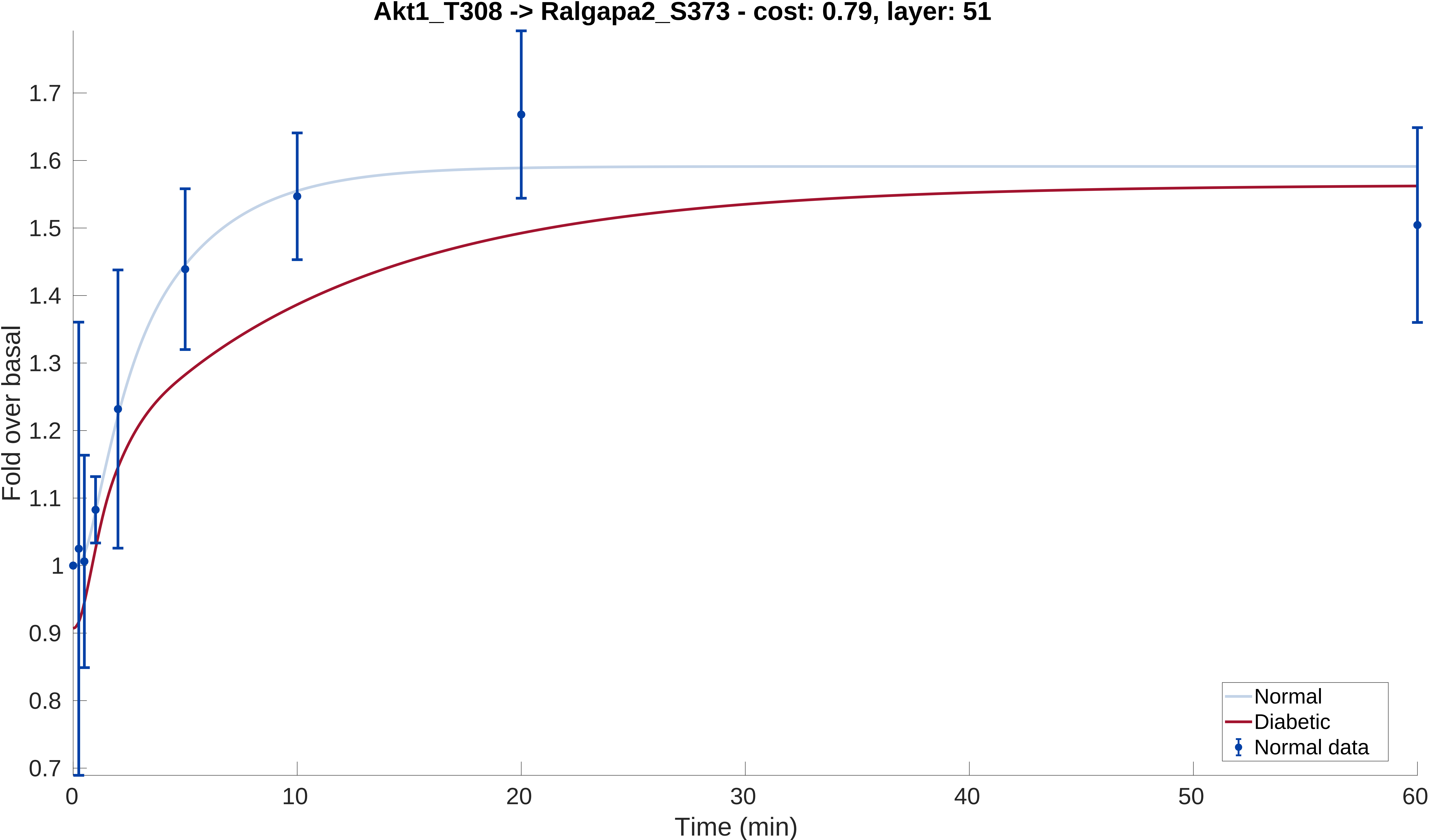

Akt1_T308 -> Ralgapa2_S373 - cost: 0.79, layer: 51
Fold over basal
1.7
1.6
1.5
1.4
1.3
1.2
1.1
1
0.9
0.8
0.7
0
10
20
30
40
50
60
Time (min)
Normal
Diabetic
Normal data

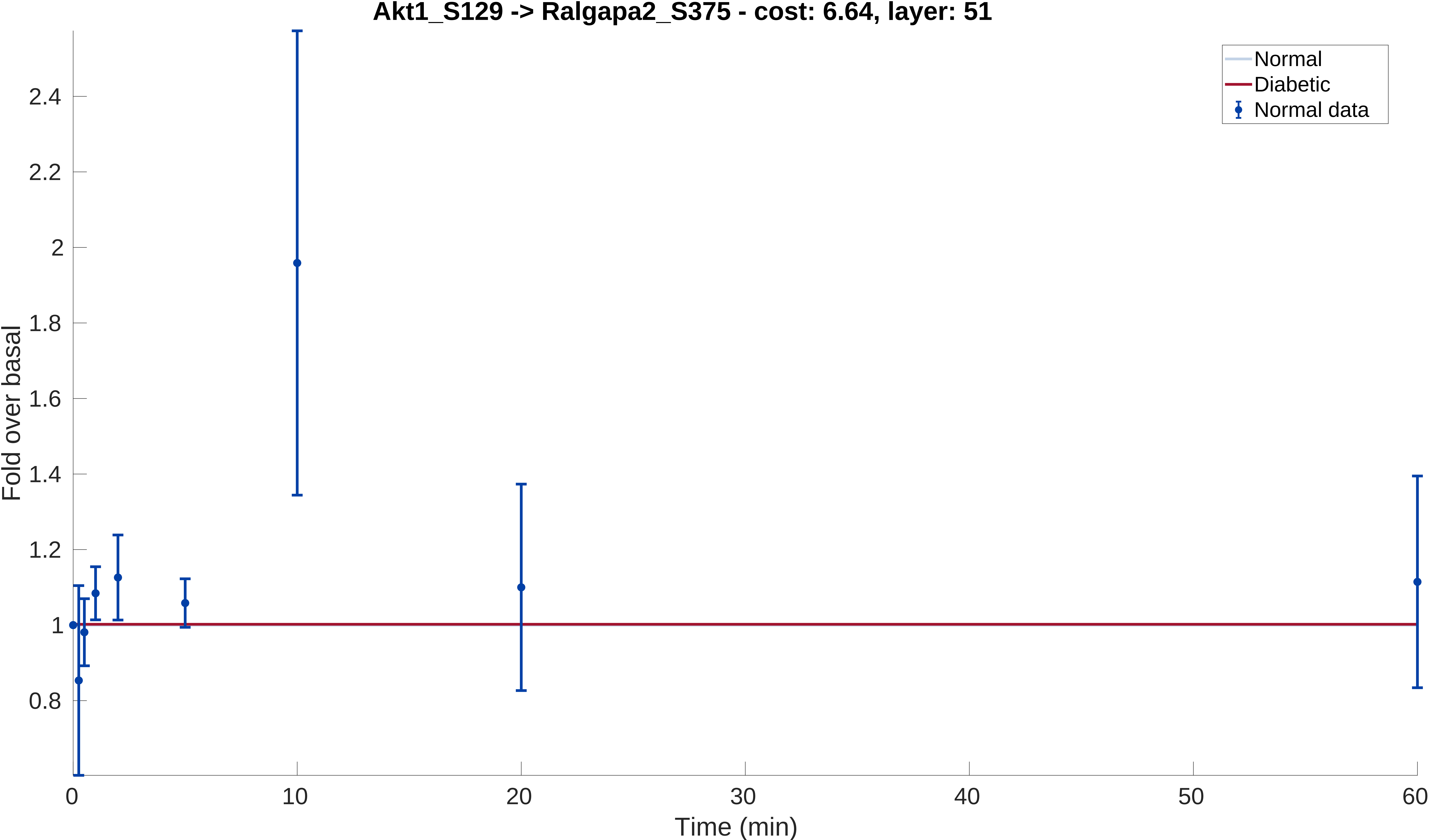

Akt1_S129 -> Ralgapa2_S375 - cost: 6.64, layer: 51
Normal
Diabetic
Normal data
Fold over basal
2.4
2.2
2
1.8
1.6
1.4
1.2
1
0.8
0
10
20
30
40
50
60
Time (min)

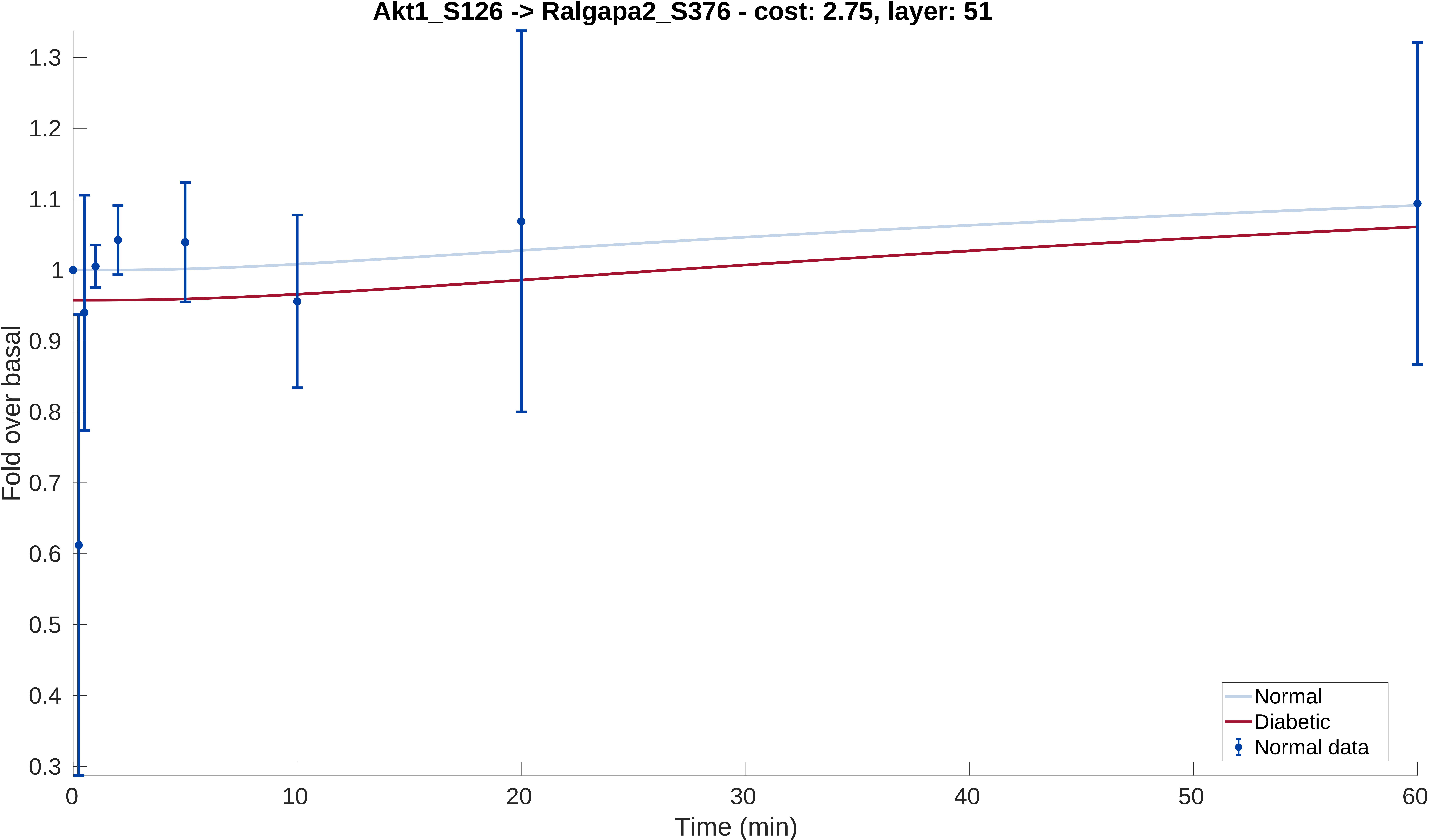

Akt1_S126 -> Ralgapa2_S376 - cost: 2.75, layer: 51
Fold over basal
Time (min)
1.3
1.2
1.1
1
0.9
0.8
0.7
0.6
0.5
0.4
0.3
0
10
20
30
40
50
60
Normal
Diabetic
Normal data

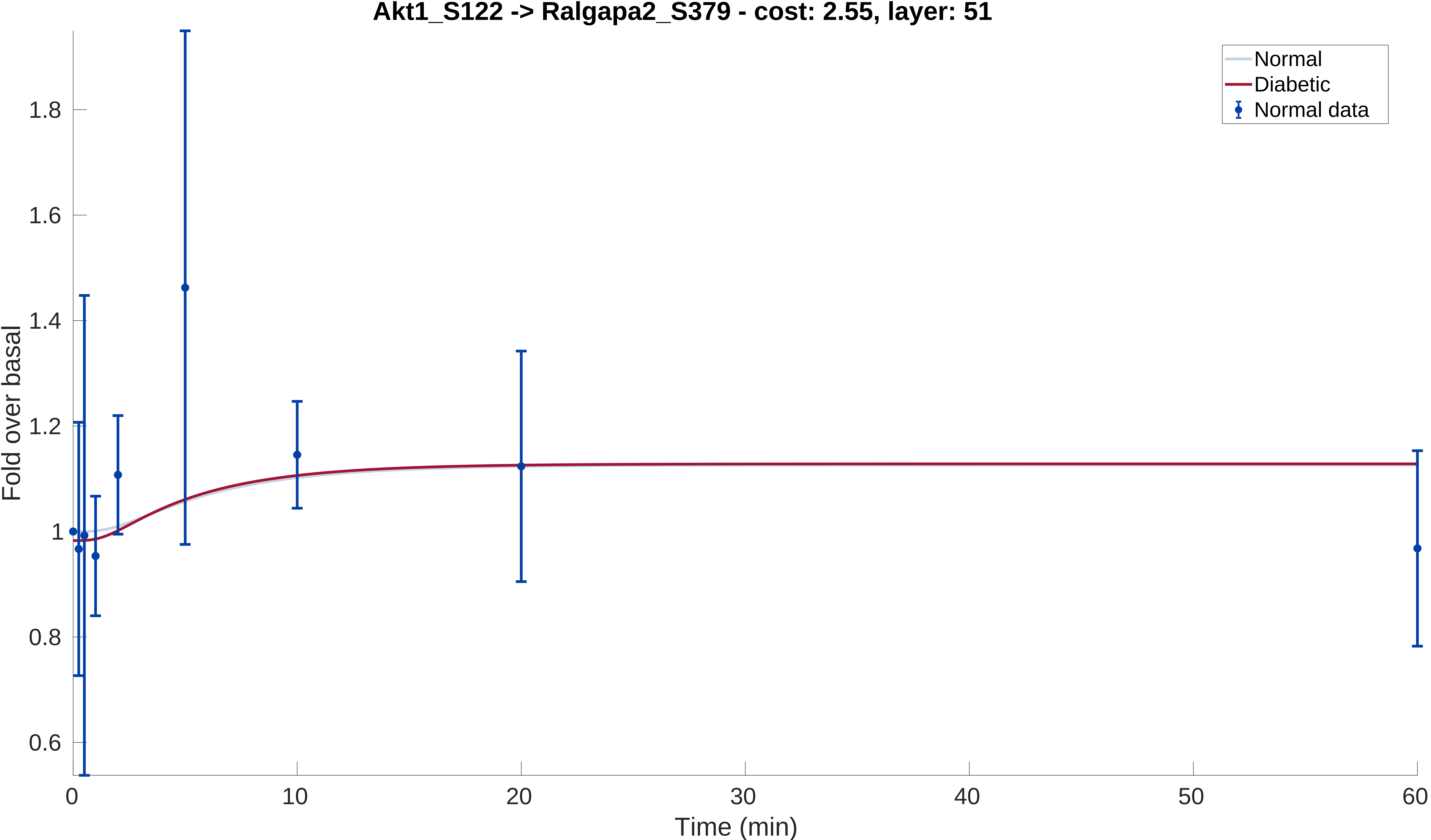
Akt1_S122 -> Ralgapa2_S379 - cost: 2.55, layer: 51
Normal
Diabetic
Normal data
Fold over basal
1.8
1.6
1.4
1.2
1
0.8
0.6
0
10
20
30
40
50
60
Time (min)

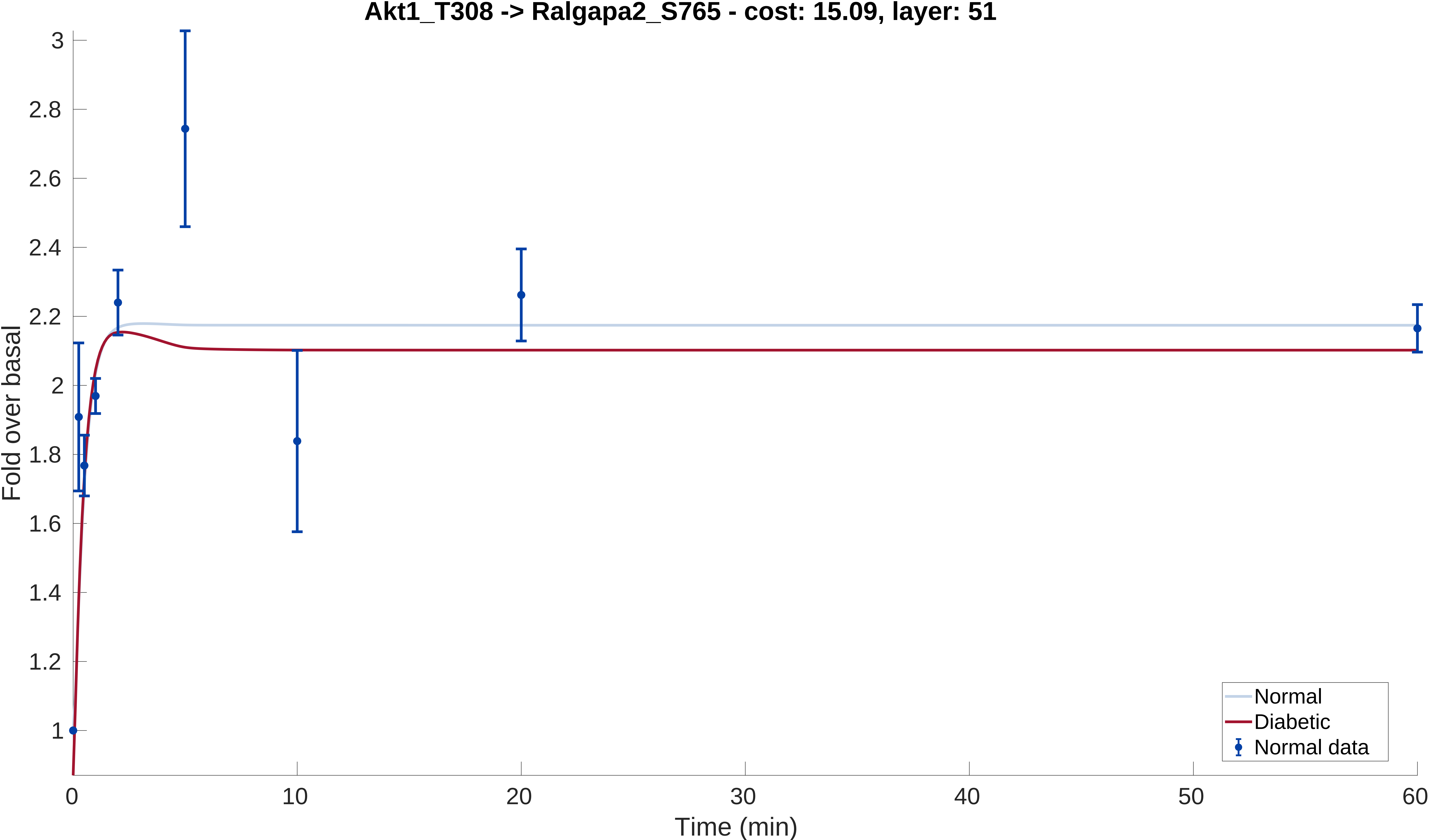
Akt1_T308 -> Ralgapa2_S765 - cost: 15.09, layer: 51
Fold over basal
Time (min)
Normal
Diabetic
Normal data
0
10
20
30
40
50
60
1
1.2
1.4
1.6
1.8
2
2.2
2.4
2.6
2.8
3

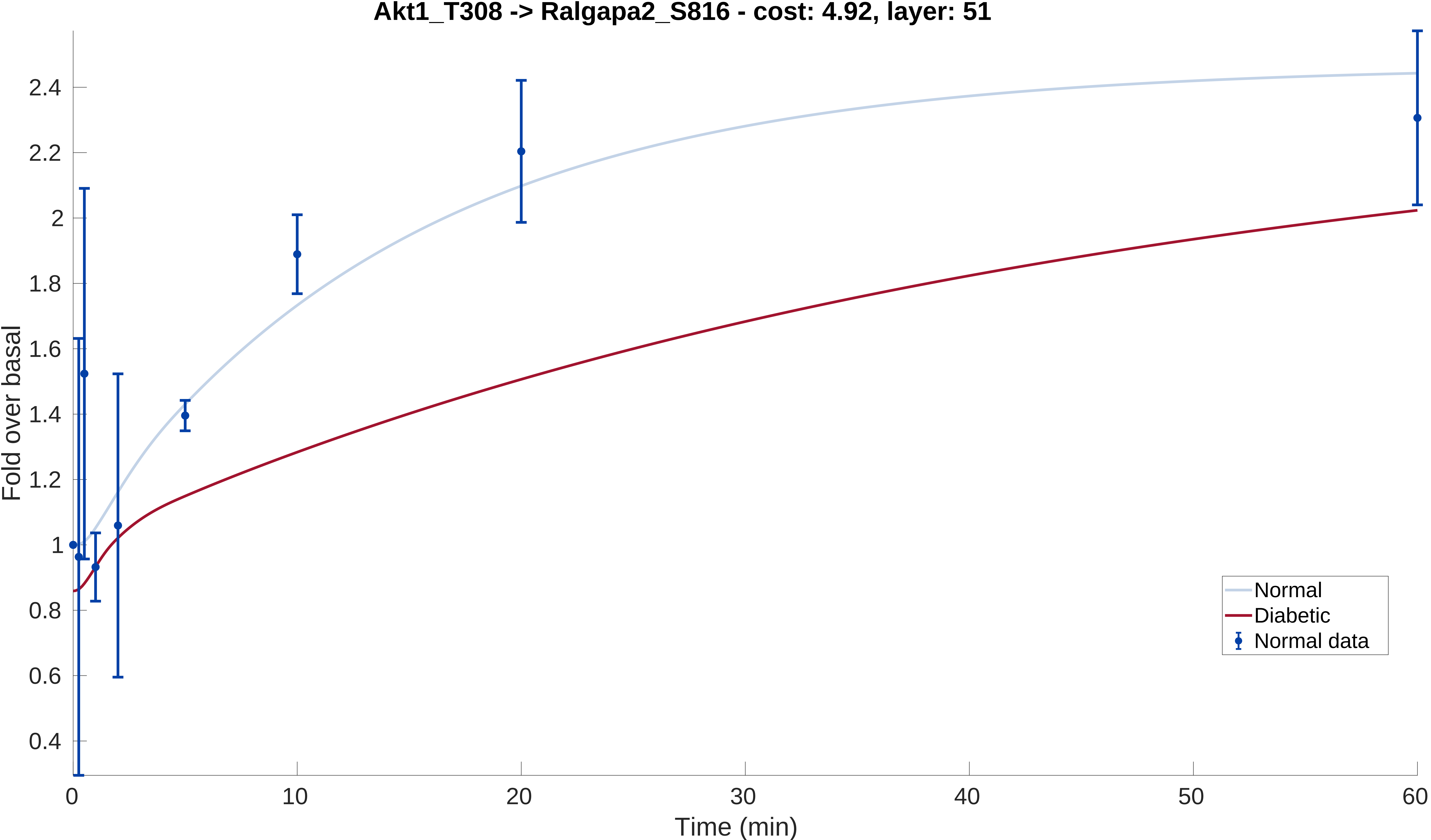
Akt1_T308 -> Ralgapa2_S816 - cost: 4.92, layer: 51
Fold over basal
2.4
2.2
2
1.8
1.6
1.4
1.2
1
0.8
0.6
0.4
0
10
20
30
40
50
60
Time (min)
Normal
Diabetic
Normal data

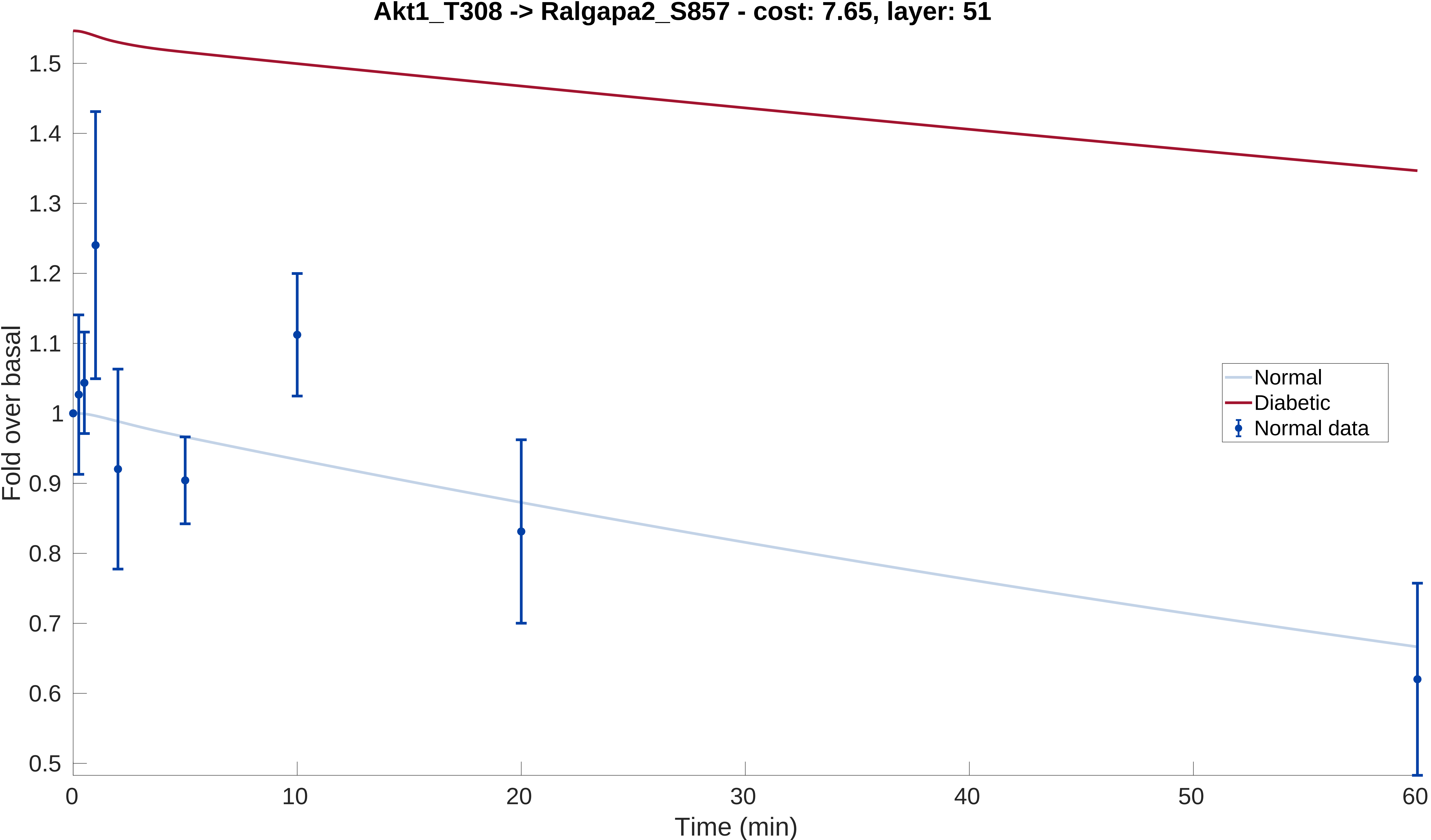
Akt1_T308 -> Ralgapa2_S857 - cost: 7.65, layer: 51
Fold over basal
Time (min)
1.5
1.4
1.3
1.2
1.1
1
0.9
0.8
0.7
0.6
0.5
0
10
20
30
40
50
60
Normal
Diabetic
Normal data

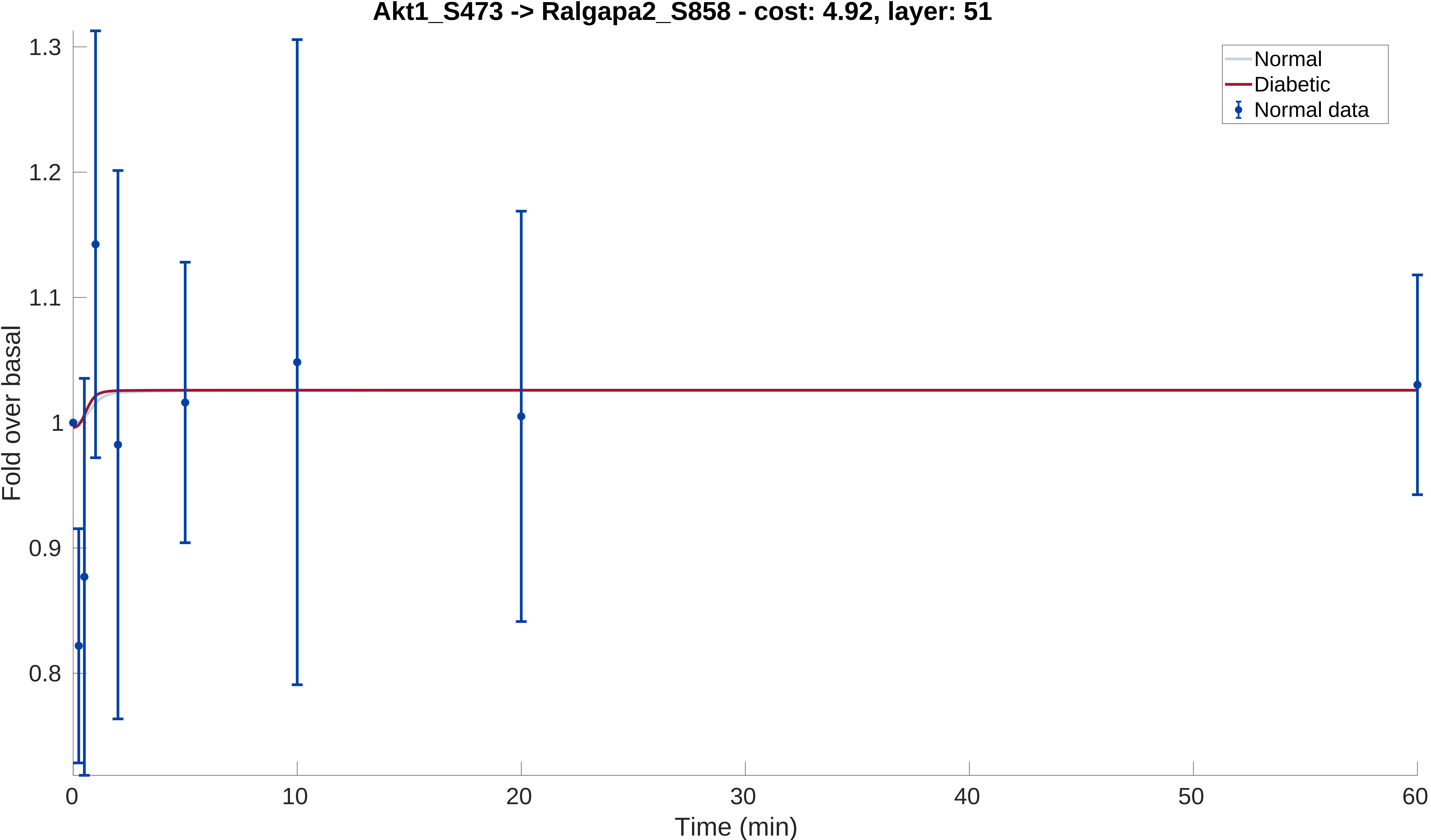

Akt1_S473 -> Ralgapa2_S858 - cost: 4.92, layer: 51
Normal
Diabetic
Normal data
Fold over basal
1.3
1.2
1.1
1
0.9
0.8
0
10
20
30
40
50
60
Time (min)

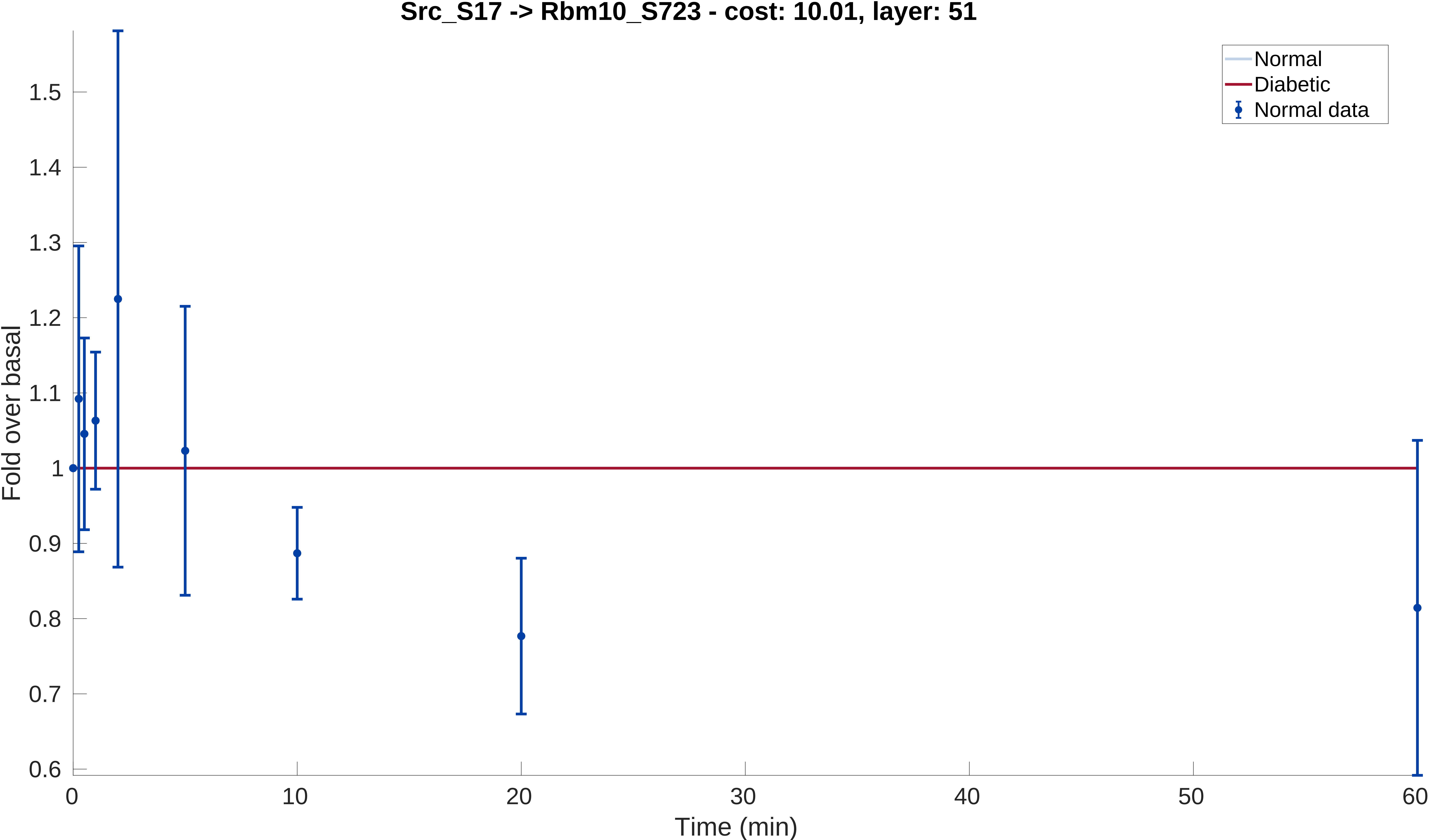

Src_S17 -> Rbm10_S723 - cost: 10.01, layer: 51
Normal
Diabetic
Normal data
Fold over basal
1.5
1.4
1.3
1.2
1.1
1
0.9
0.8
0.7
0.6
0
10
20
30
40
50
60
Time (min)

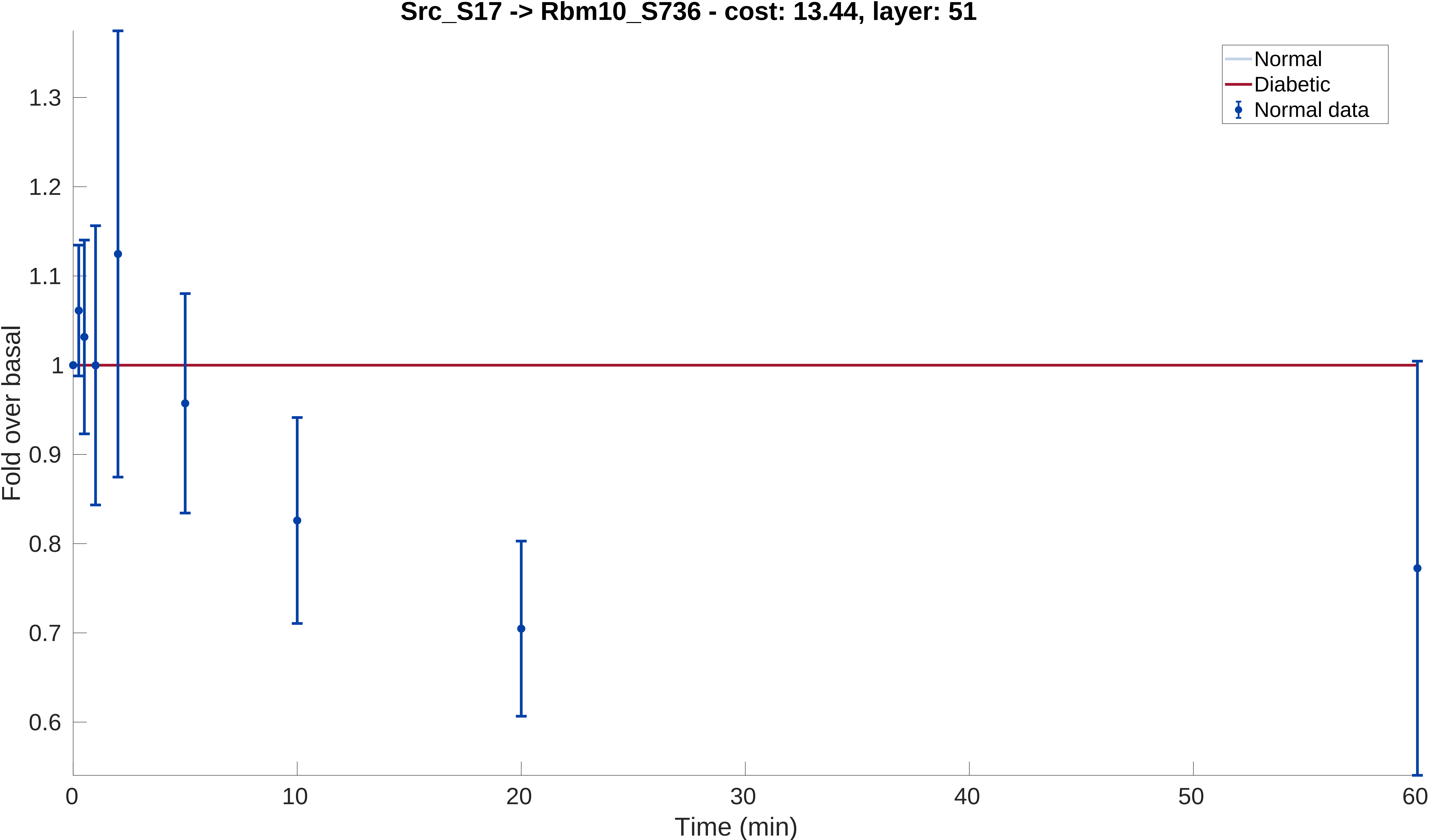

Src_S17 -> Rbm10_S736 - cost: 13.44, layer: 51
Normal
Diabetic
Normal data
Fold over basal
Time (min)
1.3
1.2
1.1
1
0.9
0.8
0.7
0.6
0
10
20
30
40
50
60

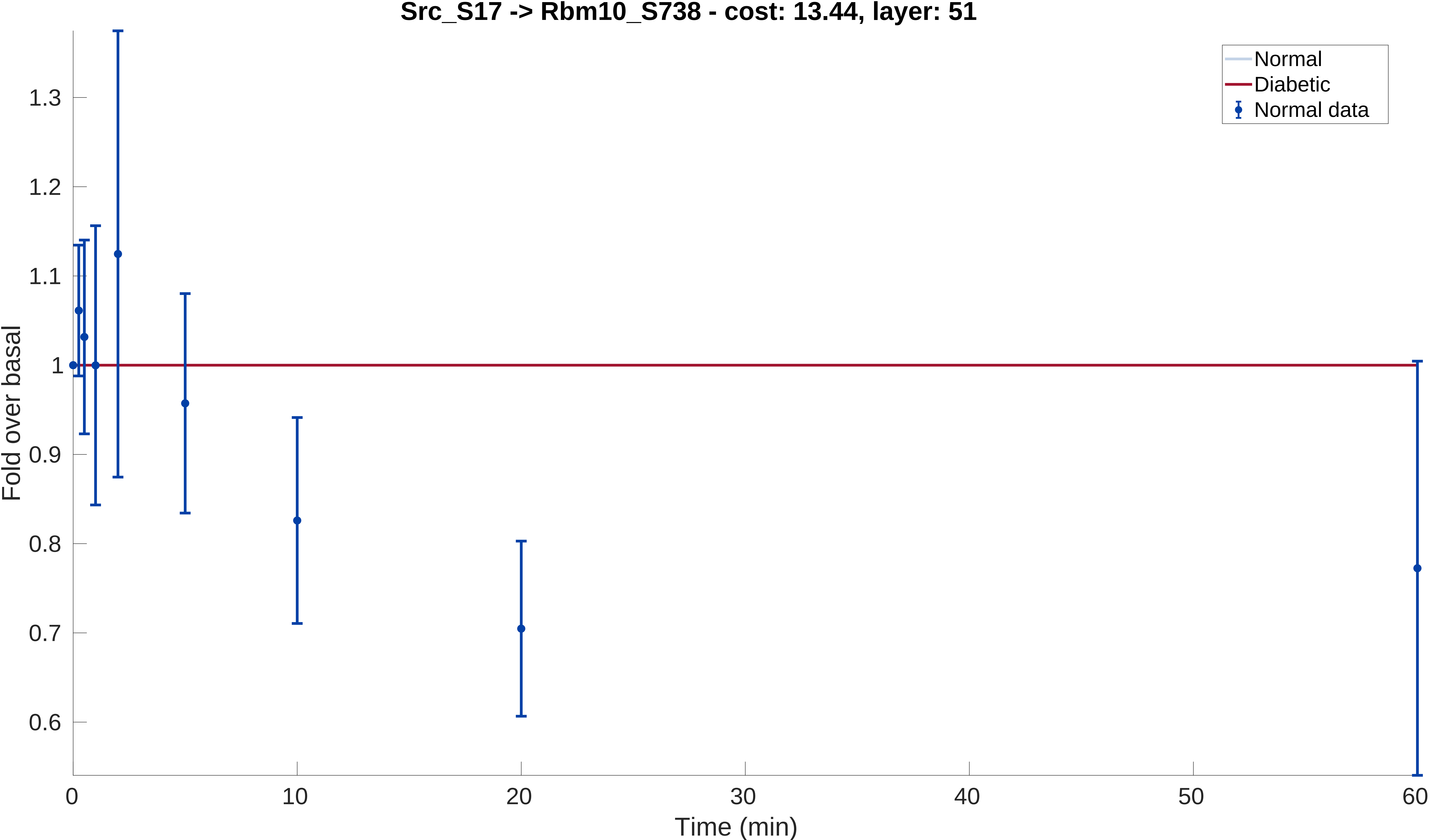

Src_S17 -> Rbm10_S738 - cost: 13.44, layer: 51
Normal
Diabetic
Normal data
Fold over basal
1.3
1.2
1.1
1
0.9
0.8
0.7
0.6
0
10
20
30
40
50
60
Time (min)

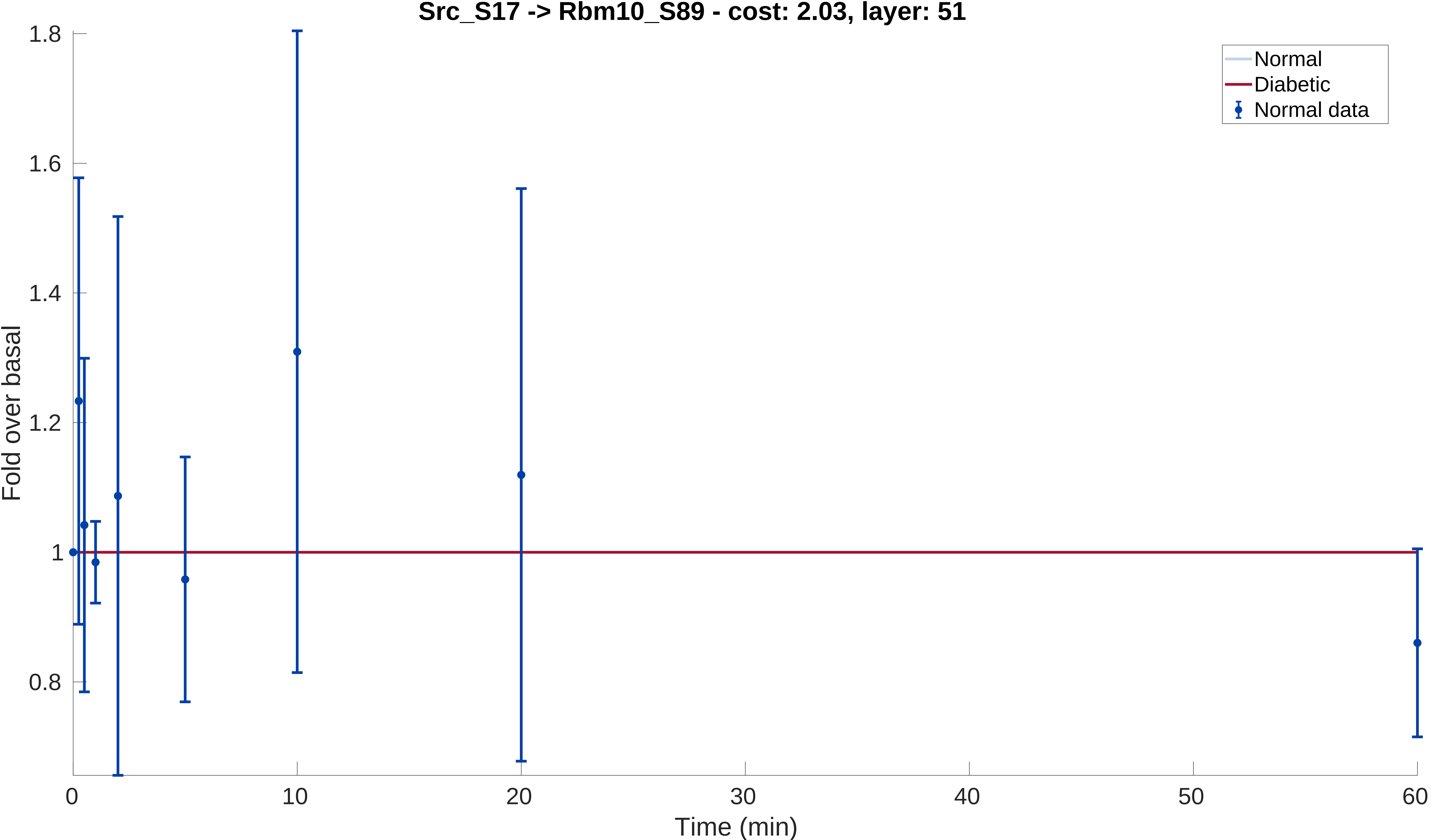

Src_S17 -> Rbm10_S89 - cost: 2.03, layer: 51
Normal
Diabetic
Normal data
Fold over basal
1.8
1.6
1.4
1.2
1
0.8
0
10
20
30
40
50
60
Time (min)

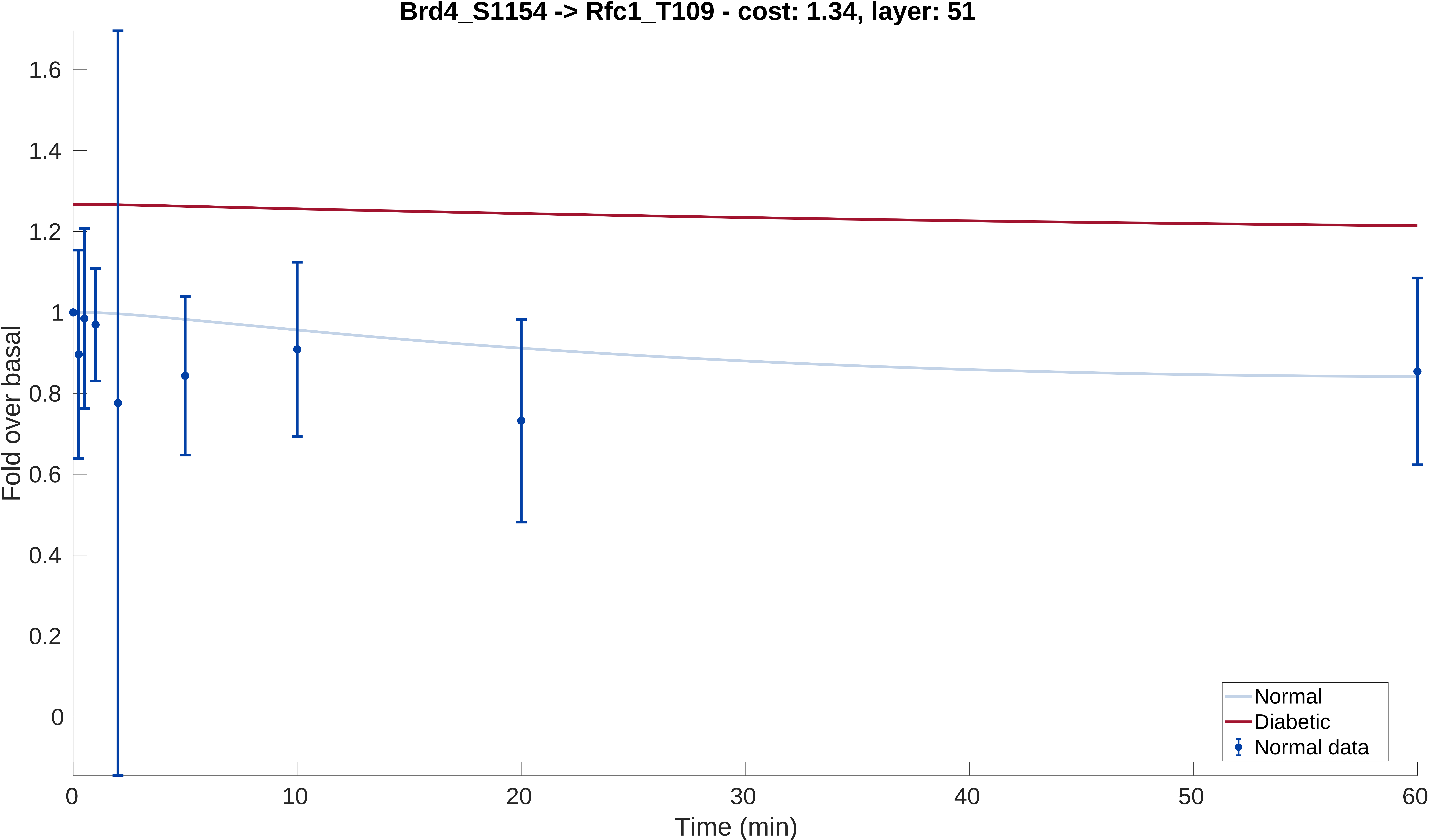
Brd4_S1154 -> Rfc1_T109 - cost: 1.34, layer: 51
Fold over basal
1.6
1.4
1.2
1
0.8
0.6
0.4
0.2
0
0
10
20
30
40
50
60
Time (min)
Normal
Diabetic
Normal data

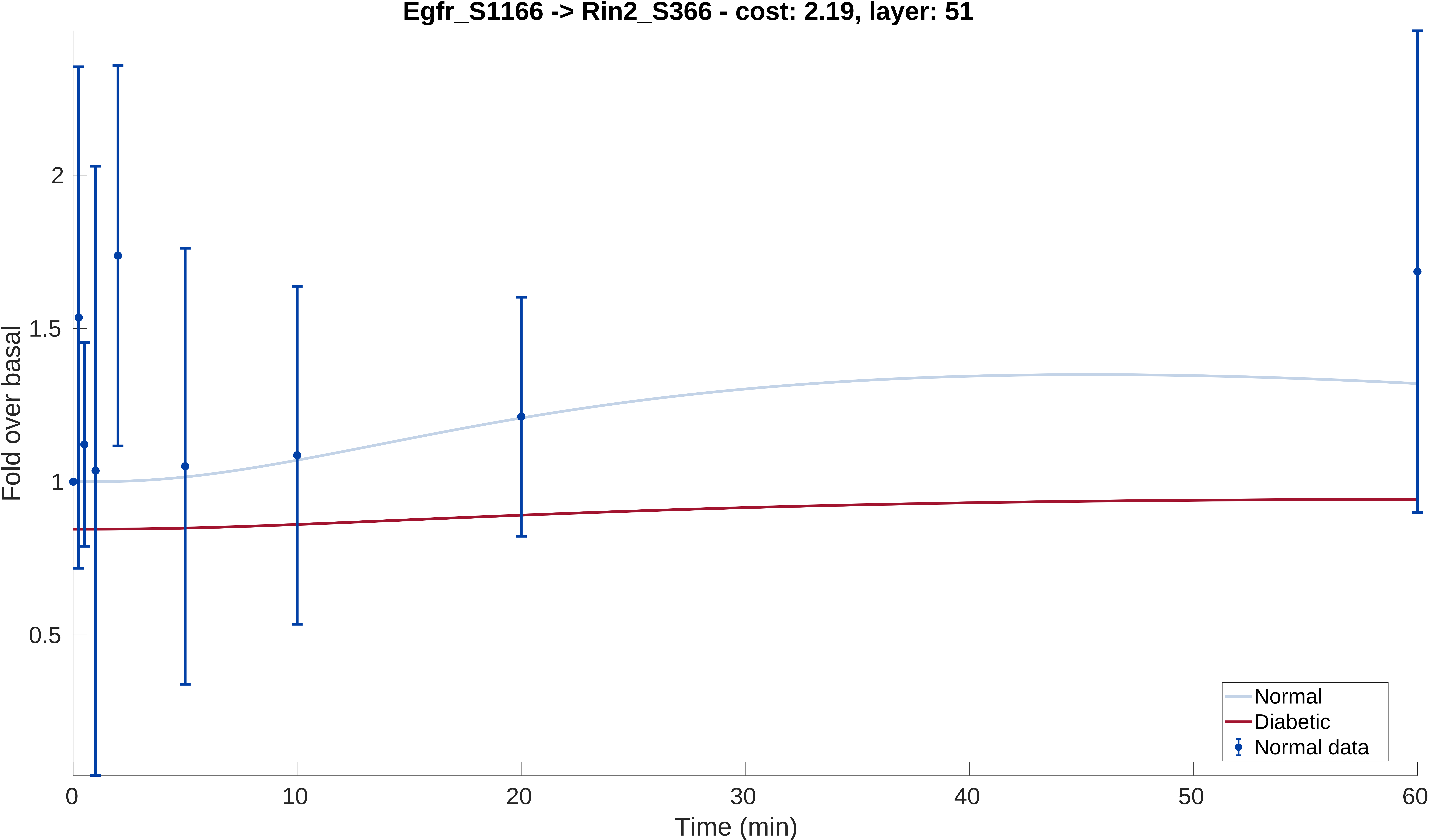
Egfr_S1166 -> Rin2_S366 - cost: 2.19, layer: 51
Fold over basal
2
1.5
1
0.5
0
10
20
30
40
50
60
Time (min)
Normal
Diabetic
Normal data

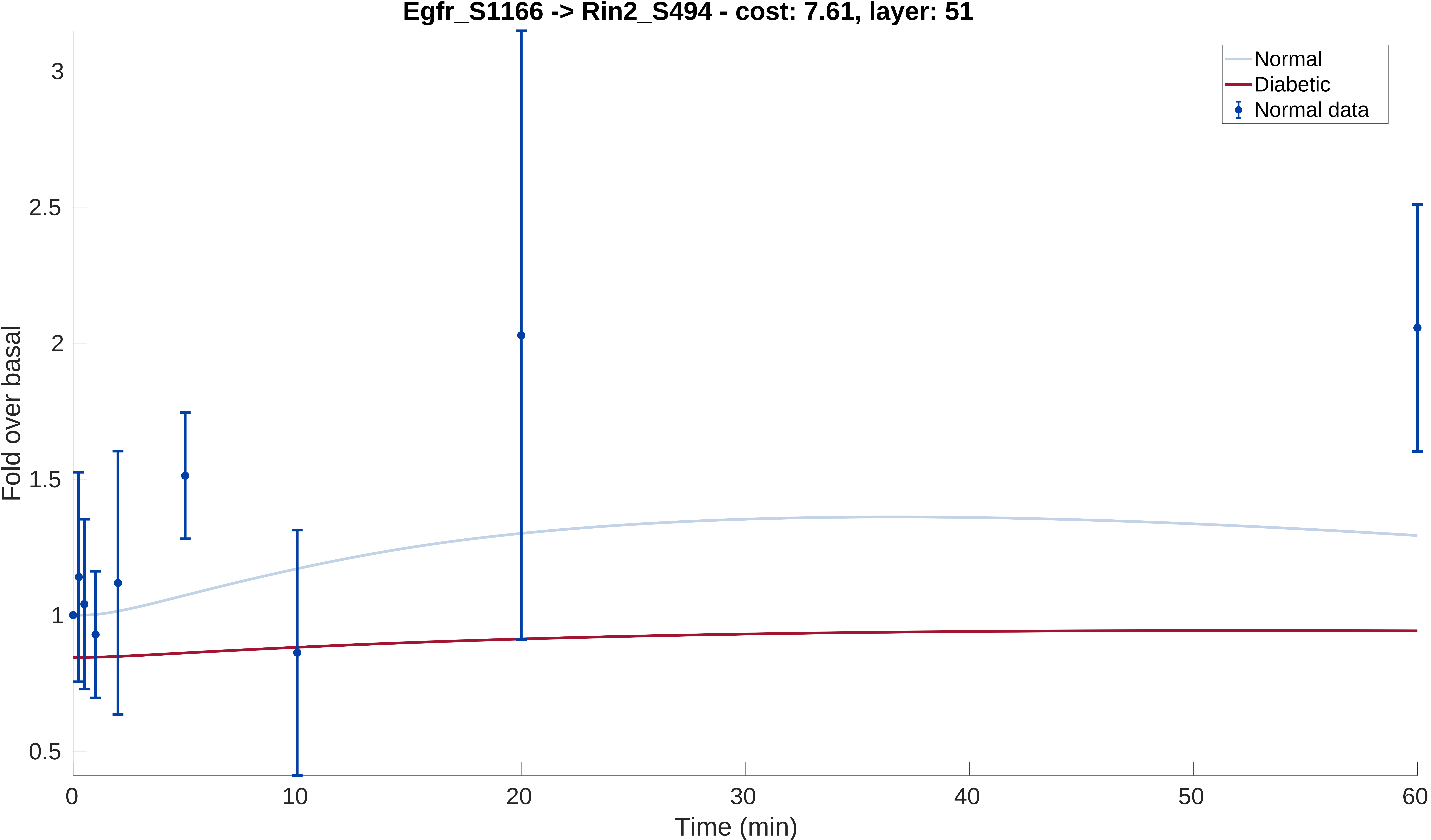
Egfr_S1166 -> Rin2_S494 - cost: 7.61, layer: 51
Normal
Diabetic
Normal data
Fold over basal
3
2.5
2
1.5
1
0.5
0
10
20
30
40
50
60
Time (min)

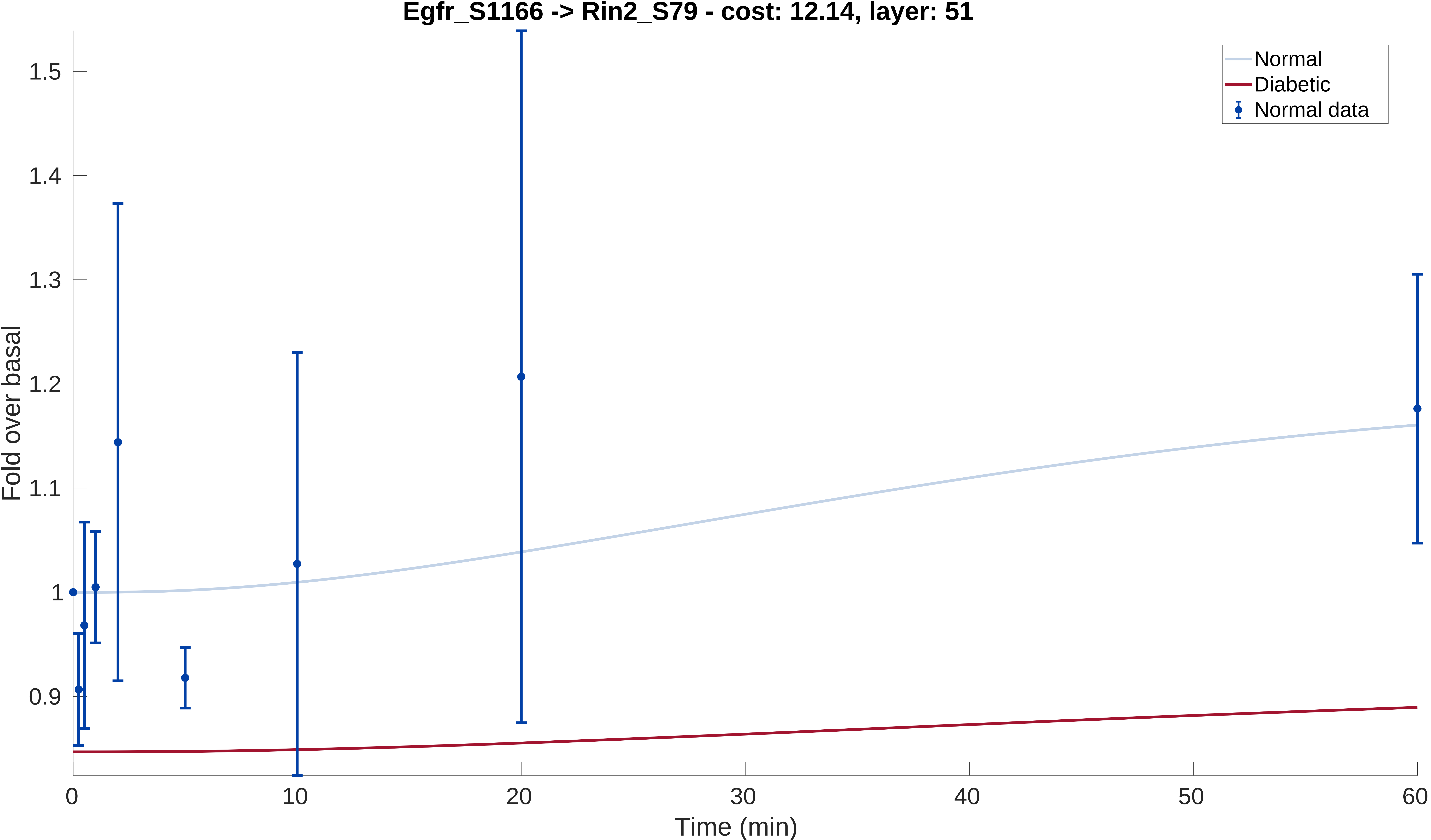

Egfr_S1166 -> Rin2_S79 - cost: 12.14, layer: 51
Normal
Diabetic
Normal data
Fold over basal
1.5
1.4
1.3
1.2
1.1
1
0.9
0
10
20
30
40
50
60
Time (min)

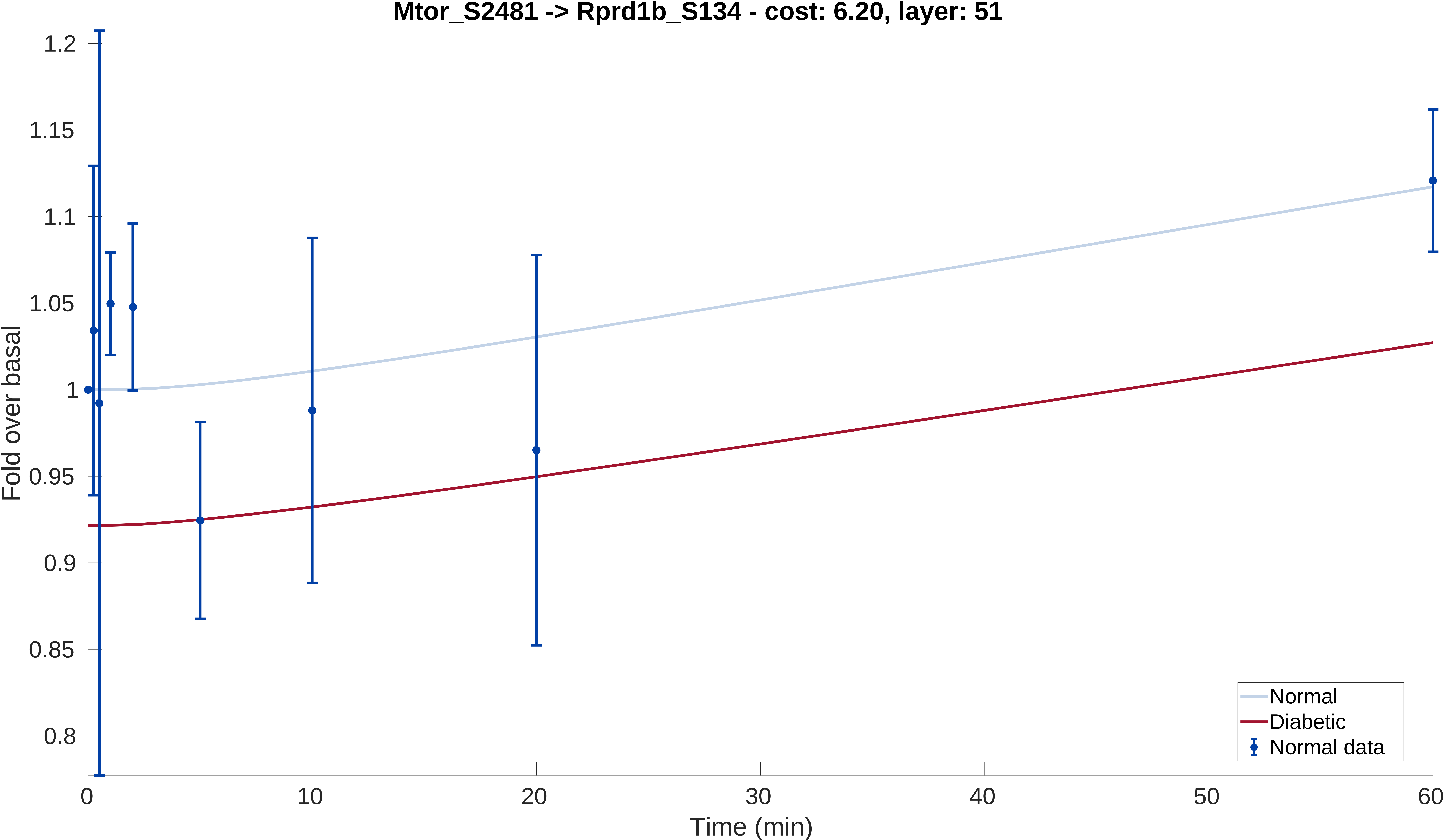

Mtor_S2481 -> Rprd1b_S134 - cost: 6.20, layer: 51
Fold over basal
Time (min)
1.2
1.15
1.1
1.05
1
0.95
0.9
0.85
0.8
0
10
20
30
40
50
60
Normal
Diabetic
Normal data

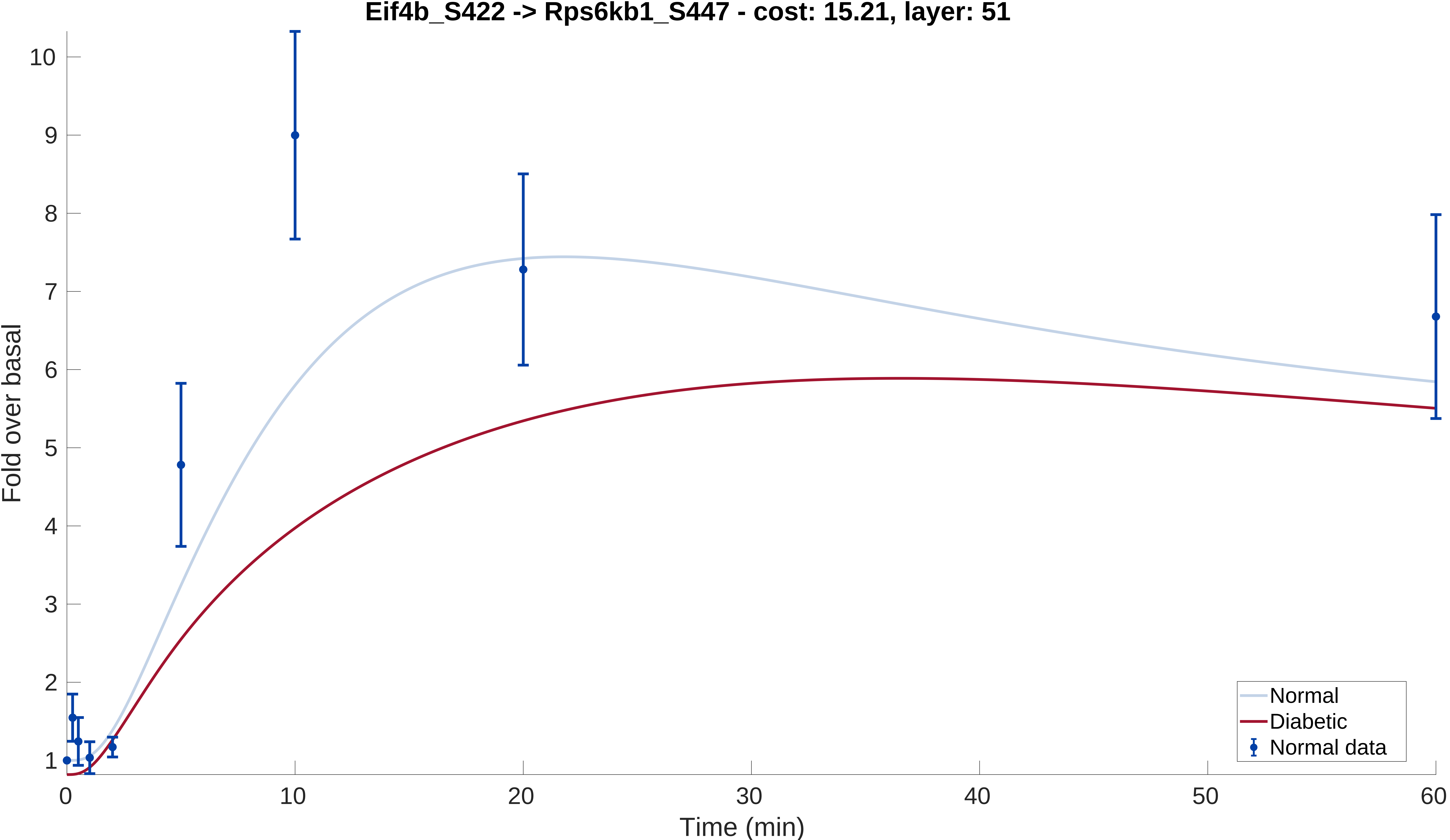

Eif4b_S422 -> Rps6kb1_S447 - cost: 15.21, layer: 51
Fold over basal
Time (min)
Normal
Diabetic
Normal data
0
10
20
30
40
50
60
1
2
3
4
5
6
7
8
9
10

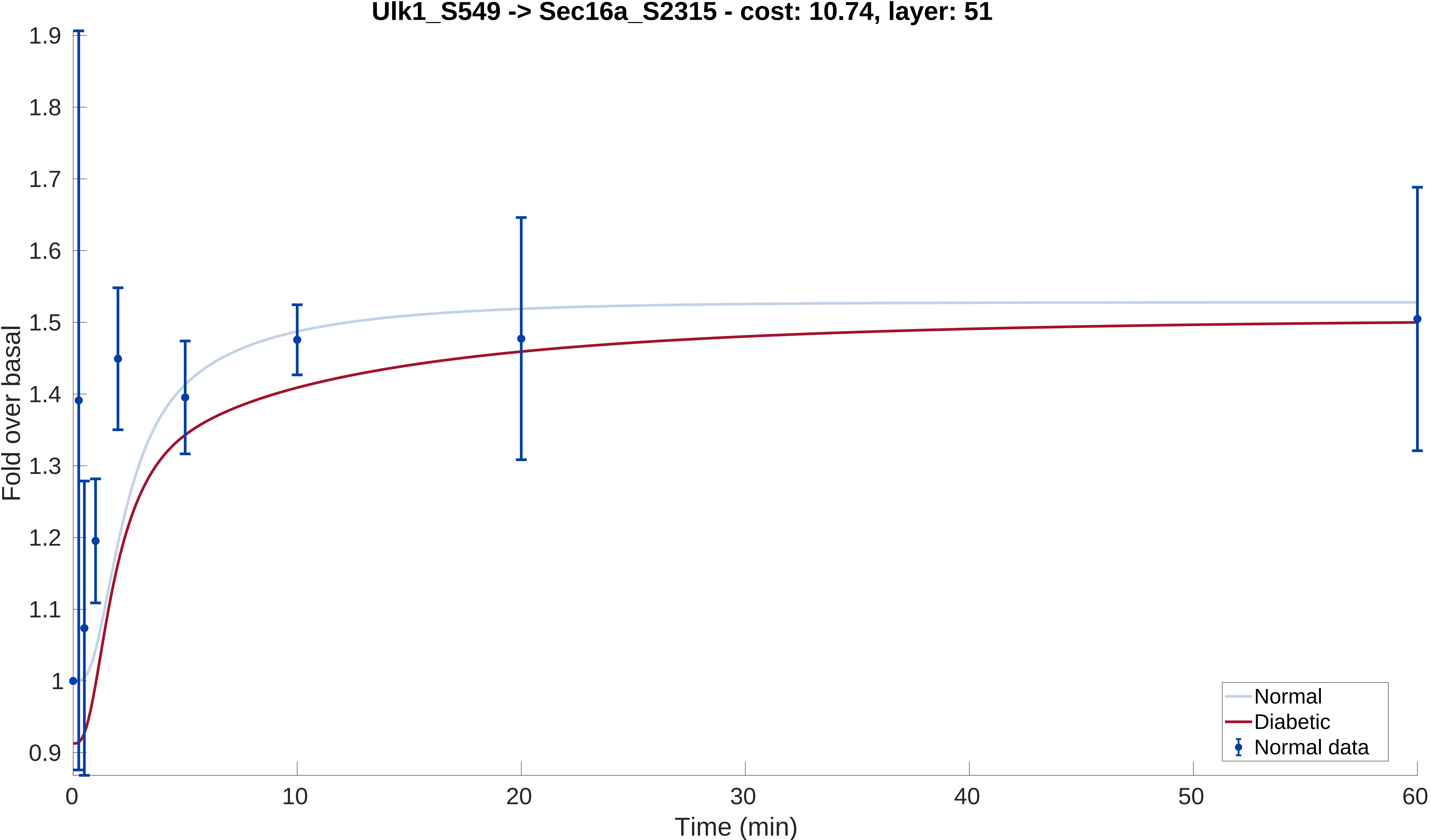
Ulk1_S549 -> Sec16a_S2315 - cost: 10.74, layer: 51
Fold over basal
1.9
1.8
1.7
1.6
1.5
1.4
1.3
1.2
1.1
1
0.9
0
10
20
30
40
50
60
Time (min)
Normal
Diabetic
Normal data

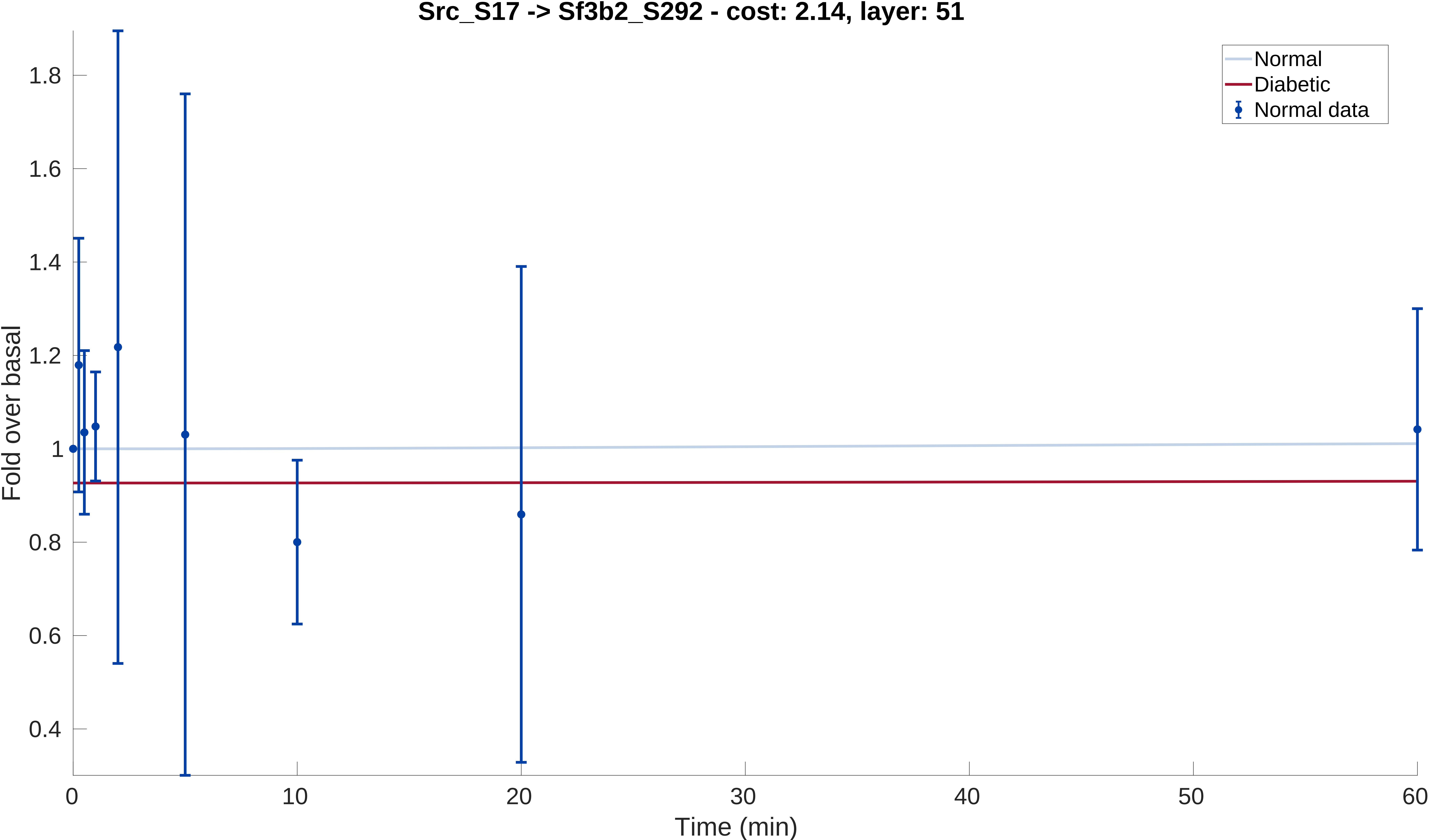

Src_S17 -> Sf3b2_S292 - cost: 2.14, layer: 51
Normal
Diabetic
Normal data
Fold over basal
1.8
1.6
1.4
1.2
1
0.8
0.6
0.4
0
10
20
30
40
50
60
Time (min)

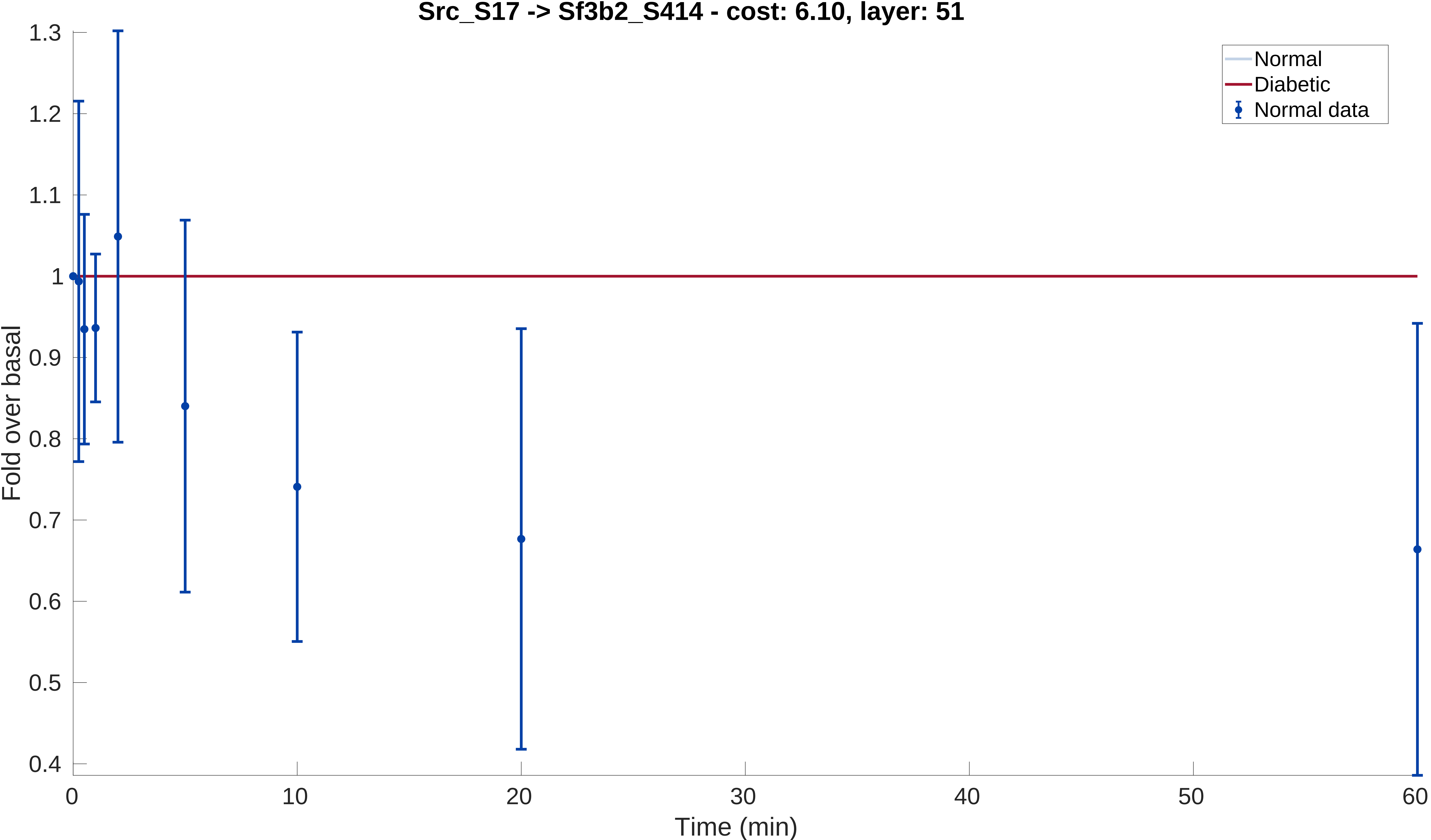

Src_S17 -> Sf3b2_S414 - cost: 6.10, layer: 51
Fold over basal
1.3
1.2
1.1
1
0.9
0.8
0.7
0.6
0.5
0.4
0
10
20
30
40
50
60
Time (min)
Normal
Diabetic
Normal data

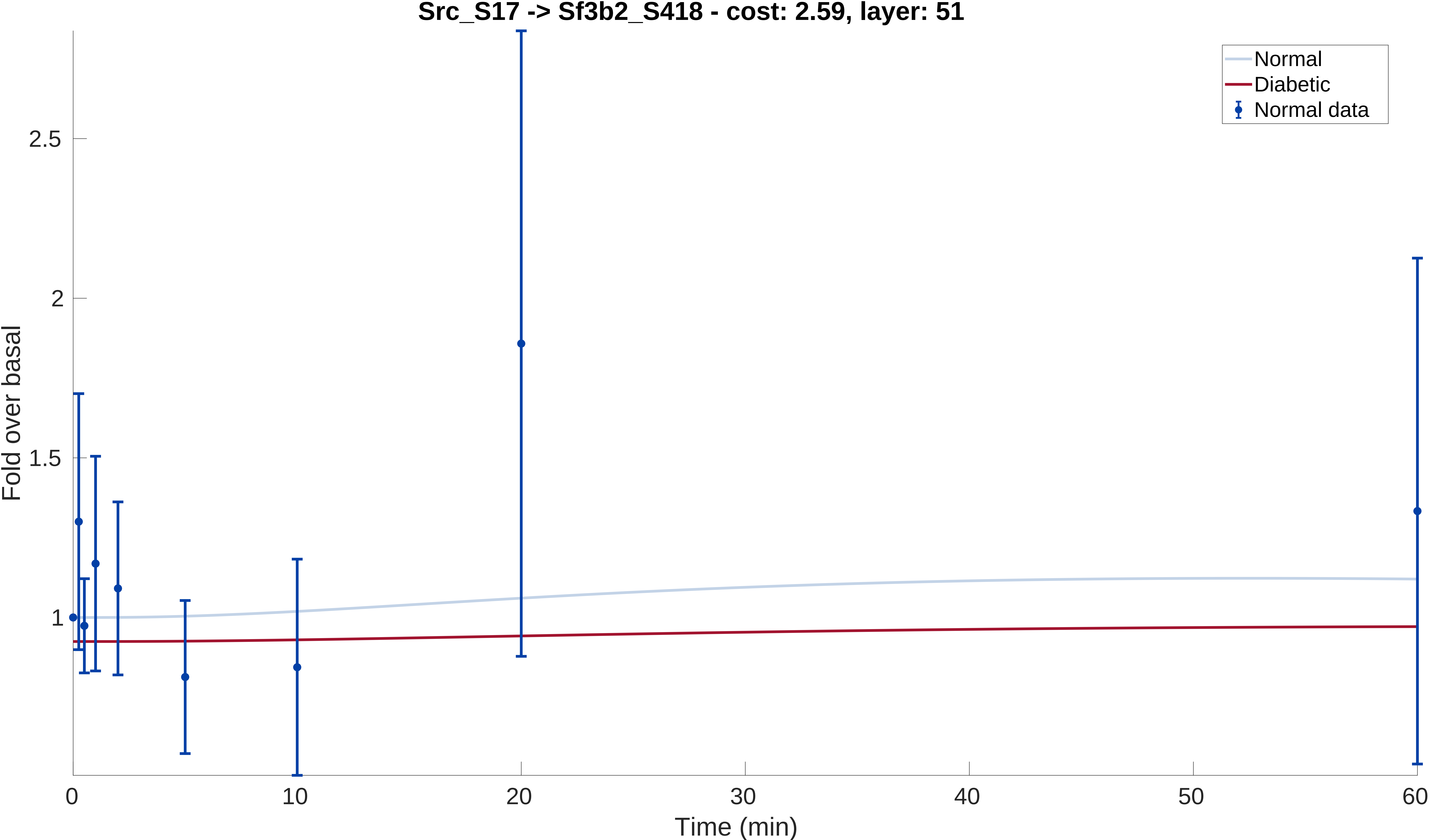

Src_S17 -> Sf3b2_S418 - cost: 2.59, layer: 51
Normal
Diabetic
Normal data
Fold over basal
2.5
2
1.5
1
0
10
20
30
40
50
60
Time (min)

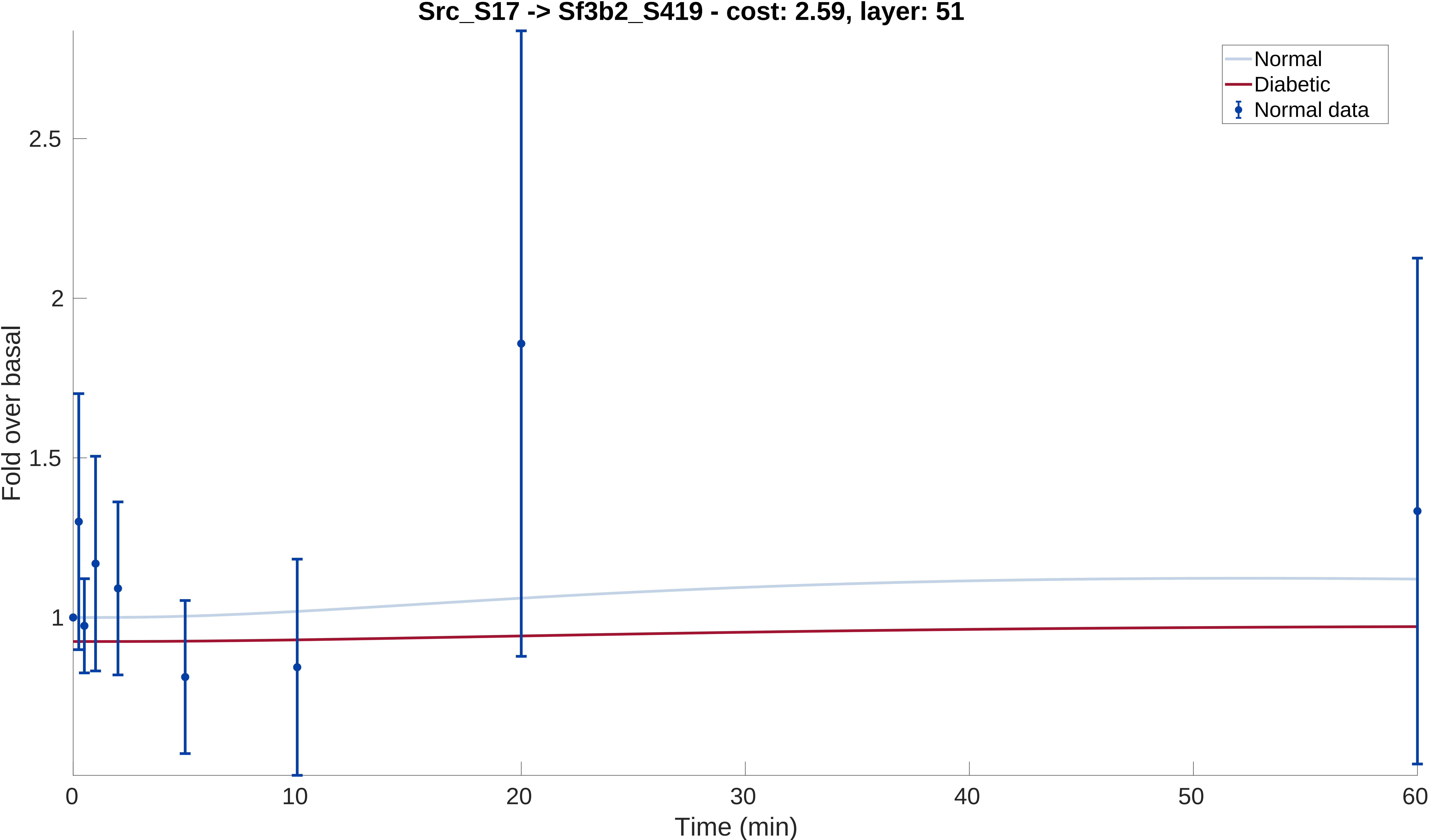

Src_S17 -> Sf3b2_S419 - cost: 2.59, layer: 51
Normal
Diabetic
Normal data
Fold over basal
2.5
2
1.5
1
0
10
20
30
40
50
60
Time (min)

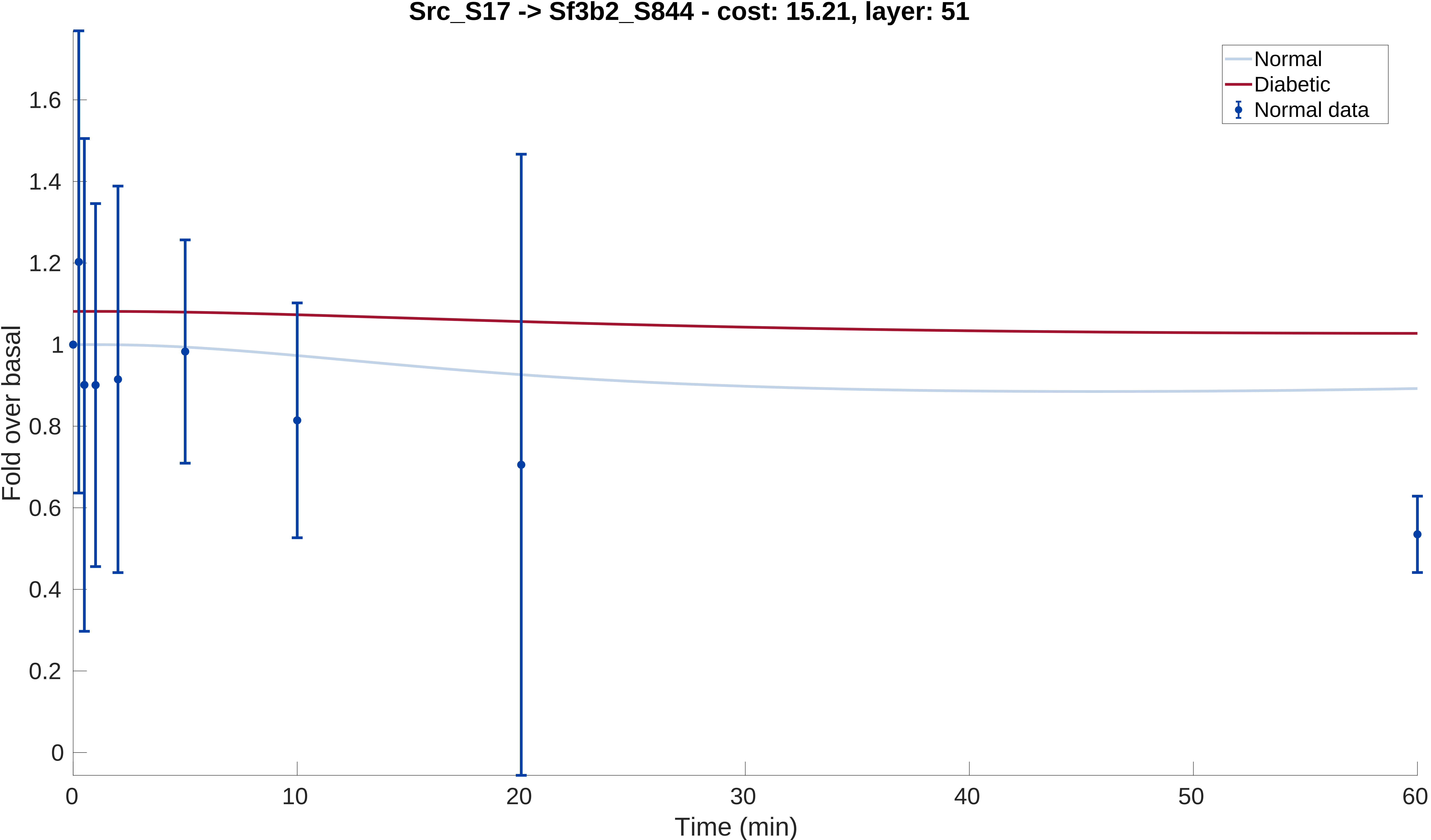

Src_S17 -> Sf3b2_S844 - cost: 15.21, layer: 51
Normal
Diabetic
Normal data
Fold over basal
Time (min)
0
0.2
0.4
0.6
0.8
1
1.2
1.4
1.6
0
10
20
30
40
50
60

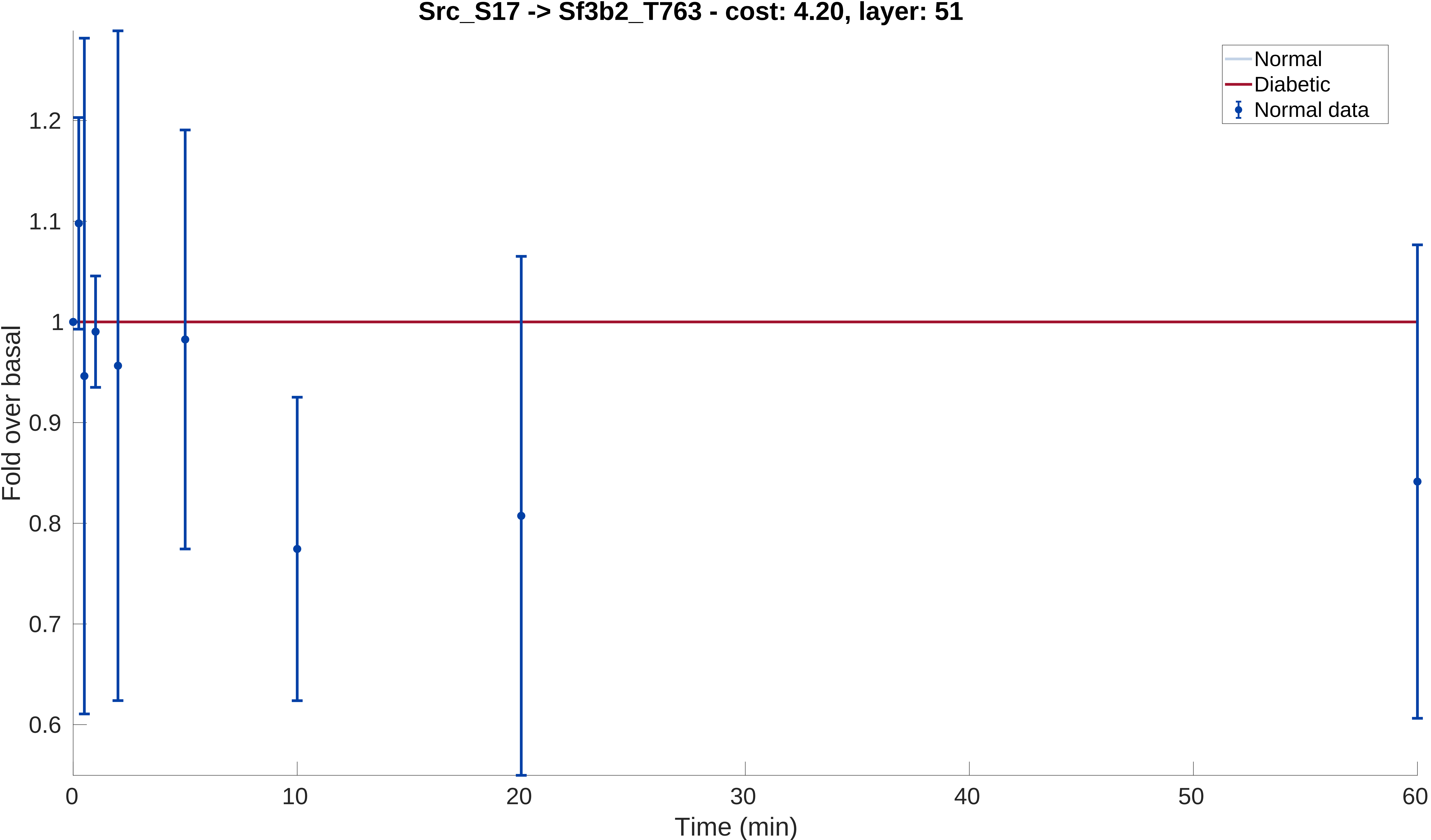

Src_S17 -> Sf3b2_T763 - cost: 4.20, layer: 51
Normal
Diabetic
Normal data
Fold over basal
1.2
1.1
1
0.9
0.8
0.7
0.6
0
10
20
30
40
50
60
Time (min)

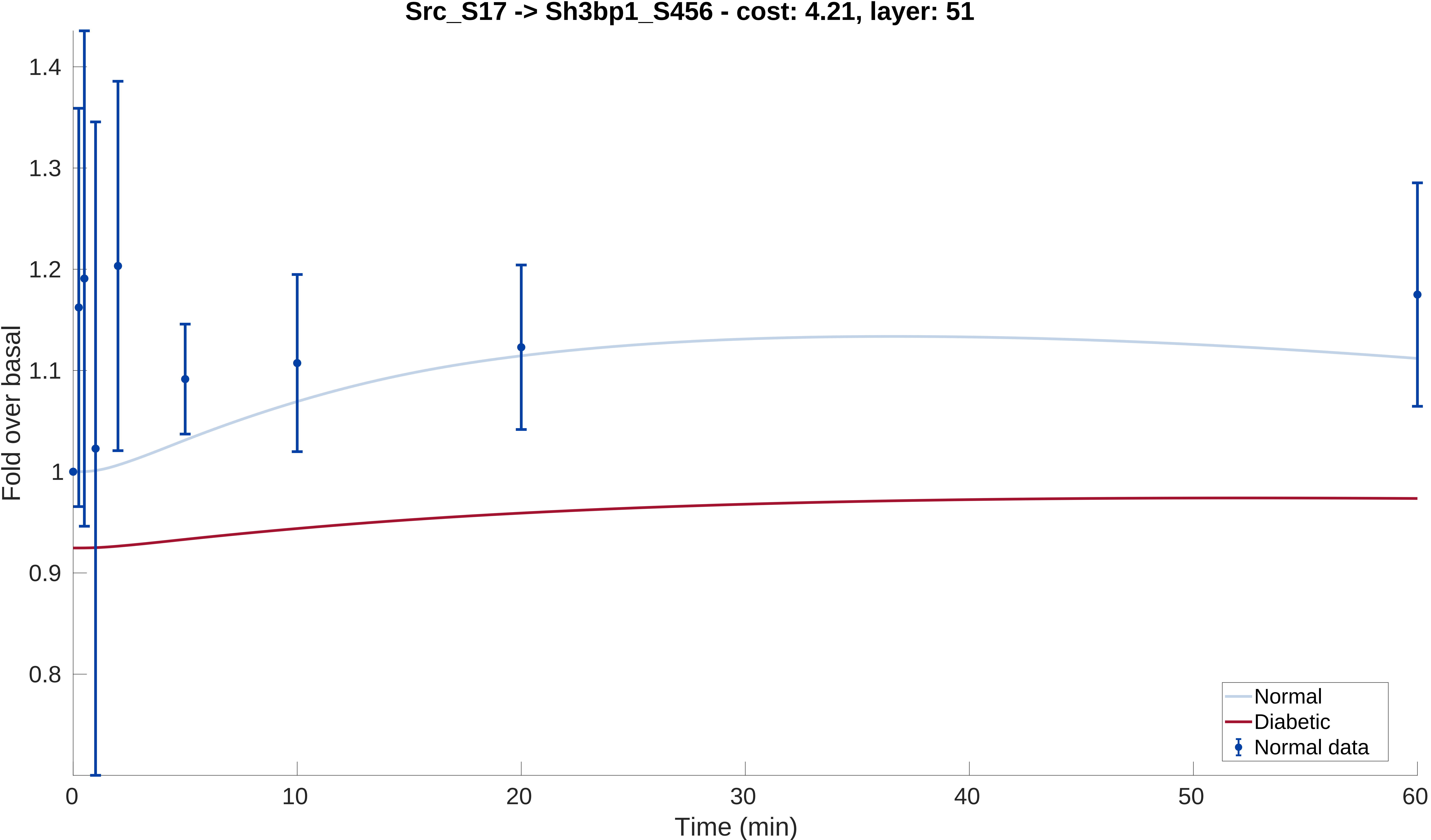

Src_S17 -> Sh3bp1_S456 - cost: 4.21, layer: 51
Fold over basal
1.4
1.3
1.2
1.1
1
0.9
0.8
0
10
20
30
40
50
60
Time (min)
Normal
Diabetic
Normal data

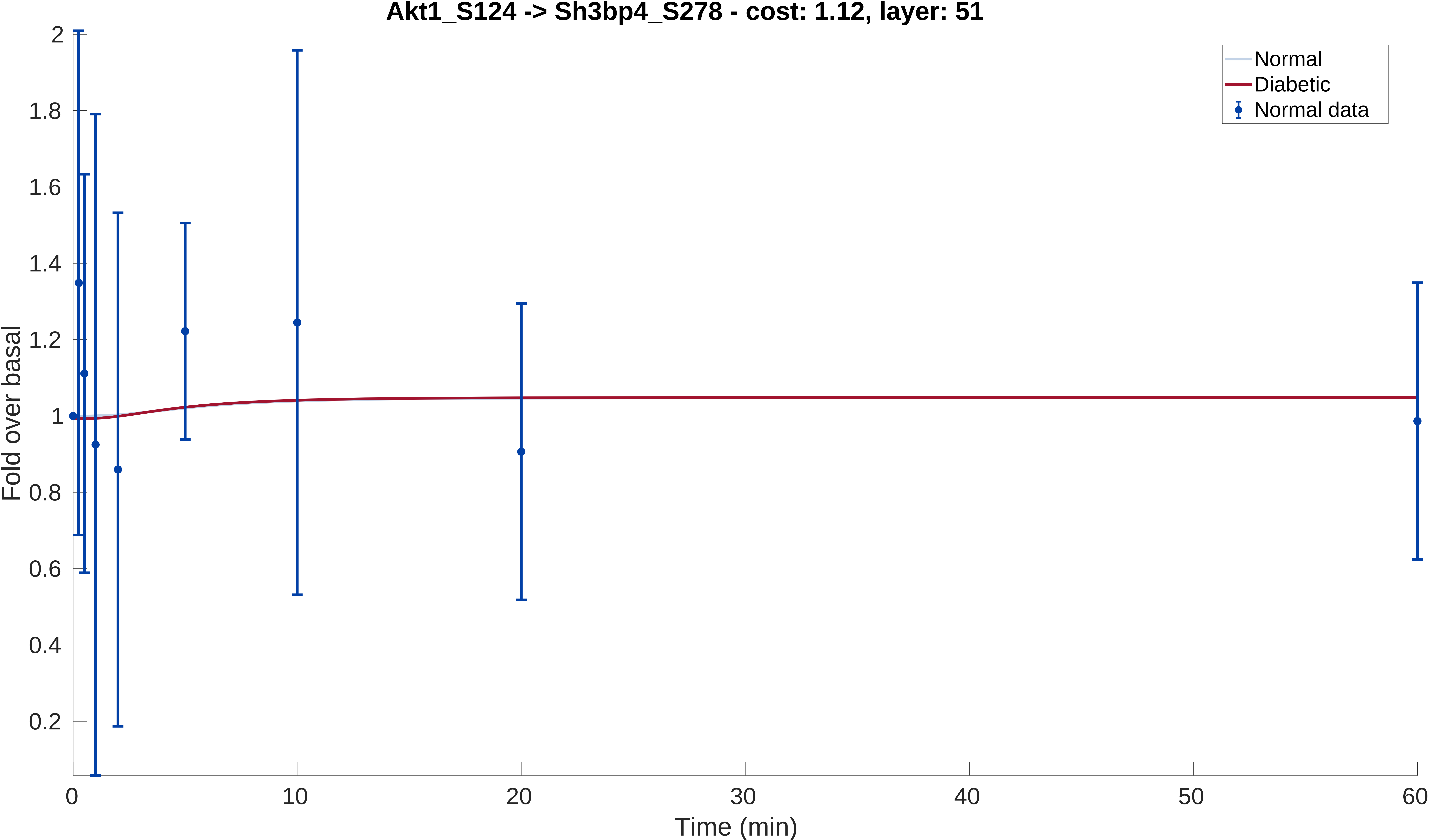
Akt1_S124 -> Sh3bp4_S278 - cost: 1.12, layer: 51
Normal
Diabetic
Normal data
Fold over basal
Time (min)
0
10
20
30
40
50
60
0.2
0.4
0.6
0.8
1
1.2
1.4
1.6
1.8
2

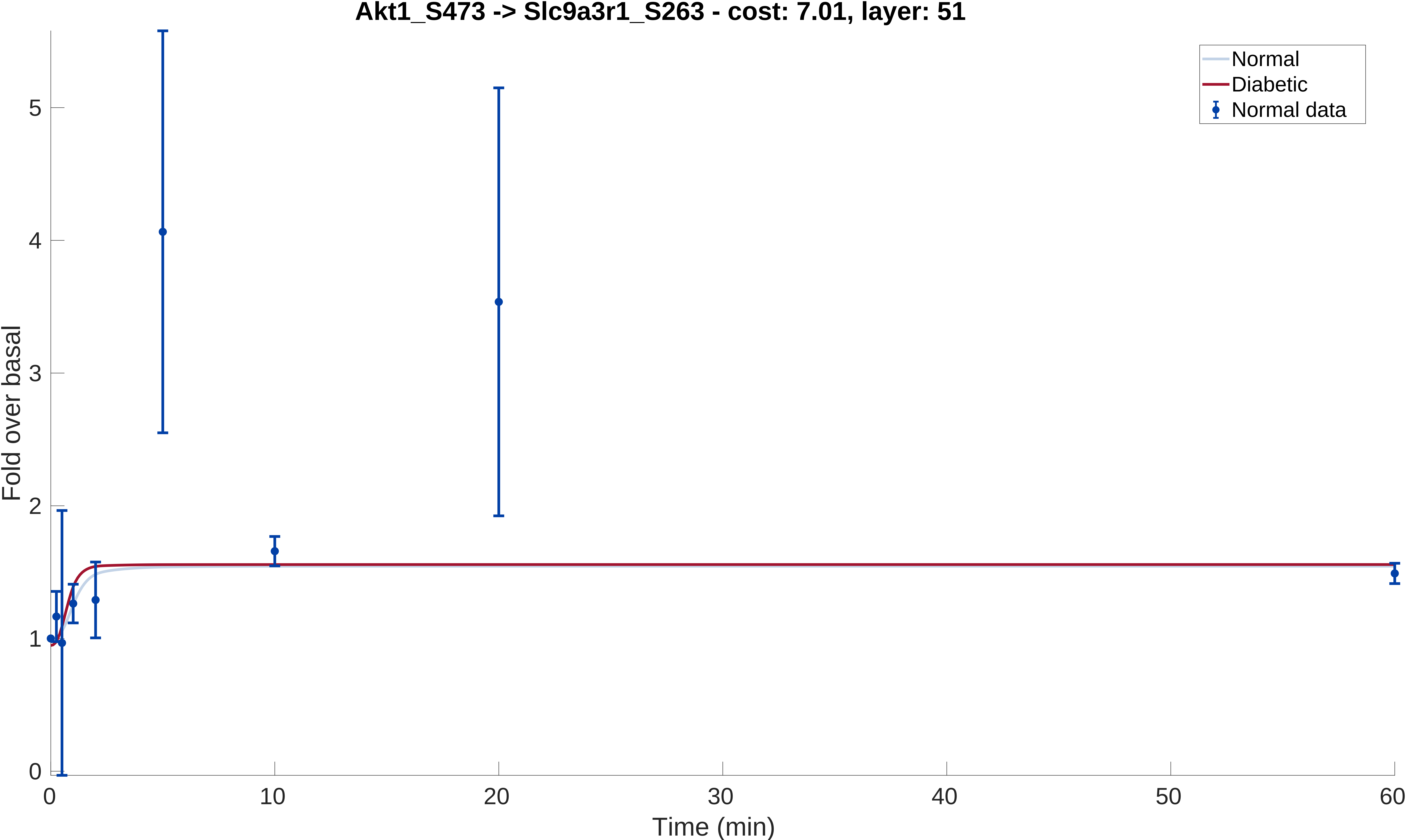

Akt1_S473 -> Slc9a3r1_S263 - cost: 7.01, layer: 51
Normal
Diabetic
Normal data
Fold over basal
0
1
2
3
4
5
0
10
20
30
40
50
60
Time (min)

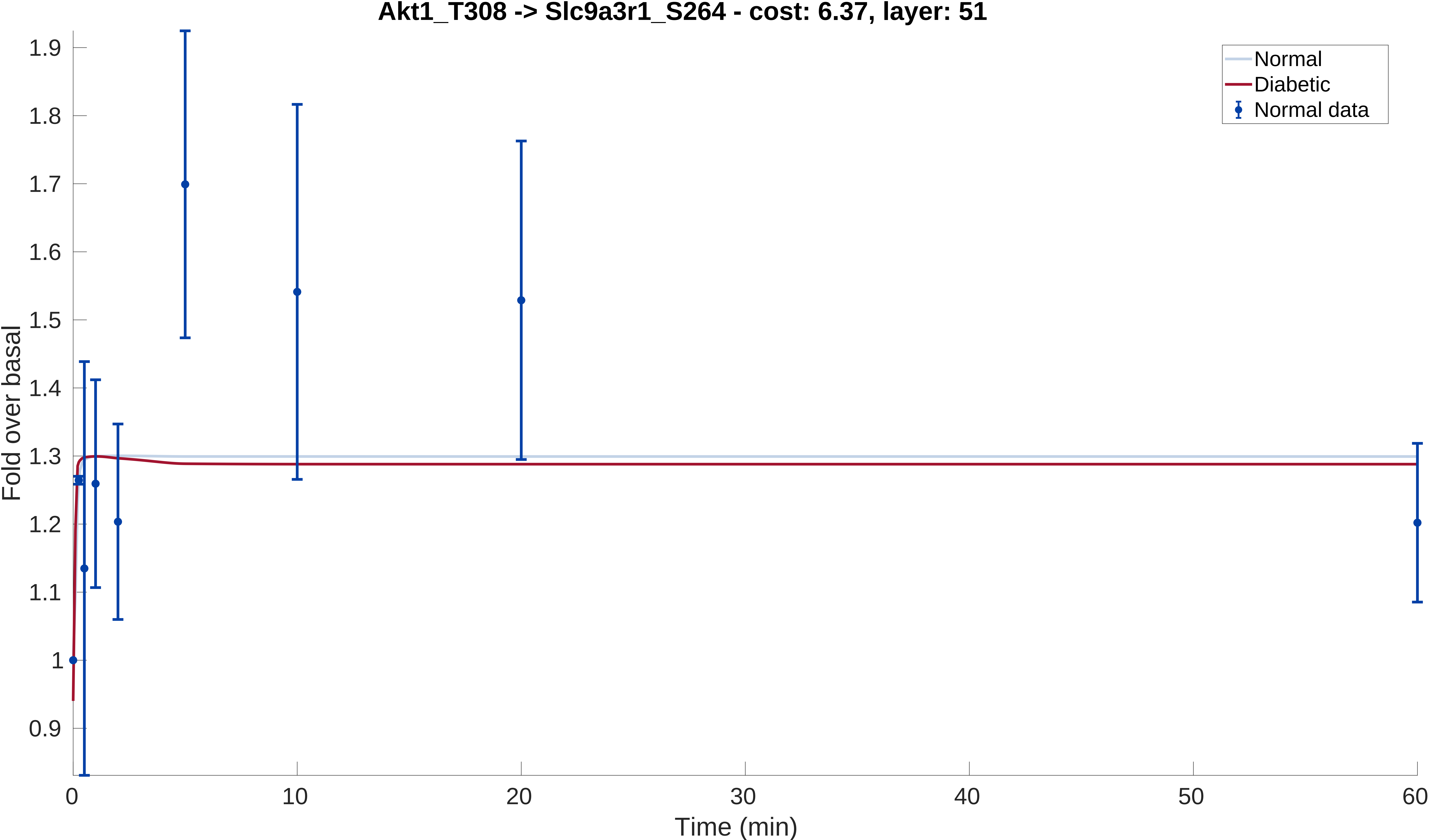

Akt1_T308 -> Slc9a3r1_S264 - cost: 6.37, layer: 51
Normal
Diabetic
Normal data
Fold over basal
1.9
1.8
1.7
1.6
1.5
1.4
1.3
1.2
1.1
1
0.9
0
10
20
30
40
50
60
Time (min)

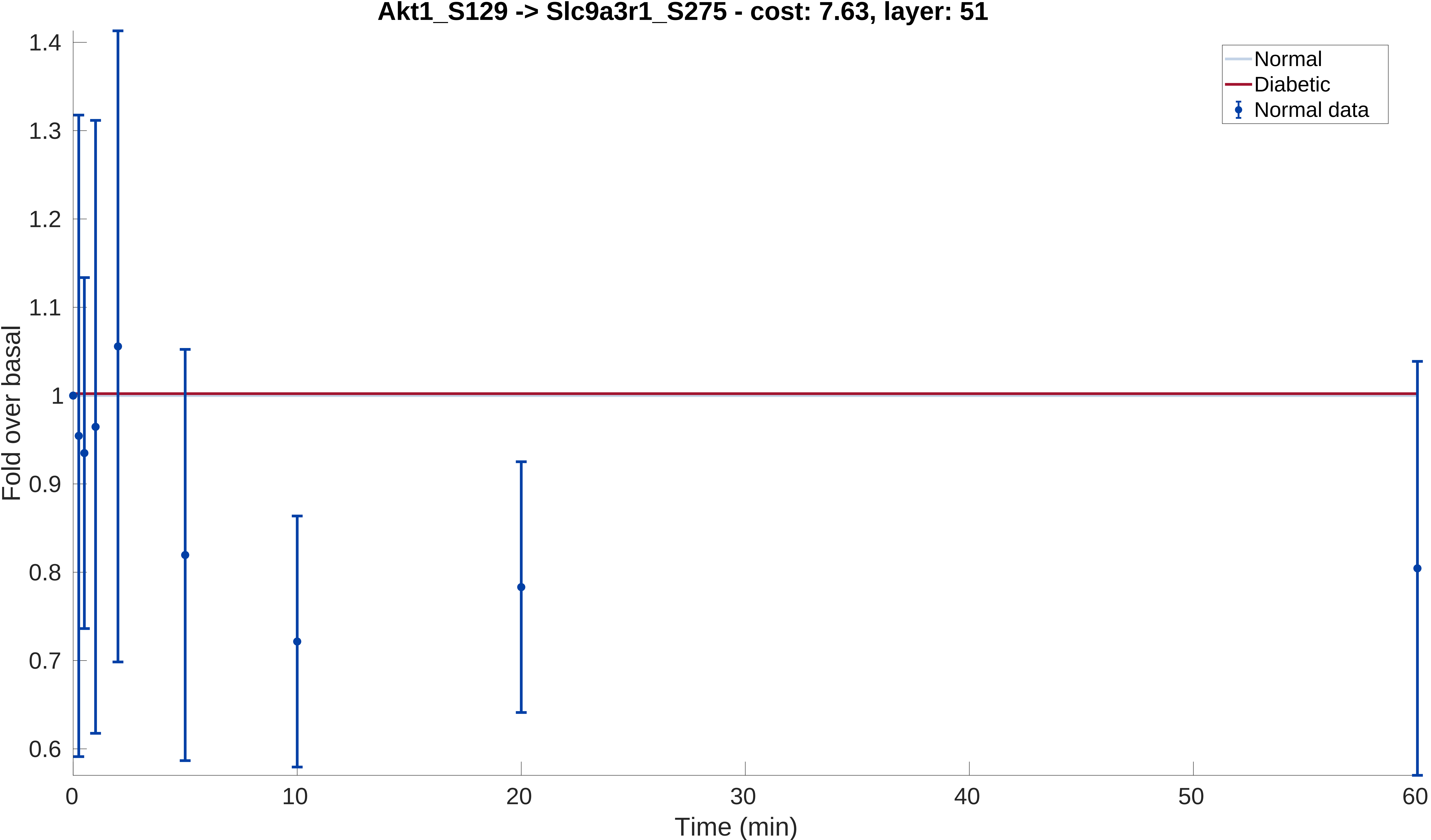

Akt1_S129 -> Slc9a3r1_S275 - cost: 7.63, layer: 51
Normal
Diabetic
Normal data
Fold over basal
1.4
1.3
1.2
1.1
1
0.9
0.8
0.7
0.6
0
10
20
30
40
50
60
Time (min)

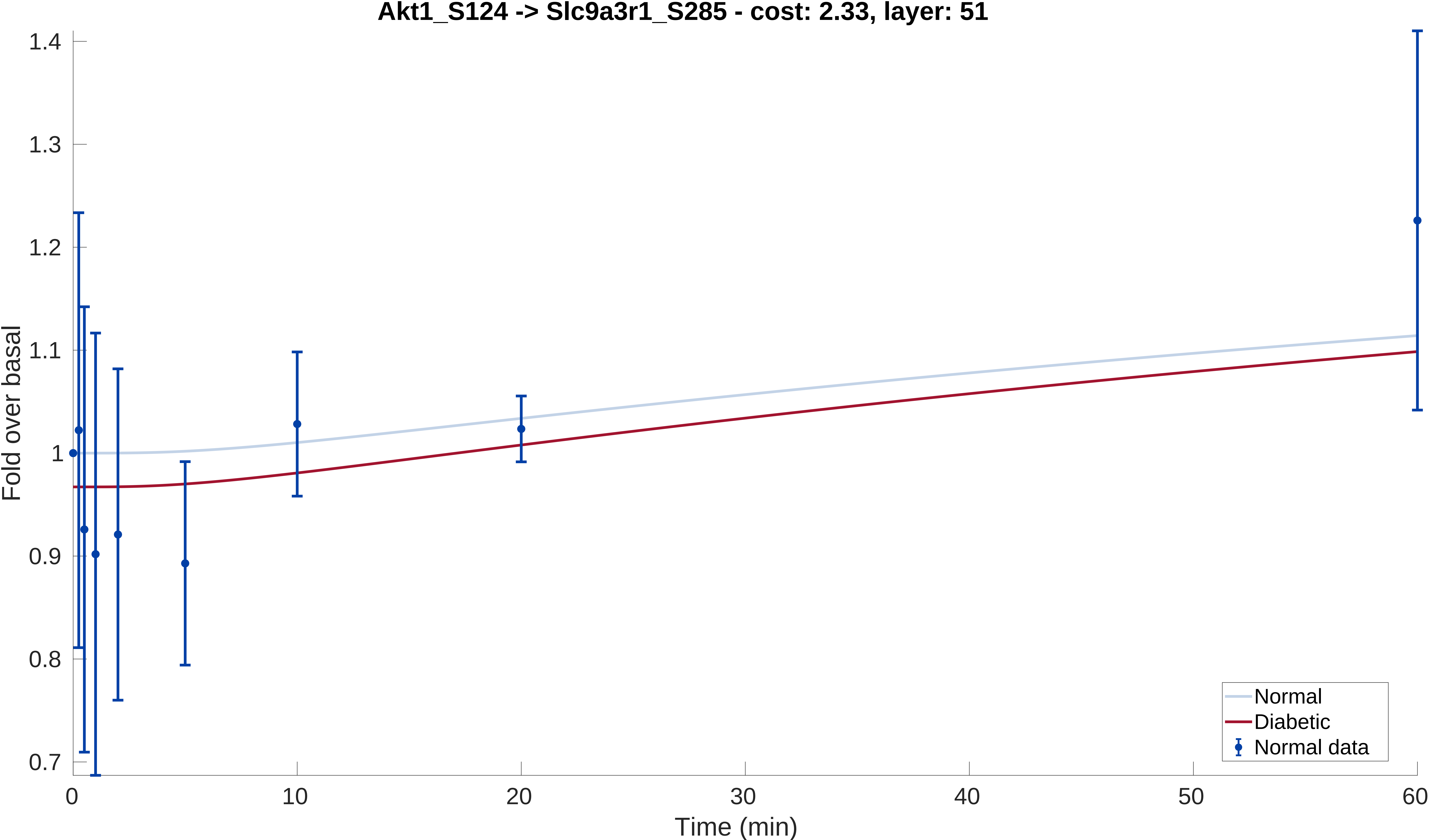

Akt1_S124 -> Slc9a3r1_S285 - cost: 2.33, layer: 51
Fold over basal
1.4
1.3
1.2
1.1
1
0.9
0.8
0.7
0
10
20
30
40
50
60
Time (min)
Normal
Diabetic
Normal data

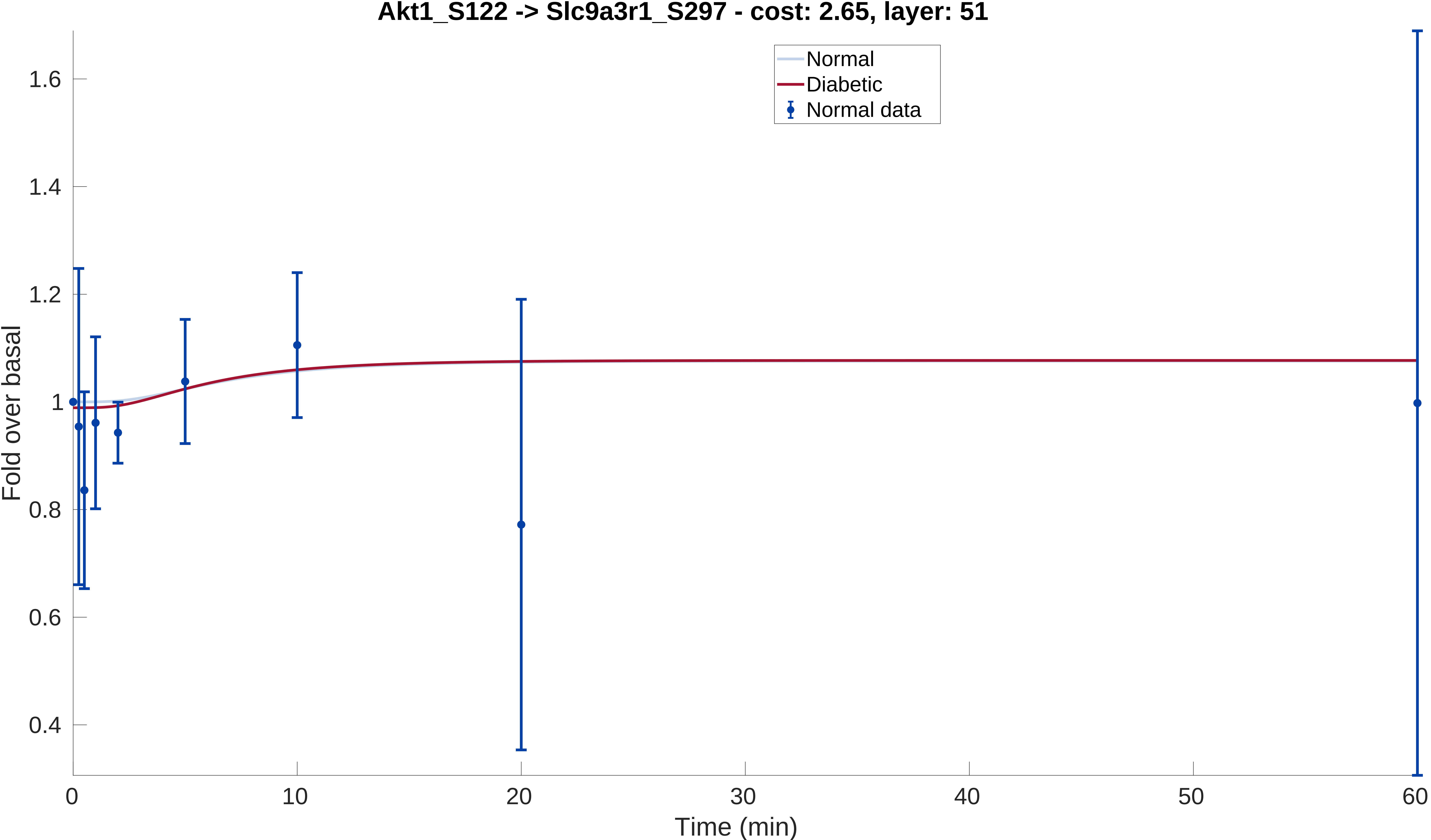

Akt1_S122 -> Slc9a3r1_S297 - cost: 2.65, layer: 51
Normal
Diabetic
Normal data
Fold over basal
1.6
1.4
1.2
1
0.8
0.6
0.4
0
10
20
30
40
50
60
Time (min)

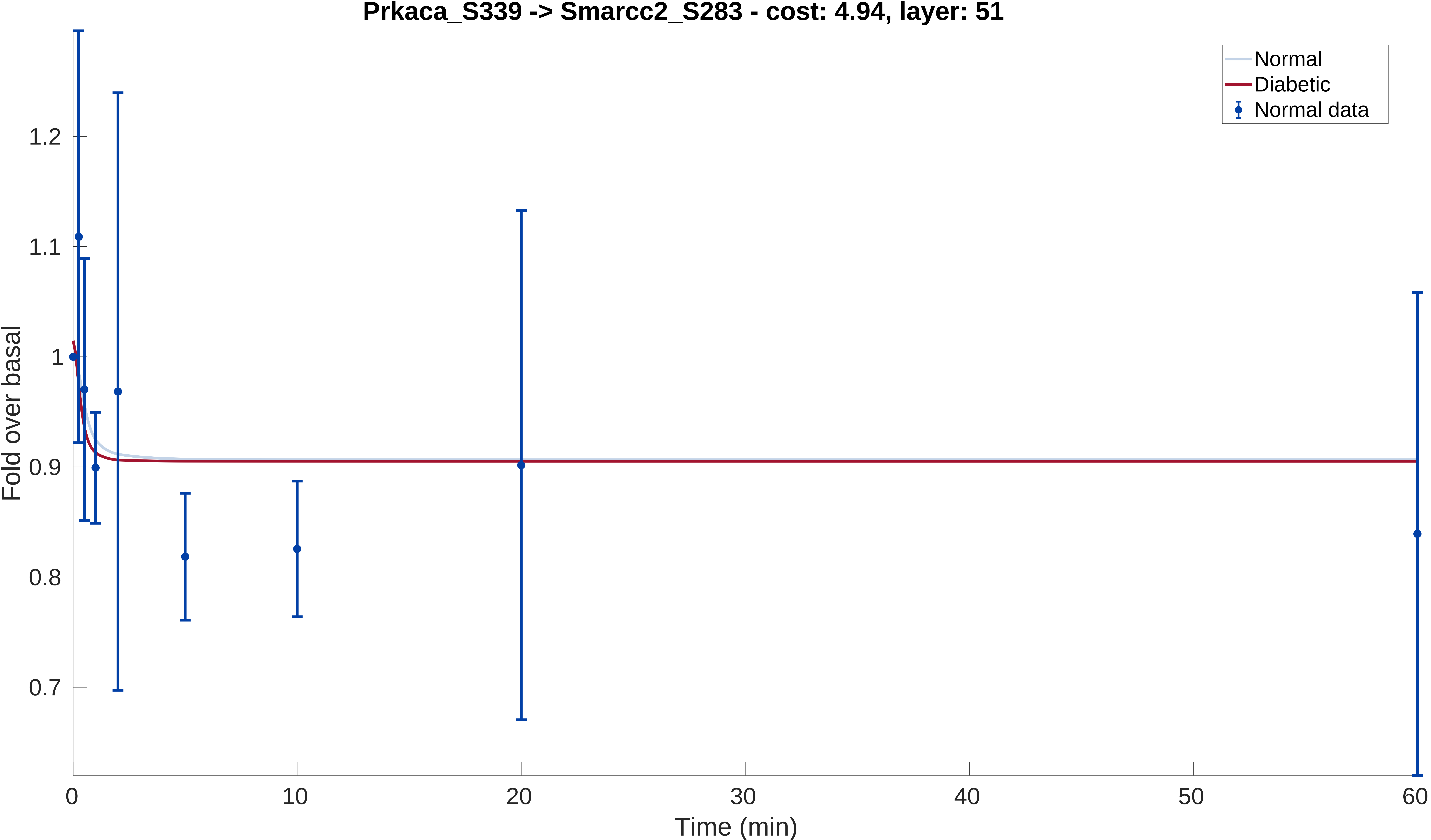

Prkaca_S339 -> Smarcc2_S283 - cost: 4.94, layer: 51
Normal
Diabetic
Normal data
Fold over basal
1.2
1.1
1
0.9
0.8
0.7
0
10
20
30
40
50
60
Time (min)

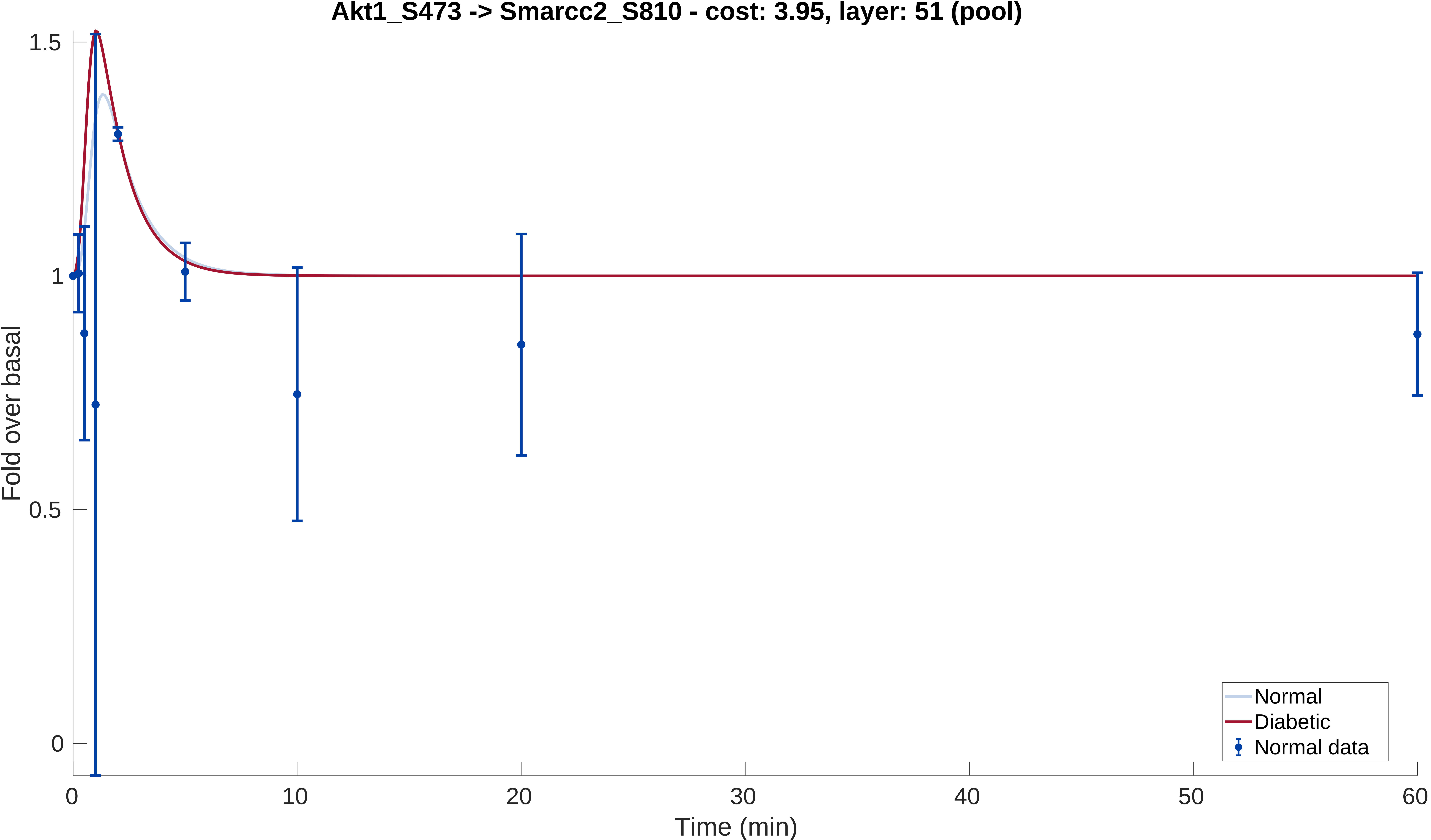
Akt1_S473 -> Smarcc2_S810 - cost: 3.95, layer: 51 (pool)
Fold over basal
1.5
1
0.5
0
0
10
20
30
40
50
60
Time (min)
Normal
Diabetic
Normal data

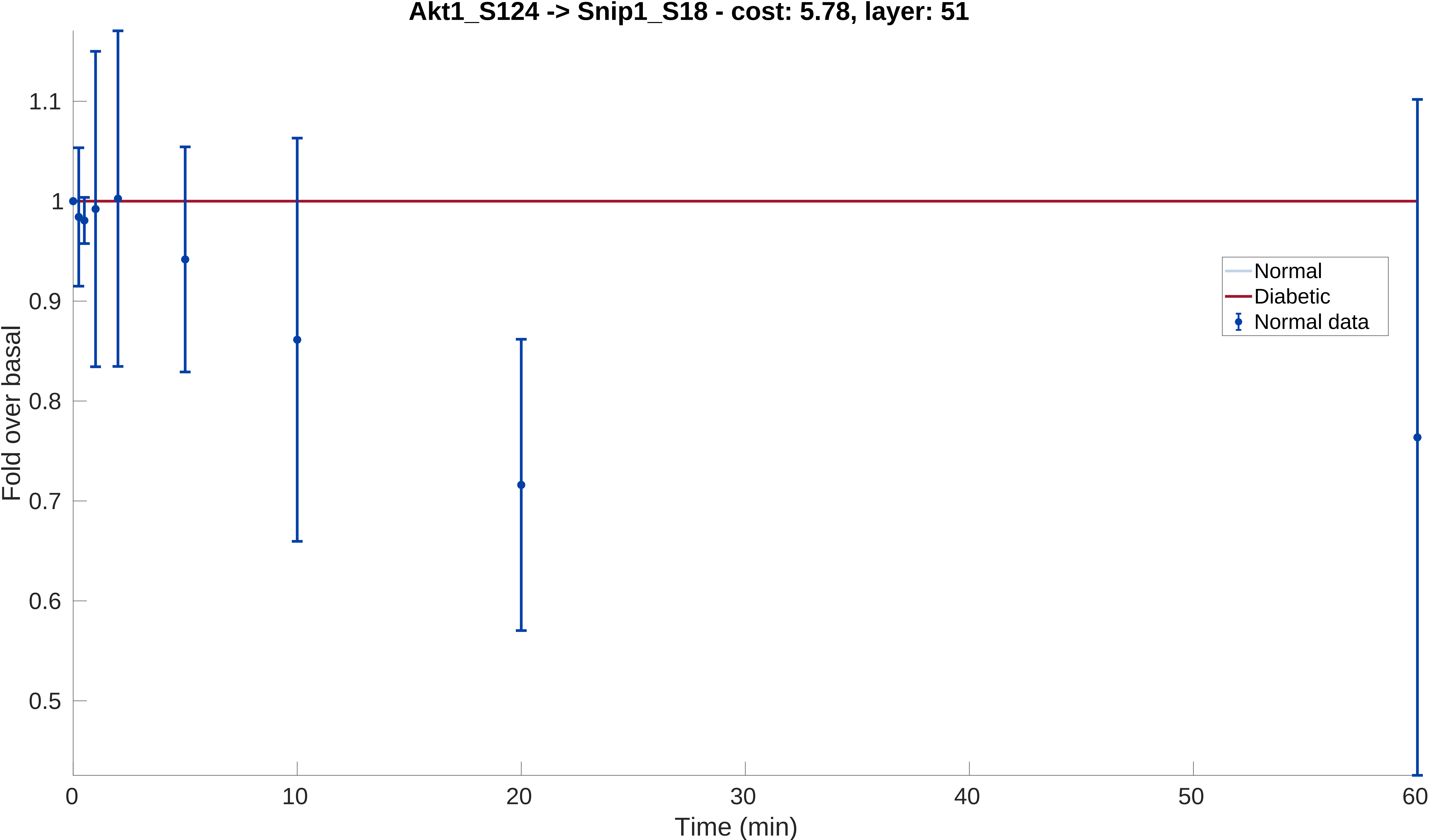
Akt1_S124 -> Snip1_S18 - cost: 5.78, layer: 51
Fold over basal
Time (min)
0.5
0.6
0.7
0.8
0.9
1
1.1
0
10
20
30
40
50
60
Normal
Diabetic
Normal data

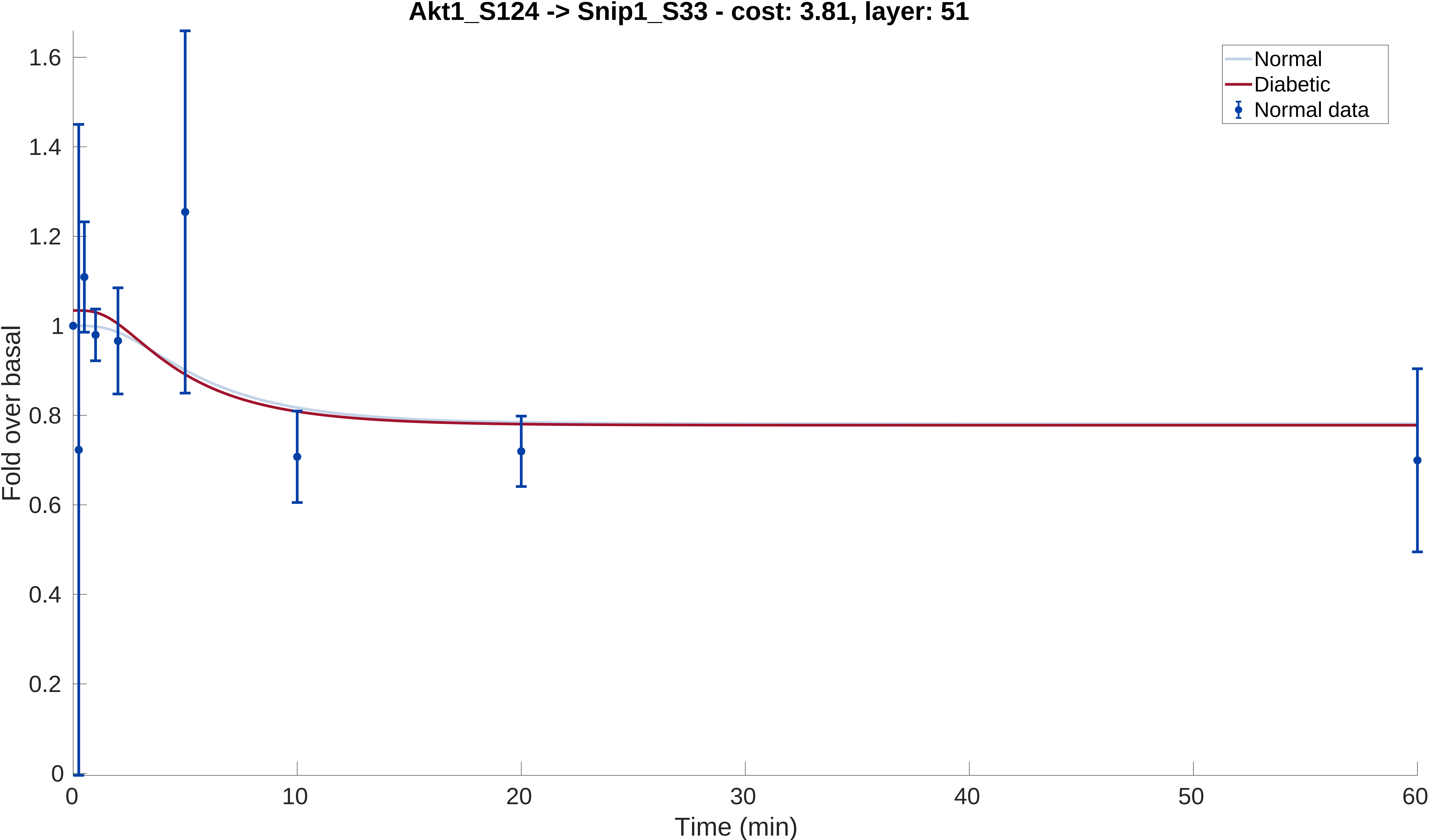

Akt1_S124 -> Snip1_S33 - cost: 3.81, layer: 51
Normal
Diabetic
Normal data
Fold over basal
Time (min)
0
0.2
0.4
0.6
0.8
1
1.2
1.4
1.6
0
10
20
30
40
50
60

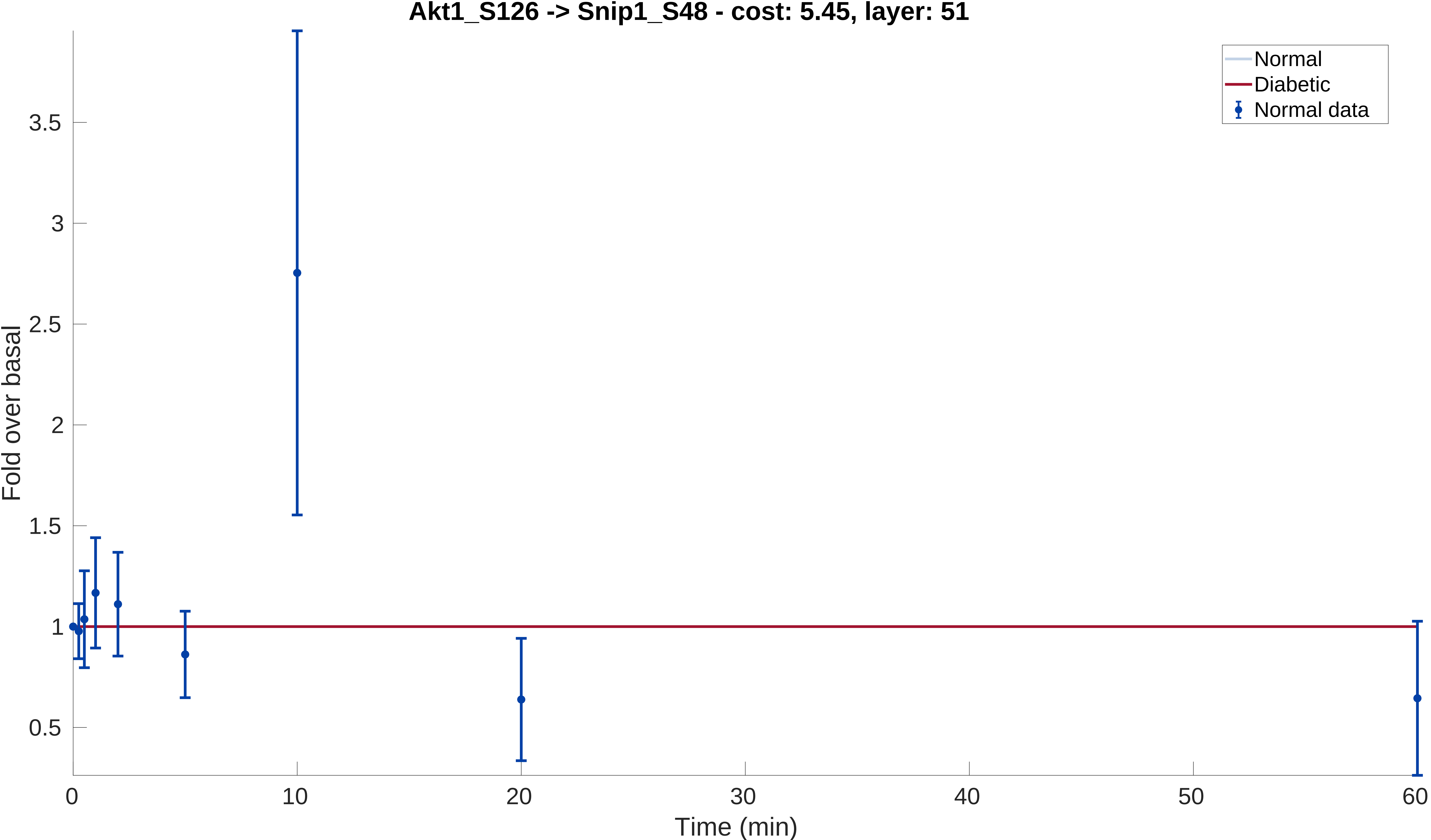
Akt1_S126 -> Snip1_S48 - cost: 5.45, layer: 51
Normal
Diabetic
Normal data
Fold over basal
3.5
3
2.5
2
1.5
1
0.5
0
10
20
30
40
50
60
Time (min)

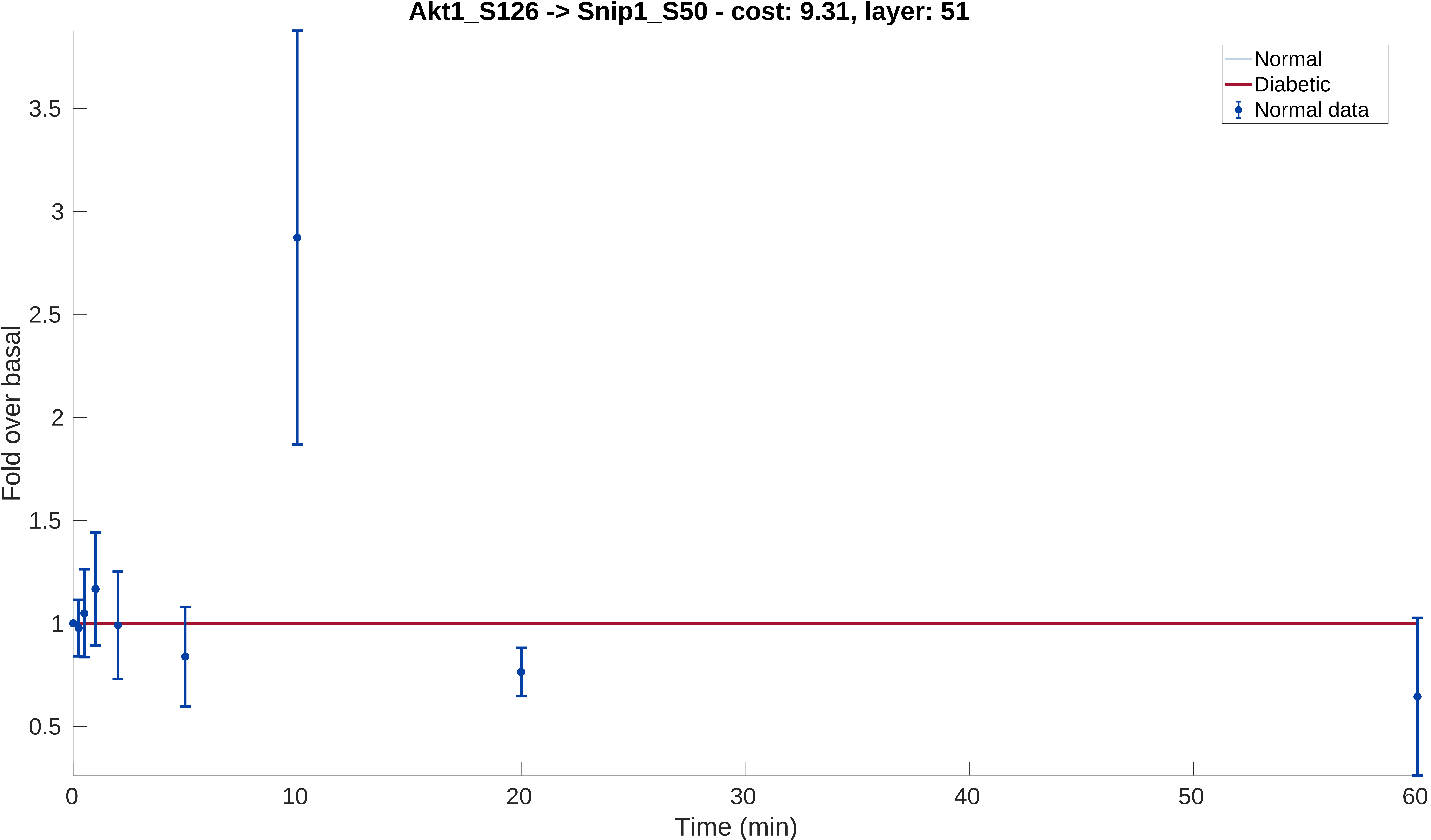

Akt1_S126 -> Snip1_S50 - cost: 9.31, layer: 51
Normal
Diabetic
Normal data
Fold over basal
3.5
3
2.5
2
1.5
1
0.5
0
10
20
30
40
50
60
Time (min)

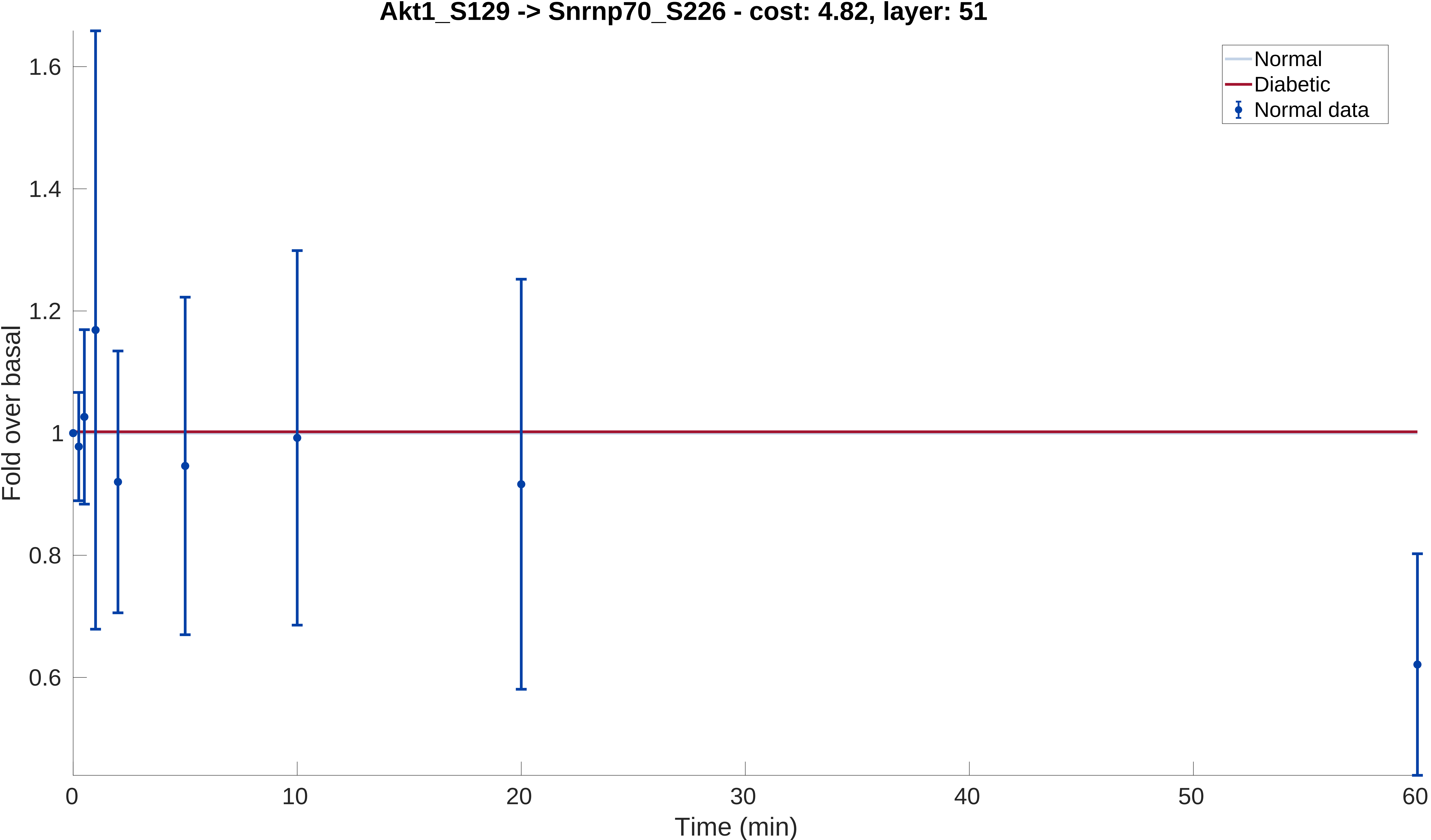
Akt1_S129 -> Snrnp70_S226 - cost: 4.82, layer: 51
Normal
Diabetic
Normal data
Fold over basal
1.6
1.4
1.2
1
0.8
0.6
0
10
20
30
40
50
60
Time (min)

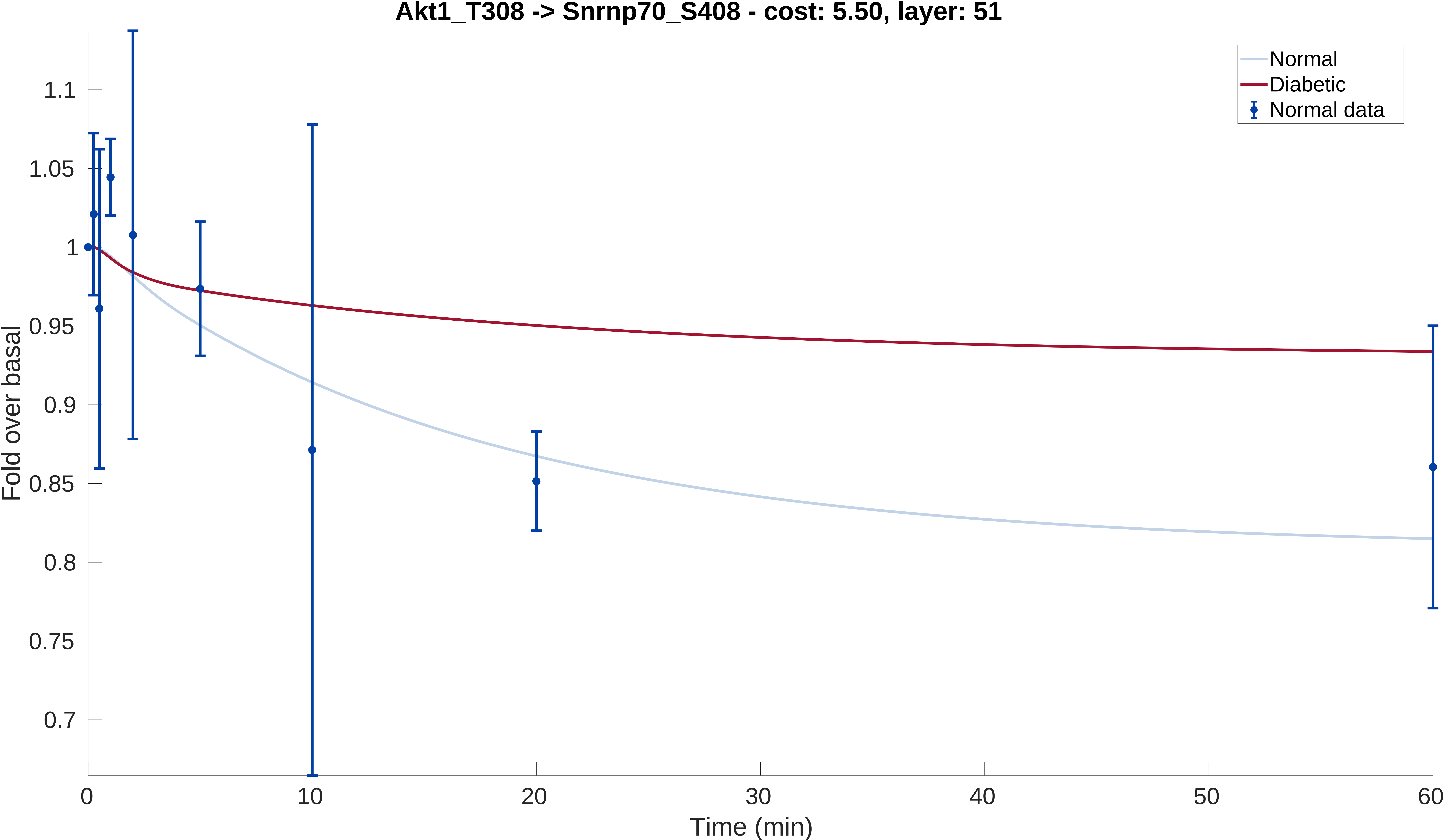
Akt1_T308 -> Snrnp70_S408 - cost: 5.50, layer: 51
Normal
Diabetic
Normal data
Fold over basal
Time (min)
1.1
1.05
1
0.95
0.9
0.85
0.8
0.75
0.7
0
10
20
30
40
50
60

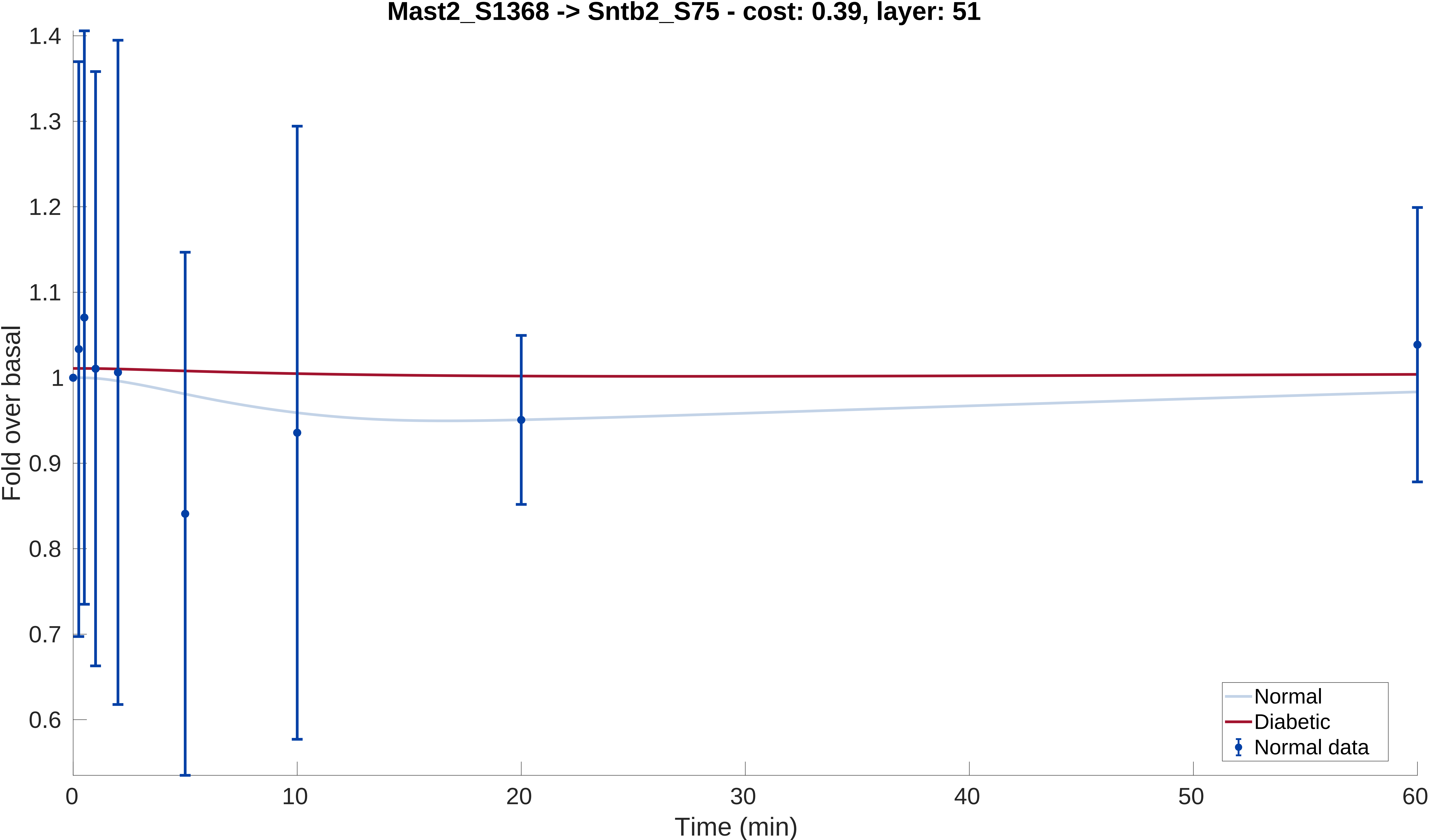

Mast2_S1368 -> Sntb2_S75 - cost: 0.39, layer: 51
Fold over basal
1.4
1.3
1.2
1.1
1
0.9
0.8
0.7
0.6
0
10
20
30
40
50
60
Time (min)
Normal
Diabetic
Normal data

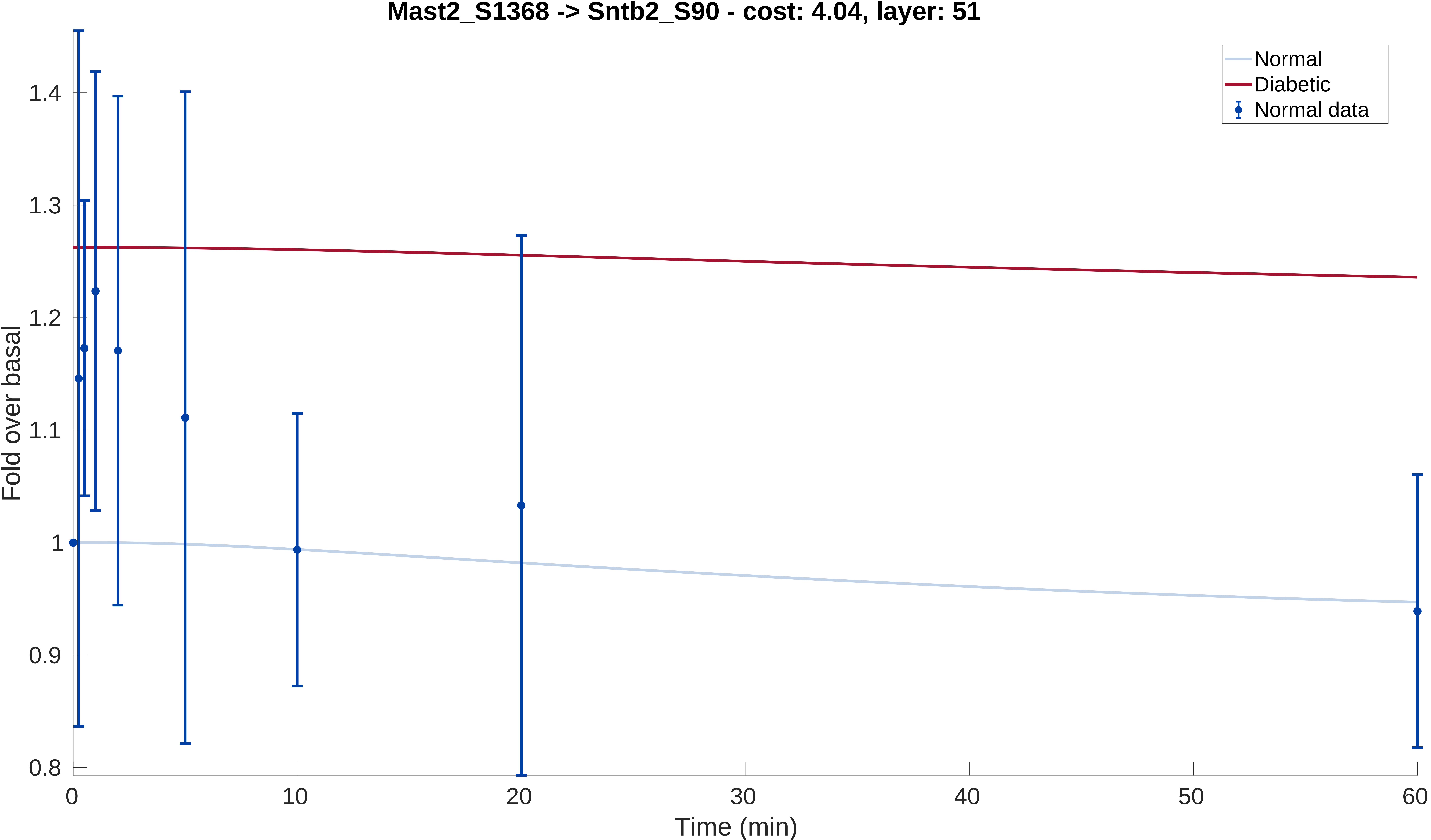

Mast2_S1368 -> Sntb2_S90 - cost: 4.04, layer: 51
Normal
Diabetic
Normal data
Fold over basal
1.4
1.3
1.2
1.1
1
0.9
0.8
0
10
20
30
40
50
60
Time (min)

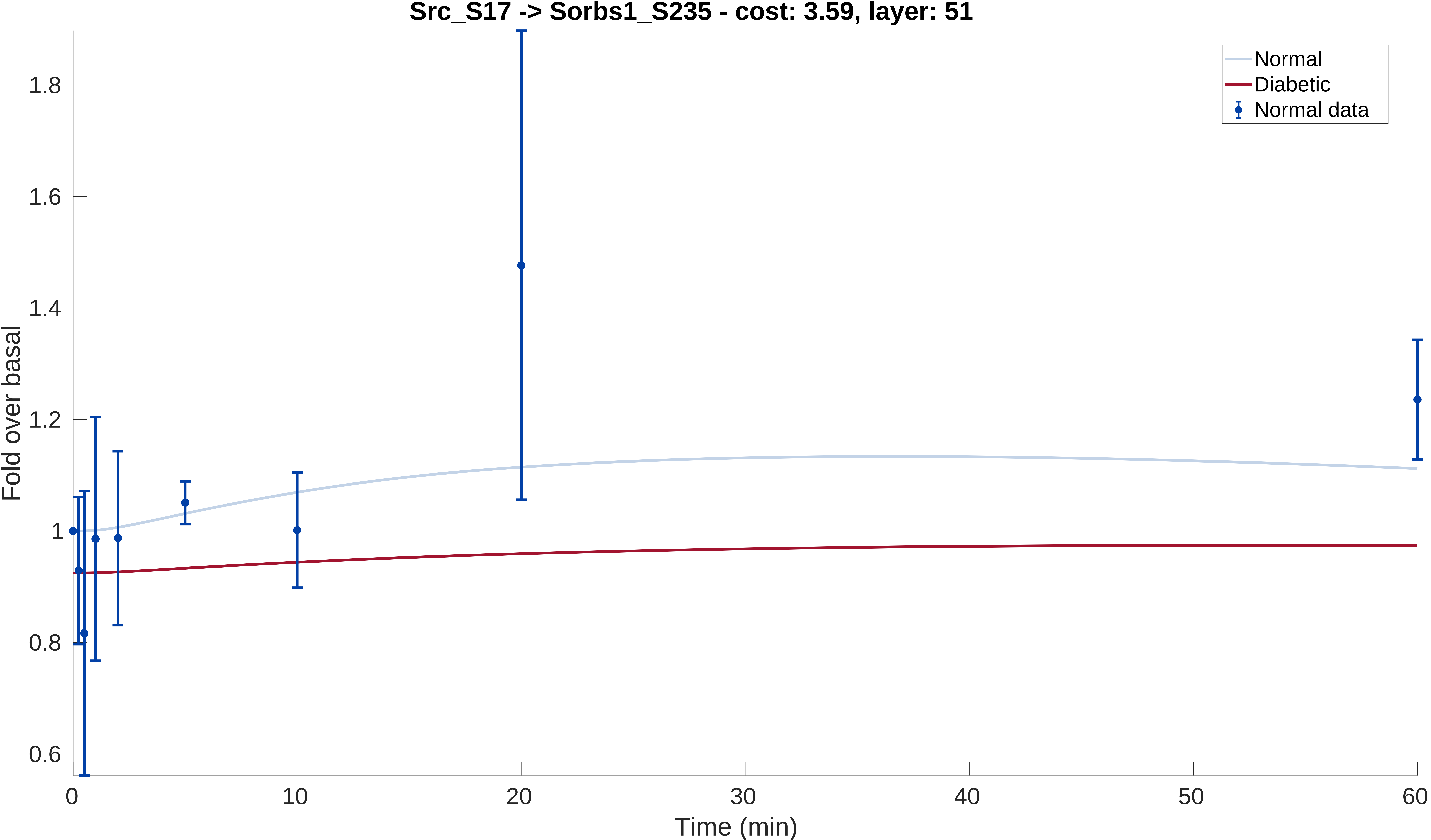

Src_S17 -> Sorbs1_S235 - cost: 3.59, layer: 51
Normal
Diabetic
Normal data
Fold over basal
1.8
1.6
1.4
1.2
1
0.8
0.6
0
10
20
30
40
50
60
Time (min)

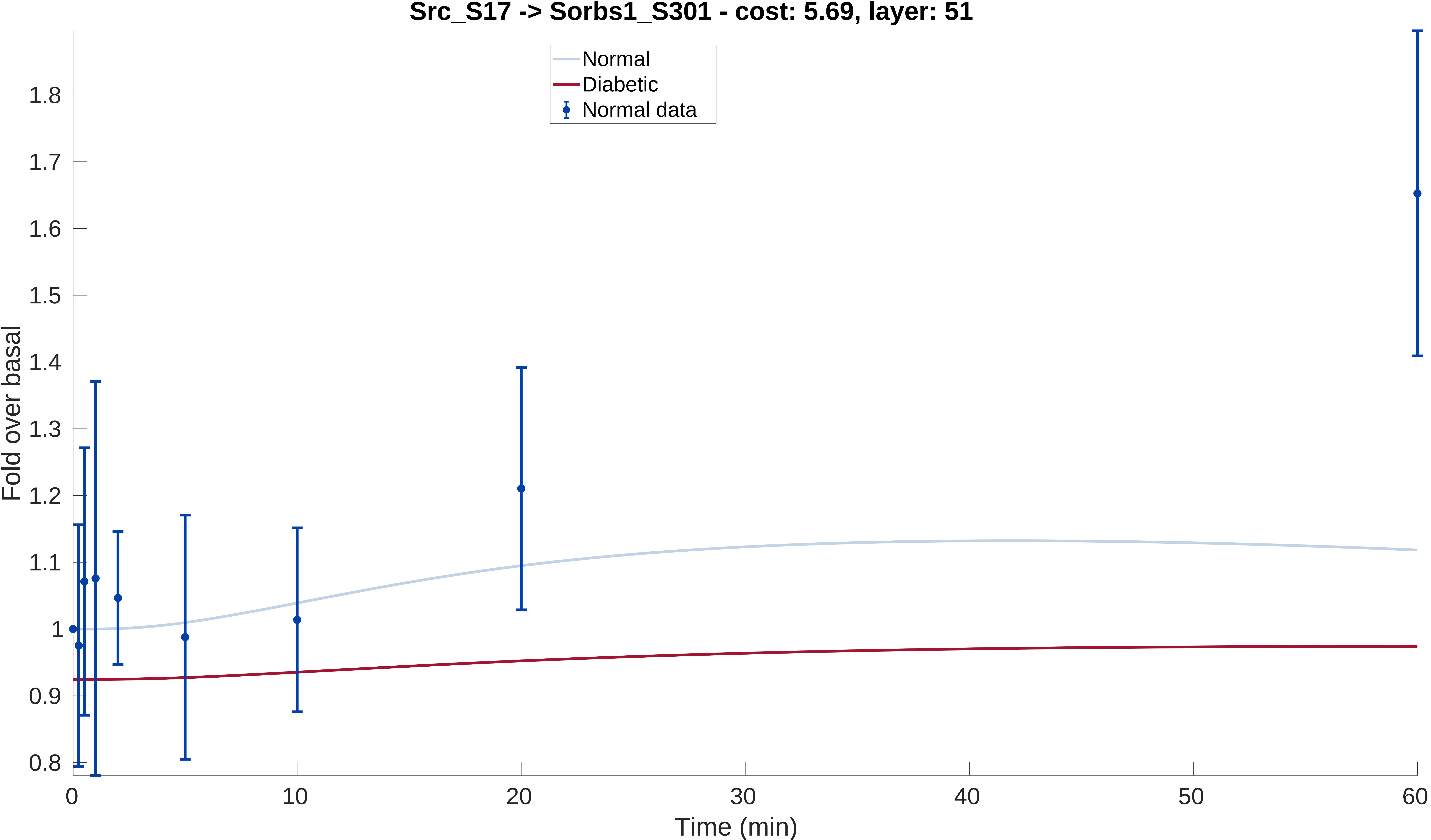
Src_S17 -> Sorbs1_S301 - cost: 5.69, layer: 51
Normal
Diabetic
Normal data
Fold over basal
1.8
1.7
1.6
1.5
1.4
1.3
1.2
1.1
1
0.9
0.8
0
10
20
30
40
50
60
Time (min)

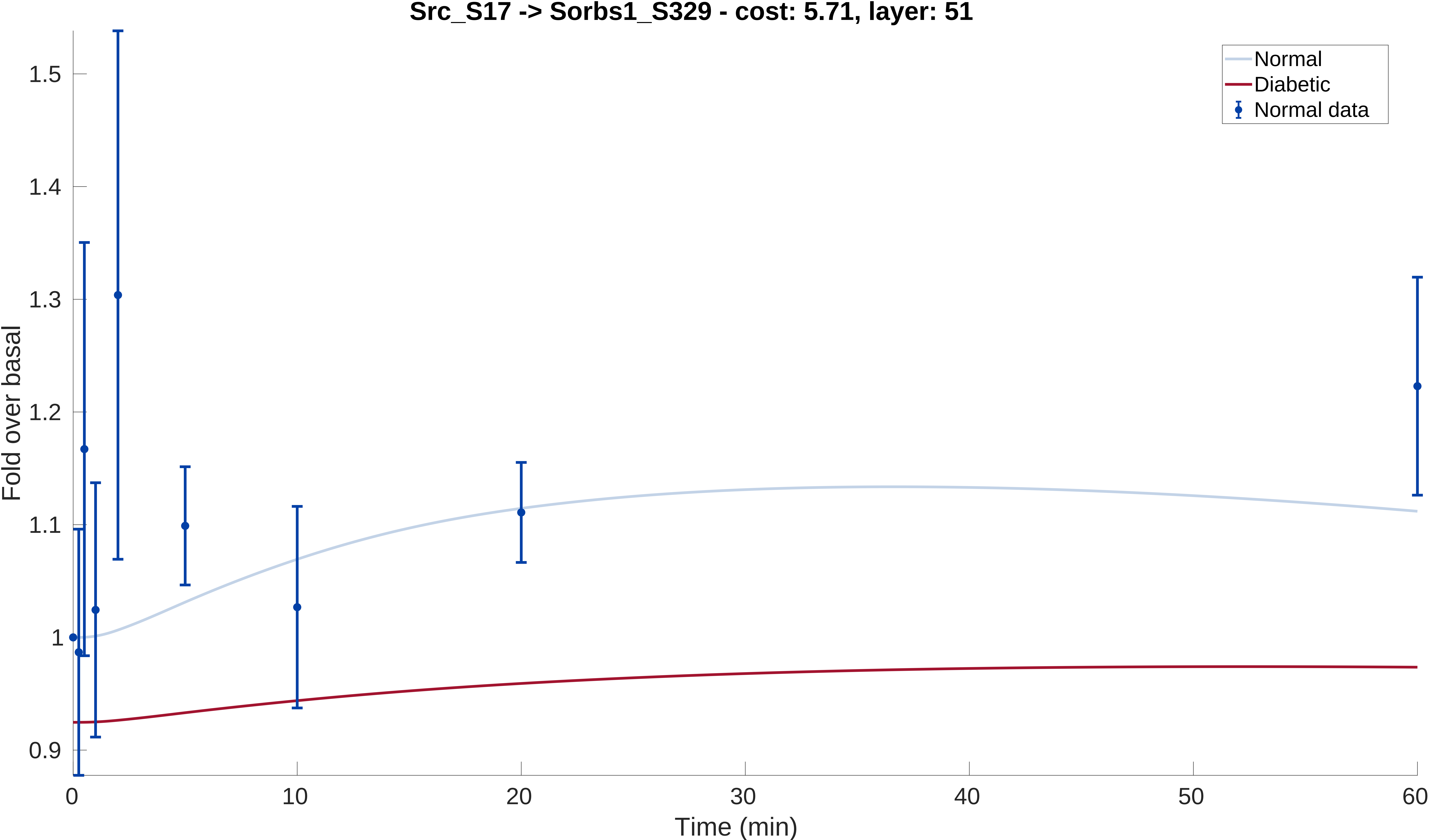

Src_S17 -> Sorbs1_S329 - cost: 5.71, layer: 51
Normal
Diabetic
Normal data
Fold over basal
Time (min)
0.9
1
1.1
1.2
1.3
1.4
1.5
0
10
20
30
40
50
60

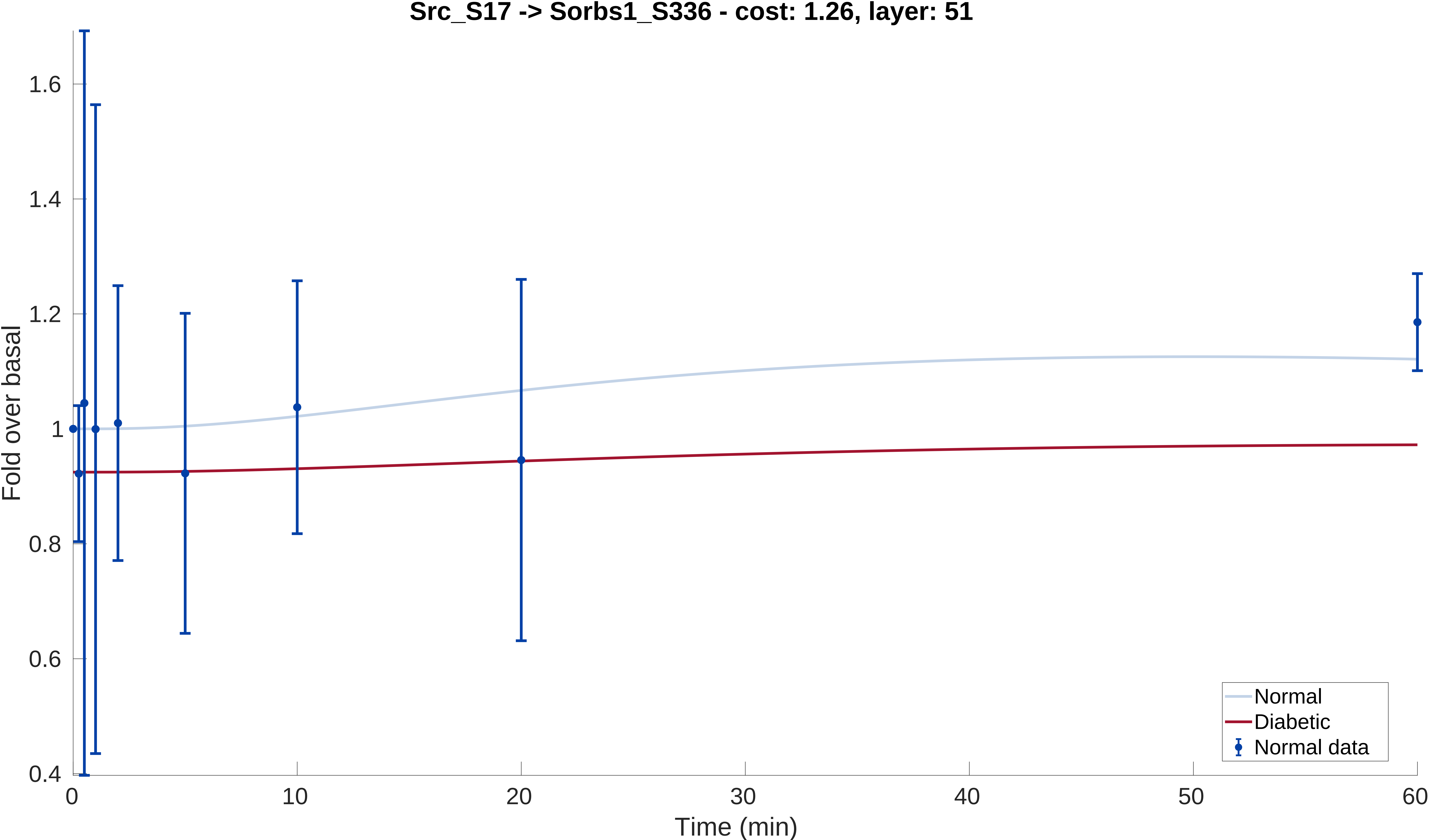

Src_S17 -> Sorbs1_S336 - cost: 1.26, layer: 51
1.6
1.4
1.2
1
0.8
0.6
0.4
Fold over basal
0
10
20
30
40
50
60
Time (min)
Normal
Diabetic
Normal data

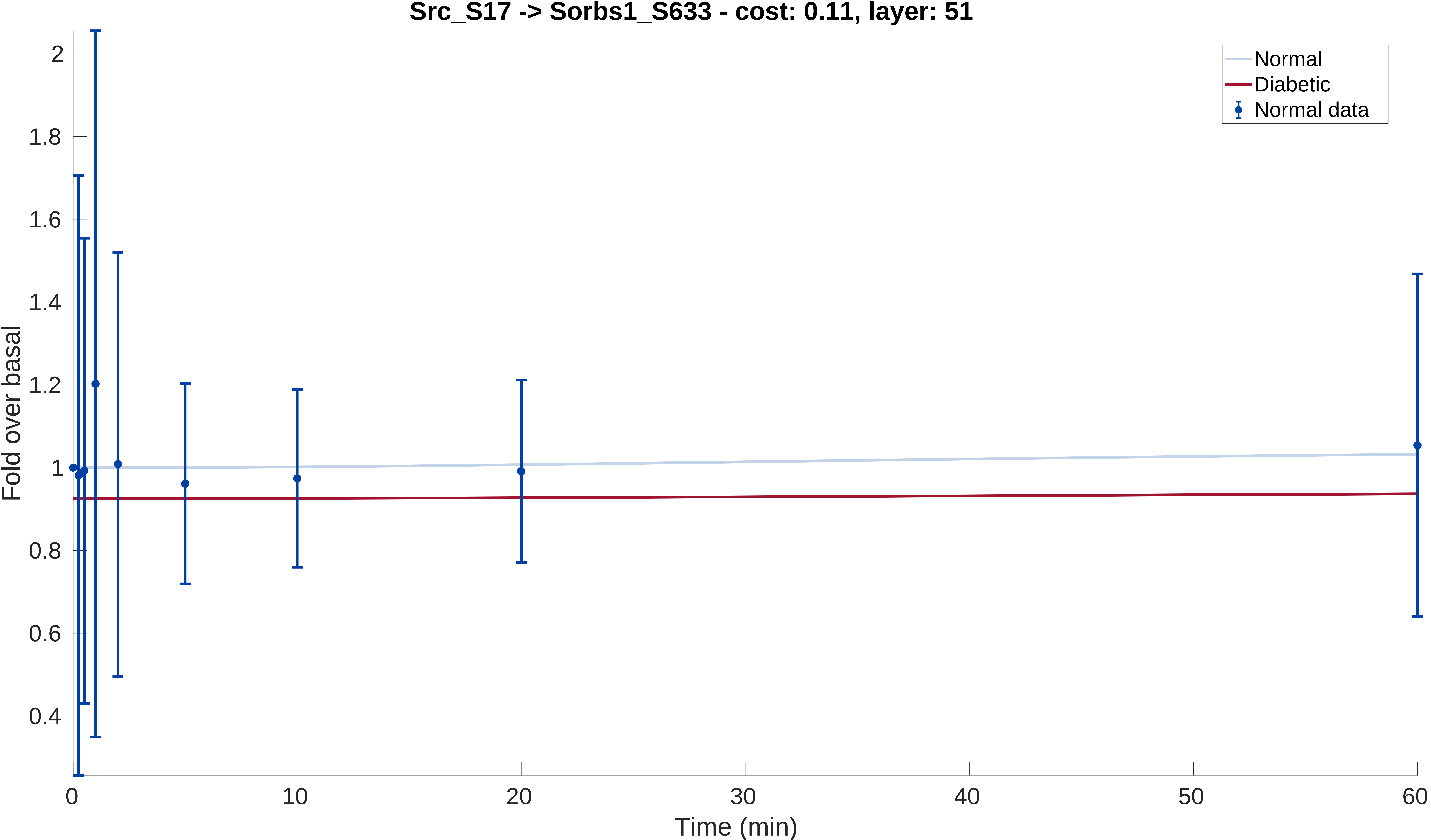

Src_S17 -> Sorbs1_S633 - cost: 0.11, layer: 51
Normal
Diabetic
Normal data
Fold over basal
2
1.8
1.6
1.4
1.2
1
0.8
0.6
0.4
0
10
20
30
40
50
60
Time (min)

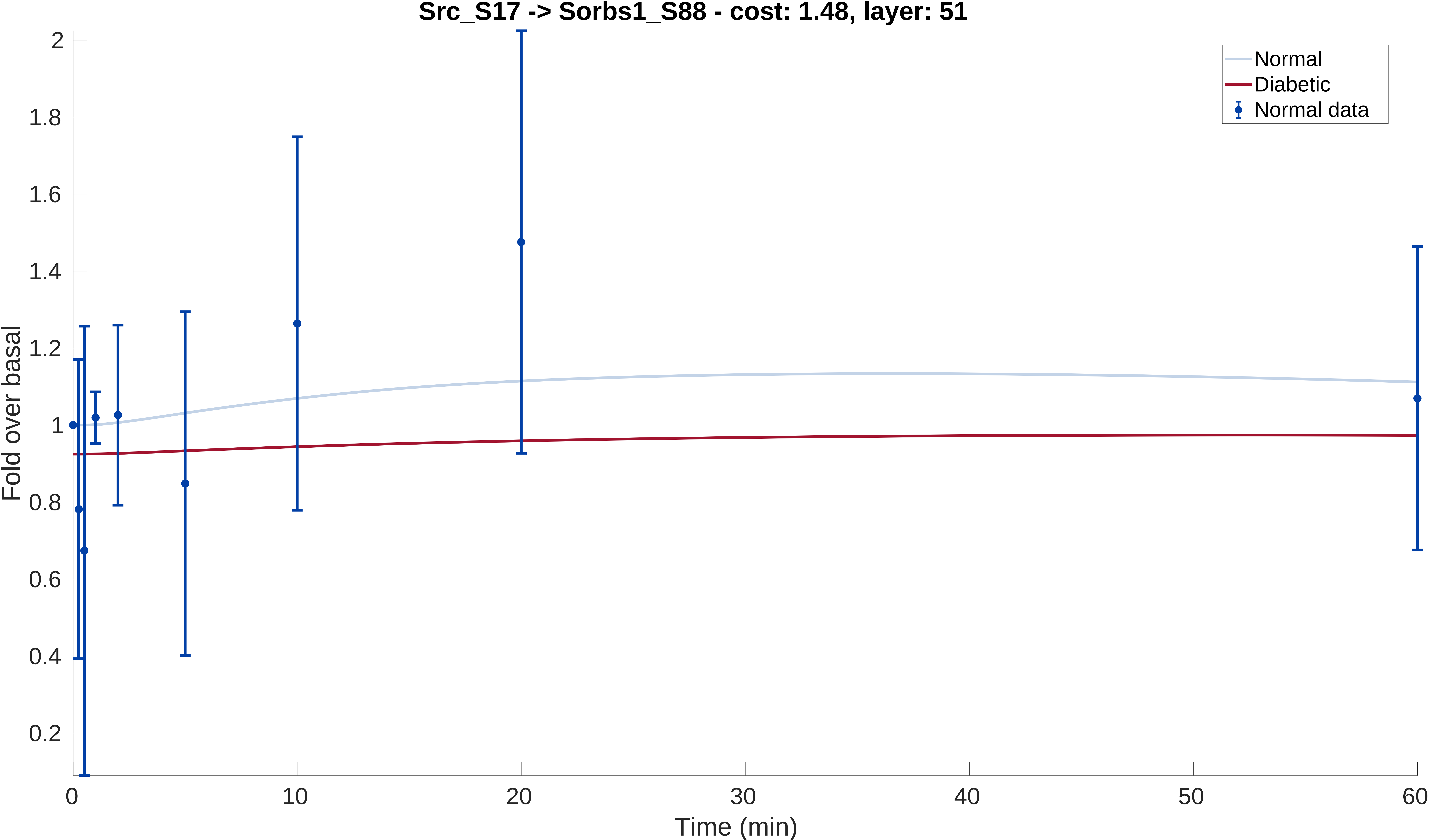

Src_S17 -> Sorbs1_S88 - cost: 1.48, layer: 51
Normal
Diabetic
Normal data
Fold over basal
2
1.8
1.6
1.4
1.2
1
0.8
0.6
0.4
0.2
0
10
20
30
40
50
60
Time (min)

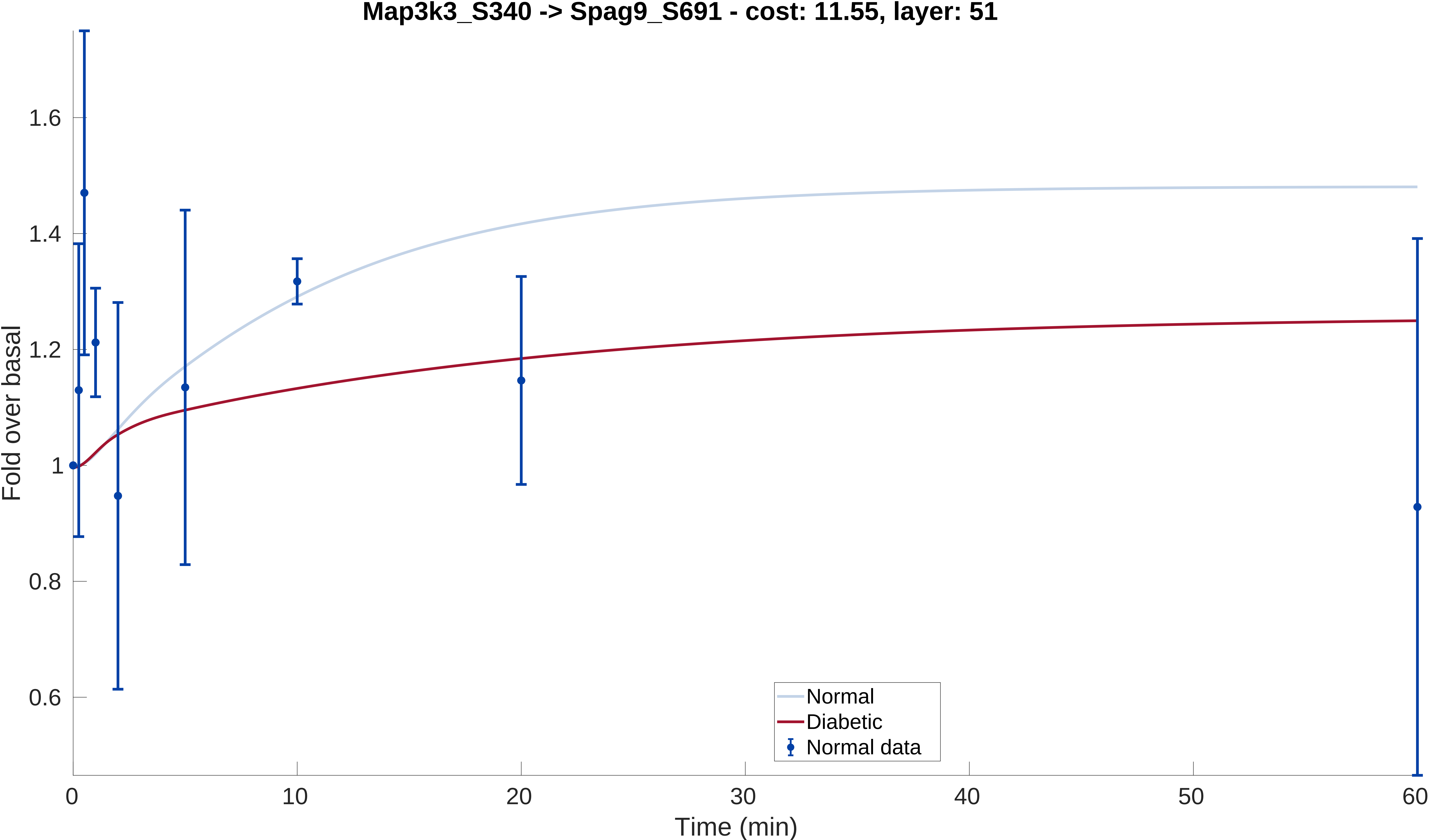
Map3k3_S340 -> Spag9_S691 - cost: 11.55, layer: 51
Fold over basal
1.6
1.4
1.2
1
0.8
0.6
0
10
20
30
40
50
60
Time (min)
Normal
Diabetic
Normal data

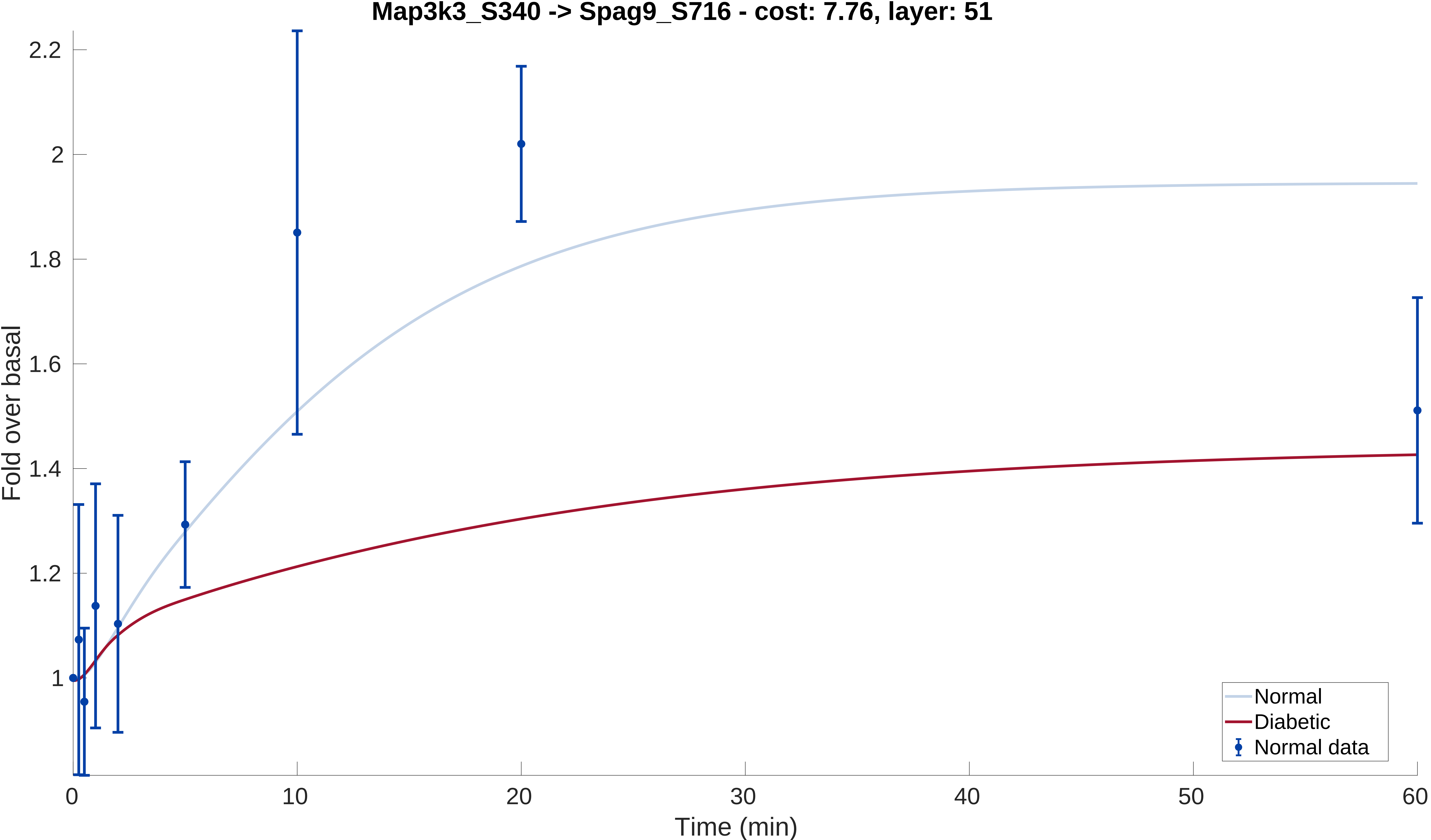
Map3k3_S340 -> Spag9_S716 - cost: 7.76, layer: 51
Fold over basal
2.2
2
1.8
1.6
1.4
1.2
1
0
10
20
30
40
50
60
Time (min)
Normal
Diabetic
Normal data

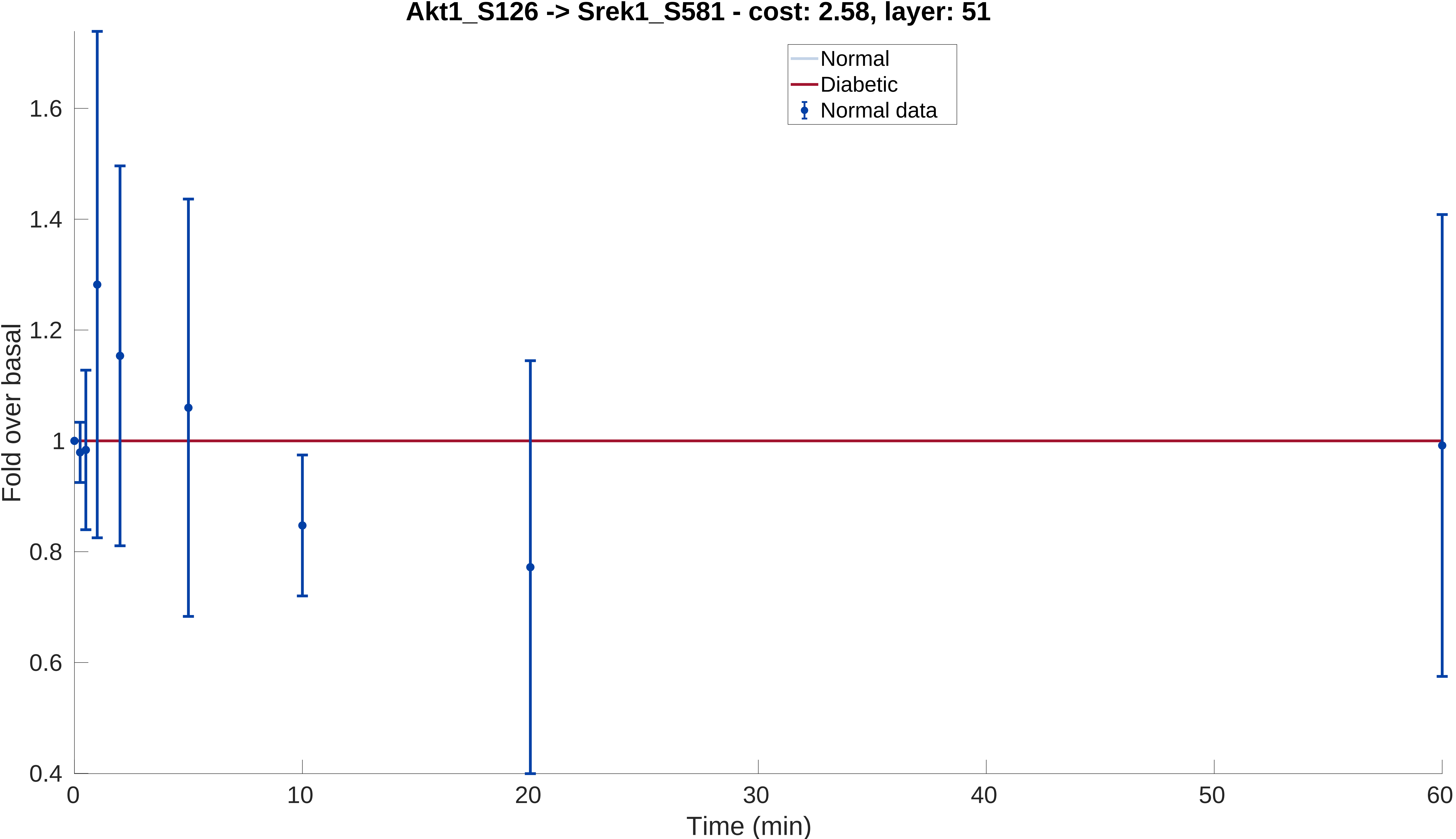

Akt1_S126 -> Srek1_S581 - cost: 2.58, layer: 51
Normal
Diabetic
Normal data
Fold over basal
1.6
1.4
1.2
1
0.8
0.6
0.4
0
10
20
30
40
50
60
Time (min)

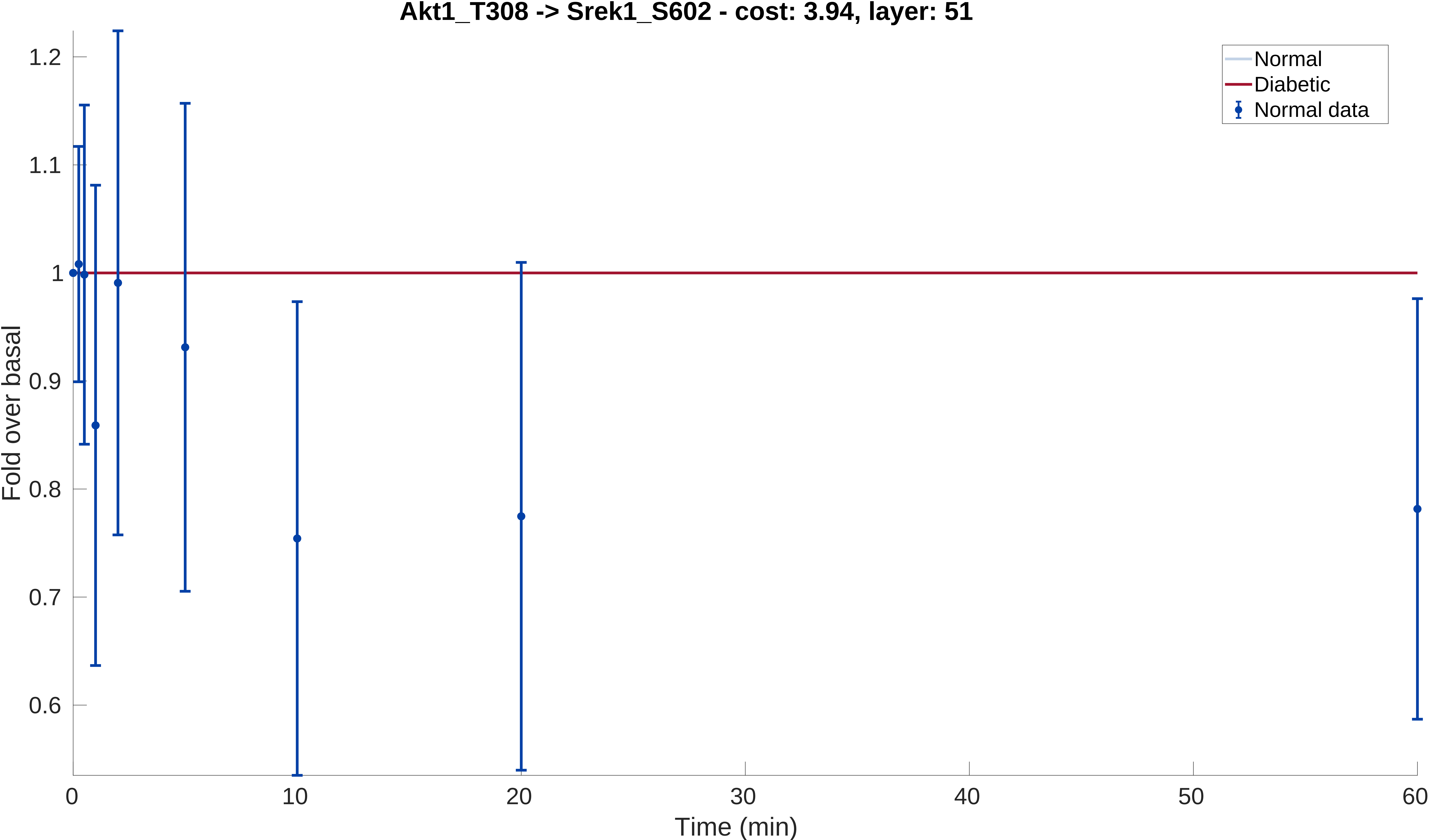

Akt1_T308 -> Srek1_S602 - cost: 3.94, layer: 51
Normal
Diabetic
Normal data
Fold over basal
1.2
1.1
1
0.9
0.8
0.7
0.6
0
10
20
30
40
50
60
Time (min)

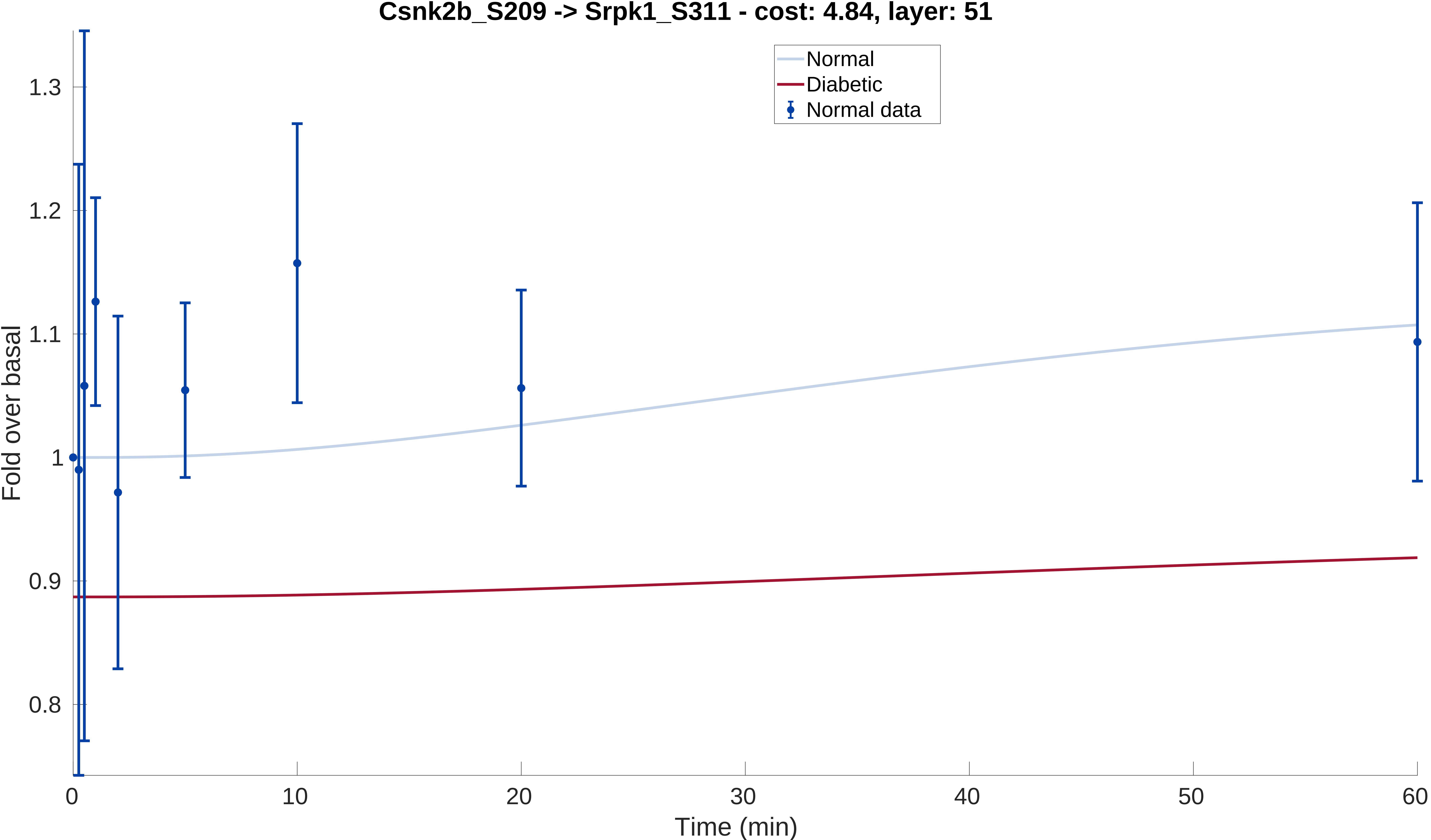

Csnk2b_S209 -> Srpk1_S311 - cost: 4.84, layer: 51
Normal
Diabetic
Normal data
Fold over basal
Time (min)
1.3
1.2
1.1
1
0.9
0.8
0
10
20
30
40
50
60

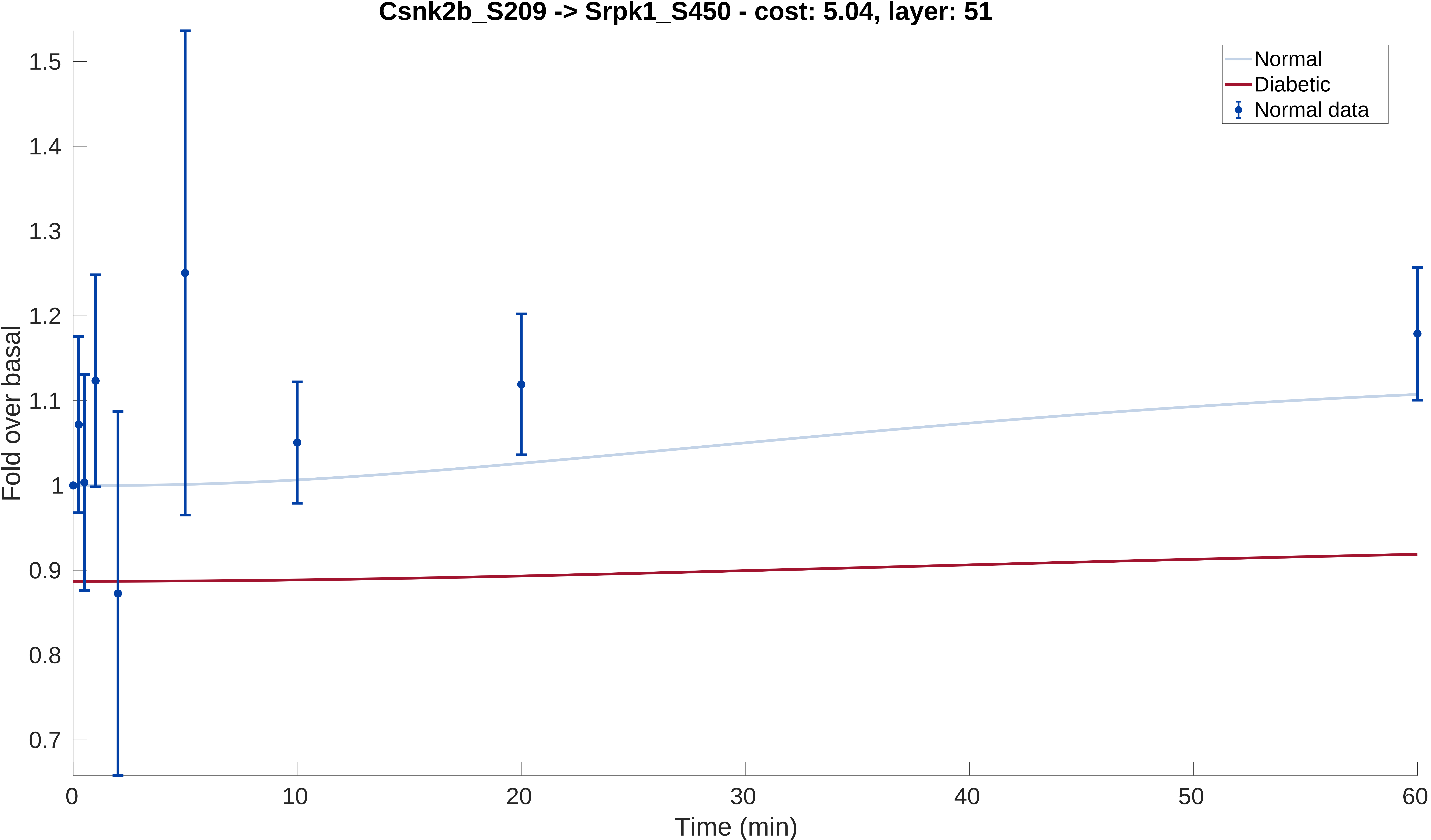

Csnk2b_S209 -> Srpk1_S450 - cost: 5.04, layer: 51
Normal
Diabetic
Normal data
Fold over basal
Time (min)
1.5
1.4
1.3
1.2
1.1
1
0.9
0.8
0.7
0
10
20
30
40
50
60

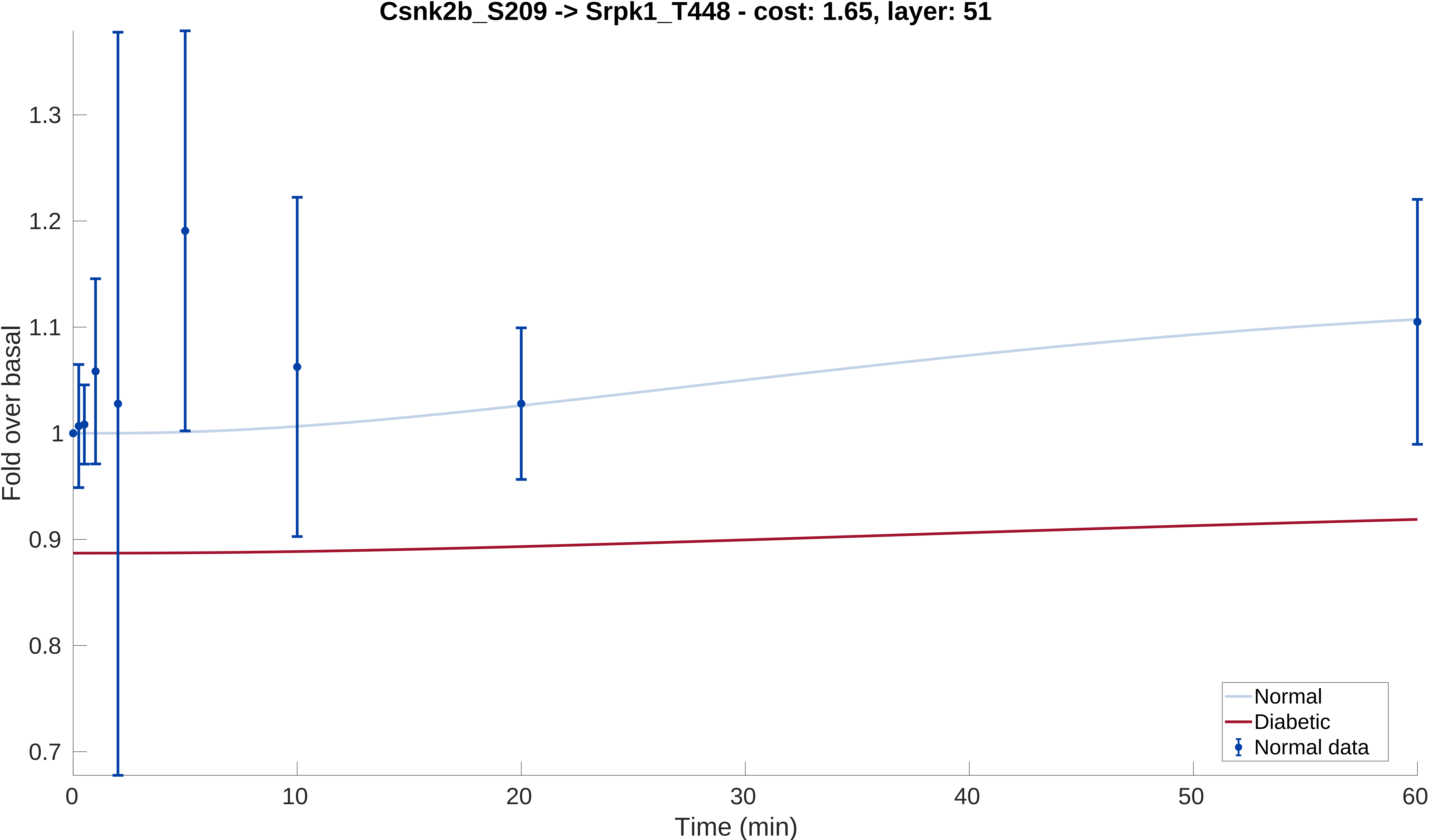

Csnk2b_S209 -> Srpk1_T448 - cost: 1.65, layer: 51
Fold over basal
1.3
1.2
1.1
1
0.9
0.8
0.7
0
10
20
30
40
50
60
Time (min)
Normal
Diabetic
Normal data

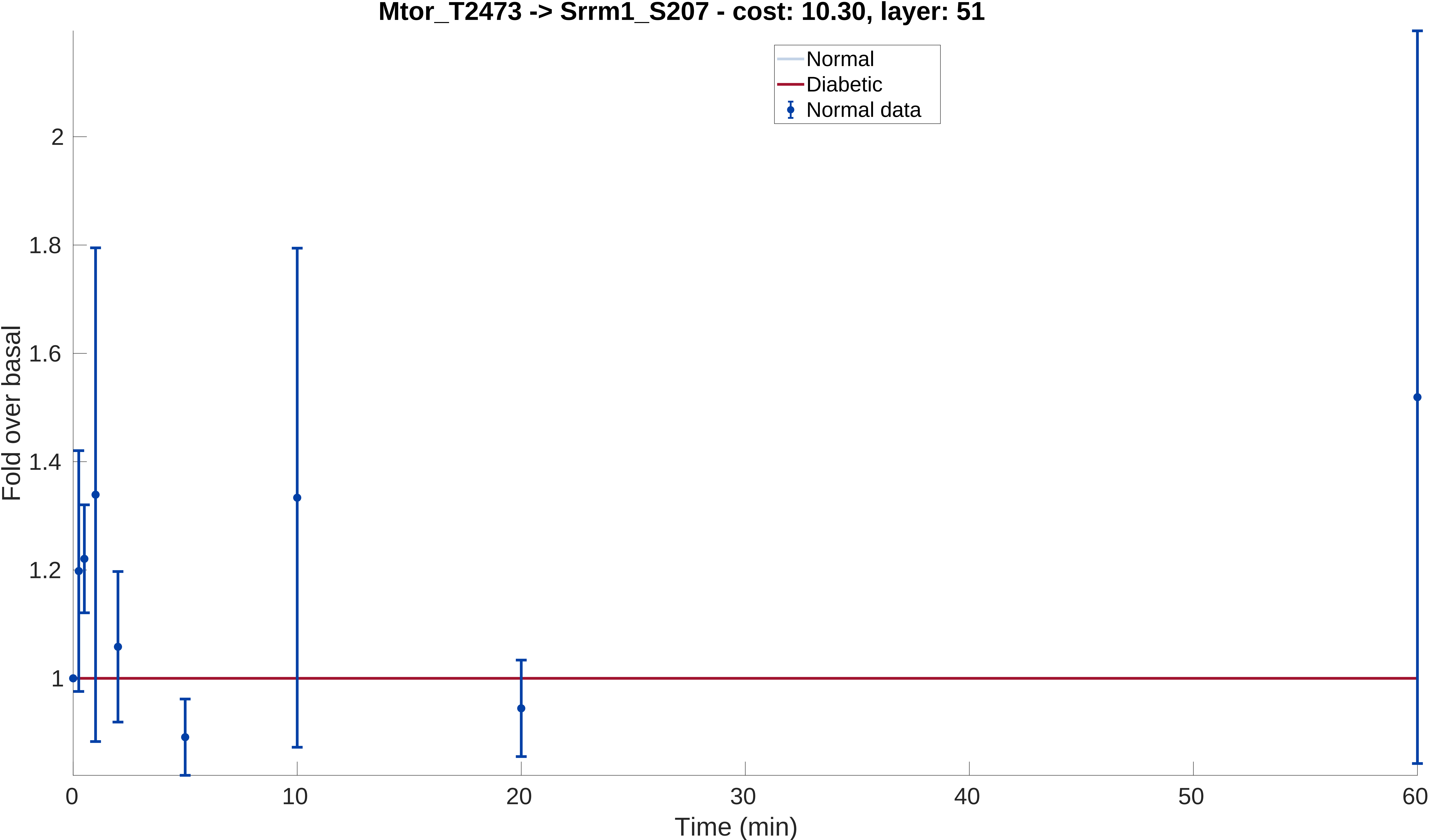

Mtor_T2473 -> Srrm1_S207 - cost: 10.30, layer: 51
Normal
Diabetic
Normal data
Fold over basal
2
1.8
1.6
1.4
1.2
1
0
10
20
30
40
50
60
Time (min)

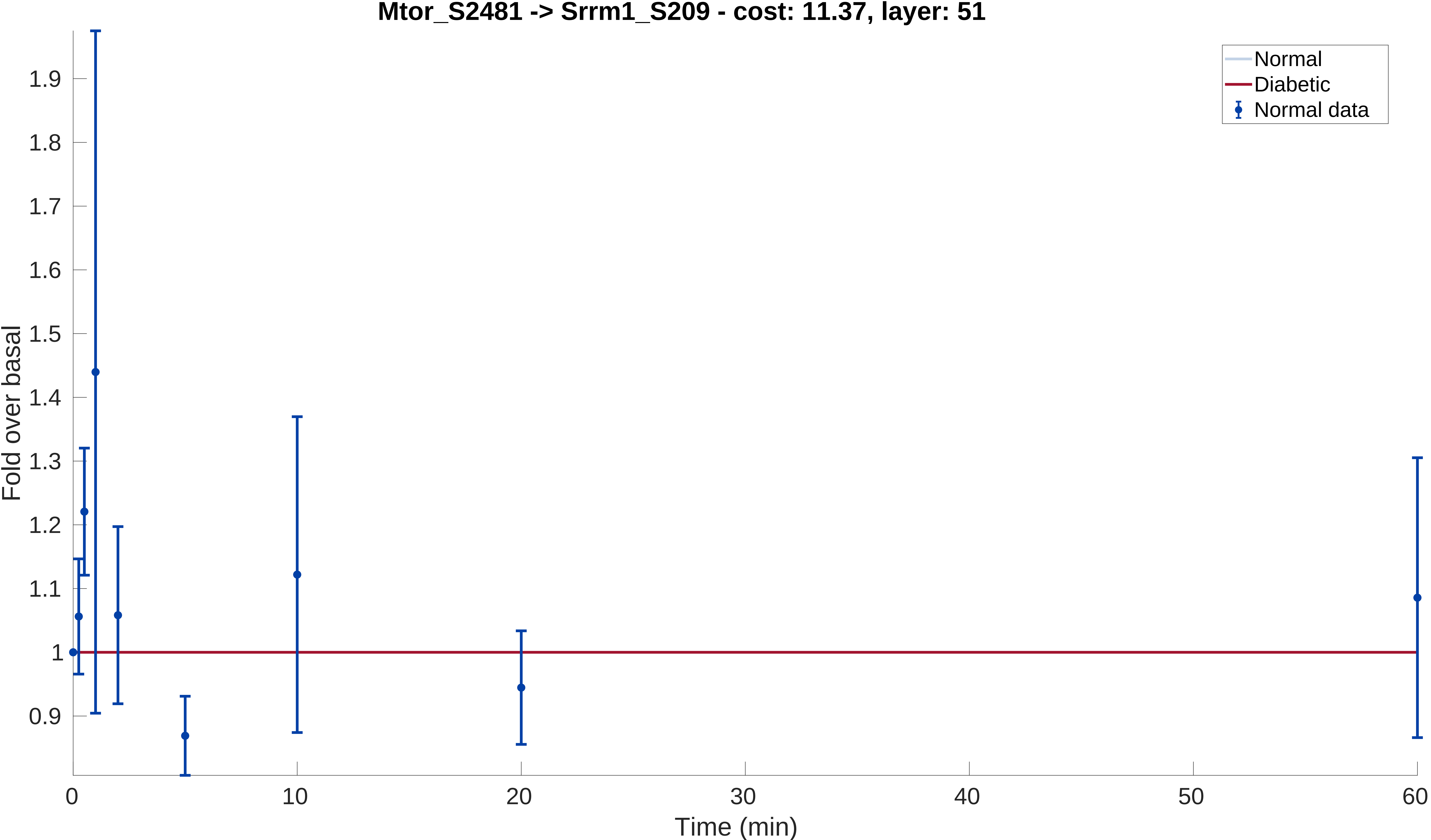

Mtor_S2481 -> Srrm1_S209 - cost: 11.37, layer: 51
Normal
Diabetic
Normal data
Fold over basal
1.9
1.8
1.7
1.6
1.5
1.4
1.3
1.2
1.1
1
0.9
0
10
20
30
40
50
60
Time (min)

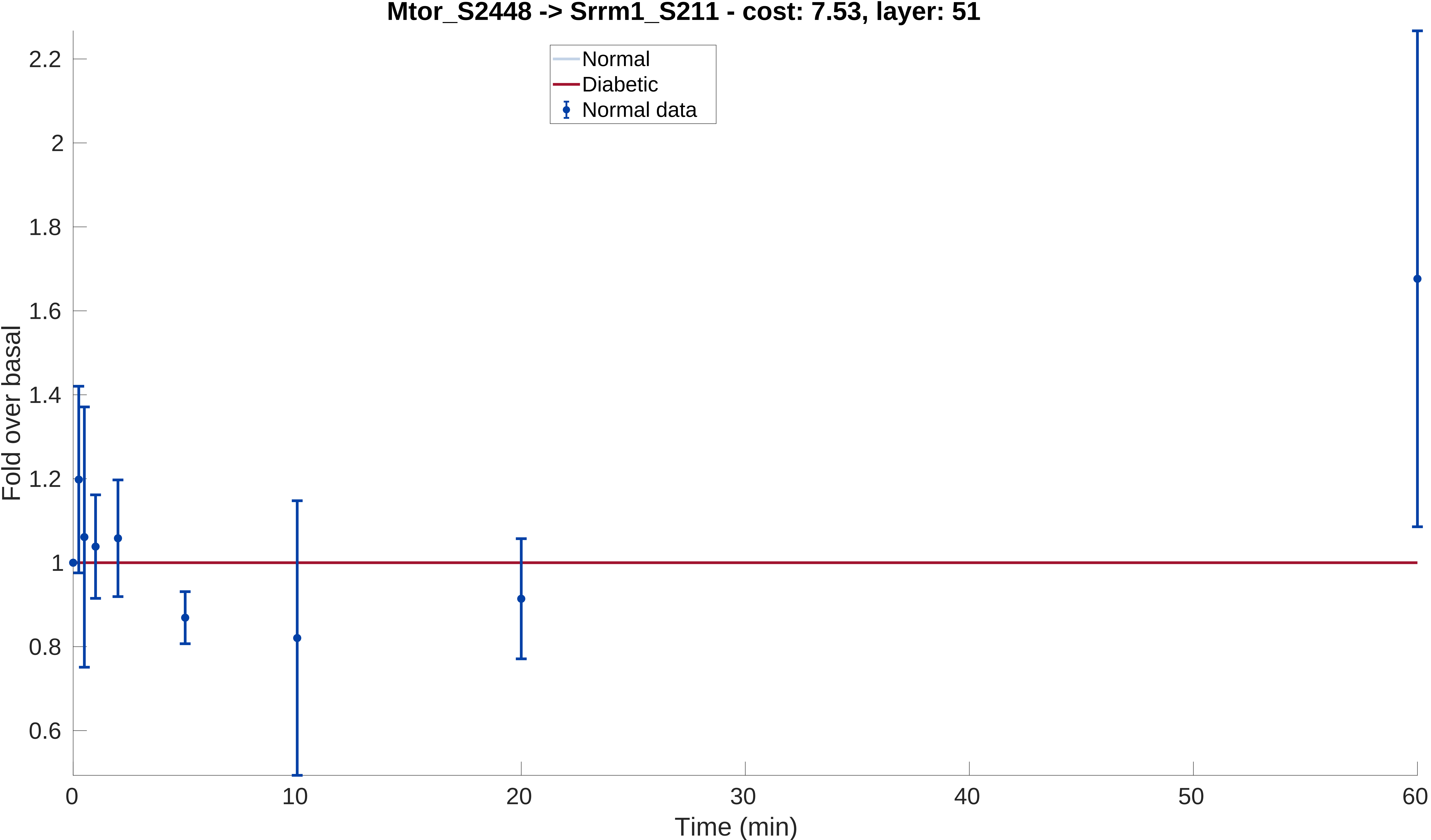

Mtor_S2448 -> Srrm1_S211 - cost: 7.53, layer: 51
Normal
Diabetic
Normal data
Fold over basal
2.2
2
1.8
1.6
1.4
1.2
1
0.8
0.6
0
10
20
30
40
50
60
Time (min)

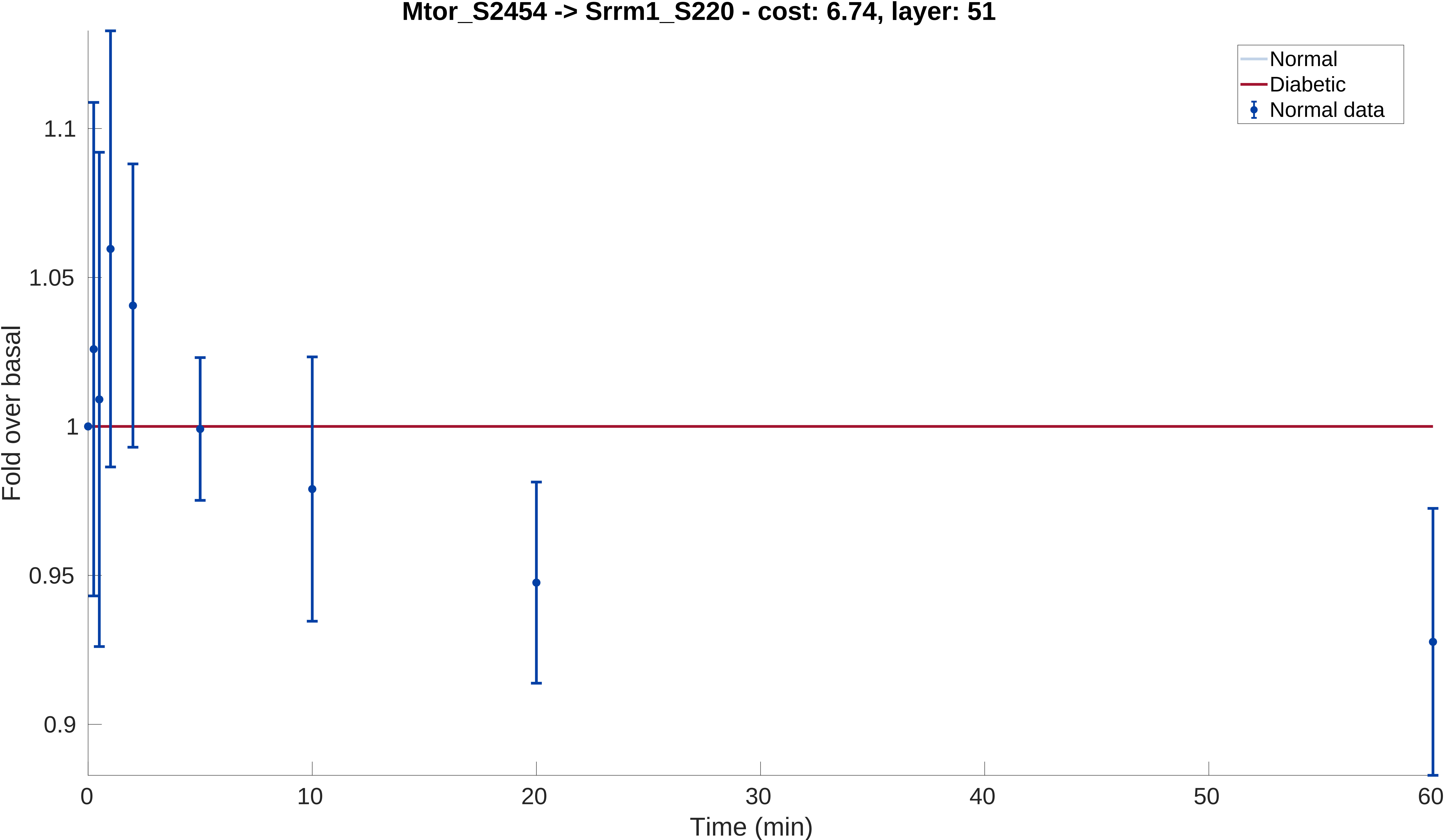

Mtor_S2454 -> Srrm1_S220 - cost: 6.74, layer: 51
Normal
Diabetic
Normal data
1.1
1.05
1
0.95
0.9
Fold over basal
0
10
20
30
40
50
60
Time (min)

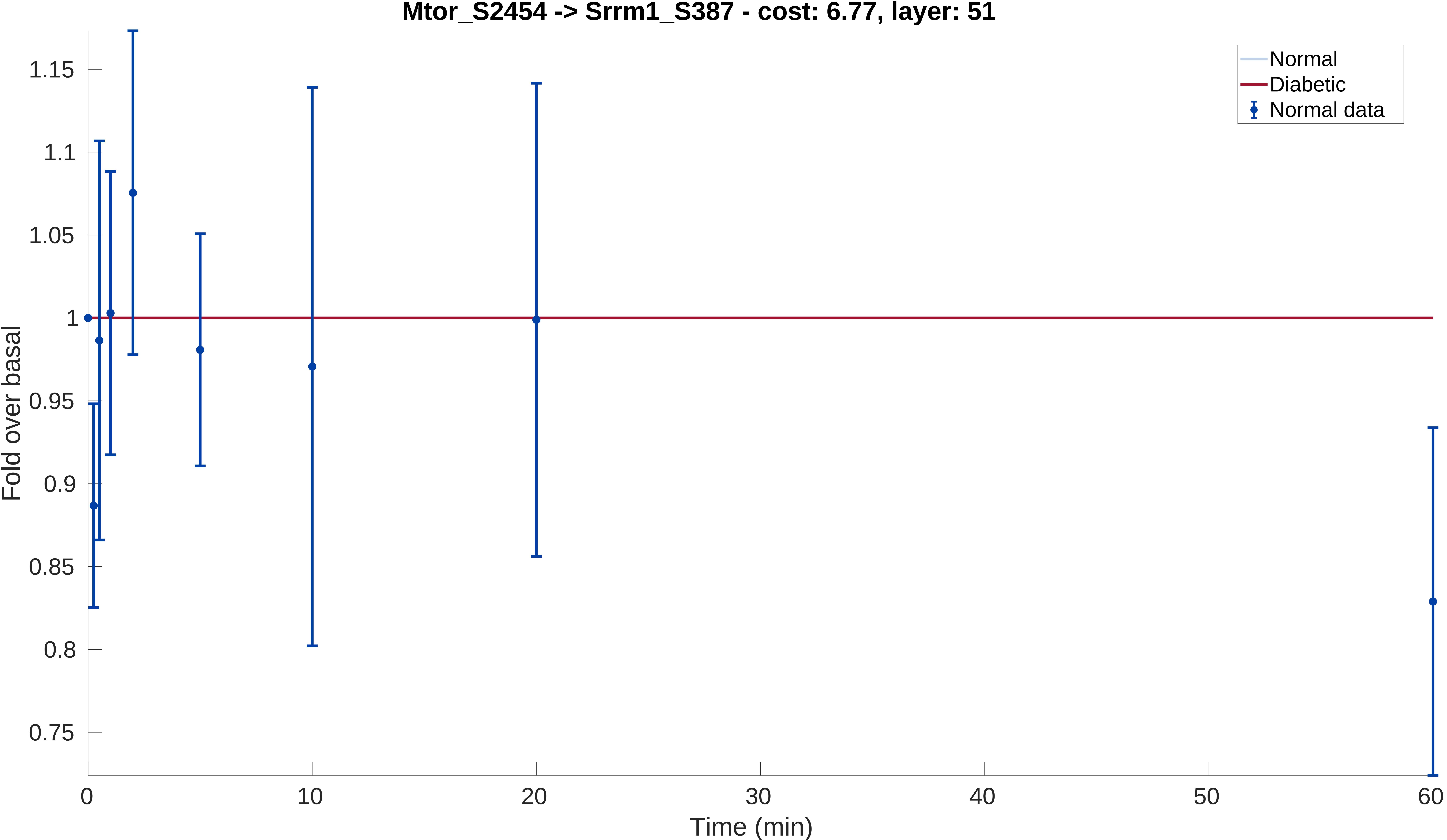
Mtor_S2454 -> Srrm1_S387 - cost: 6.77, layer: 51
Normal
Diabetic
Normal data
Fold over basal
1.15
1.1
1.05
1
0.95
0.9
0.85
0.8
0.75
0
10
20
30
40
50
60
Time (min)

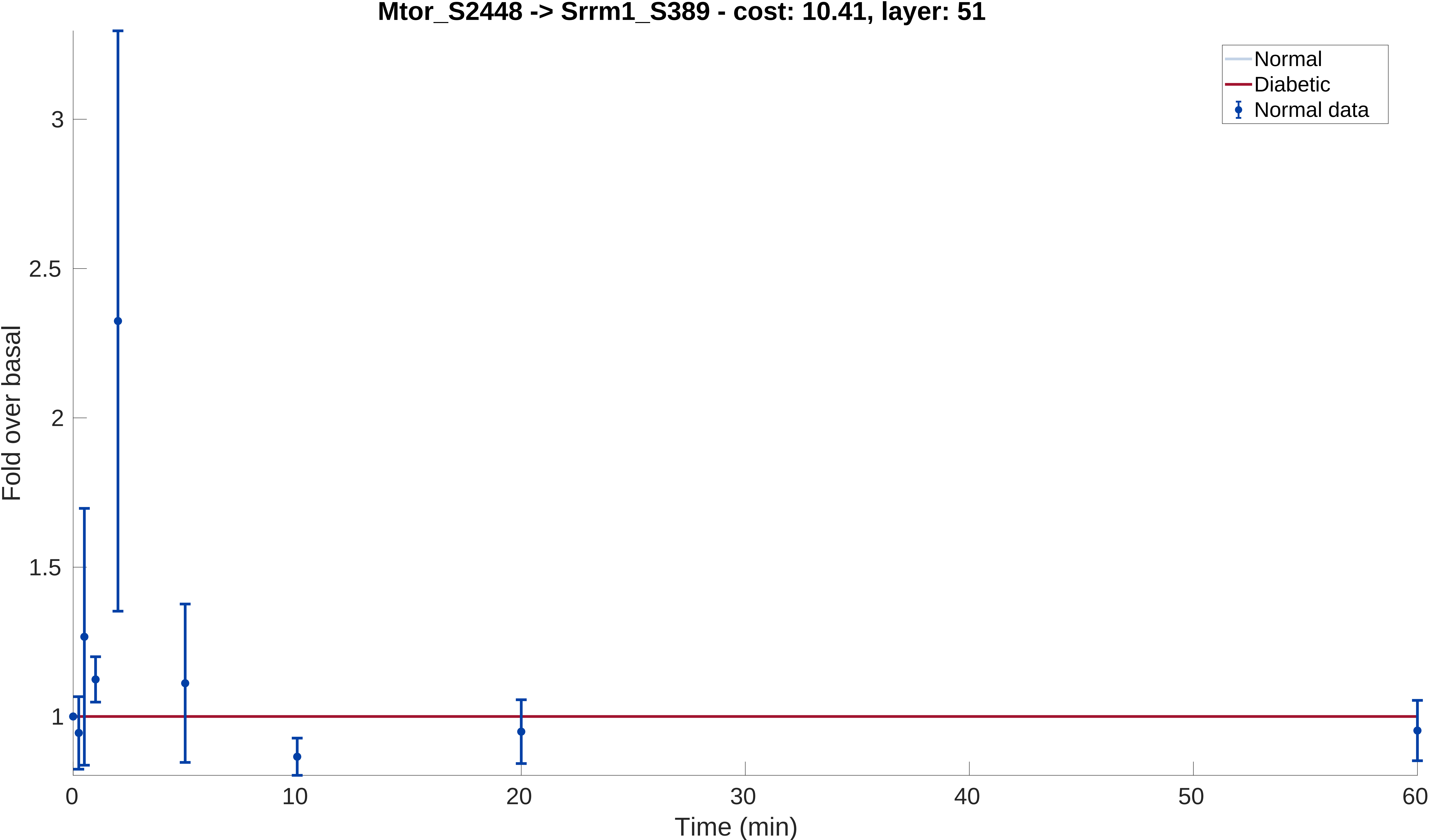
Mtor_S2448 -> Srrm1_S389 - cost: 10.41, layer: 51
Normal
Diabetic
Normal data
Fold over basal
3
2.5
2
1.5
1
0
10
20
30
40
50
60
Time (min)

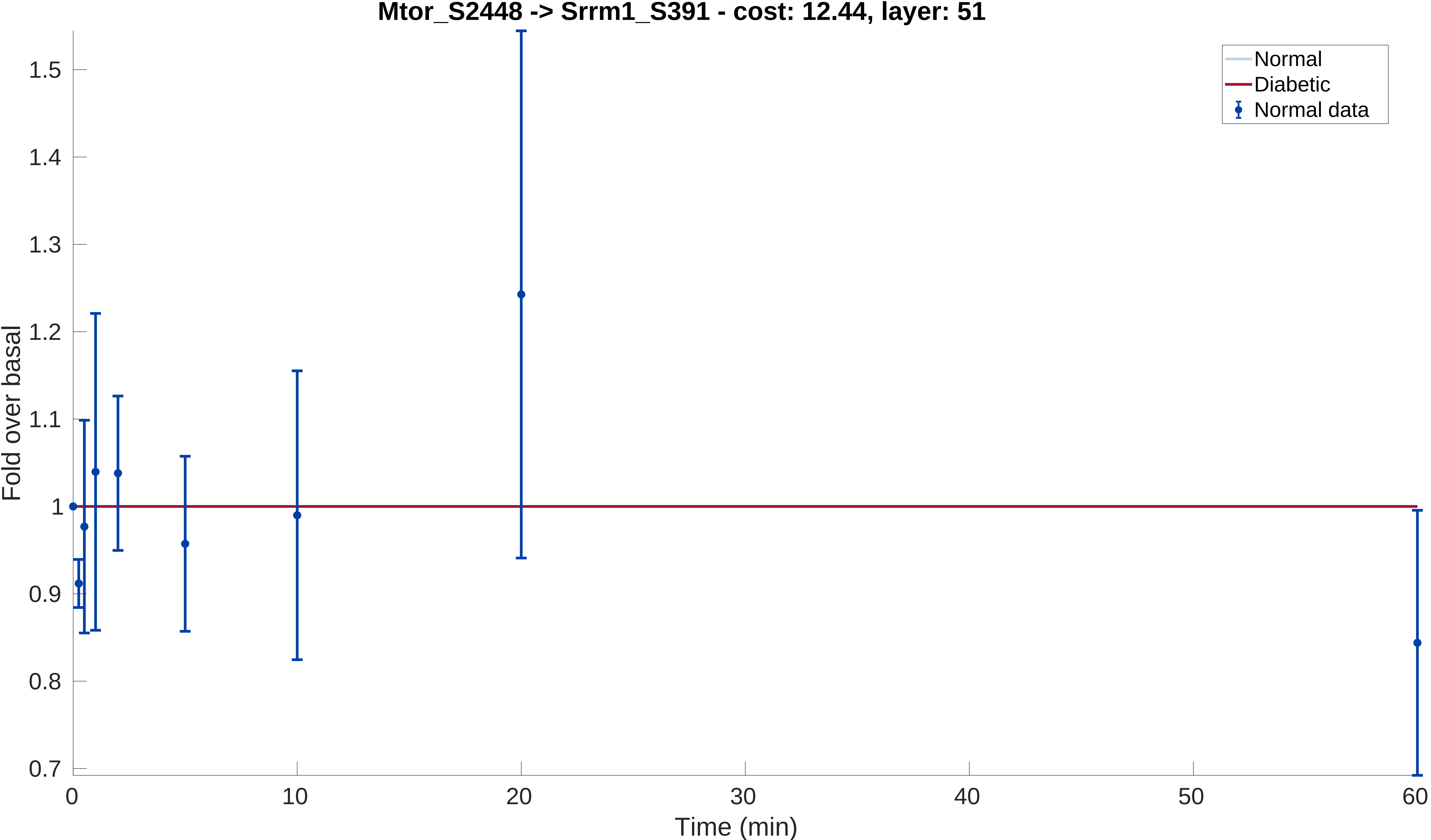

Mtor_S2448 -> Srrm1_S391 - cost: 12.44, layer: 51
Normal
Diabetic
Normal data
Fold over basal
1.5
1.4
1.3
1.2
1.1
1
0.9
0.8
0.7
0
10
20
30
40
50
60
Time (min)

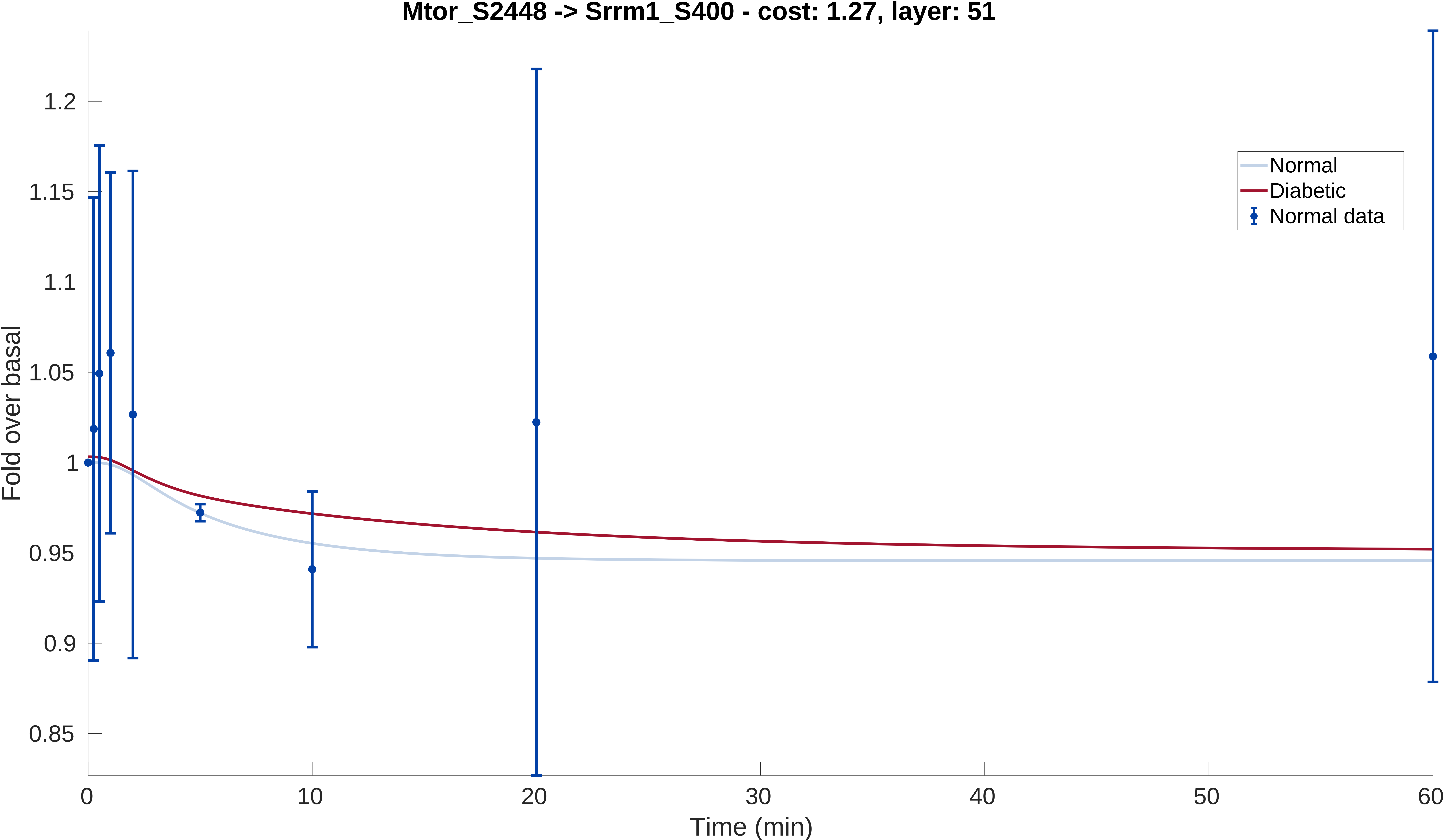

Mtor_S2448 -> Srrm1_S400 - cost: 1.27, layer: 51
Normal
Diabetic
Normal data
Fold over basal
1.2
1.15
1.1
1.05
1
0.95
0.9
0.85
0
10
20
30
40
50
60
Time (min)

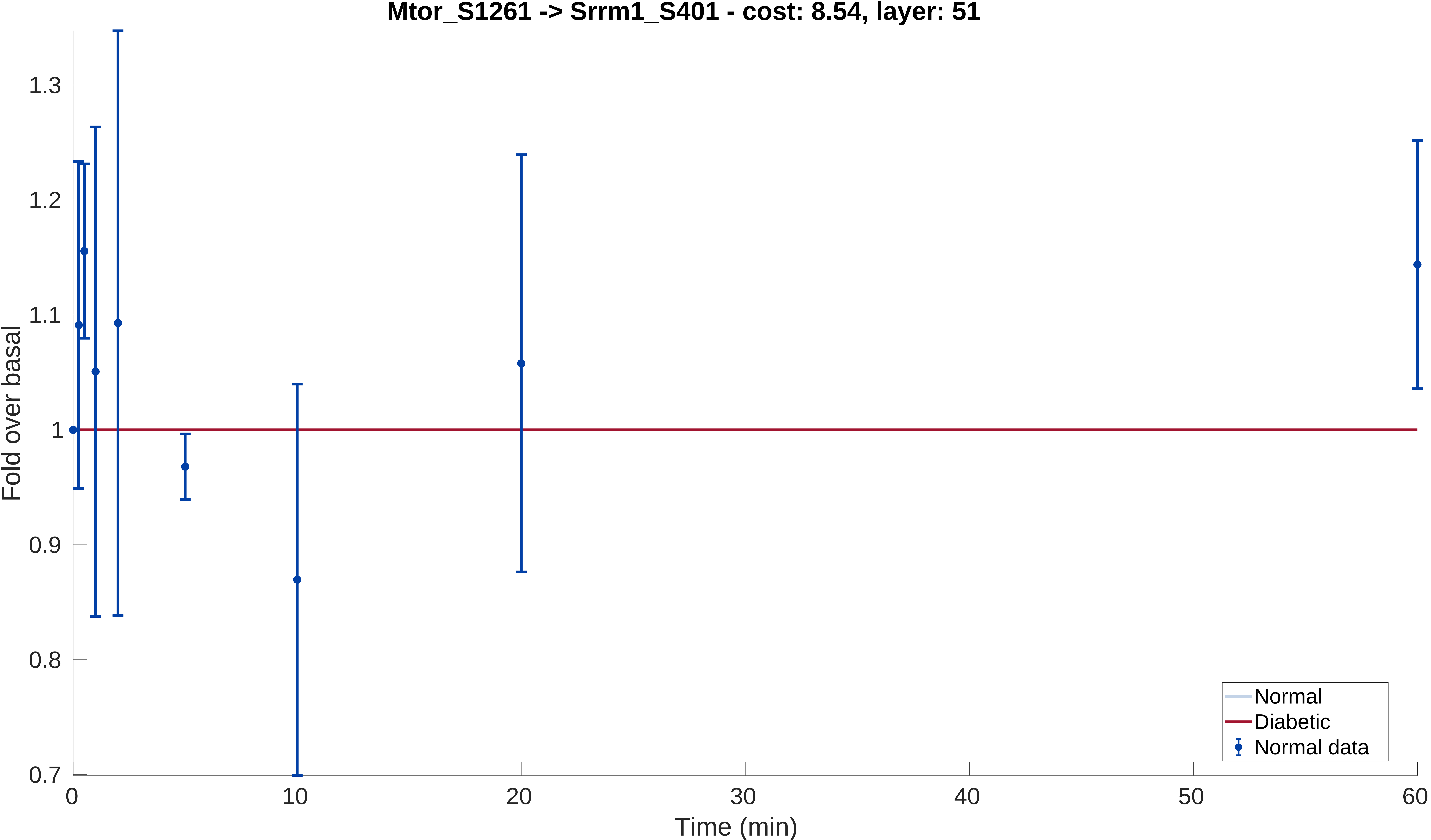

Mtor_S1261 -> Srrm1_S401 - cost: 8.54, layer: 51
Fold over basal
1.3
1.2
1.1
1
0.9
0.8
0.7
0
10
20
30
40
50
60
Time (min)
Normal
Diabetic
Normal data

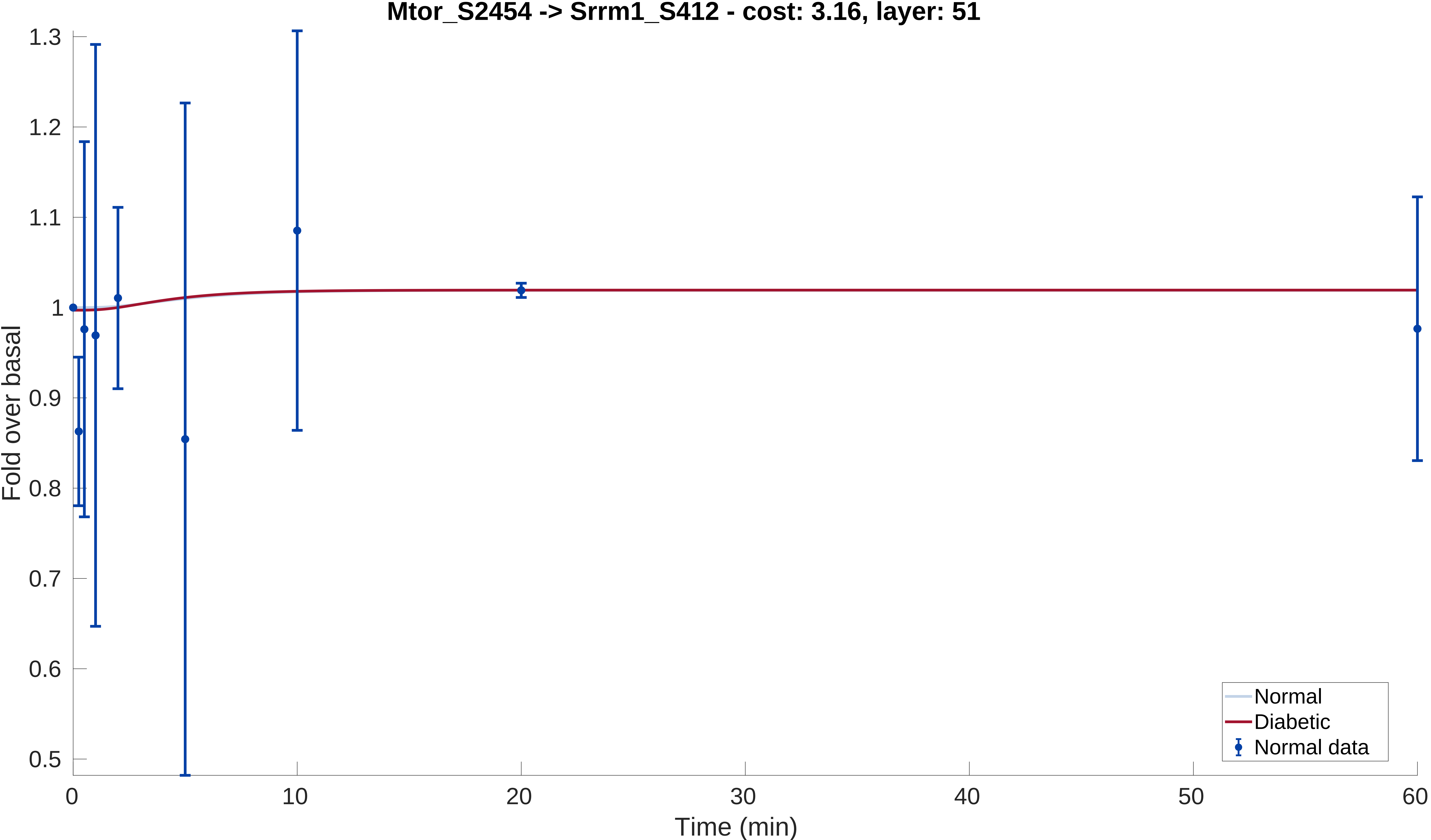
Mtor_S2454 -> Srrm1_S412 - cost: 3.16, layer: 51
Fold over basal
1.3
1.2
1.1
1
0.9
0.8
0.7
0.6
0.5
0
10
20
30
40
50
60
Time (min)
Normal
Diabetic
Normal data

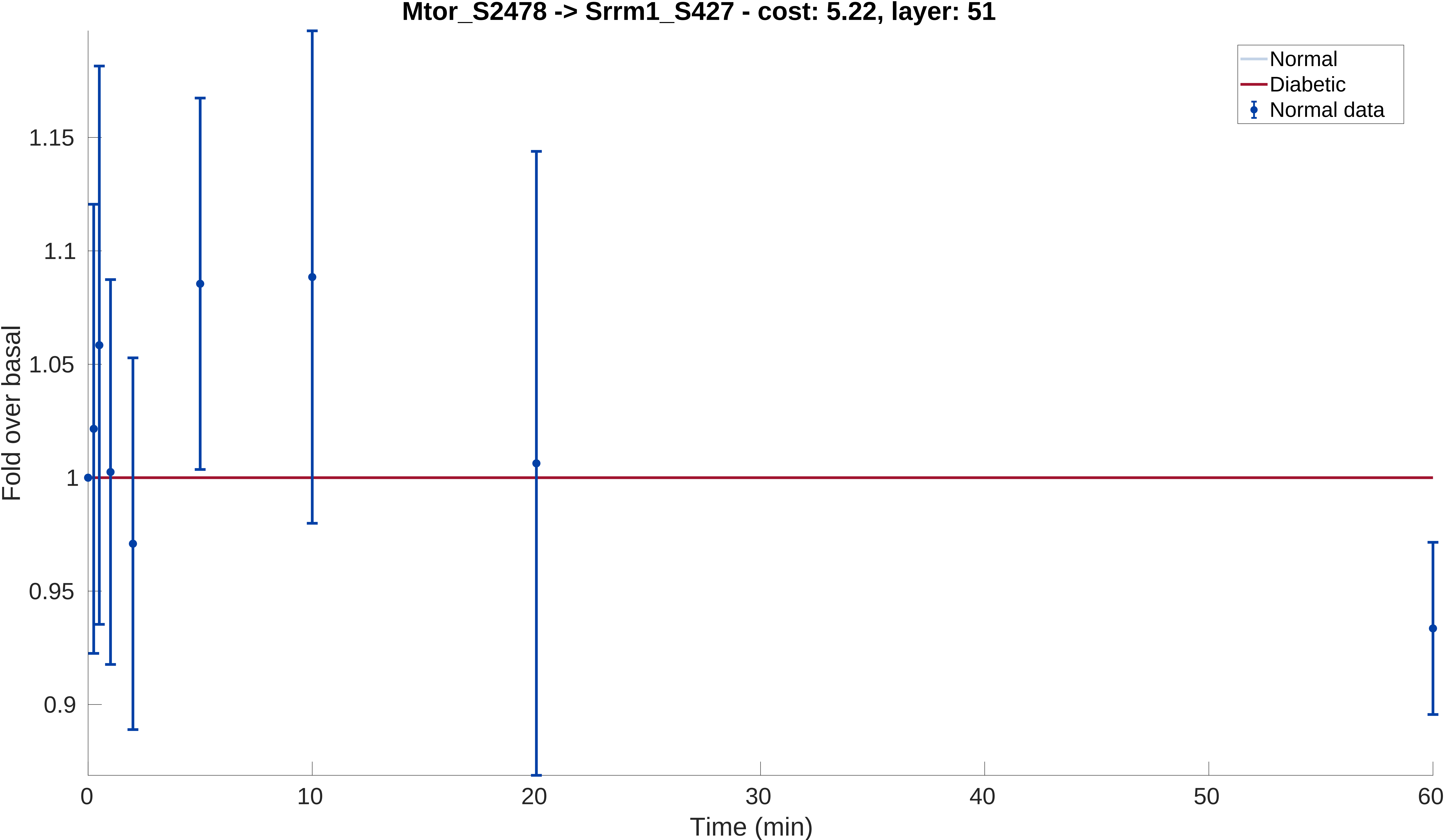

Mtor_S2478 -> Srrm1_S427 - cost: 5.22, layer: 51
Normal
Diabetic
Normal data
Fold over basal
1.15
1.1
1.05
1
0.95
0.9
0
10
20
30
40
50
60
Time (min)

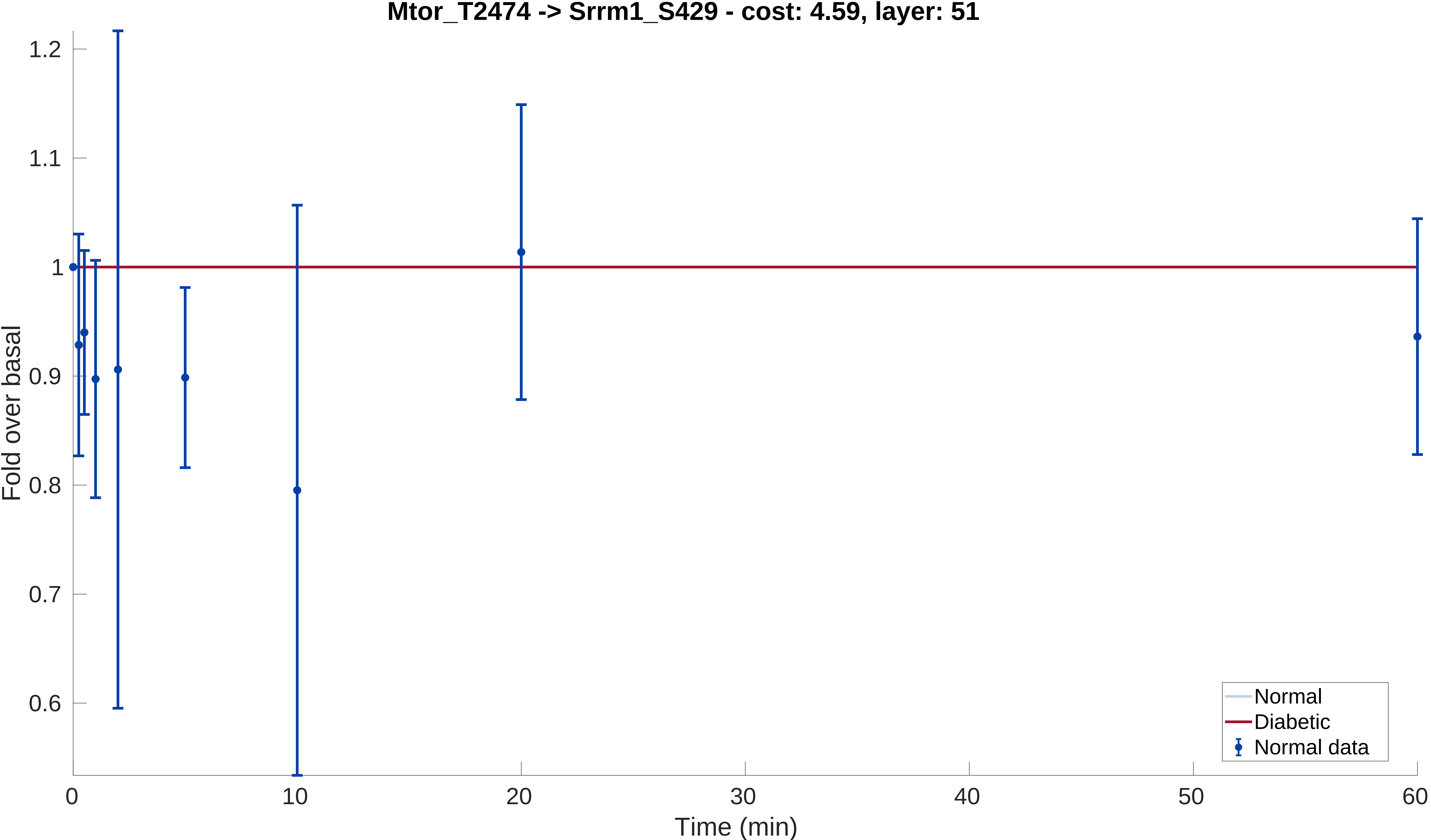
Mtor_T2474 -> Srrm1_S429 - cost: 4.59, layer: 51
Fold over basal
1.2
1.1
1
0.9
0.8
0.7
0.6
0
10
20
30
40
50
60
Time (min)
Normal
Diabetic
Normal data

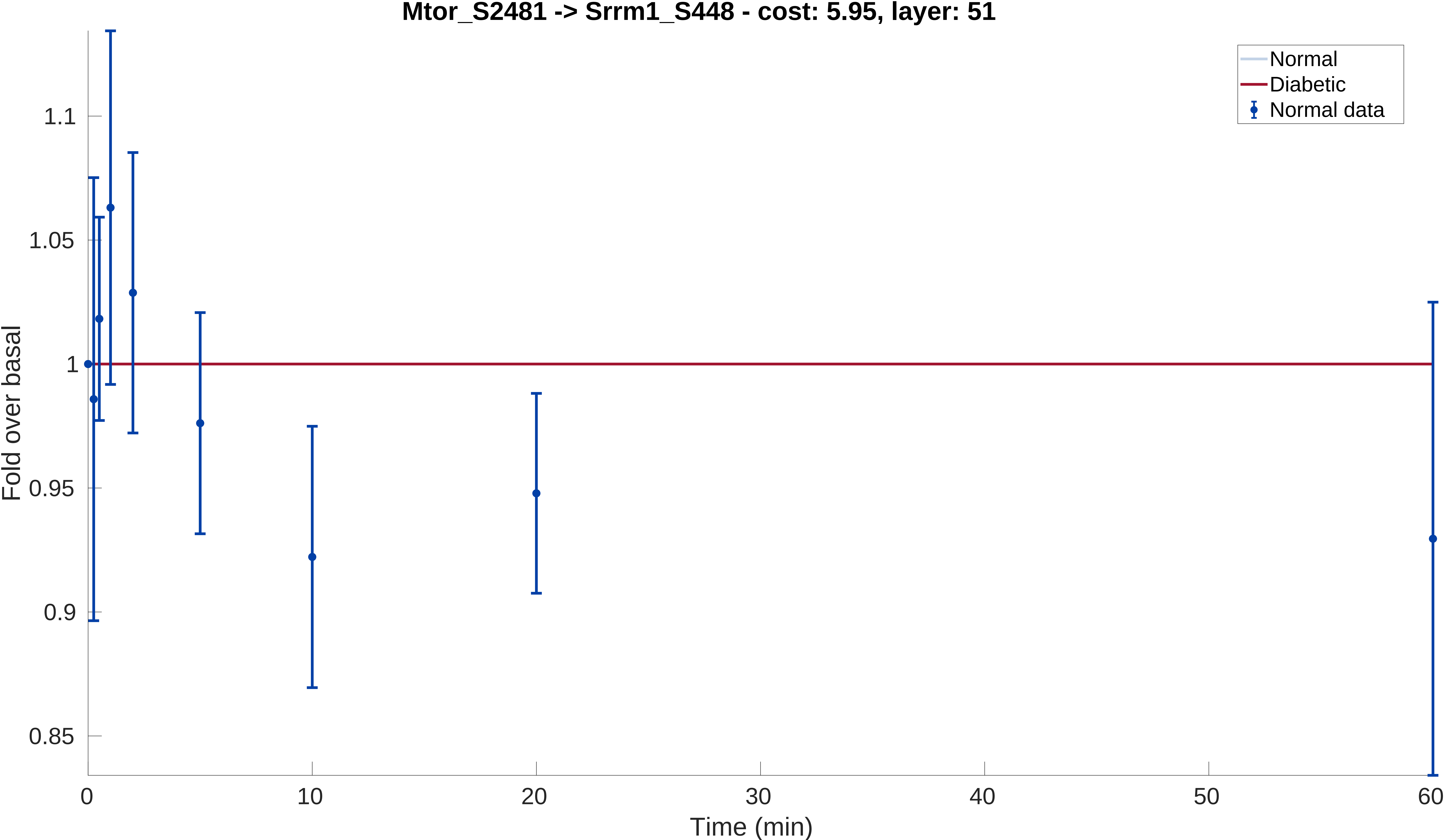
Mtor_S2481 -> Srrm1_S448 - cost: 5.95, layer: 51
Normal
Diabetic
Normal data
1.1
1.05
1
0.95
0.9
0.85
Fold over basal
0
10
20
30
40
50
60
Time (min)

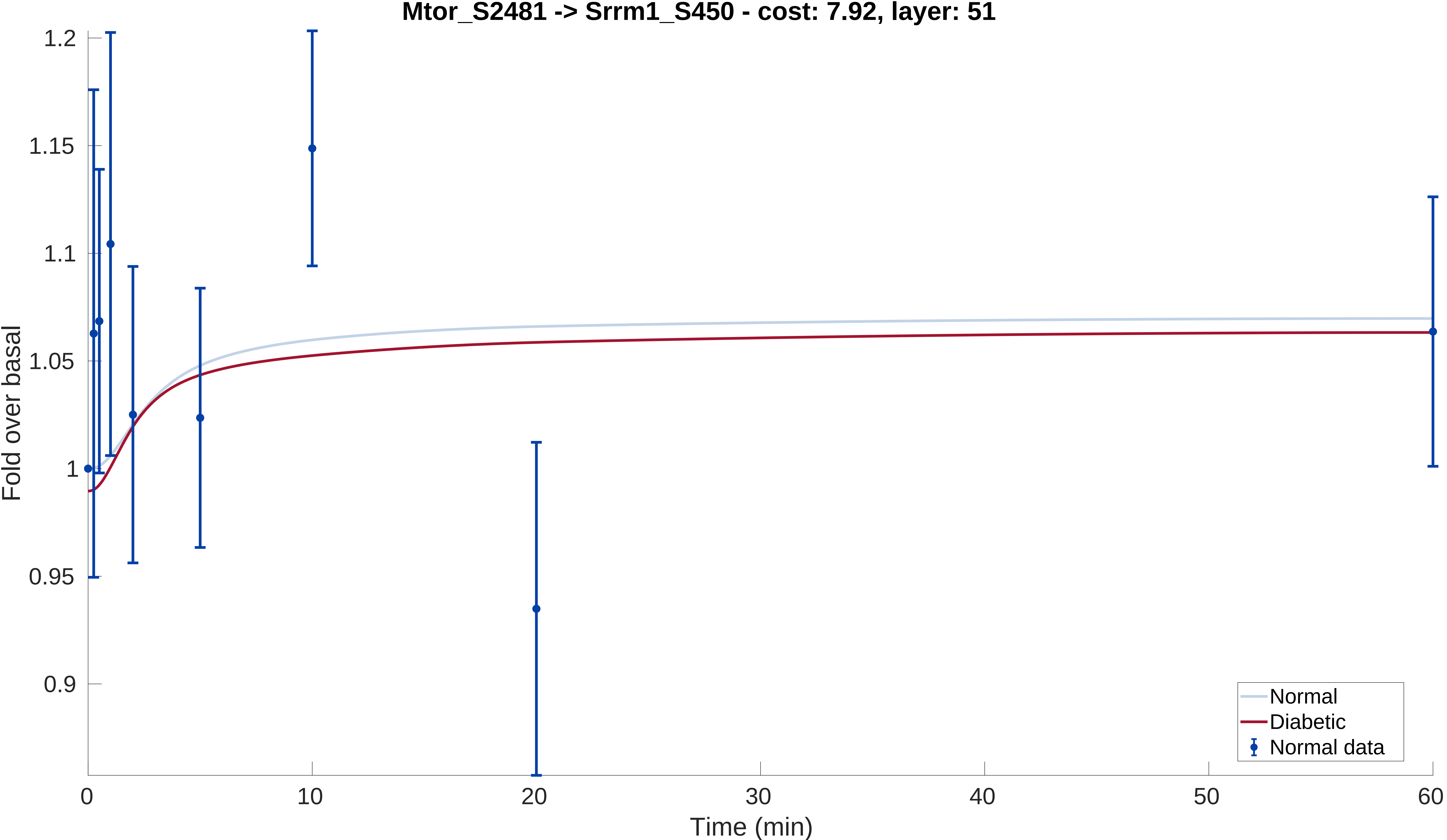
Mtor_S2481 -> Srrm1_S450 - cost: 7.92, layer: 51
Fold over basal
1.2
1.15
1.1
1.05
1
0.95
0.9
0
10
20
30
40
50
60
Time (min)
Normal
Diabetic
Normal data

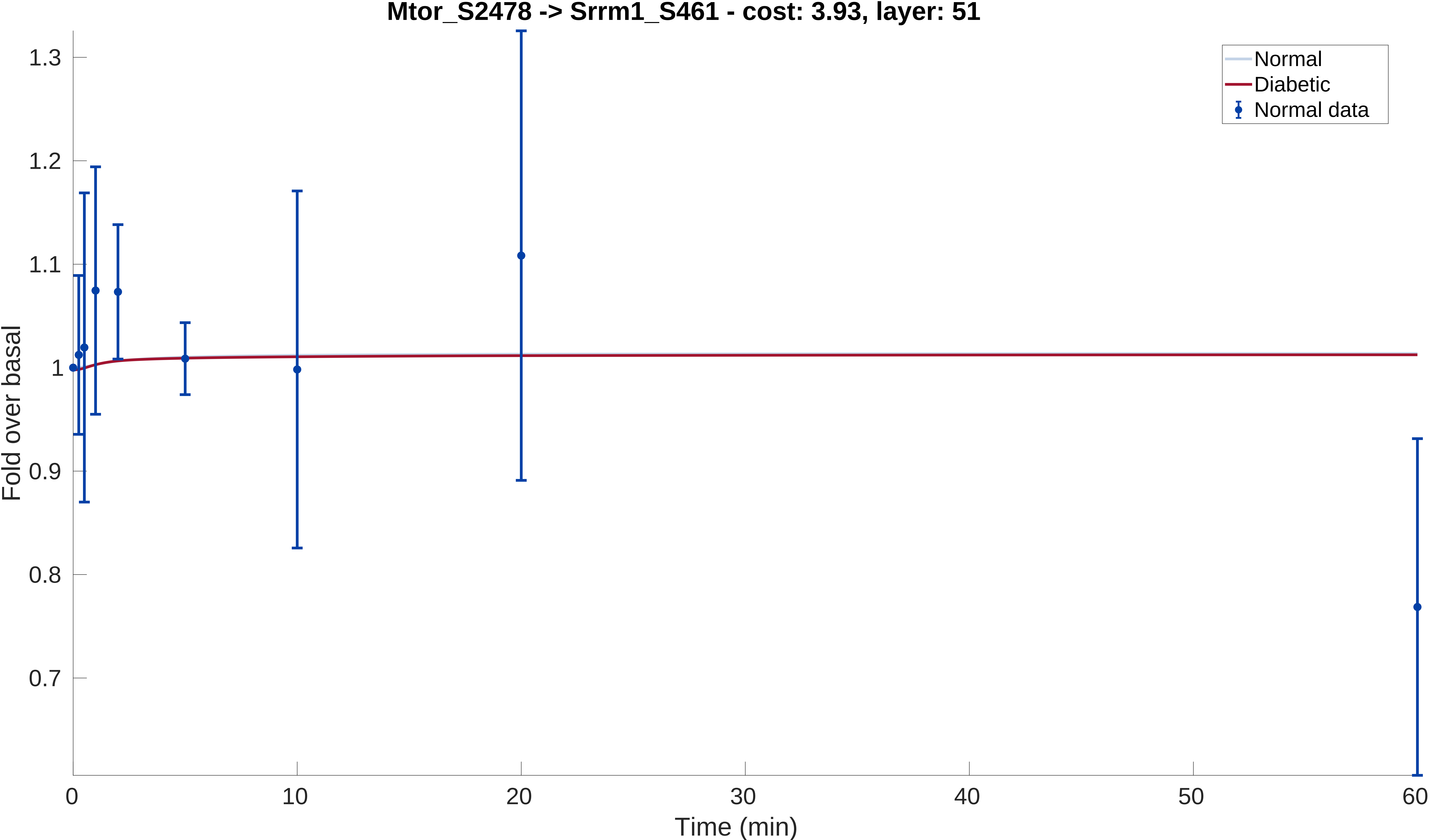

Mtor_S2478 -> Srrm1_S461 - cost: 3.93, layer: 51
Normal
Diabetic
Normal data
Fold over basal
Time (min)
0
10
20
30
40
50
60
0.7
0.8
0.9
1
1.1
1.2
1.3

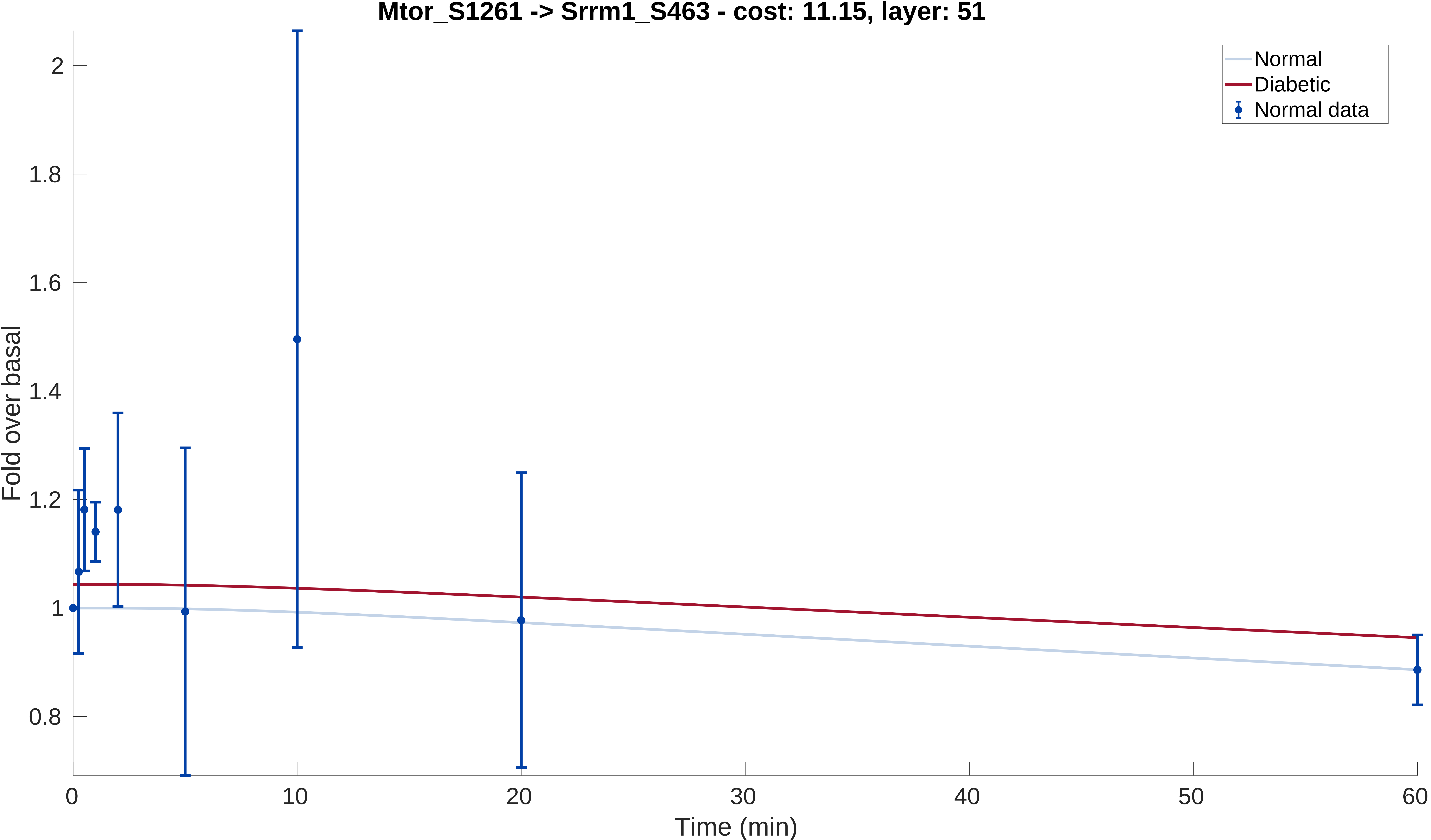
Mtor_S1261 -> Srrm1_S463 - cost: 11.15, layer: 51
Normal
Diabetic
Normal data
Fold over basal
2
1.8
1.6
1.4
1.2
1
0.8
0
10
20
30
40
50
60
Time (min)

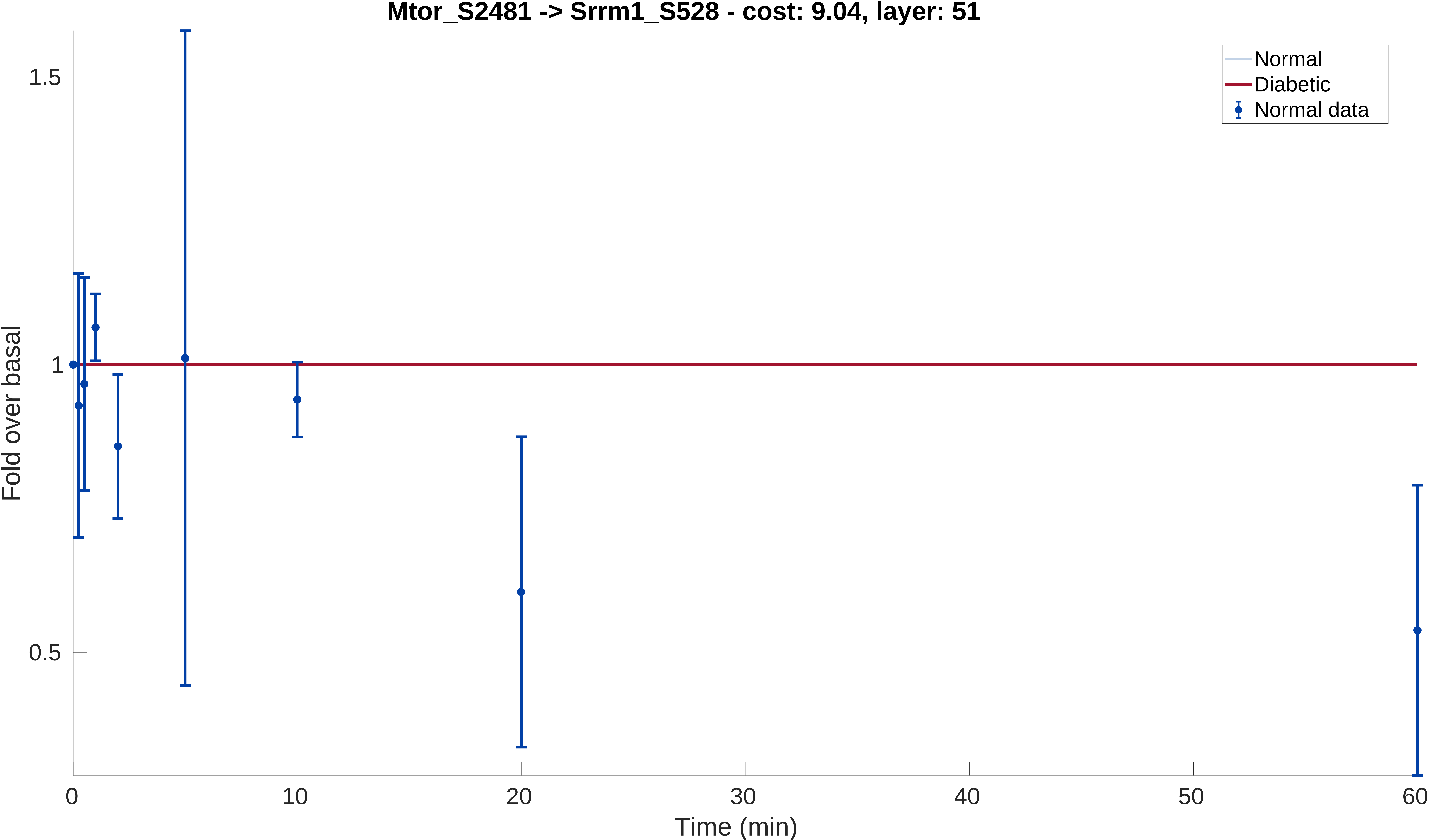
Mtor_S2481 -> Srrm1_S528 - cost: 9.04, layer: 51
Normal
Diabetic
Normal data
1.5
1
0.5
Fold over basal
0
10
20
30
40
50
60
Time (min)

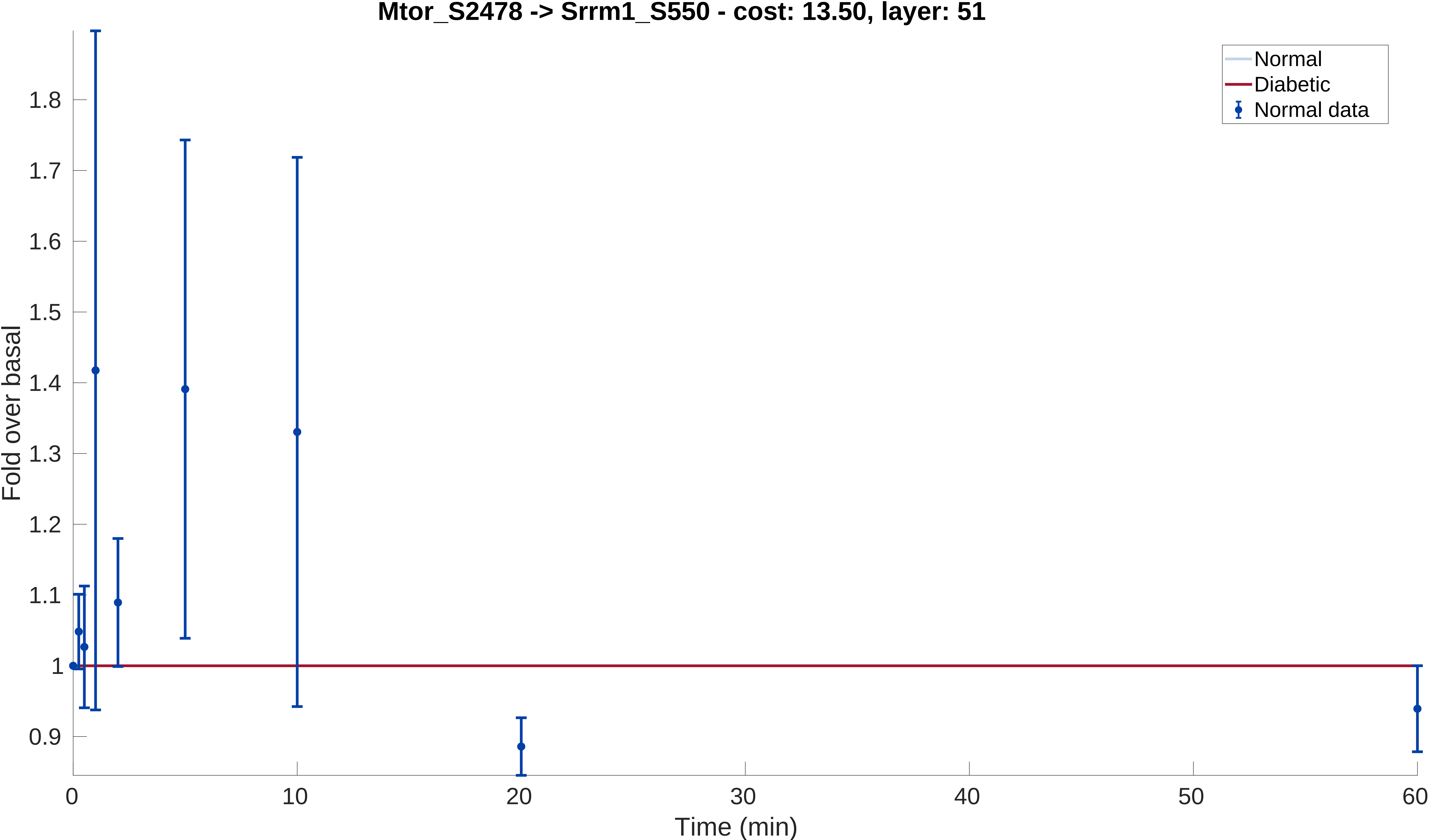

Mtor_S2478 -> Srrm1_S550 - cost: 13.50, layer: 51
Normal
Diabetic
Normal data
Fold over basal
1.8
1.7
1.6
1.5
1.4
1.3
1.2
1.1
1
0.9
0
10
20
30
40
50
60
Time (min)

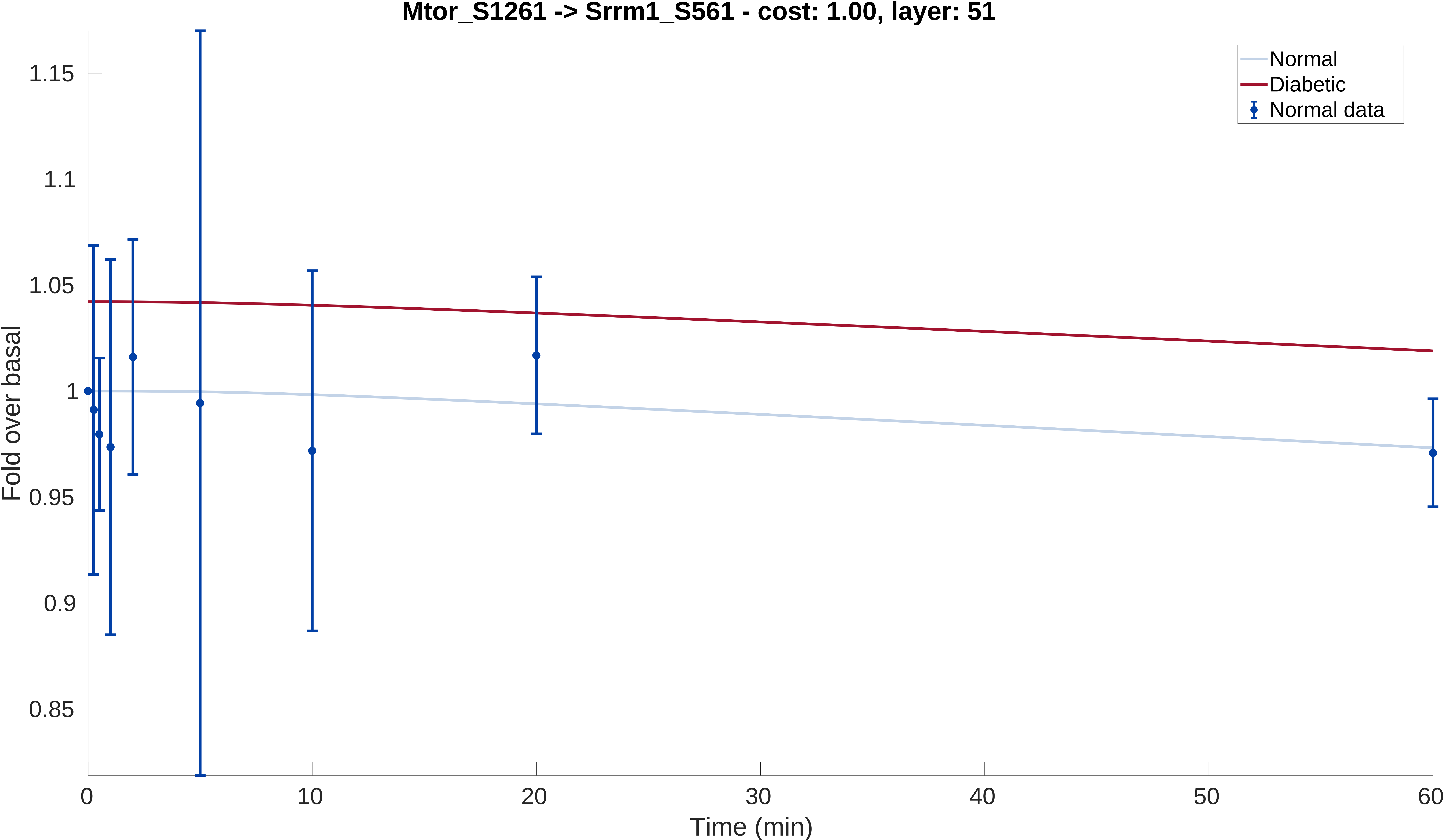
Mtor_S1261 -> Srrm1_S561 - cost: 1.00, layer: 51
Normal
Diabetic
Normal data
Fold over basal
1.15
1.1
1.05
1
0.95
0.9
0.85
0
10
20
30
40
50
60
Time (min)

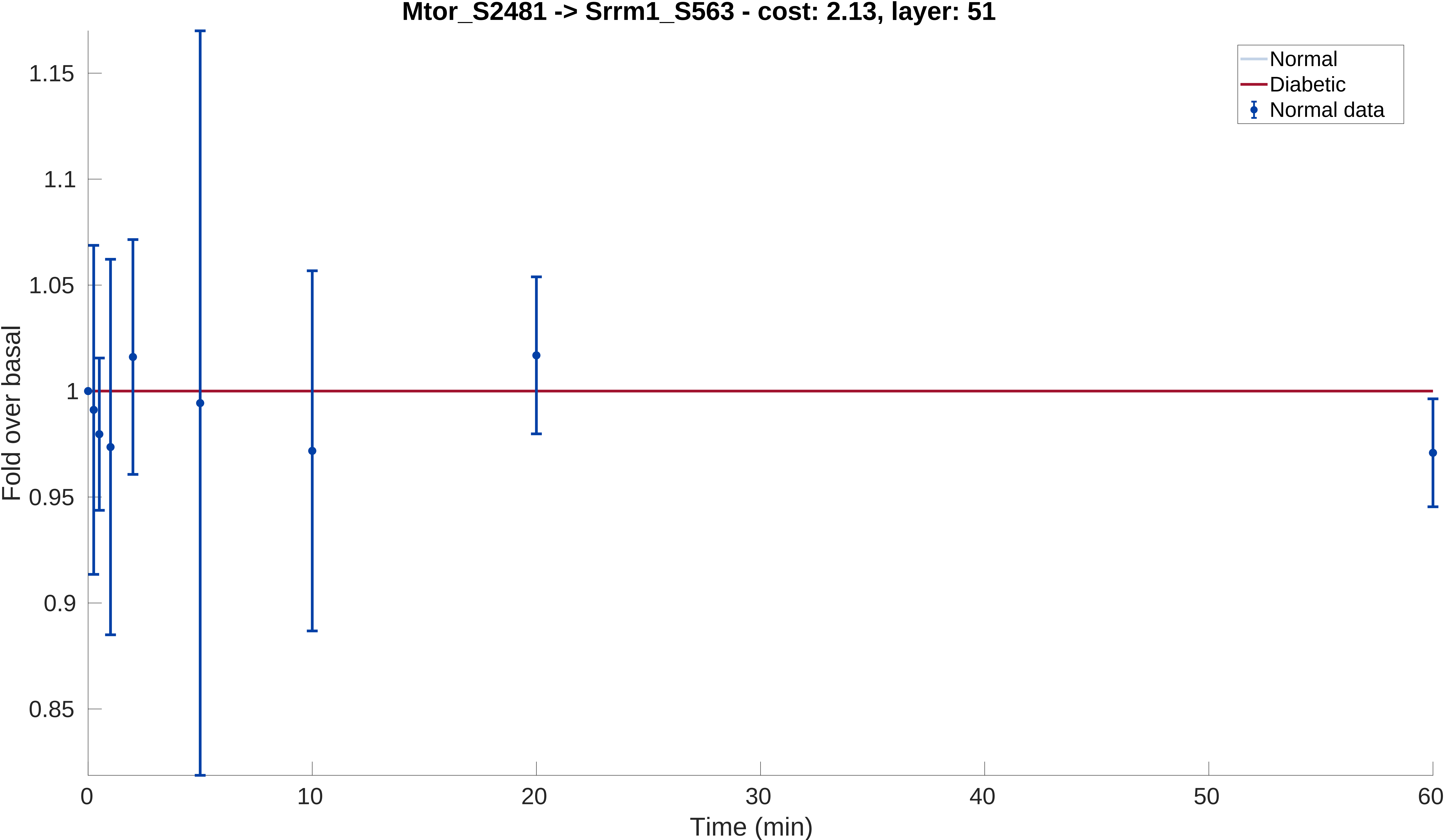

Mtor_S2481 -> Srrm1_S563 - cost: 2.13, layer: 51
Normal
Diabetic
Normal data
Fold over basal
1.15
1.1
1.05
1
0.95
0.9
0.85
0
10
20
30
40
50
60
Time (min)

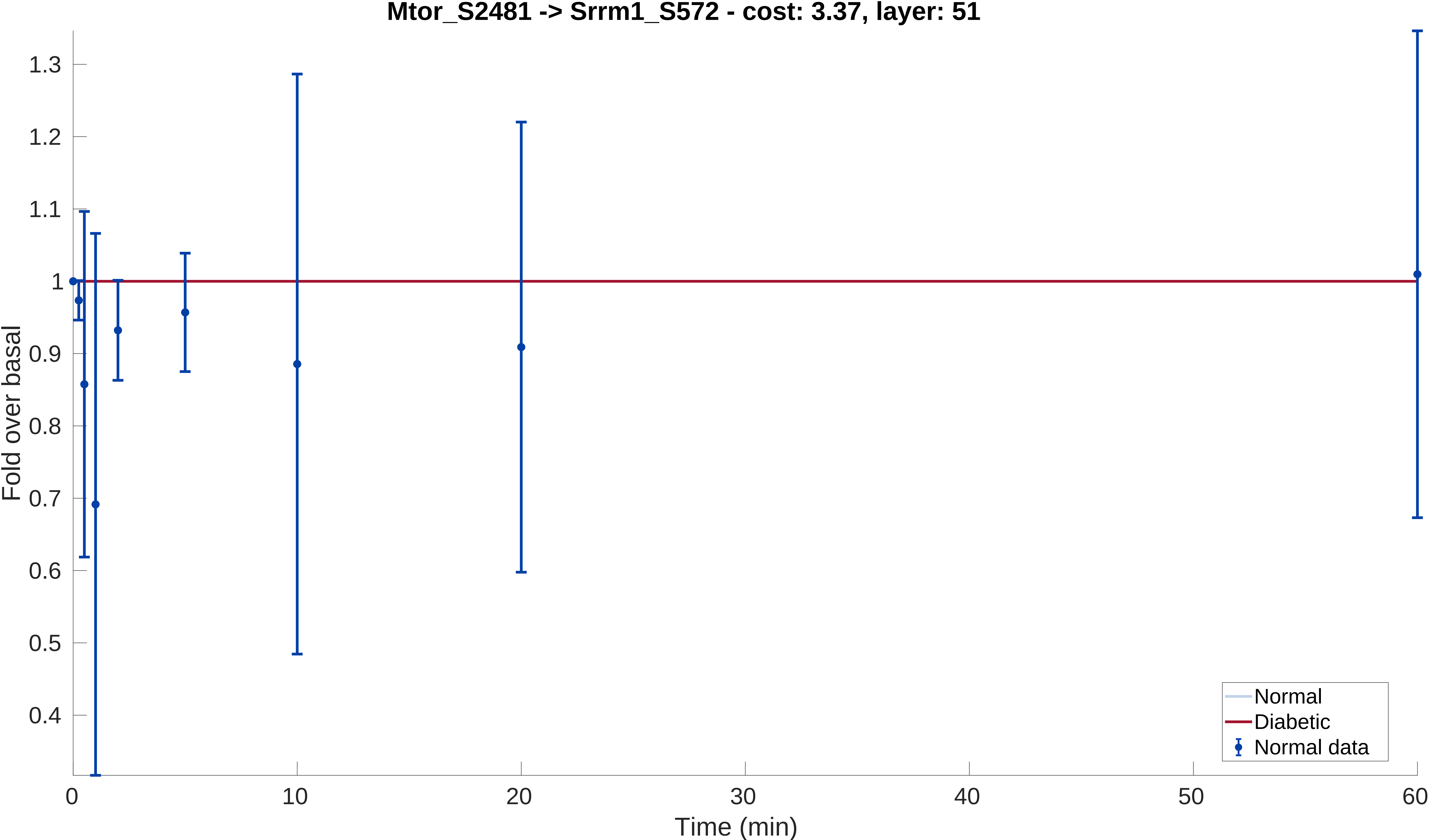

Mtor_S2481 -> Srrm1_S572 - cost: 3.37, layer: 51
Fold over basal
Time (min)
0.4
0.5
0.6
0.7
0.8
0.9
1
1.1
1.2
1.3
0
10
20
30
40
50
60
Normal
Diabetic
Normal data

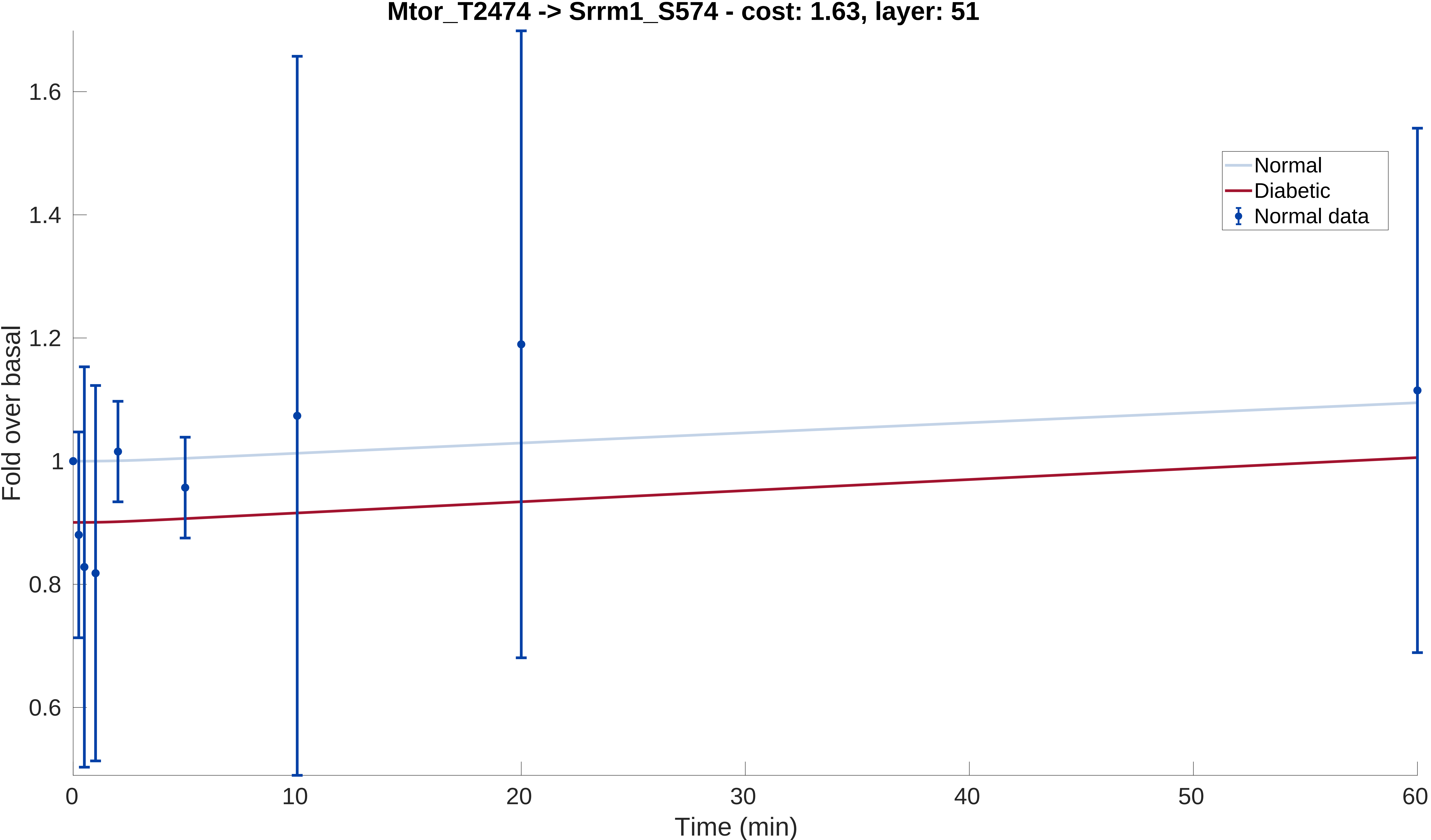

Mtor_T2474 -> Srrm1_S574 - cost: 1.63, layer: 51
Fold over basal
1.6
1.4
1.2
1
0.8
0.6
0
10
20
30
40
50
60
Time (min)
Normal
Diabetic
Normal data

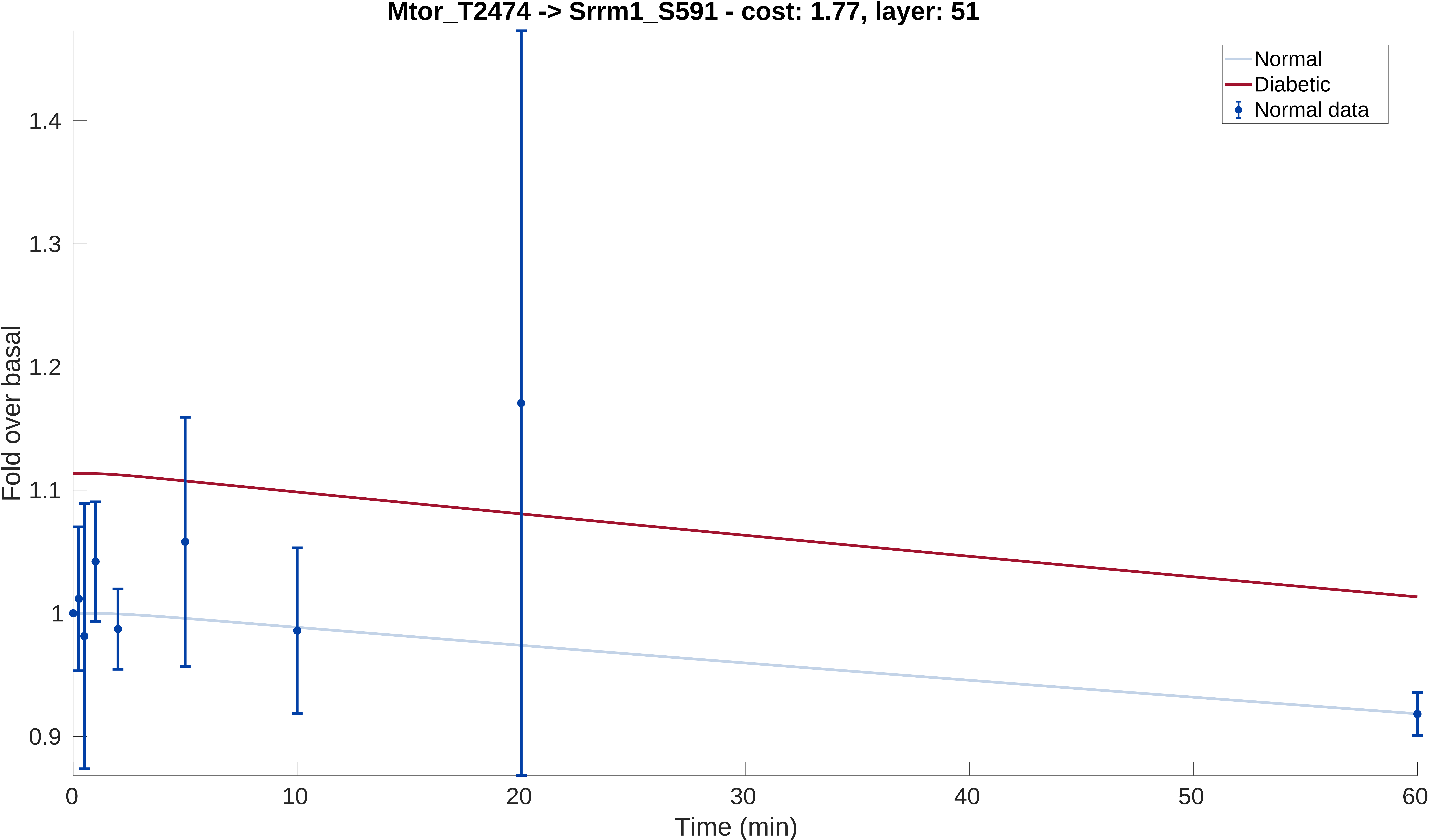

Mtor_T2474 -> Srrm1_S591 - cost: 1.77, layer: 51
Normal
Diabetic
Normal data
1.4
1.3
1.2
1.1
1
0.9
Fold over basal
0
10
20
30
40
50
60
Time (min)

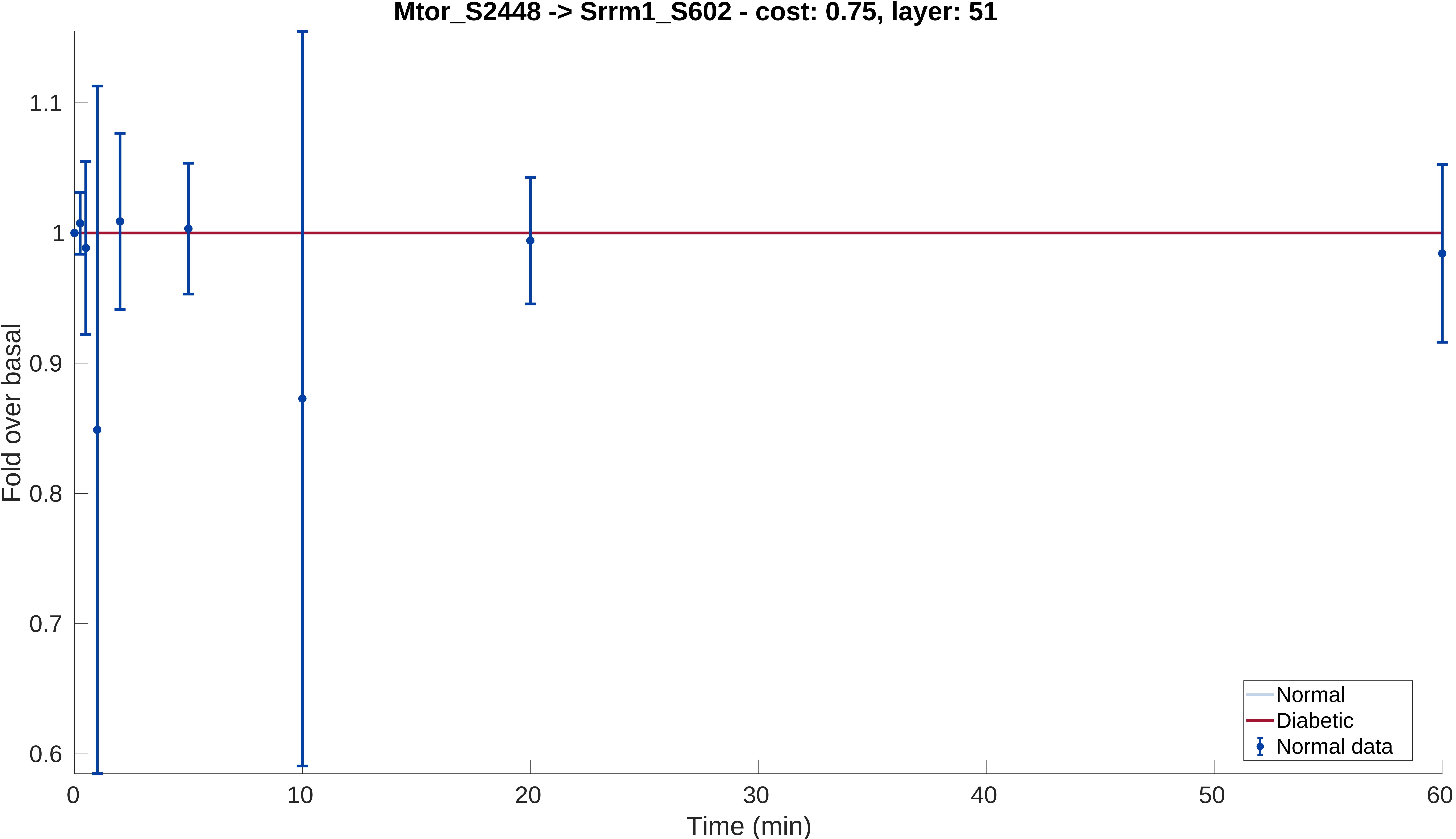
Mtor_S2448 -> Srrm1_S602 - cost: 0.75, layer: 51
Fold over basal
Time (min)
1.1
1
0.9
0.8
0.7
0.6
0
10
20
30
40
50
60
Normal
Diabetic
Normal data

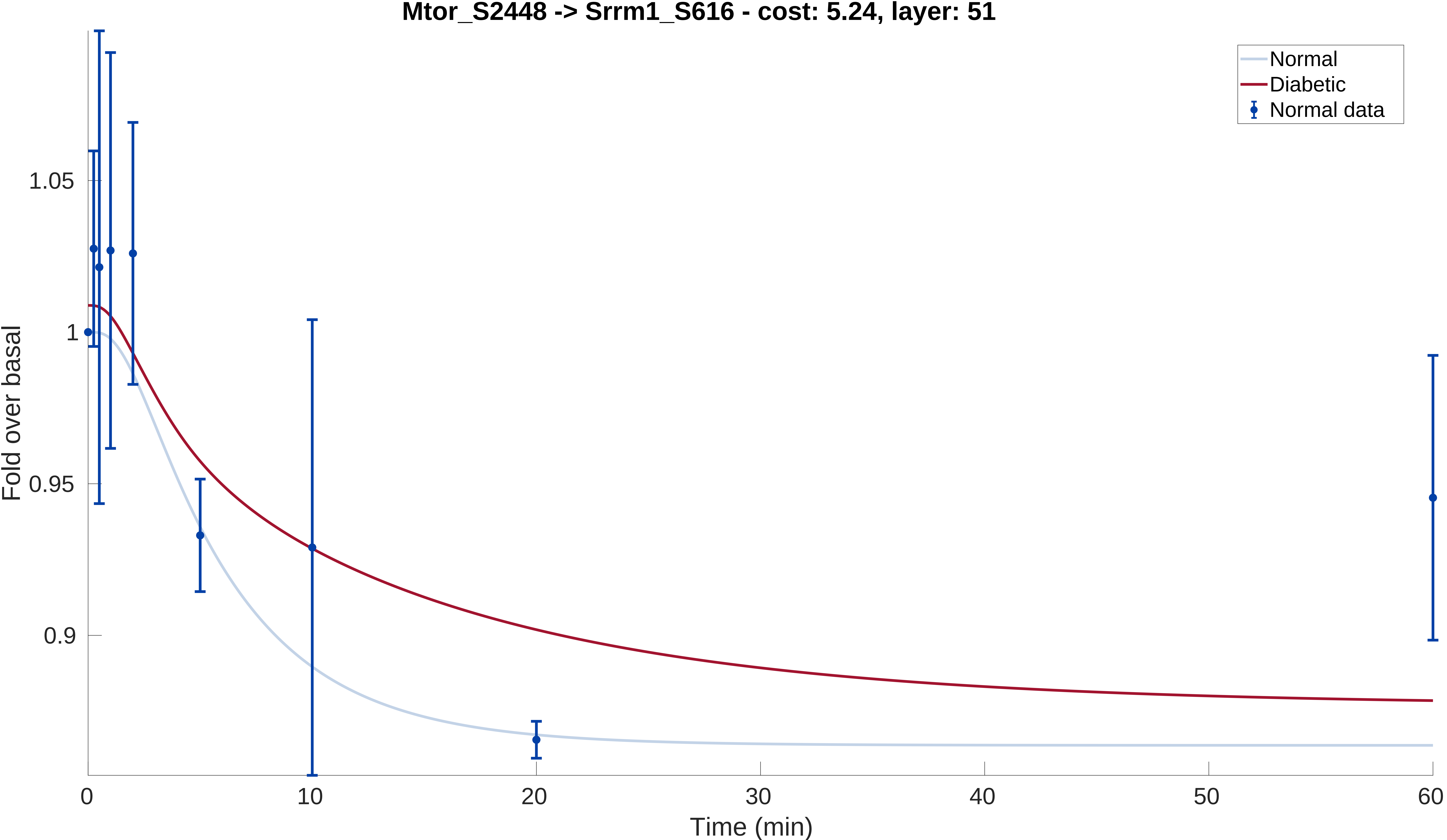

Mtor_S2448 -> Srrm1_S616 - cost: 5.24, layer: 51
Normal
Diabetic
Normal data
Fold over basal
1.05
1
0.95
0.9
0
10
20
30
40
50
60
Time (min)

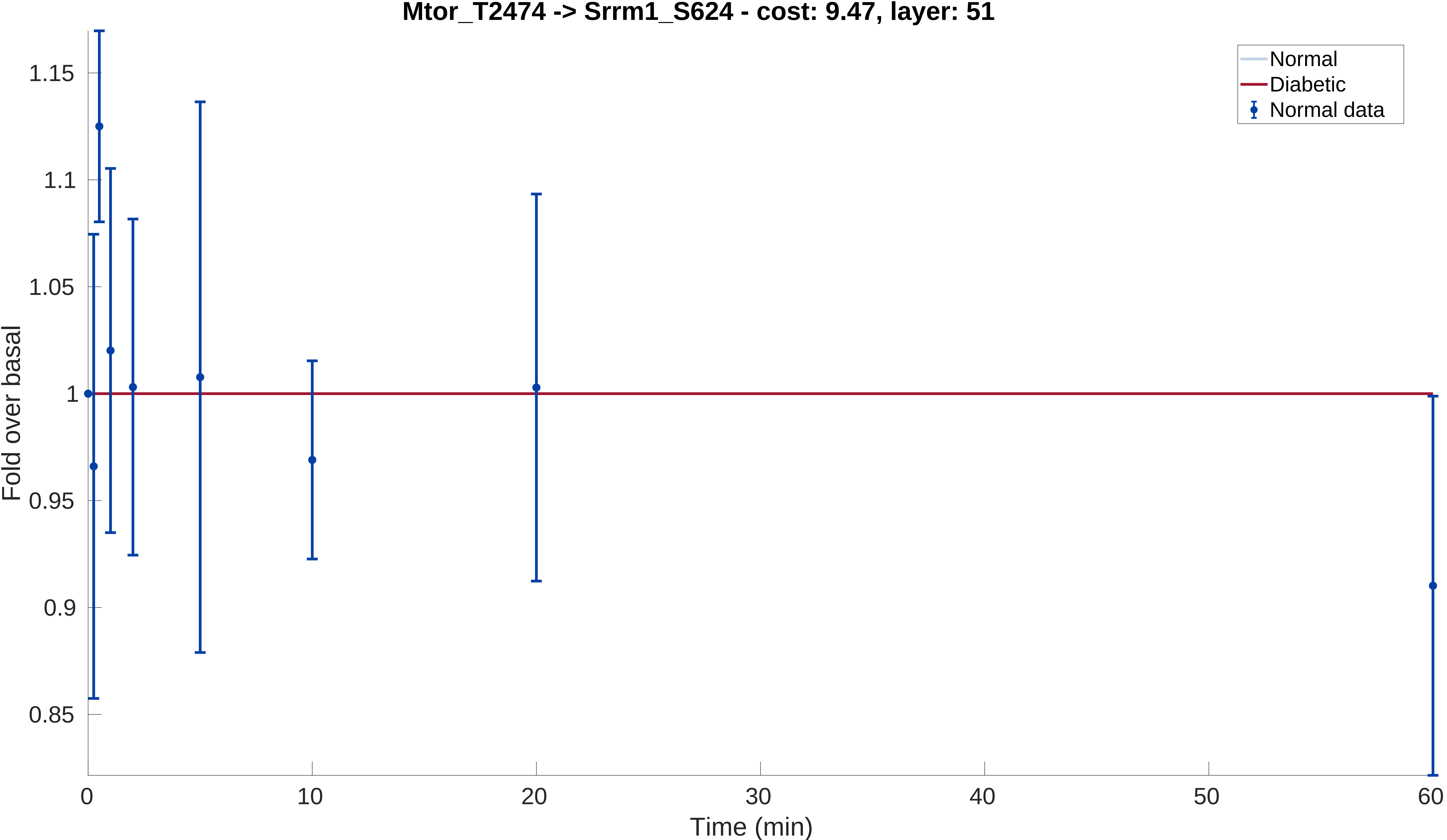

Mtor_T2474 -> Srrm1_S624 - cost: 9.47, layer: 51
Normal
Diabetic
Normal data
1.15
1.1
1.05
1
0.95
0.9
0.85
Fold over basal
0
10
20
30
40
50
60
Time (min)

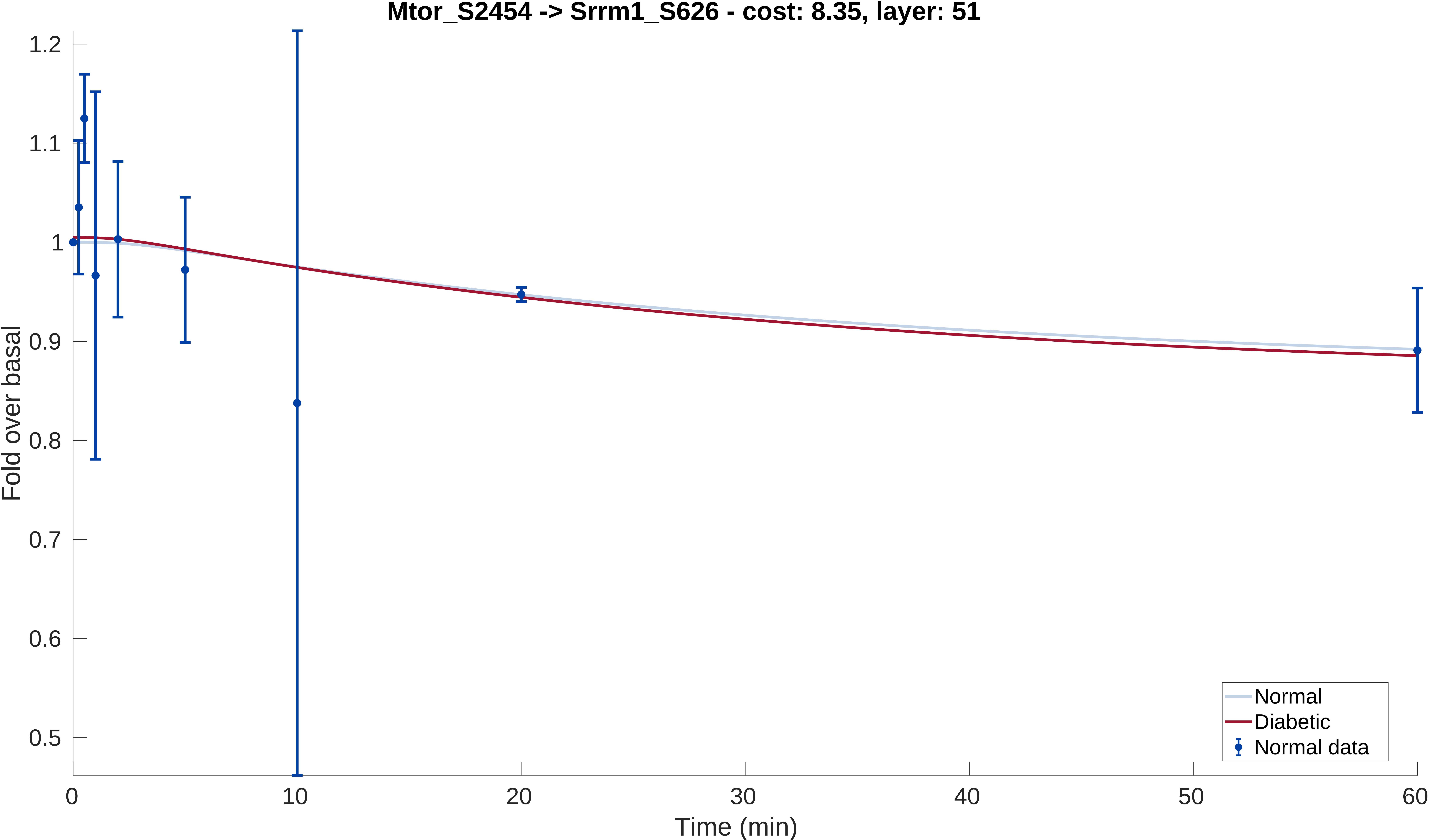
Mtor_S2454 -> Srrm1_S626 - cost: 8.35, layer: 51
Fold over basal
1.2
1.1
1
0.9
0.8
0.7
0.6
0.5
0
10
20
30
40
50
60
Time (min)
Normal
Diabetic
Normal data

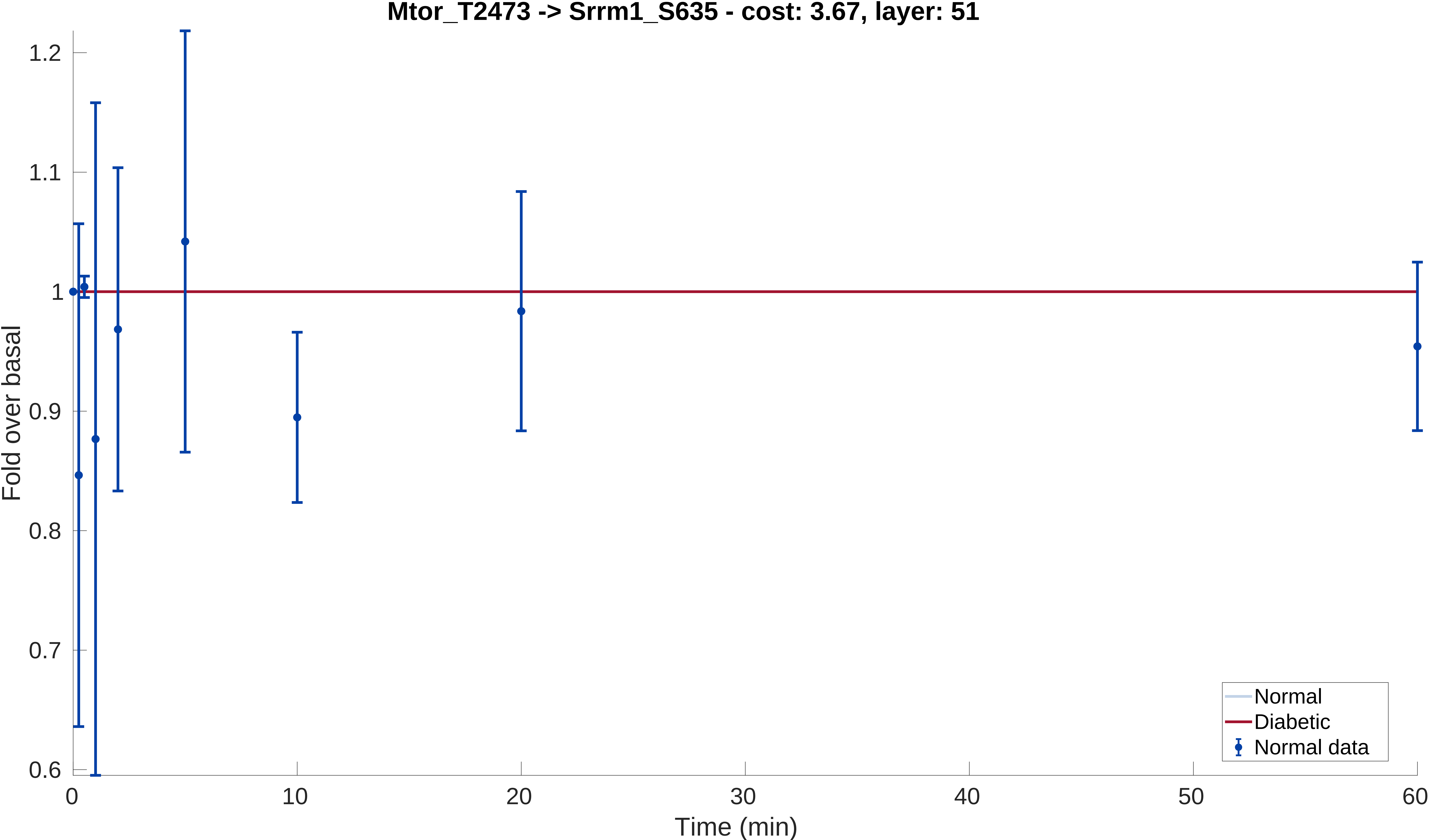
Mtor_T2473 -> Srrm1_S635 - cost: 3.67, layer: 51
Fold over basal
Time (min)
1.2
1.1
1
0.9
0.8
0.7
0.6
0
10
20
30
40
50
60
Normal
Diabetic
Normal data

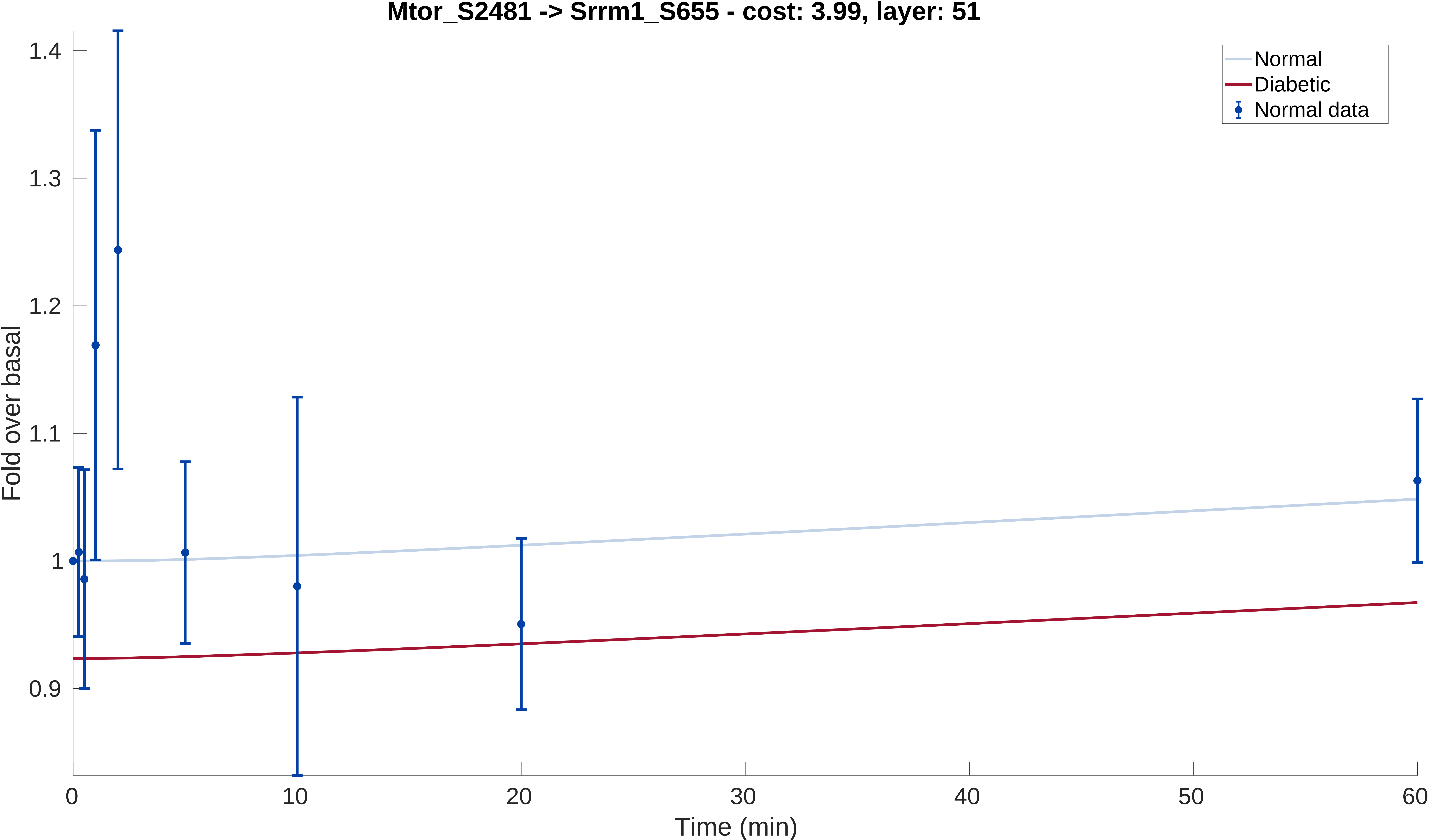

Mtor_S2481 -> Srrm1_S655 - cost: 3.99, layer: 51
Normal
Diabetic
Normal data
Fold over basal
1.4
1.3
1.2
1.1
1
0.9
0
10
20
30
40
50
60
Time (min)

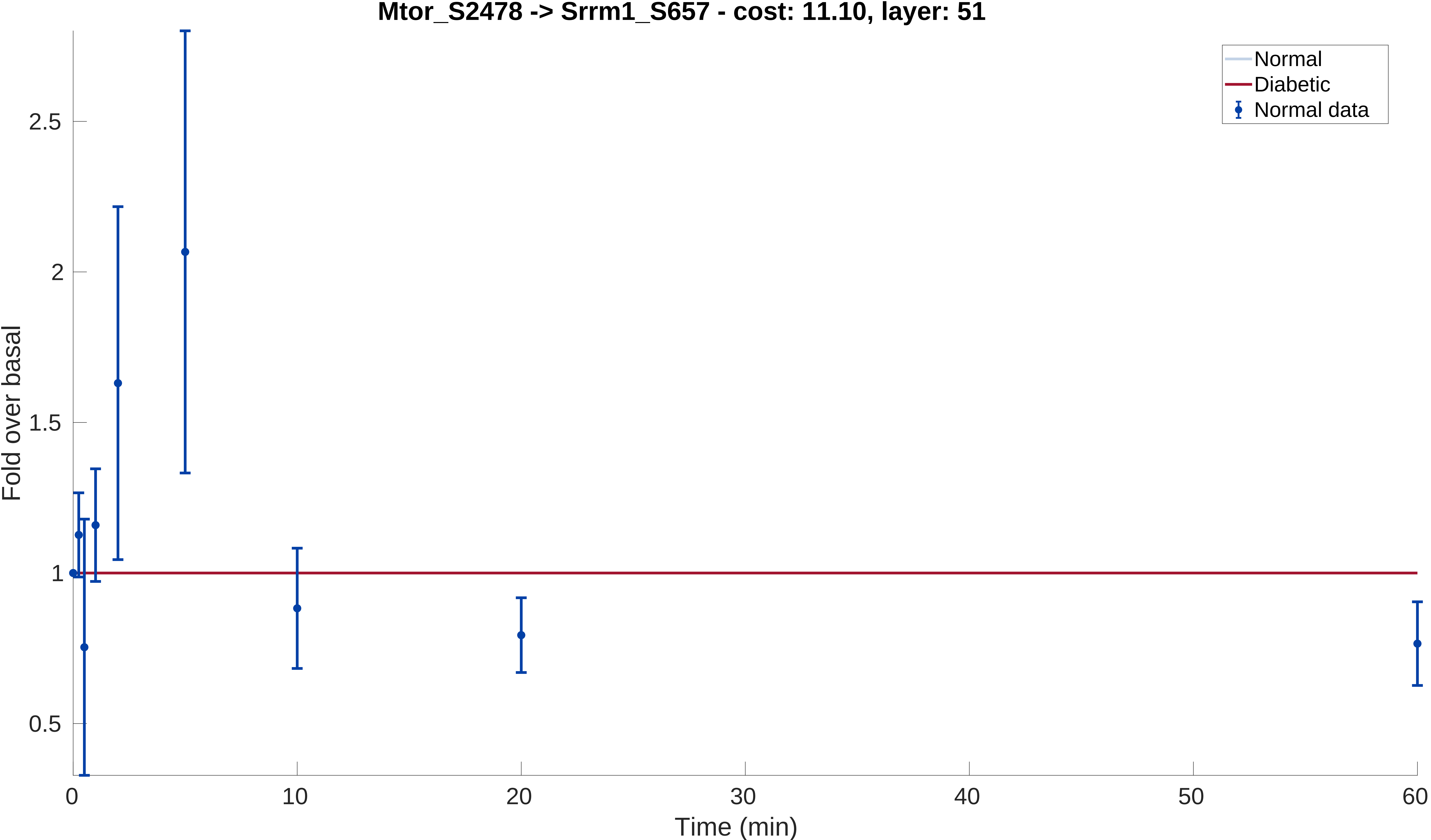

Mtor_S2478 -> Srrm1_S657 - cost: 11.10, layer: 51
Normal
Diabetic
Normal data
Fold over basal
2.5
2
1.5
1
0.5
0
10
20
30
40
50
60
Time (min)

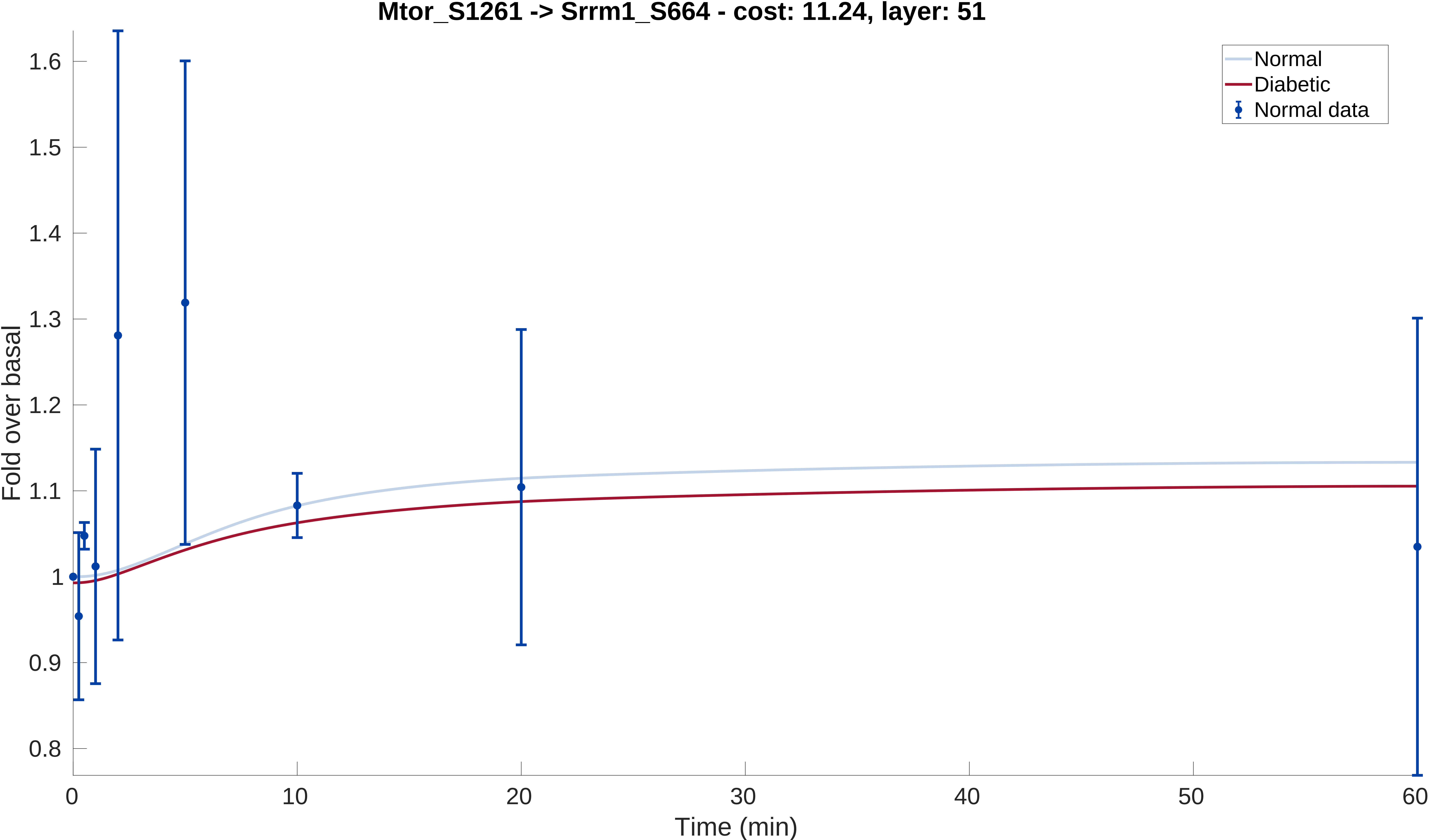
Mtor_S1261 -> Srrm1_S664 - cost: 11.24, layer: 51
Normal
Diabetic
Normal data
Fold over basal
1.6
1.5
1.4
1.3
1.2
1.1
1
0.9
0.8
0
10
20
30
40
50
60
Time (min)

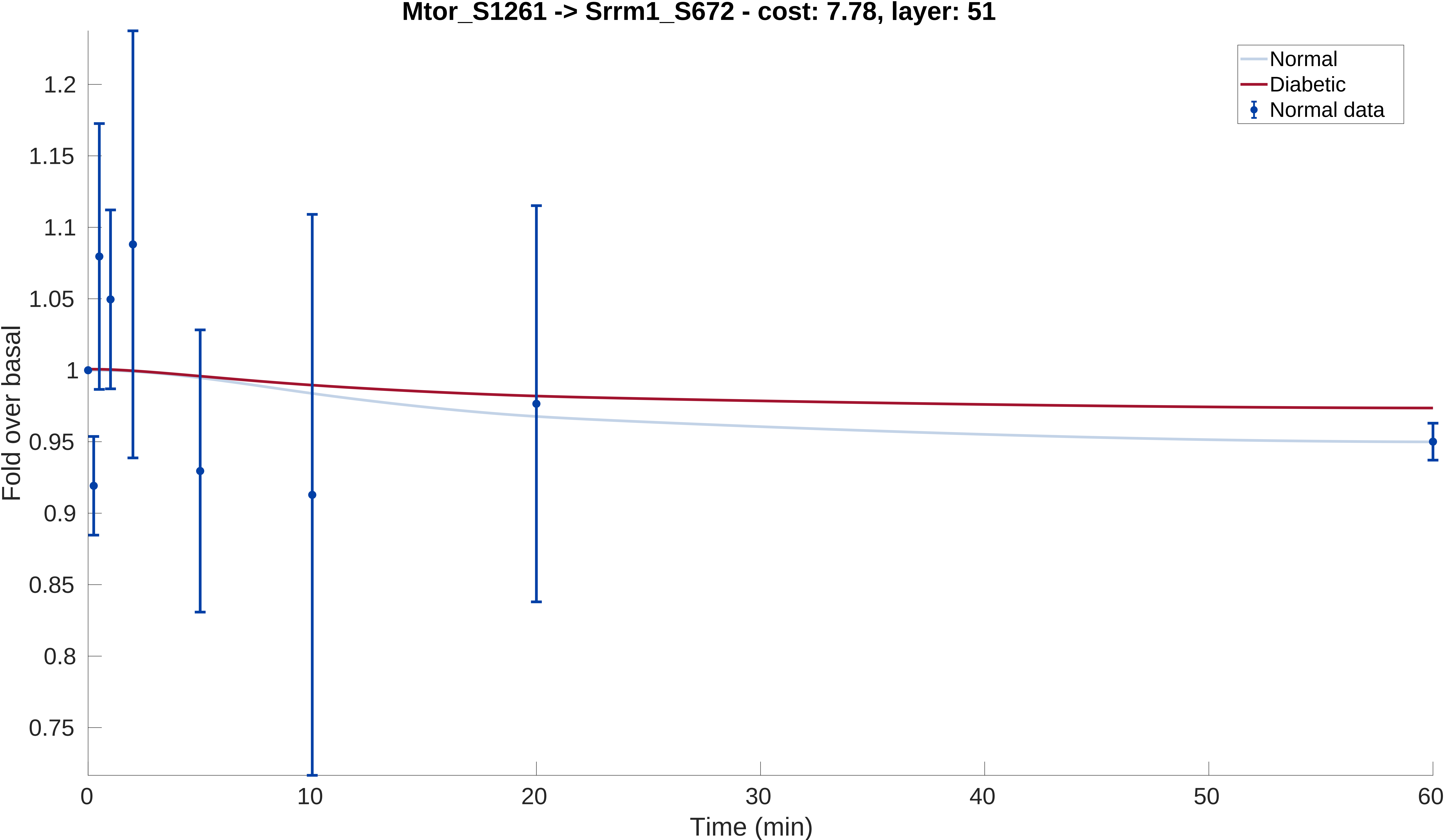

Mtor_S1261 -> Srrm1_S672 - cost: 7.78, layer: 51
Normal
Diabetic
Normal data
Fold over basal
Time (min)
1.2
1.15
1.1
1.05
1
0.95
0.9
0.85
0.8
0.75
0
10
20
30
40
50
60

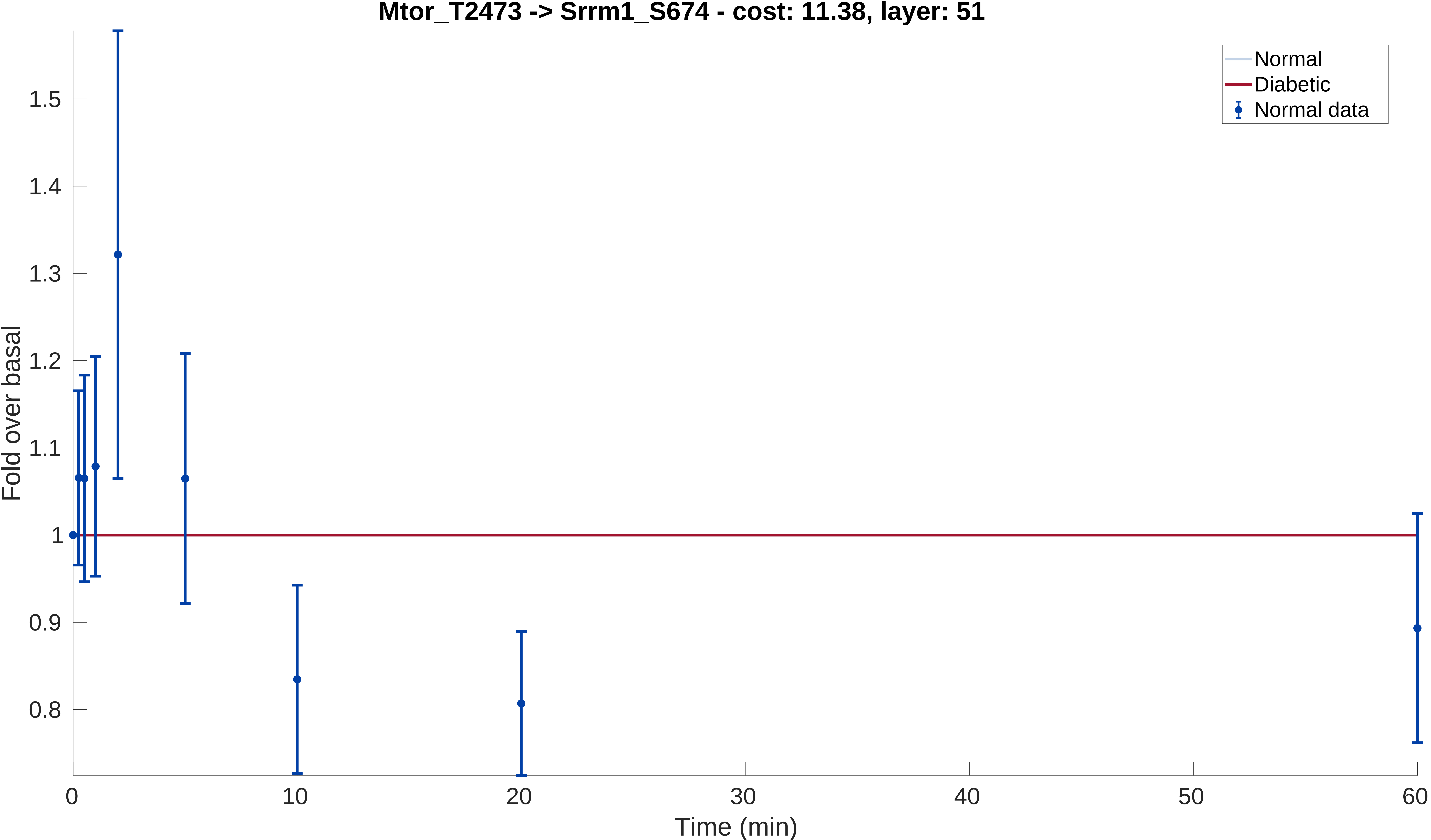
Mtor_T2473 -> Srrm1_S674 - cost: 11.38, layer: 51
Normal
Diabetic
Normal data
Fold over basal
1.5
1.4
1.3
1.2
1.1
1
0.9
0.8
0
10
20
30
40
50
60
Time (min)

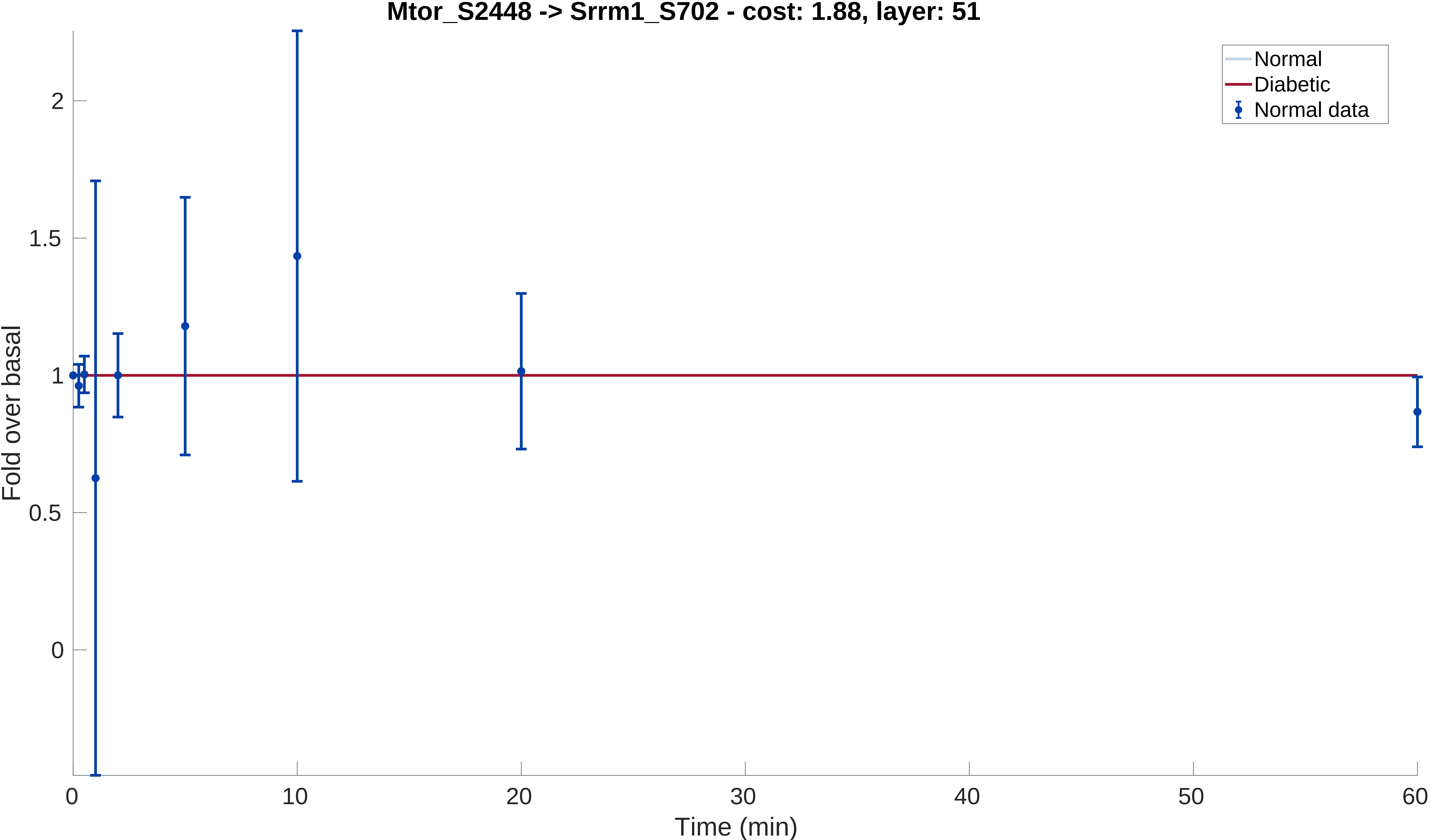

Mtor_S2448 -> Srrm1_S702 - cost: 1.88, layer: 51
Normal
Diabetic
Normal data
Fold over basal
2
1.5
1
0.5
0
0
10
20
30
40
50
60
Time (min)

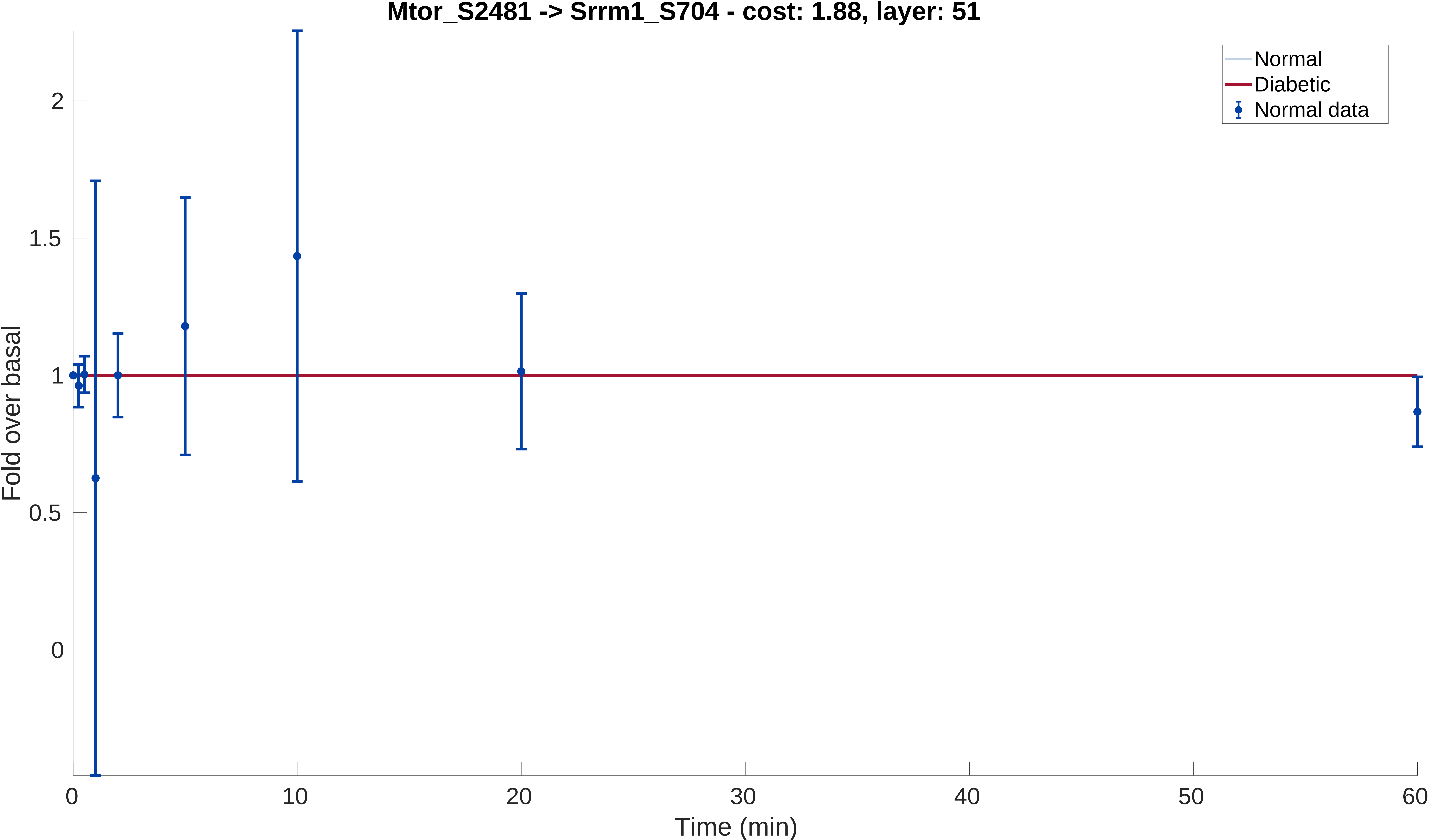

Mtor_S2481 -> Srrm1_S704 - cost: 1.88, layer: 51
Normal
Diabetic
Normal data
Fold over basal
Time (min)
0
0.5
1
1.5
2
0
10
20
30
40
50
60

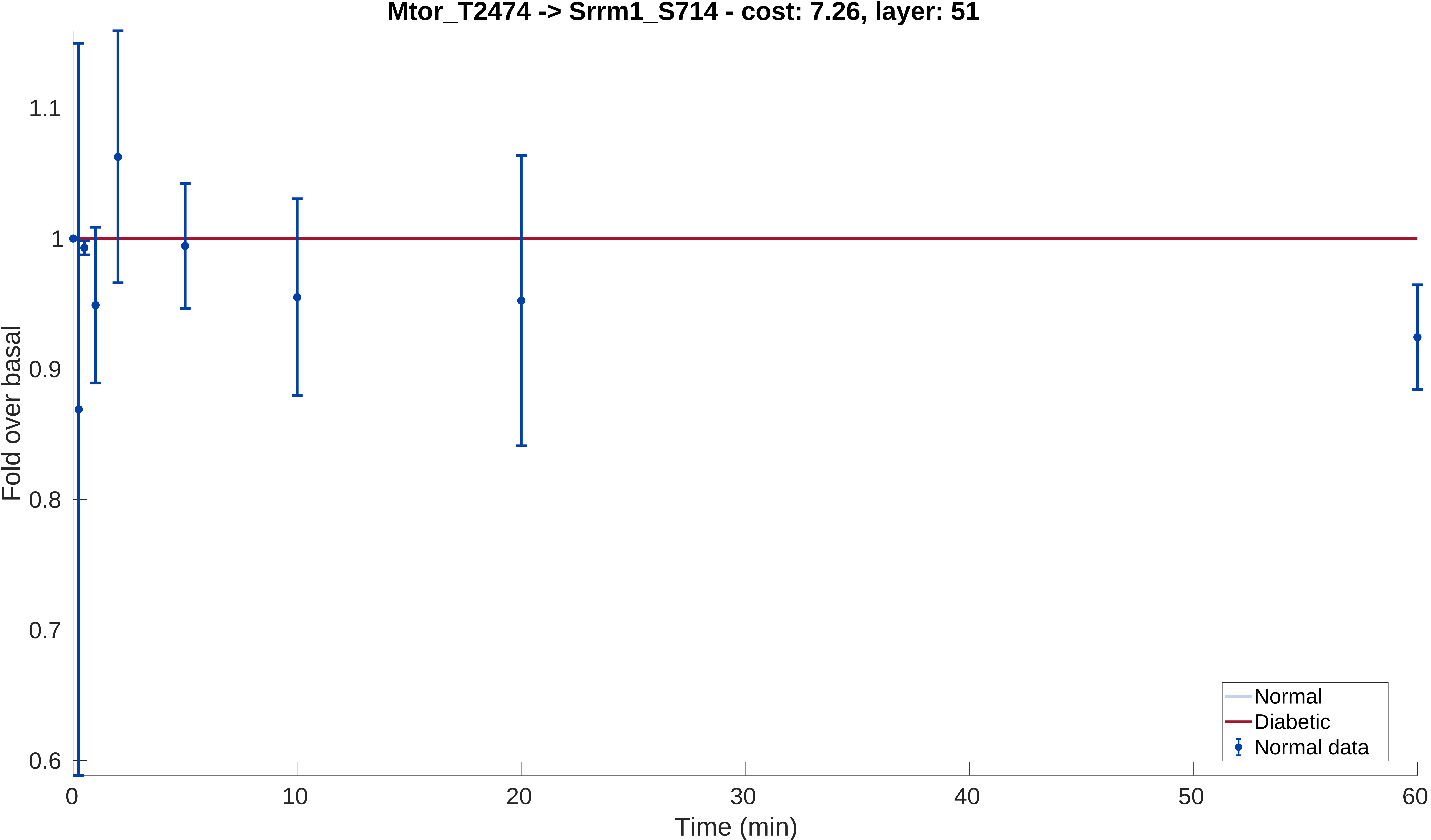

Mtor_T2474 -> Srrm1_S714 - cost: 7.26, layer: 51
Fold over basal
Time (min)
1.1
1
0.9
0.8
0.7
0.6
0
10
20
30
40
50
60
Normal
Diabetic
Normal data

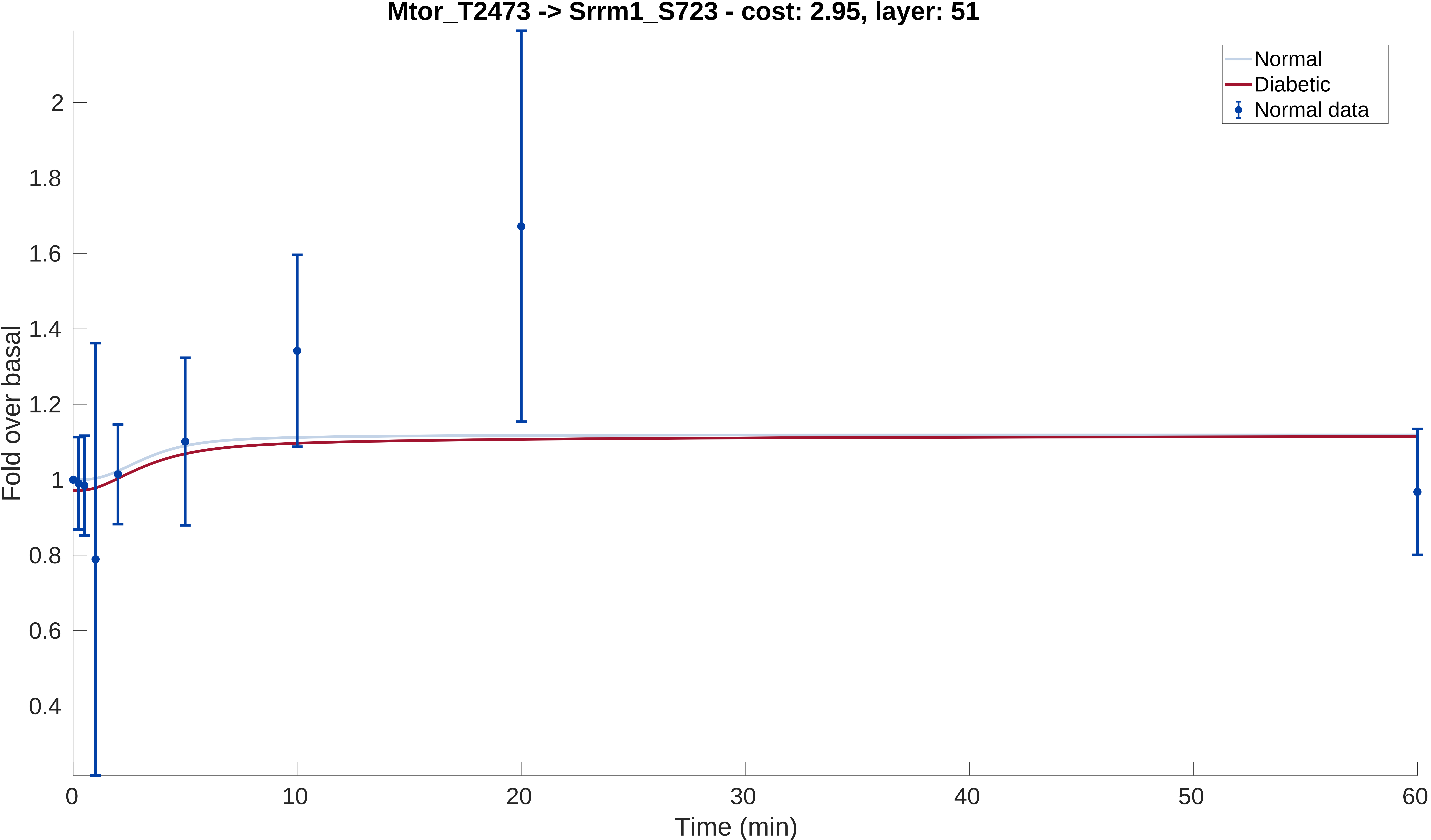

Mtor_T2473 -> Srrm1_S723 - cost: 2.95, layer: 51
Normal
Diabetic
Normal data
Fold over basal
Time (min)
0.4
0.6
0.8
1
1.2
1.4
1.6
1.8
2
0
10
20
30
40
50
60

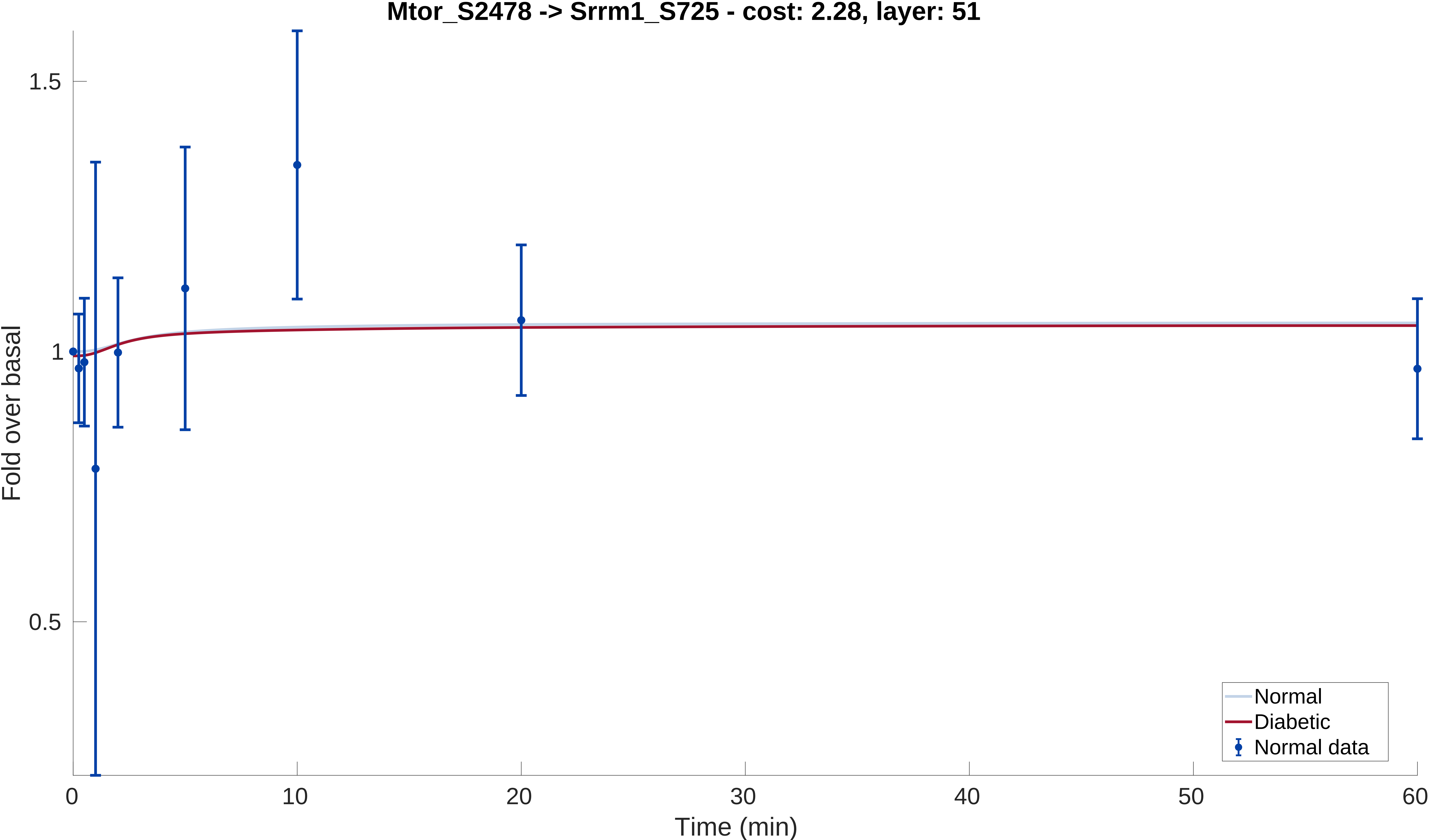

Mtor_S2478 -> Srrm1_S725 - cost: 2.28, layer: 51
1.5
1
0.5
Fold over basal
0
10
20
30
40
50
60
Time (min)
Normal
Diabetic
Normal data

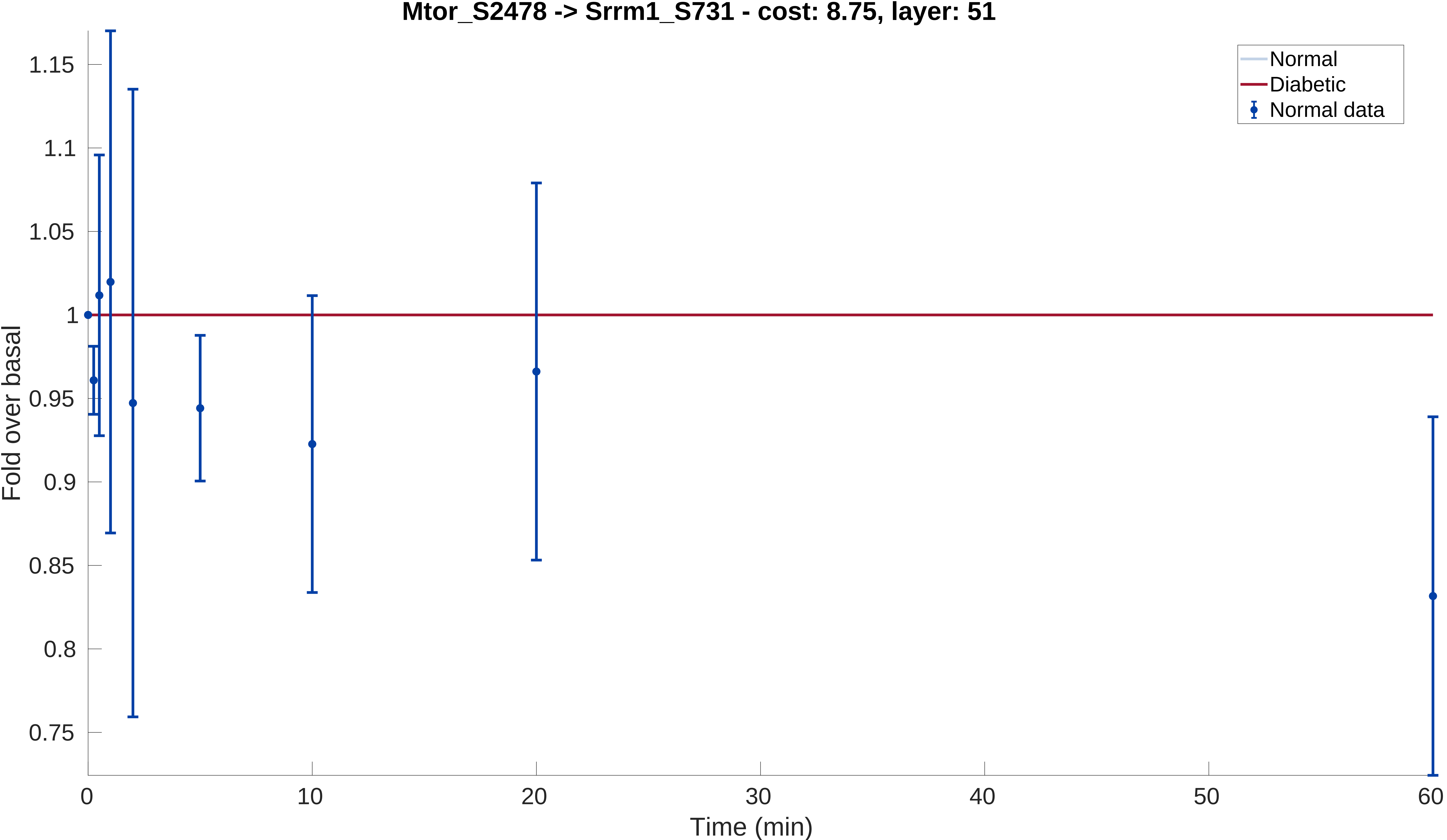
Mtor_S2478 -> Srrm1_S731 - cost: 8.75, layer: 51
Normal
Diabetic
Normal data
Fold over basal
1.15
1.1
1.05
1
0.95
0.9
0.85
0.8
0.75
0
10
20
30
40
50
60
Time (min)

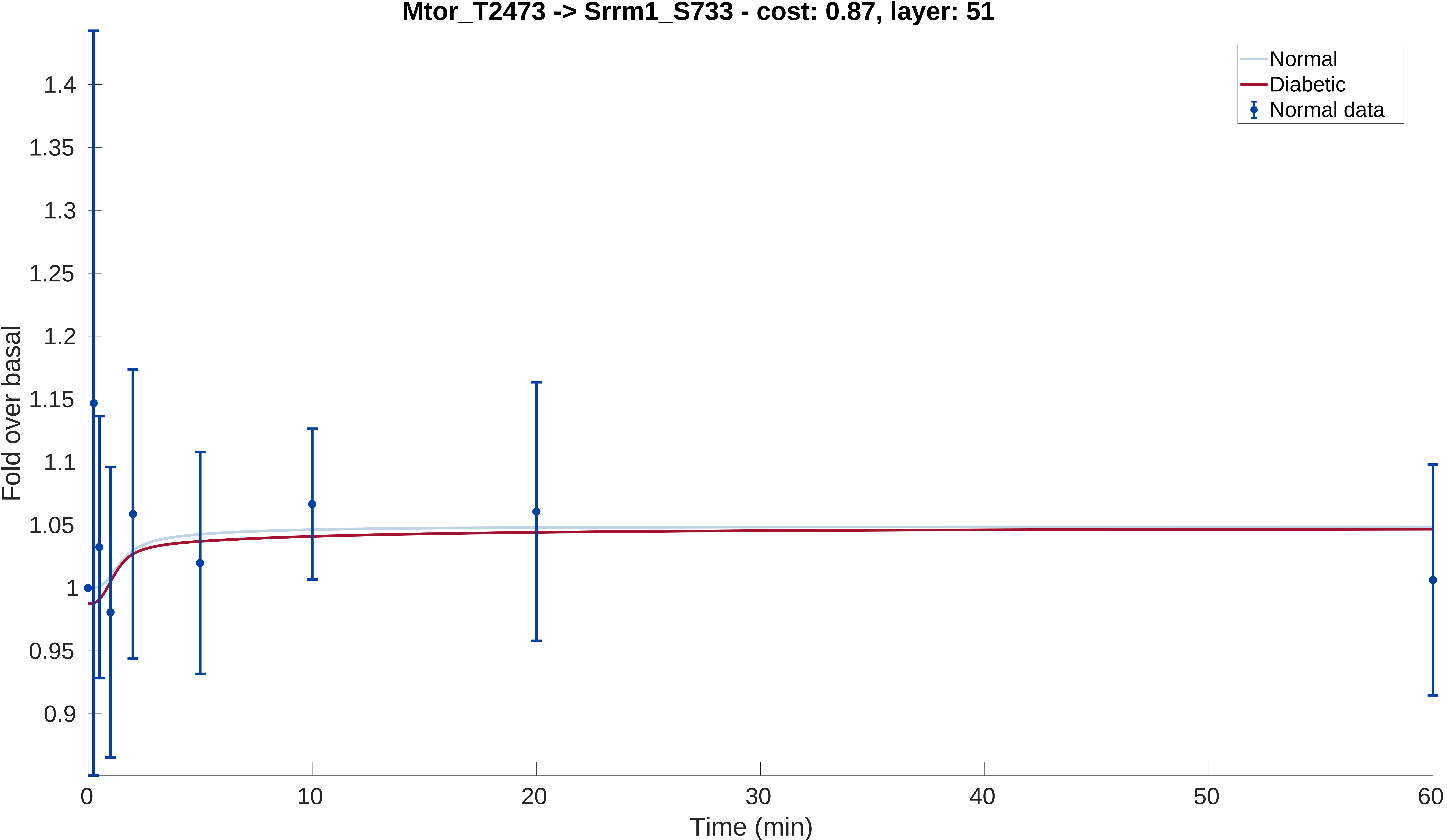

Mtor_T2473 -> Srrm1_S733 - cost: 0.87, layer: 51
Normal
Diabetic
Normal data
1.4
1.35
1.3
1.25
1.2
1.15
1.1
1.05
1
0.95
0.9
Fold over basal
0
10
20
30
40
50
60
Time (min)

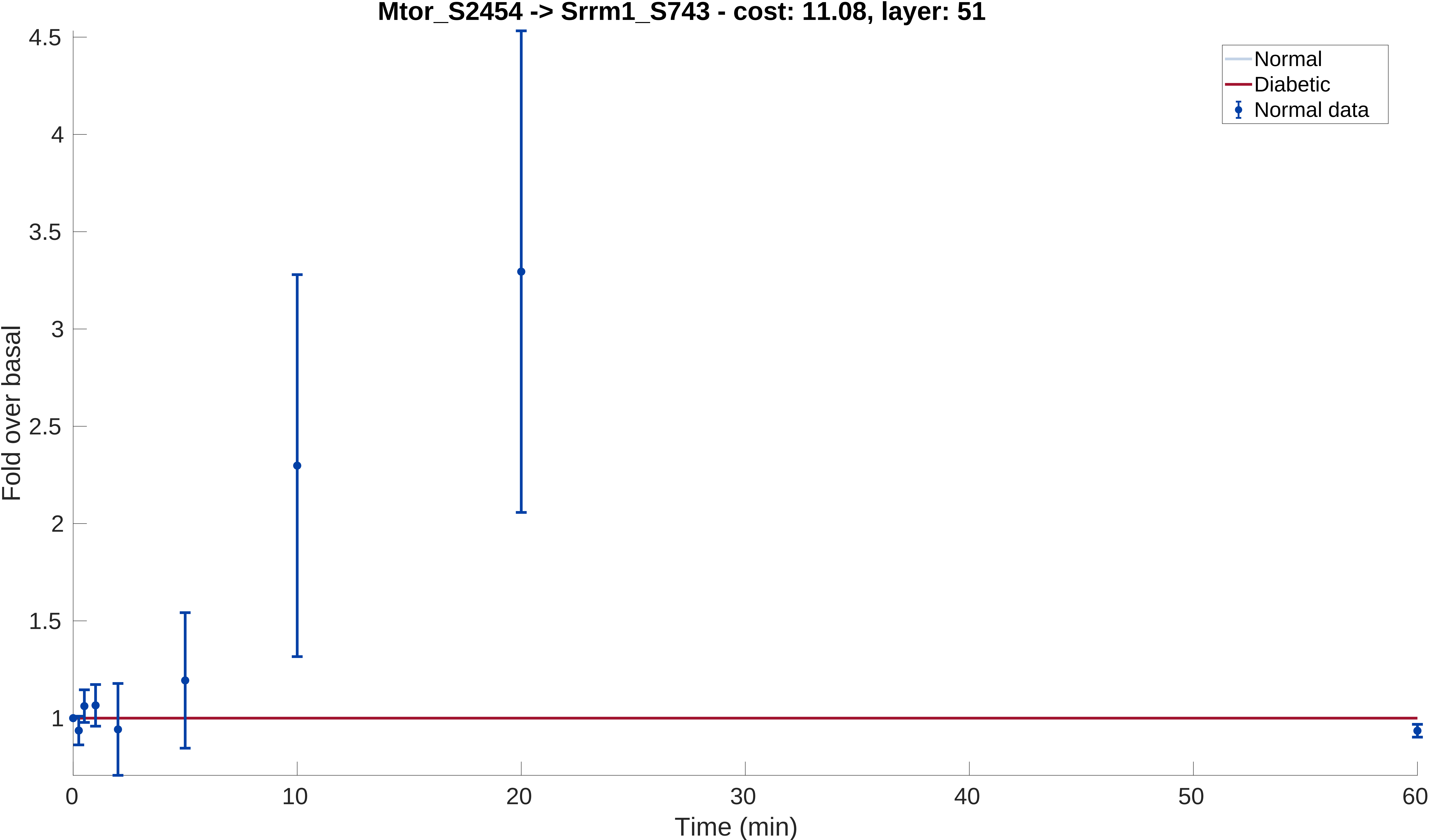
Mtor_S2454 -> Srrm1_S743 - cost: 11.08, layer: 51
Fold over basal
4.5
4
3.5
3
2.5
2
1.5
1
Time (min)
0
10
20
30
40
50
60
Normal
Diabetic
Normal data

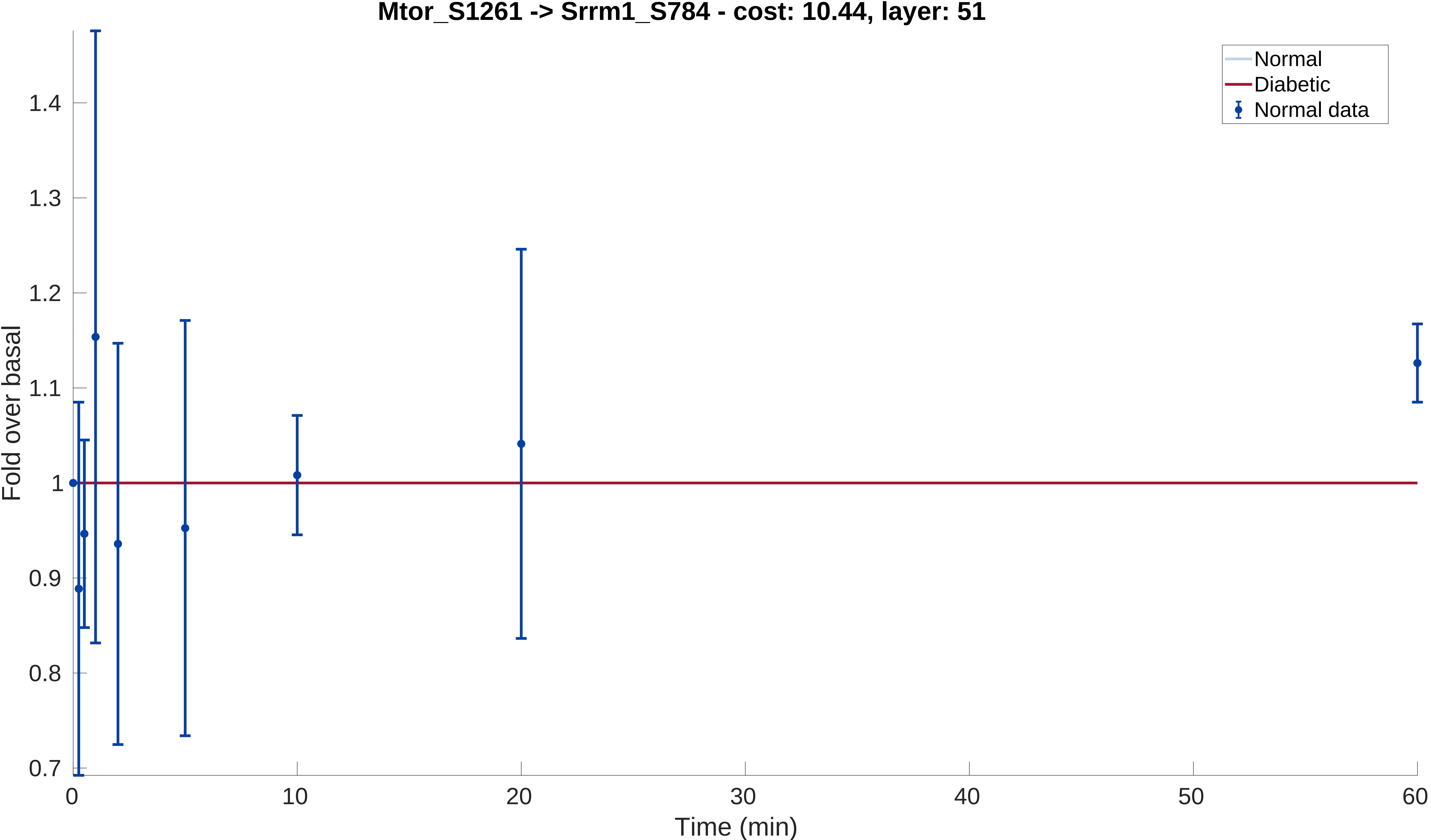
Mtor_S1261 -> Srrm1_S784 - cost: 10.44, layer: 51
Normal
Diabetic
Normal data
Fold over basal
1.4
1.3
1.2
1.1
1
0.9
0.8
0.7
0
10
20
30
40
50
60
Time (min)

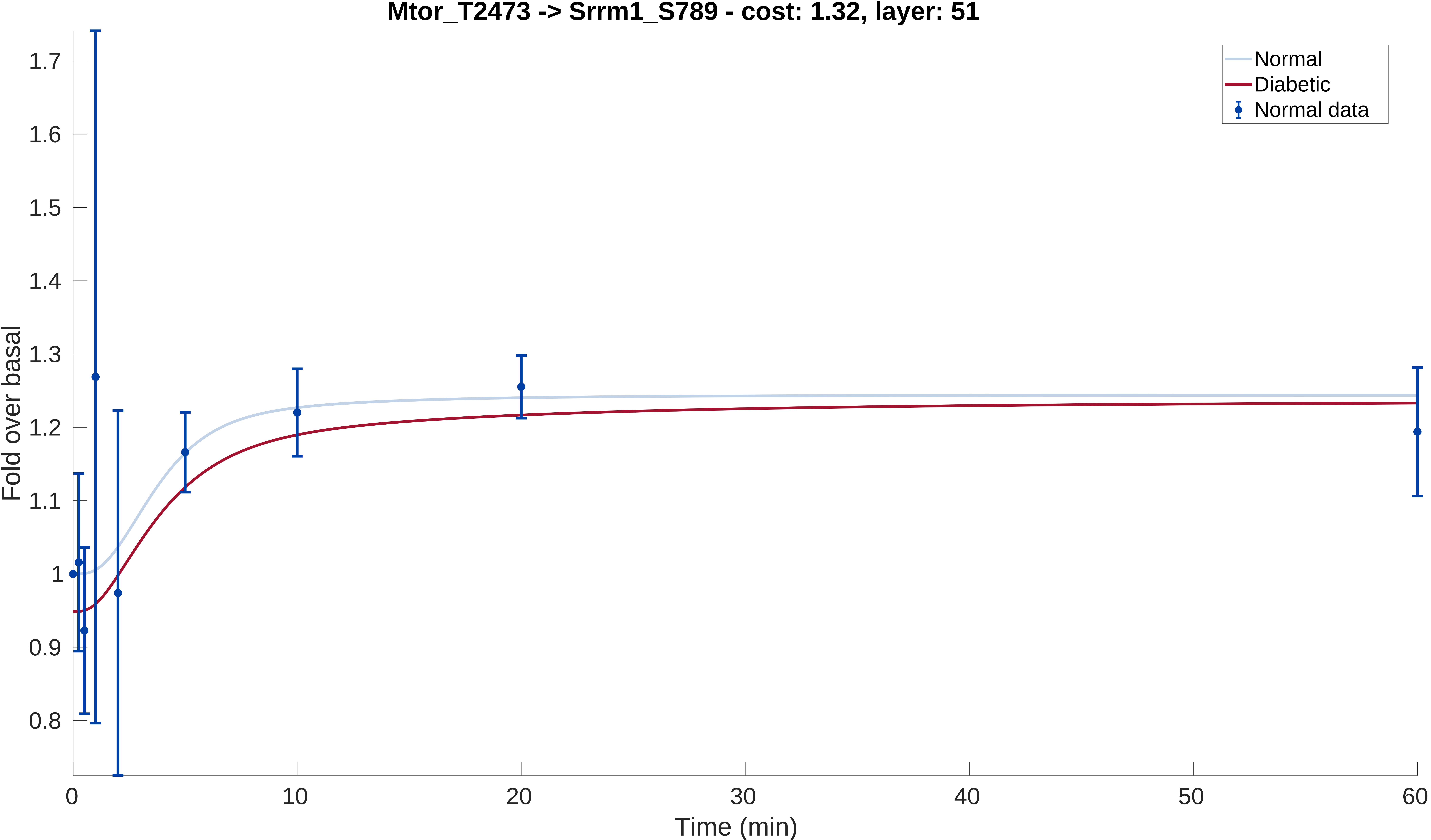

Mtor_T2473 -> Srrm1_S789 - cost: 1.32, layer: 51
Normal
Diabetic
Normal data
Fold over basal
1.7
1.6
1.5
1.4
1.3
1.2
1.1
1
0.9
0.8
0
10
20
30
40
50
60
Time (min)

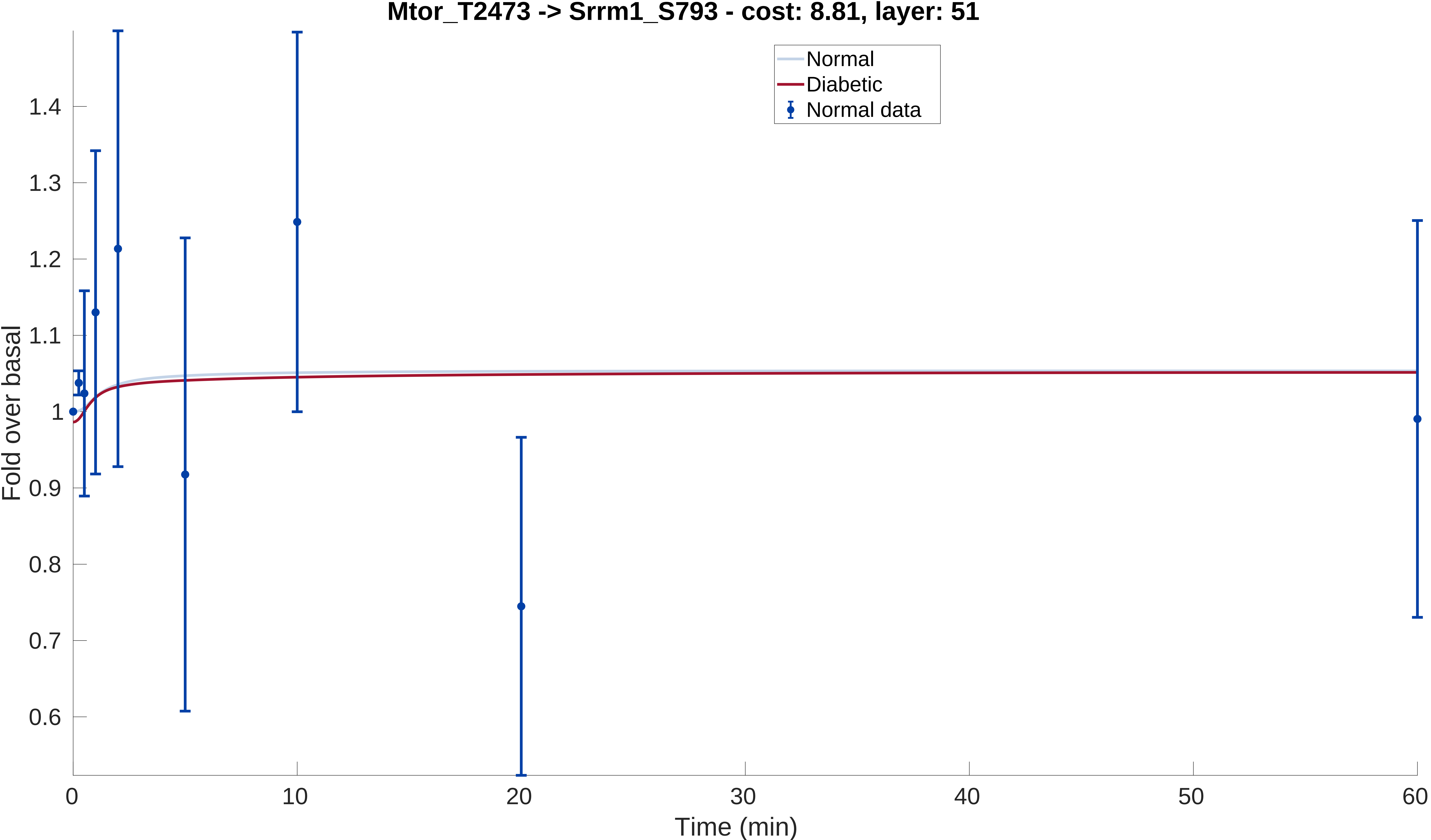

Mtor_T2473 -> Srrm1_S793 - cost: 8.81, layer: 51
Normal
Diabetic
Normal data
Fold over basal
1.4
1.3
1.2
1.1
1
0.9
0.8
0.7
0.6
0
10
20
30
40
50
60
Time (min)

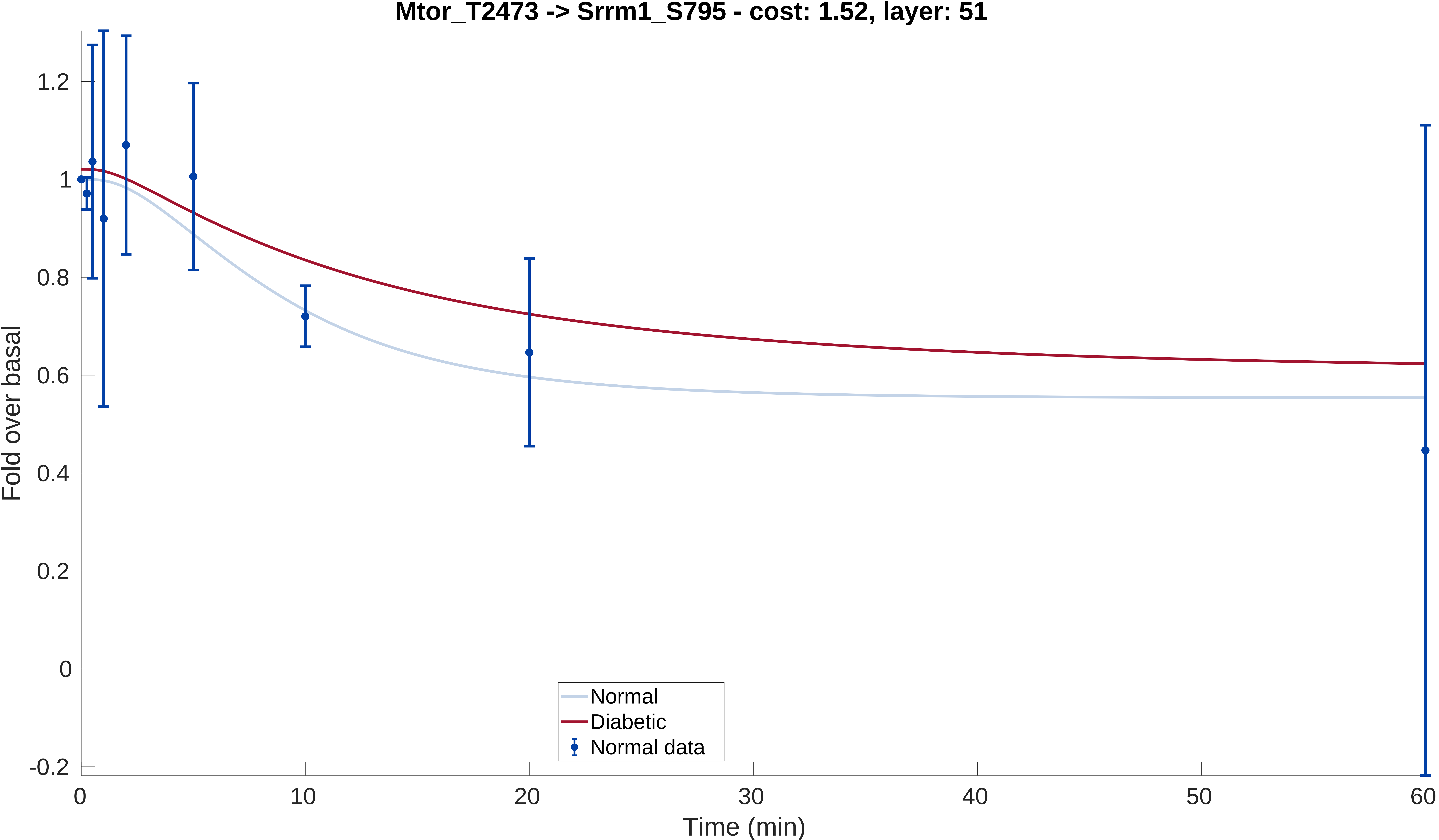

Mtor_T2473 -> Srrm1_S795 - cost: 1.52, layer: 51
Fold over basal
1.2
1
0.8
0.6
0.4
0.2
0
-0.2
0
10
20
30
40
50
60
Time (min)
Normal
Diabetic
Normal data

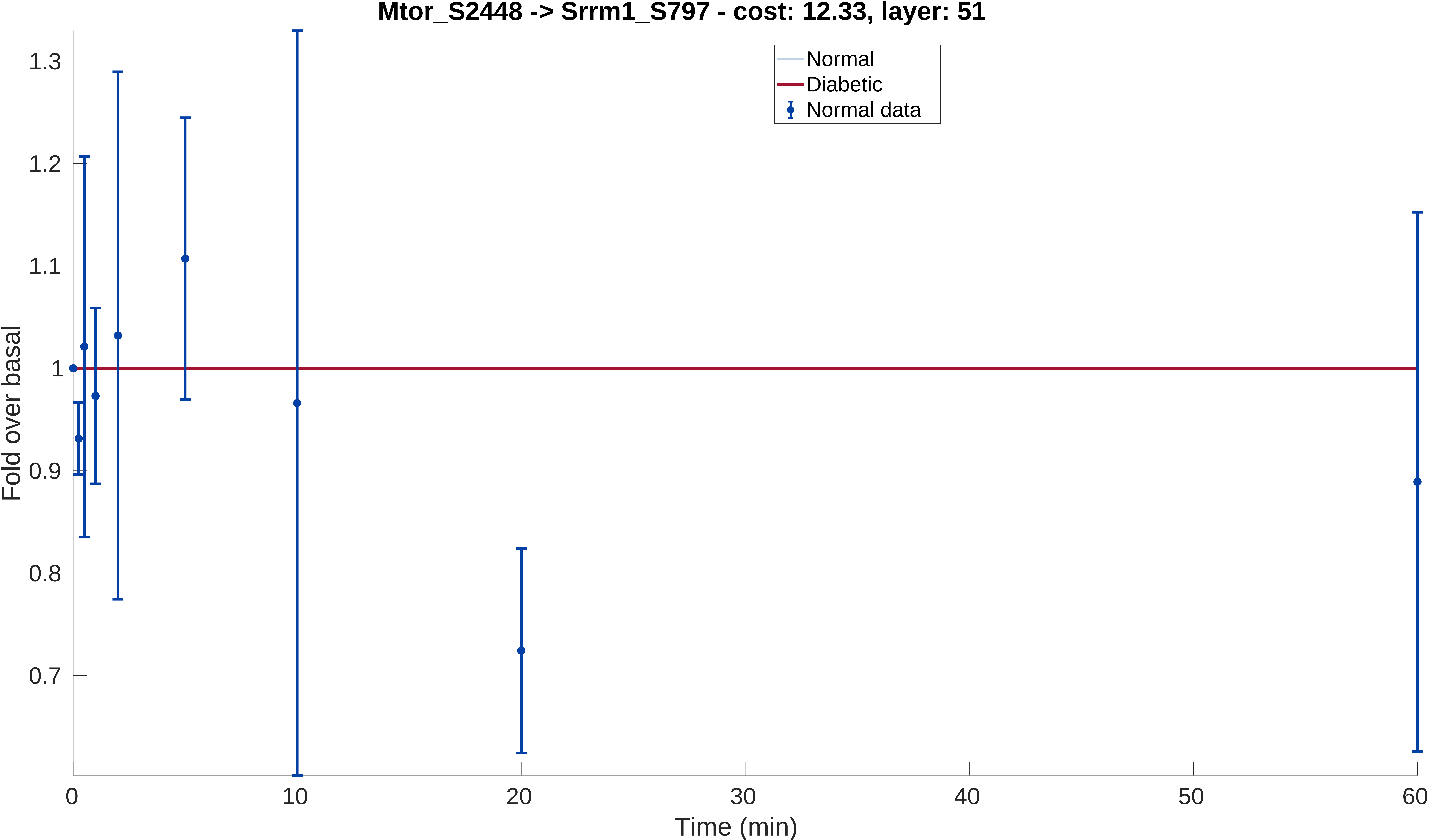
Mtor_S2448 -> Srrm1_S797 - cost: 12.33, layer: 51
Normal
Diabetic
Normal data
Fold over basal
Time (min)
0.7
0.8
0.9
1
1.1
1.2
1.3
0
10
20
30
40
50
60

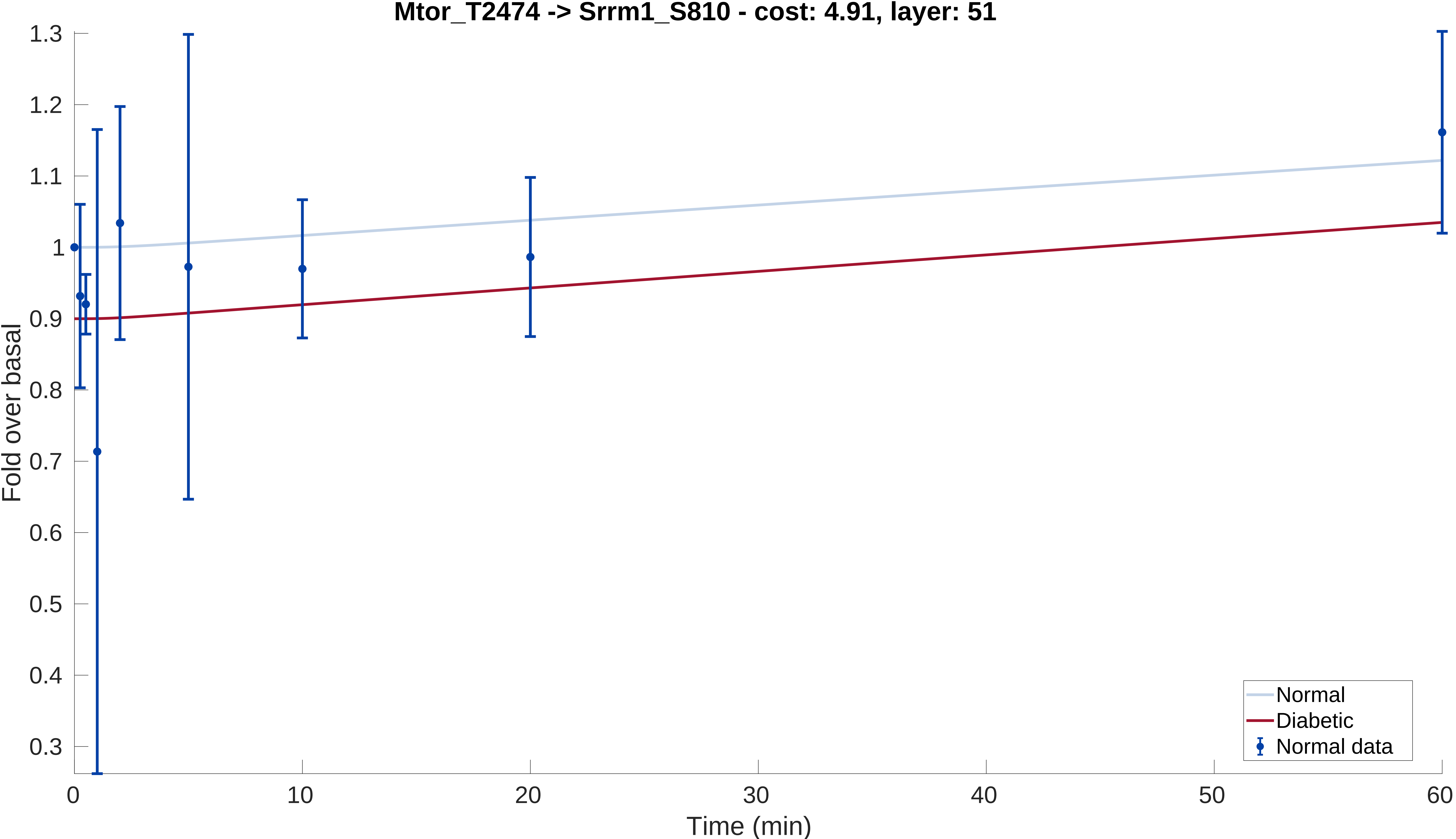

Mtor_T2474 -> Srrm1_S810 - cost: 4.91, layer: 51
Fold over basal
Time (min)
1.3
1.2
1.1
1
0.9
0.8
0.7
0.6
0.5
0.4
0.3
0
10
20
30
40
50
60
Normal
Diabetic
Normal data

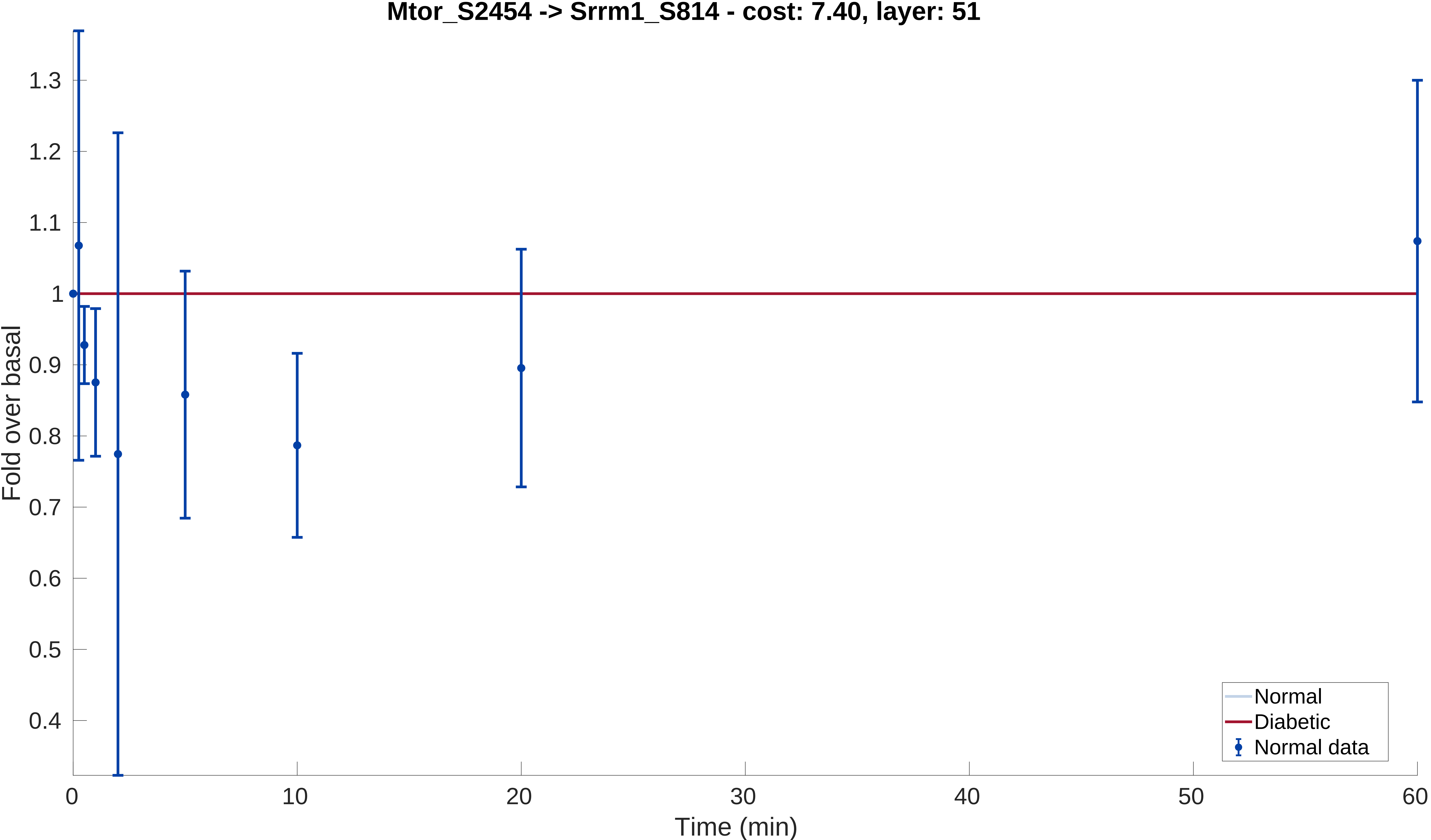

Mtor_S2454 -> Srrm1_S814 - cost: 7.40, layer: 51
1.3
1.2
1.1
1
0.9
0.8
0.7
0.6
0.5
0.4
Fold over basal
0
10
20
30
40
50
60
Time (min)
Normal
Diabetic
Normal data

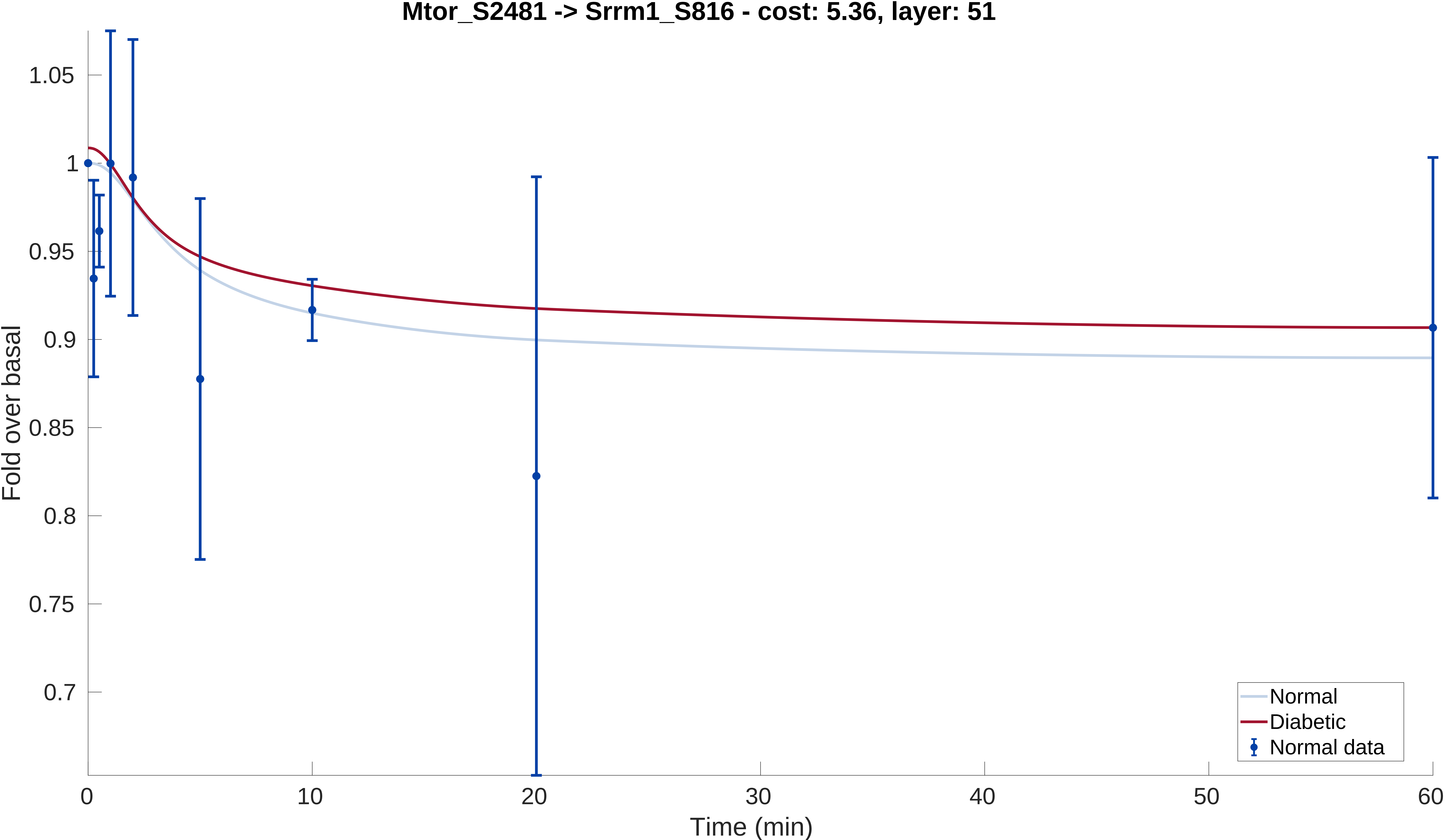
Mtor_S2481 -> Srrm1_S816 - cost: 5.36, layer: 51
Fold over basal
1.05
1
0.95
0.9
0.85
0.8
0.75
0.7
0
10
20
30
40
50
60
Time (min)
Normal
Diabetic
Normal data

#### Mtor_S2481 -> Srrm1_S822 - cost: 11.96, layer: 51

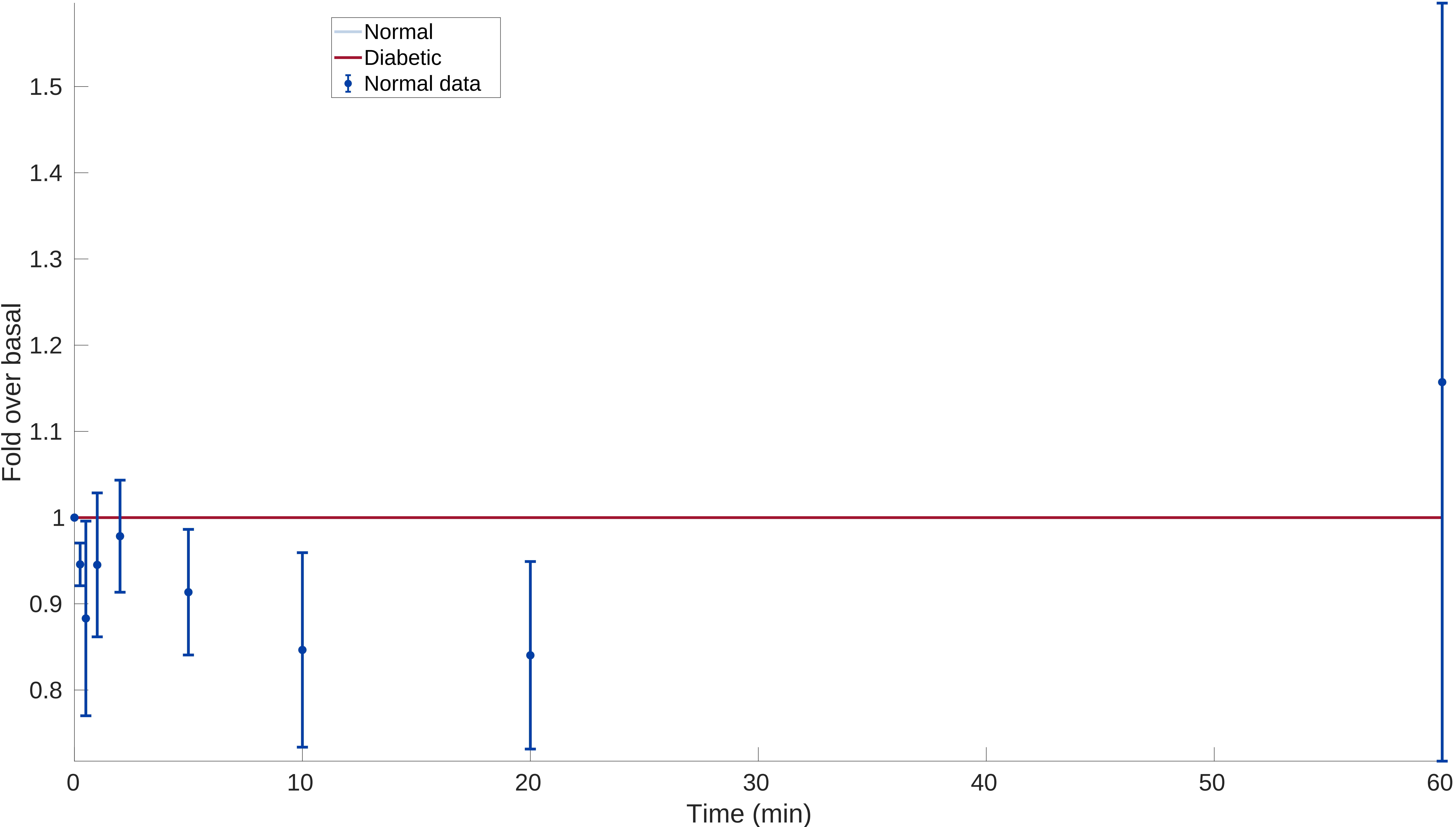

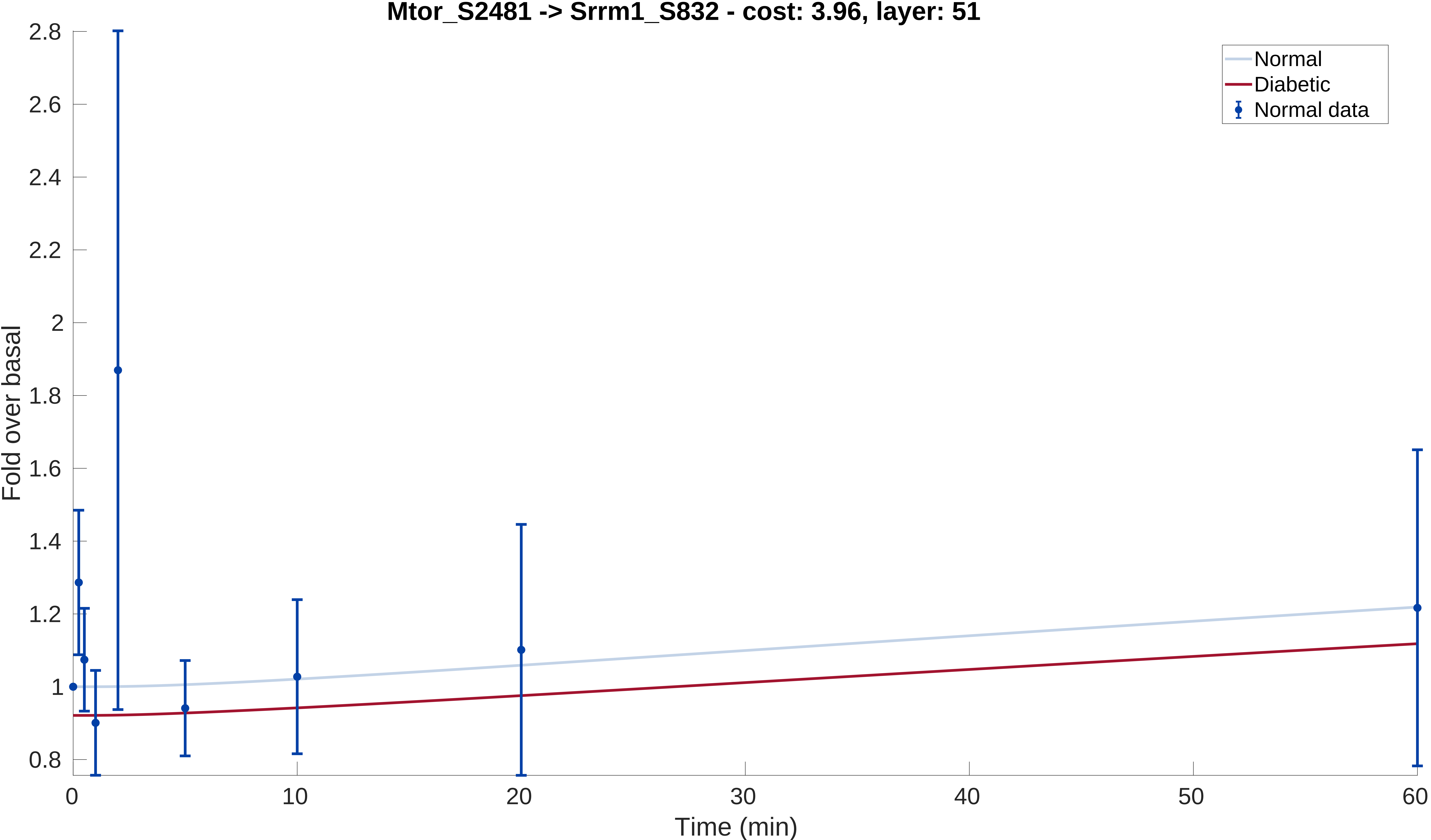

Mtor_S2481 -> Srrm1_S832 - cost: 3.96, layer: 51
Normal
Diabetic
Normal data
Fold over basal
2.8
2.6
2.4
2.2
2
1.8
1.6
1.4
1.2
1
0.8
0
10
20
30
40
50
60
Time (min)

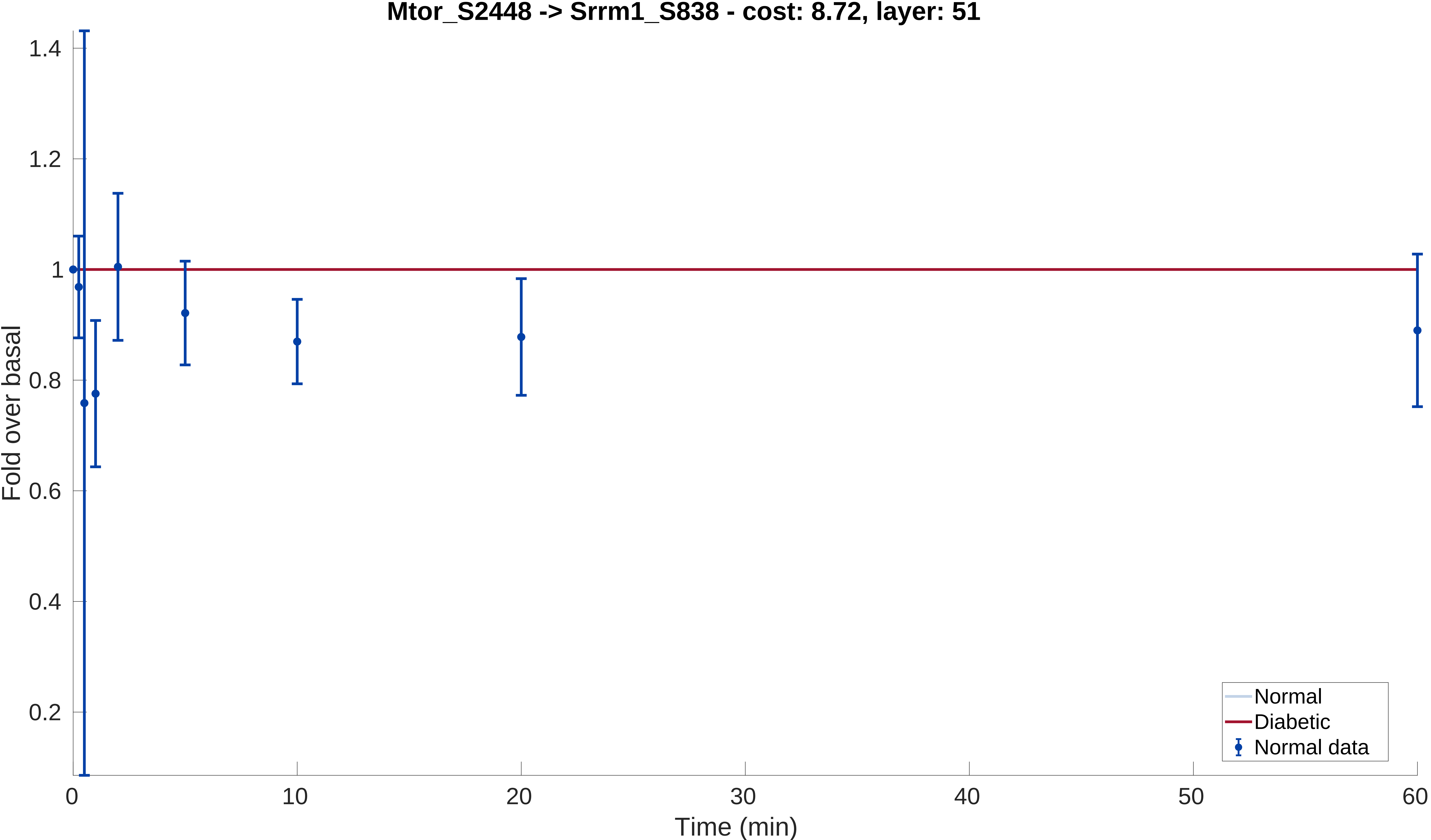

Mtor_S2448 -> Srrm1_S838 - cost: 8.72, layer: 51
Fold over basal
Time (min)
0.2
0.4
0.6
0.8
1
1.2
1.4
0
10
20
30
40
50
60
Normal
Diabetic
Normal data

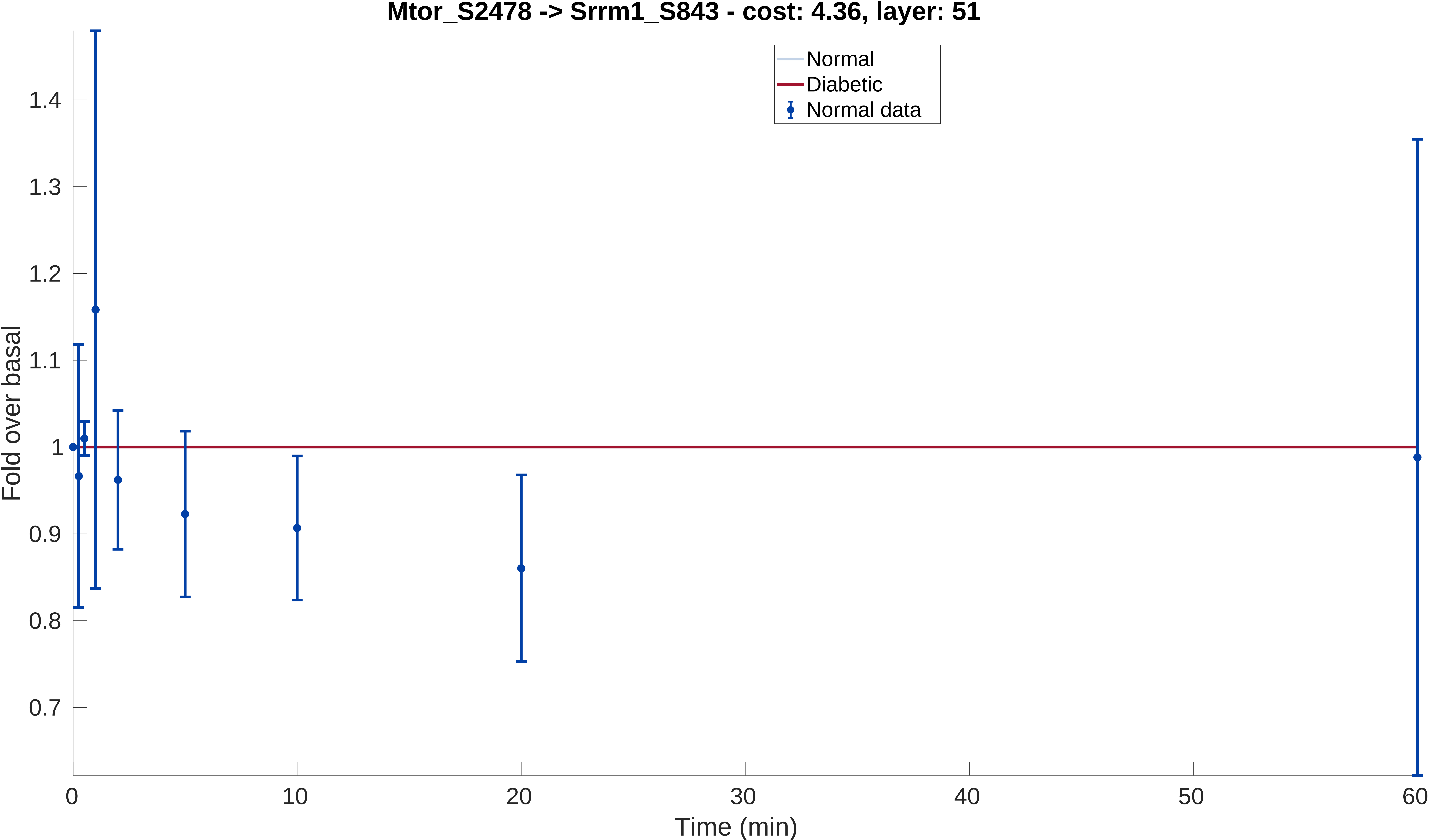
Mtor_S2478 -> Srrm1_S843 - cost: 4.36, layer: 51
Normal
Diabetic
Normal data
Fold over basal
1.4
1.3
1.2
1.1
1
0.9
0.8
0.7
0
10
20
30
40
50
60
Time (min)

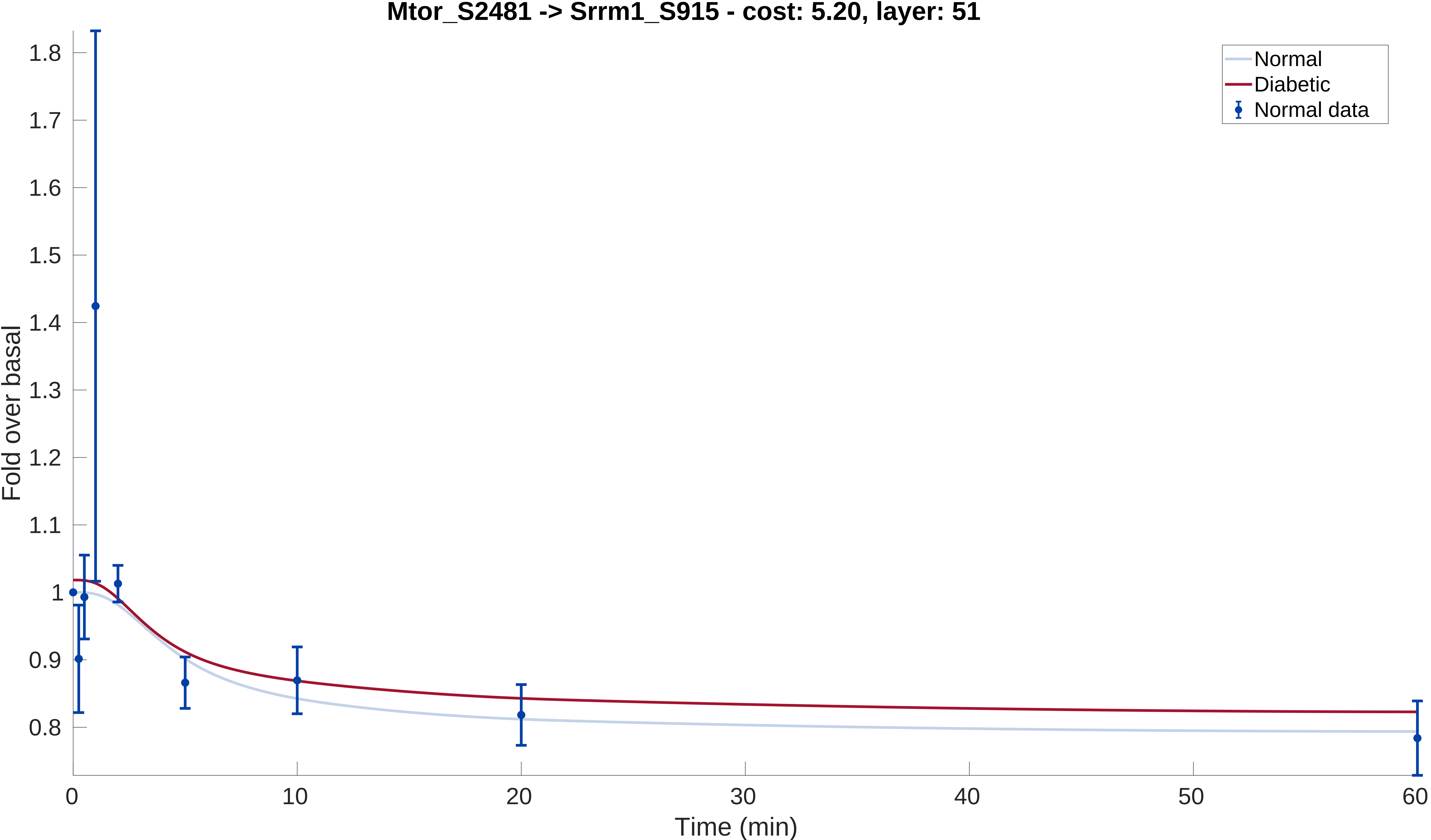

Mtor_S2481 -> Srrm1_S915 - cost: 5.20, layer: 51
Normal
Diabetic
Normal data
Fold over basal
1.8
1.7
1.6
1.5
1.4
1.3
1.2
1.1
1
0.9
0.8
0
10
20
30
40
50
60
Time (min)

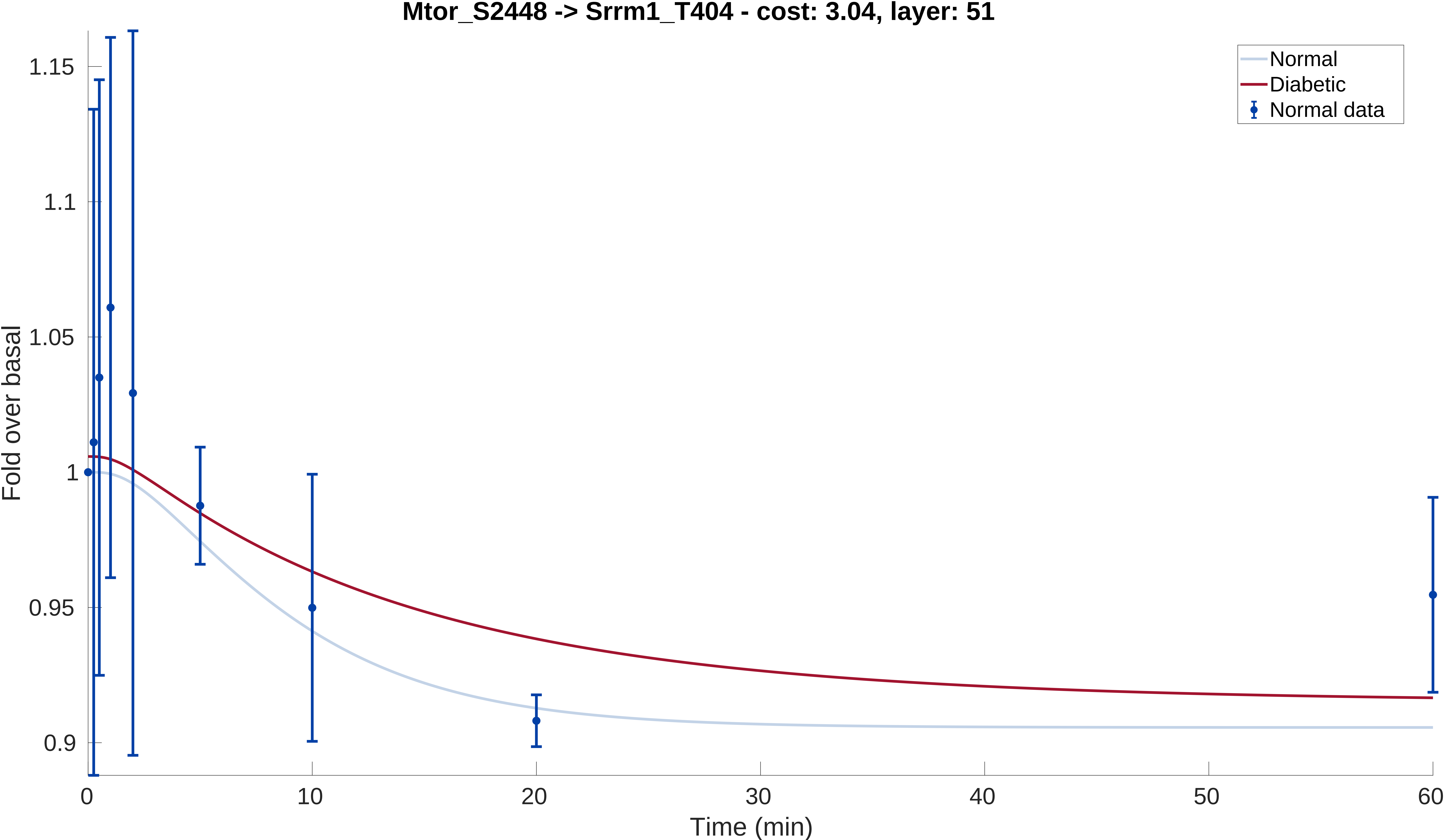

Mtor_S2448 -> Srrm1_T404 - cost: 3.04, layer: 51
Normal
Diabetic
Normal data
Fold over basal
Time (min)
1.15
1.1
1.05
1
0.95
0.9
0
10
20
30
40
50
60

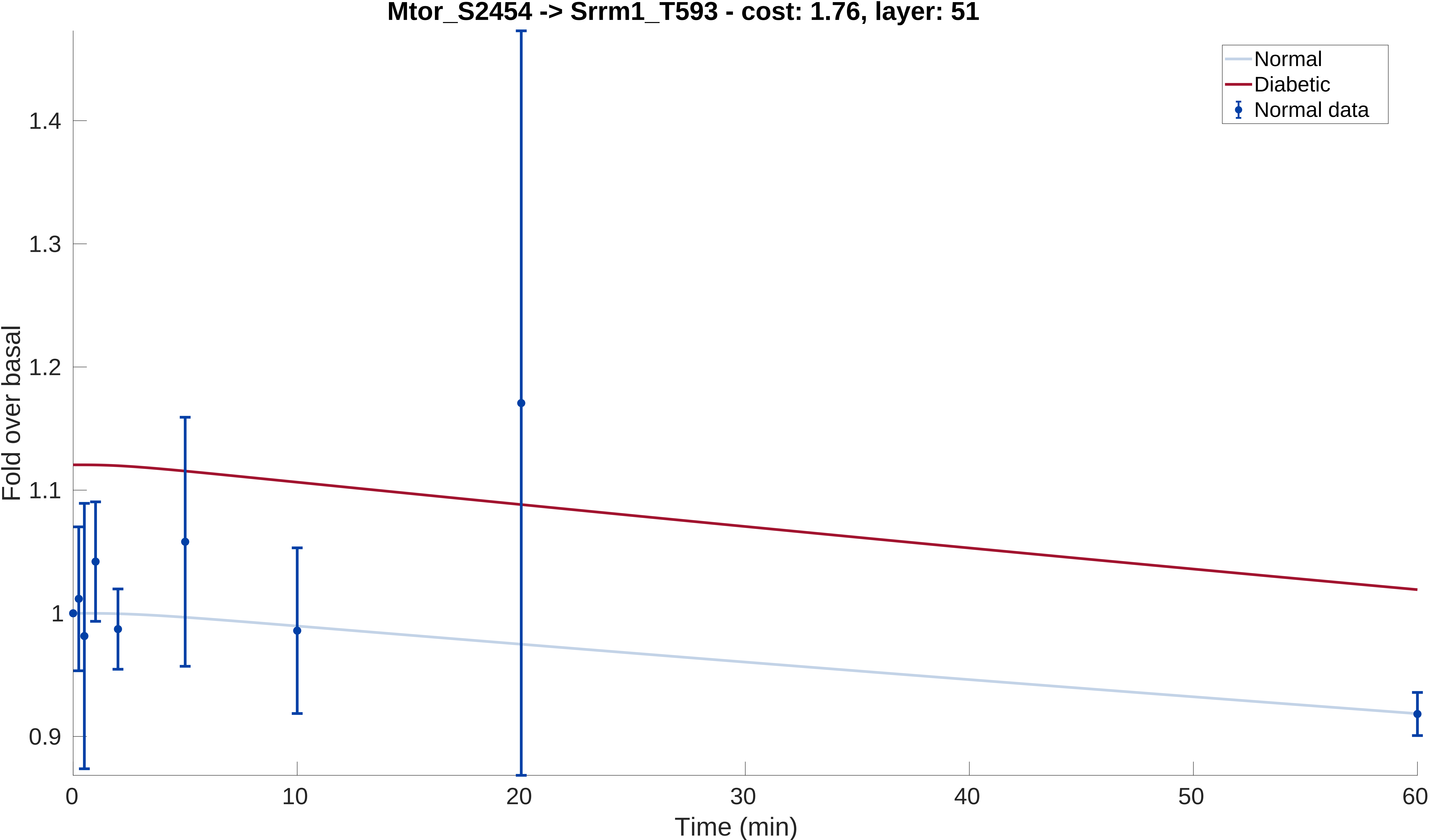

Mtor_S2454 -> Srrm1_T593 - cost: 1.76, layer: 51
Normal
Diabetic
Normal data
1.4
1.3
1.2
1.1
1
0.9
Fold over basal
0
10
20
30
40
50
60
Time (min)

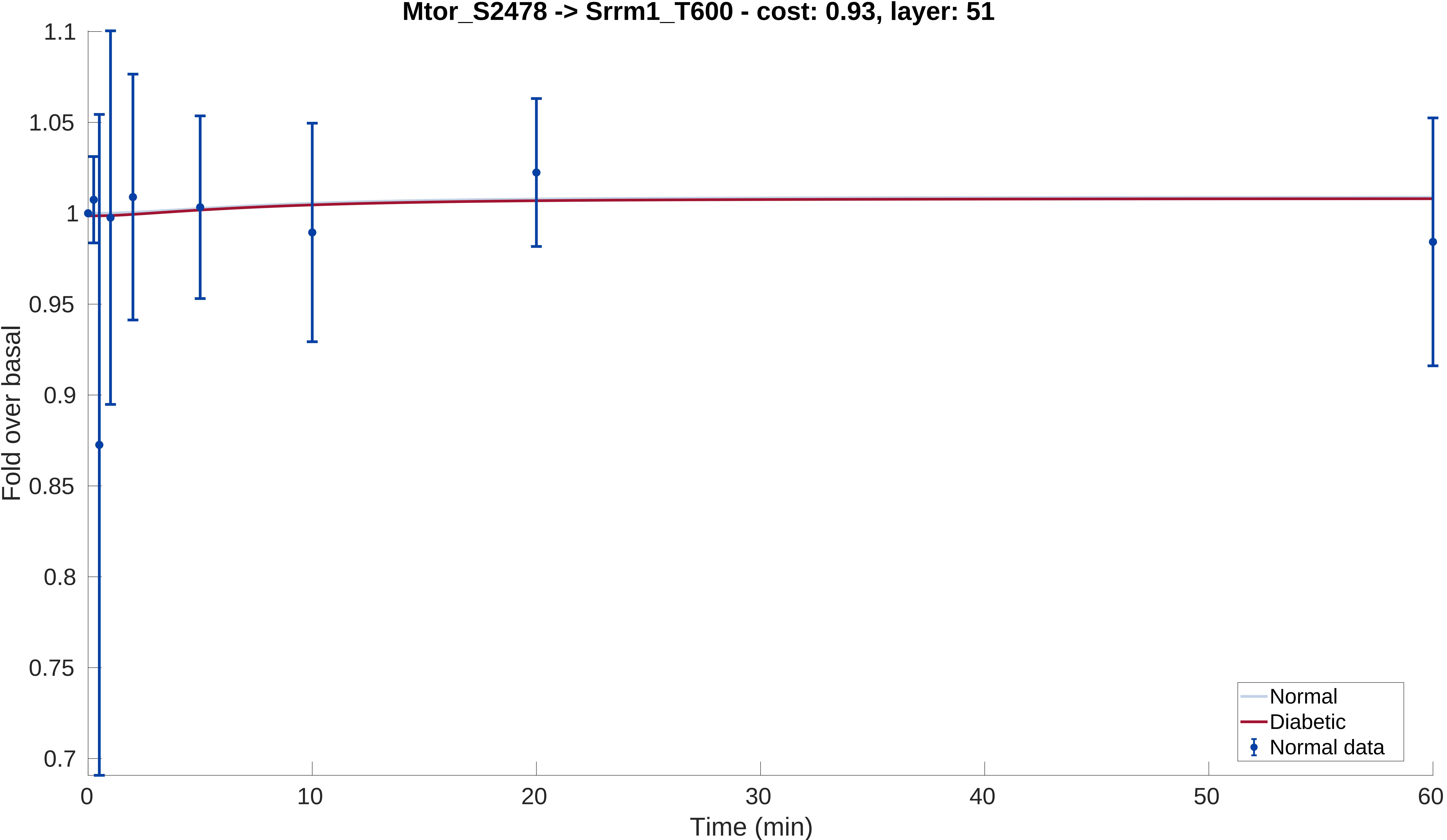
Mtor_S2478 -> Srrm1_T600 - cost: 0.93, layer: 51
1.1
1.05
1
0.95
0.9
0.85
0.8
0.75
0.7
Fold over basal
0
10
20
30
40
50
60
Time (min)
Normal
Diabetic
Normal data

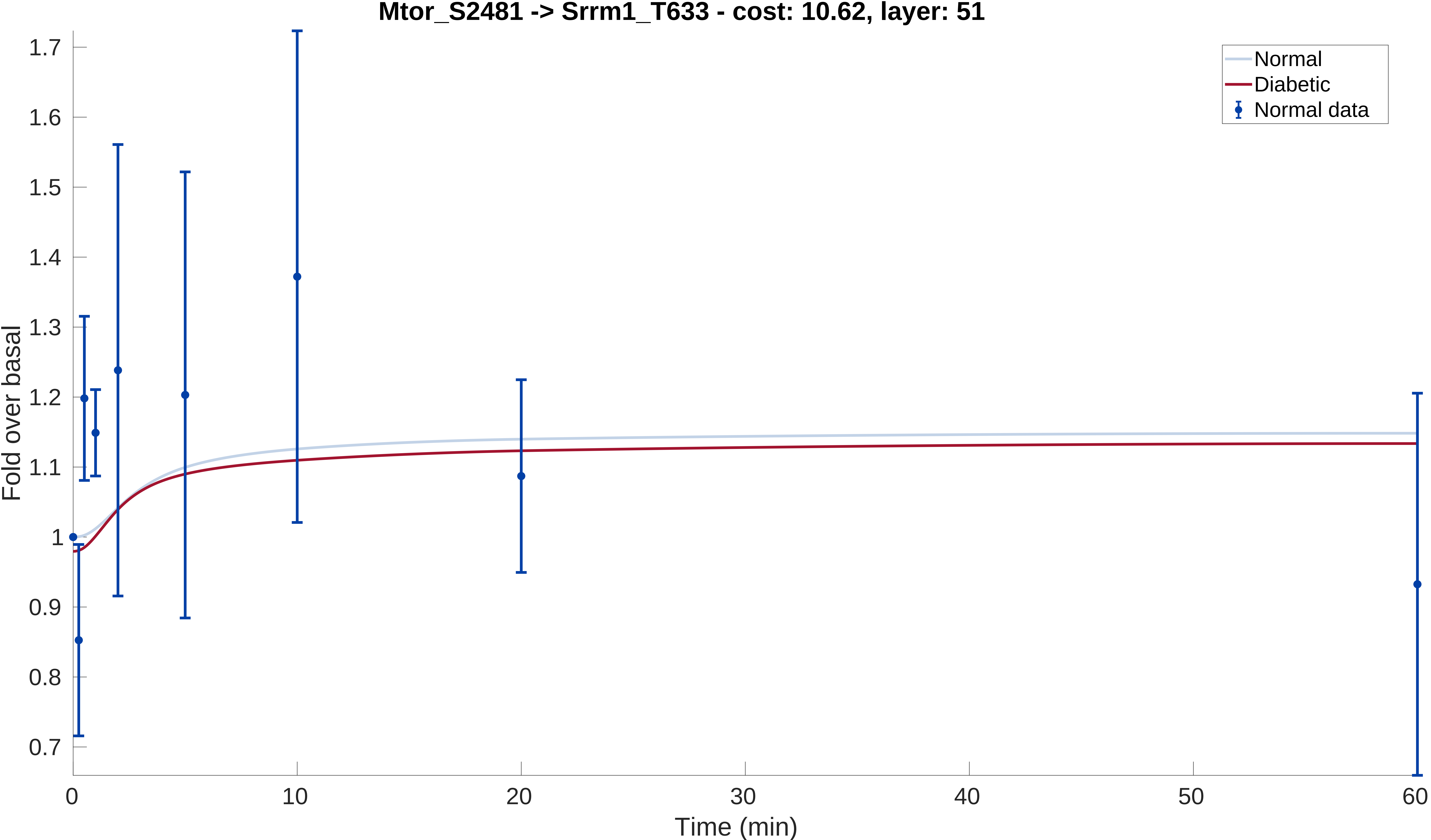

Mtor_S2481 -> Srrm1_T633 - cost: 10.62, layer: 51
Normal
Diabetic
Normal data
Fold over basal
1.7
1.6
1.5
1.4
1.3
1.2
1.1
1
0.9
0.8
0.7
0
10
20
30
40
50
60
Time (min)

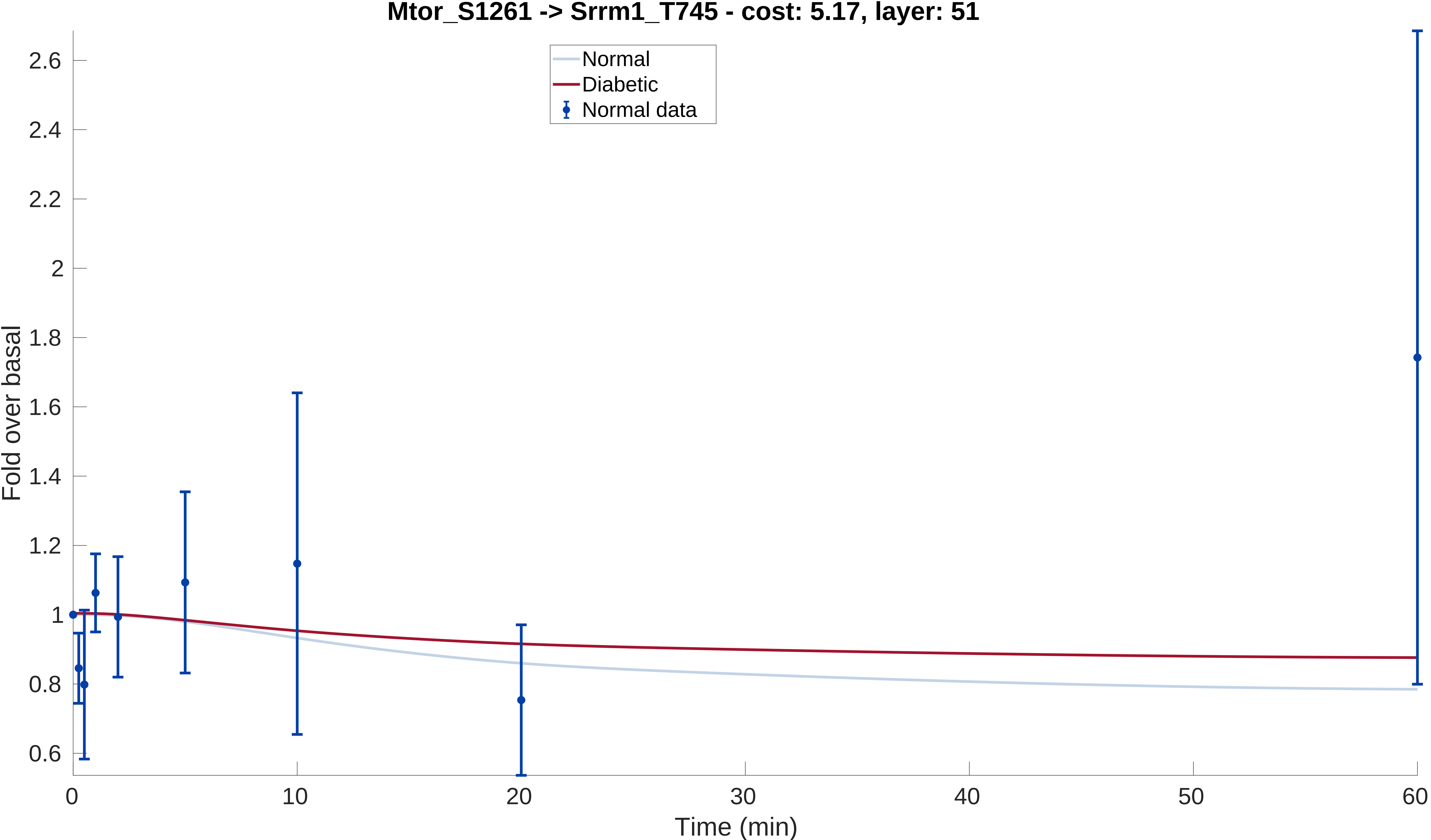

Mtor_S1261 -> Srrm1_T745 - cost: 5.17, layer: 51
Normal
Diabetic
Normal data
Fold over basal
Time (min)
0.6
0.8
1
1.2
1.4
1.6
1.8
2
2.2
2.4
2.6
0
10
20
30
40
50
60

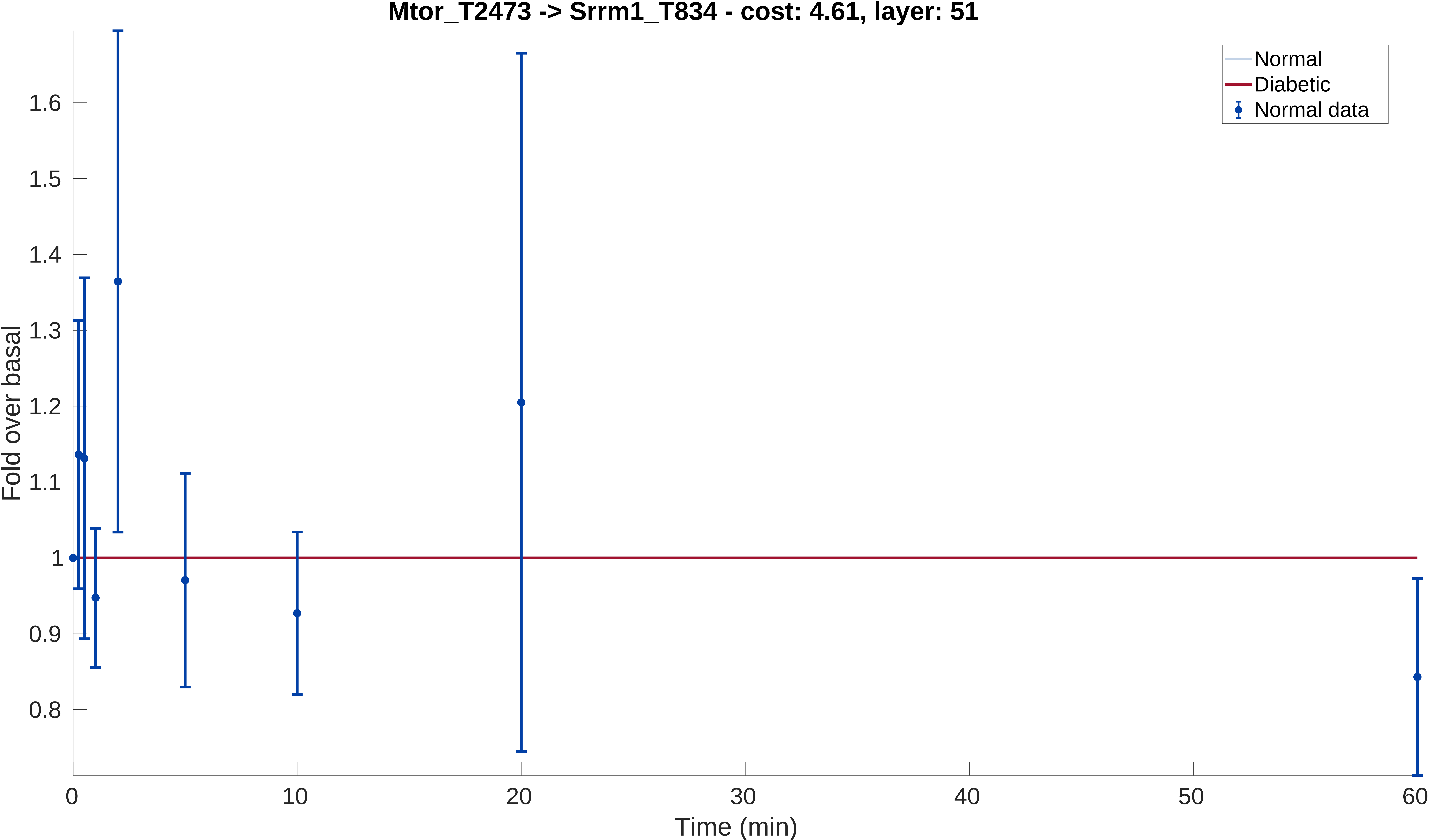

Mtor_T2473 -> Srrm1_T834 - cost: 4.61, layer: 51
Normal
Diabetic
Normal data
1.6
1.5
1.4
1.3
1.2
1.1
1
0.9
0.8
Fold over basal
0
10
20
30
40
50
60
Time (min)

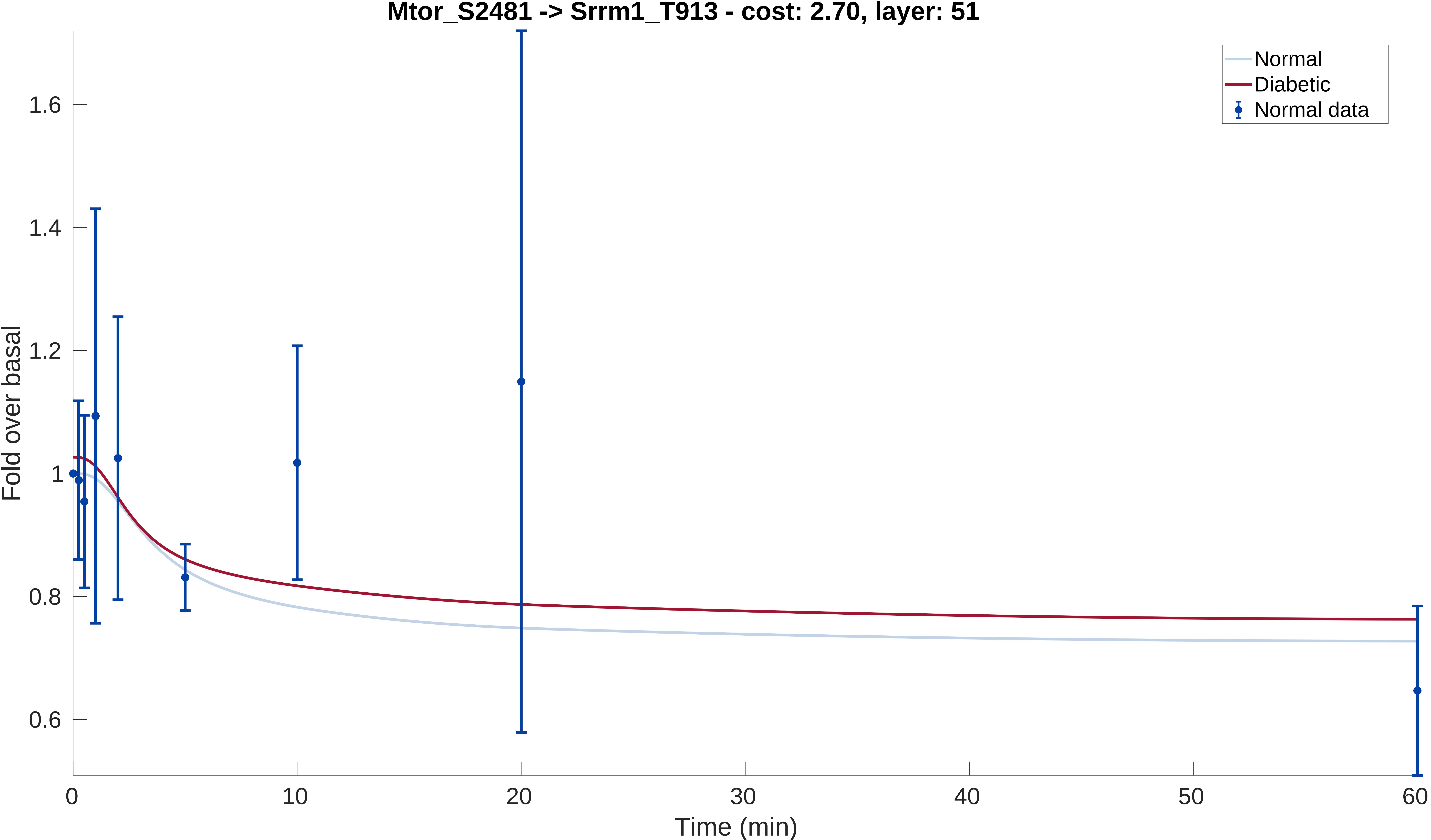

Mtor_S2481 -> Srrm1_T913 - cost: 2.70, layer: 51
Normal
Diabetic
Normal data
Fold over basal
1.6
1.4
1.2
1
0.8
0.6
0
10
20
30
40
50
60
Time (min)

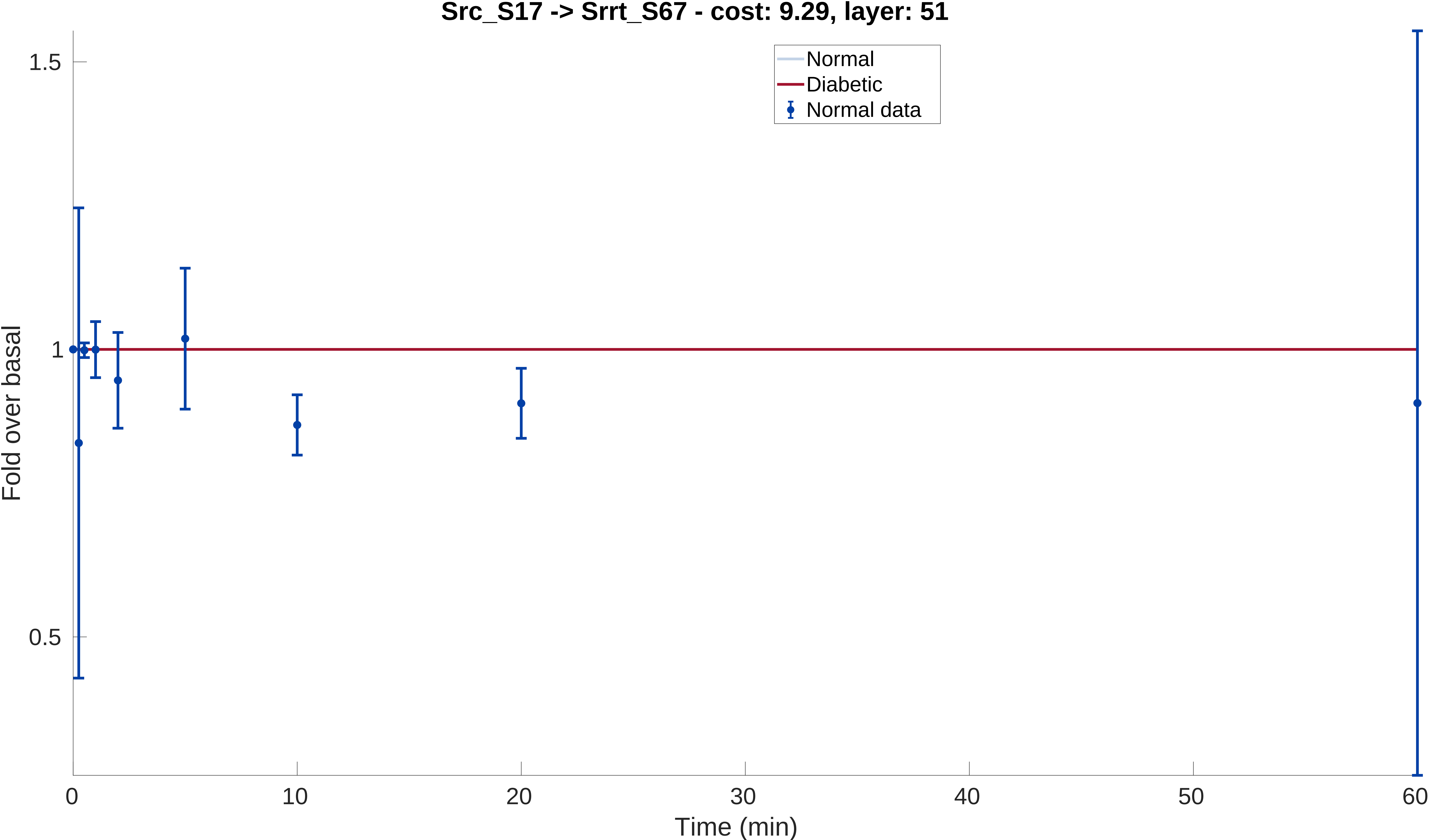

Src_S17 -> Srrt_S67 - cost: 9.29, layer: 51
Normal
Diabetic
Normal data
1.5
1
0.5
Fold over basal
0
10
20
30
40
50
60
Time (min)

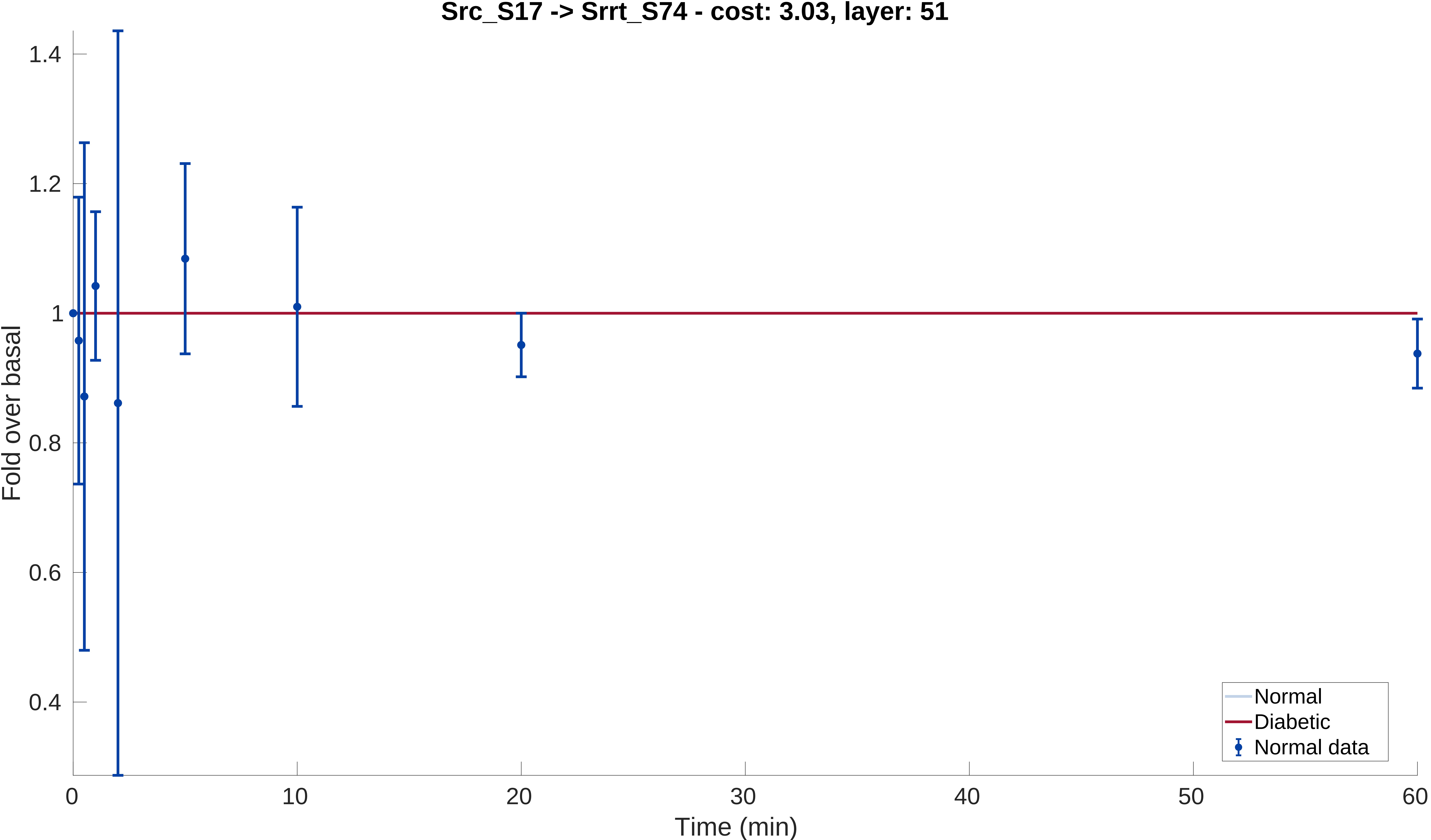

Src_S17 -> Srrt_S74 - cost: 3.03, layer: 51
Fold over basal
1.4
1.2
1
0.8
0.6
0.4
0
10
20
30
40
50
60
Time (min)
Normal
Diabetic
Normal data

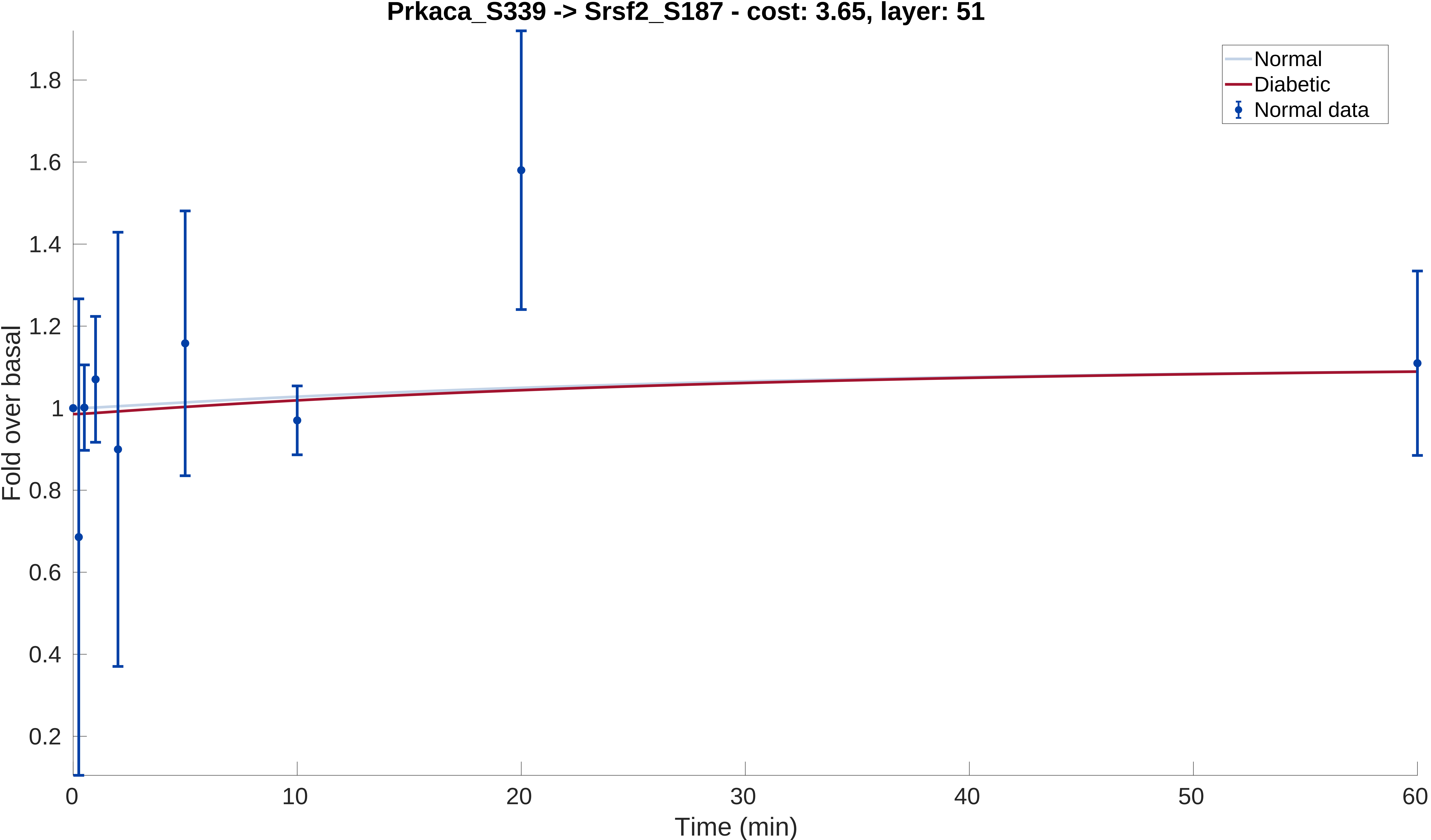
Prkaca_S339 -> Srsf2_S187 - cost: 3.65, layer: 51
Normal
Diabetic
Normal data
Fold over basal
1.8
1.6
1.4
1.2
1
0.8
0.6
0.4
0.2
0
10
20
30
40
50
60
Time (min)

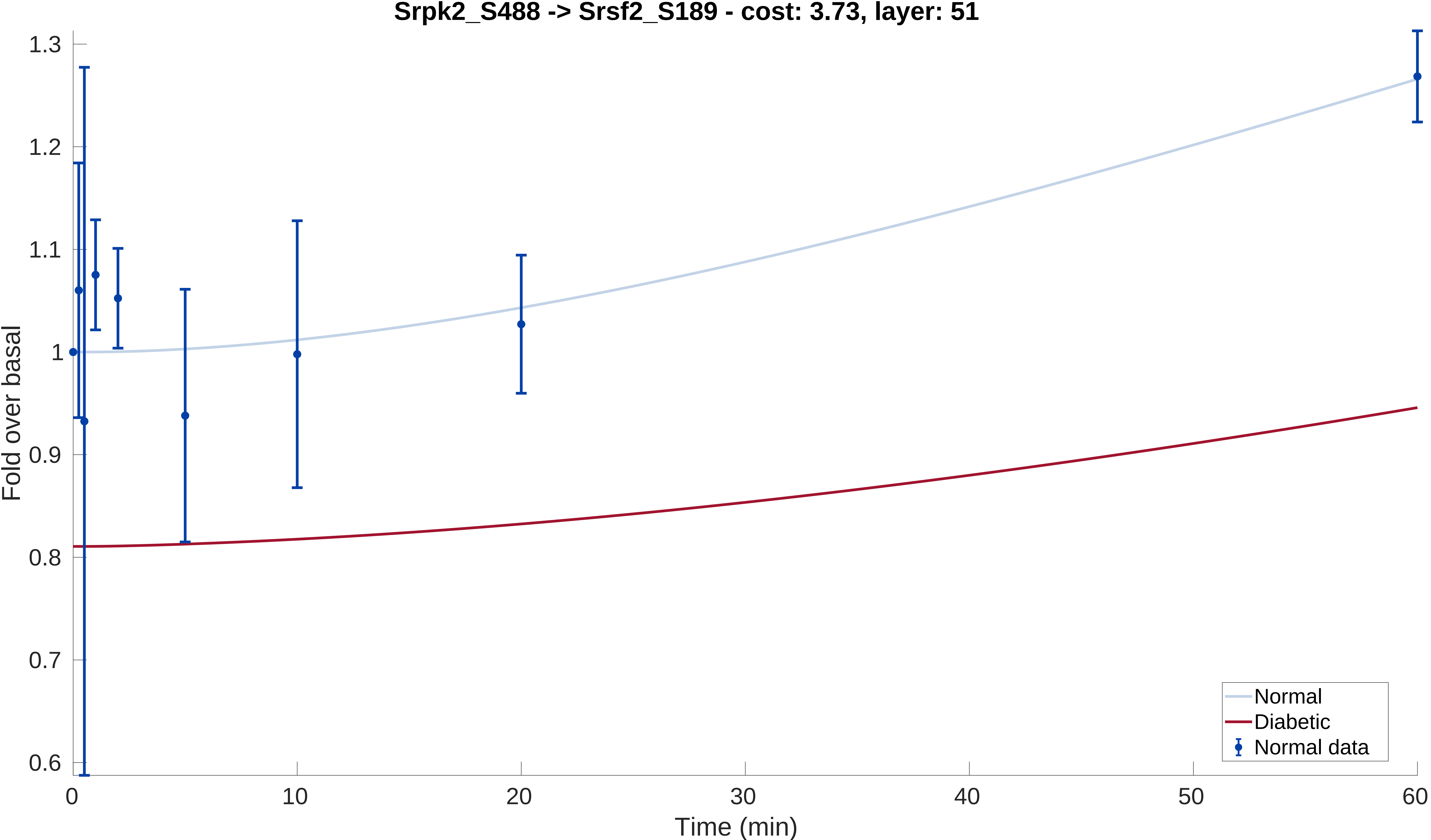

Srpk2_S488 -> Srsf2_S189 - cost: 3.73, layer: 51
1.3
1.2
1.1
1
0.9
0.8
0.7
0.6
Fold over basal
0
10
20
30
40
50
60
Time (min)
Normal
Diabetic
Normal data

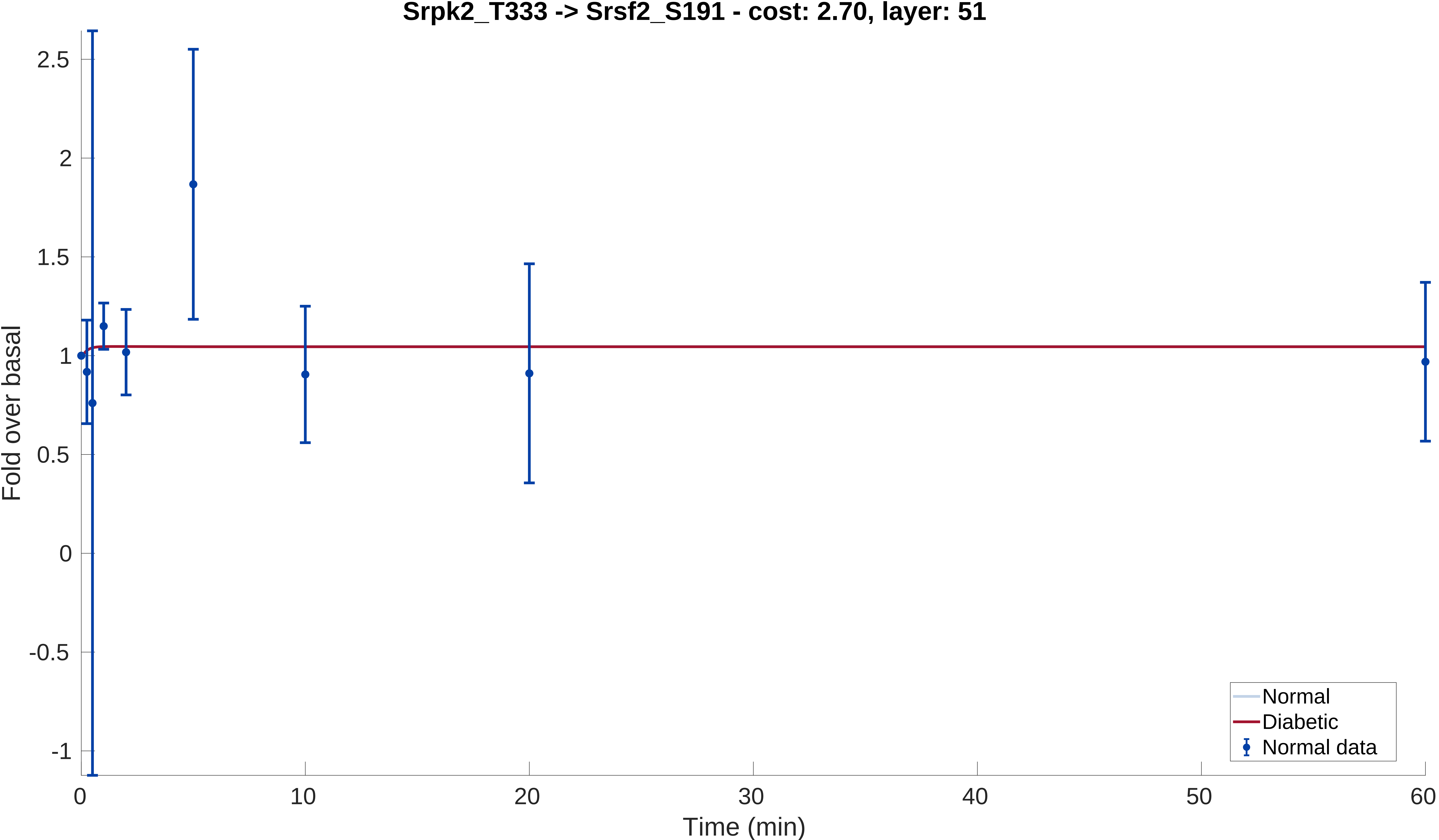

Srpk2_T333 -> Srsf2_S191 - cost: 2.70, layer: 51
Fold over basal
2.5
2
1.5
1
0.5
0
-0.5
-1
0
10
20
30
40
50
60
Time (min)
Normal
Diabetic
Normal data

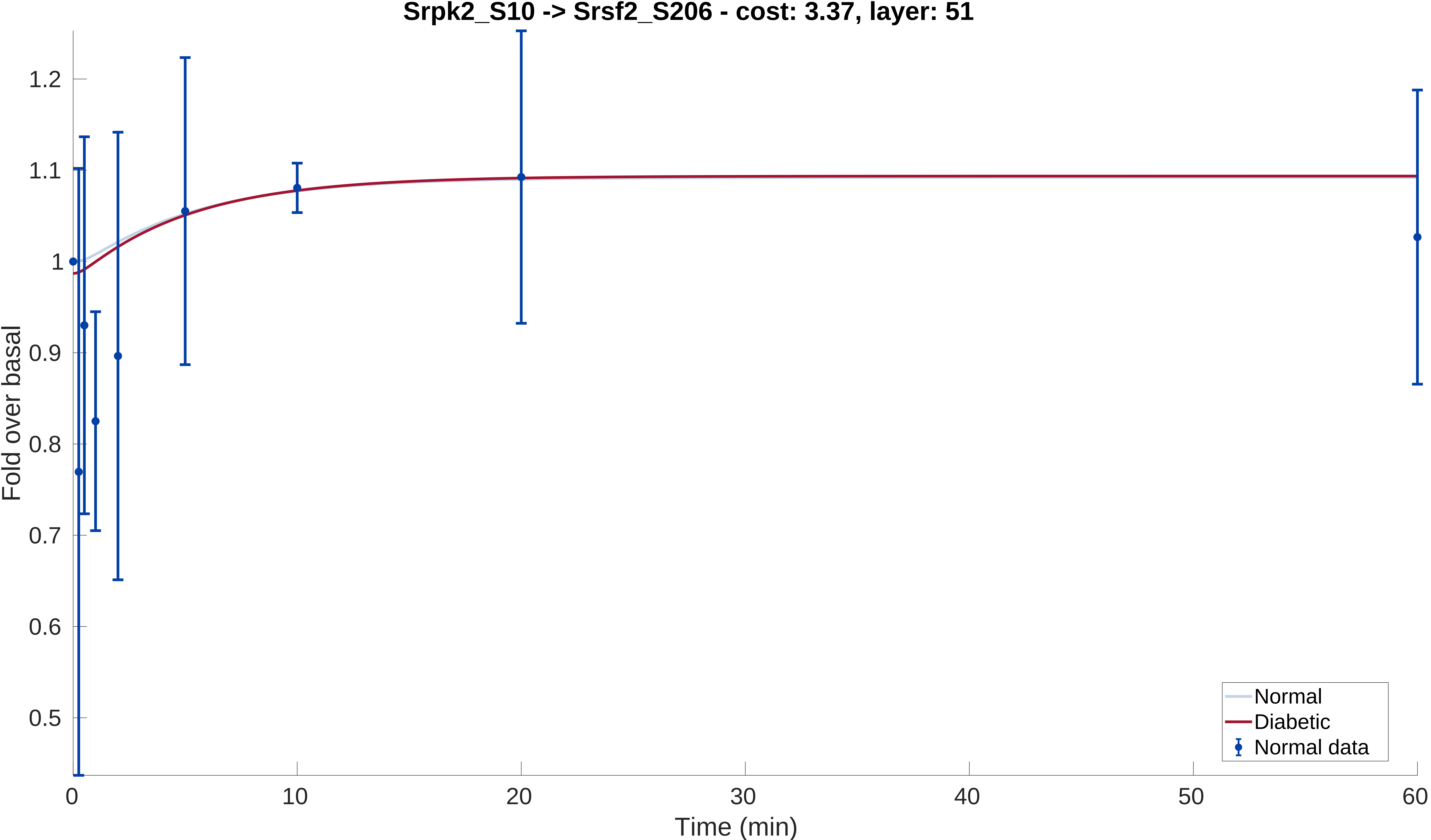

Srpk2_S10 -> Srsf2_S206 - cost: 3.37, layer: 51
Fold over basal
1.2
1.1
1
0.9
0.8
0.7
0.6
0.5
0
10
20
30
40
50
60
Time (min)
Normal
Diabetic
Normal data

Srpk2_S491 -> Srsf2_S208 - cost: 2.00, layer: 51

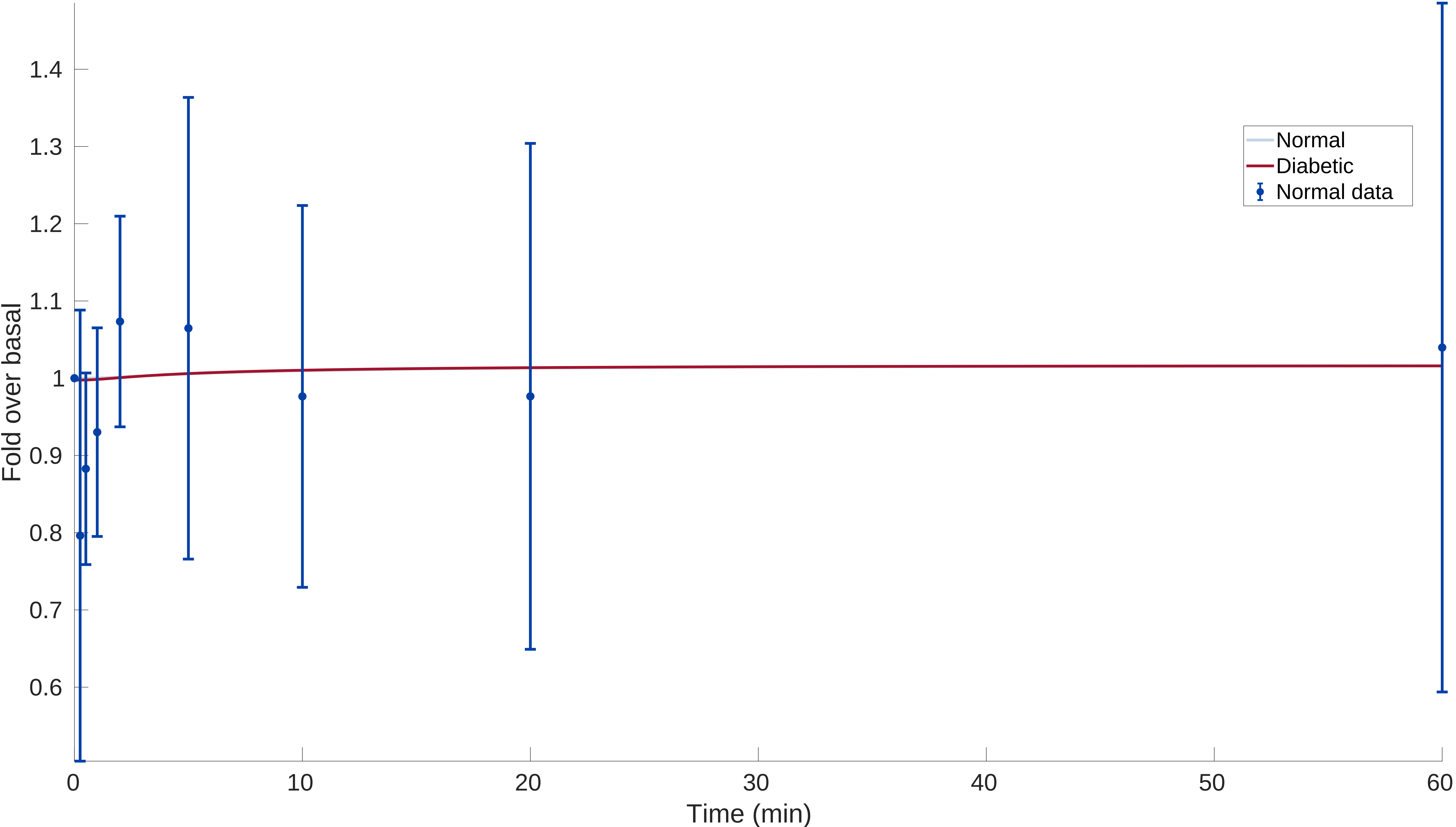

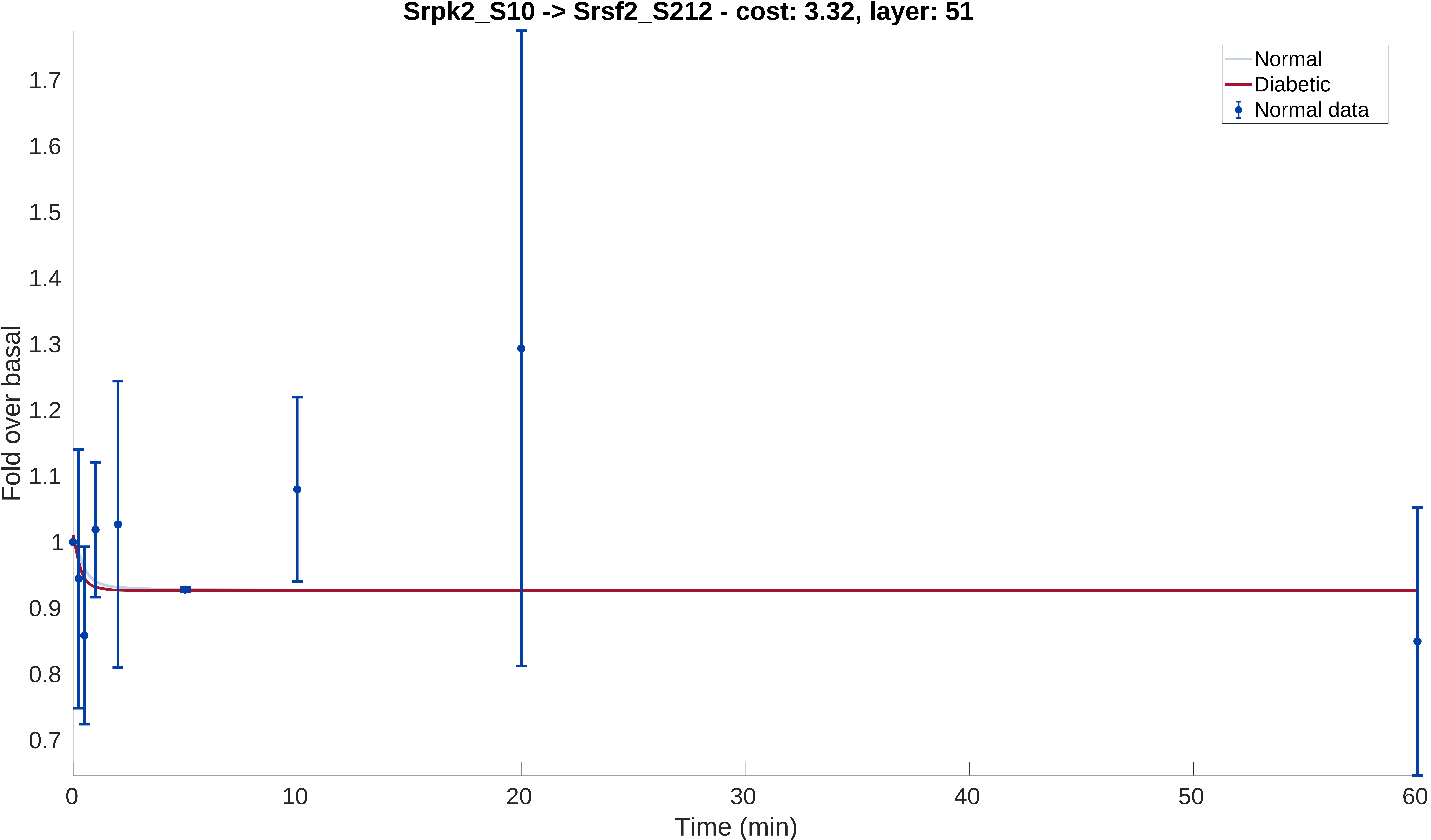
Srpk2_S10 -> Srsf2_S212 - cost: 3.32, layer: 51
Normal
Diabetic
Normal data
Fold over basal
1.7
1.6
1.5
1.4
1.3
1.2
1.1
1
0.9
0.8
0.7
0
10
20
30
40
50
60
Time (min)

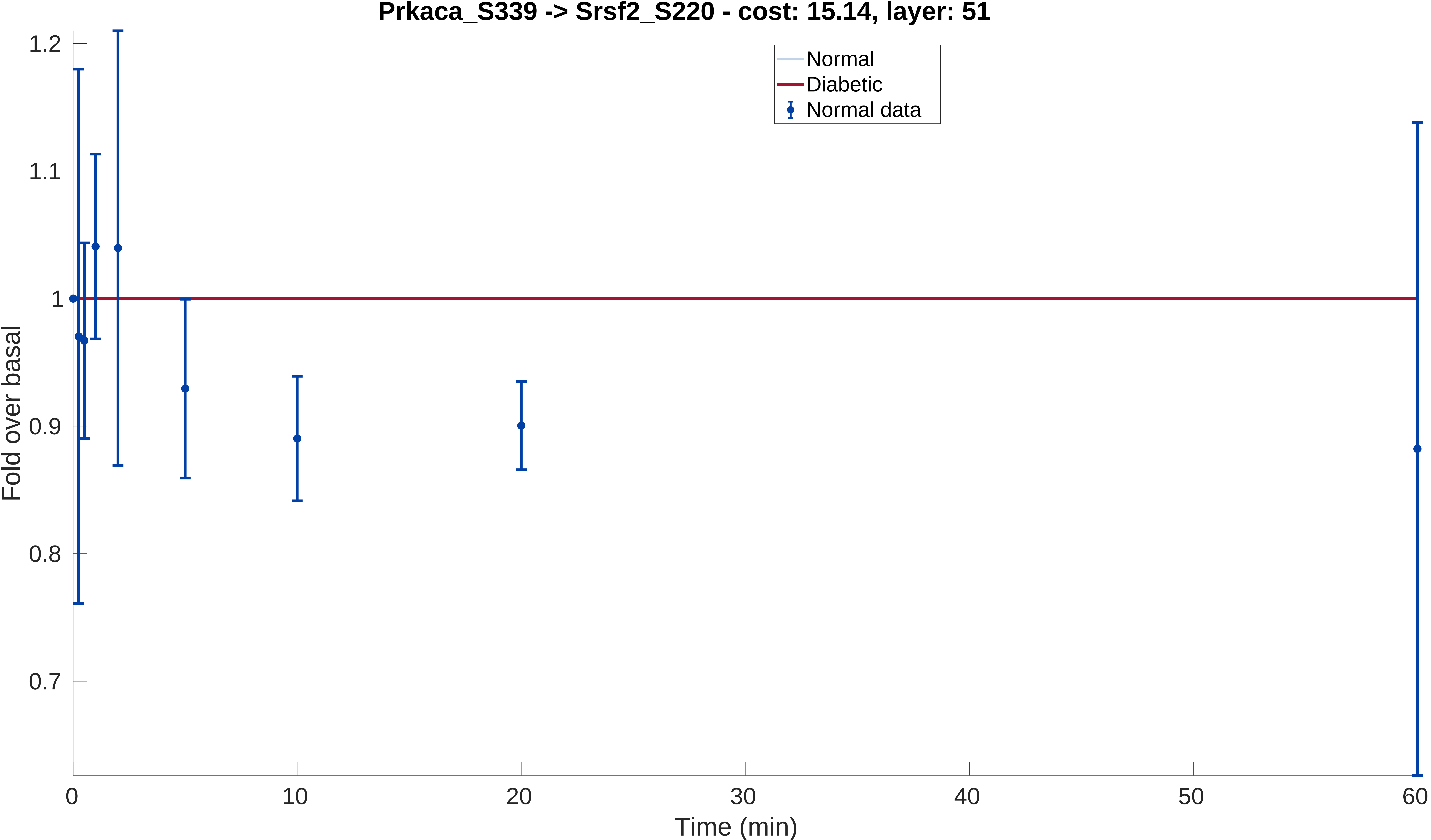
Prkaca_S339 -> Srsf2_S220 - cost: 15.14, layer: 51
Normal
Diabetic
Normal data
Fold over basal
1.2
1.1
1
0.9
0.8
0.7
0
10
20
30
40
50
60
Time (min)

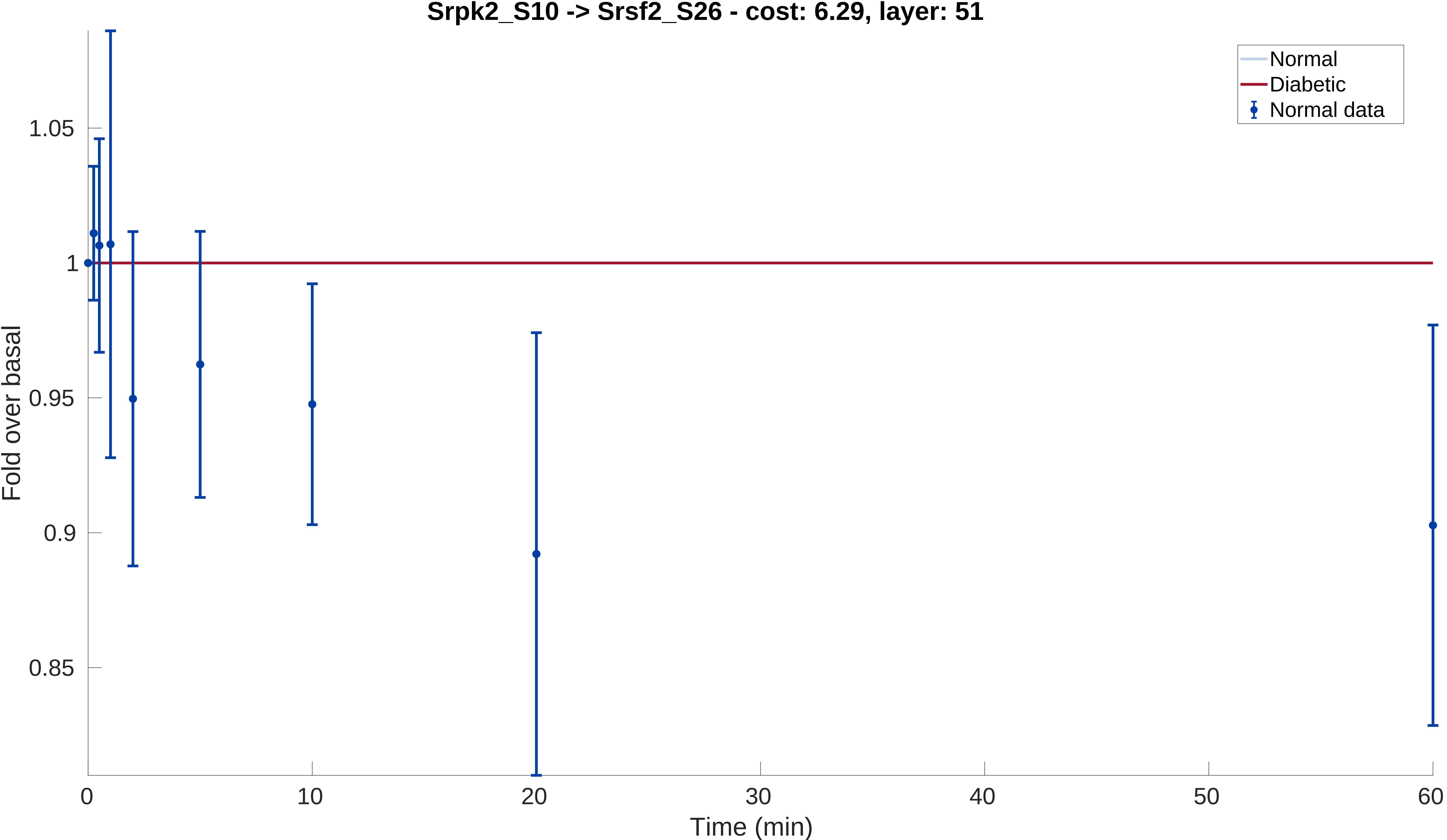

Srpk2_S10 -> Srsf2_S26 - cost: 6.29, layer: 51
Normal
Diabetic
Normal data
Fold over basal
1.05
1
0.95
0.9
0.85
0
10
20
30
40
50
60
Time (min)

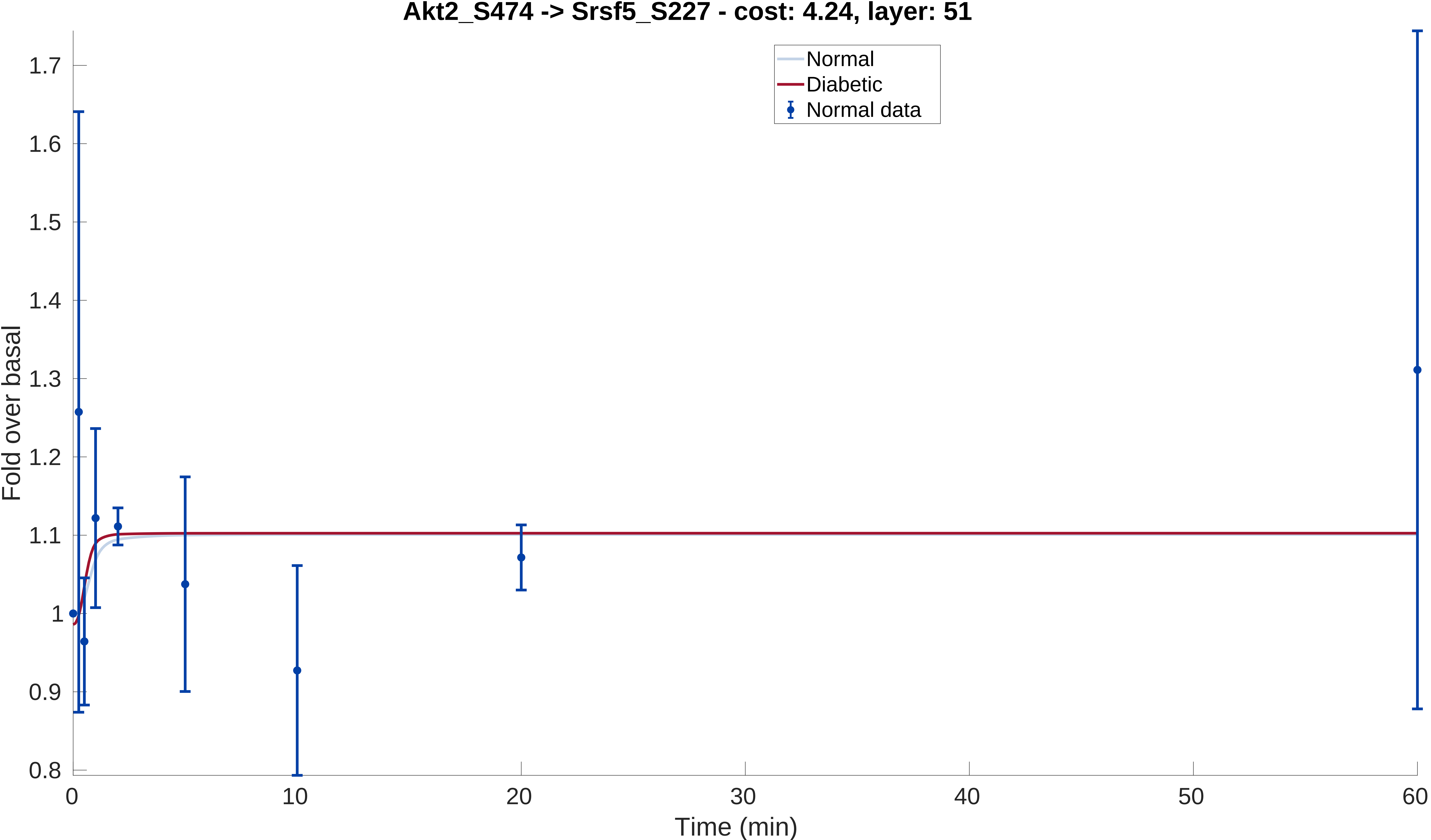
Akt2_S474 -> Srsf5_S227 - cost: 4.24, layer: 51
Normal
Diabetic
Normal data
Fold over basal
1.7
1.6
1.5
1.4
1.3
1.2
1.1
1
0.9
0.8
0
10
20
30
40
50
60
Time (min)

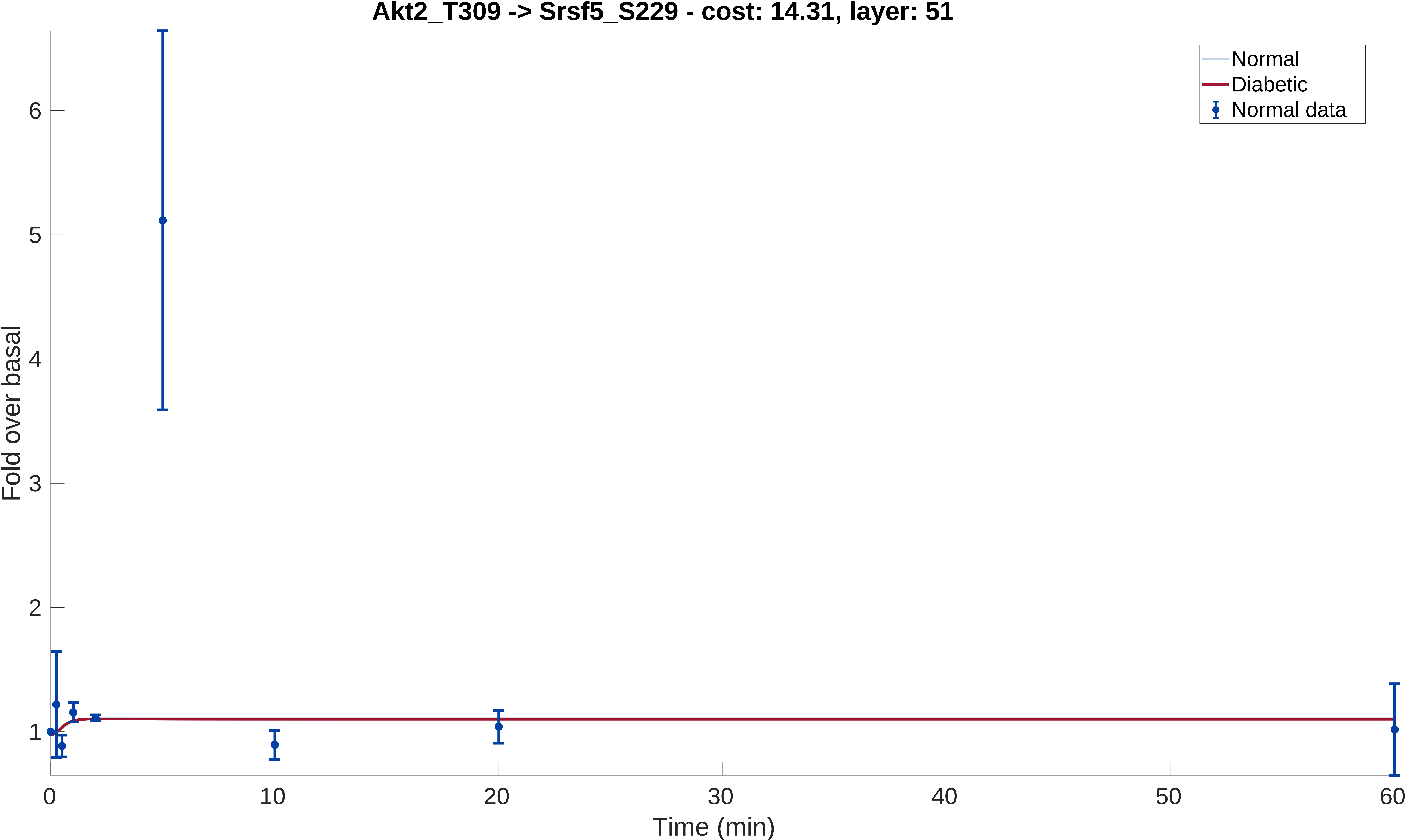
Akt2_T309 -> Srsf5_S229 - cost: 14.31, layer: 51
Normal
Diabetic
Normal data
Fold over basal
Time (min)
1
2
3
4
5
6
0
10
20
30
40
50
60

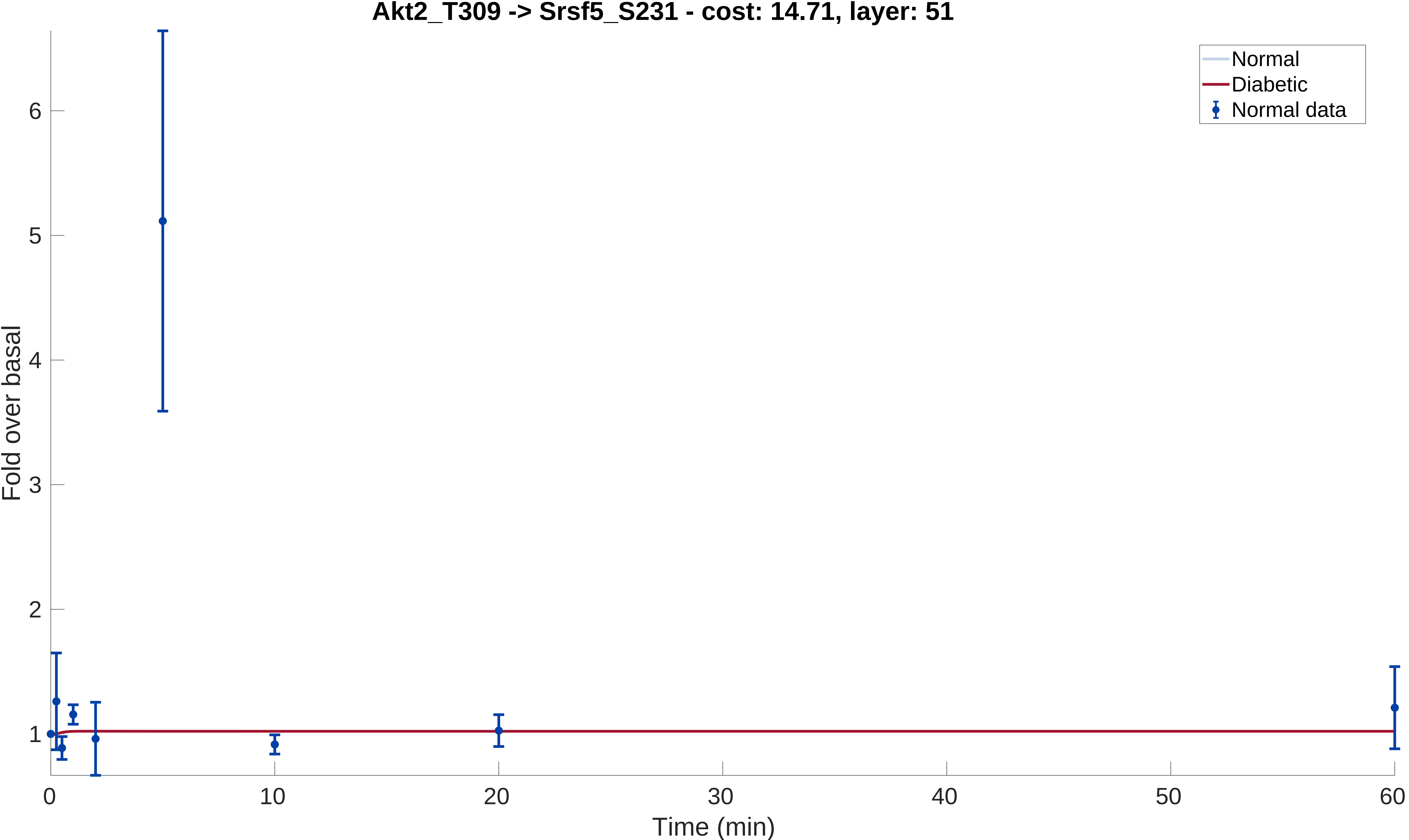
Akt2_T309 -> Srsf5_S231 - cost: 14.71, layer: 51
Normal
Diabetic
Normal data
Fold over basal
1
2
3
4
5
6
0
10
20
30
40
50
60
Time (min)

Akt2_S474 -> Srsf5_S248 - cost: 2.55, layer: 51

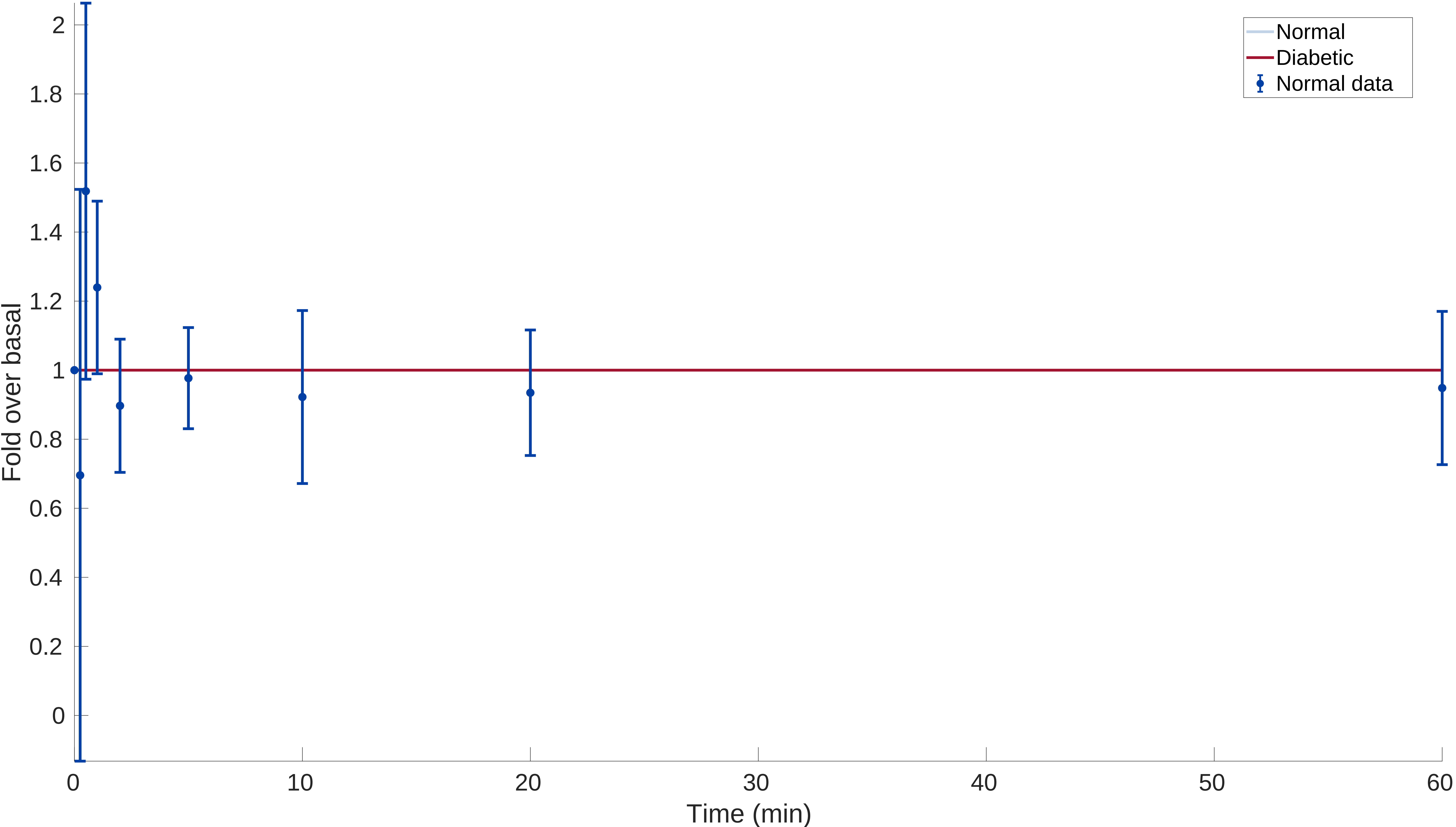

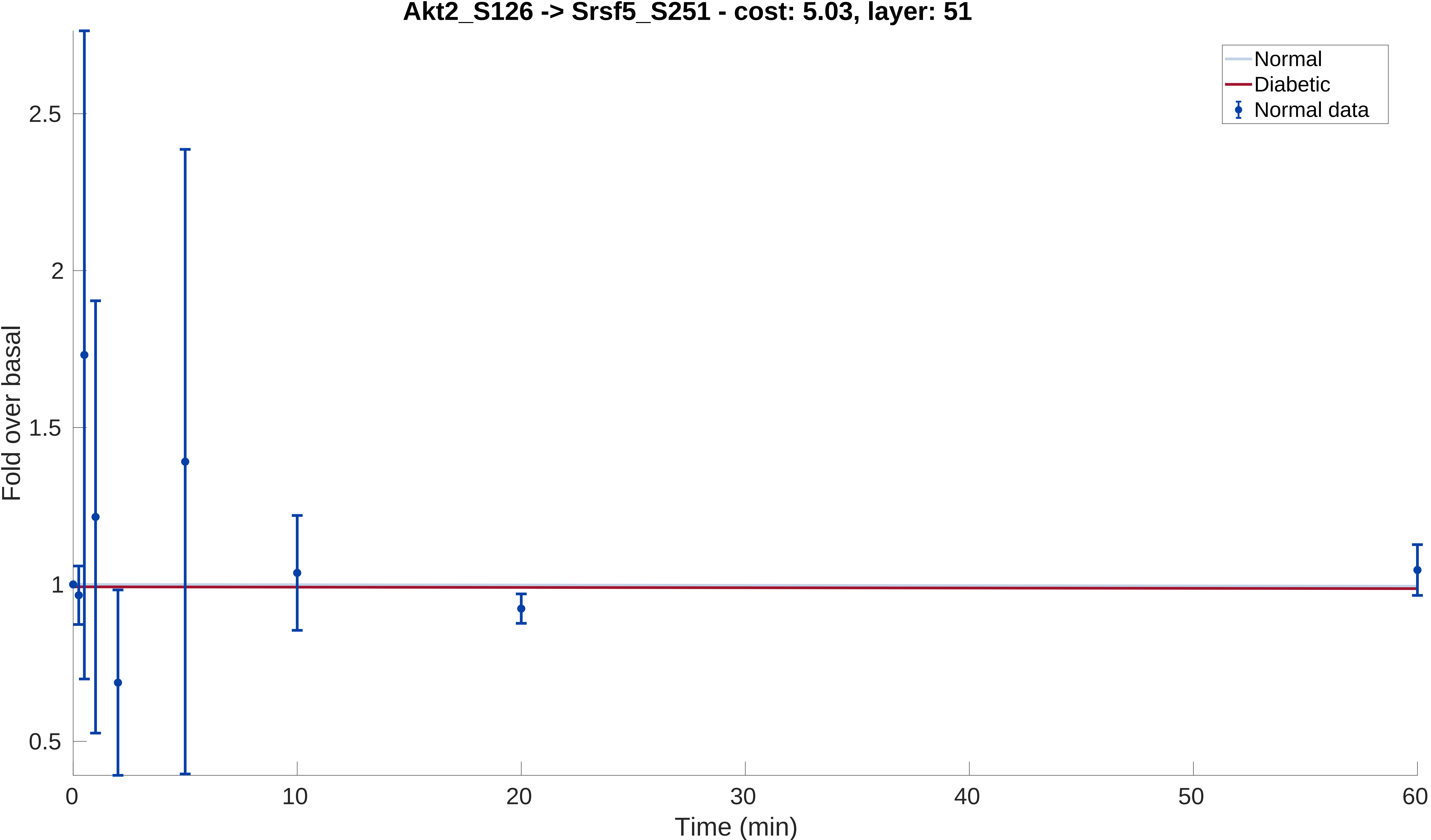
Akt2_S126 -> Srsf5_S251 - cost: 5.03, layer: 51
Normal
Diabetic
Normal data
Fold over basal
0.5
1
1.5
2
2.5
0
10
20
30
40
50
60
Time (min)

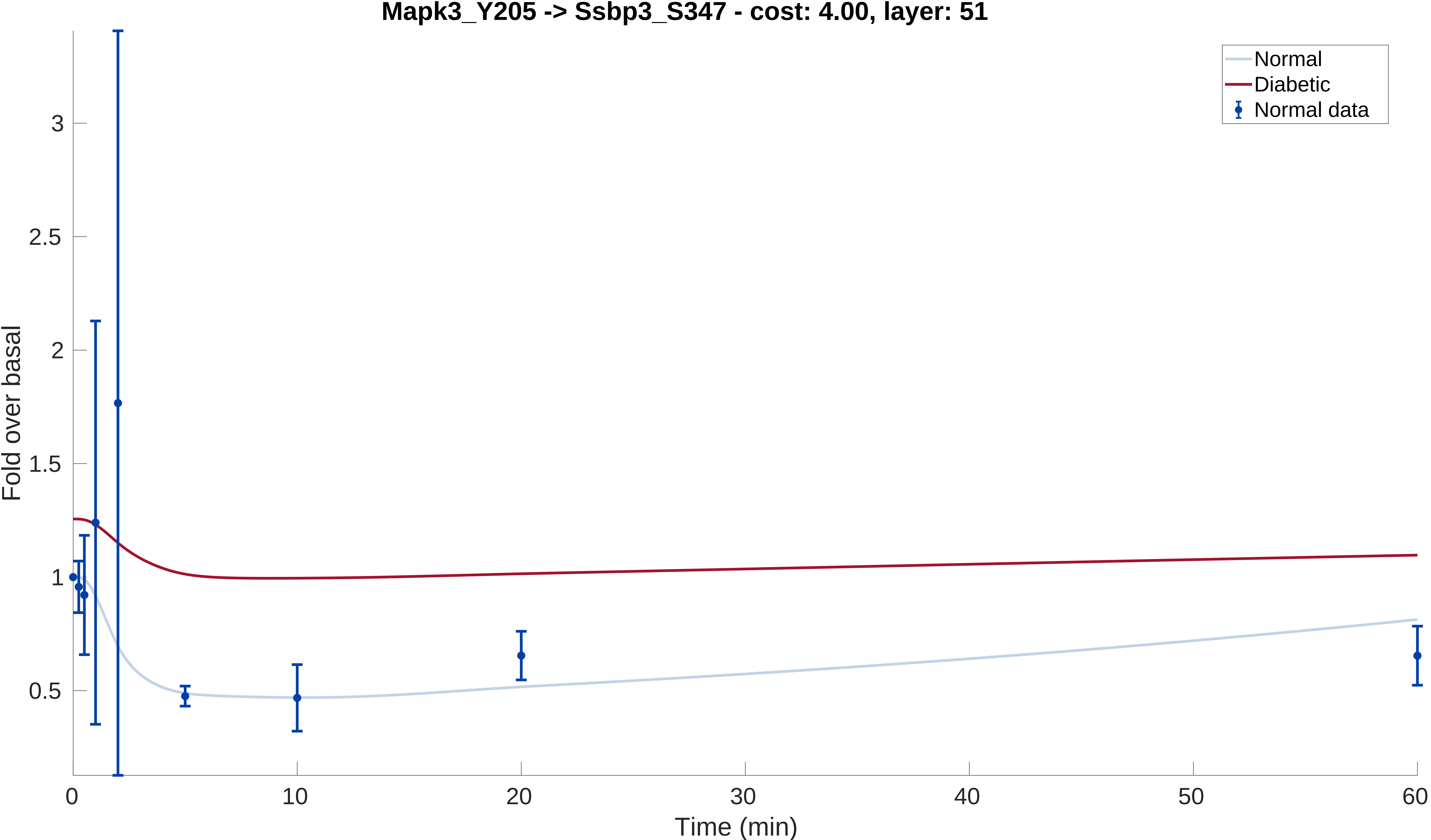

Mapk3_Y205 -> Ssbp3_S347 - cost: 4.00, layer: 51
Normal
Diabetic
Normal data
Fold over basal
3
2.5
2
1.5
1
0.5
0
10
20
30
40
50
60
Time (min)

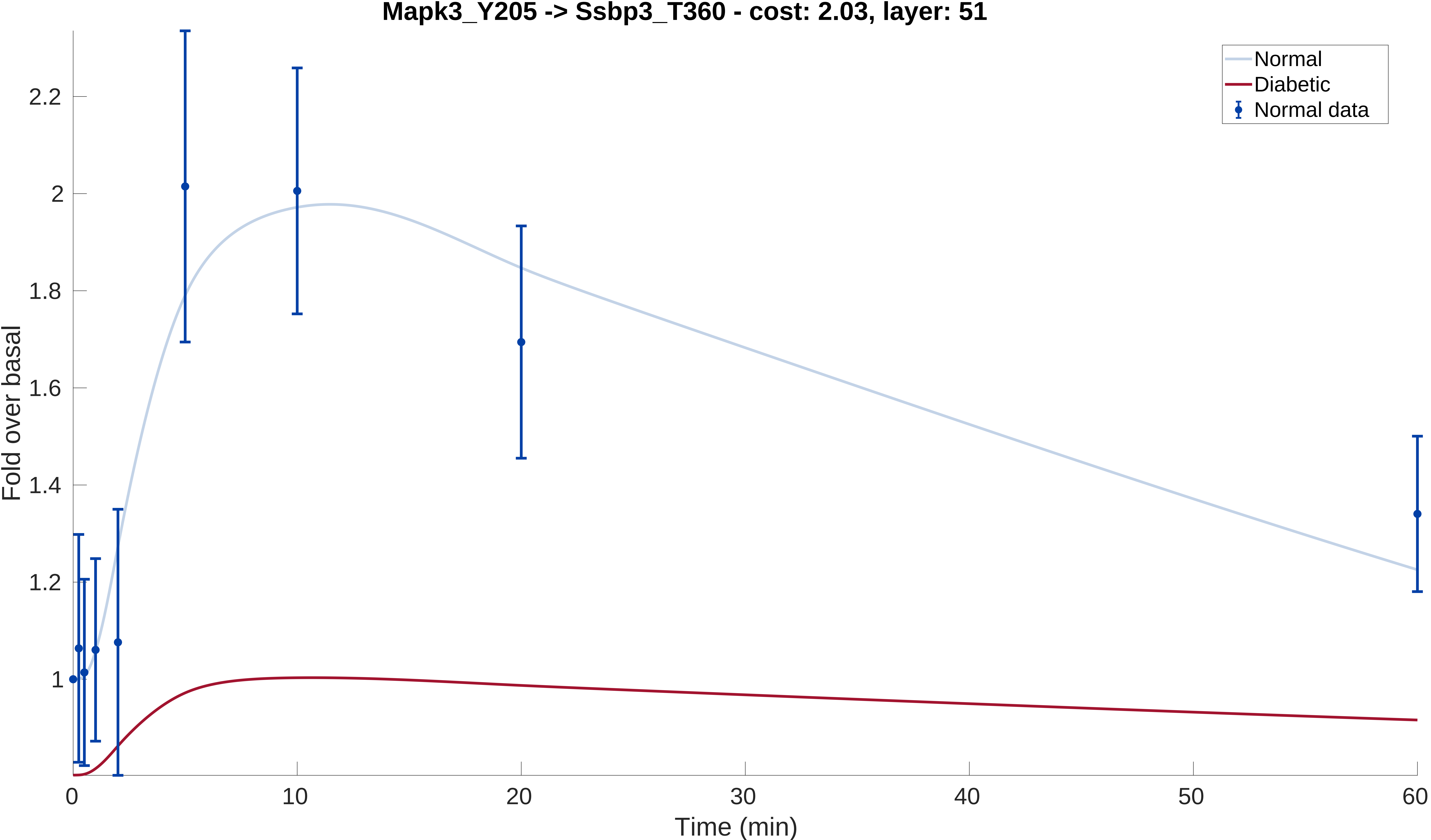

Mapk3_Y205 -> Ssbp3_T360 - cost: 2.03, layer: 51
Normal
Diabetic
Normal data
Fold over basal
2.2
2
1.8
1.6
1.4
1.2
1
0
10
20
30
40
50
60
Time (min)

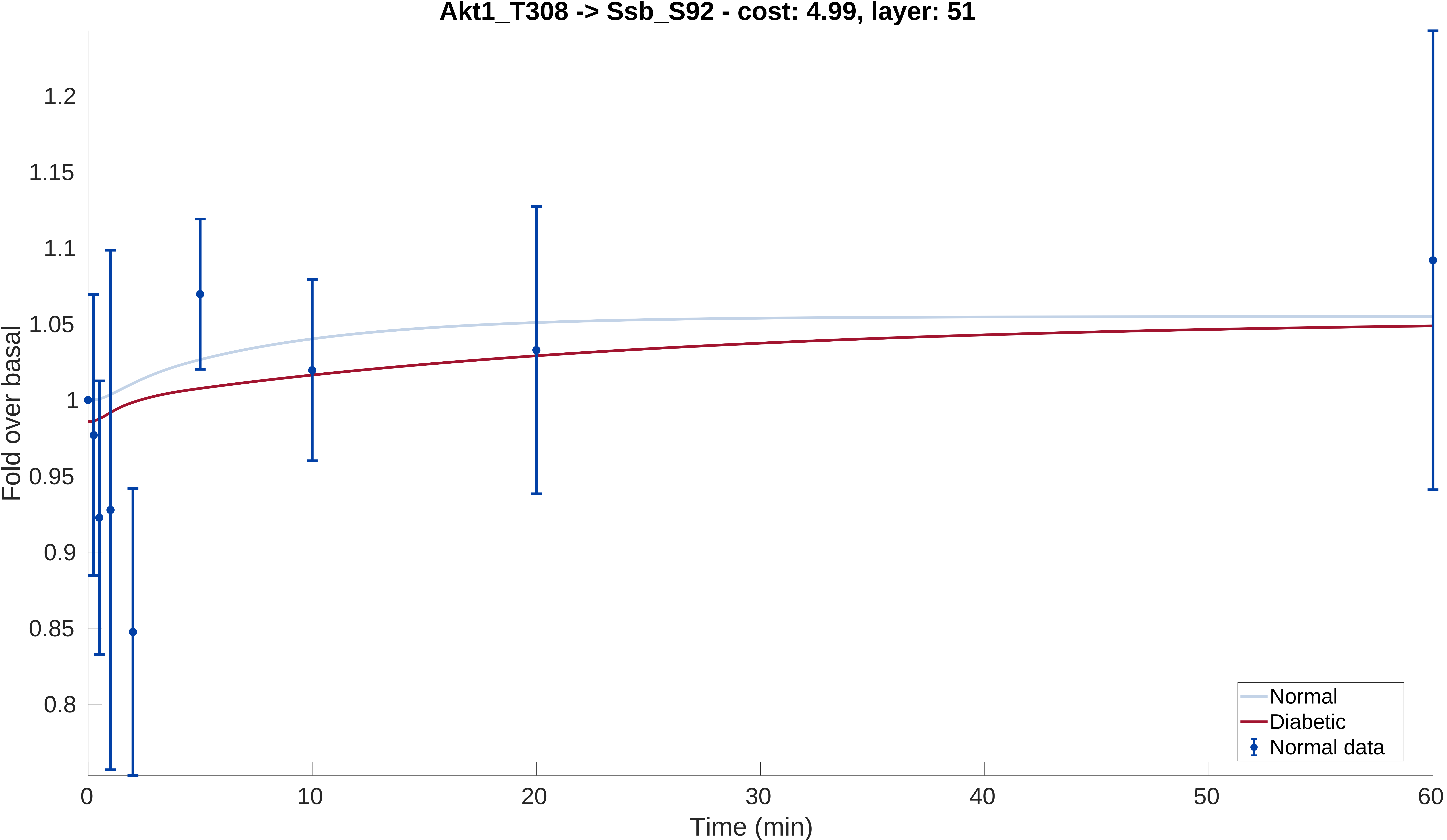

Akt1_T308 -> Ssb_S92 - cost: 4.99, layer: 51
Fold over basal
1.2
1.15
1.1
1.05
1
0.95
0.9
0.85
0.8
0
10
20
30
40
50
60
Time (min)
Normal
Diabetic
Normal data

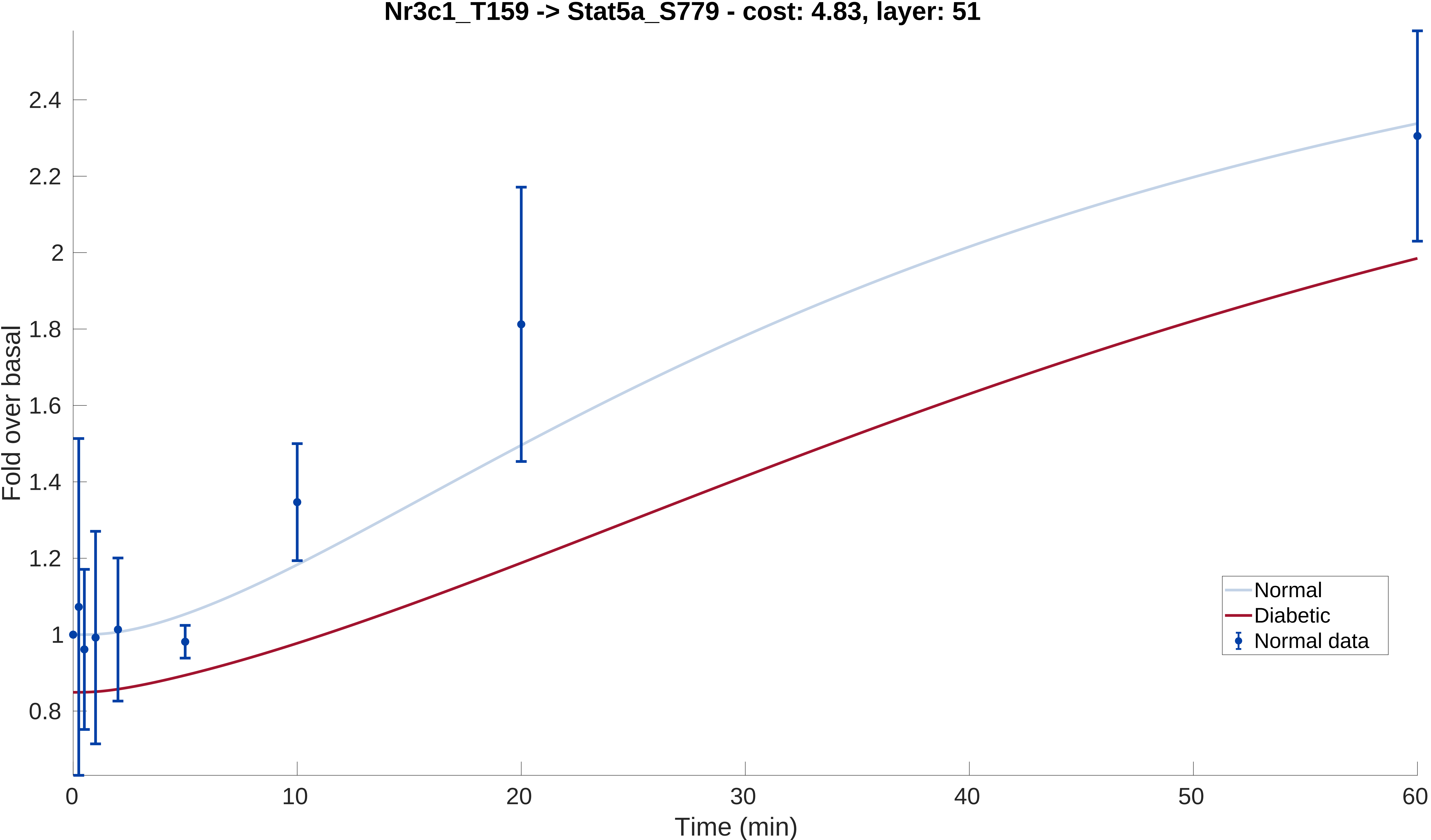

Nr3c1_T159 -> Stat5a_S779 - cost: 4.83, layer: 51
Fold over basal
2.4
2.2
2
1.8
1.6
1.4
1.2
1
0.8
0
10
20
30
40
50
60
Time (min)
Normal
Diabetic
Normal data

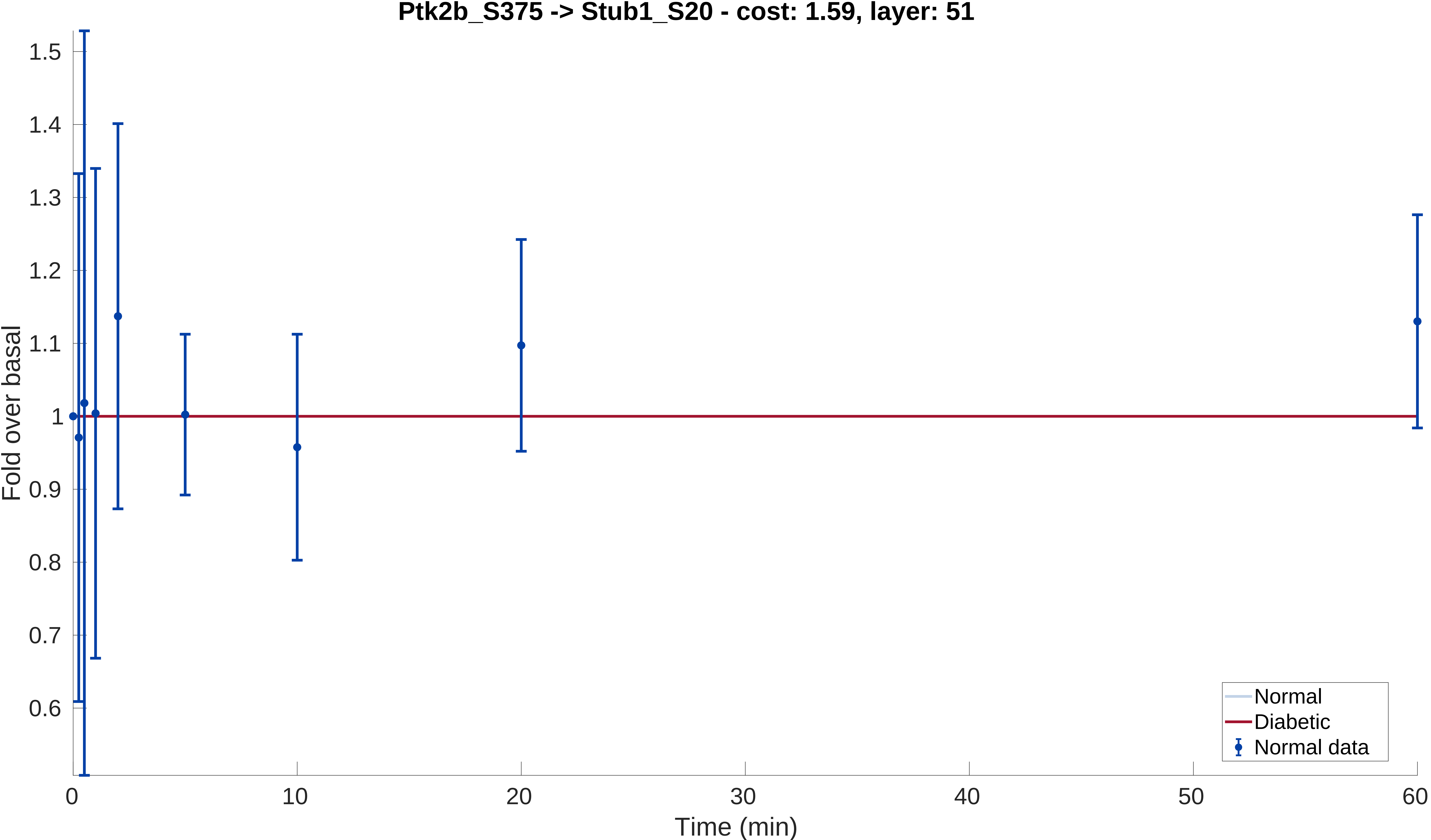

Ptk2b_S375 -> Stub1_S20 - cost: 1.59, layer: 51
Fold over basal
1.5
1.4
1.3
1.2
1.1
1
0.9
0.8
0.7
0.6
0
10
20
30
40
50
60
Time (min)
Normal
Diabetic
Normal data

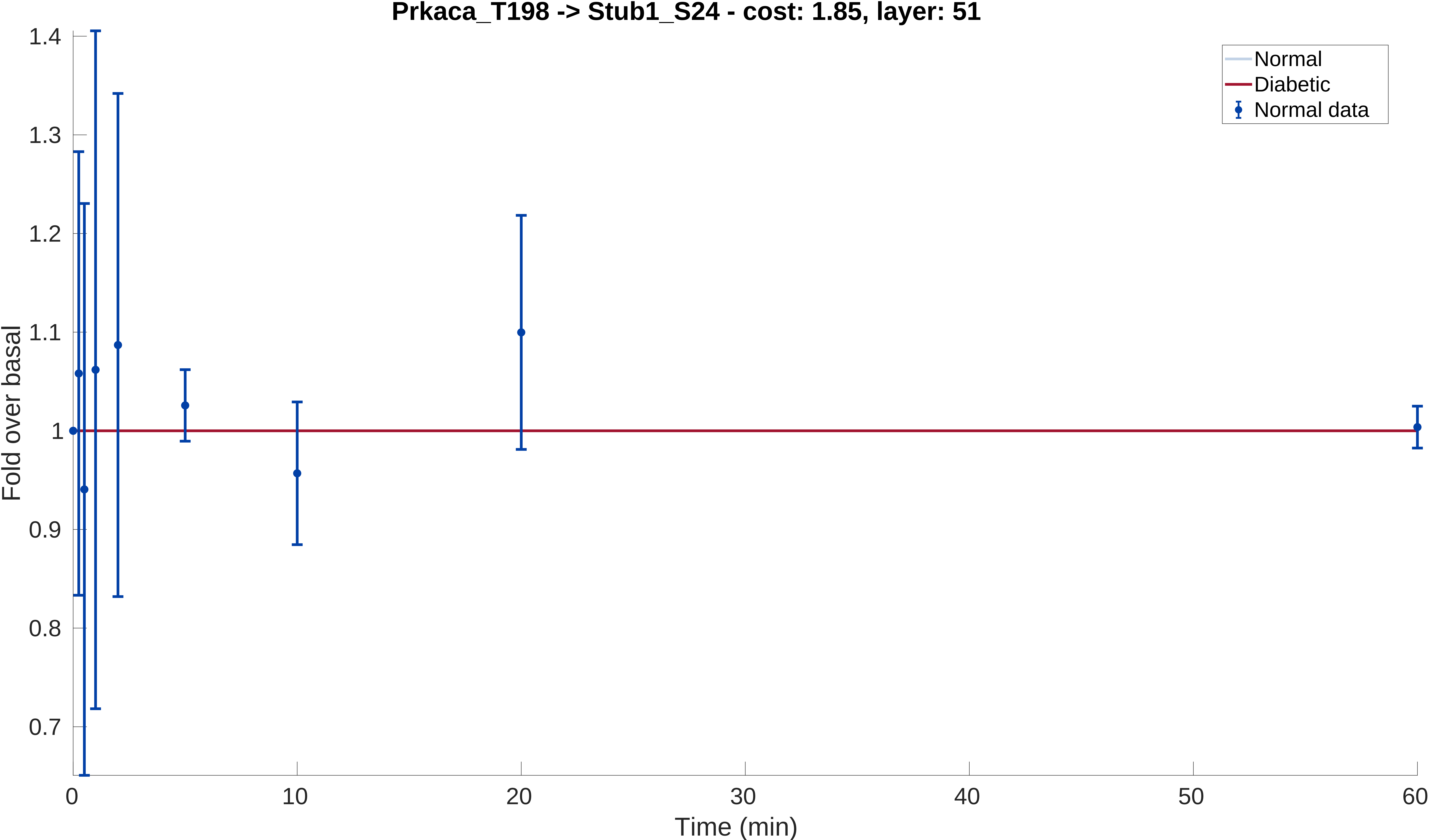

Prkaca_T198 -> Stub1_S24 - cost: 1.85, layer: 51
Normal
Diabetic
Normal data
Fold over basal
1.4
1.3
1.2
1.1
1
0.9
0.8
0.7
0
10
20
30
40
50
60
Time (min)

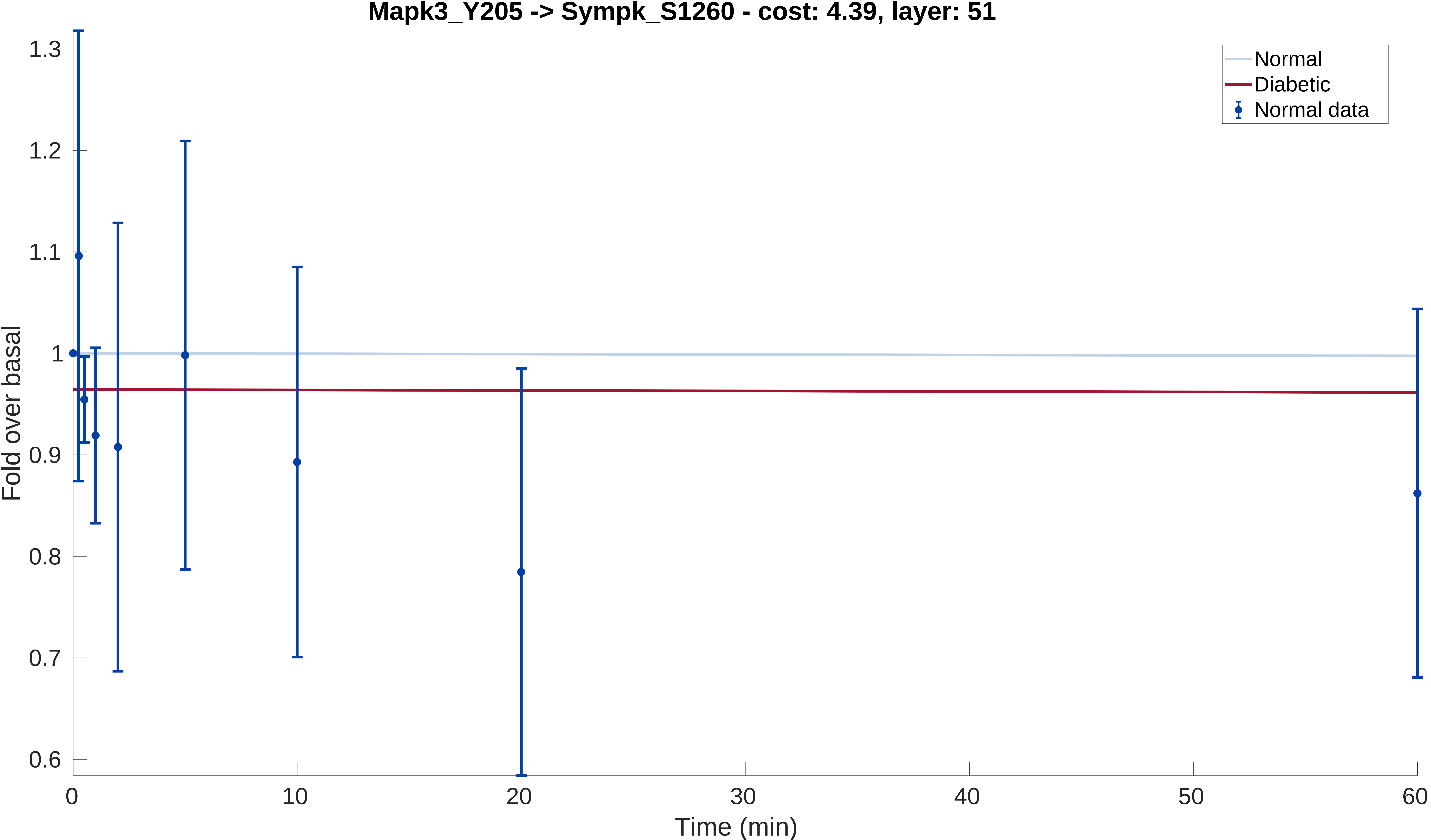

Mapk3_Y205 -> Sympk_S1260 - cost: 4.39, layer: 51
Normal
Diabetic
Normal data
Fold over basal
1.3
1.2
1.1
1
0.9
0.8
0.7
0.6
0
10
20
30
40
50
60
Time (min)

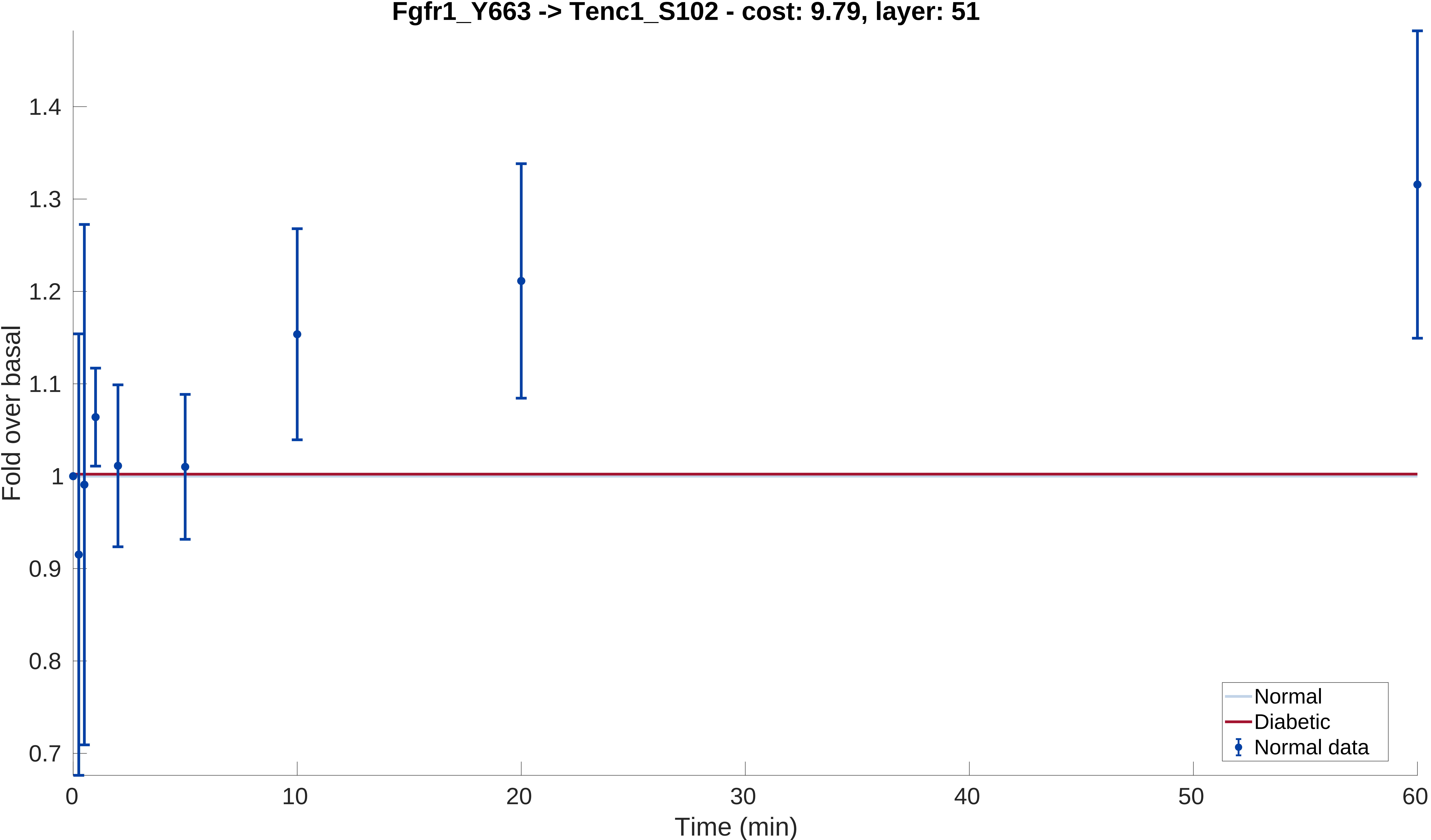

Fgfr1_Y663 -> Tenc1_S102 - cost: 9.79, layer: 51
Fold over basal
1.4
1.3
1.2
1.1
1
0.9
0.8
0.7
0
10
20
30
40
50
60
Time (min)
Normal
Diabetic
Normal data

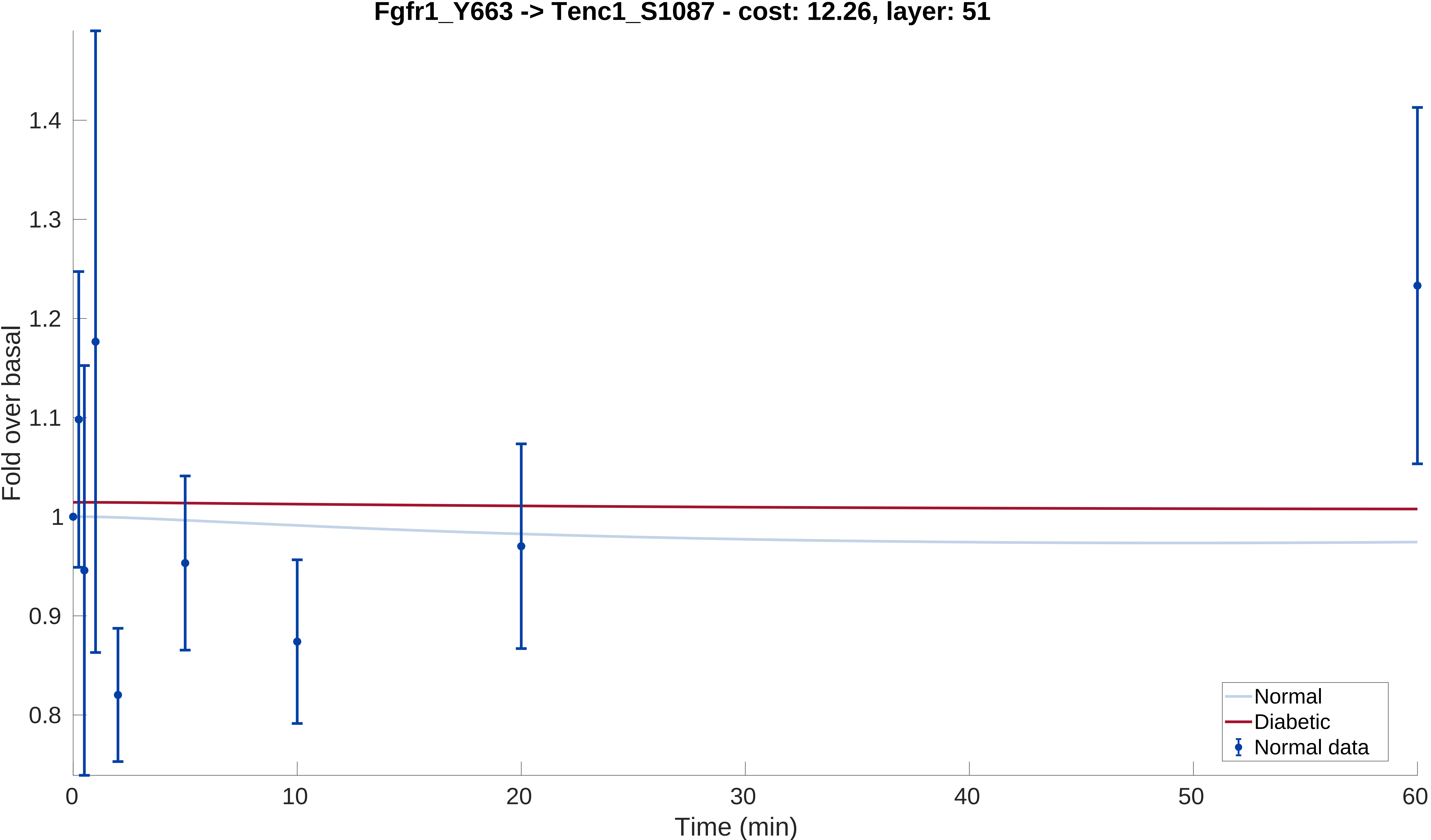

Fgfr1_Y663 -> Tenc1_S1087 - cost: 12.26, layer: 51
Fold over basal
1.4
1.3
1.2
1.1
1
0.9
0.8
0
10
20
30
40
50
60
Time (min)
Normal
Diabetic
Normal data

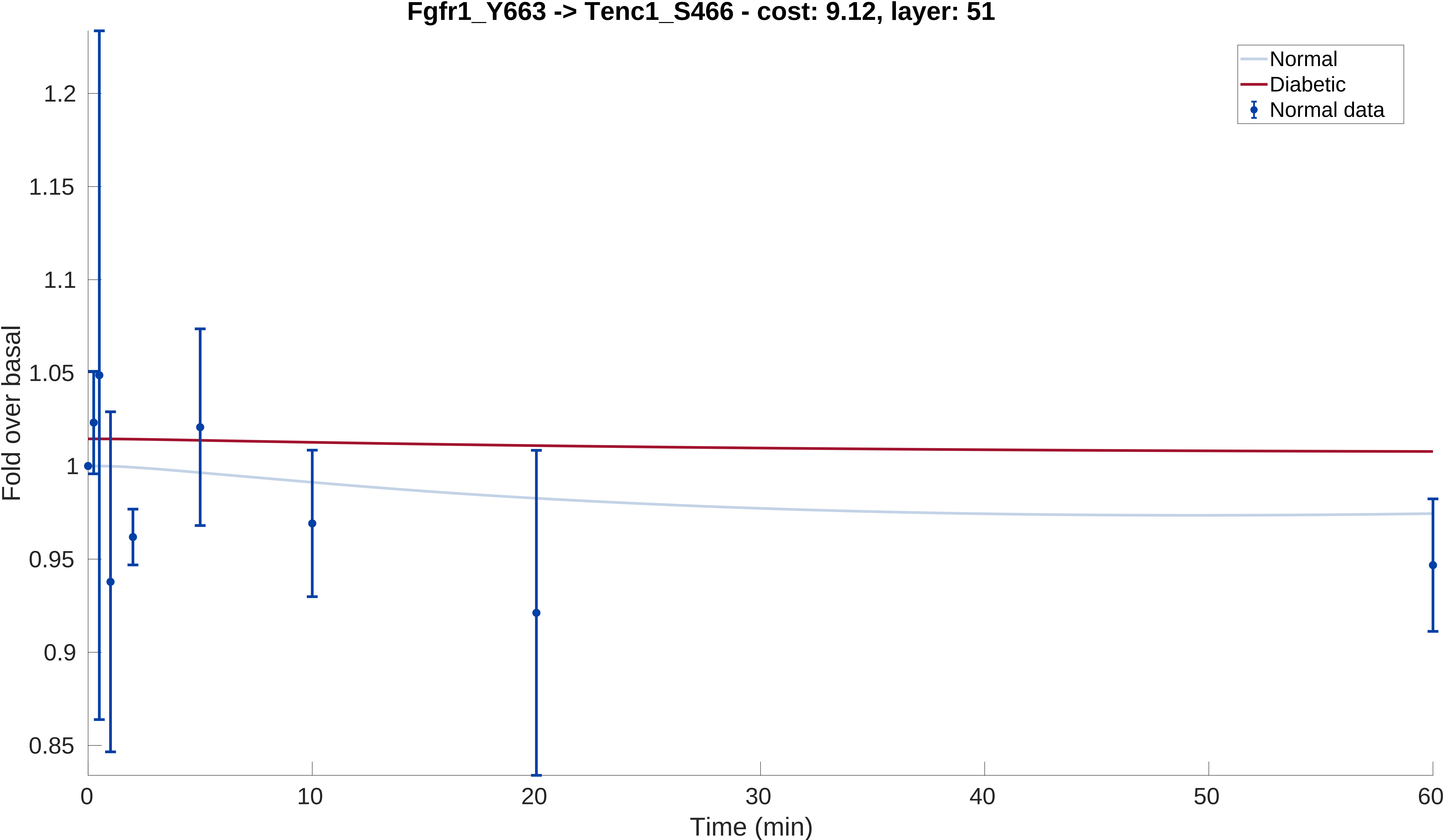

Fgfr1_Y663 -> Tenc1_S466 - cost: 9.12, layer: 51
Normal
Diabetic
Normal data
Fold over basal
1.2
1.15
1.1
1.05
1
0.95
0.9
0.85
0
10
20
30
40
50
60
Time (min)

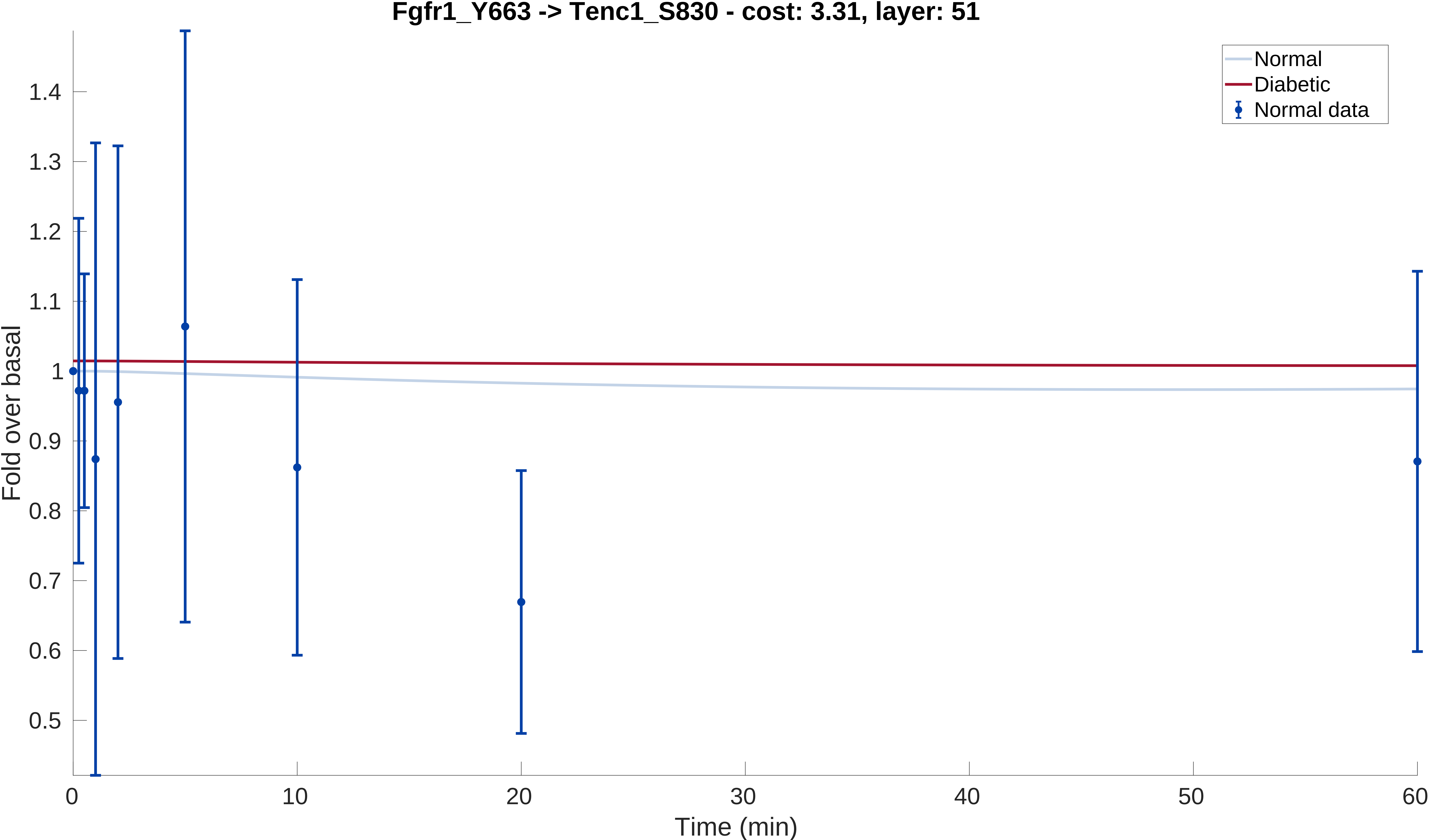

Fgfr1_Y663 -> Tenc1_S830 - cost: 3.31, layer: 51
Normal
Diabetic
Normal data
Fold over basal
Time (min)
1.4
1.3
1.2
1.1
1
0.9
0.8
0.7
0.6
0.5
0
10
20
30
40
50
60

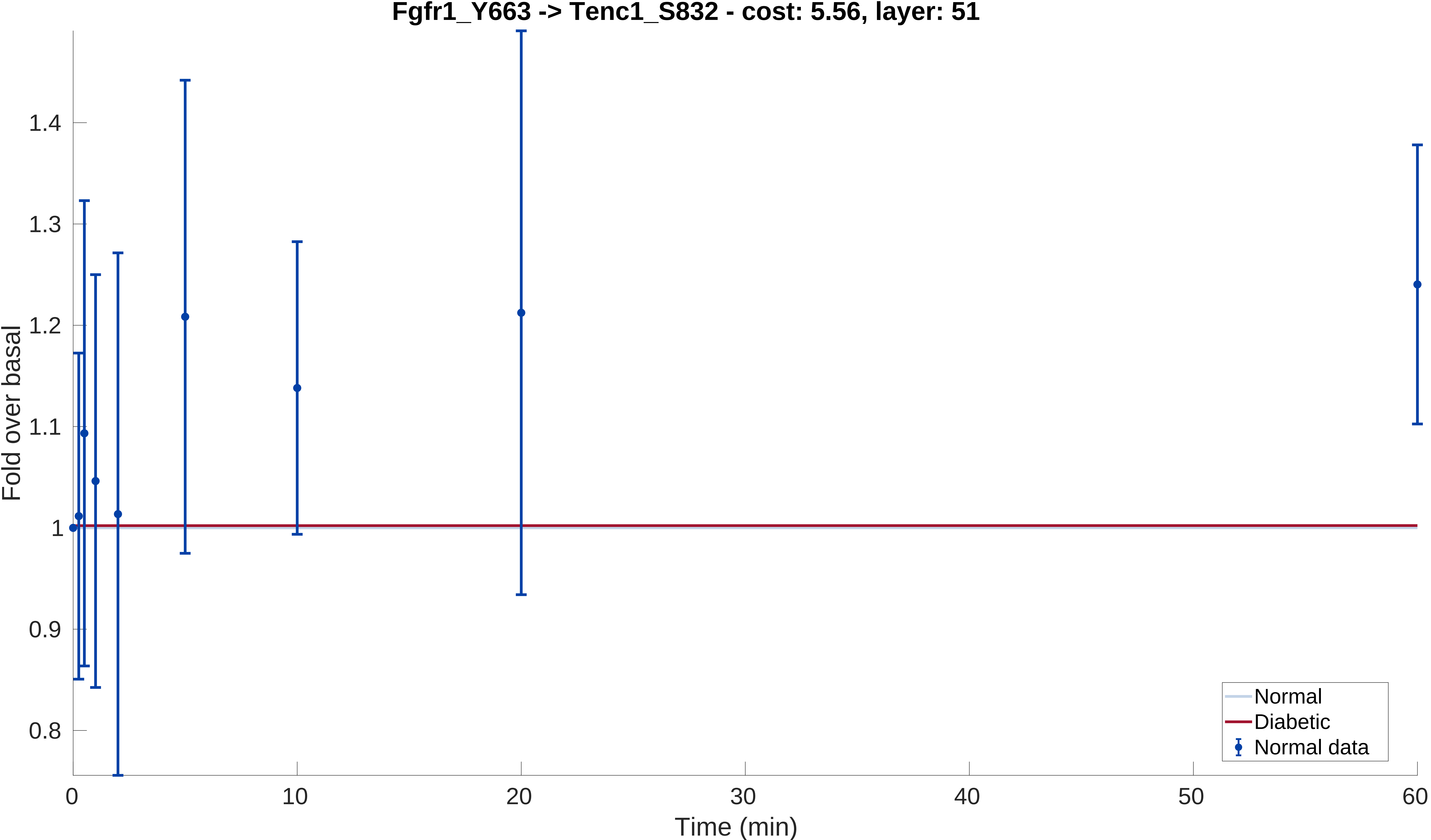

Fgfr1_Y663 -> Tenc1_S832 - cost: 5.56, layer: 51
Fold over basal
1.4
1.3
1.2
1.1
1
0.9
0.8
0
10
20
30
40
50
60
Time (min)
Normal
Diabetic
Normal data

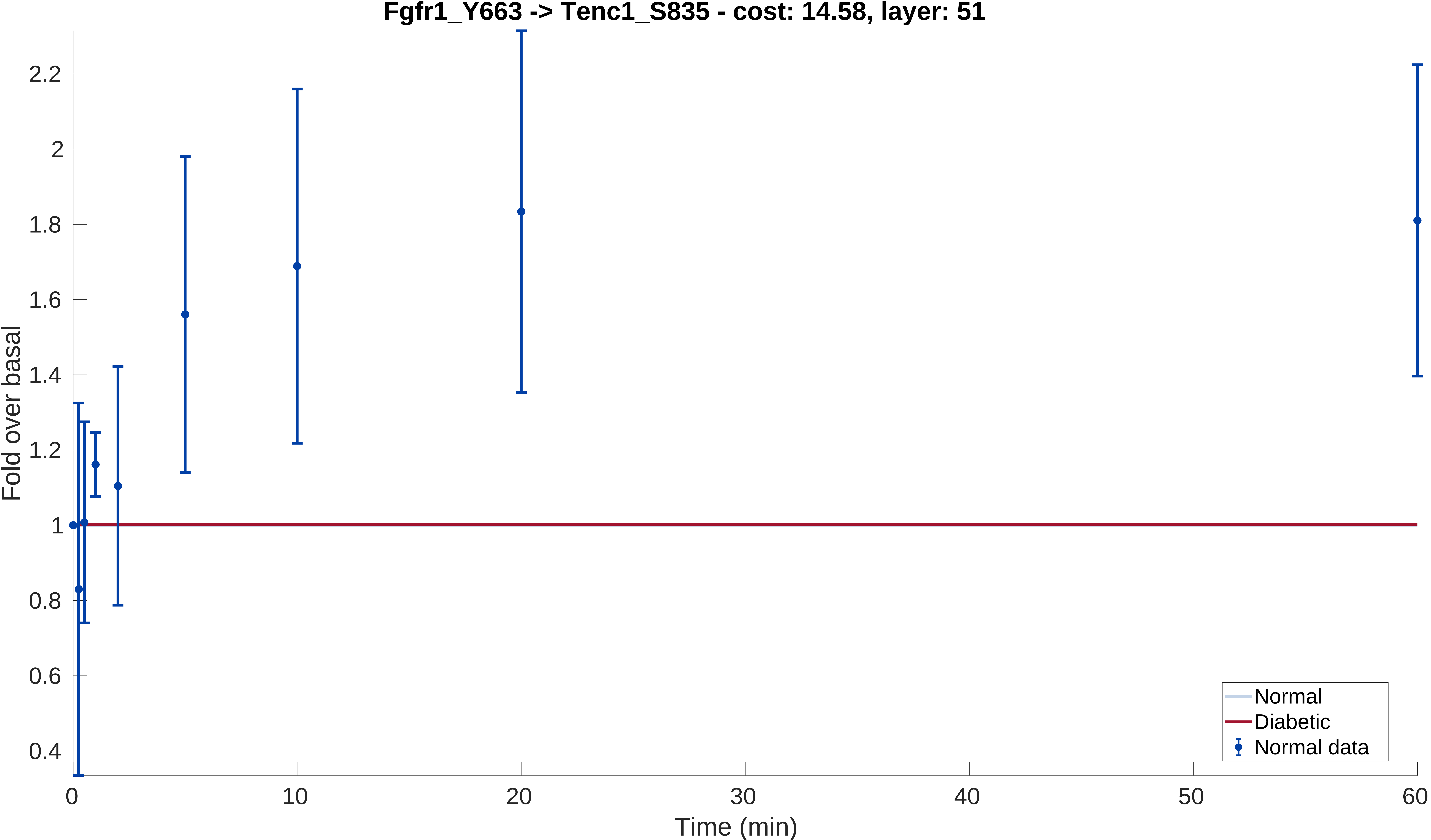

Fgfr1_Y663 -> Tenc1_S835 - cost: 14.58, layer: 51
Fold over basal
2.2
2
1.8
1.6
1.4
1.2
1
0.8
0.6
0.4
0
10
20
30
40
50
60
Time (min)
Normal
Diabetic
Normal data

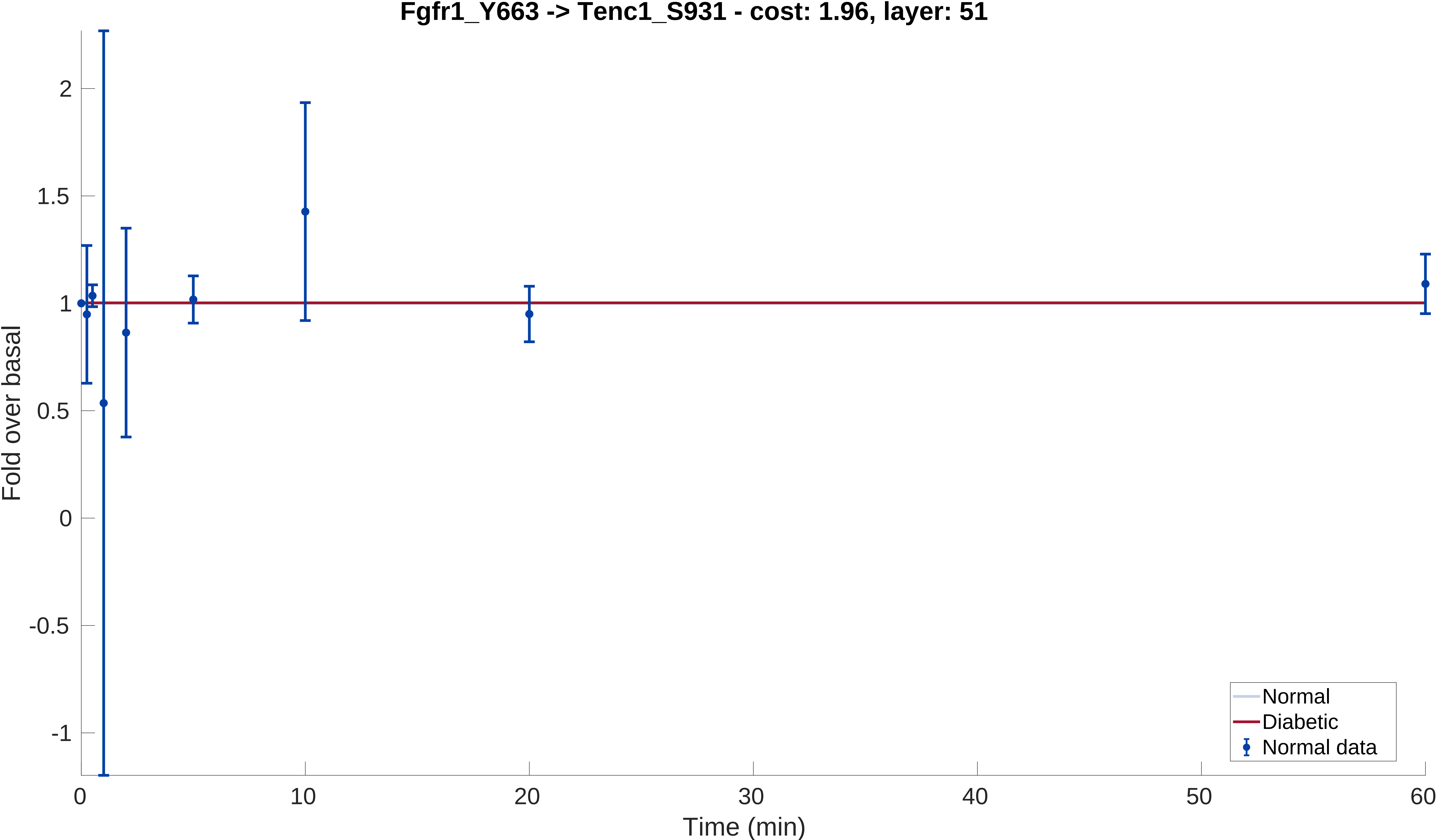

Fgfr1_Y663 -> Tenc1_S931 - cost: 1.96, layer: 51
Fold over basal
2
1.5
1
0.5
0
-0.5
-1
0
10
20
30
40
50
60
Time (min)
Normal
Diabetic
Normal data

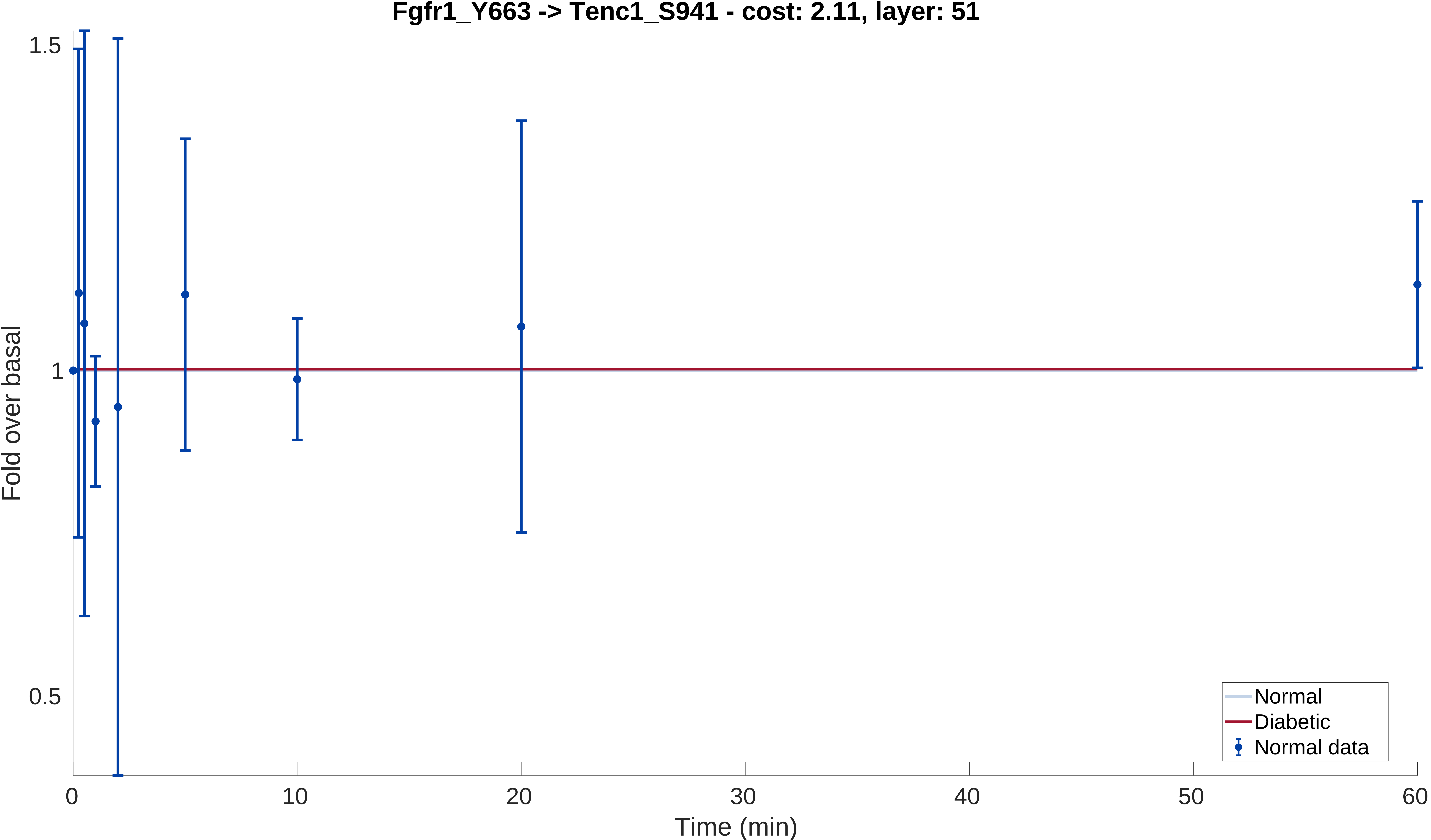

Fgfr1_Y663 -> Tenc1_S941 - cost: 2.11, layer: 51
1.5
1
0.5
Fold over basal
0
10
20
30
40
50
60
Time (min)
Normal
Diabetic
Normal data

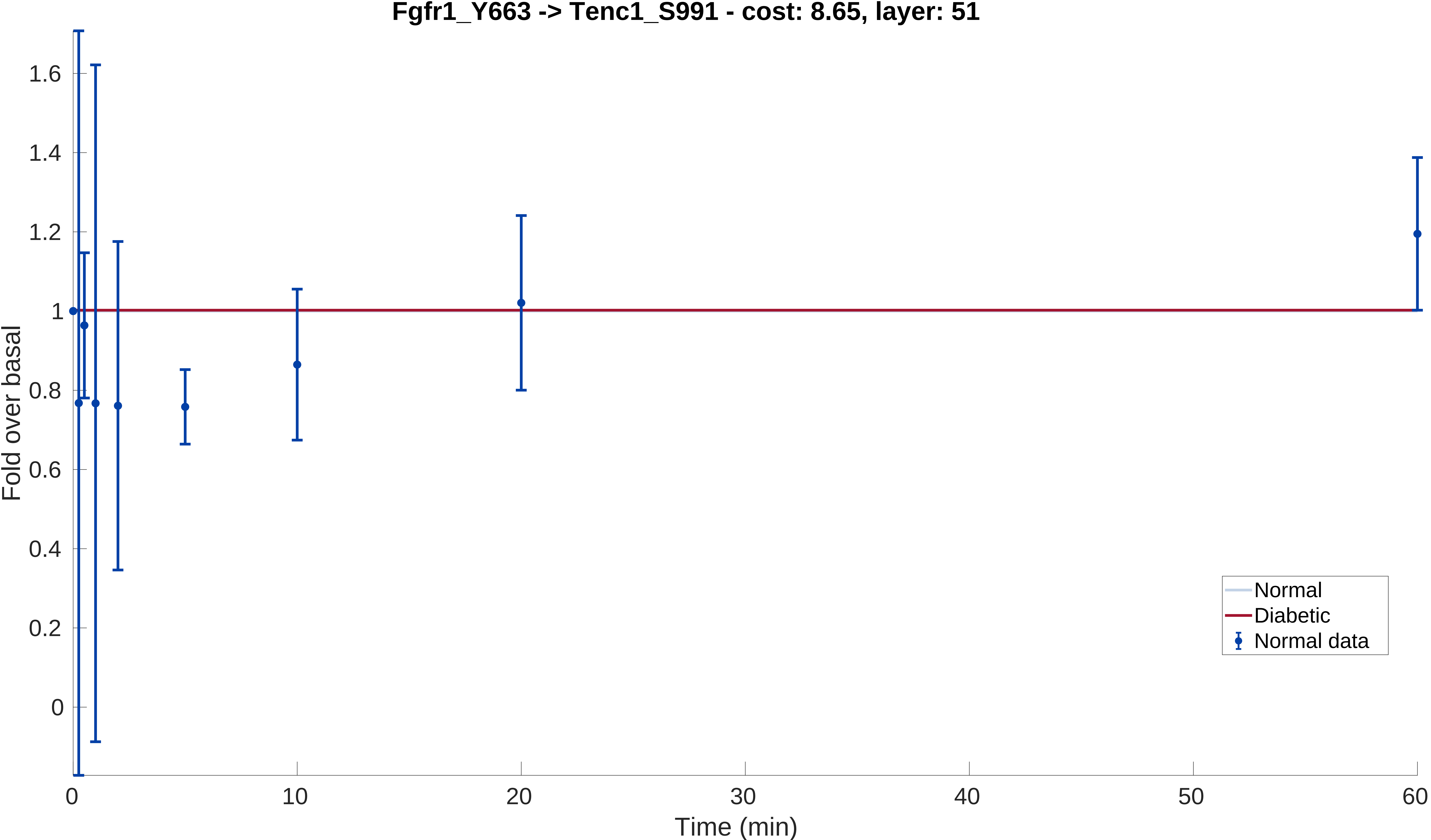
Fgfr1_Y663 -> Tenc1_S991 - cost: 8.65, layer: 51
Fold over basal
1.6
1.4
1.2
1
0.8
0.6
0.4
0.2
0
0
10
20
30
40
50
60
Time (min)
Normal
Diabetic
Normal data

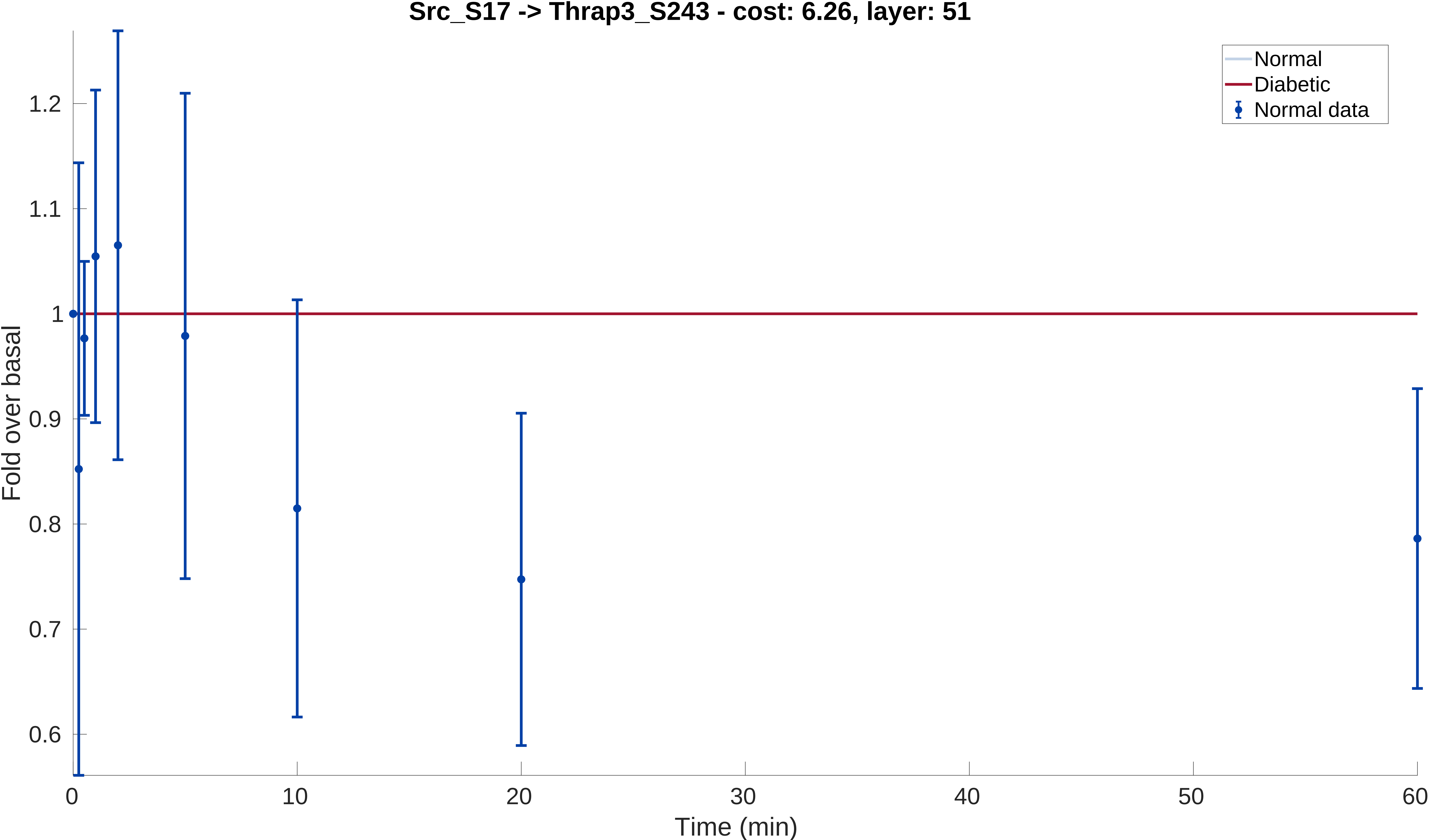

Src_S17 -> Thrap3_S243 - cost: 6.26, layer: 51
Normal
Diabetic
Normal data
Fold over basal
1.2
1.1
1
0.9
0.8
0.7
0.6
0
10
20
30
40
50
60
Time (min)

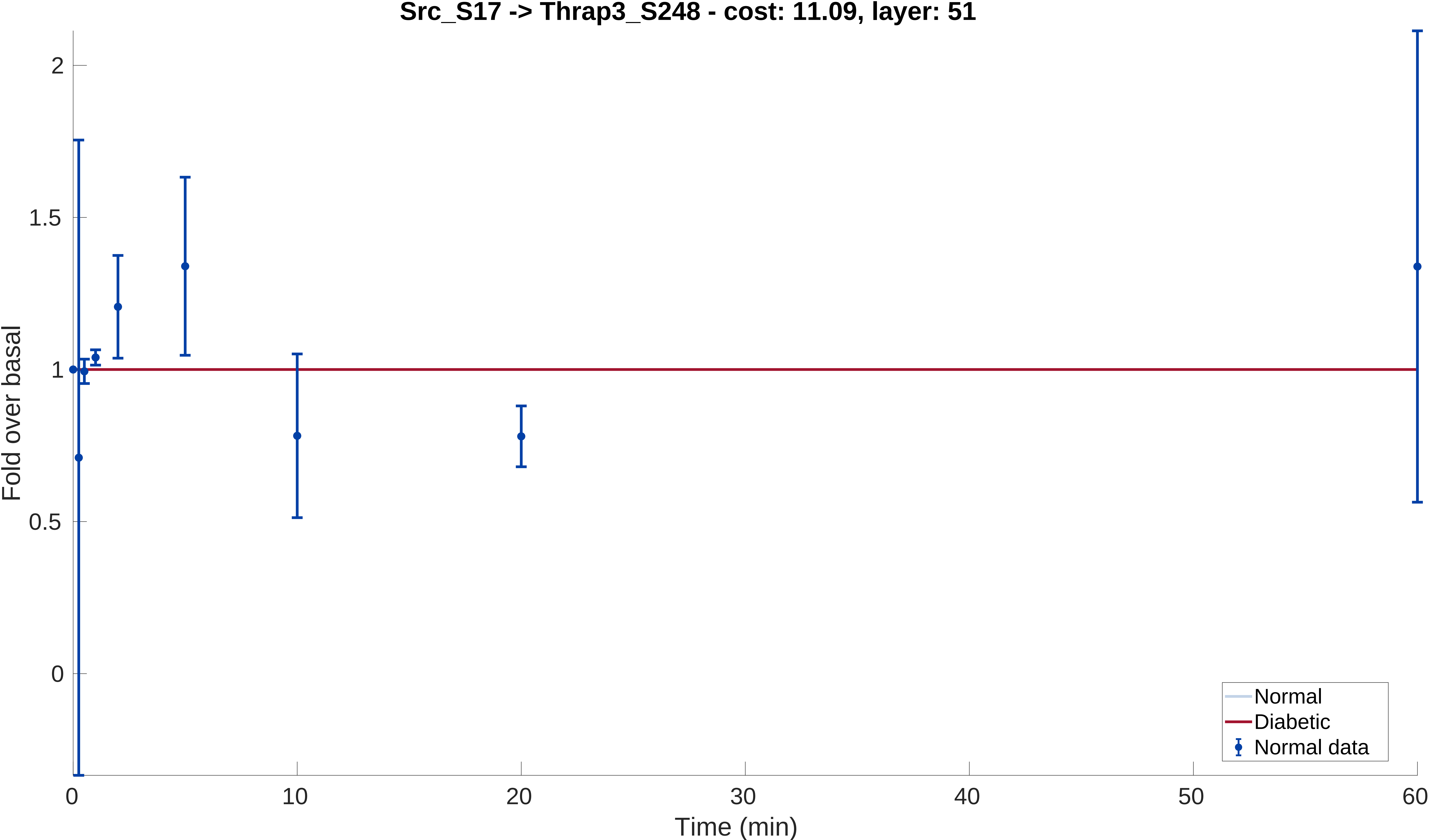

Src_S17 -> Thrap3_S248 - cost: 11.09, layer: 51
Fold over basal
2
1.5
1
0.5
0
0
10
20
30
40
50
60
Time (min)
Normal
Diabetic
Normal data

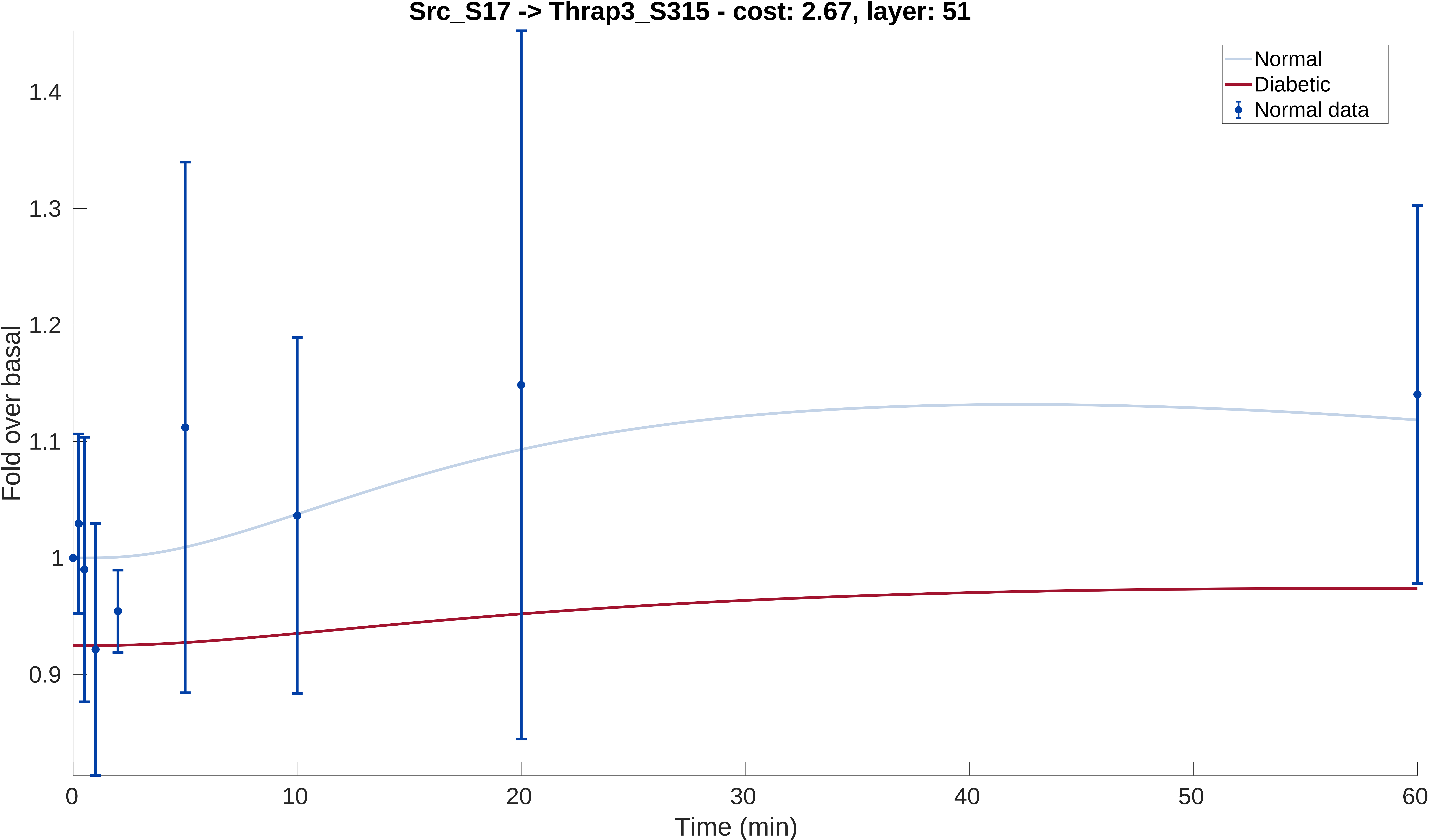
Src_S17 -> Thrap3_S315 - cost: 2.67, layer: 51
Normal
Diabetic
Normal data
1.4
1.3
1.2
1.1
1
0.9
Fold over basal
0
10
20
30
40
50
60
Time (min)

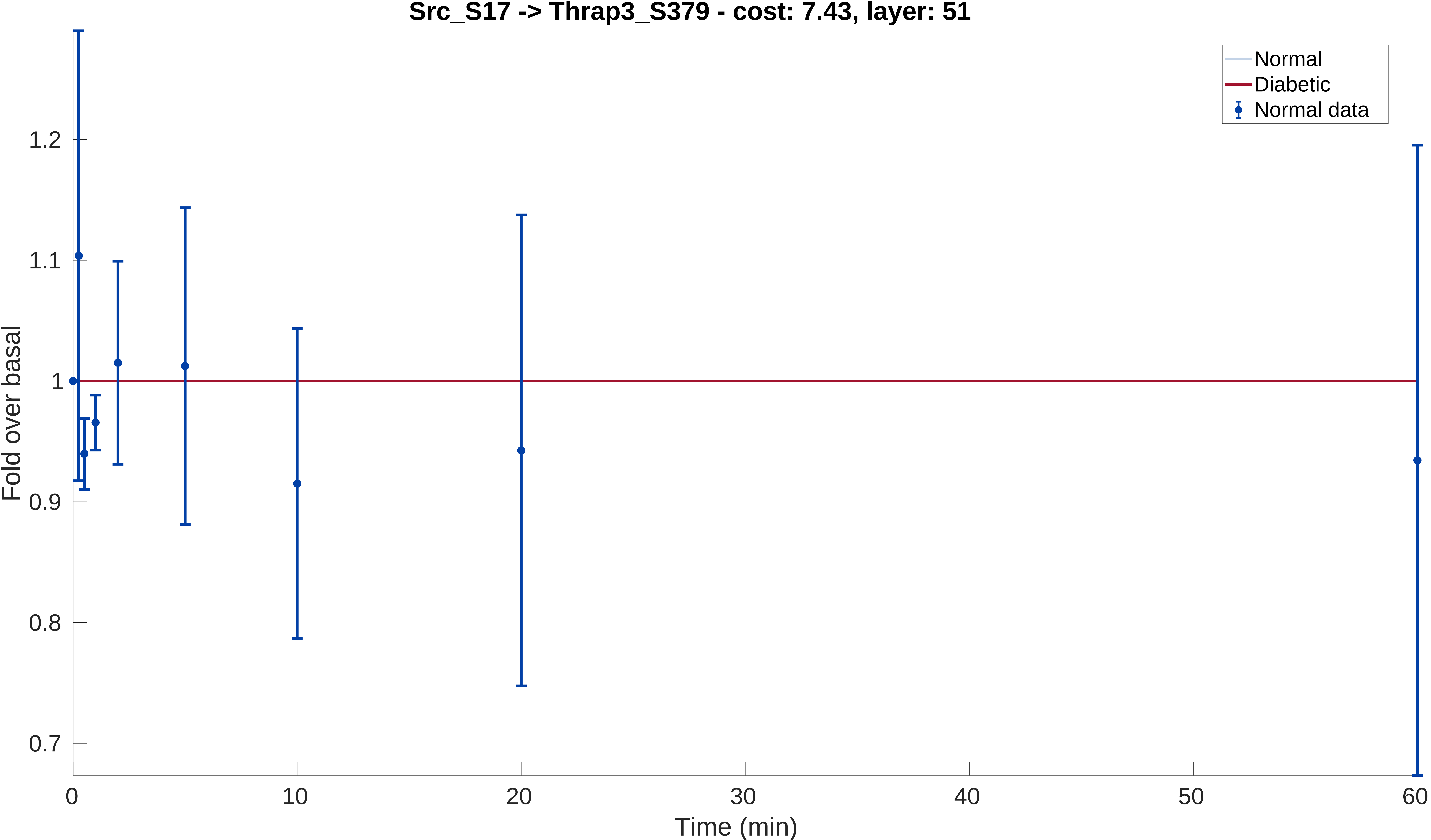

Src_S17 -> Thrap3_S379 - cost: 7.43, layer: 51
Normal
Diabetic
Normal data
Fold over basal
1.2
1.1
1
0.9
0.8
0.7
0
10
20
30
40
50
60
Time (min)

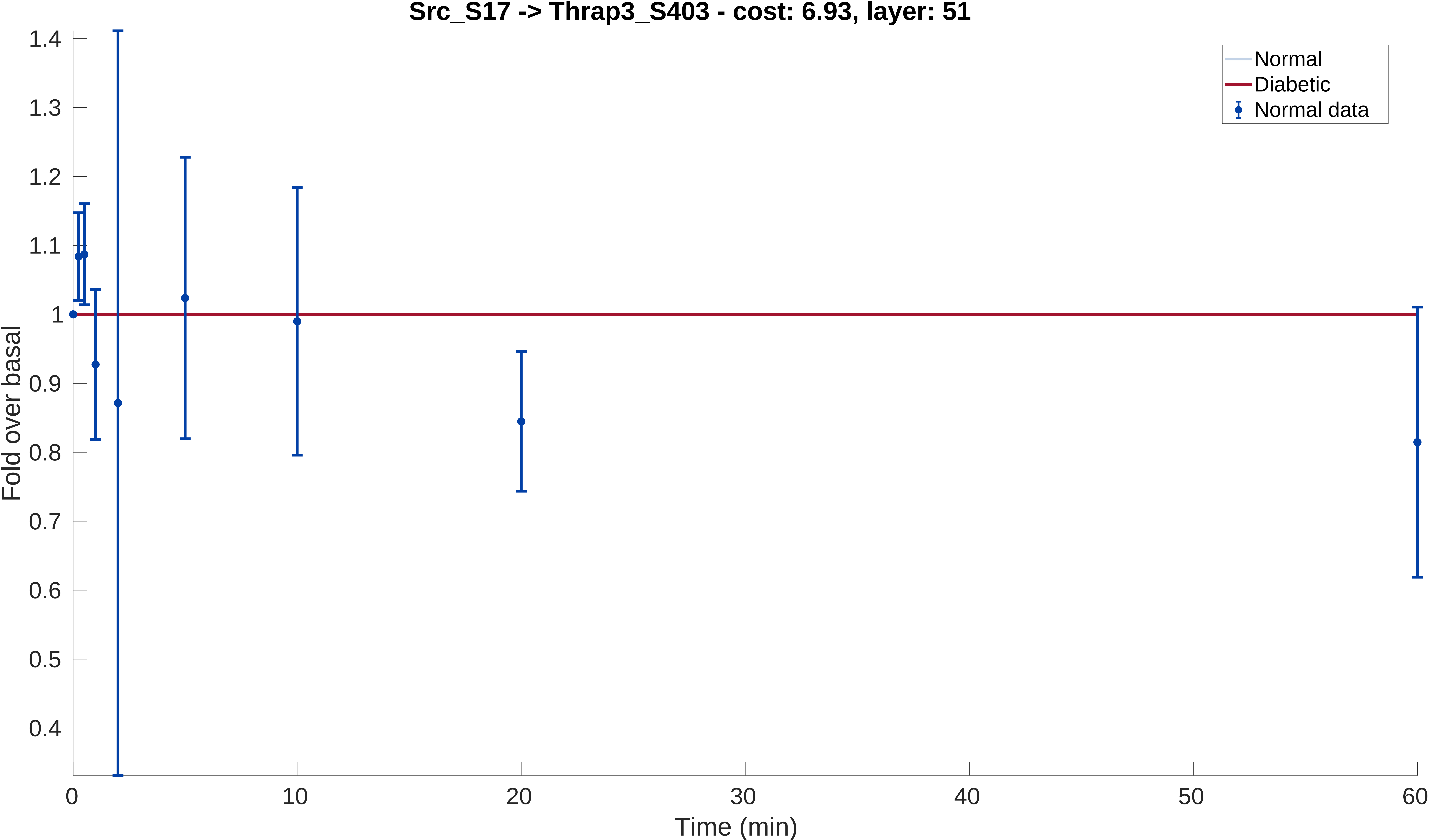

Src_S17 -> Thrap3_S403 - cost: 6.93, layer: 51
Normal
Diabetic
Normal data
Fold over basal
1.4
1.3
1.2
1.1
1
0.9
0.8
0.7
0.6
0.5
0.4
0
10
20
30
40
50
60
Time (min)

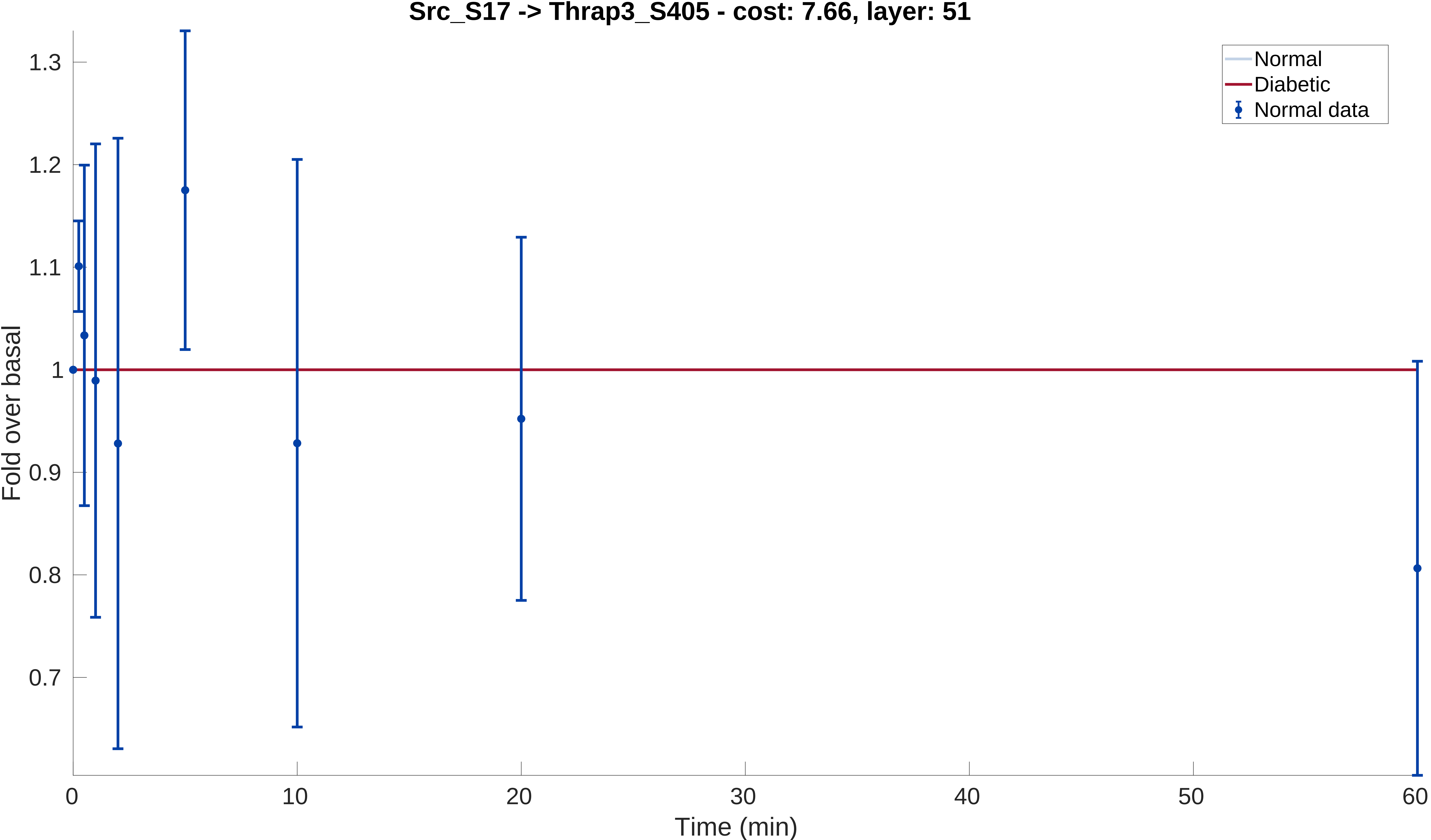

Src_S17 -> Thrap3_S405 - cost: 7.66, layer: 51
Normal
Diabetic
Normal data
Fold over basal
1.3
1.2
1.1
1
0.9
0.8
0.7
0
10
20
30
40
50
60
Time (min)

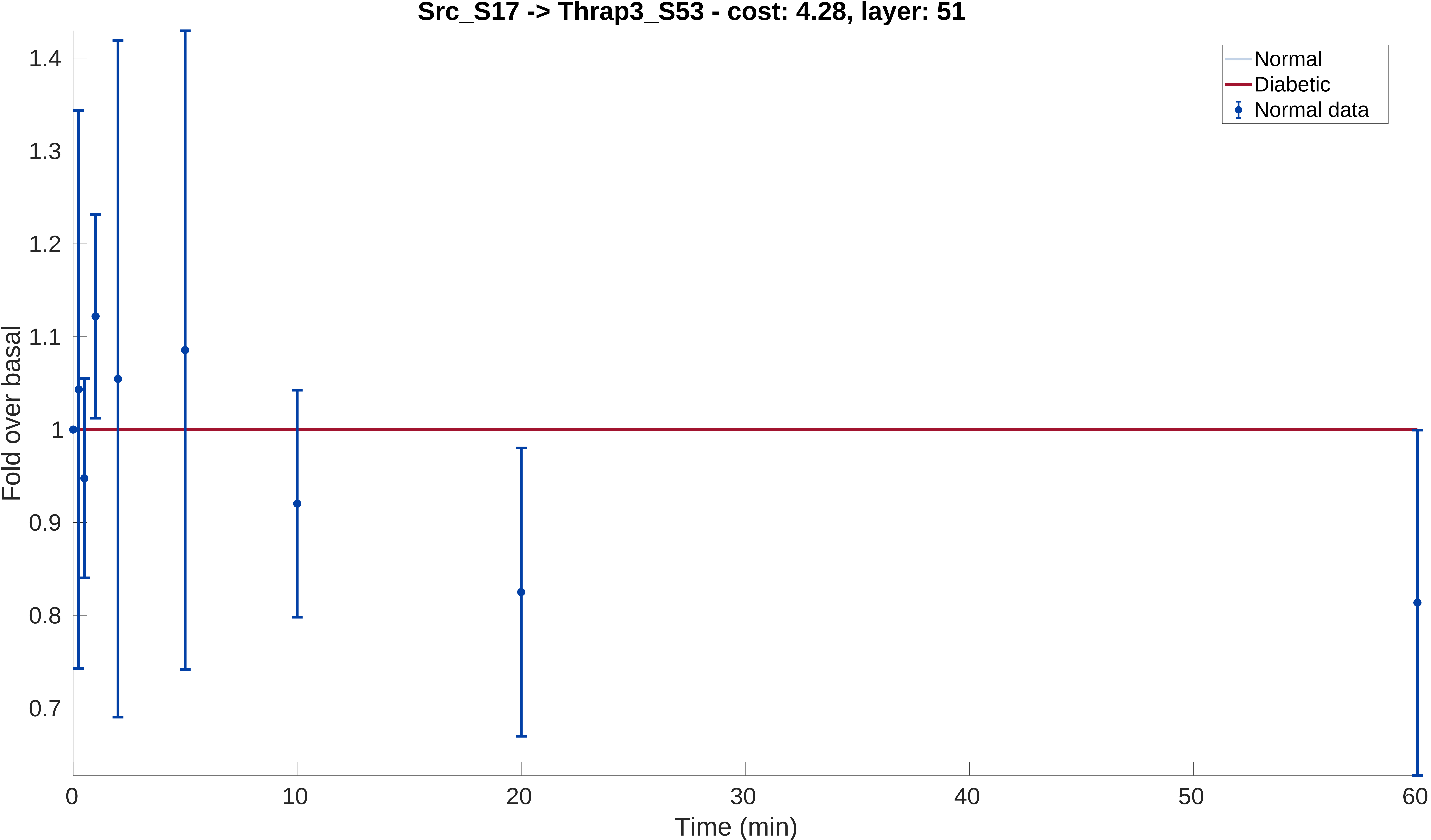

Src_S17 -> Thrap3_S53 - cost: 4.28, layer: 51
Normal
Diabetic
Normal data
Fold over basal
1.4
1.3
1.2
1.1
1
0.9
0.8
0.7
0
10
20
30
40
50
60
Time (min)

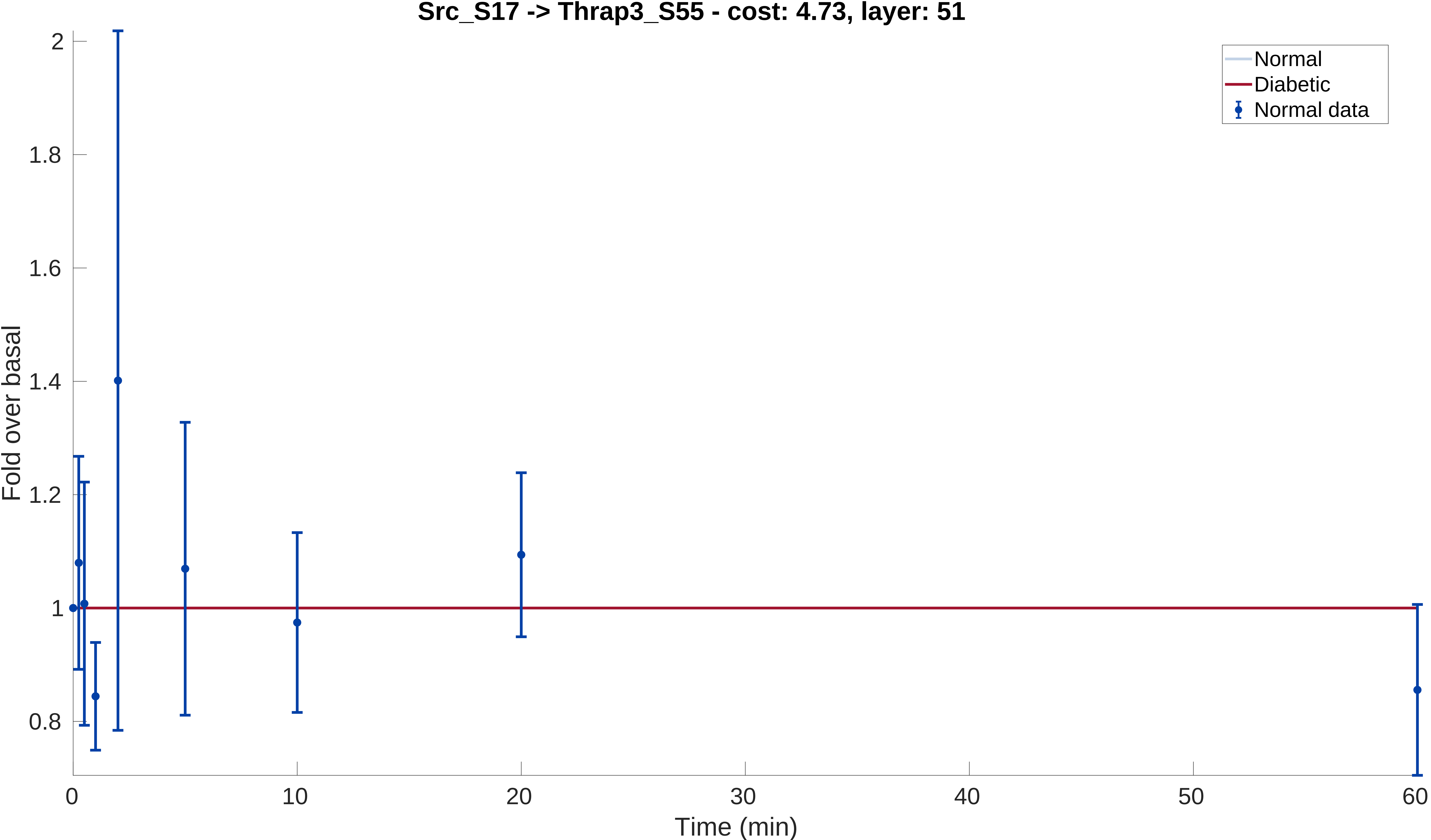
Src_S17 -> Thrap3_S55 - cost: 4.73, layer: 51
Normal
Diabetic
Normal data
Fold over basal
2
1.8
1.6
1.4
1.2
1
0.8
0
10
20
30
40
50
60
Time (min)

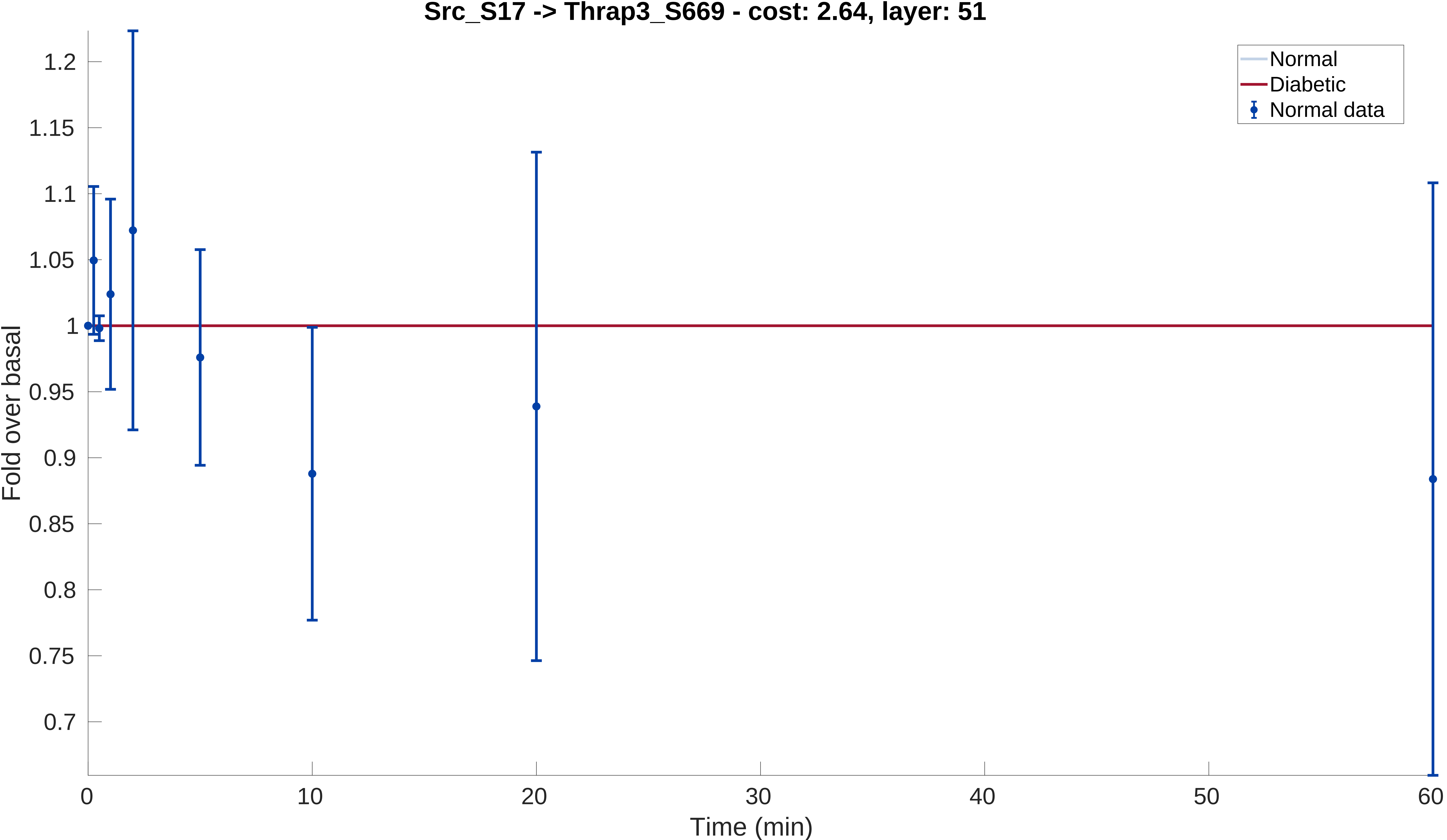

Src_S17 -> Thrap3_S669 - cost: 2.64, layer: 51
Normal
Diabetic
Normal data
1.2
1.15
1.1
1.05
1
0.95
0.9
0.85
0.8
0.75
0.7
Fold over basal
0
10
20
30
40
50
60
Time (min)

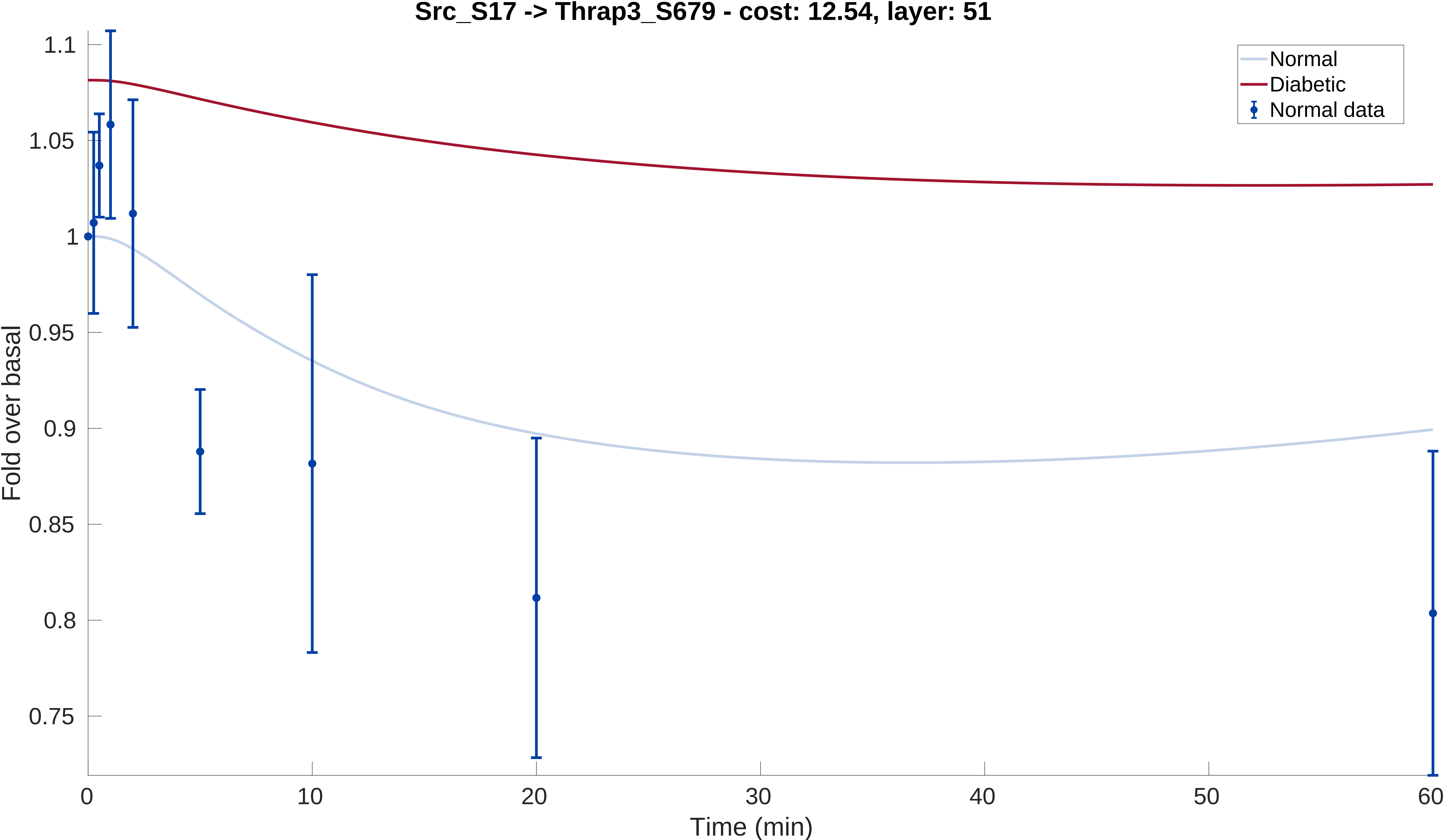
Src_S17 -> Thrap3_S679 - cost: 12.54, layer: 51
Normal
Diabetic
Normal data
Fold over basal
Time (min)
1.1
1.05
1
0.95
0.9
0.85
0.8
0.75
0
10
20
30
40
50
60

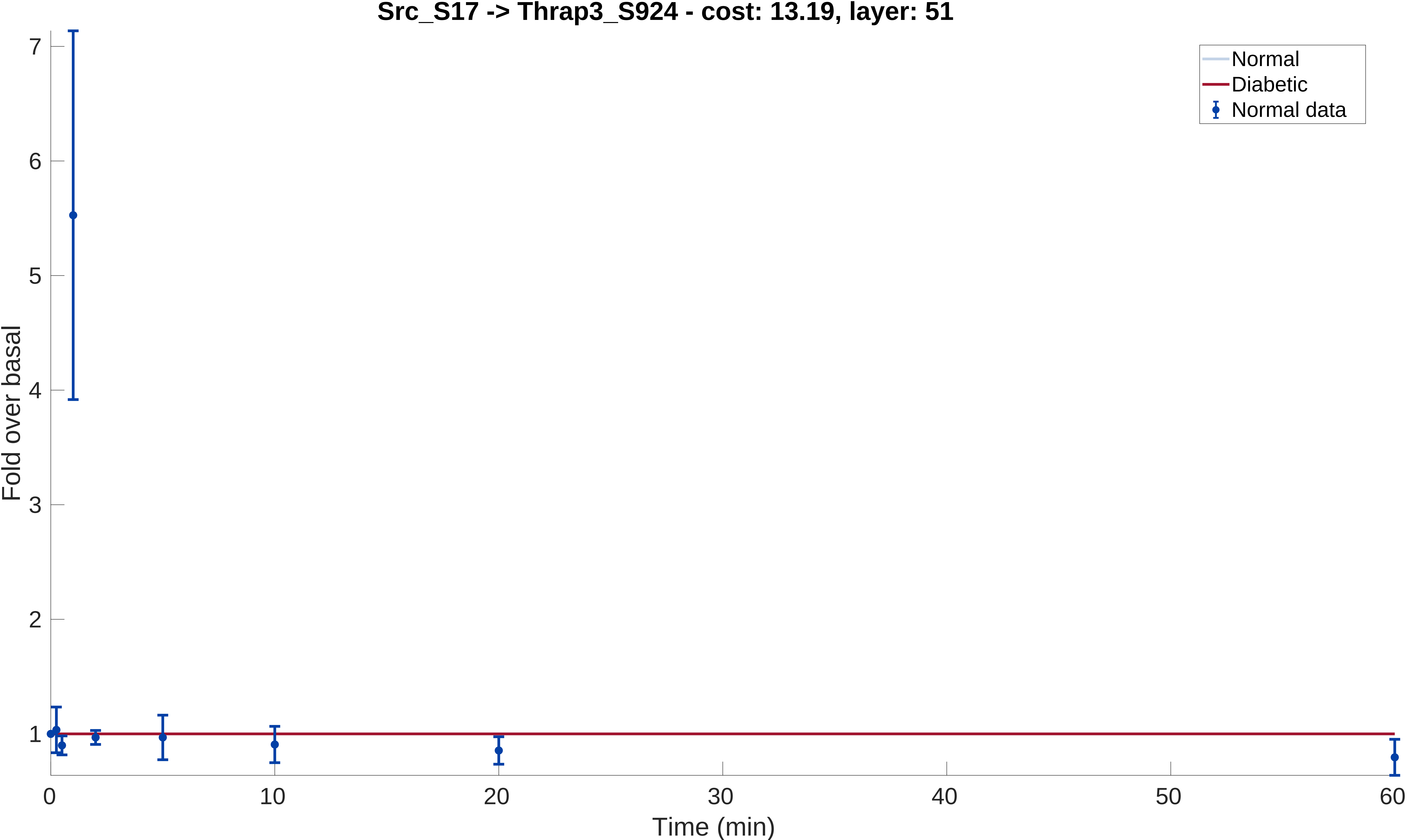

Src_S17 -> Thrap3_S924 - cost: 13.19, layer: 51
Normal
Diabetic
Normal data
Fold over basal
Time (min)
1
2
3
4
5
6
7
0
10
20
30
40
50
60

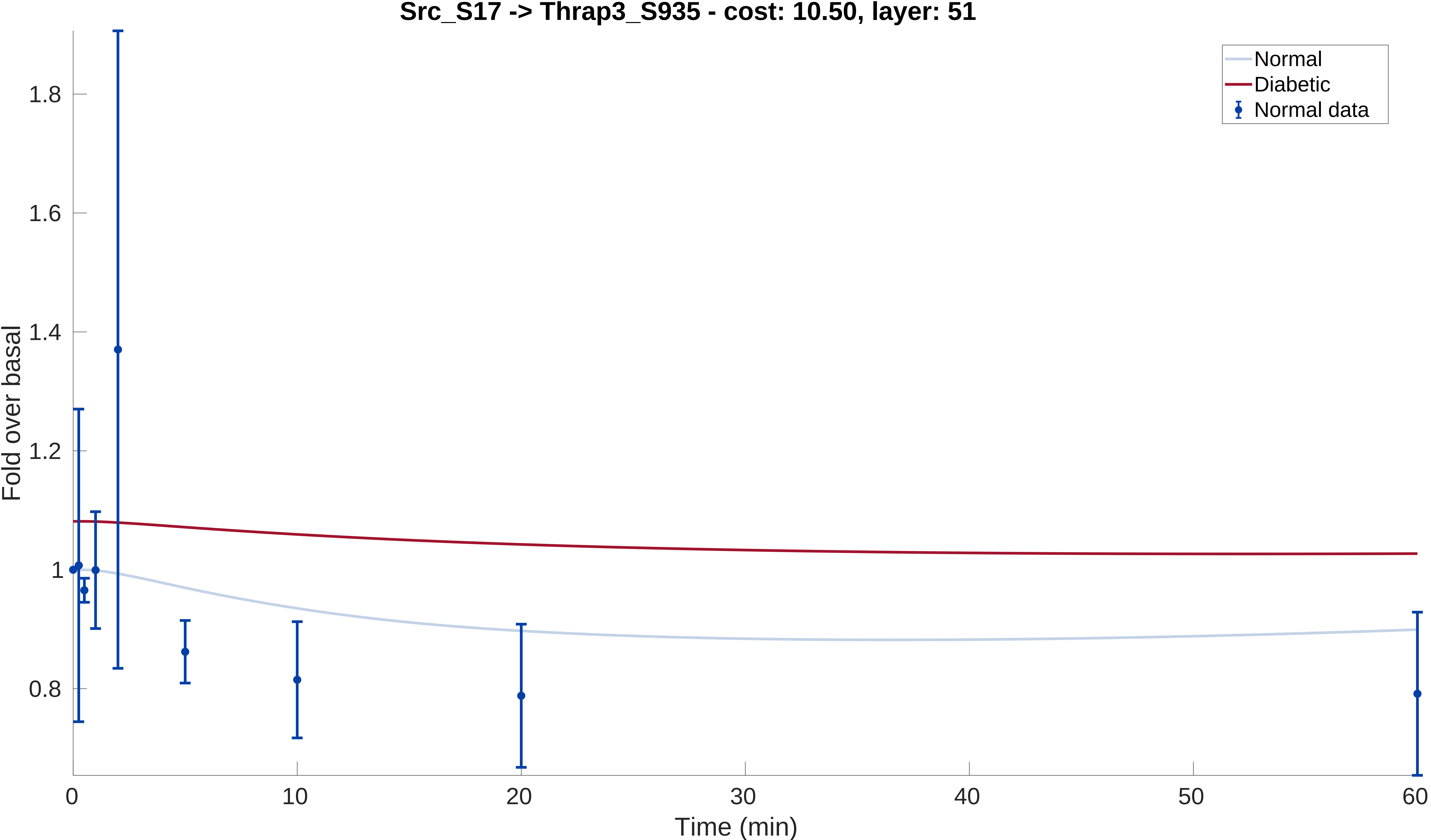
Src_S17 -> Thrap3_S935 - cost: 10.50, layer: 51
Normal
Diabetic
Normal data
Fold over basal
1.8
1.6
1.4
1.2
1
0.8
0
10
20
30
40
50
60
Time (min)

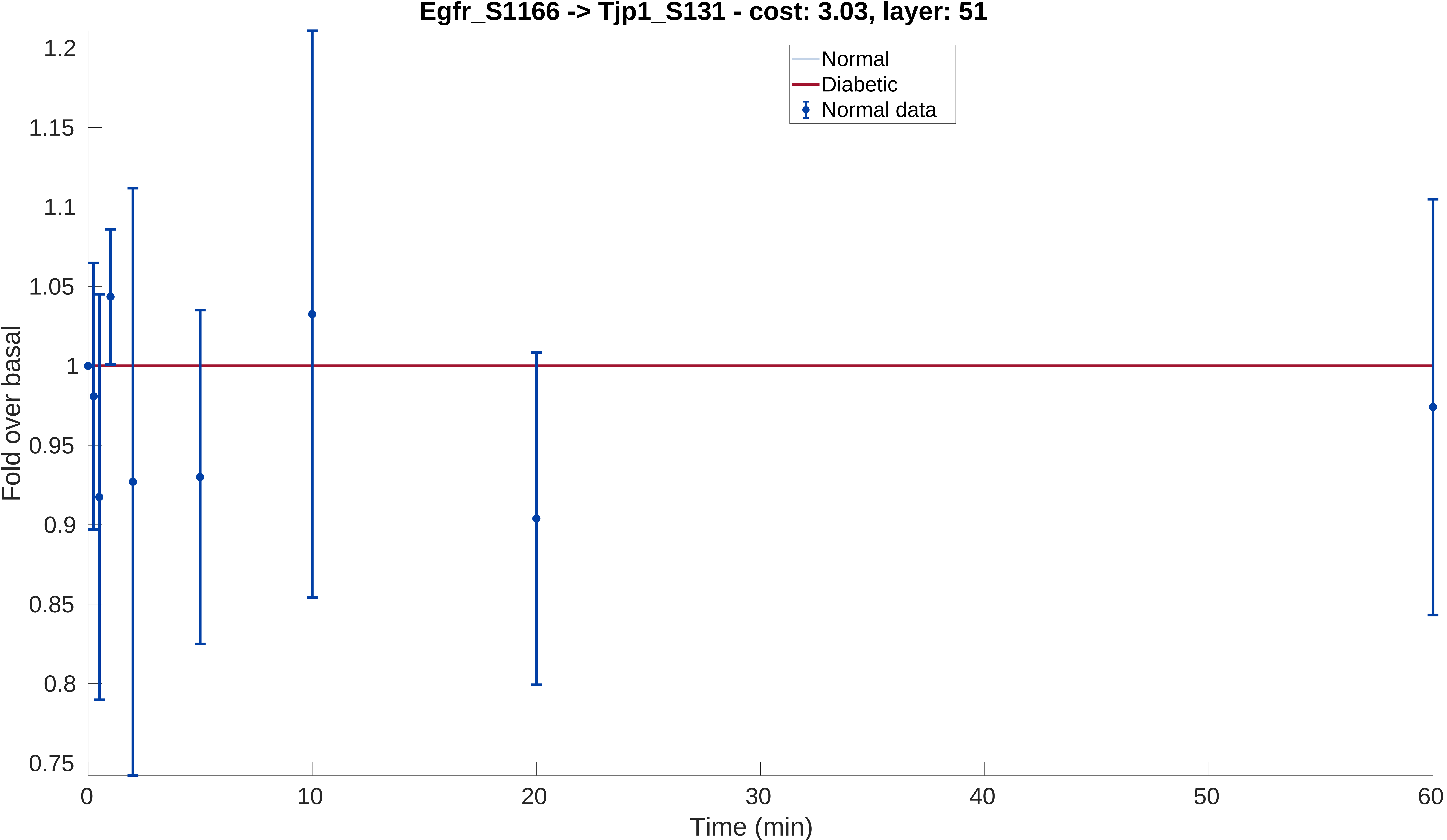

Egfr_S1166 -> Tjp1_S131 - cost: 3.03, layer: 51
Normal
Diabetic
Normal data
Fold over basal
1.2
1.15
1.1
1.05
1
0.95
0.9
0.85
0.8
0.75
0
10
20
30
40
50
60
Time (min)

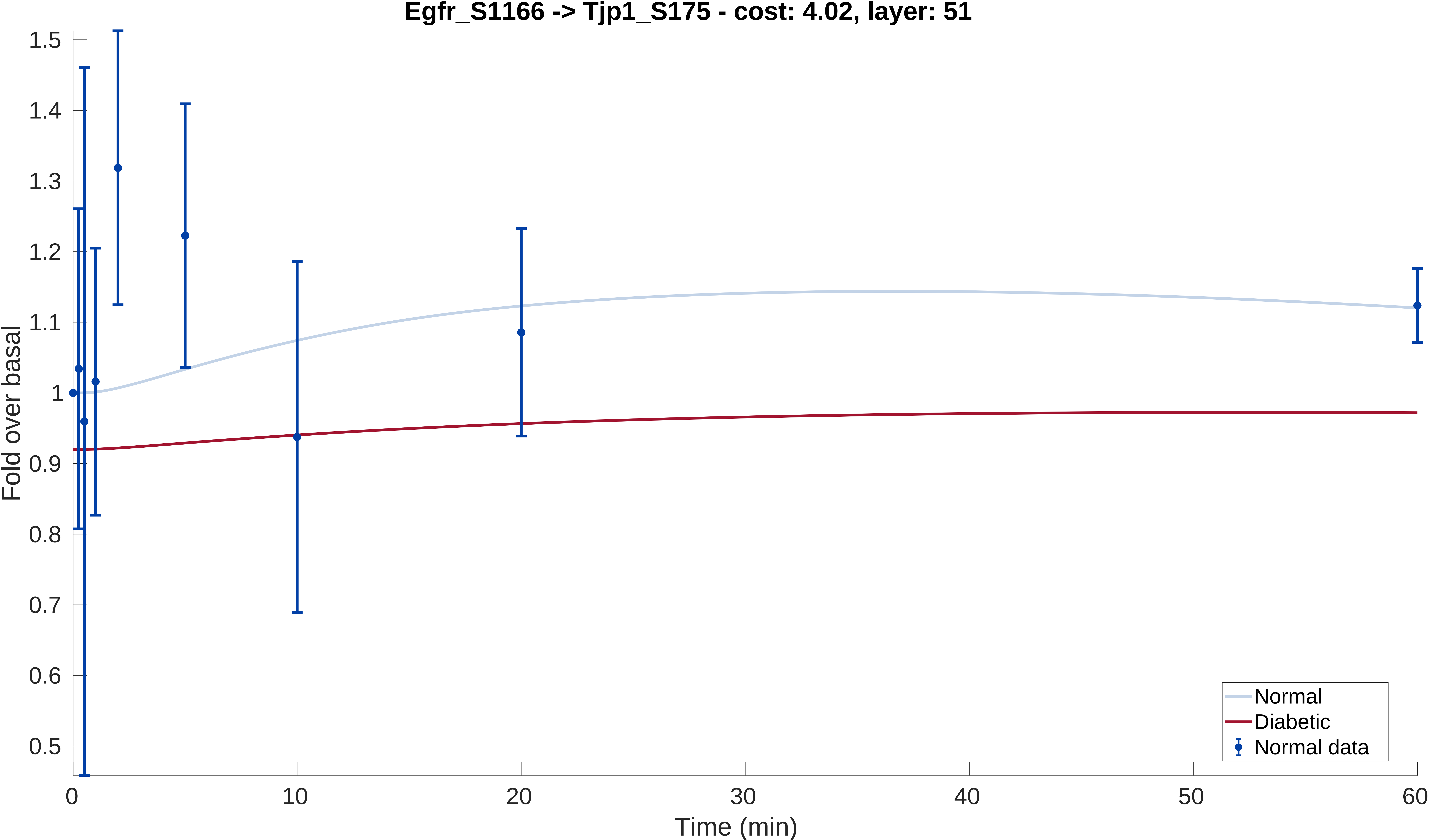

Egfr_S1166 -> Tjp1_S175 - cost: 4.02, layer: 51
Fold over basal
1.5
1.4
1.3
1.2
1.1
1
0.9
0.8
0.7
0.6
0.5
0
10
20
30
40
50
60
Time (min)
Normal
Diabetic
Normal data

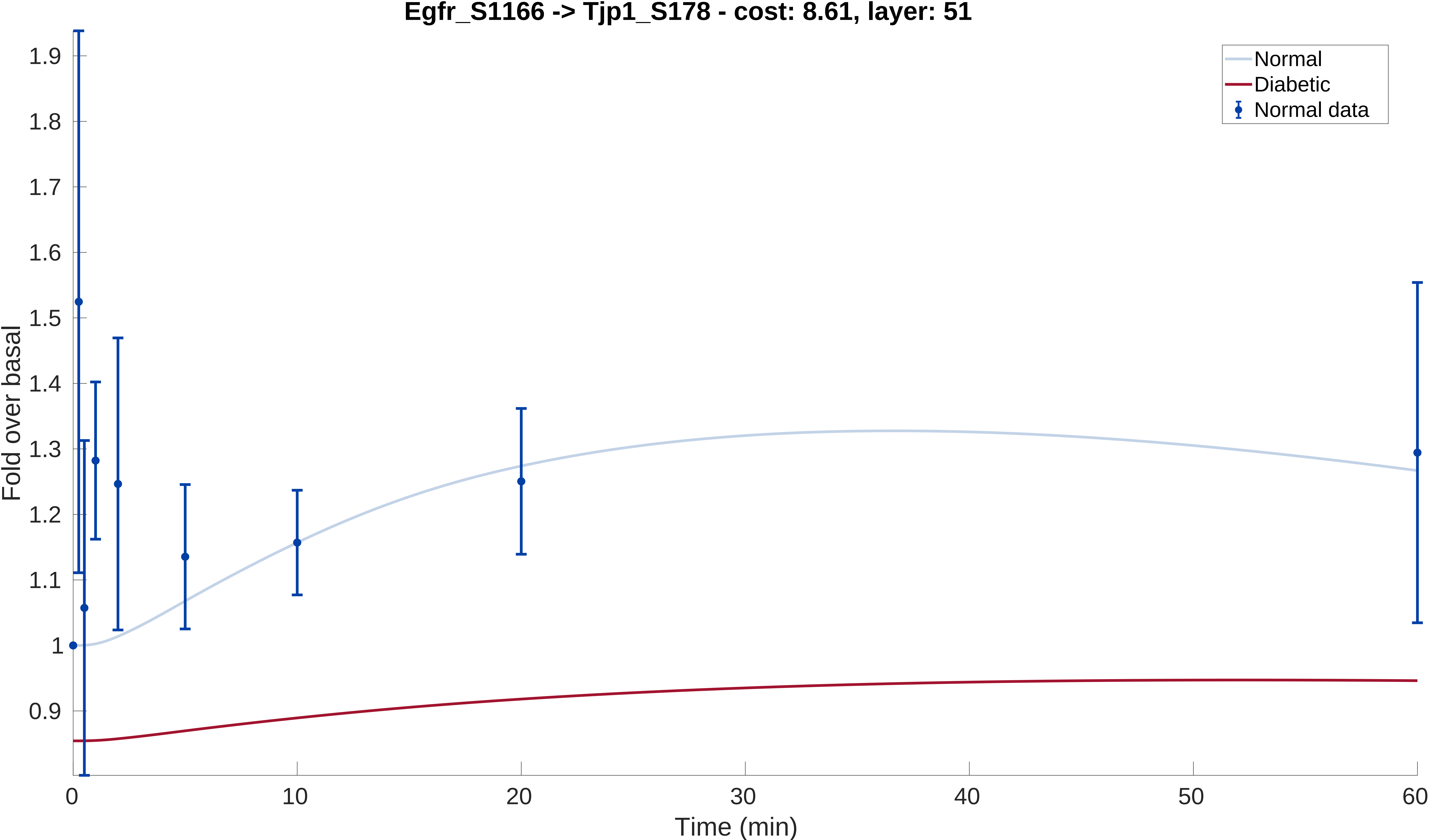

Egfr_S1166 -> Tjp1_S178 - cost: 8.61, layer: 51
Normal
Diabetic
Normal data
Fold over basal
1.9
1.8
1.7
1.6
1.5
1.4
1.3
1.2
1.1
1
0.9
0
10
20
30
40
50
60
Time (min)

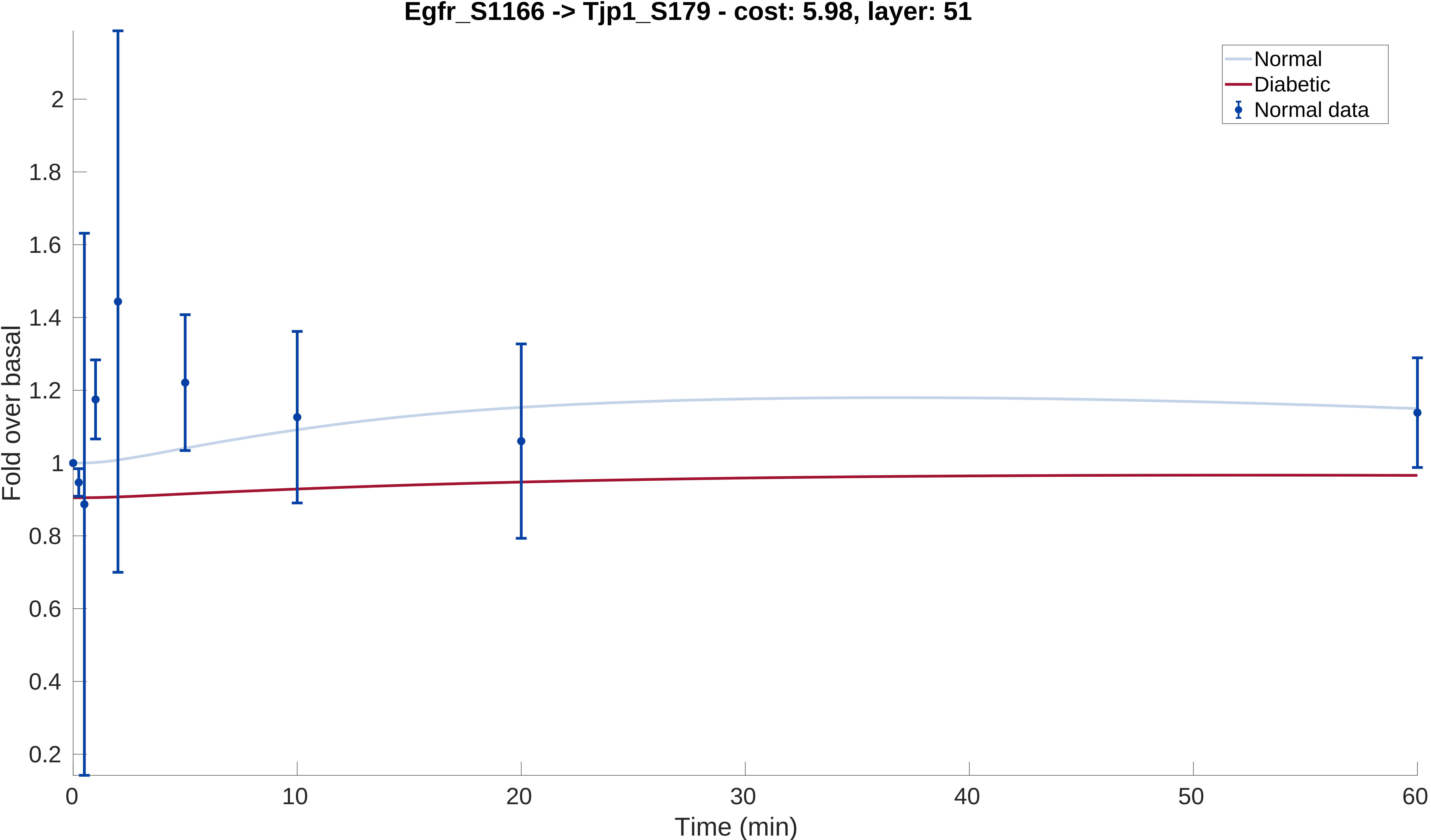
Egfr_S1166 -> Tjp1_S179 - cost: 5.98, layer: 51
Normal
Diabetic
Normal data
Fold over basal
Time (min)
0.2
0.4
0.6
0.8
1
1.2
1.4
1.6
1.8
2
0
10
20
30
40
50
60

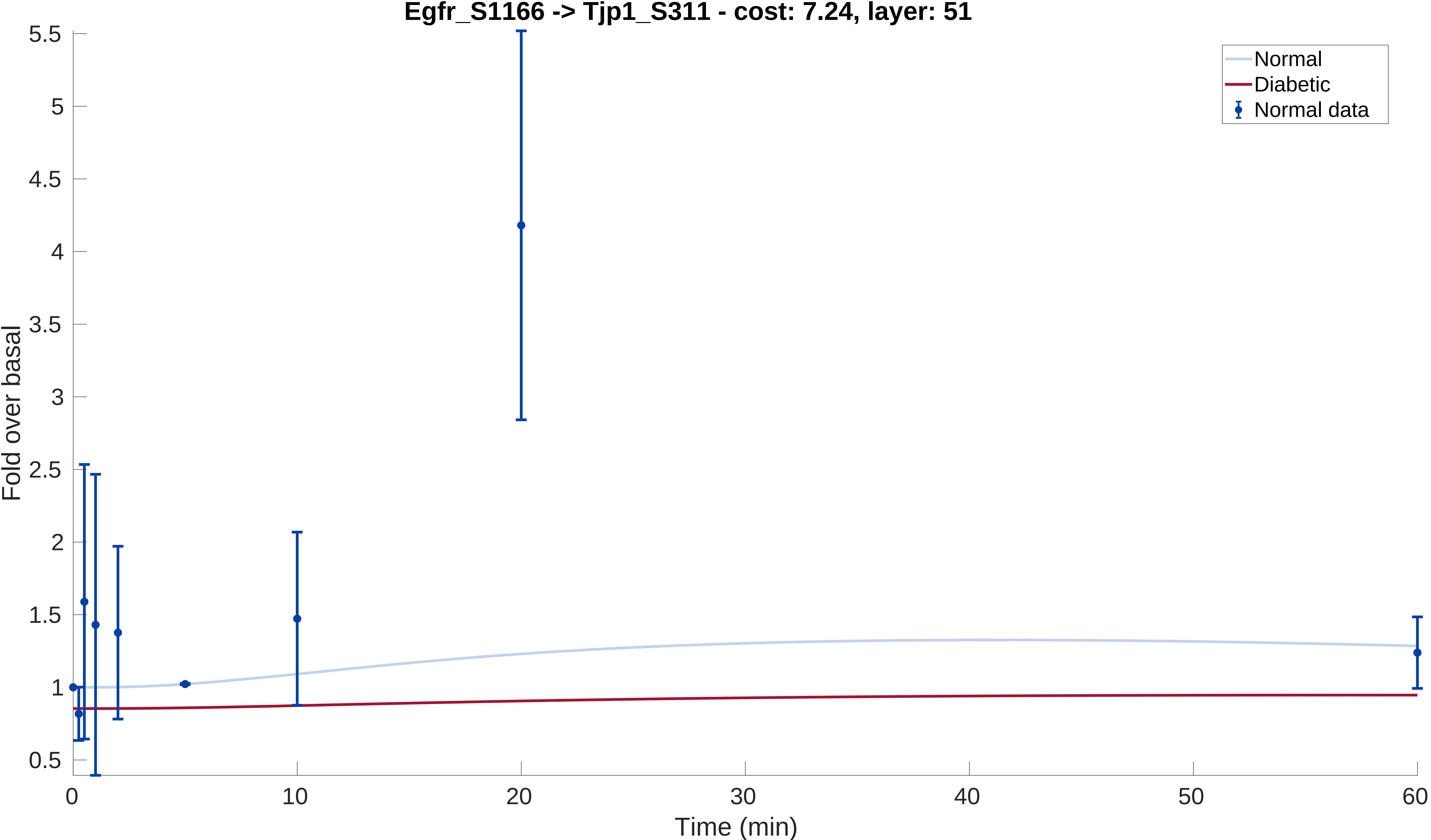
Egfr_S1166 -> Tjp1_S311 - cost: 7.24, layer: 51
Normal
Diabetic
Normal data
Fold over basal
Time (min)
0.5
1
1.5
2
2.5
3
3.5
4
4.5
5
5.5
0
10
20
30
40
50
60

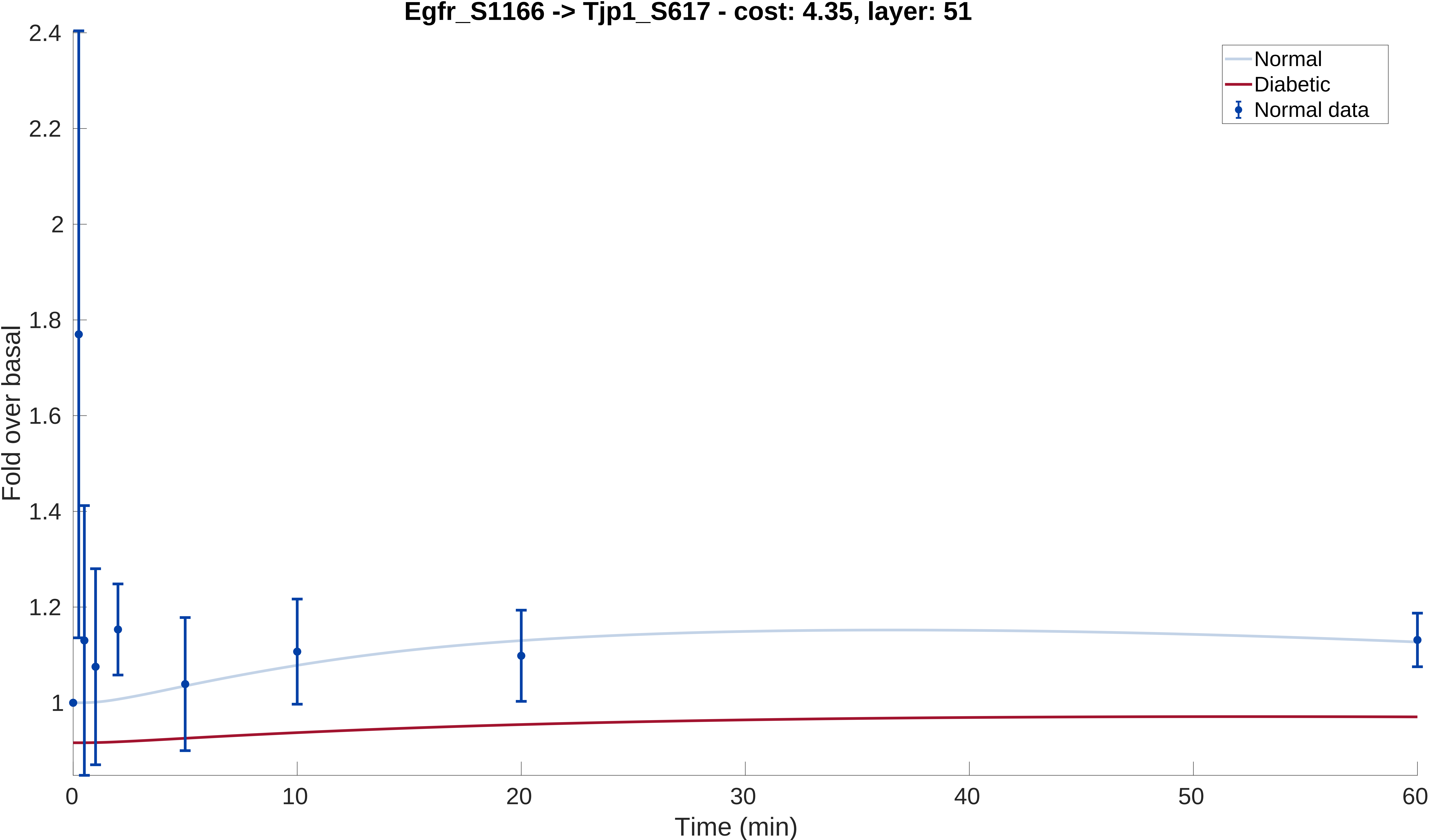

Egfr_S1166 -> Tjp1_S617 - cost: 4.35, layer: 51
Normal
Diabetic
Normal data
Fold over basal
2.4
2.2
2
1.8
1.6
1.4
1.2
1
0
10
20
30
40
50
60
Time (min)

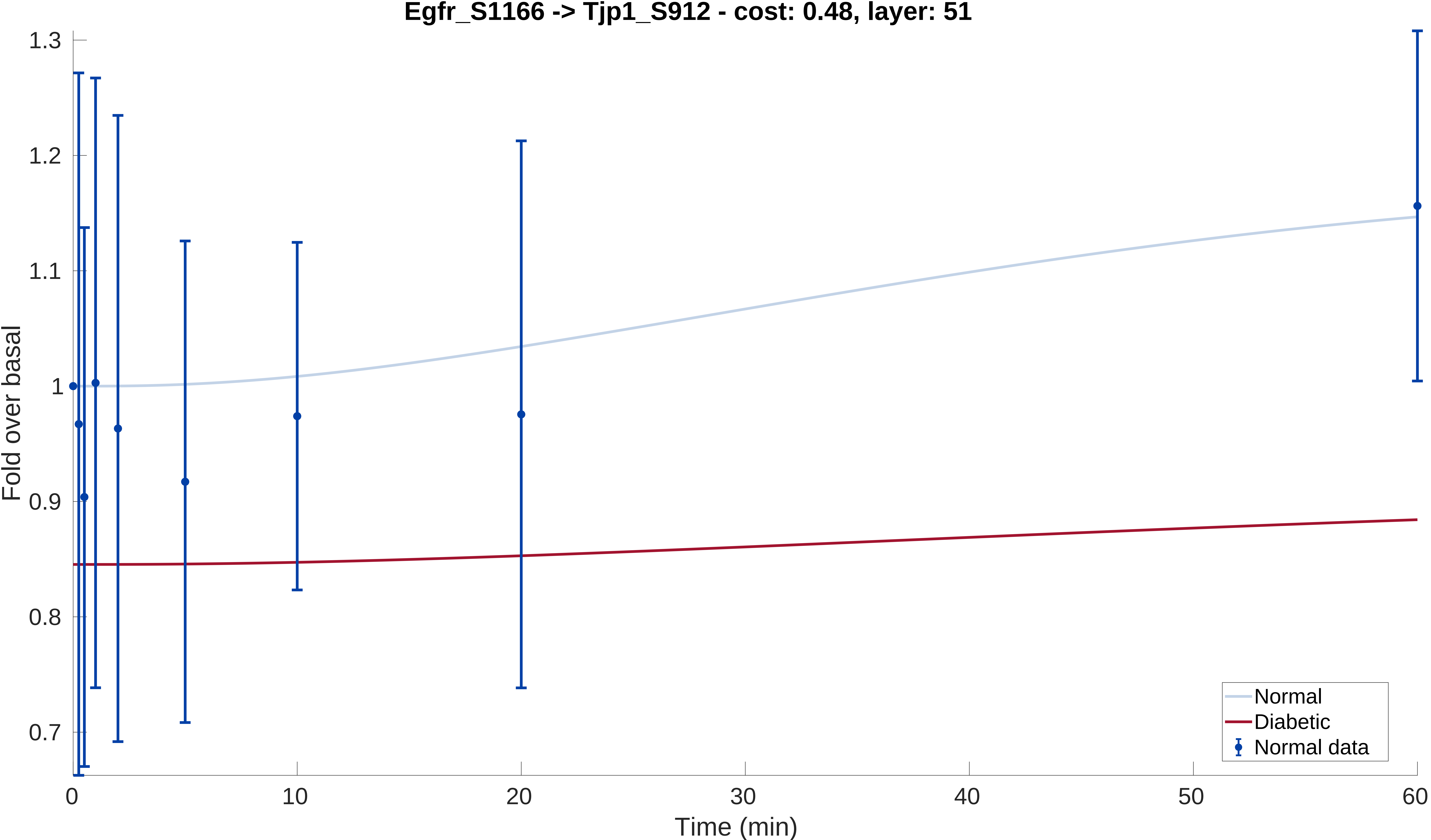

Egfr_S1166 -> Tjp1_S912 - cost: 0.48, layer: 51
Fold over basal
1.3
1.2
1.1
1
0.9
0.8
0.7
0
10
20
30
40
50
60
Time (min)
Normal
Diabetic
Normal data

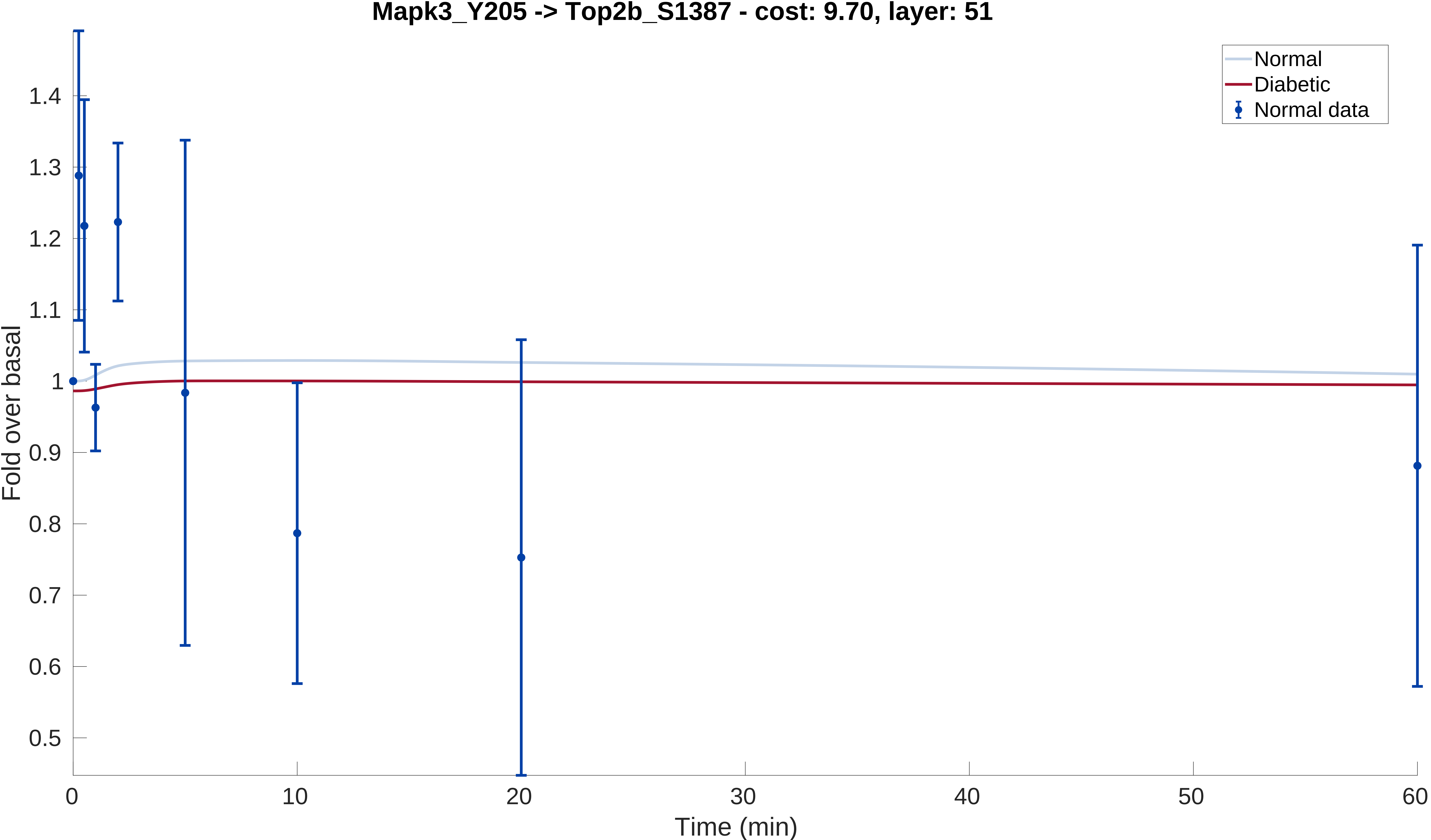

Mapk3_Y205 -> Top2b_S1387 - cost: 9.70, layer: 51
Normal
Diabetic
Normal data
Fold over basal
1.4
1.3
1.2
1.1
1
0.9
0.8
0.7
0.6
0.5
0
10
20
30
40
50
60
Time (min)

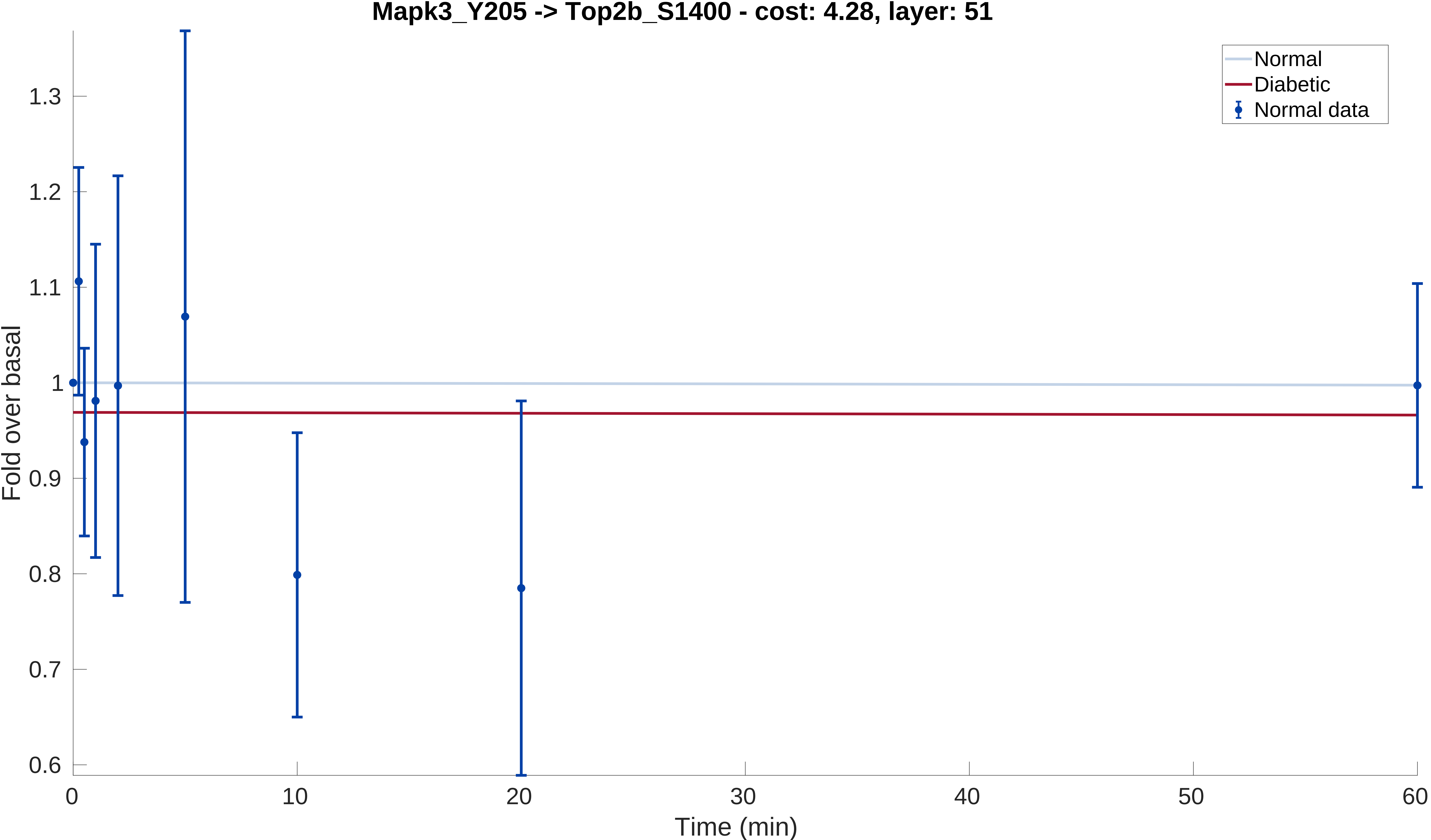
Mapk3_Y205 -> Top2b_S1400 - cost: 4.28, layer: 51
Normal
Diabetic
Normal data
Fold over basal
1.3
1.2
1.1
1
0.9
0.8
0.7
0.6
0
10
20
30
40
50
60
Time (min)

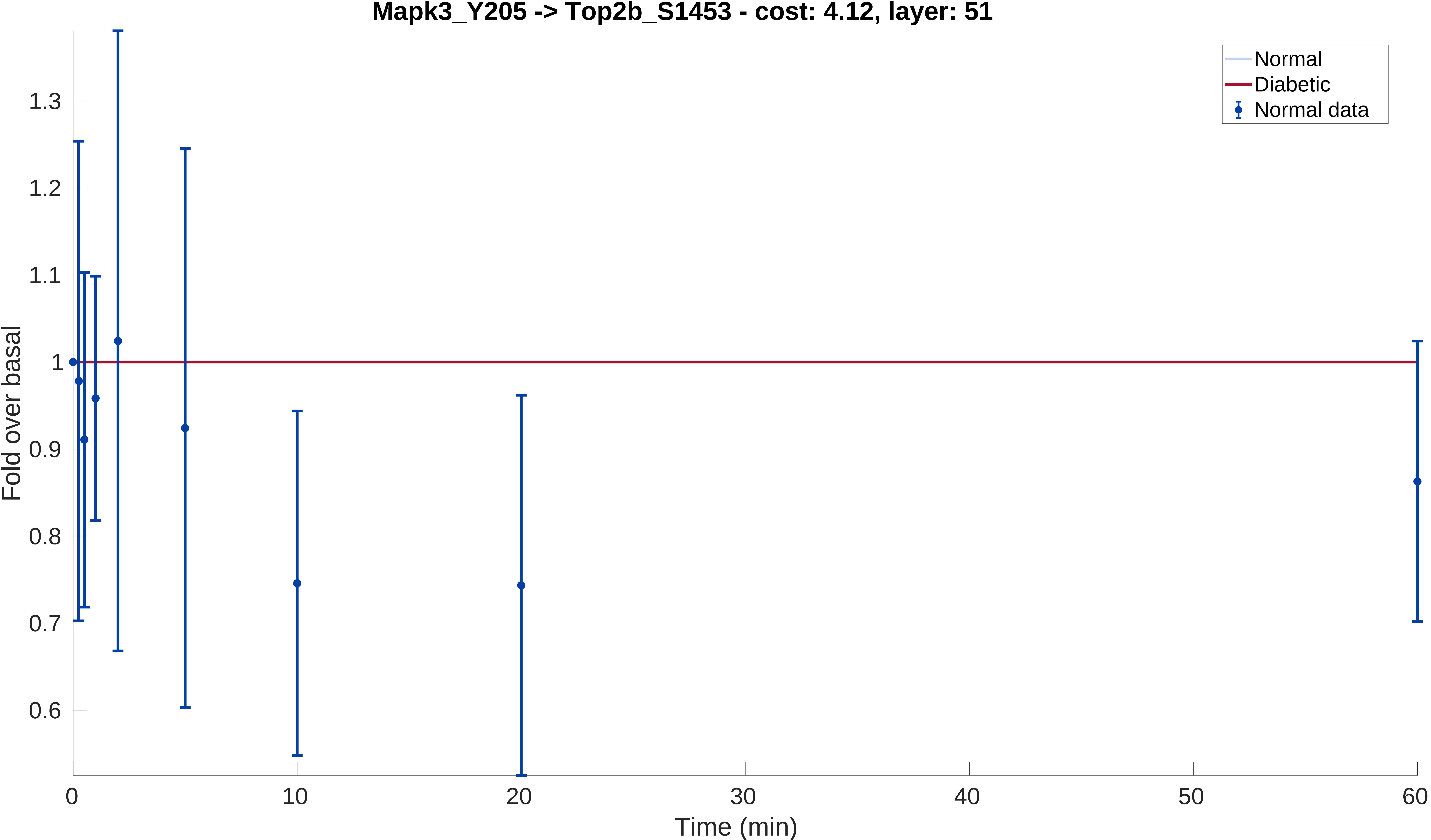
Mapk3_Y205 -> Top2b_S1453 - cost: 4.12, layer: 51
Normal
Diabetic
Normal data
Fold over basal
1.3
1.2
1.1
1
0.9
0.8
0.7
0.6
0
10
20
30
40
50
60
Time (min)

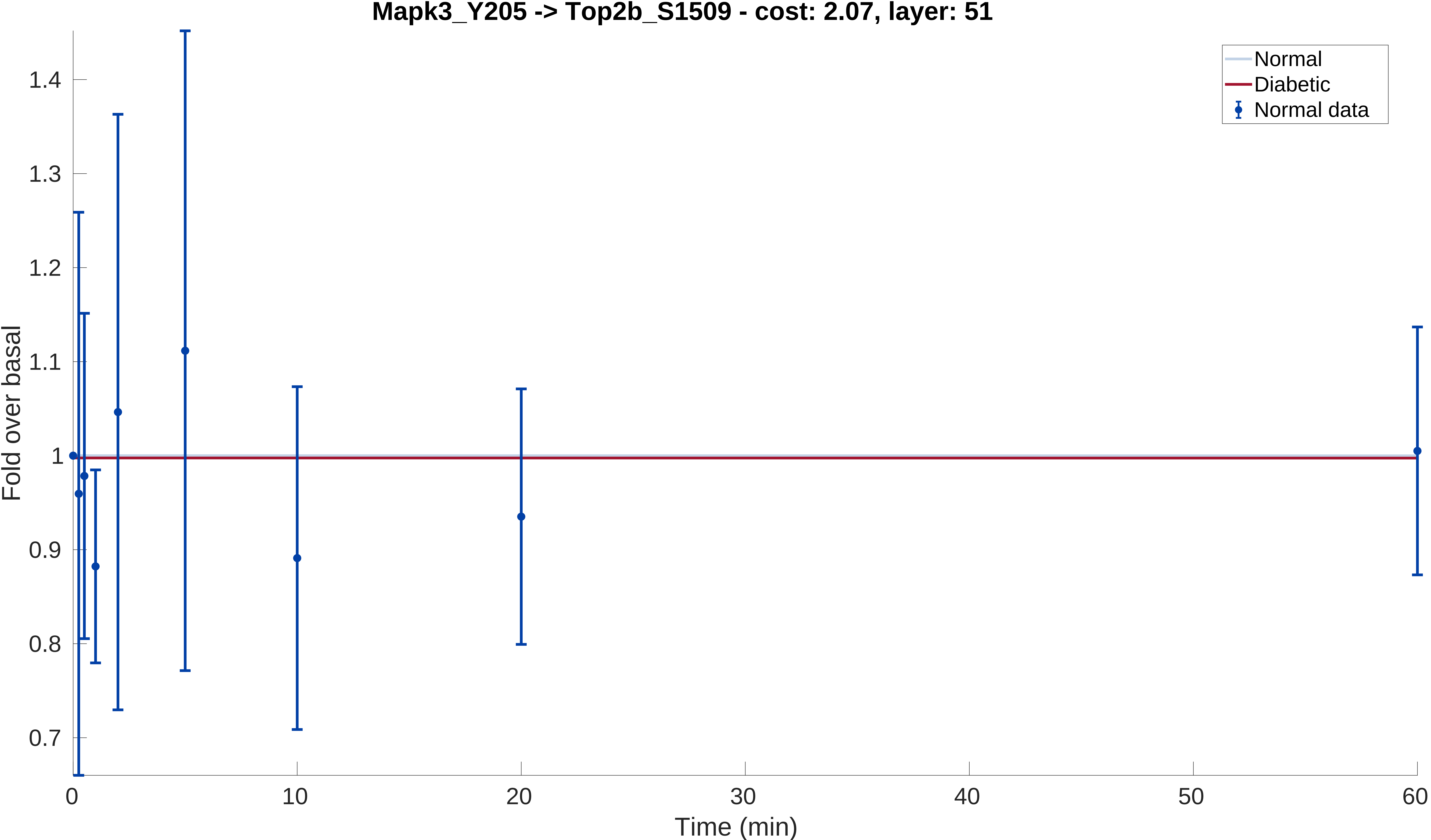

Mapk3_Y205 -> Top2b_S1509 - cost: 2.07, layer: 51
Normal
Diabetic
Normal data
Fold over basal
1.4
1.3
1.2
1.1
1
0.9
0.8
0.7
0
10
20
30
40
50
60
Time (min)

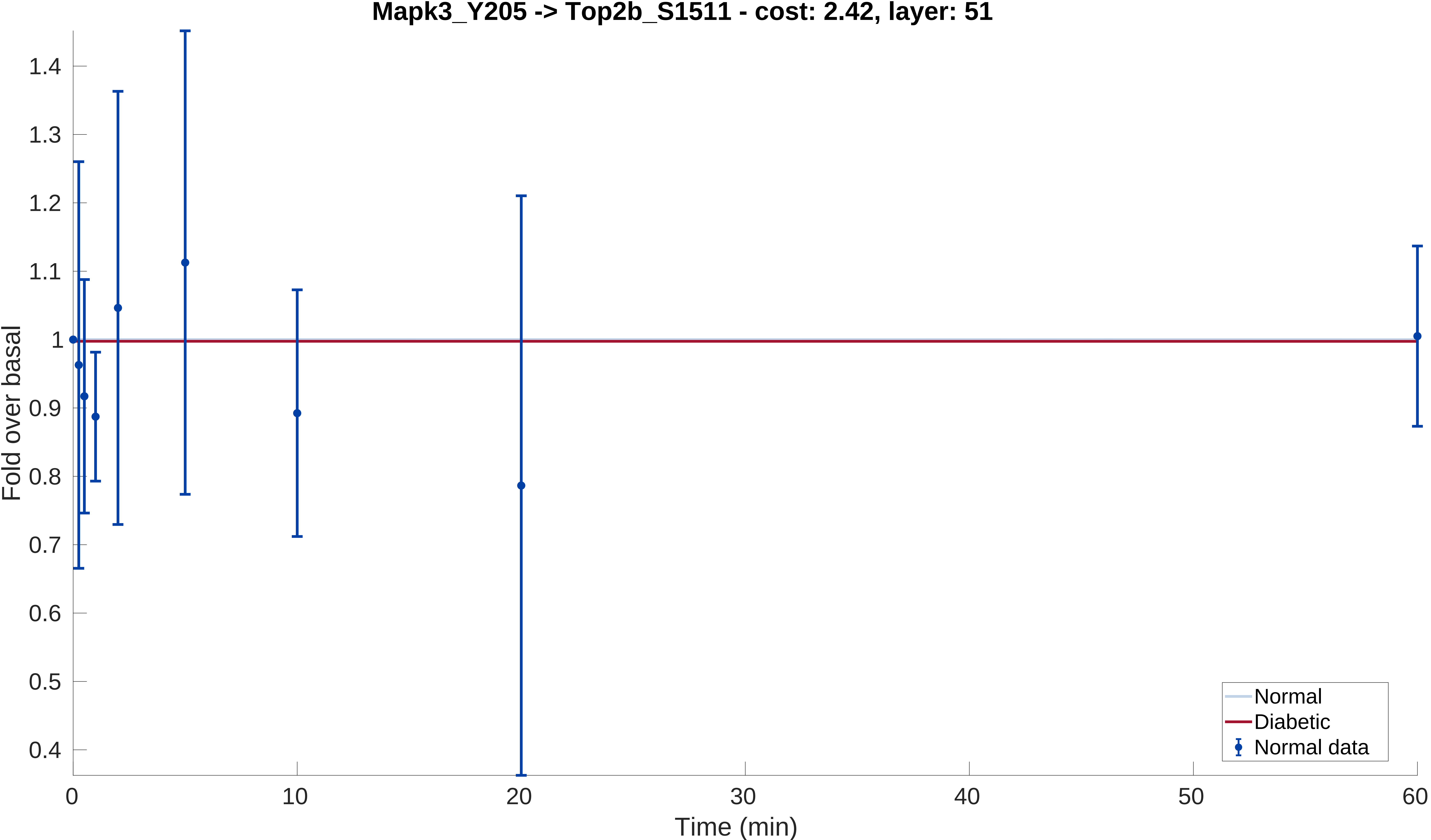

Mapk3_Y205 -> Top2b_S1511 - cost: 2.42, layer: 51
Fold over basal
1.4
1.3
1.2
1.1
1
0.9
0.8
0.7
0.6
0.5
0.4
0
10
20
30
40
50
60
Time (min)
Normal
Diabetic
Normal data

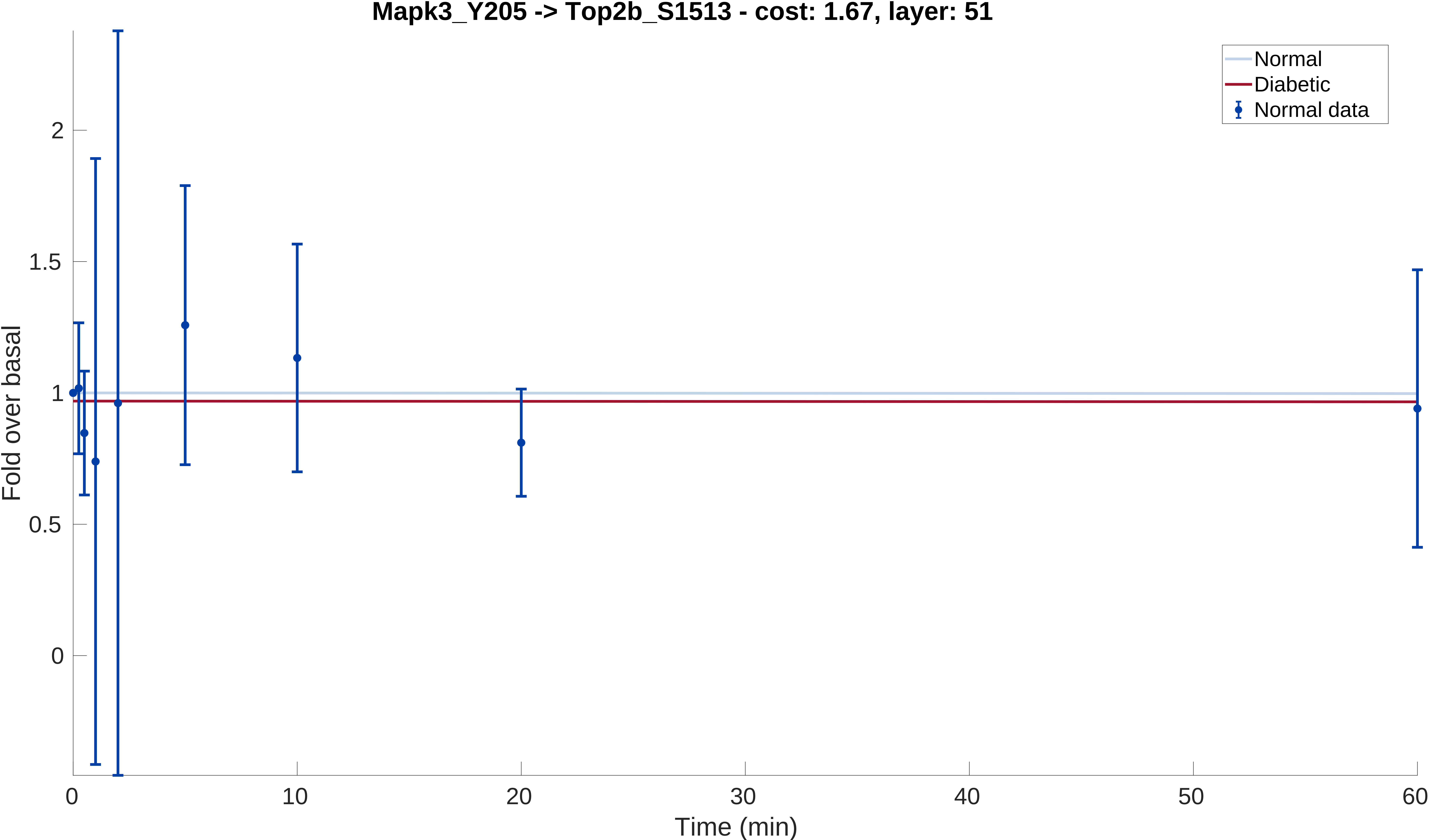

Mapk3_Y205 -> Top2b_S1513 - cost: 1.67, layer: 51
Normal
Diabetic
Normal data
Fold over basal
2
1.5
1
0.5
0
0
10
20
30
40
50
60
Time (min)

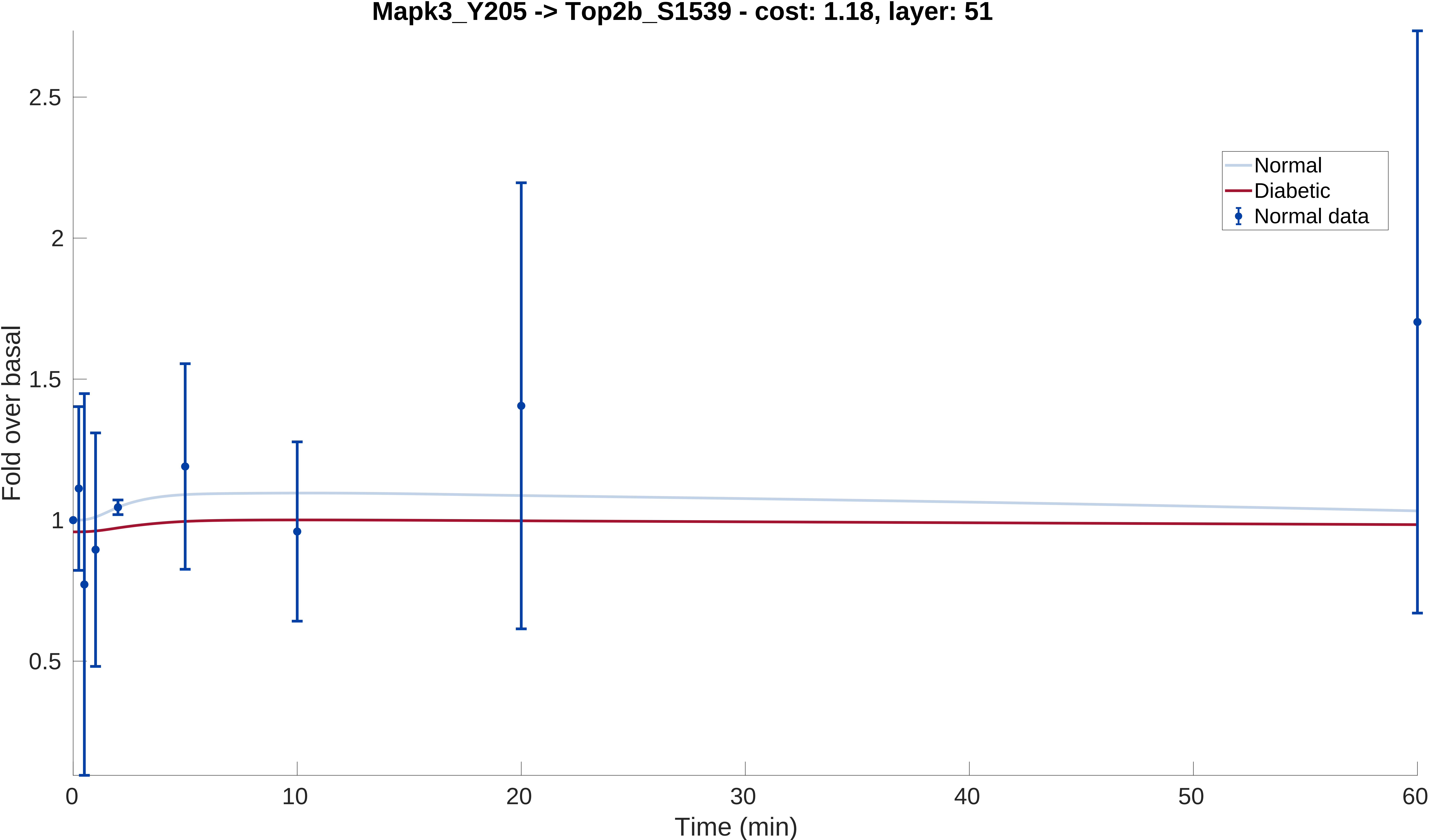
Mapk3_Y205 -> Top2b_S1539 - cost: 1.18, layer: 51
Fold over basal
Time (min)
Normal
Diabetic
Normal data
0.5
1
1.5
2
2.5
0
10
20
30
40
50
60

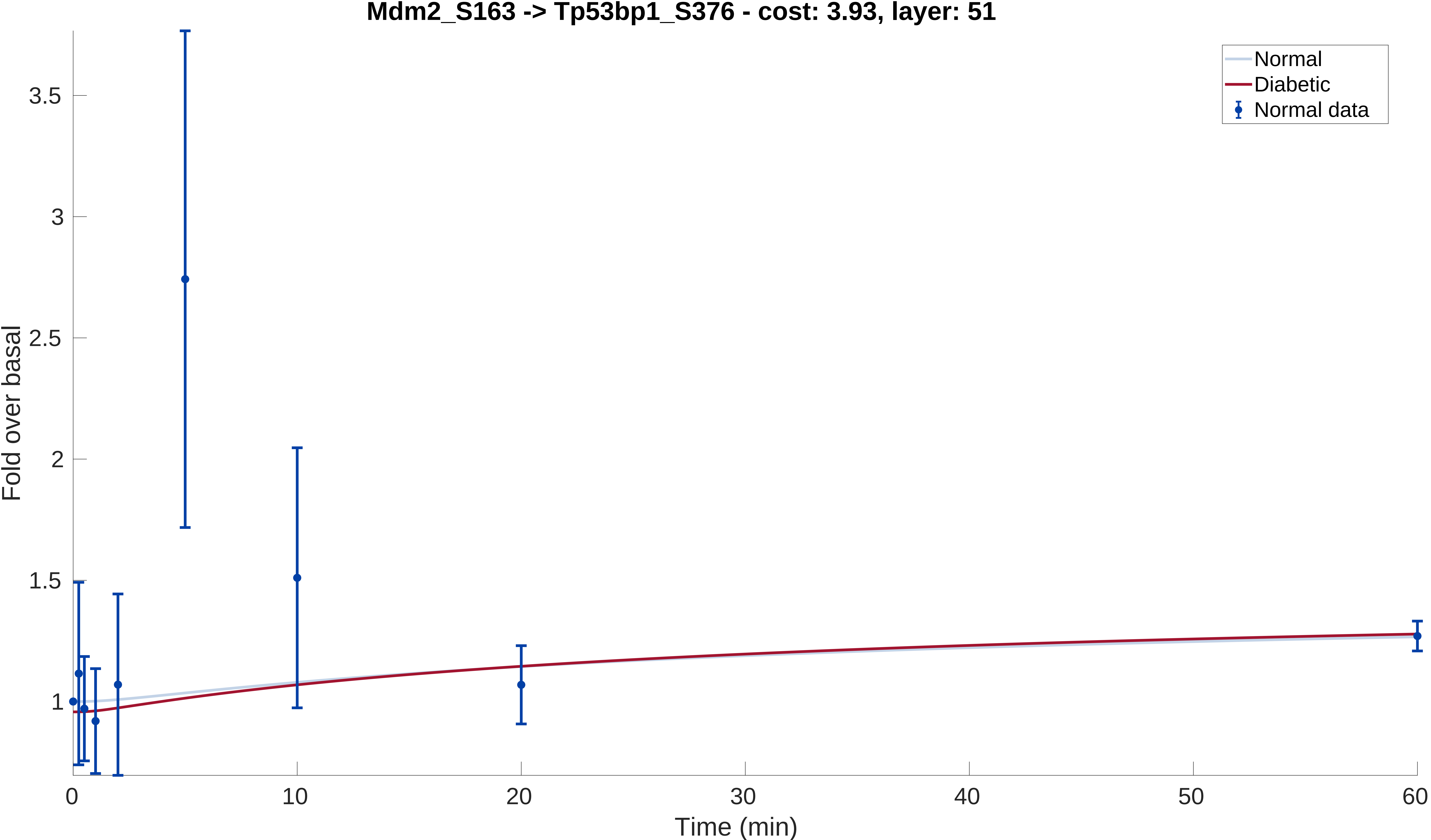

Mdm2_S163 -> Tp53bp1_S376 - cost: 3.93, layer: 51
Normal
Diabetic
Normal data
Fold over basal
Time (min)
3.5
3
2.5
2
1.5
1
0
10
20
30
40
50
60

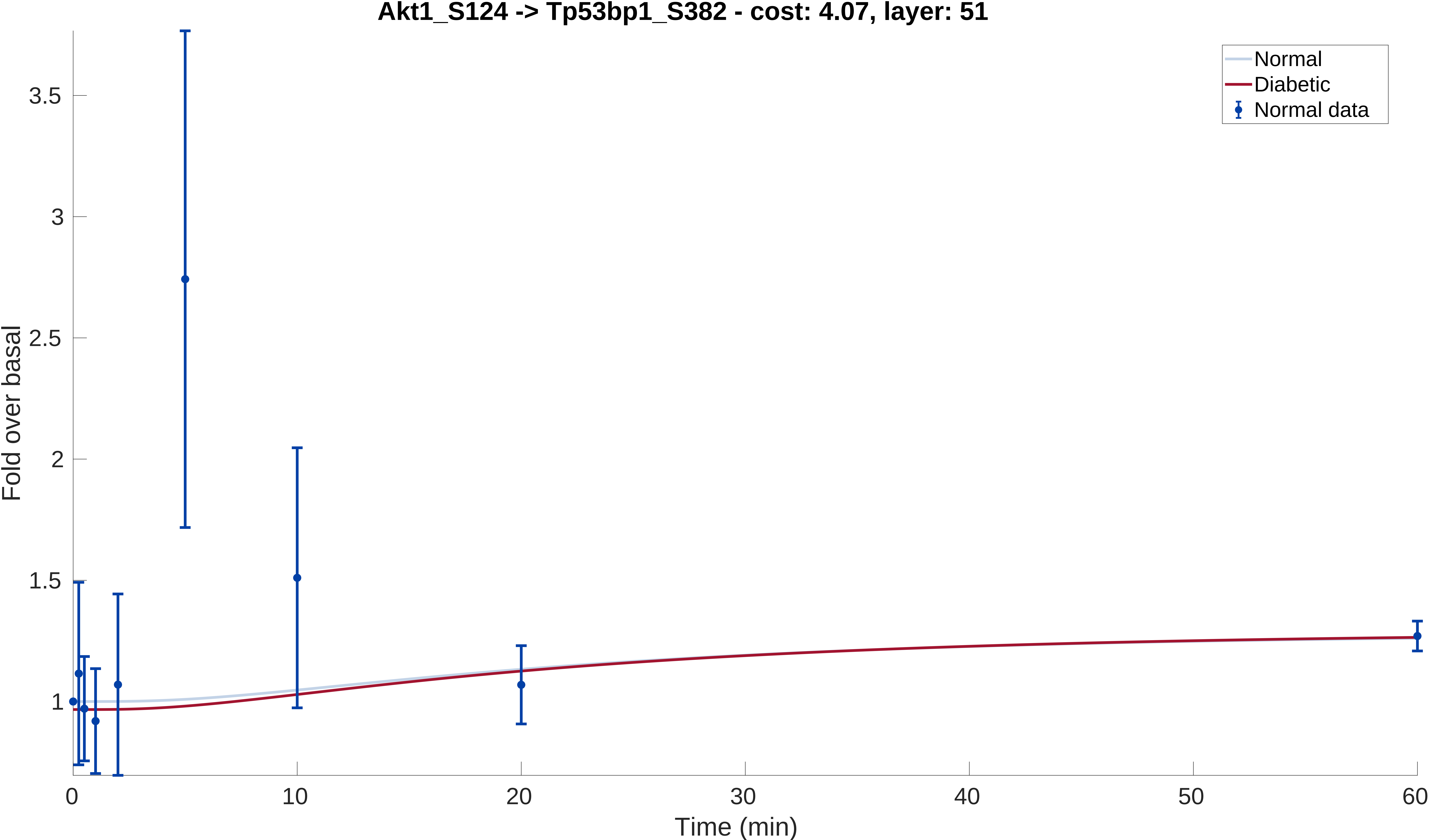
Akt1_S124 -> Tp53bp1_S382 - cost: 4.07, layer: 51
Normal
Diabetic
Normal data
3.5
3
2.5
2
1.5
1
Fold over basal
0
10
20
30
40
50
60
Time (min)

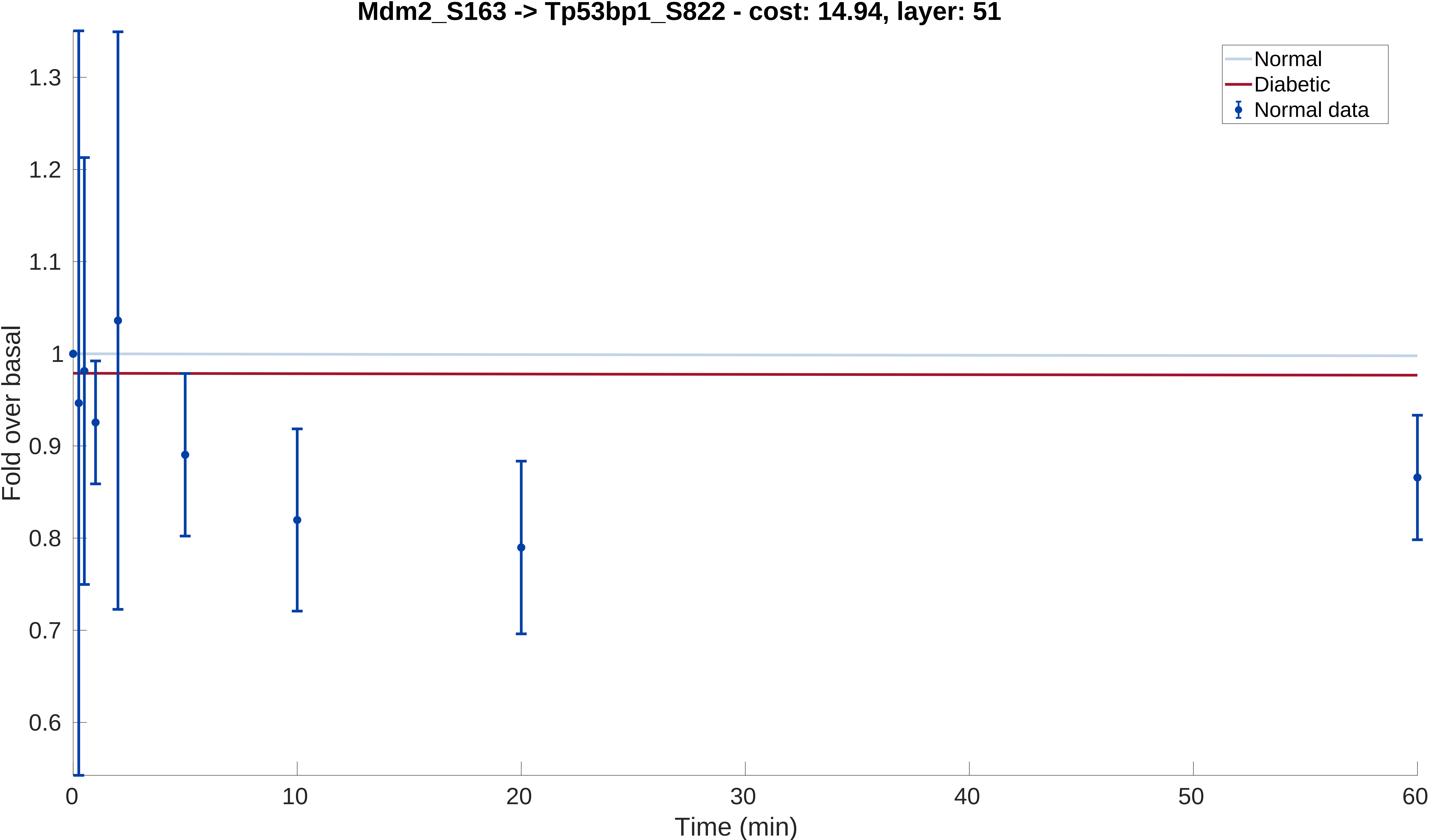

Mdm2_S163 -> Tp53bp1_S822 - cost: 14.94, layer: 51
Normal
Diabetic
Normal data
Fold over basal
1.3
1.2
1.1
1
0.9
0.8
0.7
0.6
0
10
20
30
40
50
60
Time (min)

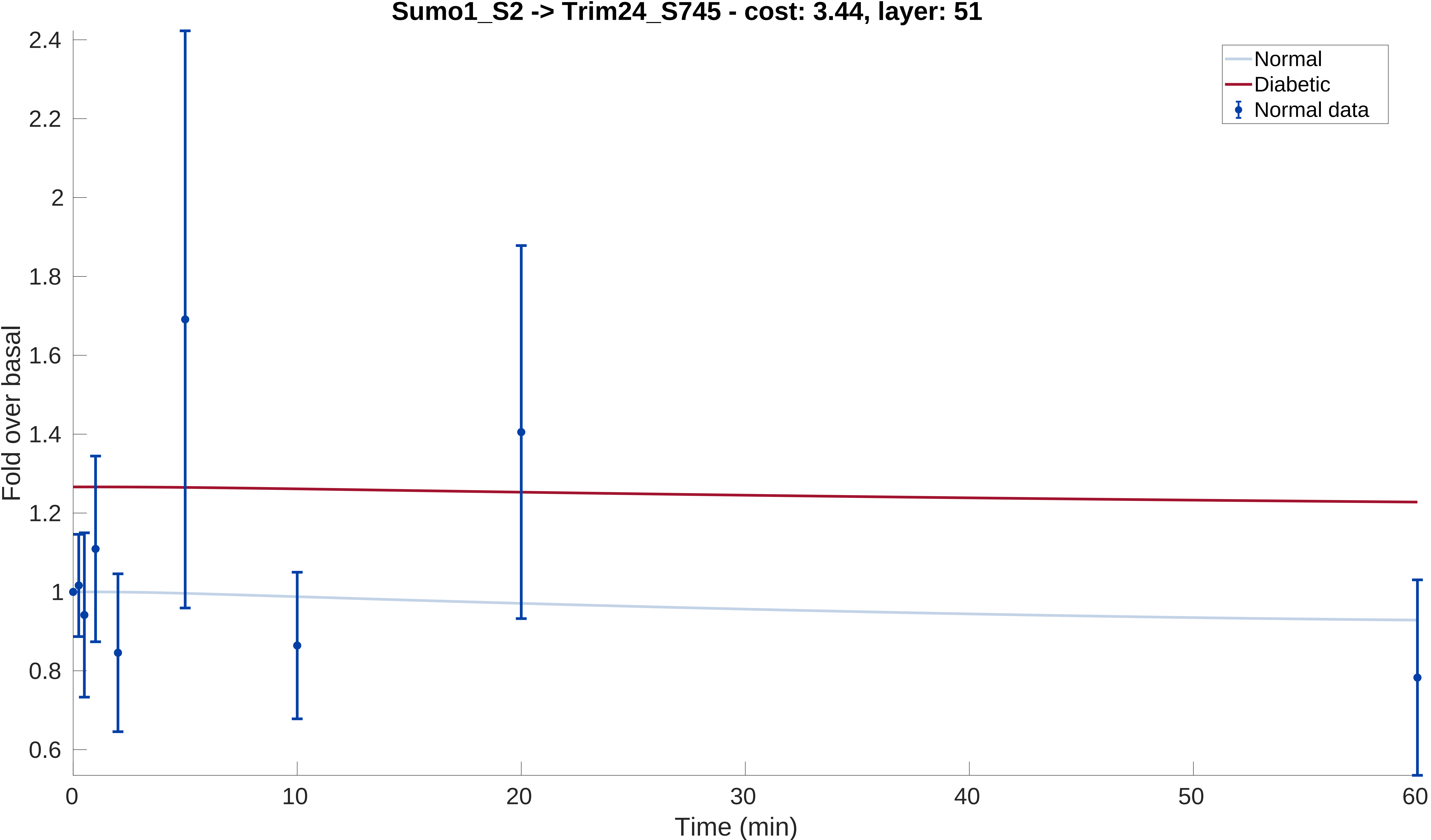

Sumo1_S2 -> Trim24_S745 - cost: 3.44, layer: 51
Normal
Diabetic
Normal data
Fold over basal
2.4
2.2
2
1.8
1.6
1.4
1.2
1
0.8
0.6
0
10
20
30
40
50
60
Time (min)

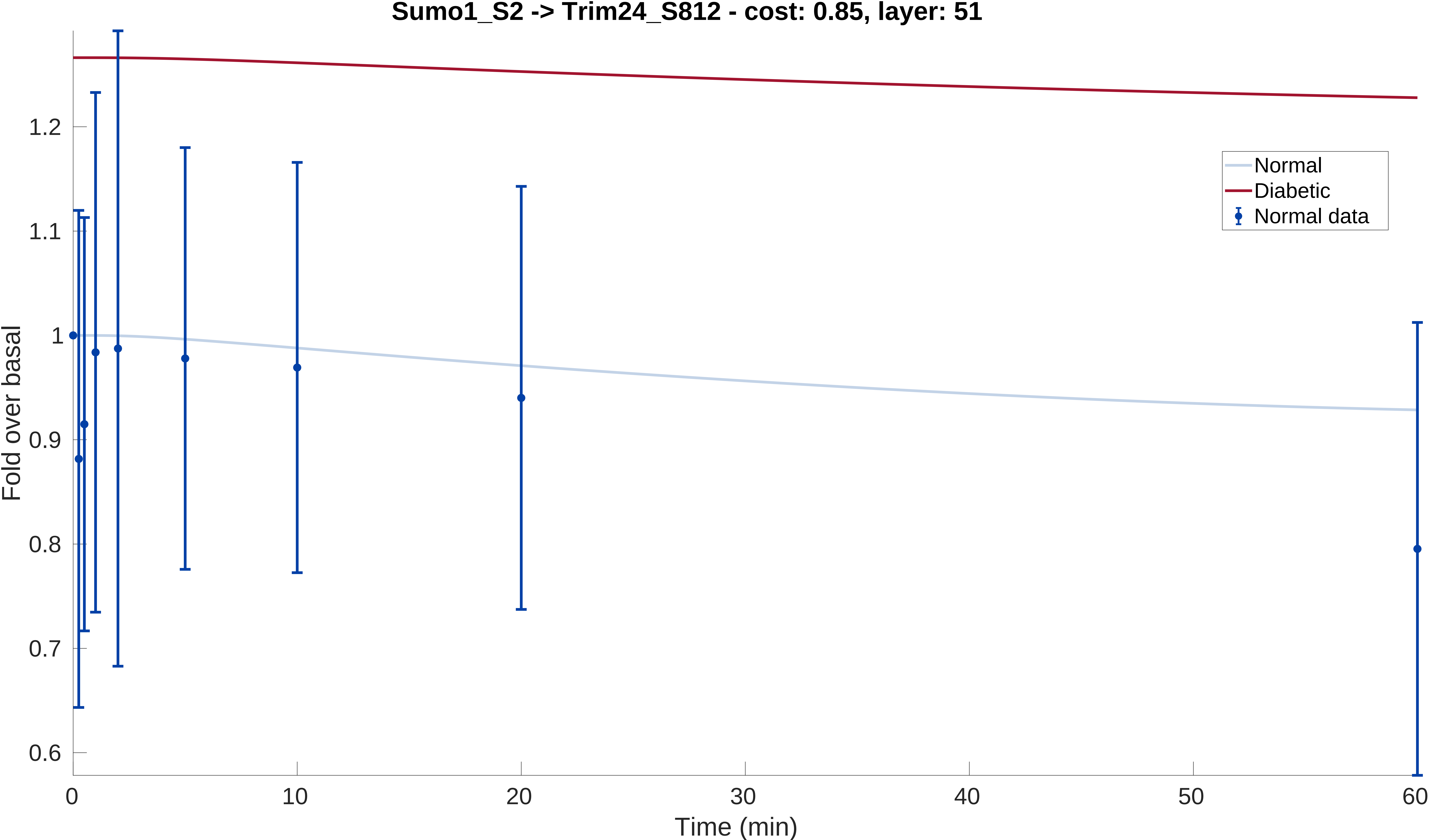

Sumo1_S2 -> Trim24_S812 - cost: 0.85, layer: 51
Normal
Diabetic
Normal data
Fold over basal
Time (min)
0.6
0.7
0.8
0.9
1
1.1
1.2
0
10
20
30
40
50
60

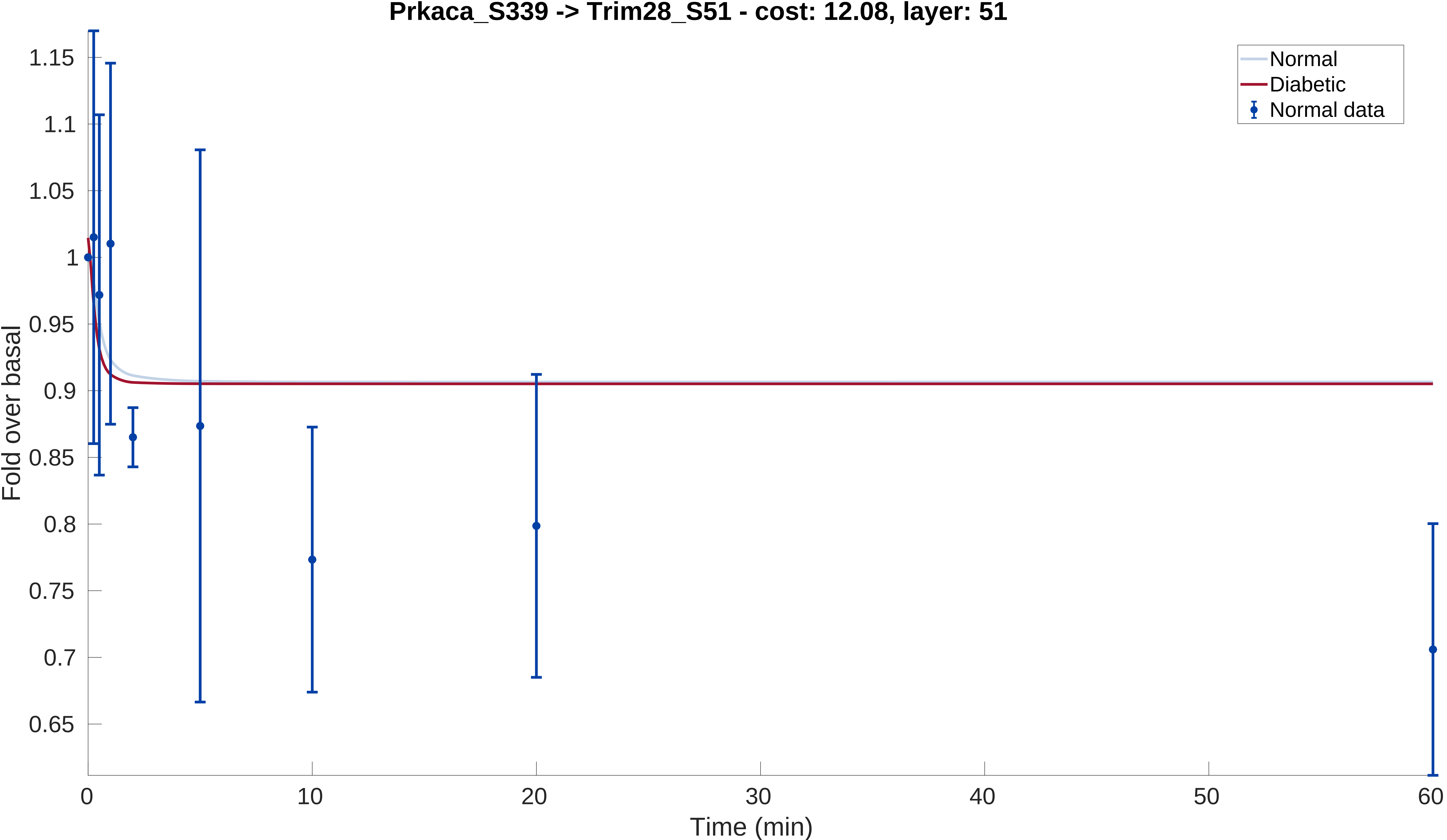

Prkaca_S339 -> Trim28_S51 - cost: 12.08, layer: 51
Normal
Diabetic
Normal data
Fold over basal
1.15
1.1
1.05
1
0.95
0.9
0.85
0.8
0.75
0.7
0.65
0
10
20
30
40
50
60
Time (min)

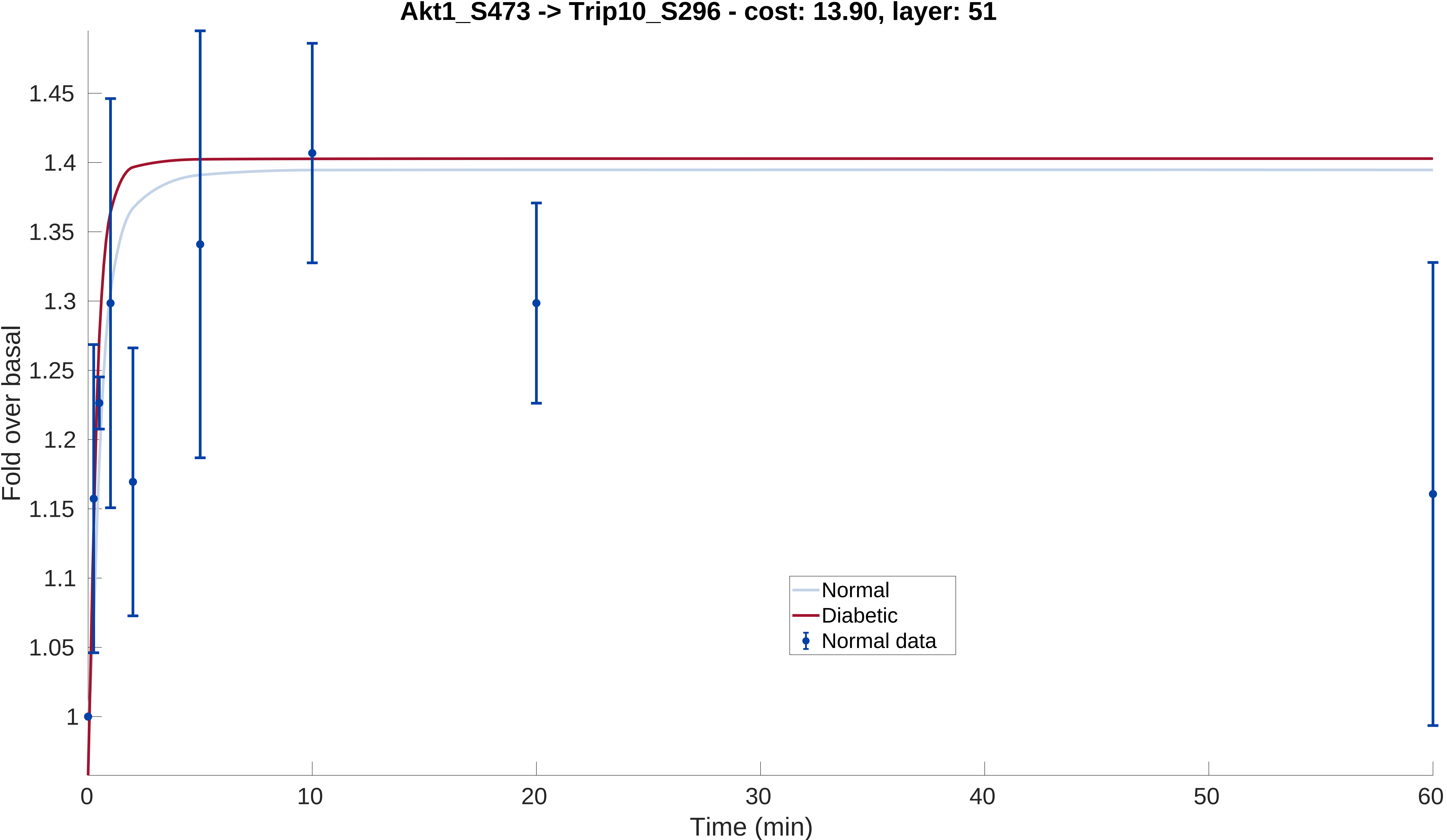

Akt1_S473 -> Trip10_S296 - cost: 13.90, layer: 51
1.45
1.4
1.35
1.3
1.25
1.2
1.15
1.1
1.05
1
Fold over basal
0
10
20
30
40
50
60
Time (min)
Normal
Diabetic
Normal data

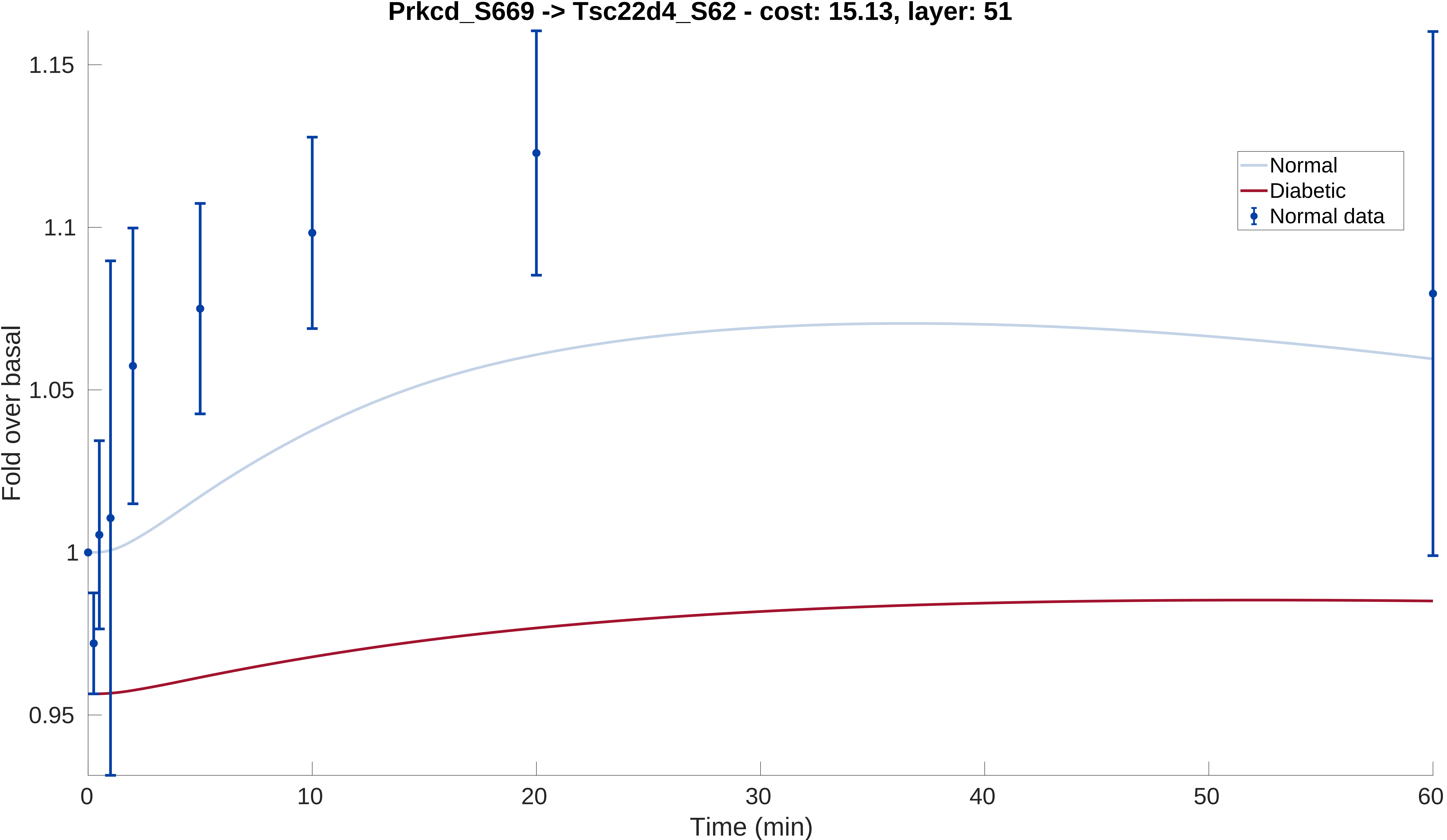

Prkcd_S669 -> Tsc22d4_S62 - cost: 15.13, layer: 51
Normal
Diabetic
Normal data
Fold over basal
1.15
1.1
1.05
1
0.95
0
10
20
30
40
50
60
Time (min)

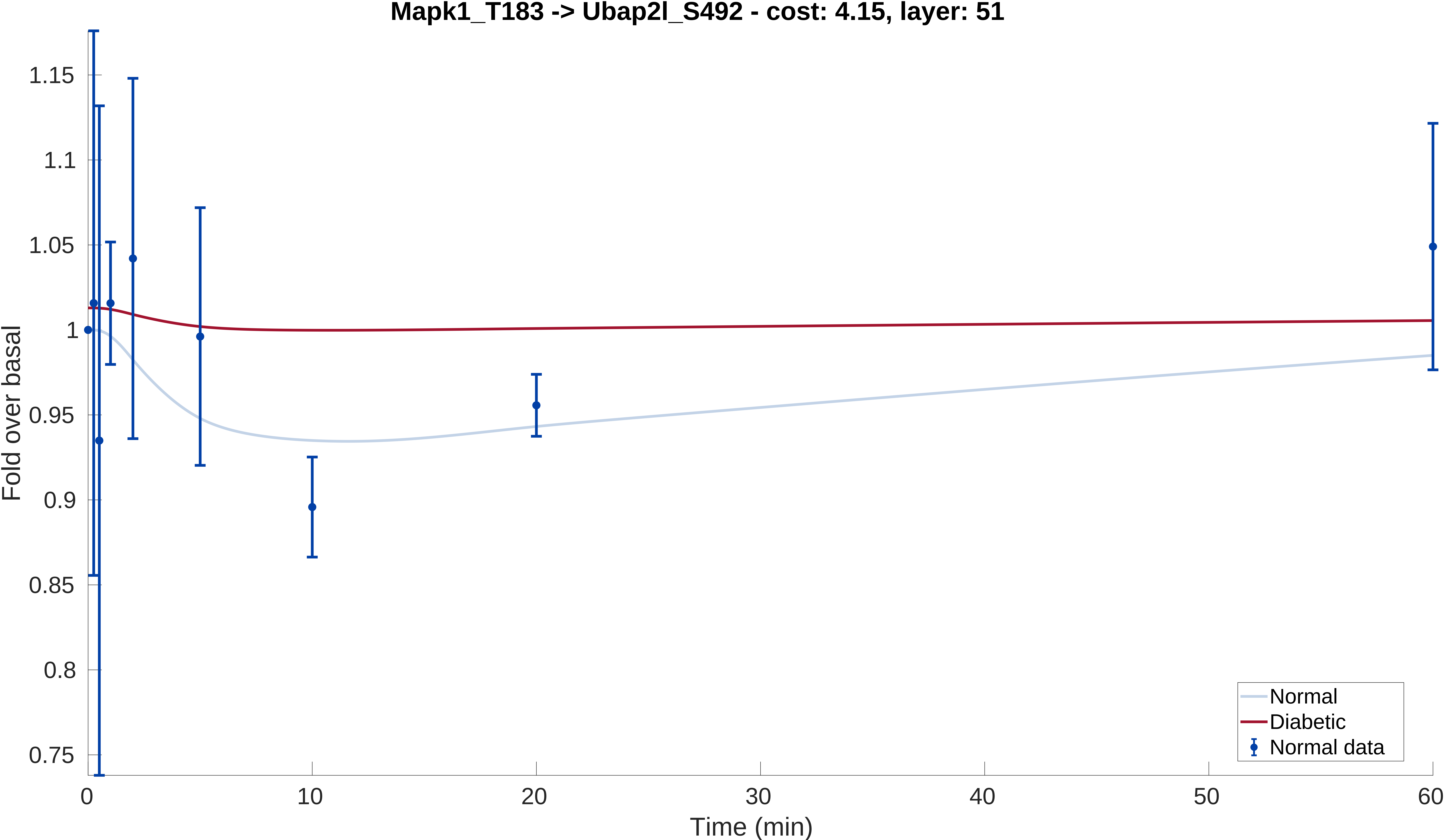

Mapk1_T183 -> Ubap2l_S492 - cost: 4.15, layer: 51
Fold over basal
1.15
1.1
1.05
1
0.95
0.9
0.85
0.8
0.75
0
10
20
30
40
50
60
Time (min)
Normal
Diabetic
Normal data

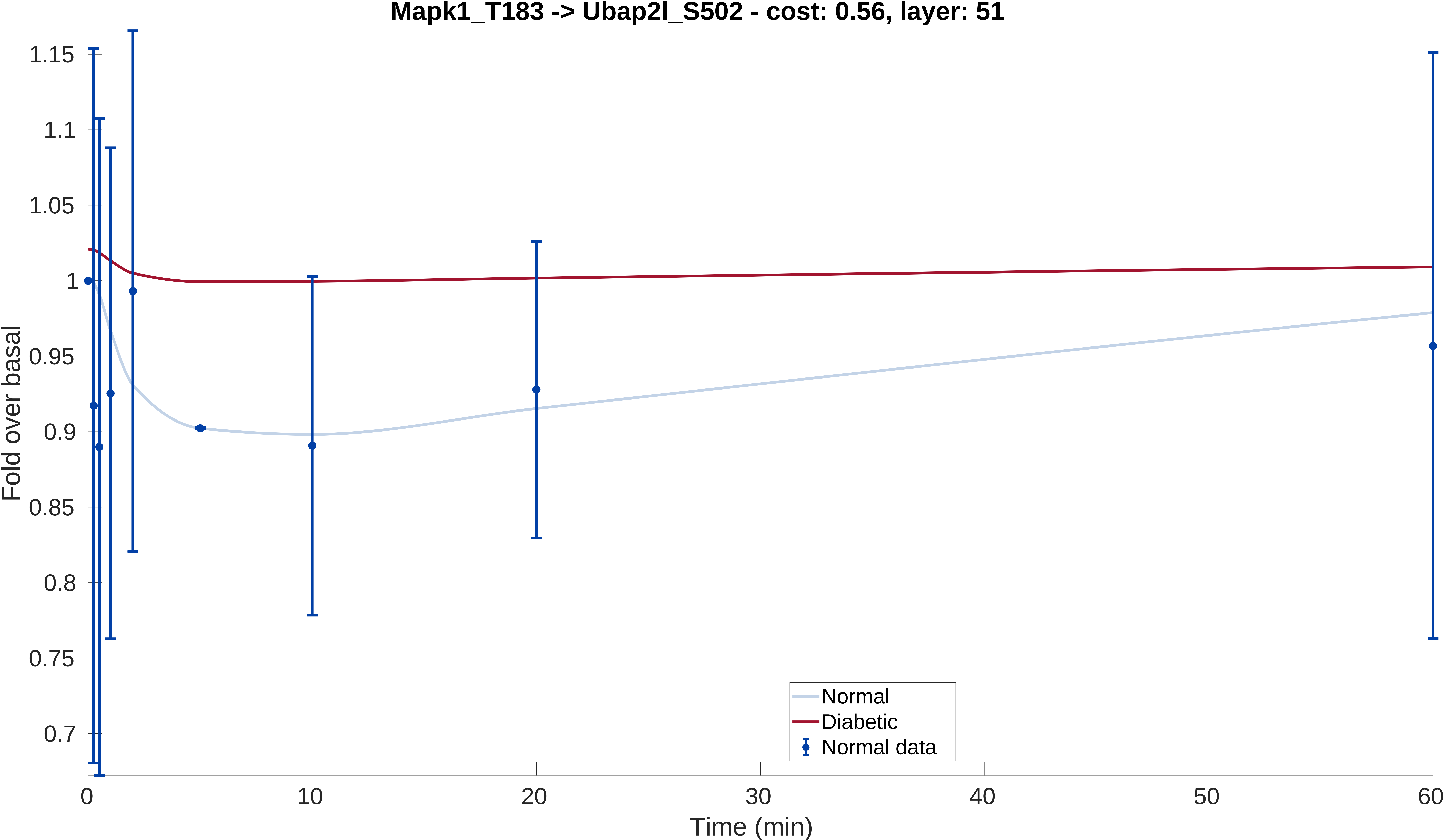

Mapk1_T183 -> Ubap2l_S502 - cost: 0.56, layer: 51
Fold over basal
1.15
1.1
1.05
1
0.95
0.9
0.85
0.8
0.75
0.7
0
10
20
30
40
50
60
Time (min)
Normal
Diabetic
Normal data

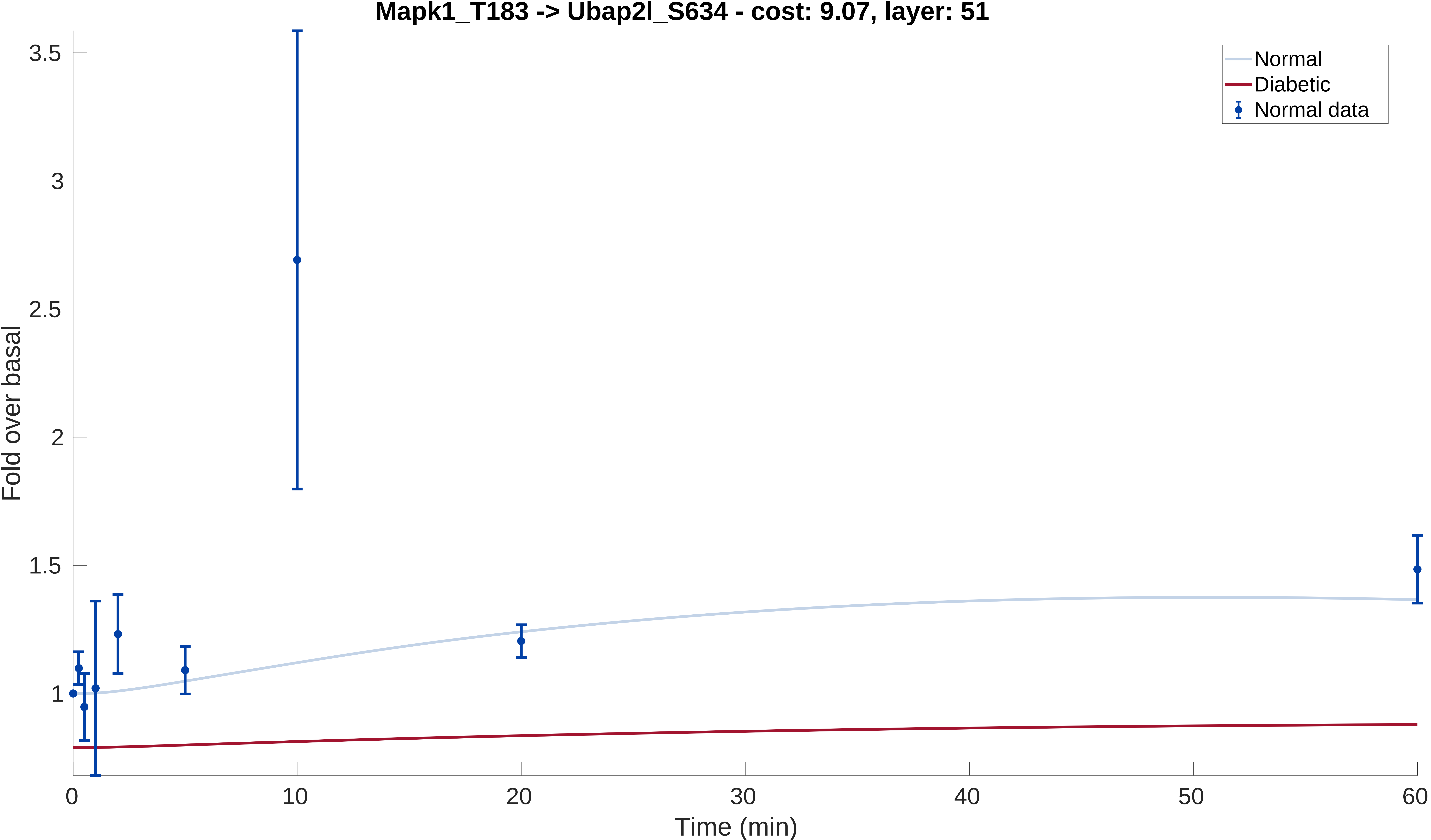

Mapk1_T183 -> Ubap2l_S634 - cost: 9.07, layer: 51
Normal
Diabetic
Normal data
Fold over basal
3.5
3
2.5
2
1.5
1
0
10
20
30
40
50
60
Time (min)

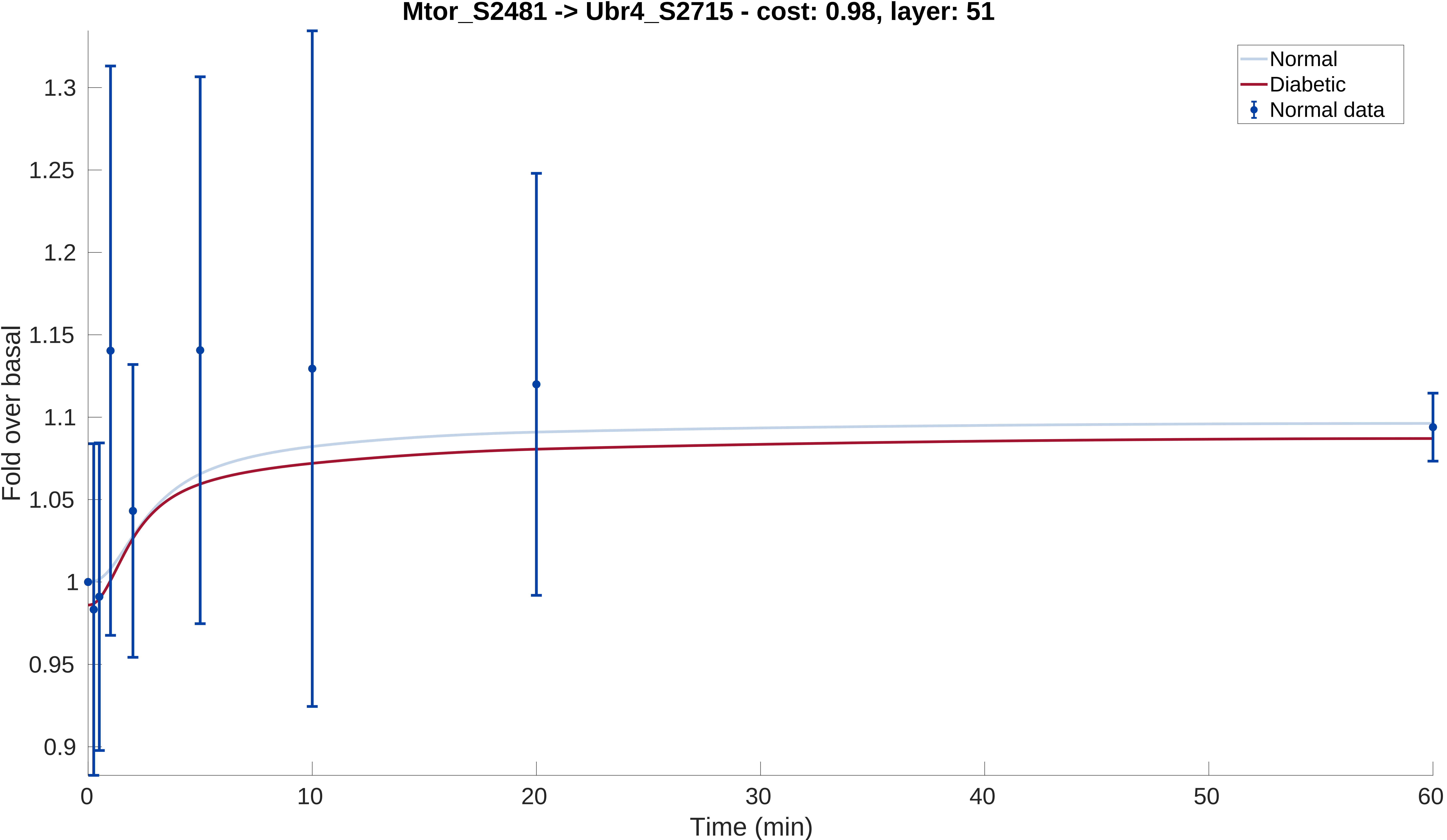

Mtor_S2481 -> Ubr4_S2715 - cost: 0.98, layer: 51
Normal
Diabetic
Normal data
Fold over basal
1.3
1.25
1.2
1.15
1.1
1.05
1
0.95
0.9
0
10
20
30
40
50
60
Time (min)

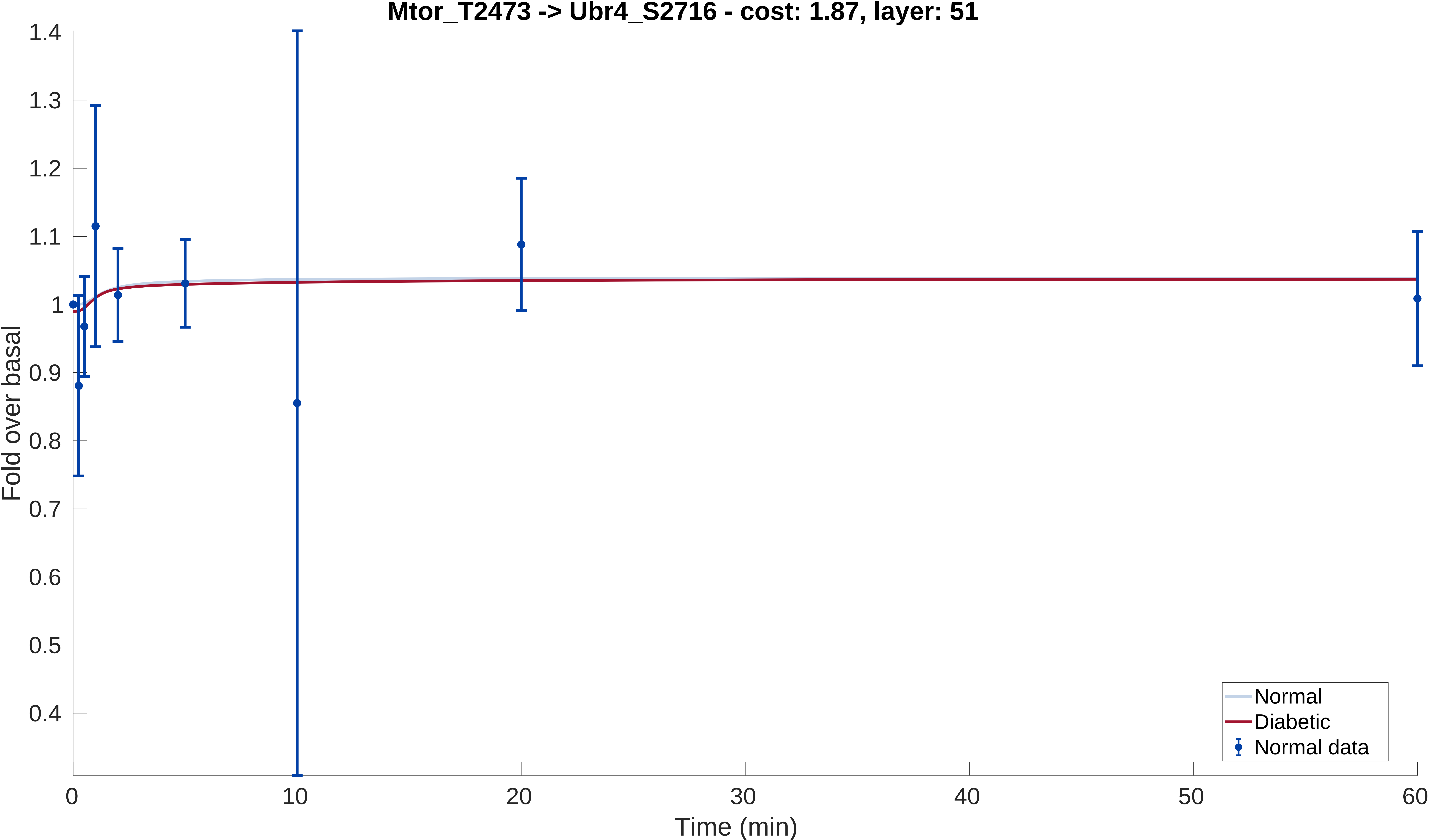

Mtor_T2473 -> Ubr4_S2716 - cost: 1.87, layer: 51
Fold over basal
Time (min)
1.4
1.3
1.2
1.1
1
0.9
0.8
0.7
0.6
0.5
0.4
0
10
20
30
40
50
60
Normal
Diabetic
Normal data

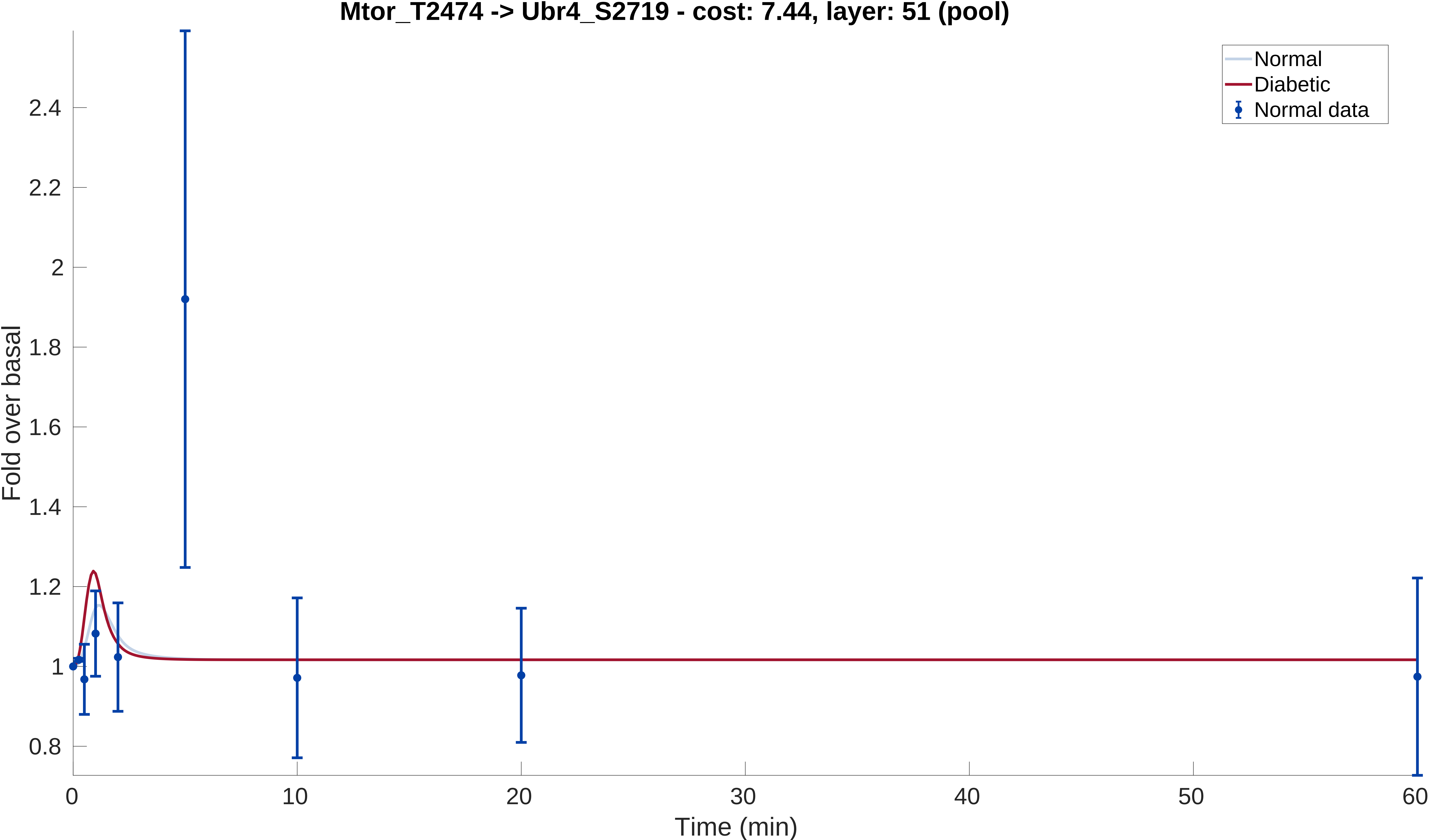
Mtor_T2474 -> Ubr4_S2719 - cost: 7.44, layer: 51 (pool)
Normal
Diabetic
Normal data
Fold over basal
2.4
2.2
2
1.8
1.6
1.4
1.2
1
0.8
0
10
20
30
40
50
60
Time (min)

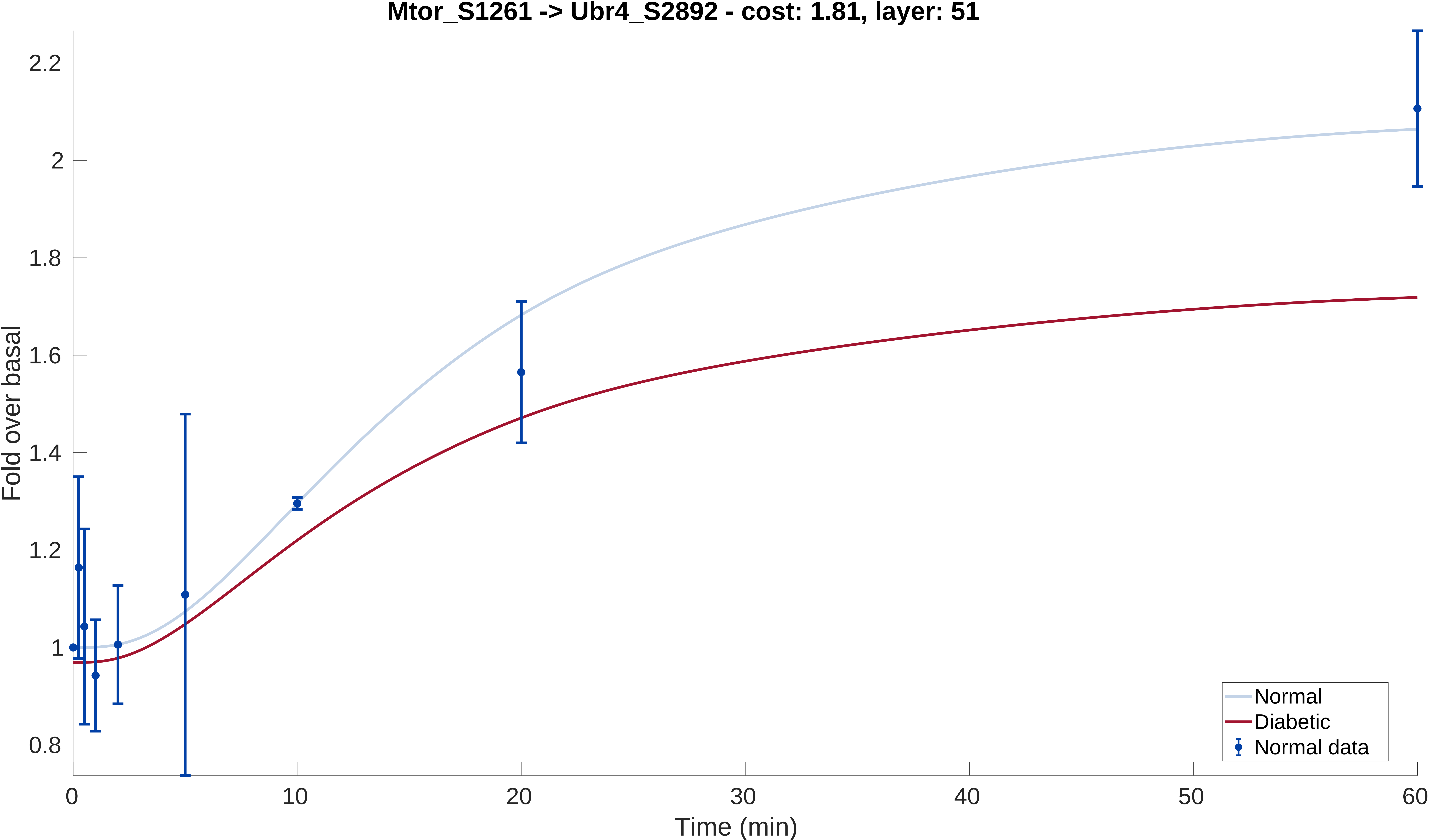
Mtor_S1261 -> Ubr4_S2892 - cost: 1.81, layer: 51
Fold over basal
Time (min)
0.8
1
1.2
1.4
1.6
1.8
2
2.2
0
10
20
30
40
50
60
Normal
Diabetic
Normal data

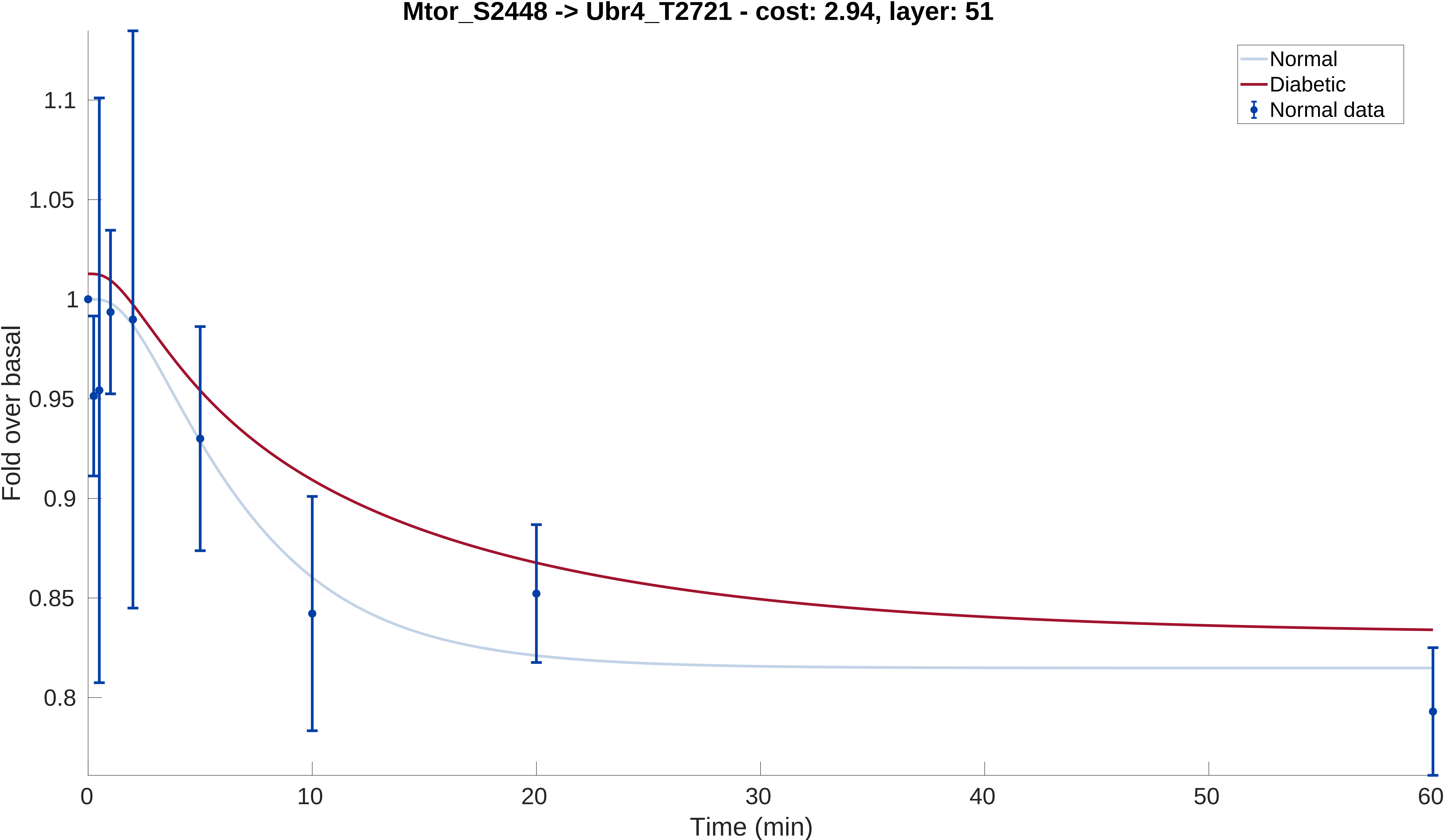
Mtor_S2448 -> Ubr4_T2721 - cost: 2.94, layer: 51
Normal
Diabetic
Normal data
Fold over basal
1.1
1.05
1
0.95
0.9
0.85
0.8
0
10
20
30
40
50
60
Time (min)

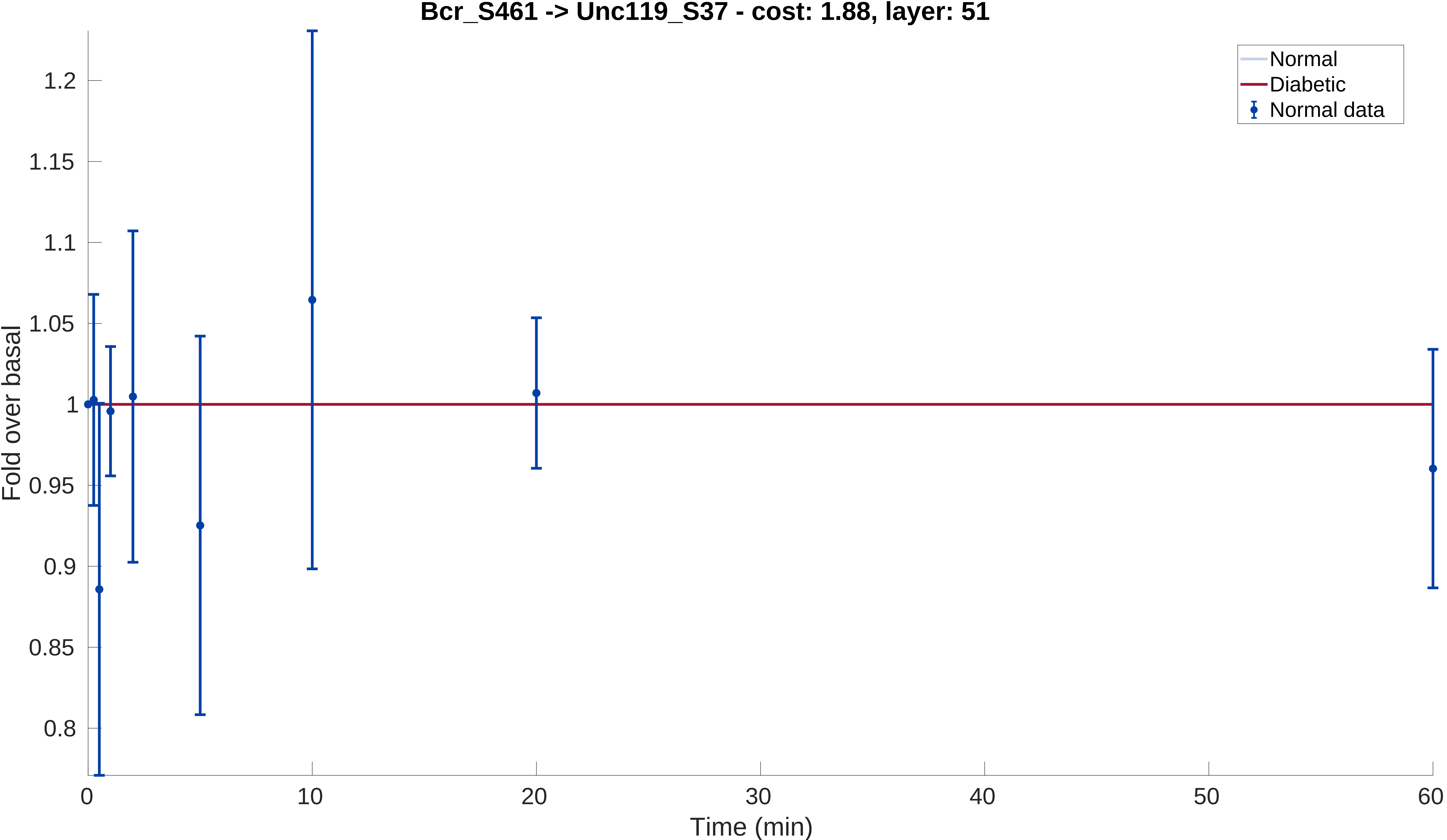
Bcr_S461 -> Unc119_S37 - cost: 1.88, layer: 51
Normal
Diabetic
Normal data
Fold over basal
1.2
1.15
1.1
1.05
1
0.95
0.9
0.85
0.8
0
10
20
30
40
50
60
Time (min)

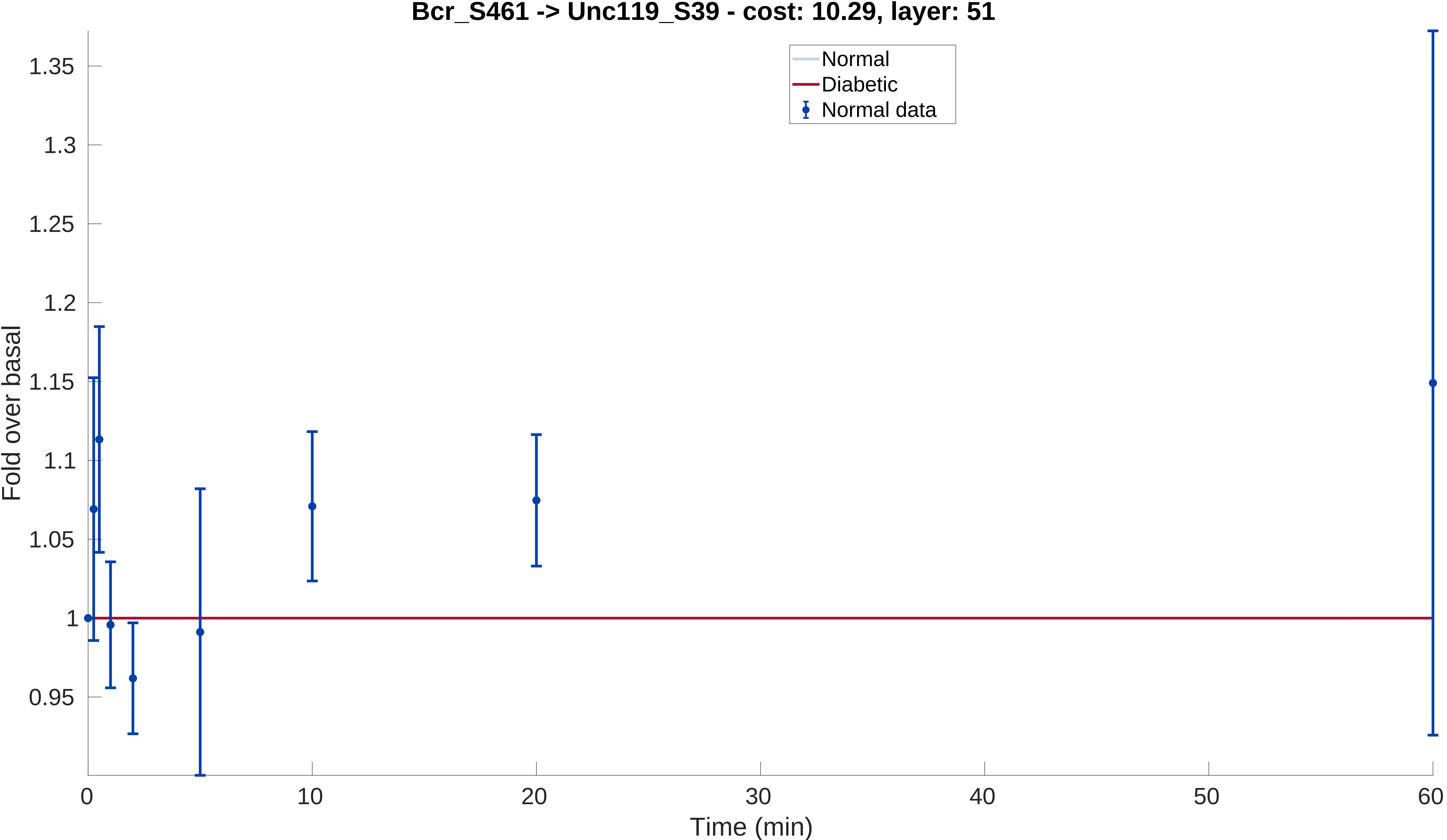

Bcr_S461 -> Unc119_S39 - cost: 10.29, layer: 51
Normal
Diabetic
Normal data
1.35
1.3
1.25
1.2
1.15
1.1
1.05
1
0.95
Fold over basal
0
10
20
30
40
50
60
Time (min)

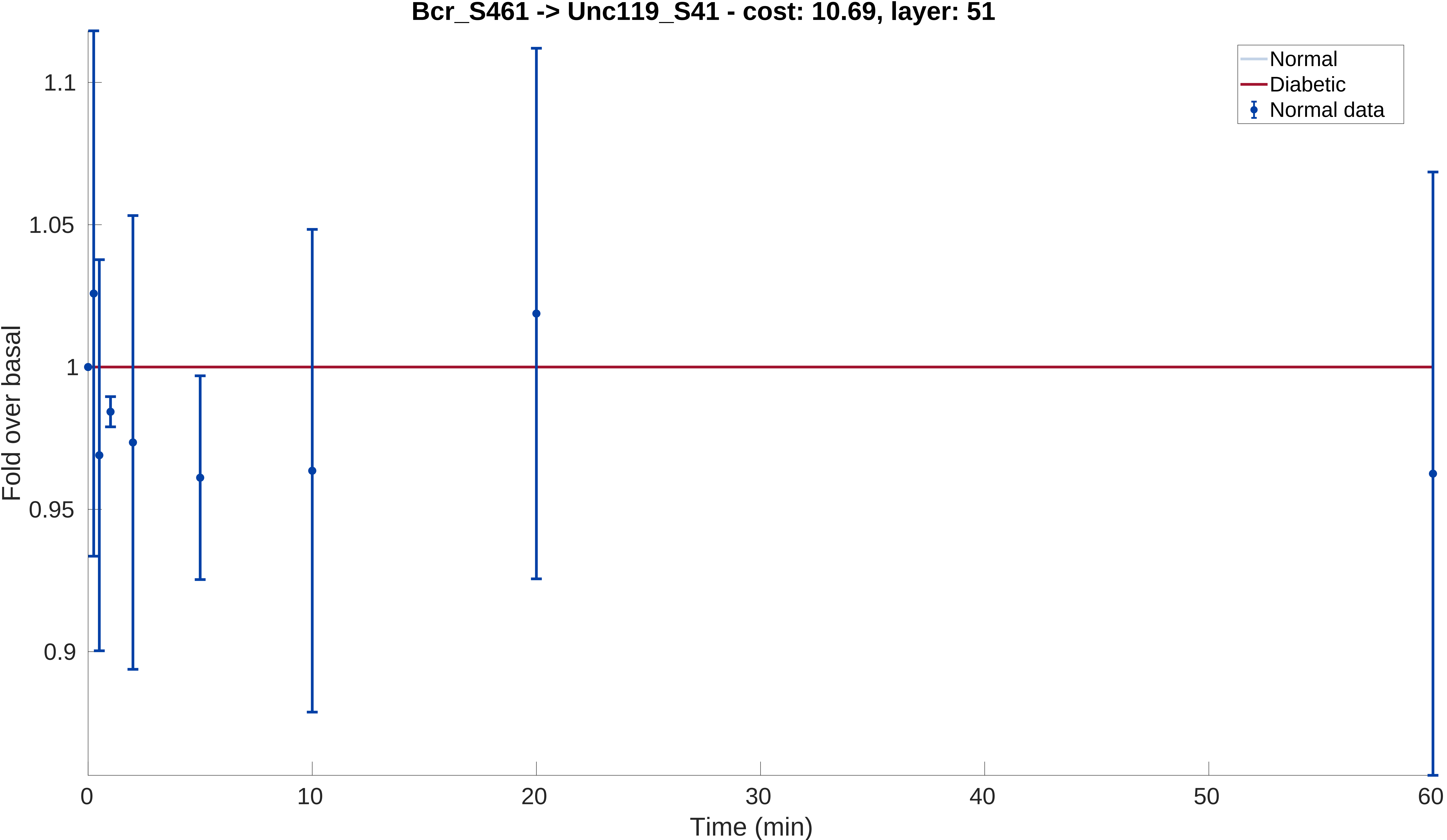

Bcr_S461 -> Unc119_S41 - cost: 10.69, layer: 51
Normal
Diabetic
Normal data
1.1
1.05
1
0.95
0.9
Fold over basal
0
10
20
30
40
50
60
Time (min)

#### Akt1_S126 -> Usp14_S393 - cost: 6.82, layer: 51

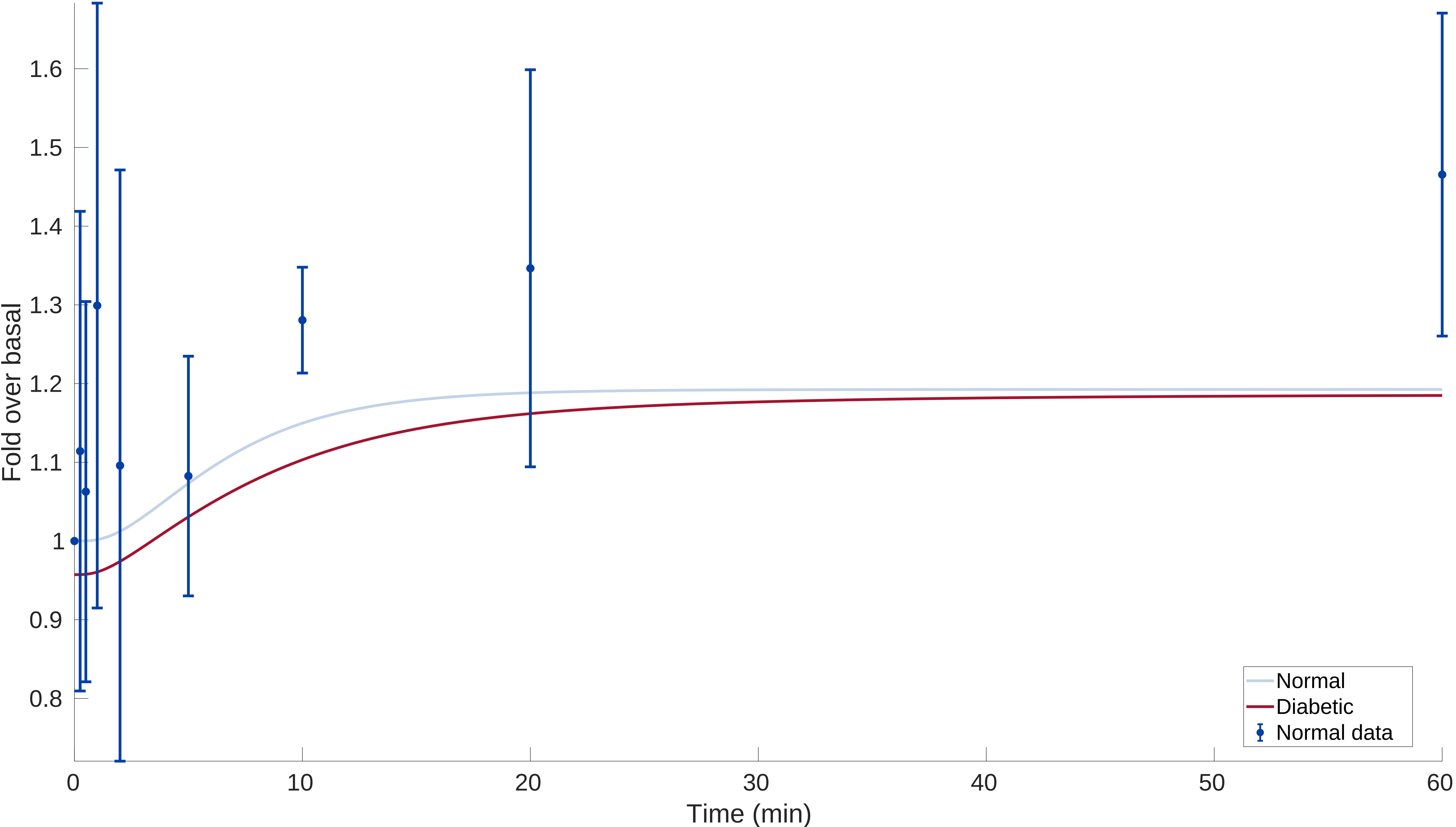

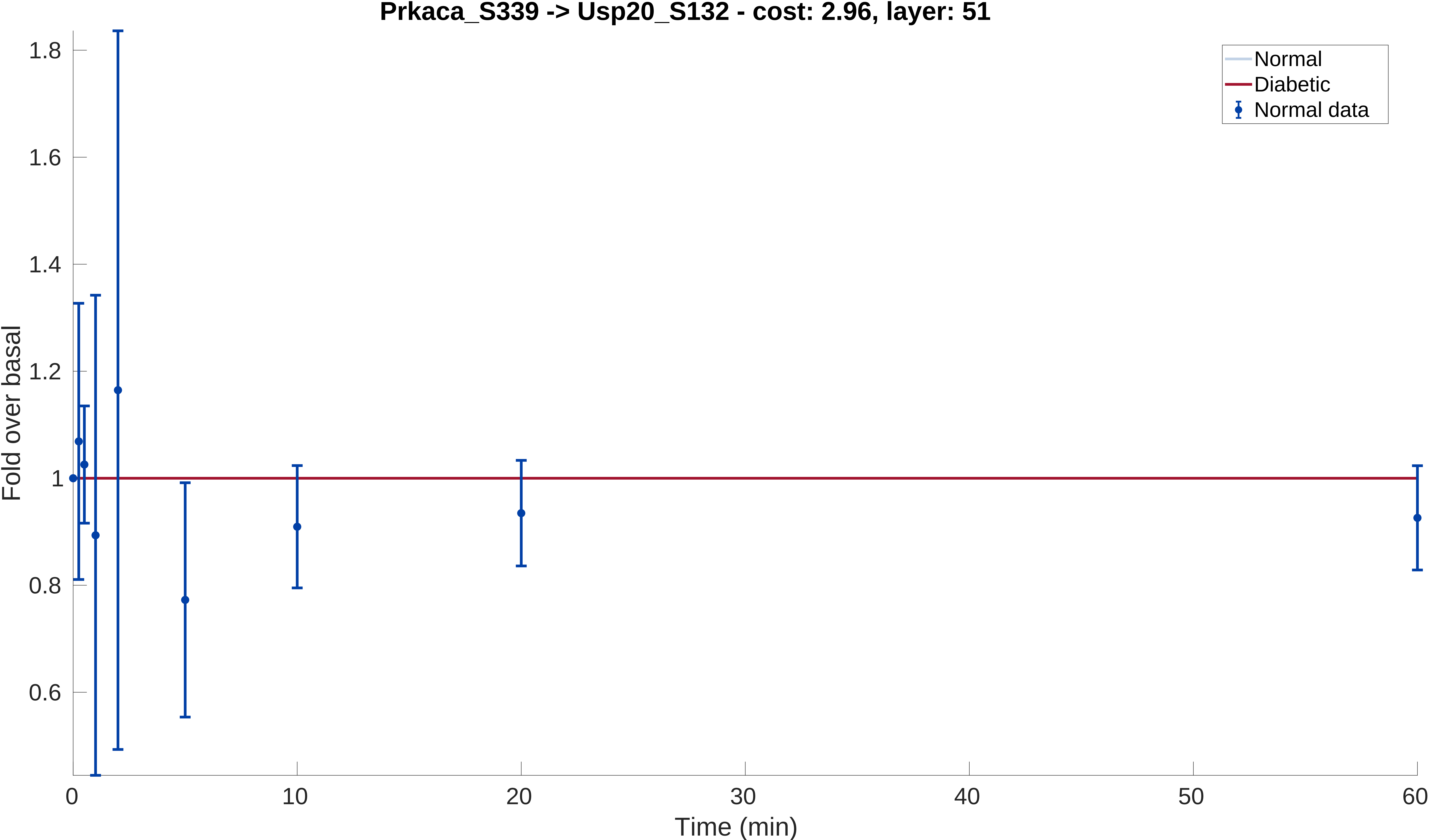

Prrkaca_S339 -> Usp20_S132 - cost: 2.96, layer: 51
Normal
Diabetic
Normal data
Fold over basal
1.8
1.6
1.4
1.2
1
0.8
0.6
0
10
20
30
40
50
60
Time (min)

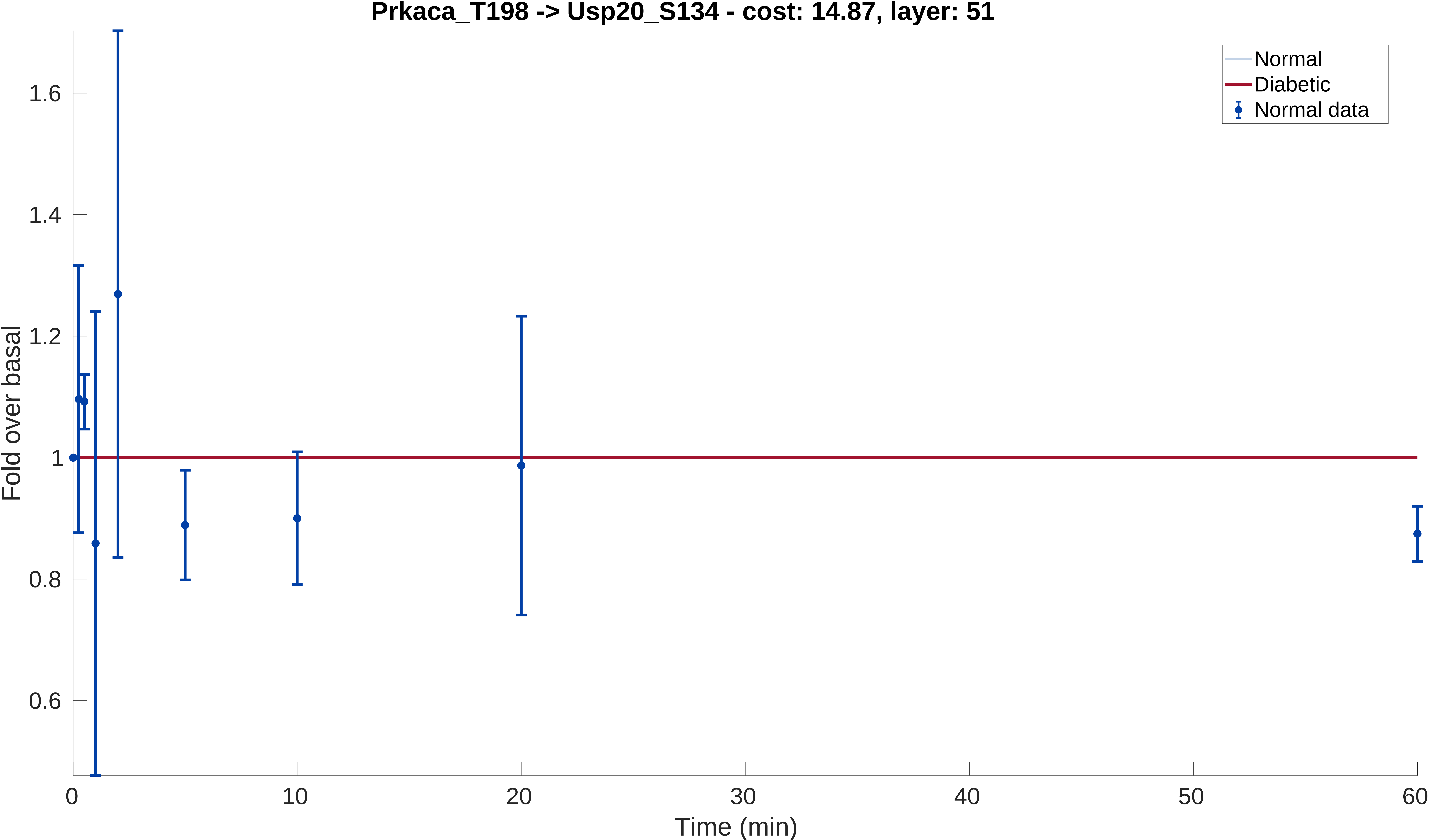

Prkaca_T198 -> Usp20_S134 - cost: 14.87, layer: 51
Normal
Diabetic
Normal data
Fold over basal
1.6
1.4
1.2
1
0.8
0.6
0
10
20
30
40
50
60
Time (min)

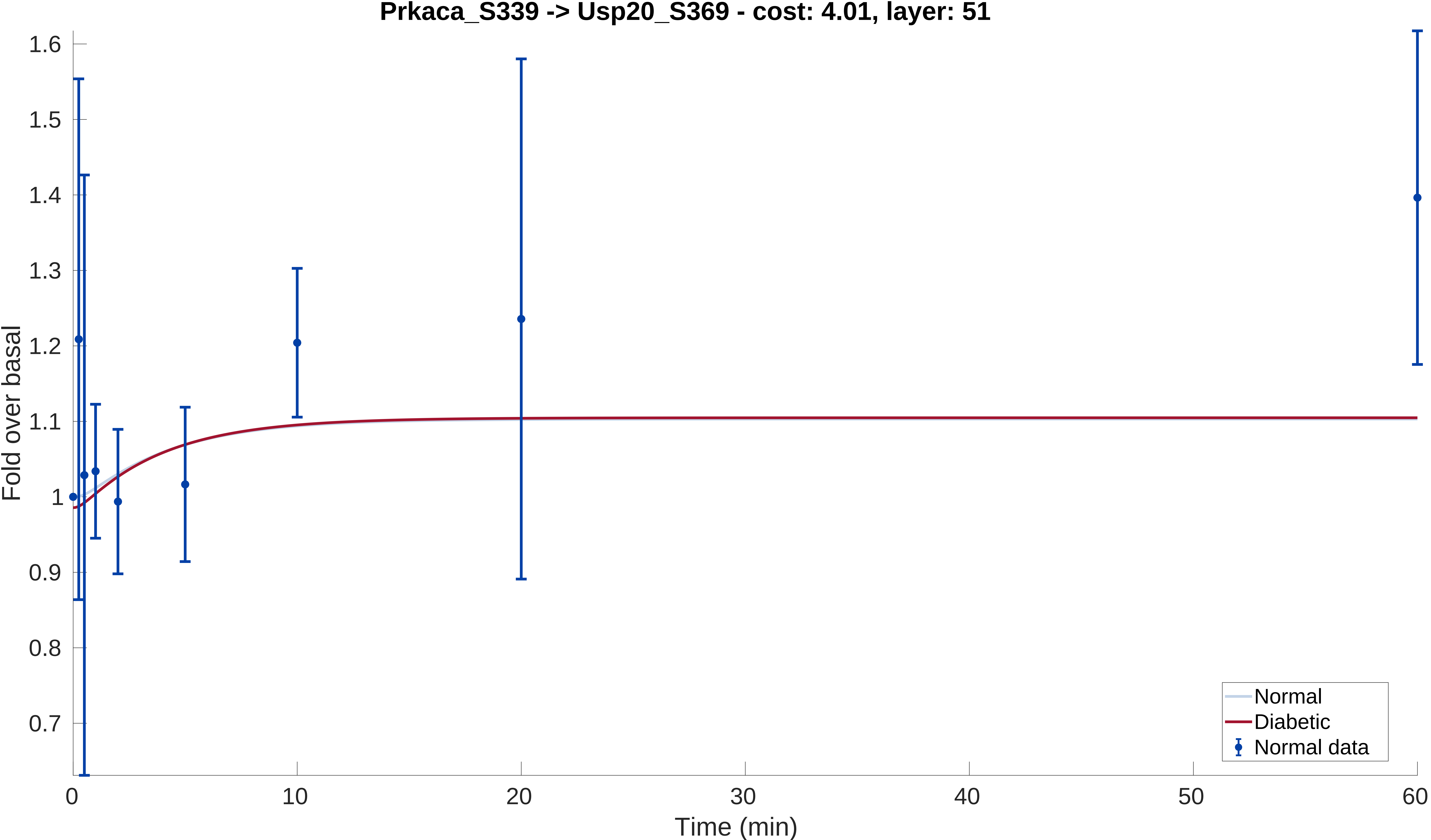

Prkaca_S339 -> Usp20_S369 - cost: 4.01, layer: 51
Fold over basal
1.6
1.5
1.4
1.3
1.2
1.1
1
0.9
0.8
0.7
0
10
20
30
40
50
60
Time (min)
Normal
Diabetic
Normal data

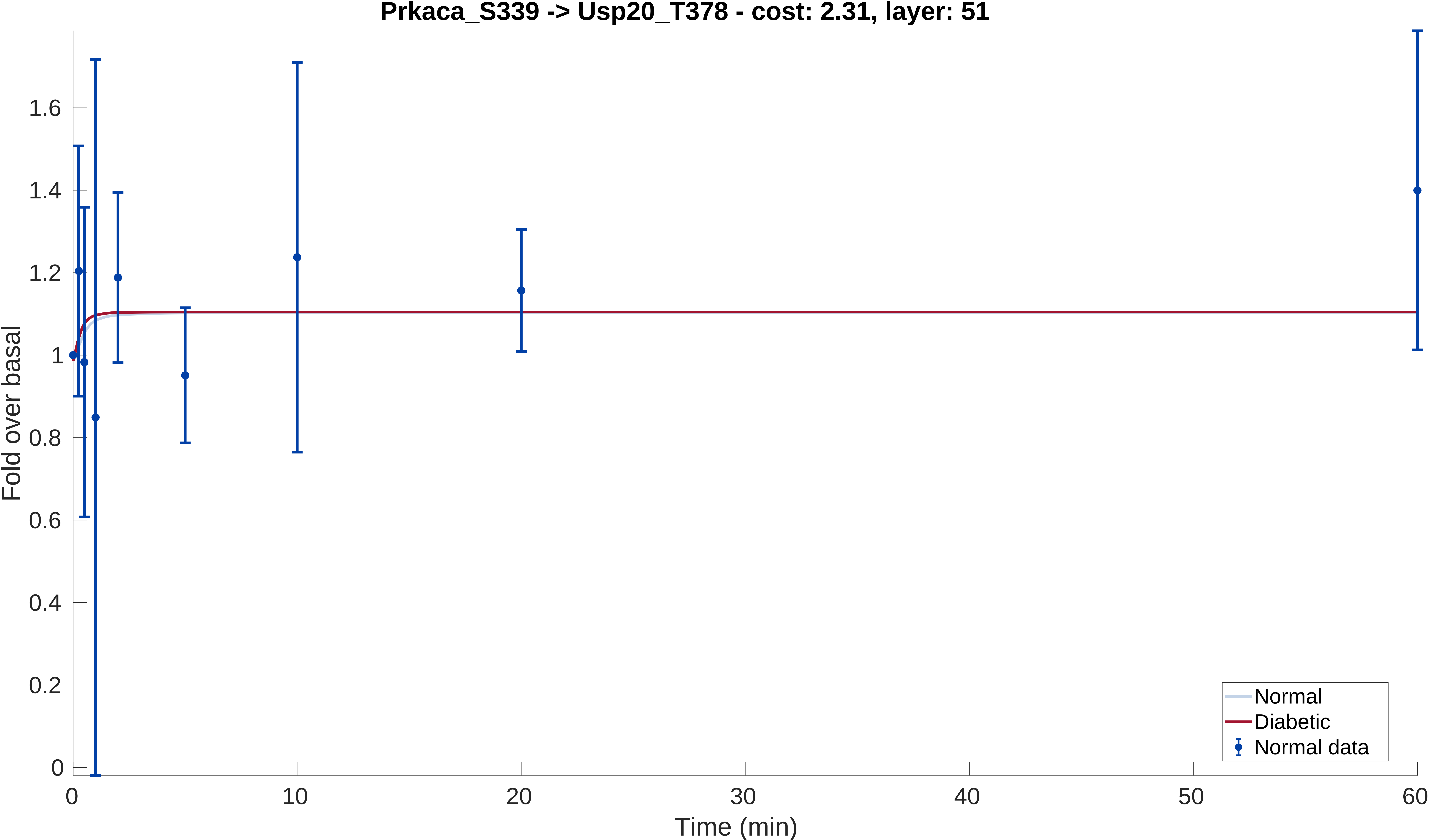

Prrkaca_S339 -> Usp20_T378 - cost: 2.31, layer: 51
Fold over basal
Time (min)
0
0.2
0.4
0.6
0.8
1
1.2
1.4
1.6
0
10
20
30
40
50
60
Normal
Diabetic
Normal data

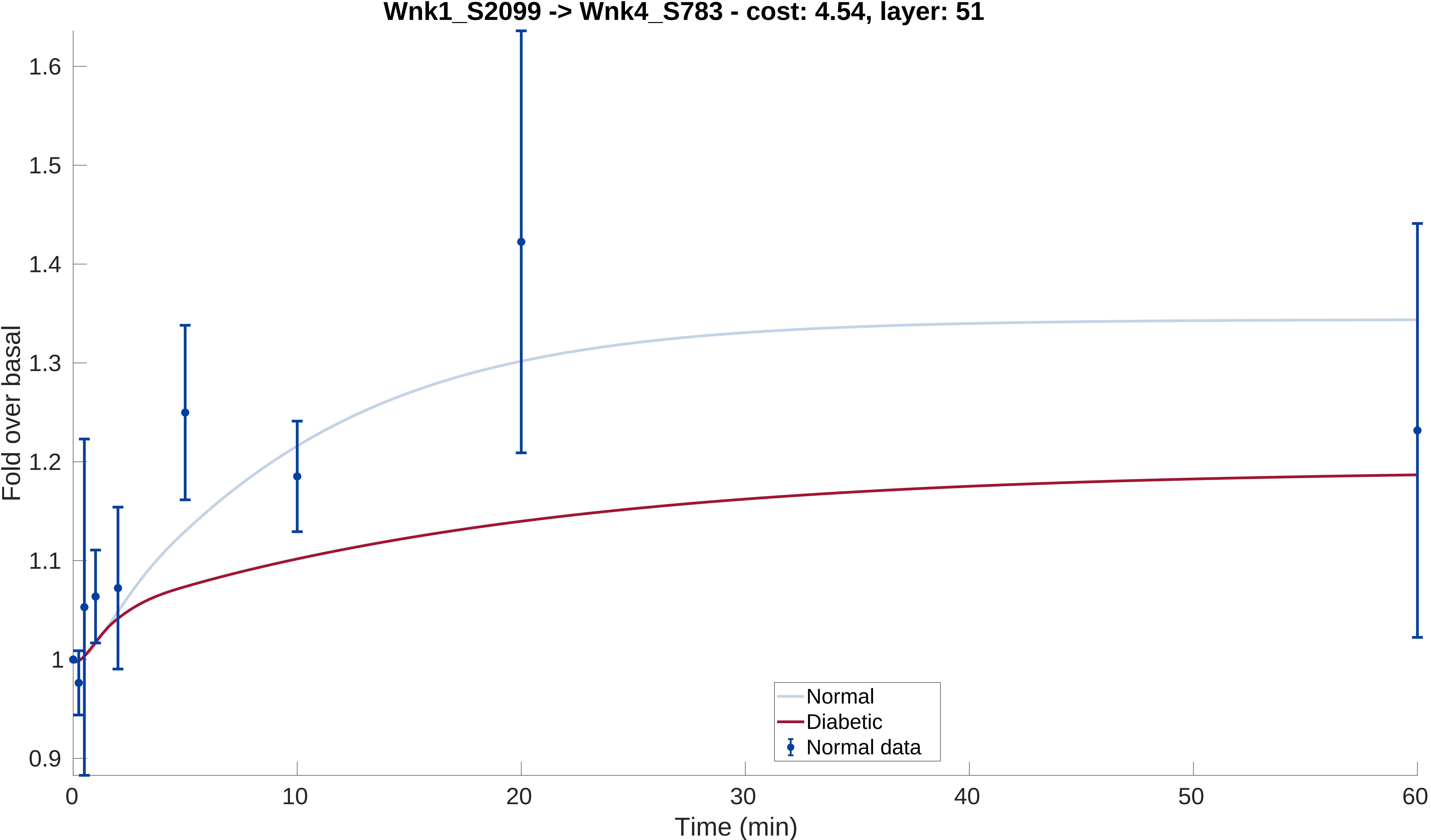

Wnk1_S2099 -> Wnk4_S783 - cost: 4.54, layer: 51
Fold over basal
1.6
1.5
1.4
1.3
1.2
1.1
1
0.9
0
10
20
30
40
50
60
Time (min)
Normal
Diabetic
Normal data

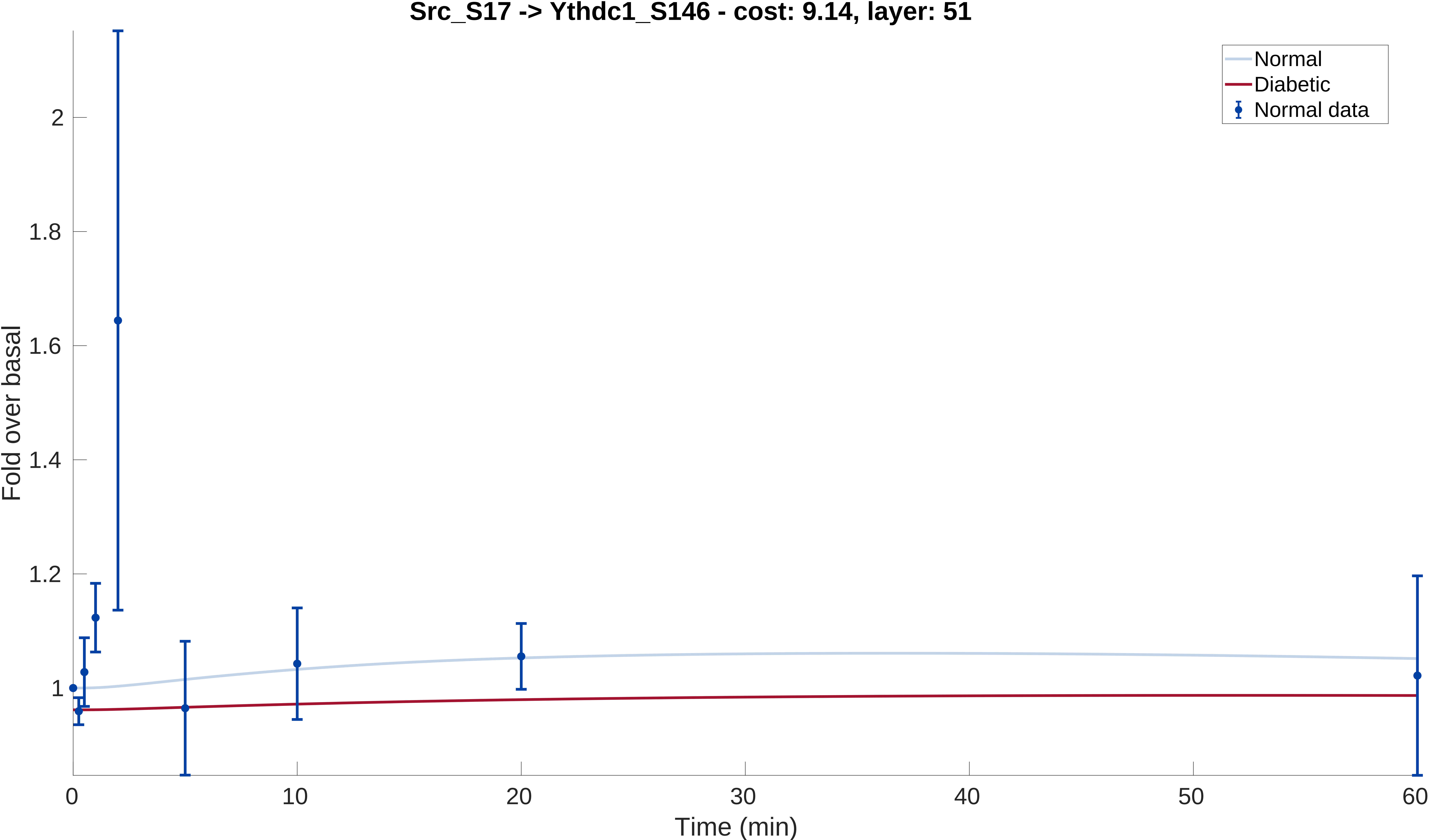
Src_S17 -> Ythdc1_S146 - cost: 9.14, layer: 51
Normal
Diabetic
Normal data
Fold over basal
2
1.8
1.6
1.4
1.2
1
0
10
20
30
40
50
60
Time (min)

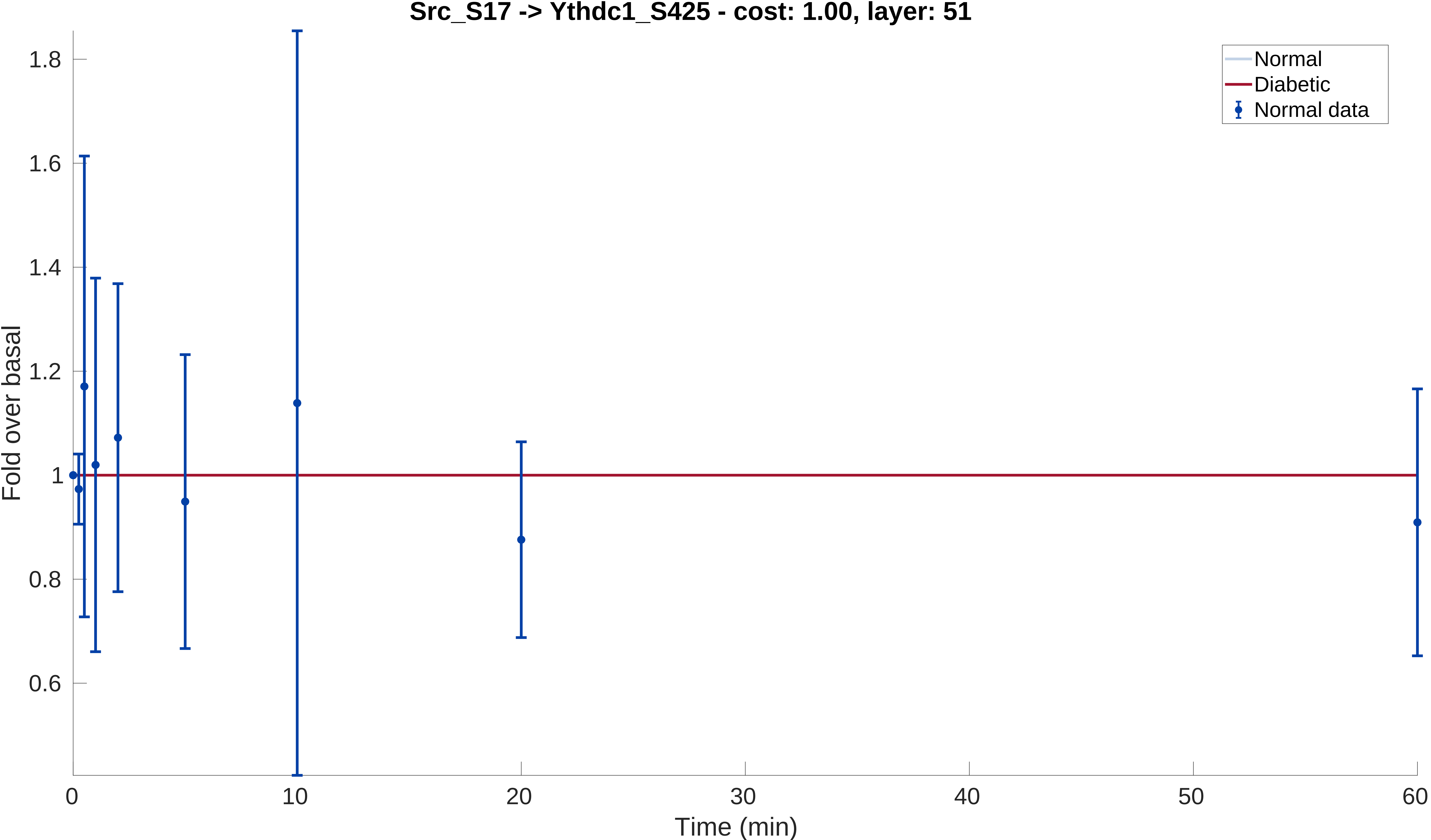

Src_S17 -> Ythdc1_S425 - cost: 1.00, layer: 51
Normal
Diabetic
Normal data
Fold over basal
1.8
1.6
1.4
1.2
1
0.8
0.6
0
10
20
30
40
50
60
Time (min)

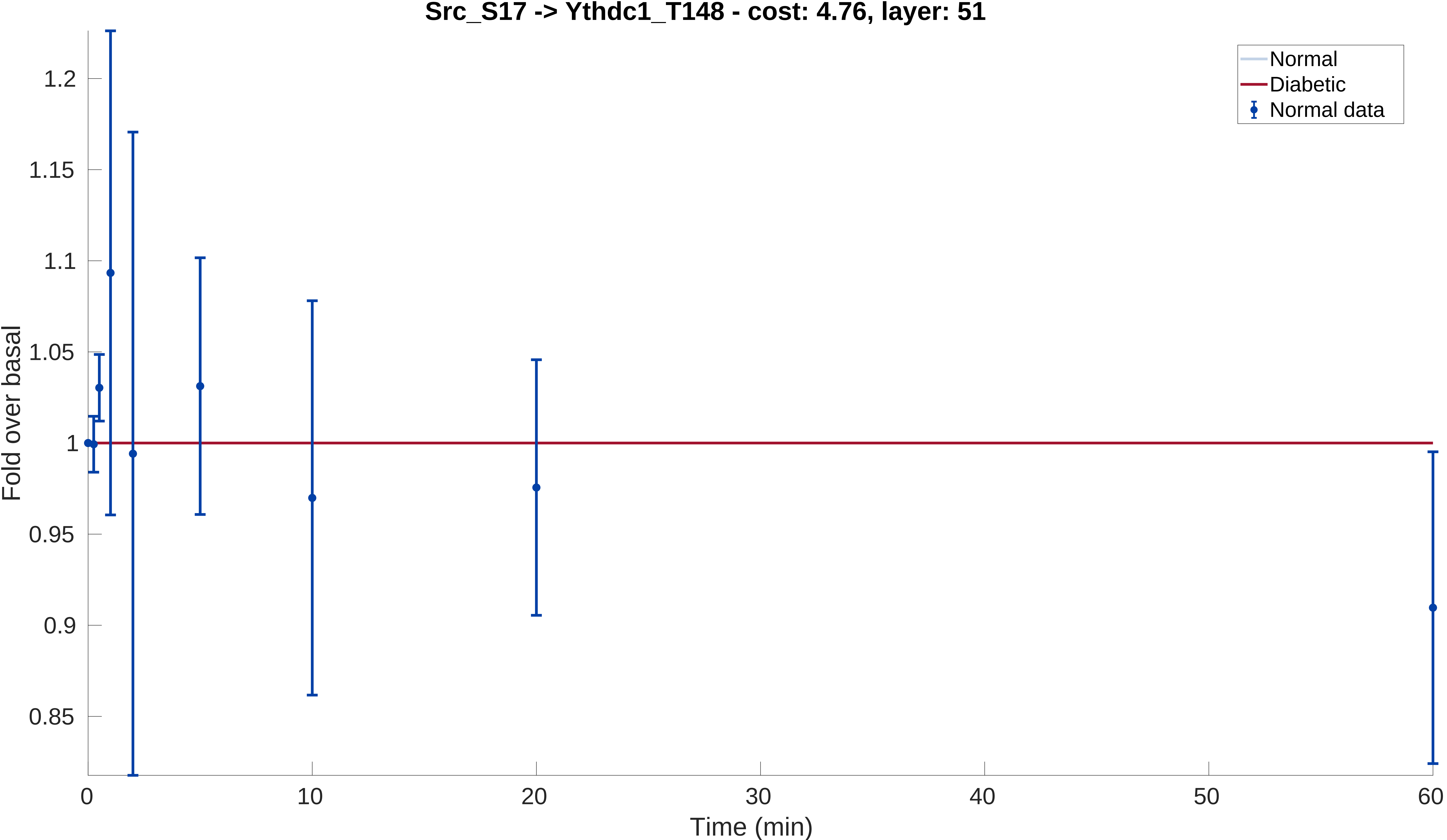

Src_S17 -> Ythdc1_T148 - cost: 4.76, layer: 51
Normal
Diabetic
Normal data
1.2
1.15
1.1
1.05
1
0.95
0.9
0.85
Fold over basal
0
10
20
30
40
50
60
Time (min)

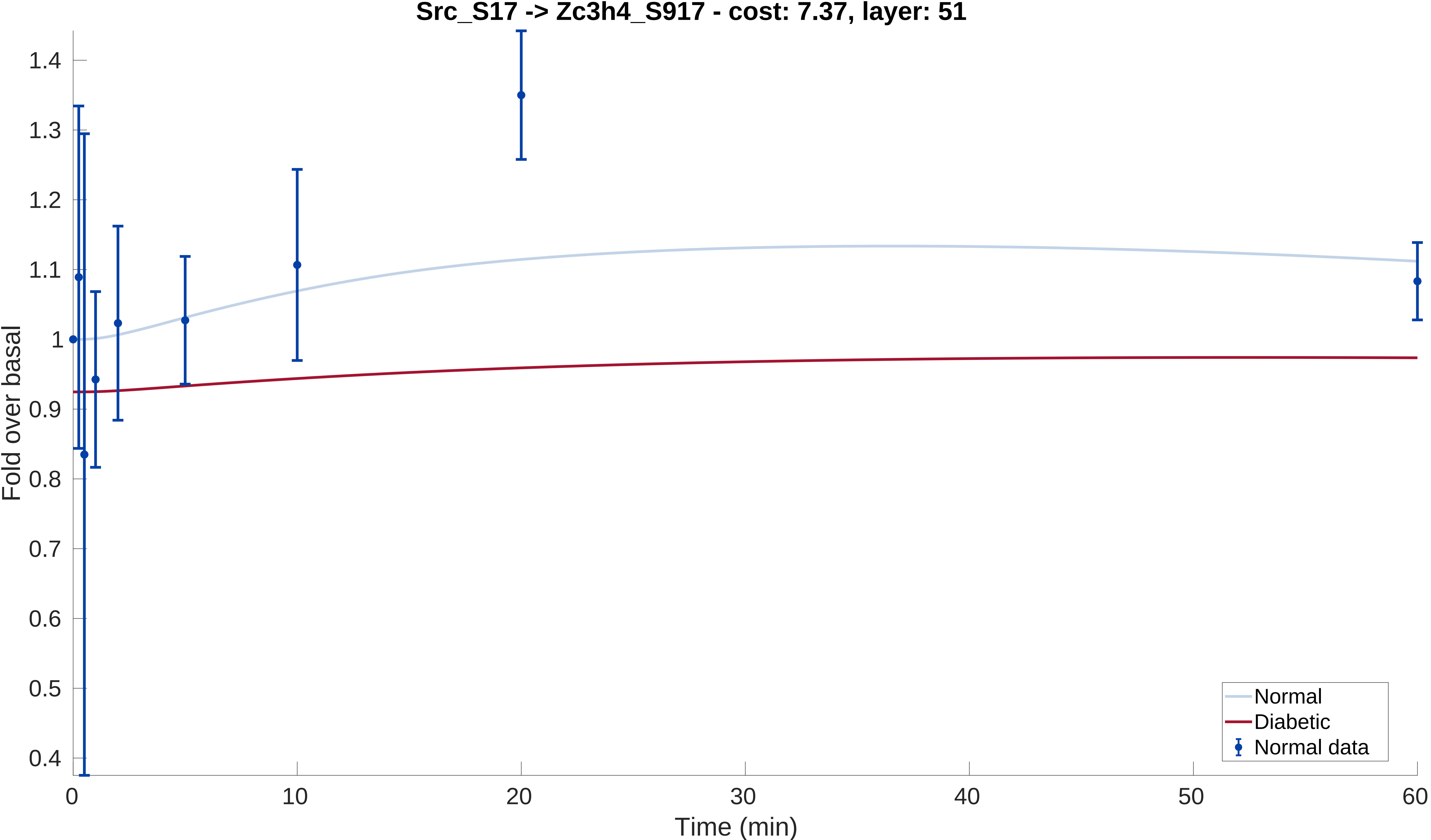
Src_S17 -> Zc3h4_S917 - cost: 7.37, layer: 51
Fold over basal
1.4
1.3
1.2
1.1
1
0.9
0.8
0.7
0.6
0.5
0.4
0
10
20
30
40
50
60
Time (min)
Normal
Diabetic
Normal data

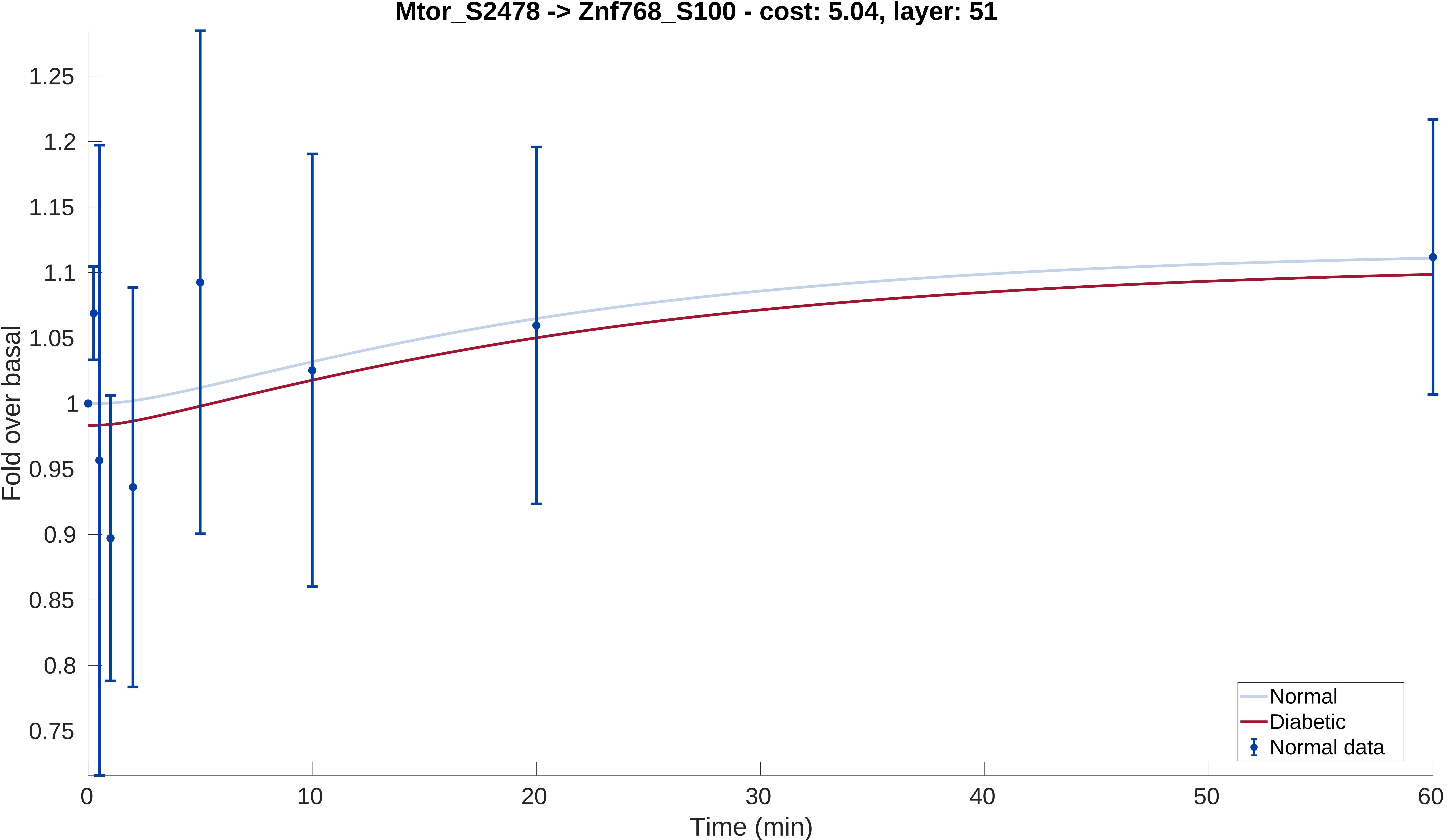
Mtor_S2478 -> Znf768_S100 - cost: 5.04, layer: 51
1.25
1.2
1.15
1.1
1.05
1
0.95
0.9
0.85
0.8
0.75
Fold over basal
0
10
20
30
40
50
60
Time (min)
Normal
Diabetic
Normal data

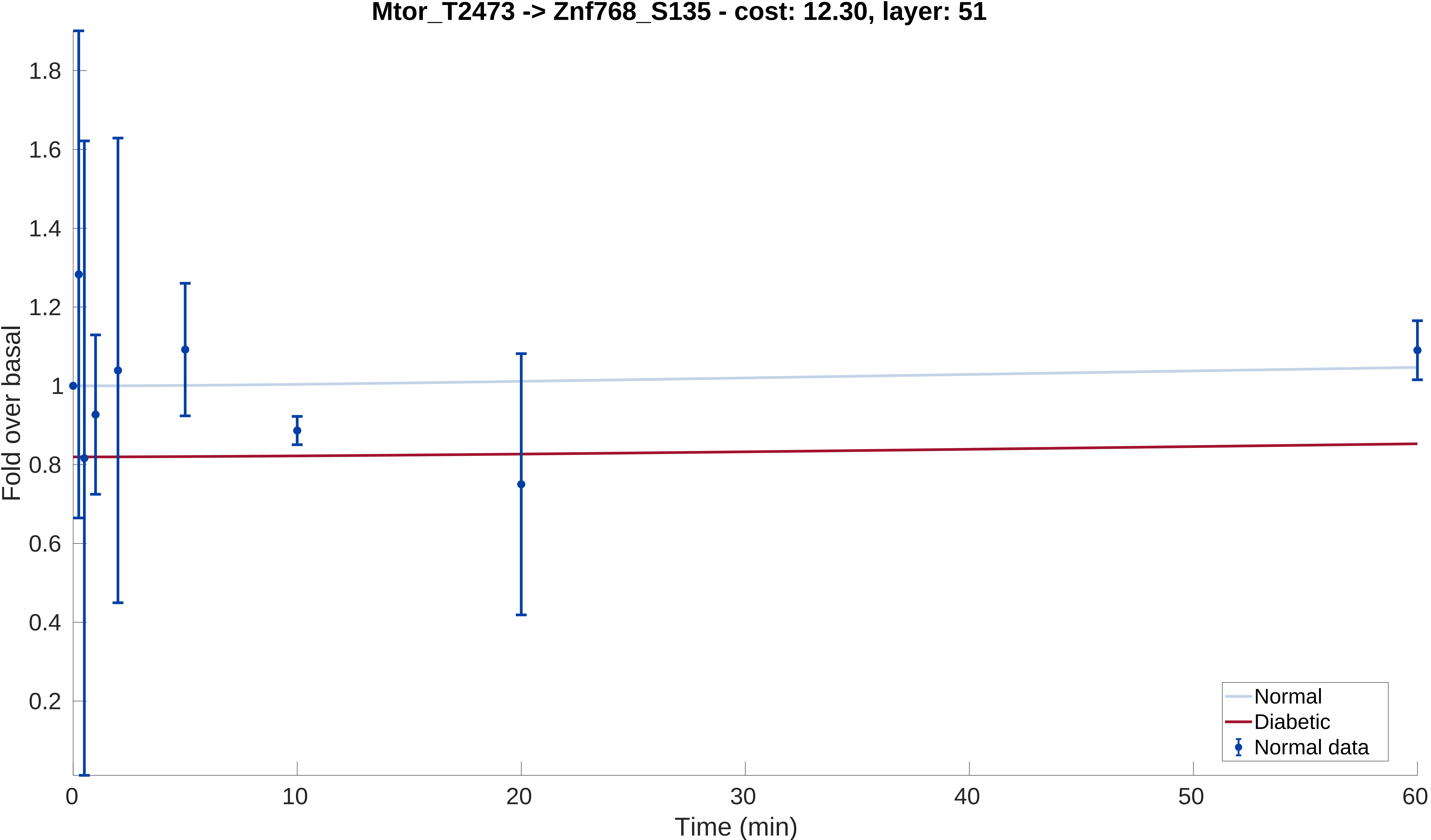

Mtor_T2473 -> Znf768_S135 - cost: 12.30, layer: 51
Fold over basal
Time (min)
1.8
1.6
1.4
1.2
1
0.8
0.6
0.4
0.2
0
10
20
30
40
50
60
Normal
Diabetic
Normal data

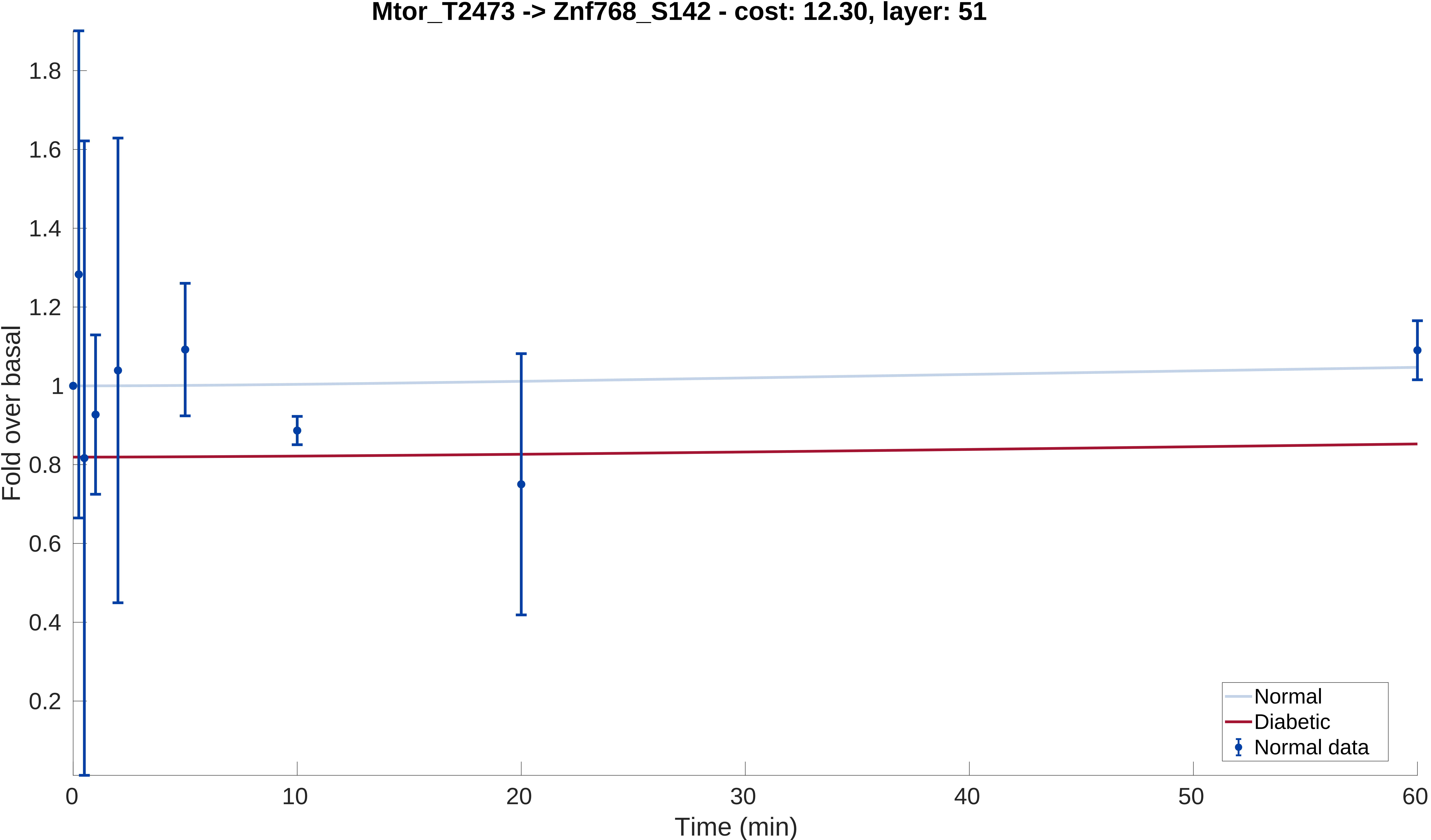
Mtor_T2473 -> Znf768_S142 - cost: 12.30, layer: 51
Fold over basal
1.8
1.6
1.4
1.2
1
0.8
0.6
0.4
0.2
0
10
20
30
40
50
60
Time (min)
Normal
Diabetic
Normal data

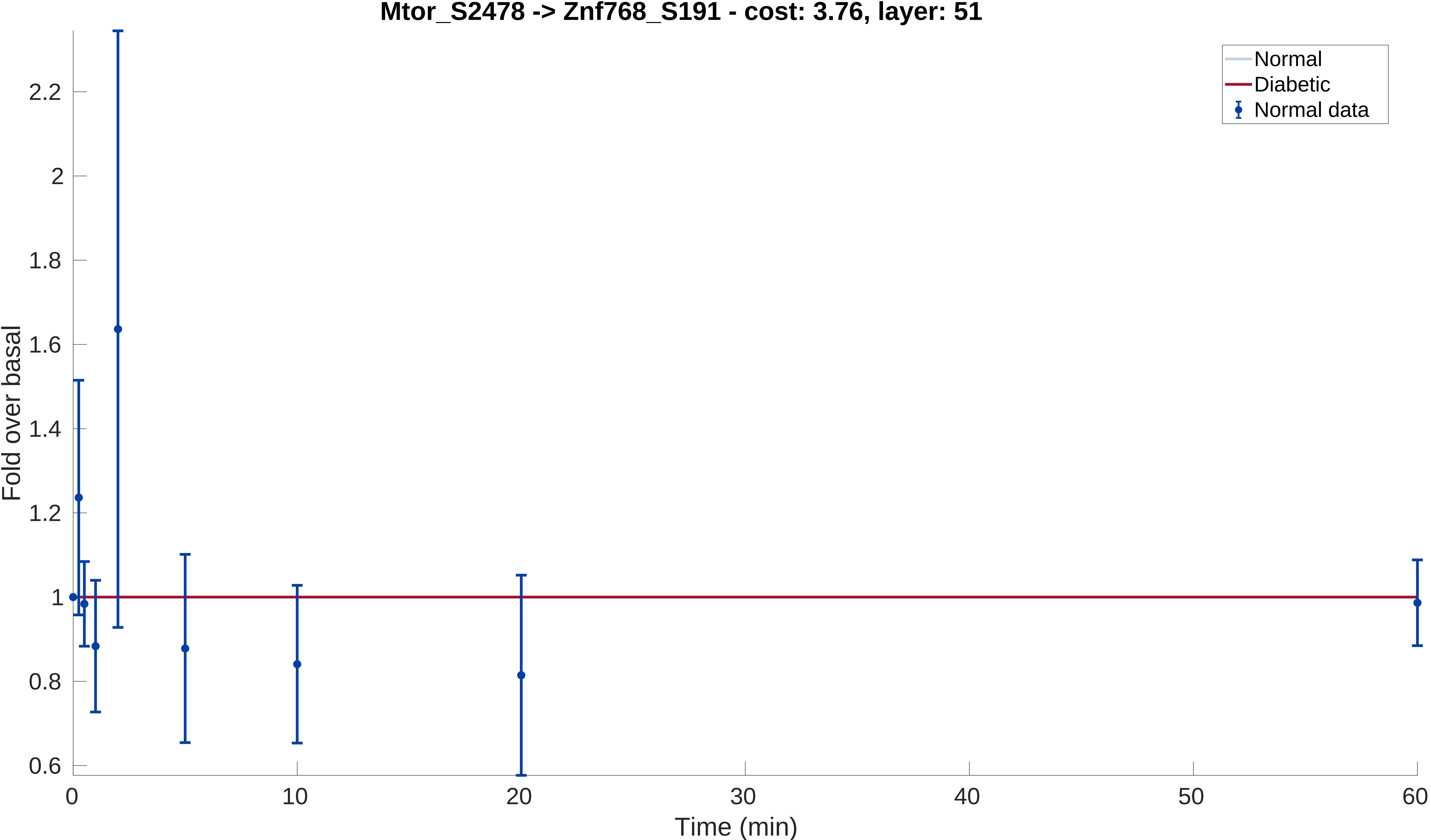

Mtor_S2478 -> Znf768_S191 - cost: 3.76, layer: 51
Normal
Diabetic
Normal data
Fold over basal
2.2
2
1.8
1.6
1.4
1.2
1
0.8
0.6
0
10
20
30
40
50
60
Time (min)

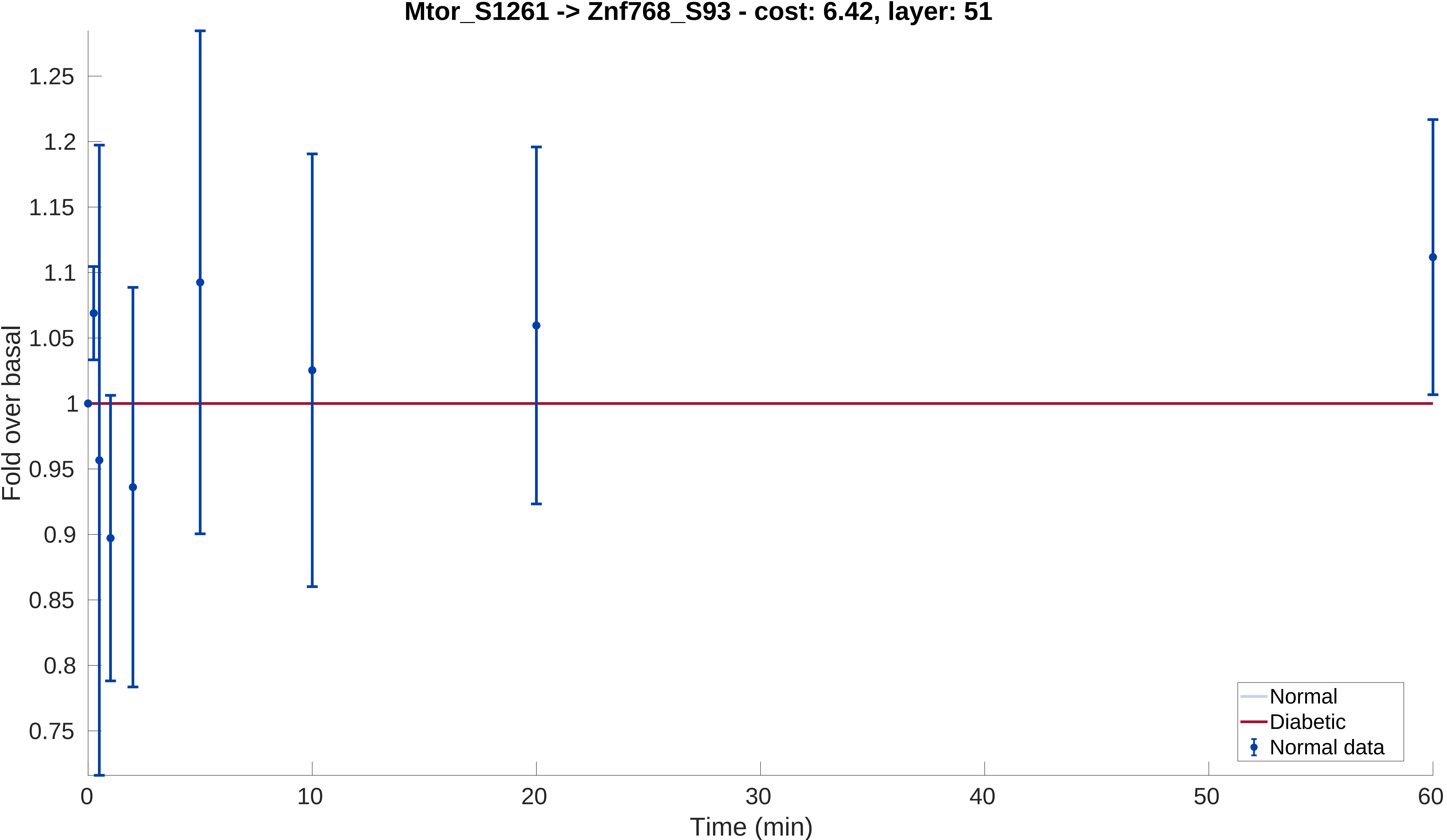

Mtor_S1261 -> Znf768_S93 - cost: 6.42, layer: 51
Fold over basal
1.25
1.2
1.15
1.1
1.05
1
0.95
0.9
0.85
0.8
0.75
0
10
20
30
40
50
60
Time (min)
Normal
Diabetic
Normal data

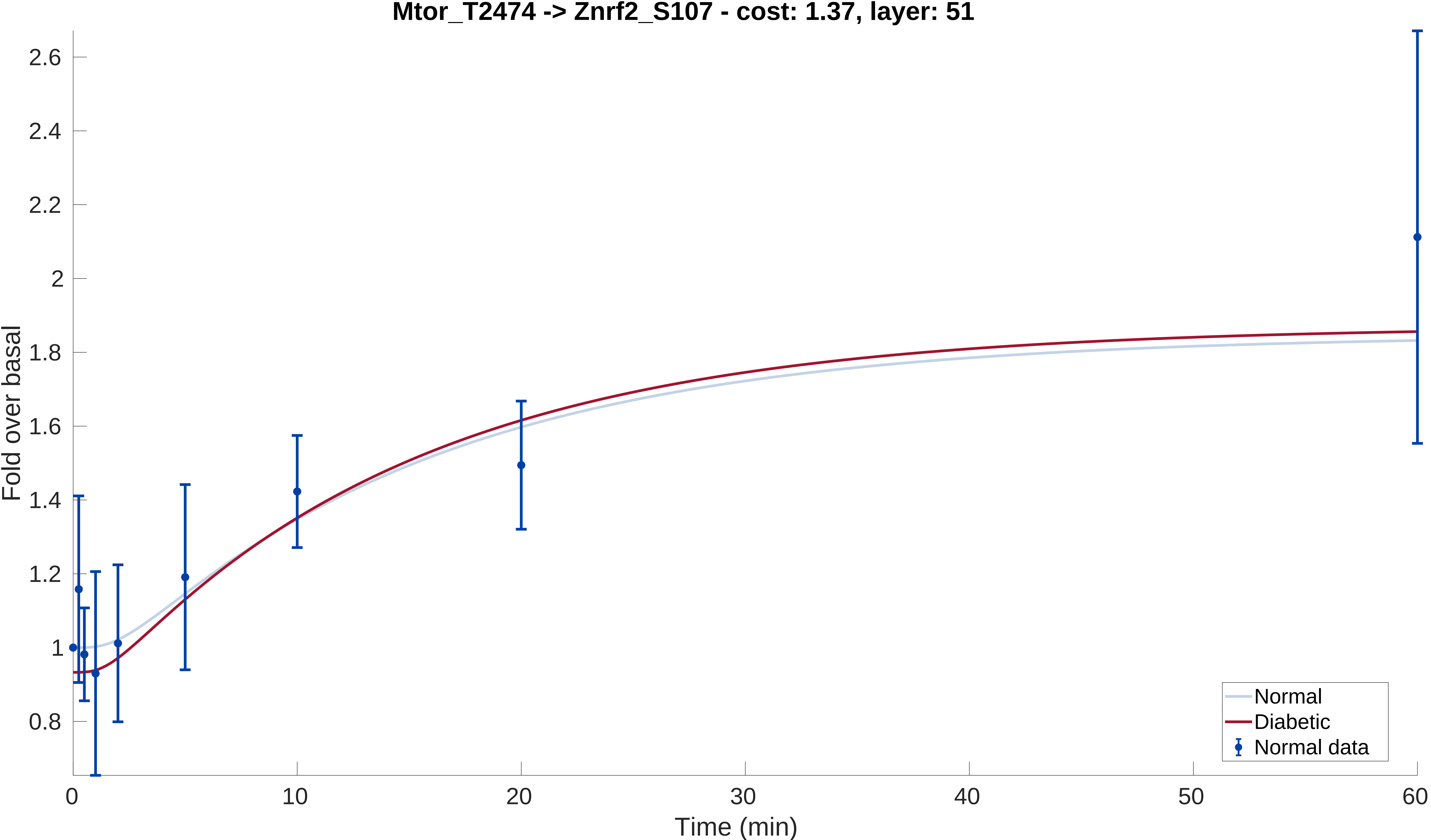
Mtor_T2474 -> Znrf2_S107 - cost: 1.37, layer: 51
Fold over basal
2.6
2.4
2.2
2
1.8
1.6
1.4
1.2
1
0.8
0
10
20
30
40
50
60
Time (min)
Normal
Diabetic
Normal data

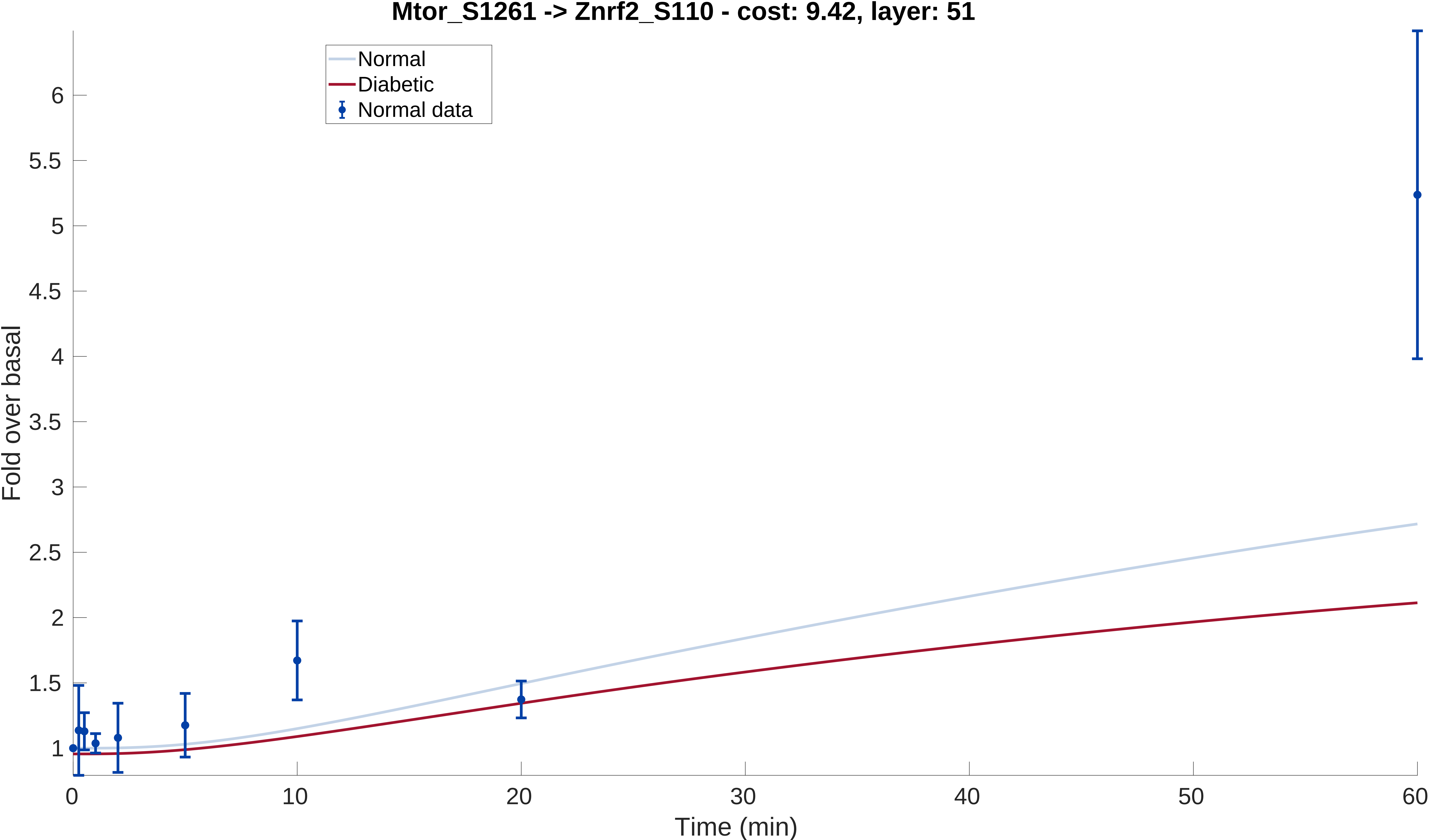

Mtor_S1261 -> Znrf2_S110 - cost: 9.42, layer: 51
Normal
Diabetic
Normal data
Fold over basal
Time (min)
1
1.5
2
2.5
3
3.5
4
4.5
5
5.5
6
0
10
20
30
40
50
60

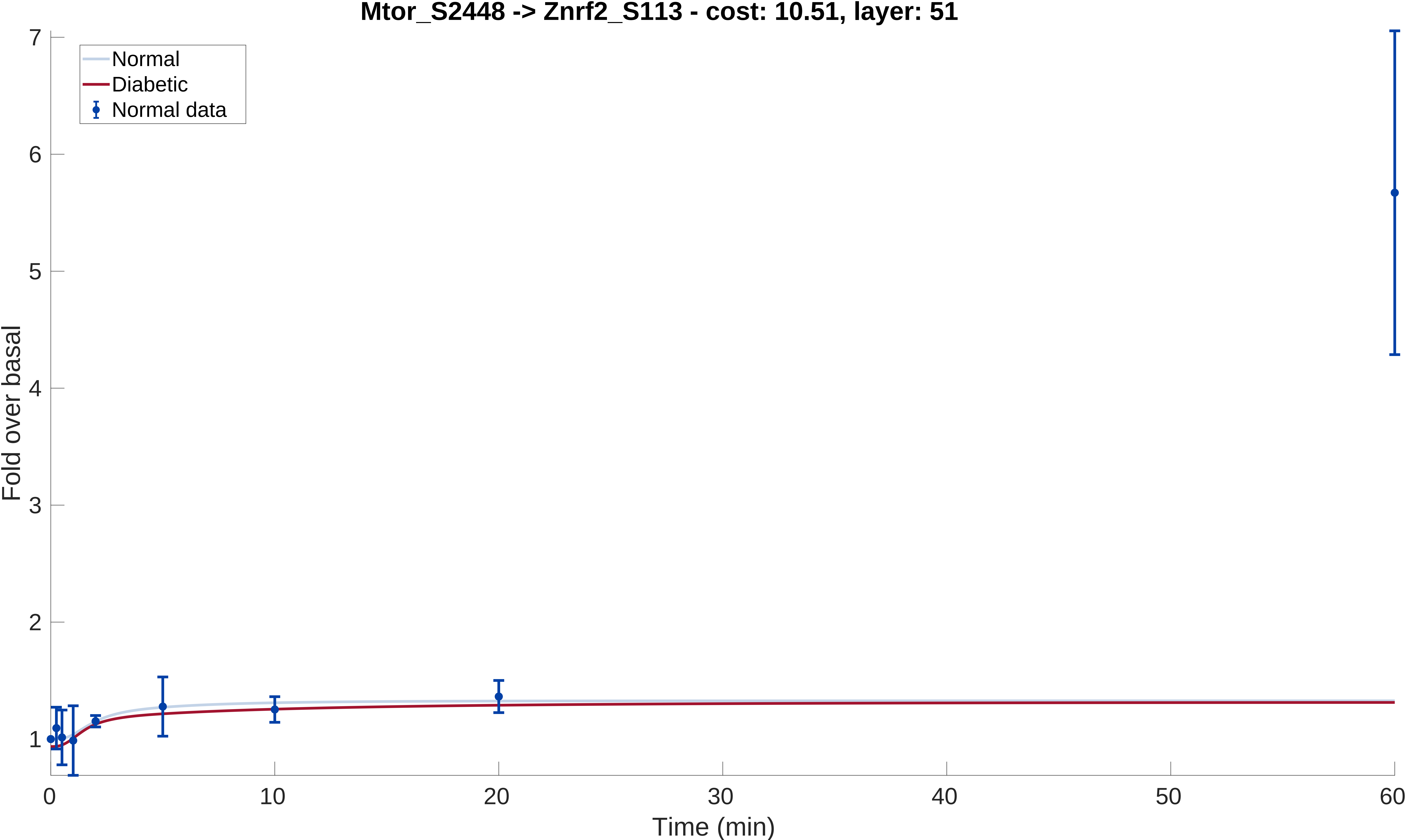

Mtor_S2448 -> Znrf2_S113 - cost: 10.51, layer: 51
Normal
Diabetic
Normal data
Fold over basal
Time (min)
1
2
3
4
5
6
7
0
10
20
30
40
50
60

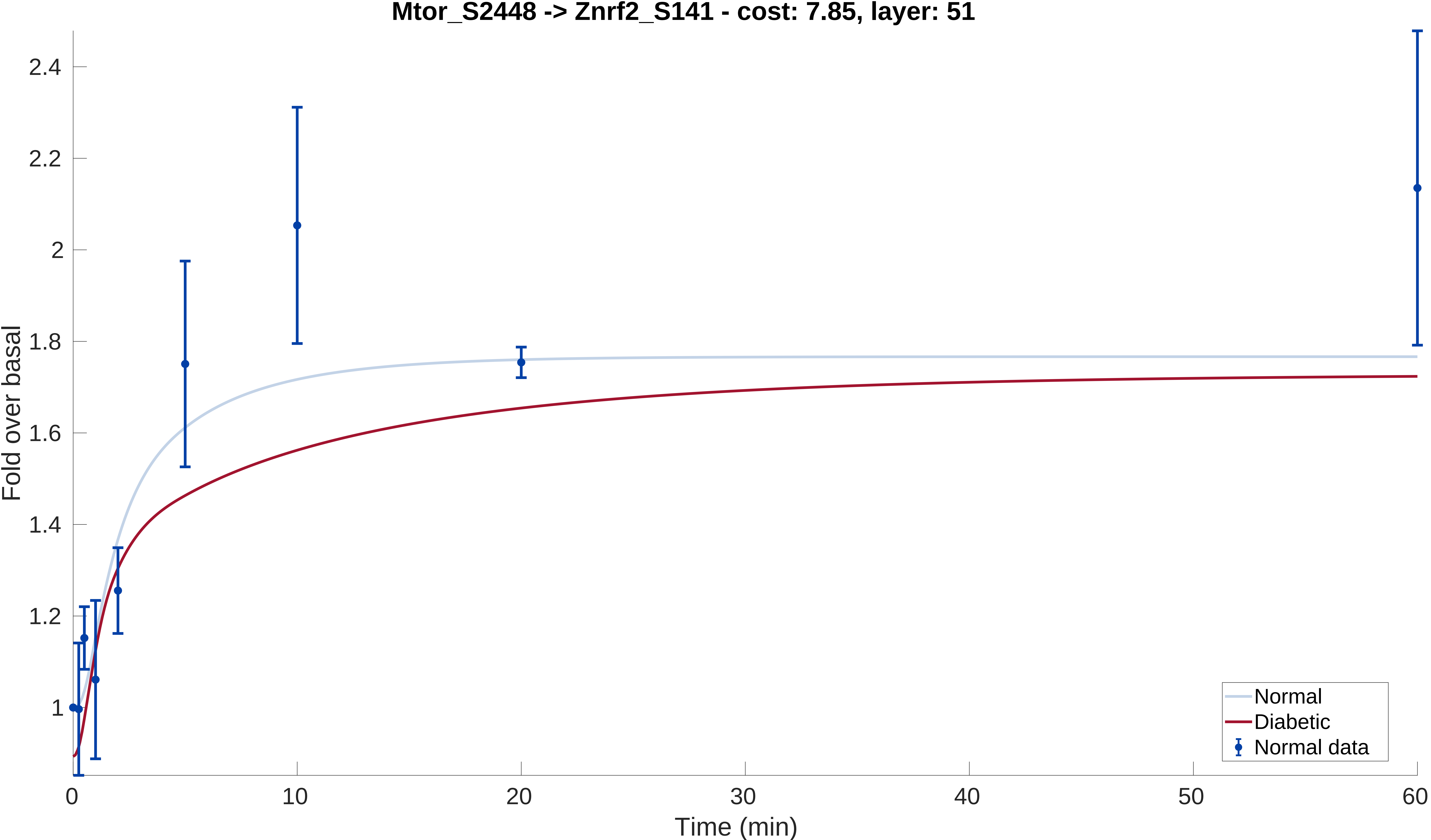

Mtor_S2448 -> Znrf2_S141 - cost: 7.85, layer: 51
Fold over basal
2.4
2.2
2
1.8
1.6
1.4
1.2
1
0
10
20
30
40
50
60
Time (min)
Normal
Diabetic
Normal data

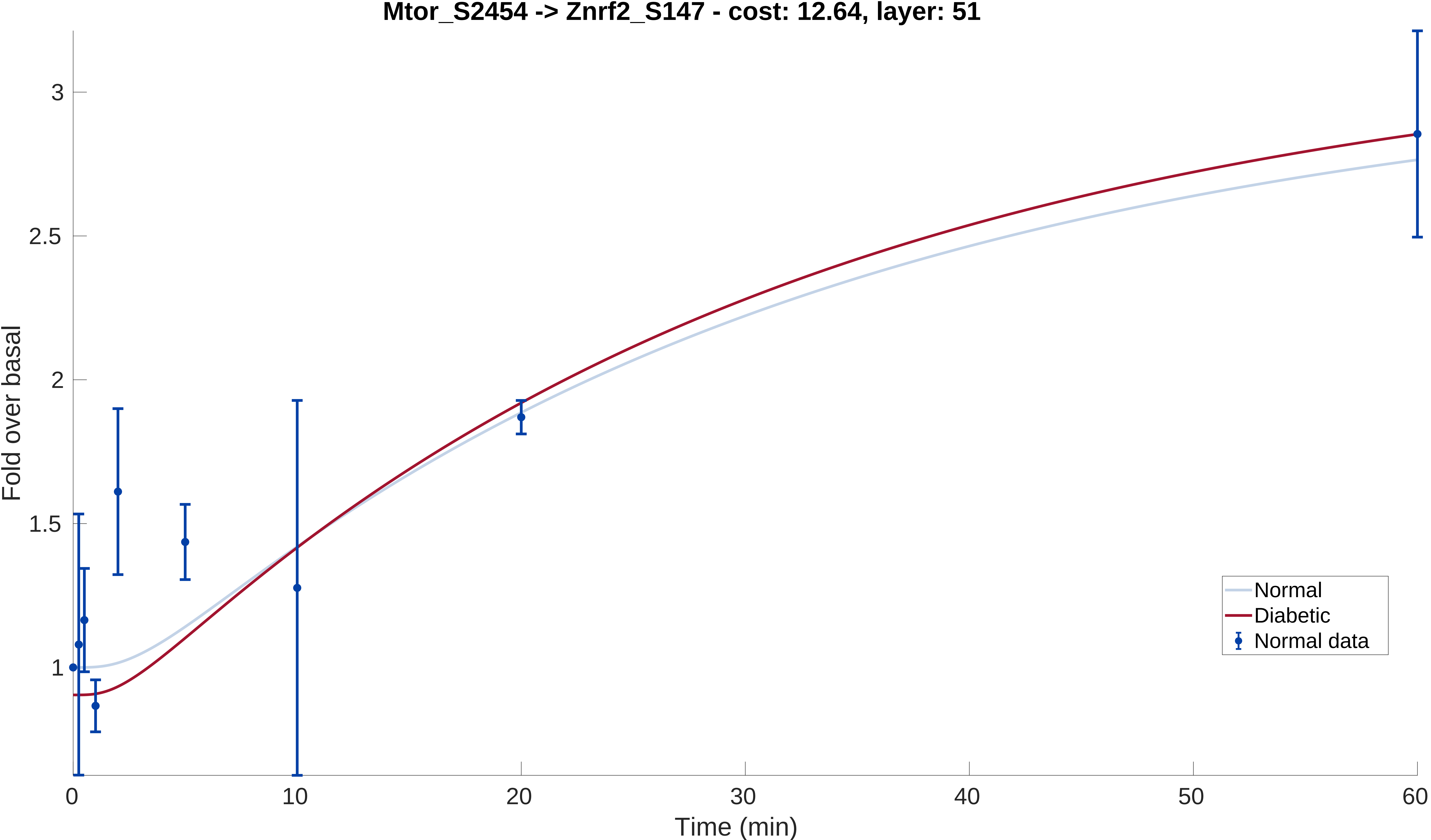

Mtor_S2454 -> Znrf2_S147 - cost: 12.64, layer: 51
Fold over basal
3
2.5
2
1.5
1
0
10
20
30
40
50
60
Time (min)
Normal
Diabetic
Normal data

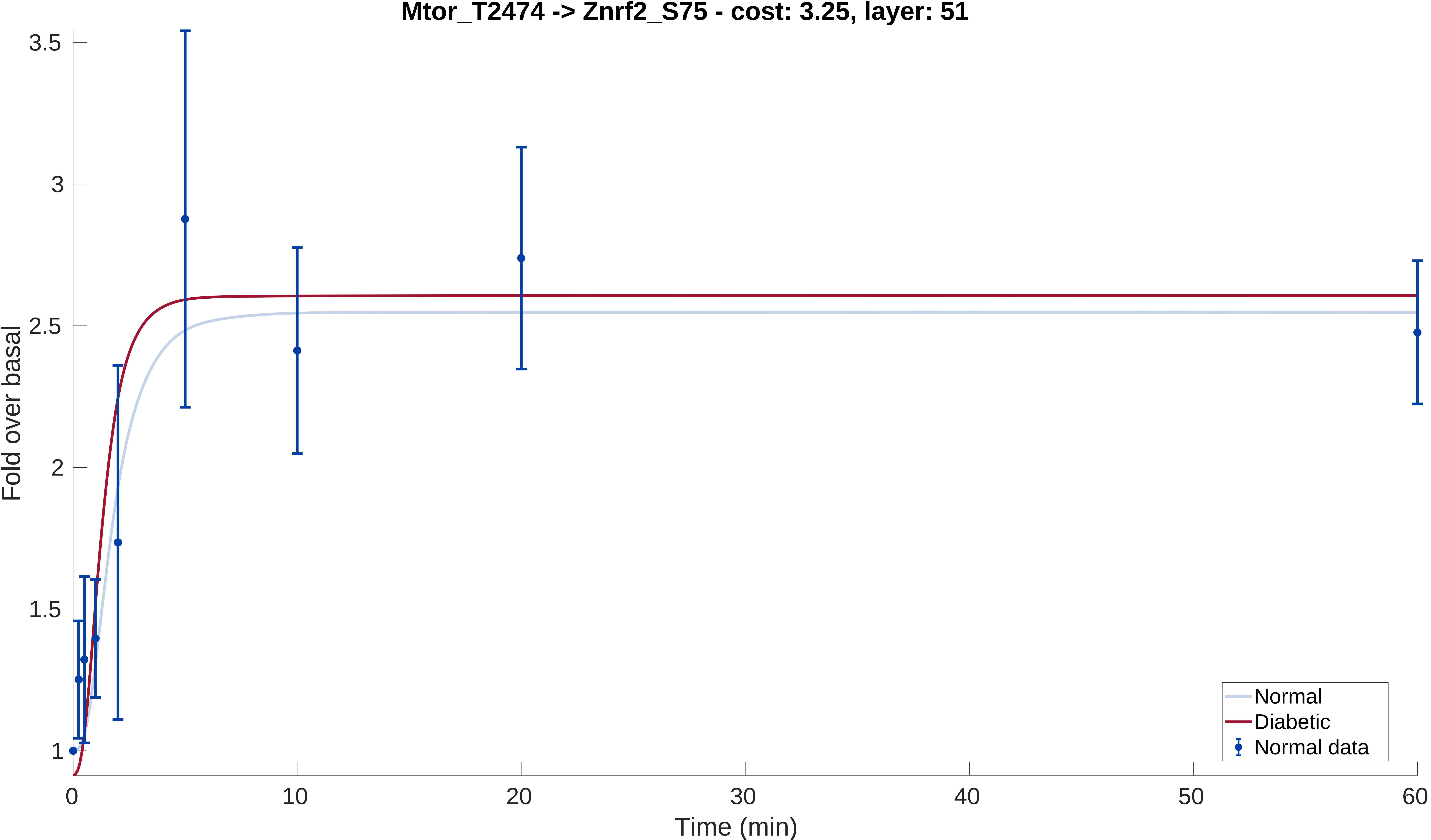

Mtor_T2474 -> Znrf2_S75 - cost: 3.25, layer: 51
3.5
3
2.5
2
1.5
1
Fold over basal
0
10
20
30
40
50
60
Time (min)
Normal
Diabetic
Normal data

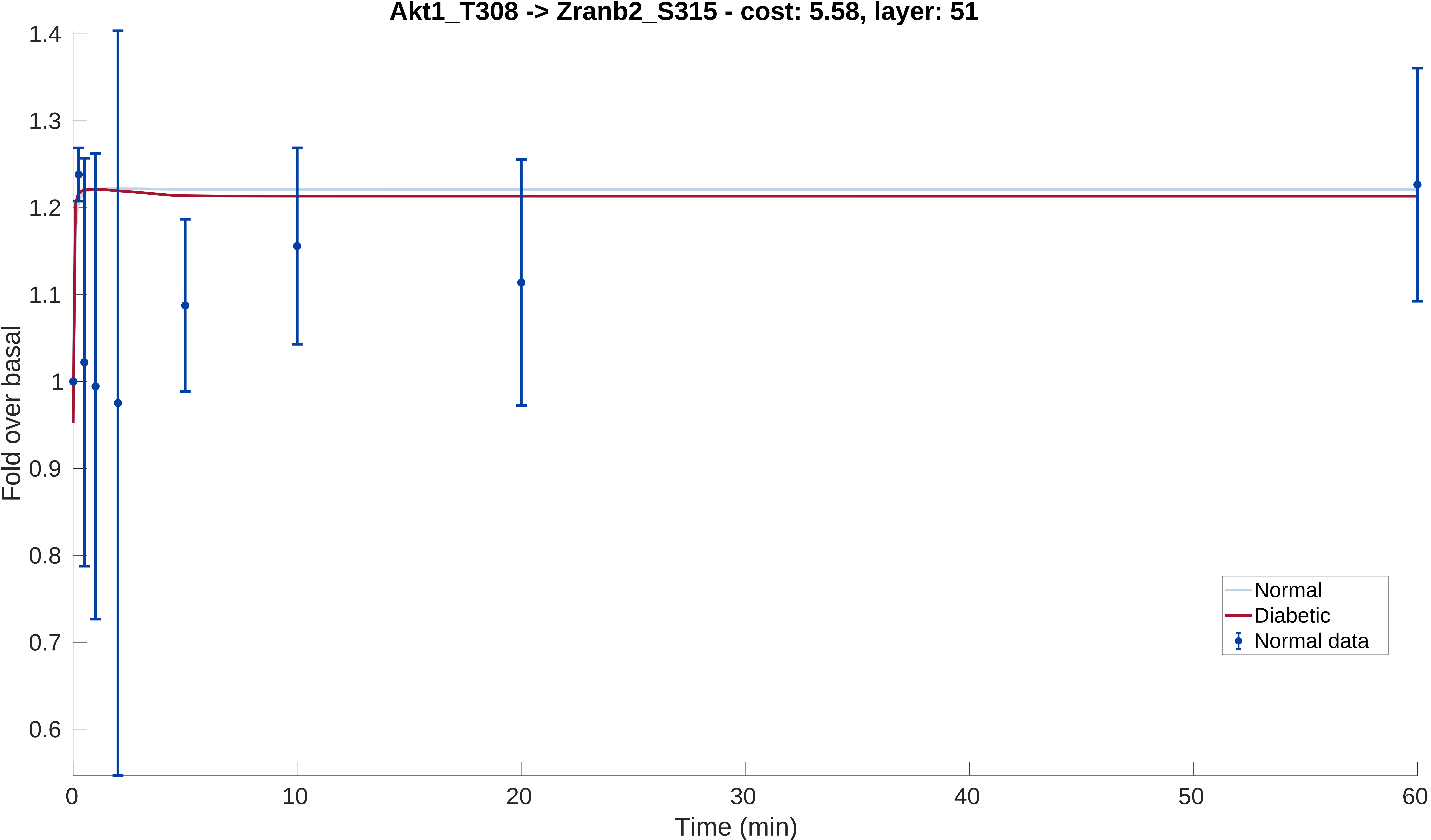

Akt1_T308 -> Zranb2_S315 - cost: 5.58, layer: 51
Fold over basal
1.4
1.3
1.2
1.1
1
0.9
0.8
0.7
0.6
0
10
20
30
40
50
60
Time (min)
Normal
Diabetic
Normal data

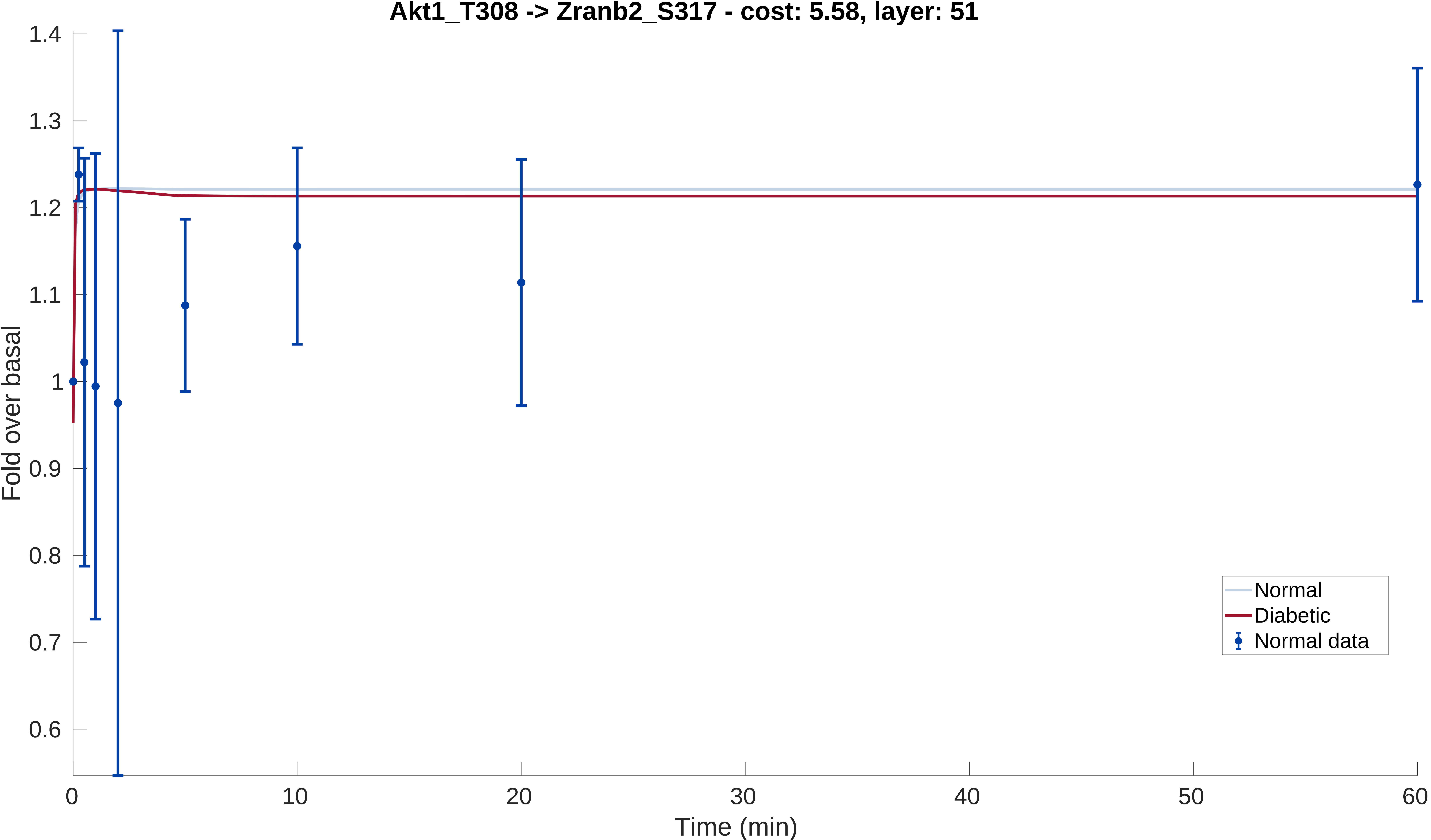

Akt1_T308 -> Zranb2_S317 - cost: 5.58, layer: 51
Fold over basal
1.4
1.3
1.2
1.1
1
0.9
0.8
0.7
0.6
0
10
20
30
40
50
60
Time (min)
Normal
Diabetic
Normal data

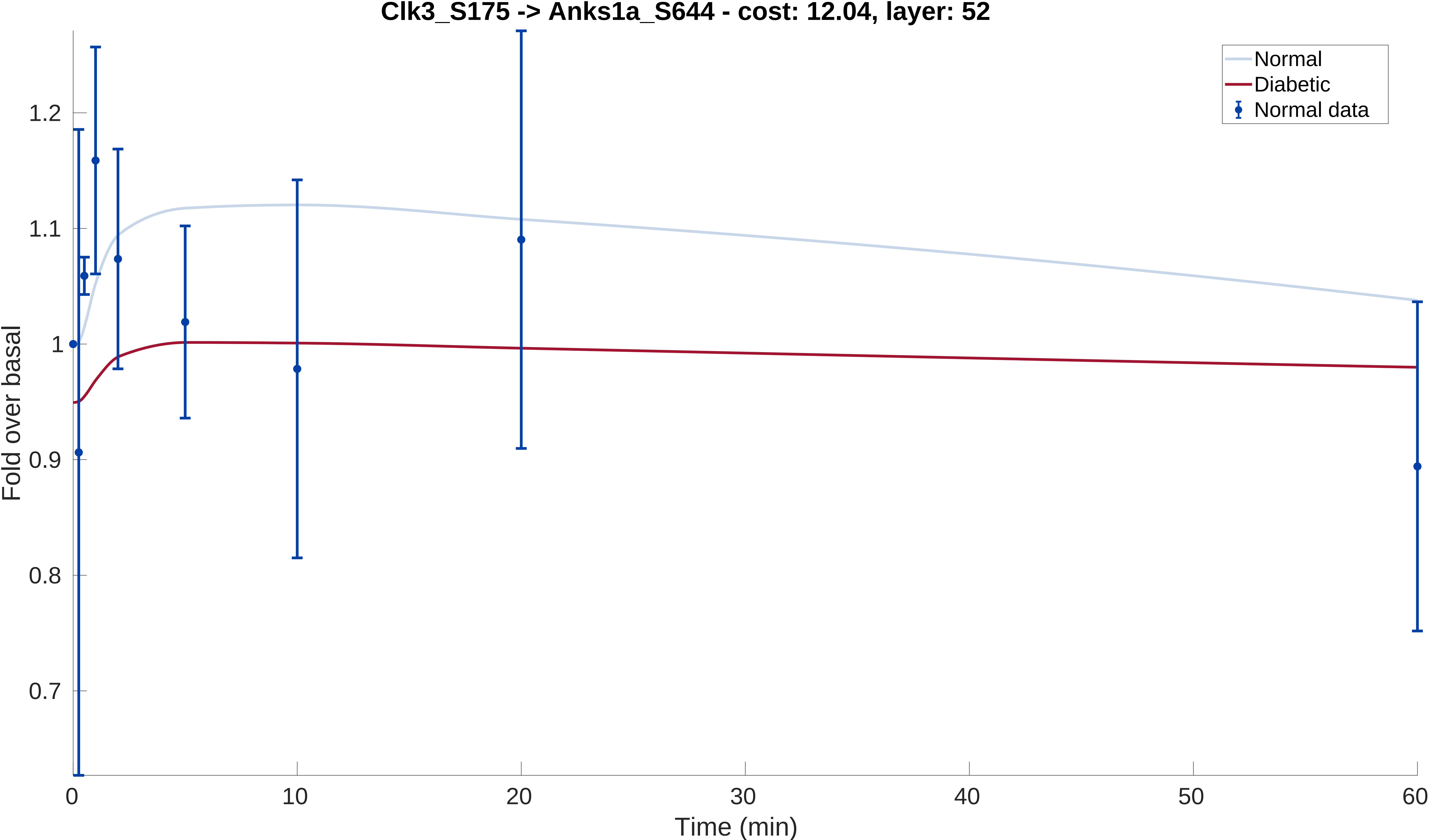

Clk3_S175 -> Anks1a_S644 - cost: 12.04, layer: 52
Normal
Diabetic
Normal data
Fold over basal
1.2
1.1
1
0.9
0.8
0.7
0
10
20
30
40
50
60
Time (min)

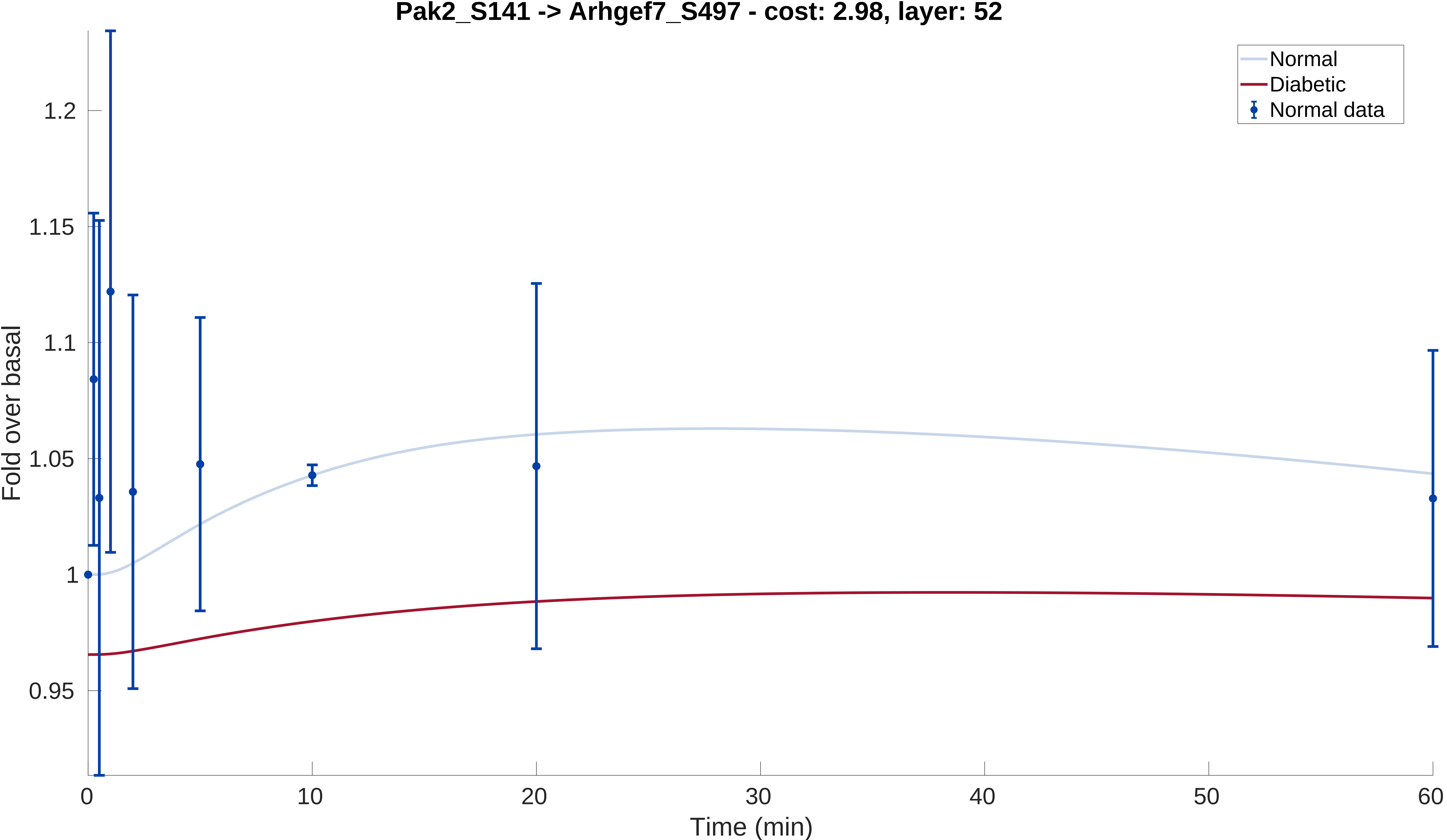

Pak2_S141 -> Arhgef7_S497 - cost: 2.98, layer: 52
Normal
Diabetic
Normal data
Fold over basal
1.2
1.15
1.1
1.05
1
0.95
0
10
20
30
40
50
60
Time (min)

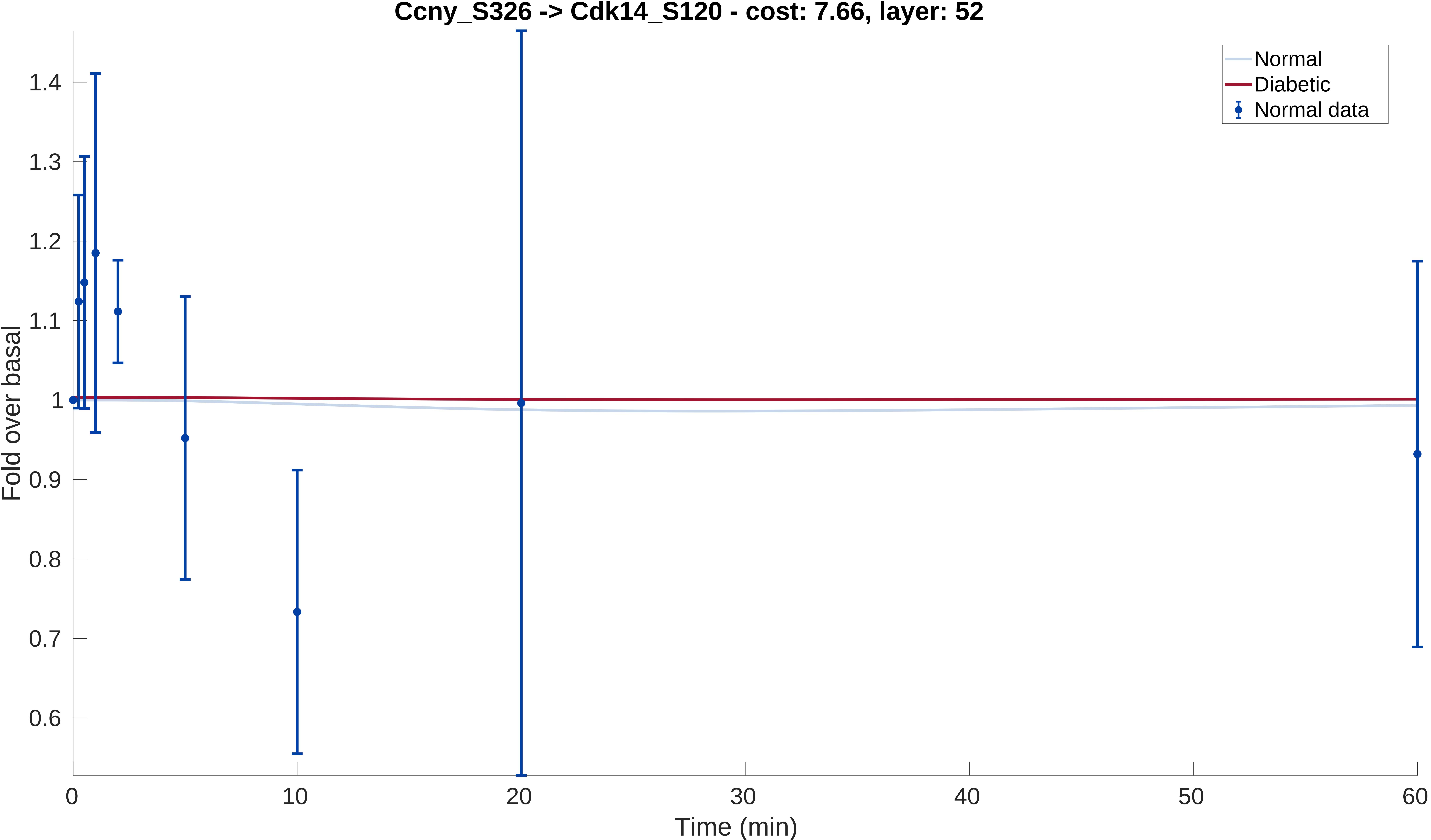

Ccny_S326 -> Cdk14_S120 - cost: 7.66, layer: 52
Normal
Diabetic
Normal data
Fold over basal
1.4
1.3
1.2
1.1
1
0.9
0.8
0.7
0.6
0
10
20
30
40
50
60
Time (min)

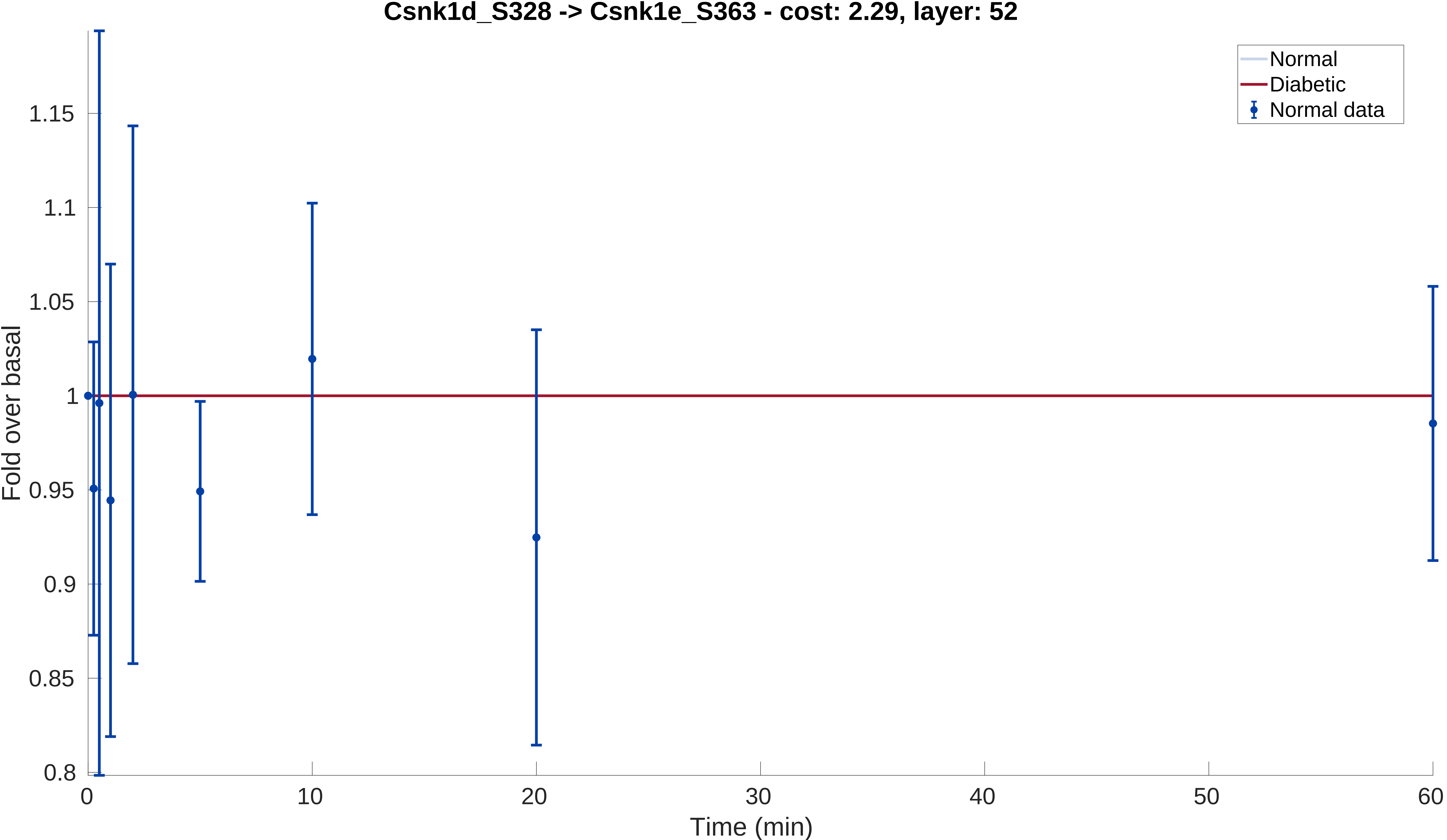

Csnk1d_S328 -> Csnk1e_S363 - cost: 2.29, layer: 52
Normal
Diabetic
Normal data
1.15
1.1
1.05
1
0.95
0.9
0.85
0.8
Fold over basal
0
10
20
30
40
50
60
Time (min)

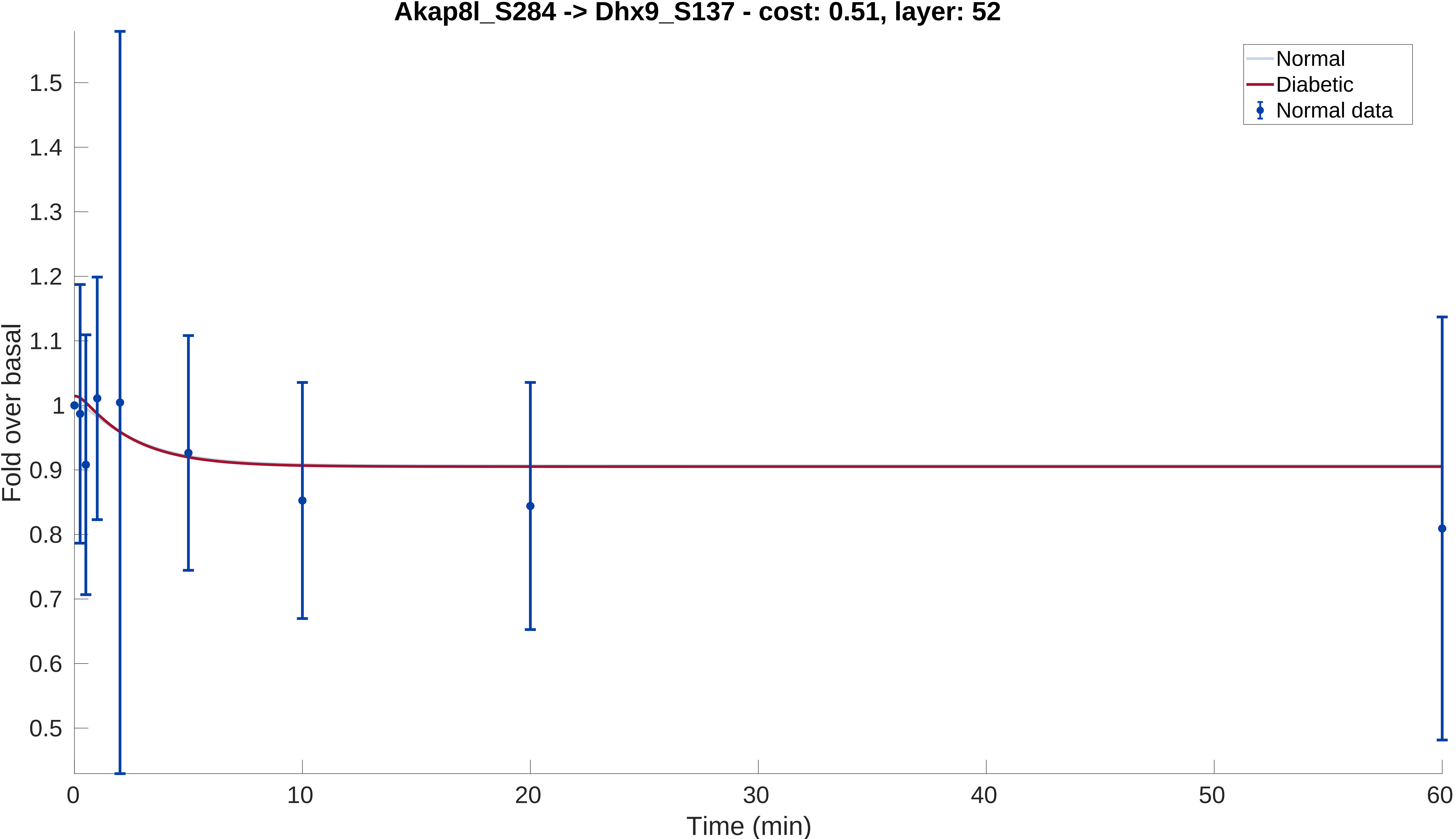

Akap8l_S284 -> Dhx9_S137 - cost: 0.51, layer: 52
Normal
Diabetic
Normal data
Fold over basal
1.5
1.4
1.3
1.2
1.1
1
0.9
0.8
0.7
0.6
0.5
0
10
20
30
40
50
60
Time (min)

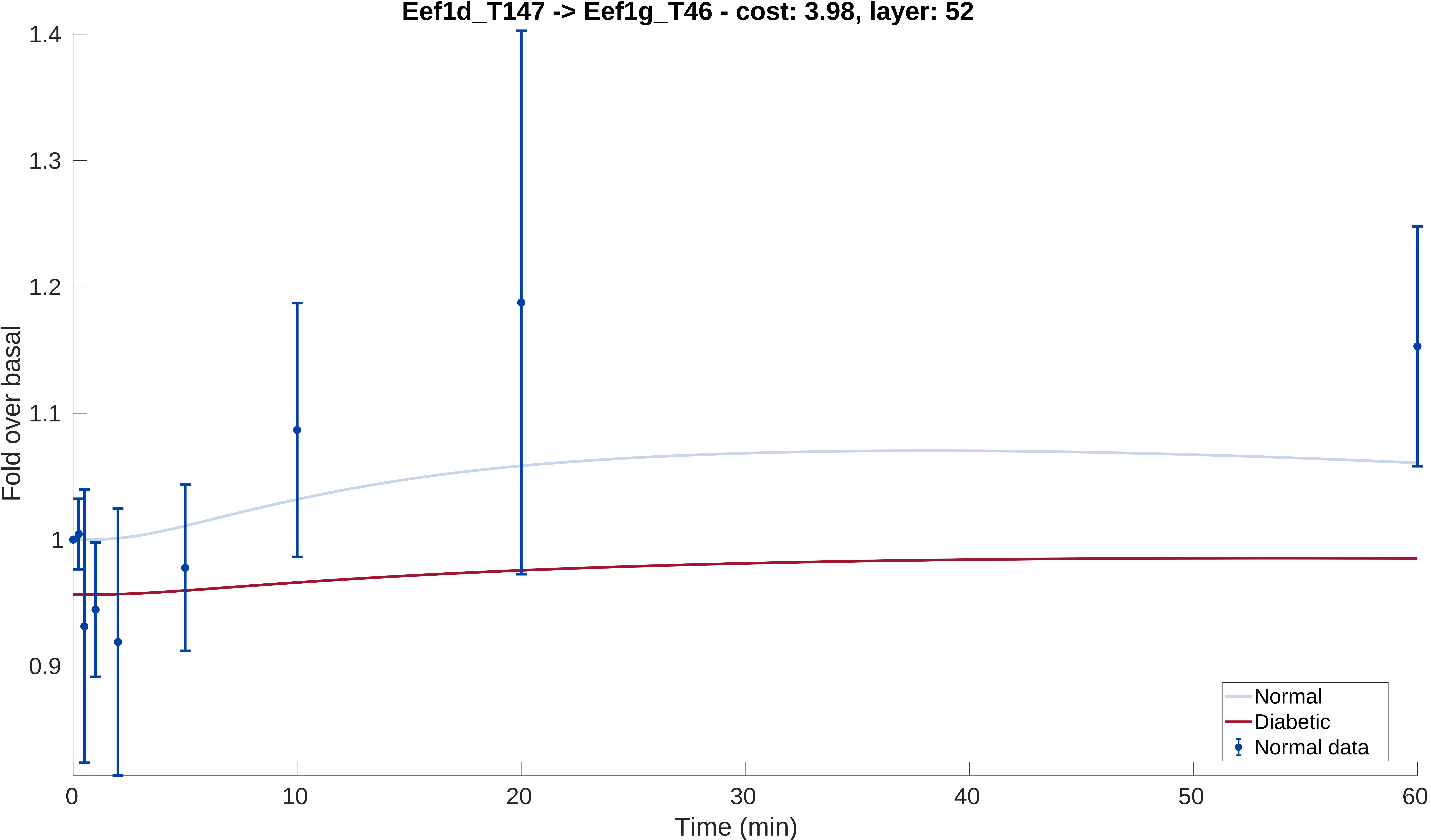

Eef1d_T147 -> Eef1g_T46 - cost: 3.98, layer: 52
1.4
1.3
1.2
1.1
1
0.9
Fold over basal
0
10
20
30
40
50
60
Time (min)
Normal
Diabetic
Normal data

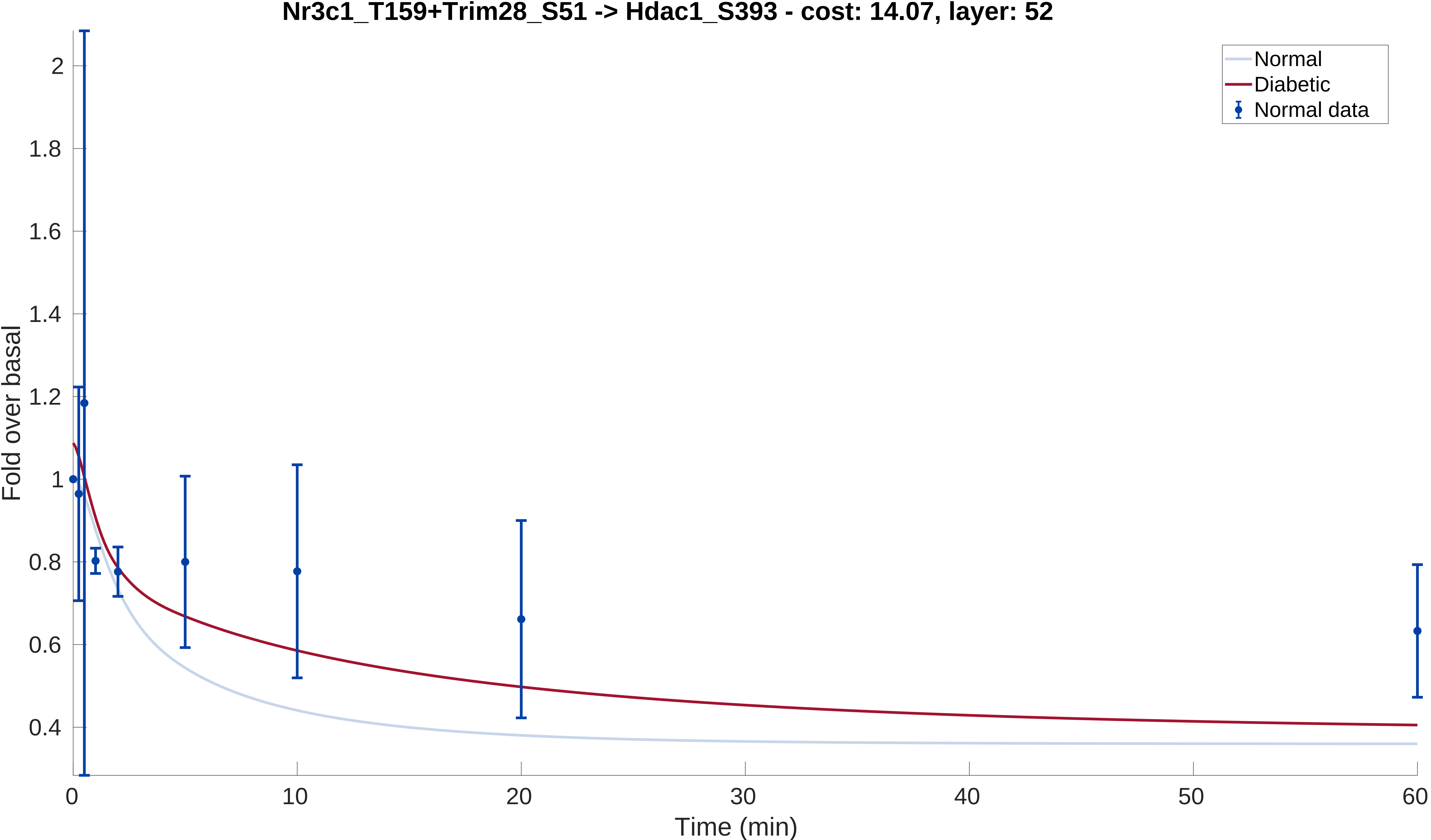

Nr3c1_T159+Trim28_S51 -> Hdac1_S393 - cost: 14.07, layer: 52
Normal
Diabetic
Normal data
Fold over basal
2
1.8
1.6
1.4
1.2
1
0.8
0.6
0.4
0
10
20
30
40
50
60
Time (min)

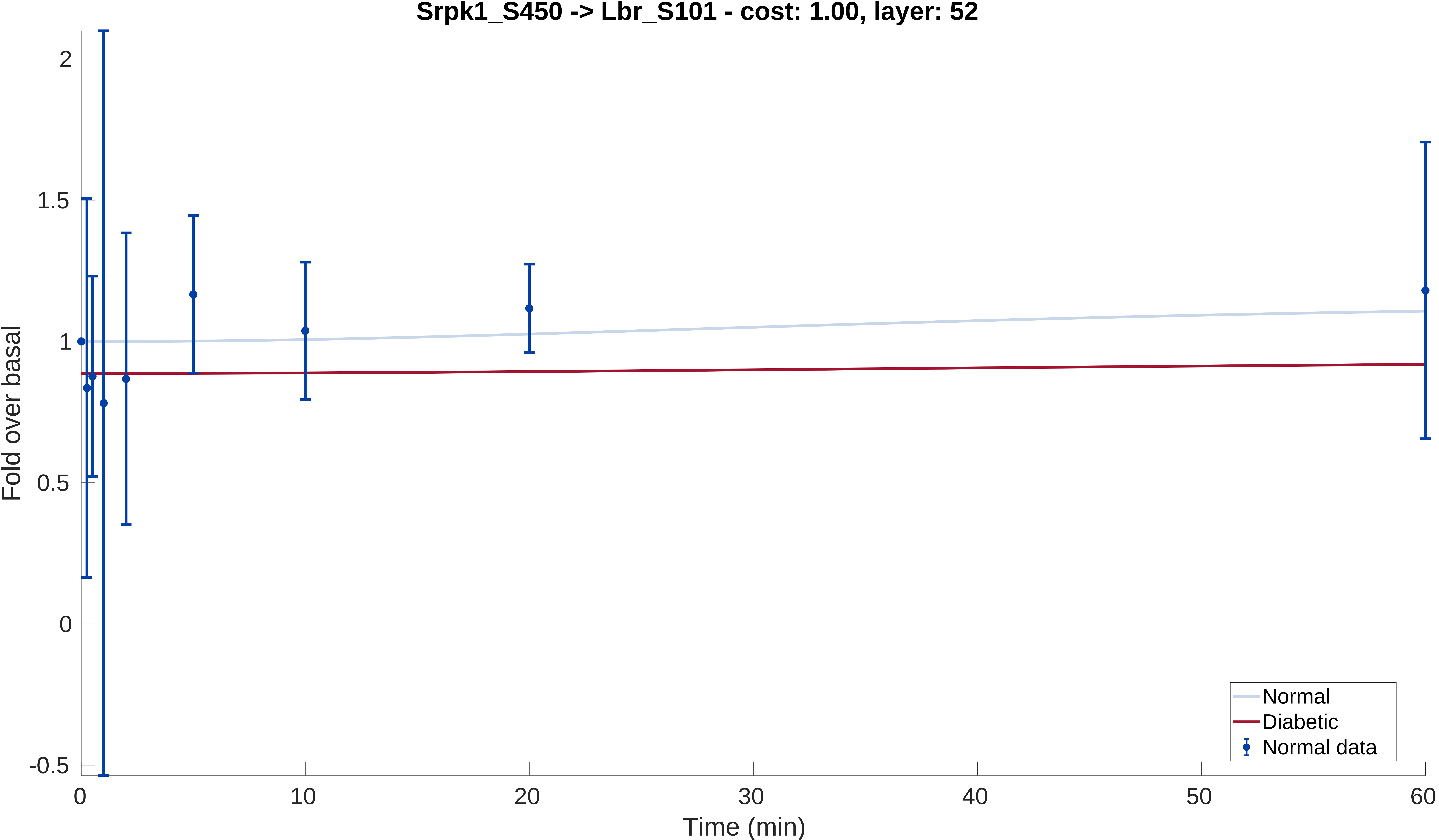
Srpk1_S450 -> Lbr_S101 - cost: 1.00, layer: 52
Fold over basal
Time (min)
2
1.5
1
0.5
0
-0.5
0
10
20
30
40
50
60
Normal
Diabetic
Normal data

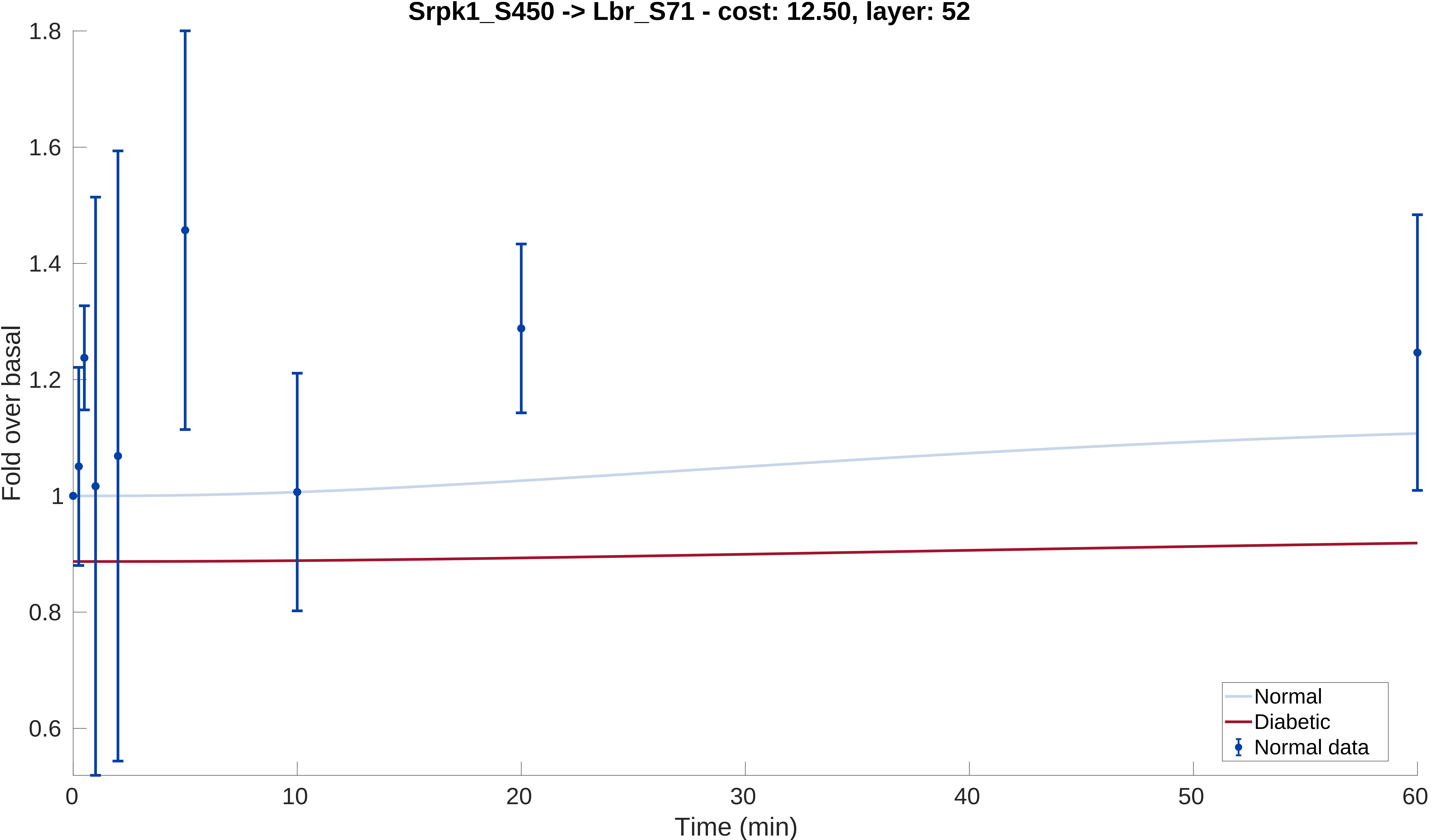

Srpk1_S450 -> Lbr_S71 - cost: 12.50, layer: 52
Fold over basal
1.8
1.6
1.4
1.2
1
0.8
0.6
0
10
20
30
40
50
60
Time (min)
Normal
Diabetic
Normal data

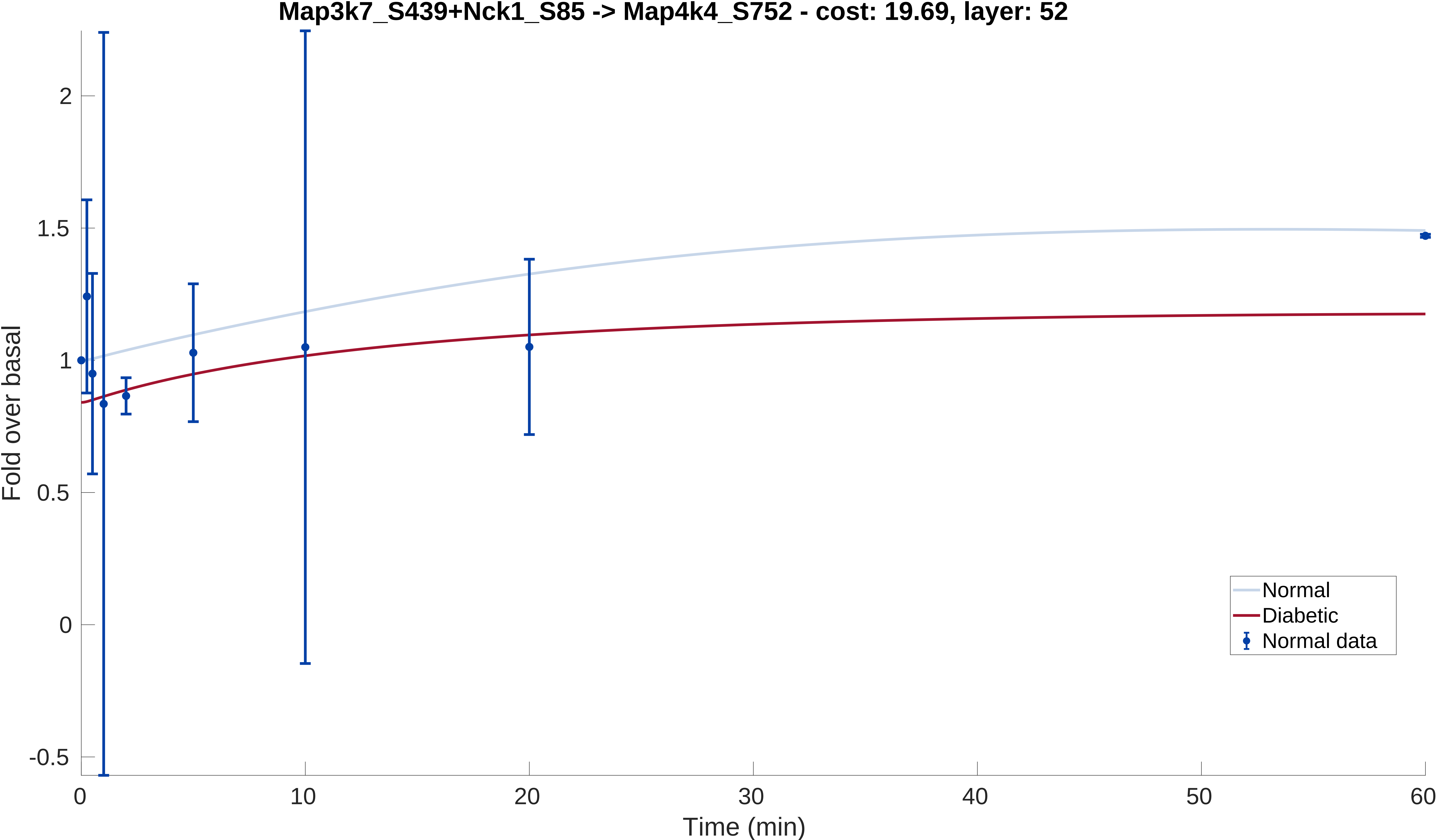

Map3k7_S439+Nck1_S85 -> Map4k4_S752 - cost: 19.69, layer: 52
Fold over basal
Time (min)
2
1.5
1
0.5
0
-0.5
0
10
20
30
40
50
60
Normal
Diabetic
Normal data

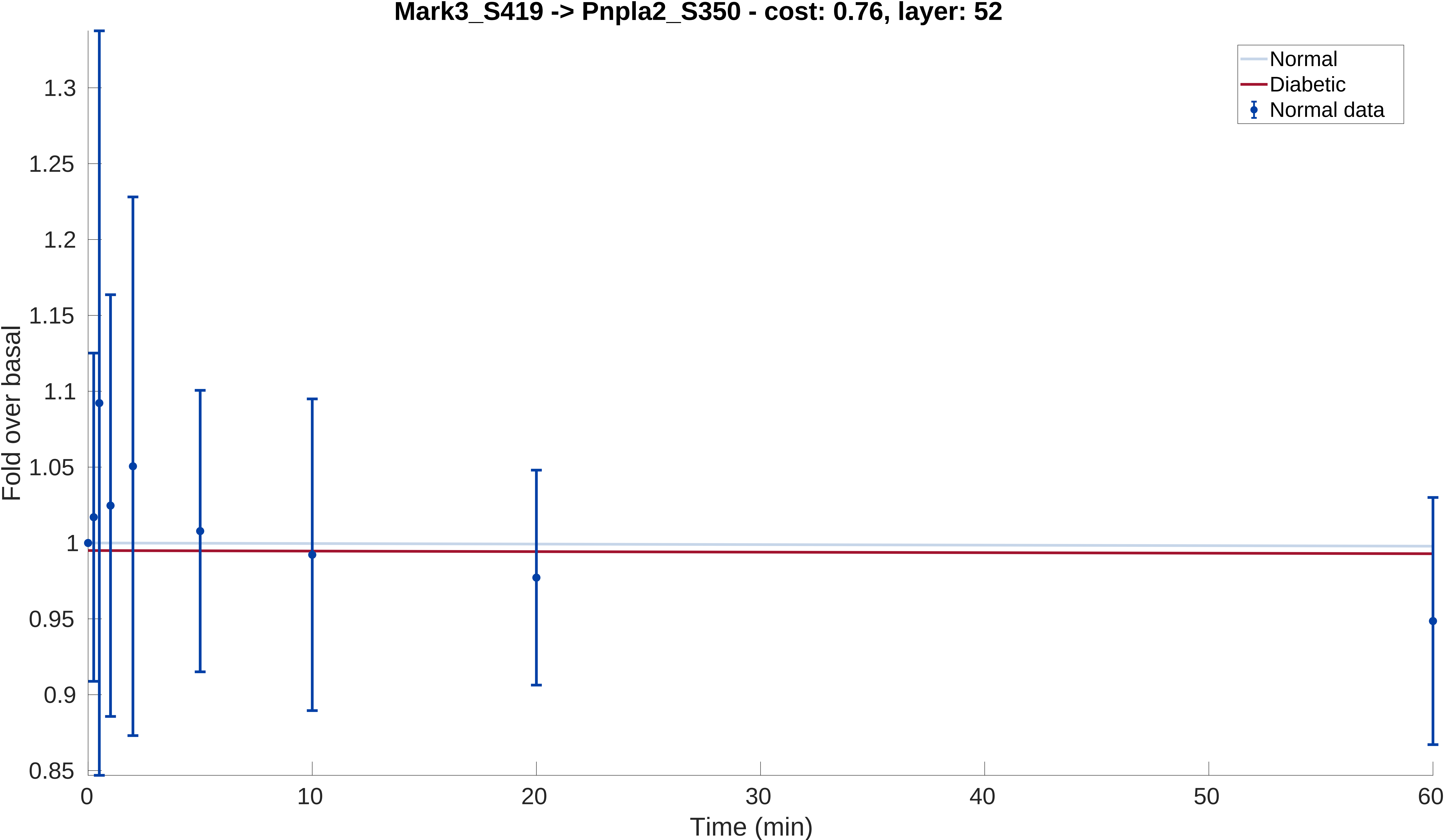

Mark3_S419 -> Pnpla2_S350 - cost: 0.76, layer: 52
Normal
Diabetic
Normal data
Fold over basal
Time (min)
1.3
1.25
1.2
1.15
1.1
1.05
1
0.95
0.9
0.85
0
10
20
30
40
50
60

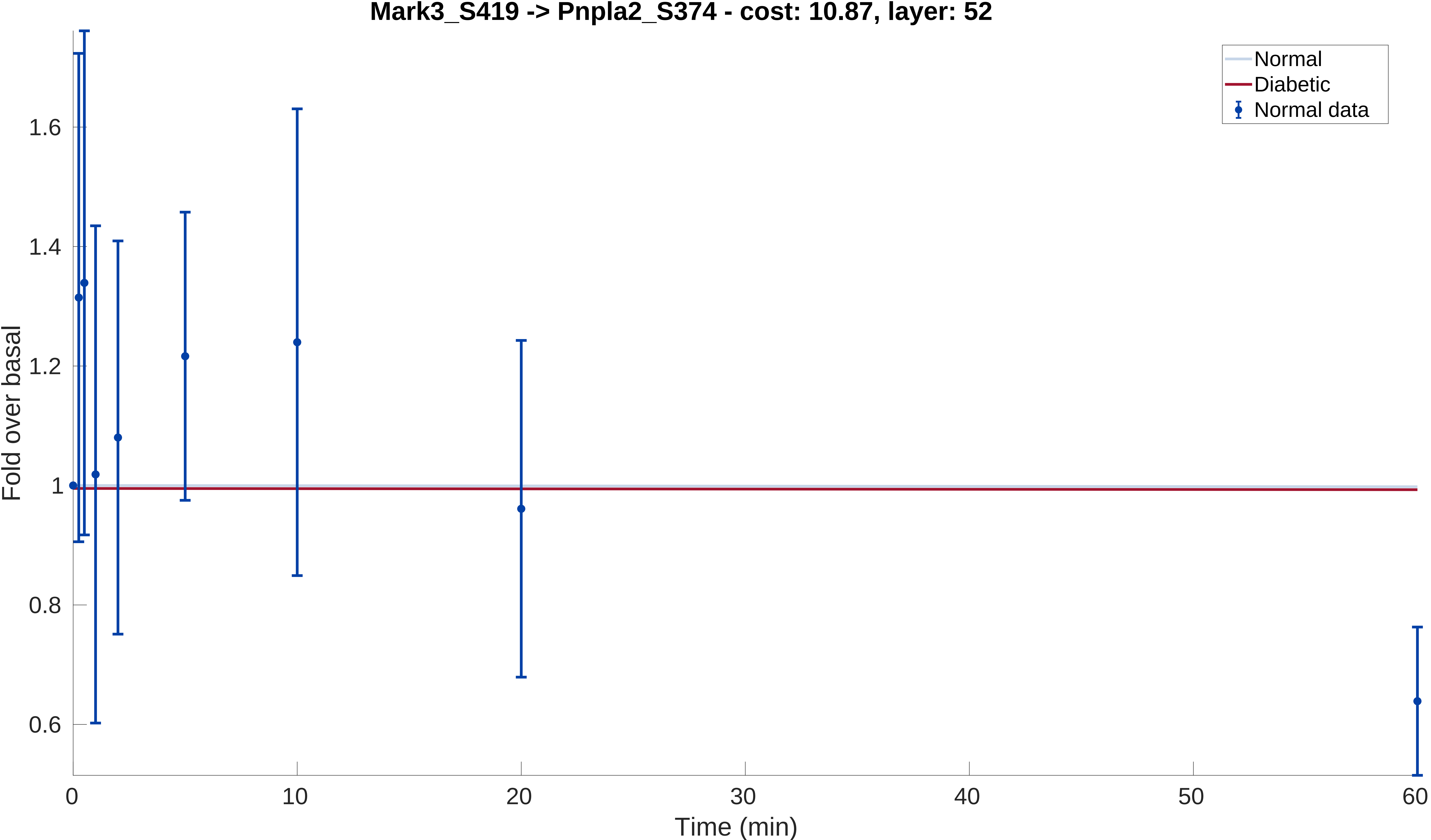

Mark3_S419 -> Pnpla2_S374 - cost: 10.87, layer: 52
Normal
Diabetic
Normal data
1.6
1.4
1.2
1
0.8
0.6
Fold over basal
0
10
20
30
40
50
60
Time (min)

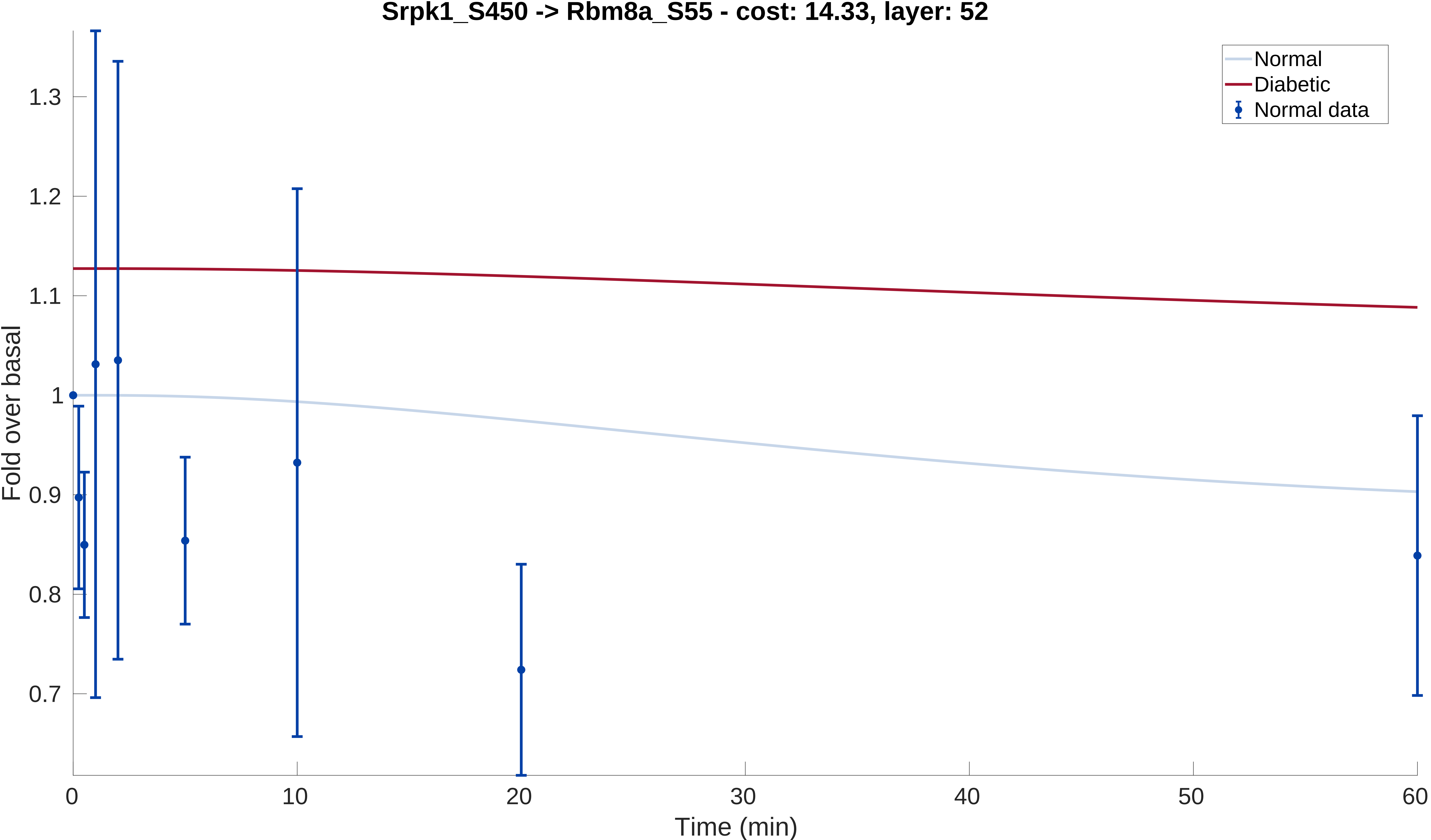
Srpk1_S450 -> Rbm8a_S55 - cost: 14.33, layer: 52
Normal
Diabetic
Normal data
Fold over basal
1.3
1.2
1.1
1
0.9
0.8
0.7
0
10
20
30
40
50
60
Time (min)

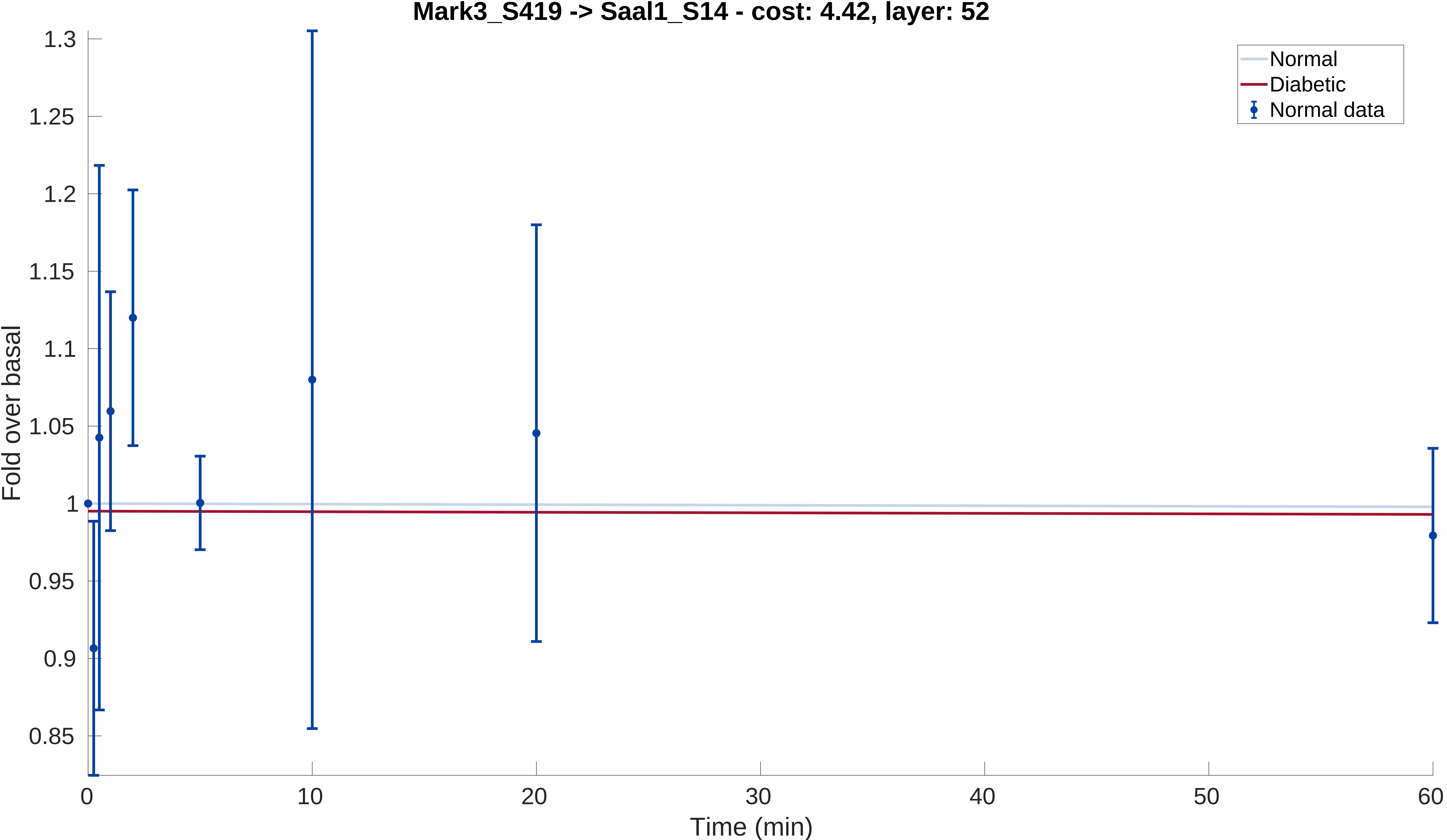
Mark3_S419 -> Saal1_S14 - cost: 4.42, layer: 52
Normal
Diabetic
Normal data
Fold over basal
1.3
1.25
1.2
1.15
1.1
1.05
1
0.95
0.9
0.85
0
10
20
30
40
50
60
Time (min)

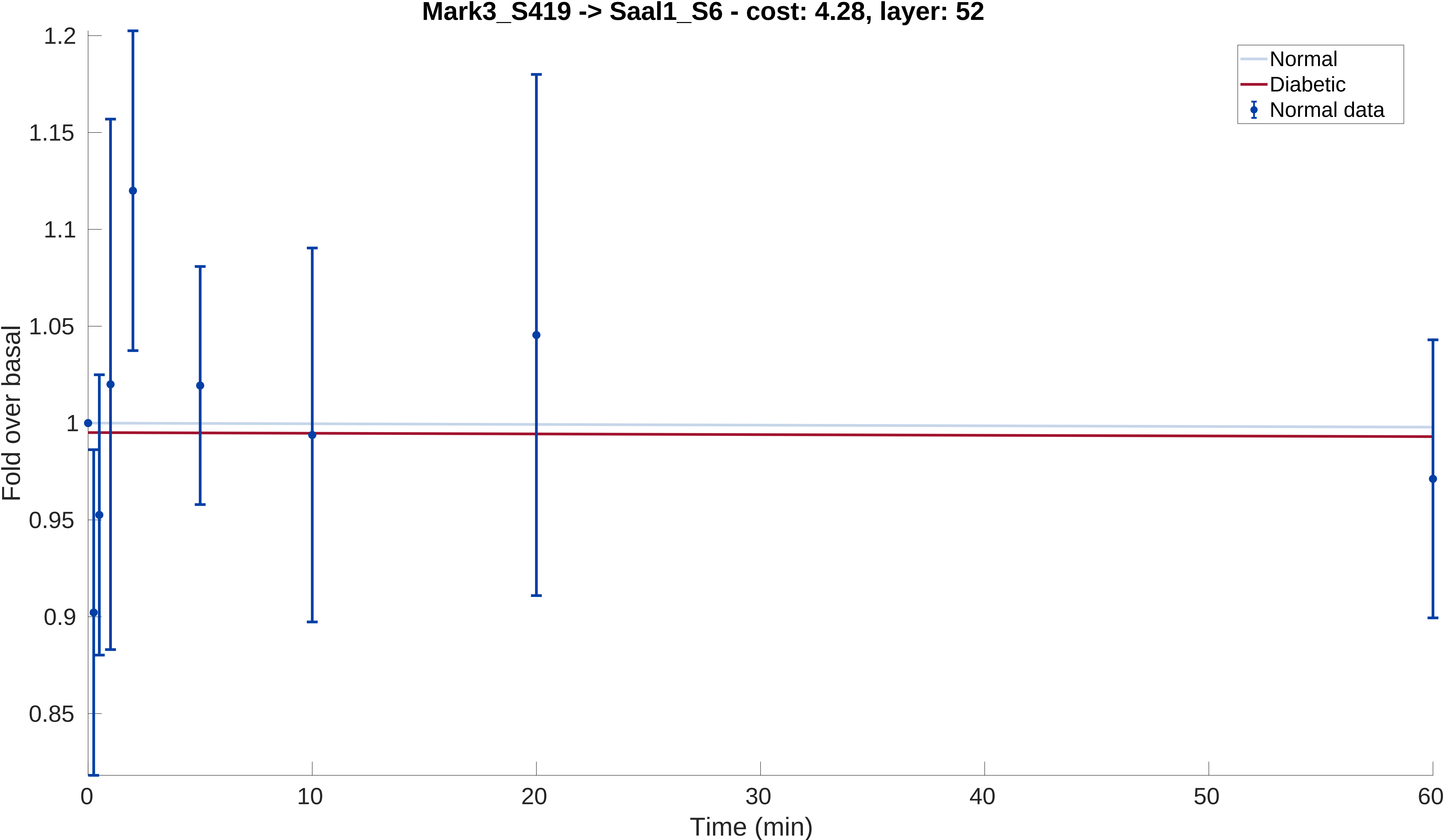

Mark3_S419 -> Saal1_S6 - cost: 4.28, layer: 52
Fold over basal
1.2
1.15
1.1
1.05
1
0.95
0.9
0.85
0
10
20
30
40
50
60
Time (min)
Normal
Diabetic
Normal data

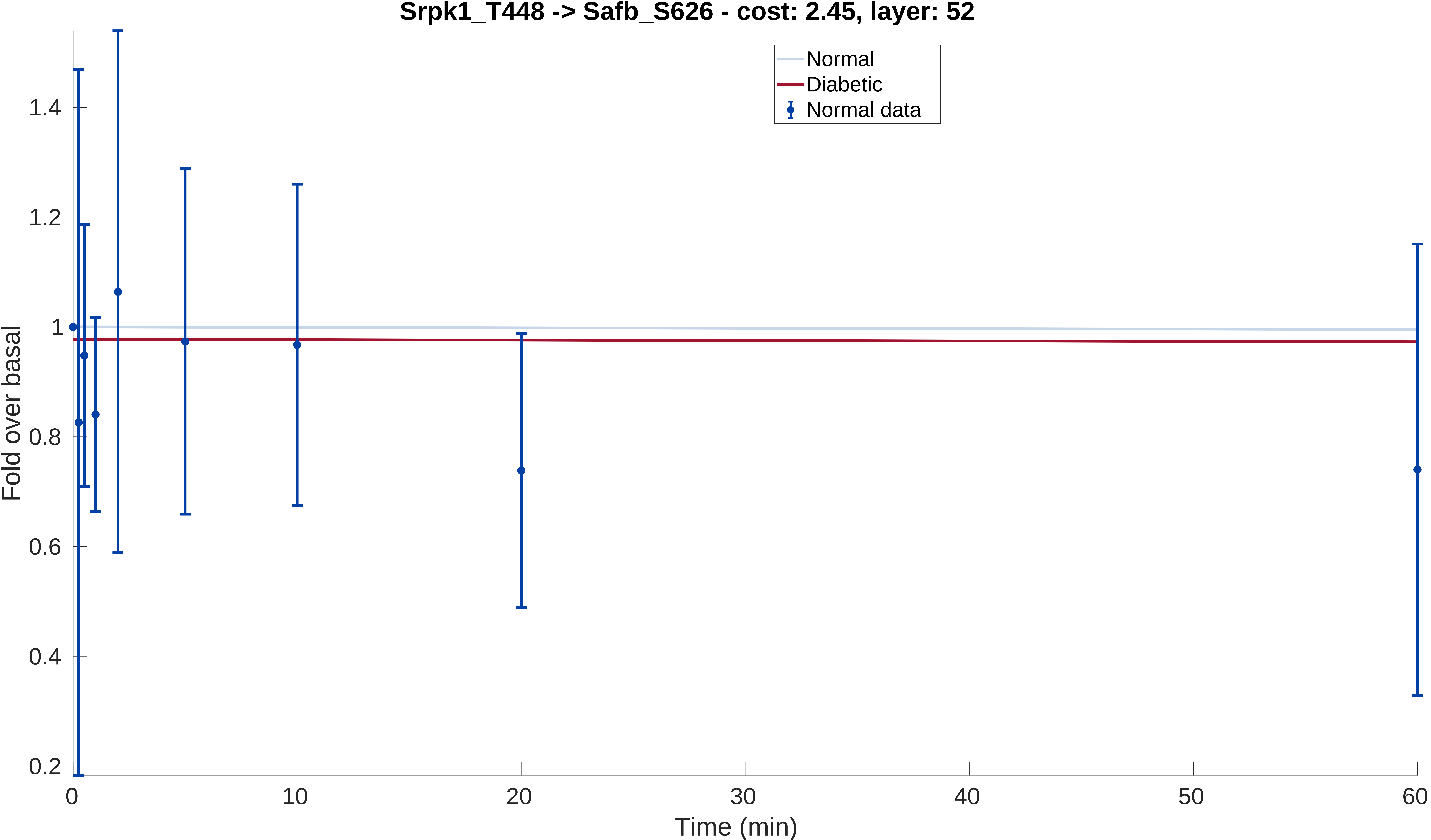
Srpk1_T448 -> Safb_S626 - cost: 2.45, layer: 52
Normal
Diabetic
Normal data
Fold over basal
1.4
1.2
1
0.8
0.6
0.4
0.2
0
10
20
30
40
50
60
Time (min)

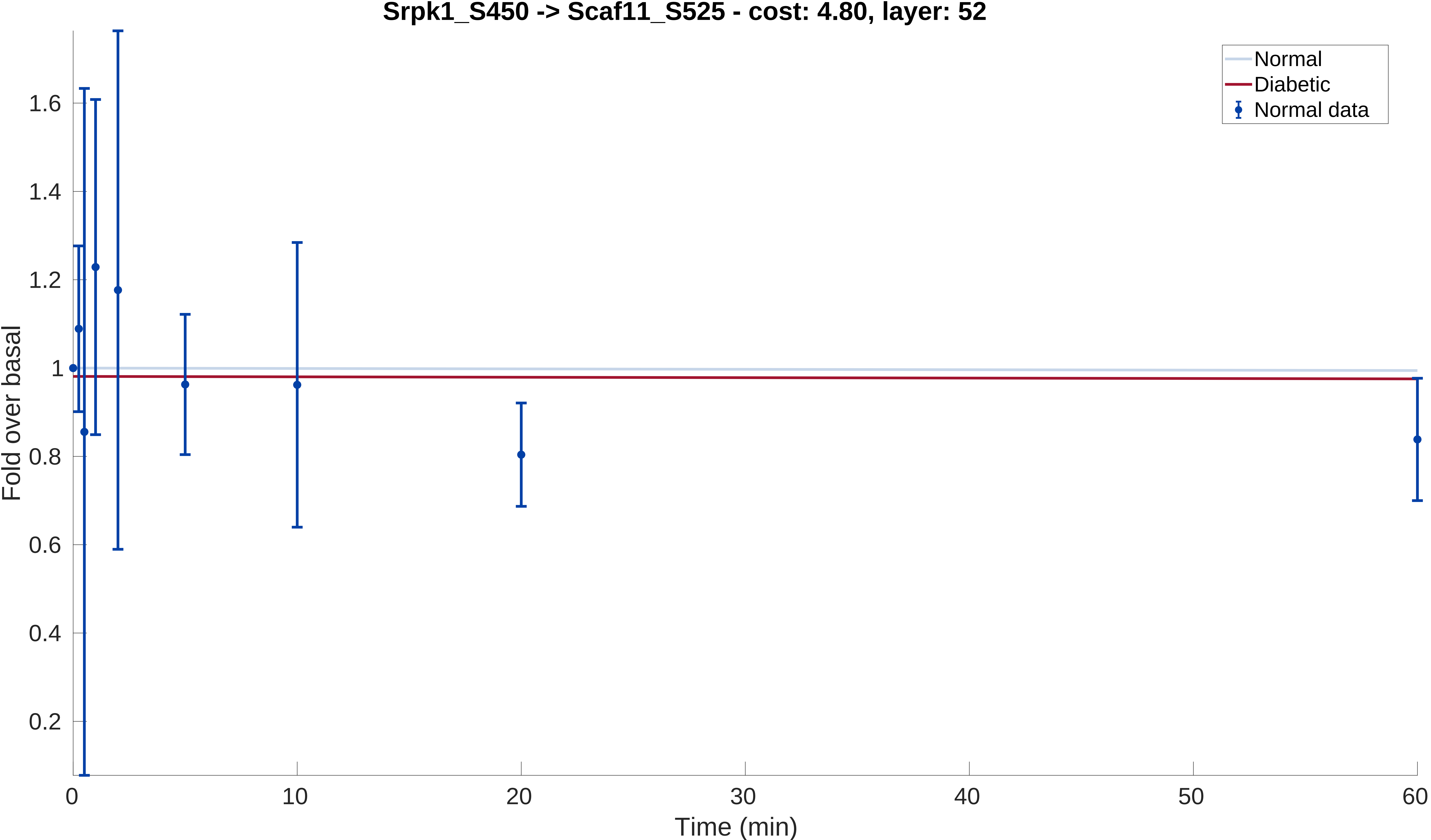
Srpk1_S450 -> Scaf11_S525 - cost: 4.80, layer: 52
Normal
Diabetic
Normal data
Fold over basal
1.6
1.4
1.2
1
0.8
0.6
0.4
0.2
0
10
20
30
40
50
60
Time (min)

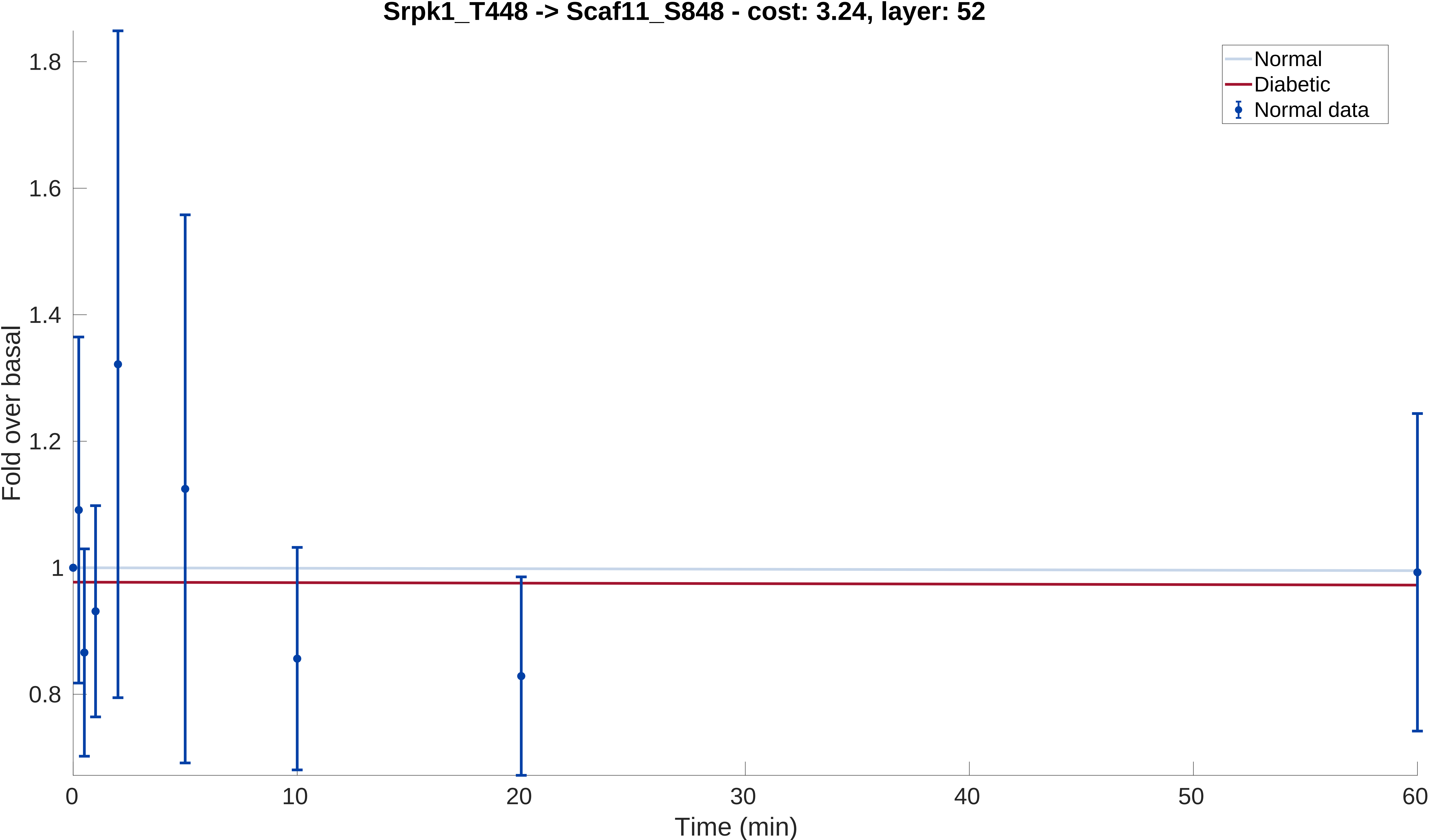

Srpk1_T448 -> Scaf11_S848 - cost: 3.24, layer: 52
Normal
Diabetic
Normal data
Fold over basal
1.8
1.6
1.4
1.2
1
0.8
0
10
20
30
40
50
60
Time (min)

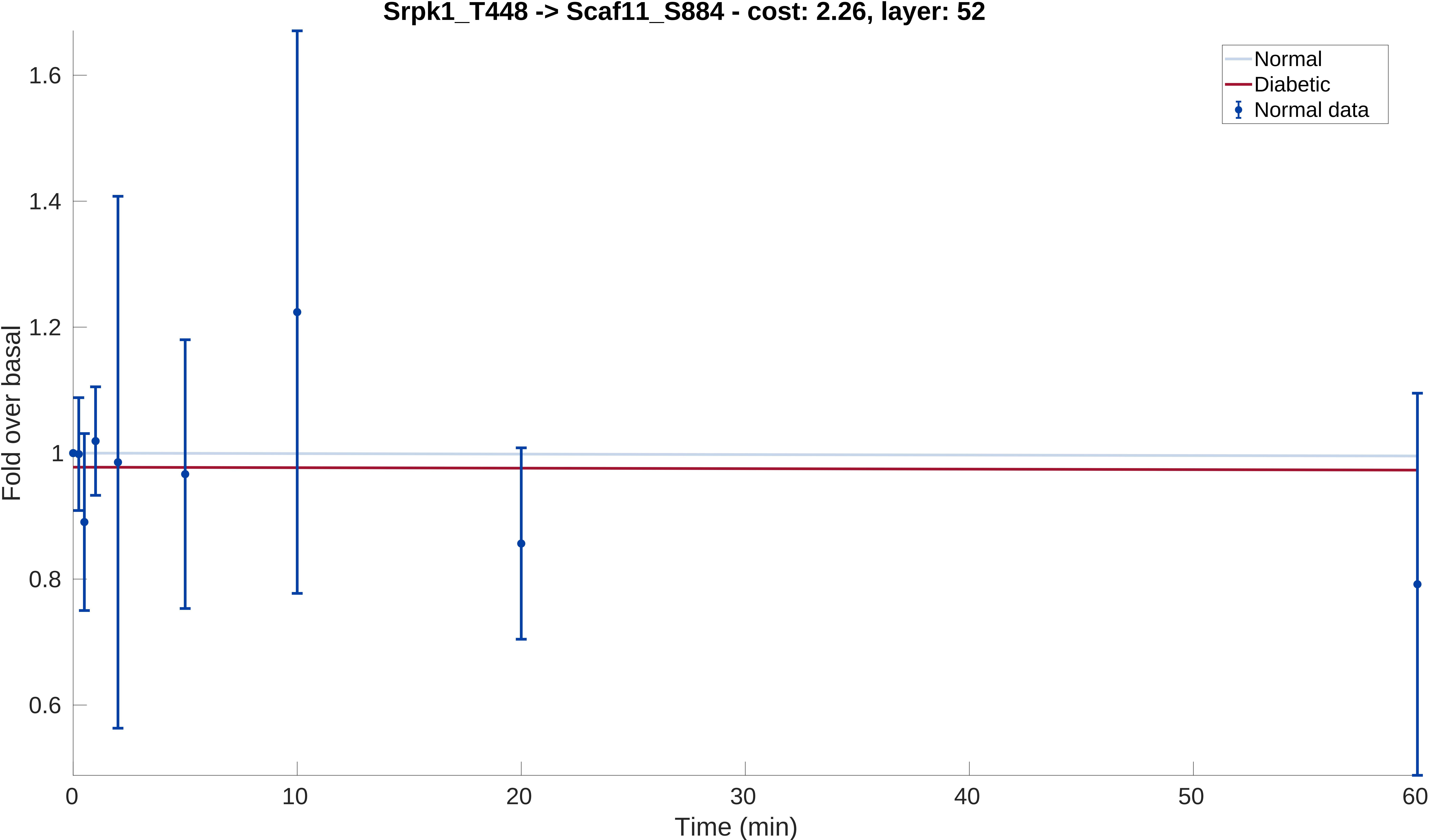

Srpk1_T448 -> Scaf11_S884 - cost: 2.26, layer: 52
Normal
Diabetic
Normal data
Fold over basal
Time (min)
1.6
1.4
1.2
1
0.8
0.6
0
10
20
30
40
50
60

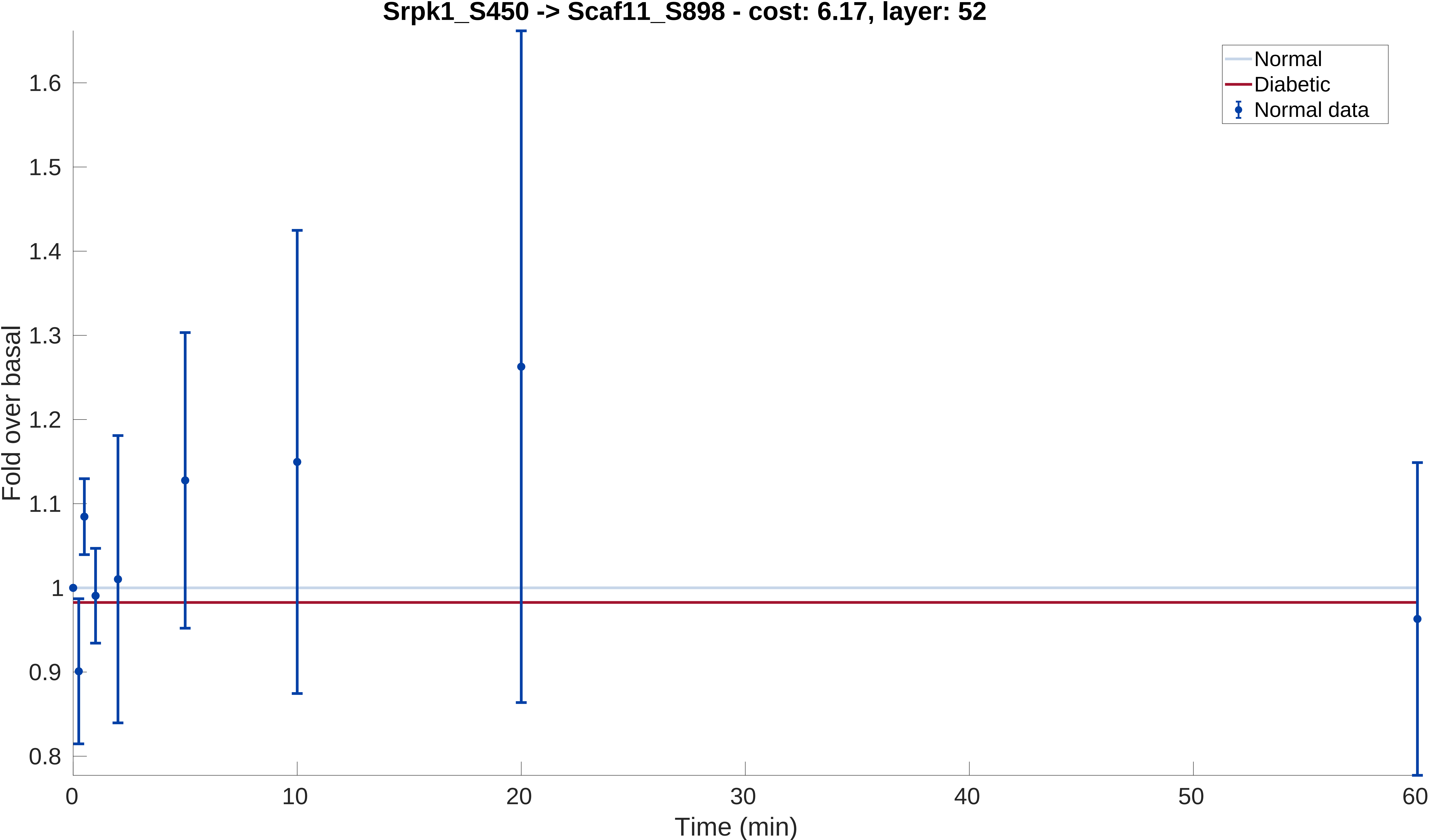
Srpk1_S450 -> Scaf11_S898 - cost: 6.17, layer: 52
Normal
Diabetic
Normal data
Fold over basal
1.6
1.5
1.4
1.3
1.2
1.1
1
0.9
0.8
0
10
20
30
40
50
60
Time (min)

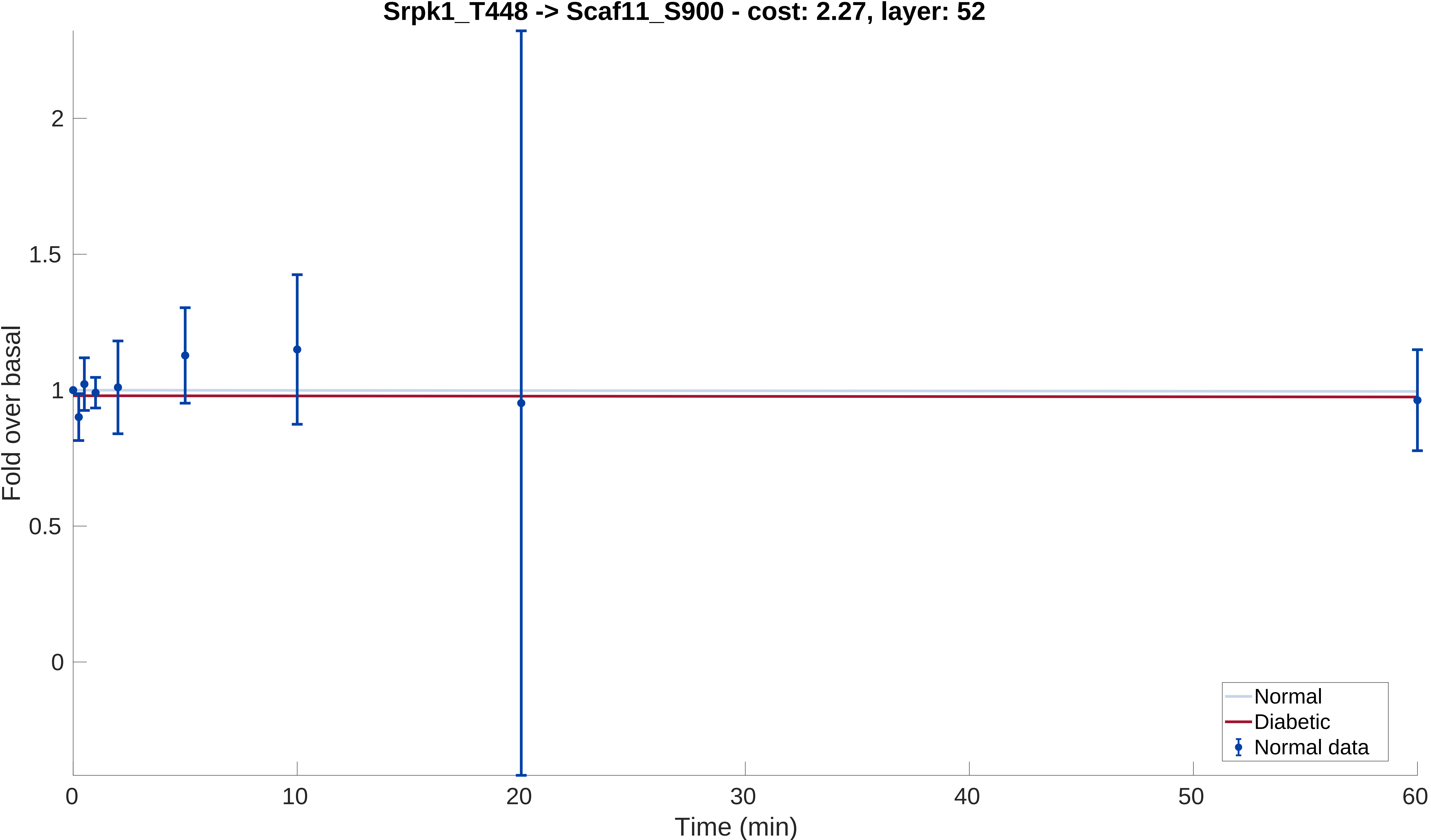

Srpk1_T448 -> Scaf11_S900 - cost: 2.27, layer: 52
Fold over basal
Time (min)
0
0.5
1
1.5
2
0
10
20
30
40
50
60
Normal
Diabetic
Normal data

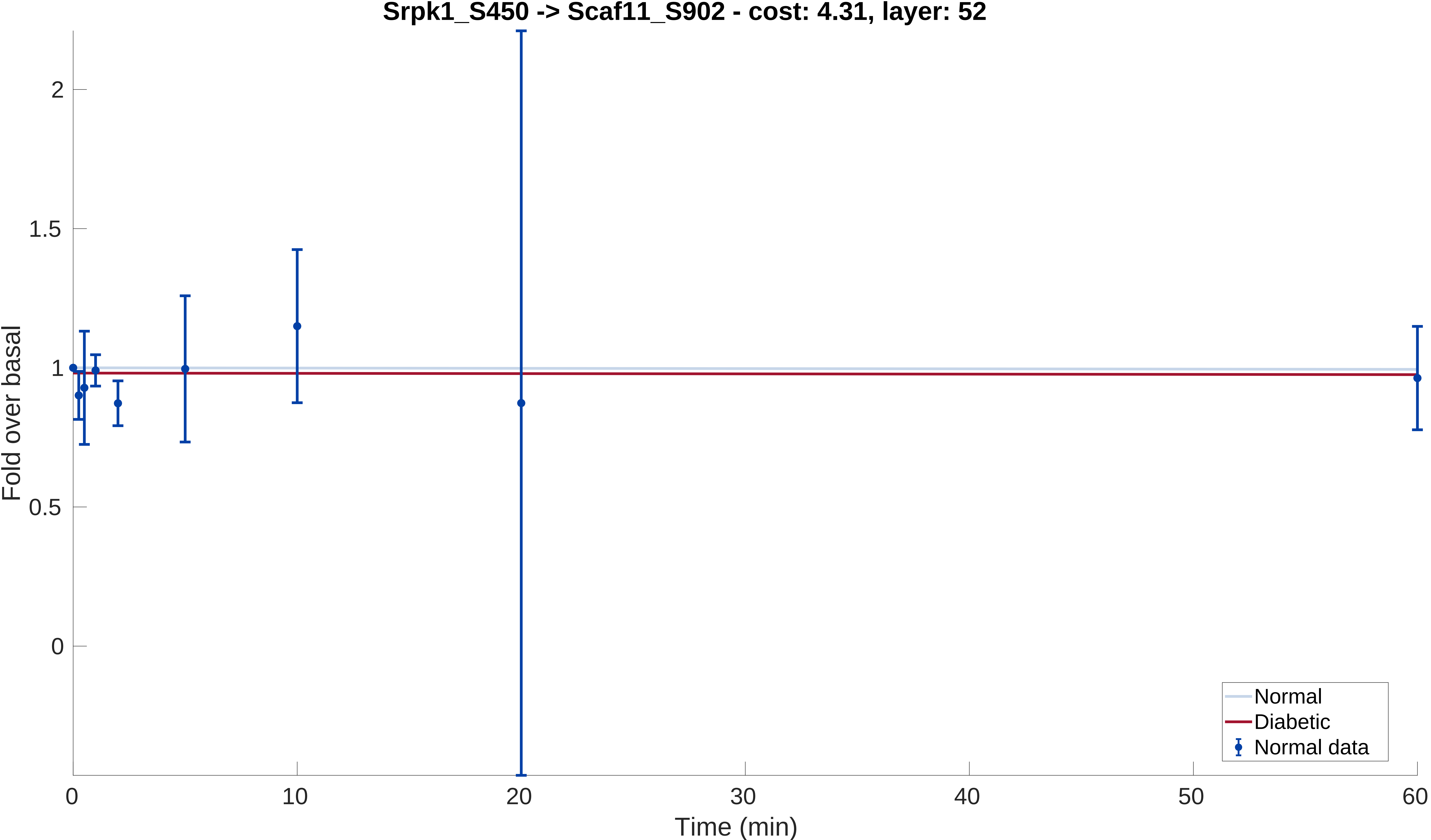

Srpk1_S450 -> Scaf11_S902 - cost: 4.31, layer: 52
Fold over basal
2
1.5
1
0.5
0
0
10
20
30
40
50
60
Time (min)
Normal
Diabetic
Normal data

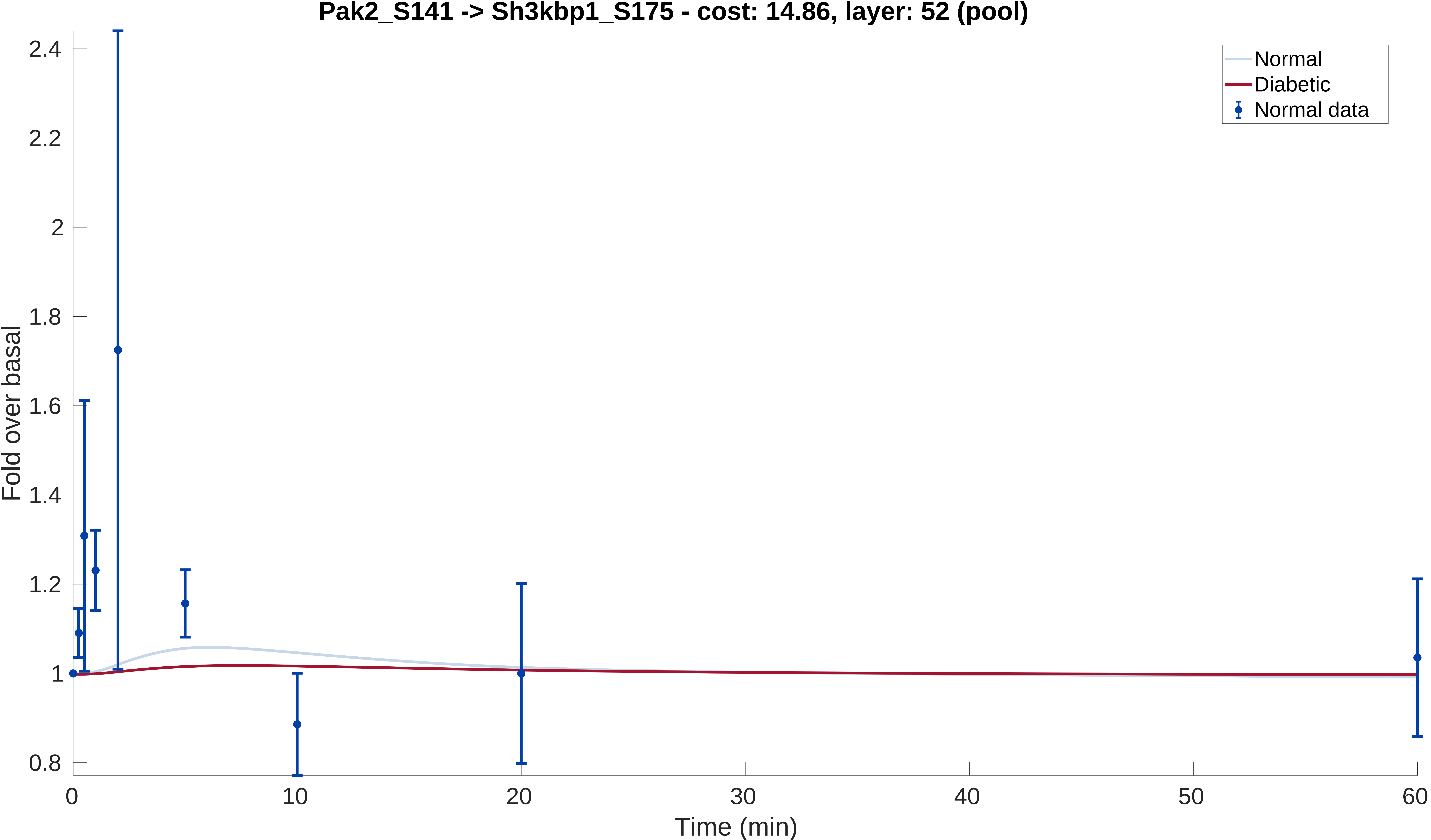

Pak2_S141 -> Sh3kbp1_S175 - cost: 14.86, layer: 52 (pool)
Normal
Diabetic
Normal data
Fold over basal
Time (min)
2.4
2.2
2
1.8
1.6
1.4
1.2
1
0.8
0
10
20
30
40
50
60

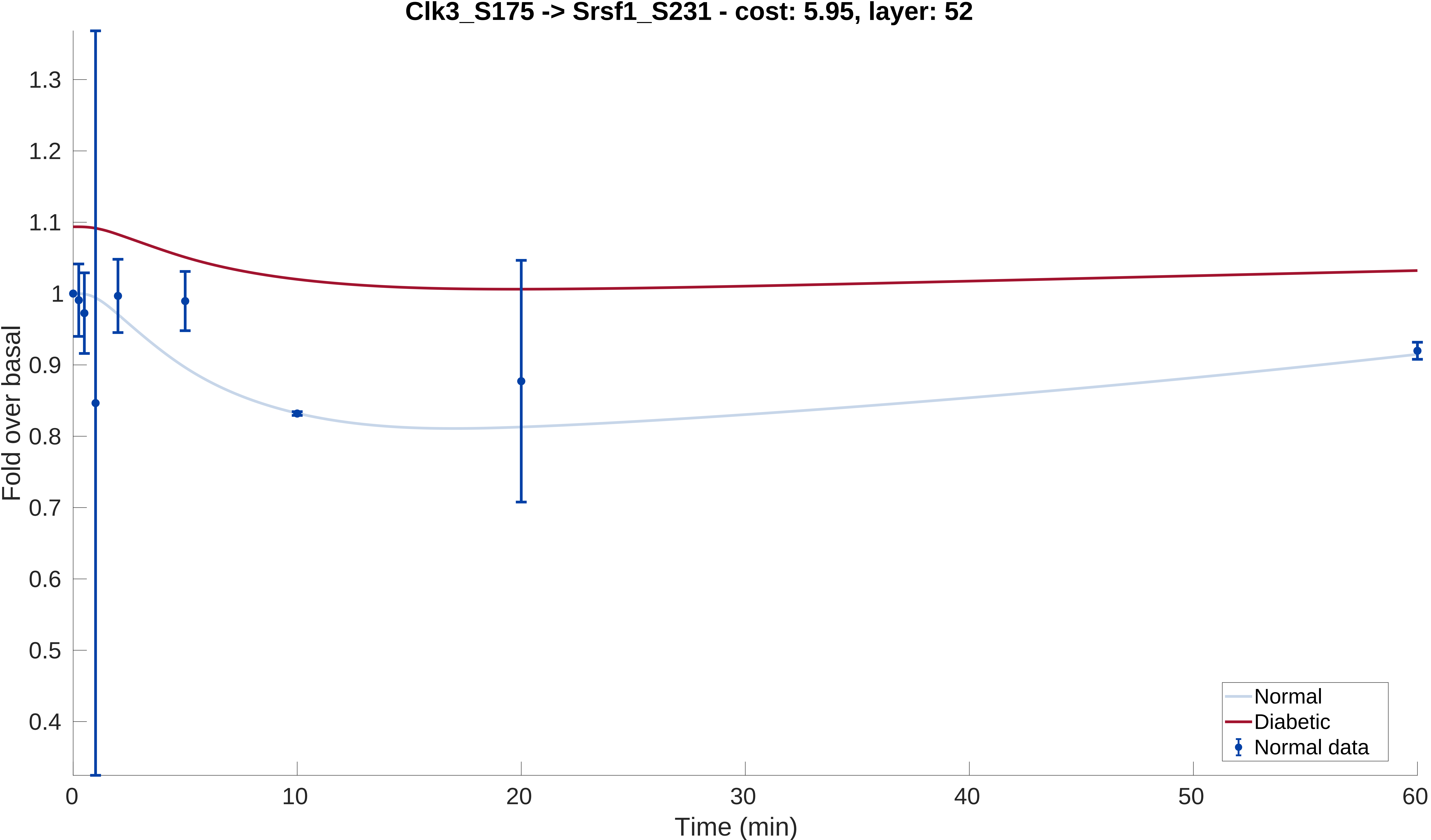

Clk3_S175 -> Srsf1_S231 - cost: 5.95, layer: 52
Fold over basal
1.3
1.2
1.1
1
0.9
0.8
0.7
0.6
0.5
0.4
0
10
20
30
40
50
60
Time (min)
Normal
Diabetic
Normal data

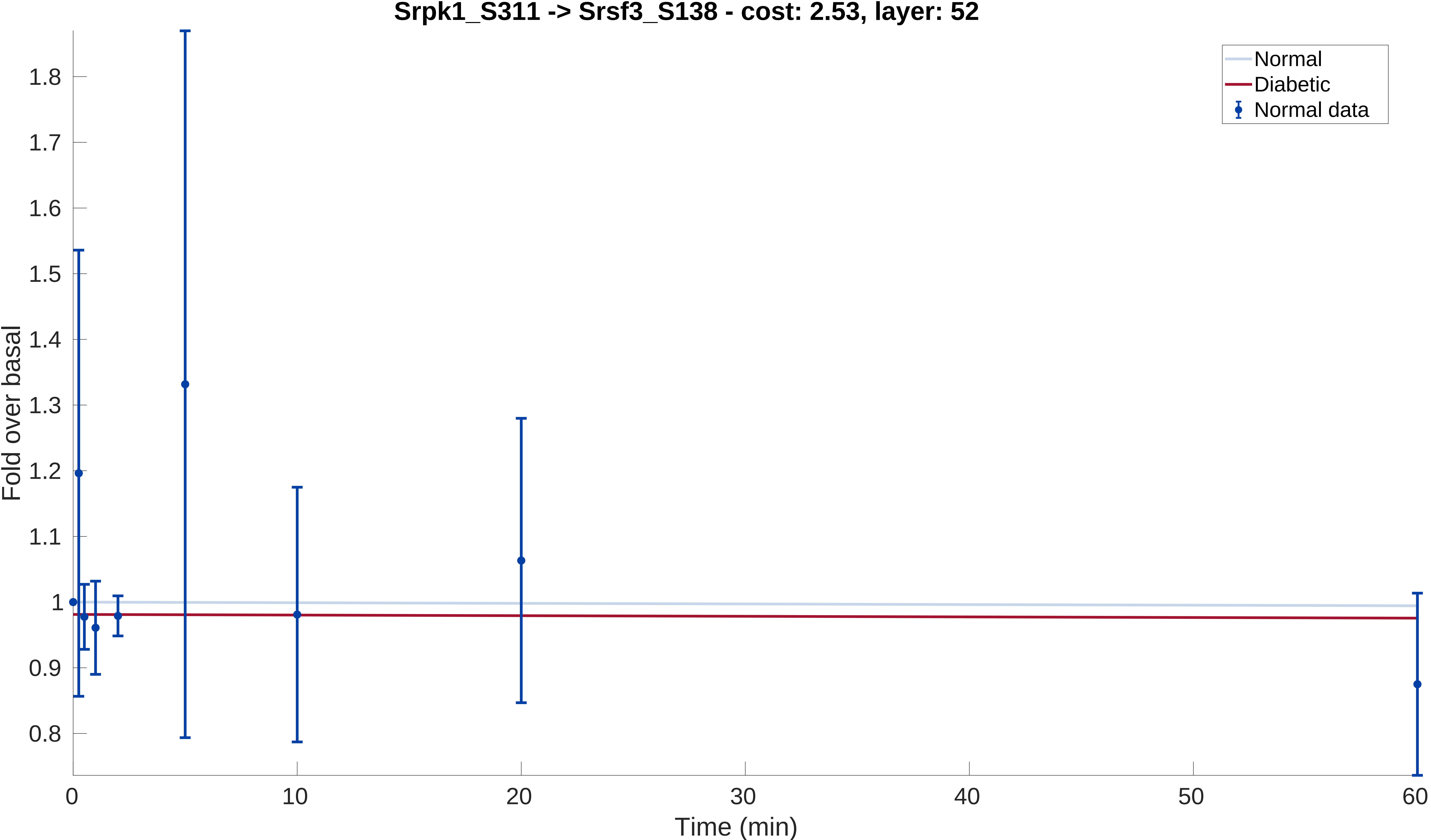

Srpk1_S311 -> Srsf3_S138 - cost: 2.53, layer: 52
Normal
Diabetic
Normal data
Fold over basal
1.8
1.7
1.6
1.5
1.4
1.3
1.2
1.1
1
0.9
0.8
0
10
20
30
40
50
60
Time (min)

#### Srpk1_S450 -> Srsf3_S140 - cost: 6.45, layer: 52

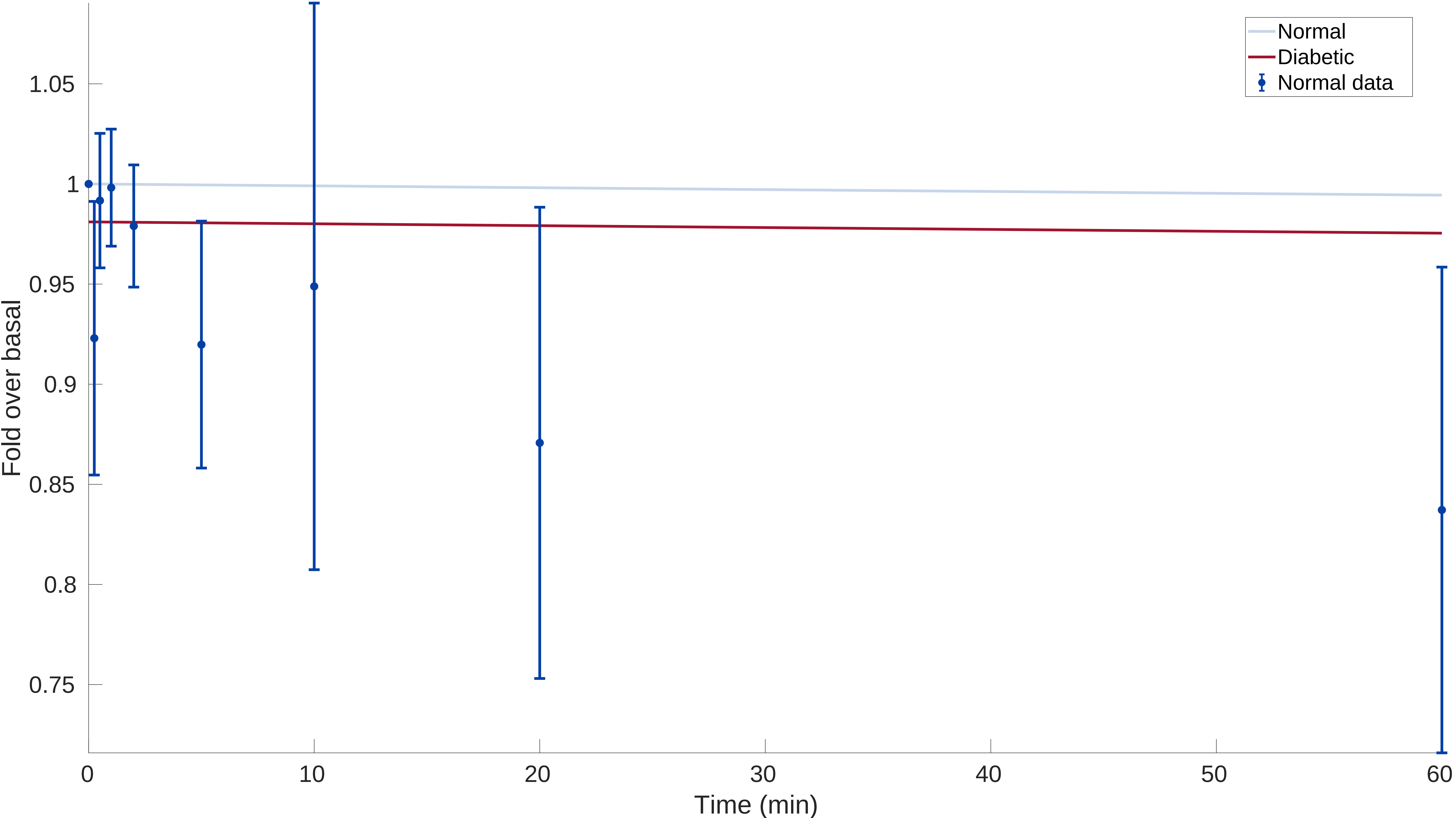

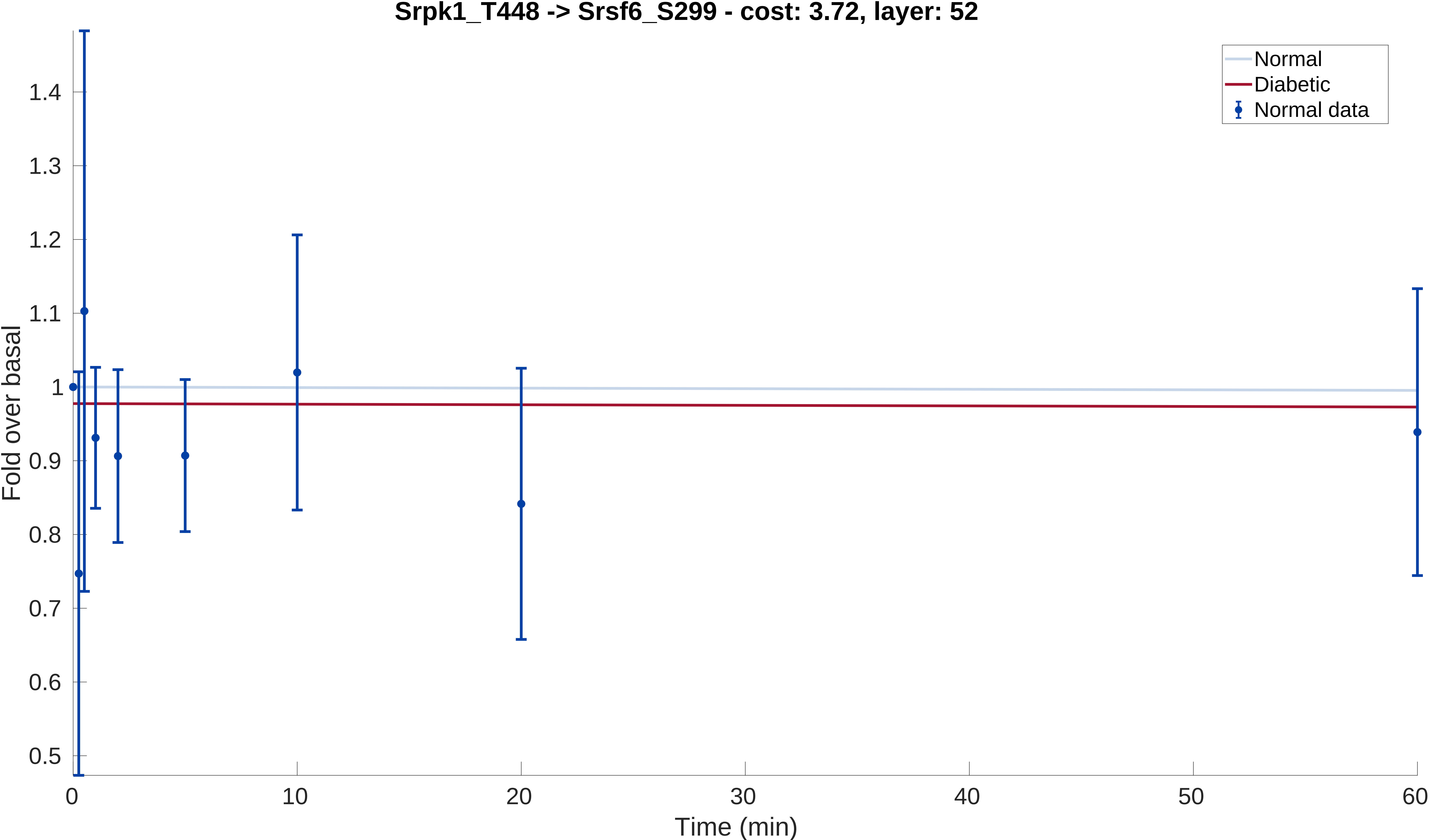

Srpk1_T448 -> Srsf6_S299 - cost: 3.72, layer: 52
Normal
Diabetic
Normal data
Fold over basal
1.4
1.3
1.2
1.1
1
0.9
0.8
0.7
0.6
0.5
0
10
20
30
40
50
60
Time (min)

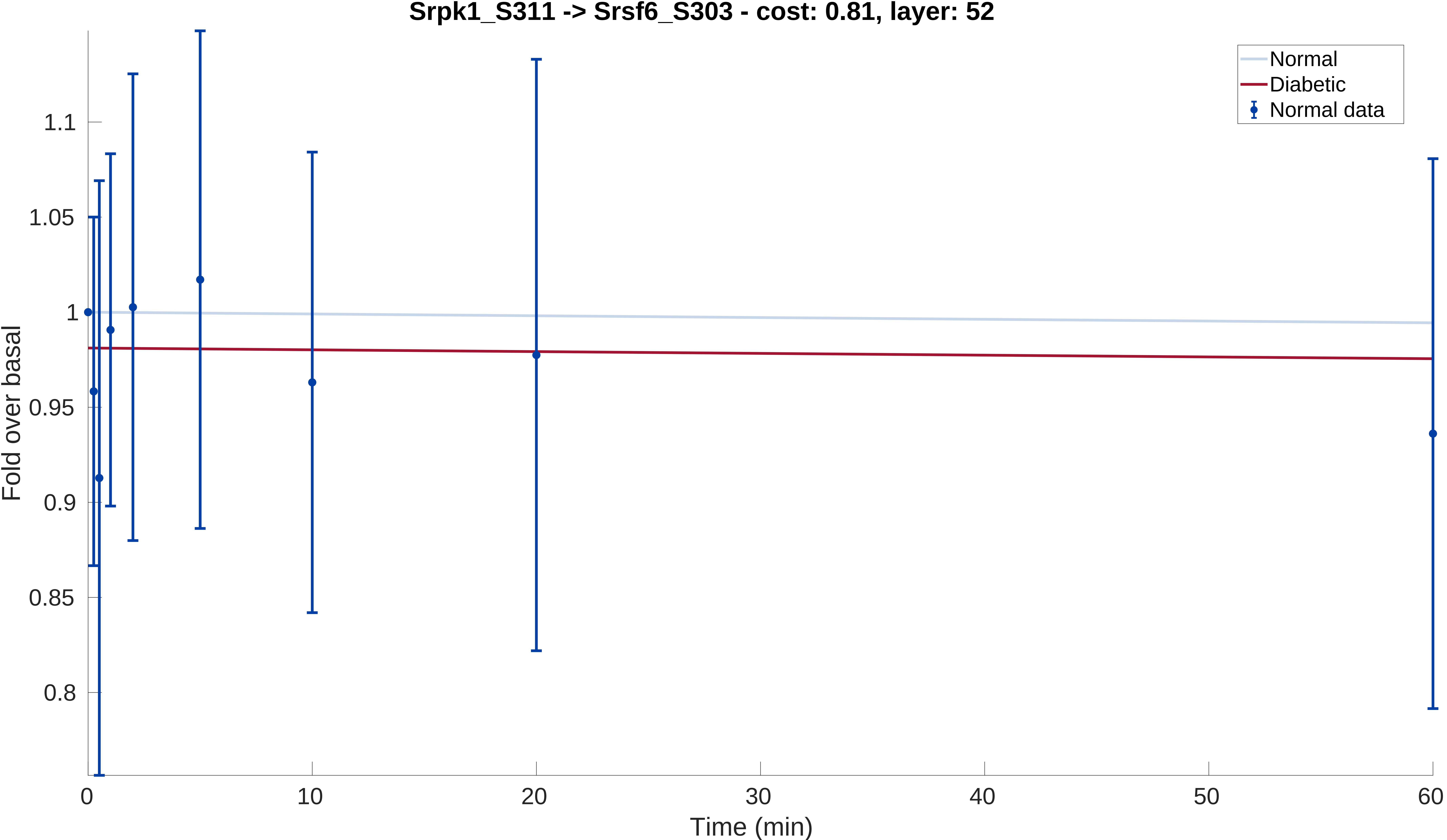

Srpk1_S311 -> Srsf6_S303 - cost: 0.81, layer: 52
Normal
Diabetic
Normal data
1.1
1.05
1
0.95
0.9
0.85
0.8
Fold over basal
0
10
20
30
40
50
60
Time (min)

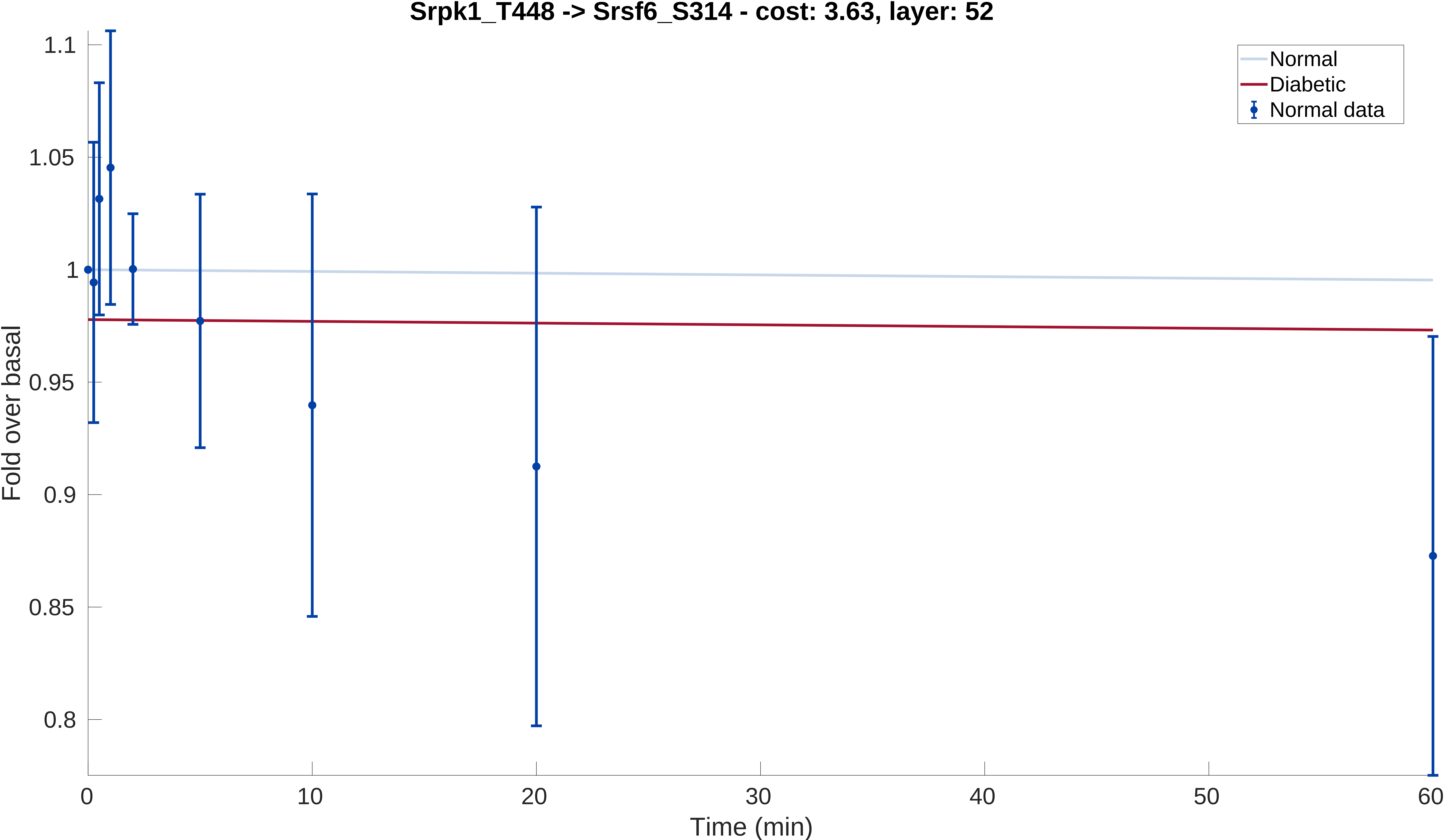
Srpk1_T448 -> Srsf6_S314 - cost: 3.63, layer: 52
Normal
Diabetic
Normal data
Fold over basal
1.1
1.05
1
0.95
0.9
0.85
0.8
0
10
20
30
40
50
60
Time (min)

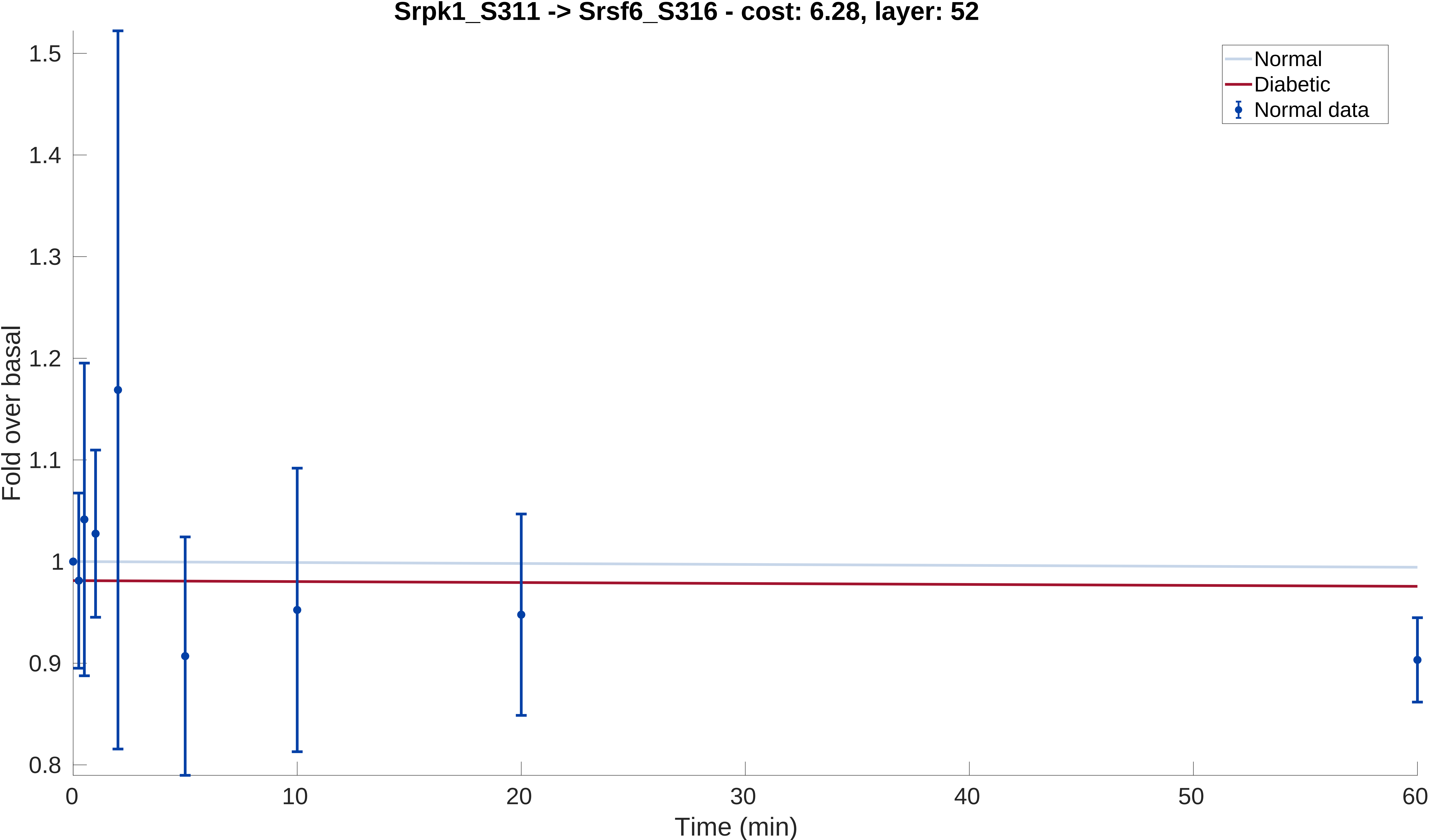

Srpk1_S311 -> Srsf6_S316 - cost: 6.28, layer: 52
Normal
Diabetic
Normal data
Fold over basal
1.5
1.4
1.3
1.2
1.1
1
0.9
0.8
0
10
20
30
40
50
60
Time (min)

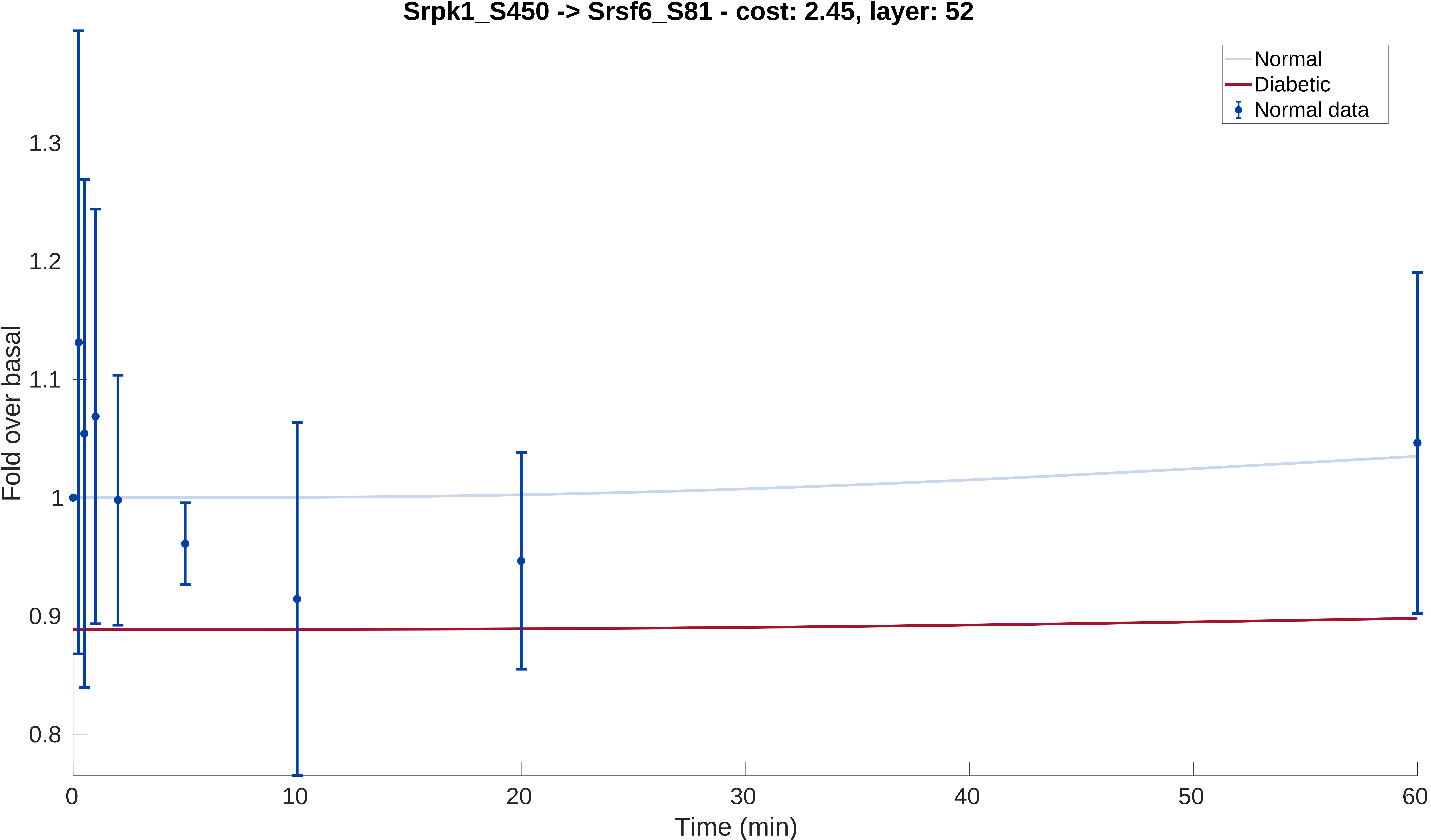
Srpk1_S450 -> Srsf6_S81 - cost: 2.45, layer: 52
Normal
Diabetic
Normal data
Fold over basal
1.3
1.2
1.1
1
0.9
0.8
0
10
20
30
40
50
60
Time (min)

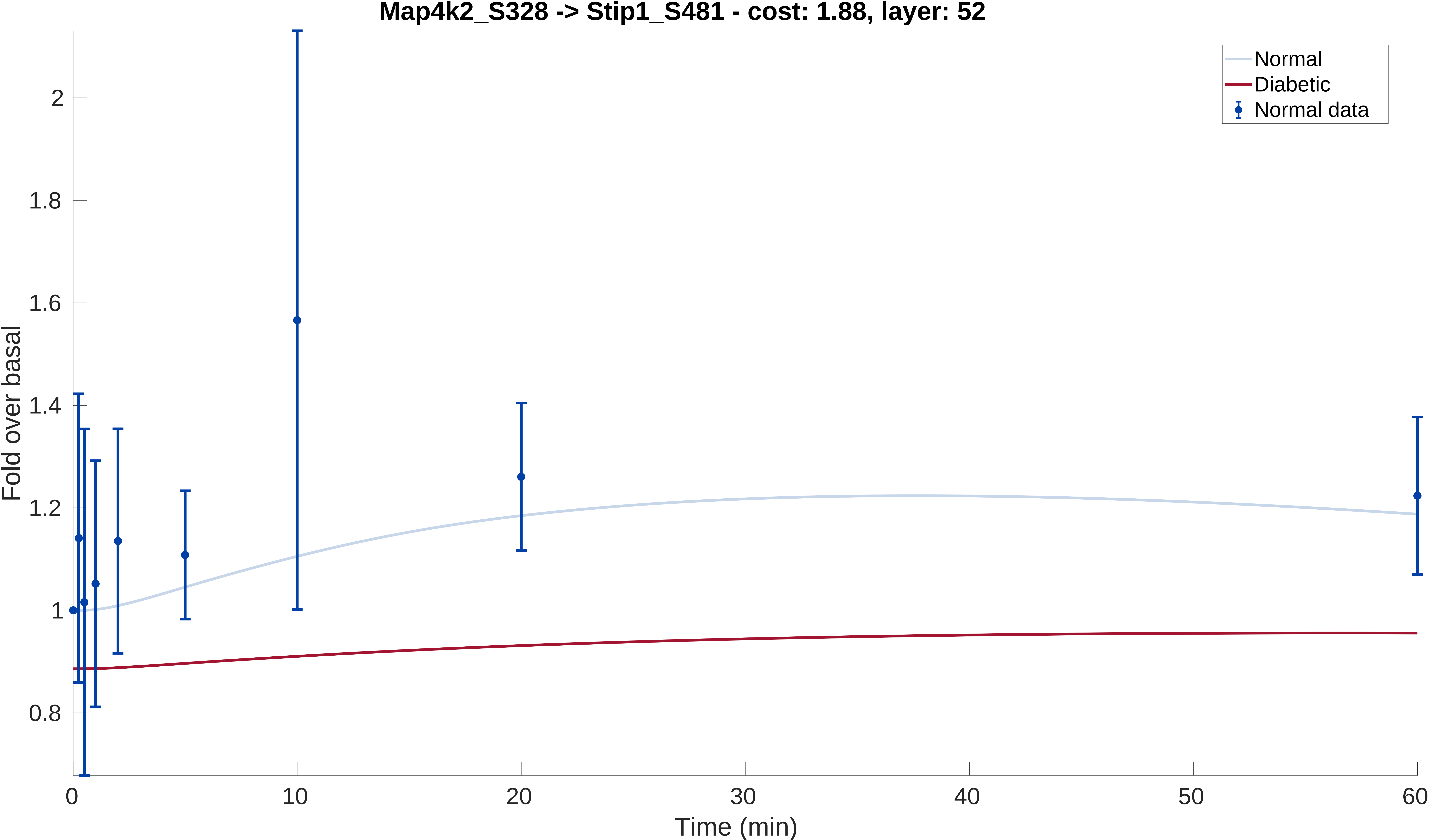

Map4k2_S328 -> Stip1_S481 - cost: 1.88, layer: 52
Normal
Diabetic
Normal data
Fold over basal
2
1.8
1.6
1.4
1.2
1
0.8
0
10
20
30
40
50
60
Time (min)

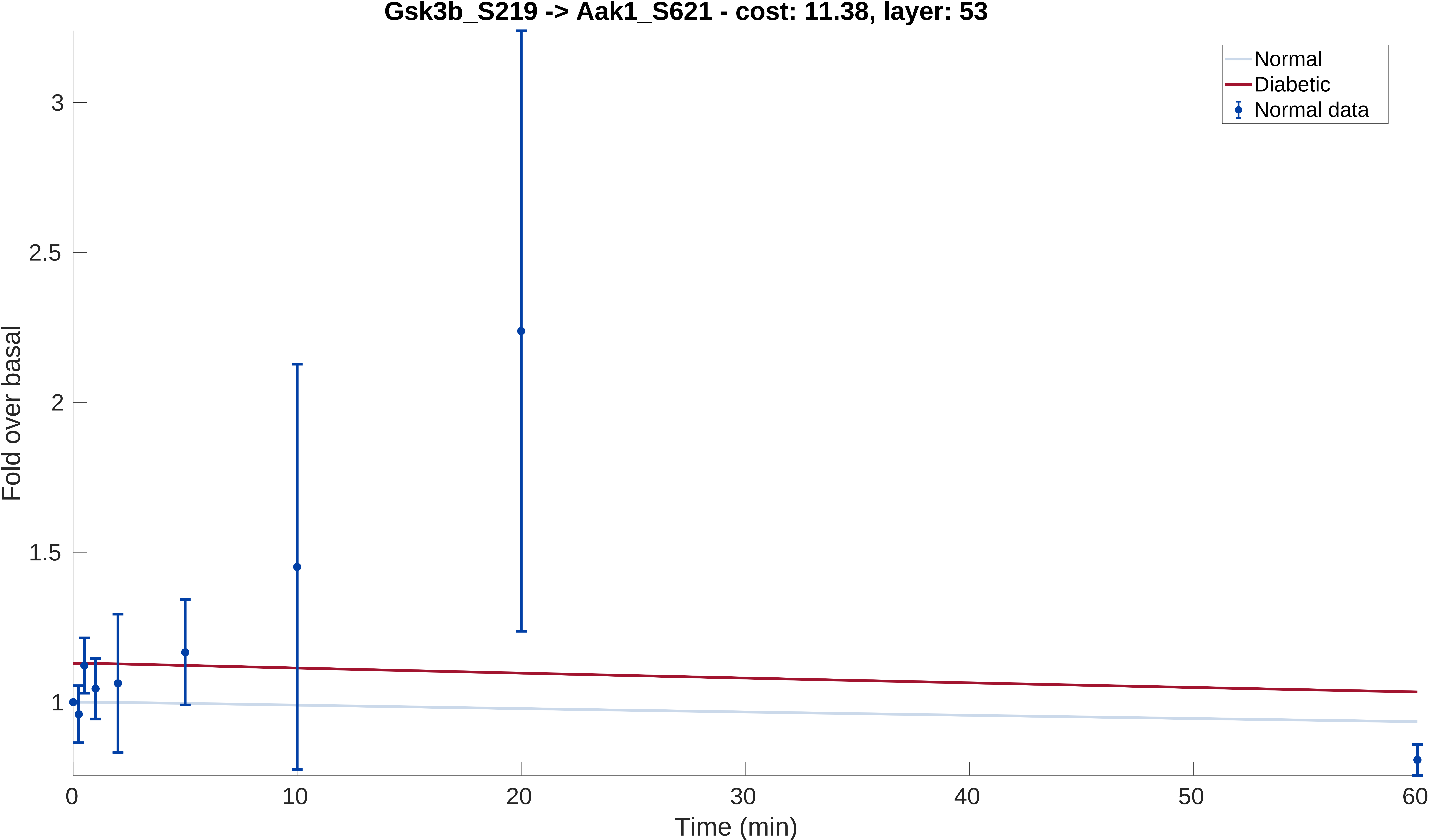

Gsk3b_S219 -> Aak1_S621 - cost: 11.38, layer: 53
Normal
Diabetic
Normal data
Fold over basal
3
2.5
2
1.5
1
0
10
20
30
40
50
60
Time (min)

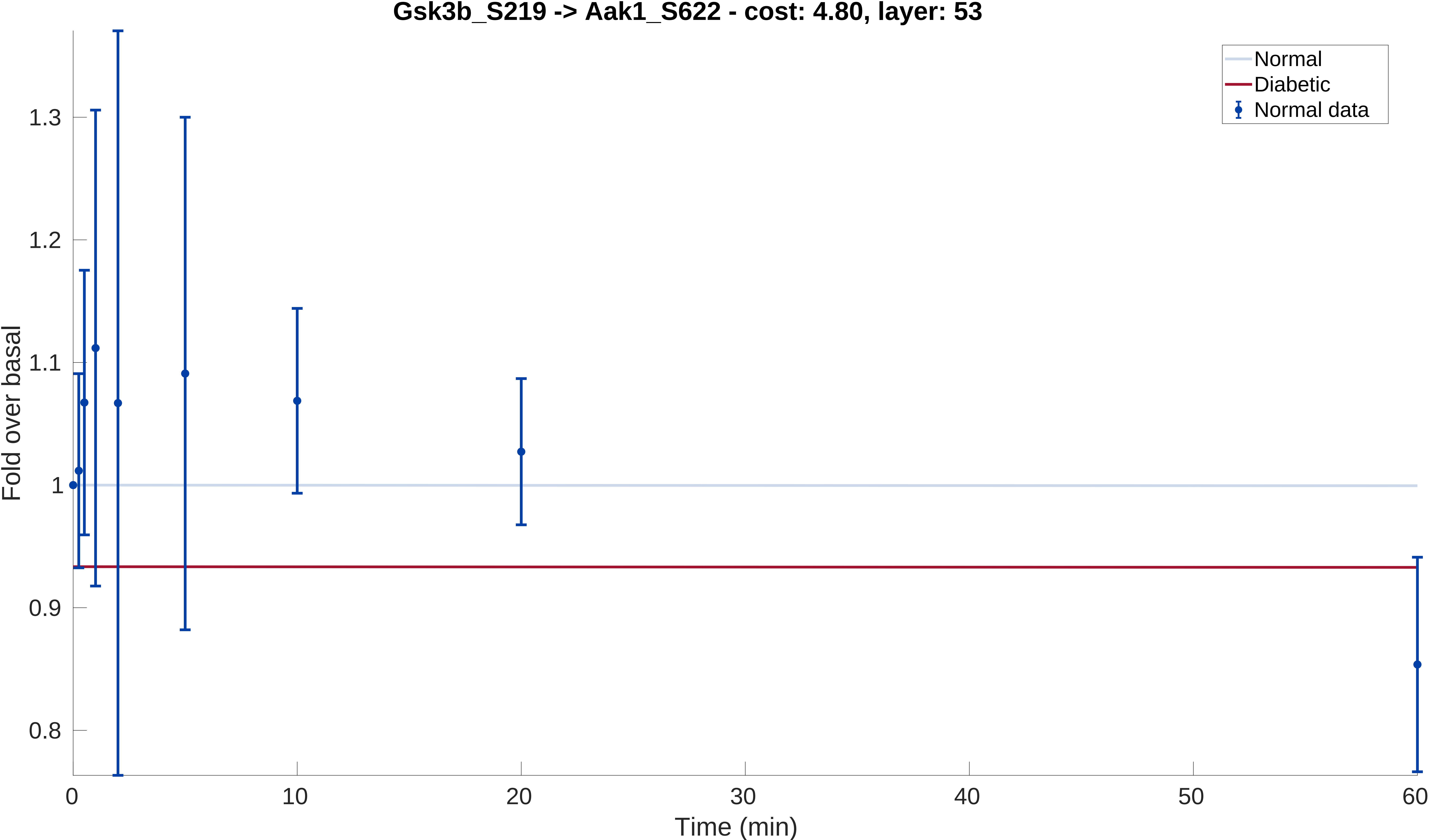

Gsk3b_S219 -> Aak1_S622 - cost: 4.80, layer: 53
Normal
Diabetic
Normal data
Fold over basal
1.3
1.2
1.1
1
0.9
0.8
0
10
20
30
40
50
60
Time (min)

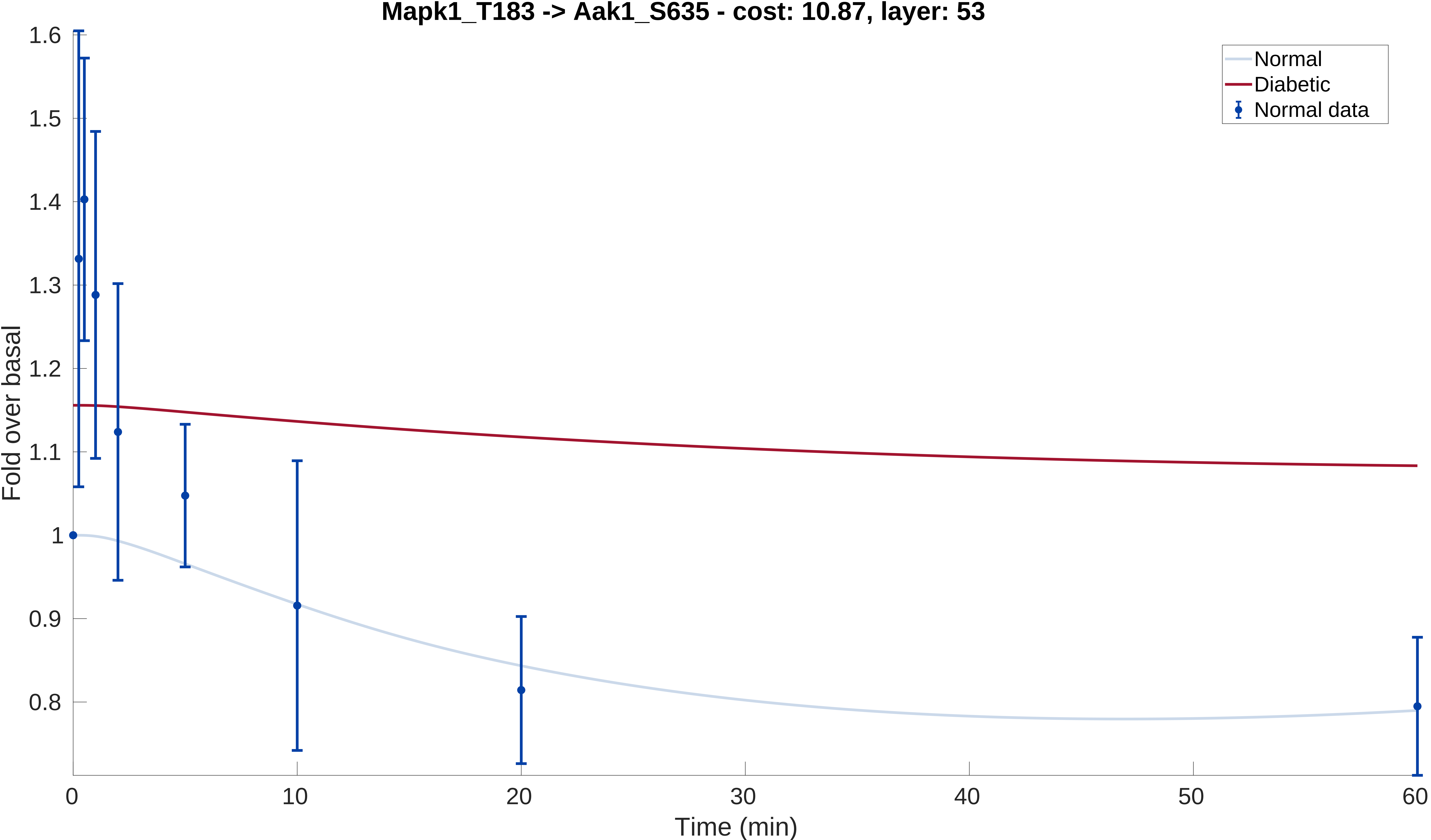

Mapk1_T183 -> Aak1_S635 - cost: 10.87, layer: 53
Normal
Diabetic
Normal data
Fold over basal
1.6
1.5
1.4
1.3
1.2
1.1
1
0.9
0.8
0
10
20
30
40
50
60
Time (min)

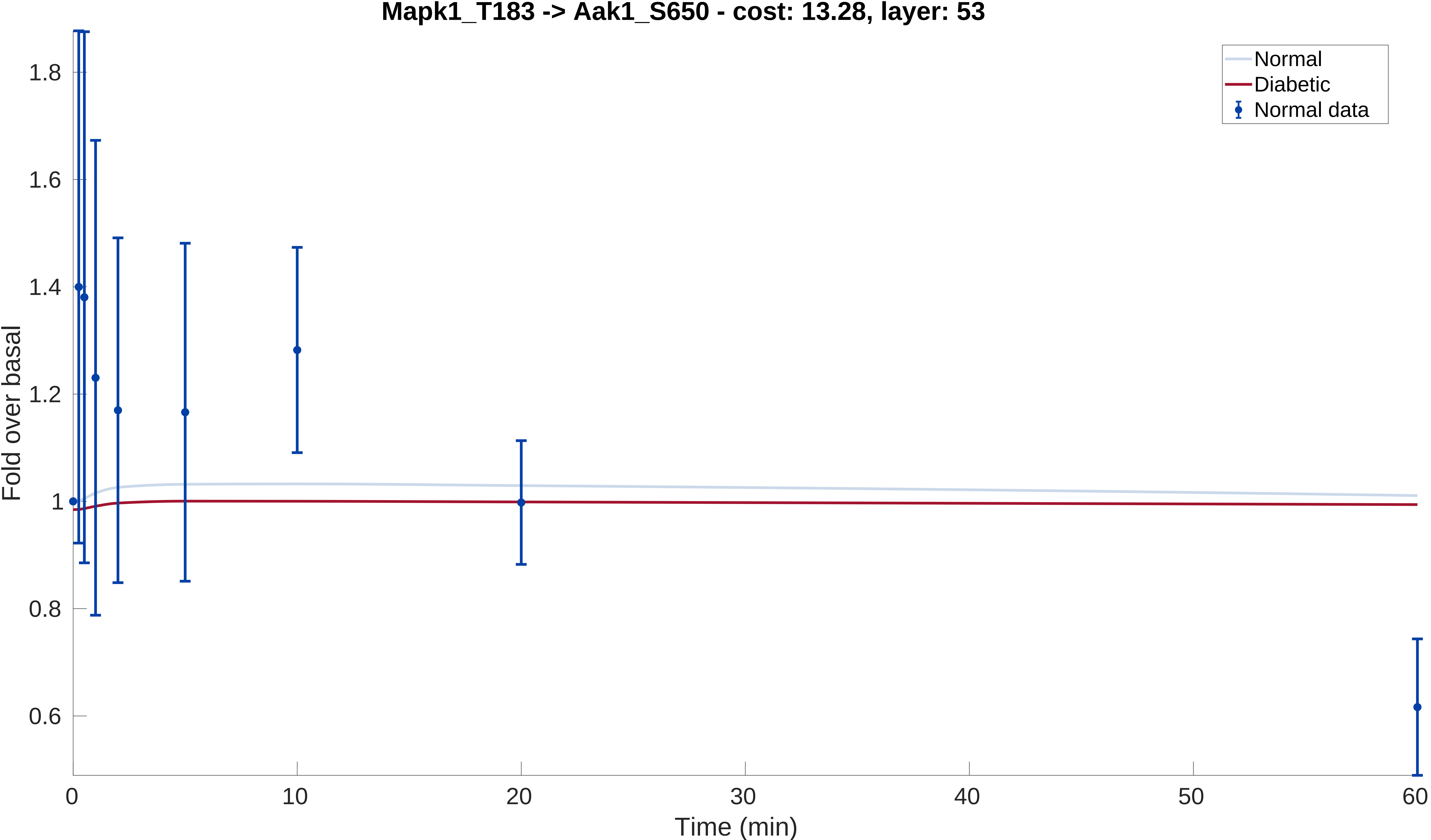

Mapk1_T183 -> Aak1_S650 - cost: 13.28, layer: 53
Normal
Diabetic
Normal data
Fold over basal
1.8
1.6
1.4
1.2
1
0.8
0.6
0
10
20
30
40
50
60
Time (min)

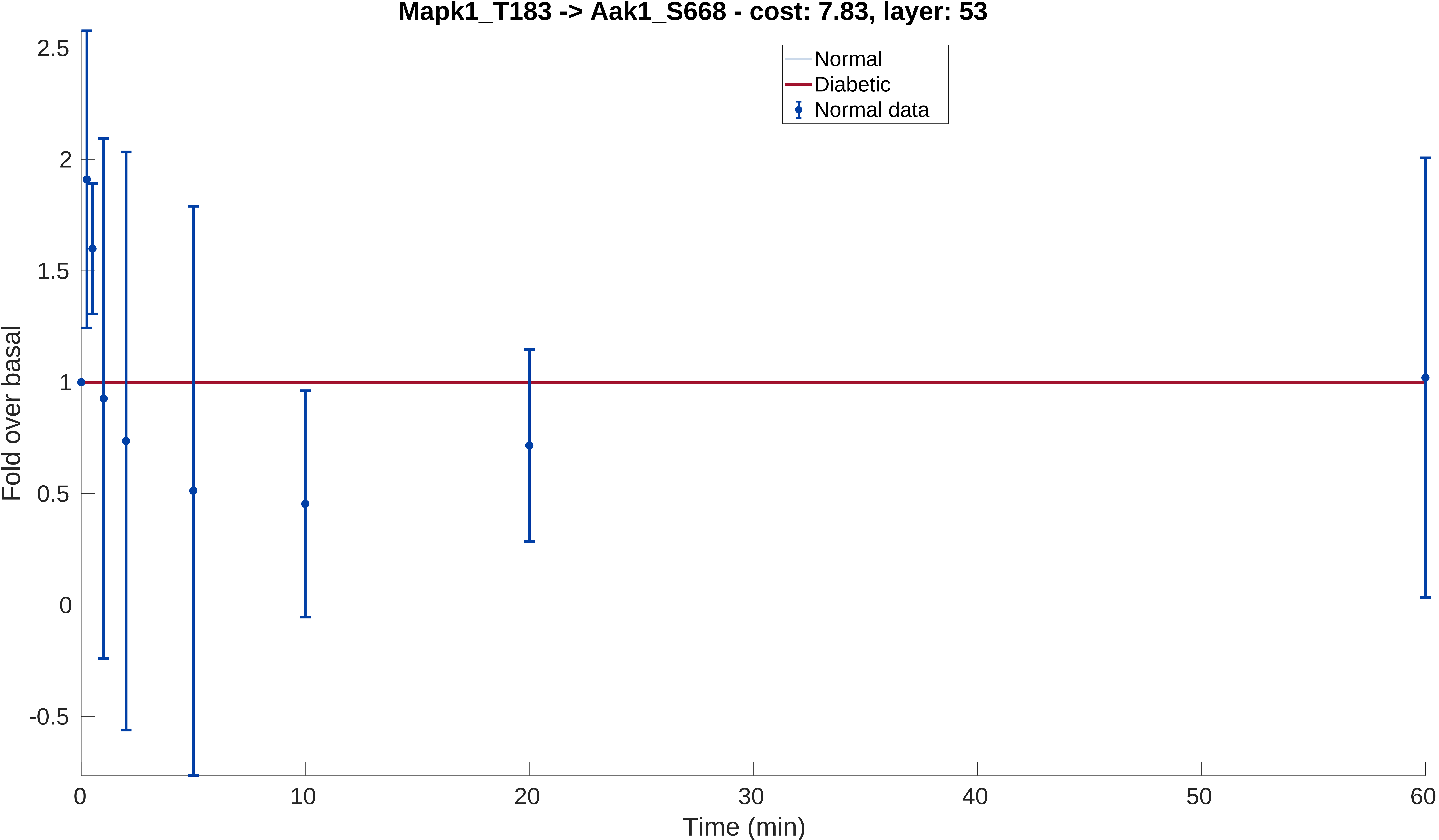

Mapk1_T183 -> Aak1_S668 - cost: 7.83, layer: 53
Normal
Diabetic
Normal data
Fold over basal
Time (min)
2.5
2
1.5
1
0.5
0
-0.5
0
10
20
30
40
50
60

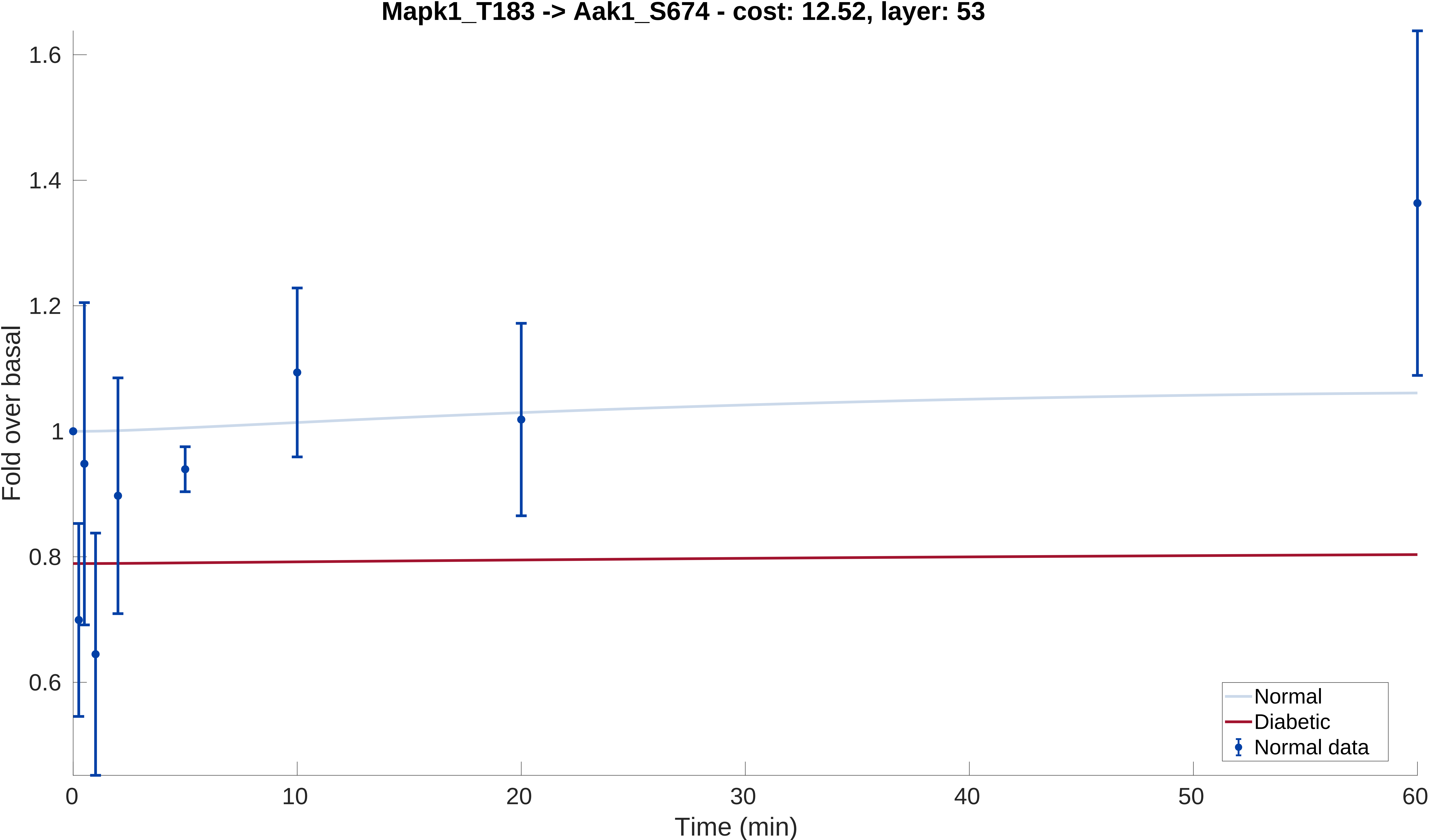

Mapk1_T183 -> Aak1_S674 - cost: 12.52, layer: 53
1.6
1.4
1.2
1
0.8
0.6
Fold over basal
0
10
20
30
40
50
60
Time (min)
Normal
Diabetic
Normal data

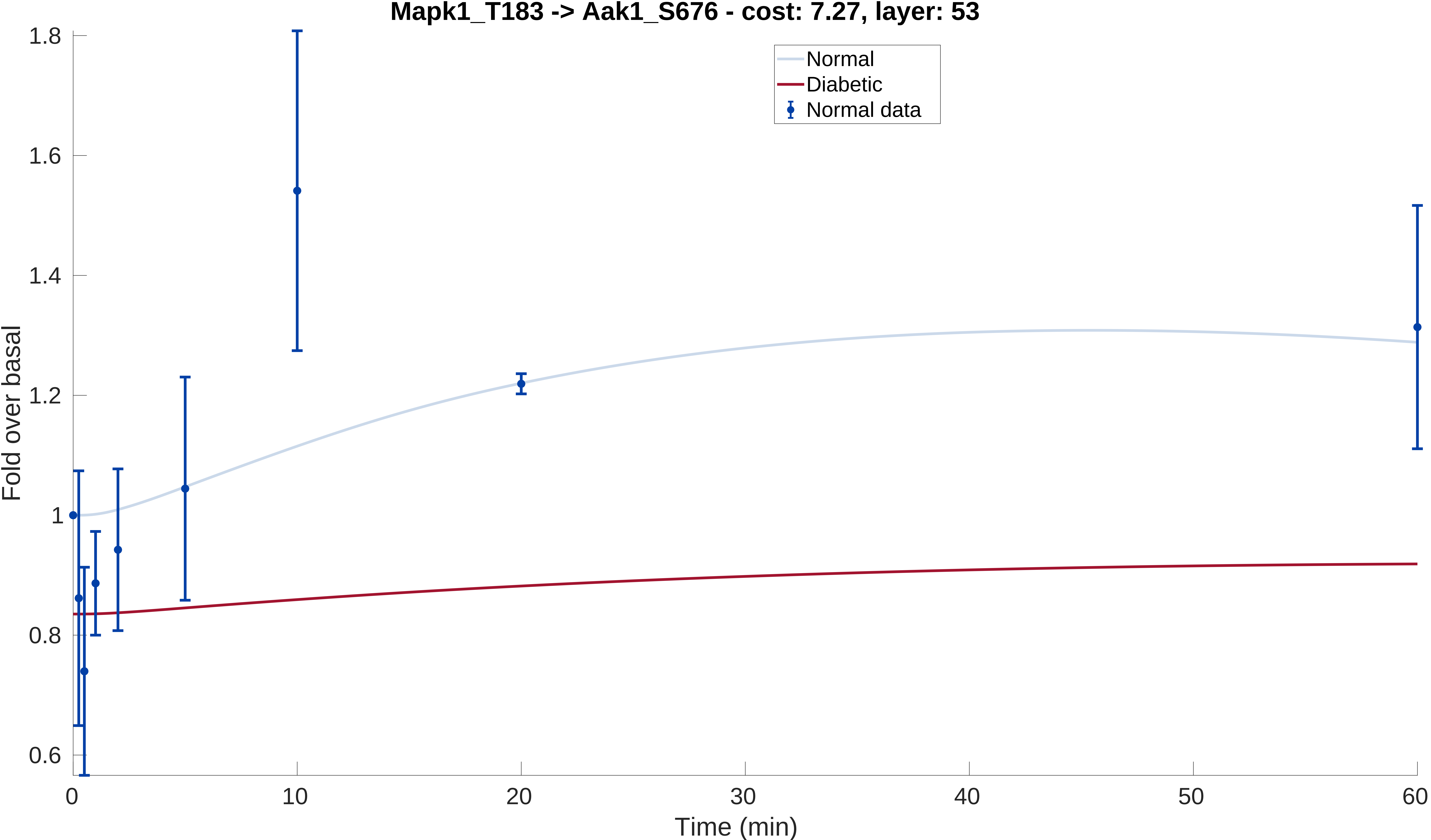

Mapk1_T183 -> Aak1_S676 - cost: 7.27, layer: 53
Normal
Diabetic
Normal data
Fold over basal
Time (min)
1.8
1.6
1.4
1.2
1
0.8
0.6
0
10
20
30
40
50
60

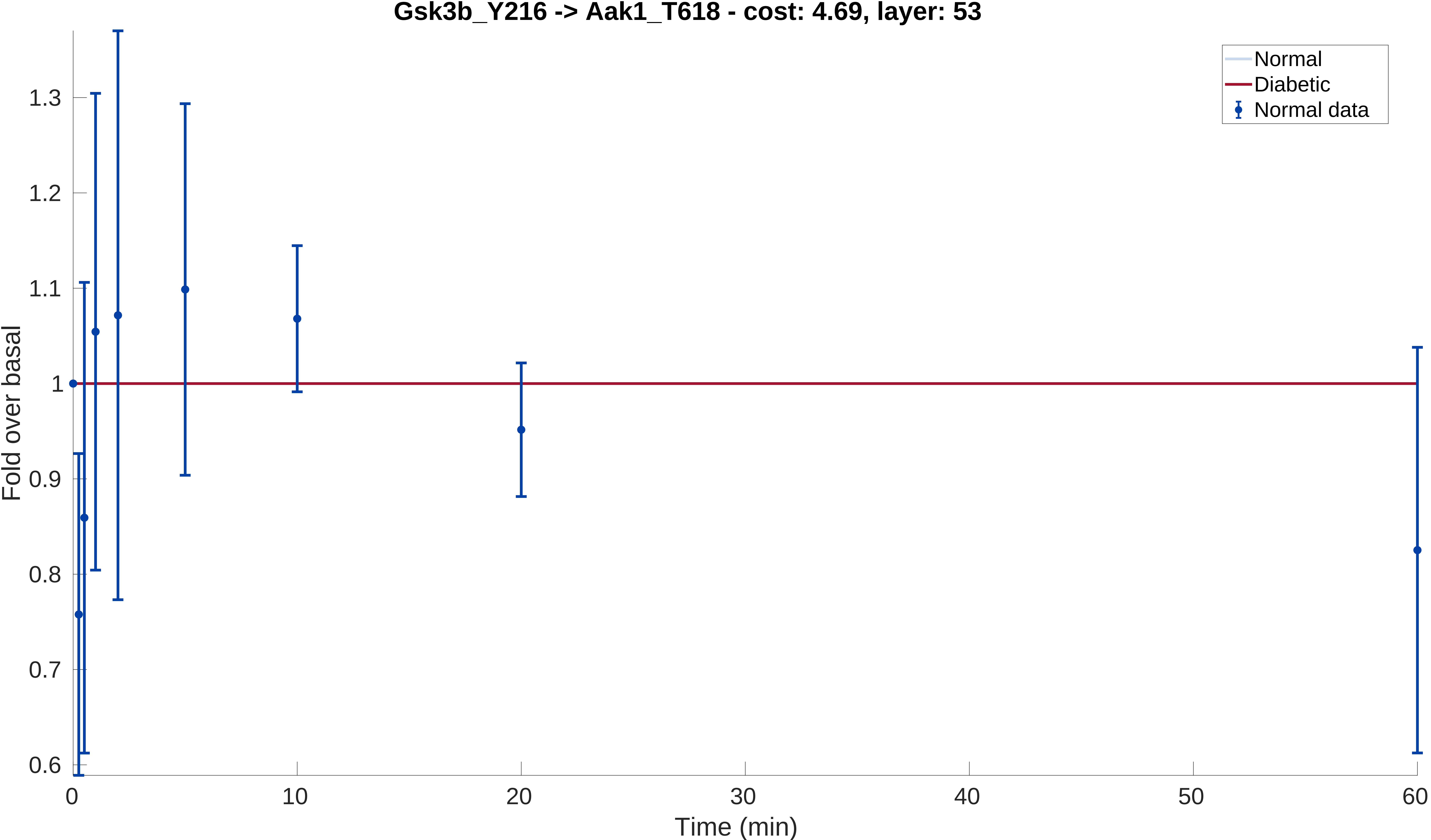

Gsk3b_Y216 -> Aak1_T618 - cost: 4.69, layer: 53
Normal
Diabetic
Normal data
Fold over basal
Time (min)
1.3
1.2
1.1
1
0.9
0.8
0.7
0.6
0
10
20
30
40
50
60

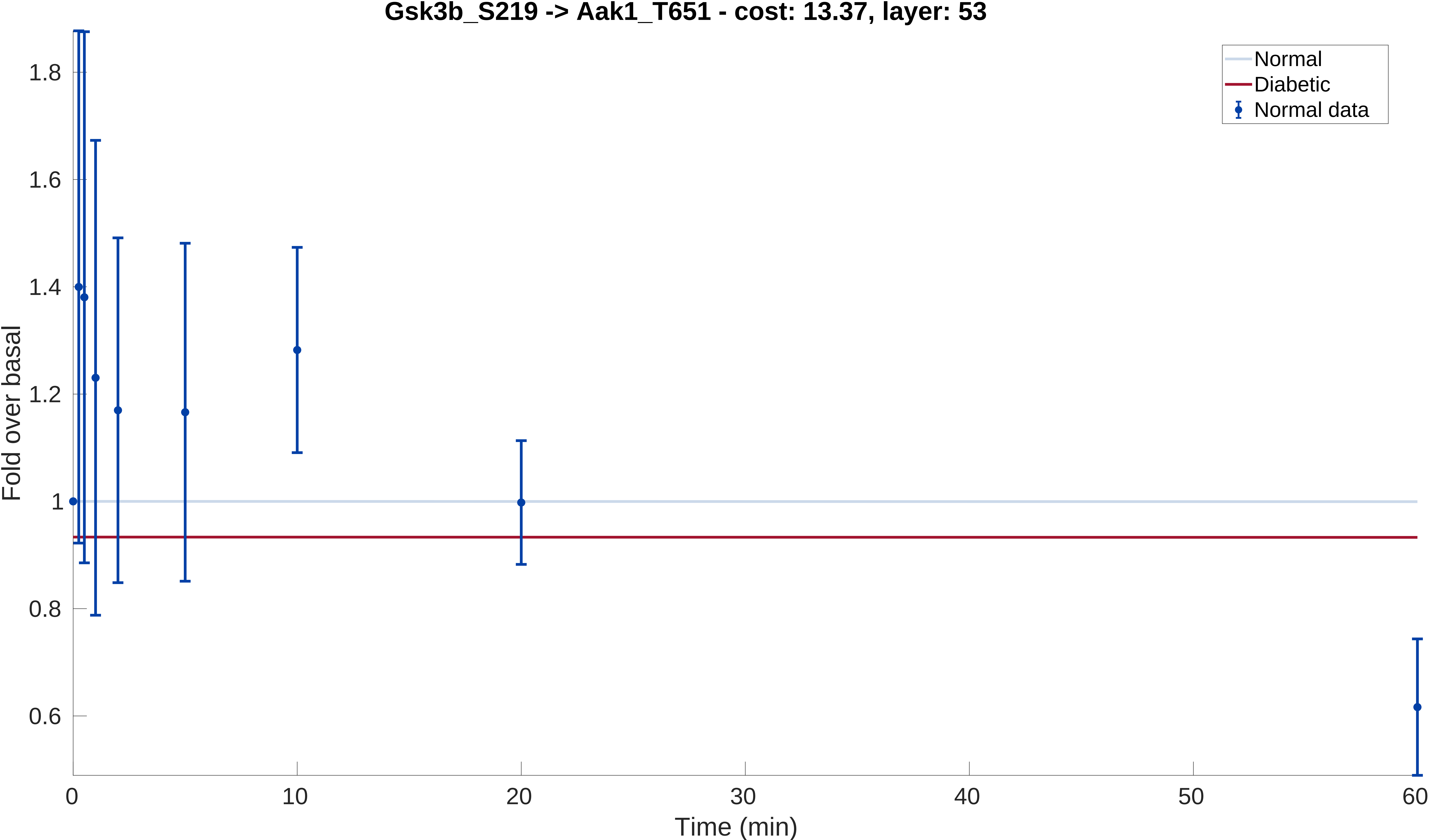

Gsk3b_S219 -> Aak1_T651 - cost: 13.37, layer: 53
Normal
Diabetic
Normal data
Fold over basal
1.8
1.6
1.4
1.2
1
0.8
0.6
0
10
20
30
40
50
60
Time (min)

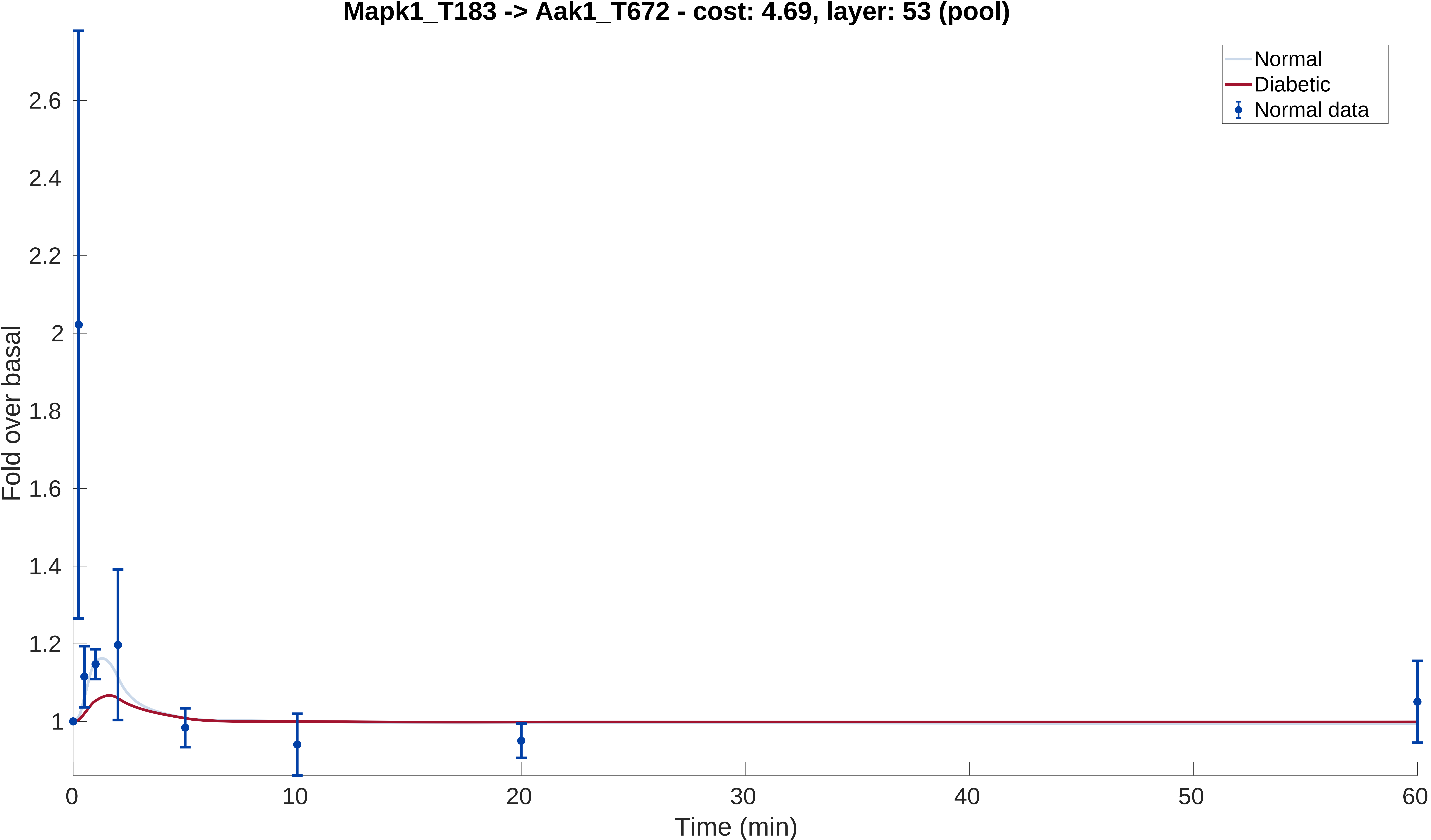

Mapk1_T183 -> Aak1_T672 - cost: 4.69, layer: 53 (pool)
Normal
Diabetic
Normal data
Fold over basal
2.6
2.4
2.2
2
1.8
1.6
1.4
1.2
1
0
10
20
30
40
50
60
Time (min)

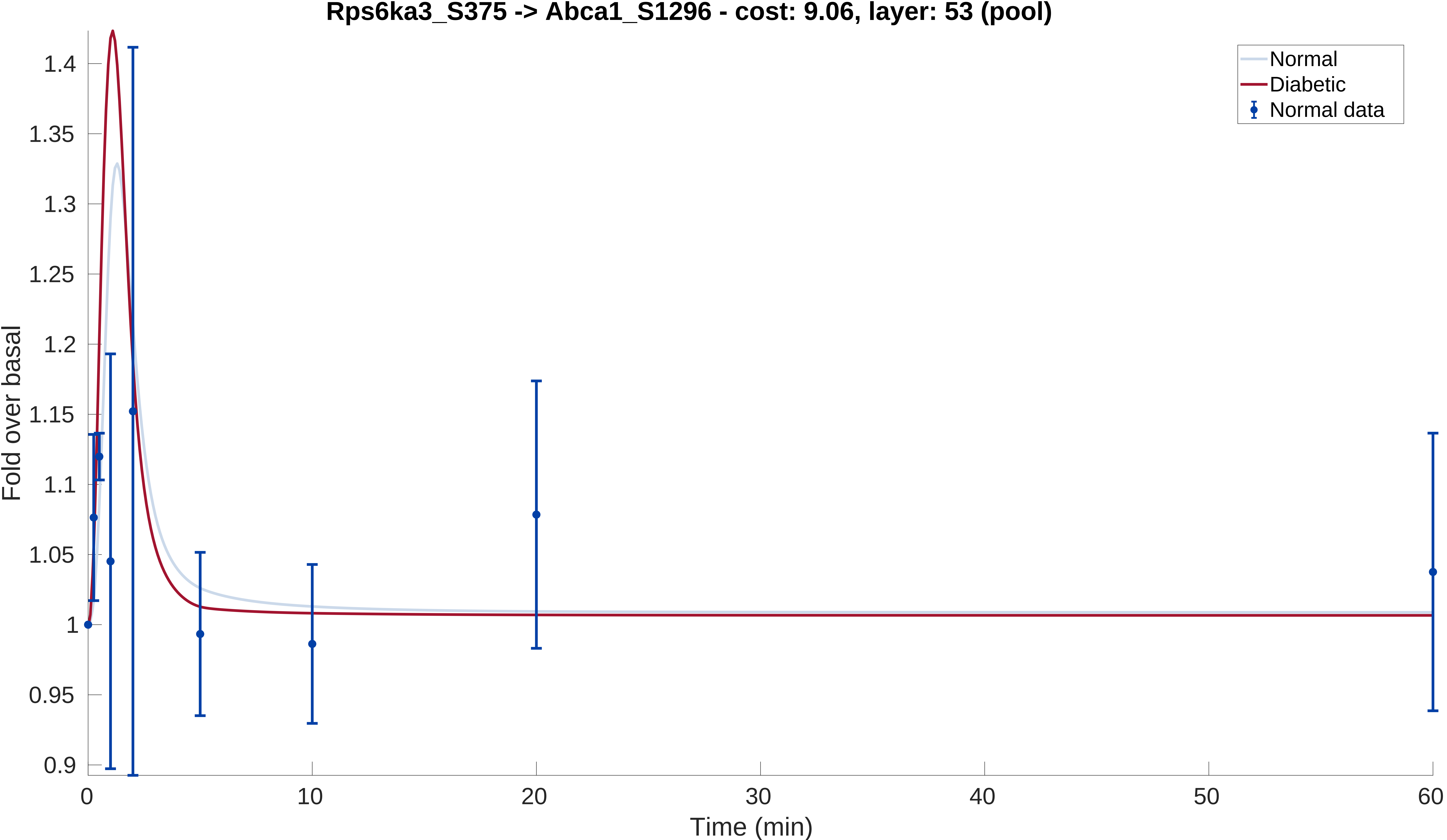

Rps6ka3_S375 -> Abca1_S1296 - cost: 9.06, layer: 53 (pool)
Normal
Diabetic
Normal data
Fold over basal
1.4
1.35
1.3
1.25
1.2
1.15
1.1
1.05
1
0.95
0.9
0
10
20
30
40
50
60
Time (min)

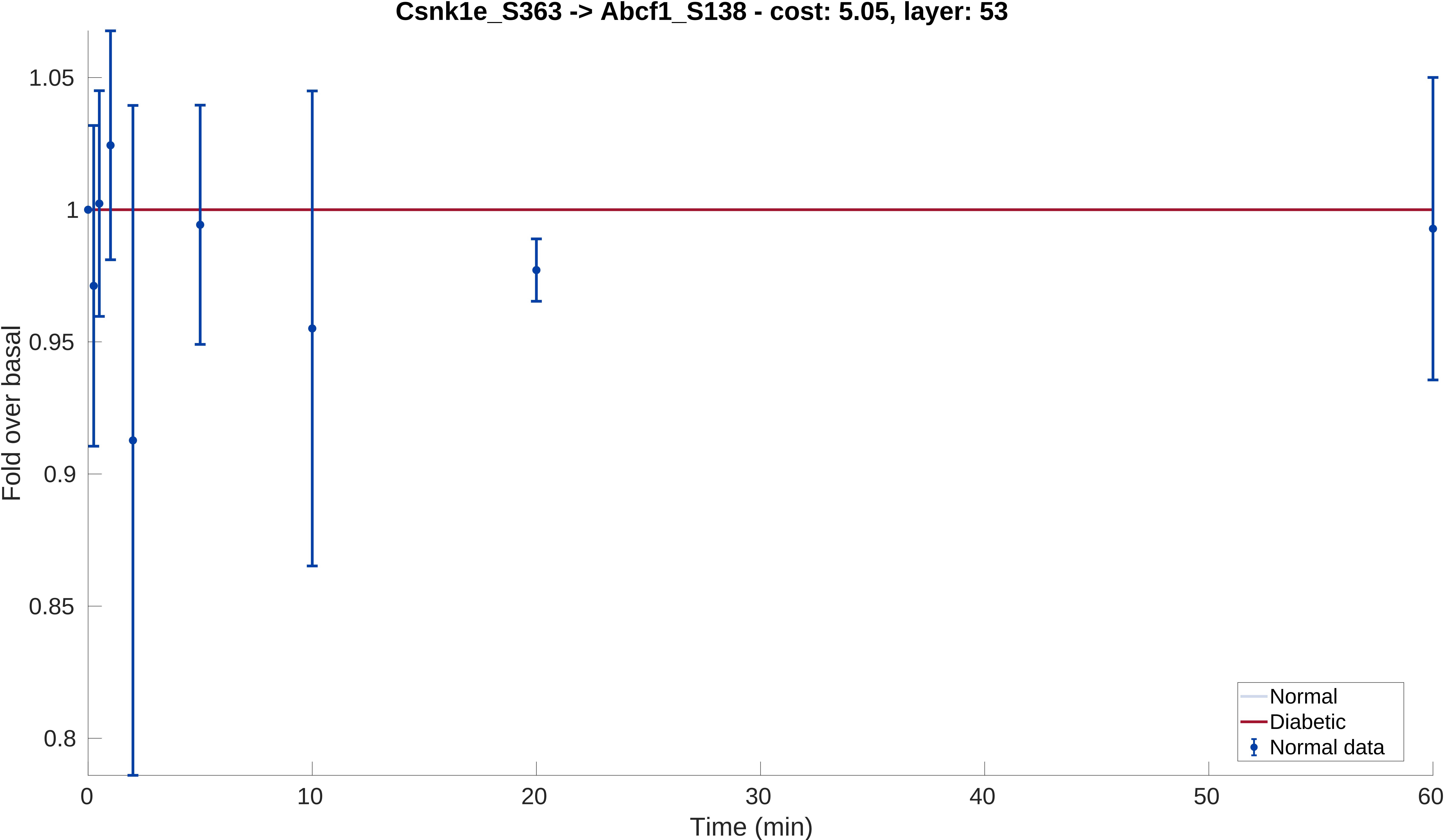

Csnk1e_S363 -> Abcf1_S138 - cost: 5.05, layer: 53
Fold over basal
1.05
1
0.95
0.9
0.85
0.8
0
10
20
30
40
50
60
Time (min)
Normal
Diabetic
Normal data

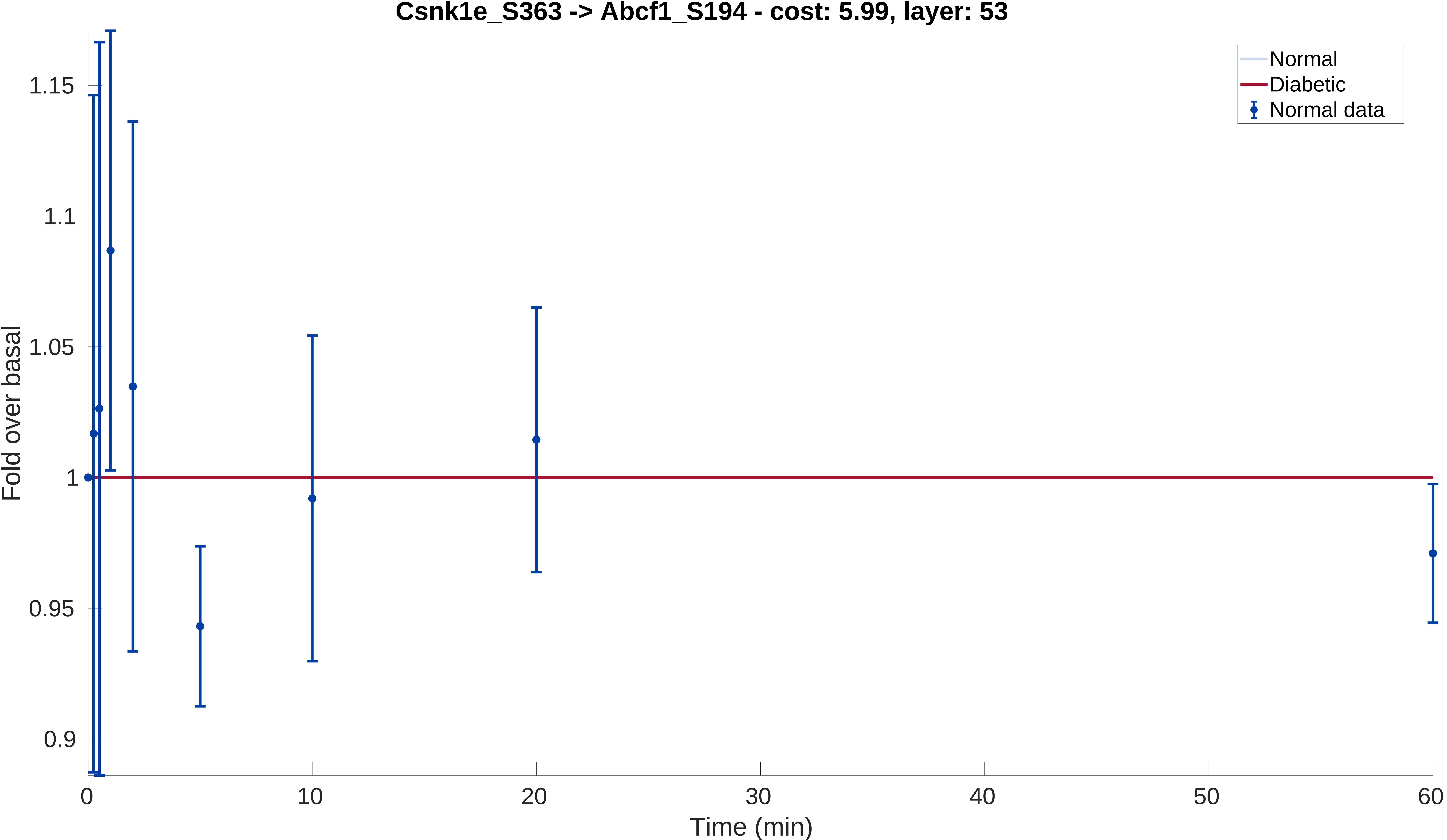
Csnk1e_S363 -> Abcf1_S194 - cost: 5.99, layer: 53
Normal
Diabetic
Normal data
Fold over basal
1.15
1.1
1.05
1
0.95
0.9
0
10
20
30
40
50
60
Time (min)

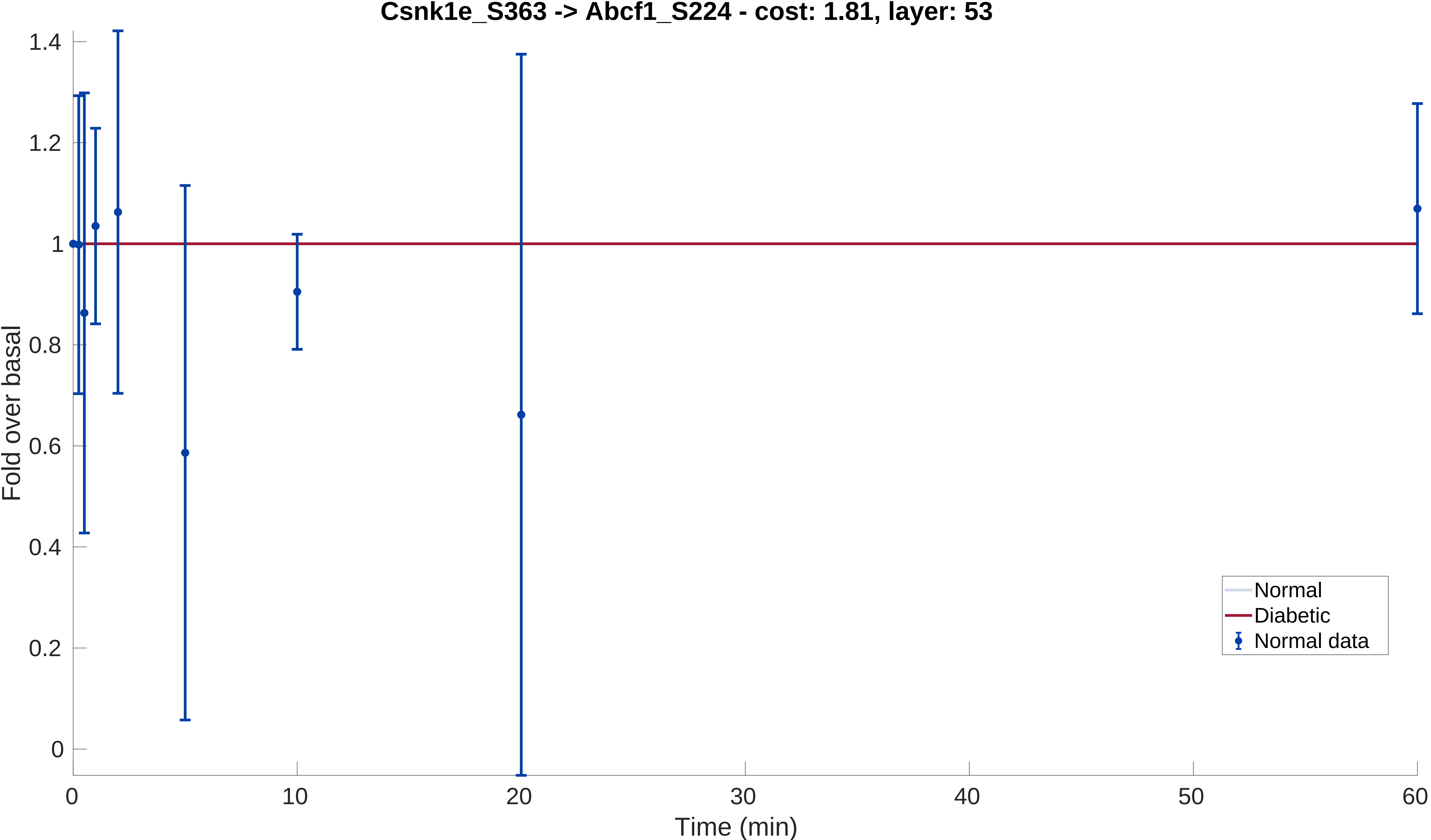
Csnk1e_S363 -> Abcf1_S224 - cost: 1.81, layer: 53
Fold over basal
Time (min)
0
0.2
0.4
0.6
0.8
1
1.2
1.4
0
10
20
30
40
50
60
Normal
Diabetic
Normal data

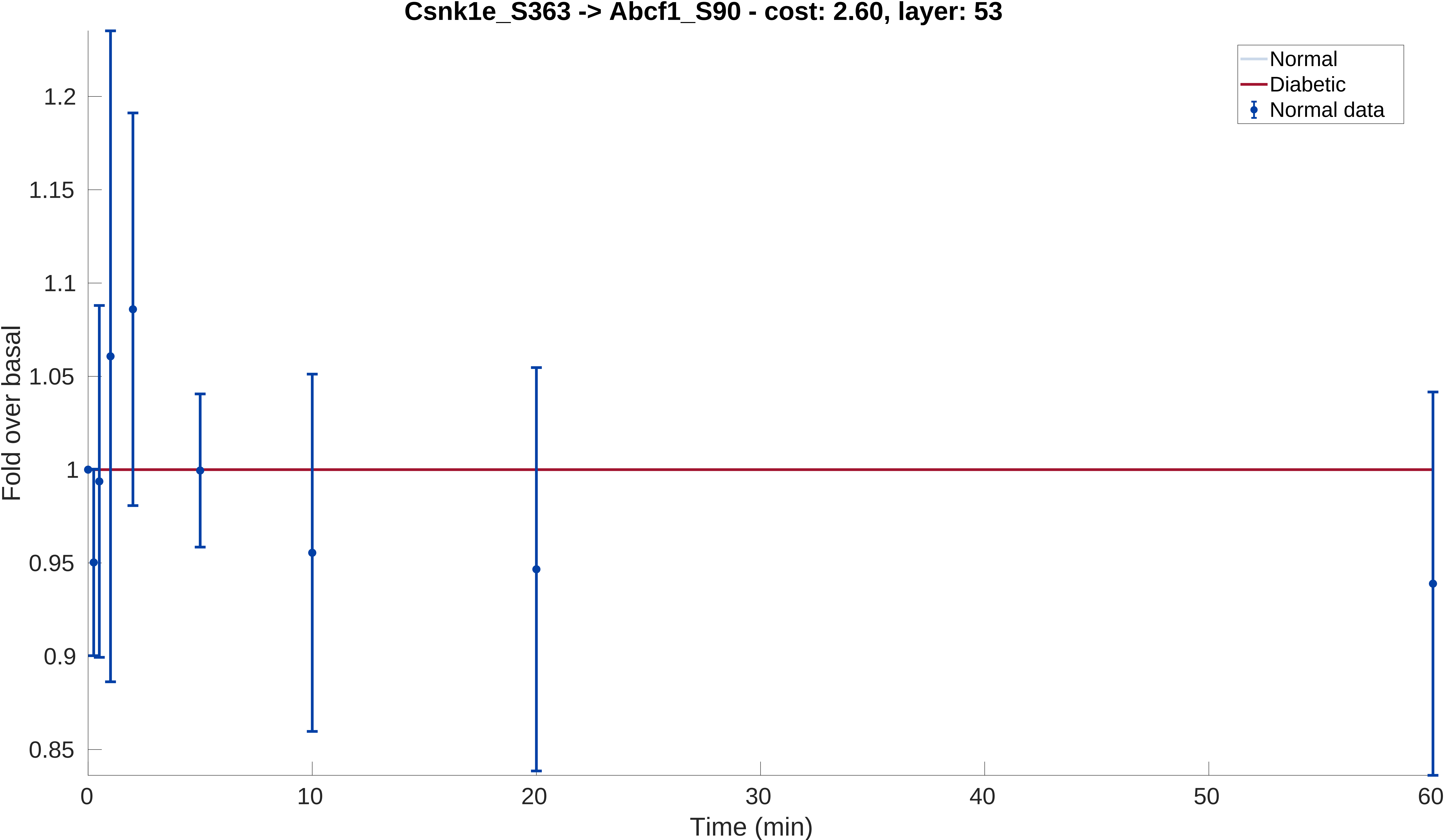

Csnk1e_S363 -> Abcf1_S90 - cost: 2.60, layer: 53
Normal
Diabetic
Normal data
Fold over basal
1.2
1.15
1.1
1.05
1
0.95
0.9
0.85
0
10
20
30
40
50
60
Time (min)

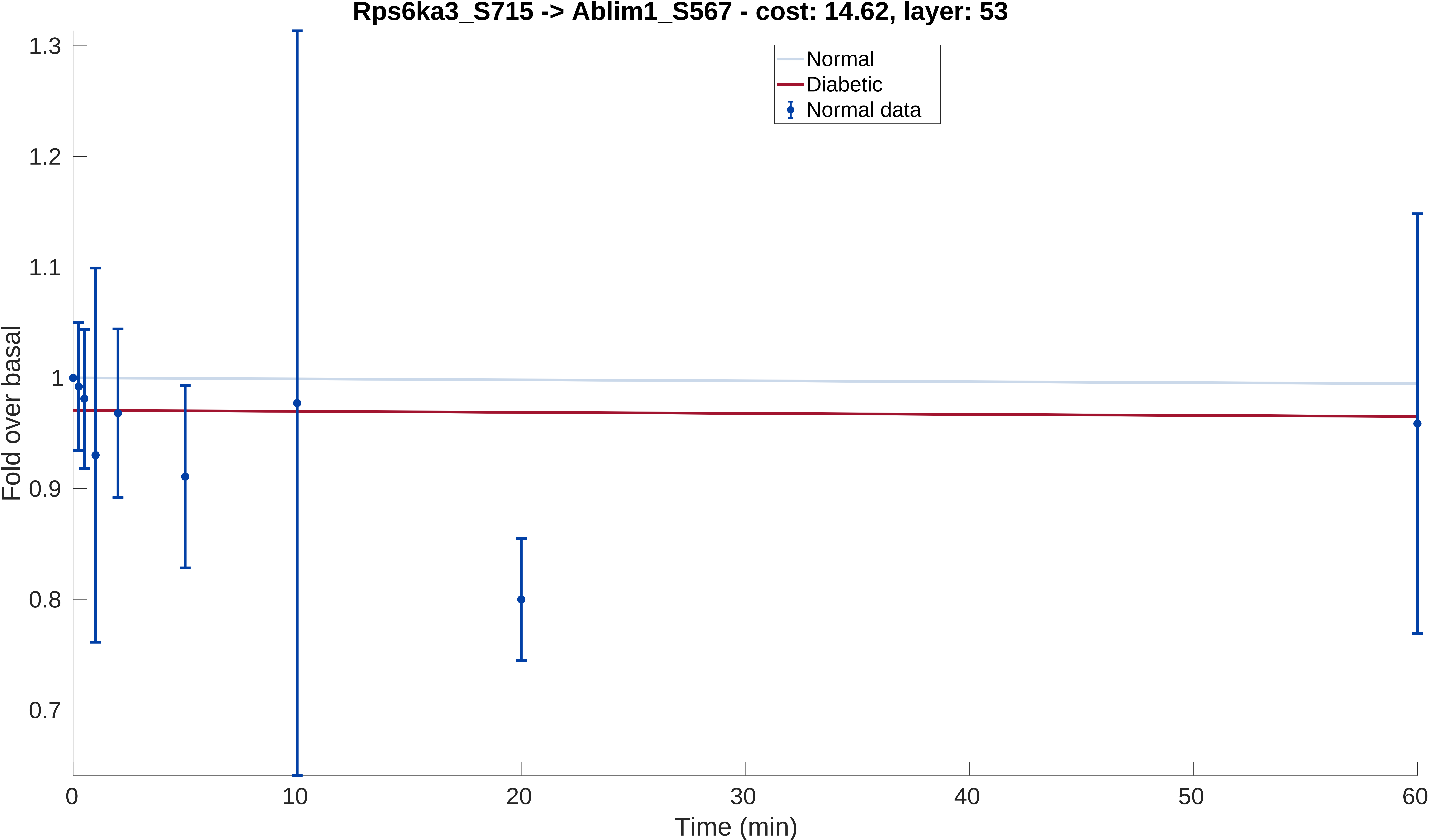

Rps6ka3_S715 -> Ablim1_S567 - cost: 14.62, layer: 53
Normal
Diabetic
Normal data
Fold over basal
1.3
1.2
1.1
1
0.9
0.8
0.7
0
10
20
30
40
50
60
Time (min)

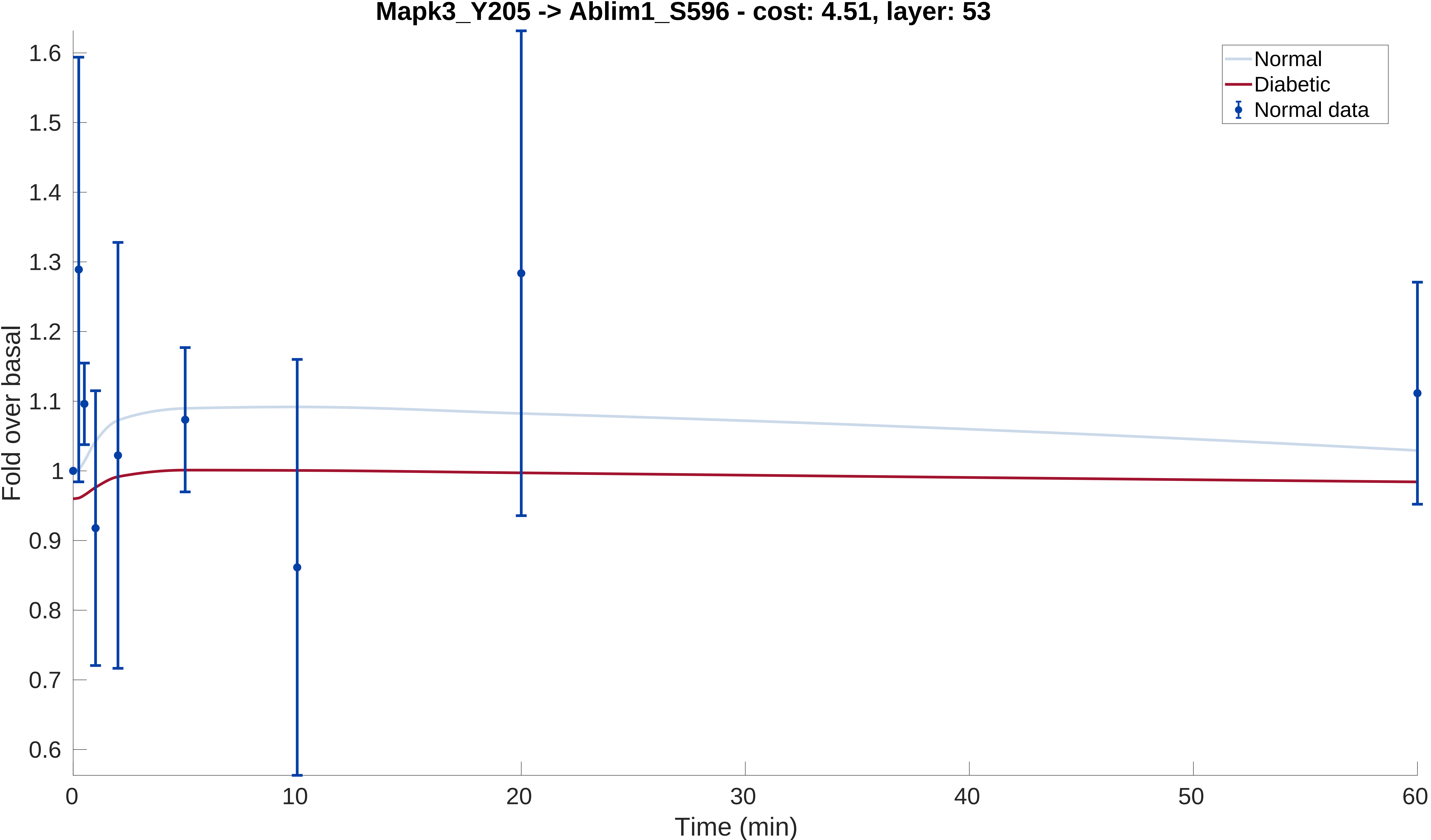
Mapk3_Y205 -> Ablim1_S596 - cost: 4.51, layer: 53
Normal
Diabetic
Normal data
Fold over basal
1.6
1.5
1.4
1.3
1.2
1.1
1
0.9
0.8
0.7
0.6
0
10
20
30
40
50
60
Time (min)

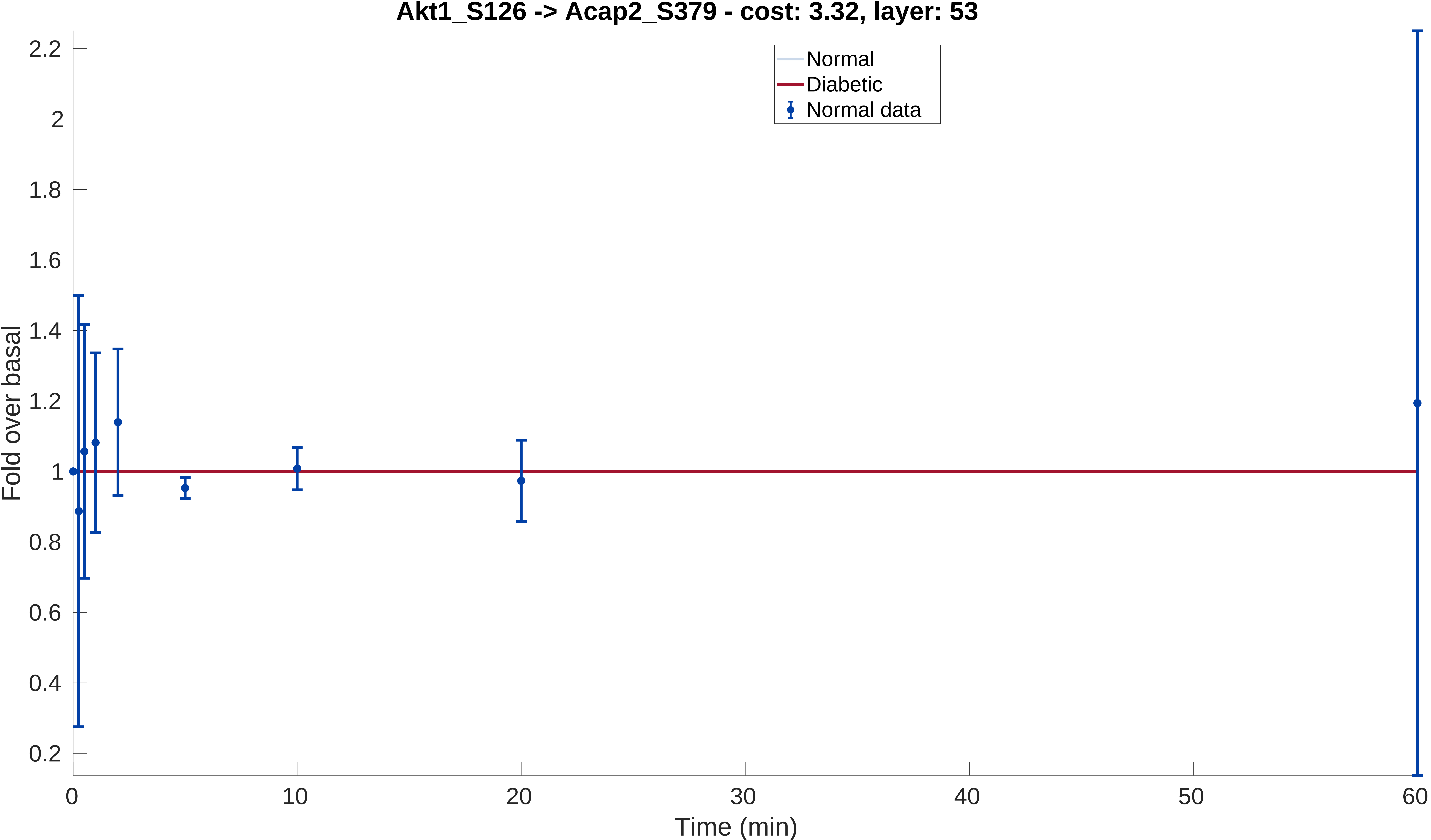
Akt1_S126 -> Acap2_S379 - cost: 3.32, layer: 53
Normal
Diabetic
Normal data
Fold over basal
Time (min)
0.2
0.4
0.6
0.8
1
1.2
1.4
1.6
1.8
2
2.2
0
10
20
30
40
50
60

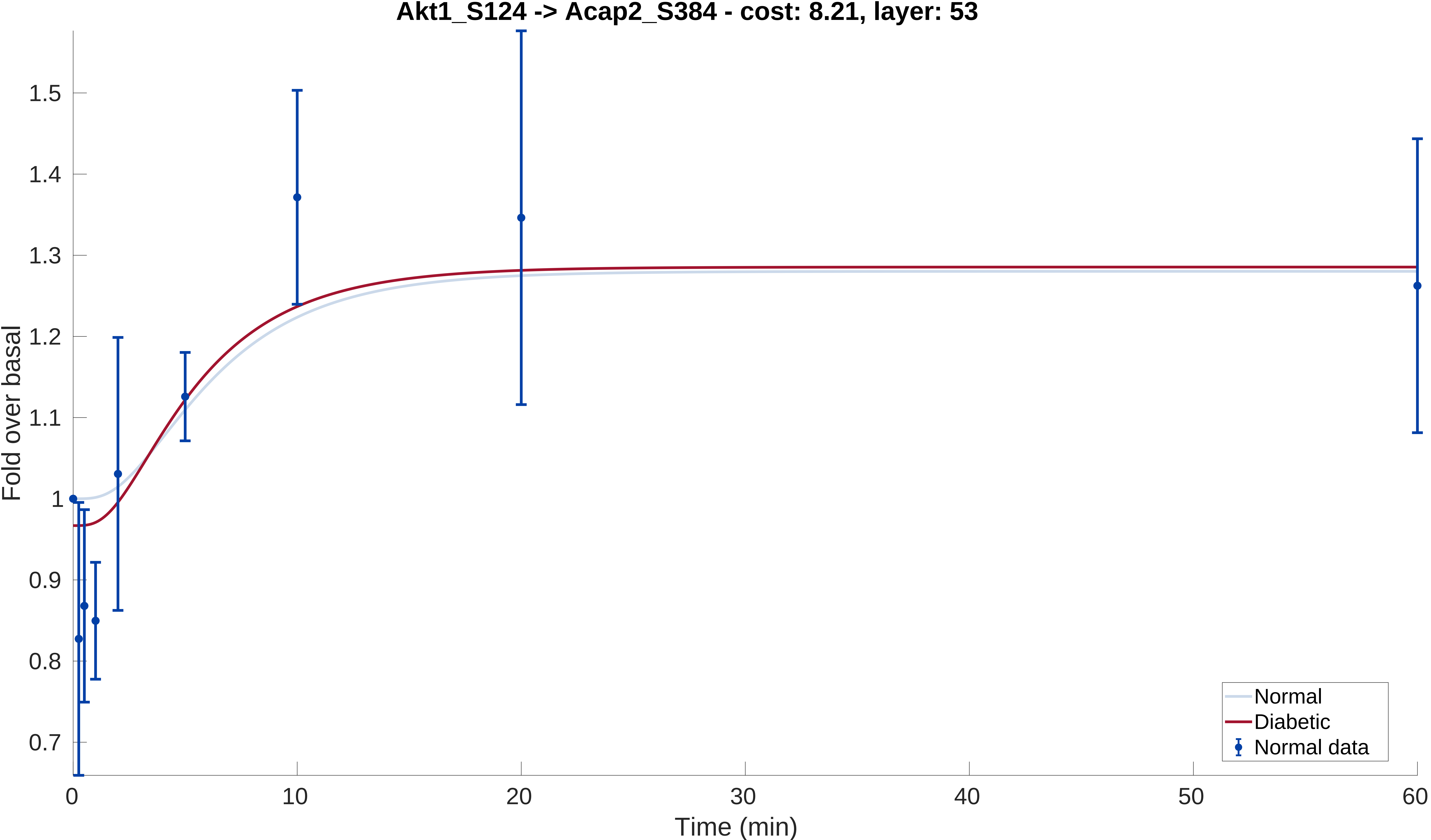

Akt1_S124 -> Acap2_S384 - cost: 8.21, layer: 53
Fold over basal
1.5
1.4
1.3
1.2
1.1
1
0.9
0.8
0.7
0
10
20
30
40
50
60
Time (min)
Normal
Diabetic
Normal data

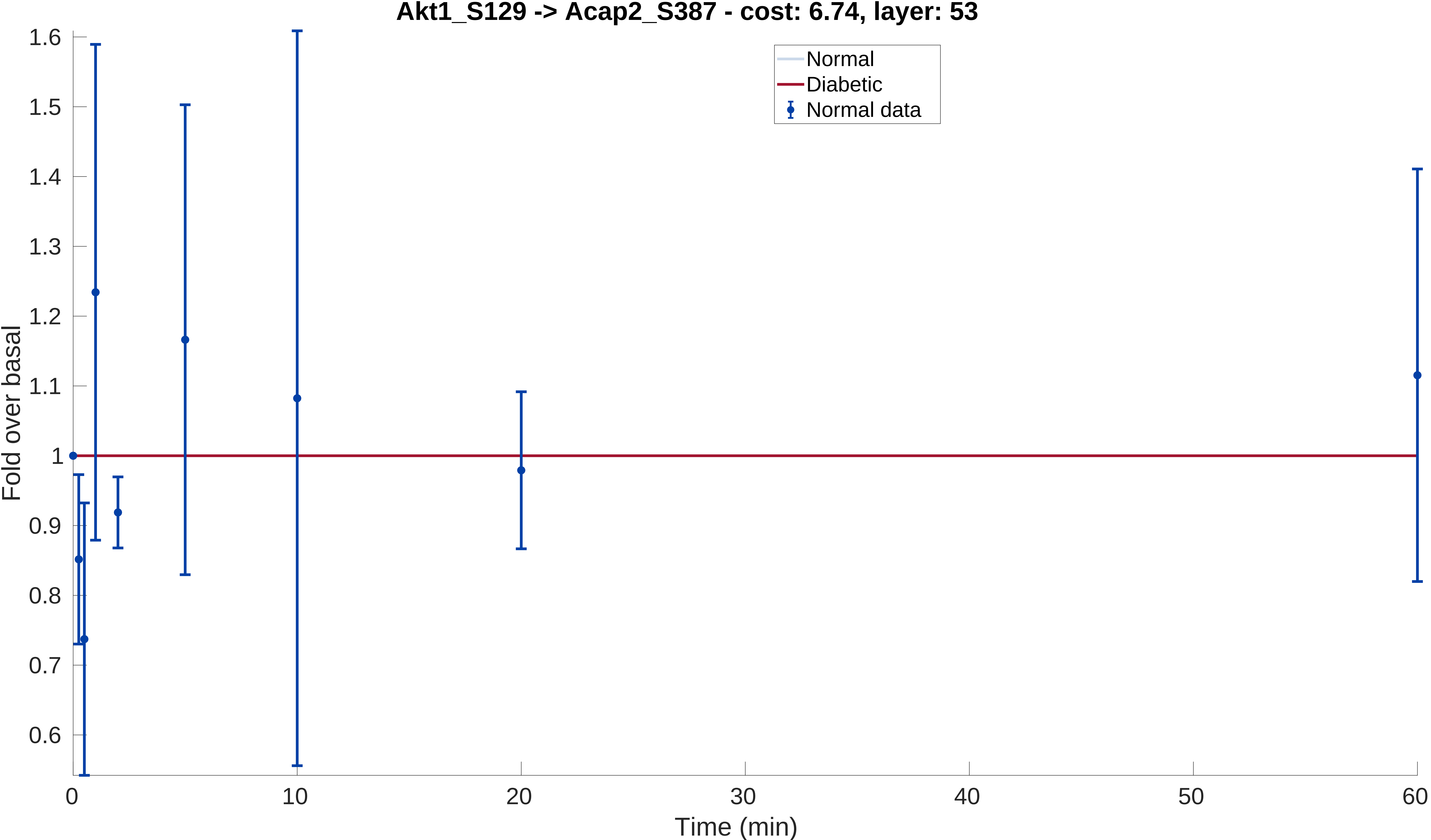

Akt1_S129 -> Acap2_S387 - cost: 6.74, layer: 53
Normal
Diabetic
Normal data
Fold over basal
1.6
1.5
1.4
1.3
1.2
1.1
1
0.9
0.8
0.7
0.6
0
10
20
30
40
50
60
Time (min)

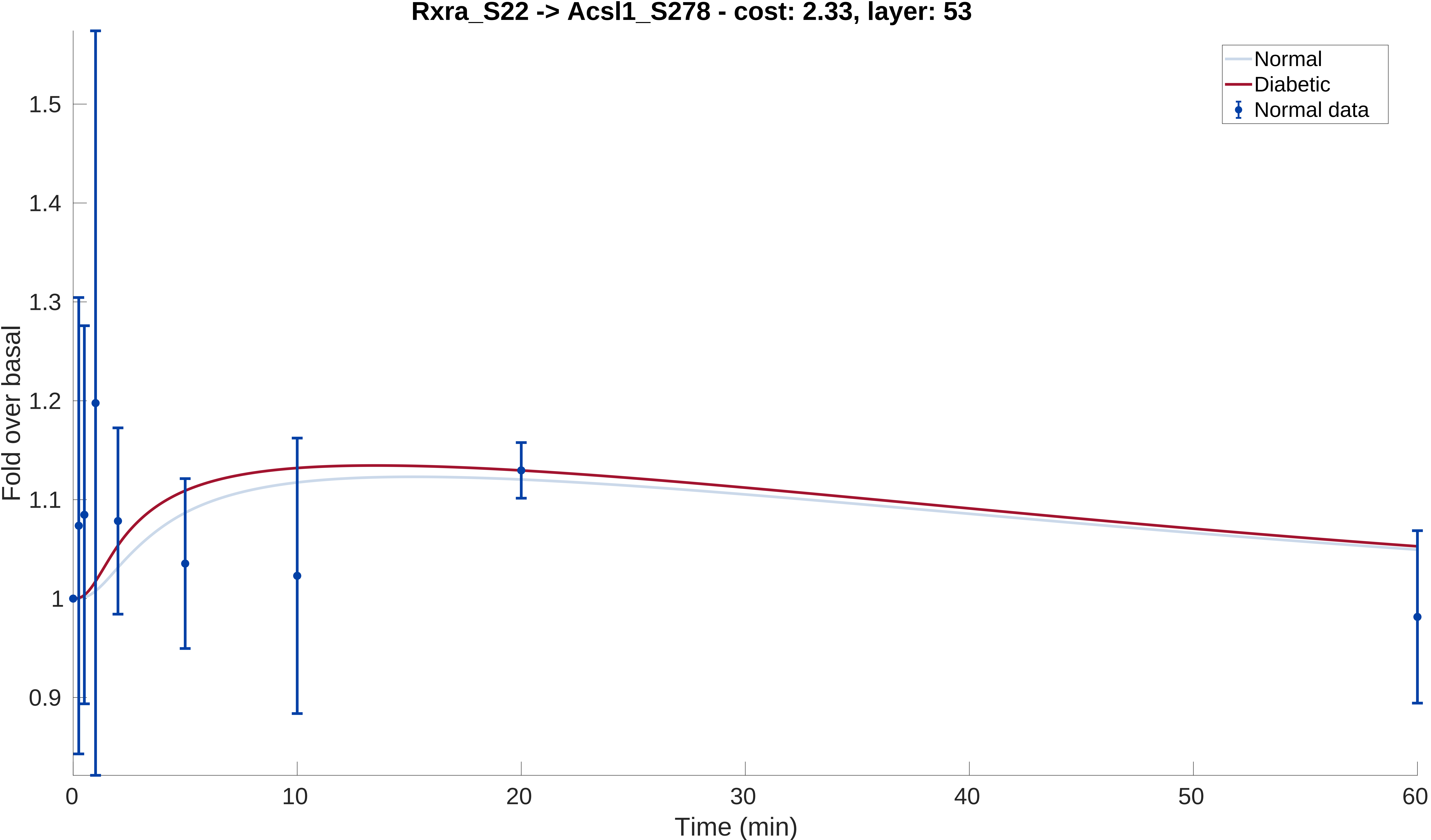
Rxra_S22 -> Acsl1_S278 - cost: 2.33, layer: 53
Normal
Diabetic
Normal data
Fold over basal
1.5
1.4
1.3
1.2
1.1
1
0.9
0
10
20
30
40
50
60
Time (min)

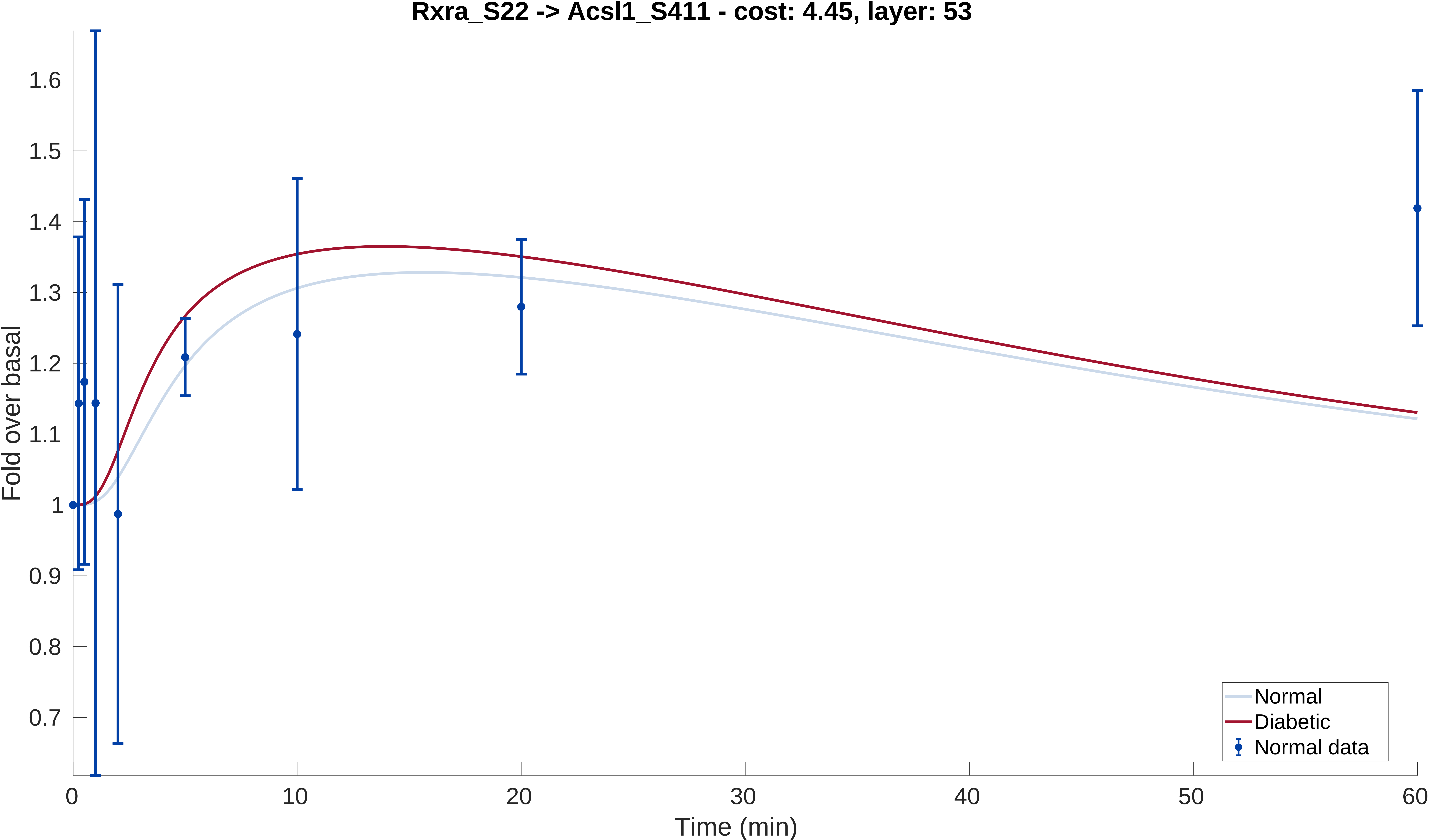

Rxra_S22 -> Acsl1_S411 - cost: 4.45, layer: 53
Fold over basal
1.6
1.5
1.4
1.3
1.2
1.1
1
0.9
0.8
0.7
0
10
20
30
40
50
60
Time (min)
Normal
Diabetic
Normal data

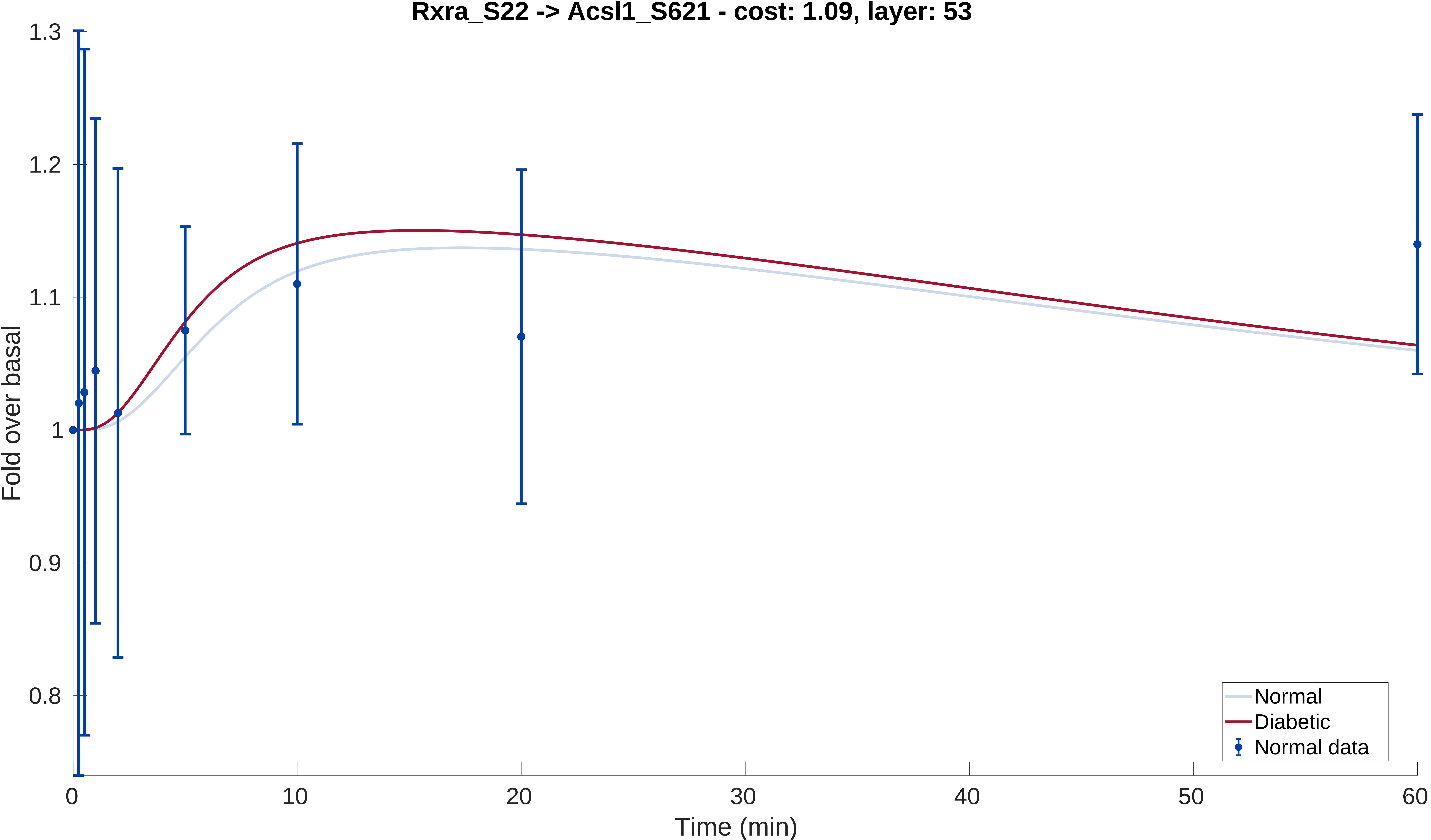

Rxra_S22 -> Acsl1_S621 - cost: 1.09, layer: 53
Fold over basal
1.3
1.2
1.1
1
0.9
0.8
0
10
20
30
40
50
60
Time (min)
Normal
Diabetic
Normal data

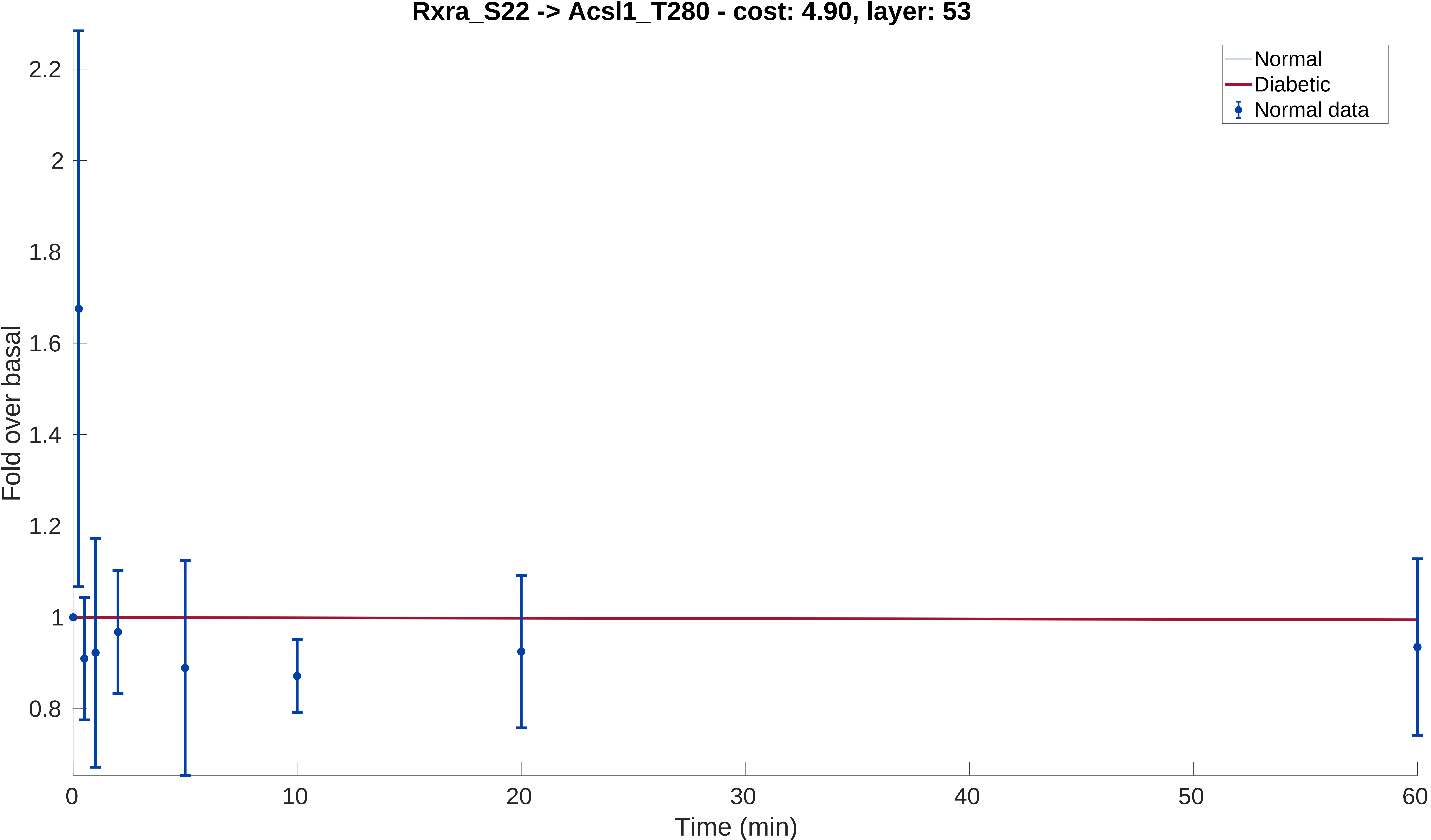

Rxra_S22 -> Acsl1_T280 - cost: 4.90, layer: 53
Normal
Diabetic
Normal data
Fold over basal
Time (min)
0.8
1
1.2
1.4
1.6
1.8
2
2.2
0
10
20
30
40
50
60

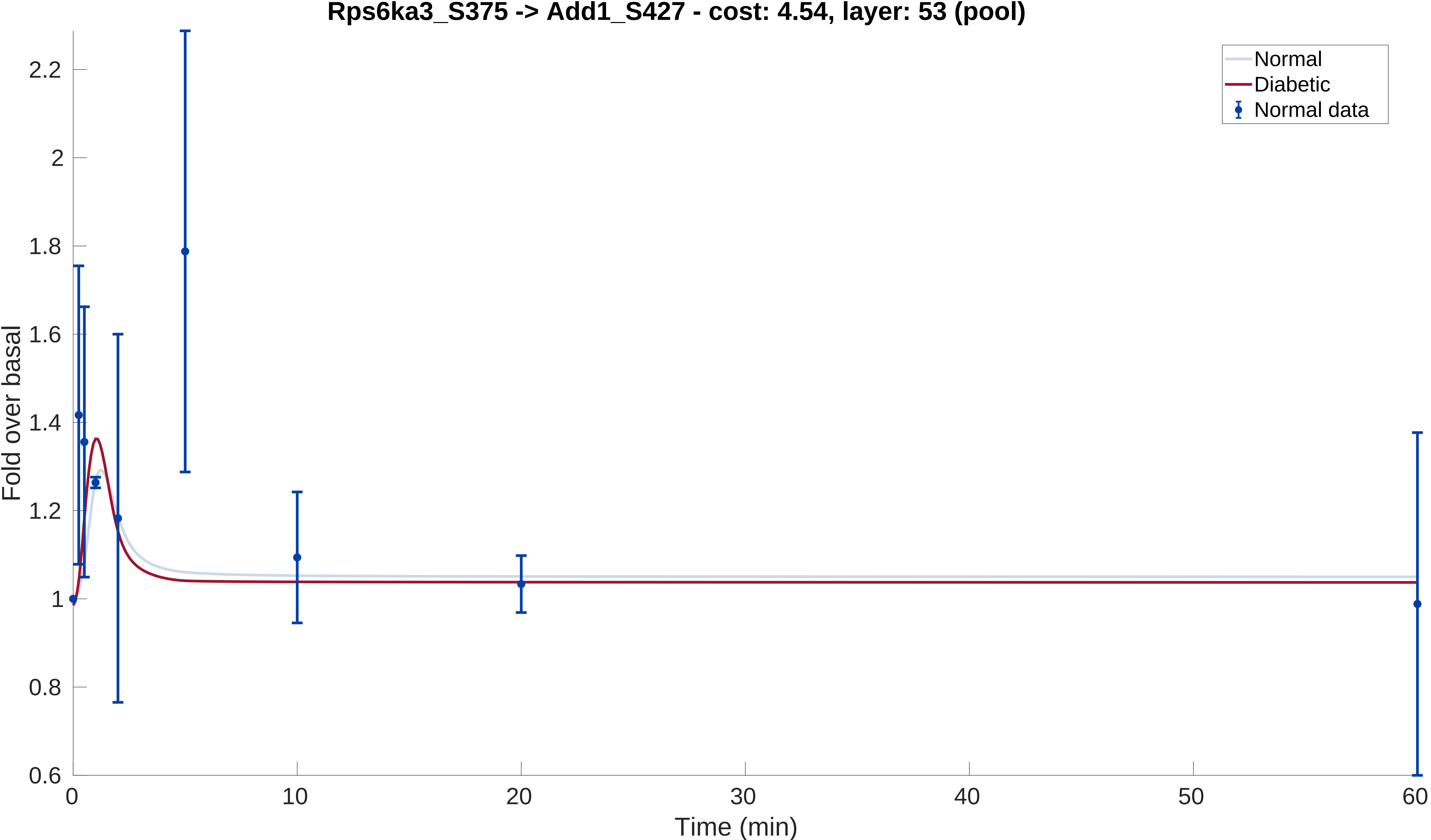
Rps6ka3_S375 -> Add1_S427 - cost: 4.54, layer: 53 (pool)
Normal
Diabetic
Normal data
Fold over basal
2.2
2
1.8
1.6
1.4
1.2
1
0.8
0.6
0
10
20
30
40
50
60
Time (min)

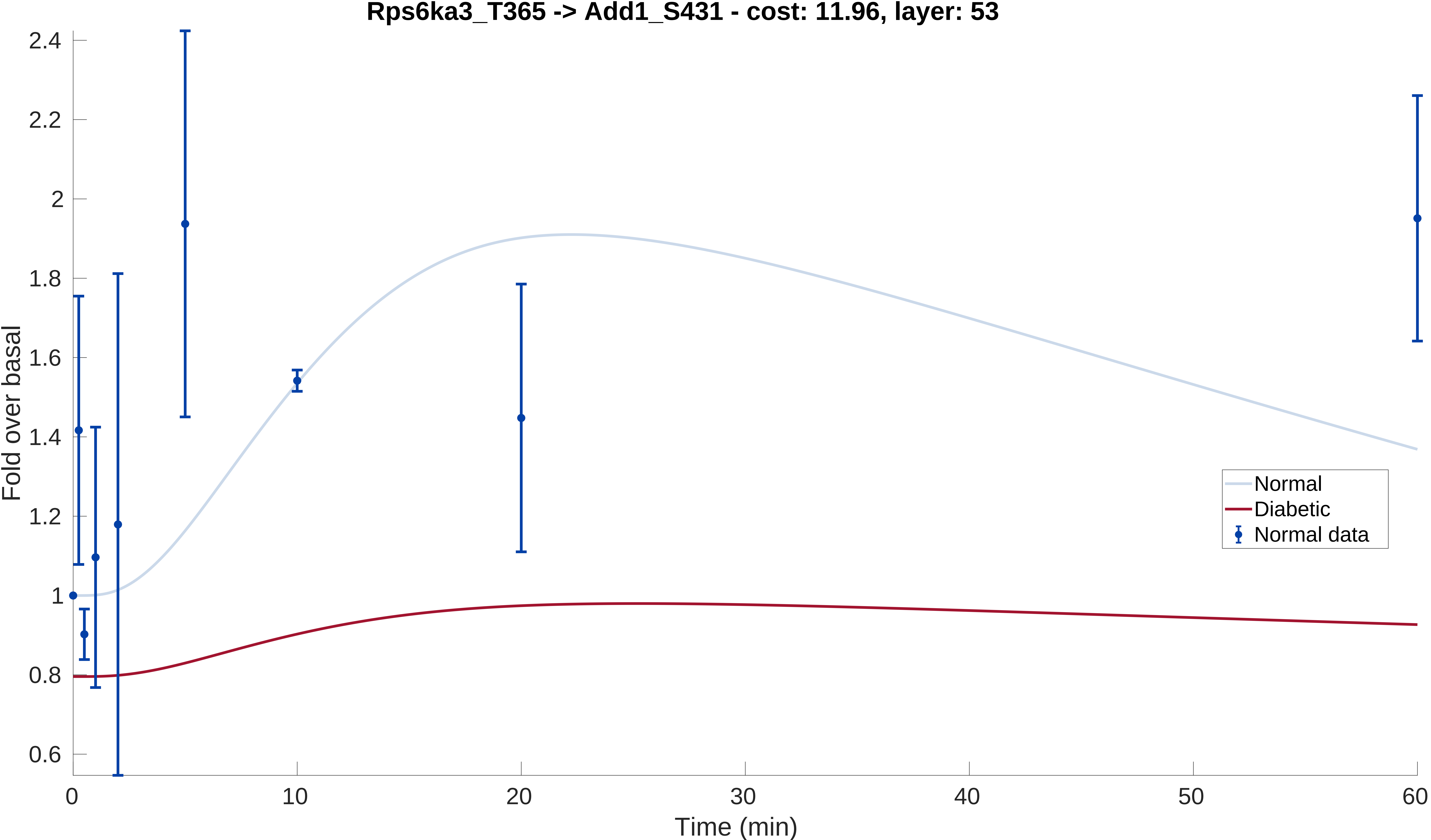

Rps6ka3_T365 -> Add1_S431 - cost: 11.96, layer: 53
Fold over basal
2.4
2.2
2
1.8
1.6
1.4
1.2
1
0.8
0.6
0
10
20
30
40
50
60
Time (min)
Normal
Diabetic
Normal data

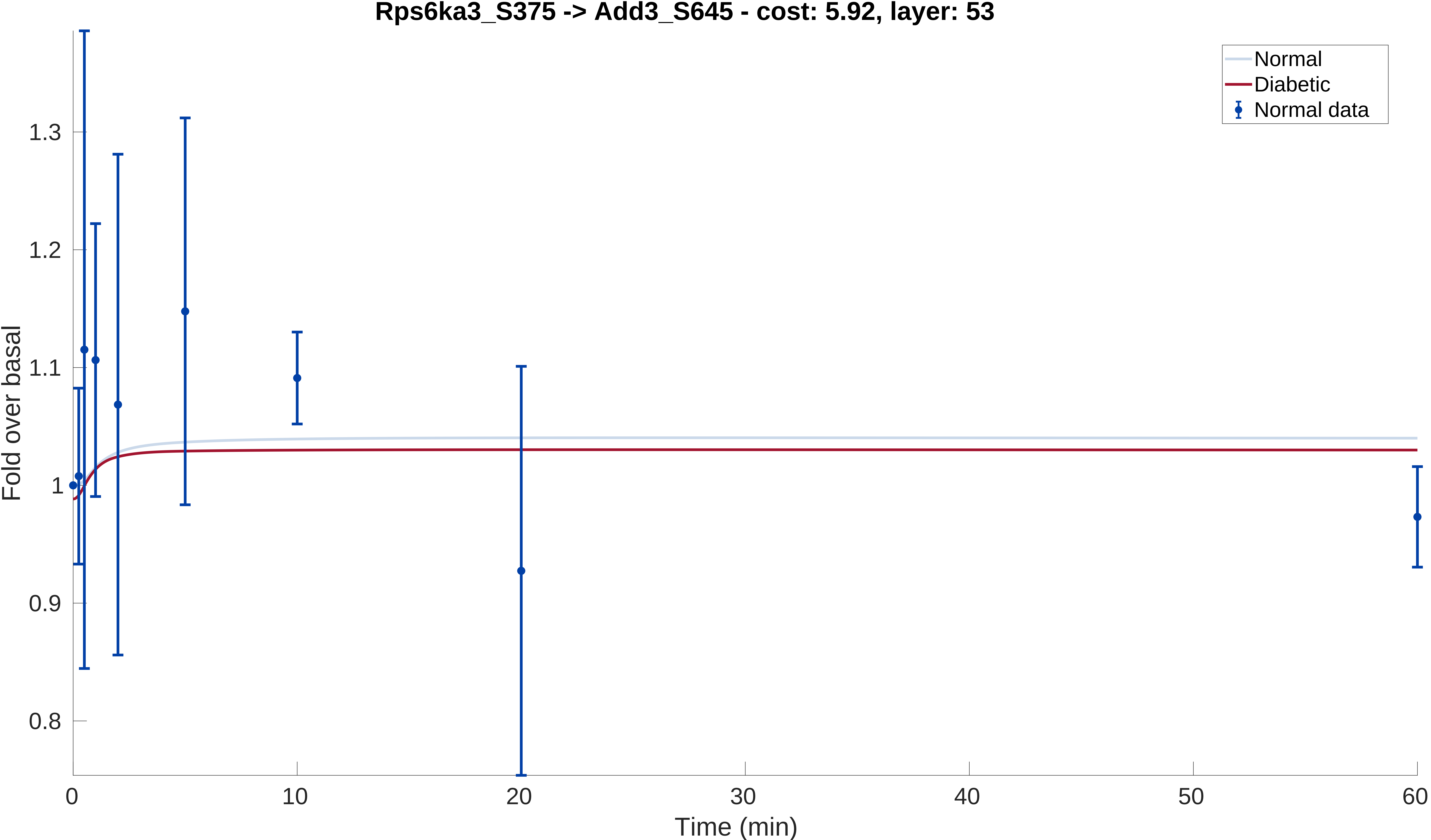

Rps6ka3_S375 -> Add3_S645 - cost: 5.92, layer: 53
Normal
Diabetic
Normal data
Fold over basal
1.3
1.2
1.1
1
0.9
0.8
0
10
20
30
40
50
60
Time (min)

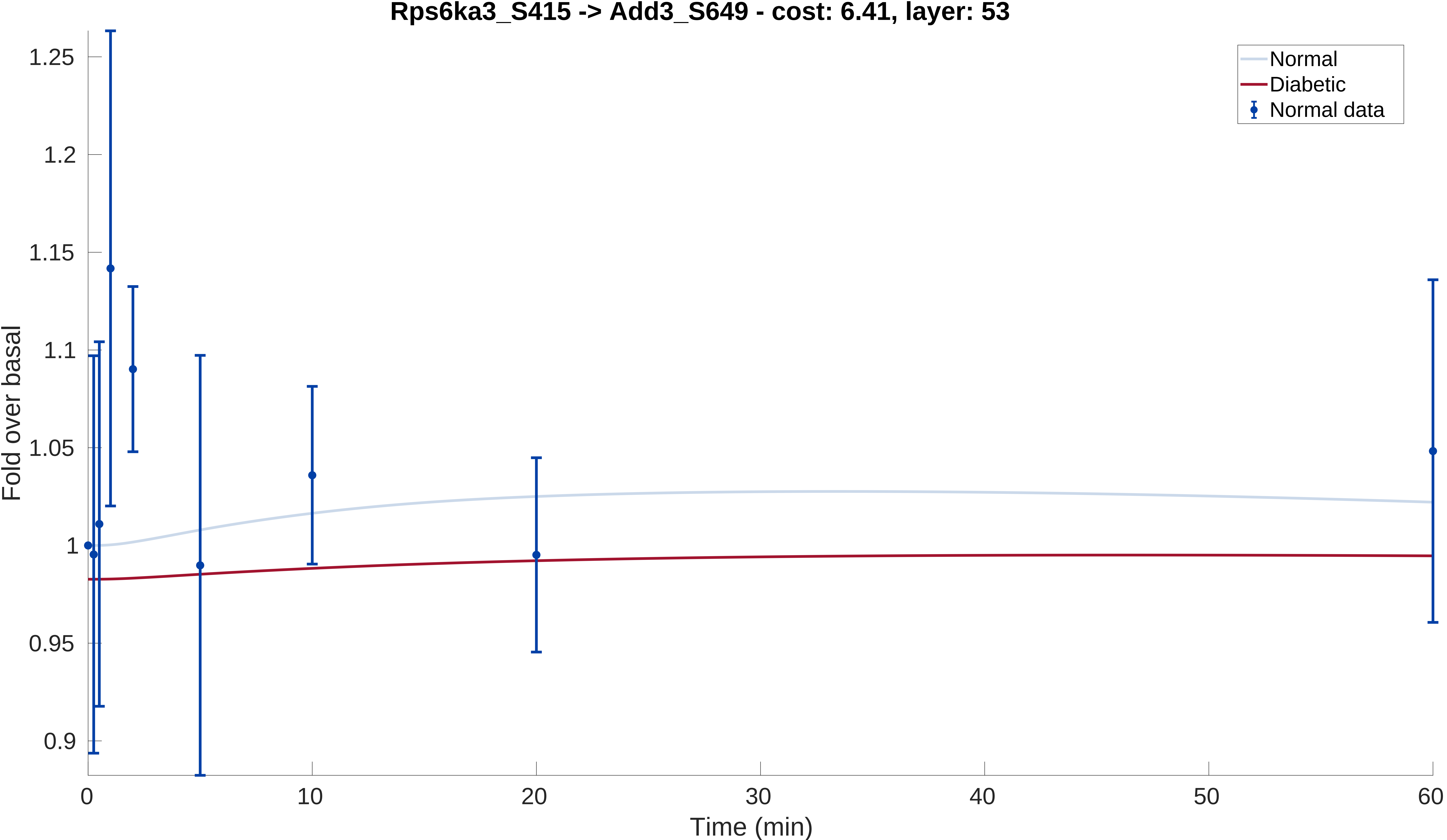

Rps6ka3_S415 -> Add3_S649 - cost: 6.41, layer: 53
Normal
Diabetic
Normal data
Fold over basal
1.25
1.2
1.15
1.1
1.05
1
0.95
0.9
0
10
20
30
40
50
60
Time (min)

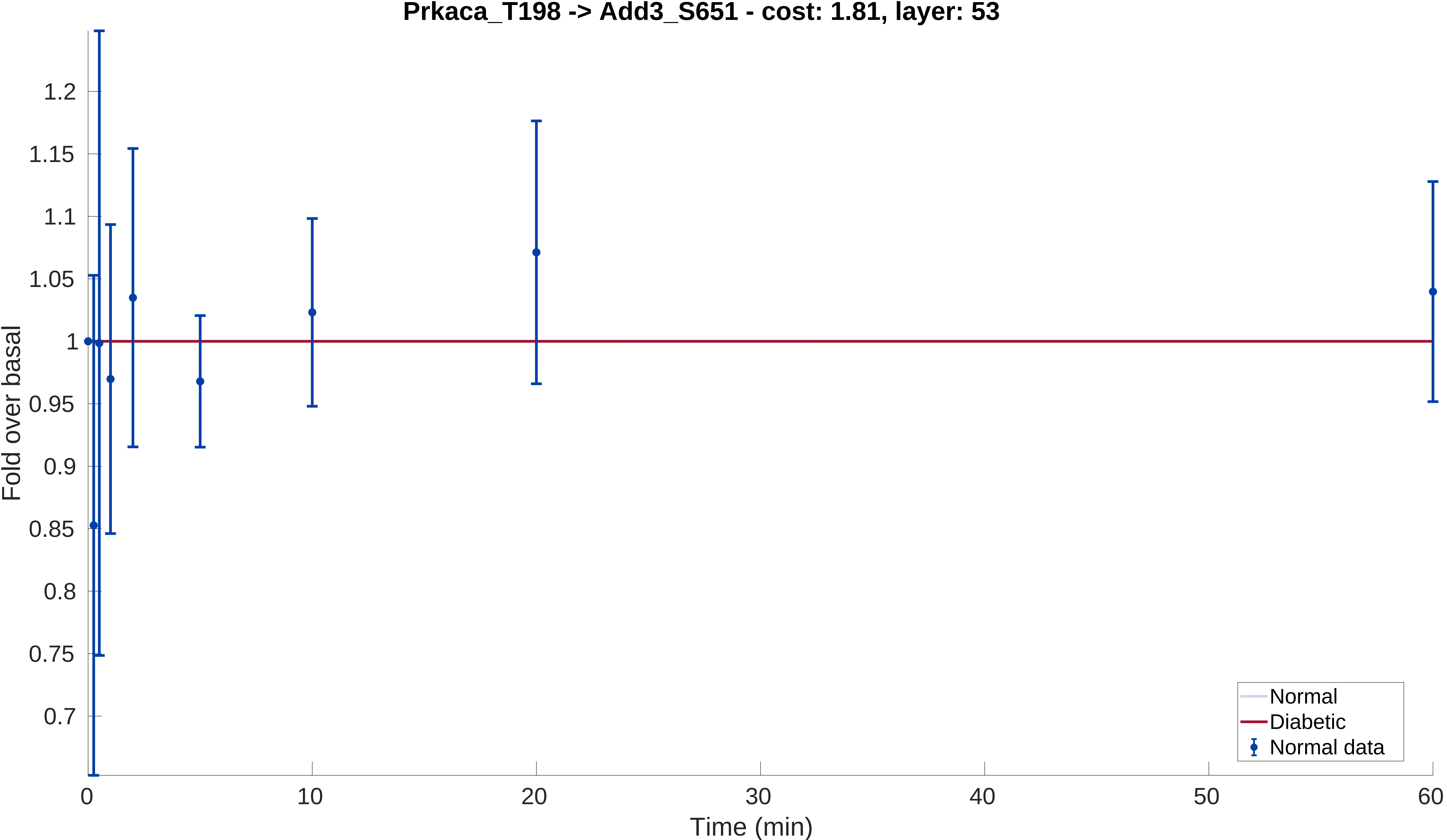

Prkaca_T198 -> Add3_S651 - cost: 1.81, layer: 53
Fold over basal
1.2
1.15
1.1
1.05
1
0.95
0.9
0.85
0.8
0.75
0.7
0
10
20
30
40
50
60
Time (min)
Normal
Diabetic
Normal data

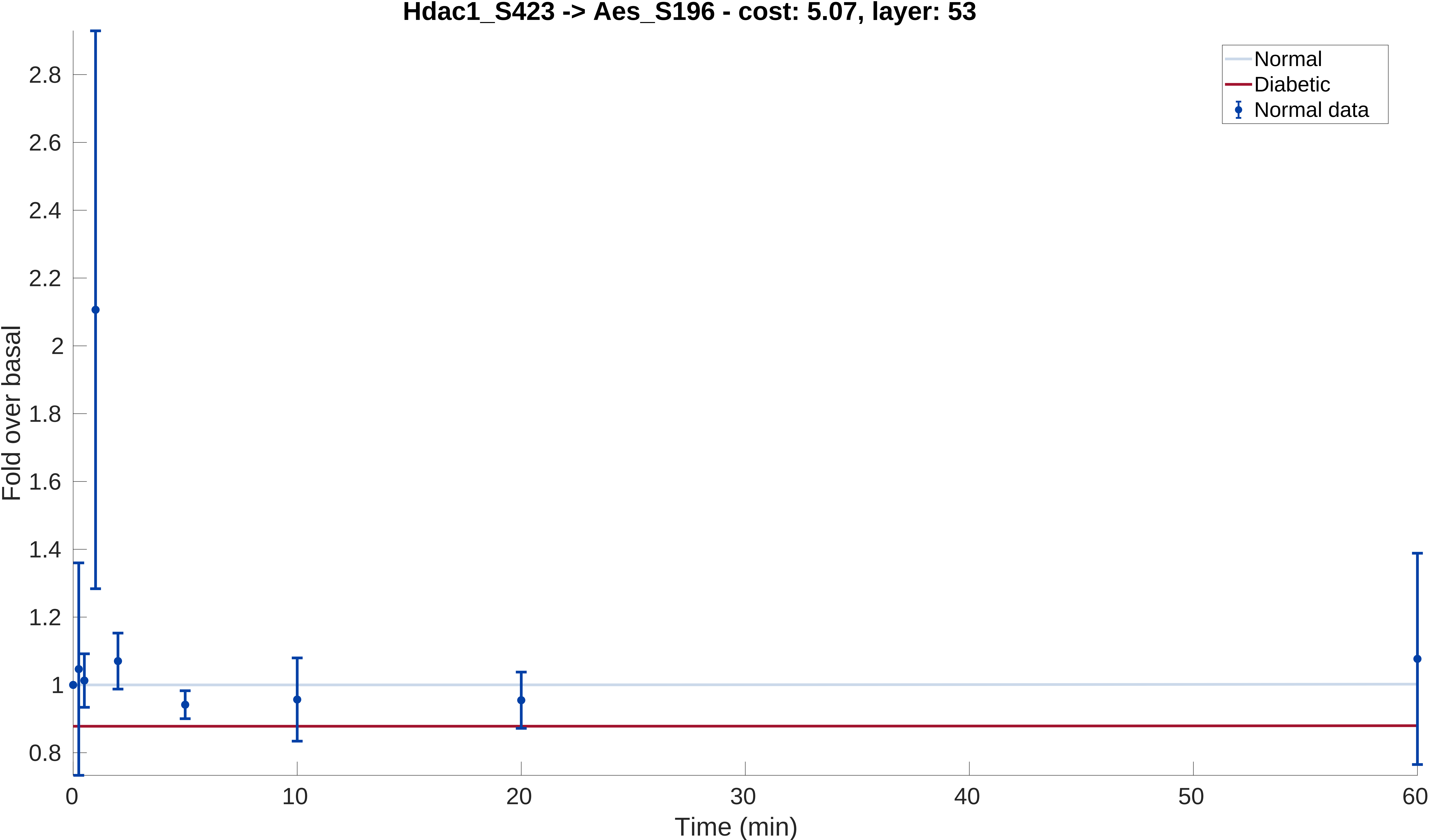

Hdac1_S423 -> Aes_S196 - cost: 5.07, layer: 53
Normal
Diabetic
Normal data
Fold over basal
2.8
2.6
2.4
2.2
2
1.8
1.6
1.4
1.2
1
0.8
0
10
20
30
40
50
60
Time (min)

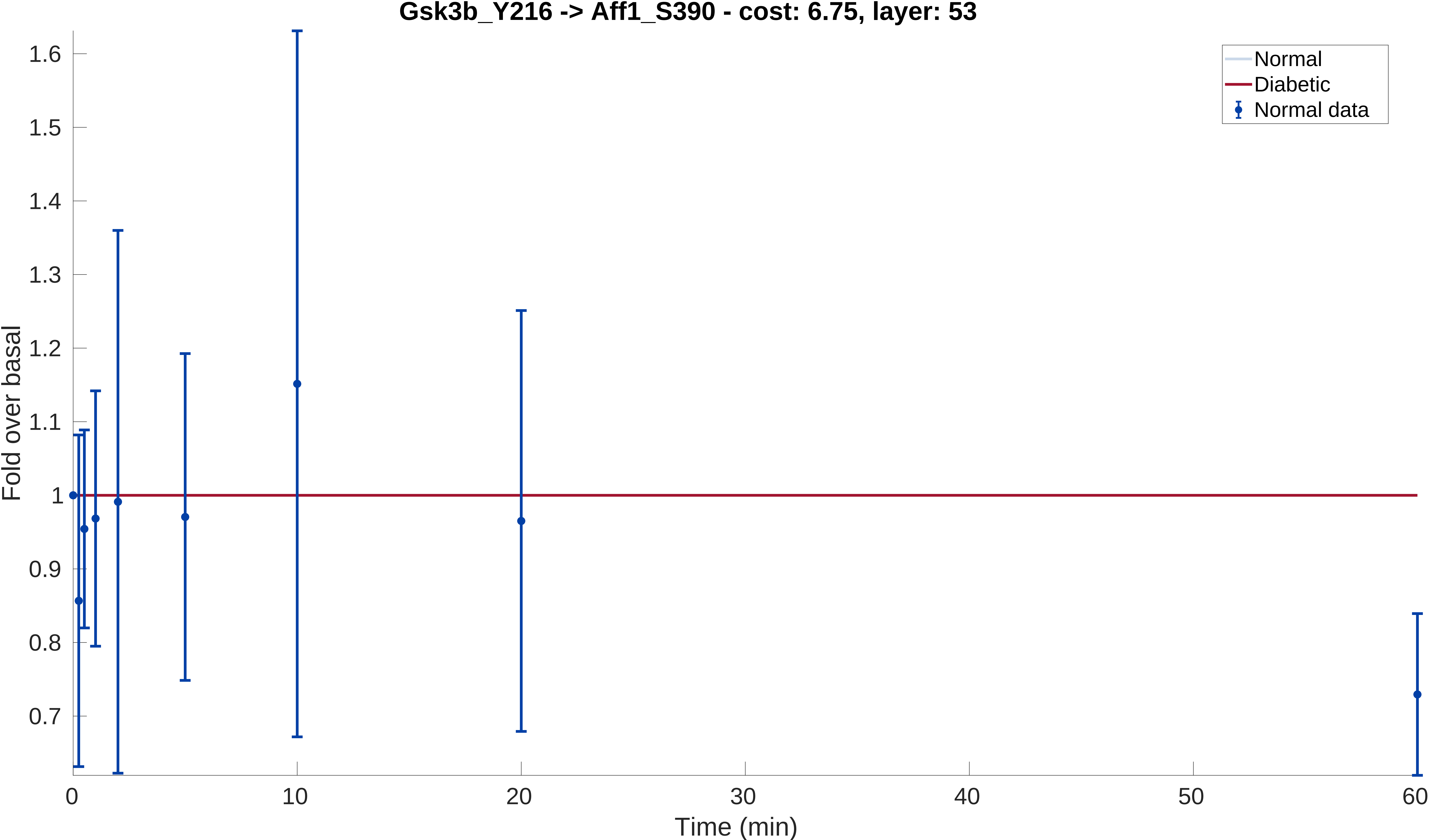
Gsk3b_Y216 -> Aff1_S390 - cost: 6.75, layer: 53
Normal
Diabetic
Normal data
Fold over basal
1.6
1.5
1.4
1.3
1.2
1.1
1
0.9
0.8
0.7
0
10
20
30
40
50
60
Time (min)

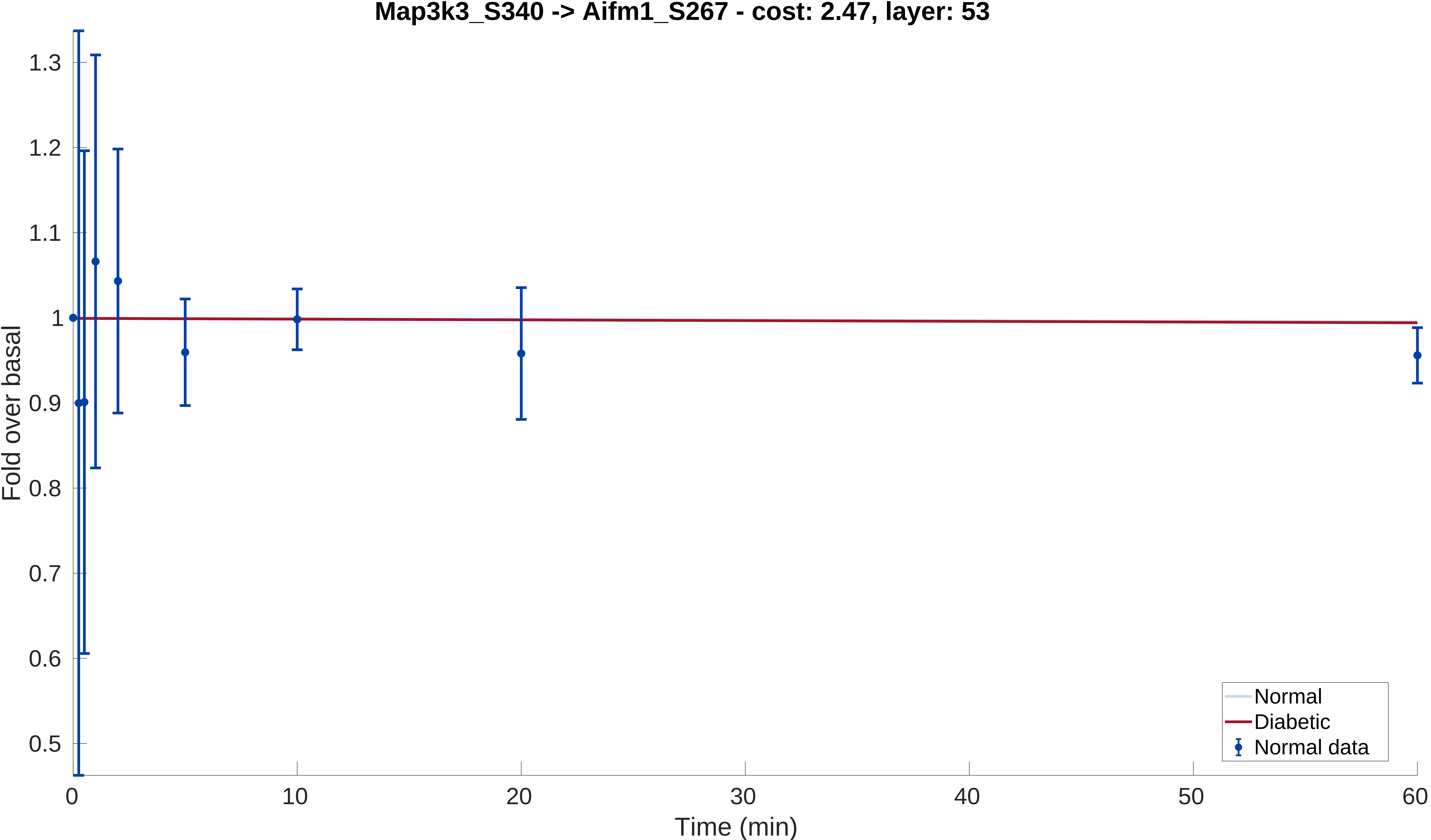

Map3k3_S340 -> Aifm1_S267 - cost: 2.47, layer: 53
Fold over basal
1.3
1.2
1.1
1
0.9
0.8
0.7
0.6
0.5
0
10
20
30
40
50
60
Time (min)
Normal
Diabetic
Normal data

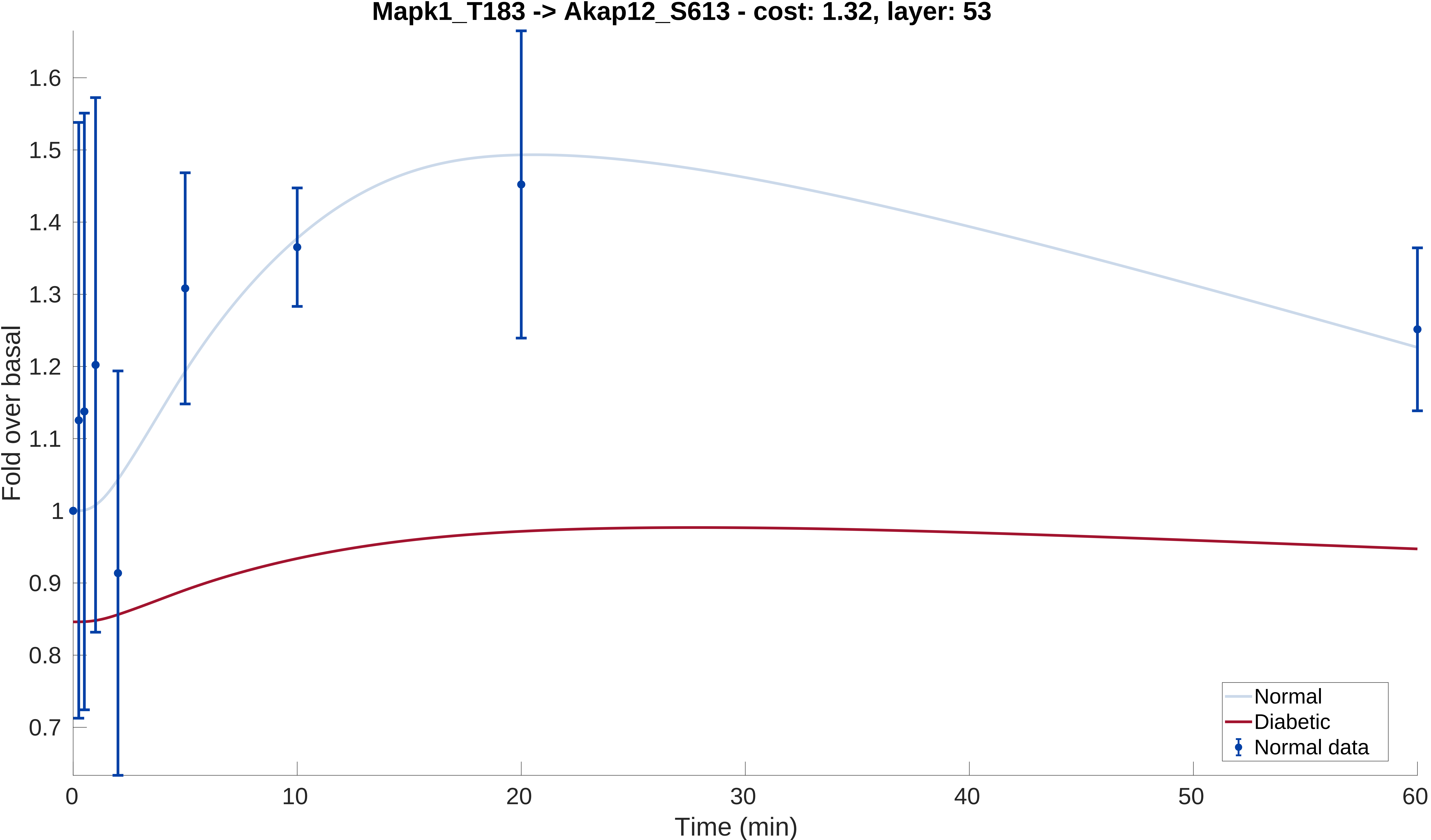

Mapk1_T183 -> Akap12_S613 - cost: 1.32, layer: 53
Fold over basal
1.6
1.5
1.4
1.3
1.2
1.1
1
0.9
0.8
0.7
0
10
20
30
40
50
60
Time (min)
Normal
Diabetic
Normal data

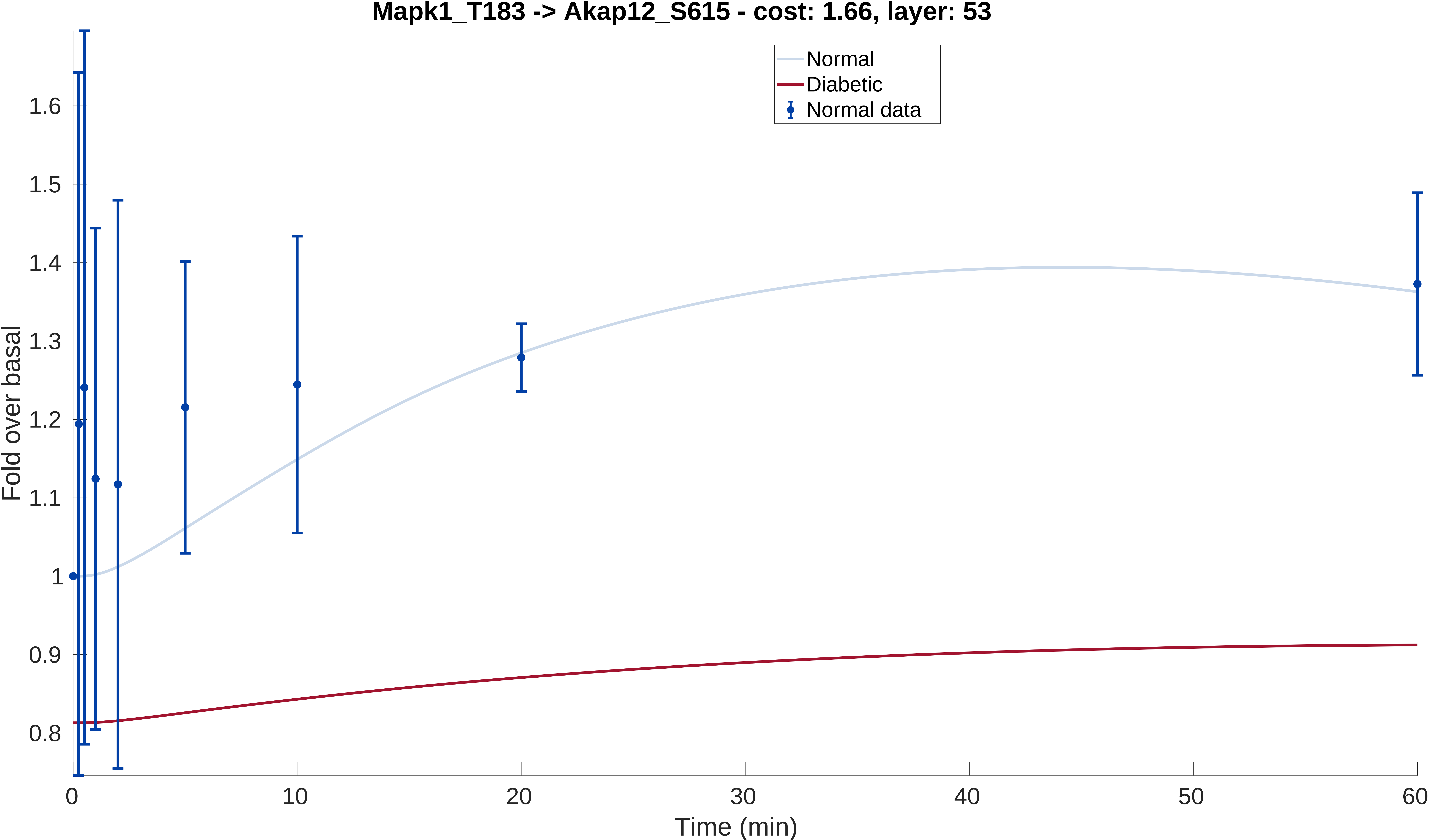
Mapk1_T183 -> Akap12_S615 - cost: 1.66, layer: 53
Normal
Diabetic
Normal data
Fold over basal
1.6
1.5
1.4
1.3
1.2
1.1
1
0.9
0.8
0
10
20
30
40
50
60
Time (min)

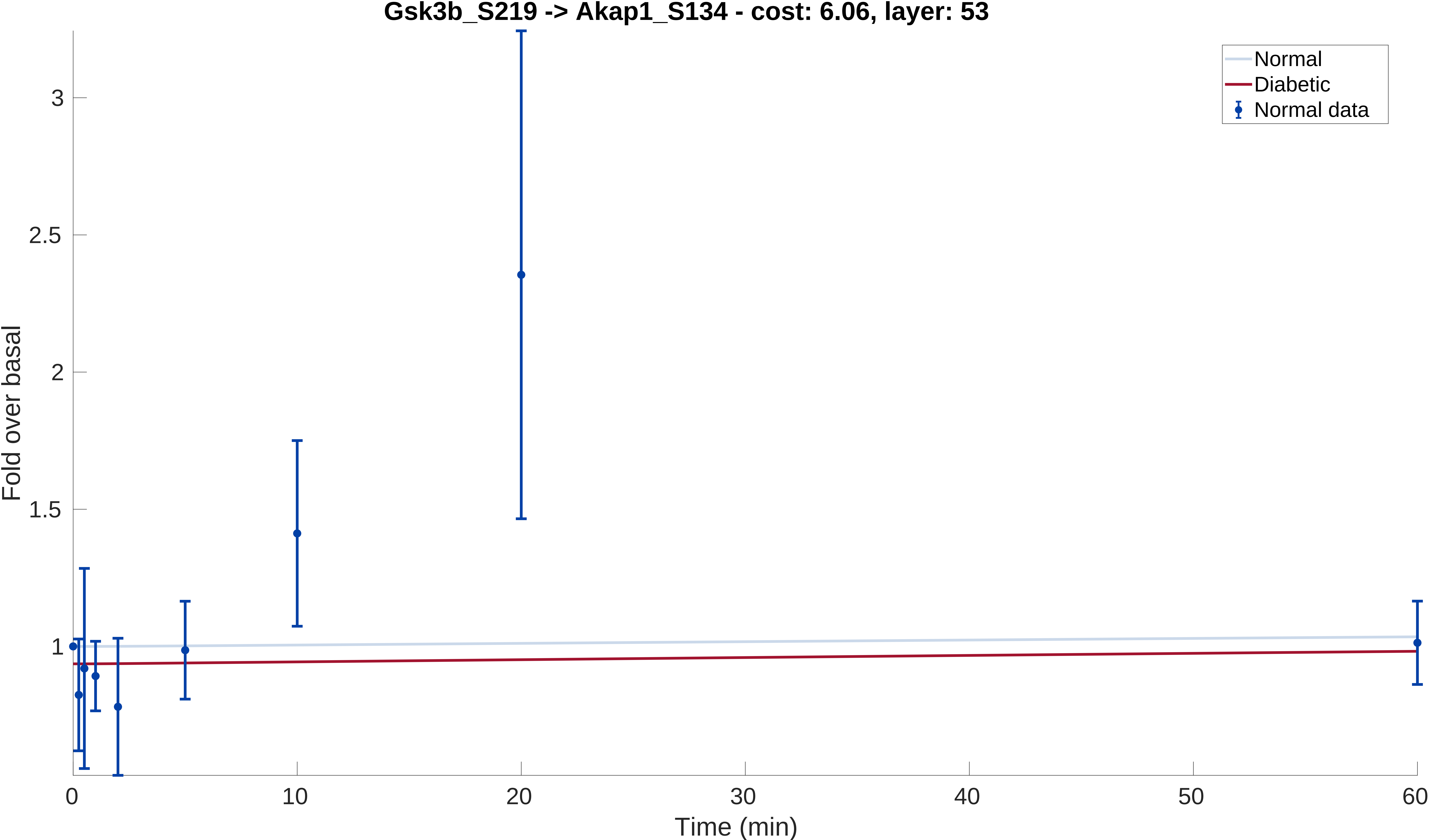

Gsk3b_S219 -> Akap1_S134 - cost: 6.06, layer: 53
Normal
Diabetic
Normal data
Fold over basal
3
2.5
2
1.5
1
0
10
20
30
40
50
60
Time (min)

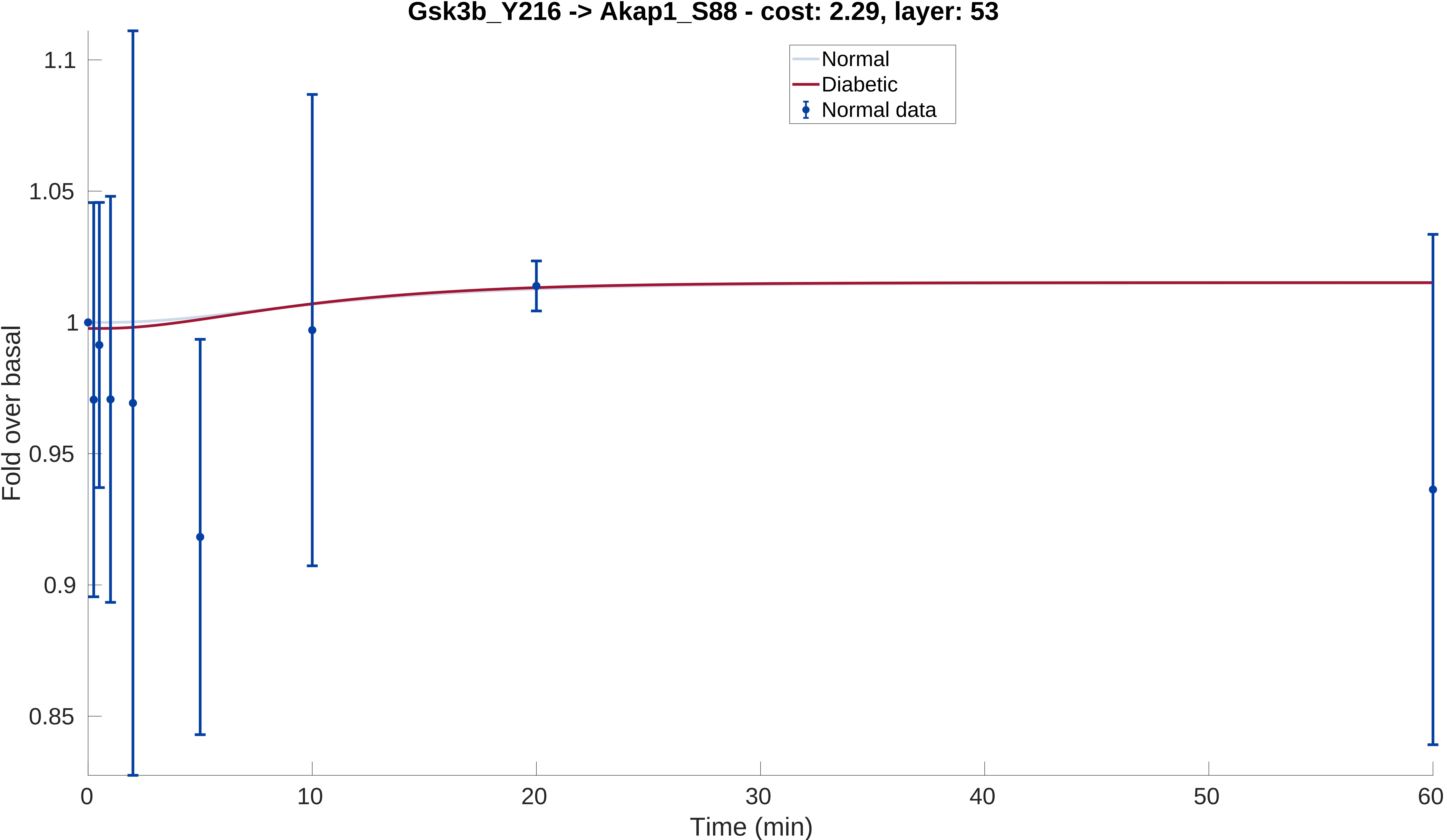

Gsk3b_Y216 -> Akap1_S88 - cost: 2.29, layer: 53
Normal
Diabetic
Normal data
Fold over basal
1.1
1.05
1
0.95
0.9
0.85
0
10
20
30
40
50
60
Time (min)

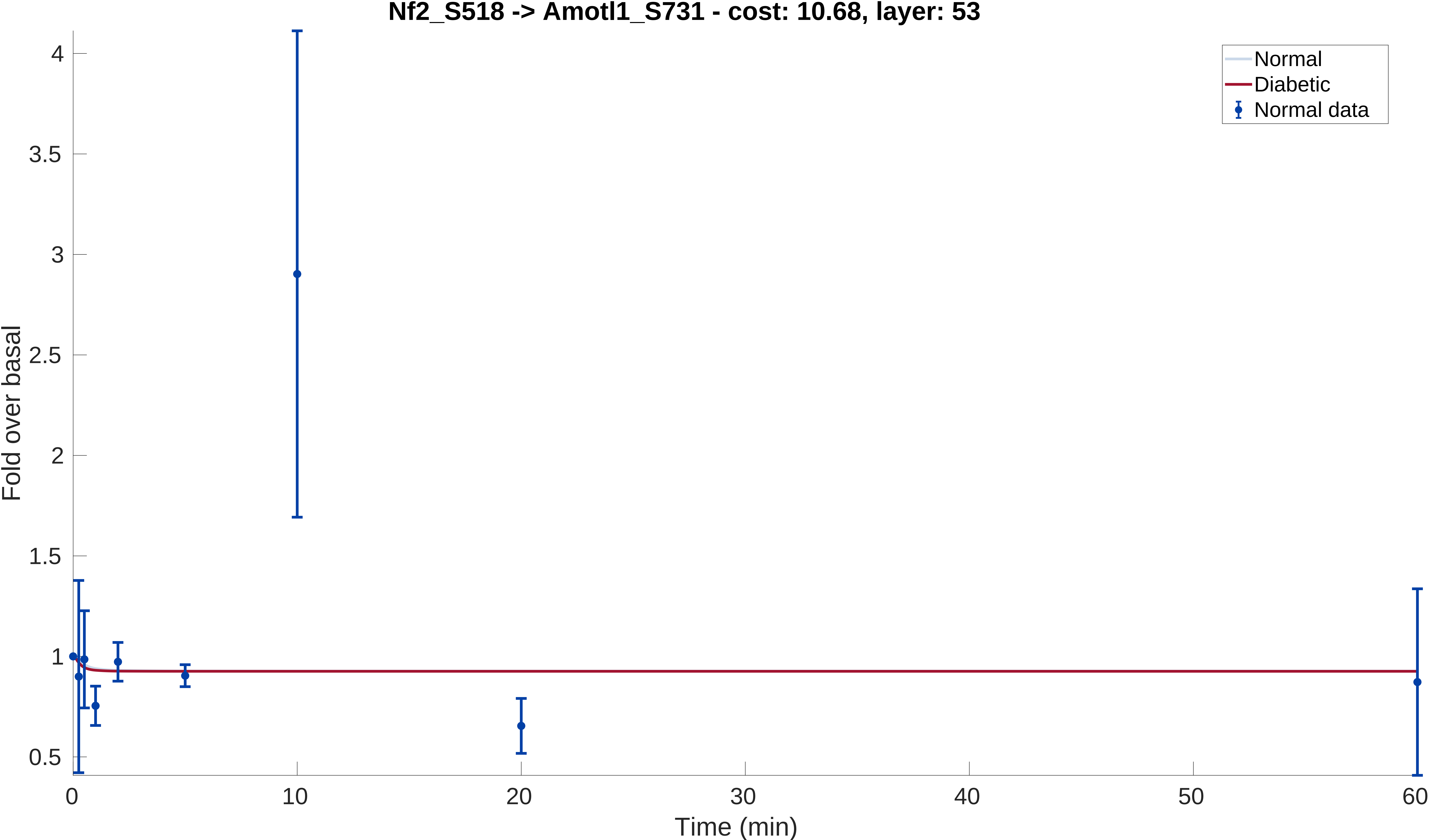
Nf2_S518 -> Amotl1_S731 - cost: 10.68, layer: 53
Normal
Diabetic
Normal data
Fold over basal
Time (min)
0.5
1
1.5
2
2.5
3
3.5
4
0
10
20
30
40
50
60

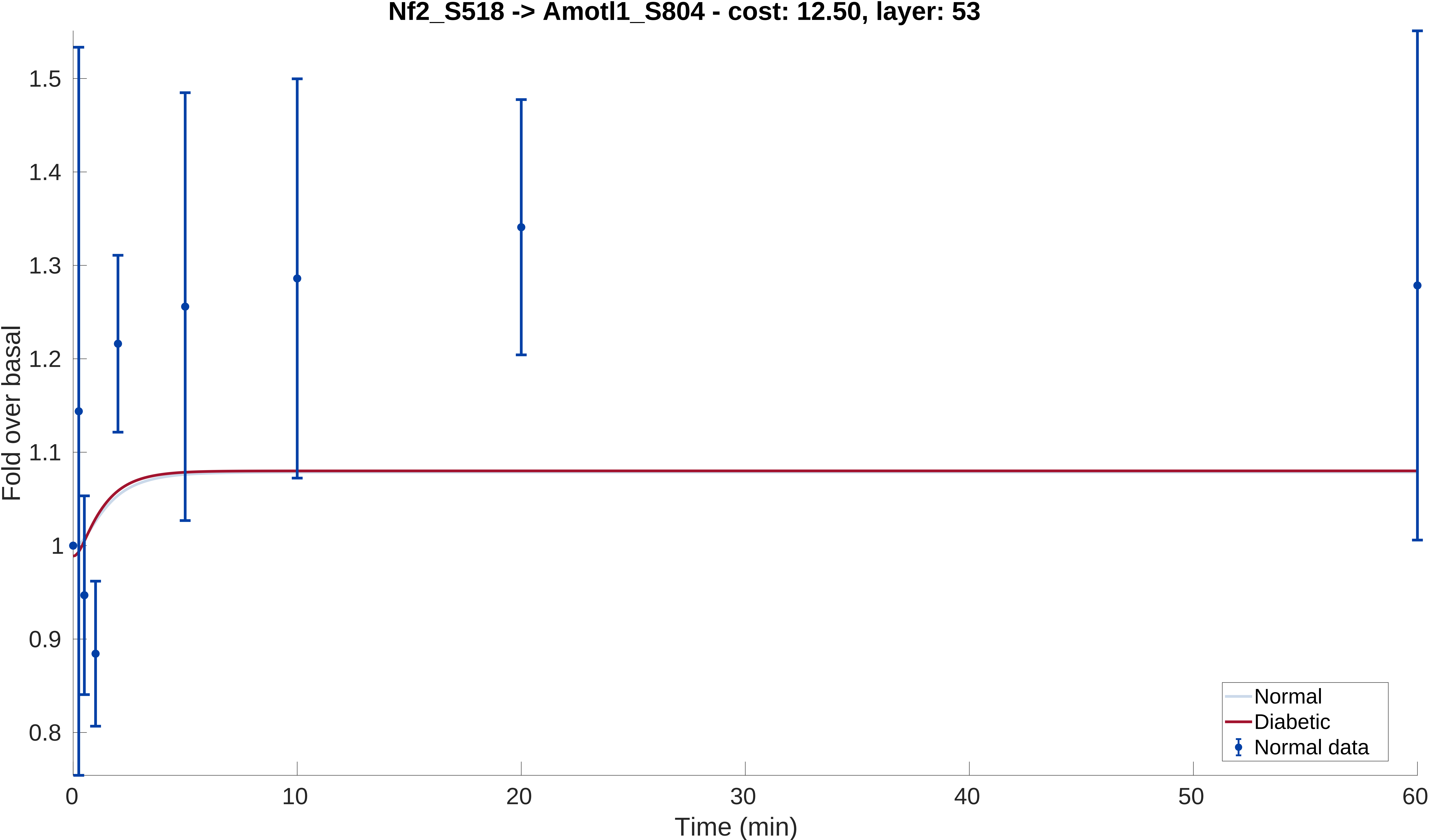

Nf2_S518 -> Amotl1_S804 - cost: 12.50, layer: 53
Fold over basal
1.5
1.4
1.3
1.2
1.1
1
0.9
0.8
0
10
20
30
40
50
60
Time (min)
Normal
Diabetic
Normal data

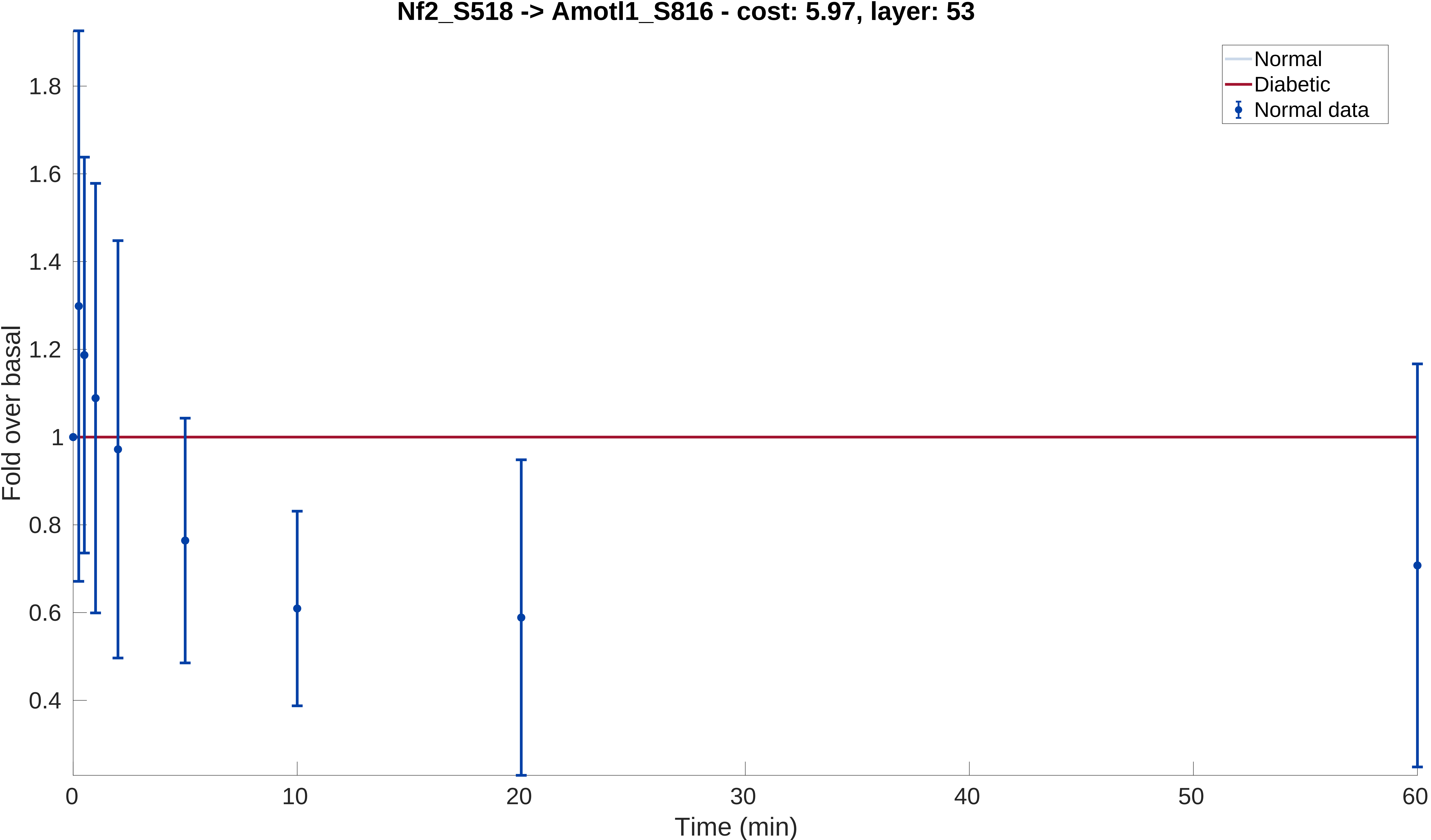
Nf2_S518 -> Amotl1_S816 - cost: 5.97, layer: 53
Normal
Diabetic
Normal data
Fold over basal
1.8
1.6
1.4
1.2
1
0.8
0.6
0.4
0
10
20
30
40
50
60
Time (min)

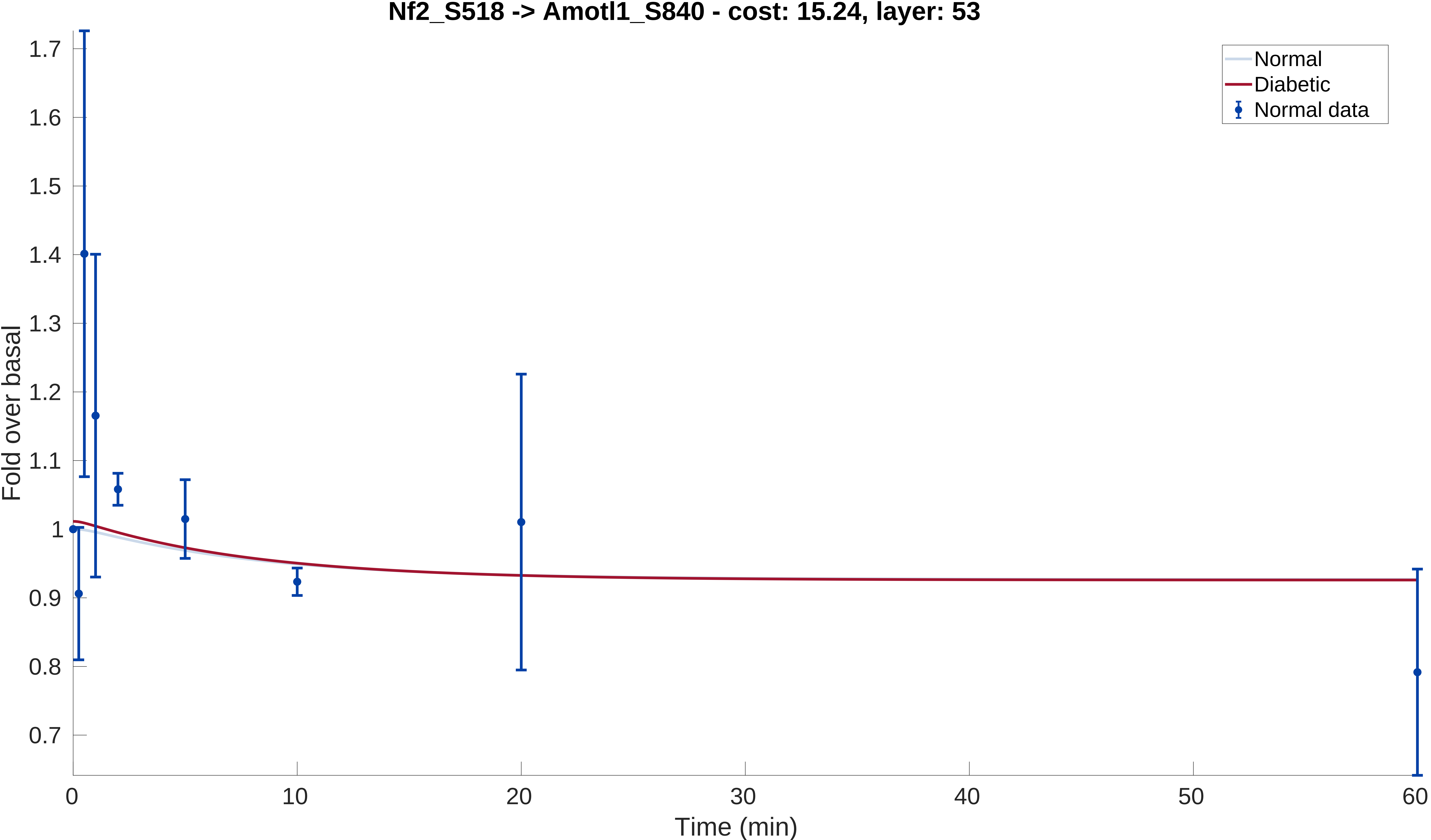

Nf2_S518 -> Amotl1_S840 - cost: 15.24, layer: 53
Normal
Diabetic
Normal data
Fold over basal
Time (min)
1.7
1.6
1.5
1.4
1.3
1.2
1.1
1
0.9
0.8
0.7
0
10
20
30
40
50
60

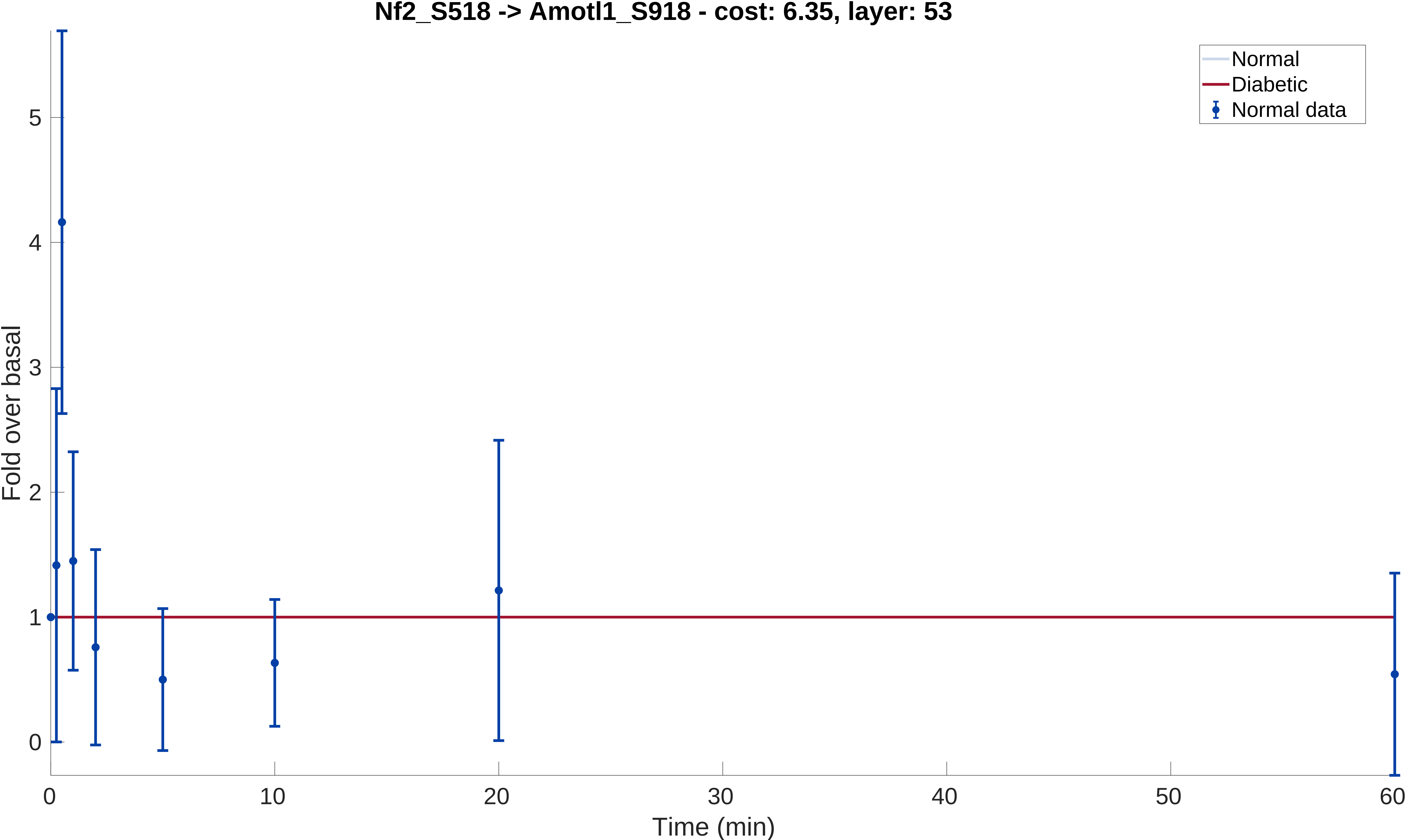

Nf2_S518 -> Amotl1_S918 - cost: 6.35, layer: 53
Normal
Diabetic
Normal data
Fold over basal
0
1
2
3
4
5
0
10
20
30
40
50
60
Time (min)

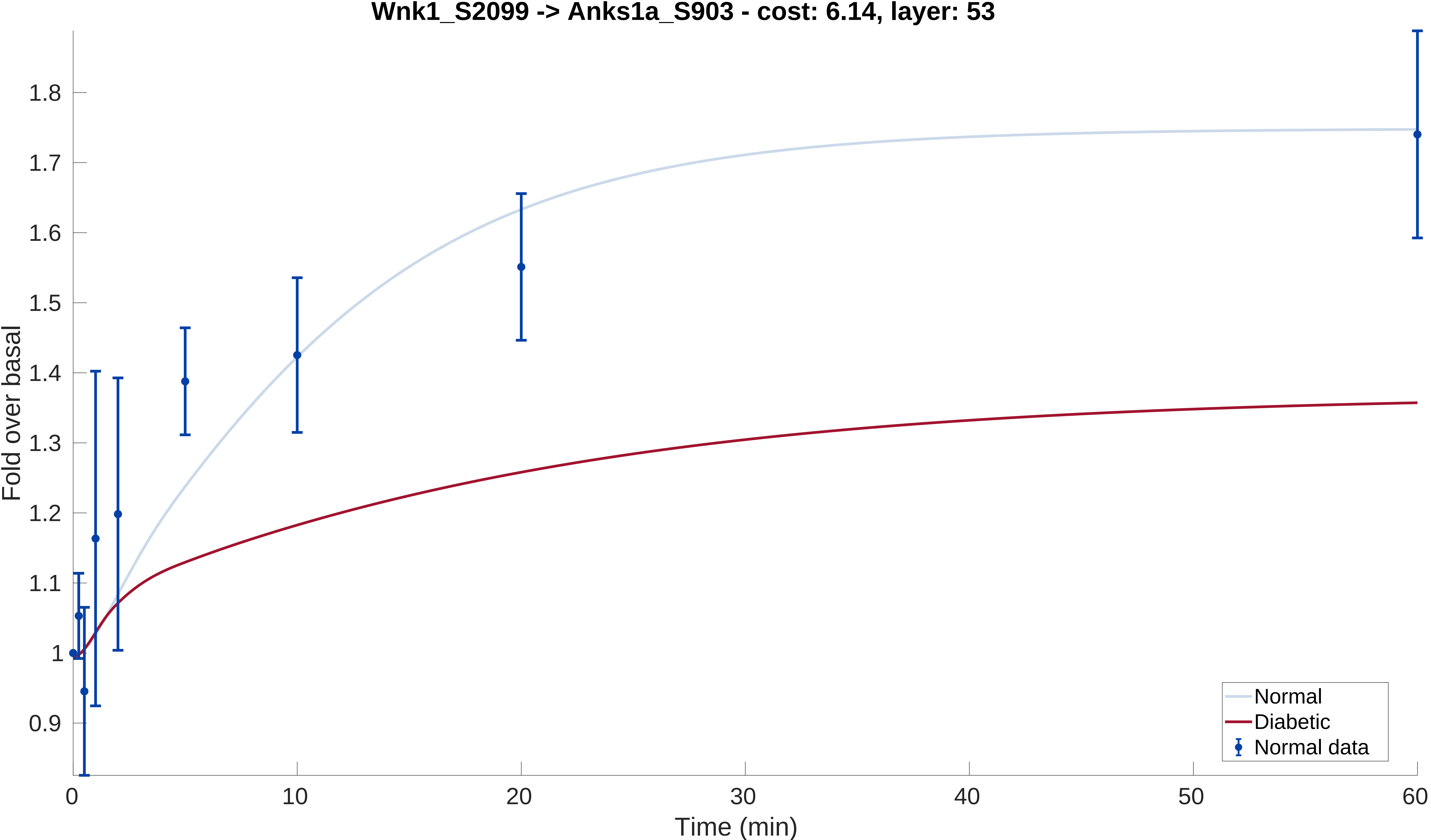

Wnk1_S2099 -> Anks1a_S903 - cost: 6.14, layer: 53
Fold over basal
1.8
1.7
1.6
1.5
1.4
1.3
1.2
1.1
1
0.9
0
10
20
30
40
50
60
Time (min)
Normal
Diabetic
Normal data

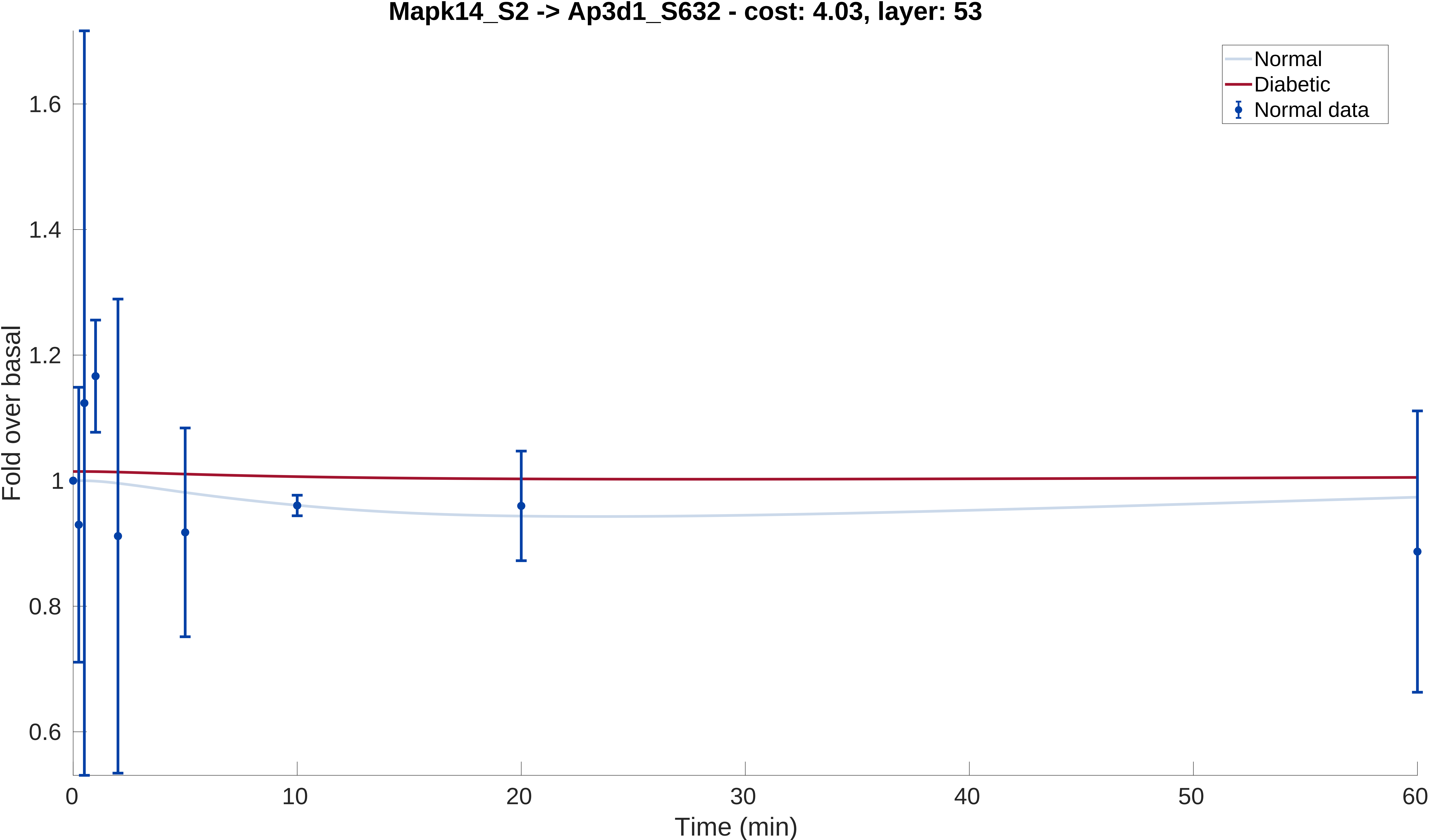
Mapk14_S2 -> Ap3d1_S632 - cost: 4.03, layer: 53
Normal
Diabetic
Normal data
Fold over basal
1.6
1.4
1.2
1
0.8
0.6
0
10
20
30
40
50
60
Time (min)

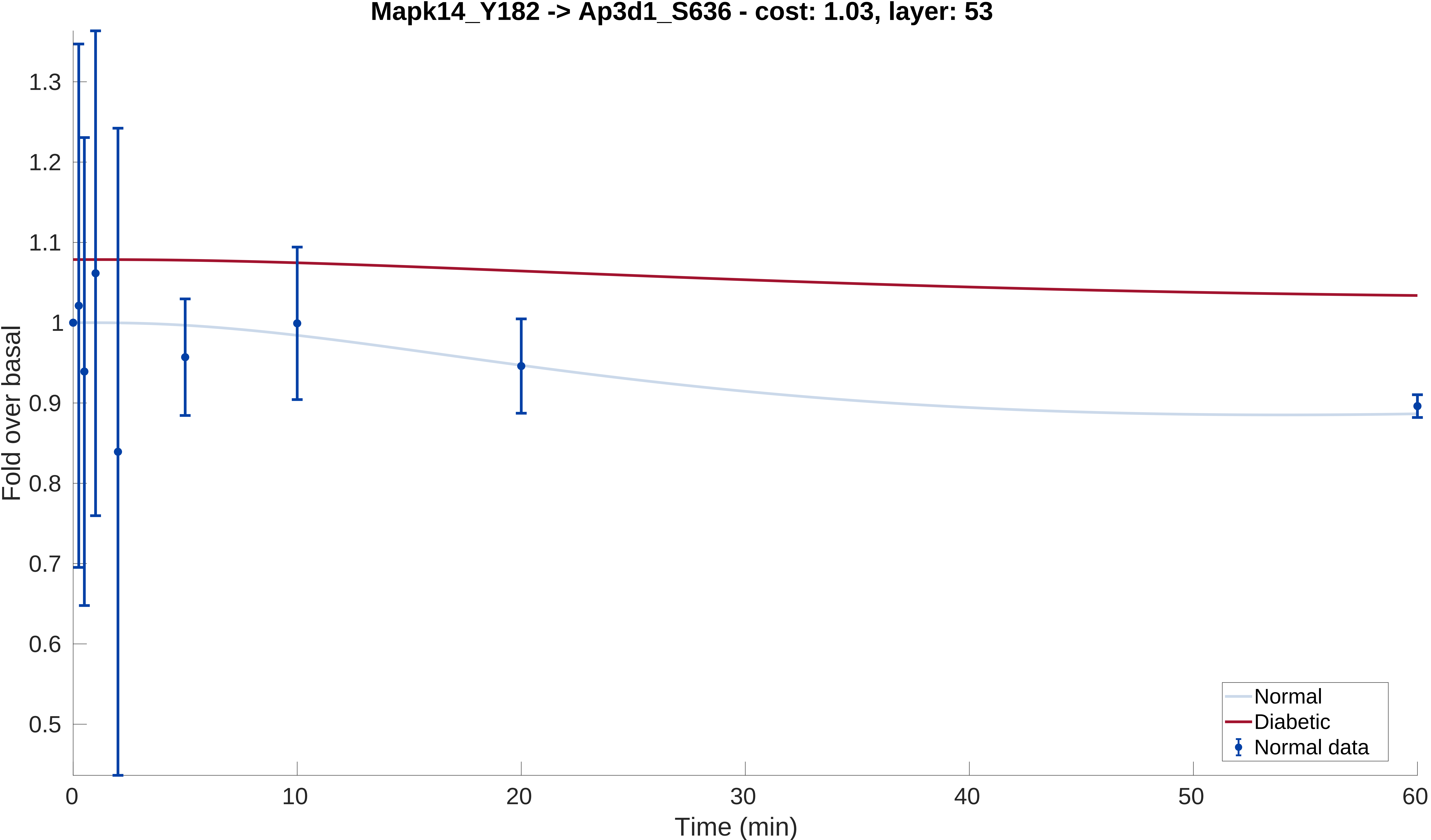
Mapk14_Y182 -> Ap3d1_S636 - cost: 1.03, layer: 53
Fold over basal
1.3
1.2
1.1
1
0.9
0.8
0.7
0.6
0.5
0
10
20
30
40
50
60
Time (min)
Normal
Diabetic
Normal data

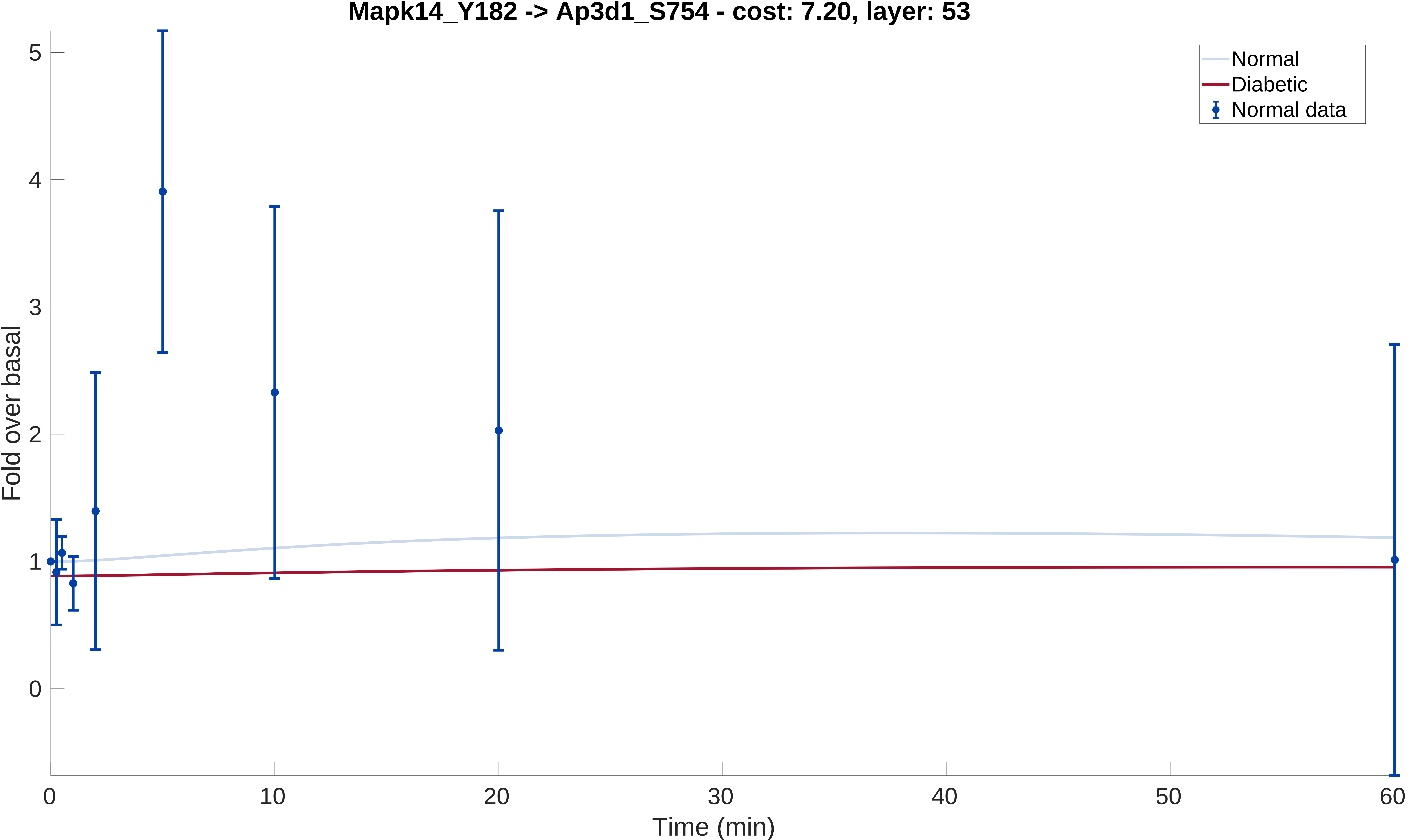

Mapk14_Y182 -> Ap3d1_S754 - cost: 7.20, layer: 53
Normal
Diabetic
Normal data
Fold over basal
Time (min)
0
1
2
3
4
5
0
10
20
30
40
50
60

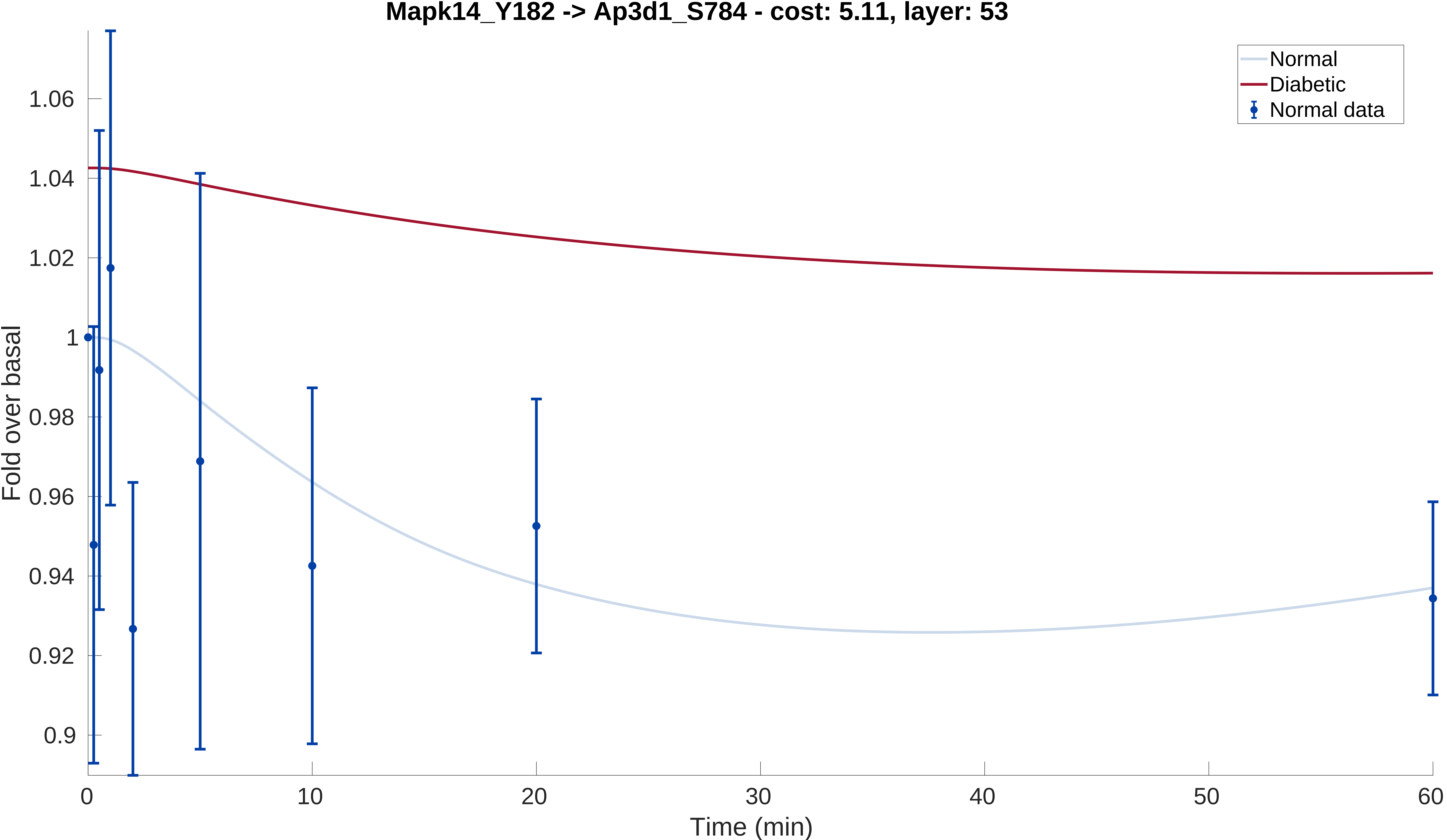
Mapk14_Y182 -> Ap3d1_S784 - cost: 5.11, layer: 53
Normal
Diabetic
Normal data
Fold over basal
1.06
1.04
1.02
1
0.98
0.96
0.94
0.92
0.9
0
10
20
30
40
50
60
Time (min)

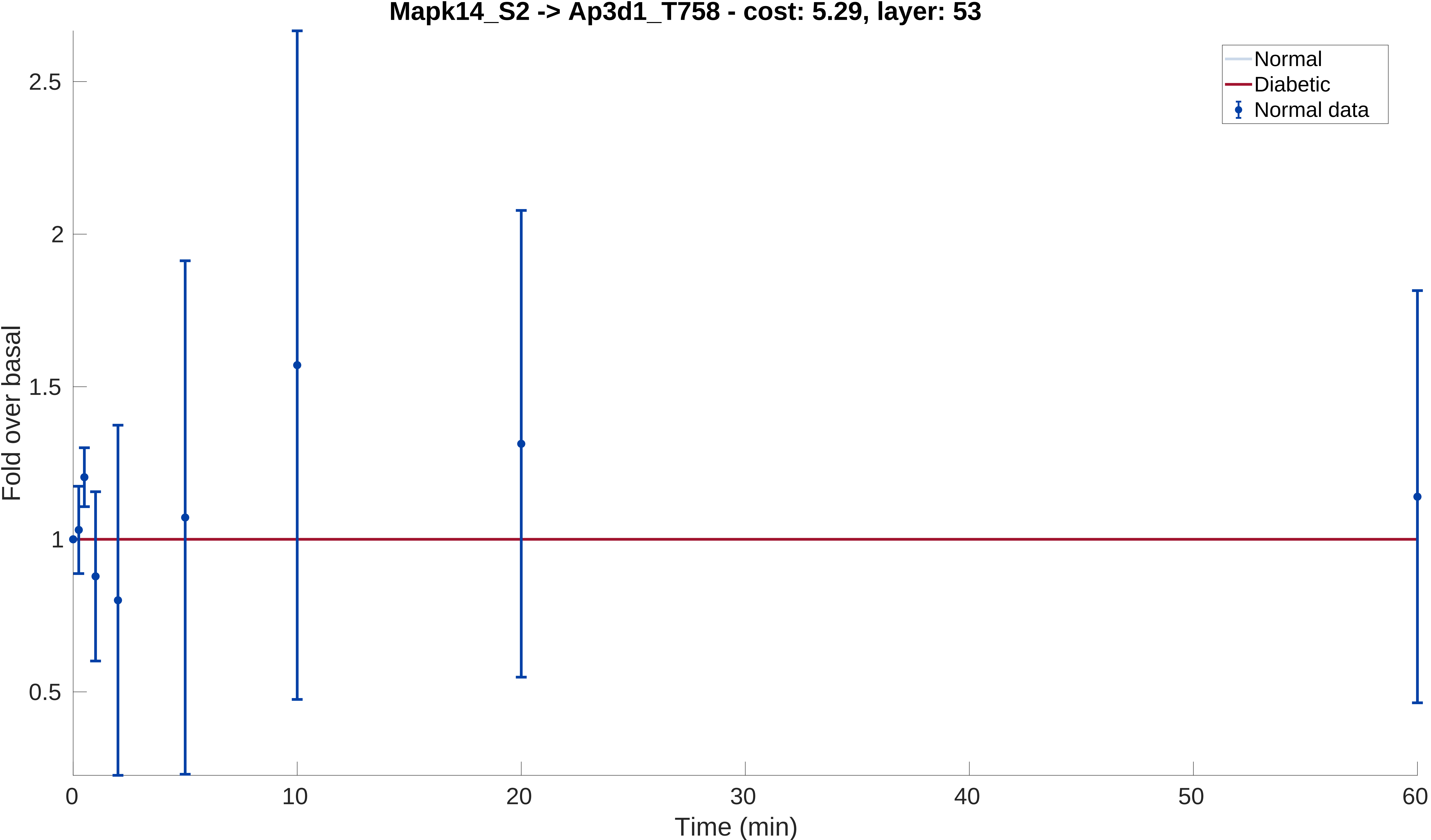
Mapk14_S2 -> Ap3d1_T758 - cost: 5.29, layer: 53
Normal
Diabetic
Normal data
Fold over basal
2.5
2
1.5
1
0.5
0
10
20
30
40
50
60
Time (min)

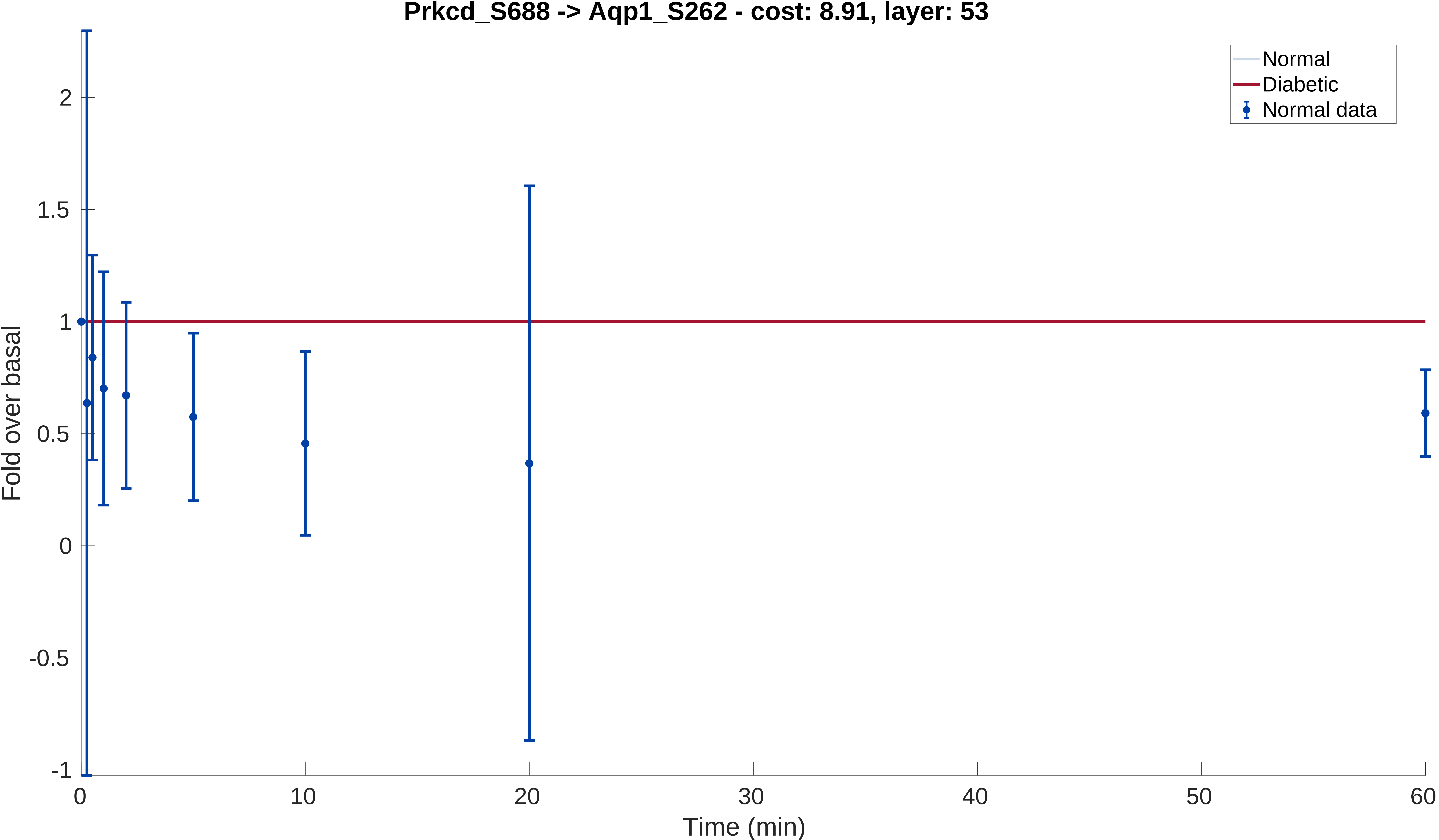

Prkcd_S688 -> Aqp1_S262 - cost: 8.91, layer: 53
Normal
Diabetic
Normal data
Fold over basal
Time (min)
2
1.5
1
0.5
0
-0.5
-1
0
10
20
30
40
50
60

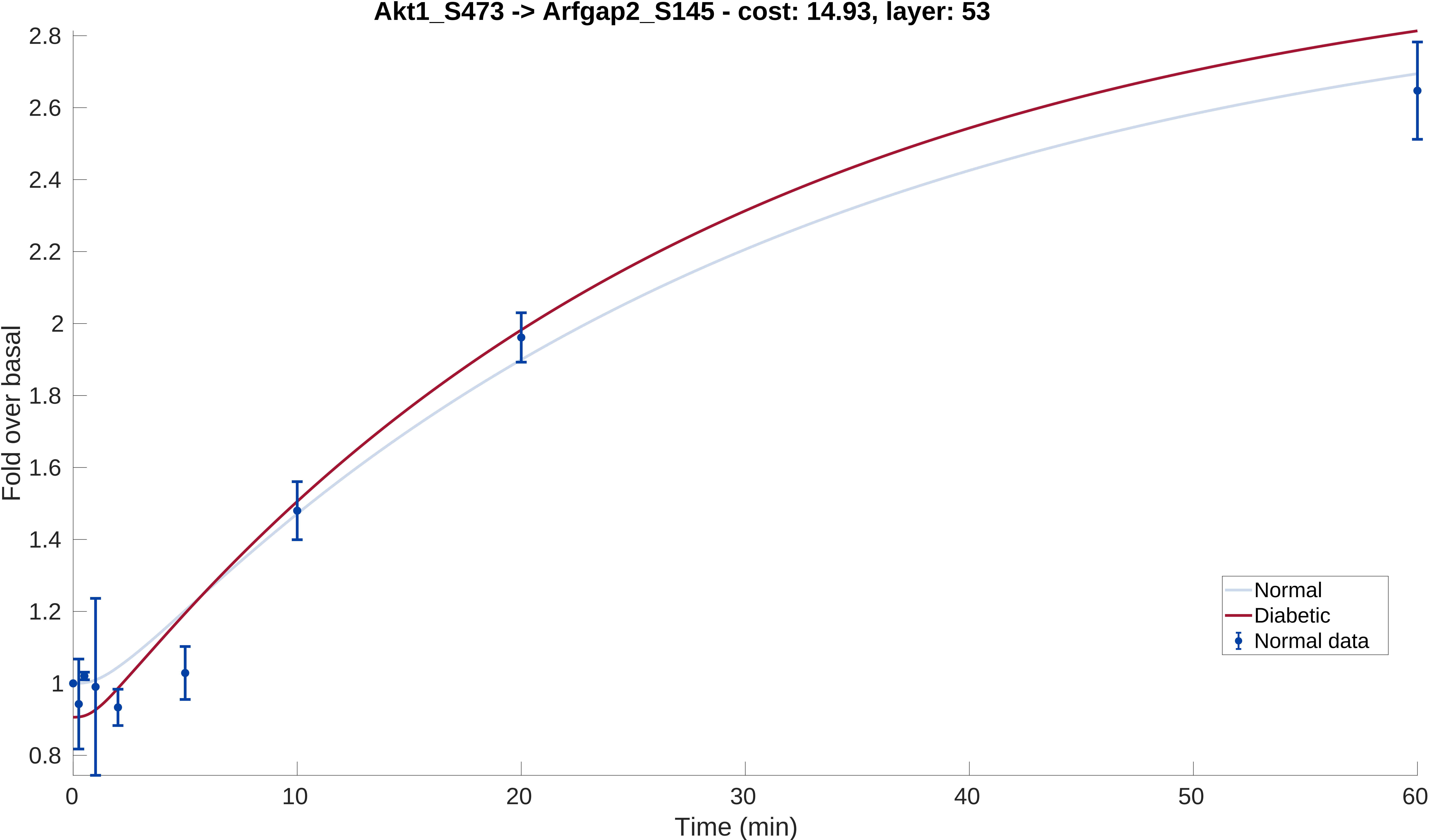
Akt1_S473 -> Arfgap2_S145 - cost: 14.93, layer: 53
Fold over basal
2.8
2.6
2.4
2.2
2
1.8
1.6
1.4
1.2
1
0.8
0
10
20
30
40
50
60
Time (min)
Normal
Diabetic
Normal data

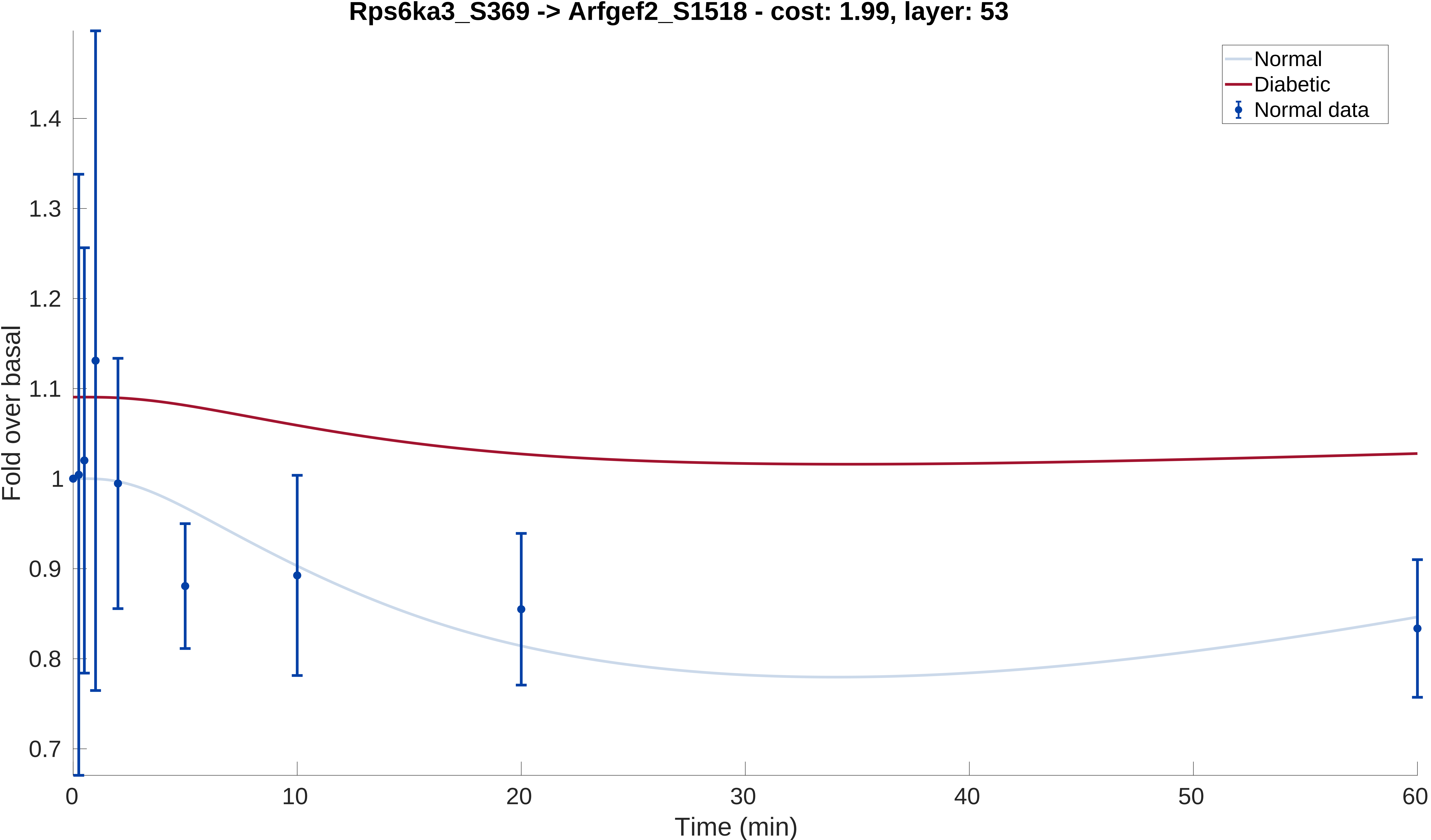

Rps6ka3_S369 -> Arfgef2_S1518 - cost: 1.99, layer: 53
Normal
Diabetic
Normal data
1.4
1.3
1.2
1.1
1
0.9
0.8
0.7
Fold over basal
0
10
20
30
40
50
60
Time (min)

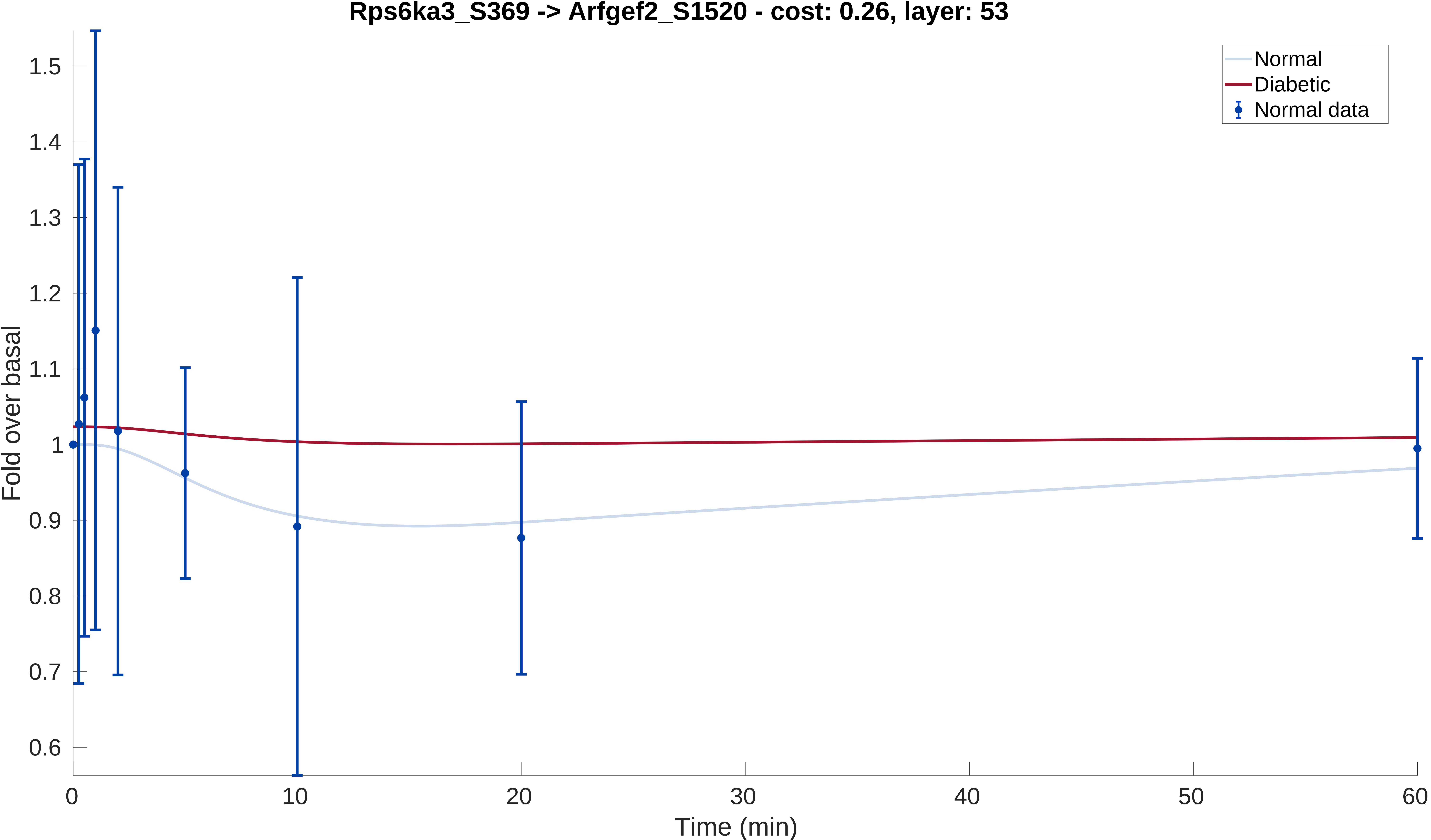

Rps6ka3_S369 -> Arfgef2_S1520 - cost: 0.26, layer: 53
Normal
Diabetic
Normal data
Fold over basal
Time (min)
1.5
1.4
1.3
1.2
1.1
1
0.9
0.8
0.7
0.6
0
10
20
30
40
50
60

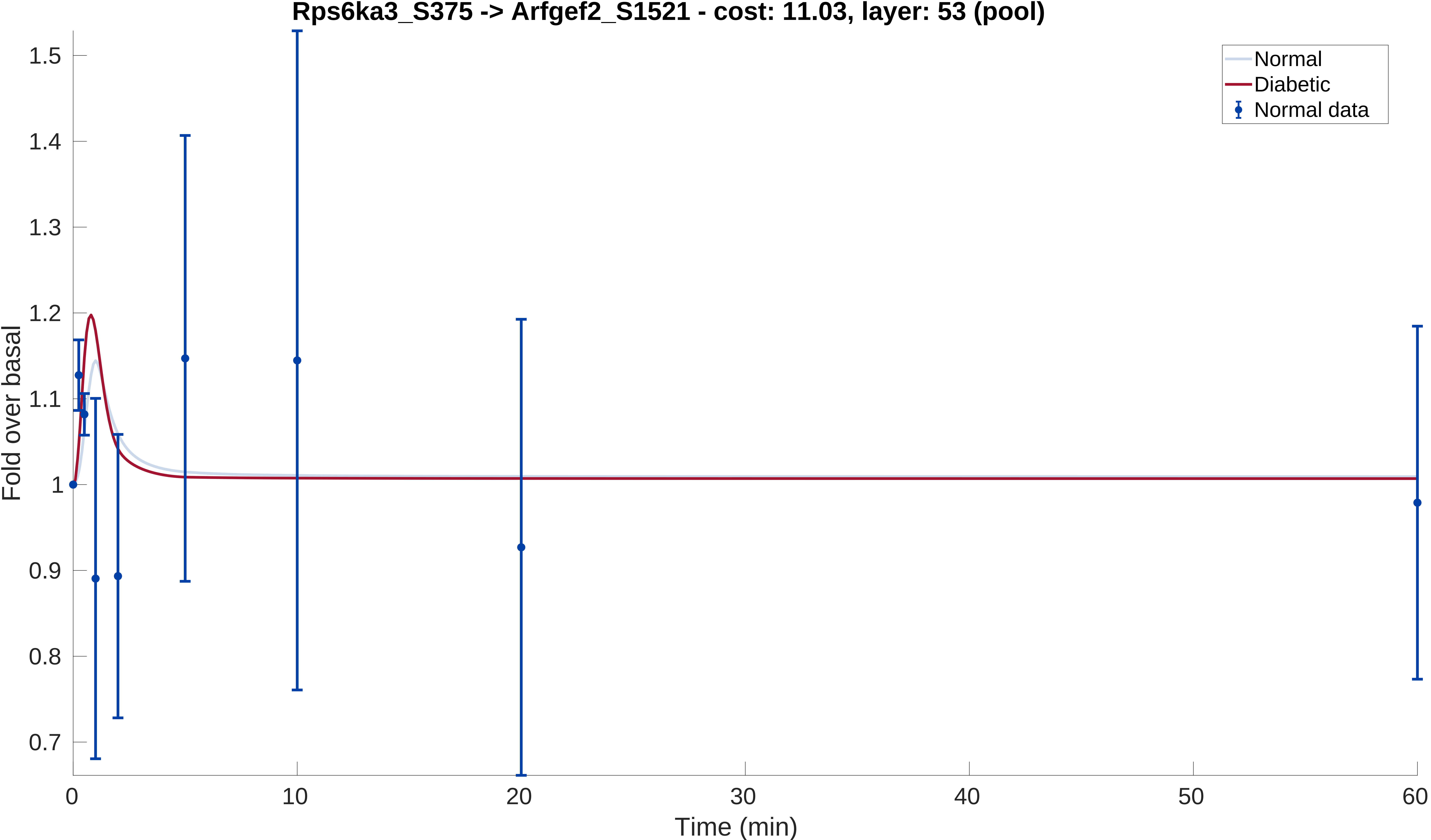

Rps6ka3_S375 -> Arfgef2_S1521 - cost: 11.03, layer: 53 (pool)
Normal
Diabetic
Normal data
Fold over basal
1.5
1.4
1.3
1.2
1.1
1
0.9
0.8
0.7
0
10
20
30
40
50
60
Time (min)

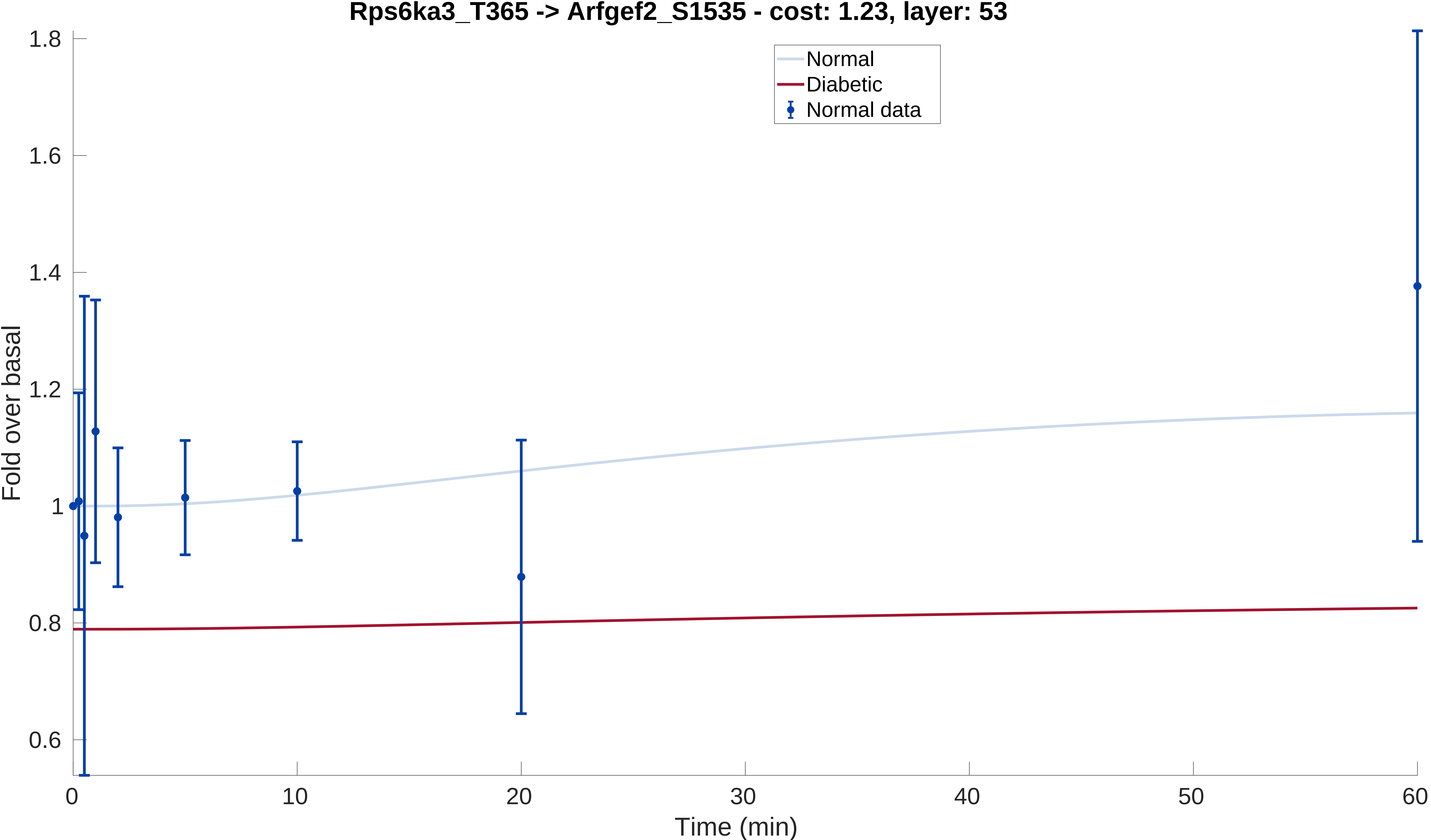

Rps6ka3_T365 -> Arfgef2_S1535 - cost: 1.23, layer: 53
Normal
Diabetic
Normal data
Fold over basal
1.8
1.6
1.4
1.2
1
0.8
0.6
0
10
20
30
40
50
60
Time (min)

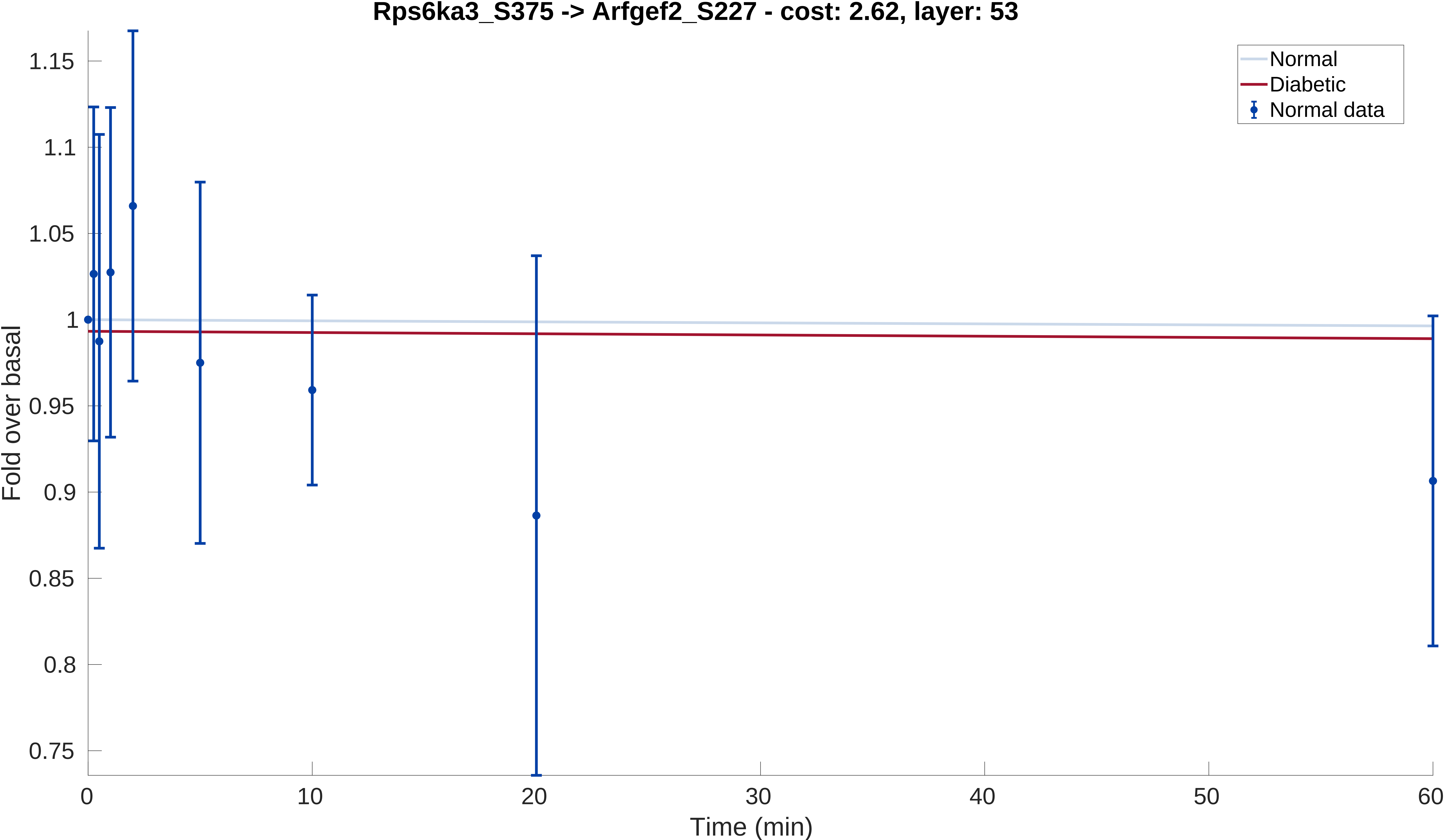
Rps6ka3_S375 -> Arfgef2_S227 - cost: 2.62, layer: 53
Normal
Diabetic
Normal data
Fold over basal
1.15
1.1
1.05
1
0.95
0.9
0.85
0.8
0.75
0
10
20
30
40
50
60
Time (min)

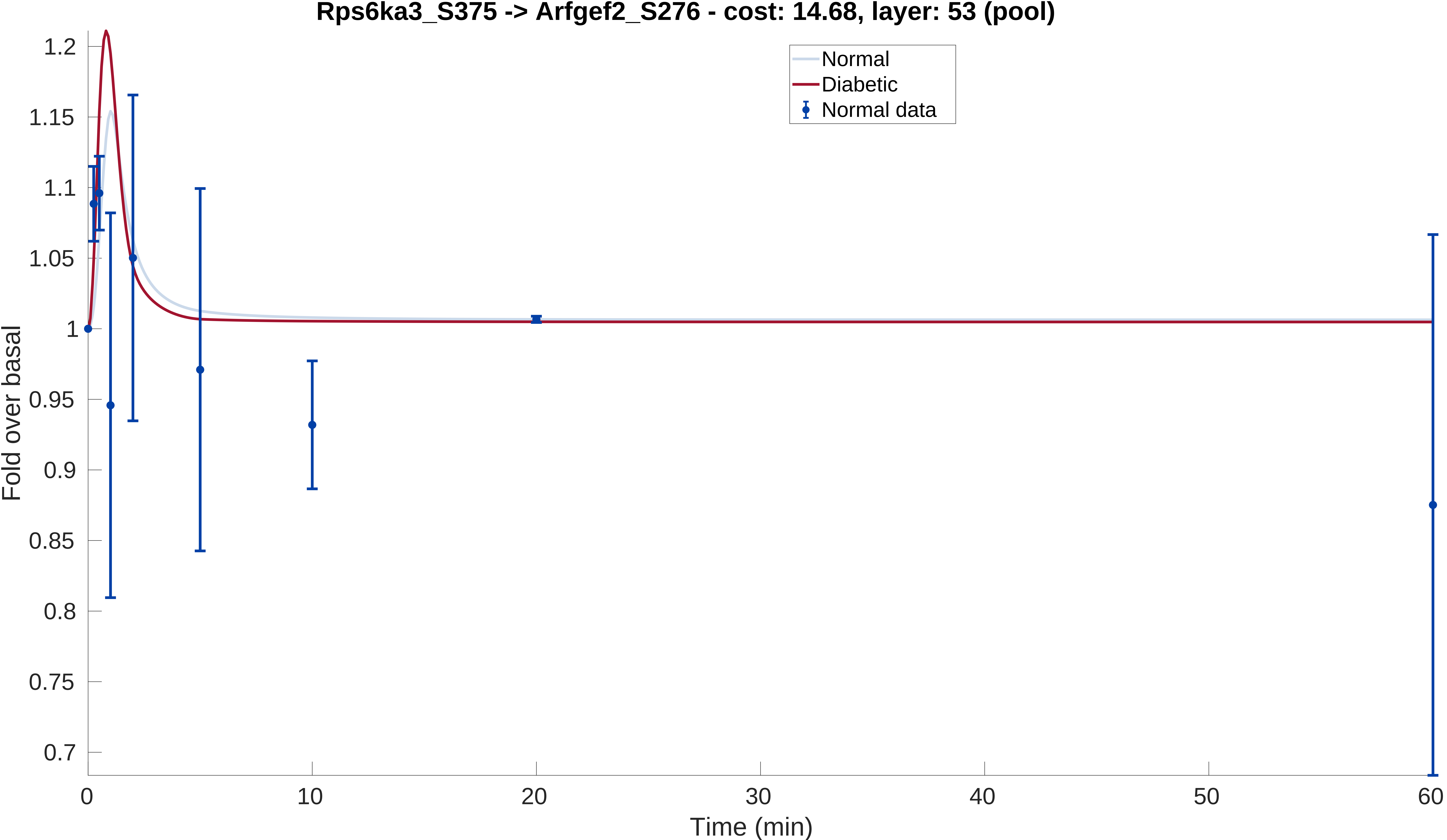

Rps6ka3_S375 -> Arfgef2_S276 - cost: 14.68, layer: 53 (pool)
Normal
Diabetic
Normal data
Fold over basal
1.2
1.15
1.1
1.05
1
0.95
0.9
0.85
0.8
0.75
0.7
0
10
20
30
40
50
60
Time (min)

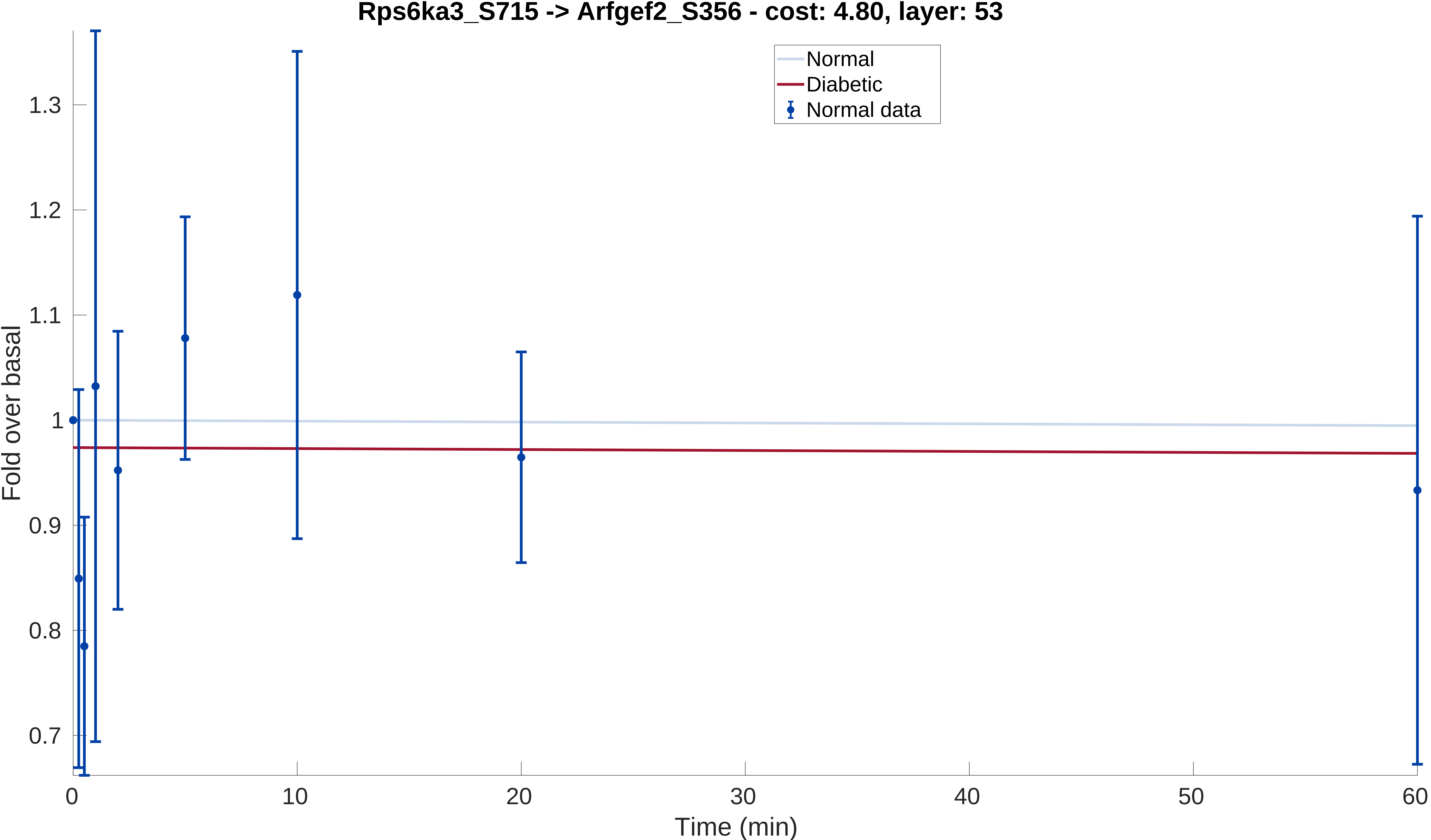

Rps6ka3_S715 -> Arfgef2_S356 - cost: 4.80, layer: 53
Normal
Diabetic
Normal data
Fold over basal
Time (min)
0.7
0.8
0.9
1
1.1
1.2
1.3
0
10
20
30
40
50
60

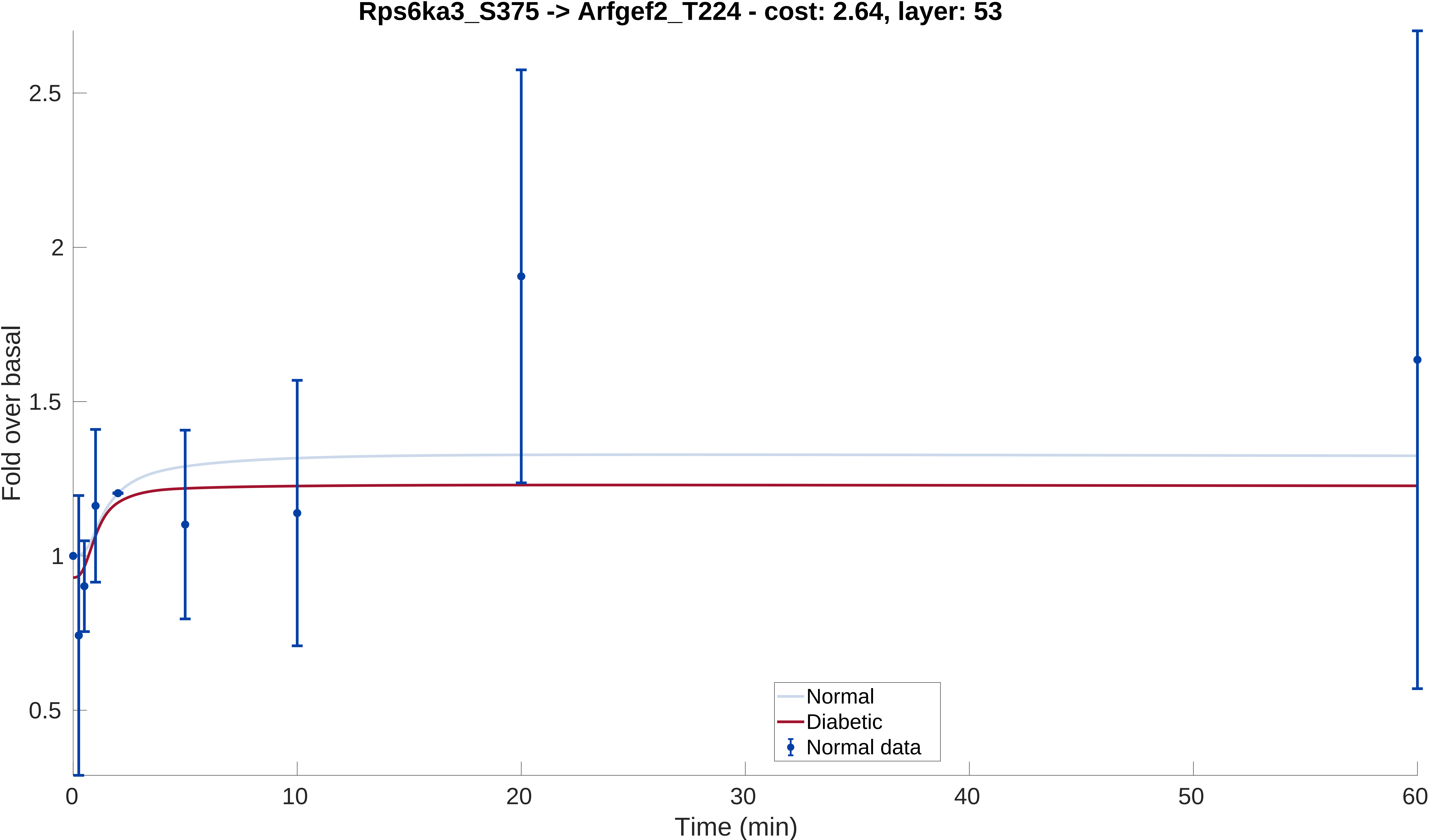

Rps6ka3_S375 -> Arfgef2_T224 - cost: 2.64, layer: 53
Fold over basal
2.5
2
1.5
1
0.5
0
10
20
30
40
50
60
Time (min)
Normal
Diabetic
Normal data

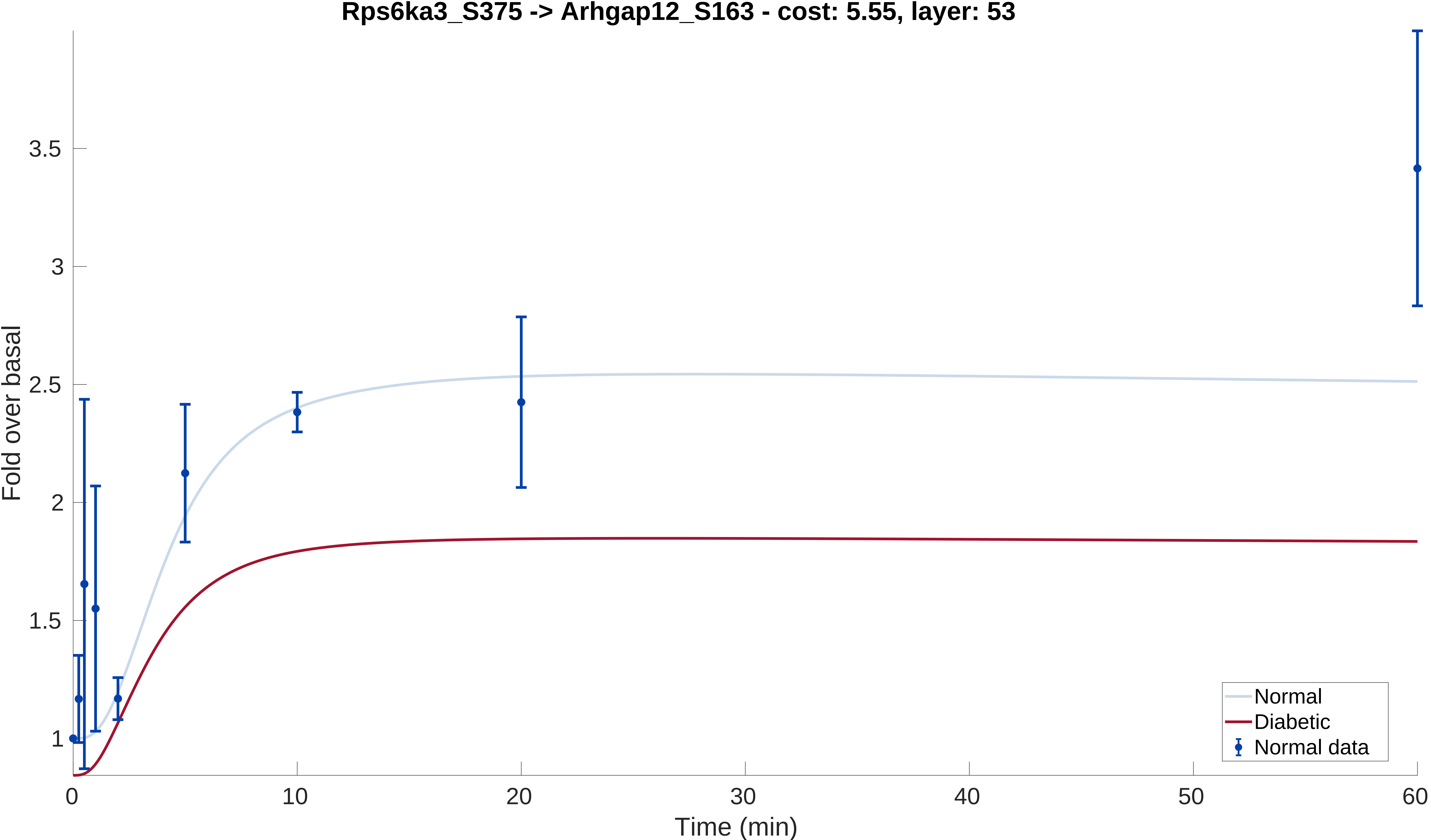

Rps6ka3_S375 -> Arhgap12_S163 - cost: 5.55, layer: 53
Fold over basal
Time (min)
3.5
3
2.5
2
1.5
1
0
10
20
30
40
50
60
Normal
Diabetic
Normal data

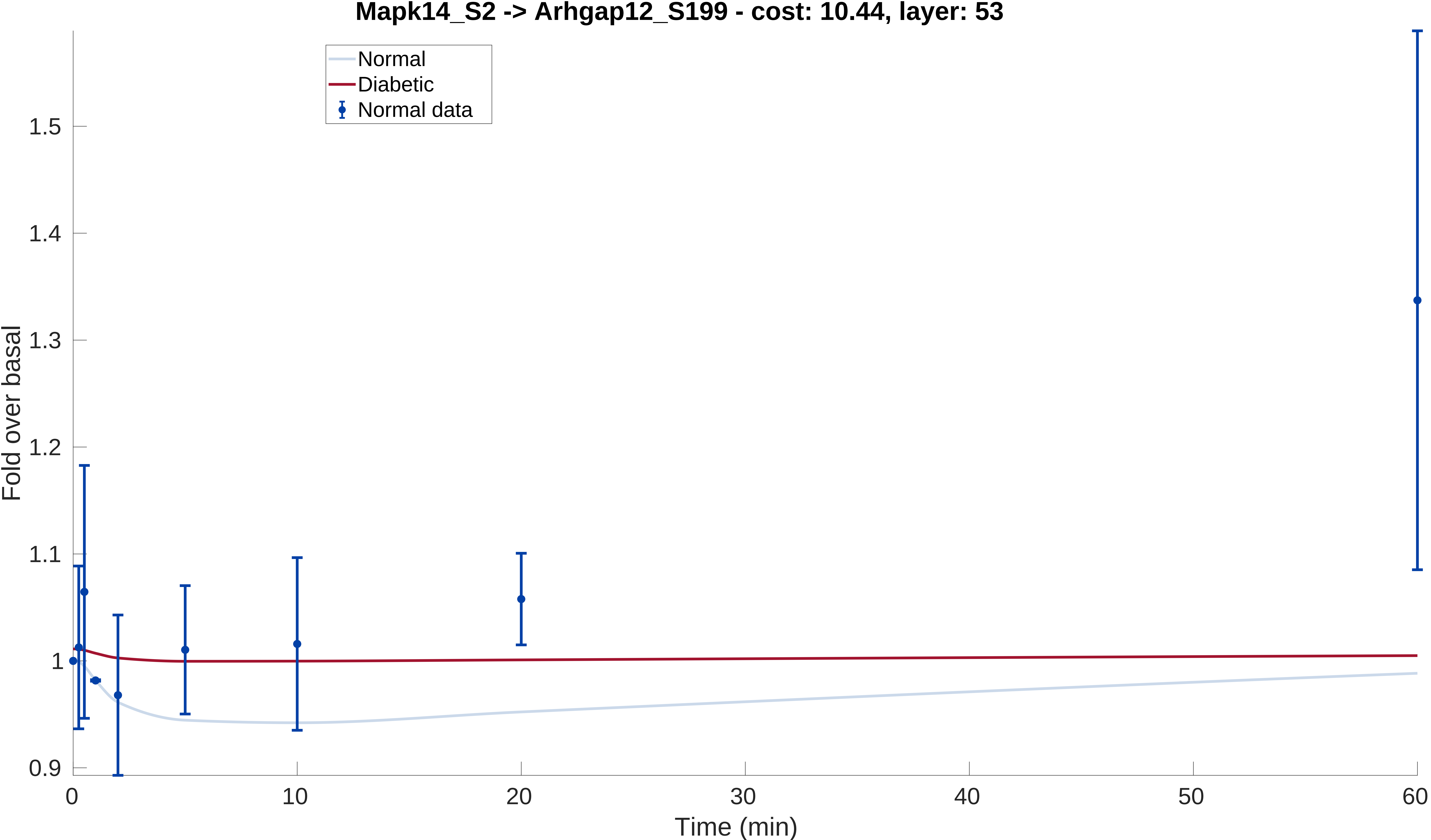

Mapk14_S2 -> Arhgap12_S199 - cost: 10.44, layer: 53
Normal
Diabetic
Normal data
Fold over basal
1.5
1.4
1.3
1.2
1.1
1
0.9
0
10
20
30
40
50
60
Time (min)

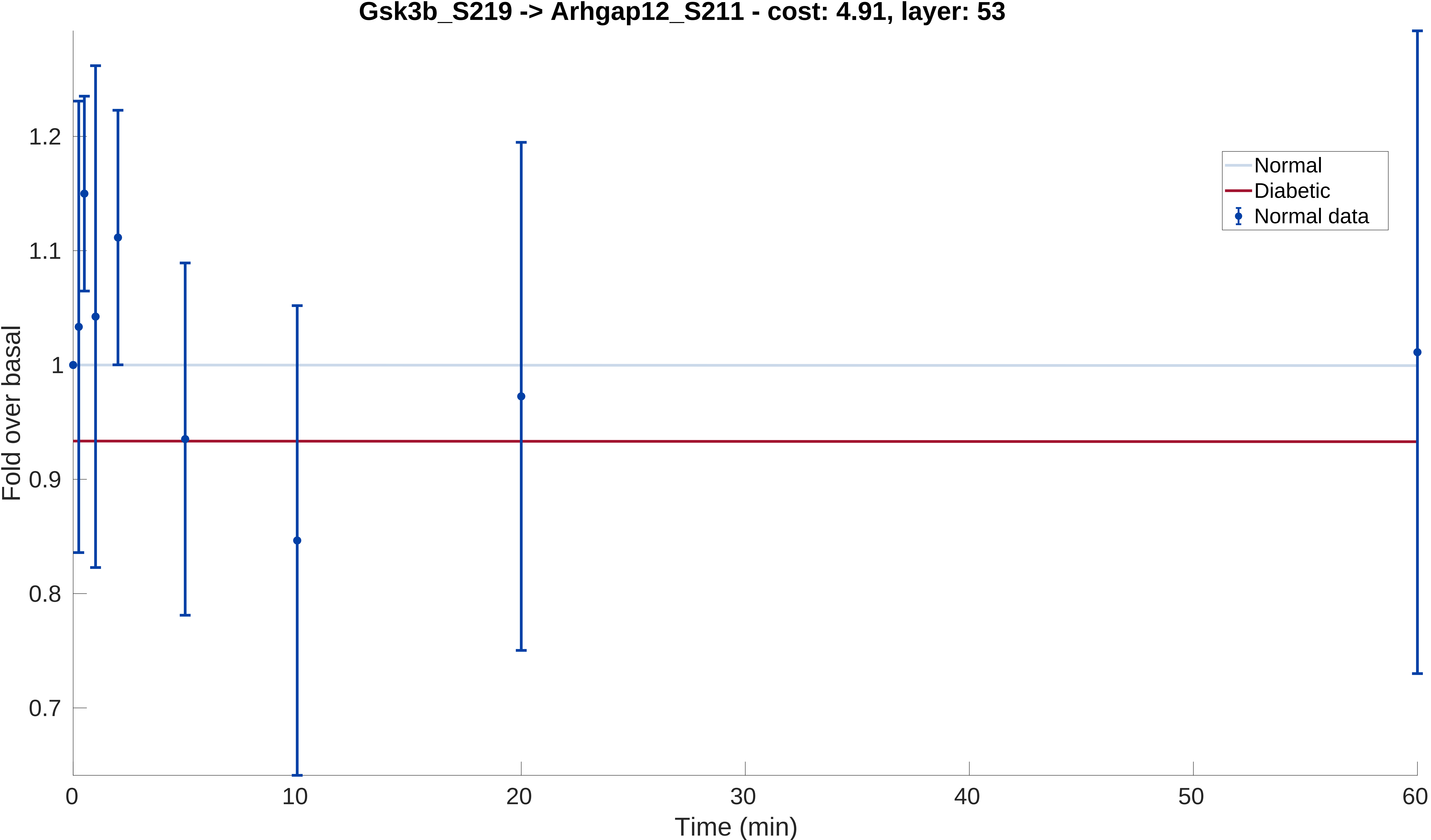
Gsk3b_S219 -> Arhgap12_S211 - cost: 4.91, layer: 53
Normal
Diabetic
Normal data
Fold over basal
0.7
0.8
0.9
1
1.1
1.2
0
10
20
30
40
50
60
Time (min)

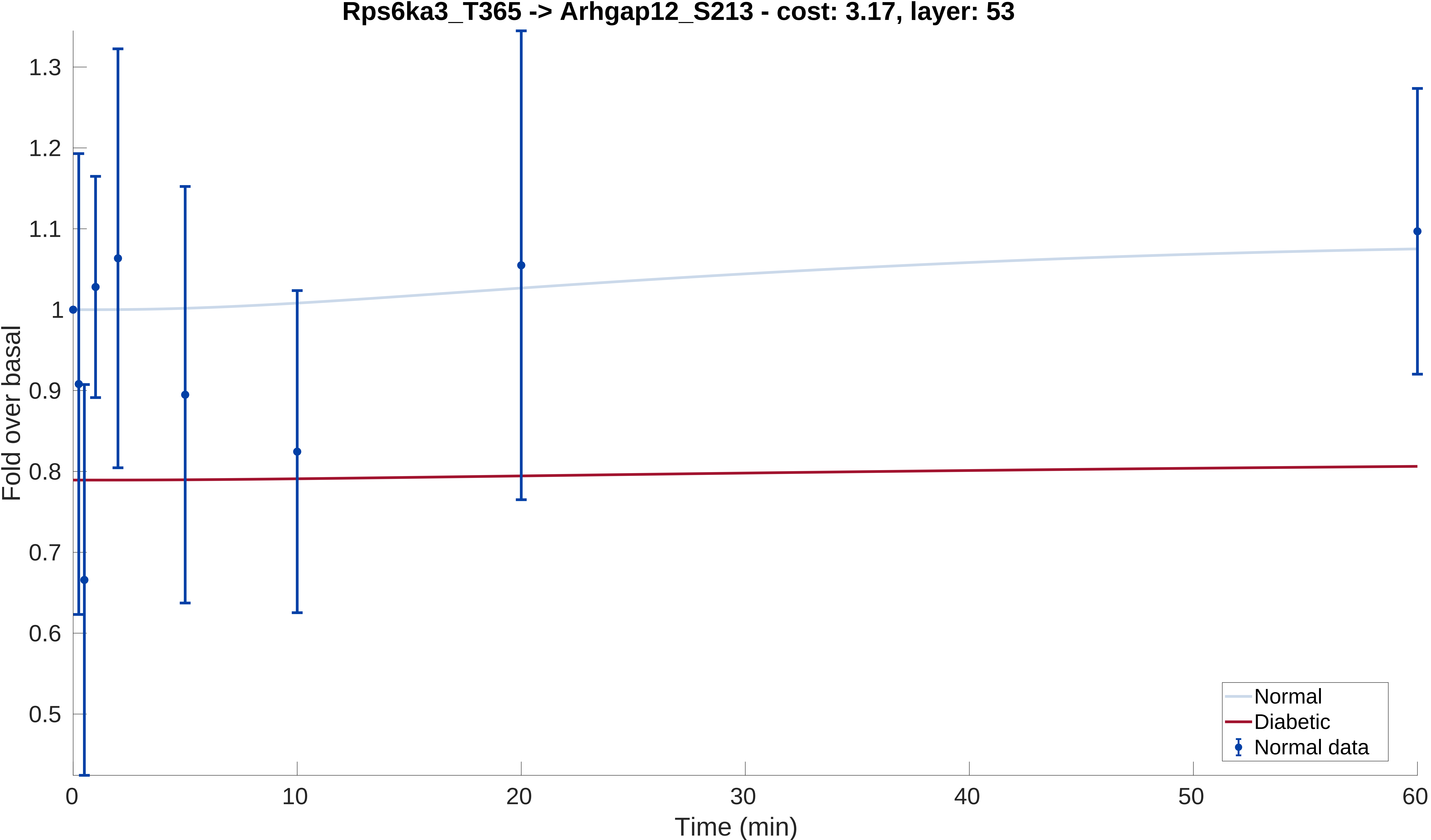

Rps6ka3_T365 -> Arhgap12_S213 - cost: 3.17, layer: 53
Fold over basal
1.3
1.2
1.1
1
0.9
0.8
0.7
0.6
0.5
0
10
20
30
40
50
60
Time (min)
Normal
Diabetic
Normal data

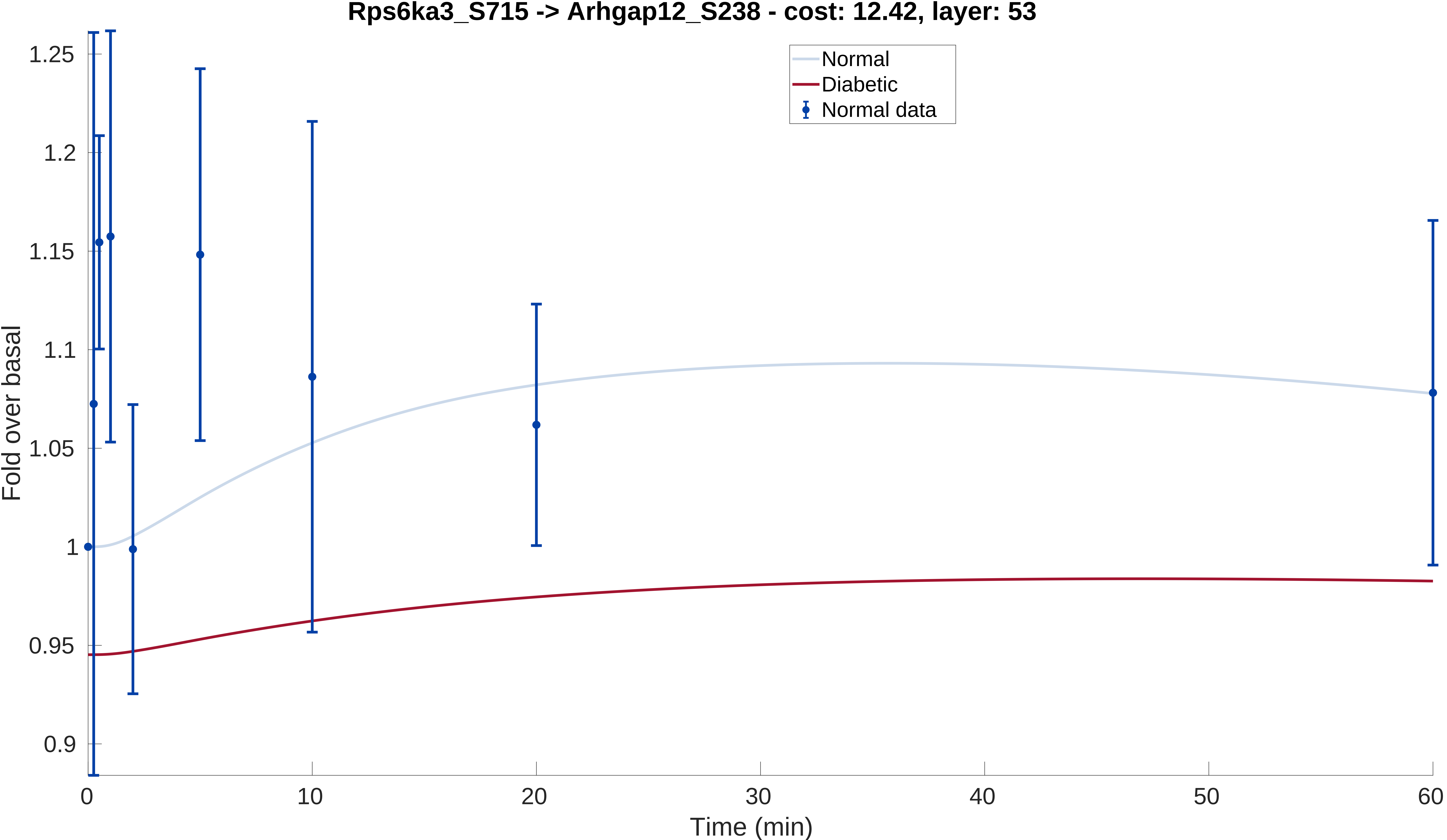

Rps6ka3_S715 -> Arhgap12_S238 - cost: 12.42, layer: 53
Normal
Diabetic
Normal data
Fold over basal
1.25
1.2
1.15
1.1
1.05
1
0.95
0.9
0
10
20
30
40
50
60
Time (min)

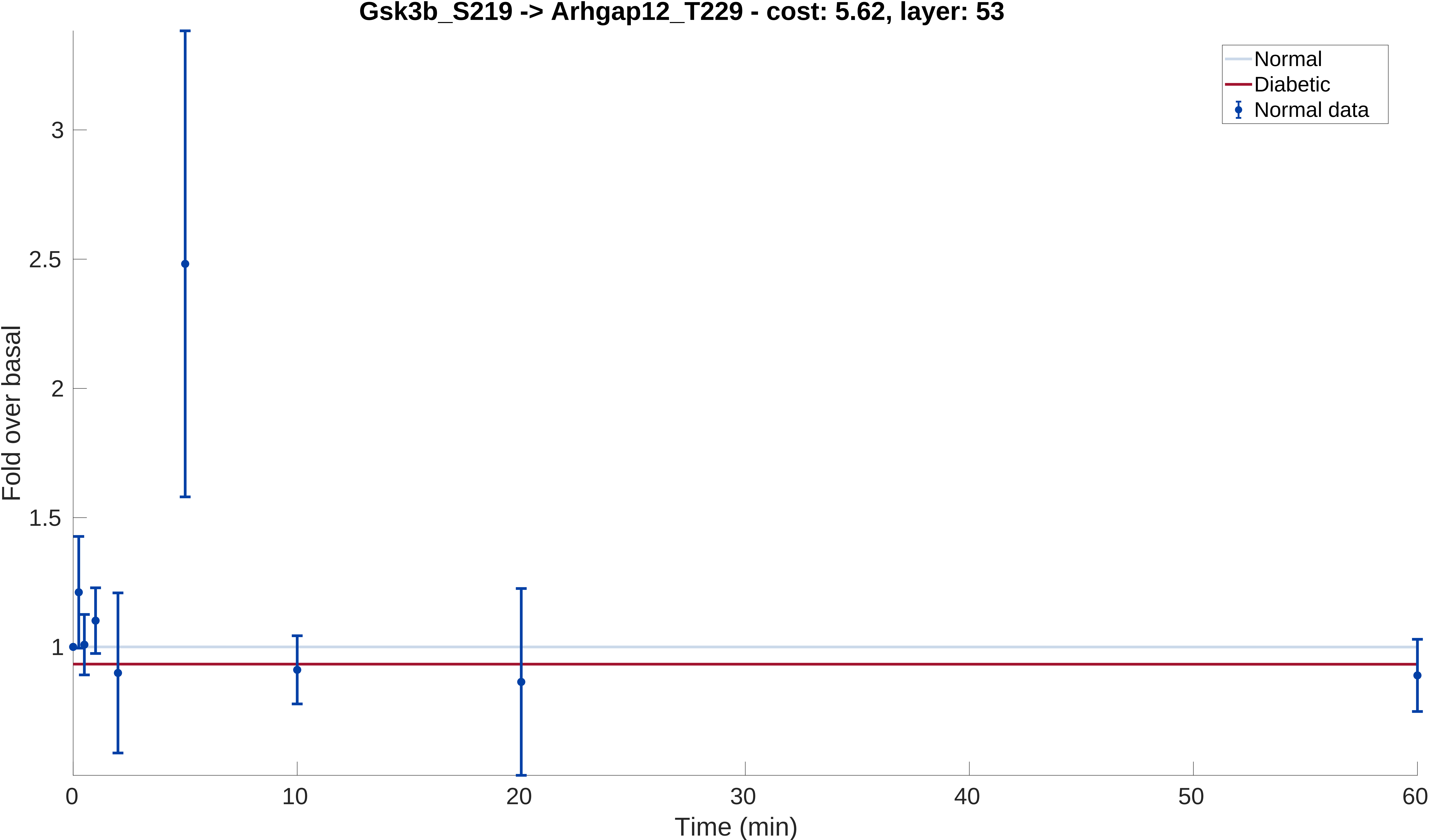

Gsk3b_S219 -> Arhgap12_T229 - cost: 5.62, layer: 53
Normal
Diabetic
Normal data
Fold over basal
3
2.5
2
1.5
1
0
10
20
30
40
50
60
Time (min)

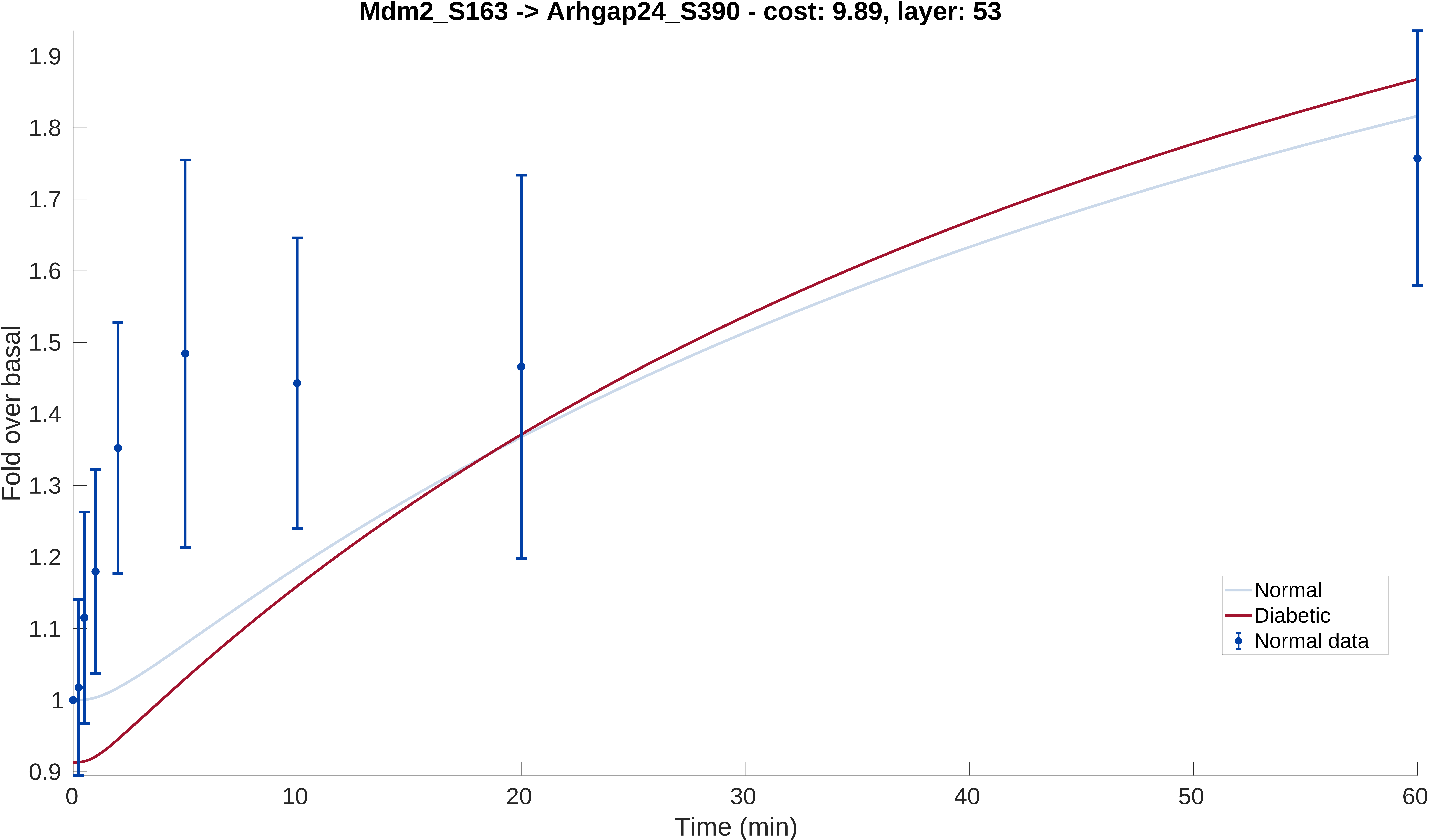

Mdm2_S163 -> Arhgap24_S390 - cost: 9.89, layer: 53
Fold over basal
Time (min)
0.9
1
1.1
1.2
1.3
1.4
1.5
1.6
1.7
1.8
1.9
0
10
20
30
40
50
60
Normal
Diabetic
Normal data

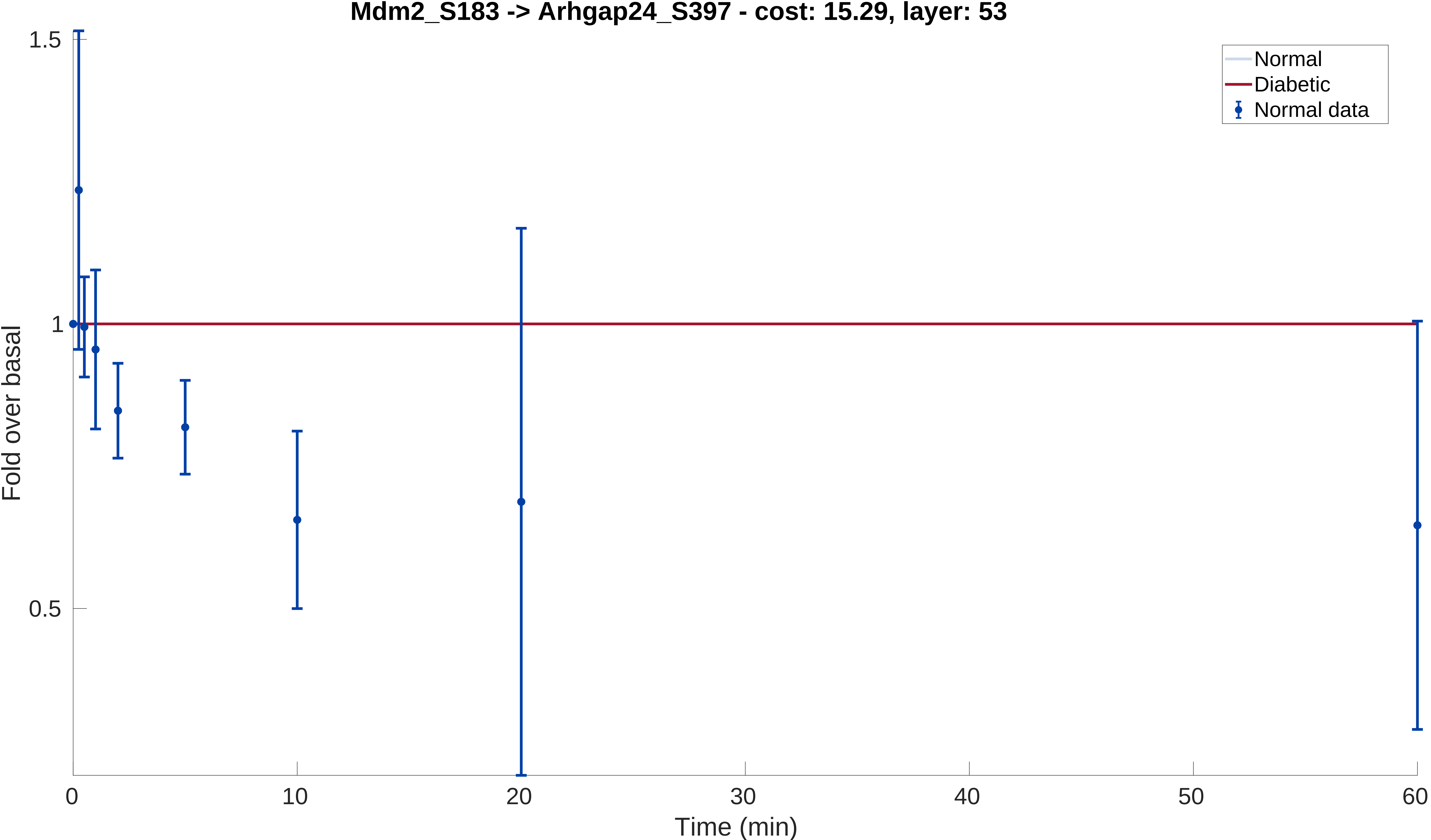

Mdm2_S183 -> Arhgap24_S397 - cost: 15.29, layer: 53
Fold over basal
1.5
1
0.5
0
10
20
30
40
50
60
Time (min)
Normal
Diabetic
Normal data

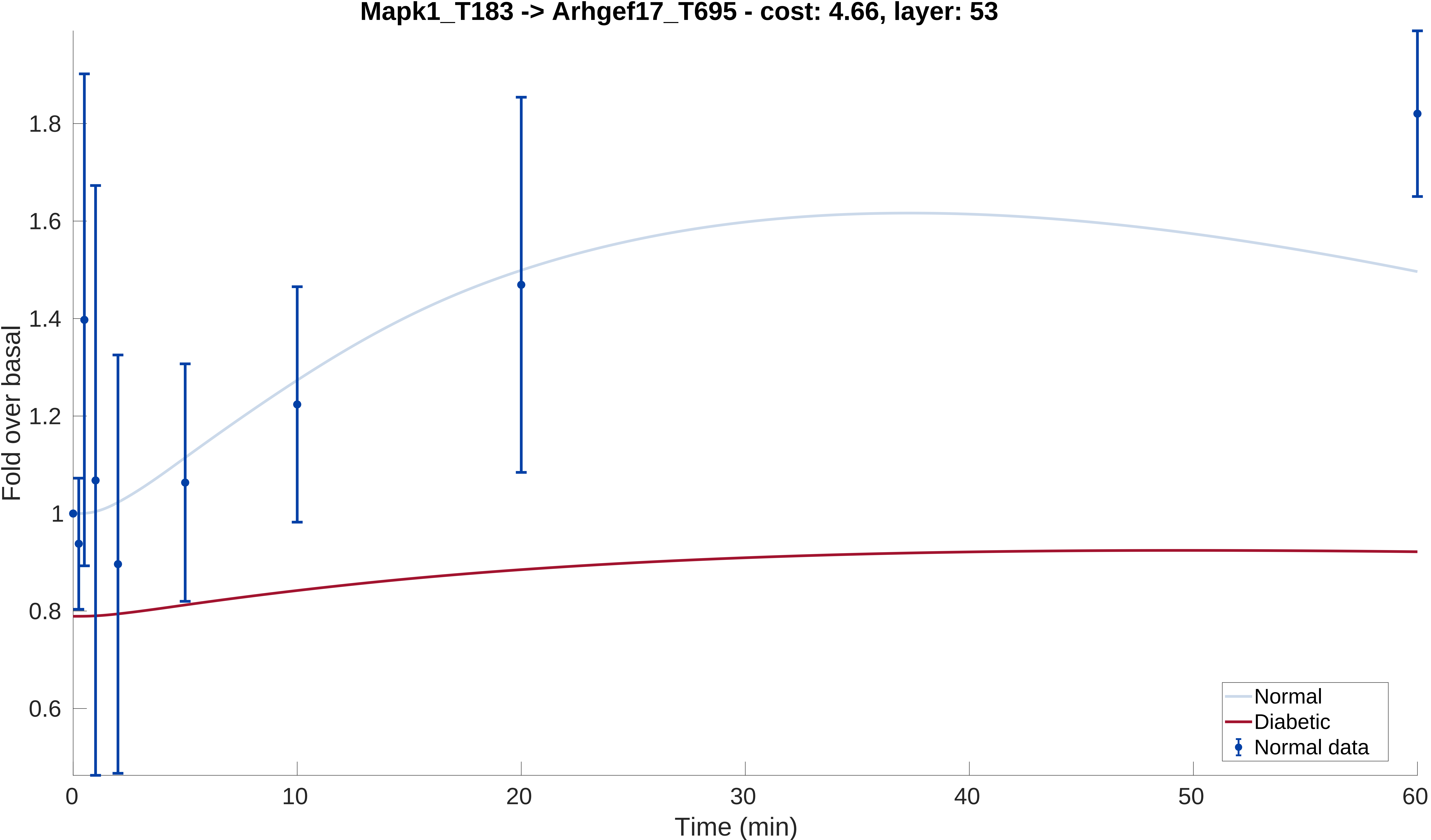

Mapk1_T183 -> Arhgef17_T695 - cost: 4.66, layer: 53
Fold over basal
1.8
1.6
1.4
1.2
1
0.8
0.6
0
10
20
30
40
50
60
Time (min)
Normal
Diabetic
Normal data

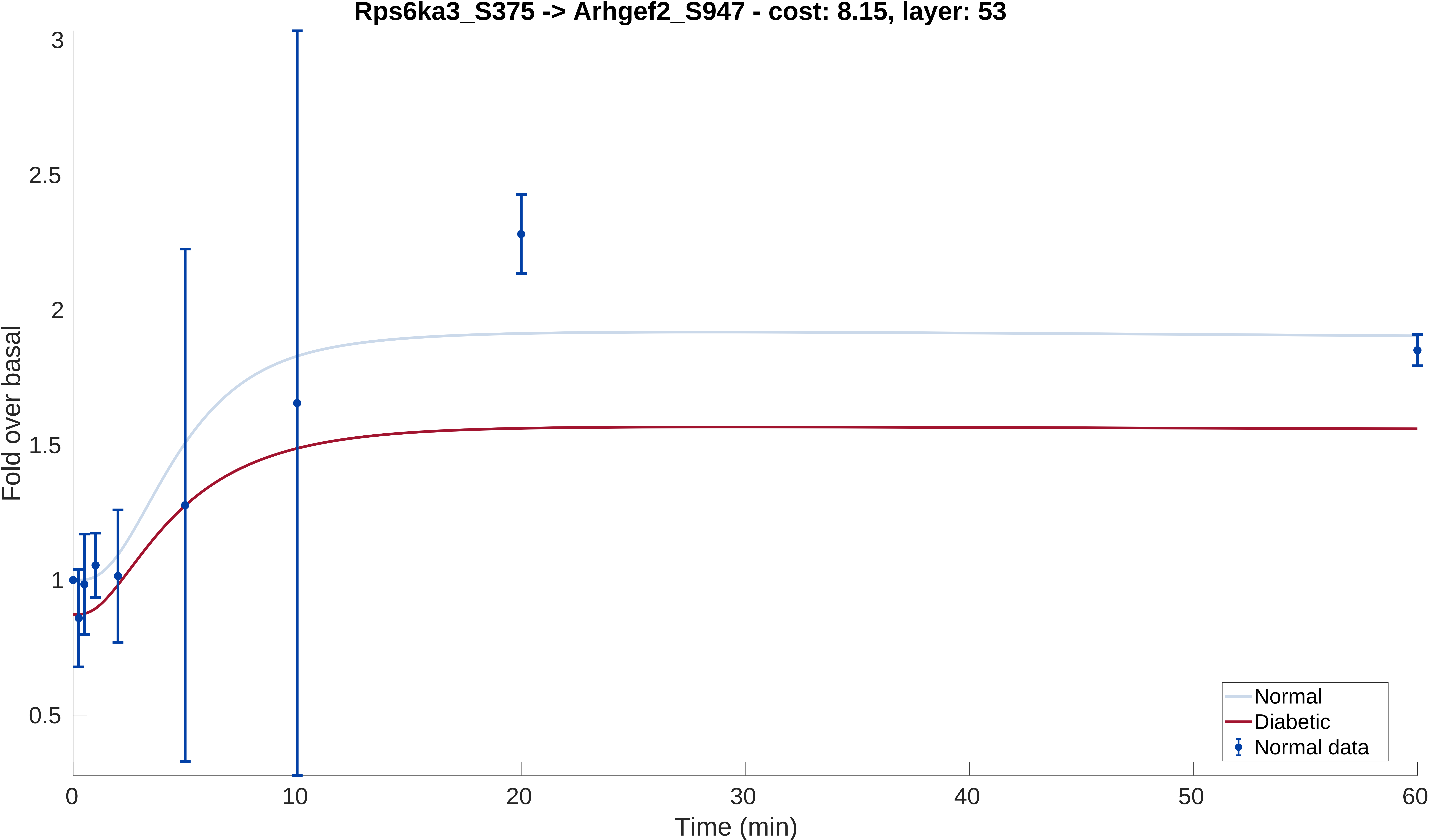
Rps6ka3_S375 -> Arhgef2_S947 - cost: 8.15, layer: 53
Fold over basal
3
2.5
2
1.5
1
0.5
0
10
20
30
40
50
60
Time (min)
Normal
Diabetic
Normal data

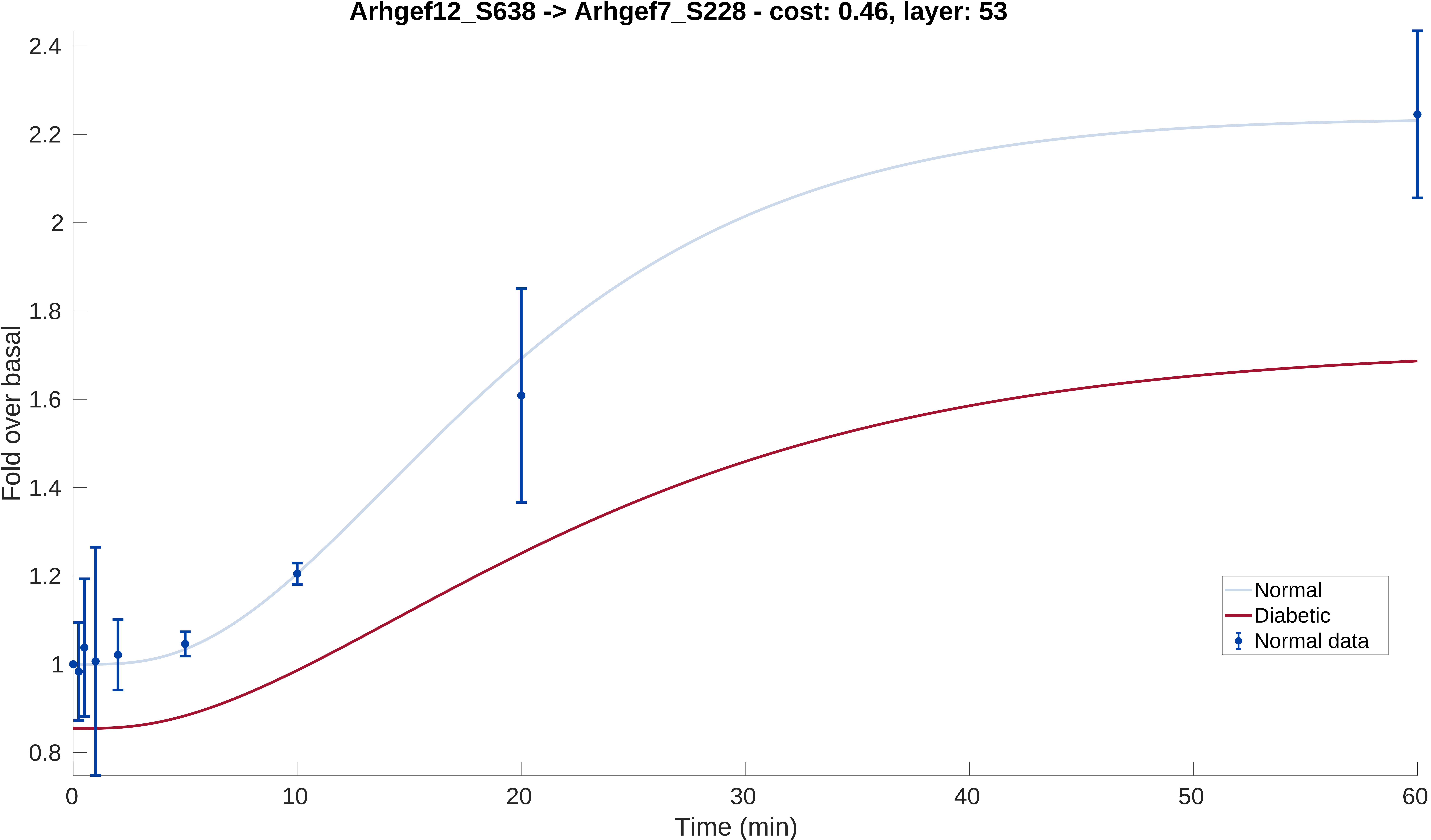

Arhgef12_S638 -> Arhgef7_S228 - cost: 0.46, layer: 53
Fold over basal
2.4
2.2
2
1.8
1.6
1.4
1.2
1
0.8
0
10
20
30
40
50
60
Time (min)
Normal
Diabetic
Normal data

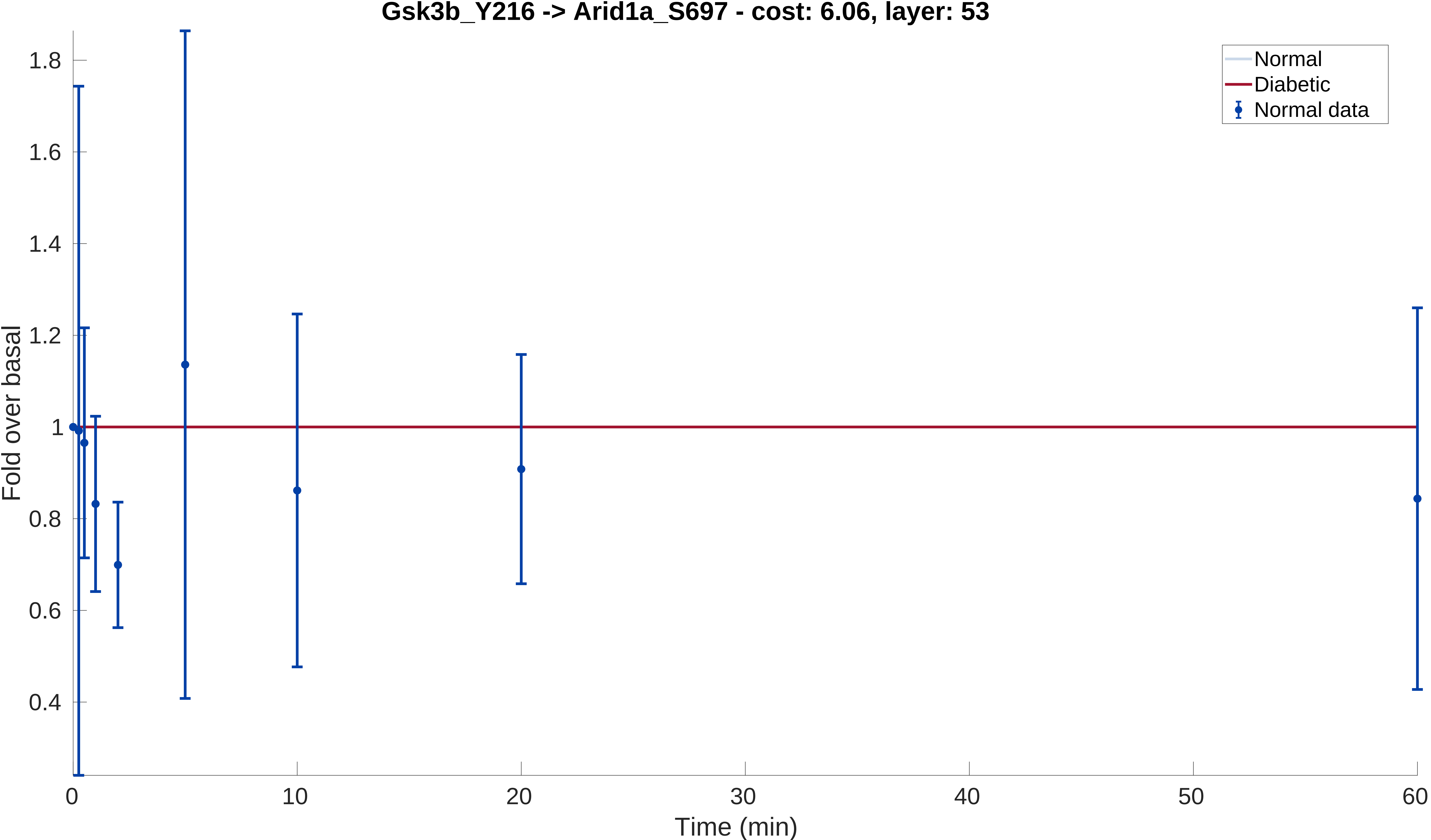

Gsk3b_Y216 -> Arid1a_S697 - cost: 6.06, layer: 53
Normal
Diabetic
Normal data
1.8
1.6
1.4
1.2
1
0.8
0.6
0.4
Fold over basal
0
10
20
30
40
50
60
Time (min)

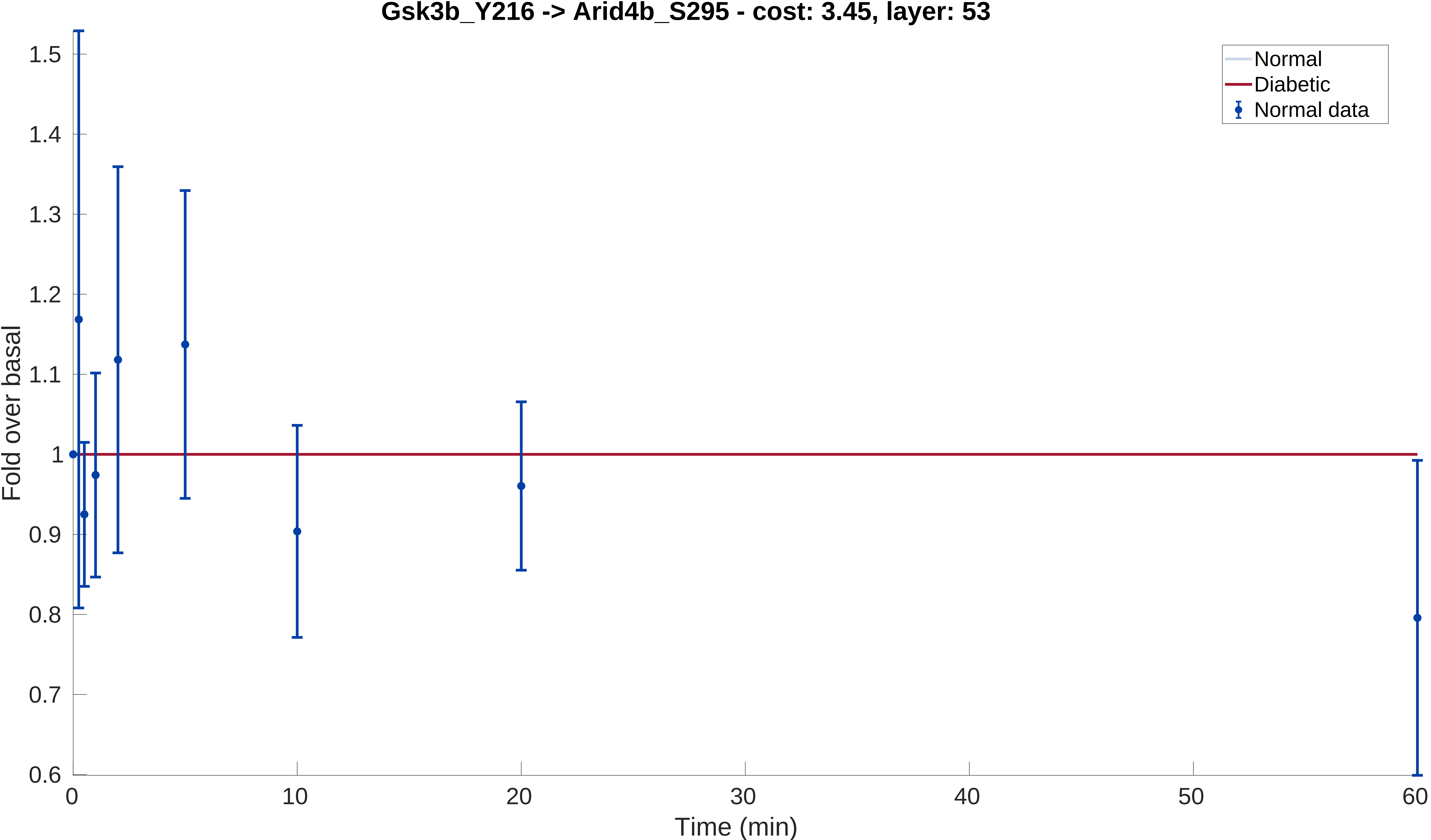

Gsk3b_Y216 -> Arid4b_S295 - cost: 3.45, layer: 53
Normal
Diabetic
Normal data
Fold over basal
1.5
1.4
1.3
1.2
1.1
1
0.9
0.8
0.7
0.6
0
10
20
30
40
50
60
Time (min)

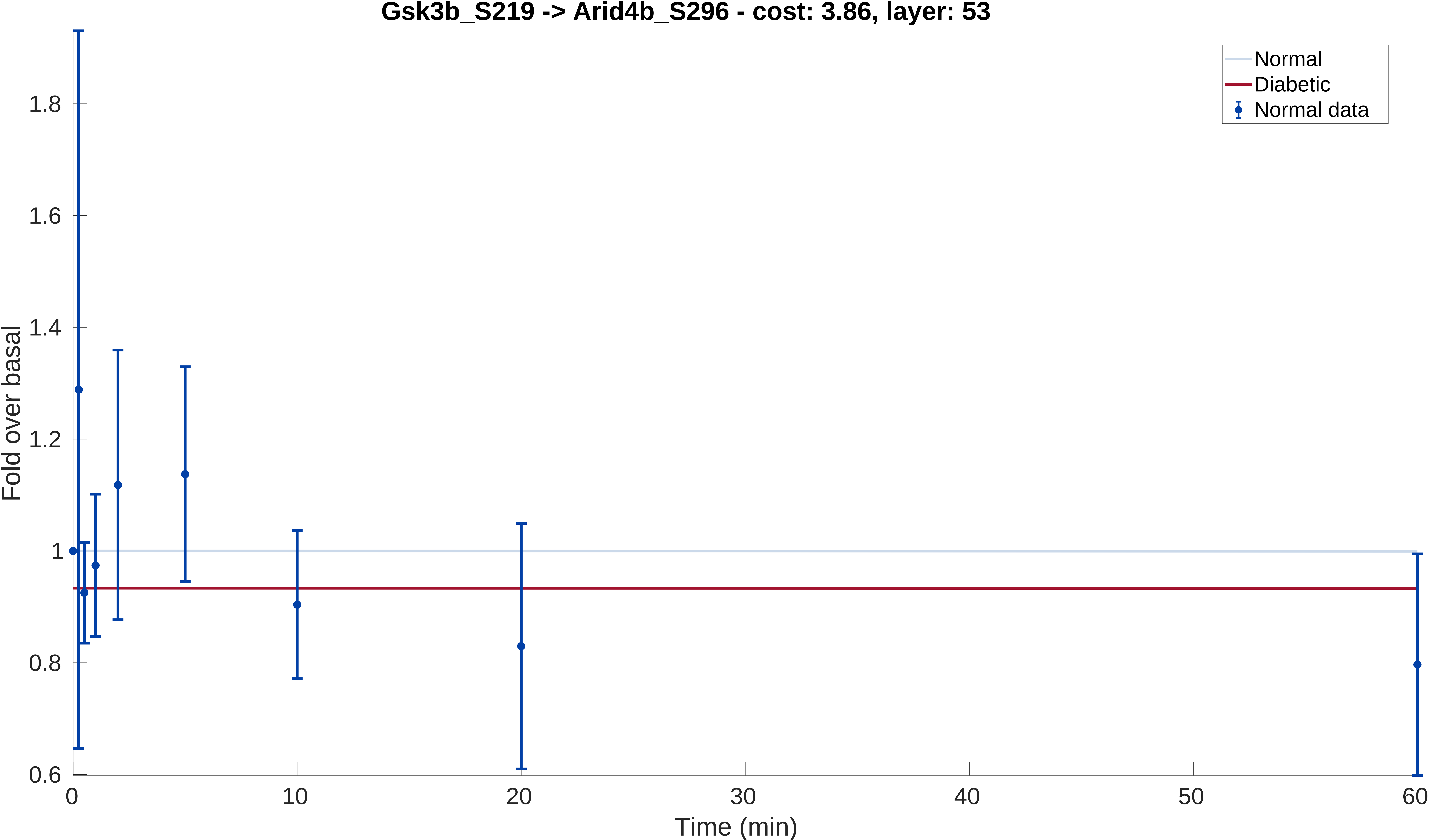

Gsk3b_S219 -> Arid4b_S296 - cost: 3.86, layer: 53
Fold over basal
Time (min)
Normal
Diabetic
Normal data
0.6
0.8
1
1.2
1.4
1.6
1.8
0
10
20
30
40
50
60

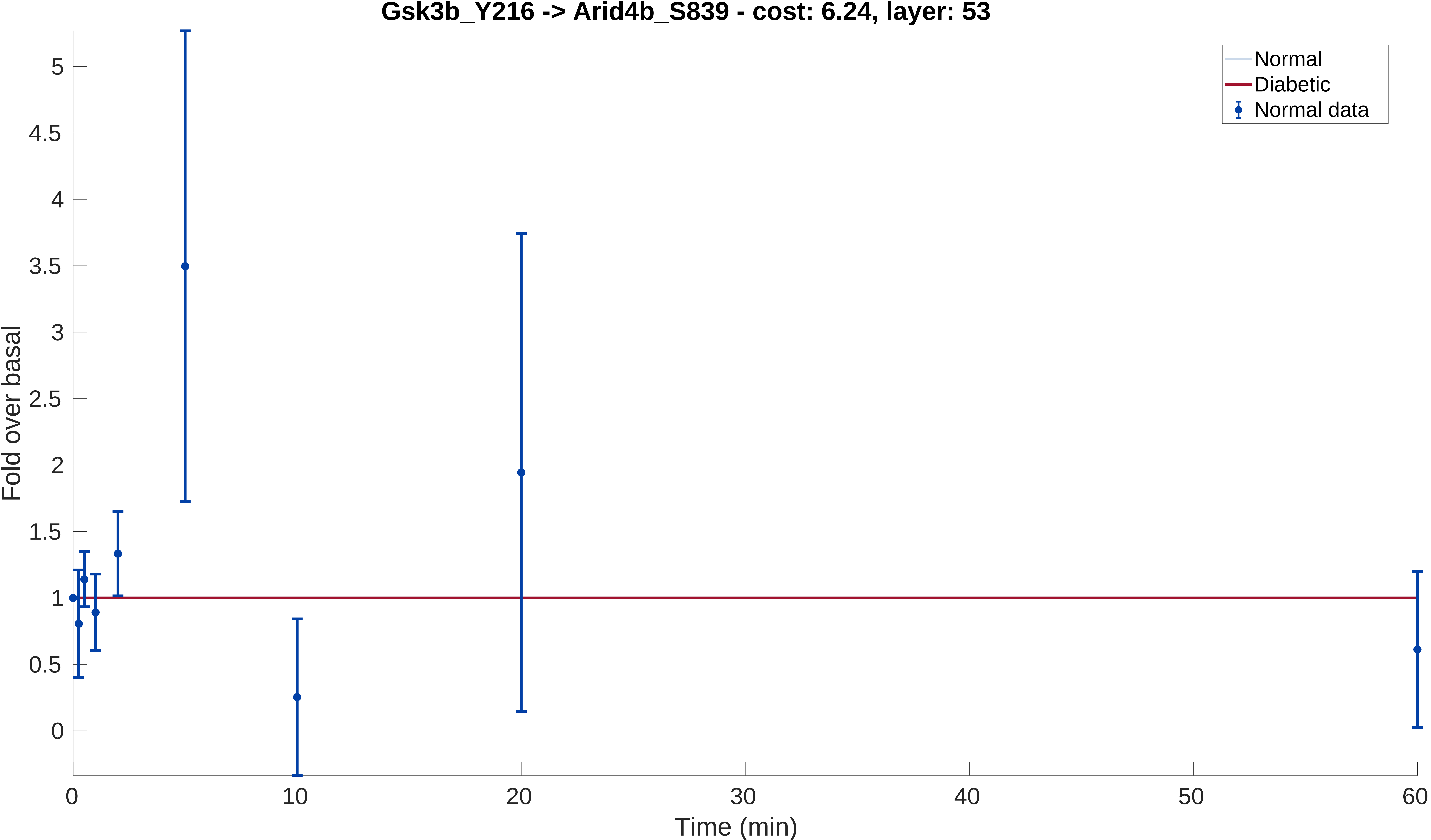
Gsk3b_Y216 -> Arid4b_S839 - cost: 6.24, layer: 53
Normal
Diabetic
Normal data
Fold over basal
Time (min)
0
0.5
1
1.5
2
2.5
3
3.5
4
4.5
5
0
10
20
30
40
50
60

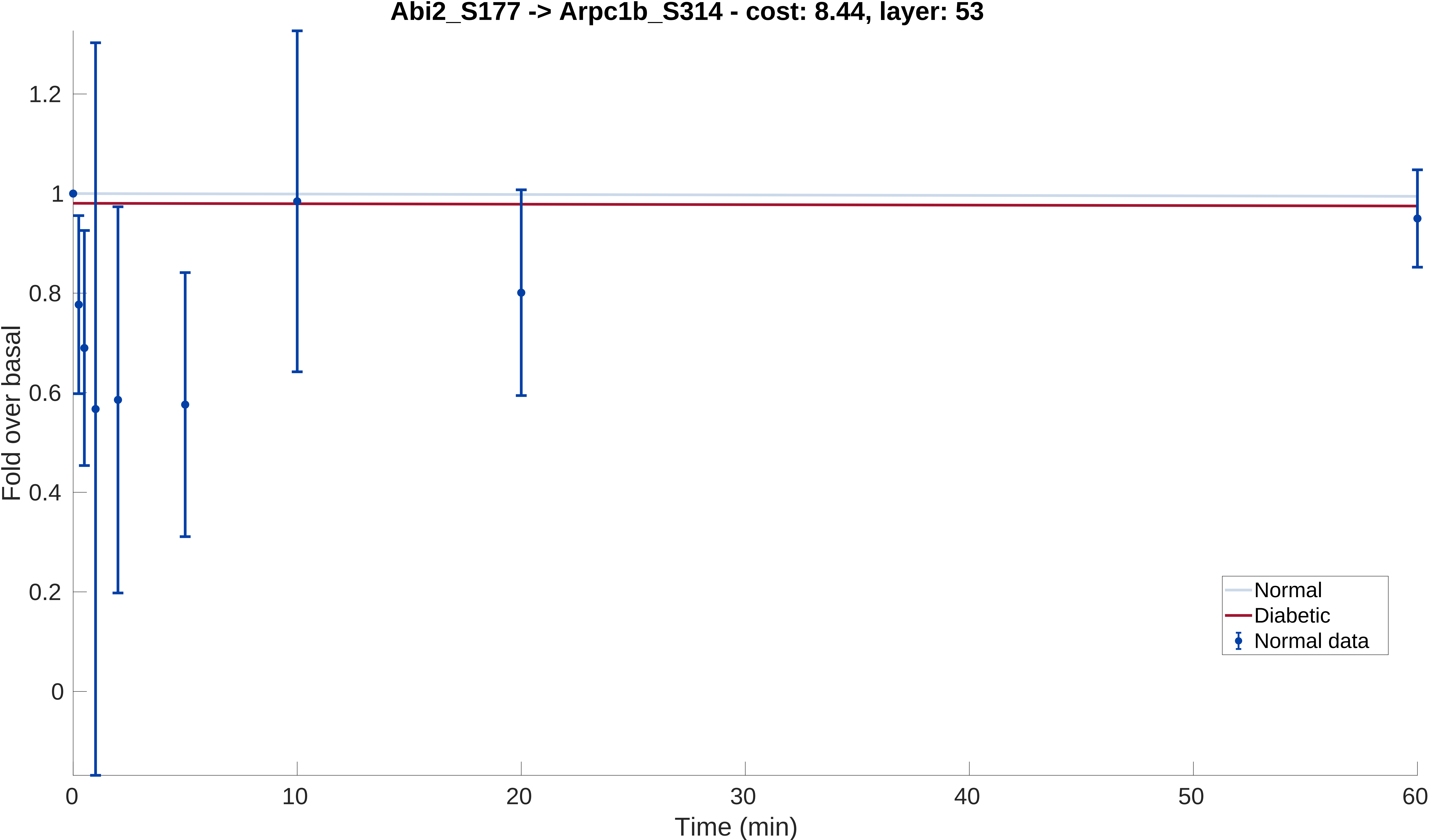
Abi2_S177 -> Arpc1b_S314 - cost: 8.44, layer: 53
Fold over basal
1.2
1
0.8
0.6
0.4
0.2
0
Time (min)
0
10
20
30
40
50
60
Normal
Diabetic
Normal data

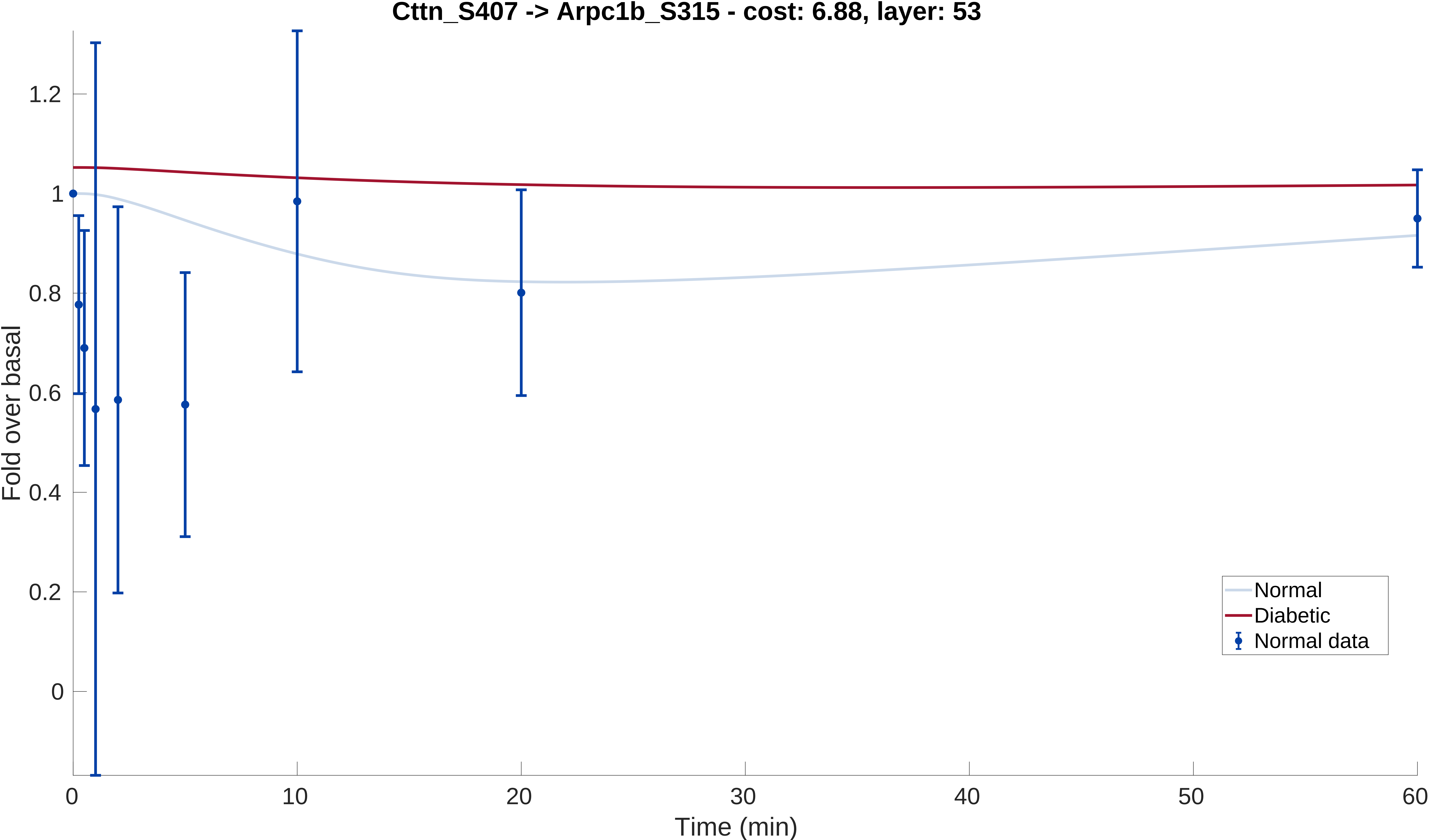

Cttn_S407 -> Arpc1b_S315 - cost: 6.88, layer: 53
Fold over basal
1.2
1
0.8
0.6
0.4
0.2
0
0
10
20
30
40
50
60
Time (min)
Normal
Diabetic
Normal data

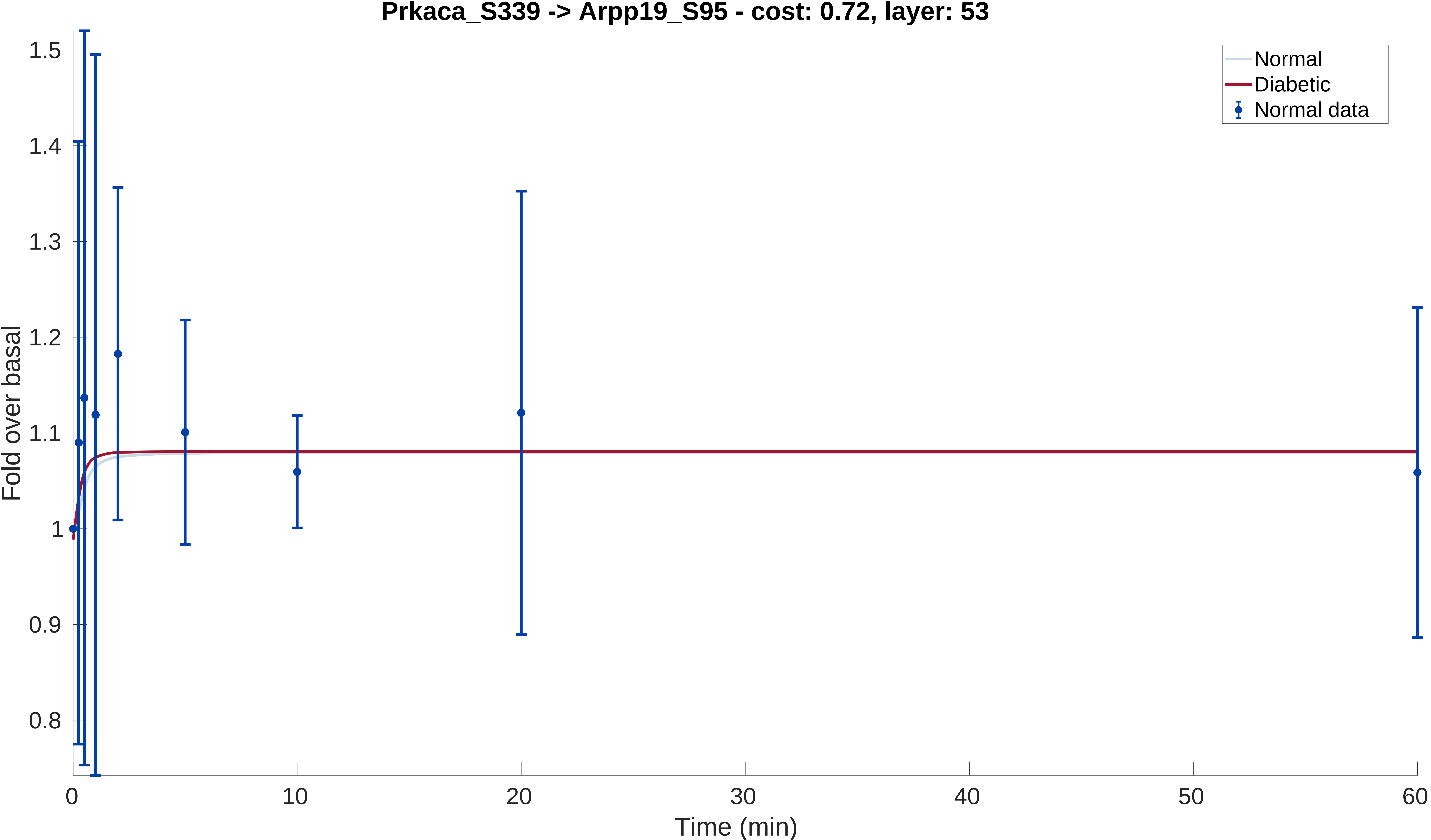
Prkaca_S339 -> Arpp19_S95 - cost: 0.72, layer: 53
Normal
Diabetic
Normal data
Fold over basal
1.5
1.4
1.3
1.2
1.1
1
0.9
0.8
0
10
20
30
40
50
60
Time (min)

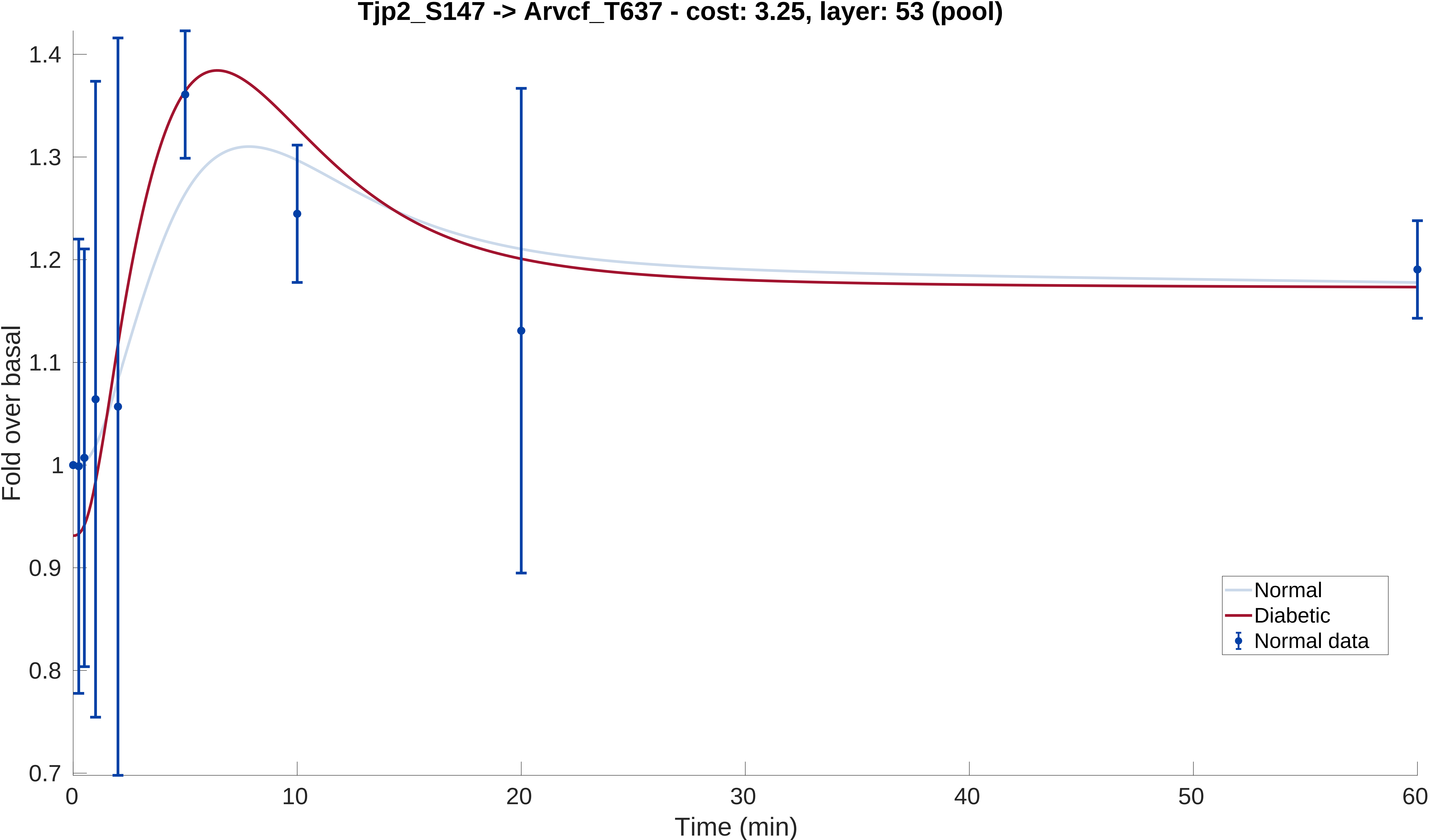

Tjp2_S147 -> Arvcf_T637 - cost: 3.25, layer: 53 (pool)
Fold over basal
1.4
1.3
1.2
1.1
1
0.9
0.8
0.7
0
10
20
30
40
50
60
Time (min)
Normal
Diabetic
Normal data

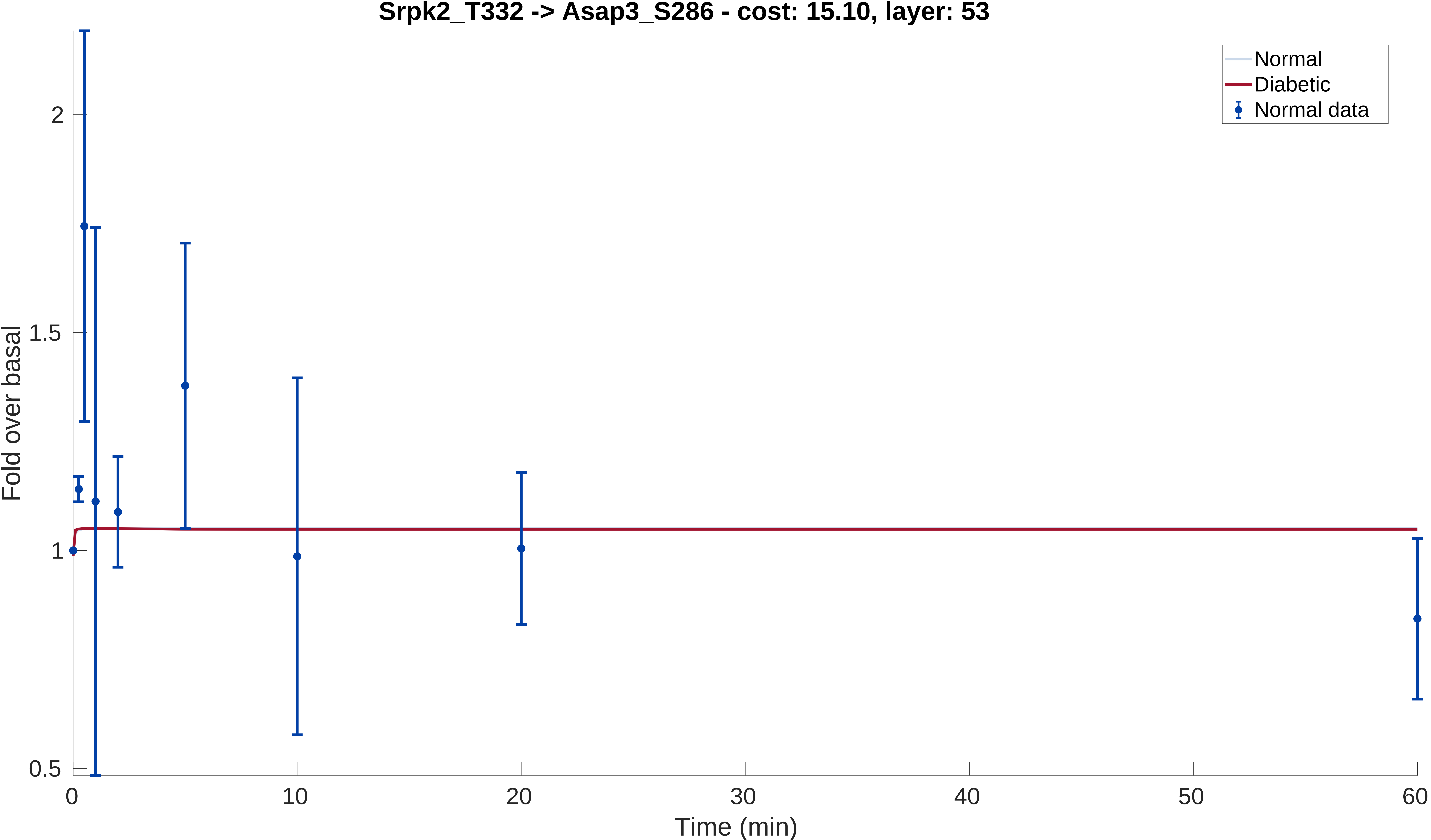

Srpk2_T332 -> Asap3_S286 - cost: 15.10, layer: 53
Normal
Diabetic
Normal data
Fold over basal
0.5
1
1.5
2
0
10
20
30
40
50
60
Time (min)

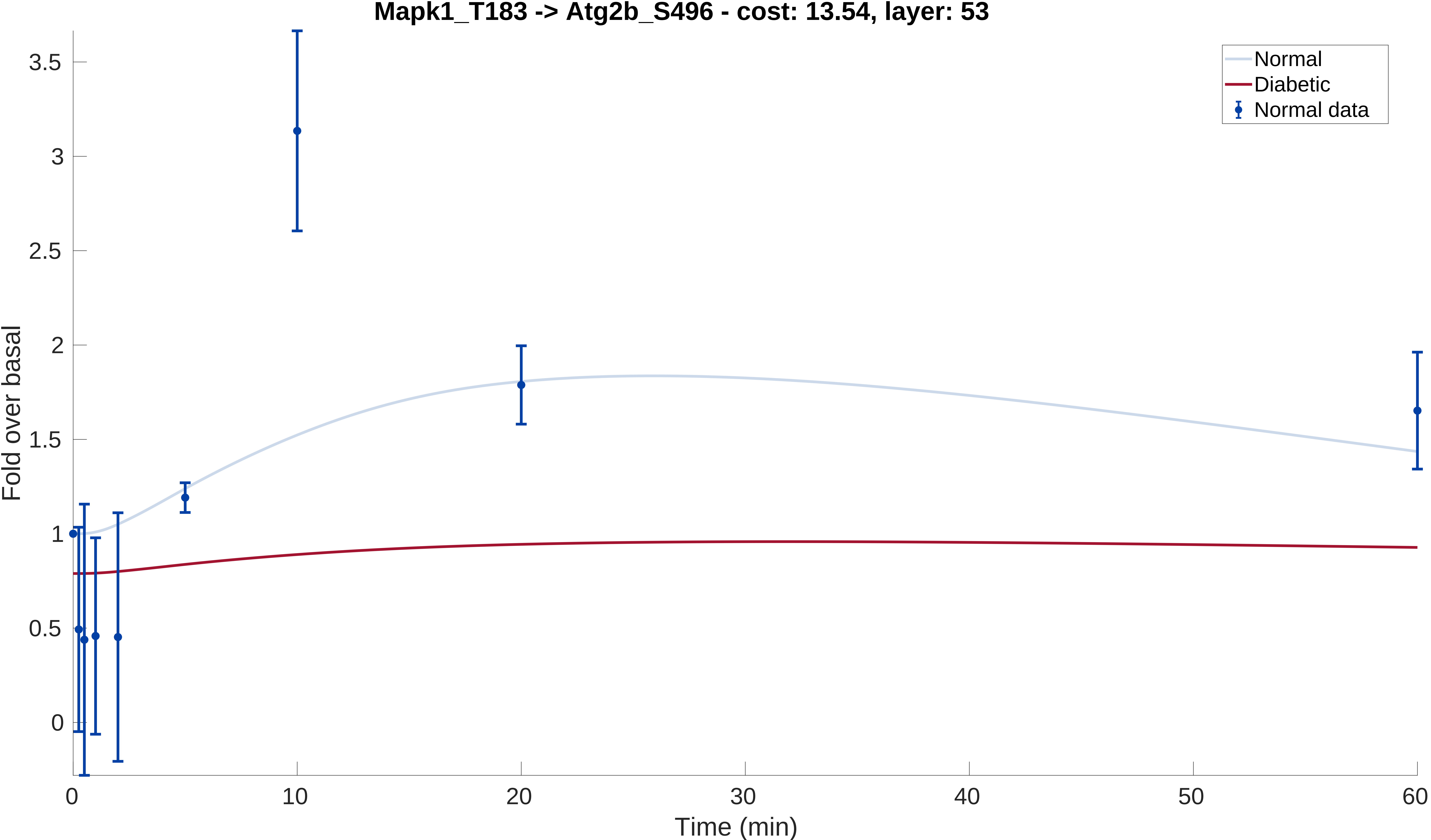

Mapk1_T183 -> Atg2b_S496 - cost: 13.54, layer: 53
Normal
Diabetic
Normal data
Fold over basal
Time (min)
3.5
3
2.5
2
1.5
1
0.5
0
0
10
20
30
40
50
60

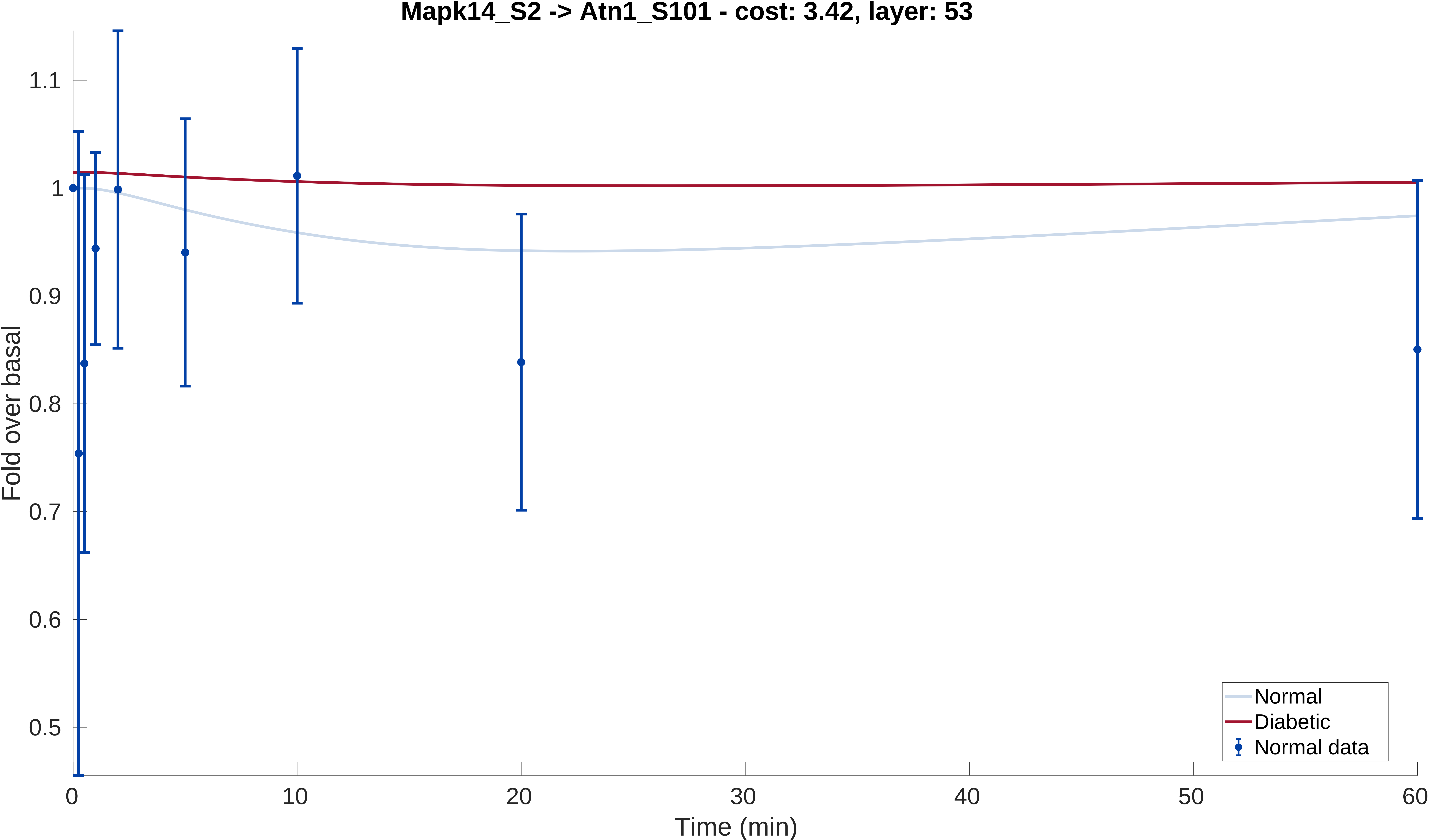

Mapk14_S2 -> Atn1_S101 - cost: 3.42, layer: 53
1.1
1
0.9
0.8
0.7
0.6
0.5
Fold over basal
0
10
20
30
40
50
60
Time (min)
Normal
Diabetic
Normal data

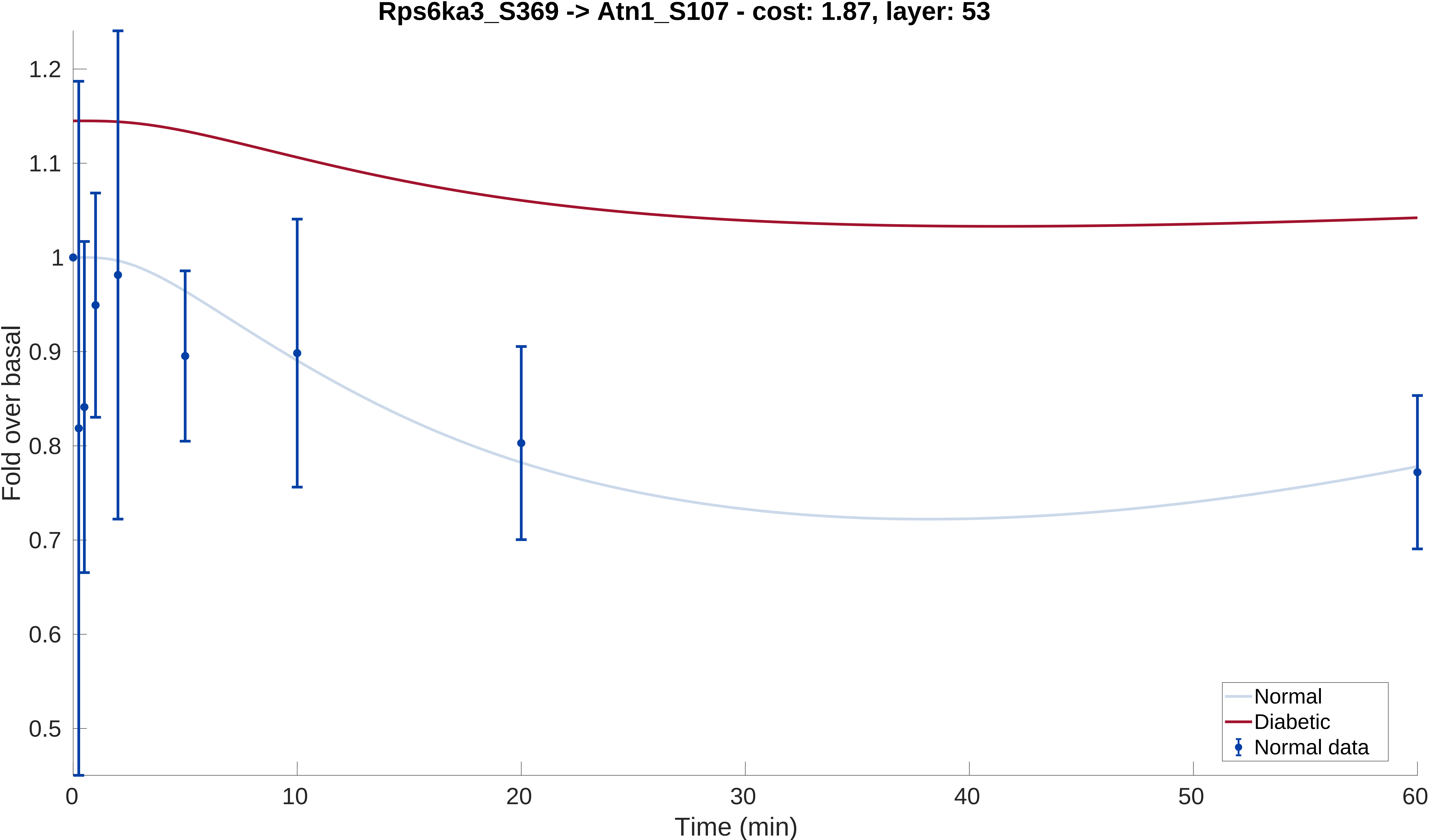

Rps6ka3_S369 -> Atn1_S107 - cost: 1.87, layer: 53
Fold over basal
1.2
1.1
1
0.9
0.8
0.7
0.6
0.5
0
10
20
30
40
50
60
Time (min)
Normal
Diabetic
Normal data

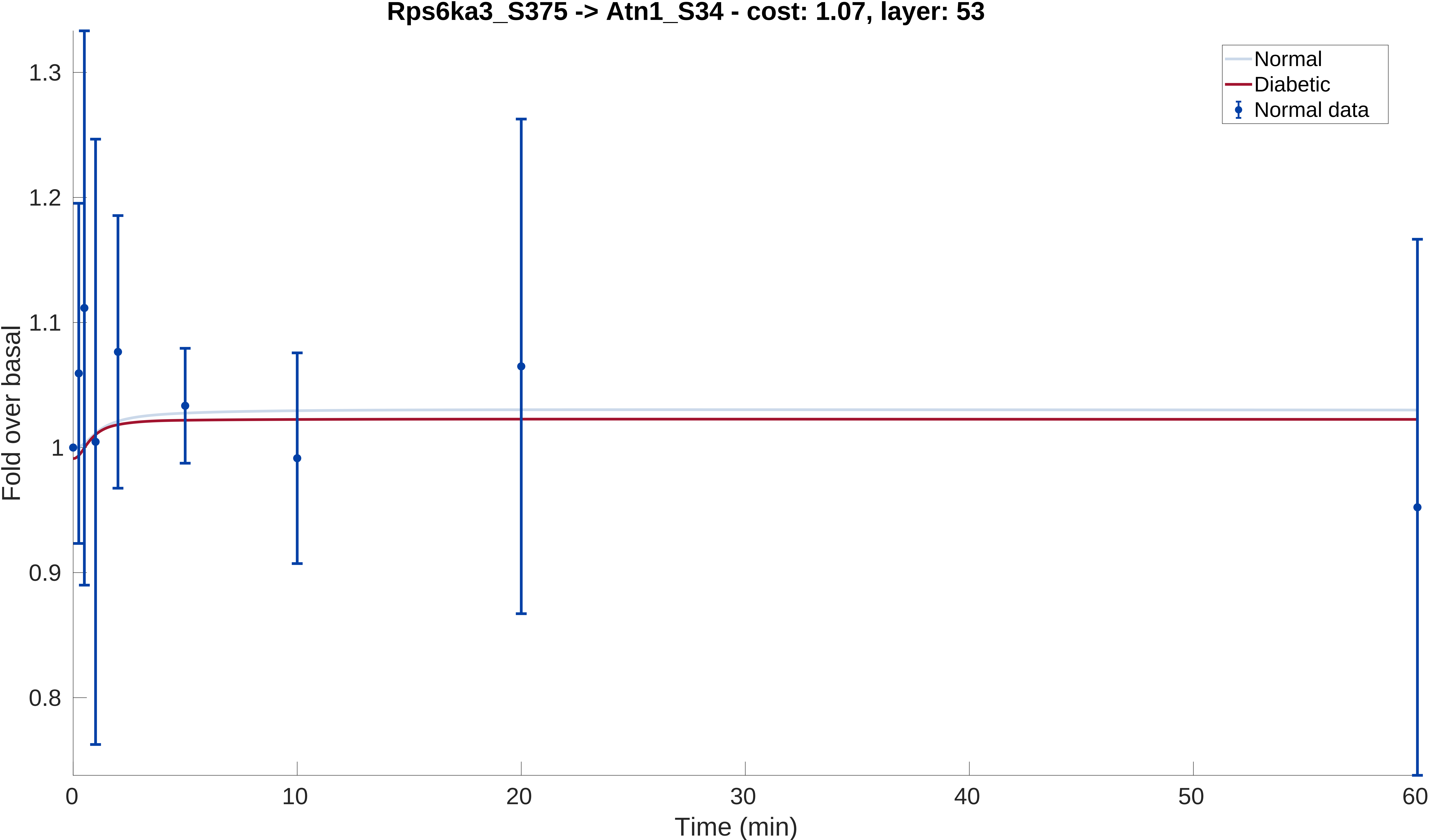
Rps6ka3_S375 -> Atn1_S34 - cost: 1.07, layer: 53
Normal
Diabetic
Normal data
Fold over basal
1.3
1.2
1.1
1
0.9
0.8
0
10
20
30
40
50
60
Time (min)

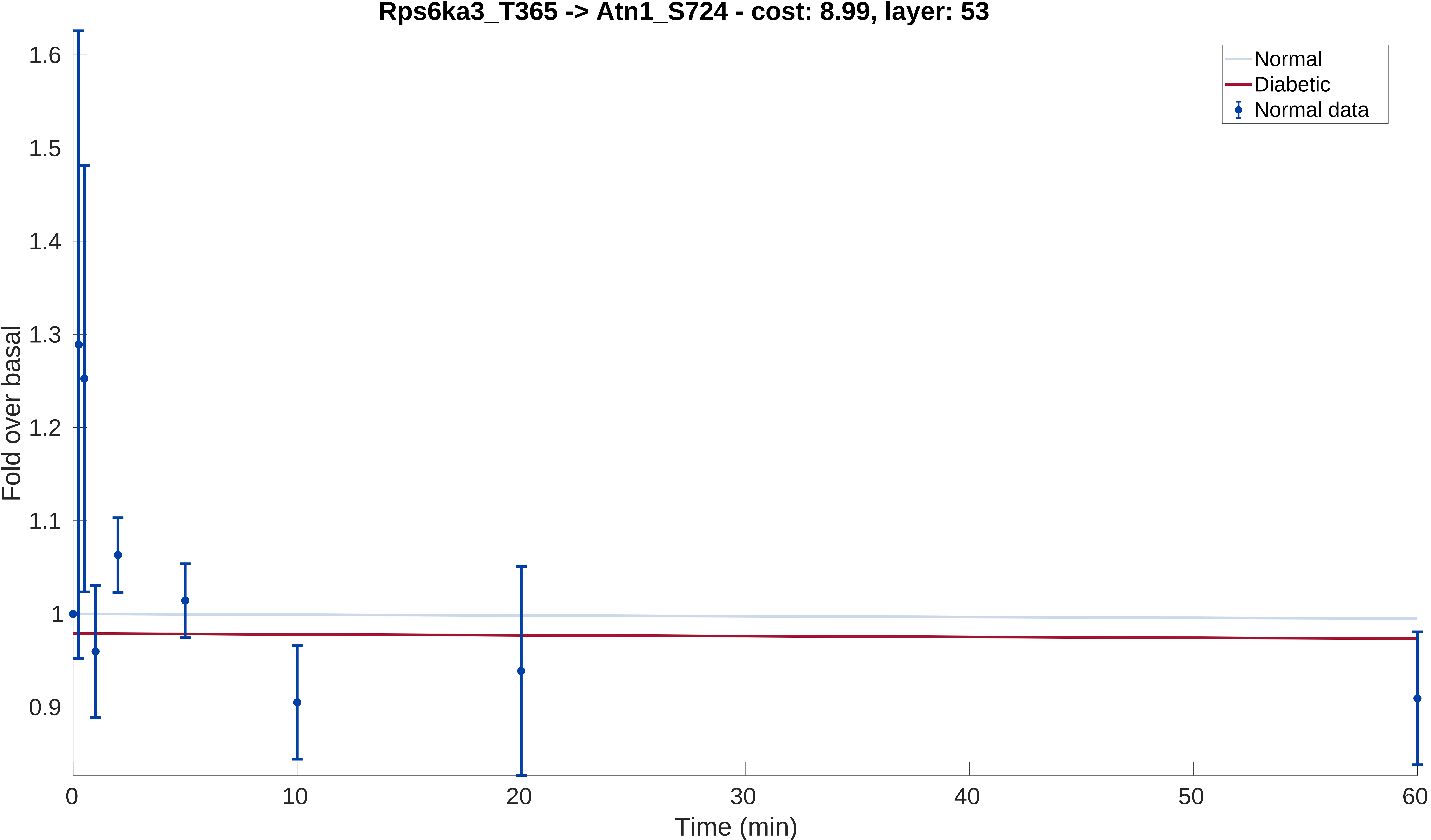

Rps6ka3_T365 -> Atn1_S724 - cost: 8.99, layer: 53
Normal
Diabetic
Normal data
Fold over basal
1.6
1.5
1.4
1.3
1.2
1.1
1
0.9
0
10
20
30
40
50
60
Time (min)

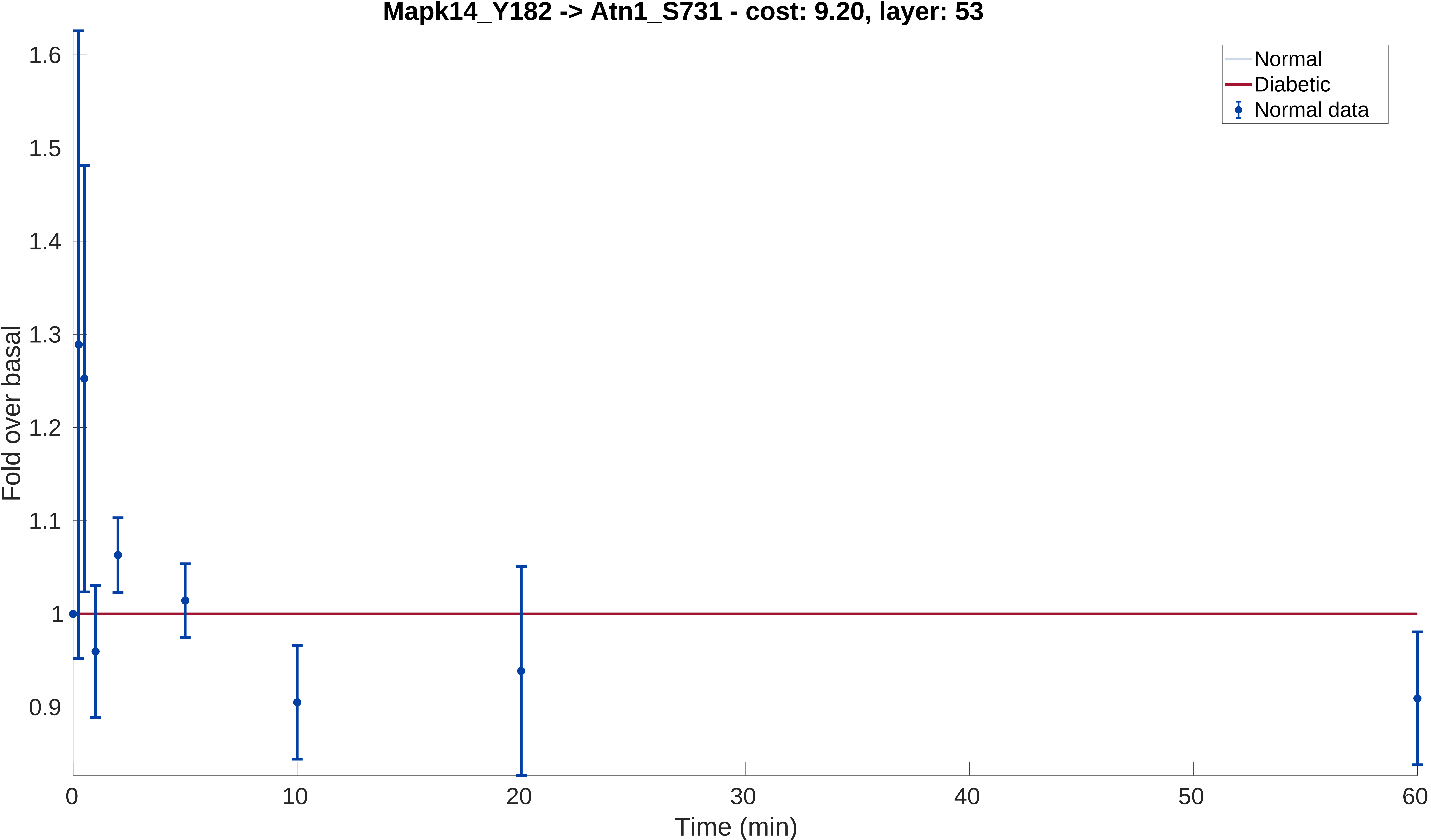

Mapk14_Y182 -> Atn1_S731 - cost: 9.20, layer: 53
Normal
Diabetic
Normal data
Fold over basal
1.6
1.5
1.4
1.3
1.2
1.1
1
0.9
0
10
20
30
40
50
60
Time (min)

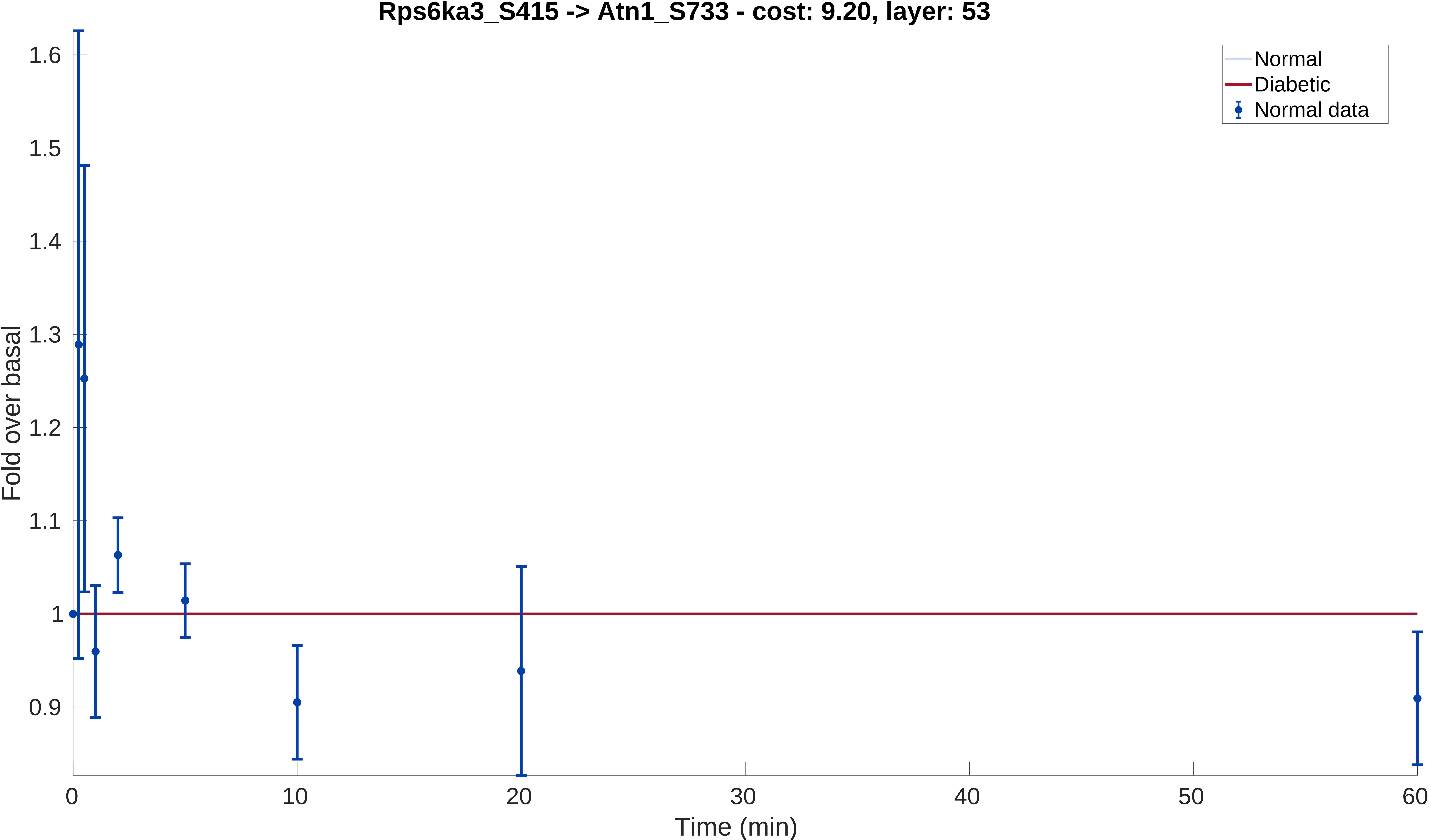

Rps6ka3_S415 -> Atn1_S733 - cost: 9.20, layer: 53
Fold over basal
1.6
1.5
1.4
1.3
1.2
1.1
1
0.9
0
10
20
30
40
50
60
Time (min)
Normal
Diabetic
Normal data

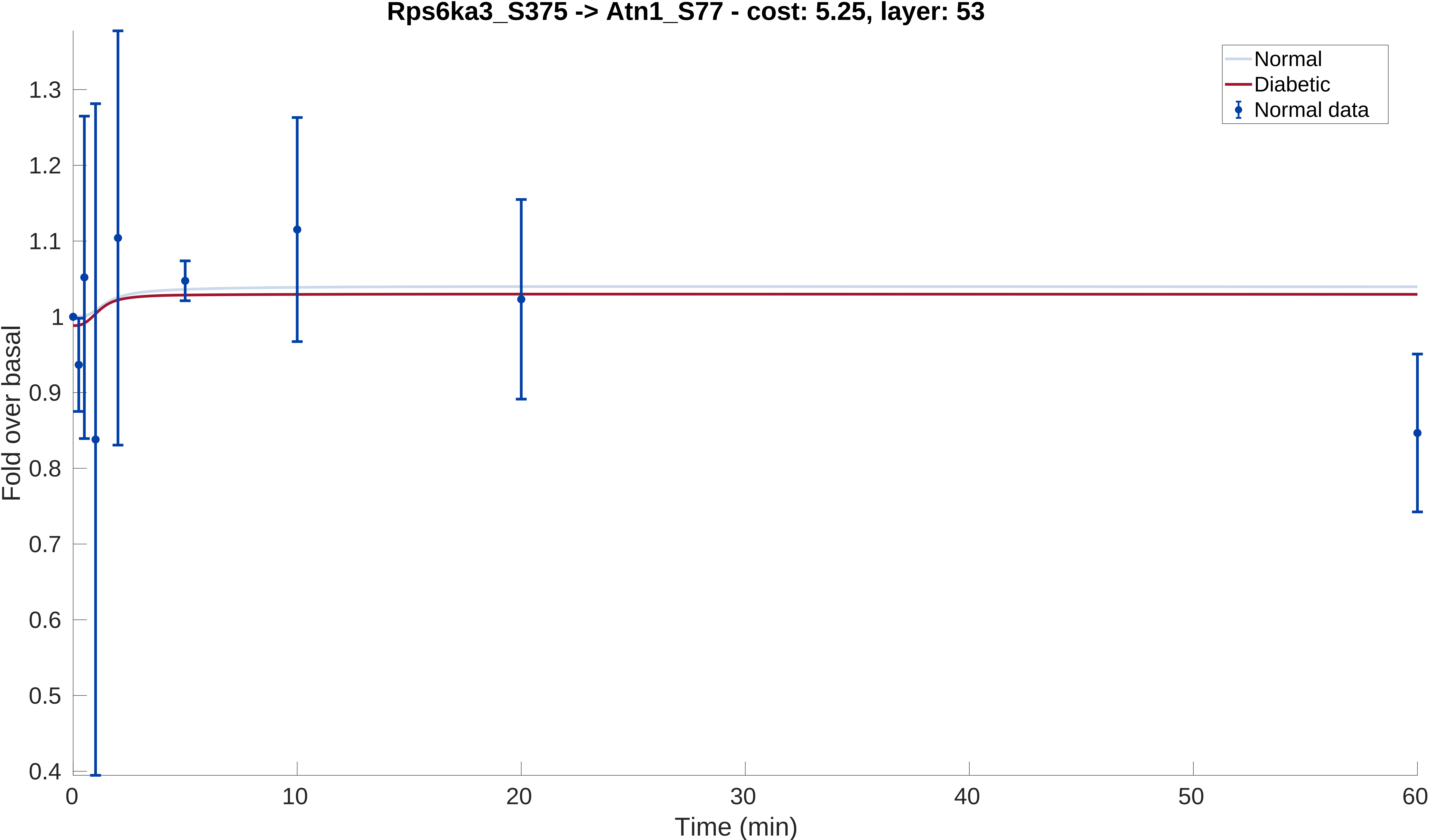

Rps6ka3_S375 -> Atn1_S77 - cost: 5.25, layer: 53
Normal
Diabetic
Normal data
Fold over basal
1.3
1.2
1.1
1
0.9
0.8
0.7
0.6
0.5
0.4
0
10
20
30
40
50
60
Time (min)

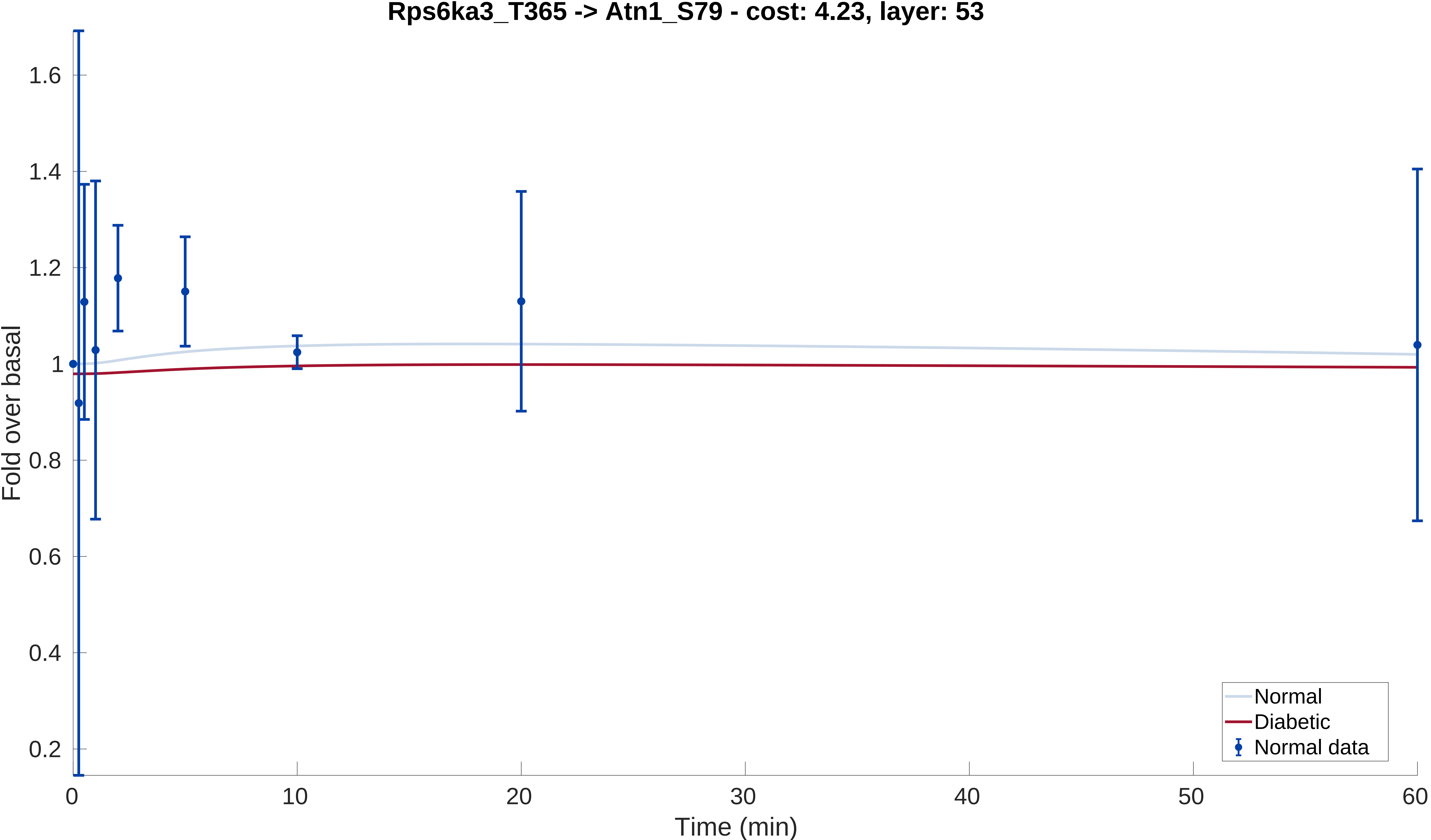
Rps6ka3_T365 -> Atn1_S79 - cost: 4.23, layer: 53
Fold over basal
1.6
1.4
1.2
1
0.8
0.6
0.4
0.2
0
10
20
30
40
50
60
Time (min)
Normal
Diabetic
Normal data

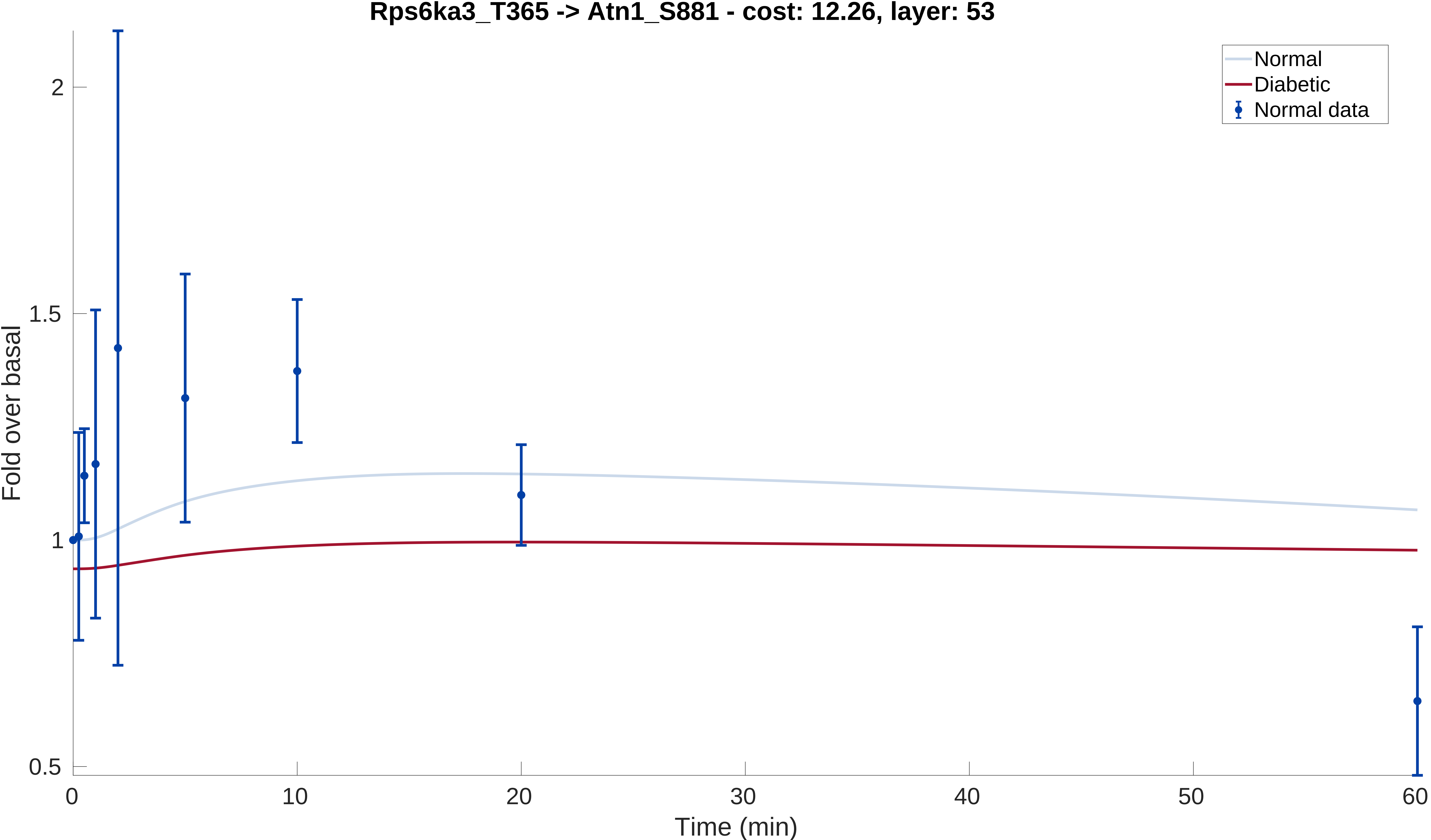

Rps6ka3_T365 -> Atn1_S881 - cost: 12.26, layer: 53
Normal
Diabetic
Normal data
Fold over basal
Time (min)
0.5
1
1.5
2
0
10
20
30
40
50
60

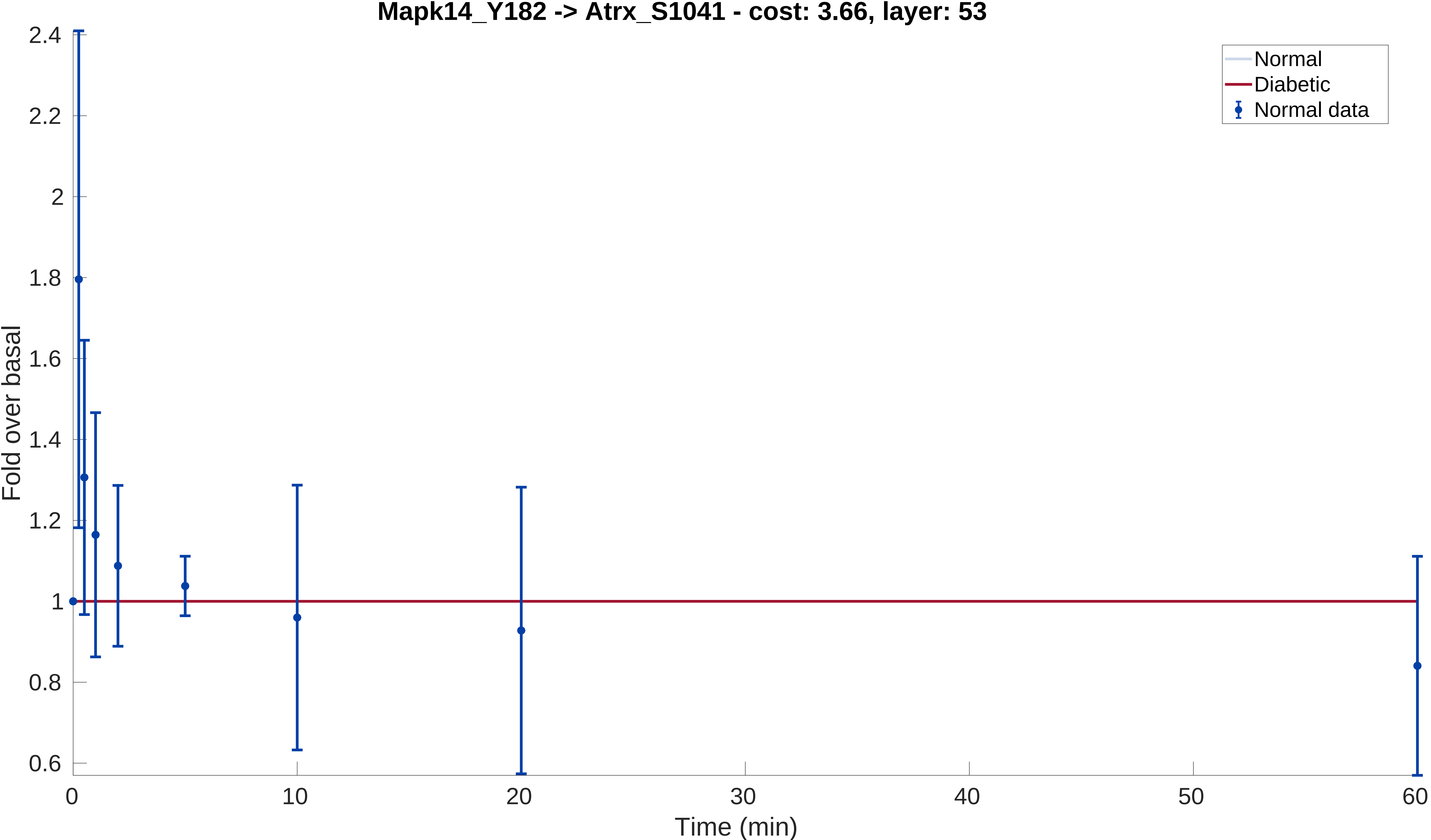

Mapk14_Y182 -> Atrx_S1041 - cost: 3.66, layer: 53
Normal
Diabetic
Normal data
Fold over basal
Time (min)
2.4
2.2
2
1.8
1.6
1.4
1.2
1
0.8
0.6
0
10
20
30
40
50
60

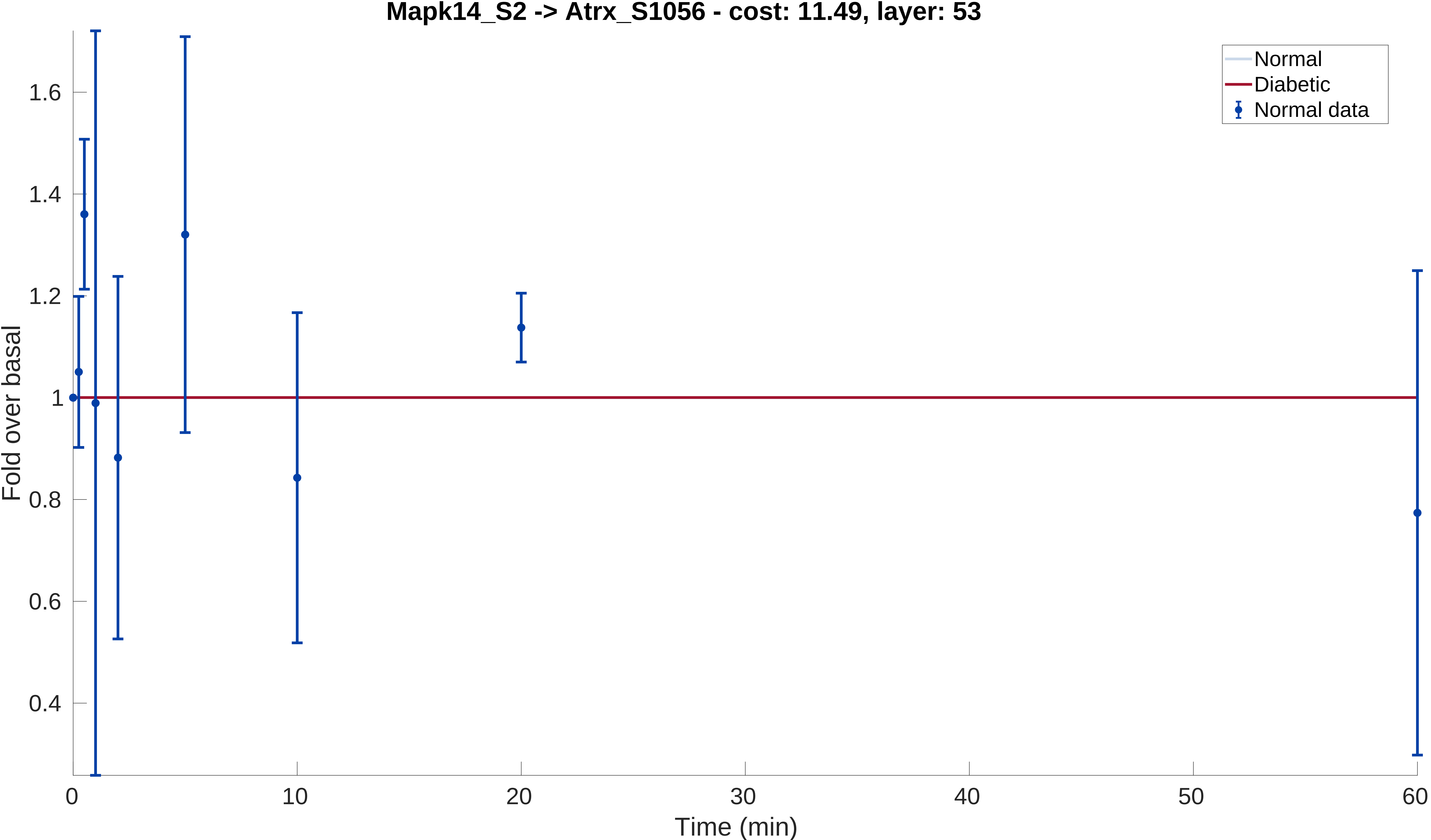

Mapk14_S2 -> Atrx_S1056 - cost: 11.49, layer: 53
Normal
Diabetic
Normal data
Fold over basal
1.6
1.4
1.2
1
0.8
0.6
0.4
0
10
20
30
40
50
60
Time (min)

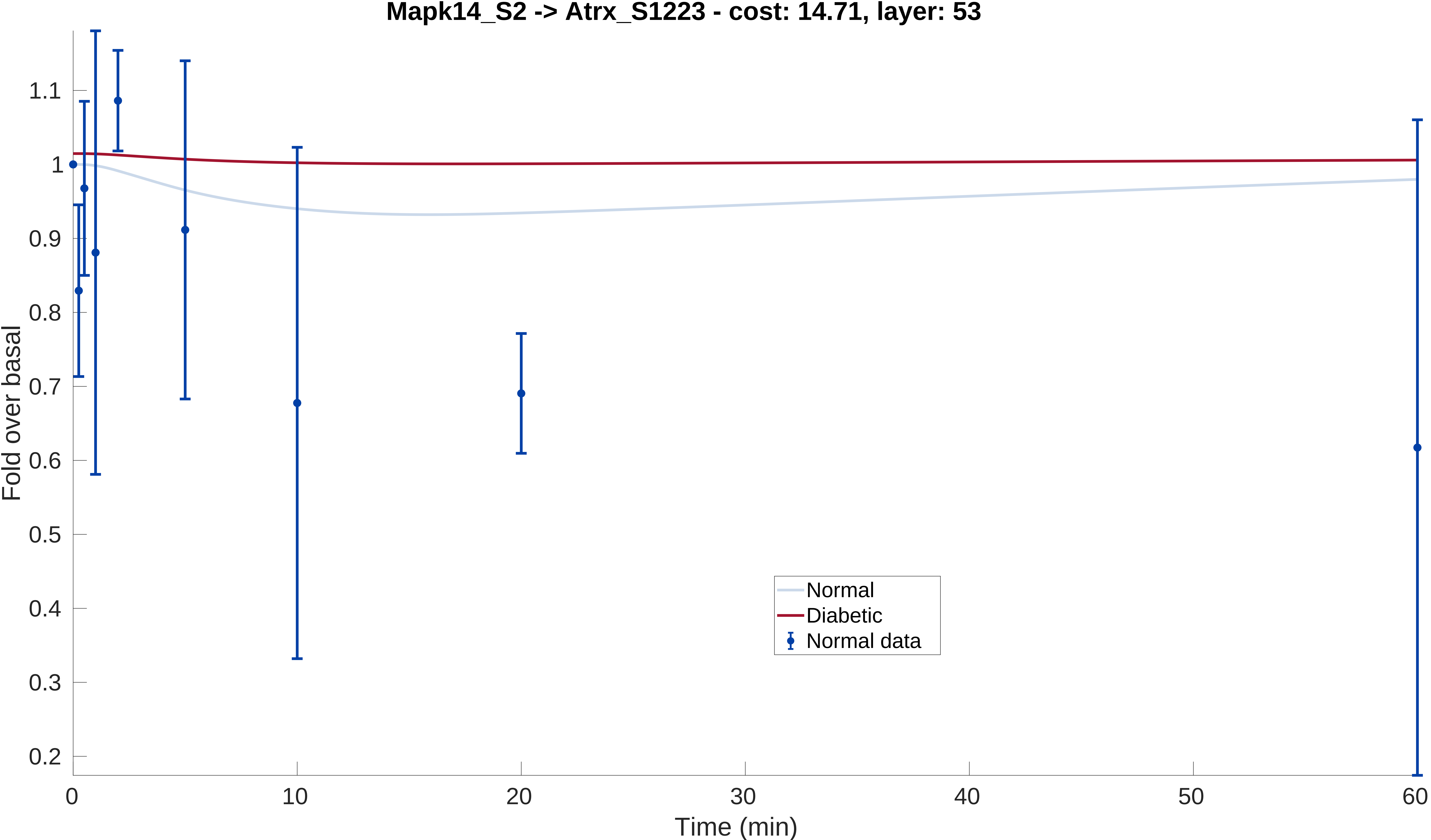
Mapk14_S2 -> Atrx_S1223 - cost: 14.71, layer: 53
Fold over basal
1.1
1
0.9
0.8
0.7
0.6
0.5
0.4
0.3
0.2
0
10
20
30
40
50
60
Time (min)
Normal
Diabetic
Normal data

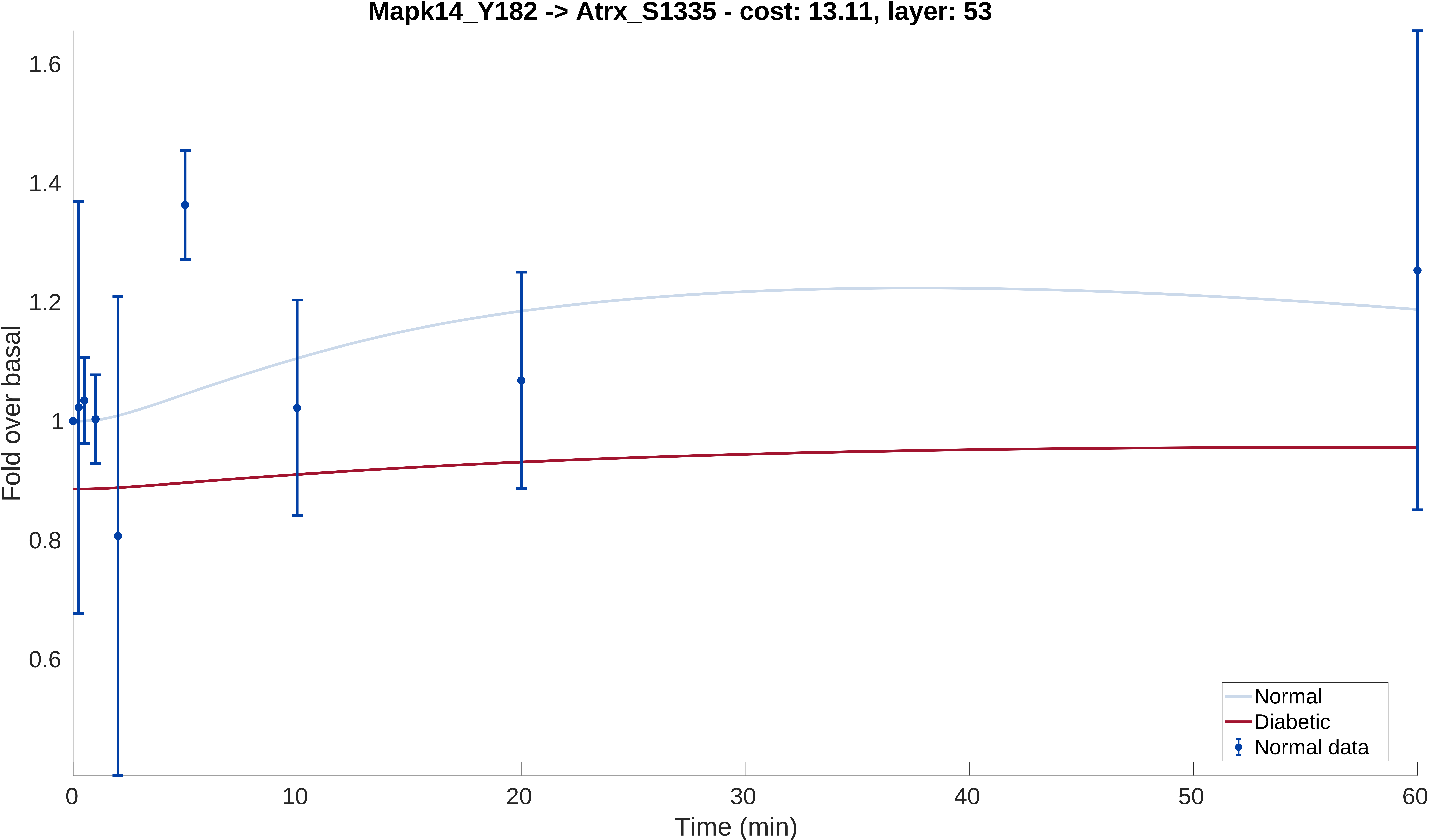

Mapk14_Y182 -> Atrx_S1335 - cost: 13.11, layer: 53
Fold over basal
1.6
1.4
1.2
1
0.8
0.6
0
10
20
30
40
50
60
Time (min)
Normal
Diabetic
Normal data

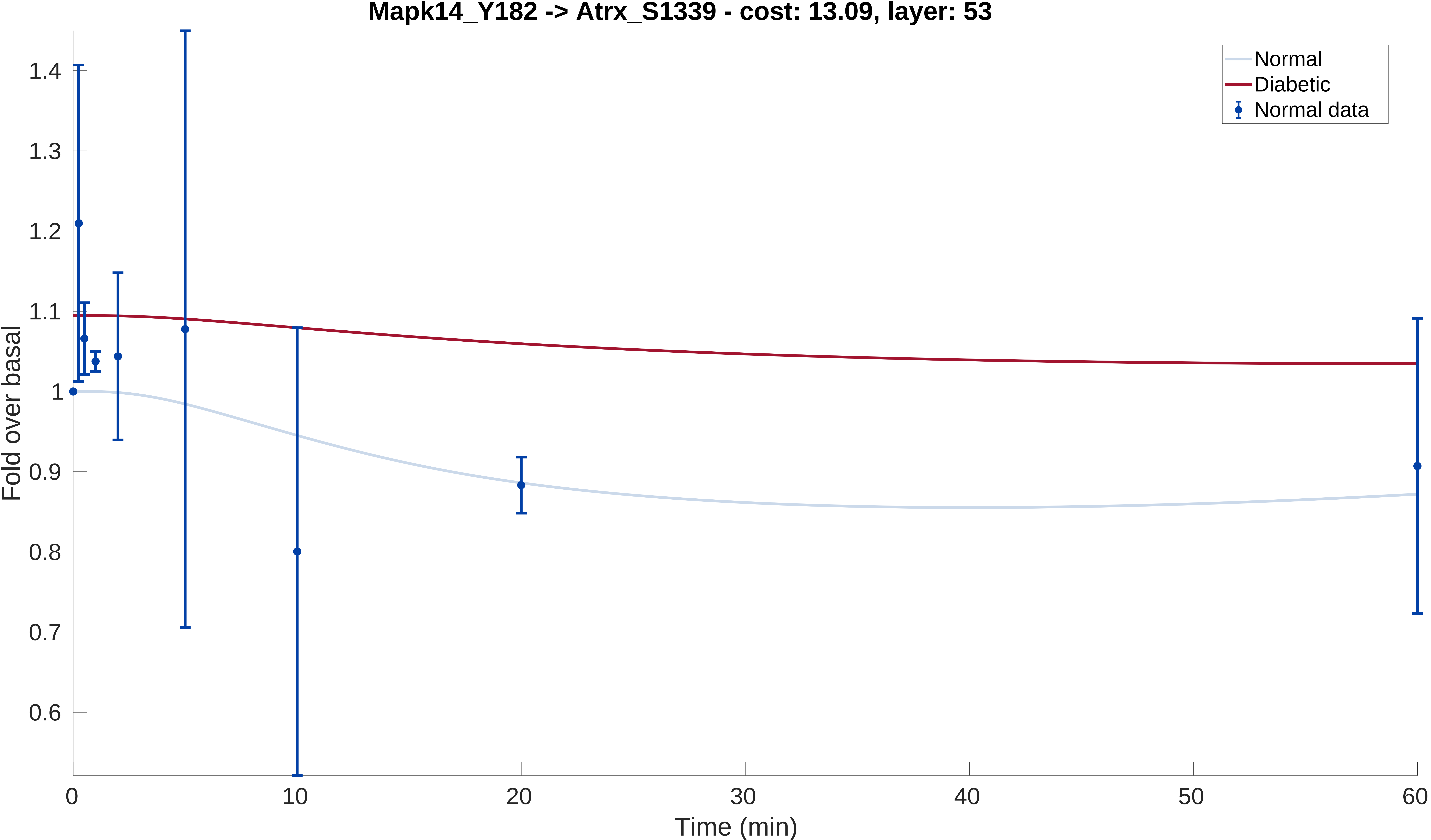

Mapk14_Y182 -> Atrx_S1339 - cost: 13.09, layer: 53
Normal
Diabetic
Normal data
Fold over basal
1.4
1.3
1.2
1.1
1
0.9
0.8
0.7
0.6
0
10
20
30
40
50
60
Time (min)

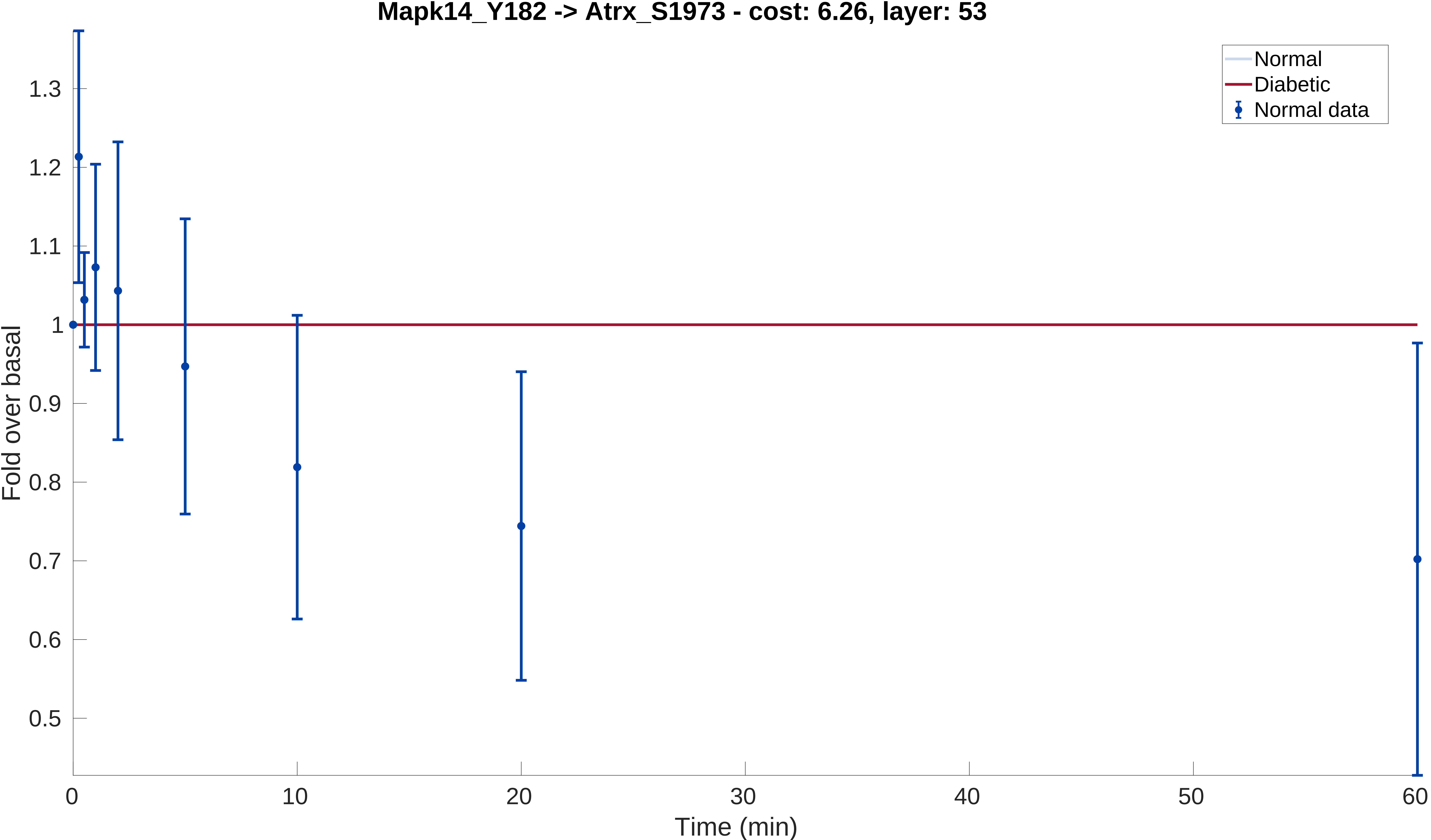

Mapk14_Y182 -> Atrx_S1973 - cost: 6.26, layer: 53
Normal
Diabetic
Normal data
Fold over basal
1.3
1.2
1.1
1
0.9
0.8
0.7
0.6
0.5
0
10
20
30
40
50
60
Time (min)

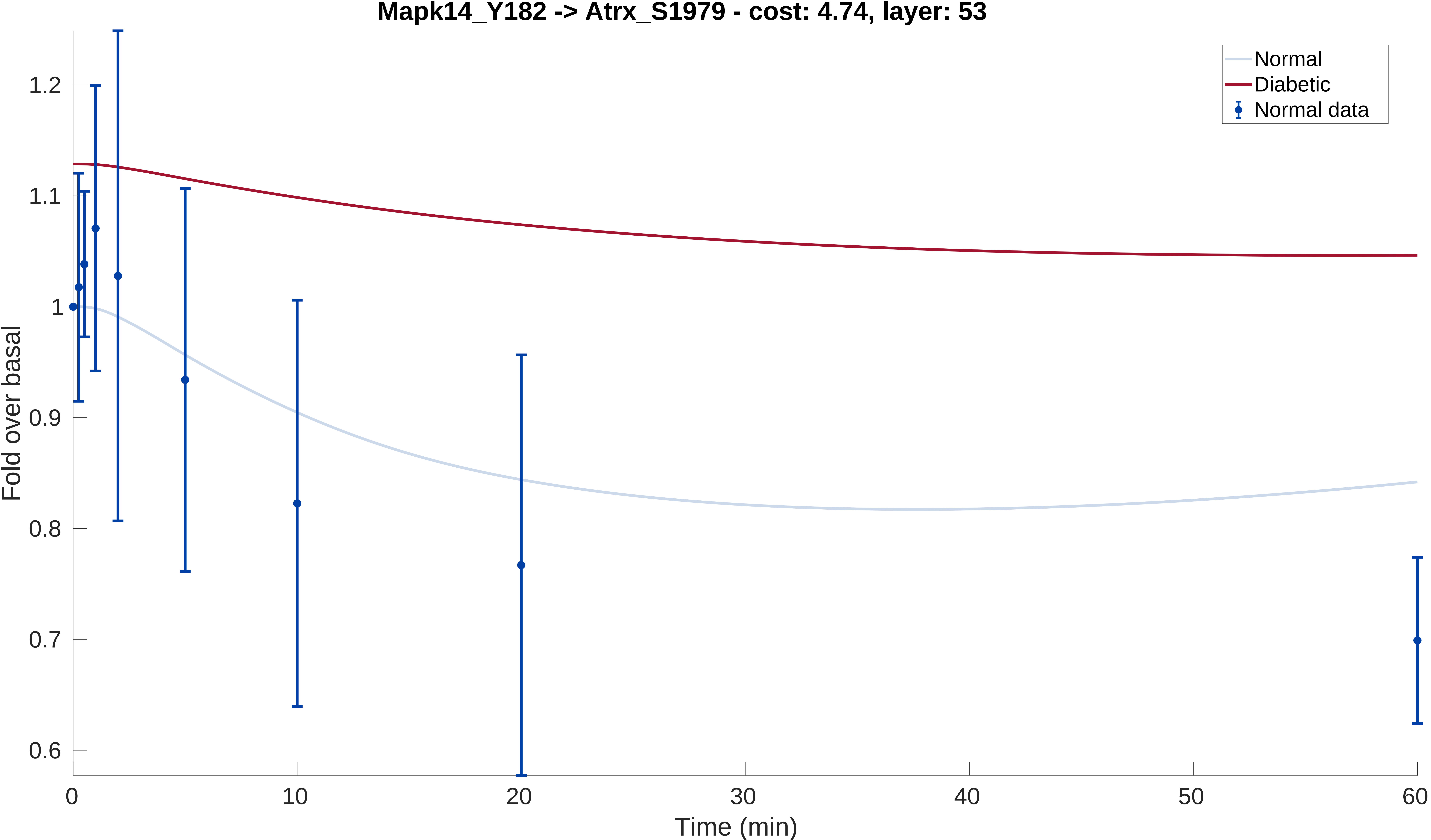
Mapk14_Y182 -> Atrx_S1979 - cost: 4.74, layer: 53
Normal
Diabetic
Normal data
Fold over basal
1.2
1.1
1
0.9
0.8
0.7
0.6
0
10
20
30
40
50
60
Time (min)

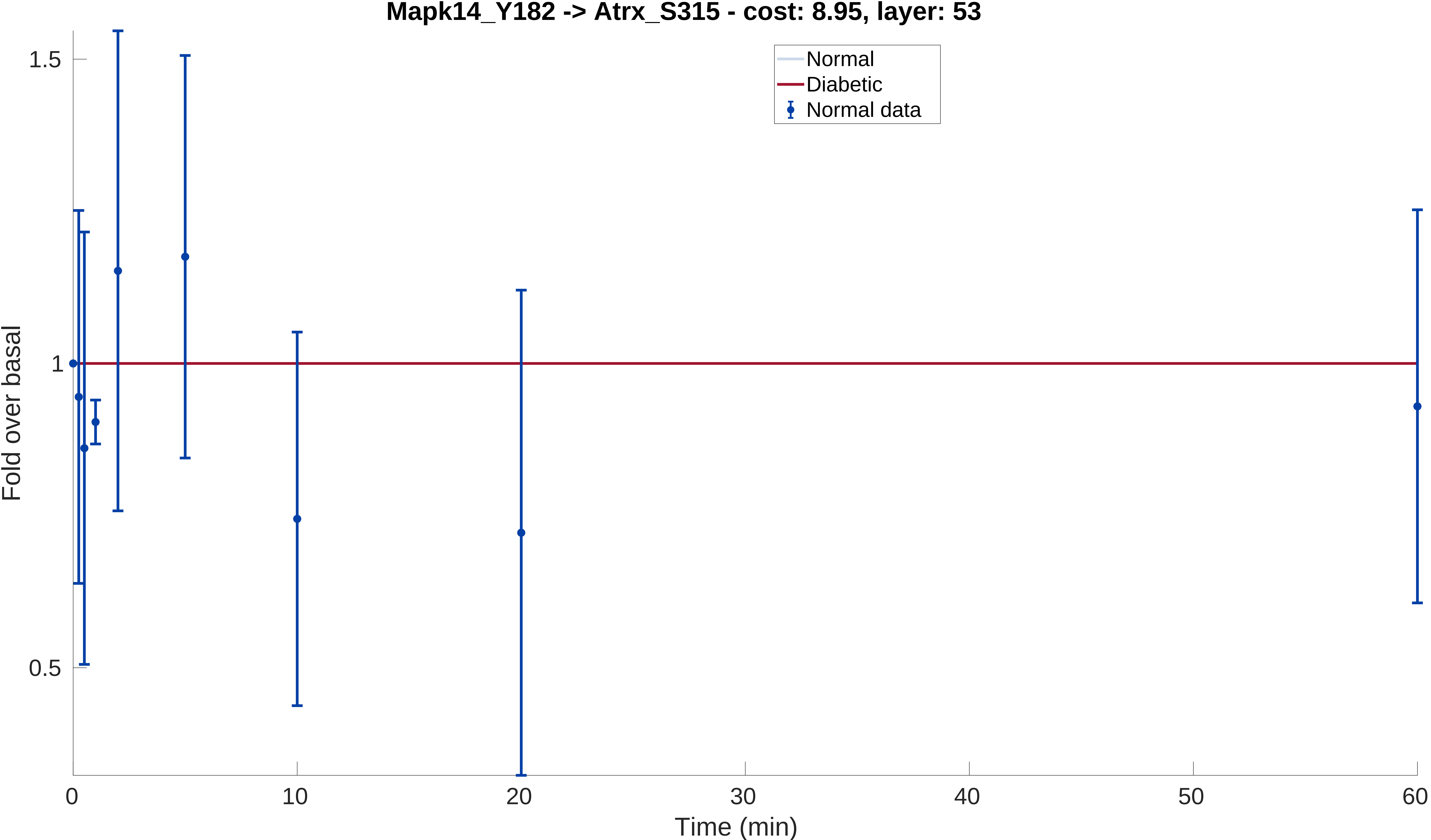

Mapk14_Y182 -> Atrx_S315 - cost: 8.95, layer: 53
Normal
Diabetic
Normal data
1.5
1
0.5
Fold over basal
0
10
20
30
40
50
60
Time (min)

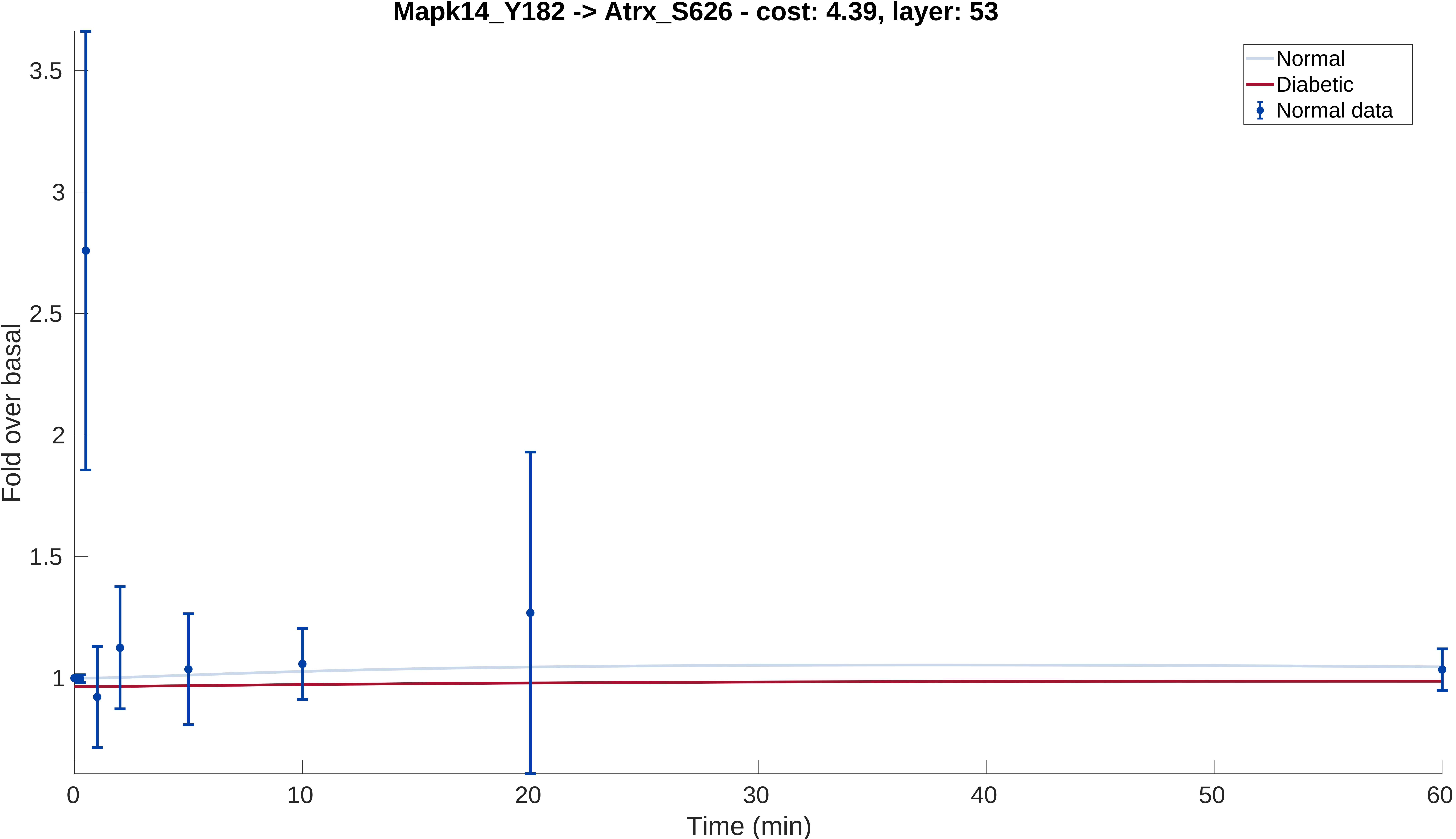

Mapk14_Y182 -> Atrx_S626 - cost: 4.39, layer: 53
Normal
Diabetic
Normal data
Fold over basal
3.5
3
2.5
2
1.5
1
0
10
20
30
40
50
60
Time (min)

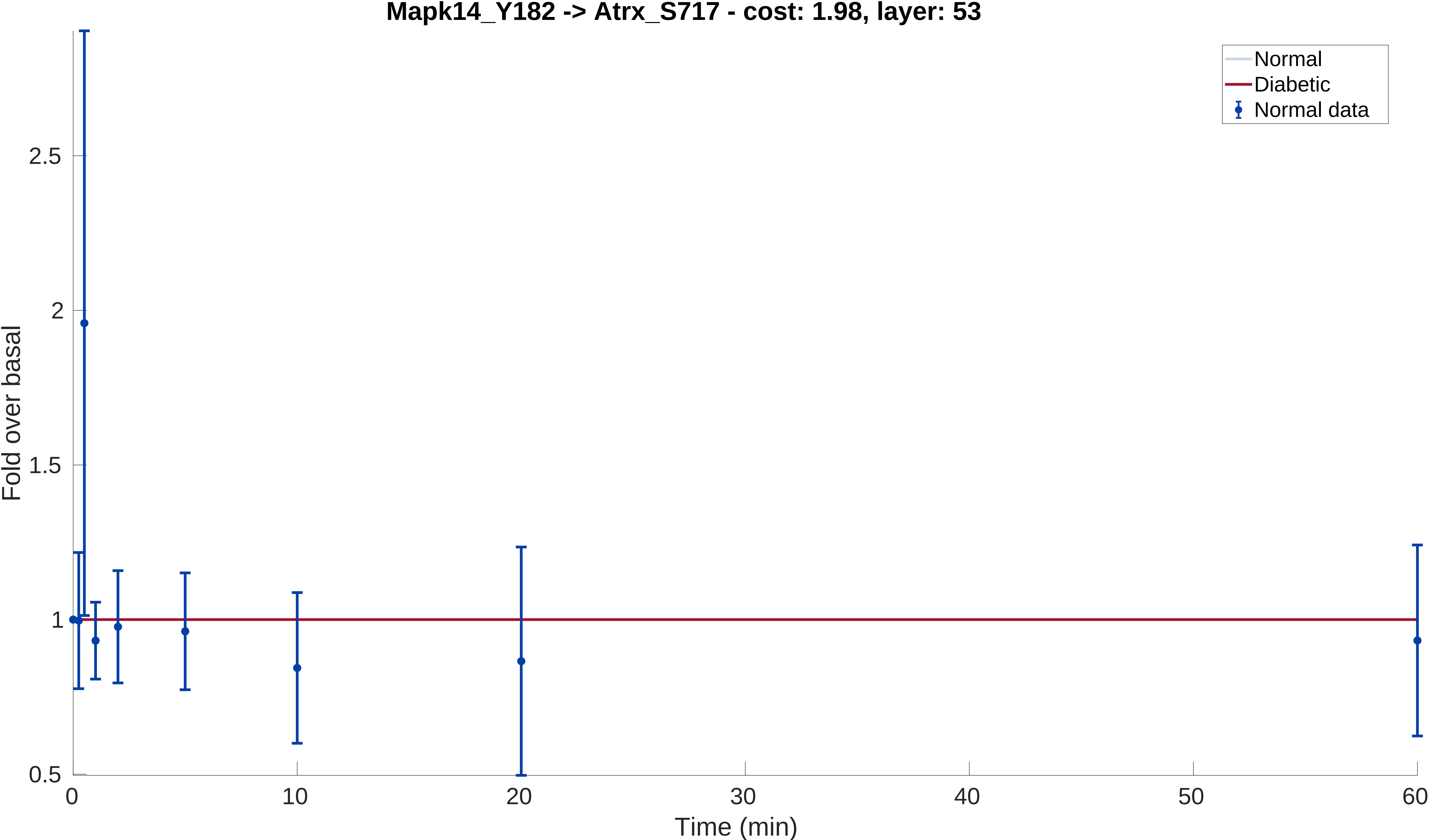

Mapk14_Y182 -> Atrx_S717 - cost: 1.98, layer: 53
Normal
Diabetic
Normal data
Fold over basal
2.5
2
1.5
1
0.5
0
10
20
30
40
50
60
Time (min)

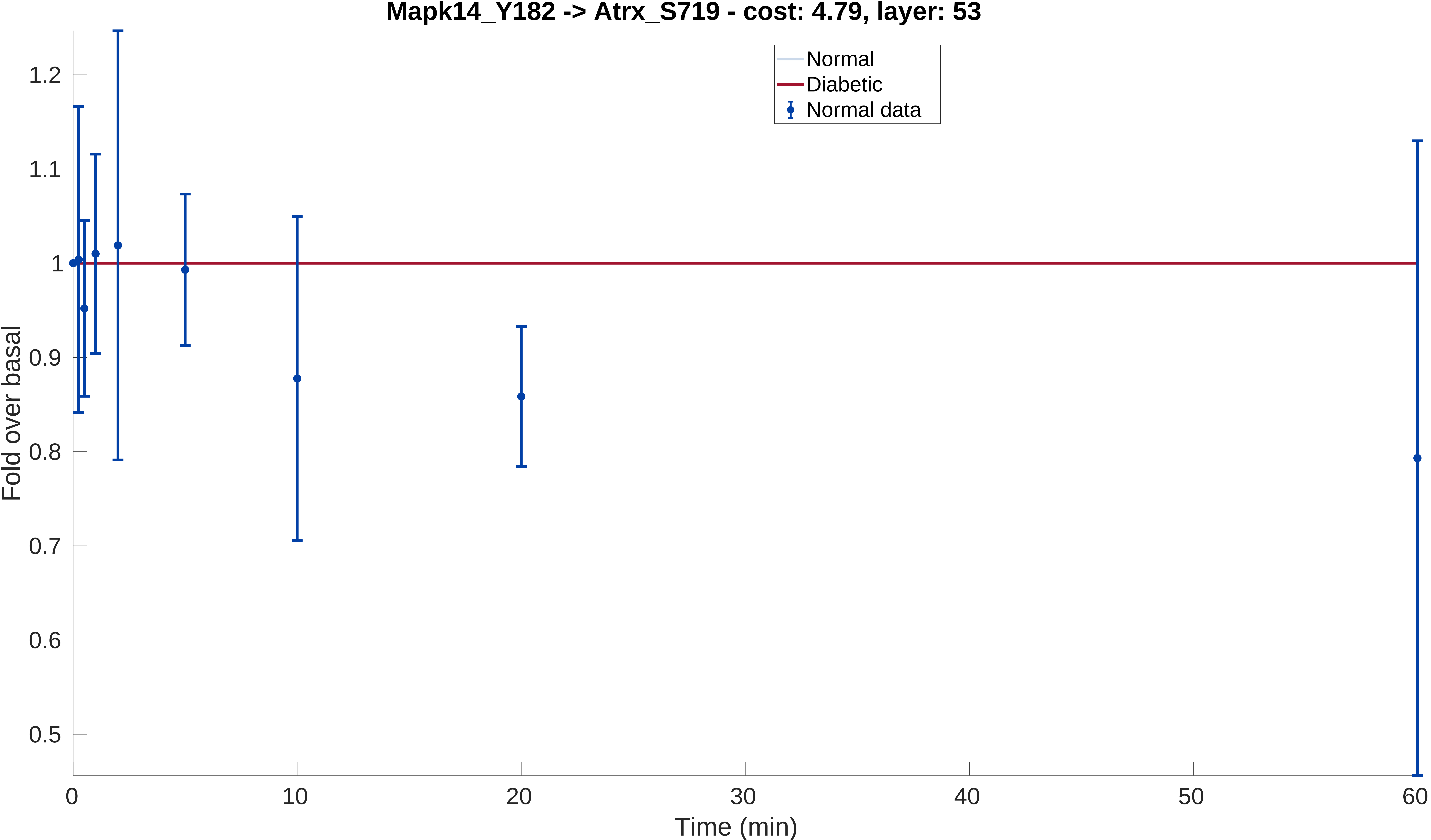

Mapk14_Y182 -> Atrx_S719 - cost: 4.79, layer: 53
Normal
Diabetic
Normal data
Fold over basal
1.2
1.1
1
0.9
0.8
0.7
0.6
0.5
0
10
20
30
40
50
60
Time (min)

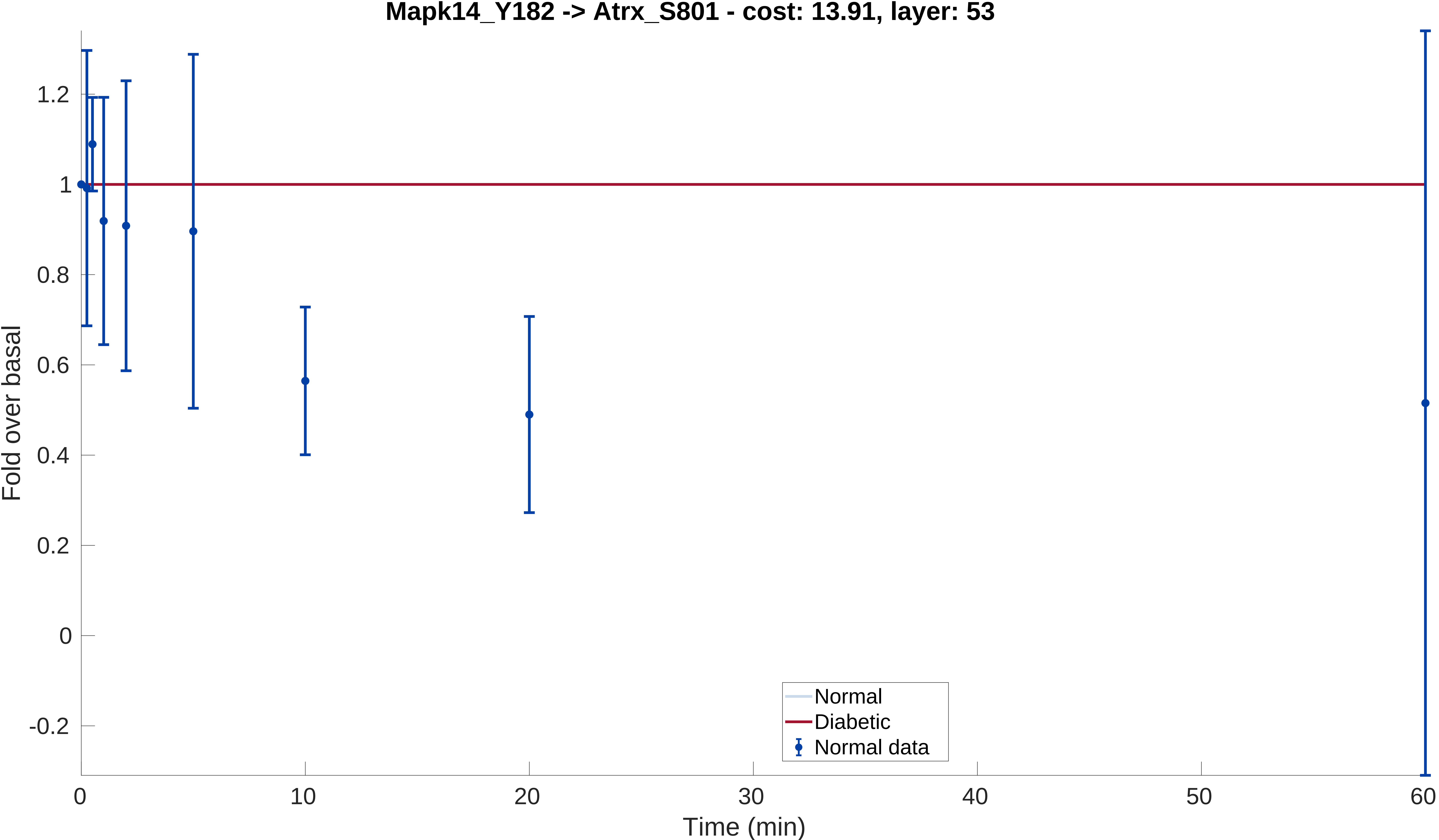

Mapk14_Y182 -> Atrx_S801 - cost: 13.91, layer: 53
Fold over basal
1.2
1
0.8
0.6
0.4
0.2
0
-0.2
0
10
20
30
40
50
60
Time (min)
Normal
Diabetic
Normal data

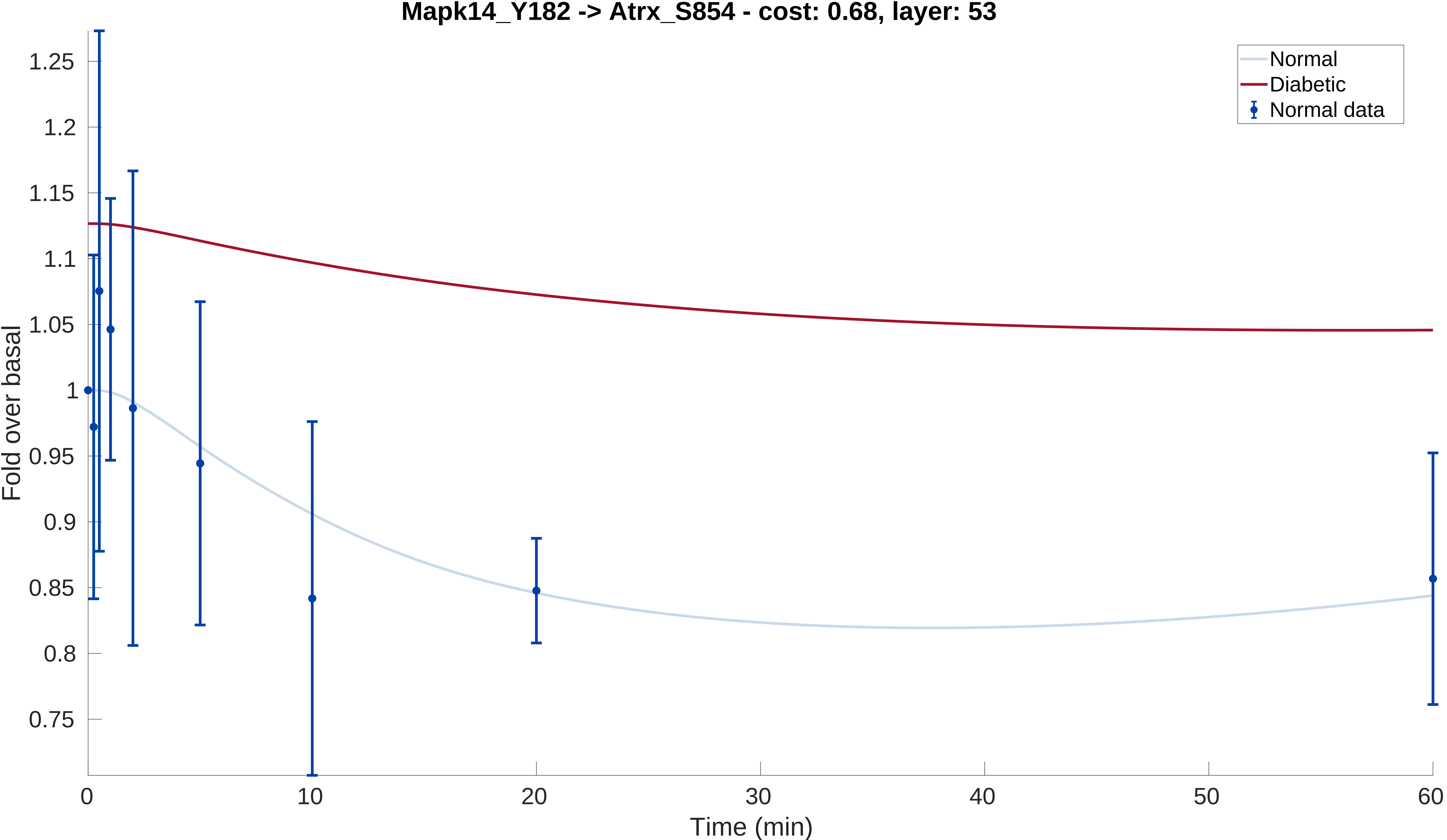

Mapk14_Y182 -> Atrx_S854 - cost: 0.68, layer: 53
Normal
Diabetic
Normal data
Fold over basal
1.25
1.2
1.15
1.1
1.05
1
0.95
0.9
0.85
0.8
0.75
0
10
20
30
40
50
60
Time (min)

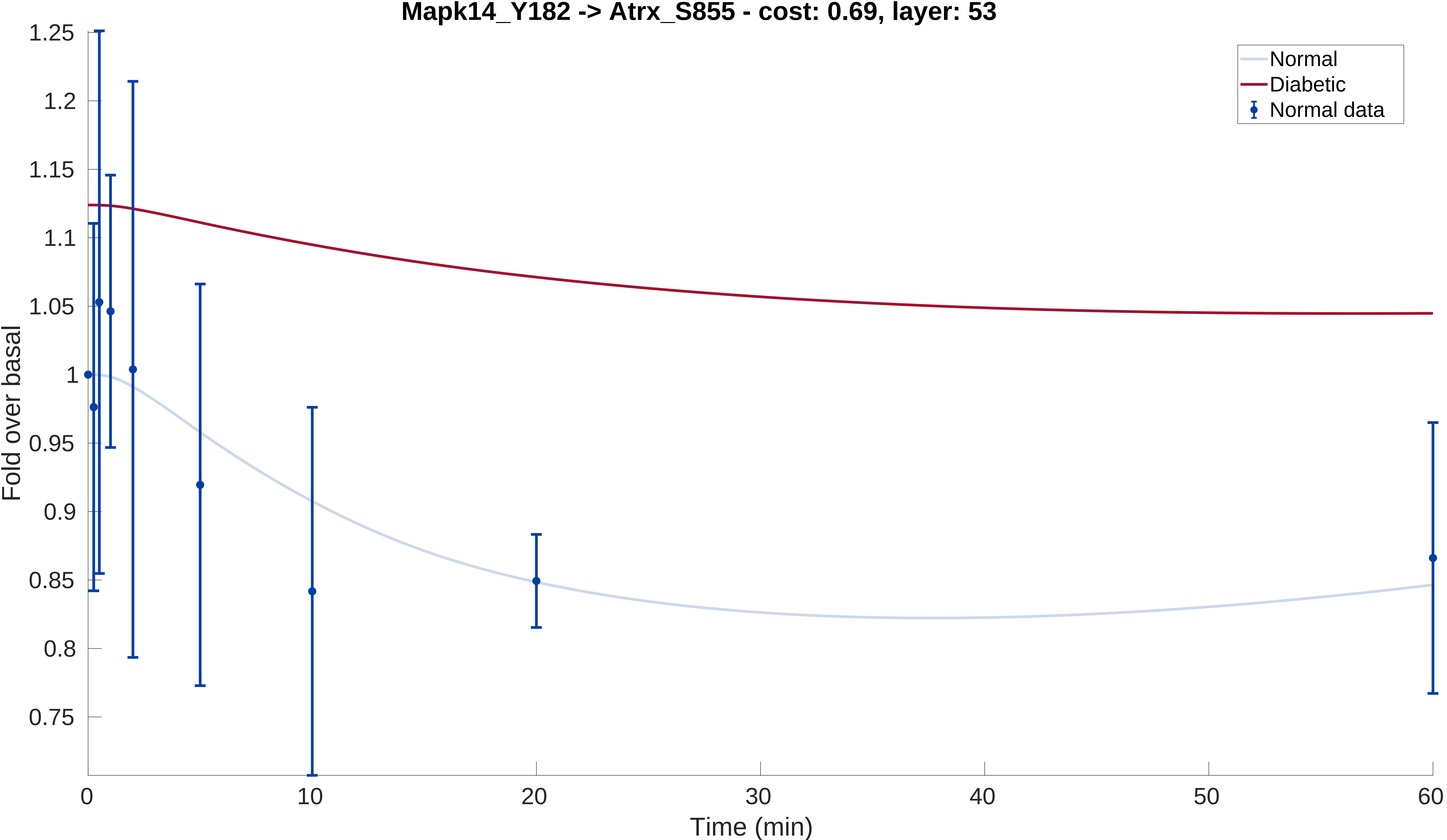

Mapk14_Y182 -> Atrx_S855 - cost: 0.69, layer: 53
Normal
Diabetic
Normal data
Fold over basal
1.25
1.2
1.15
1.1
1.05
1
0.95
0.9
0.85
0.8
0.75
0
10
20
30
40
50
60
Time (min)

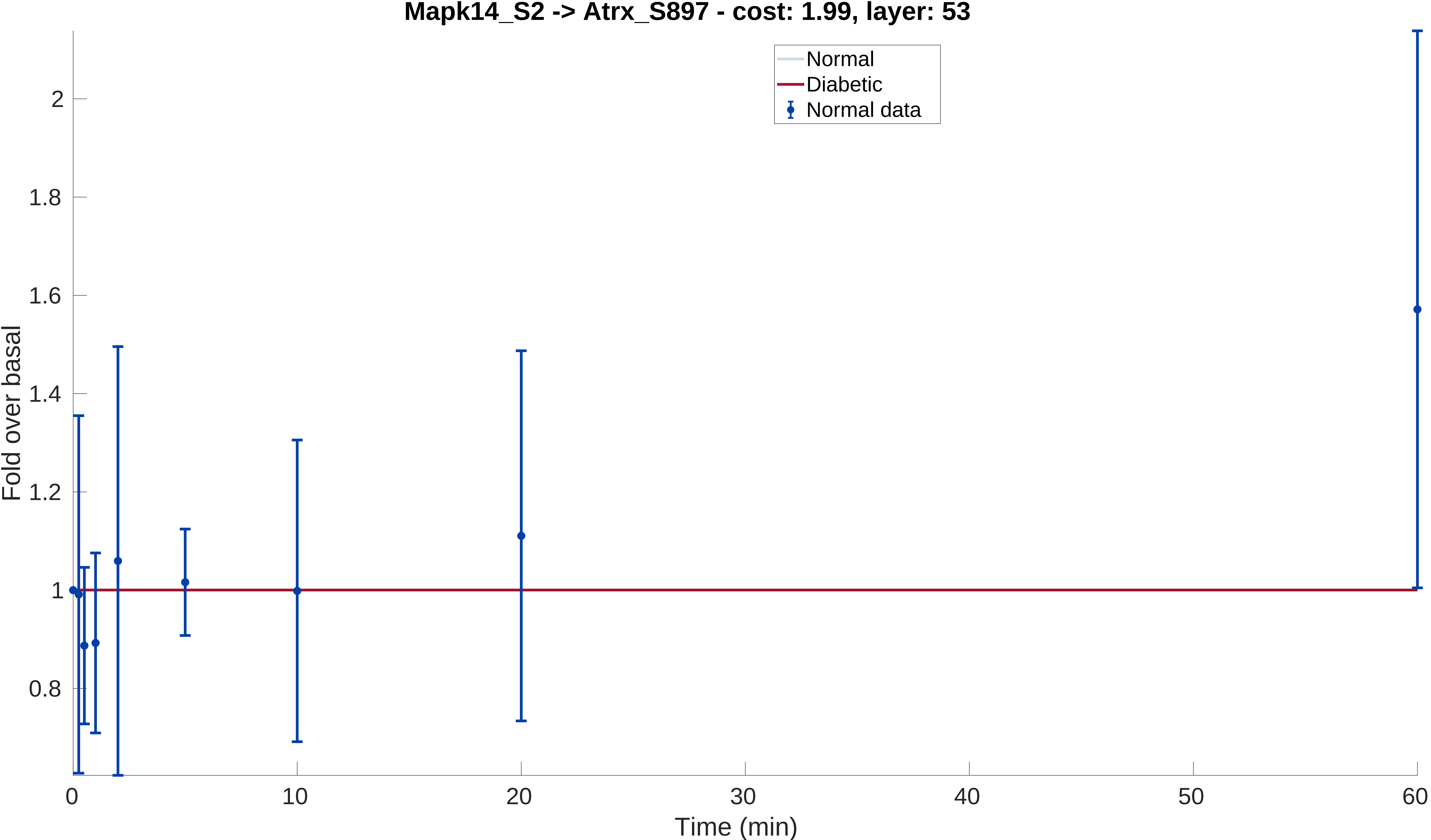
Mapk14_S2 -> Atrx_S897 - cost: 1.99, layer: 53
Normal
Diabetic
Normal data
Fold over basal
2
1.8
1.6
1.4
1.2
1
0.8
0
10
20
30
40
50
60
Time (min)

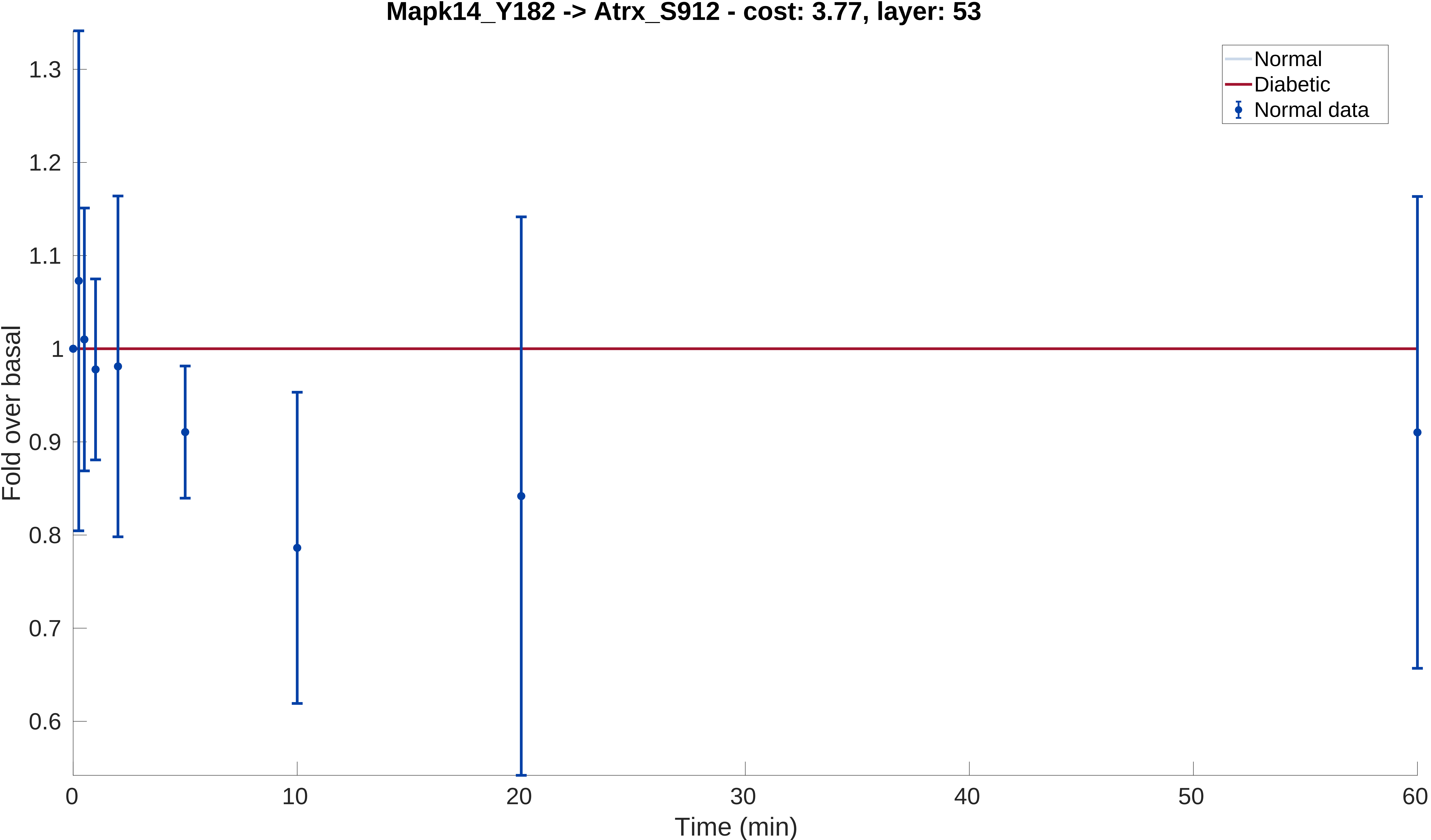

Mapk14_Y182 -> Atrx_S912 - cost: 3.77, layer: 53
Normal
Diabetic
Normal data
Fold over basal
1.3
1.2
1.1
1
0.9
0.8
0.7
0.6
0
10
20
30
40
50
60
Time (min)

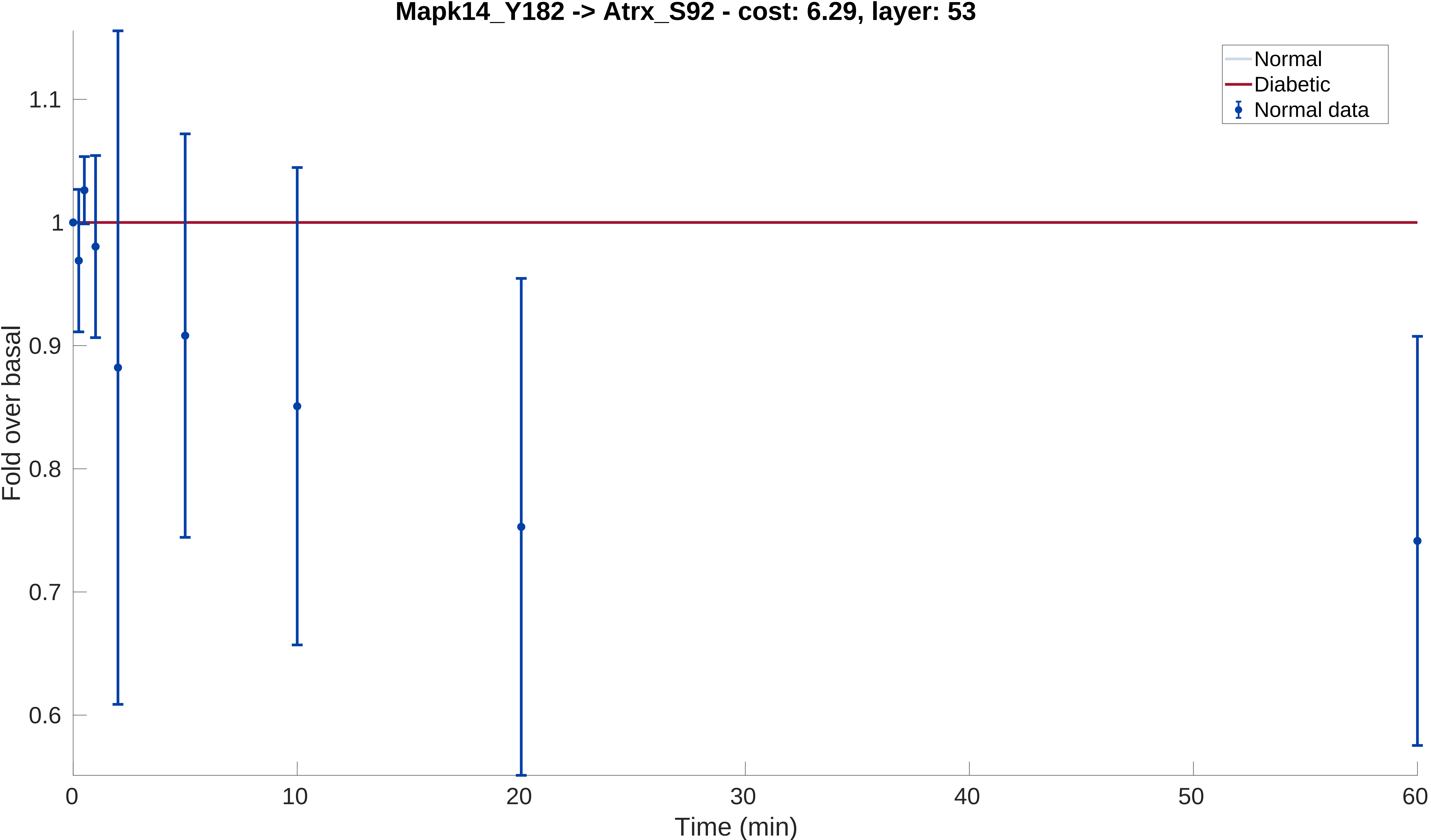
Mapk14_Y182 -> Atrx_S92 - cost: 6.29, layer: 53
Normal
Diabetic
Normal data
1.1
1
0.9
0.8
0.7
0.6
Fold over basal
0
10
20
30
40
50
60
Time (min)

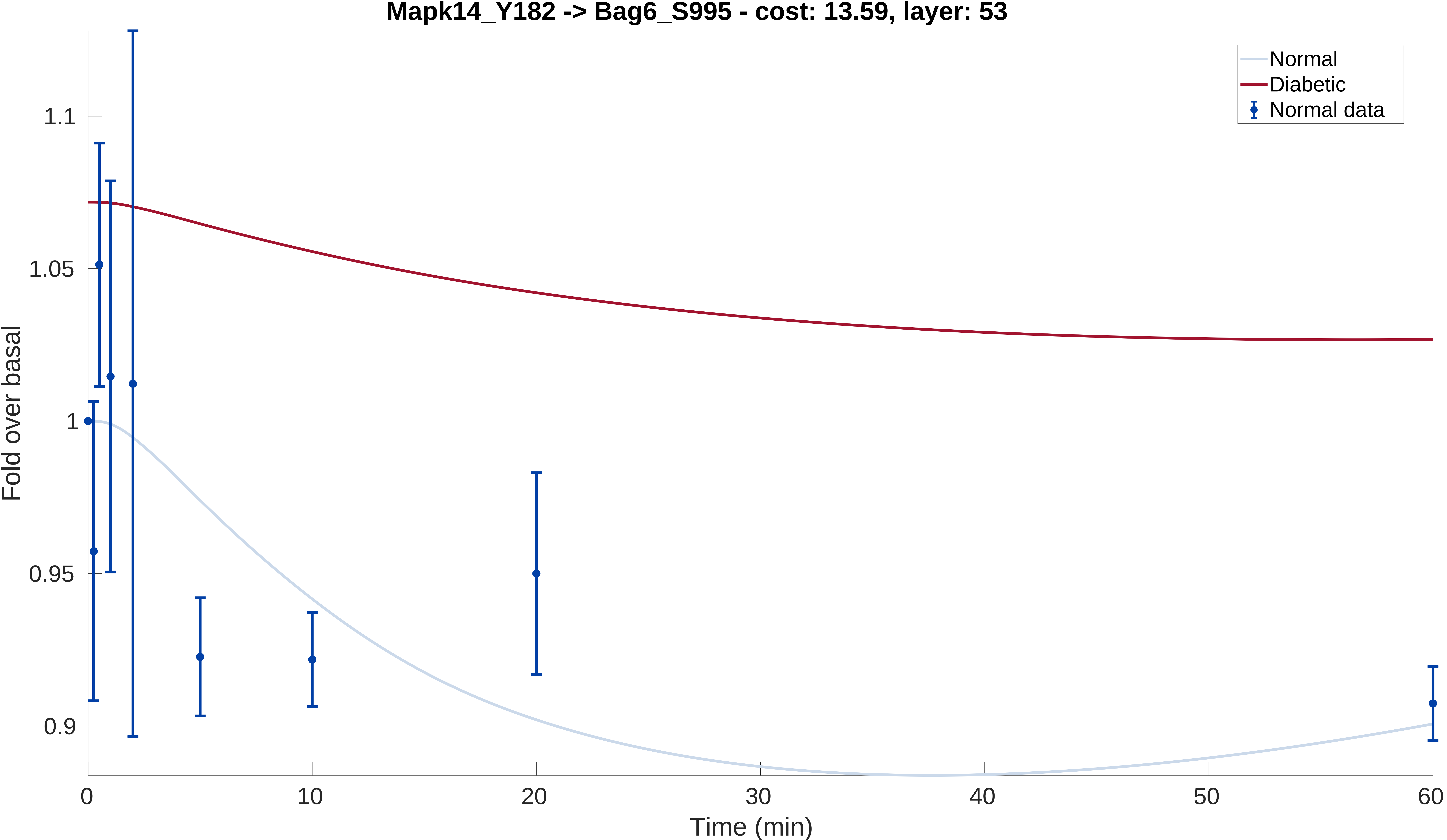

Mapk14_Y182 -> Bag6_S995 - cost: 13.59, layer: 53
Normal
Diabetic
Normal data
Fold over basal
1.1
1.05
1
0.95
0.9
0
10
20
30
40
50
60
Time (min)

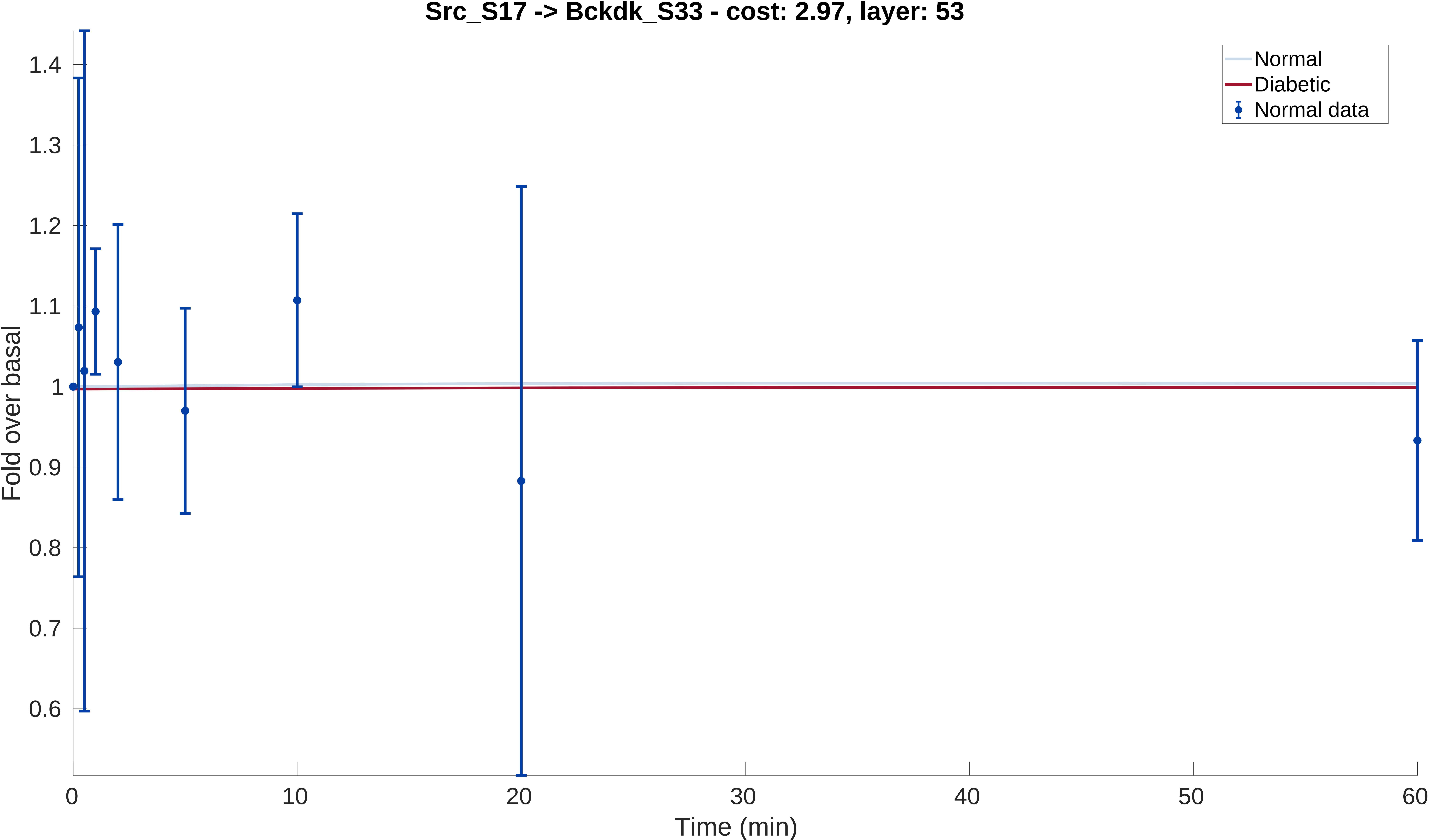
Src_S17 -> Bckdk_S33 - cost: 2.97, layer: 53
Normal
Diabetic
Normal data
Fold over basal
1.4
1.3
1.2
1.1
1
0.9
0.8
0.7
0.6
0
10
20
30
40
50
60
Time (min)

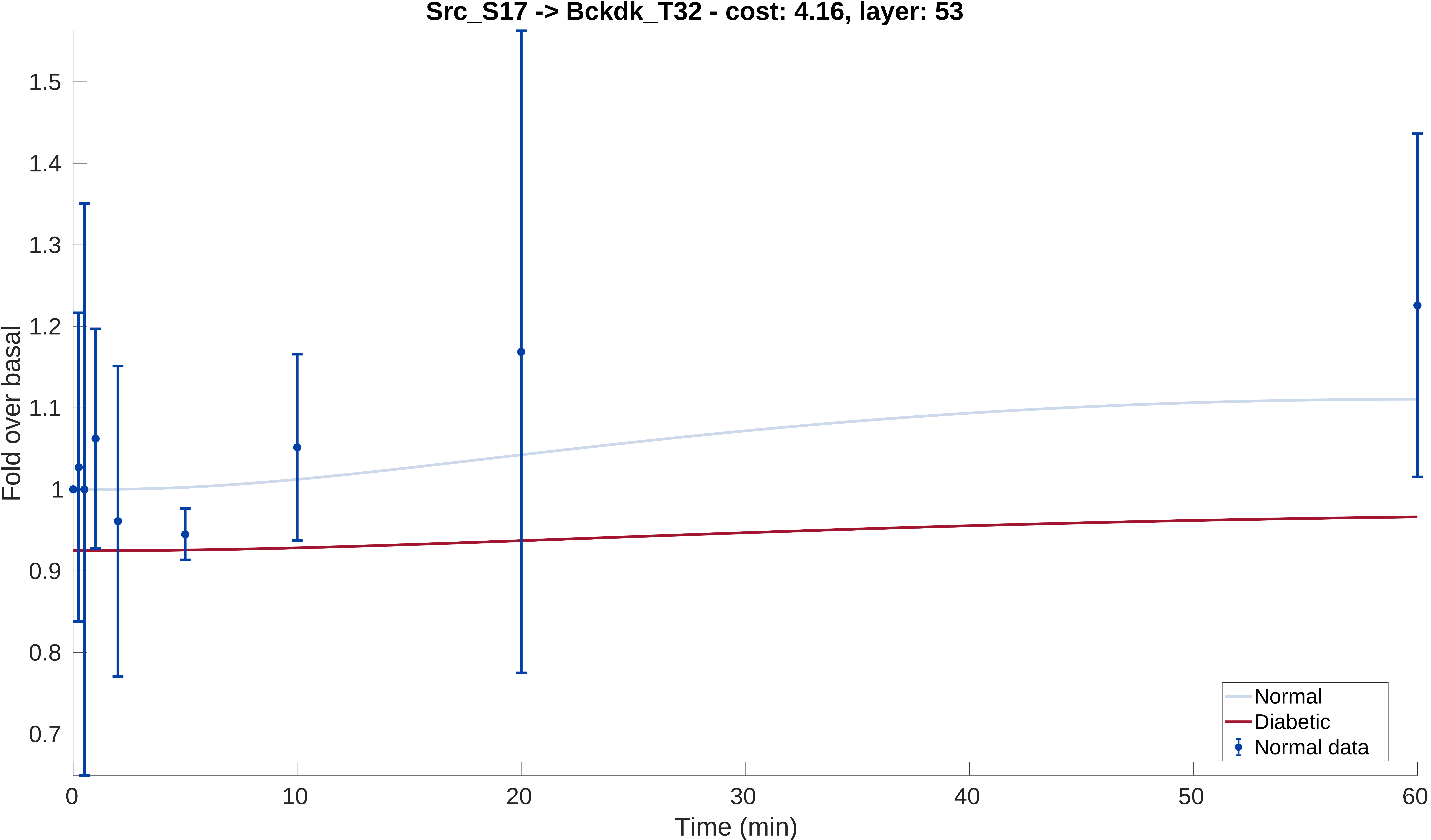

Src_S17 -> Bckdk_T32 - cost: 4.16, layer: 53
Fold over basal
1.5
1.4
1.3
1.2
1.1
1
0.9
0.8
0.7
0
10
20
30
40
50
60
Time (min)
Normal
Diabetic
Normal data

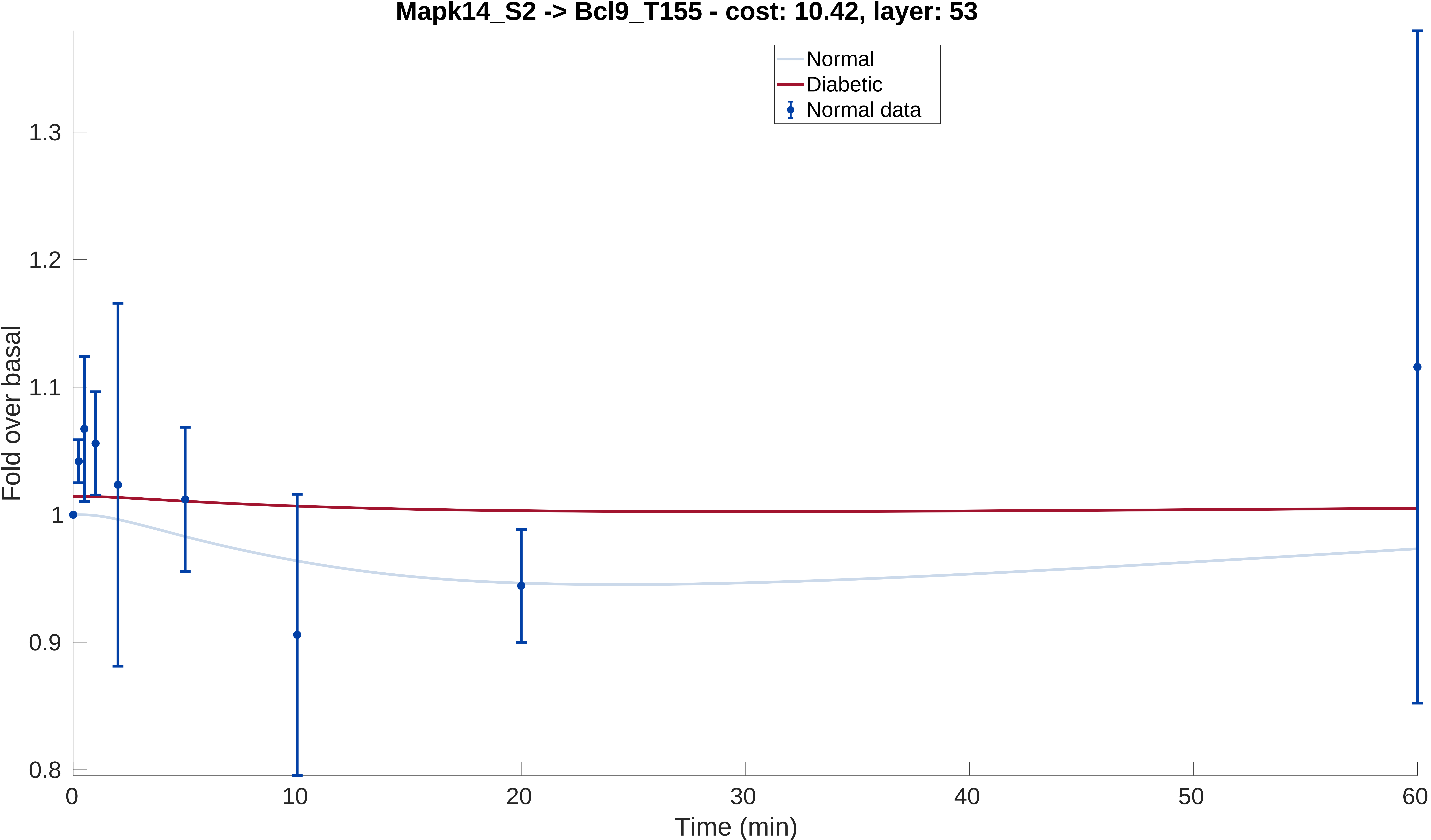

Mapk14_S2 -> Bcl9_T155 - cost: 10.42, layer: 53
Normal
Diabetic
Normal data
Fold over basal
1.3
1.2
1.1
1
0.9
0.8
0
10
20
30
40
50
60
Time (min)

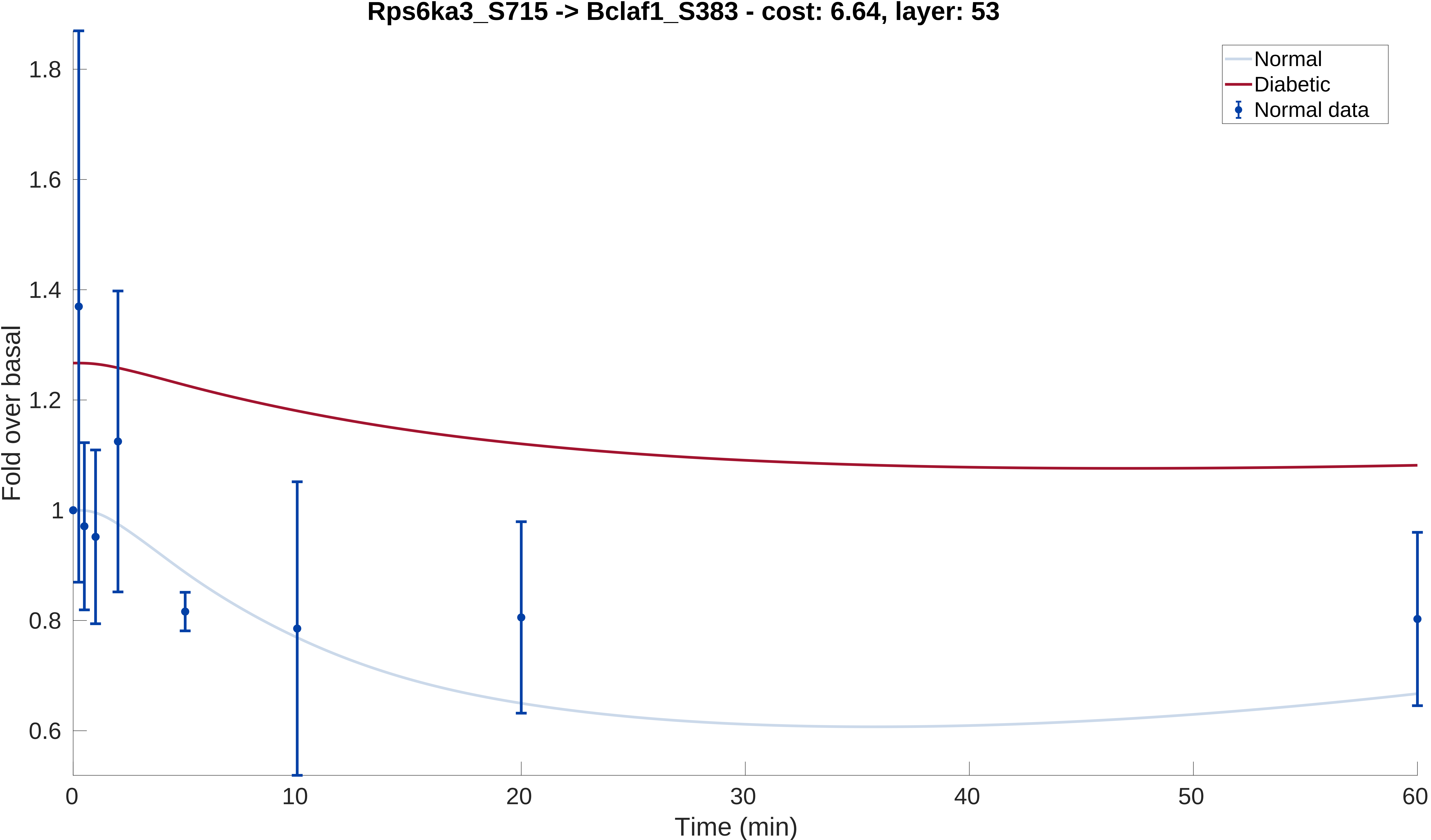

Rps6ka3_S715 -> Bclaf1_S383 - cost: 6.64, layer: 53
Normal
Diabetic
Normal data
Fold over basal
Time (min)
0.6
0.8
1
1.2
1.4
1.6
1.8
0
10
20
30
40
50
60

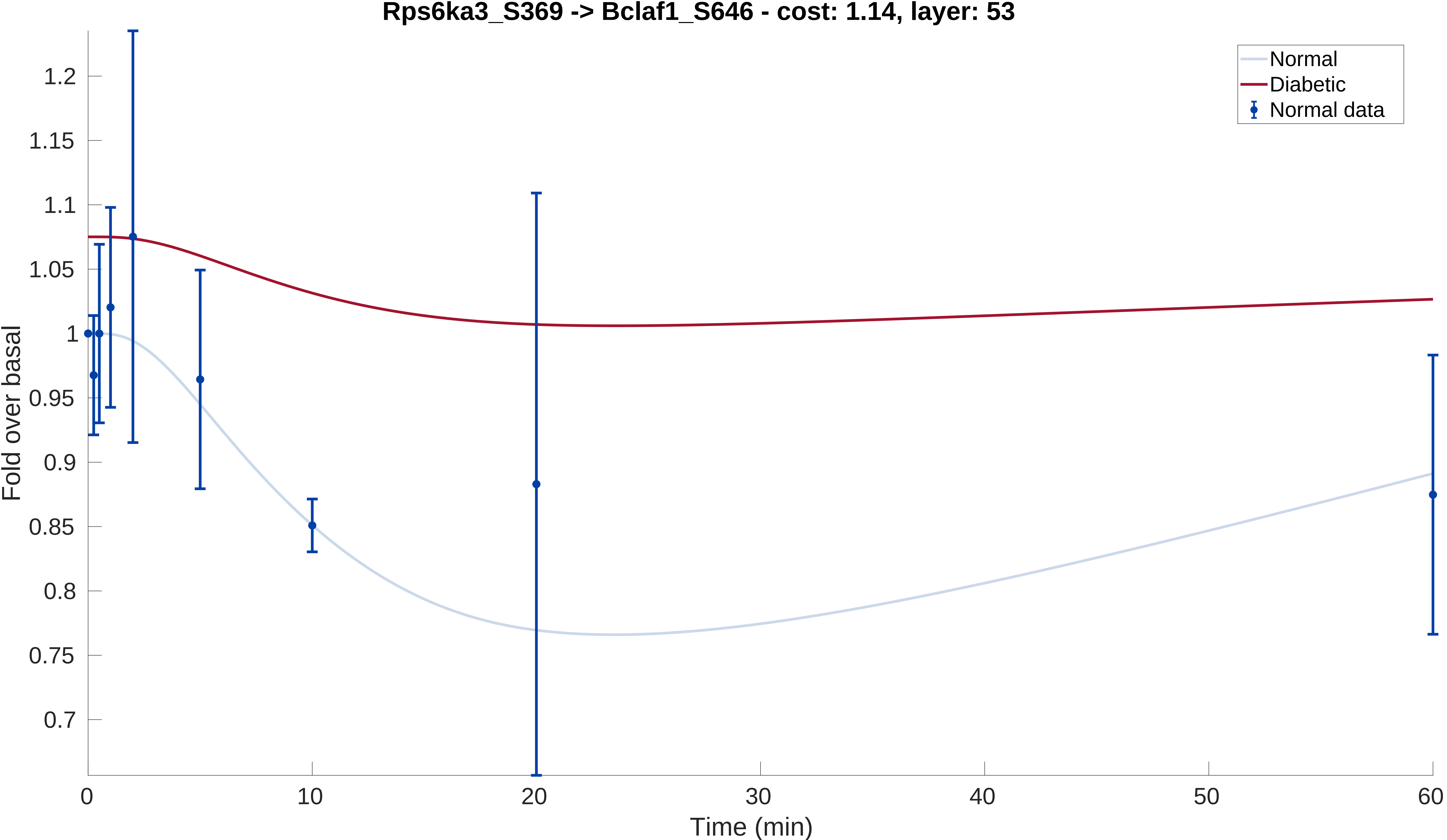
Rps6ka3_S369 -> Bclaf1_S646 - cost: 1.14, layer: 53
Normal
Diabetic
Normal data
Fold over basal
1.2
1.15
1.1
1.05
1
0.95
0.9
0.85
0.8
0.75
0.7
0
10
20
30
40
50
60
Time (min)

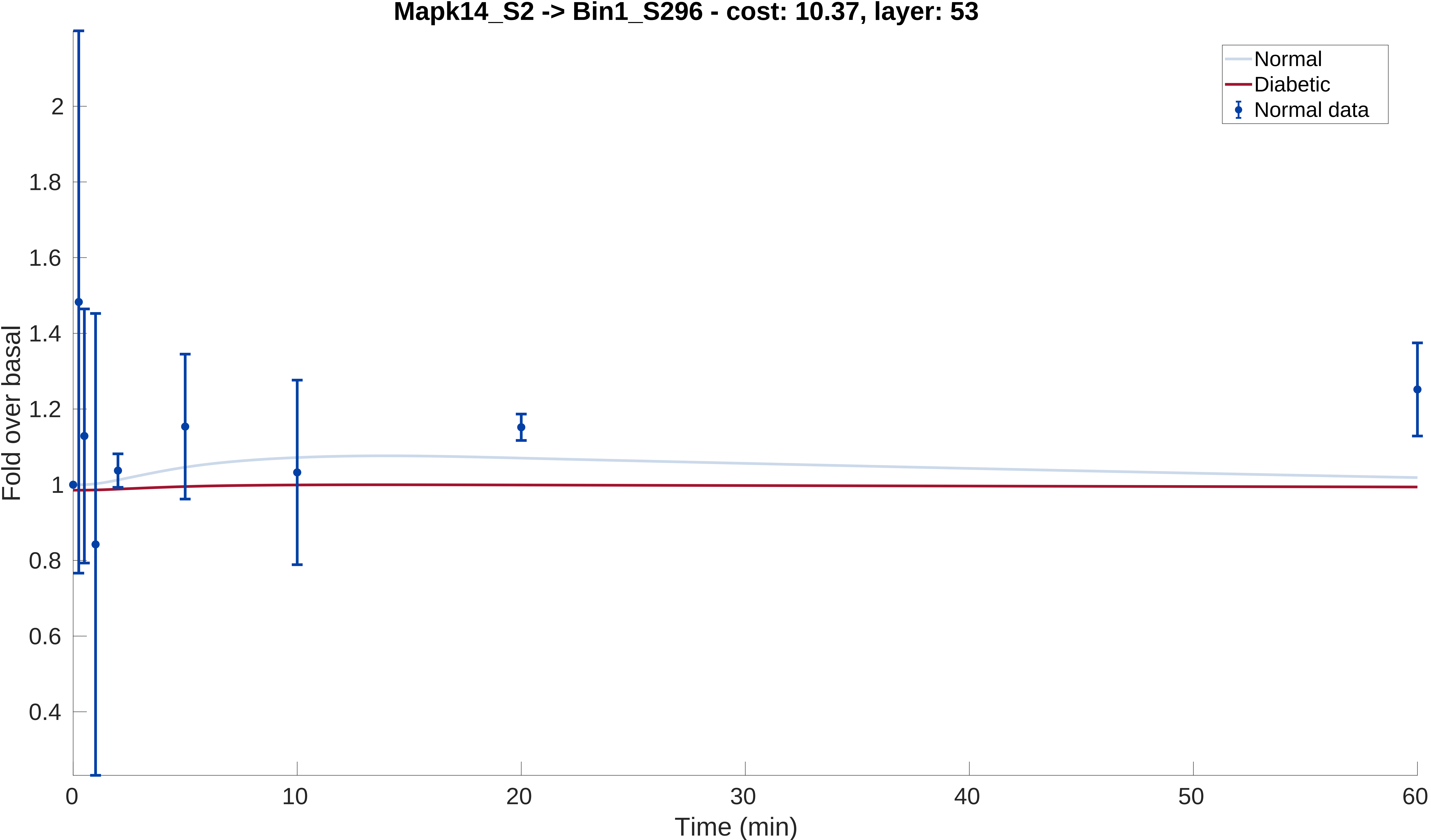
Mapk14_S2 -> Bin1_S296 - cost: 10.37, layer: 53
Normal
Diabetic
Normal data
Fold over basal
2
1.8
1.6
1.4
1.2
1
0.8
0.6
0.4
0
10
20
30
40
50
60
Time (min)

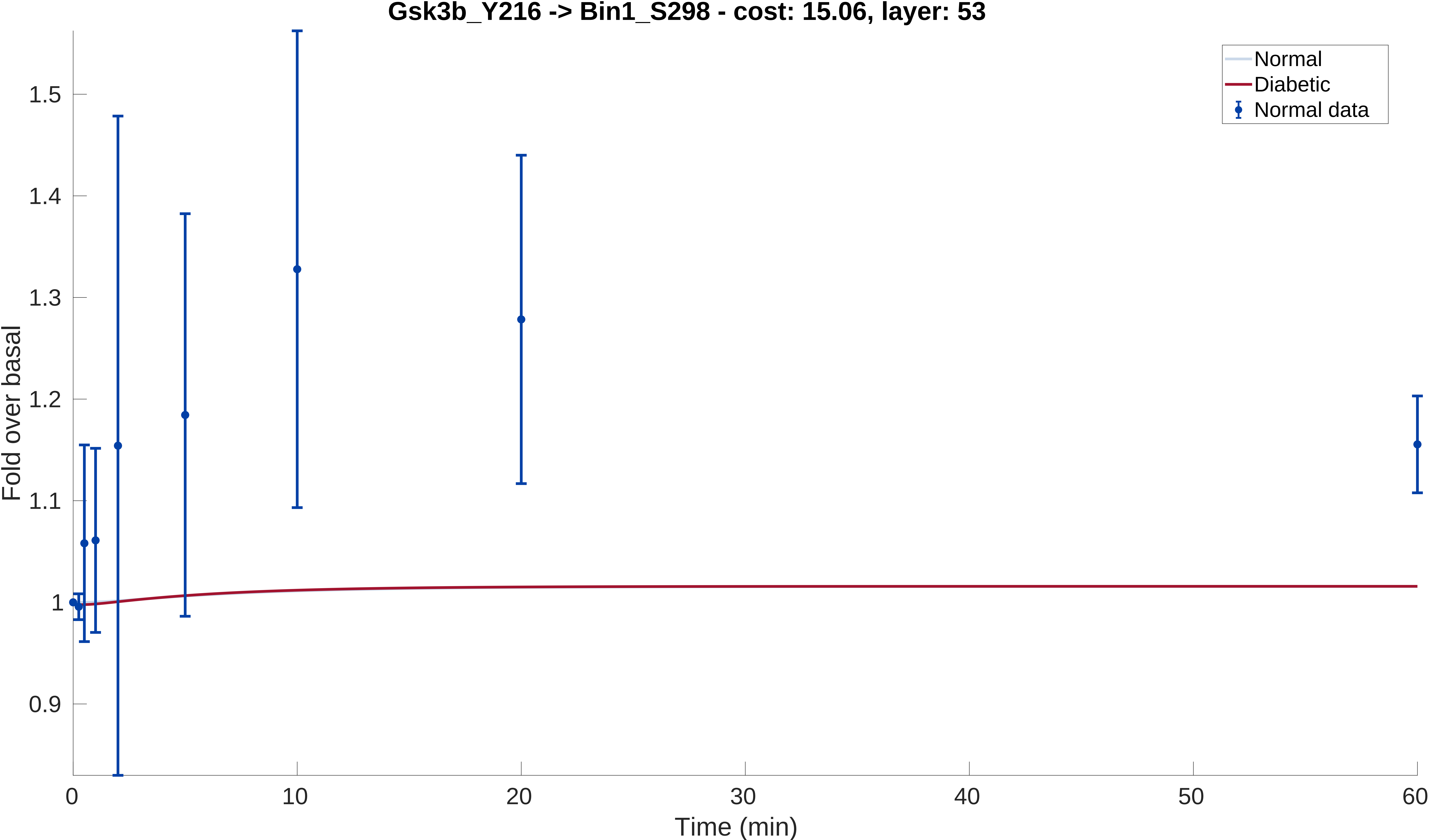

Gsk3b_Y216 -> Bin1_S298 - cost: 15.06, layer: 53
Normal
Diabetic
Normal data
Fold over basal
1.5
1.4
1.3
1.2
1.1
1
0.9
0
10
20
30
40
50
60
Time (min)

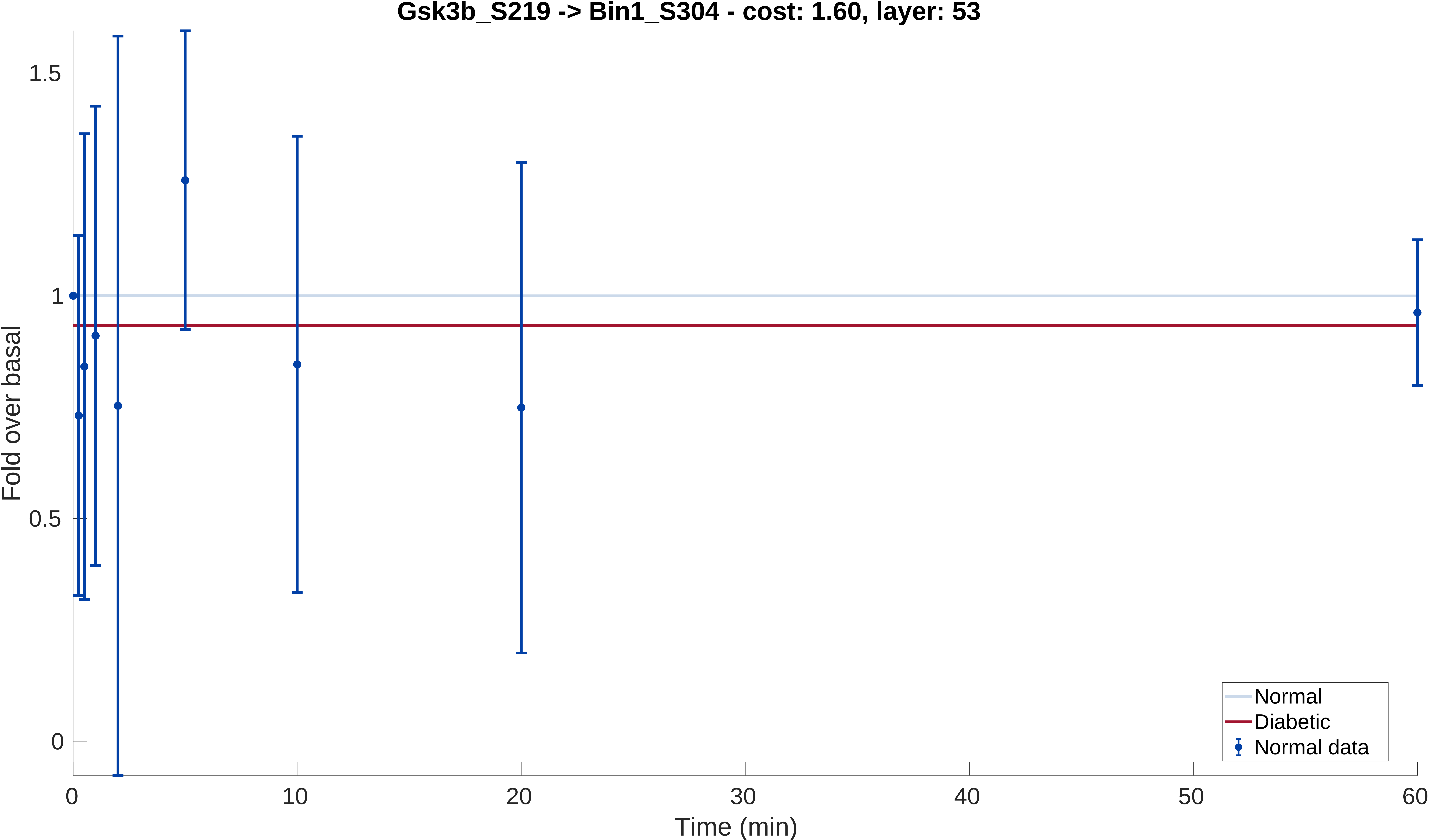

Gsk3b_S219 -> Bin1_S304 - cost: 1.60, layer: 53
Fold over basal
1.5
1
0.5
0
Time (min)
0
10
20
30
40
50
60
Normal
Diabetic
Normal data

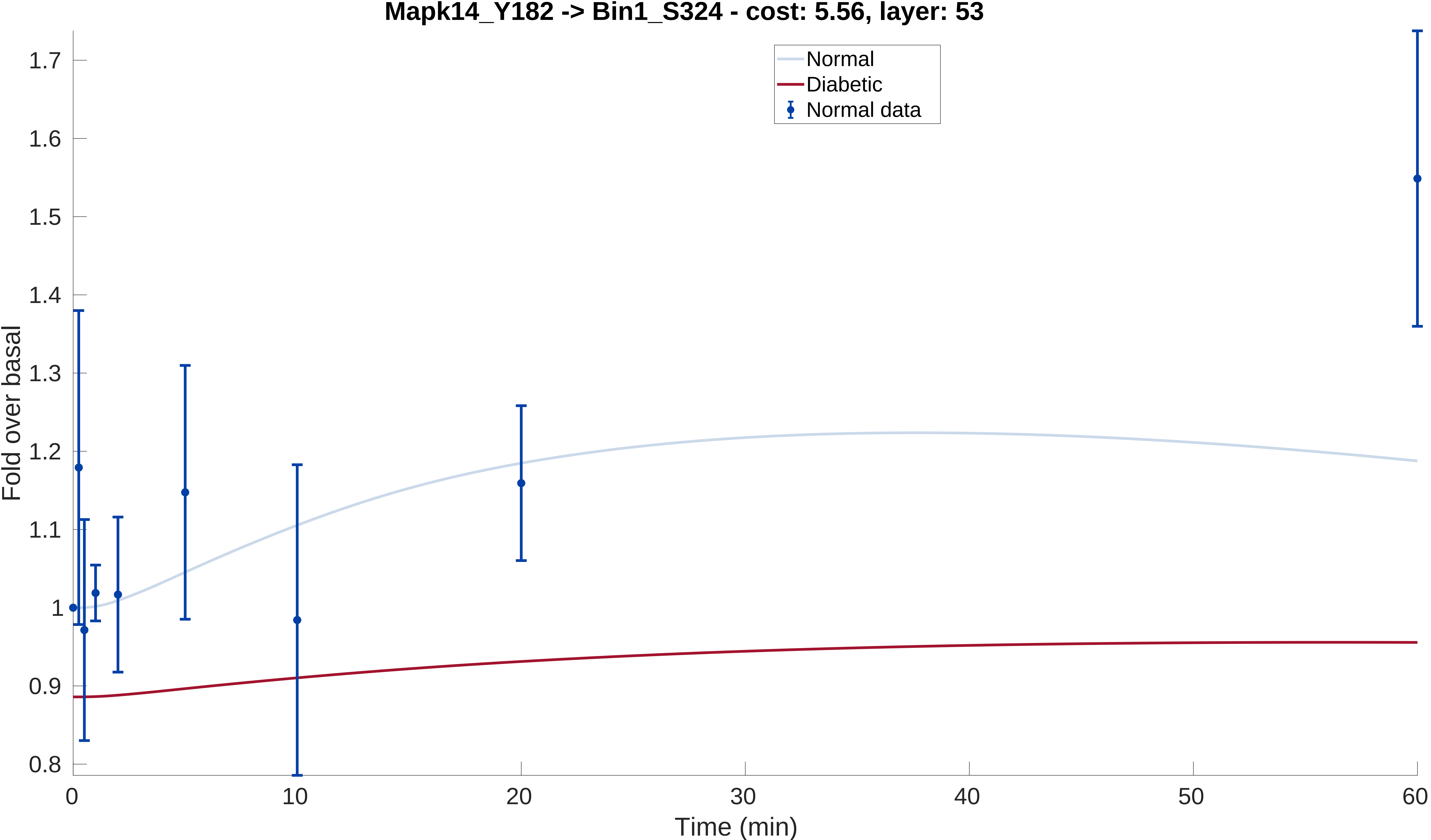

Mapk14_Y182 -> Bin1_S324 - cost: 5.56, layer: 53
Normal
Diabetic
Normal data
Fold over basal
1.7
1.6
1.5
1.4
1.3
1.2
1.1
1
0.9
0.8
0
10
20
30
40
50
60
Time (min)

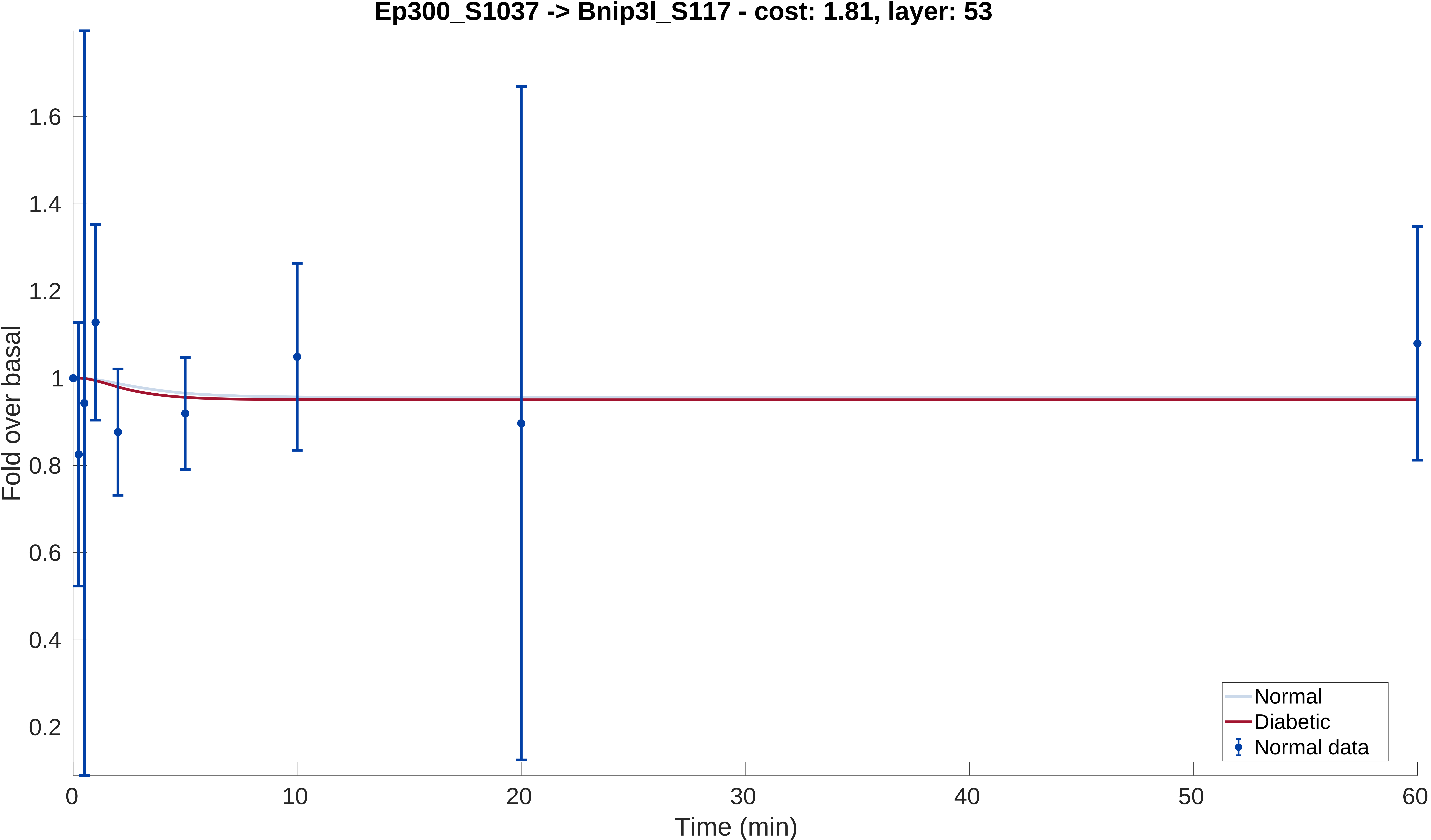

Ep300_S1037 -> Bnip3l_S117 - cost: 1.81, layer: 53
Fold over basal
Time (min)
0.2
0.4
0.6
0.8
1
1.2
1.4
1.6
0
10
20
30
40
50
60
Normal
Diabetic
Normal data

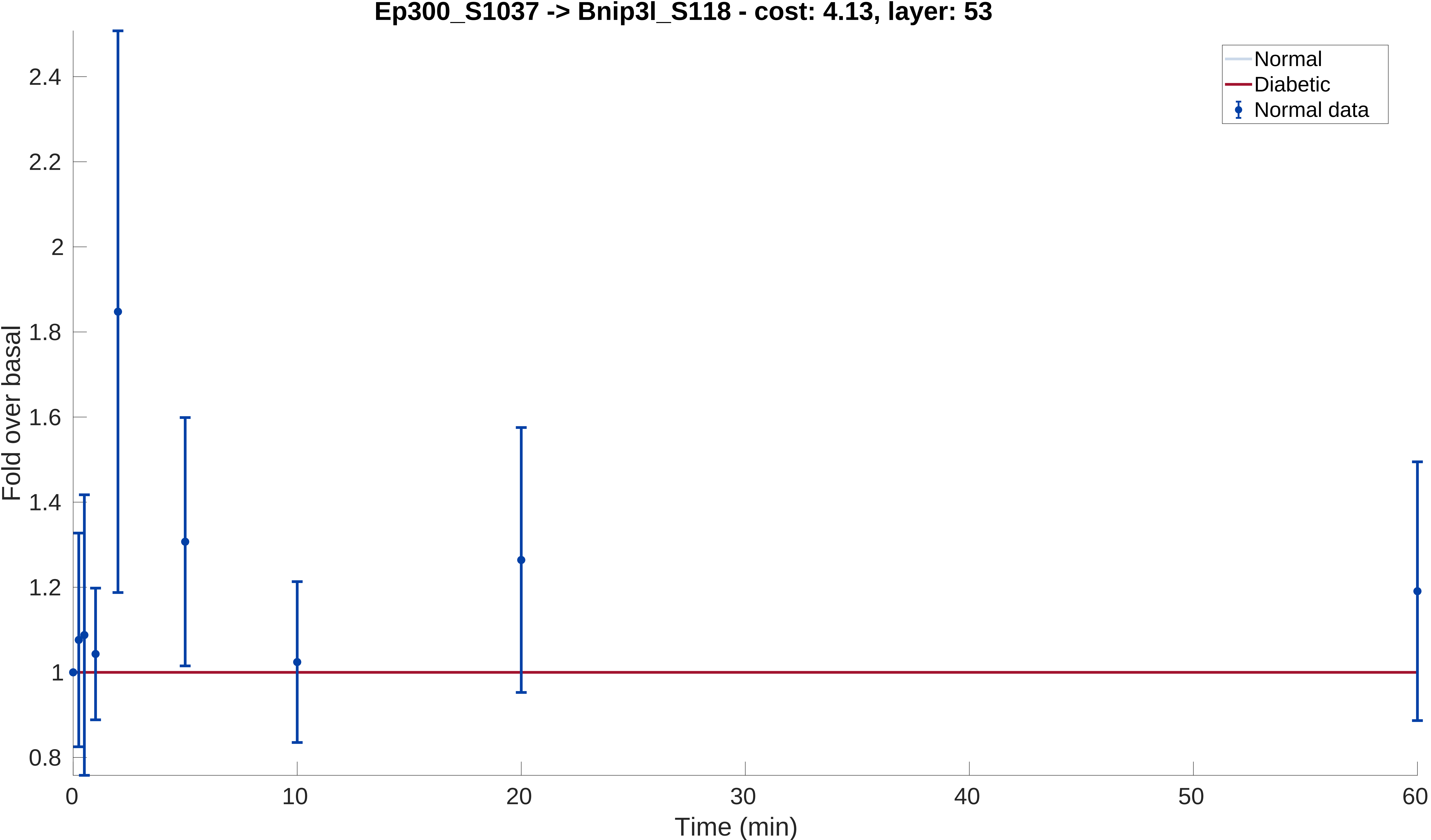

Ep300_S1037 -> Bnip3l_S118 - cost: 4.13, layer: 53
Normal
Diabetic
Normal data
Fold over basal
Time (min)
2.4
2.2
2
1.8
1.6
1.4
1.2
1
0.8
0
10
20
30
40
50
60

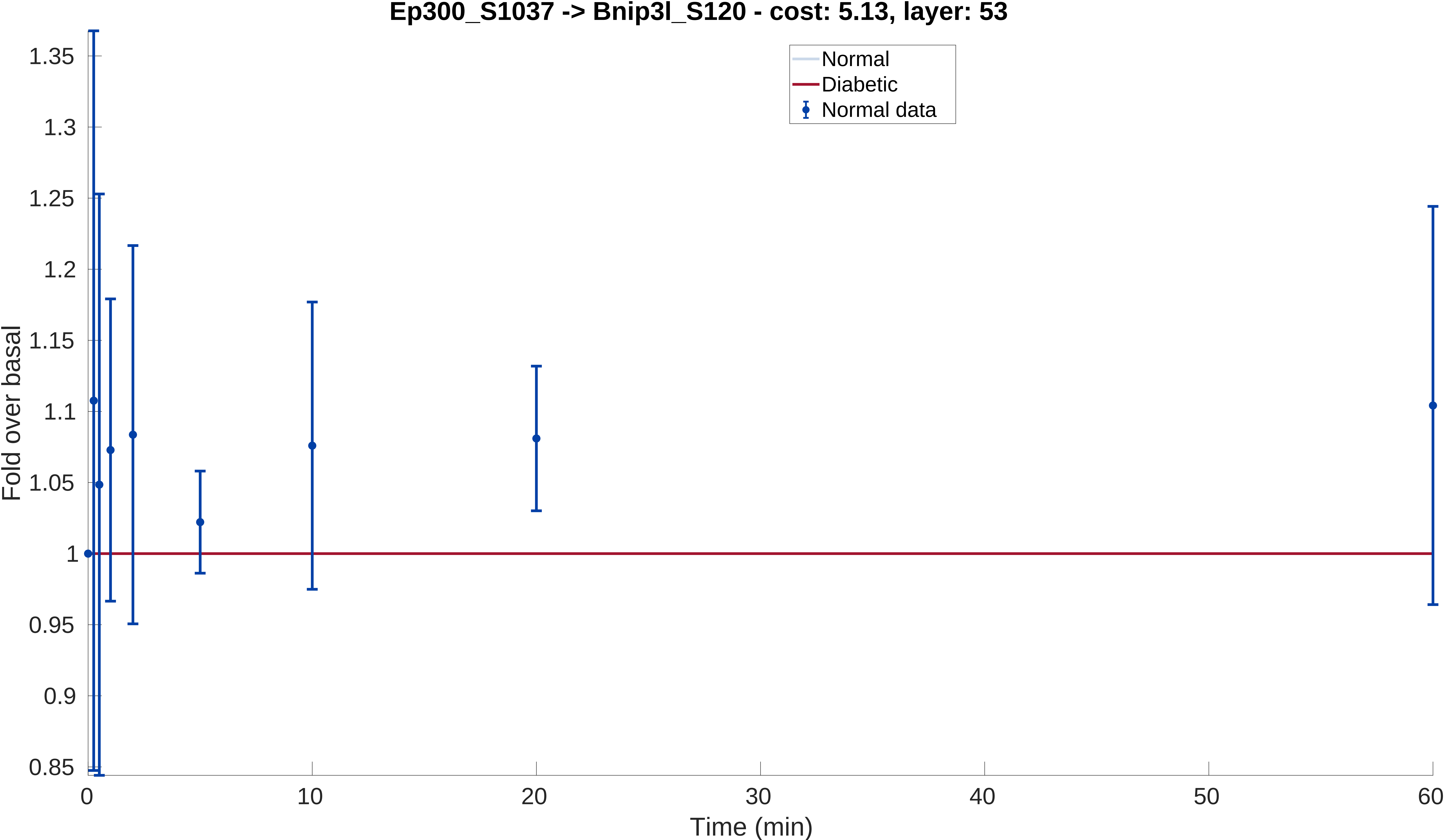

Ep300_S1037 -> Bnip3l_S120 - cost: 5.13, layer: 53
Normal
Diabetic
Normal data
Fold over basal
1.35
1.3
1.25
1.2
1.15
1.1
1.05
1
0.95
0.9
0.85
0
10
20
30
40
50
60
Time (min)

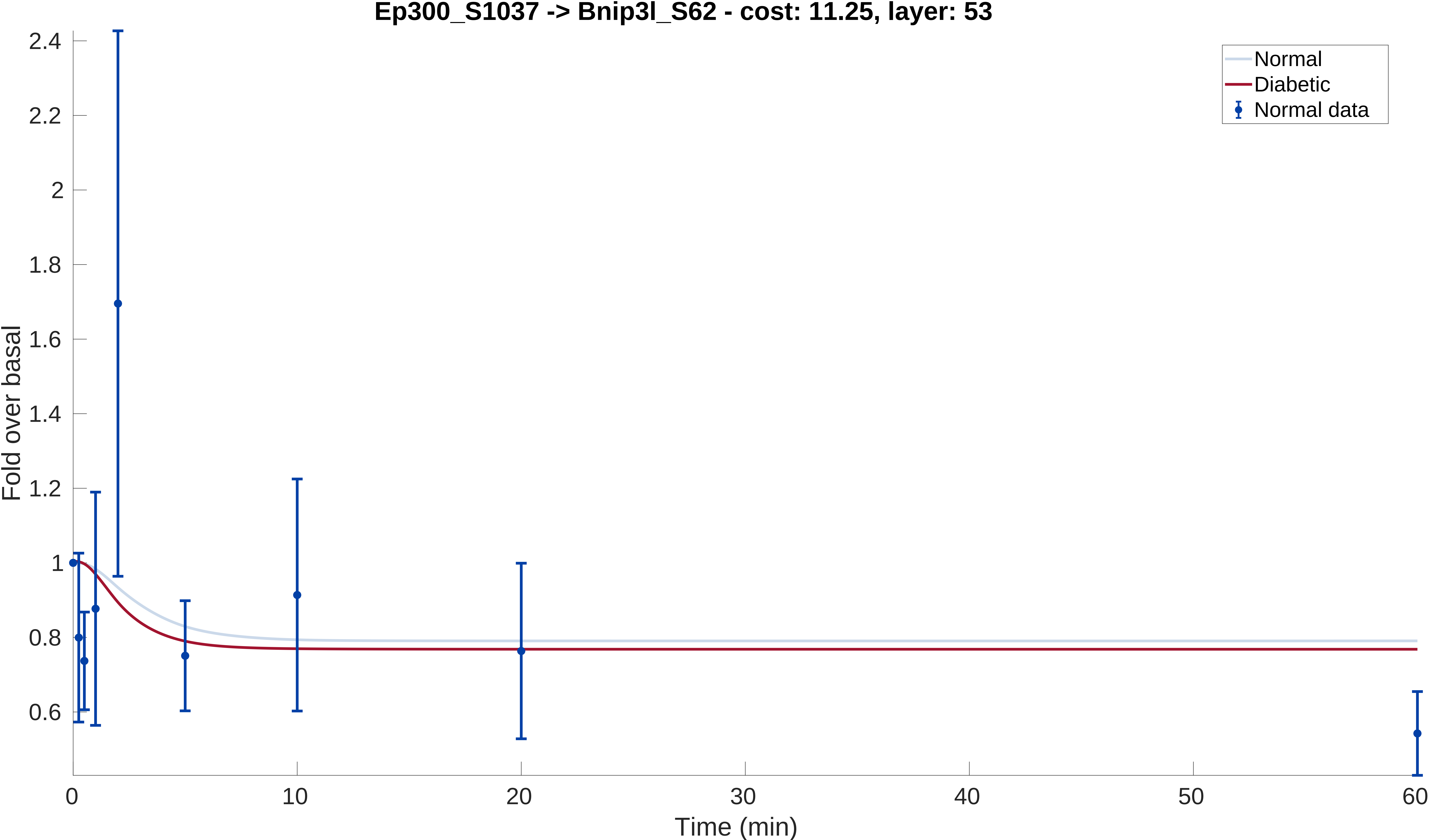

Ep300_S1037 -> Bnip3l_S62 - cost: 11.25, layer: 53
Fold over basal
2.4
2.2
2
1.8
1.6
1.4
1.2
1
0.8
0.6
0
10
20
30
40
50
60
Time (min)
Normal
Diabetic
Normal data

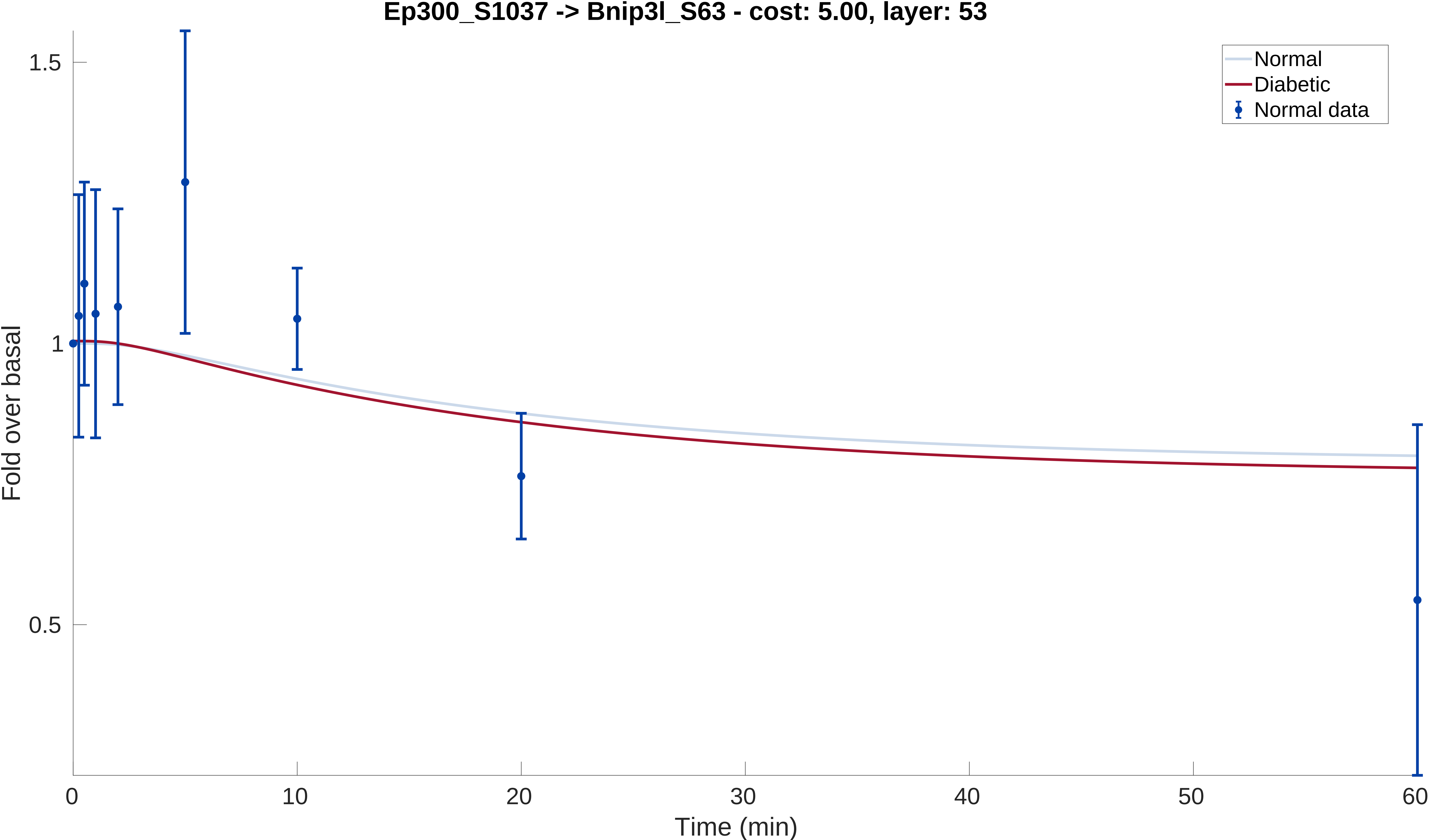

Ep300_S1037 -> Bnip3l_S63 - cost: 5.00, layer: 53
Normal
Diabetic
Normal data
Fold over basal
1.5
1
0.5
0
10
20
30
40
50
60
Time (min)

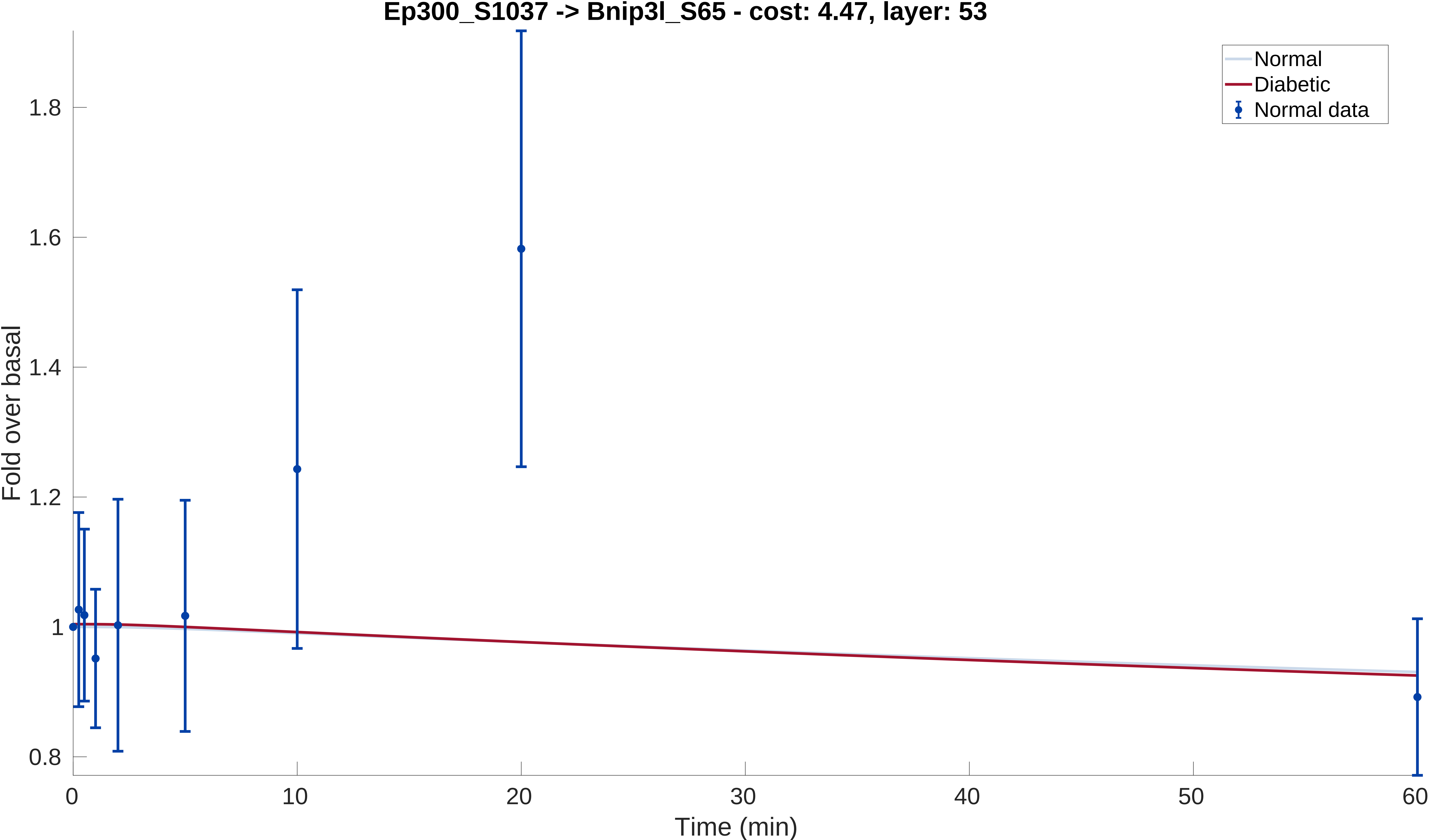

Ep300_S1037 -> Bnip3l_S65 - cost: 4.47, layer: 53
Normal
Diabetic
Normal data
Fold over basal
Time (min)
0.8
1
1.2
1.4
1.6
1.8
0
10
20
30
40
50
60

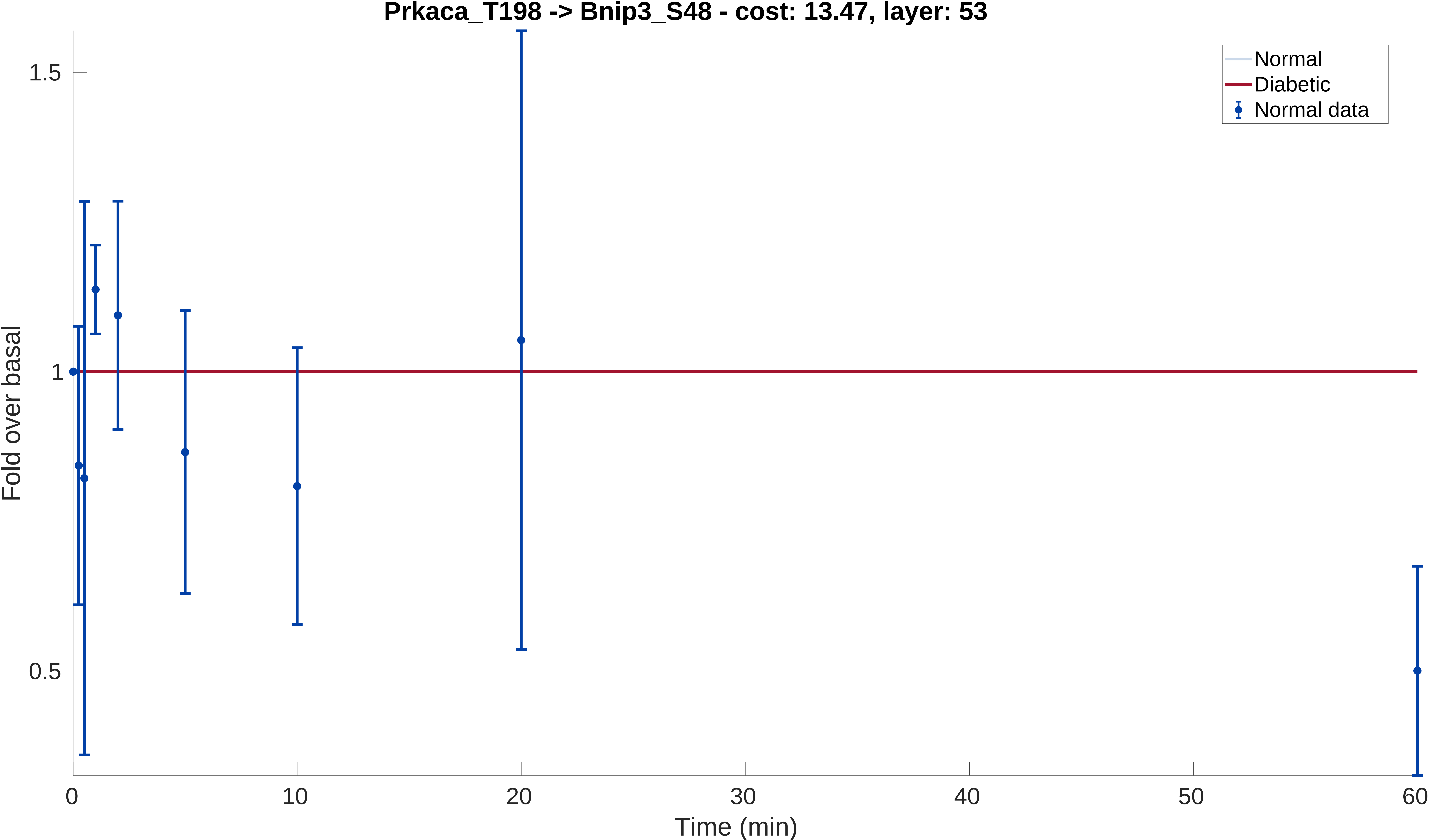

Prkaca_T198 -> Bnip3_S48 - cost: 13.47, layer: 53
Normal
Diabetic
Normal data
1.5
1
0.5
Fold over basal
0
10
20
30
40
50
60
Time (min)

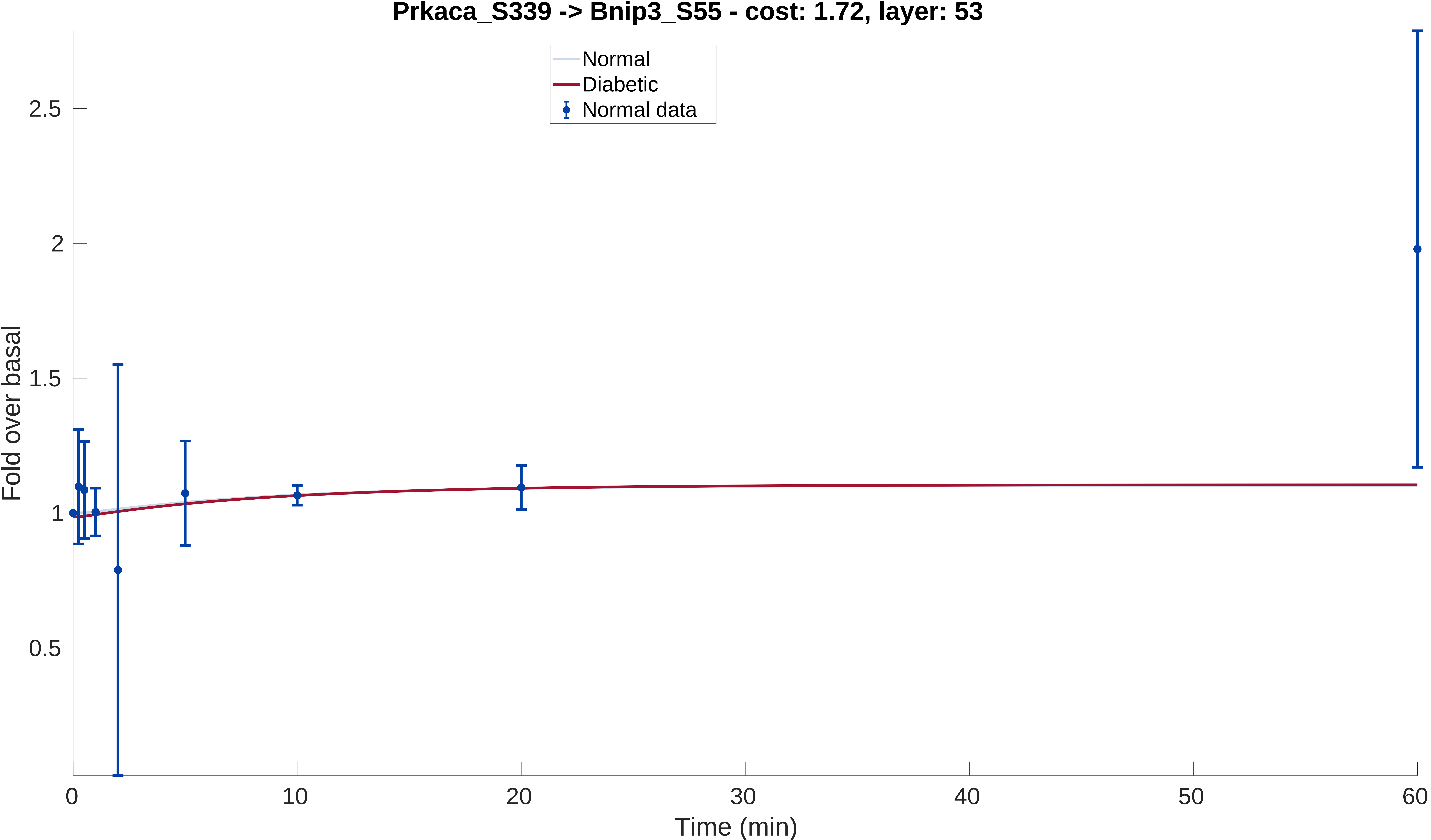
Prkaca_S339 -> Bnip3_S55 - cost: 1.72, layer: 53
Normal
Diabetic
Normal data
Fold over basal
Time (min)
0.5
1
1.5
2
2.5
0
10
20
30
40
50
60

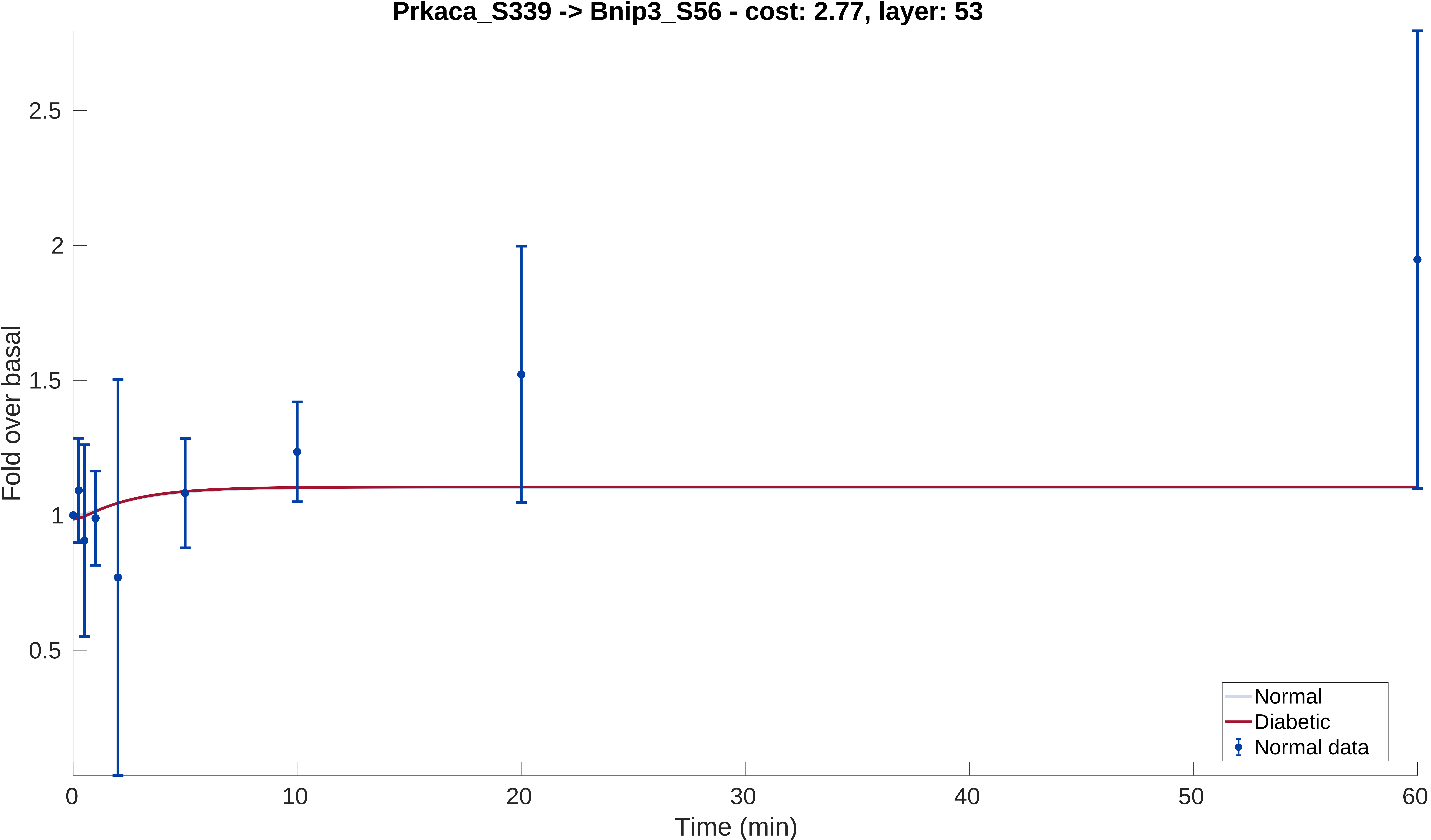

Prkaca_S339 -> Bnip3_S56 - cost: 2.77, layer: 53
Fold over basal
Time (min)
0.5
1
1.5
2
2.5
0
10
20
30
40
50
60
Normal
Diabetic
Normal data

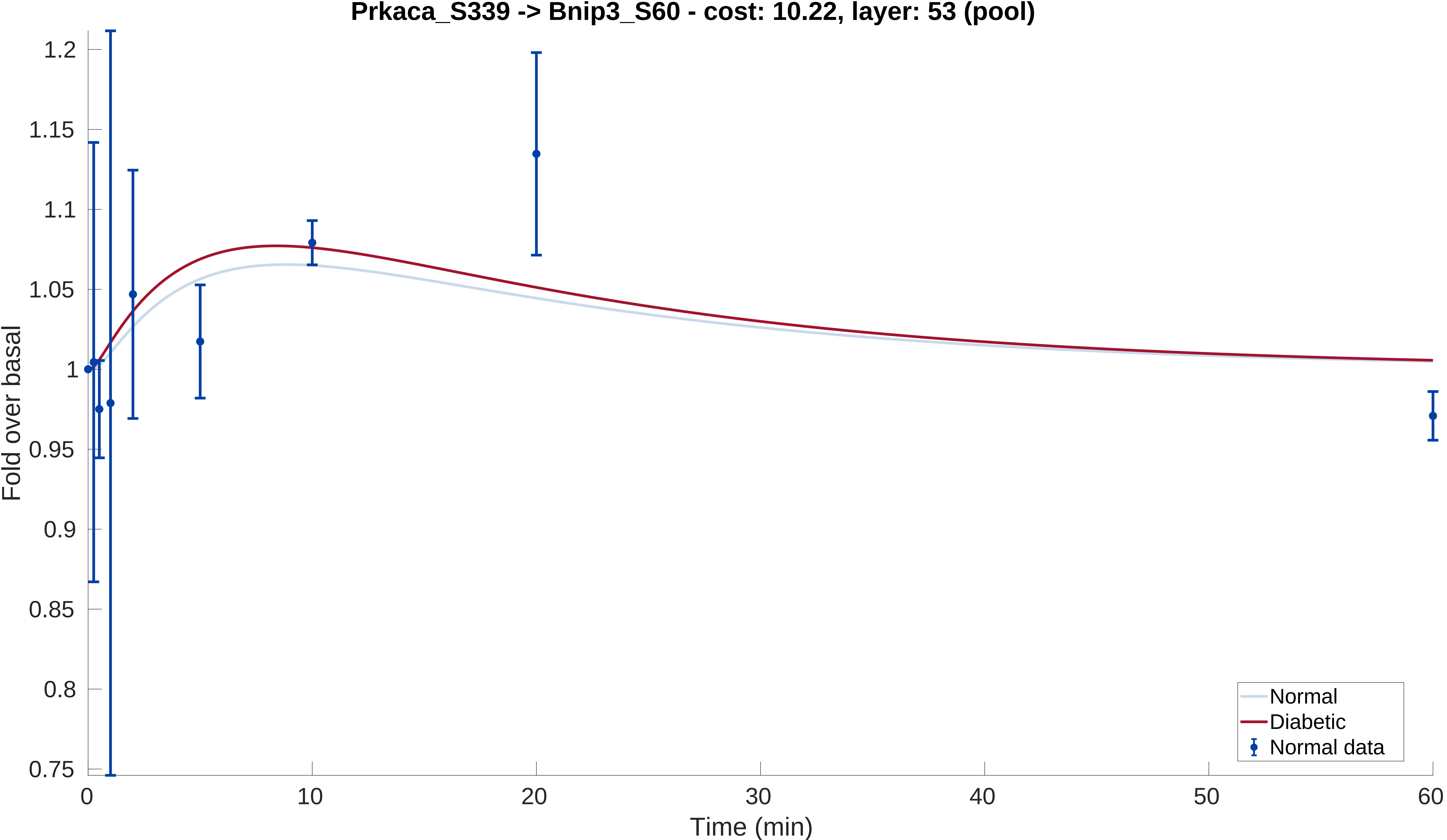

Prkaca_S339 -> Bnip3_S60 - cost: 10.22, layer: 53 (pool)
Fold over basal
1.2
1.15
1.1
1.05
1
0.95
0.9
0.85
0.8
0.75
0
10
20
30
40
50
60
Time (min)
Normal
Diabetic
Normal data

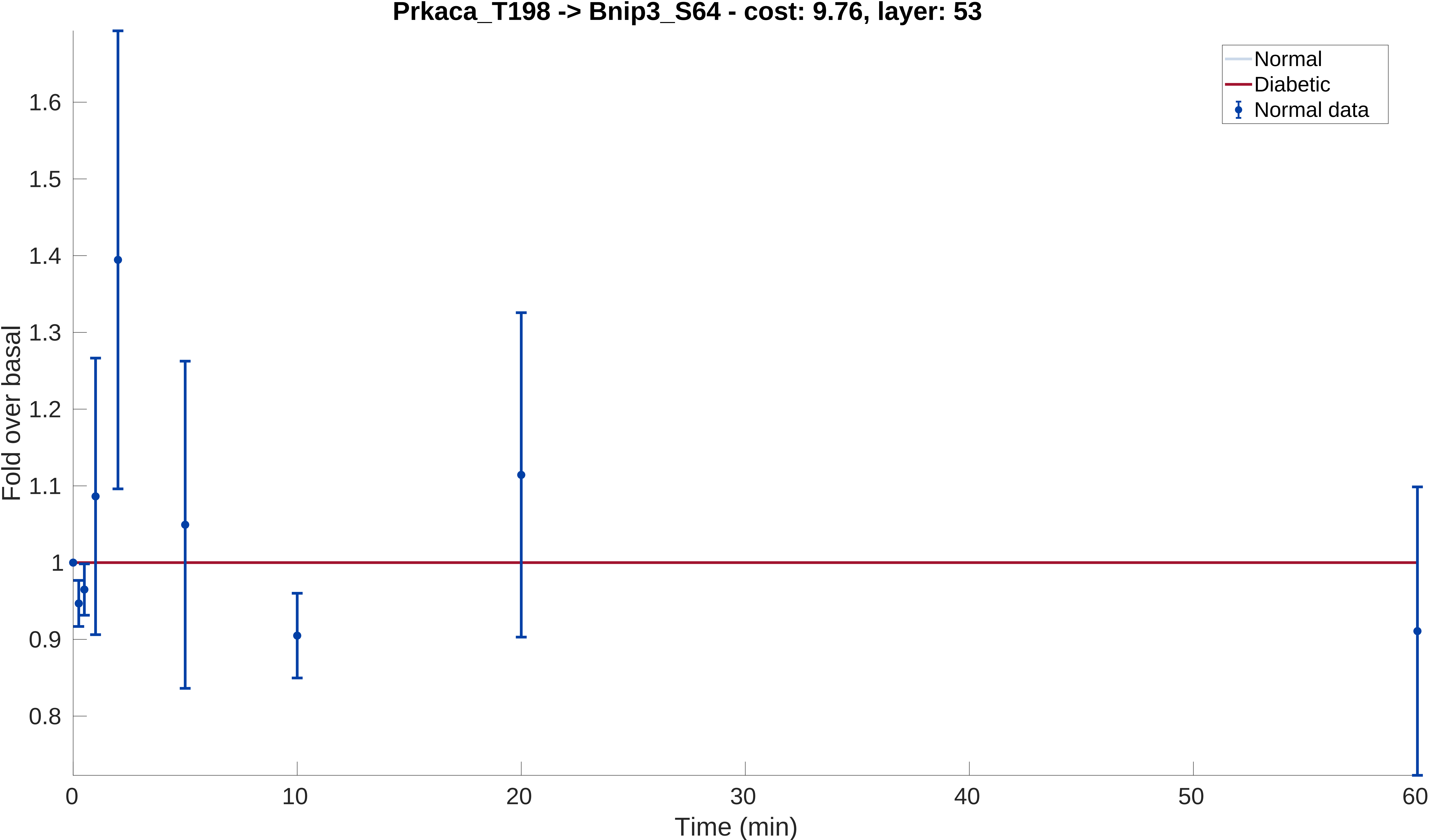

Prkaca_T198 -> Bnip3_S64 - cost: 9.76, layer: 53
Normal
Diabetic
Normal data
Fold over basal
1.6
1.5
1.4
1.3
1.2
1.1
1
0.9
0.8
0
10
20
30
40
50
60
Time (min)

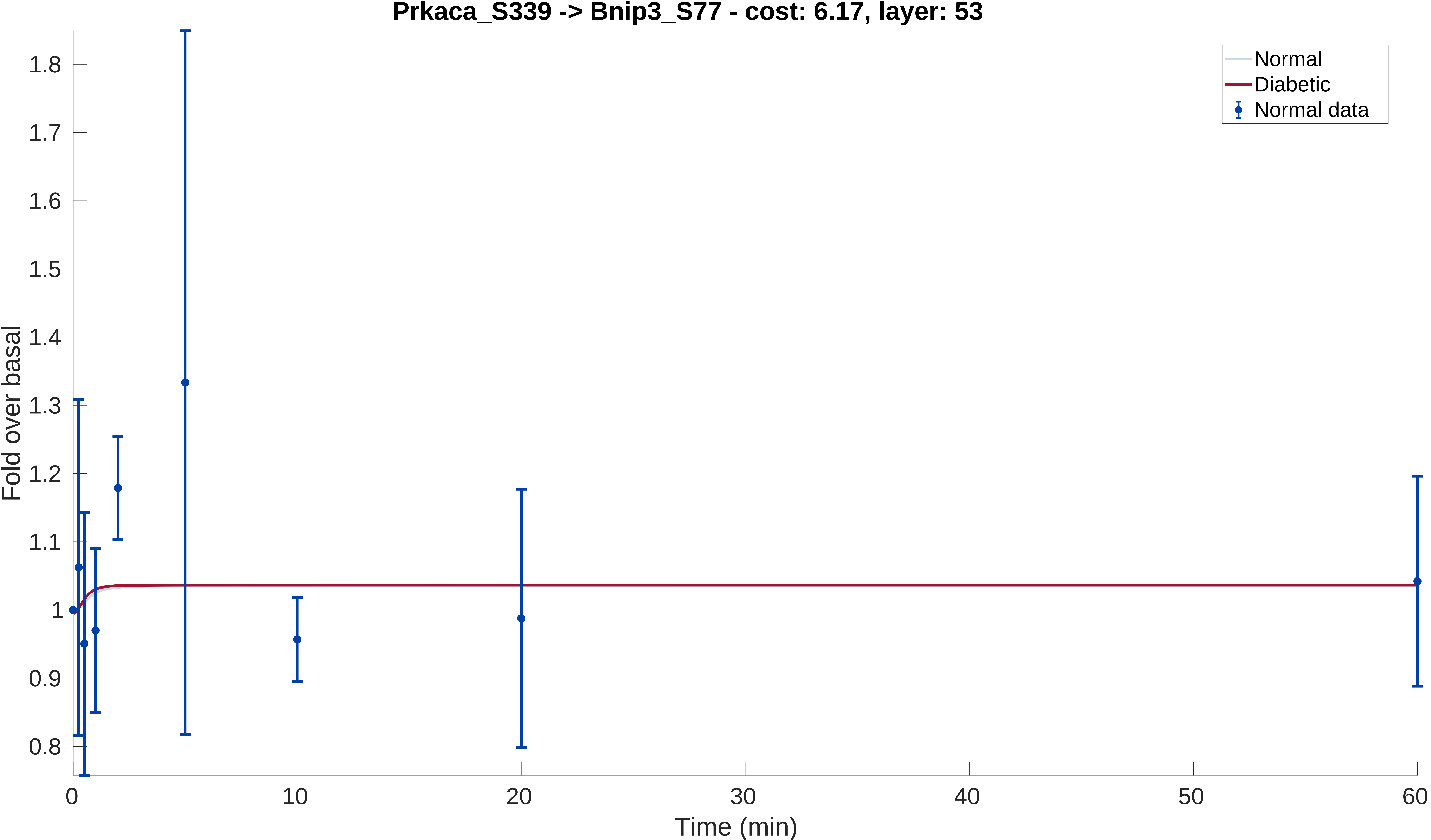

Prrkaca_S339 -> Bnip3_S77 - cost: 6.17, layer: 53
Normal
Diabetic
Normal data
Fold over basal
1.8
1.7
1.6
1.5
1.4
1.3
1.2
1.1
1
0.9
0.8
0
10
20
30
40
50
60
Time (min)

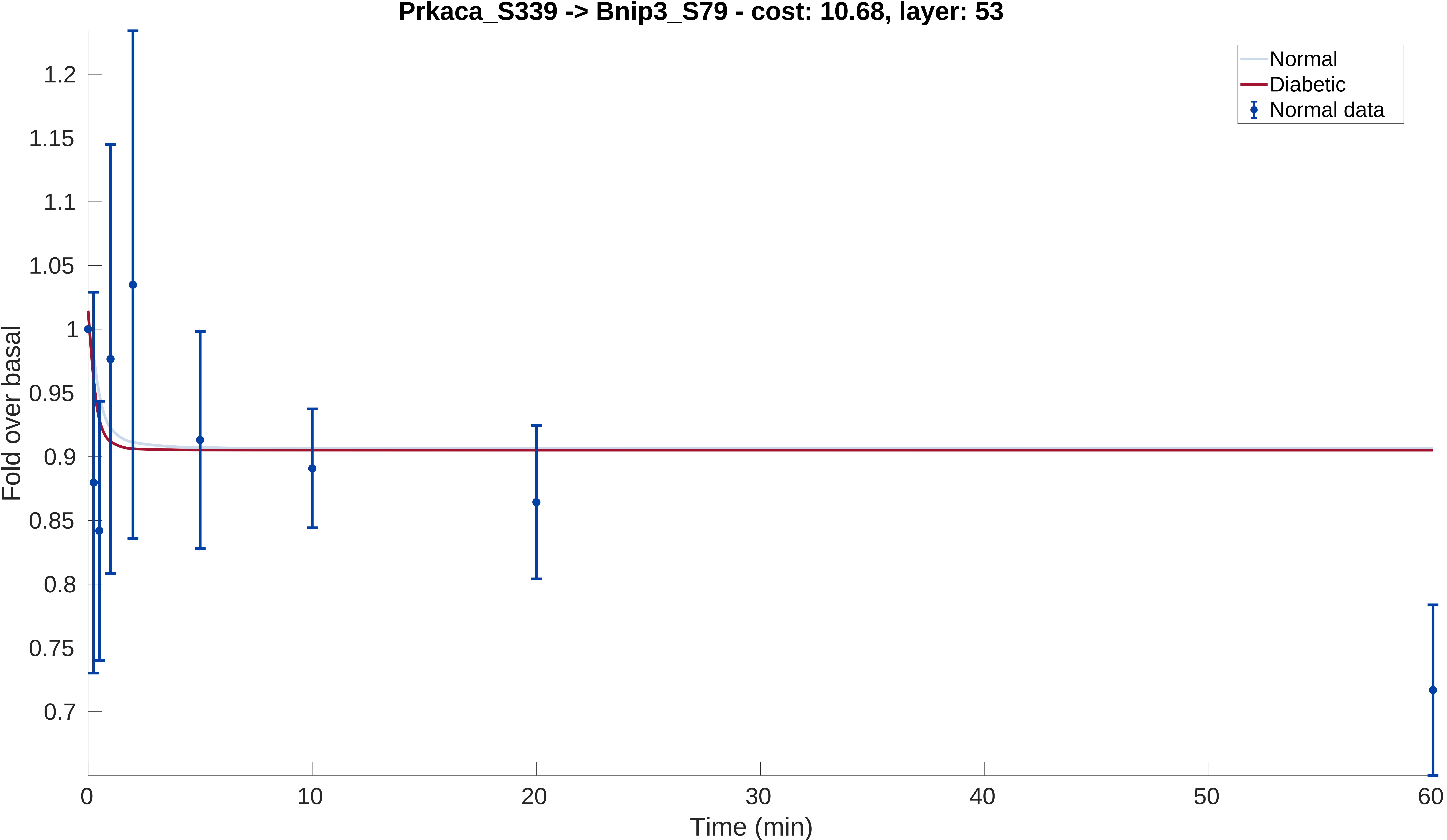
Prkaca_S339 -> Bnip3_S79 - cost: 10.68, layer: 53
Normal
Diabetic
Normal data
Fold over basal
1.2
1.15
1.1
1.05
1
0.95
0.9
0.85
0.8
0.75
0.7
0
10
20
30
40
50
60
Time (min)

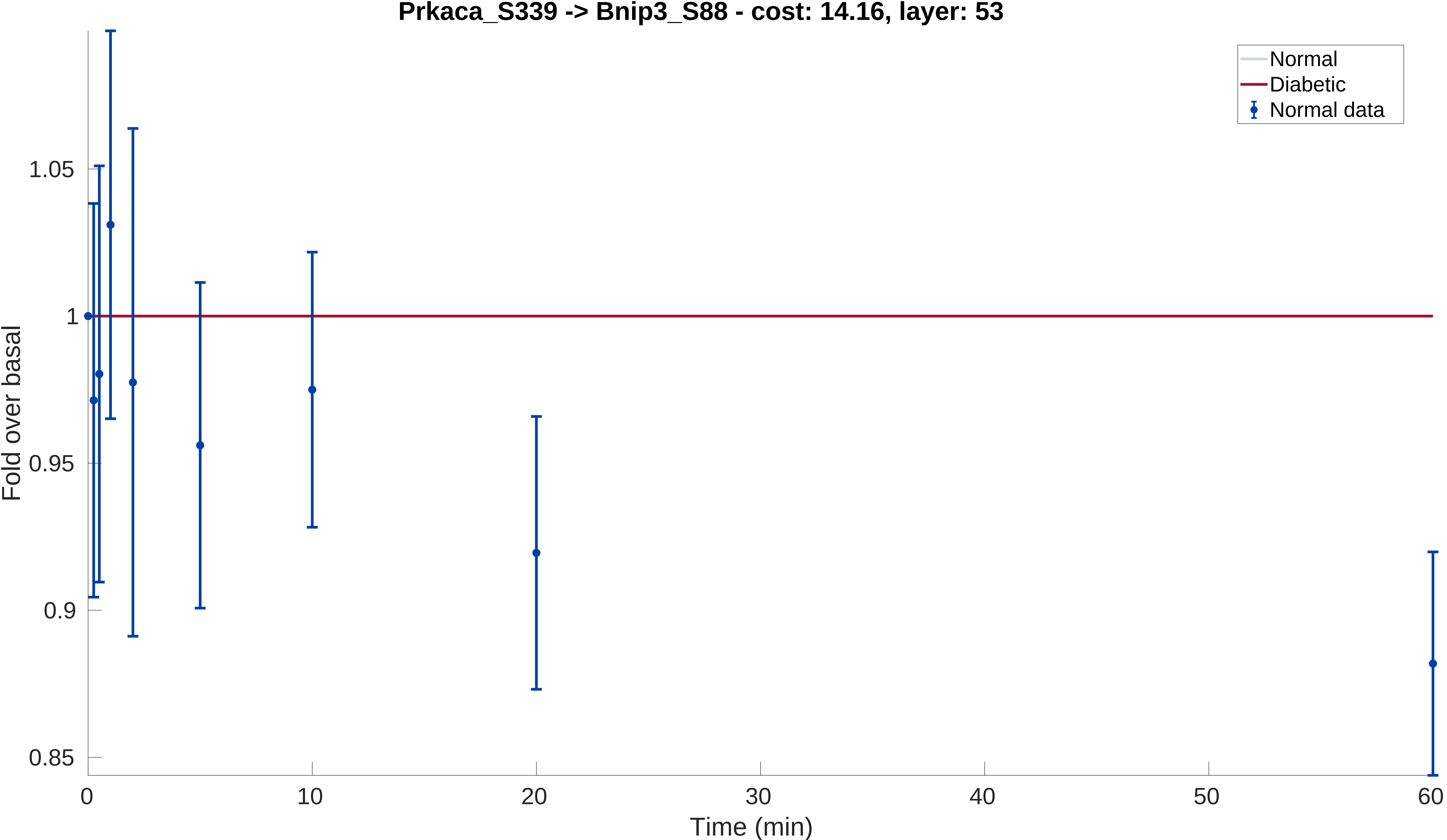

Prkaca_S339 -> Bnip3_S88 - cost: 14.16, layer: 53
Normal
Diabetic
Normal data
1.05
1
0.95
0.9
0.85
Fold over basal
0
10
20
30
40
50
60
Time (min)

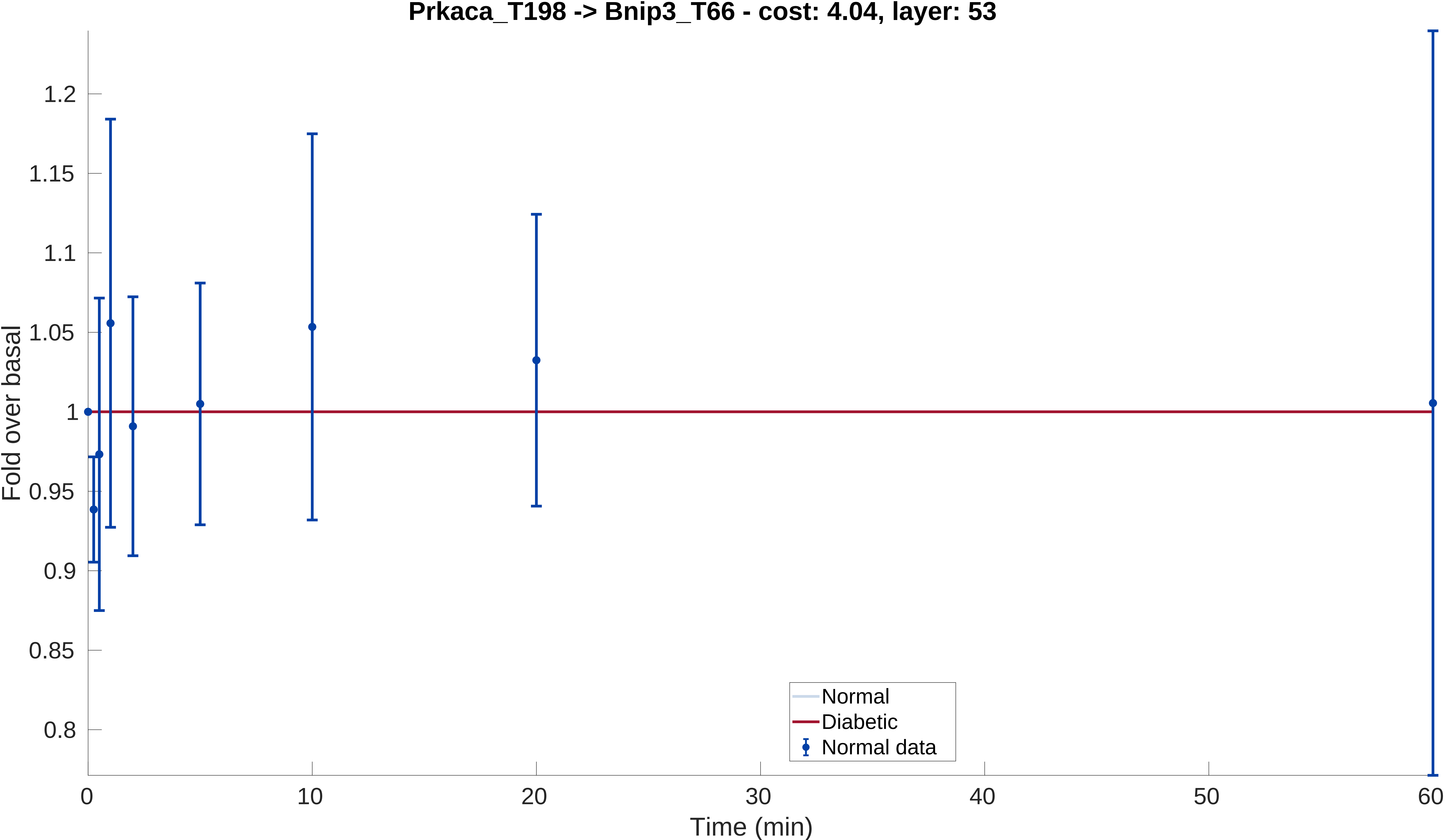

Prkaca_T198 -> Bnip3_T66 - cost: 4.04, layer: 53
Fold over basal
1.2
1.15
1.1
1.05
1
0.95
0.9
0.85
0.8
0
10
20
30
40
50
60
Time (min)
Normal
Diabetic
Normal data

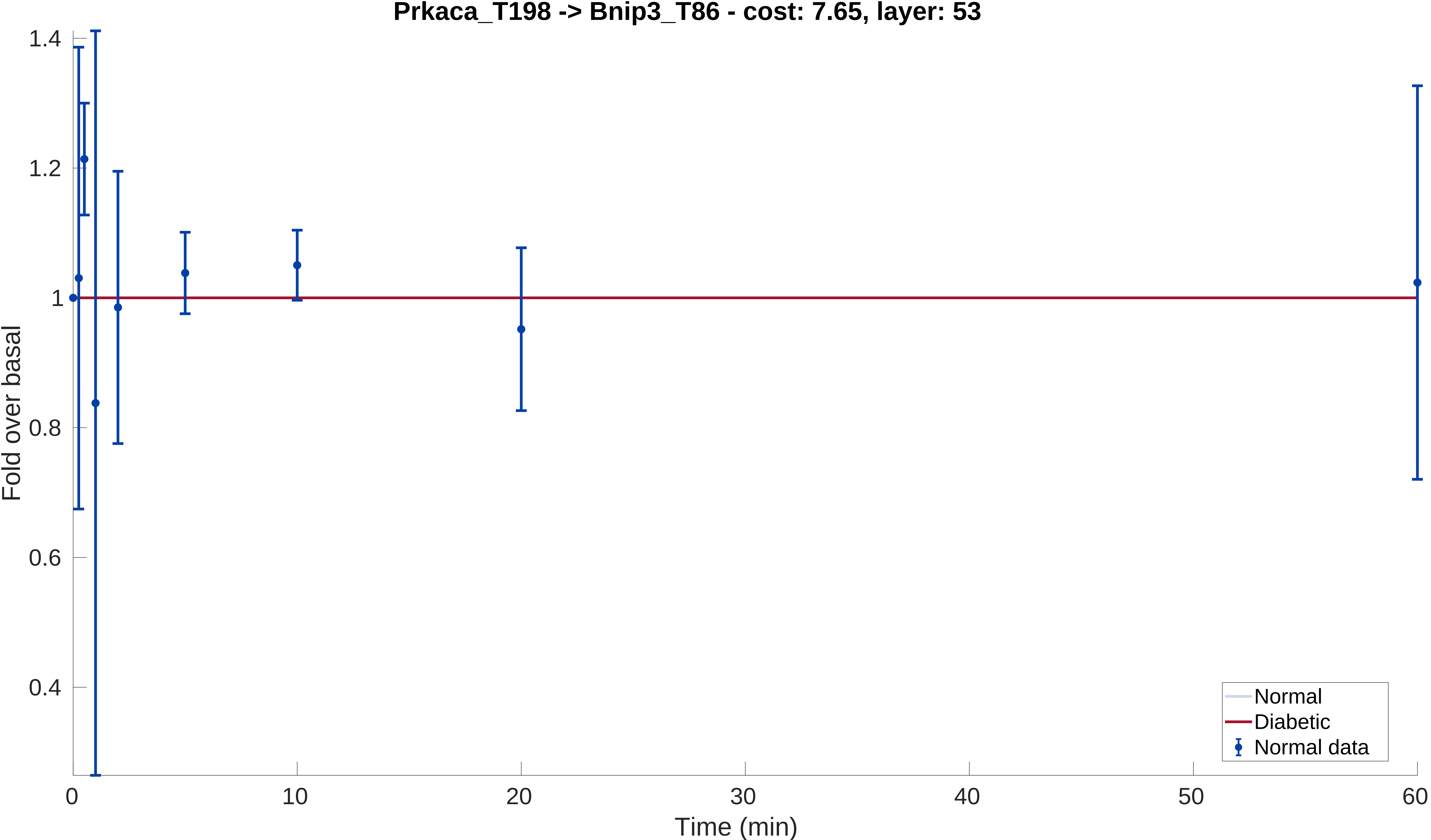

Prkaca_T198 -> Bnip3_T86 - cost: 7.65, layer: 53
1.4
1.2
1
0.8
0.6
0.4
Fold over basal
0
10
20
30
40
50
60
Time (min)
Normal
Diabetic
Normal data

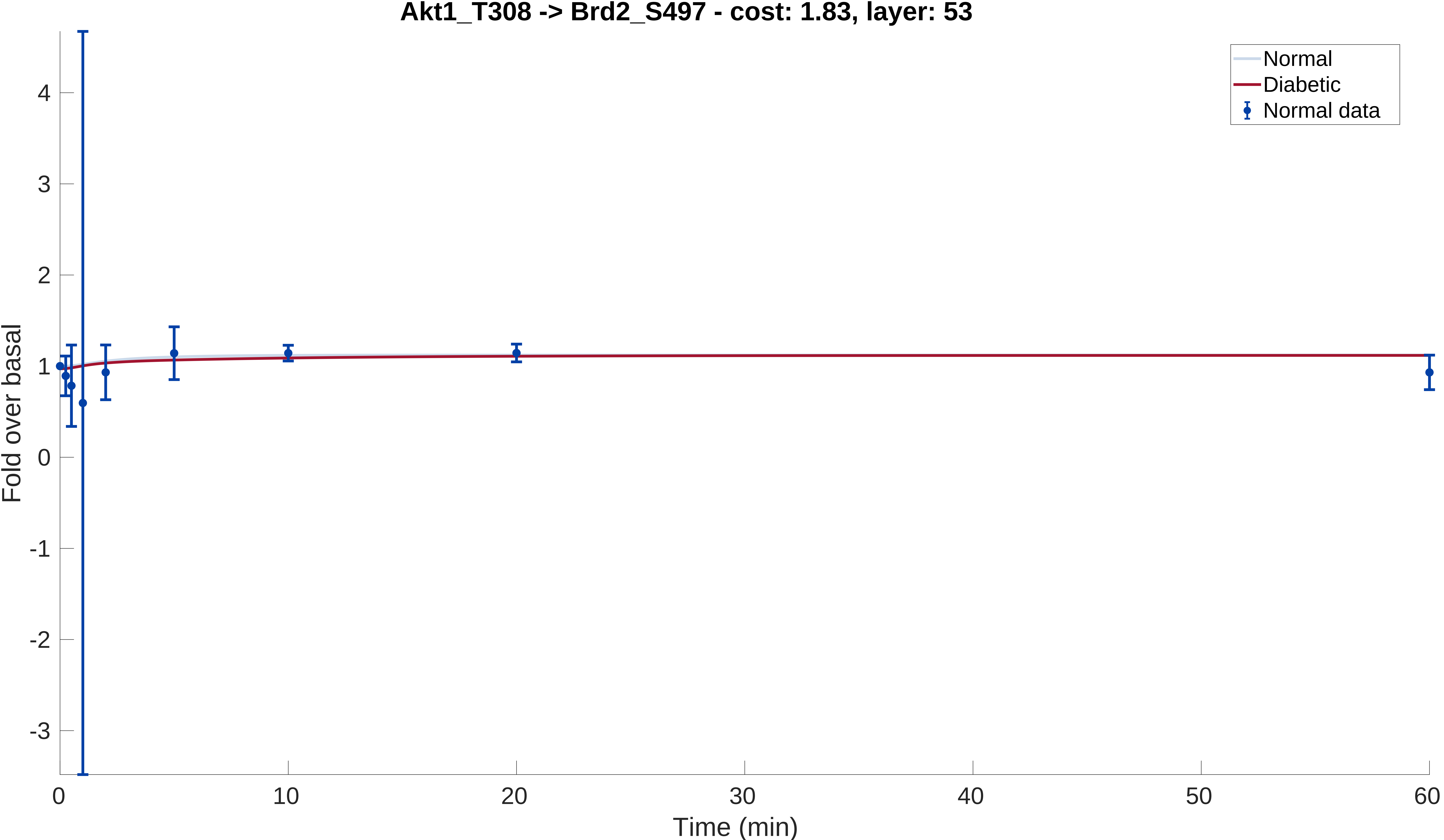

Akt1_T308 -> Brd2_S497 - cost: 1.83, layer: 53
Normal
Diabetic
Normal data
Fold over basal
4
3
2
1
0
-1
-2
-3
0
10
20
30
40
50
60
Time (min)

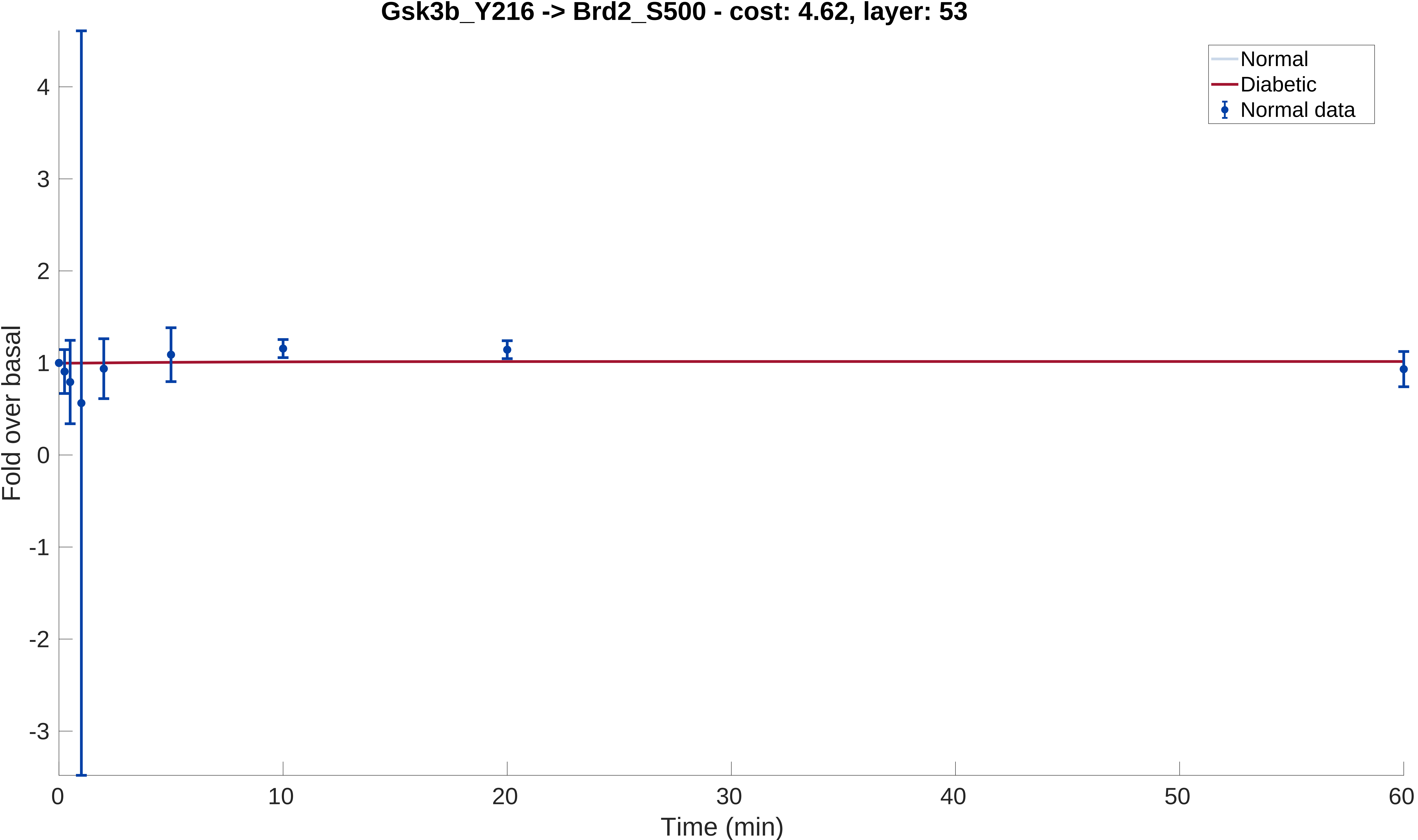

Gsk3b_Y216 -> Brd2_S500 - cost: 4.62, layer: 53
Normal
Diabetic
Normal data
Fold over basal
Time (min)
4
3
2
1
0
-1
-2
-3
0
10
20
30
40
50
60

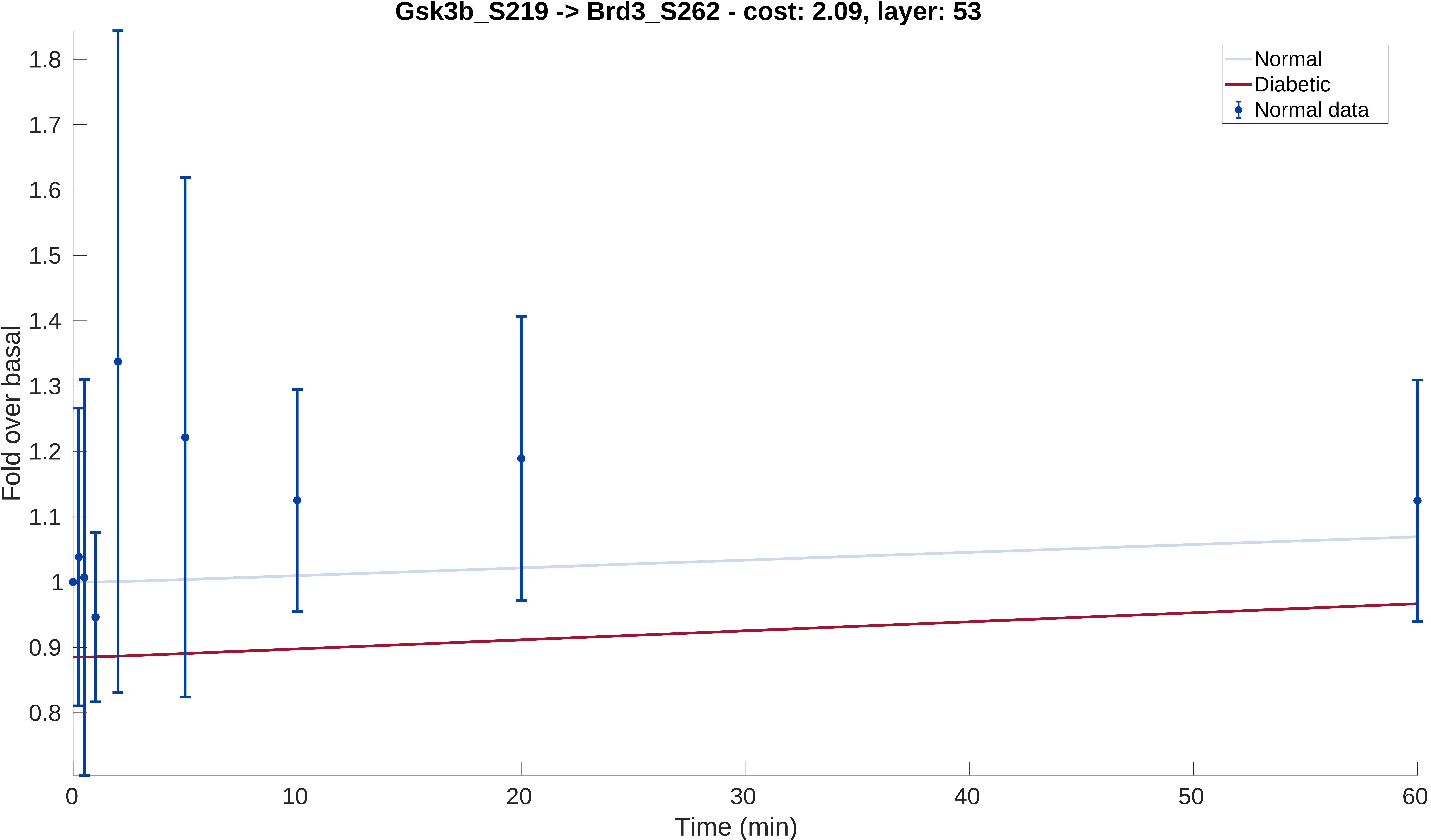

Gsk3b_S219 -> Brd3_S262 - cost: 2.09, layer: 53
Normal
Diabetic
Normal data
Fold over basal
1.8
1.7
1.6
1.5
1.4
1.3
1.2
1.1
1
0.9
0.8
0
10
20
30
40
50
60
Time (min)

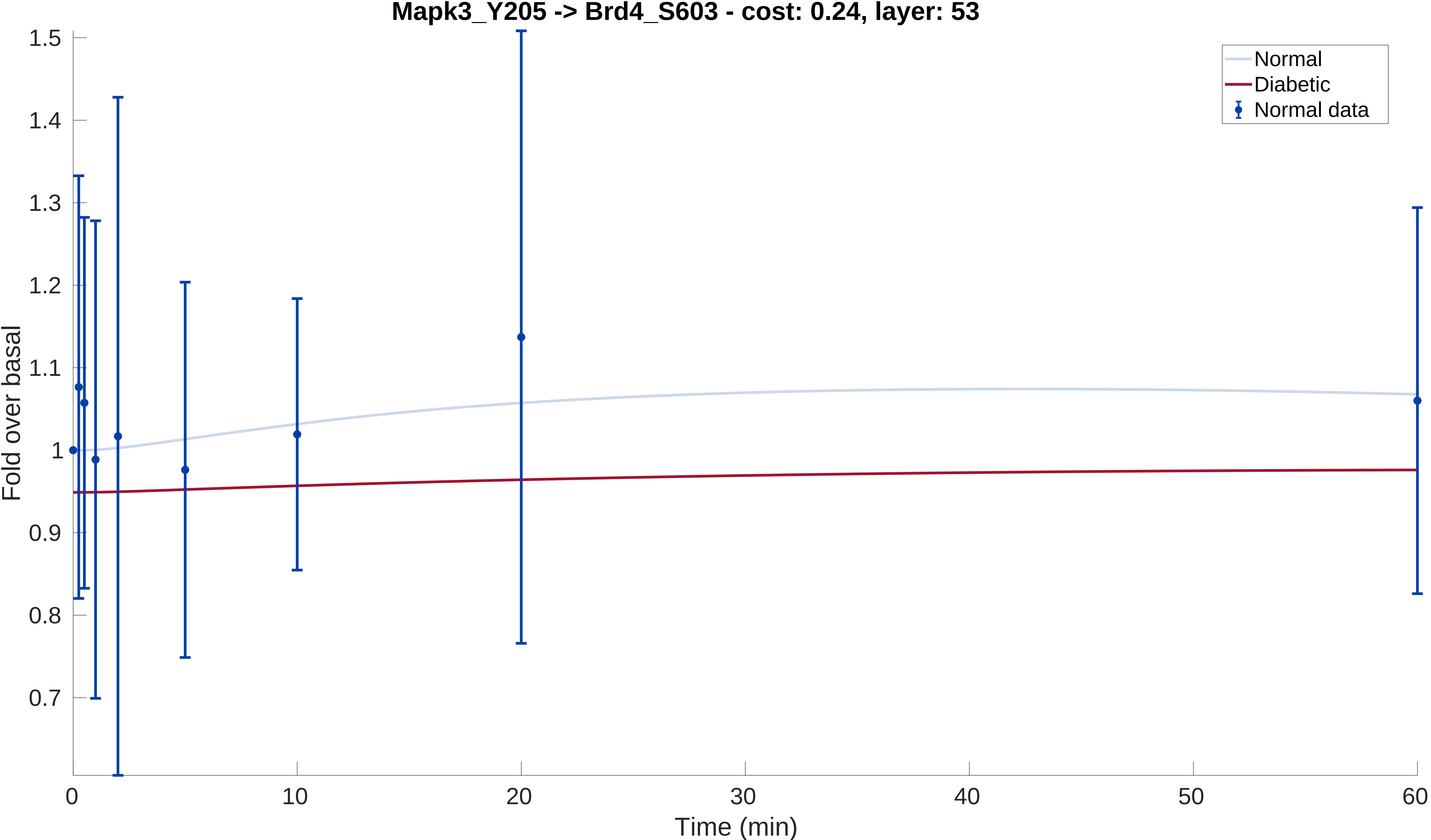
Mapk3_Y205 -> Brd4_S603 - cost: 0.24, layer: 53
Normal
Diabetic
Normal data
Fold over basal
Time (min)
1.5
1.4
1.3
1.2
1.1
1
0.9
0.8
0.7
0
10
20
30
40
50
60

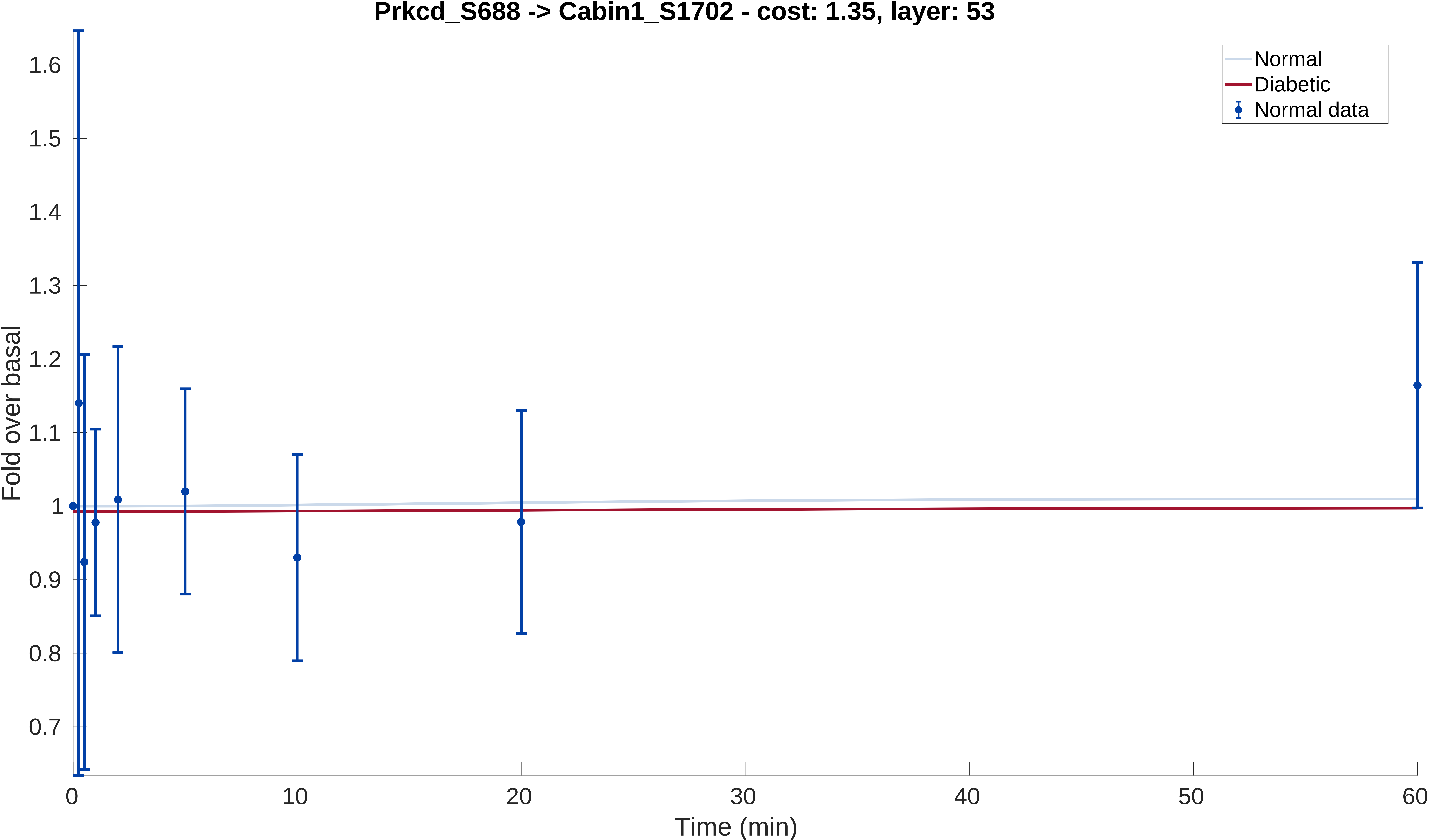

Prkcd_S688 -> Cabin1_S1702 - cost: 1.35, layer: 53
Normal
Diabetic
Normal data
Fold over basal
1.6
1.5
1.4
1.3
1.2
1.1
1
0.9
0.8
0.7
0
10
20
30
40
50
60
Time (min)

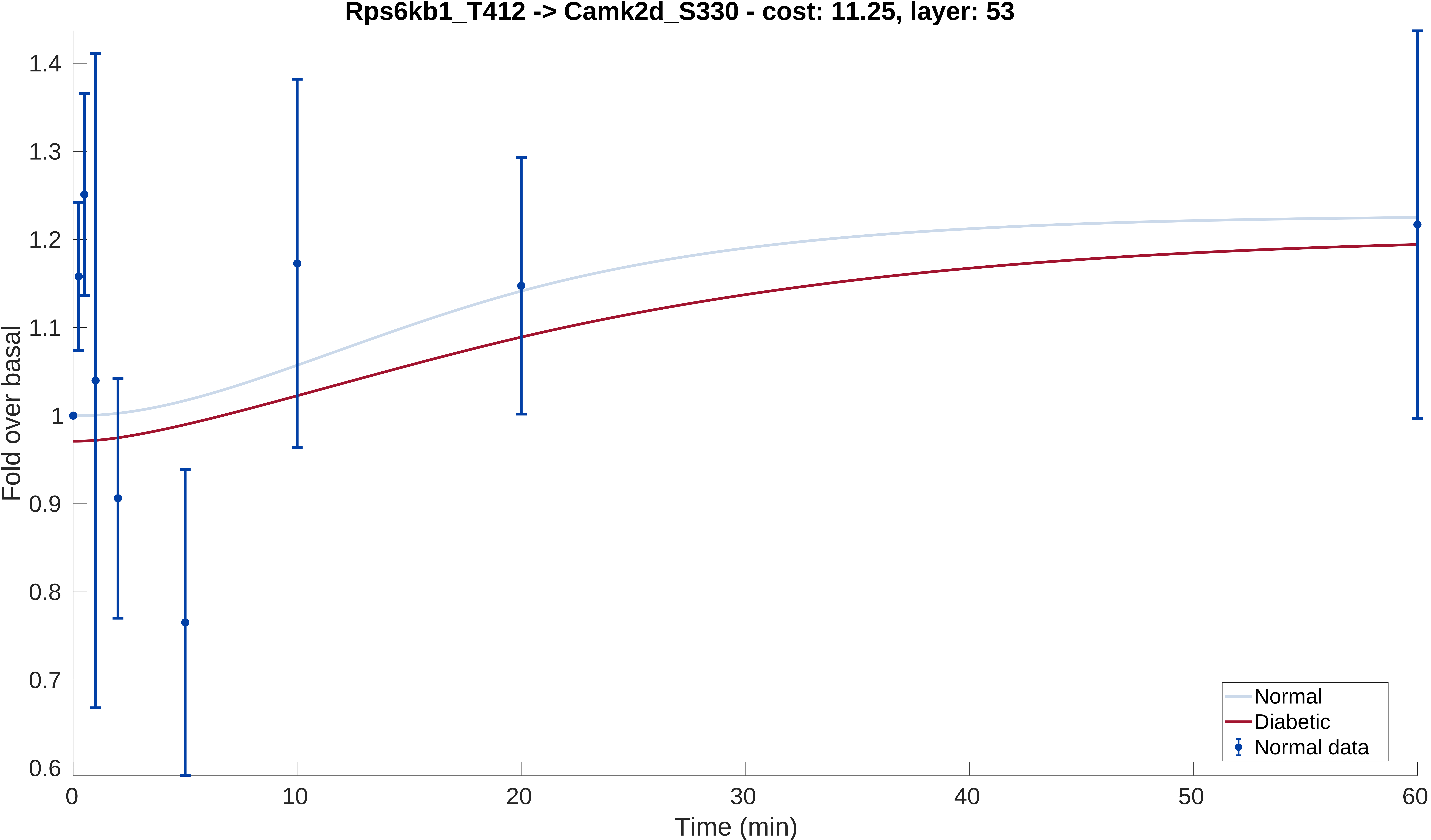

Rps6kb1_T412 -> Camk2d_S330 - cost: 11.25, layer: 53
Fold over basal
1.4
1.3
1.2
1.1
1
0.9
0.8
0.7
0.6
0
10
20
30
40
50
60
Time (min)
Normal
Diabetic
Normal data

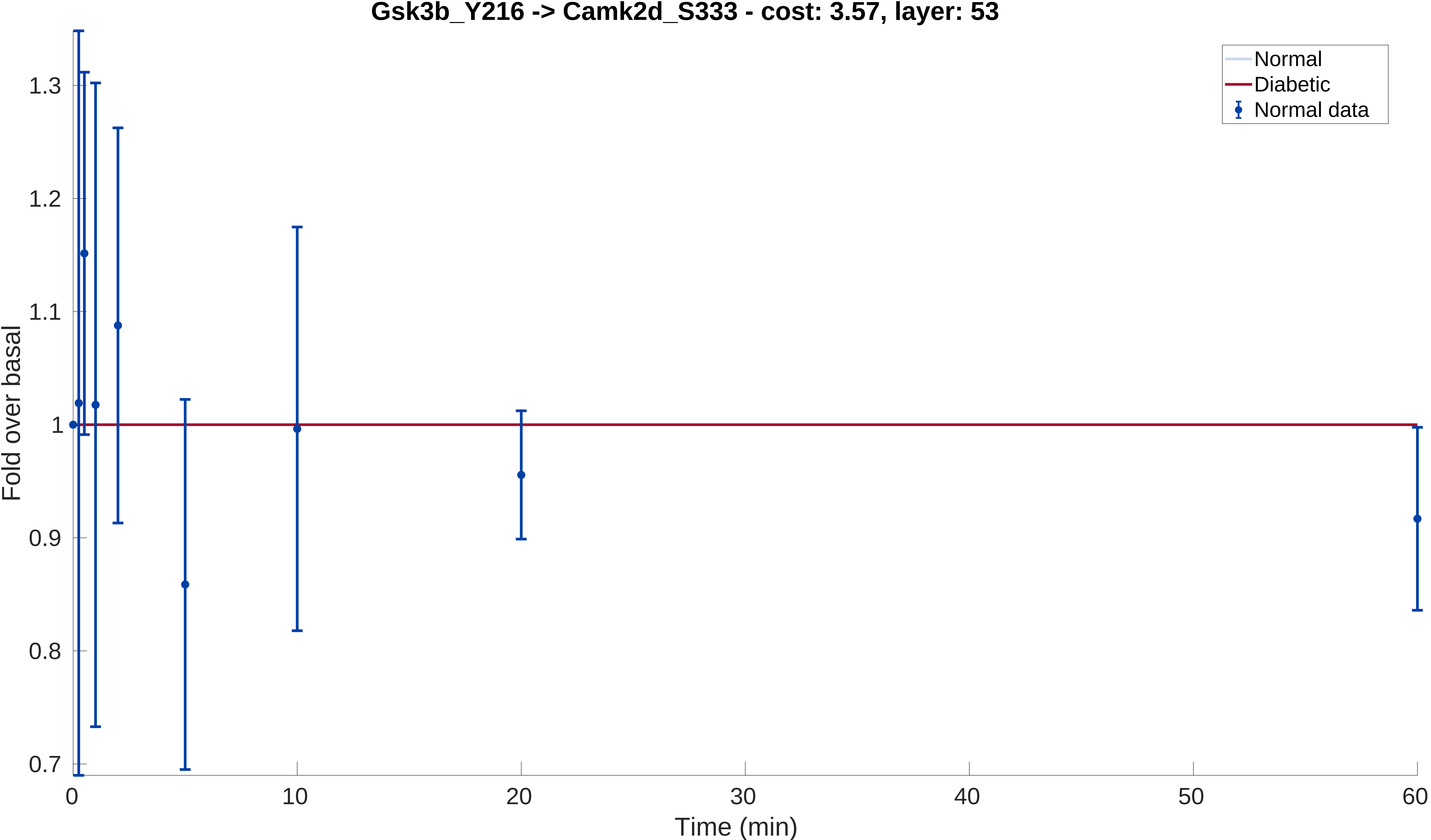

Gsk3b_Y216 -> Camk2d_S333 - cost: 3.57, layer: 53
Normal
Diabetic
Normal data
1.3
1.2
1.1
1
0.9
0.8
0.7
Fold over basal
0
10
20
30
40
50
60
Time (min)

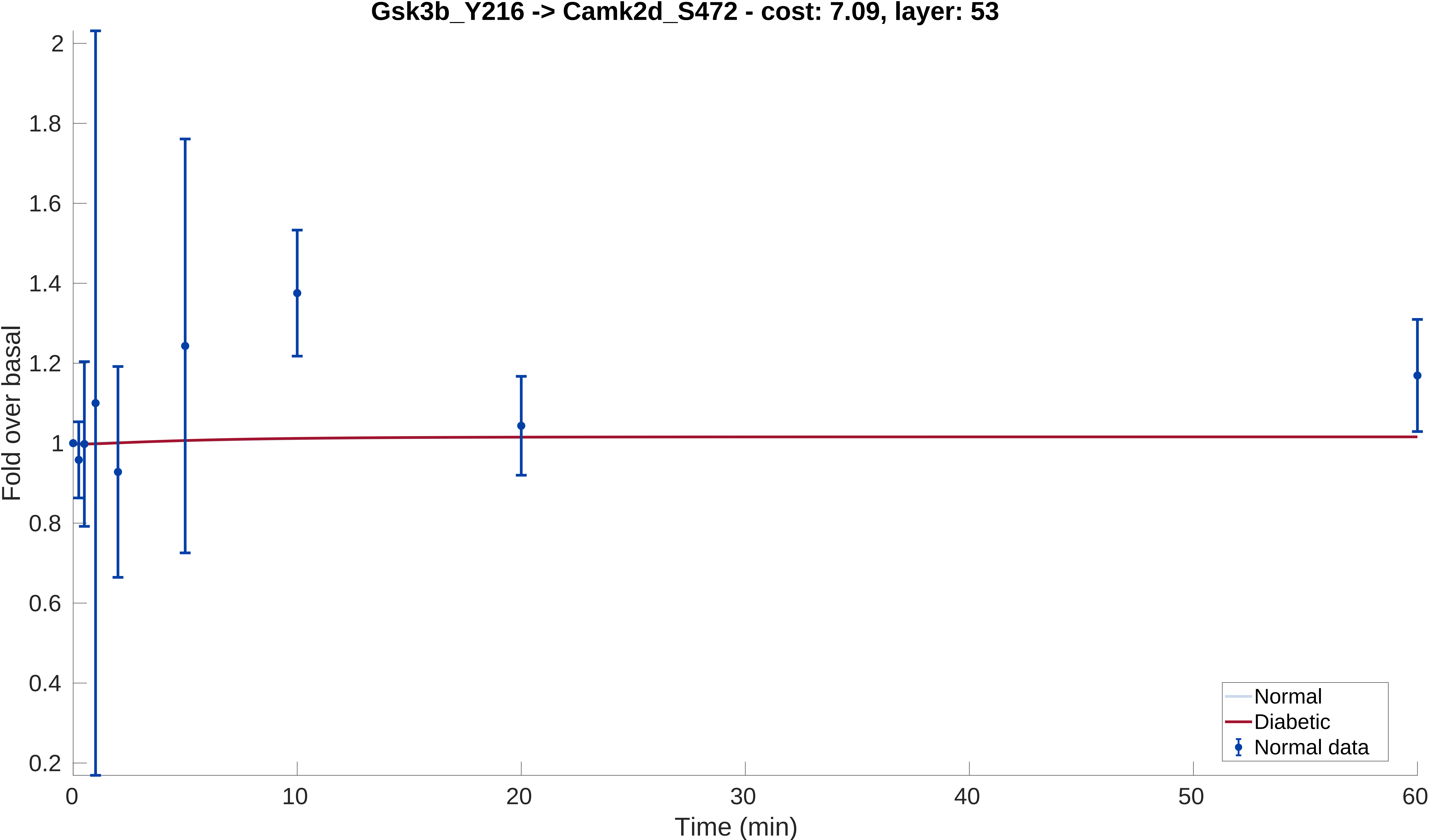

Gsk3b_Y216 -> Camk2d_S472 - cost: 7.09, layer: 53
Fold over basal
2
1.8
1.6
1.4
1.2
1
0.8
0.6
0.4
0.2
0
10
20
30
40
50
60
Time (min)
Normal
Diabetic
Normal data

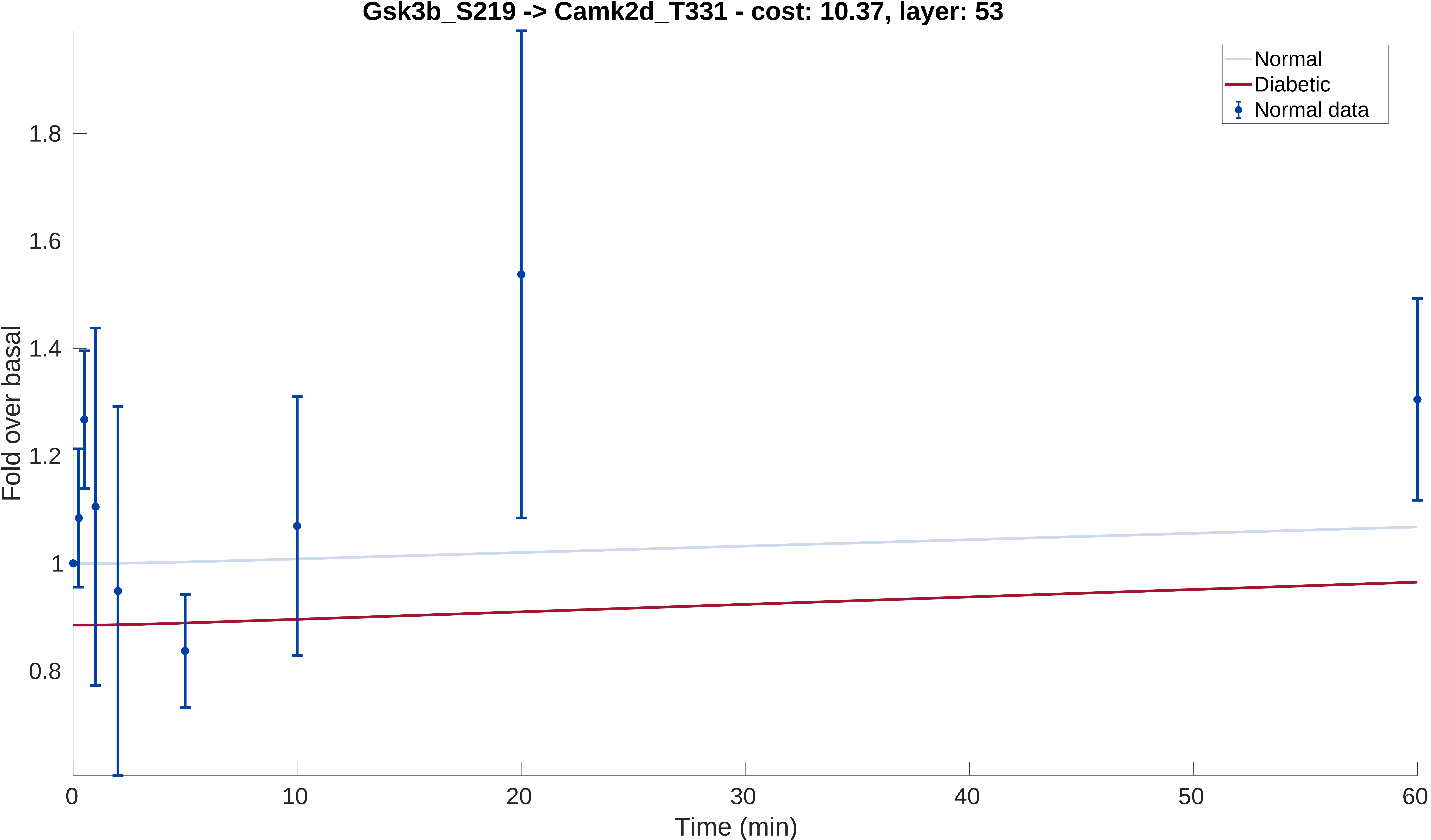

Gsk3b_S219 -> Camk2d_T331 - cost: 10.37, layer: 53
Normal
Diabetic
Normal data
Fold over basal
1.8
1.6
1.4
1.2
1
0.8
0
10
20
30
40
50
60
Time (min)

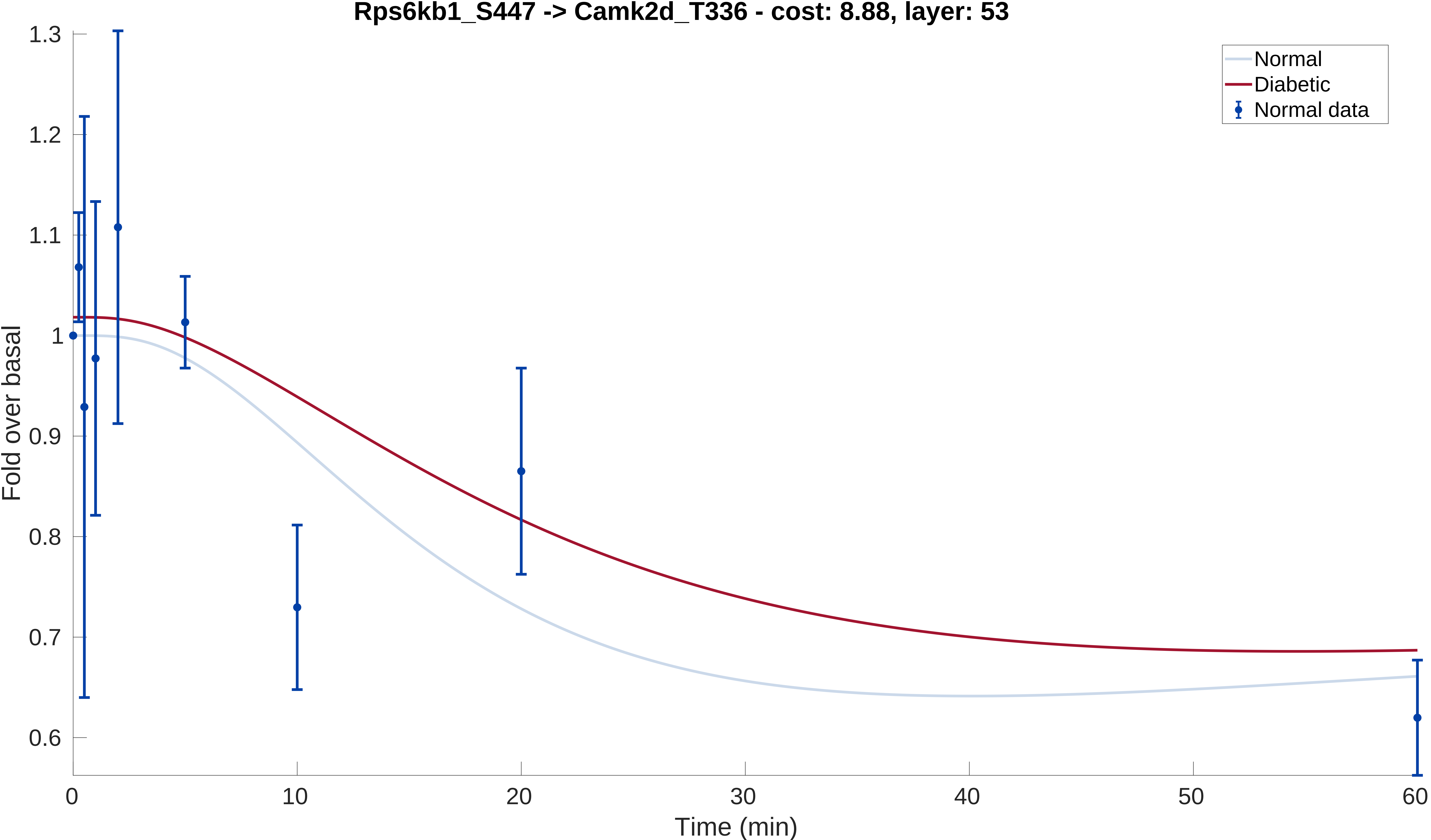

Rps6kb1_S447 -> Camk2d_T336 - cost: 8.88, layer: 53
Fold over basal
Time (min)
Normal
Diabetic
Normal data
1.3
1.2
1.1
1
0.9
0.8
0.7
0.6
0
10
20
30
40
50
60

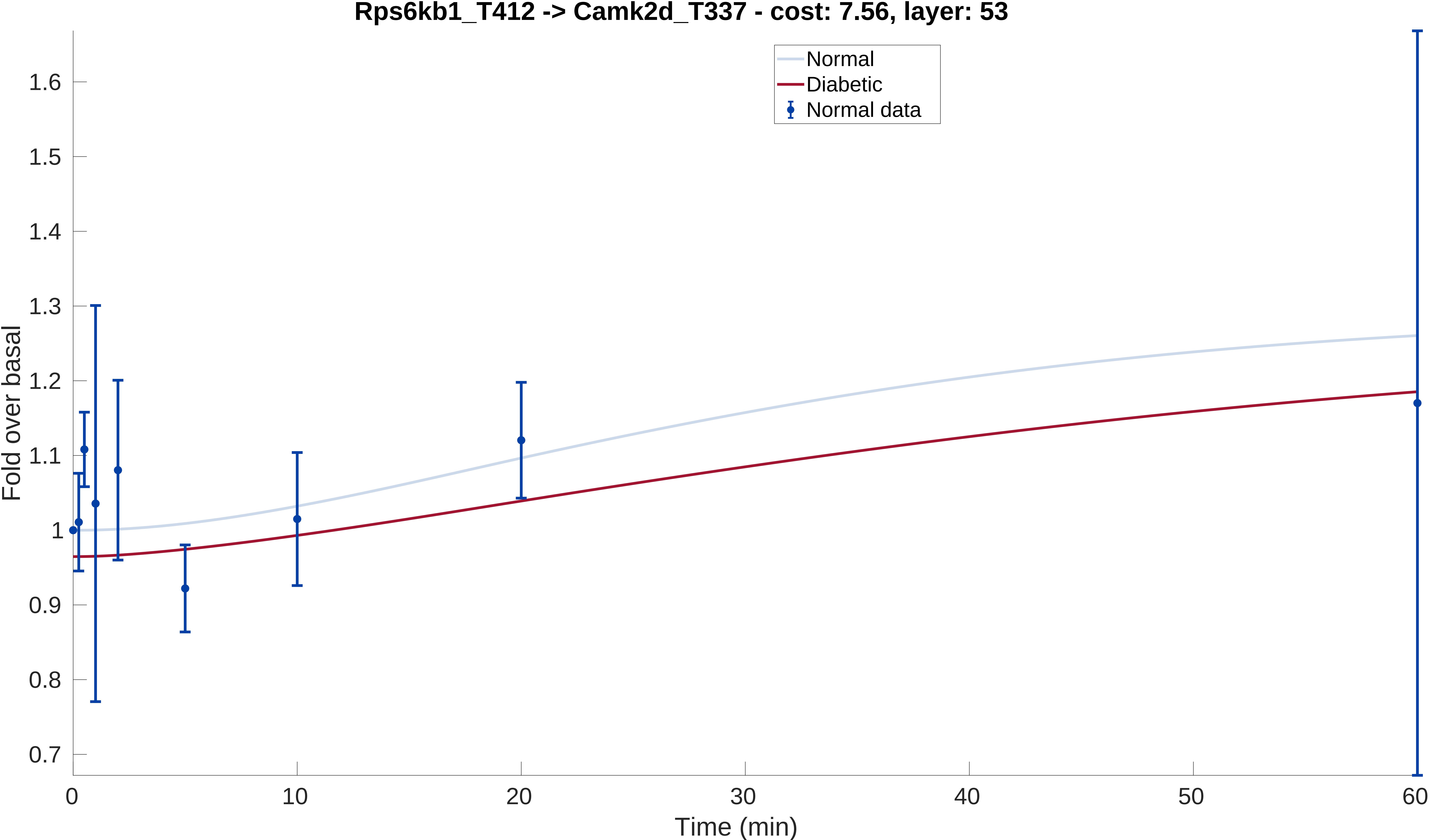
Rps6kb1_T412 -> Camk2d_T337 - cost: 7.56, layer: 53
Normal
Diabetic
Normal data
Fold over basal
Time (min)
0.7
0.8
0.9
1
1.1
1.2
1.3
1.4
1.5
1.6
0
10
20
30
40
50
60

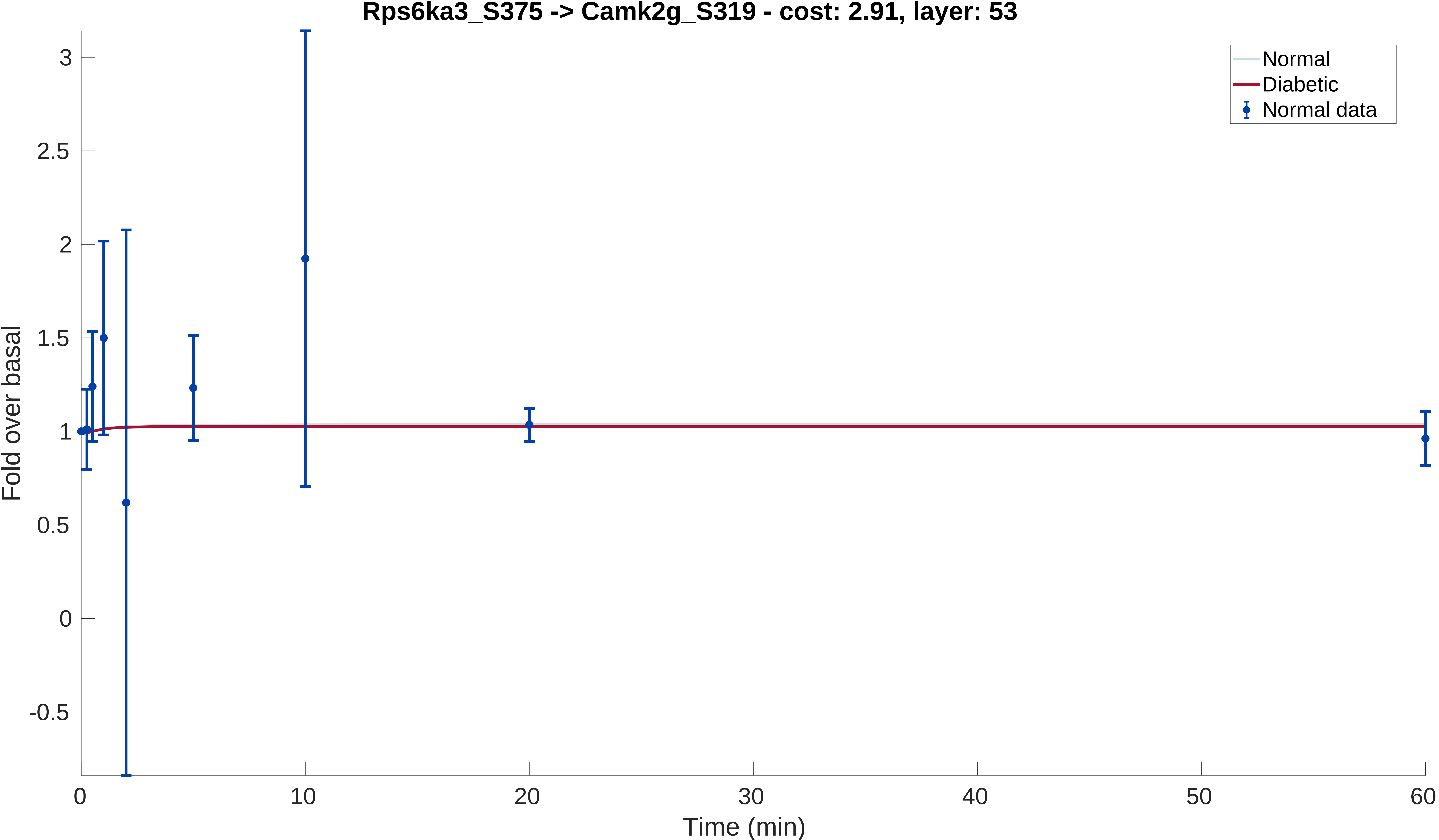
Rps6ka3_S375 -> Camk2g_S319 - cost: 2.91, layer: 53
Normal
Diabetic
Normal data
Fold over basal
Time (min)
3
2.5
2
1.5
1
0.5
0
-0.5
0
10
20
30
40
50
60

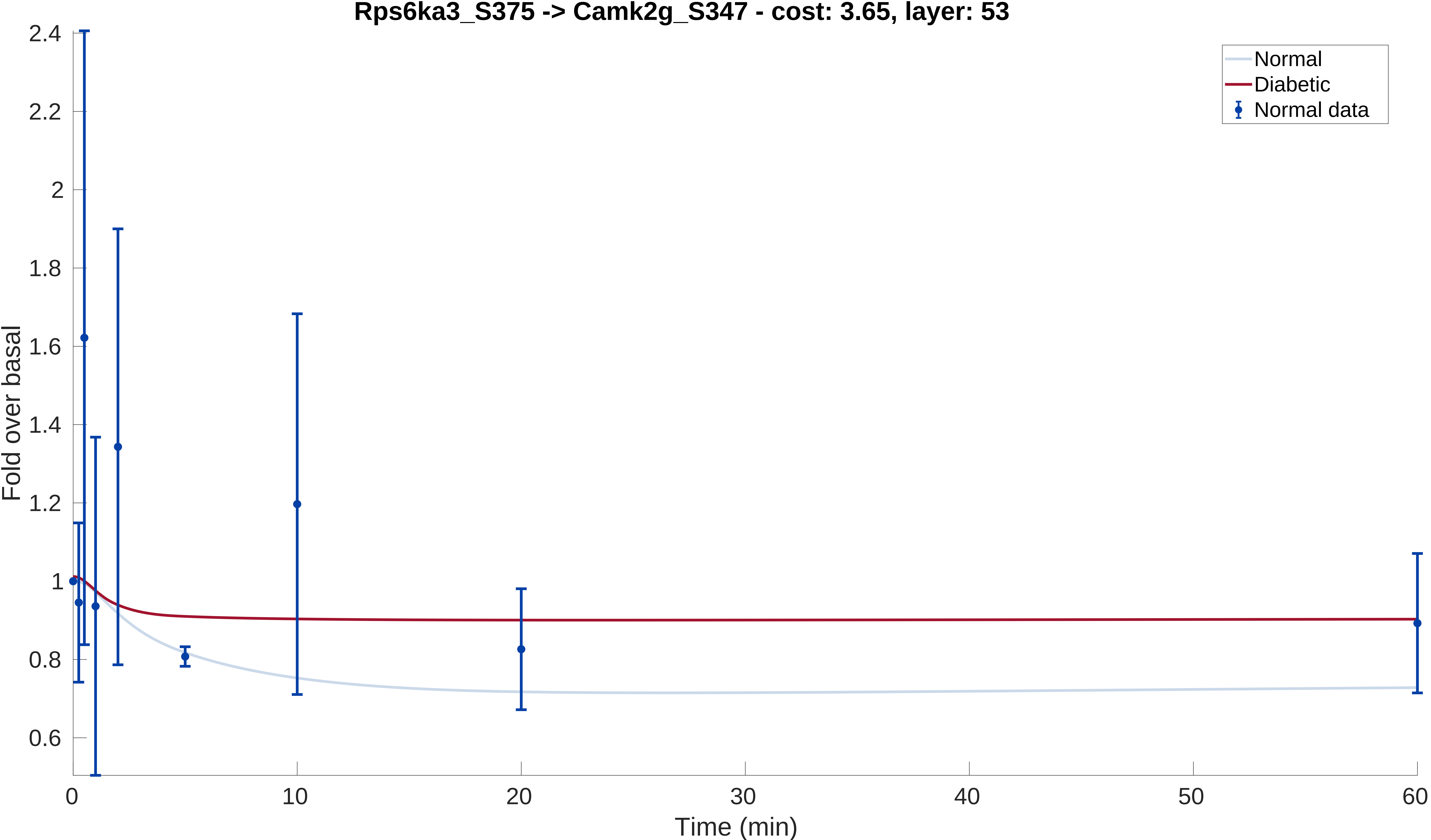
Rps6ka3_S375 -> Camk2g_S347 - cost: 3.65, layer: 53
Normal
Diabetic
Normal data
Fold over basal
Time (min)
2.4
2.2
2
1.8
1.6
1.4
1.2
1
0.8
0.6
0
10
20
30
40
50
60

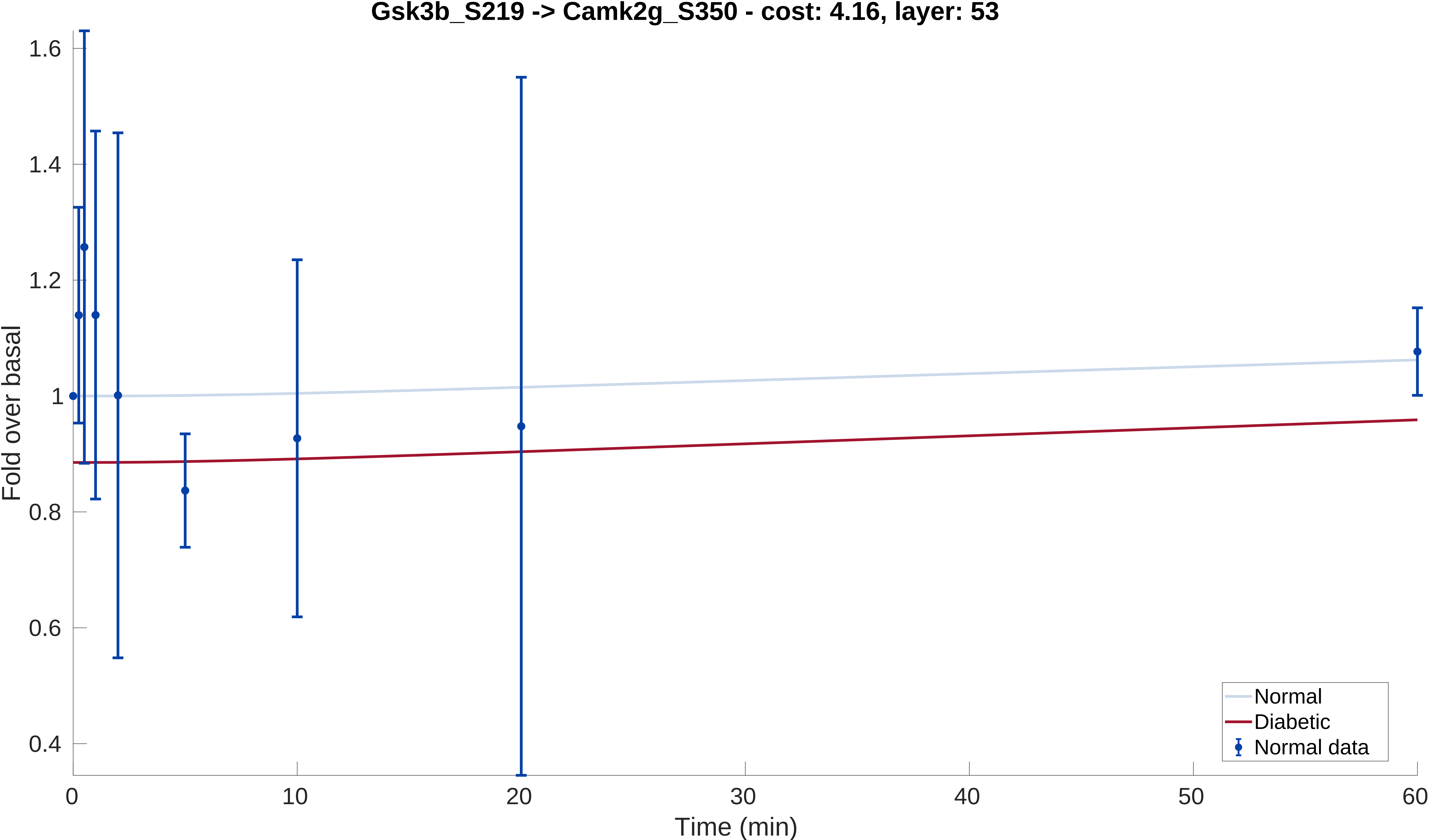

Gsk3b_S219 -> Camk2g_S350 - cost: 4.16, layer: 53
Fold over basal
1.6
1.4
1.2
1
0.8
0.6
0.4
0
10
20
30
40
50
60
Time (min)
Normal
Diabetic
Normal data

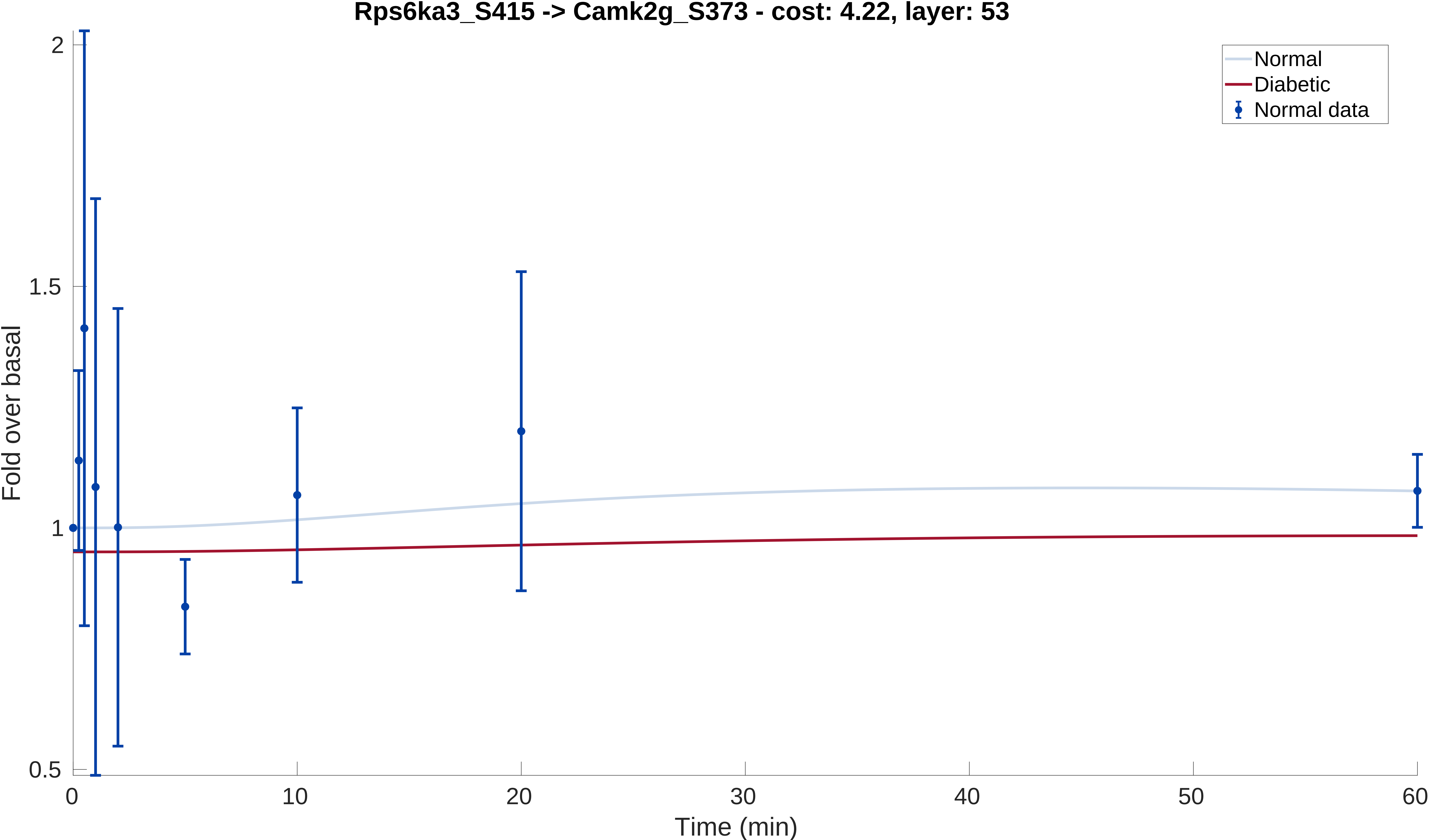

Rps6ka3_S415 -> Camk2g_S373 - cost: 4.22, layer: 53
Normal
Diabetic
Normal data
Fold over basal
Time (min)
2
1.5
1
0.5
0
10
20
30
40
50
60

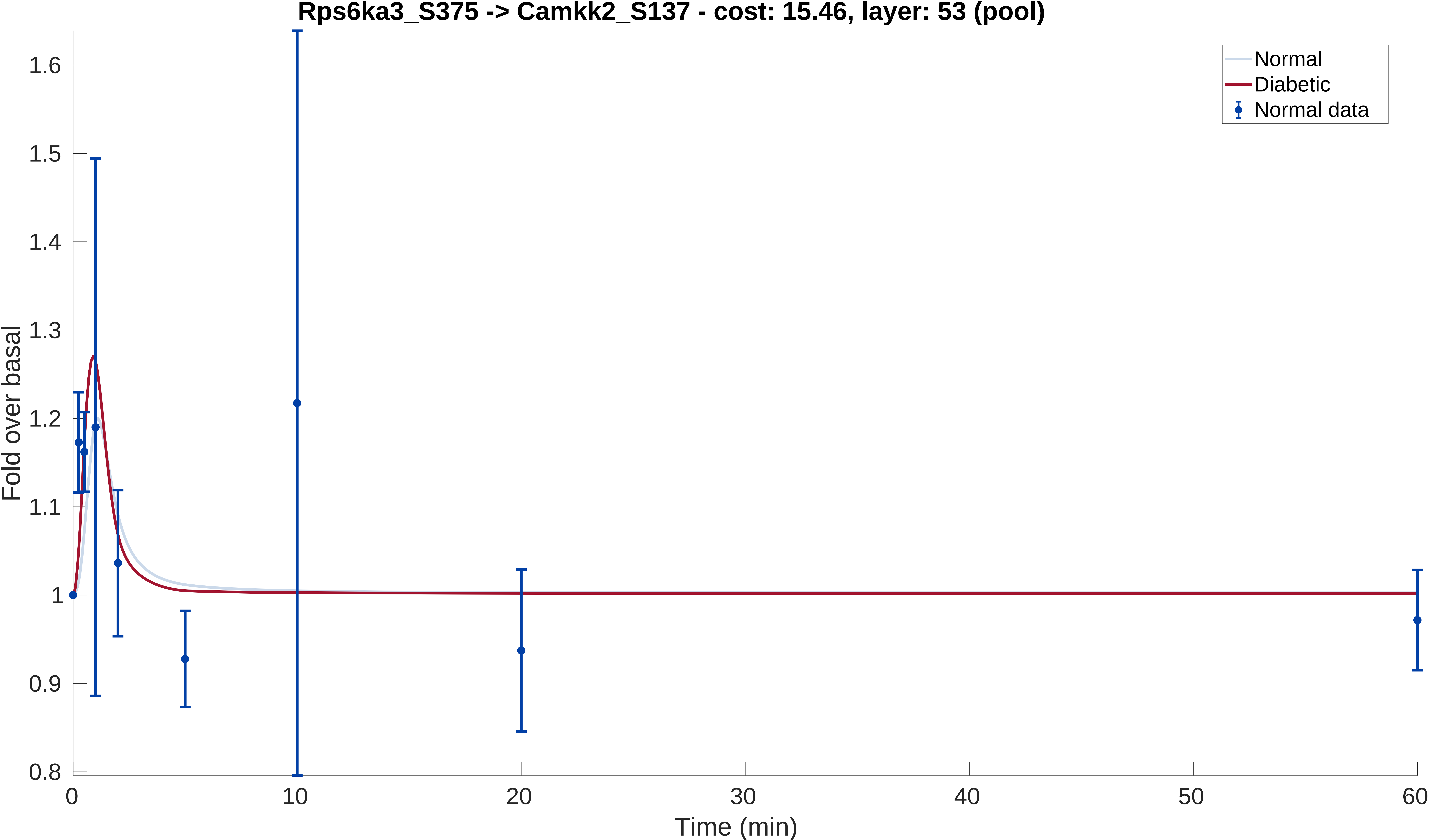

Rps6ka3_S375 -> Camkk2_S137 - cost: 15.46, layer: 53 (pool)
Normal
Diabetic
Normal data
Fold over basal
Time (min)
0.8
0.9
1
1.1
1.2
1.3
1.4
1.5
1.6
0
10
20
30
40
50
60

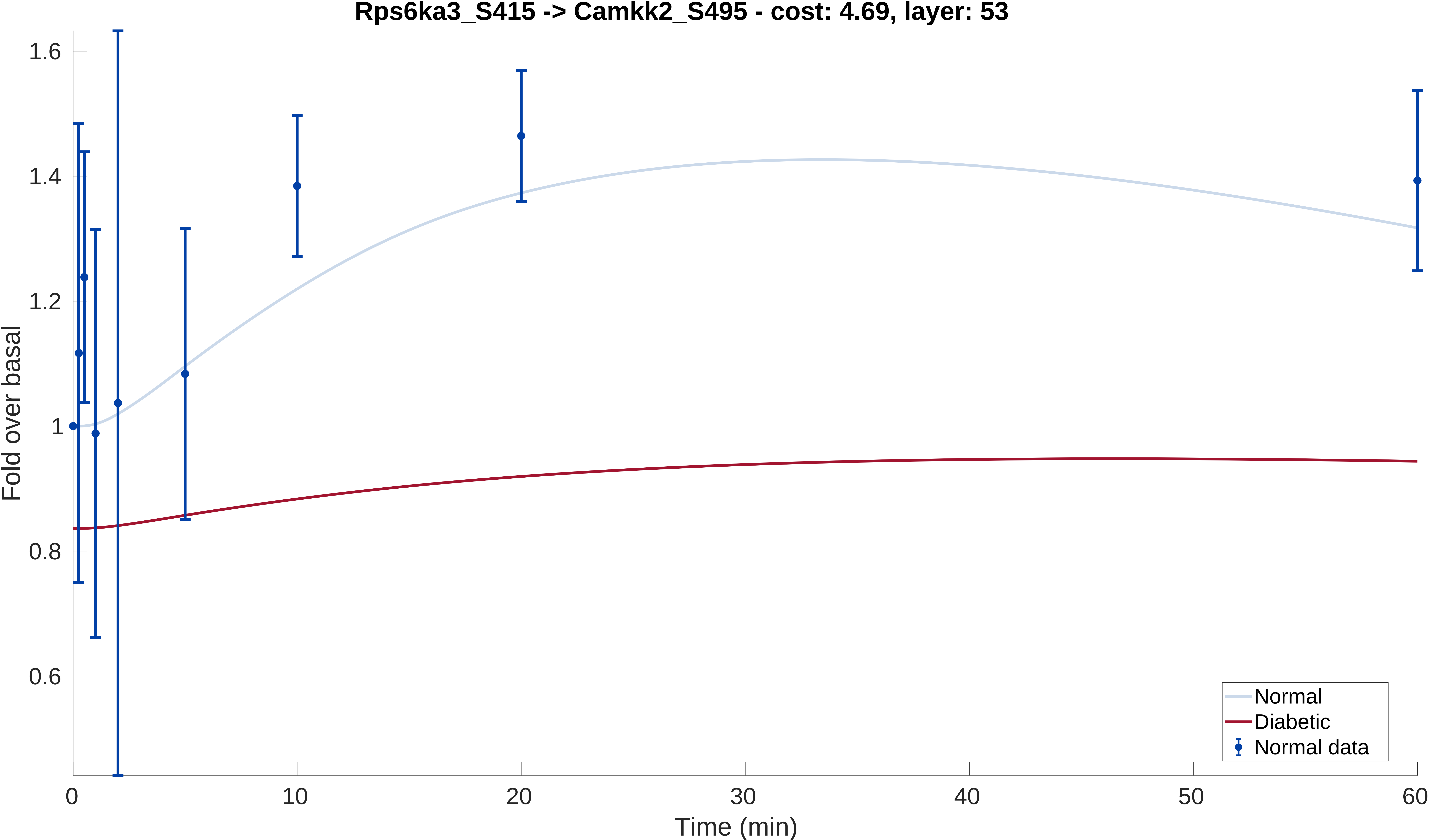

Rps6ka3_S415 -> Camkk2_S495 - cost: 4.69, layer: 53
Fold over basal
1.6
1.4
1.2
1
0.8
0.6
0
10
20
30
40
50
60
Time (min)
Normal
Diabetic
Normal data

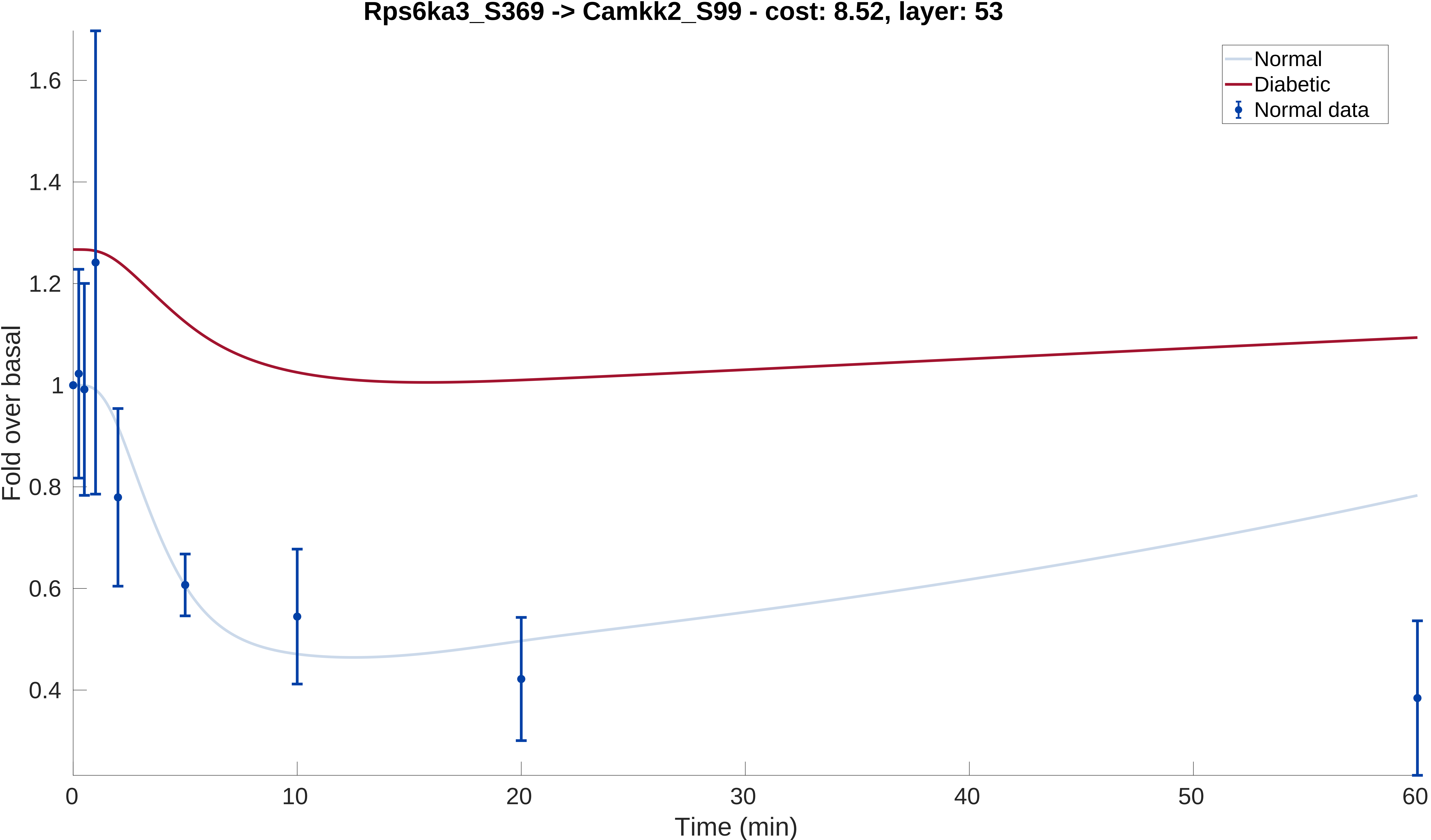

Rps6ka3_S369 -> Camkk2_S99 - cost: 8.52, layer: 53
Normal
Diabetic
Normal data
Fold over basal
1.6
1.4
1.2
1
0.8
0.6
0.4
0
10
20
30
40
50
60
Time (min)

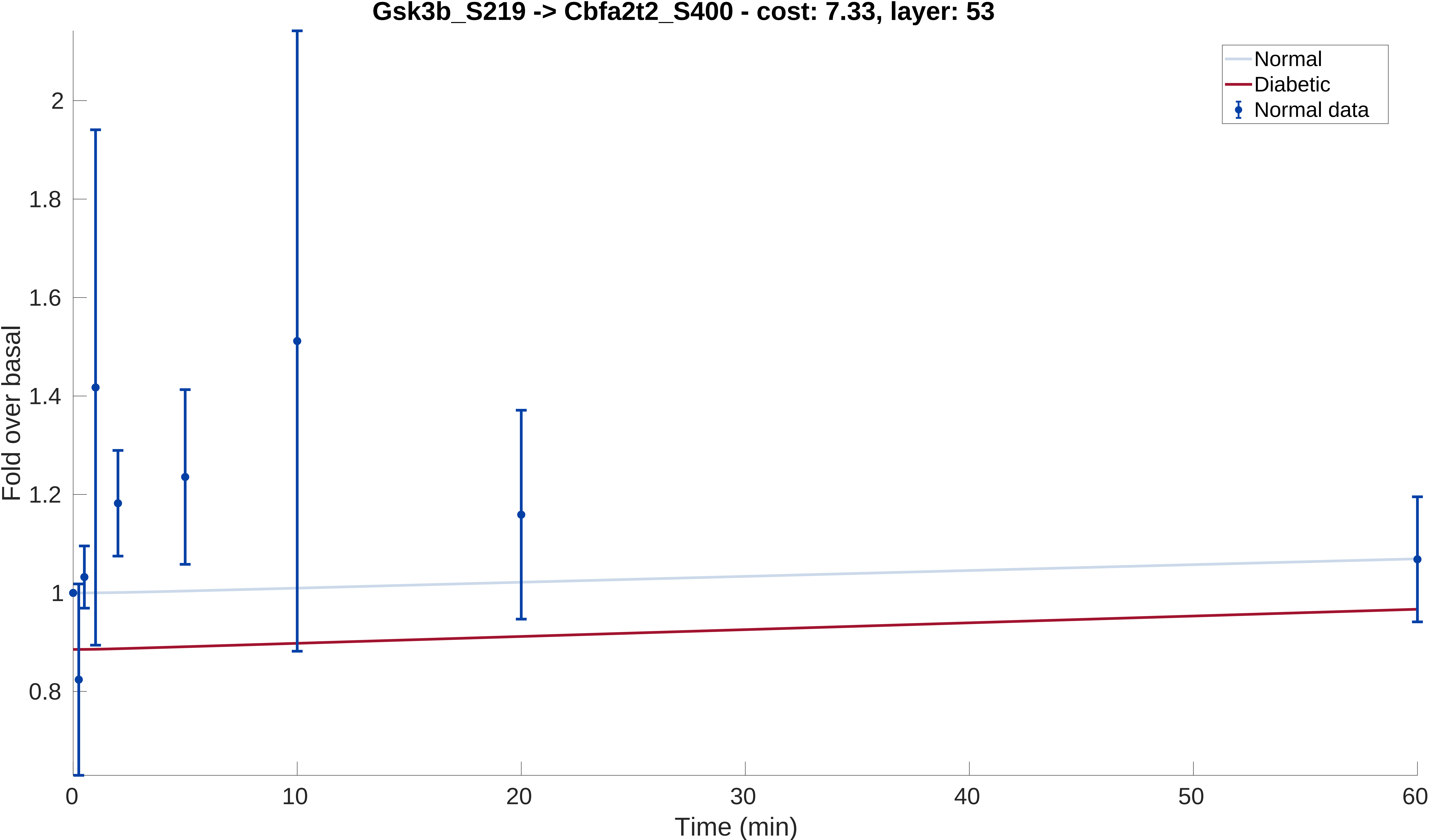

Gsk3b_S219 -> Cbfa2t2_S400 - cost: 7.33, layer: 53
Normal
Diabetic
Normal data
Fold over basal
2
1.8
1.6
1.4
1.2
1
0.8
0
10
20
30
40
50
60
Time (min)

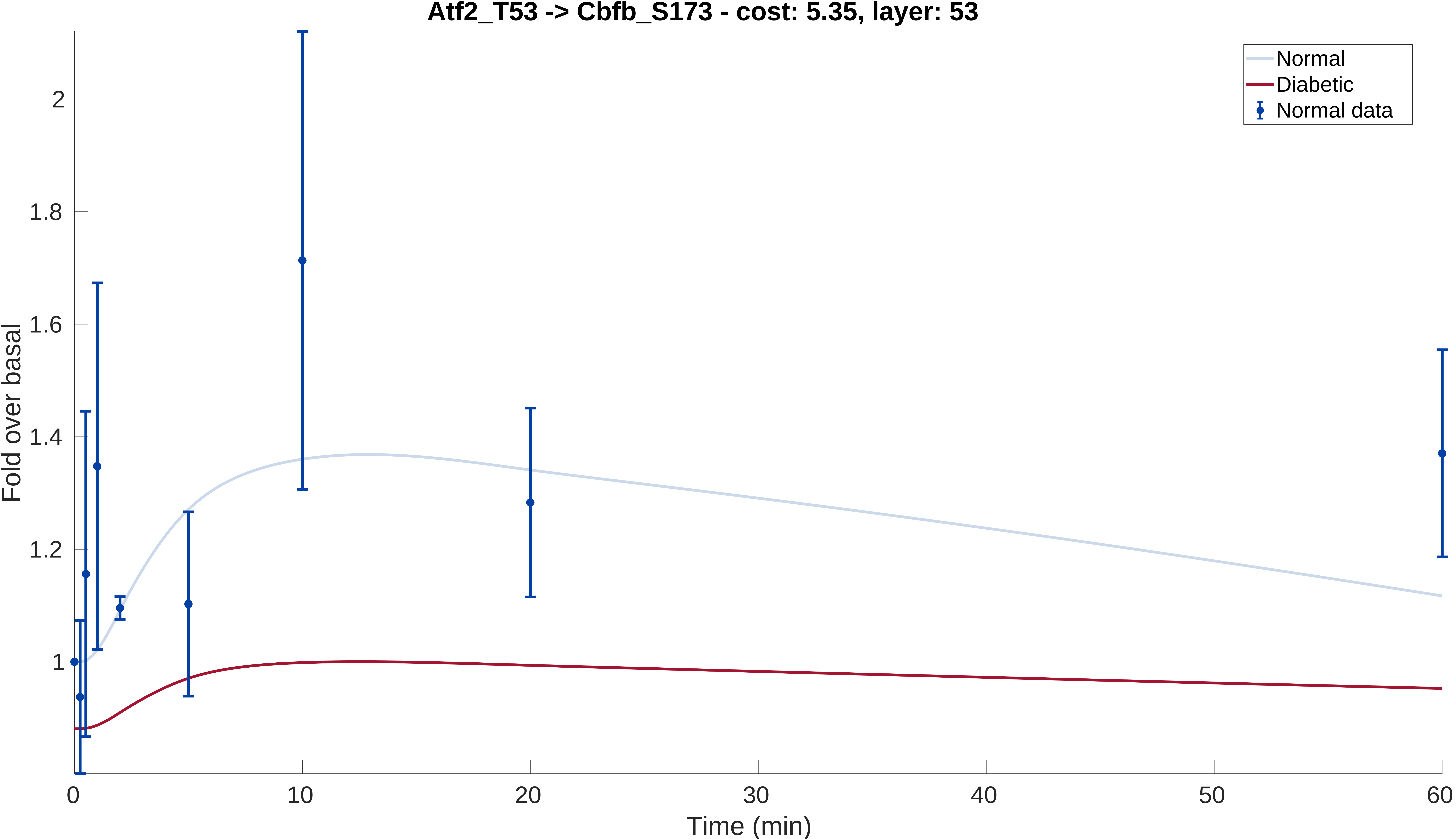

Atf2_T53 -> Cbfb_S173 - cost: 5.35, layer: 53
Normal
Diabetic
Normal data
Fold over basal
Time (min)
2
1.8
1.6
1.4
1.2
1
0
10
20
30
40
50
60

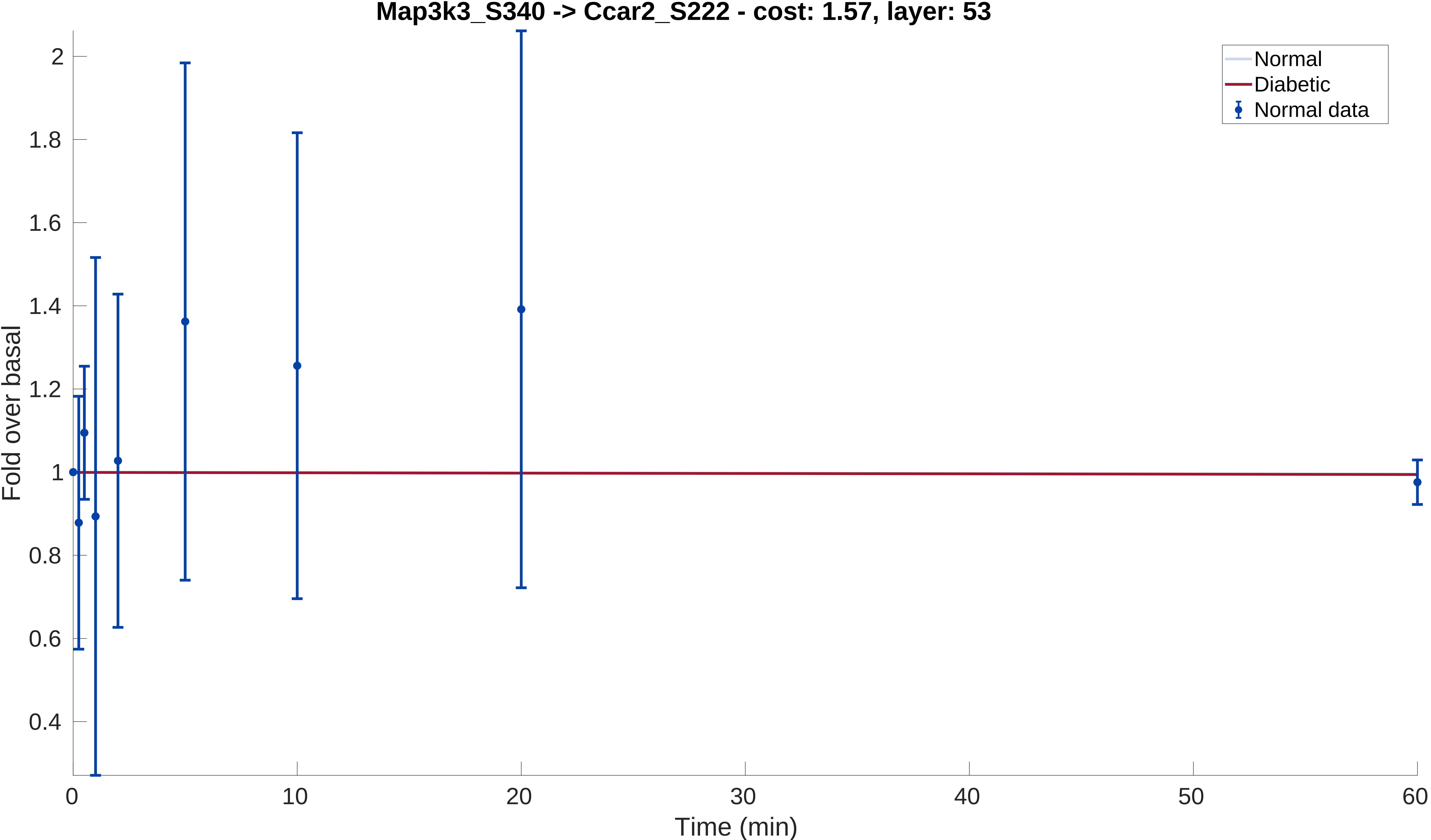
Map3k3_S340 -> Ccar2_S222 - cost: 1.57, layer: 53
Normal
Diabetic
Normal data
Fold over basal
Time (min)
0.4
0.6
0.8
1
1.2
1.4
1.6
1.8
2
0
10
20
30
40
50
60

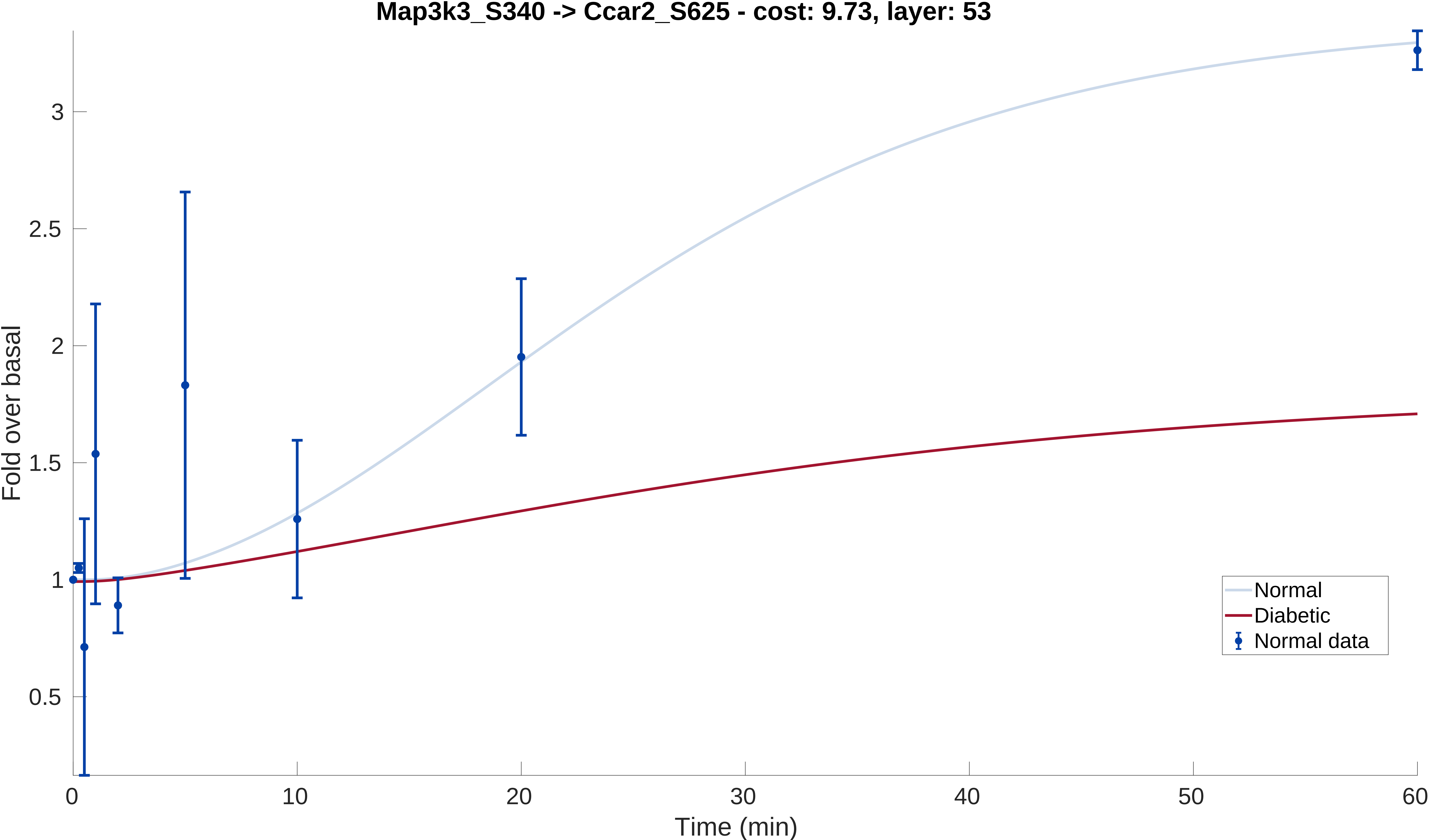

Map3k3_S340 -> Ccar2_S625 - cost: 9.73, layer: 53
Fold over basal
0.5
1
1.5
2
2.5
3
0
10
20
30
40
50
60
Time (min)
Normal
Diabetic
Normal data

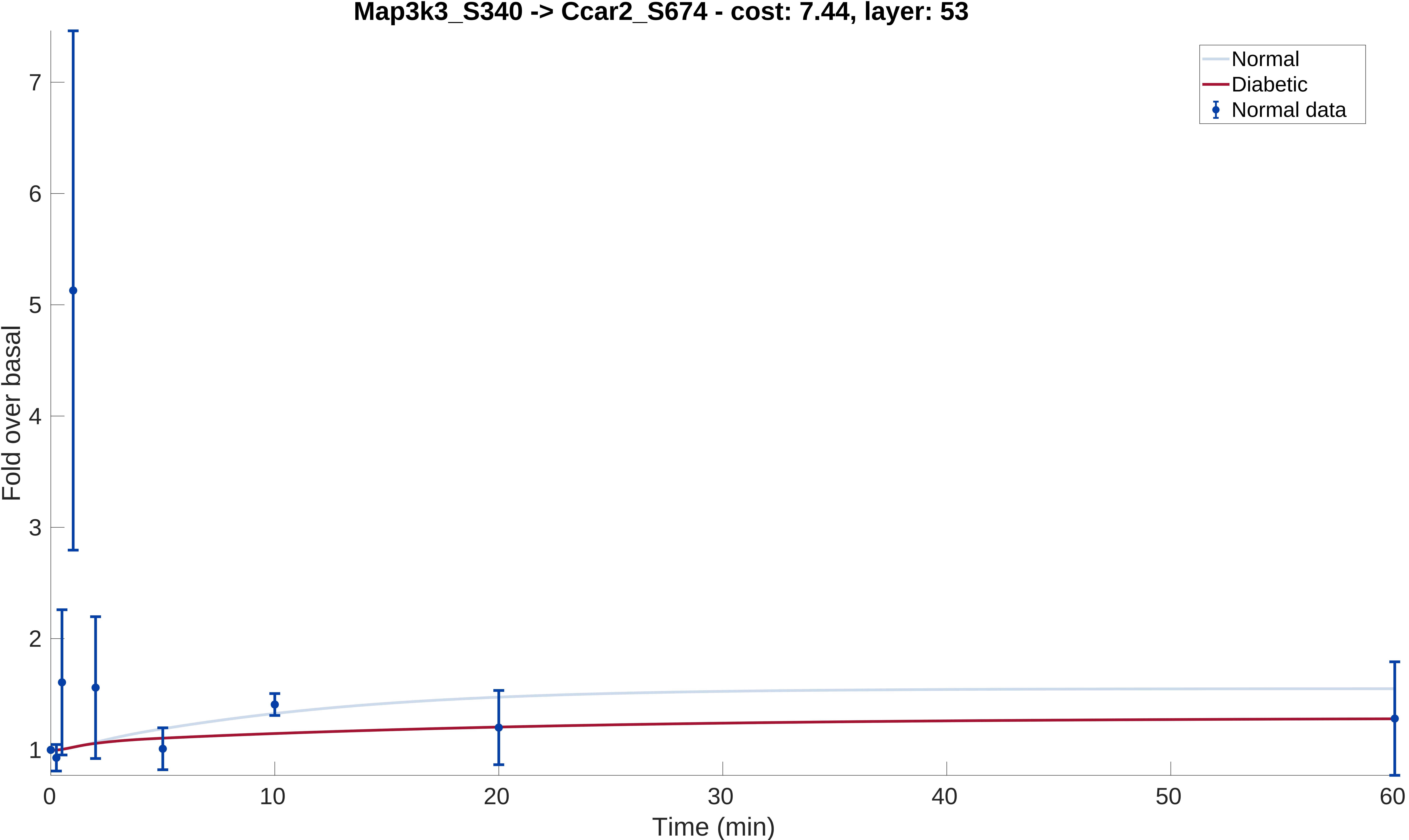

Map3k3_S340 -> Ccar2_S674 - cost: 7.44, layer: 53
Fold over basal
Time (min)
Normal
Diabetic
Normal data
0
10
20
30
40
50
60
1
2
3
4
5
6
7

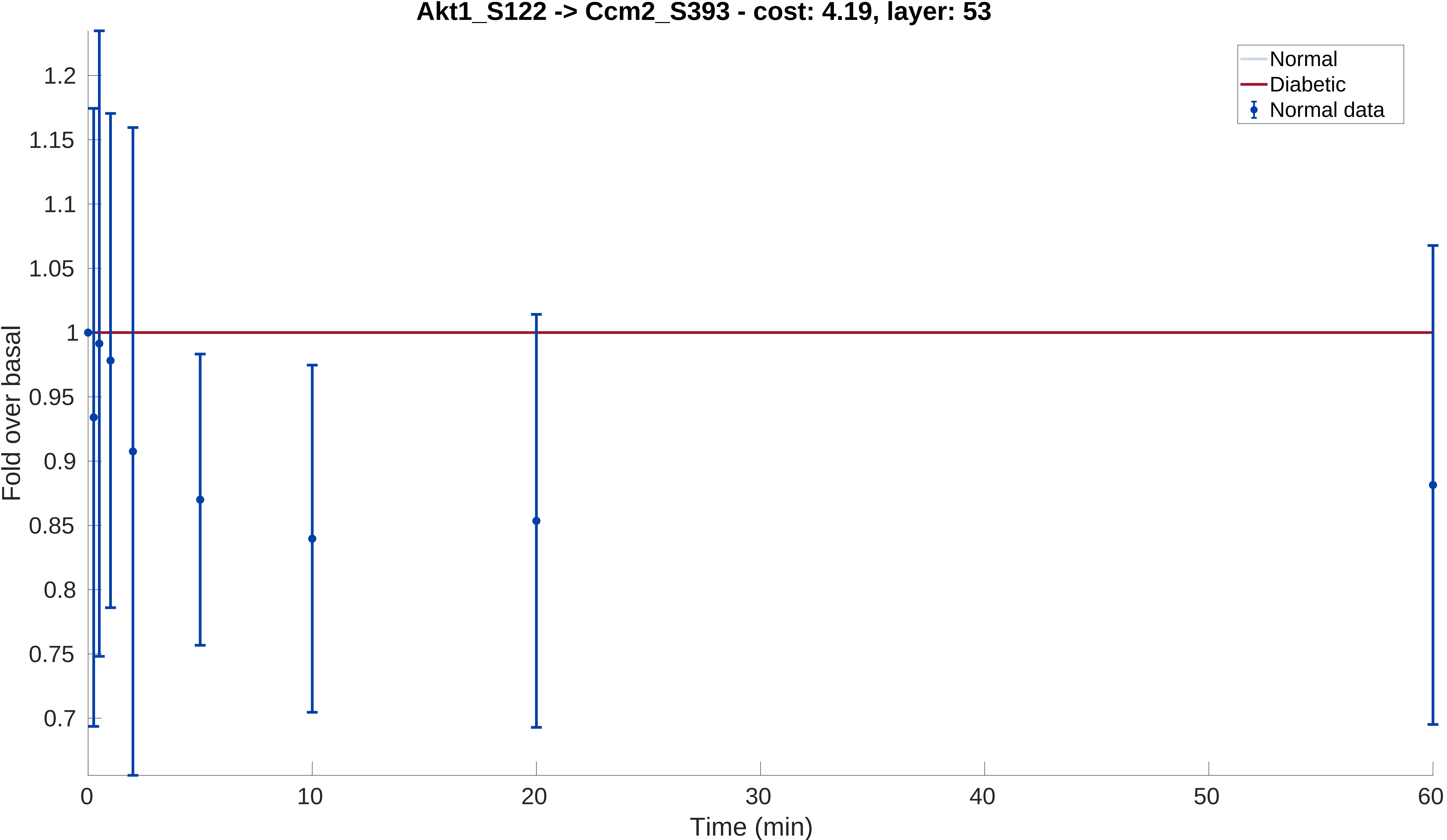

Akt1_S122 -> Ccm2_S393 - cost: 4.19, layer: 53
Normal
Diabetic
Normal data
Fold over basal
1.2
1.15
1.1
1.05
1
0.95
0.9
0.85
0.8
0.75
0.7
0
10
20
30
40
50
60
Time (min)

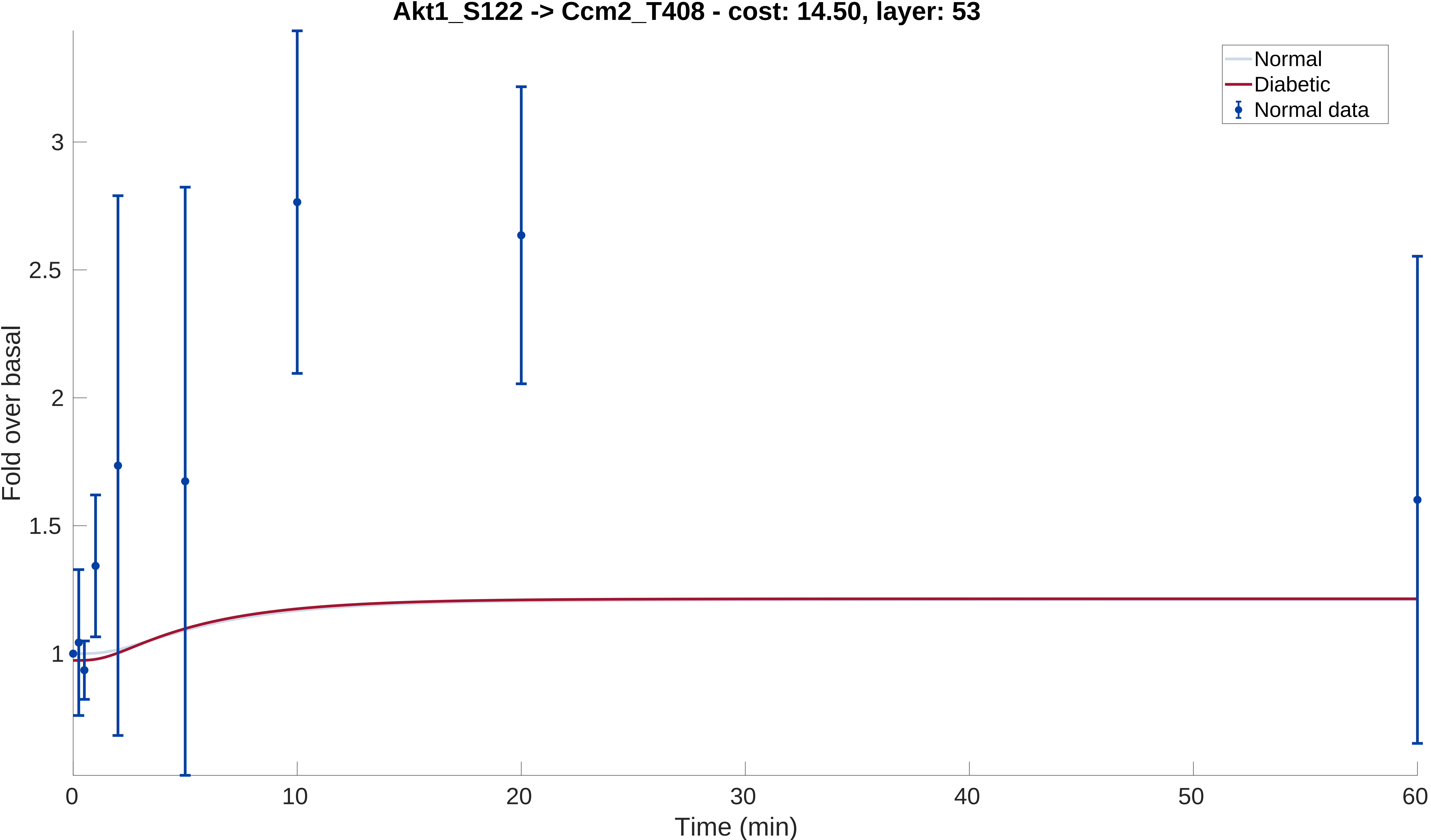

Akt1_S122 -> Ccm2_T408 - cost: 14.50, layer: 53
Normal
Diabetic
Normal data
Fold over basal
3
2.5
2
1.5
1
0
10
20
30
40
50
60
Time (min)

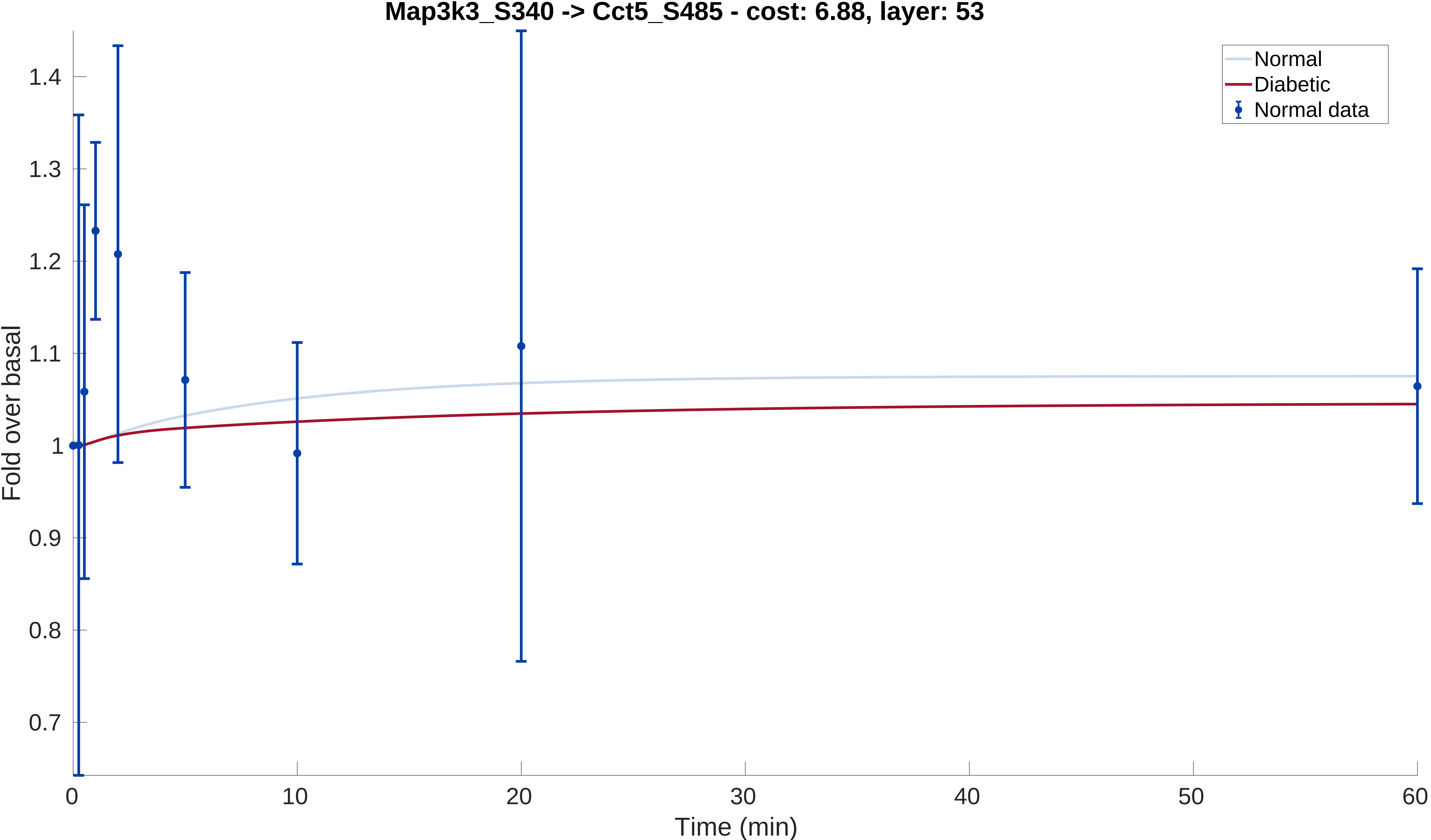

Map3k3_S340 -> Cct5_S485 - cost: 6.88, layer: 53
Normal
Diabetic
Normal data
Fold over basal
1.4
1.3
1.2
1.1
1
0.9
0.8
0.7
0
10
20
30
40
50
60
Time (min)

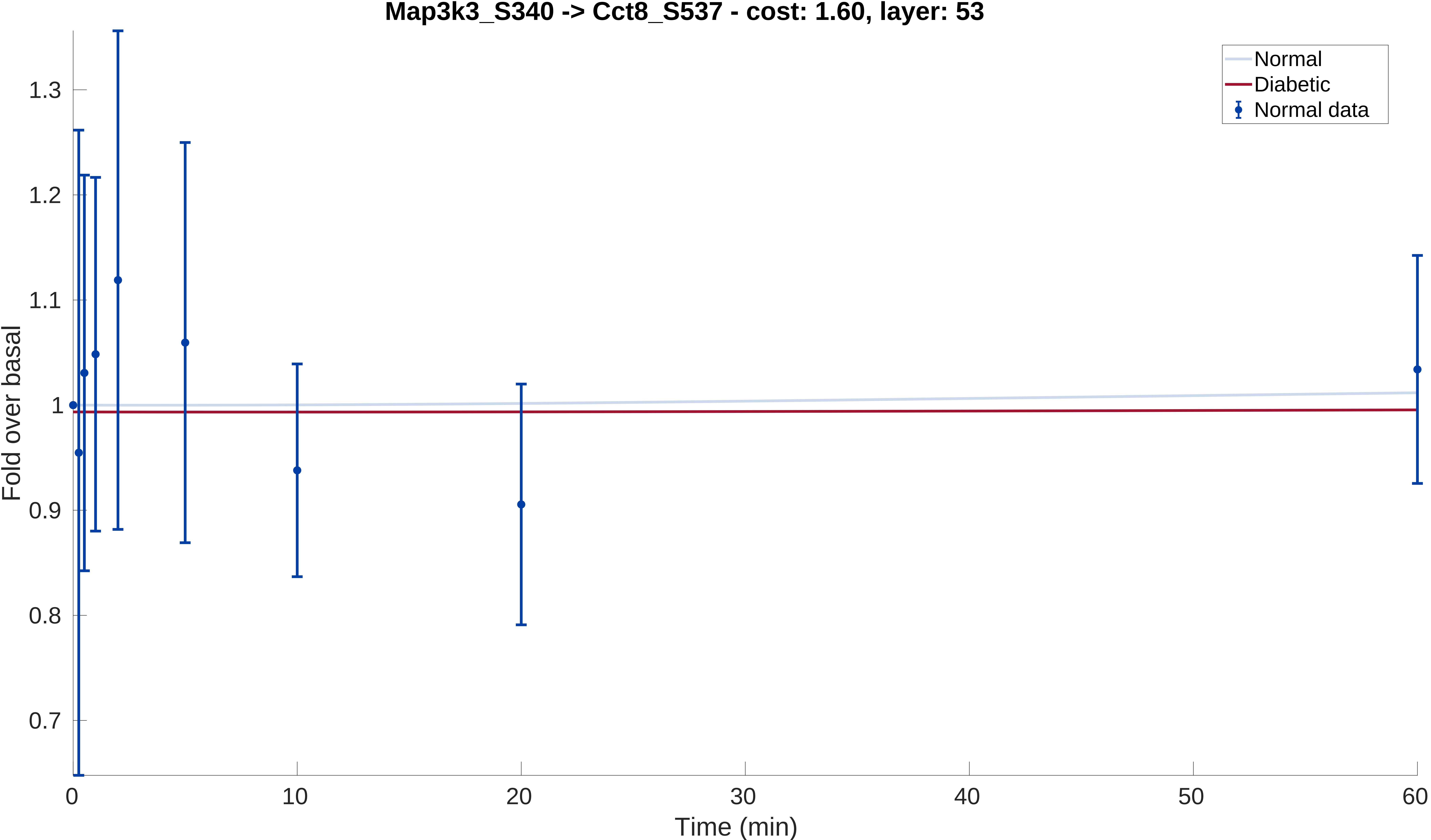
Map3k3_S340 -> Cct8_S537 - cost: 1.60, layer: 53
Normal
Diabetic
Normal data
Fold over basal
1.3
1.2
1.1
1
0.9
0.8
0.7
0
10
20
30
40
50
60
Time (min)

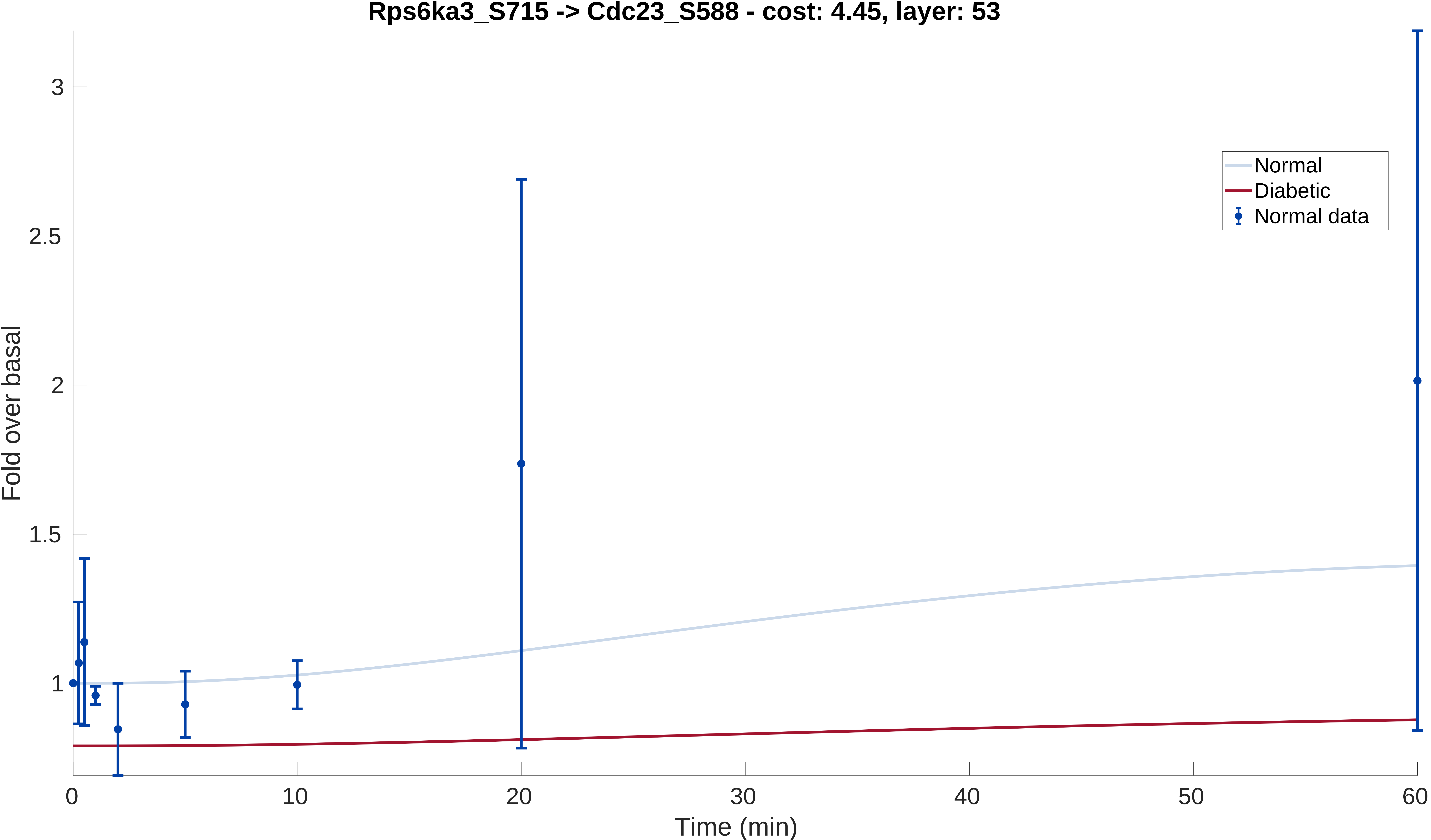

Rps6ka3_S715 -> Cdc23_S588 - cost: 4.45, layer: 53
Normal
Diabetic
Normal data
Fold over basal
Time (min)
3
2.5
2
1.5
1
0
10
20
30
40
50
60

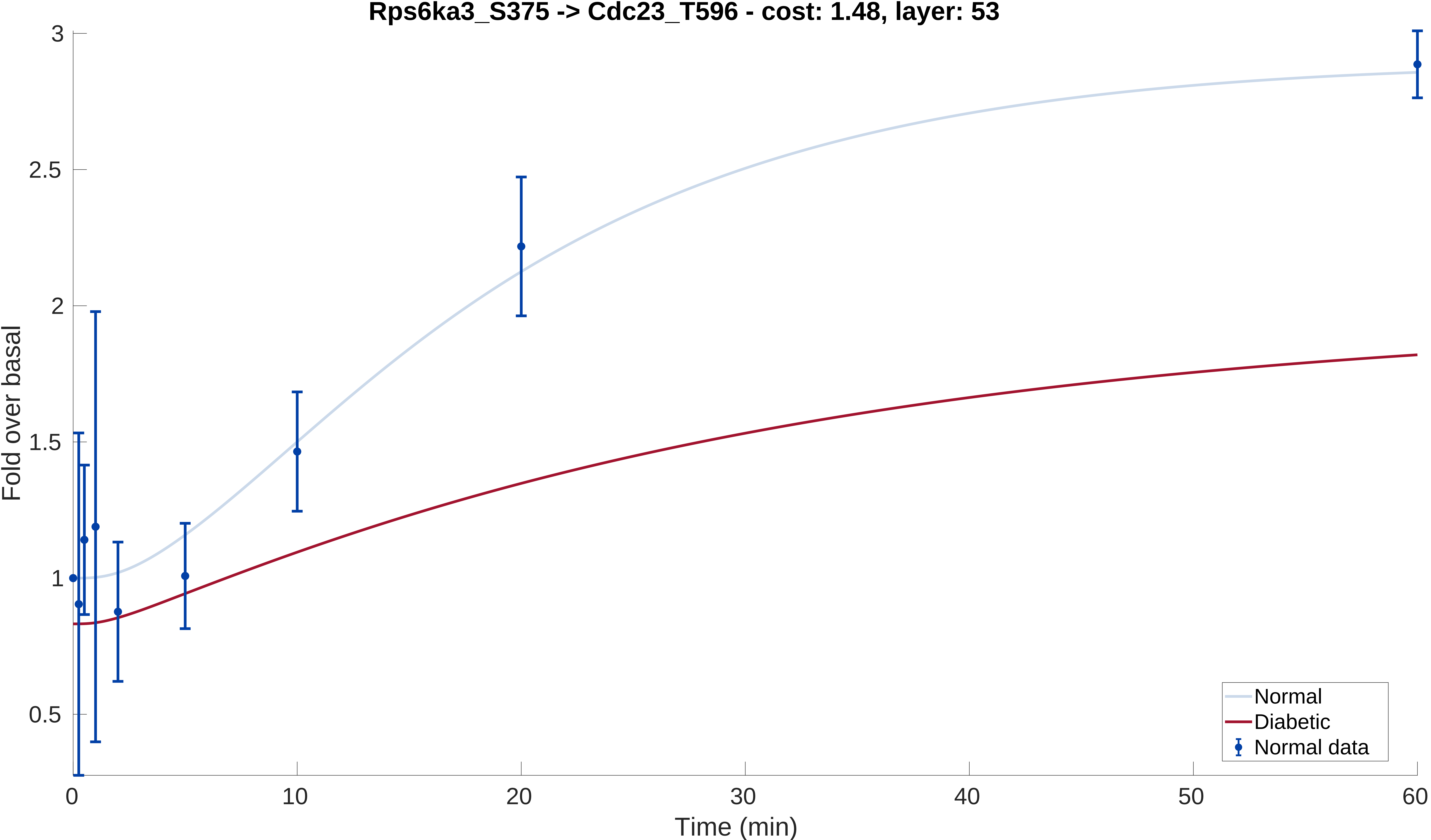

Rps6ka3_S375 -> Cdc23_T596 - cost: 1.48, layer: 53
Fold over basal
Time (min)
0
10
20
30
40
50
60
0.5
1
1.5
2
2.5
3
Normal
Diabetic
Normal data

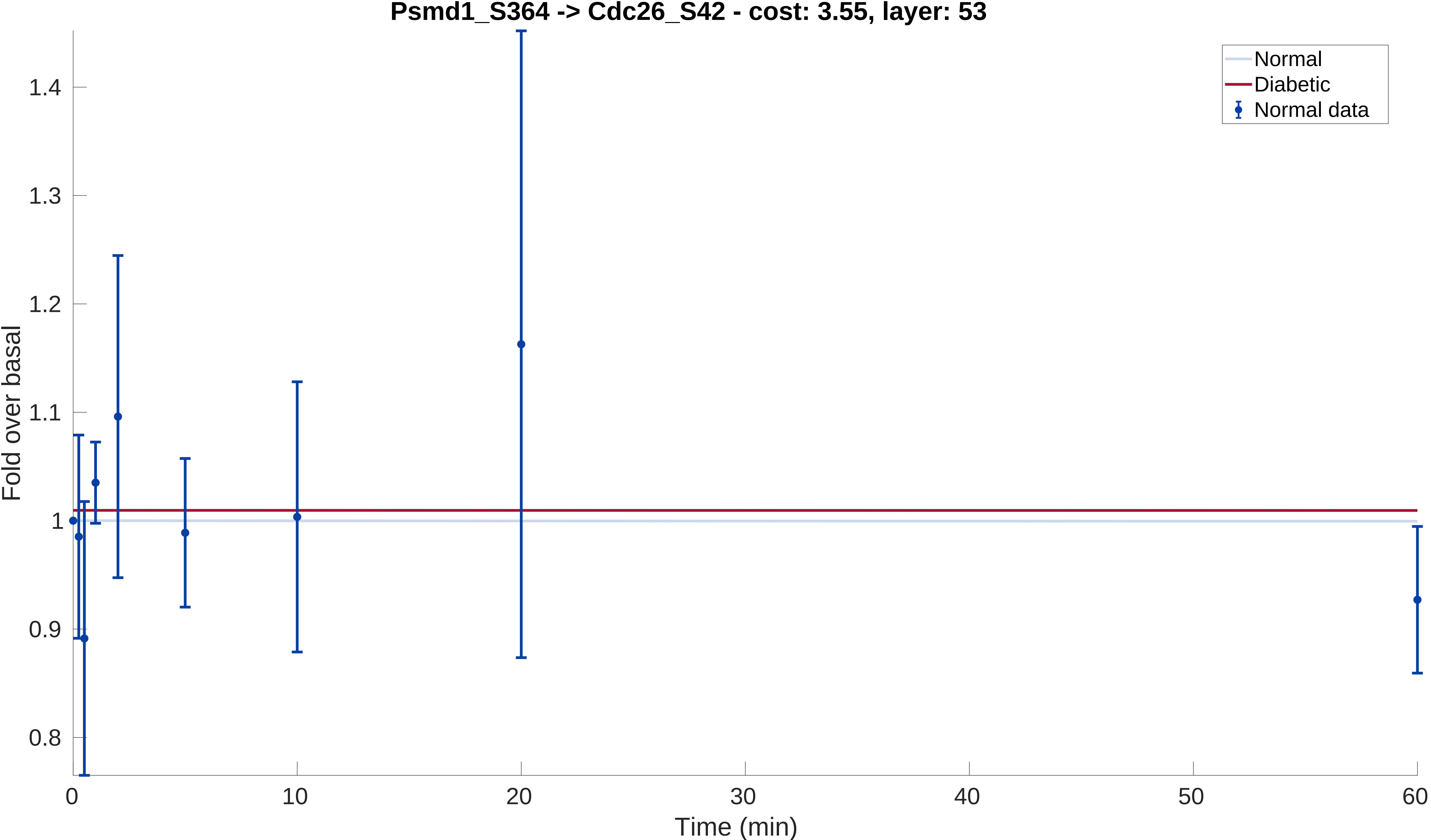
Psmd1_S364 -> Cdc26_S42 - cost: 3.55, layer: 53
Normal
Diabetic
Normal data
Fold over basal
Time (min)
0.8
0.9
1
1.1
1.2
1.3
1.4
0
10
20
30
40
50
60

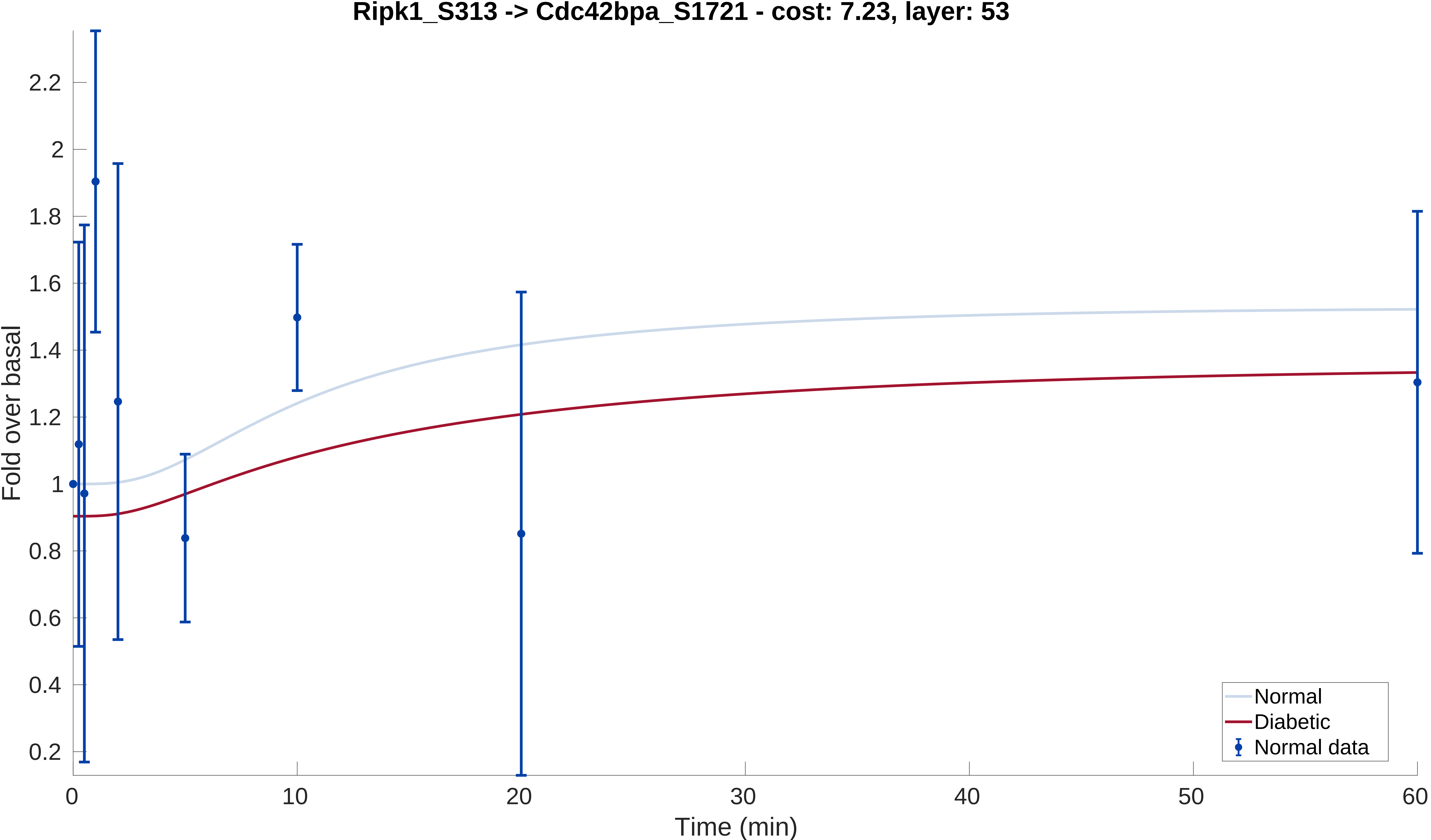

Ripk1_S313 -> Cdc42bpa_S1721 - cost: 7.23, layer: 53
Fold over basal
Time (min)
0.2
0.4
0.6
0.8
1
1.2
1.4
1.6
1.8
2
2.2
0
10
20
30
40
50
60
Normal
Diabetic
Normal data

#### Ripk1_S313 -> Cdc42bpa_S1724 - cost: 2.25, layer: 53

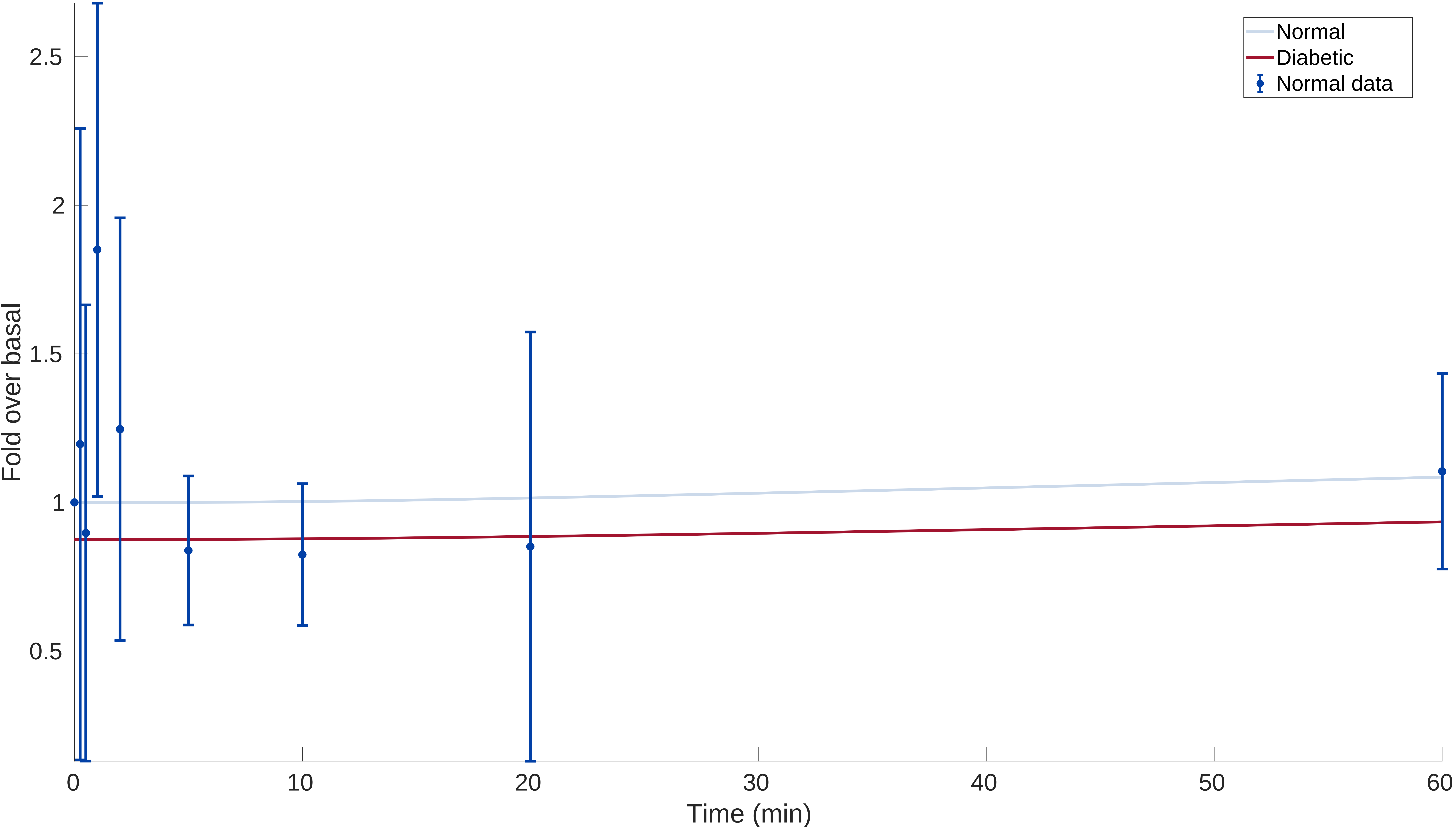

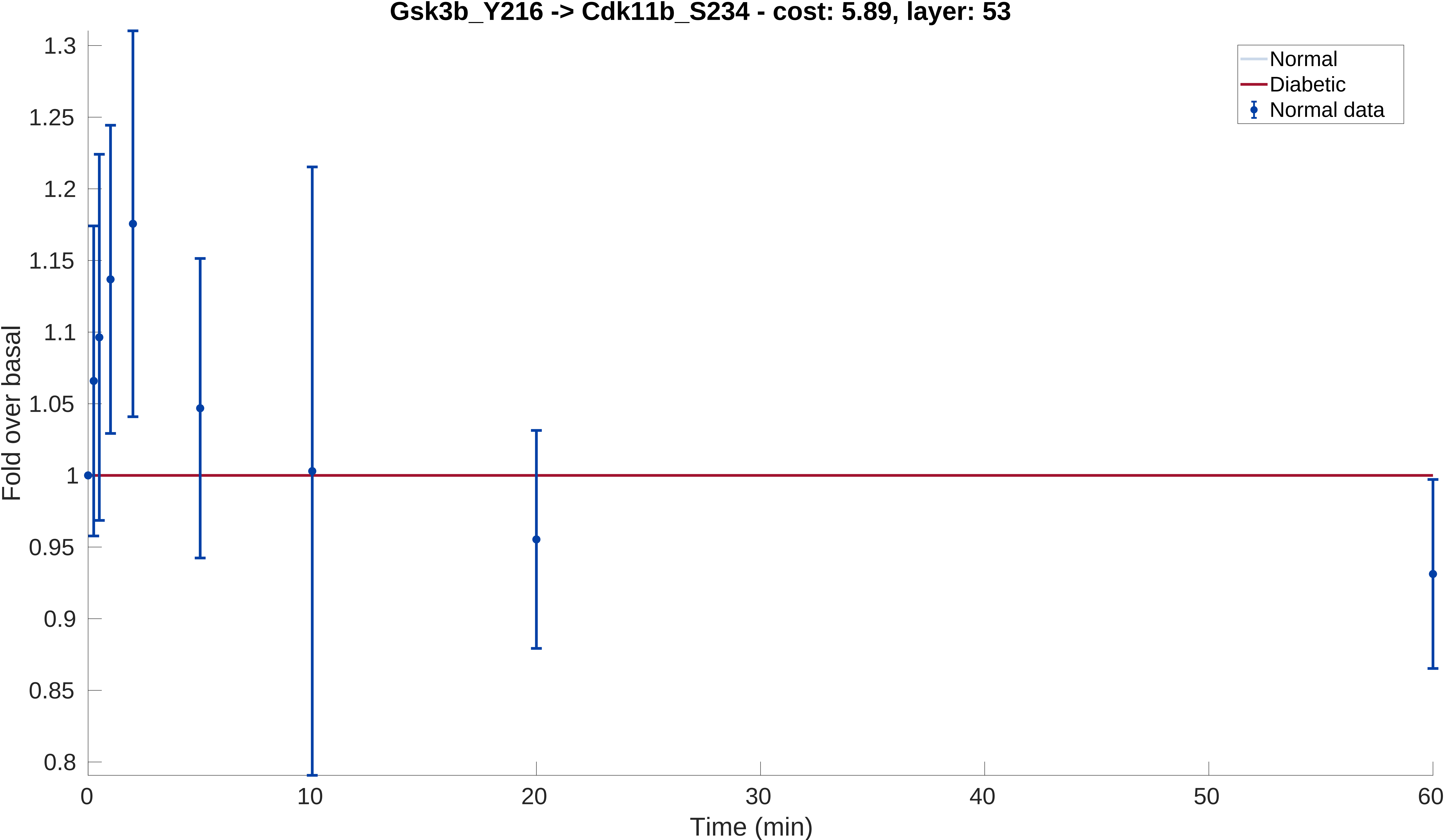

Gsk3b_Y216 -> Cdk11b_S234 - cost: 5.89, layer: 53
Normal
Diabetic
Normal data
Fold over basal
Time (min)
1.3
1.25
1.2
1.15
1.1
1.05
1
0.95
0.9
0.85
0.8
0
10
20
30
40
50
60

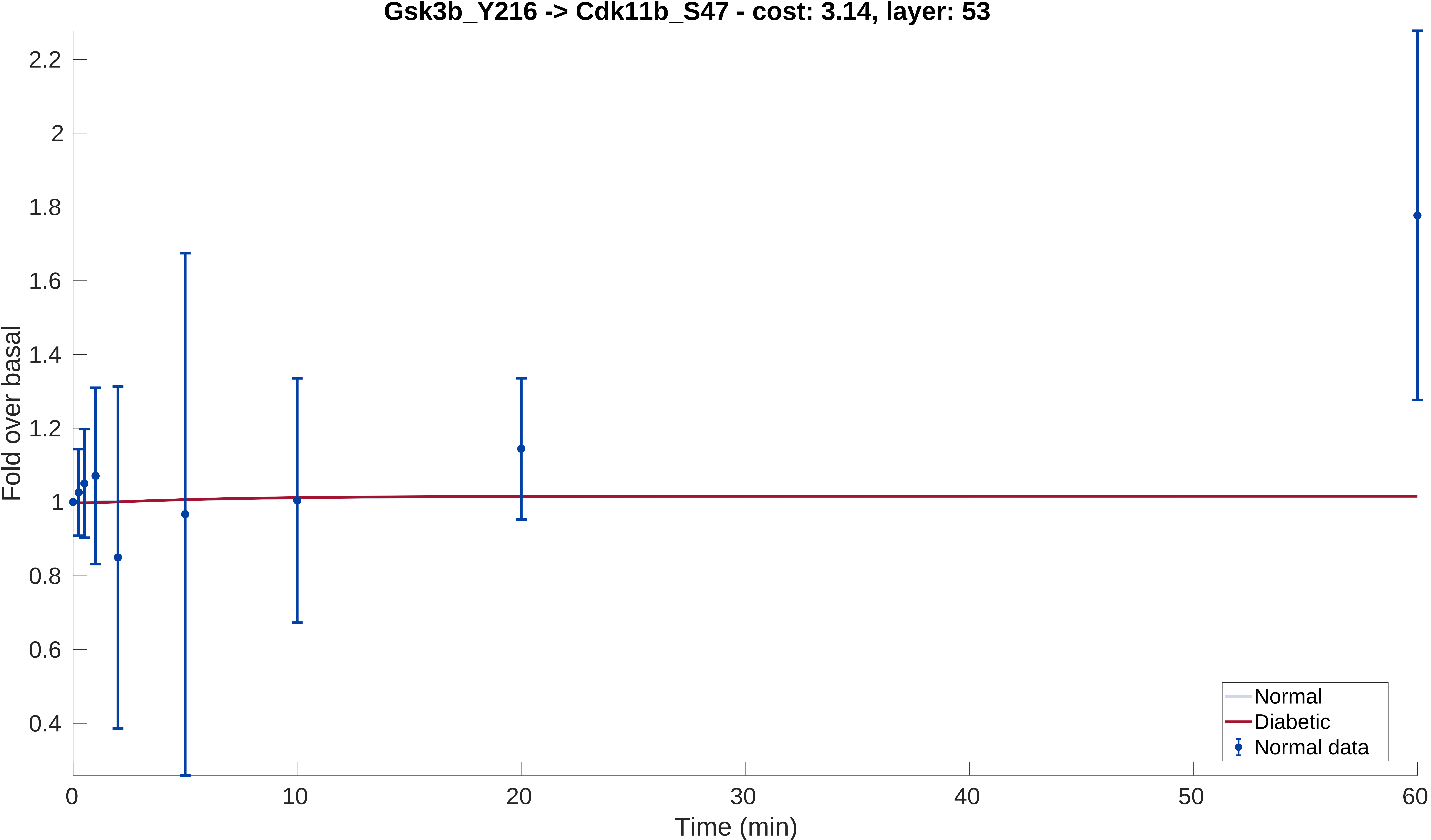

Gsk3b_Y216 -> Cdk11b_S47 - cost: 3.14, layer: 53
Fold over basal
2.2
2
1.8
1.6
1.4
1.2
1
0.8
0.6
0.4
0
10
20
30
40
50
60
Time (min)
Normal
Diabetic
Normal data

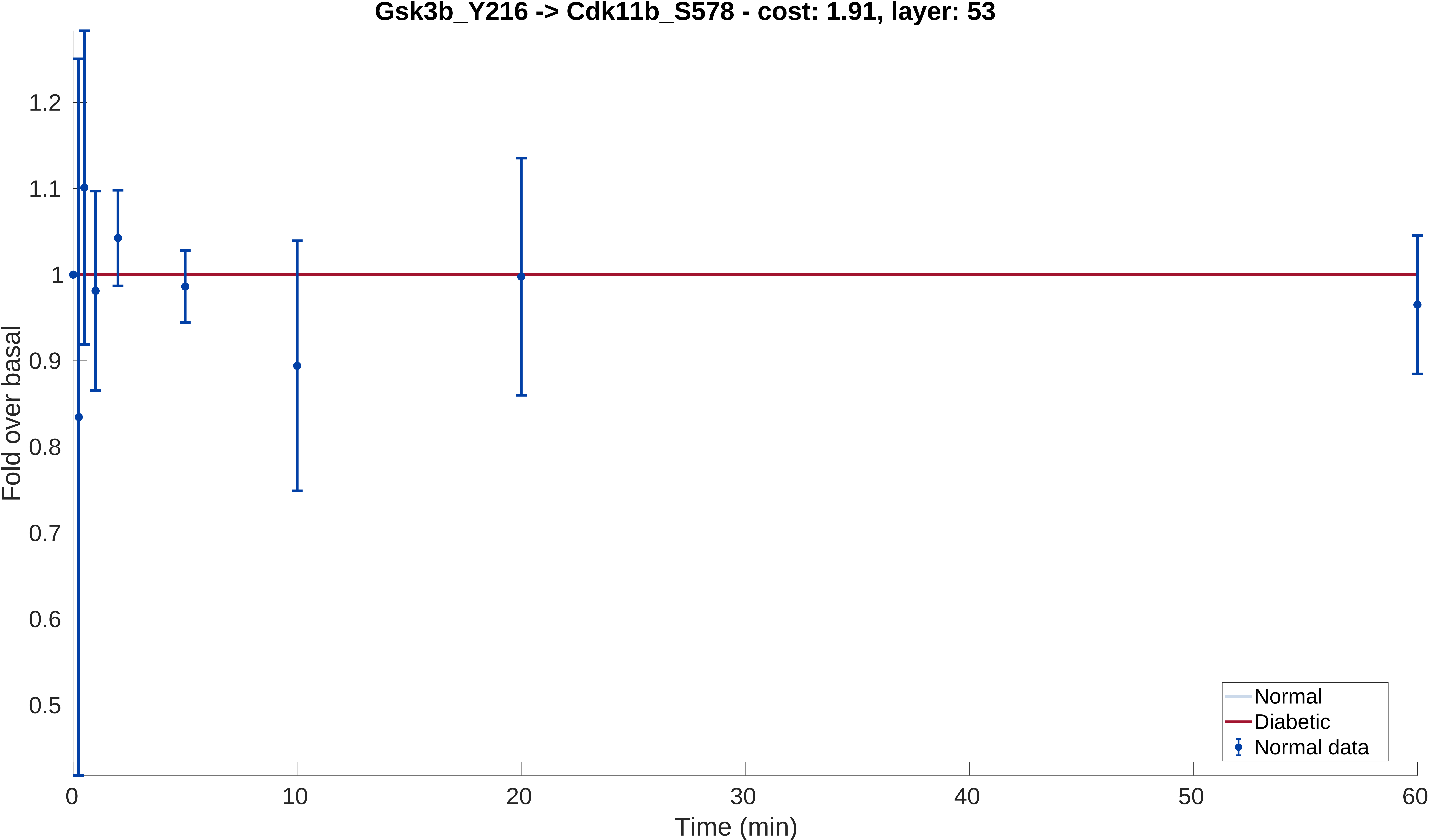

Gsk3b_Y216 -> Cdk11b_S578 - cost: 1.91, layer: 53
Fold over basal
1.2
1.1
1
0.9
0.8
0.7
0.6
0.5
0
10
20
30
40
50
60
Time (min)
Normal
Diabetic
Normal data

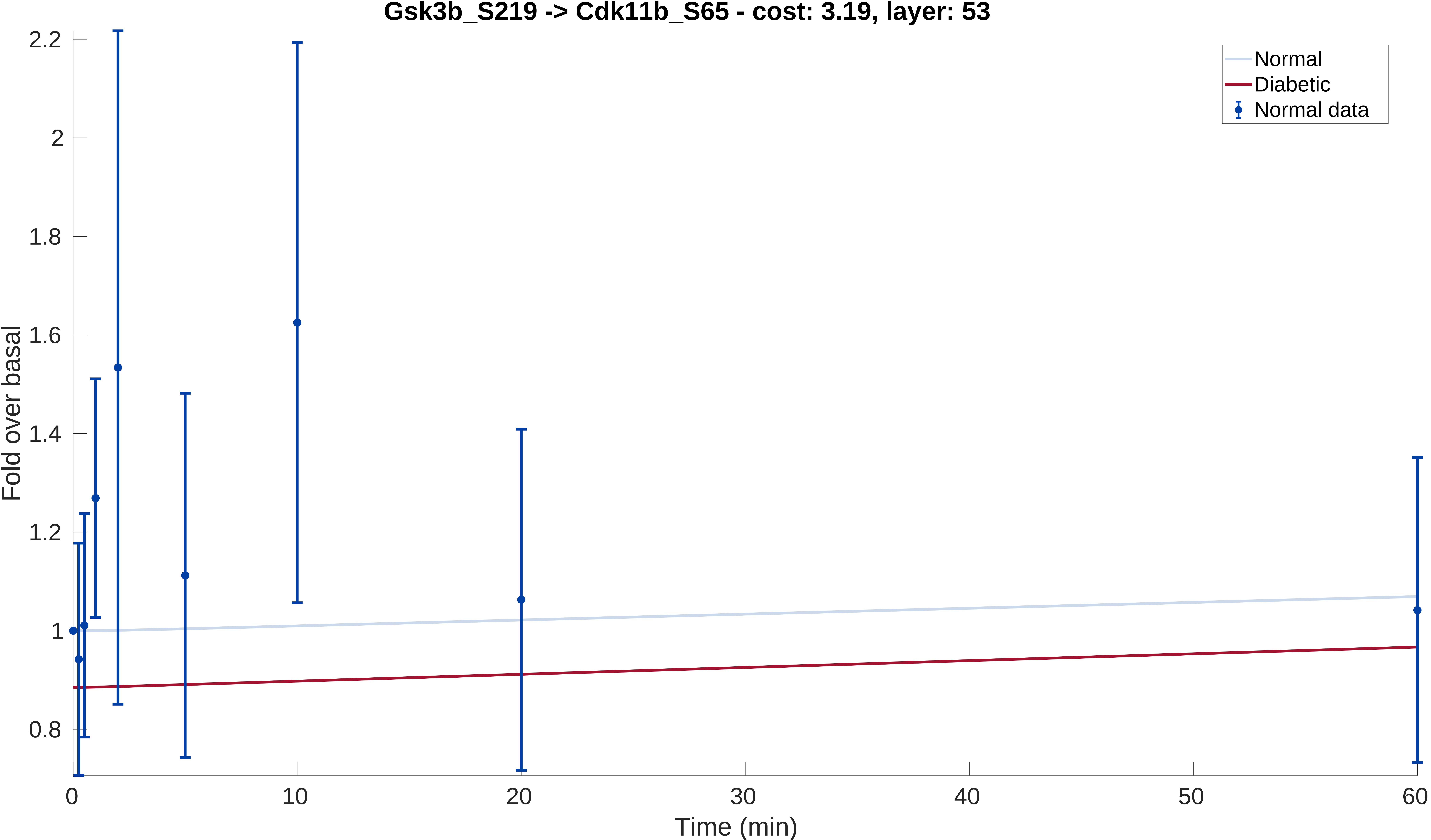

Gsk3b_S219 -> Cdk11b_S65 - cost: 3.19, layer: 53
Normal
Diabetic
Normal data
Fold over basal
Time (min)
0.8
1
1.2
1.4
1.6
1.8
2
2.2
0
10
20
30
40
50
60

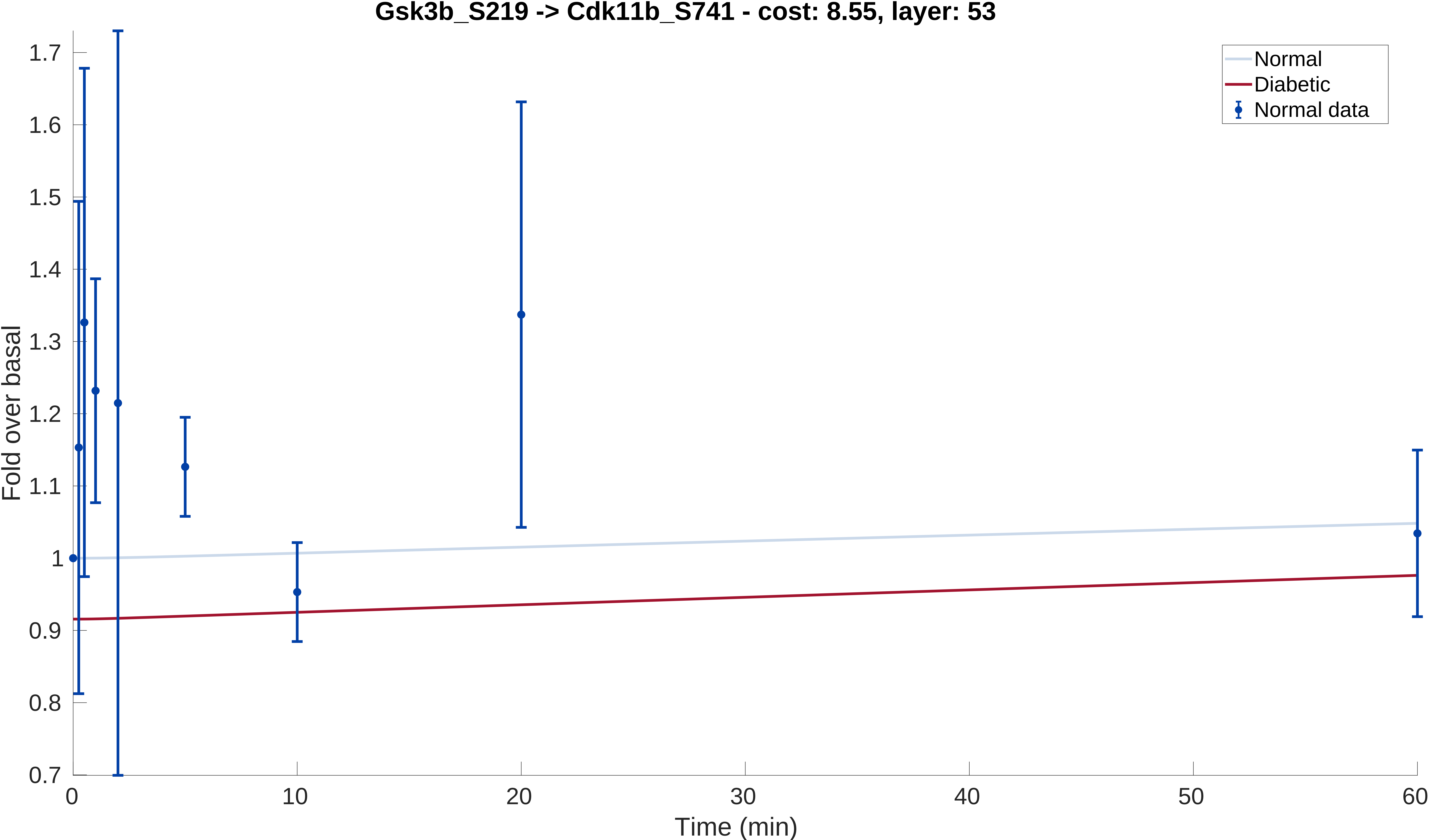

Gsk3b_S219 -> Cdk11b_S741 - cost: 8.55, layer: 53
Normal
Diabetic
Normal data
Fold over basal
1.7
1.6
1.5
1.4
1.3
1.2
1.1
1
0.9
0.8
0.7
0
10
20
30
40
50
60
Time (min)

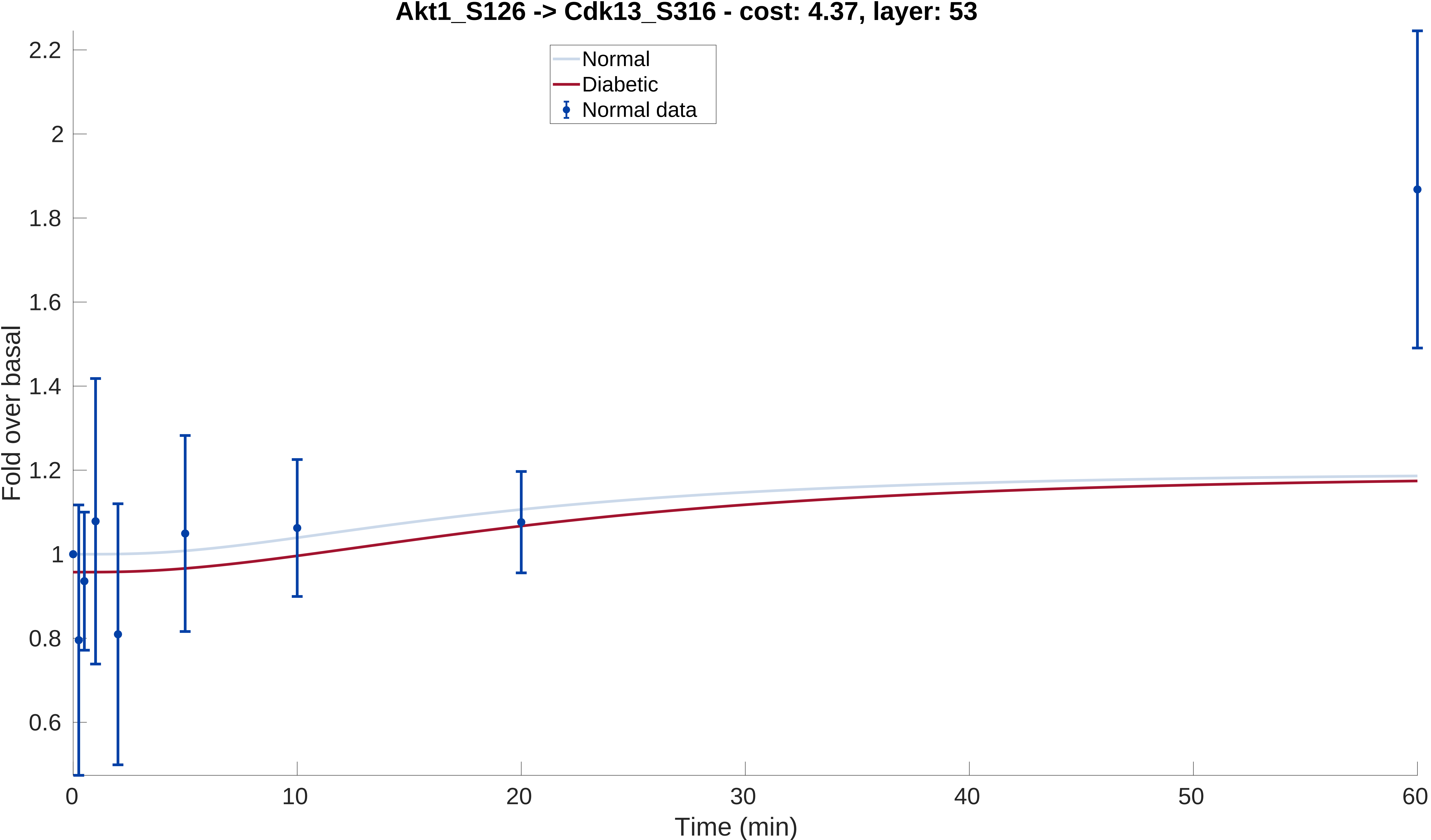

Akt1_S126 -> Cdk13_S316 - cost: 4.37, layer: 53
Normal
Diabetic
Normal data
Fold over basal
Time (min)
0.6
0.8
1
1.2
1.4
1.6
1.8
2
2.2
0
10
20
30
40
50
60

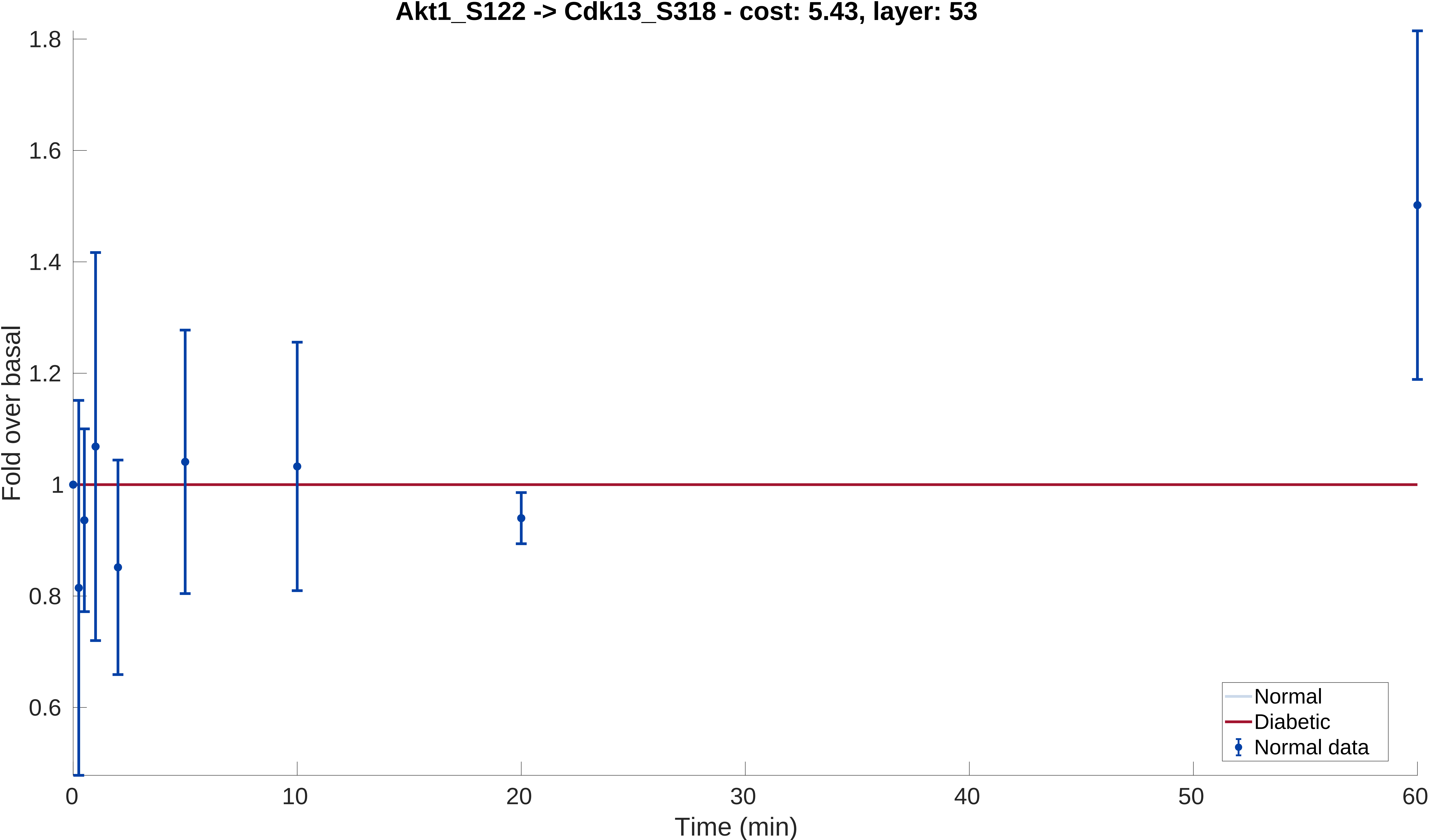

Akt1_S122 -> Cdk13_S318 - cost: 5.43, layer: 53
Fold over basal
1.8
1.6
1.4
1.2
1
0.8
0.6
0
10
20
30
40
50
60
Time (min)
Normal
Diabetic
Normal data

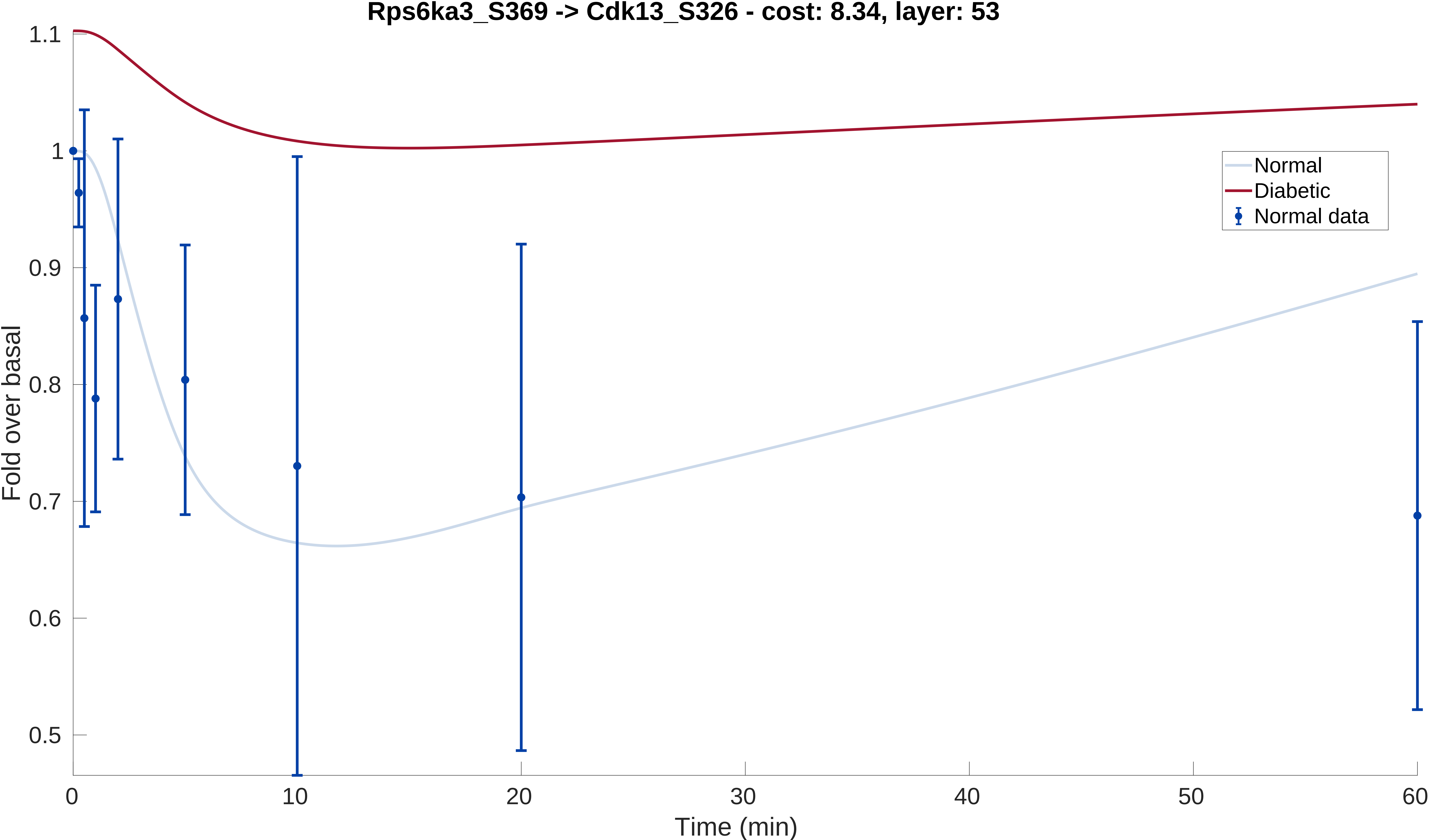
Rps6ka3_S369 -> Cdk13_S326 - cost: 8.34, layer: 53
Fold over basal
1.1
1
0.9
0.8
0.7
0.6
0.5
0
10
20
30
40
50
60
Time (min)
Normal
Diabetic
Normal data

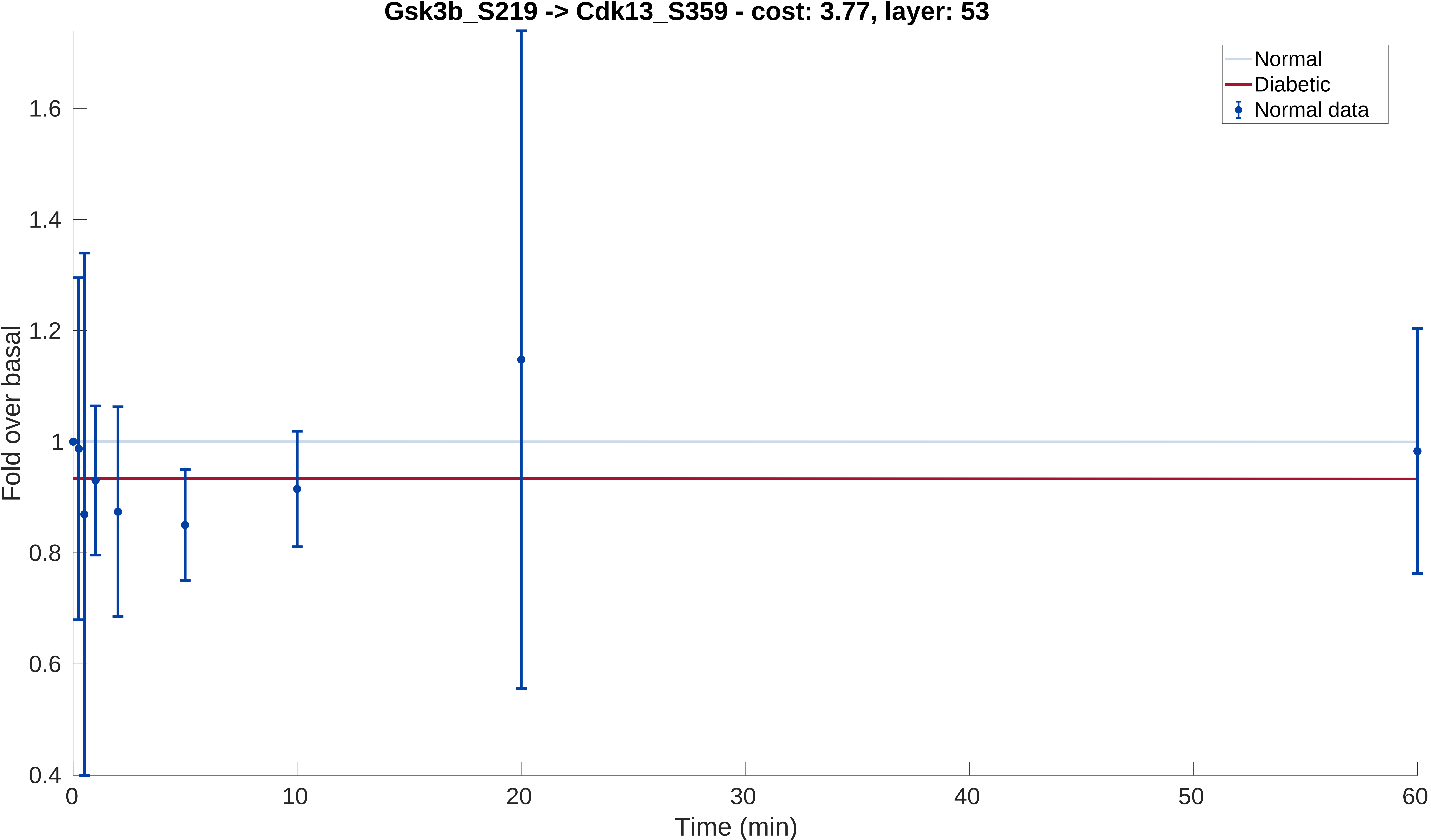

Gsk3b_S219 -> Cdk13_S359 - cost: 3.77, layer: 53
Normal
Diabetic
Normal data
Fold over basal
1.6
1.4
1.2
1
0.8
0.6
0.4
0
10
20
30
40
50
60
Time (min)

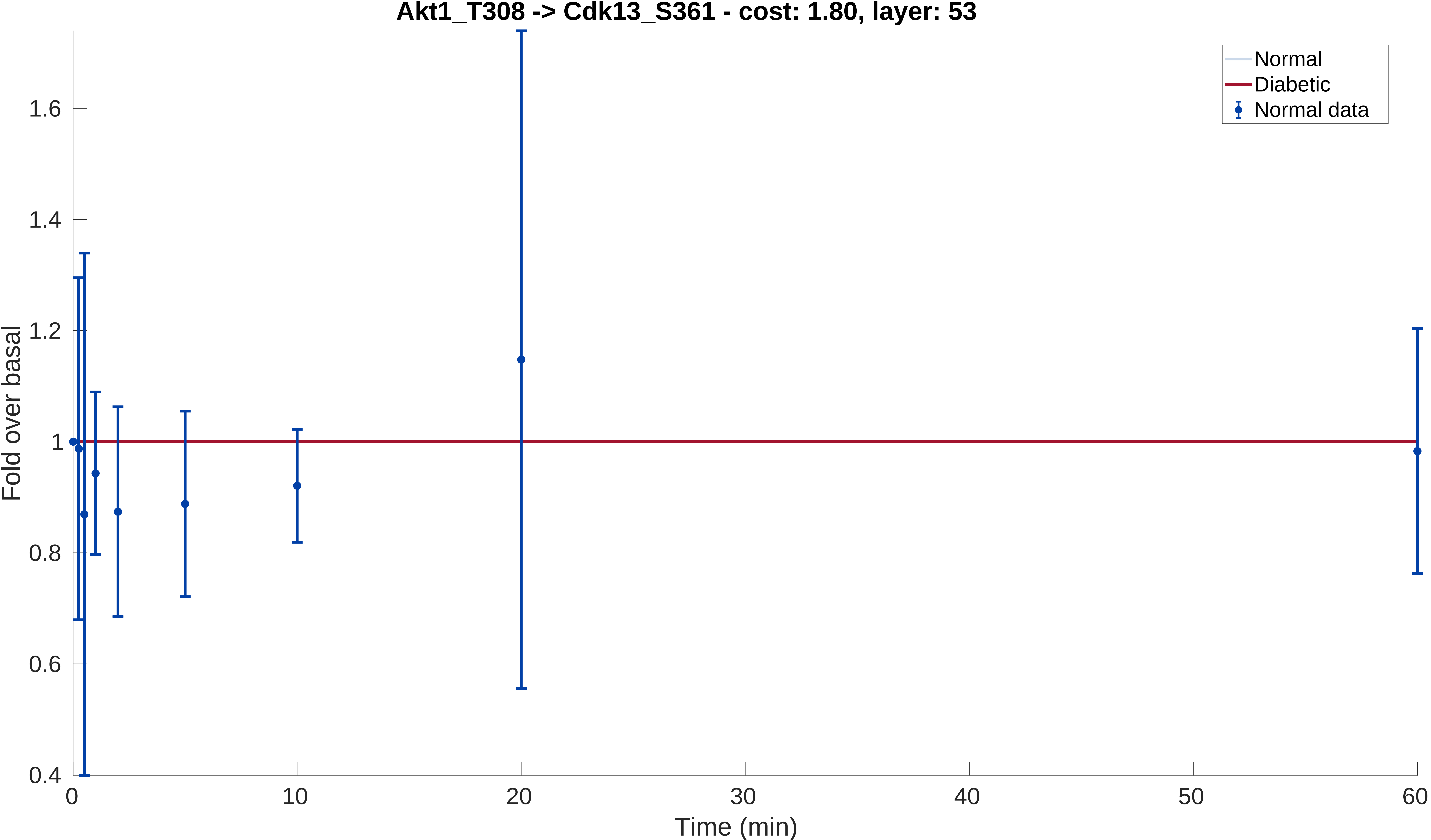

Akt1_T308 -> Cdk13_S361 - cost: 1.80, layer: 53
Normal
Diabetic
Normal data
Fold over basal
1.6
1.4
1.2
1
0.8
0.6
0.4
0
10
20
30
40
50
60
Time (min)

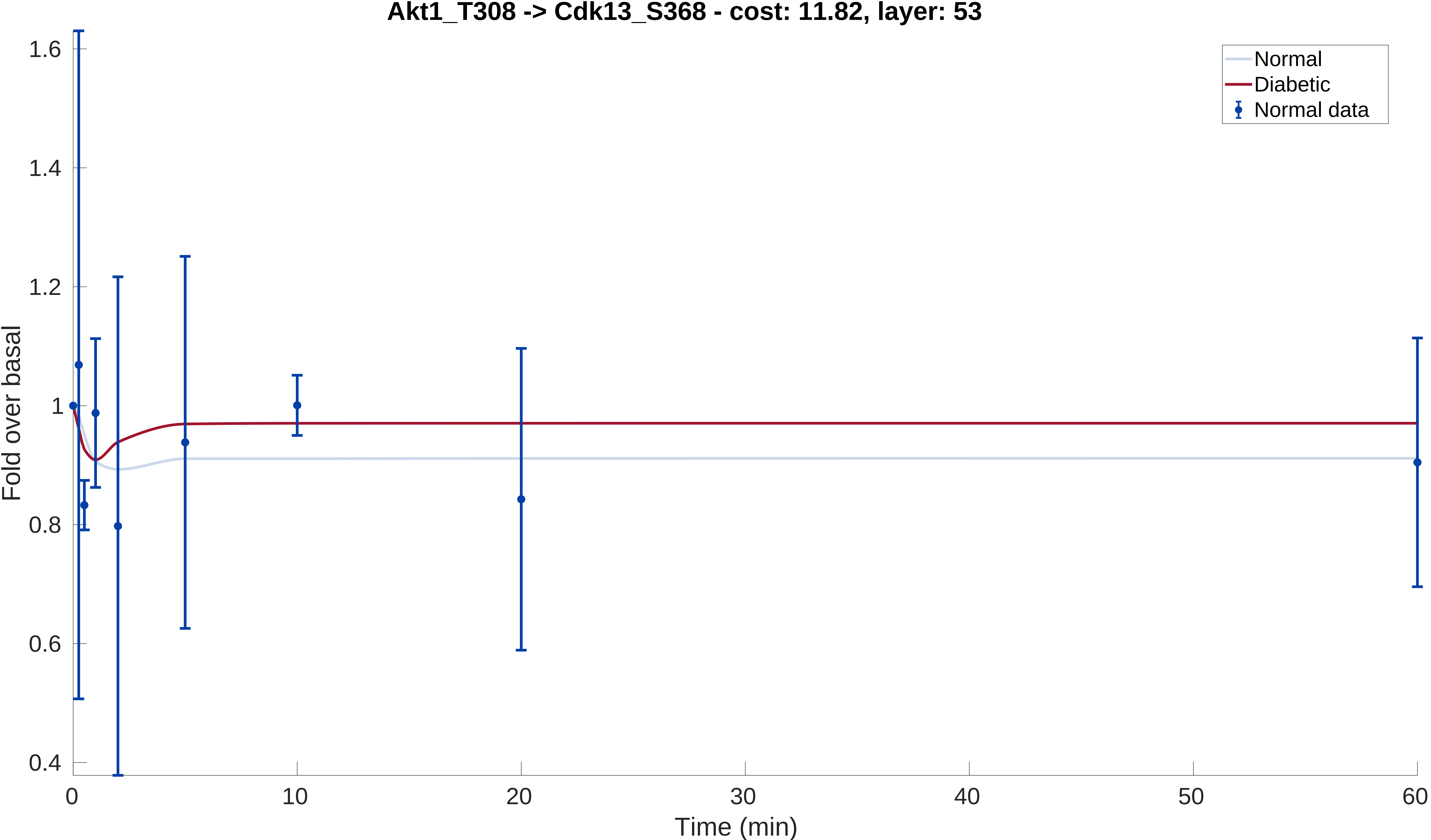
Akt1_T308 -> Cdk13_S368 - cost: 11.82, layer: 53
Normal
Diabetic
Normal data
Fold over basal
1.6
1.4
1.2
1
0.8
0.6
0.4
0
10
20
30
40
50
60
Time (min)

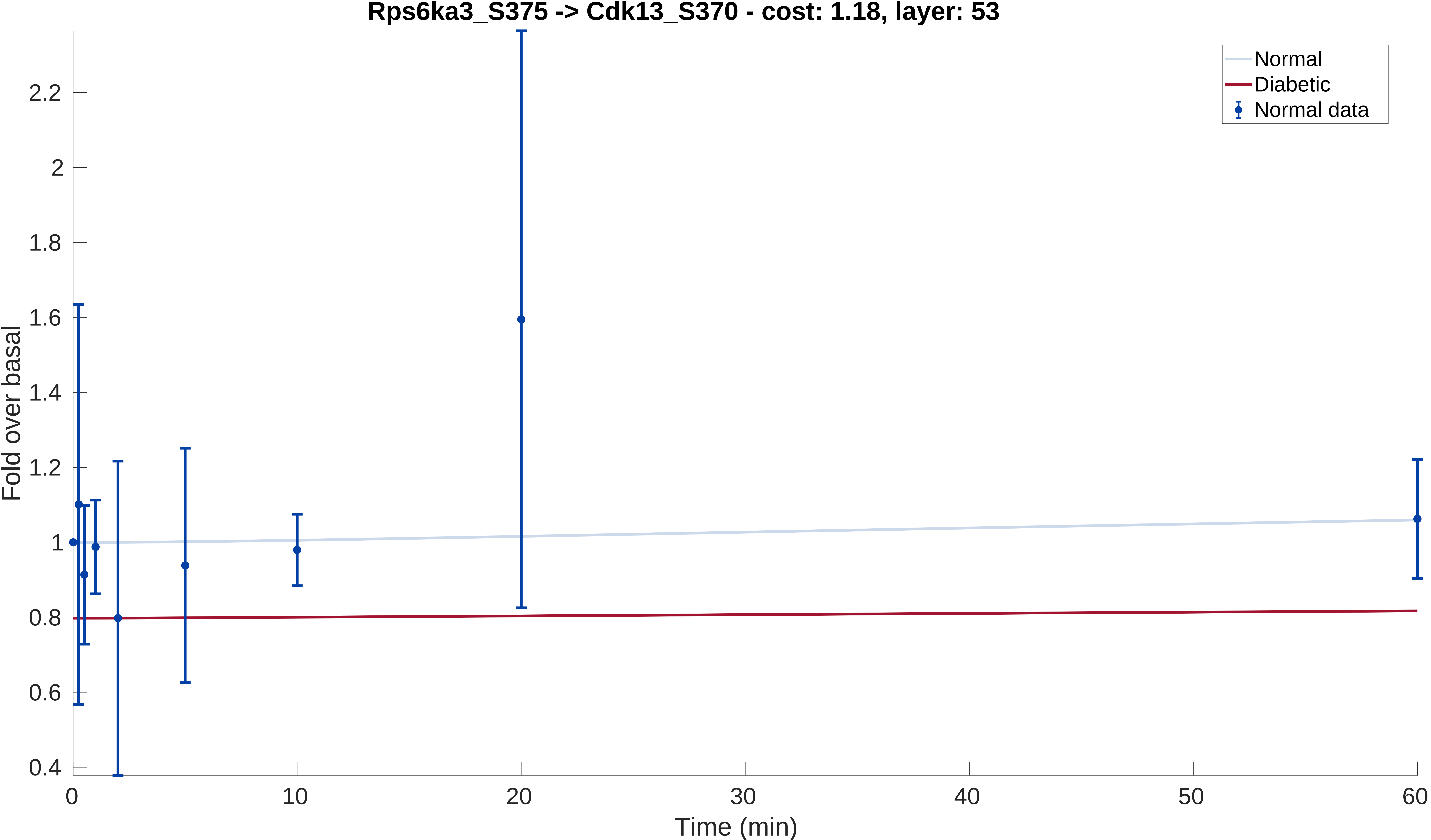

Rps6ka3_S375 -> Cdk13_S370 - cost: 1.18, layer: 53
Normal
Diabetic
Normal data
Fold over basal
Time (min)
0.4
0.6
0.8
1
1.2
1.4
1.6
1.8
2
2.2
0
10
20
30
40
50
60

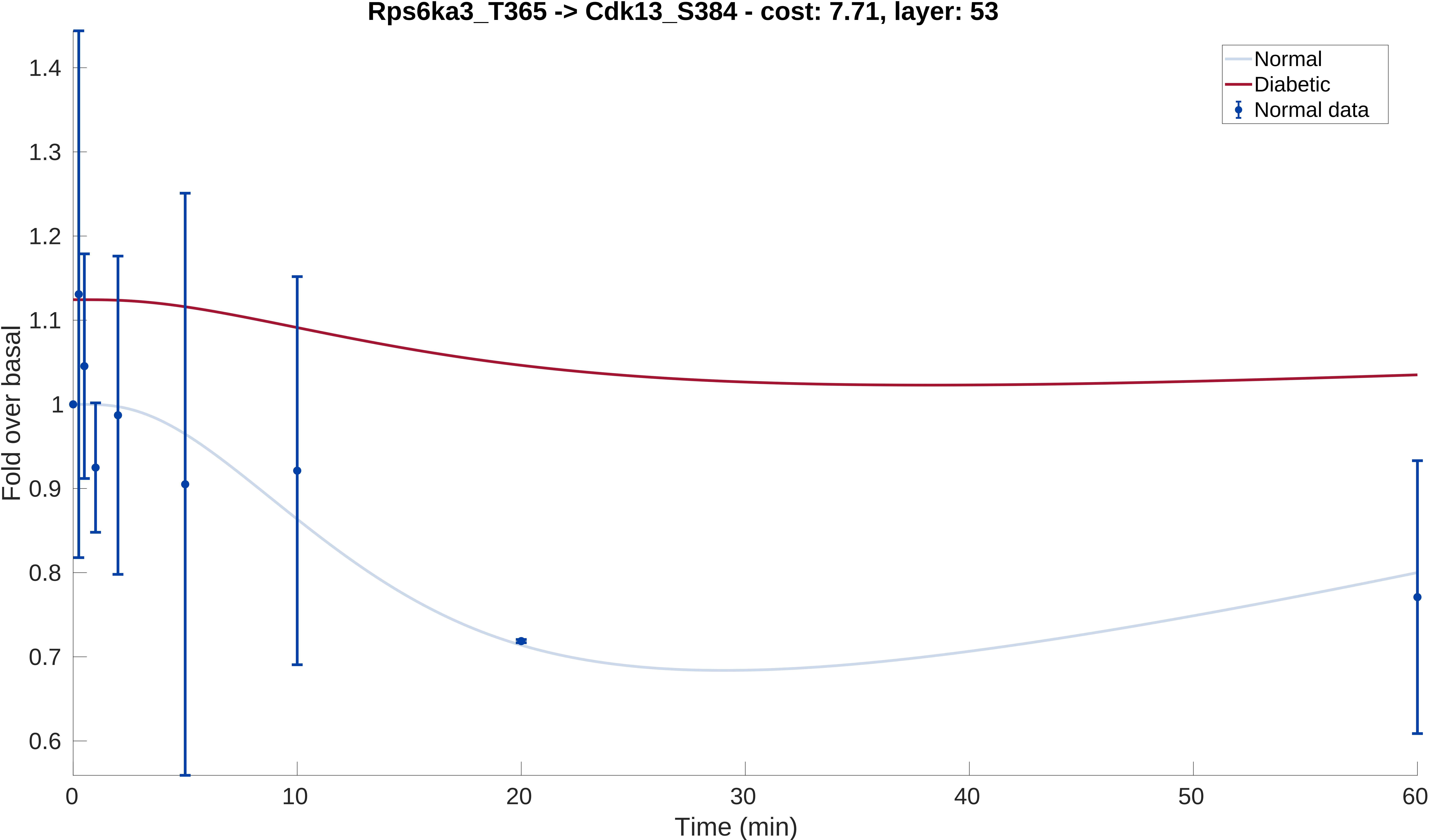

Rps6ka3_T365 -> Cdk13_S384 - cost: 7.71, layer: 53
Normal
Diabetic
Normal data
Fold over basal
1.4
1.3
1.2
1.1
1
0.9
0.8
0.7
0.6
0
10
20
30
40
50
60
Time (min)

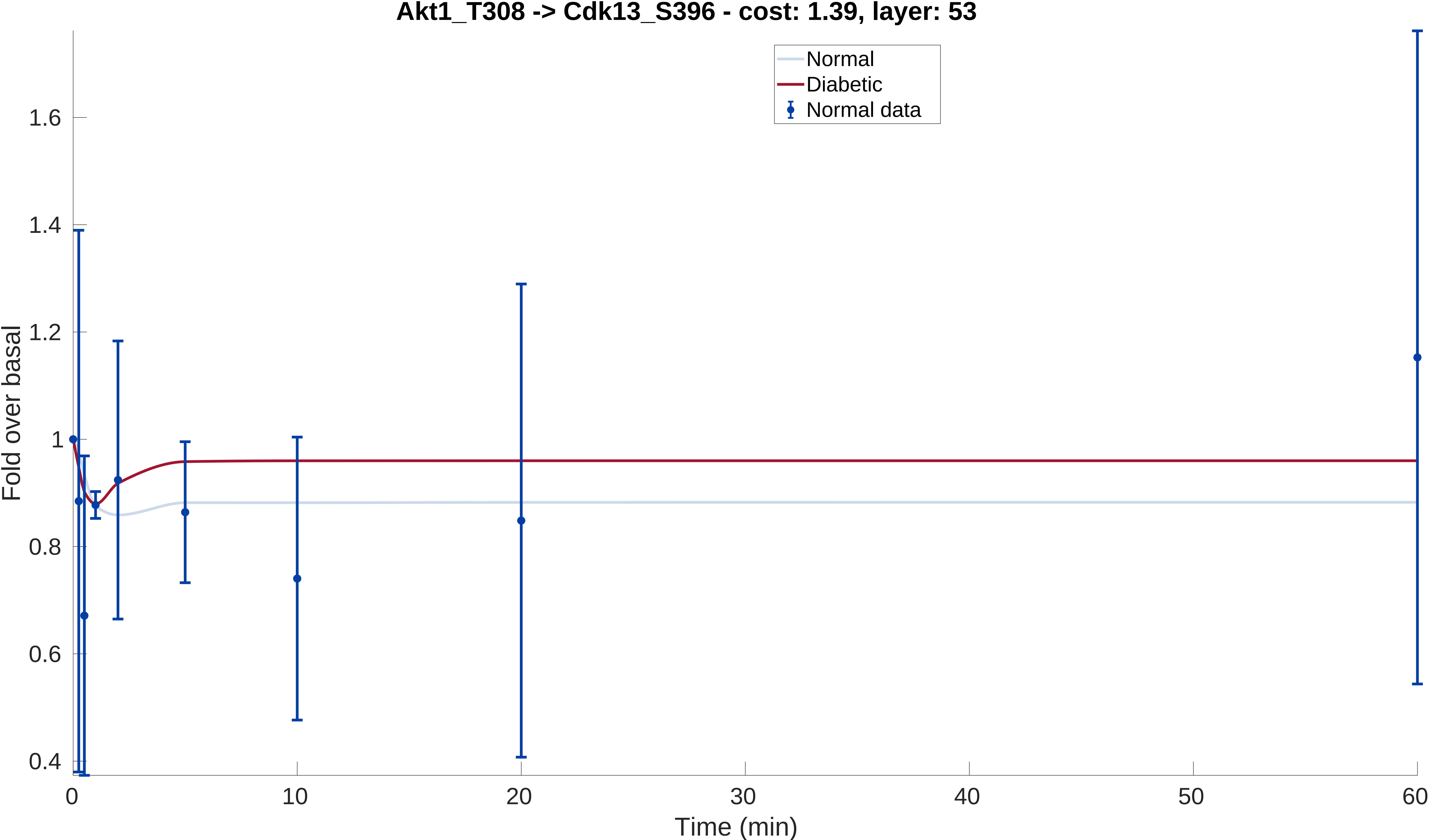

Akt1_T308 -> Cdk13_S396 - cost: 1.39, layer: 53
Normal
Diabetic
Normal data
Fold over basal
1.6
1.4
1.2
1
0.8
0.6
0.4
0
10
20
30
40
50
60
Time (min)

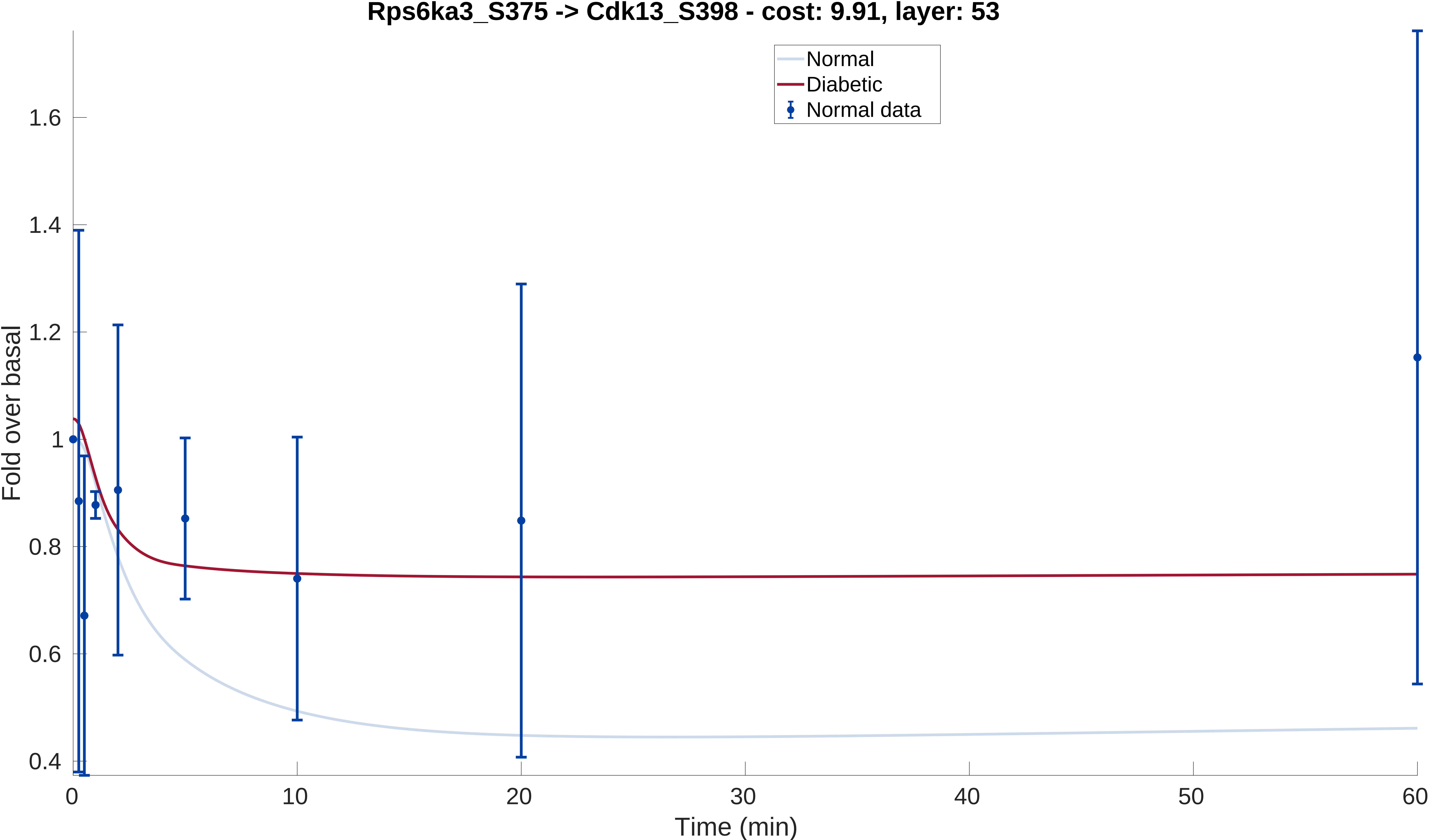

Rps6ka3_S375 -> Cdk13_S398 - cost: 9.91, layer: 53
Normal
Diabetic
Normal data
Fold over basal
1.6
1.4
1.2
1
0.8
0.6
0.4
0
10
20
30
40
50
60
Time (min)

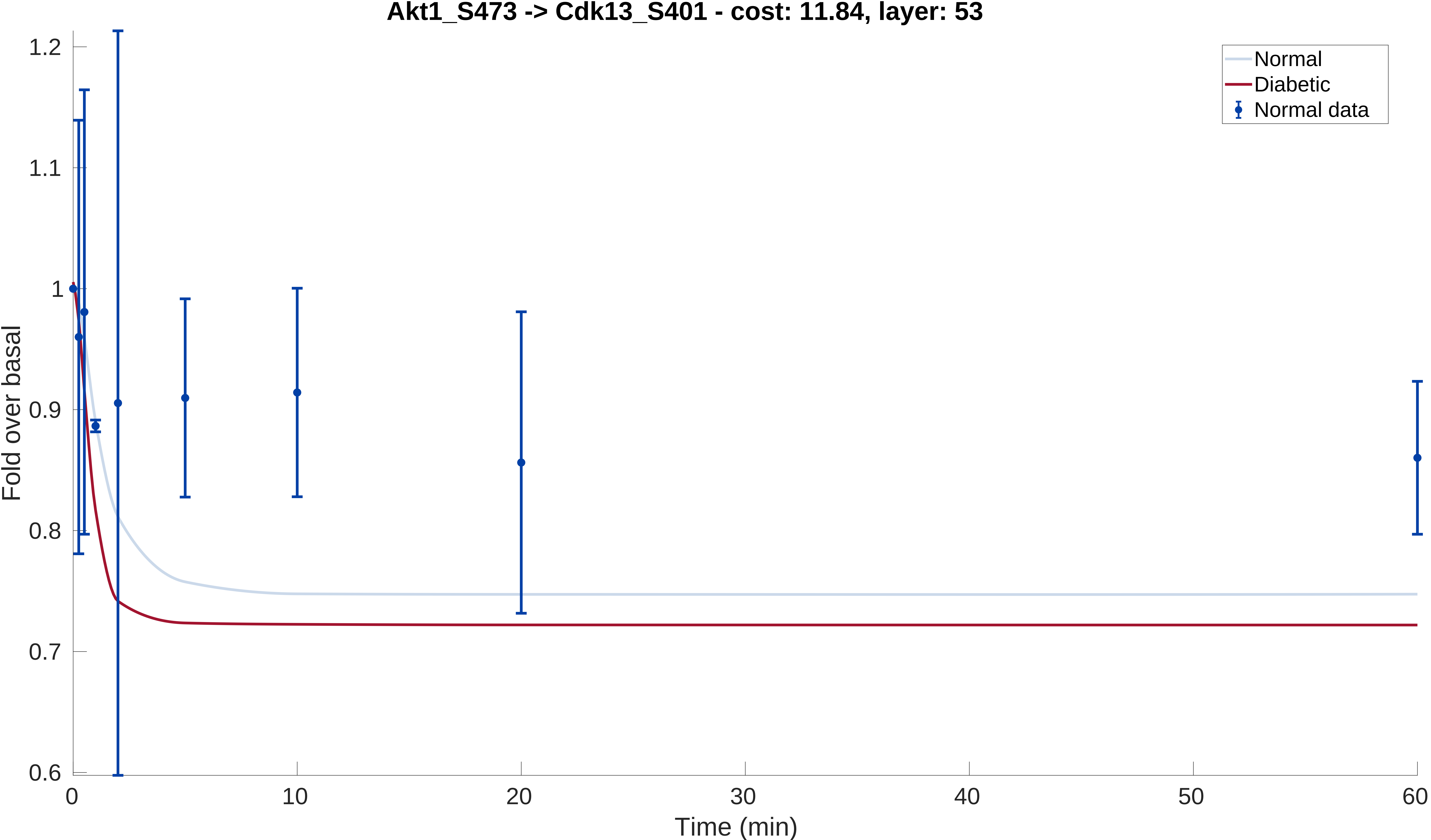

Akt1_S473 -> Cdk13_S401 - cost: 11.84, layer: 53
Normal
Diabetic
Normal data
Fold over basal
1.2
1.1
1
0.9
0.8
0.7
0.6
0
10
20
30
40
50
60
Time (min)

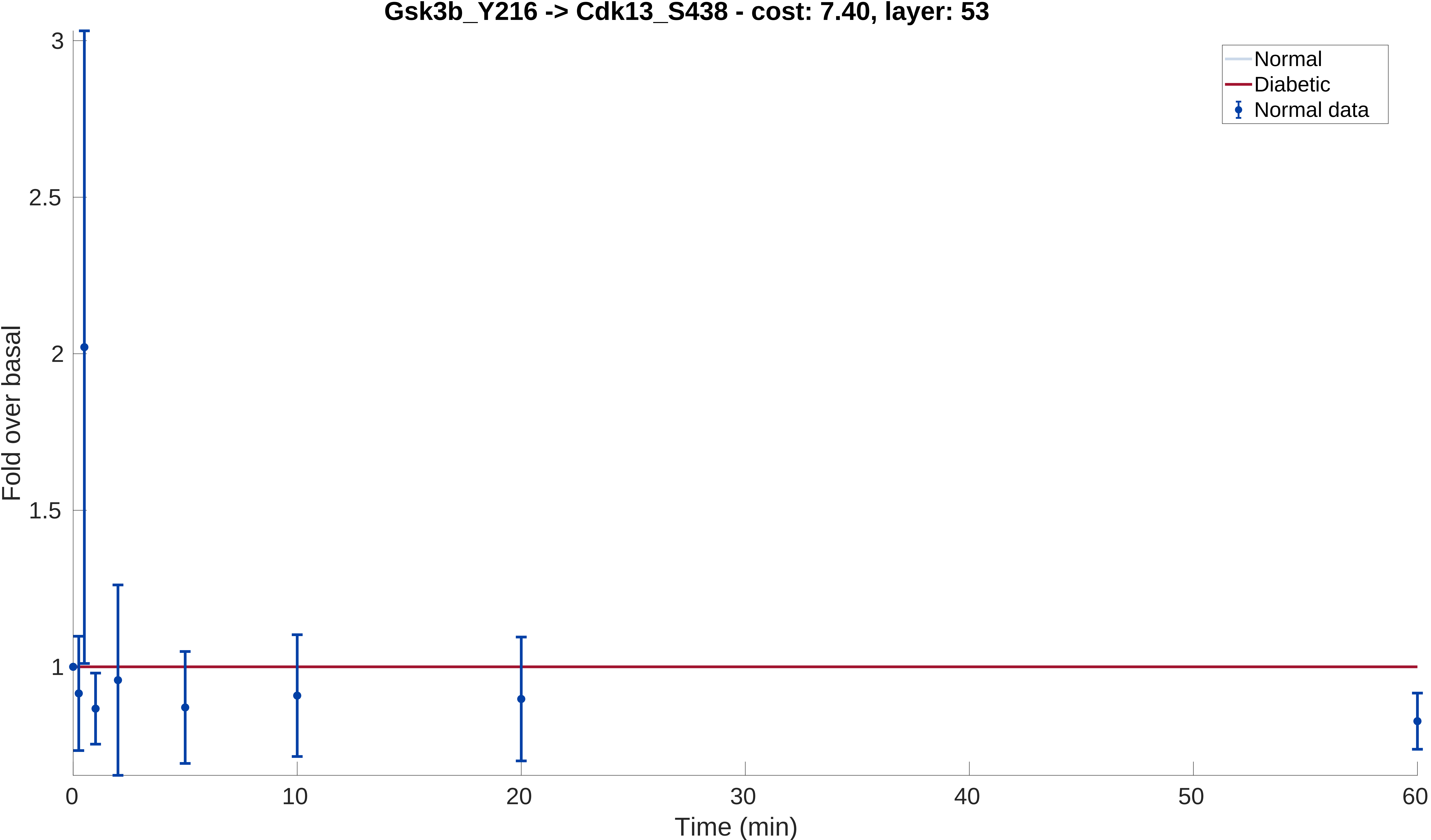

Gsk3b_Y216 -> Cdk13_S438 - cost: 7.40, layer: 53
Normal
Diabetic
Normal data
Fold over basal
3
2.5
2
1.5
1
0
10
20
30
40
50
60
Time (min)

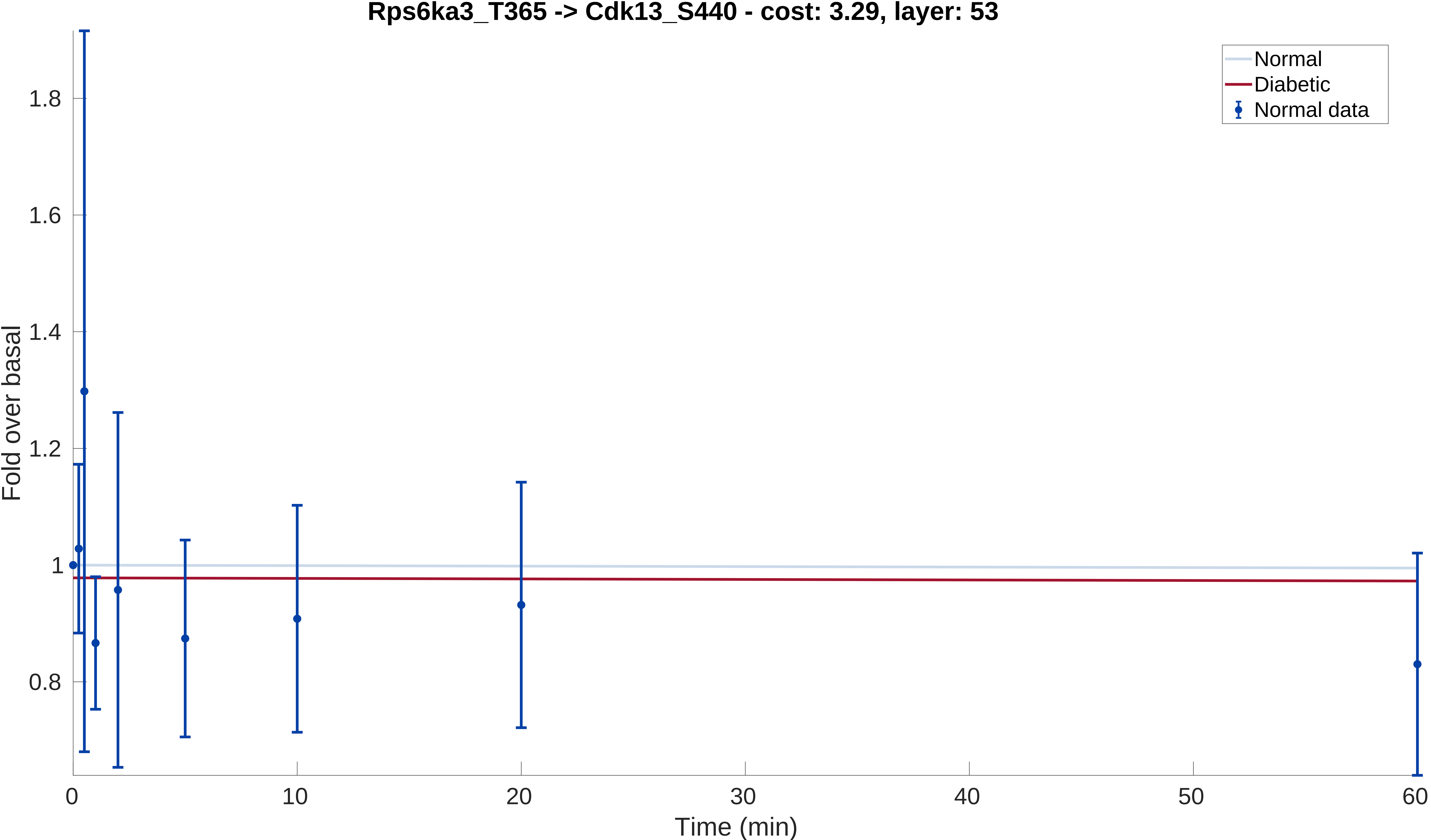

Rps6ka3_T365 -> Cdk13_S440 - cost: 3.29, layer: 53
Normal
Diabetic
Normal data
Fold over basal
1.8
1.6
1.4
1.2
1
0.8
0
10
20
30
40
50
60
Time (min)

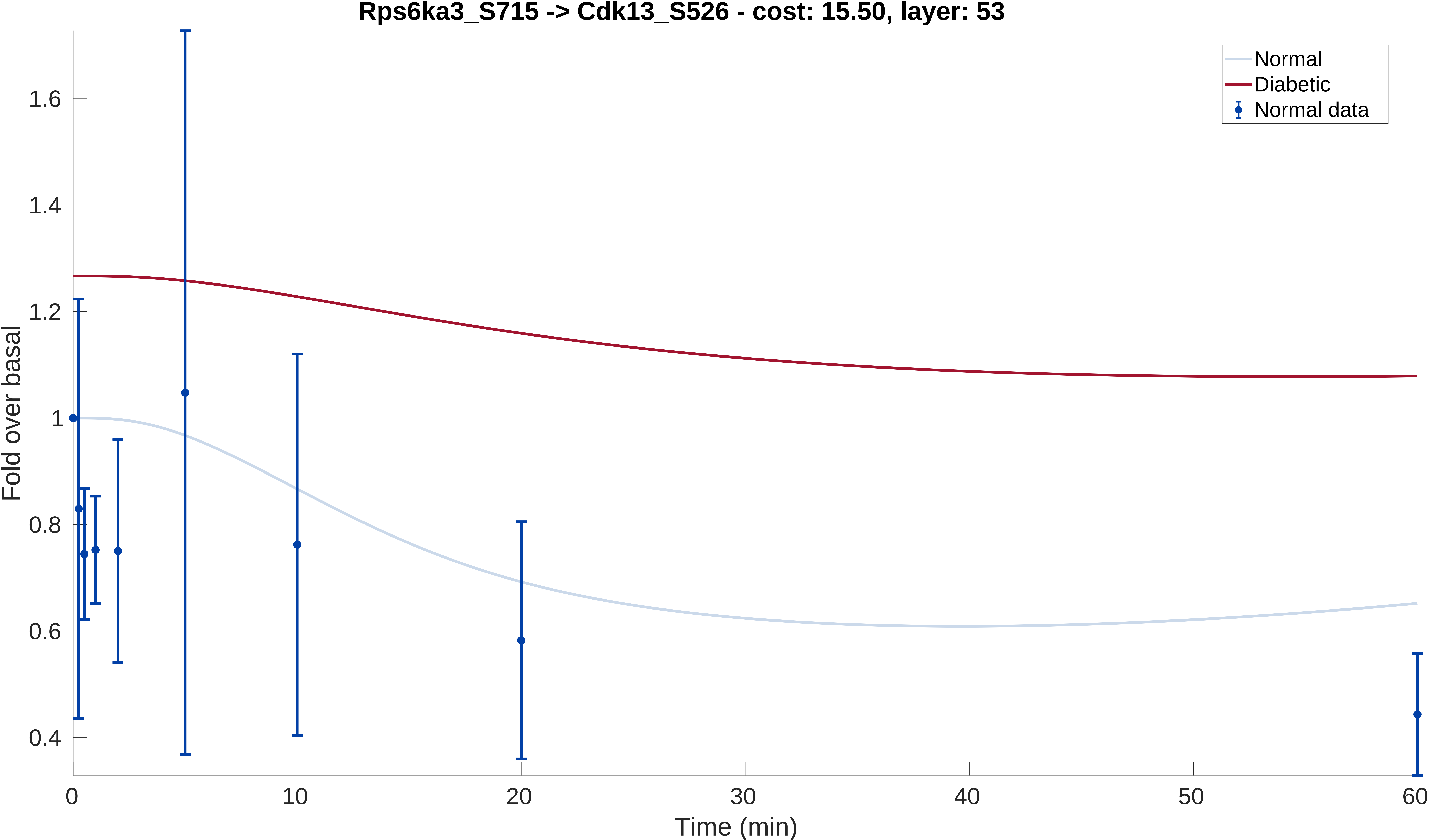

Rps6ka3_S715 -> Cdk13_S526 - cost: 15.50, layer: 53
Normal
Diabetic
Normal data
Fold over basal
Time (min)
1.6
1.4
1.2
1
0.8
0.6
0.4
0
10
20
30
40
50
60

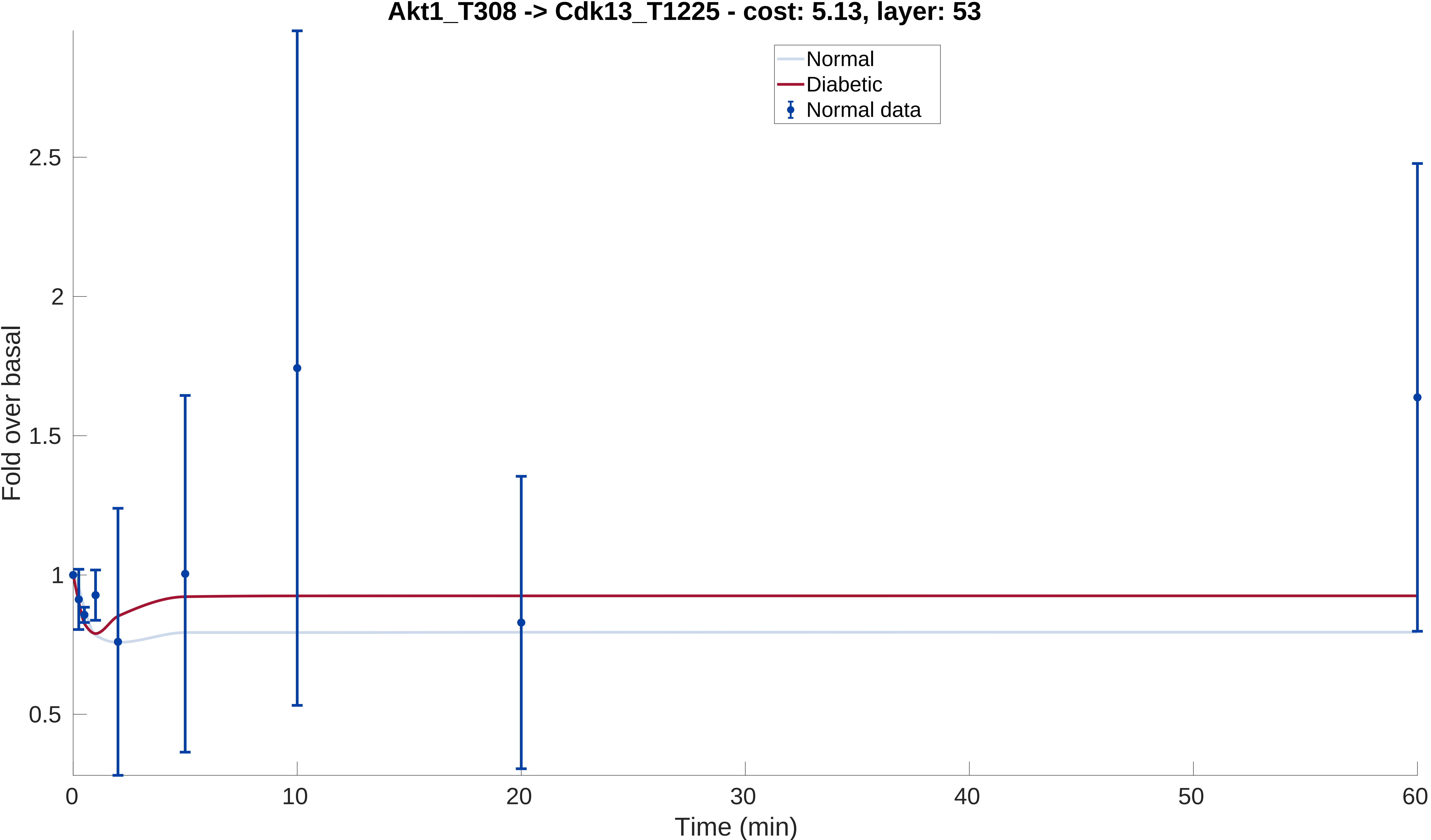

Akt1_T308 -> Cdk13_T1225 - cost: 5.13, layer: 53
Normal
Diabetic
Normal data
Fold over basal
2.5
2
1.5
1
0.5
0
10
20
30
40
50
60
Time (min)

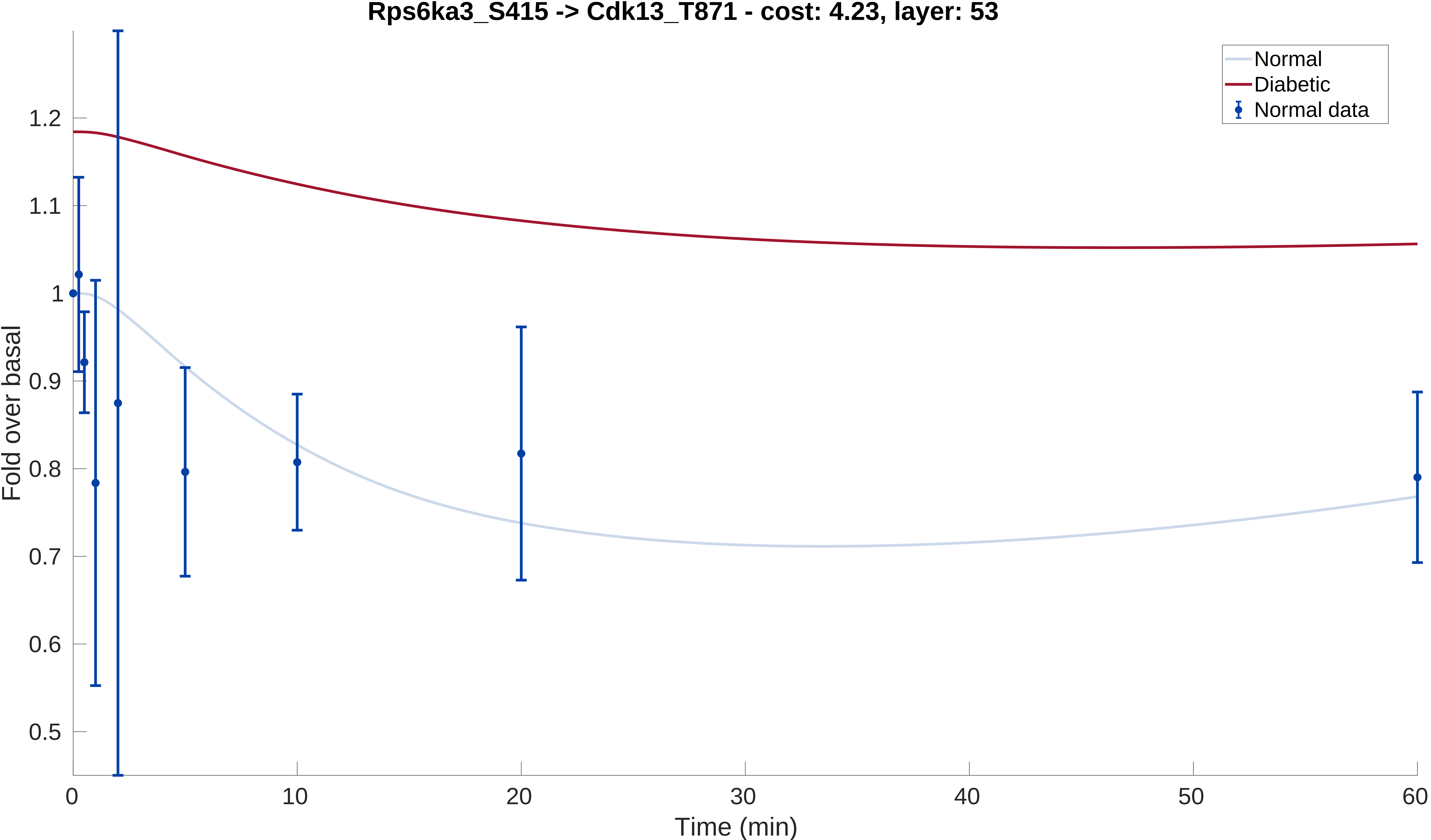

Rps6ka3_S415 -> Cdk13_T871 - cost: 4.23, layer: 53
Normal
Diabetic
Normal data
Fold over basal
1.2
1.1
1
0.9
0.8
0.7
0.6
0.5
0
10
20
30
40
50
60
Time (min)

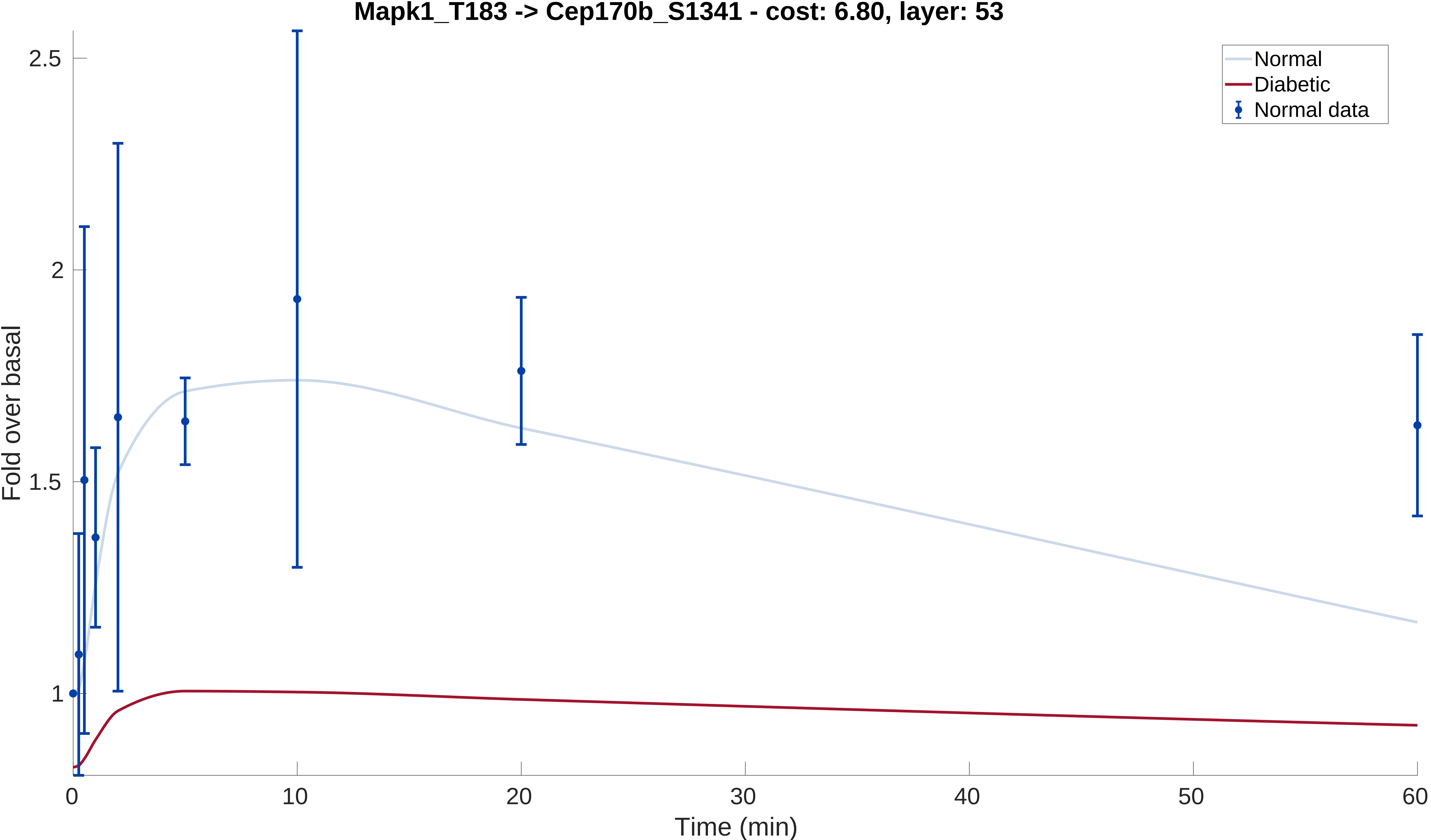
Mapk1_T183 -> Cep170b_S1341 - cost: 6.80, layer: 53
Normal
Diabetic
Normal data
Fold over basal
2.5
2
1.5
1
0
10
20
30
40
50
60
Time (min)

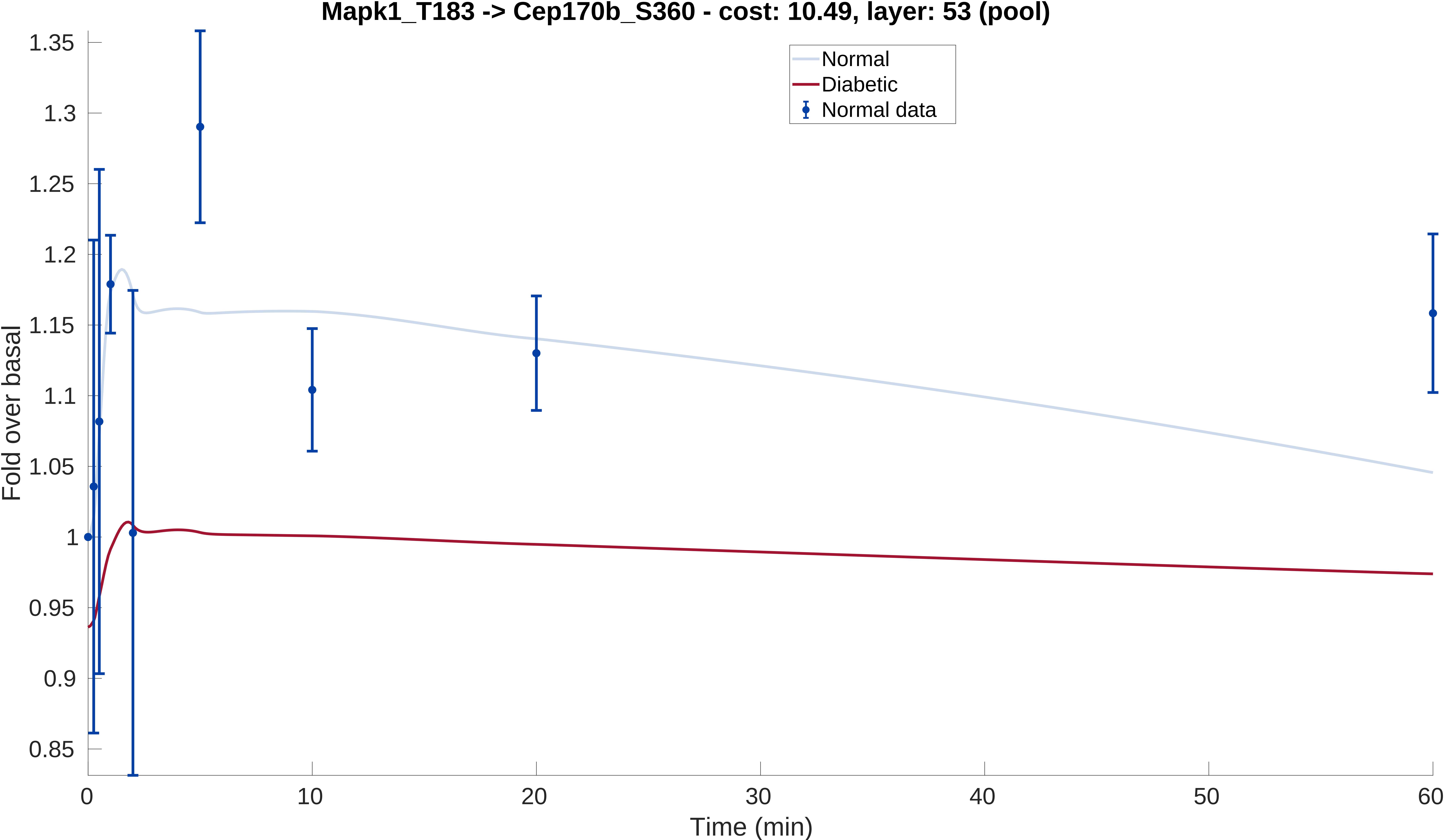
Mapk1_T183 -> Cep170b_S360 - cost: 10.49, layer: 53 (pool)
Normal
Diabetic
Normal data
Fold over basal
Time (min)
1.35
1.3
1.25
1.2
1.15
1.1
1.05
1
0.95
0.9
0.85
0
10
20
30
40
50
60

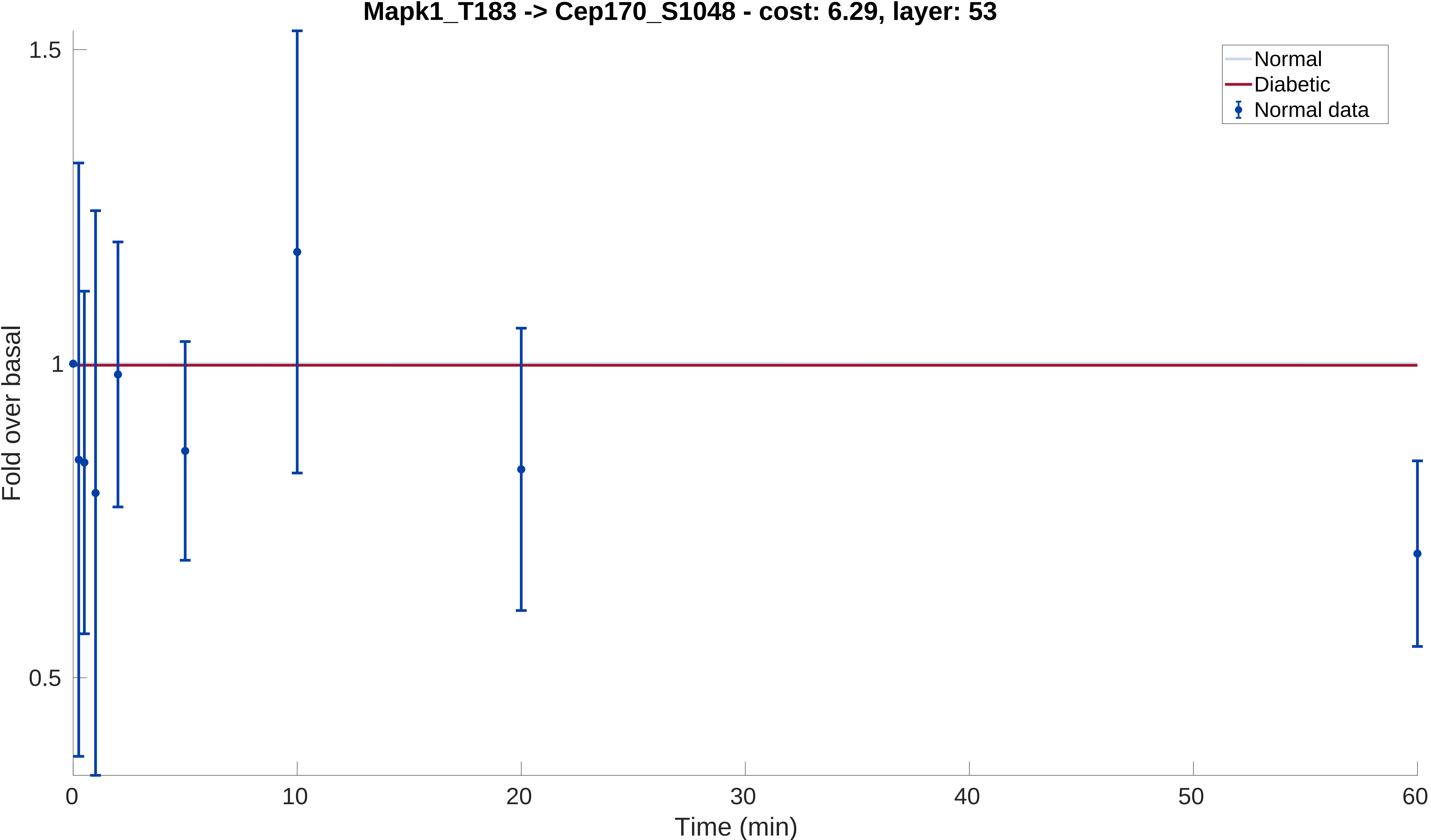

Mapk1_T183 -> Cep170_S1048 - cost: 6.29, layer: 53
Normal
Diabetic
Normal data
Fold over basal
Time (min)
1.5
1
0.5
0
10
20
30
40
50
60

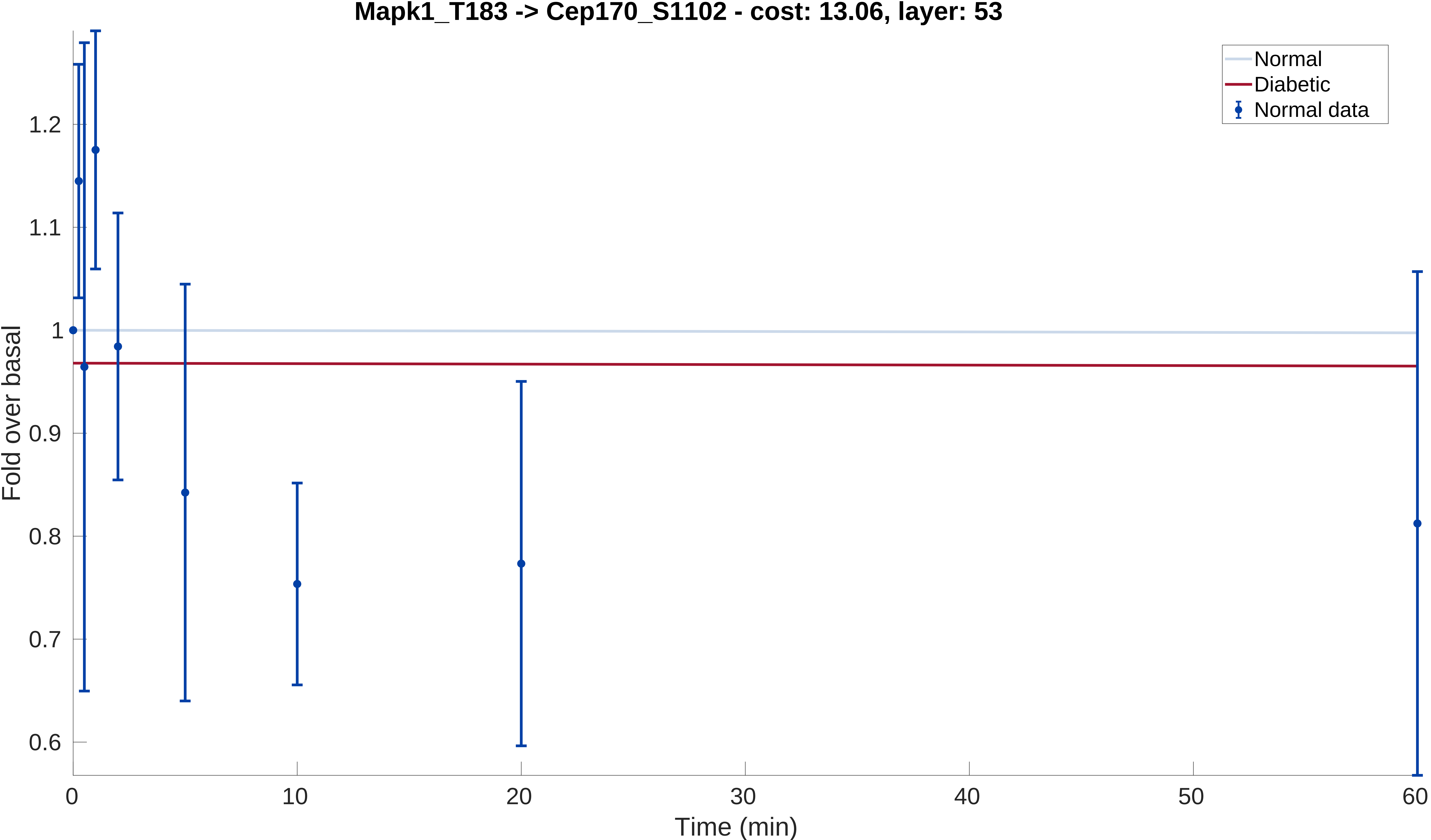

Mapk1_T183 -> Cep170_S1102 - cost: 13.06, layer: 53
Normal
Diabetic
Normal data
Fold over basal
Time (min)
0.6
0.7
0.8
0.9
1
1.1
1.2
0
10
20
30
40
50
60

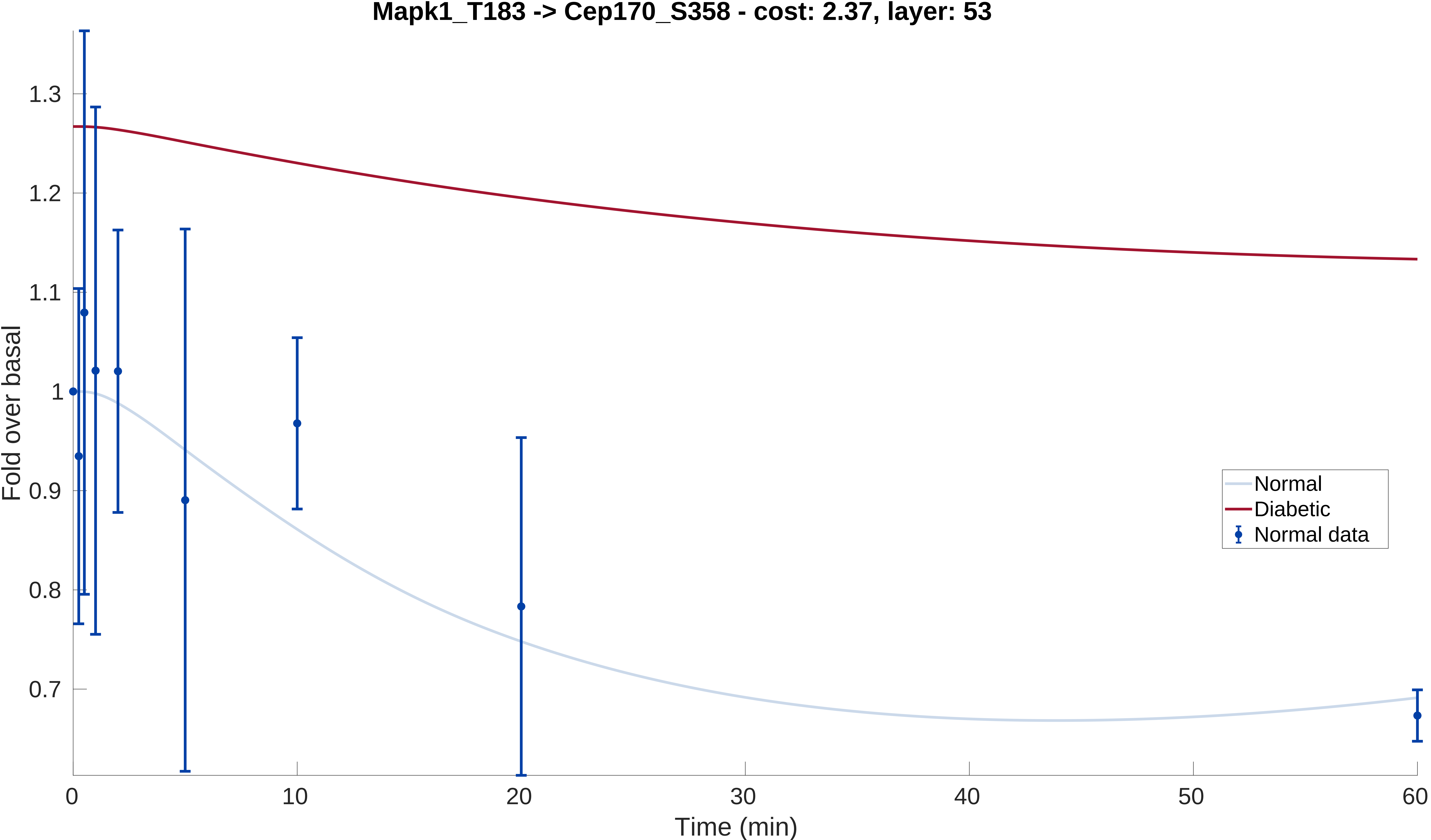

Mapk1_T183 -> Cep170_S358 - cost: 2.37, layer: 53
Fold over basal
1.3
1.2
1.1
1
0.9
0.8
0.7
0
10
20
30
40
50
60
Time (min)
Normal
Diabetic
Normal data

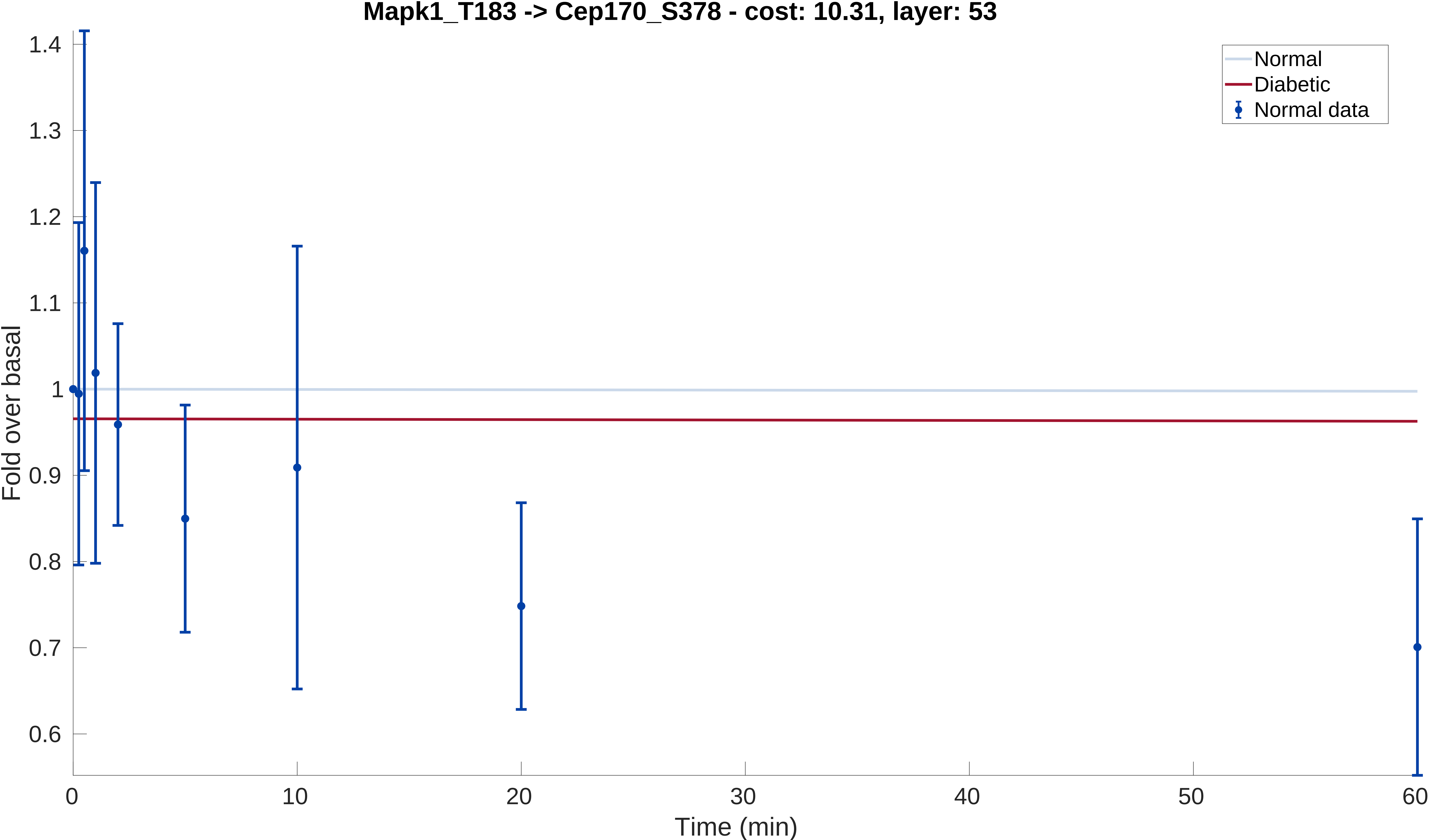

Mapk1_T183 -> Cep170_S378 - cost: 10.31, layer: 53
Normal
Diabetic
Normal data
Fold over basal
1.4
1.3
1.2
1.1
1
0.9
0.8
0.7
0.6
0
10
20
30
40
50
60
Time (min)

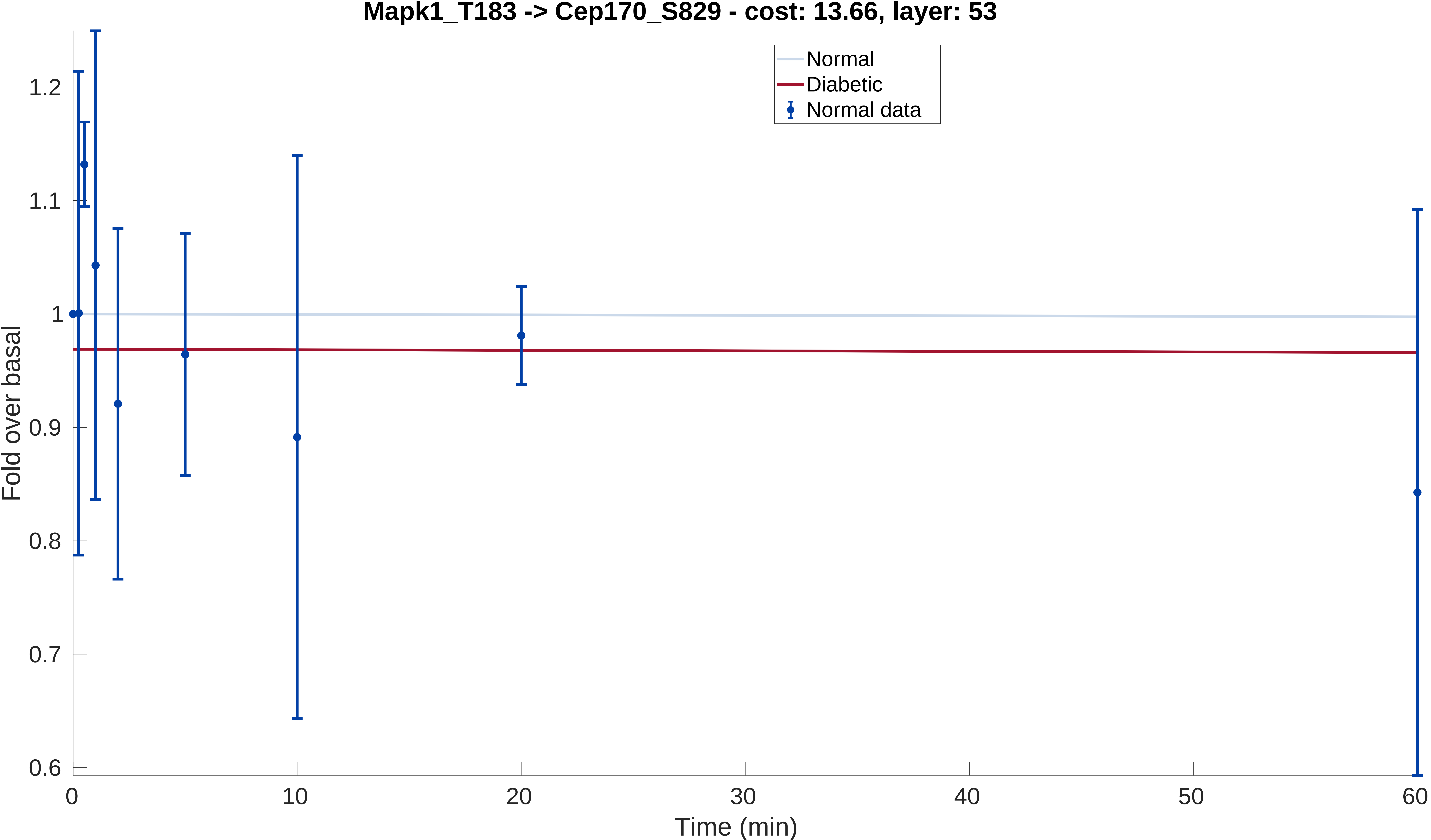

Mapk1_T183 -> Cep170_S829 - cost: 13.66, layer: 53
Normal
Diabetic
Normal data
Fold over basal
1.2
1.1
1
0.9
0.8
0.7
0.6
0
10
20
30
40
50
60
Time (min)

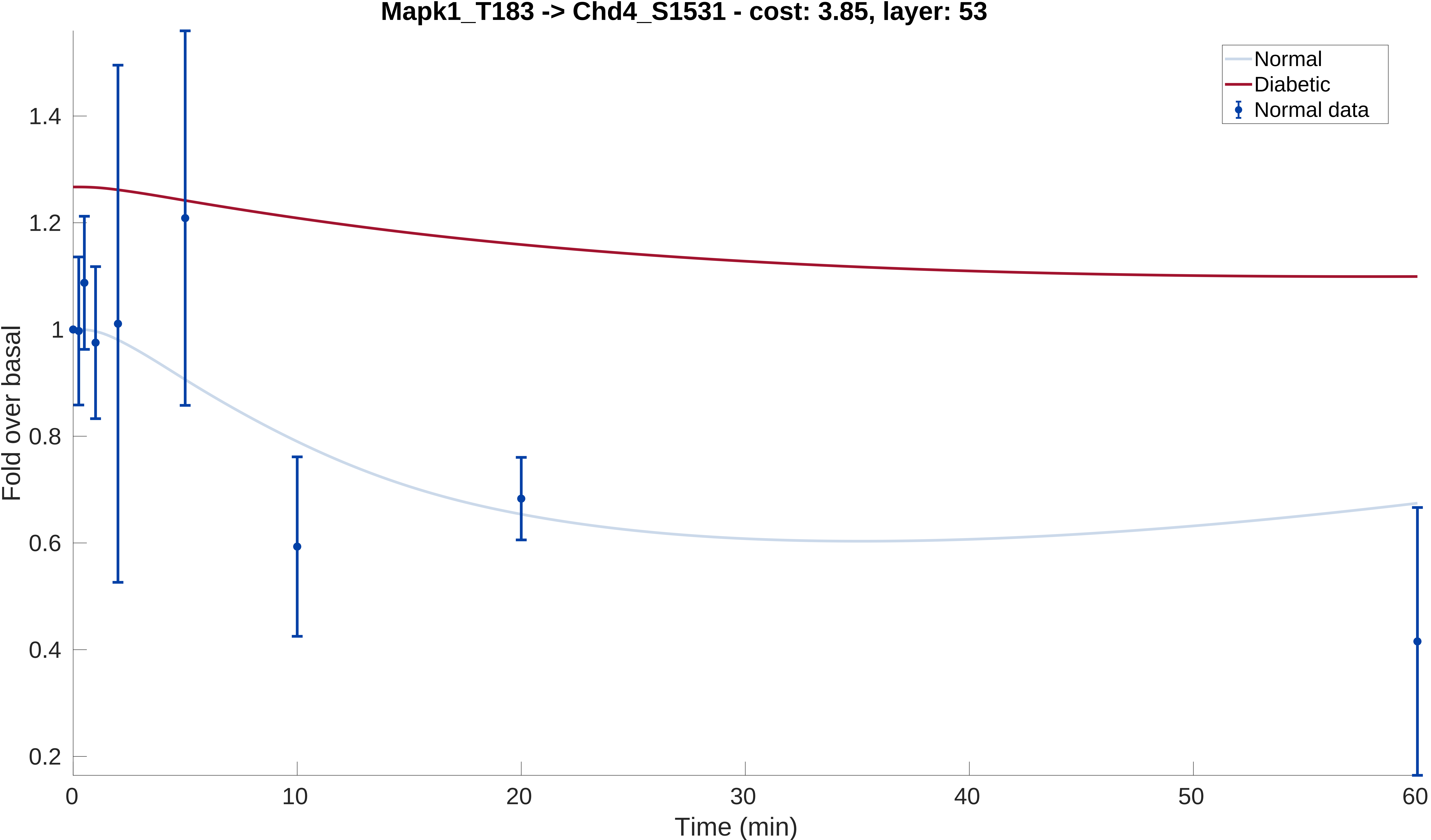

Mapk1_T183 -> Chd4_S1531 - cost: 3.85, layer: 53
Normal
Diabetic
Normal data
Fold over basal
1.4
1.2
1
0.8
0.6
0.4
0.2
0
10
20
30
40
50
60
Time (min)

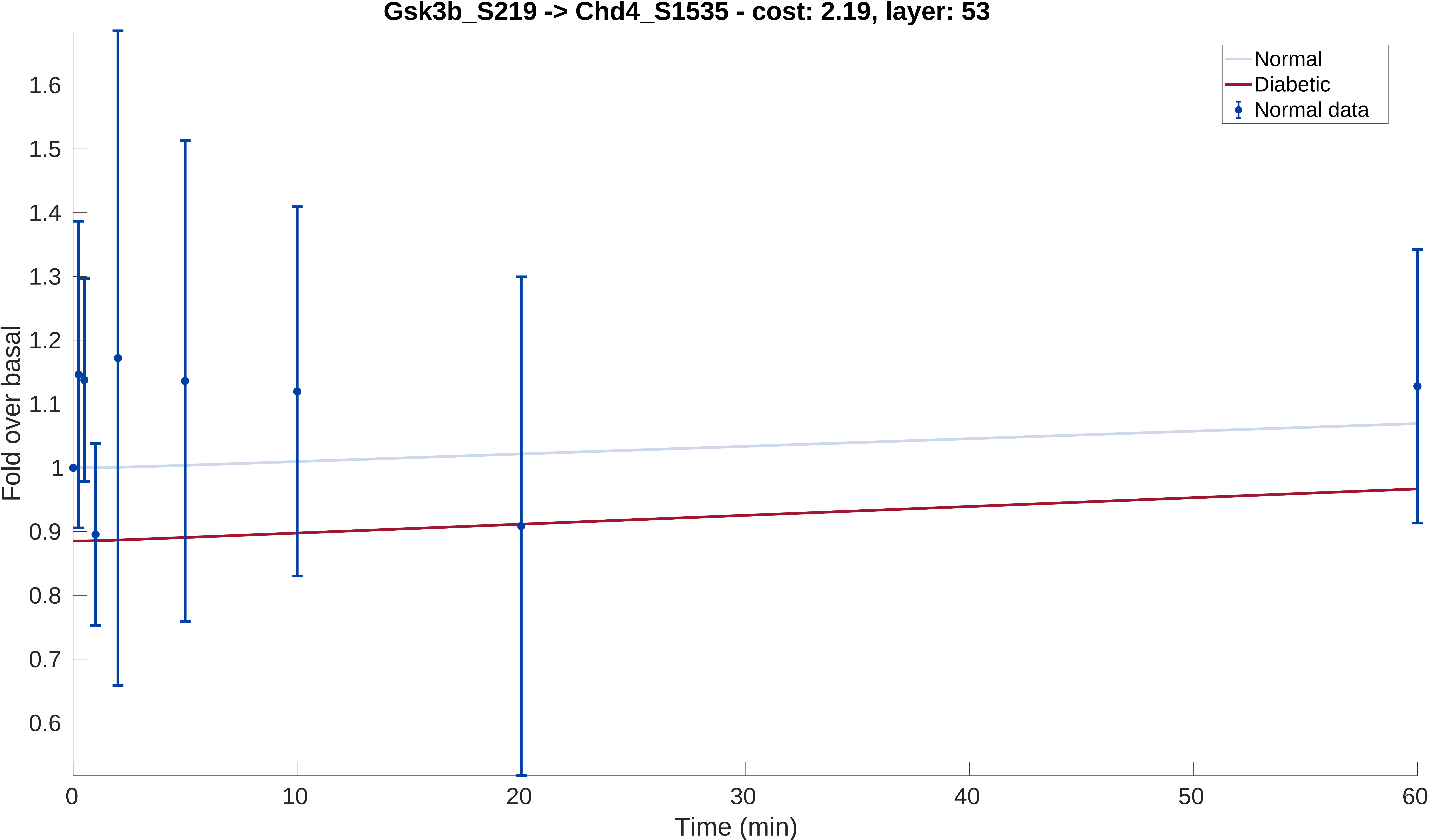

Gsk3b_S219 -> Chd4_S1535 - cost: 2.19, layer: 53
Normal
Diabetic
Normal data
Fold over basal
1.6
1.5
1.4
1.3
1.2
1.1
1
0.9
0.8
0.7
0.6
0
10
20
30
40
50
60
Time (min)

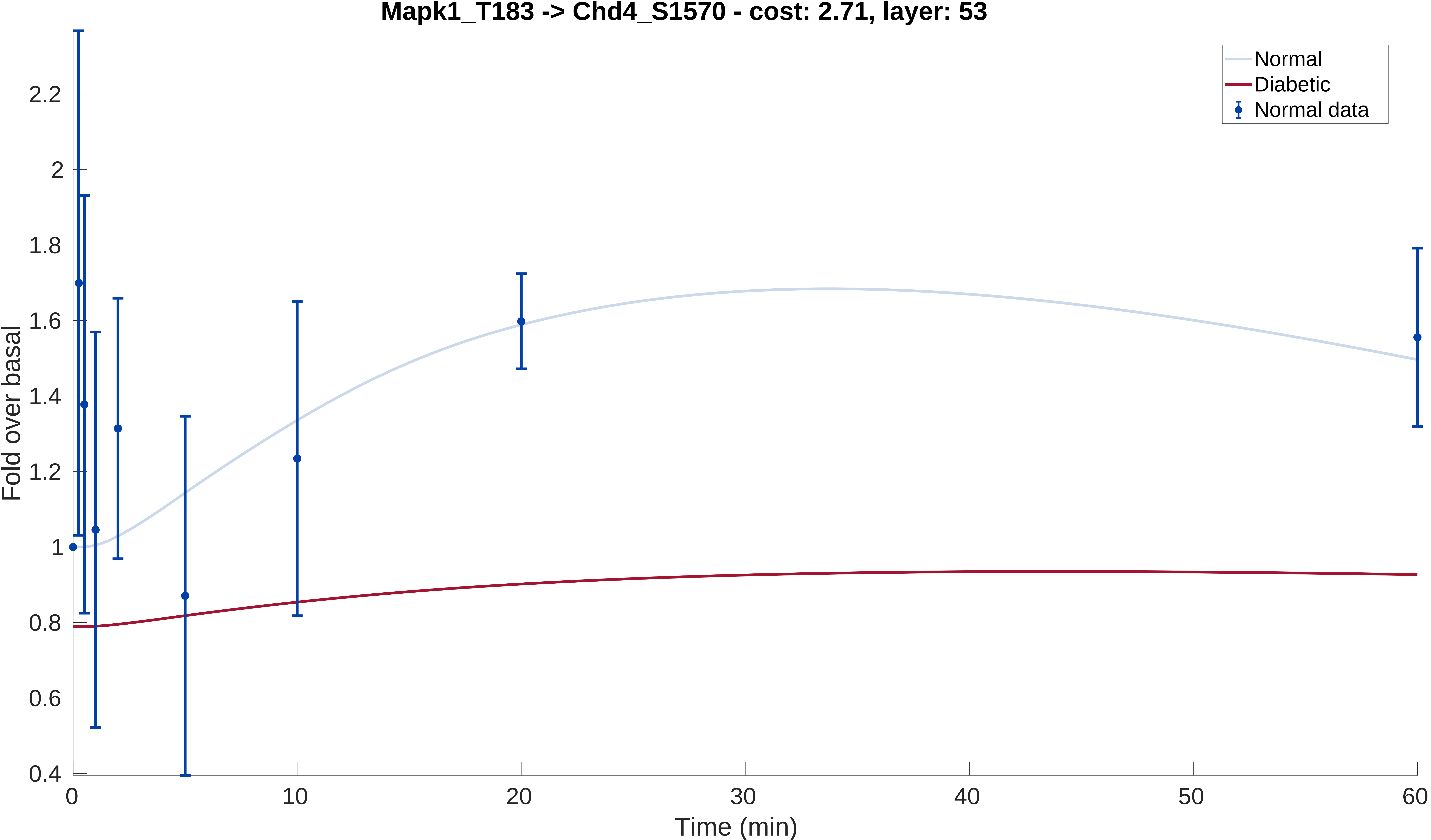

Mapk1_T183 -> Chd4_S1570 - cost: 2.71, layer: 53
Normal
Diabetic
Normal data
Fold over basal
Time (min)
0.4
0.6
0.8
1
1.2
1.4
1.6
1.8
2
2.2
0
10
20
30
40
50
60

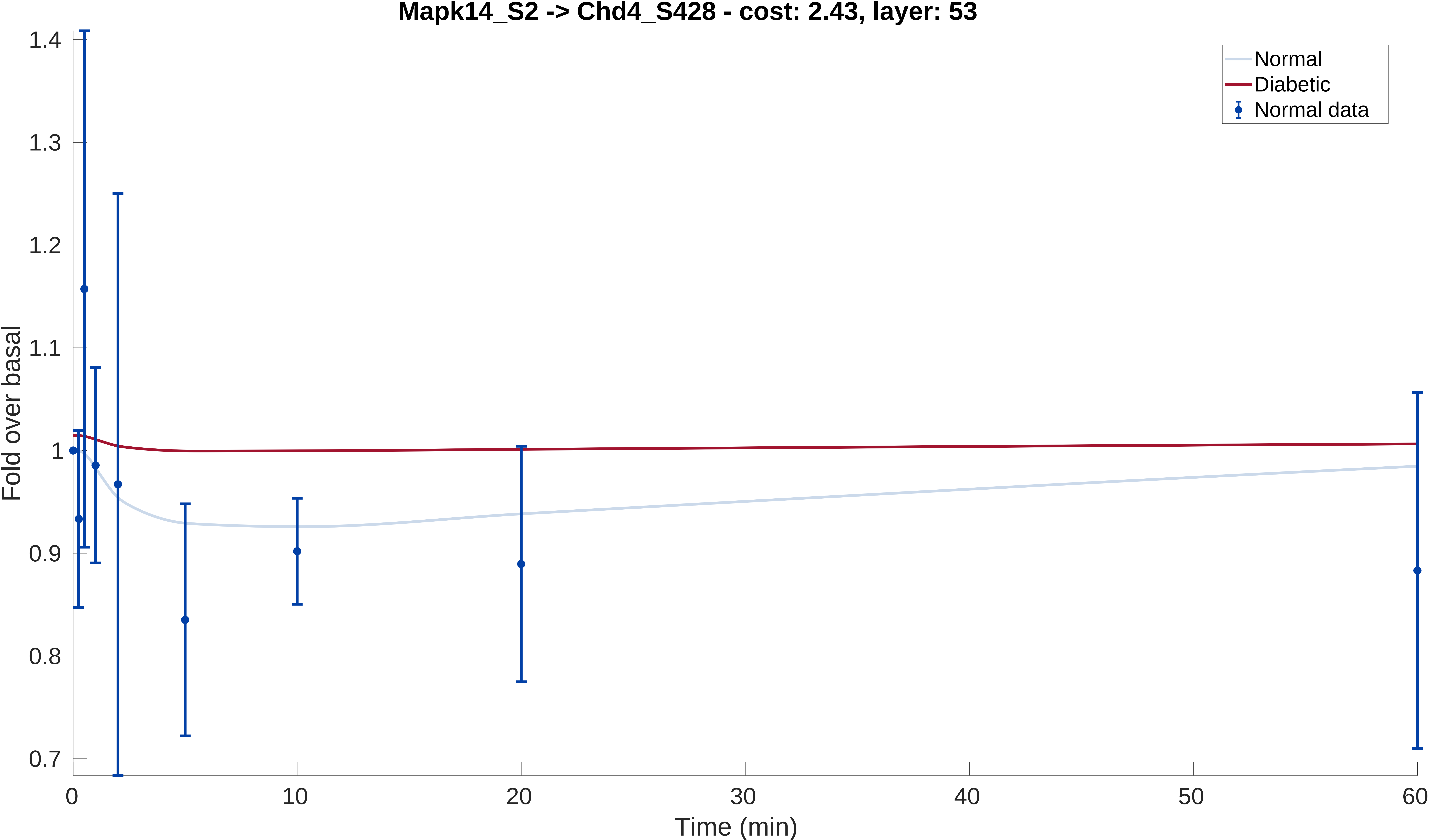

Mapk14_S2 -> Chd4_S428 - cost: 2.43, layer: 53
Normal
Diabetic
Normal data
Fold over basal
1.4
1.3
1.2
1.1
1
0.9
0.8
0.7
0
10
20
30
40
50
60
Time (min)

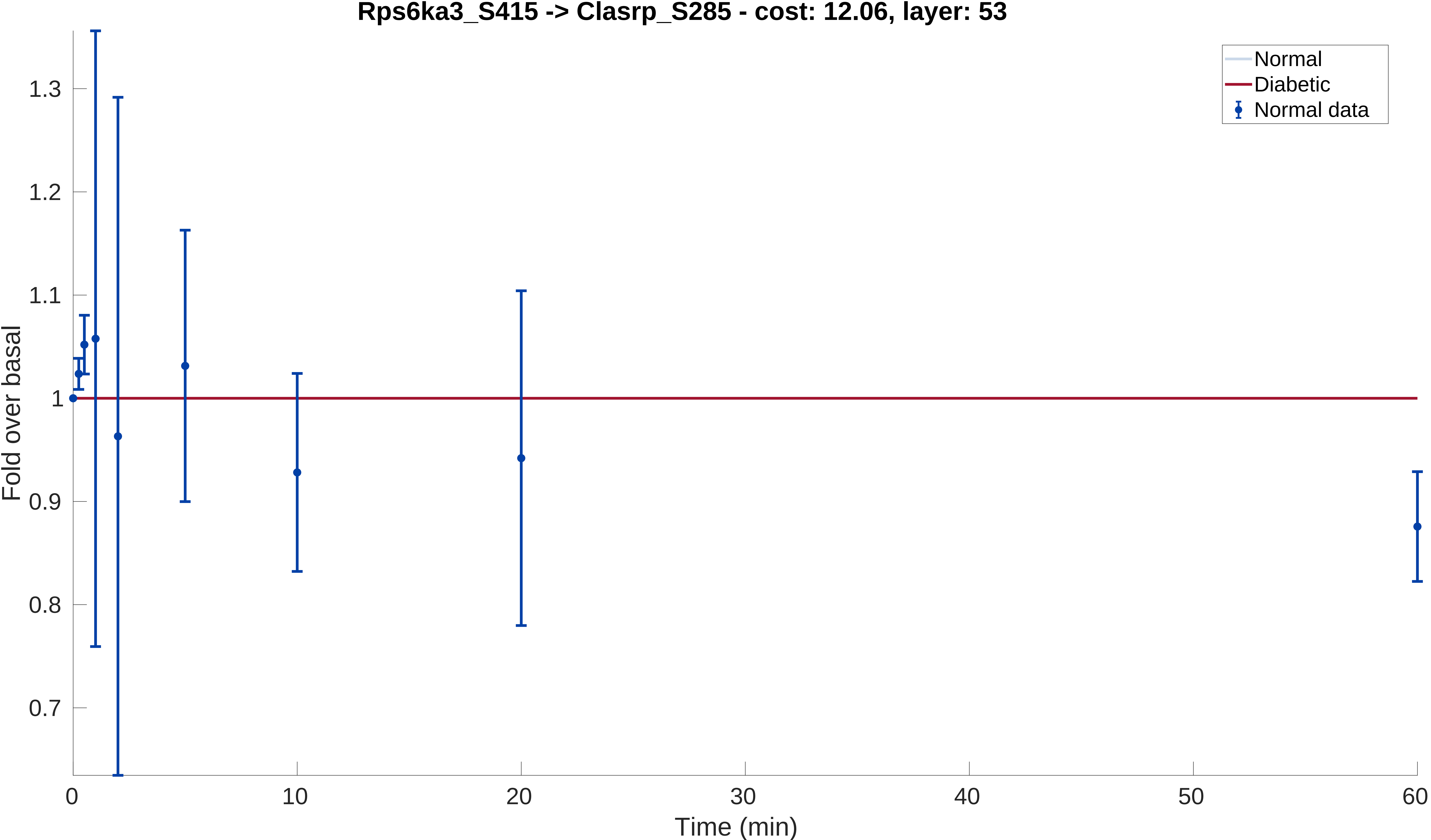

Rps6ka3_S415 -> Clasrp_S285 - cost: 12.06, layer: 53
Normal
Diabetic
Normal data
Fold over basal
1.3
1.2
1.1
1
0.9
0.8
0.7
0
10
20
30
40
50
60
Time (min)

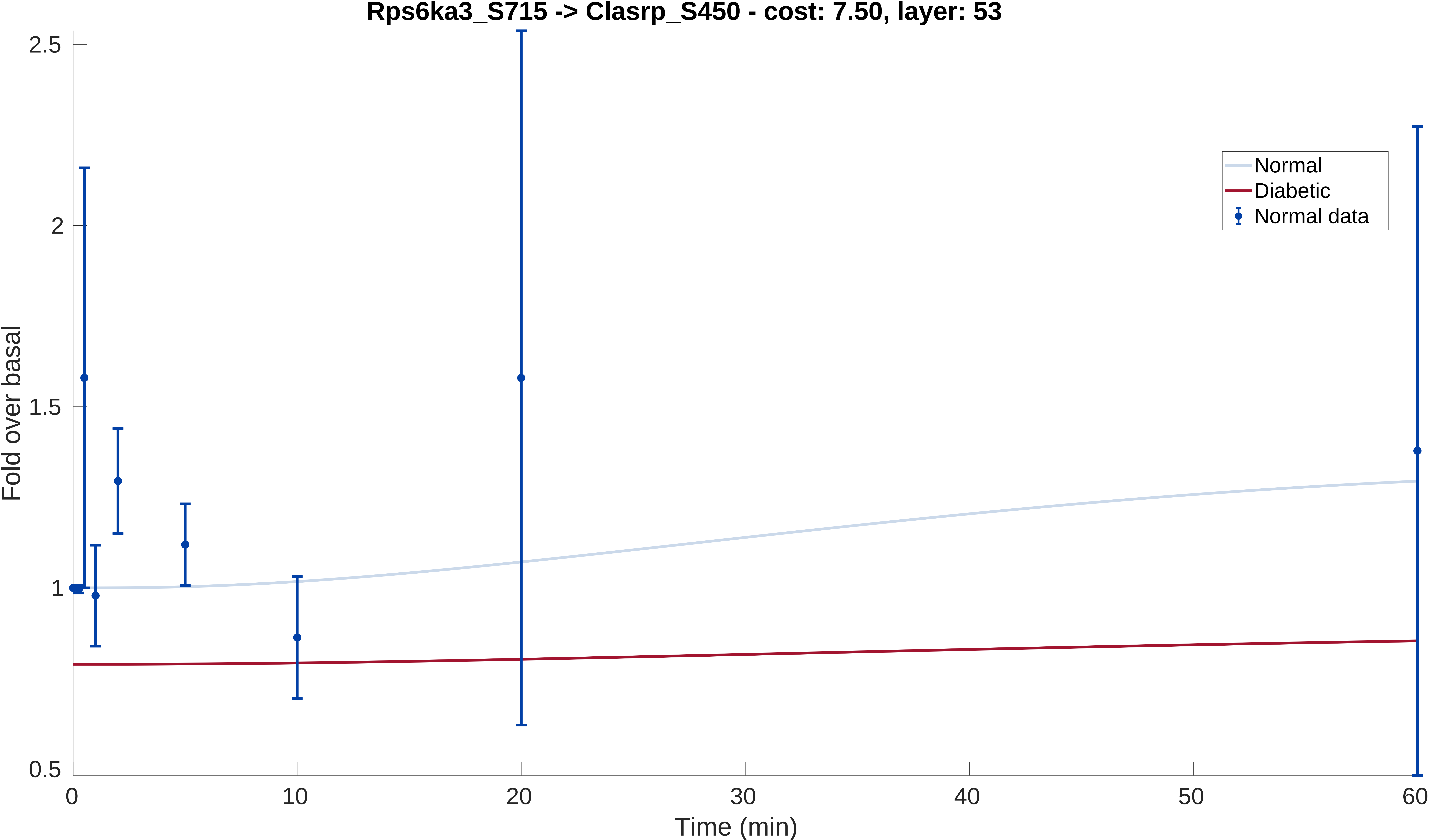

Rps6ka3_S715 -> Clasrp_S450 - cost: 7.50, layer: 53
Fold over basal
Time (min)
Normal
Diabetic
Normal data
0.5
1
1.5
2
2.5
0
10
20
30
40
50
60

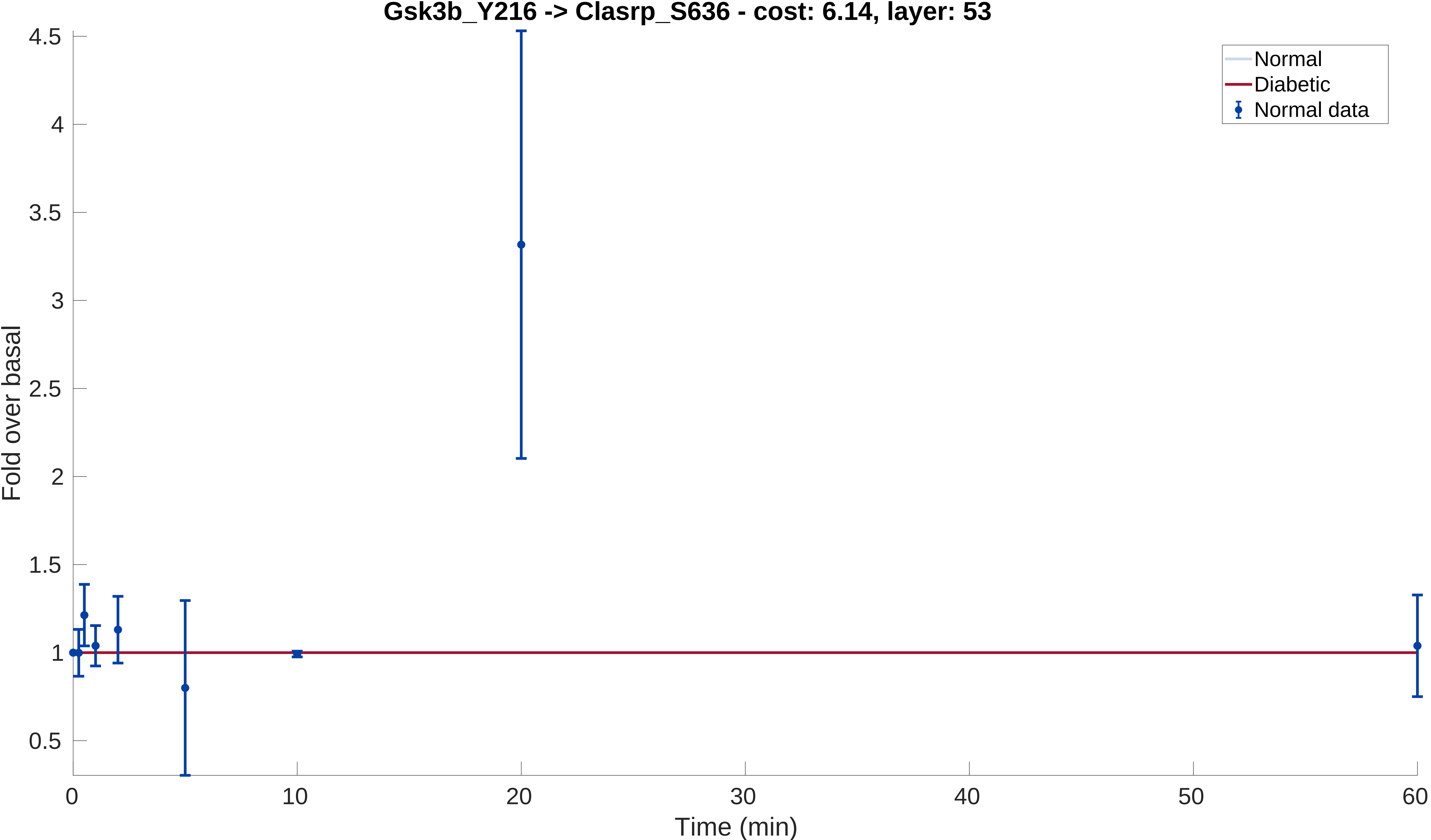

Gsk3b_Y216 -> Clasrp_S636 - cost: 6.14, layer: 53
Normal
Diabetic
Normal data
Fold over basal
Time (min)
0.5
1
1.5
2
2.5
3
3.5
4
4.5
0
10
20
30
40
50
60

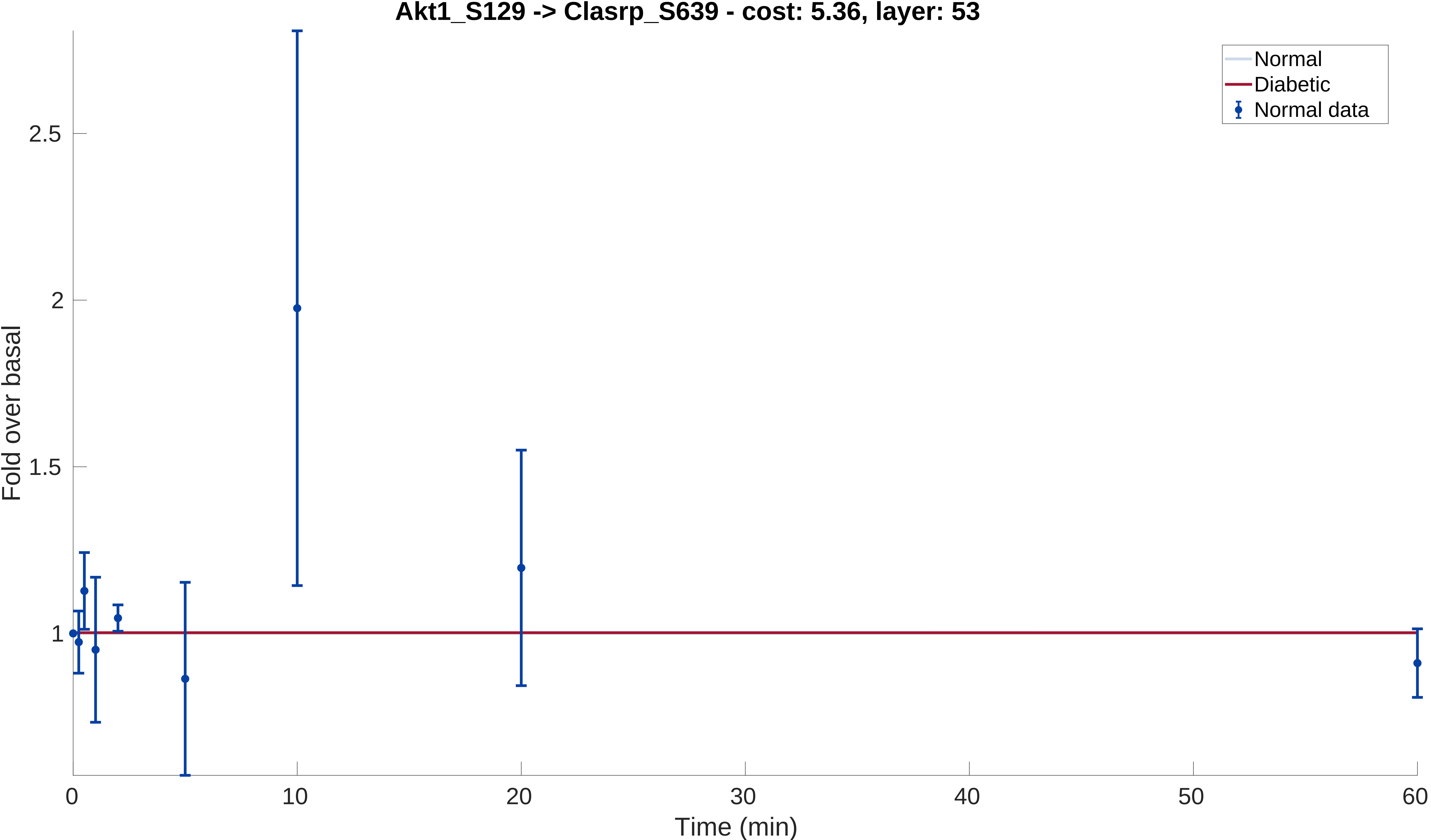

Akt1_S129 -> Clasrp_S639 - cost: 5.36, layer: 53
Normal
Diabetic
Normal data
Fold over basal
2.5
2
1.5
1
0
10
20
30
40
50
60
Time (min)

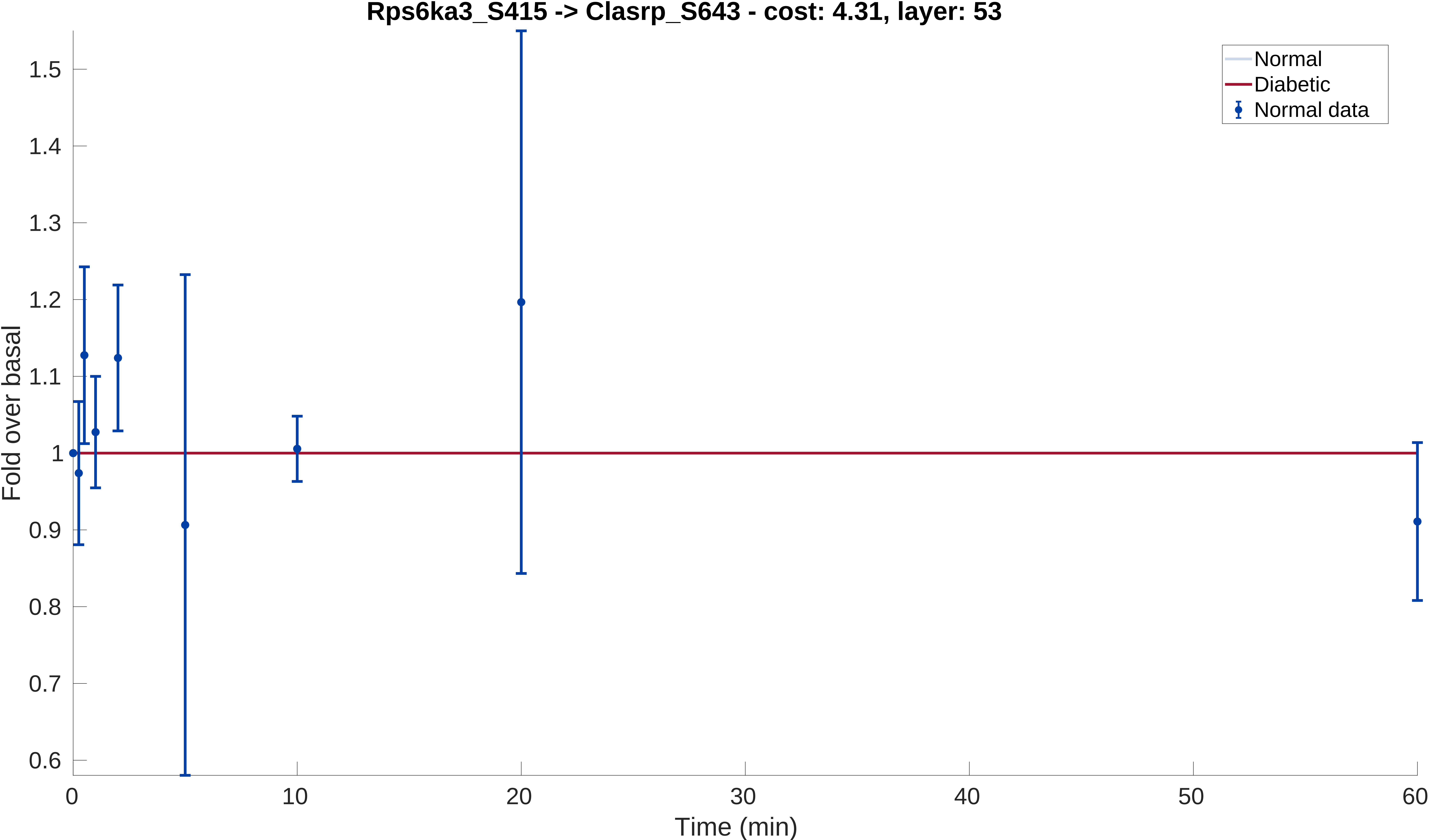

Rps6ka3_S415 -> Clasrp_S643 - cost: 4.31, layer: 53
Normal
Diabetic
Normal data
Fold over basal
Time (min)
1.5
1.4
1.3
1.2
1.1
1
0.9
0.8
0.7
0.6
0
10
20
30
40
50
60

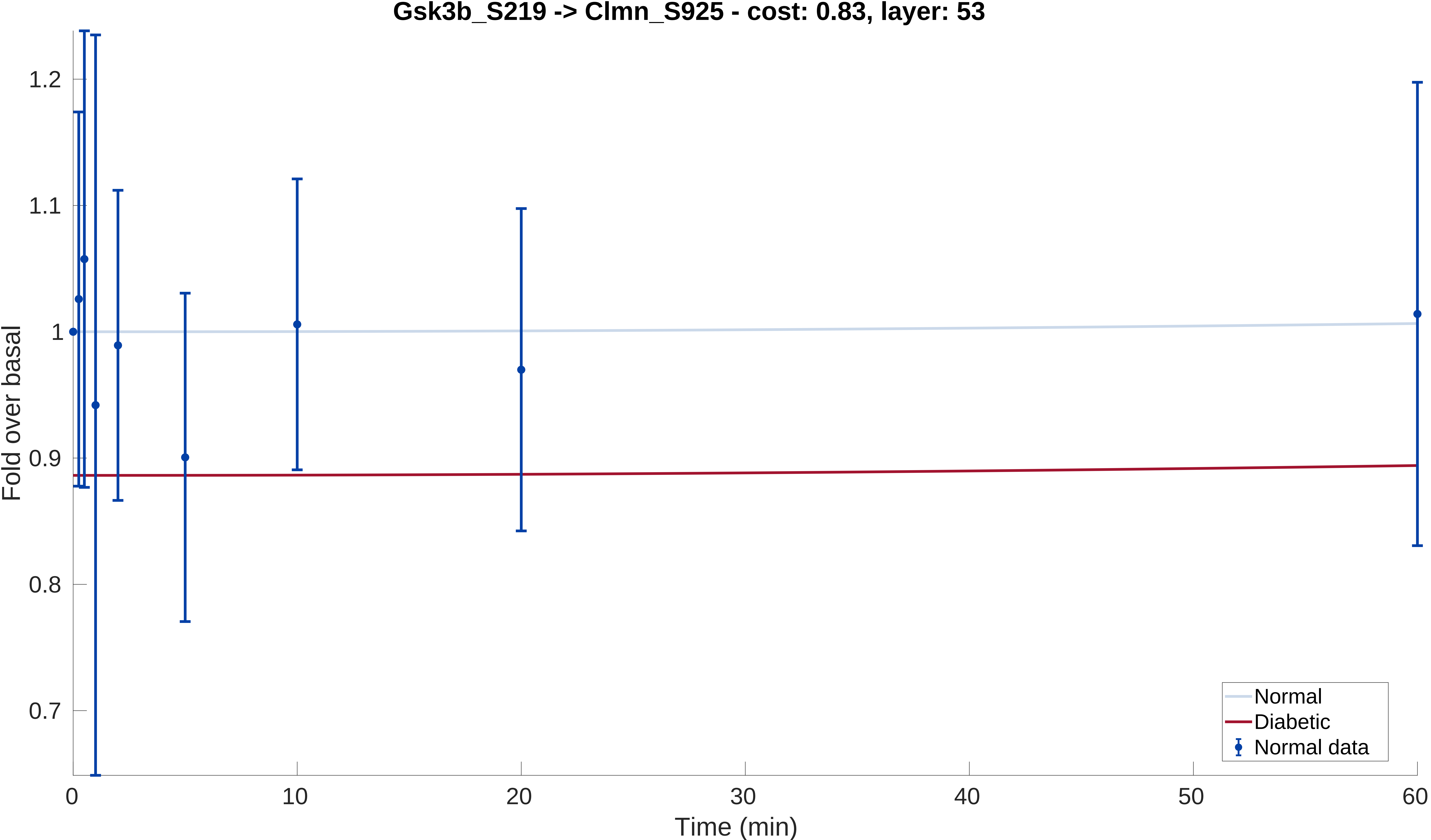

Gsk3b_S219 -> Clmn_S925 - cost: 0.83, layer: 53
Fold over basal
1.2
1.1
1
0.9
0.8
0.7
0
10
20
30
40
50
60
Time (min)
Normal
Diabetic
Normal data

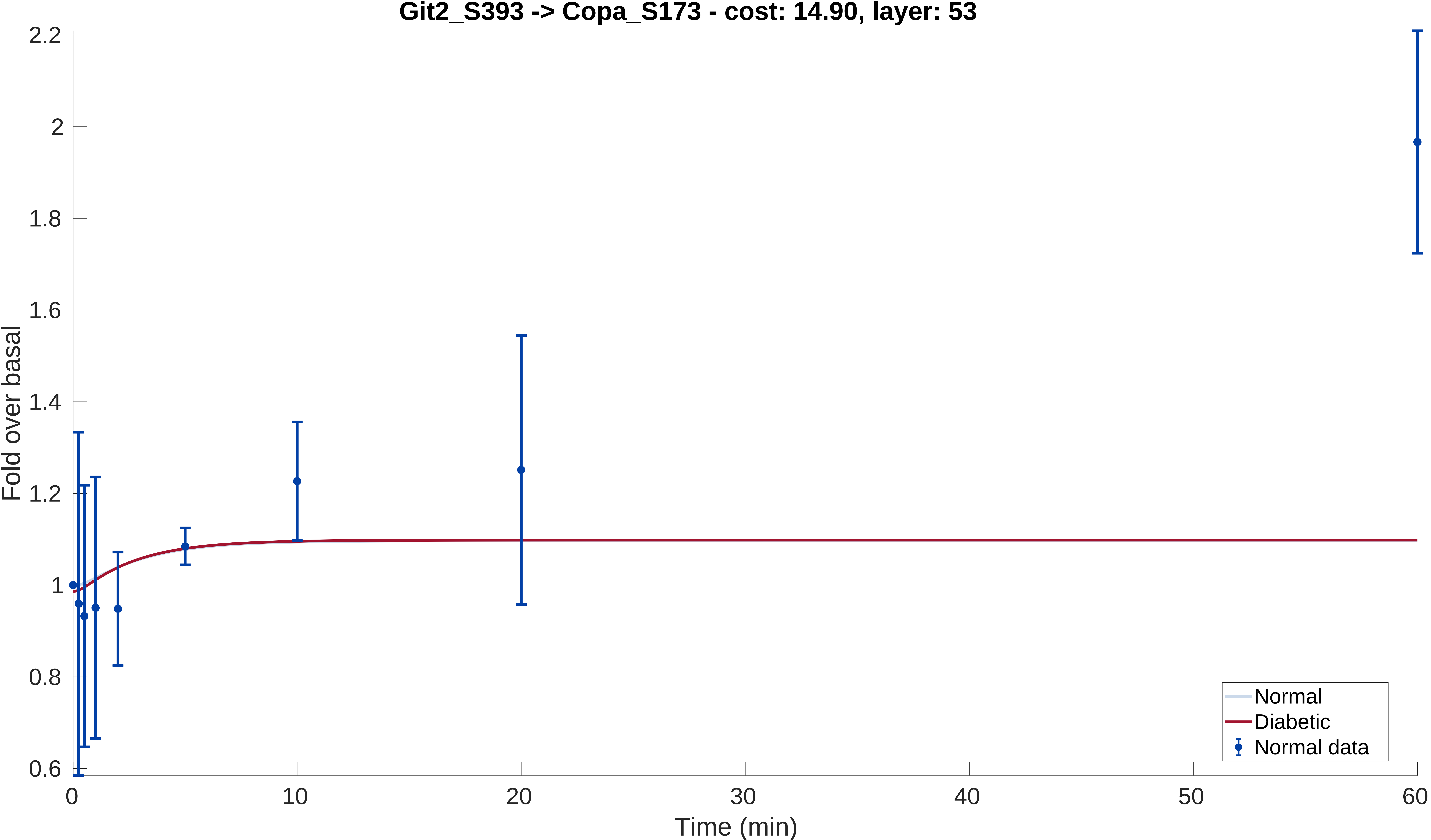

Git2_S393 -> Copa_S173 - cost: 14.90, layer: 53
Fold over basal
Time (min)
0.6
0.8
1
1.2
1.4
1.6
1.8
2
2.2
0
10
20
30
40
50
60
Normal
Diabetic
Normal data

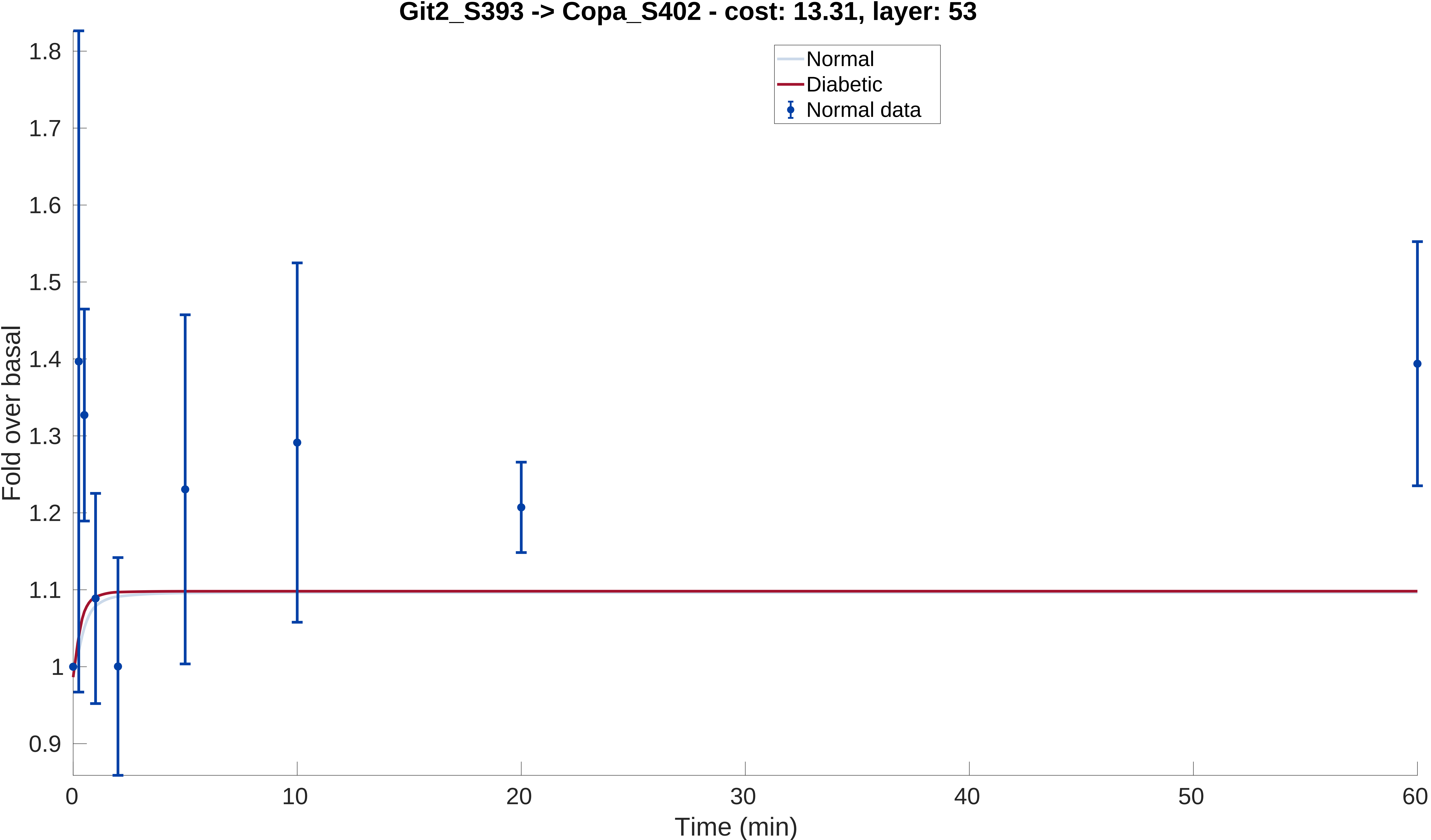

Git2_S393 -> Copa_S402 - cost: 13.31, layer: 53
Normal
Diabetic
Normal data
1.8
1.7
1.6
1.5
1.4
1.3
1.2
1.1
1
0.9
Fold over basal
0
10
20
30
40
50
60
Time (min)

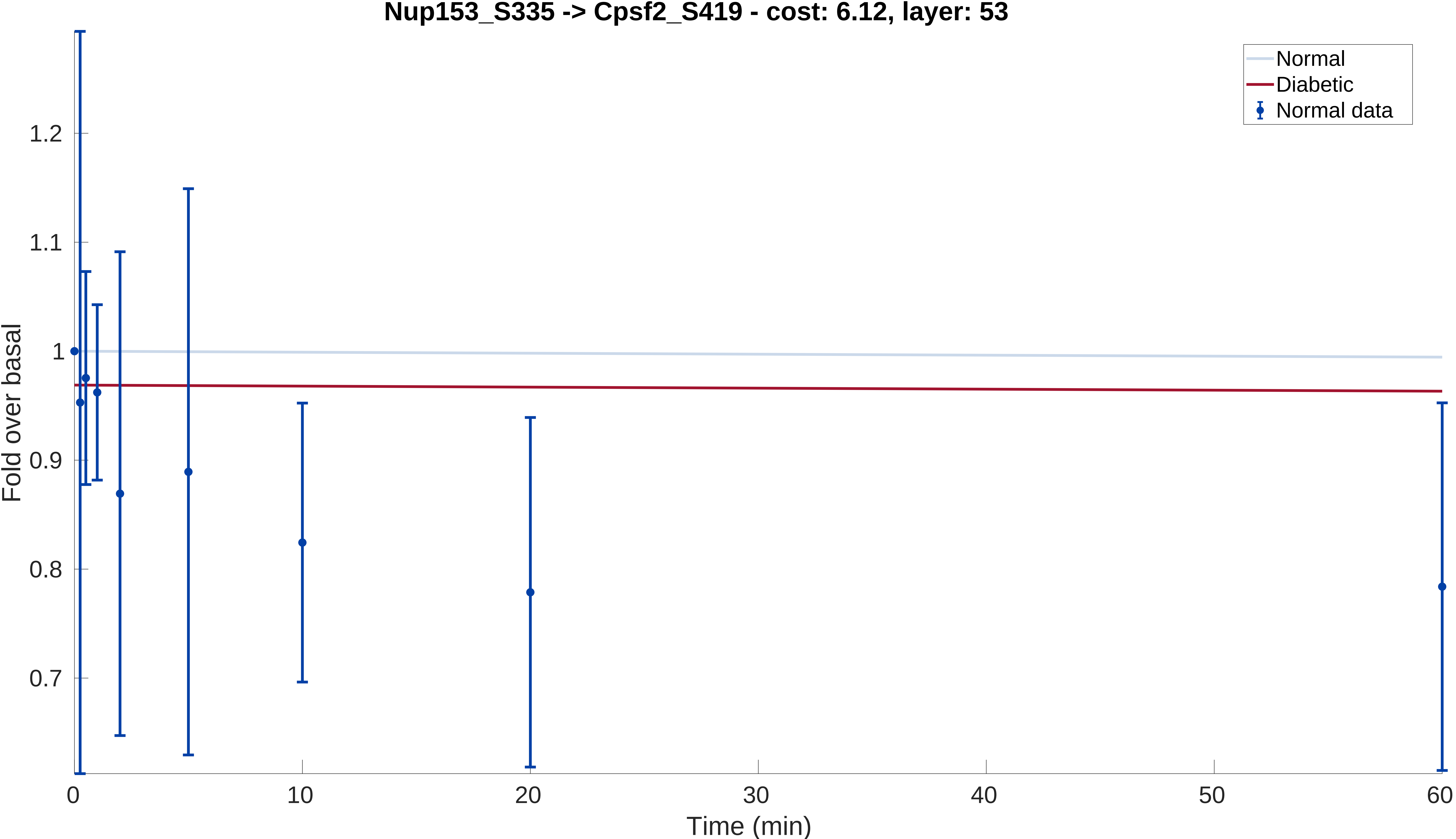

Nup153_S335 -> Cpsf2_S419 - cost: 6.12, layer: 53
Normal
Diabetic
Normal data
1.2
1.1
1
0.9
0.8
0.7
Fold over basal
0
10
20
30
40
50
60
Time (min)

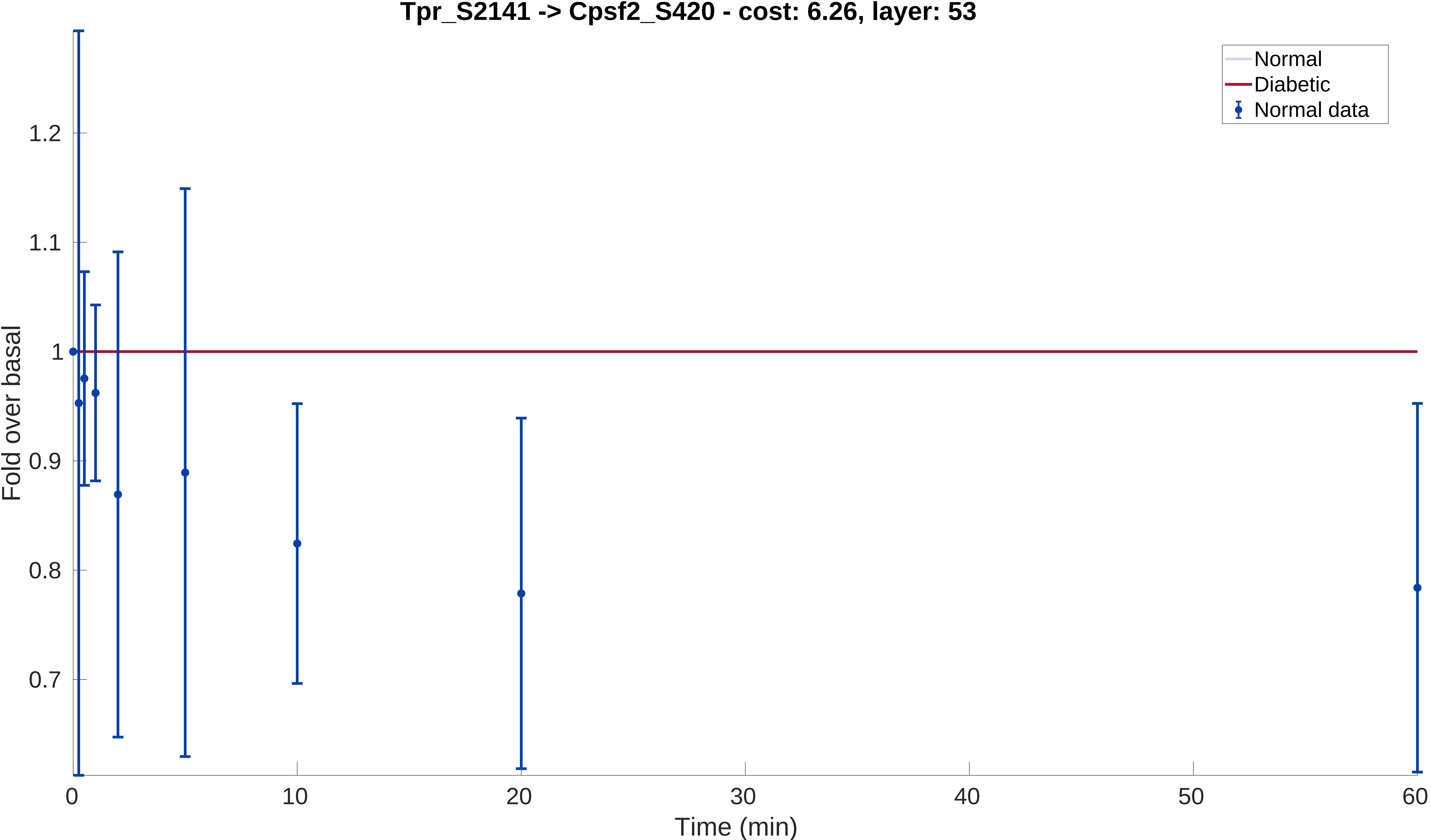
Tpr_S2141 -> Cpsf2_S420 - cost: 6.26, layer: 53
Normal
Diabetic
Normal data
Fold over basal
1.2
1.1
1
0.9
0.8
0.7
0
10
20
30
40
50
60
Time (min)

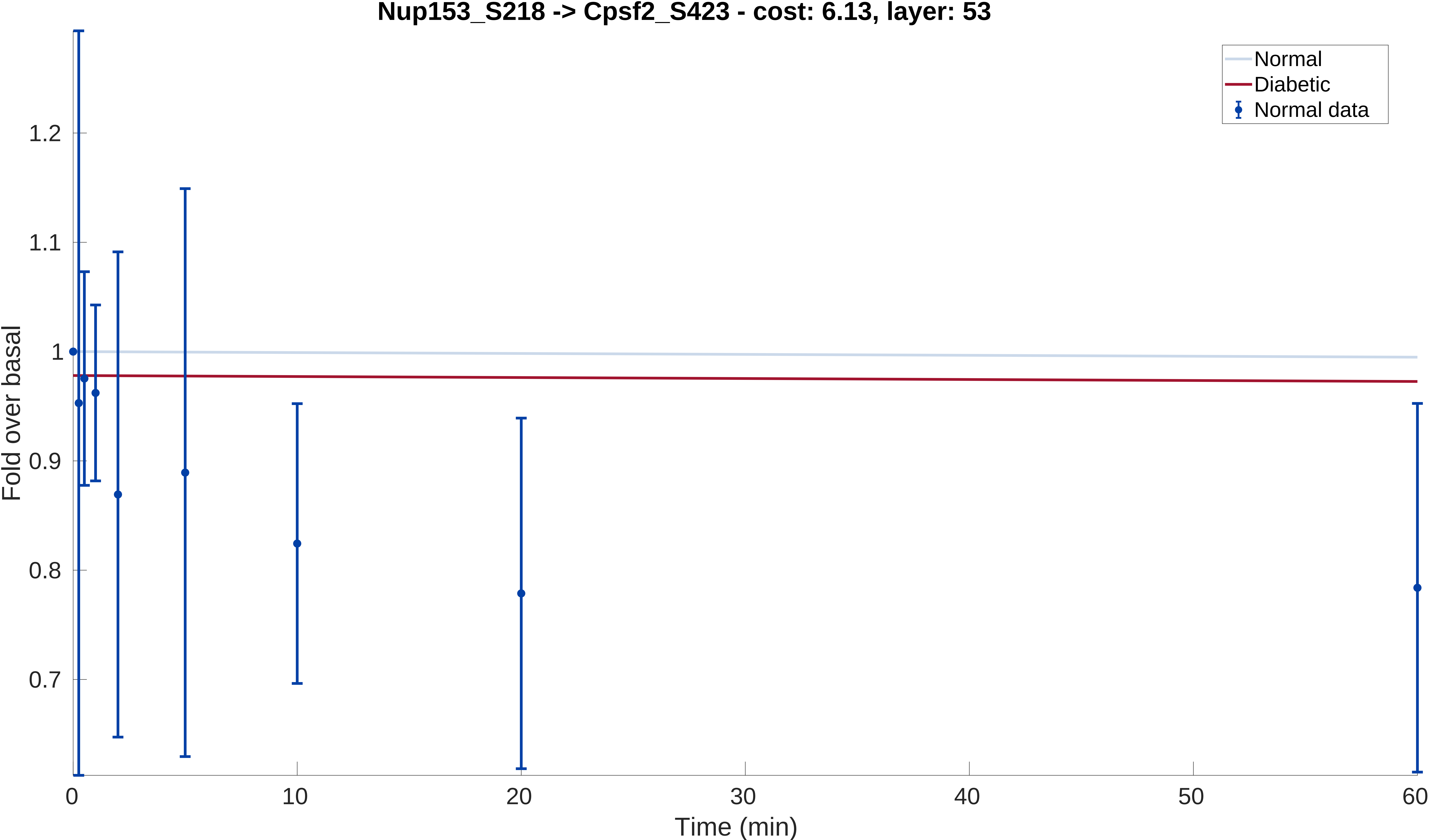

Nup153_S218 -> Cpsf2_S423 - cost: 6.13, layer: 53
Normal
Diabetic
Normal data
Fold over basal
1.2
1.1
1
0.9
0.8
0.7
0
10
20
30
40
50
60
Time (min)

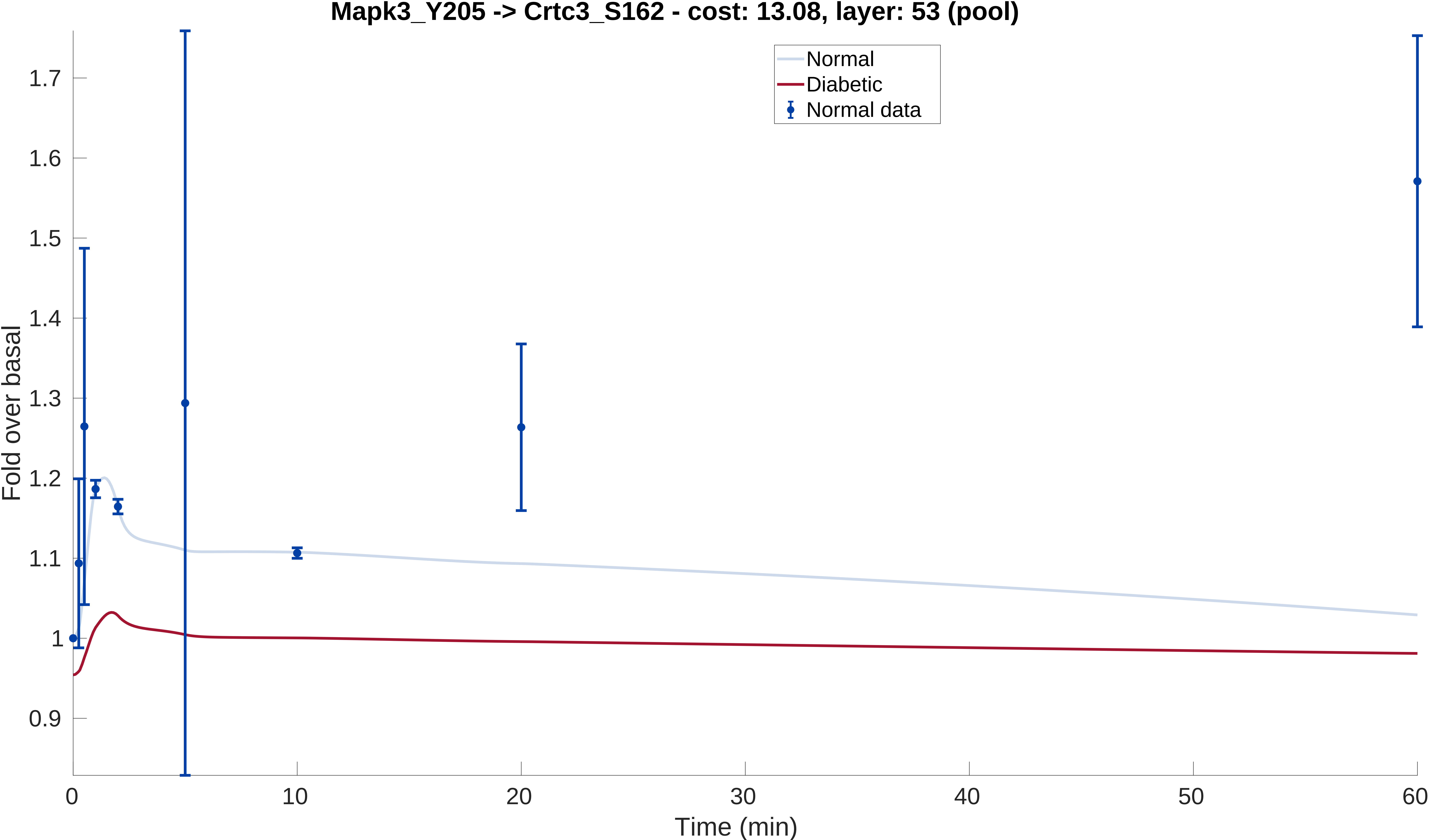

Mapk3_Y205 -> Crtc3_S162 - cost: 13.08, layer: 53 (pool)
Normal
Diabetic
Normal data
Fold over basal
1.7
1.6
1.5
1.4
1.3
1.2
1.1
1
0.9
0
10
20
30
40
50
60
Time (min)

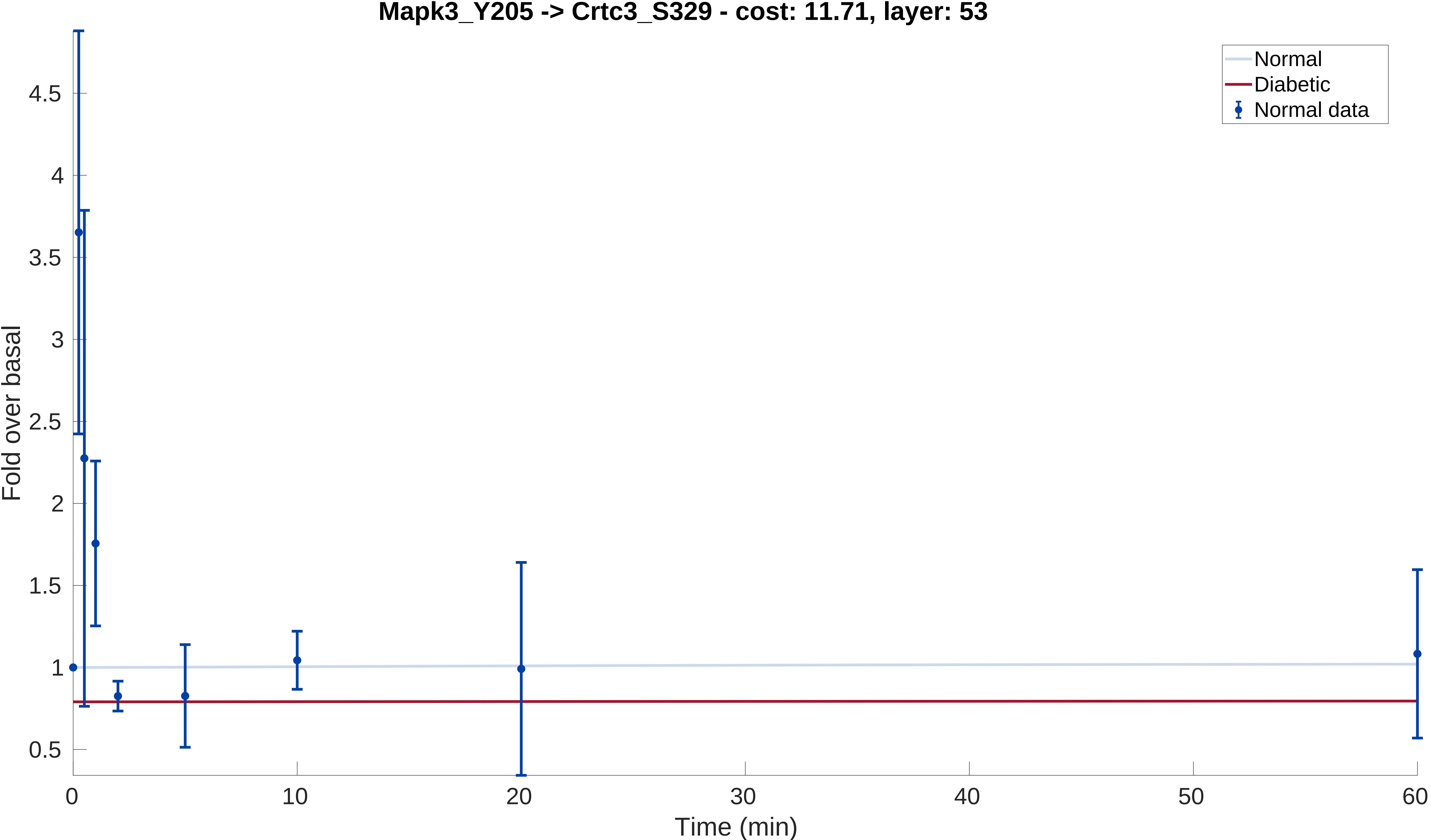

Mapk3_Y205 -> Crtc3_S329 - cost: 11.71, layer: 53
Normal
Diabetic
Normal data
Fold over basal
4.5
4
3.5
3
2.5
2
1.5
1
0.5
0
10
20
30
40
50
60
Time (min)

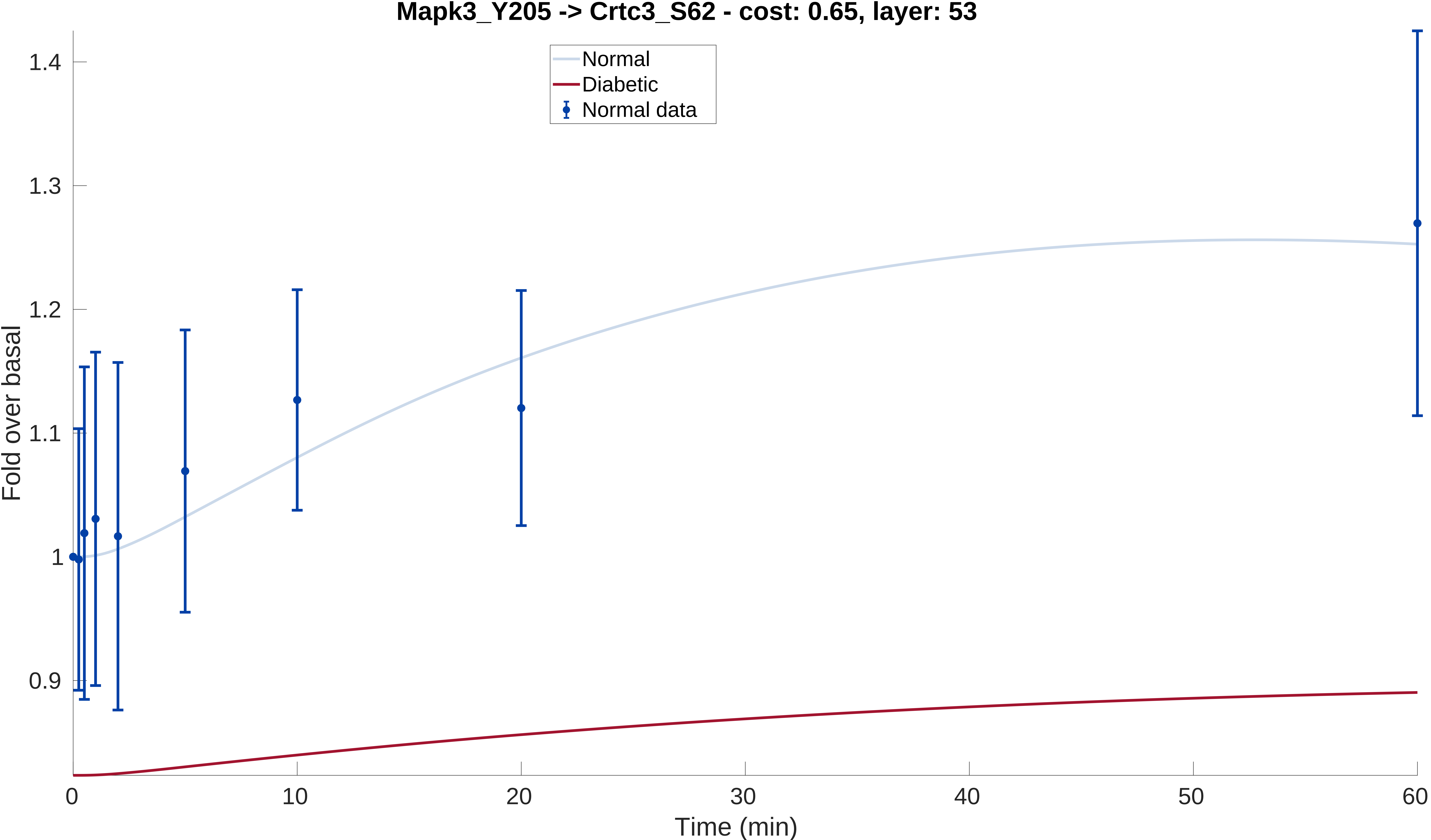

Mapk3_Y205 -> Crtc3_S62 - cost: 0.65, layer: 53
Normal
Diabetic
Normal data
Fold over basal
1.4
1.3
1.2
1.1
1
0.9
0
10
20
30
40
50
60
Time (min)

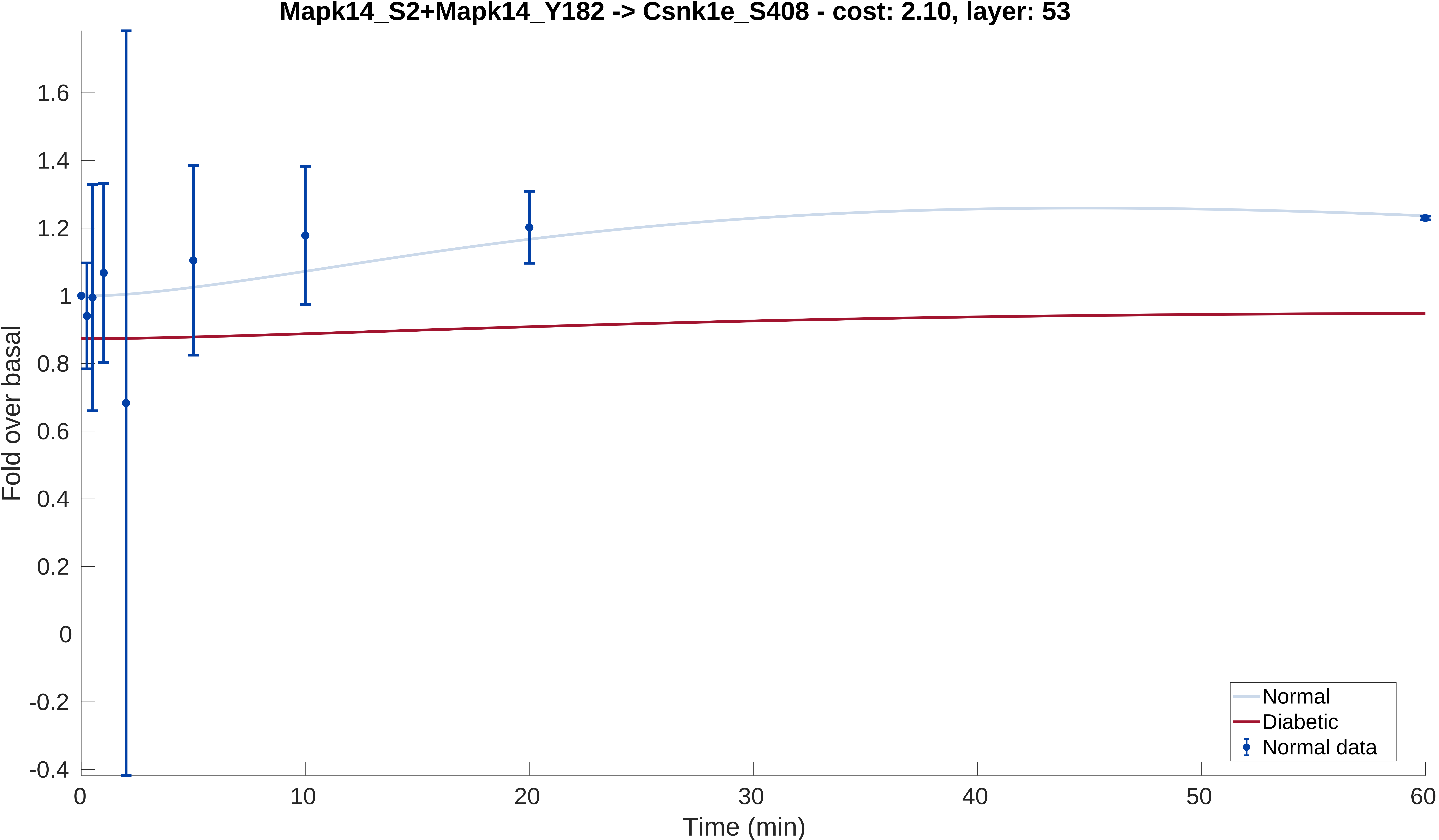

Mapk14_S2+Mapk14_Y182 -> Csnk1e_S408 - cost: 2.10, layer: 53
Fold over basal
1.6
1.4
1.2
1
0.8
0.6
0.4
0.2
0
-0.2
-0.4
0
10
20
30
40
50
60
Time (min)
Normal
Diabetic
Normal data

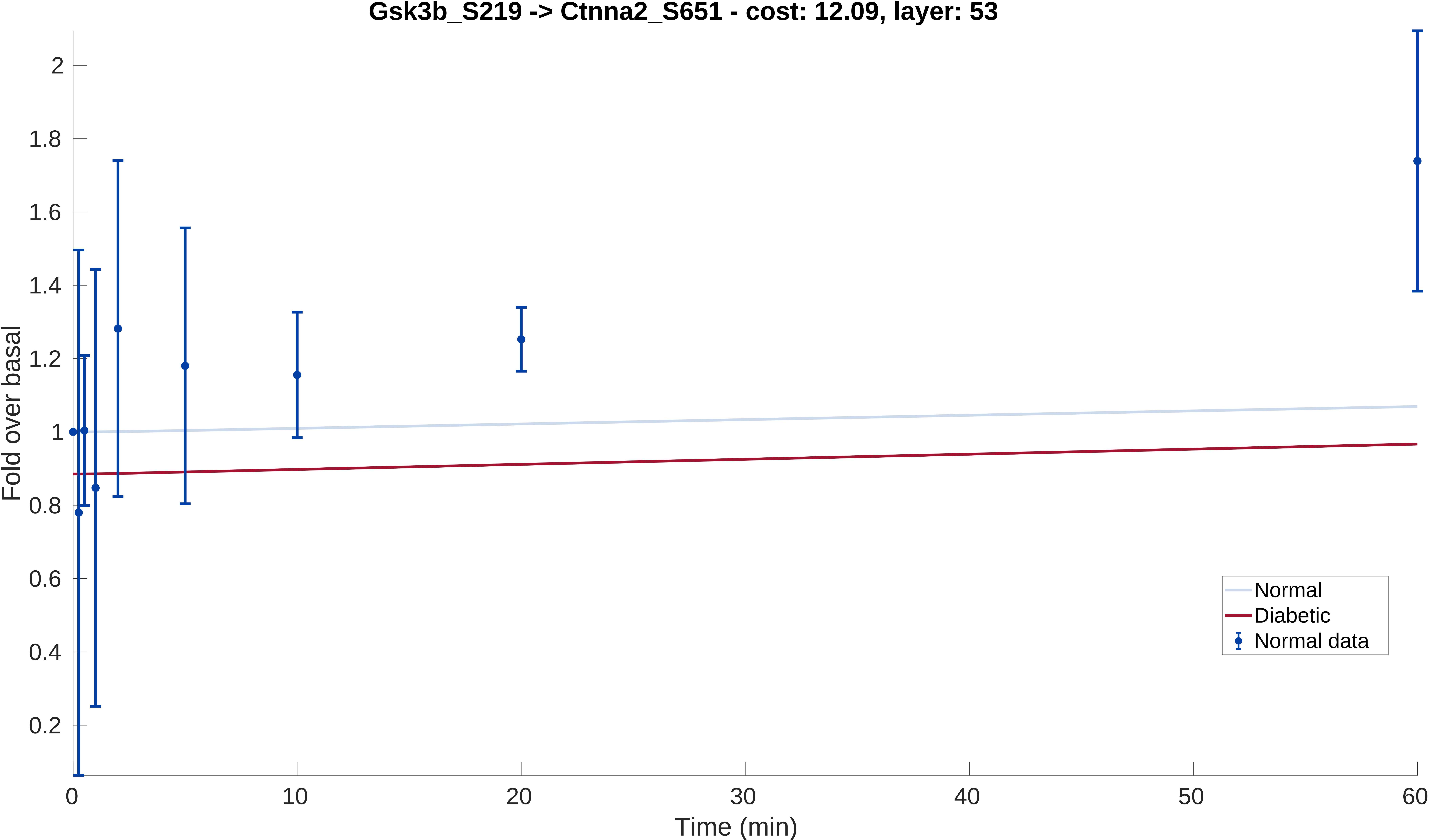

Gsk3b_S219 -> Ctnna2_S651 - cost: 12.09, layer: 53
Fold over basal
Time (min)
2
1.8
1.6
1.4
1.2
1
0.8
0.6
0.4
0.2
0
10
20
30
40
50
60
Normal
Diabetic
Normal data

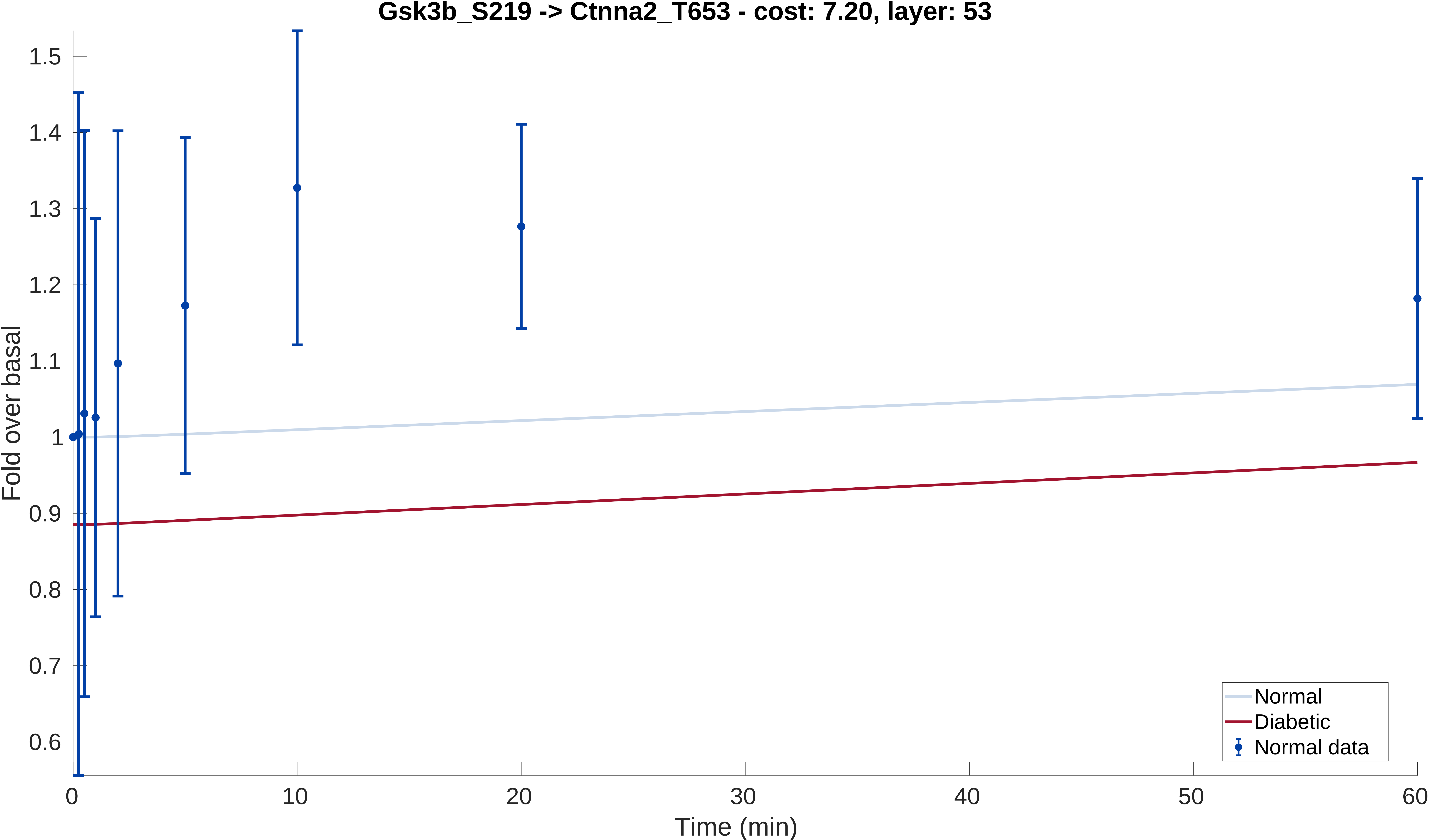

Gsk3b_S219 -> Ctnna2_T653 - cost: 7.20, layer: 53
Fold over basal
Time (min)
1.5
1.4
1.3
1.2
1.1
1
0.9
0.8
0.7
0.6
0
10
20
30
40
50
60
Normal
Diabetic
Normal data

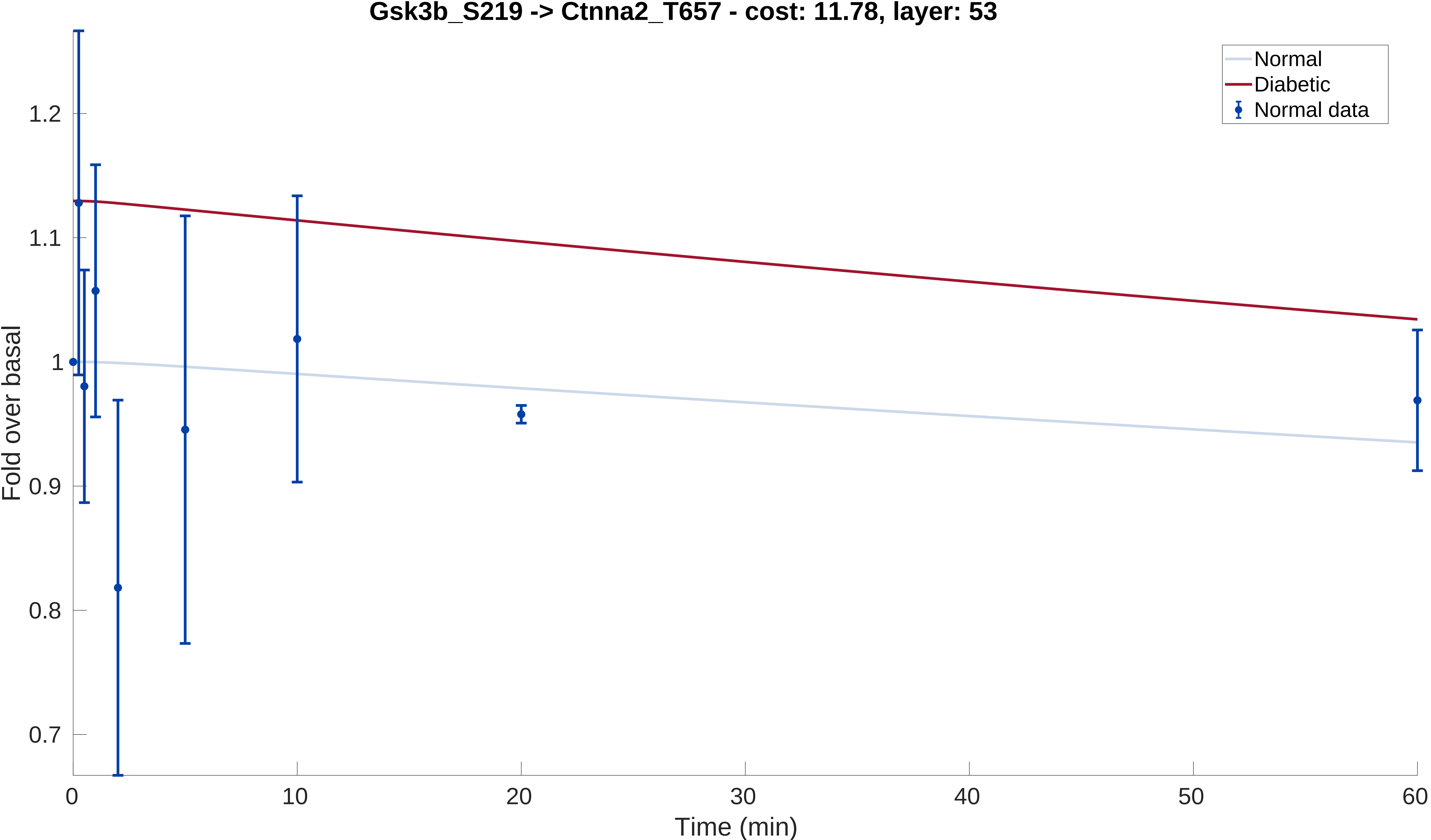

Gsk3b_S219 -> Ctnna2_T657 - cost: 11.78, layer: 53
Normal
Diabetic
Normal data
1.2
1.1
1
0.9
0.8
0.7
Fold over basal
0
10
20
30
40
50
60
Time (min)

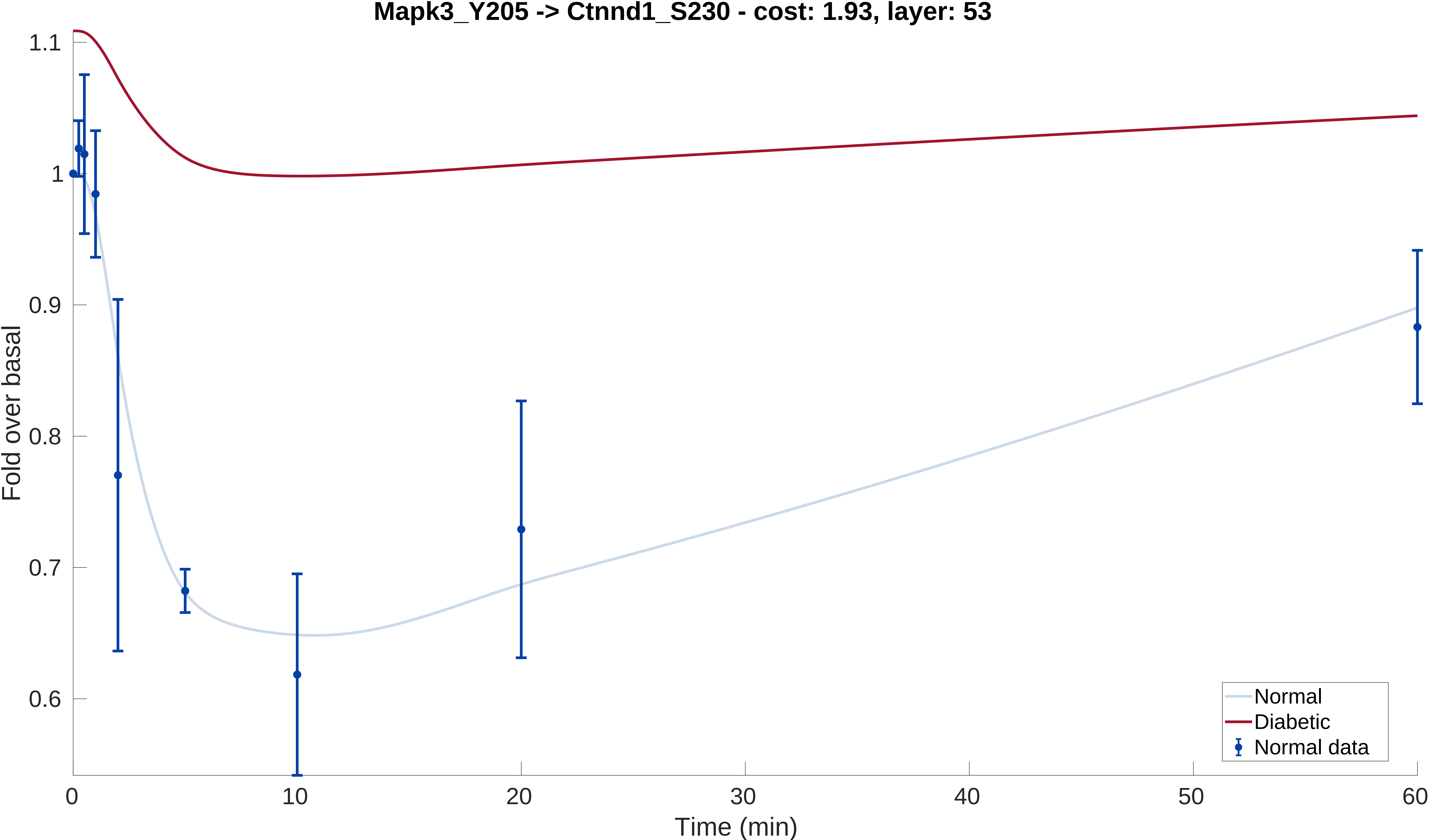

Mapk3_Y205 -> Ctnnd1_S230 - cost: 1.93, layer: 53
1.1
1
0.9
0.8
0.7
0.6
Fold over basal
0
10
20
30
40
50
60
Time (min)
Normal
Diabetic
Normal data

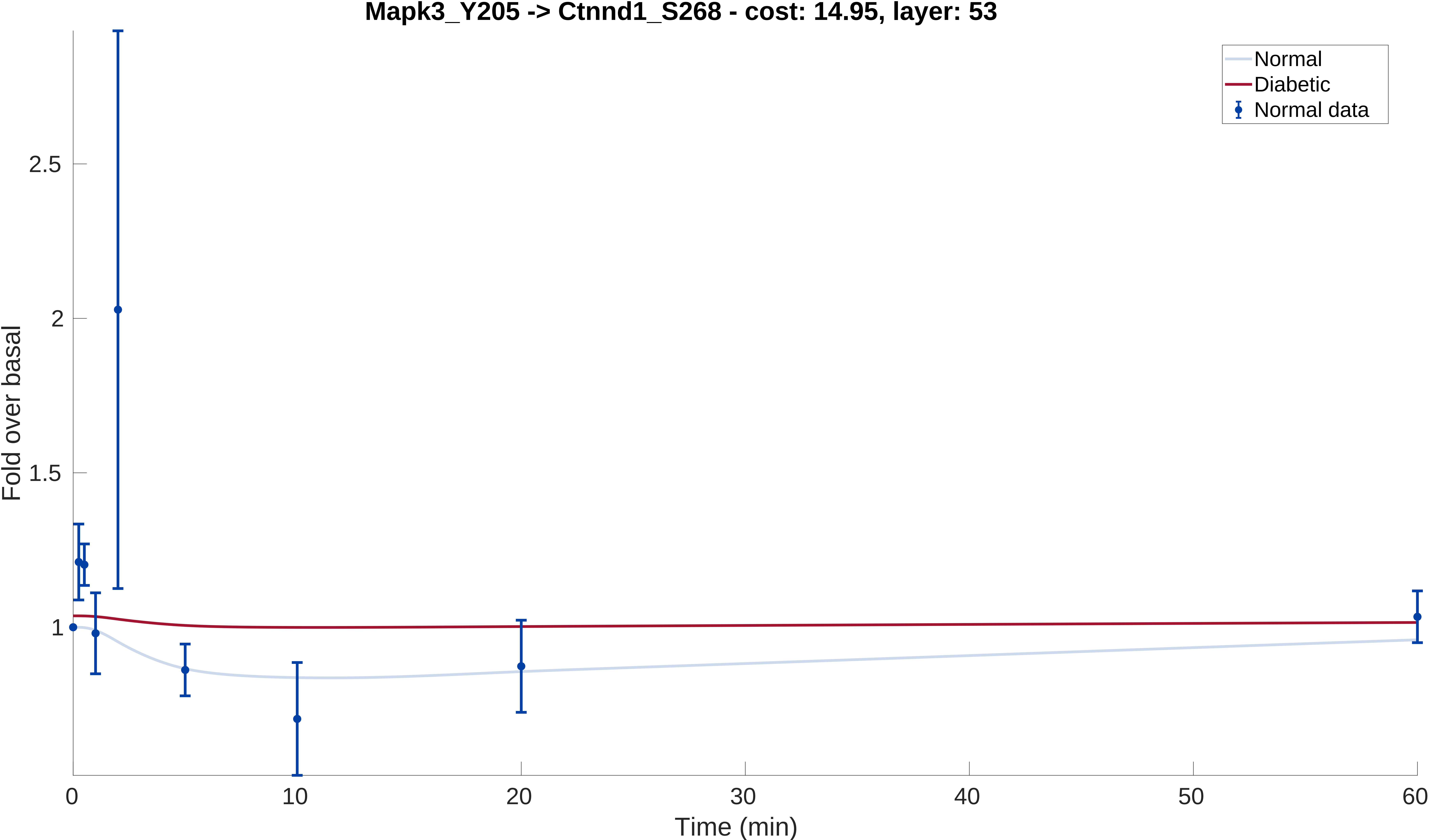
Mapk3_Y205 -> Ctnnd1_S268 - cost: 14.95, layer: 53
Normal
Diabetic
Normal data
Fold over basal
2.5
2
1.5
1
0
10
20
30
40
50
60
Time (min)

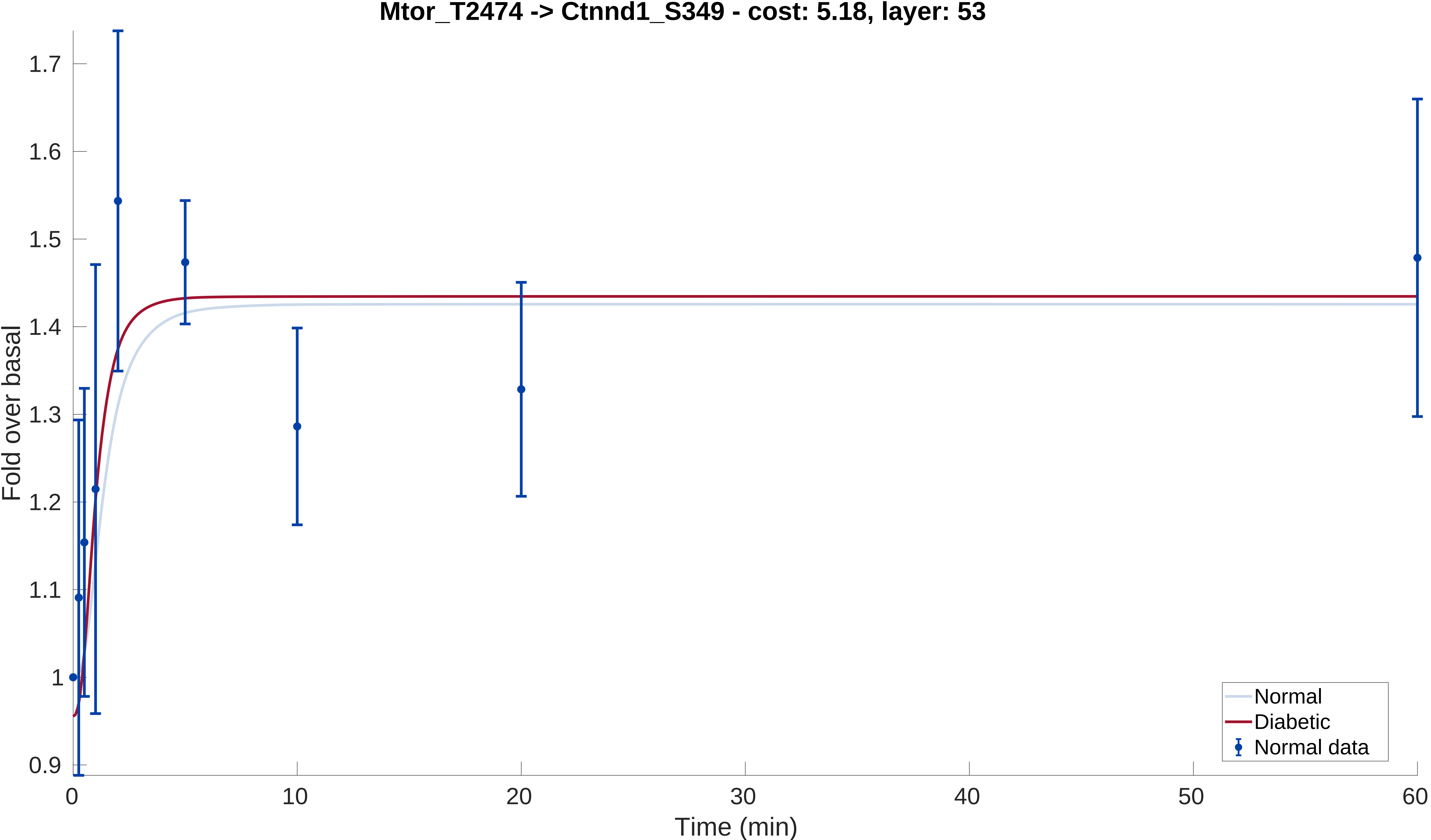

Mtor_T2474 -> Ctnnd1_S349 - cost: 5.18, layer: 53
Fold over basal
1.7
1.6
1.5
1.4
1.3
1.2
1.1
1
0.9
0
10
20
30
40
50
60
Time (min)
Normal
Diabetic
Normal data

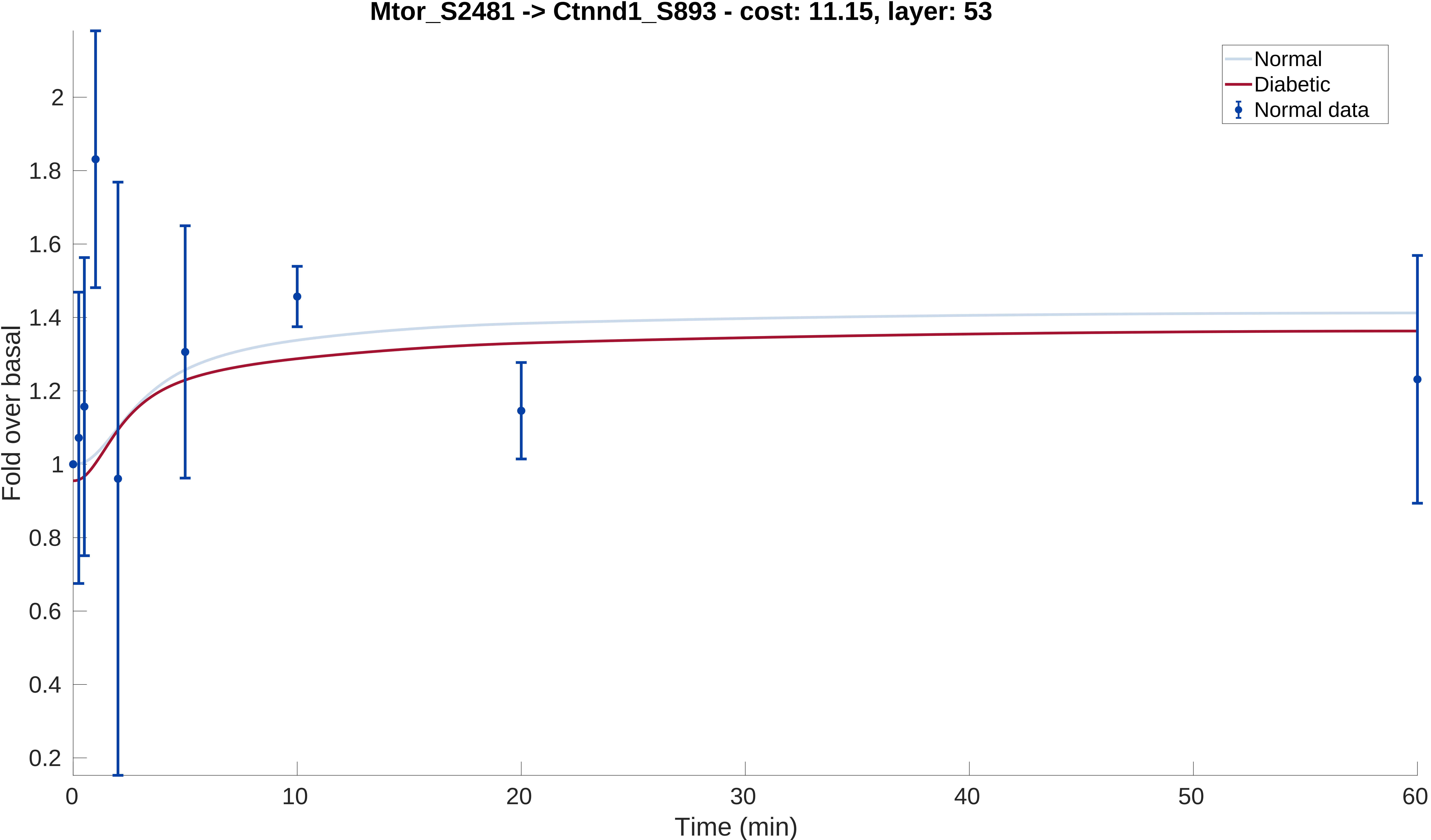
Mtor_S2481 -> Ctnnd1_S893 - cost: 11.15, layer: 53
Normal
Diabetic
Normal data
Fold over basal
2
1.8
1.6
1.4
1.2
1
0.8
0.6
0.4
0.2
0
10
20
30
40
50
60
Time (min)

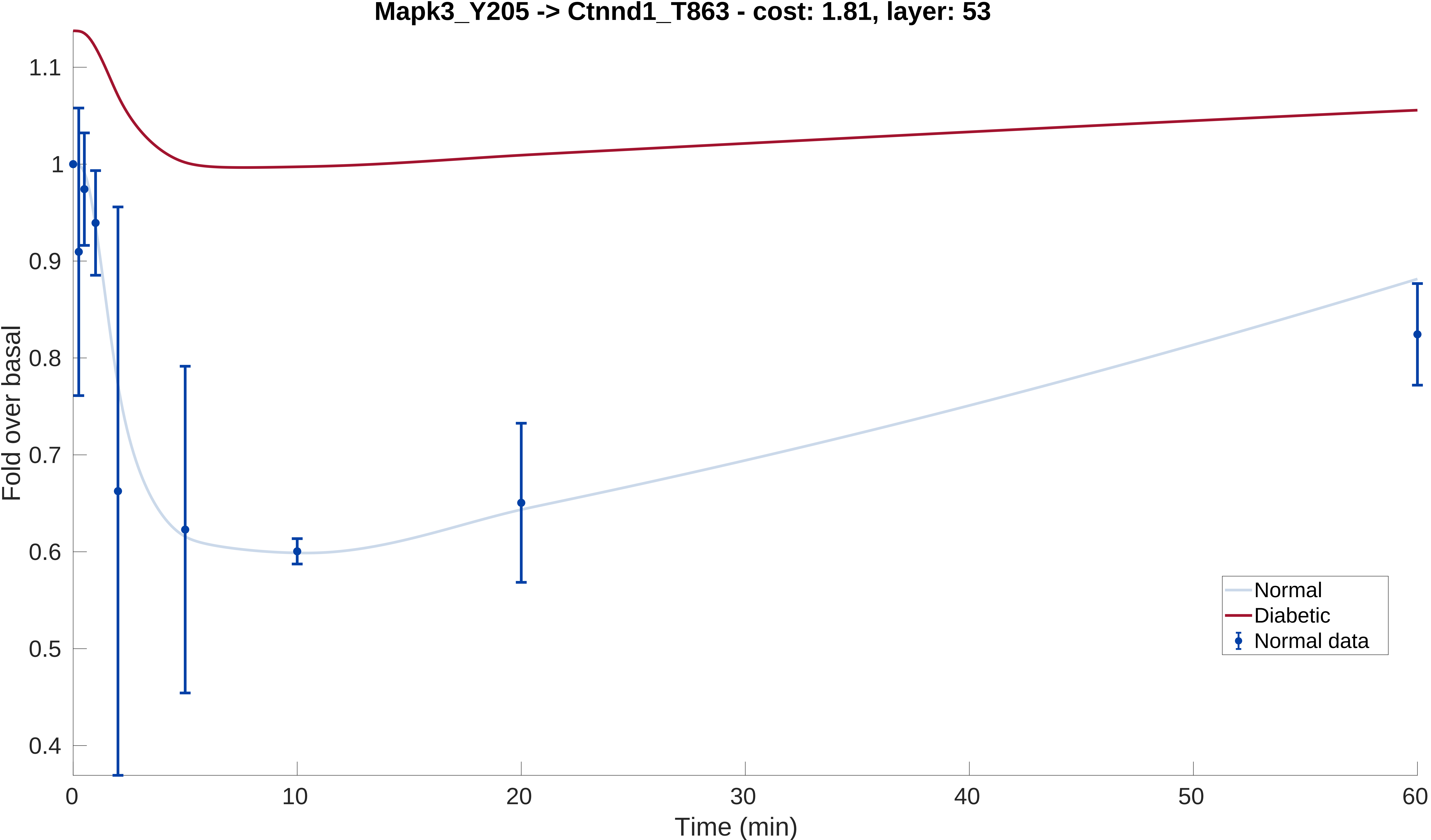

Mapk3_Y205 -> Ctnnd1_T863 - cost: 1.81, layer: 53
Fold over basal
1.1
1
0.9
0.8
0.7
0.6
0.5
0.4
0
10
20
30
40
50
60
Time (min)
Normal
Diabetic
Normal data

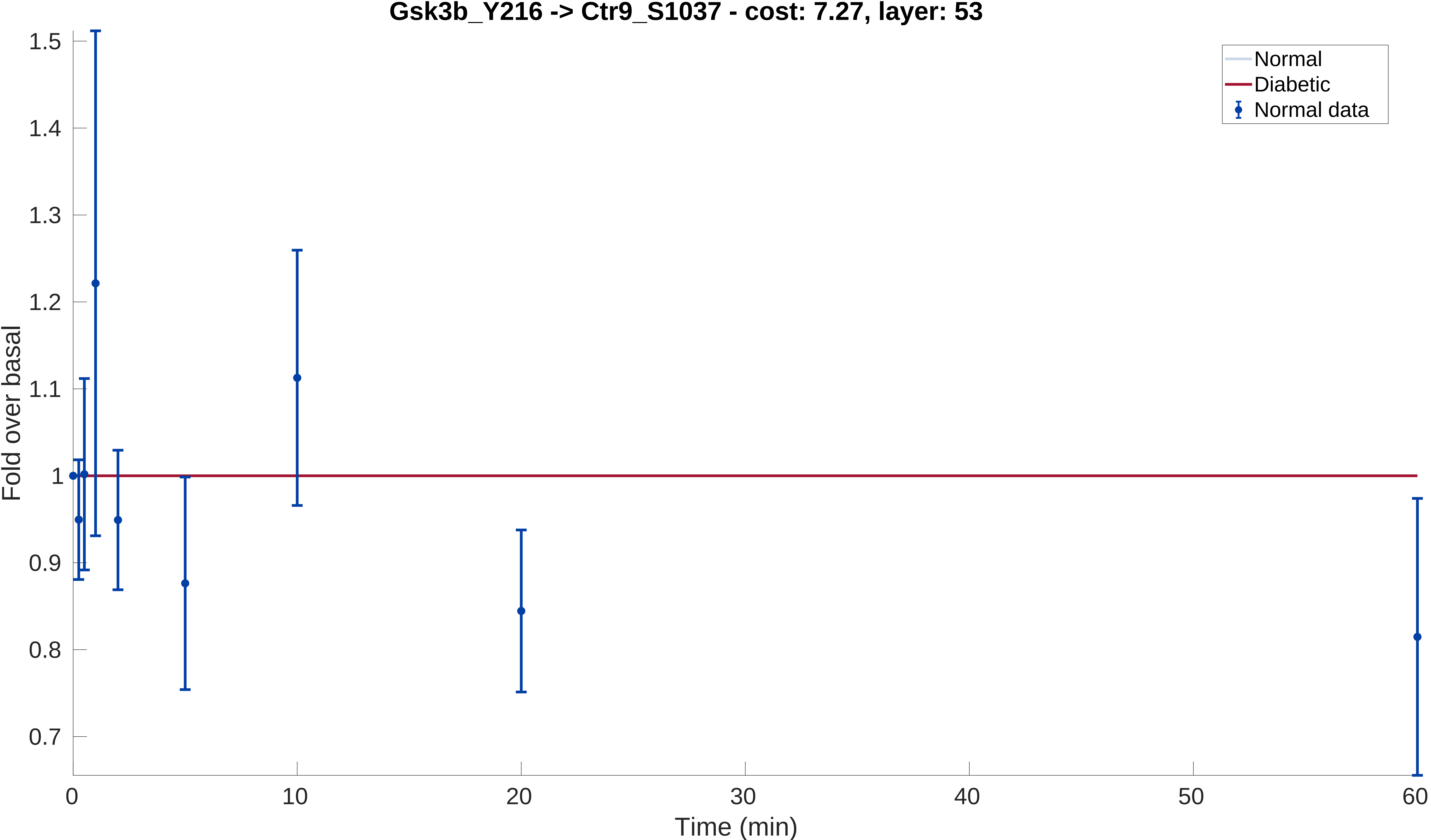

Gsk3b_Y216 -> Ctr9_S1037 - cost: 7.27, layer: 53
Normal
Diabetic
Normal data
Fold over basal
Time (min)
1.5
1.4
1.3
1.2
1.1
1
0.9
0.8
0.7
0
10
20
30
40
50
60

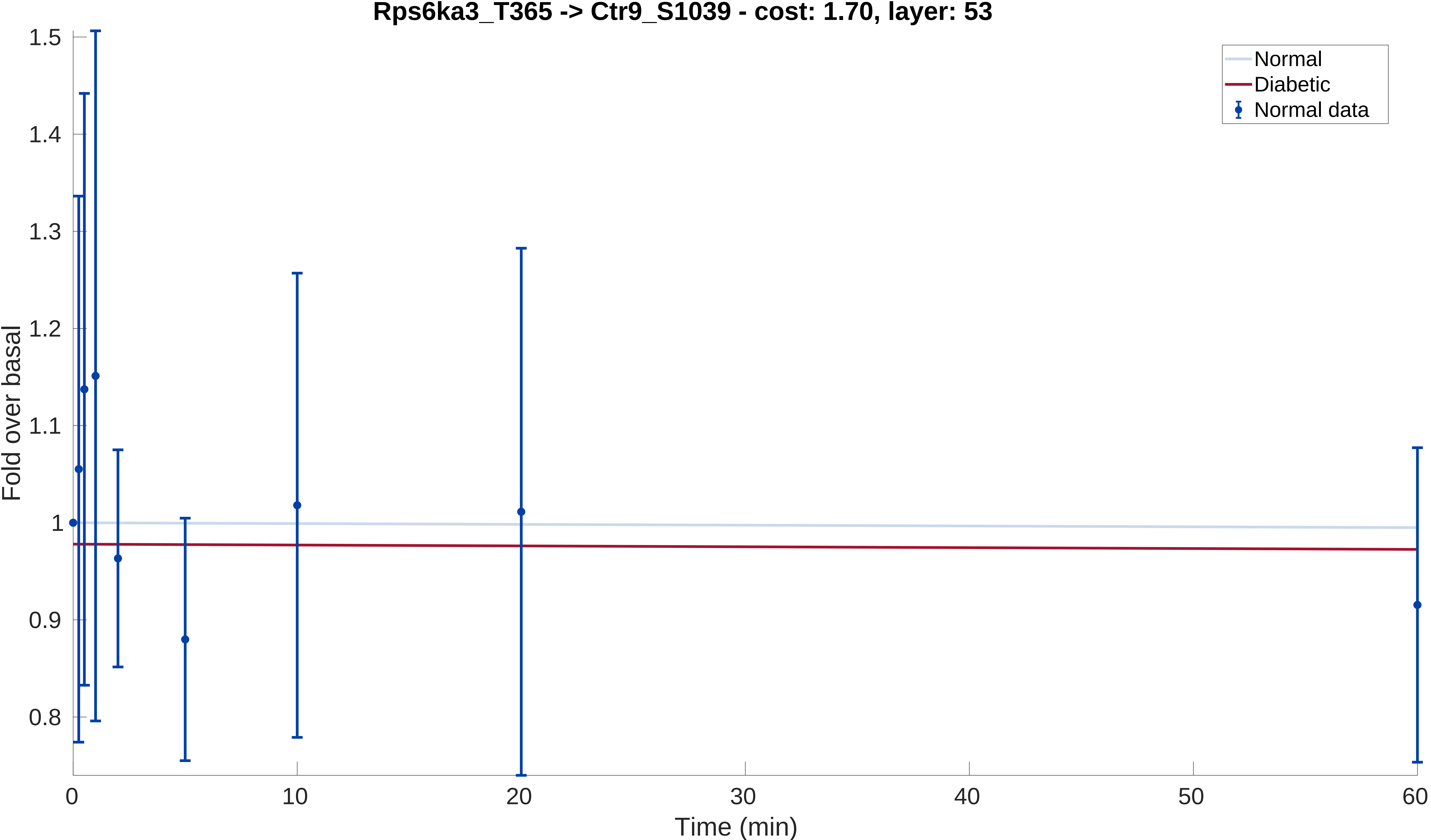

Rps6ka3_T365 -> Ctr9_S1039 - cost: 1.70, layer: 53
Normal
Diabetic
Normal data
Fold over basal
Time (min)
1.5
1.4
1.3
1.2
1.1
1
0.9
0.8
0
10
20
30
40
50
60

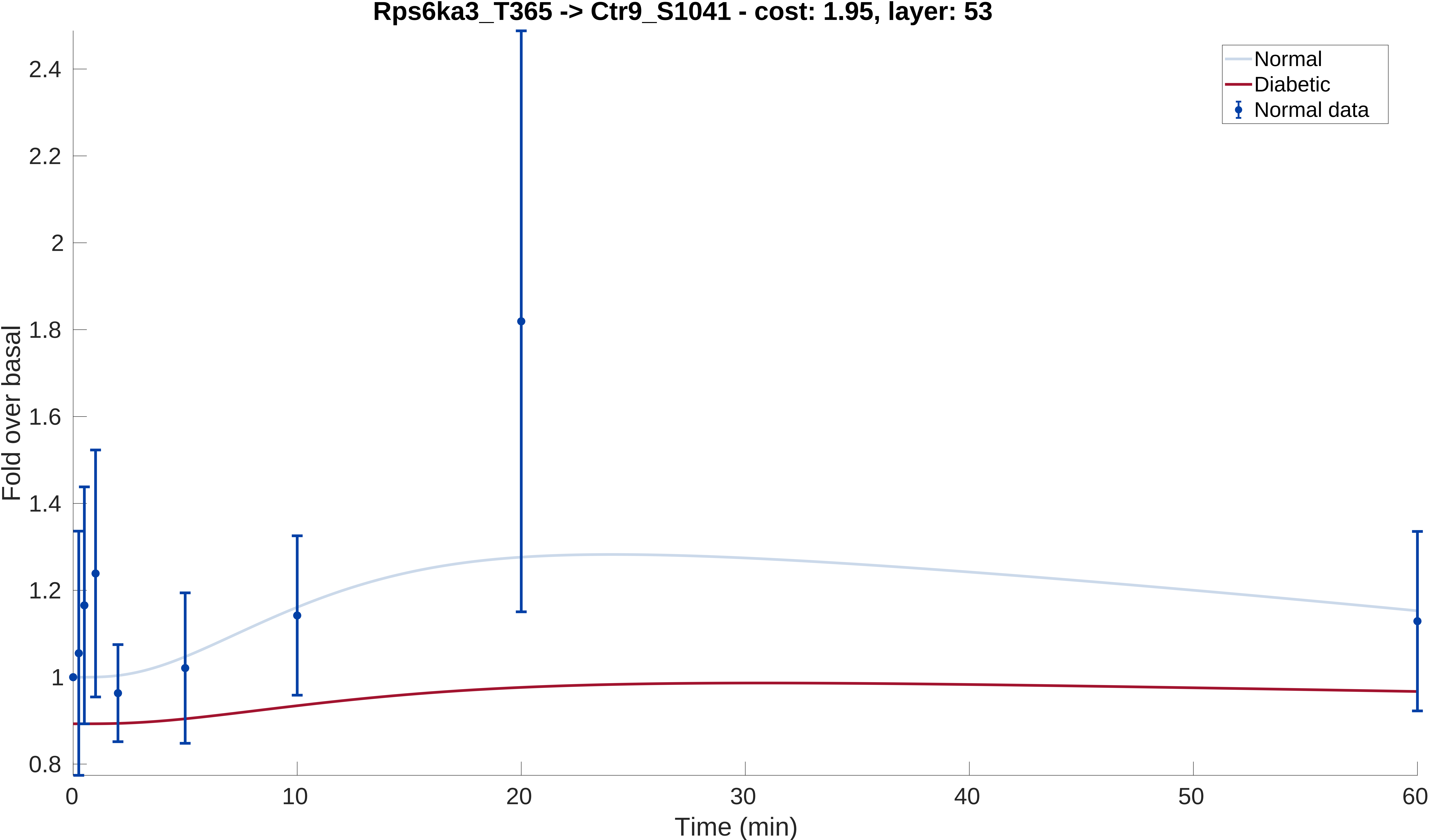

Rps6ka3_T365 -> Ctr9_S1041 - cost: 1.95, layer: 53
Normal
Diabetic
Normal data
Fold over basal
2.4
2.2
2
1.8
1.6
1.4
1.2
1
0.8
0
10
20
30
40
50
60
Time (min)

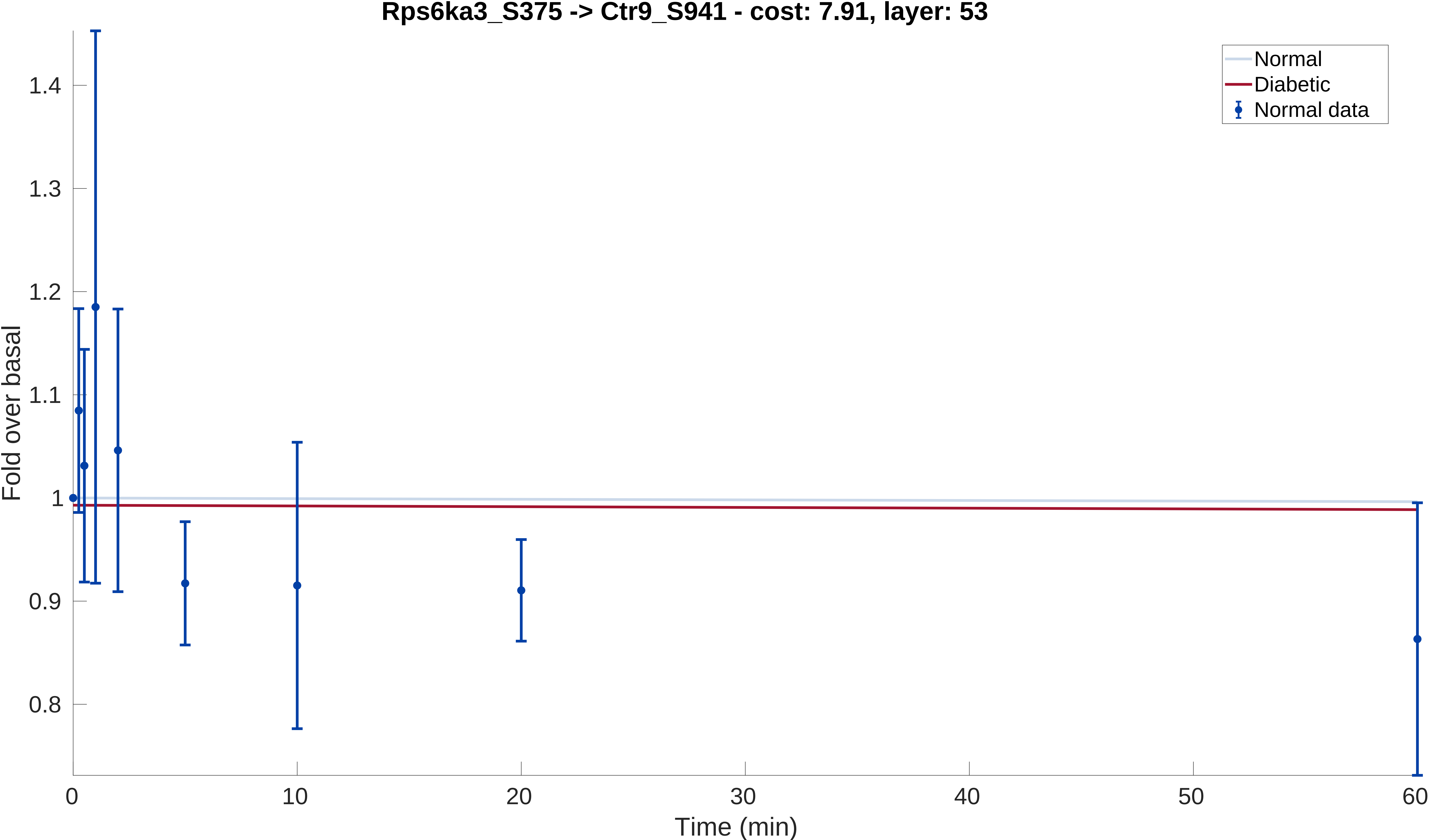
Rps6ka3_S375 -> Ctr9_S941 - cost: 7.91, layer: 53
Normal
Diabetic
Normal data
Fold over basal
1.4
1.3
1.2
1.1
1
0.9
0.8
0
10
20
30
40
50
60
Time (min)

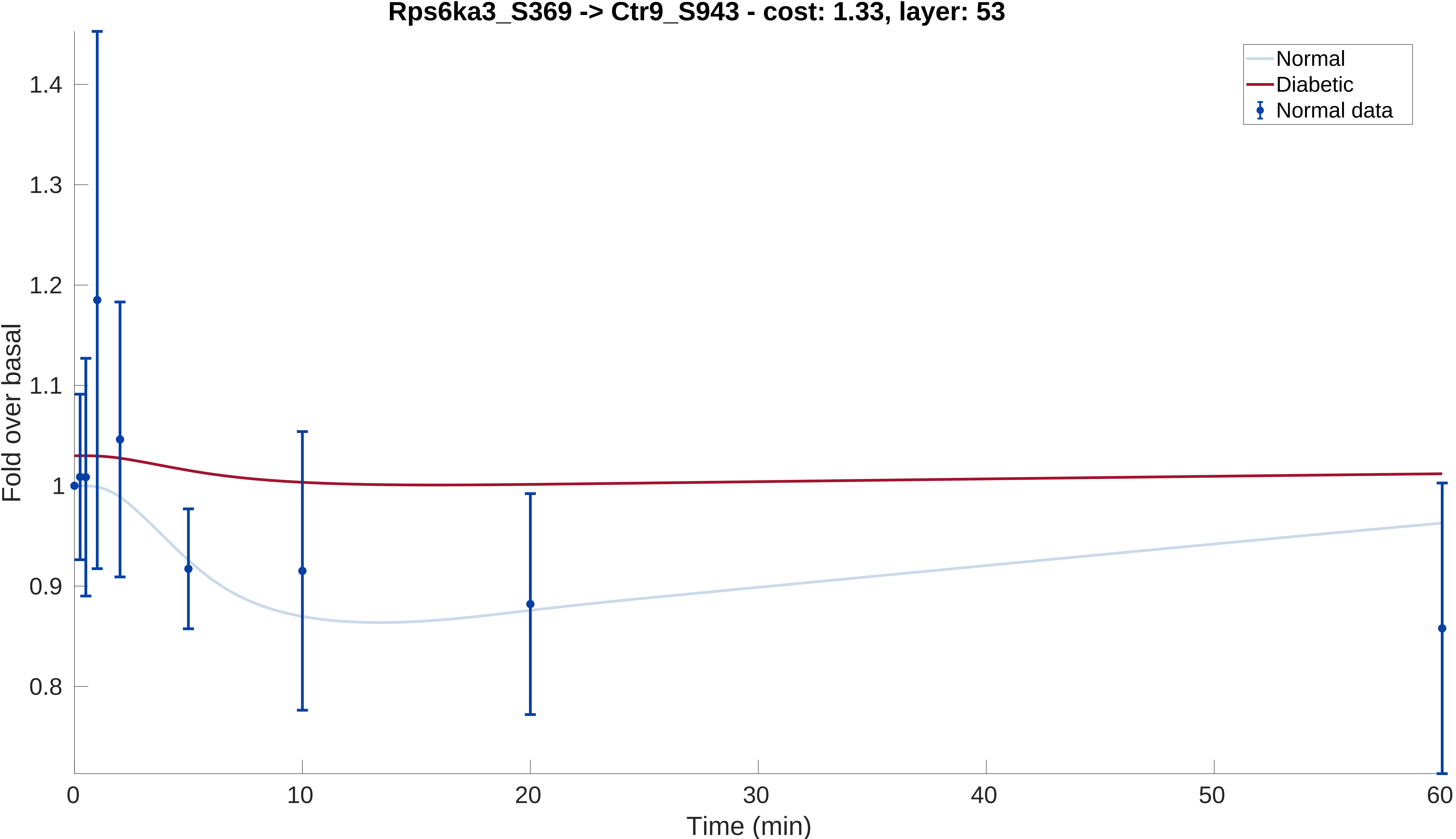

Rps6ka3_S369 -> Ctr9_S943 - cost: 1.33, layer: 53
Normal
Diabetic
Normal data
Fold over basal
1.4
1.3
1.2
1.1
1
0.9
0.8
0
10
20
30
40
50
60
Time (min)

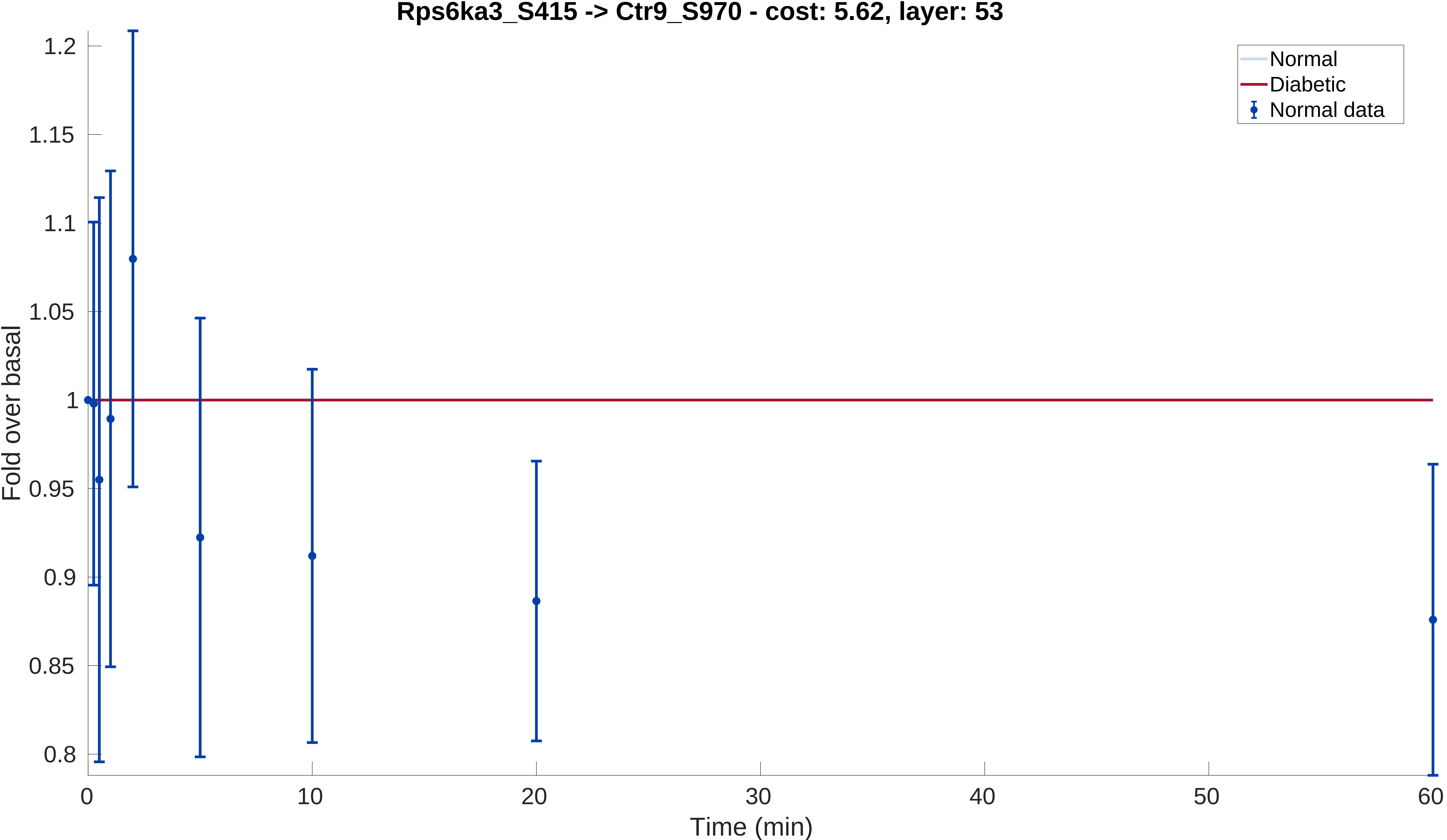
Rps6ka3_S415 -> Ctr9_S970 - cost: 5.62, layer: 53
Normal
Diabetic
Normal data
Fold over basal
1.2
1.15
1.1
1.05
1
0.95
0.9
0.85
0.8
0
10
20
30
40
50
60
Time (min)

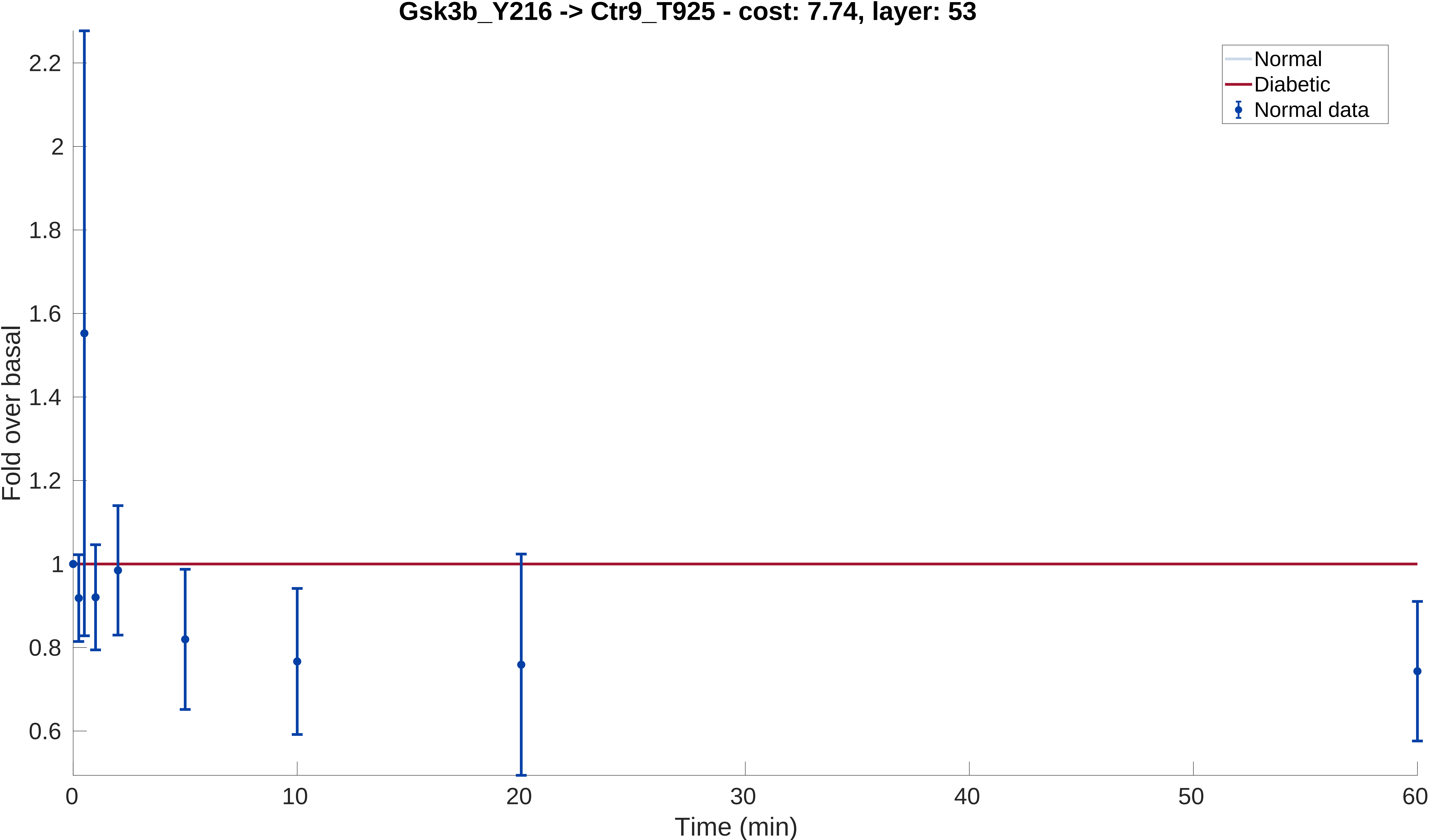

Gsk3b_Y216 -> Ctr9_T925 - cost: 7.74, layer: 53
Fold over basal
2.2
2
1.8
1.6
1.4
1.2
1
0.8
0.6
0
10
20
30
40
50
60
Time (min)
Normal
Diabetic
Normal data

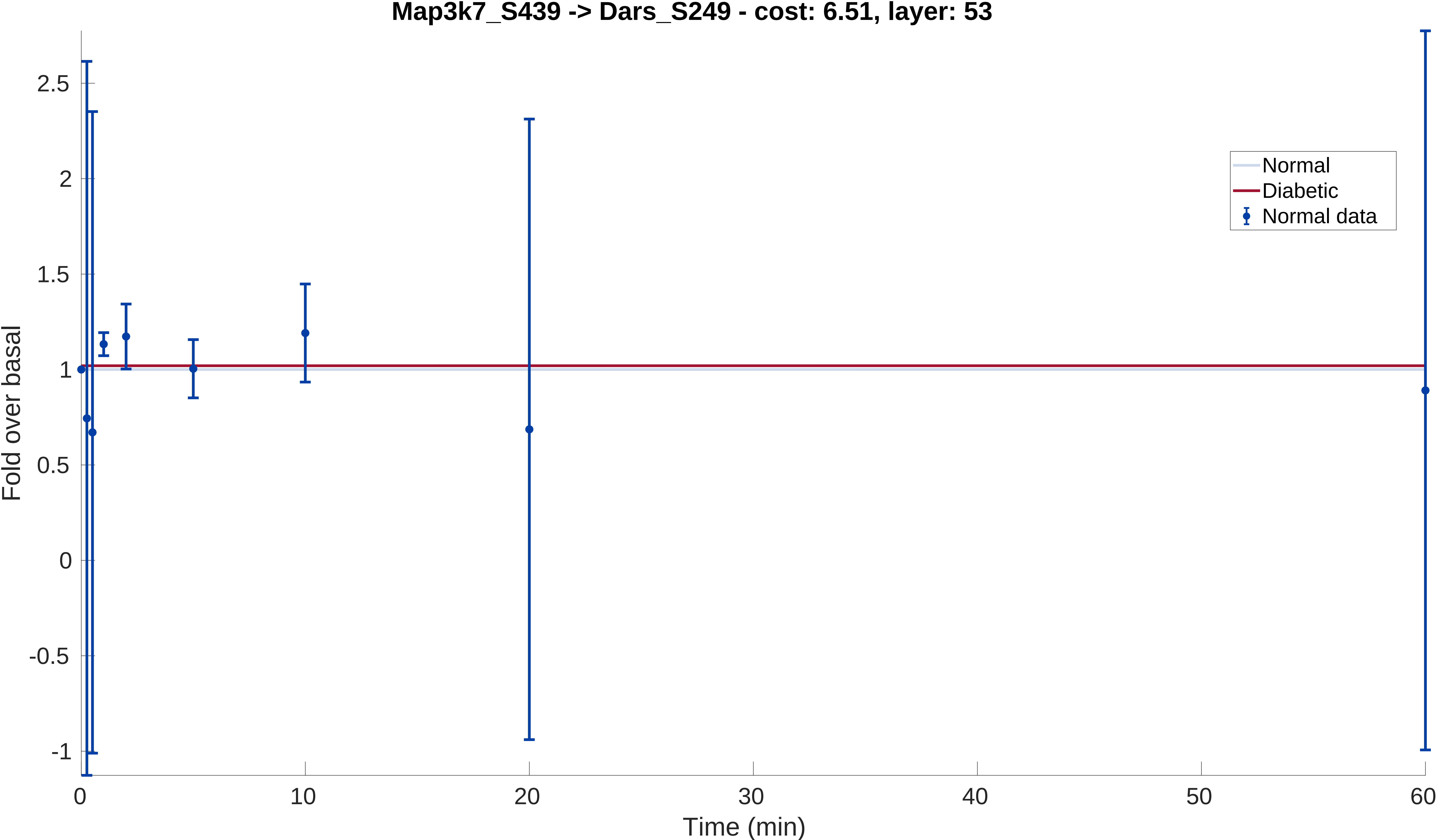
Map3k7_S439 -> Dars_S249 - cost: 6.51, layer: 53
Fold over basal
Time (min)
Normal
Diabetic
Normal data
2.5
2
1.5
1
0.5
0
-0.5
-1
0
10
20
30
40
50
60

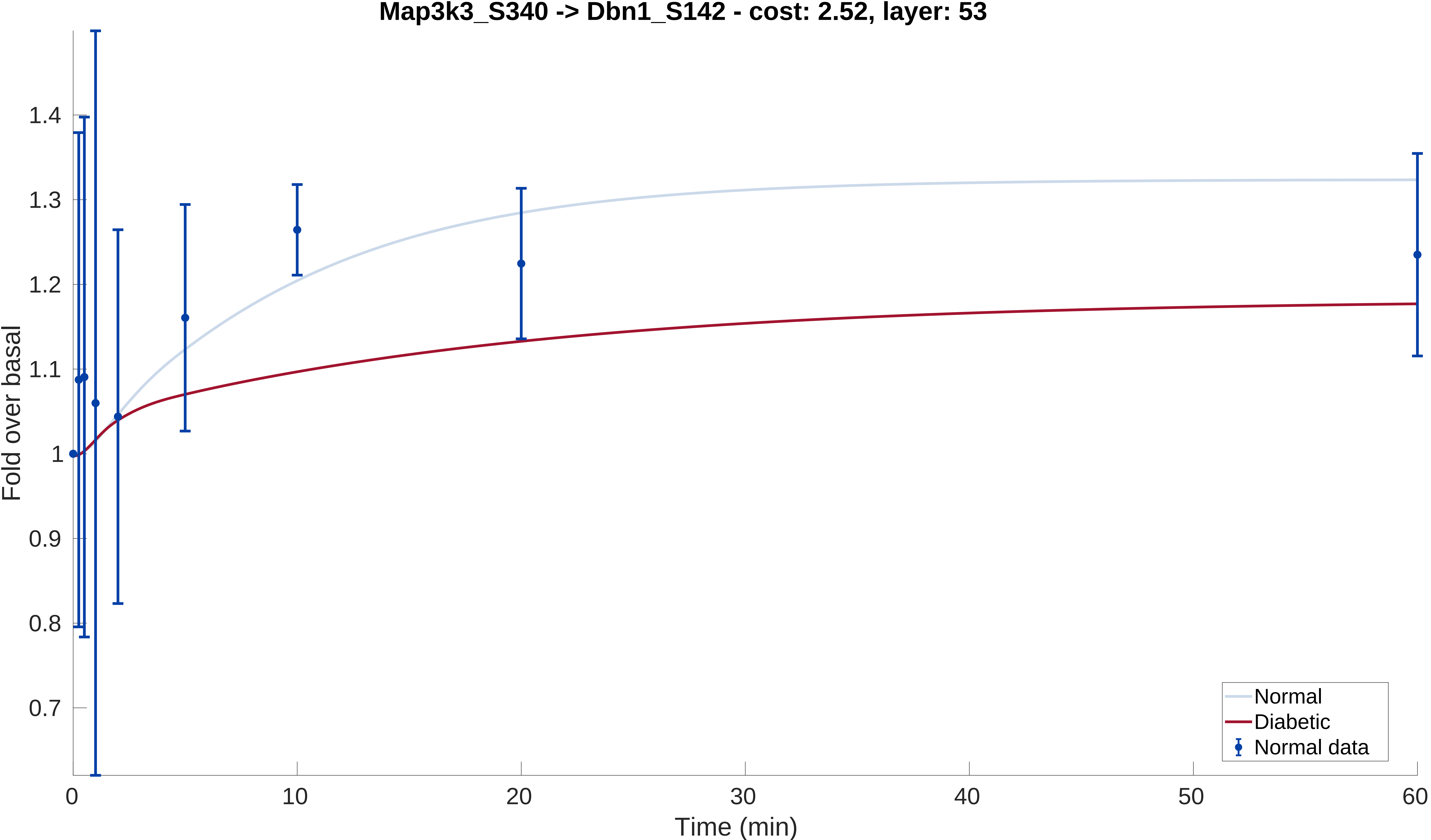

Map3k3_S340 -> Dbn1_S142 - cost: 2.52, layer: 53
Fold over basal
1.4
1.3
1.2
1.1
1
0.9
0.8
0.7
0
10
20
30
40
50
60
Time (min)
Normal
Diabetic
Normal data

Rps6ka3_S369 -> Dclk1_S307 - cost: 4.03, layer: 53

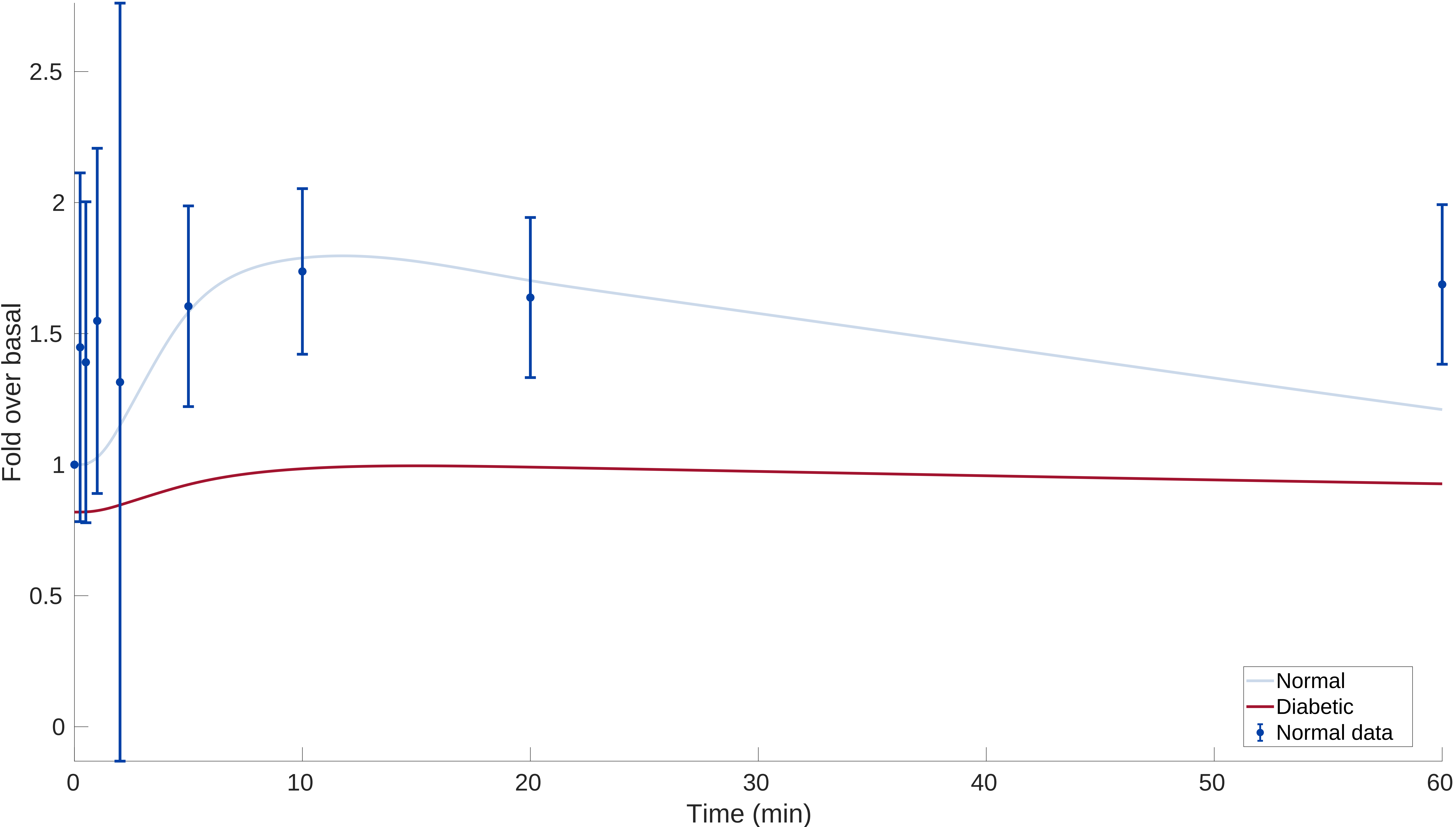

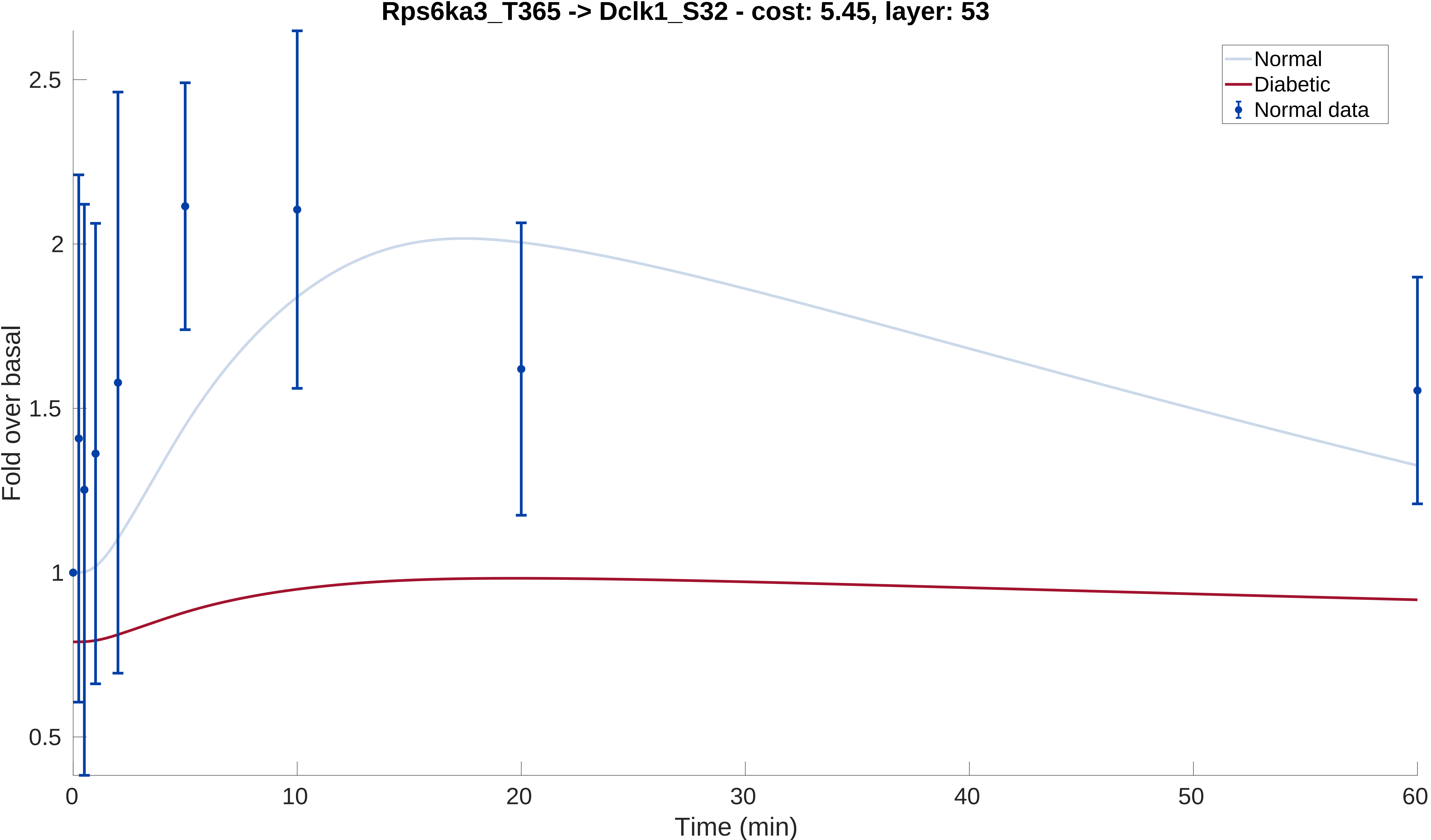

Rps6ka3_T365 -> Dclk1_S32 - cost: 5.45, layer: 53
Normal
Diabetic
Normal data
Fold over basal
2.5
2
1.5
1
0.5
0
10
20
30
40
50
60
Time (min)

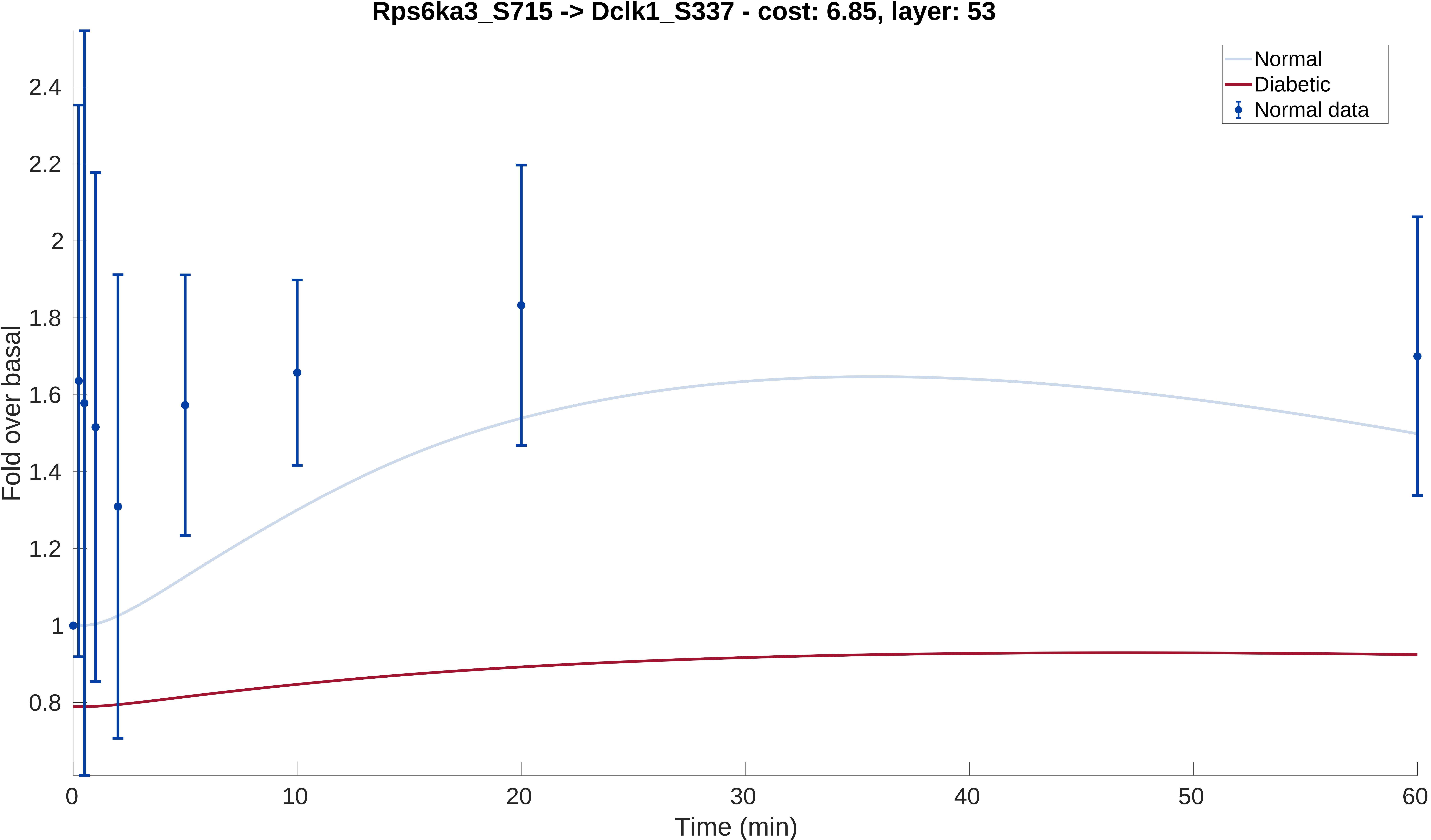

Rps6ka3_S715 -> Dclk1_S337 - cost: 6.85, layer: 53
Normal
Diabetic
Normal data
Fold over basal
2.4
2.2
2
1.8
1.6
1.4
1.2
1
0.8
0
10
20
30
40
50
60
Time (min)

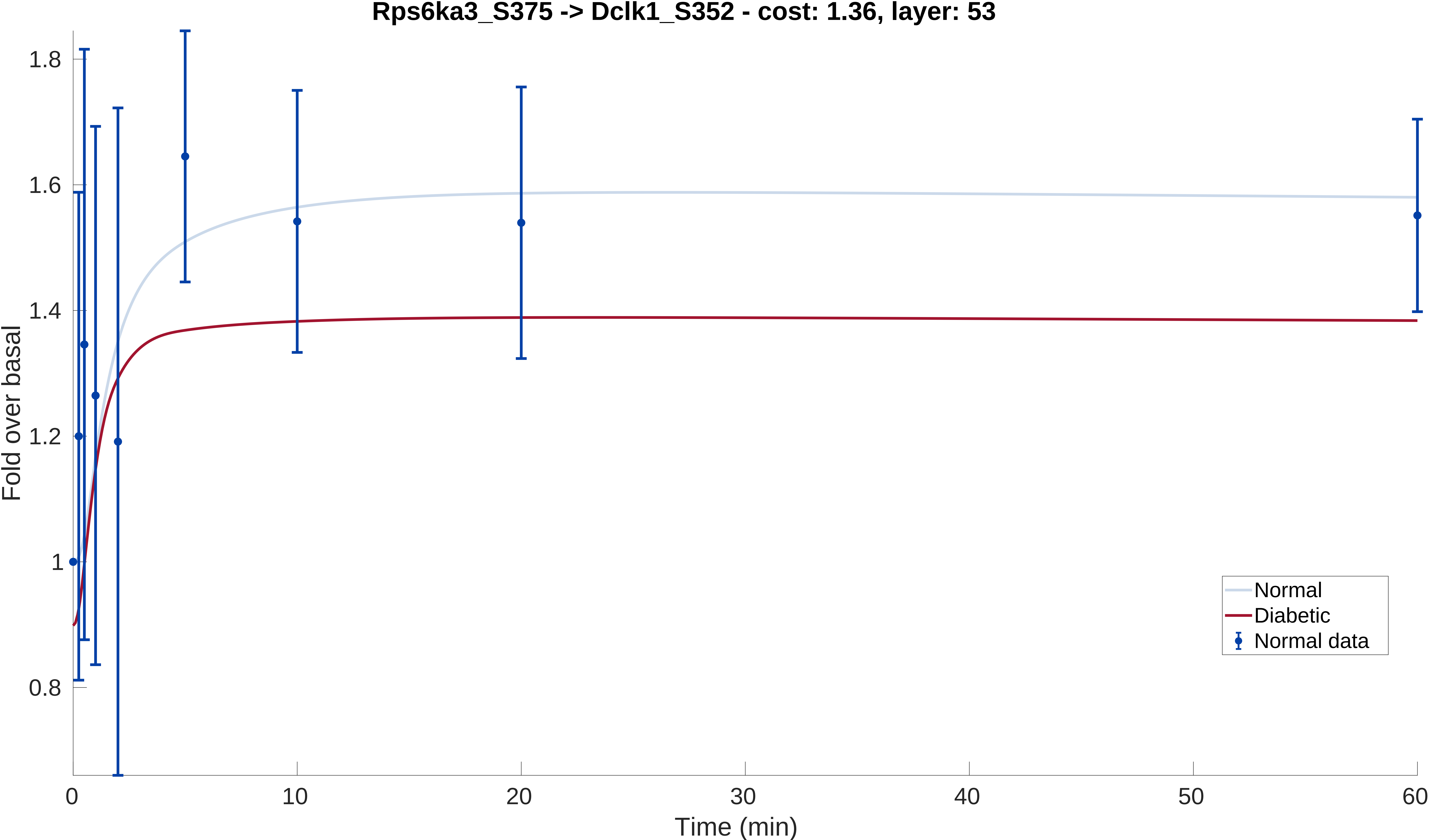

Rps6ka3_S375 -> Dclk1_S352 - cost: 1.36, layer: 53
Fold over basal
1.8
1.6
1.4
1.2
1
0.8
0
10
20
30
40
50
60
Time (min)
Normal
Diabetic
Normal data

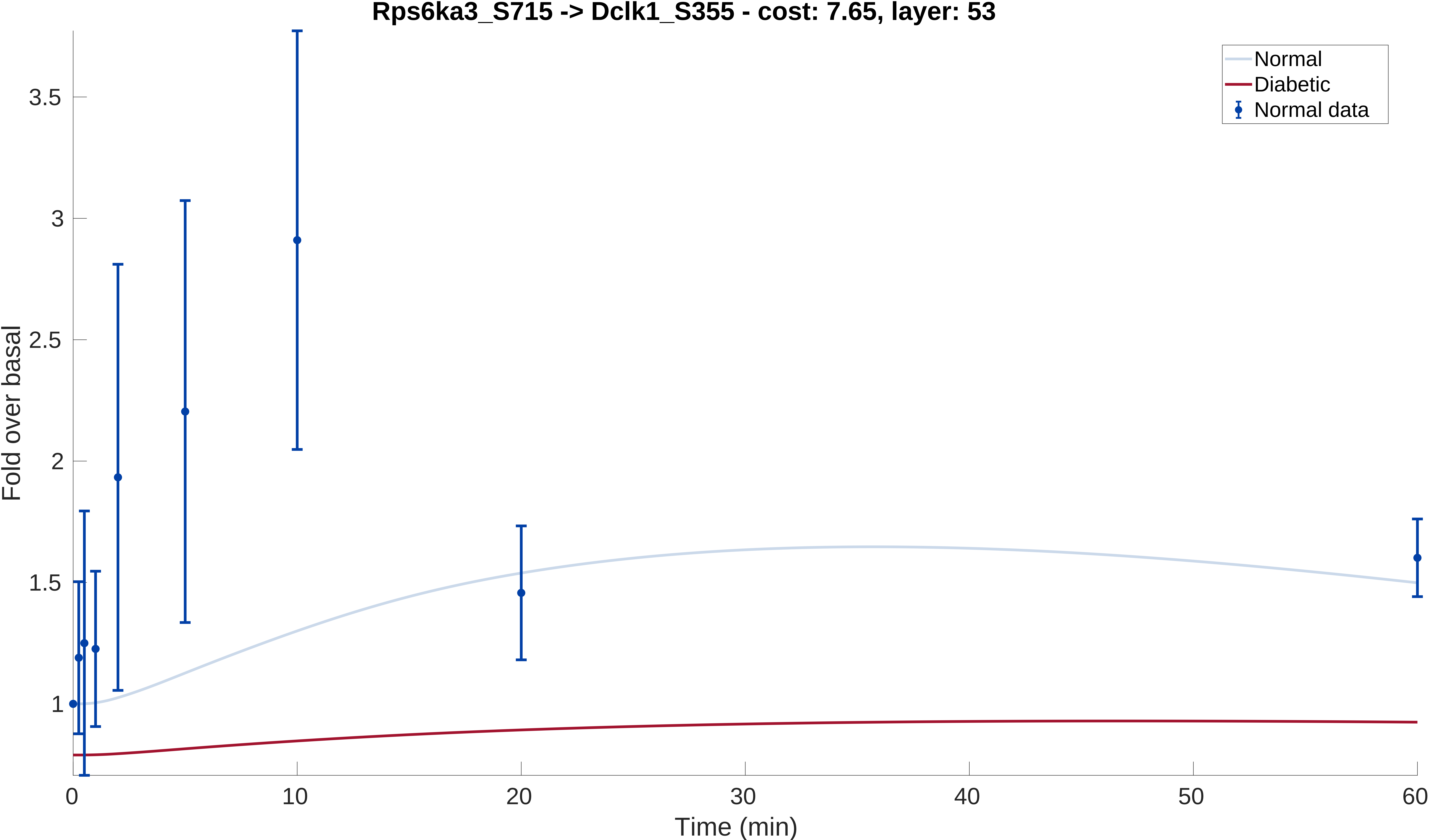
Rps6ka3_S715 -> Dclk1_S355 - cost: 7.65, layer: 53
Normal
Diabetic
Normal data
Fold over basal
Time (min)
3.5
3
2.5
2
1.5
1
0
10
20
30
40
50
60

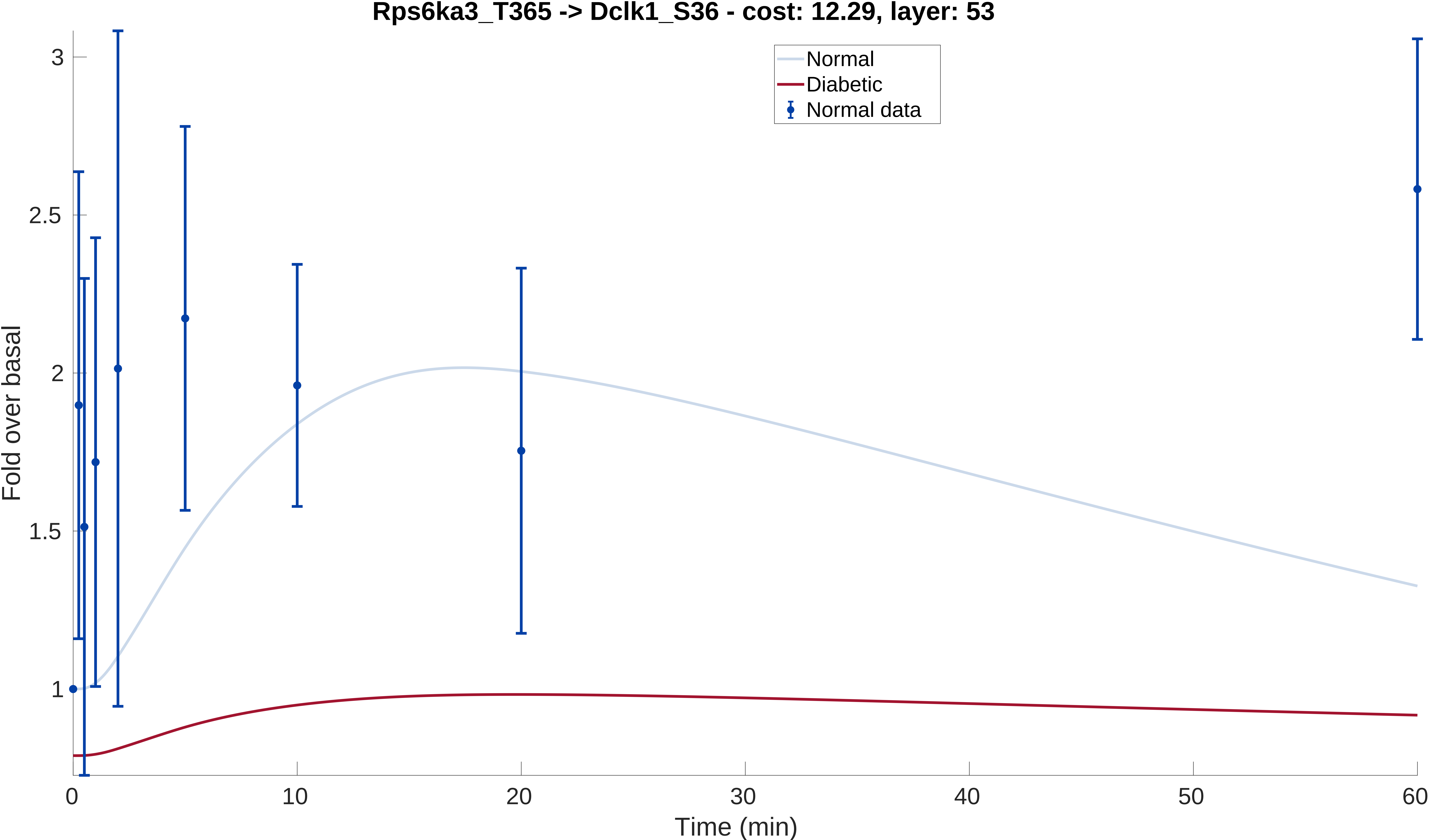
Rps6ka3_T365 -> Dclk1_S36 - cost: 12.29, layer: 53
Normal
Diabetic
Normal data
Fold over basal
Time (min)
3
2.5
2
1.5
1
0
10
20
30
40
50
60

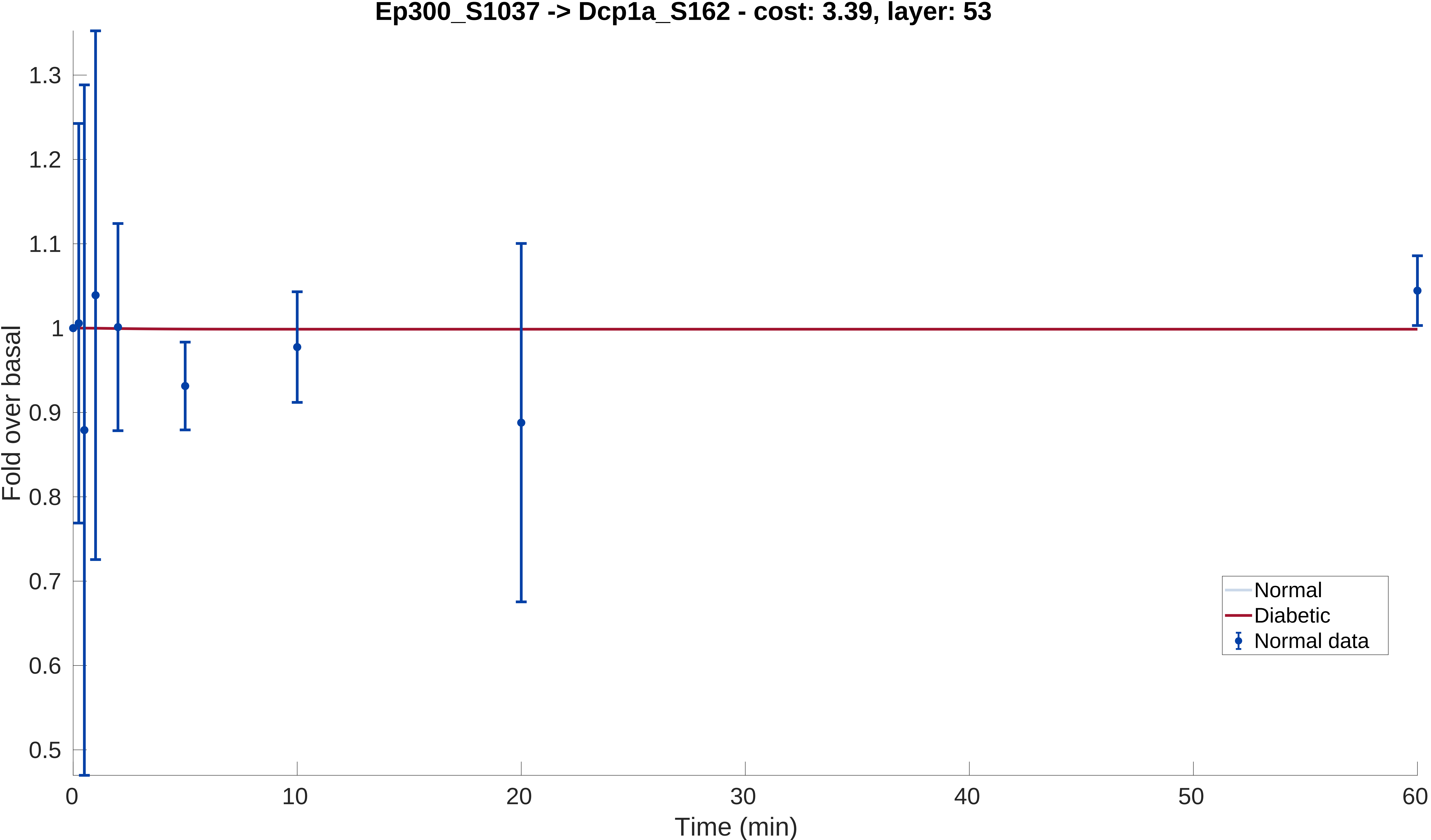

Ep300_S1037 -> Dcp1a_S162 - cost: 3.39, layer: 53
Fold over basal
1.3
1.2
1.1
1
0.9
0.8
0.7
0.6
0.5
0
10
20
30
40
50
60
Time (min)
Normal
Diabetic
Normal data

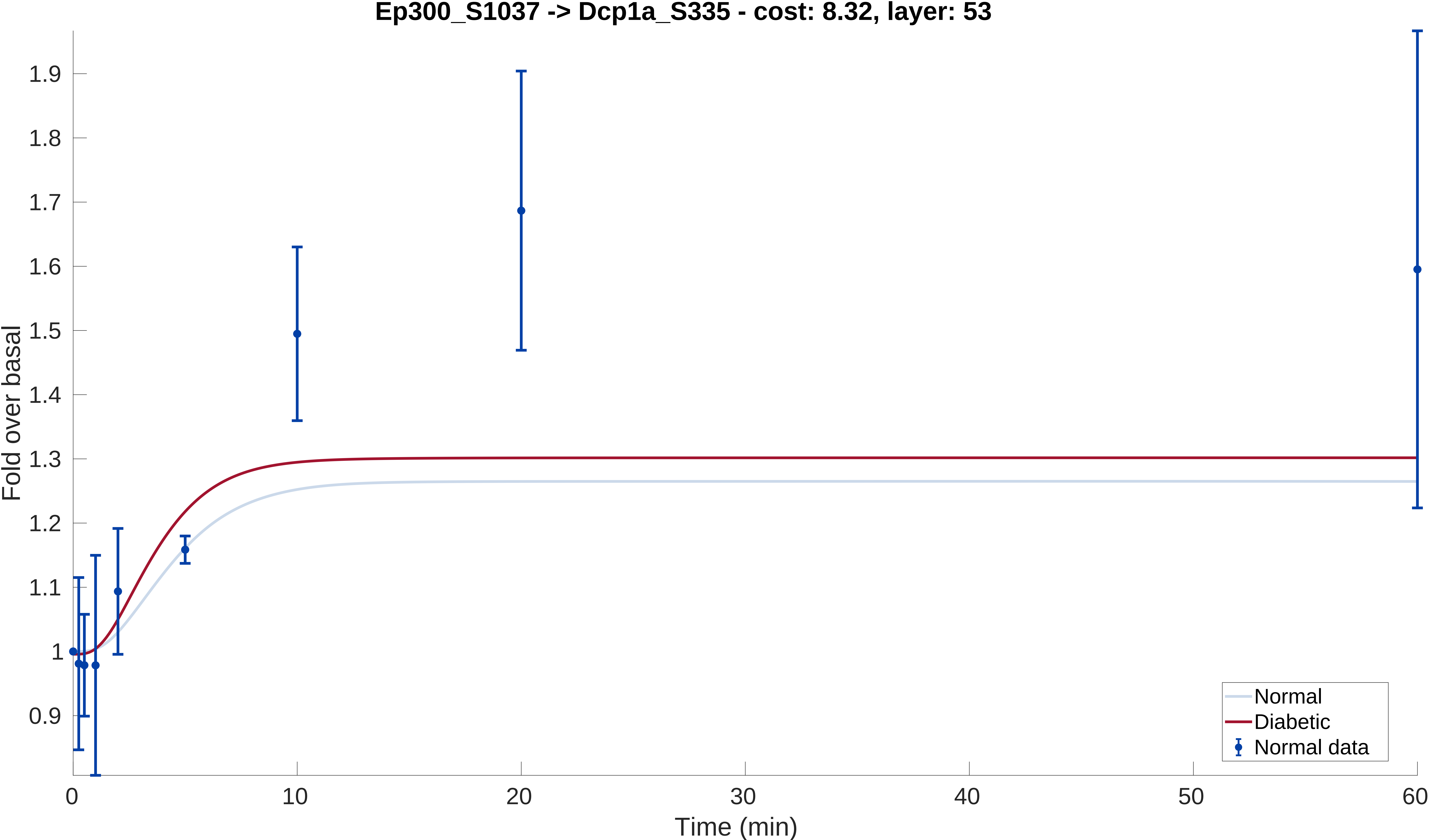

Ep300_S1037 -> Dcp1a_S335 - cost: 8.32, layer: 53
Fold over basal
1.9
1.8
1.7
1.6
1.5
1.4
1.3
1.2
1.1
1
0.9
0
10
20
30
40
50
60
Time (min)
Normal
Diabetic
Normal data

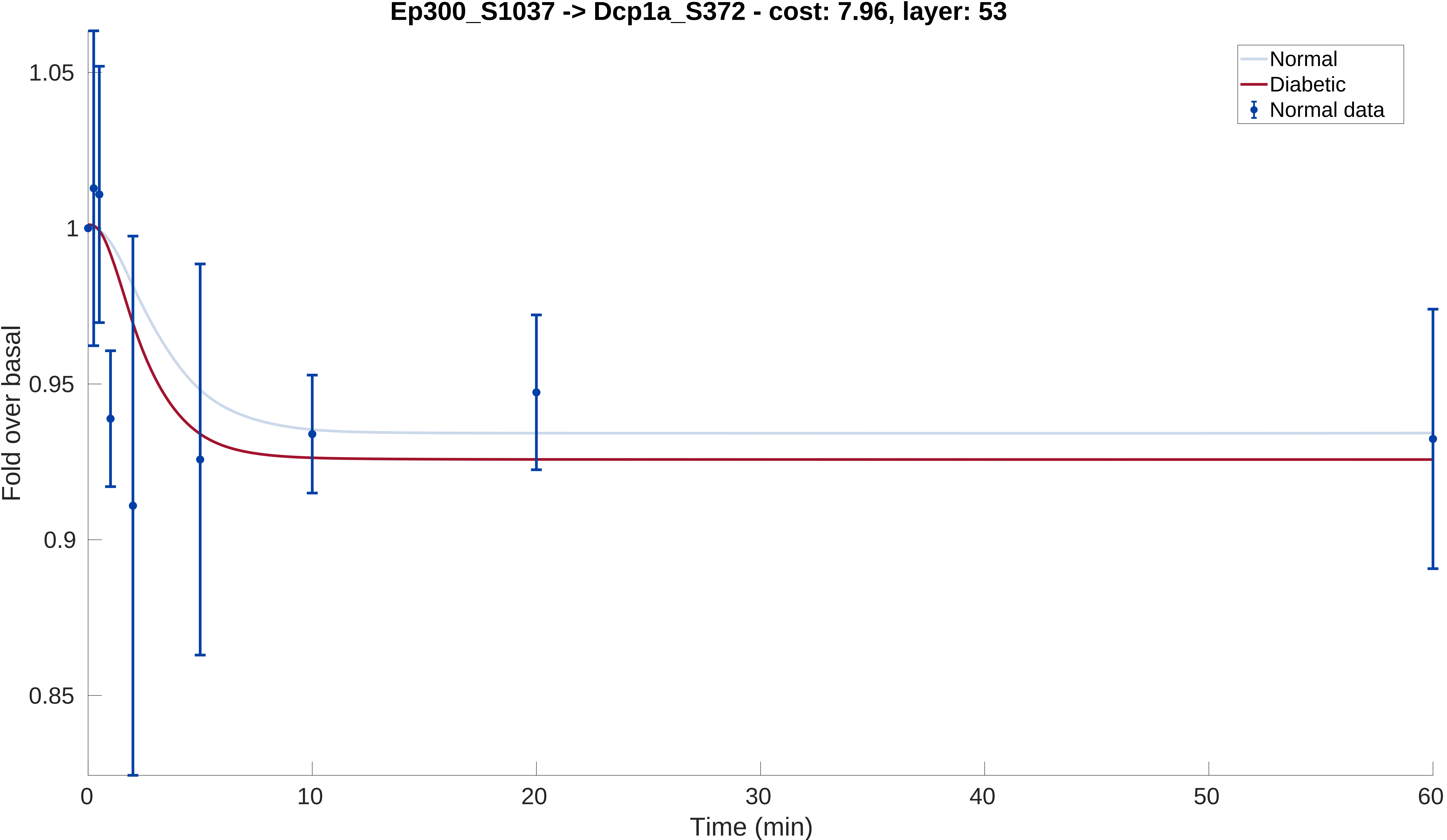

Ep300_S1037 -> Dcp1a_S372 - cost: 7.96, layer: 53
Normal
Diabetic
Normal data
1.05
1
0.95
0.9
0.85
Fold over basal
0
10
20
30
40
50
60
Time (min)

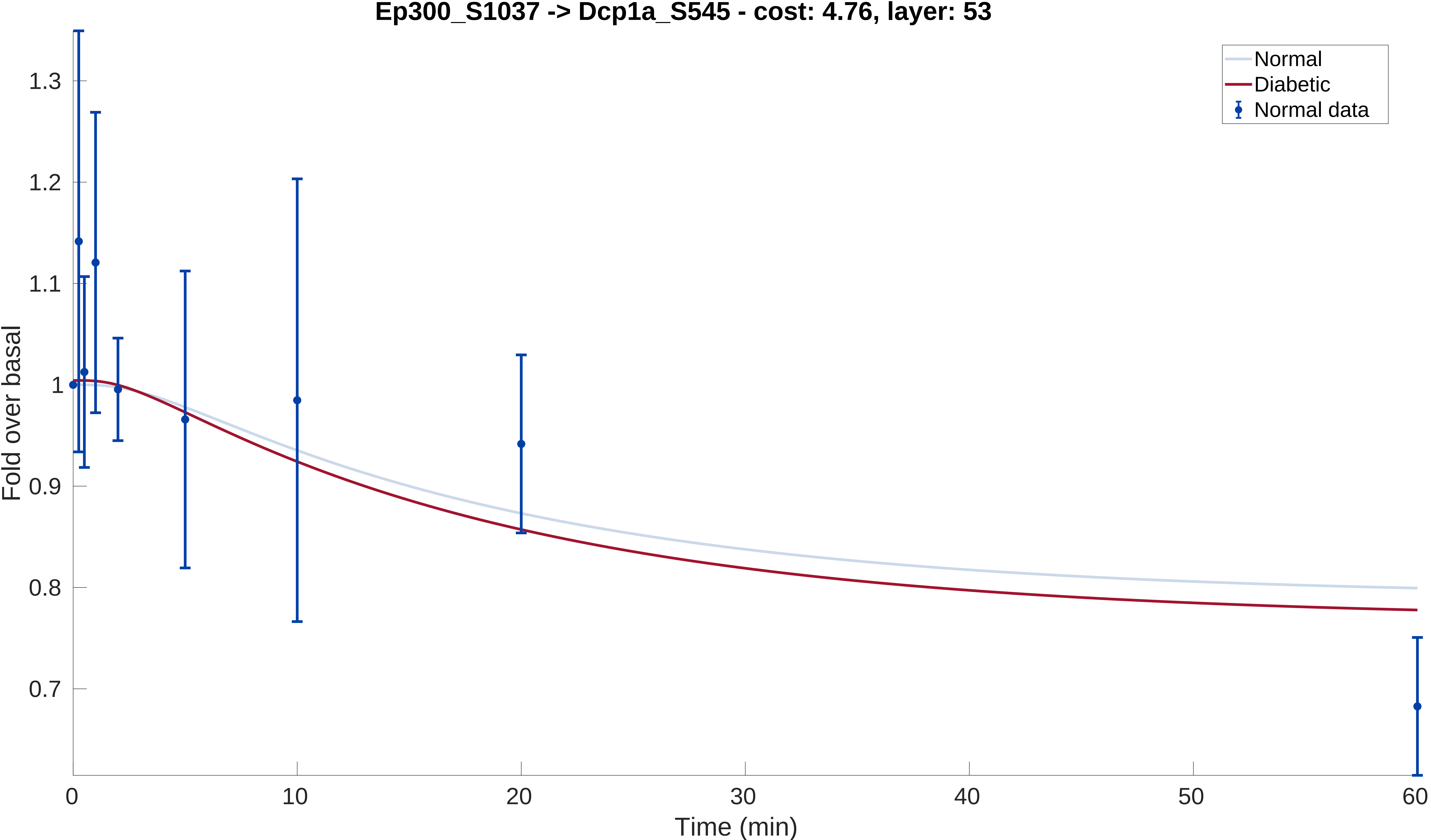

Ep300_S1037 -> Dcp1a_S545 - cost: 4.76, layer: 53
Normal
Diabetic
Normal data
Fold over basal
1.3
1.2
1.1
1
0.9
0.8
0.7
0
10
20
30
40
50
60
Time (min)

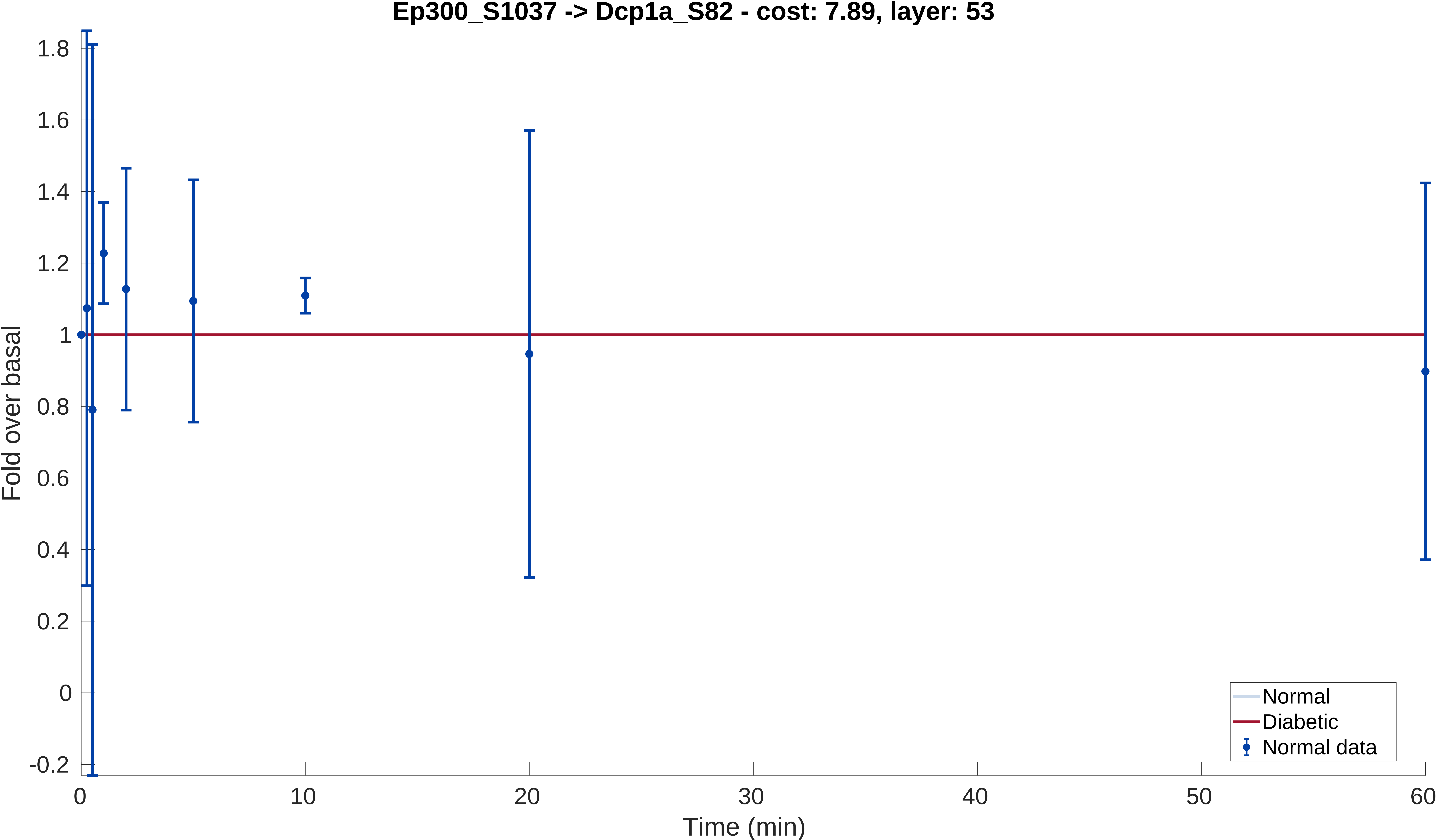

Ep300_S1037 -> Dcp1a_S82 - cost: 7.89, layer: 53
Fold over basal
1.8
1.6
1.4
1.2
1
0.8
0.6
0.4
0.2
0
-0.2
0
10
20
30
40
50
60
Time (min)
Normal
Diabetic
Normal data

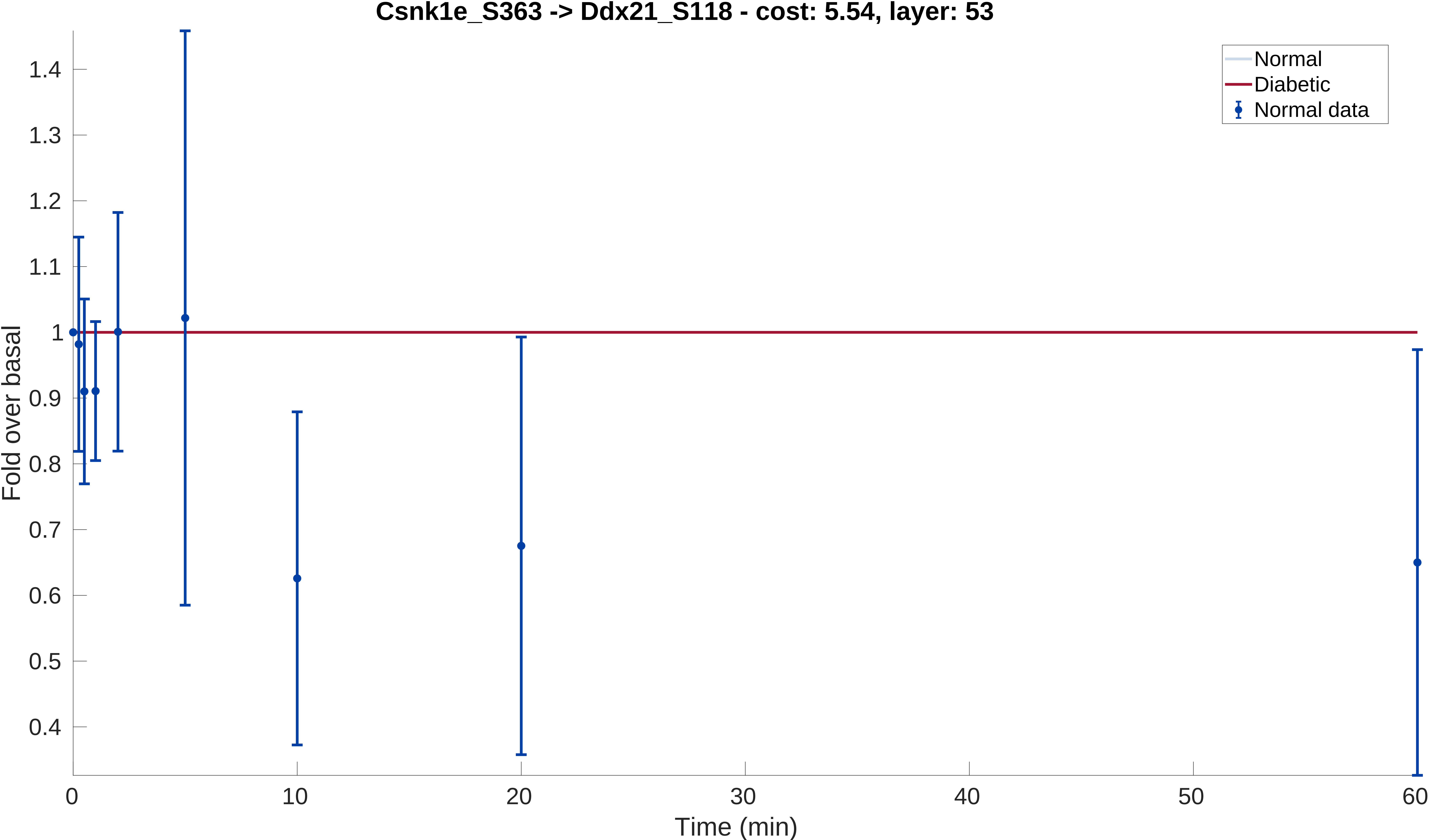

Csnk1e_S363 -> Ddx21_S118 - cost: 5.54, layer: 53
Normal
Diabetic
Normal data
Fold over basal
1.4
1.3
1.2
1.1
1
0.9
0.8
0.7
0.6
0.5
0.4
0
10
20
30
40
50
60
Time (min)

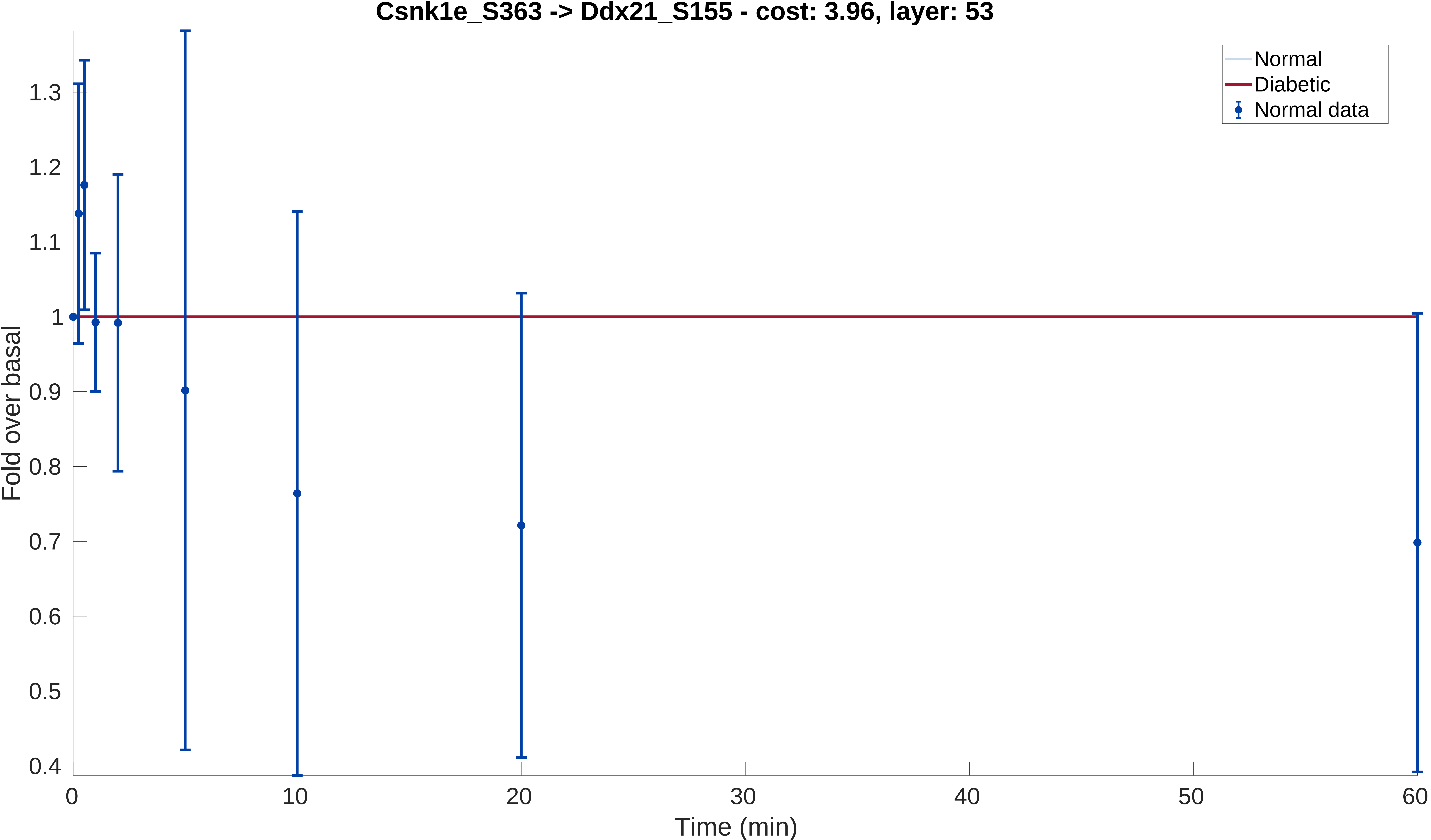
Csnk1e_S363 -> Ddx21_S155 - cost: 3.96, layer: 53
Normal
Diabetic
Normal data
Fold over basal
1.3
1.2
1.1
1
0.9
0.8
0.7
0.6
0.5
0.4
0
10
20
30
40
50
60
Time (min)

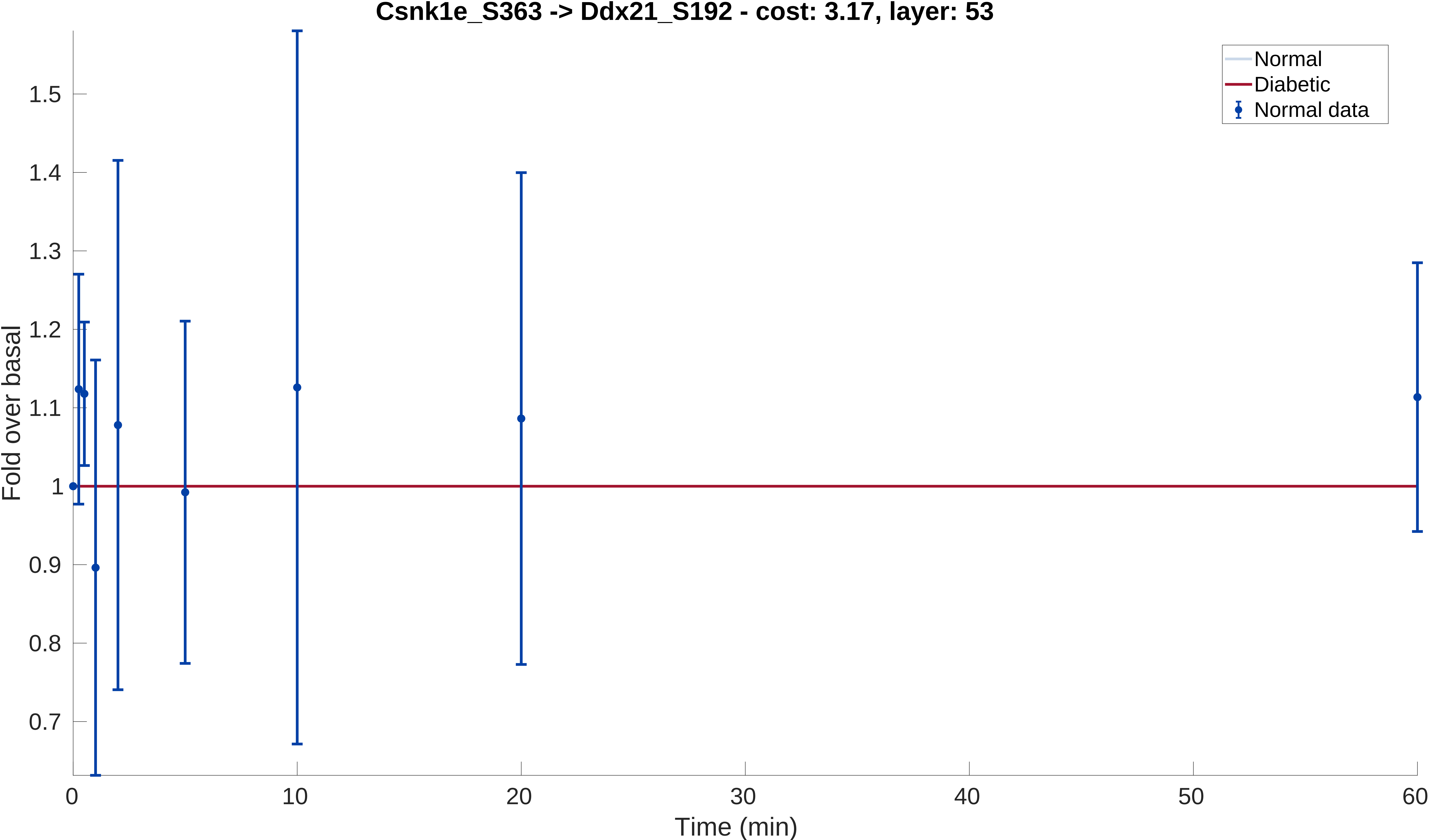
Csnk1e_S363 -> Ddx21_S192 - cost: 3.17, layer: 53
Normal
Diabetic
Normal data
Fold over basal
1.5
1.4
1.3
1.2
1.1
1
0.9
0.8
0.7
0
10
20
30
40
50
60
Time (min)

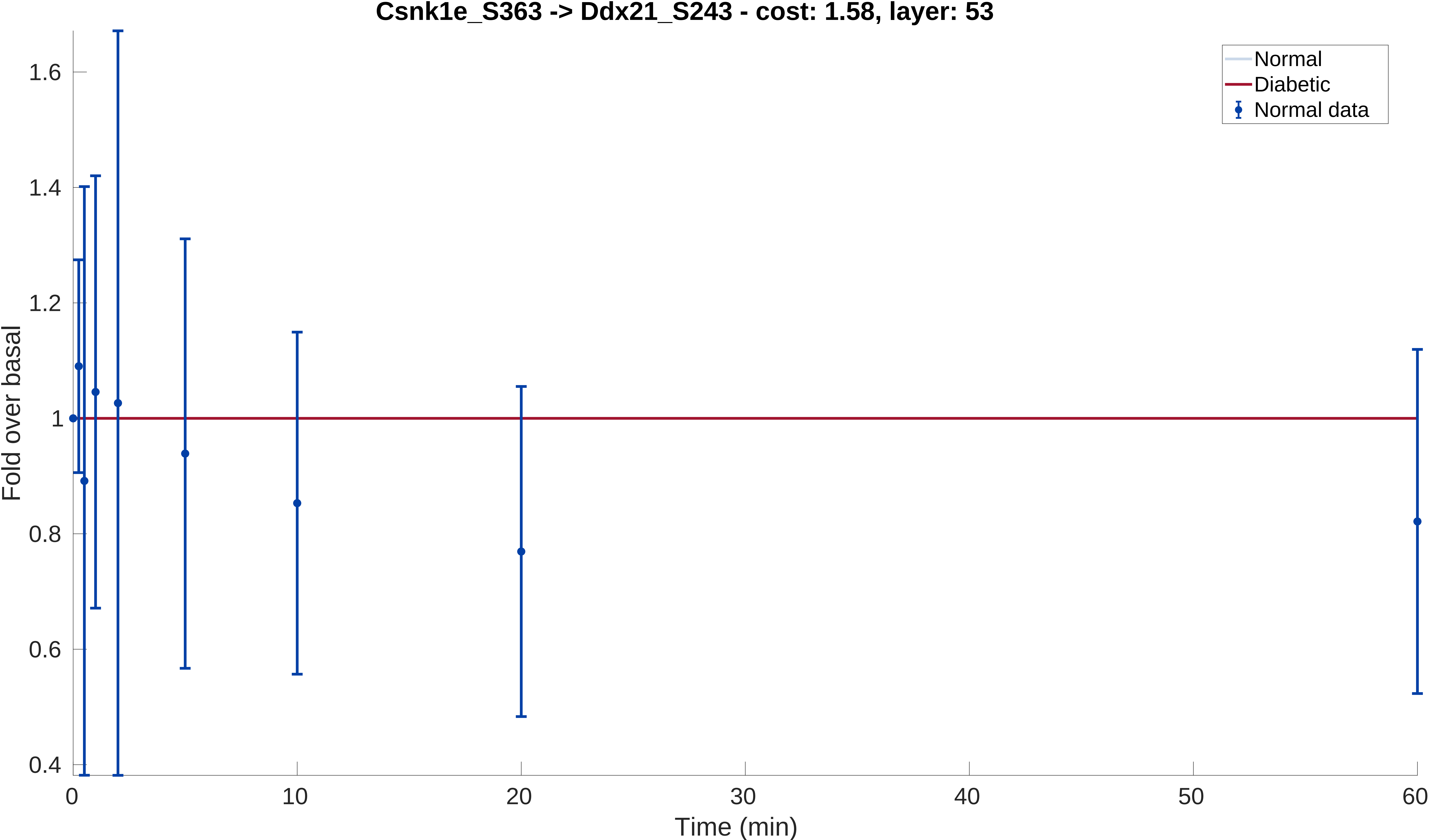

Csnk1e_S363 -> Ddx21_S243 - cost: 1.58, layer: 53
Normal
Diabetic
Normal data
Fold over basal
Time (min)
0.4
0.6
0.8
1
1.2
1.4
1.6
0
10
20
30
40
50
60

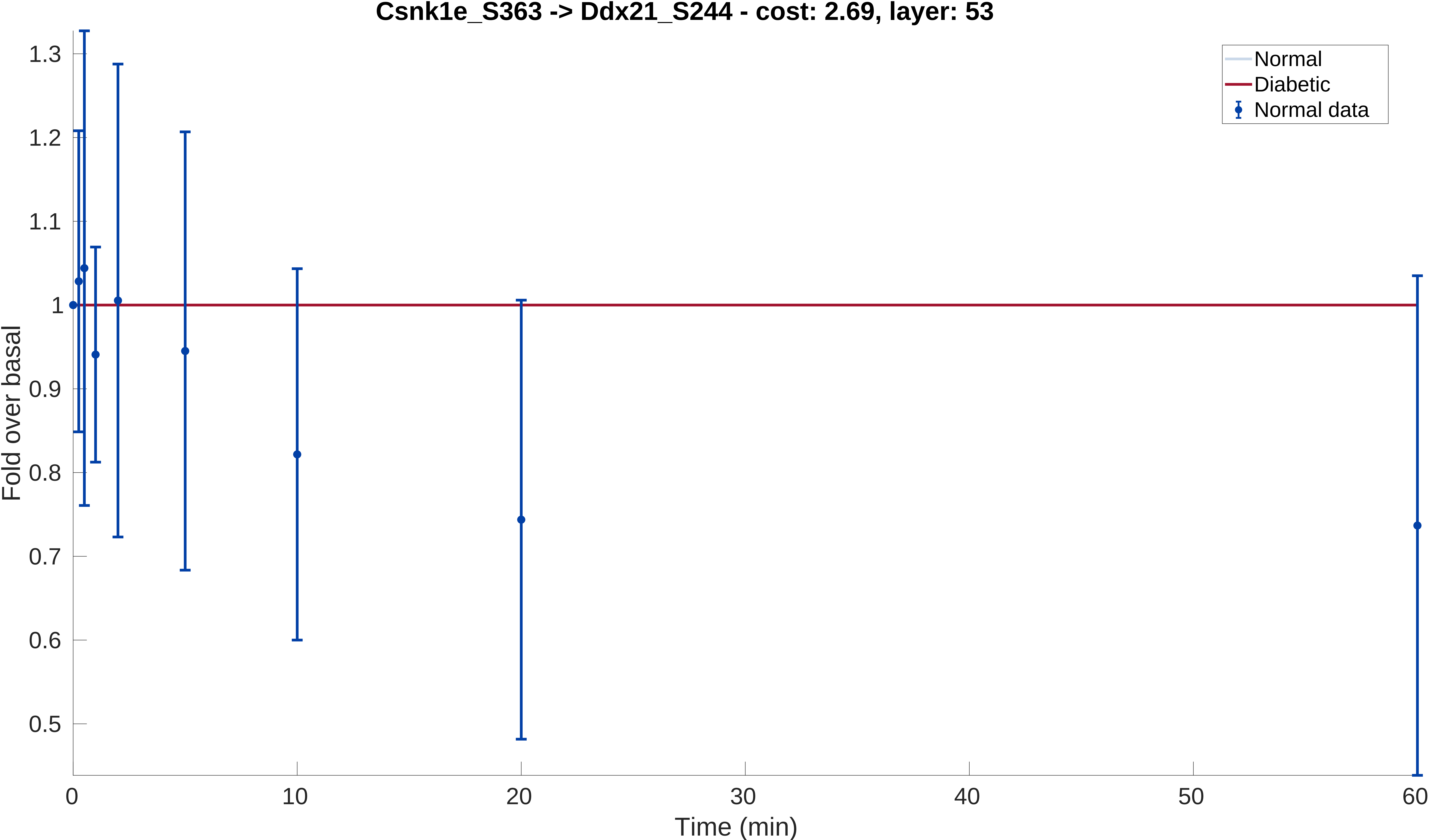
Csnk1e_S363 -> Ddx21_S244 - cost: 2.69, layer: 53
Normal
Diabetic
Normal data
Fold over basal
1.3
1.2
1.1
1
0.9
0.8
0.7
0.6
0.5
0
10
20
30
40
50
60
Time (min)

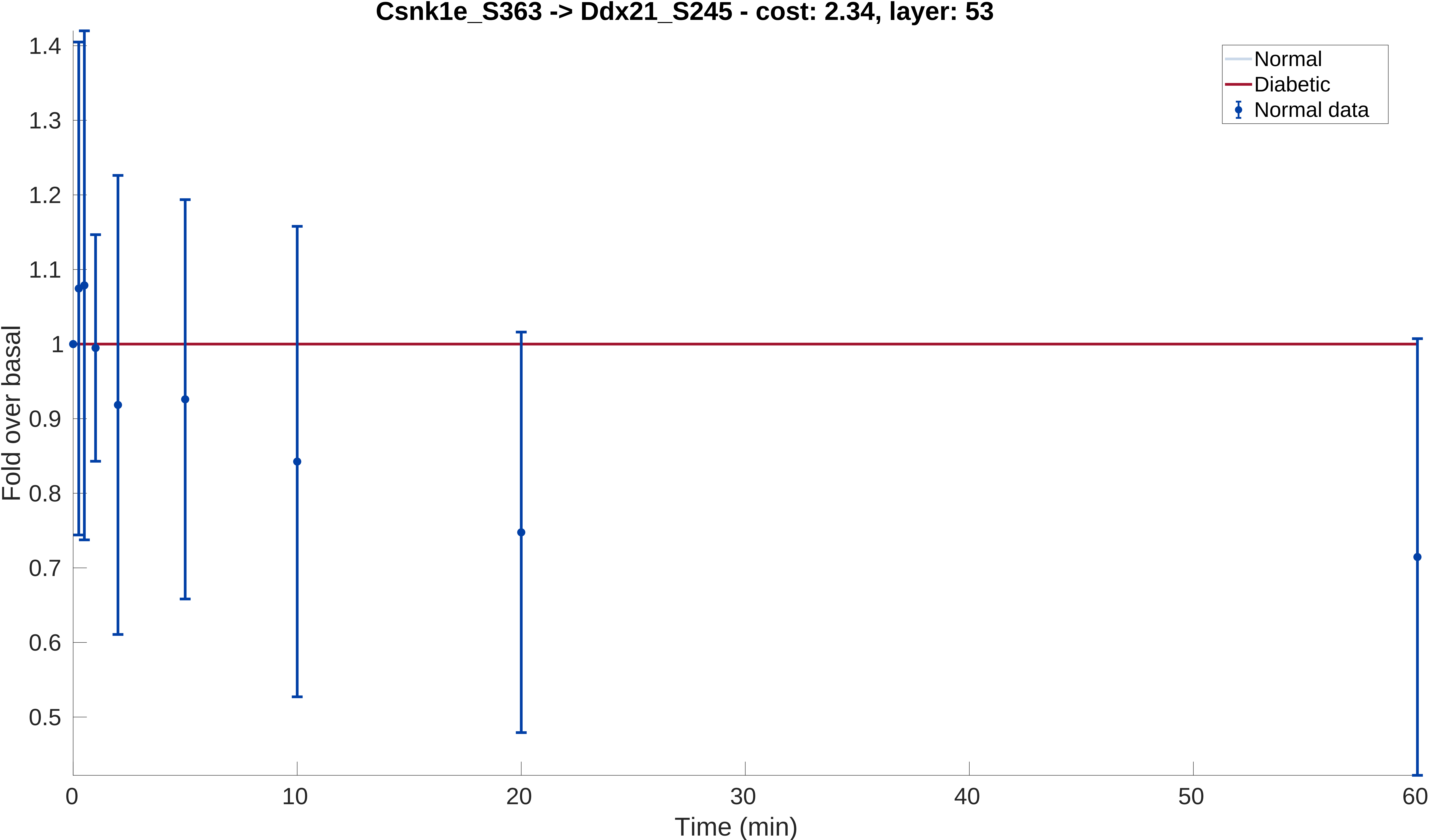

Csnk1e_S363 -> Ddx21_S245 - cost: 2.34, layer: 53
Normal
Diabetic
Normal data
Fold over basal
Time (min)
0.5
0.6
0.7
0.8
0.9
1
1.1
1.2
1.3
1.4
0
10
20
30
40
50
60

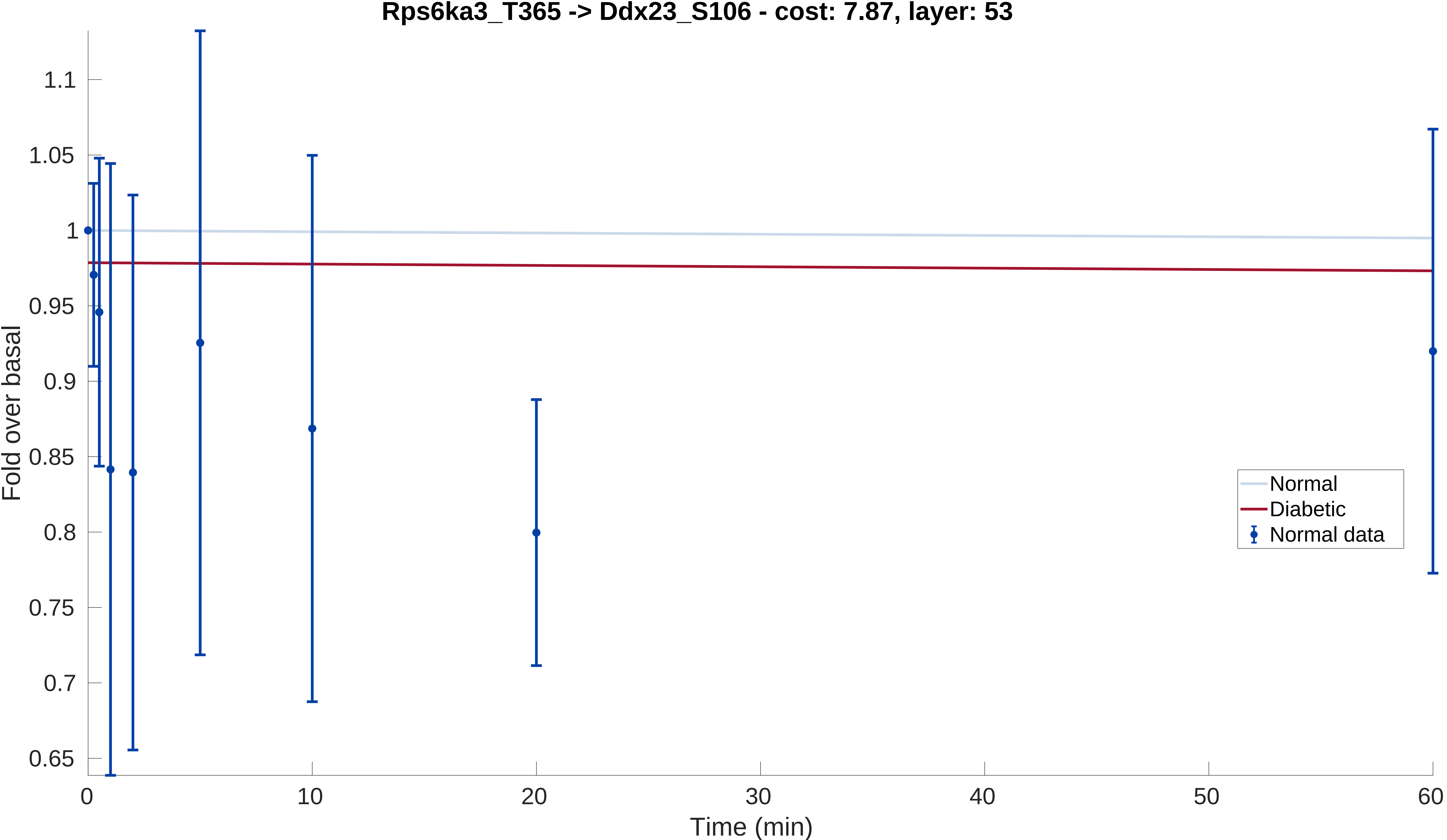

Rps6ka3_T365 -> Ddx23_S106 - cost: 7.87, layer: 53
Fold over basal
1.1
1.05
1
0.95
0.9
0.85
0.8
0.75
0.7
0.65
0
10
20
30
40
50
60
Time (min)
Normal
Diabetic
Normal data

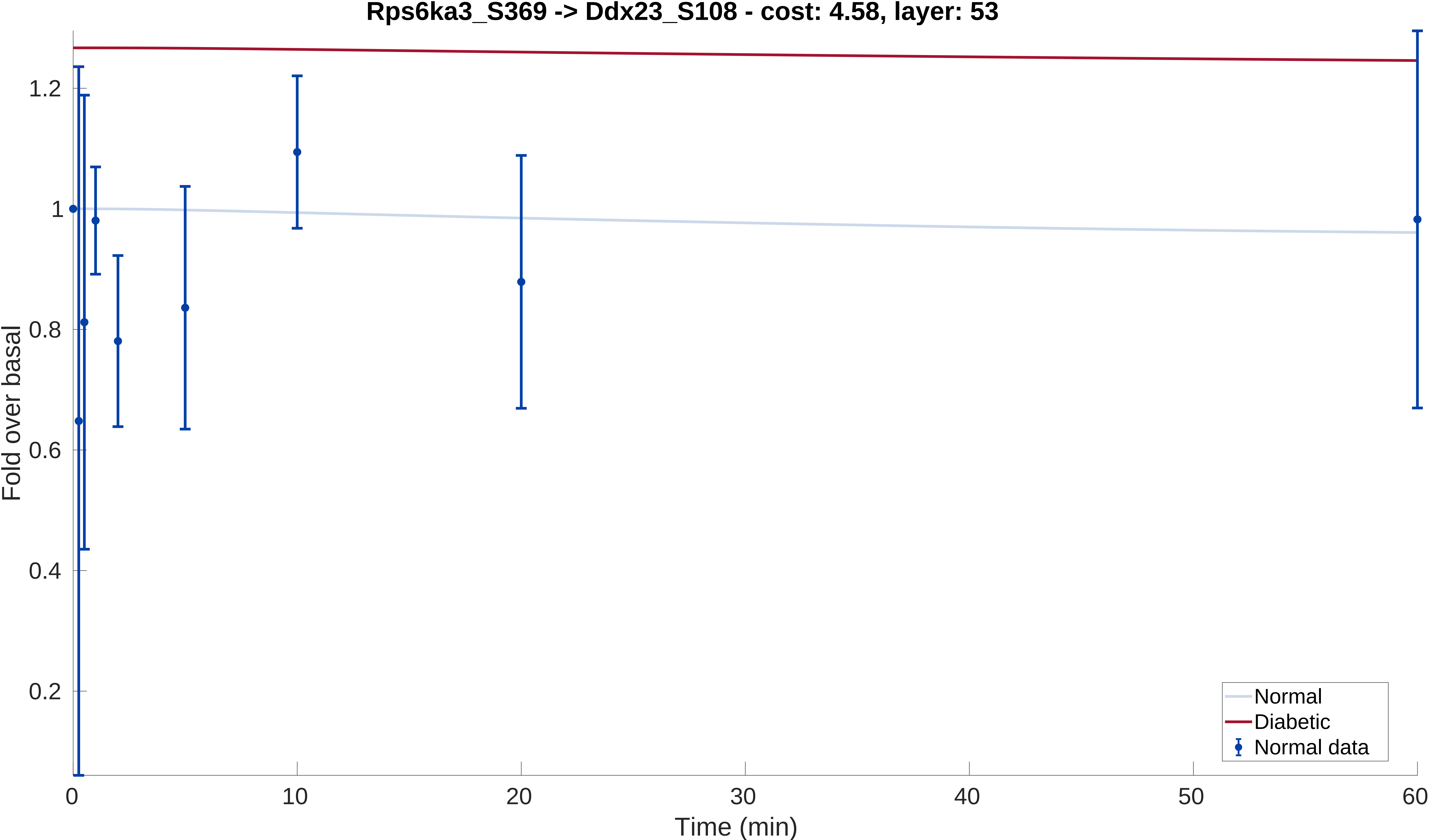

Rps6ka3_S369 -> Ddx23_S108 - cost: 4.58, layer: 53
Fold over basal
1.2
1
0.8
0.6
0.4
0.2
0
10
20
30
40
50
60
Time (min)
Normal
Diabetic
Normal data

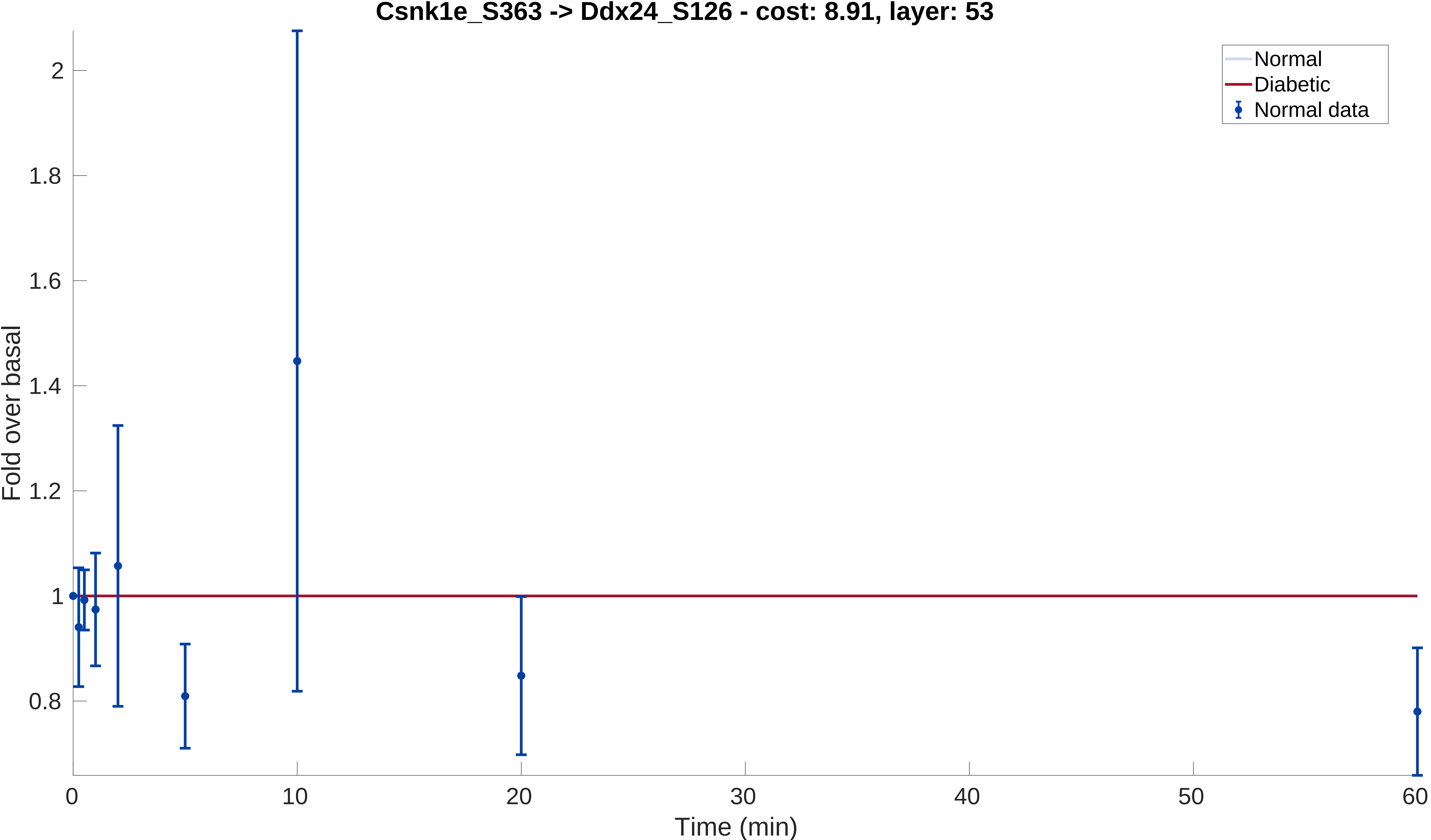
Csnk1e_S363 -> Ddx24_S126 - cost: 8.91, layer: 53
Normal
Diabetic
Normal data
Fold over basal
Time (min)
0.8
1
1.2
1.4
1.6
1.8
2
0
10
20
30
40
50
60

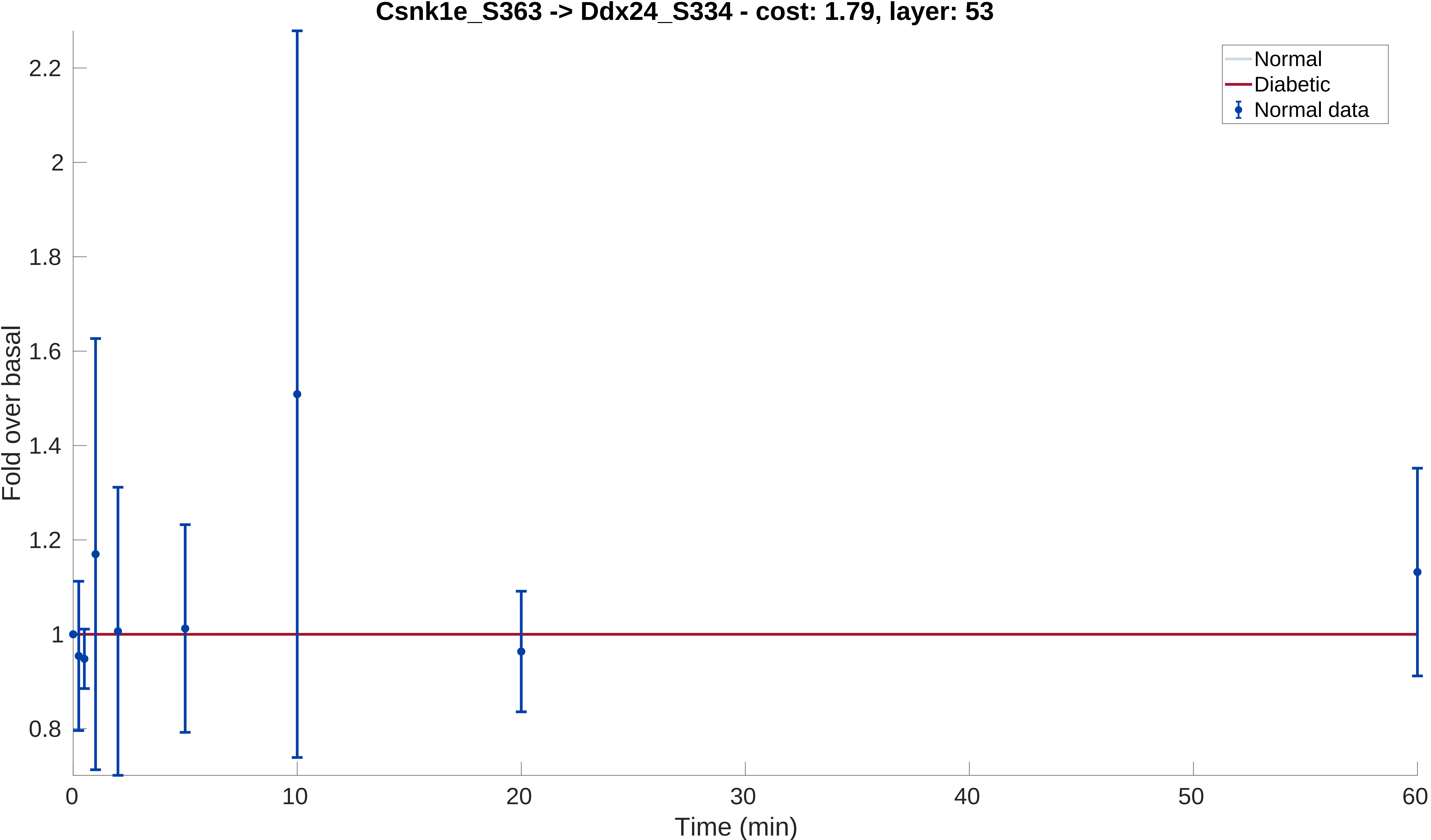
Csnk1e_S363 -> Ddx24_S334 - cost: 1.79, layer: 53
Normal
Diabetic
Normal data
Fold over basal
2.2
2
1.8
1.6
1.4
1.2
1
0.8
0
10
20
30
40
50
60
Time (min)

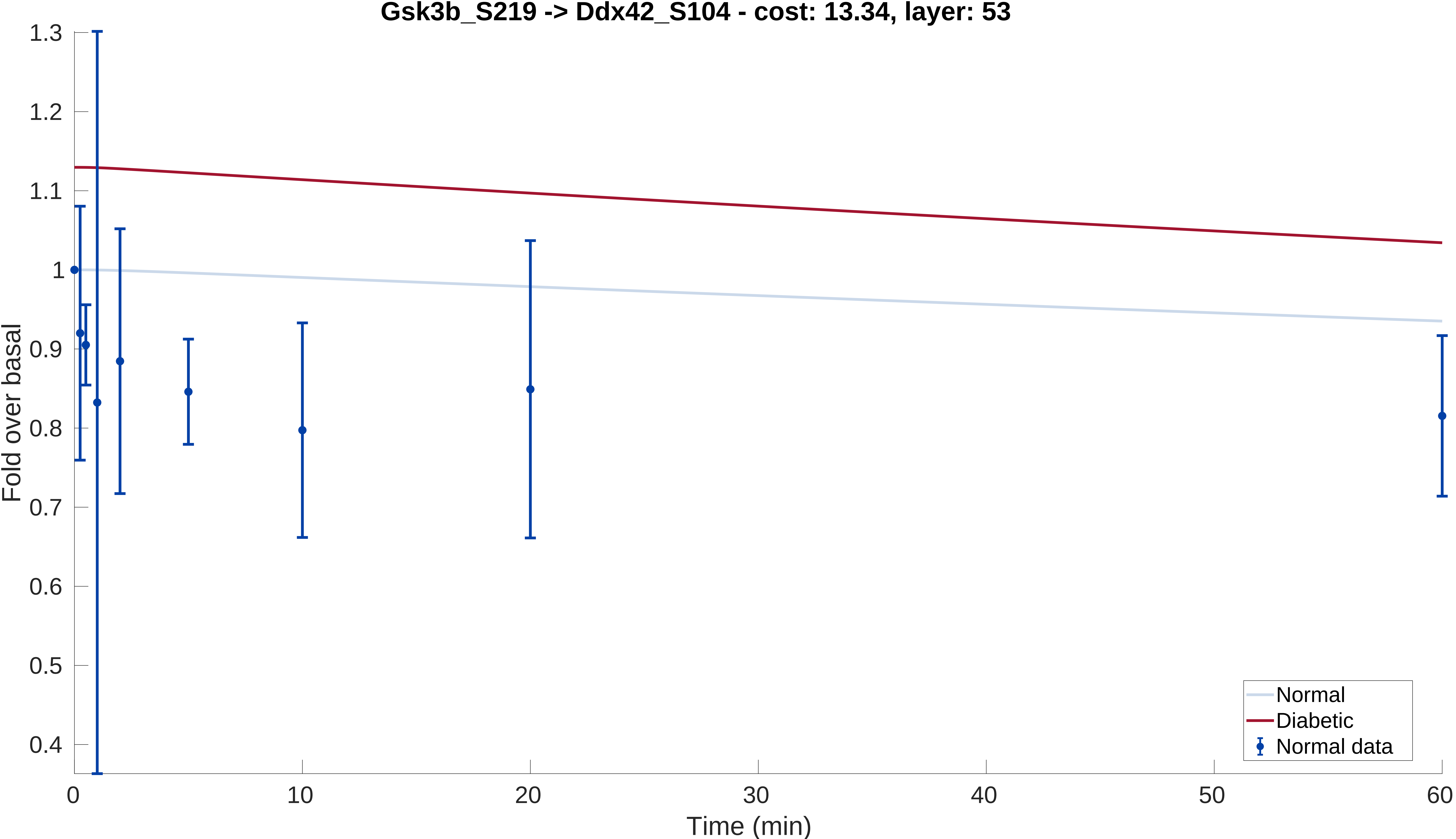

Gsk3b_S219 -> Ddx42_S104 - cost: 13.34, layer: 53
Fold over basal
1.3
1.2
1.1
1
0.9
0.8
0.7
0.6
0.5
0.4
0
10
20
30
40
50
60
Time (min)
Normal
Diabetic
Normal data

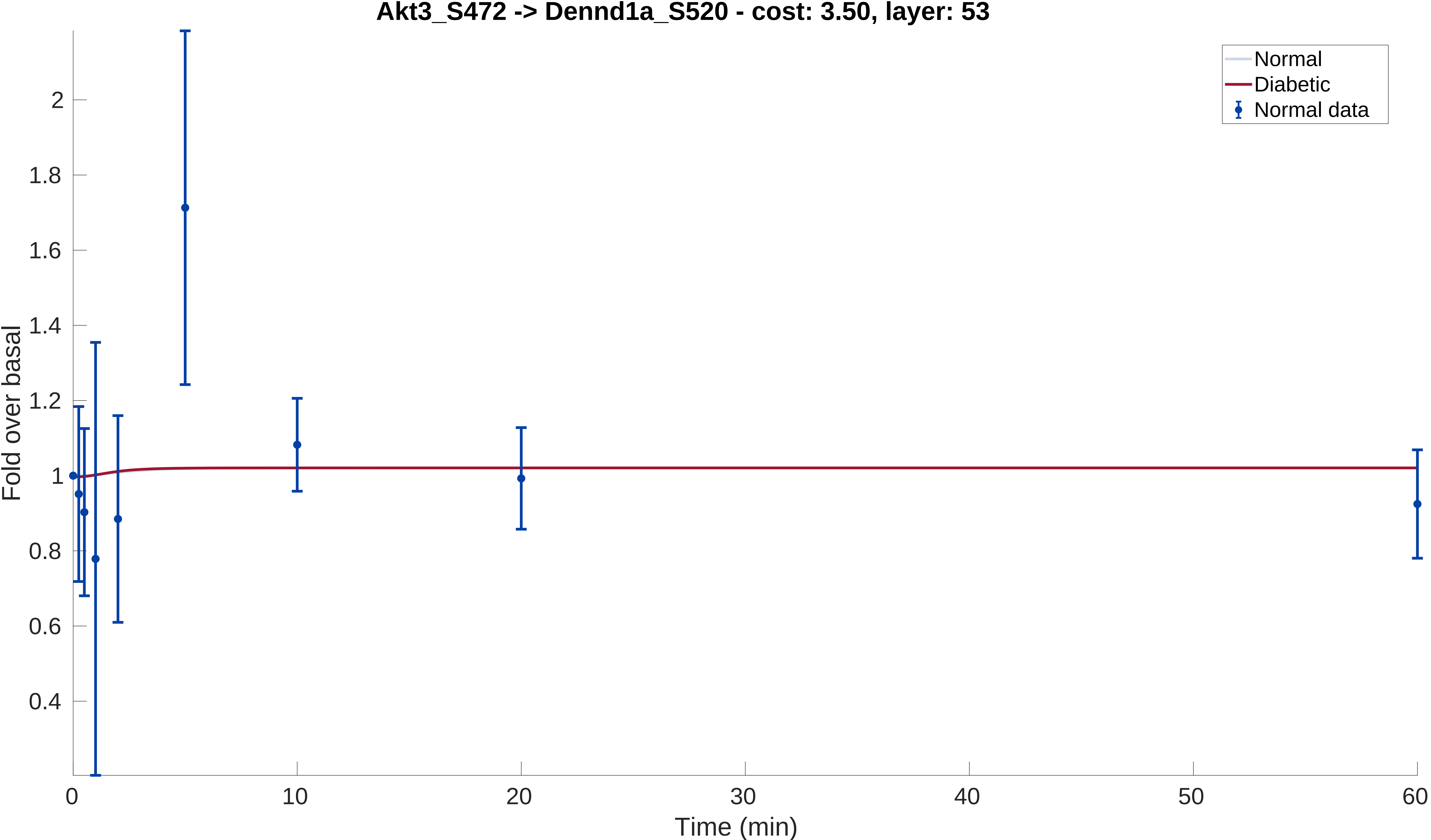

Akt3_S472 -> Dennd1a_S520 - cost: 3.50, layer: 53
Normal
Diabetic
Normal data
Fold over basal
0.4
0.6
0.8
1
1.2
1.4
1.6
1.8
2
0
10
20
30
40
50
60
Time (min)

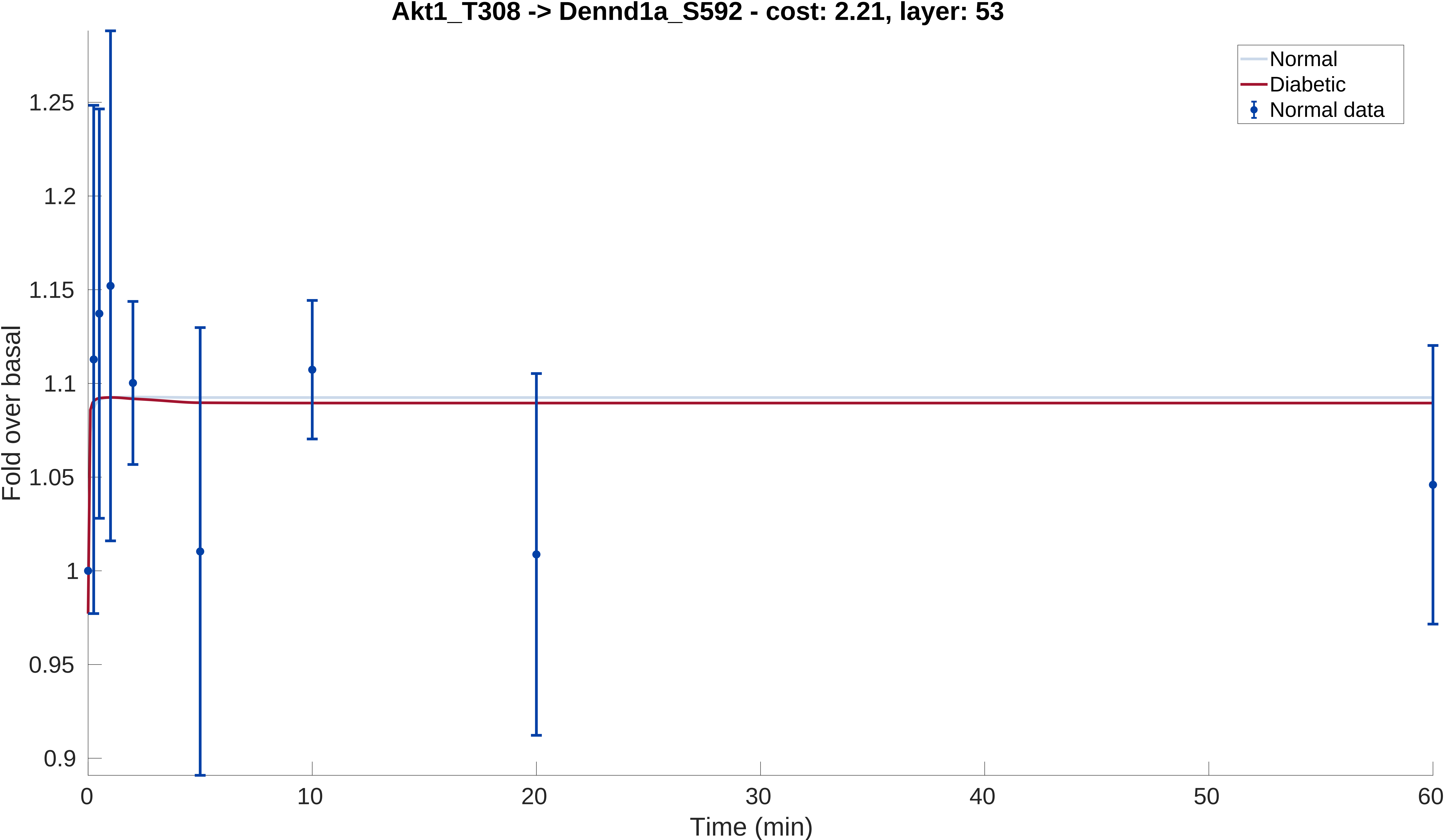

Akt1_T308 -> Dennd1a_S592 - cost: 2.21, layer: 53
Normal
Diabetic
Normal data
Fold over basal
1.25
1.2
1.15
1.1
1.05
1
0.95
0.9
0
10
20
30
40
50
60
Time (min)

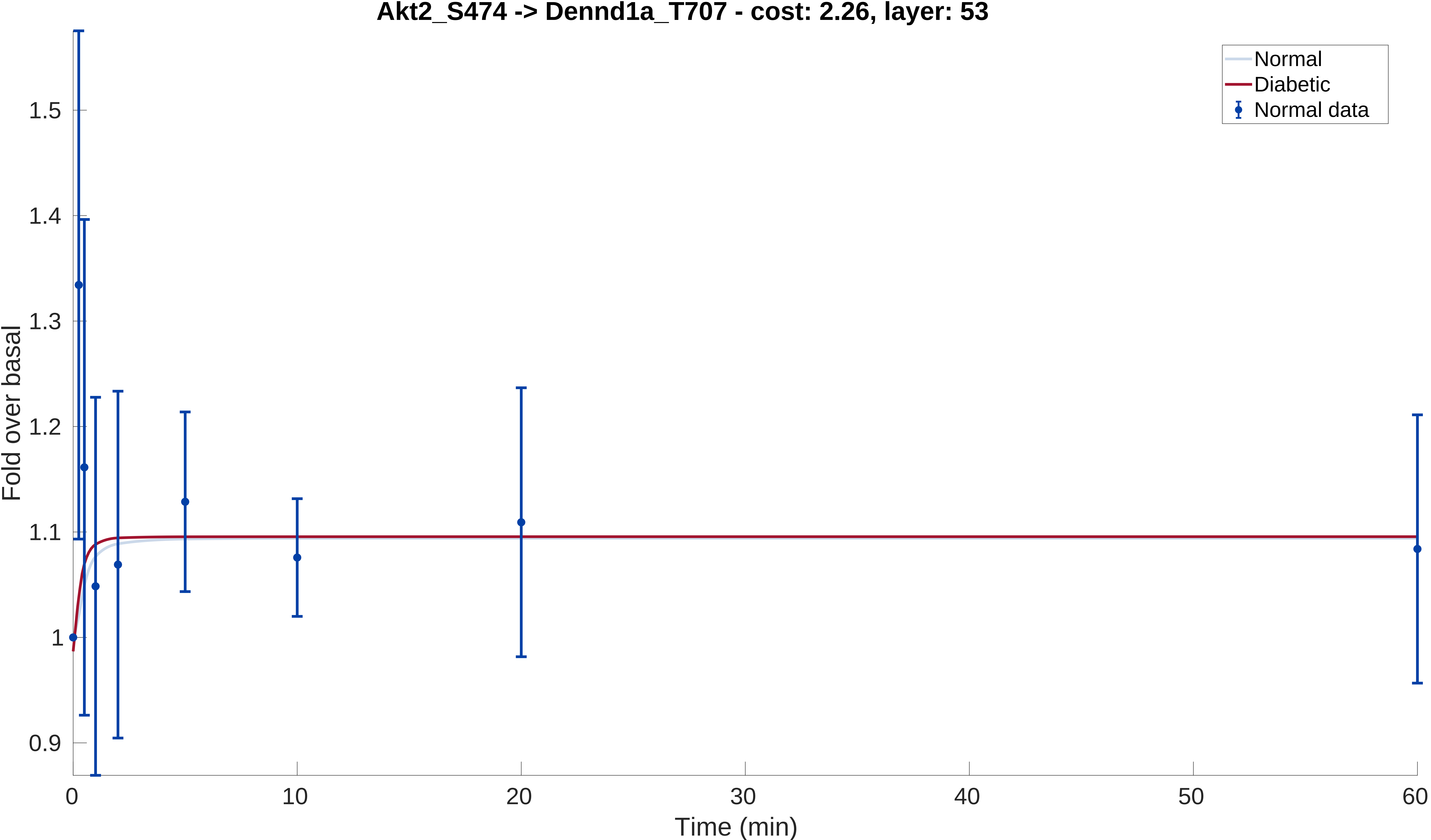

Akt2_S474 -> Dennd1a_T707 - cost: 2.26, layer: 53
Normal
Diabetic
Normal data
Fold over basal
0.9
1
1.1
1.2
1.3
1.4
1.5
0
10
20
30
40
50
60
Time (min)

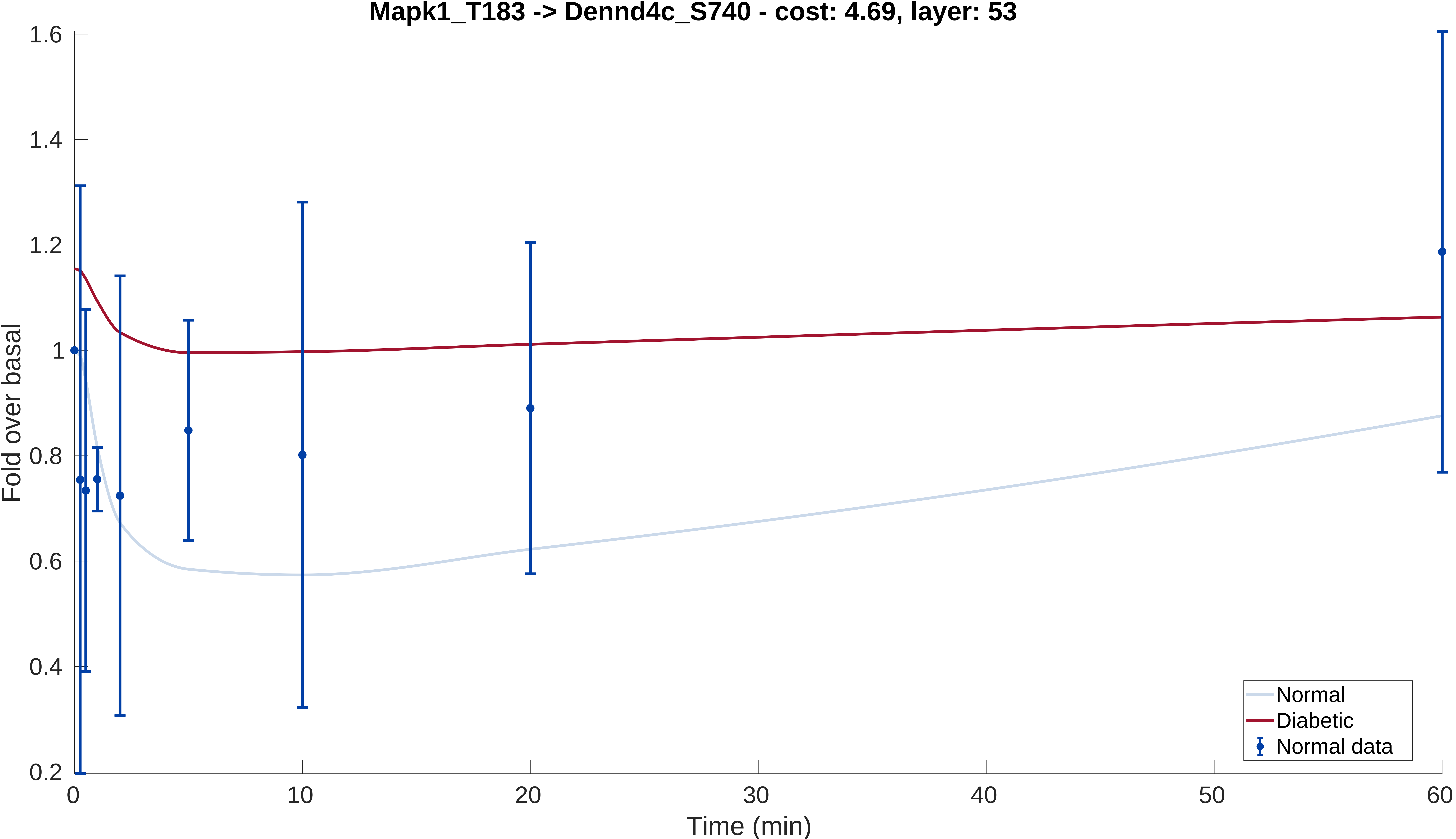

Mapk1_T183 -> Dennd4c_S740 - cost: 4.69, layer: 53
1.6
1.4
1.2
1
0.8
0.6
0.4
0.2
Fold over basal
0
10
20
30
40
50
60
Time (min)
Normal
Diabetic
Normal data

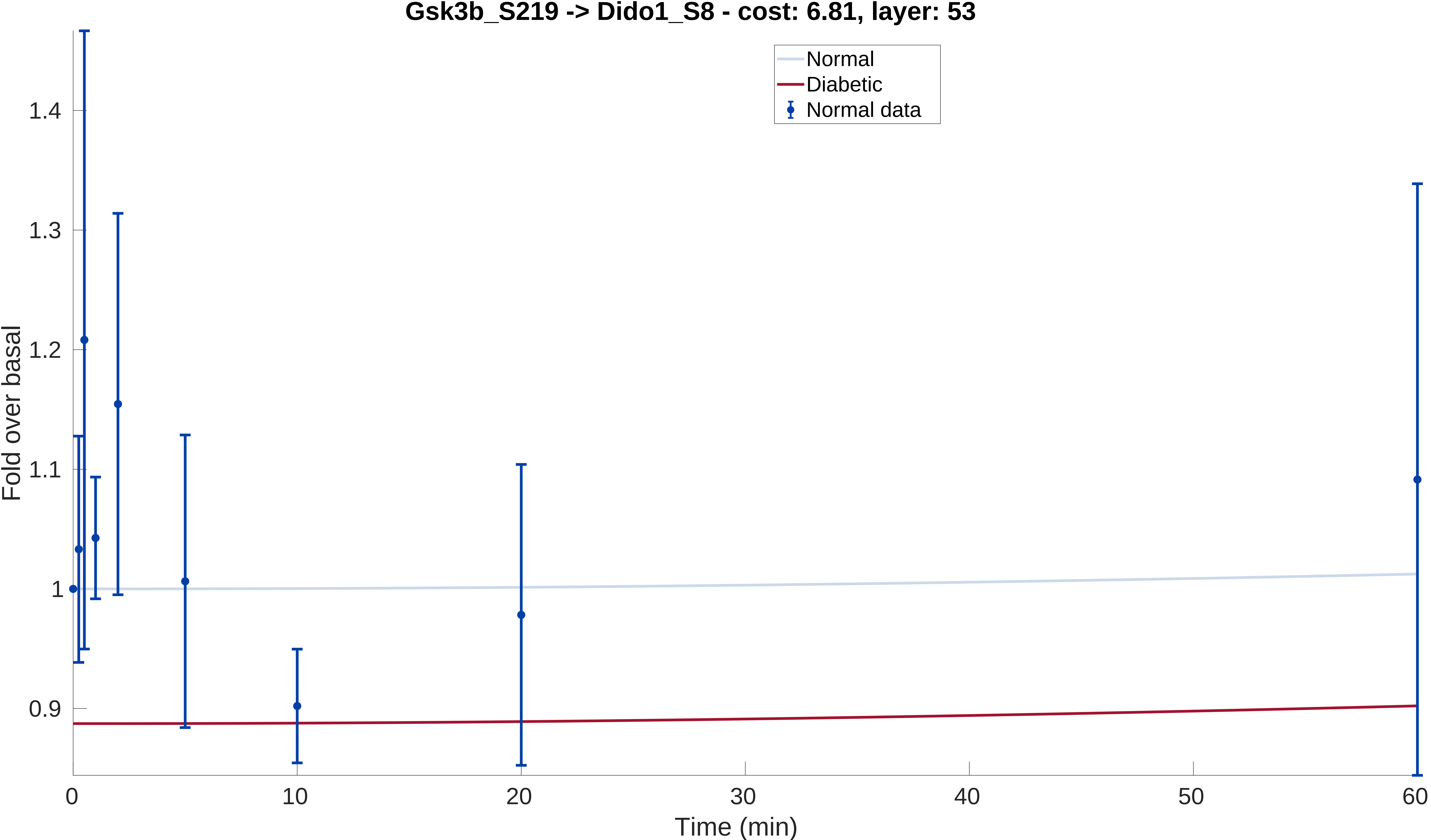

Gsk3b_S219 -> Dido1_S8 - cost: 6.81, layer: 53
Normal
Diabetic
Normal data
Fold over basal
1.4
1.3
1.2
1.1
1
0.9
0
10
20
30
40
50
60
Time (min)

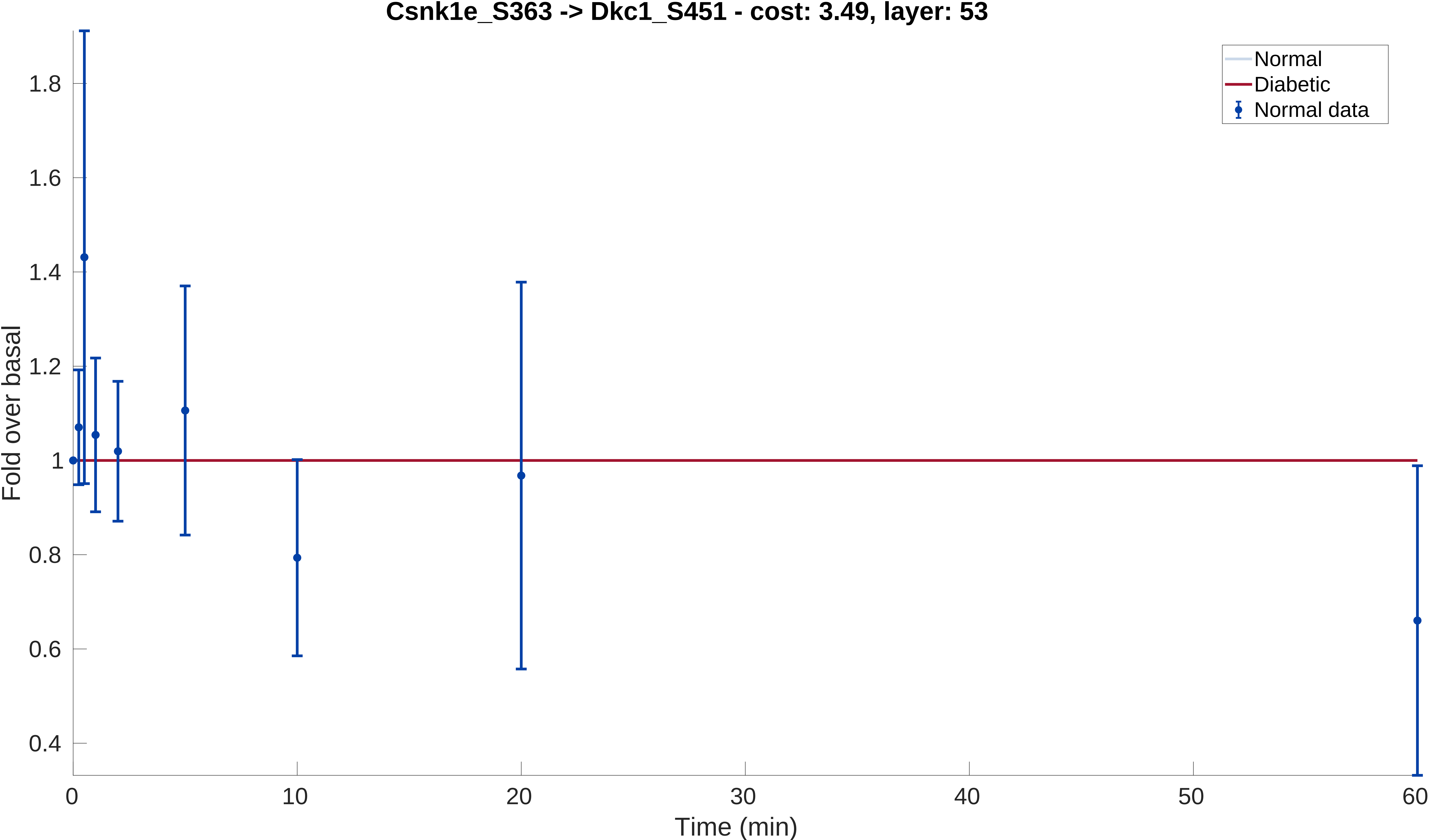

Csnk1e_S363 -> Dkc1_S451 - cost: 3.49, layer: 53
Normal
Diabetic
Normal data
Fold over basal
Time (min)
1.8
1.6
1.4
1.2
1
0.8
0.6
0.4
0
10
20
30
40
50
60

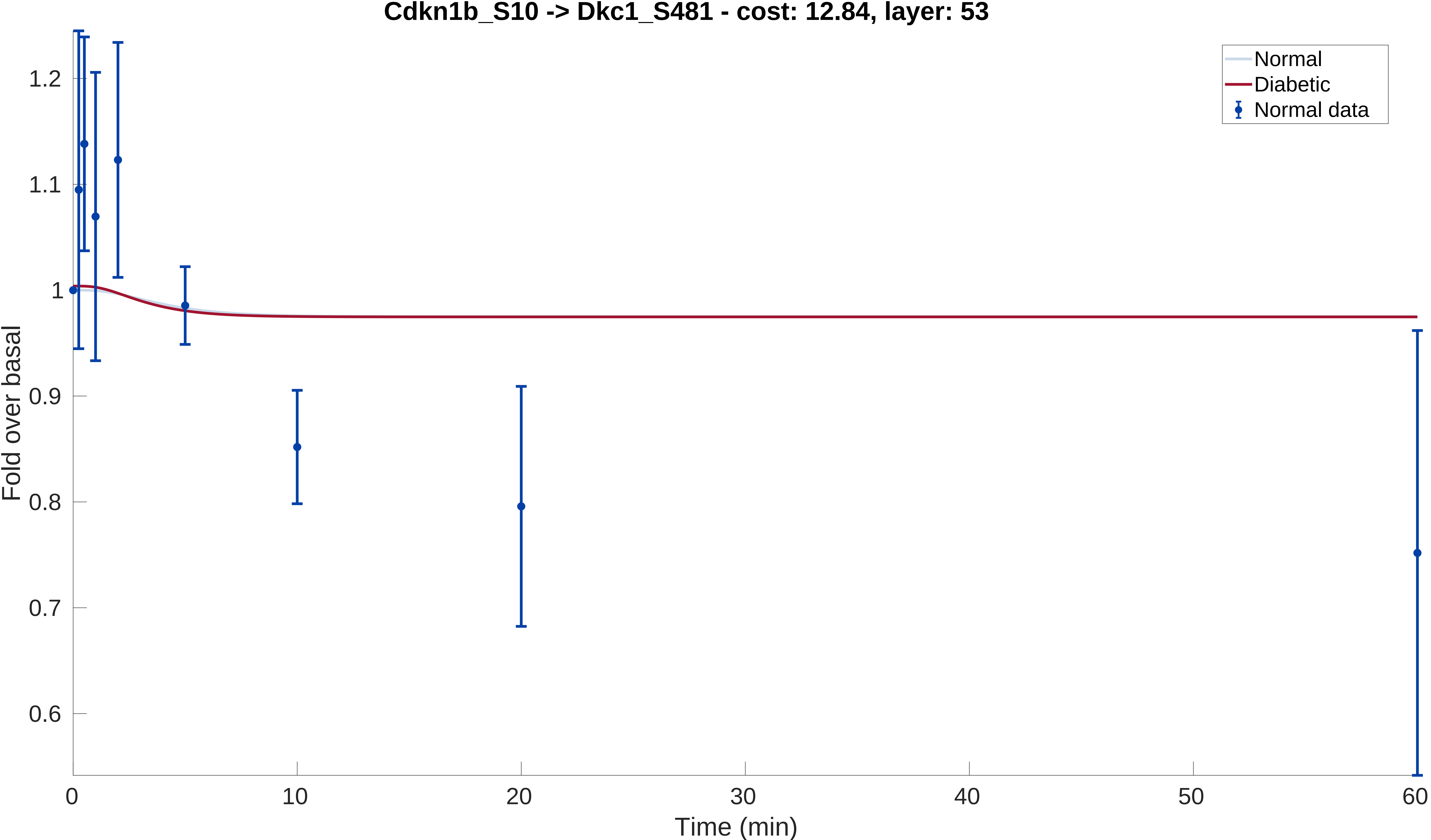
Cdkn1b_S10 -> Dkc1_S481 - cost: 12.84, layer: 53
Normal
Diabetic
Normal data
Fold over basal
1.2
1.1
1
0.9
0.8
0.7
0.6
0
10
20
30
40
50
60
Time (min)

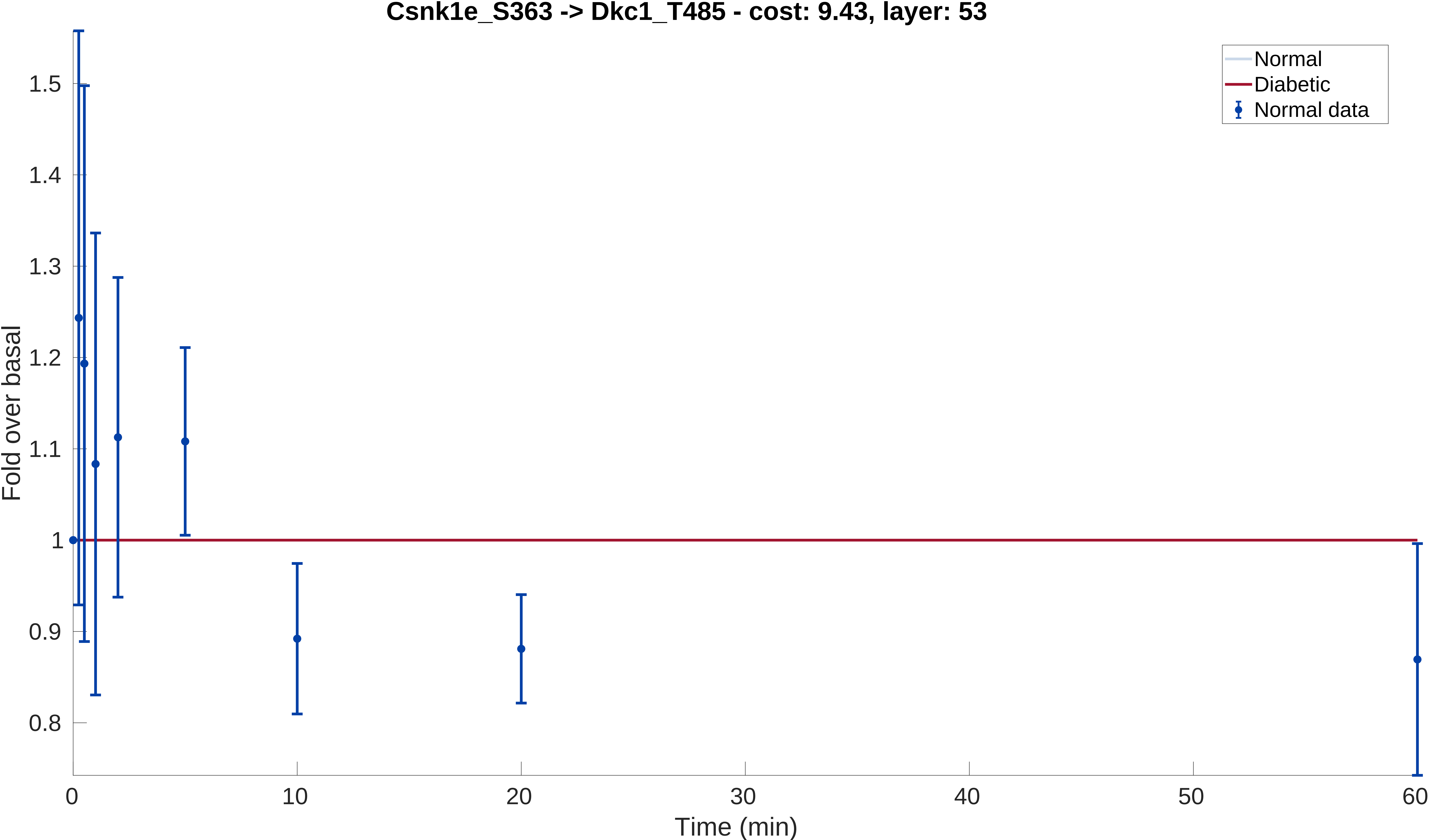

Csnk1e_S363 -> Dkc1_T485 - cost: 9.43, layer: 53
Normal
Diabetic
Normal data
Fold over basal
1.5
1.4
1.3
1.2
1.1
1
0.9
0.8
0
10
20
30
40
50
60
Time (min)

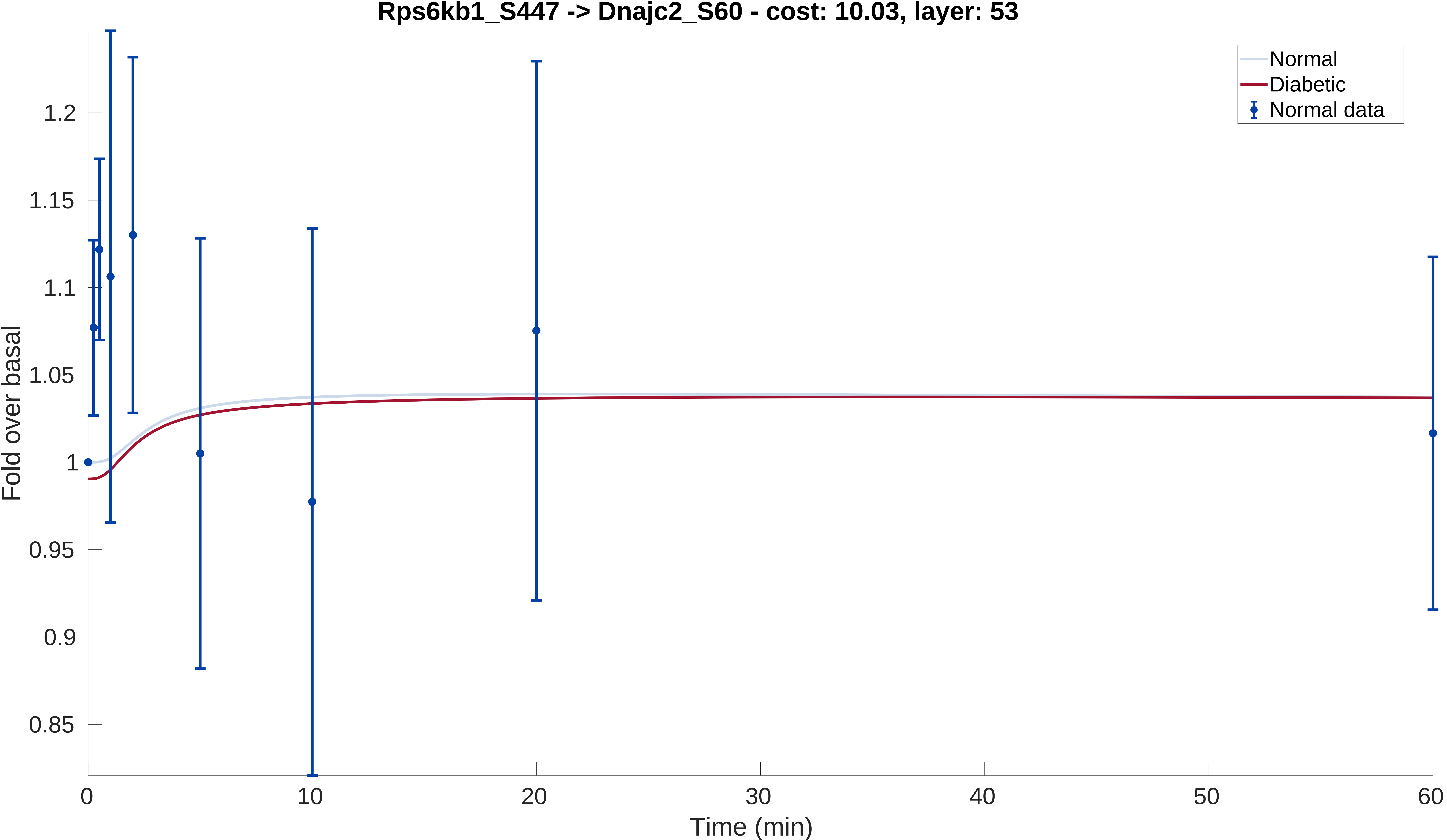

Rps6kb1_S447 -> Dnajc2_S60 - cost: 10.03, layer: 53
Normal
Diabetic
Normal data
Fold over basal
1.2
1.15
1.1
1.05
1
0.95
0.9
0.85
0
10
20
30
40
50
60
Time (min)

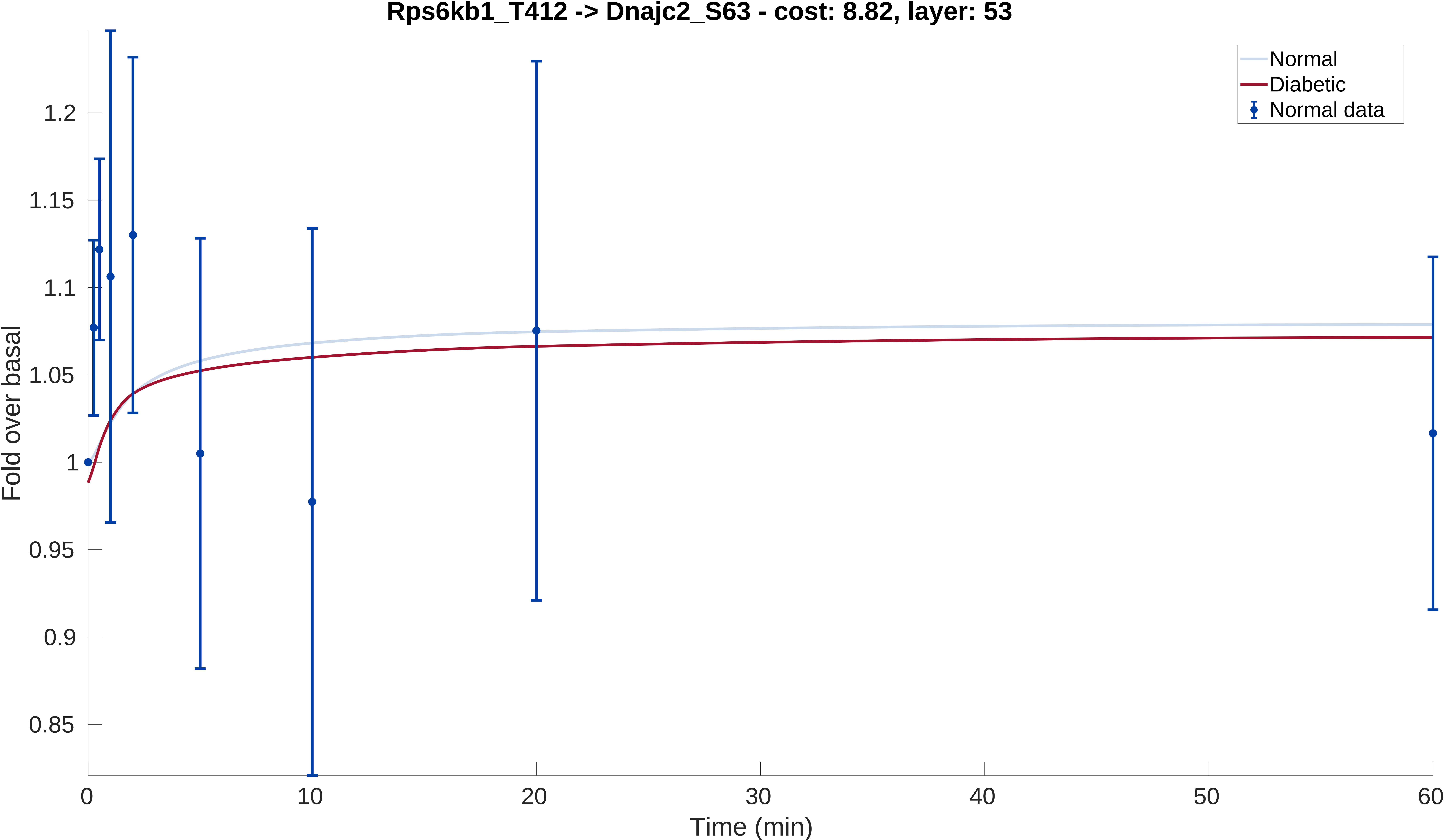

Rps6kb1_T412 -> Dnajc2_S63 - cost: 8.82, layer: 53
Normal
Diabetic
Normal data
1.2
1.15
1.1
1.05
1
0.95
0.9
0.85
Fold over basal
0
10
20
30
40
50
60
Time (min)

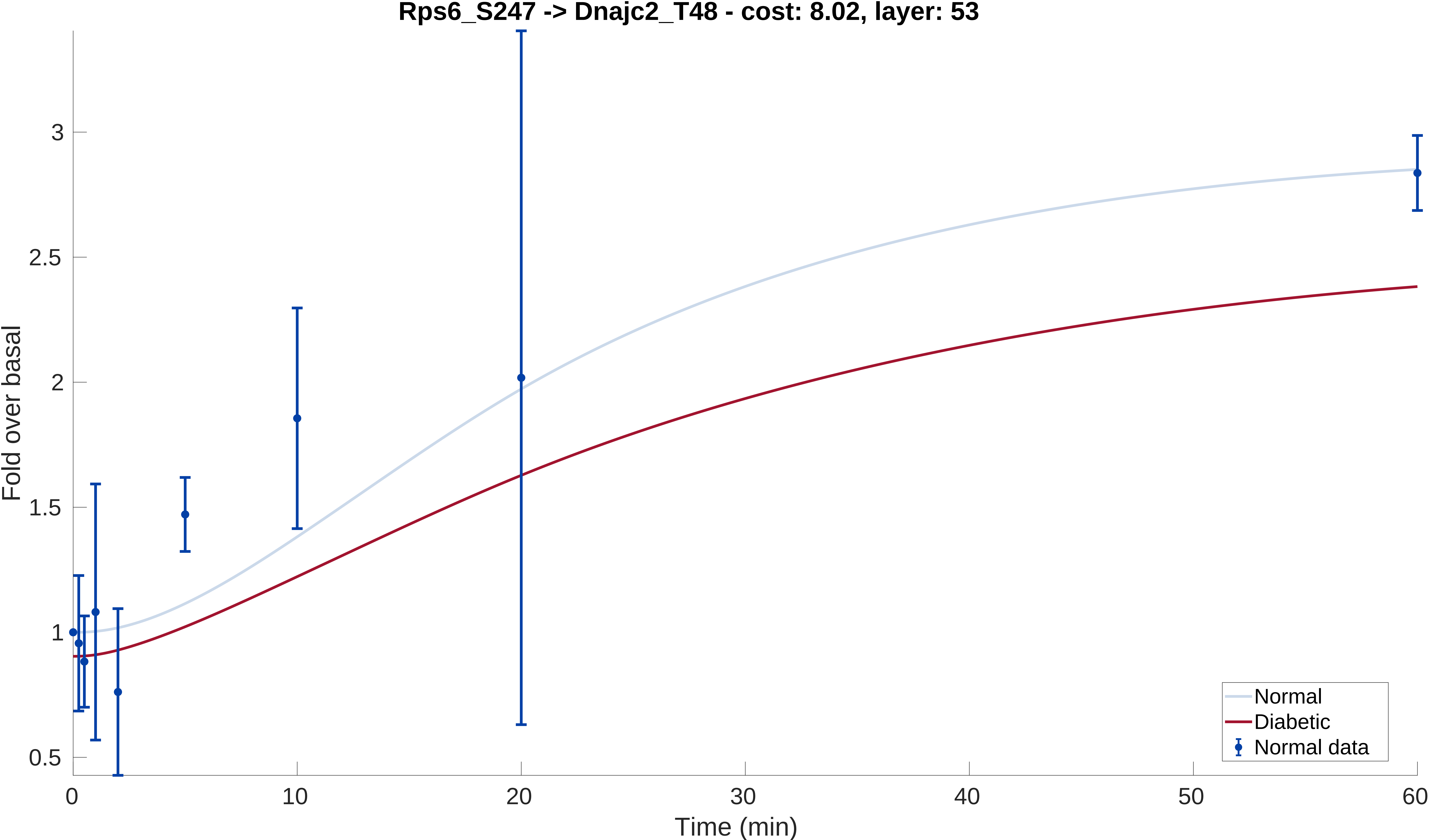

Rps6_S247 -> Dnajc2_T48 - cost: 8.02, layer: 53
Fold over basal
Time (min)
0.5
1
1.5
2
2.5
3
0
10
20
30
40
50
60
Normal
Diabetic
Normal data

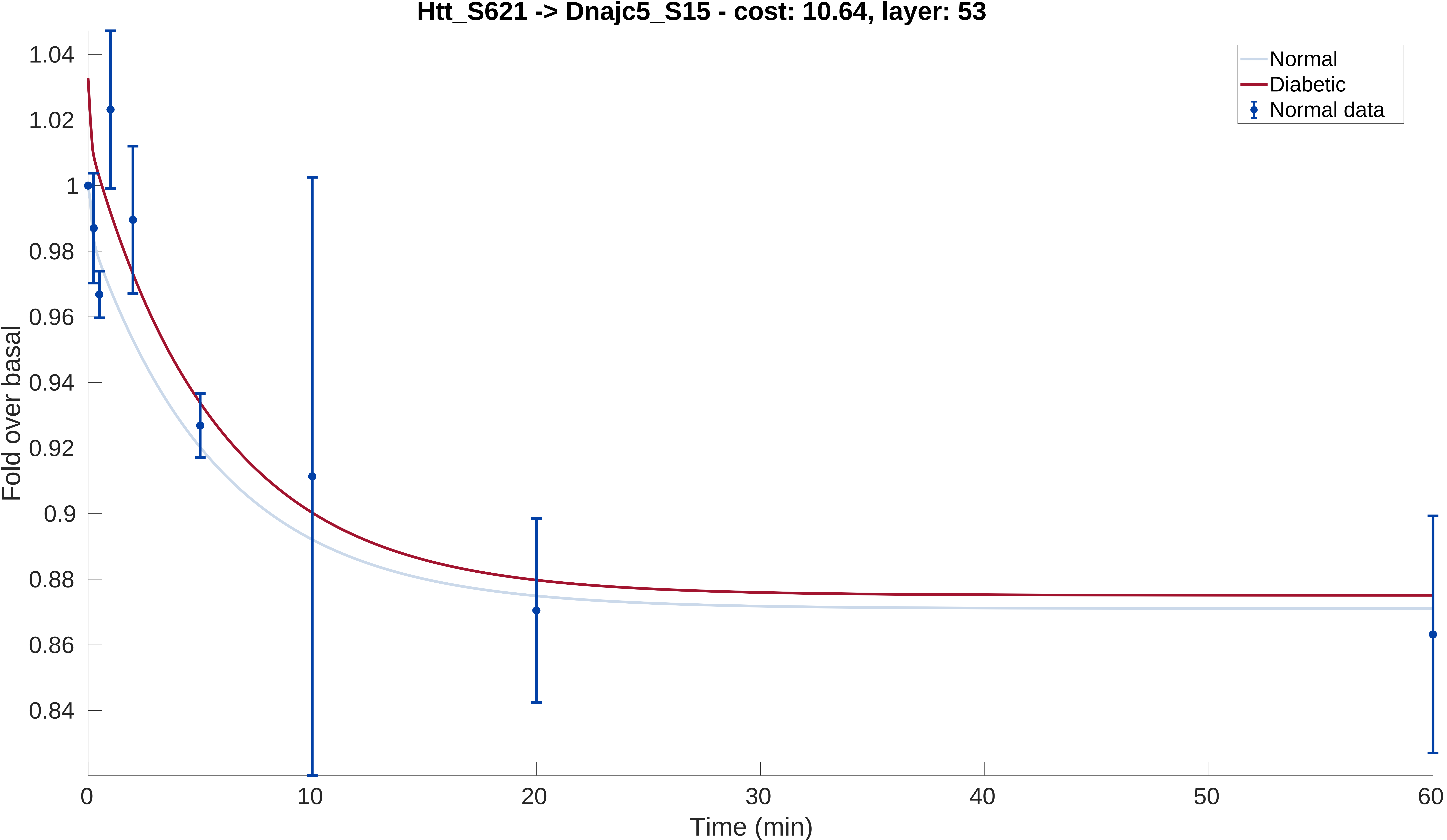

Htt_S621 -> Dnajc5_S15 - cost: 10.64, layer: 53
Normal
Diabetic
Normal data
Fold over basal
Time (min)
1.04
1.02
1
0.98
0.96
0.94
0.92
0.9
0.88
0.86
0.84
0
10
20
30
40
50
60

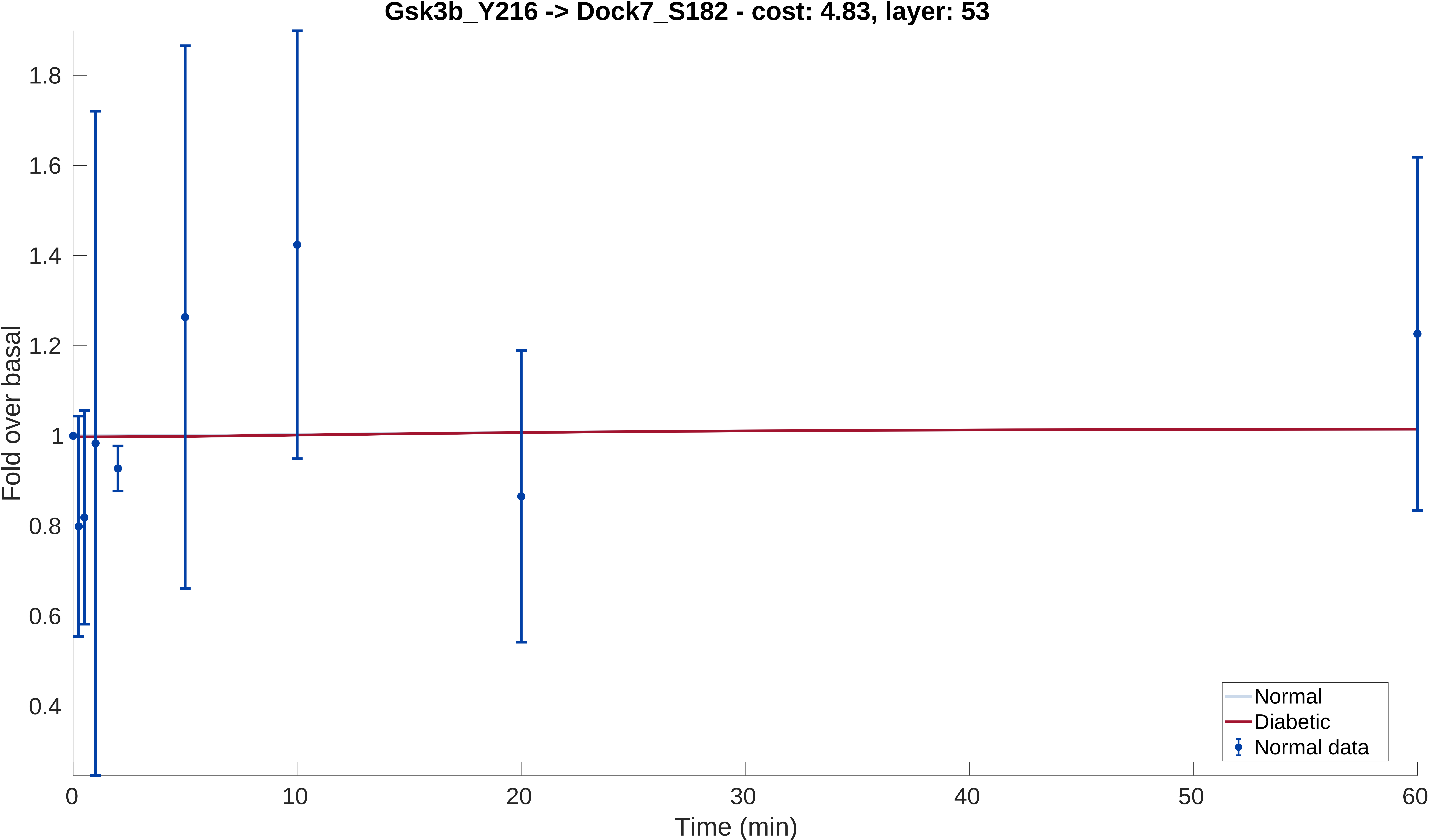

Gsk3b_Y216 -> Dock7_S182 - cost: 4.83, layer: 53
Fold over basal
1.8
1.6
1.4
1.2
1
0.8
0.6
0.4
0
10
20
30
40
50
60
Time (min)
Normal
Diabetic
Normal data

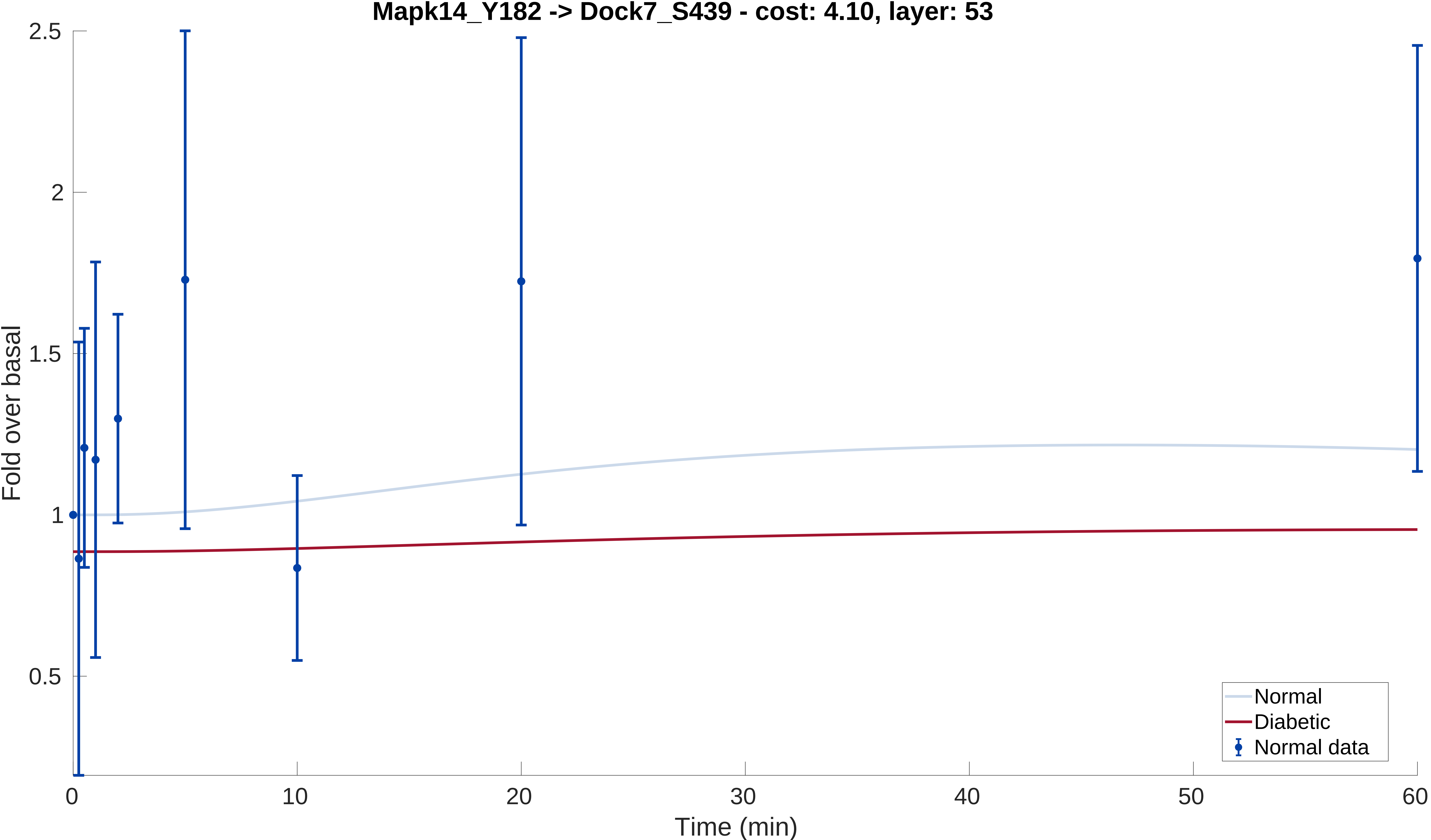

Mapk14_Y182 -> Dock7_S439 - cost: 4.10, layer: 53
Fold over basal
2.5
2
1.5
1
0.5
0
10
20
30
40
50
60
Time (min)
Normal
Diabetic
Normal data

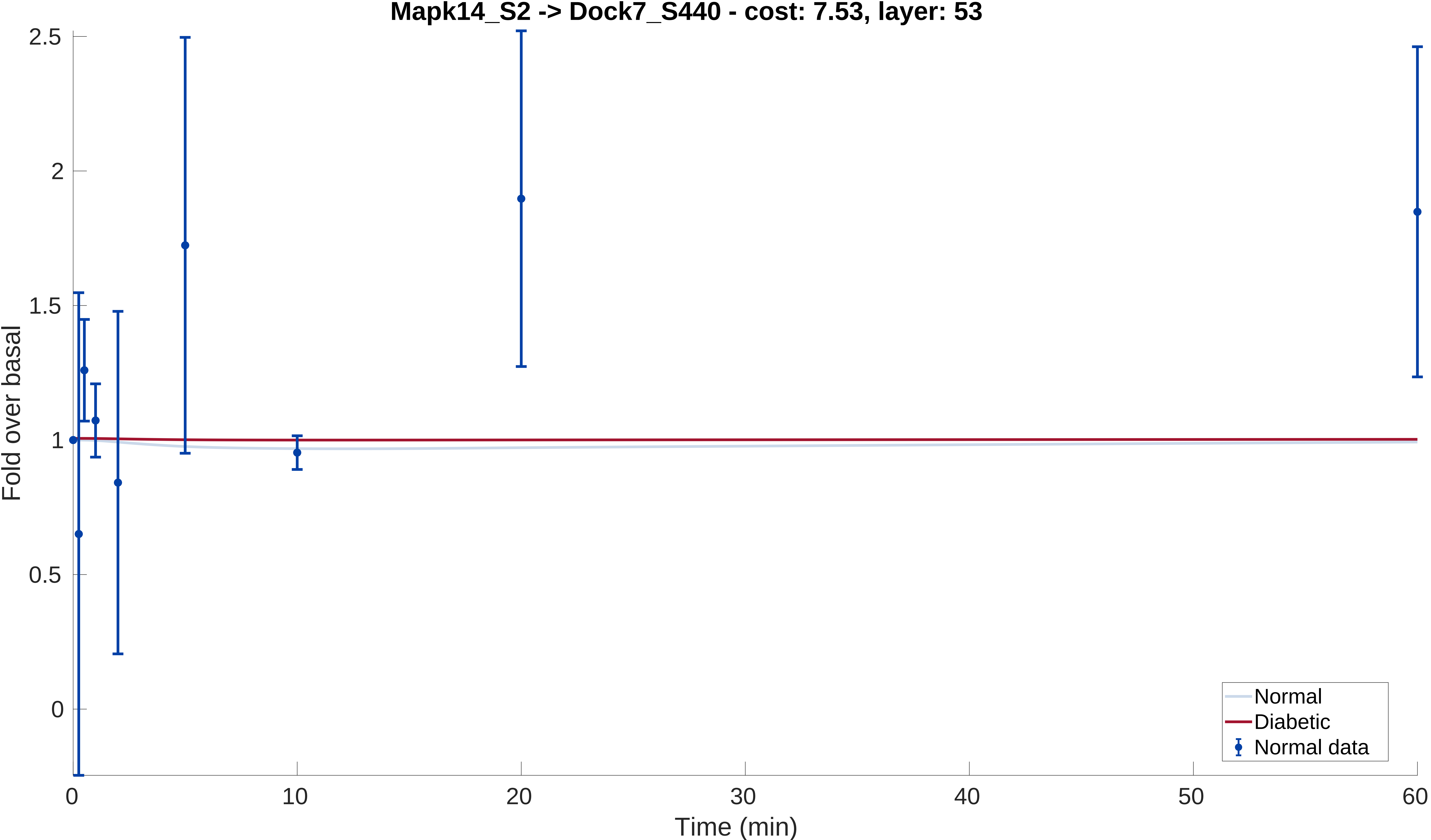

Mapk14_S2 -> Dock7_S440 - cost: 7.53, layer: 53
Fold over basal
Time (min)
2.5
2
1.5
1
0.5
0
0
10
20
30
40
50
60
Normal
Diabetic
Normal data

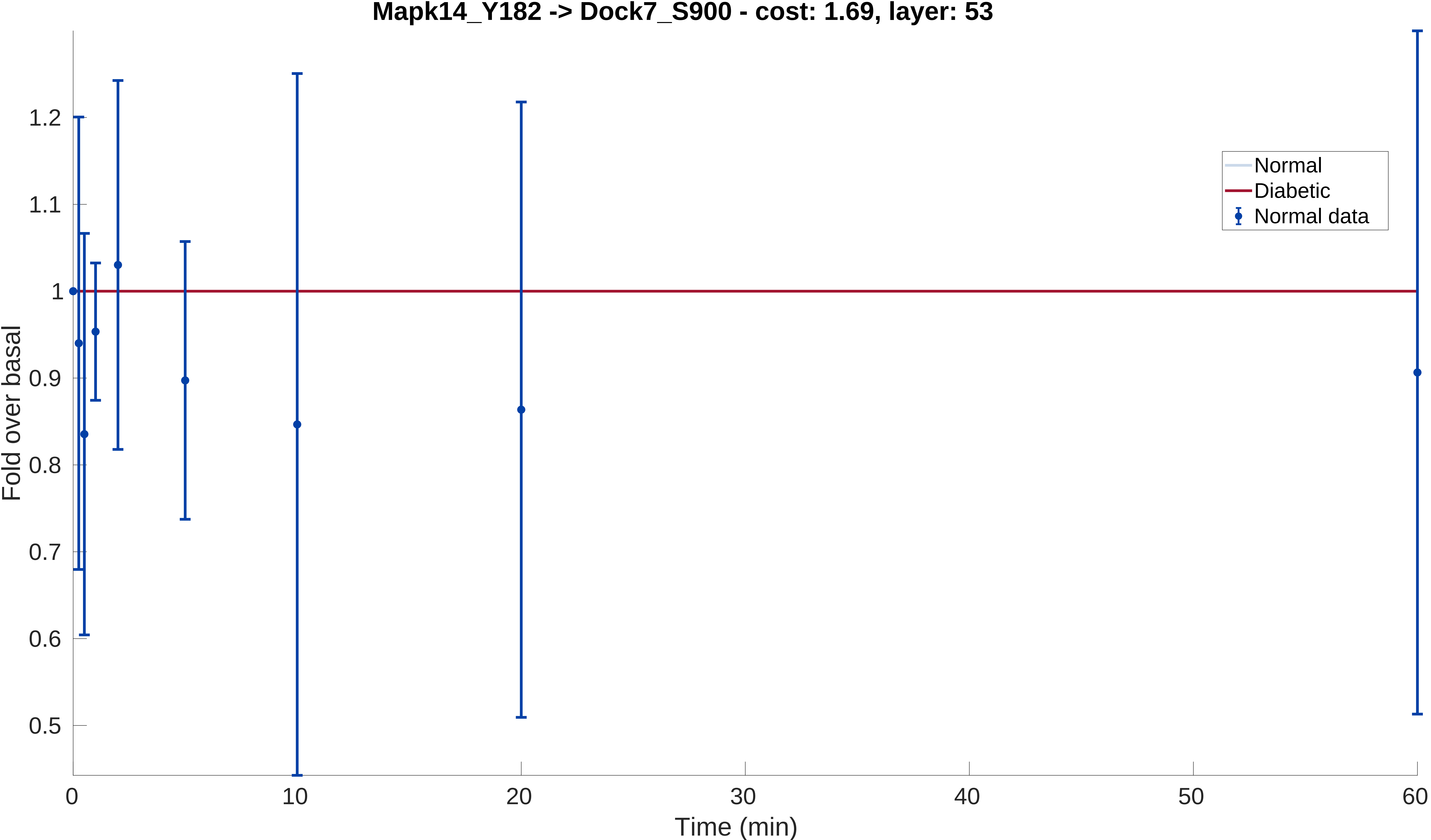

Mapk14_Y182 -> Dock7_S900 - cost: 1.69, layer: 53
Fold over basal
Time (min)
0
10
20
30
40
50
60
0.5
0.6
0.7
0.8
0.9
1
1.1
1.2
Normal
Diabetic
Normal data

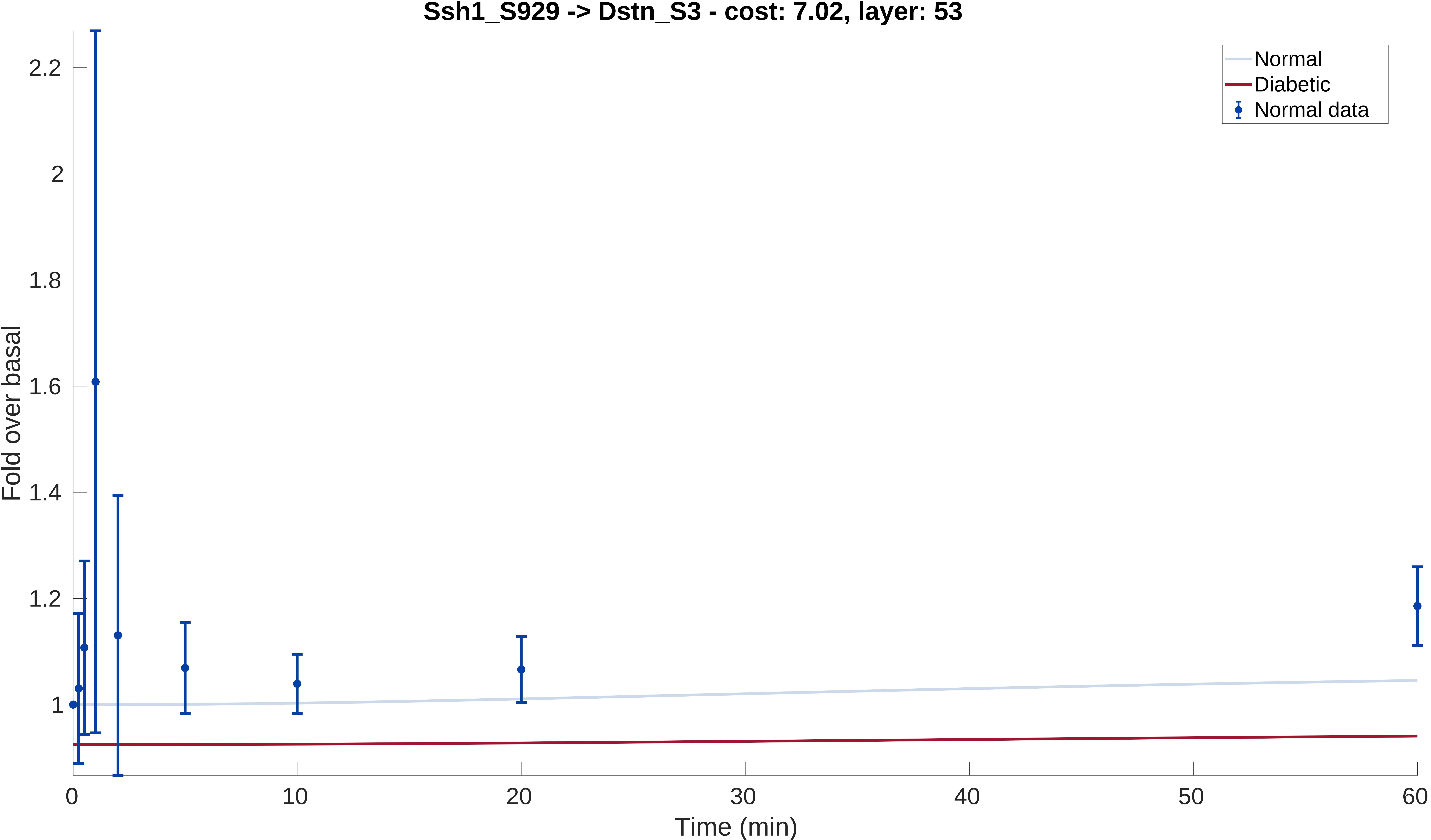

Ssh1_S929 -> Dstn_S3 - cost: 7.02, layer: 53
Fold over basal
Time (min)
Normal
Diabetic
Normal data
2.2
2
1.8
1.6
1.4
1.2
1
0
10
20
30
40
50
60

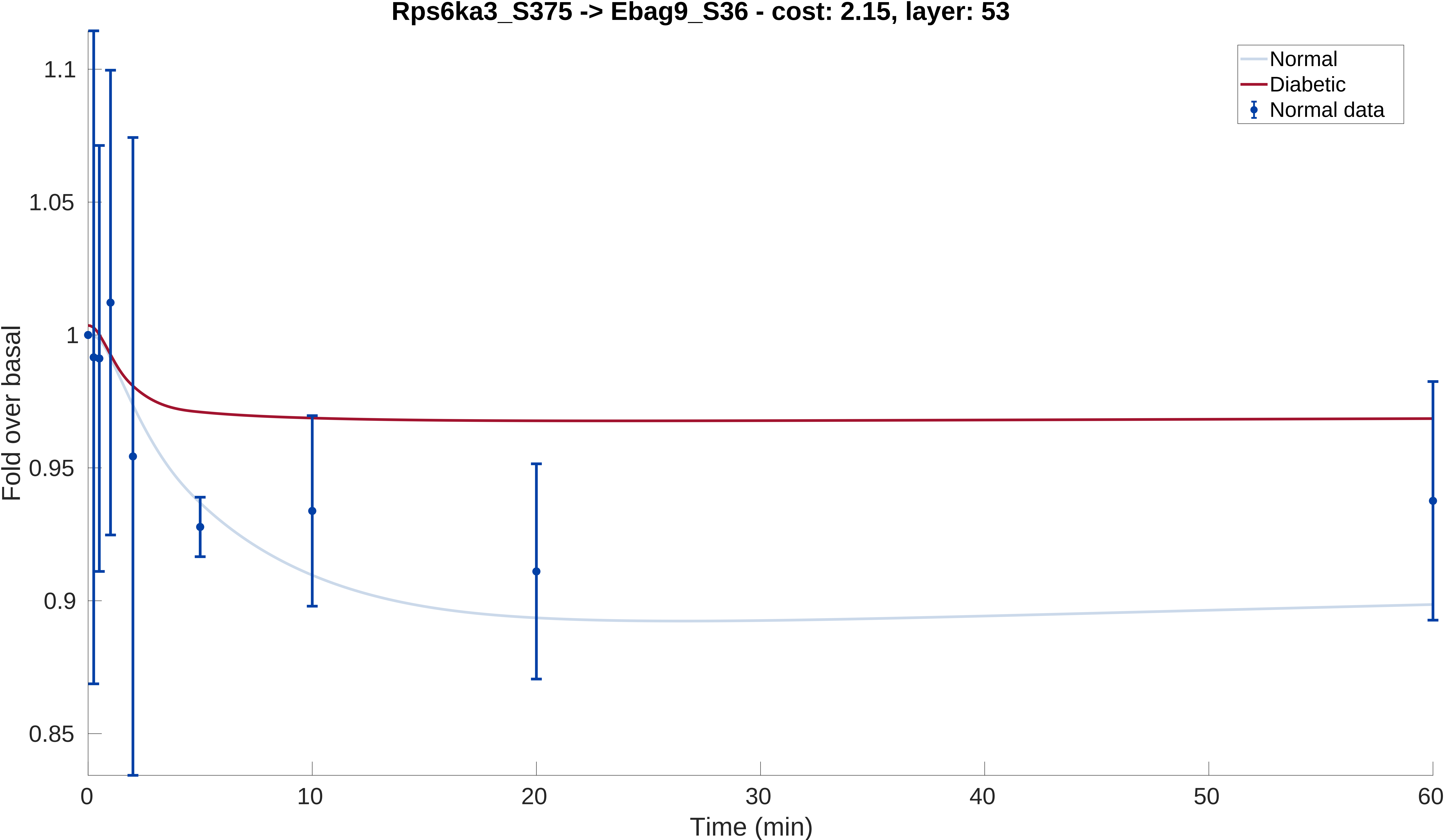

Rps6ka3_S375 -> Ebag9_S36 - cost: 2.15, layer: 53
Normal
Diabetic
Normal data
Fold over basal
1.1
1.05
1
0.95
0.9
0.85
0
10
20
30
40
50
60
Time (min)

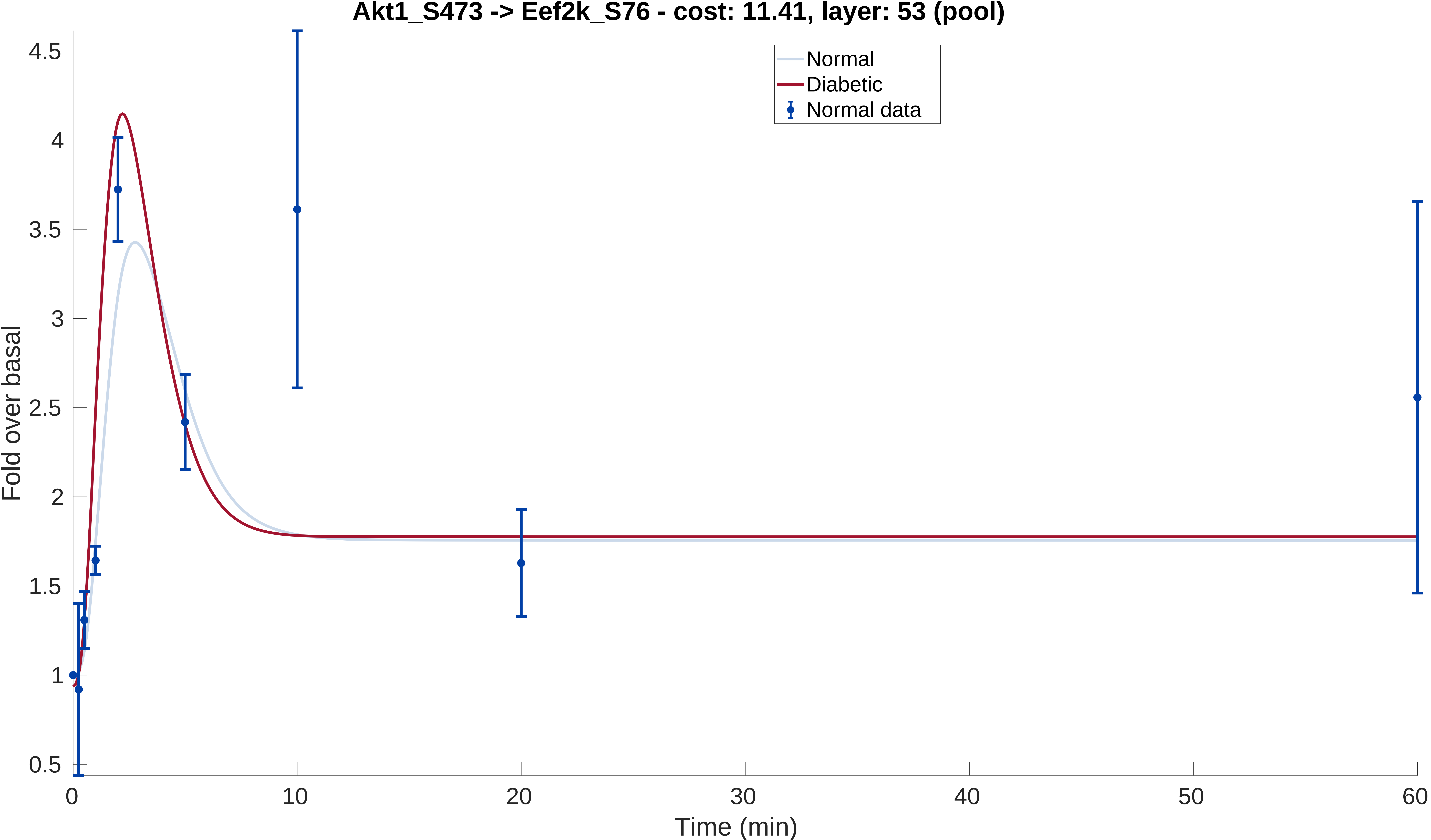

Akt1_S473 -> Eef2k_S76 - cost: 11.41, layer: 53 (pool)
Normal
Diabetic
Normal data
Fold over basal
4.5
4
3.5
3
2.5
2
1.5
1
0.5
0
10
20
30
40
50
60
Time (min)

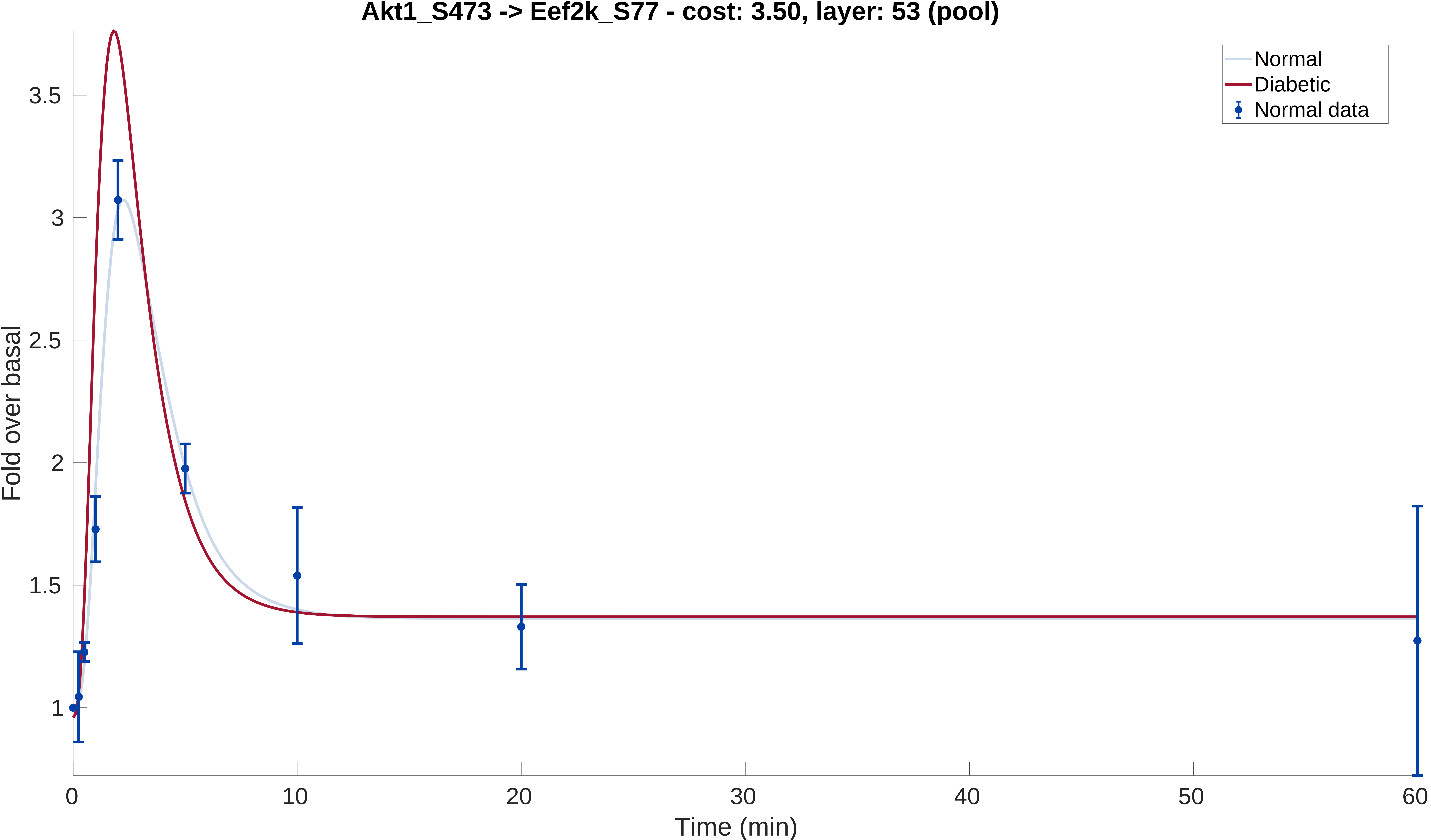

Akt1_S473 -> Eef2k_S77 - cost: 3.50, layer: 53 (pool)
Normal
Diabetic
Normal data
Fold over basal
1
1.5
2
2.5
3
3.5
0
10
20
30
40
50
60
Time (min)

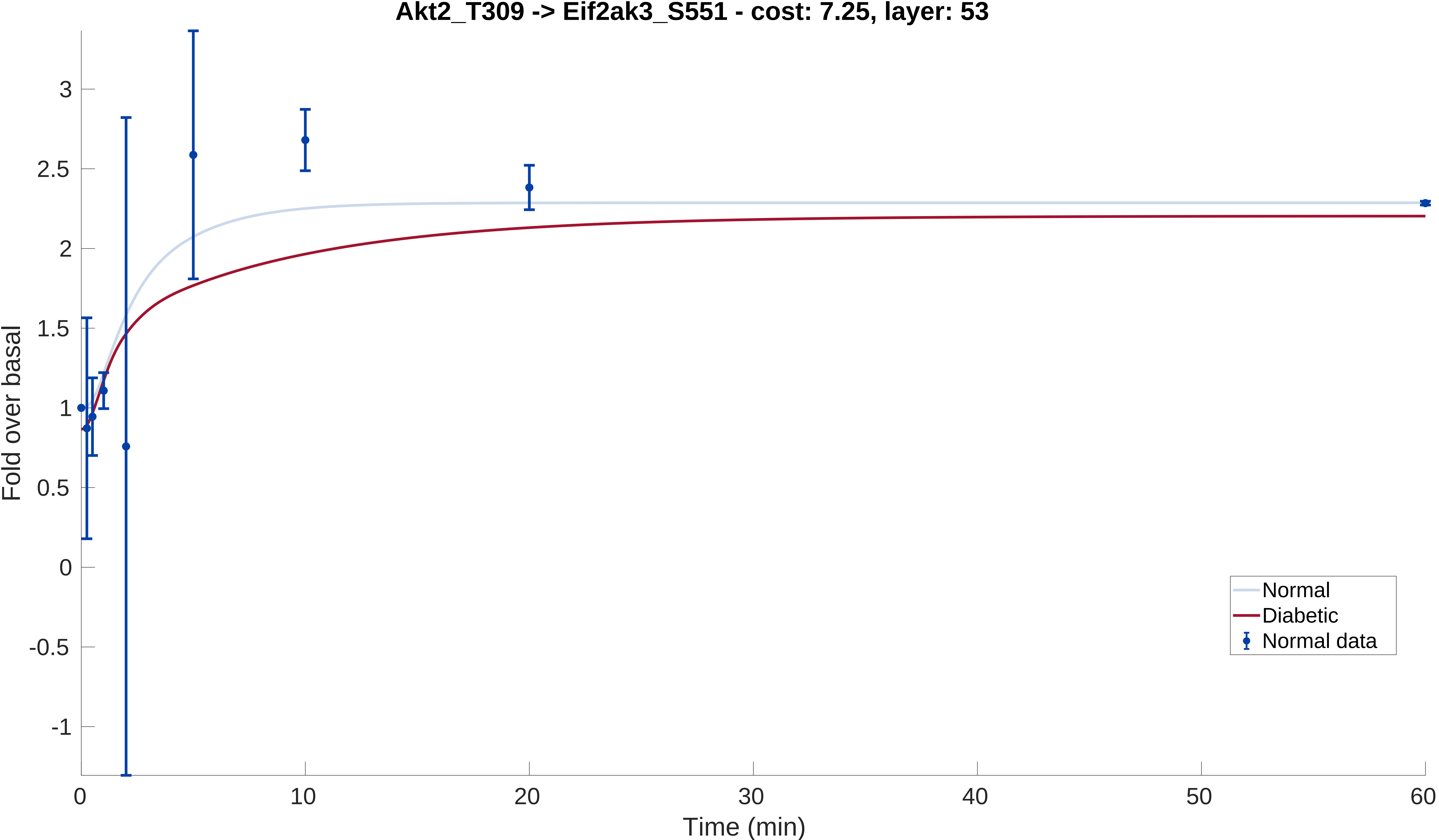

Akt2_T309 -> Eif2ak3_S551 - cost: 7.25, layer: 53
Fold over basal
Time (min)
3
2.5
2
1.5
1
0.5
0
-0.5
-1
0
10
20
30
40
50
60
Normal
Diabetic
Normal data

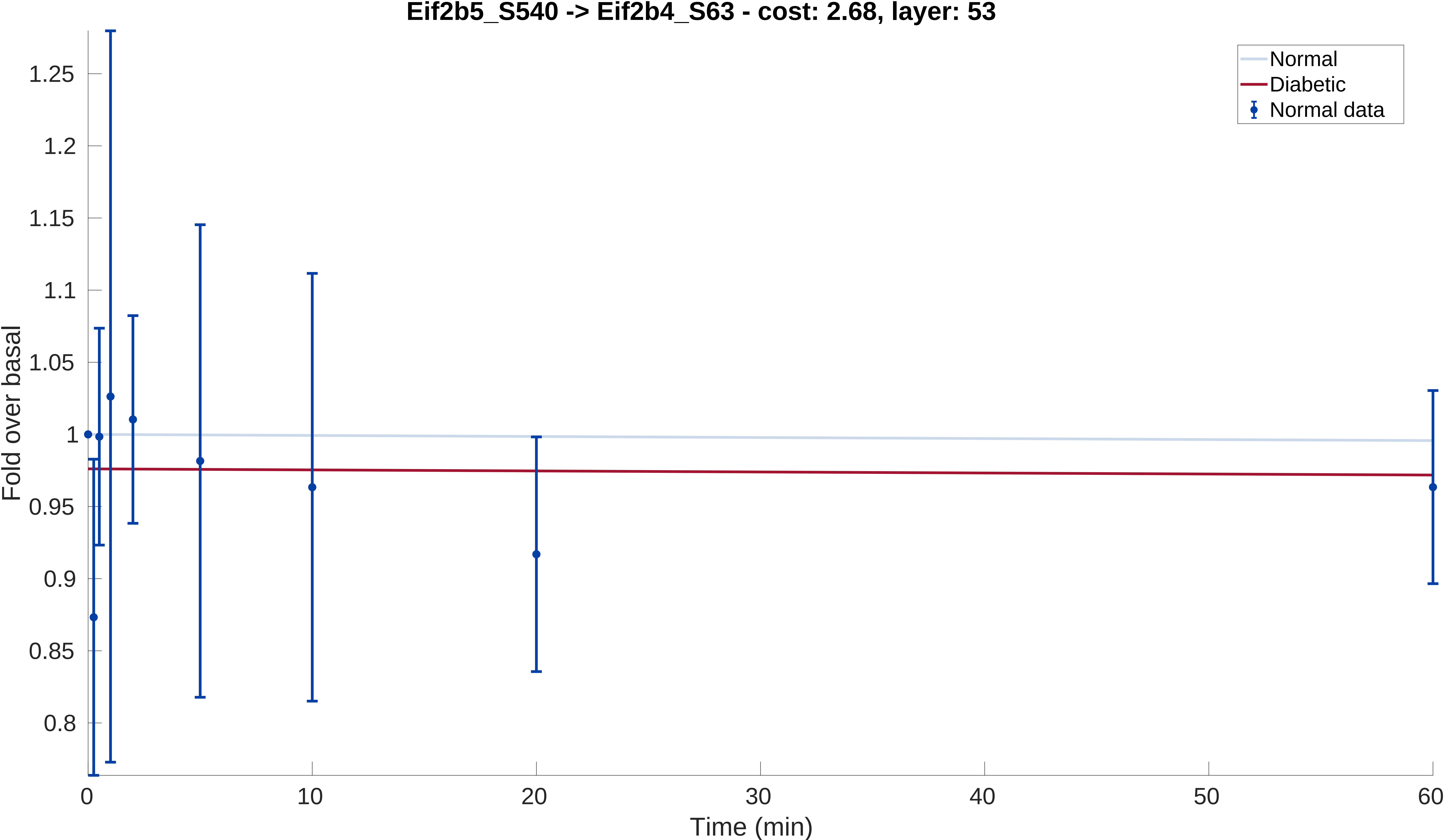

Eif2b5_S540 -> Eif2b4_S63 - cost: 2.68, layer: 53
Normal
Diabetic
Normal data
Fold over basal
1.25
1.2
1.15
1.1
1.05
1
0.95
0.9
0.85
0.8
0
10
20
30
40
50
60
Time (min)

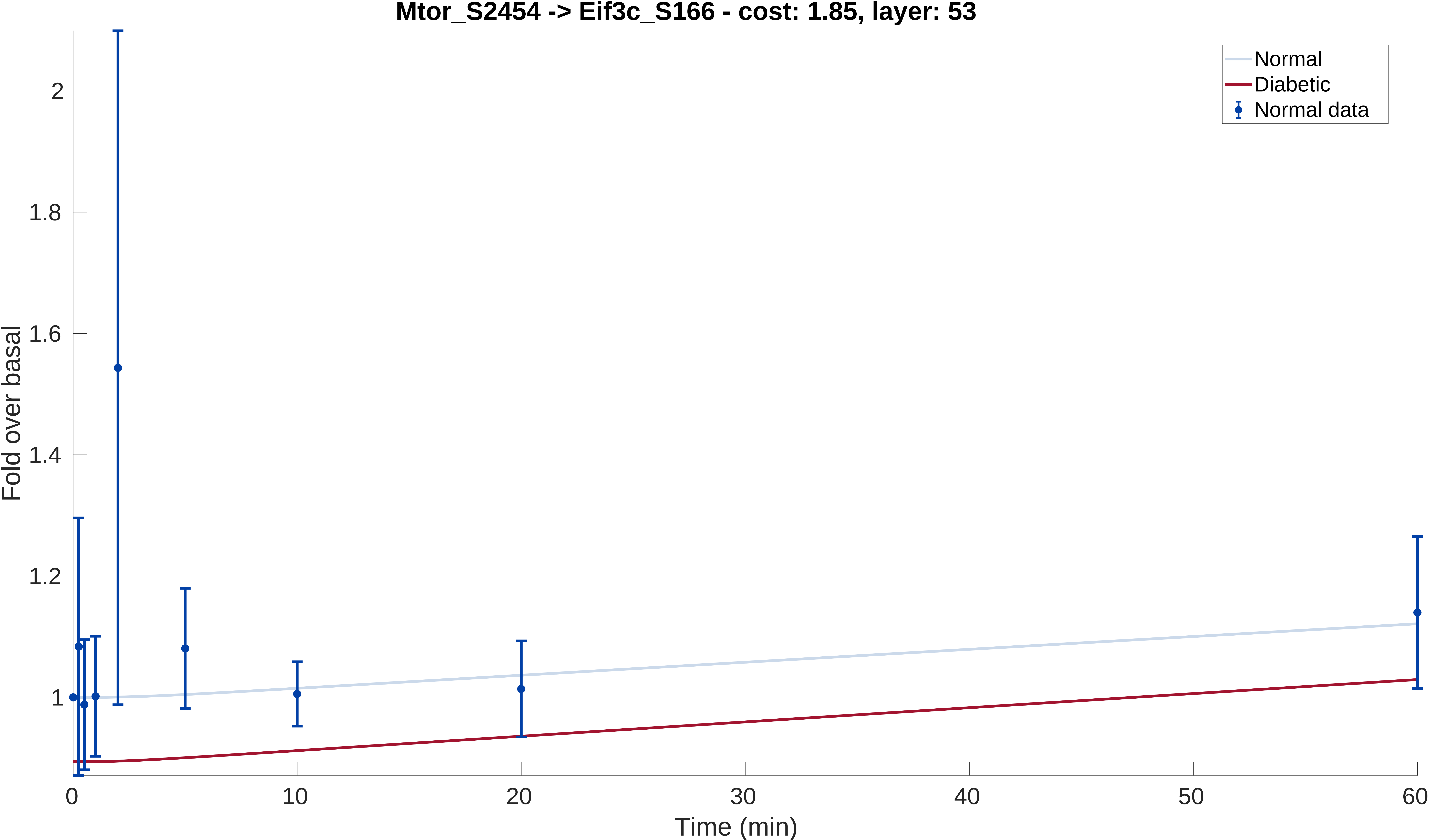
Mtor_S2454 -> Eif3c_S166 - cost: 1.85, layer: 53
Normal
Diabetic
Normal data
Fold over basal
2
1.8
1.6
1.4
1.2
1
0
10
20
30
40
50
60
Time (min)

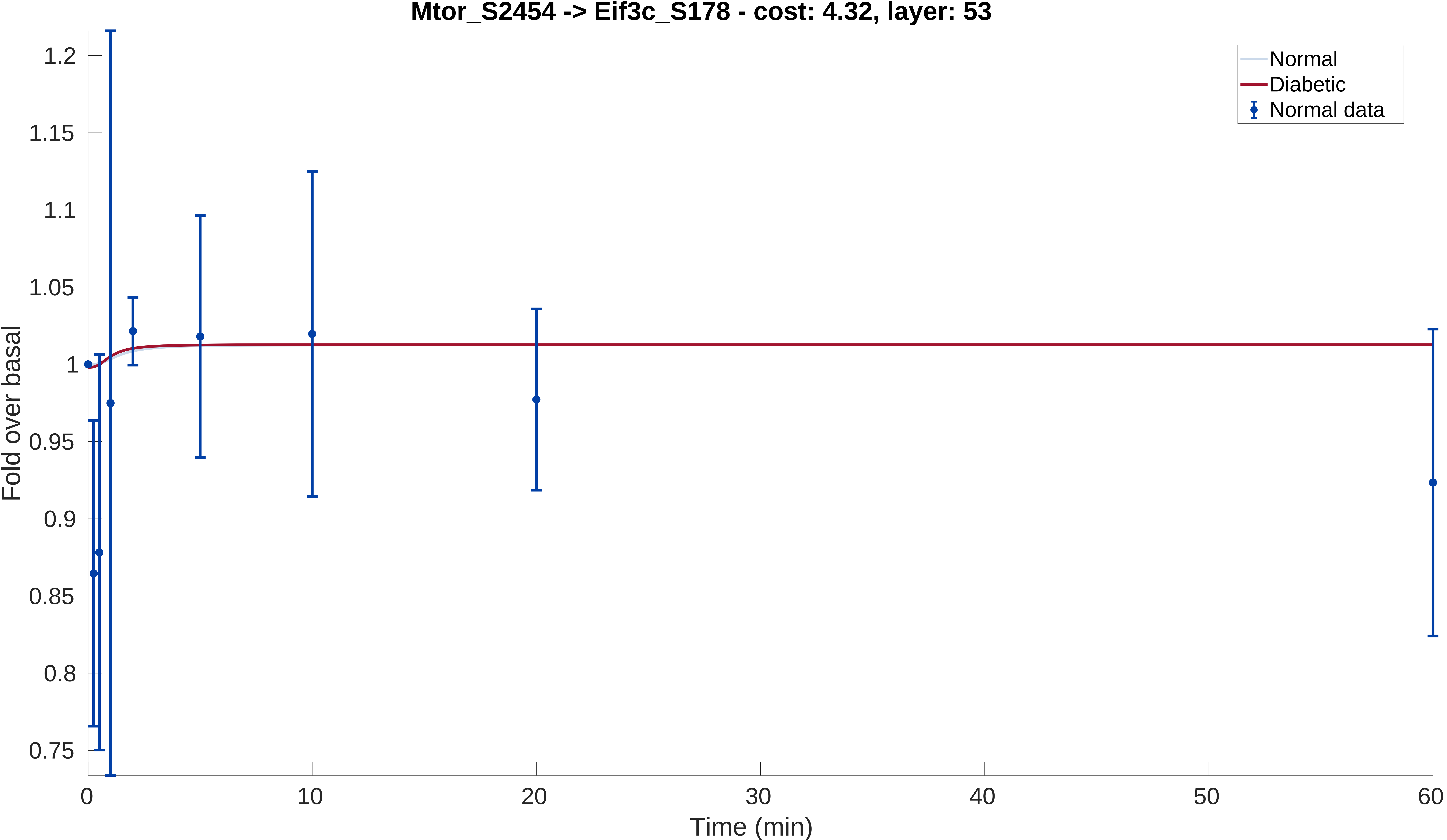

Mtor_S2454 -> Eif3c_S178 - cost: 4.32, layer: 53
Normal
Diabetic
Normal data
Fold over basal
Time (min)
1.2
1.15
1.1
1.05
1
0.95
0.9
0.85
0.8
0.75
0
10
20
30
40
50
60

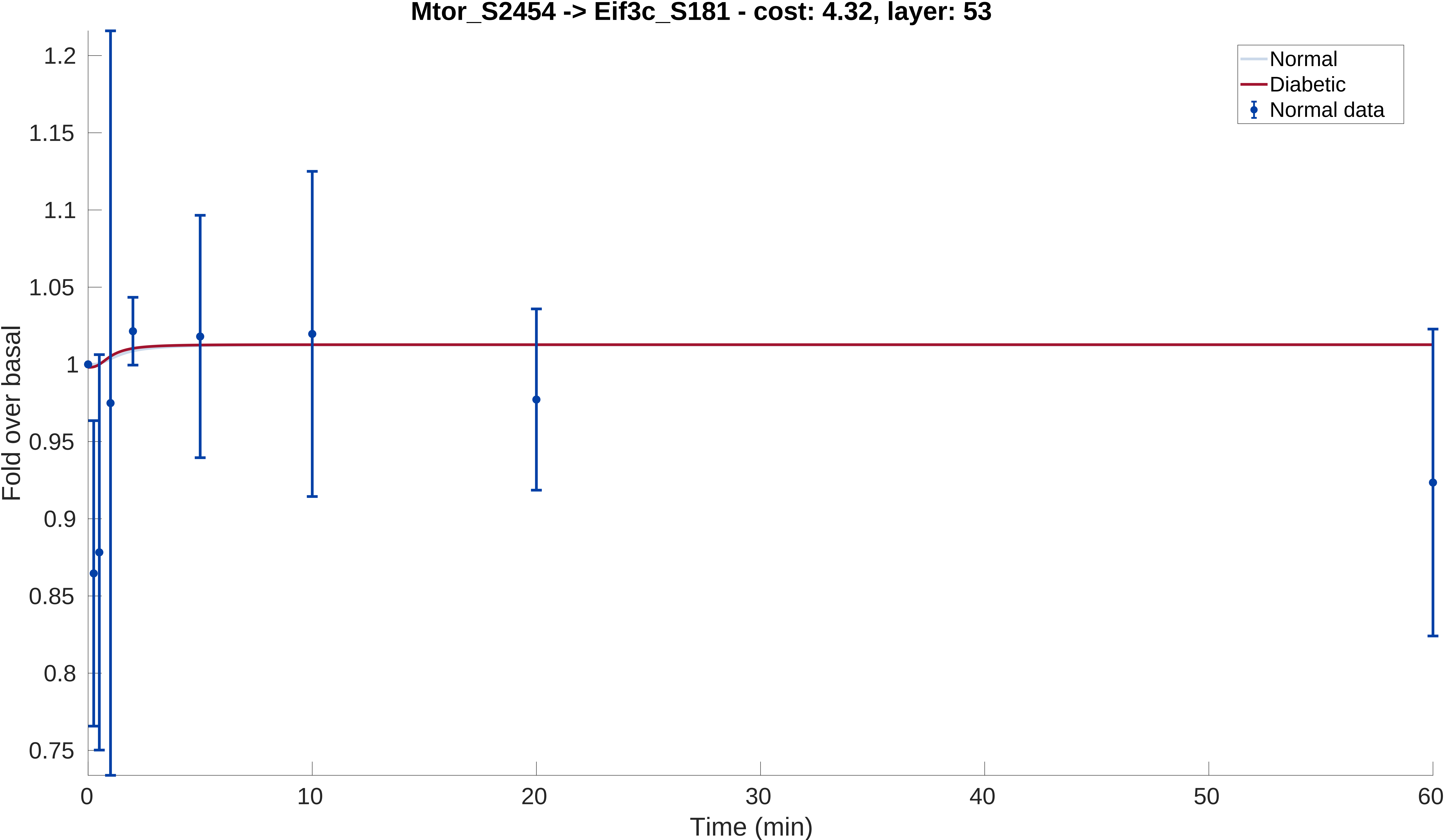

Mtor_S2454 -> Eif3c_S181 - cost: 4.32, layer: 53
Normal
Diabetic
Normal data
Fold over basal
Time (min)
1.2
1.15
1.1
1.05
1
0.95
0.9
0.85
0.8
0.75
0
10
20
30
40
50
60

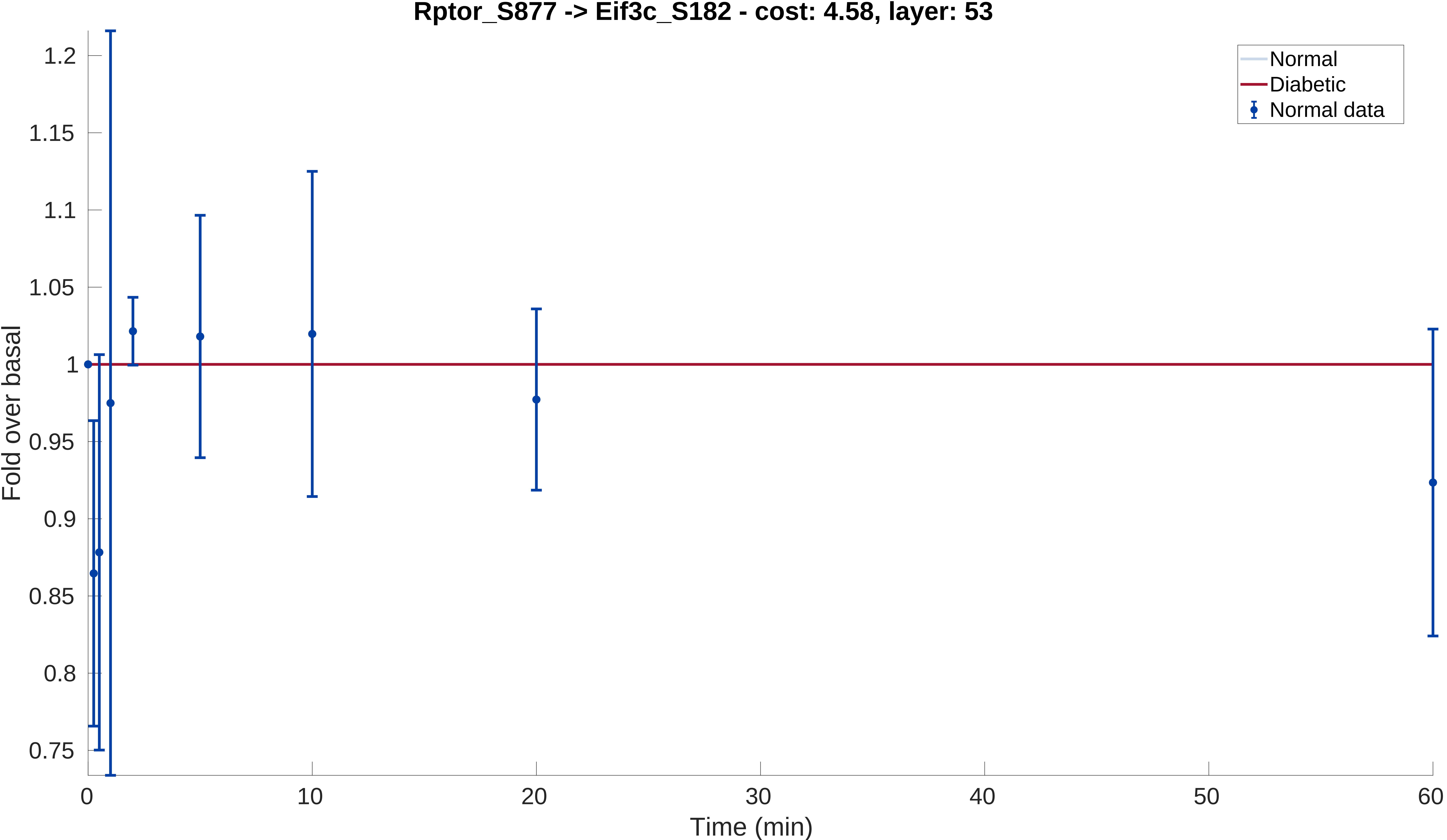

Rptor_S877 -> Eif3c_S182 - cost: 4.58, layer: 53
Normal
Diabetic
Normal data
Fold over basal
1.2
1.15
1.1
1.05
1
0.95
0.9
0.85
0.8
0.75
0
10
20
30
40
50
60
Time (min)

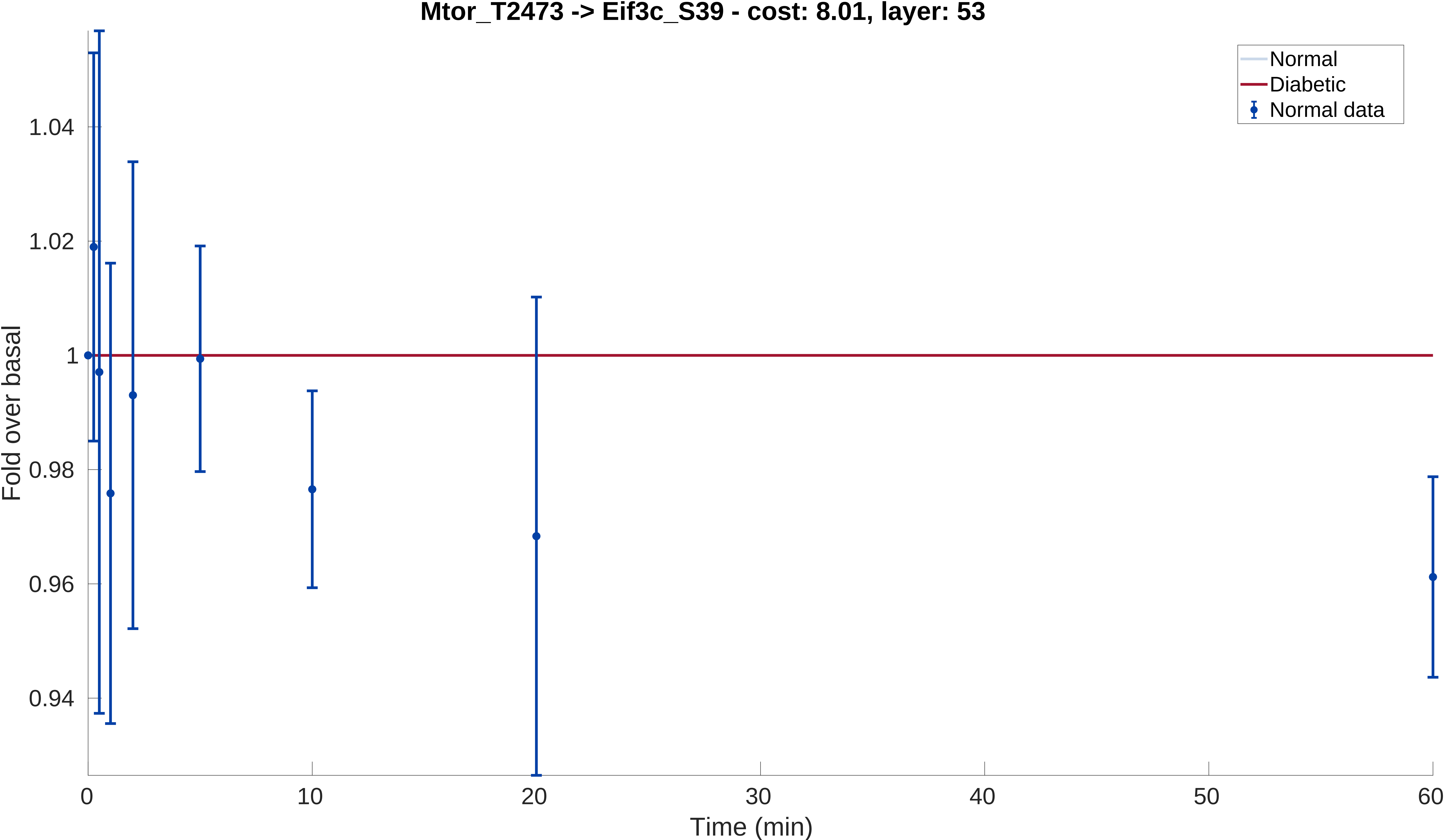

Mtor_T2473 -> Eif3c_S39 - cost: 8.01, layer: 53
Normal
Diabetic
Normal data
Fold over basal
1.04
1.02
1
0.98
0.96
0.94
0
10
20
30
40
50
60
Time (min)

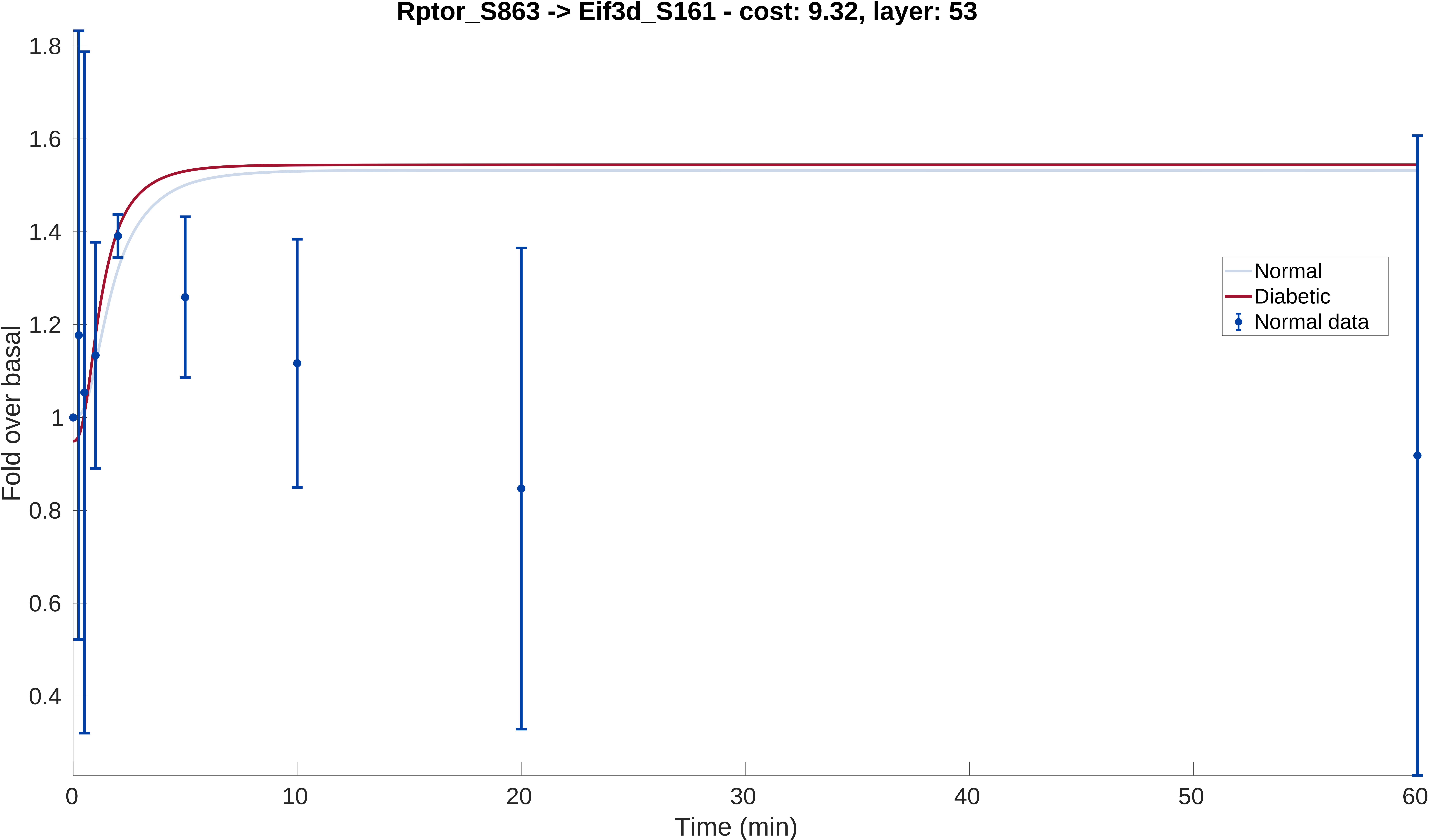

Rptor_S863 -> Eif3d_S161 - cost: 9.32, layer: 53
Fold over basal
1.8
1.6
1.4
1.2
1
0.8
0.6
0.4
0
10
20
30
40
50
60
Time (min)
Normal
Diabetic
Normal data

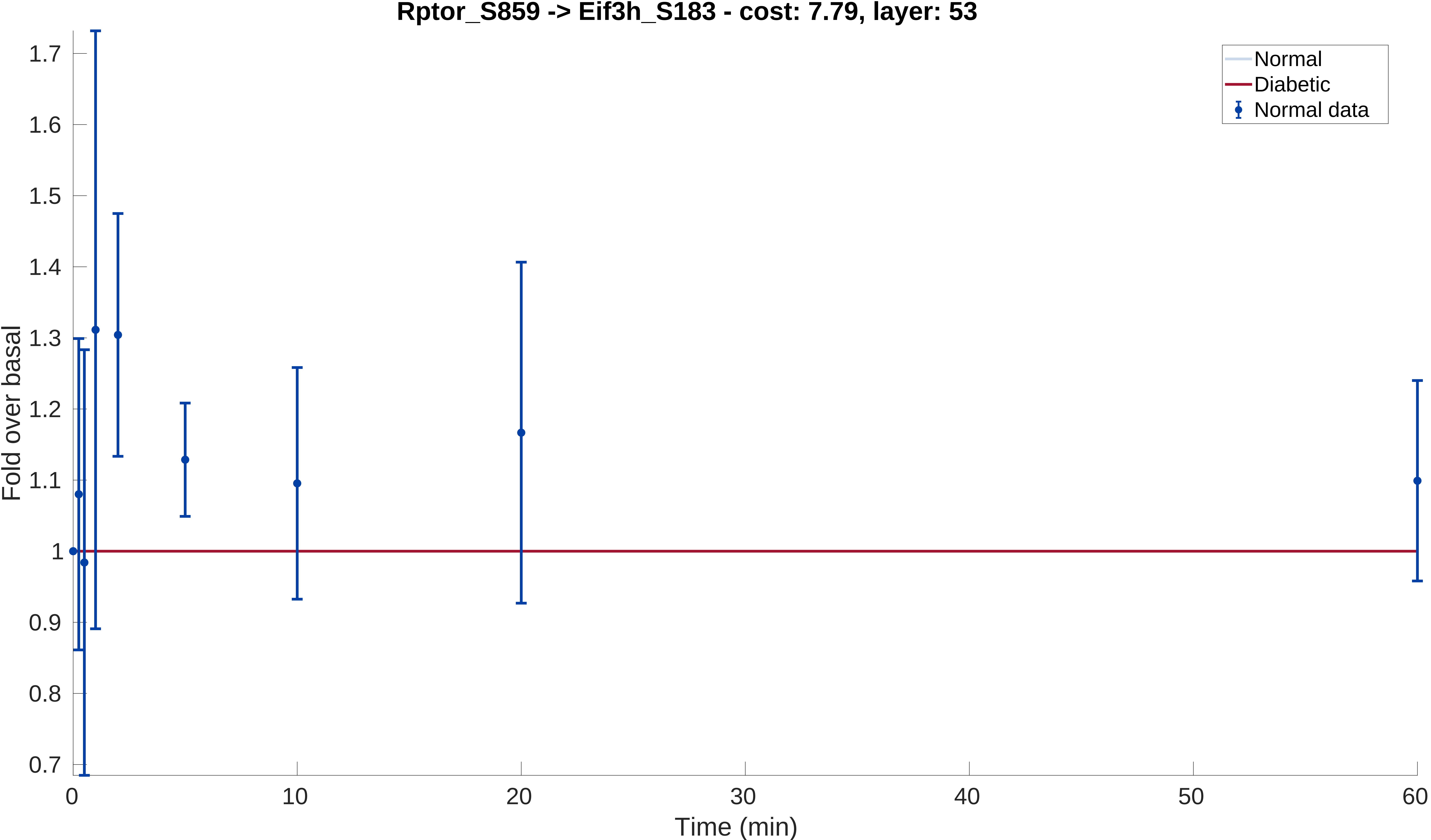

Rptor_S859 -> Eif3h_S183 - cost: 7.79, layer: 53
Normal
Diabetic
Normal data
Fold over basal
1.7
1.6
1.5
1.4
1.3
1.2
1.1
1
0.9
0.8
0.7
0
10
20
30
40
50
60
Time (min)

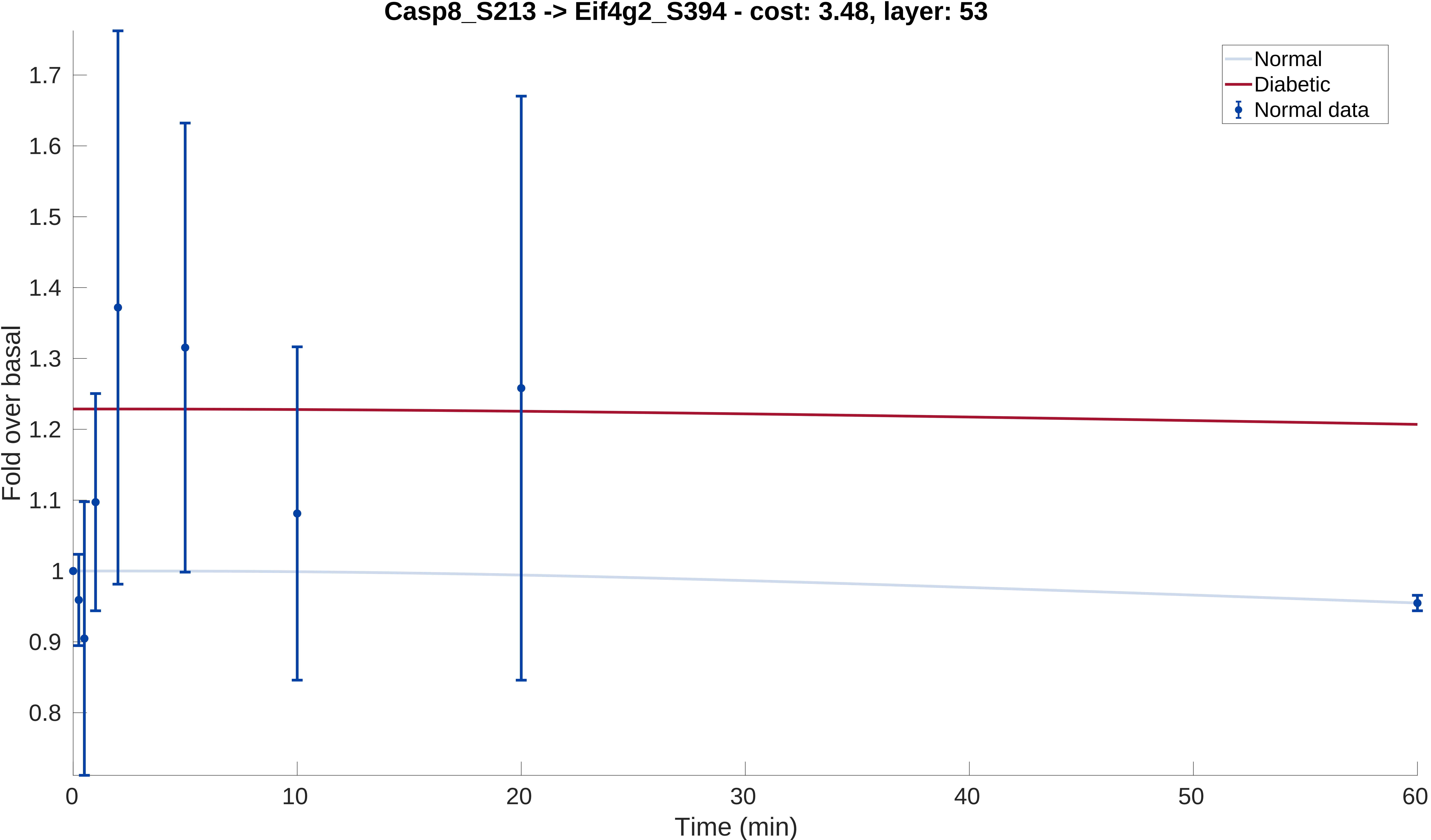

Casp8_S213 -> Eif4g2_S394 - cost: 3.48, layer: 53
Normal
Diabetic
Normal data
1.7
1.6
1.5
1.4
1.3
1.2
1.1
1
0.9
0.8
Fold over basal
0
10
20
30
40
50
60
Time (min)

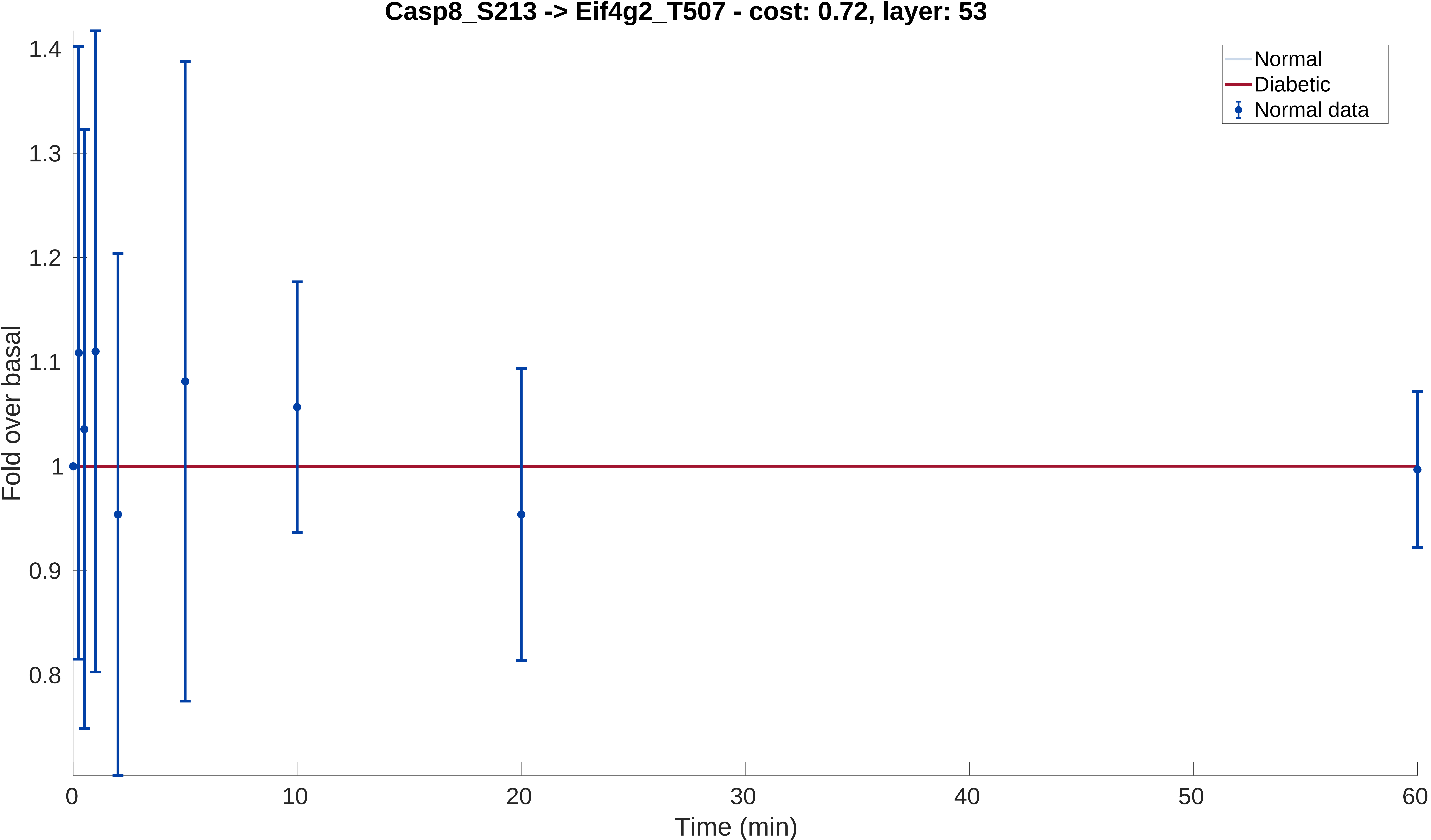

Casp8_S213 -> Eif4g2_T507 - cost: 0.72, layer: 53
Normal
Diabetic
Normal data
Fold over basal
Time (min)
1.4
1.3
1.2
1.1
1
0.9
0.8
0
10
20
30
40
50
60

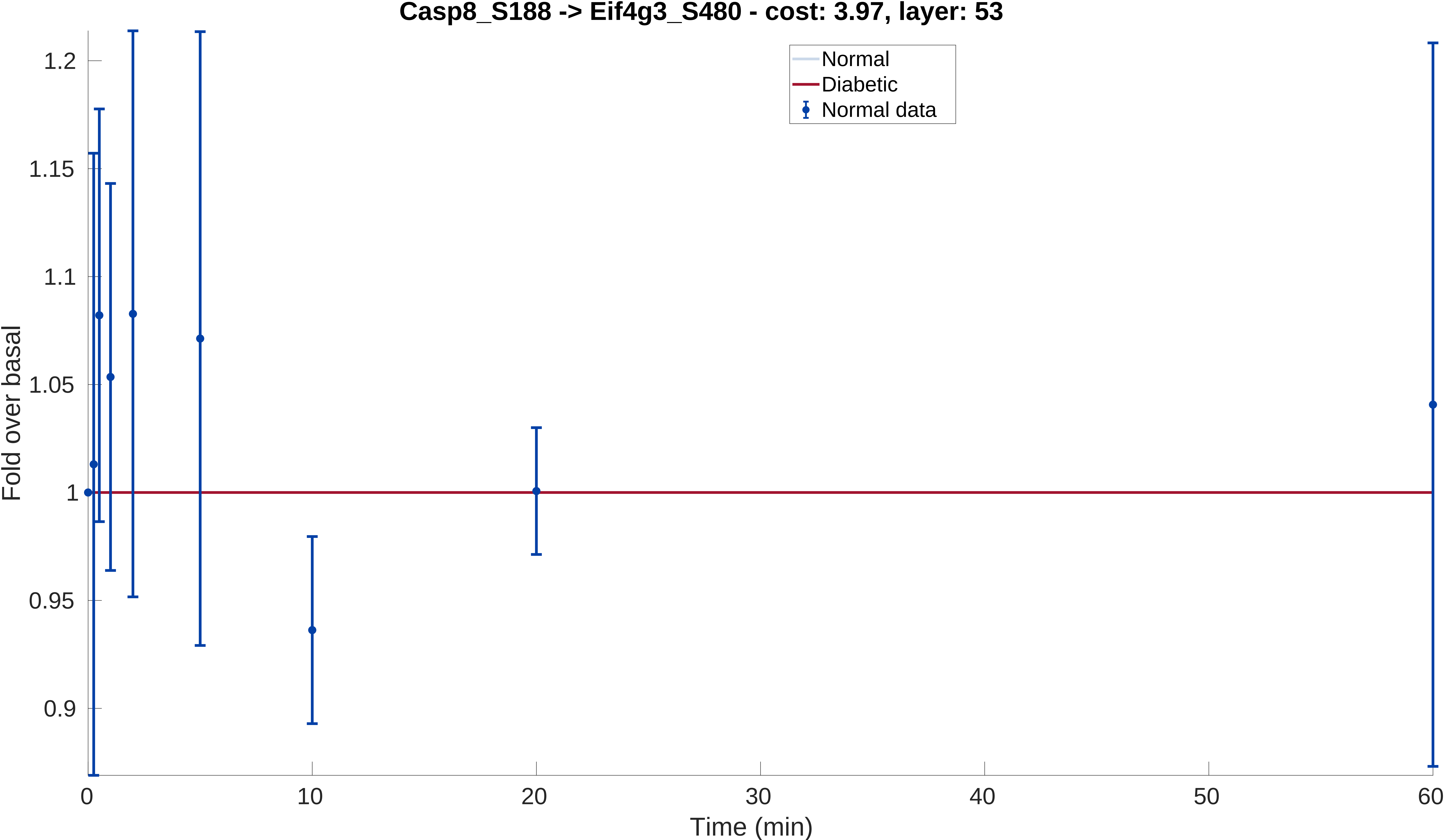
Casp8_S188 -> Eif4g3_S480 - cost: 3.97, layer: 53
Normal
Diabetic
Normal data
Fold over basal
1.2
1.15
1.1
1.05
1
0.95
0.9
0
10
20
30
40
50
60
Time (min)

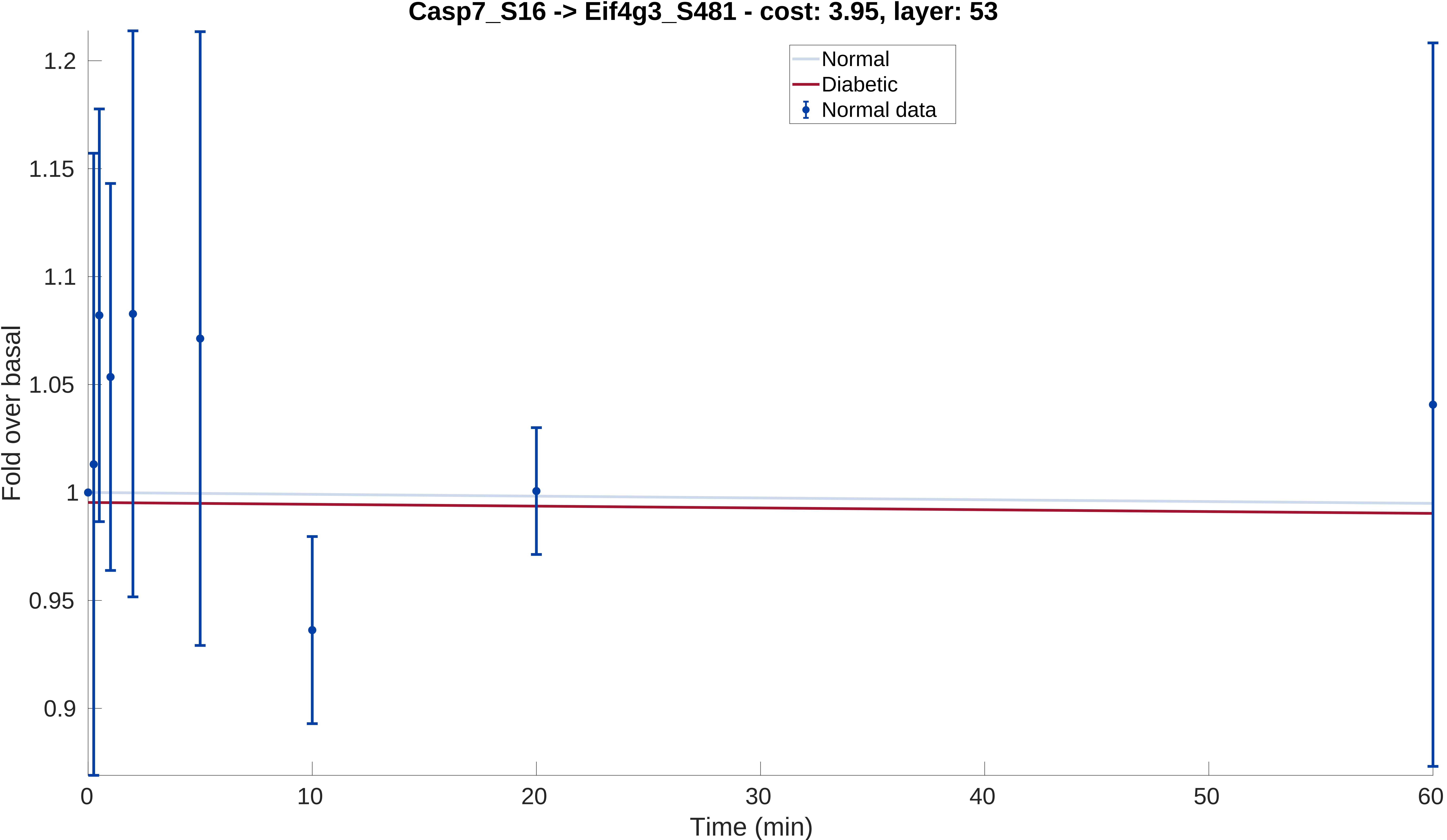

Casp7_S16 -> Eif4g3_S481 - cost: 3.95, layer: 53
Normal
Diabetic
Normal data
Fold over basal
Time (min)
1.2
1.15
1.1
1.05
1
0.95
0.9
0
10
20
30
40
50
60

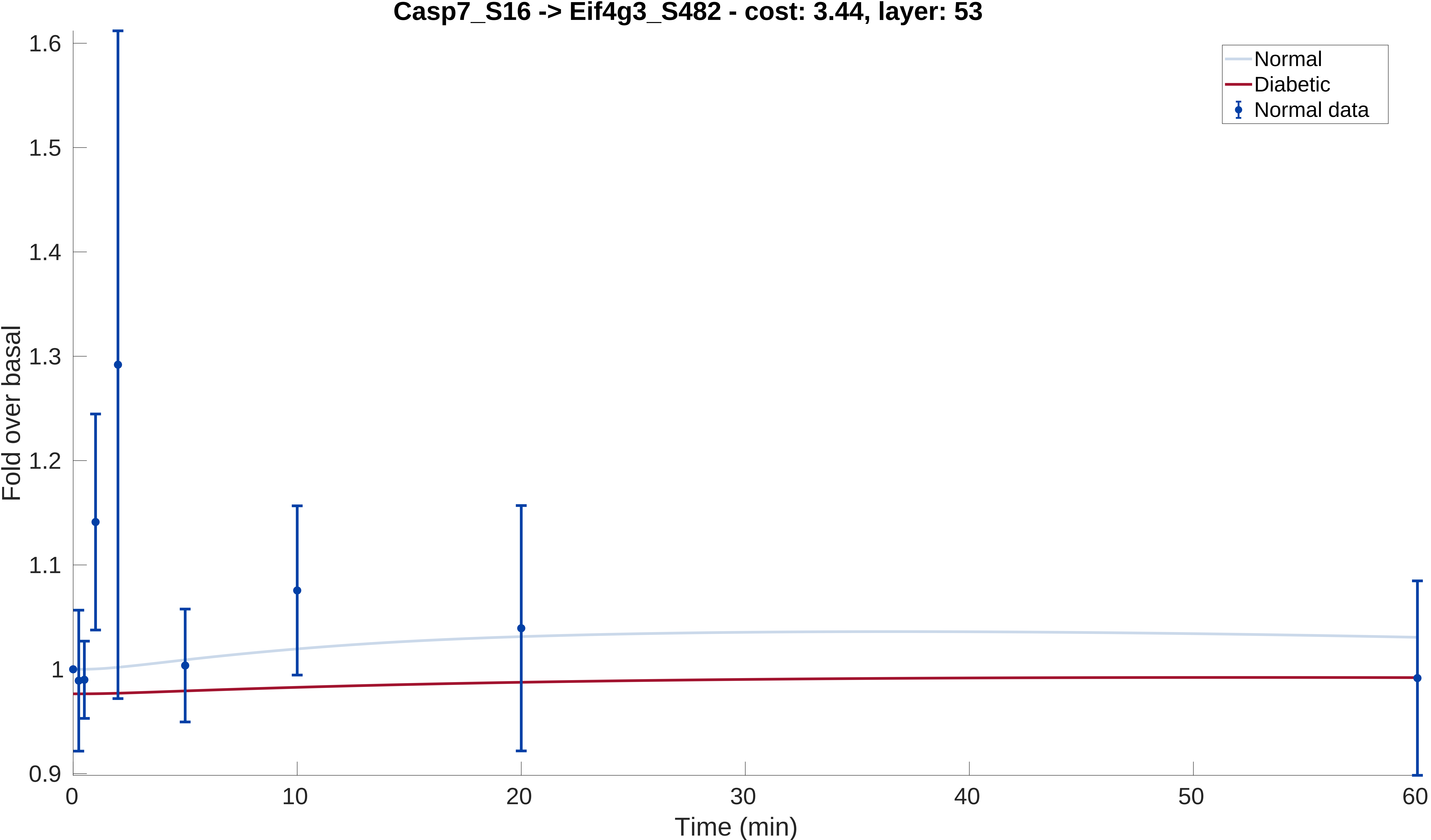

Casp7_S16 -> Eif4g3_S482 - cost: 3.44, layer: 53
Normal
Diabetic
Normal data
Fold over basal
1.6
1.5
1.4
1.3
1.2
1.1
1
0.9
0
10
20
30
40
50
60
Time (min)

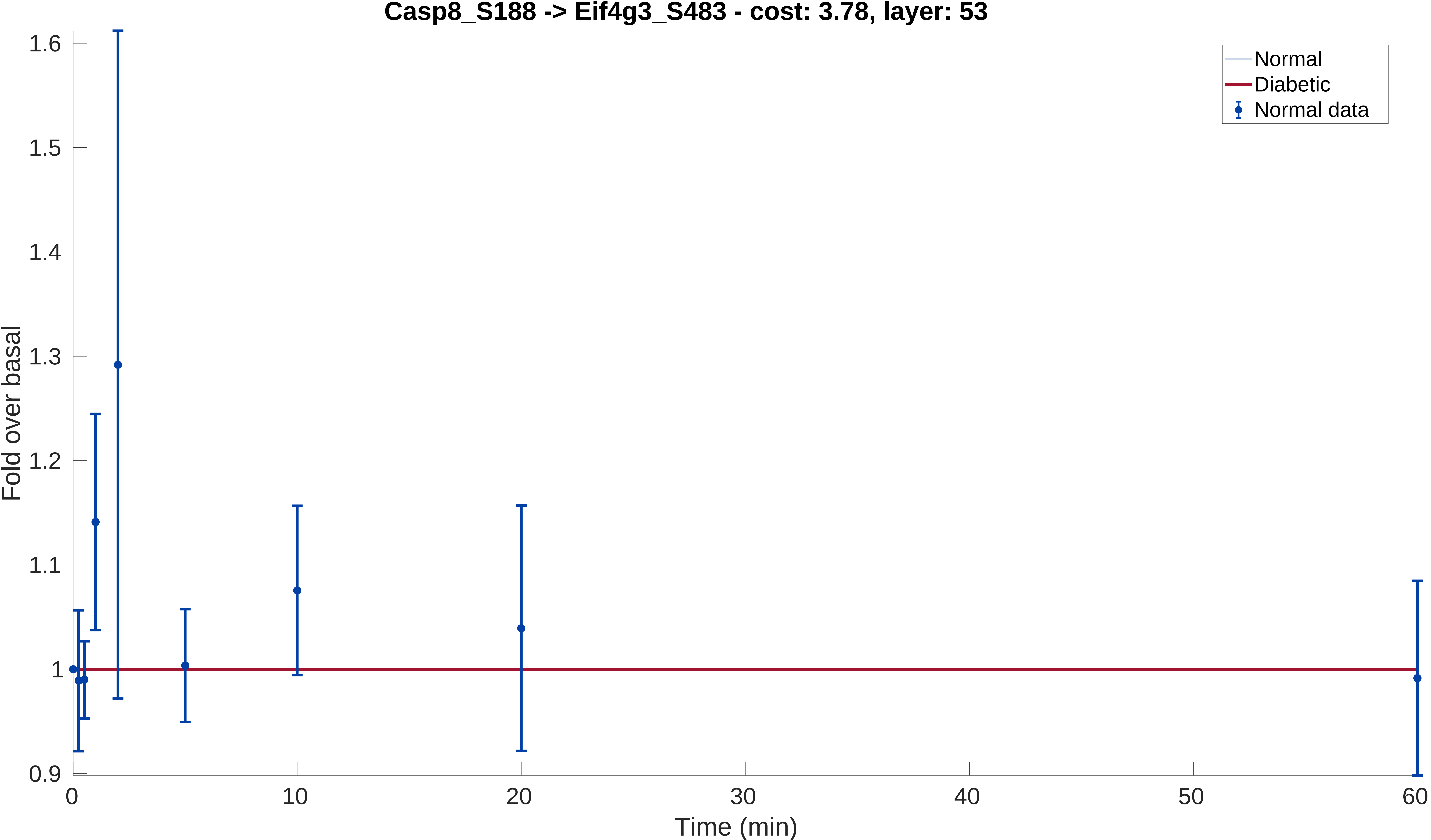

Casp8_S188 -> Eif4g3_S483 - cost: 3.78, layer: 53
Fold over basal
Time (min)
Normal
Diabetic
Normal data
1.6
1.5
1.4
1.3
1.2
1.1
1
0.9
0
10
20
30
40
50
60

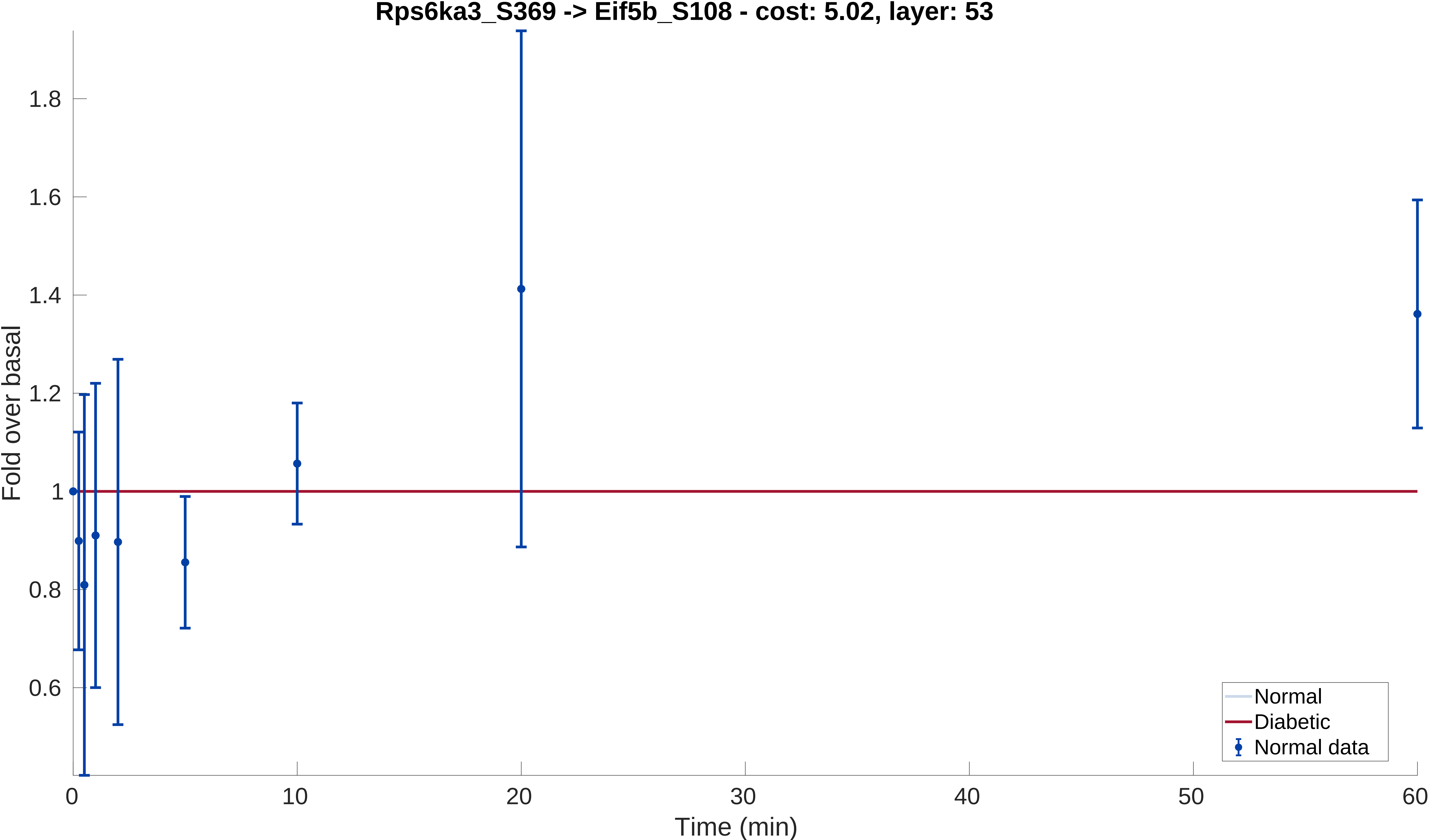
Rps6ka3_S369 -> Eif5b_S108 - cost: 5.02, layer: 53
Fold over basal
1.8
1.6
1.4
1.2
1
0.8
0.6
0
10
20
30
40
50
60
Time (min)
Normal
Diabetic
Normal data

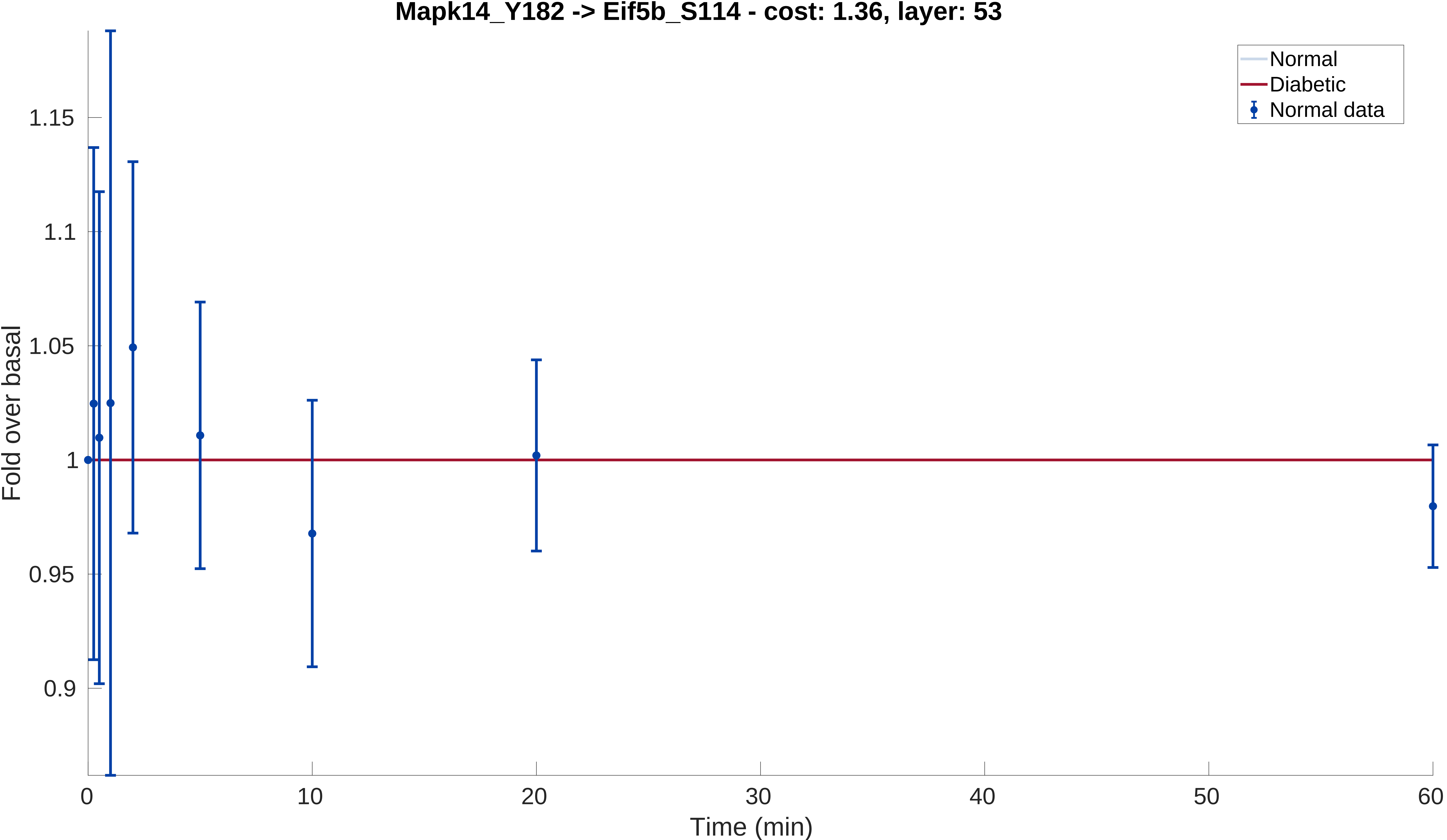

Mapk14_Y182 -> Eif5b_S114 - cost: 1.36, layer: 53
Normal
Diabetic
Normal data
Fold over basal
1.15
1.1
1.05
1
0.95
0.9
0
10
20
30
40
50
60
Time (min)

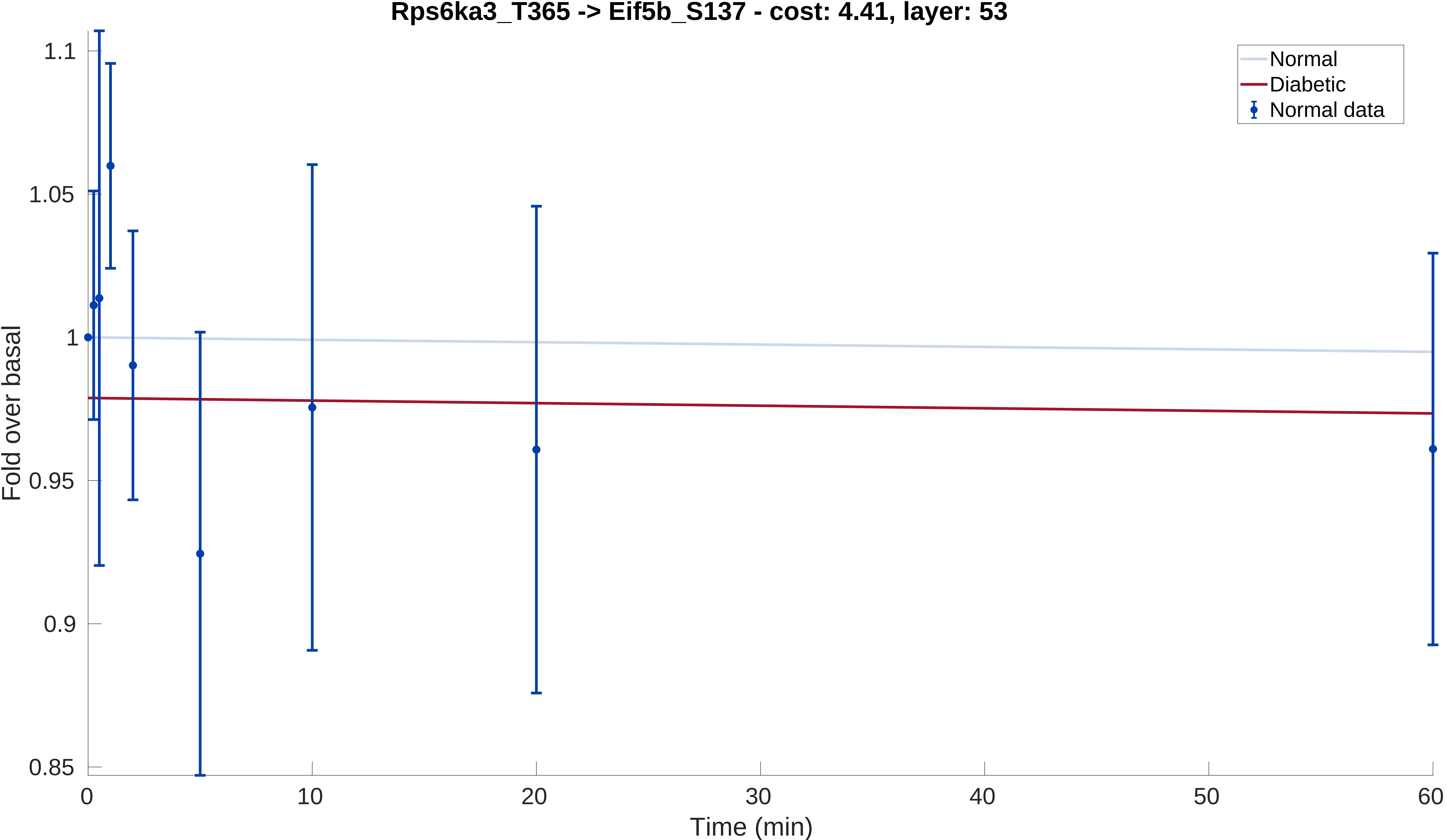

Rps6ka3_T365 -> Eif5b_S137 - cost: 4.41, layer: 53
Normal
Diabetic
Normal data
Fold over basal
1.1
1.05
1
0.95
0.9
0.85
0
10
20
30
40
50
60
Time (min)

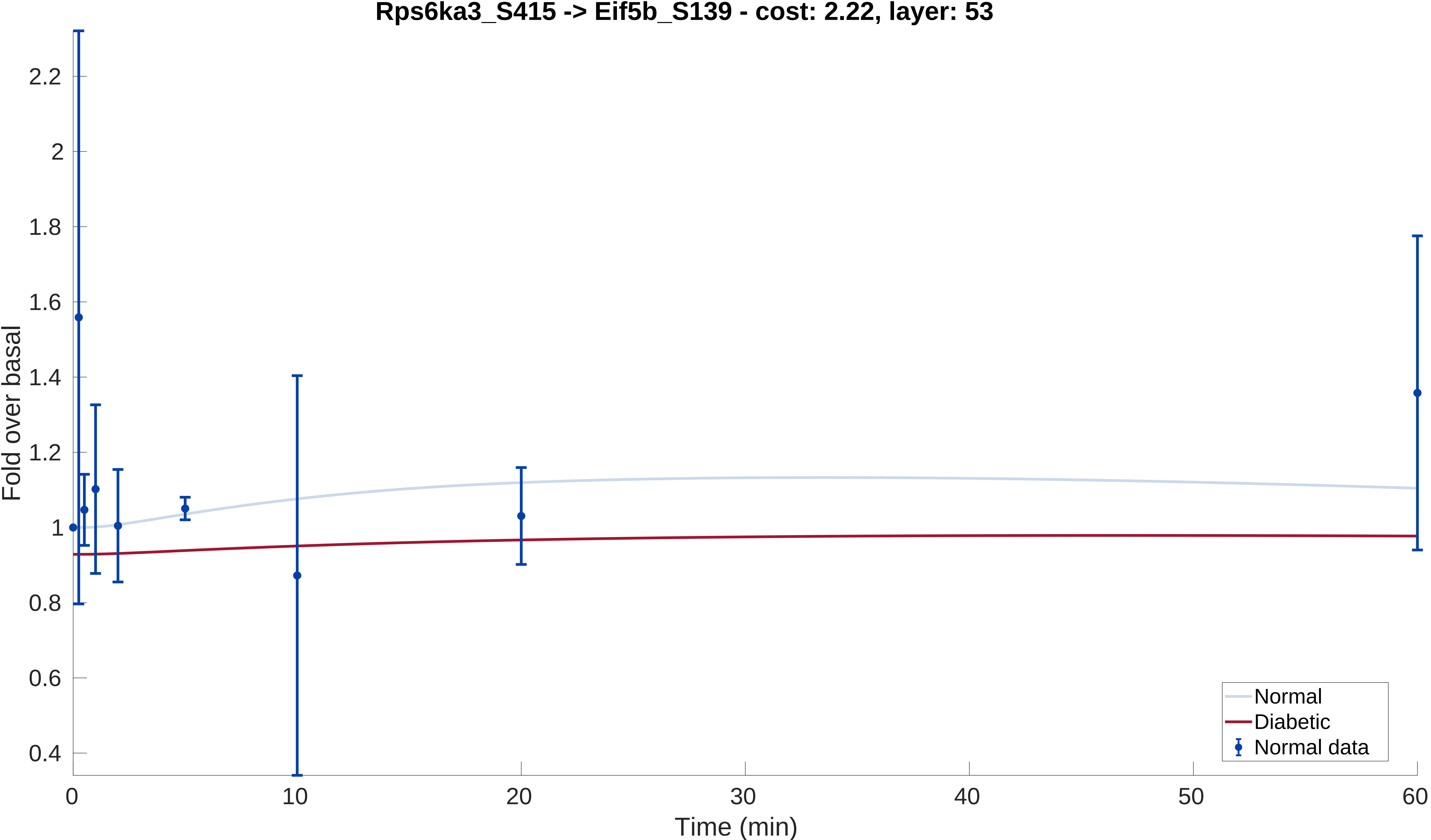

Rps6ka3_S415 -> Eif5b_S139 - cost: 2.22, layer: 53
Fold over basal
2.2
2
1.8
1.6
1.4
1.2
1
0.8
0.6
0.4
0
10
20
30
40
50
60
Time (min)
Normal
Diabetic
Normal data

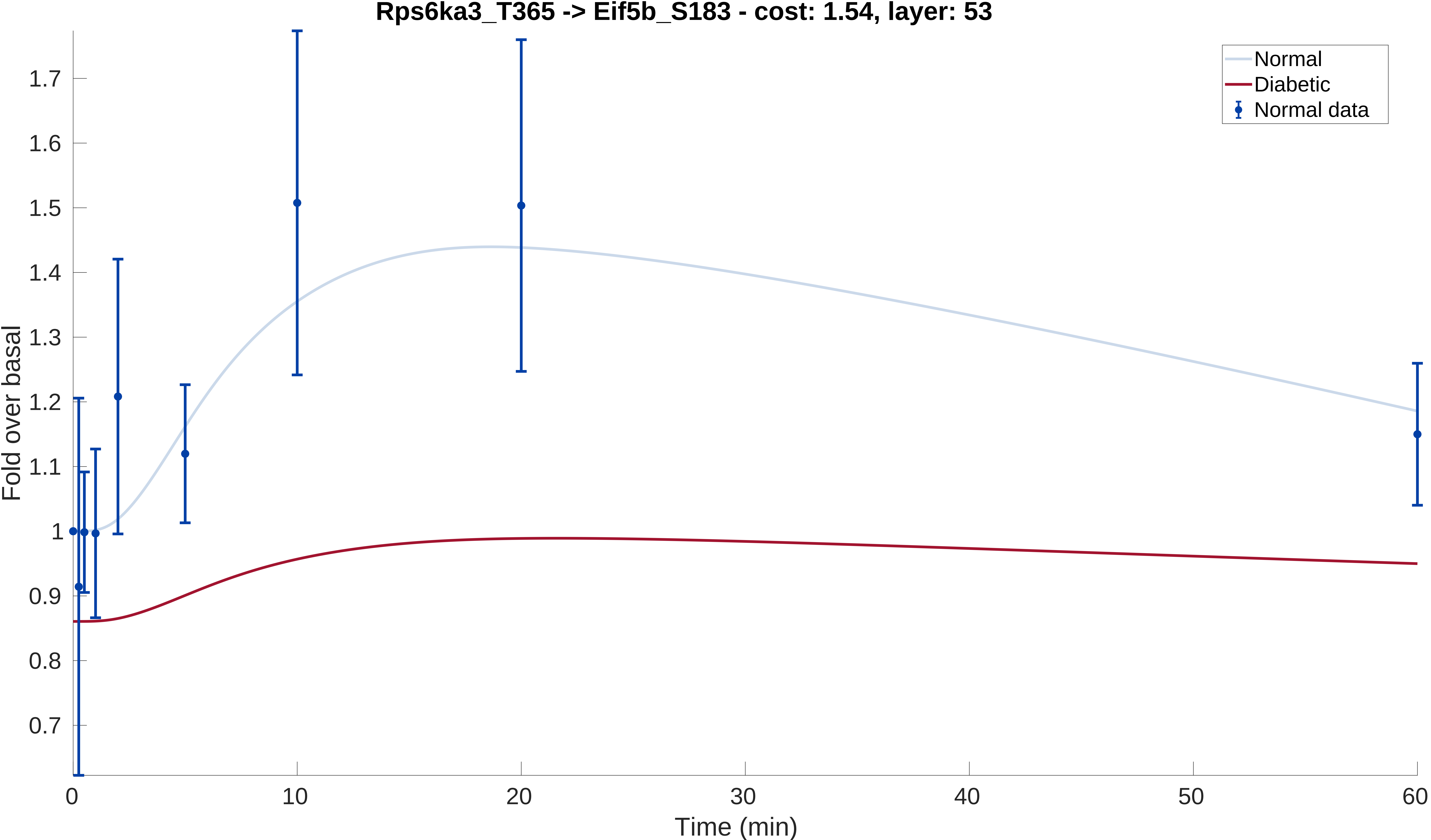

Rps6ka3_T365 -> Eif5b_S183 - cost: 1.54, layer: 53
Normal
Diabetic
Normal data
Fold over basal
1.7
1.6
1.5
1.4
1.3
1.2
1.1
1
0.9
0.8
0.7
0
10
20
30
40
50
60
Time (min)

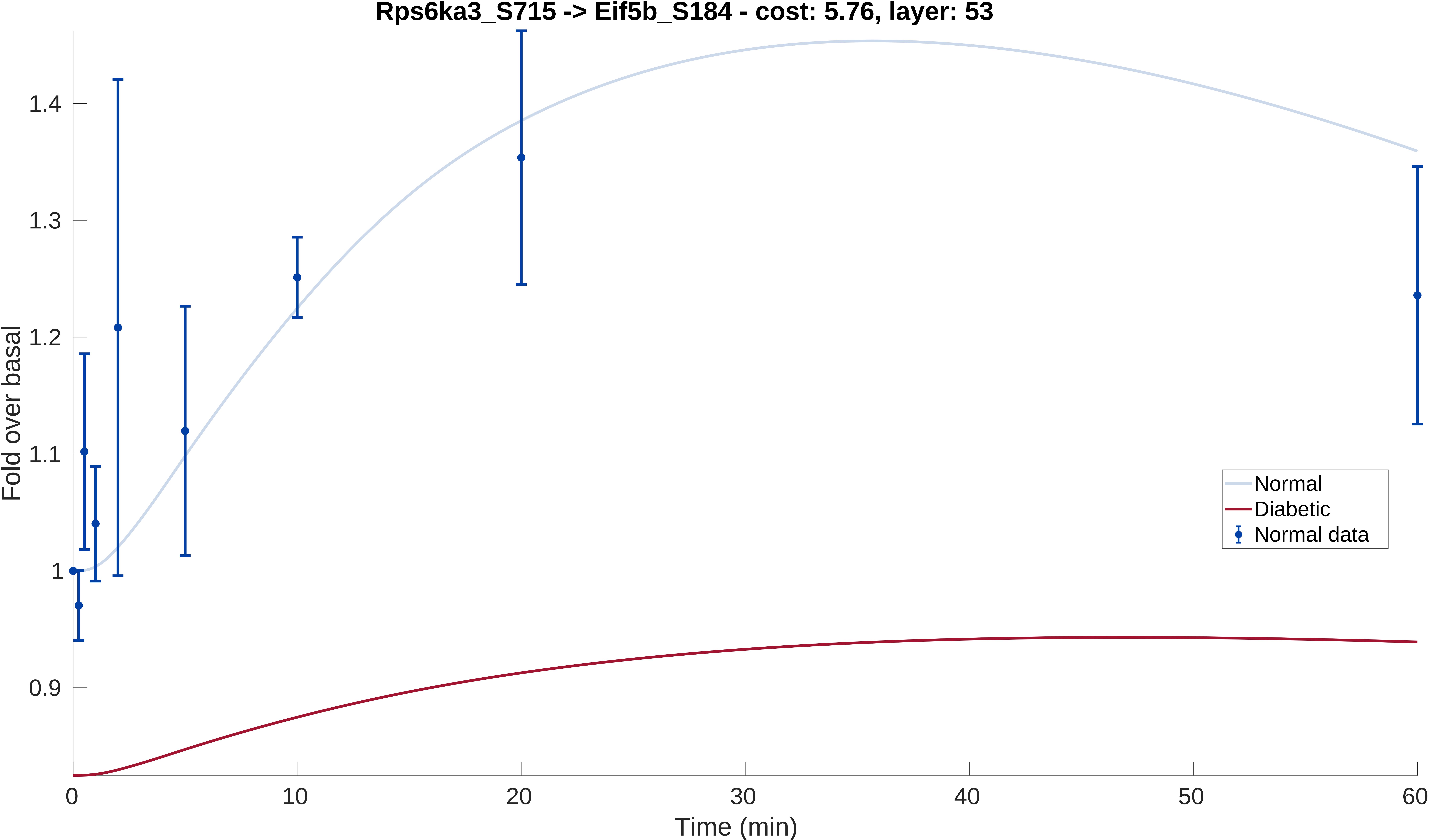

Rps6ka3_S715 -> Eif5b_S184 - cost: 5.76, layer: 53
Fold over basal
Time (min)
1.4
1.3
1.2
1.1
1
0.9
0
10
20
30
40
50
60
Normal
Diabetic
Normal data

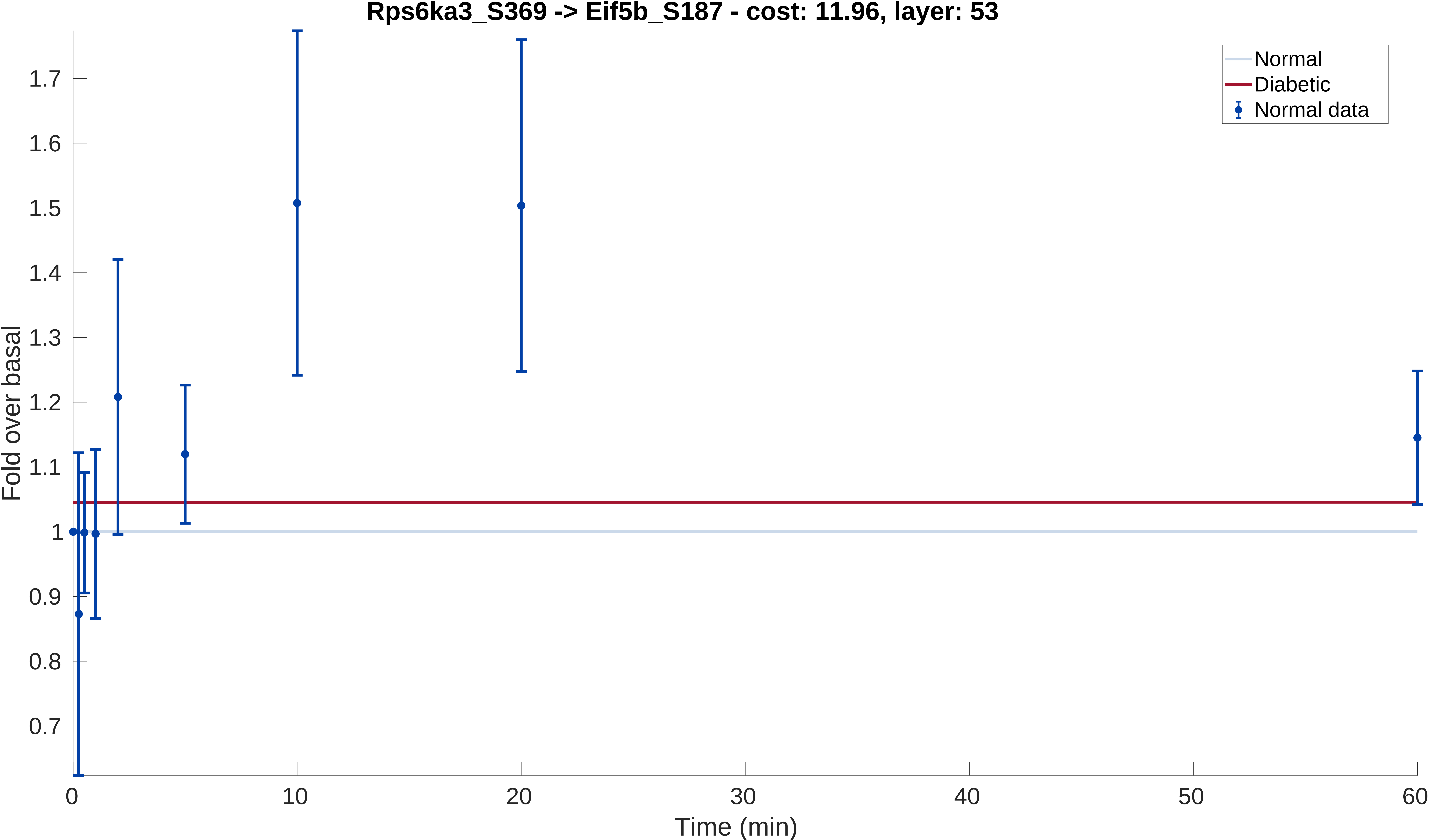

Rps6ka3_S369 -> Eif5b_S187 - cost: 11.96, layer: 53
Normal
Diabetic
Normal data
Fold over basal
1.7
1.6
1.5
1.4
1.3
1.2
1.1
1
0.9
0.8
0.7
0
10
20
30
40
50
60
Time (min)

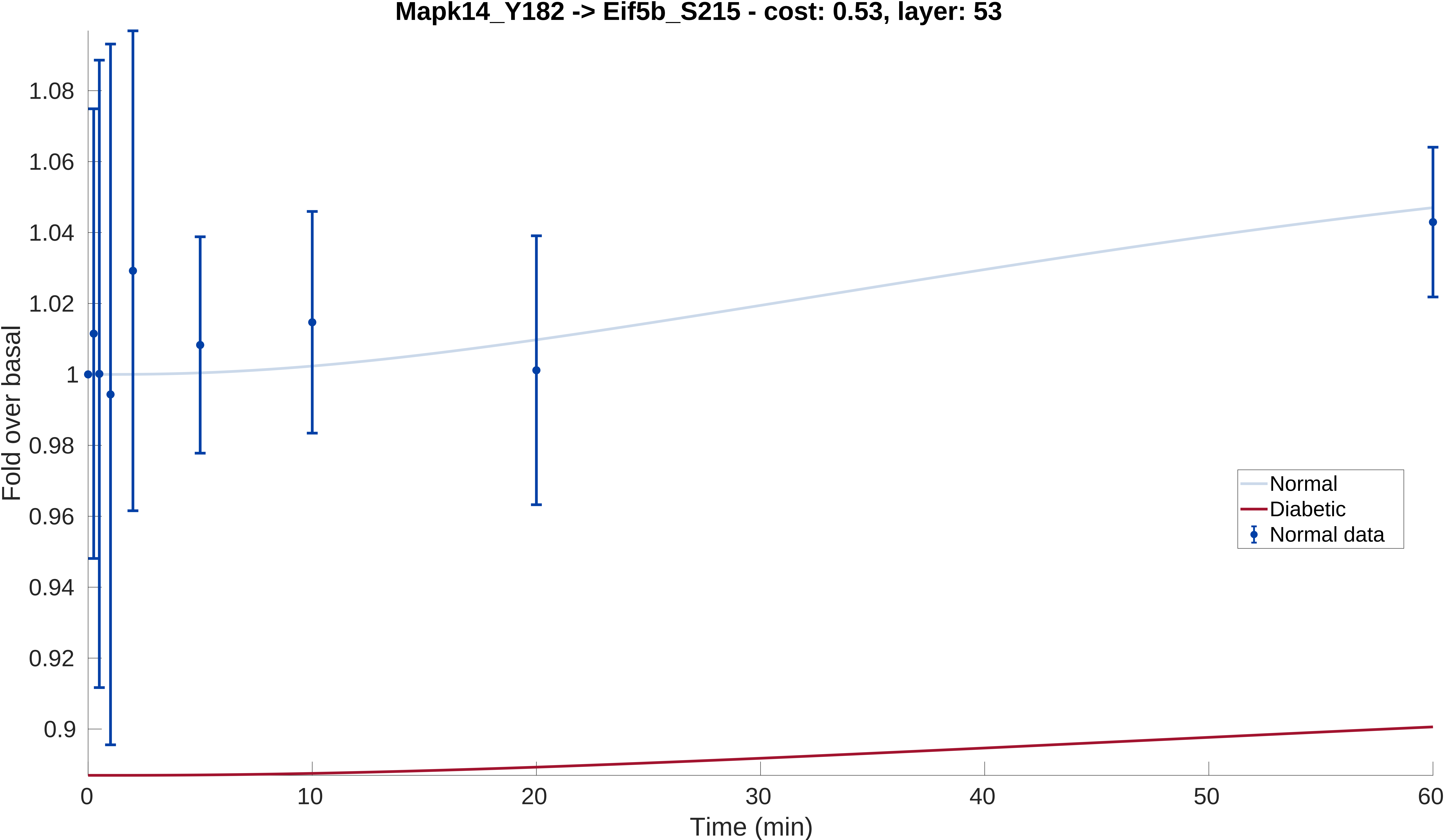

Mapk14_Y182 -> Eif5b_S215 - cost: 0.53, layer: 53
Fold over basal
Time (min)
1.08
1.06
1.04
1.02
1
0.98
0.96
0.94
0.92
0.9
0
10
20
30
40
50
60
Normal
Diabetic
Normal data

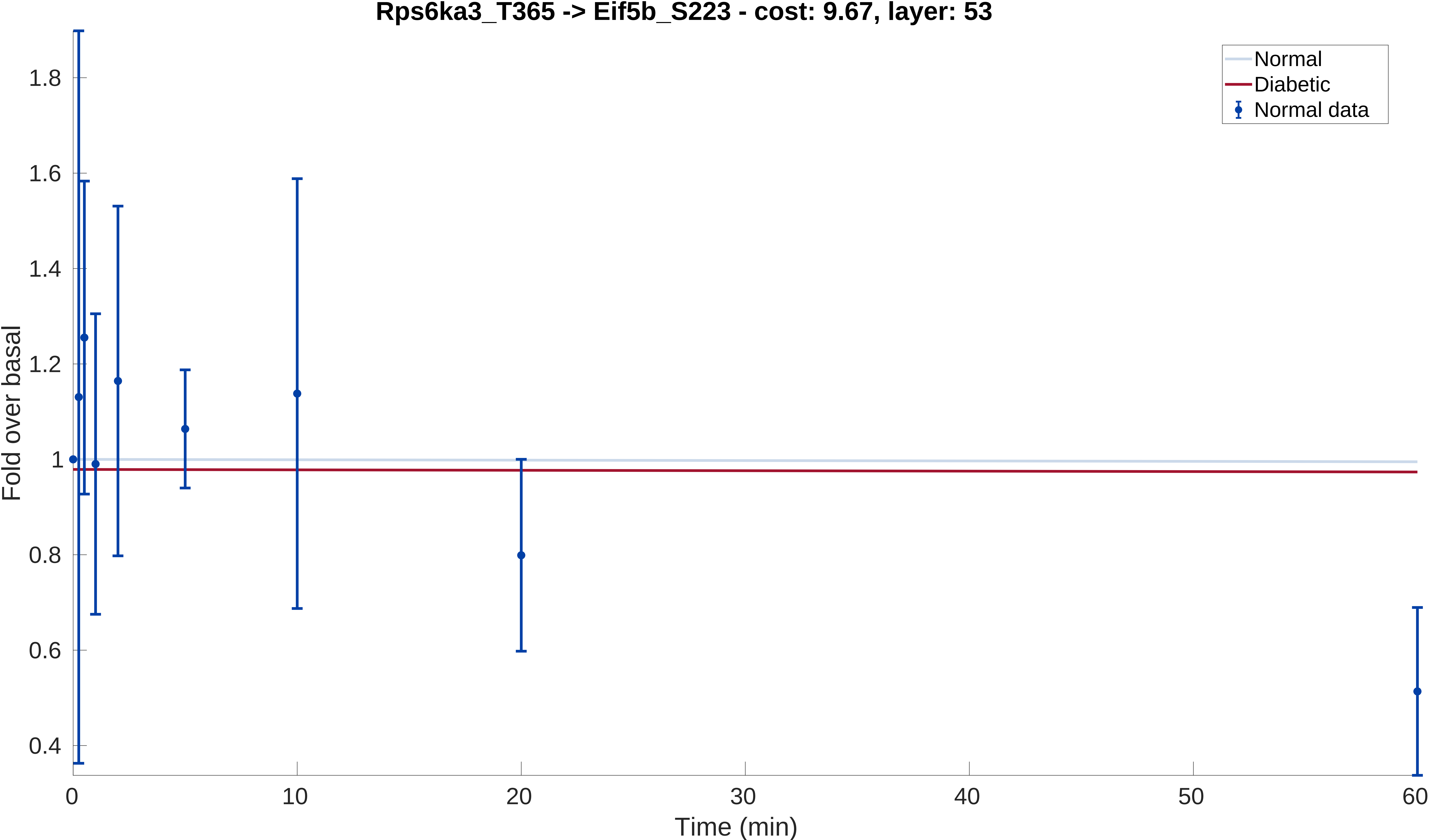

Rps6ka3_T365 -> Eif5b_S223 - cost: 9.67, layer: 53
Normal
Diabetic
Normal data
Fold over basal
1.8
1.6
1.4
1.2
1
0.8
0.6
0.4
0
10
20
30
40
50
60
Time (min)

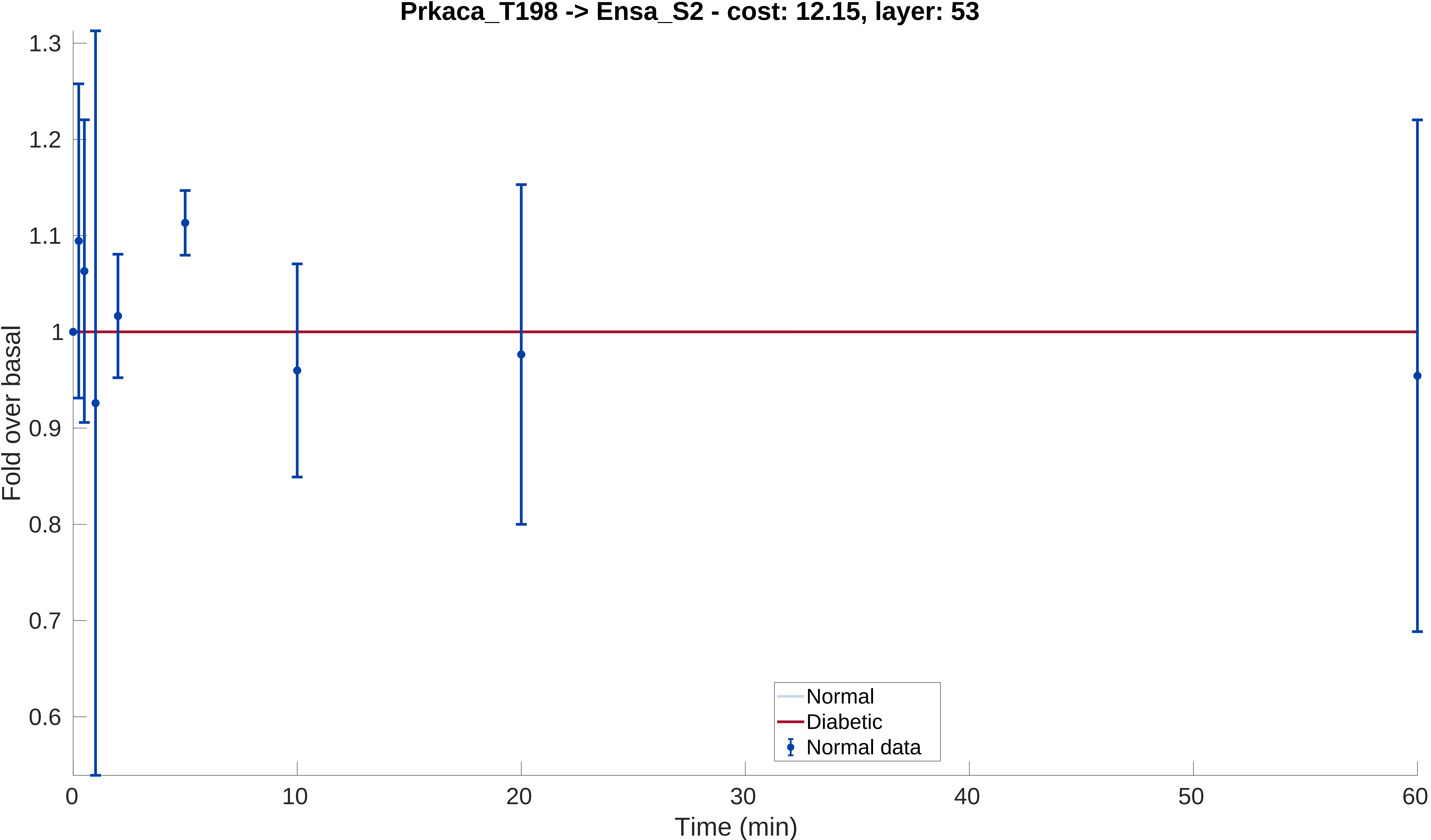
Prkaca_T198 -> Ensa_S2 - cost: 12.15, layer: 53
Fold over basal
Time (min)
1.3
1.2
1.1
1
0.9
0.8
0.7
0.6
0
10
20
30
40
50
60
Normal
Diabetic
Normal data

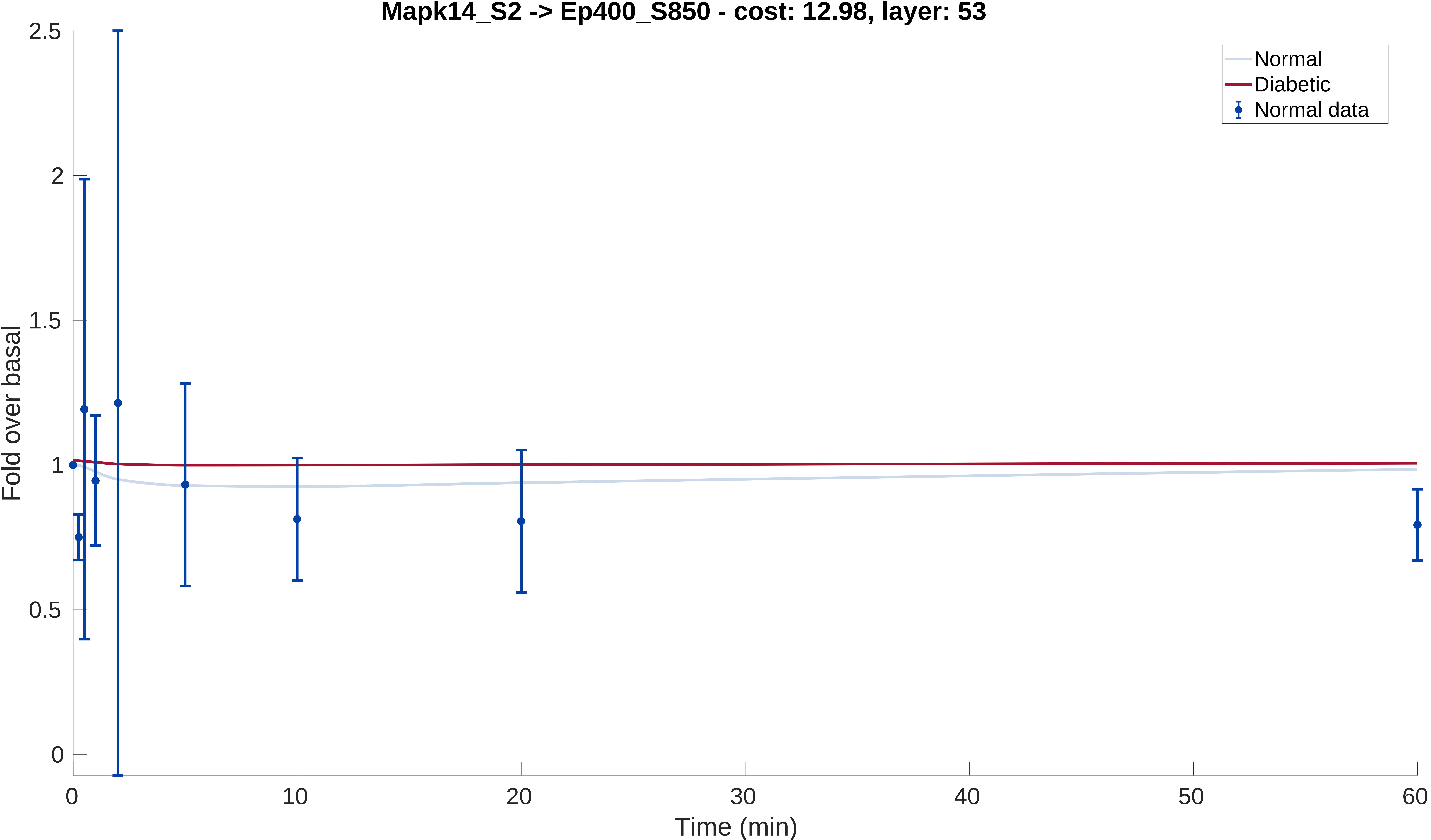

Mapk14_S2 -> Ep400_S850 - cost: 12.98, layer: 53
Normal
Diabetic
Normal data
Fold over basal
2.5
2
1.5
1
0.5
0
0
10
20
30
40
50
60
Time (min)

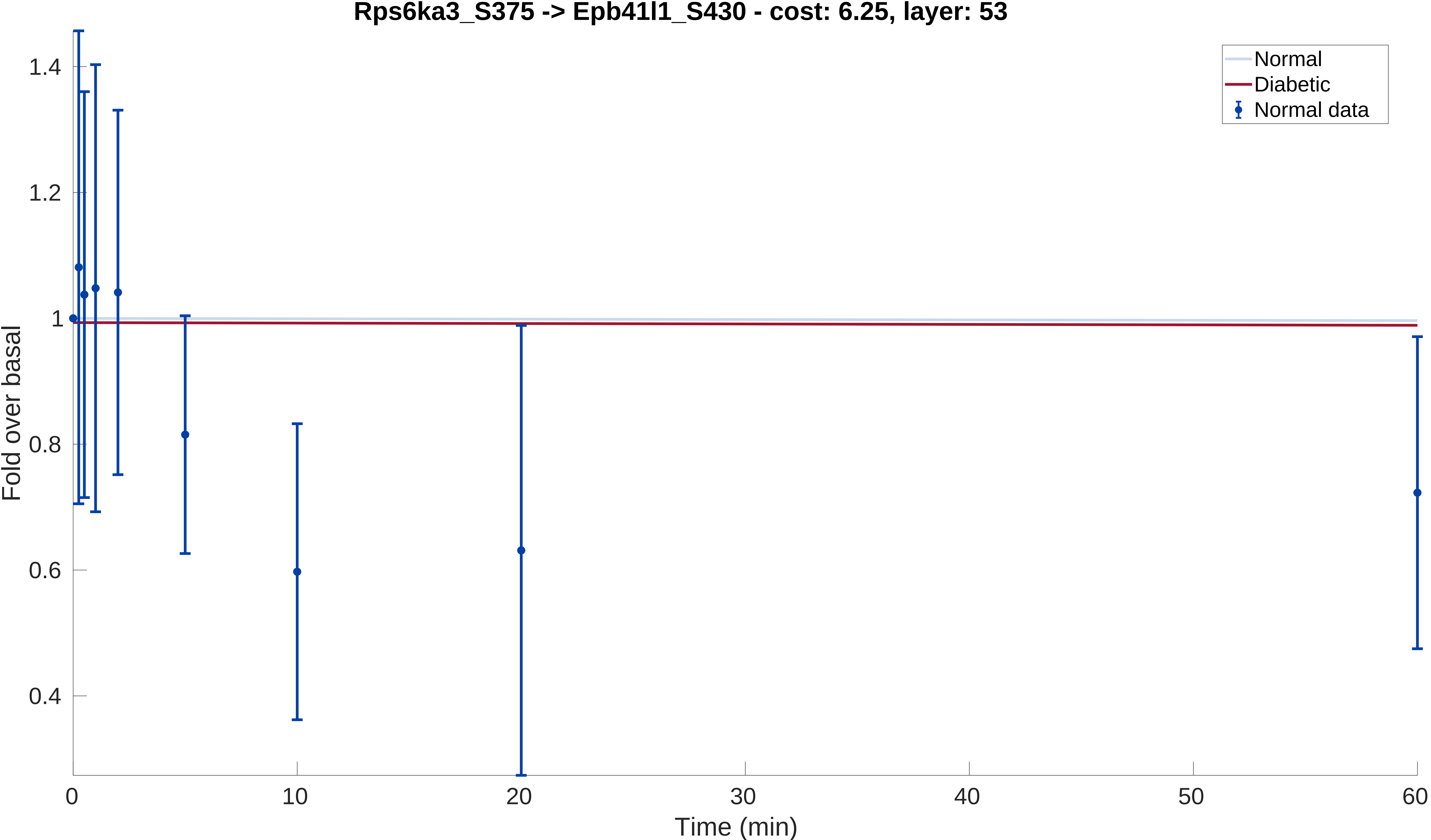

Rps6ka3_S375 -> Epb41l1_S430 - cost: 6.25, layer: 53
Normal
Diabetic
Normal data
Fold over basal
1.4
1.2
1
0.8
0.6
0.4
0
10
20
30
40
50
60
Time (min)

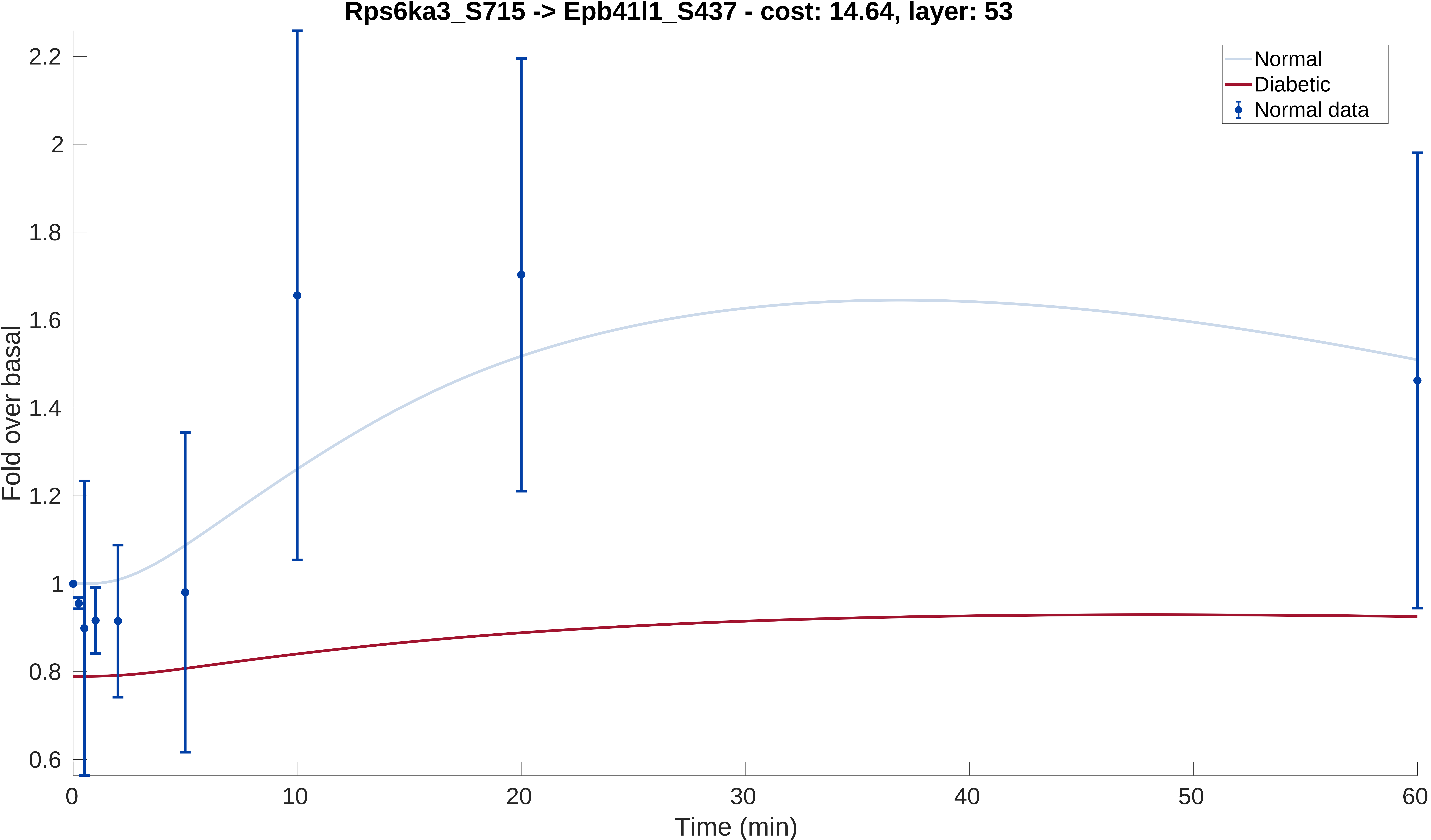
Rps6ka3_S715 -> Epb41l1_S437 - cost: 14.64, layer: 53
Normal
Diabetic
Normal data
Fold over basal
Time (min)
0.6
0.8
1
1.2
1.4
1.6
1.8
2
2.2
0
10
20
30
40
50
60

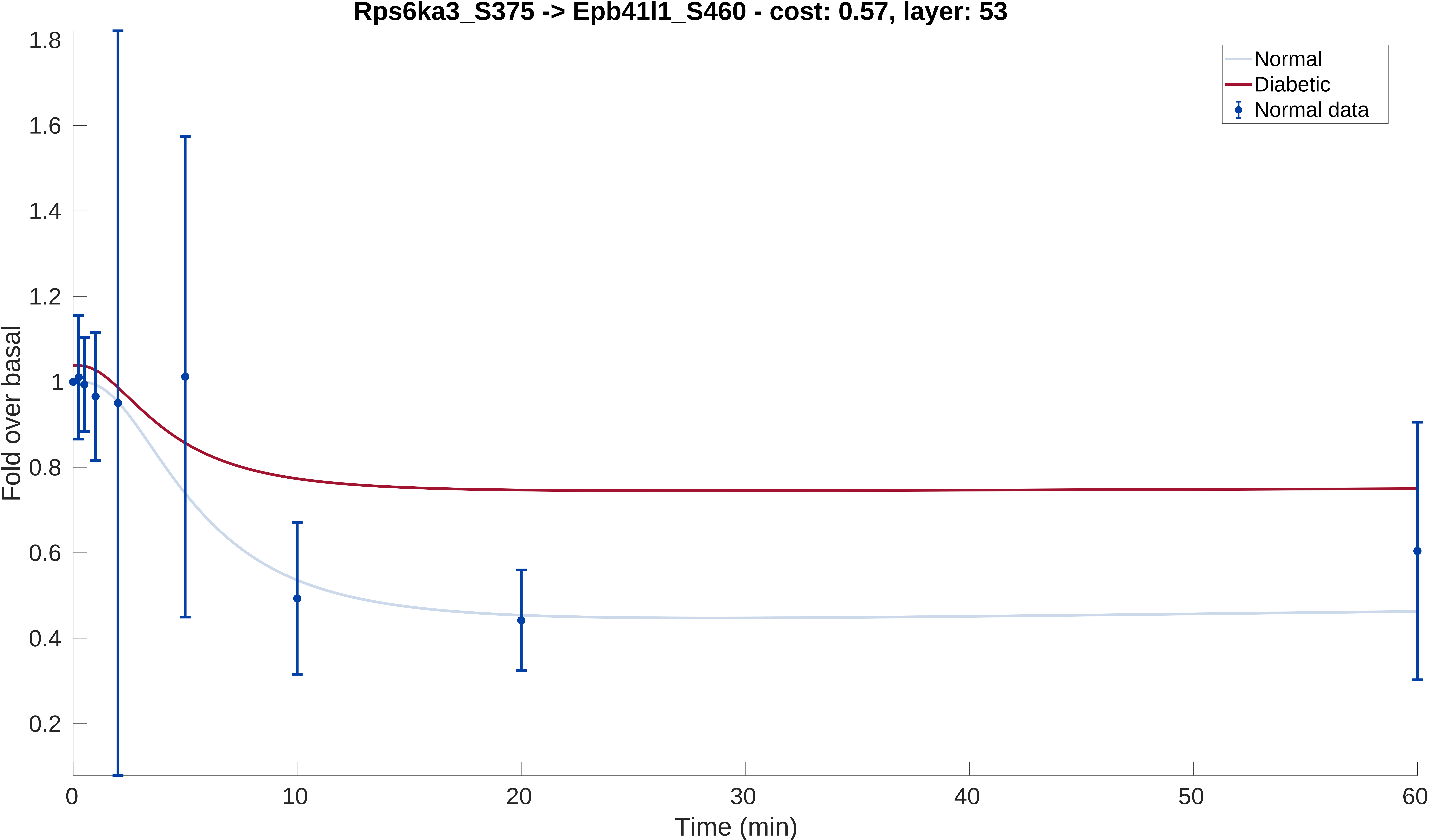
Rps6ka3_S375 -> Epb41l1_S460 - cost: 0.57, layer: 53
Normal
Diabetic
Normal data
Fold over basal
1.8
1.6
1.4
1.2
1
0.8
0.6
0.4
0.2
0
10
20
30
40
50
60
Time (min)

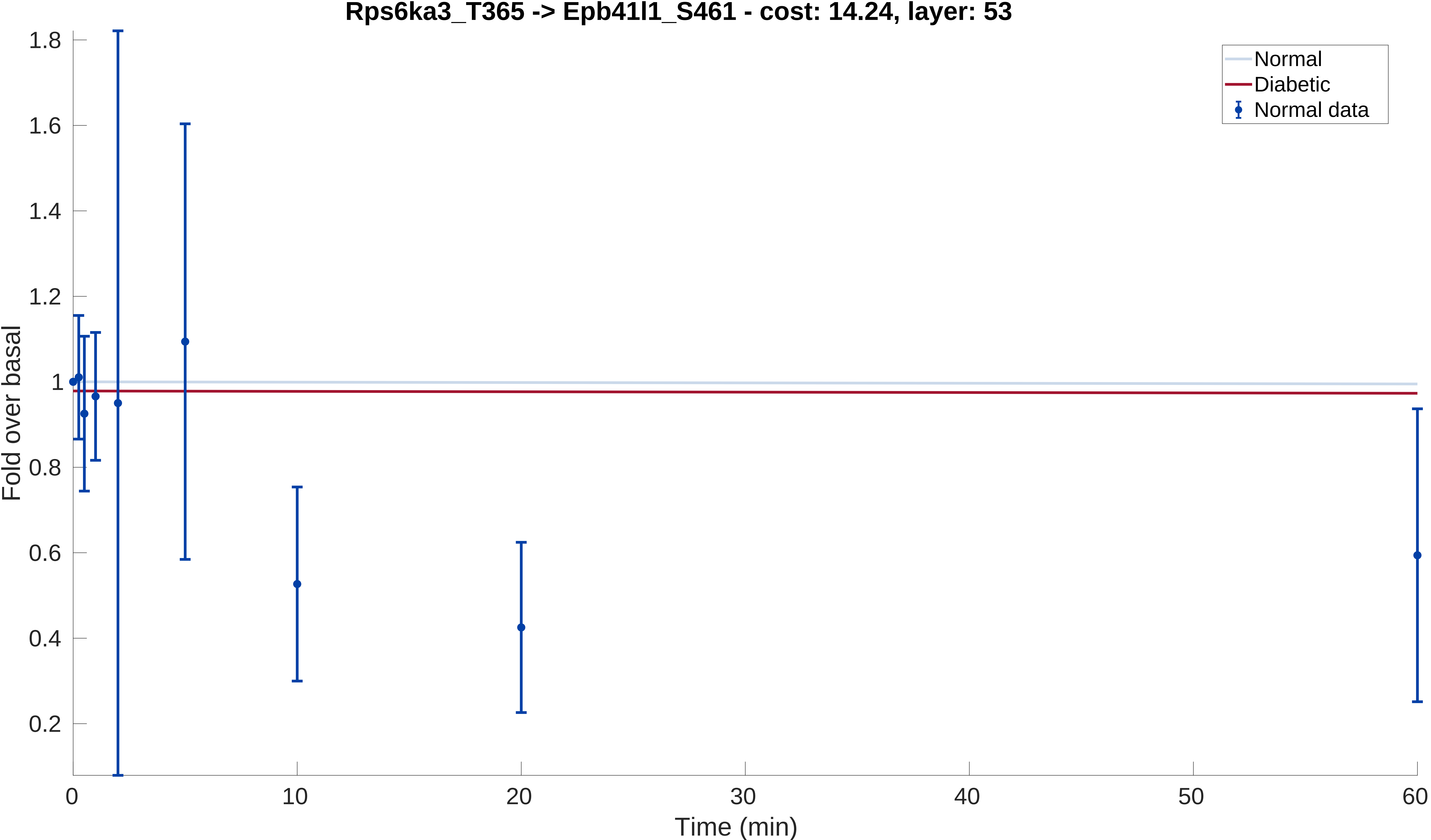

Rps6ka3_T365 -> Epb41l1_S461 - cost: 14.24, layer: 53
Normal
Diabetic
Normal data
Fold over basal
1.8
1.6
1.4
1.2
1
0.8
0.6
0.4
0.2
0
10
20
30
40
50
60
Time (min)

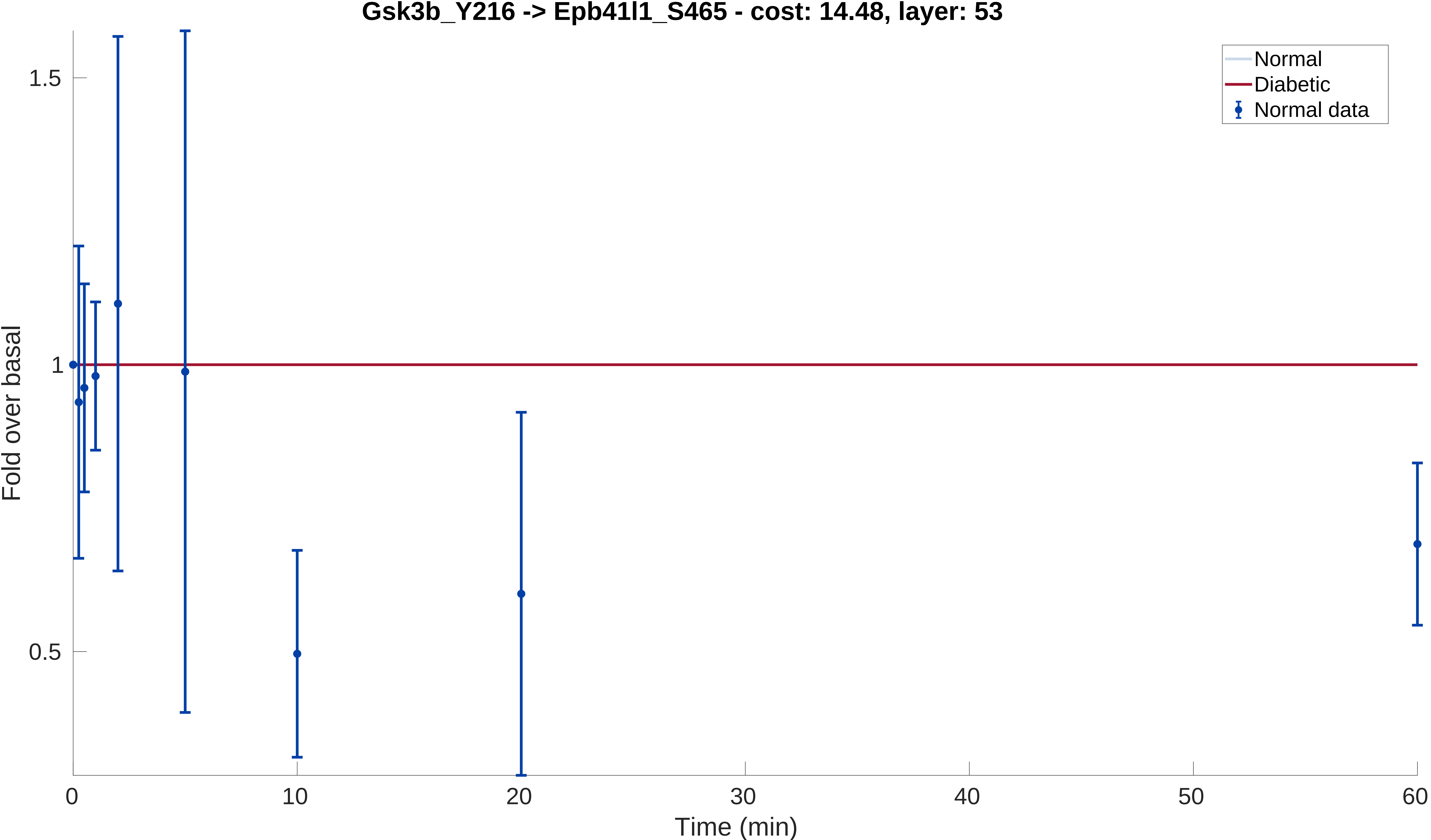

Gsk3b_Y216 -> Epb41l1_S465 - cost: 14.48, layer: 53
Normal
Diabetic
Normal data
Fold over basal
1.5
1
0.5
0
10
20
30
40
50
60
Time (min)

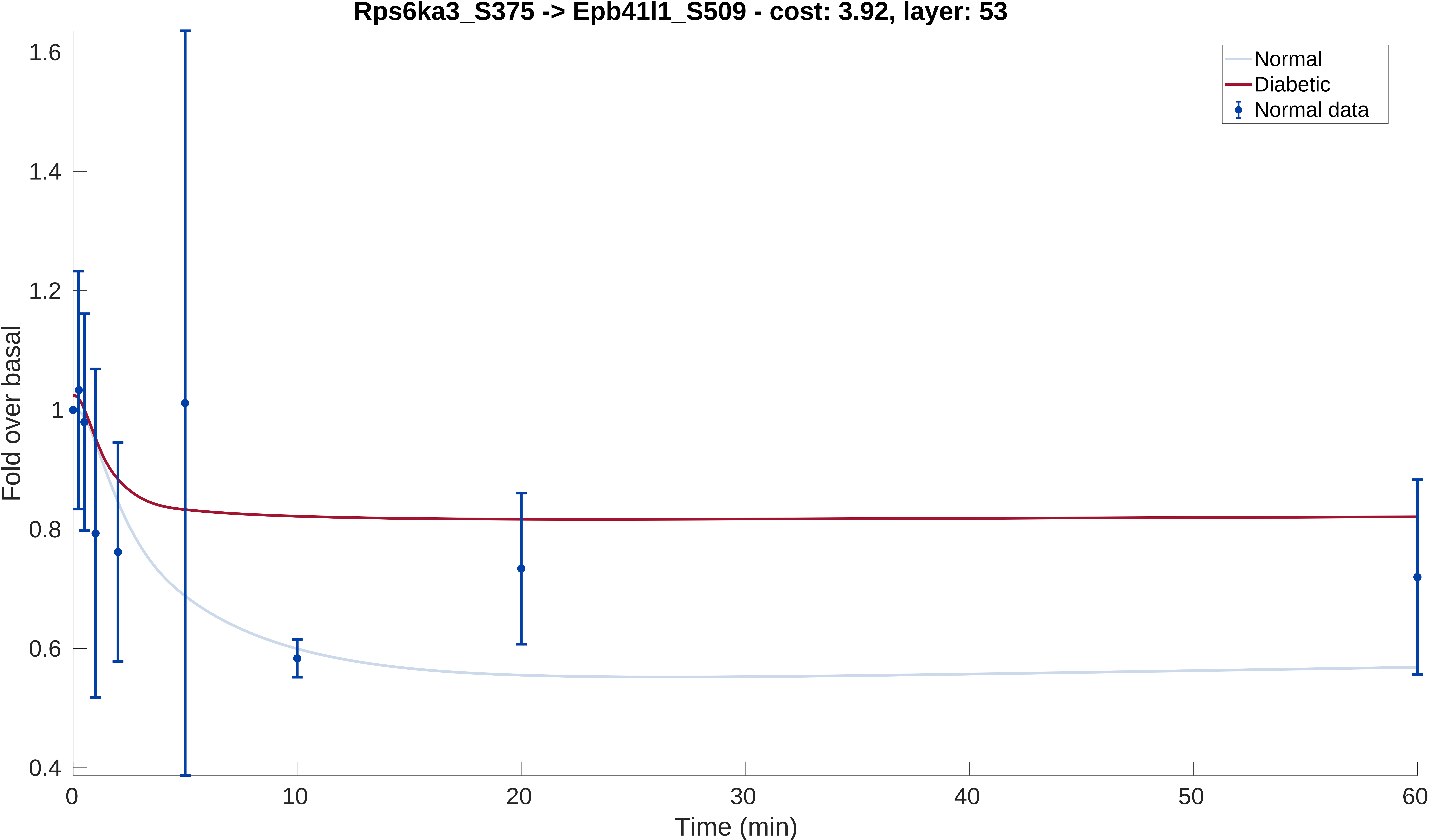

Rps6ka3_S375 -> Epb41l1_S509 - cost: 3.92, layer: 53
Normal
Diabetic
Normal data
Fold over basal
1.6
1.4
1.2
1
0.8
0.6
0.4
0
10
20
30
40
50
60
Time (min)

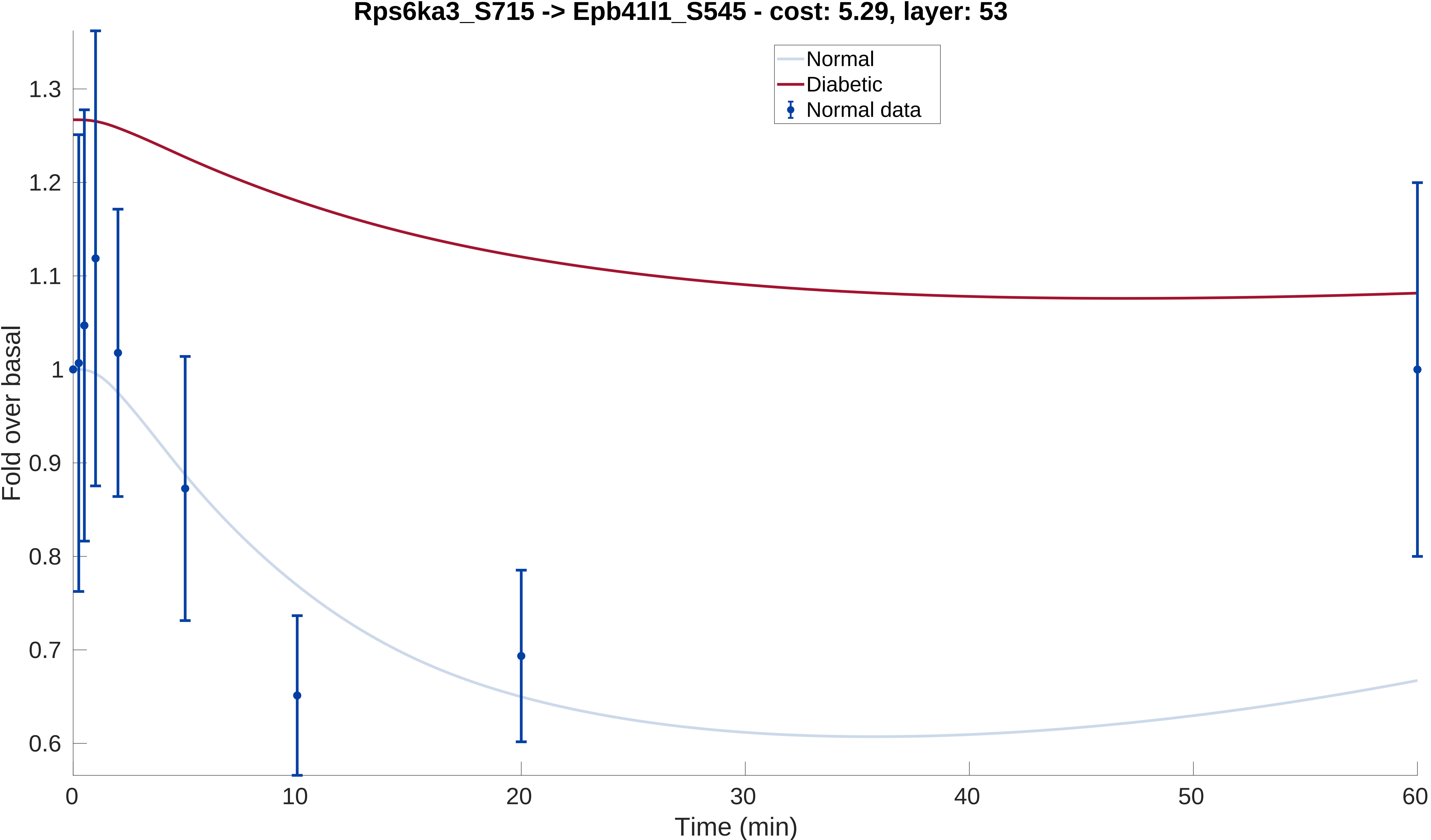

Rps6ka3_S715 -> Epb41l1_S545 - cost: 5.29, layer: 53
Normal
Diabetic
Normal data
1.3
1.2
1.1
1
0.9
0.8
0.7
0.6
Fold over basal
0
10
20
30
40
50
60
Time (min)

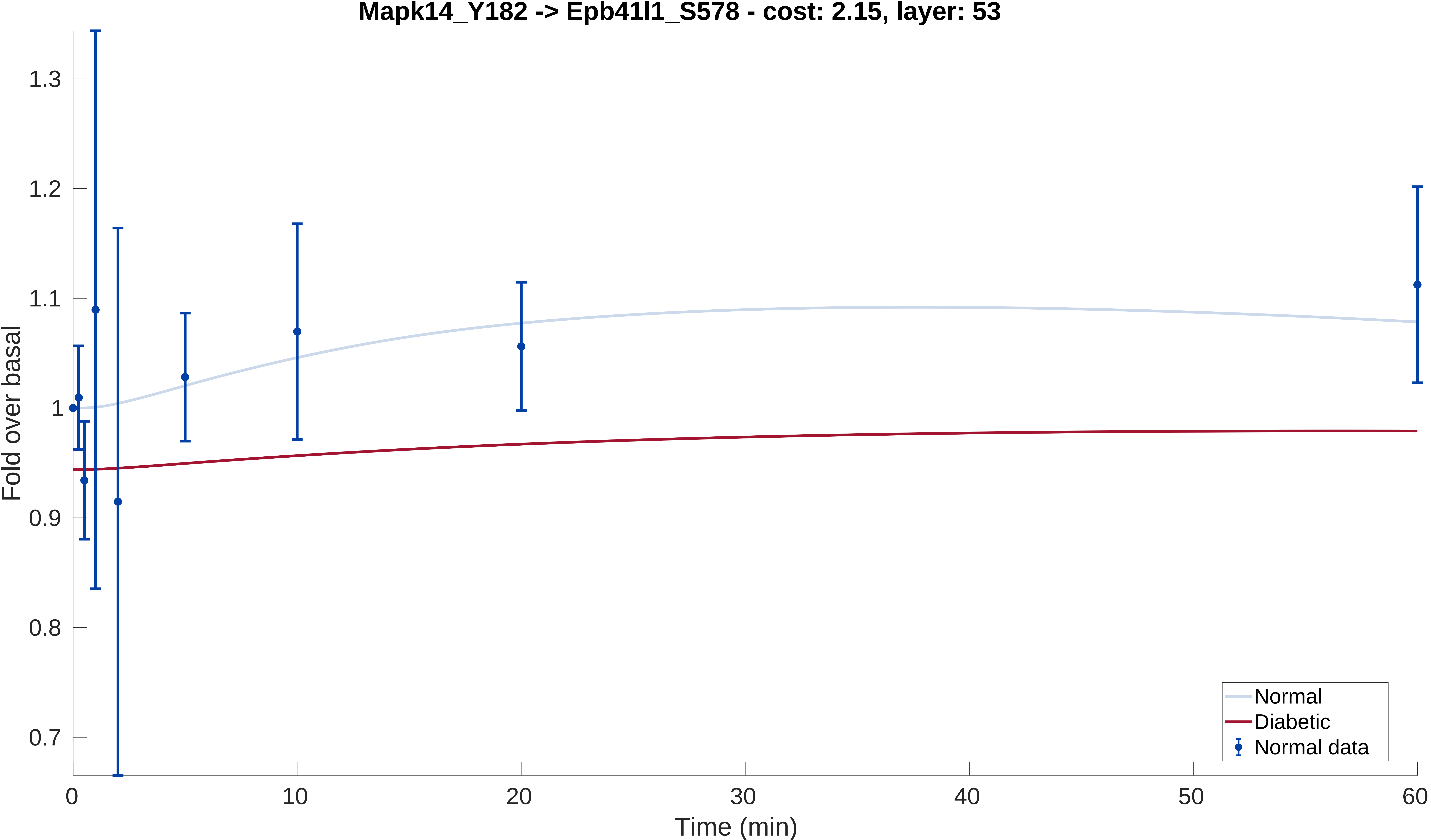

Mapk14_Y182 -> Epb41l1_S578 - cost: 2.15, layer: 53
Fold over basal
1.3
1.2
1.1
1
0.9
0.8
0.7
0
10
20
30
40
50
60
Time (min)
Normal
Diabetic
Normal data

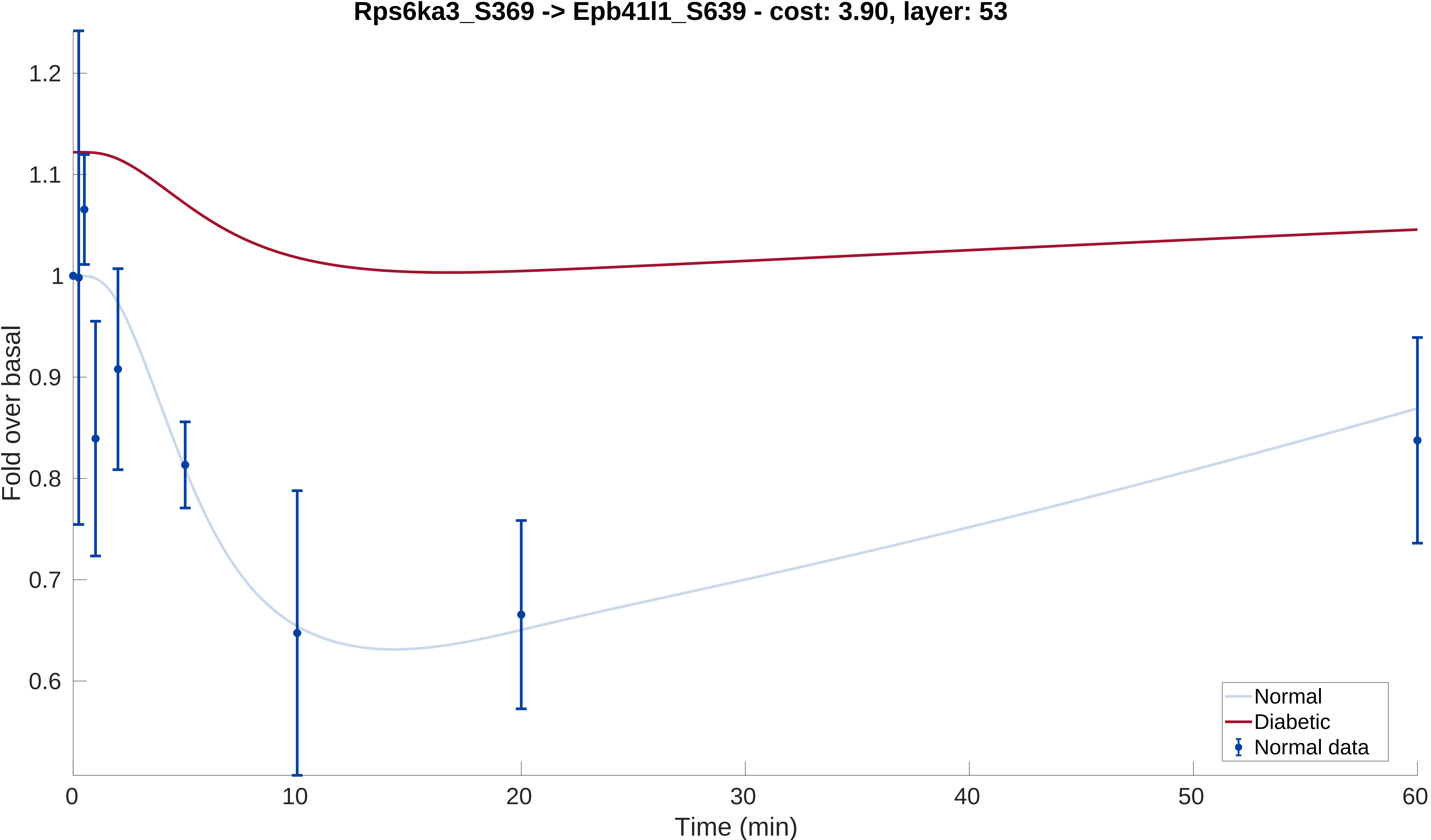

Rps6ka3_S369 -> Epb41l1_S639 - cost: 3.90, layer: 53
Fold over basal
Time (min)
1.2
1.1
1
0.9
0.8
0.7
0.6
0
10
20
30
40
50
60
Normal
Diabetic
Normal data

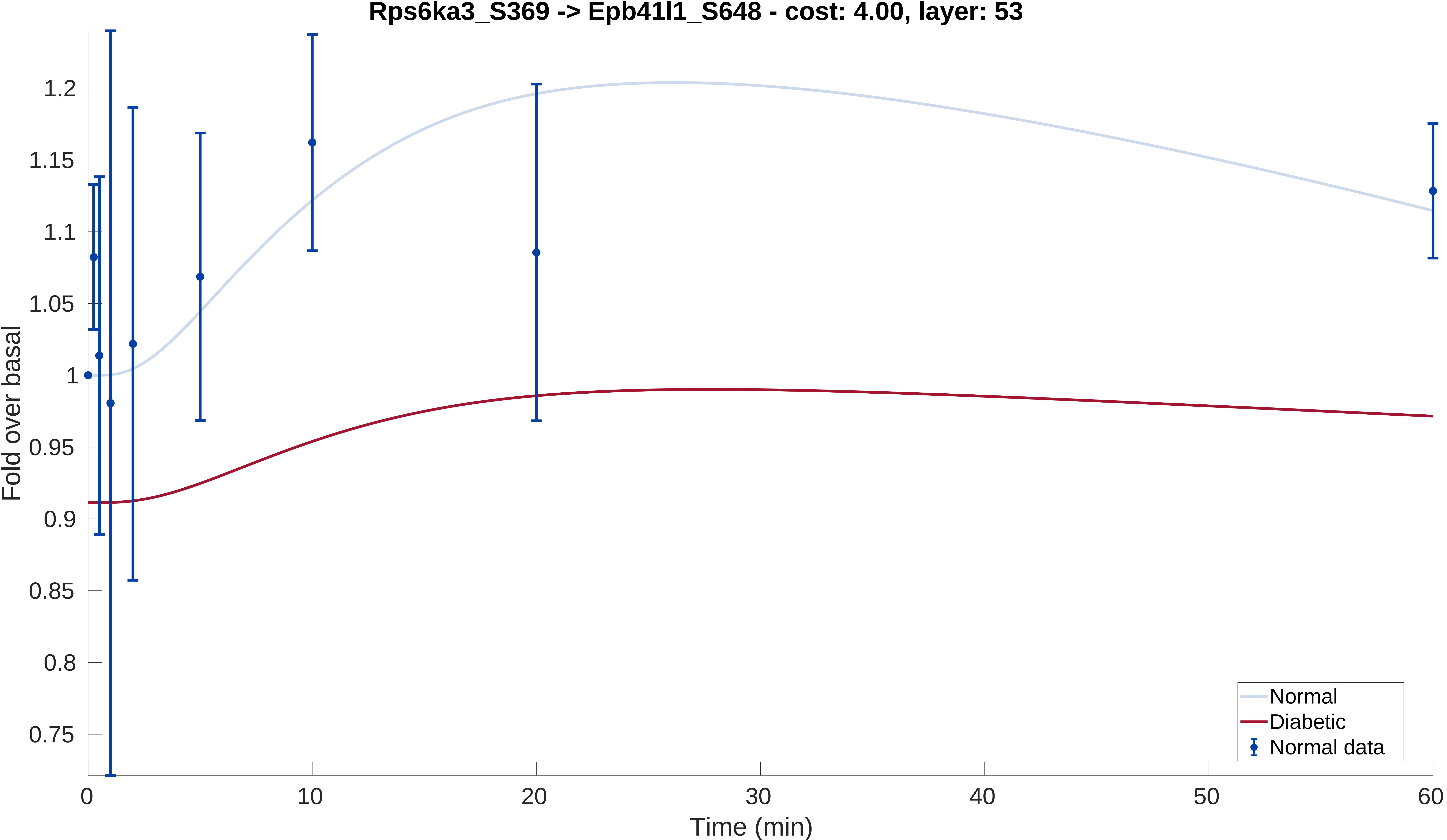

Rps6ka3_S369 -> Epb41l1_S648 - cost: 4.00, layer: 53
Fold over basal
1.2
1.15
1.1
1.05
1
0.95
0.9
0.85
0.8
0.75
0
10
20
30
40
50
60
Time (min)
Normal
Diabetic
Normal data

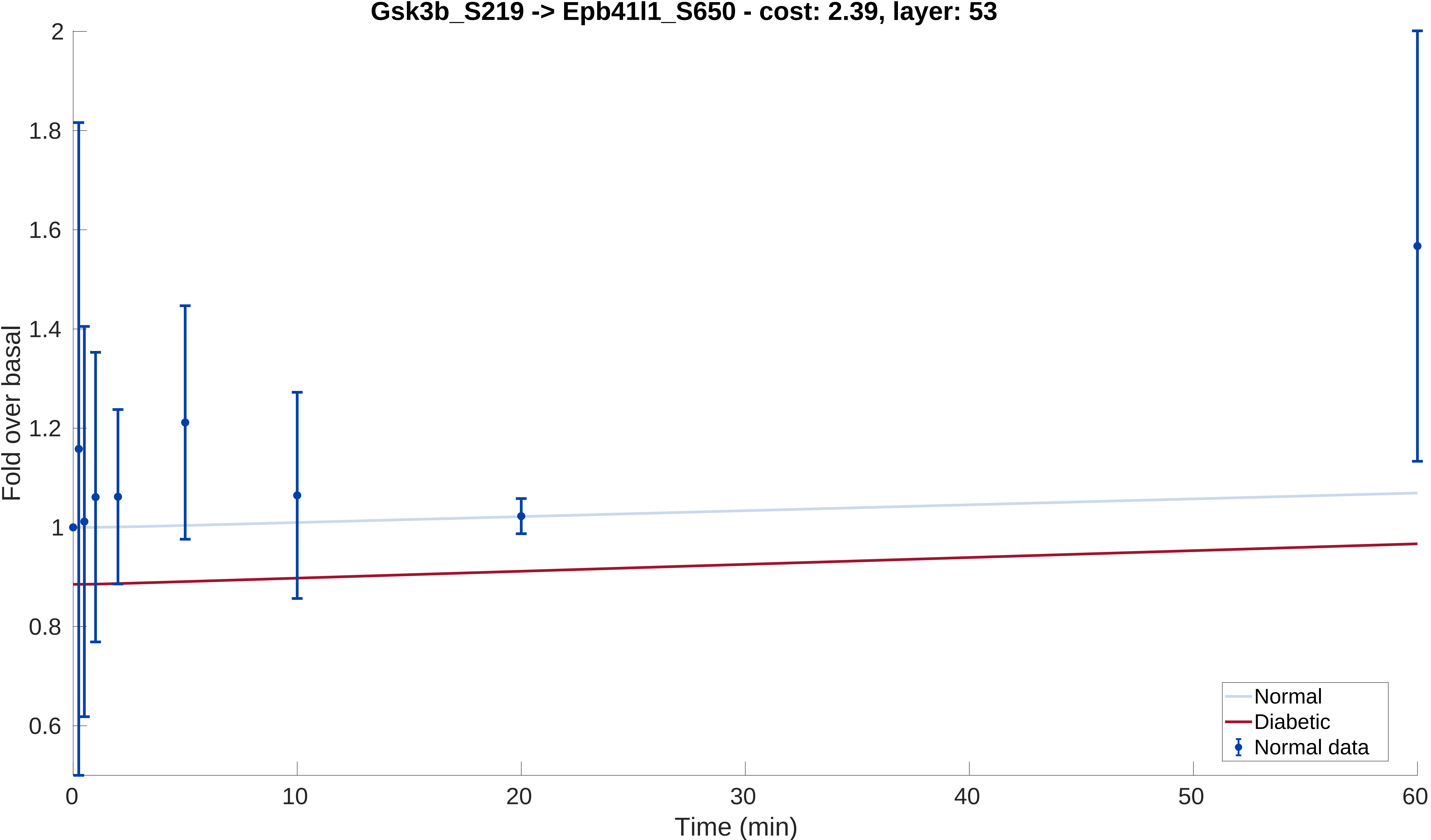

Gsk3b_S219 -> Epb41l1_S650 - cost: 2.39, layer: 53
Fold over basal
Time (min)
2
1.8
1.6
1.4
1.2
1
0.8
0.6
0
10
20
30
40
50
60
Normal
Diabetic
Normal data

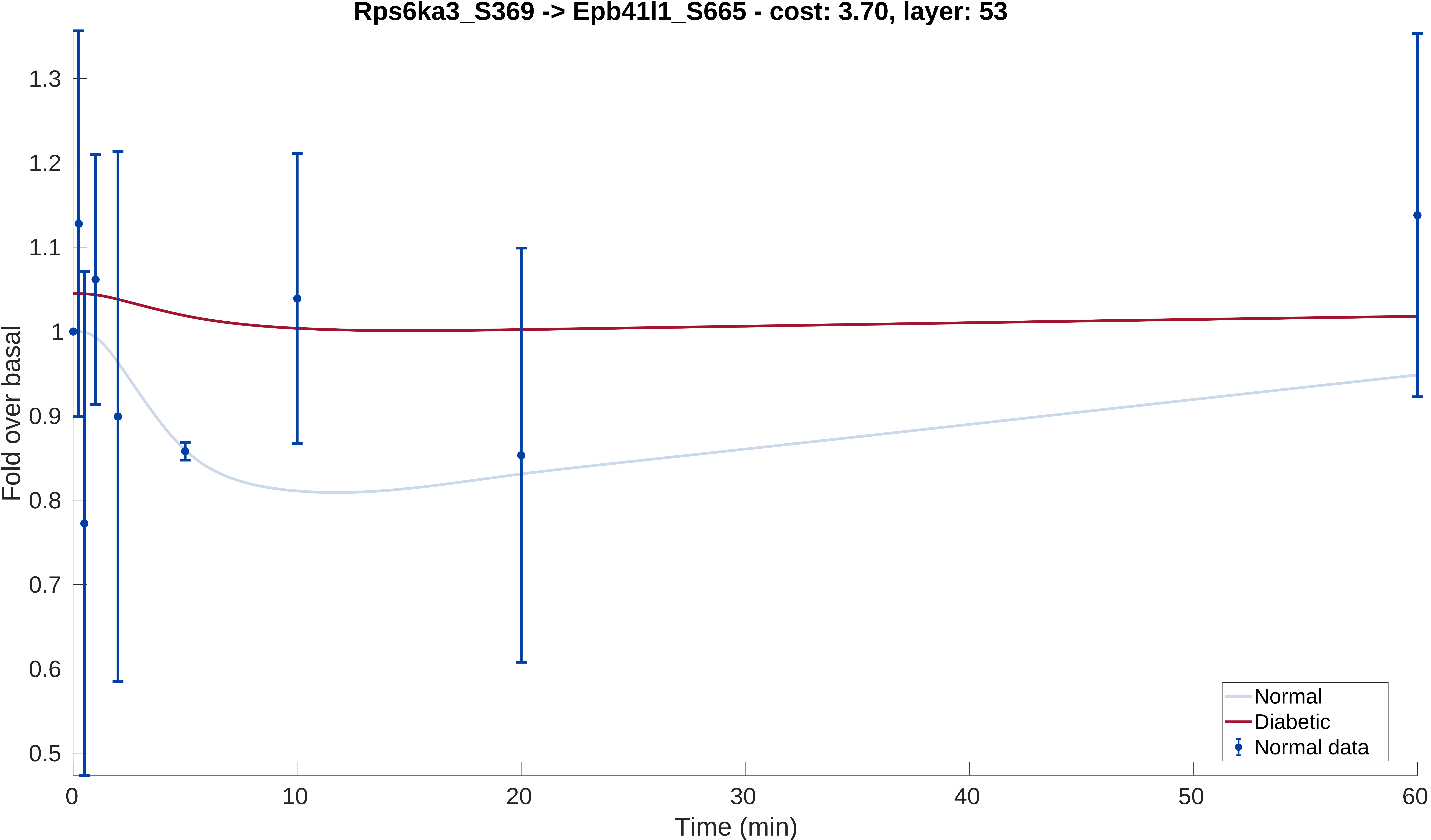
Rps6ka3_S369 -> Epb41l1_S665 - cost: 3.70, layer: 53
Fold over basal
1.3
1.2
1.1
1
0.9
0.8
0.7
0.6
0.5
0
10
20
30
40
50
60
Time (min)
Normal
Diabetic
Normal data

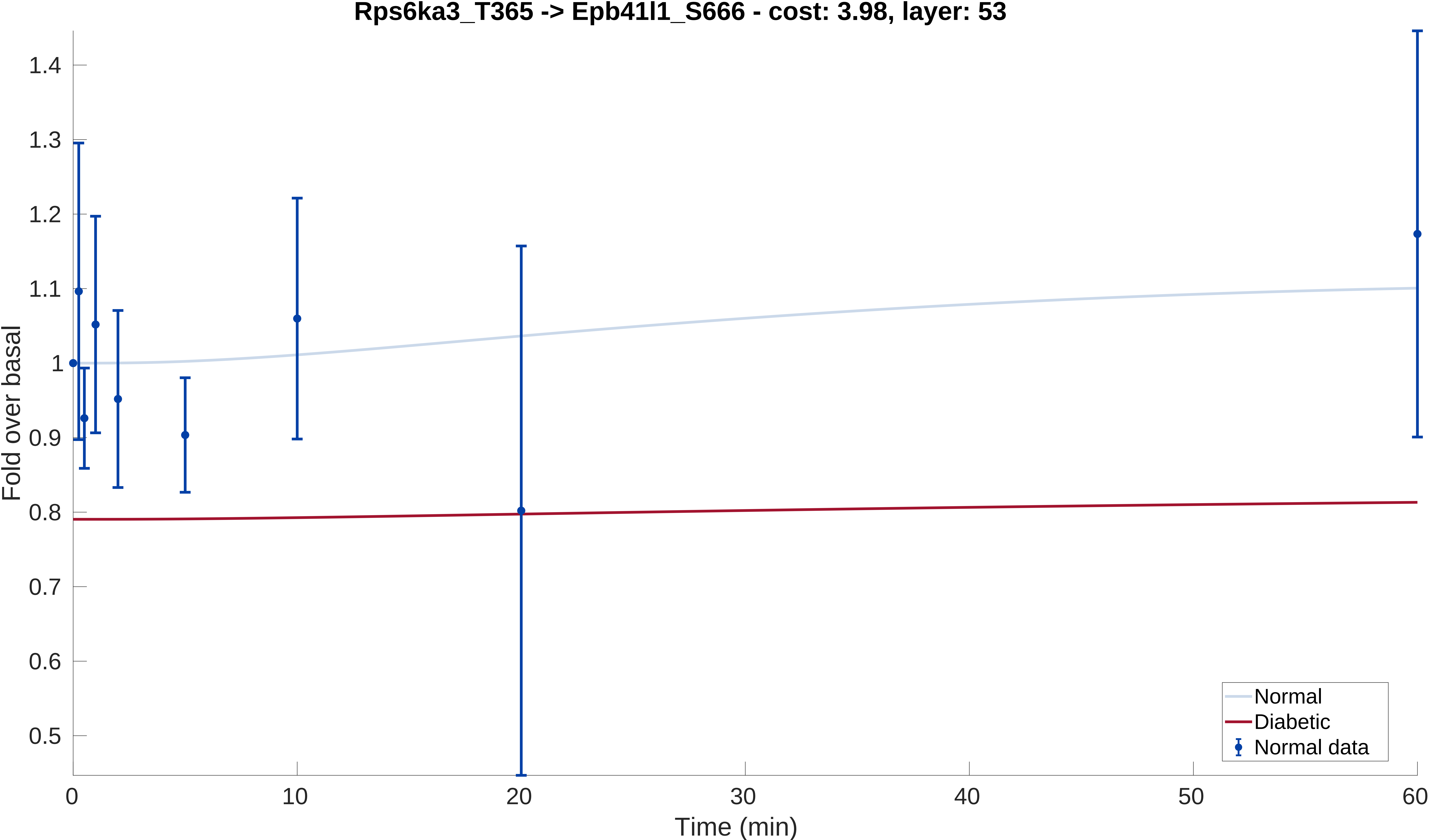

Rps6ka3_T365 -> Epb41l1_S666 - cost: 3.98, layer: 53
Fold over basal
Time (min)
1.4
1.3
1.2
1.1
1
0.9
0.8
0.7
0.6
0.5
0
10
20
30
40
50
60
Normal
Diabetic
Normal data

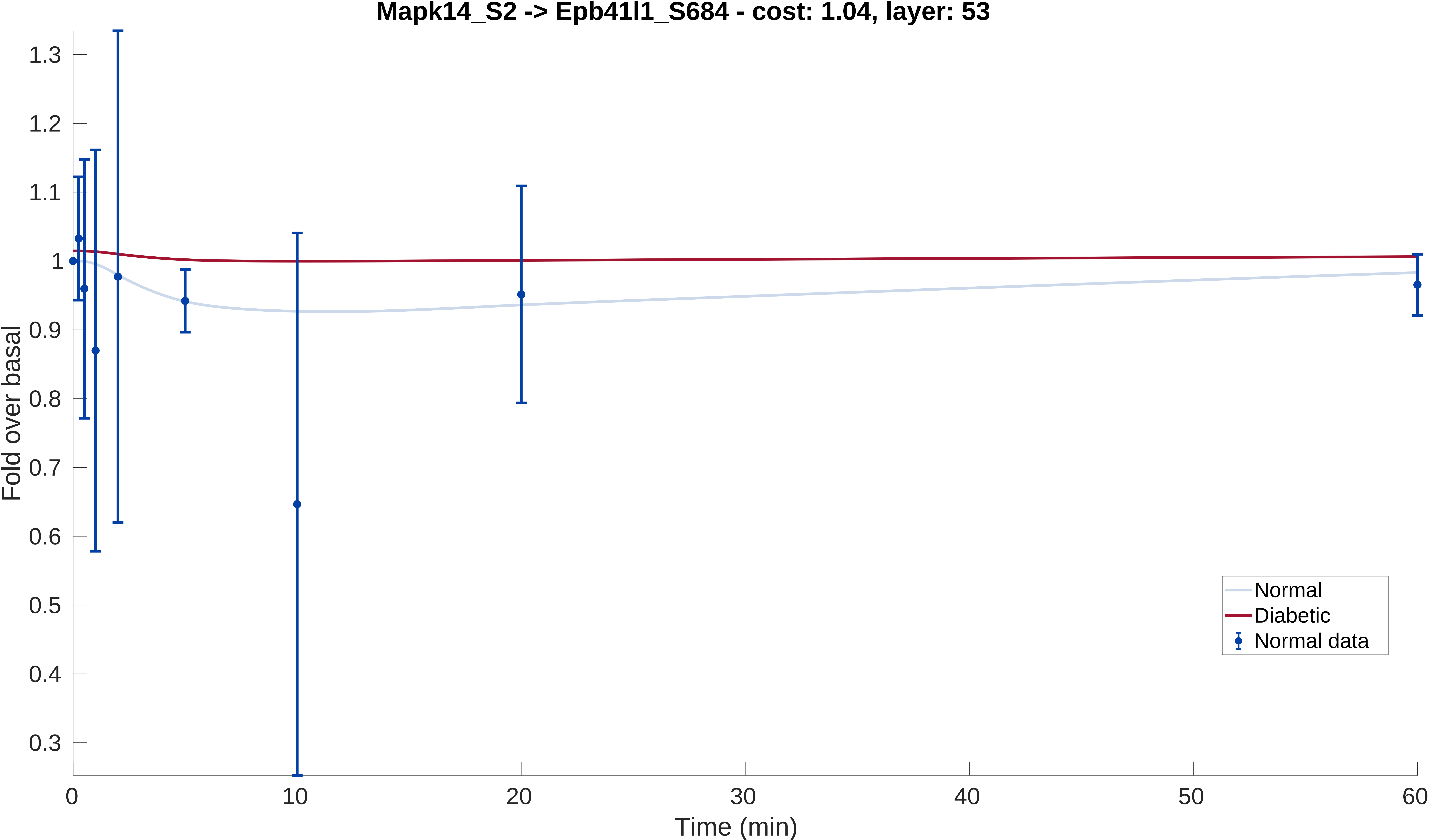

Mapk14_S2 -> Epb41l1_S684 - cost: 1.04, layer: 53
Fold over basal
Time (min)
1.3
1.2
1.1
1
0.9
0.8
0.7
0.6
0.5
0.4
0.3
0
10
20
30
40
50
60
Normal
Diabetic
Normal data

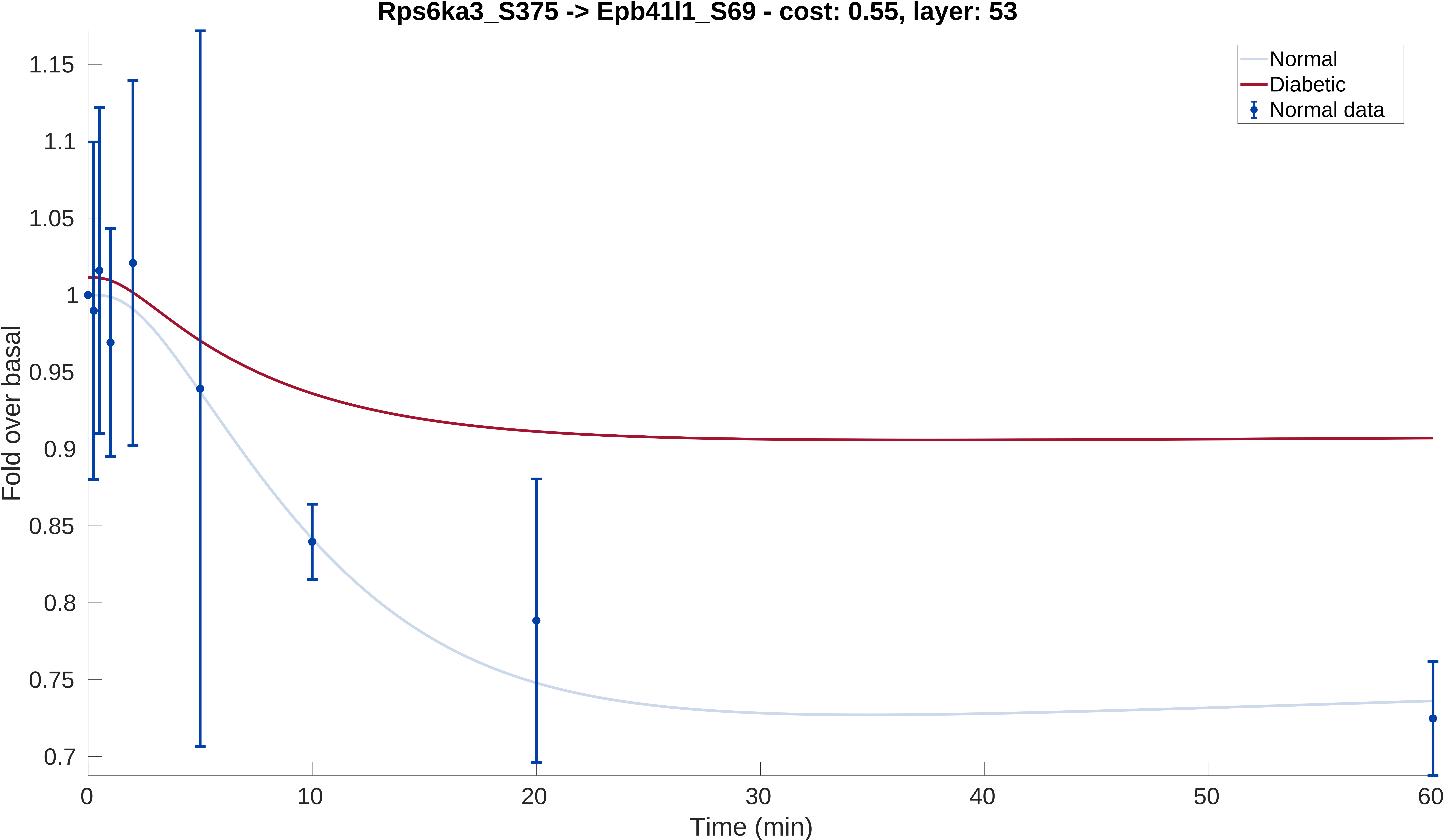

Rps6ka3_S375 -> Epb41l1_S69 - cost: 0.55, layer: 53
Normal
Diabetic
Normal data
Fold over basal
1.15
1.1
1.05
1
0.95
0.9
0.85
0.8
0.75
0.7
0
10
20
30
40
50
60
Time (min)

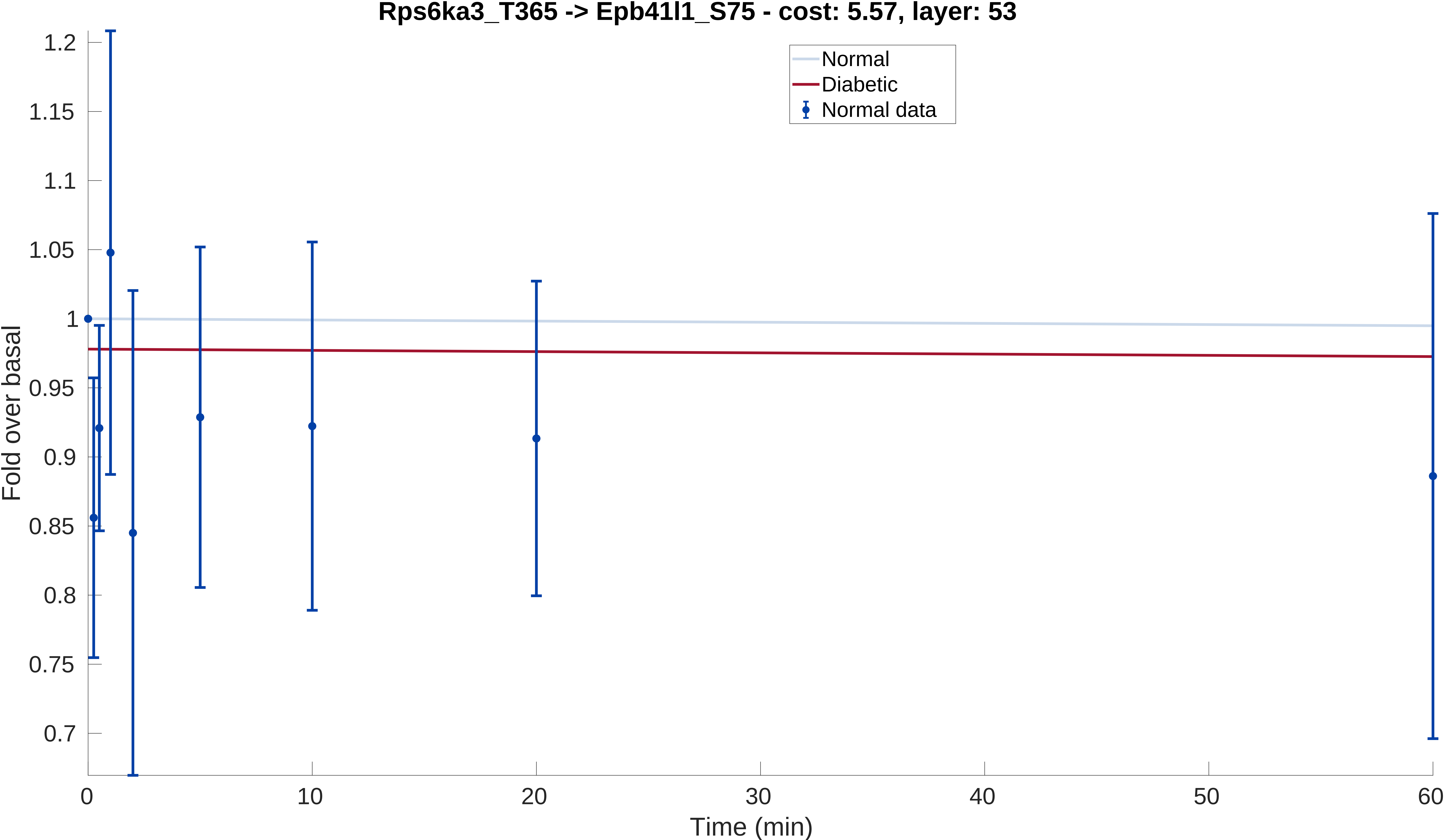

Rps6ka3_T365 -> Epb41l1_S75 - cost: 5.57, layer: 53
Normal
Diabetic
Normal data
Fold over basal
Time (min)
1.2
1.15
1.1
1.05
1
0.95
0.9
0.85
0.8
0.75
0.7
0
10
20
30
40
50
60

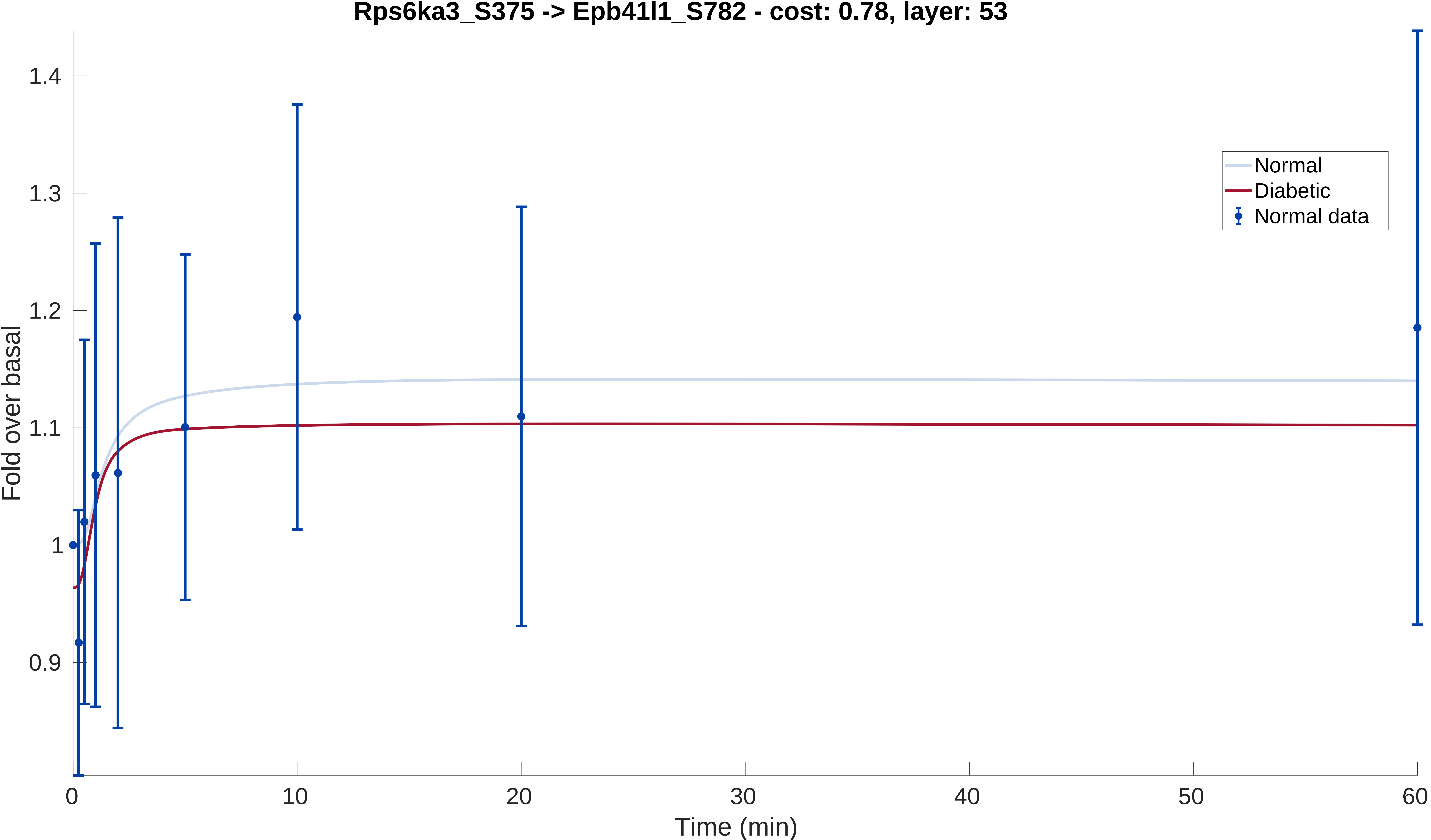

Rps6ka3_S375 -> Epb41l1_S782 - cost: 0.78, layer: 53
Fold over basal
Time (min)
Normal
Diabetic
Normal data
1.4
1.3
1.2
1.1
1
0.9
0
10
20
30
40
50
60

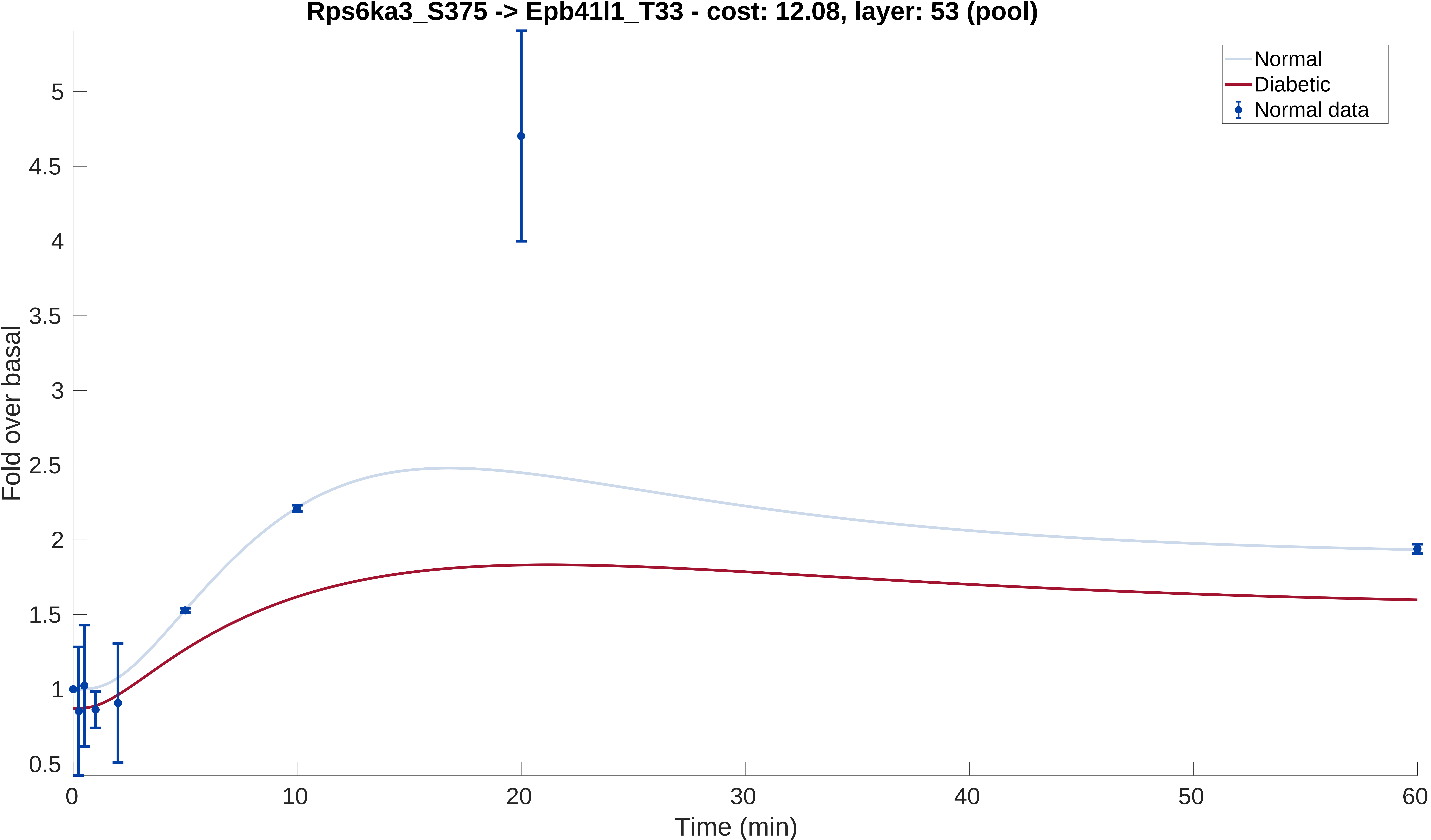

Rps6ka3_S375 -> Epb41l1_T33 - cost: 12.08, layer: 53 (pool)
Normal
Diabetic
Normal data
Fold over basal
Time (min)
5
4.5
4
3.5
3
2.5
2
1.5
1
0.5
0
10
20
30
40
50
60

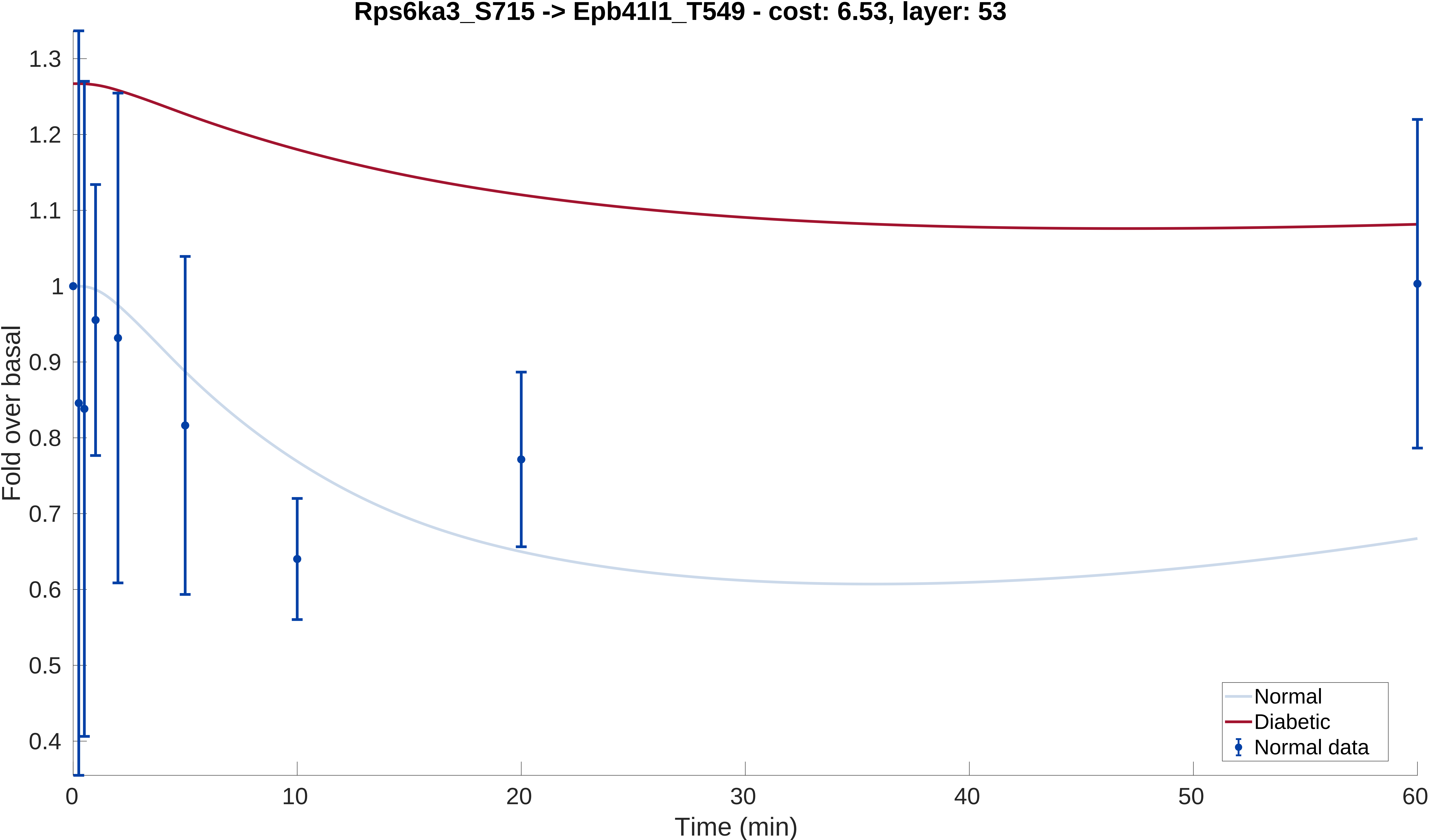

Rps6ka3_S715 -> Epb41l1_T549 - cost: 6.53, layer: 53
Fold over basal
Time (min)
1.3
1.2
1.1
1
0.9
0.8
0.7
0.6
0.5
0.4
0
10
20
30
40
50
60
Normal
Diabetic
Normal data

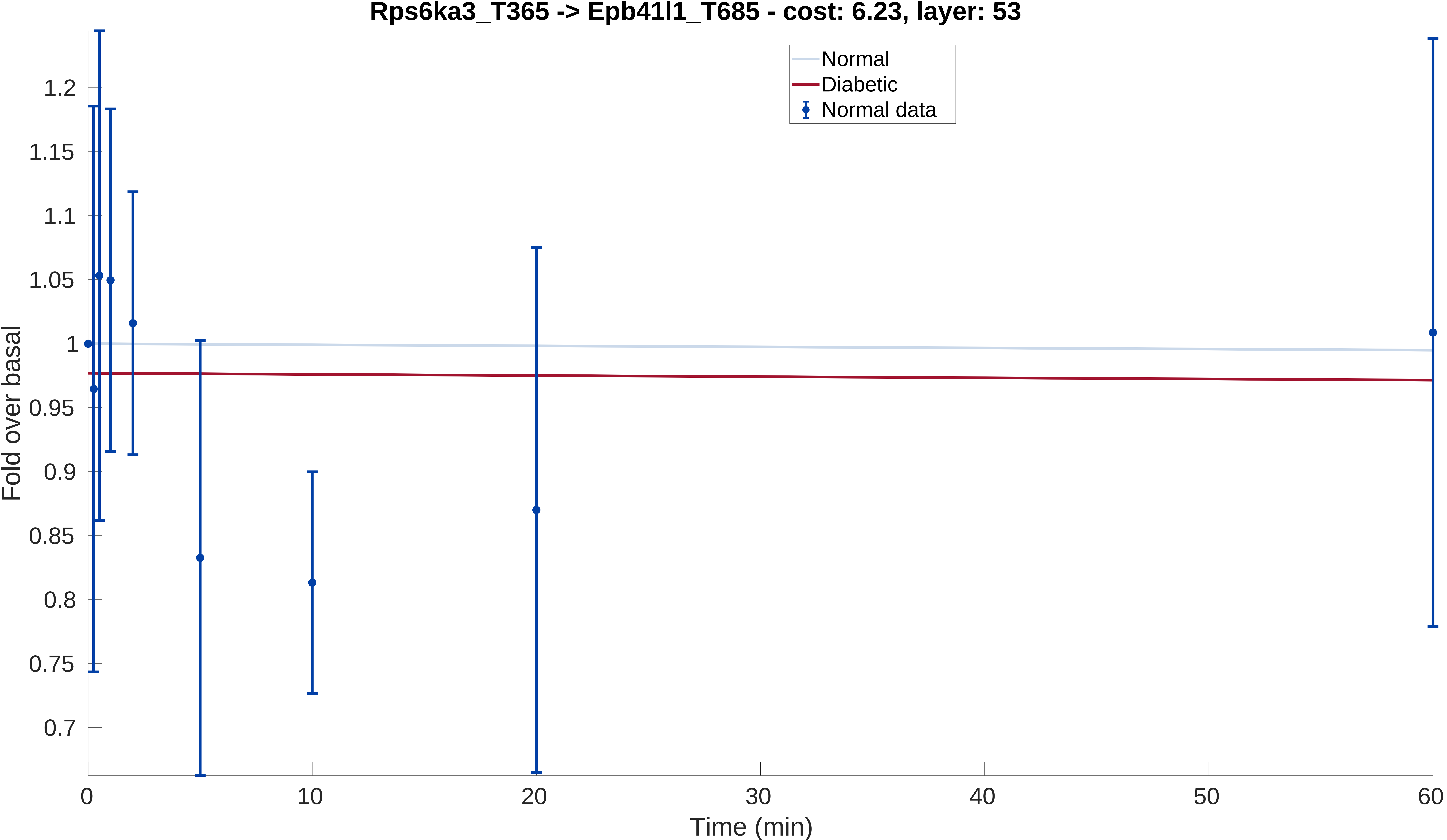

Rps6ka3_T365 -> Epb41l1_T685 - cost: 6.23, layer: 53
Normal
Diabetic
Normal data
Fold over basal
1.2
1.15
1.1
1.05
1
0.95
0.9
0.85
0.8
0.75
0.7
0
10
20
30
40
50
60
Time (min)

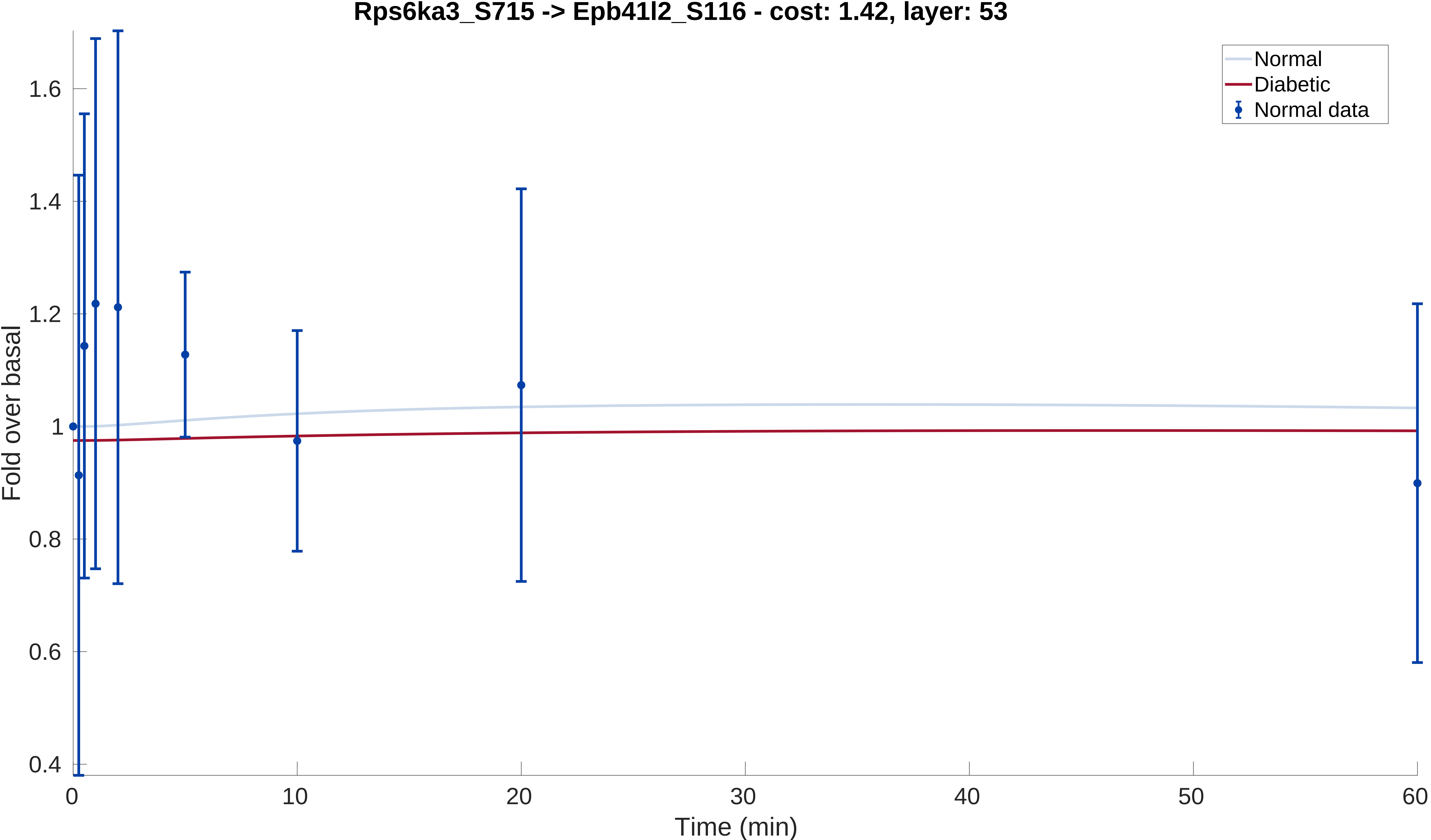

Rps6ka3_S715 -> Epb41l2_S116 - cost: 1.42, layer: 53
Normal
Diabetic
Normal data
Fold over basal
1.6
1.4
1.2
1
0.8
0.6
0.4
0
10
20
30
40
50
60
Time (min)

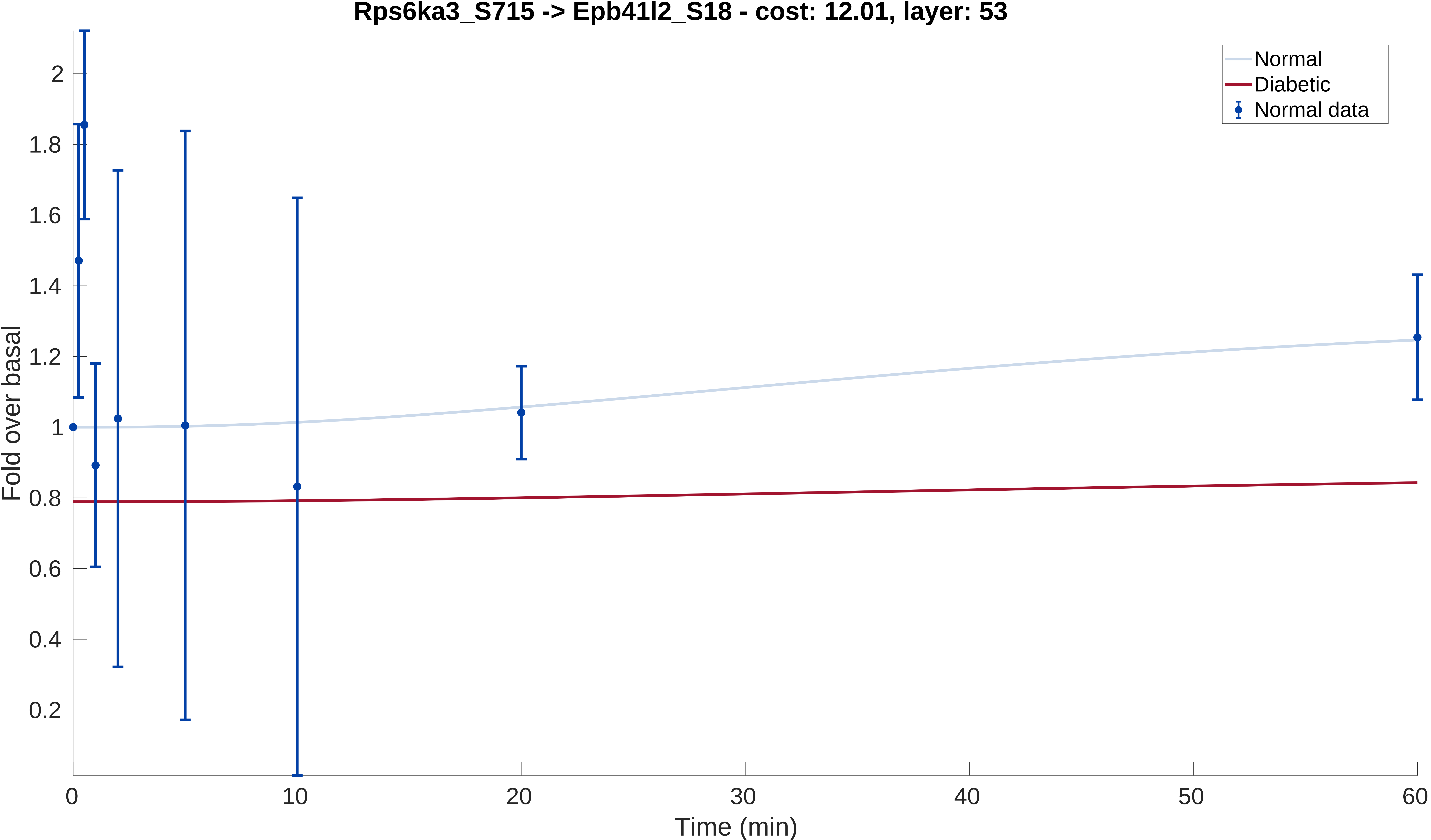

Rps6ka3_S715 -> Epb41l2_S18 - cost: 12.01, layer: 53
Normal
Diabetic
Normal data
Fold over basal
2
1.8
1.6
1.4
1.2
1
0.8
0.6
0.4
0.2
0
10
20
30
40
50
60
Time (min)

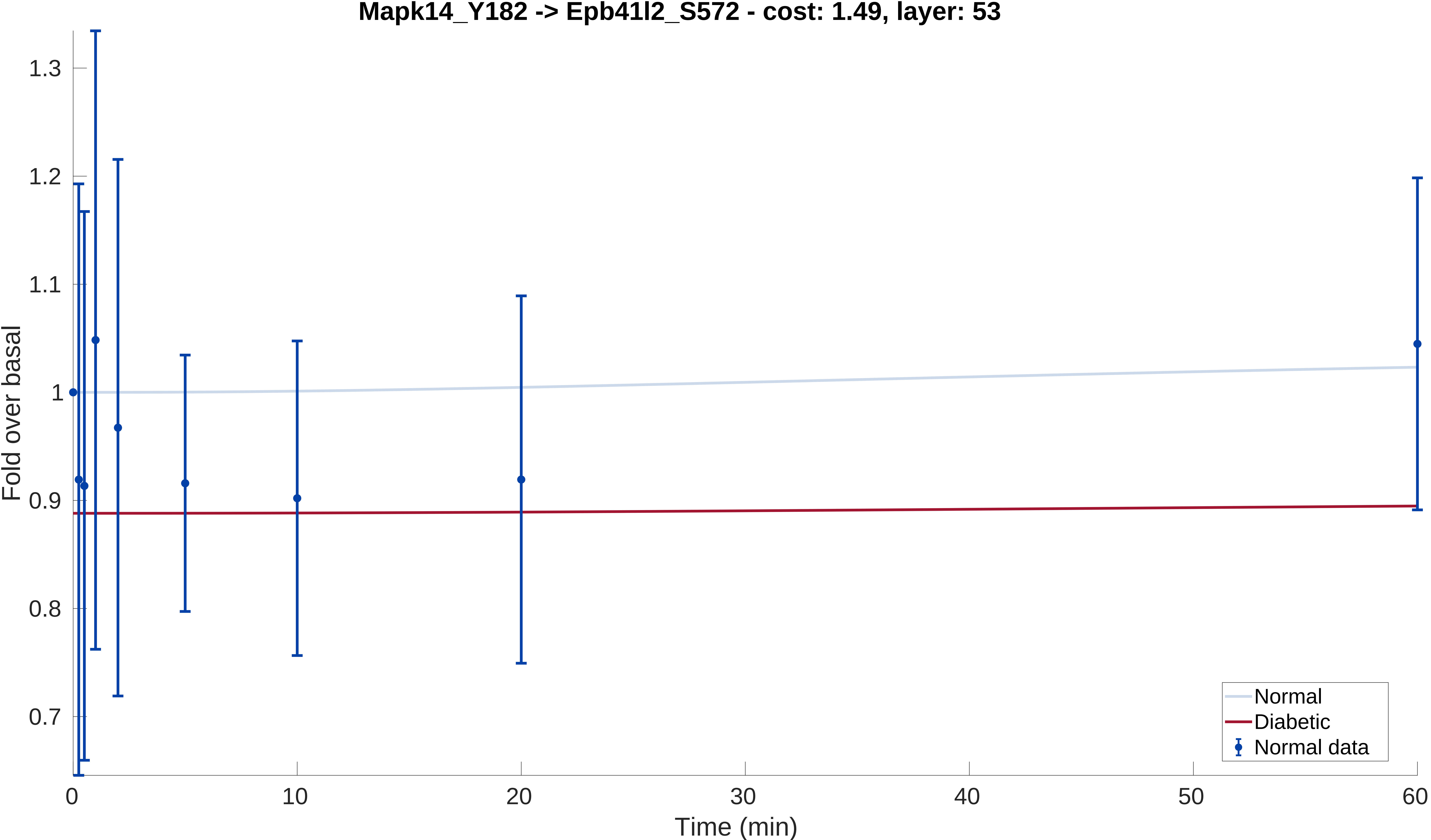

Mapk14_Y182 -> Epb41l2_S572 - cost: 1.49, layer: 53
Fold over basal
1.3
1.2
1.1
1
0.9
0.8
0.7
0
10
20
30
40
50
60
Time (min)
Normal
Diabetic
Normal data

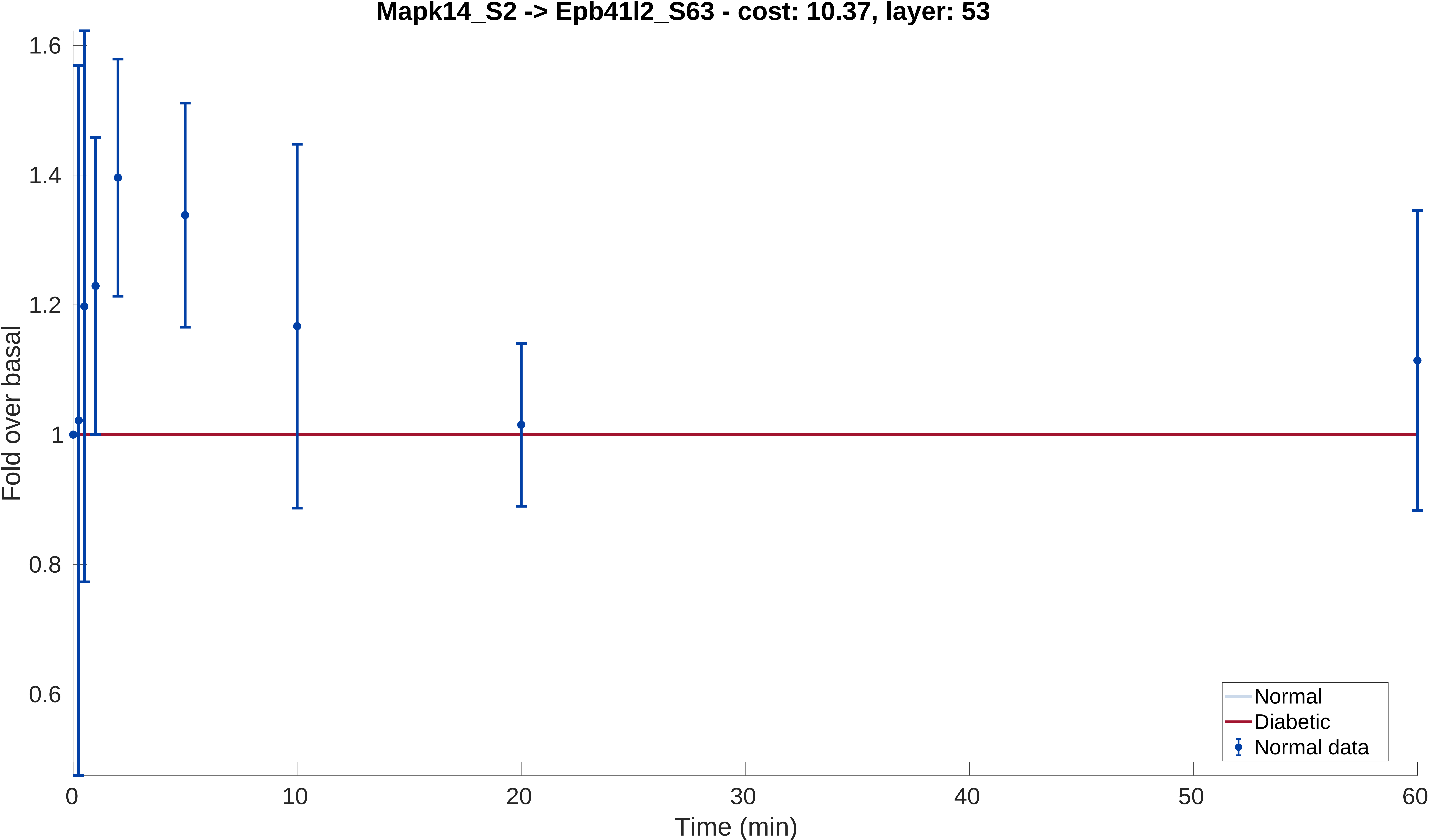
Mapk14_S2 -> Epb41l2_S63 - cost: 10.37, layer: 53
Fold over basal
1.6
1.4
1.2
1
0.8
0.6
0
10
20
30
40
50
60
Time (min)
Normal
Diabetic
Normal data

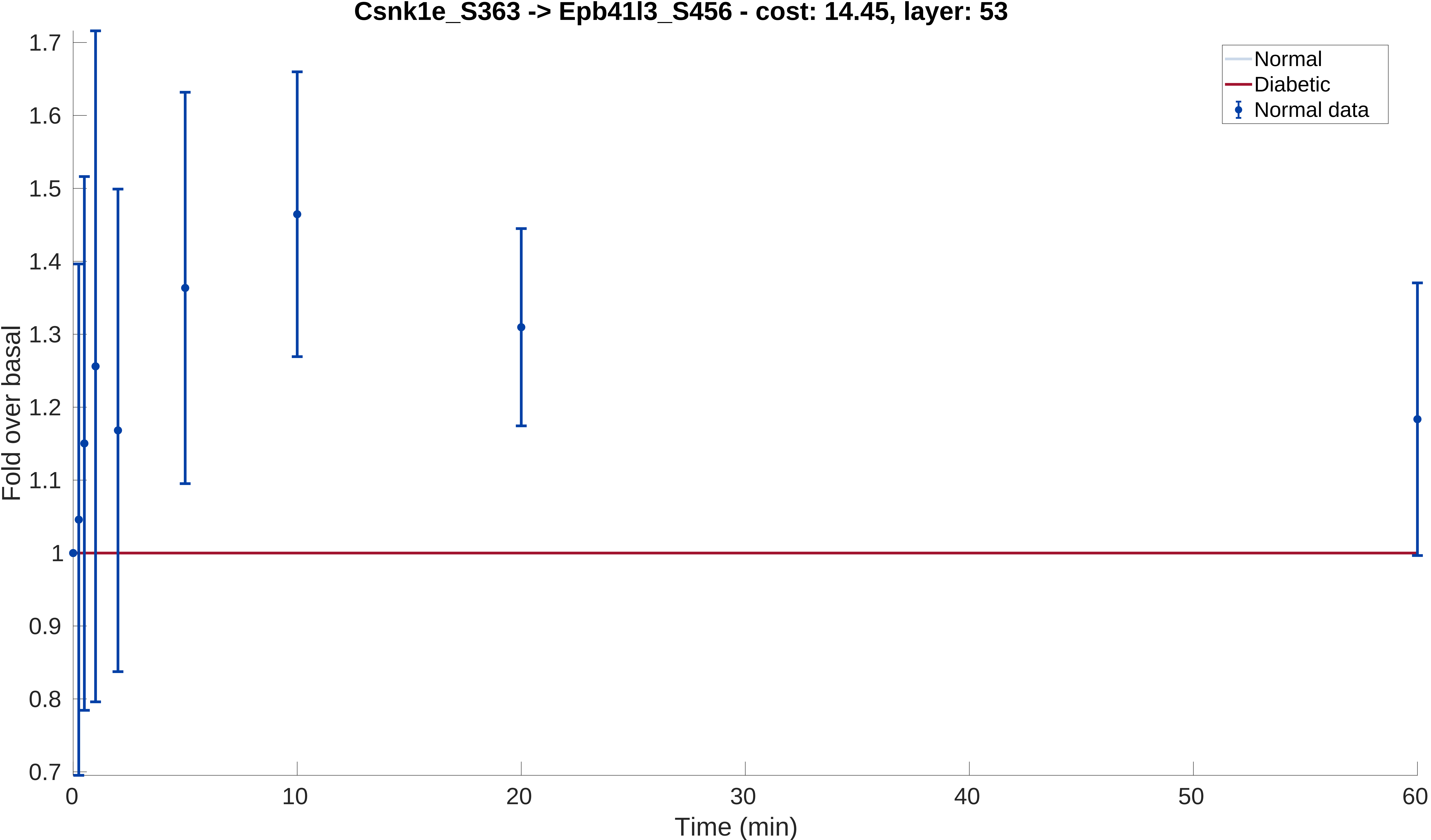

Csnk1e_S363 -> Epb41l3_S456 - cost: 14.45, layer: 53
Normal
Diabetic
Normal data
Fold over basal
1.7
1.6
1.5
1.4
1.3
1.2
1.1
1
0.9
0.8
0.7
0
10
20
30
40
50
60
Time (min)

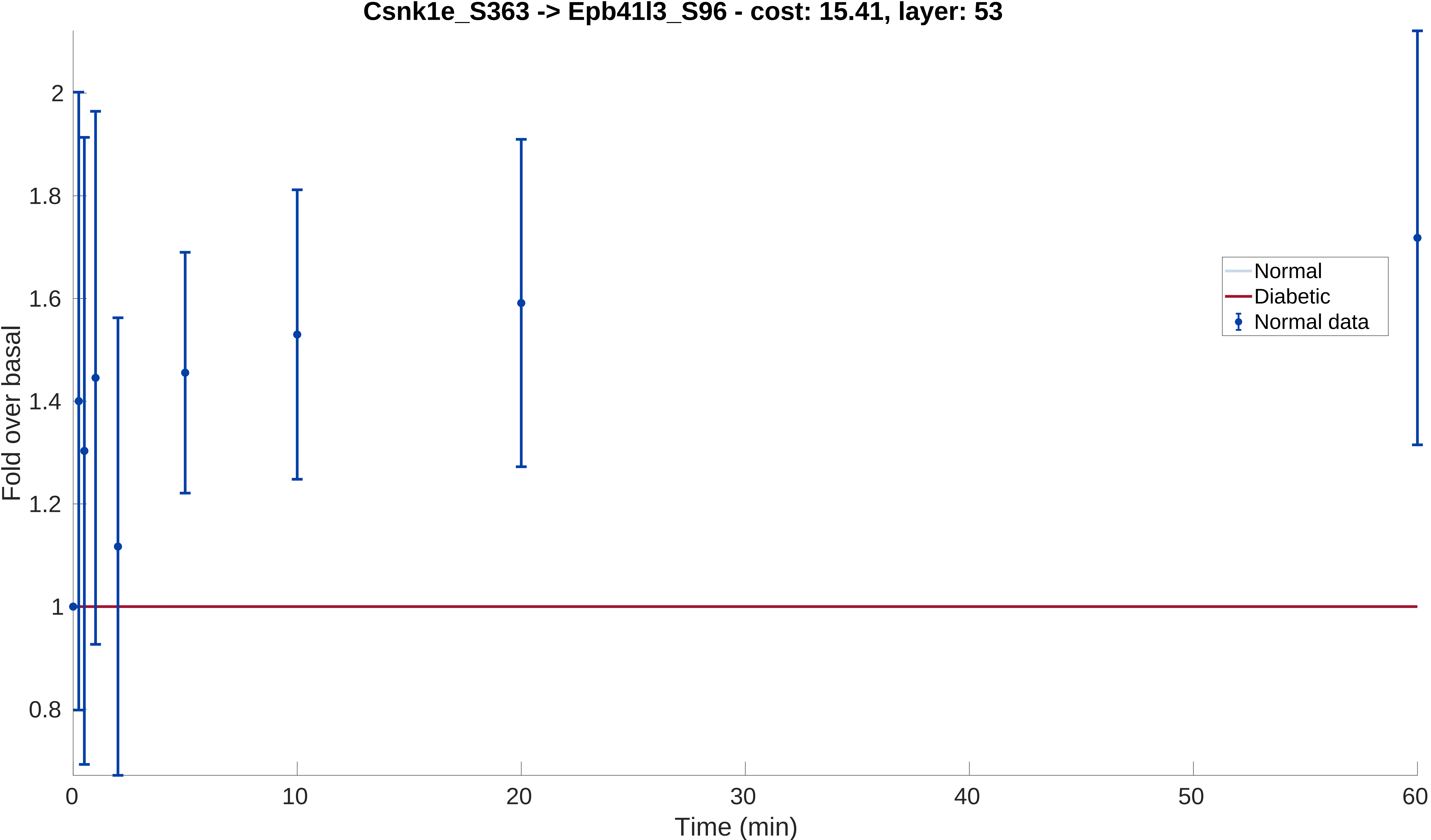
Csnk1e_S363 -> Epb41l3_S96 - cost: 15.41, layer: 53
Fold over basal
2
1.8
1.6
1.4
1.2
1
0.8
0
10
20
30
40
50
60
Time (min)
Normal
Diabetic
Normal data

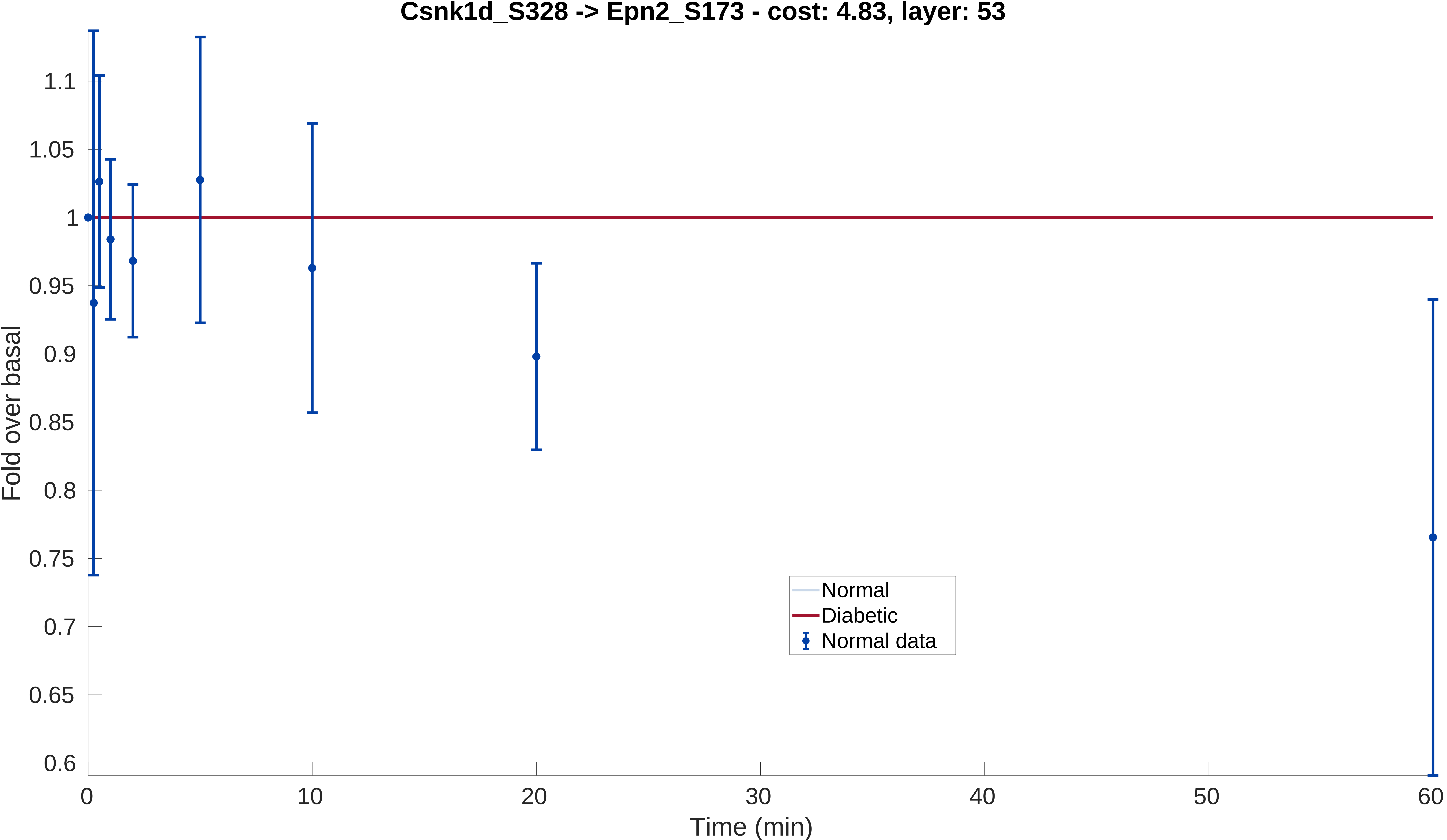

Csnk1d_S328 -> Epn2_S173 - cost: 4.83, layer: 53
Fold over basal
1.1
1.05
1
0.95
0.9
0.85
0.8
0.75
0.7
0.65
0.6
0
10
20
30
40
50
60
Time (min)
Normal
Diabetic
Normal data

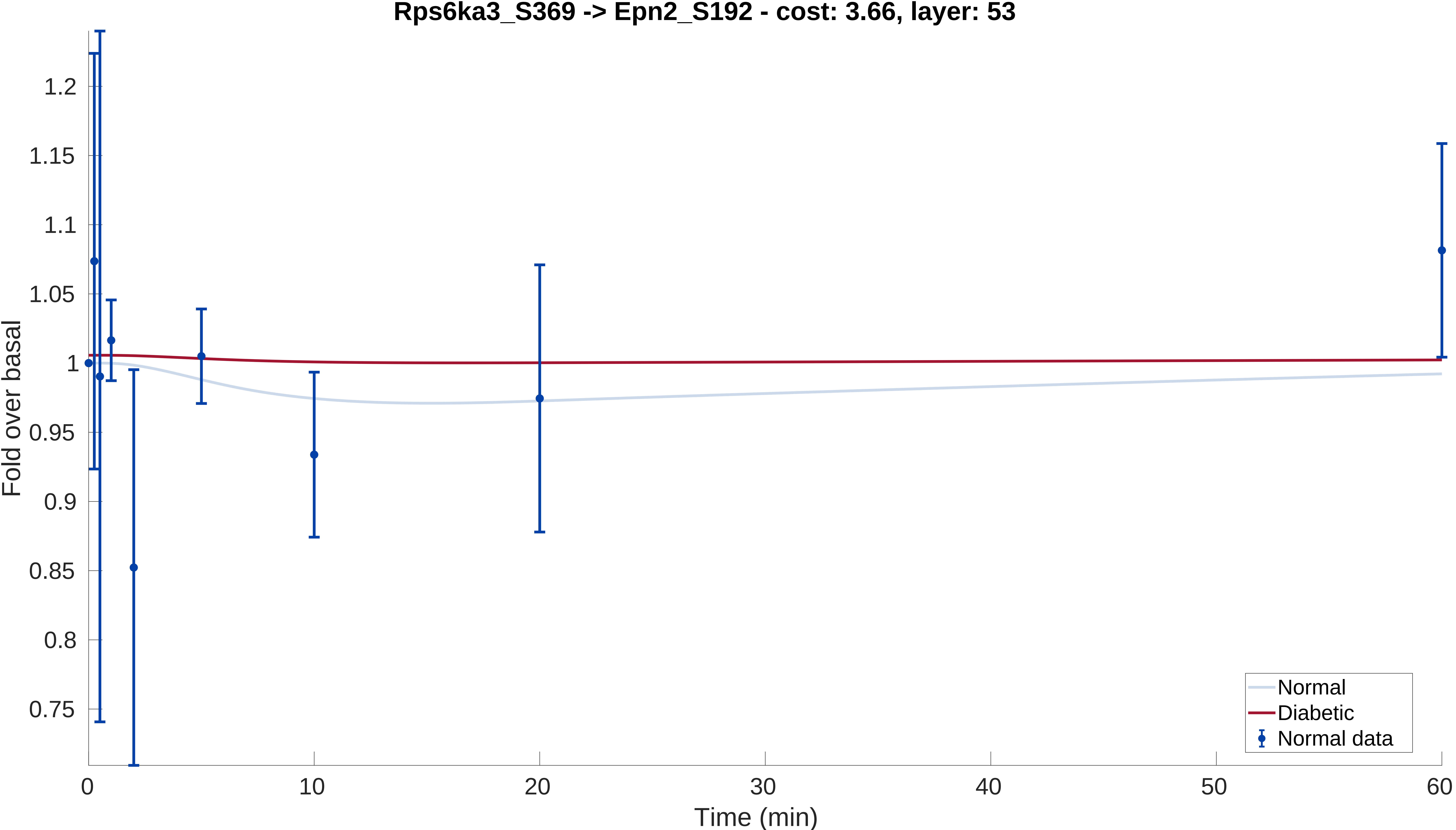

Rps6ka3_S369 -> Epn2_S192 - cost: 3.66, layer: 53
Fold over basal
1.2
1.15
1.1
1.05
1
0.95
0.9
0.85
0.8
0.75
0
10
20
30
40
50
60
Time (min)
Normal
Diabetic
Normal data

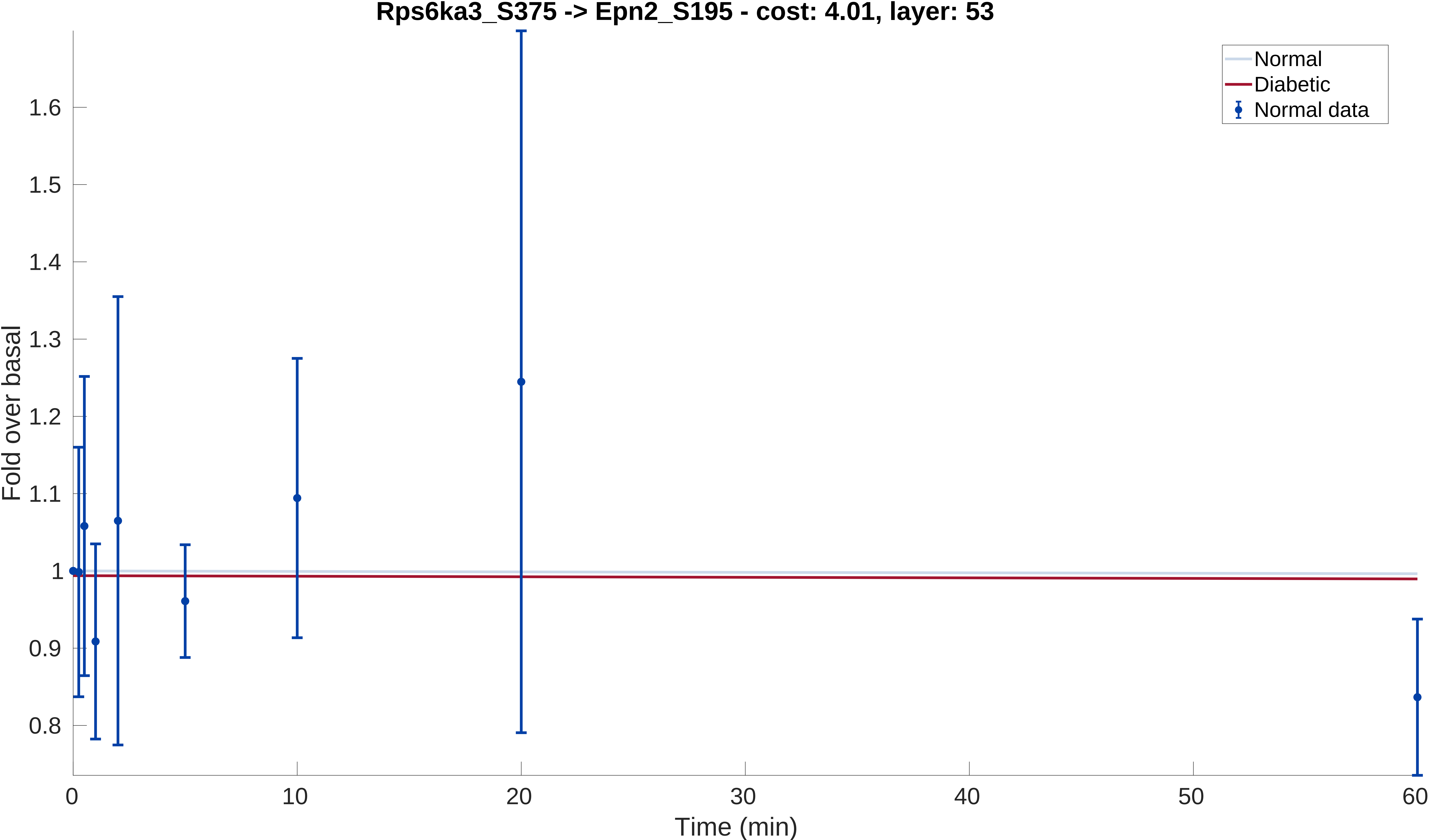

Rps6ka3_S375 -> Epn2_S195 - cost: 4.01, layer: 53
Normal
Diabetic
Normal data
Fold over basal
1.6
1.5
1.4
1.3
1.2
1.1
1
0.9
0.8
0
10
20
30
40
50
60
Time (min)

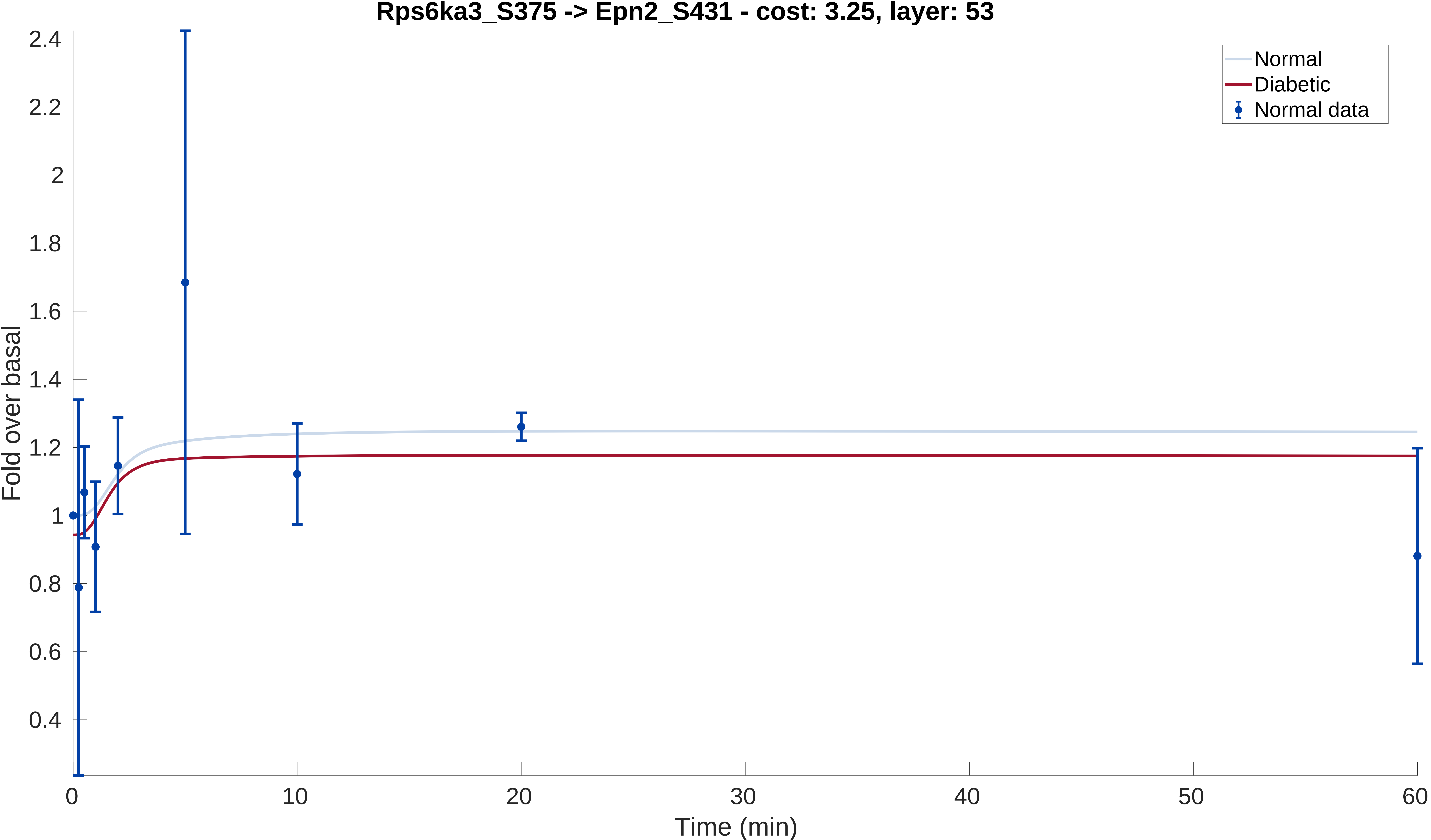

Rps6ka3_S375 -> Epn2_S431 - cost: 3.25, layer: 53
Fold over basal
2.4
2.2
2
1.8
1.6
1.4
1.2
1
0.8
0.6
0.4
0
10
20
30
40
50
60
Time (min)
Normal
Diabetic
Normal data

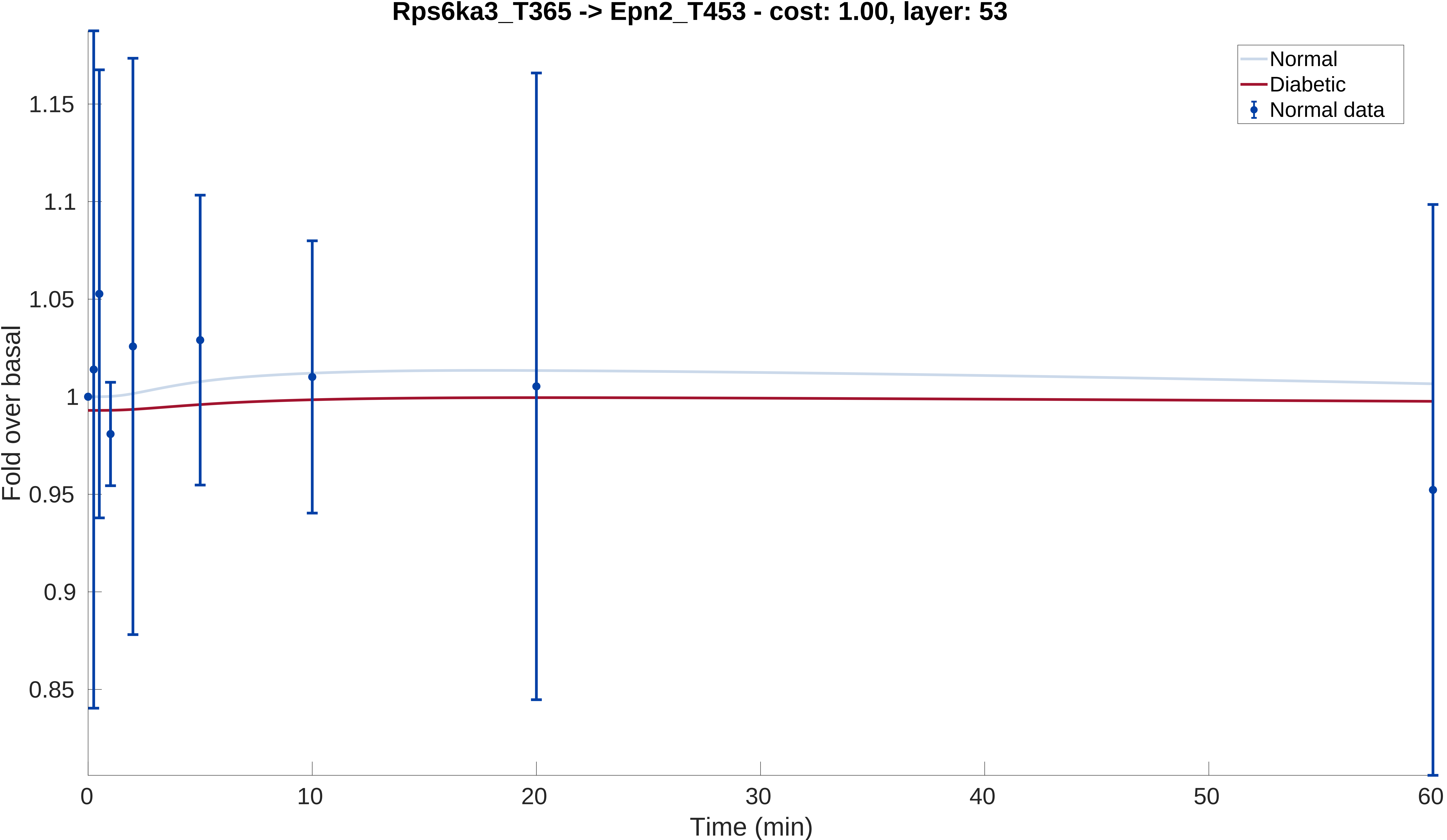

Rps6ka3_T365 -> Epn2_T453 - cost: 1.00, layer: 53
Normal
Diabetic
Normal data
Fold over basal
1.15
1.1
1.05
1
0.95
0.9
0.85
0
10
20
30
40
50
60
Time (min)

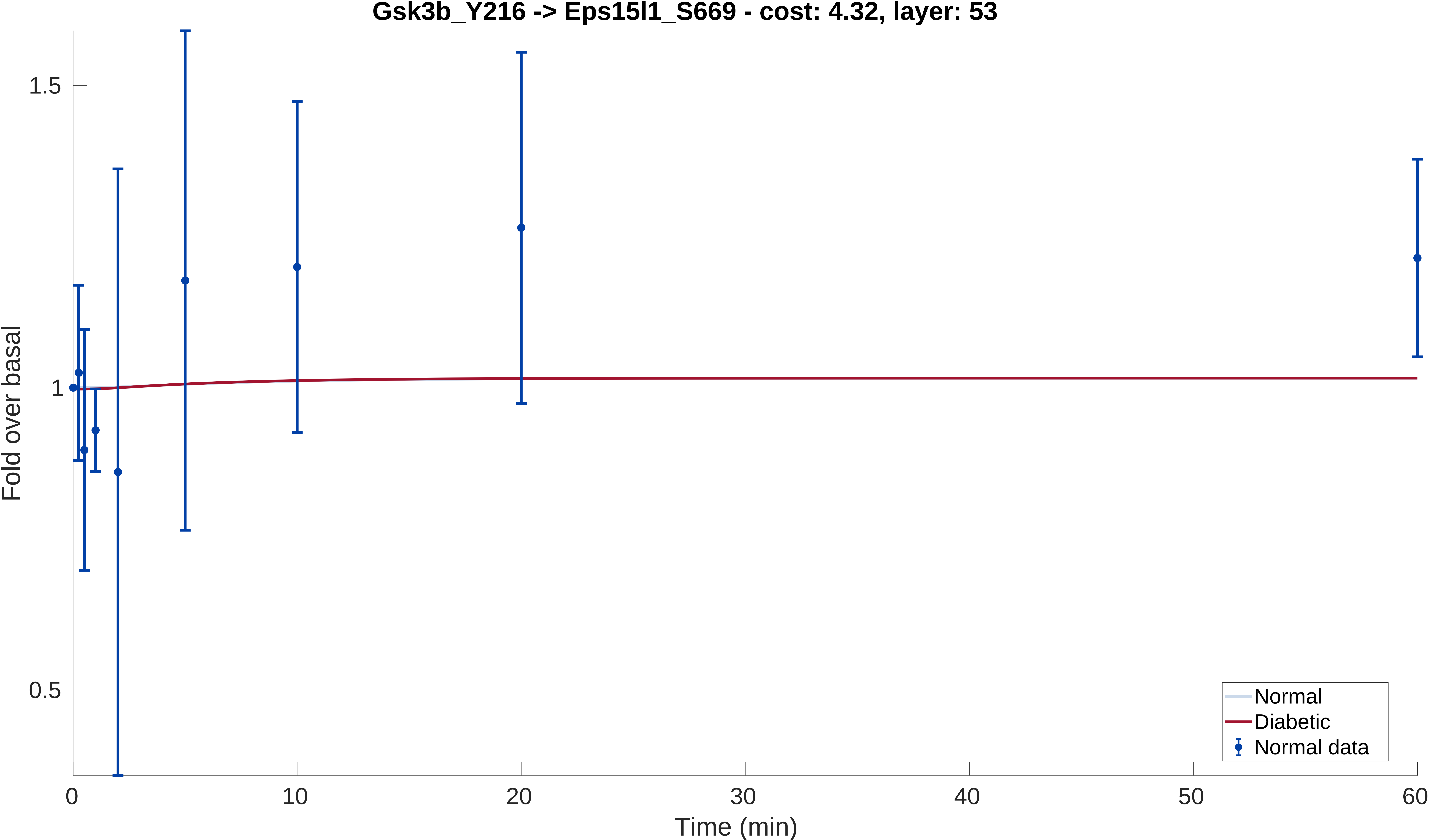

Gsk3b_Y216 -> Eps15l1_S669 - cost: 4.32, layer: 53
1.5
1
0.5
Fold over basal
0
10
20
30
40
50
60
Time (min)
Normal
Diabetic
Normal data

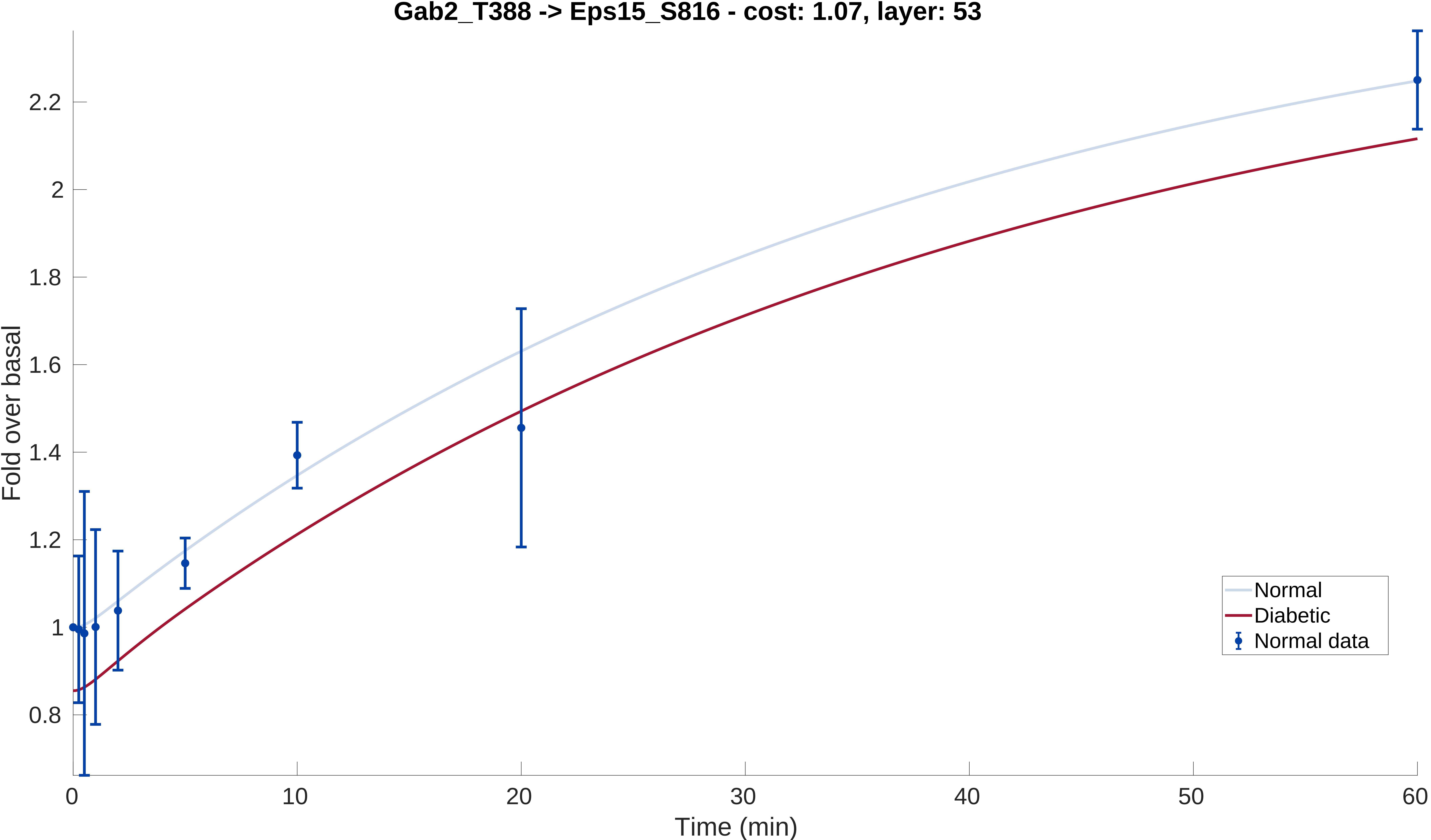

Gab2_T388 -> Eps15_S816 - cost: 1.07, layer: 53
Fold over basal
2.2
2
1.8
1.6
1.4
1.2
1
0.8
0
10
20
30
40
50
60
Time (min)
Normal
Diabetic
Normal data

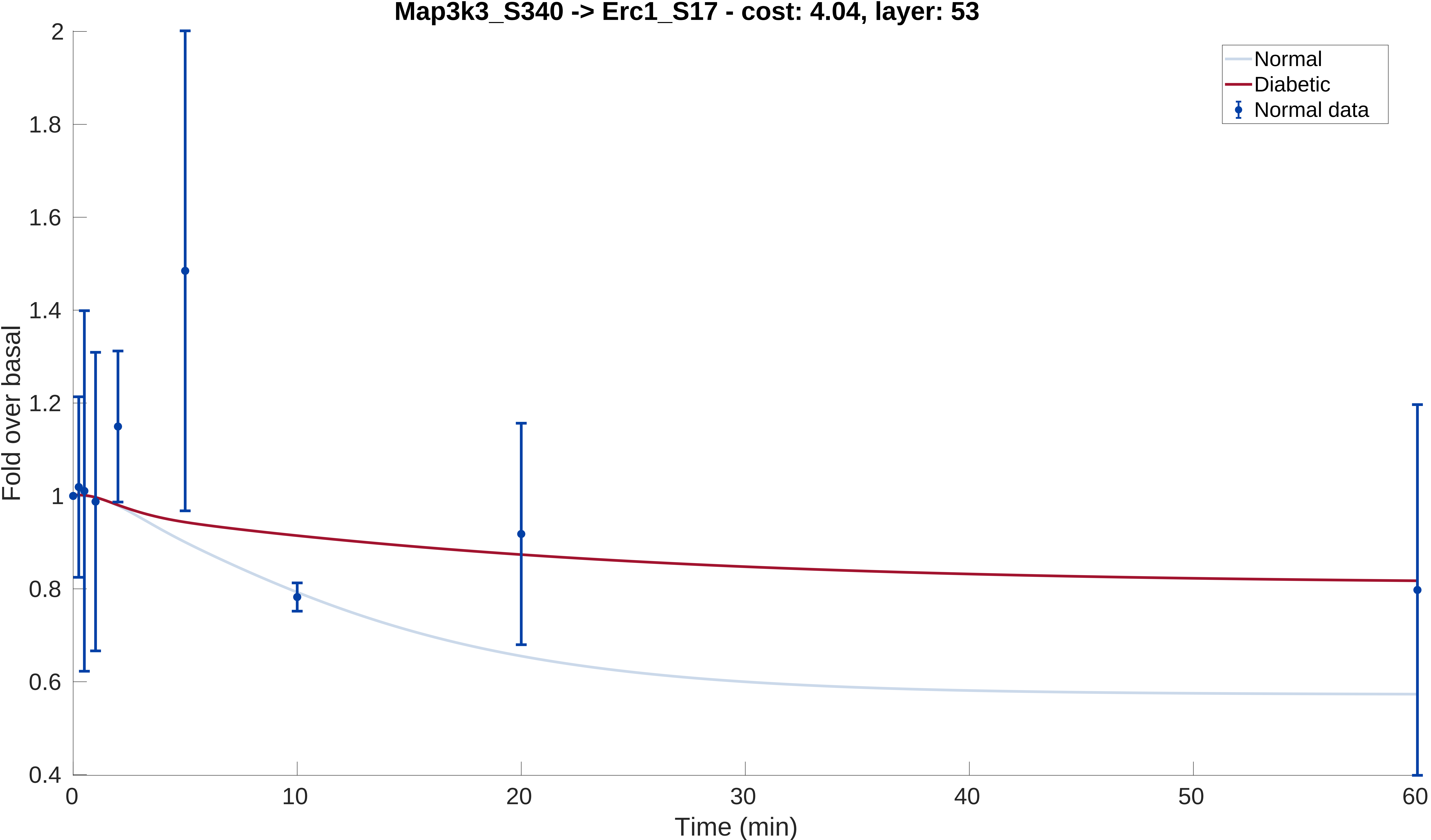
Map3k3_S340 -> Erc1_S17 - cost: 4.04, layer: 53
Normal
Diabetic
Normal data
Fold over basal
2
1.8
1.6
1.4
1.2
1
0.8
0.6
0.4
0
10
20
30
40
50
60
Time (min)

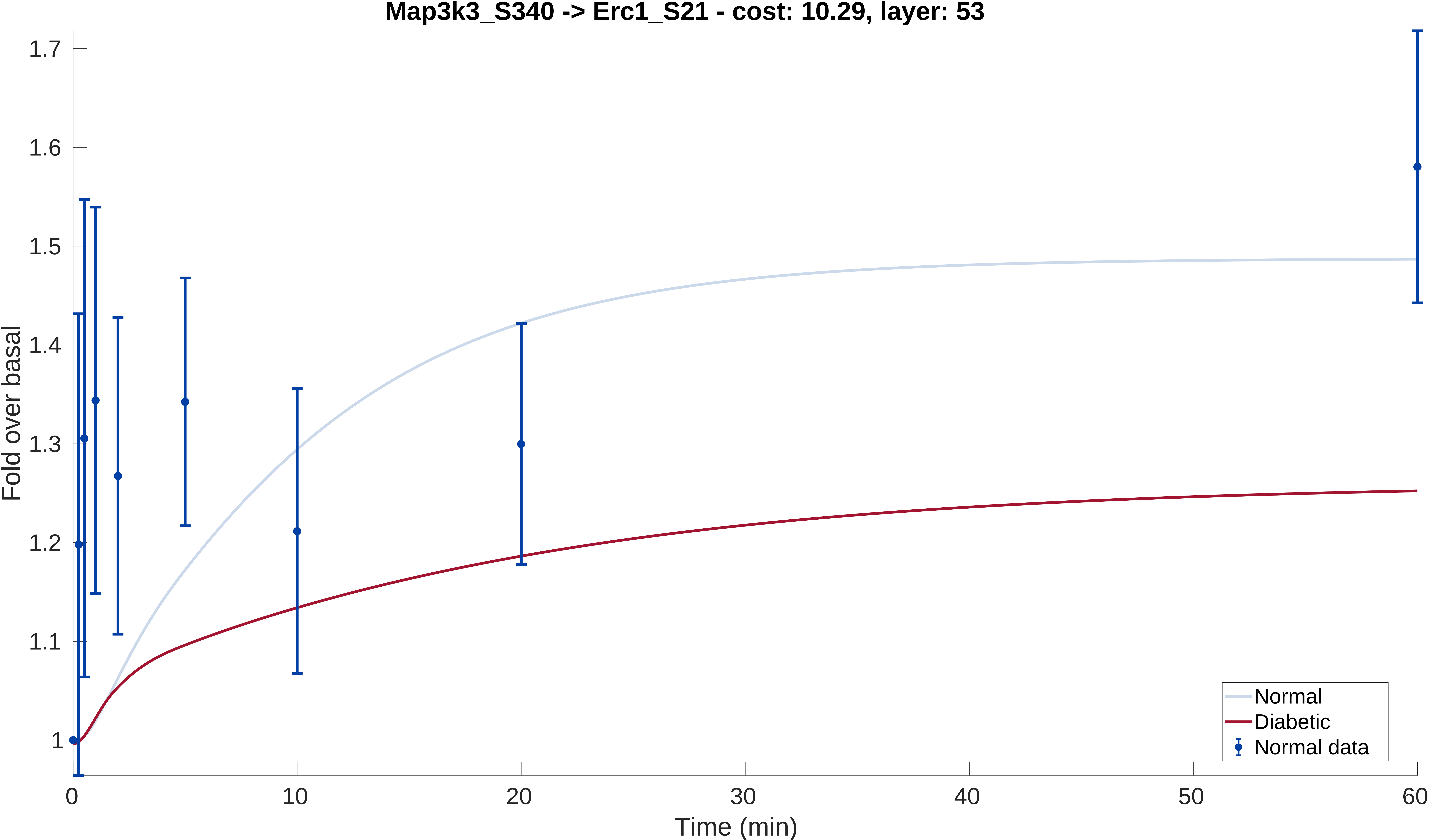

Map3k3_S340 -> Erc1_S21 - cost: 10.29, layer: 53
Fold over basal
1.7
1.6
1.5
1.4
1.3
1.2
1.1
1
0
10
20
30
40
50
60
Time (min)
Normal
Diabetic
Normal data

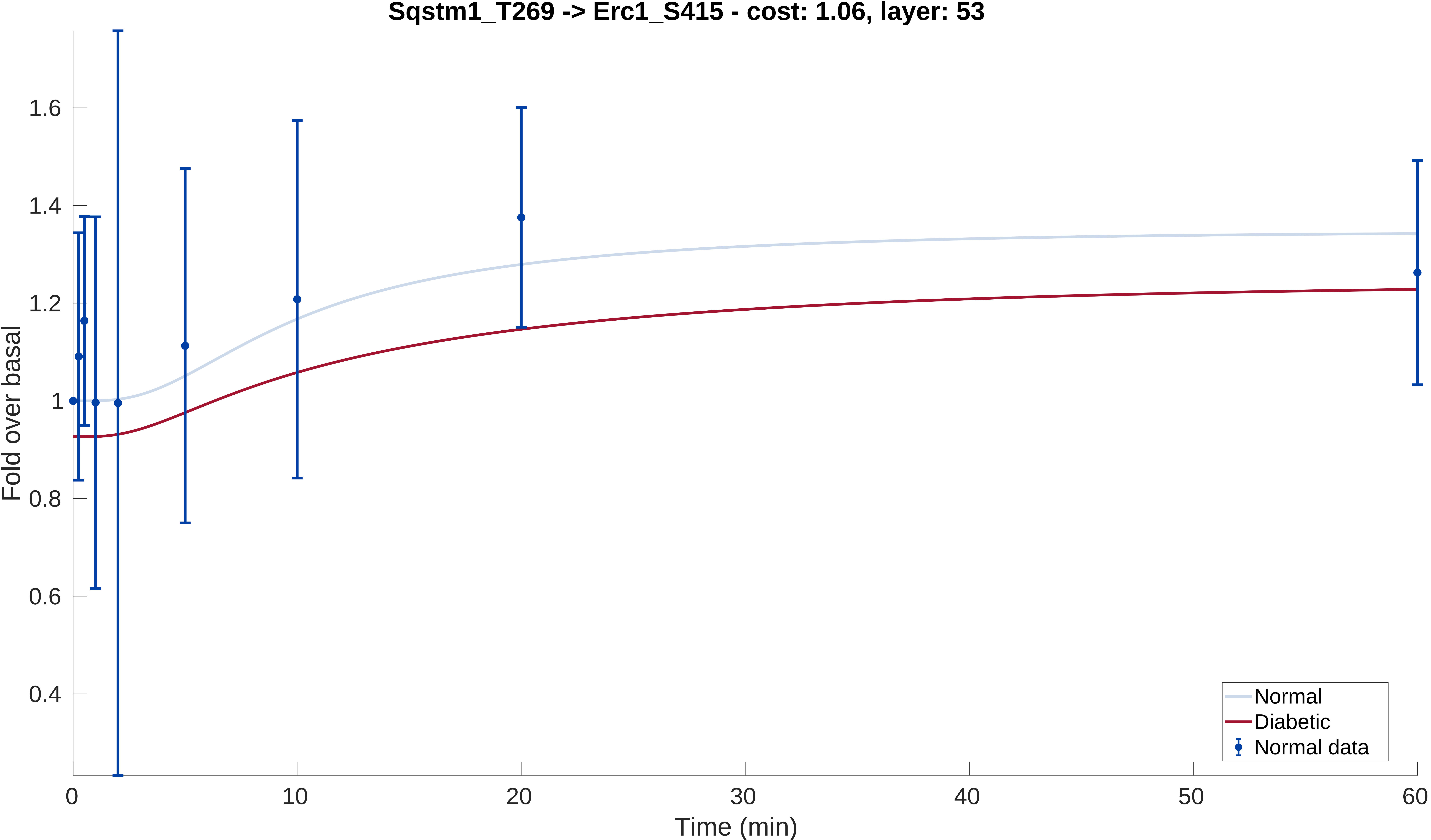

Sqstm1_T269 -> Erc1_S415 - cost: 1.06, layer: 53
Fold over basal
1.6
1.4
1.2
1
0.8
0.6
0.4
0
10
20
30
40
50
60
Time (min)
Normal
Diabetic
Normal data

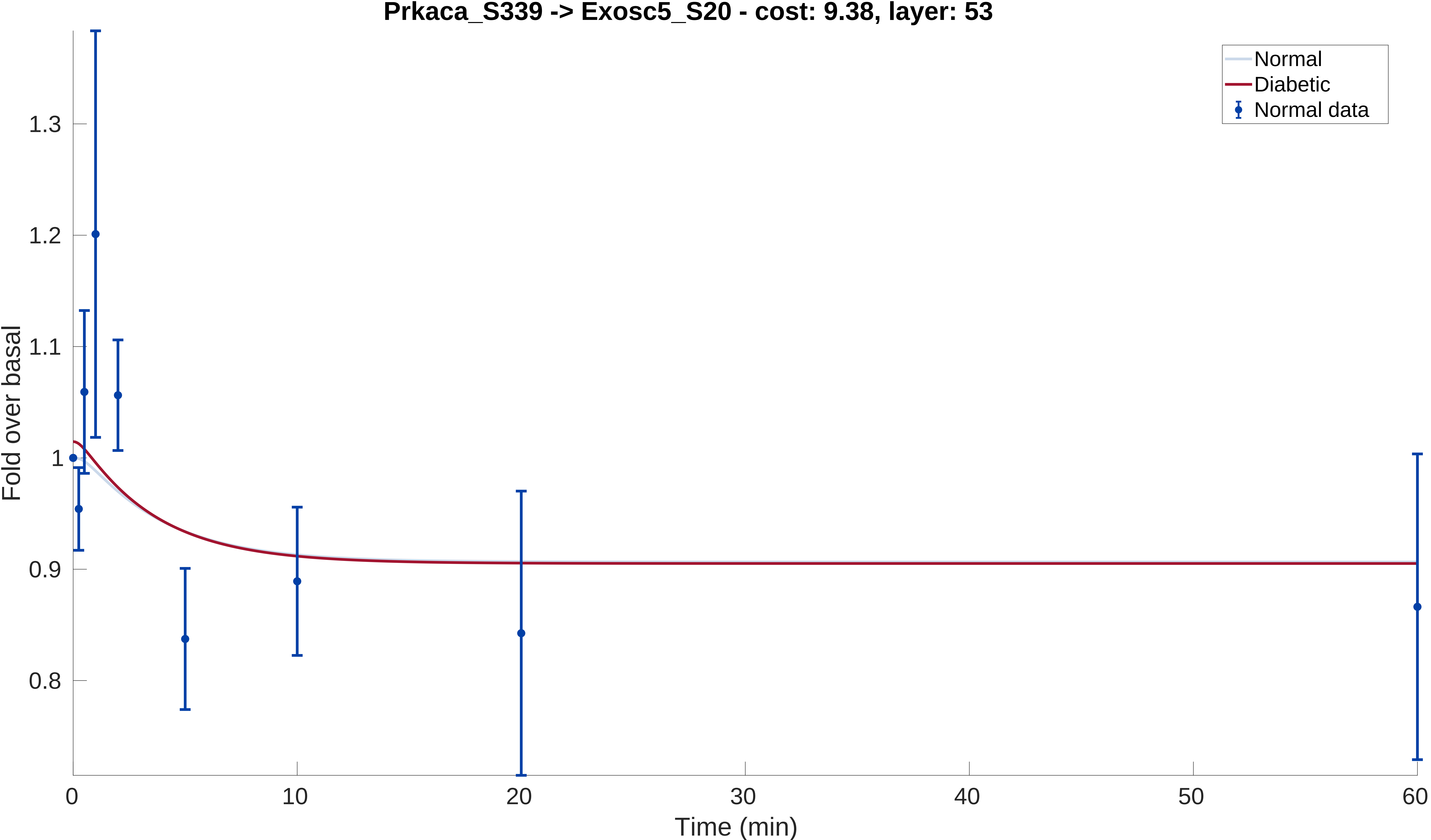

Prkaca_S339 -> Exosc5_S20 - cost: 9.38, layer: 53
Normal
Diabetic
Normal data
Fold over basal
1.3
1.2
1.1
1
0.9
0.8
0
10
20
30
40
50
60
Time (min)

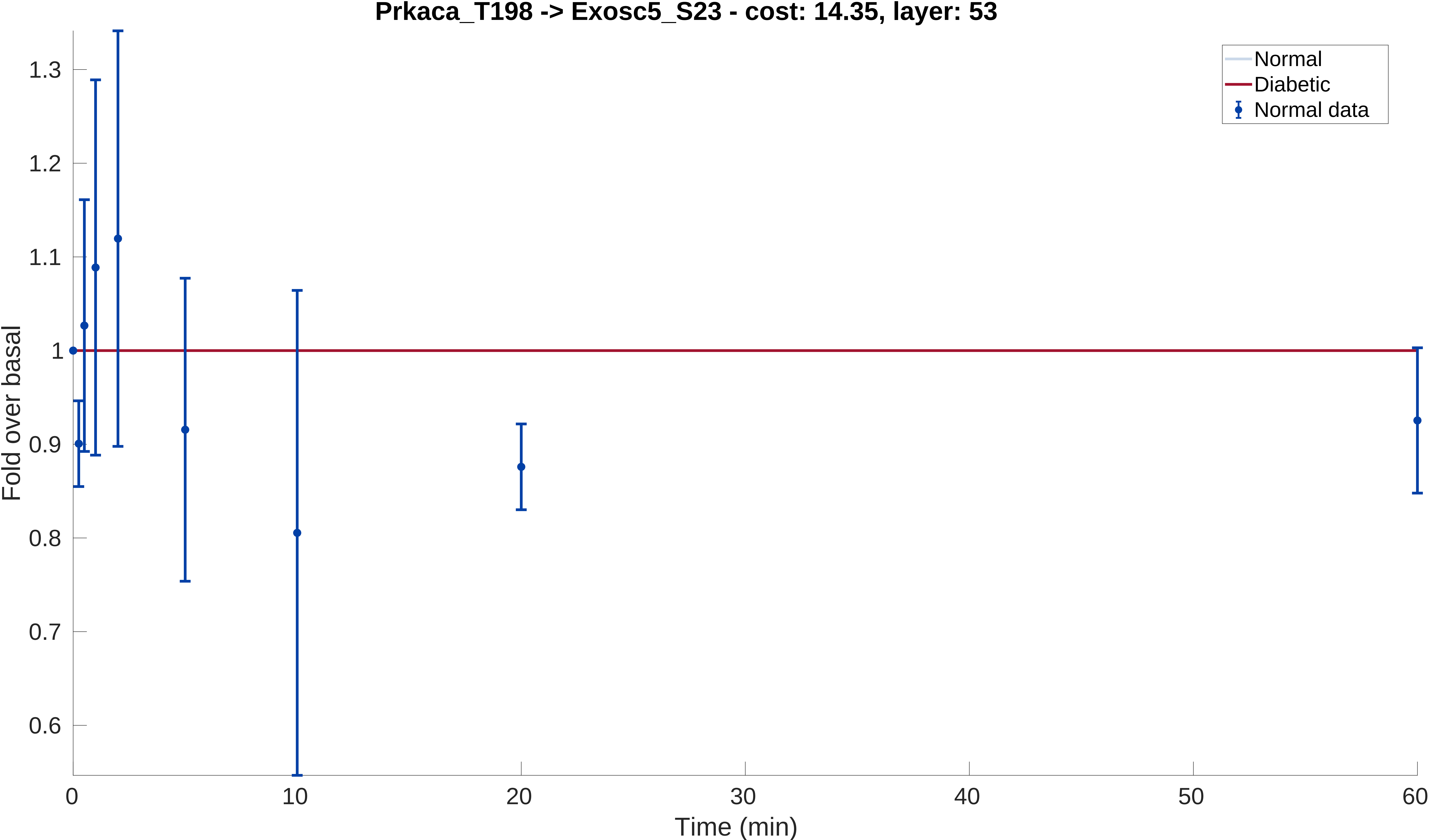

Prkaca_T198 -> Exosc5_S23 - cost: 14.35, layer: 53
Normal
Diabetic
Normal data
Fold over basal
1.3
1.2
1.1
1
0.9
0.8
0.7
0.6
0
10
20
30
40
50
60
Time (min)

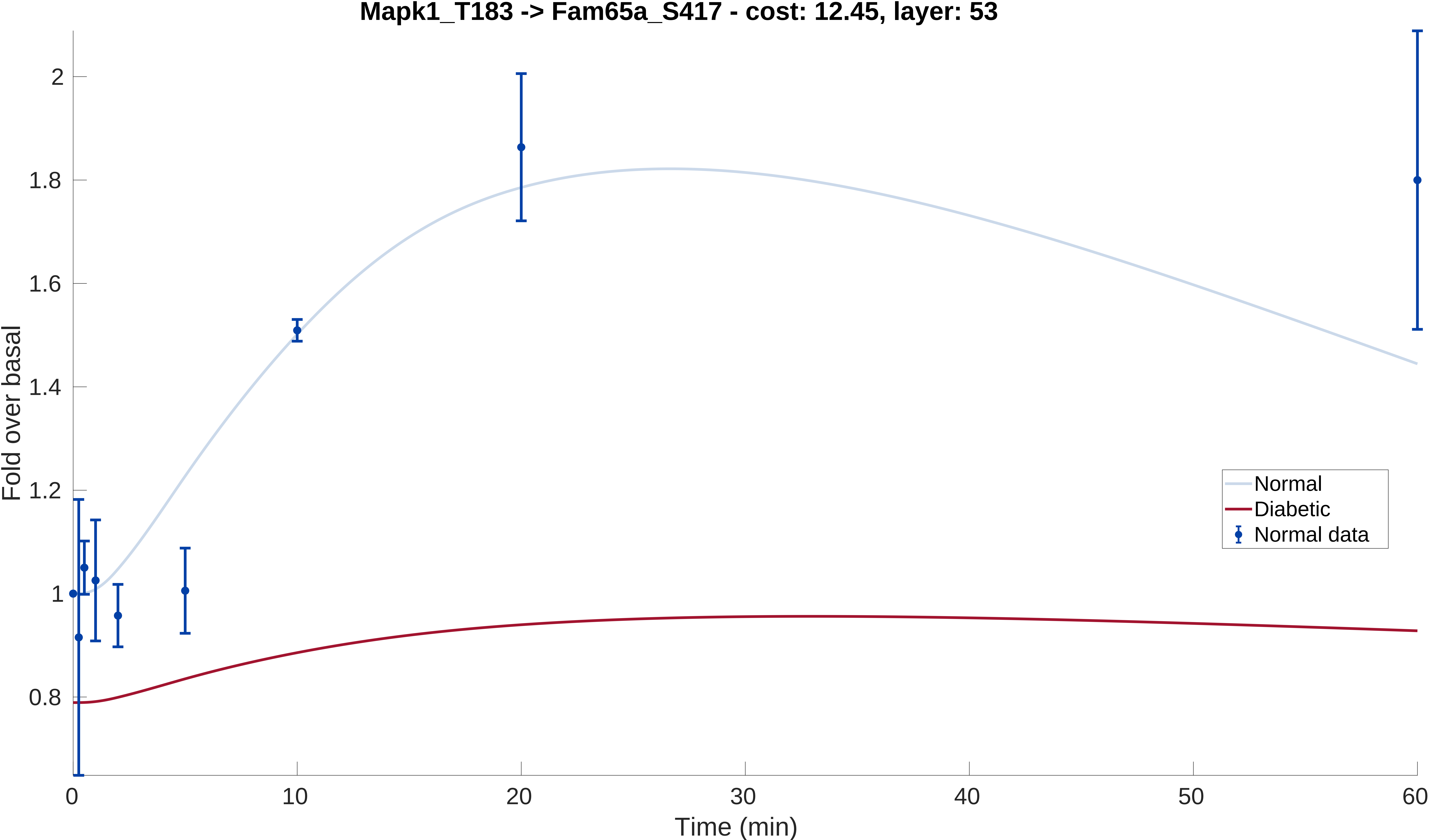

Mapk1_T183 -> Fam65a_S417 - cost: 12.45, layer: 53
Fold over basal
2
1.8
1.6
1.4
1.2
1
0.8
0
10
20
30
40
50
60
Time (min)
Normal
Diabetic
Normal data

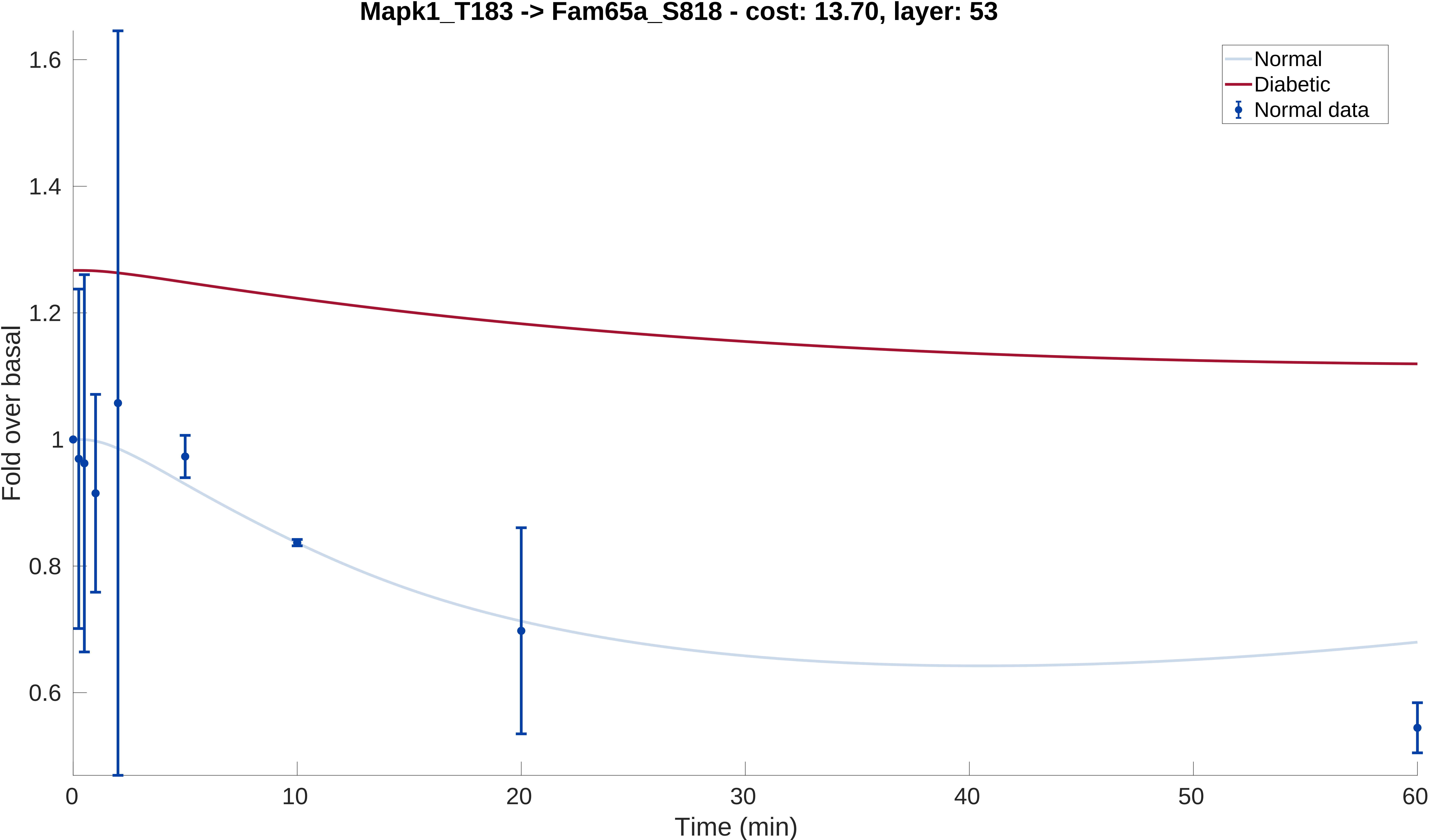

Mapk1_T183 -> Fam65a_S818 - cost: 13.70, layer: 53
Normal
Diabetic
Normal data
Fold over basal
1.6
1.4
1.2
1
0.8
0.6
0
10
20
30
40
50
60
Time (min)

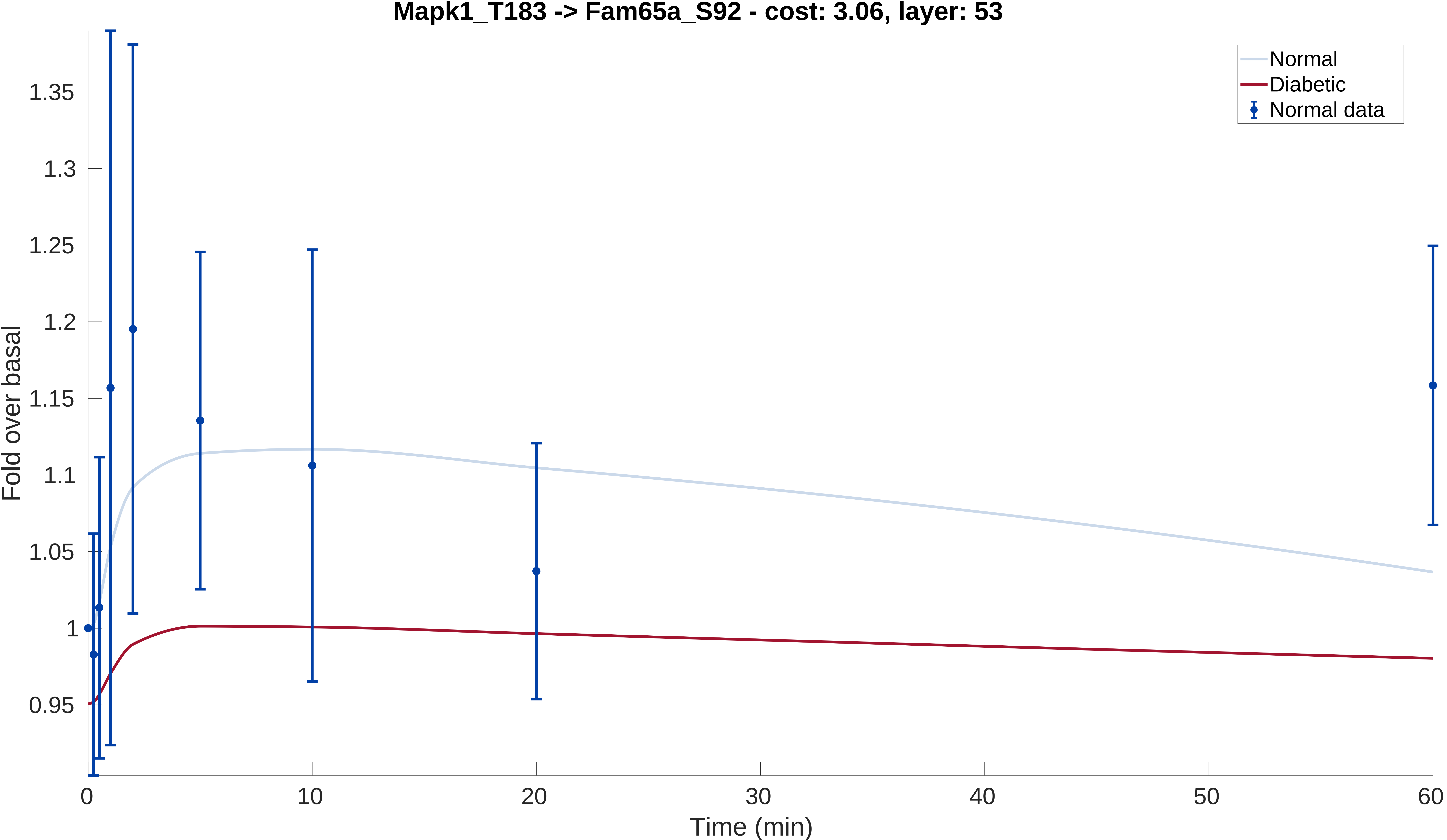
Mapk1_T183 -> Fam65a_S92 - cost: 3.06, layer: 53
Normal
Diabetic
Normal data
Fold over basal
1.35
1.3
1.25
1.2
1.15
1.1
1.05
1
0.95
0
10
20
30
40
50
60
Time (min)

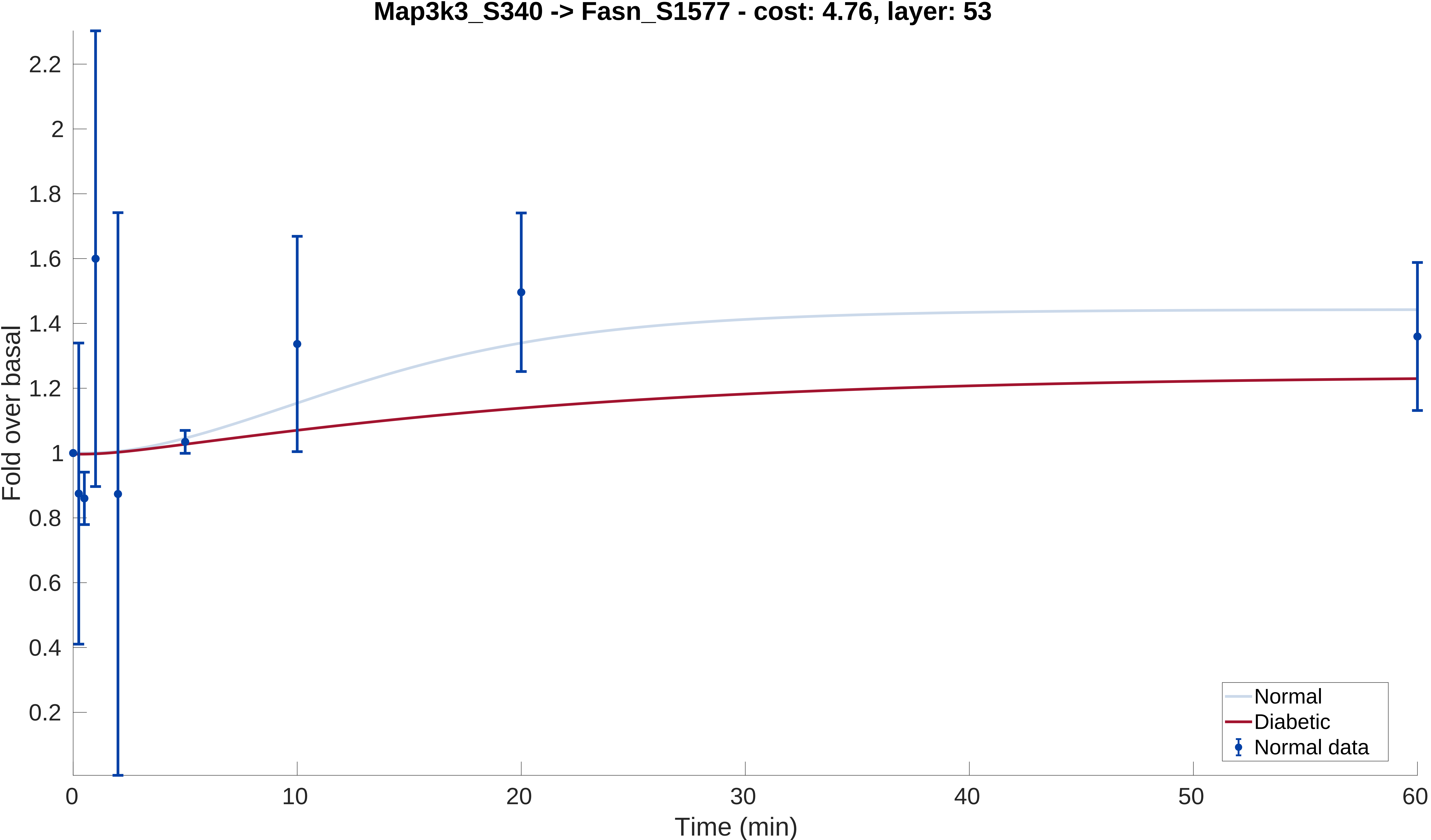
Map3k3_S340 -> Fasn_S1577 - cost: 4.76, layer: 53
Fold over basal
2.2
2
1.8
1.6
1.4
1.2
1
0.8
0.6
0.4
0.2
0
10
20
30
40
50
60
Time (min)
Normal
Diabetic
Normal data

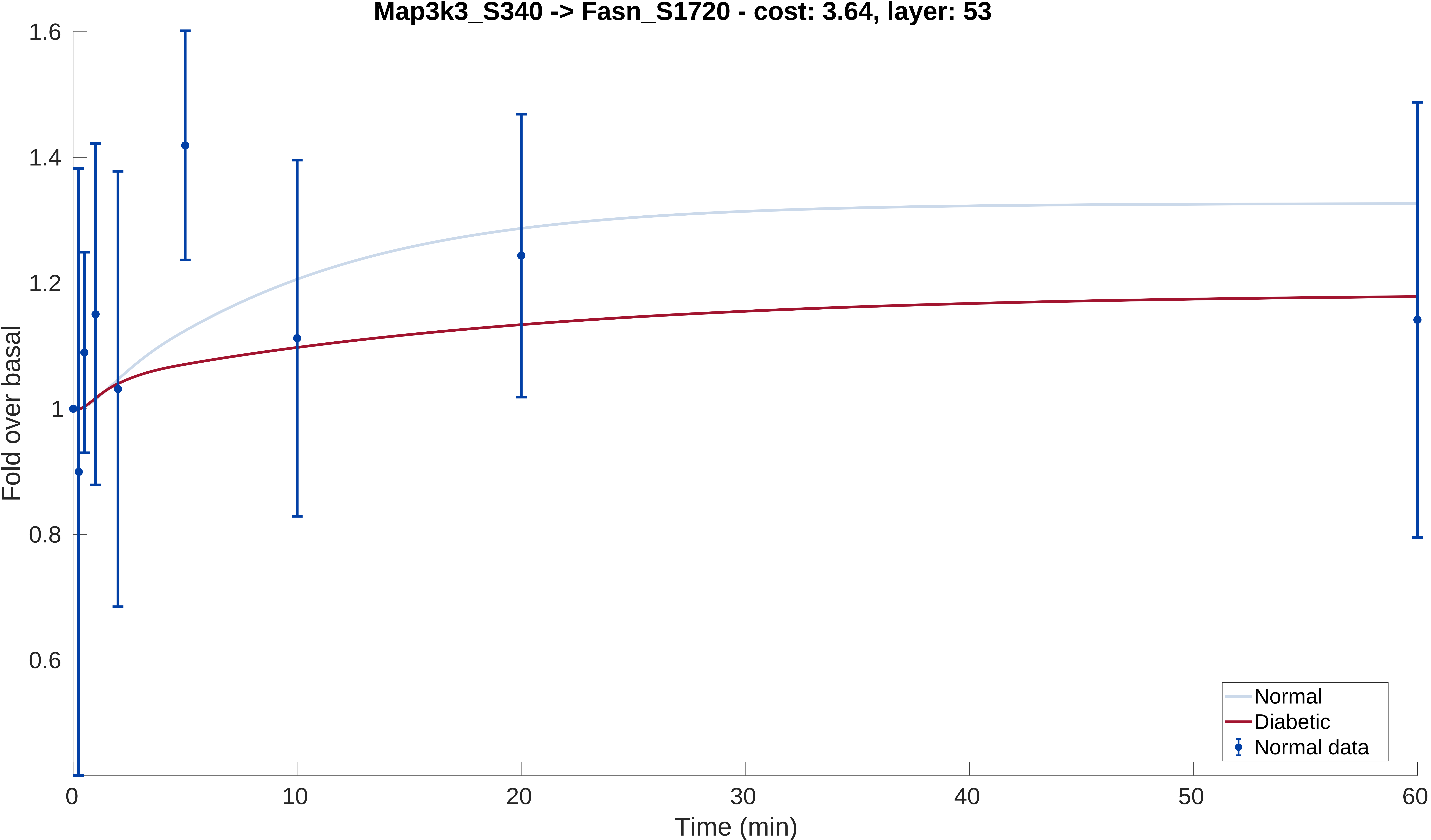
Map3k3_S340 -> Fasn_S1720 - cost: 3.64, layer: 53
1.6
1.4
1.2
1
0.8
0.6
Fold over basal
0
10
20
30
40
50
60
Time (min)
Normal
Diabetic
Normal data

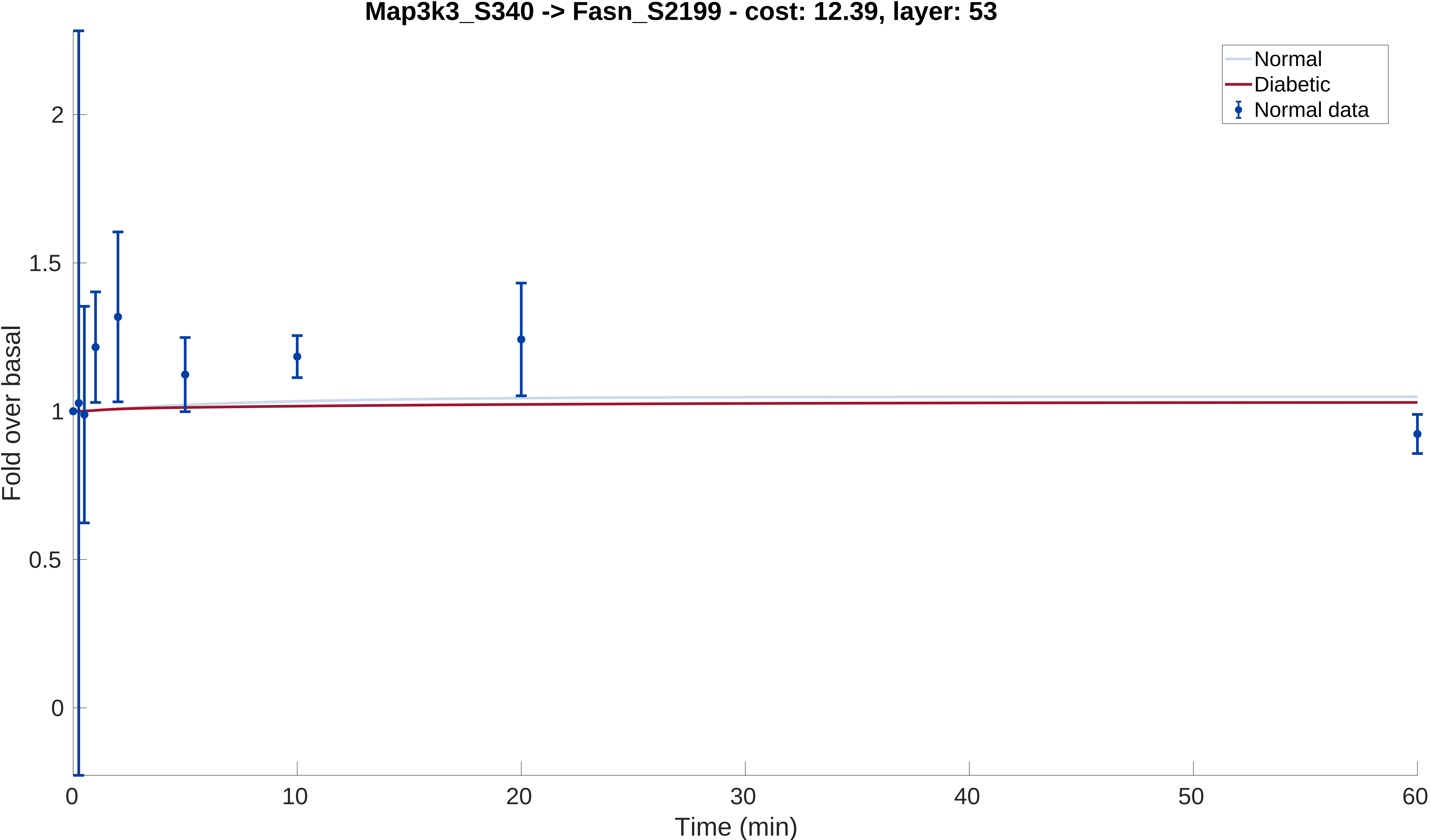
Map3k3_S340 -> Fasn_S2199 - cost: 12.39, layer: 53
Normal
Diabetic
Normal data
Fold over basal
2
1.5
1
0.5
0
0
10
20
30
40
50
60
Time (min)

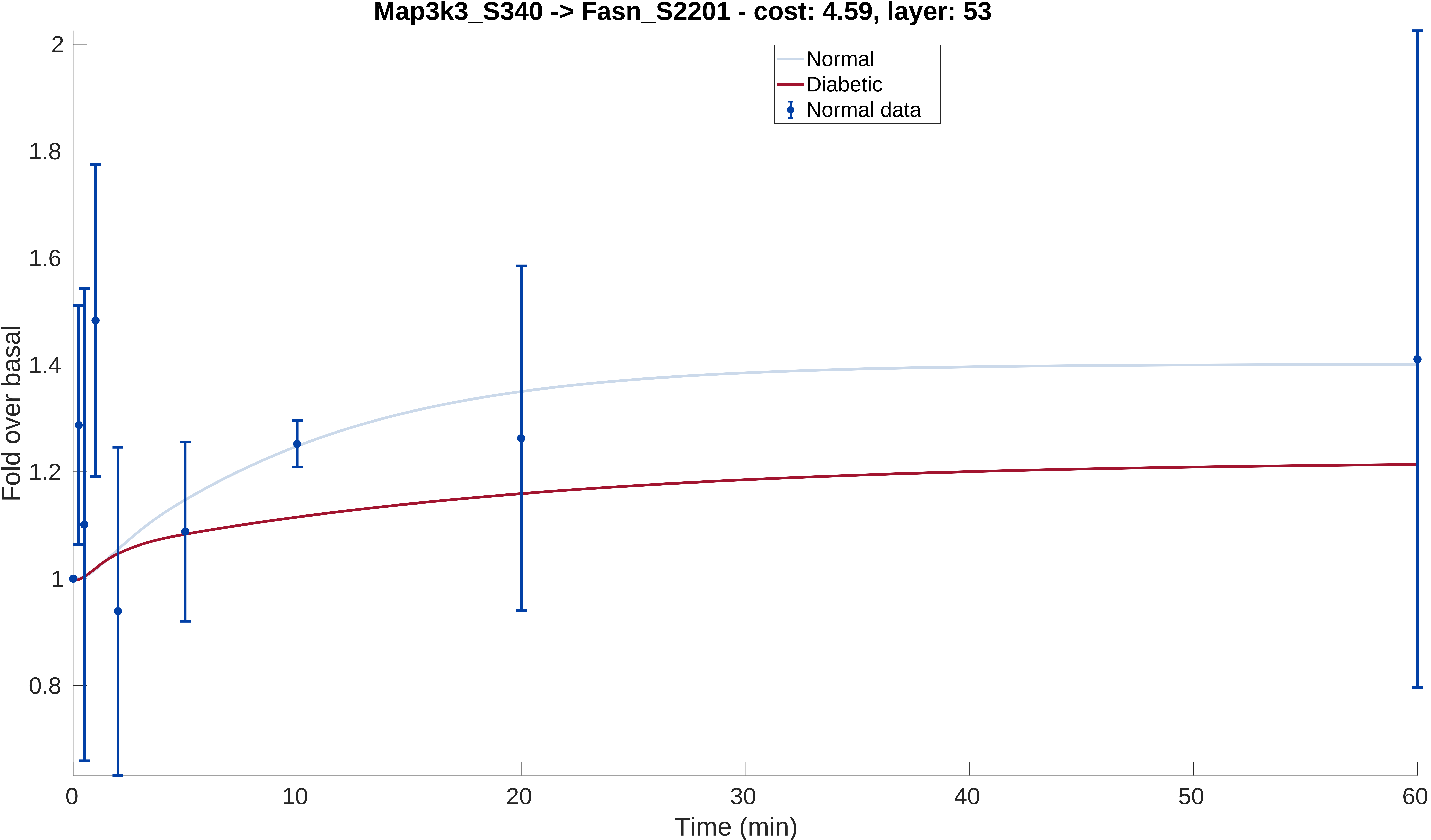
Map3k3_S340 -> Fasn_S2201 - cost: 4.59, layer: 53
Normal
Diabetic
Normal data
Fold over basal
2
1.8
1.6
1.4
1.2
1
0.8
0
10
20
30
40
50
60
Time (min)

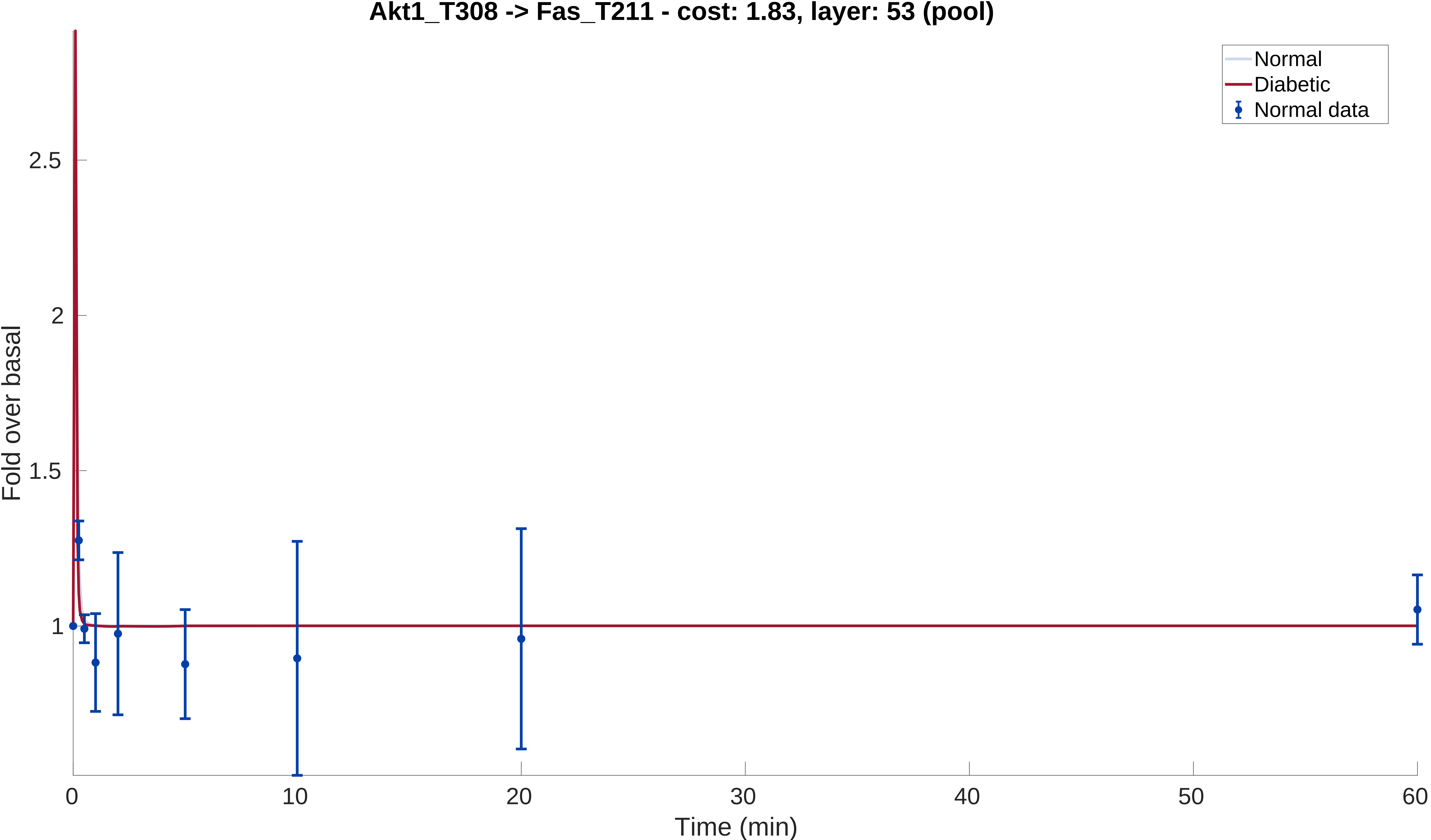

Akt1_T308 -> Fas_T211 - cost: 1.83, layer: 53 (pool)
Normal
Diabetic
Normal data
Fold over basal
2.5
2
1.5
1
0
10
20
30
40
50
60
Time (min)

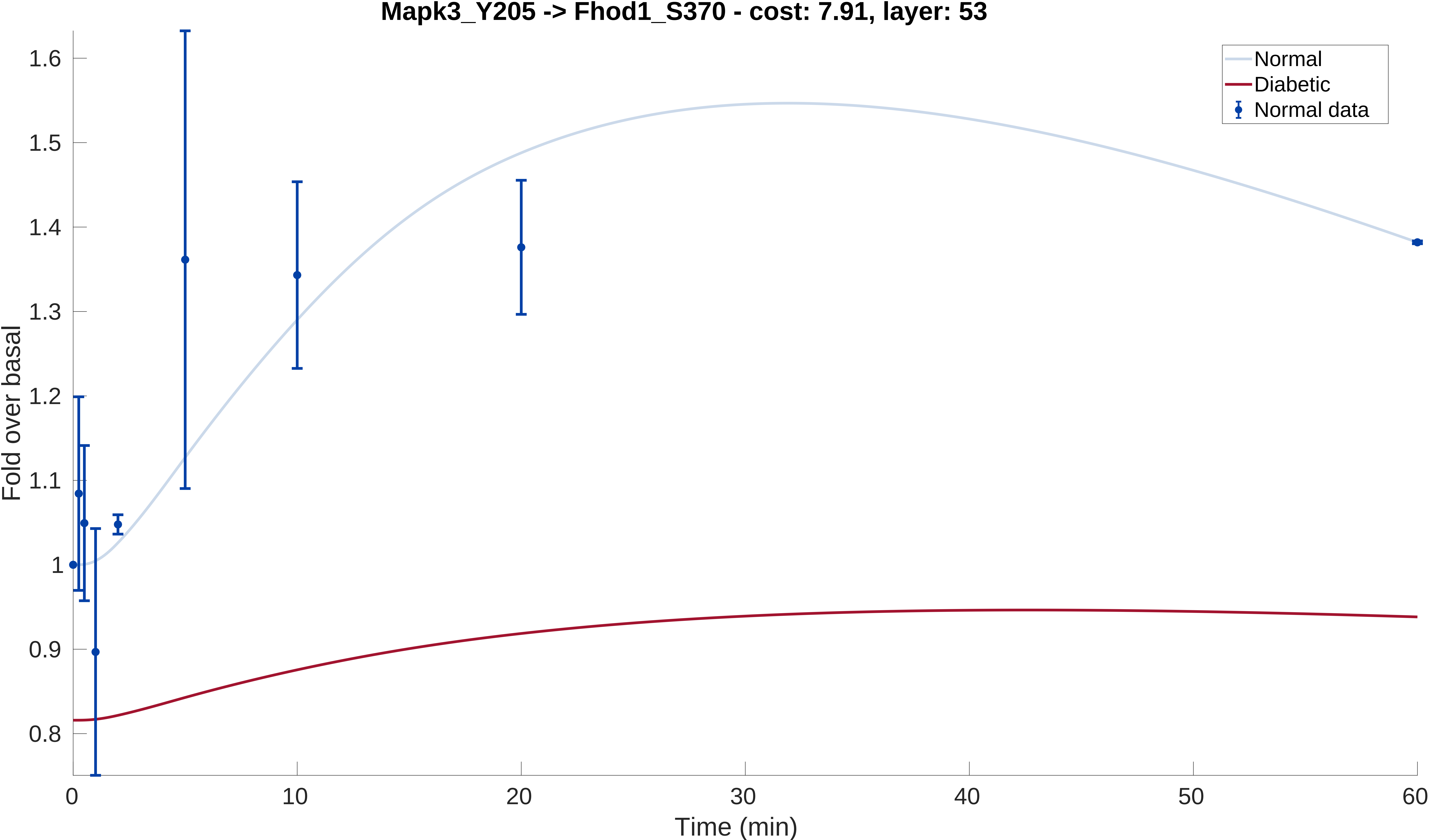

Mapk3_Y205 -> Fhod1_S370 - cost: 7.91, layer: 53
Normal
Diabetic
Normal data
Fold over basal
1.6
1.5
1.4
1.3
1.2
1.1
1
0.9
0.8
0
10
20
30
40
50
60
Time (min)

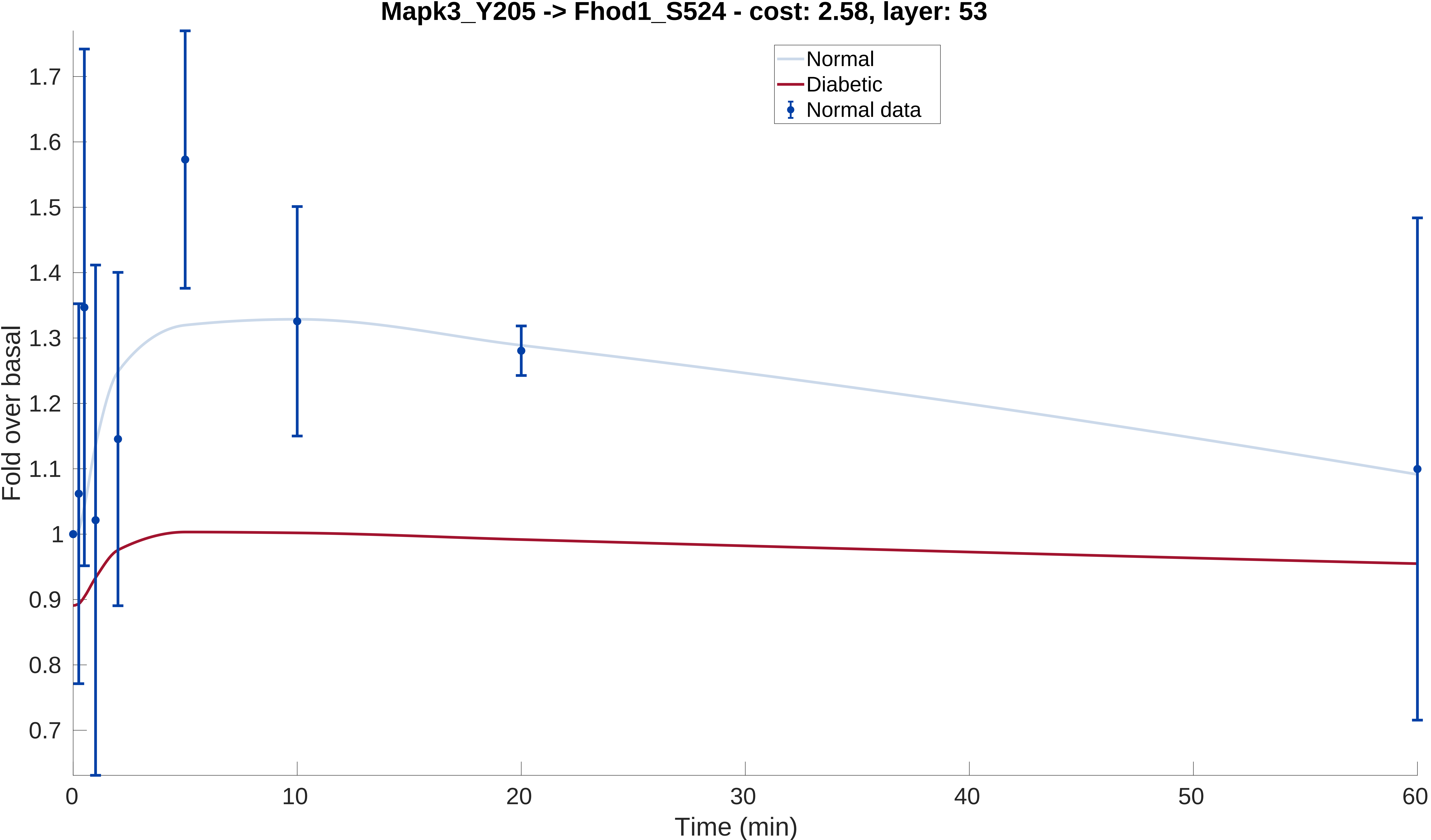

Mapk3_Y205 -> Fhod1_S524 - cost: 2.58, layer: 53
Normal
Diabetic
Normal data
Fold over basal
Time (min)
0.7
0.8
0.9
1
1.1
1.2
1.3
1.4
1.5
1.6
1.7
0
10
20
30
40
50
60

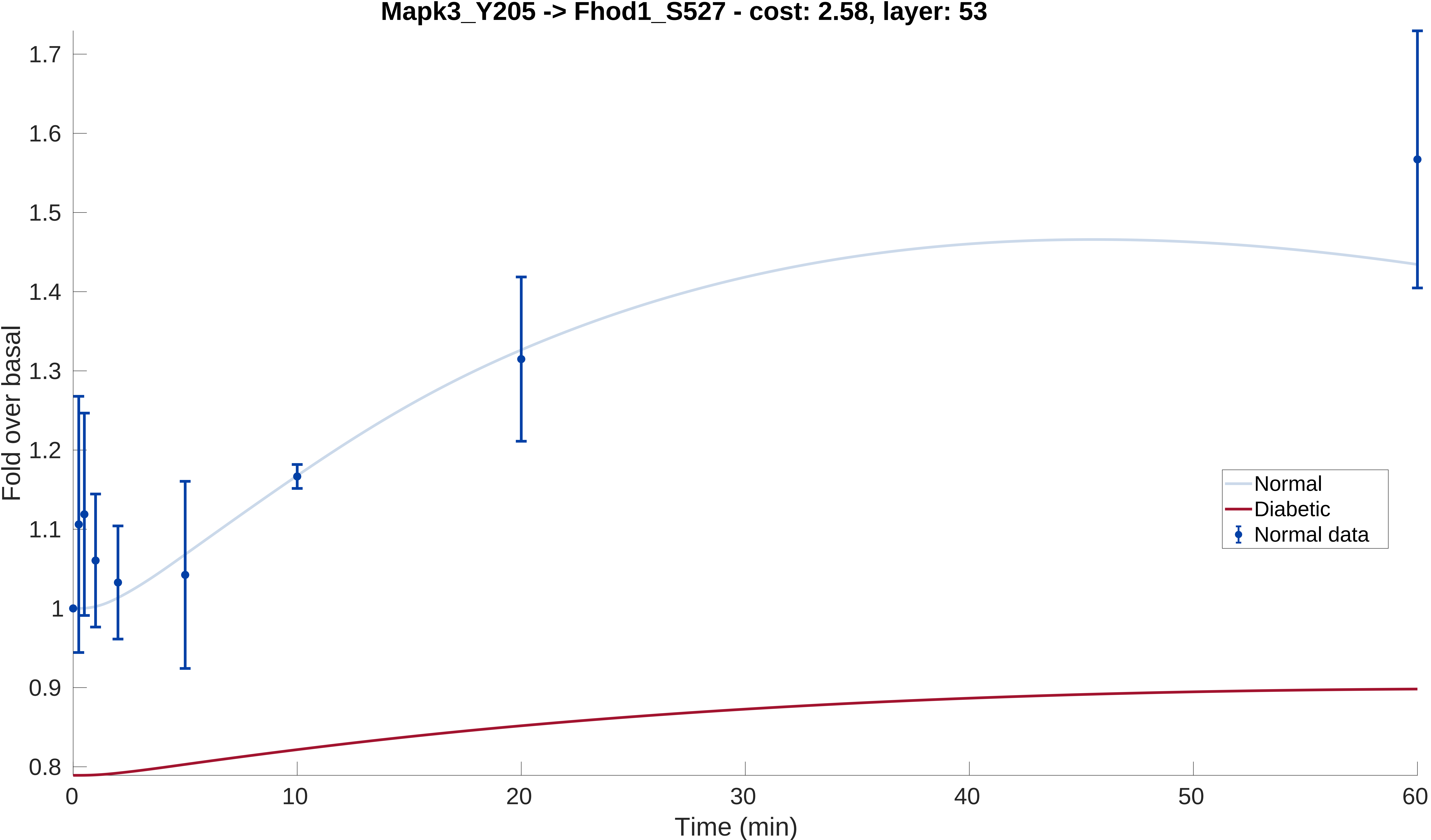

Mapk3_Y205 -> Fhod1_S527 - cost: 2.58, layer: 53
Fold over basal
1.7
1.6
1.5
1.4
1.3
1.2
1.1
1
0.9
0.8
0
10
20
30
40
50
60
Time (min)
Normal
Diabetic
Normal data

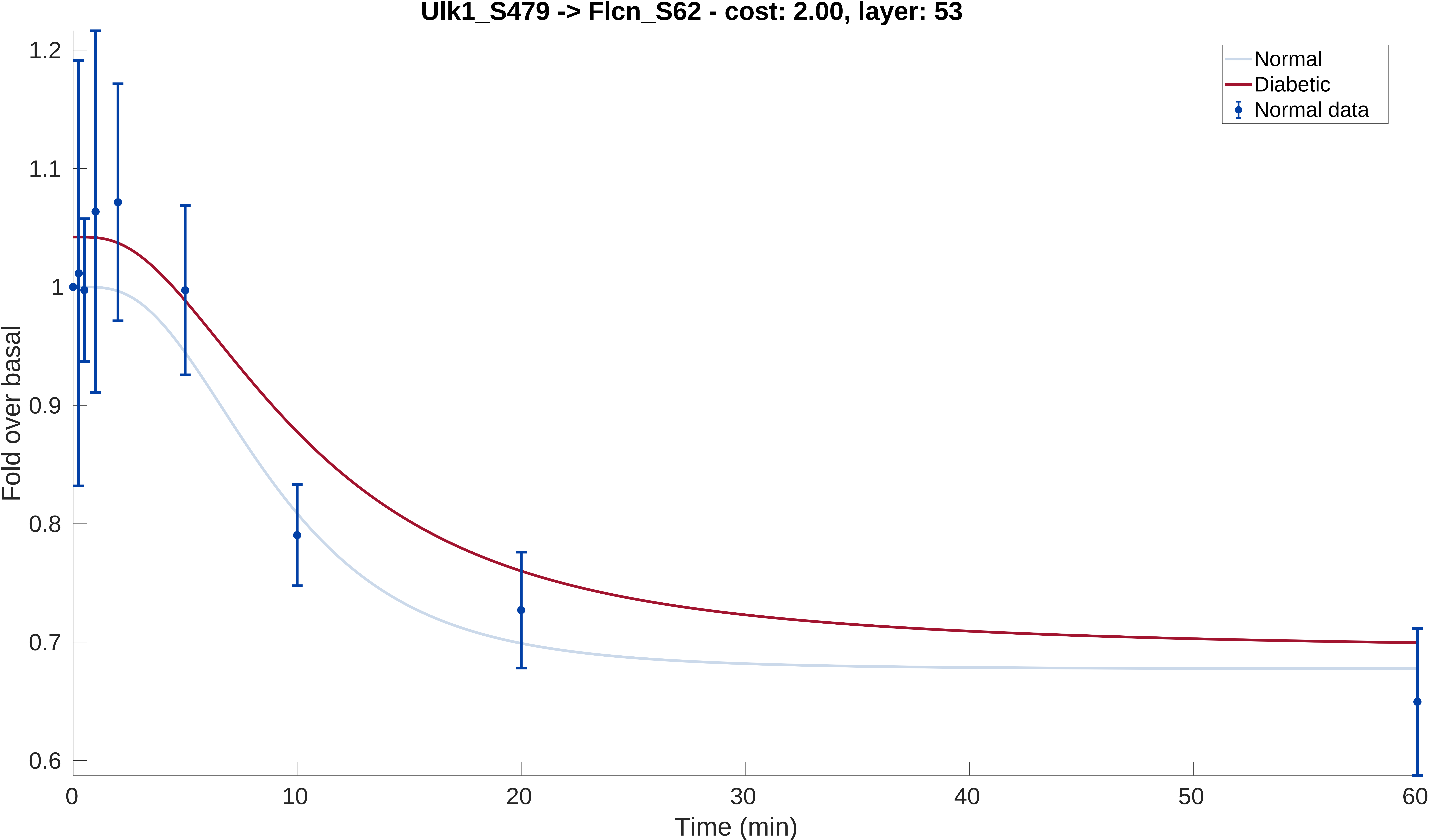

Ulk1_S479 -> Flcn_S62 - cost: 2.00, layer: 53
Normal
Diabetic
Normal data
Fold over basal
1.2
1.1
1
0.9
0.8
0.7
0.6
0
10
20
30
40
50
60
Time (min)

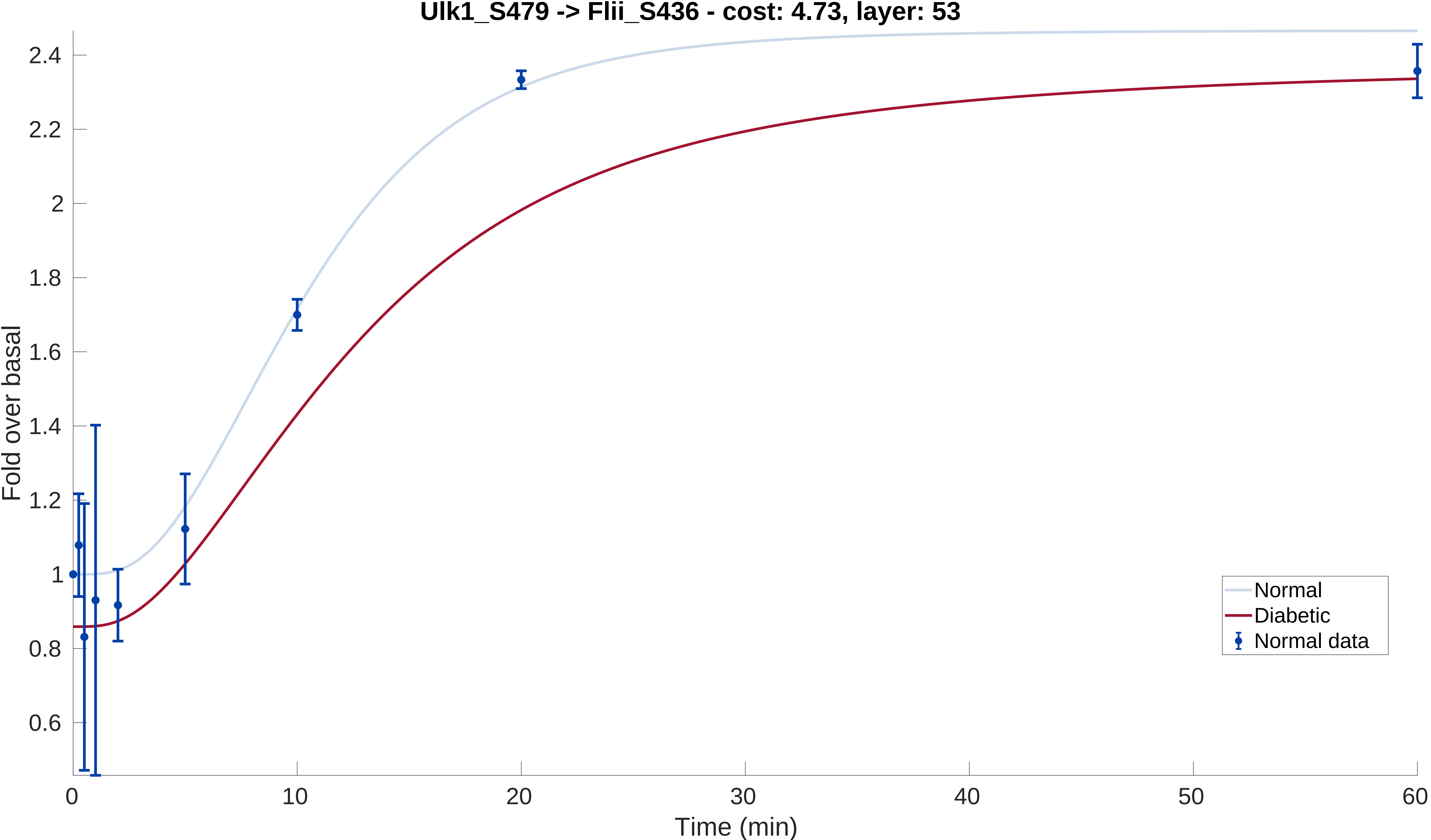
Ulk1_S479 -> Flii_S436 - cost: 4.73, layer: 53
Fold over basal
2.4
2.2
2
1.8
1.6
1.4
1.2
1
0.8
0.6
0
10
20
30
40
50
60
Time (min)
Normal
Diabetic
Normal data

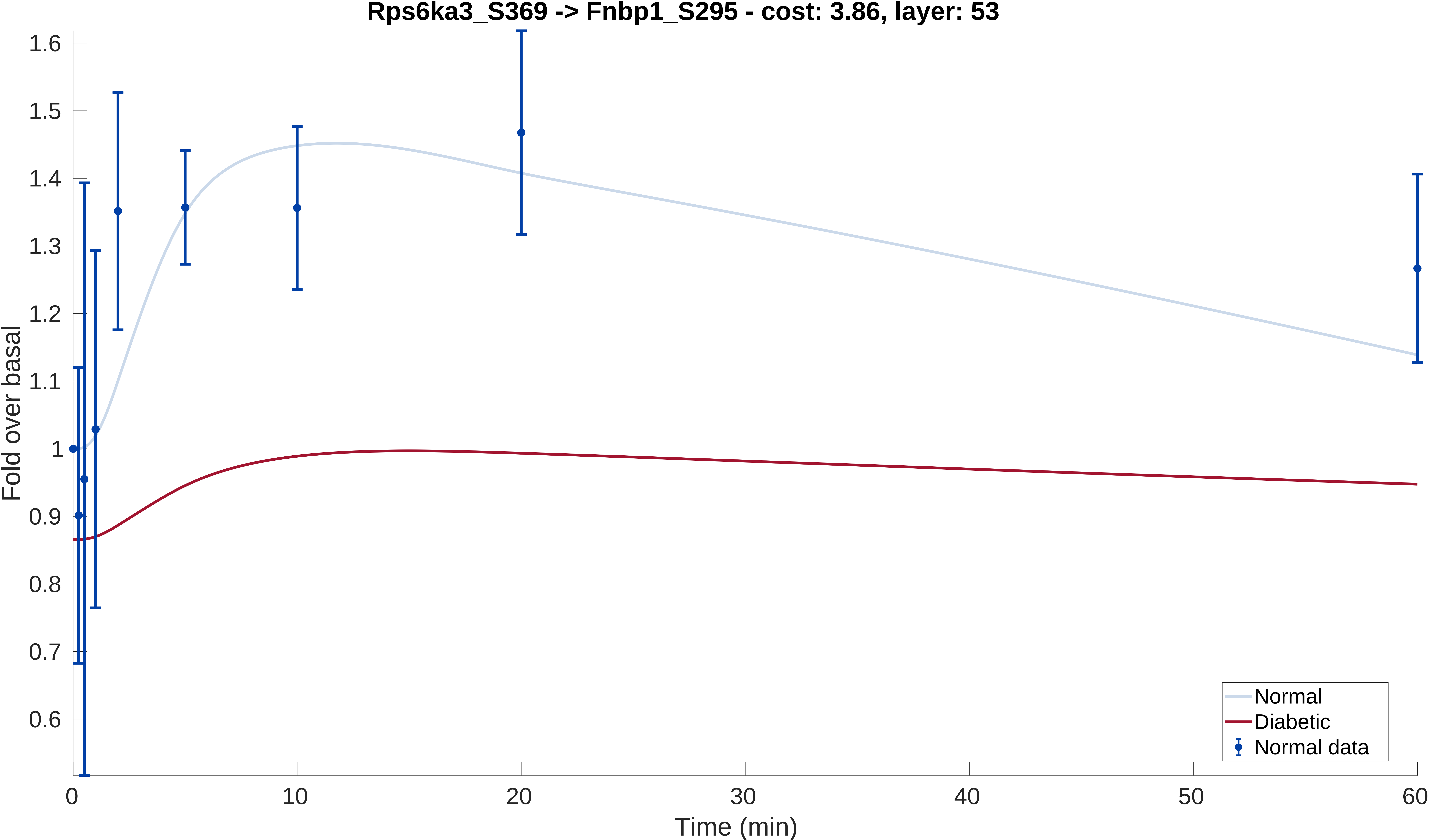

Rps6ka3_S369 -> Fnbp1_S295 - cost: 3.86, layer: 53
Fold over basal
1.6
1.5
1.4
1.3
1.2
1.1
1
0.9
0.8
0.7
0.6
0
10
20
30
40
50
60
Time (min)
Normal
Diabetic
Normal data

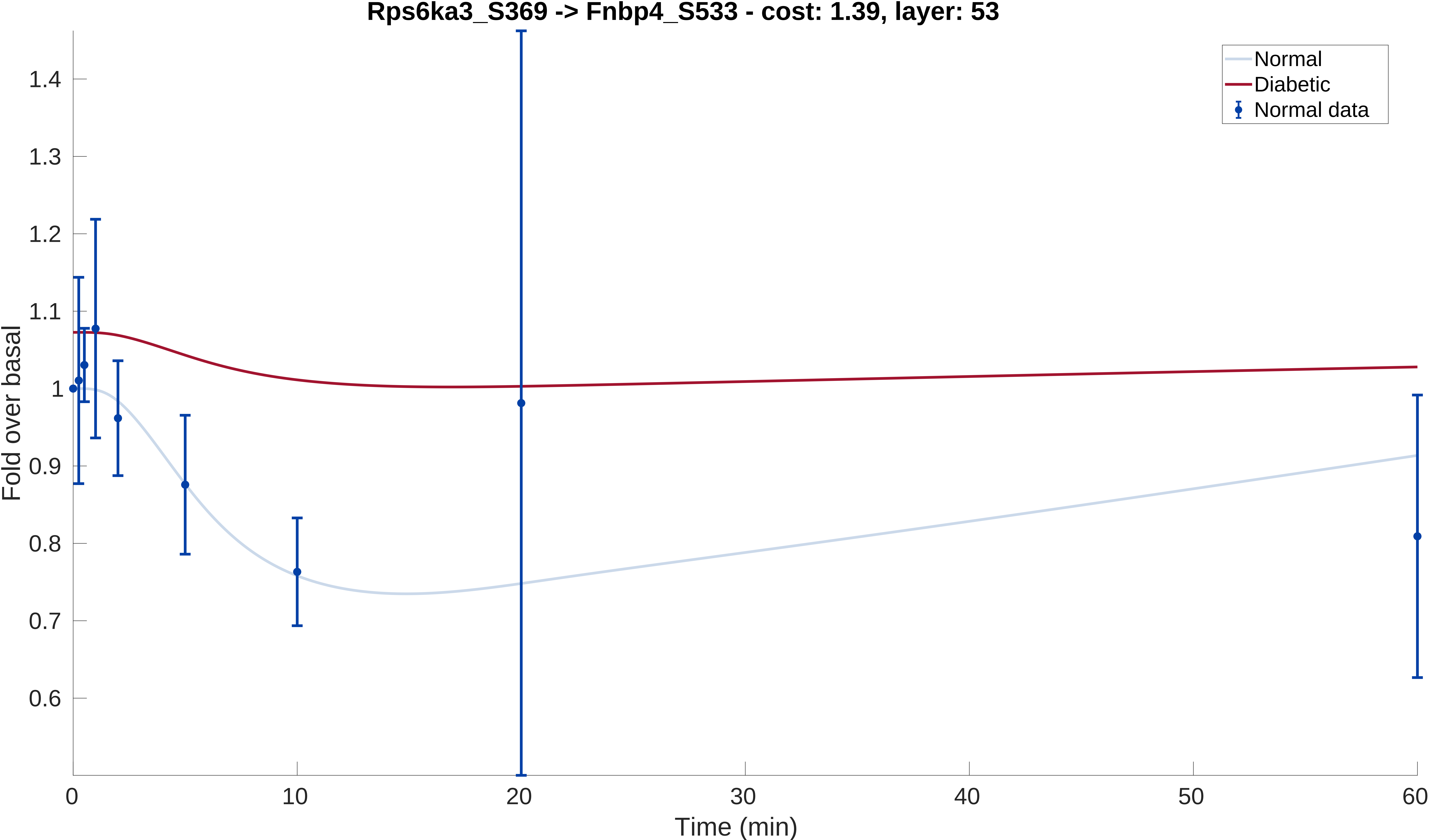
Rps6ka3_S369 -> Fnbp4_S533 - cost: 1.39, layer: 53
Normal
Diabetic
Normal data
Fold over basal
Time (min)
0.6
0.7
0.8
0.9
1
1.1
1.2
1.3
1.4
0
10
20
30
40
50
60

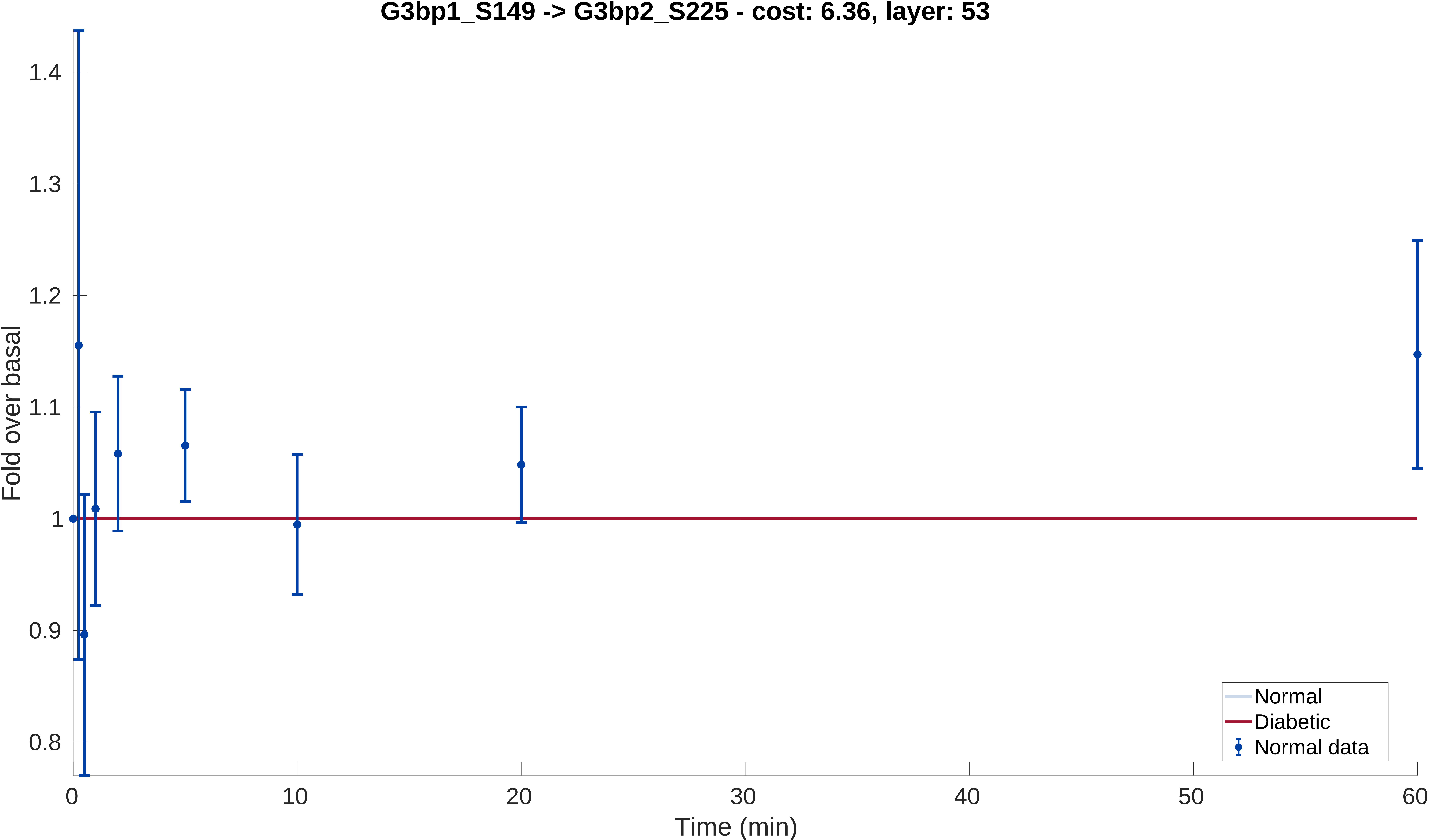

G3bp1_S149 -> G3bp2_S225 - cost: 6.36, layer: 53
Fold over basal
1.4
1.3
1.2
1.1
1
0.9
0.8
0
10
20
30
40
50
60
Time (min)
Normal
Diabetic
Normal data

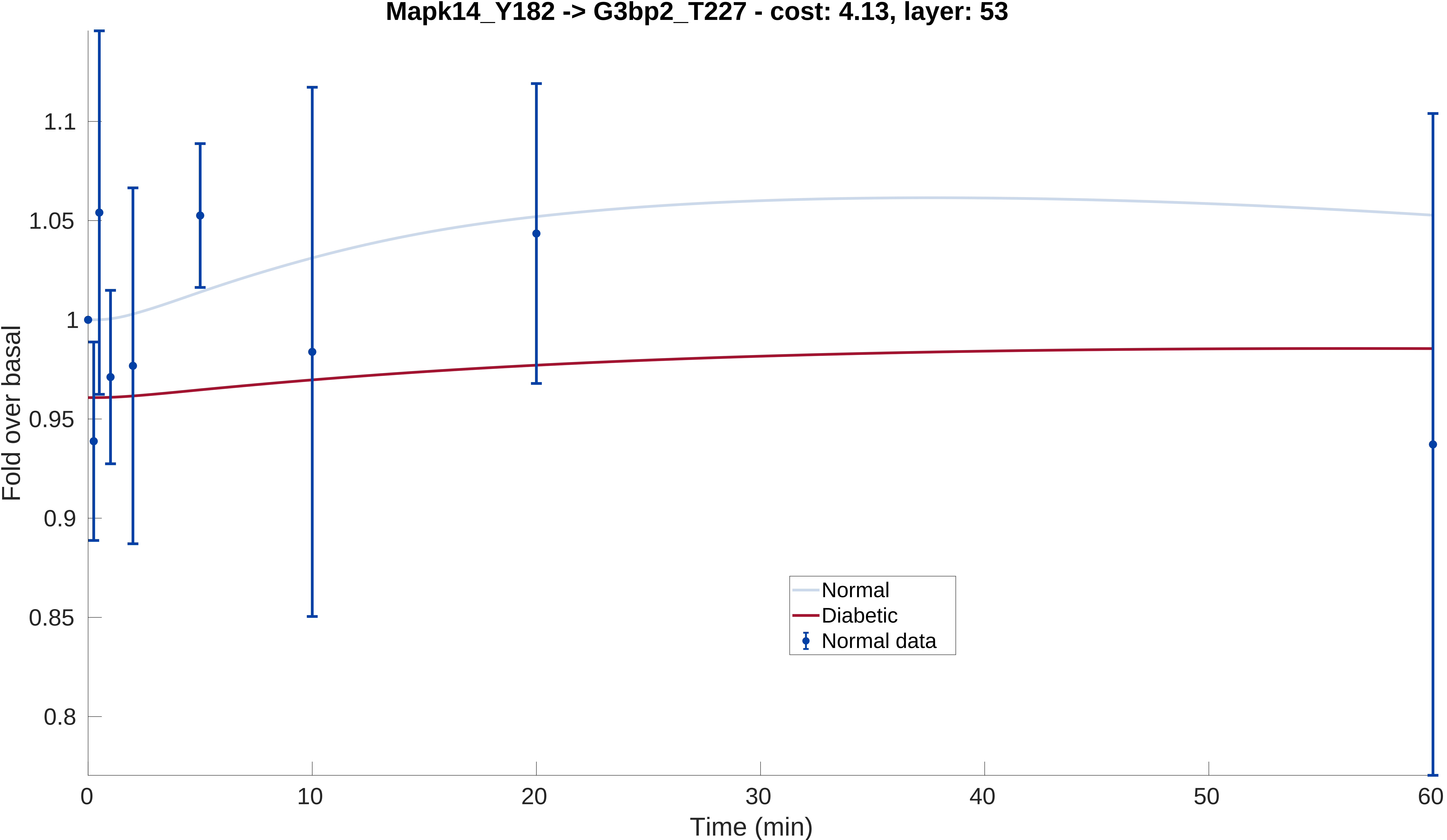

Mapk14_Y182 -> G3bp2_T227 - cost: 4.13, layer: 53
Fold over basal
1.1
1.05
1
0.95
0.9
0.85
0.8
0
10
20
30
40
50
60
Time (min)
Normal
Diabetic
Normal data

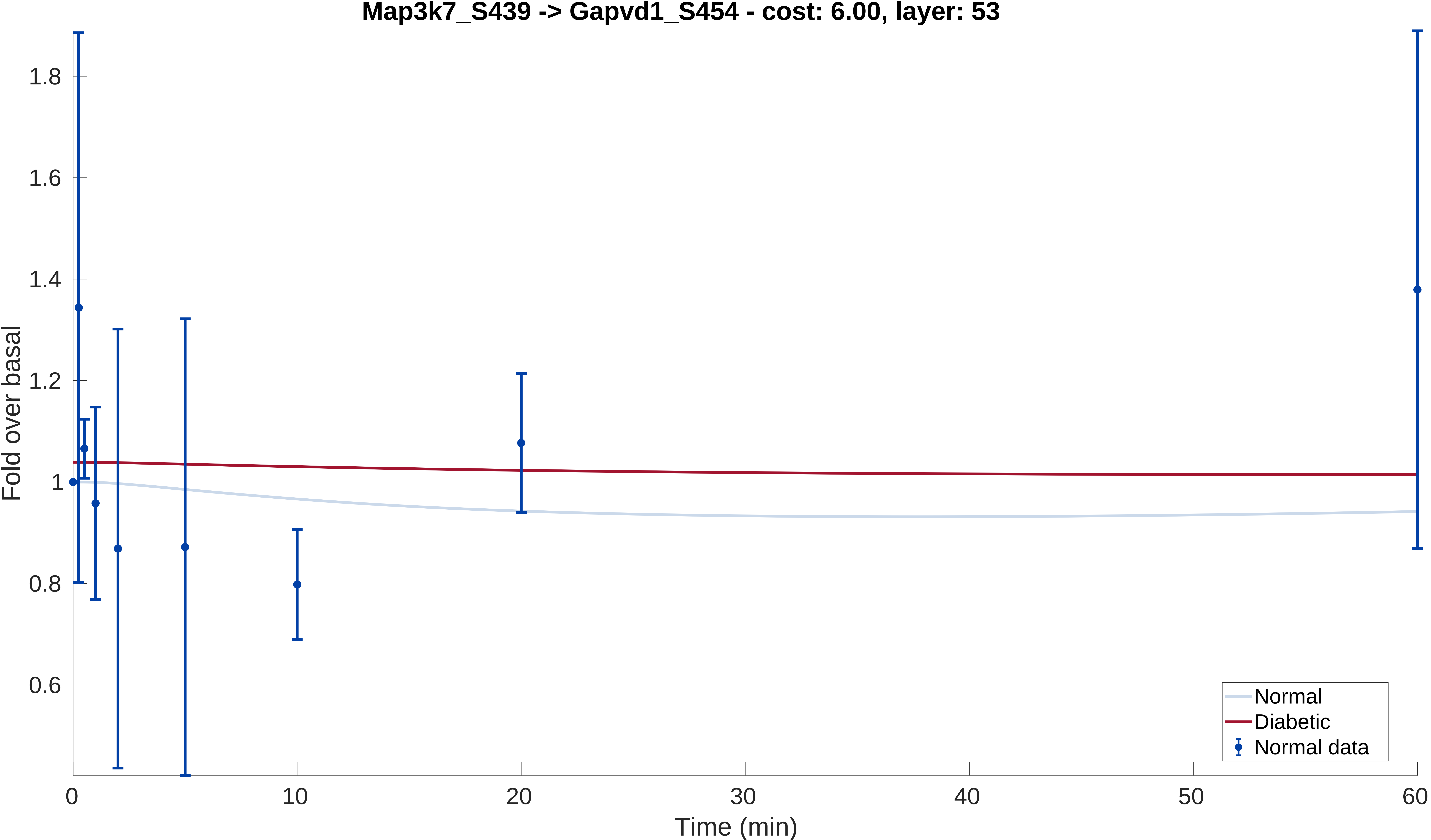

Map3k7_S439 -> Gapvd1_S454 - cost: 6.00, layer: 53
Fold over basal
1.8
1.6
1.4
1.2
1
0.8
0.6
0
10
20
30
40
50
60
Time (min)
Normal
Diabetic
Normal data

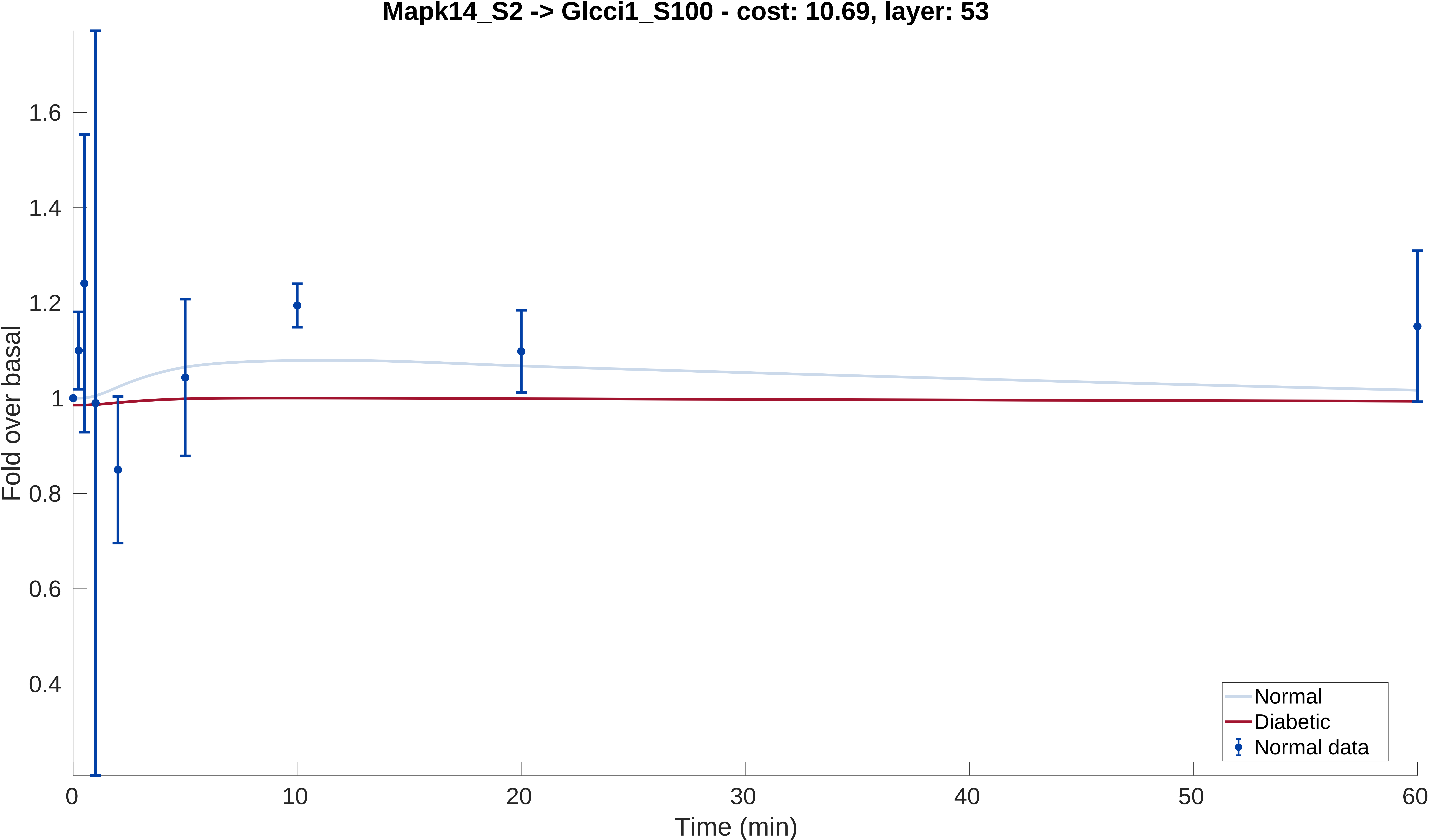

Mapk14_S2 -> Glcci1_S100 - cost: 10.69, layer: 53
Fold over basal
1.6
1.4
1.2
1
0.8
0.6
0.4
0
10
20
30
40
50
60
Time (min)
Normal
Diabetic
Normal data

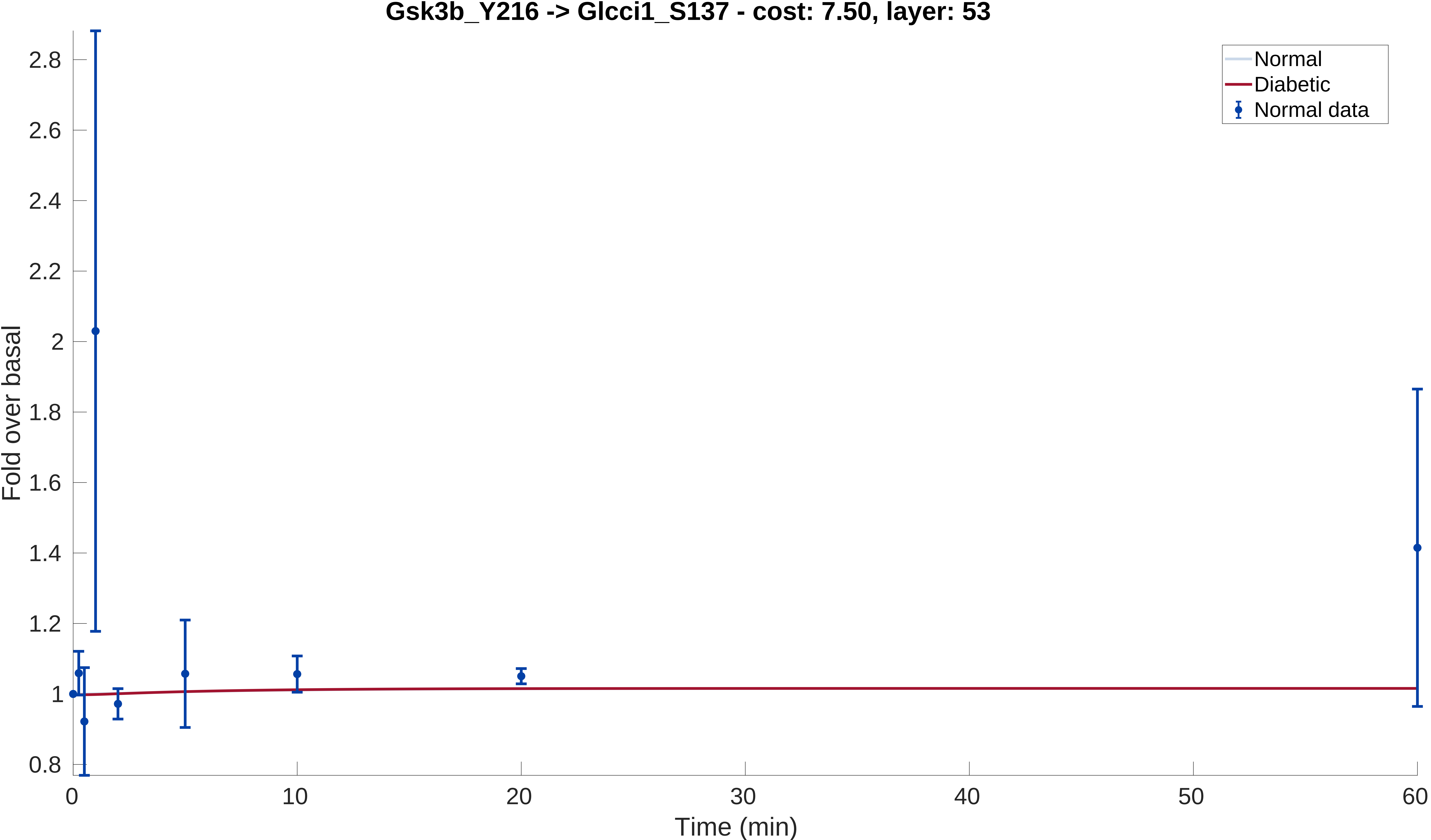

Gsk3b_Y216 -> Glcci1_S137 - cost: 7.50, layer: 53
Normal
Diabetic
Normal data
Fold over basal
2.8
2.6
2.4
2.2
2
1.8
1.6
1.4
1.2
1
0.8
0
10
20
30
40
50
60
Time (min)

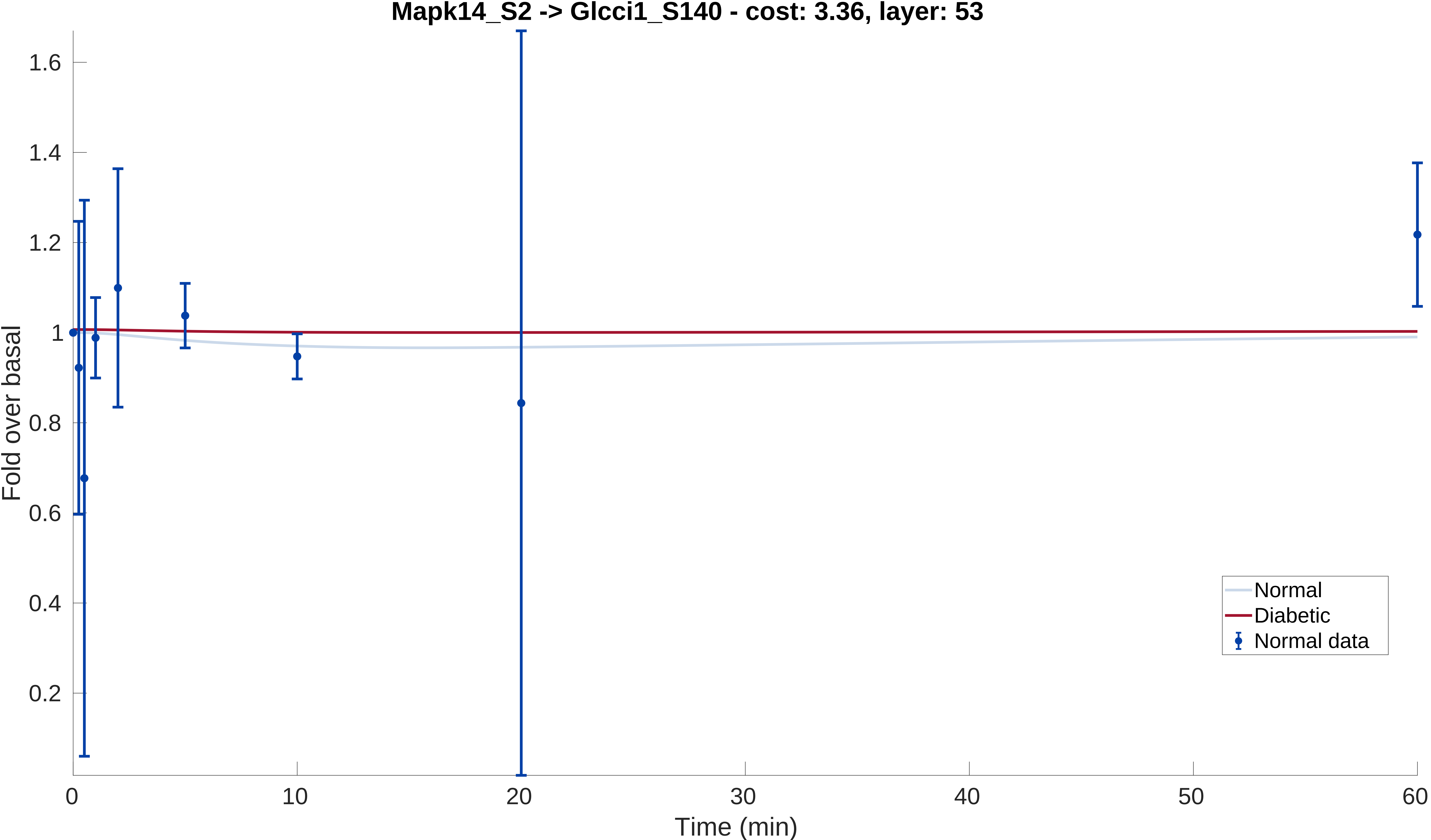

Mapk14_S2 -> Glcci1_S140 - cost: 3.36, layer: 53
Fold over basal
Time (min)
1.6
1.4
1.2
1
0.8
0.6
0.4
0.2
0
10
20
30
40
50
60
Normal
Diabetic
Normal data

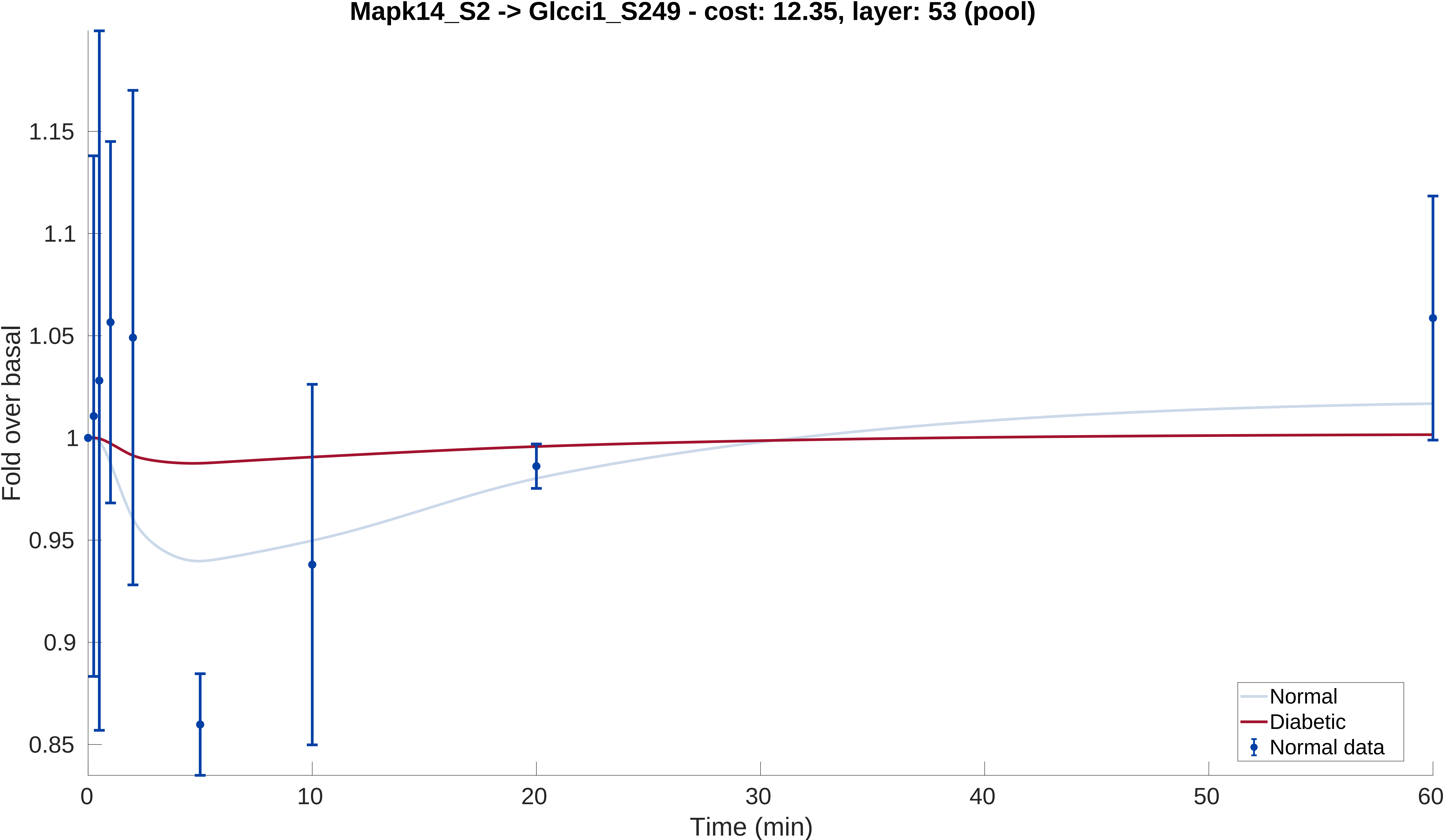
Mapk14_S2 -> Glcci1_S249 - cost: 12.35, layer: 53 (pool)
Fold over basal
1.15
1.1
1.05
1
0.95
0.9
0.85
0
10
20
30
40
50
60
Time (min)
Normal
Diabetic
Normal data

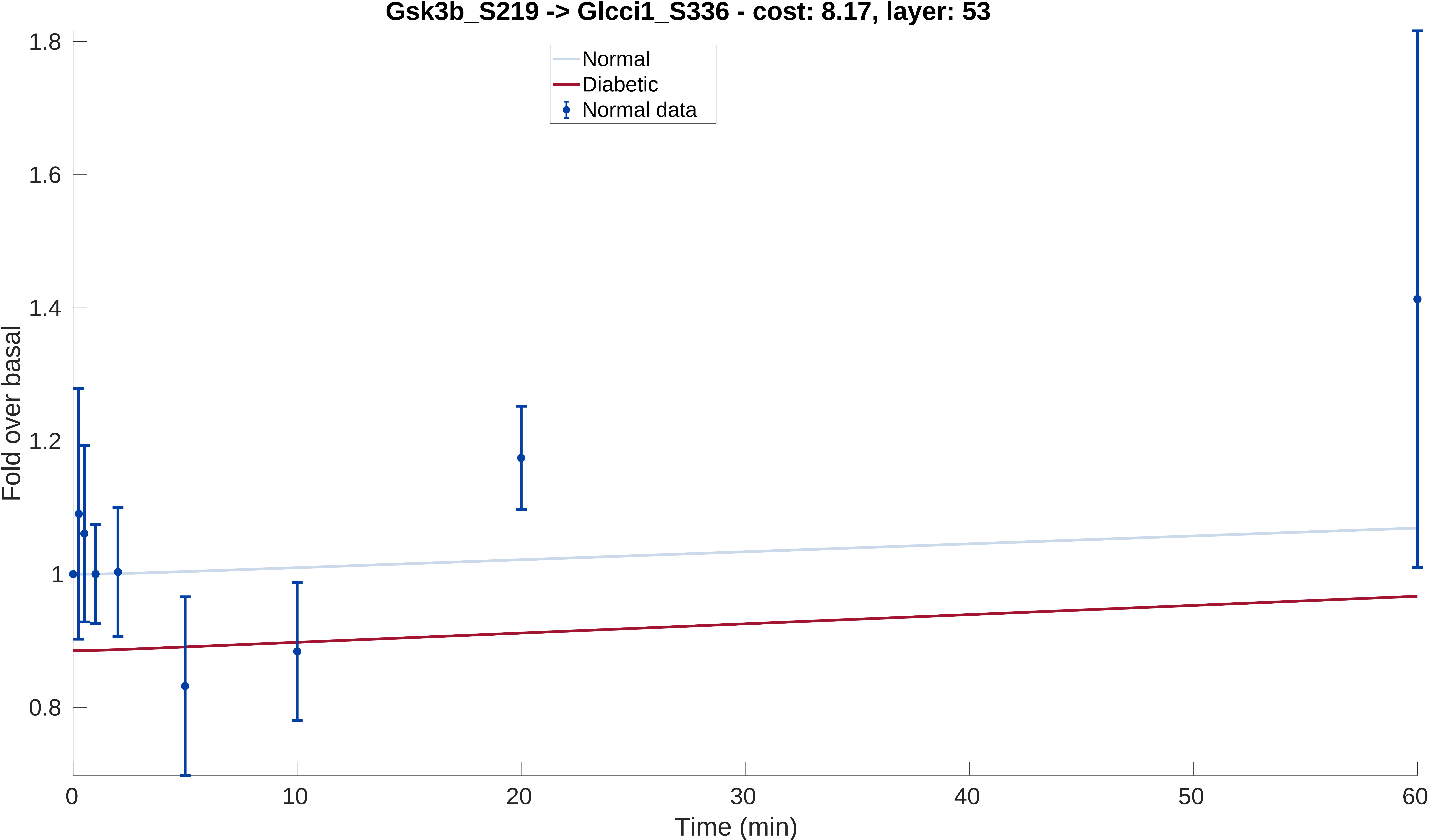

Gsk3b_S219 -> Glcci1_S336 - cost: 8.17, layer: 53
Normal
Diabetic
Normal data
Fold over basal
1.8
1.6
1.4
1.2
1
0.8
0
10
20
30
40
50
60
Time (min)

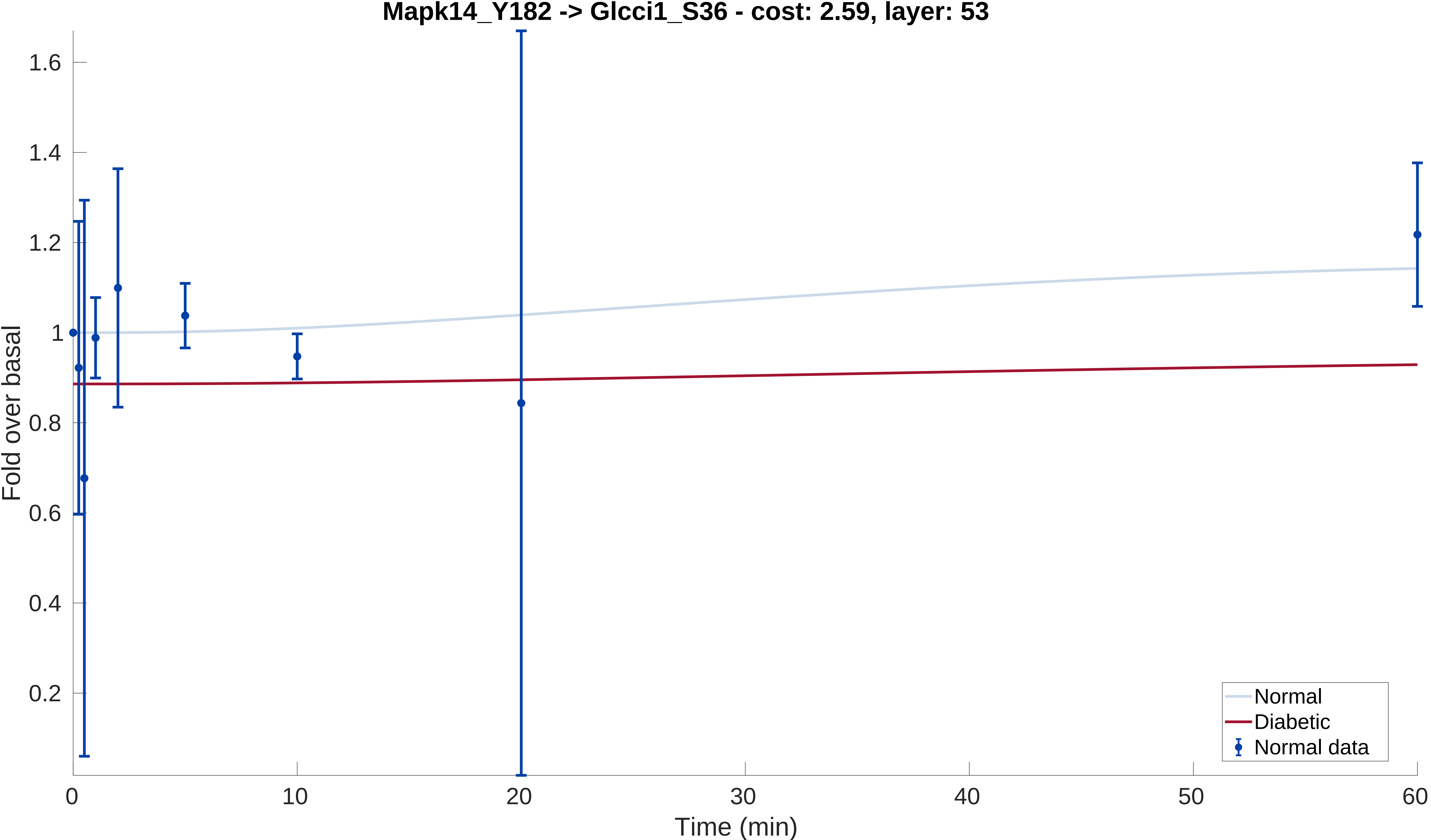
Mapk14_Y182 -> Glcci1_S36 - cost: 2.59, layer: 53
Fold over basal
Time (min)
1.6
1.4
1.2
1
0.8
0.6
0.4
0.2
0
10
20
30
40
50
60
Normal
Diabetic
Normal data

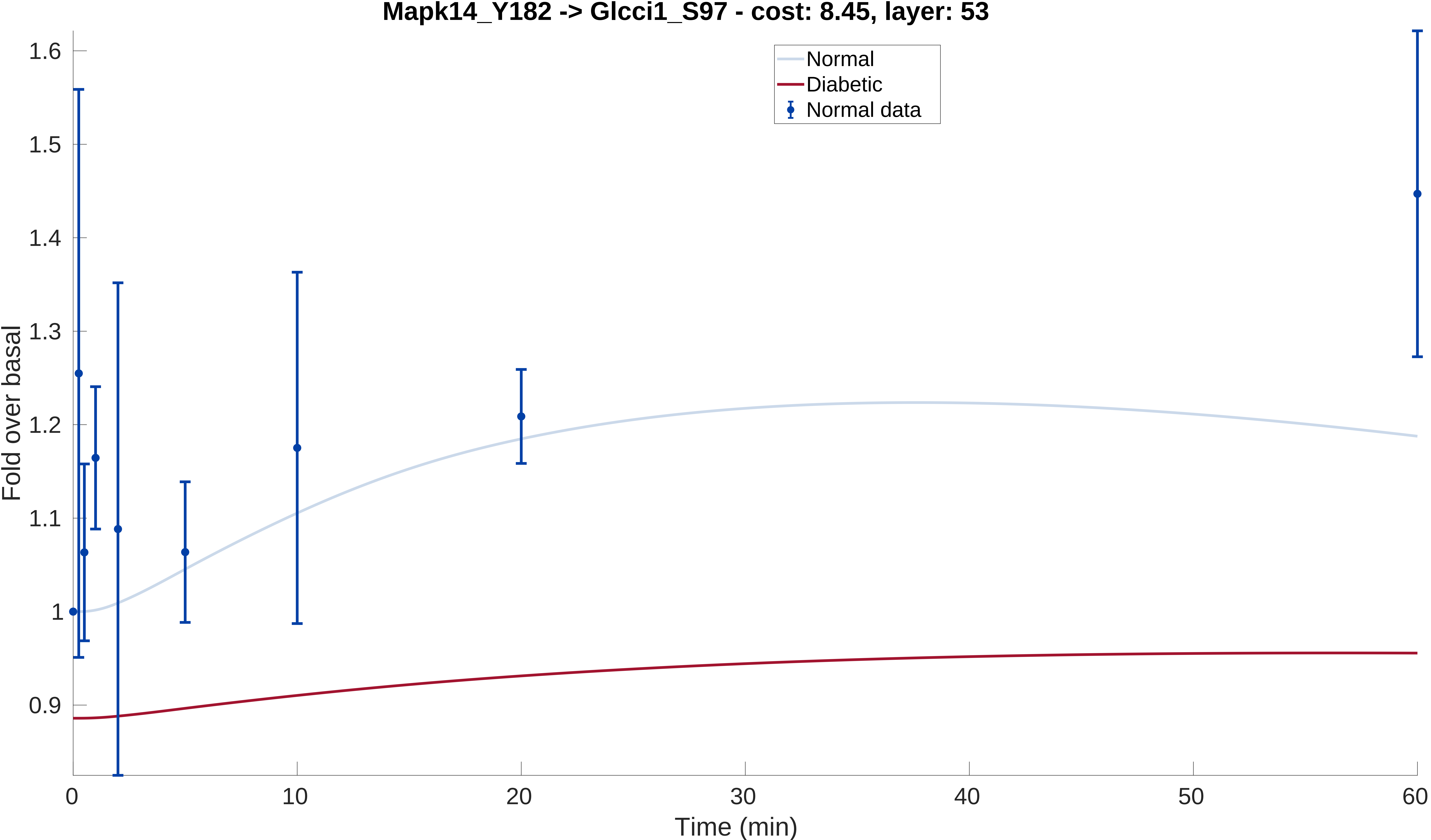

Mapk14_Y182 -> Glcci1_S97 - cost: 8.45, layer: 53
Normal
Diabetic
Normal data
Fold over basal
1.6
1.5
1.4
1.3
1.2
1.1
1
0.9
0
10
20
30
40
50
60
Time (min)

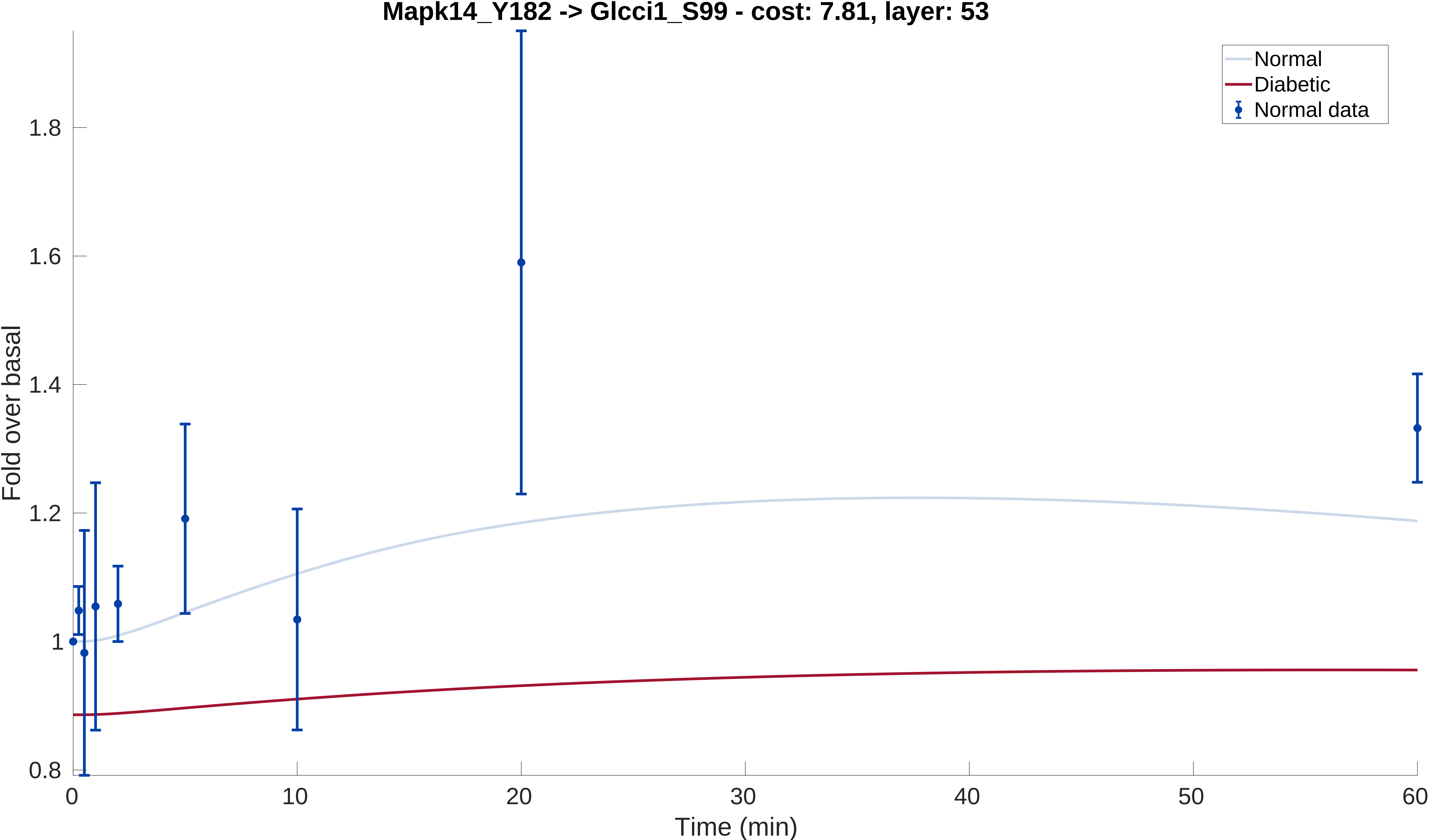

Mapk14_Y182 -> Glcci1_S99 - cost: 7.81, layer: 53
Normal
Diabetic
Normal data
Fold over basal
1.8
1.6
1.4
1.2
1
0.8
0
10
20
30
40
50
60
Time (min)

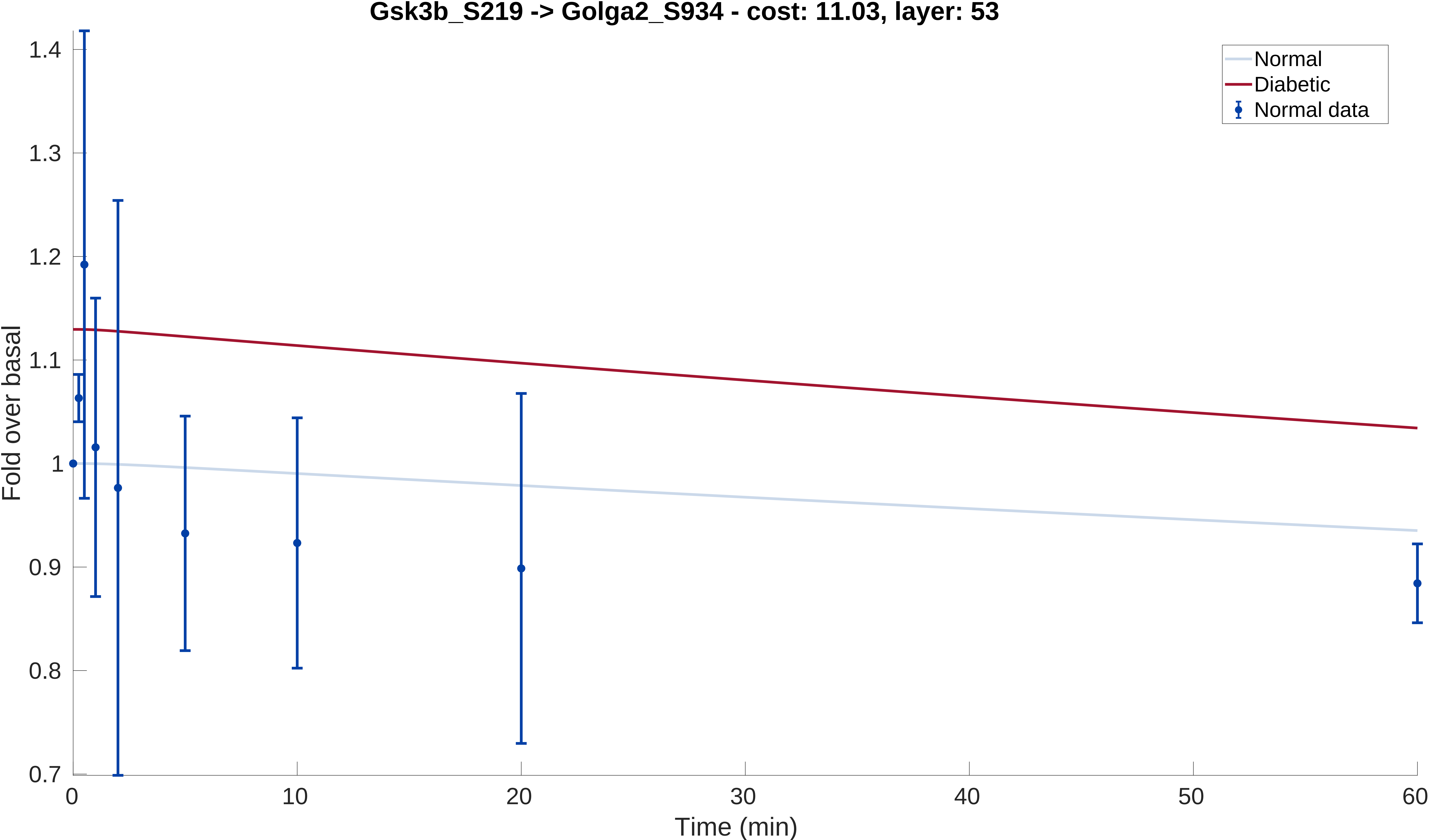

Gsk3b_S219 -> Golga2_S934 - cost: 11.03, layer: 53
Normal
Diabetic
Normal data
Fold over basal
1.4
1.3
1.2
1.1
1
0.9
0.8
0.7
0
10
20
30
40
50
60
Time (min)

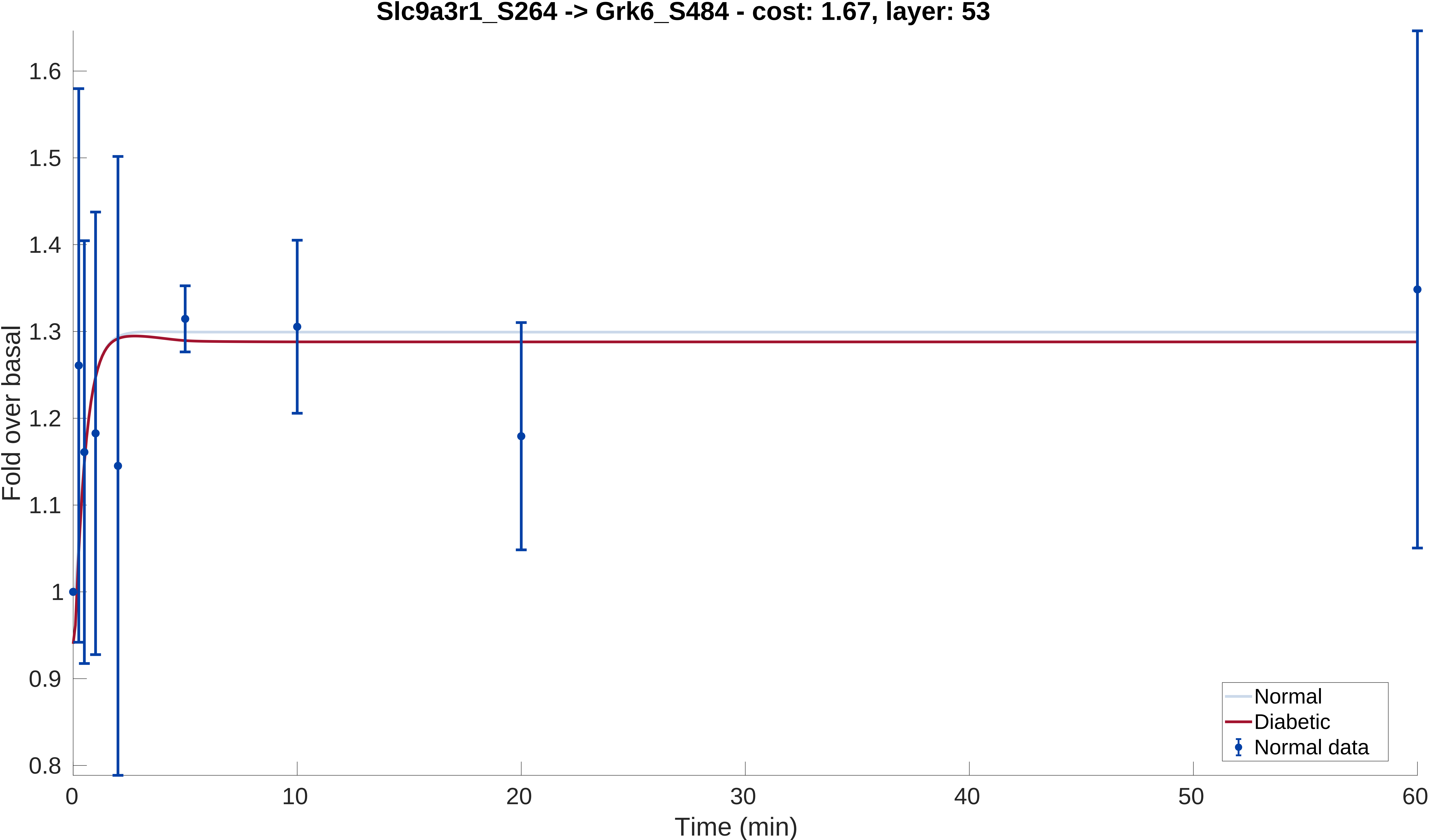

Slc9a3r1_S264 -> Grk6_S484 - cost: 1.67, layer: 53
Fold over basal
1.6
1.5
1.4
1.3
1.2
1.1
1
0.9
0.8
0
10
20
30
40
50
60
Time (min)
Normal
Diabetic
Normal data

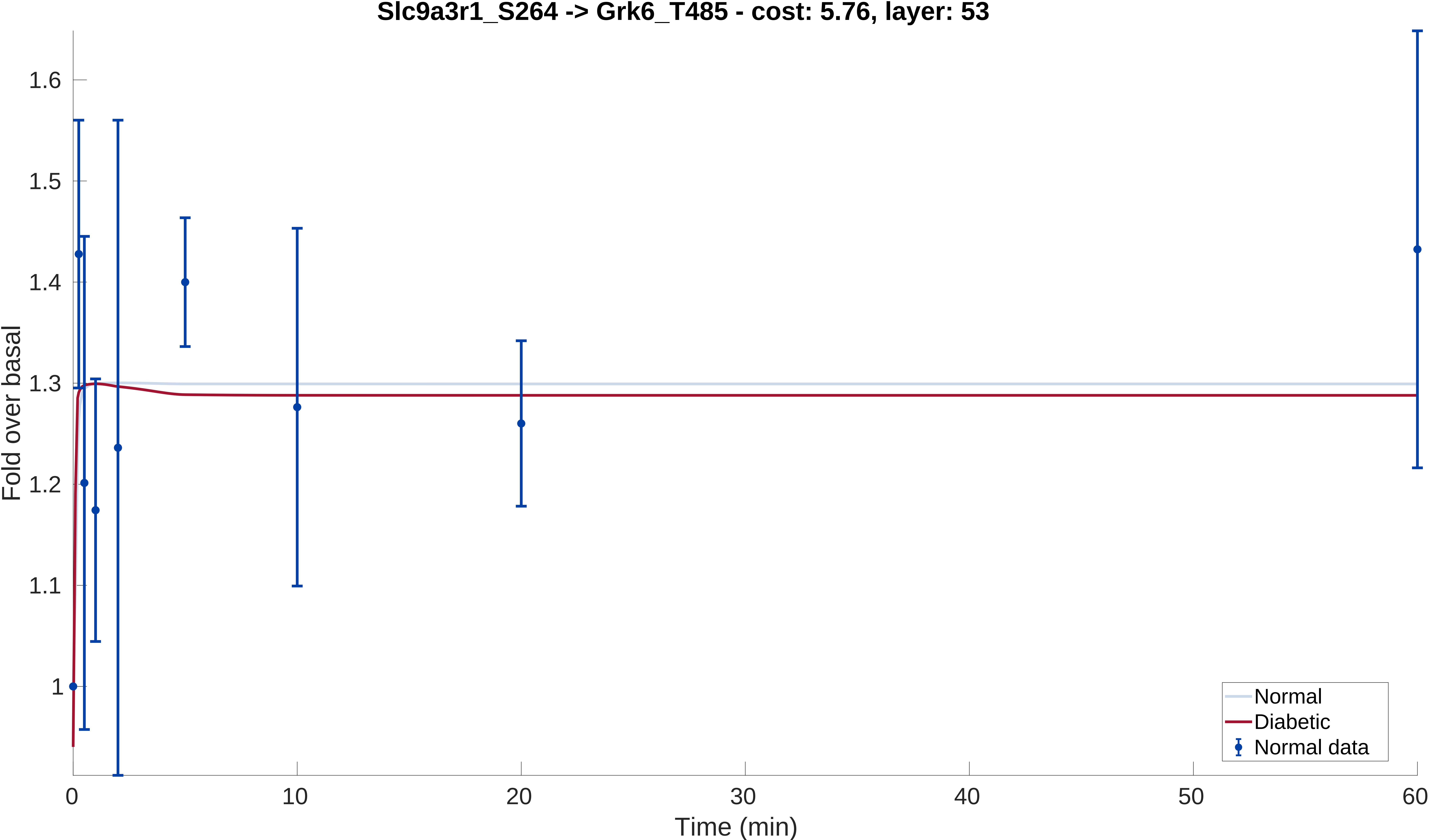

Slc9a3r1_S264 -> Grk6_T485 - cost: 5.76, layer: 53
Fold over basal
1.6
1.5
1.4
1.3
1.2
1.1
1
0
10
20
30
40
50
60
Time (min)
Normal
Diabetic
Normal data

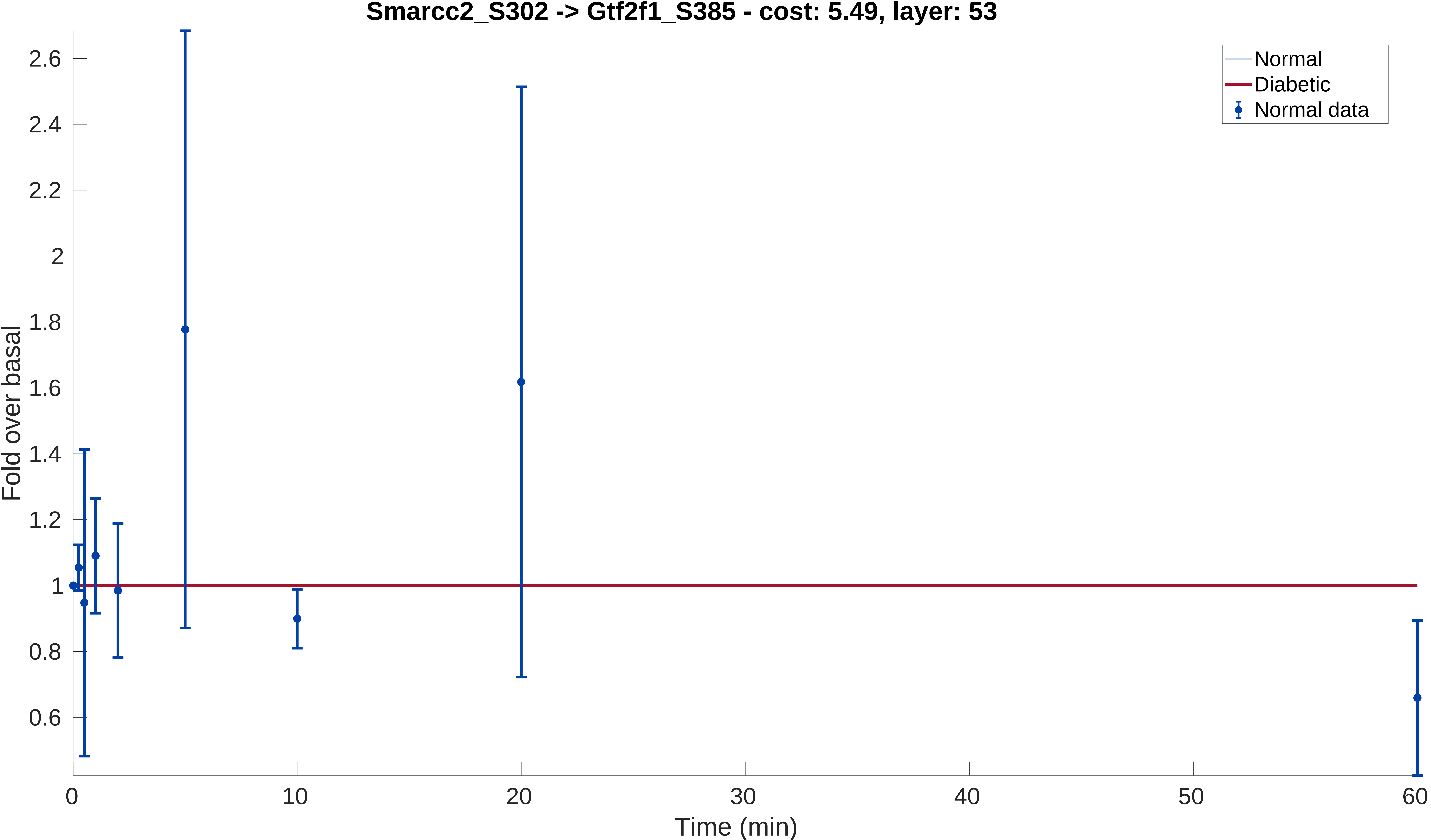
Smarcc2_S302 -> Gtf2f1_S385 - cost: 5.49, layer: 53
Normal
Diabetic
Normal data
Fold over basal
Time (min)
0.6
0.8
1
1.2
1.4
1.6
1.8
2
2.2
2.4
2.6
0
10
20
30
40
50
60

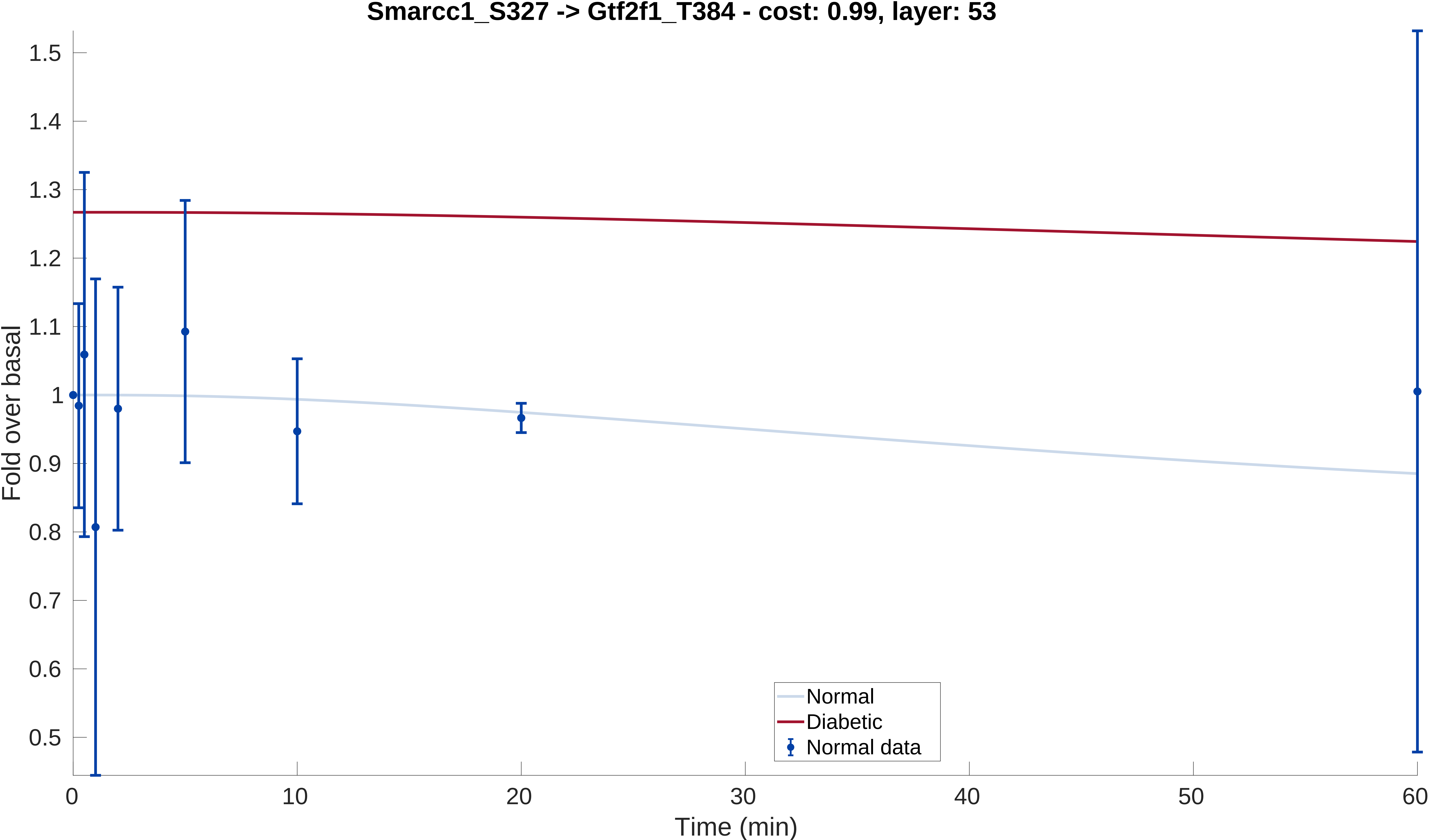

Smarcc1_S327 -> Gtf2f1_T384 - cost: 0.99, layer: 53
Fold over basal
Time (min)
1.5
1.4
1.3
1.2
1.1
1
0.9
0.8
0.7
0.6
0.5
0
10
20
30
40
50
60
Normal
Diabetic
Normal data

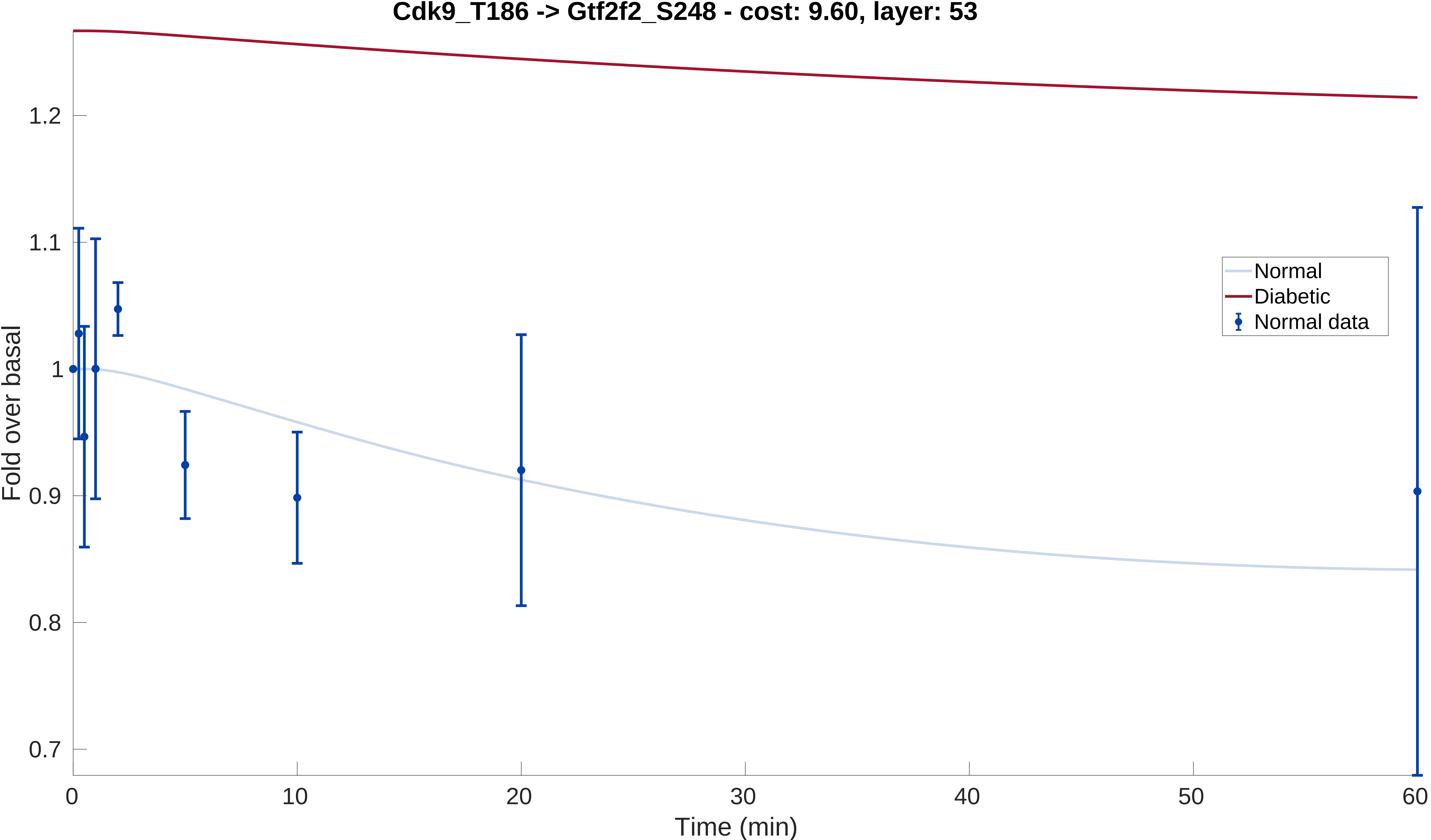
Cdk9_T186 -> Gtf2f2_S248 - cost: 9.60, layer: 53
Normal
Diabetic
Normal data
Fold over basal
1.2
1.1
1
0.9
0.8
0.7
0
10
20
30
40
50
60
Time (min)

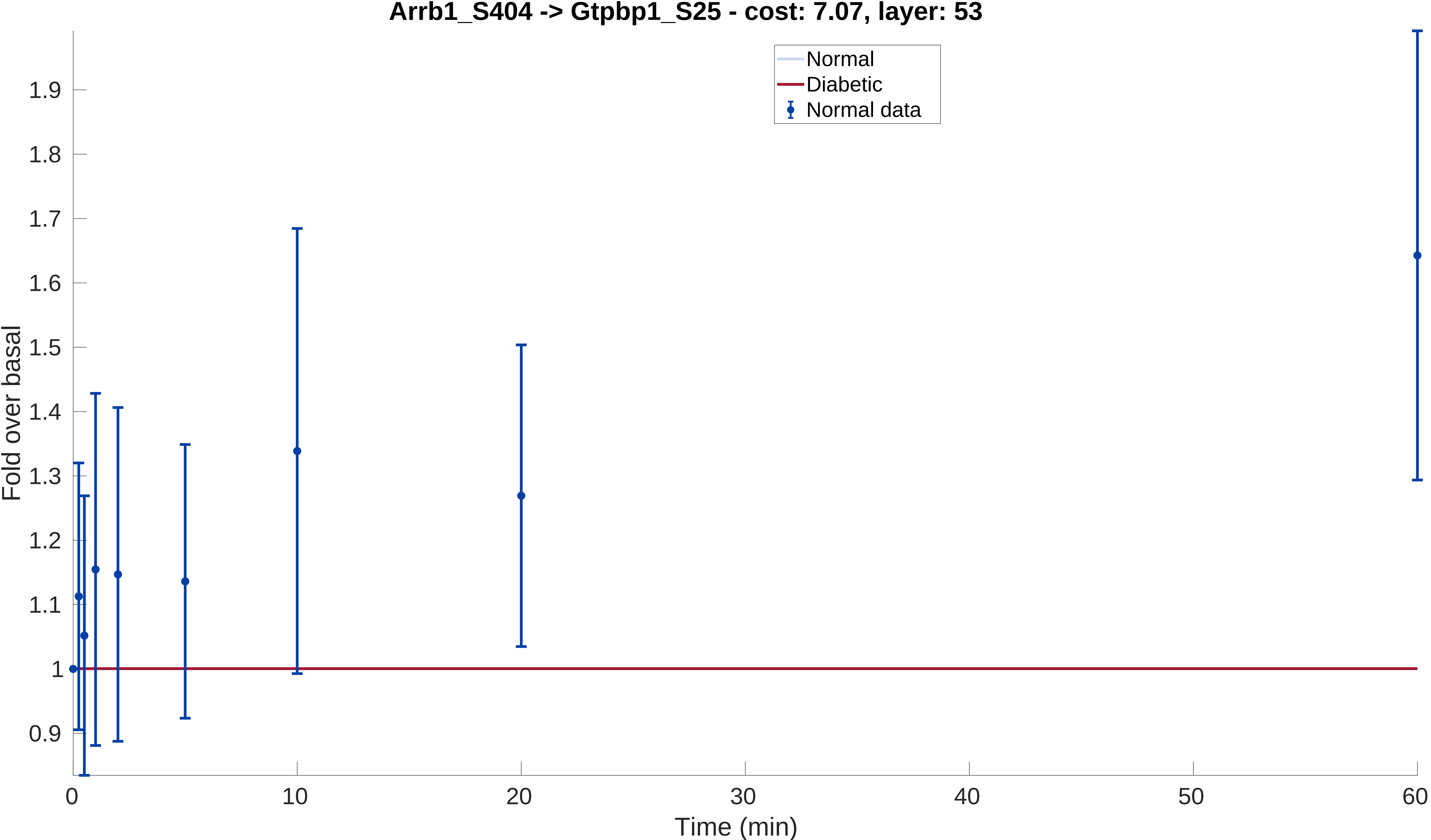

Arrb1_S404 -> Gtpbp1_S25 - cost: 7.07, layer: 53
Normal
Diabetic
Normal data
Fold over basal
1.9
1.8
1.7
1.6
1.5
1.4
1.3
1.2
1.1
1
0.9
0
10
20
30
40
50
60
Time (min)

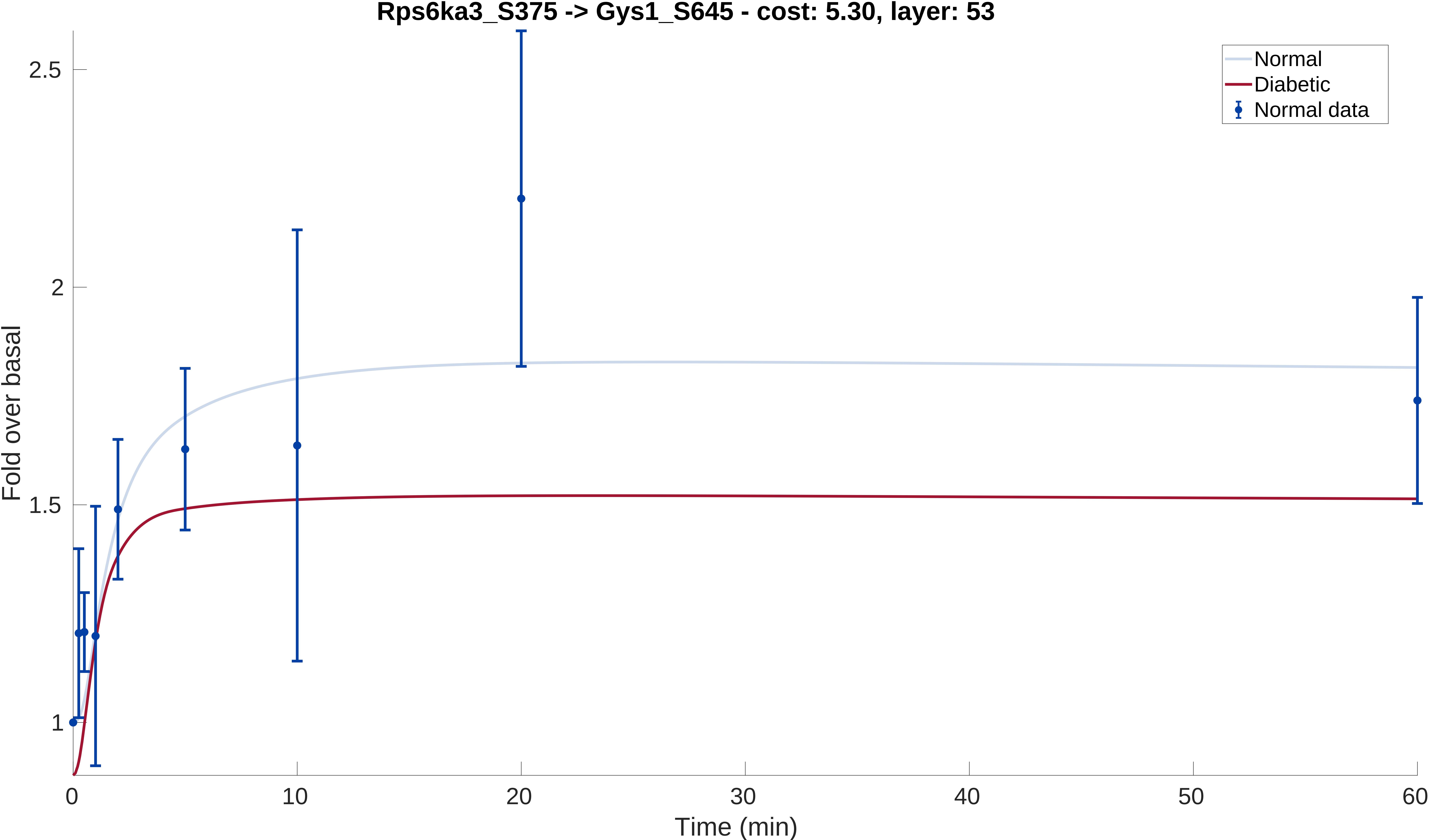
Rps6ka3_S375 -> Gys1_S645 - cost: 5.30, layer: 53
Normal
Diabetic
Normal data
Fold over basal
2.5
2
1.5
1
0
10
20
30
40
50
60
Time (min)

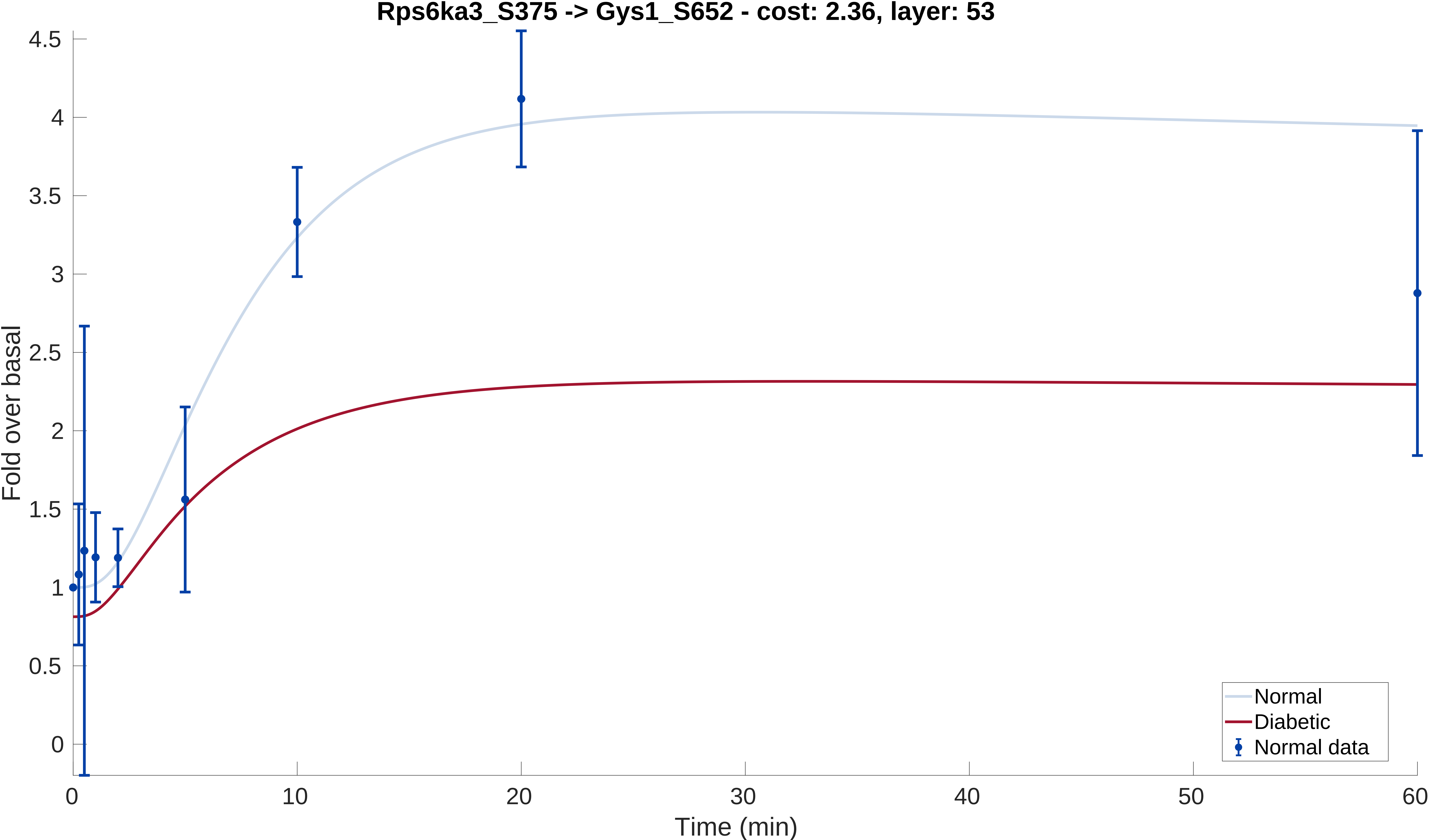

Rps6ka3_S375 -> Gys1_S652 - cost: 2.36, layer: 53
Fold over basal
Time (min)
0
0.5
1
1.5
2
2.5
3
3.5
4
4.5
0
10
20
30
40
50
60
Normal
Diabetic
Normal data

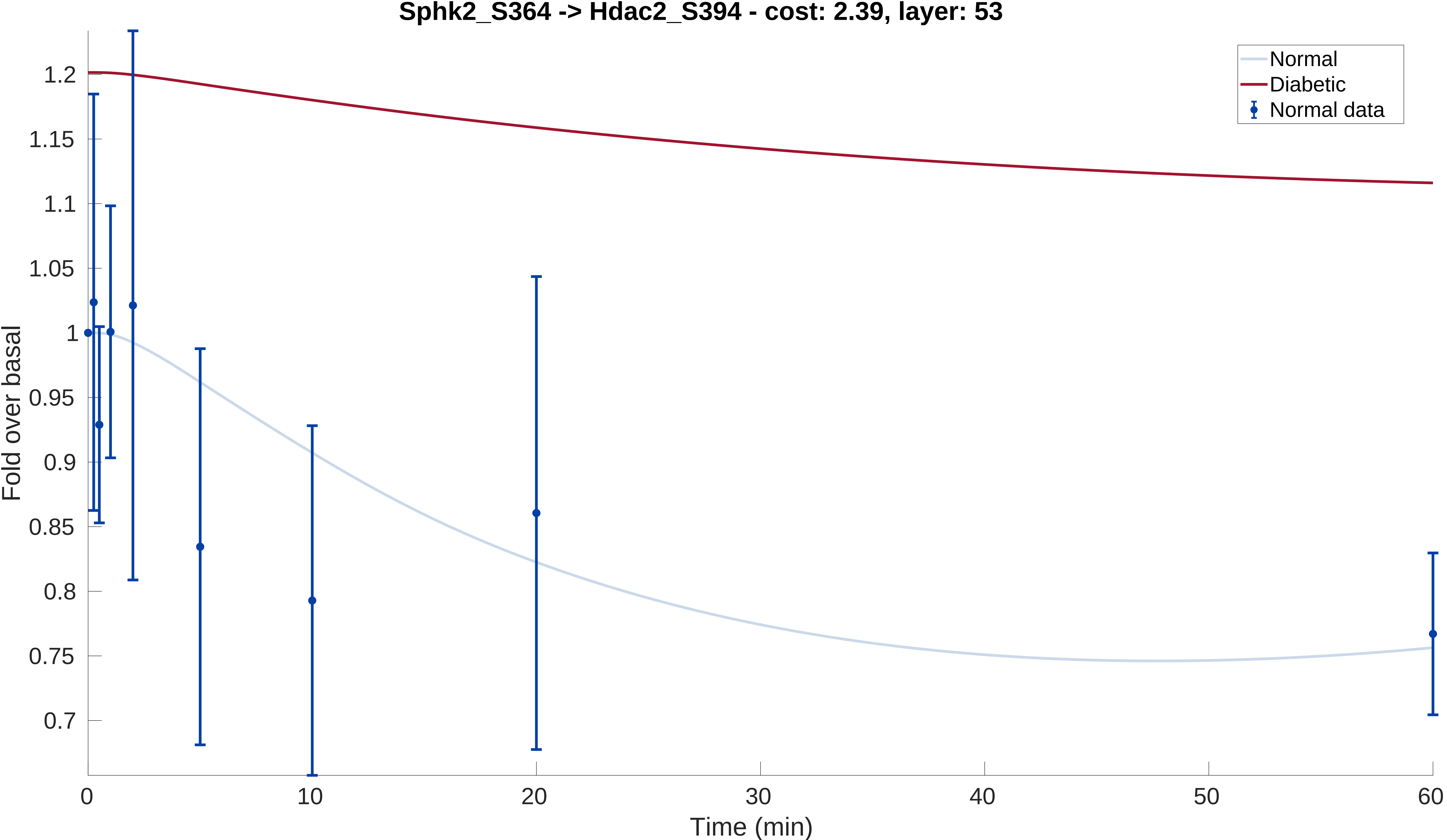

Sphk2_S364 -> Hdac2_S394 - cost: 2.39, layer: 53
Normal
Diabetic
Normal data
Fold over basal
Time (min)
1.2
1.15
1.1
1.05
1
0.95
0.9
0.85
0.8
0.75
0.7
0
10
20
30
40
50
60

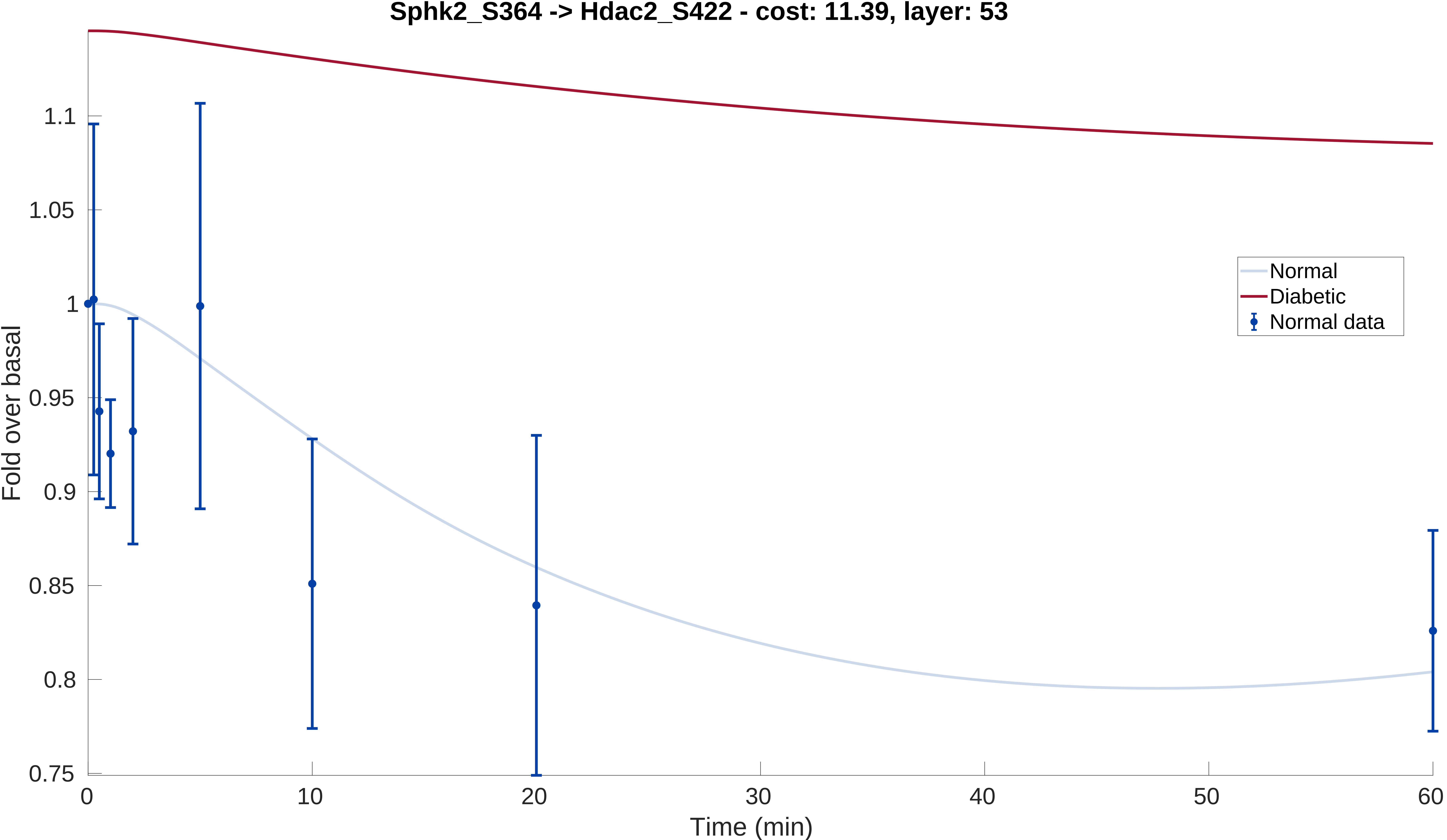

Sphk2_S364 -> Hdac2_S422 - cost: 11.39, layer: 53
Fold over basal
1.1
1.05
1
0.95
0.9
0.85
0.8
0.75
0
10
20
30
40
50
60
Time (min)
Normal
Diabetic
Normal data

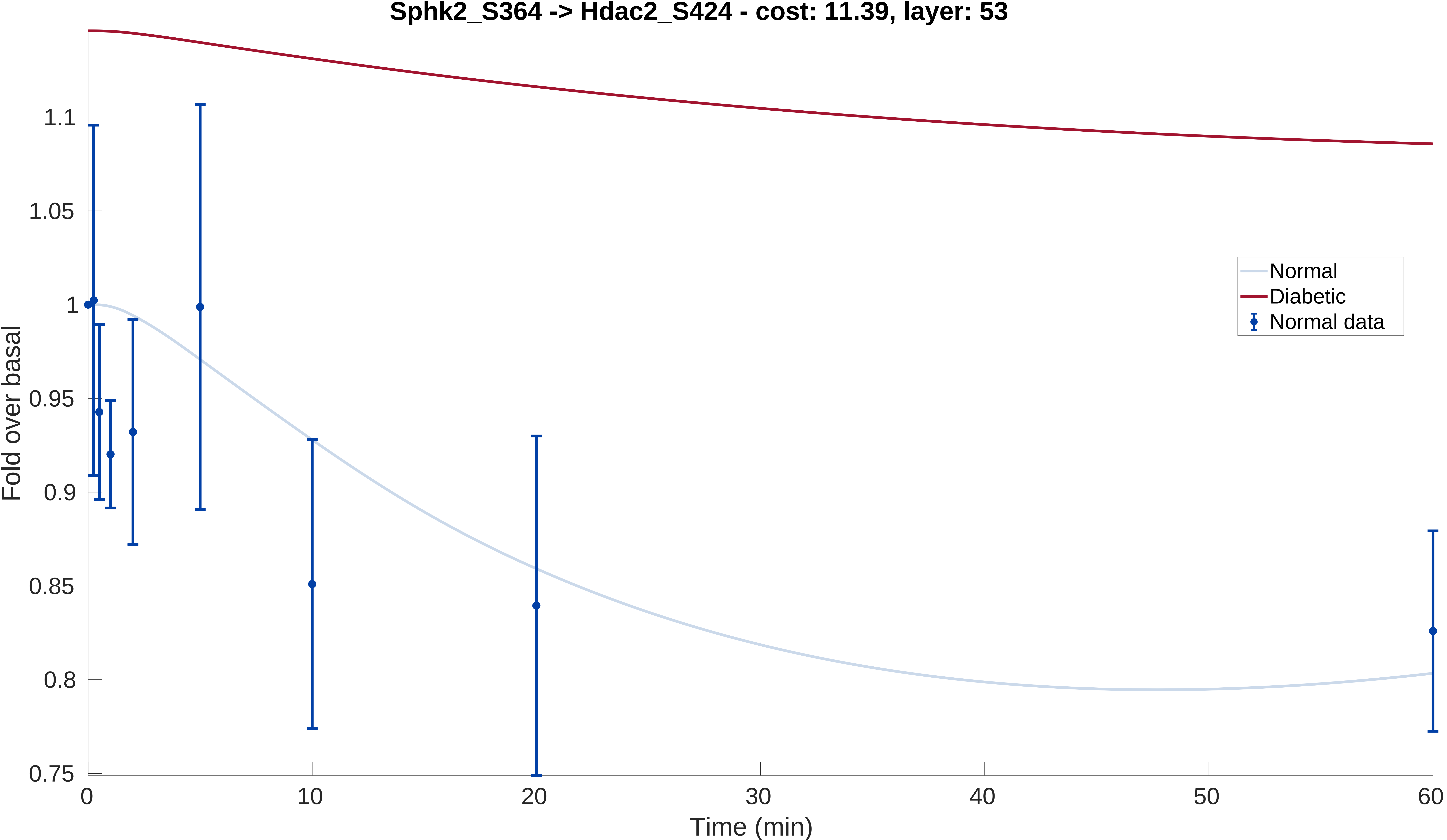

Sphk2_S364 -> Hdac2_S424 - cost: 11.39, layer: 53
Fold over basal
1.1
1.05
1
0.95
0.9
0.85
0.8
0.75
0
10
20
30
40
50
60
Time (min)
Normal
Diabetic
Normal data

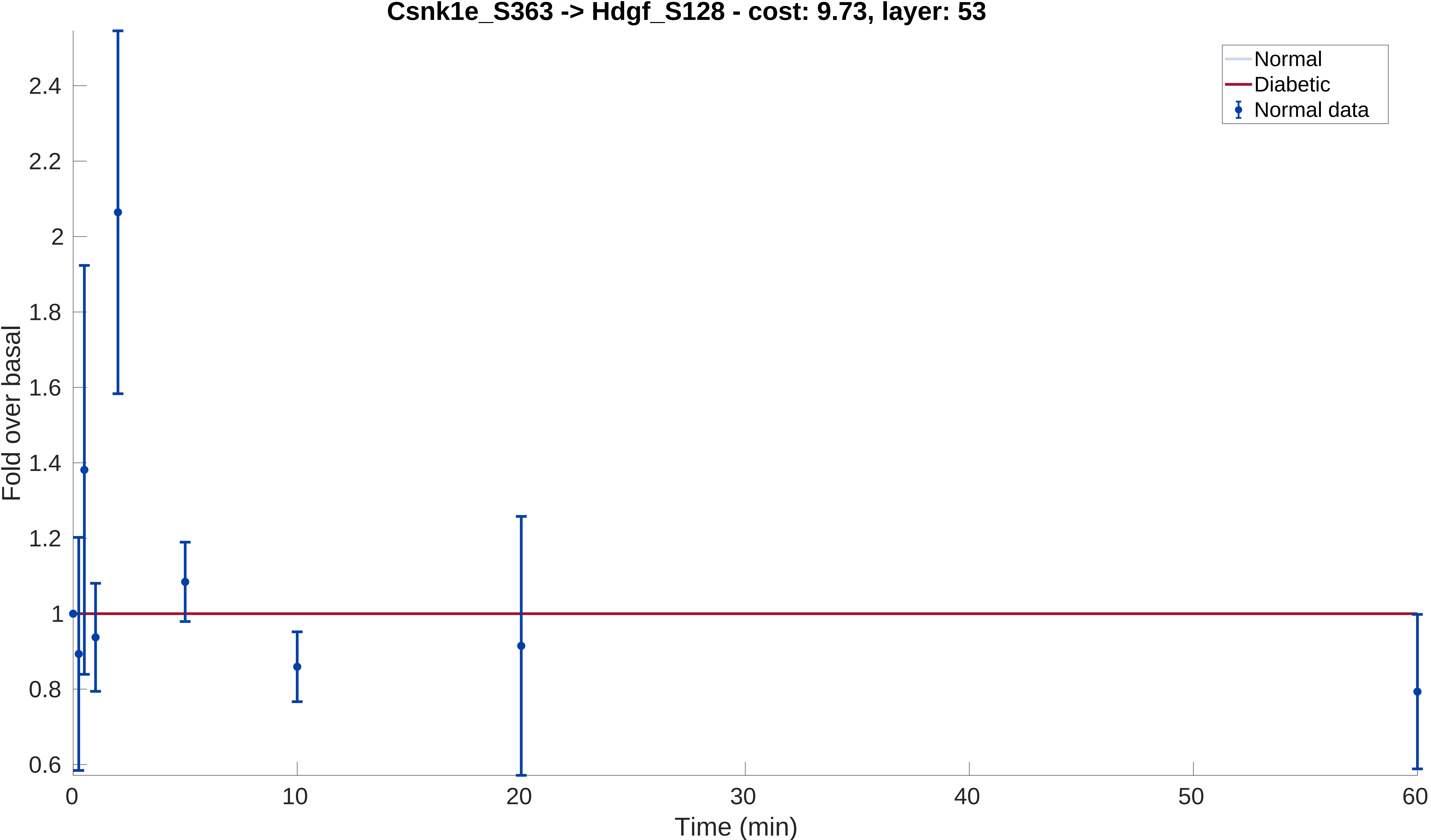

Csnk1e_S363 -> Hdgf_S128 - cost: 9.73, layer: 53
Normal
Diabetic
Normal data
Fold over basal
Time (min)
0
10
20
30
40
50
60
0.6
0.8
1
1.2
1.4
1.6
1.8
2
2.2
2.4

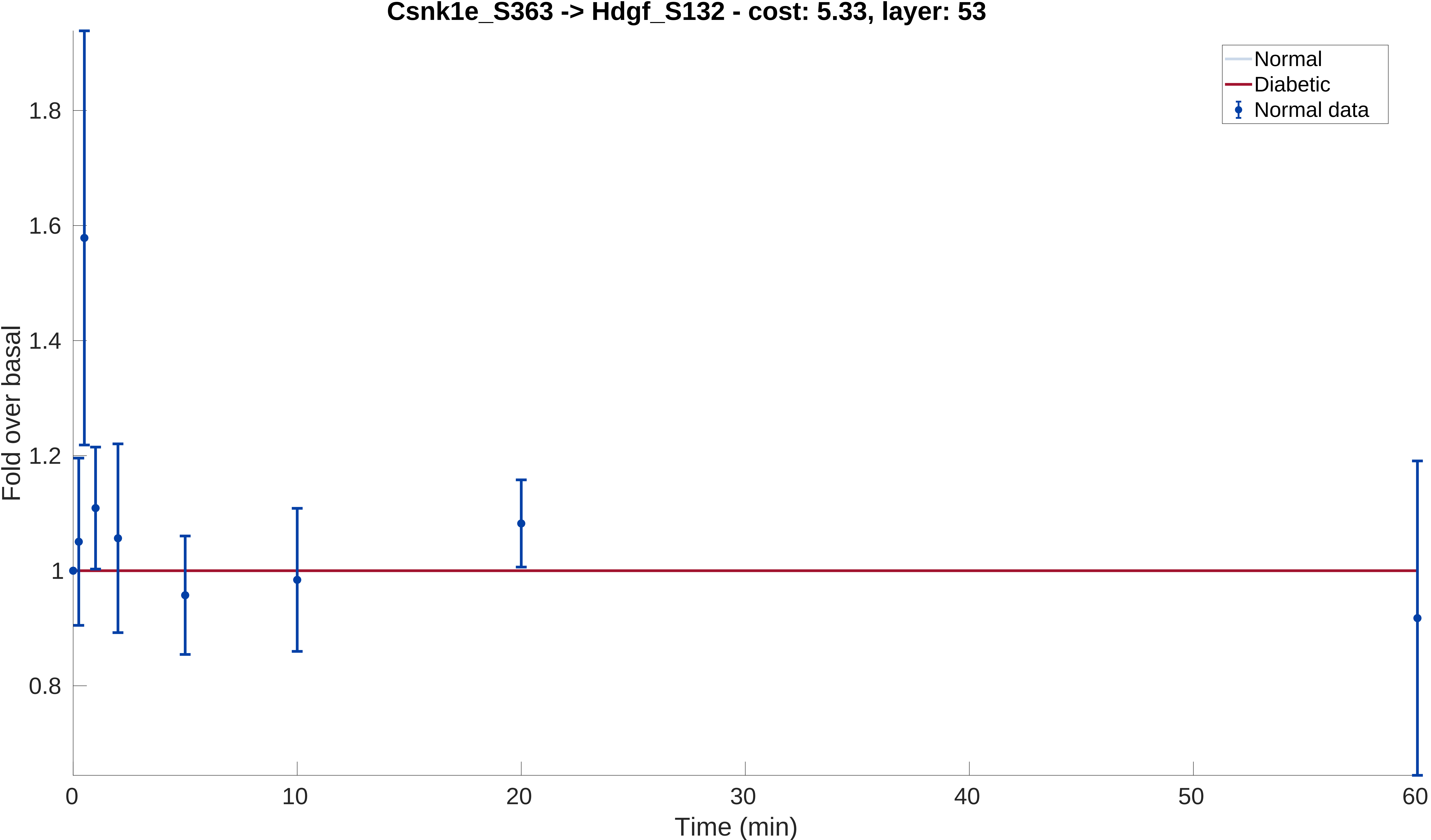
Csnk1e_S363 -> Hdgf_S132 - cost: 5.33, layer: 53
Normal
Diabetic
Normal data
Fold over basal
1.8
1.6
1.4
1.2
1
0.8
0
10
20
30
40
50
60
Time (min)

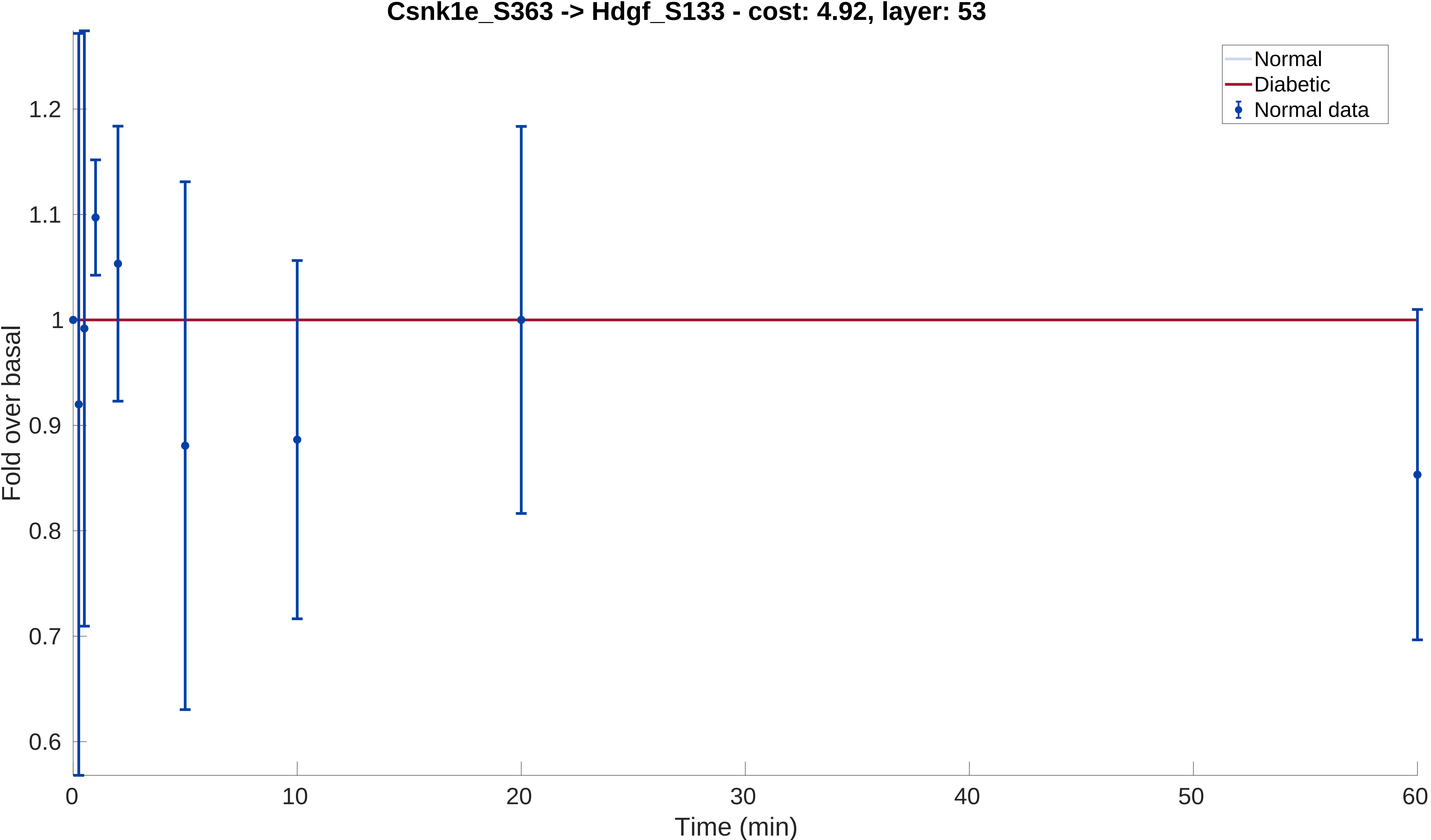

Csnk1e_S363 -> Hdgf_S133 - cost: 4.92, layer: 53
Normal
Diabetic
Normal data
Fold over basal
1.2
1.1
1
0.9
0.8
0.7
0.6
0
10
20
30
40
50
60
Time (min)

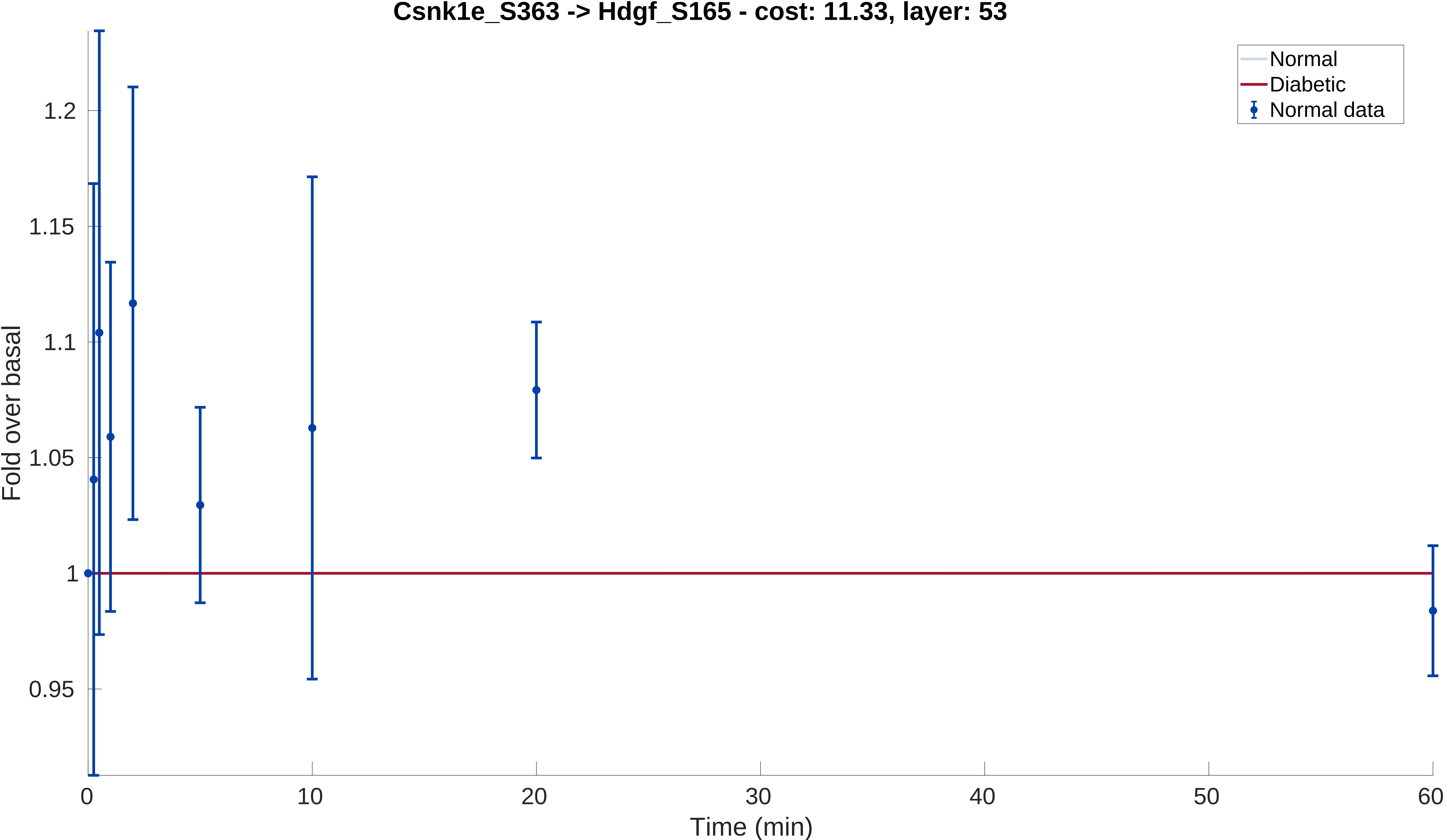
Csnk1e_S363 -> Hdgf_S165 - cost: 11.33, layer: 53
Normal
Diabetic
Normal data
Fold over basal
1.2
1.15
1.1
1.05
1
0.95
0
10
20
30
40
50
60
Time (min)

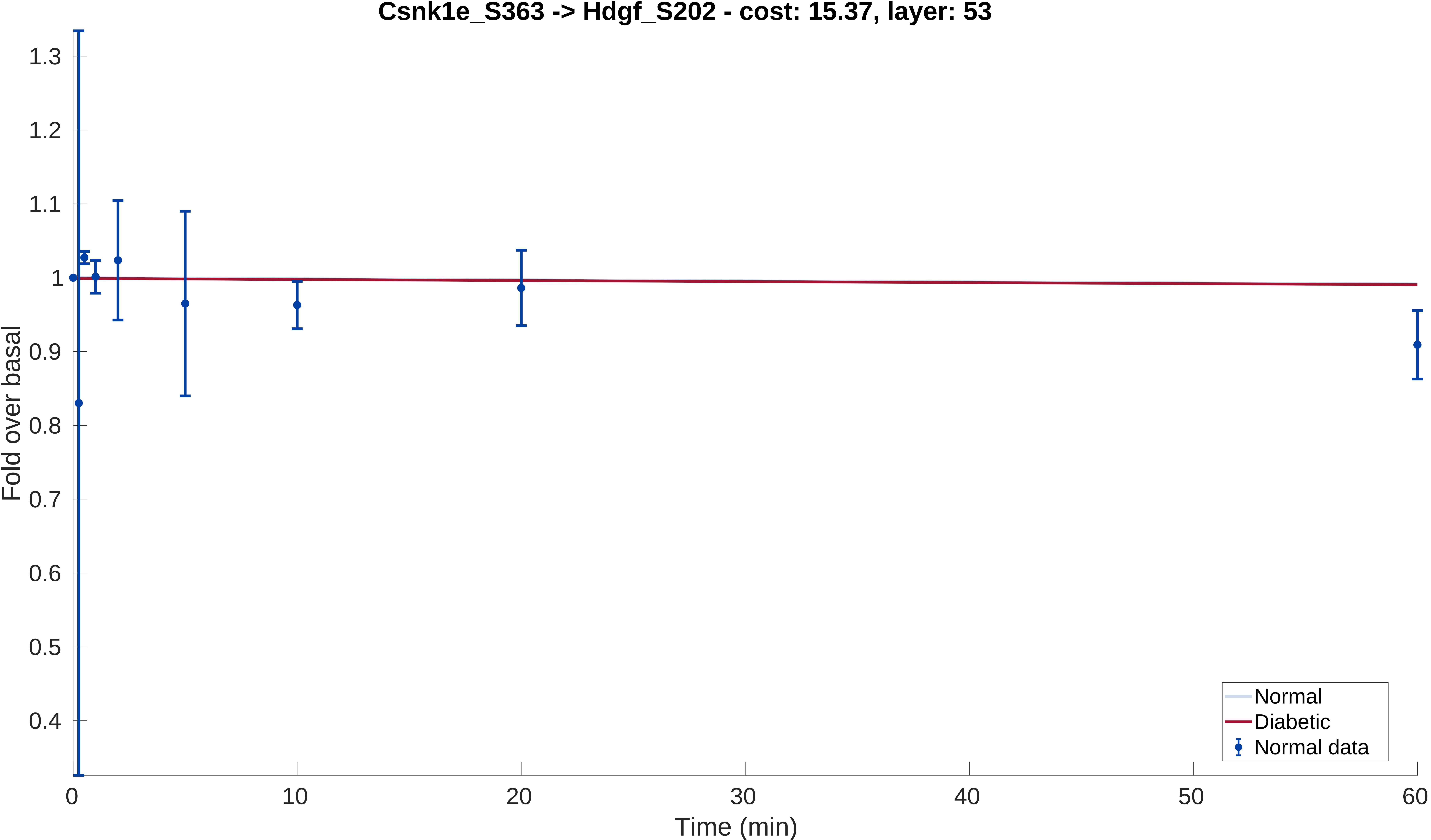

Csnk1e_S363 -> Hdgf_S202 - cost: 15.37, layer: 53
Fold over basal
1.3
1.2
1.1
1
0.9
0.8
0.7
0.6
0.5
0.4
0
10
20
30
40
50
60
Time (min)
Normal
Diabetic
Normal data

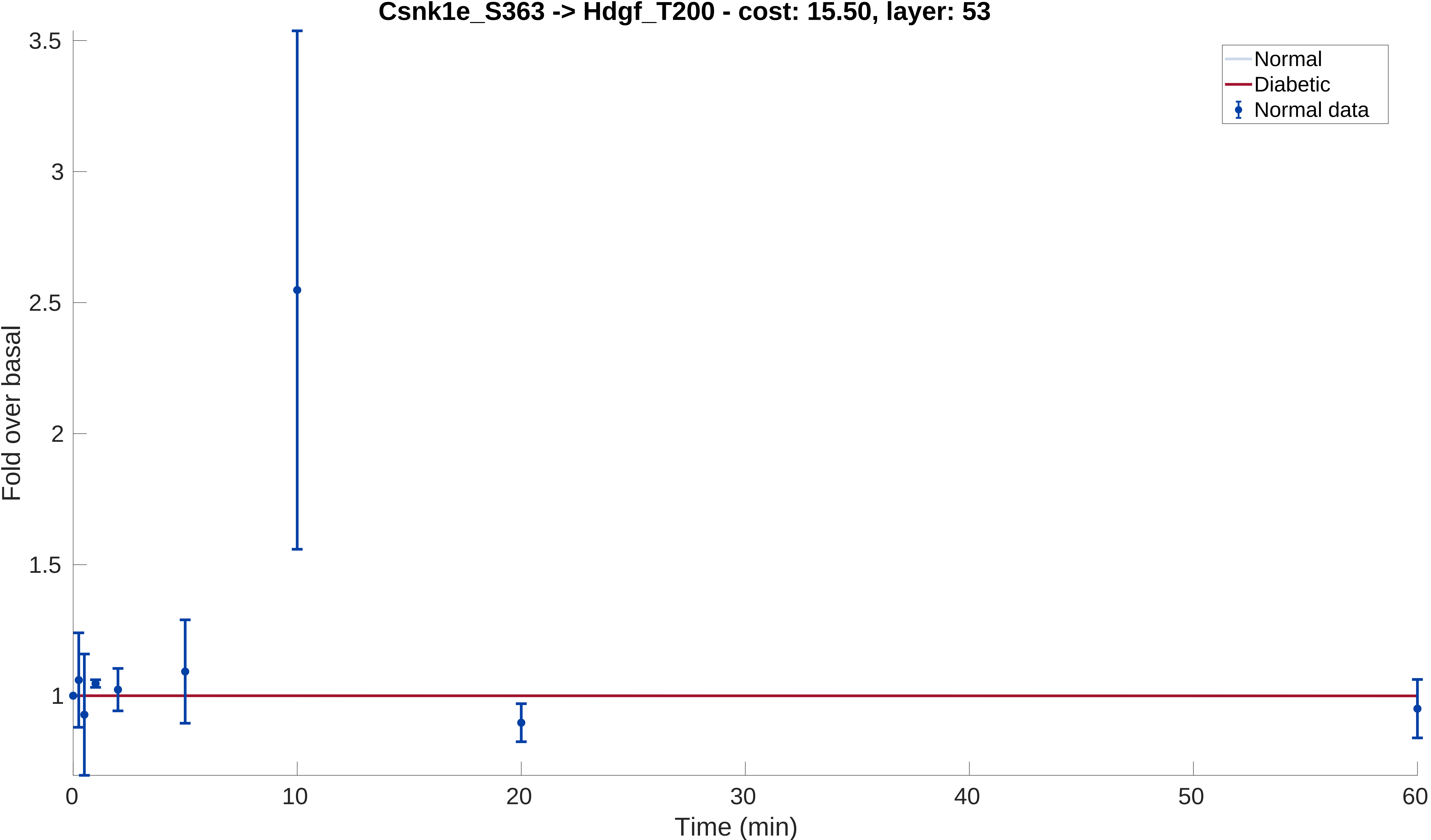

Csnk1e_S363 -> Hdgf_T200 - cost: 15.50, layer: 53
Normal
Diabetic
Normal data
Fold over basal
3.5
3
2.5
2
1.5
1
0
10
20
30
40
50
60
Time (min)

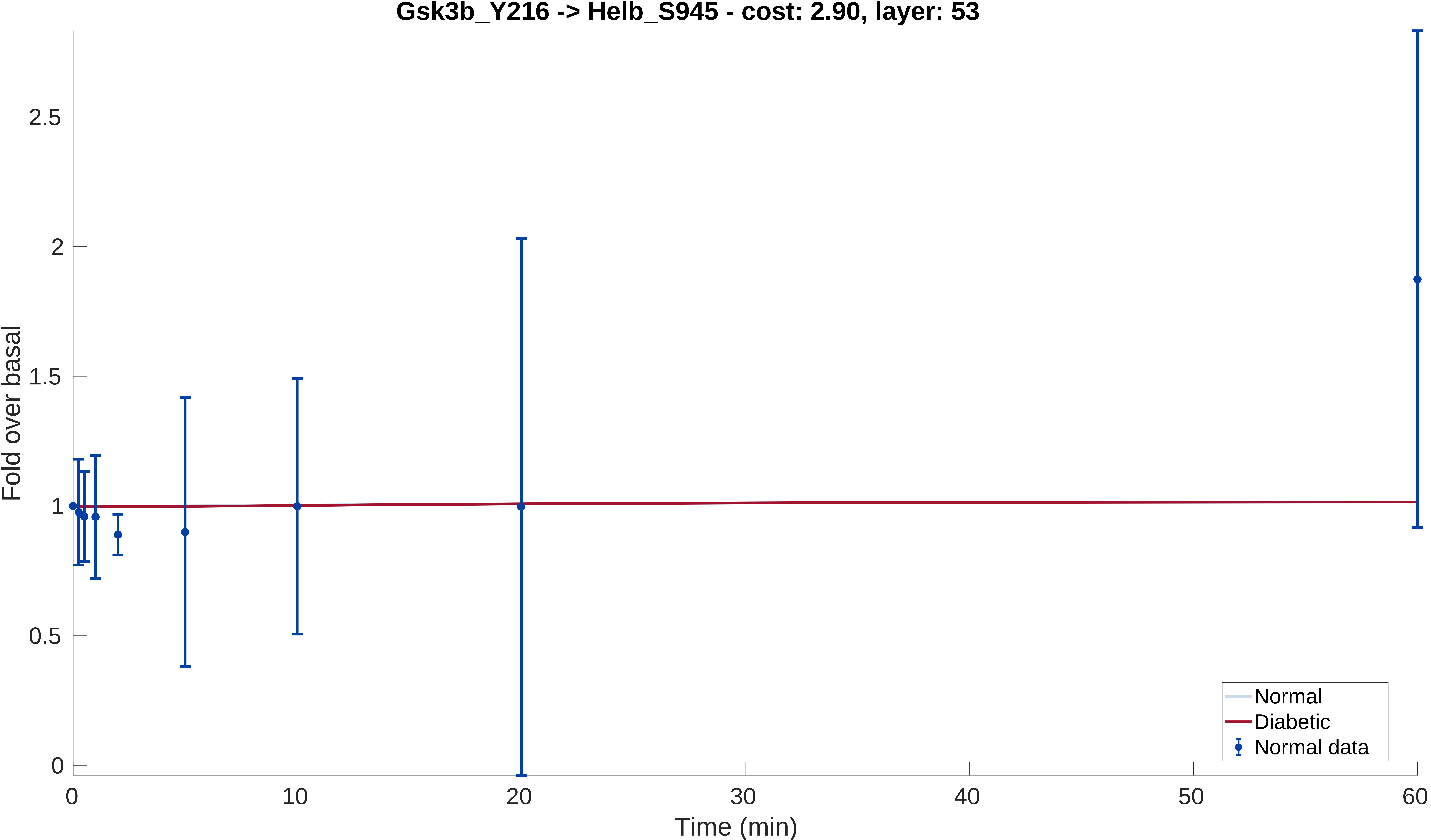

Gsk3b_Y216 -> Helb_S945 - cost: 2.90, layer: 53
Fold over basal
Time (min)
0
0.5
1
1.5
2
2.5
0
10
20
30
40
50
60
Normal
Diabetic
Normal data

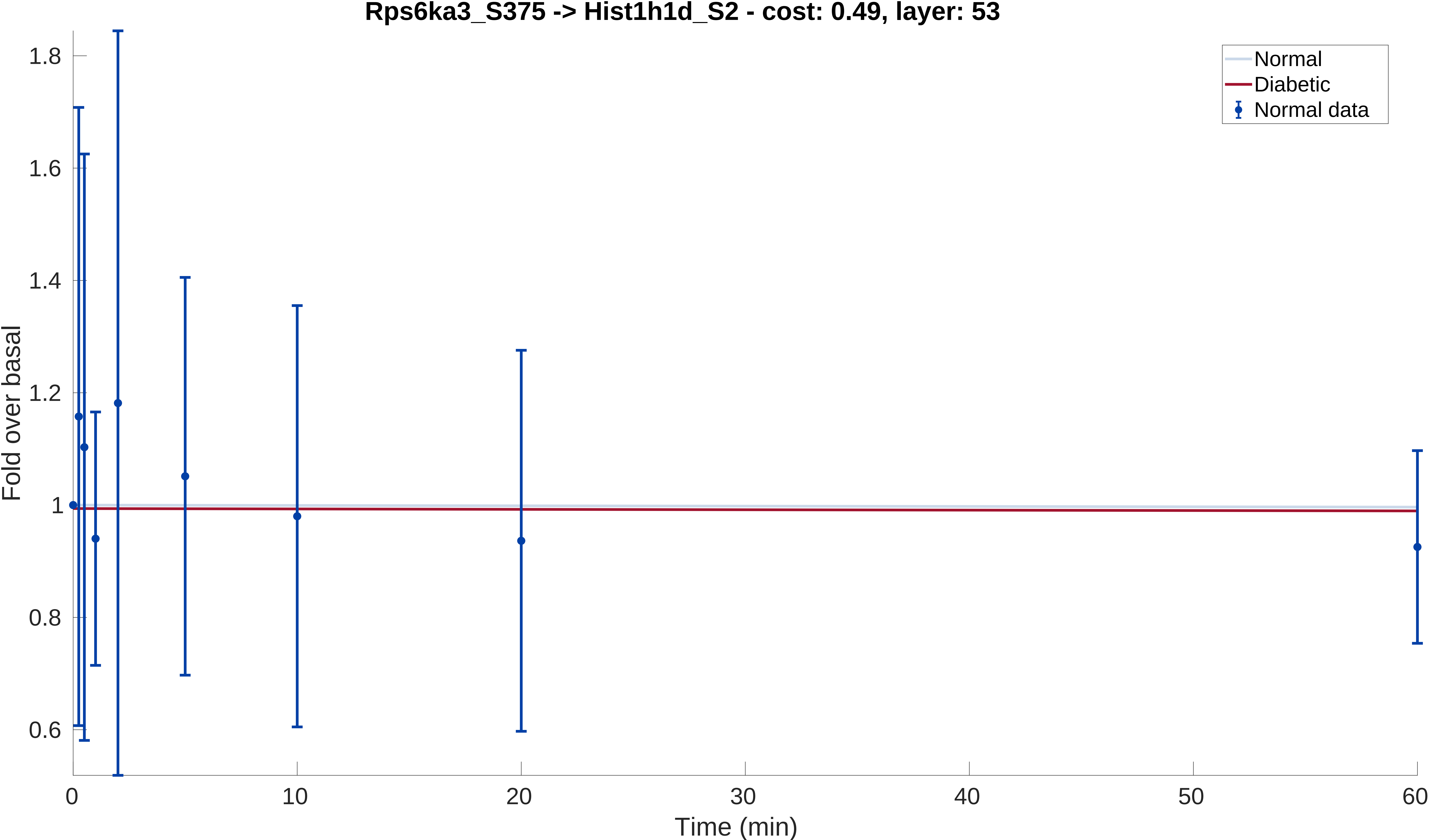

Rps6ka3_S375 -> Hist1h1d_S2 - cost: 0.49, layer: 53
Normal
Diabetic
Normal data
Fold over basal
Time (min)
1.8
1.6
1.4
1.2
1
0.8
0.6
0
10
20
30
40
50
60

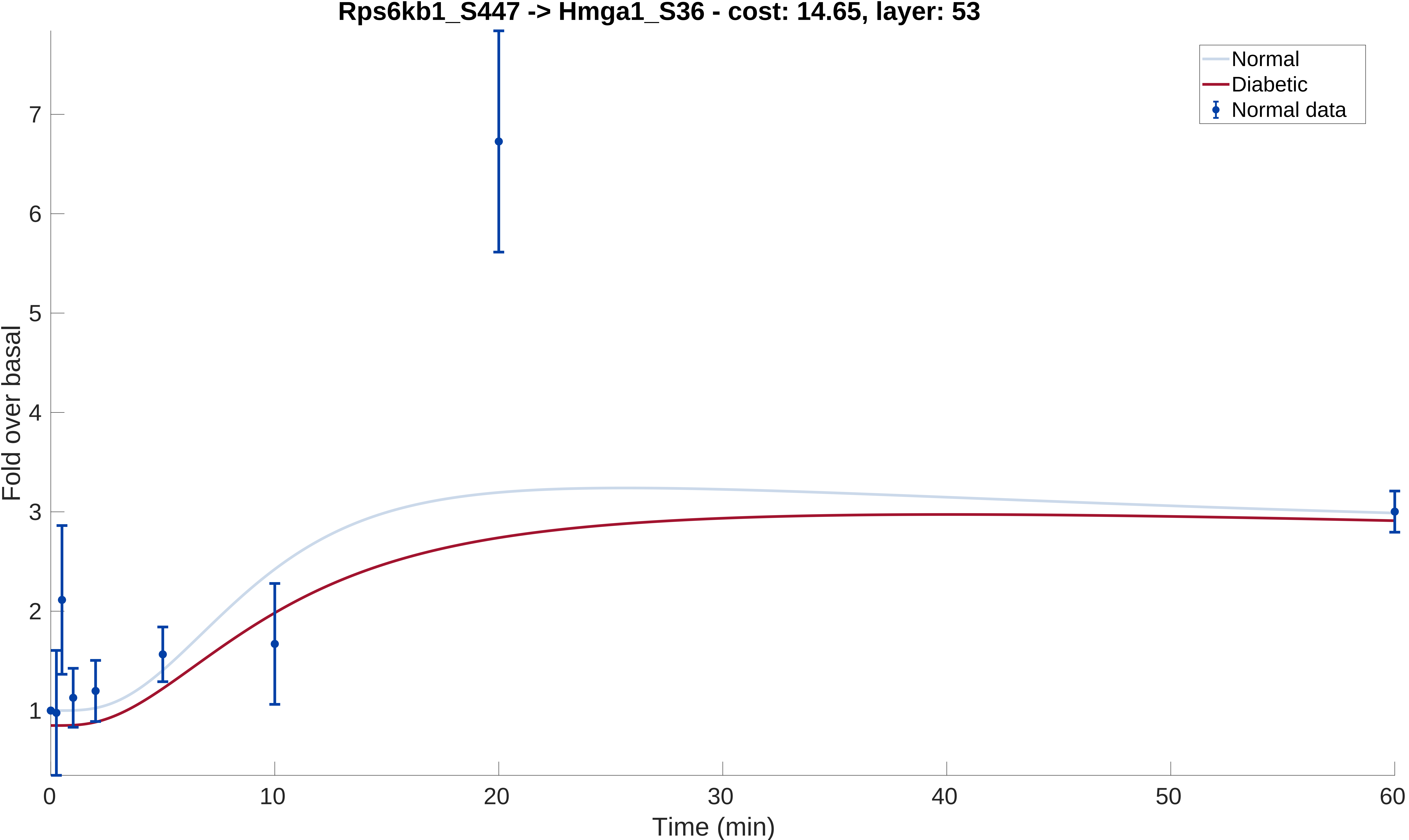

Rps6kb1_S447 -> Hmga1_S36 - cost: 14.65, layer: 53
Normal
Diabetic
Normal data
Fold over basal
Time (min)
1
2
3
4
5
6
7
0
10
20
30
40
50
60

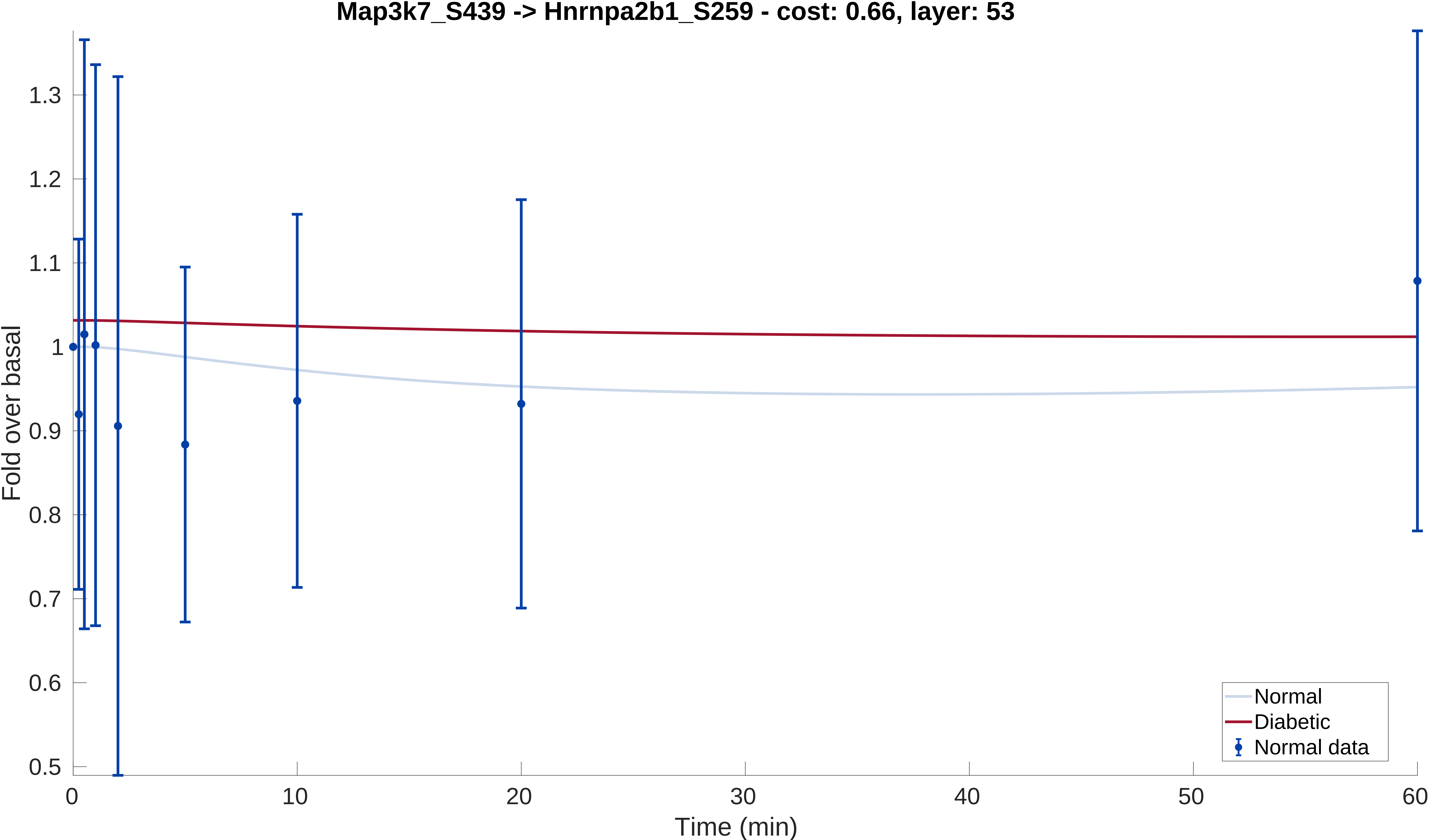

Map3k7_S439 -> Hnrnpa2b1_S259 - cost: 0.66, layer: 53
1.3
1.2
1.1
1
0.9
0.8
0.7
0.6
0.5
Fold over basal
0
10
20
30
40
50
60
Time (min)
Normal
Diabetic
Normal data

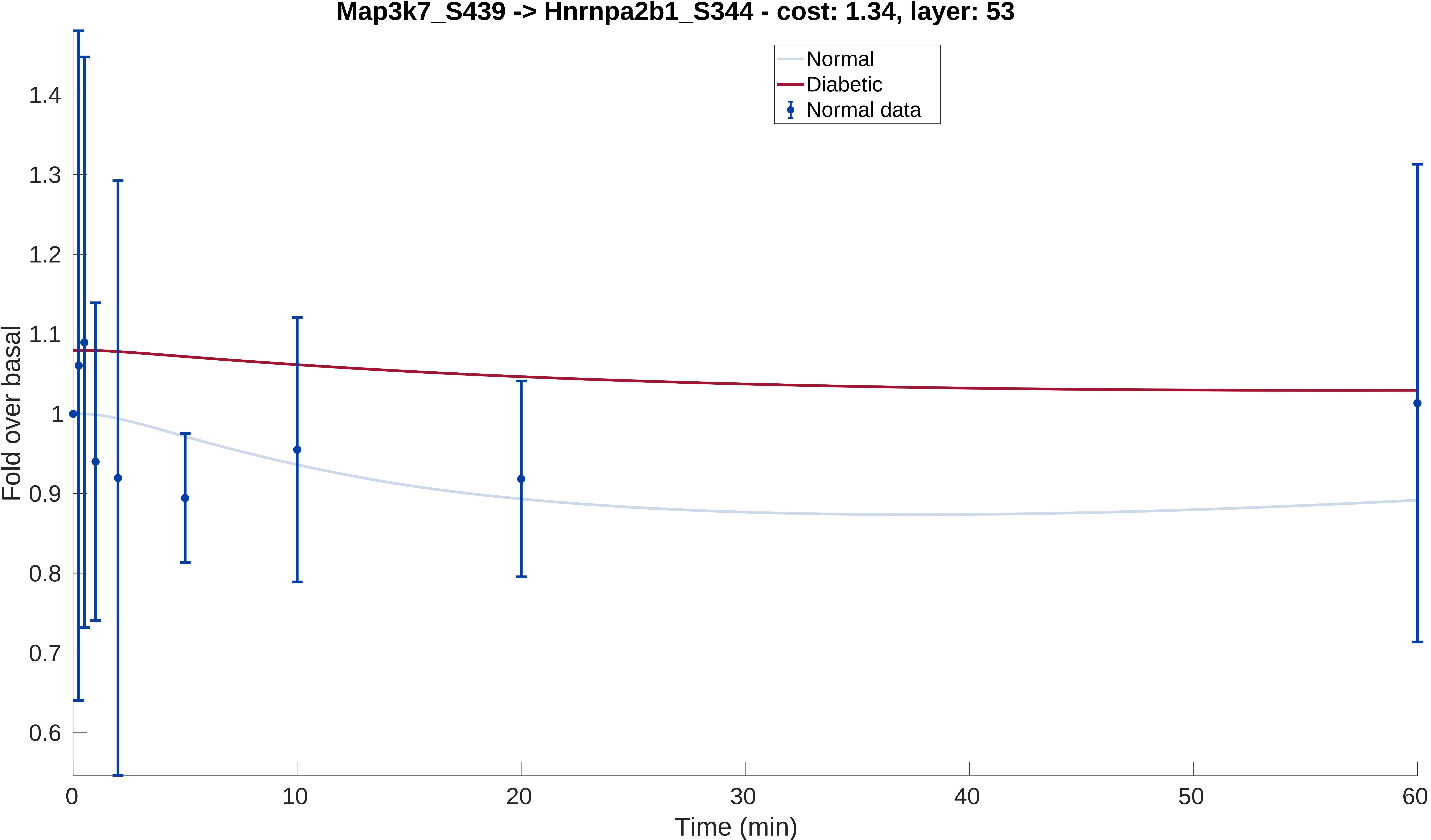

Map3k7_S439 -> Hnrnpa2b1_S344 - cost: 1.34, layer: 53
Normal
Diabetic
Normal data
Fold over basal
Time (min)
1.4
1.3
1.2
1.1
1
0.9
0.8
0.7
0.6
0
10
20
30
40
50
60

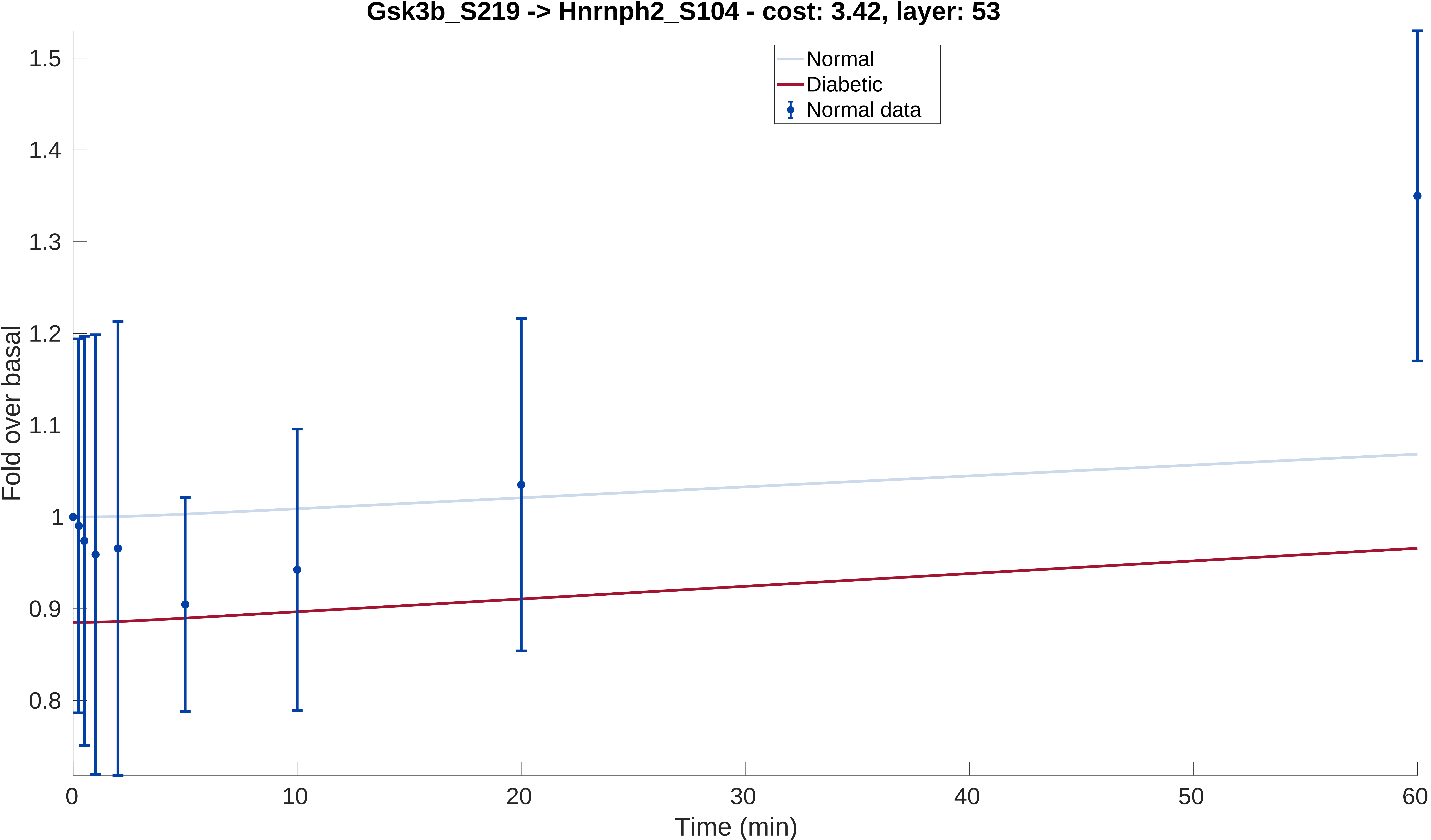
Gsk3b_S219 -> Hnrnph2_S104 - cost: 3.42, layer: 53
Normal
Diabetic
Normal data
Fold over basal
1.5
1.4
1.3
1.2
1.1
1
0.9
0.8
0
10
20
30
40
50
60
Time (min)

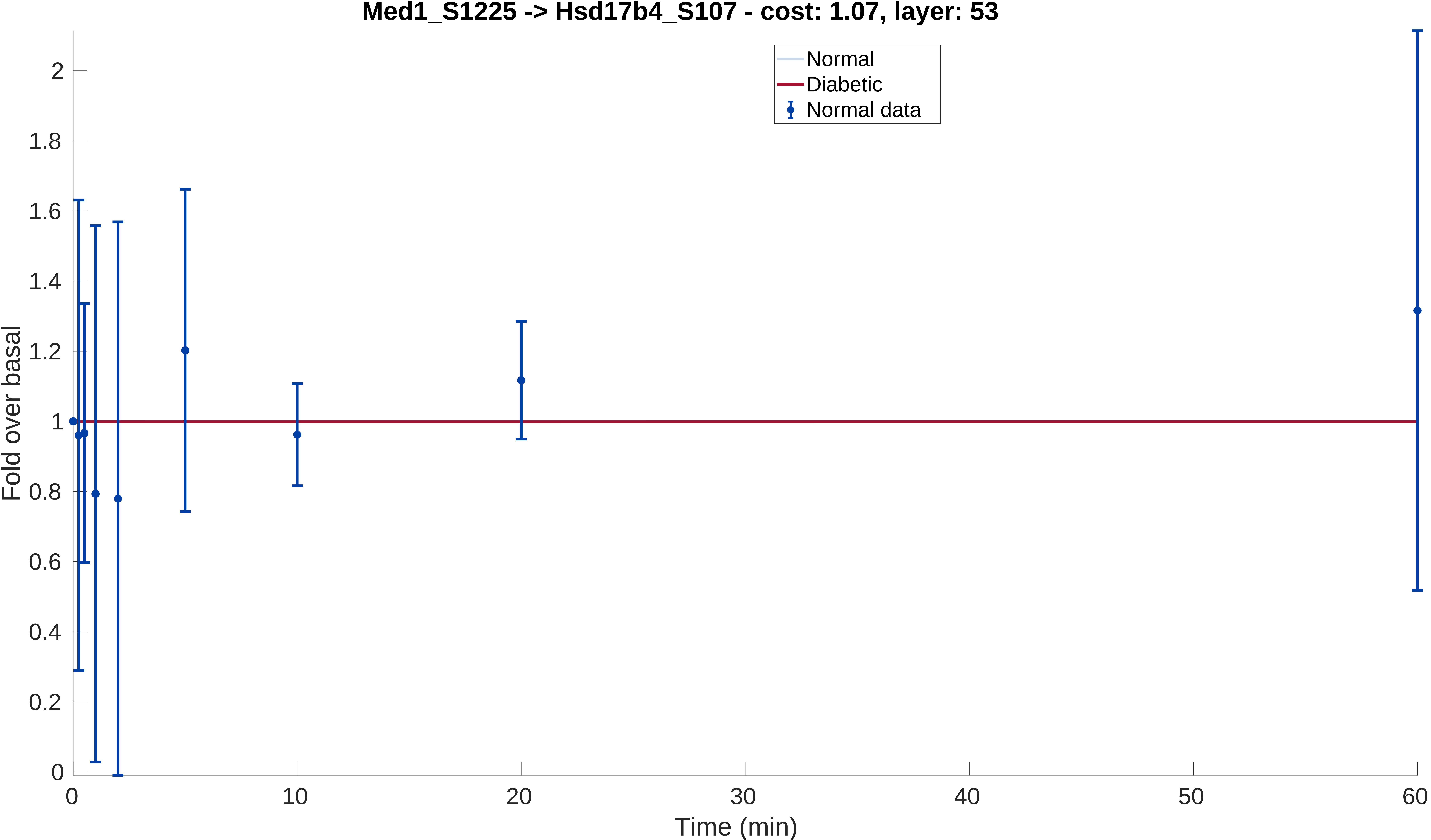

Med1_S1225 -> Hsd17b4_S107 - cost: 1.07, layer: 53
Normal
Diabetic
Normal data
Fold over basal
Time (min)
0
0.2
0.4
0.6
0.8
1
1.2
1.4
1.6
1.8
2
0
10
20
30
40
50
60

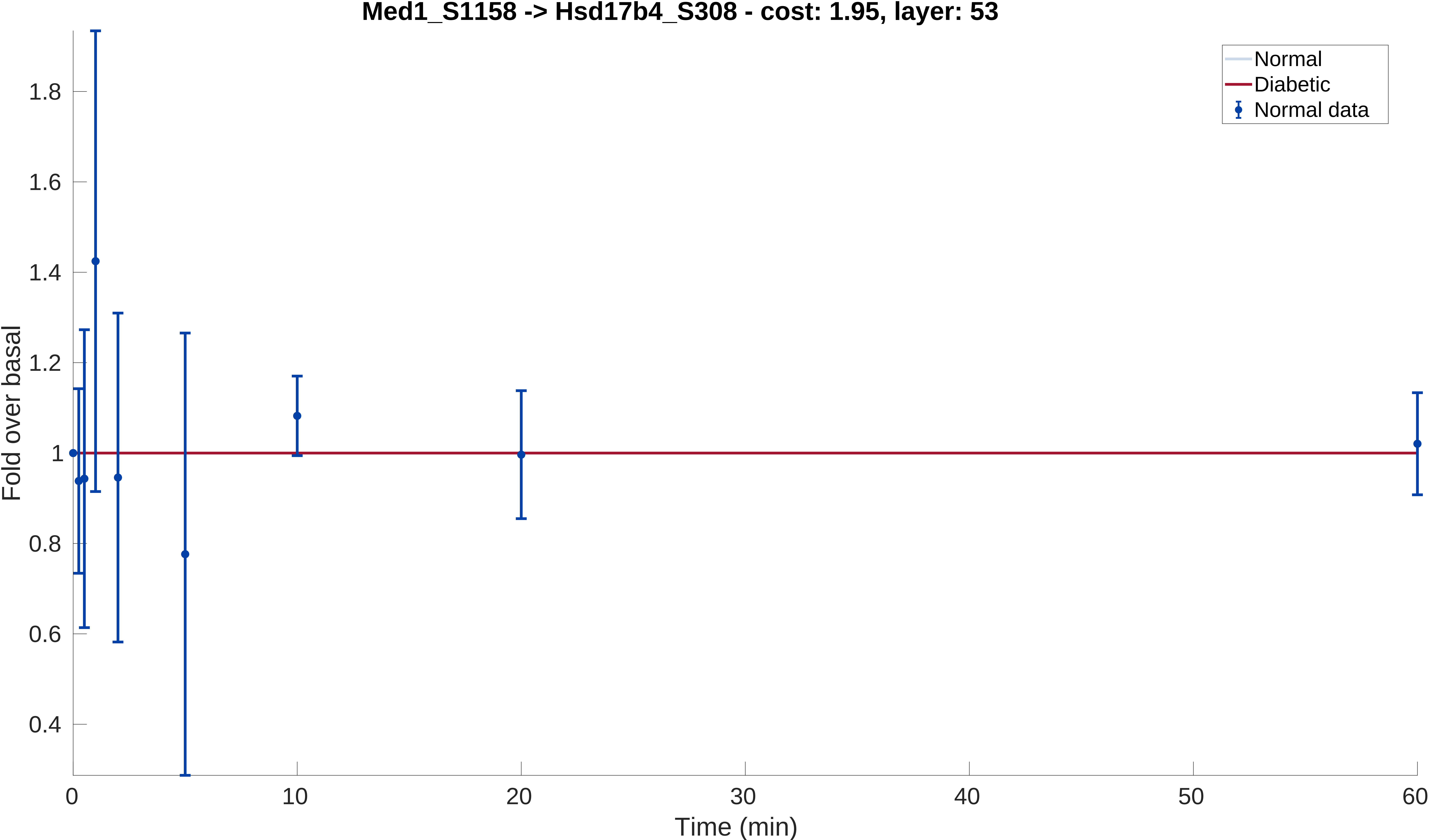

Med1_S1158 -> Hsd17b4_S308 - cost: 1.95, layer: 53
Normal
Diabetic
Normal data
Fold over basal
1.8
1.6
1.4
1.2
1
0.8
0.6
0.4
0
10
20
30
40
50
60
Time (min)

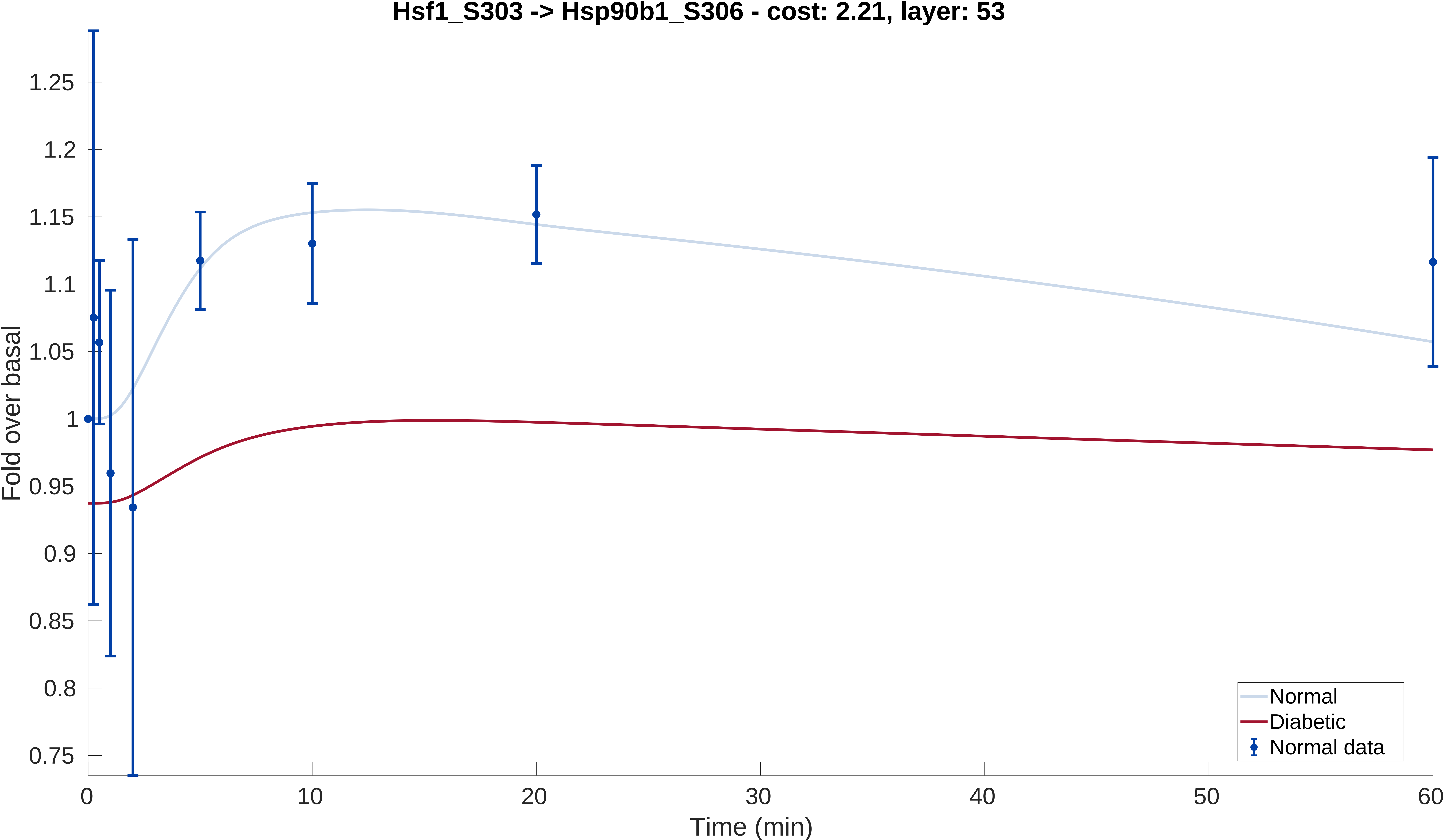

Hsf1_S303 -> Hsp90b1_S306 - cost: 2.21, layer: 53
Fold over basal
1.25
1.2
1.15
1.1
1.05
1
0.95
0.9
0.85
0.8
0.75
0
10
20
30
40
50
60
Time (min)
Normal
Diabetic
Normal data

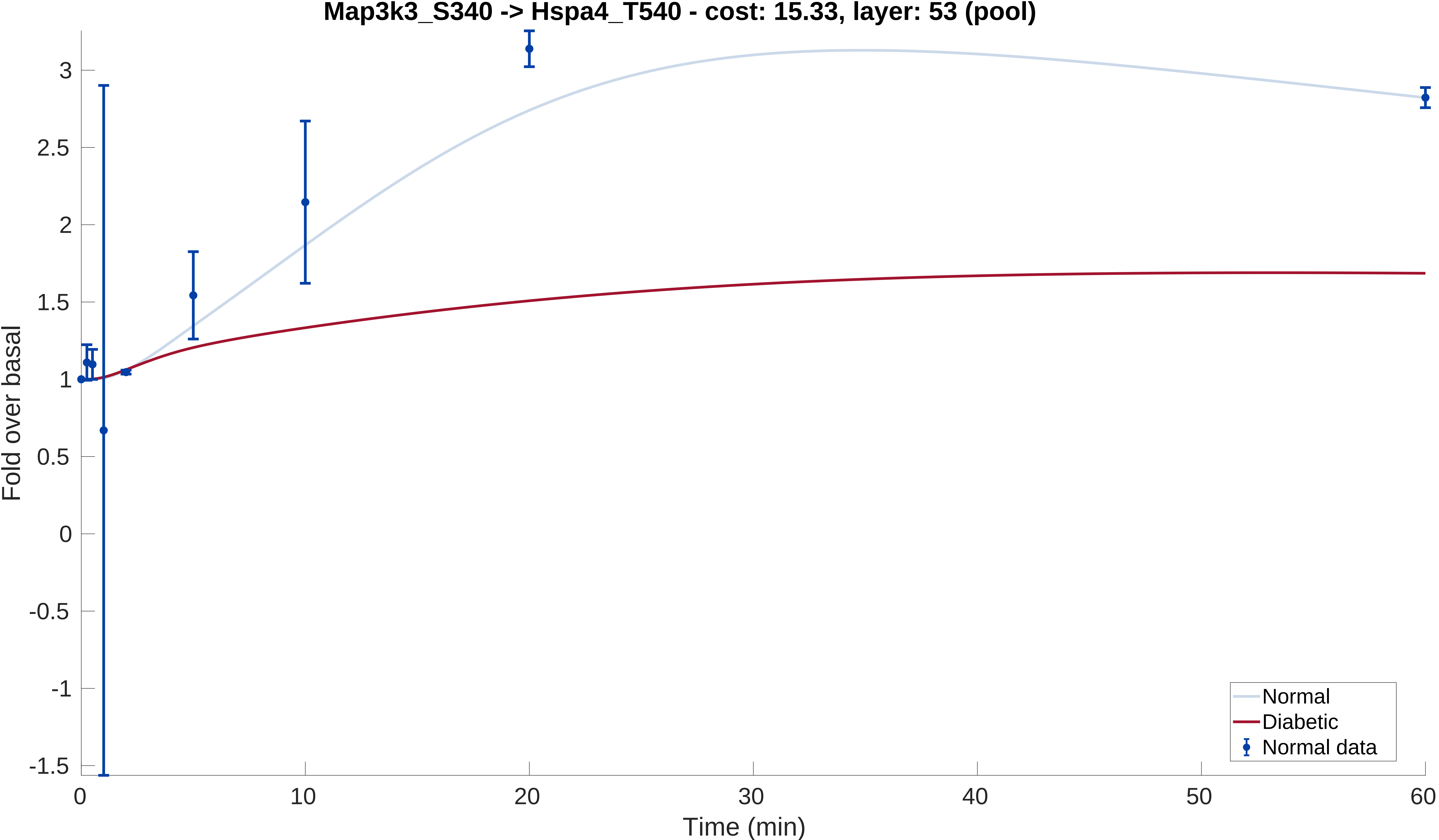

Map3k3_S340 -> Hspa4_T540 - cost: 15.33, layer: 53 (pool)
Fold over basal
3
2.5
2
1.5
1
0.5
0
-0.5
-1
-1.5
0
10
20
30
40
50
60
Time (min)
Normal
Diabetic
Normal data

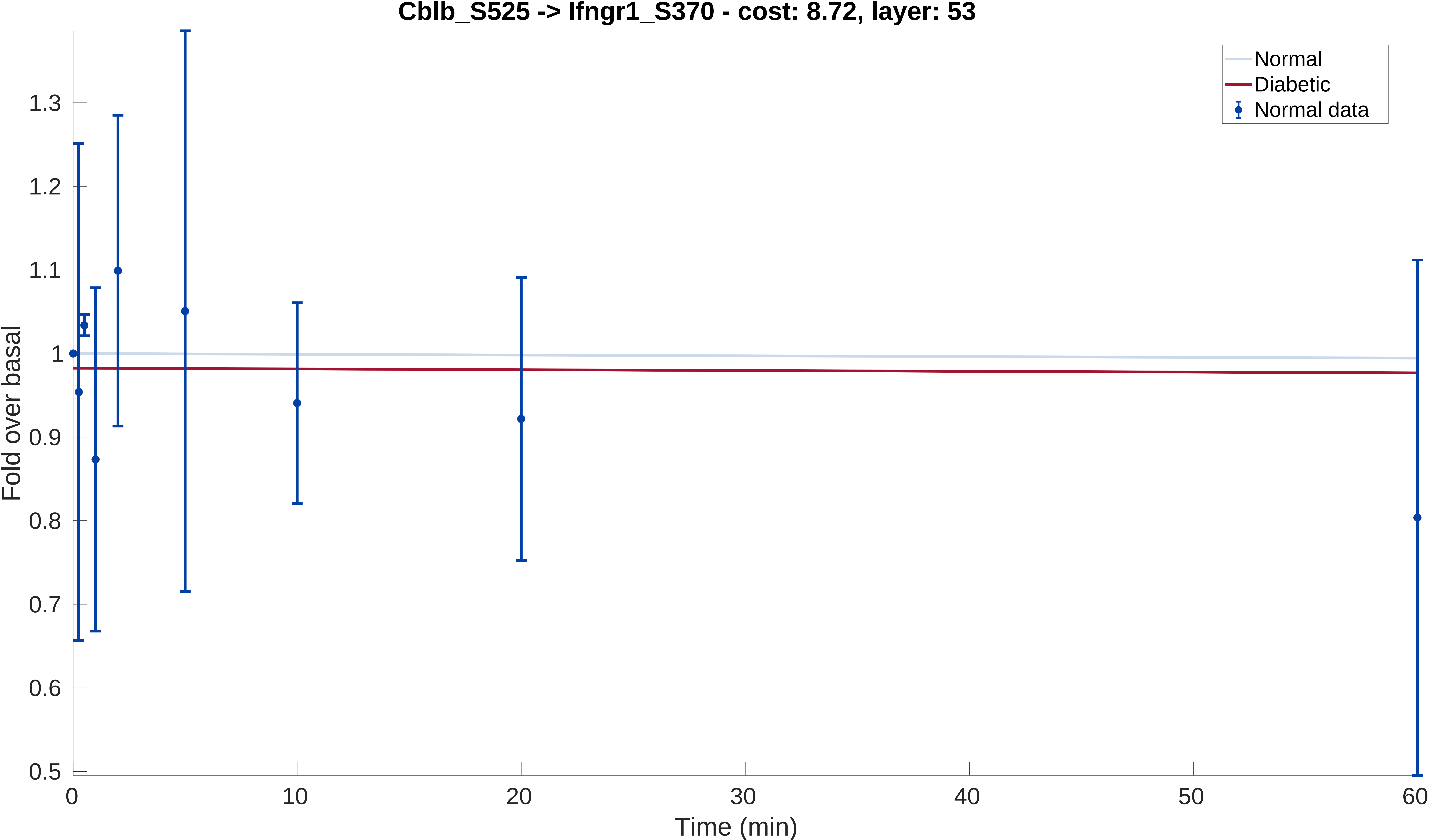

Cblb_S525 -> Ifngr1_S370 - cost: 8.72, layer: 53
Normal
Diabetic
Normal data
Fold over basal
Time (min)
0.5
0.6
0.7
0.8
0.9
1
1.1
1.2
1.3
0
10
20
30
40
50
60

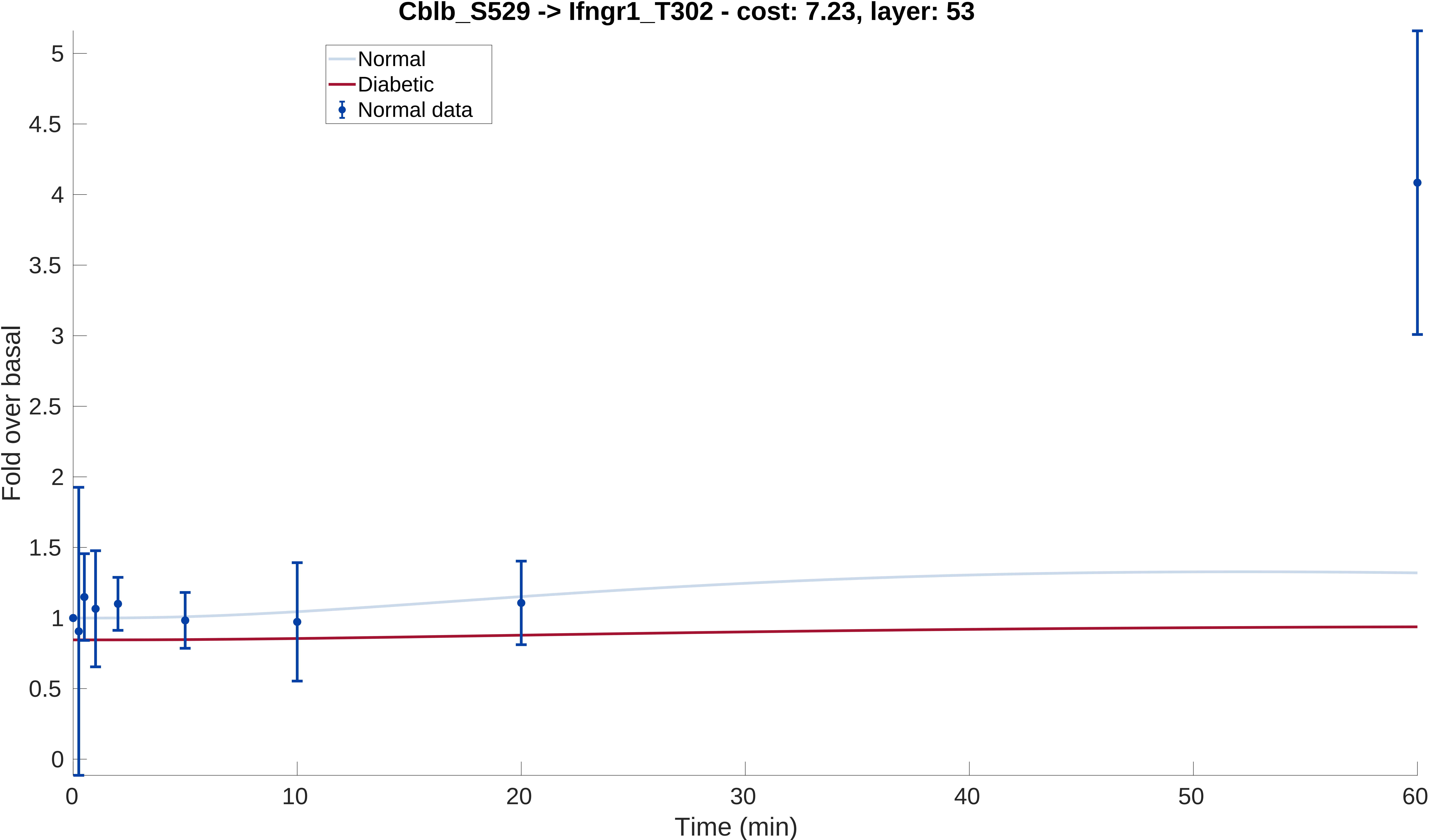

Cblb_S529 -> Ifngr1_T302 - cost: 7.23, layer: 53
Normal
Diabetic
Normal data
Fold over basal
Time (min)
0
0.5
1
1.5
2
2.5
3
3.5
4
4.5
5
0
10
20
30
40
50
60

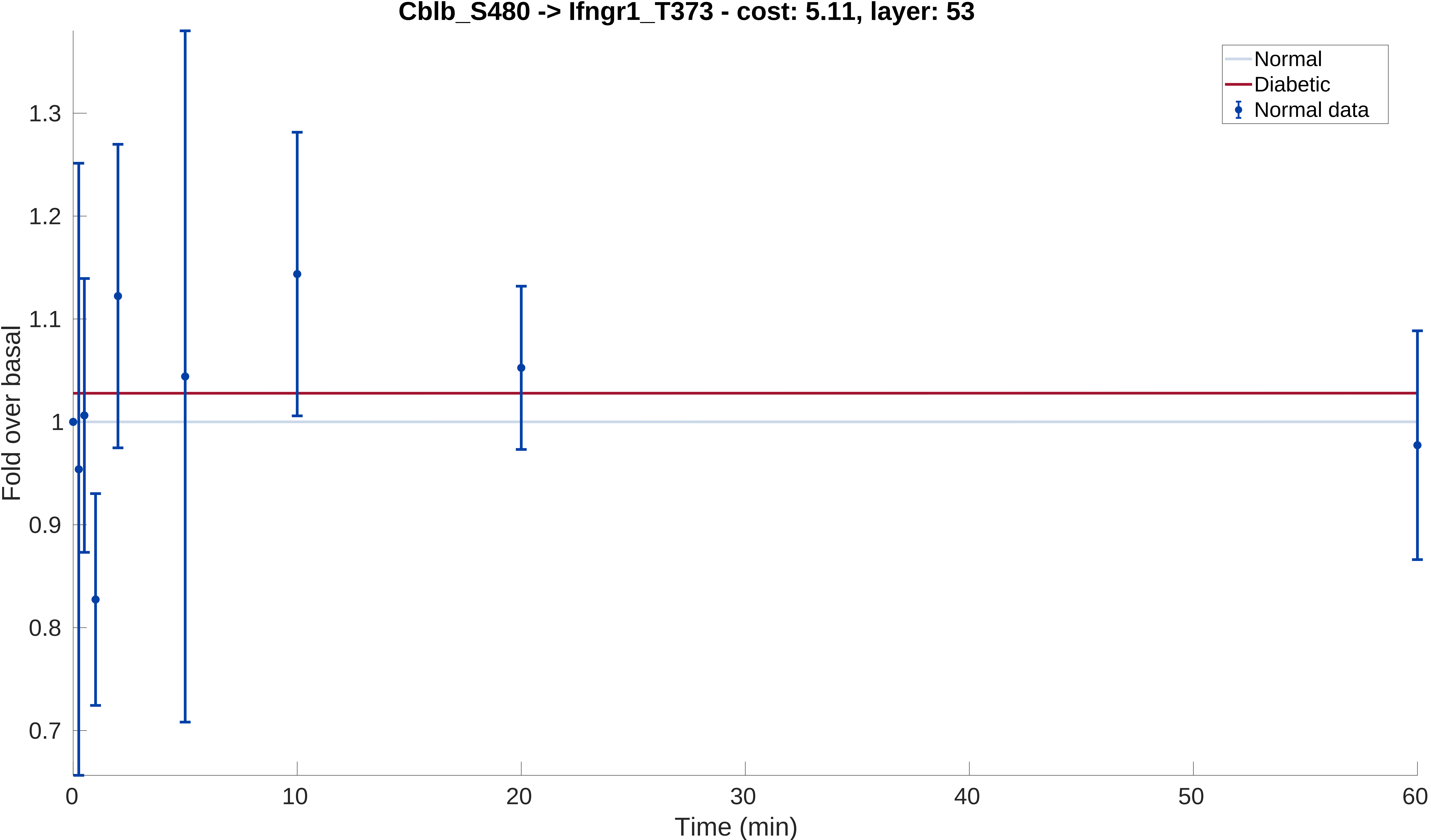

Cblb_S480 -> Ifngr1_T373 - cost: 5.11, layer: 53
Normal
Diabetic
Normal data
Fold over basal
1.3
1.2
1.1
1
0.9
0.8
0.7
0
10
20
30
40
50
60
Time (min)

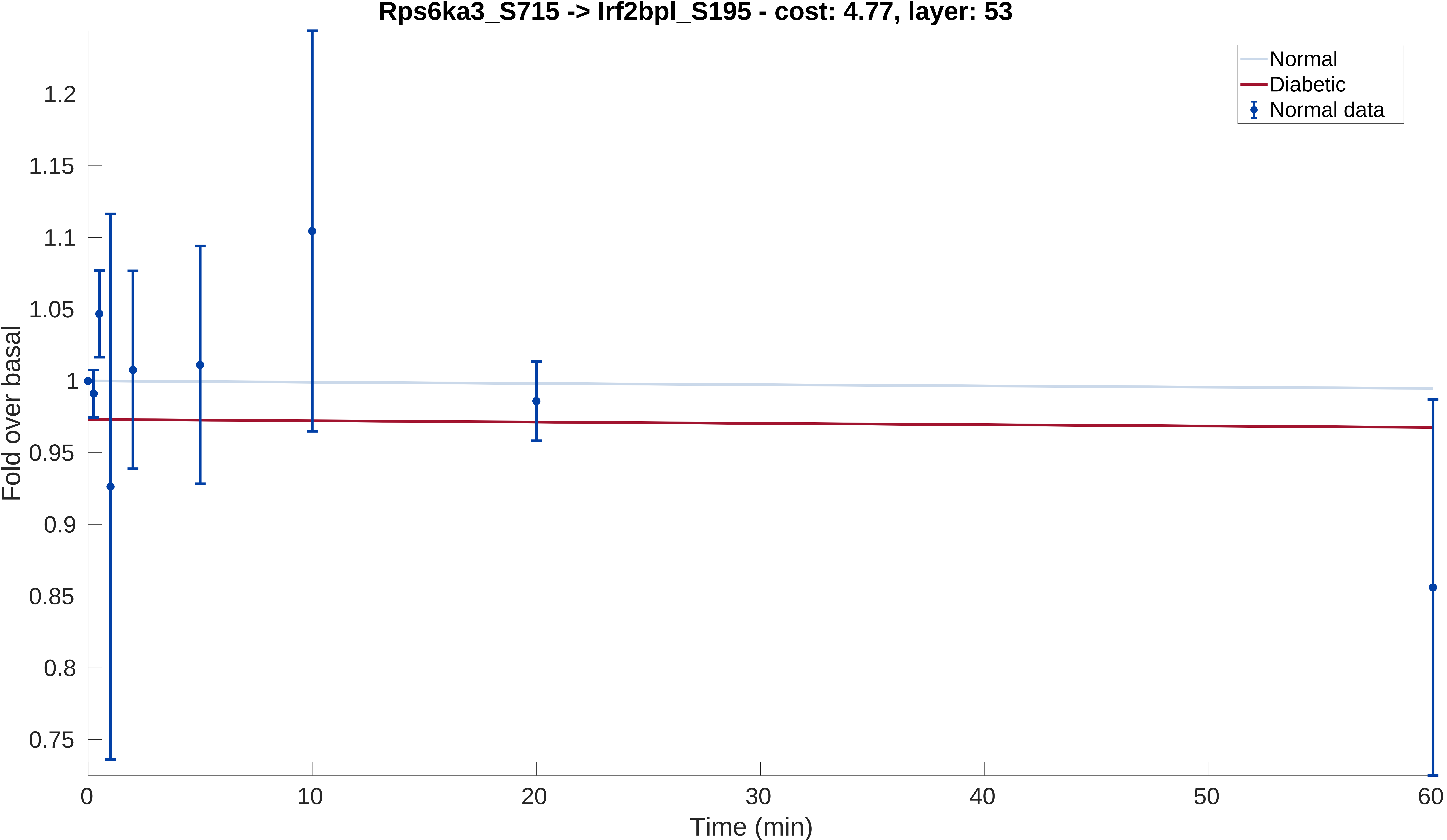

Rps6ka3_S715 -> Irf2bpl_S195 - cost: 4.77, layer: 53
Normal
Diabetic
Normal data
Fold over basal
Time (min)
1.2
1.15
1.1
1.05
1
0.95
0.9
0.85
0.8
0.75
0
10
20
30
40
50
60

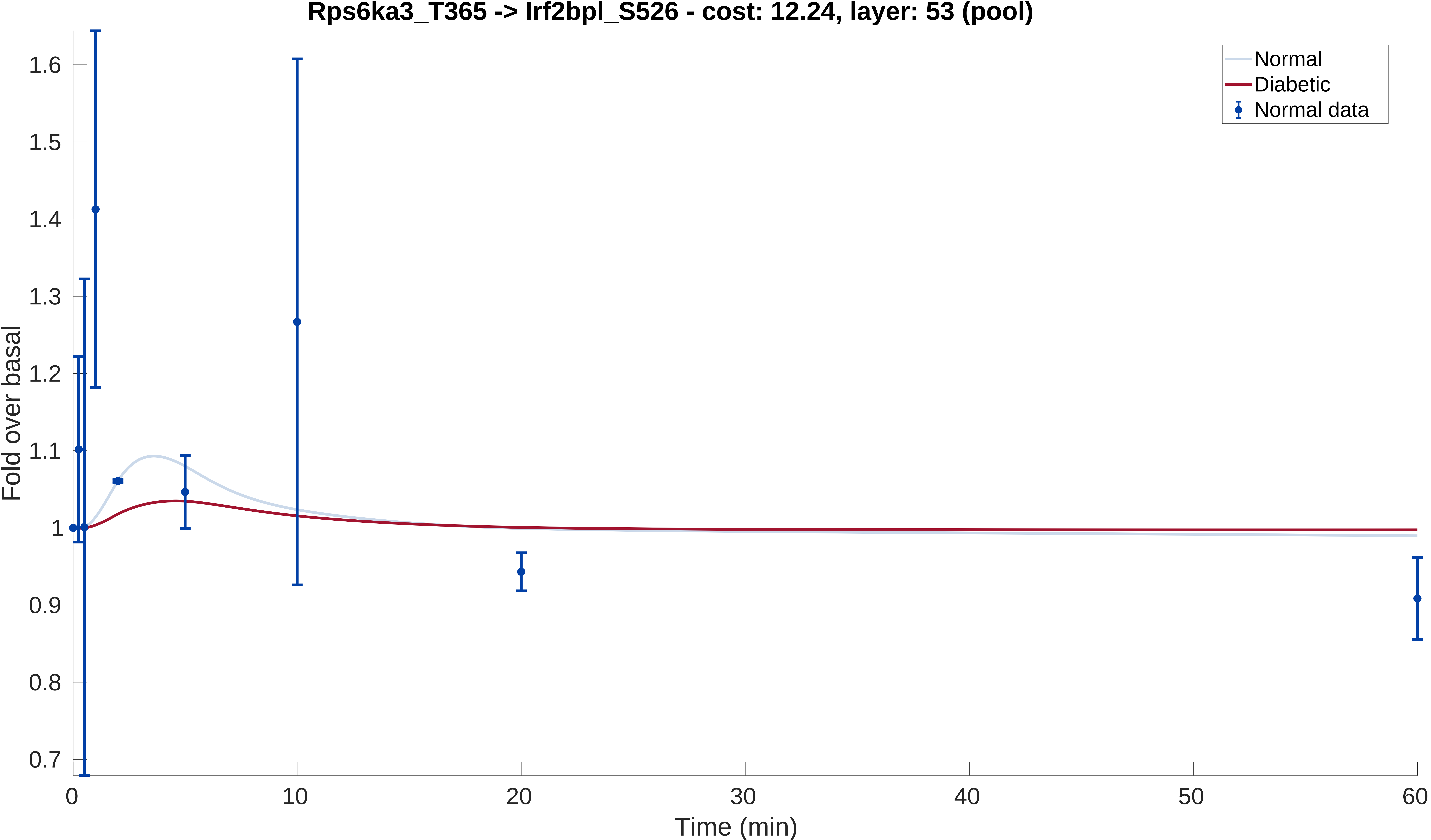

Rps6ka3_T365 -> Irf2bpl_S526 - cost: 12.24, layer: 53 (pool)
Normal
Diabetic
Normal data
Fold over basal
1.6
1.5
1.4
1.3
1.2
1.1
1
0.9
0.8
0.7
0
10
20
30
40
50
60
Time (min)

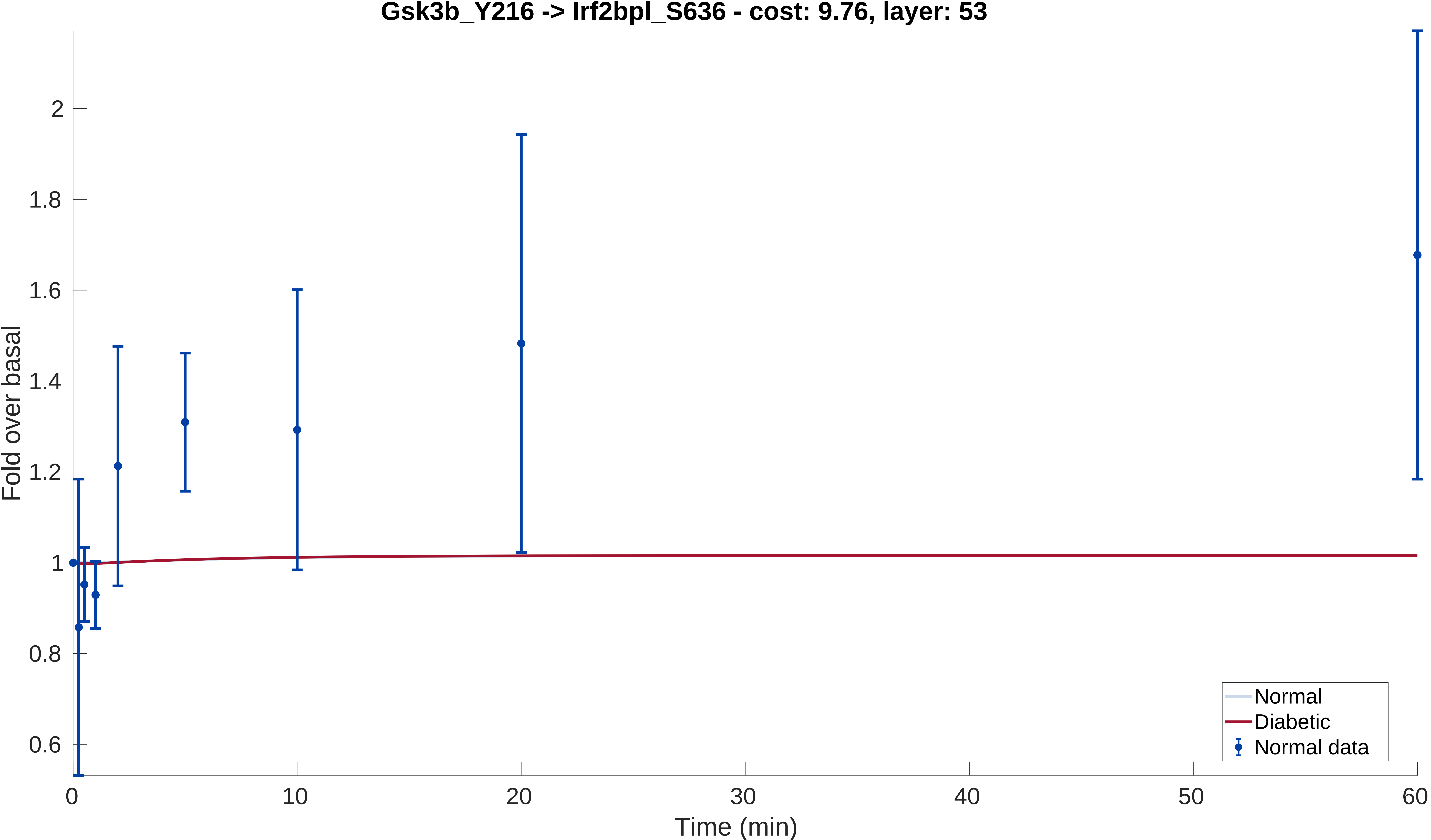

Gsk3b_Y216 -> Irf2bpl_S636 - cost: 9.76, layer: 53
Fold over basal
2
1.8
1.6
1.4
1.2
1
0.8
0.6
0
10
20
30
40
50
60
Time (min)
Normal
Diabetic
Normal data

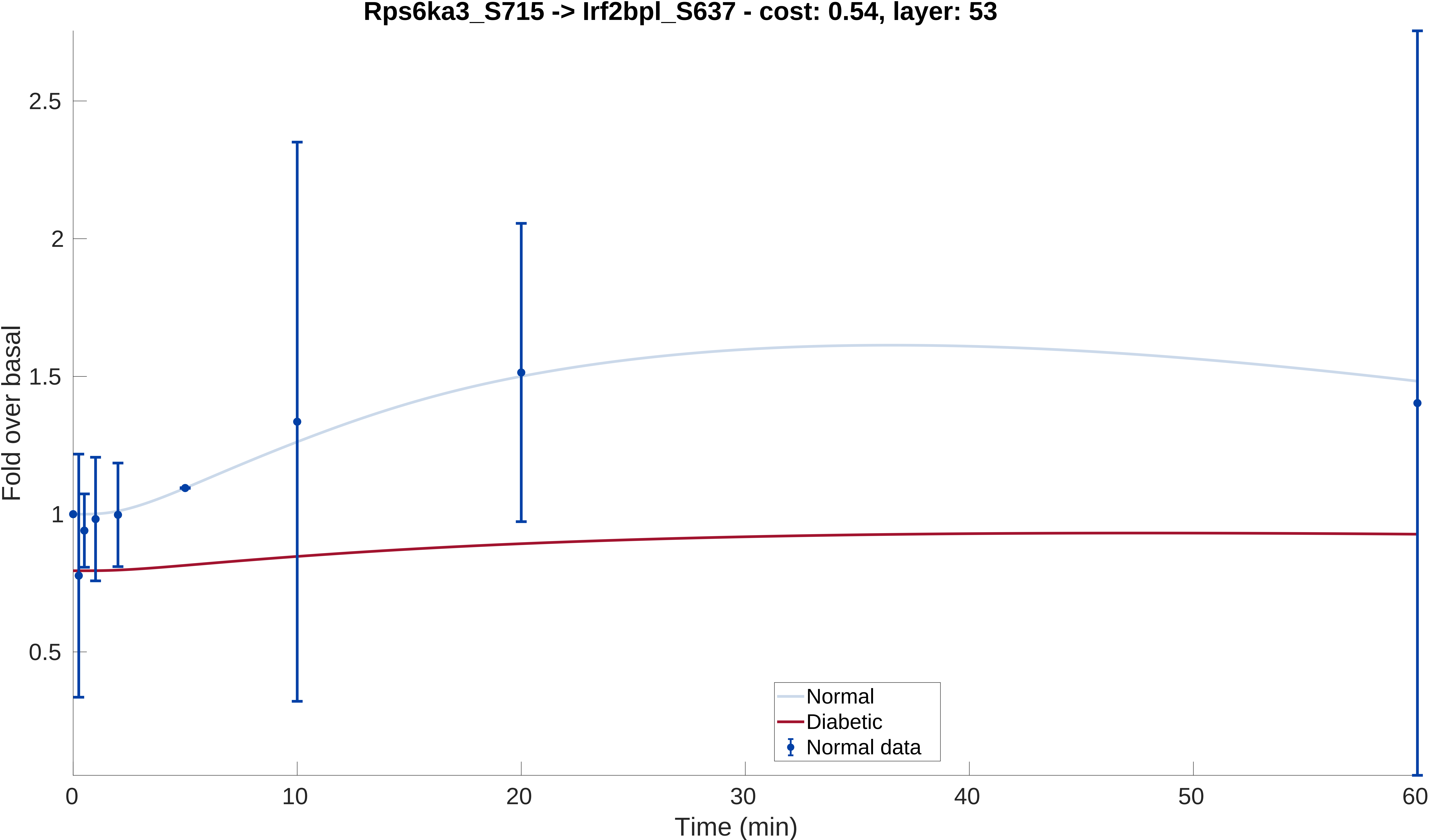
Rps6ka3_S715 -> Irf2bpl_S637 - cost: 0.54, layer: 53
Fold over basal
2.5
2
1.5
1
0.5
0
10
20
30
40
50
60
Time (min)
Normal
Diabetic
Normal data

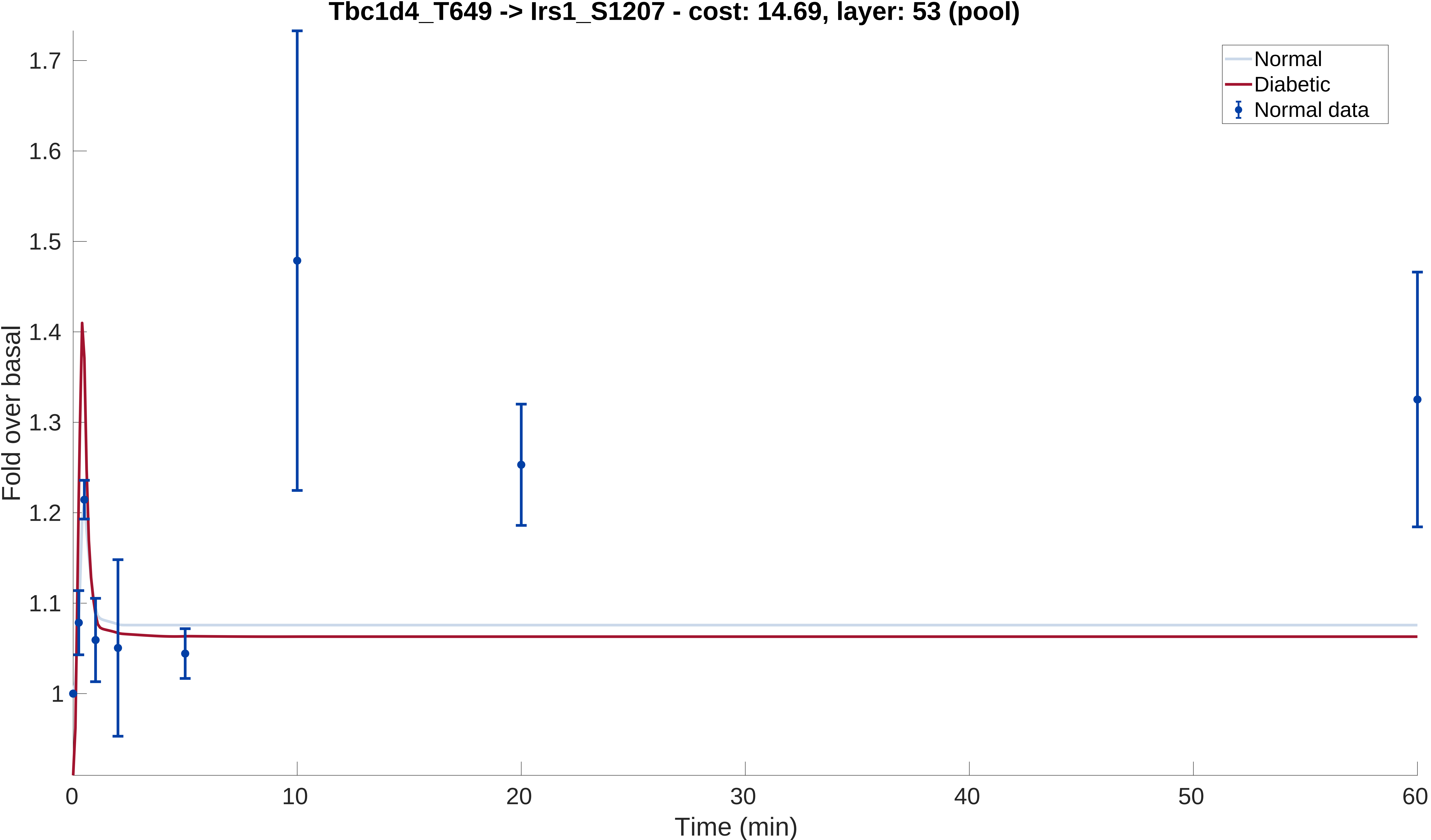

Tbc1d4_T649 -> Irs1_S1207 - cost: 14.69, layer: 53 (pool)
Normal
Diabetic
Normal data
Fold over basal
1.7
1.6
1.5
1.4
1.3
1.2
1.1
1
0
10
20
30
40
50
60
Time (min)

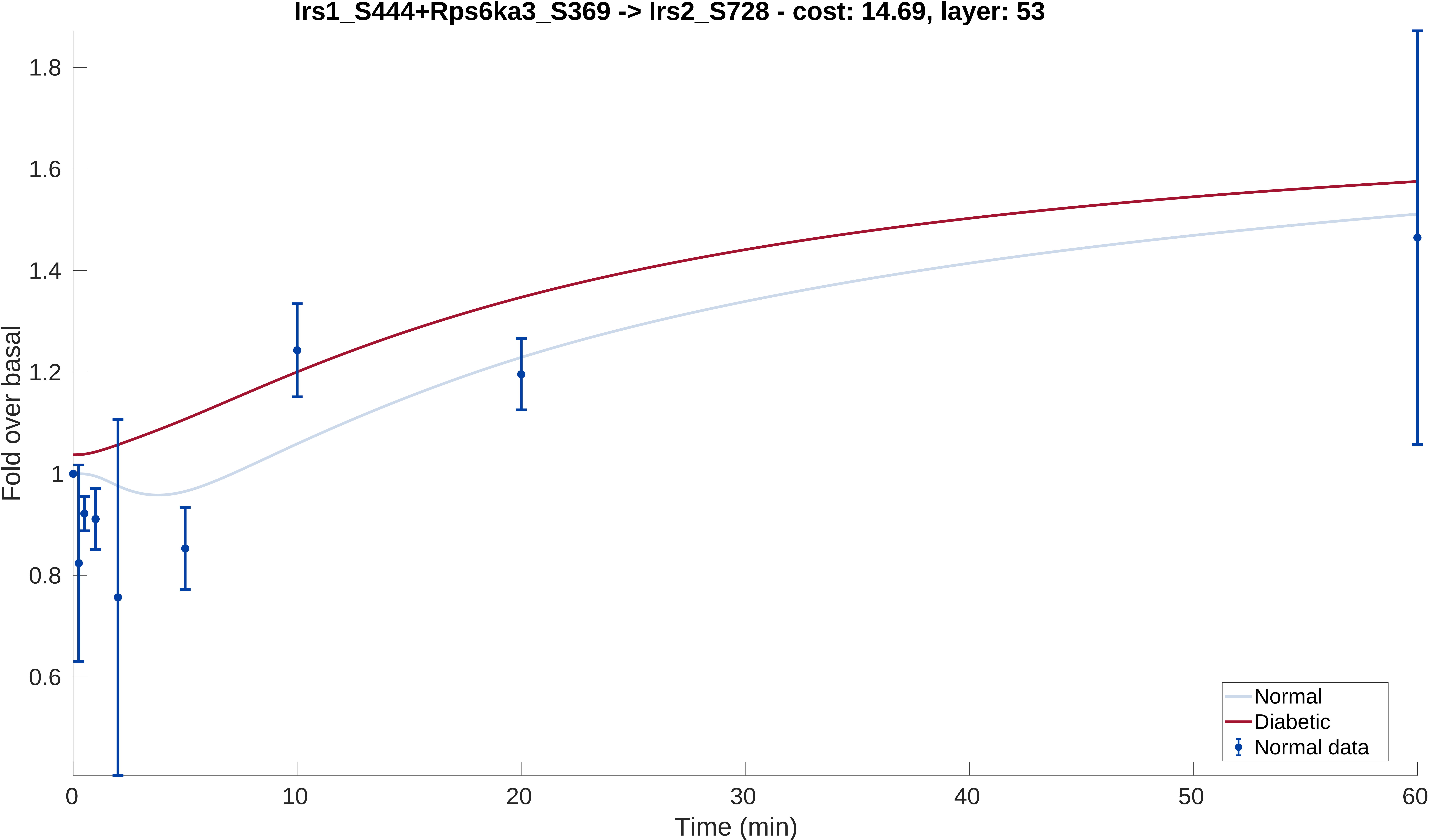

Irs1_S444+Rps6ka3_S369 -> Irs2_S728 - cost: 14.69, layer: 53
Fold over basal
1.8
1.6
1.4
1.2
1
0.8
0.6
0
10
20
30
40
50
60
Time (min)
Normal
Diabetic
Normal data

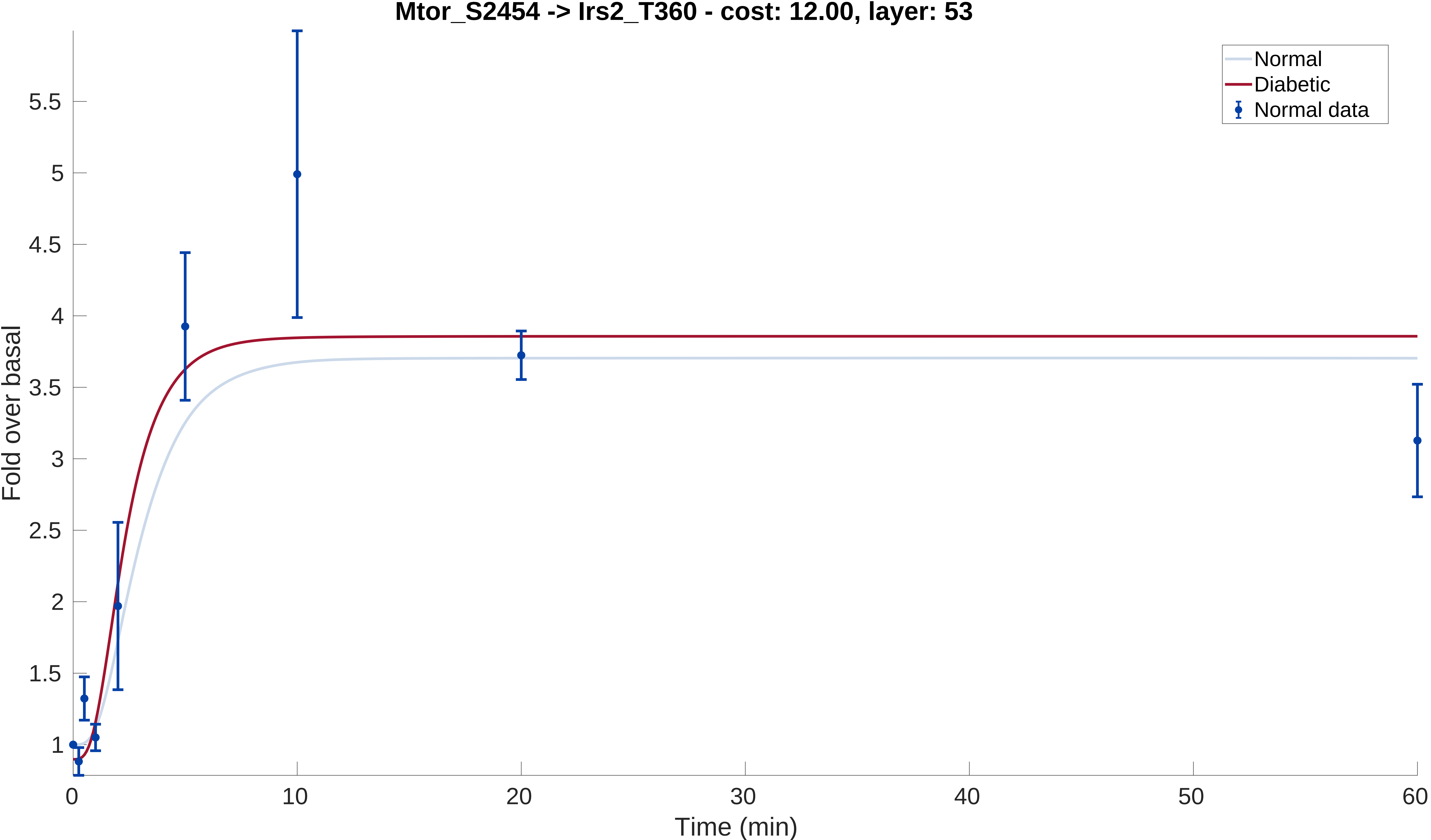

Mtor_S2454 -> Irs2_T360 - cost: 12.00, layer: 53
Normal
Diabetic
Normal data
Fold over basal
5.5
5
4.5
4
3.5
3
2.5
2
1.5
1
0
10
20
30
40
50
60
Time (min)

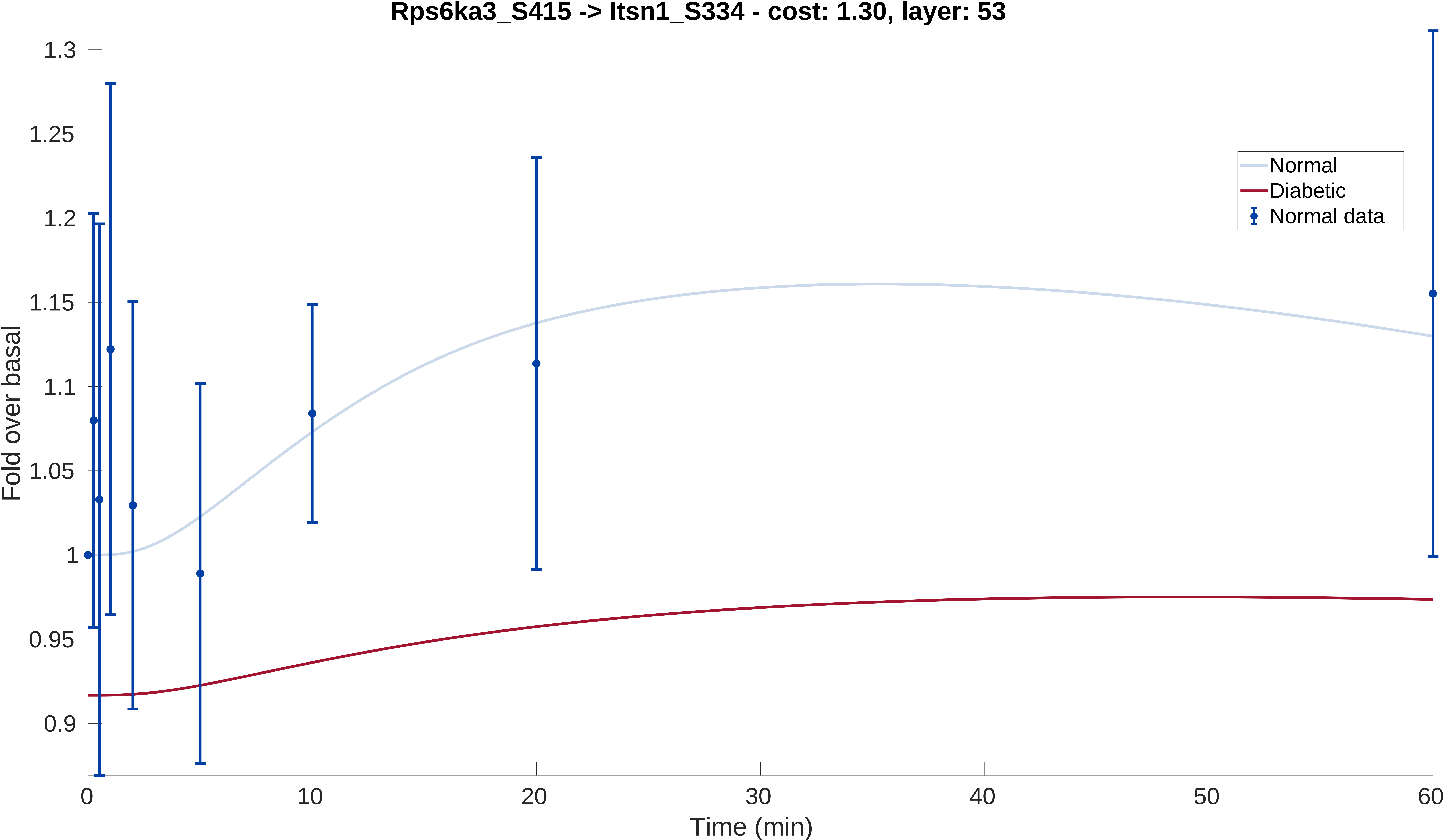

Rps6ka3_S415 -> Itsn1_S334 - cost: 1.30, layer: 53
Fold over basal
1.3
1.25
1.2
1.15
1.1
1.05
1
0.95
0.9
0
10
20
30
40
50
60
Time (min)
Normal
Diabetic
Normal data

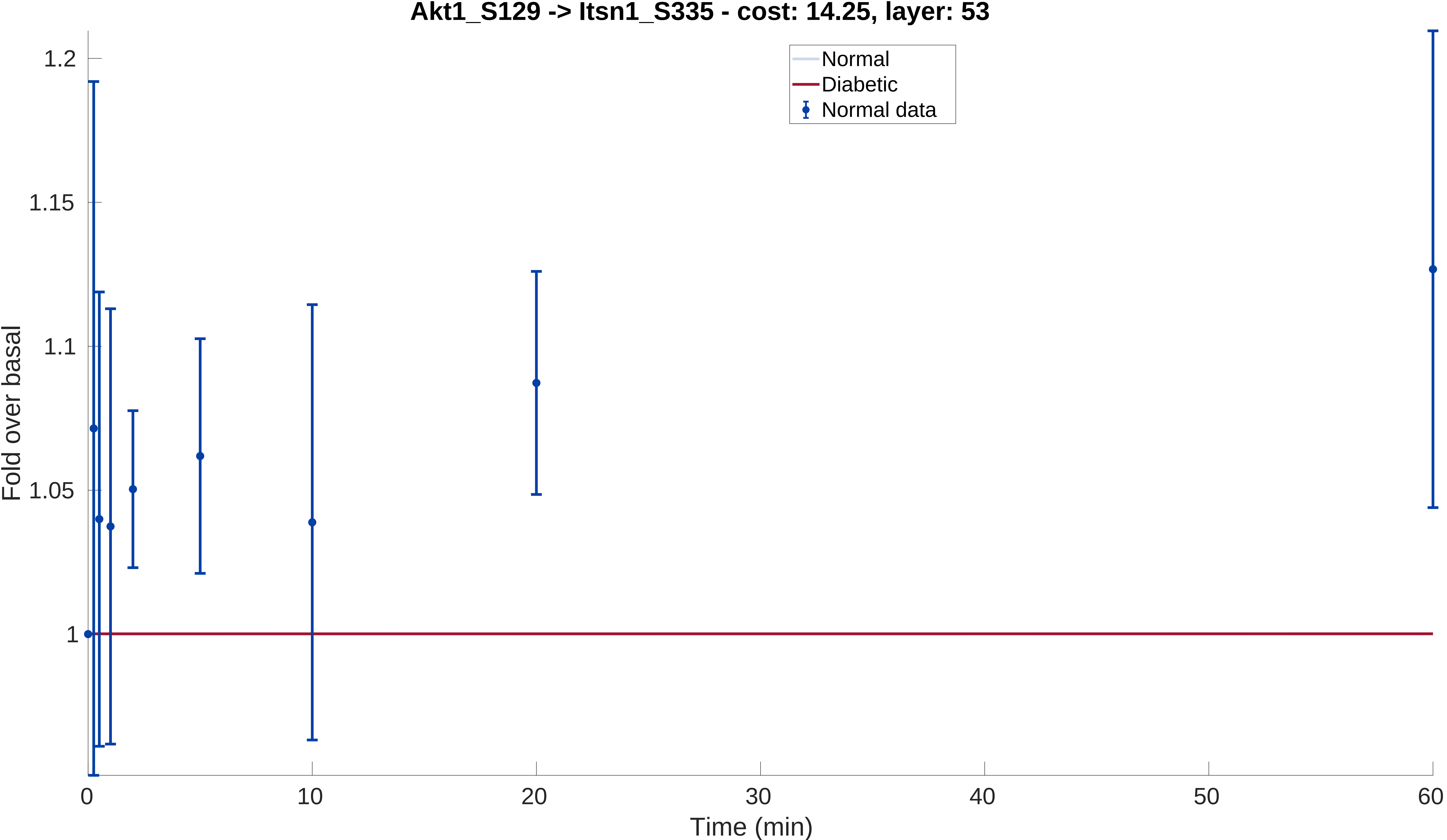

Akt1_S129 -> Itsn1_S335 - cost: 14.25, layer: 53
Normal
Diabetic
Normal data
1.2
1.15
1.1
1.05
1
Fold over basal
0
10
20
30
40
50
60
Time (min)

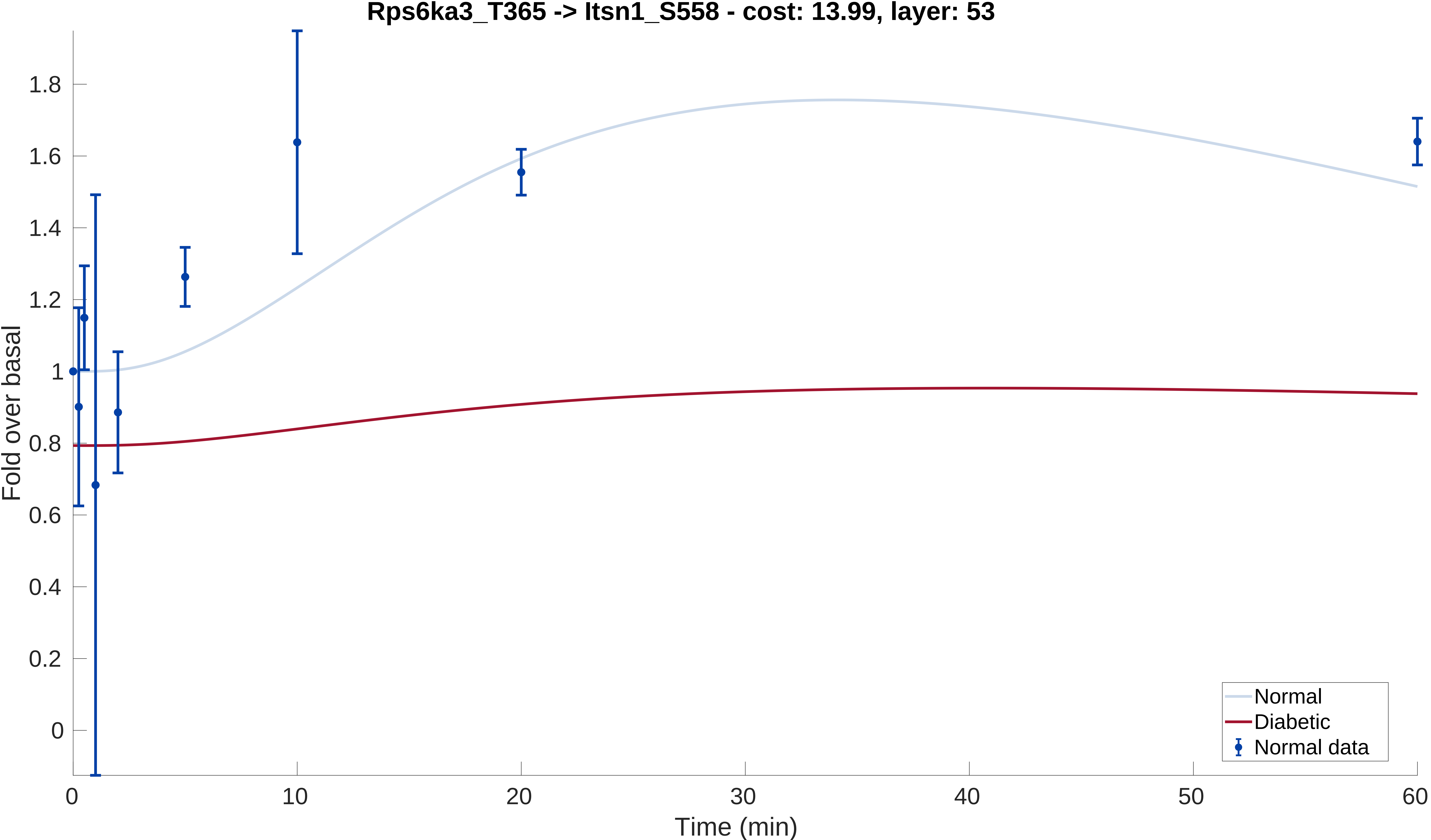

Rps6ka3_T365 -> Itsn1_S558 - cost: 13.99, layer: 53
Fold over basal
1.8
1.6
1.4
1.2
1
0.8
0.6
0.4
0.2
0
0
10
20
30
40
50
60
Time (min)
Normal
Diabetic
Normal data

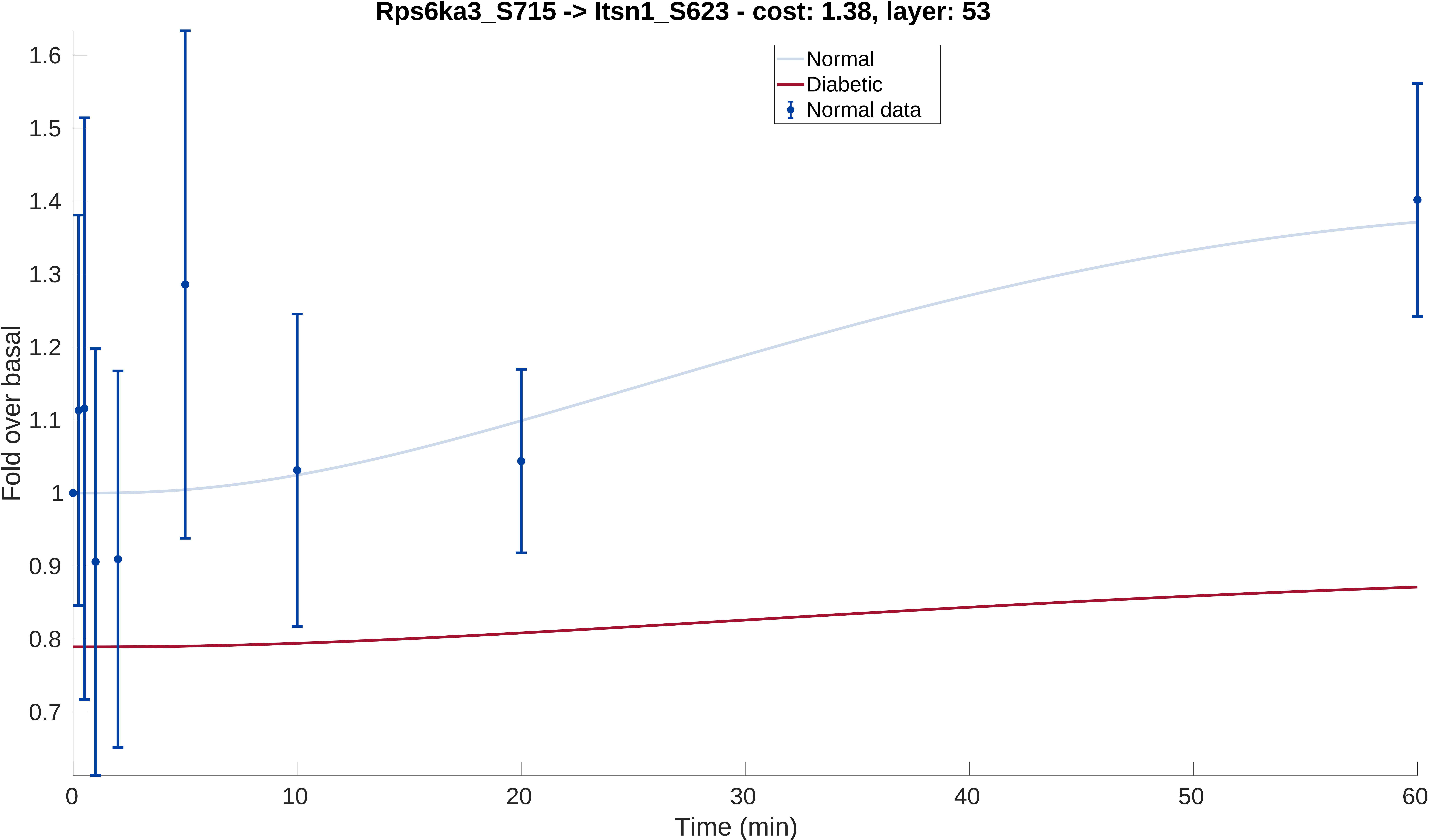

Rps6ka3_S715 -> Itsn1_S623 - cost: 1.38, layer: 53
Normal
Diabetic
Normal data
Fold over basal
Time (min)
1.6
1.5
1.4
1.3
1.2
1.1
1
0.9
0.8
0.7
0
10
20
30
40
50
60

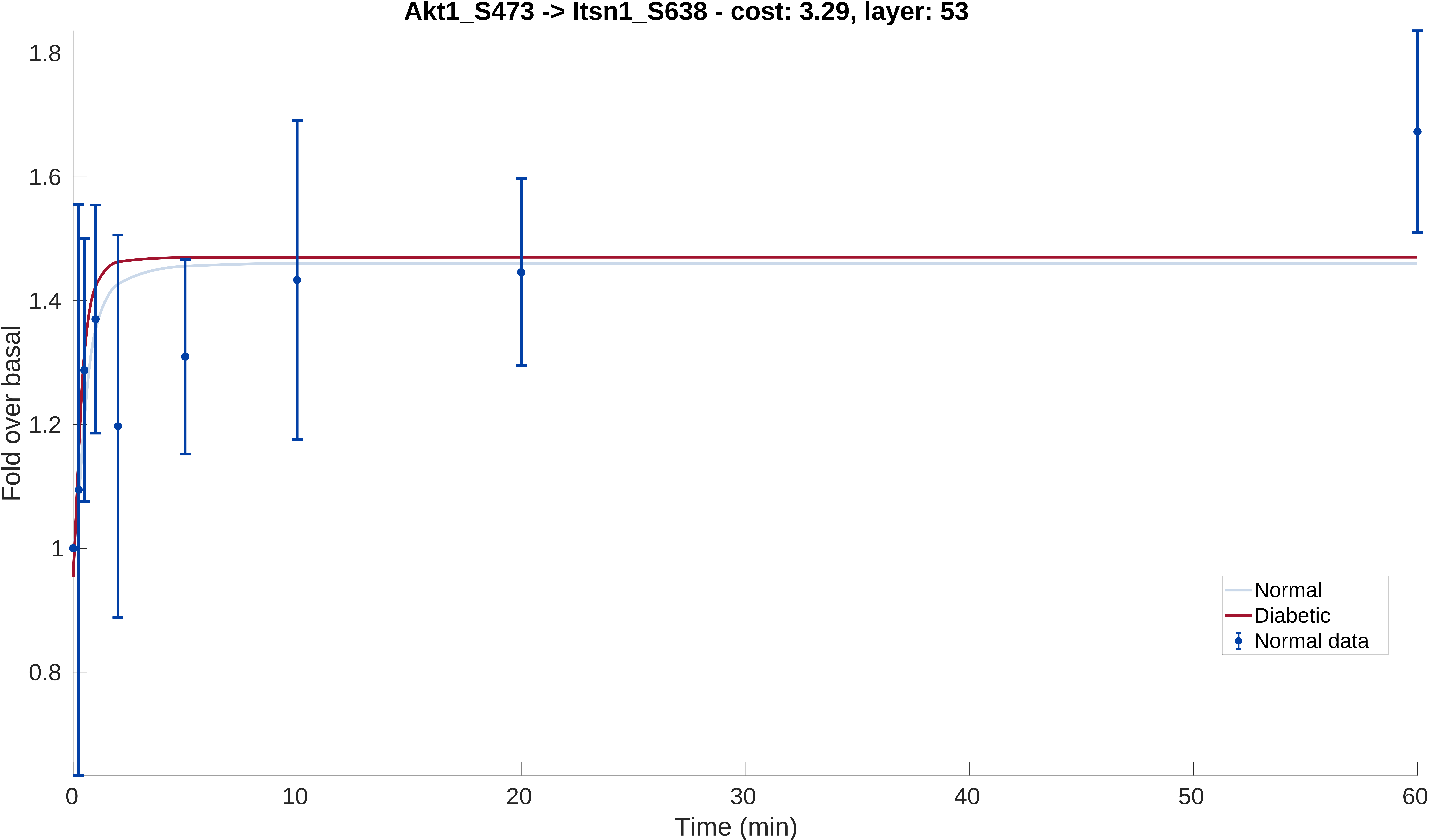

Akt1_S473 -> Itsn1_S638 - cost: 3.29, layer: 53
Fold over basal
1.8
1.6
1.4
1.2
1
0.8
0
10
20
30
40
50
60
Time (min)
Normal
Diabetic
Normal data

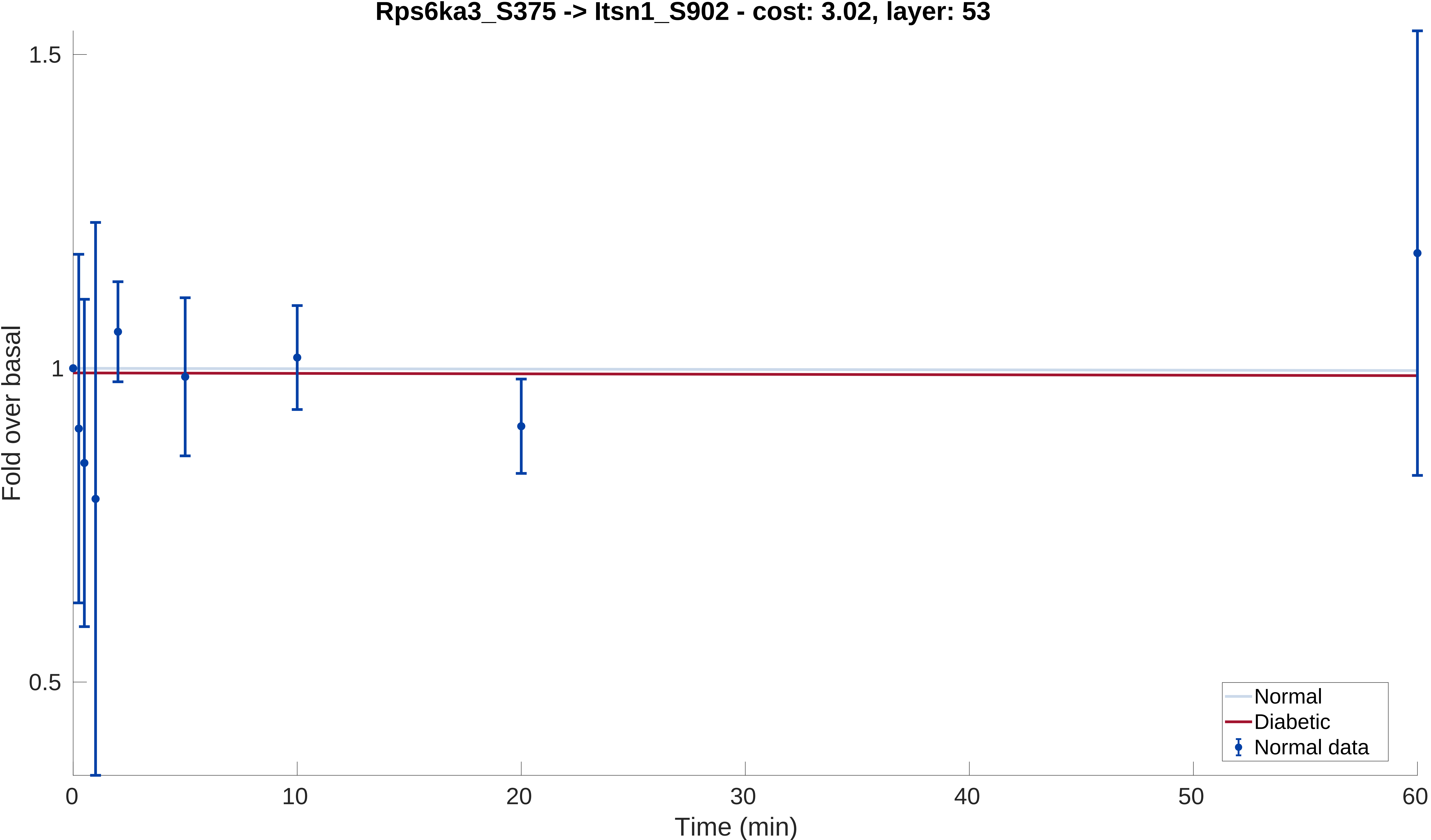

Rps6ka3_S375 -> Itsn1_S902 - cost: 3.02, layer: 53
1.5
1
0.5
Fold over basal
0
10
20
30
40
50
60
Time (min)
Normal
Diabetic
Normal data

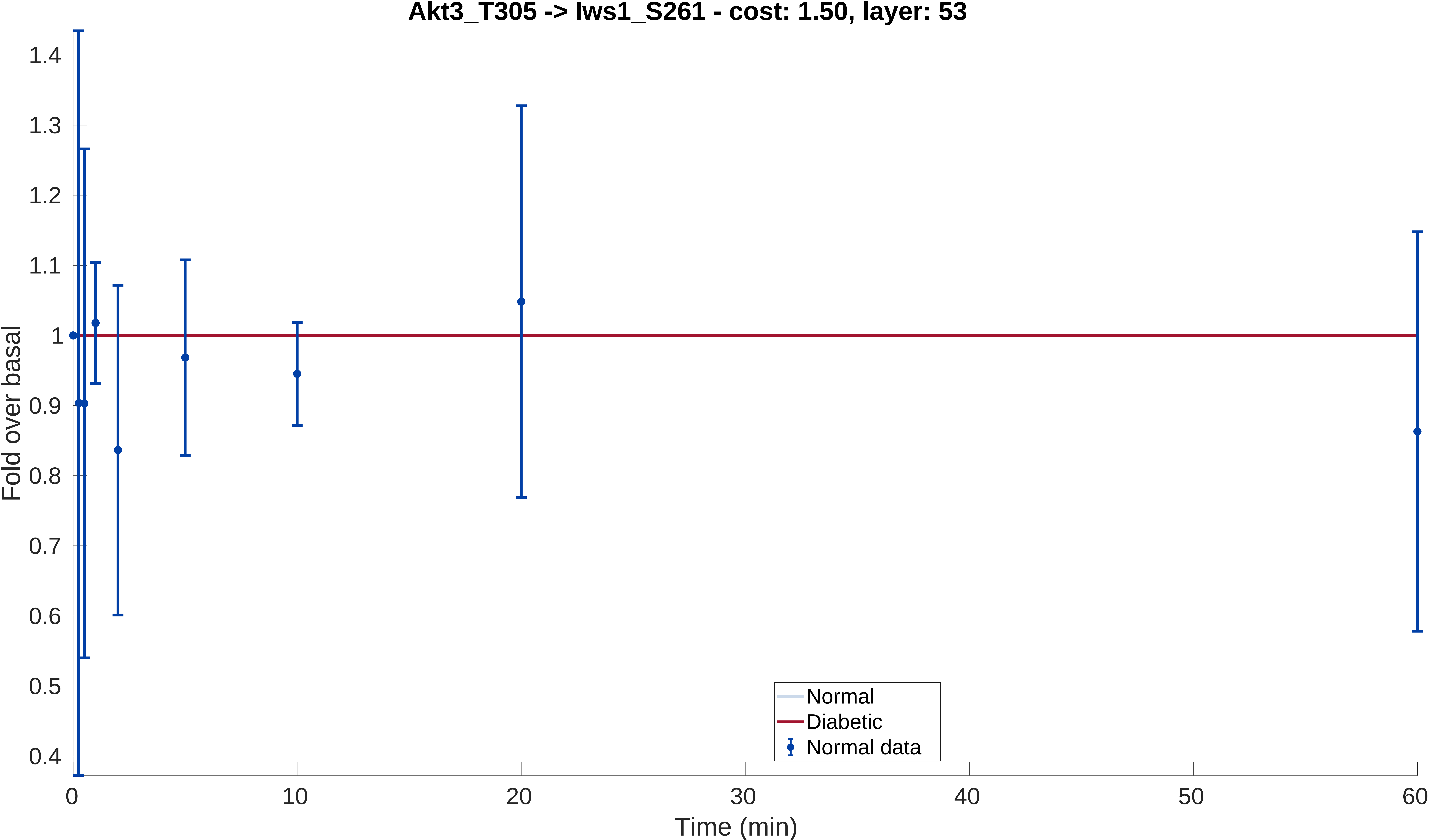

Akt3_T305 -> Iws1_S261 - cost: 1.50, layer: 53
Fold over basal
1.4
1.3
1.2
1.1
1
0.9
0.8
0.7
0.6
0.5
0.4
0
10
20
30
40
50
60
Time (min)
Normal
Diabetic
Normal data

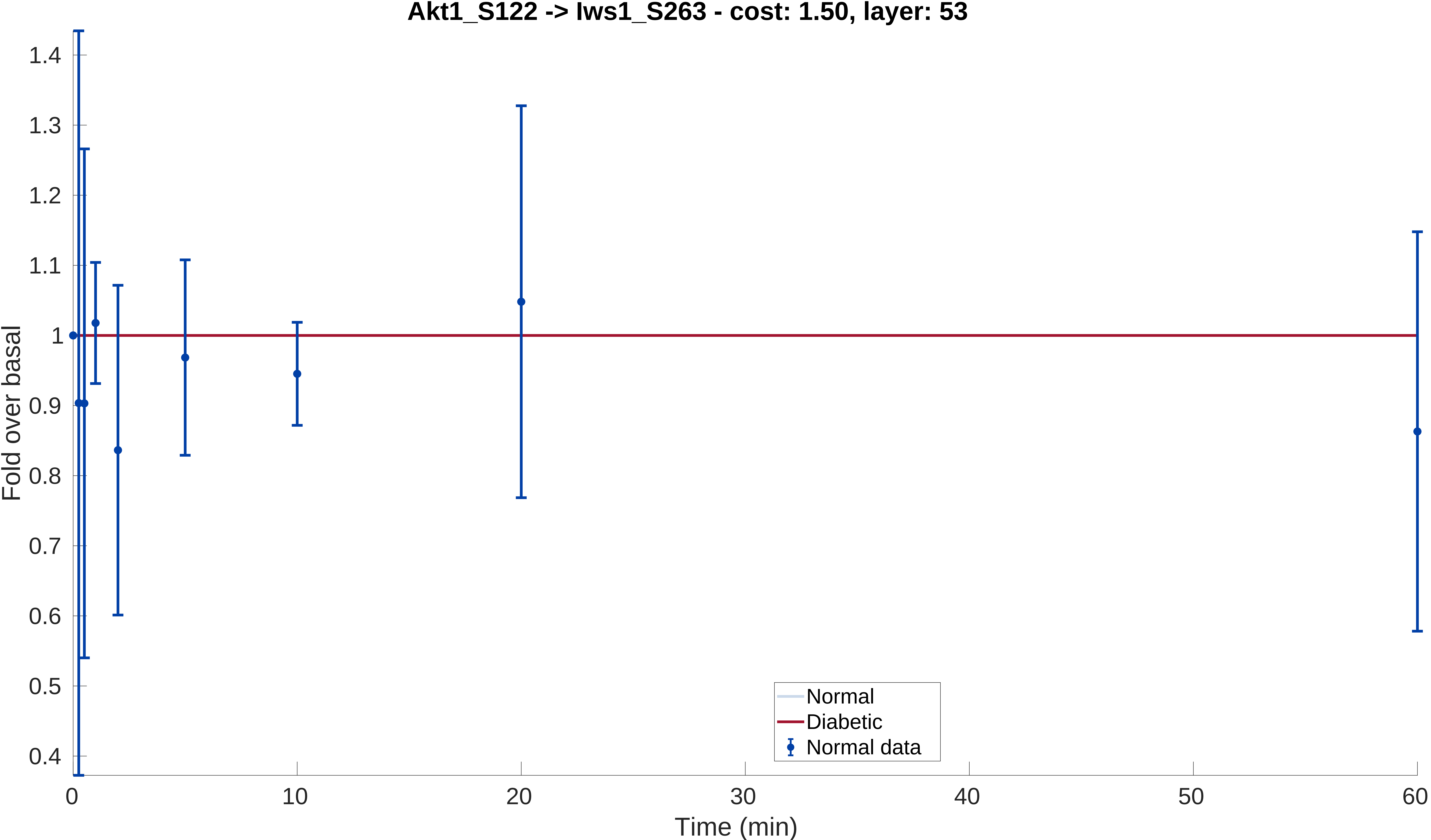

Akt1_S122 -> Iws1_S263 - cost: 1.50, layer: 53
Fold over basal
1.4
1.3
1.2
1.1
1
0.9
0.8
0.7
0.6
0.5
0.4
0
10
20
30
40
50
60
Time (min)
Normal
Diabetic
Normal data

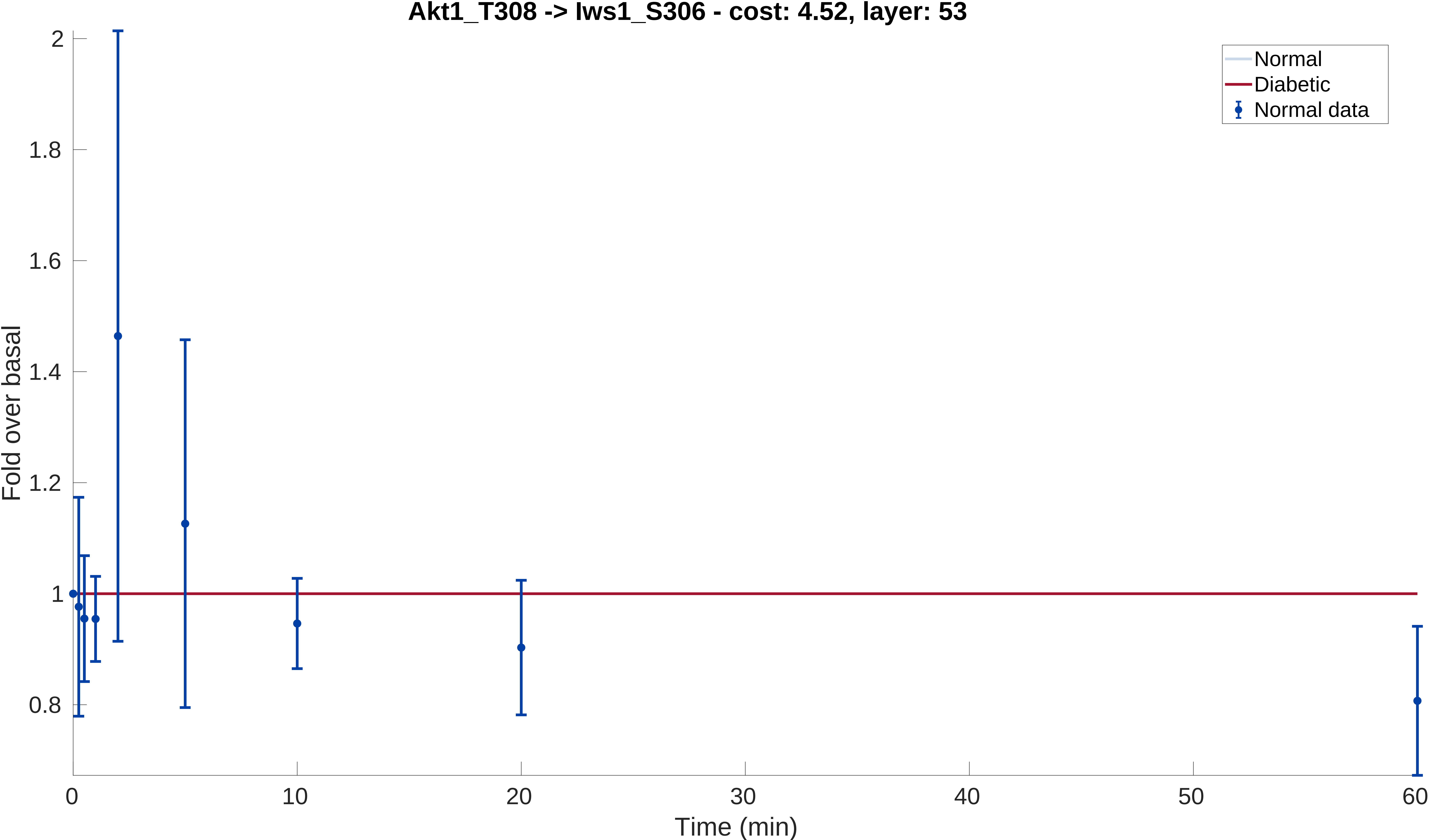

Akt1_T308 -> Iws1_S306 - cost: 4.52, layer: 53
Normal
Diabetic
Normal data
Fold over basal
2
1.8
1.6
1.4
1.2
1
0.8
0
10
20
30
40
50
60
Time (min)

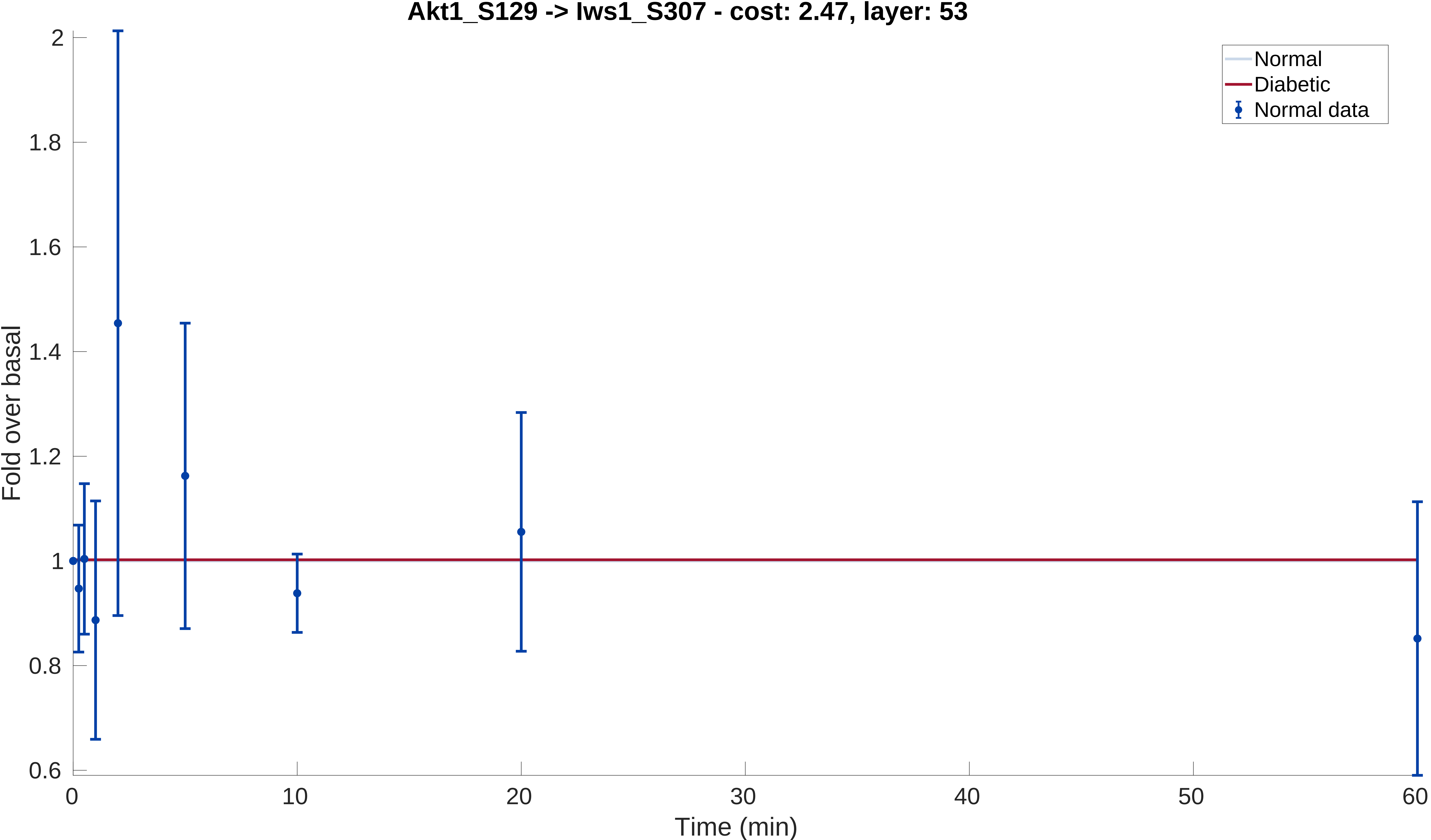

Akt1_S129 -> Iws1_S307 - cost: 2.47, layer: 53
Normal
Diabetic
Normal data
Fold over basal
Time (min)
0.6
0.8
1
1.2
1.4
1.6
1.8
2
0
10
20
30
40
50
60

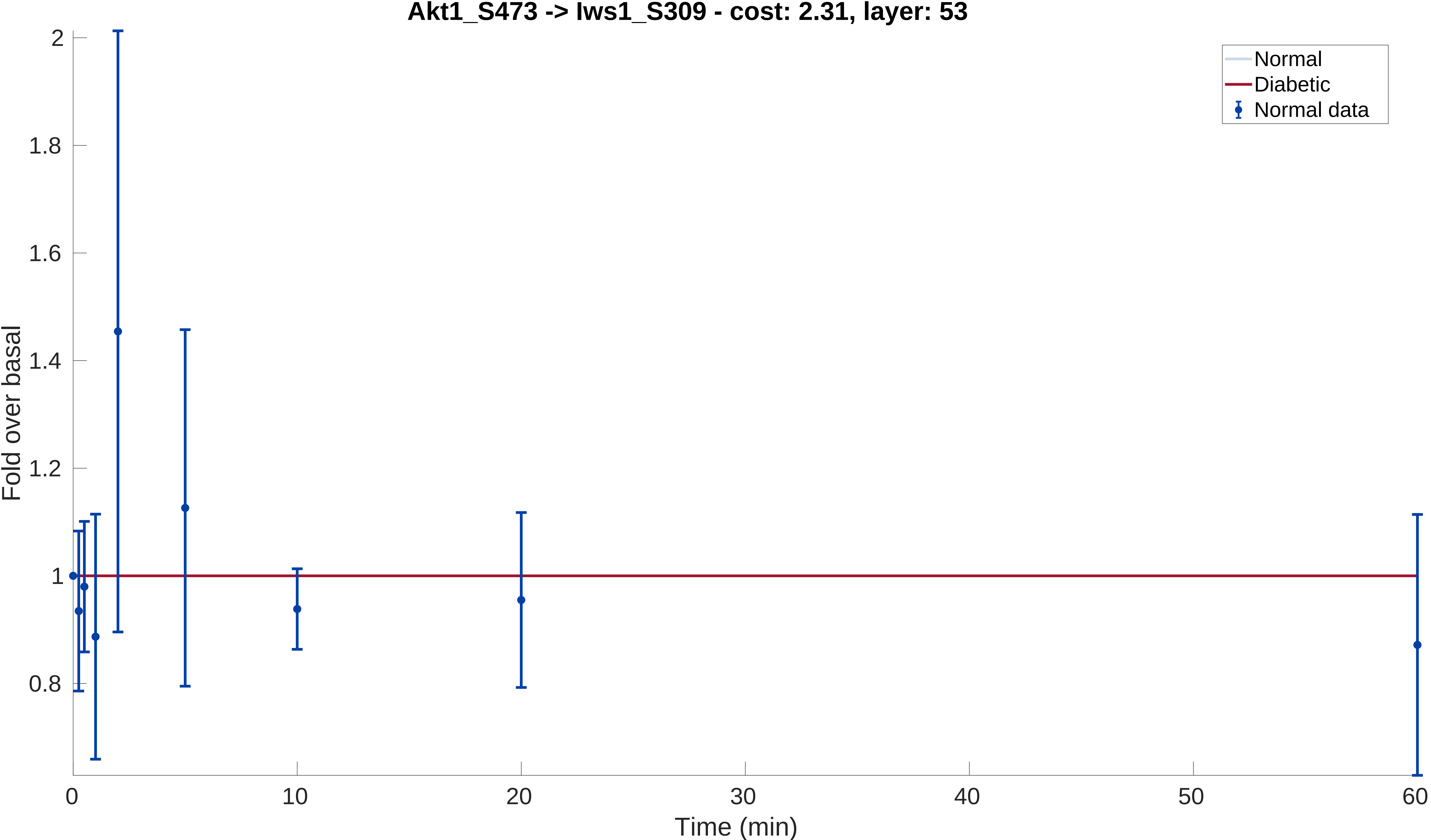

Akt1_S473 -> Iws1_S309 - cost: 2.31, layer: 53
Normal
Diabetic
Normal data
Fold over basal
2
1.8
1.6
1.4
1.2
1
0.8
0
10
20
30
40
50
60
Time (min)

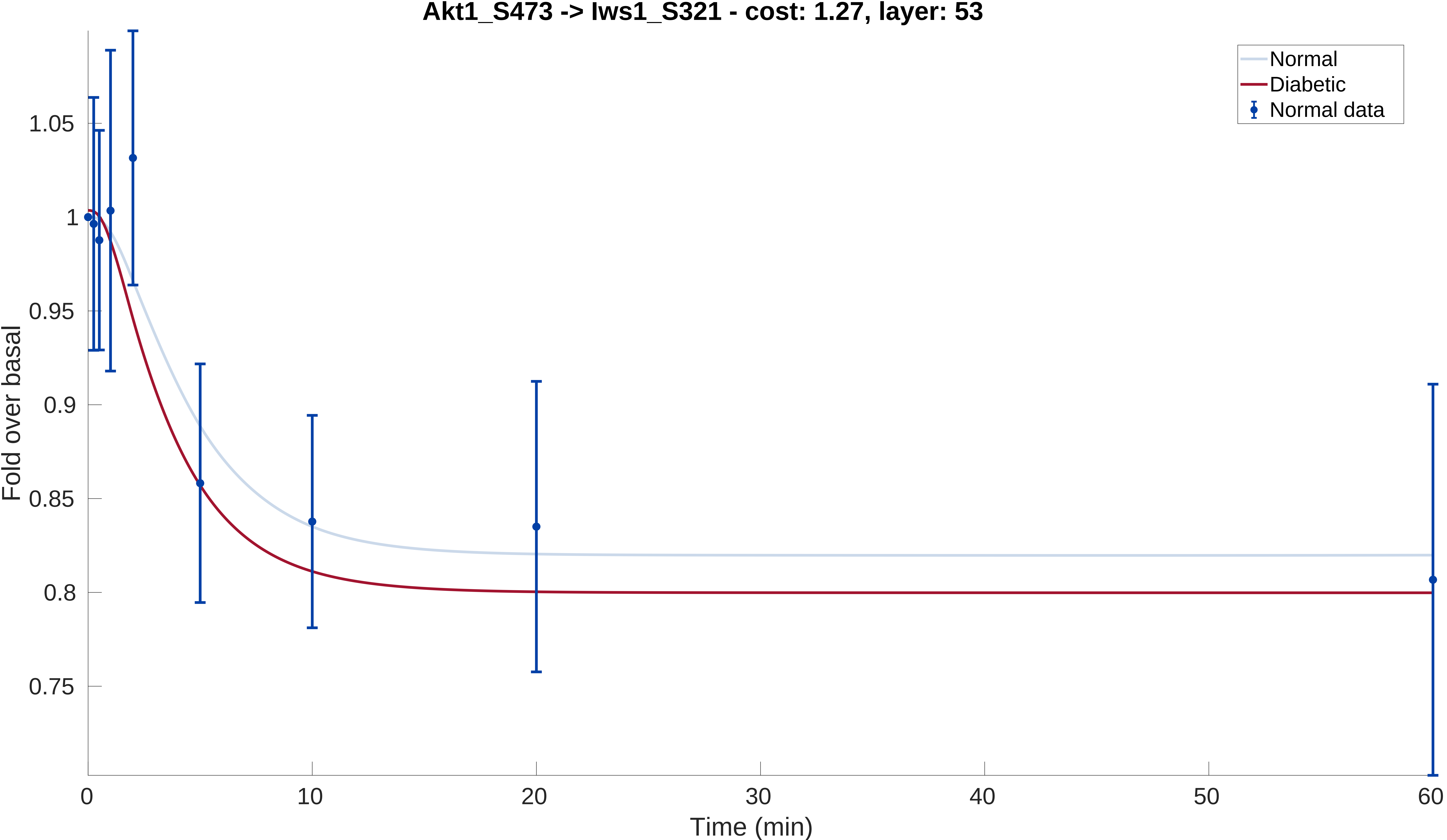
Akt1_S473 -> Iws1_S321 - cost: 1.27, layer: 53
Normal
Diabetic
Normal data
1.05
1
0.95
0.9
0.85
0.8
0.75
Fold over basal
0
10
20
30
40
50
60
Time (min)

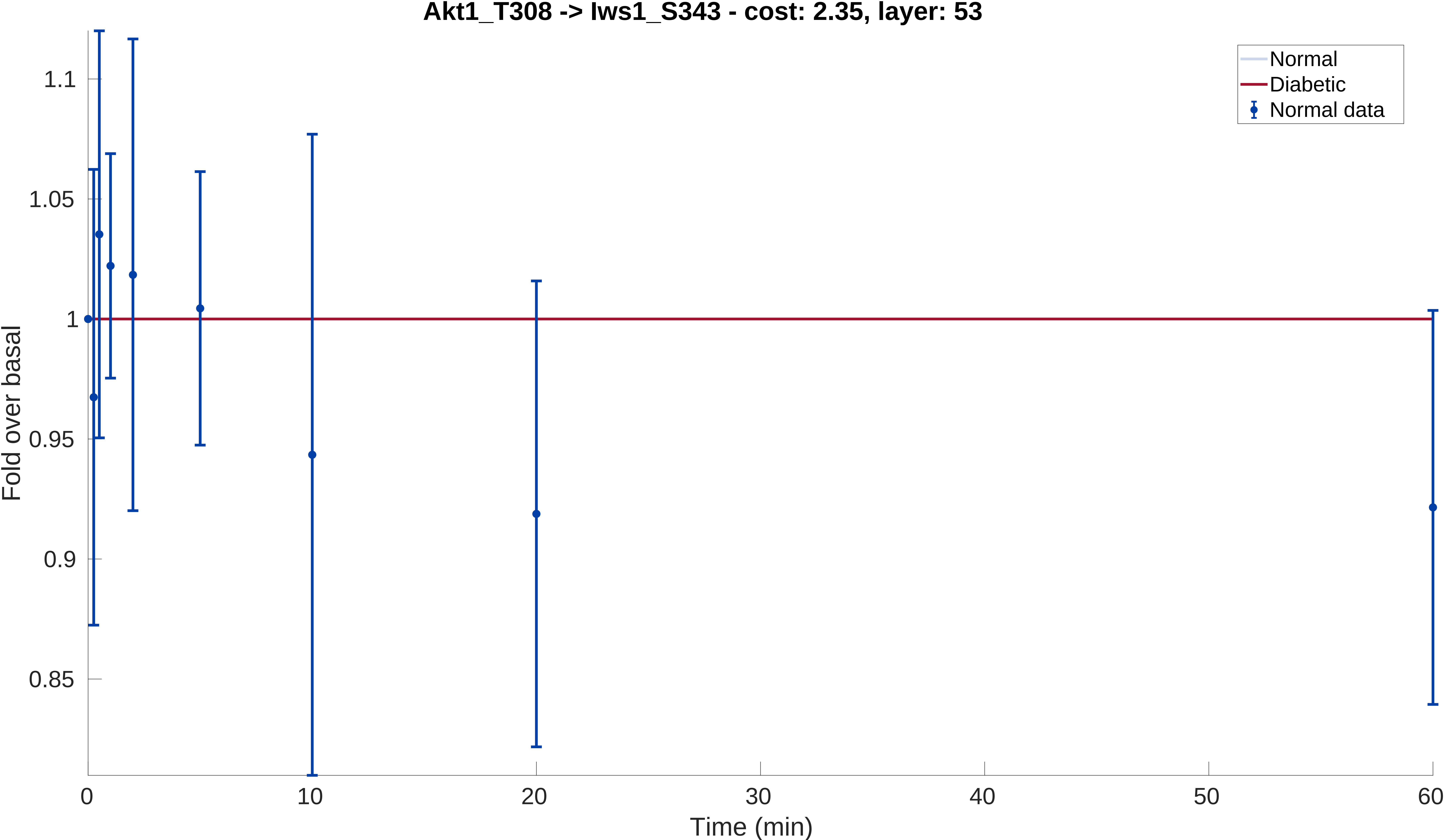

Akt1_T308 -> Iws1_S343 - cost: 2.35, layer: 53
Normal
Diabetic
Normal data
Fold over basal
1.1
1.05
1
0.95
0.9
0.85
0
10
20
30
40
50
60
Time (min)

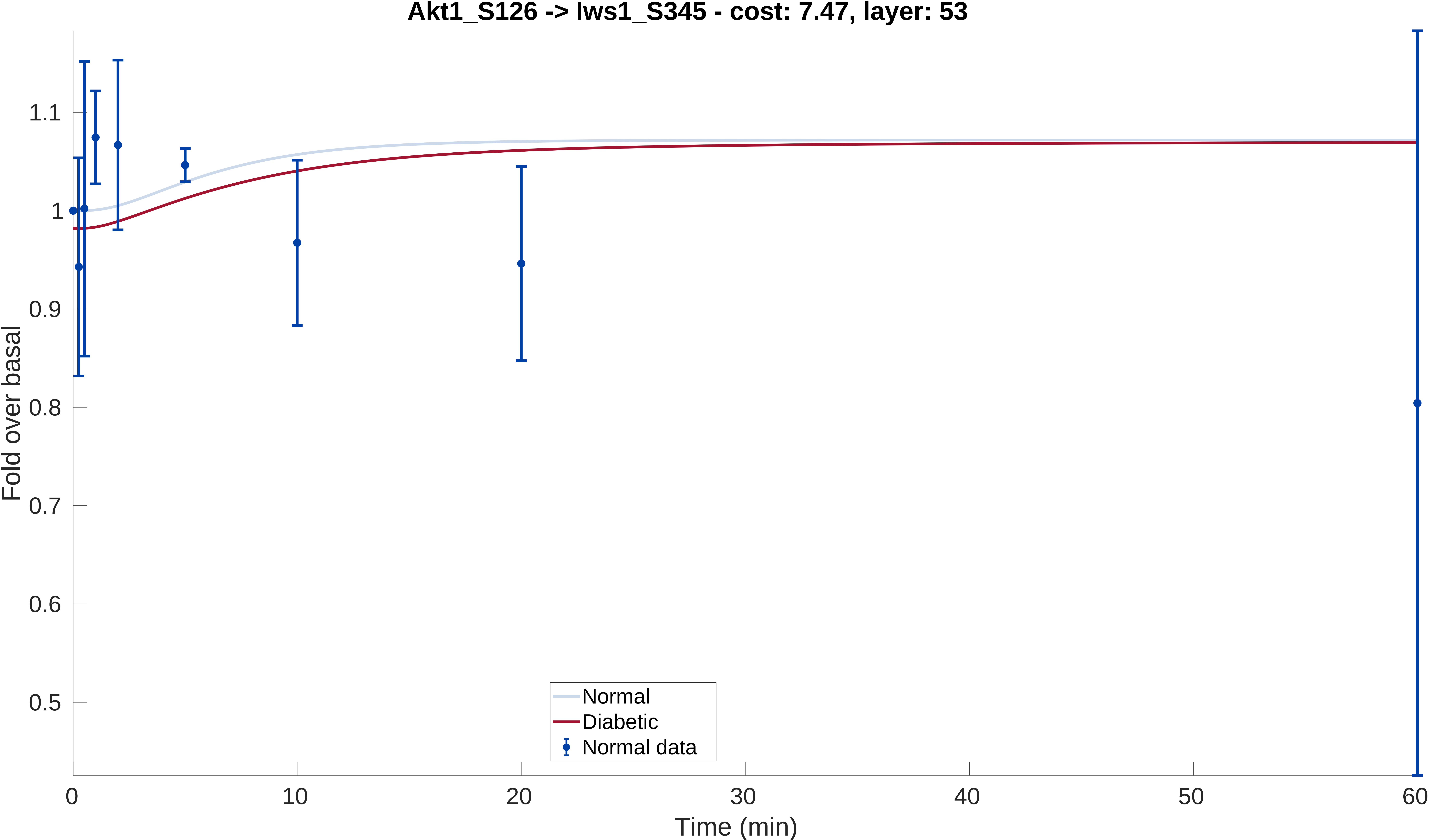

Akt1_S126 -> Iws1_S345 - cost: 7.47, layer: 53
Fold over basal
1.1
1
0.9
0.8
0.7
0.6
0.5
0
10
20
30
40
50
60
Time (min)
Normal
Diabetic
Normal data

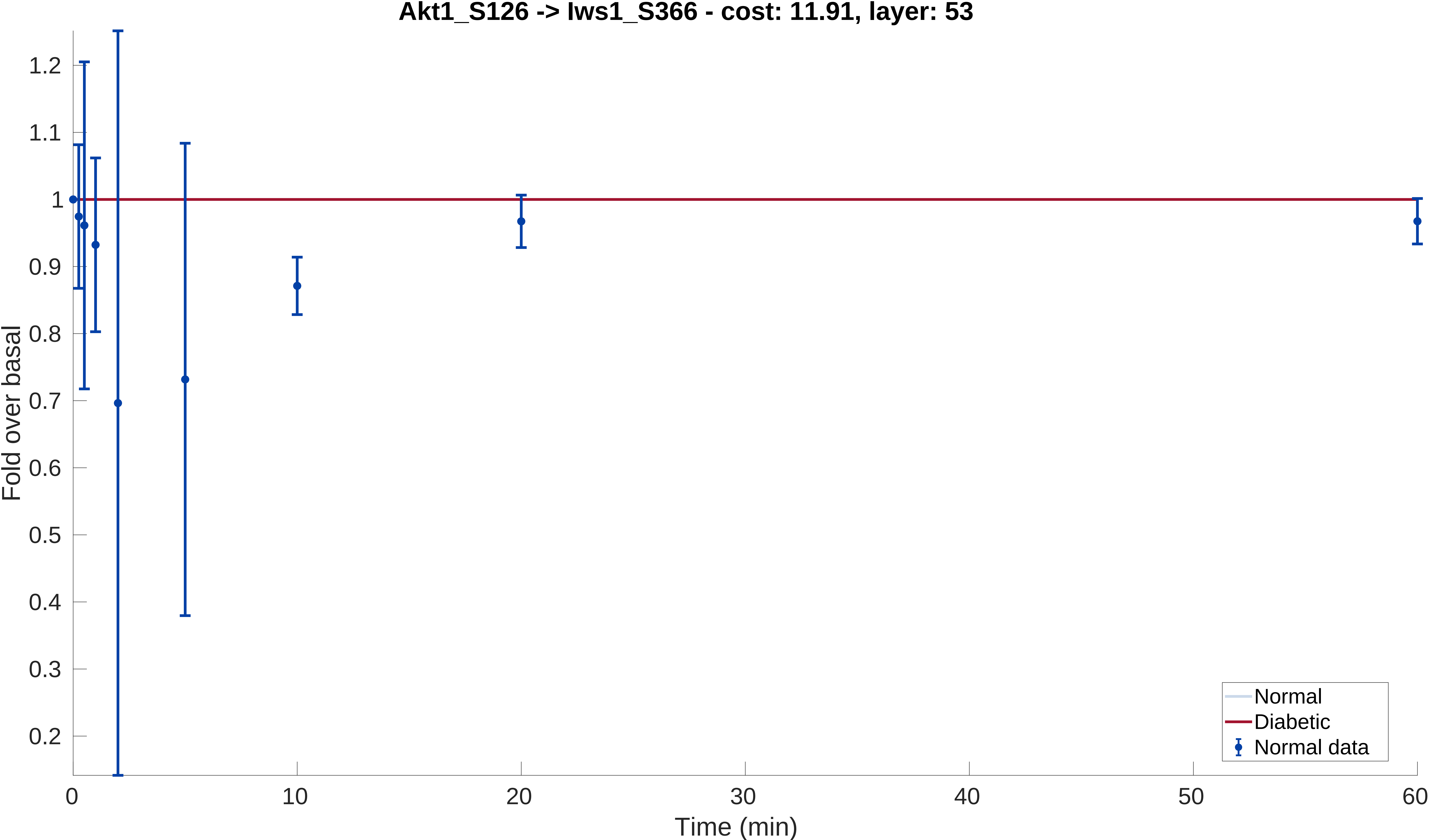

Akt1_S126 -> Iws1_S366 - cost: 11.91, layer: 53
Fold over basal
1.2
1.1
1
0.9
0.8
0.7
0.6
0.5
0.4
0.3
0.2
0
10
20
30
40
50
60
Time (min)
Normal
Diabetic
Normal data

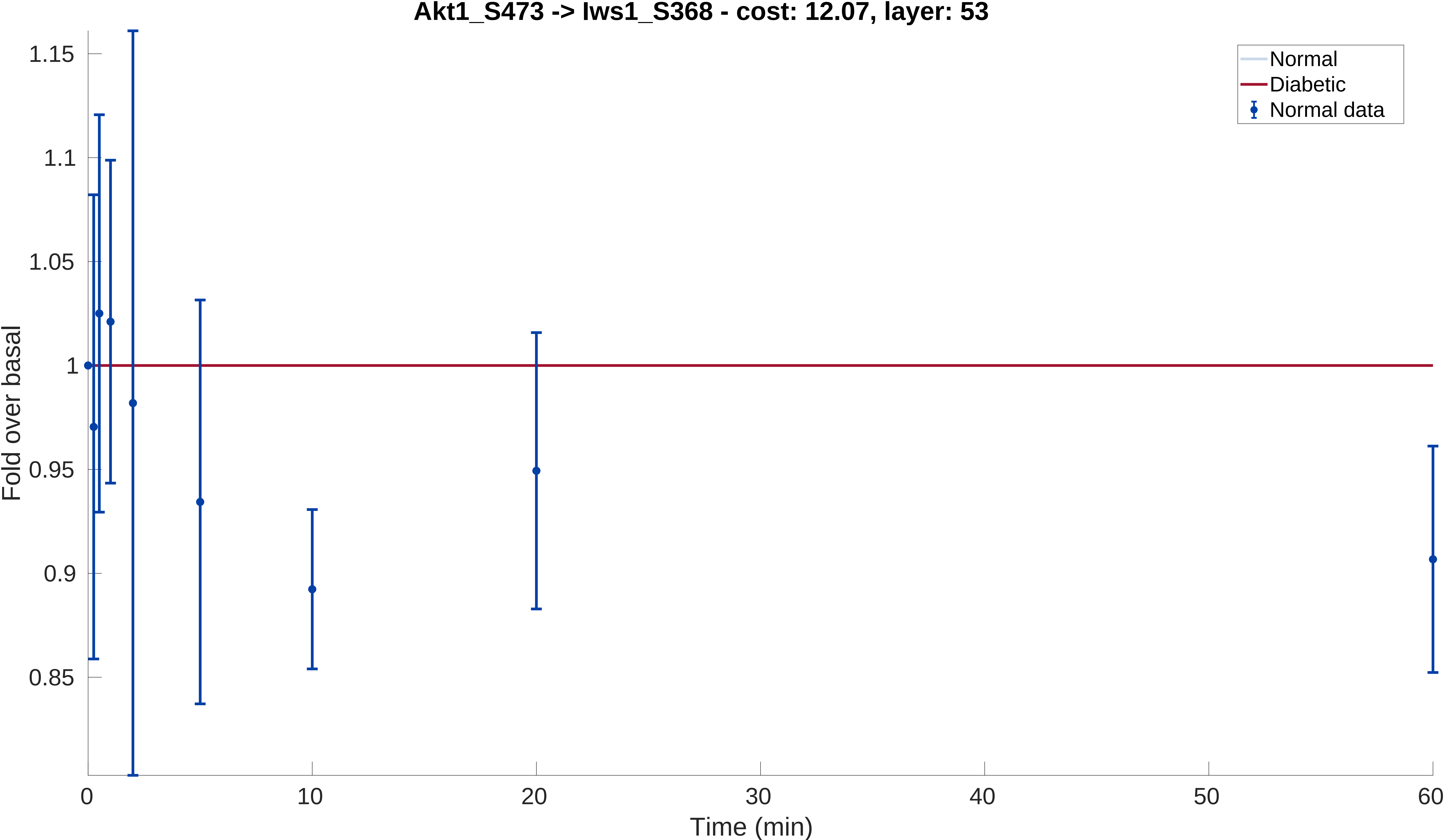

Akt1_S473 -> Iws1_S368 - cost: 12.07, layer: 53
Normal
Diabetic
Normal data
Fold over basal
1.15
1.1
1.05
1
0.95
0.9
0.85
0
10
20
30
40
50
60
Time (min)

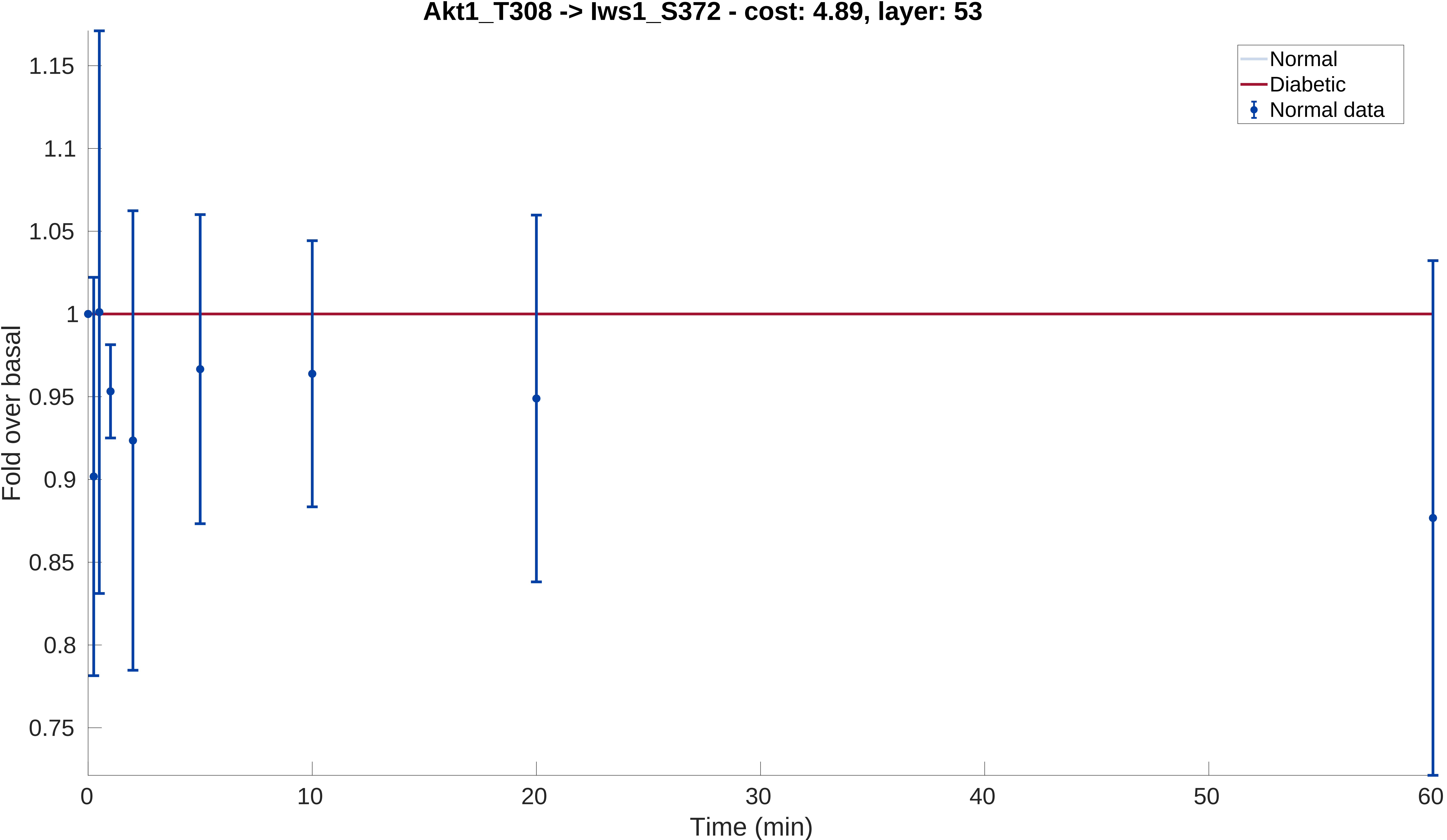

Akt1_T308 -> Iws1_S372 - cost: 4.89, layer: 53
Normal
Diabetic
Normal data
Fold over basal
1.15
1.1
1.05
1
0.95
0.9
0.85
0.8
0.75
0
10
20
30
40
50
60
Time (min)

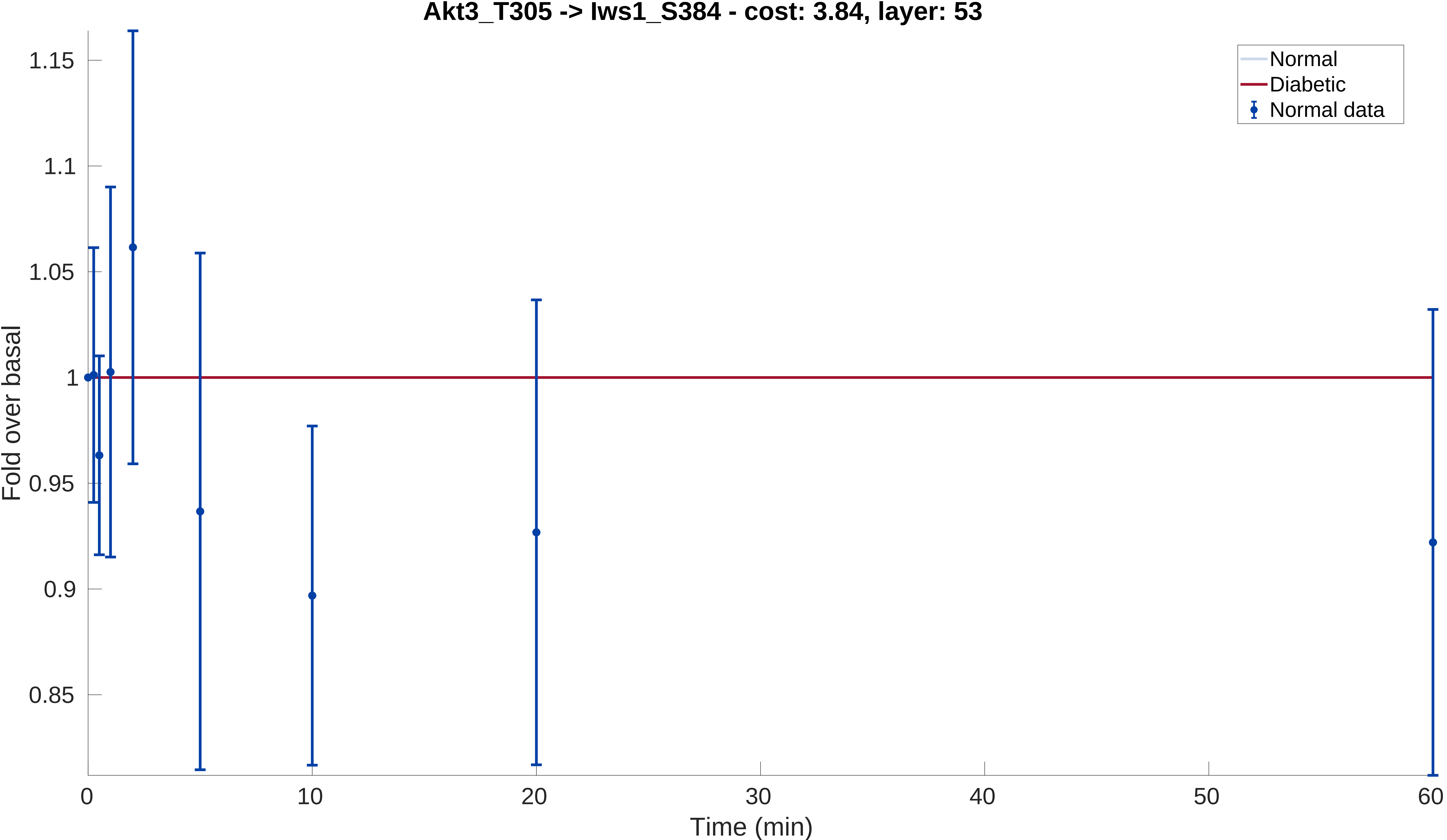

Akt3_T305 -> Iws1_S384 - cost: 3.84, layer: 53
Normal
Diabetic
Normal data
Fold over basal
1.15
1.1
1.05
1
0.95
0.9
0.85
0
10
20
30
40
50
60
Time (min)

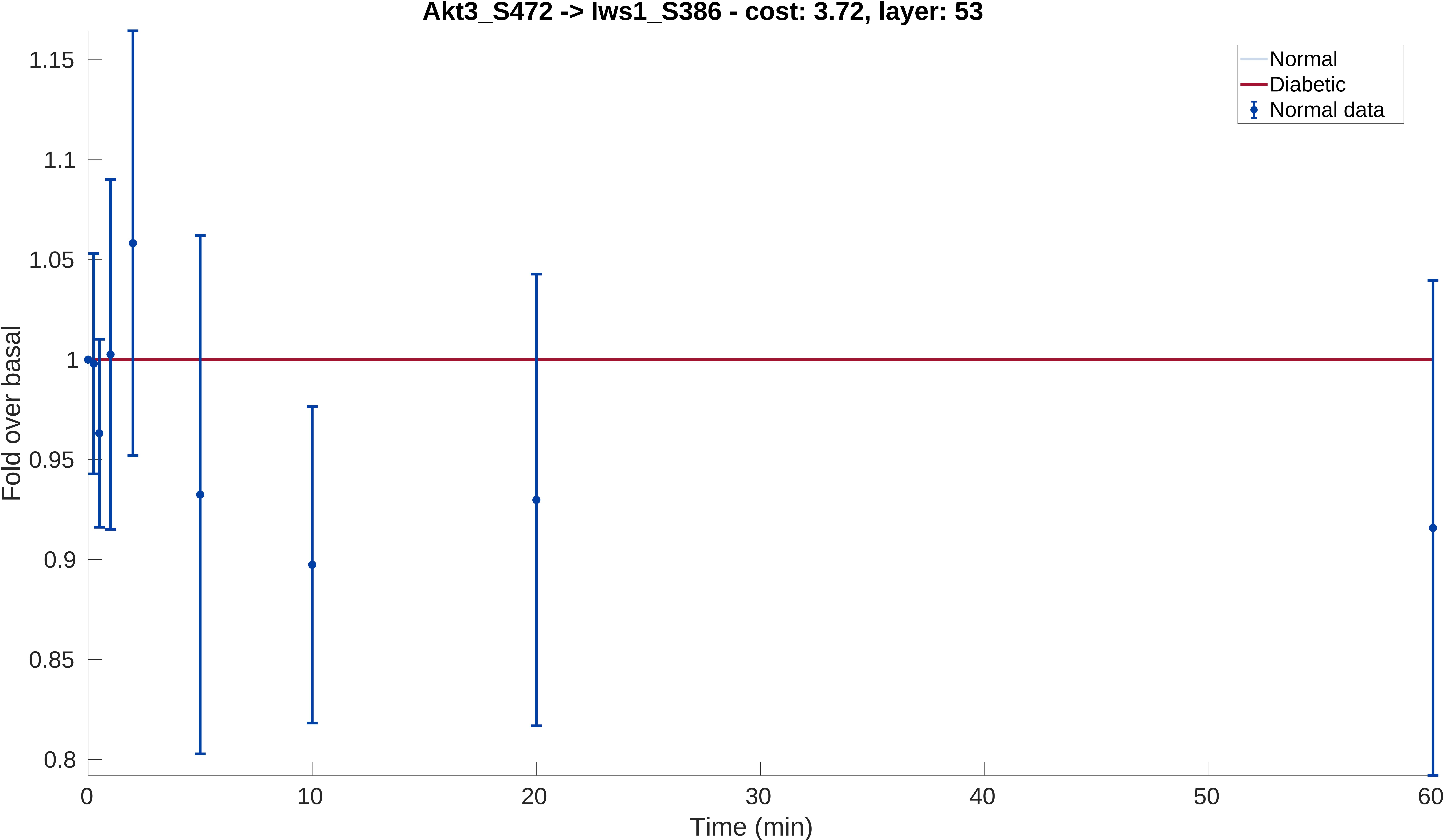
Akt3_S472 -> Iws1_S386 - cost: 3.72, layer: 53
Normal
Diabetic
Normal data
Fold over basal
1.15
1.1
1.05
1
0.95
0.9
0.85
0.8
0
10
20
30
40
50
60
Time (min)

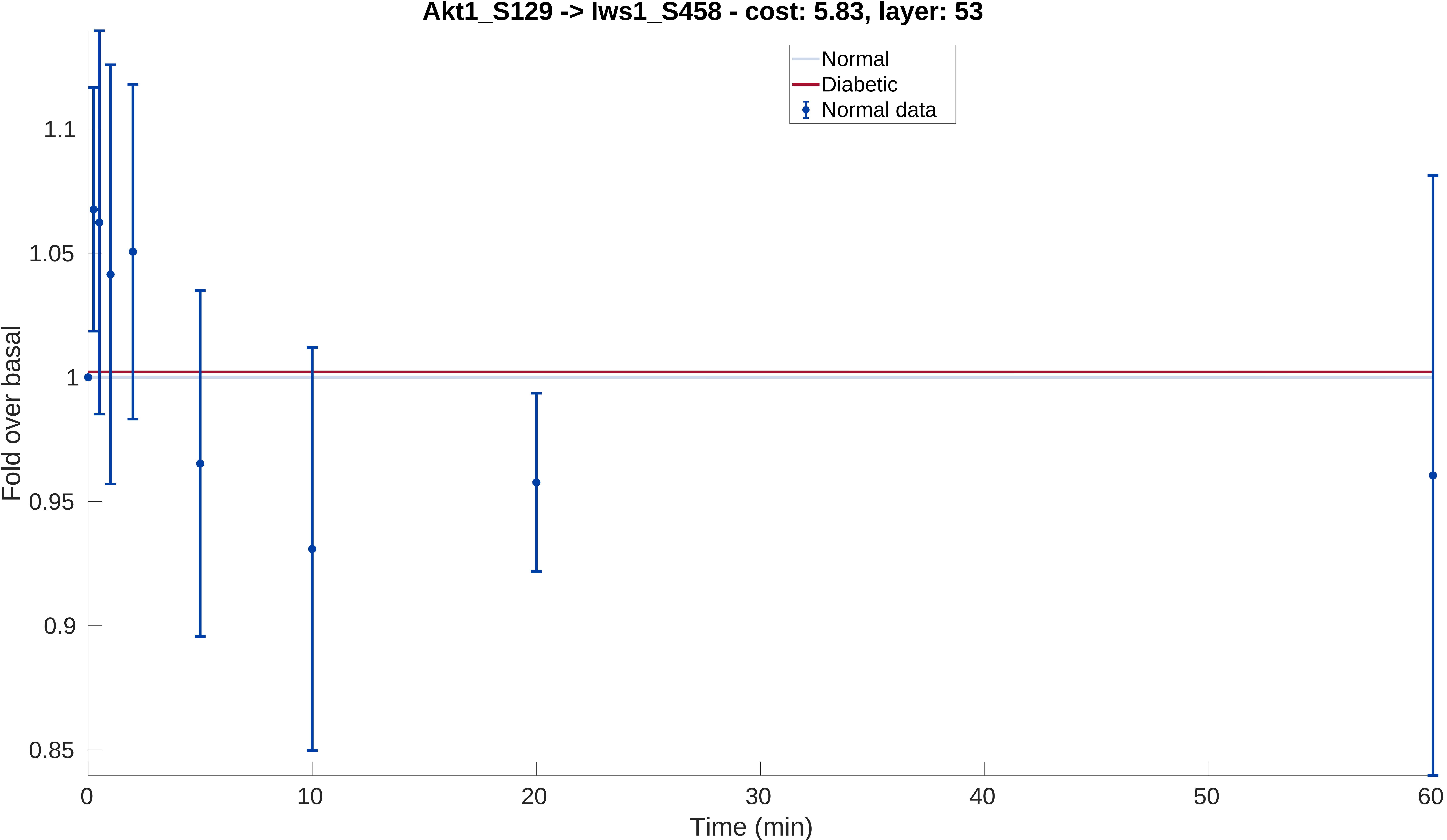

Akt1_S129 -> Iws1_S458 - cost: 5.83, layer: 53
Normal
Diabetic
Normal data
Fold over basal
1.1
1.05
1
0.95
0.9
0.85
0
10
20
30
40
50
60
Time (min)

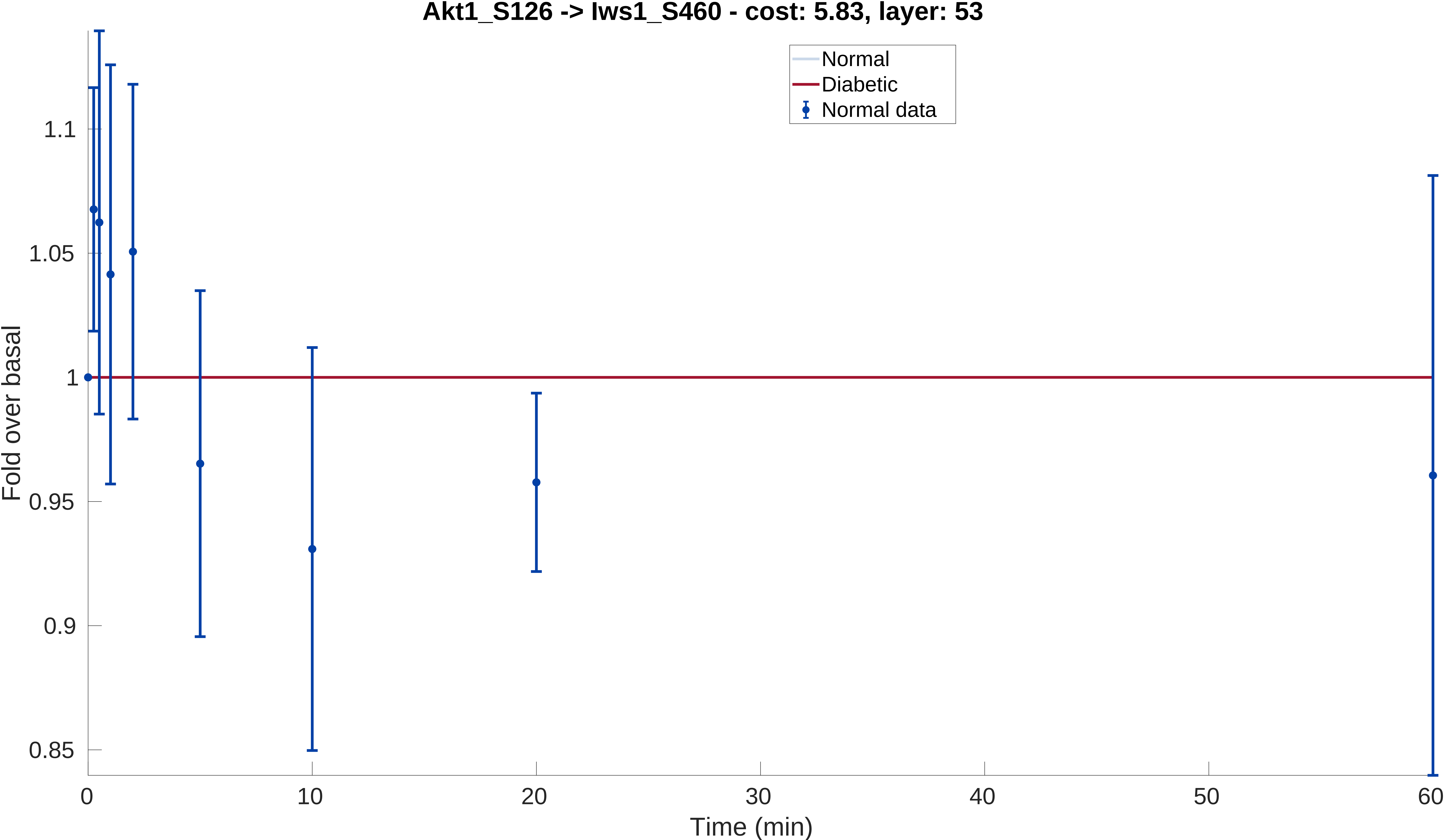

Akt1_S126 -> Iws1_S460 - cost: 5.83, layer: 53
Normal
Diabetic
Normal data
Fold over basal
1.1
1.05
1
0.95
0.9
0.85
0
10
20
30
40
50
60
Time (min)

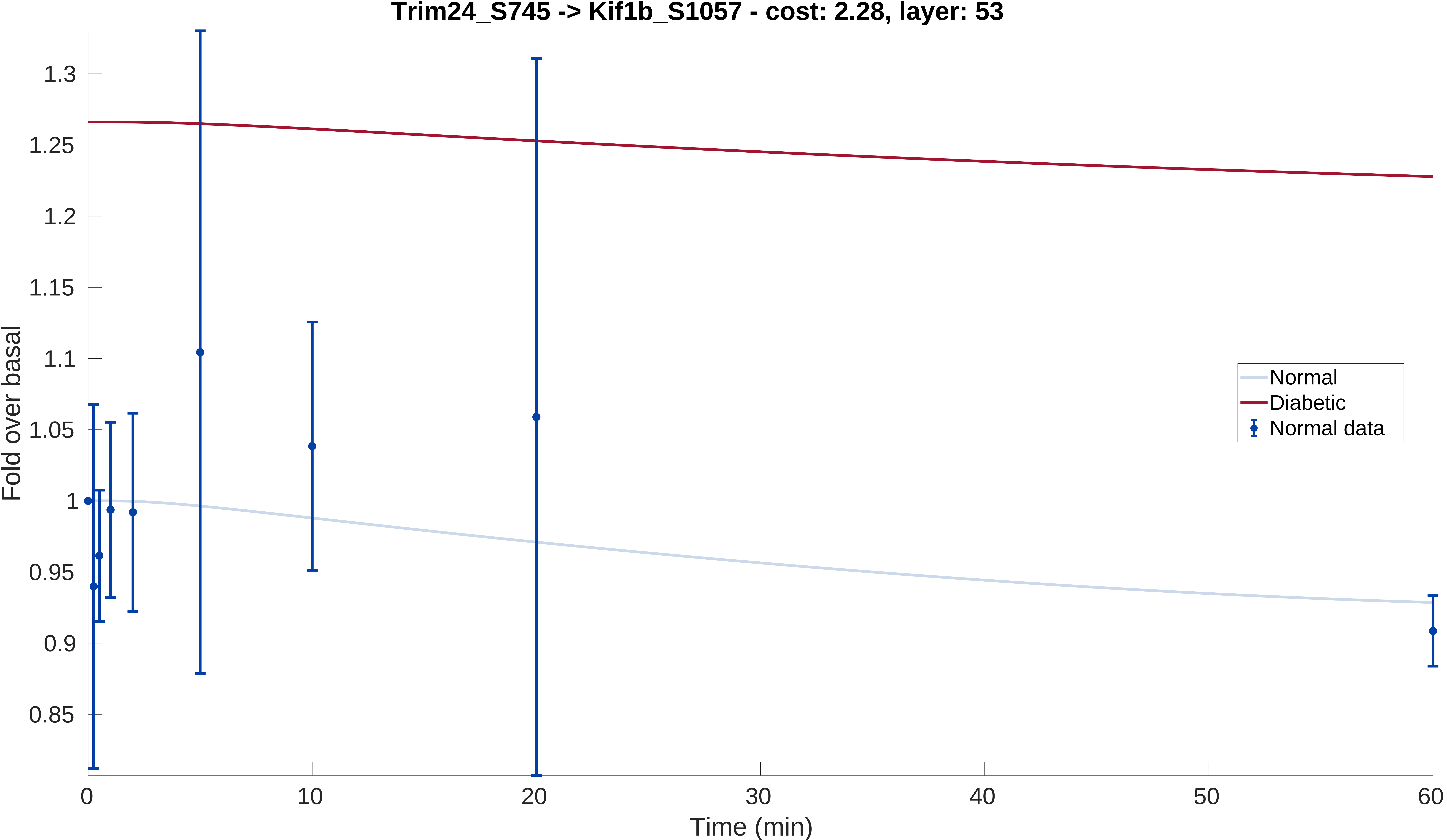

Trim24_S745 -> Kif1b_S1057 - cost: 2.28, layer: 53
Fold over basal
1.3
1.25
1.2
1.15
1.1
1.05
1
0.95
0.9
0.85
0
10
20
30
40
50
60
Time (min)
Normal
Diabetic
Normal data

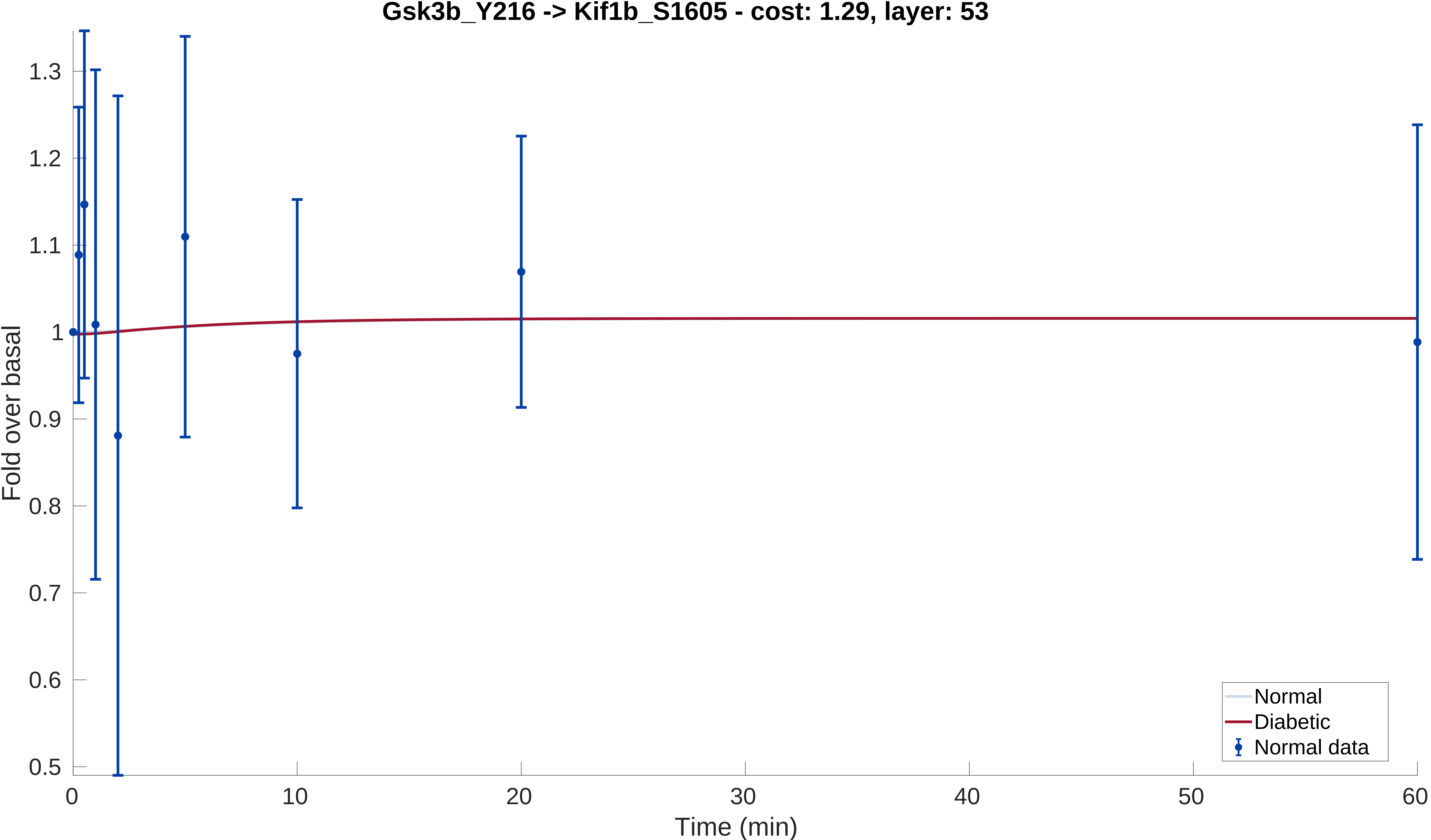

Gsk3b_Y216 -> Kif1b_S1605 - cost: 1.29, layer: 53
Fold over basal
1.3
1.2
1.1
1
0.9
0.8
0.7
0.6
0.5
0
10
20
30
40
50
60
Time (min)
Normal
Diabetic
Normal data

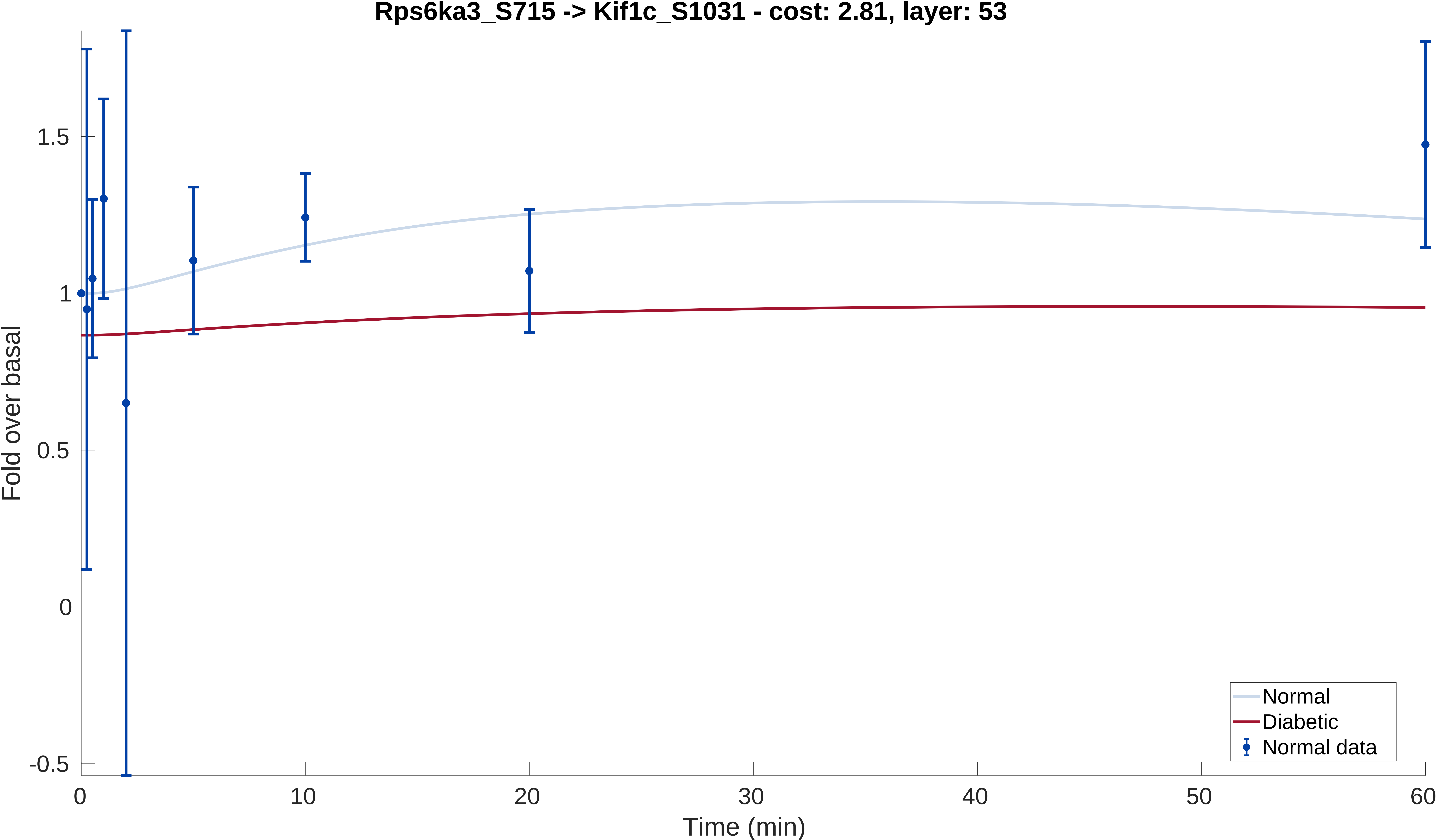
Rps6ka3_S715 -> Kif1c_S1031 - cost: 2.81, layer: 53
Fold over basal
1.5
1
0.5
0
-0.5
0
10
20
30
40
50
60
Time (min)
Normal
Diabetic
Normal data

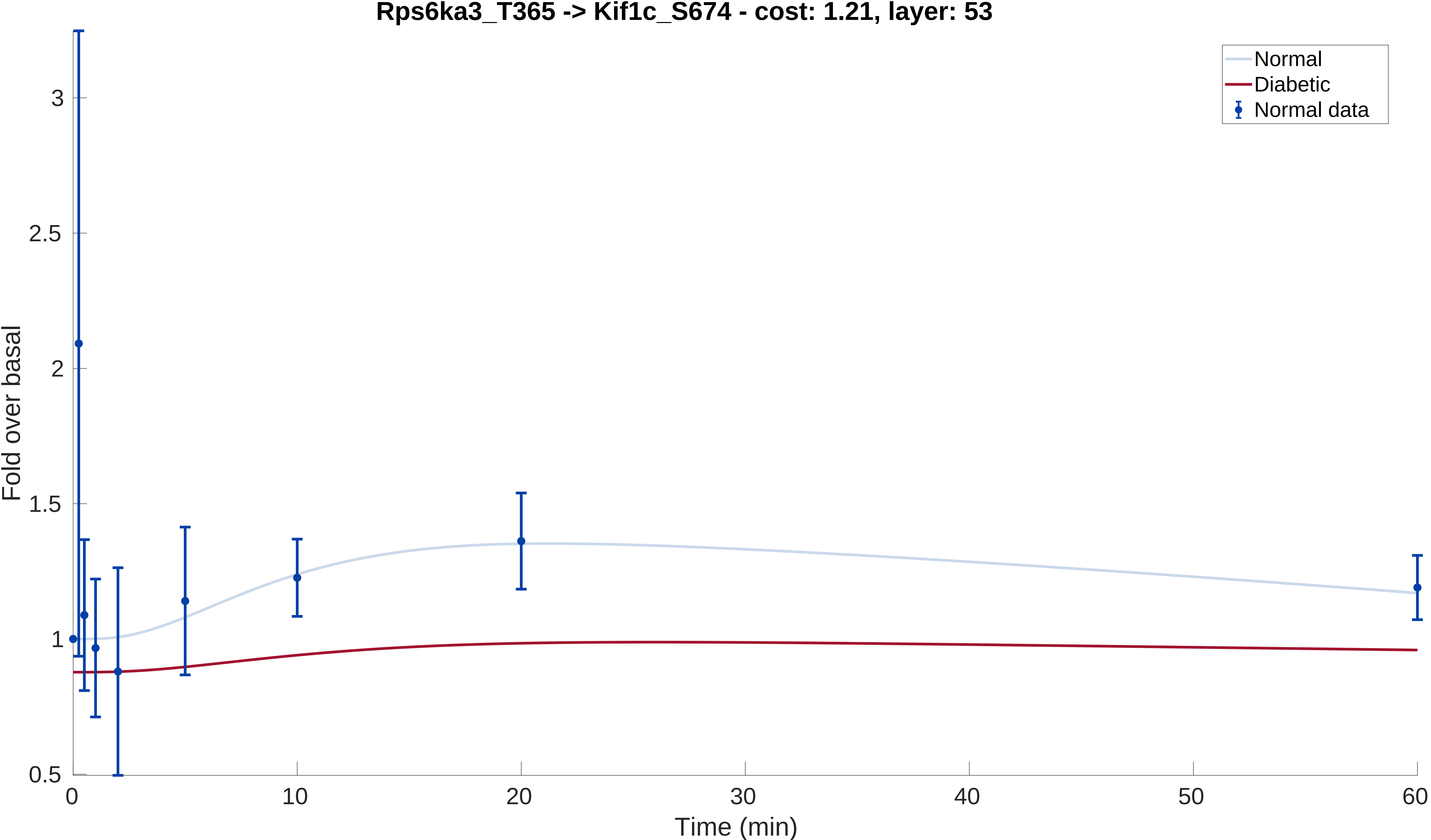

Rps6ka3_T365 -> Kif1c_S674 - cost: 1.21, layer: 53
Normal
Diabetic
Normal data
Fold over basal
3
2.5
2
1.5
1
0.5
0
10
20
30
40
50
60
Time (min)

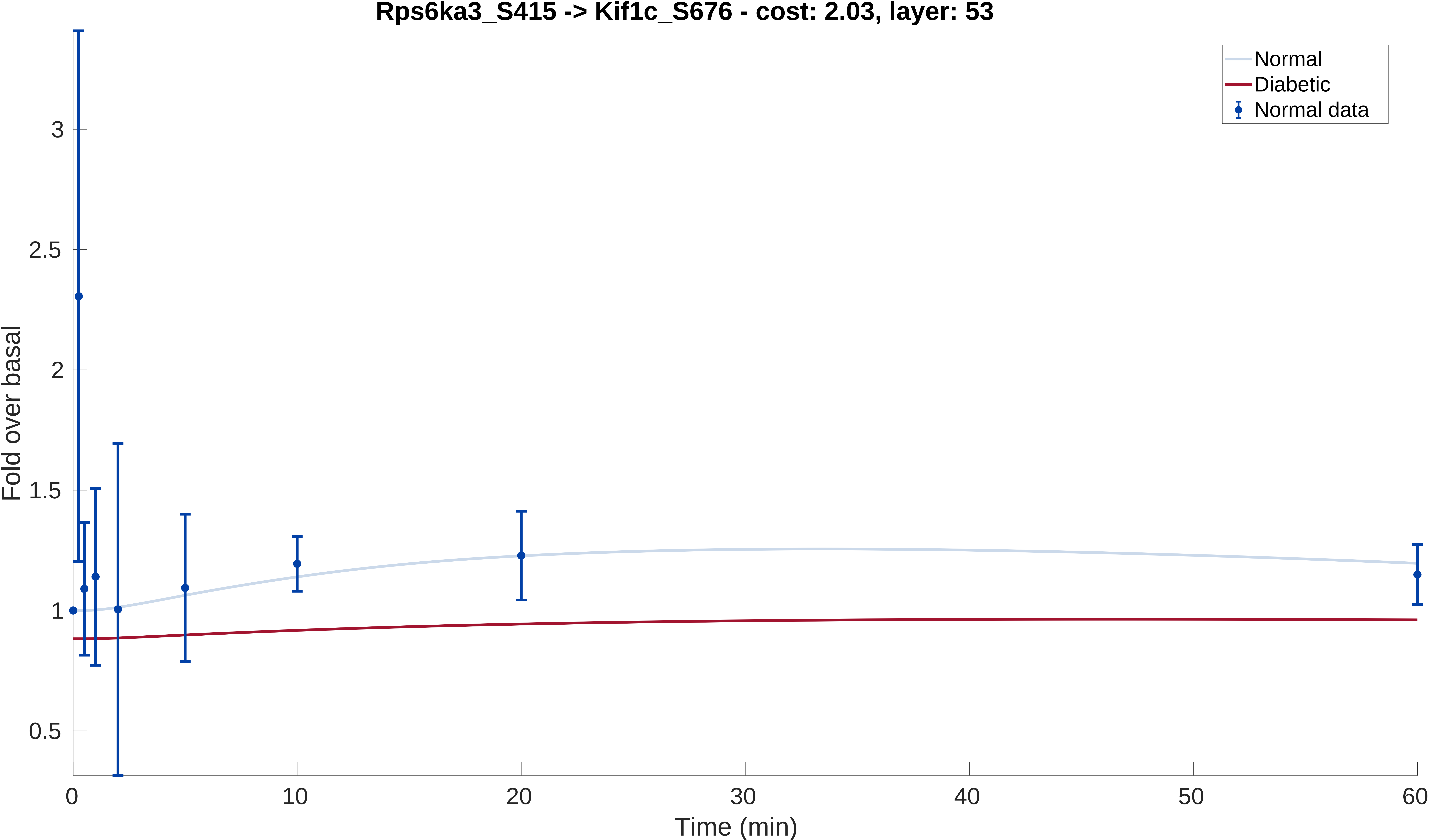

Rps6ka3_S415 -> Kif1c_S676 - cost: 2.03, layer: 53
Normal
Diabetic
Normal data
Fold over basal
0.5
1
1.5
2
2.5
3
0
10
20
30
40
50
60
Time (min)

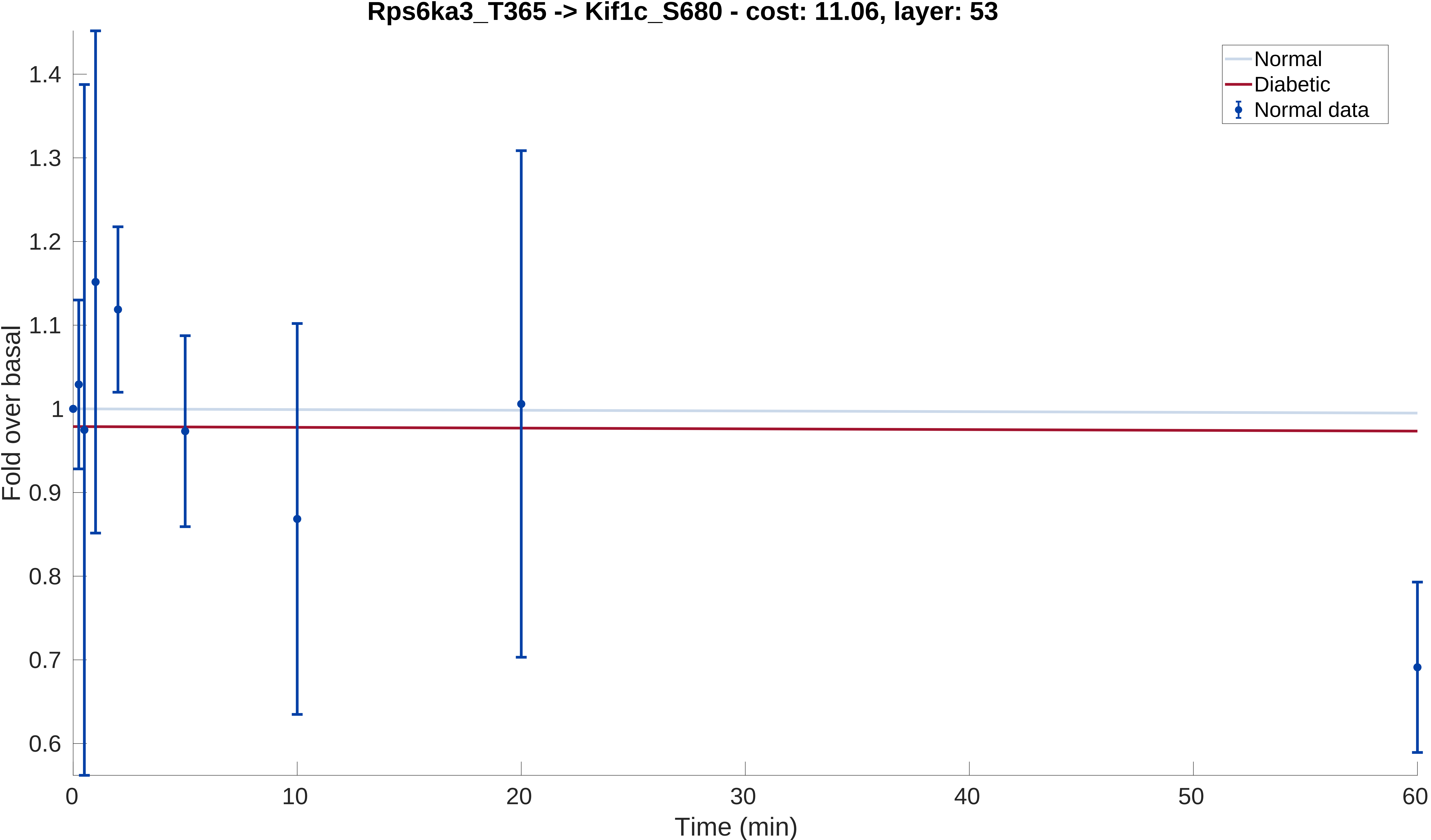

Rps6ka3_T365 -> Kif1c_S680 - cost: 11.06, layer: 53
Normal
Diabetic
Normal data
Fold over basal
Time (min)
1.4
1.3
1.2
1.1
1
0.9
0.8
0.7
0.6
0
10
20
30
40
50
60

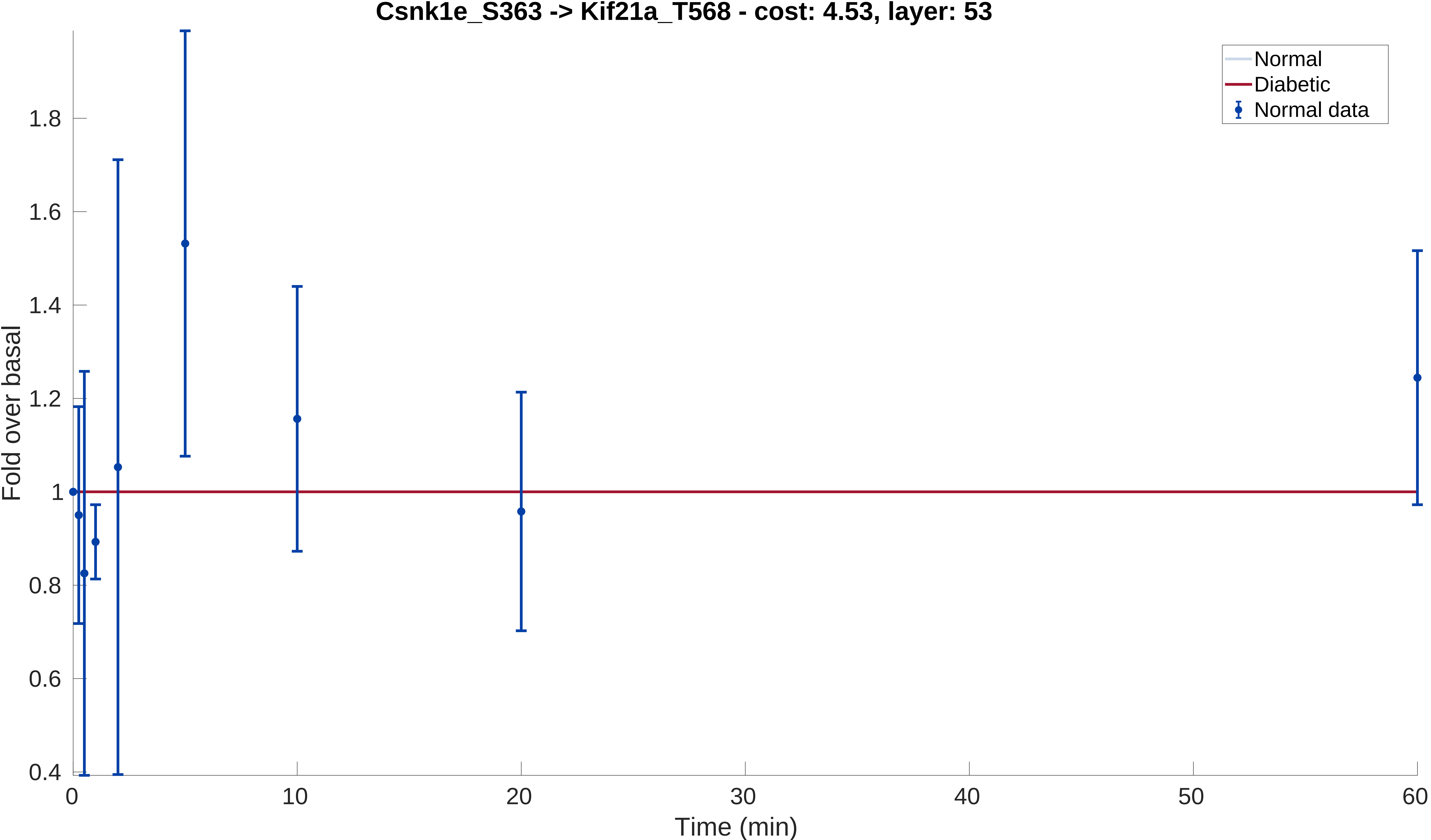

Csnk1e_S363 -> Kif21a_T568 - cost: 4.53, layer: 53
Normal
Diabetic
Normal data
Fold over basal
1.8
1.6
1.4
1.2
1
0.8
0.6
0.4
0
10
20
30
40
50
60
Time (min)

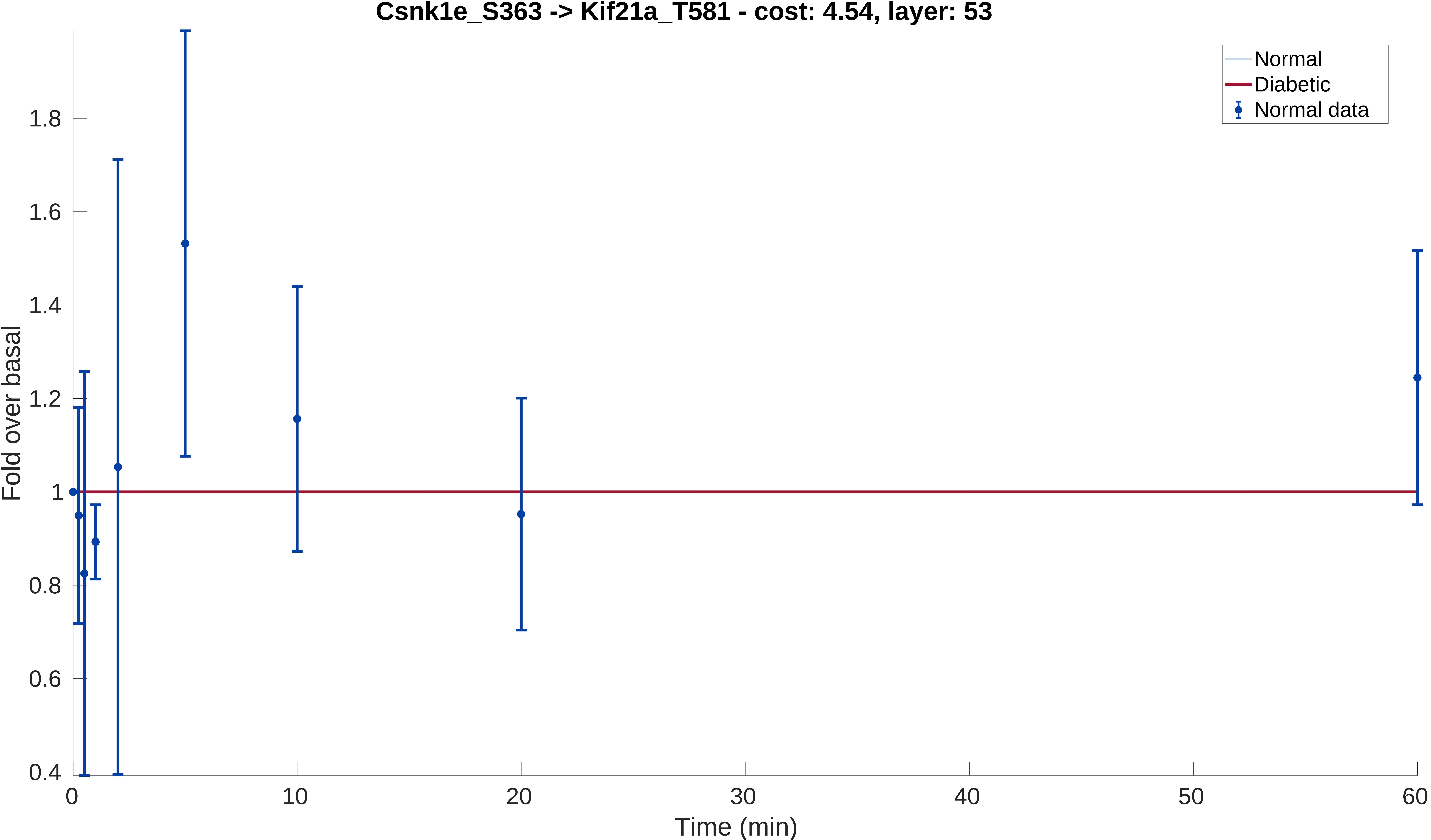
Csnk1e_S363 -> Kif21a_T581 - cost: 4.54, layer: 53
Normal
Diabetic
Normal data
Fold over basal
Time (min)
0.4
0.6
0.8
1
1.2
1.4
1.6
1.8
0
10
20
30
40
50
60

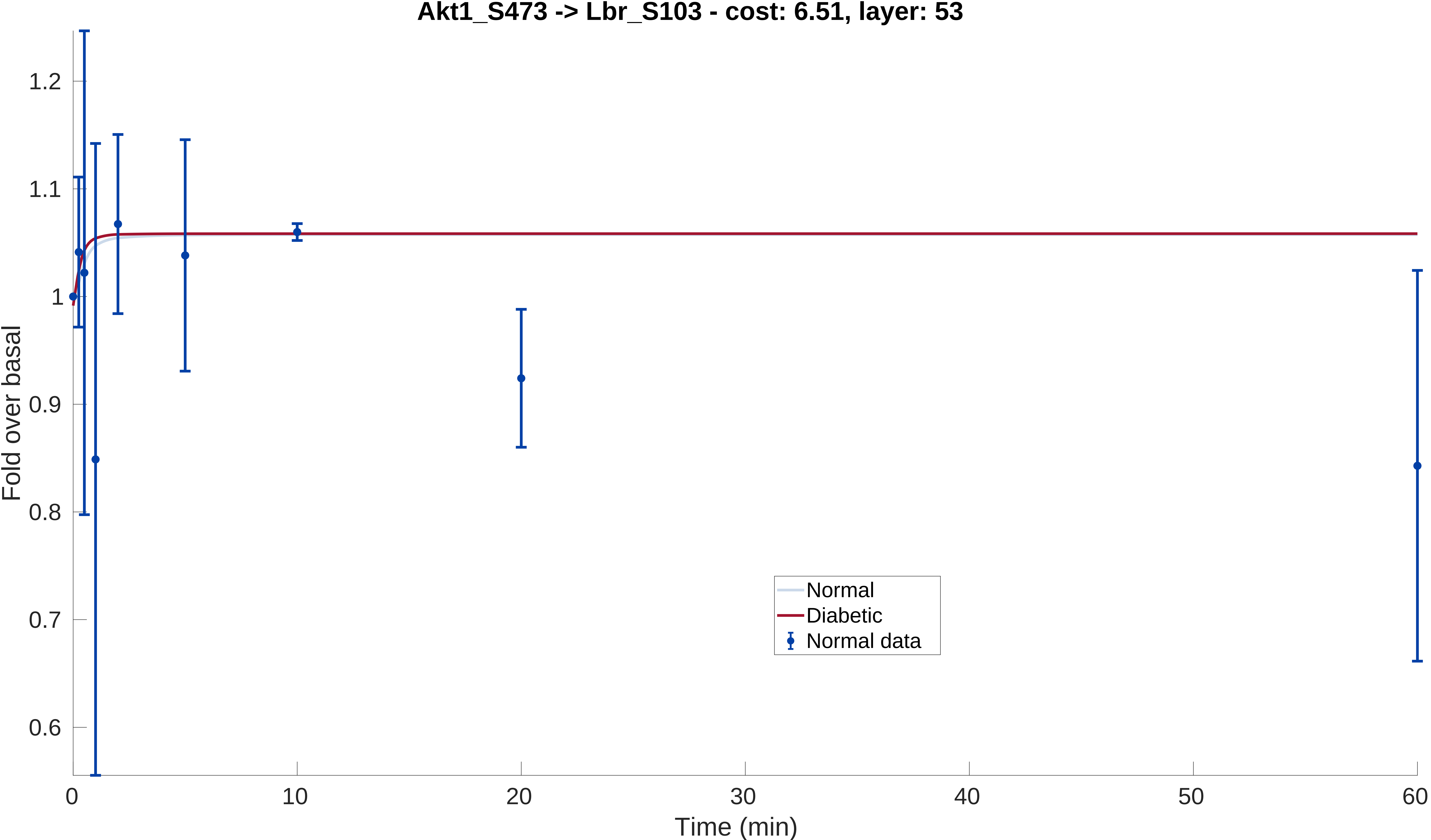

Akt1_S473 -> Lbr_S103 - cost: 6.51, layer: 53
Fold over basal
1.2
1.1
1
0.9
0.8
0.7
0.6
0
10
20
30
40
50
60
Time (min)
Normal
Diabetic
Normal data

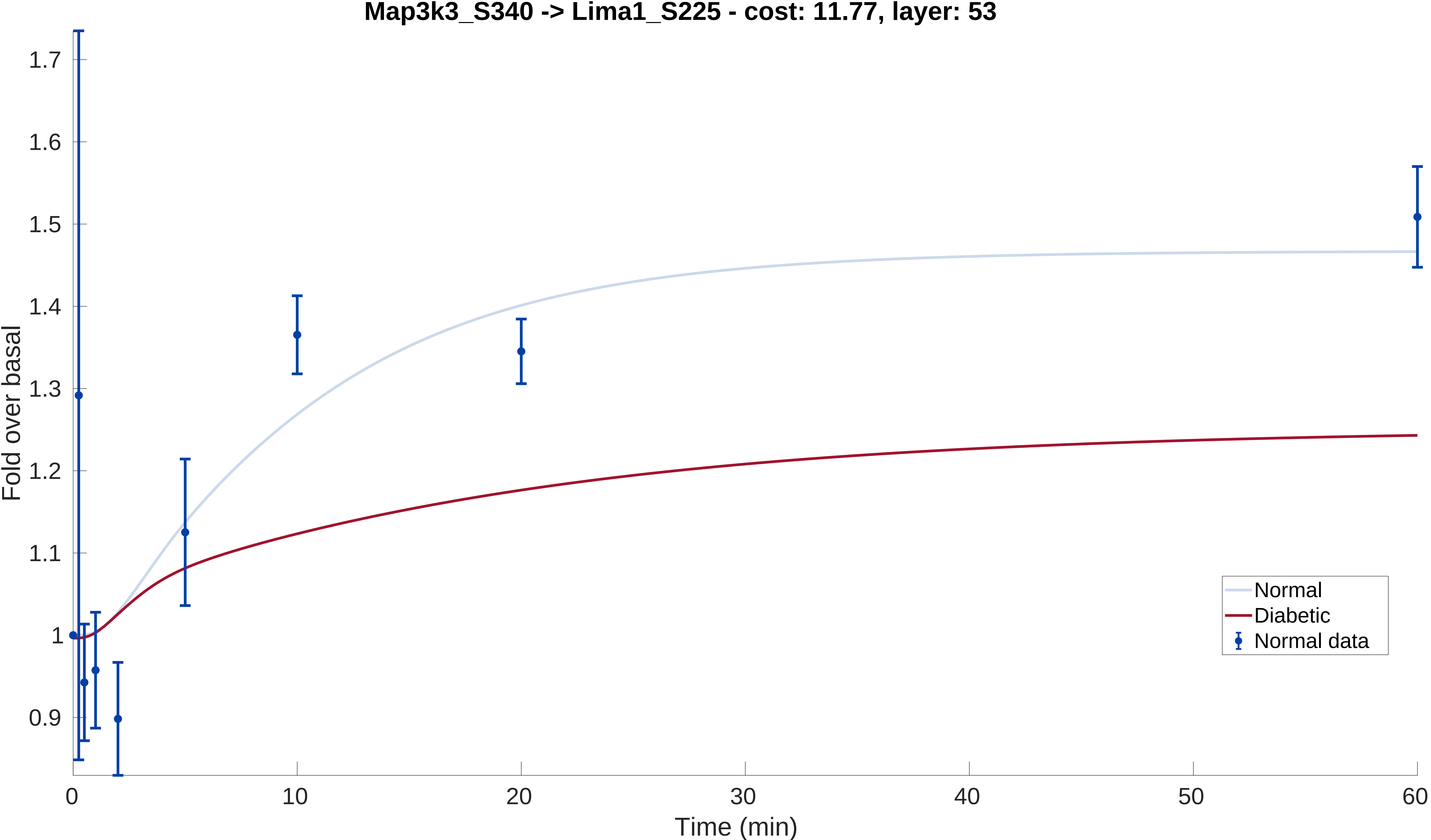

Map3k3_S340 -> Lima1_S225 - cost: 11.77, layer: 53
Fold over basal
Time (min)
0.9
1
1.1
1.2
1.3
1.4
1.5
1.6
1.7
0
10
20
30
40
50
60
Normal
Diabetic
Normal data

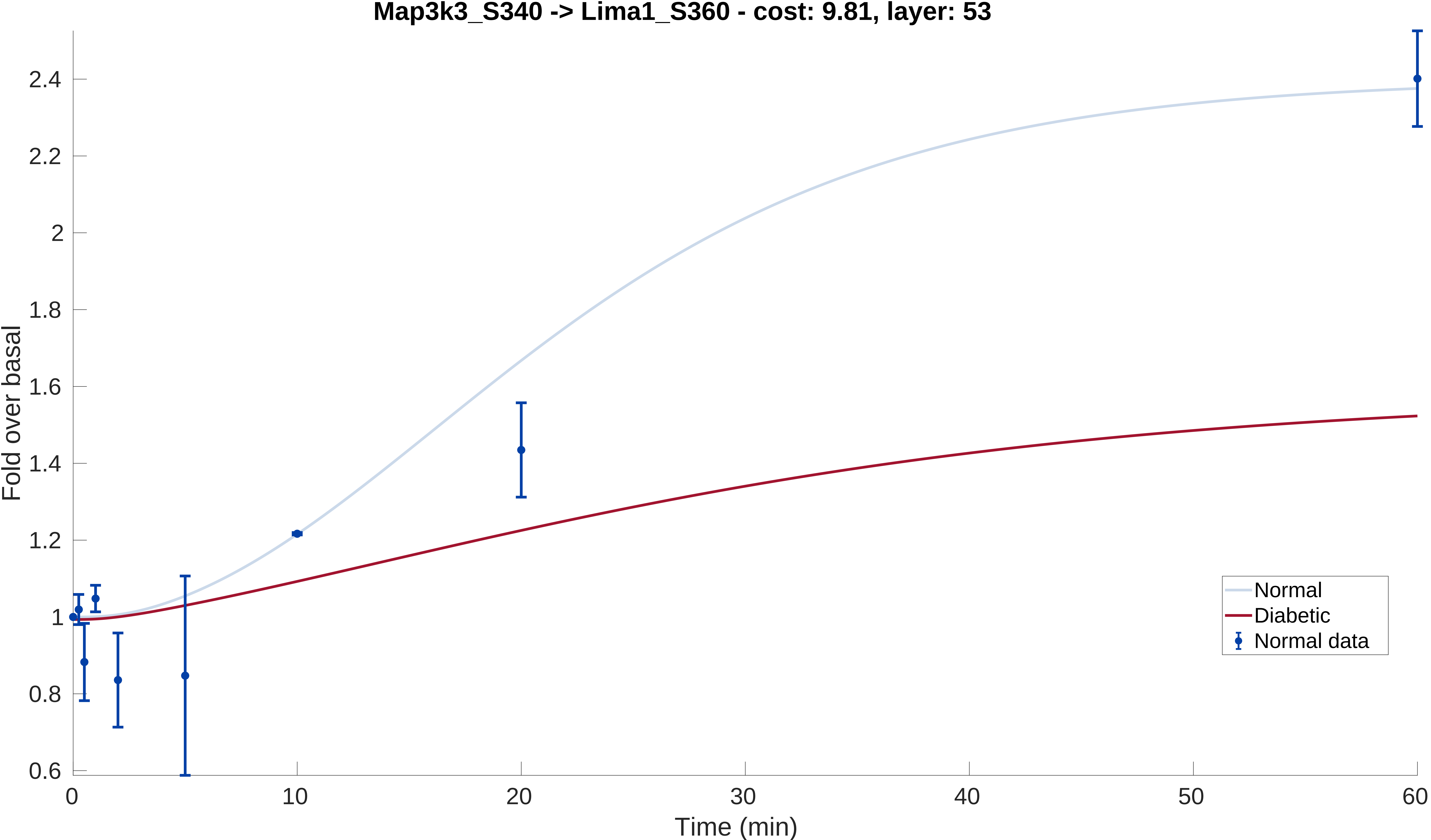
Map3k3_S340 -> Lima1_S360 - cost: 9.81, layer: 53
Fold over basal
Time (min)
0.6
0.8
1
1.2
1.4
1.6
1.8
2
2.2
2.4
0
10
20
30
40
50
60
Normal
Diabetic
Normal data

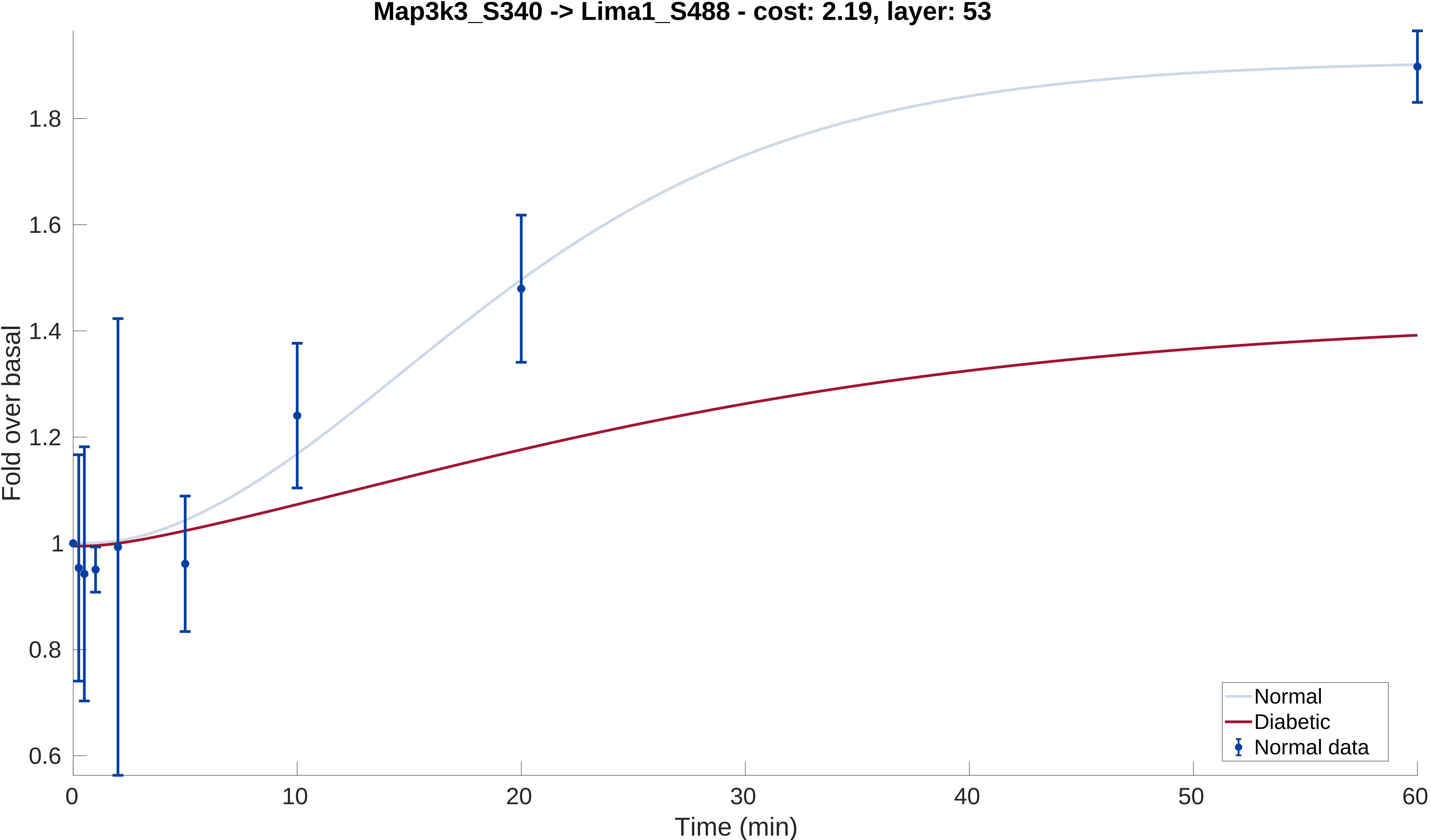
Map3k3_S340 -> Lima1_S488 - cost: 2.19, layer: 53
Fold over basal
1.8
1.6
1.4
1.2
1
0.8
0.6
0
10
20
30
40
50
60
Time (min)
Normal
Diabetic
Normal data

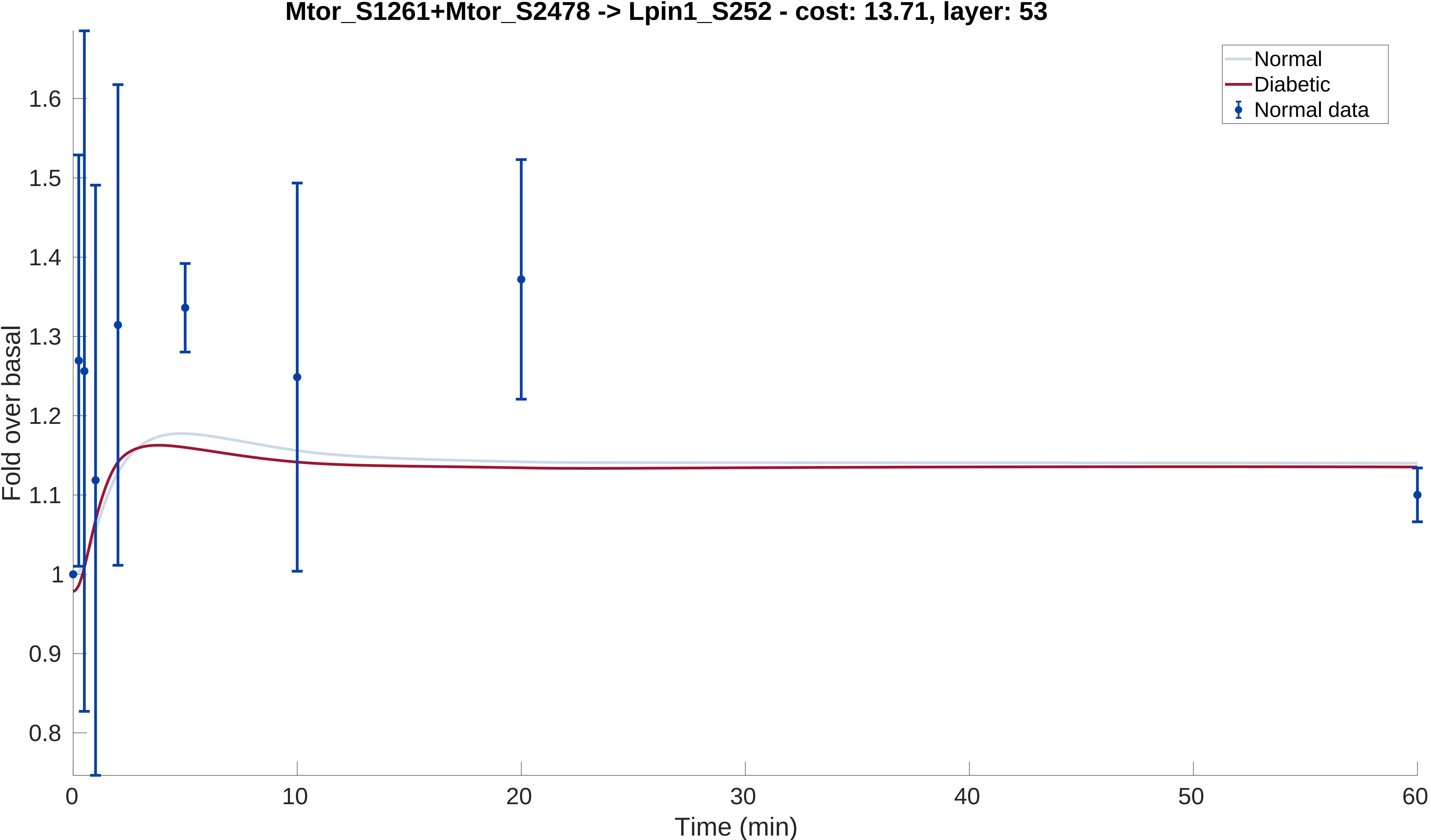
Mtor_S1261+Mtor_S2478 -> Lpin1_S252 - cost: 13.71, layer: 53
Normal
Diabetic
Normal data
Fold over basal
1.6
1.5
1.4
1.3
1.2
1.1
1
0.9
0.8
0
10
20
30
40
50
60
Time (min)

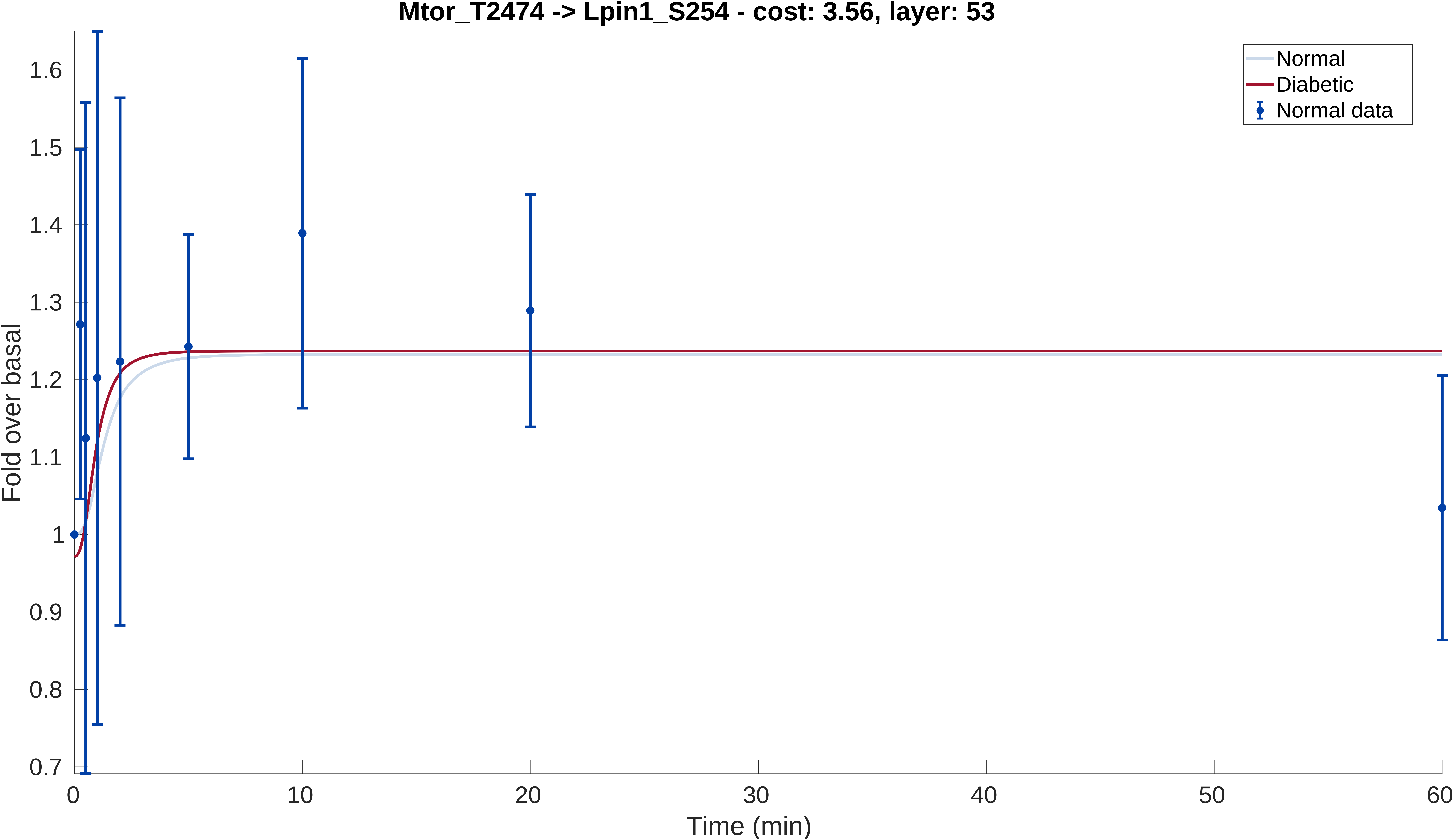

Mtor_T2474 -> Lpin1_S254 - cost: 3.56, layer: 53
Normal
Diabetic
Normal data
Fold over basal
Time (min)
1.6
1.5
1.4
1.3
1.2
1.1
1
0.9
0.8
0.7
0
10
20
30
40
50
60

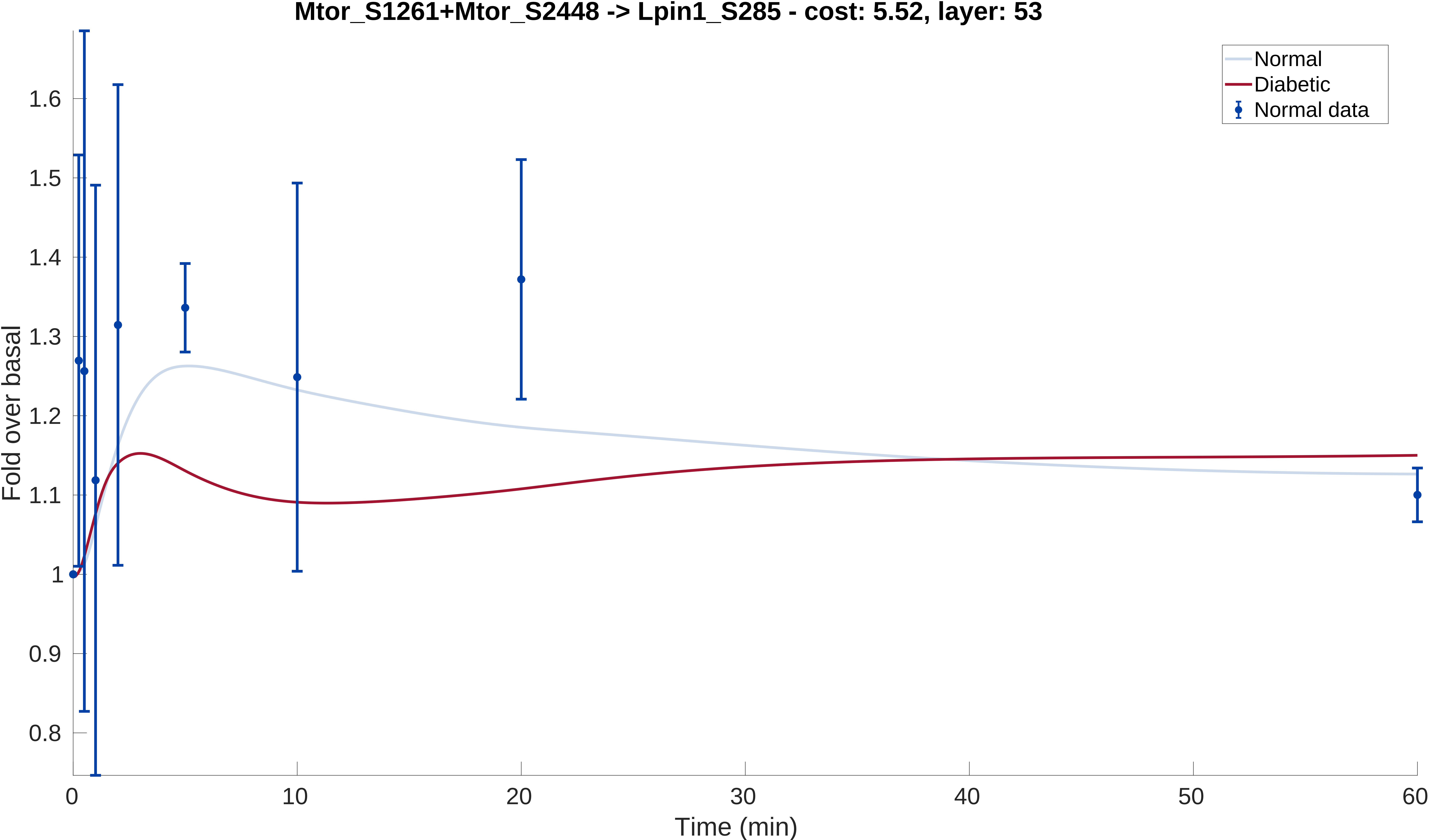
Mtor_S1261+Mtor_S2448 -> Lpin1_S285 - cost: 5.52, layer: 53
Normal
Diabetic
Normal data
Fold over basal
Time (min)
0.8
0.9
1
1.1
1.2
1.3
1.4
1.5
1.6
0
10
20
30
40
50
60

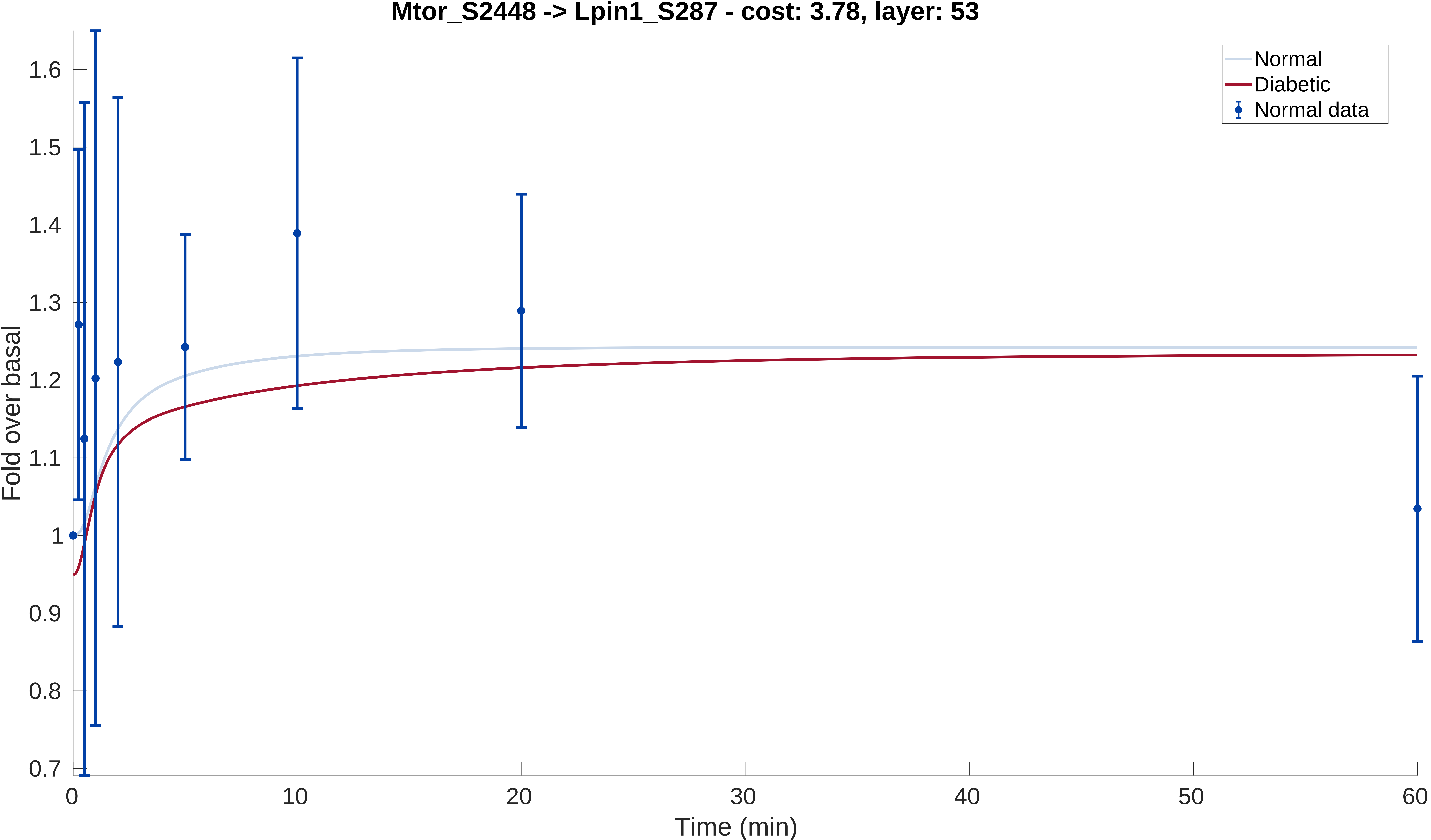
Mtor_S2448 -> Lpin1_S287 - cost: 3.78, layer: 53
Normal
Diabetic
Normal data
Fold over basal
1.6
1.5
1.4
1.3
1.2
1.1
1
0.9
0.8
0.7
0
10
20
30
40
50
60
Time (min)

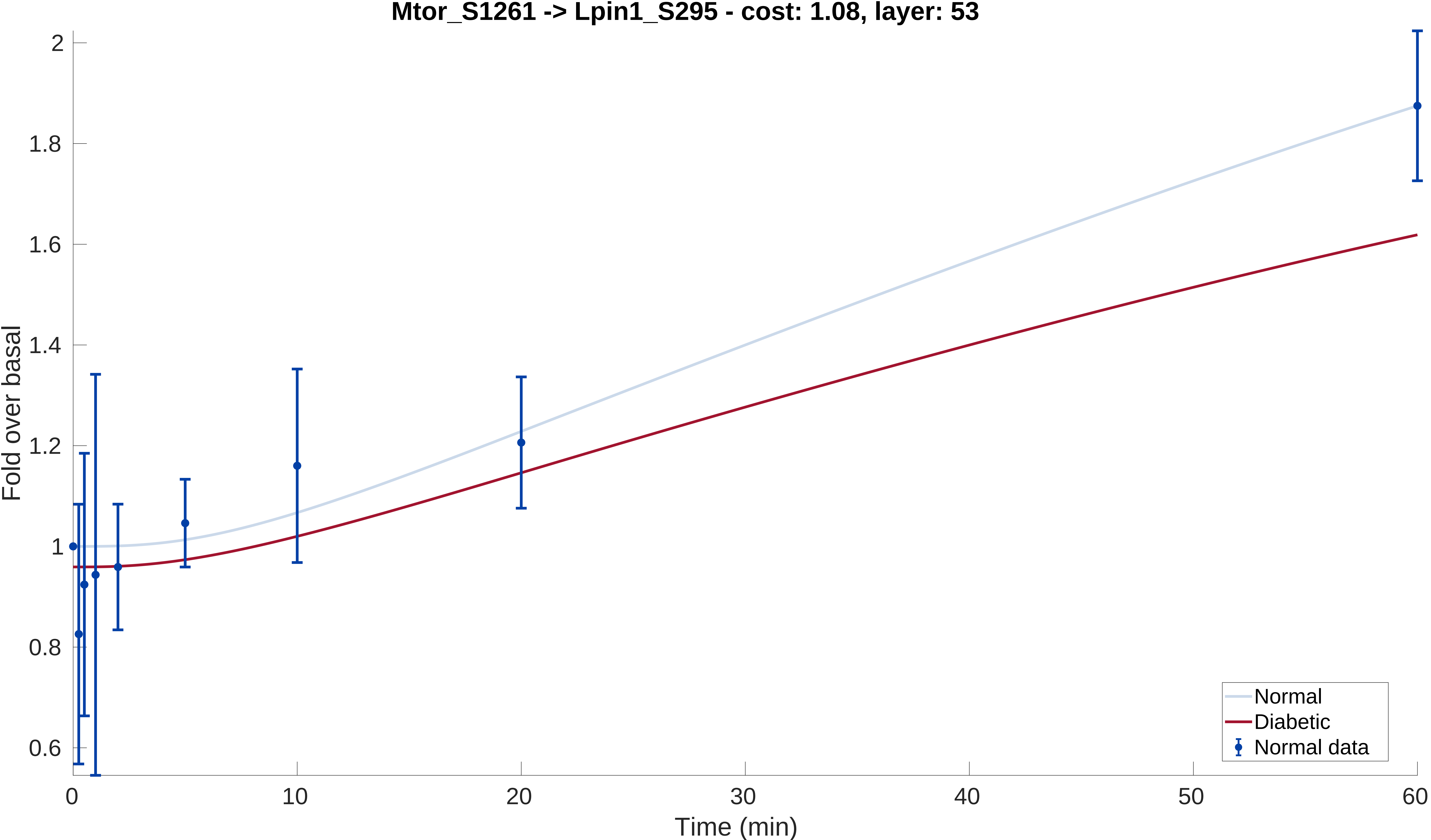
Mtor_S1261 -> Lpin1_S295 - cost: 1.08, layer: 53
Fold over basal
2
1.8
1.6
1.4
1.2
1
0.8
0.6
0
10
20
30
40
50
60
Time (min)
Normal
Diabetic
Normal data

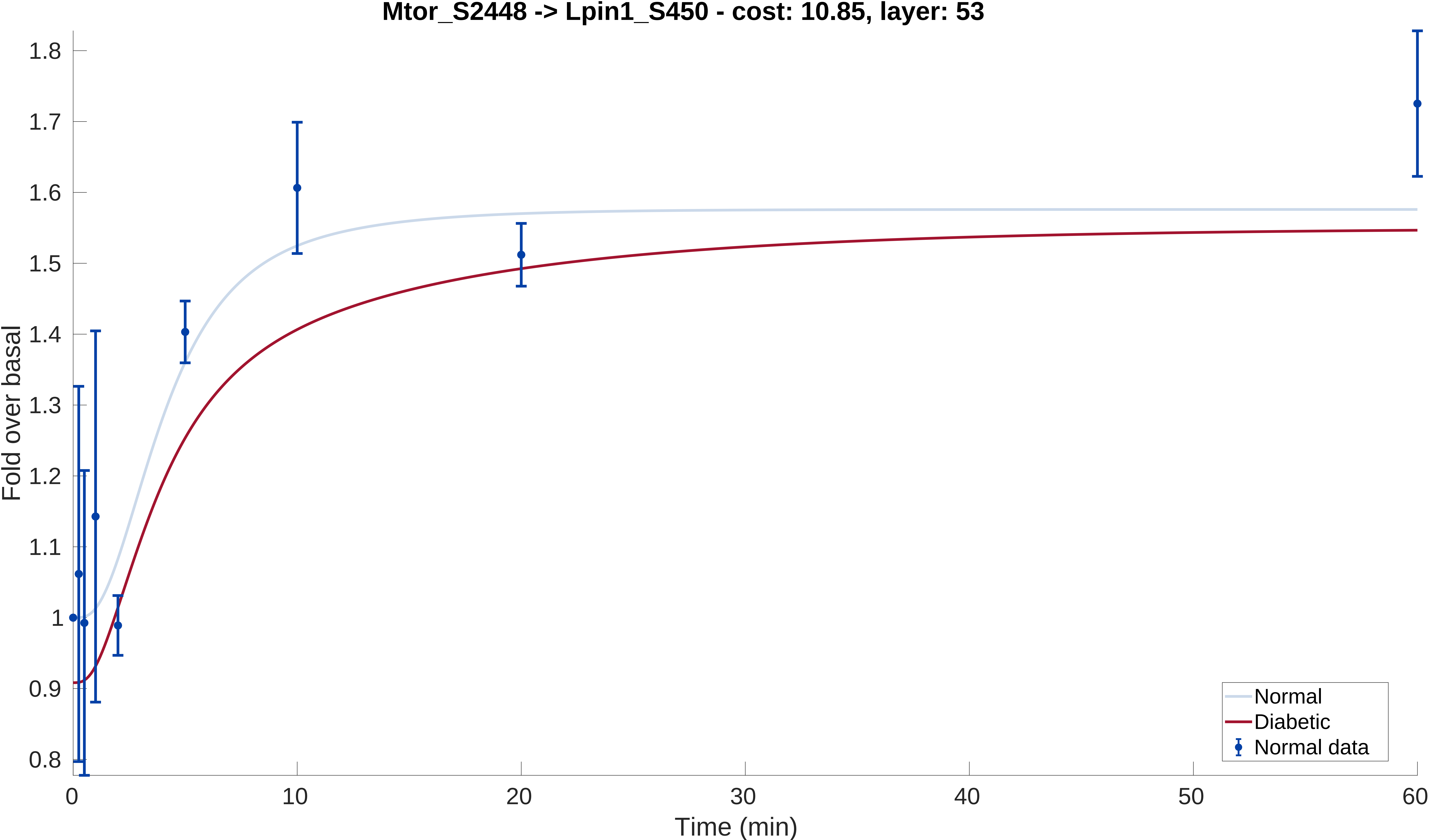

Mtor_S2448 -> Lpin1_S450 - cost: 10.85, layer: 53
Fold over basal
1.8
1.7
1.6
1.5
1.4
1.3
1.2
1.1
1
0.9
0.8
0
10
20
30
40
50
60
Time (min)
Normal
Diabetic
Normal data

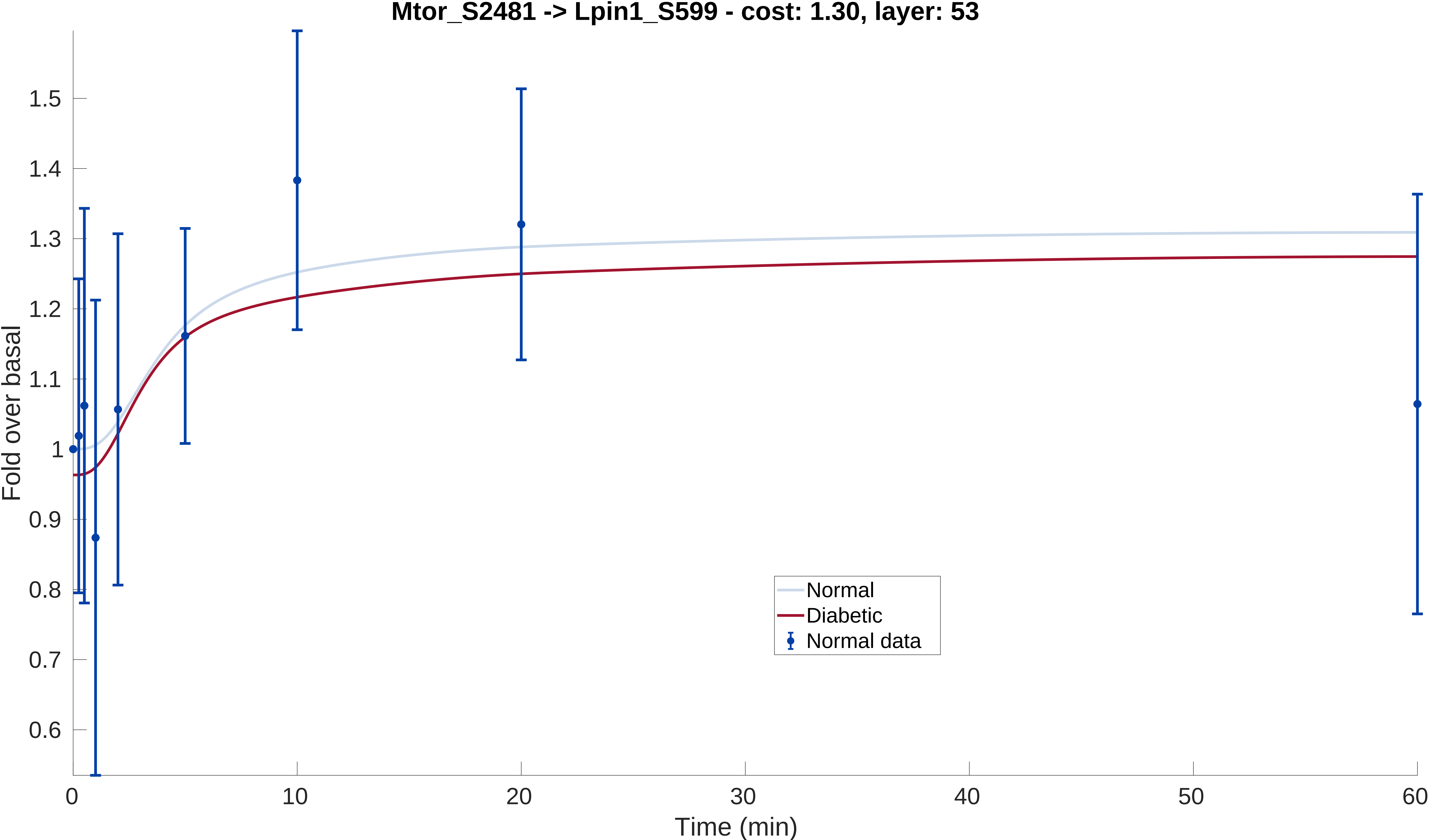

Mtor_S2481 -> Lpin1_S599 - cost: 1.30, layer: 53
Fold over basal
1.5
1.4
1.3
1.2
1.1
1
0.9
0.8
0.7
0.6
0
10
20
30
40
50
60
Time (min)
Normal
Diabetic
Normal data

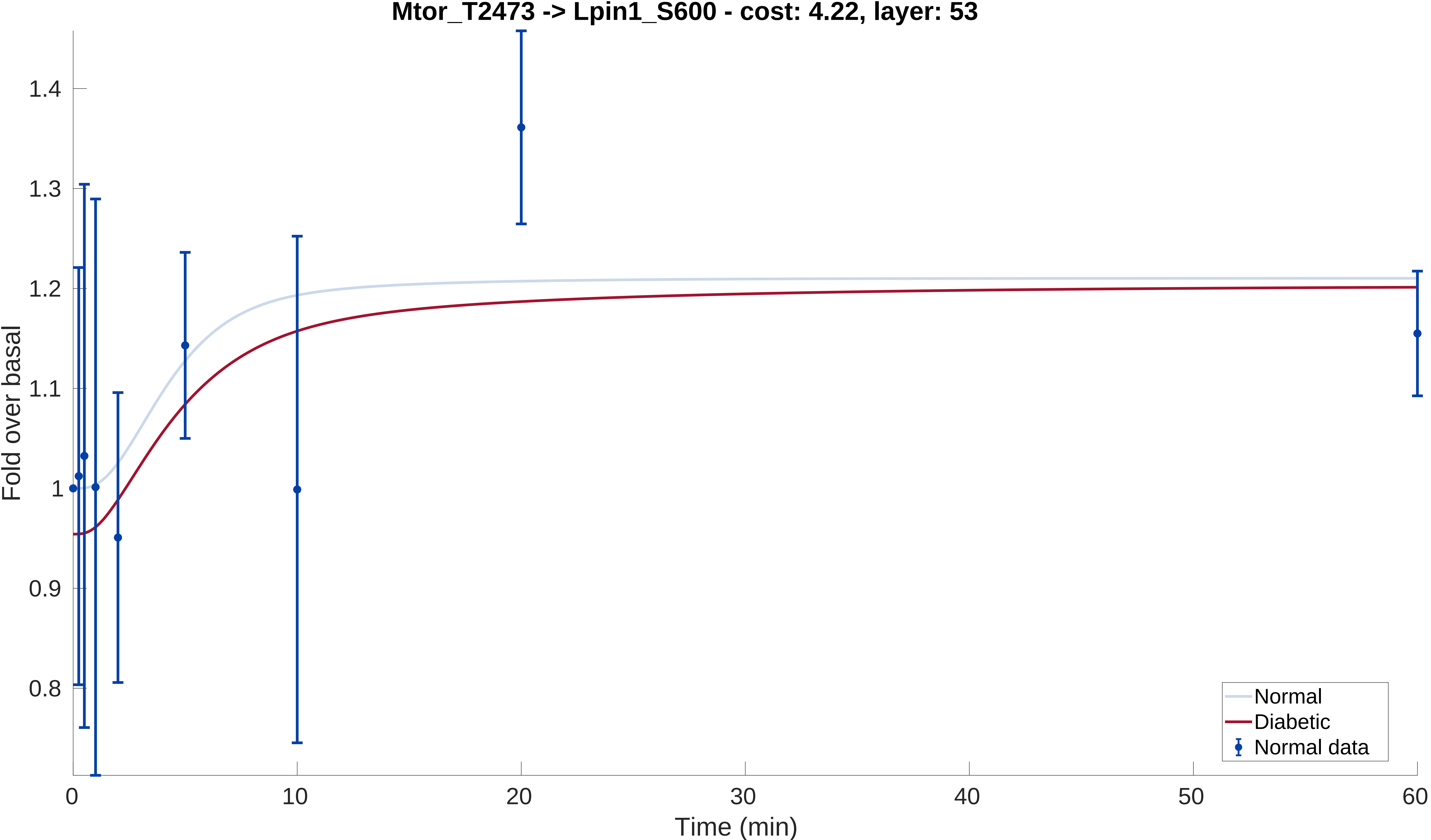
Mtor_T2473 -> Lpin1_S600 - cost: 4.22, layer: 53
Fold over basal
1.4
1.3
1.2
1.1
1
0.9
0.8
0
10
20
30
40
50
60
Time (min)
Normal
Diabetic
Normal data

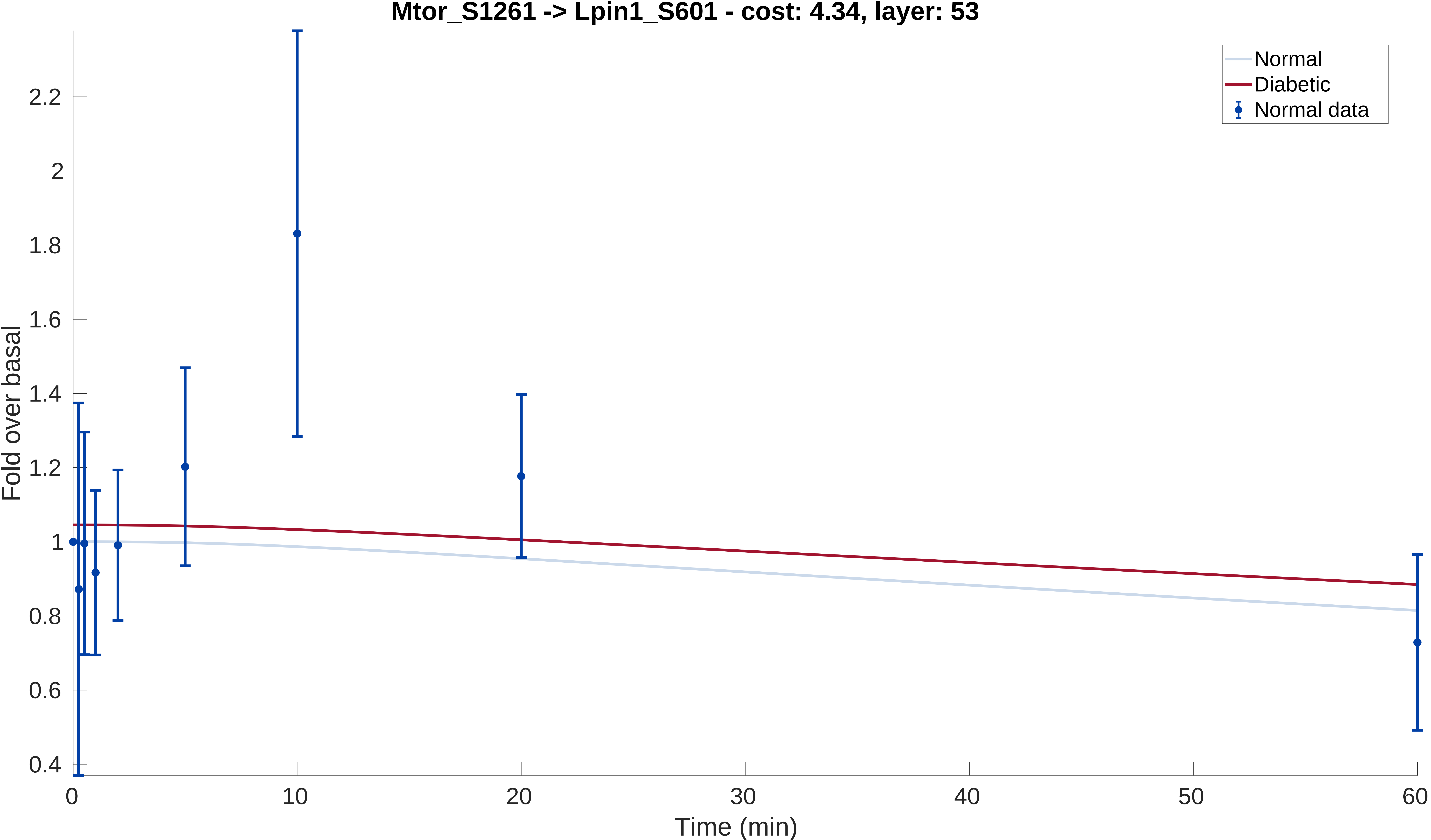

Mtor_S1261 -> Lpin1_S601 - cost: 4.34, layer: 53
Normal
Diabetic
Normal data
Fold over basal
2.2
2
1.8
1.6
1.4
1.2
1
0.8
0.6
0.4
0
10
20
30
40
50
60
Time (min)

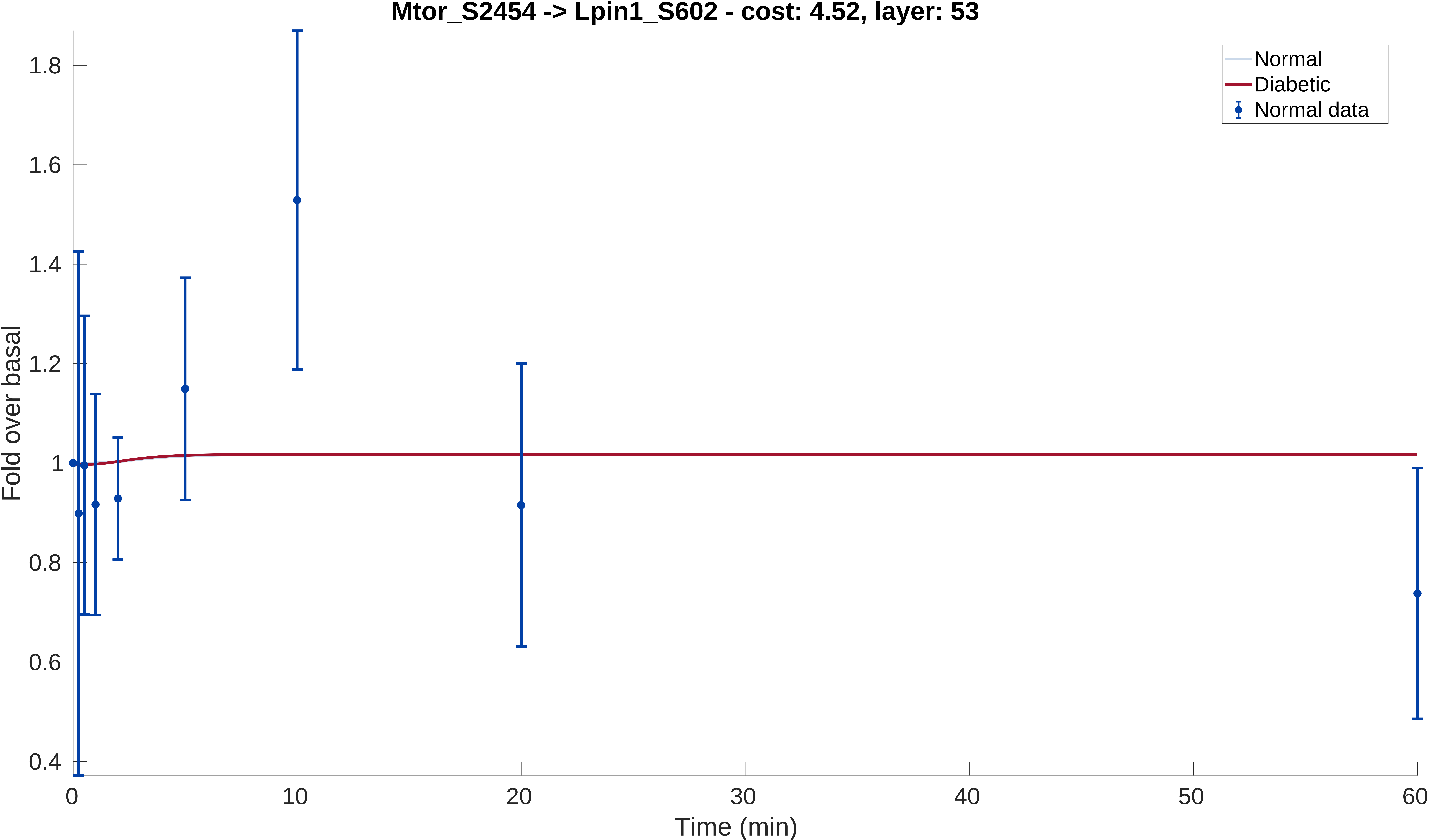

Mtor_S2454 -> Lpin1_S602 - cost: 4.52, layer: 53
Normal
Diabetic
Normal data
Fold over basal
1.8
1.6
1.4
1.2
1
0.8
0.6
0.4
0
10
20
30
40
50
60
Time (min)

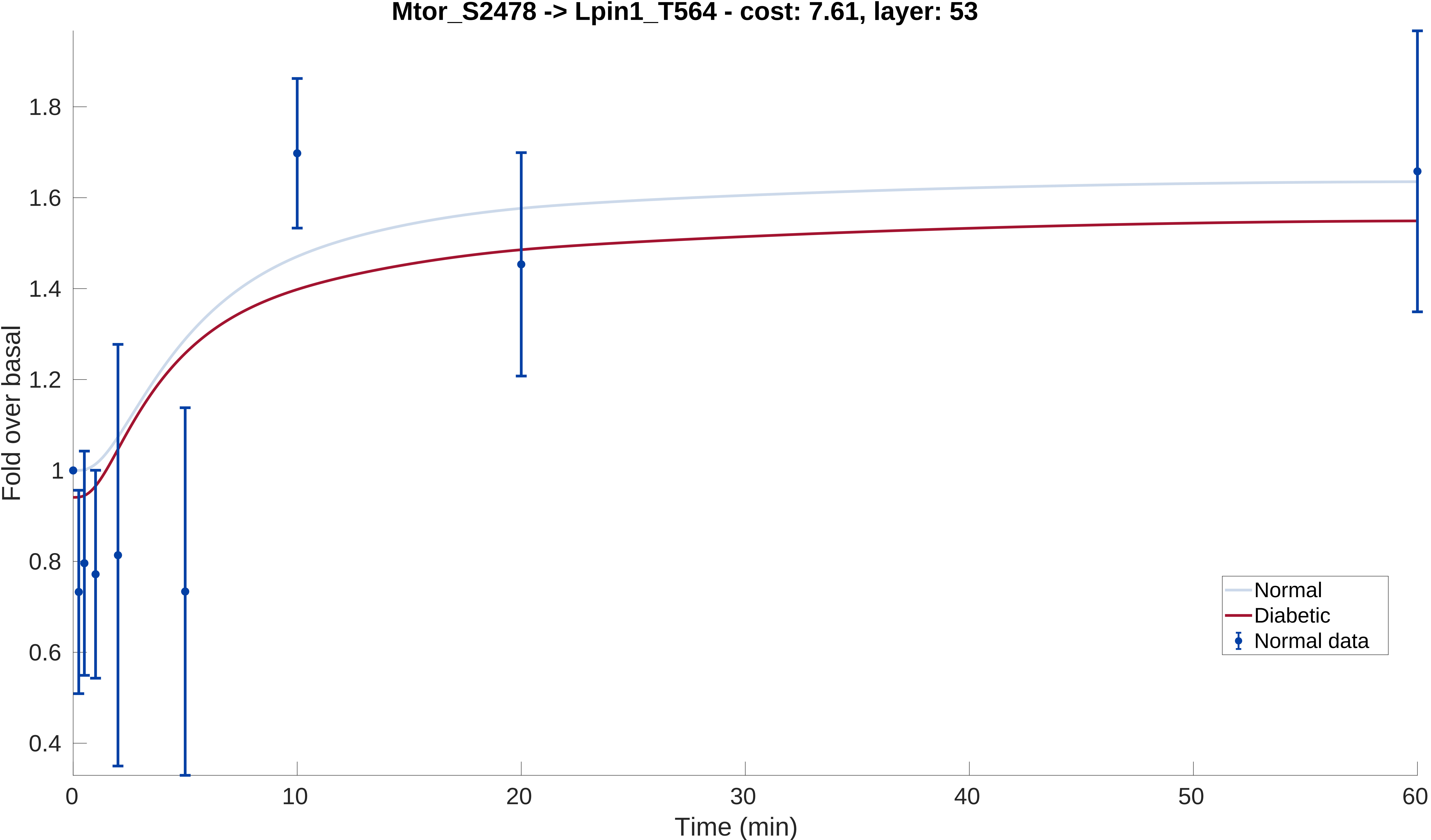

Mtor_S2478 -> Lpin1_T564 - cost: 7.61, layer: 53
Fold over basal
1.8
1.6
1.4
1.2
1
0.8
0.6
0.4
0
10
20
30
40
50
60
Time (min)
Normal
Diabetic
Normal data

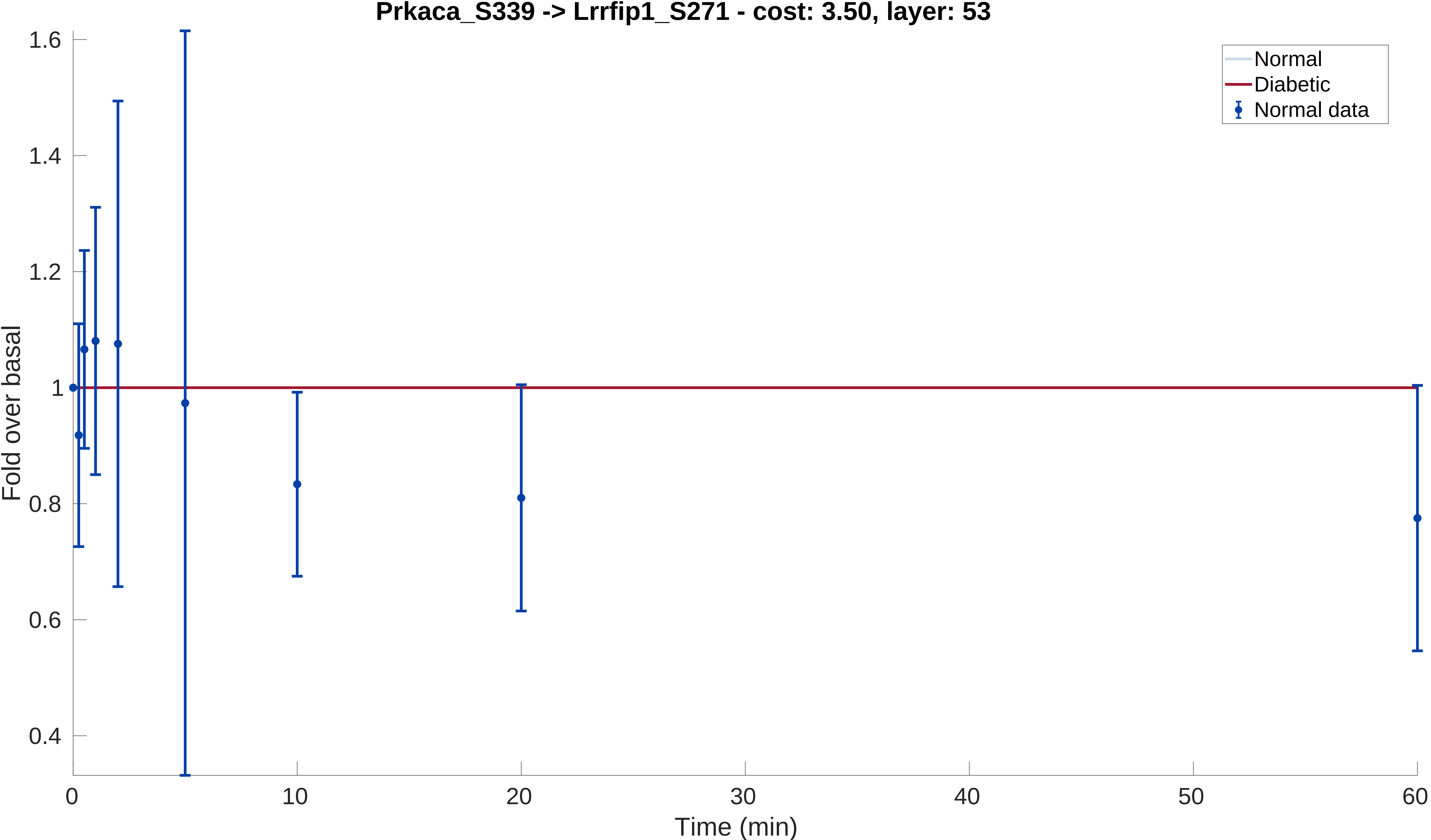
Prkaca_S339 -> Lrrfip1_S271 - cost: 3.50, layer: 53
Normal
Diabetic
Normal data
Fold over basal
Time (min)
1.6
1.4
1.2
1
0.8
0.6
0.4
0
10
20
30
40
50
60

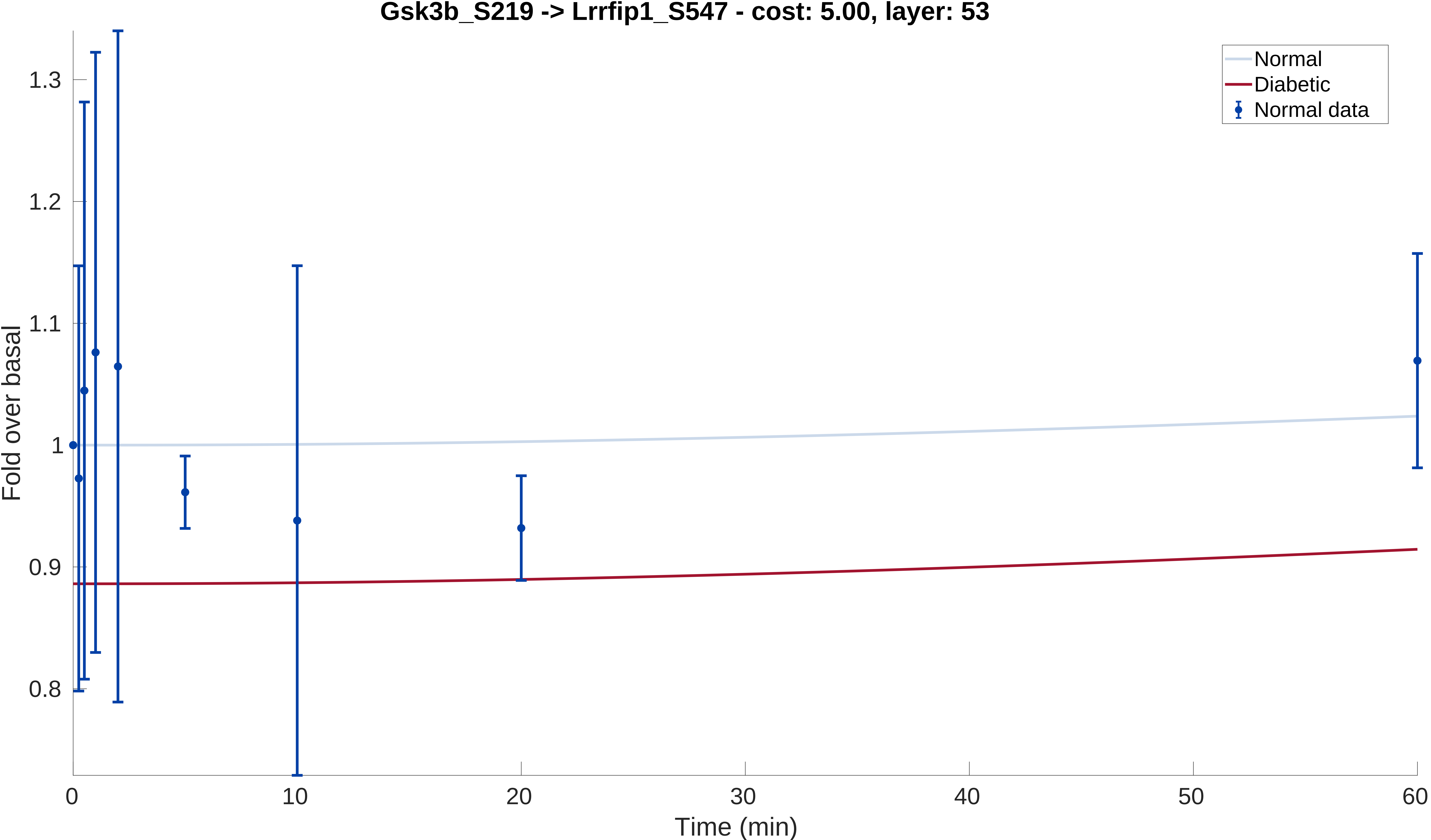

Gsk3b_S219 -> Lrrfip1_S547 - cost: 5.00, layer: 53
Normal
Diabetic
Normal data
Fold over basal
Time (min)
1.3
1.2
1.1
1
0.9
0.8
0
10
20
30
40
50
60

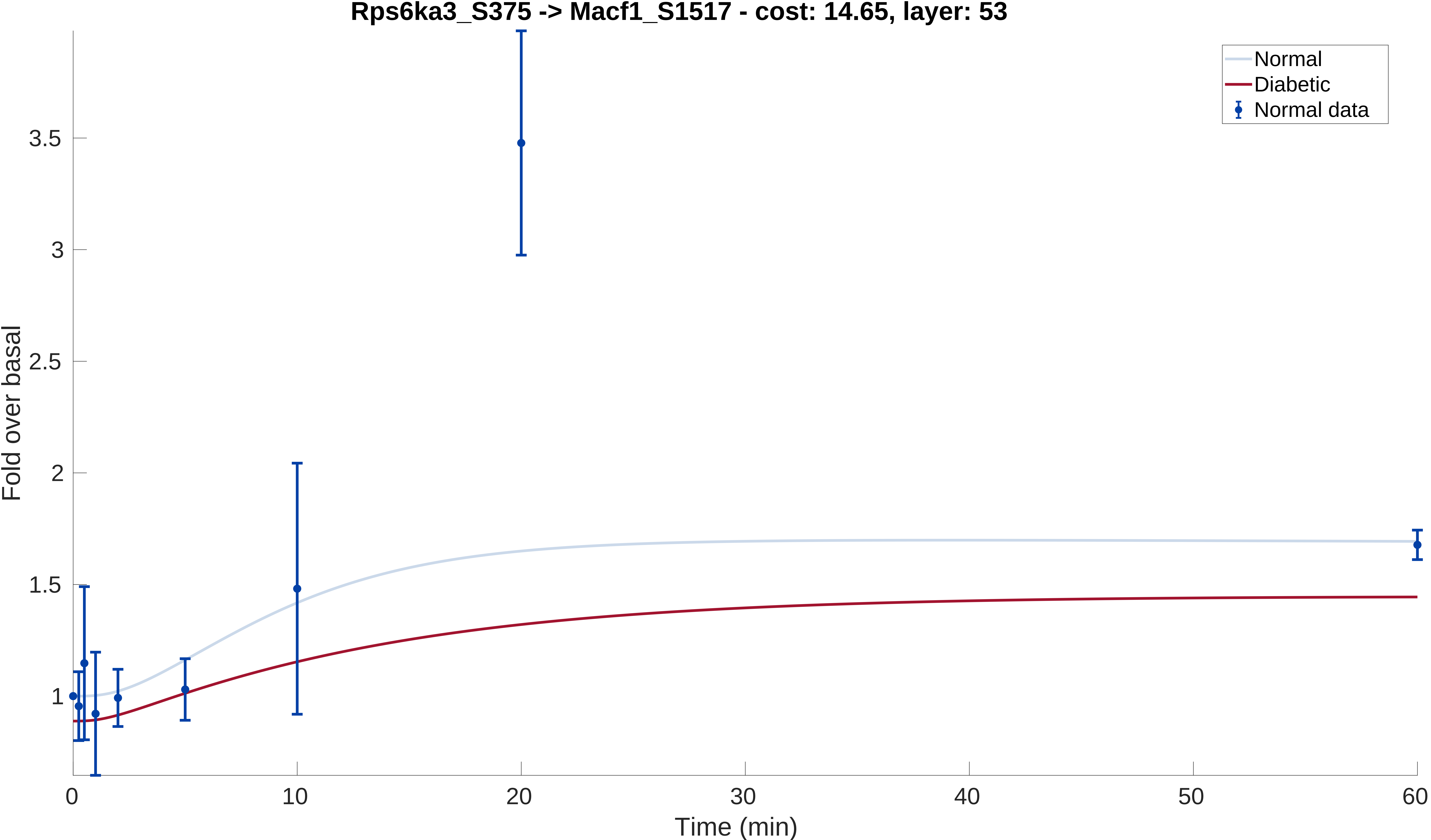

Rps6ka3_S375 -> Macf1_S1517 - cost: 14.65, layer: 53
Normal
Diabetic
Normal data
Fold over basal
3.5
3
2.5
2
1.5
1
0
10
20
30
40
50
60
Time (min)

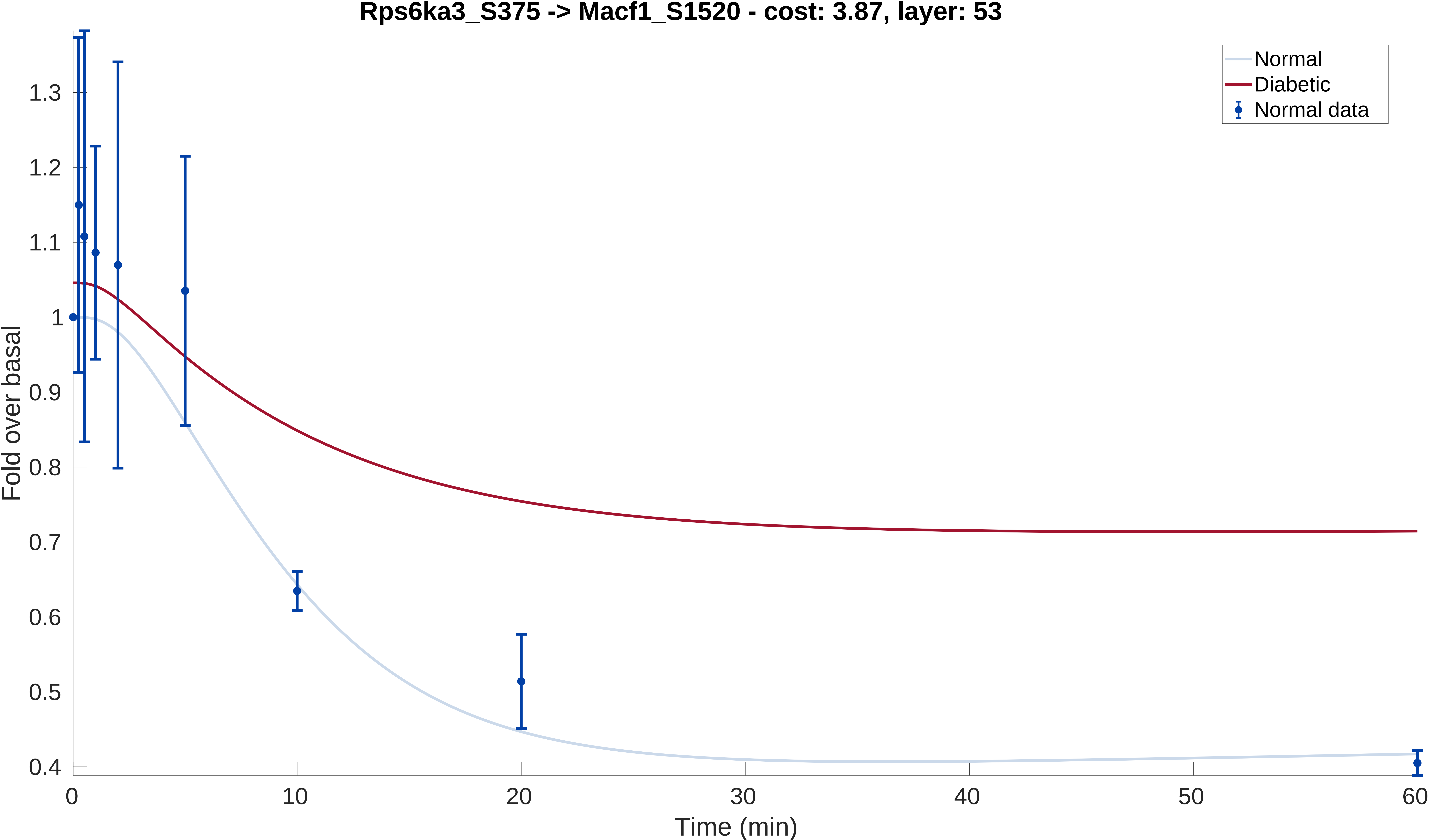

Rps6ka3_S375 -> Macf1_S1520 - cost: 3.87, layer: 53
Normal
Diabetic
Normal data
Fold over basal
Time (min)
0
10
20
30
40
50
60
0.4
0.5
0.6
0.7
0.8
0.9
1
1.1
1.2
1.3

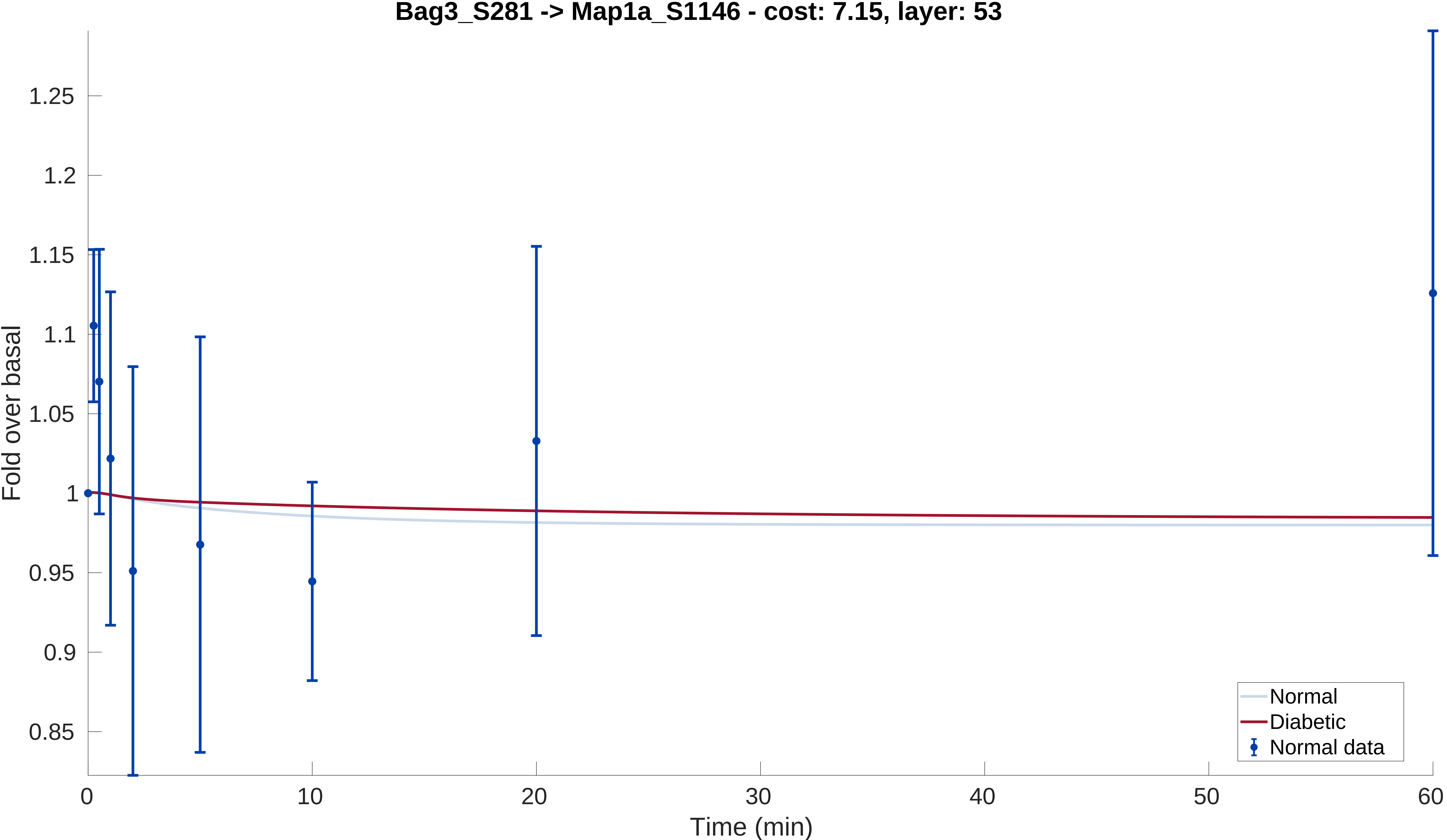

Bag3_S281 -> Map1a_S1146 - cost: 7.15, layer: 53
Fold over basal
1.25
1.2
1.15
1.1
1.05
1
0.95
0.9
0.85
0
10
20
30
40
50
60
Time (min)
Normal
Diabetic
Normal data

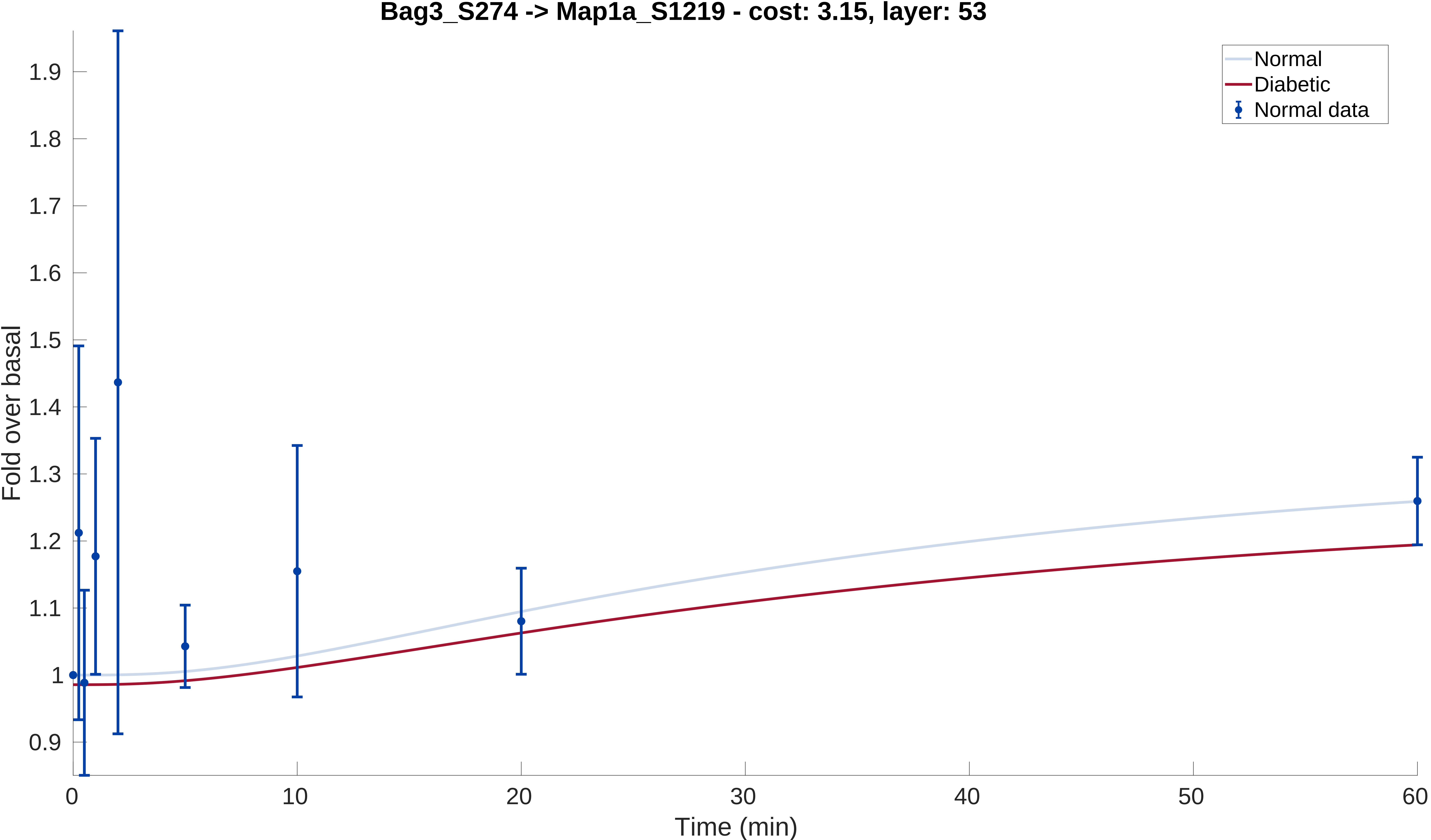

Bag3_S274 -> Map1a_S1219 - cost: 3.15, layer: 53
Normal
Diabetic
Normal data
Fold over basal
1.9
1.8
1.7
1.6
1.5
1.4
1.3
1.2
1.1
1
0.9
0
10
20
30
40
50
60
Time (min)

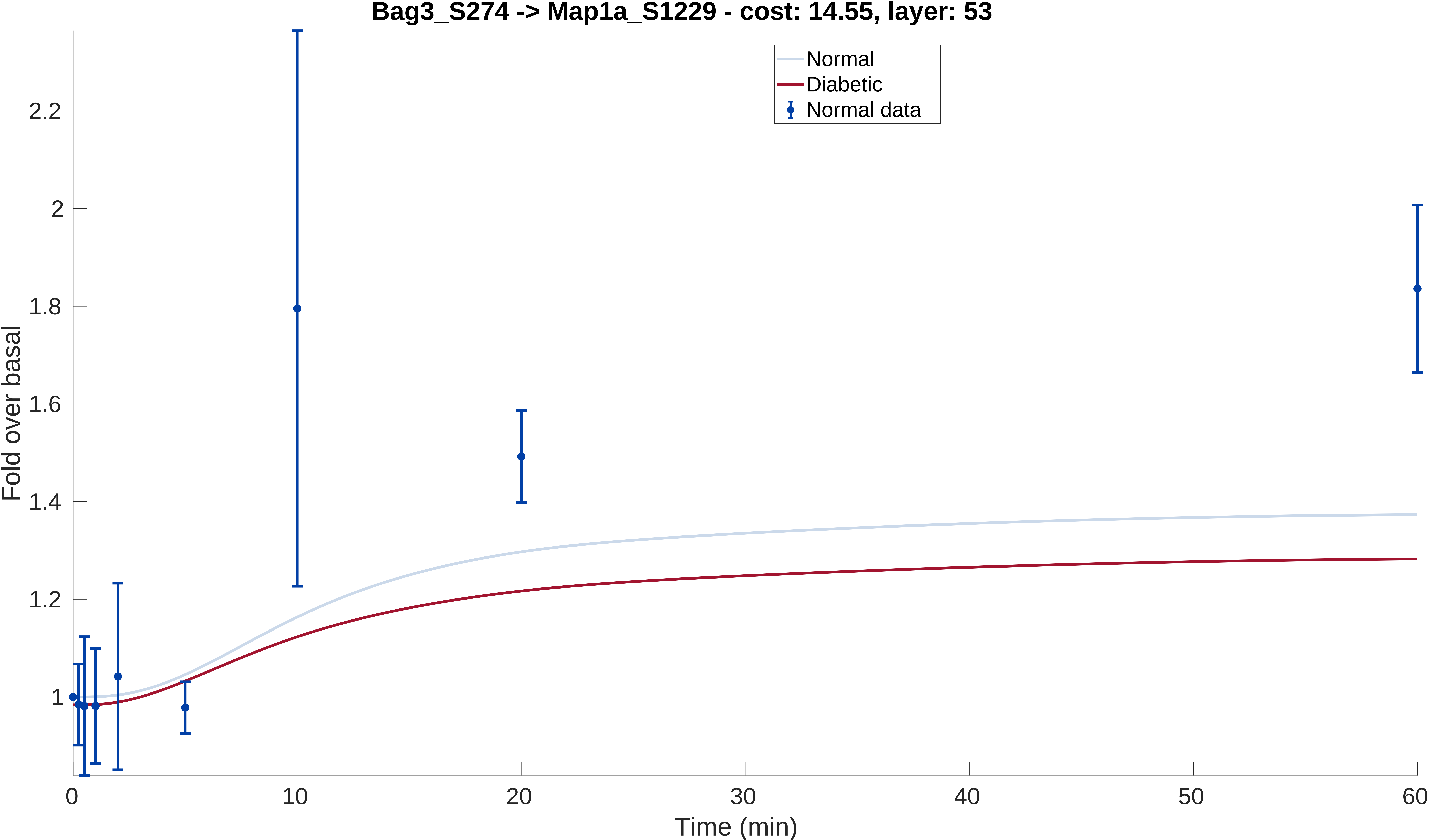

Bag3_S274 -> Map1a_S1229 - cost: 14.55, layer: 53
Normal
Diabetic
Normal data
Fold over basal
Time (min)
2.2
2
1.8
1.6
1.4
1.2
1
0
10
20
30
40
50
60

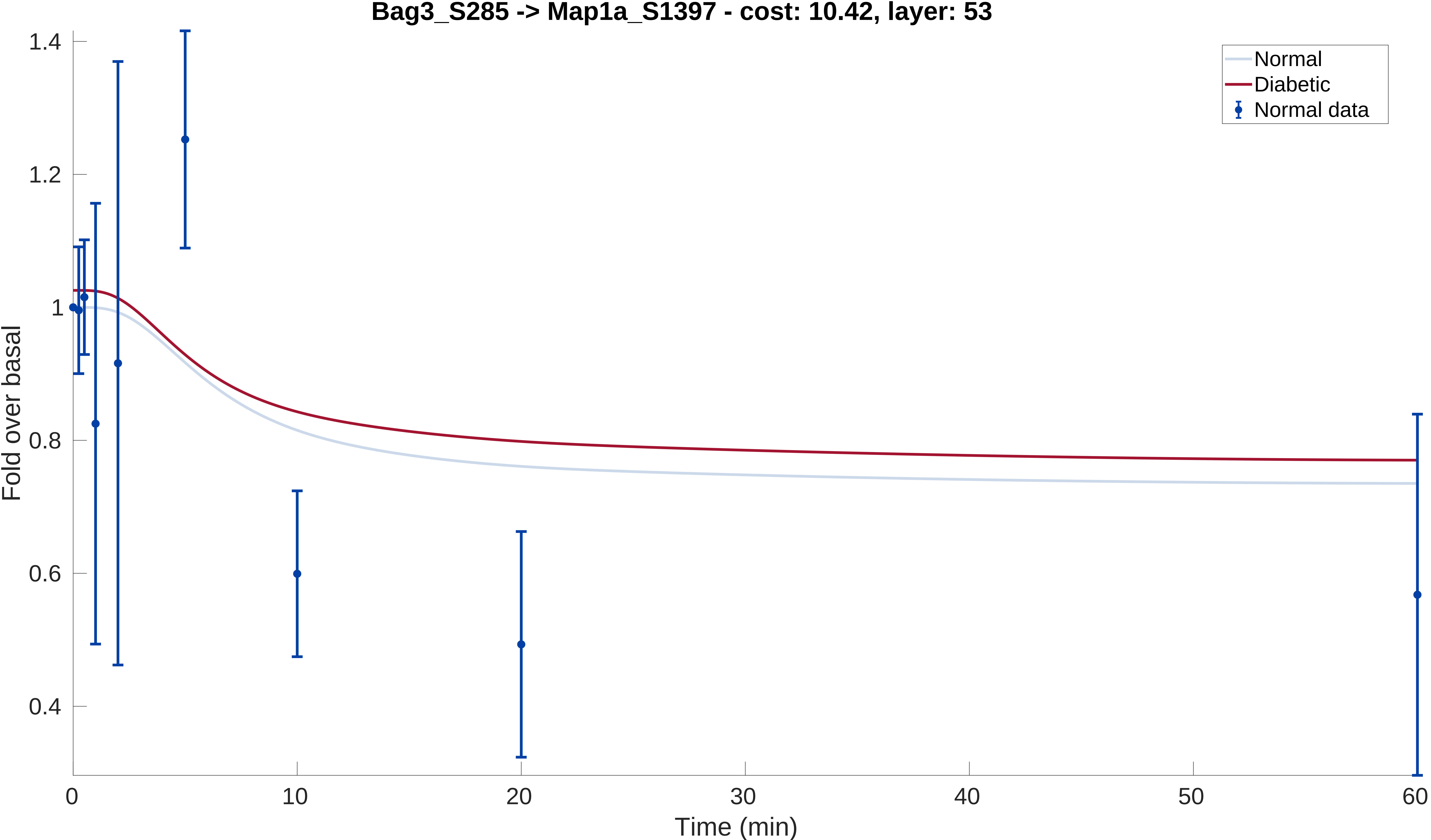

Bag3_S285 -> Map1a_S1397 - cost: 10.42, layer: 53
Normal
Diabetic
Normal data
Fold over basal
1.4
1.2
1
0.8
0.6
0.4
0
10
20
30
40
50
60
Time (min)

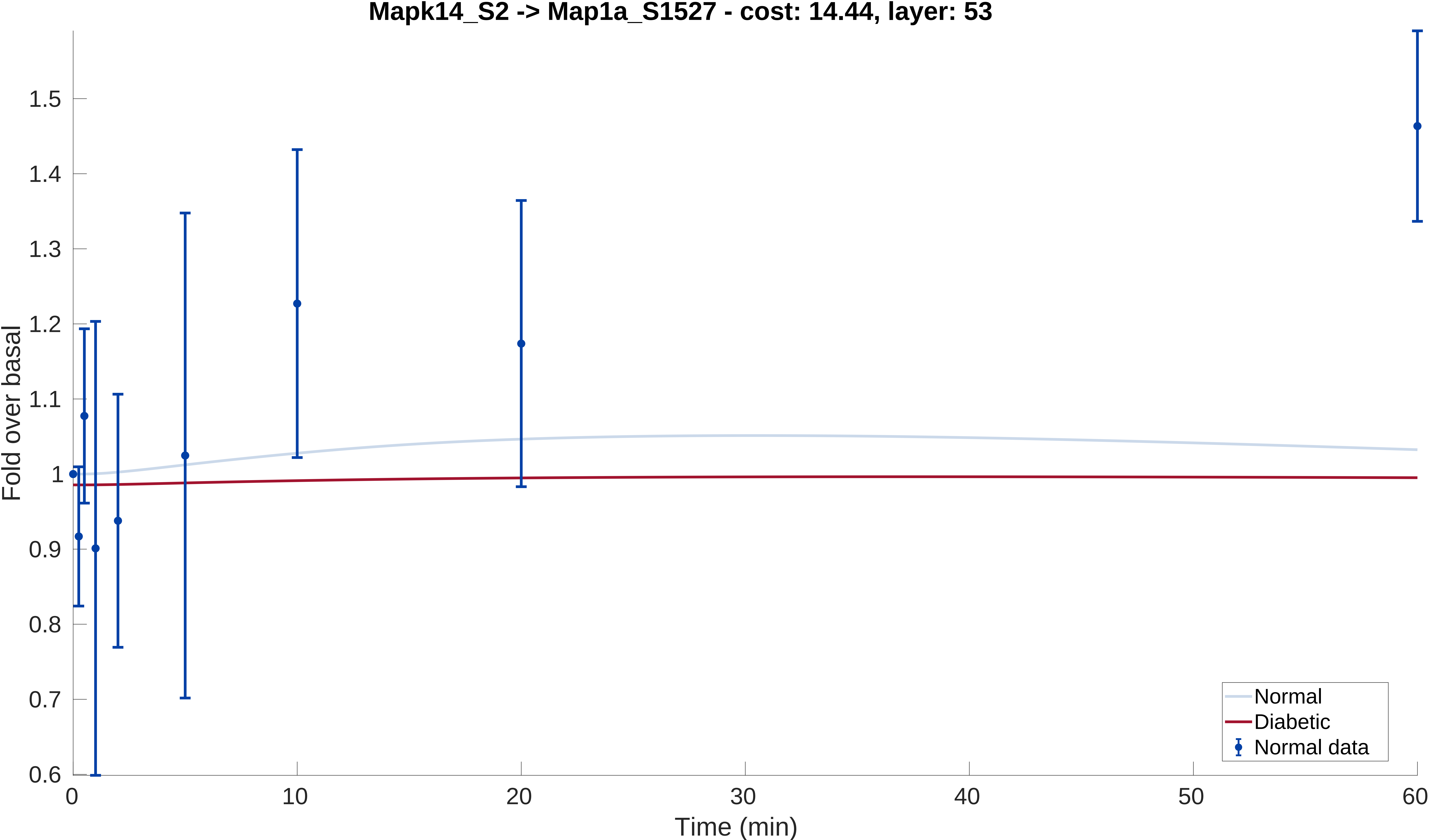

Mapk14_S2 -> Map1a_S1527 - cost: 14.44, layer: 53
Fold over basal
1.5
1.4
1.3
1.2
1.1
1
0.9
0.8
0.7
0.6
0
10
20
30
40
50
60
Time (min)
Normal
Diabetic
Normal data

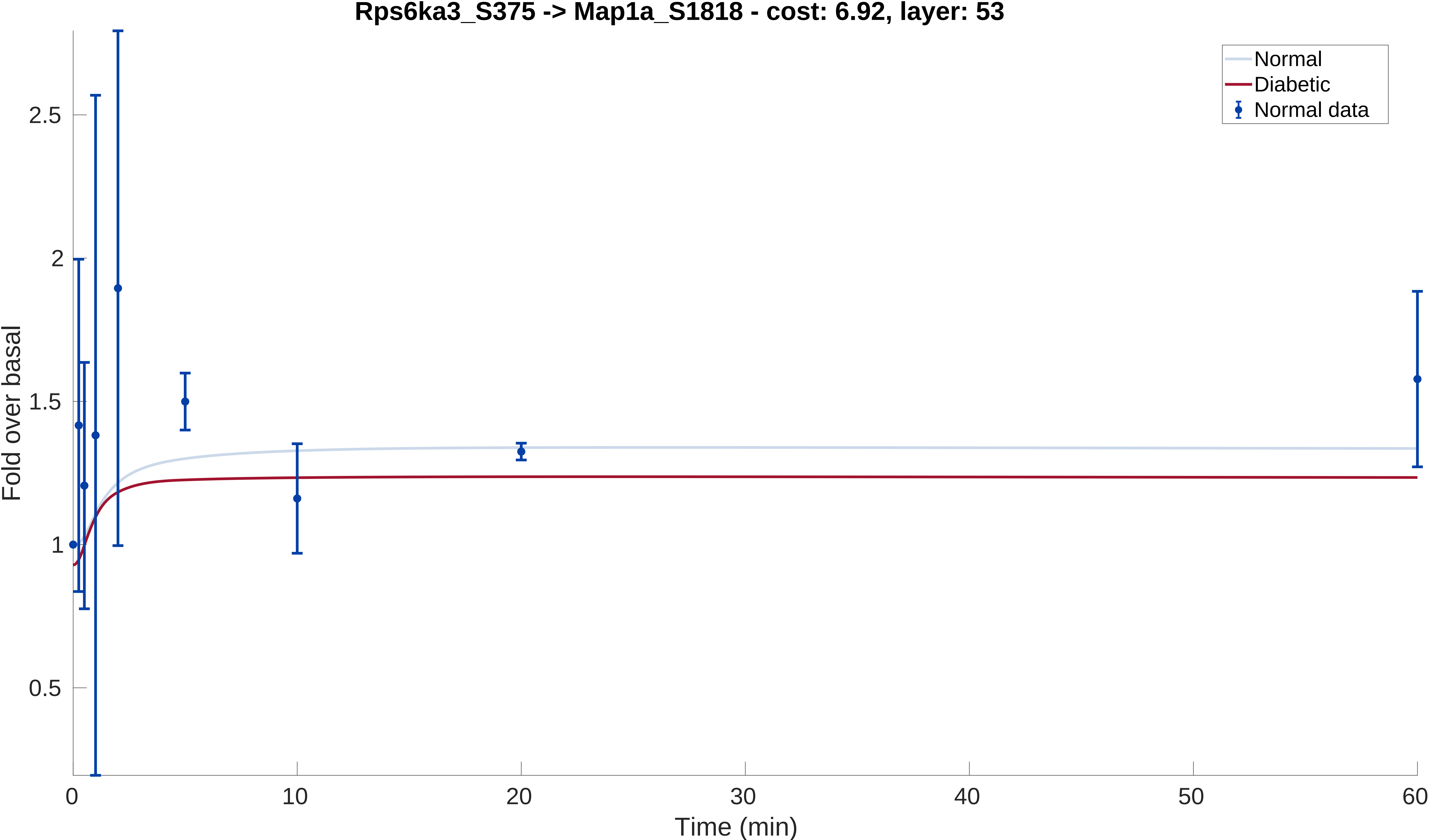
Rps6ka3_S375 -> Map1a_S1818 - cost: 6.92, layer: 53
Normal
Diabetic
Normal data
Fold over basal
2.5
2
1.5
1
0.5
0
10
20
30
40
50
60
Time (min)

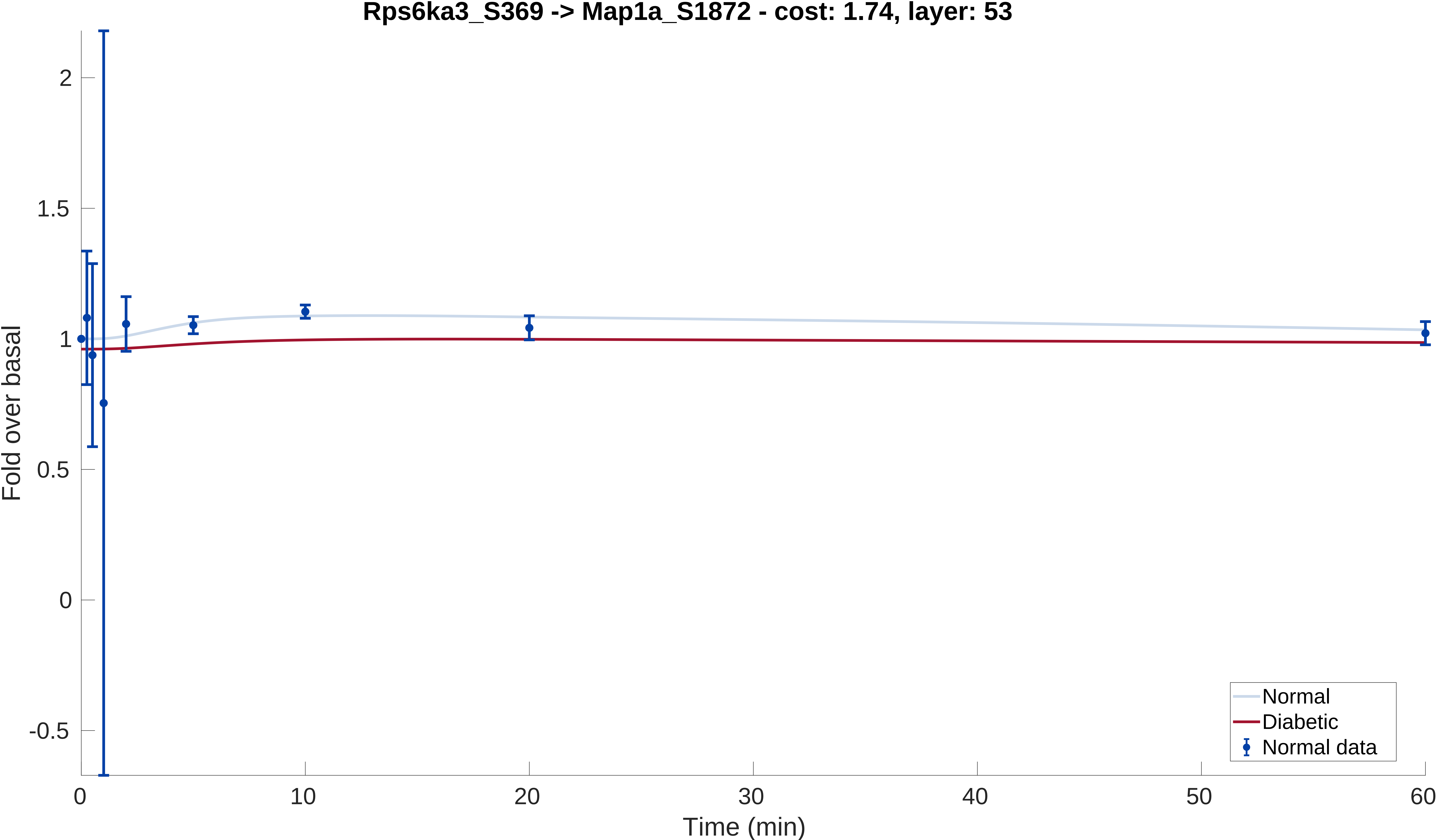

Rps6ka3_S369 -> Map1a_S1872 - cost: 1.74, layer: 53
Fold over basal
Time (min)
2
1.5
1
0.5
0
-0.5
0
10
20
30
40
50
60
Normal
Diabetic
Normal data

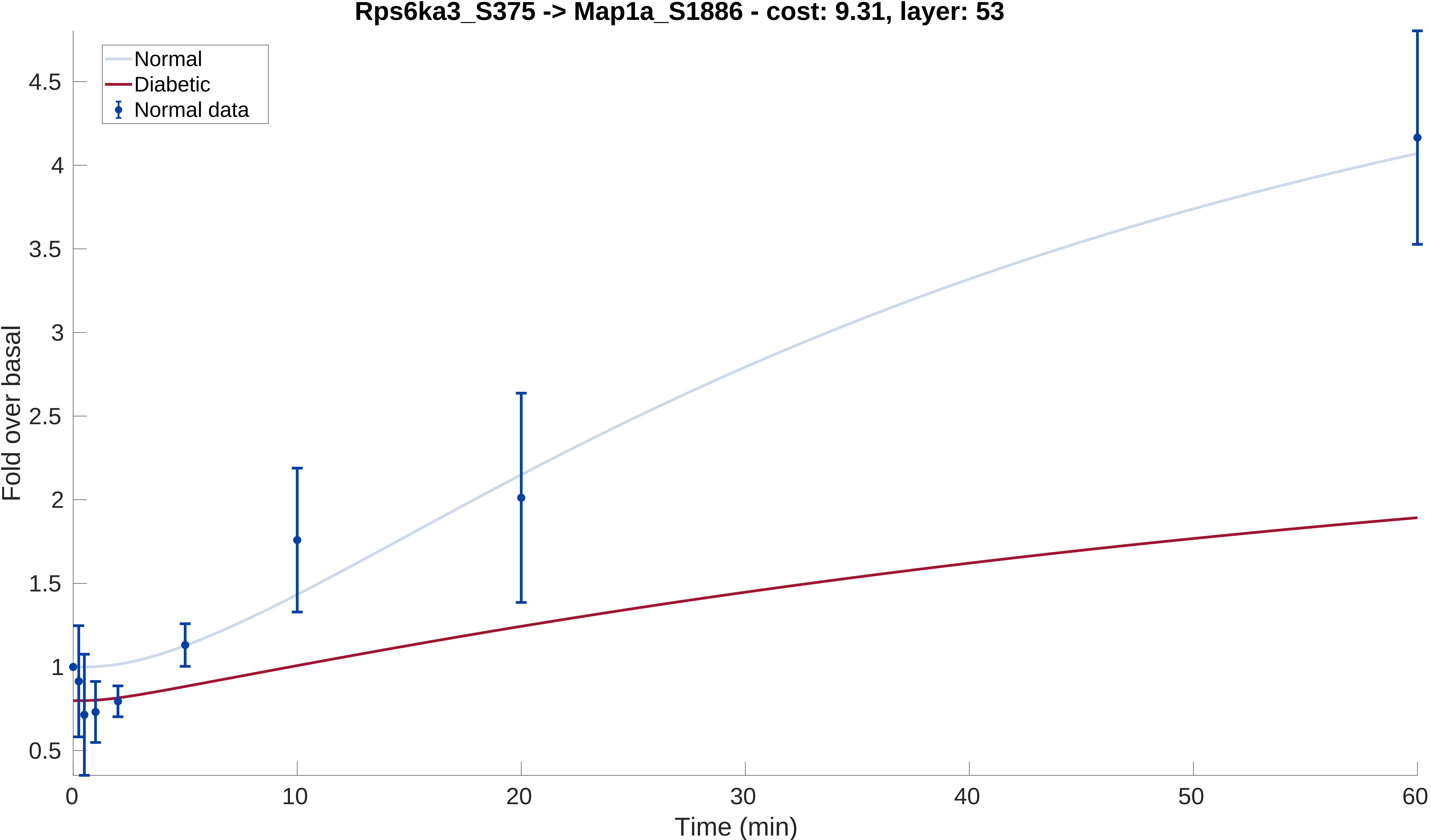
Rps6ka3_S375 -> Map1a_S1886 - cost: 9.31, layer: 53
Normal
Diabetic
Normal data
Fold over basal
4.5
4
3.5
3
2.5
2
1.5
1
0.5
0
10
20
30
40
50
60
Time (min)

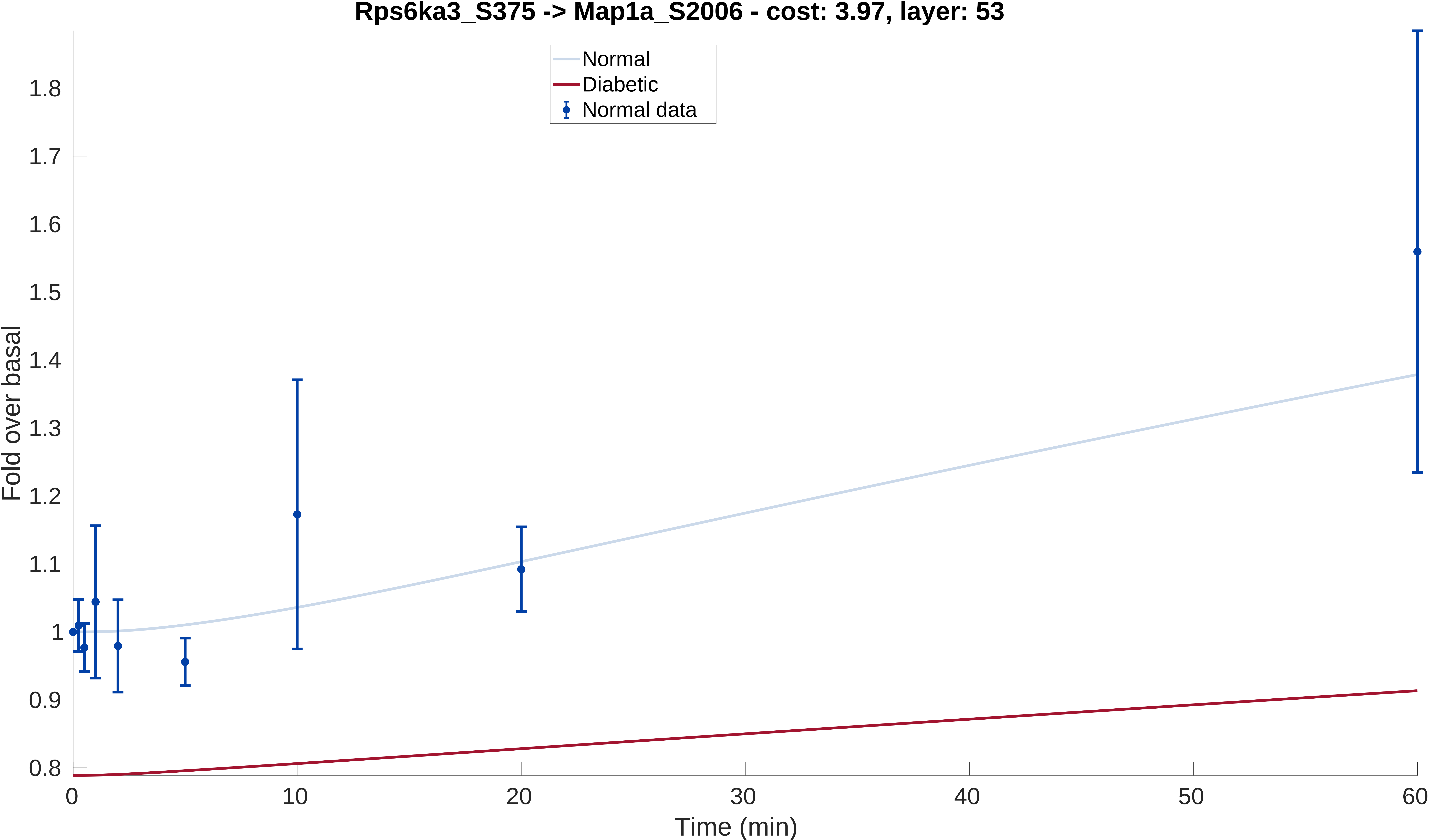

Rps6ka3_S375 -> Map1a_S2006 - cost: 3.97, layer: 53
Normal
Diabetic
Normal data
Fold over basal
1.8
1.7
1.6
1.5
1.4
1.3
1.2
1.1
1
0.9
0.8
0
10
20
30
40
50
60
Time (min)

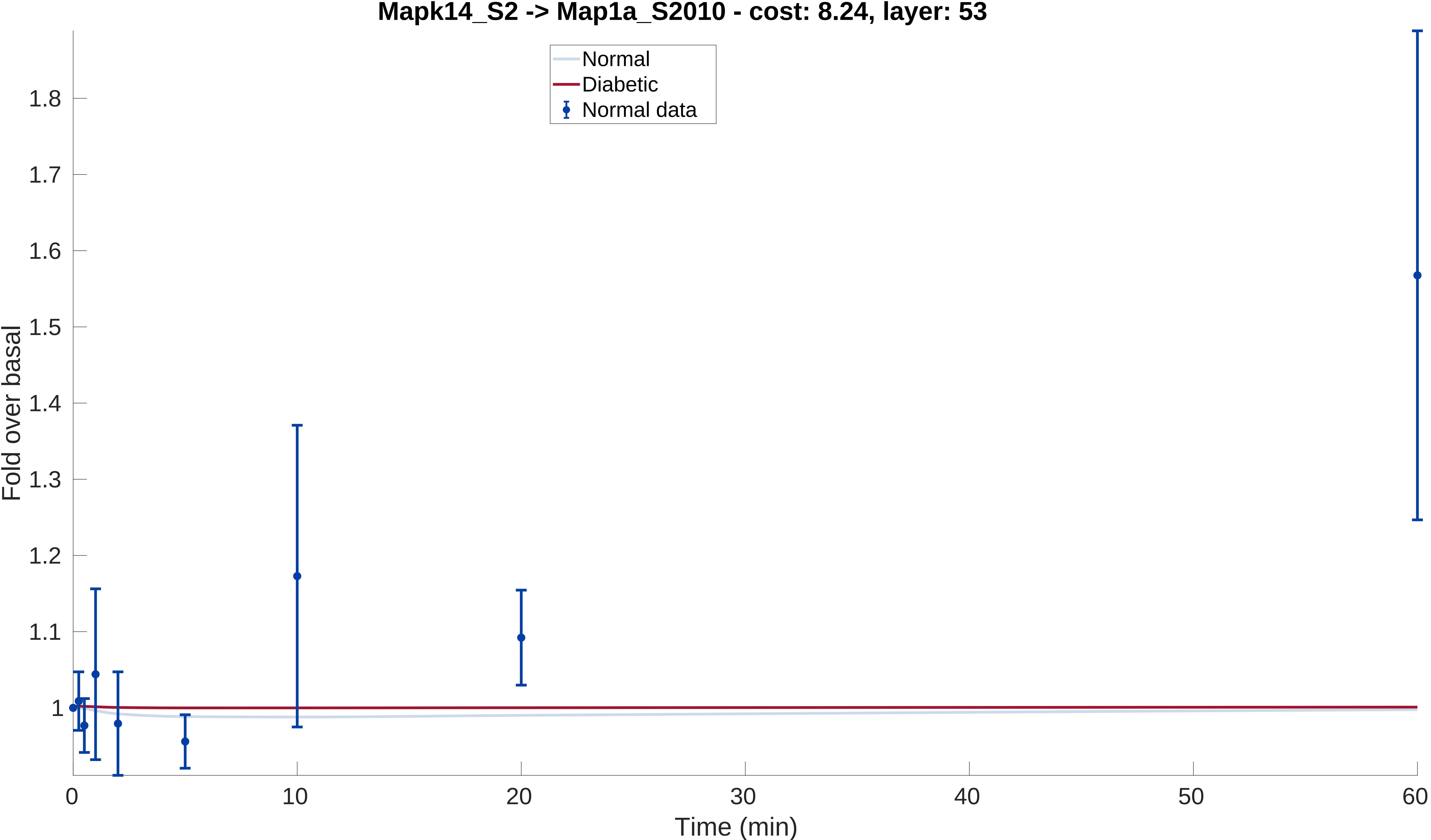
Mapk14_S2 -> Map1a_S2010 - cost: 8.24, layer: 53
Normal
Diabetic
Normal data
Fold over basal
1.8
1.7
1.6
1.5
1.4
1.3
1.2
1.1
1
0
10
20
30
40
50
60
Time (min)

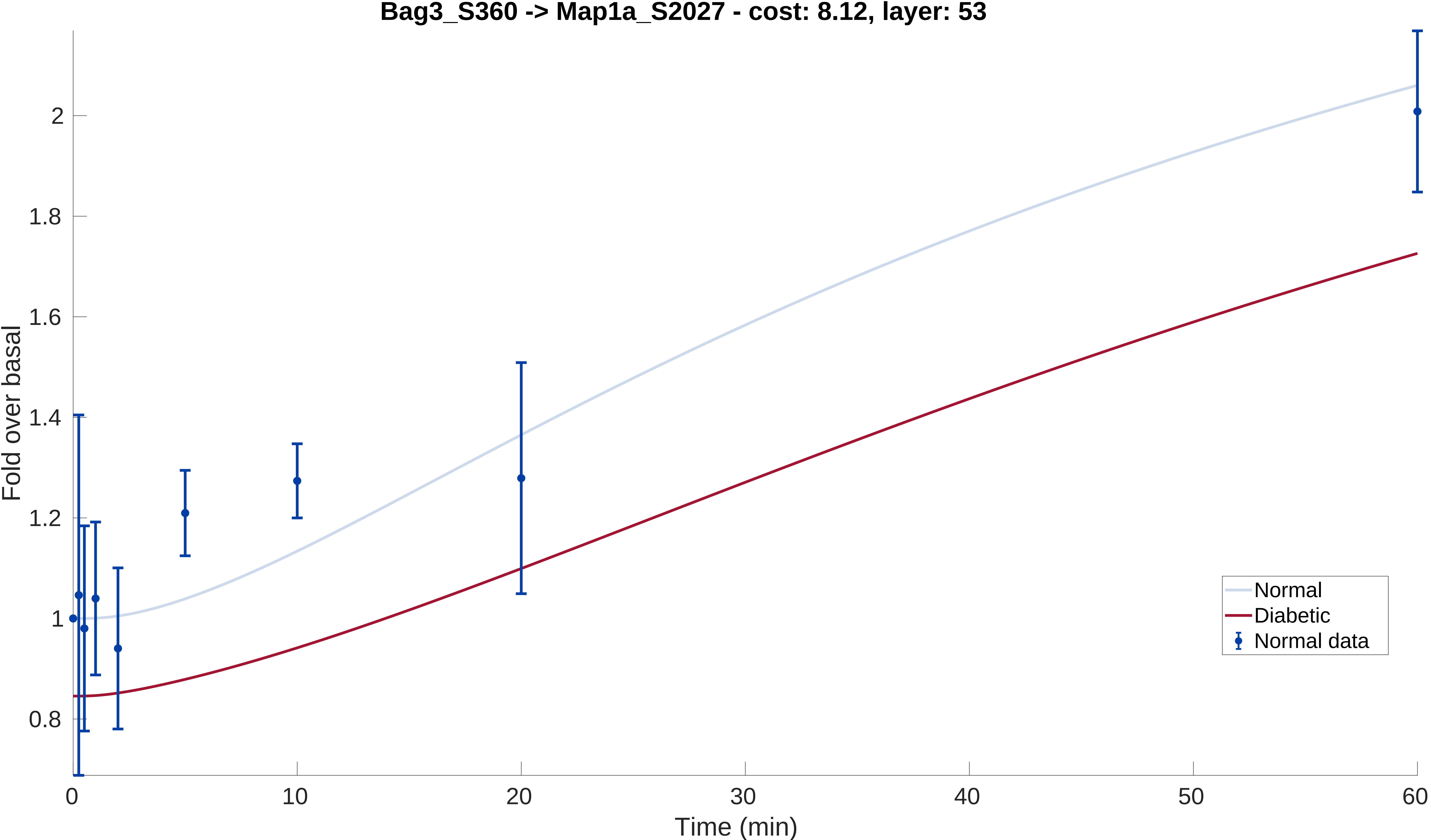

Bag3_S360 -> Map1a_S2027 - cost: 8.12, layer: 53
Fold over basal
Time (min)
0.8
1
1.2
1.4
1.6
1.8
2
0
10
20
30
40
50
60
Normal
Diabetic
Normal data

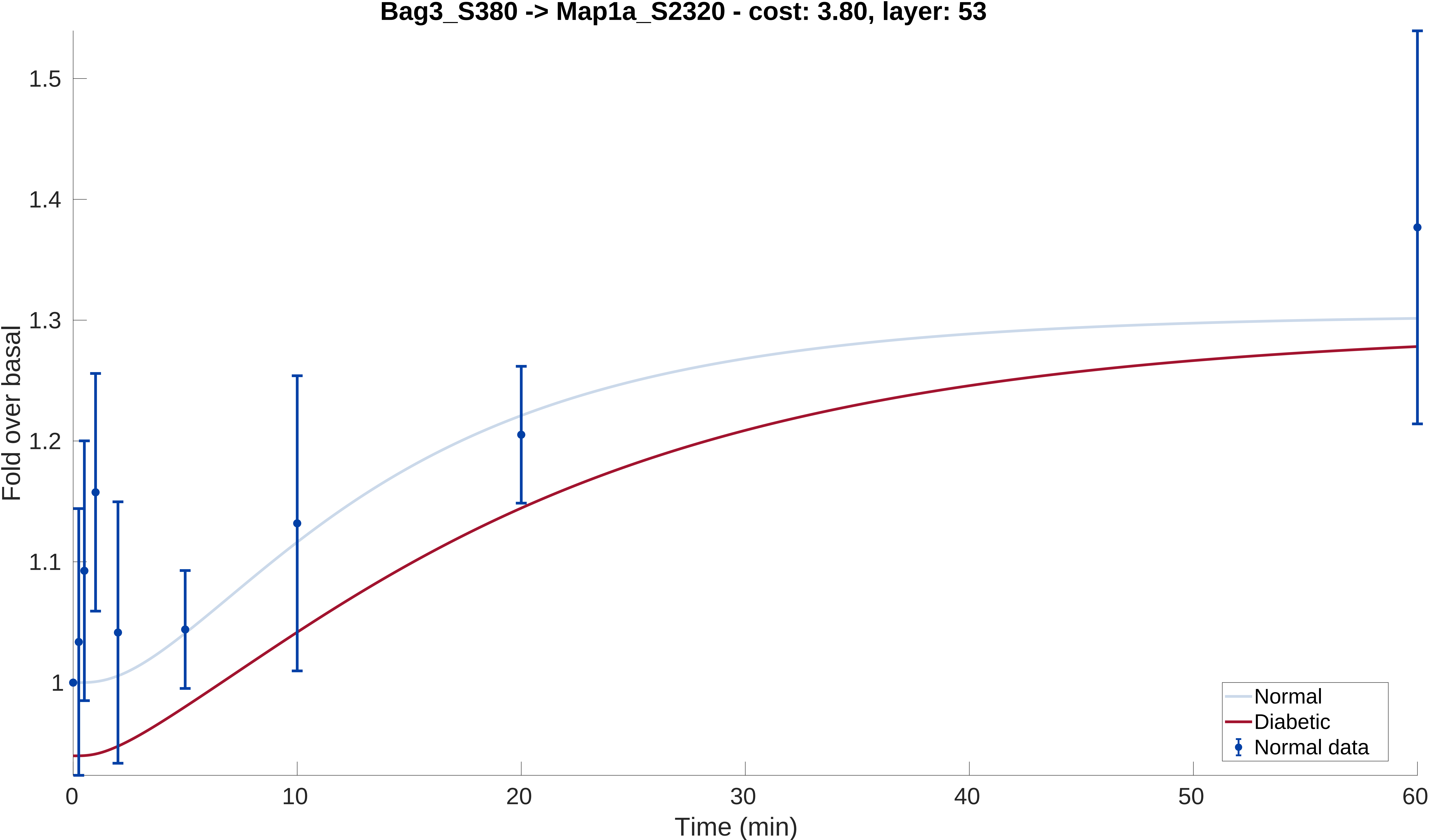
Bag3_S380 -> Map1a_S2320 - cost: 3.80, layer: 53
Fold over basal
1.5
1.4
1.3
1.2
1.1
1
0
10
20
30
40
50
60
Time (min)
Normal
Diabetic
Normal data

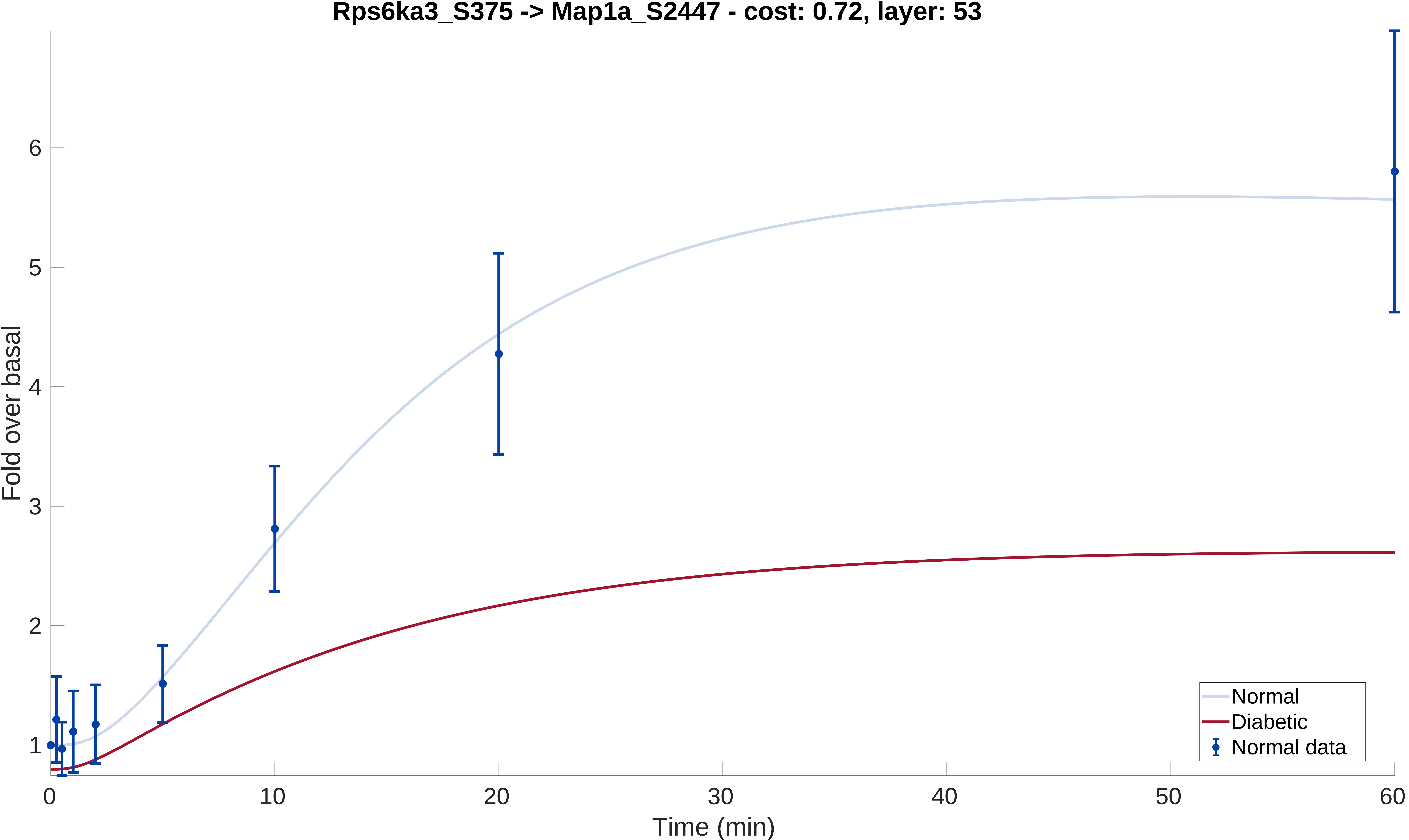

Rps6ka3_S375 -> Map1a_S2447 - cost: 0.72, layer: 53
Fold over basal
1
2
3
4
5
6
0
10
20
30
40
50
60
Time (min)
Normal
Diabetic
Normal data

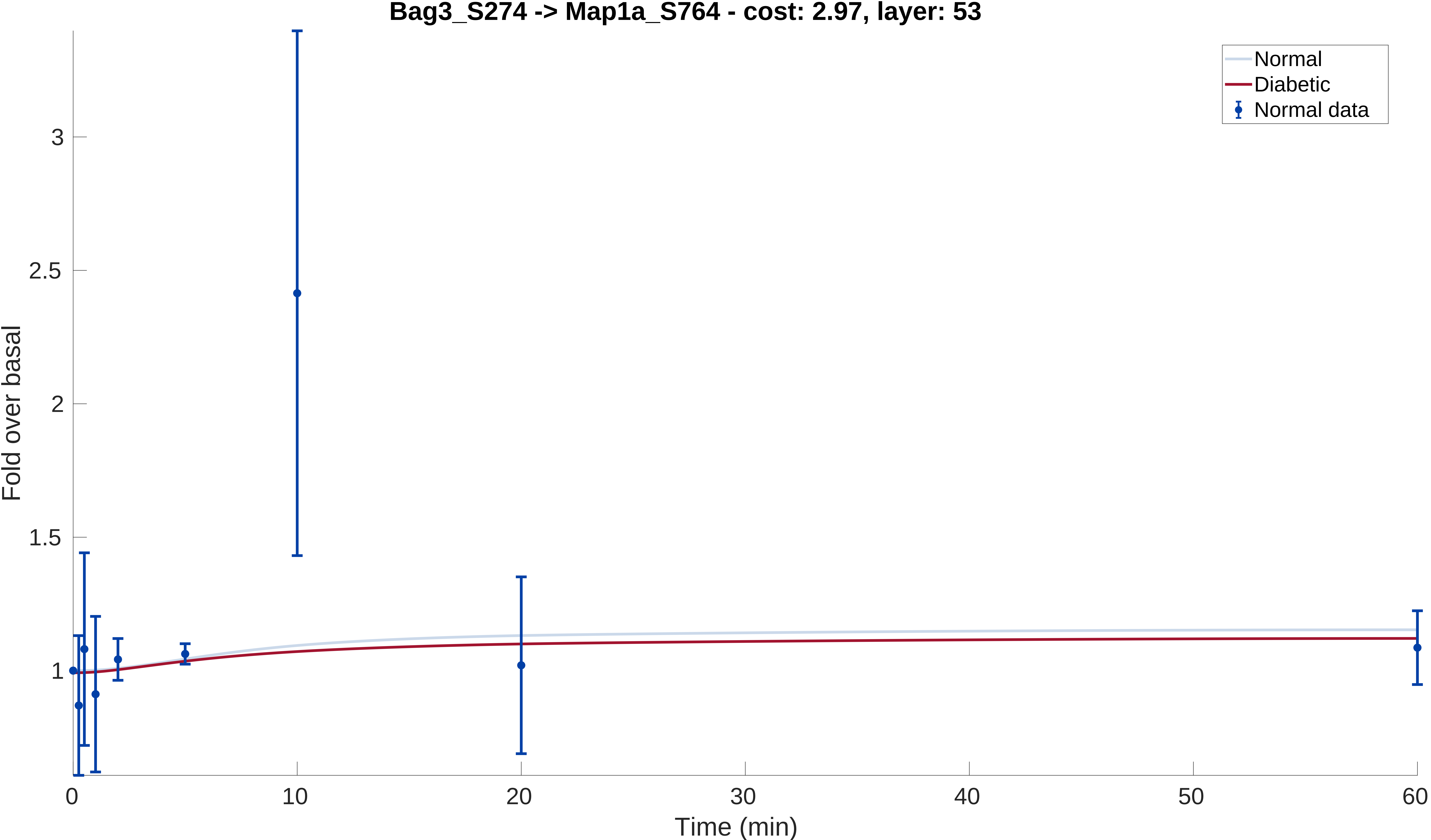
Bag3_S274 -> Map1a_S764 - cost: 2.97, layer: 53
Normal
Diabetic
Normal data
Fold over basal
3
2.5
2
1.5
1
0
10
20
30
40
50
60
Time (min)

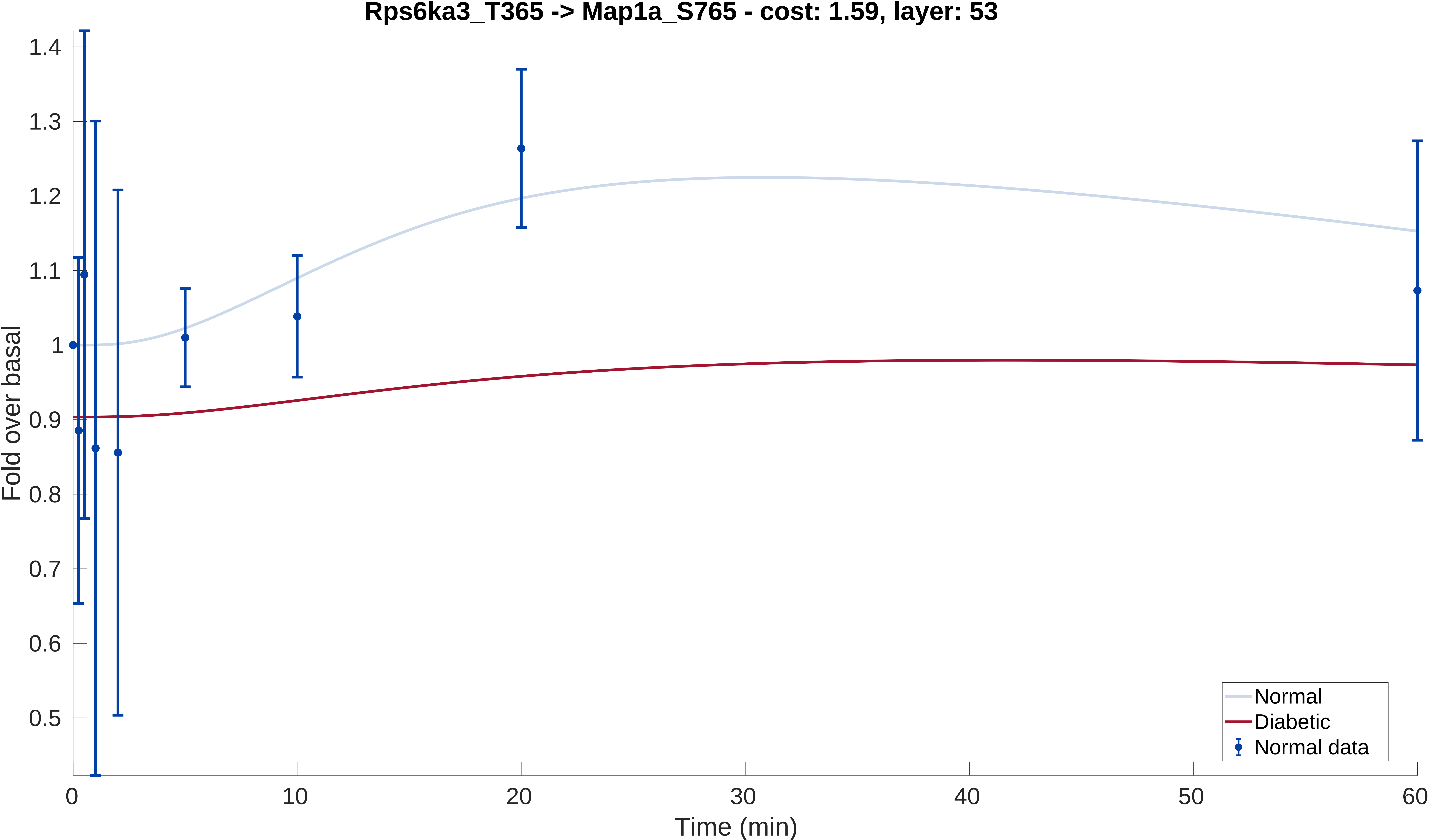
Rps6ka3_T365 -> Map1a_S765 - cost: 1.59, layer: 53
Fold over basal
1.4
1.3
1.2
1.1
1
0.9
0.8
0.7
0.6
0.5
0
10
20
30
40
50
60
Time (min)
Normal
Diabetic
Normal data

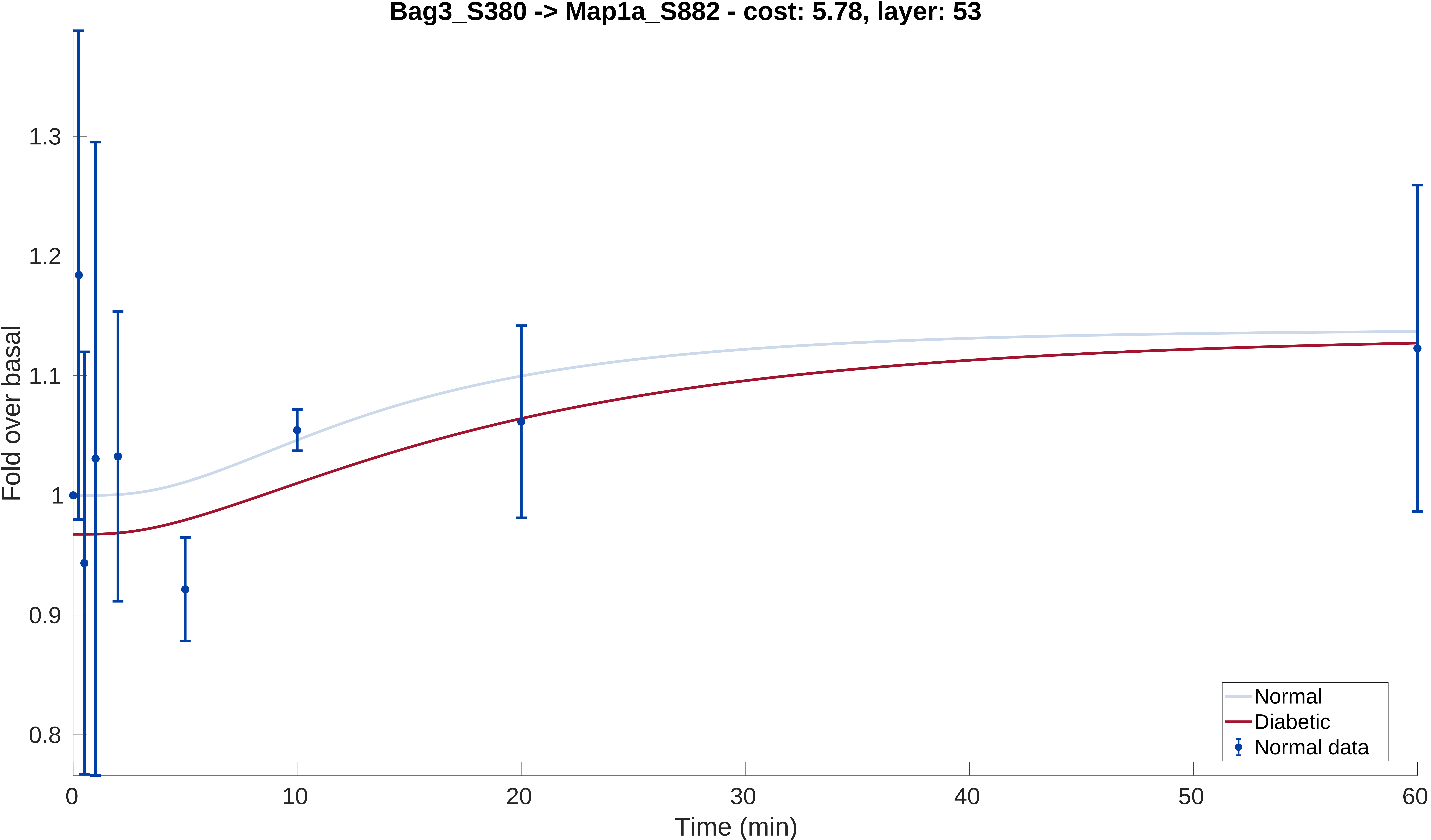

Bag3_S380 -> Map1a_S882 - cost: 5.78, layer: 53
Fold over basal
1.3
1.2
1.1
1
0.9
0.8
0
10
20
30
40
50
60
Time (min)
Normal
Diabetic
Normal data

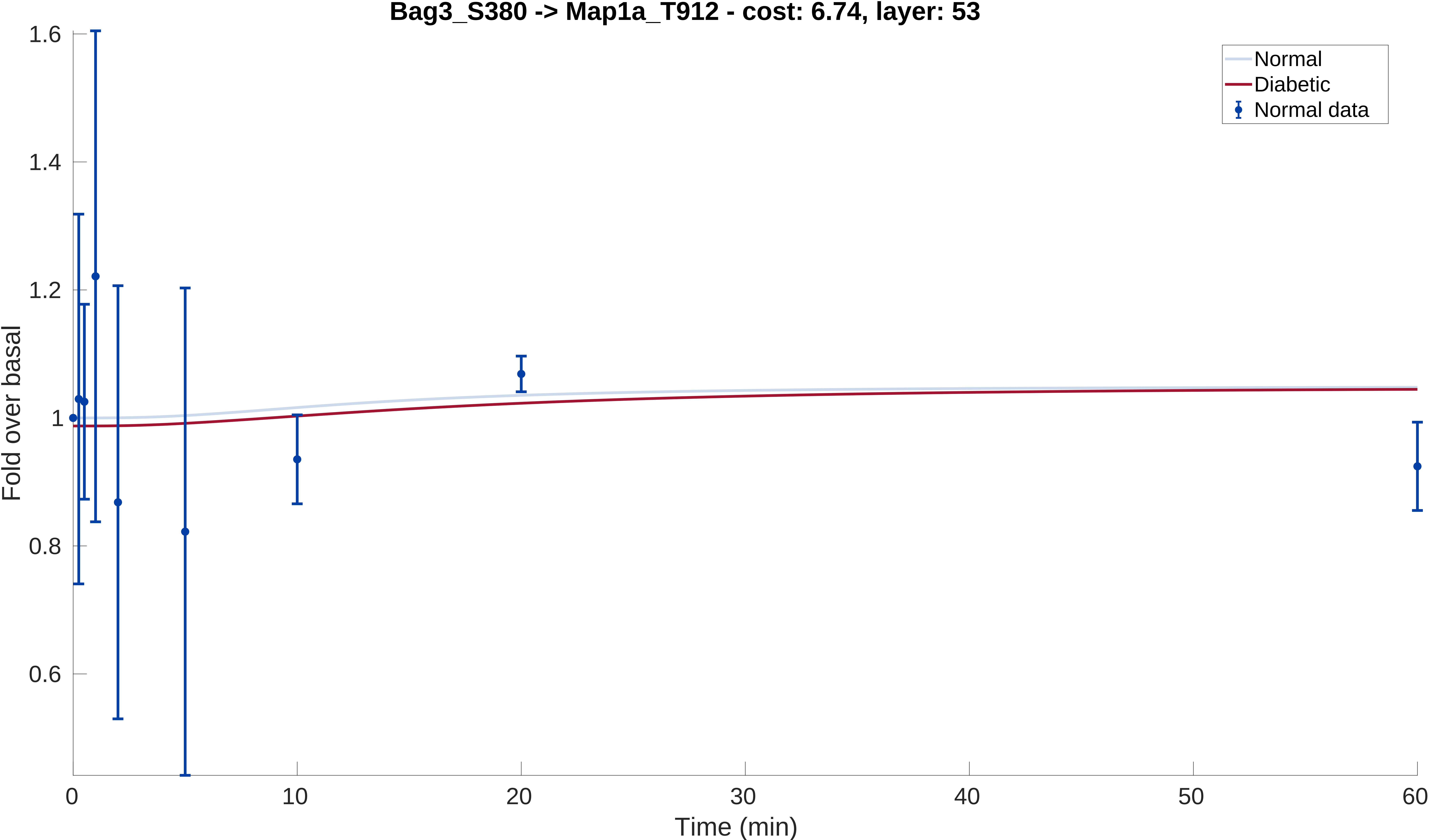
Bag3_S380 -> Map1a_T912 - cost: 6.74, layer: 53
Normal
Diabetic
Normal data
Fold over basal
1.6
1.4
1.2
1
0.8
0.6
0
10
20
30
40
50
60
Time (min)

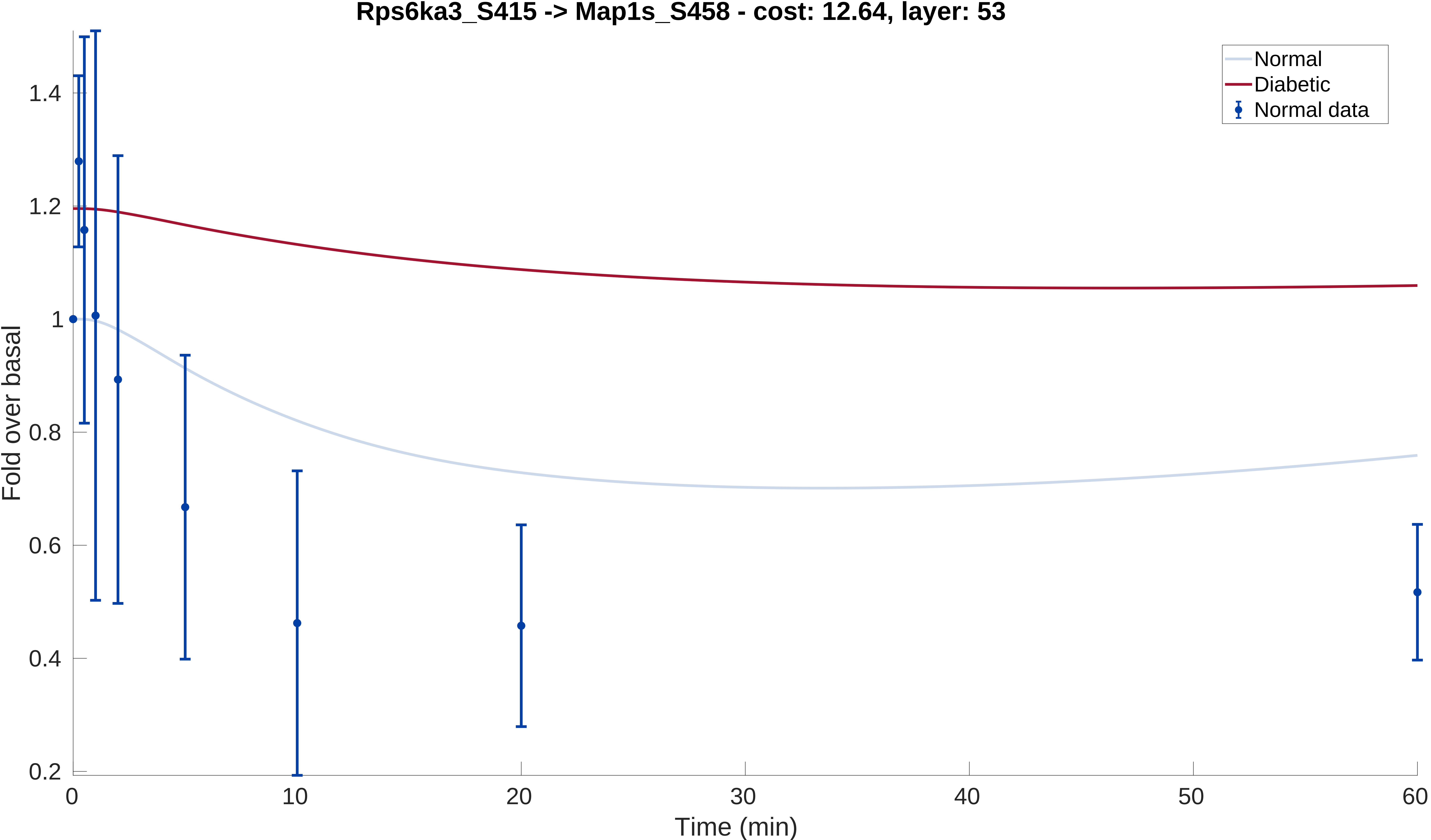

Rps6ka3_S415 -> Map1s_S458 - cost: 12.64, layer: 53
Normal
Diabetic
Normal data
Fold over basal
1.4
1.2
1
0.8
0.6
0.4
0.2
0
10
20
30
40
50
60
Time (min)

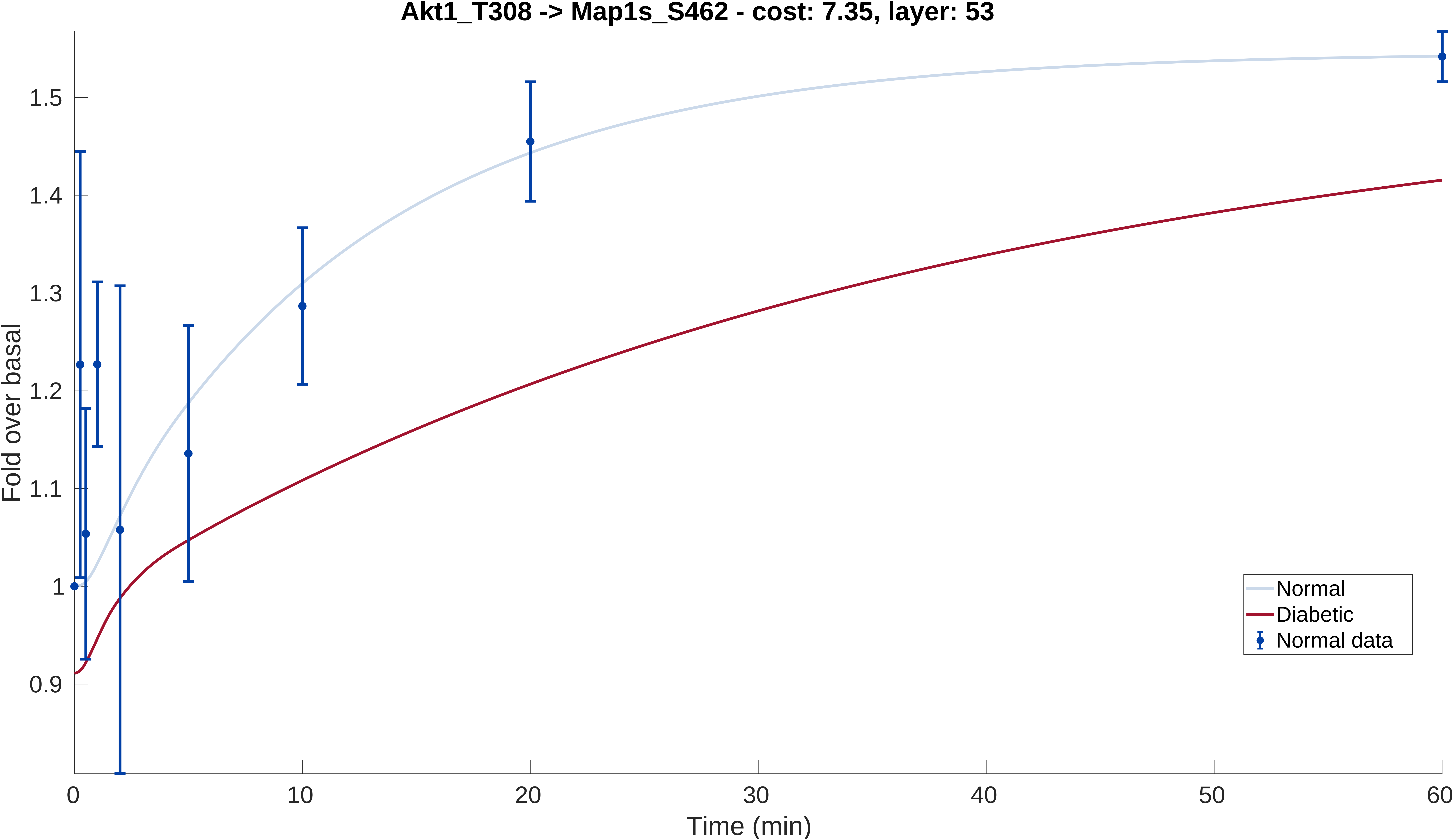

Akt1_T308 -> Map1s_S462 - cost: 7.35, layer: 53
Fold over basal
1.5
1.4
1.3
1.2
1.1
1
0.9
0
10
20
30
40
50
60
Time (min)
Normal
Diabetic
Normal data

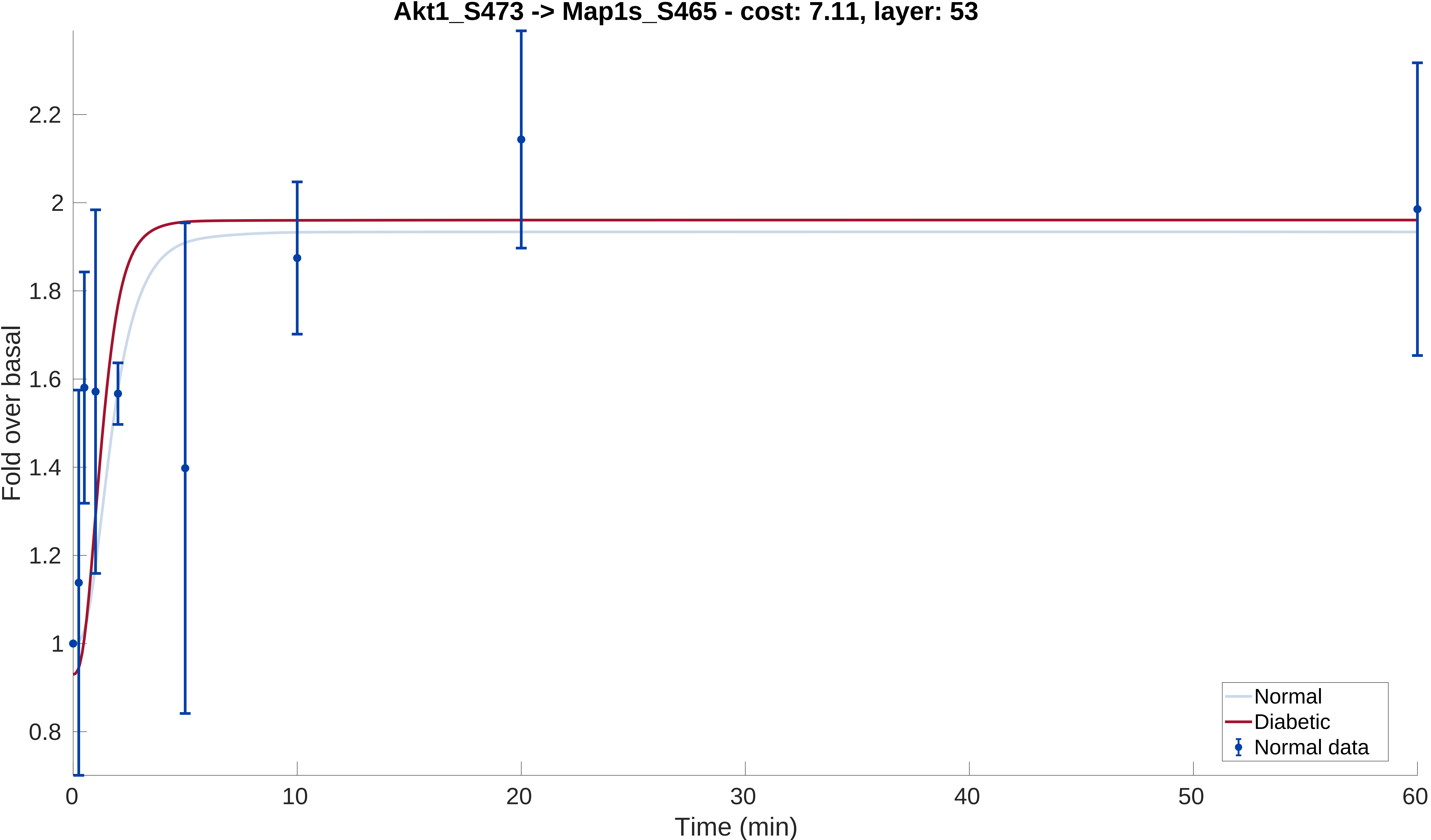

Akt1_S473 -> Map1s_S465 - cost: 7.11, layer: 53
Fold over basal
2.2
2
1.8
1.6
1.4
1.2
1
0.8
0
10
20
30
40
50
60
Time (min)
Normal
Diabetic
Normal data

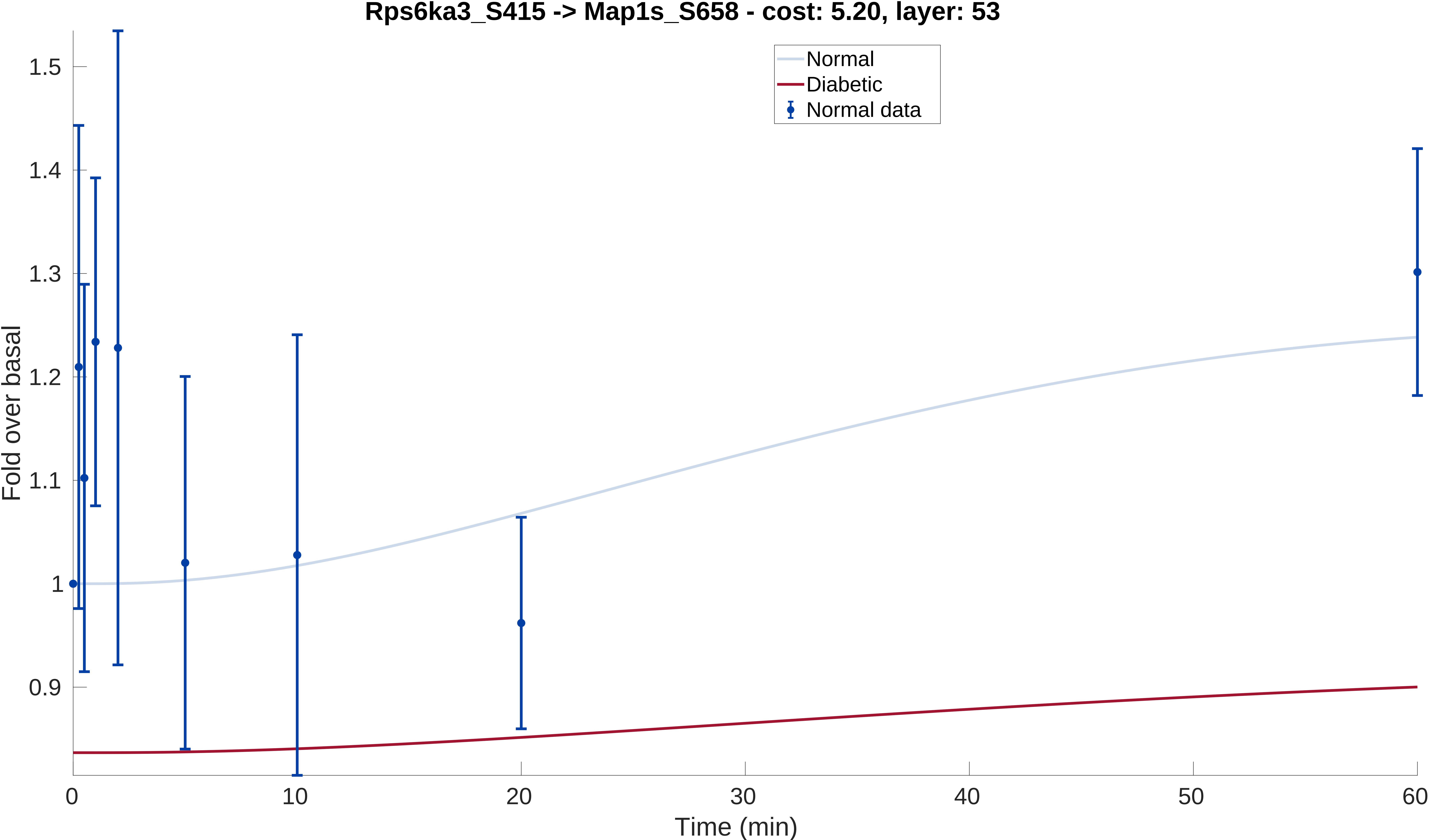

Rps6ka3_S415 -> Map1s_S658 - cost: 5.20, layer: 53
Normal
Diabetic
Normal data
Fold over basal
1.5
1.4
1.3
1.2
1.1
1
0.9
0
10
20
30
40
50
60
Time (min)

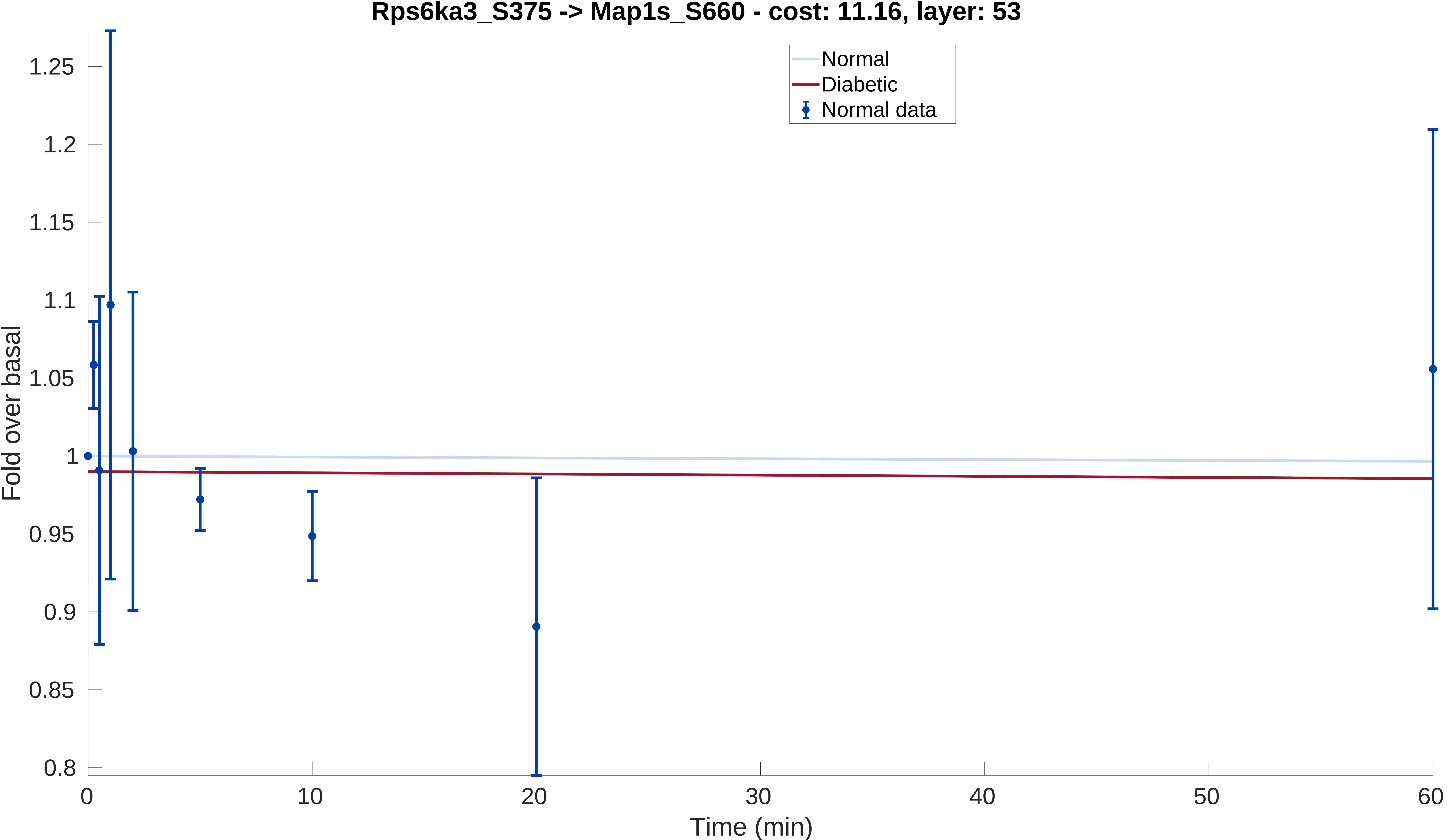

Rps6ka3_S375 -> Map1s_S660 - cost: 11.16, layer: 53
Normal
Diabetic
Normal data
Fold over basal
1.25
1.2
1.15
1.1
1.05
1
0.95
0.9
0.85
0.8
0
10
20
30
40
50
60
Time (min)

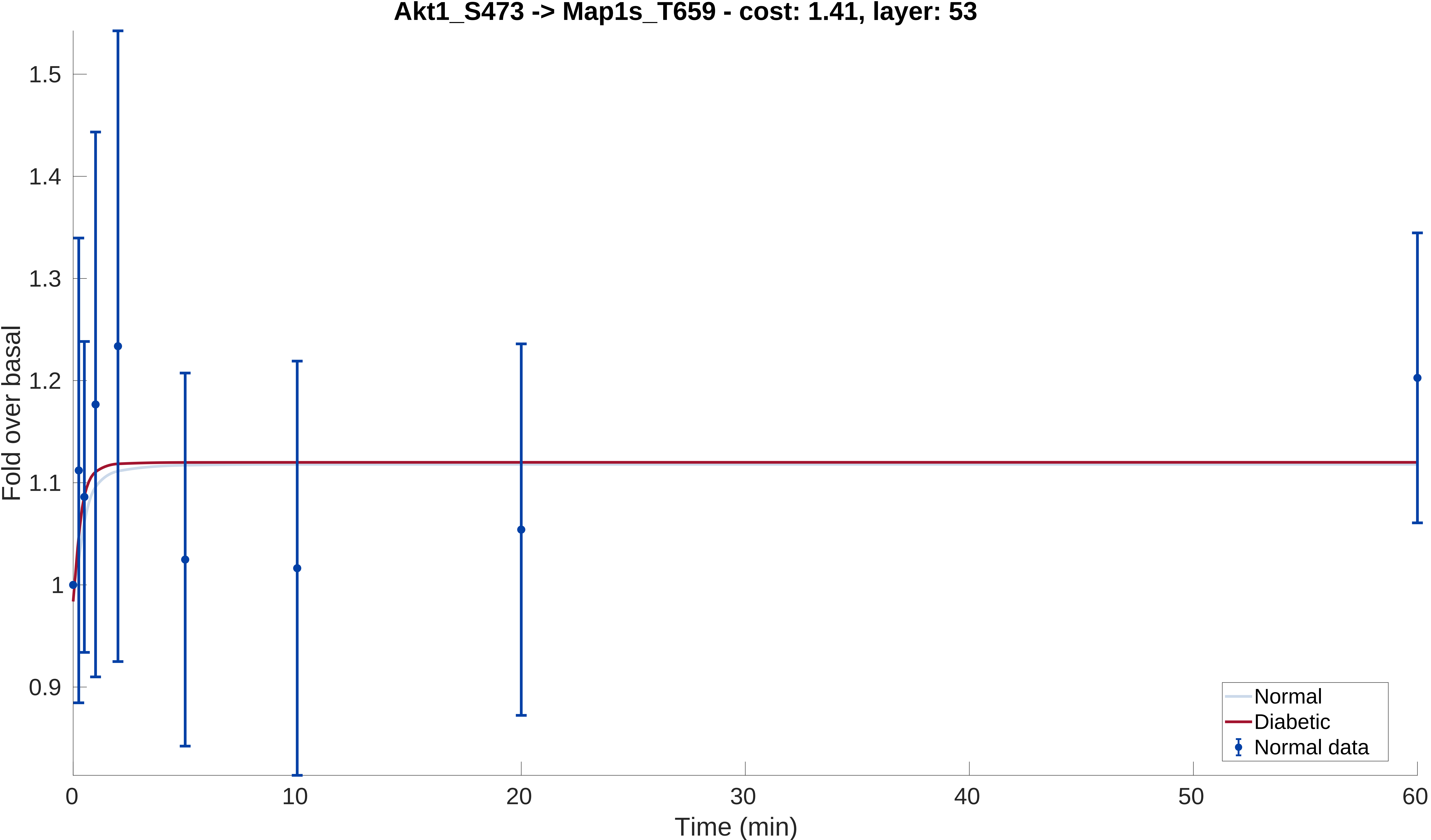
Akt1_S473 -> Map1s_T659 - cost: 1.41, layer: 53
Fold over basal
1.5
1.4
1.3
1.2
1.1
1
0.9
0
10
20
30
40
50
60
Time (min)
Normal
Diabetic
Normal data

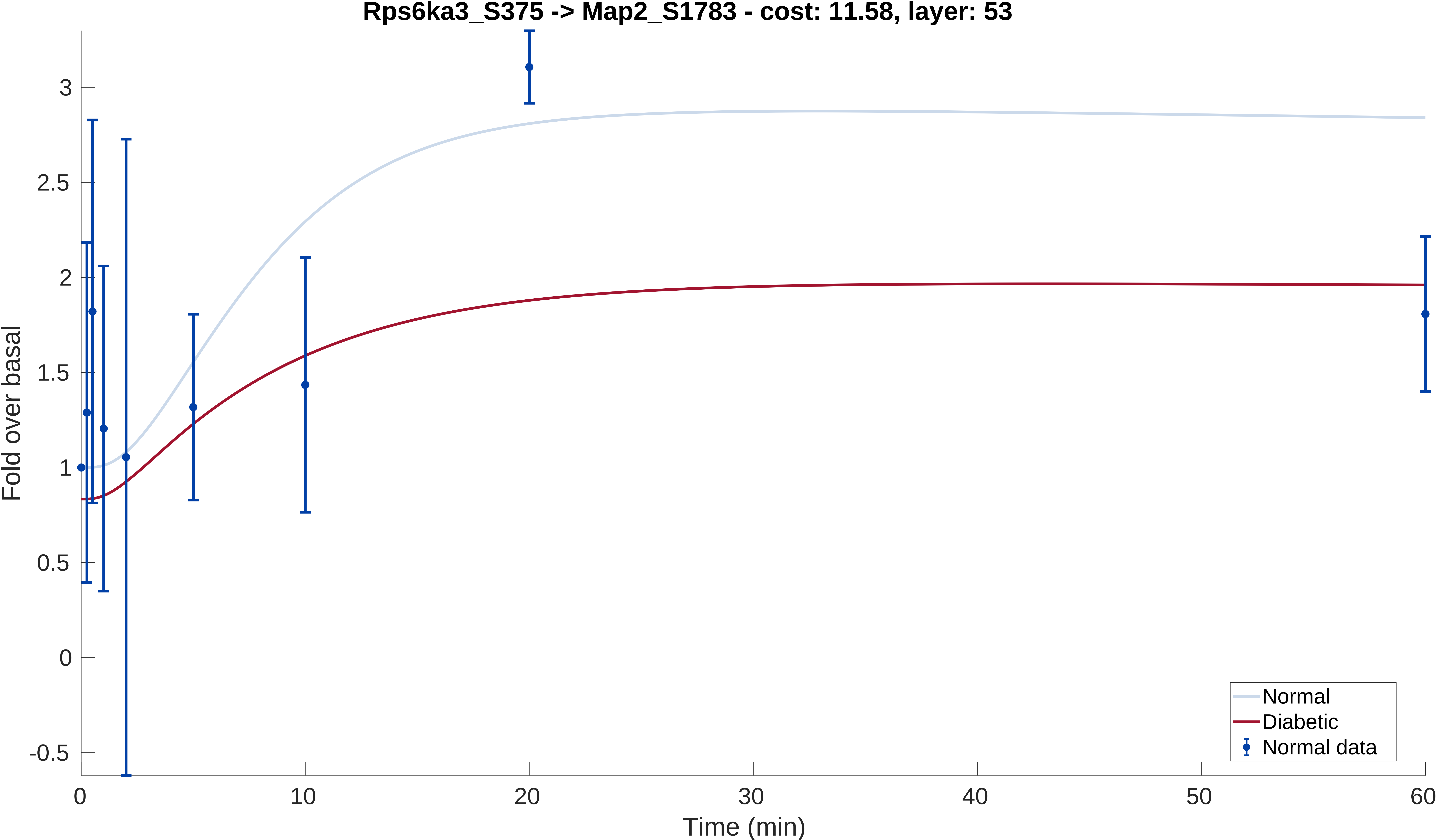

Rps6ka3_S375 -> Map2_S1783 - cost: 11.58, layer: 53
Fold over basal
3
2.5
2
1.5
1
0.5
0
-0.5
0
10
20
30
40
50
60
Time (min)
Normal
Diabetic
Normal data

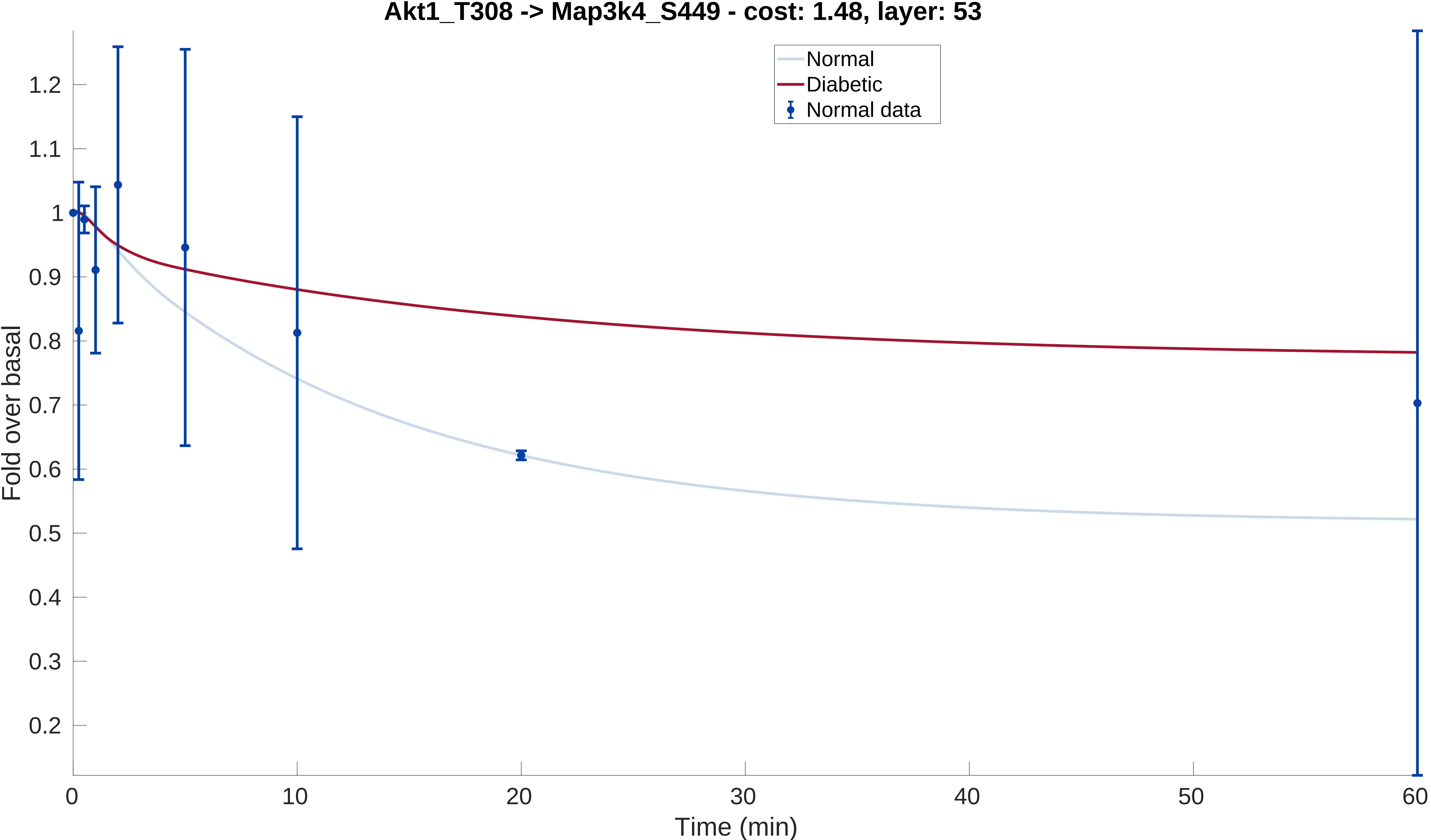

Akt1_T308 -> Map3k4_S449 - cost: 1.48, layer: 53
Normal
Diabetic
Normal data
Fold over basal
Time (min)
0.2
0.3
0.4
0.5
0.6
0.7
0.8
0.9
1
1.1
1.2
0
10
20
30
40
50
60

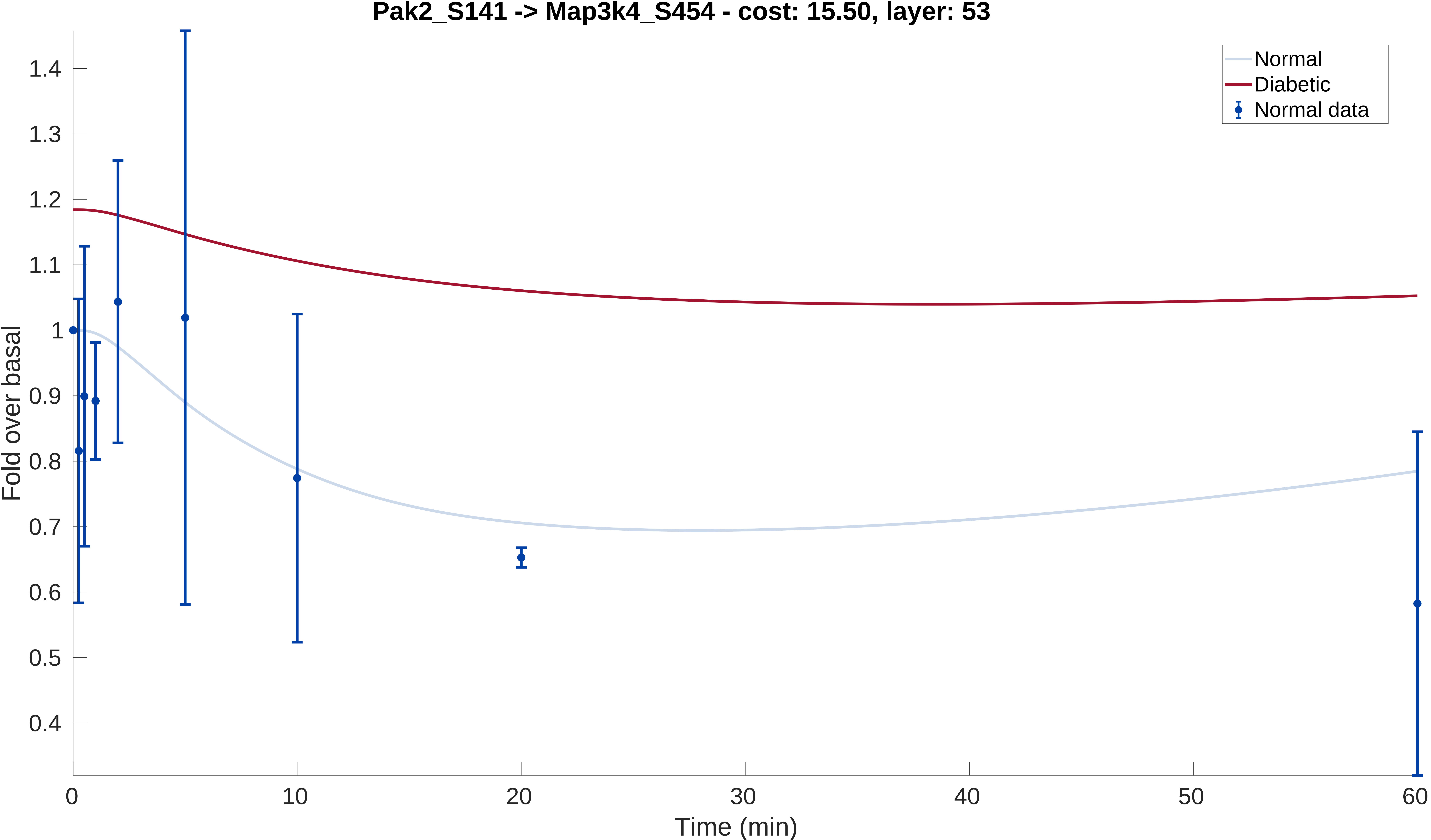

Pak2_S141 -> Map3k4_S454 - cost: 15.50, layer: 53
Normal
Diabetic
Normal data
Fold over basal
1.4
1.3
1.2
1.1
1
0.9
0.8
0.7
0.6
0.5
0.4
0
10
20
30
40
50
60
Time (min)

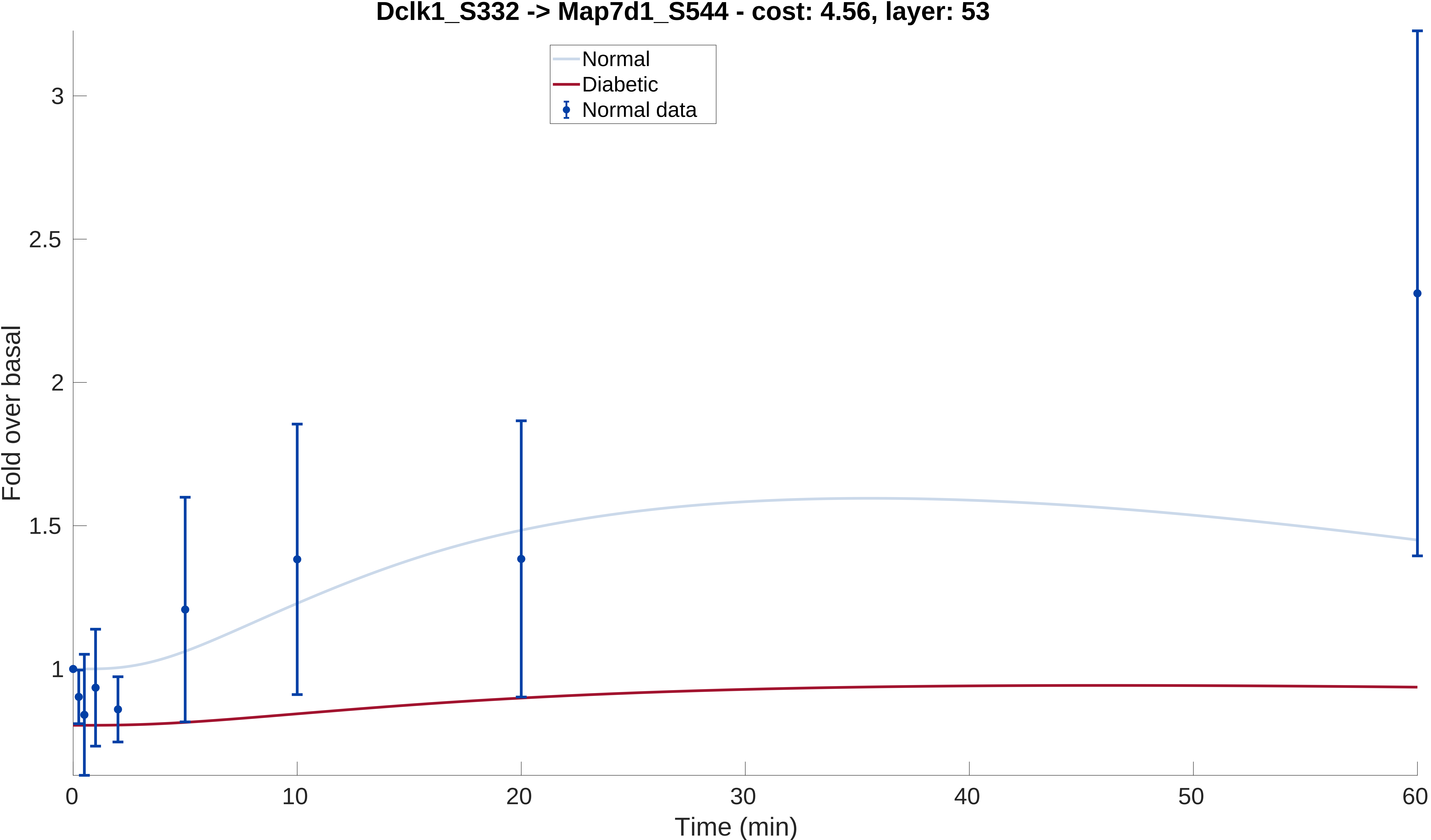

Dclk1_S332 -> Map7d1_S544 - cost: 4.56, layer: 53
Normal
Diabetic
Normal data
Fold over basal
3
2.5
2
1.5
1
0
10
20
30
40
50
60
Time (min)

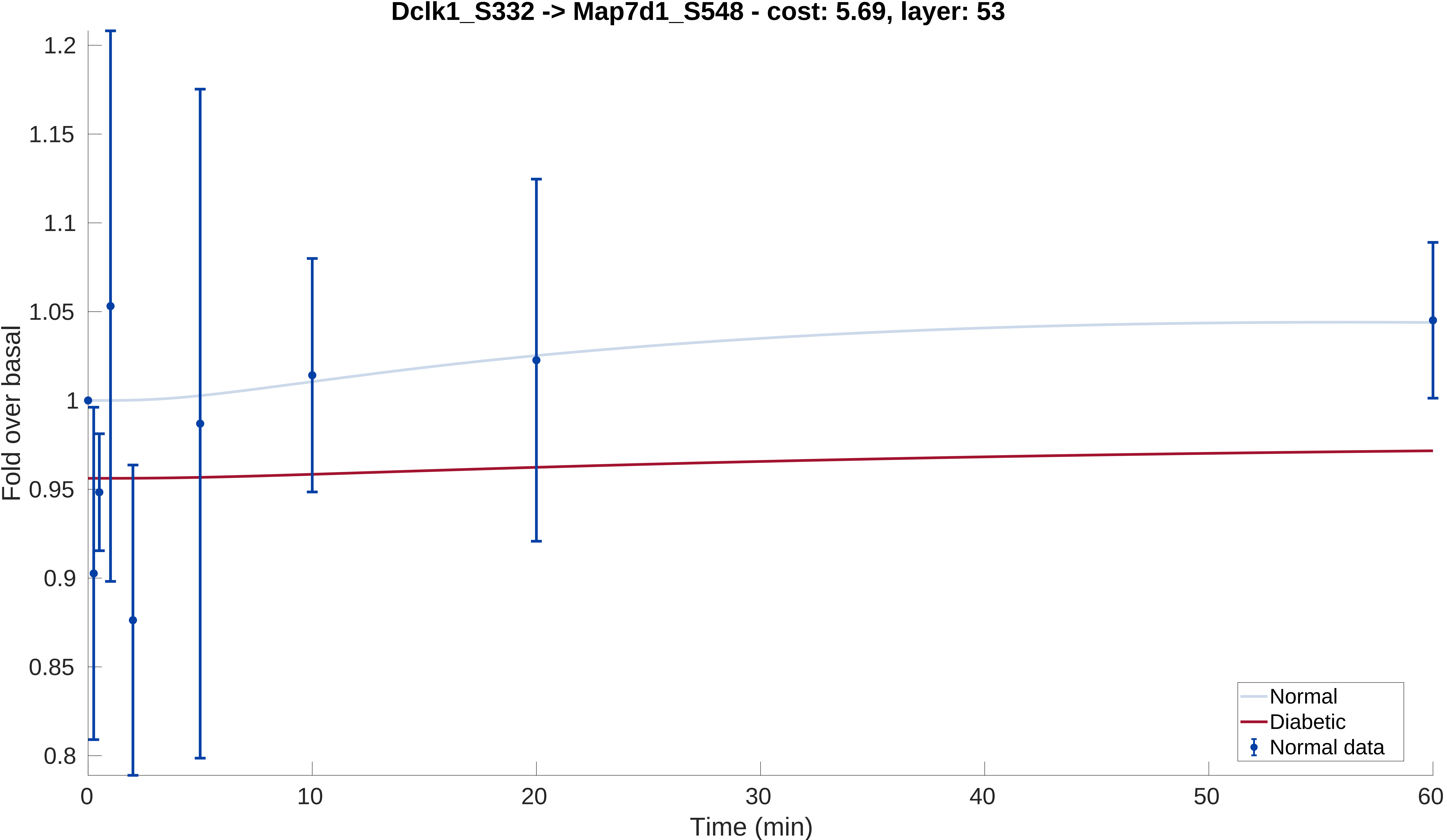

Dclk1_S332 -> Map7d1_S548 - cost: 5.69, layer: 53
Fold over basal
Time (min)
1.2
1.15
1.1
1.05
1
0.95
0.9
0.85
0.8
0
10
20
30
40
50
60
Normal
Diabetic
Normal data

#### Dclk1_S332 -> Map7d1_S552 - cost: 1.92, layer: 53

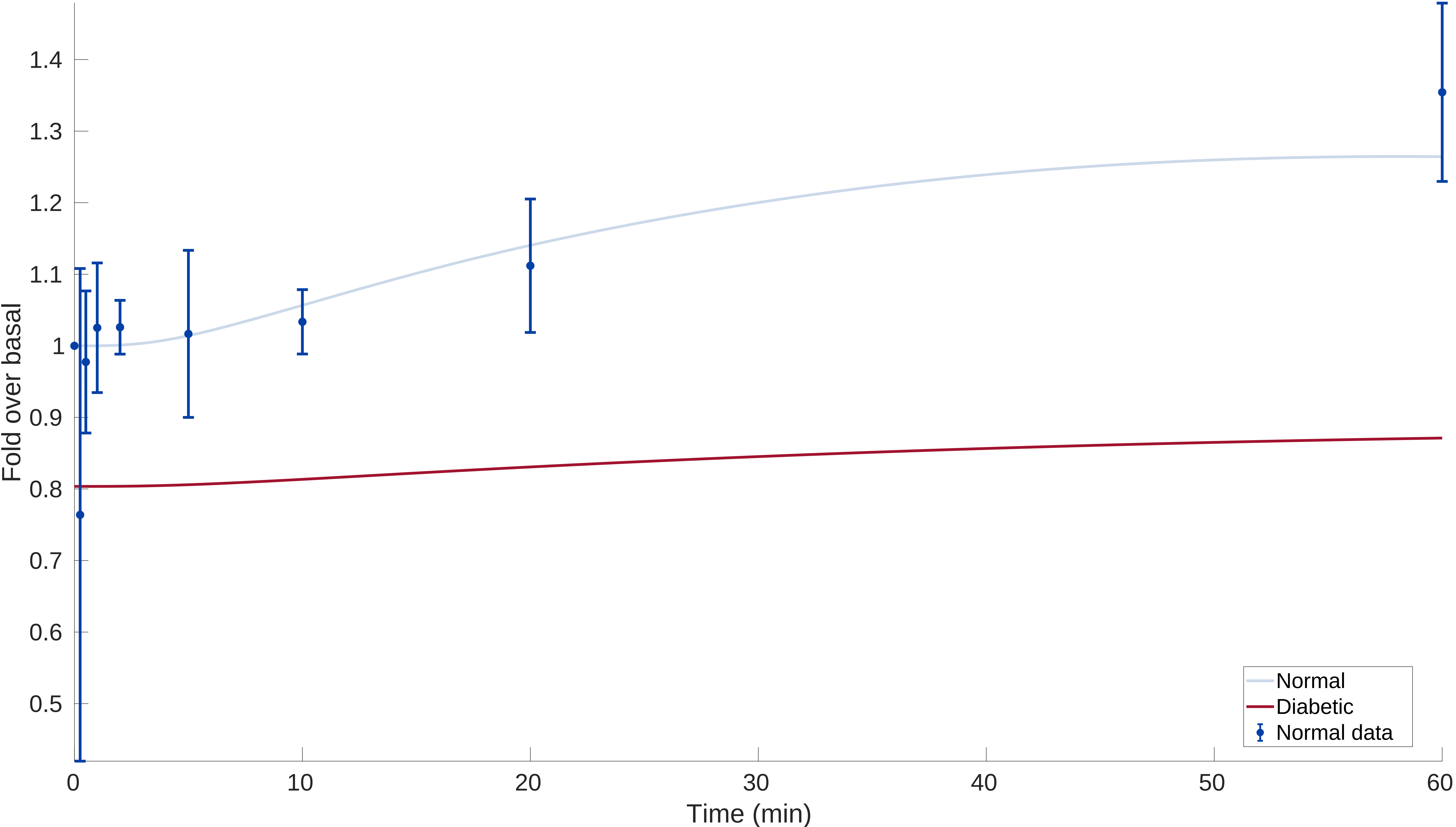

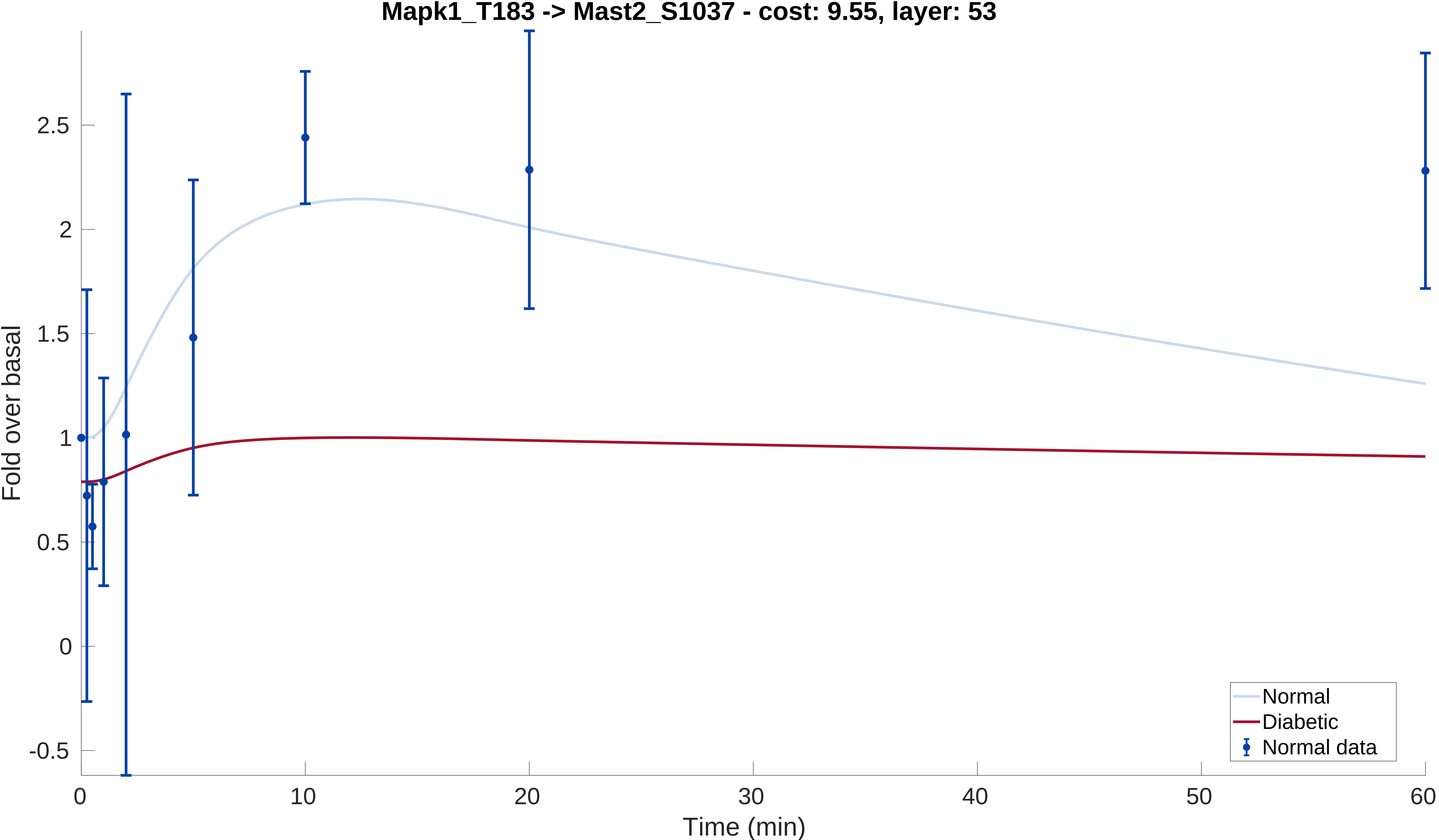
Mapk1_T183 -> Mast2_S1037 - cost: 9.55, layer: 53
Fold over basal
2.5
2
1.5
1
0.5
0
-0.5
0
10
20
30
40
50
60
Time (min)
Normal
Diabetic
Normal data

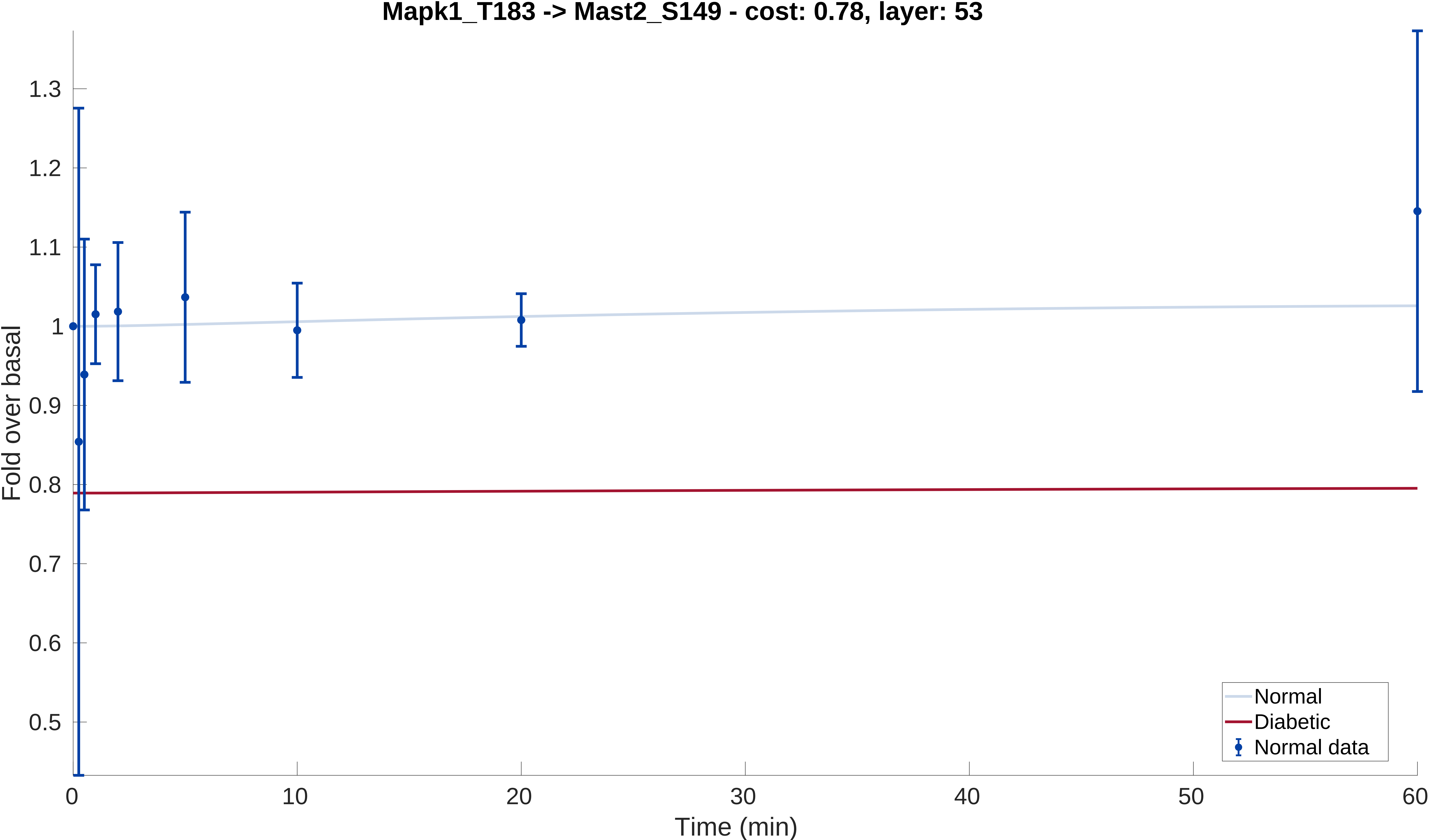
Mapk1_T183 -> Mast2_S149 - cost: 0.78, layer: 53
Fold over basal
1.3
1.2
1.1
1
0.9
0.8
0.7
0.6
0.5
0
10
20
30
40
50
60
Time (min)
Normal
Diabetic
Normal data

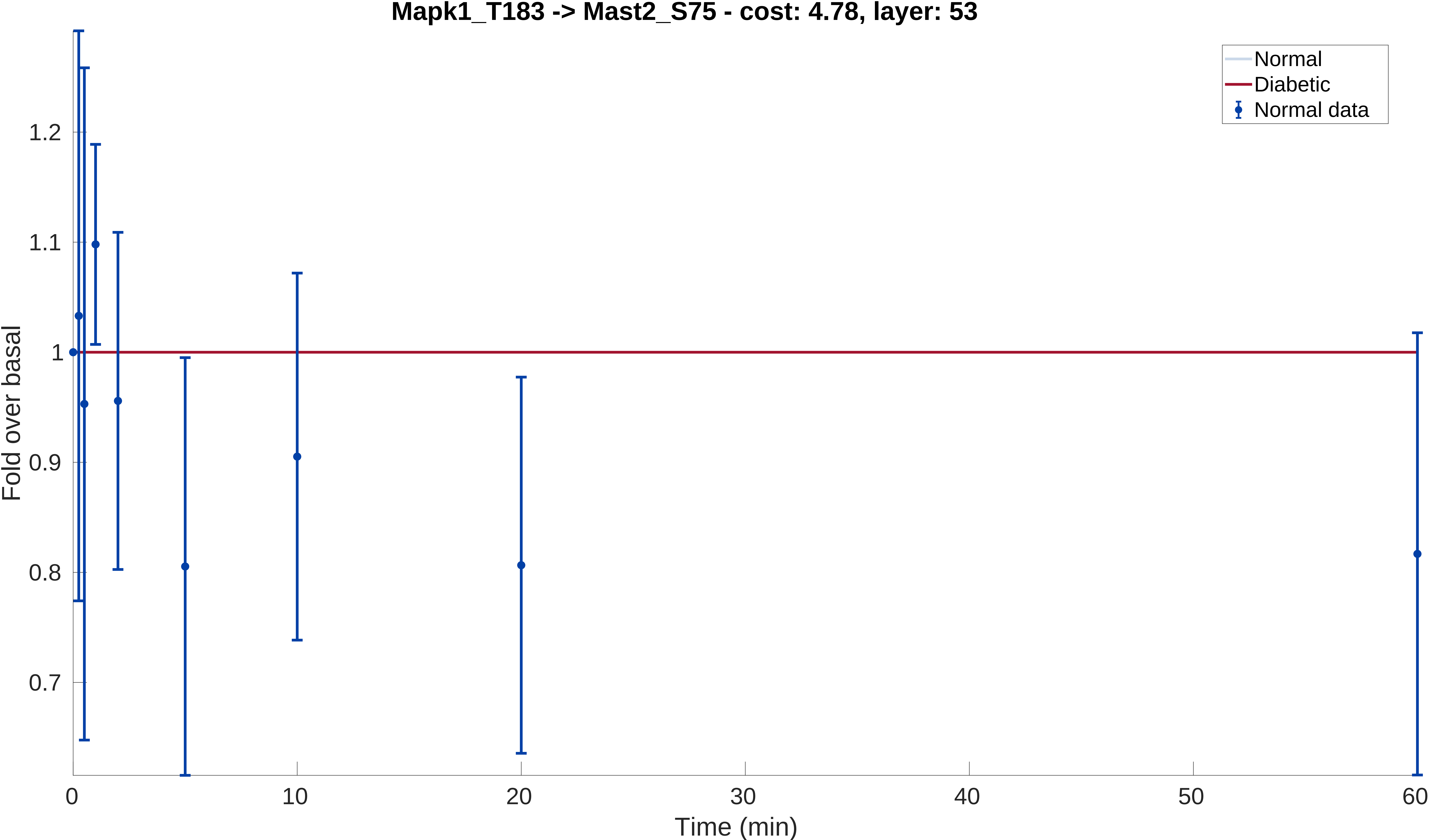

Mapk1_T183 -> Mast2_S75 - cost: 4.78, layer: 53
Normal
Diabetic
Normal data
Fold over basal
1.2
1.1
1
0.9
0.8
0.7
0
10
20
30
40
50
60
Time (min)

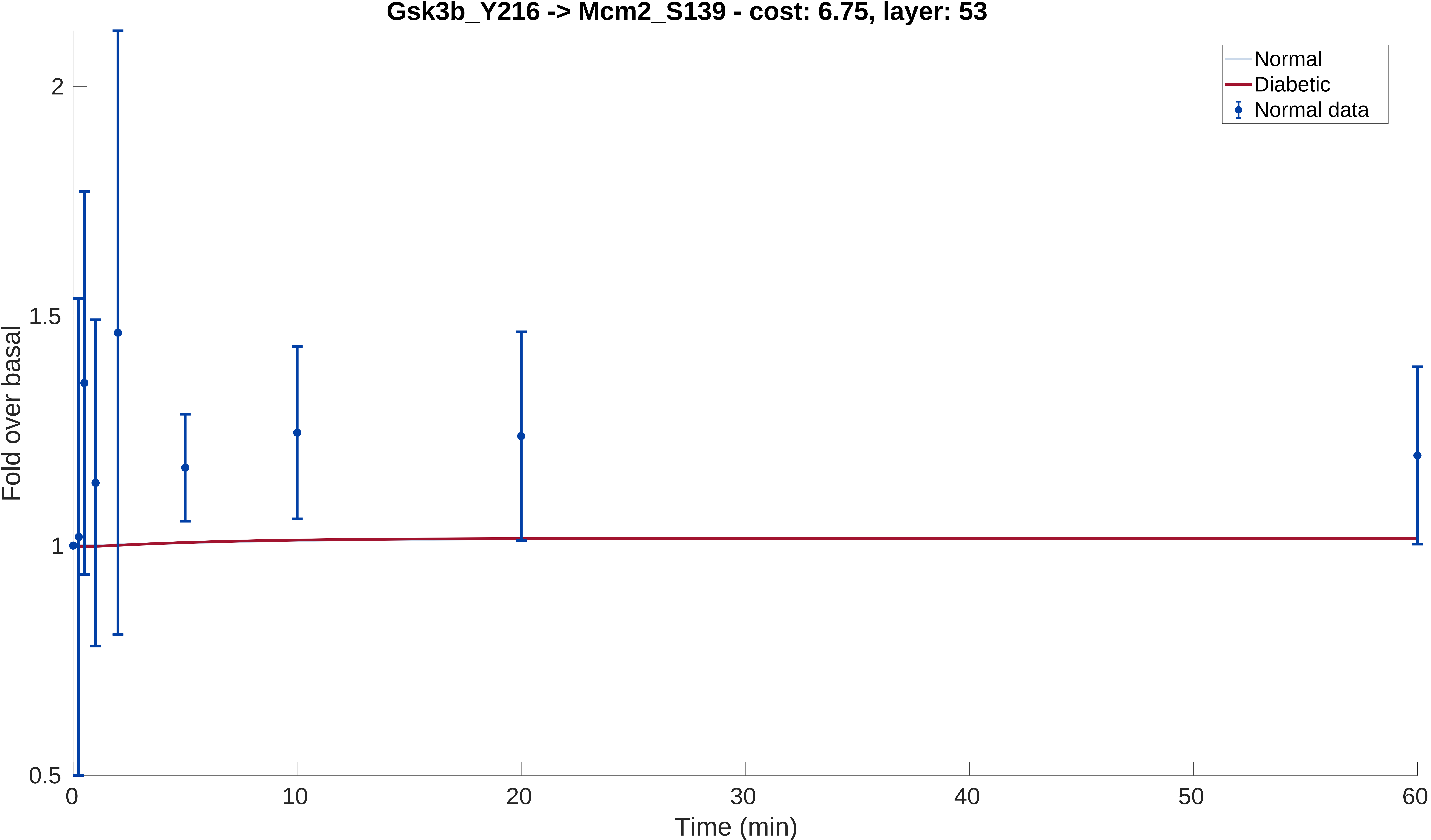
Gsk3b_Y216 -> Mcm2_S139 - cost: 6.75, layer: 53
Normal
Diabetic
Normal data
2
1.5
1
0.5
Fold over basal
0
10
20
30
40
50
60
Time (min)

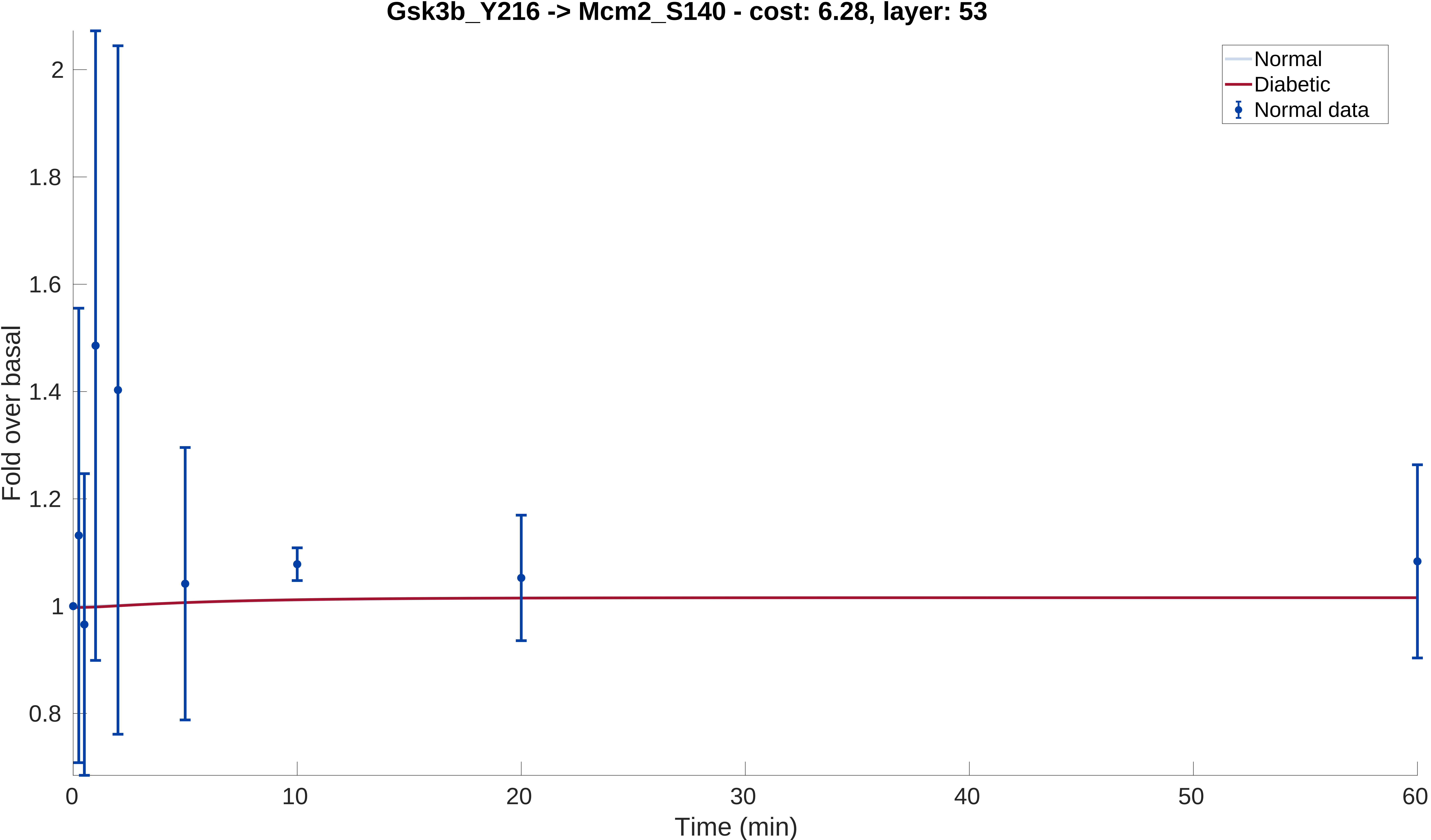

Gsk3b_Y216 -> Mcm2_S140 - cost: 6.28, layer: 53
Normal
Diabetic
Normal data
Fold over basal
2
1.8
1.6
1.4
1.2
1
0.8
0
10
20
30
40
50
60
Time (min)

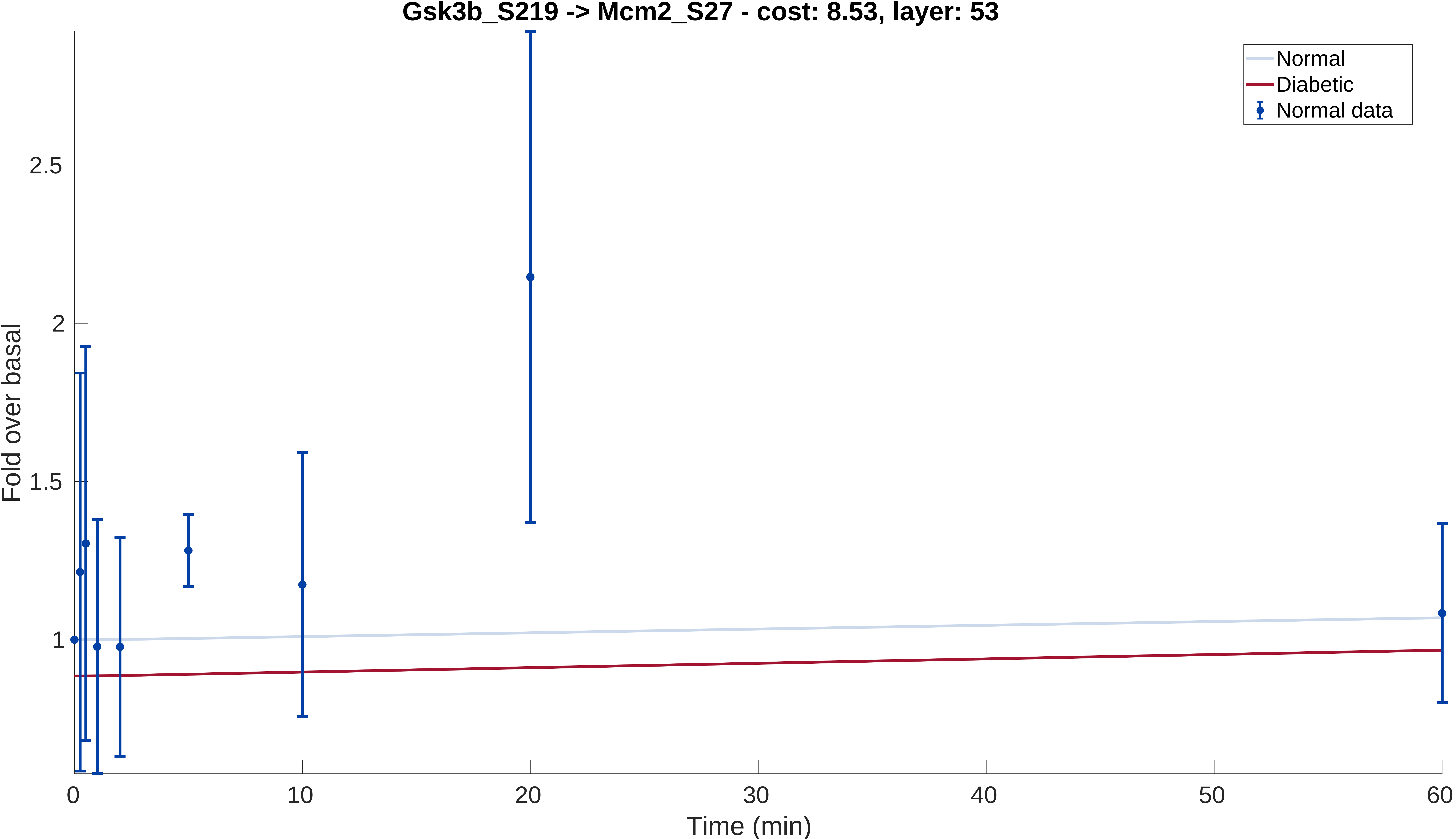
Gsk3b_S219 -> Mcm2_S27 - cost: 8.53, layer: 53
Normal
Diabetic
Normal data
Fold over basal
2.5
2
1.5
1
0
10
20
30
40
50
60
Time (min)

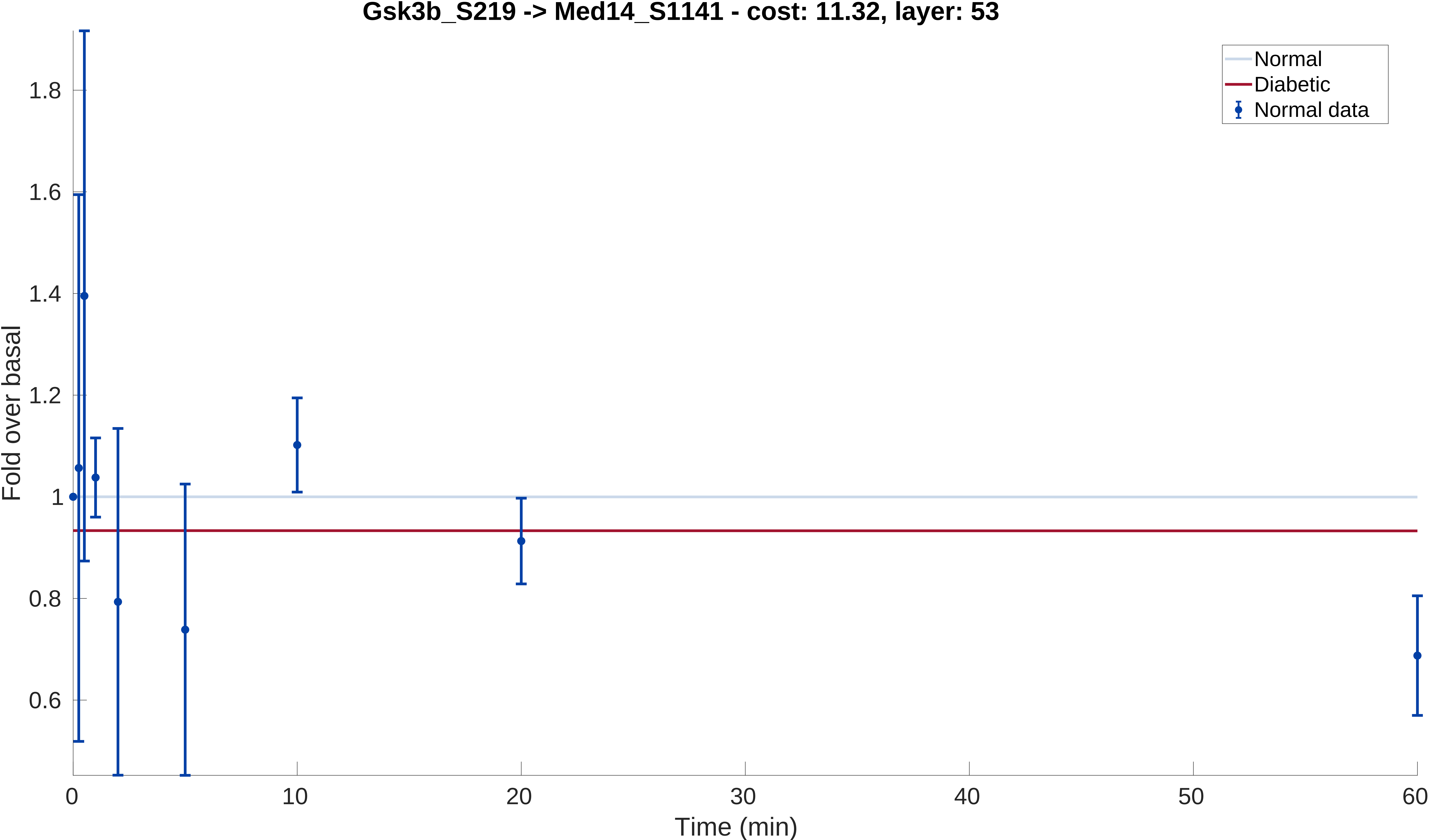

Gsk3b_S219 -> Med14_S1141 - cost: 11.32, layer: 53
Normal
Diabetic
Normal data
Fold over basal
1.8
1.6
1.4
1.2
1
0.8
0.6
0
10
20
30
40
50
60
Time (min)

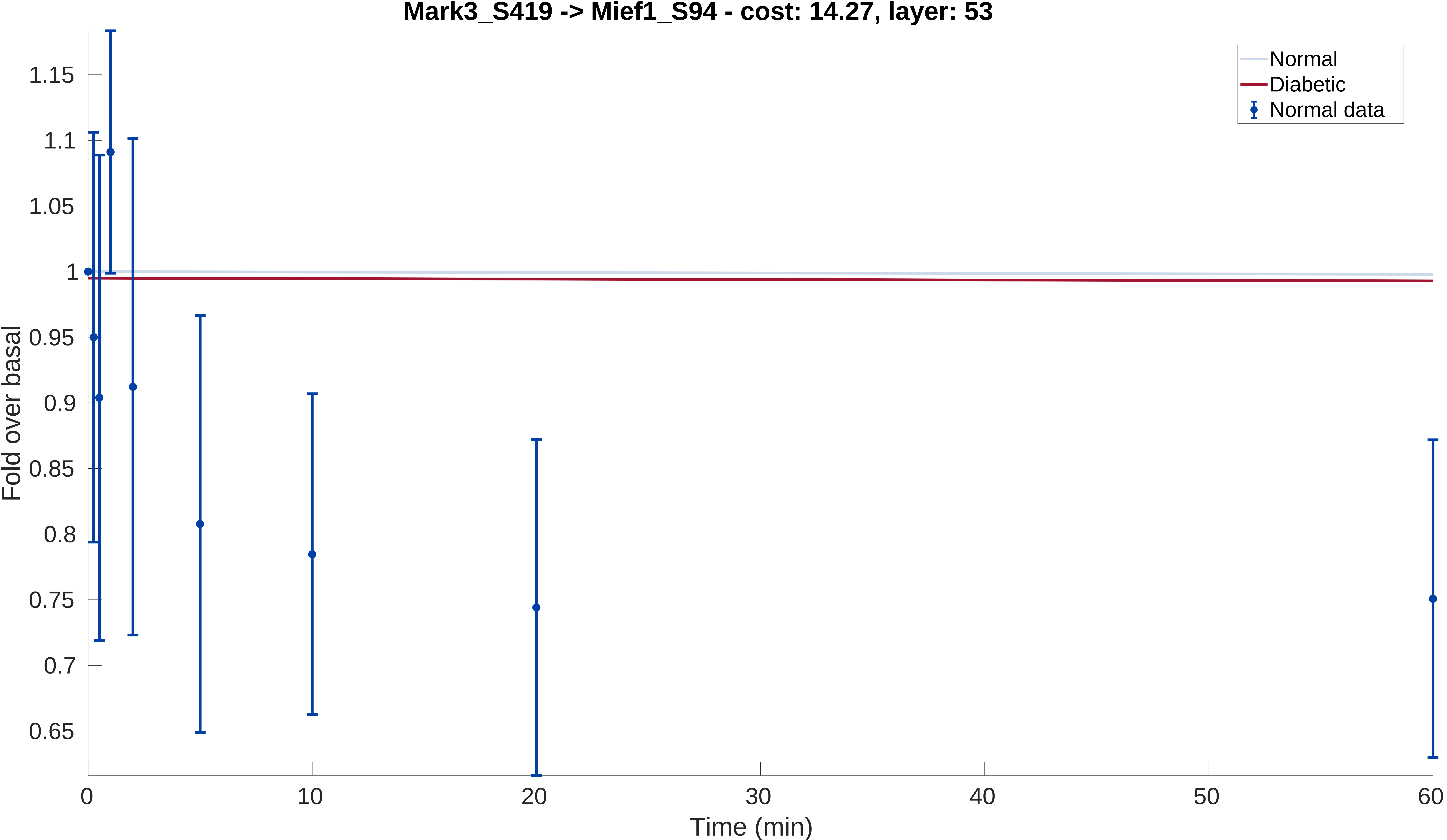
Mark3_S419 -> Mief1_S94 - cost: 14.27, layer: 53
Normal
Diabetic
Normal data
Fold over basal
Time (min)
1.15
1.1
1.05
1
0.95
0.9
0.85
0.8
0.75
0.7
0.65
0
10
20
30
40
50
60

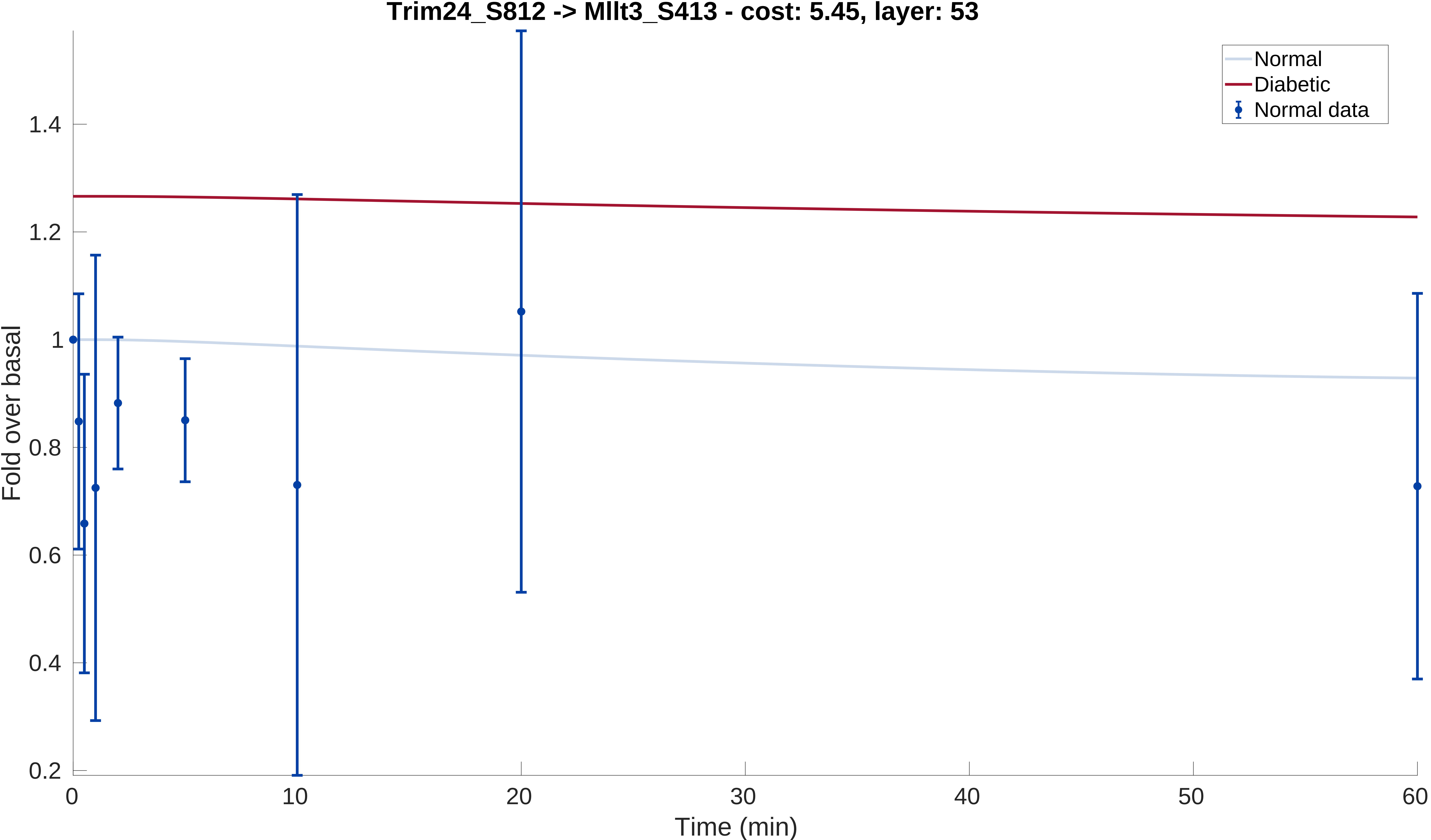

Trim24_S812 -> Mllt3_S413 - cost: 5.45, layer: 53
Normal
Diabetic
Normal data
Fold over basal
1.4
1.2
1
0.8
0.6
0.4
0.2
0
10
20
30
40
50
60
Time (min)

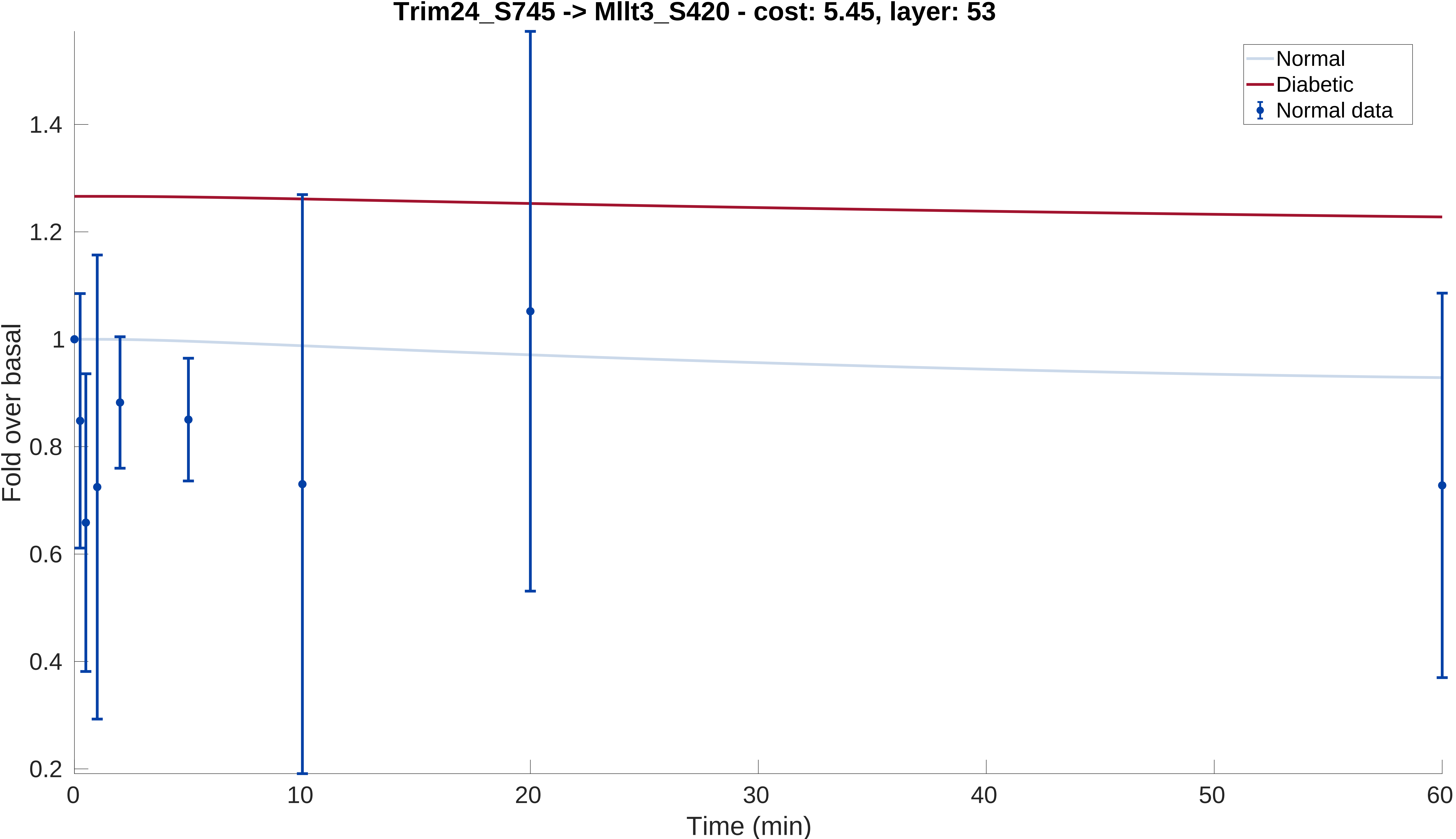

Trim24_S745 -> Mllt3_S420 - cost: 5.45, layer: 53
Normal
Diabetic
Normal data
Fold over basal
1.4
1.2
1
0.8
0.6
0.4
0.2
0
10
20
30
40
50
60
Time (min)

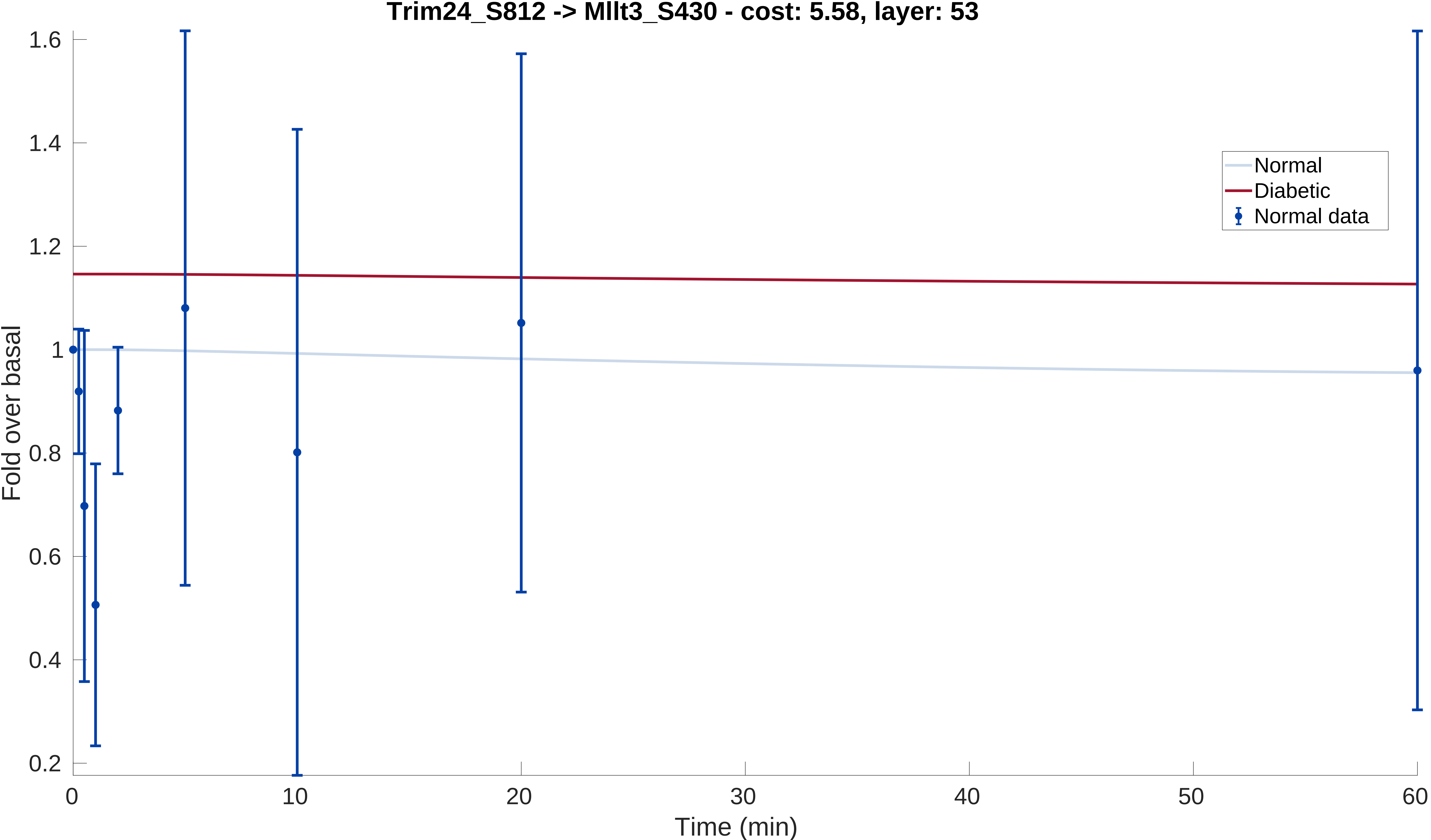
Trim24_S812 -> Mllt3_S430 - cost: 5.58, layer: 53
Fold over basal
Time (min)
Normal
Diabetic
Normal data
0.2
0.4
0.6
0.8
1
1.2
1.4
1.6
0
10
20
30
40
50
60

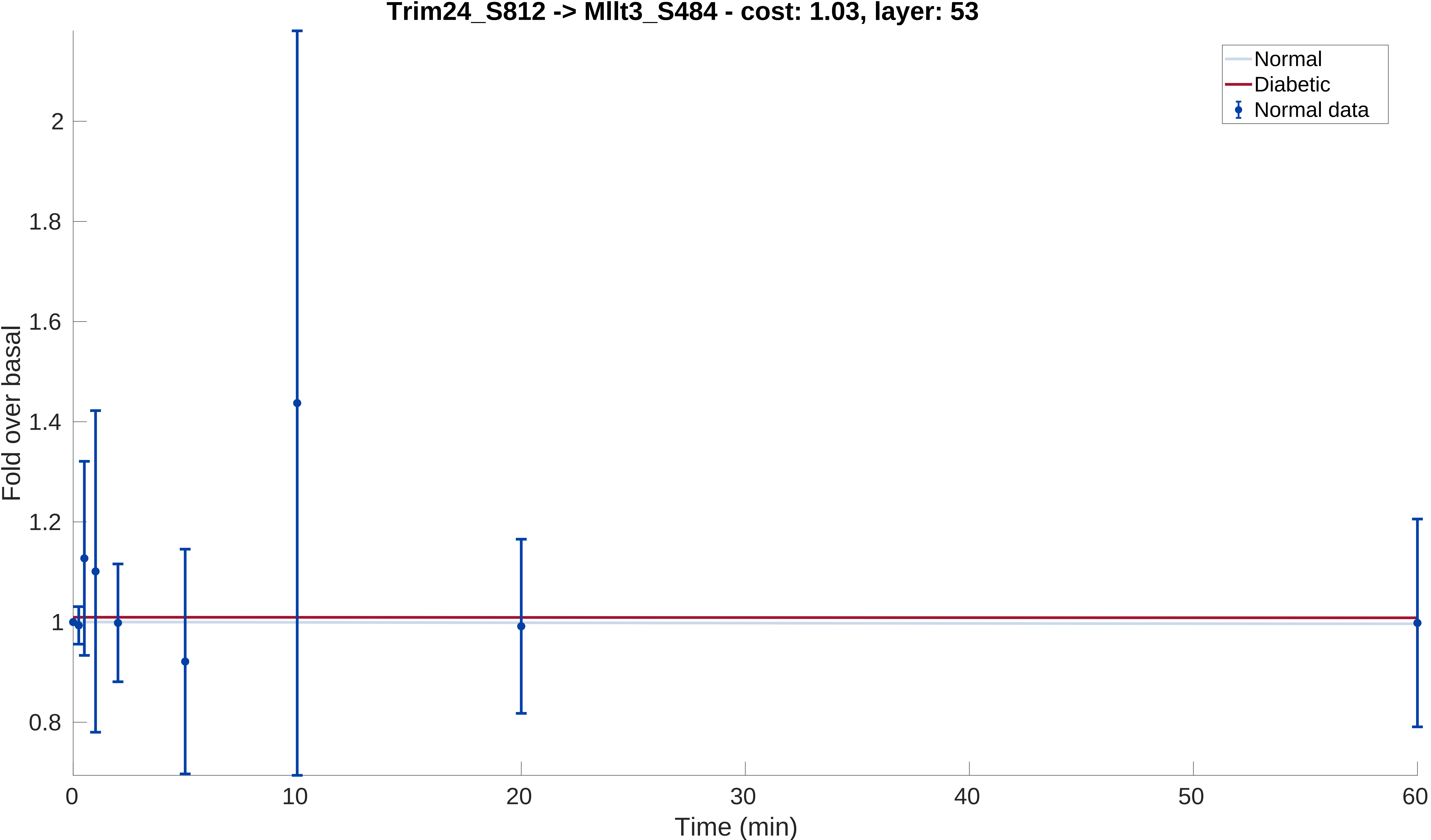

Trim24_S812 -> Mllt3_S484 - cost: 1.03, layer: 53
Normal
Diabetic
Normal data
Fold over basal
2
1.8
1.6
1.4
1.2
1
0.8
0
10
20
30
40
50
60
Time (min)

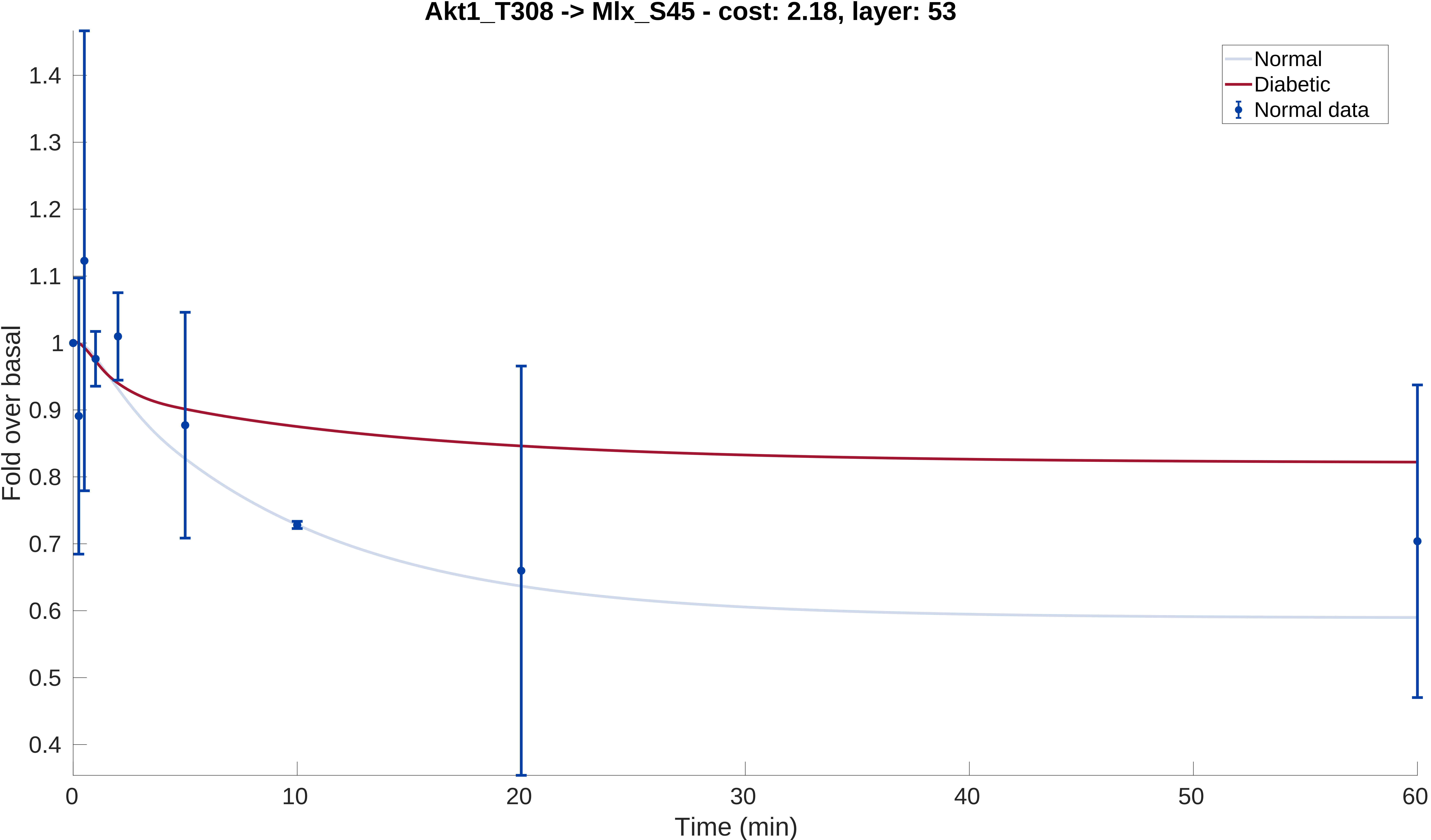

Akt1_T308 -> Mlx_S45 - cost: 2.18, layer: 53
Normal
Diabetic
Normal data
Fold over basal
Time (min)
1.4
1.3
1.2
1.1
1
0.9
0.8
0.7
0.6
0.5
0.4
0
10
20
30
40
50
60

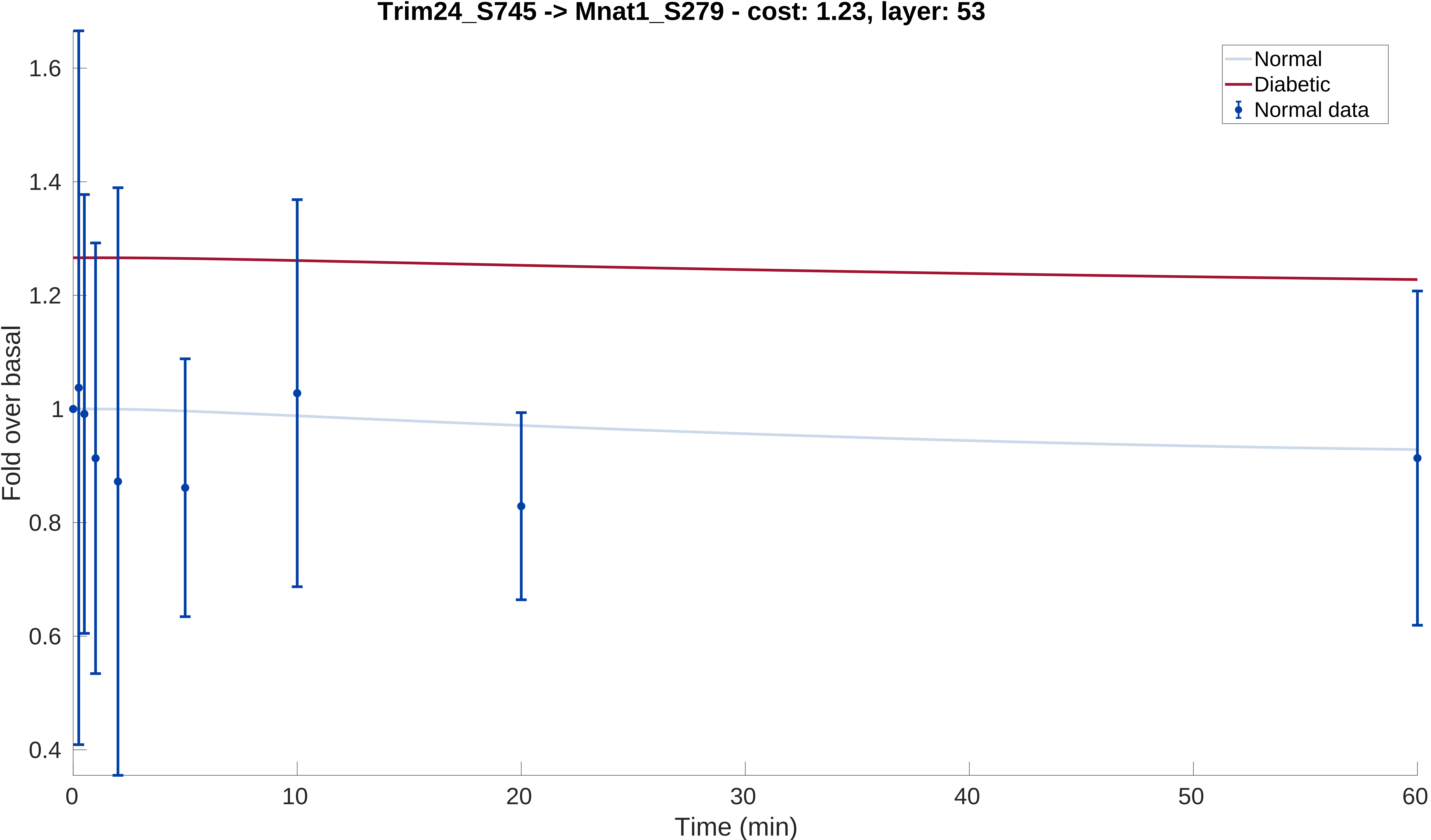

Trim24_S745 -> Mnat1_S279 - cost: 1.23, layer: 53
Normal
Diabetic
Normal data
Fold over basal
1.6
1.4
1.2
1
0.8
0.6
0.4
0
10
20
30
40
50
60
Time (min)

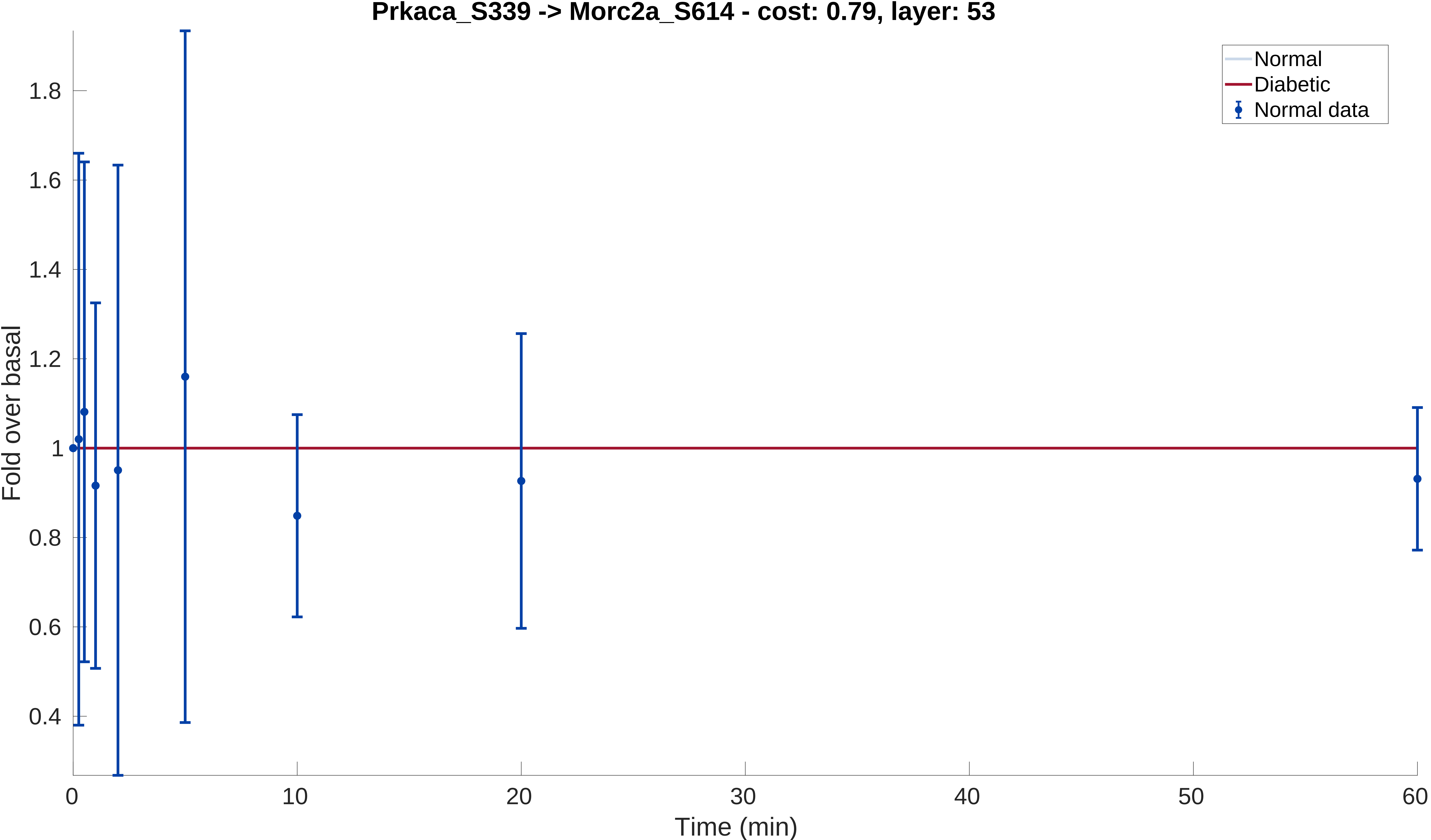

Prkaca_S339 -> Morc2a_S614 - cost: 0.79, layer: 53
Normal
Diabetic
Normal data
Fold over basal
Time (min)
1.8
1.6
1.4
1.2
1
0.8
0.6
0.4
0
10
20
30
40
50
60

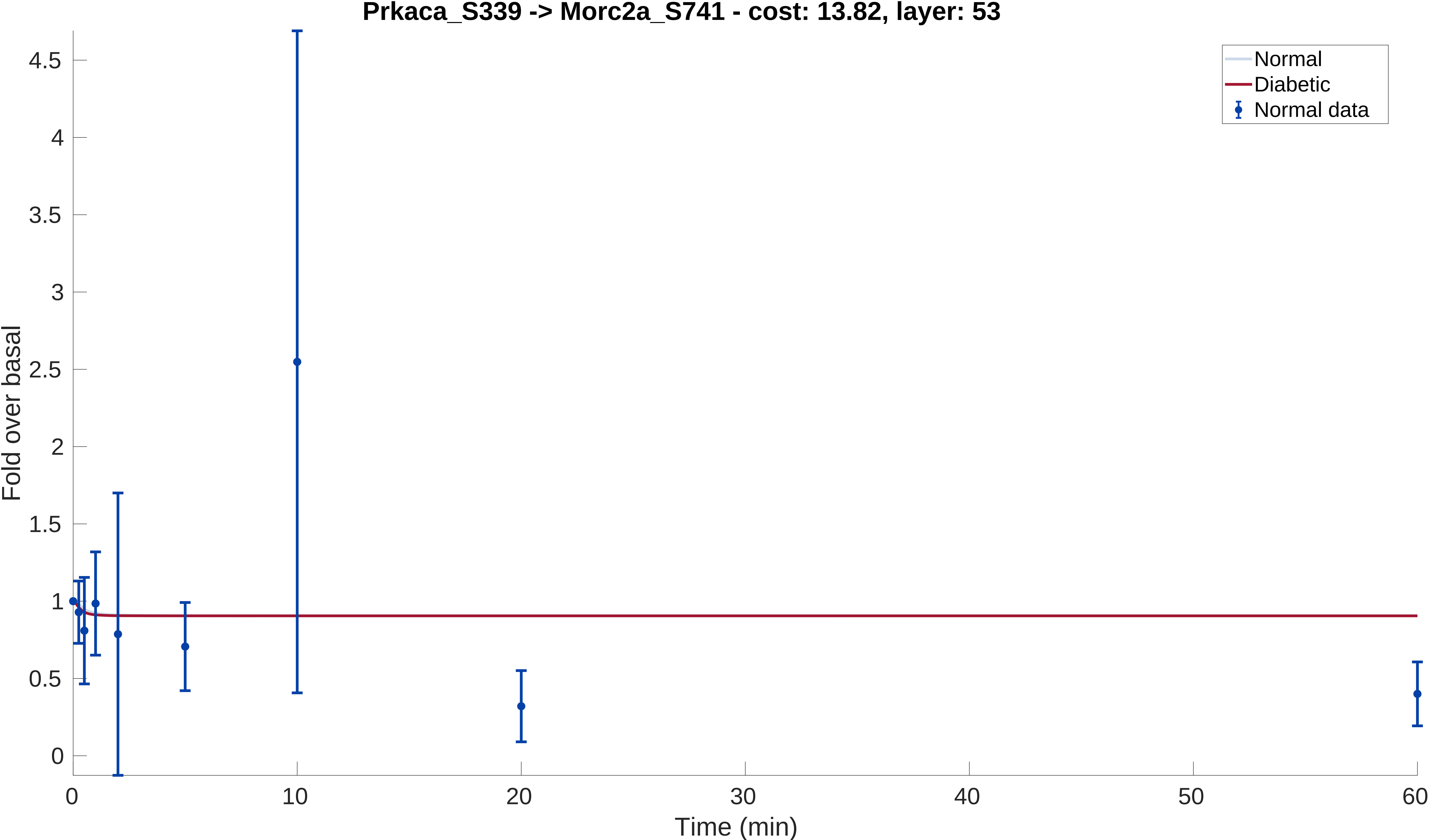

Prkaca_S339 -> Morc2a_S741 - cost: 13.82, layer: 53
Normal
Diabetic
Normal data
Fold over basal
Time (min)
0
0.5
1
1.5
2
2.5
3
3.5
4
4.5
0
10
20
30
40
50
60

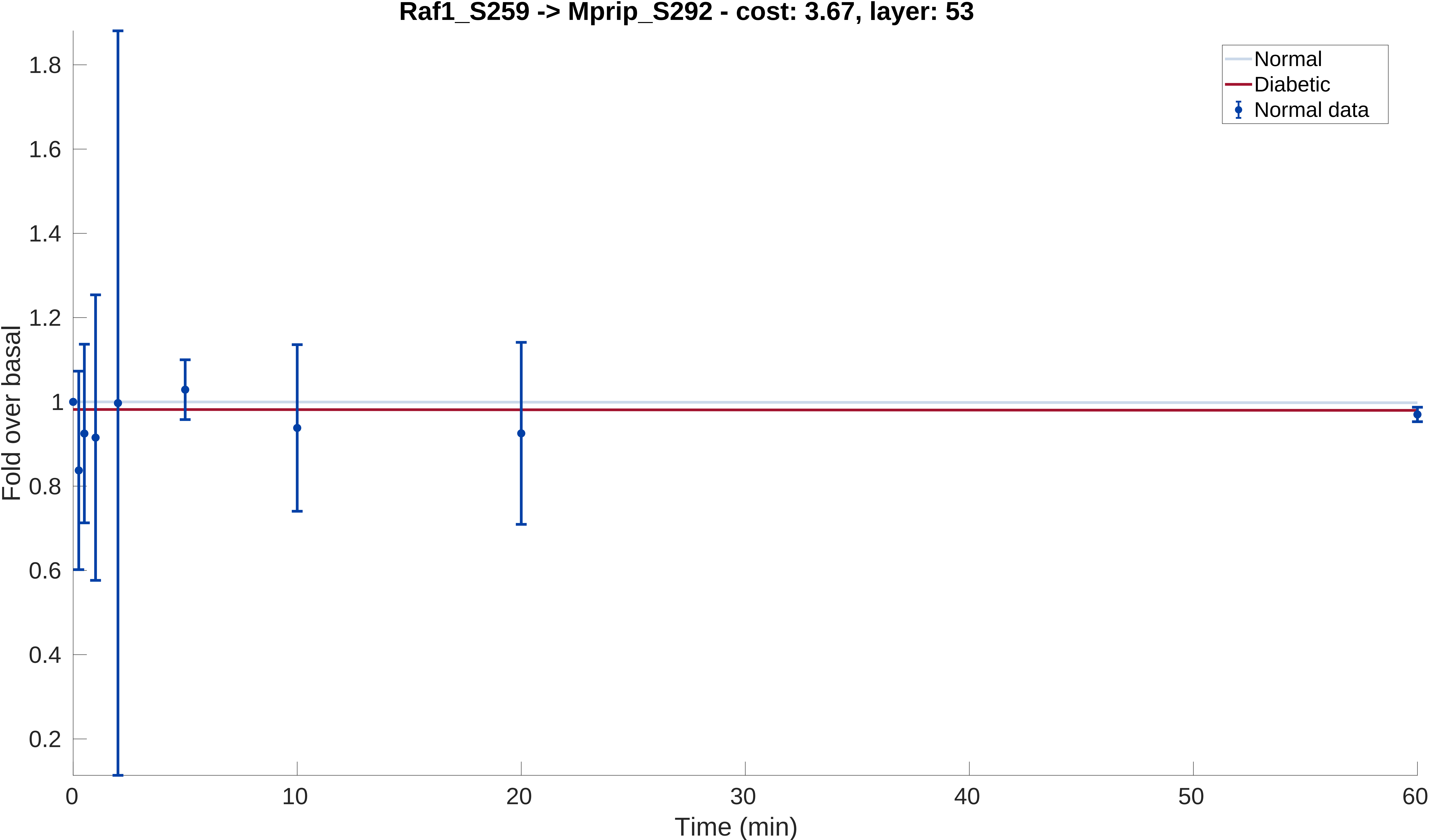

Raf1_S259 -> Mprip_S292 - cost: 3.67, layer: 53
Normal
Diabetic
Normal data
Fold over basal
1.8
1.6
1.4
1.2
1
0.8
0.6
0.4
0.2
0
10
20
30
40
50
60
Time (min)

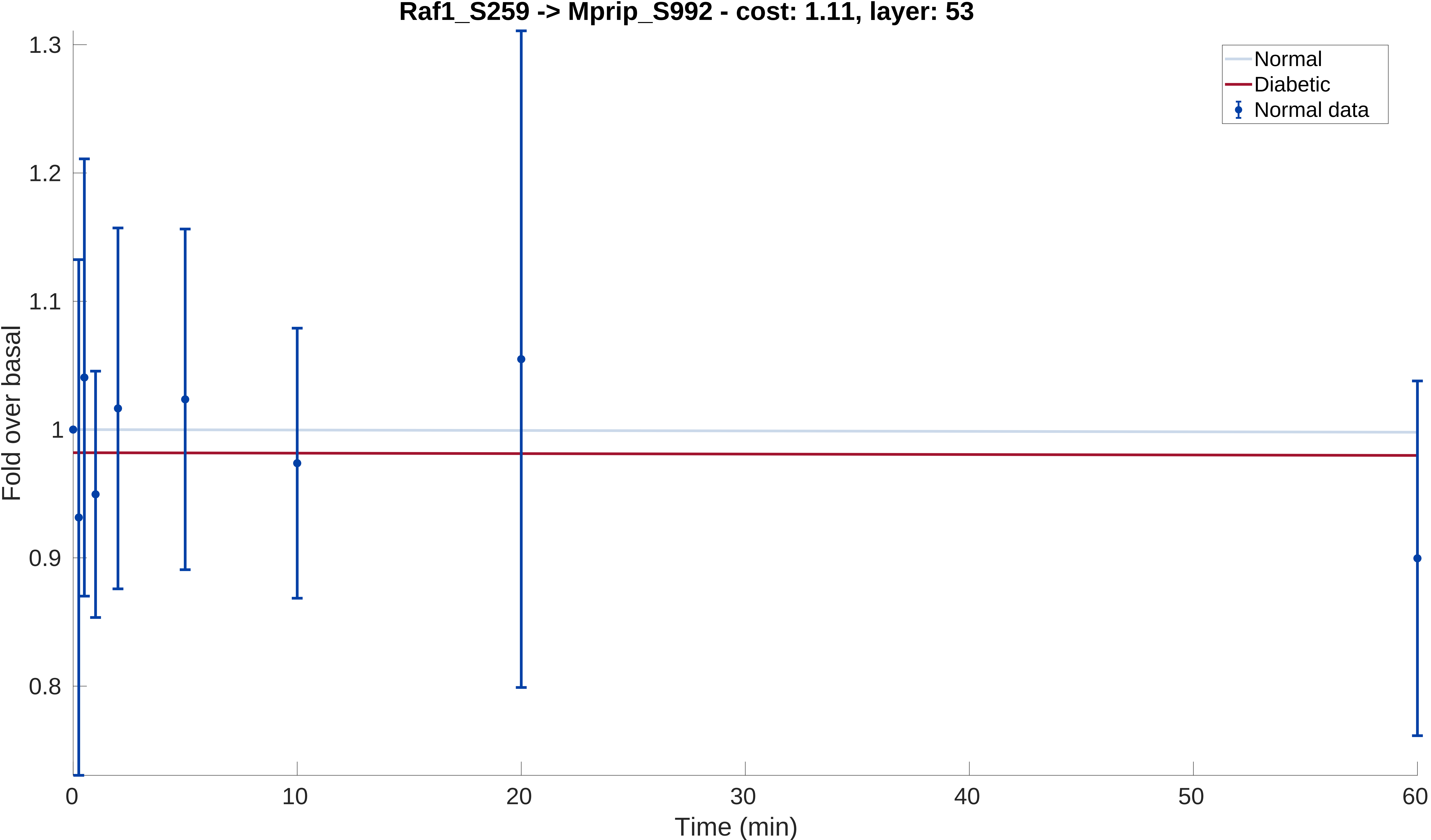

Raf1_S259 -> Mprip_S992 - cost: 1.11, layer: 53
Normal
Diabetic
Normal data
Fold over basal
1.3
1.2
1.1
1
0.9
0.8
0
10
20
30
40
50
60
Time (min)

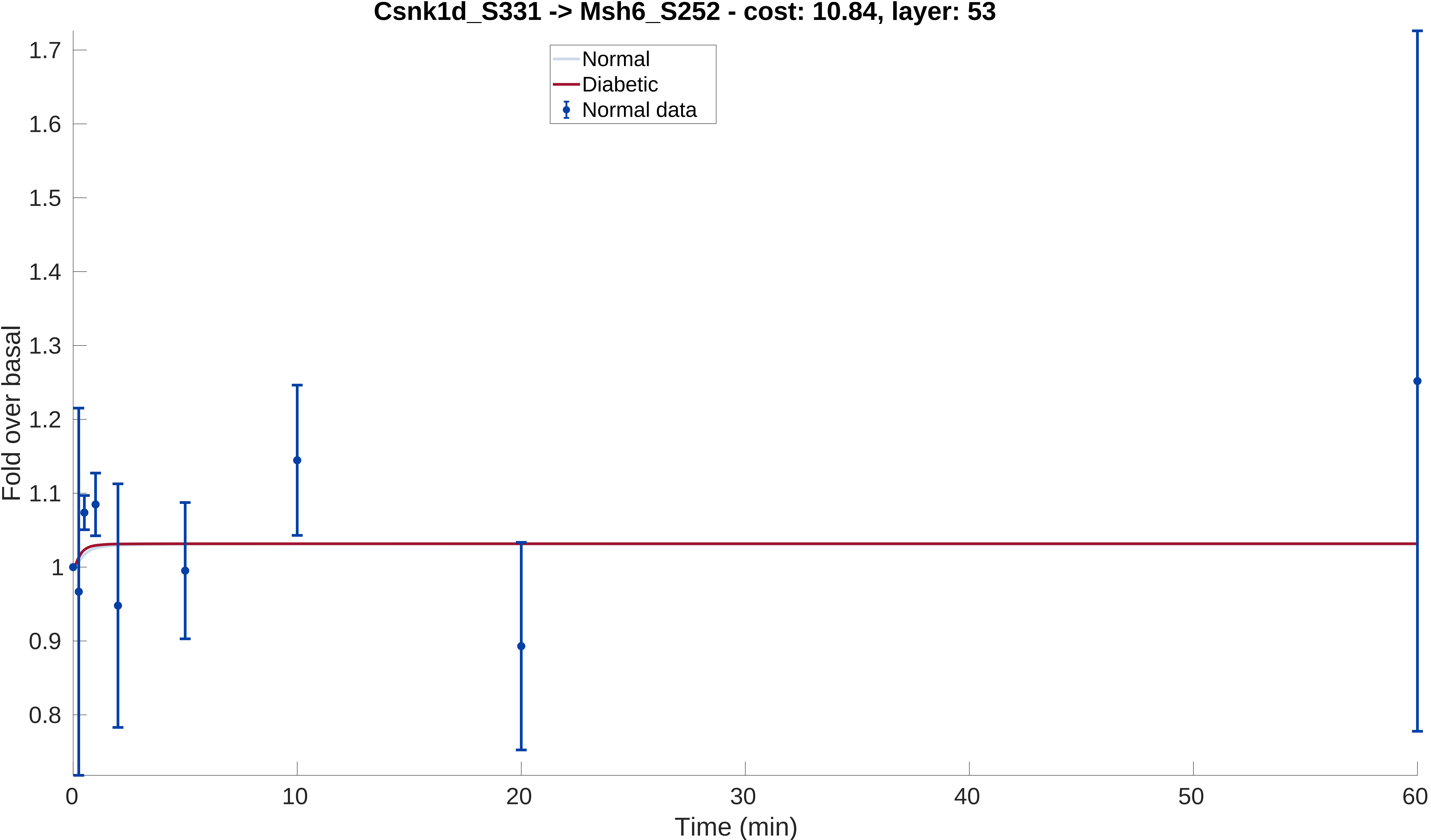

Csnk1d_S331 -> Msh6_S252 - cost: 10.84, layer: 53
Normal
Diabetic
Normal data
Fold over basal
1.7
1.6
1.5
1.4
1.3
1.2
1.1
1
0.9
0.8
0
10
20
30
40
50
60
Time (min)

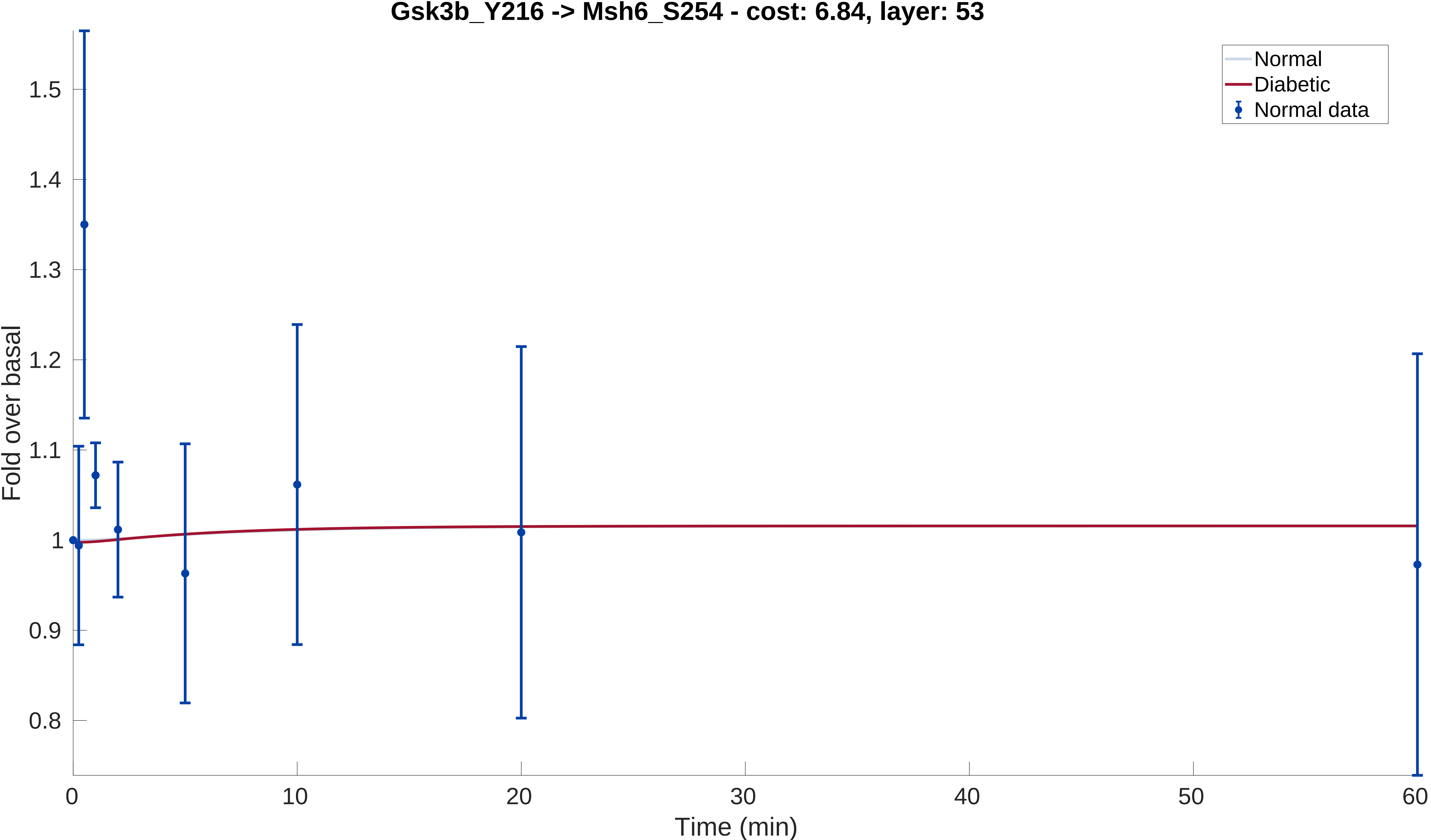

Gsk3b_Y216 -> Msh6_S254 - cost: 6.84, layer: 53
Normal
Diabetic
Normal data
Fold over basal
1.5
1.4
1.3
1.2
1.1
1
0.9
0.8
0
10
20
30
40
50
60
Time (min)

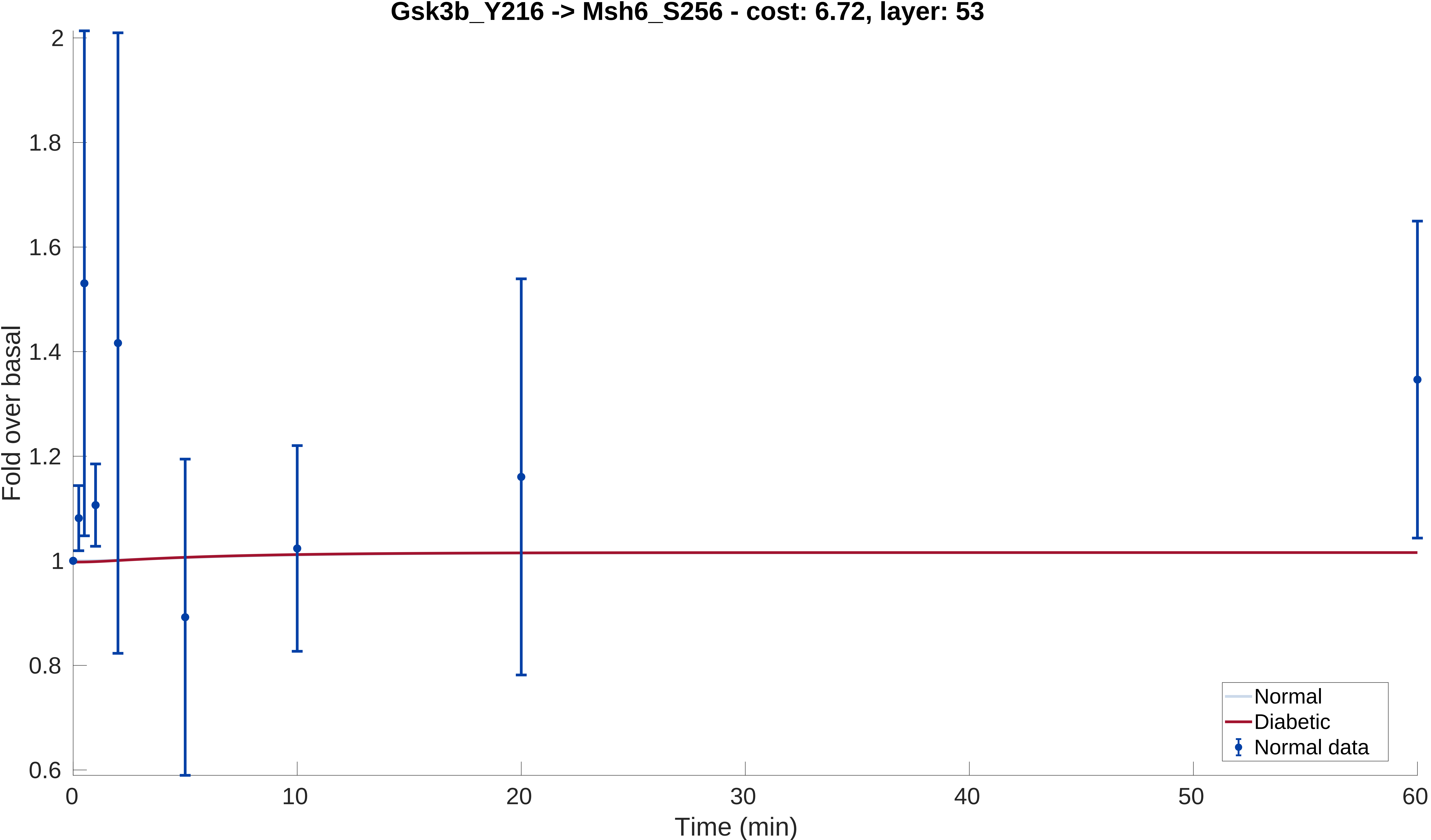

Gsk3b_Y216 -> Msh6_S256 - cost: 6.72, layer: 53
Fold over basal
2
1.8
1.6
1.4
1.2
1
0.8
0.6
0
10
20
30
40
50
60
Time (min)
Normal
Diabetic
Normal data

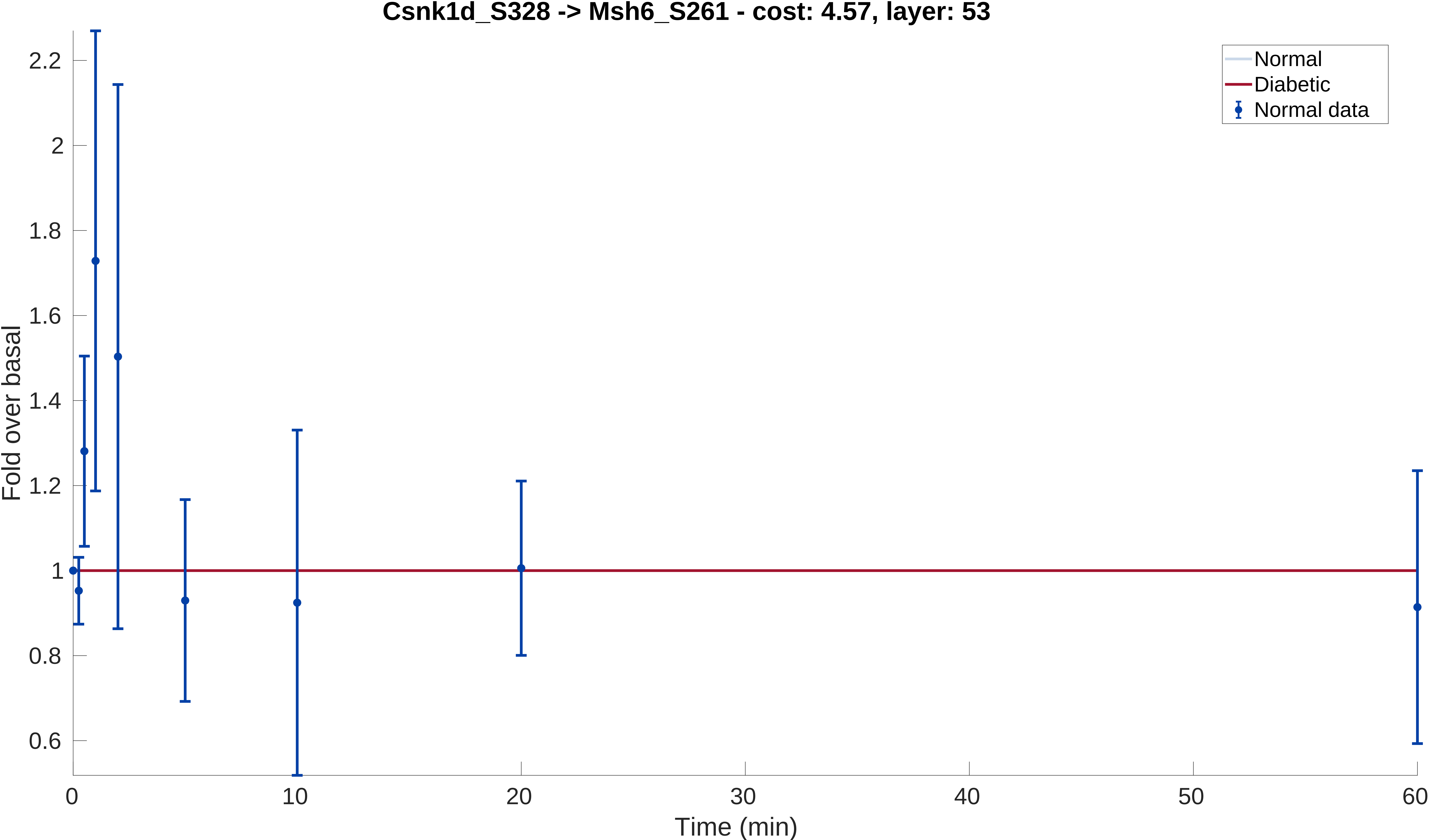

Csnk1d_S328 -> Msh6_S261 - cost: 4.57, layer: 53
Normal
Diabetic
Normal data
Fold over basal
2.2
2
1.8
1.6
1.4
1.2
1
0.8
0.6
0
10
20
30
40
50
60
Time (min)

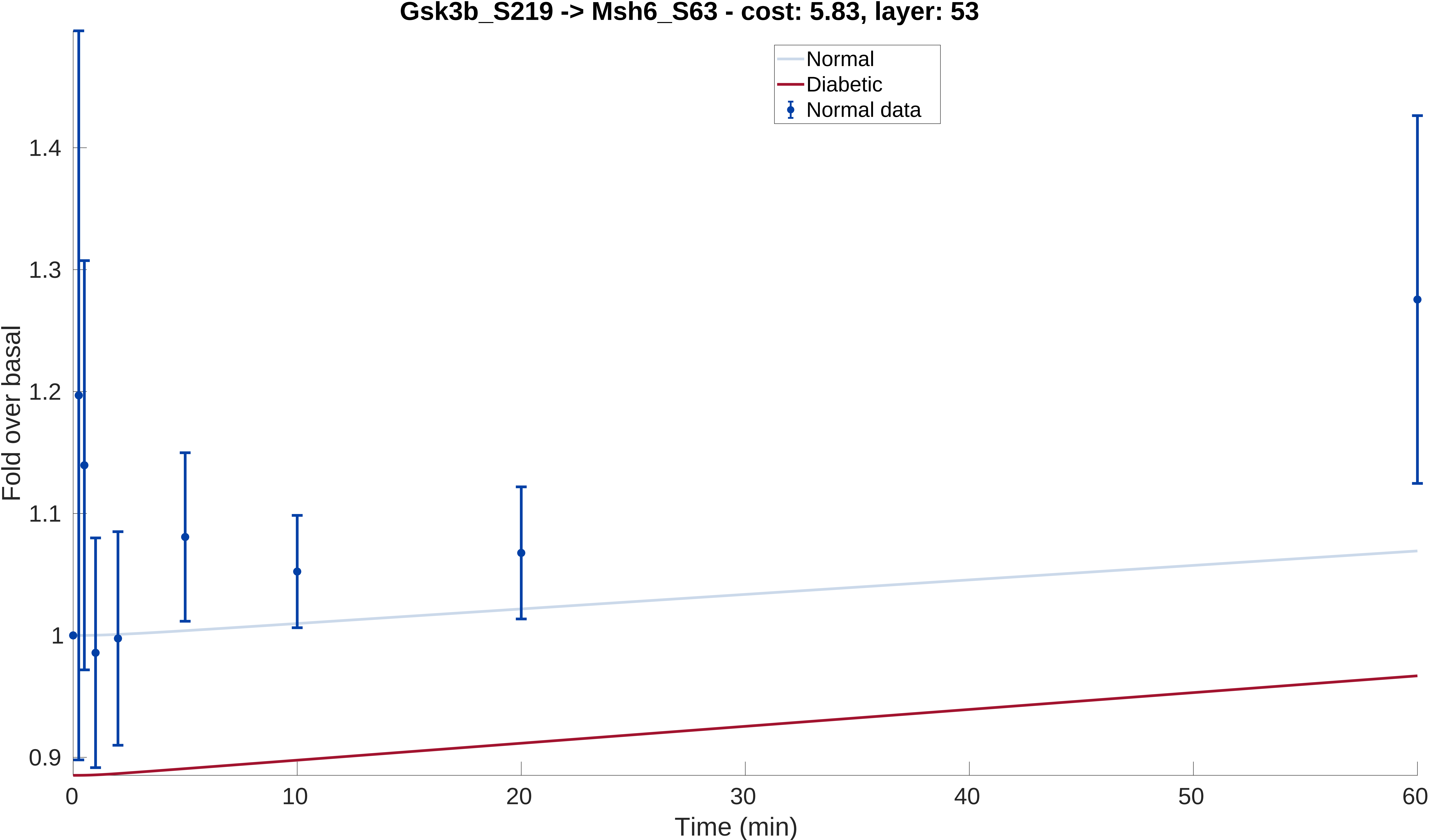

Gsk3b_S219 -> Msh6_S63 - cost: 5.83, layer: 53
Normal
Diabetic
Normal data
Fold over basal
1.4
1.3
1.2
1.1
1
0.9
0
10
20
30
40
50
60
Time (min)

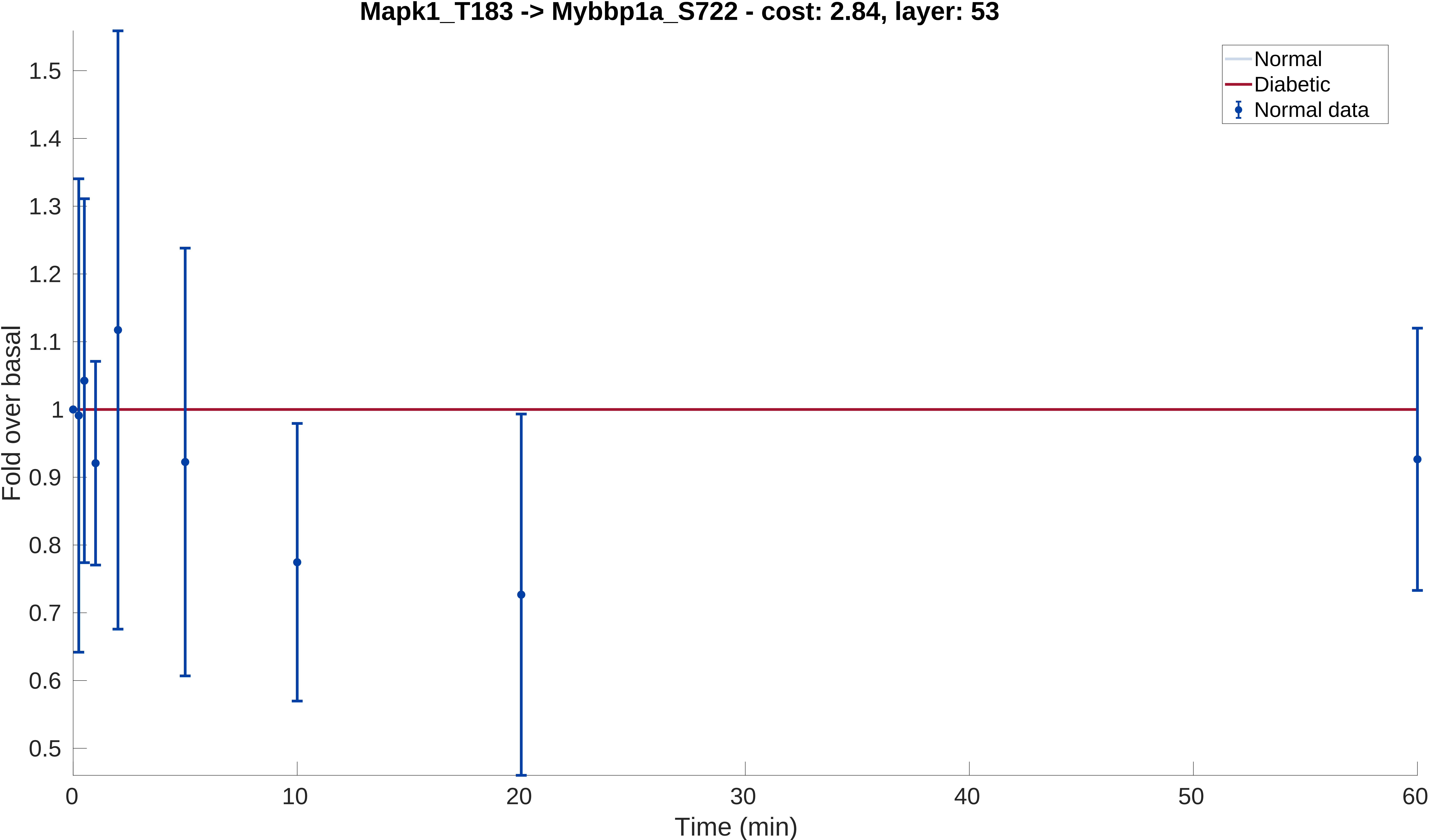

Mapk1_T183 -> Mybbp1a_S722 - cost: 2.84, layer: 53
Normal
Diabetic
Normal data
Fold over basal
Time (min)
1.5
1.4
1.3
1.2
1.1
1
0.9
0.8
0.7
0.6
0.5
0
10
20
30
40
50
60

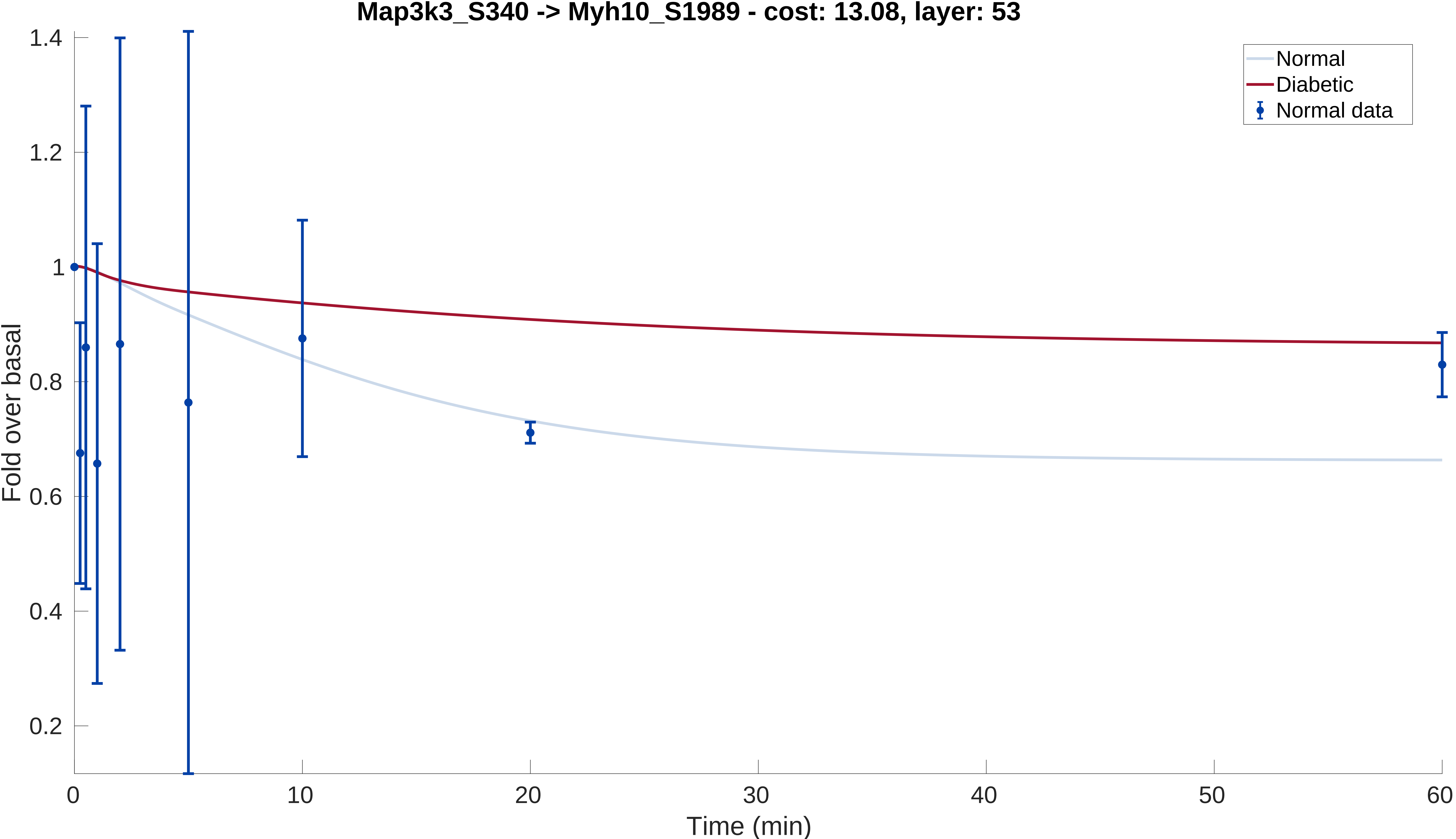
Map3k3_S340 -> Myh10_S1989 - cost: 13.08, layer: 53
Normal
Diabetic
Normal data
Fold over basal
1.4
1.2
1
0.8
0.6
0.4
0.2
0
10
20
30
40
50
60
Time (min)

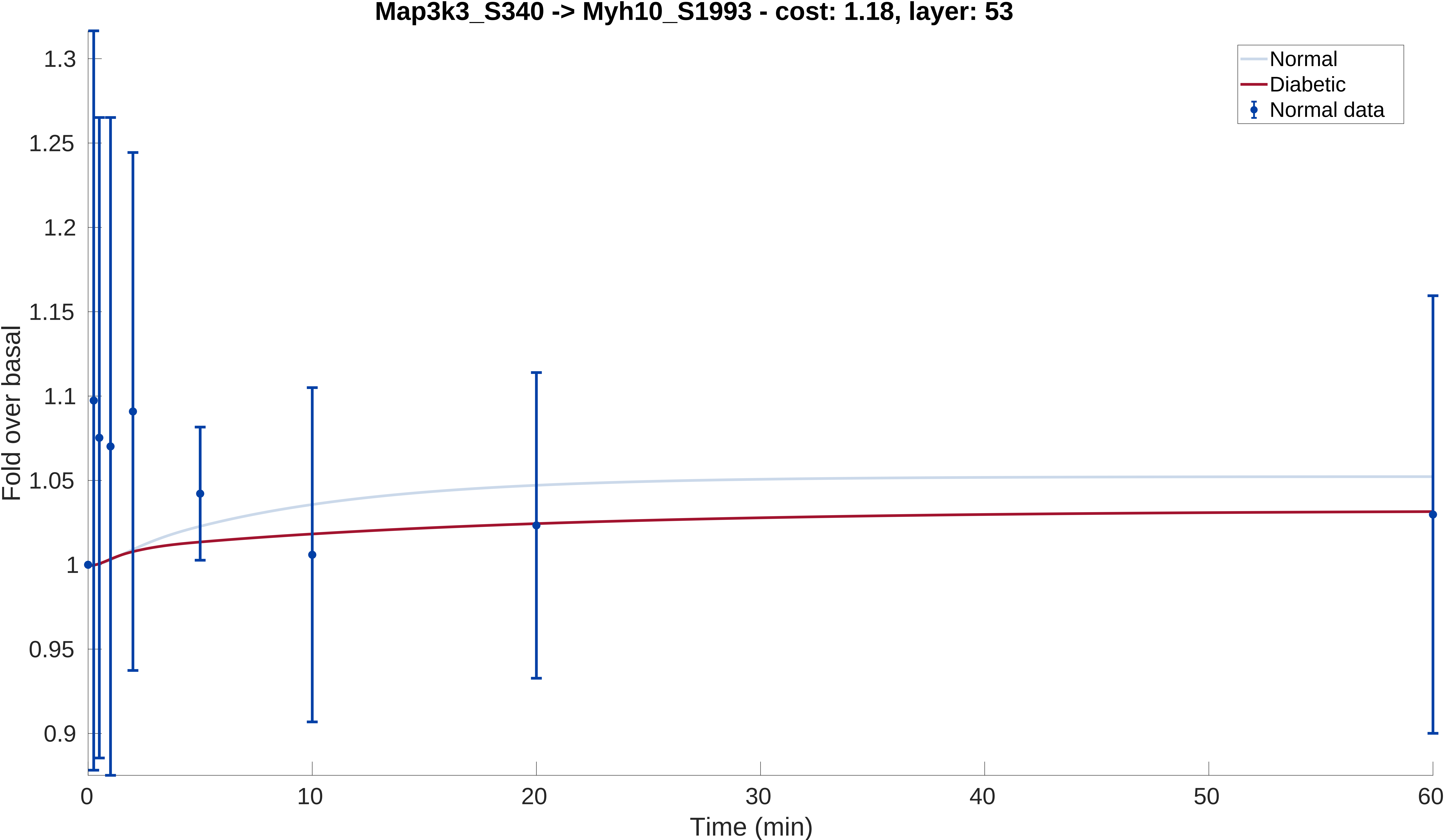
Map3k3_S340 -> Myh10_S1993 - cost: 1.18, layer: 53
Fold over basal
1.3
1.25
1.2
1.15
1.1
1.05
1
0.95
0.9
0
10
20
30
40
50
60
Time (min)
Normal
Diabetic
Normal data

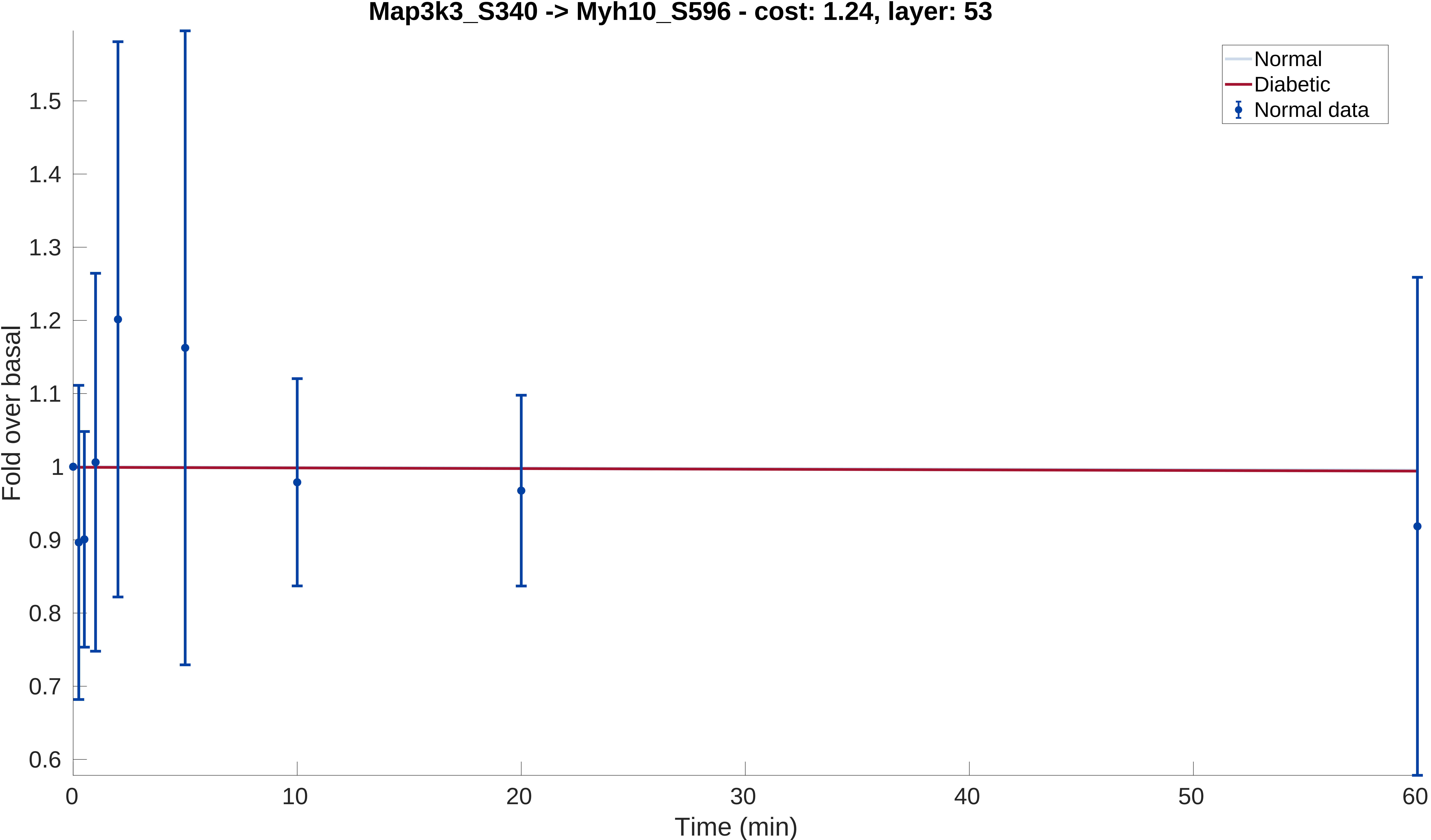

Map3k3_S340 -> Myh10_S596 - cost: 1.24, layer: 53
Normal
Diabetic
Normal data
Fold over basal
1.5
1.4
1.3
1.2
1.1
1
0.9
0.8
0.7
0.6
0
10
20
30
40
50
60
Time (min)

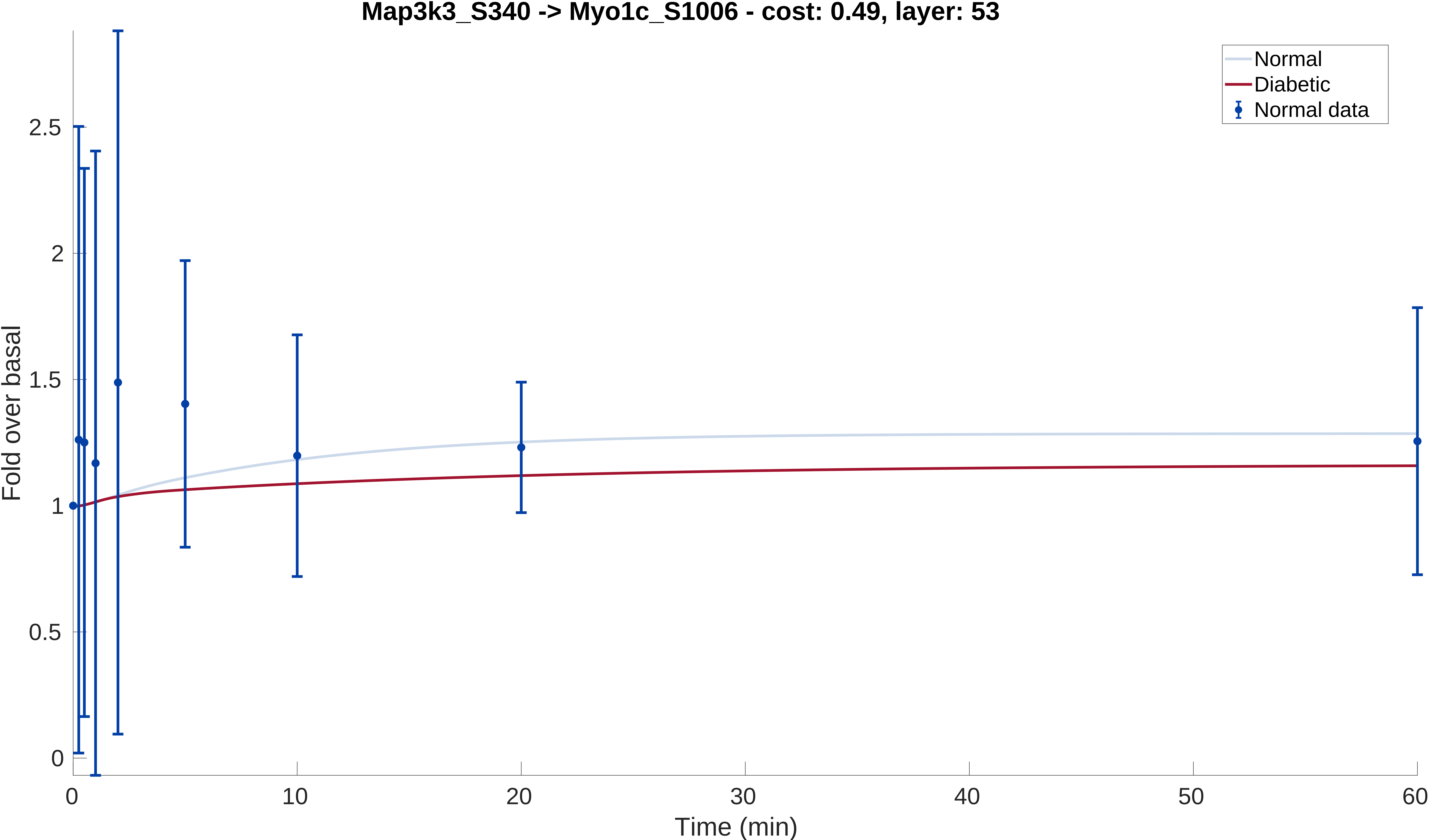

Map3k3_S340 -> Myo1c_S1006 - cost: 0.49, layer: 53
Normal
Diabetic
Normal data
Fold over basal
0
0.5
1
1.5
2
2.5
0
10
20
30
40
50
60
Time (min)

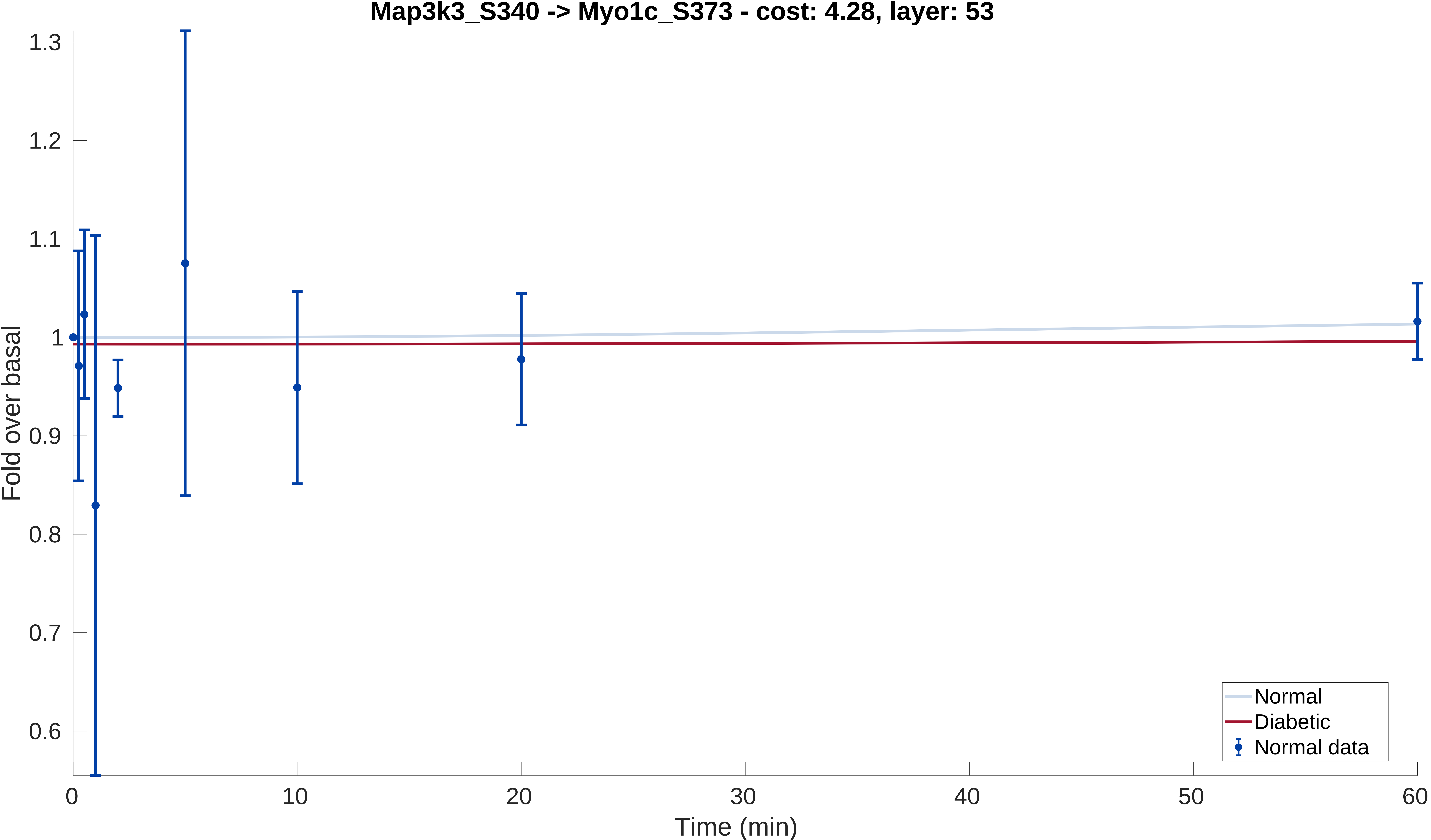
Map3k3_S340 -> Myo1c_S373 - cost: 4.28, layer: 53
Fold over basal
1.3
1.2
1.1
1
0.9
0.8
0.7
0.6
0
10
20
30
40
50
60
Time (min)
Normal
Diabetic
Normal data

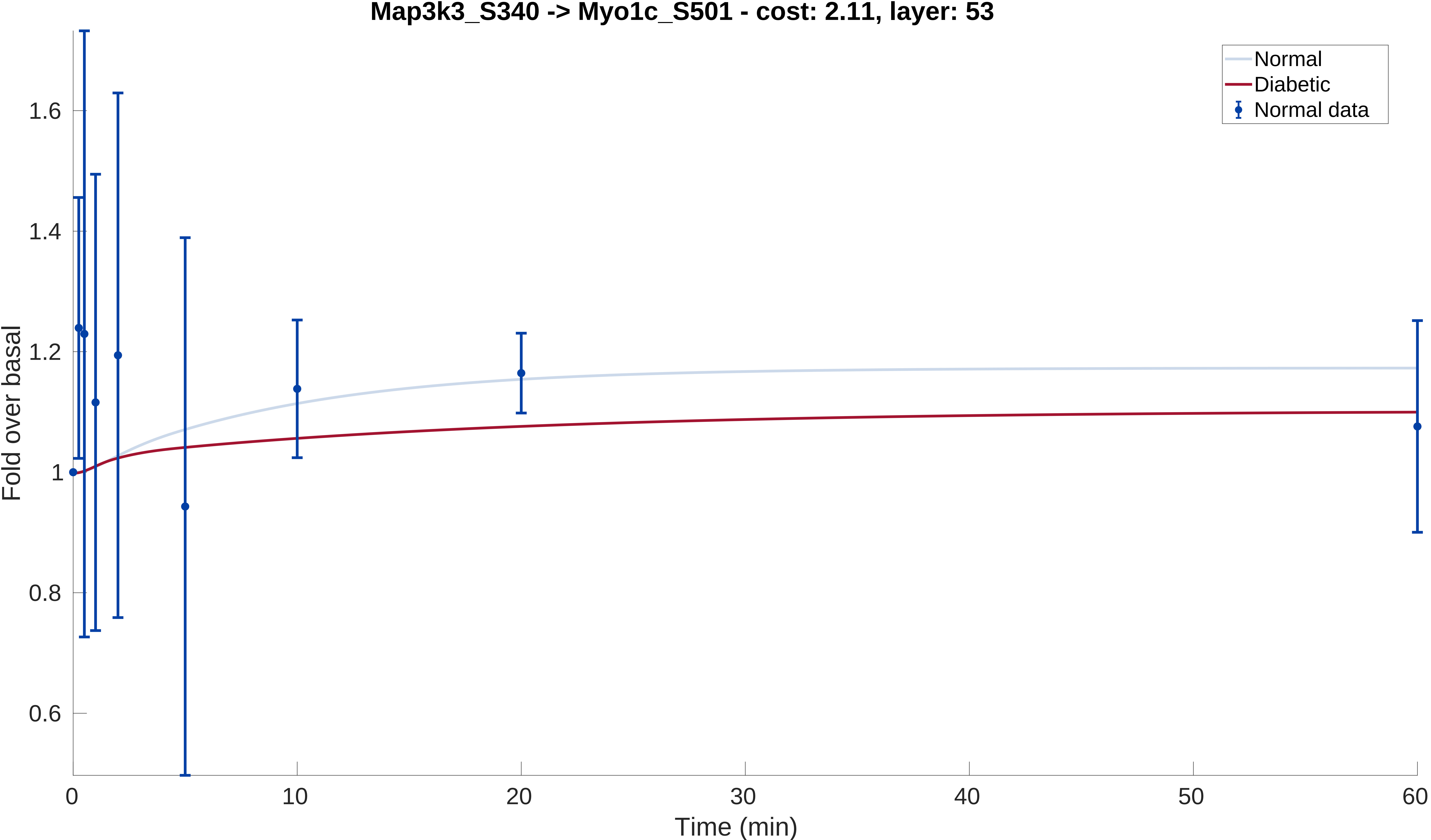

Map3k3_S340 -> Myo1c_S501 - cost: 2.11, layer: 53
Normal
Diabetic
Normal data
Fold over basal
1.6
1.4
1.2
1
0.8
0.6
0
10
20
30
40
50
60
Time (min)

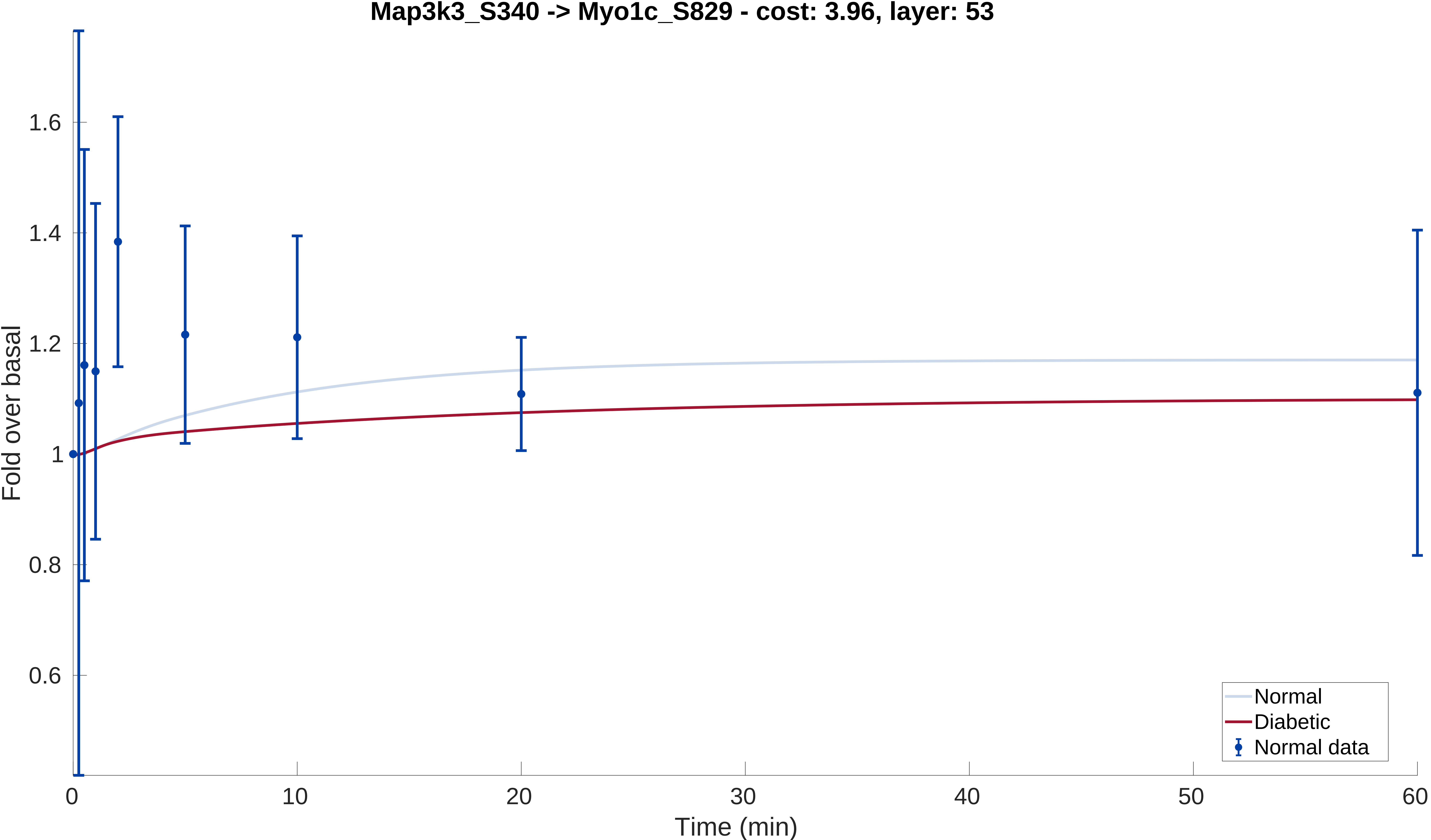

Map3k3_S340 -> Myo1c_S829 - cost: 3.96, layer: 53
Fold over basal
1.6
1.4
1.2
1
0.8
0.6
0
10
20
30
40
50
60
Time (min)
Normal
Diabetic
Normal data

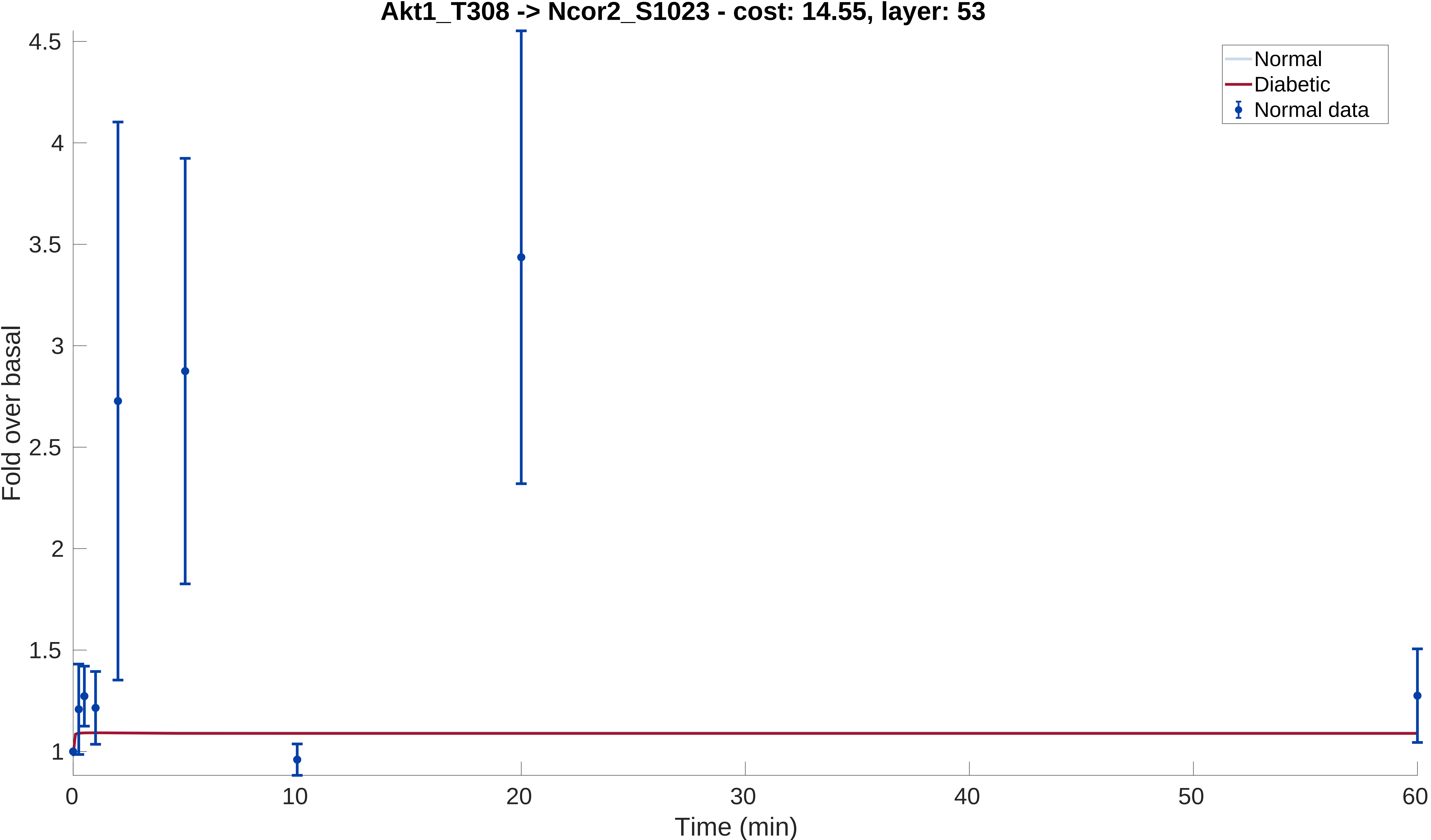

Akt1_T308 -> Ncor2_S1023 - cost: 14.55, layer: 53
Normal
Diabetic
Normal data
Fold over basal
4.5
4
3.5
3
2.5
2
1.5
1
0
10
20
30
40
50
60
Time (min)

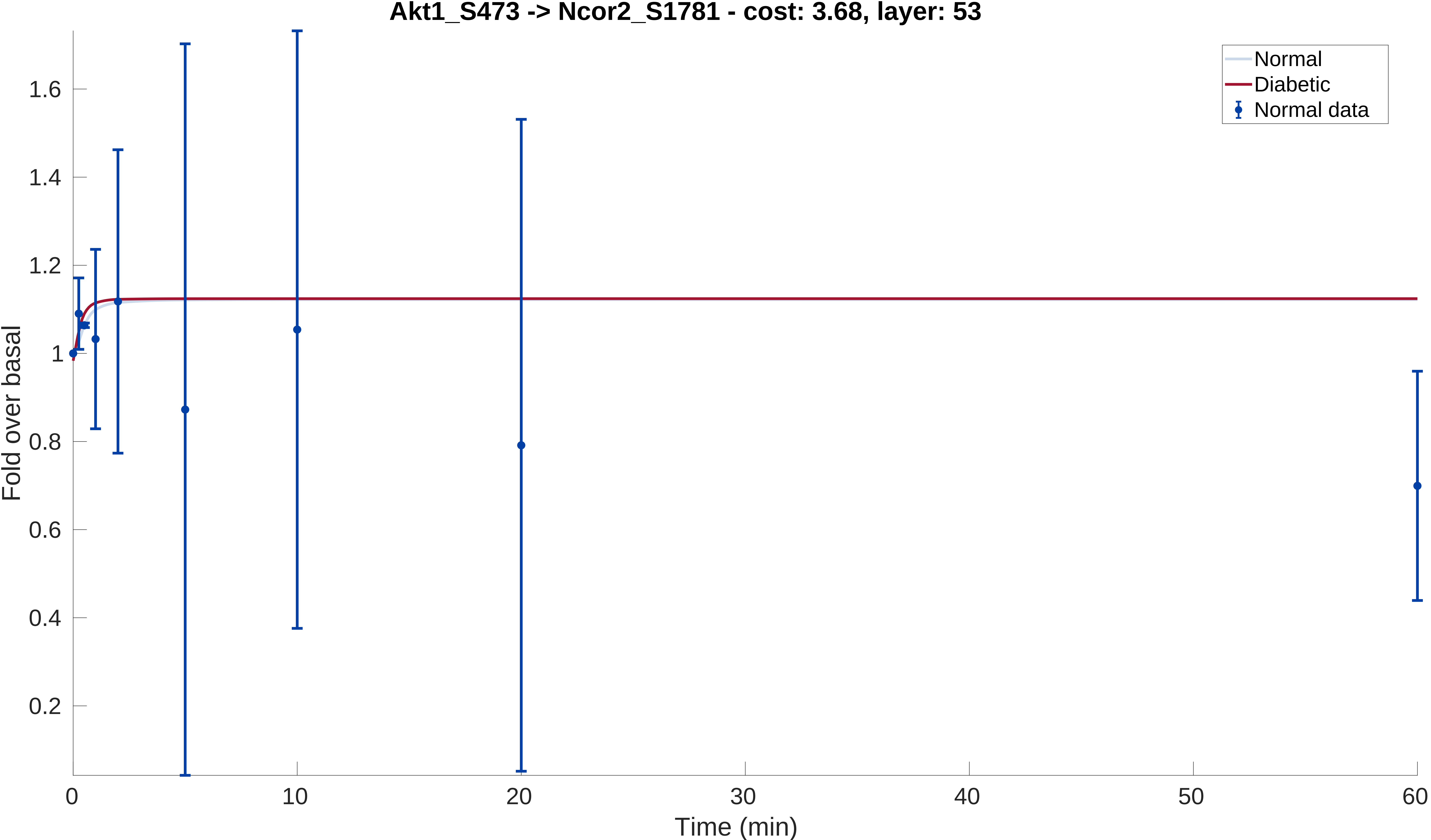

Akt1_S473 -> Ncor2_S1781 - cost: 3.68, layer: 53
Normal
Diabetic
Normal data
Fold over basal
1.6
1.4
1.2
1
0.8
0.6
0.4
0.2
0
10
20
30
40
50
60
Time (min)

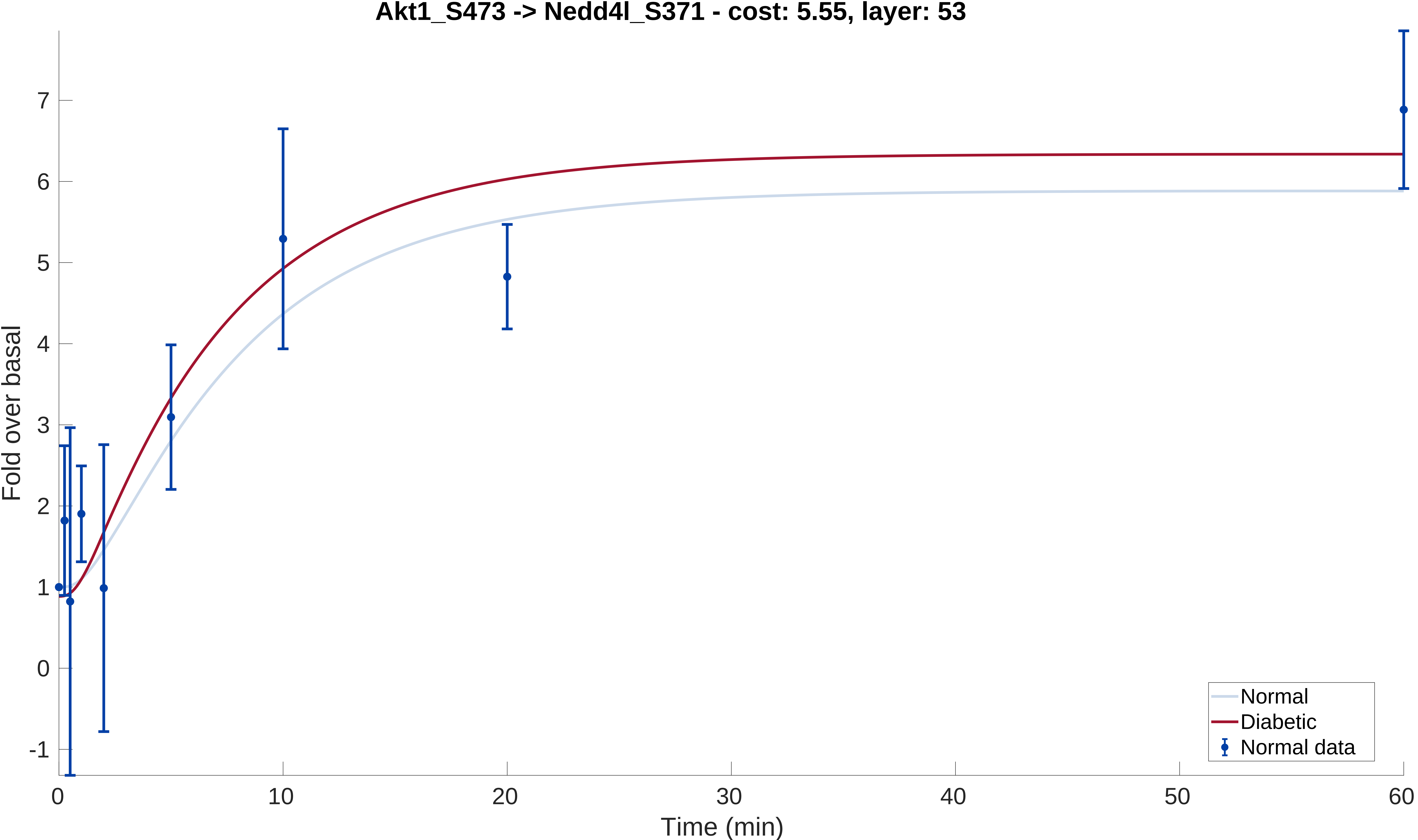
Akt1_S473 -> Nedd4l_S371 - cost: 5.55, layer: 53
Fold over basal
Time (min)
7
6
5
4
3
2
1
0
-1
0
10
20
30
40
50
60
Normal
Diabetic
Normal data

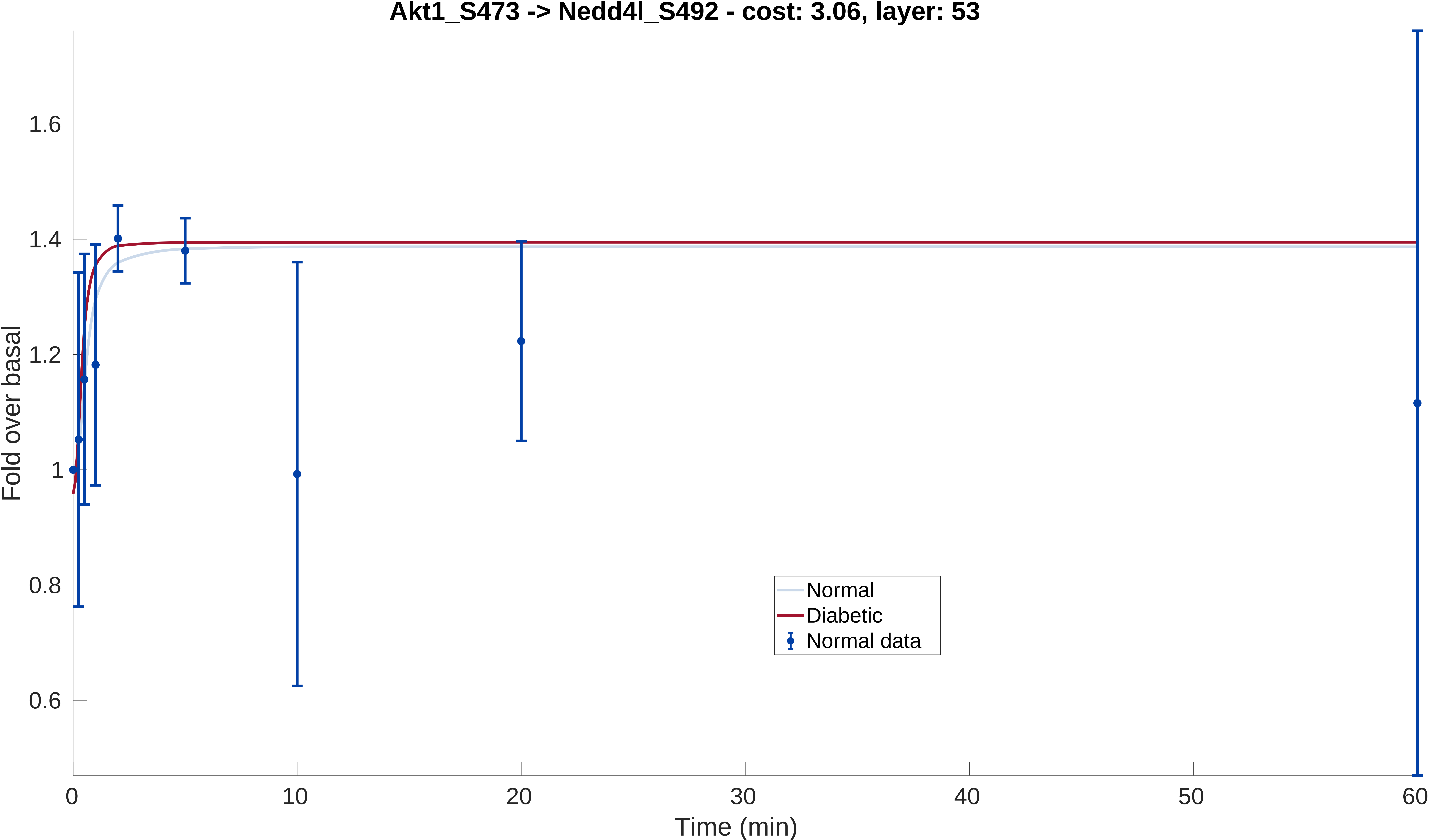

Akt1_S473 -> Nedd4l_S492 - cost: 3.06, layer: 53
Fold over basal
1.6
1.4
1.2
1
0.8
0.6
0
10
20
30
40
50
60
Time (min)
Normal
Diabetic
Normal data

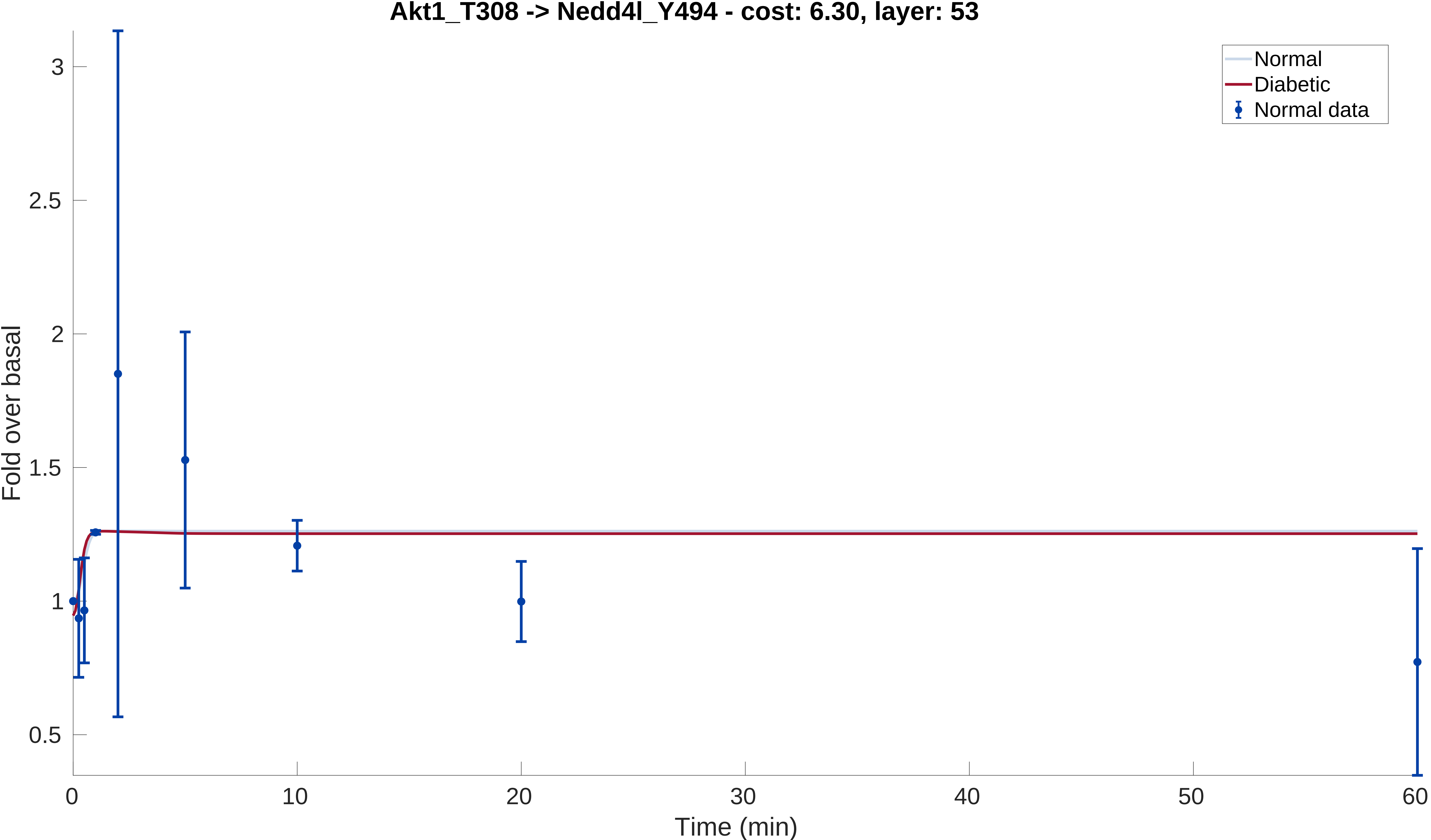

Akt1_T308 -> Nedd4l_Y494 - cost: 6.30, layer: 53
Normal
Diabetic
Normal data
Fold over basal
Time (min)
0.5
1
1.5
2
2.5
3
0
10
20
30
40
50
60

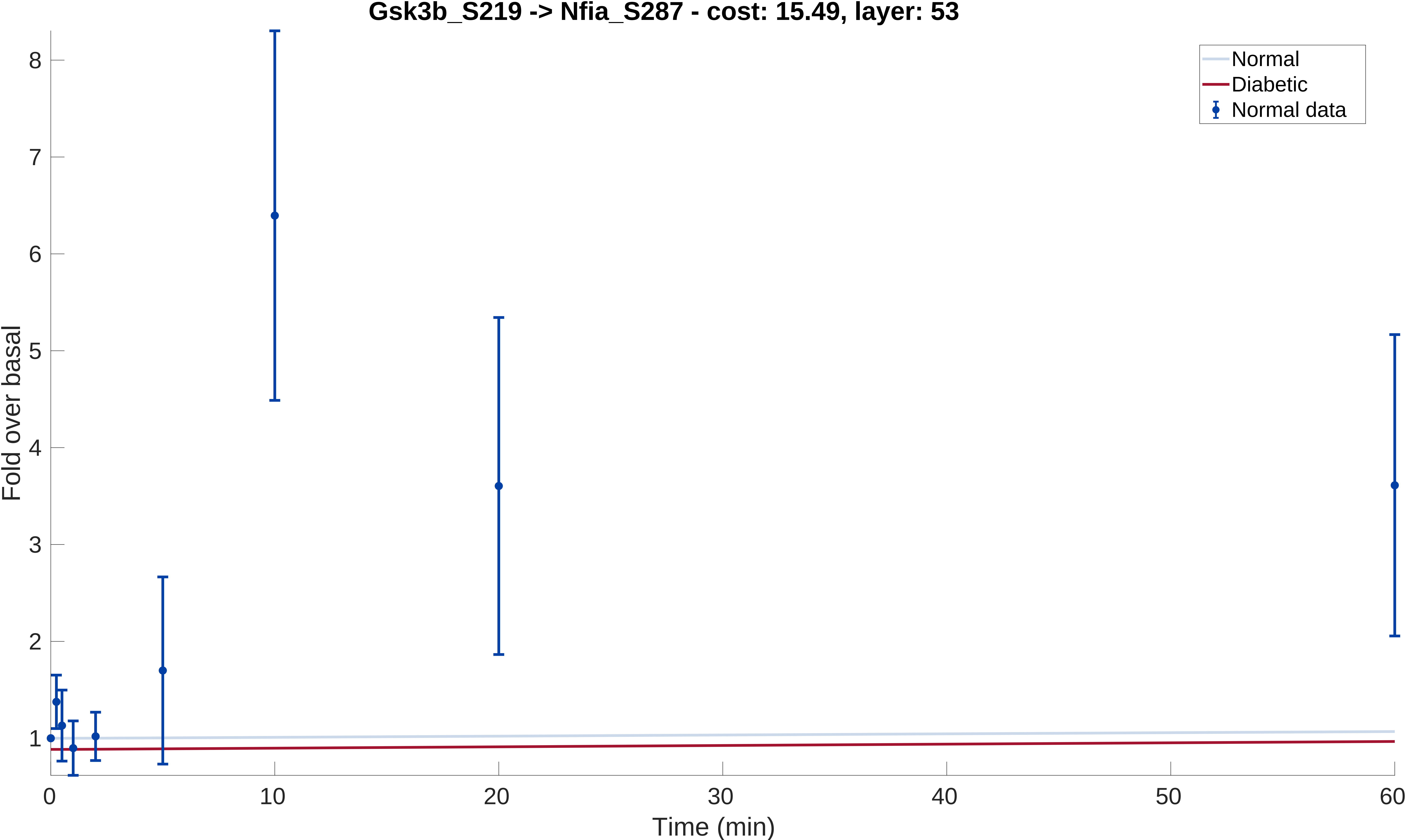

Gsk3b_S219 -> Nfia_S287 - cost: 15.49, layer: 53
Normal
Diabetic
Normal data
Fold over basal
Time (min)
0
10
20
30
40
50
60
1
2
3
4
5
6
7
8

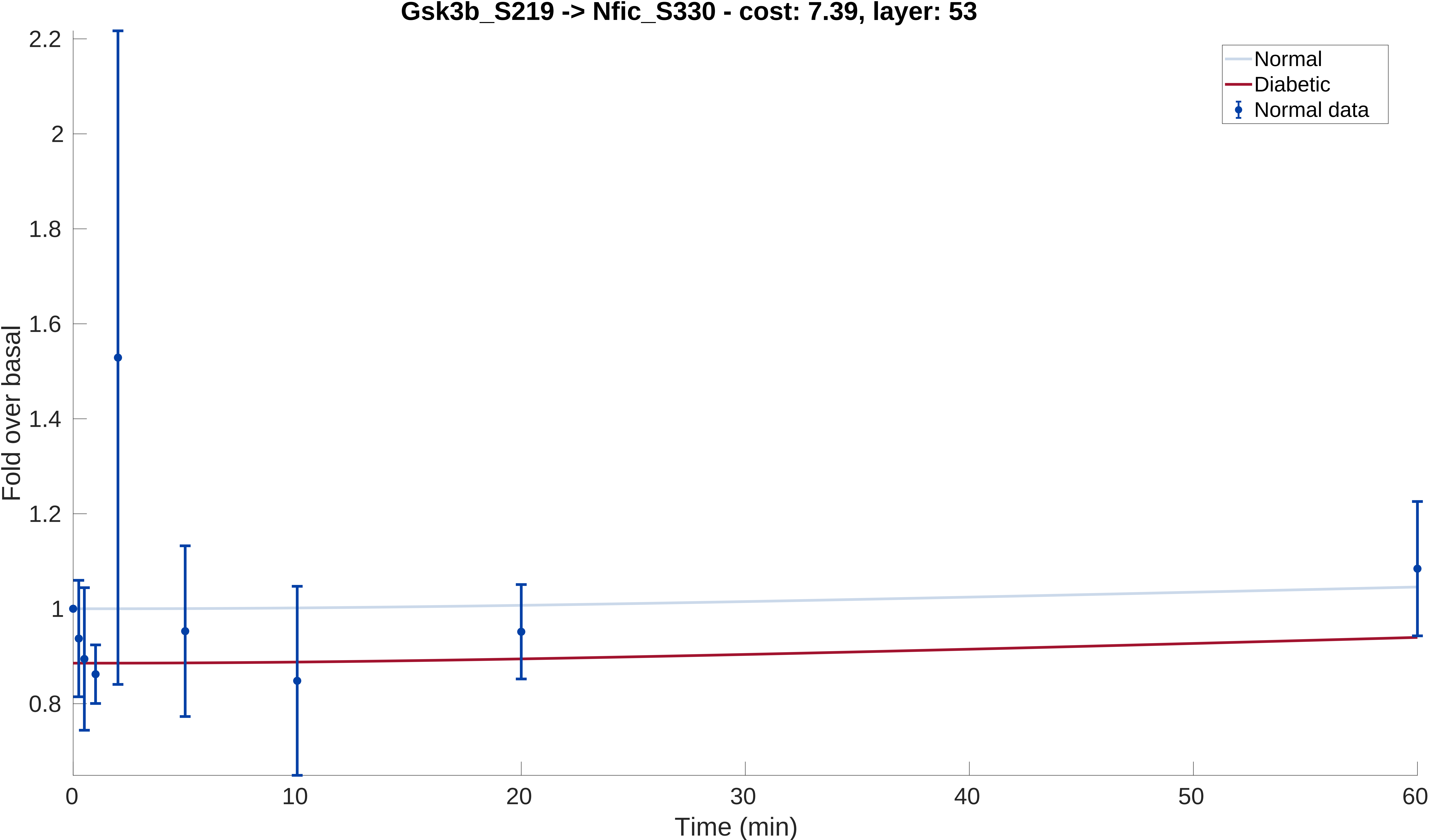
Gsk3b_S219 -> Nfic_S330 - cost: 7.39, layer: 53
Fold over basal
Time (min)
Normal
Diabetic
Normal data
0.8
1
1.2
1.4
1.6
1.8
2
2.2
0
10
20
30
40
50
60

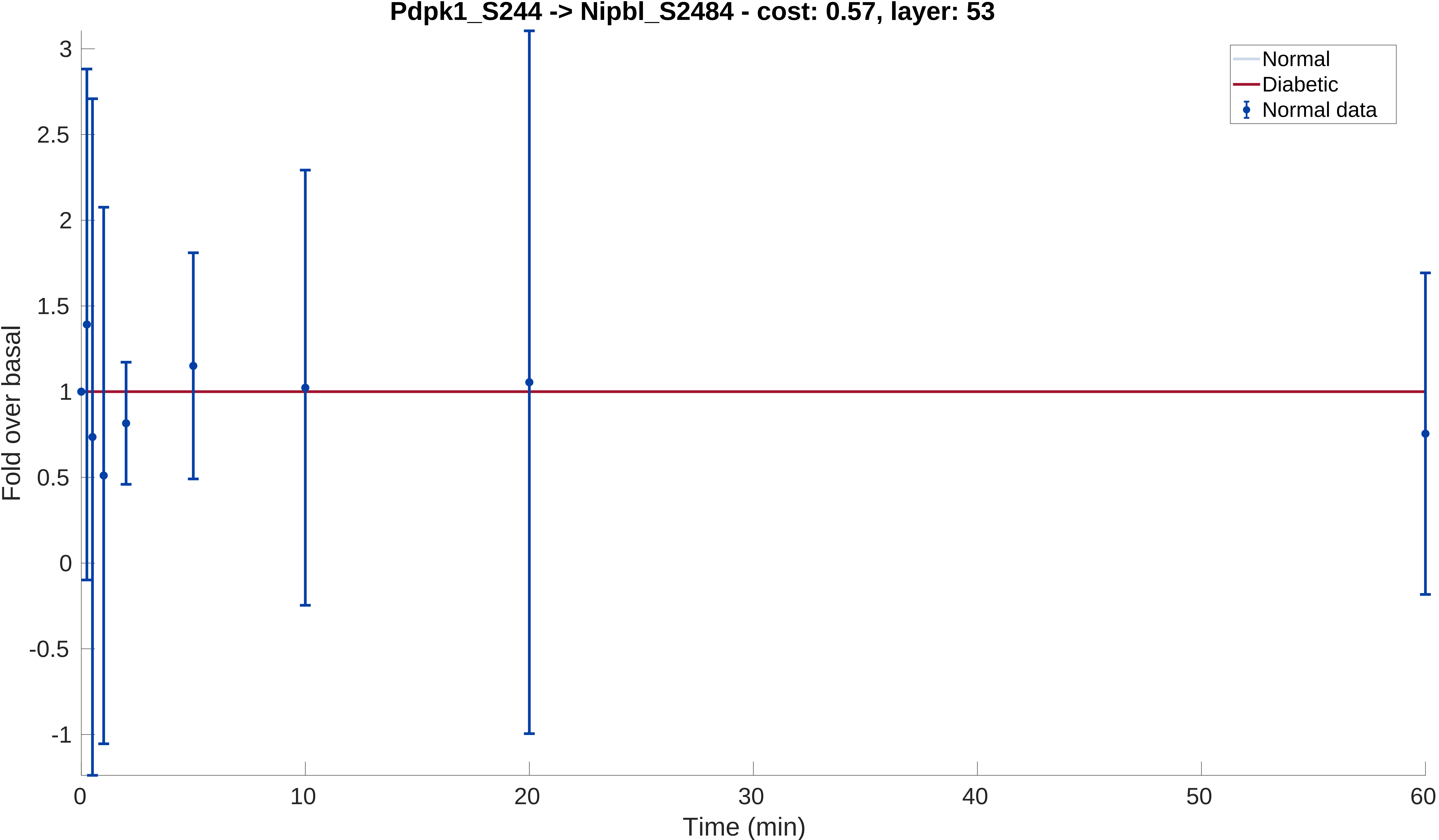

Pdpk1_S244 -> Nipbl_S2484 - cost: 0.57, layer: 53
Normal
Diabetic
Normal data
Fold over basal
3
2.5
2
1.5
1
0.5
0
-0.5
-1
0
10
20
30
40
50
60
Time (min)

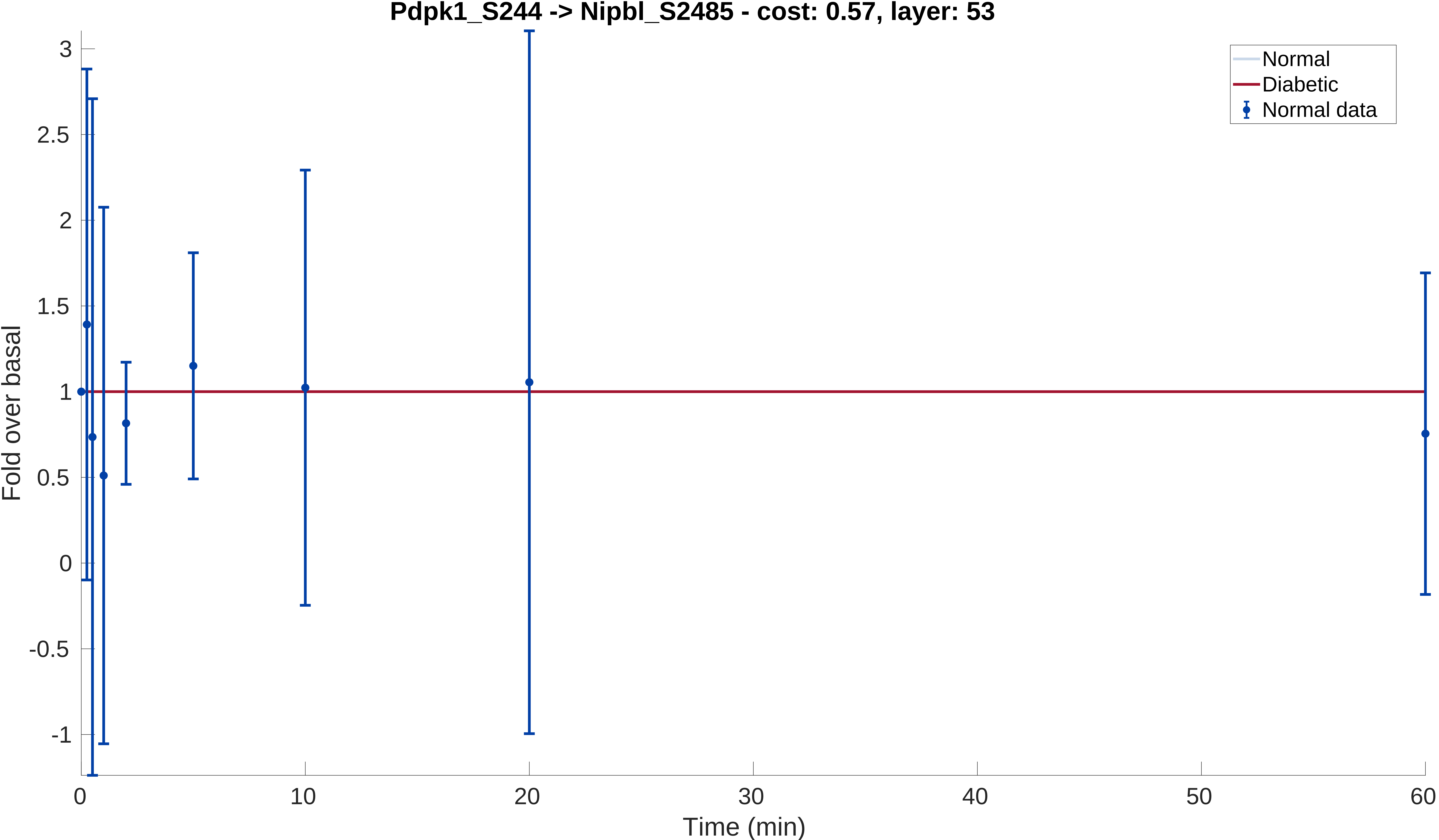

Pdpk1_S244 -> Nipbl_S2485 - cost: 0.57, layer: 53
Normal
Diabetic
Normal data
Fold over basal
Time (min)
3
2.5
2
1.5
1
0.5
0
-0.5
-1
0
10
20
30
40
50
60

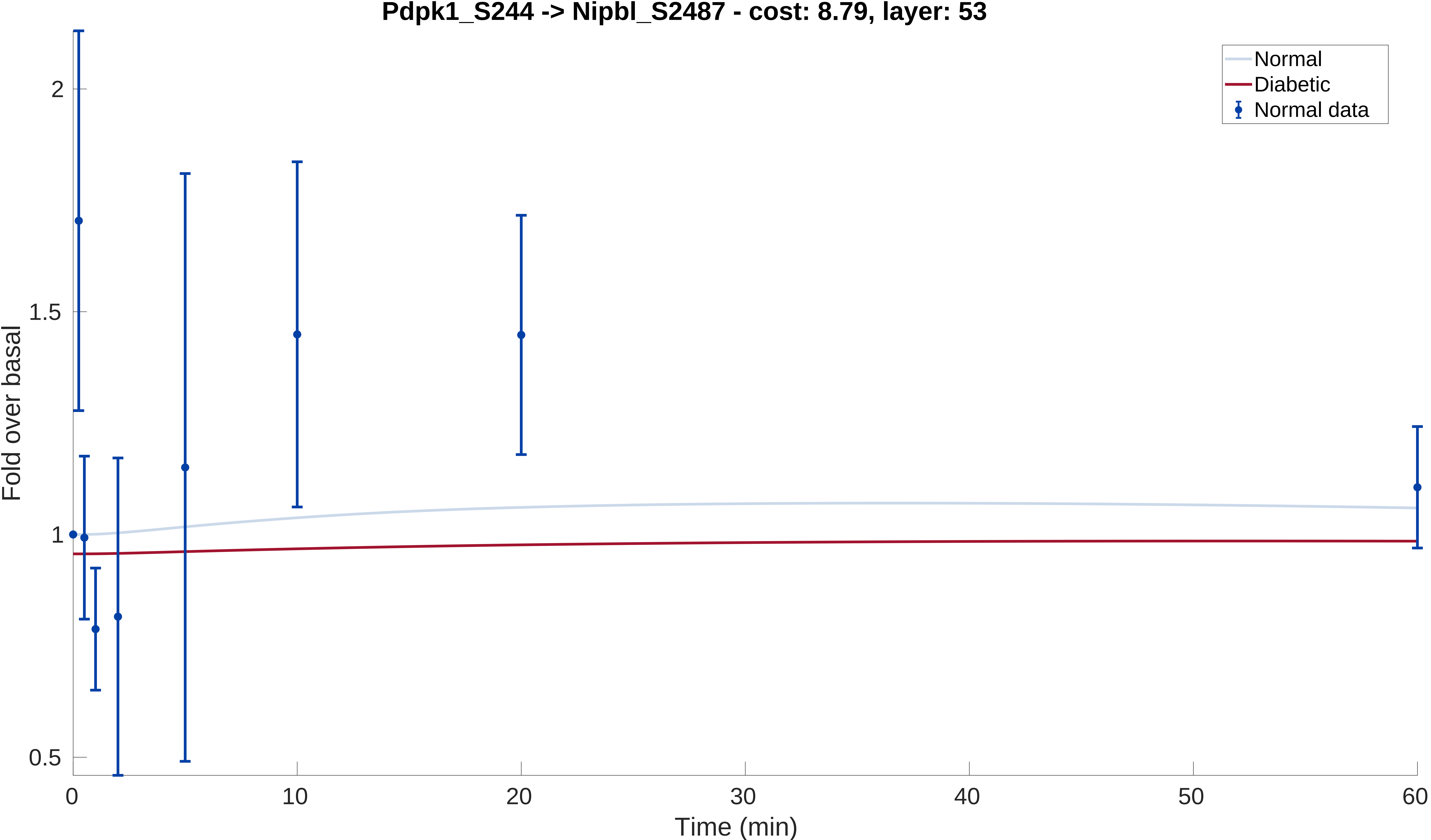

Pdpk1_S244 -> Nipbl_S2487 - cost: 8.79, layer: 53
Normal
Diabetic
Normal data
Fold over basal
2
1.5
1
0.5
0
10
20
30
40
50
60
Time (min)

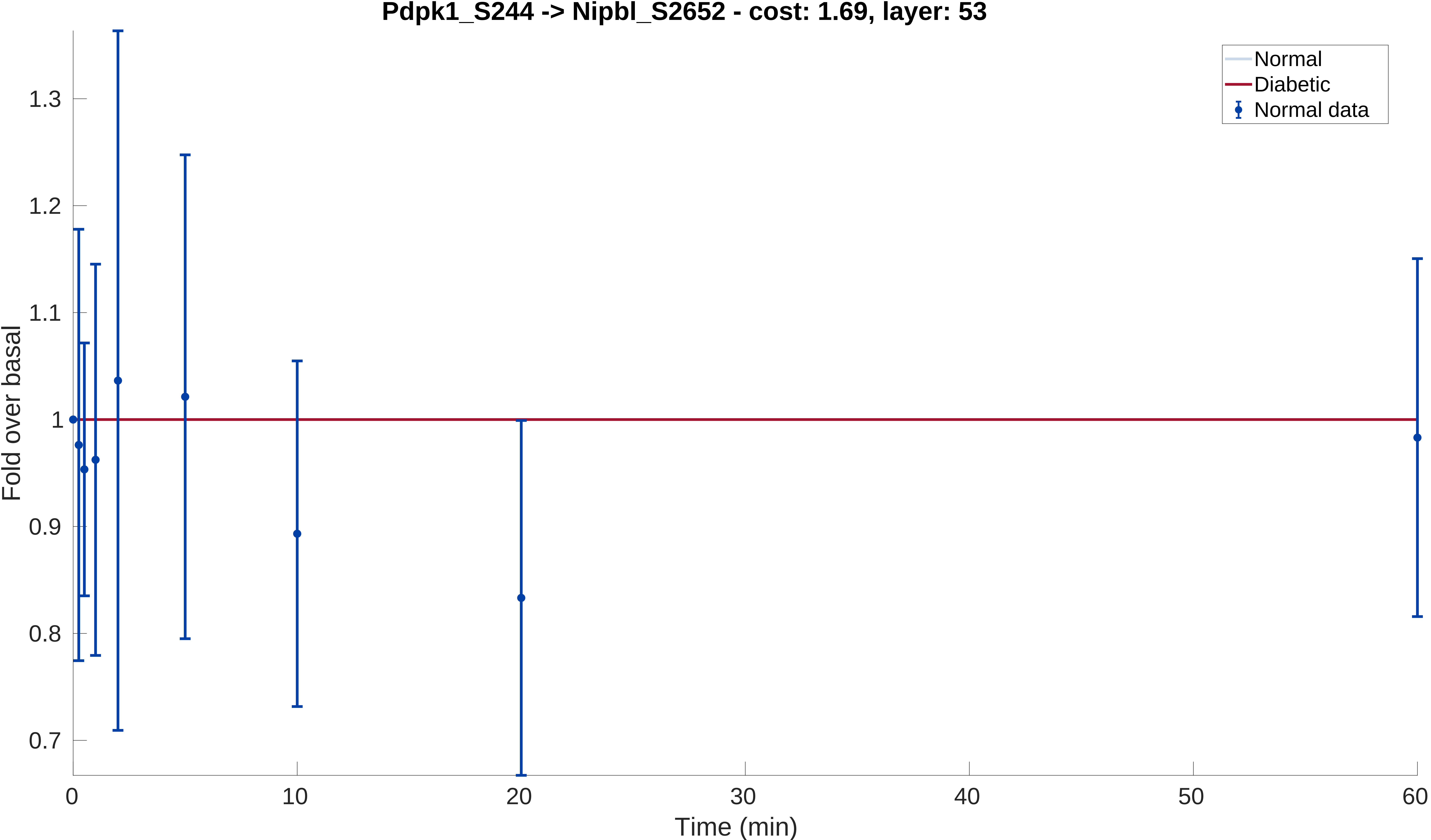

Pdpk1_S244 -> Nipbl_S2652 - cost: 1.69, layer: 53
Normal
Diabetic
Normal data
Fold over basal
1.3
1.2
1.1
1
0.9
0.8
0.7
0
10
20
30
40
50
60
Time (min)

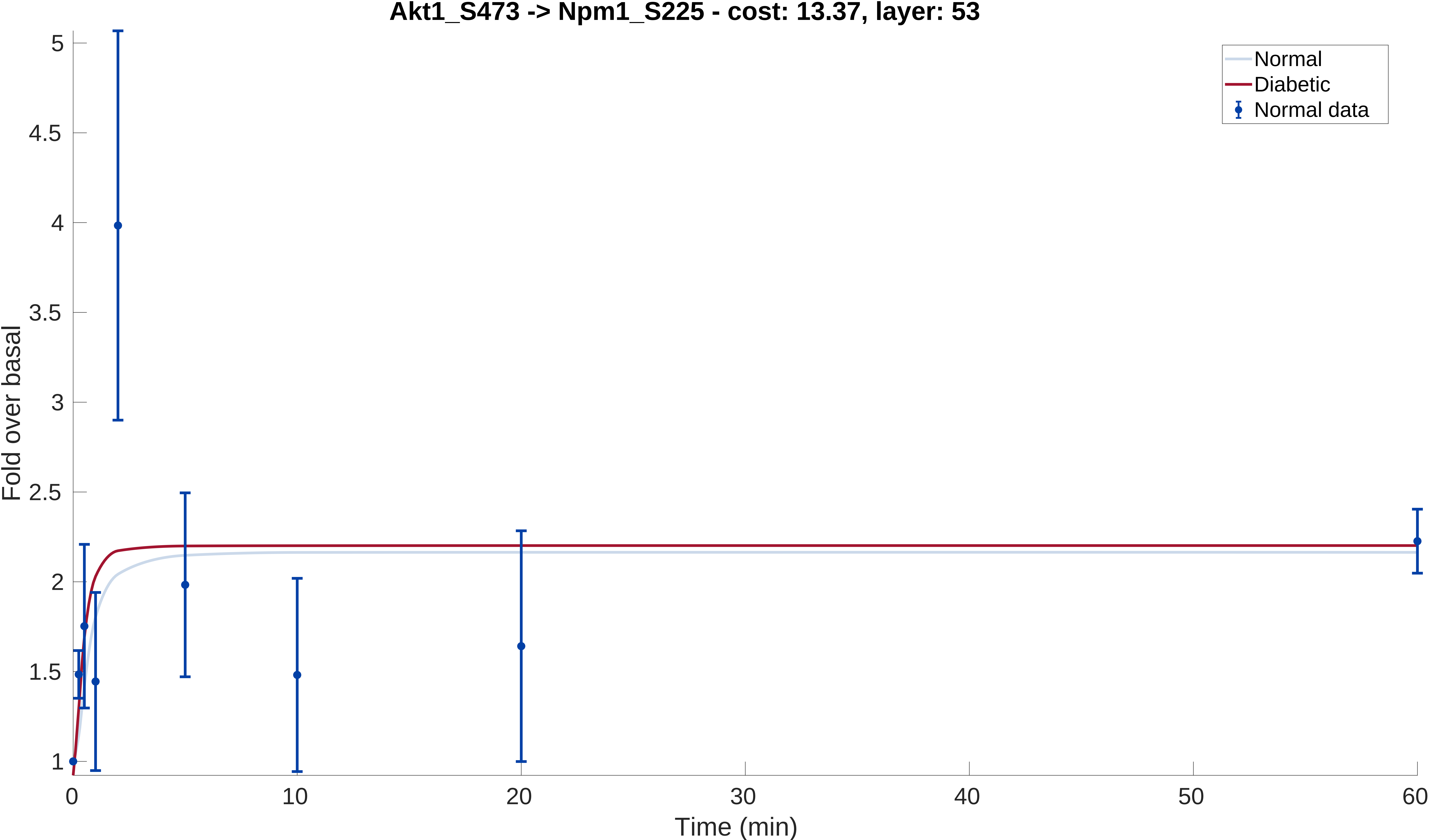

Akt1_S473 -> Npm1_S225 - cost: 13.37, layer: 53
Normal
Diabetic
Normal data
Fold over basal
5
4.5
4
3.5
3
2.5
2
1.5
1
0
10
20
30
40
50
60
Time (min)

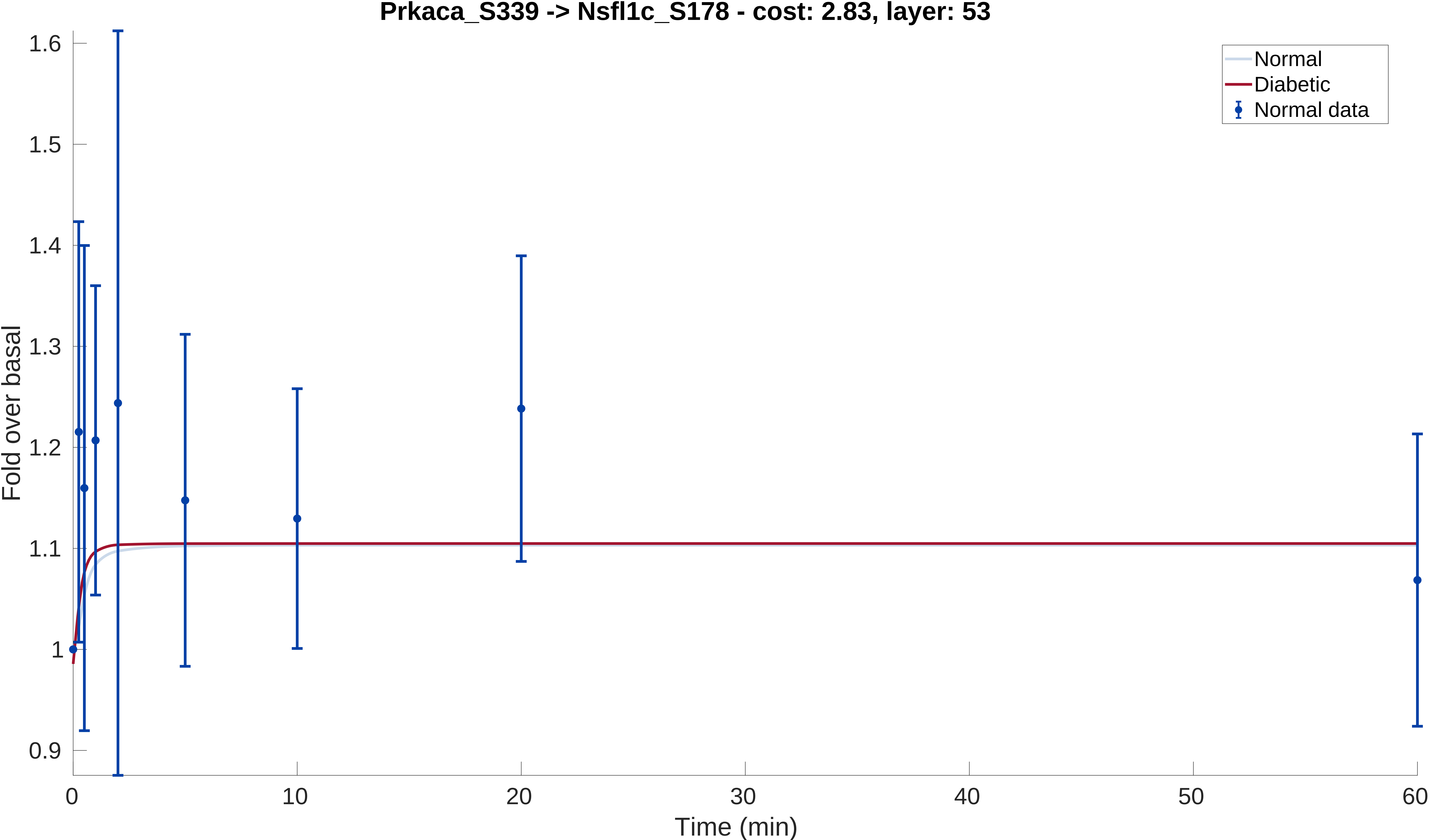

Prkaca_S339 -> Nsfl1c_S178 - cost: 2.83, layer: 53
Normal
Diabetic
Normal data
Fold over basal
Time (min)
0.9
1
1.1
1.2
1.3
1.4
1.5
1.6
0
10
20
30
40
50
60

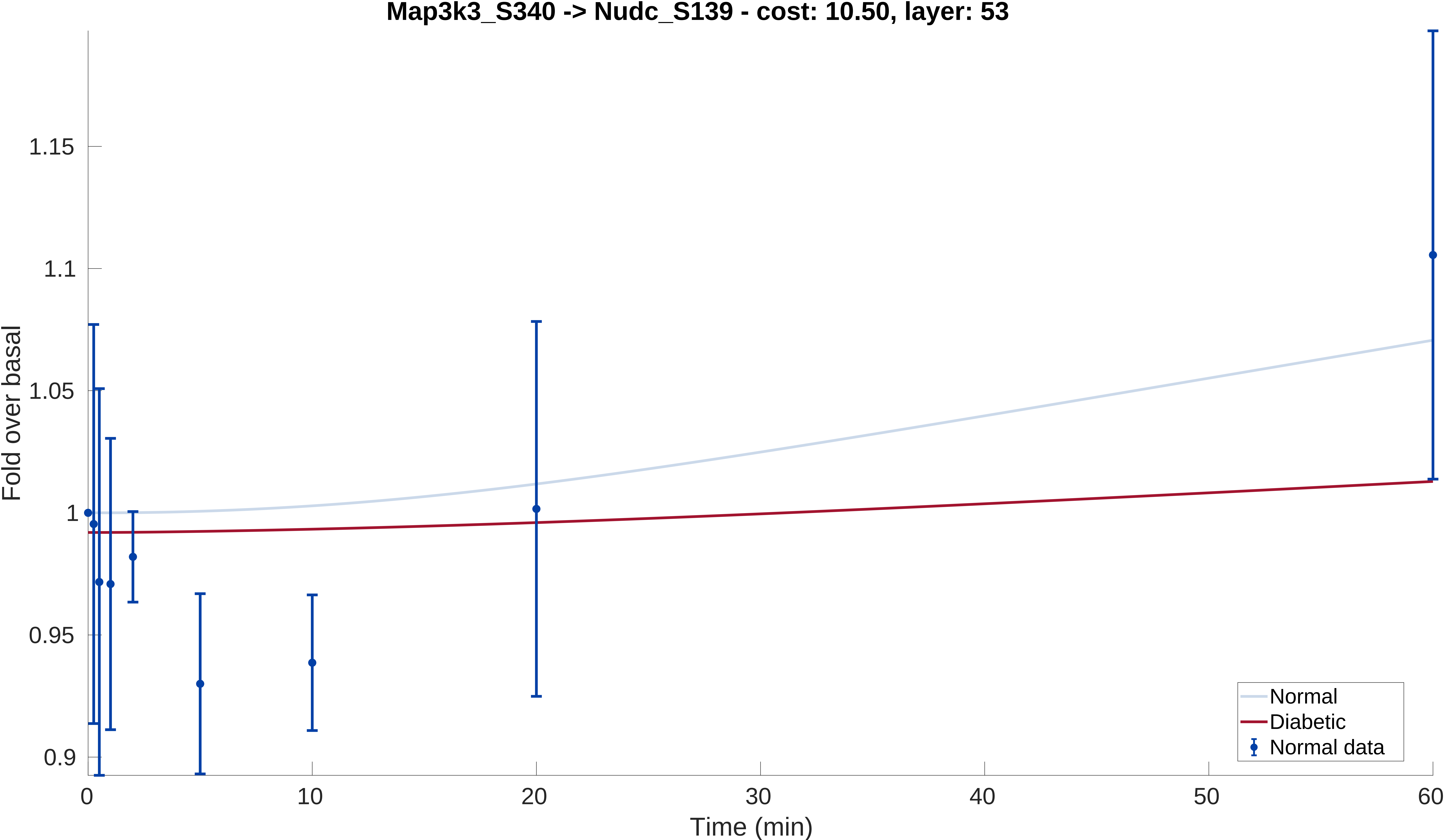

Map3k3_S340 -> Nudc_S139 - cost: 10.50, layer: 53
Fold over basal
1.15
1.1
1.05
1
0.95
0.9
0
10
20
30
40
50
60
Time (min)
Normal
Diabetic
Normal data

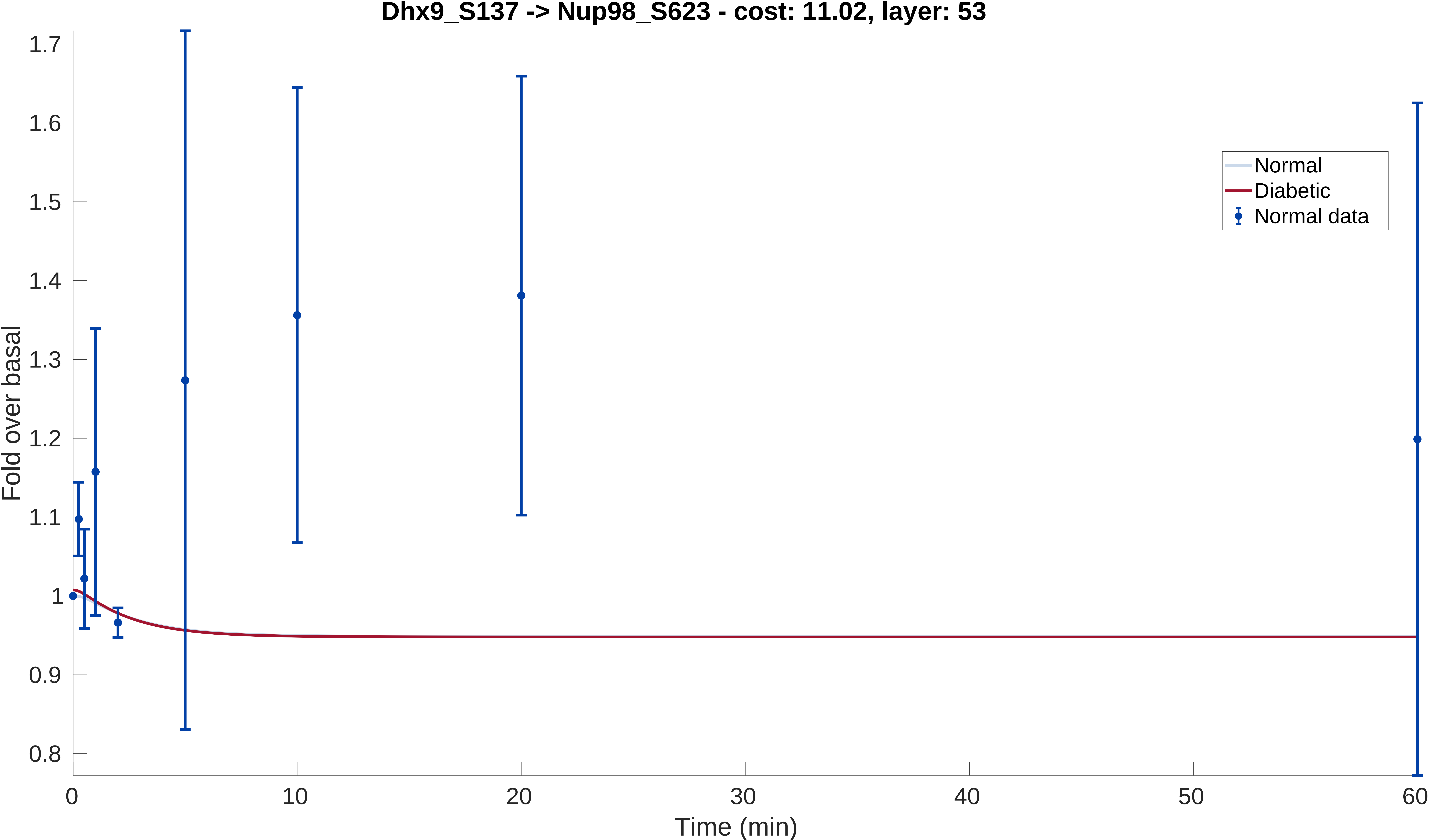
Dhx9_S137 -> Nup98_S623 - cost: 11.02, layer: 53
Fold over basal
Time (min)
Normal
Diabetic
Normal data
0.8
0.9
1
1.1
1.2
1.3
1.4
1.5
1.6
1.7
0
10
20
30
40
50
60

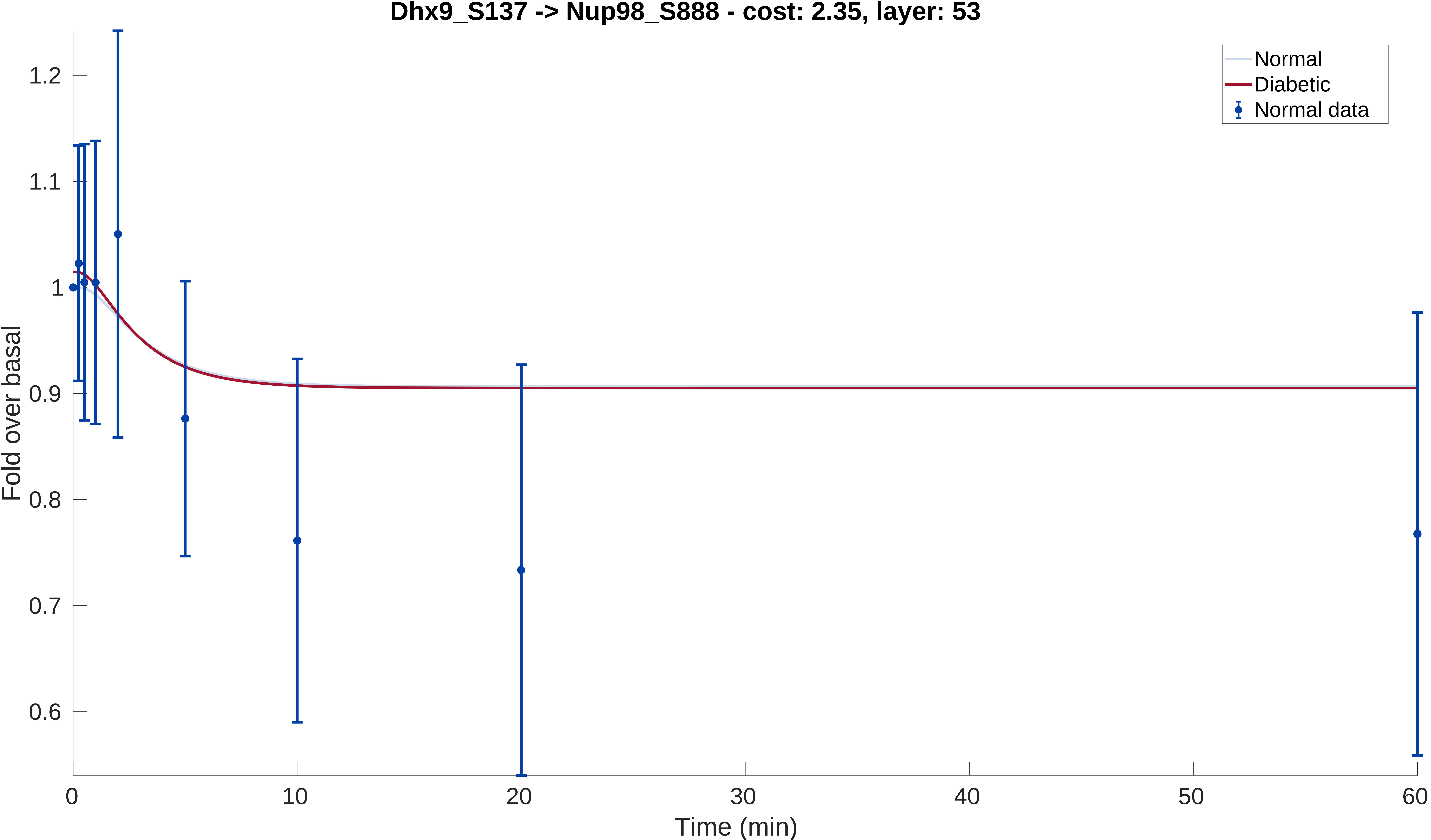

Dhx9_S137 -> Nup98_S888 - cost: 2.35, layer: 53
Normal
Diabetic
Normal data
Fold over basal
1.2
1.1
1
0.9
0.8
0.7
0.6
0
10
20
30
40
50
60
Time (min)

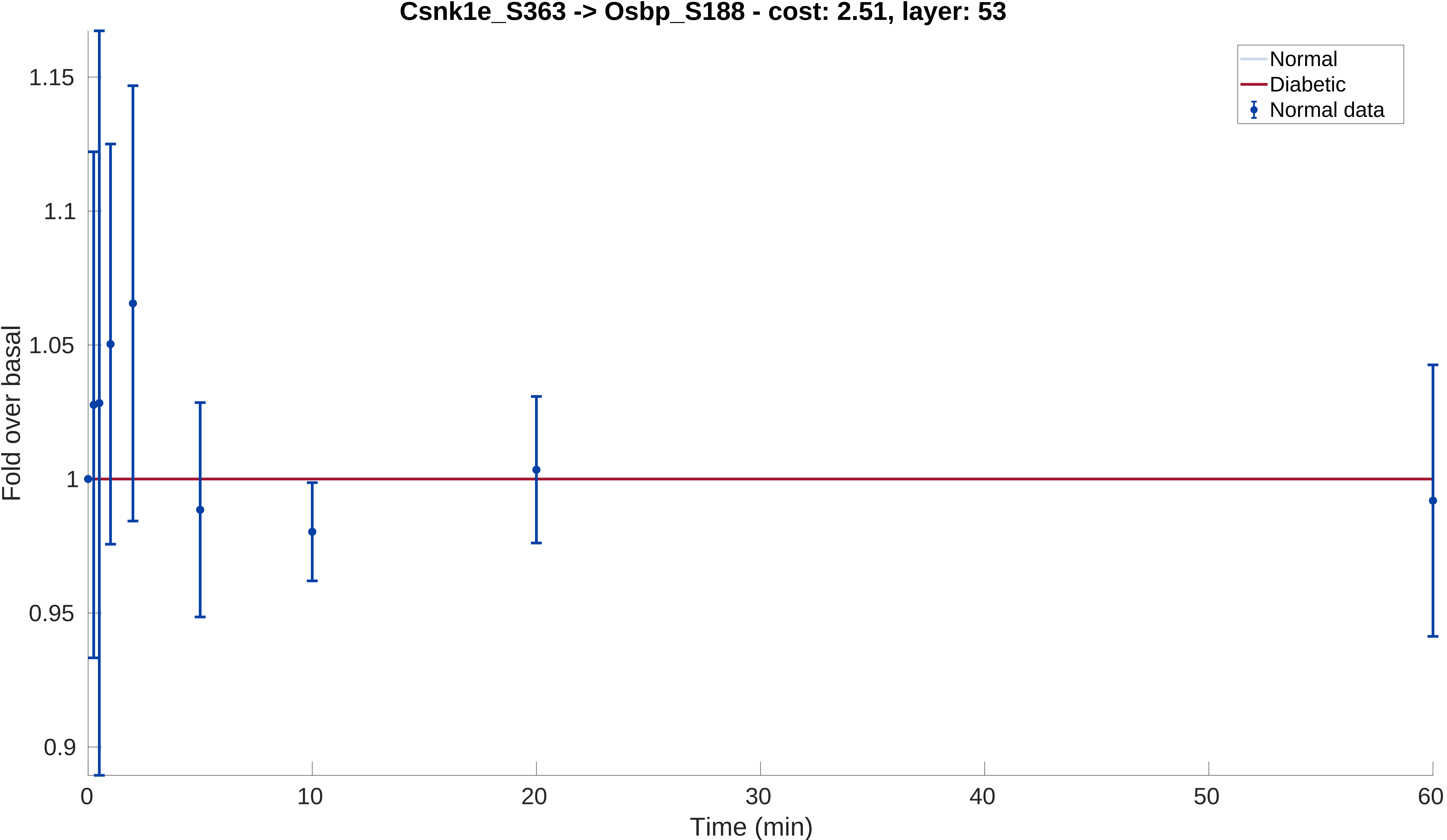

Csnk1e_S363 -> Osbp_S188 - cost: 2.51, layer: 53
Normal
Diabetic
Normal data
Fold over basal
1.15
1.1
1.05
1
0.95
0.9
0
10
20
30
40
50
60
Time (min)

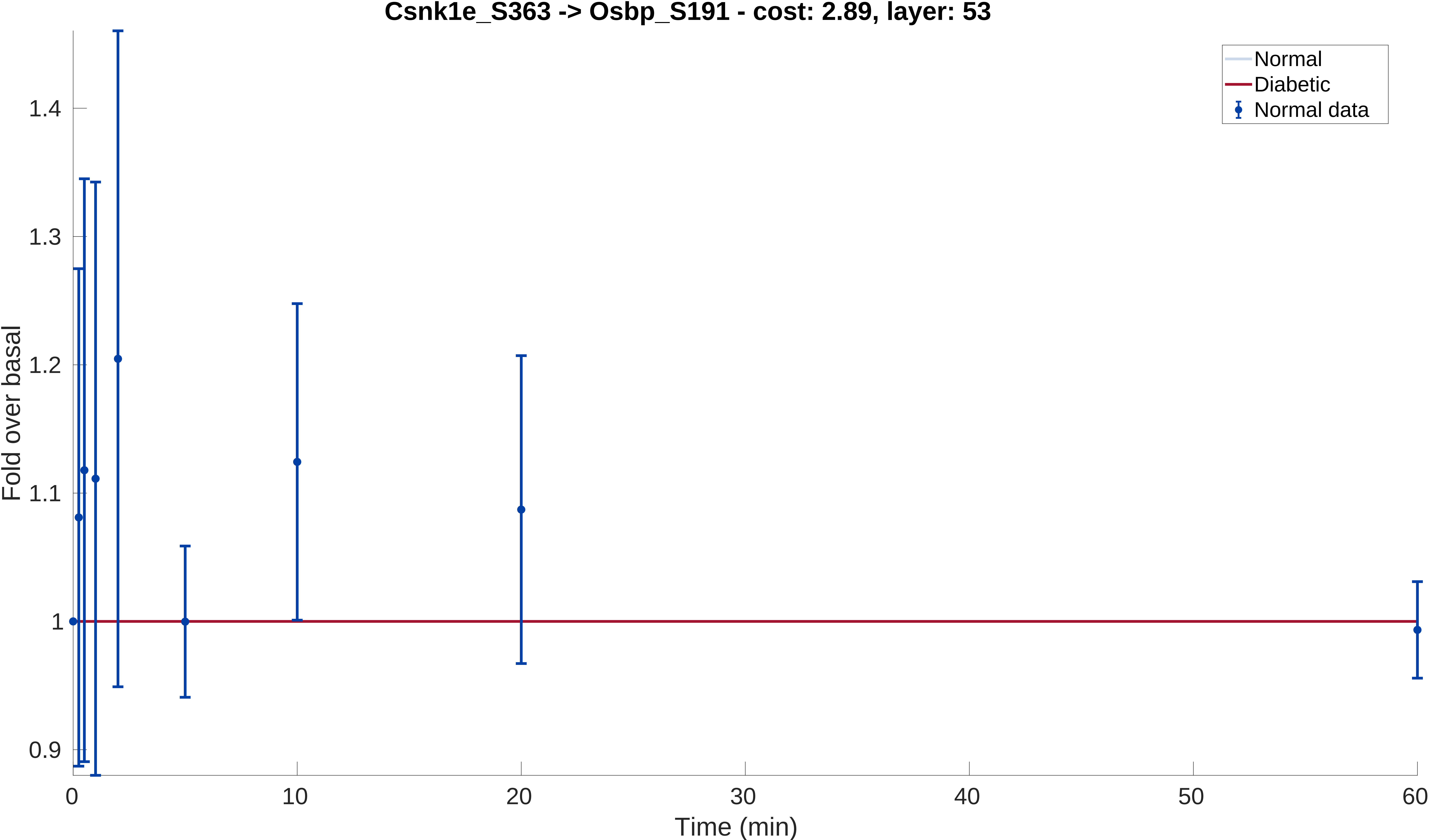

Csnk1e_S363 -> Osbp_S191 - cost: 2.89, layer: 53
Normal
Diabetic
Normal data
Fold over basal
Time (min)
1.4
1.3
1.2
1.1
1
0.9
0
10
20
30
40
50
60

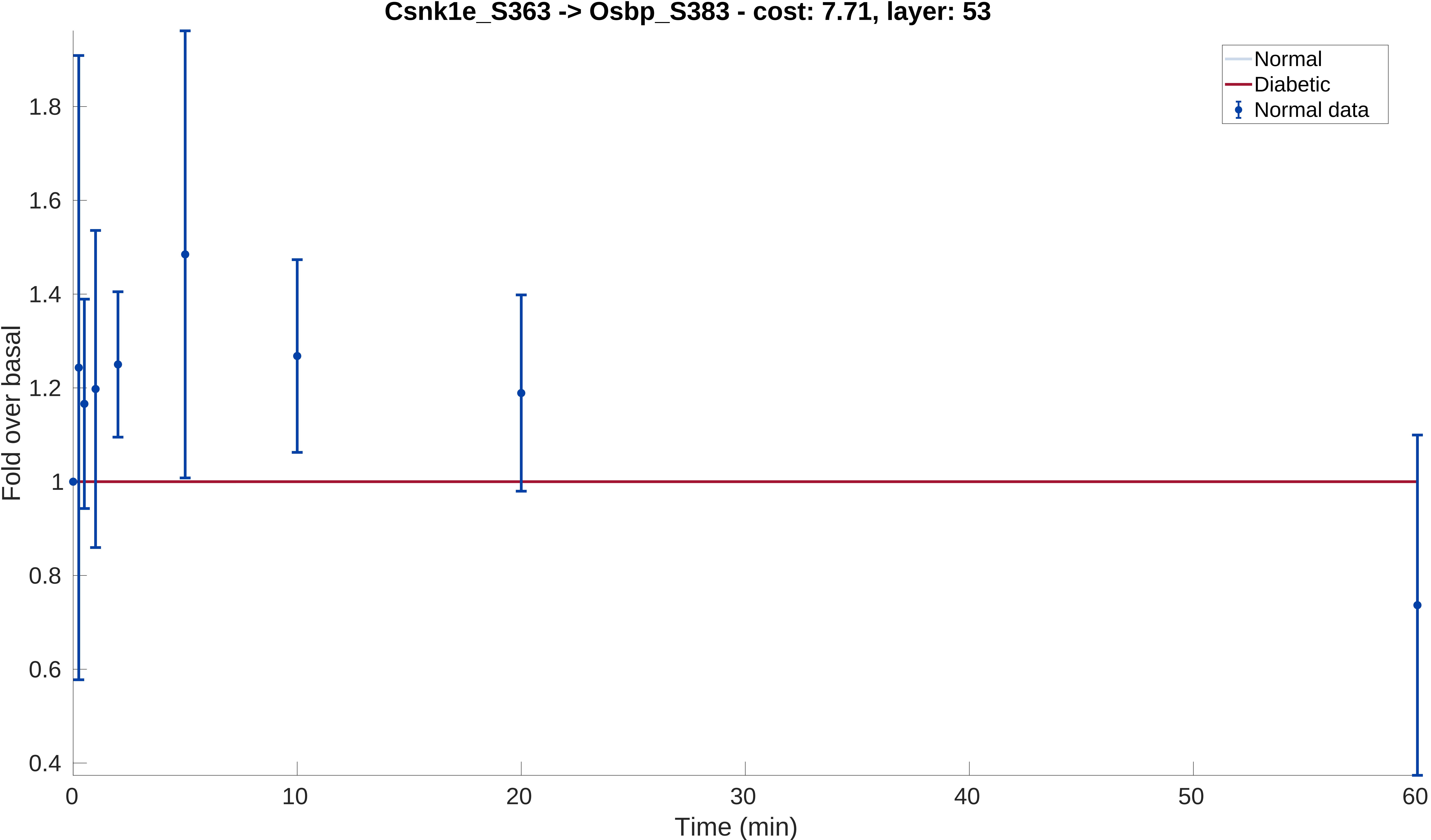

Csnk1e_S363 -> Osbp_S383 - cost: 7.71, layer: 53
Normal
Diabetic
Normal data
Fold over basal
0.4
0.6
0.8
1
1.2
1.4
1.6
1.8
0
10
20
30
40
50
60
Time (min)

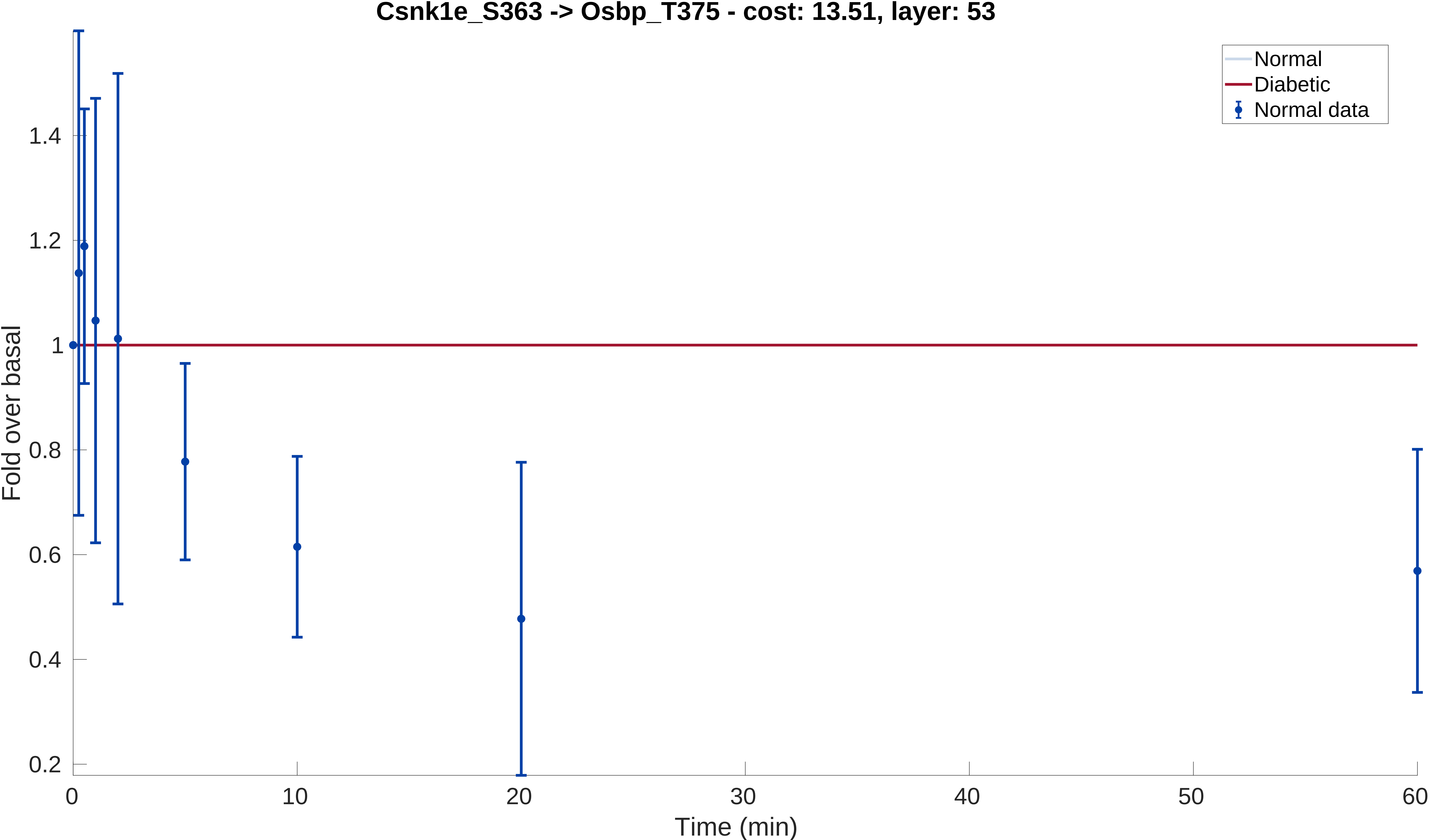
Csnk1e_S363 -> Osbp_T375 - cost: 13.51, layer: 53
Normal
Diabetic
Normal data
Fold over basal
1.4
1.2
1
0.8
0.6
0.4
0.2
0
10
20
30
40
50
60
Time (min)

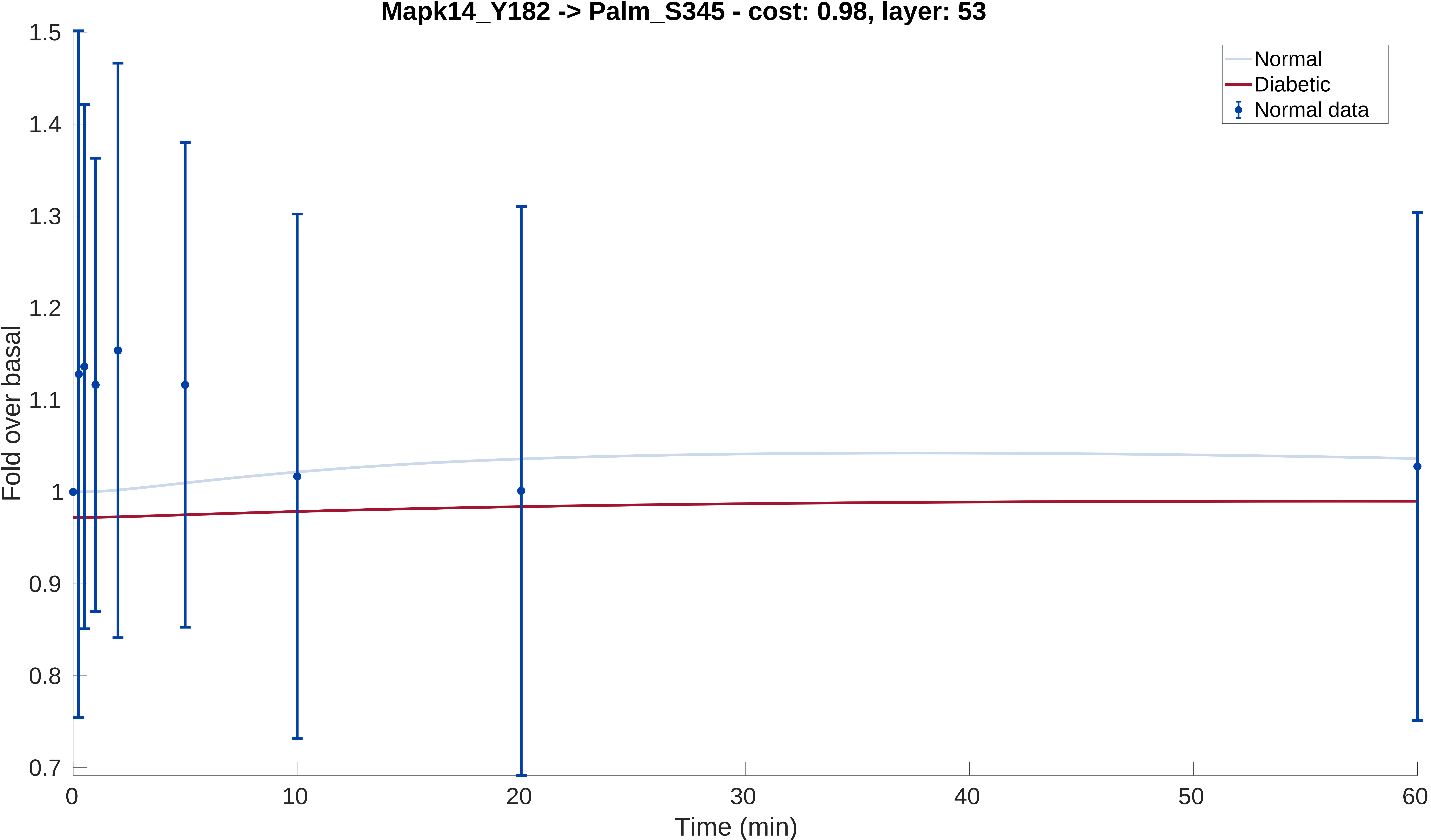

Mapk14_Y182 -> Palm_S345 - cost: 0.98, layer: 53
Normal
Diabetic
Normal data
Fold over basal
1.5
1.4
1.3
1.2
1.1
1
0.9
0.8
0.7
0
10
20
30
40
50
60
Time (min)

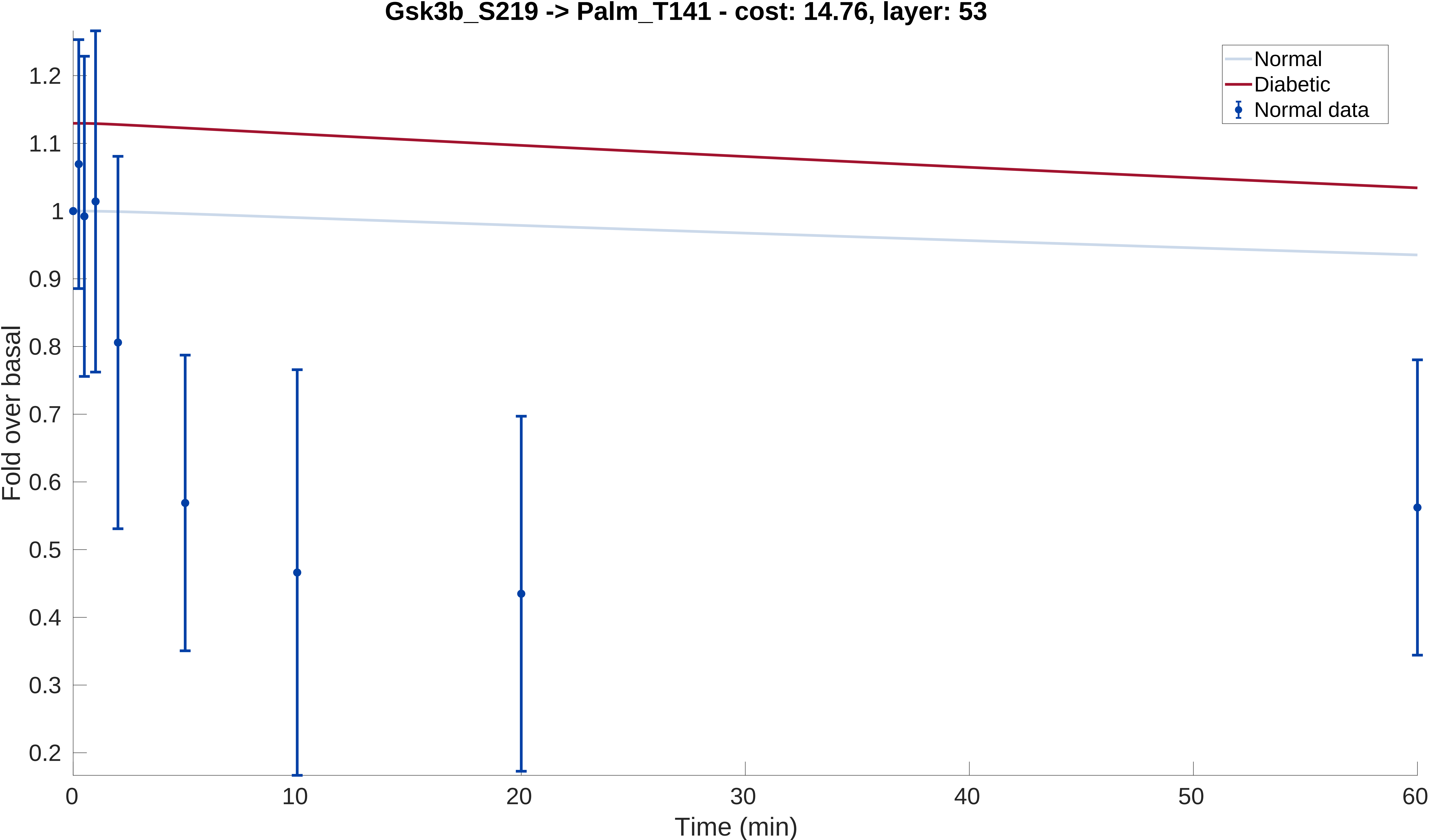
Gsk3b_S219 -> Palm_T141 - cost: 14.76, layer: 53
Normal
Diabetic
Normal data
Fold over basal
Time (min)
1.2
1.1
1
0.9
0.8
0.7
0.6
0.5
0.4
0.3
0.2
0
10
20
30
40
50
60

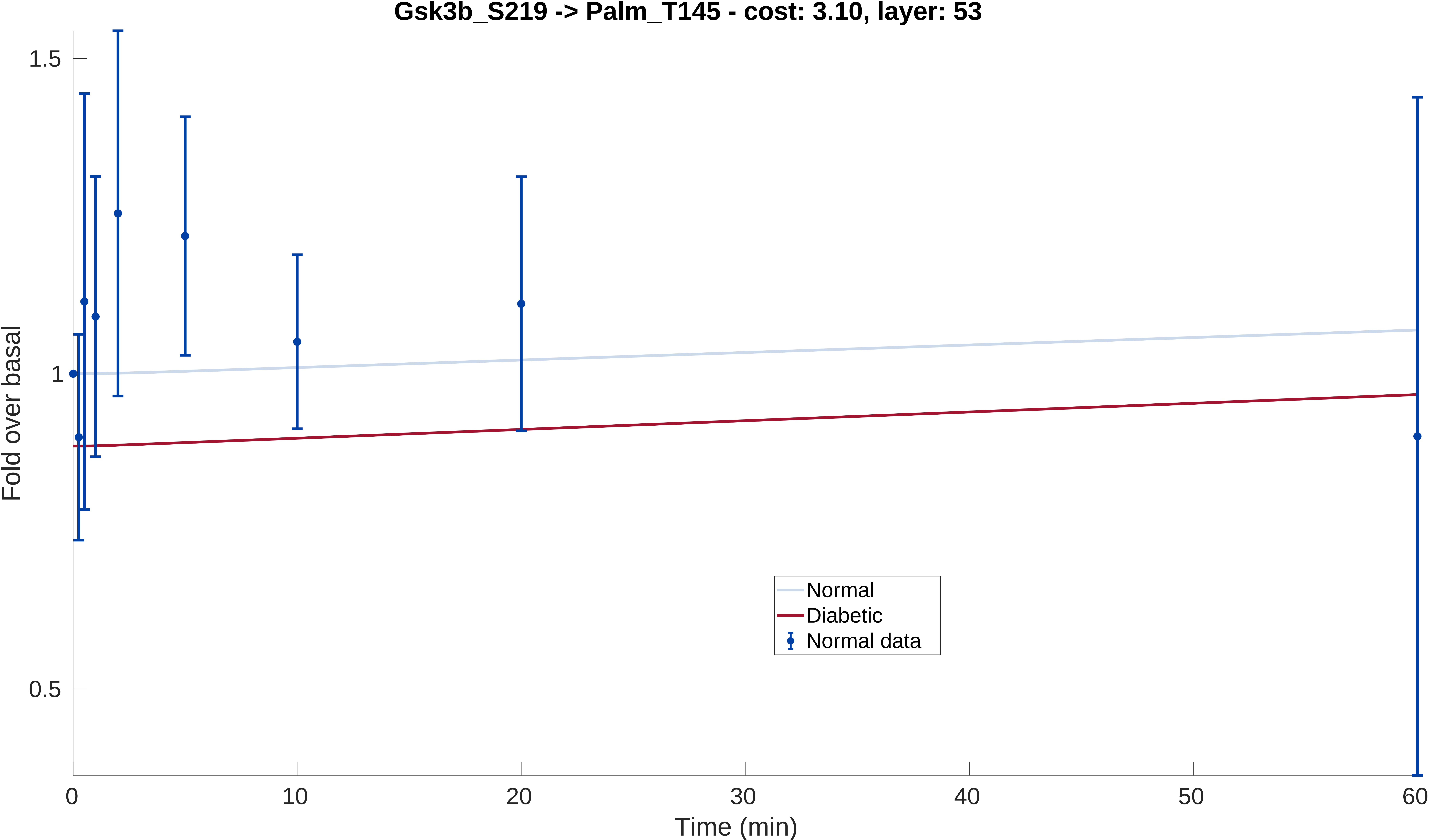
Gsk3b_S219 -> Palm_T145 - cost: 3.10, layer: 53
1.5
1
0.5
Fold over basal
0
10
20
30
40
50
60
Time (min)
Normal
Diabetic
Normal data

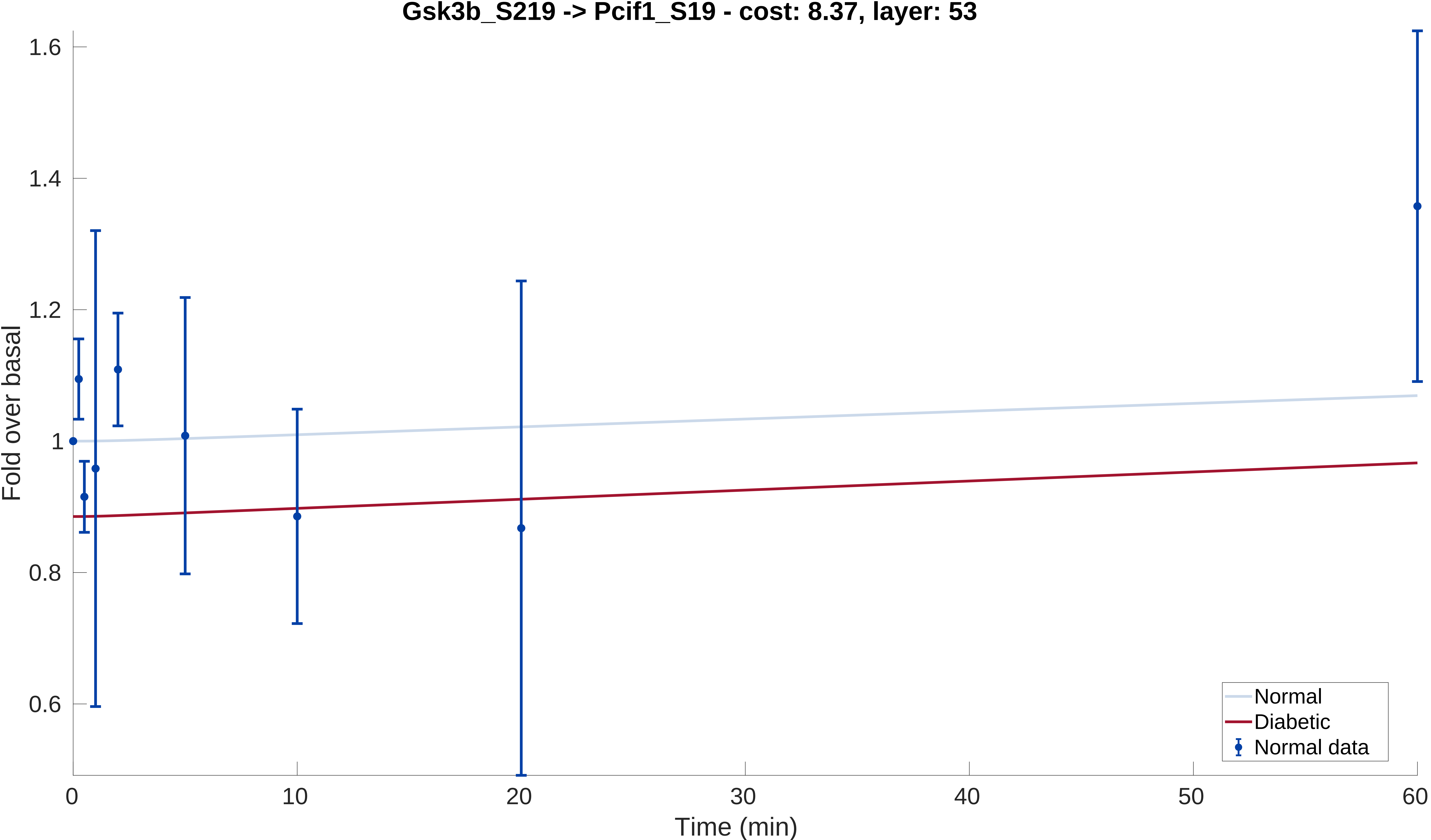

Gsk3b_S219 -> Pcif1_S19 - cost: 8.37, layer: 53
1.6
1.4
1.2
1
0.8
0.6
Fold over basal
0
10
20
30
40
50
60
Time (min)
Normal
Diabetic
Normal data

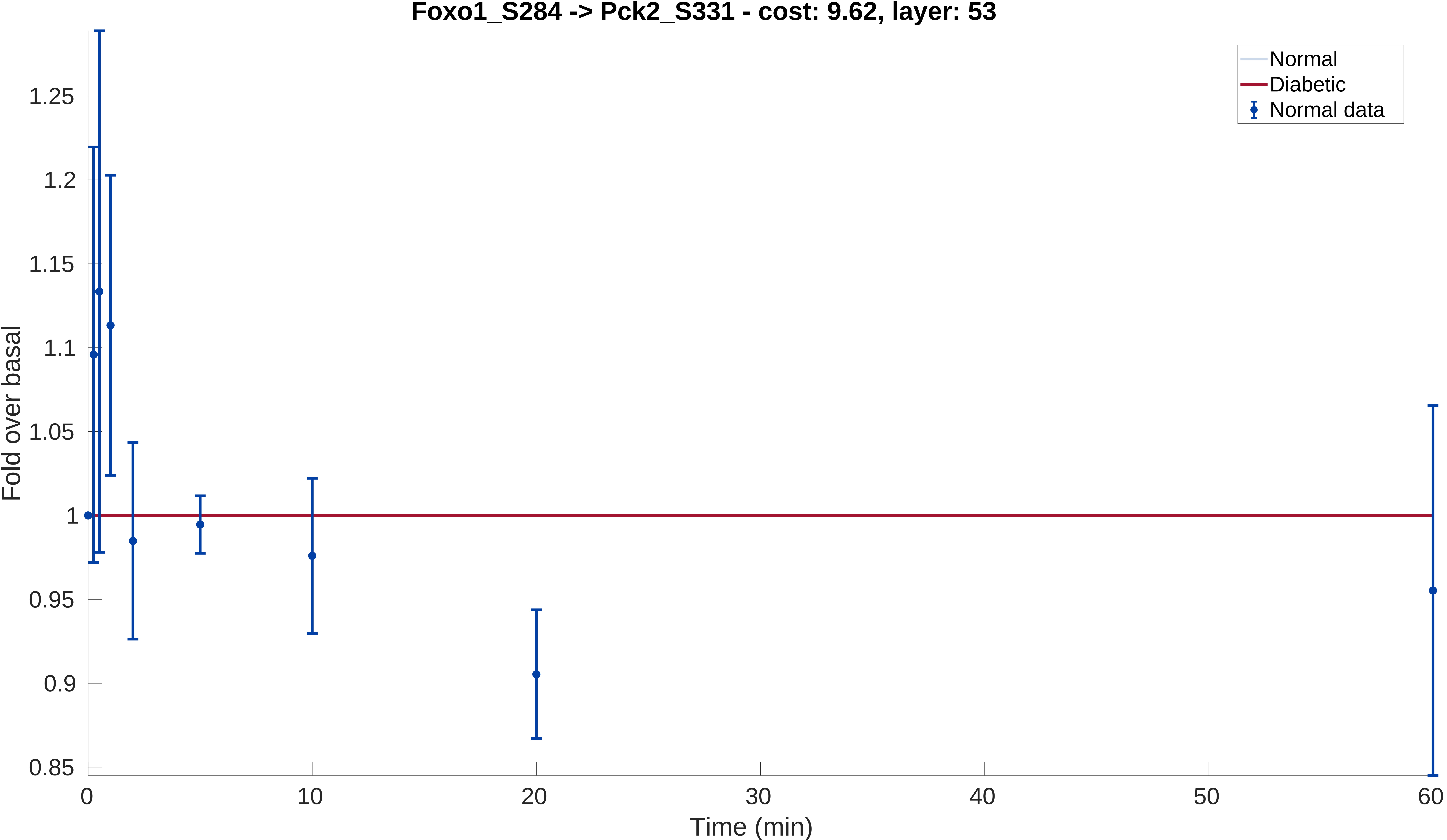
Foxo1_S284 -> Pck2_S331 - cost: 9.62, layer: 53
Normal
Diabetic
Normal data
Fold over basal
1.25
1.2
1.15
1.1
1.05
1
0.95
0.9
0.85
0
10
20
30
40
50
60
Time (min)

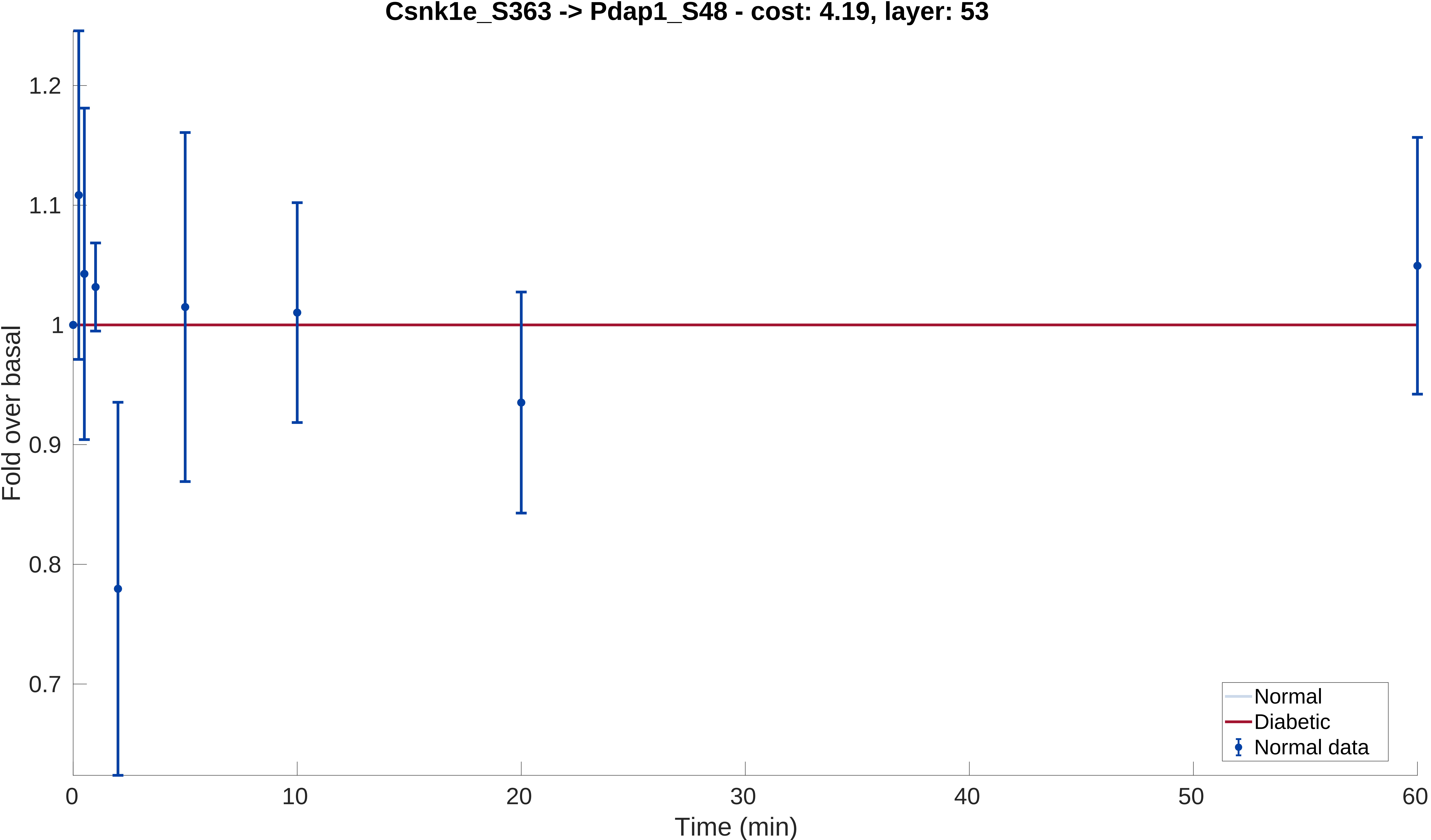
Csnk1e_S363 -> Pdap1_S48 - cost: 4.19, layer: 53
Fold over basal
1.2
1.1
1
0.9
0.8
0.7
0
10
20
30
40
50
60
Time (min)
Normal
Diabetic
Normal data

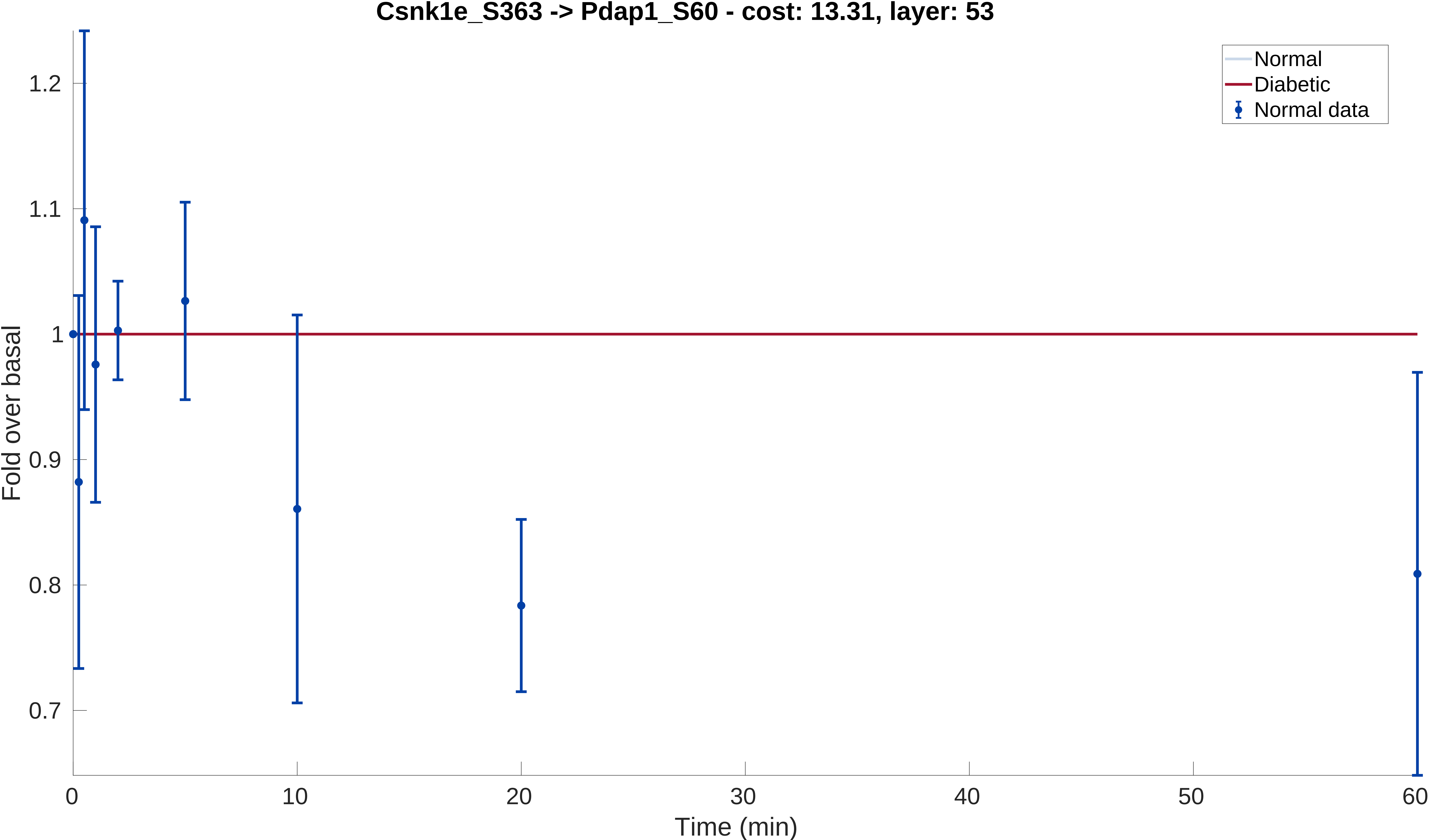
Csnk1e_S363 -> Pdap1_S60 - cost: 13.31, layer: 53
Normal
Diabetic
Normal data
Fold over basal
1.2
1.1
1
0.9
0.8
0.7
0
10
20
30
40
50
60
Time (min)

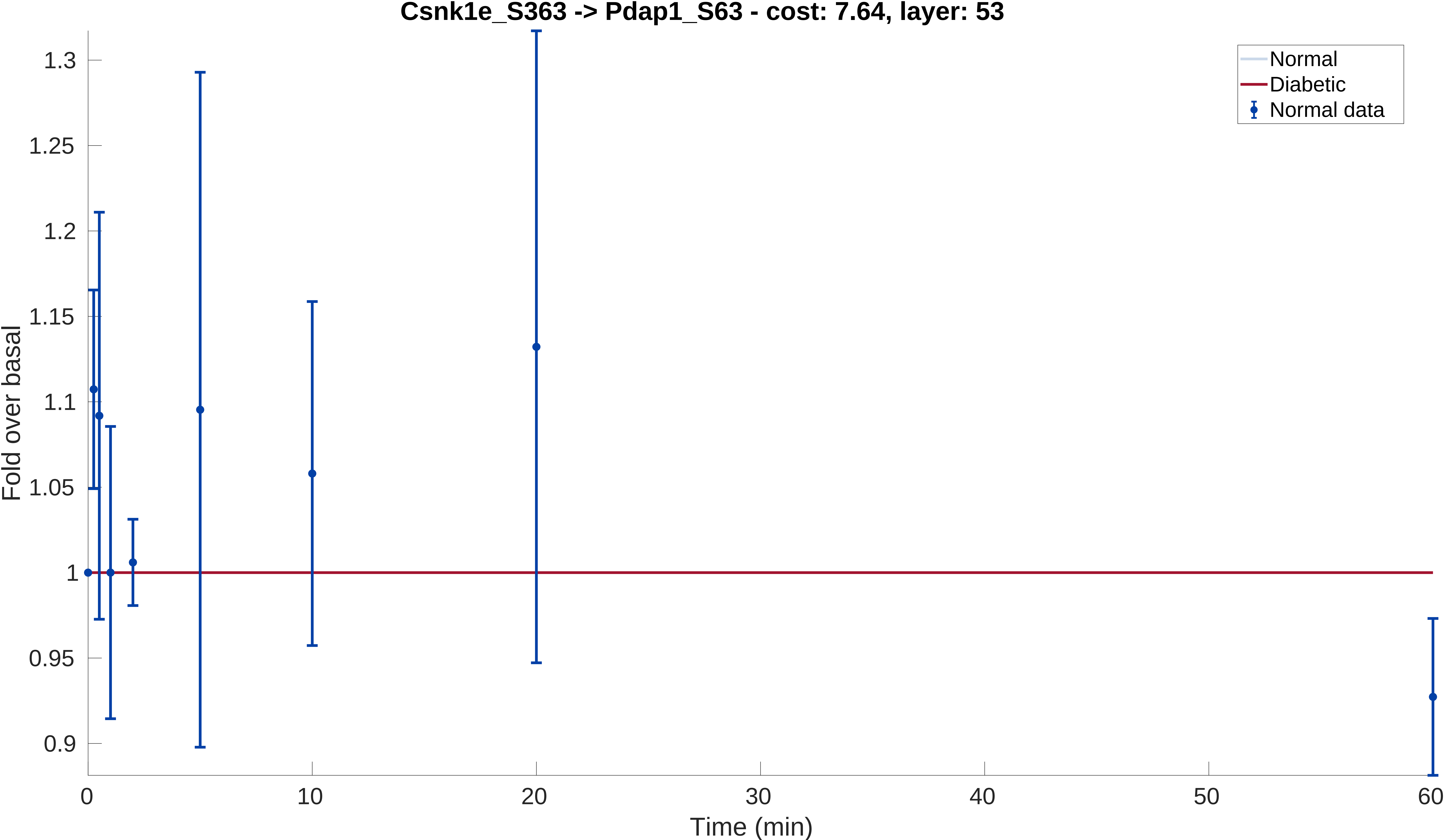
Csnk1e_S363 -> Pdap1_S63 - cost: 7.64, layer: 53
Normal
Diabetic
Normal data
Fold over basal
1.3
1.25
1.2
1.15
1.1
1.05
1
0.95
0.9
0
10
20
30
40
50
60
Time (min)

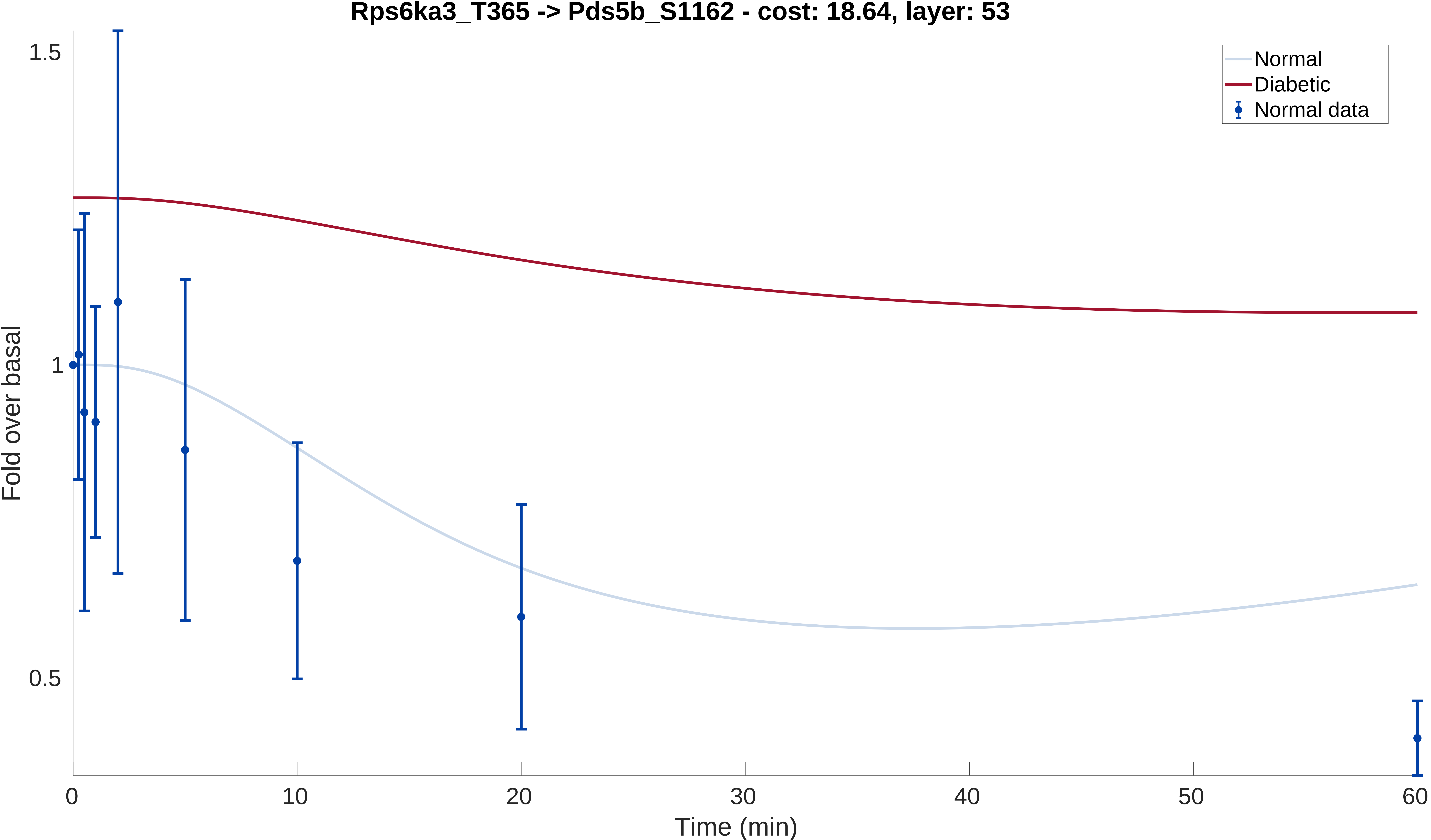

Rps6ka3_T365 -> Pds5b_S1162 - cost: 18.64, layer: 53
Normal
Diabetic
Normal data
1.5
1
0.5
Fold over basal
0
10
20
30
40
50
60
Time (min)

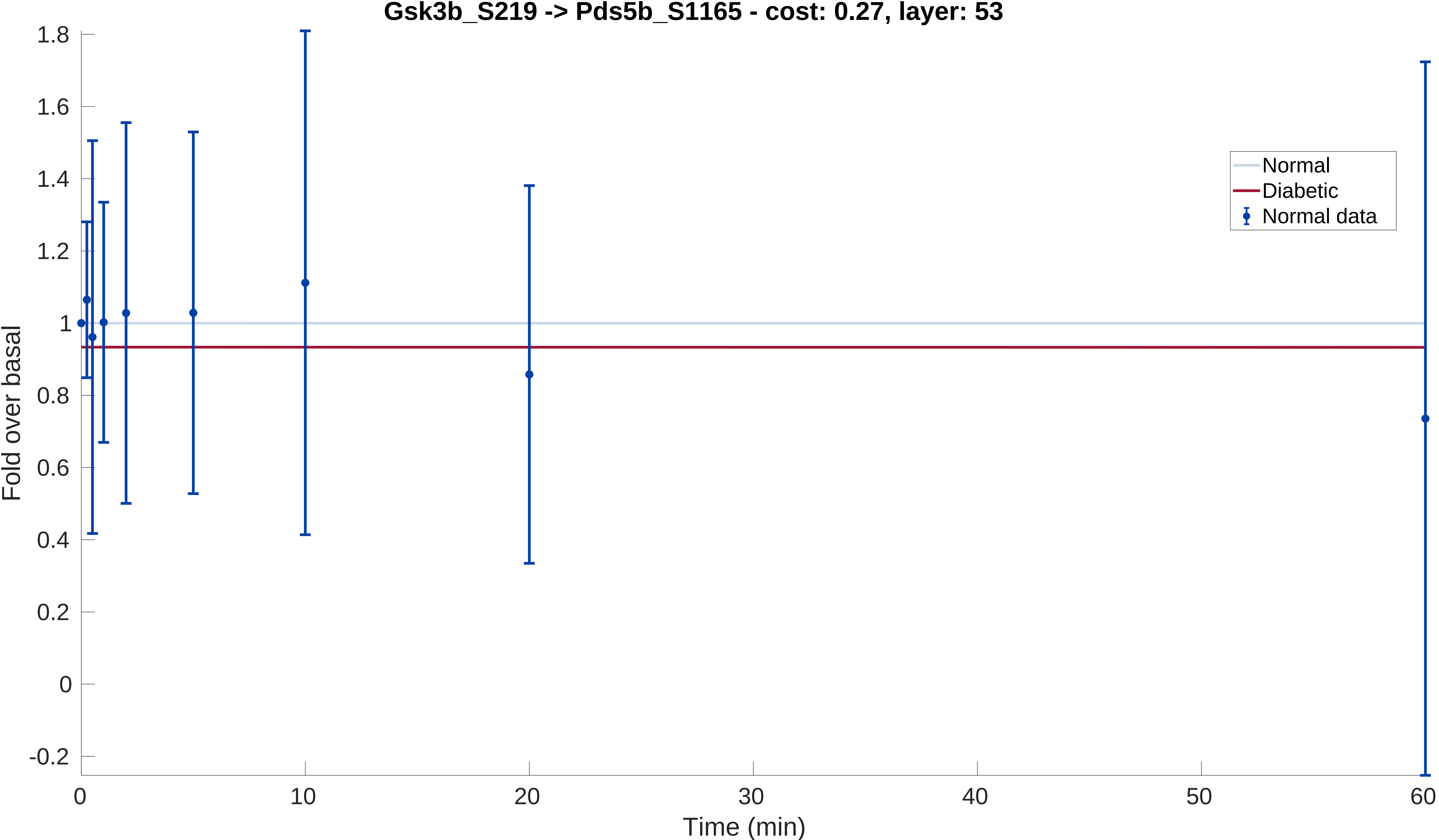

Gsk3b_S219 -> Pds5b_S1165 - cost: 0.27, layer: 53
1.8
1.6
1.4
1.2
1
0.8
0.6
0.4
0.2
0
-0.2
Fold over basal
0
10
20
30
40
50
60
Time (min)
Normal
Diabetic
Normal data

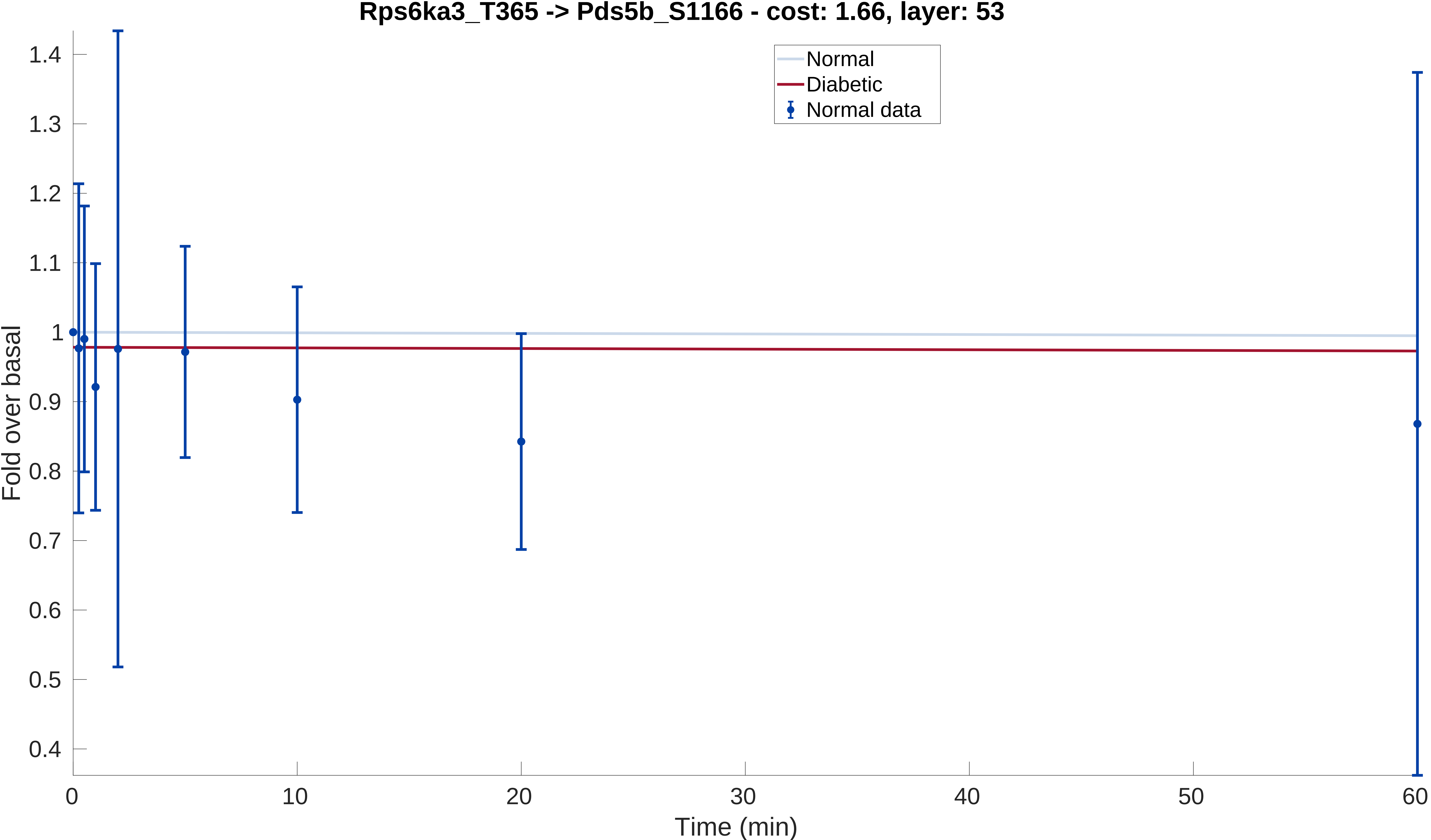

Rps6ka3_T365 -> Pds5b_S1166 - cost: 1.66, layer: 53
Normal
Diabetic
Normal data
Fold over basal
1.4
1.3
1.2
1.1
1
0.9
0.8
0.7
0.6
0.5
0.4
0
10
20
30
40
50
60
Time (min)

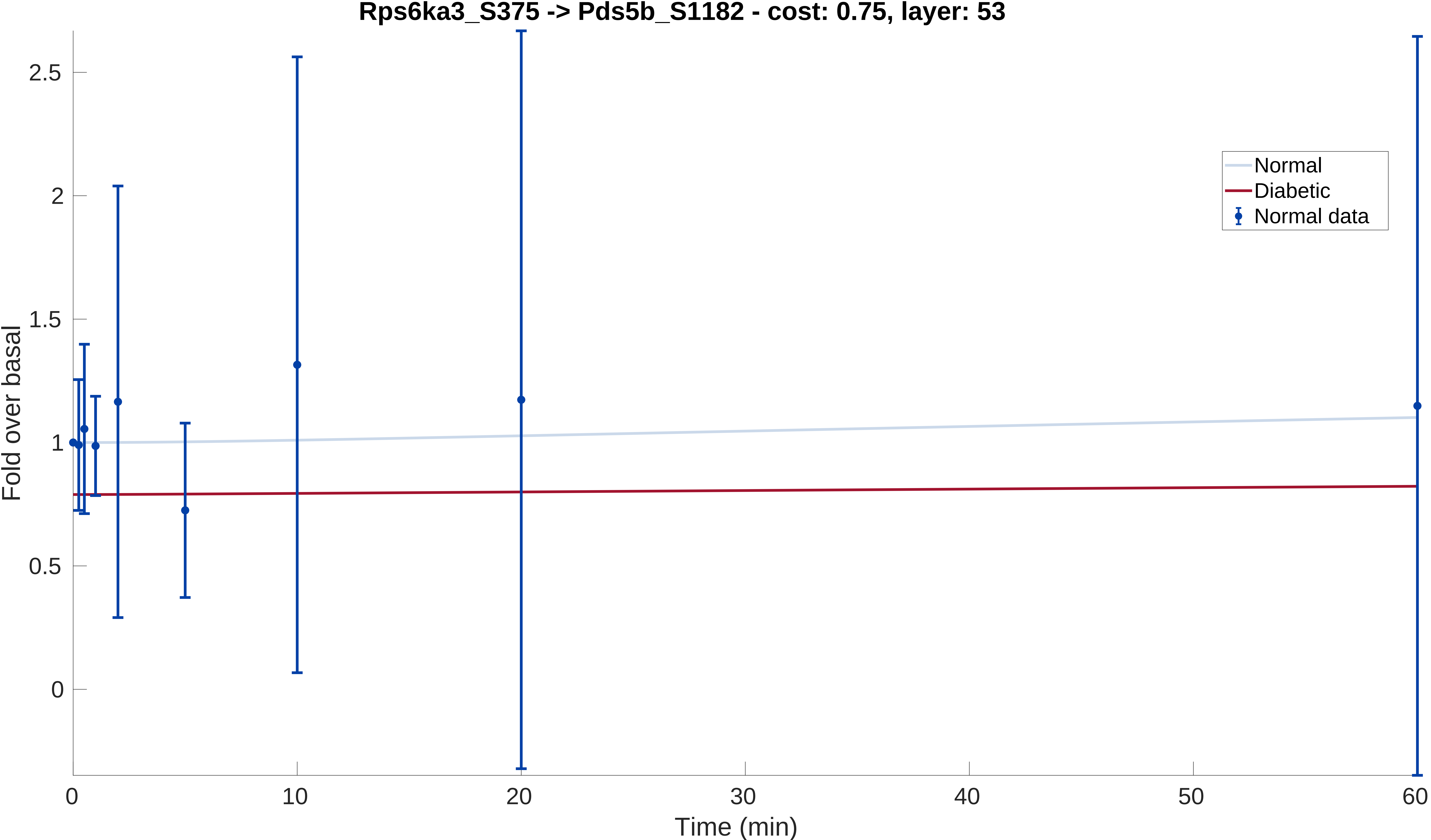
Rps6ka3_S375 -> Pds5b_S1182 - cost: 0.75, layer: 53
Normal
Diabetic
Normal data
Fold over basal
Time (min)
2.5
2
1.5
1
0.5
0
0
10
20
30
40
50
60

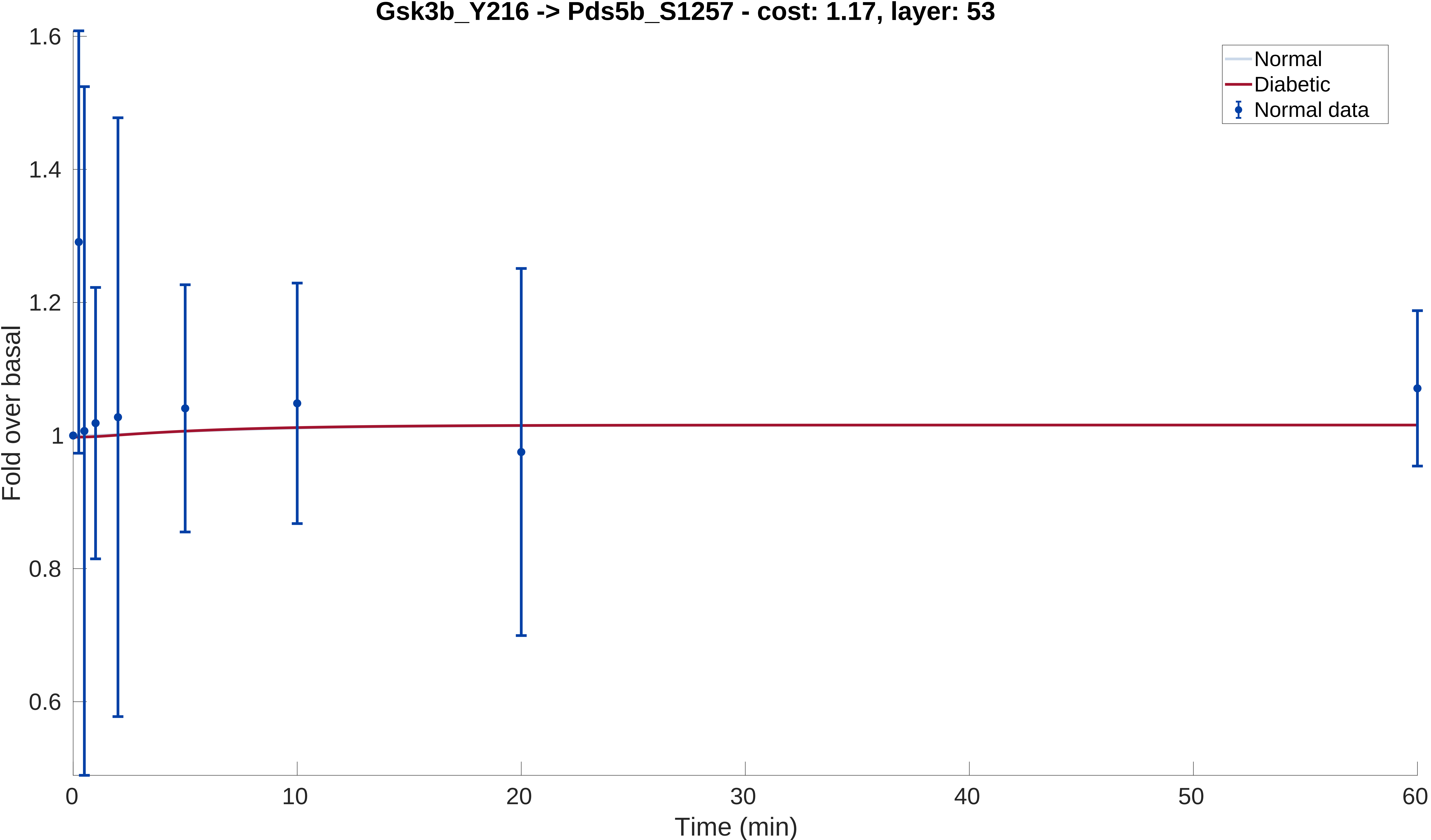

Gsk3b_Y216 -> Pds5b_S1257 - cost: 1.17, layer: 53
Normal
Diabetic
Normal data
Fold over basal
1.6
1.4
1.2
1
0.8
0.6
0
10
20
30
40
50
60
Time (min)

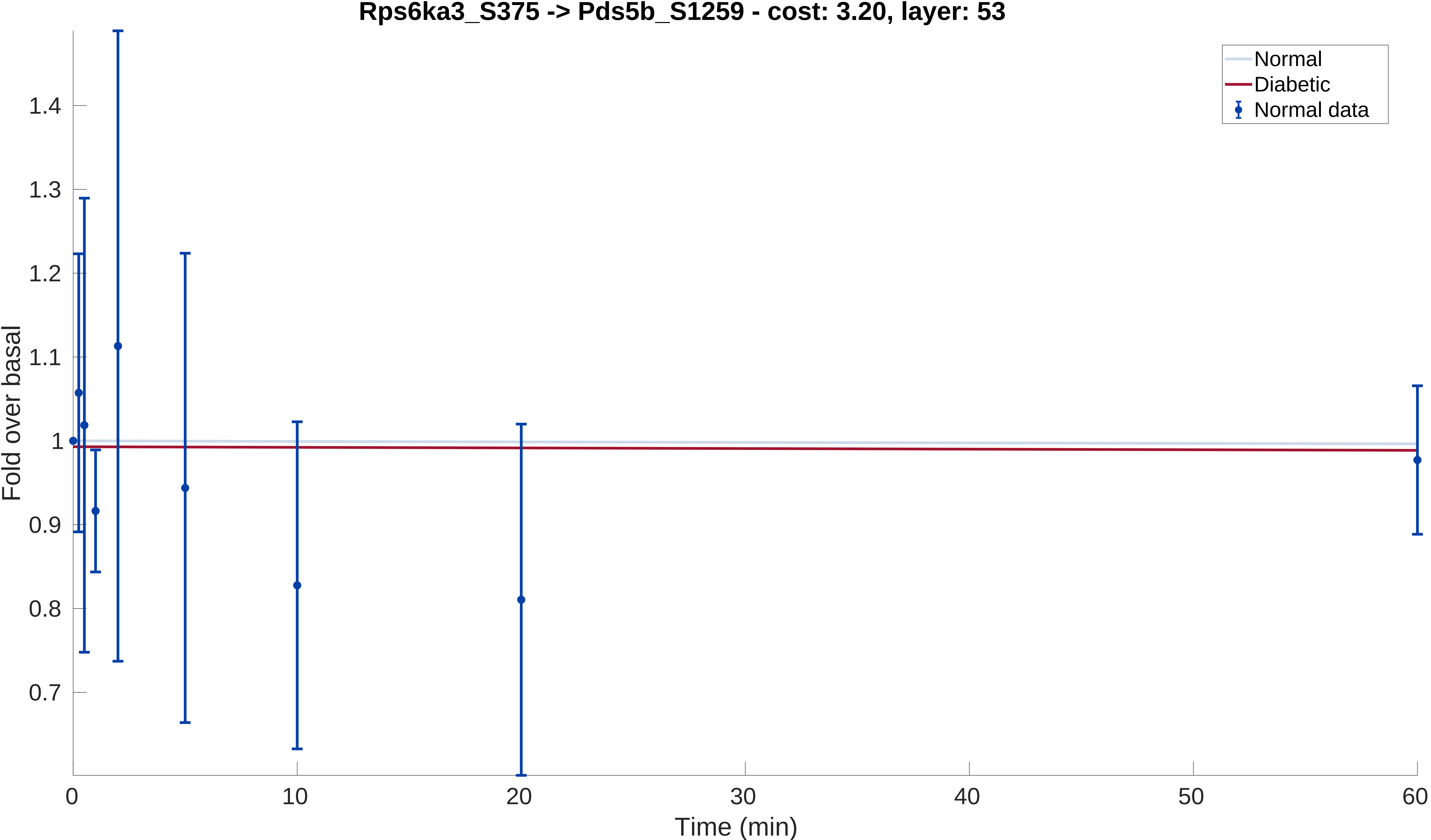

Rps6ka3_S375 -> Pds5b_S1259 - cost: 3.20, layer: 53
Normal
Diabetic
Normal data
Fold over basal
1.4
1.3
1.2
1.1
1
0.9
0.8
0.7
0
10
20
30
40
50
60
Time (min)

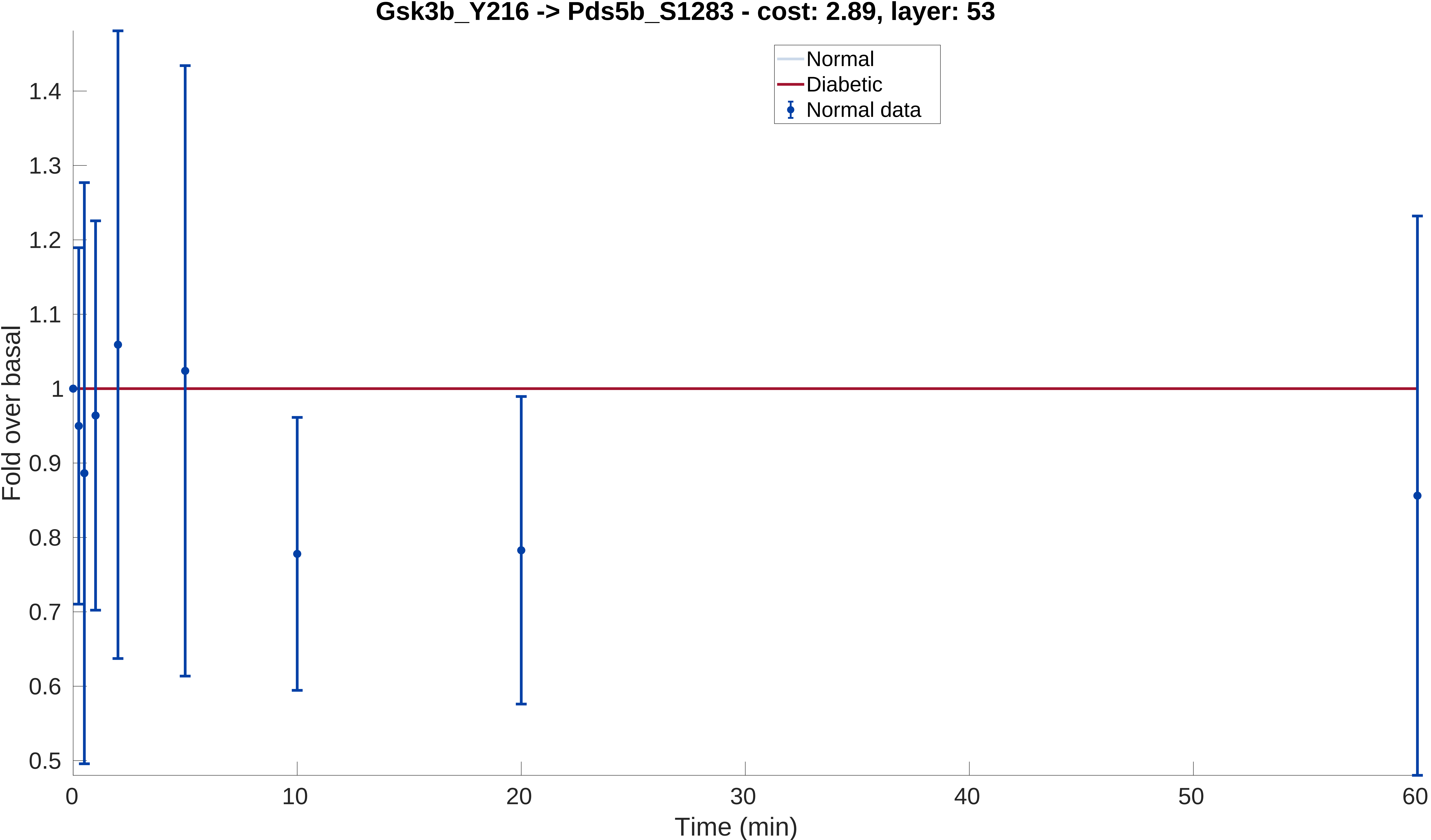

Gsk3b_Y216 -> Pds5b_S1283 - cost: 2.89, layer: 53
Normal
Diabetic
Normal data
Fold over basal
1.4
1.3
1.2
1.1
1
0.9
0.8
0.7
0.6
0.5
0
10
20
30
40
50
60
Time (min)

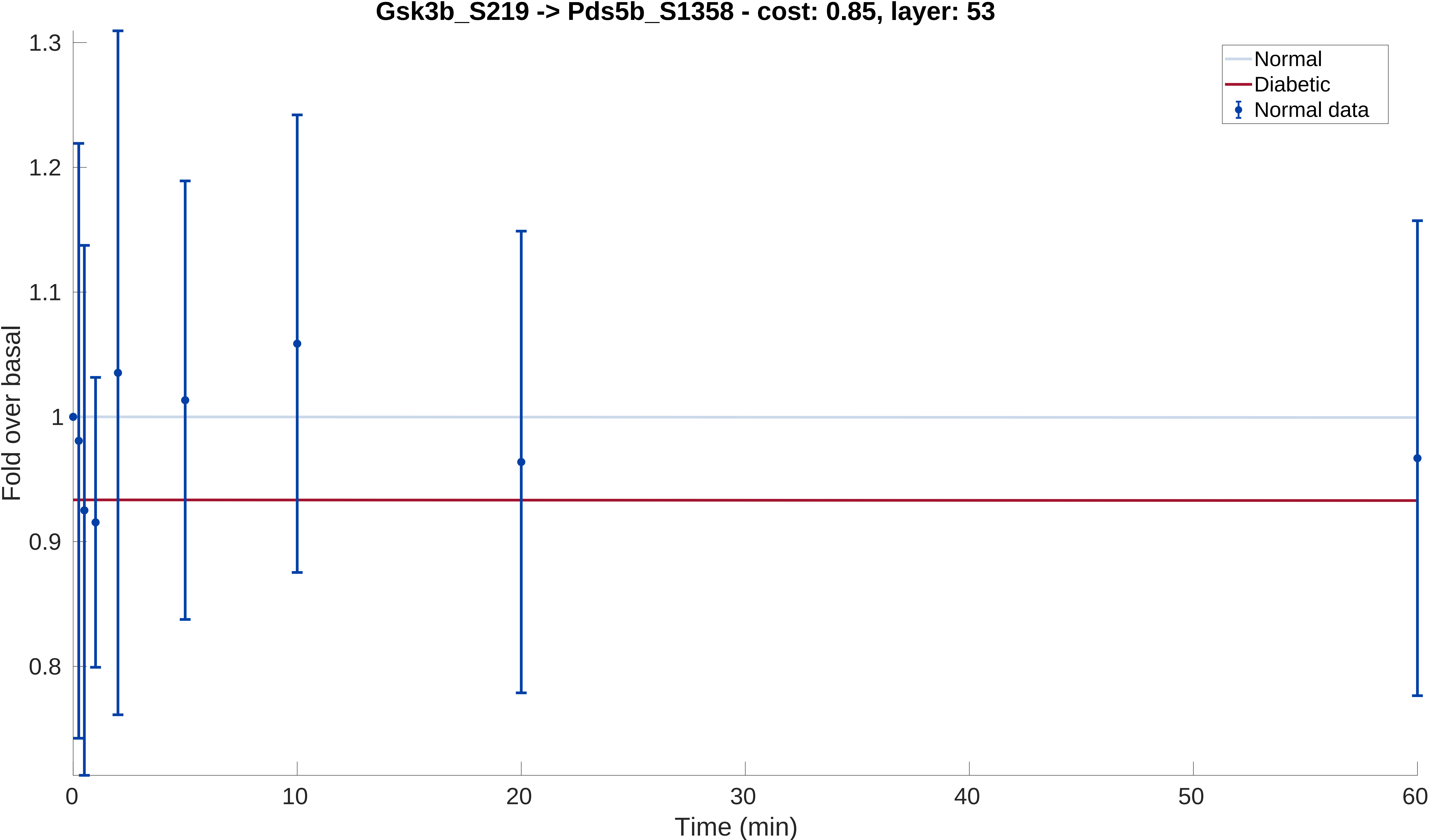
Gsk3b_S219 -> Pds5b_S1358 - cost: 0.85, layer: 53
Normal
Diabetic
Normal data
Fold over basal
1.3
1.2
1.1
1
0.9
0.8
0
10
20
30
40
50
60
Time (min)

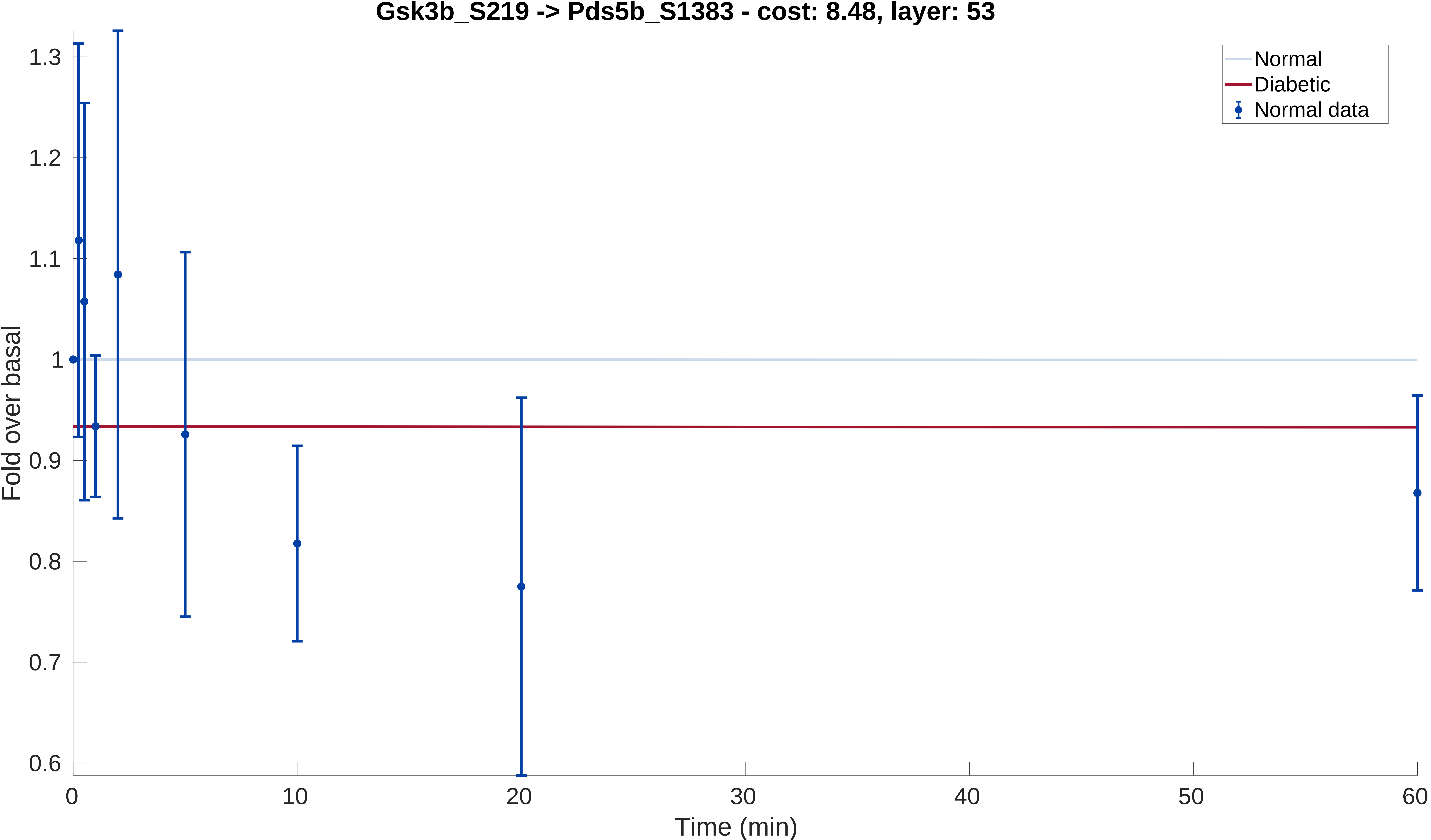
Gsk3b_S219 -> Pds5b_S1383 - cost: 8.48, layer: 53
Normal
Diabetic
Normal data
Fold over basal
Time (min)
1.3
1.2
1.1
1
0.9
0.8
0.7
0.6
0
10
20
30
40
50
60

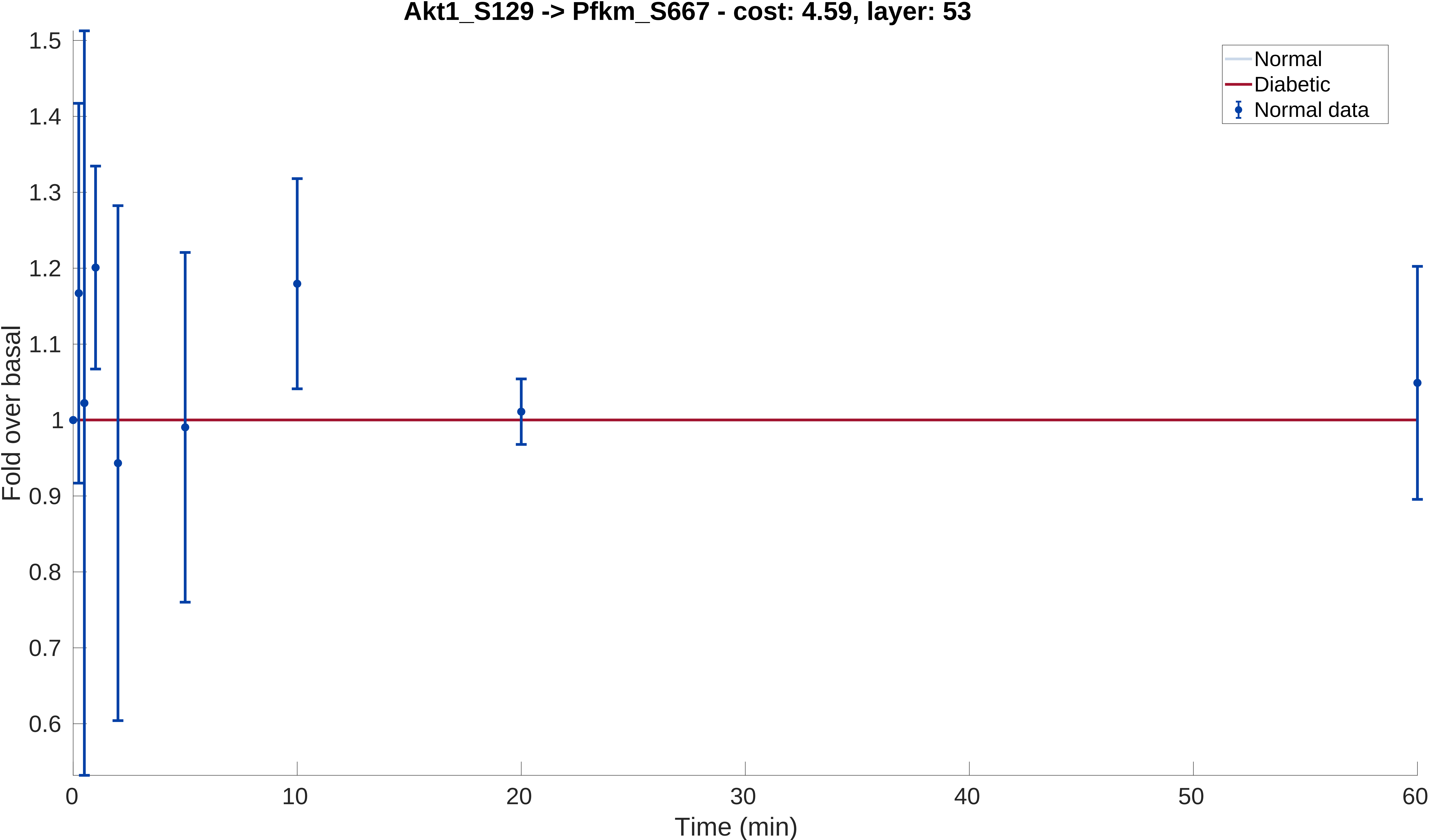

Akt1_S129 -> Pfkm_S667 - cost: 4.59, layer: 53
Normal
Diabetic
Normal data
Fold over basal
1.5
1.4
1.3
1.2
1.1
1
0.9
0.8
0.7
0.6
0
10
20
30
40
50
60
Time (min)

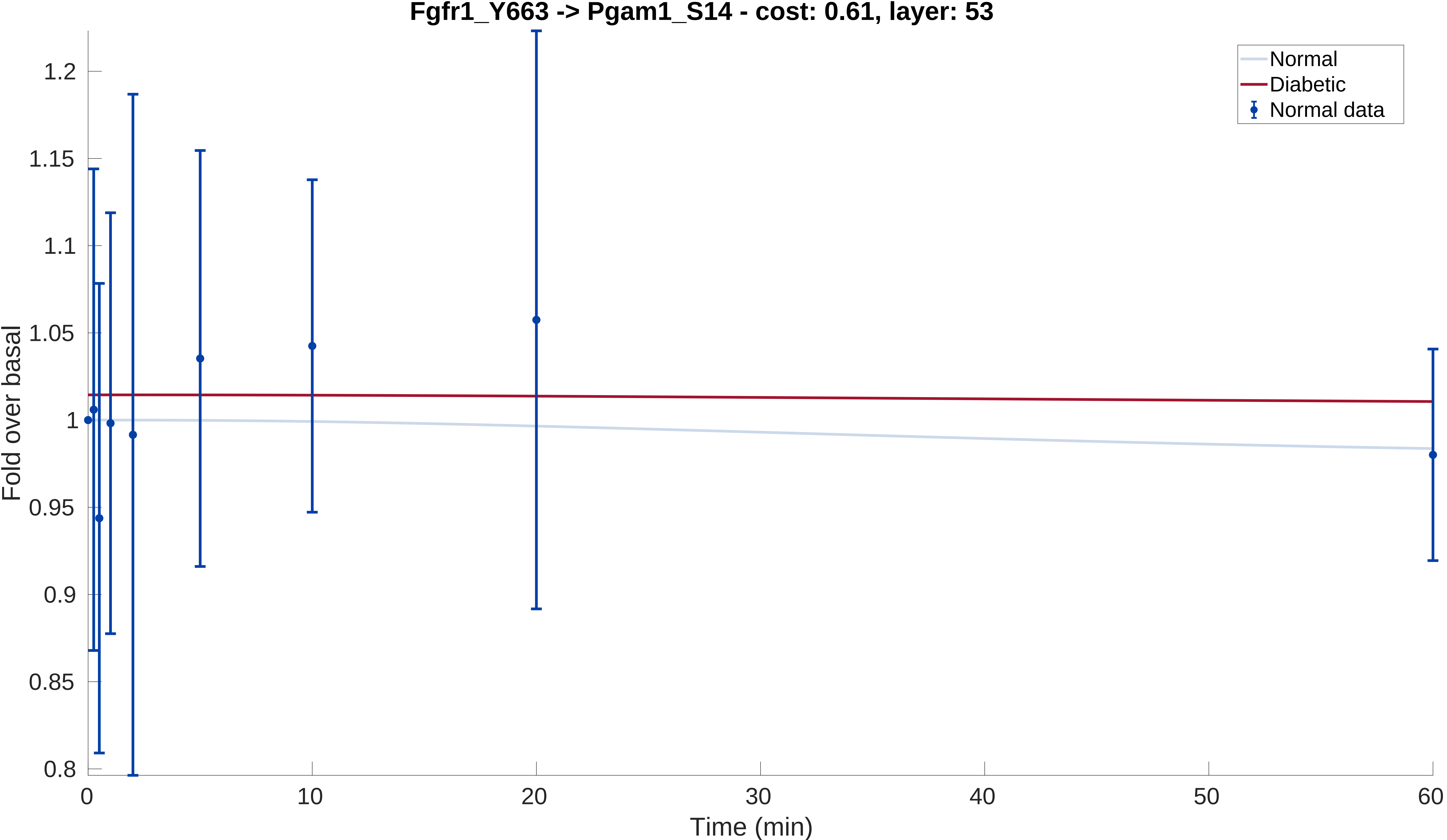

Fgfr1_Y663 -> Pgam1_S14 - cost: 0.61, layer: 53
Normal
Diabetic
Normal data
Fold over basal
Time (min)
0.8
0.85
0.9
0.95
1
1.05
1.1
1.15
1.2
0
10
20
30
40
50
60

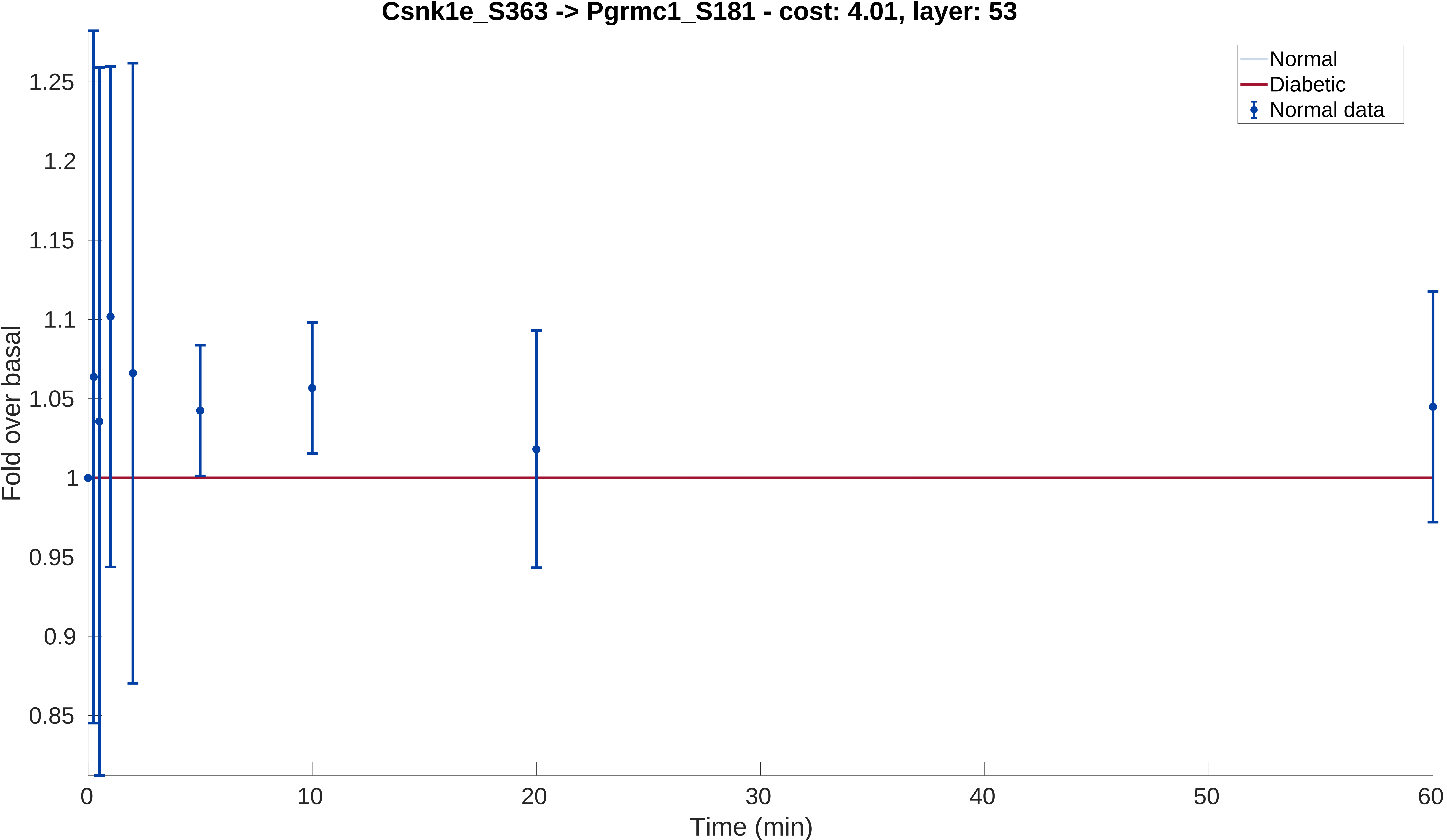

Csnk1e_S363 -> Pgrmc1_S181 - cost: 4.01, layer: 53
Normal
Diabetic
Normal data
Fold over basal
1.25
1.2
1.15
1.1
1.05
1
0.95
0.9
0.85
0
10
20
30
40
50
60
Time (min)

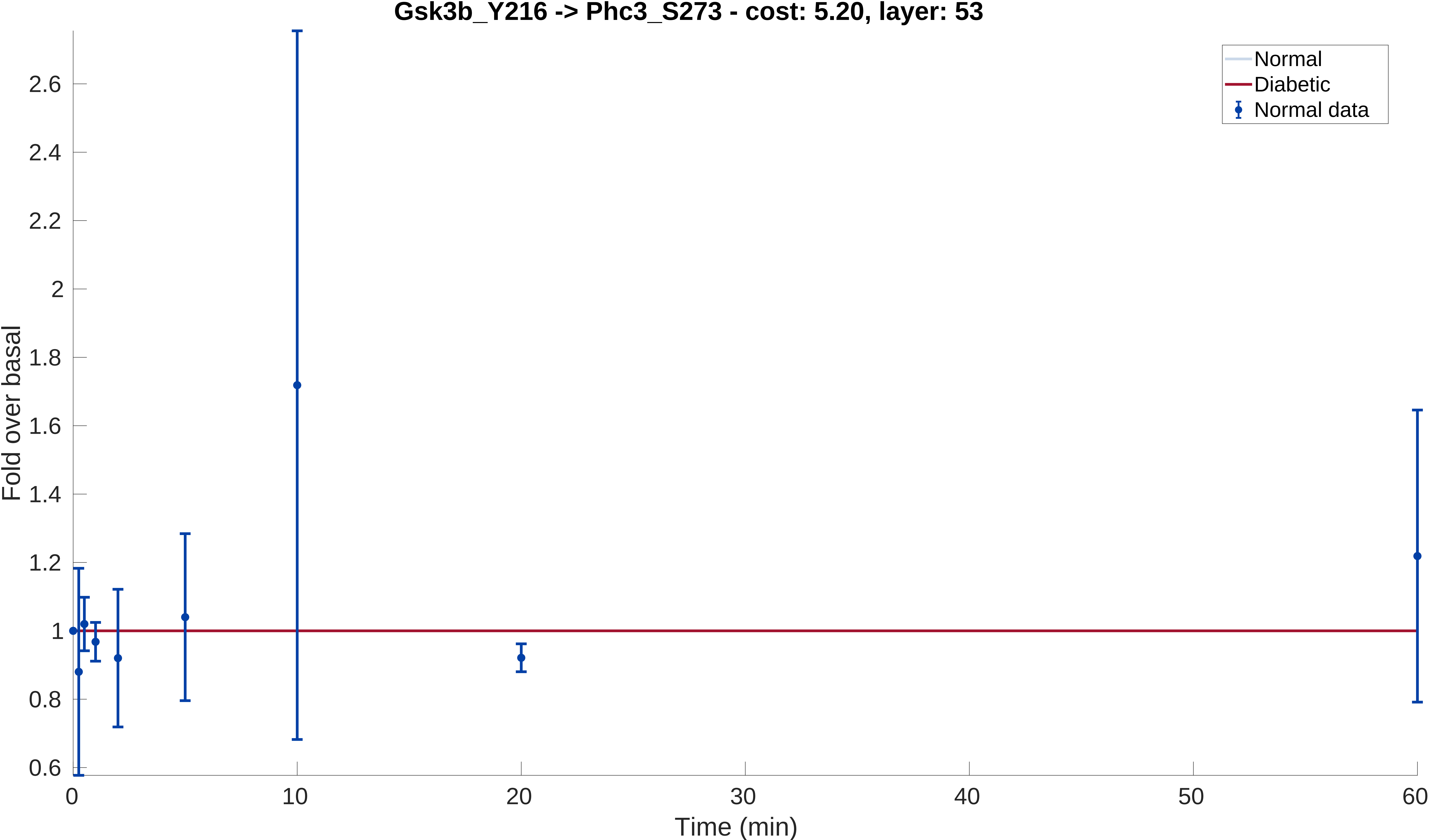

Gsk3b_Y216 -> Phc3_S273 - cost: 5.20, layer: 53
Normal
Diabetic
Normal data
Fold over basal
2.6
2.4
2.2
2
1.8
1.6
1.4
1.2
1
0.8
0.6
0
10
20
30
40
50
60
Time (min)

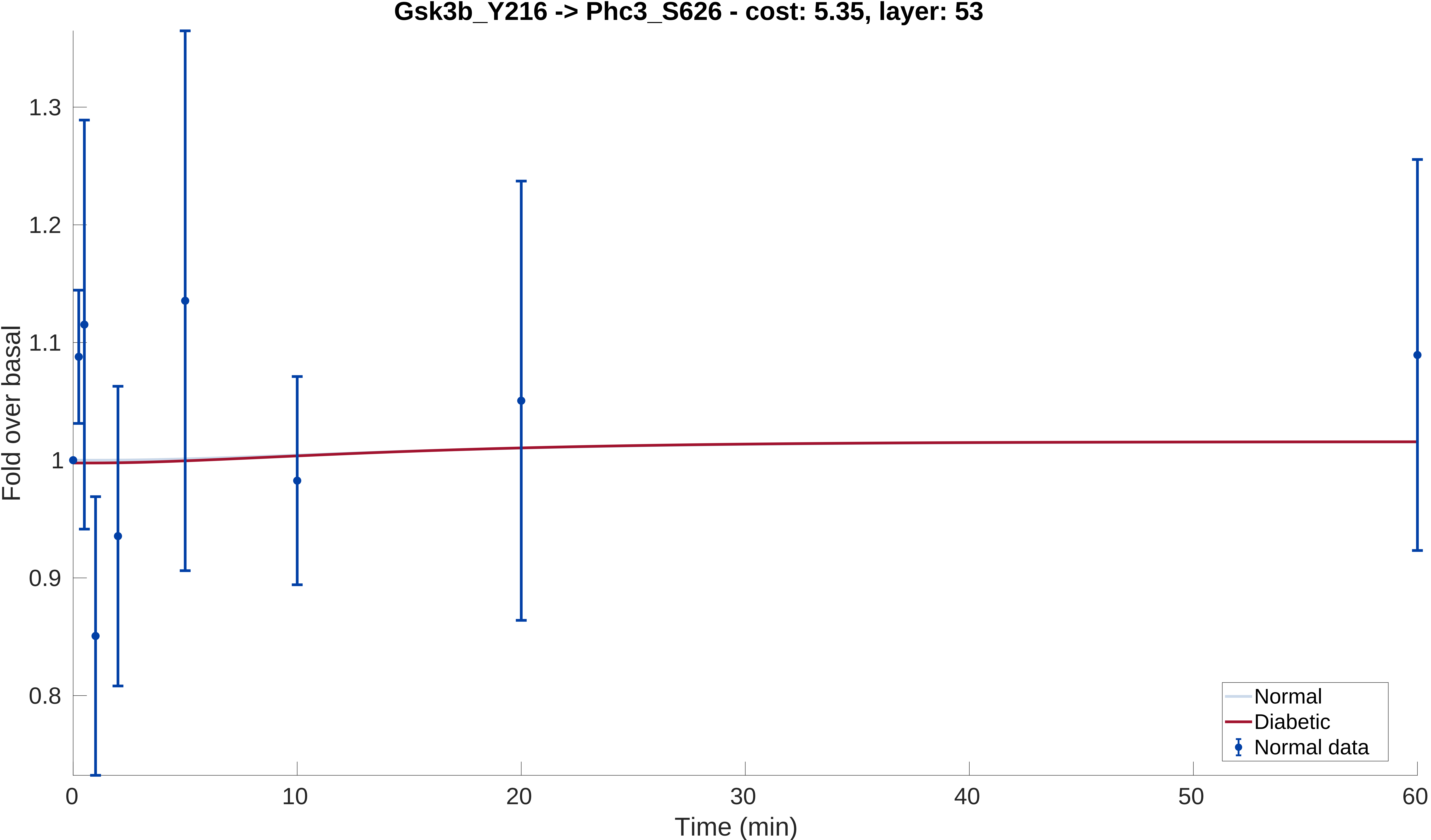

Gsk3b_Y216 -> Phc3_S626 - cost: 5.35, layer: 53
Fold over basal
1.3
1.2
1.1
1
0.9
0.8
0
10
20
30
40
50
60
Time (min)
Normal
Diabetic
Normal data

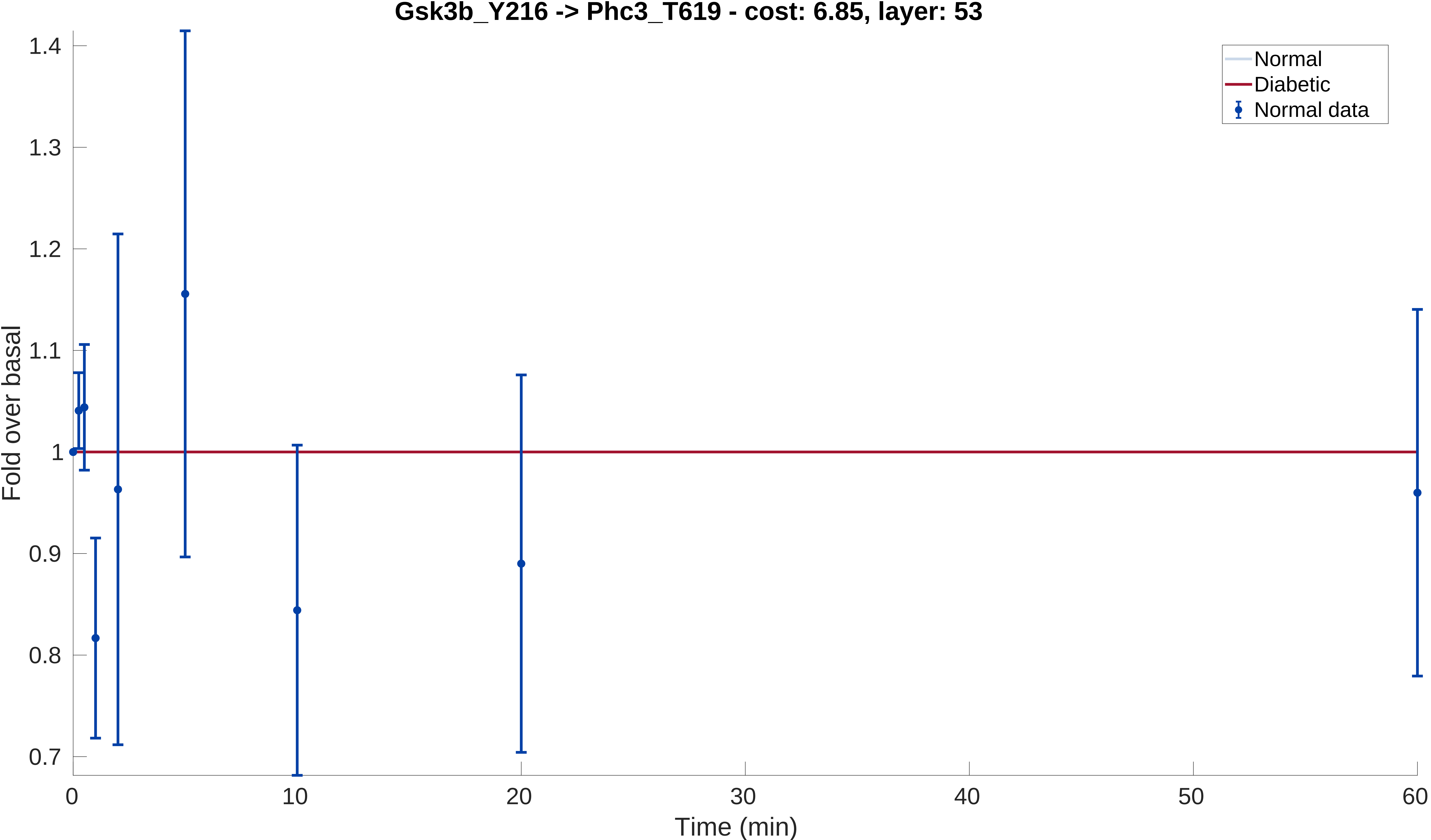

Gsk3b_Y216 -> Phc3_T619 - cost: 6.85, layer: 53
Normal
Diabetic
Normal data
Fold over basal
1.4
1.3
1.2
1.1
1
0.9
0.8
0.7
0
10
20
30
40
50
60
Time (min)

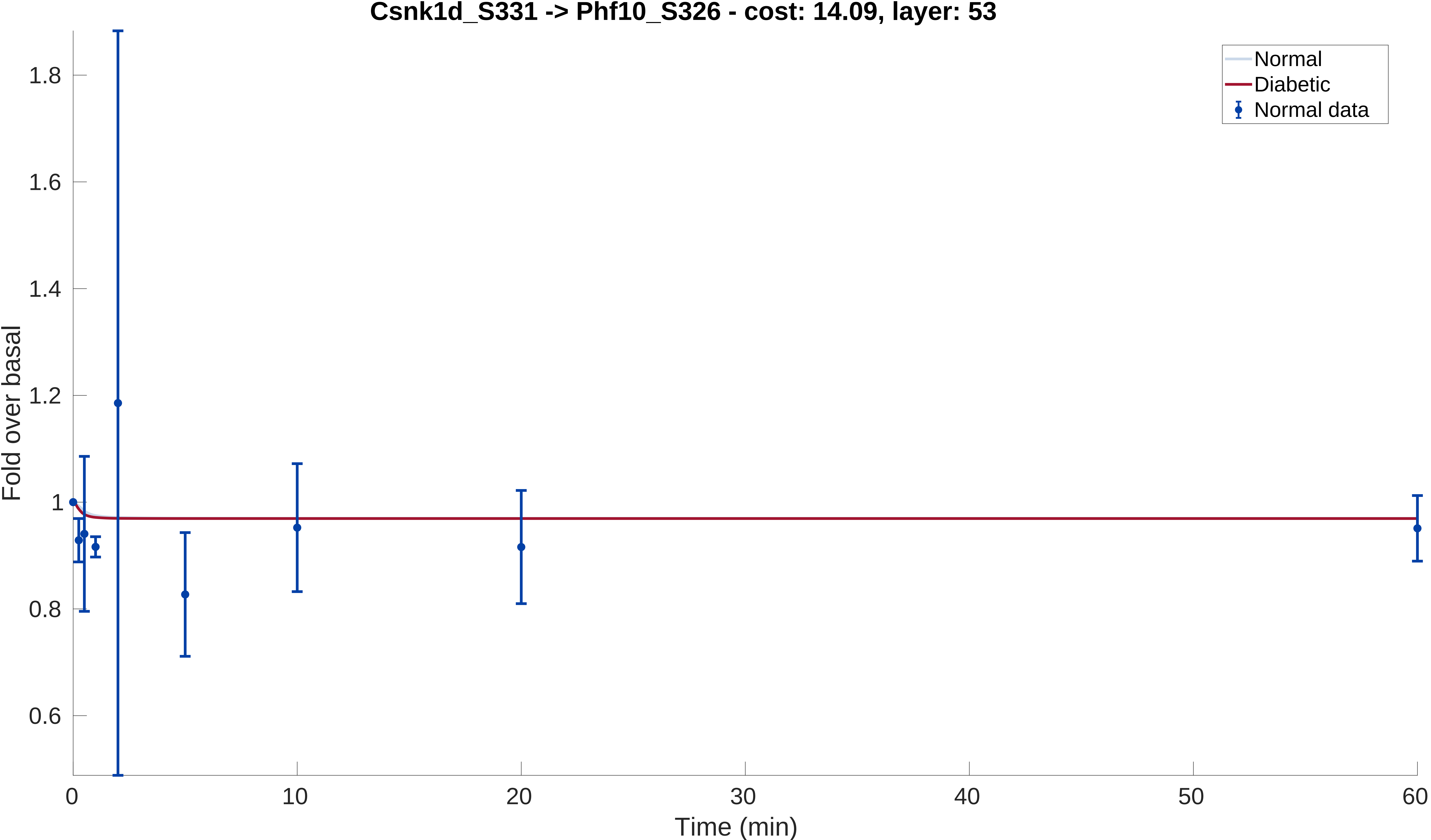
Csnk1d_S331 -> Phf10_S326 - cost: 14.09, layer: 53
Normal
Diabetic
Normal data
Fold over basal
Time (min)
1.8
1.6
1.4
1.2
1
0.8
0.6
0
10
20
30
40
50
60

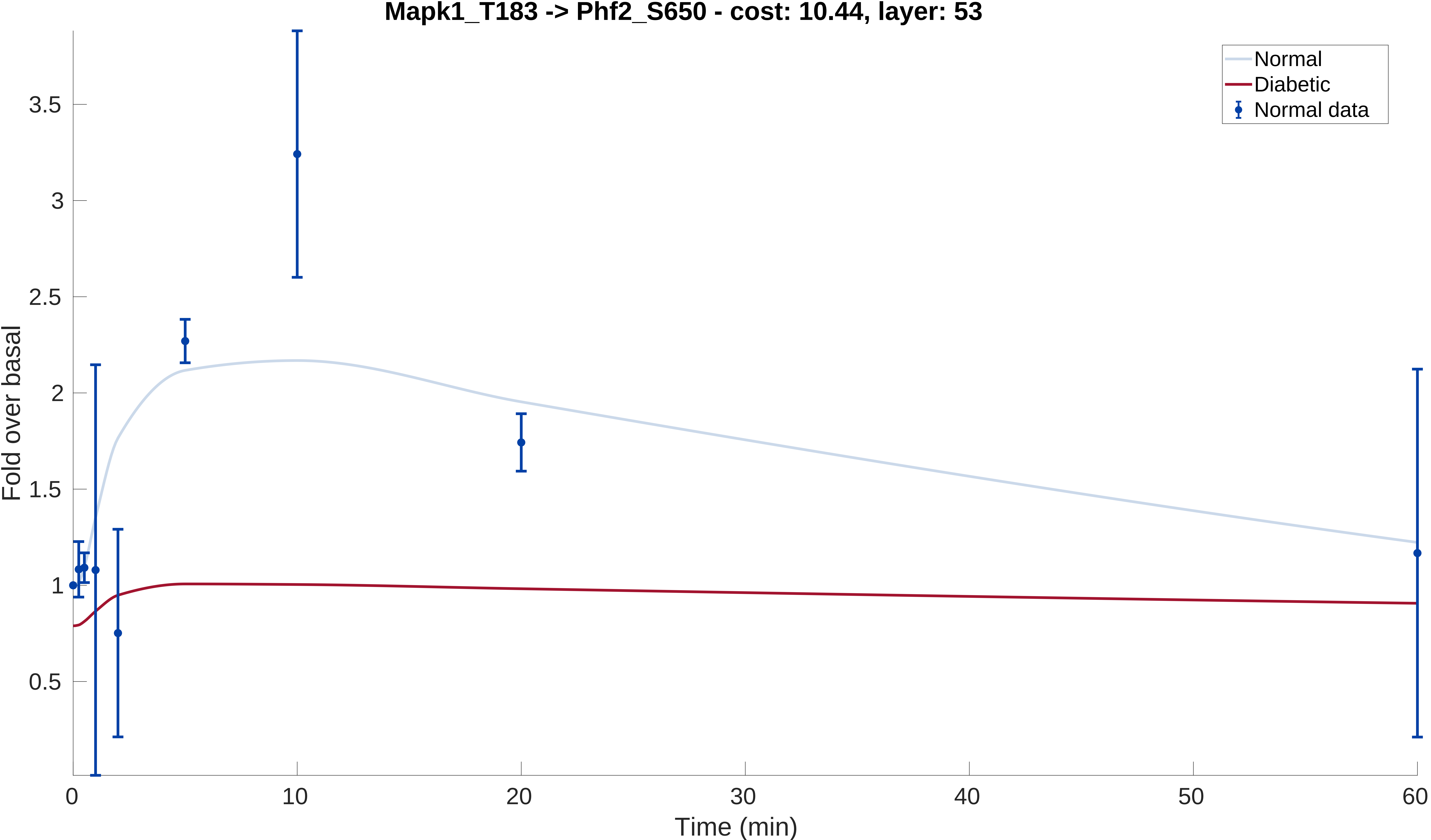

Mapk1_T183 -> Phf2_S650 - cost: 10.44, layer: 53
Normal
Diabetic
Normal data
Fold over basal
Time (min)
0.5
1
1.5
2
2.5
3
3.5
0
10
20
30
40
50
60

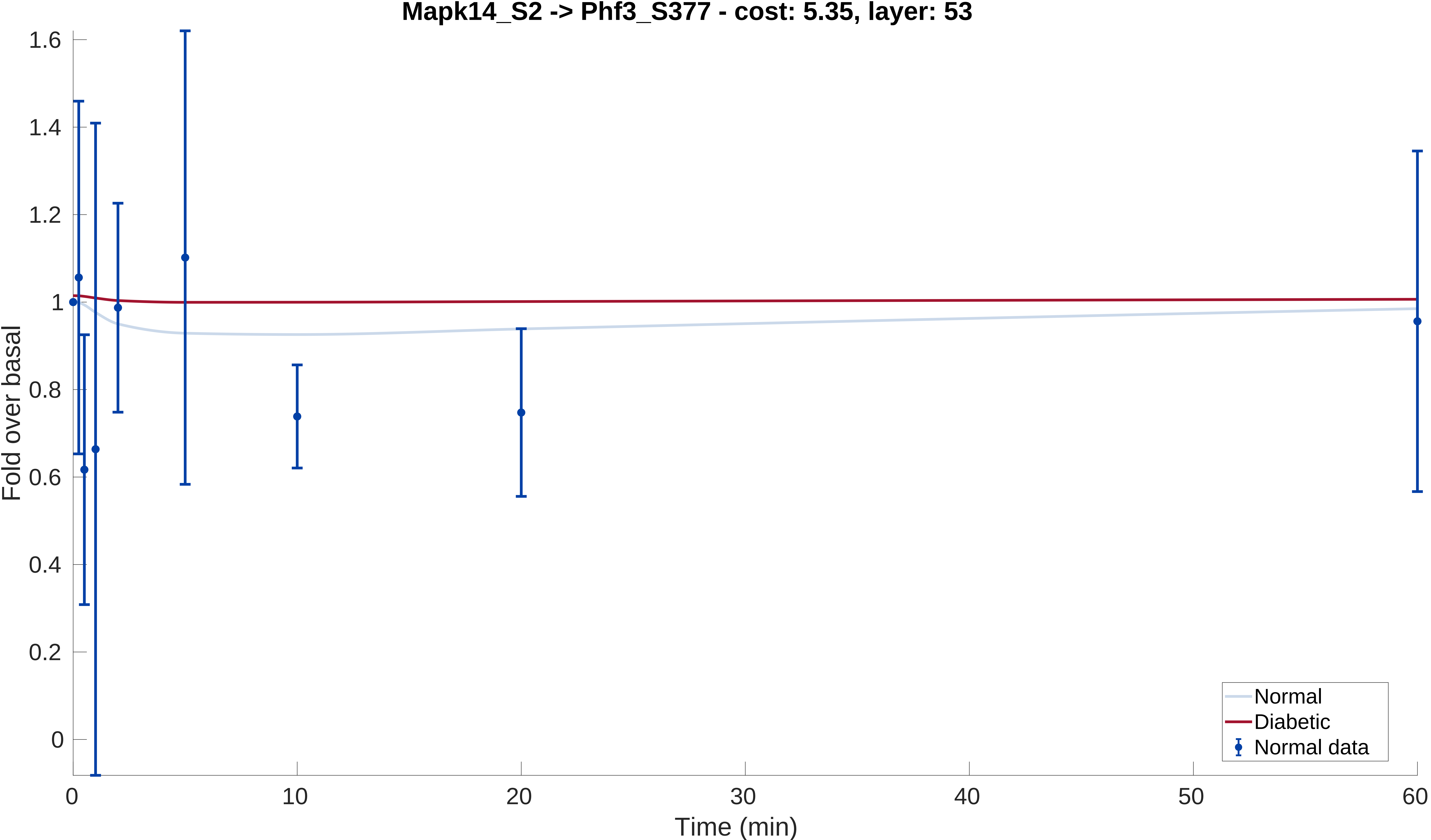

Mapk14_S2 -> Phf3_S377 - cost: 5.35, layer: 53
Fold over basal
Time (min)
0
0.2
0.4
0.6
0.8
1
1.2
1.4
1.6
0
10
20
30
40
50
60
Normal
Diabetic
Normal data

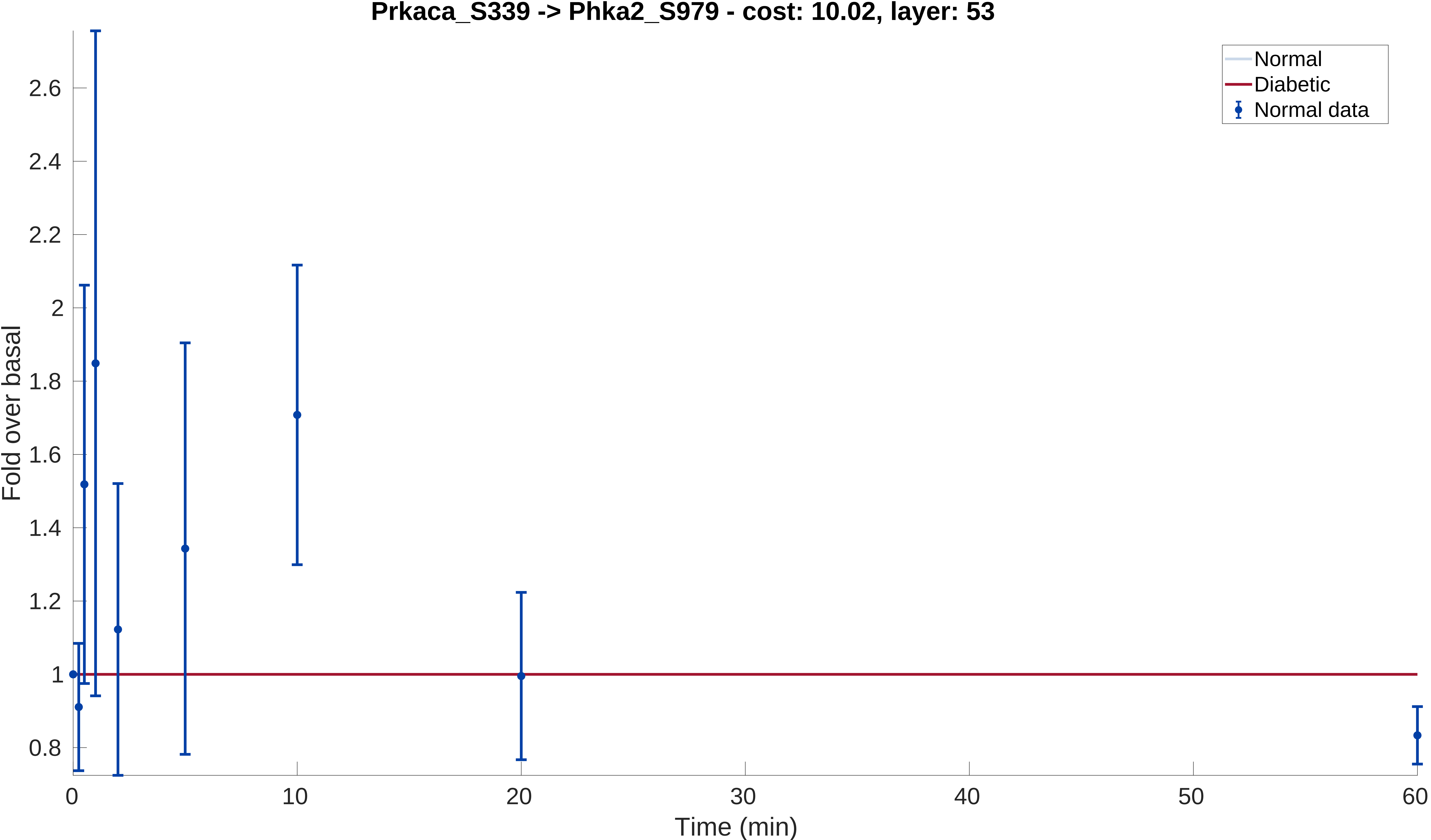
Prkaca_S339 -> Phka2_S979 - cost: 10.02, layer: 53
Normal
Diabetic
Normal data
Fold over basal
2.6
2.4
2.2
2
1.8
1.6
1.4
1.2
1
0.8
0
10
20
30
40
50
60
Time (min)

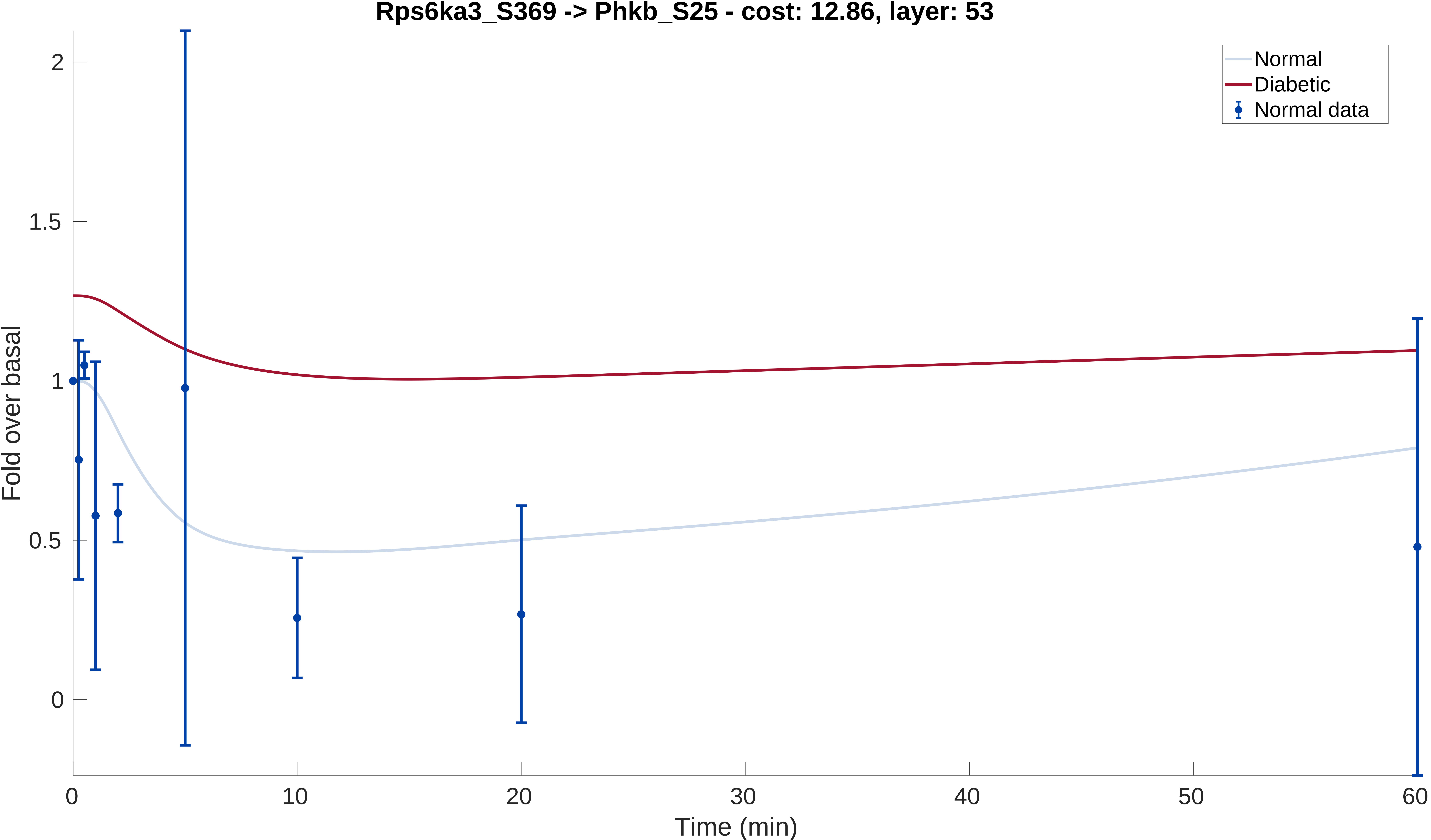

Rps6ka3_S369 -> Phkb_S25 - cost: 12.86, layer: 53
Normal
Diabetic
Normal data
Fold over basal
2
1.5
1
0.5
0
0
10
20
30
40
50
60
Time (min)

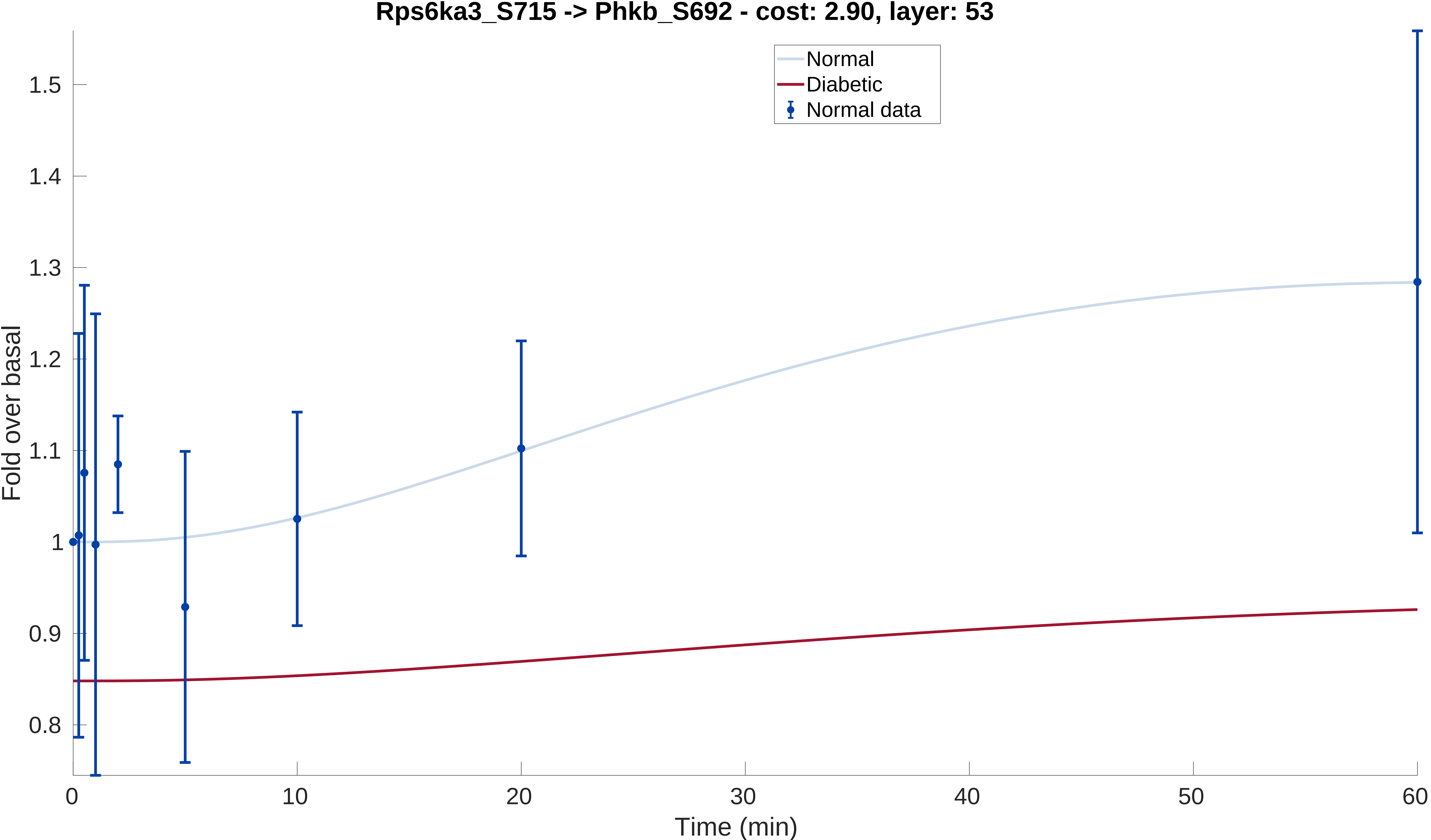

Rps6ka3_S715 -> Phkb_S692 - cost: 2.90, layer: 53
Normal
Diabetic
Normal data
Fold over basal
1.5
1.4
1.3
1.2
1.1
1
0.9
0.8
0
10
20
30
40
50
60
Time (min)

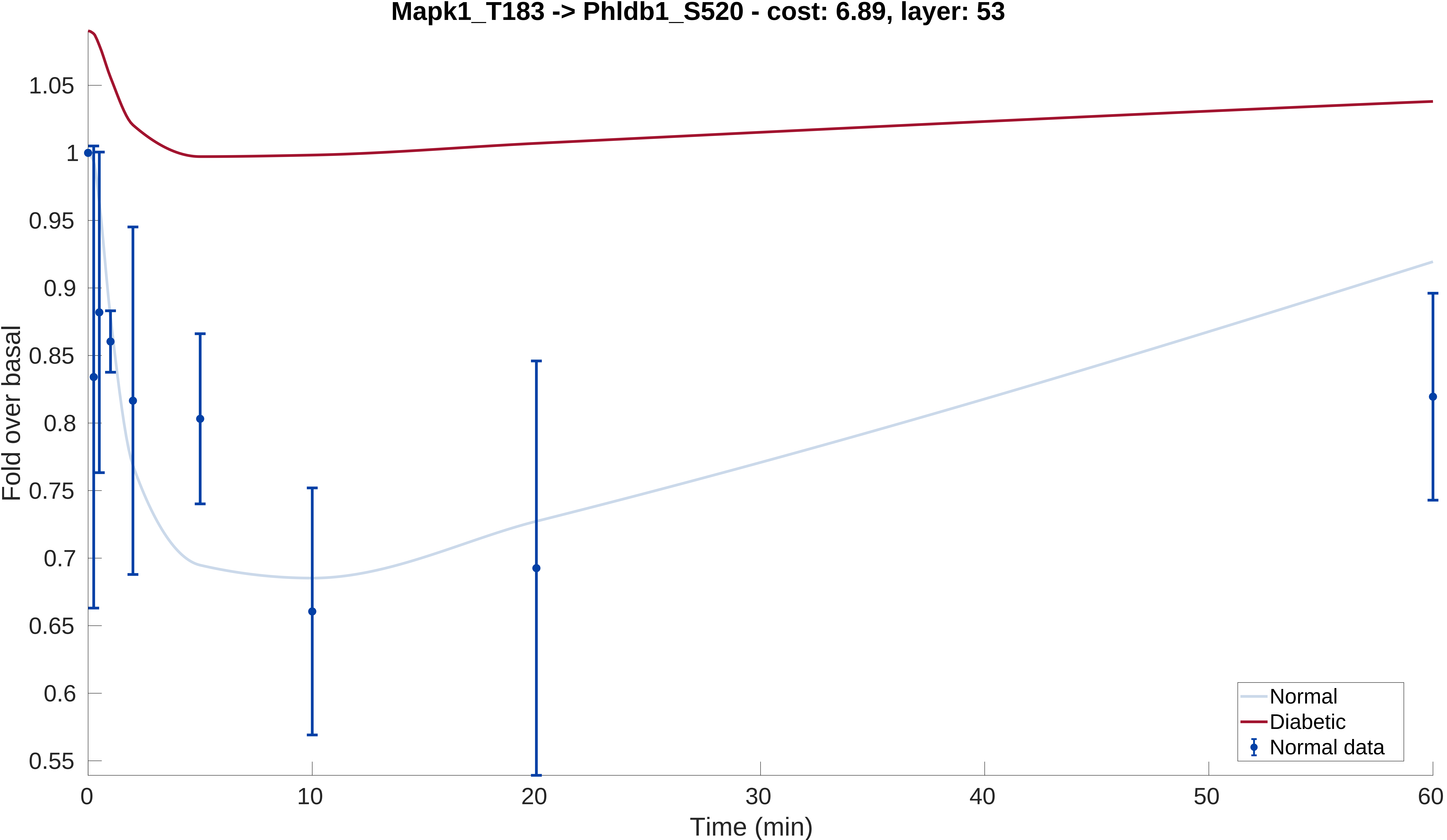
Mapk1_T183 -> Phldb1_S520 - cost: 6.89, layer: 53
Fold over basal
1.05
1
0.95
0.9
0.85
0.8
0.75
0.7
0.65
0.6
0.55
0
10
20
30
40
50
60
Time (min)
Normal
Diabetic
Normal data

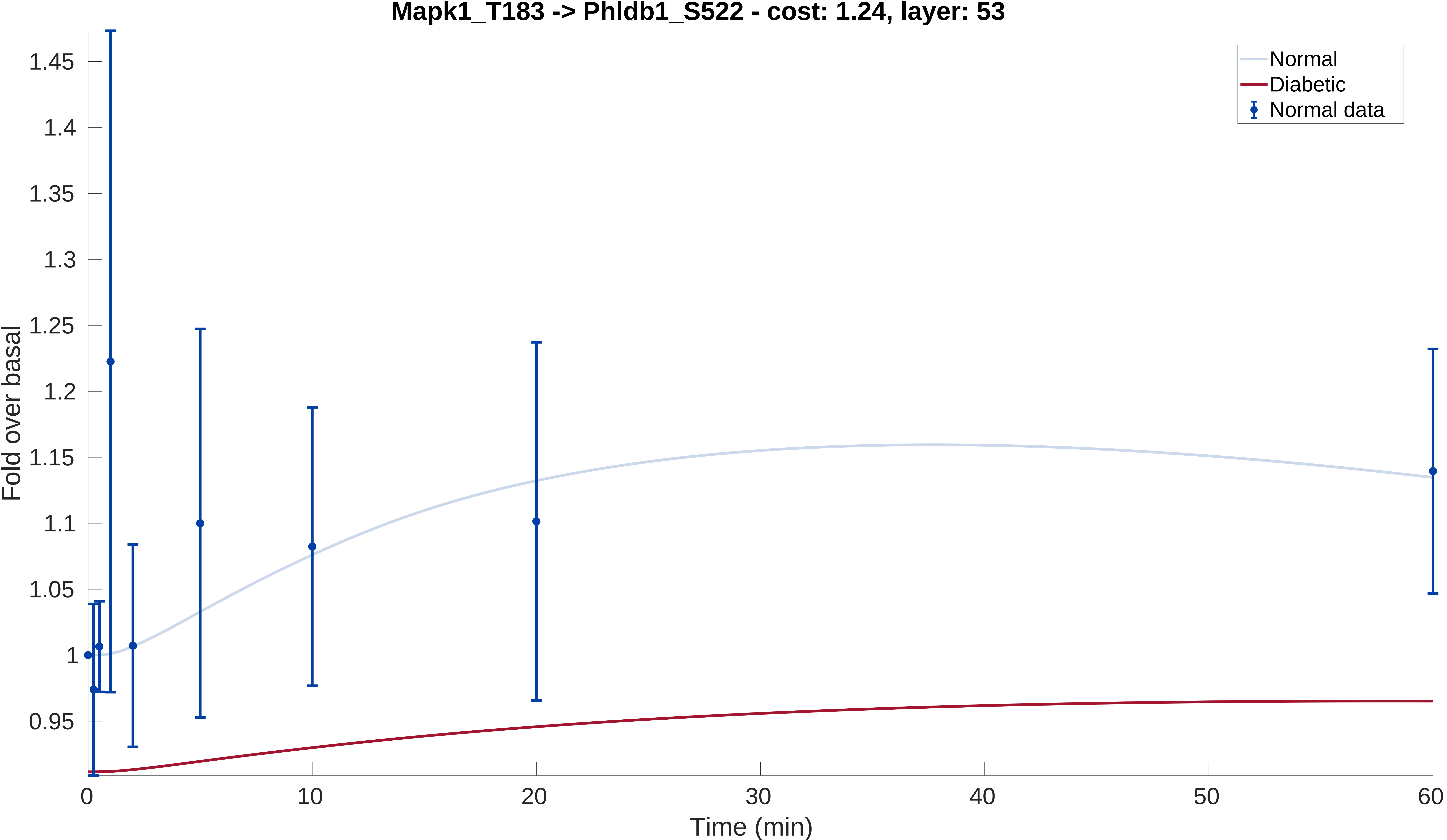

Mapk1_T183 -> Phldb1_S522 - cost: 1.24, layer: 53
Normal
Diabetic
Normal data
Fold over basal
1.45
1.4
1.35
1.3
1.25
1.2
1.15
1.1
1.05
1
0.95
0
10
20
30
40
50
60
Time (min)

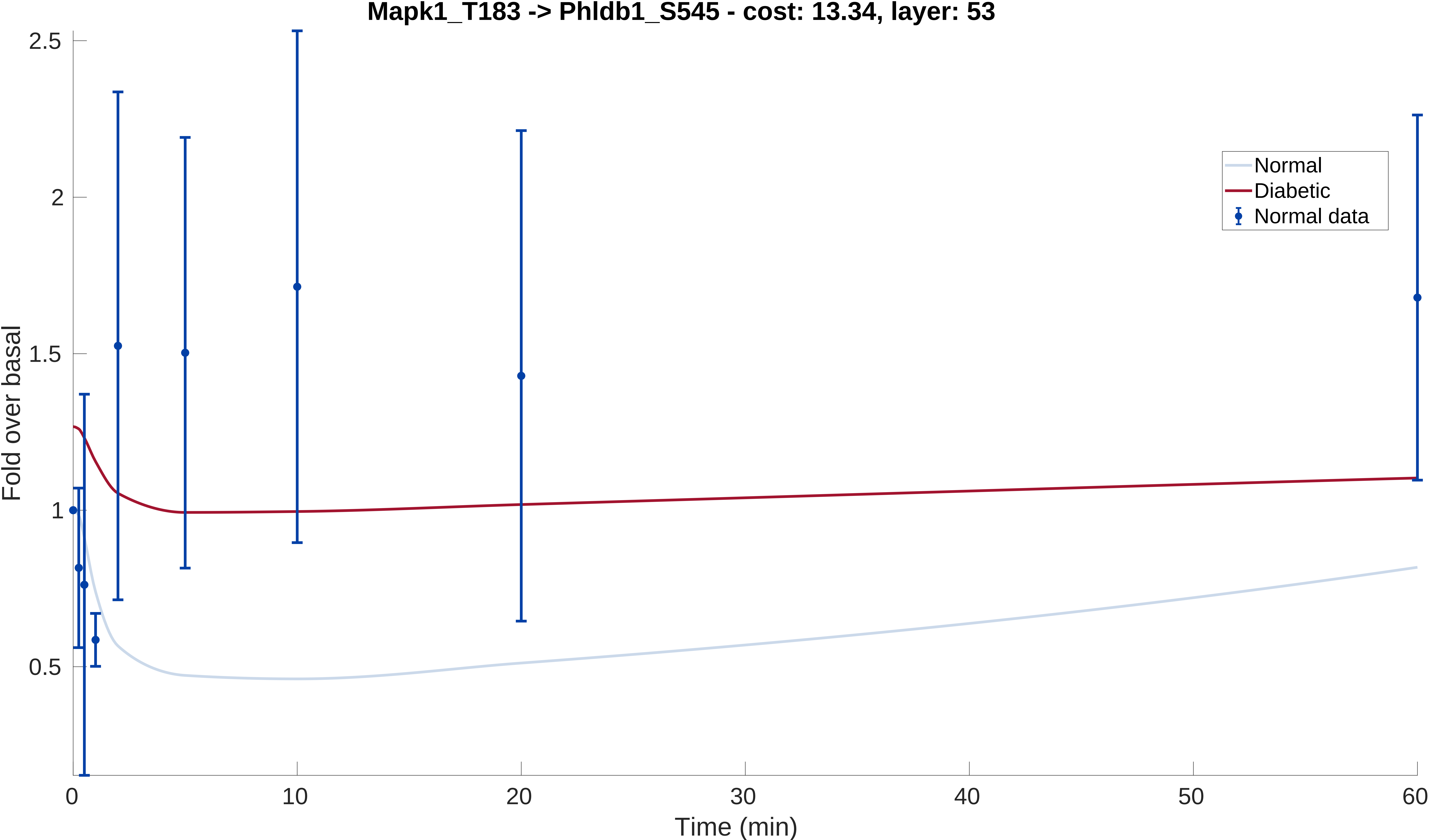

Mapk1_T183 -> Phldb1_S545 - cost: 13.34, layer: 53
Normal
Diabetic
Normal data
Fold over basal
Time (min)
0.5
1
1.5
2
2.5
0
10
20
30
40
50
60

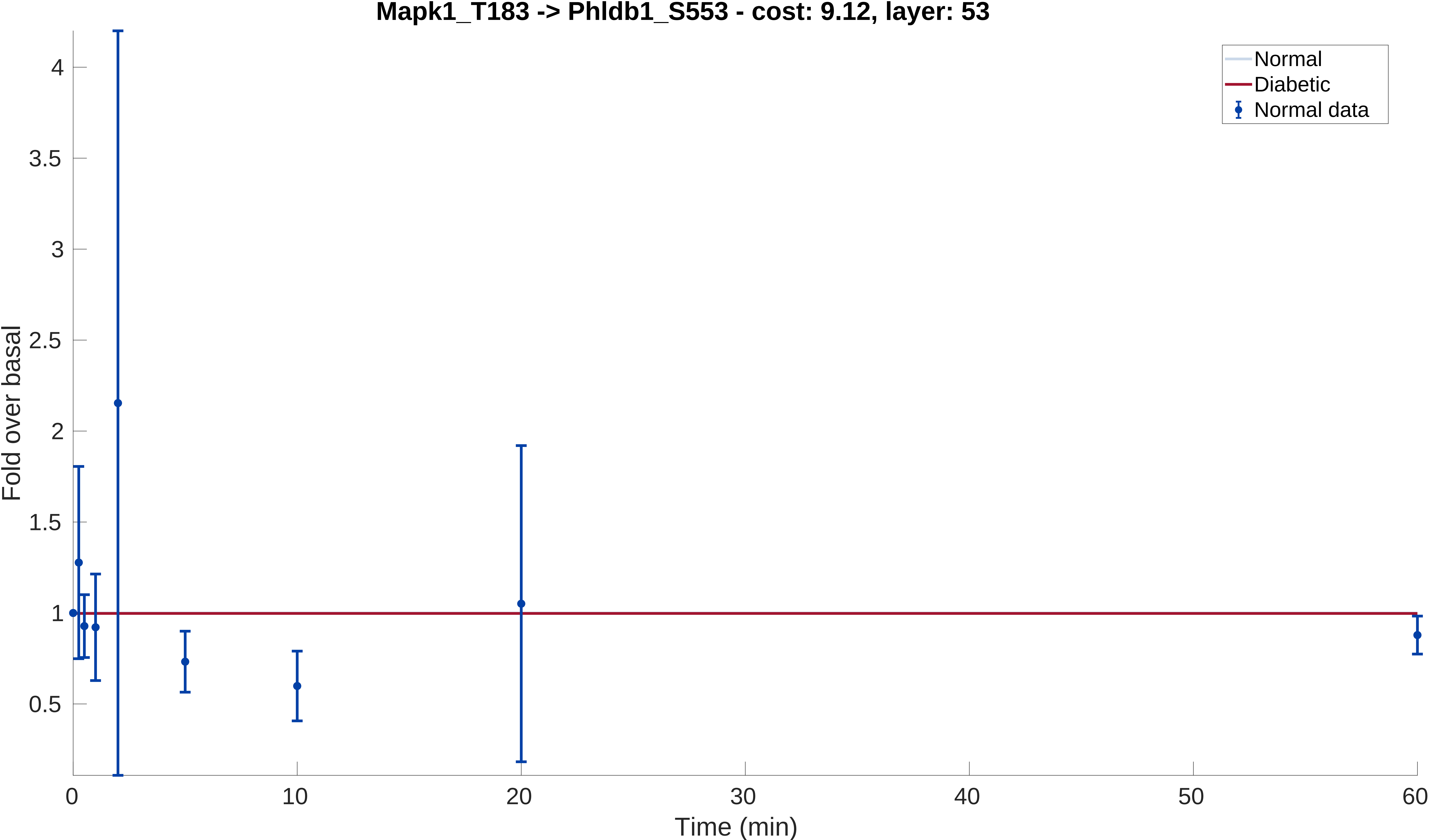

Mapk1_T183 -> Phldb1_S553 - cost: 9.12, layer: 53
Normal
Diabetic
Normal data
Fold over basal
4
3.5
3
2.5
2
1.5
1
0.5
0
10
20
30
40
50
60
Time (min)

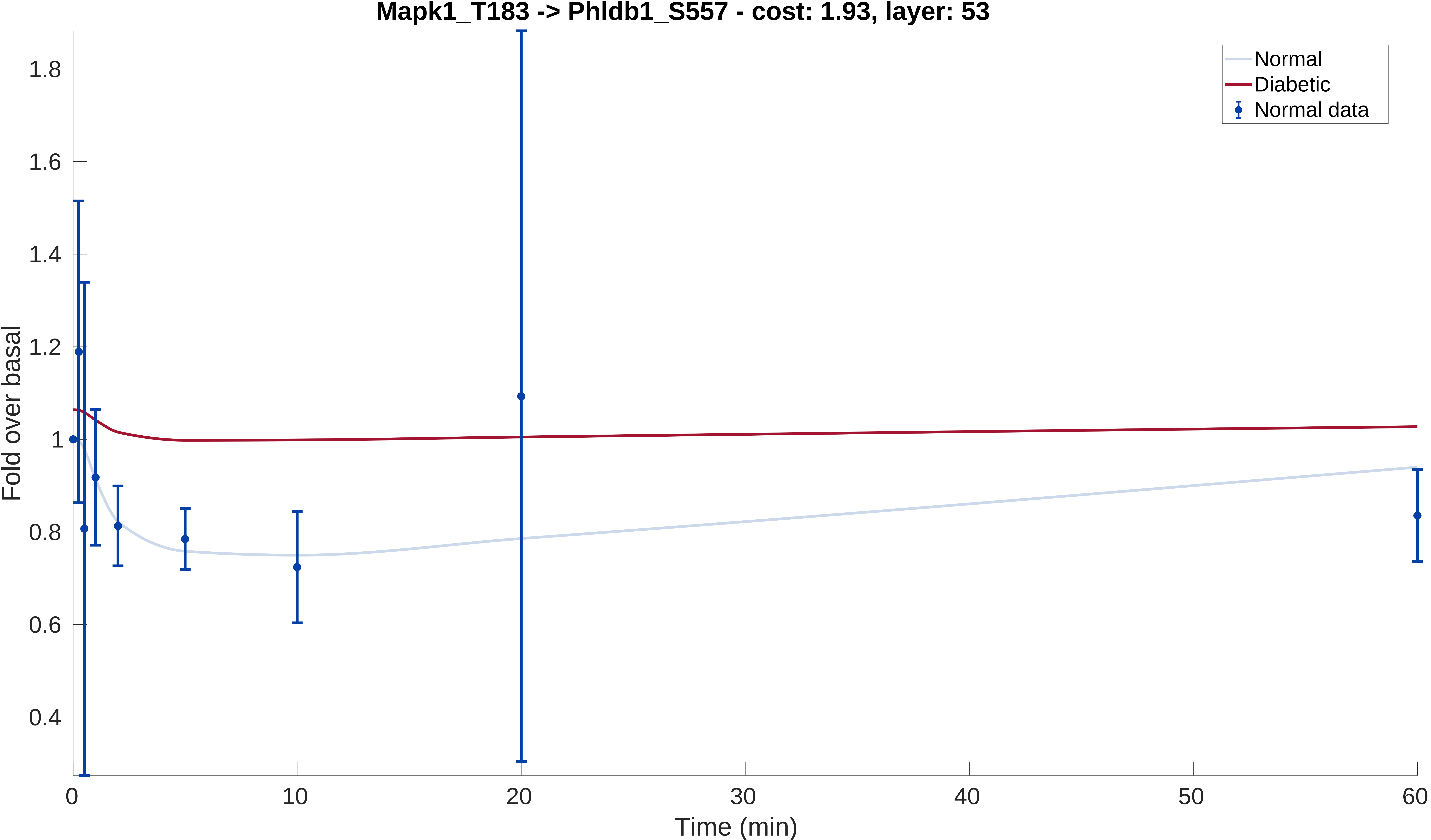
Mapk1_T183 -> Phldb1_S557 - cost: 1.93, layer: 53
Normal
Diabetic
Normal data
Fold over basal
1.8
1.6
1.4
1.2
1
0.8
0.6
0.4
0
10
20
30
40
50
60
Time (min)

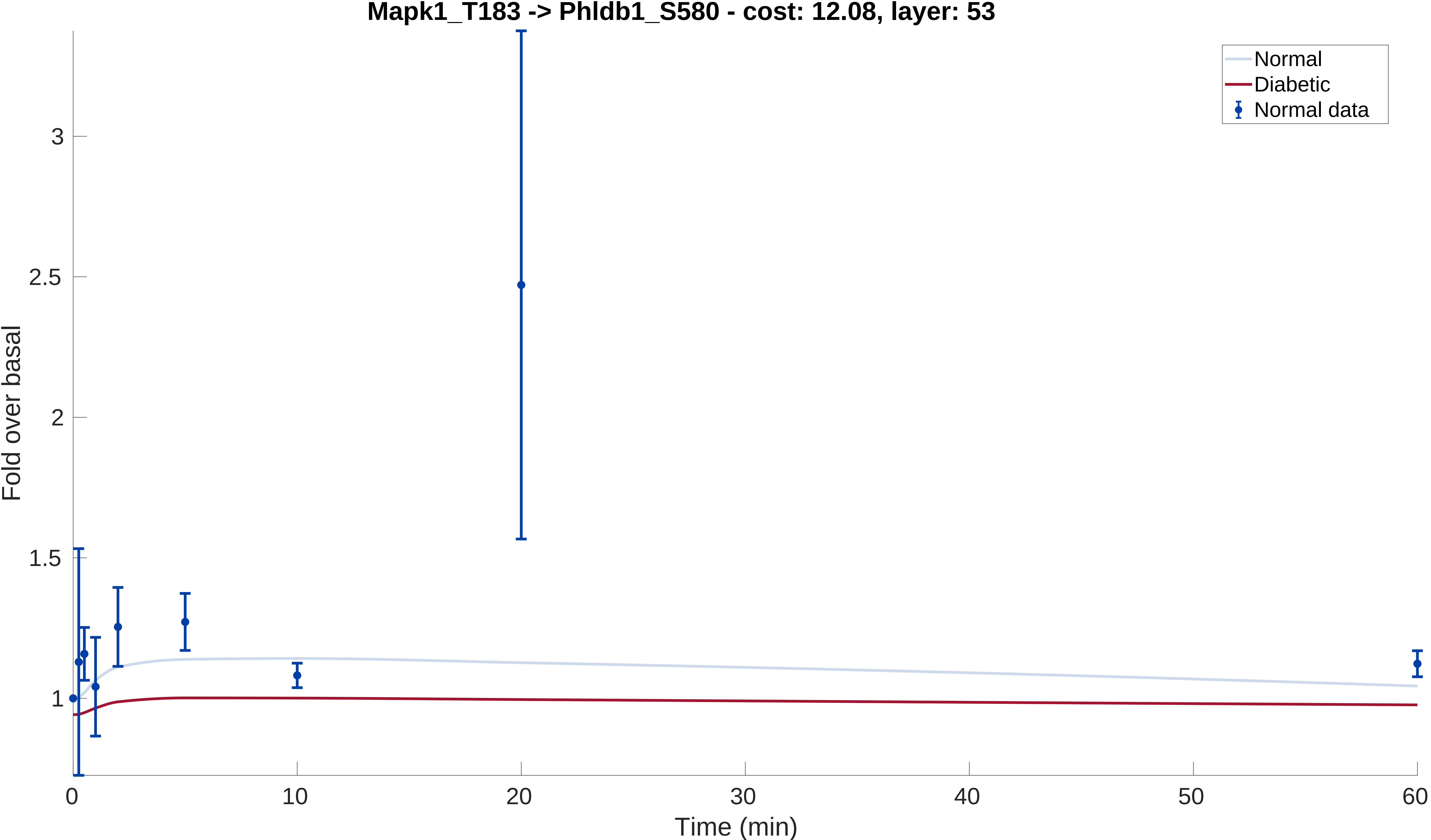

Mapk1_T183 -> Phldb1_S580 - cost: 12.08, layer: 53
Normal
Diabetic
Normal data
Fold over basal
3
2.5
2
1.5
1
0
10
20
30
40
50
60
Time (min)

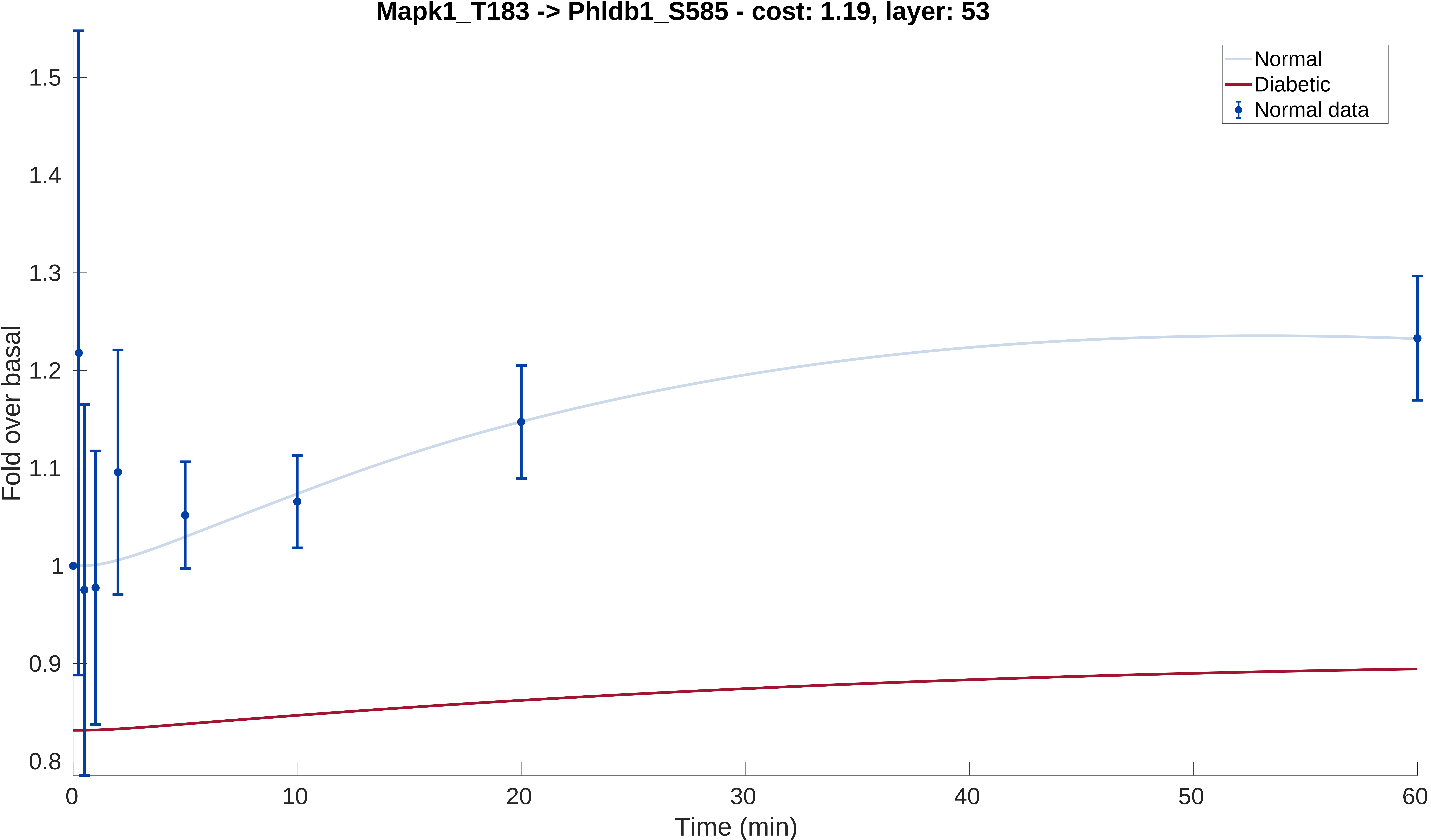
Mapk1_T183 -> Phldb1_S585 - cost: 1.19, layer: 53
Normal
Diabetic
Normal data
Fold over basal
1.5
1.4
1.3
1.2
1.1
1
0.9
0.8
0
10
20
30
40
50
60
Time (min)

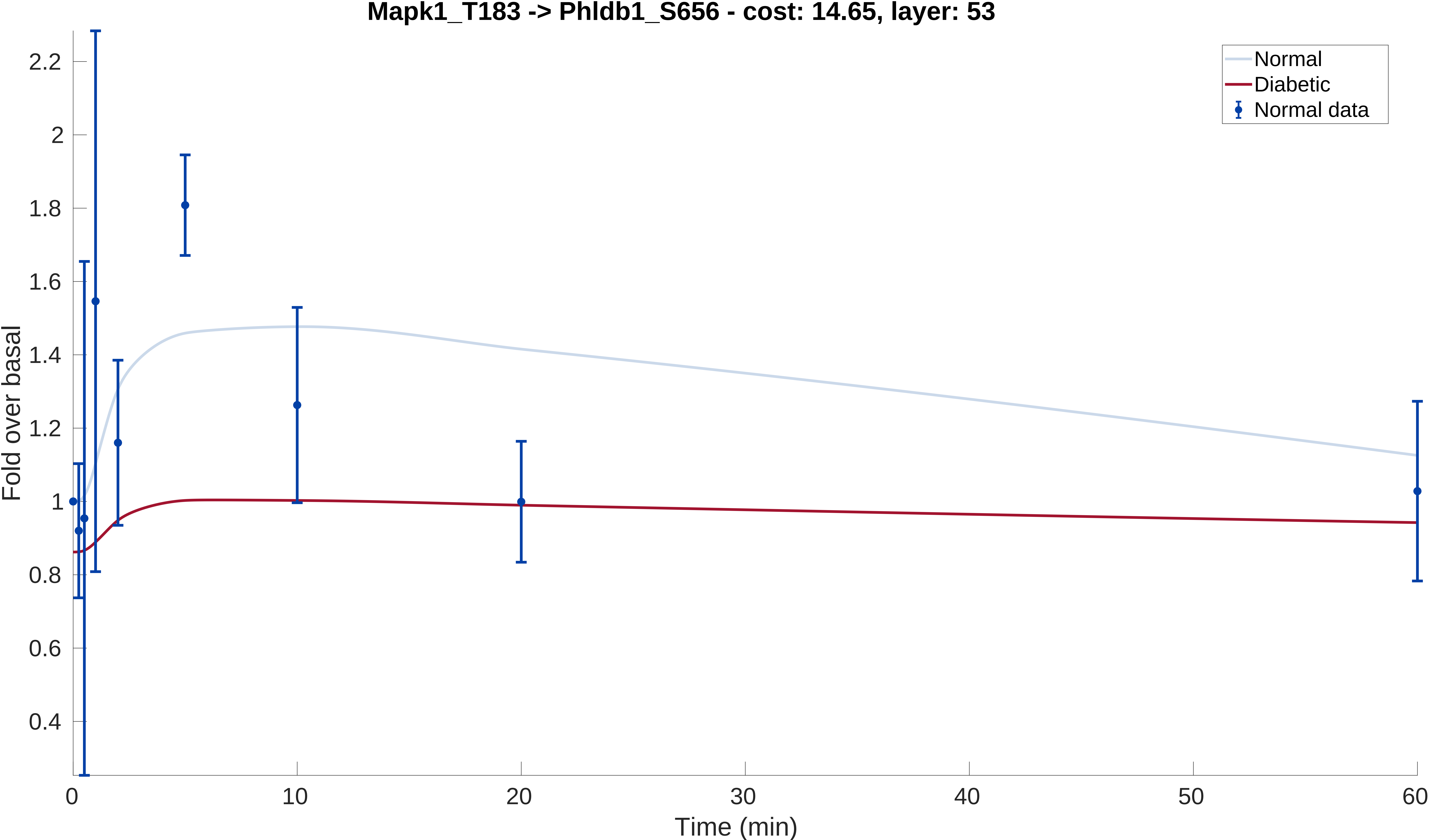

Mapk1_T183 -> Phldb1_S656 - cost: 14.65, layer: 53
Normal
Diabetic
Normal data
Fold over basal
2.2
2
1.8
1.6
1.4
1.2
1
0.8
0.6
0.4
0
10
20
30
40
50
60
Time (min)

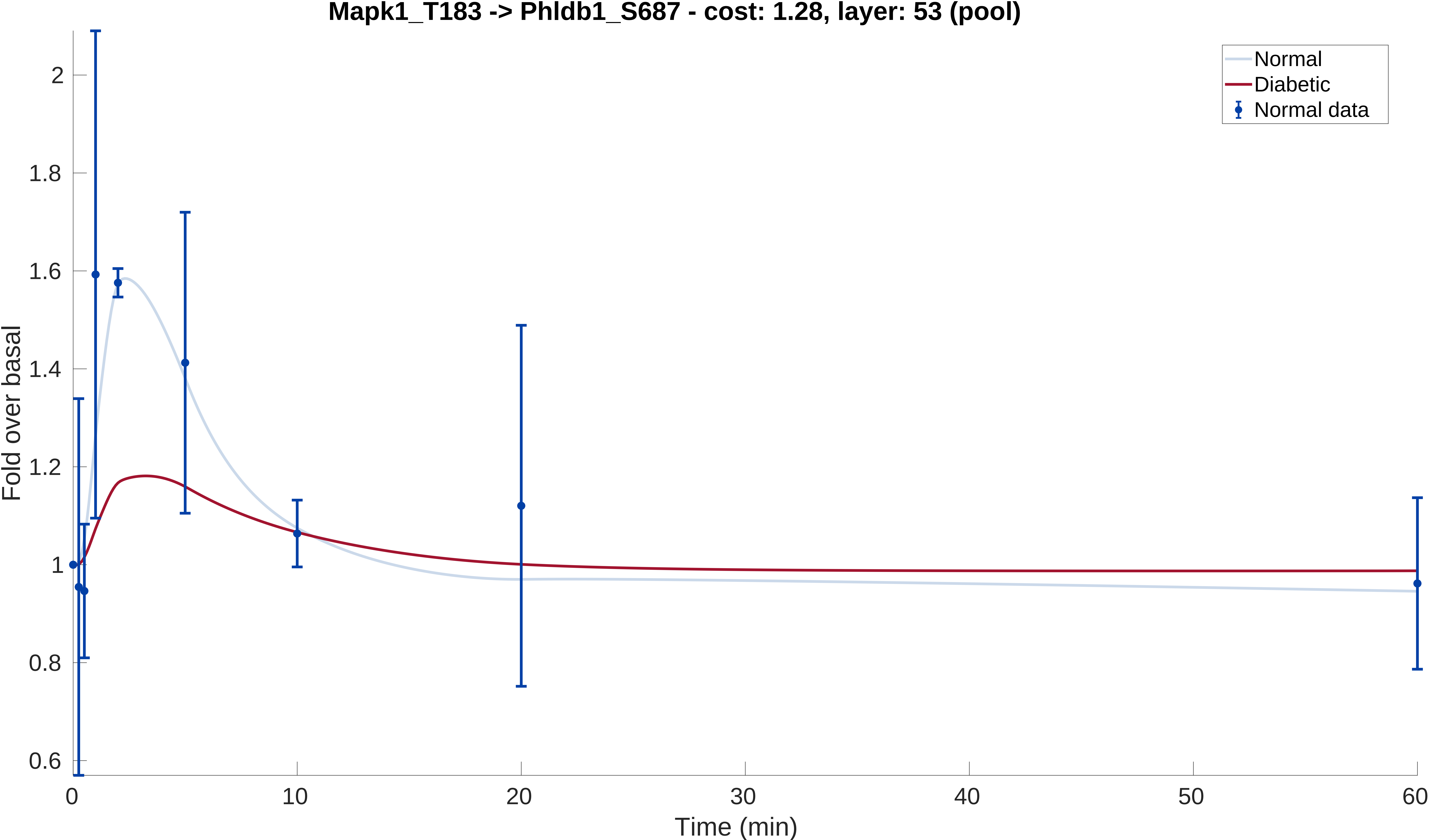
Mapk1_T183 -> Phldb1_S687 - cost: 1.28, layer: 53 (pool)
Normal
Diabetic
Normal data
Fold over basal
Time (min)
0.6
0.8
1
1.2
1.4
1.6
1.8
2
0
10
20
30
40
50
60

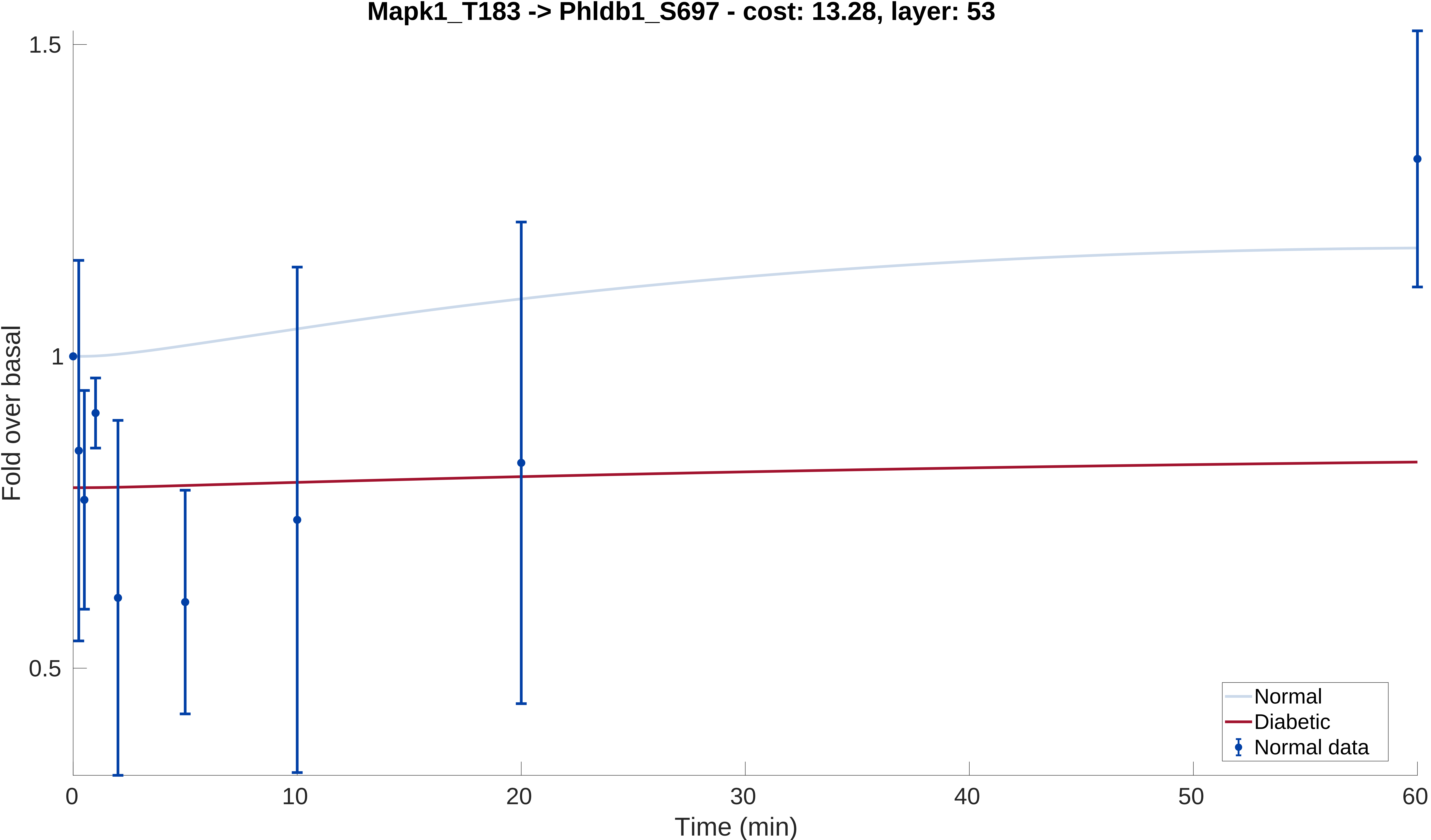
Mapk1_T183 -> Phldb1_S697 - cost: 13.28, layer: 53
1.5
1
0.5
Fold over basal
0
10
20
30
40
50
60
Time (min)
Normal
Diabetic
Normal data

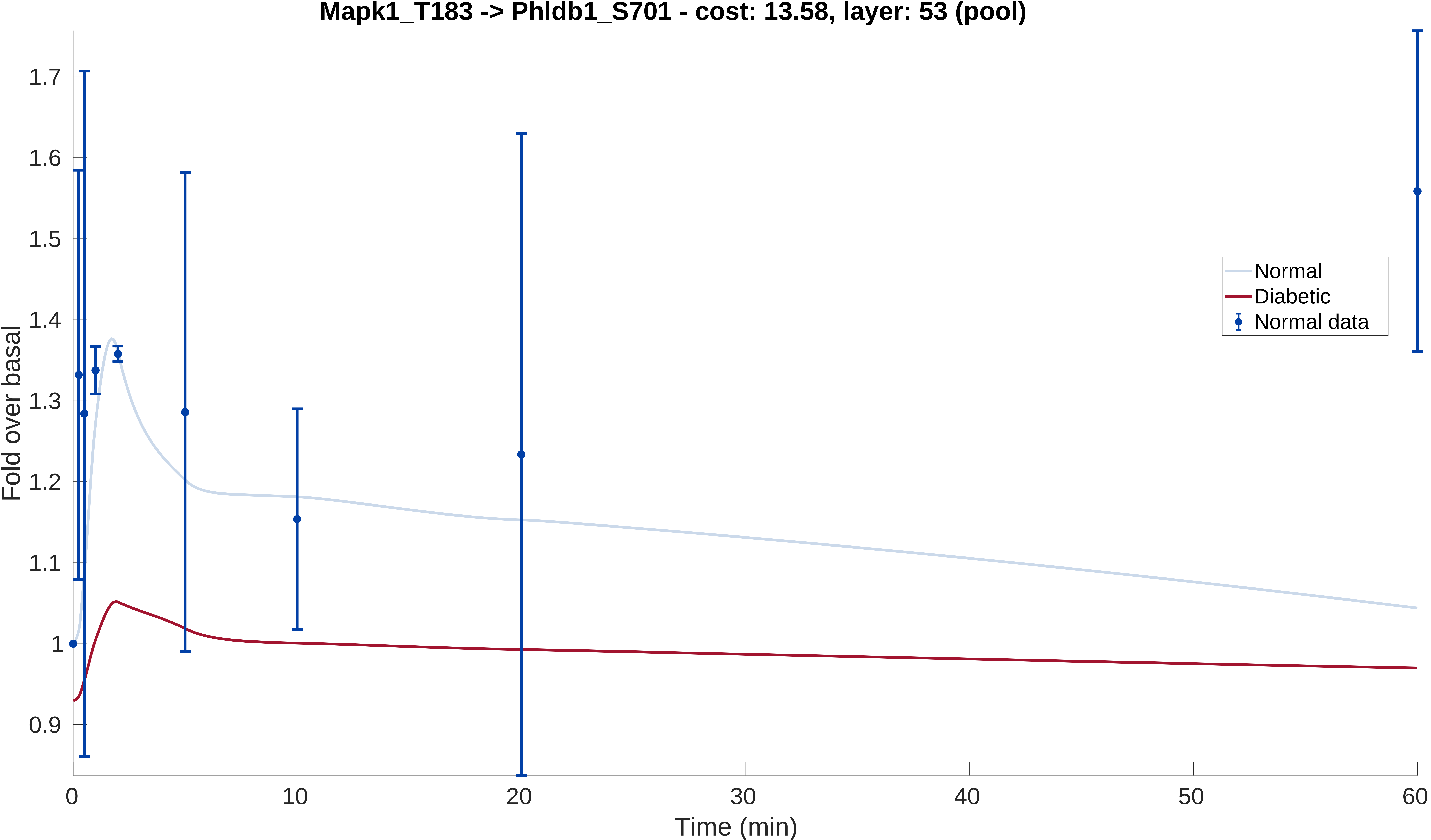

Mapk1_T183 -> Phldb1_S701 - cost: 13.58, layer: 53 (pool)
Fold over basal
1.7
1.6
1.5
1.4
1.3
1.2
1.1
1
0.9
0
10
20
30
40
50
60
Time (min)
Normal
Diabetic
Normal data

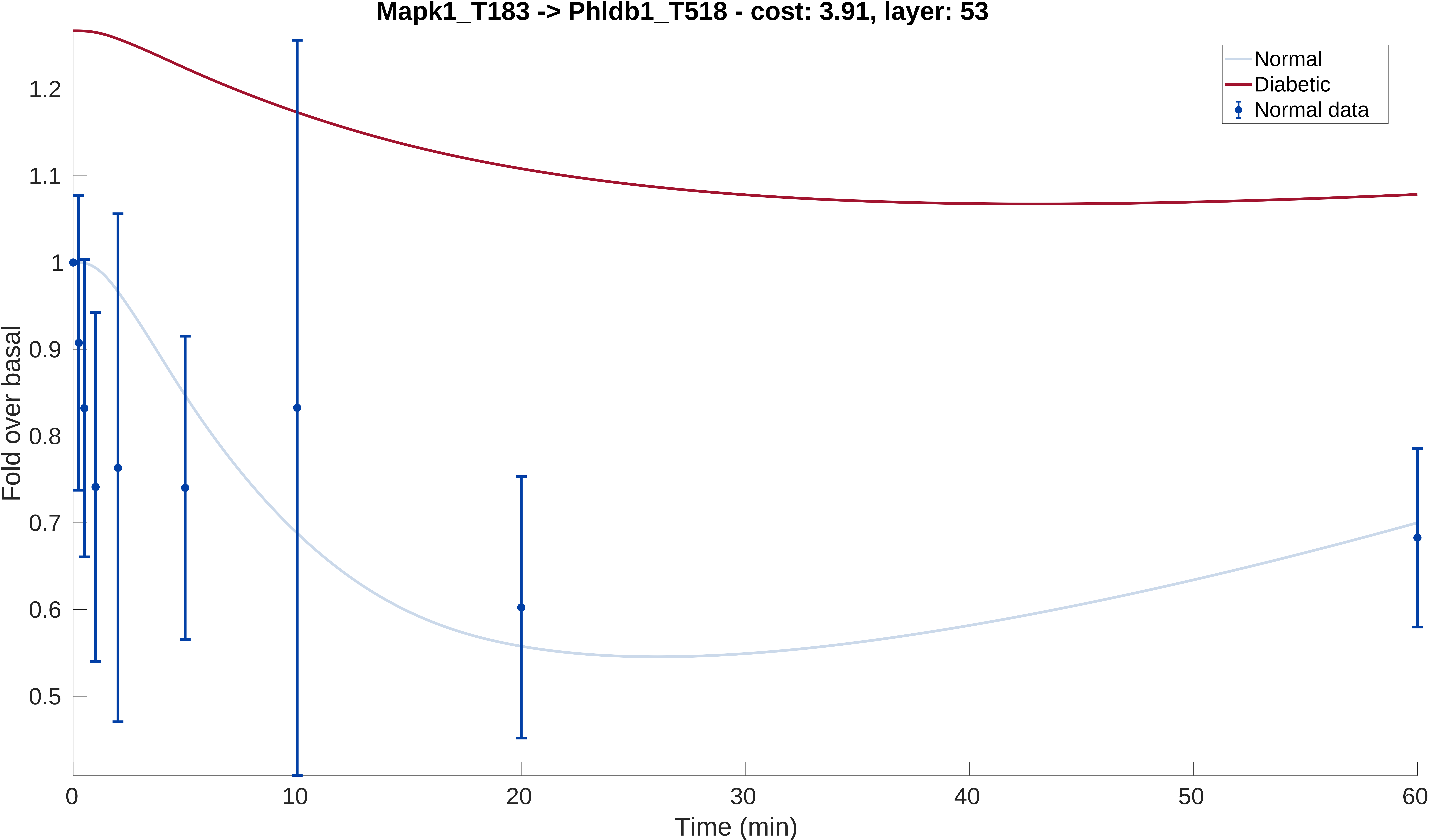

Mapk1_T183 -> Phldb1_T518 - cost: 3.91, layer: 53
Normal
Diabetic
Normal data
Fold over basal
Time (min)
1.2
1.1
1
0.9
0.8
0.7
0.6
0.5
0
10
20
30
40
50
60

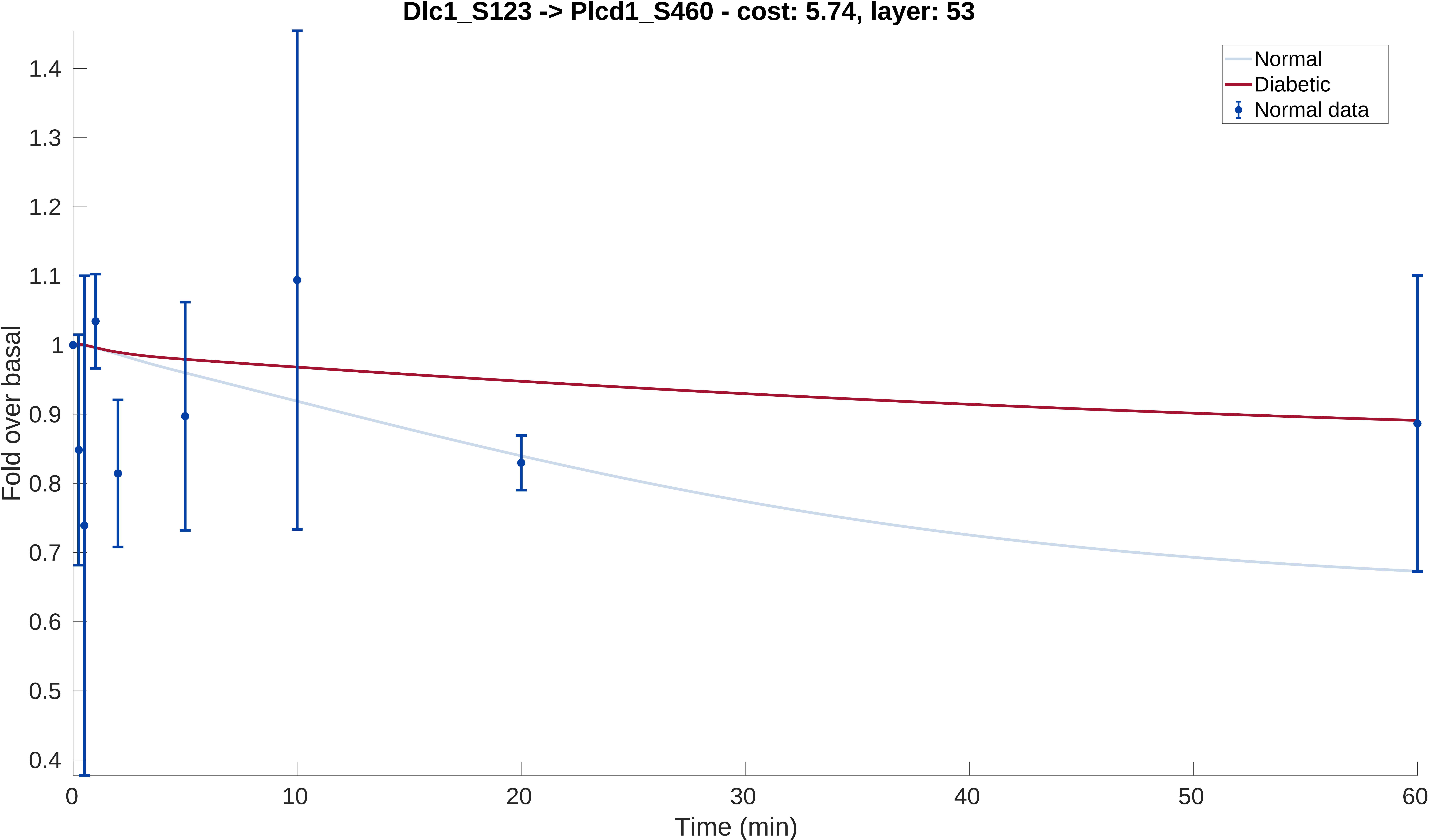
Dlc1_S123 -> Plcd1_S460 - cost: 5.74, layer: 53
Normal
Diabetic
Normal data
Fold over basal
Time (min)
1.4
1.3
1.2
1.1
1
0.9
0.8
0.7
0.6
0.5
0.4
0
10
20
30
40
50
60

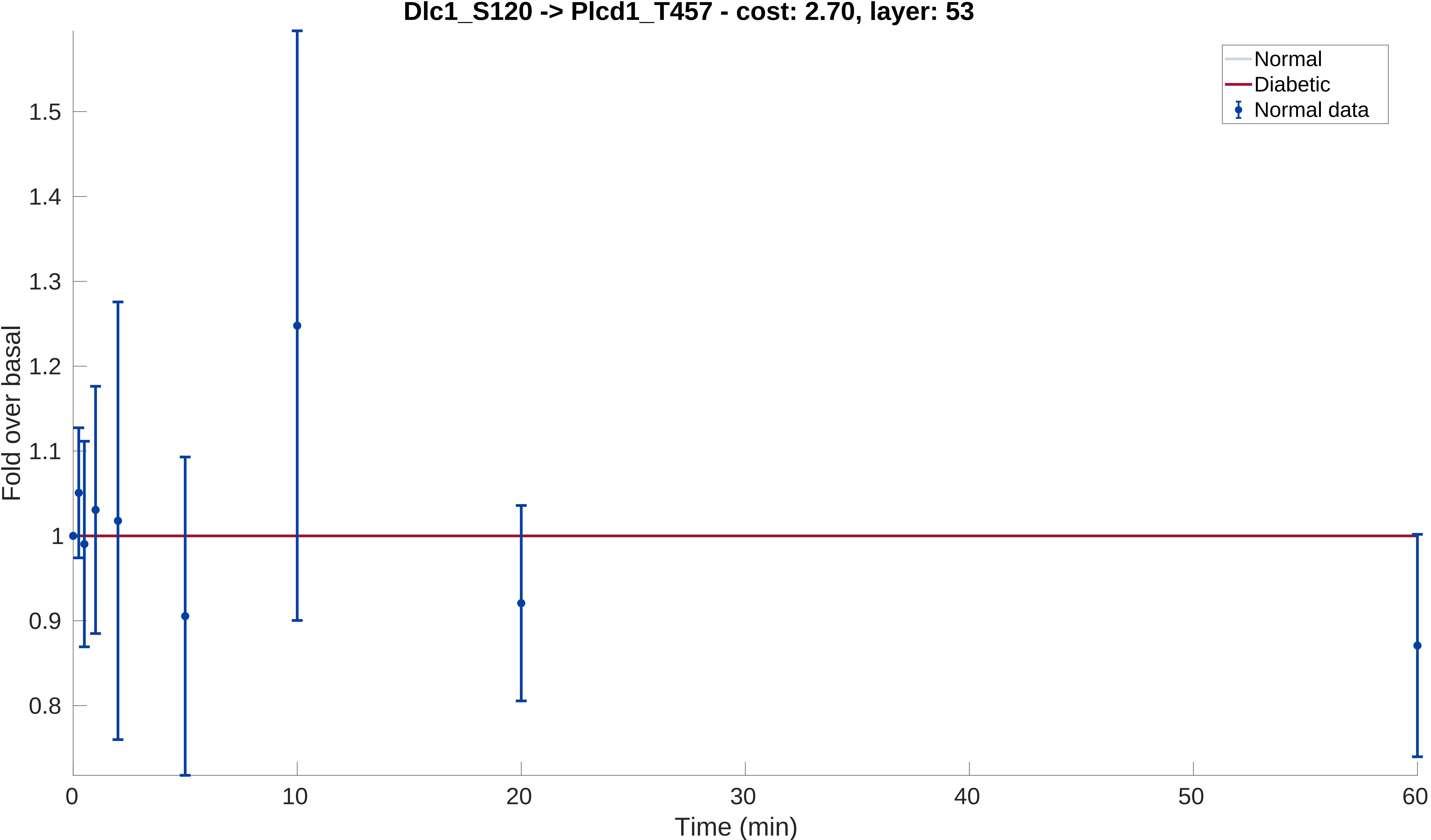

Dlc1_S120 -> Plcd1_T457 - cost: 2.70, layer: 53
Normal
Diabetic
Normal data
Fold over basal
1.5
1.4
1.3
1.2
1.1
1
0.9
0.8
0
10
20
30
40
50
60
Time (min)

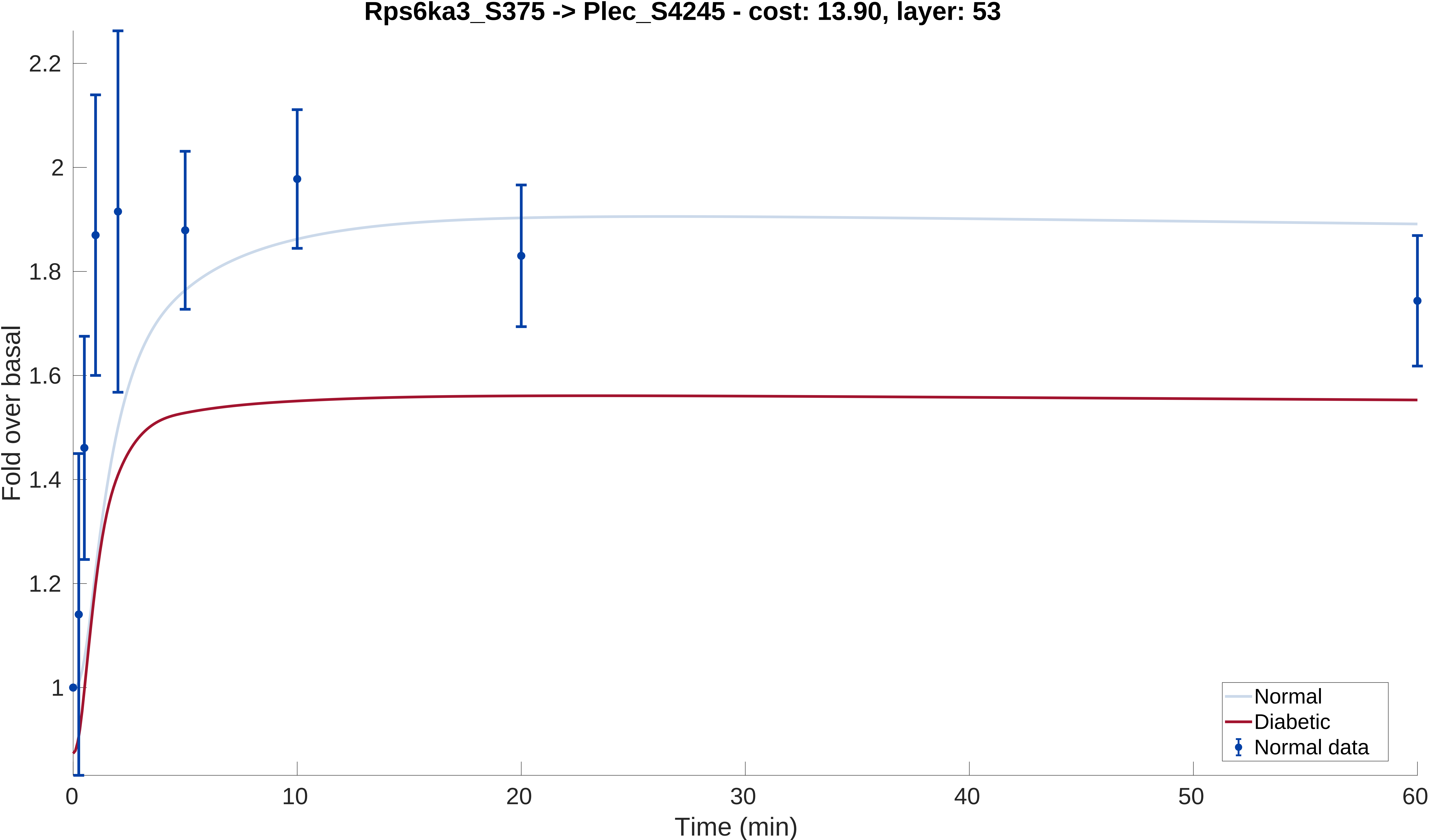
Rps6ka3_S375 -> Plec_S4245 - cost: 13.90, layer: 53
Fold over basal
2.2
2
1.8
1.6
1.4
1.2
1
0
10
20
30
40
50
60
Time (min)
Normal
Diabetic
Normal data

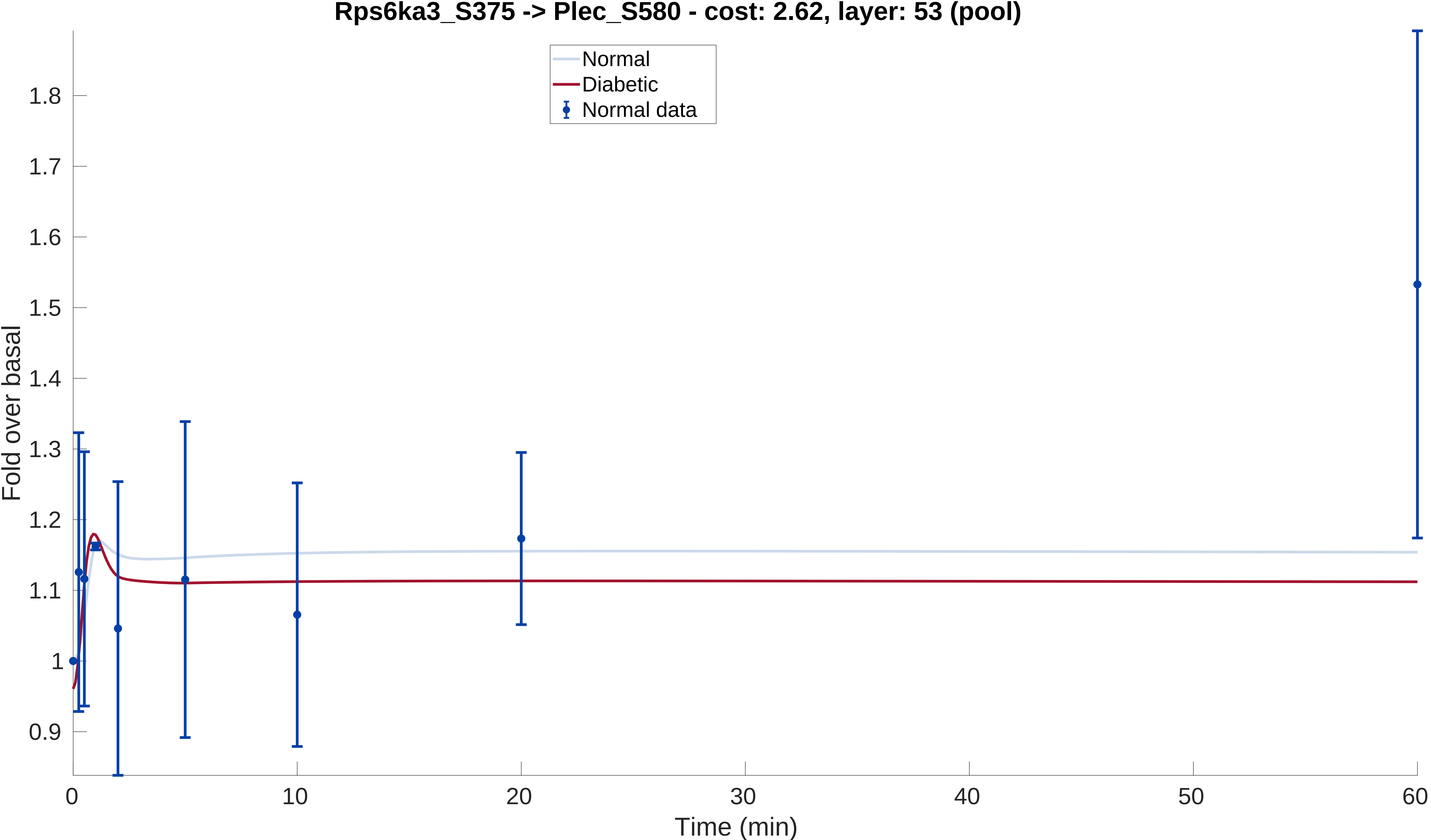

Rps6ka3_S375 -> Plec_S580 - cost: 2.62, layer: 53 (pool)
Normal
Diabetic
Normal data
Fold over basal
1.8
1.7
1.6
1.5
1.4
1.3
1.2
1.1
1
0.9
0
10
20
30
40
50
60
Time (min)

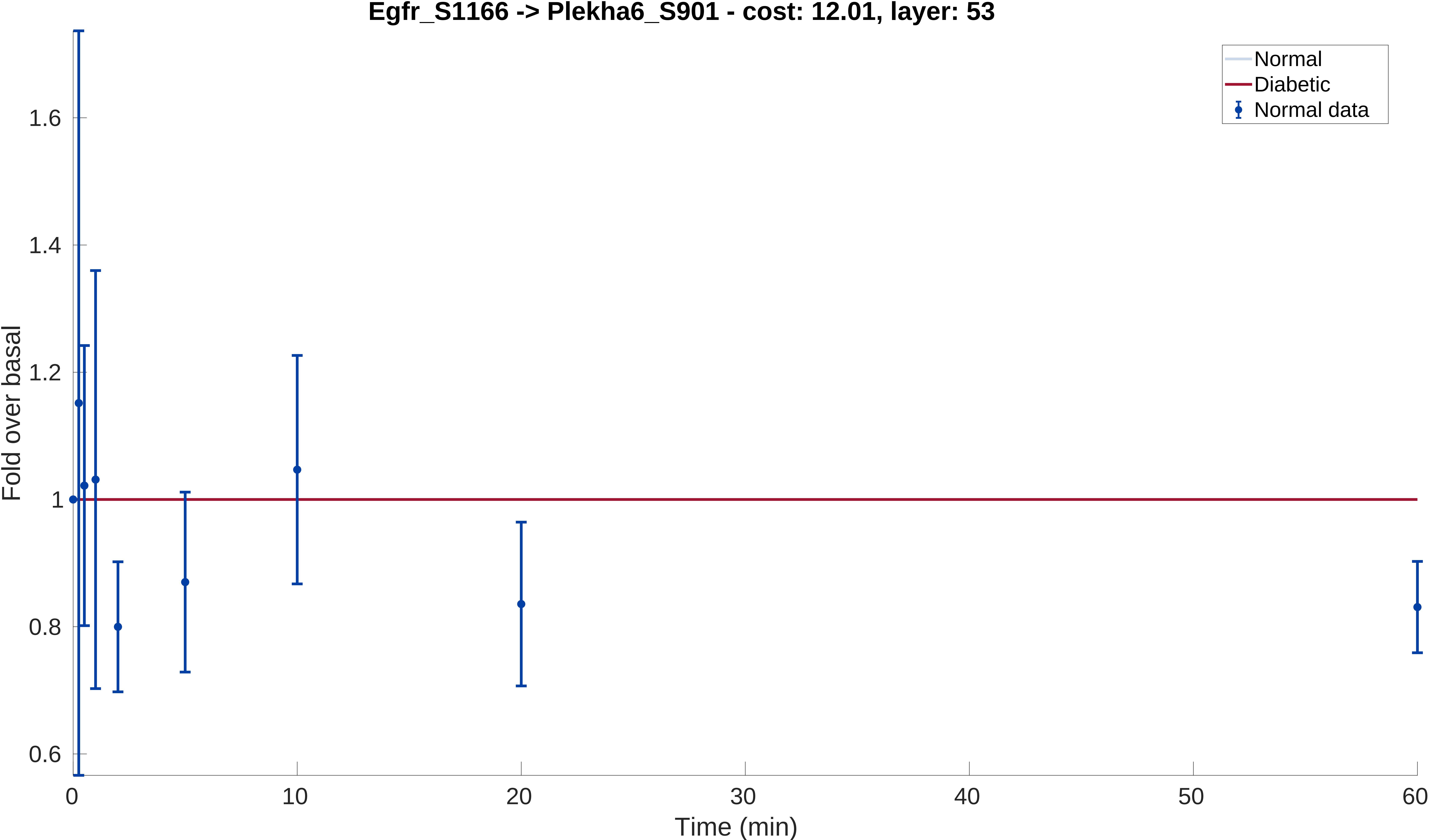

Egfr_S1166 -> Plekha6_S901 - cost: 12.01, layer: 53
Normal
Diabetic
Normal data
Fold over basal
1.6
1.4
1.2
1
0.8
0.6
0
10
20
30
40
50
60
Time (min)

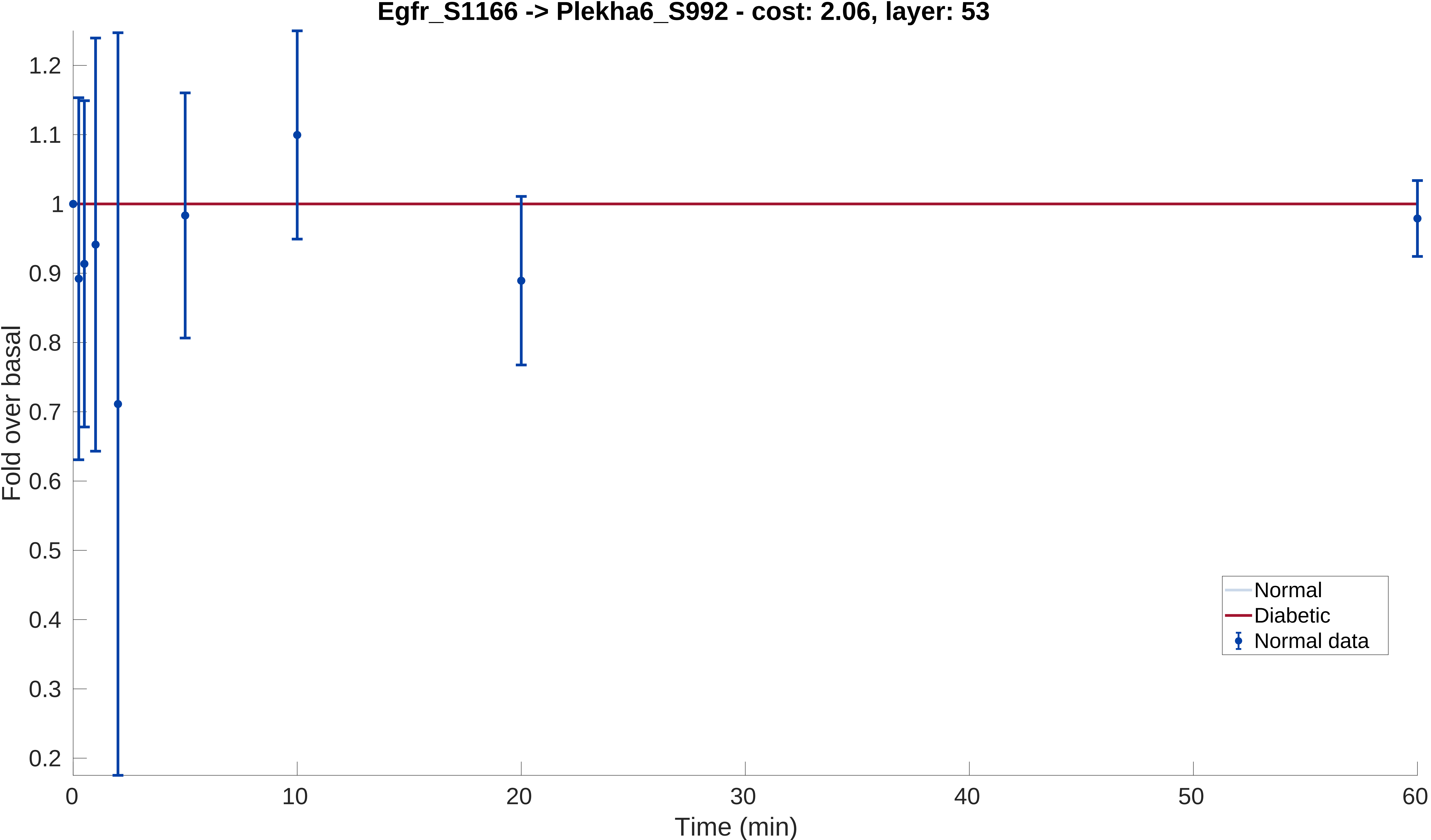

Egfr_S1166 -> Plekha6_S992 - cost: 2.06, layer: 53
Fold over basal
1.2
1.1
1
0.9
0.8
0.7
0.6
0.5
0.4
0.3
0.2
0
10
20
30
40
50
60
Time (min)
Normal
Diabetic
Normal data

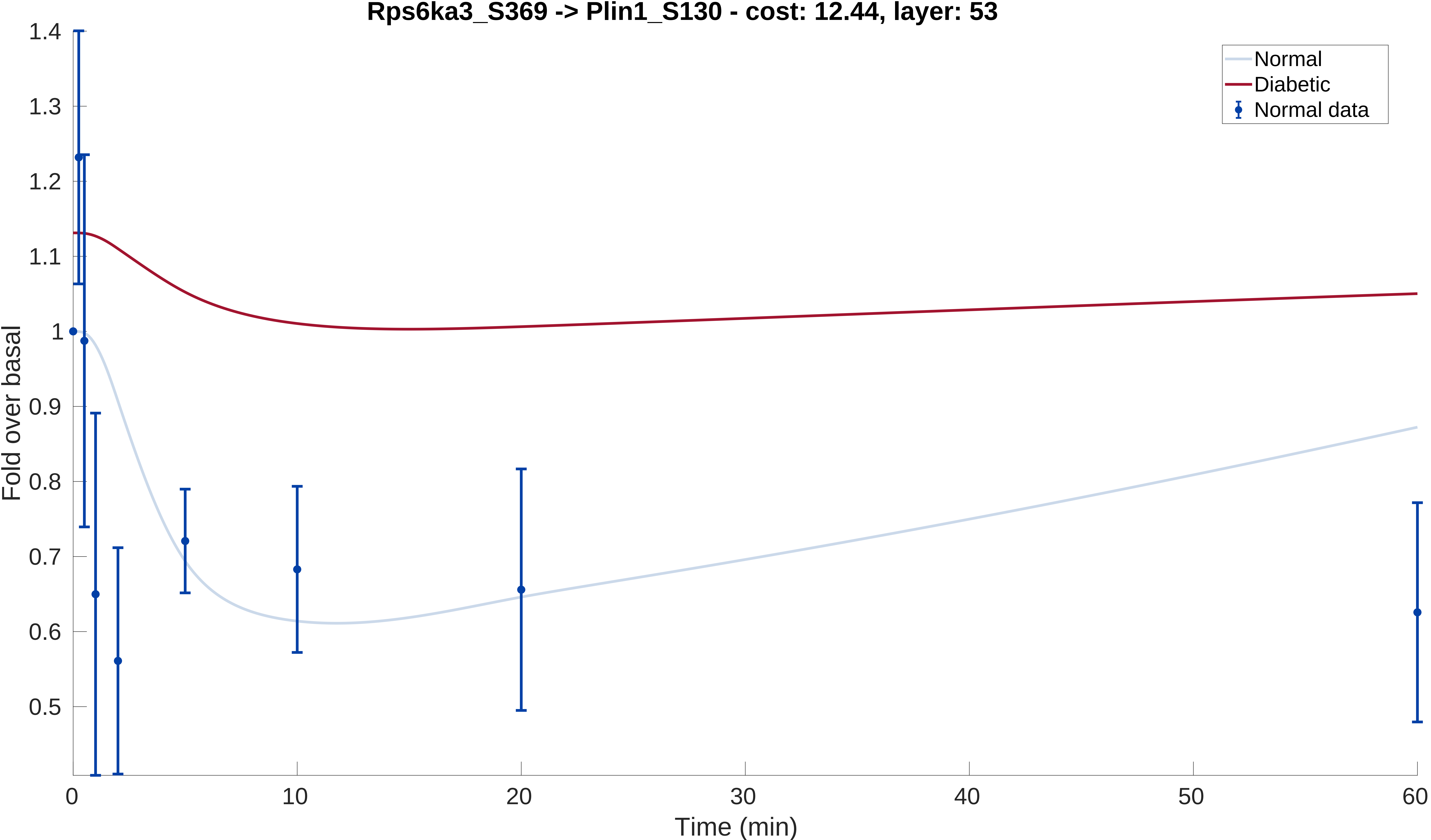

Rps6ka3_S369 -> Plin1_S130 - cost: 12.44, layer: 53
Normal
Diabetic
Normal data
Fold over basal
Time (min)
1.4
1.3
1.2
1.1
1
0.9
0.8
0.7
0.6
0.5
0
10
20
30
40
50
60

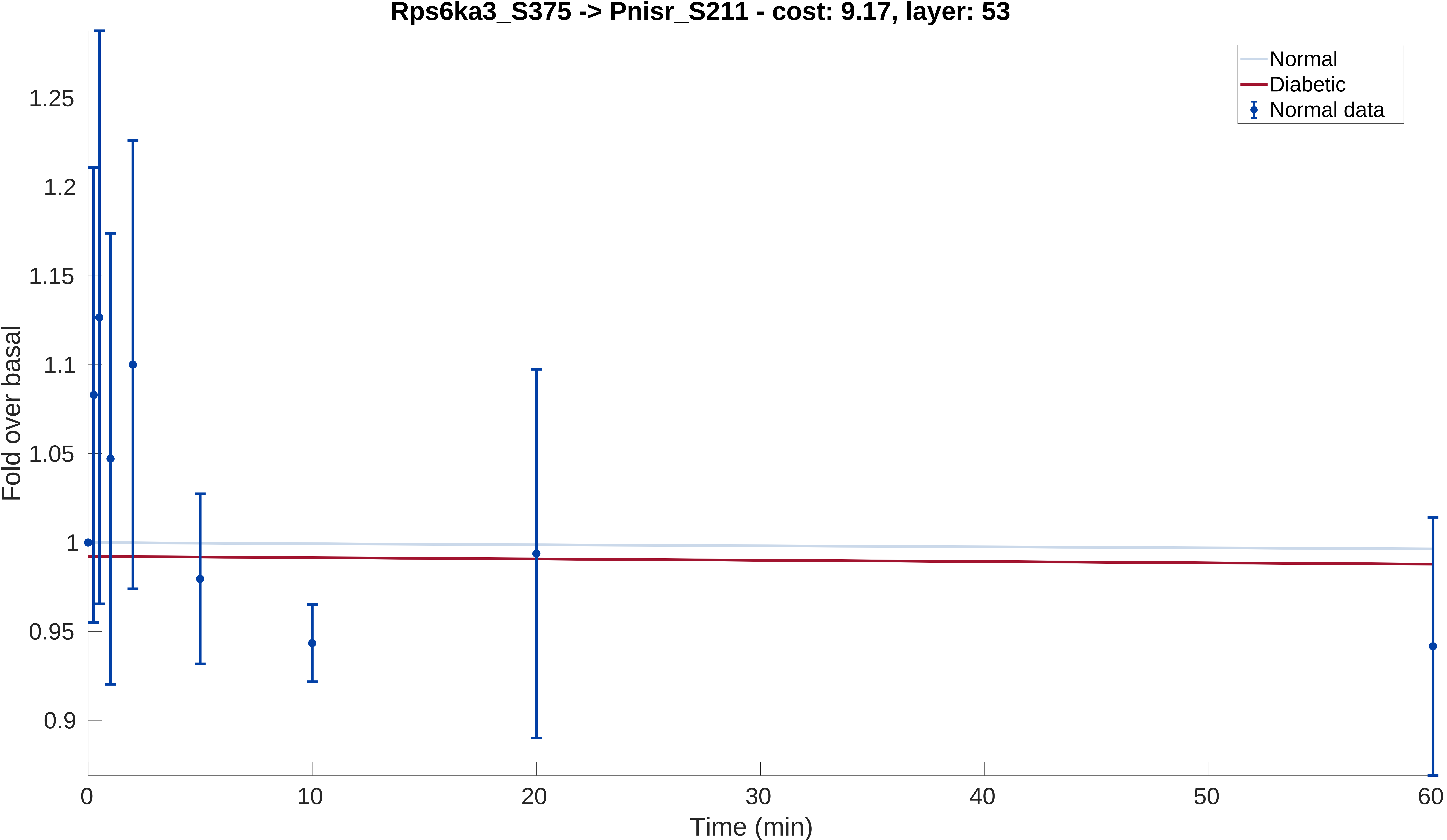

Rps6ka3_S375 -> Pnisr_S211 - cost: 9.17, layer: 53
Normal
Diabetic
Normal data
Fold over basal
1.25
1.2
1.15
1.1
1.05
1
0.95
0.9
0
10
20
30
40
50
60
Time (min)

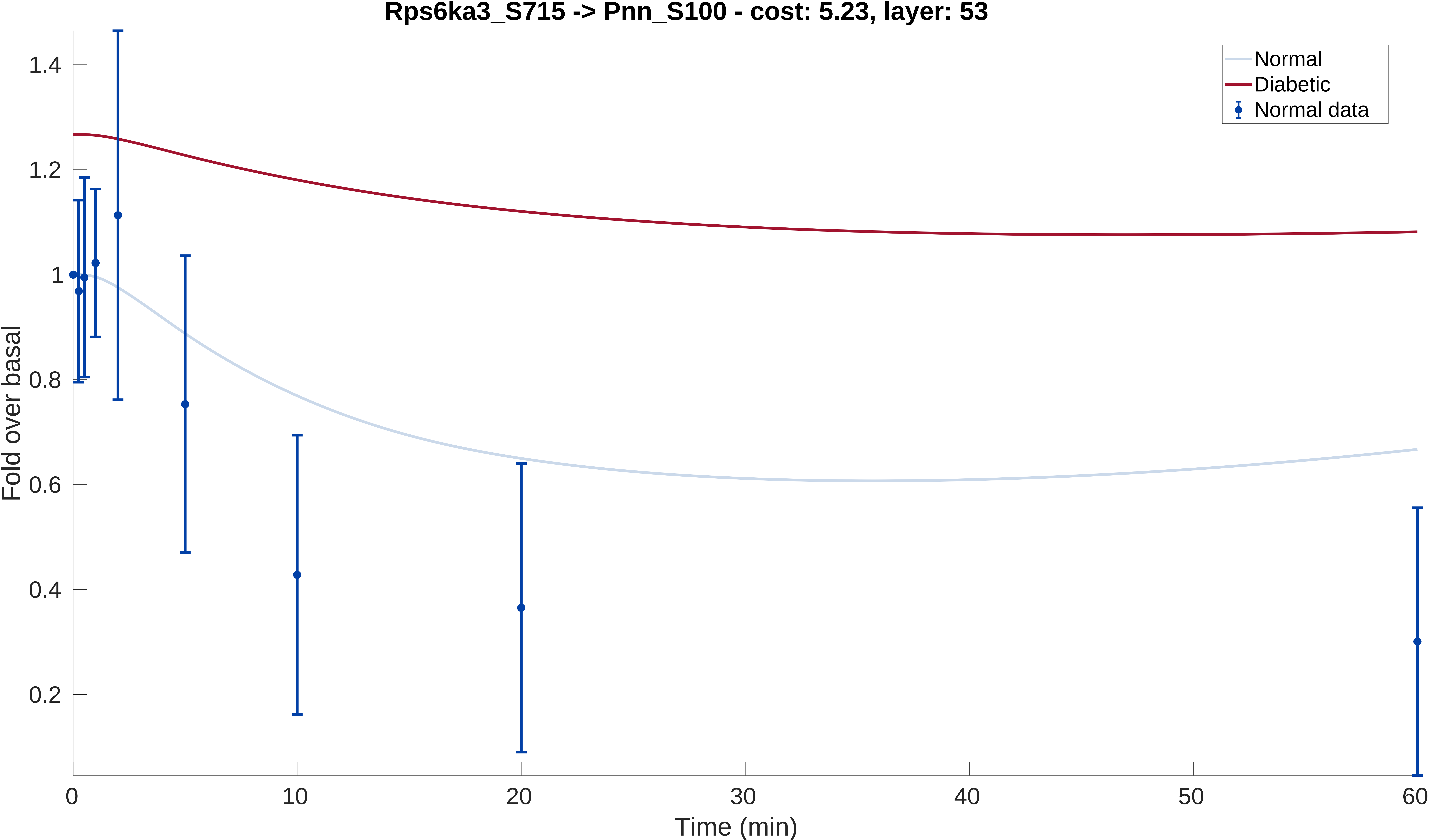
Rps6ka3_S715 -> Pnn_S100 - cost: 5.23, layer: 53
Normal
Diabetic
Normal data
Fold over basal
1.4
1.2
1
0.8
0.6
0.4
0.2
0
10
20
30
40
50
60
Time (min)

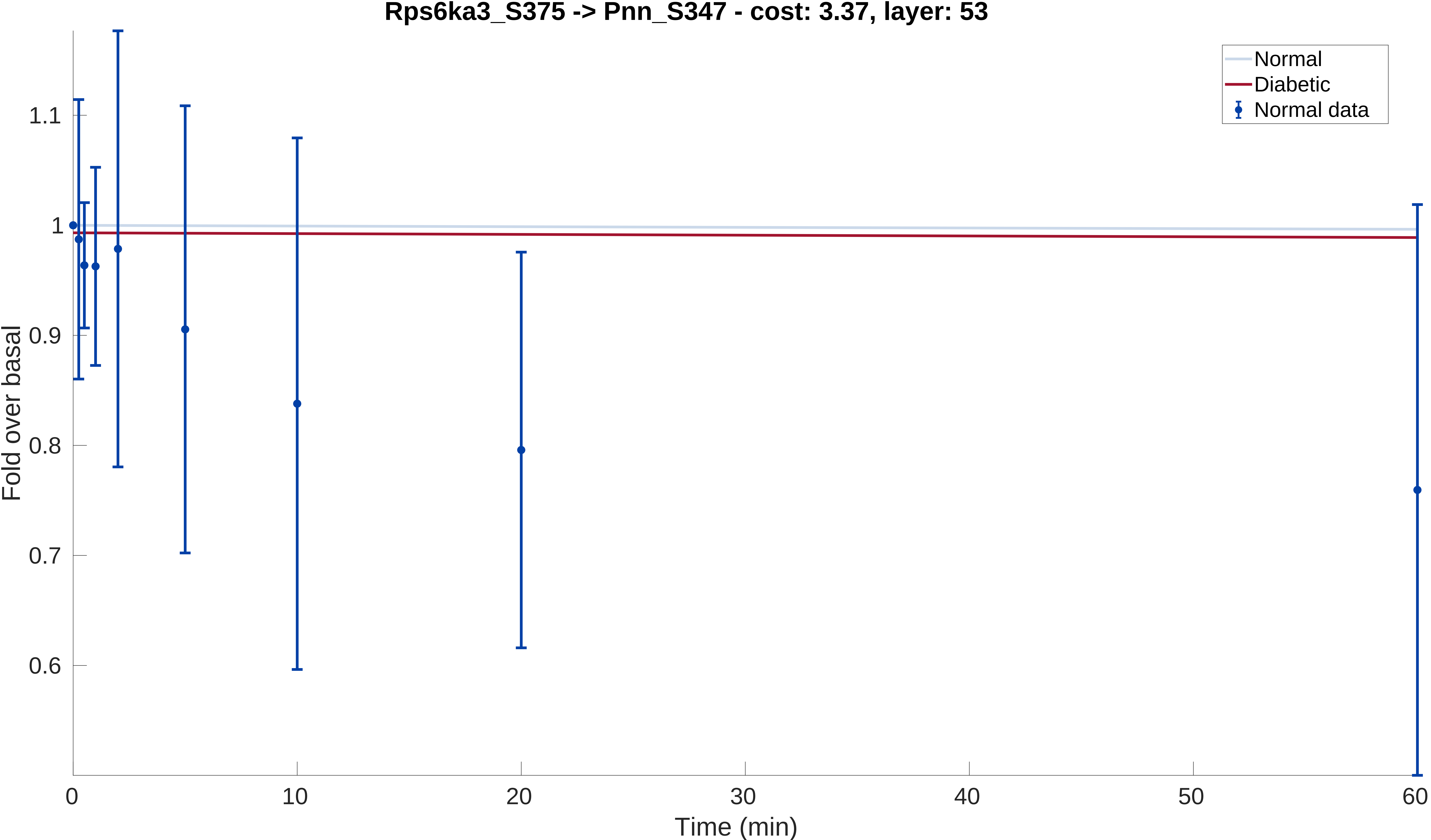

Rps6ka3_S375 -> Pnn_S347 - cost: 3.37, layer: 53
Normal
Diabetic
Normal data
Fold over basal
1.1
1
0.9
0.8
0.7
0.6
0
10
20
30
40
50
60
Time (min)

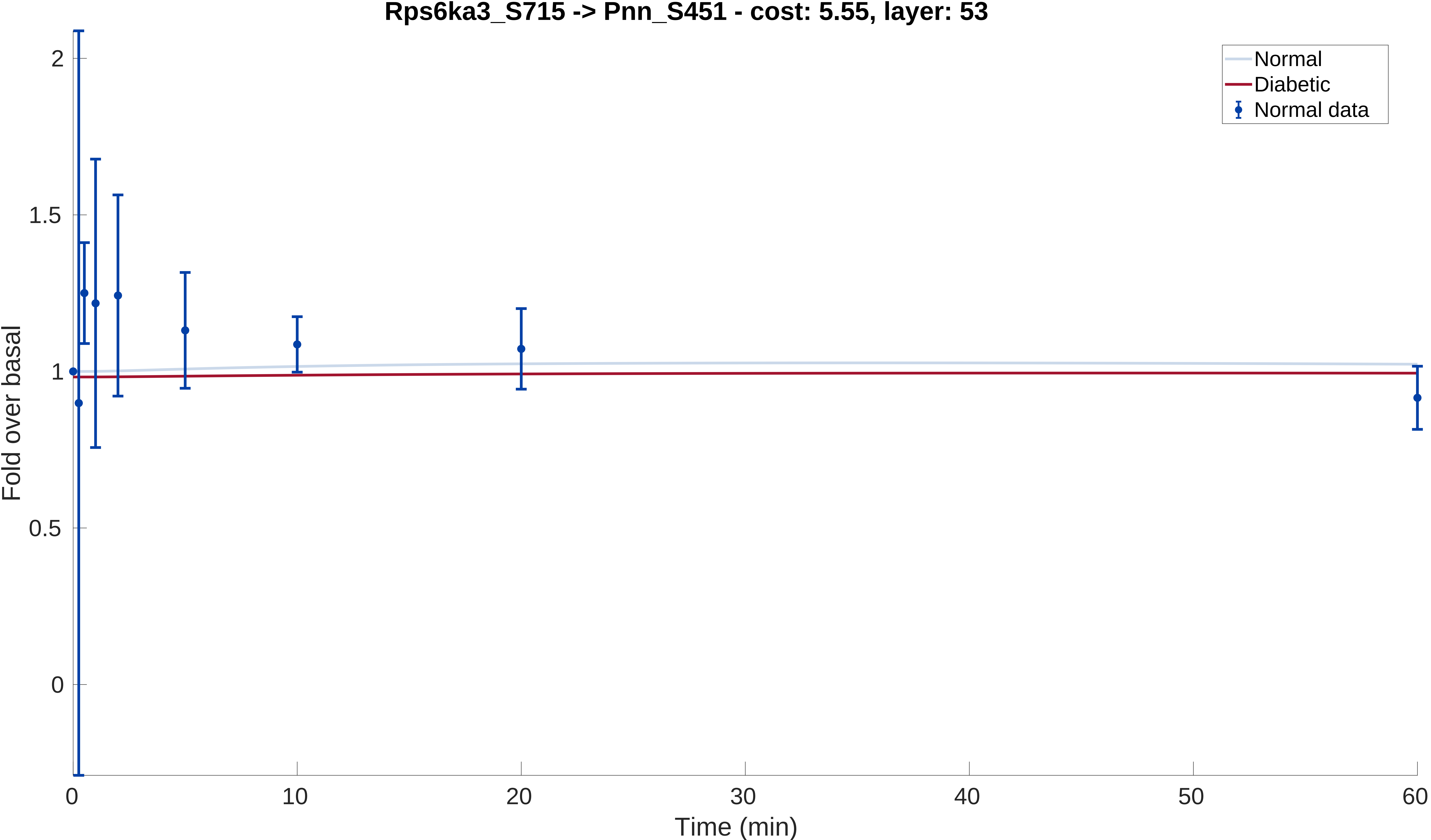

Rps6ka3_S715 -> Pnn_S451 - cost: 5.55, layer: 53
Normal
Diabetic
Normal data
Fold over basal
2
1.5
1
0.5
0
0
10
20
30
40
50
60
Time (min)

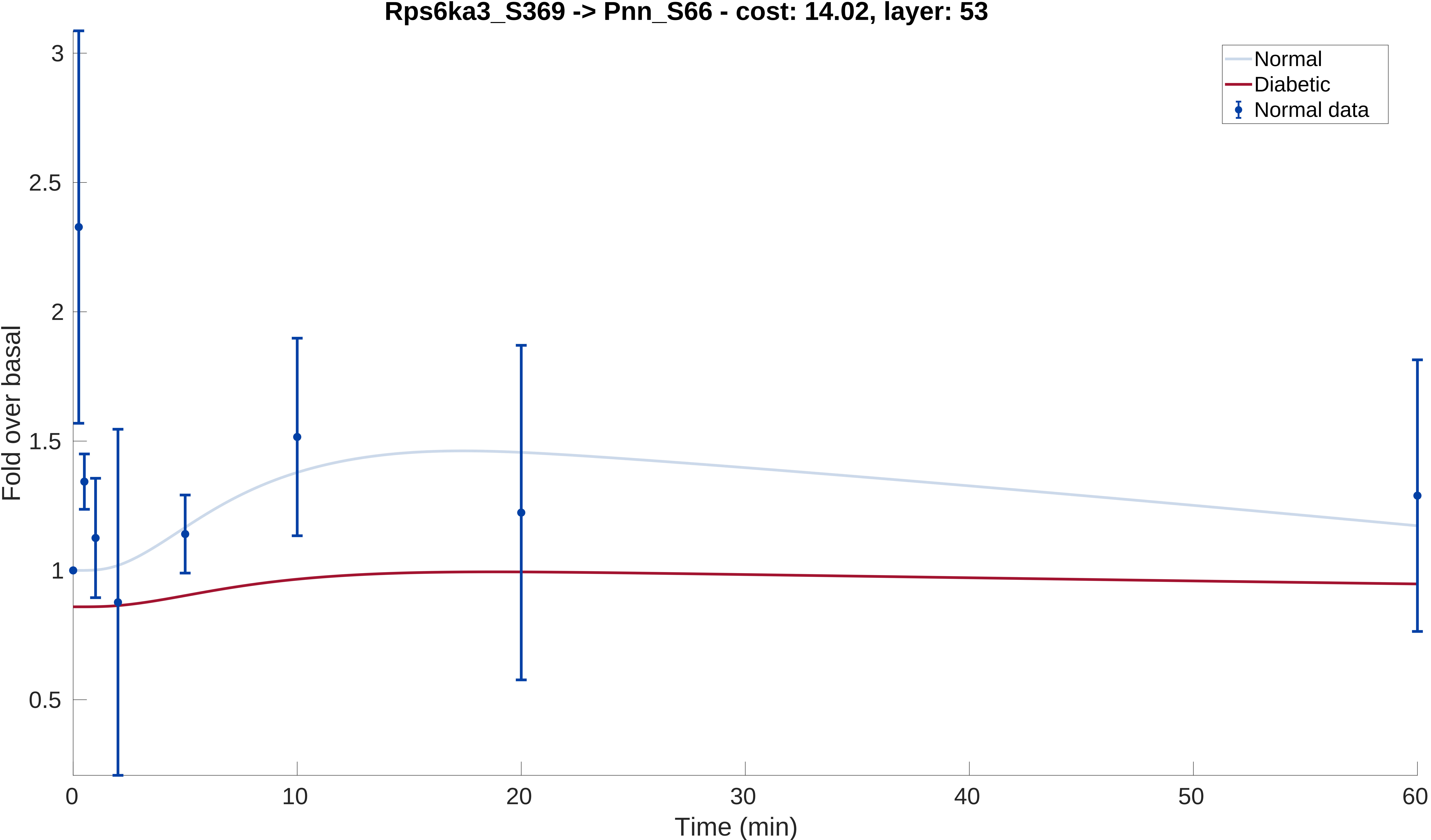

Rps6ka3_S369 -> Pnn_S66 - cost: 14.02, layer: 53
Normal
Diabetic
Normal data
Fold over basal
0.5
1
1.5
2
2.5
3
0
10
20
30
40
50
60
Time (min)

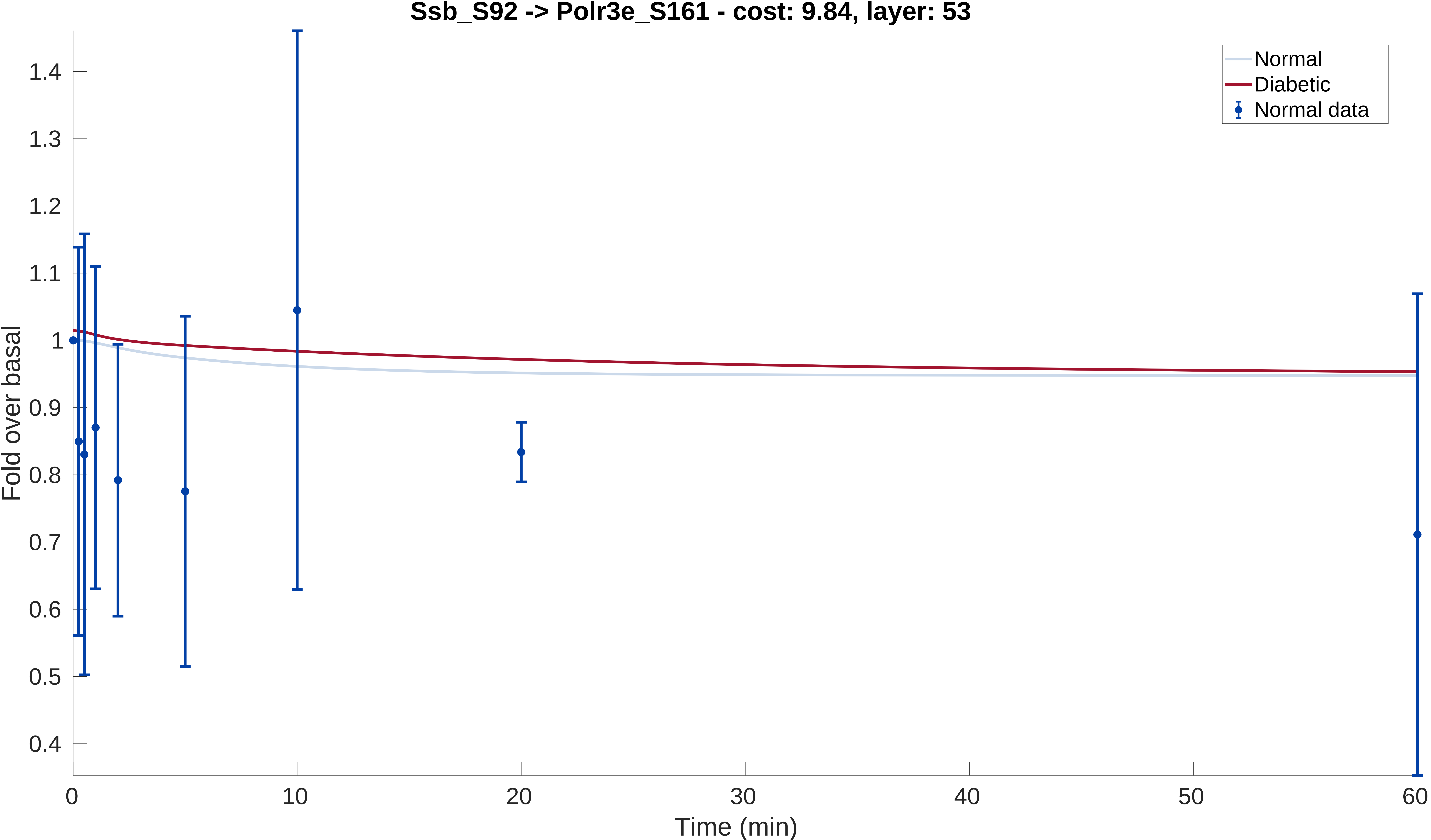
Ssb_S92 -> Polr3e_S161 - cost: 9.84, layer: 53
Normal
Diabetic
Normal data
Fold over basal
Time (min)
1.4
1.3
1.2
1.1
1
0.9
0.8
0.7
0.6
0.5
0.4
0
10
20
30
40
50
60

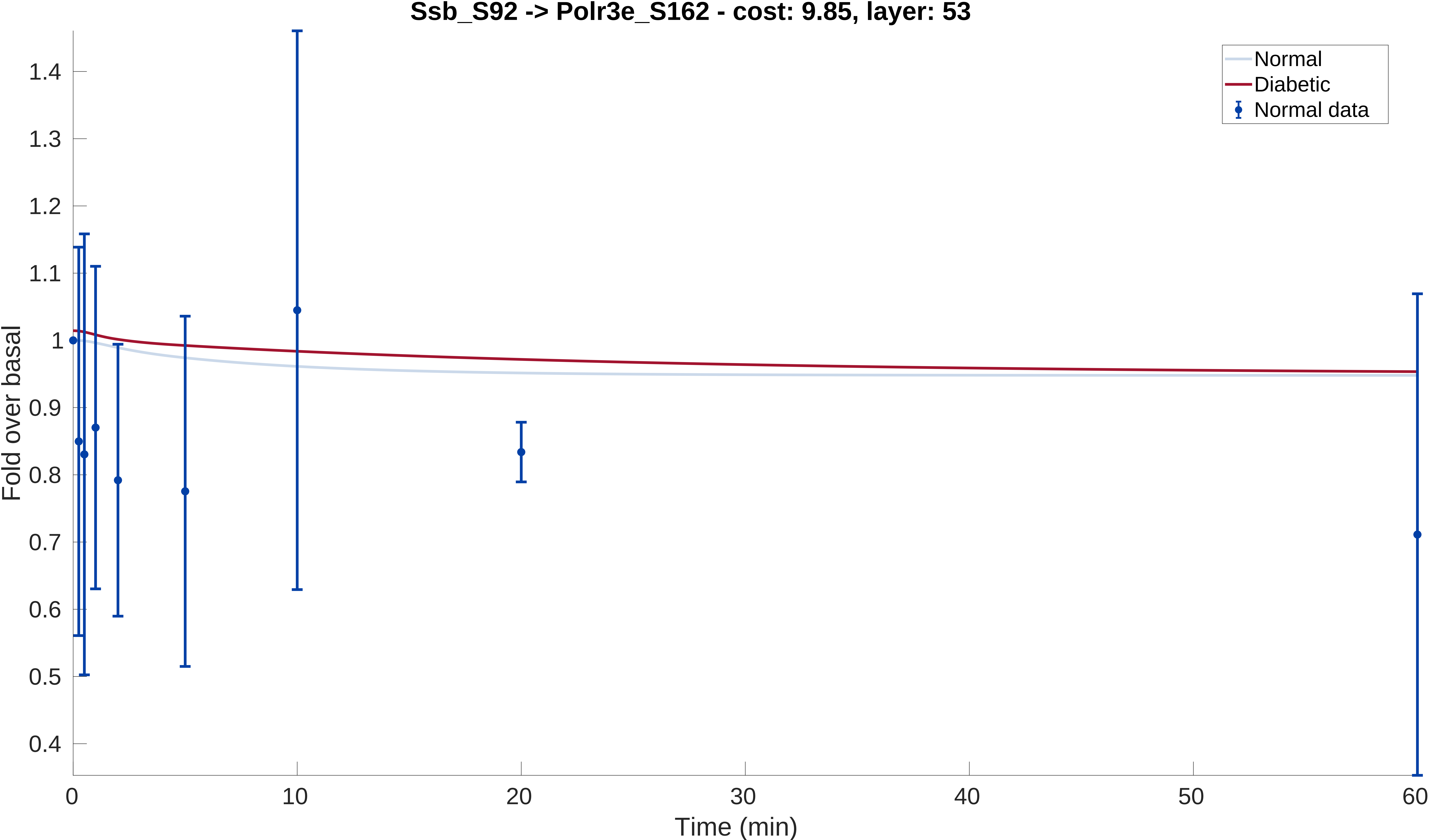
Ssb_S92 -> Polr3e_S162 - cost: 9.85, layer: 53
Normal
Diabetic
Normal data
Fold over basal
1.4
1.3
1.2
1.1
1
0.9
0.8
0.7
0.6
0.5
0.4
0
10
20
30
40
50
60
Time (min)

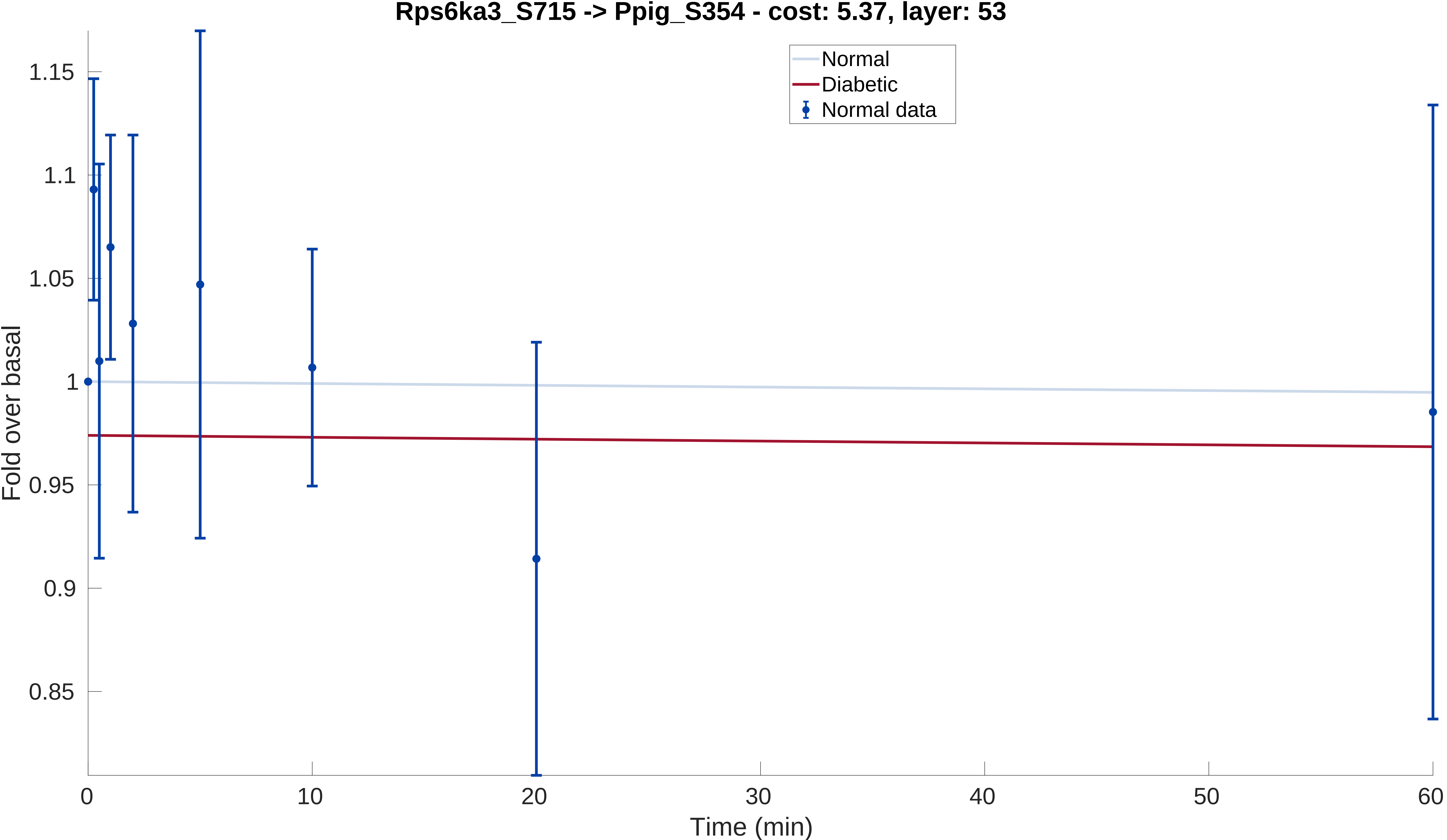

Rps6ka3_S715 -> Ppig_S354 - cost: 5.37, layer: 53
Normal
Diabetic
Normal data
Fold over basal
1.15
1.1
1.05
1
0.95
0.9
0.85
0
10
20
30
40
50
60
Time (min)

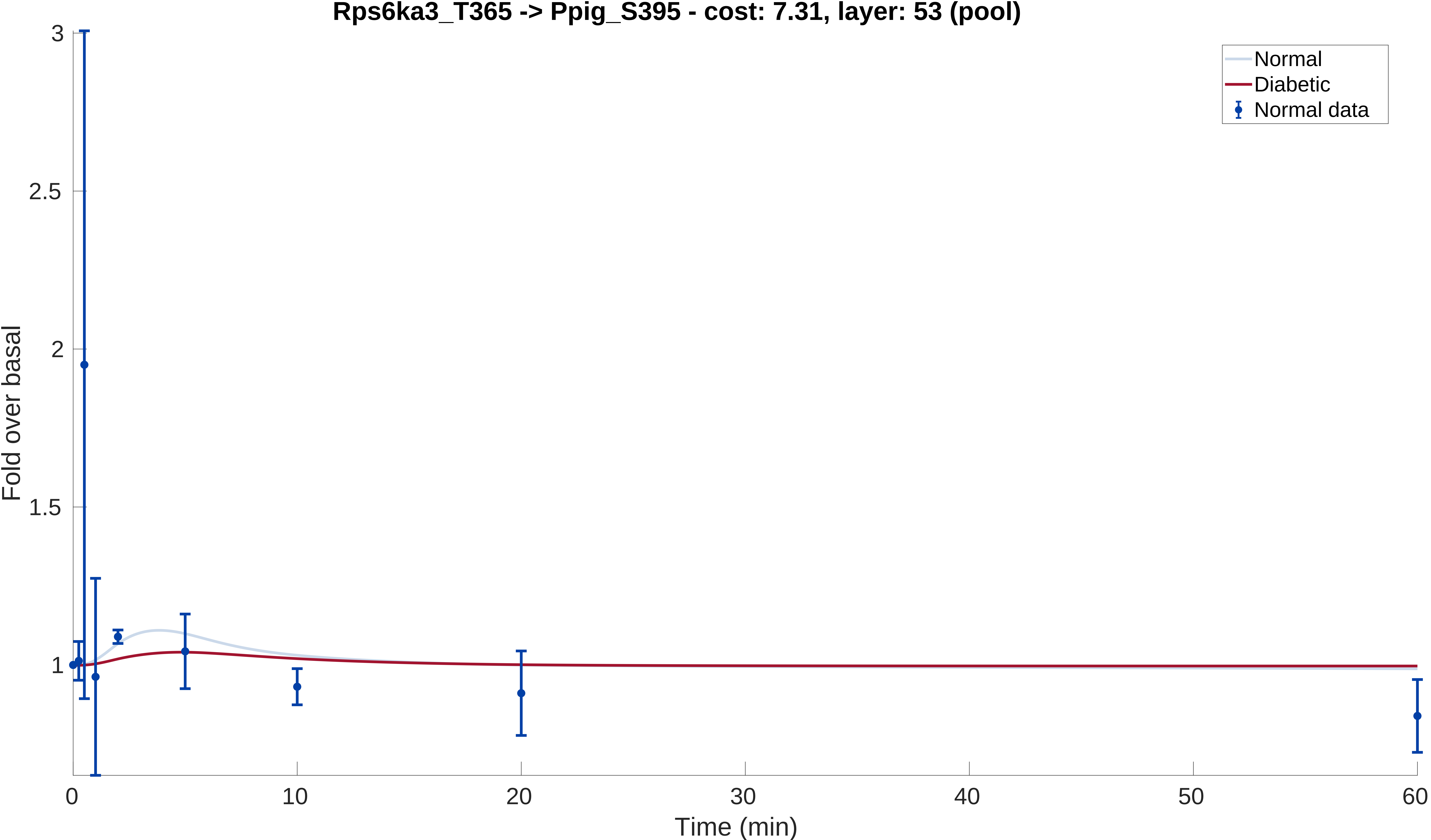

Rps6ka3_T365 -> Ppig_S395 - cost: 7.31, layer: 53 (pool)
Normal
Diabetic
Normal data
Fold over basal
3
2.5
2
1.5
1
0
10
20
30
40
50
60
Time (min)

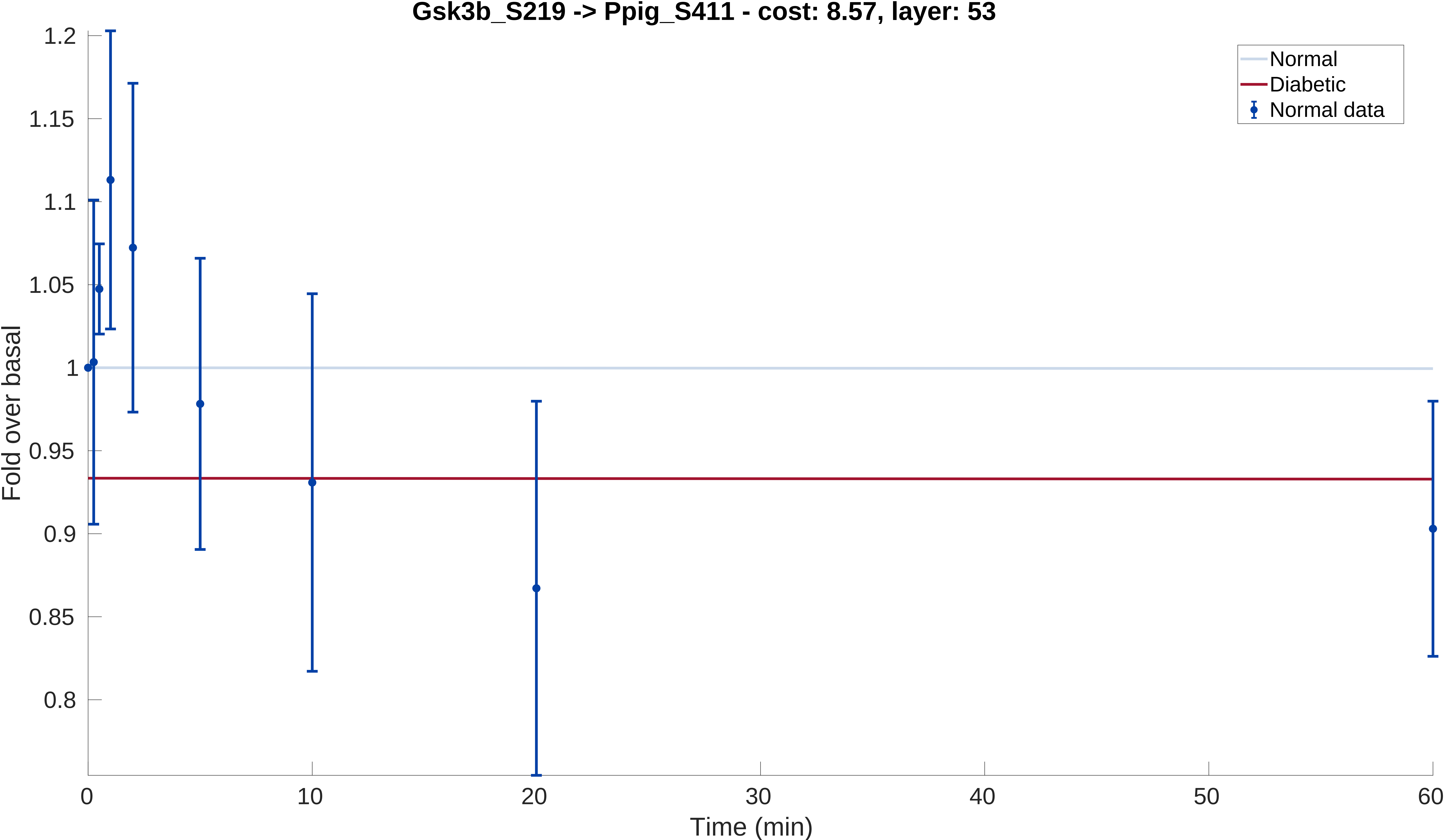

Gsk3b_S219 -> Ppig_S411 - cost: 8.57, layer: 53
Normal
Diabetic
Normal data
Fold over basal
Time (min)
1.2
1.15
1.1
1.05
1
0.95
0.9
0.85
0.8
0
10
20
30
40
50
60

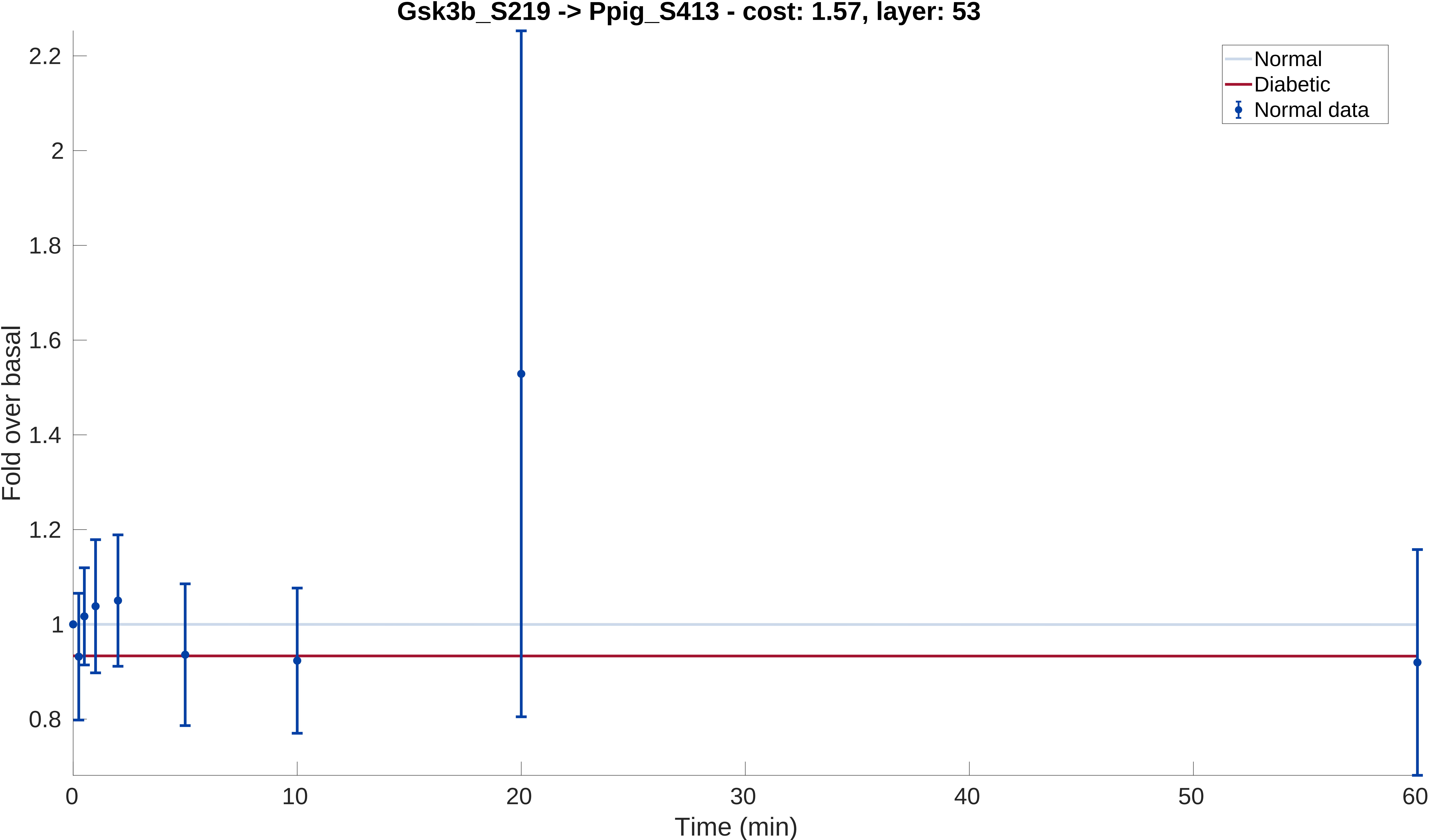

Gsk3b_S219 -> Ppig_S413 - cost: 1.57, layer: 53
Normal
Diabetic
Normal data
Fold over basal
2.2
2
1.8
1.6
1.4
1.2
1
0.8
0
10
20
30
40
50
60
Time (min)

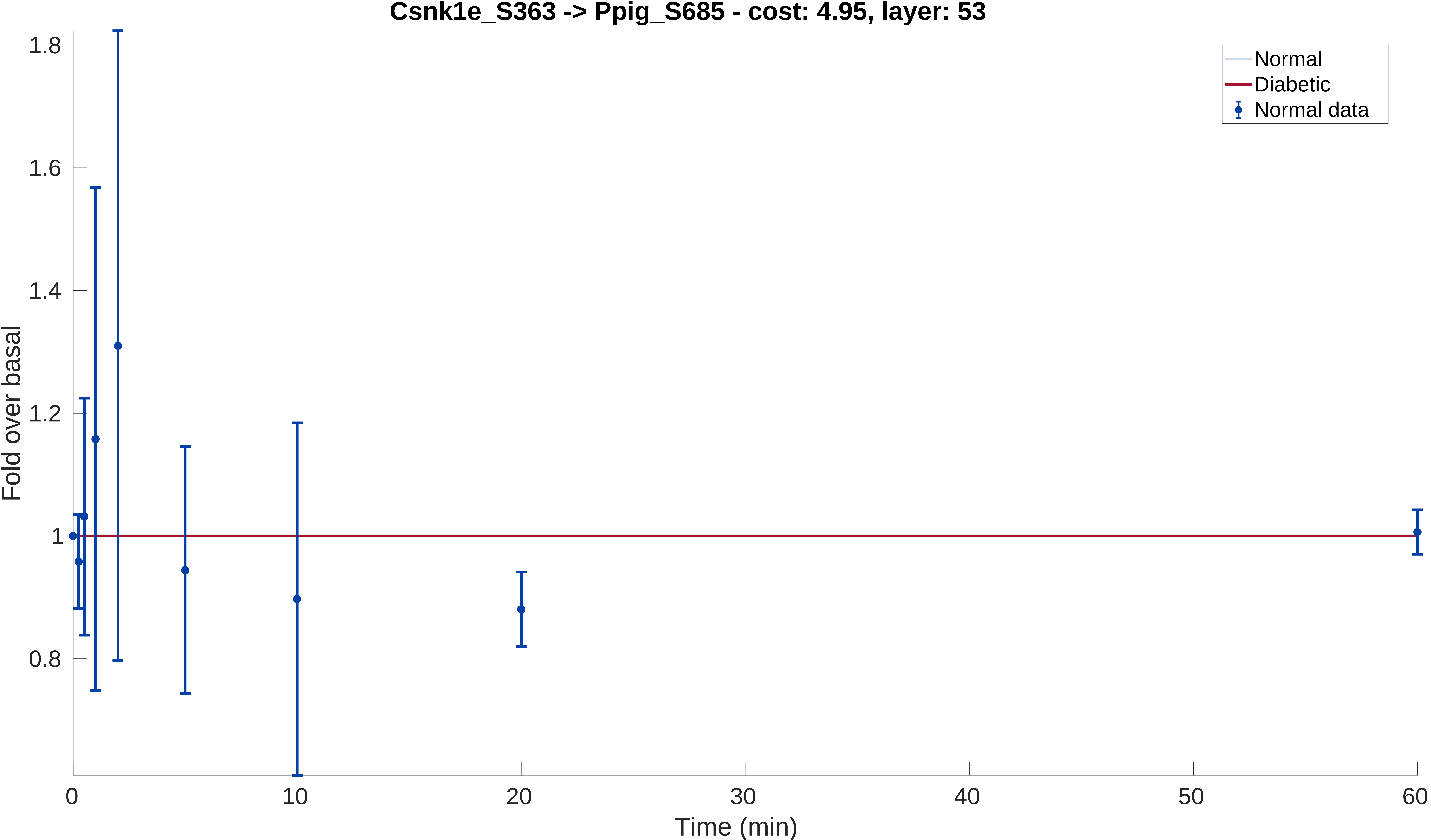

Csnk1e_S363 -> Ppig_S685 - cost: 4.95, layer: 53
Normal
Diabetic
Normal data
Fold over basal
1.8
1.6
1.4
1.2
1
0.8
0
10
20
30
40
50
60
Time (min)

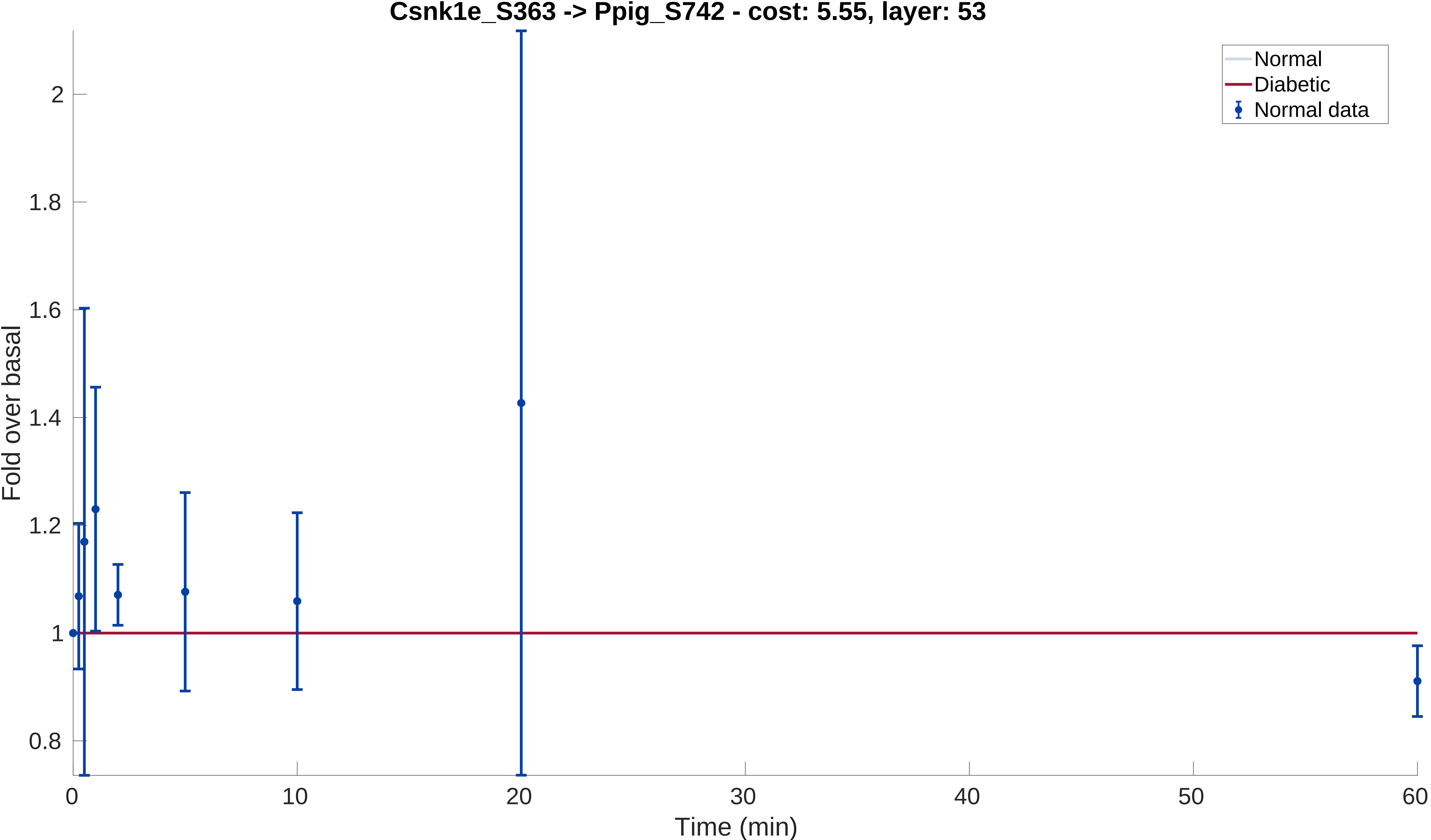

Csnk1e_S363 -> Ppig_S742 - cost: 5.55, layer: 53
Normal
Diabetic
Normal data
Fold over basal
2
1.8
1.6
1.4
1.2
1
0.8
0
10
20
30
40
50
60
Time (min)

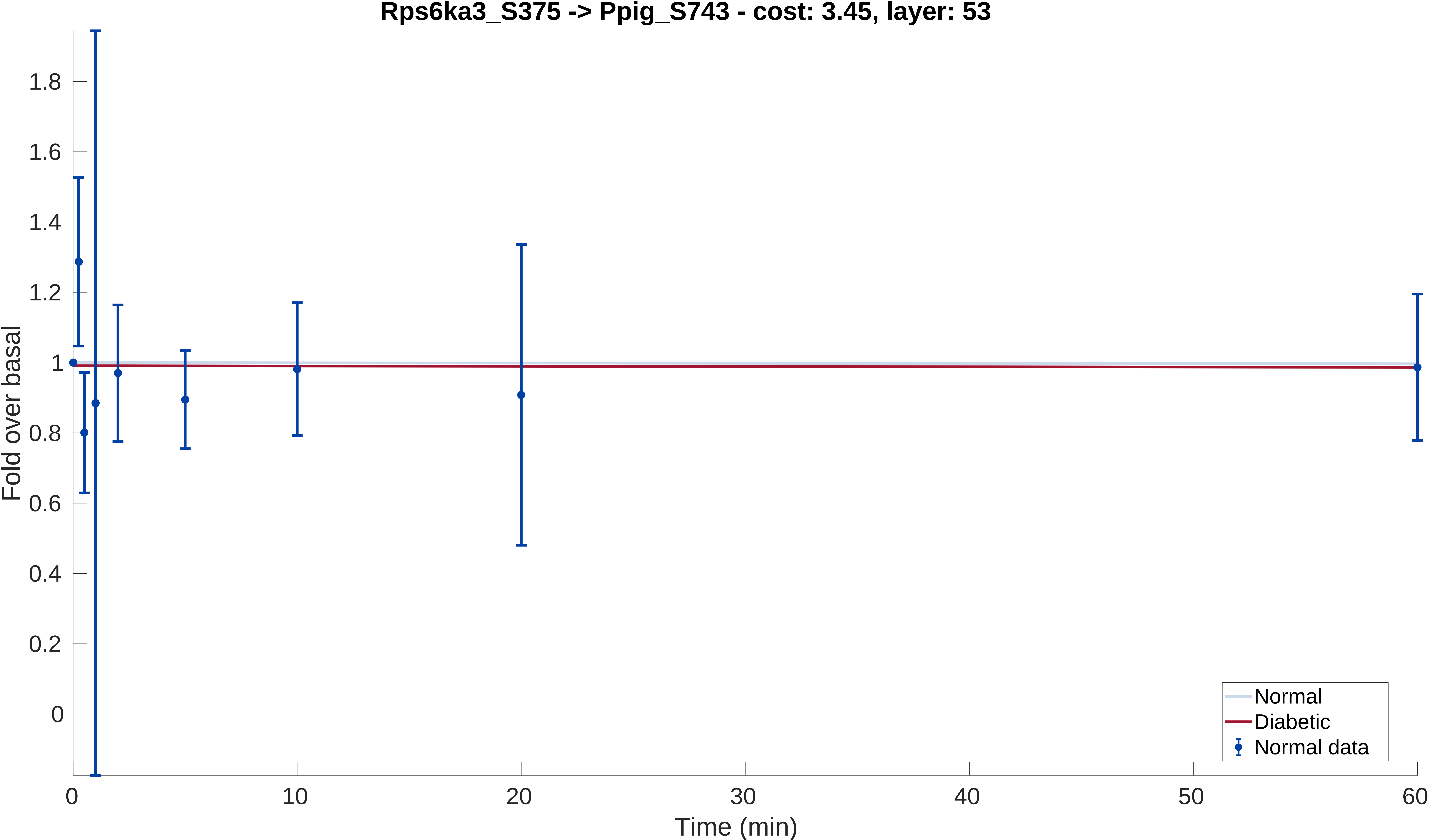

Rps6ka3_S375 -> Ppig_S743 - cost: 3.45, layer: 53
Fold over basal
1.8
1.6
1.4
1.2
1
0.8
0.6
0.4
0.2
0
Normal
Diabetic
Normal data
0
10
20
30
40
50
60
Time (min)

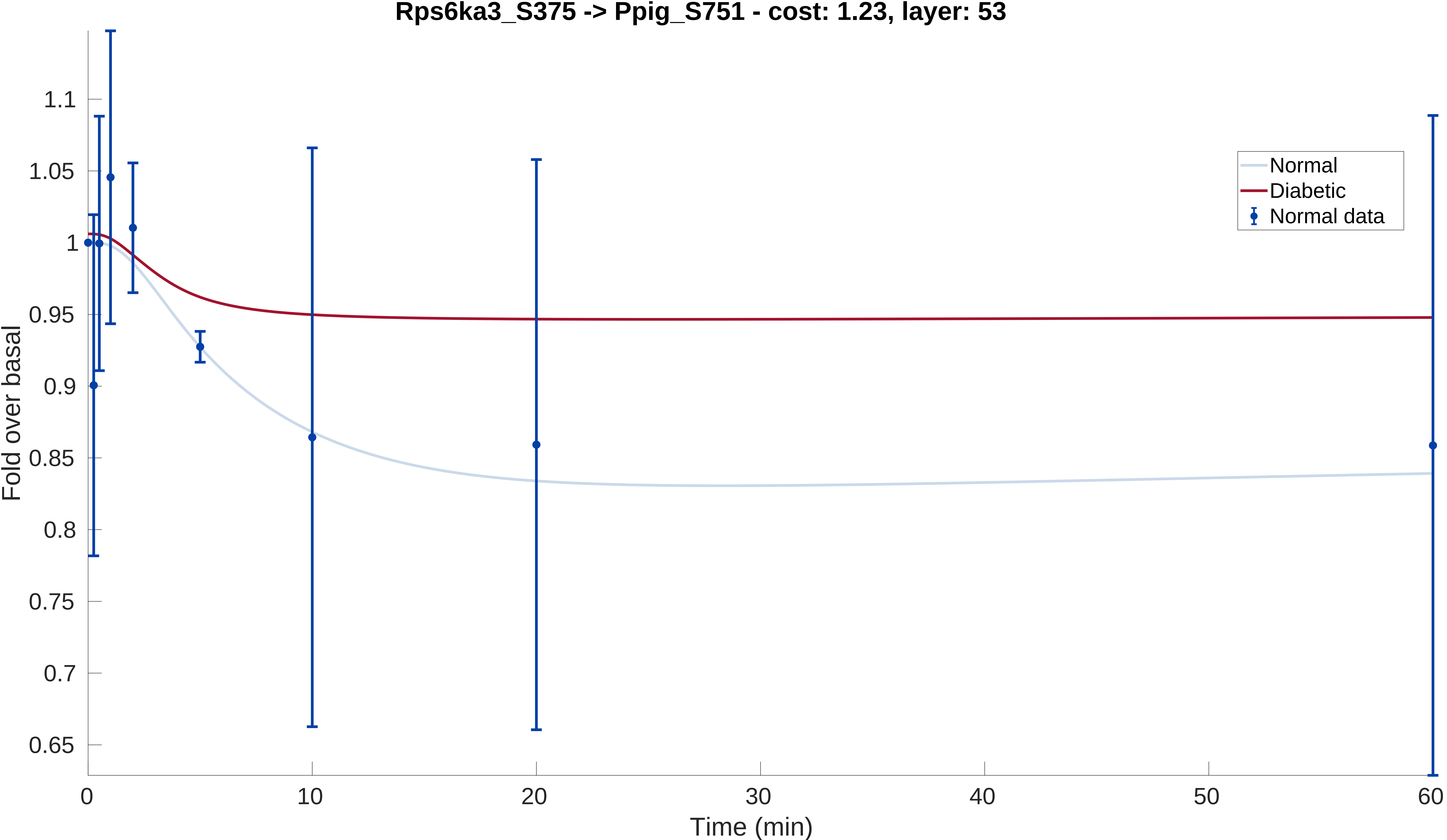

Rps6ka3_S375 -> Ppig_S751 - cost: 1.23, layer: 53
Fold over basal
1.1
1.05
1
0.95
0.9
0.85
0.8
0.75
0.7
0.65
0
10
20
30
40
50
60
Time (min)
Normal
Diabetic
Normal data

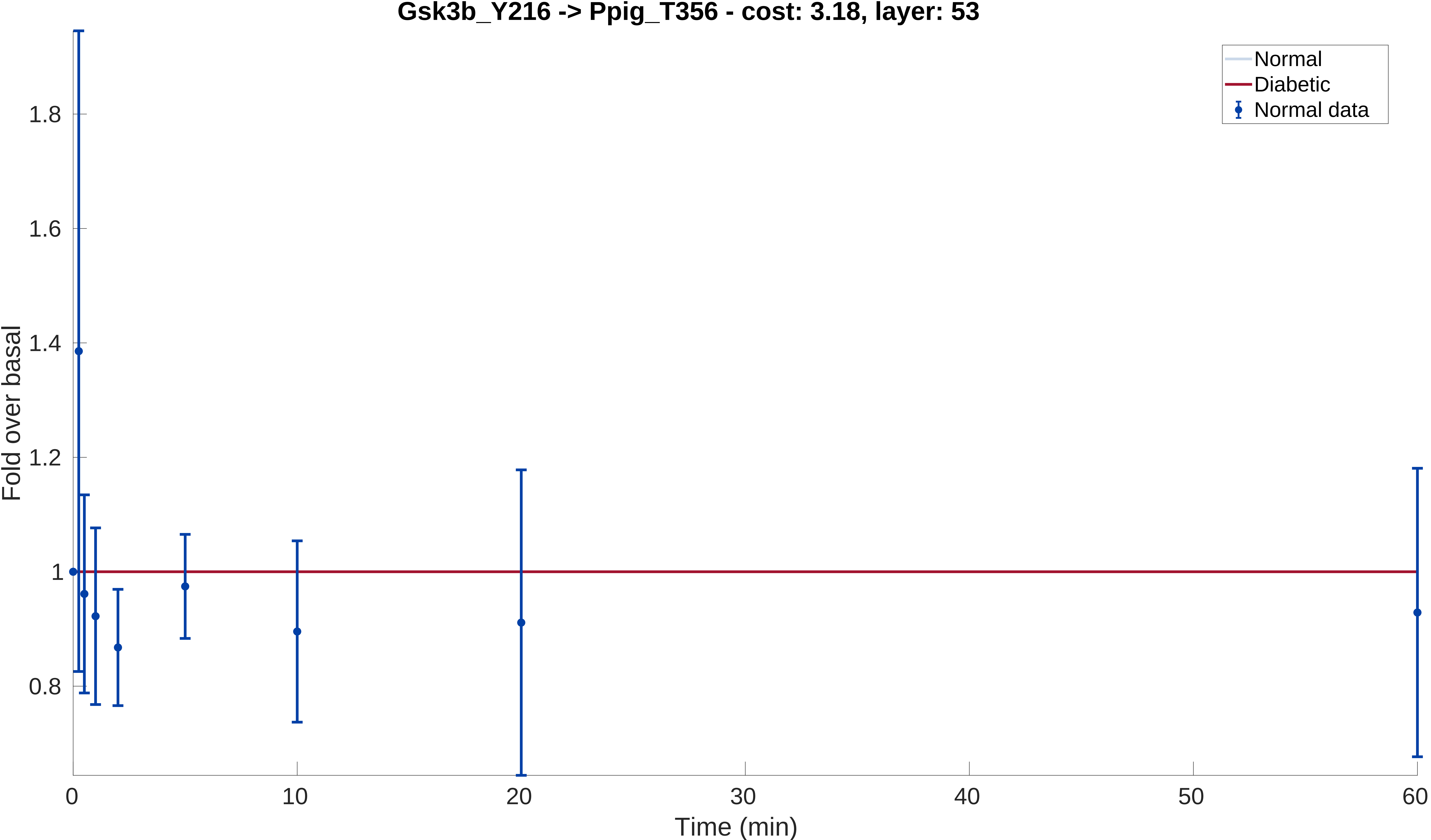
Gsk3b_Y216 -> Ppig_T356 - cost: 3.18, layer: 53
Normal
Diabetic
Normal data
Fold over basal
1.8
1.6
1.4
1.2
1
0.8
0
10
20
30
40
50
60
Time (min)

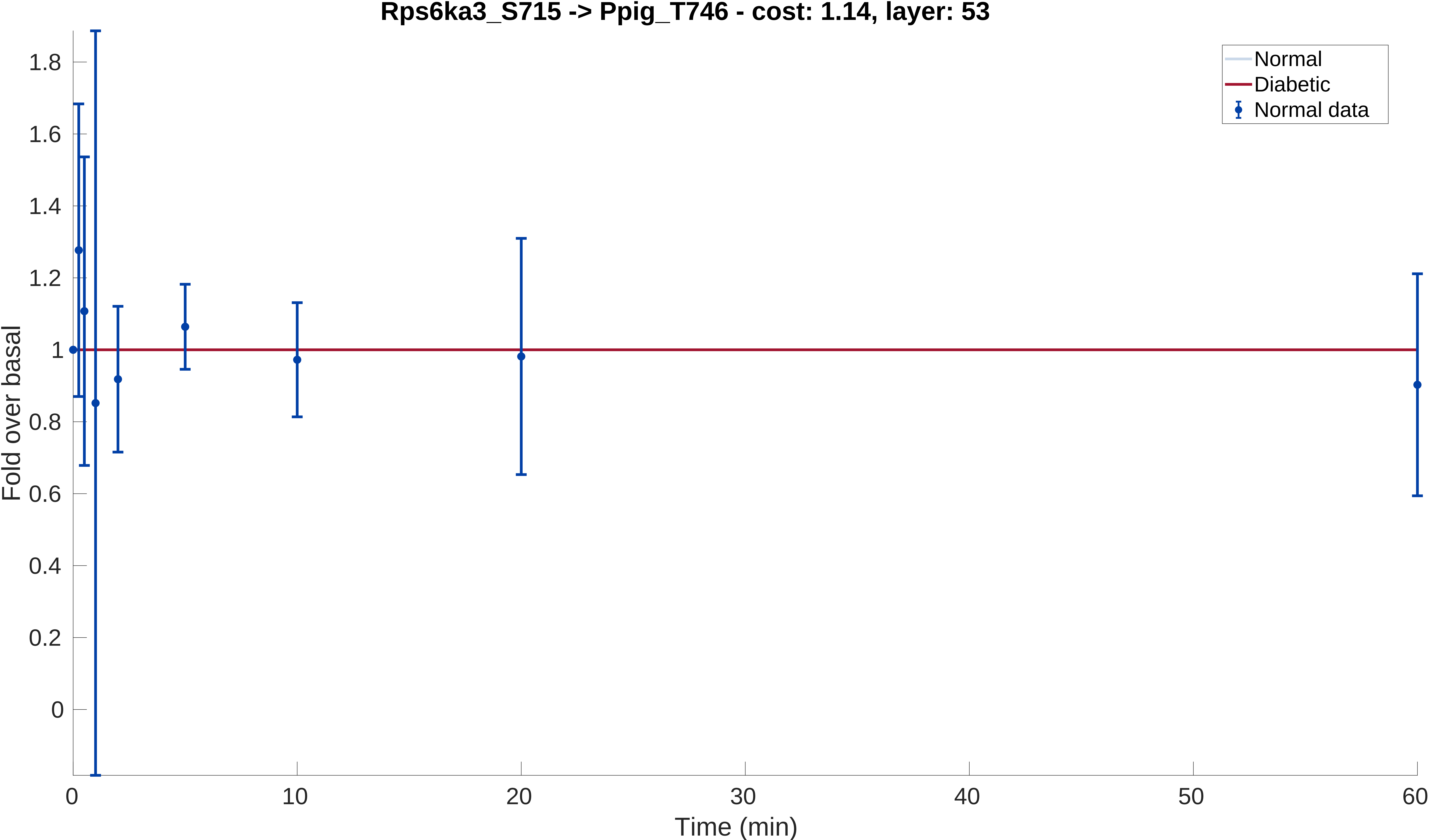
Rps6ka3_S715 -> Ppig_T746 - cost: 1.14, layer: 53
Normal
Diabetic
Normal data
1.8
1.6
1.4
1.2
1
0.8
0.6
0.4
0.2
0
Fold over basal
0
10
20
30
40
50
60
Time (min)

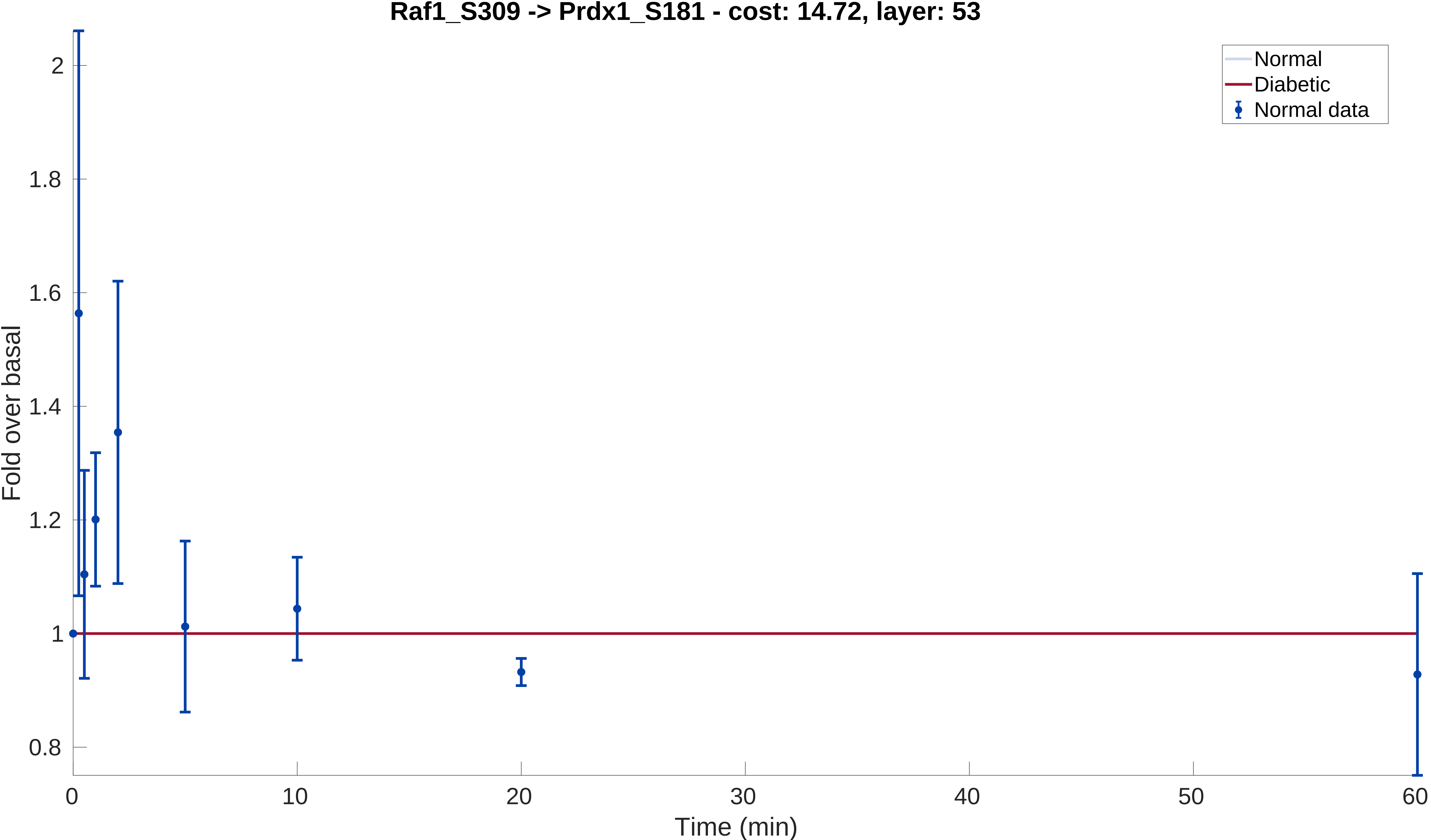

Raf1_S309 -> Prdx1_S181 - cost: 14.72, layer: 53
Normal
Diabetic
Normal data
Fold over basal
2
1.8
1.6
1.4
1.2
1
0.8
0
10
20
30
40
50
60
Time (min)

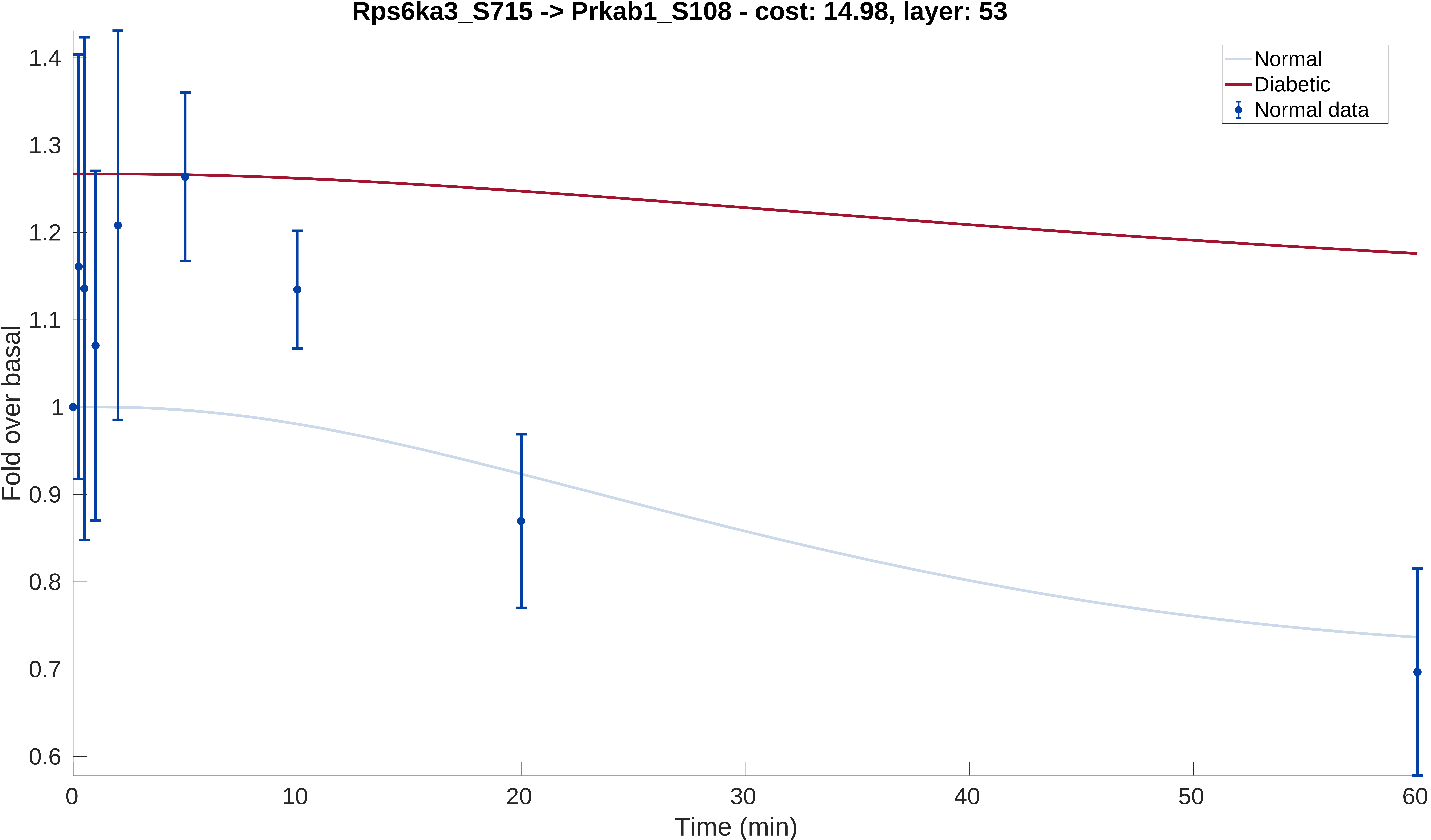

Rps6ka3_S715 -> Prkab1_S108 - cost: 14.98, layer: 53
Normal
Diabetic
Normal data
Fold over basal
Time (min)
1.4
1.3
1.2
1.1
1
0.9
0.8
0.7
0.6
0
10
20
30
40
50
60

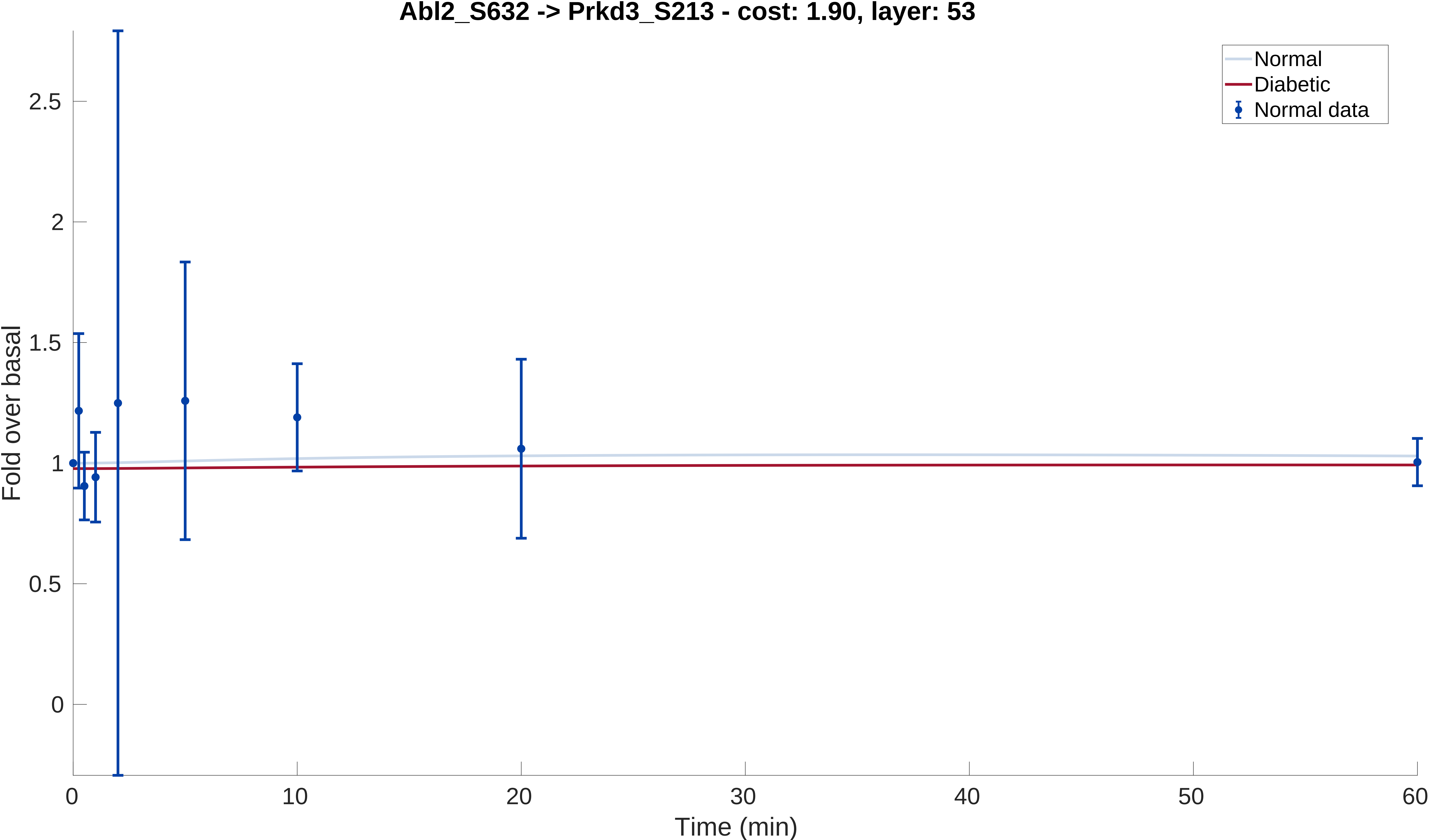

Abl2_S632 -> Prkd3_S213 - cost: 1.90, layer: 53
Normal
Diabetic
Normal data
Fold over basal
Time (min)
0
0.5
1
1.5
2
2.5
0
10
20
30
40
50
60

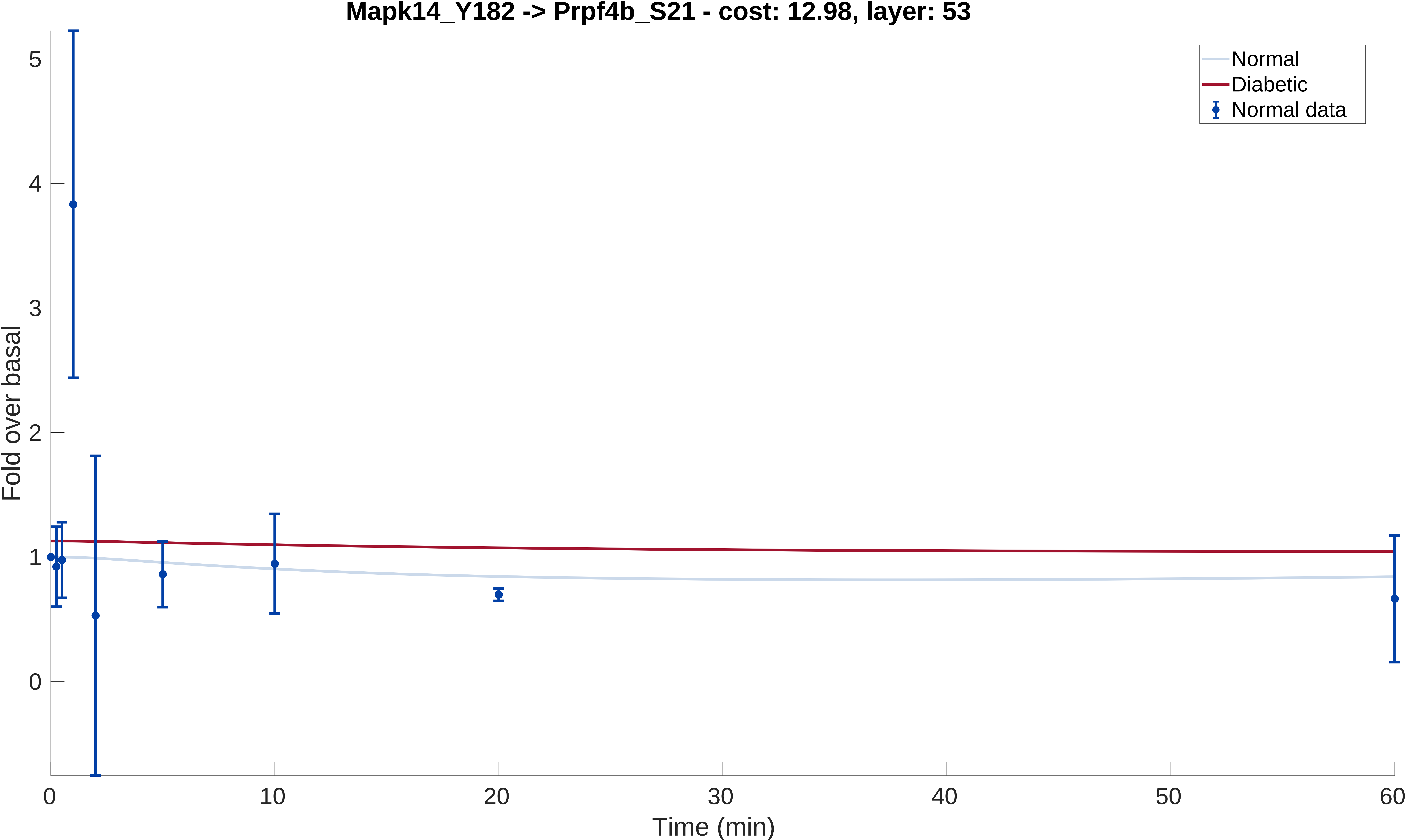

Mapk14_Y182 -> Prpf4b_S21 - cost: 12.98, layer: 53
Normal
Diabetic
Normal data
Fold over basal
0
1
2
3
4
5
0
10
20
30
40
50
60
Time (min)

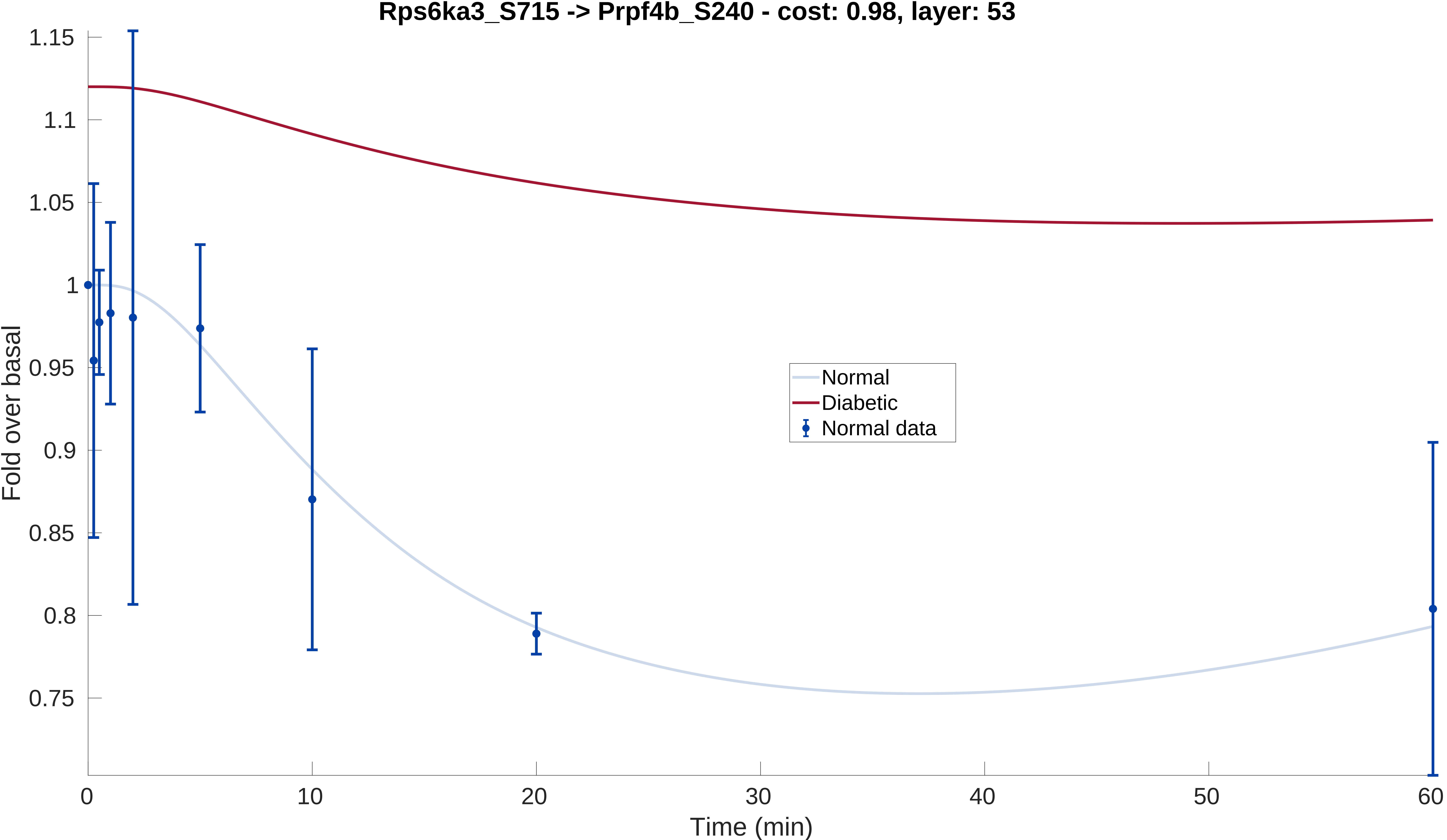
Rps6ka3_S715 -> Prpf4b_S240 - cost: 0.98, layer: 53
Fold over basal
1.15
1.1
1.05
1
0.95
0.9
0.85
0.8
0.75
0
10
20
30
40
50
60
Time (min)
Normal
Diabetic
Normal data

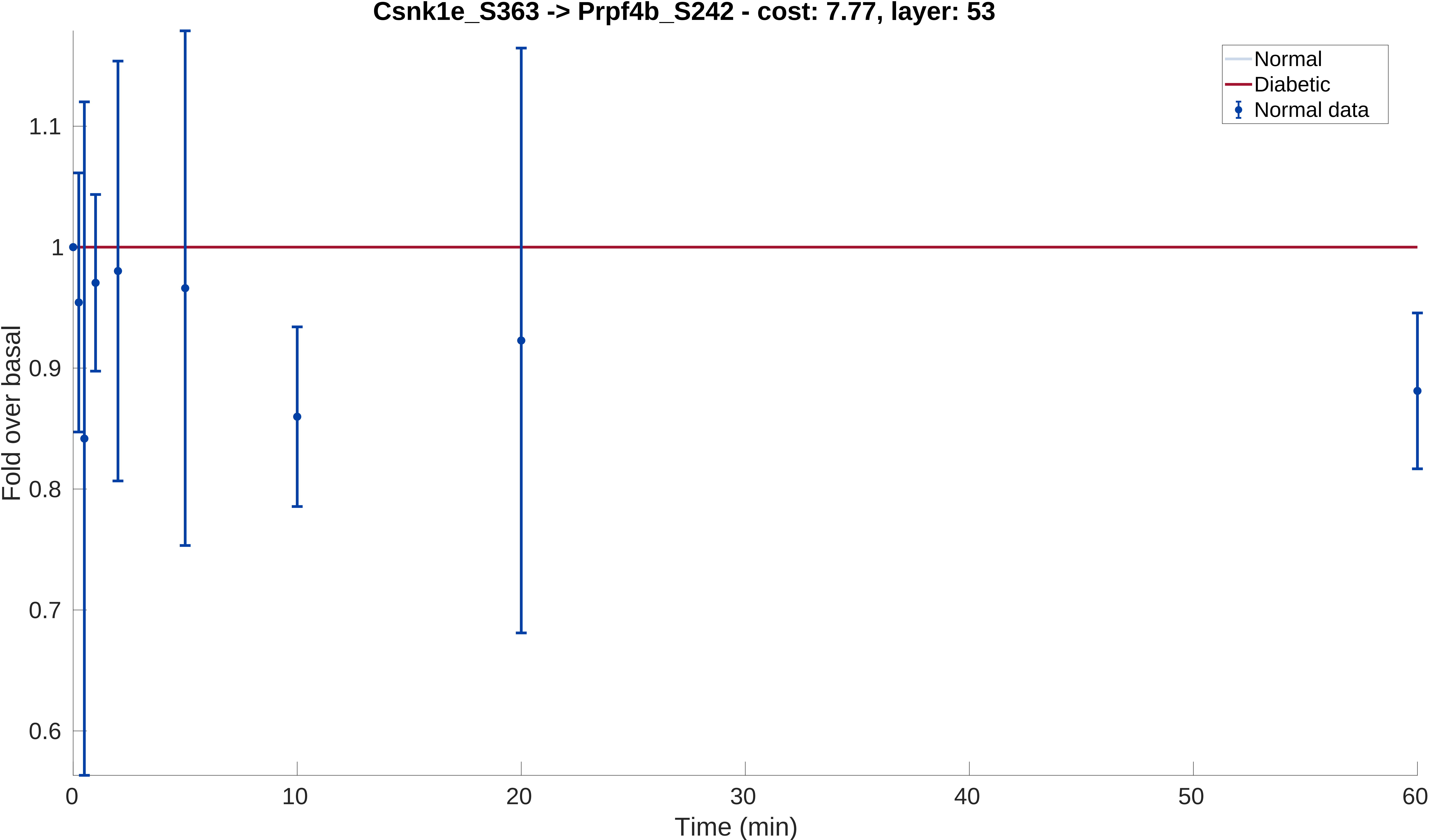

Csnk1e_S363 -> Prpf4b_S242 - cost: 7.77, layer: 53
Normal
Diabetic
Normal data
Fold over basal
Time (min)
0.6
0.7
0.8
0.9
1
1.1
0
10
20
30
40
50
60

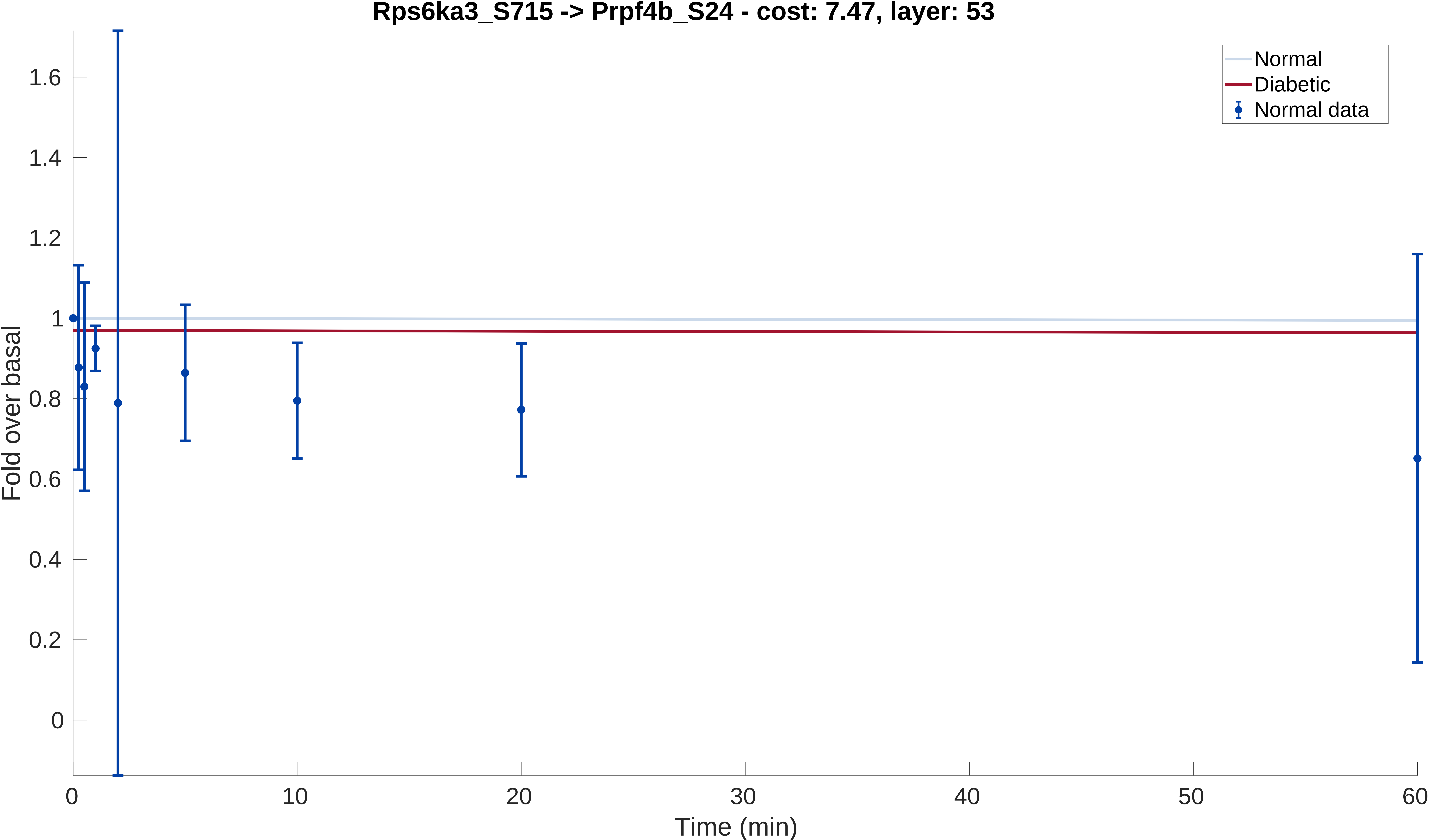

Rps6ka3_S715 -> Prpf4b_S24 - cost: 7.47, layer: 53
Normal
Diabetic
Normal data
Fold over basal
1.6
1.4
1.2
1
0.8
0.6
0.4
0.2
0
0
10
20
30
40
50
60
Time (min)

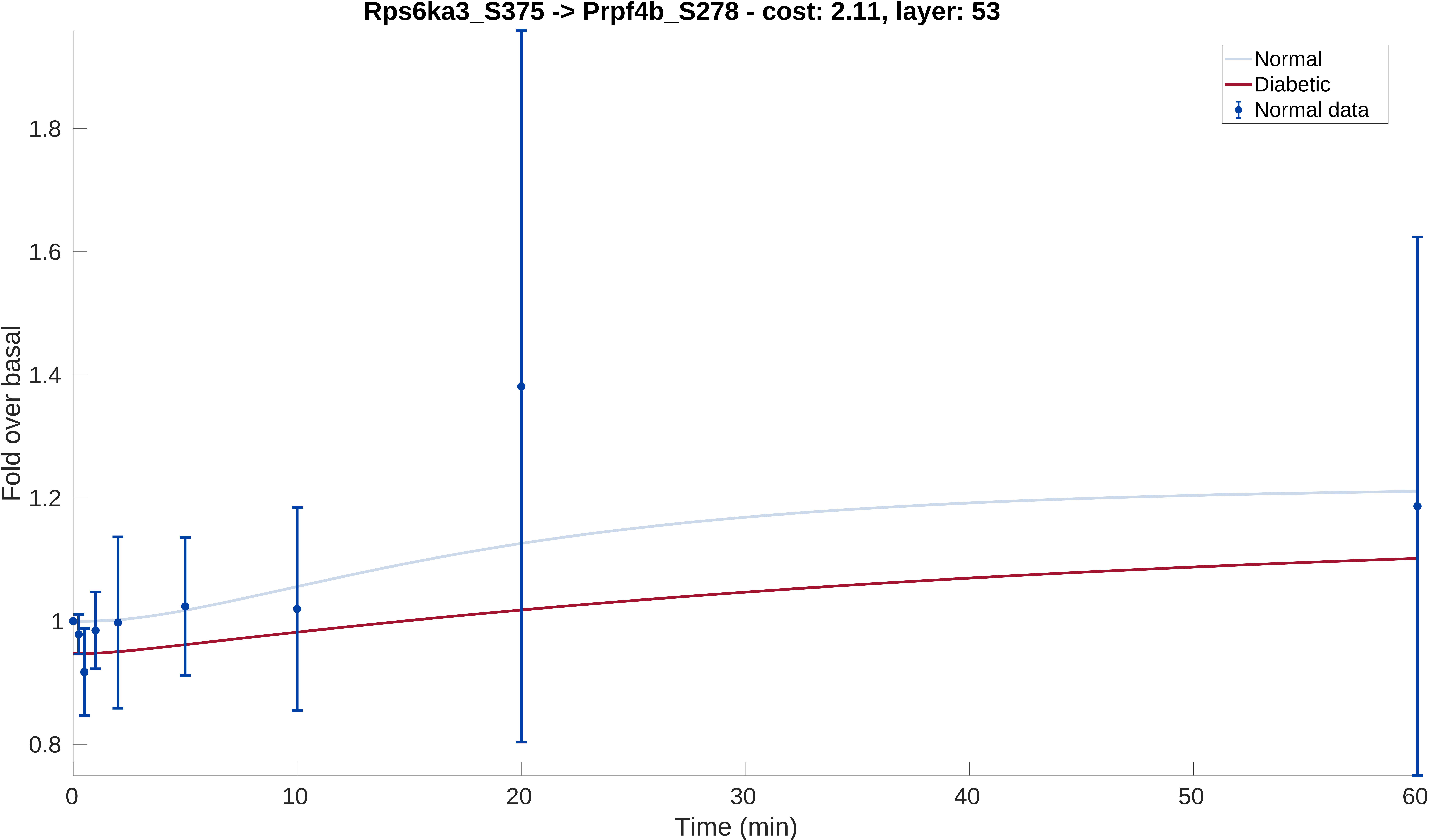

Rps6ka3_S375 -> Prpf4b_S278 - cost: 2.11, layer: 53
Normal
Diabetic
Normal data
Fold over basal
1.8
1.6
1.4
1.2
1
0.8
0
10
20
30
40
50
60
Time (min)

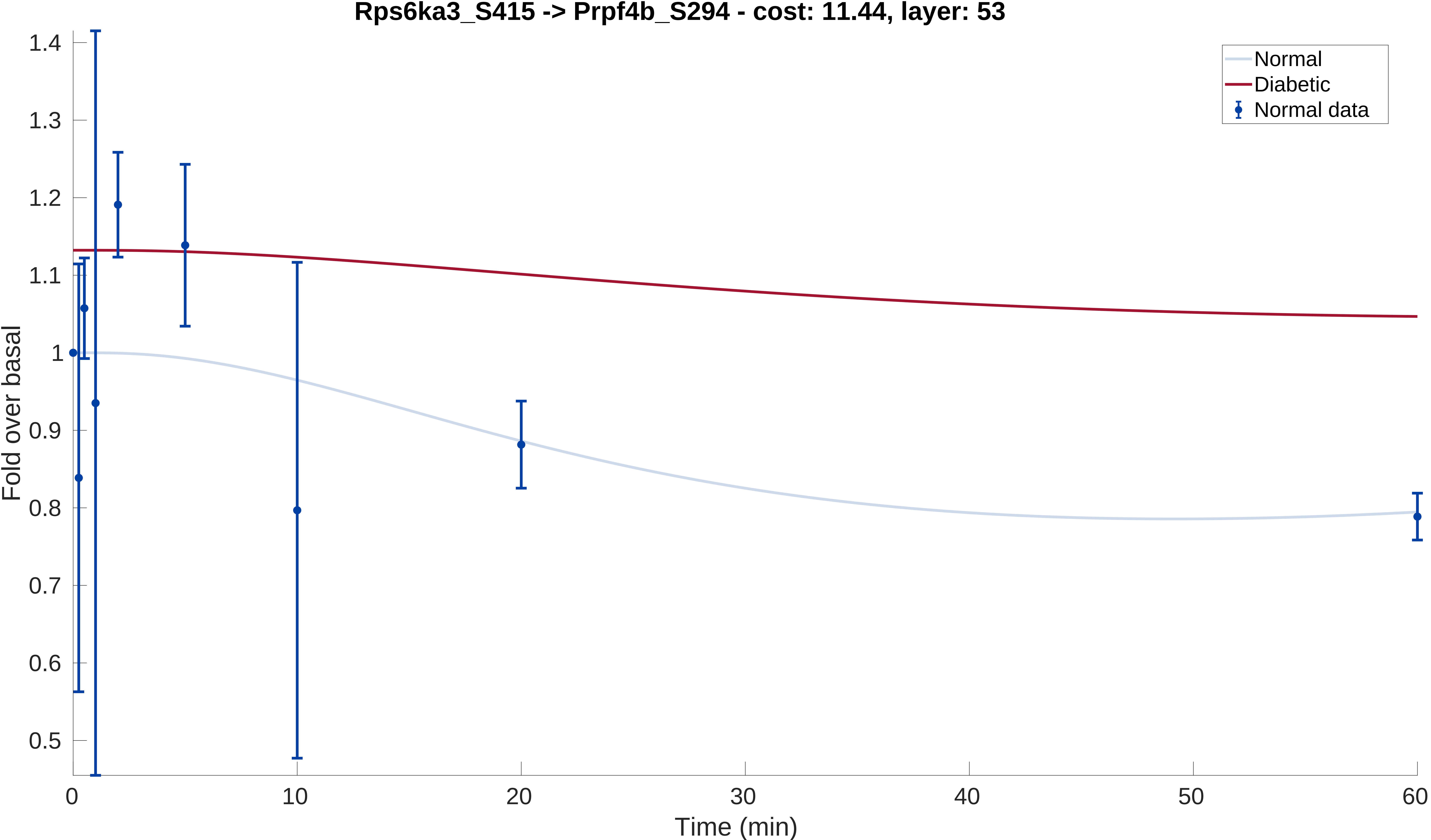
Rps6ka3_S415 -> Prpf4b_S294 - cost: 11.44, layer: 53
Normal
Diabetic
Normal data
Fold over basal
Time (min)
1.4
1.3
1.2
1.1
1
0.9
0.8
0.7
0.6
0.5
0
10
20
30
40
50
60

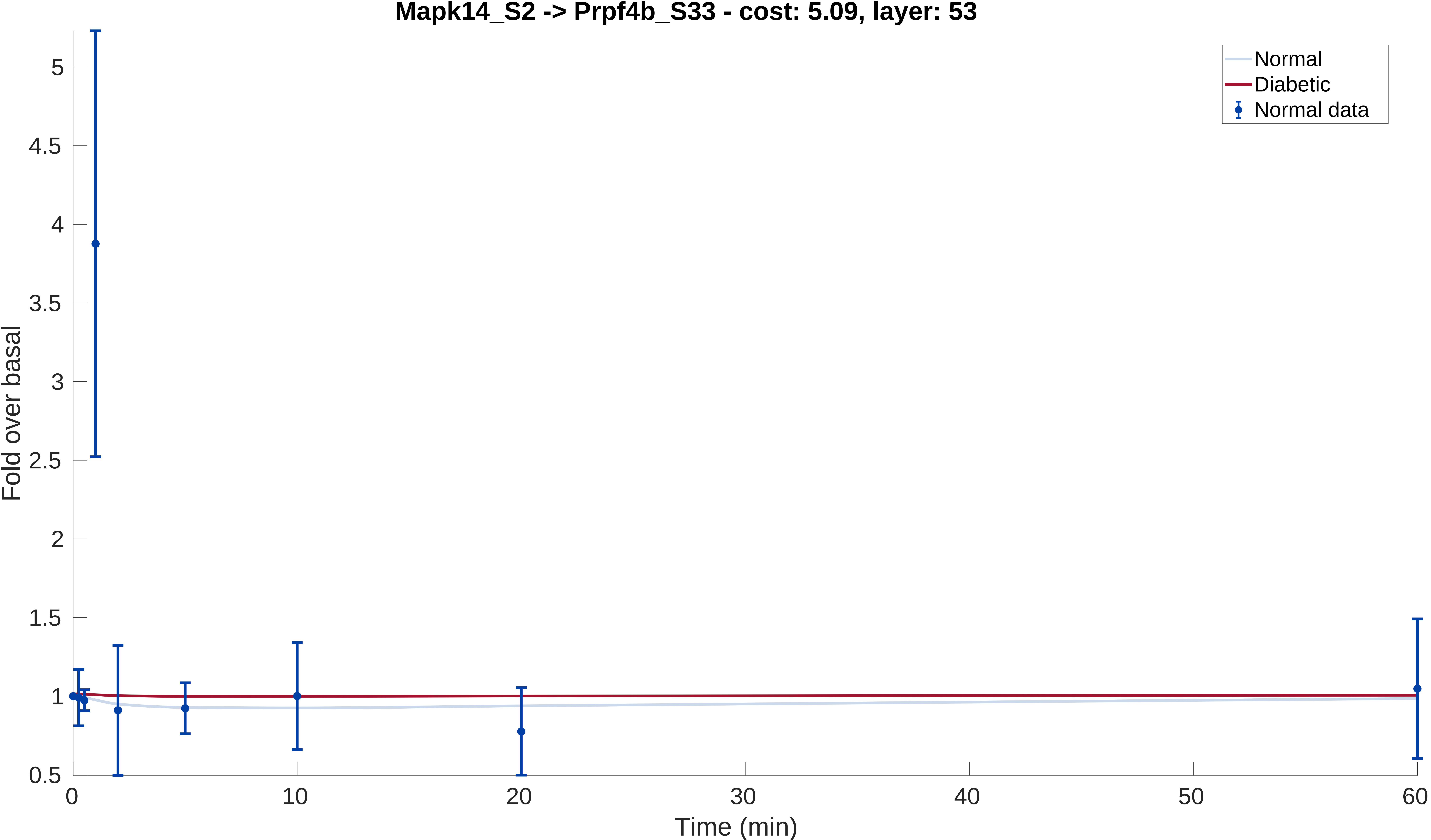
Mapk14_S2 -> Prpf4b_S33 - cost: 5.09, layer: 53
Fold over basal
Time (min)
0.5
1
1.5
2
2.5
3
3.5
4
4.5
5
0
10
20
30
40
50
60
Normal
Diabetic
Normal data

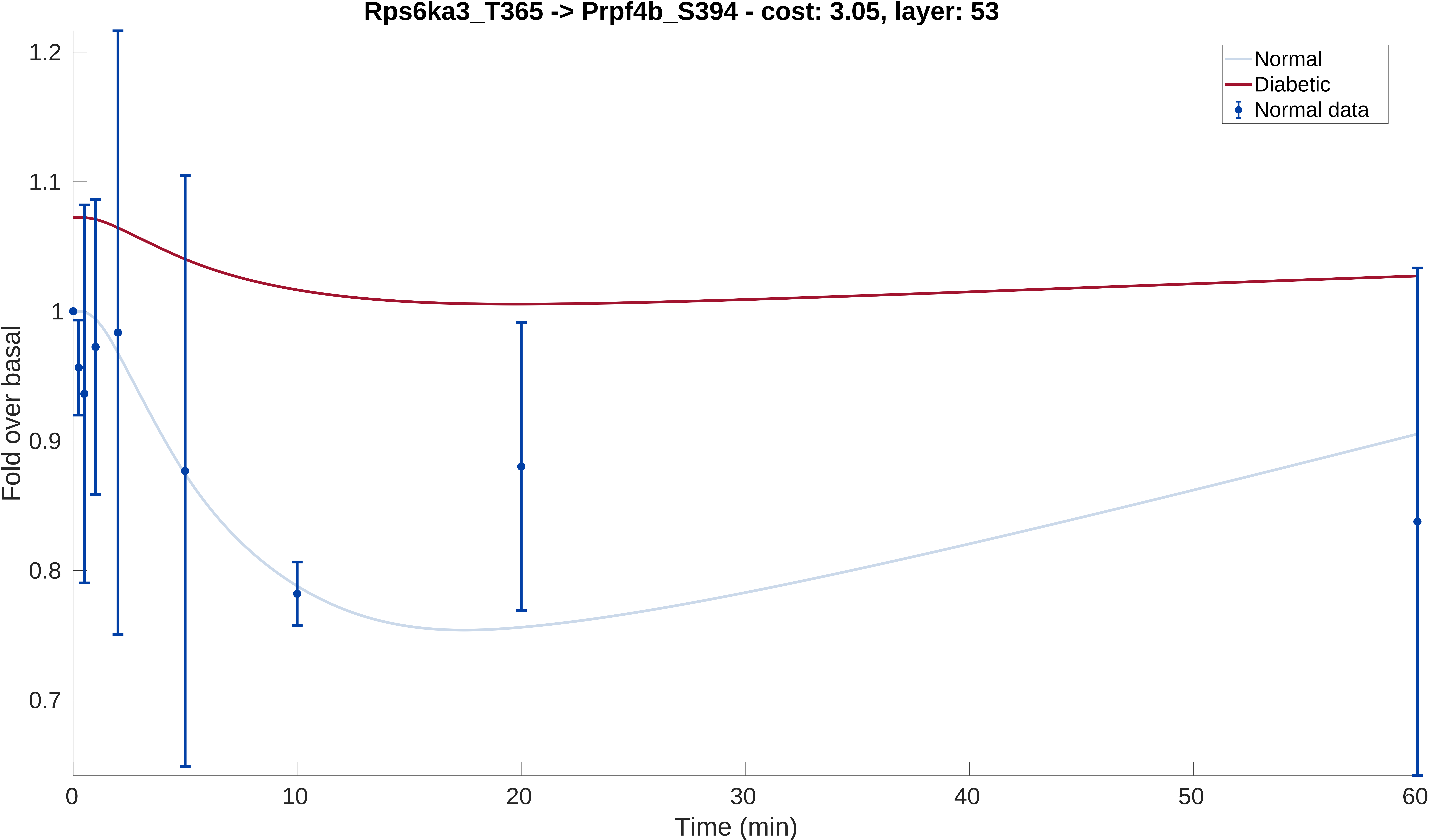

Rps6ka3_T365 -> Prpf4b_S394 - cost: 3.05, layer: 53
Normal
Diabetic
Normal data
Fold over basal
1.2
1.1
1
0.9
0.8
0.7
0
10
20
30
40
50
60
Time (min)

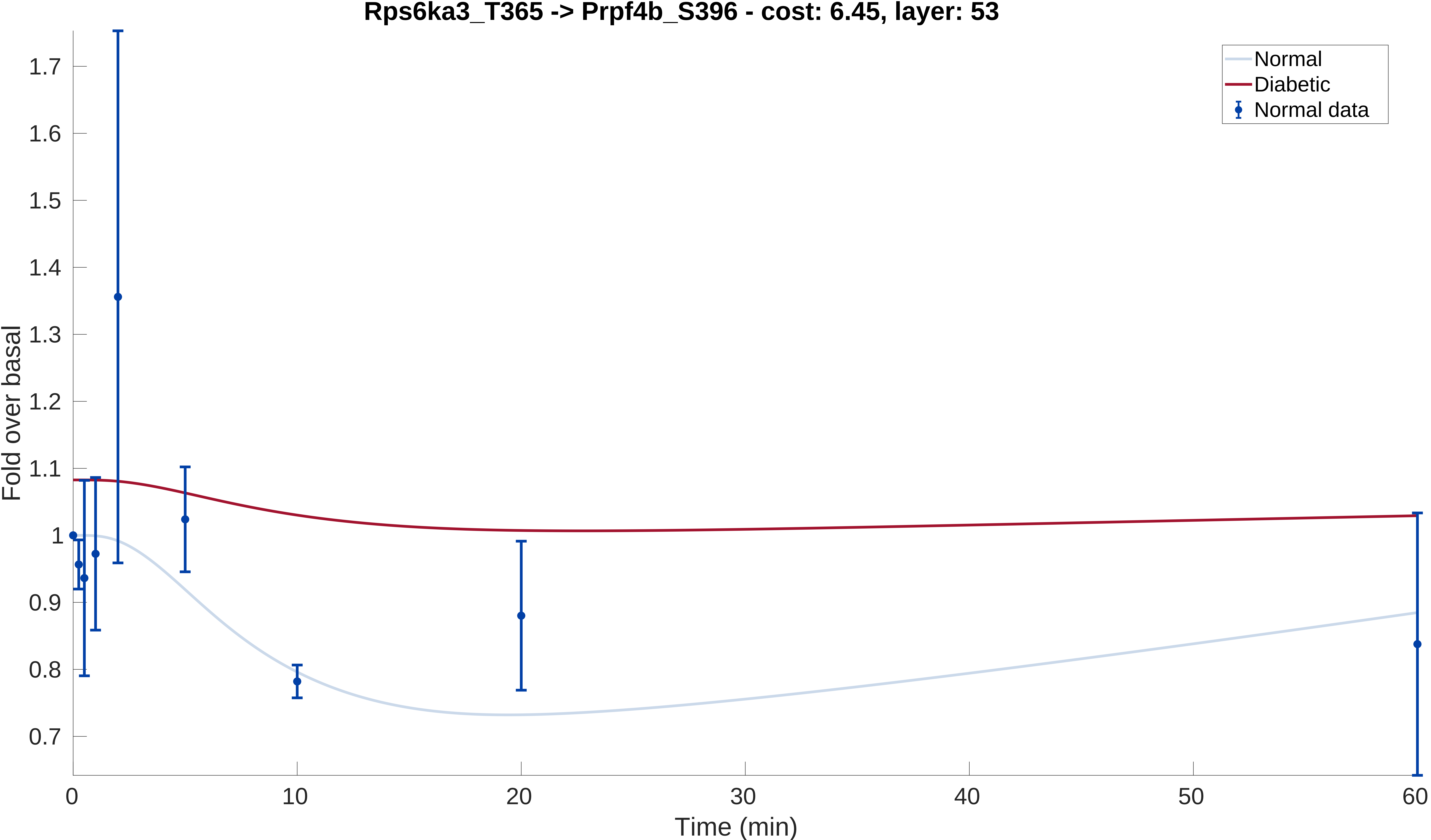

Rps6ka3_T365 -> Prpf4b_S396 - cost: 6.45, layer: 53
Normal
Diabetic
Normal data
Fold over basal
1.7
1.6
1.5
1.4
1.3
1.2
1.1
1
0.9
0.8
0.7
0
10
20
30
40
50
60
Time (min)

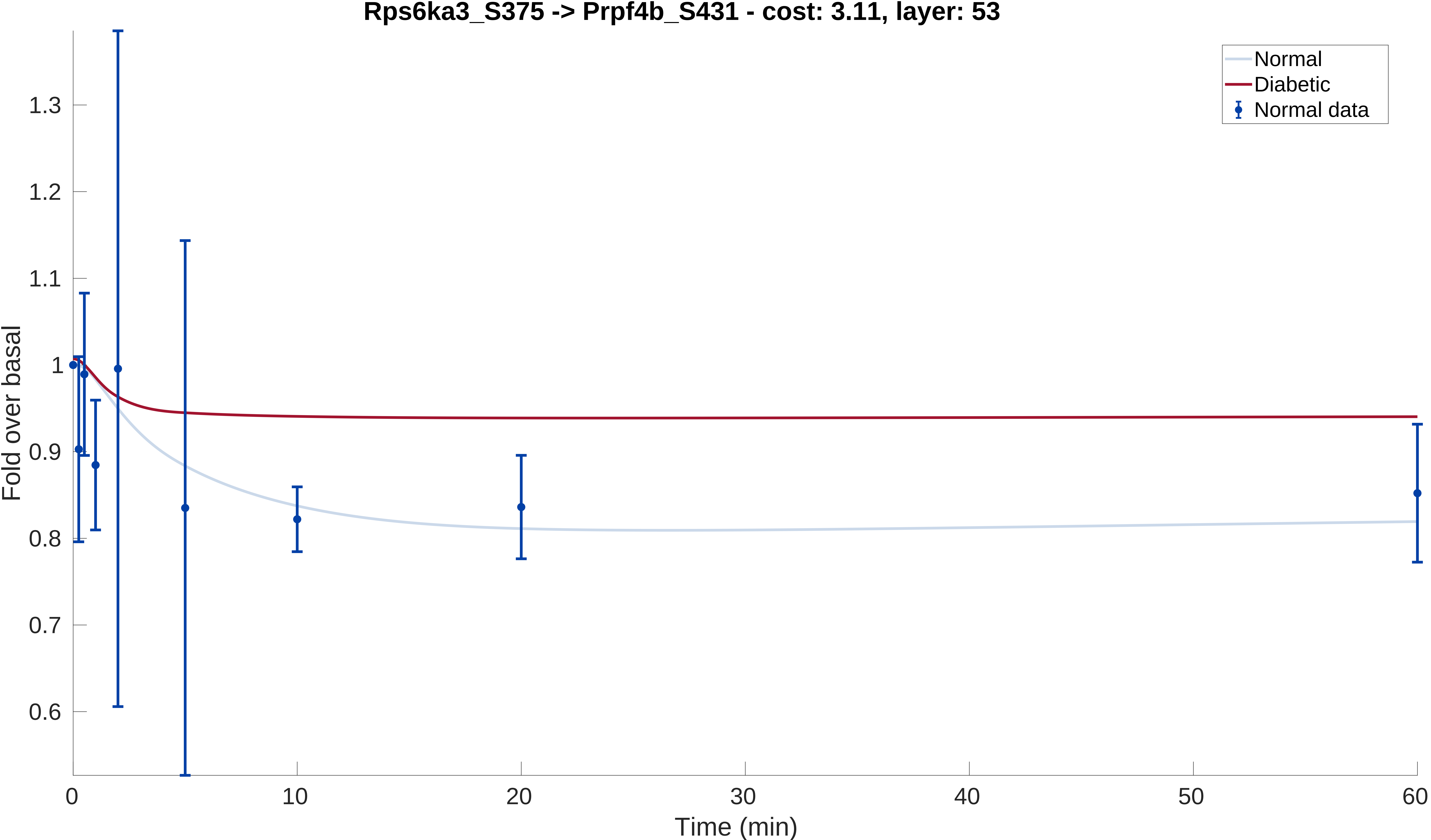

Rps6ka3_S375 -> Prpf4b_S431 - cost: 3.11, layer: 53
Normal
Diabetic
Normal data
Fold over basal
1.3
1.2
1.1
1
0.9
0.8
0.7
0.6
0
10
20
30
40
50
60
Time (min)

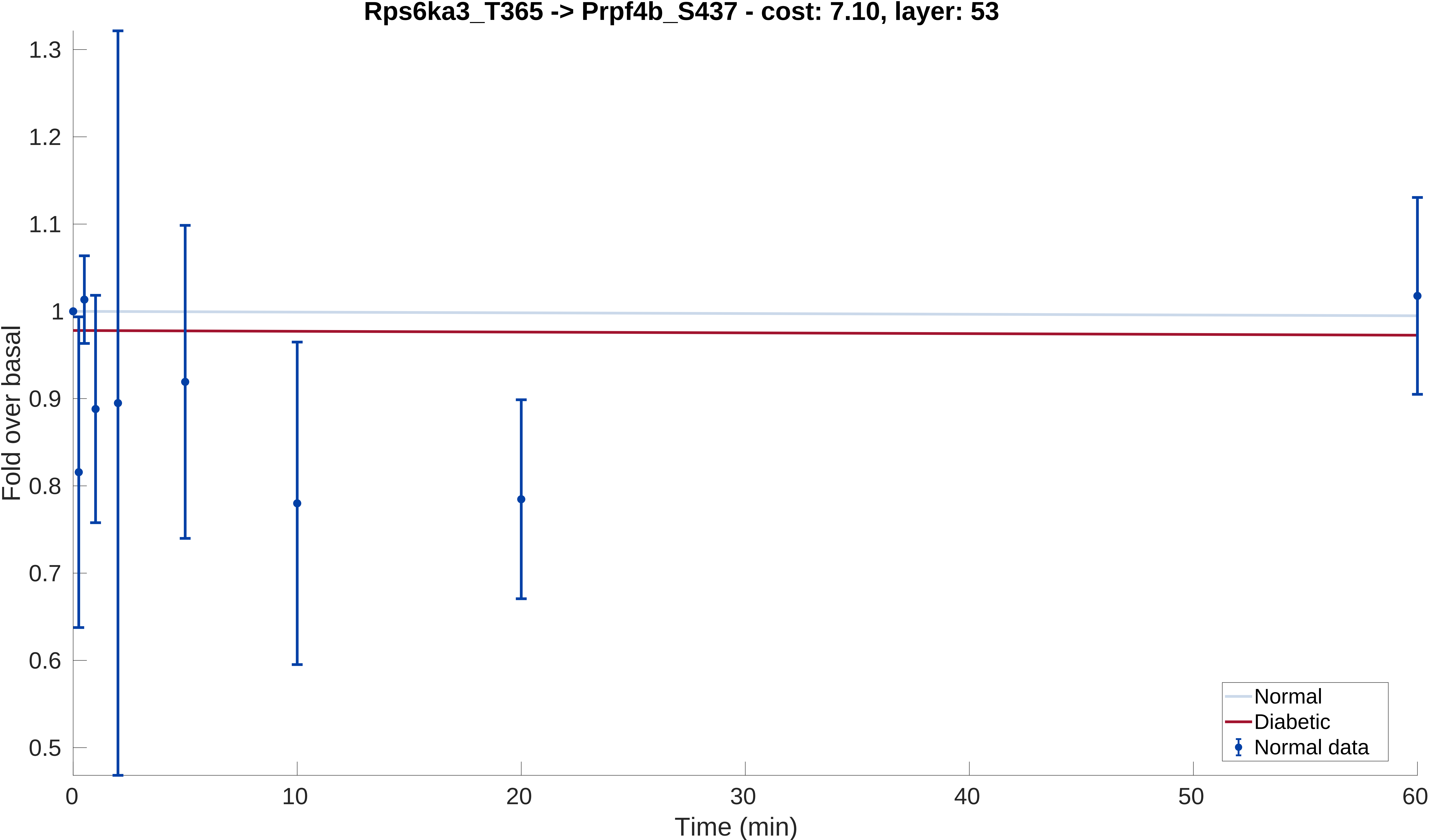
Rps6ka3_T365 -> Prpf4b_S437 - cost: 7.10, layer: 53
Fold over basal
1.3
1.2
1.1
1
0.9
0.8
0.7
0.6
0.5
0
10
20
30
40
50
60
Time (min)
Normal
Diabetic
Normal data

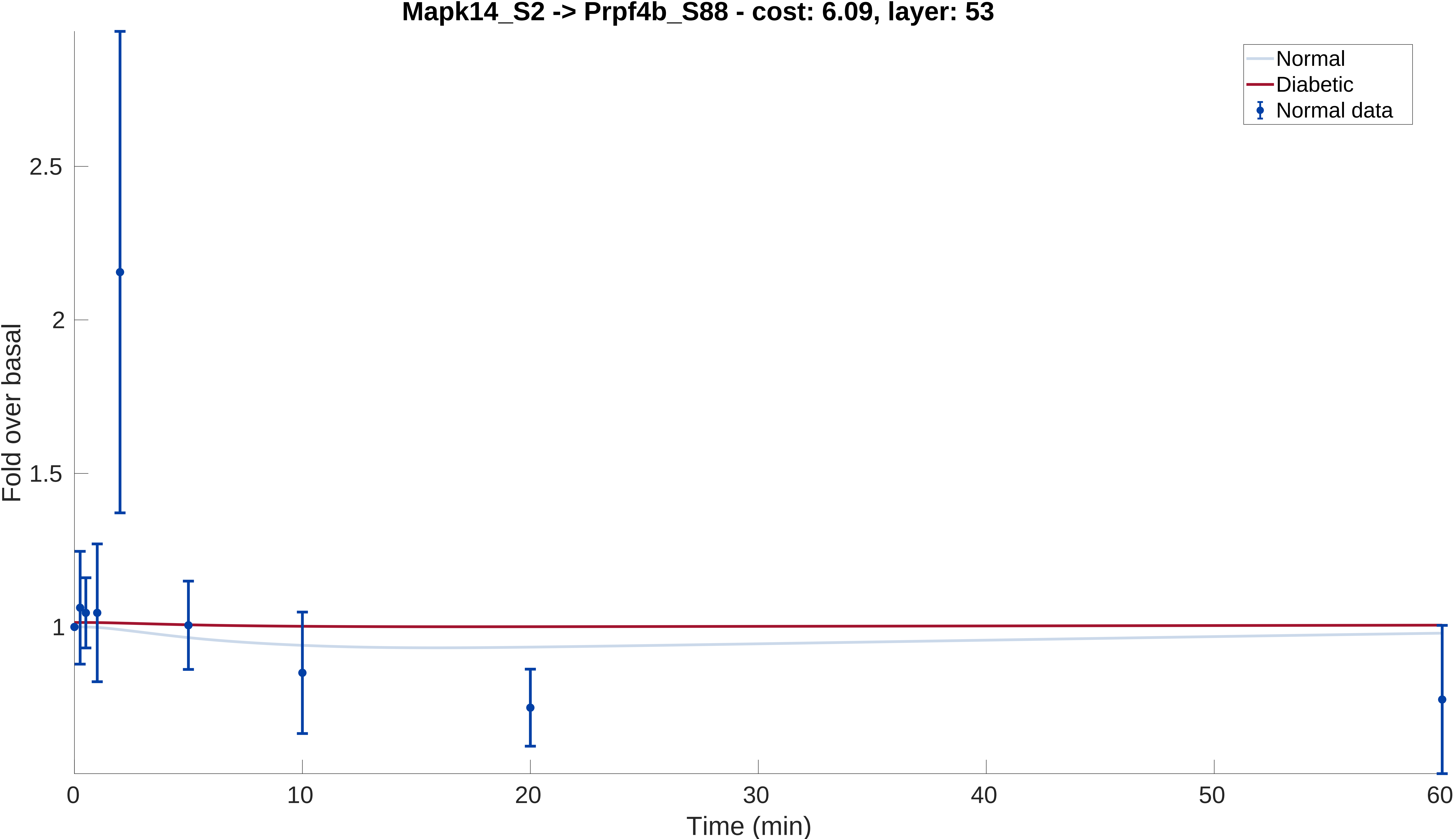
Mapk14_S2 -> Prpf4b_S88 - cost: 6.09, layer: 53
Normal
Diabetic
Normal data
Fold over basal
2.5
2
1.5
1
0
10
20
30
40
50
60
Time (min)

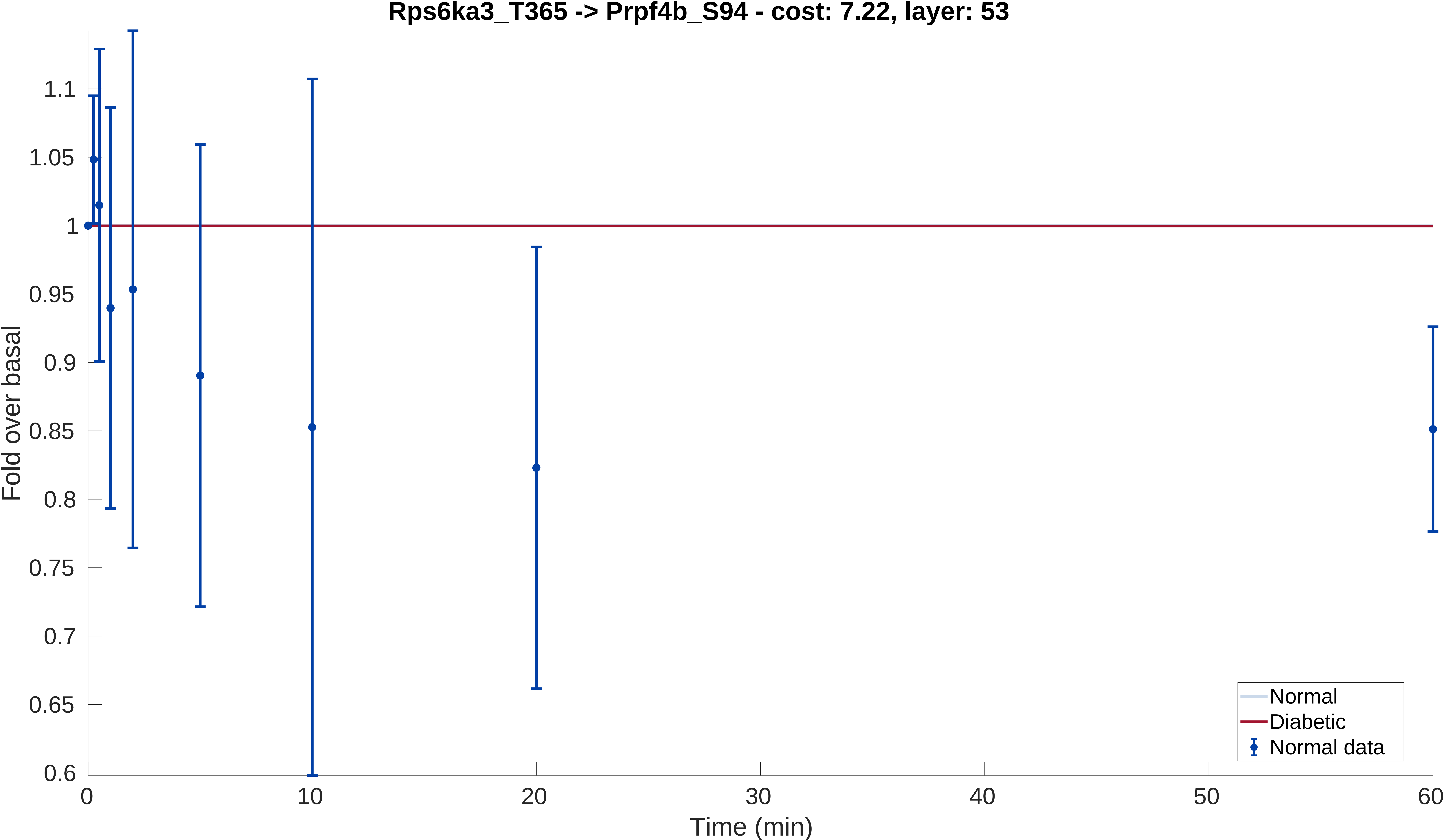

Rps6ka3_T365 -> Prpf4b_S94 - cost: 7.22, layer: 53
Fold over basal
Time (min)
1.1
1.05
1
0.95
0.9
0.85
0.8
0.75
0.7
0.65
0.6
0
10
20
30
40
50
60
Normal
Diabetic
Normal data

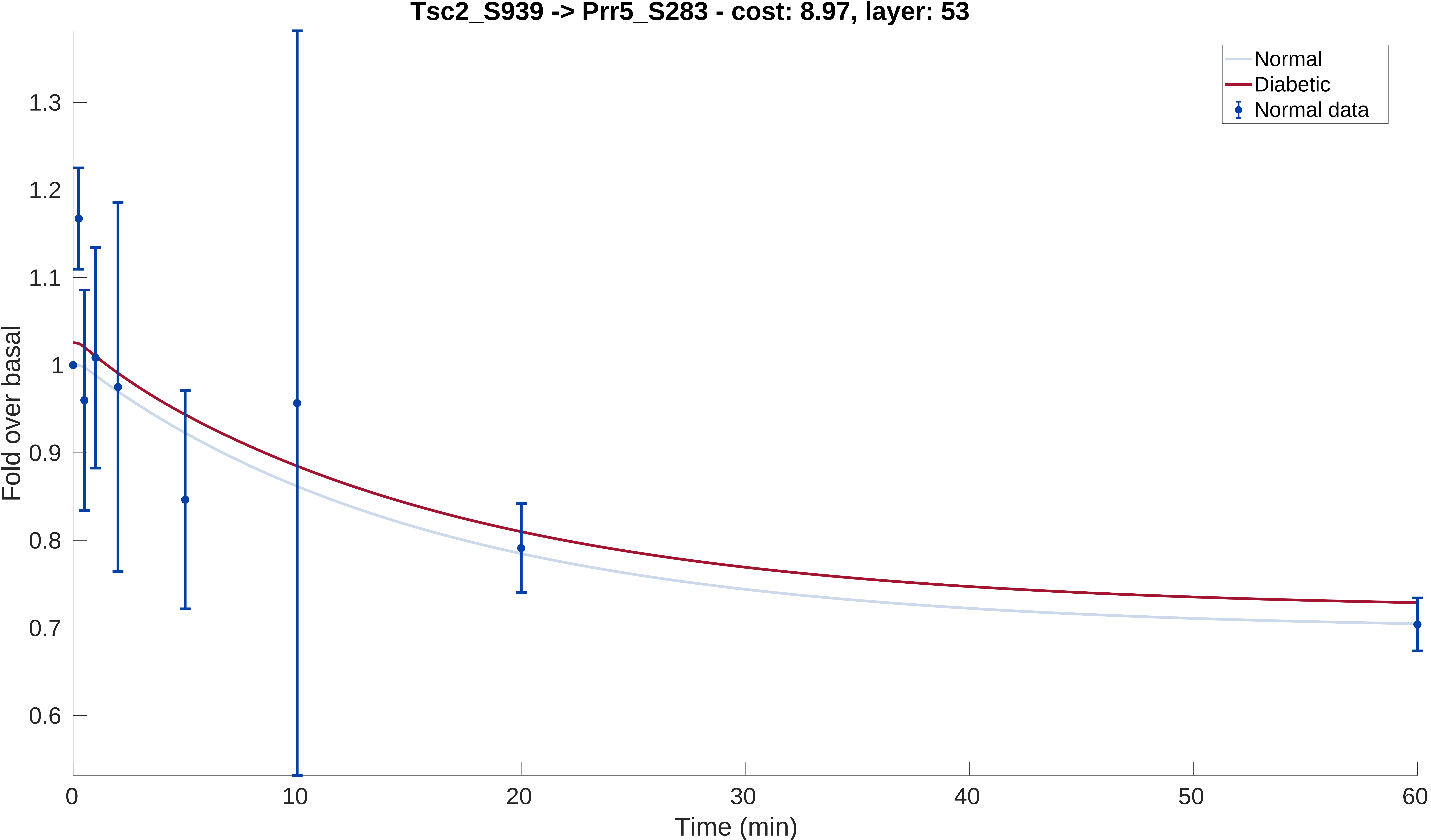

Tsc2_S939 -> Prr5_S283 - cost: 8.97, layer: 53
Normal
Diabetic
Normal data
Fold over basal
1.3
1.2
1.1
1
0.9
0.8
0.7
0.6
0
10
20
30
40
50
60
Time (min)

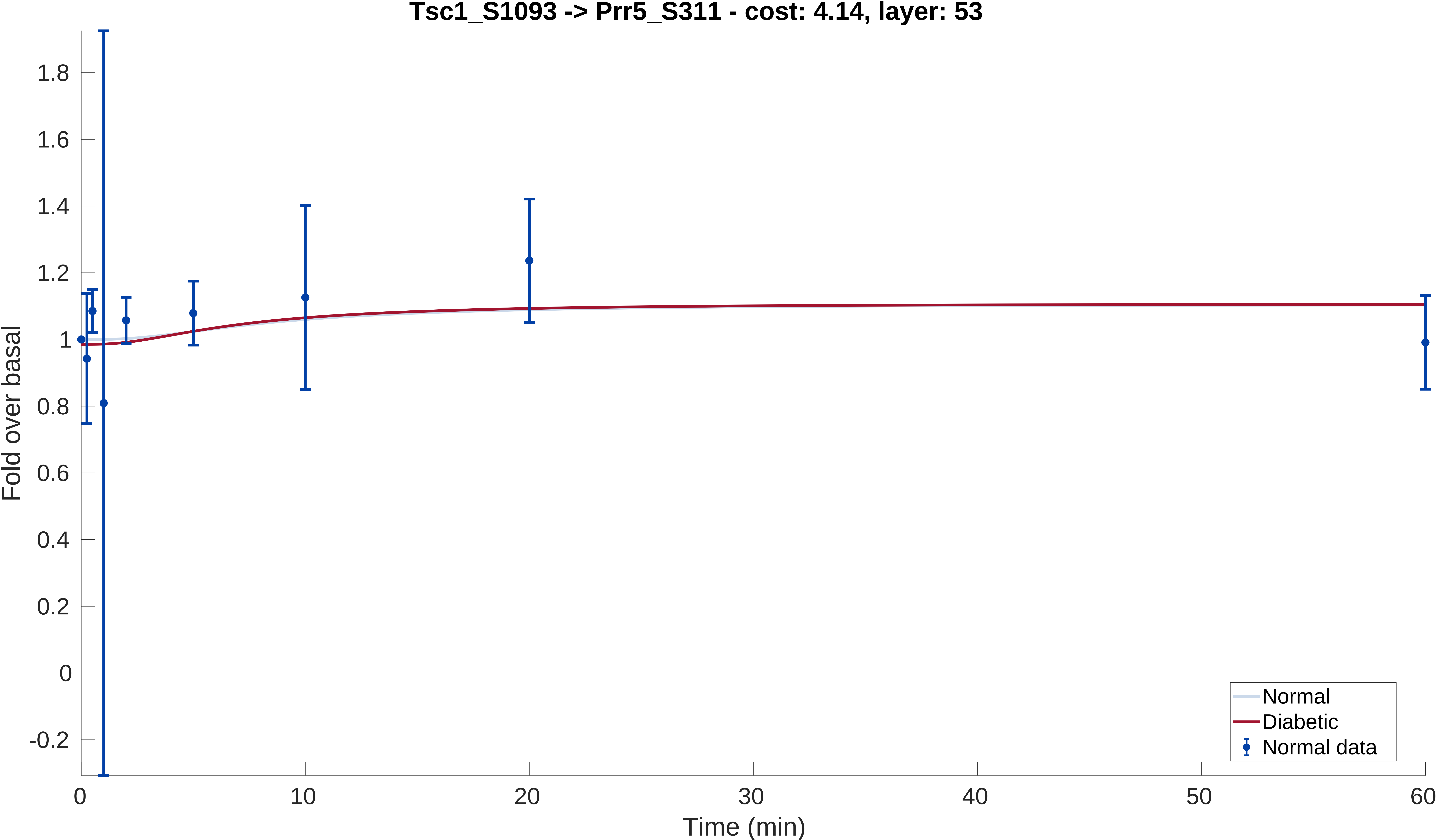

Tsc1_S1093 -> Prr5_S311 - cost: 4.14, layer: 53
Fold over basal
1.8
1.6
1.4
1.2
1
0.8
0.6
0.4
0.2
0
-0.2
0
10
20
30
40
50
60
Time (min)
Normal
Diabetic
Normal data

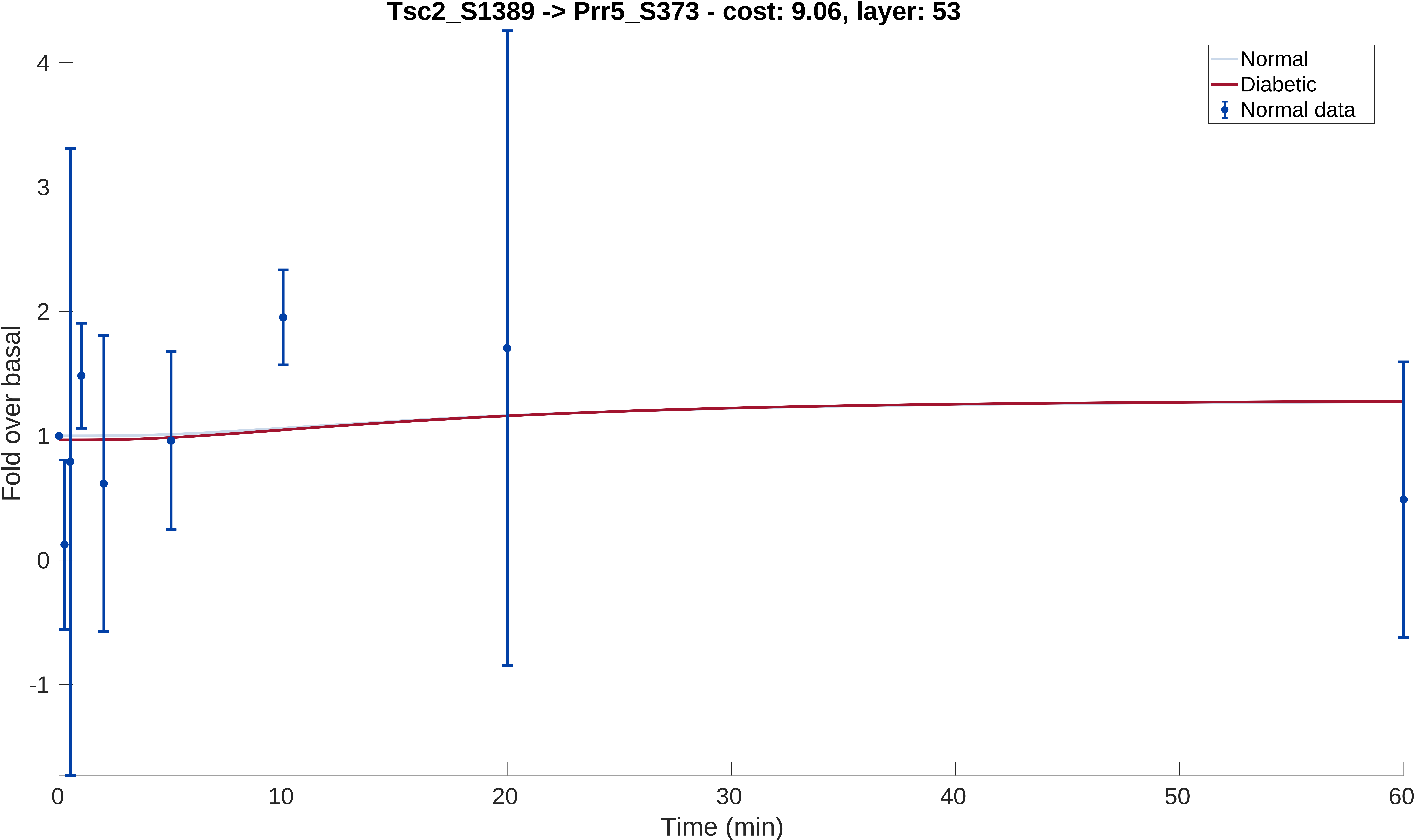

Tsc2_S1389 -> Prr5_S373 - cost: 9.06, layer: 53
Normal
Diabetic
Normal data
Fold over basal
Time (min)
4
3
2
1
0
-1
0
10
20
30
40
50
60

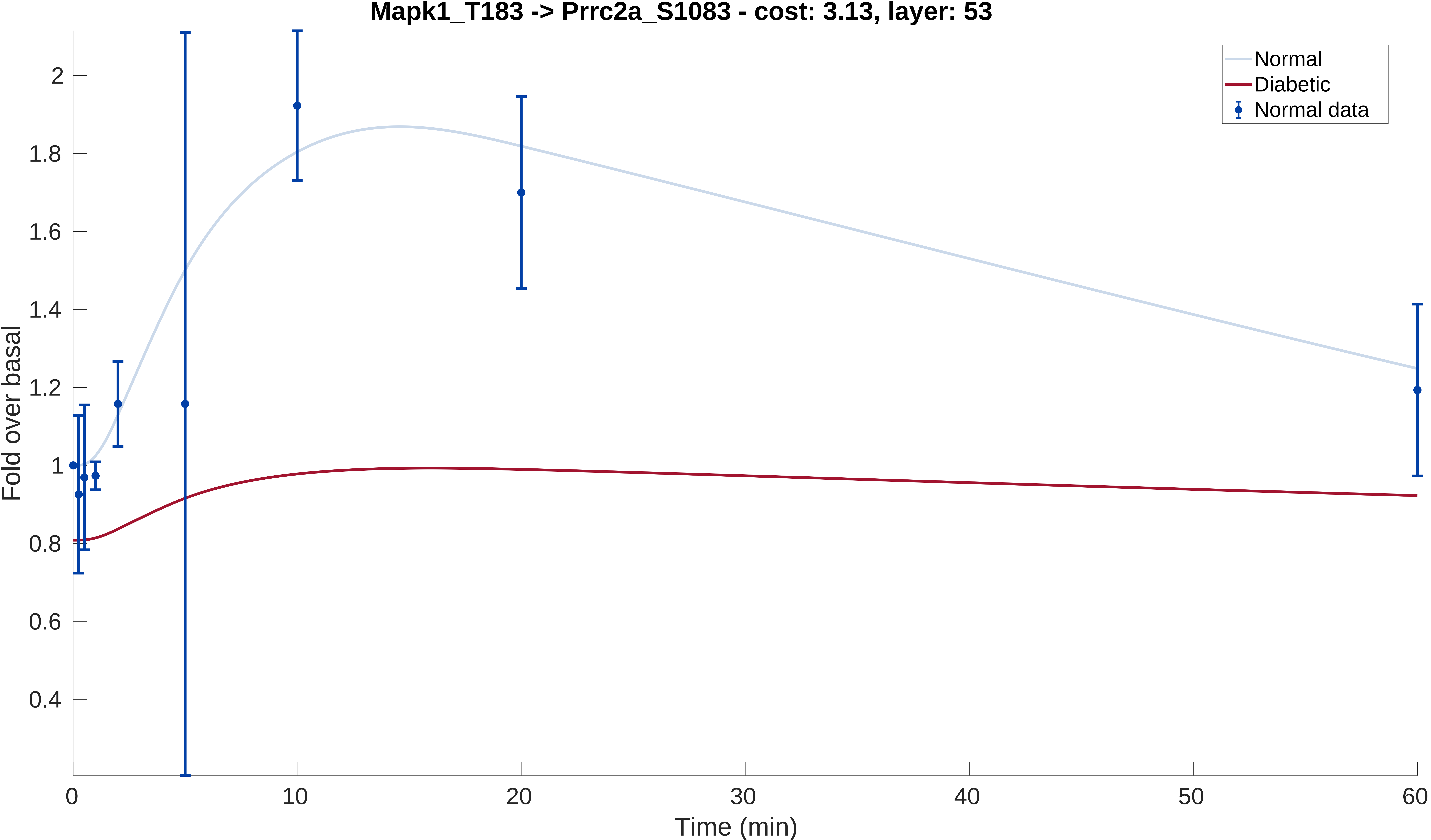
Mapk1_T183 -> Prrc2a_S1083 - cost: 3.13, layer: 53
Normal
Diabetic
Normal data
Fold over basal
2
1.8
1.6
1.4
1.2
1
0.8
0.6
0.4
0
10
20
30
40
50
60
Time (min)

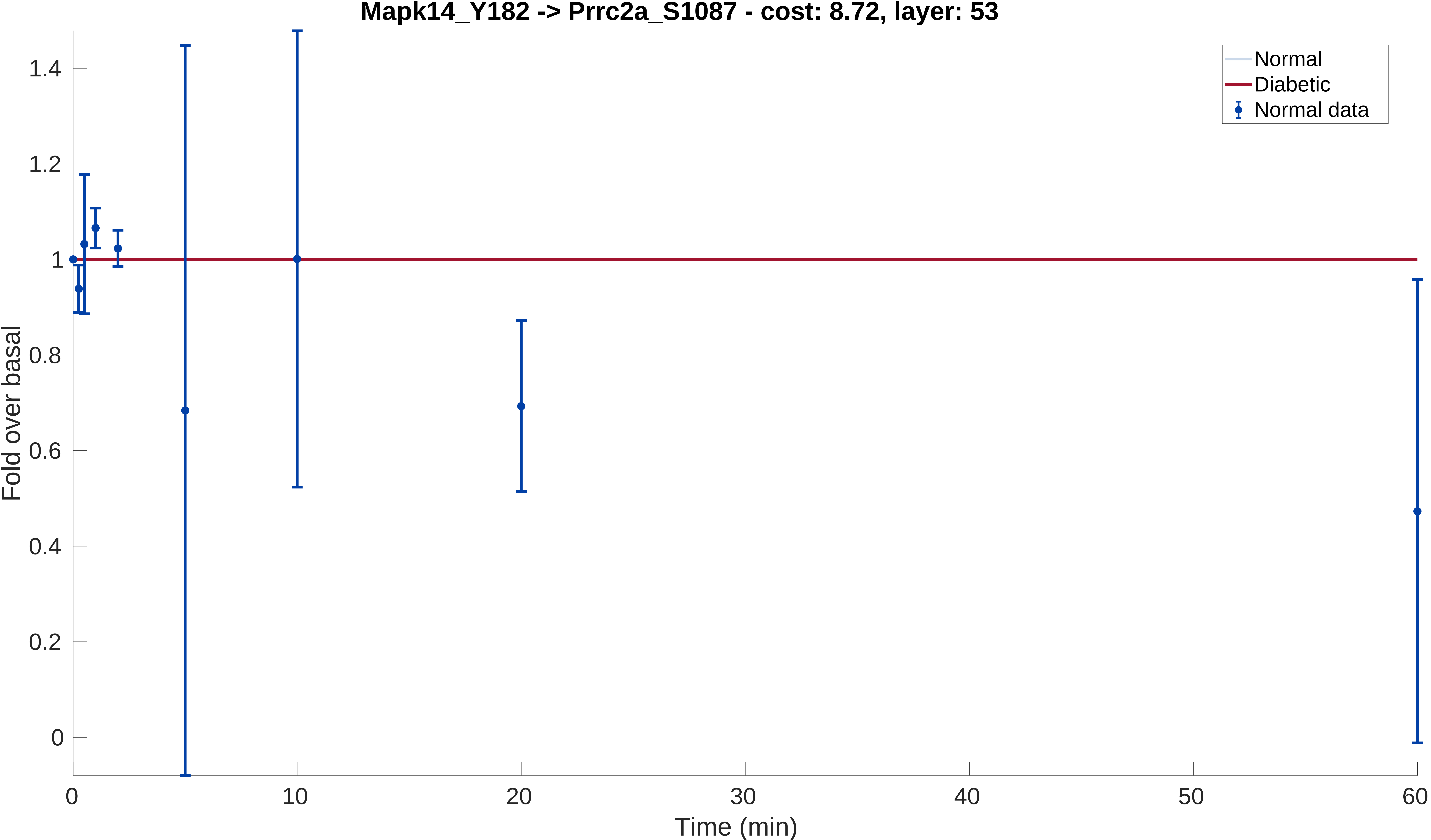

Mapk14_Y182 -> Prrc2a_S1087 - cost: 8.72, layer: 53
Normal
Diabetic
Normal data
Fold over basal
Time (min)
0
0.2
0.4
0.6
0.8
1
1.2
1.4
0
10
20
30
40
50
60

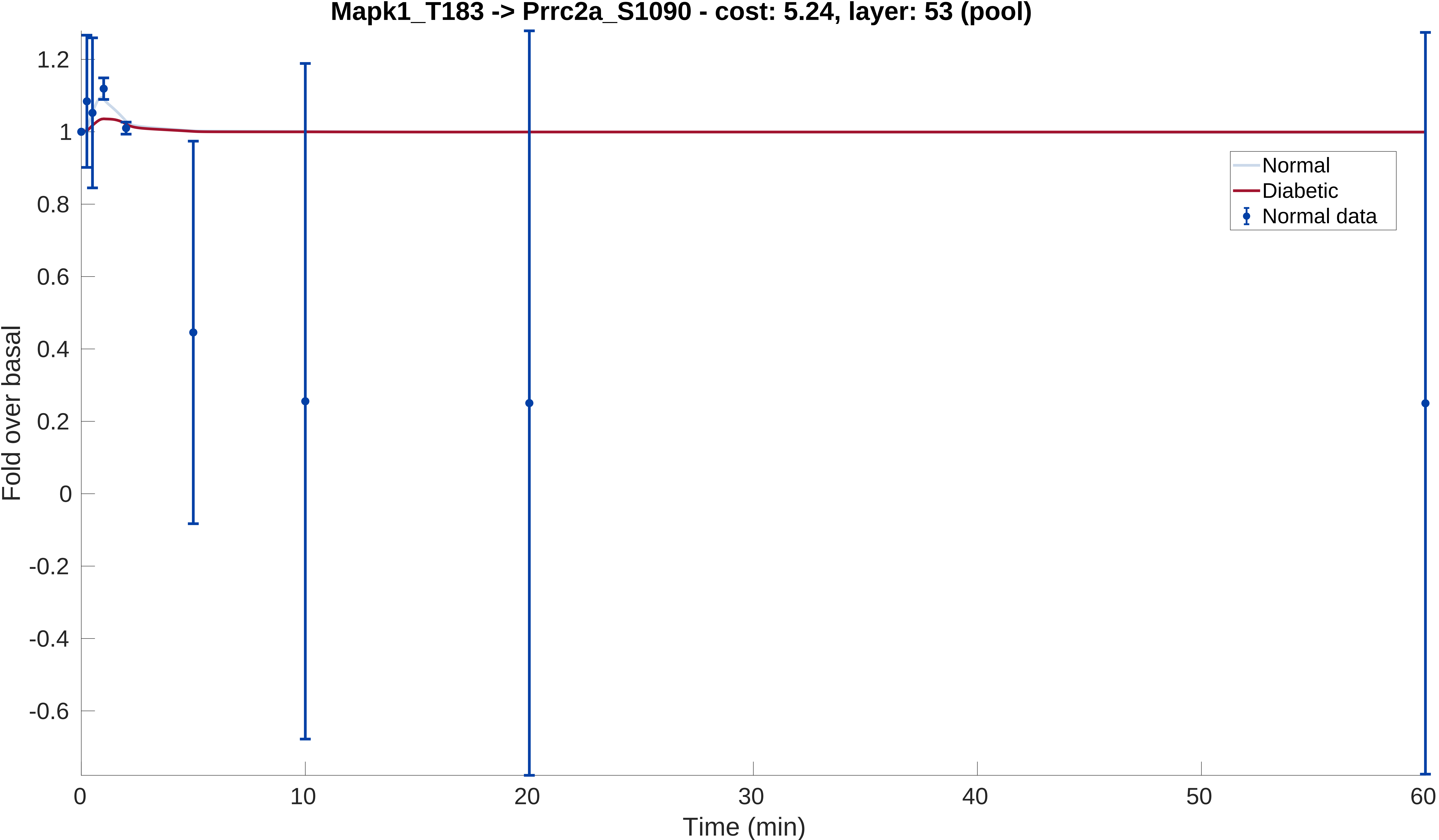
Mapk1_T183 -> Prrc2a_S1090 - cost: 5.24, layer: 53 (pool)
Fold over basal
Time (min)
Normal
Diabetic
Normal data
1.2
1
0.8
0.6
0.4
0.2
0
-0.2
-0.4
-0.6
0
10
20
30
40
50
60

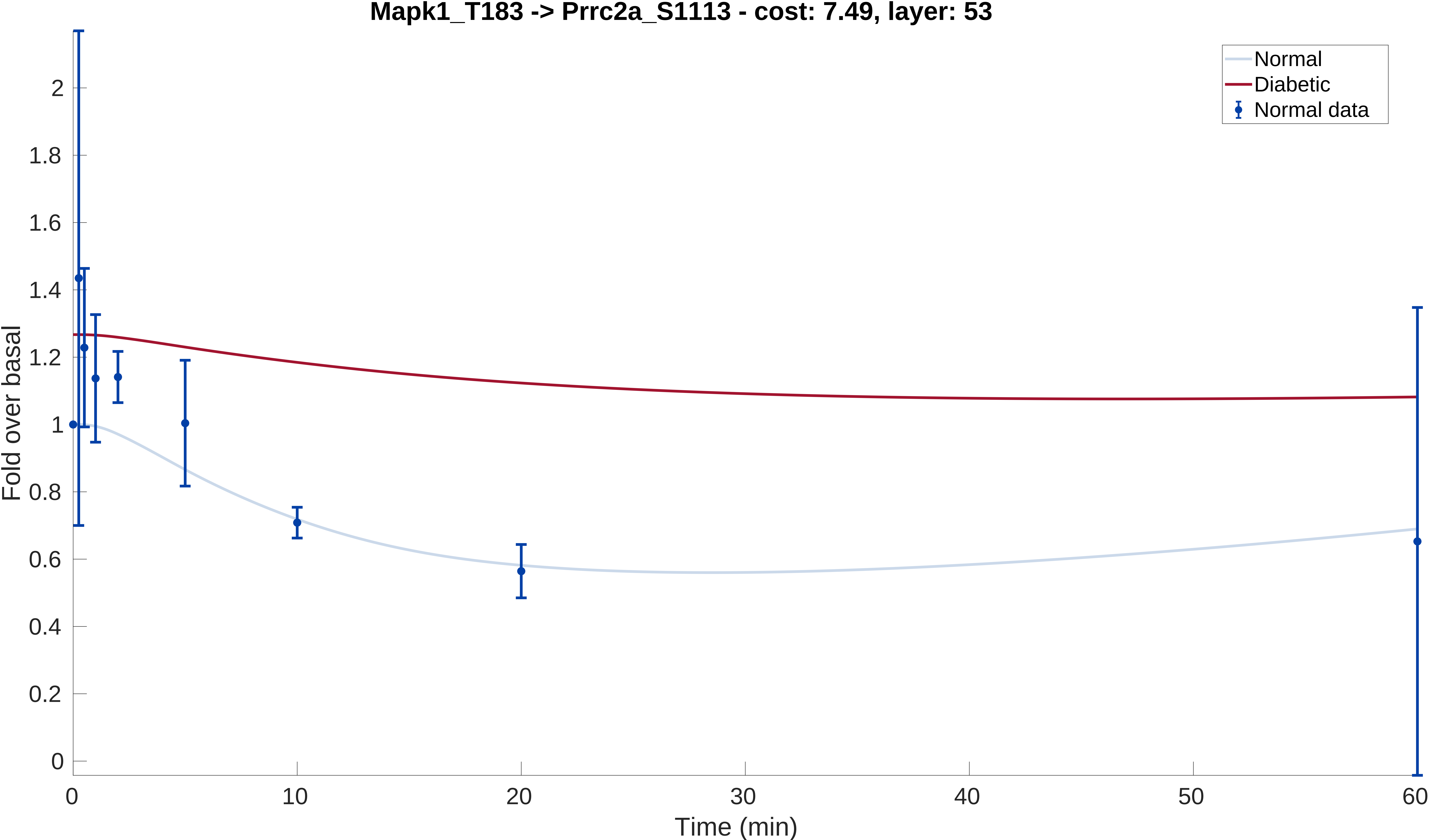

Mapk1_T183 -> Prrc2a_S1113 - cost: 7.49, layer: 53
Normal
Diabetic
Normal data
Fold over basal
Time (min)
0
0.2
0.4
0.6
0.8
1
1.2
1.4
1.6
1.8
2
0
10
20
30
40
50
60

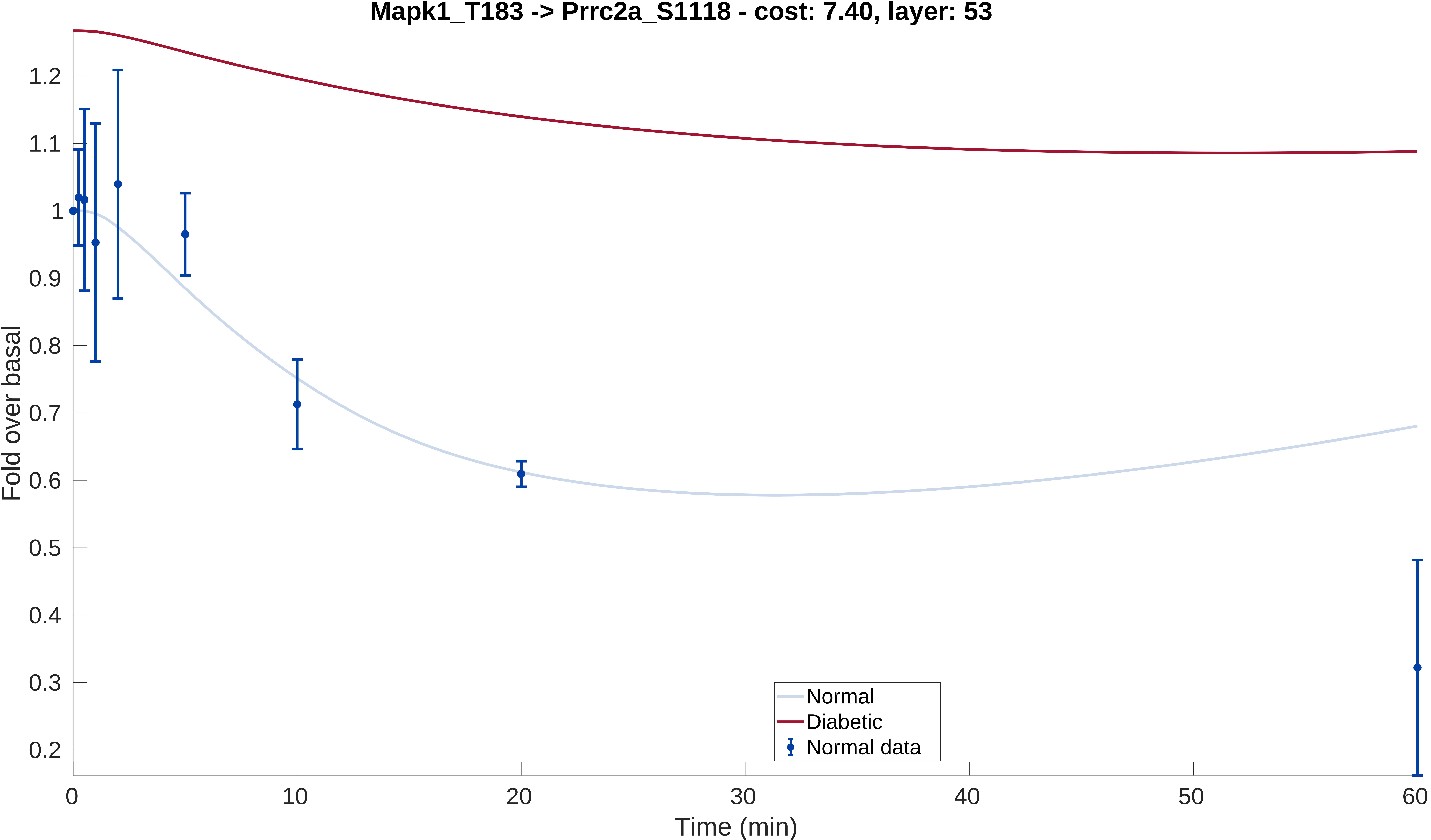

Mapk1_T183 -> Prrc2a_S1118 - cost: 7.40, layer: 53
Fold over basal
1.2
1.1
1
0.9
0.8
0.7
0.6
0.5
0.4
0.3
0.2
0
10
20
30
40
50
60
Time (min)
Normal
Diabetic
Normal data

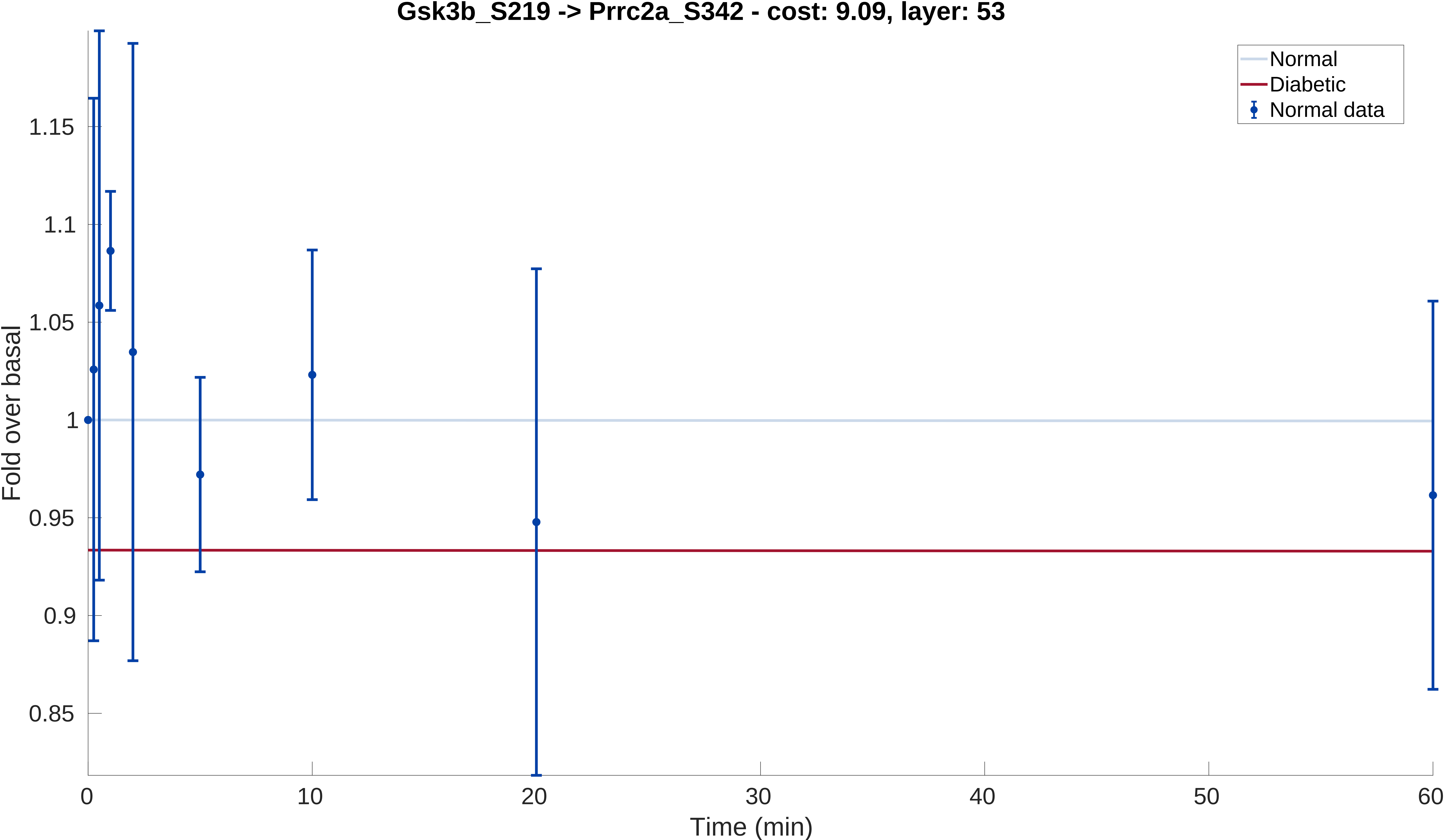
Gsk3b_S219 -> Prrc2a_S342 - cost: 9.09, layer: 53
Normal
Diabetic
Normal data
1.15
1.1
1.05
1
0.95
0.9
0.85
Fold over basal
0
10
20
30
40
50
60
Time (min)

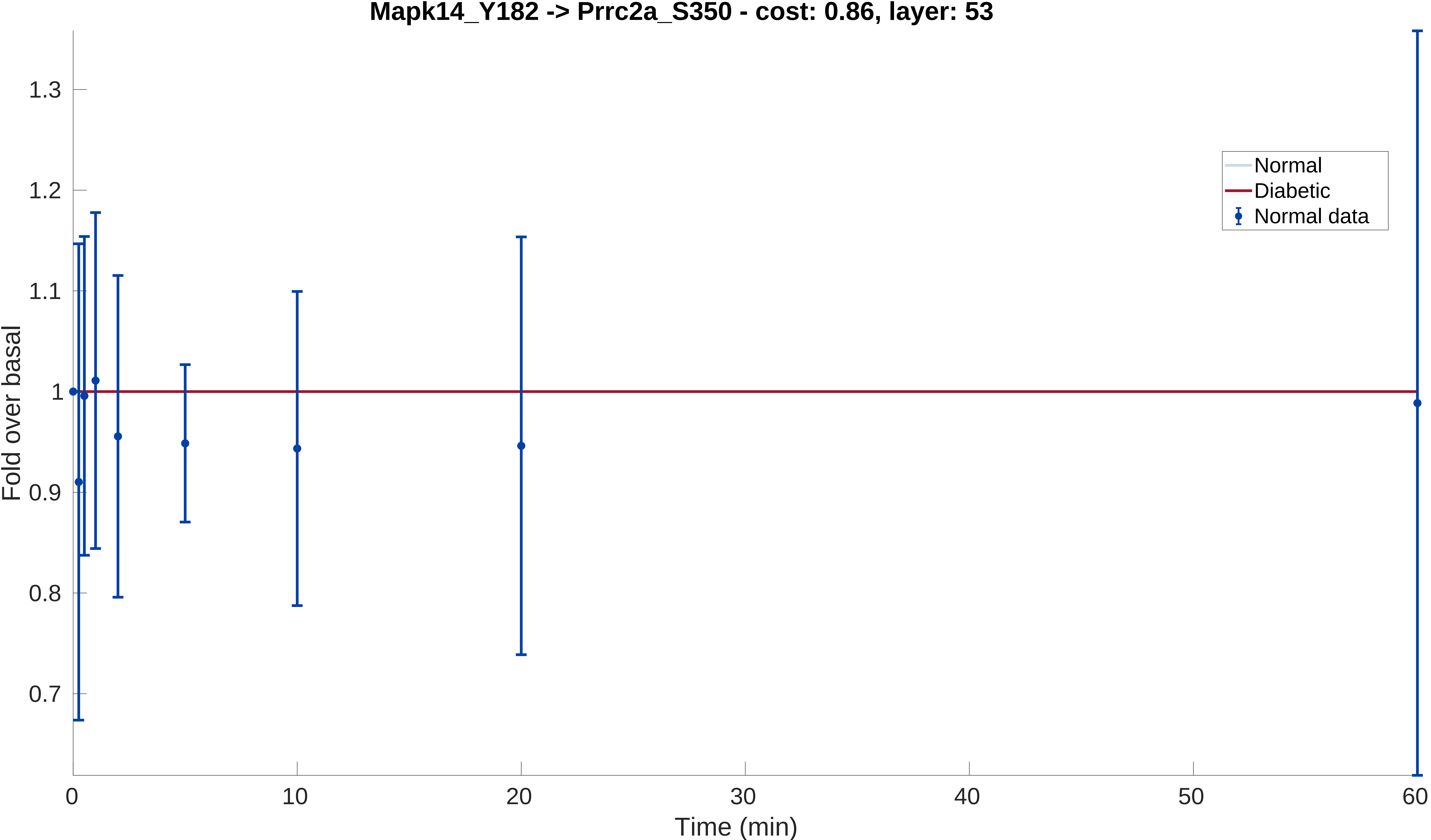

Mapk14_Y182 -> Prrc2a_S350 - cost: 0.86, layer: 53
Fold over basal
1.3
1.2
1.1
1
0.9
0.8
0.7
0
10
20
30
40
50
60
Time (min)
Normal
Diabetic
Normal data

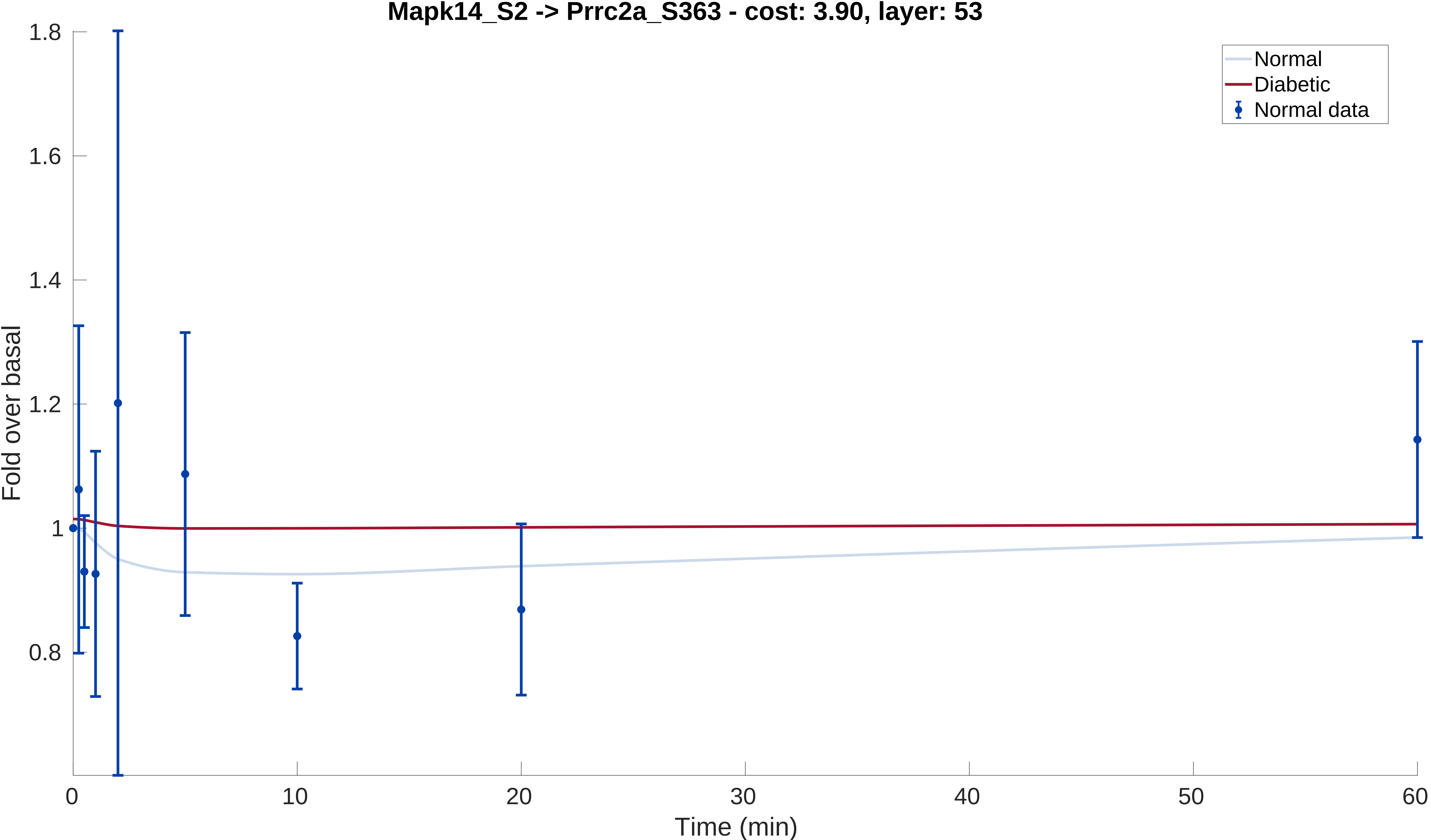

Mapk14_S2 -> Prrc2a_S363 - cost: 3.90, layer: 53
Normal
Diabetic
Normal data
Fold over basal
Time (min)
1.8
1.6
1.4
1.2
1
0.8
0
10
20
30
40
50
60

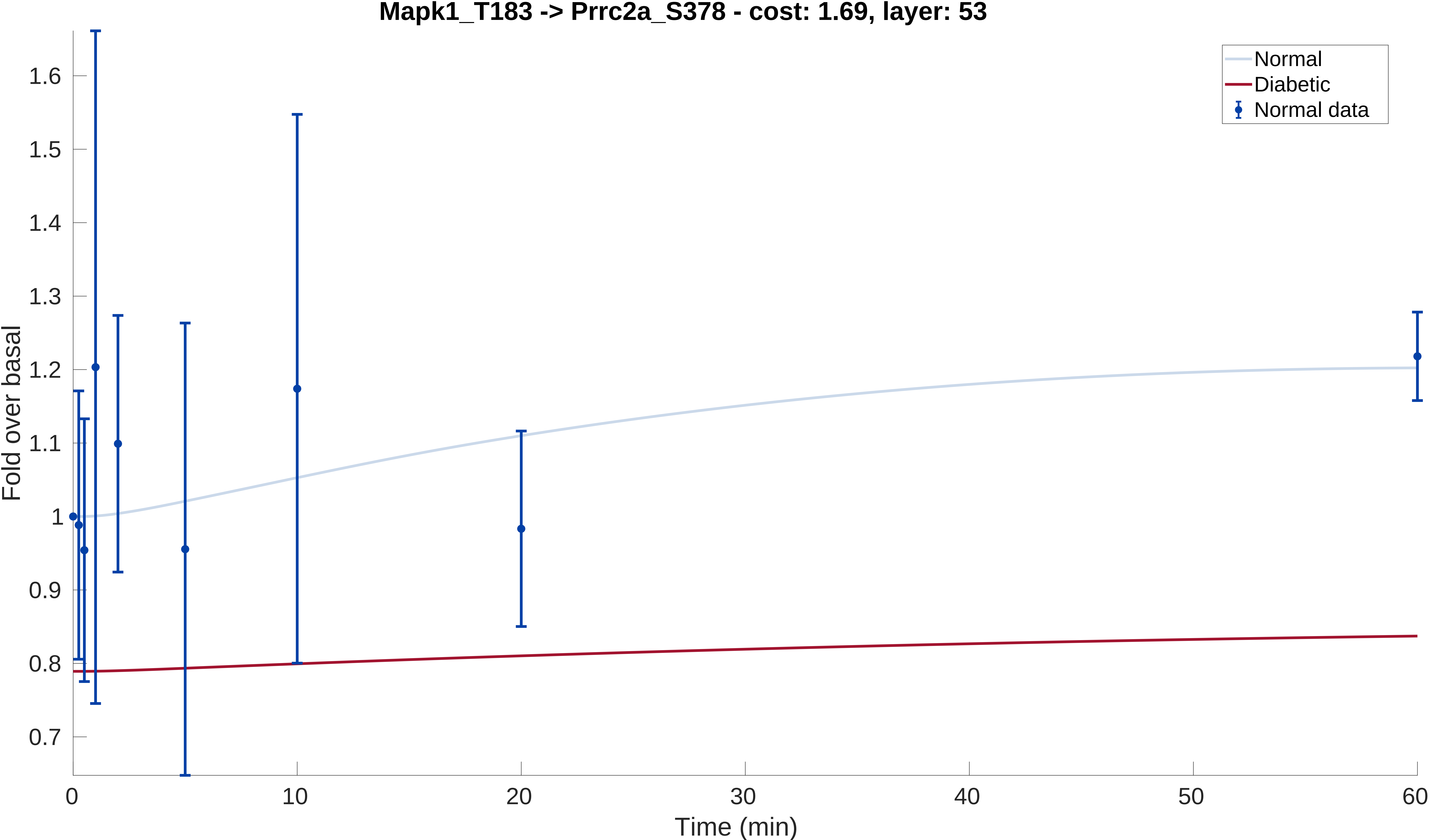

Mapk1_T183 -> Prrc2a_S378 - cost: 1.69, layer: 53
Normal
Diabetic
Normal data
Fold over basal
Time (min)
0.7
0.8
0.9
1
1.1
1.2
1.3
1.4
1.5
1.6
0
10
20
30
40
50
60

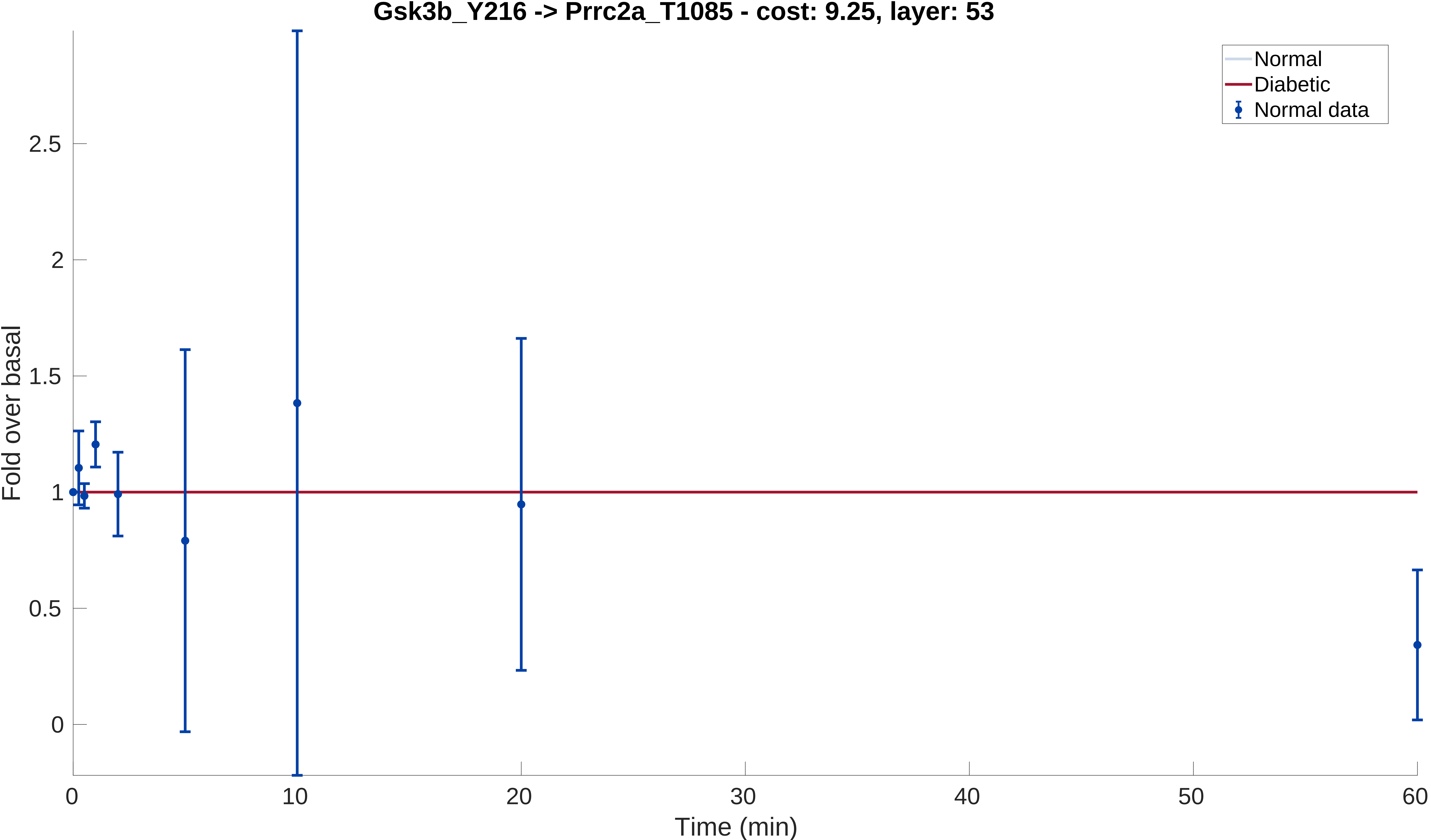

Gsk3b_Y216 -> Prrc2a_T1085 - cost: 9.25, layer: 53
Normal
Diabetic
Normal data
Fold over basal
Time (min)
0
0.5
1
1.5
2
2.5
0
10
20
30
40
50
60

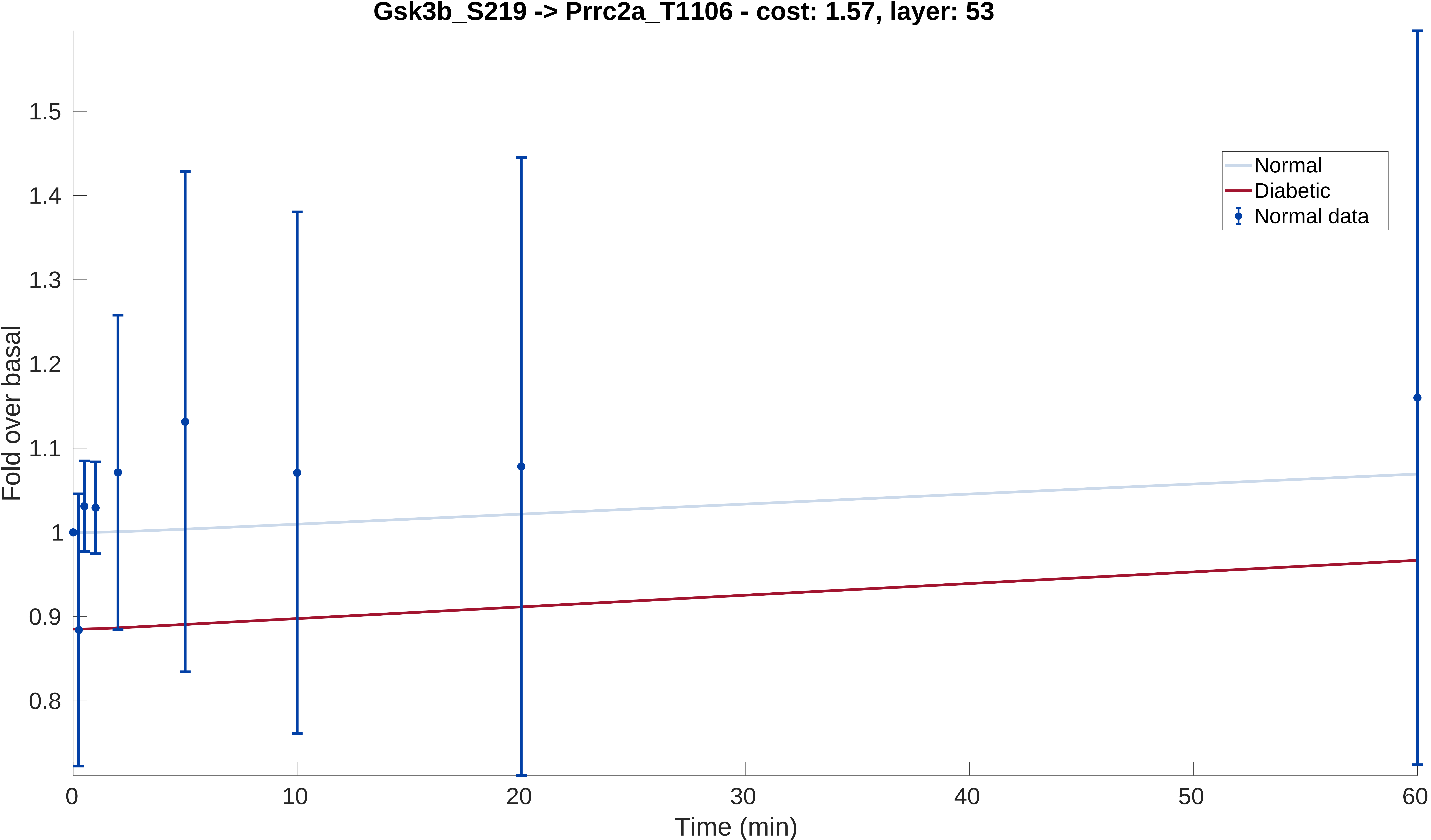

Gsk3b_S219 -> Prrc2a_T1106 - cost: 1.57, layer: 53
Fold over basal
1.5
1.4
1.3
1.2
1.1
1
0.9
0.8
0
10
20
30
40
50
60
Time (min)
Normal
Diabetic
Normal data

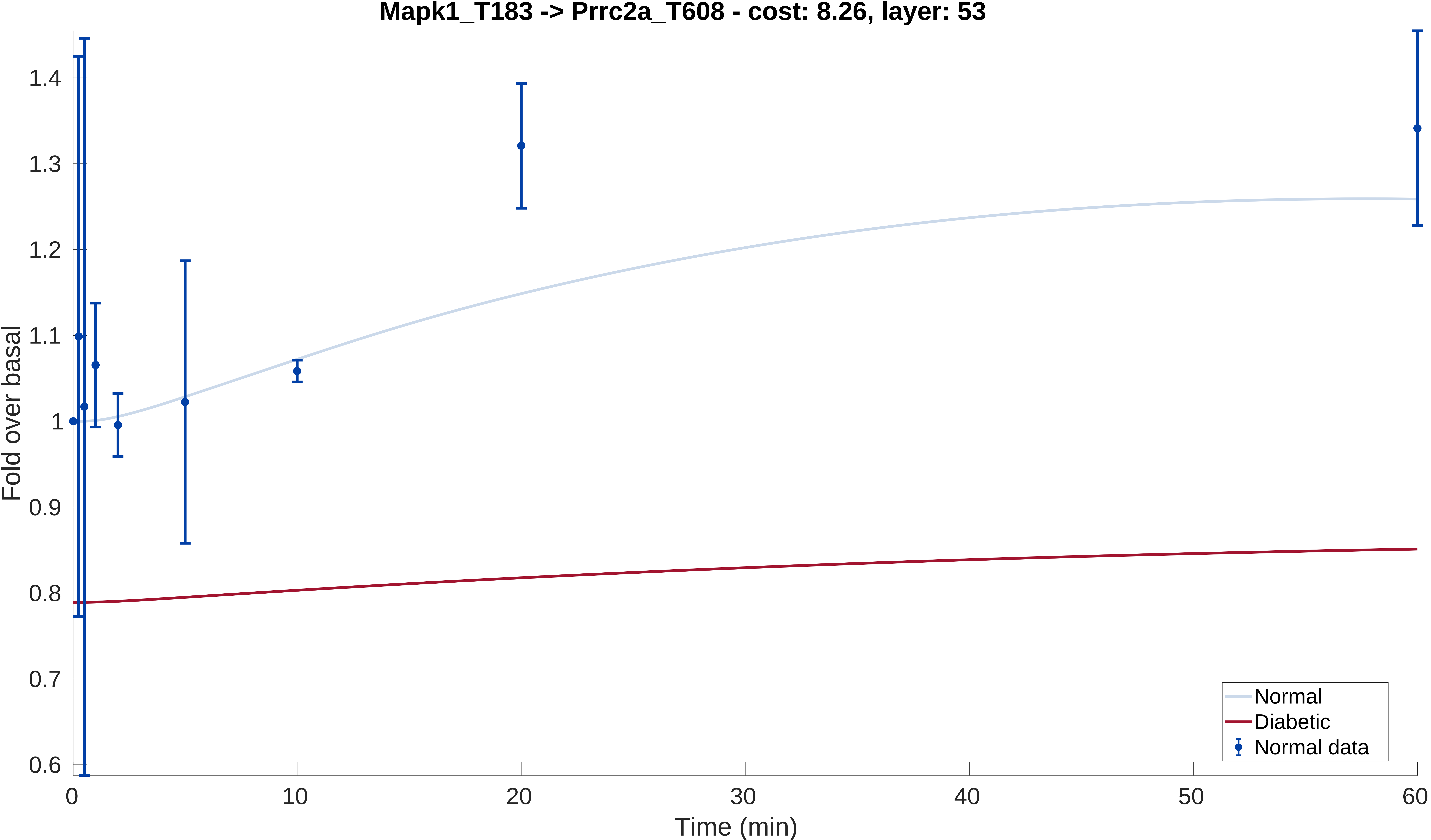
Mapk1_T183 -> Prrc2a_T608 - cost: 8.26, layer: 53
Fold over basal
1.4
1.3
1.2
1.1
1
0.9
0.8
0.7
0.6
0
10
20
30
40
50
60
Time (min)
Normal
Diabetic
Normal data

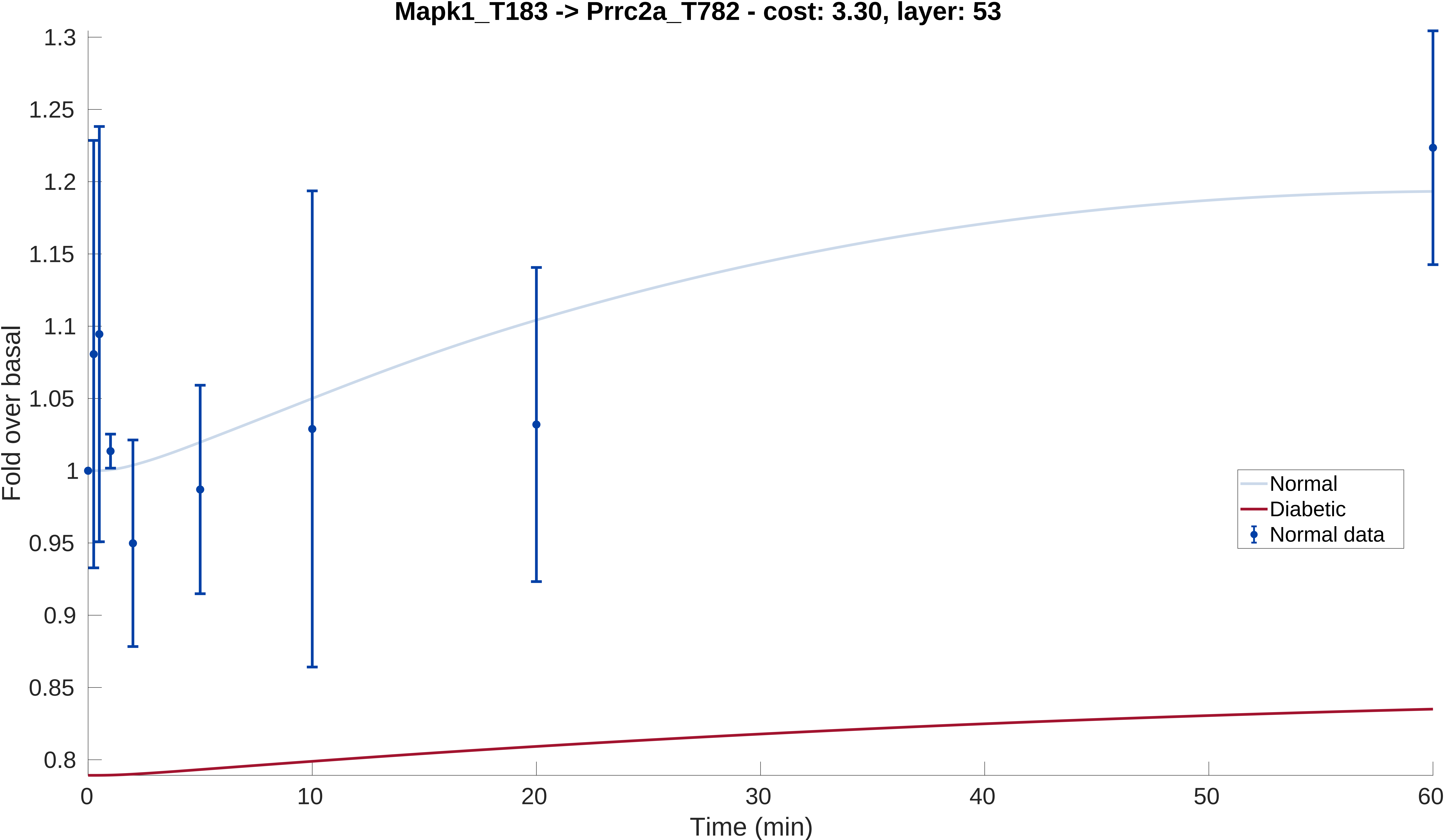
Mapk1_T183 -> Prrc2a_T782 - cost: 3.30, layer: 53
Fold over basal
1.3
1.25
1.2
1.15
1.1
1.05
1
0.95
0.9
0.85
0.8
0
10
20
30
40
50
60
Time (min)
Normal
Diabetic
Normal data

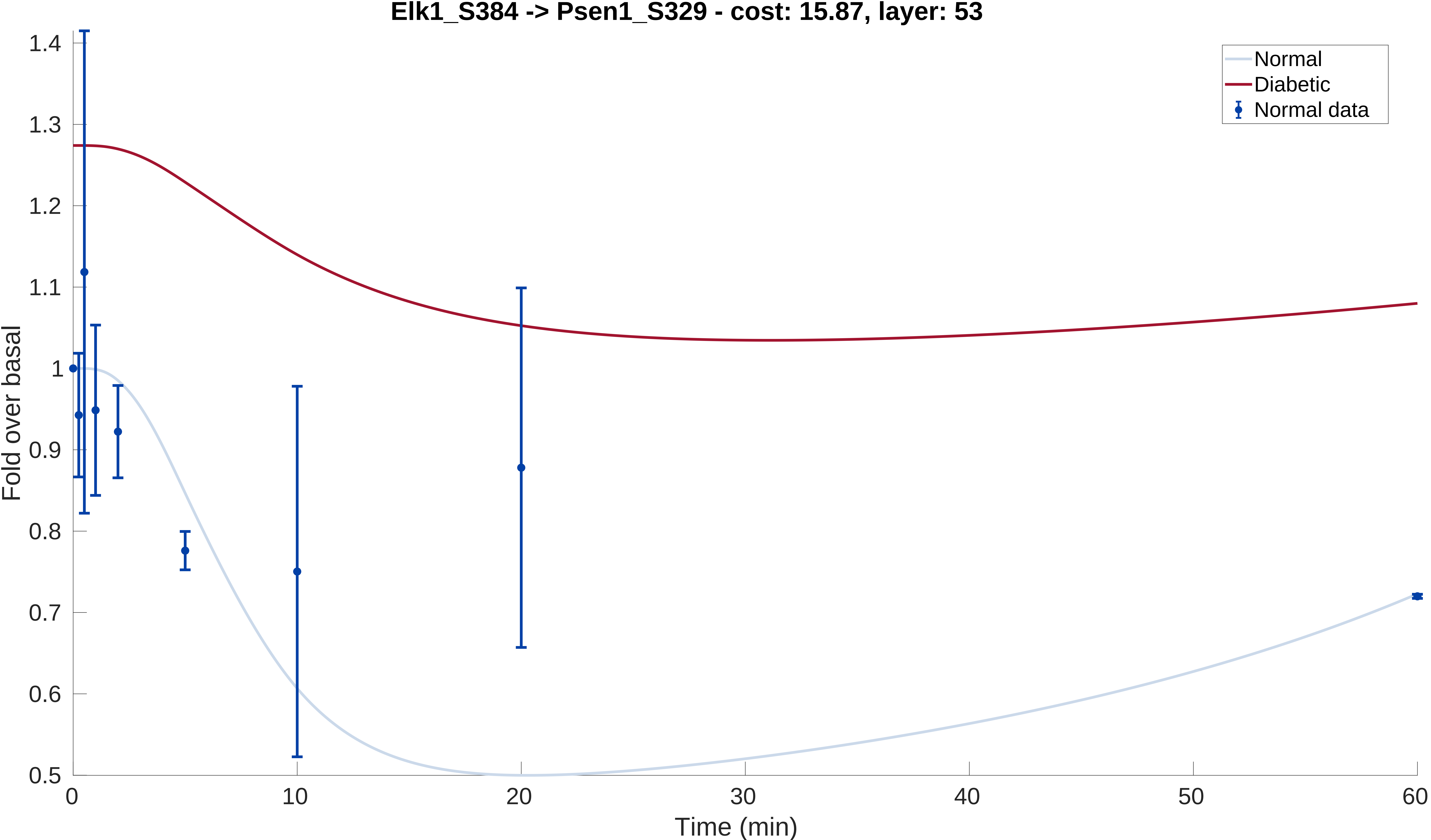

Elk1_S384 -> Psen1_S329 - cost: 15.87, layer: 53
Normal
Diabetic
Normal data
Fold over basal
1.4
1.3
1.2
1.1
1
0.9
0.8
0.7
0.6
0.5
0
10
20
30
40
50
60
Time (min)

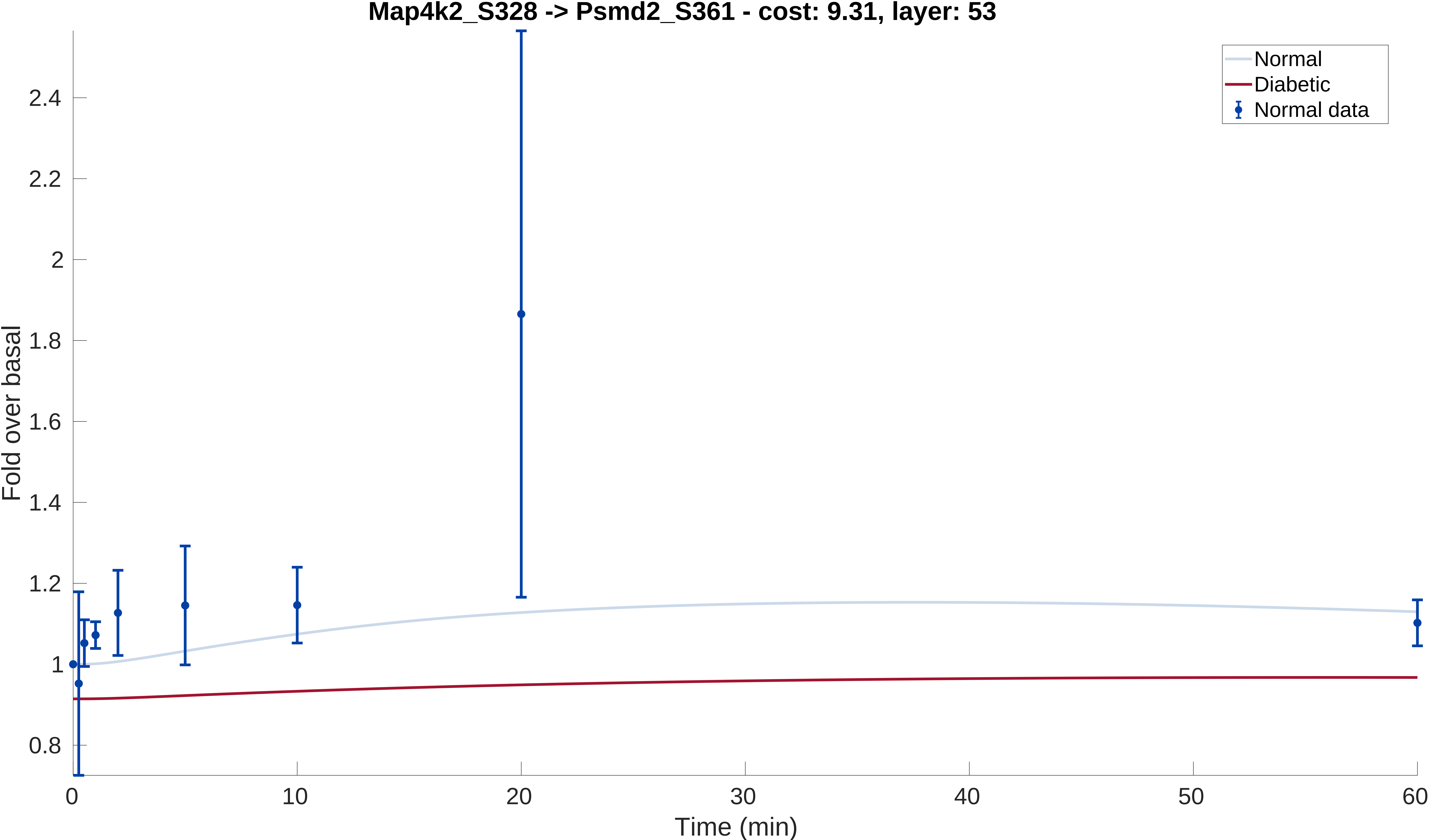

Map4k2_S328 -> Psmd2_S361 - cost: 9.31, layer: 53
Normal
Diabetic
Normal data
Fold over basal
2.4
2.2
2
1.8
1.6
1.4
1.2
1
0.8
0
10
20
30
40
50
60
Time (min)

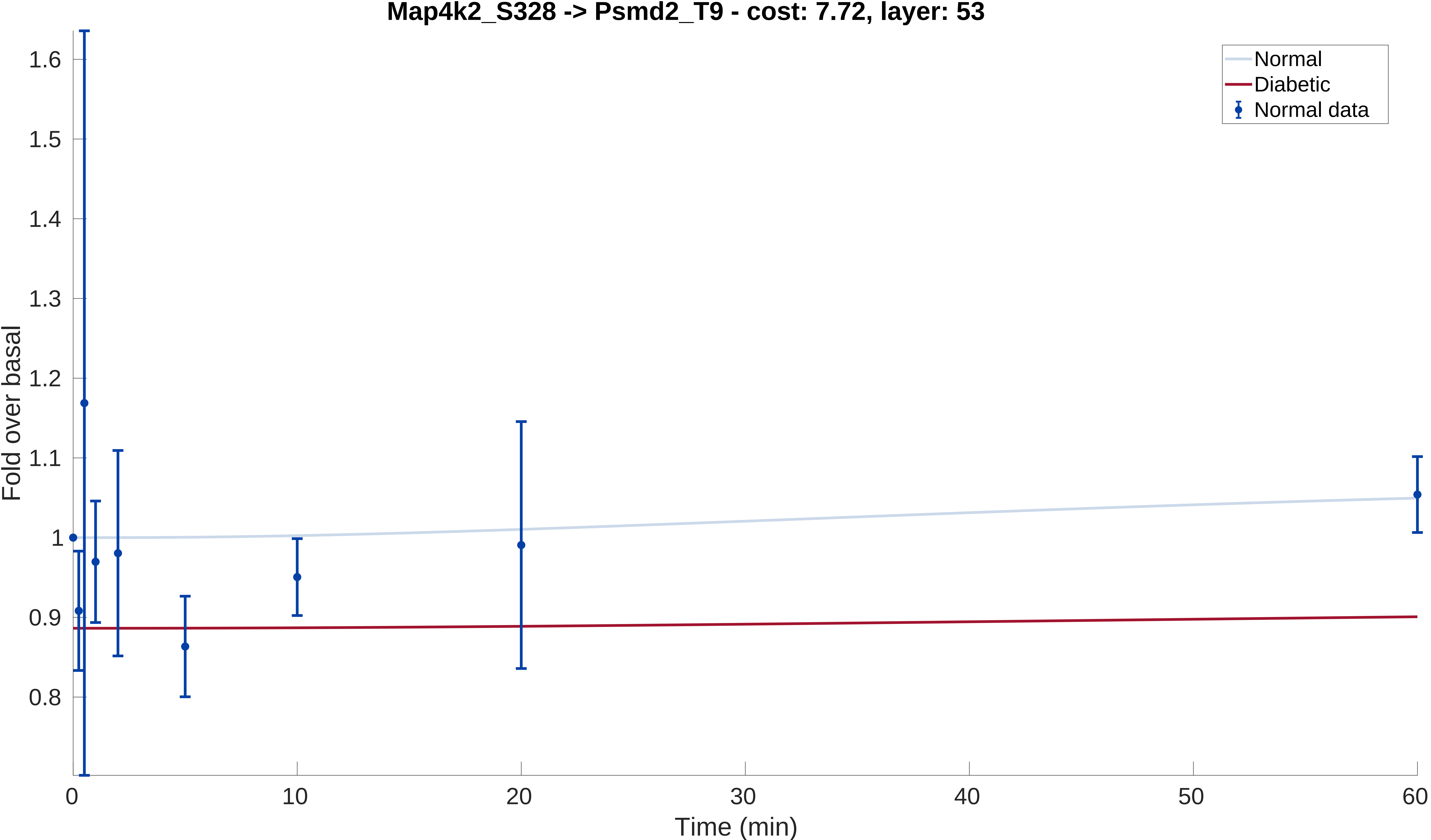

Map4k2_S328 -> Psmd2_T9 - cost: 7.72, layer: 53
Normal
Diabetic
Normal data
Fold over basal
1.6
1.5
1.4
1.3
1.2
1.1
1
0.9
0.8
0
10
20
30
40
50
60
Time (min)

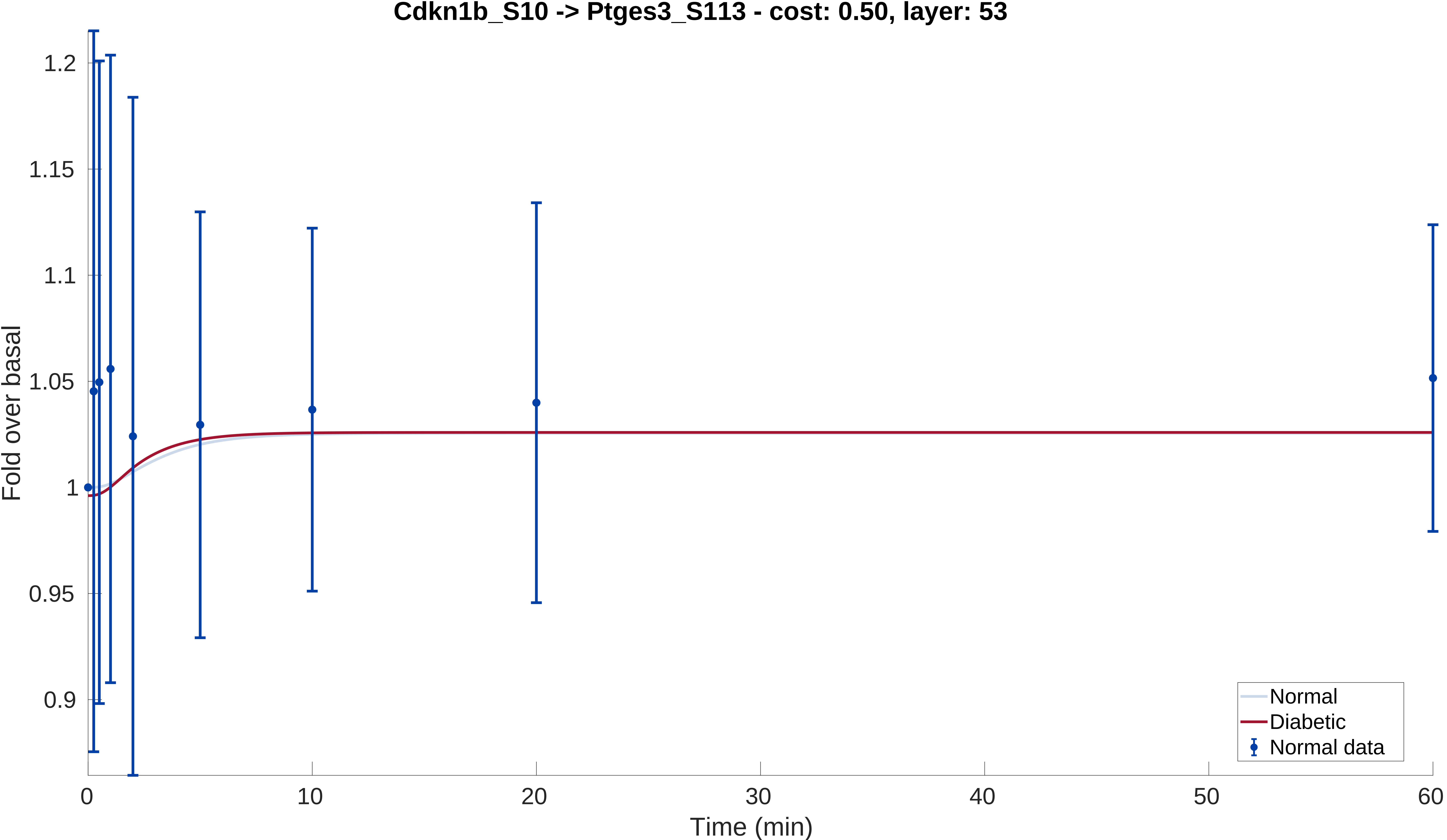

Cdkn1b_S10 -> Ptges3_S113 - cost: 0.50, layer: 53
Fold over basal
1.2
1.15
1.1
1.05
1
0.95
0.9
0
10
20
30
40
50
60
Time (min)
Normal
Diabetic
Normal data

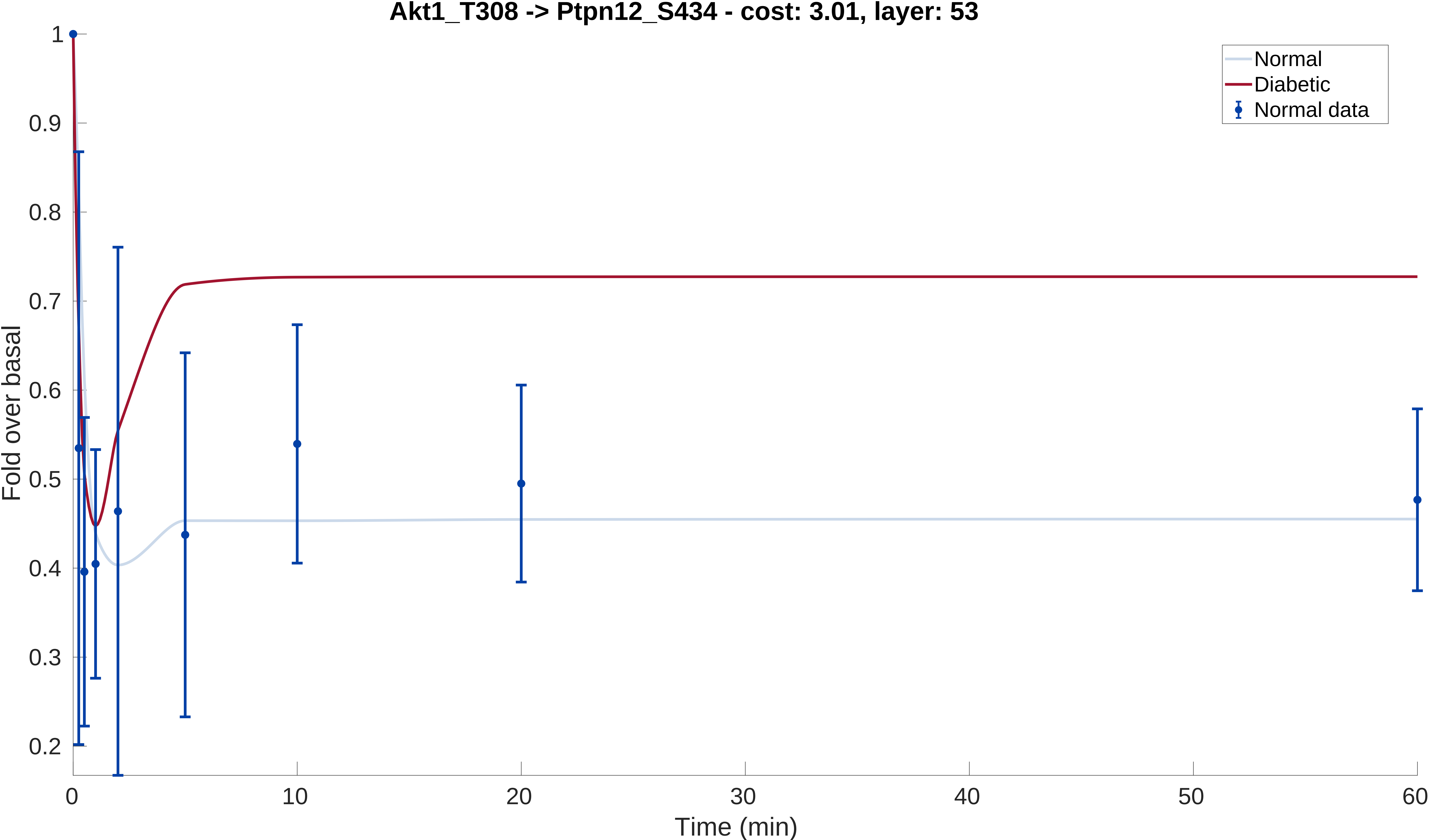
Akt1_T308 -> Ptpn12_S434 - cost: 3.01, layer: 53
Normal
Diabetic
Normal data
Fold over basal
Time (min)
1
0.9
0.8
0.7
0.6
0.5
0.4
0.3
0.2
0
10
20
30
40
50
60

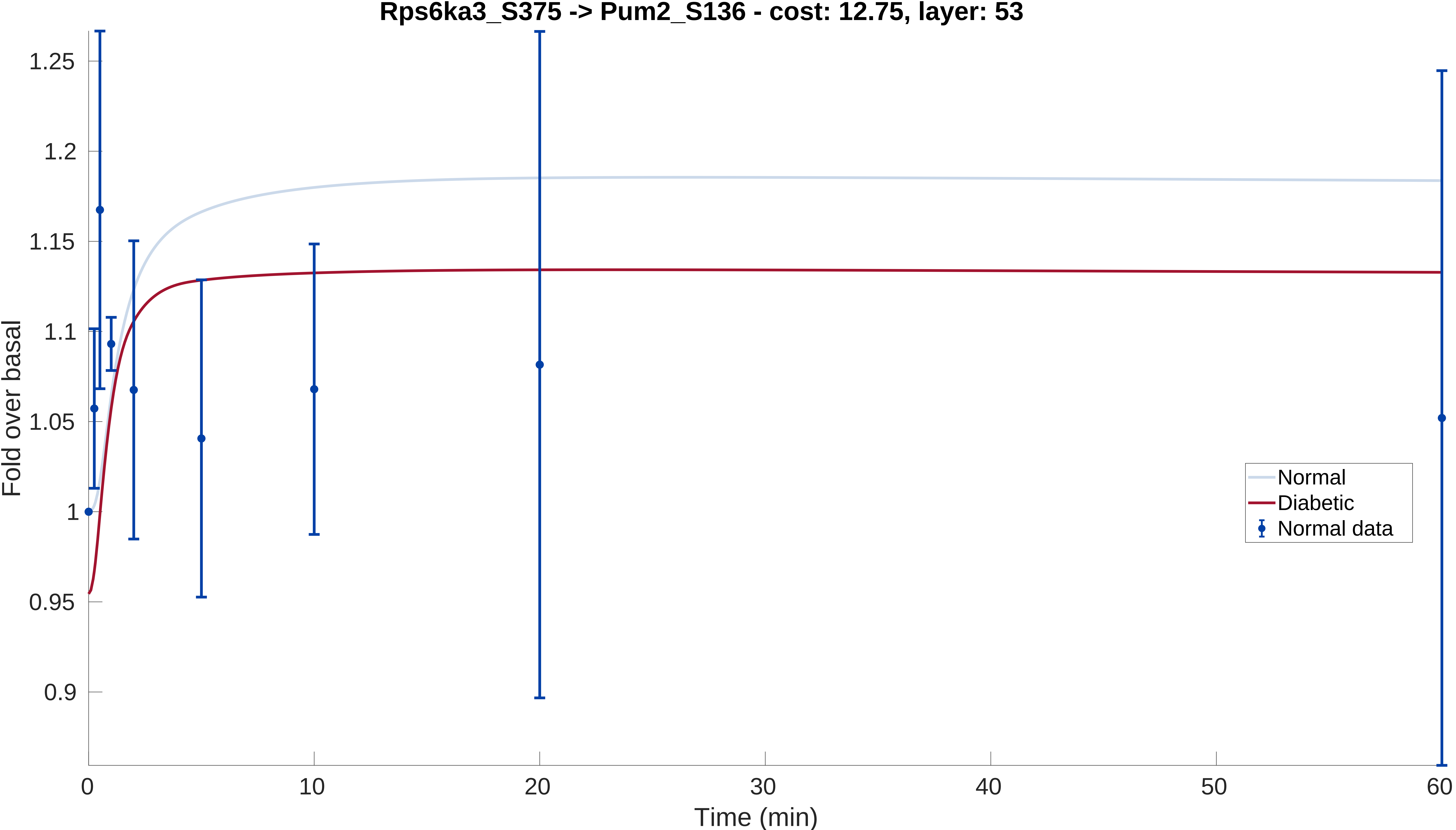
Rps6ka3_S375 -> Pum2_S136 - cost: 12.75, layer: 53
Fold over basal
1.25
1.2
1.15
1.1
1.05
1
0.95
0.9
0
10
20
30
40
50
60
Time (min)
Normal
Diabetic
Normal data

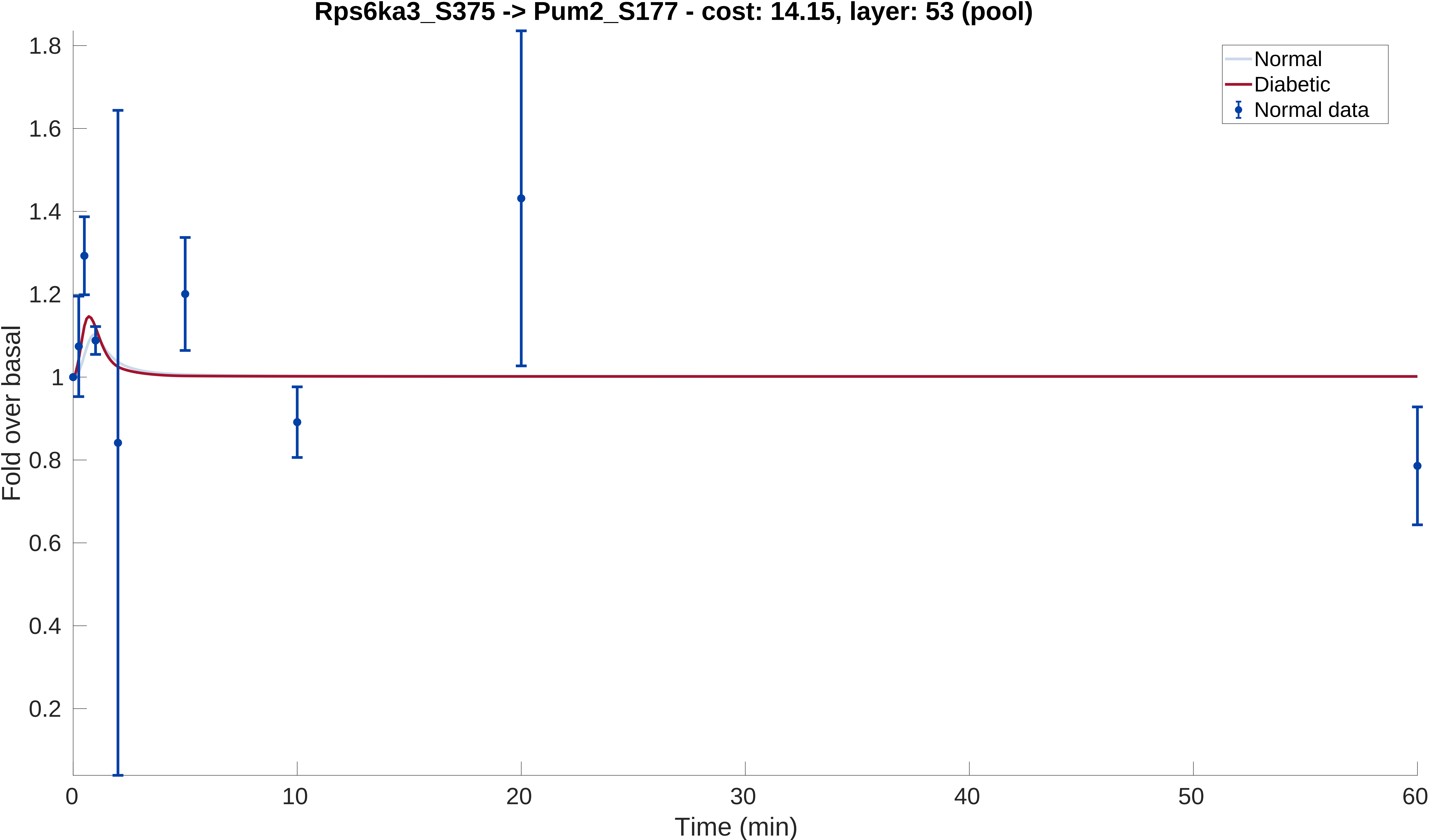

Rps6ka3_S375 -> Pum2_S177 - cost: 14.15, layer: 53 (pool)
Normal
Diabetic
Normal data
Fold over basal
1.8
1.6
1.4
1.2
1
0.8
0.6
0.4
0.2
0
10
20
30
40
50
60
Time (min)

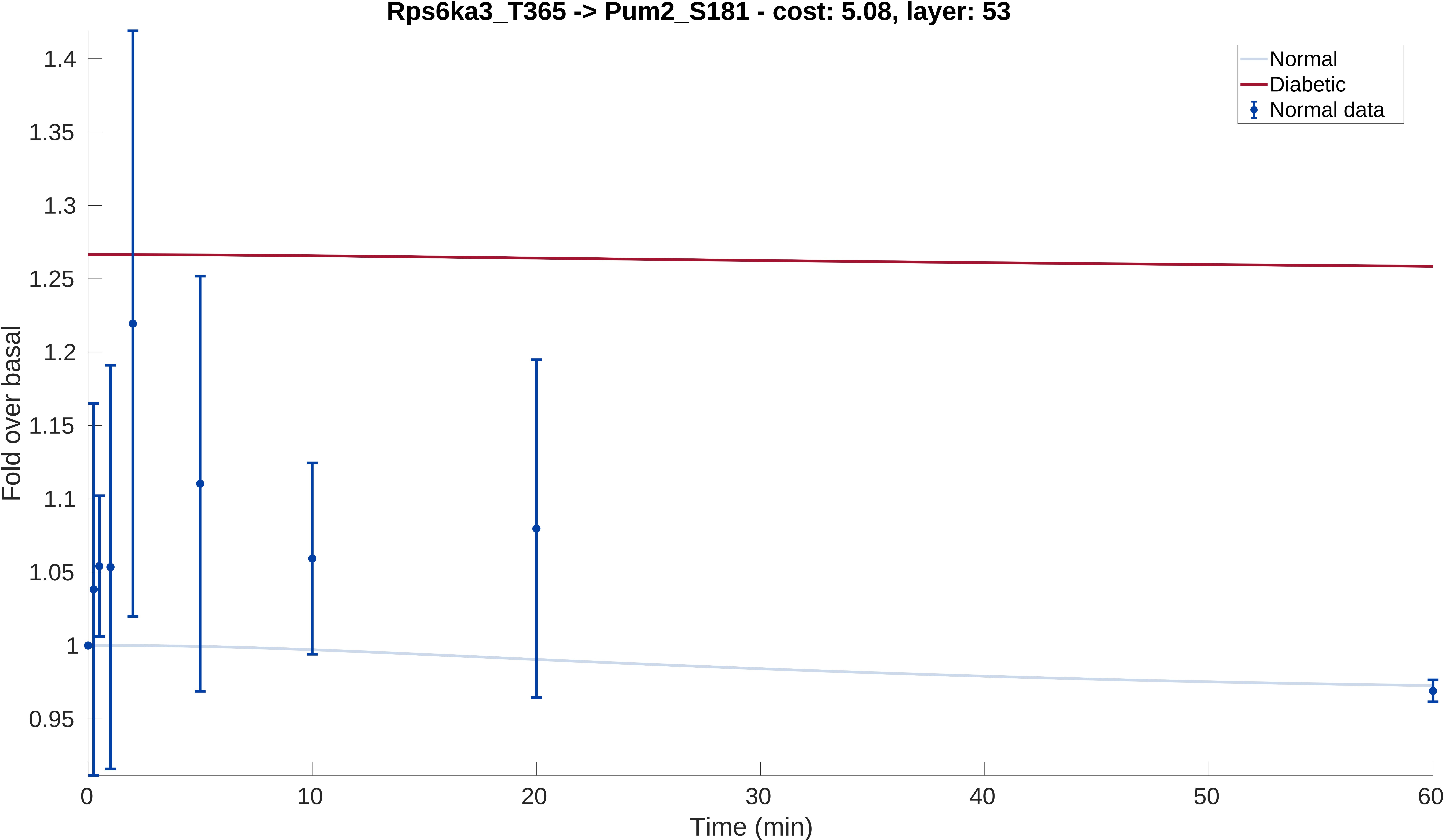
Rps6ka3_T365 -> Pum2_S181 - cost: 5.08, layer: 53
Normal
Diabetic
Normal data
Fold over basal
1.4
1.35
1.3
1.25
1.2
1.15
1.1
1.05
1
0.95
0
10
20
30
40
50
60
Time (min)

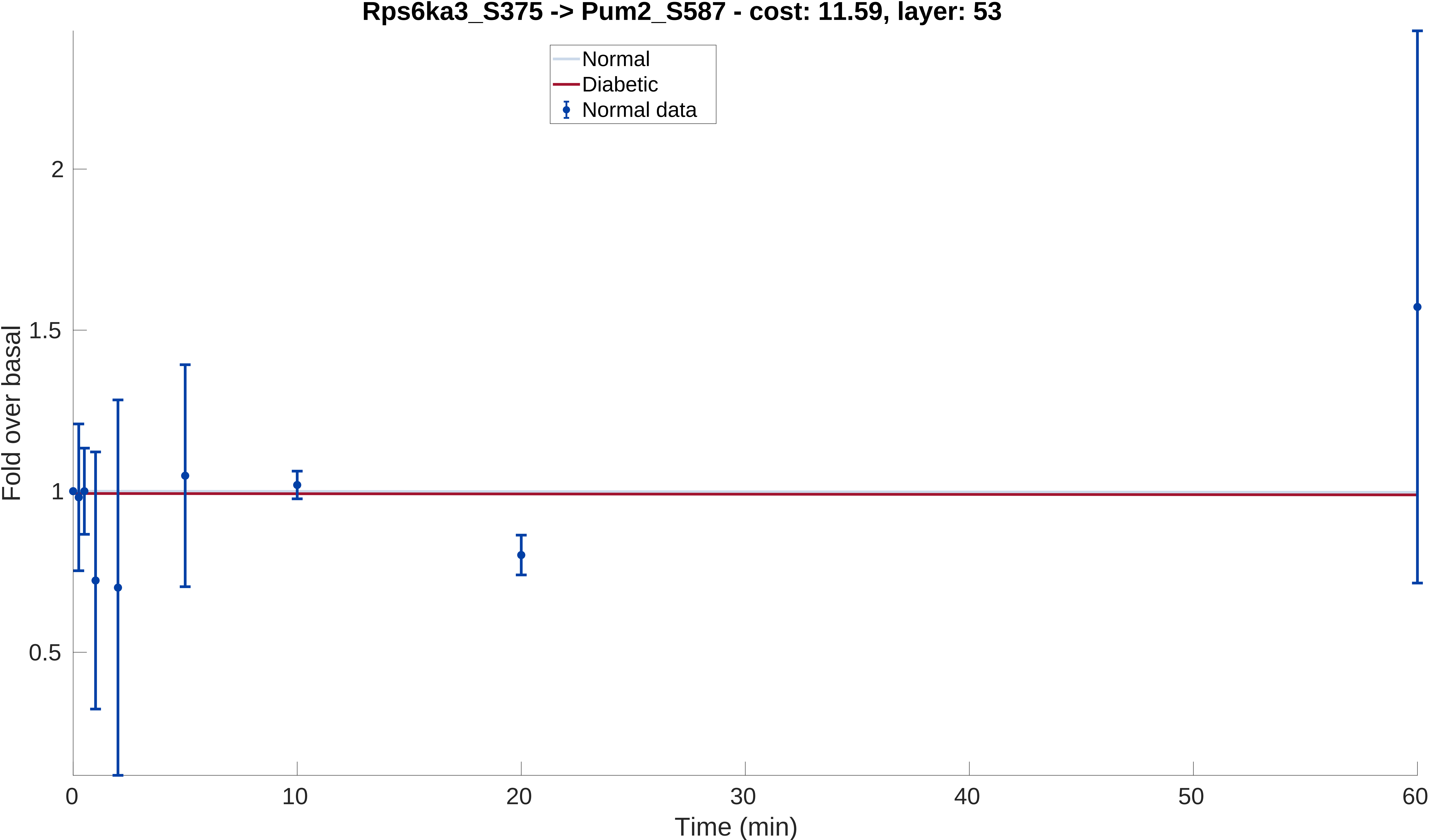

Rps6ka3_S375 -> Pum2_S587 - cost: 11.59, layer: 53
Normal
Diabetic
Normal data
Fold over basal
2
1.5
1
0.5
0
10
20
30
40
50
60
Time (min)

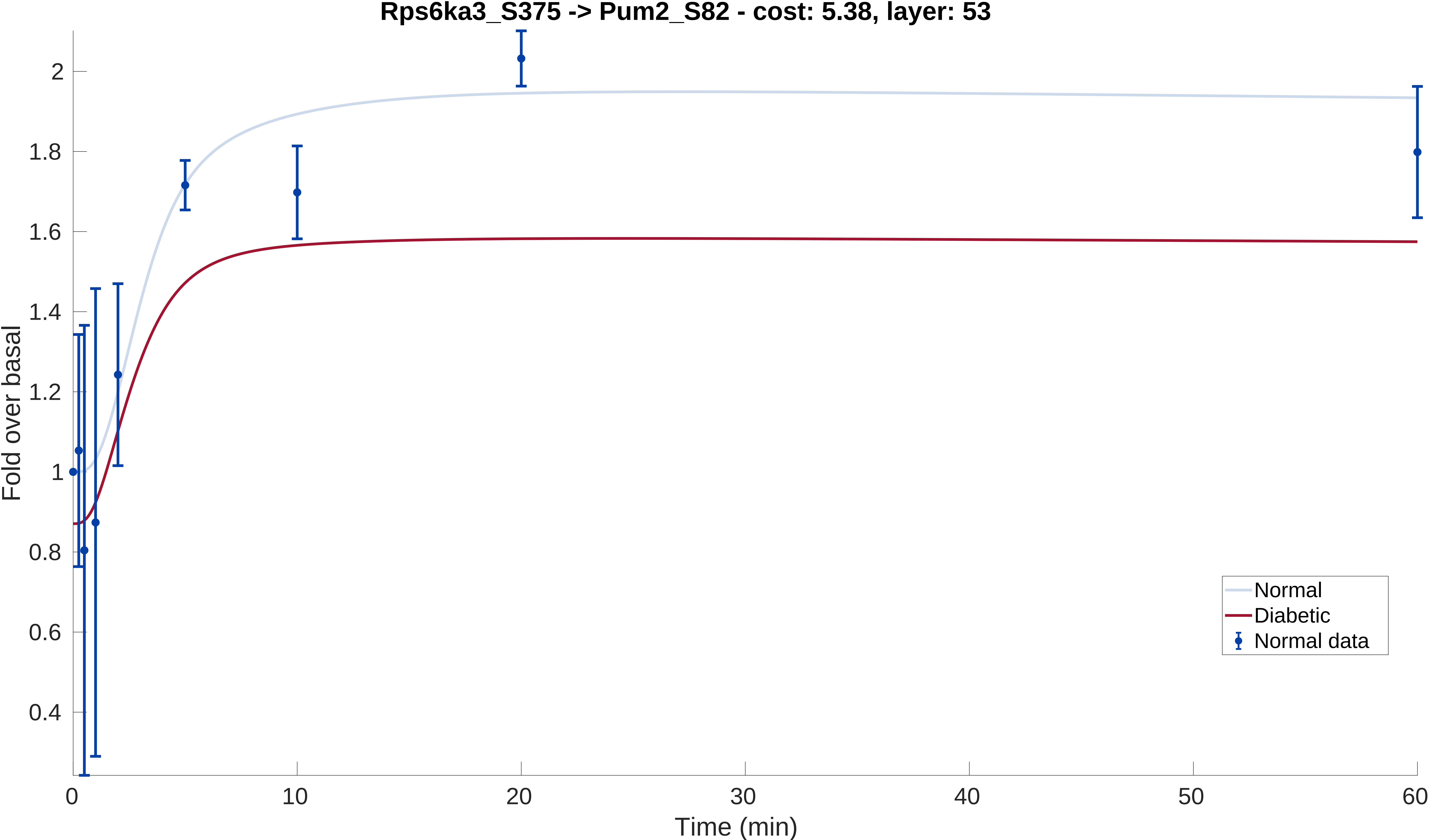

Rps6ka3_S375 -> Pum2_S82 - cost: 5.38, layer: 53
Fold over basal
2
1.8
1.6
1.4
1.2
1
0.8
0.6
0.4
0
10
20
30
40
50
60
Time (min)
Normal
Diabetic
Normal data

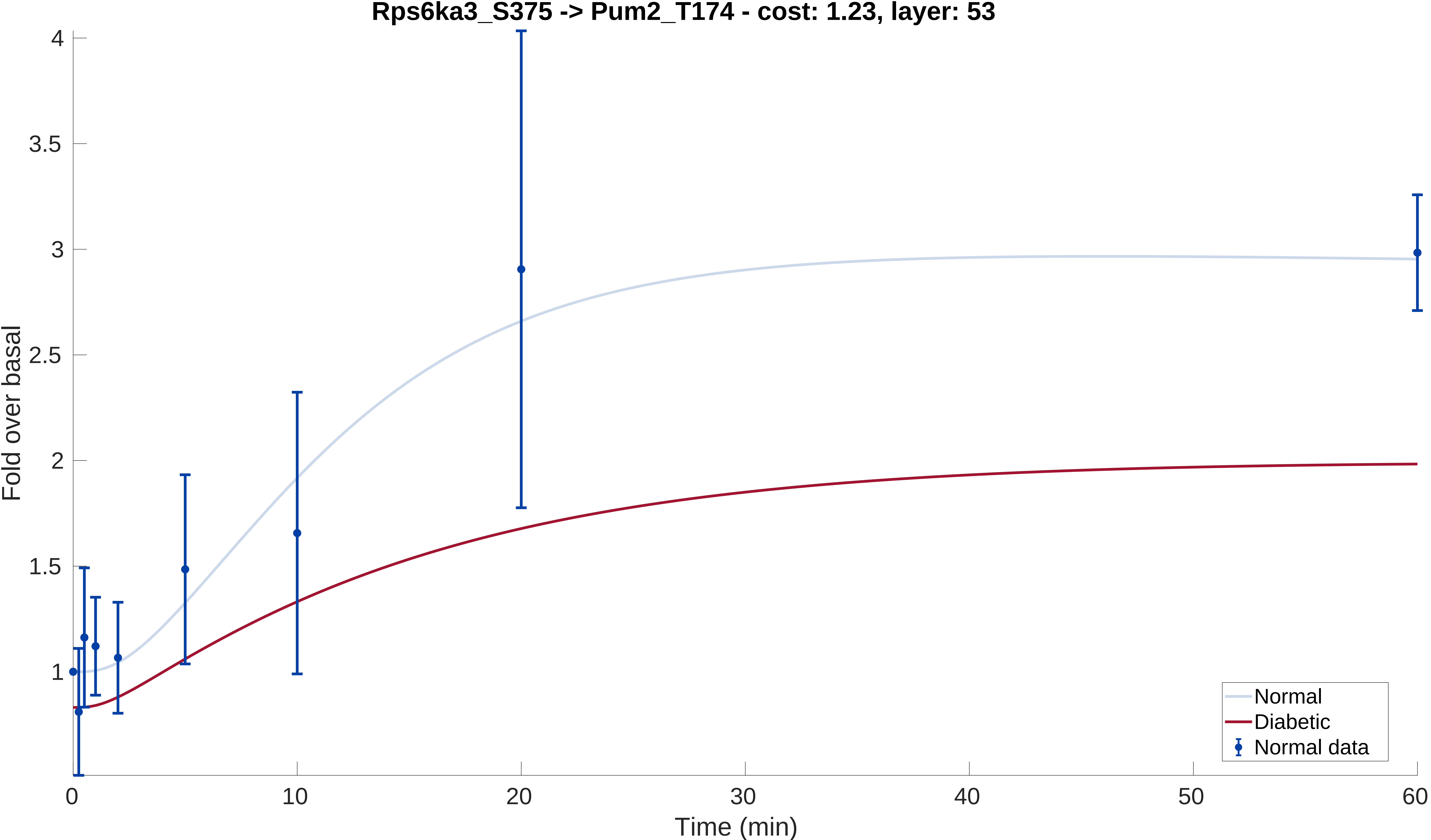
Rps6ka3_S375 -> Pum2_T174 - cost: 1.23, layer: 53
Fold over basal
4
3.5
3
2.5
2
1.5
1
0
10
20
30
40
50
60
Time (min)
Normal
Diabetic
Normal data

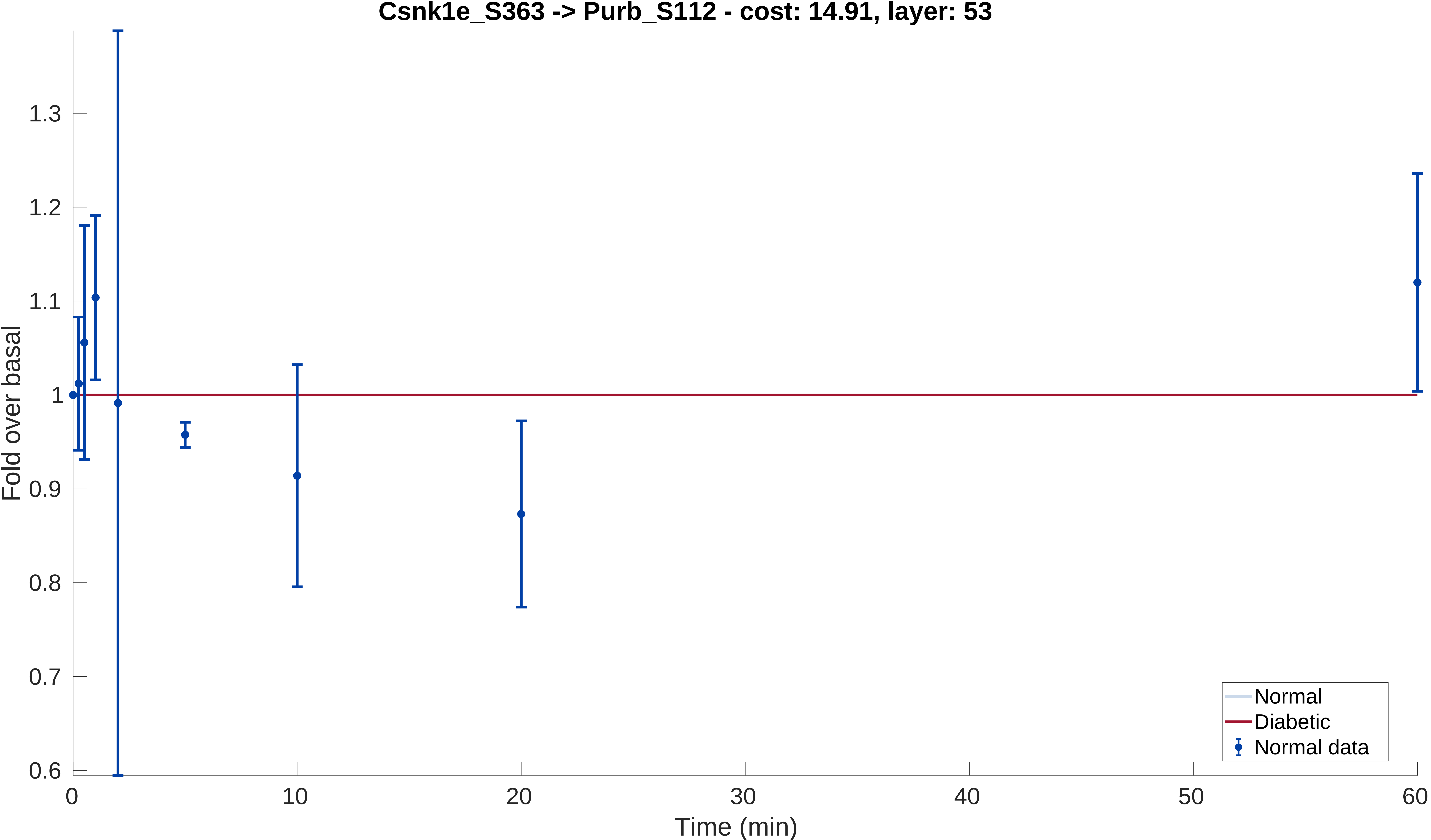
Csnk1e_S363 -> Purb_S112 - cost: 14.91, layer: 53
Fold over basal
1.3
1.2
1.1
1
0.9
0.8
0.7
0.6
0
10
20
30
40
50
60
Time (min)
Normal
Diabetic
Normal data

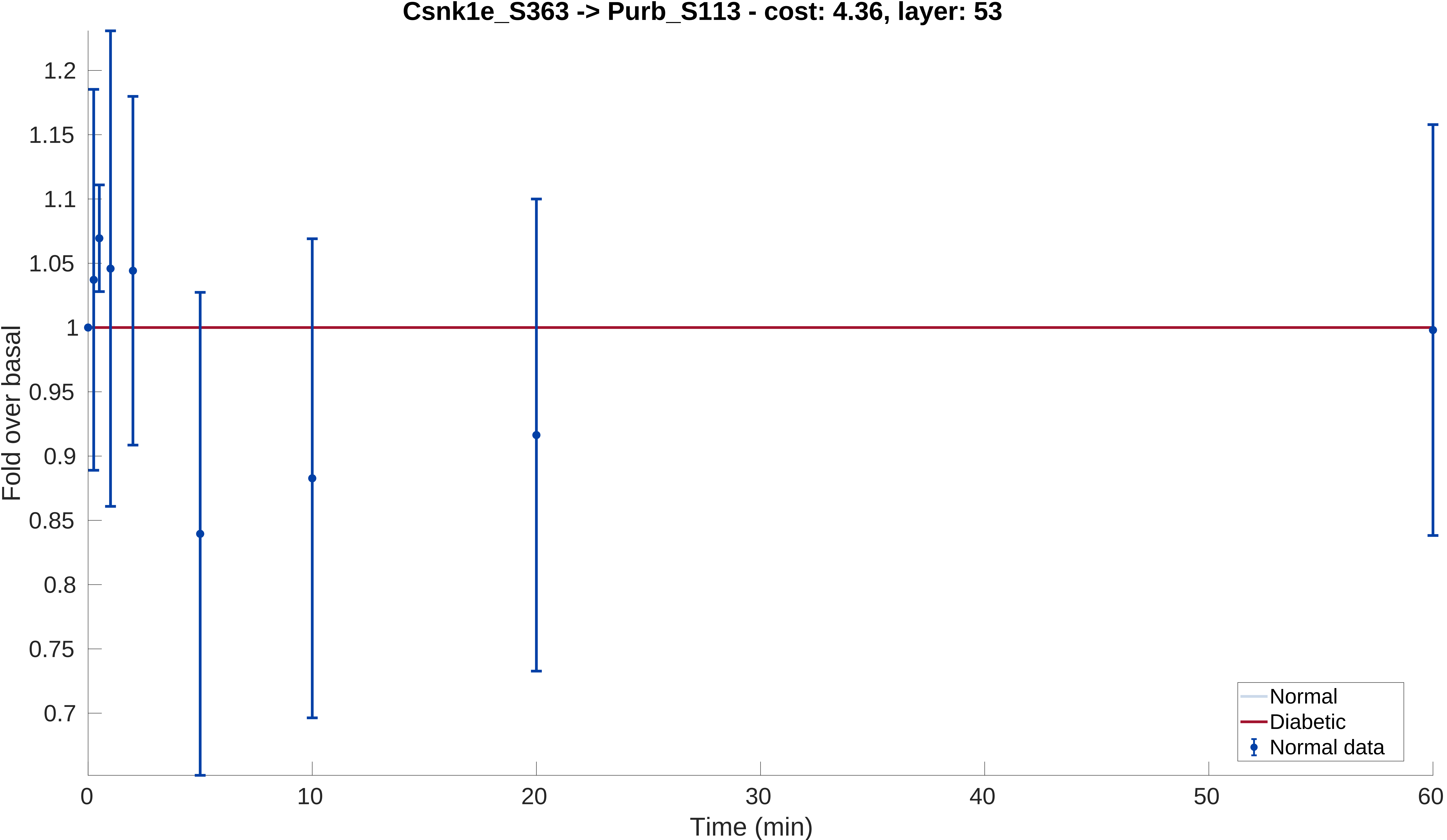

Csnk1e_S363 -> Purb_S113 - cost: 4.36, layer: 53
Fold over basal
1.2
1.15
1.1
1.05
1
0.95
0.9
0.85
0.8
0.75
0.7
0
10
20
30
40
50
60
Time (min)
Normal
Diabetic
Normal data

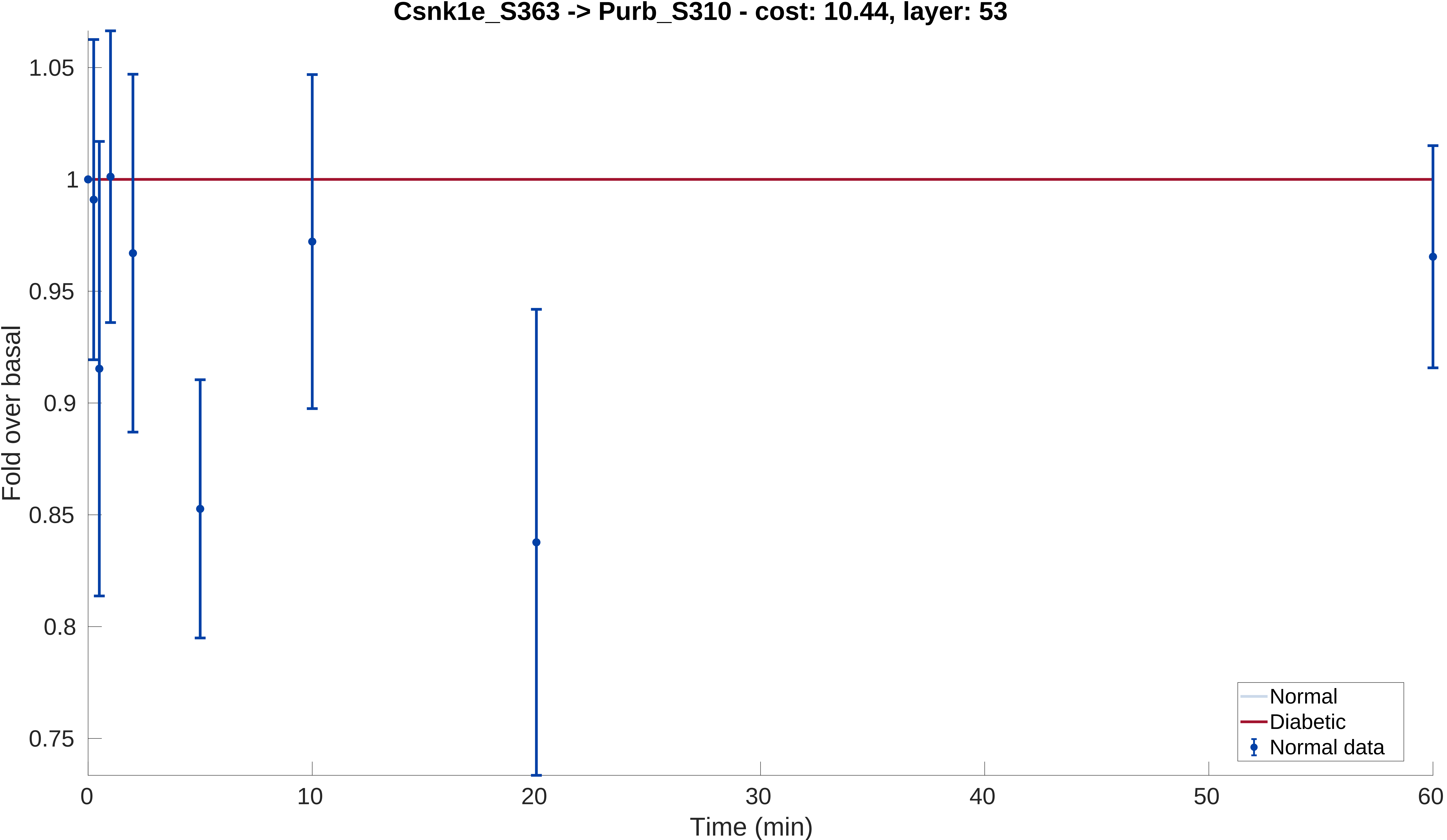

Csnk1e_S363 -> Purb_S310 - cost: 10.44, layer: 53
Fold over basal
1.05
1
0.95
0.9
0.85
0.8
0.75
0
10
20
30
40
50
60
Time (min)
Normal
Diabetic
Normal data

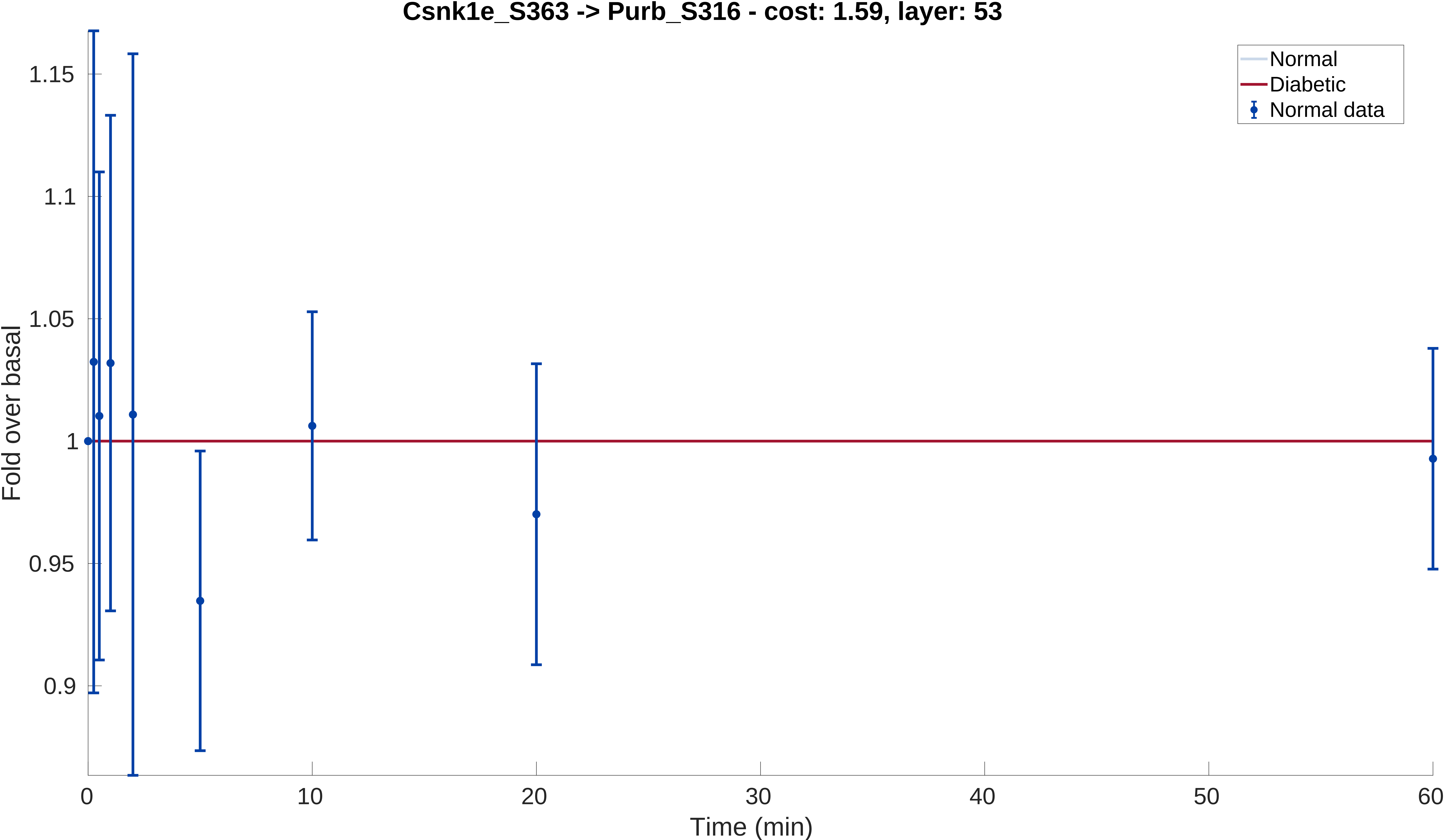

Csnk1e_S363 -> Purb_S316 - cost: 1.59, layer: 53
Normal
Diabetic
Normal data
Fold over basal
1.15
1.1
1.05
1
0.95
0.9
0
10
20
30
40
50
60
Time (min)

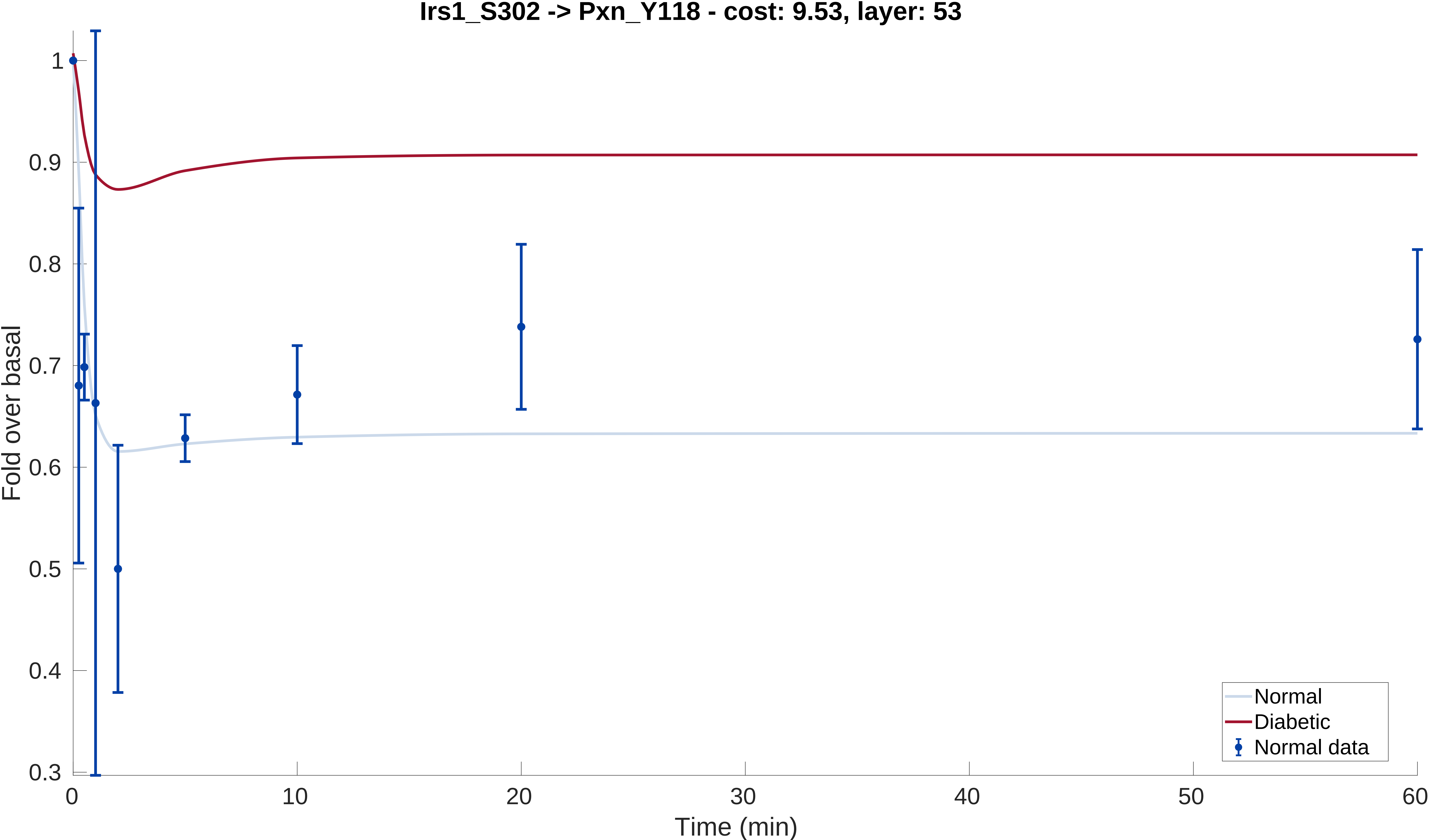

Irs1_S302 -> Pxn_Y118 - cost: 9.53, layer: 53
Fold over basal
1
0.9
0.8
0.7
0.6
0.5
0.4
0.3
0
10
20
30
40
50
60
Time (min)
Normal
Diabetic
Normal data

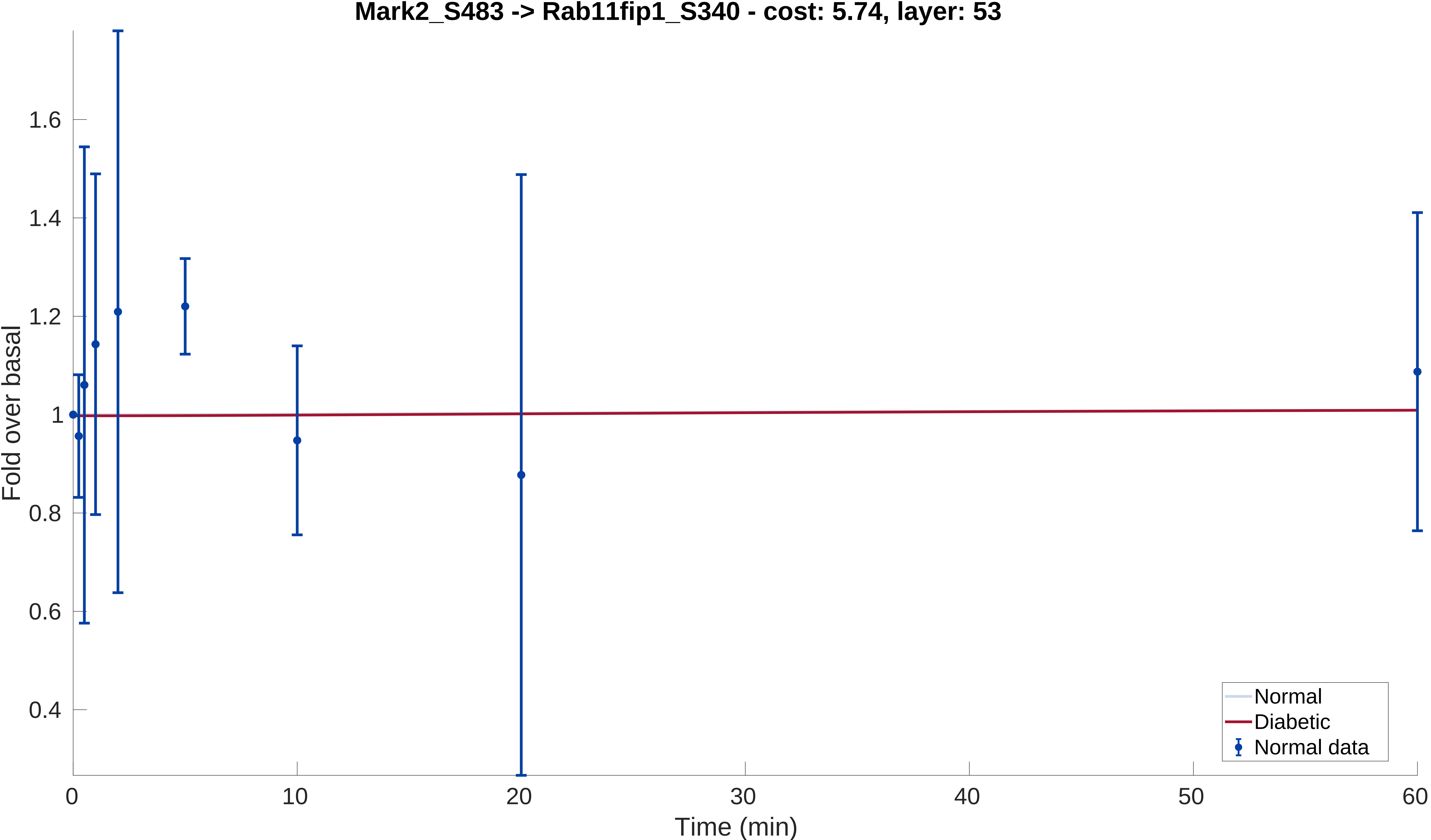

Mark2_S483 -> Rab11fip1_S340 - cost: 5.74, layer: 53
Fold over basal
1.6
1.4
1.2
1
0.8
0.6
0.4
0
10
20
30
40
50
60
Time (min)
Normal
Diabetic
Normal data

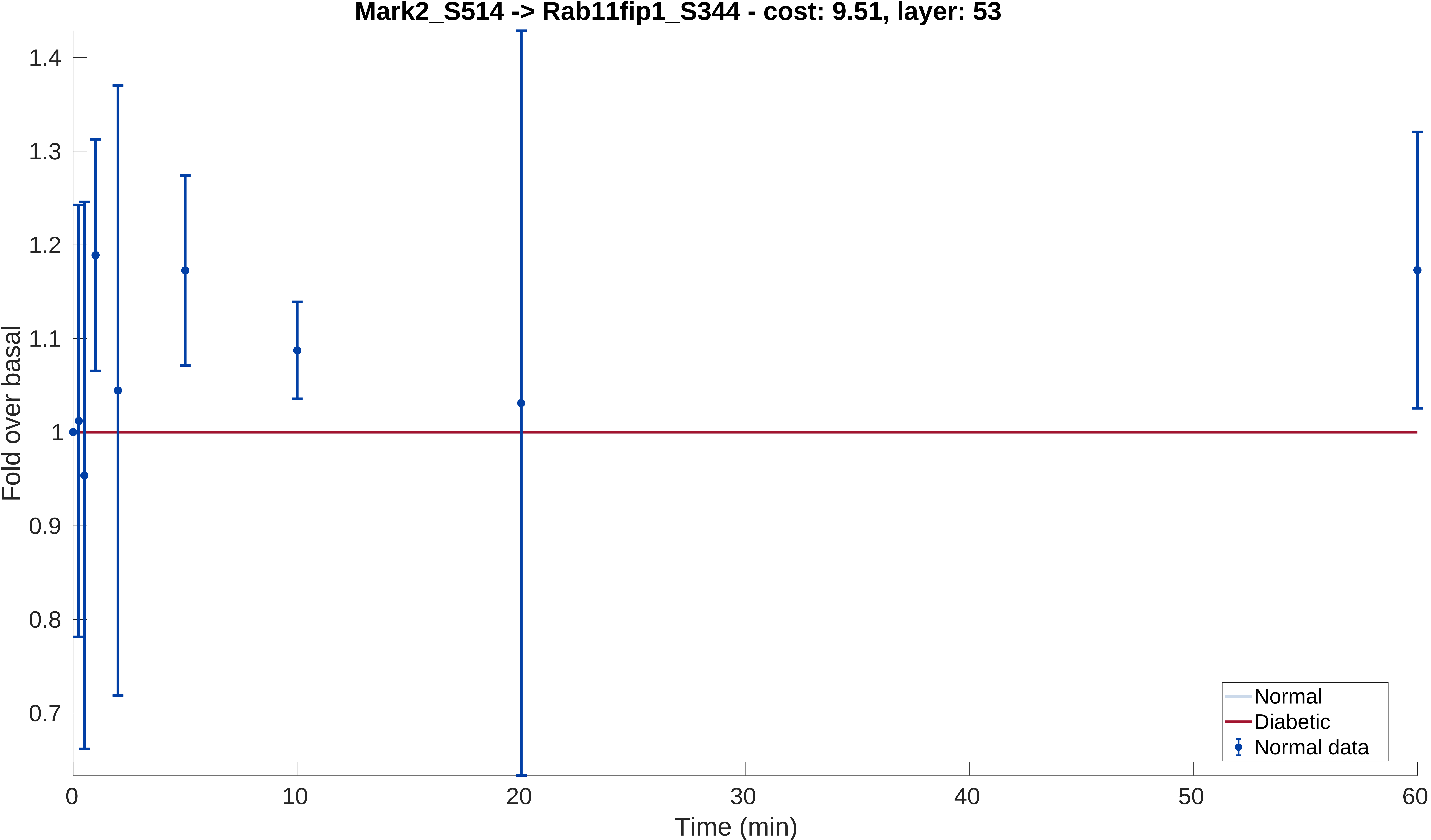

Mark2_S514 -> Rab11fip1_S344 - cost: 9.51, layer: 53
1.4
1.3
1.2
1.1
1
0.9
0.8
0.7
Fold over basal
0
10
20
30
40
50
60
Time (min)
Normal
Diabetic
Normal data

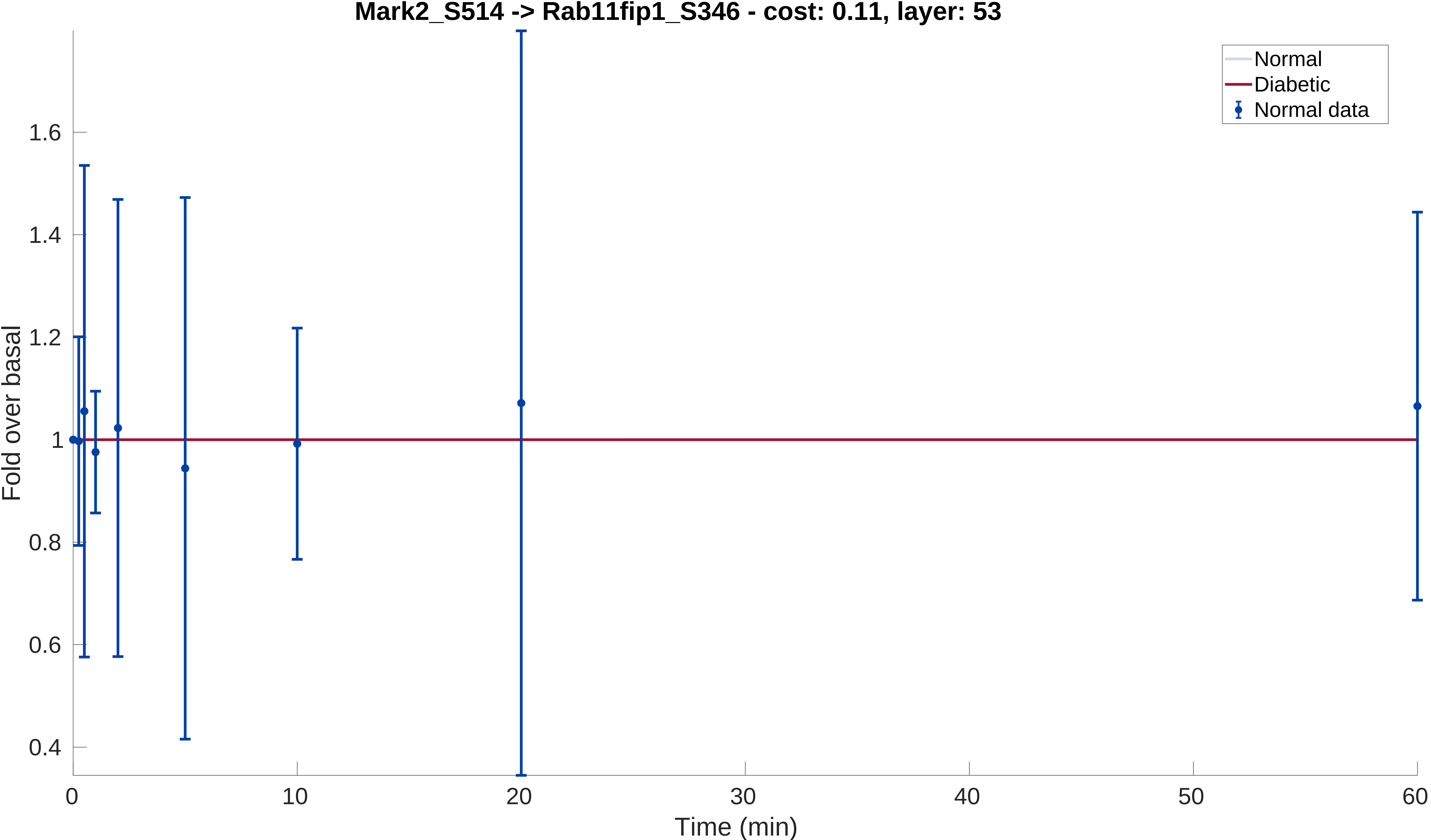

Mark2_S514 -> Rab11fip1_S346 - cost: 0.11, layer: 53
Normal
Diabetic
Normal data
Fold over basal
1.6
1.4
1.2
1
0.8
0.6
0.4
0
10
20
30
40
50
60
Time (min)

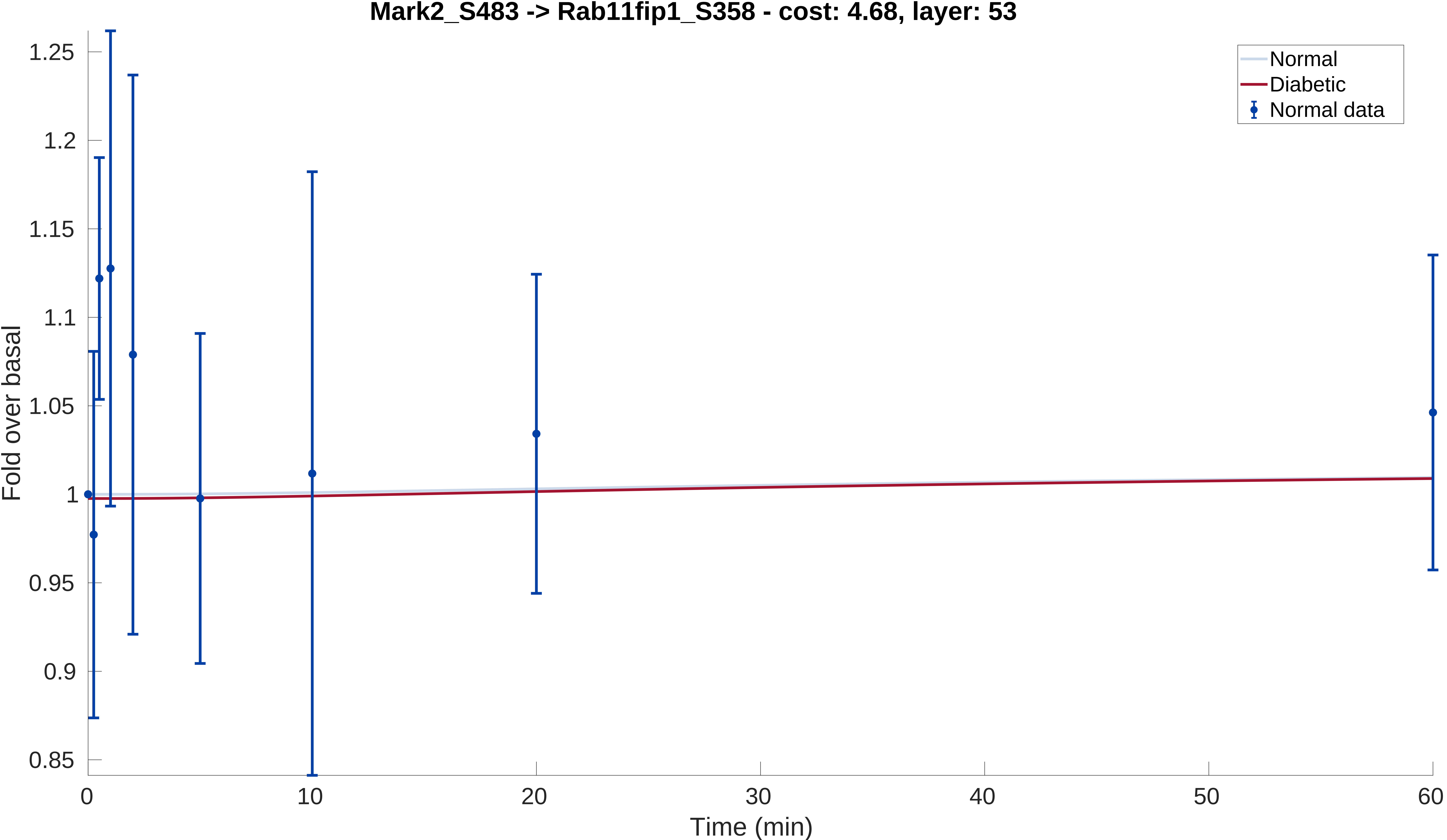

Mark2_S483 -> Rab11fip1_S358 - cost: 4.68, layer: 53
Normal
Diabetic
Normal data
Fold over basal
Time (min)
1.25
1.2
1.15
1.1
1.05
1
0.95
0.9
0.85
0
10
20
30
40
50
60

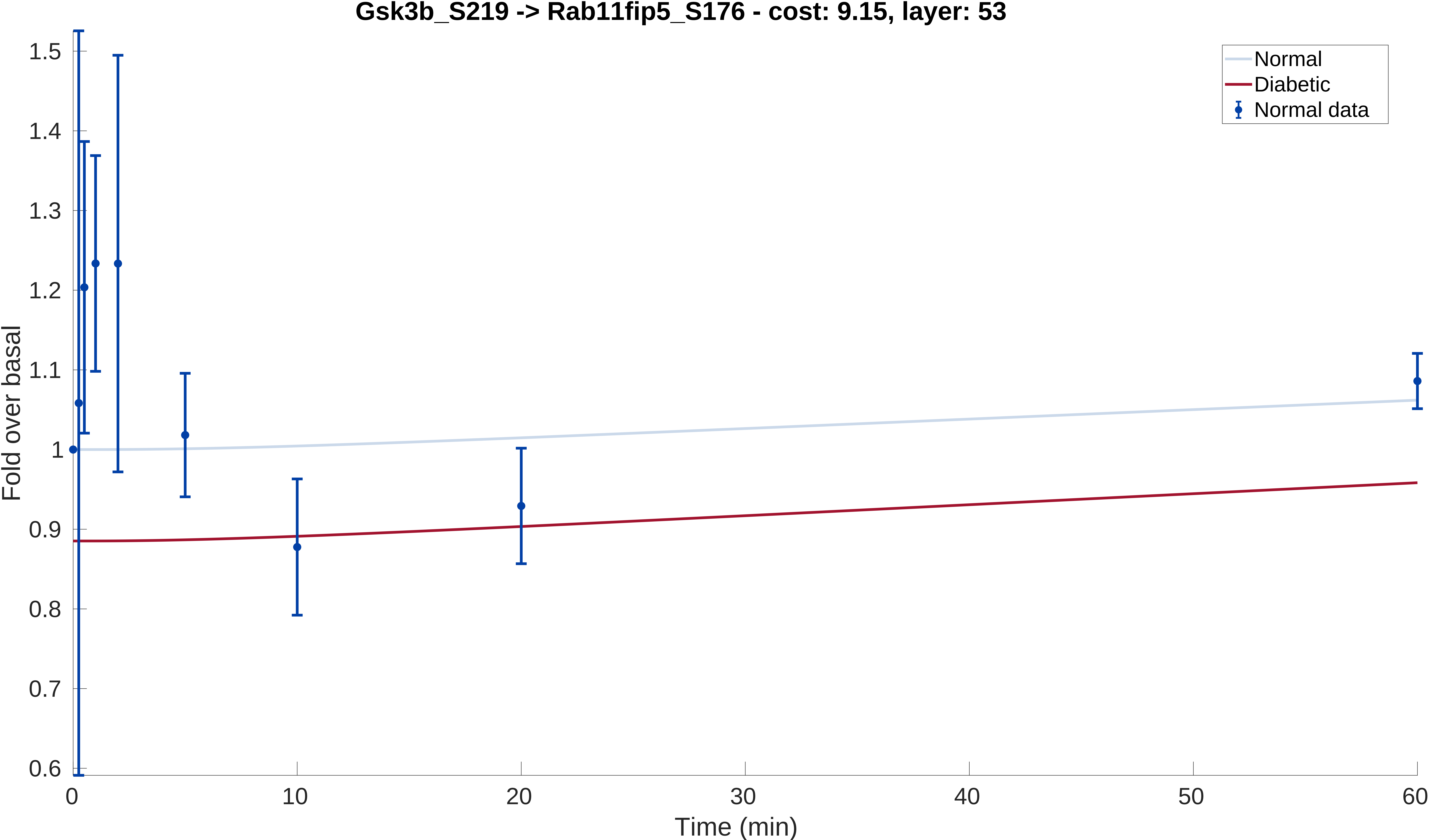

Gsk3b_S219 -> Rab11fip5_S176 - cost: 9.15, layer: 53
Normal
Diabetic
Normal data
Fold over basal
Time (min)
0.6
0.7
0.8
0.9
1
1.1
1.2
1.3
1.4
1.5
0
10
20
30
40
50
60

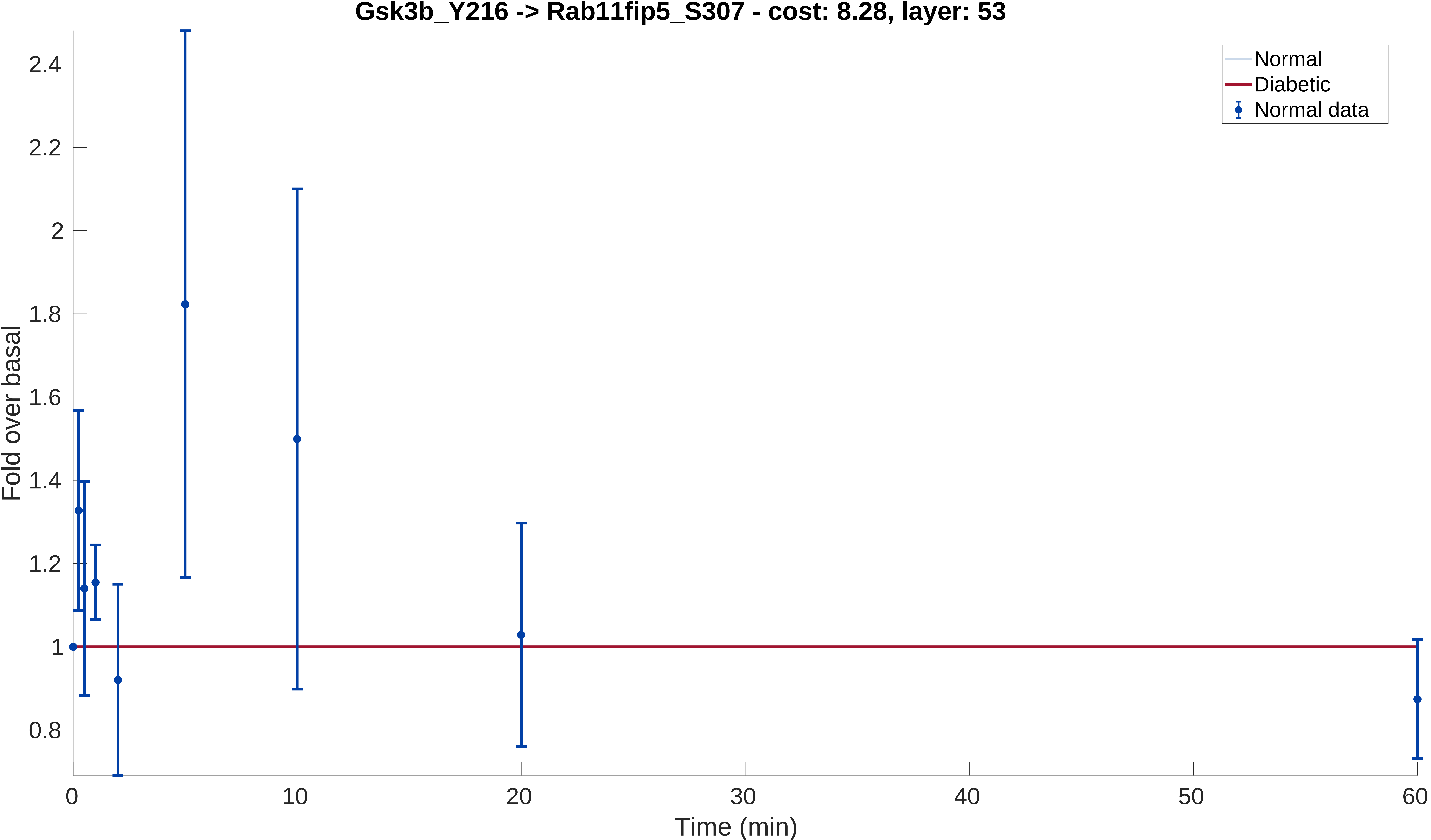

Gsk3b_Y216 -> Rab11fip5_S307 - cost: 8.28, layer: 53
Normal
Diabetic
Normal data
Fold over basal
2.4
2.2
2
1.8
1.6
1.4
1.2
1
0.8
0
10
20
30
40
50
60
Time (min)

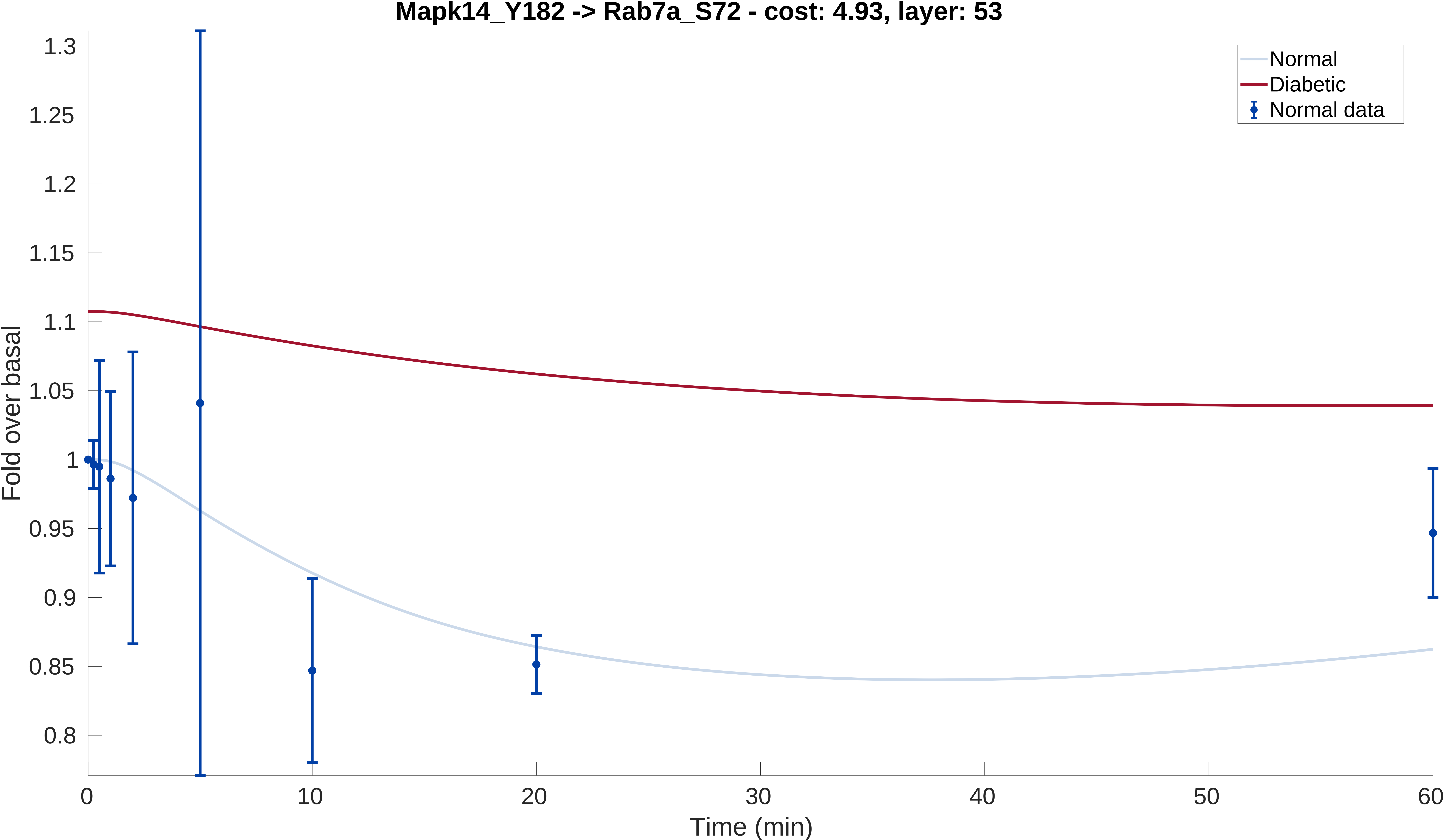
Mapk14_Y182 -> Rab7a_S72 - cost: 4.93, layer: 53
Normal
Diabetic
Normal data
Fold over basal
Time (min)
1.3
1.25
1.2
1.15
1.1
1.05
1
0.95
0.9
0.85
0.8
0
10
20
30
40
50
60

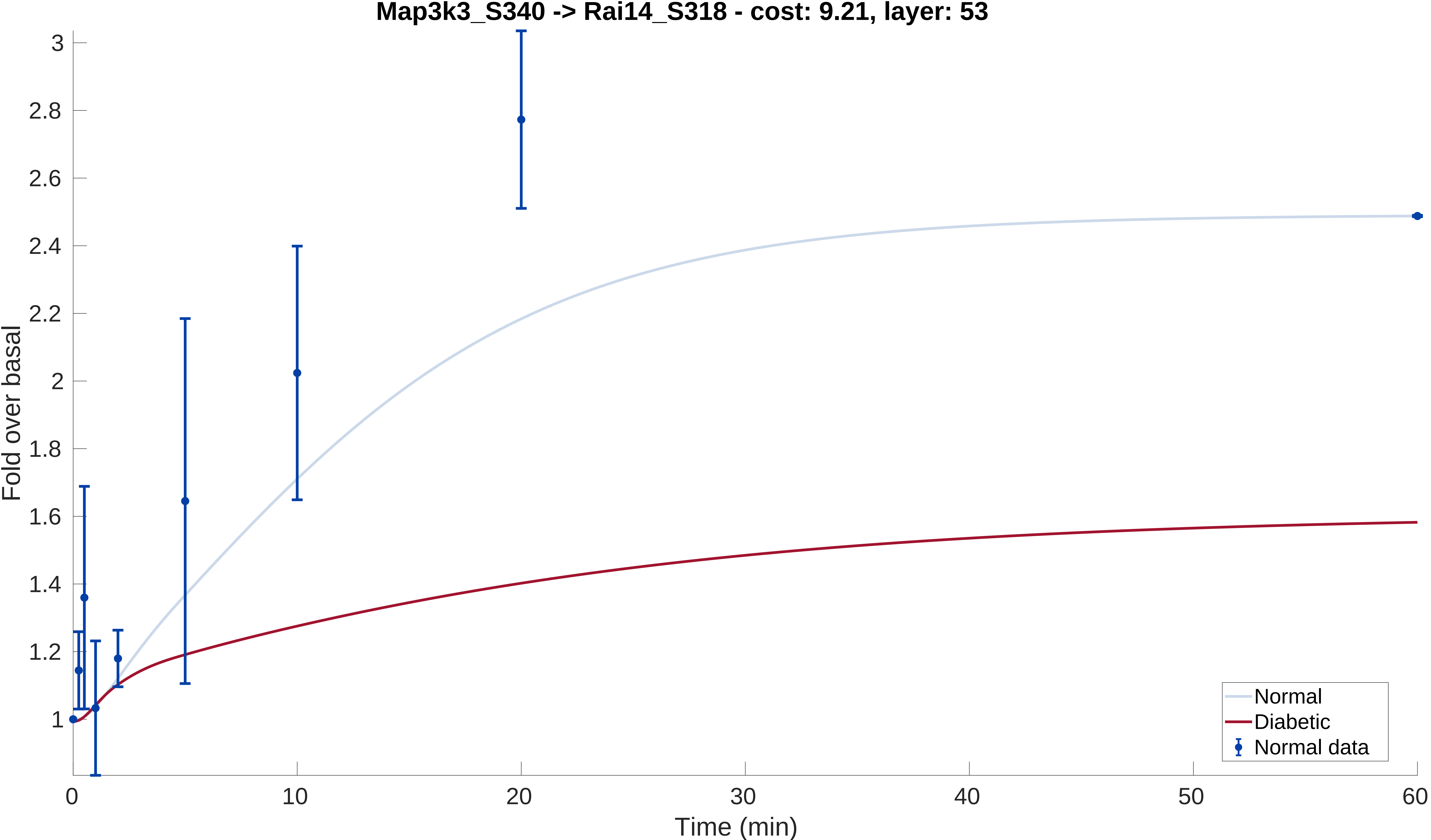

Map3k3_S340 -> Rai14_S318 - cost: 9.21, layer: 53
Fold over basal
Time (min)
3
2.8
2.6
2.4
2.2
2
1.8
1.6
1.4
1.2
1
0
10
20
30
40
50
60
Normal
Diabetic
Normal data

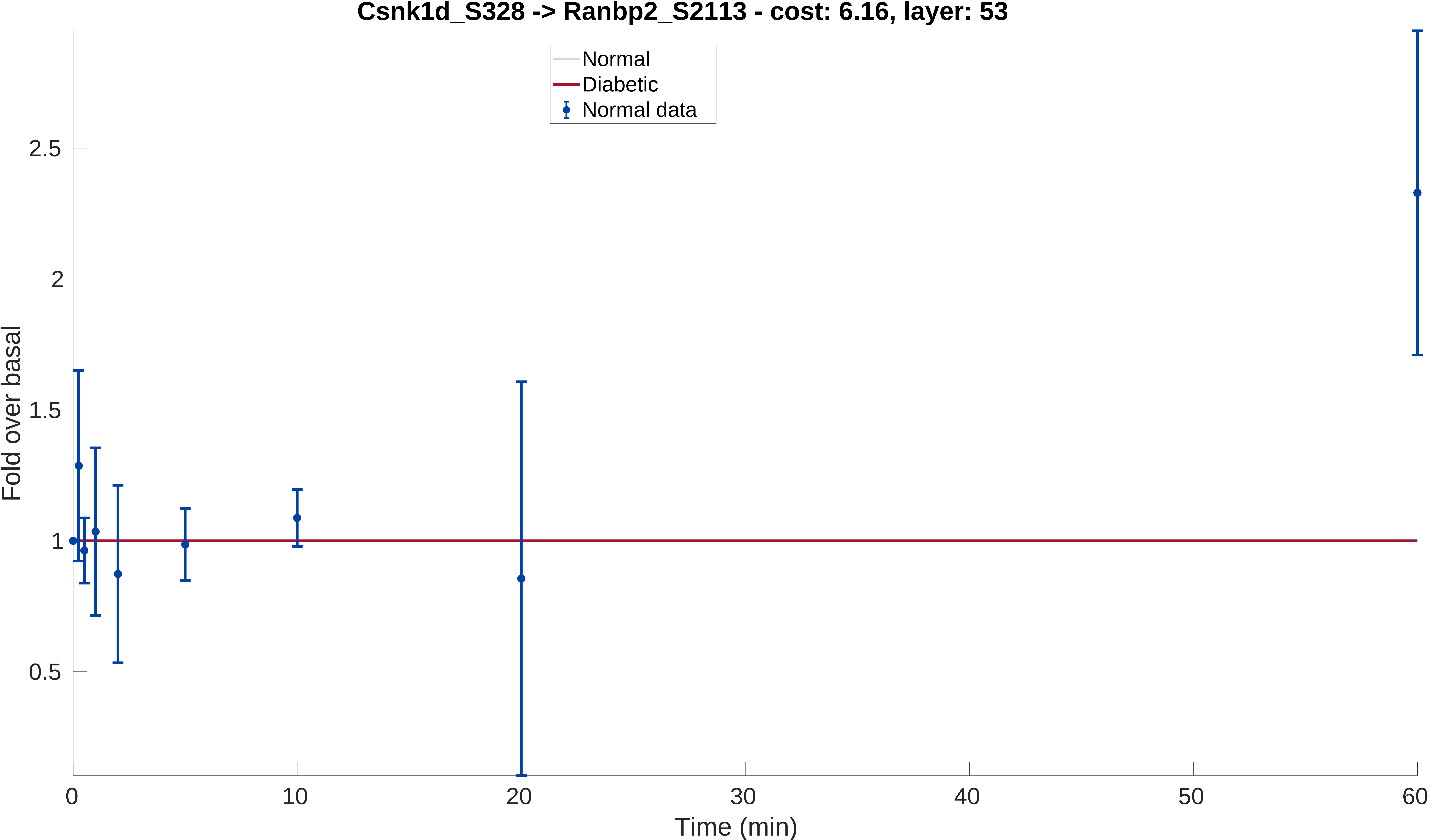

Csnk1d_S328 -> Ranbp2_S2113 - cost: 6.16, layer: 53
Normal
Diabetic
Normal data
Fold over basal
2.5
2
1.5
1
0.5
0
10
20
30
40
50
60
Time (min)

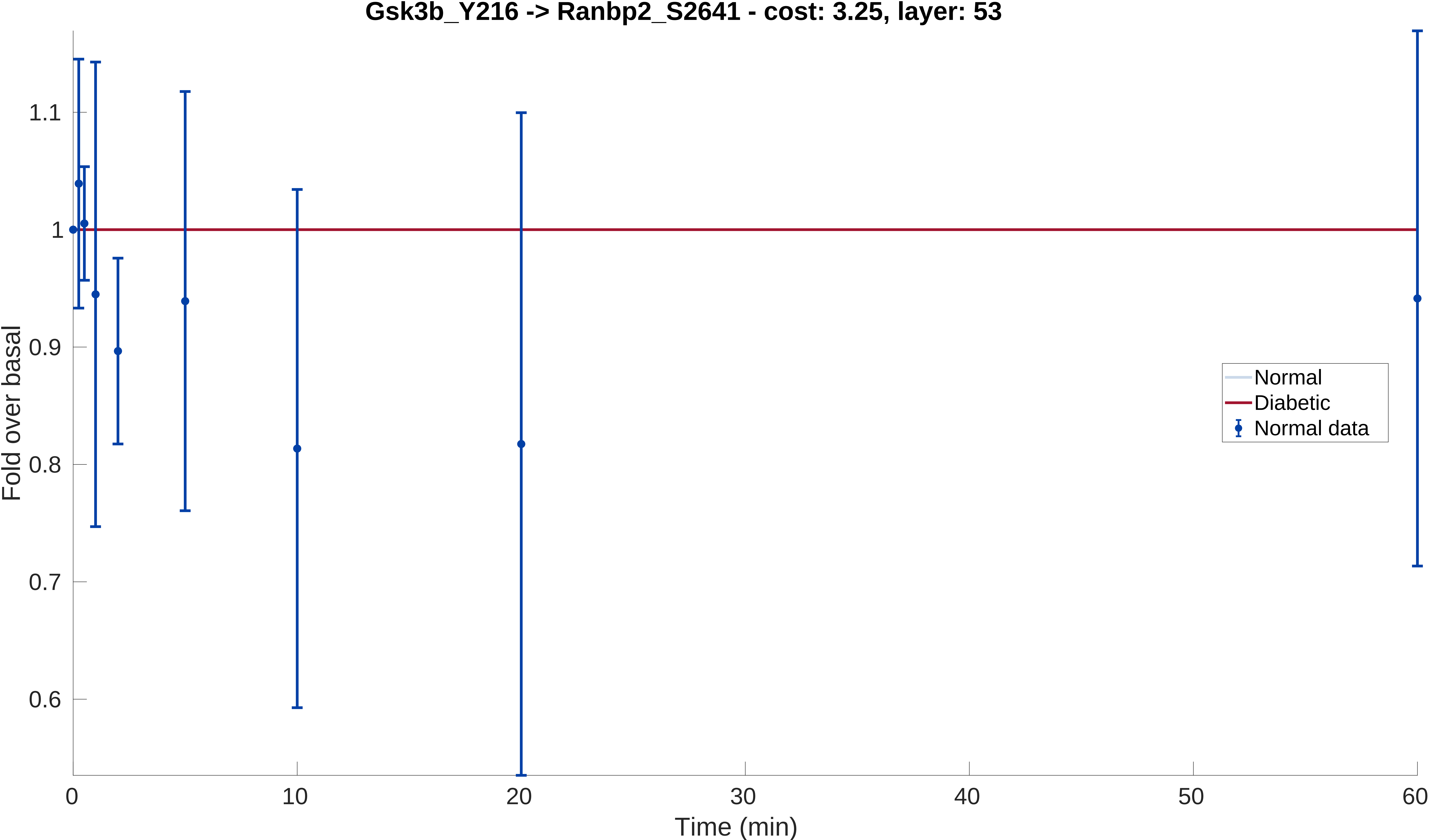

Gsk3b_Y216 -> Ranbp2_S2641 - cost: 3.25, layer: 53
Fold over basal
1.1
1
0.9
0.8
0.7
0.6
0
10
20
30
40
50
60
Time (min)
Normal
Diabetic
Normal data

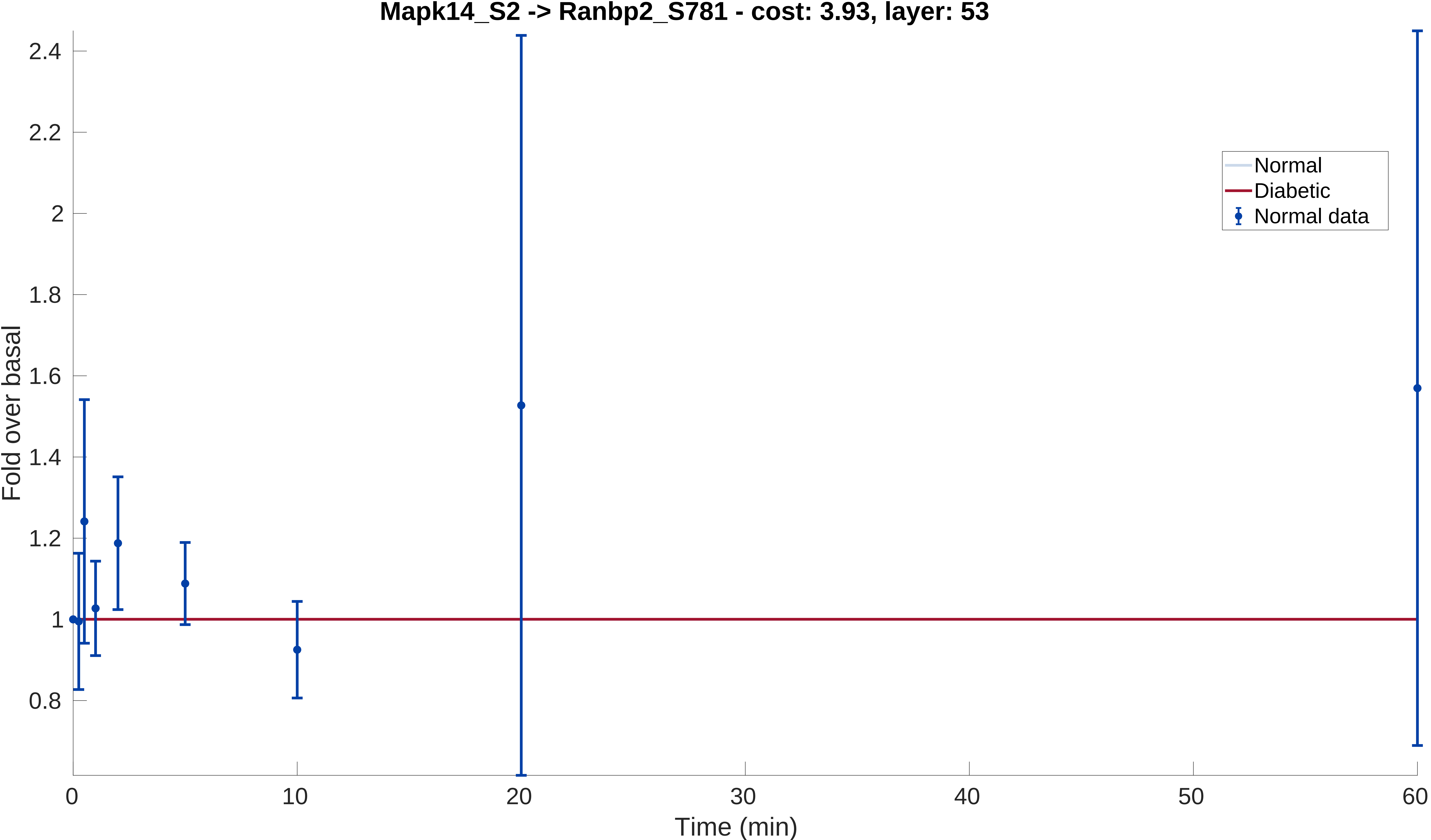

Mapk14_S2 -> Ranbp2_S781 - cost: 3.93, layer: 53
Fold over basal
Time (min)
Normal
Diabetic
Normal data
2.4
2.2
2
1.8
1.6
1.4
1.2
1
0.8
0
10
20
30
40
50
60

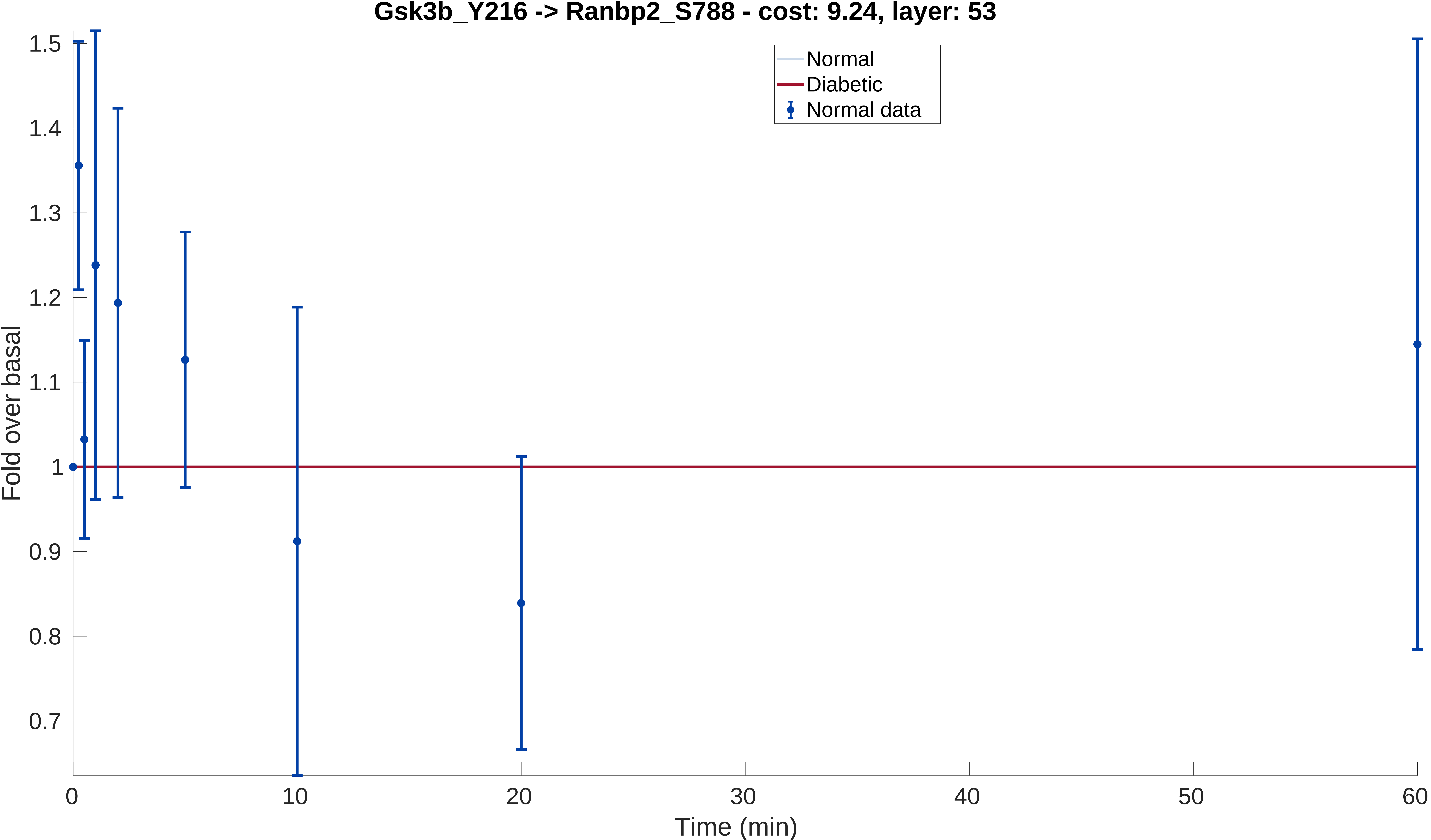

Gsk3b_Y216 -> Ranbp2_S788 - cost: 9.24, layer: 53
Normal
Diabetic
Normal data
Fold over basal
1.5
1.4
1.3
1.2
1.1
1
0.9
0.8
0.7
0
10
20
30
40
50
60
Time (min)

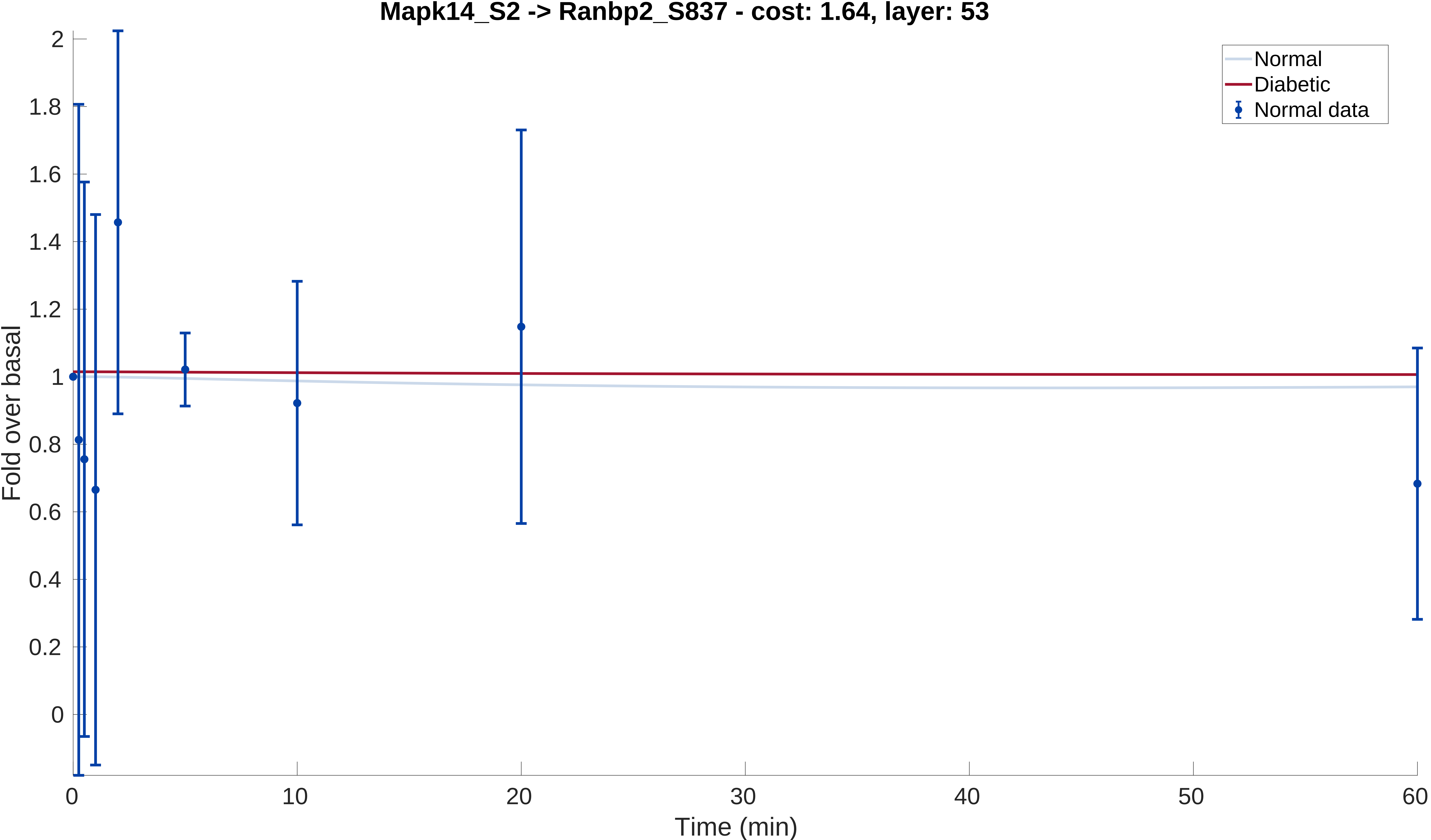
Mapk14_S2 -> Ranbp2_S837 - cost: 1.64, layer: 53
Normal
Diabetic
Normal data
Fold over basal
2
1.8
1.6
1.4
1.2
1
0.8
0.6
0.4
0.2
0
0
10
20
30
40
50
60
Time (min)

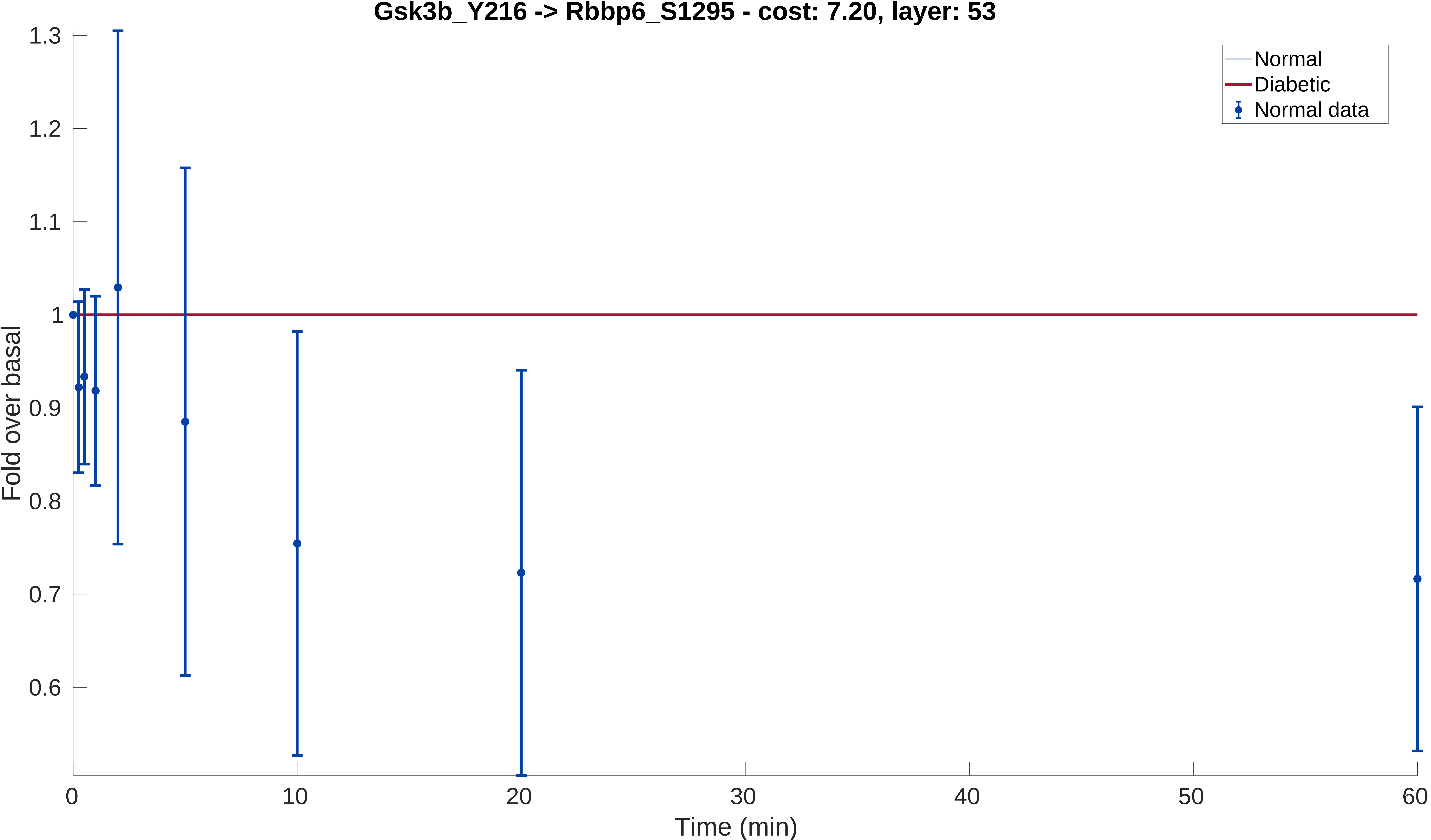

Gsk3b_Y216 -> Rbbp6_S1295 - cost: 7.20, layer: 53
Normal
Diabetic
Normal data
Fold over basal
Time (min)
1.3
1.2
1.1
1
0.9
0.8
0.7
0.6
0
10
20
30
40
50
60

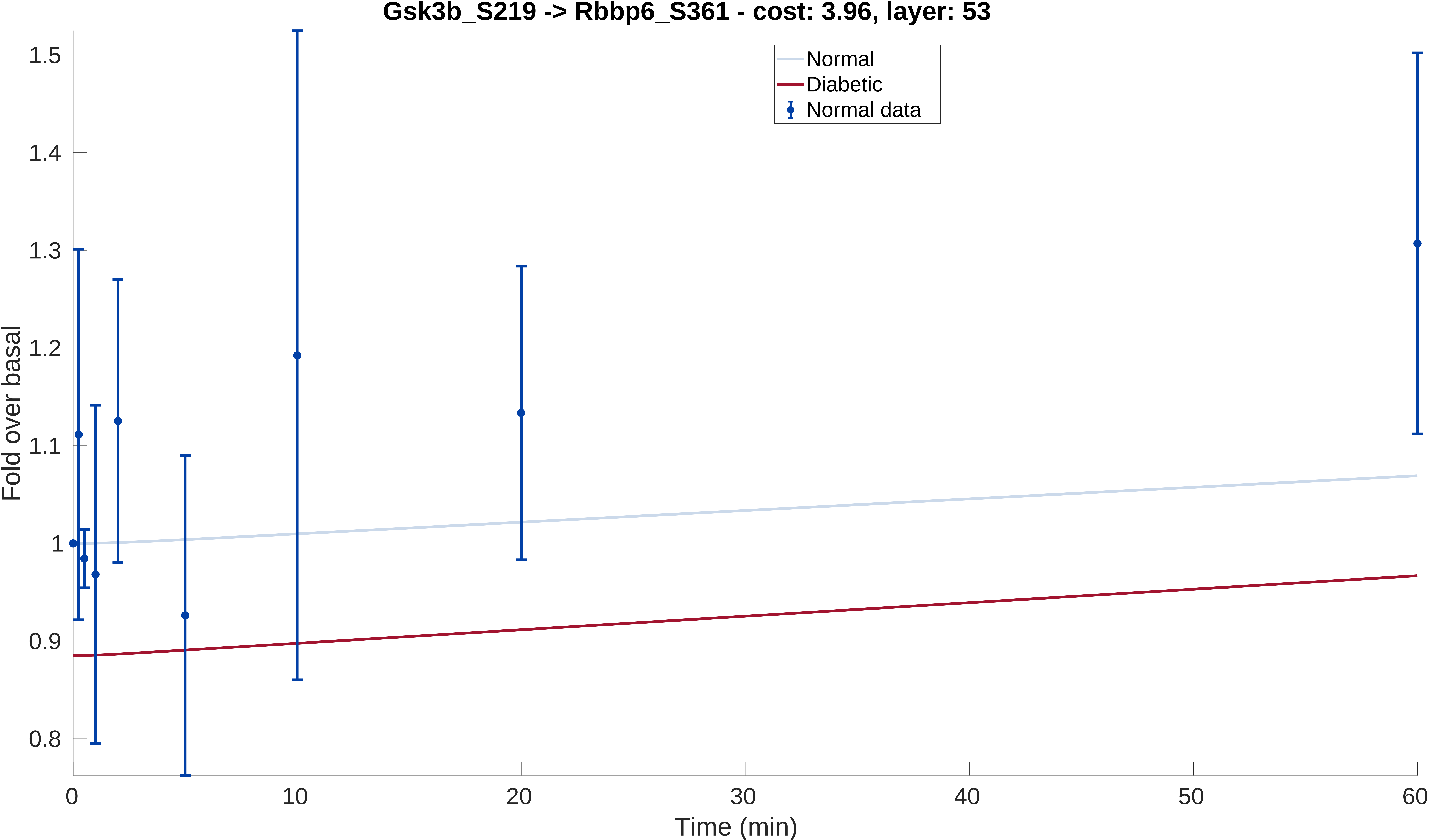

Gsk3b_S219 -> Rbbp6_S361 - cost: 3.96, layer: 53
Normal
Diabetic
Normal data
Fold over basal
0.8
0.9
1
1.1
1.2
1.3
1.4
1.5
0
10
20
30
40
50
60
Time (min)

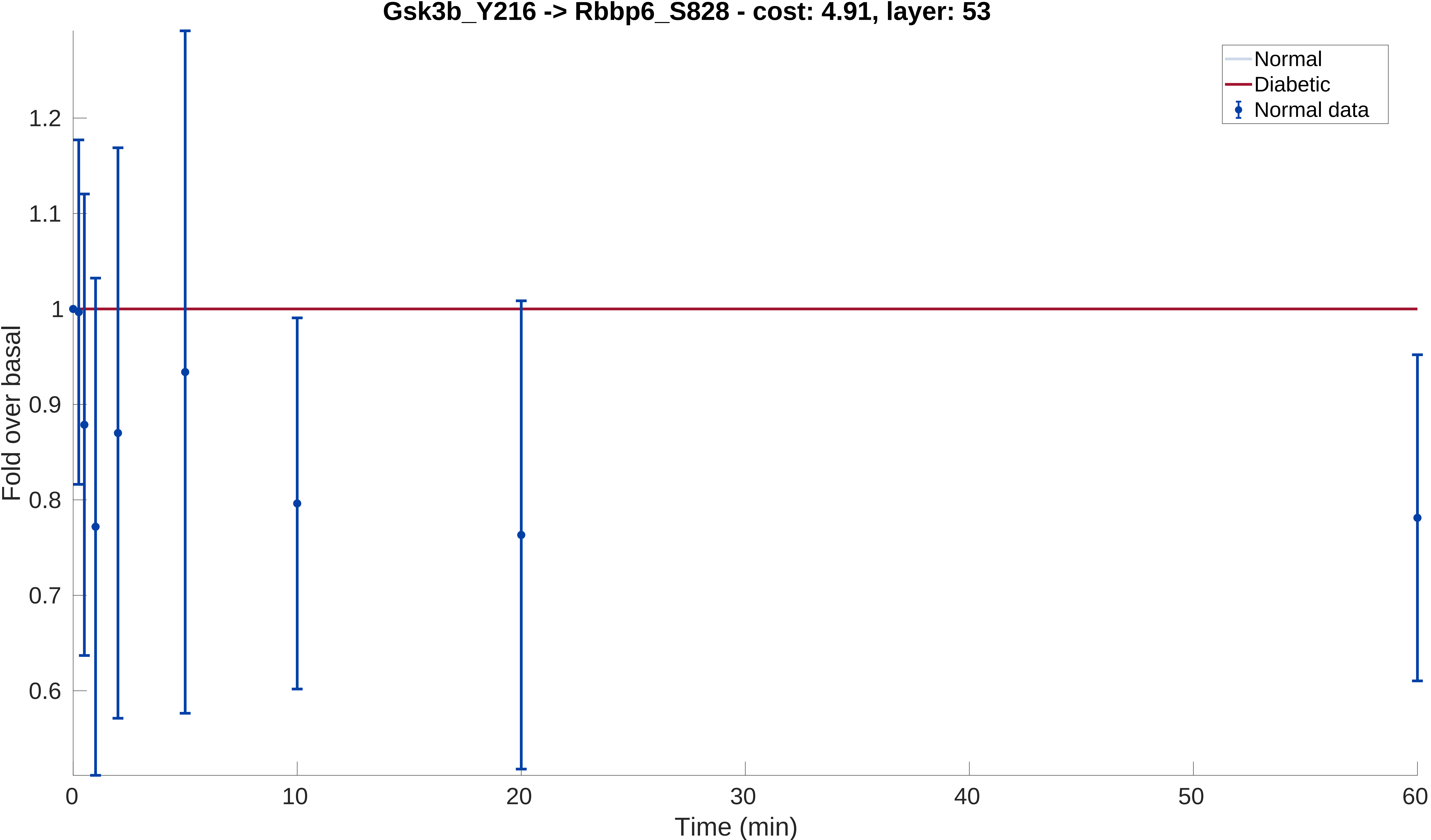
Gsk3b_Y216 -> Rbbp6_S828 - cost: 4.91, layer: 53
Normal
Diabetic
Normal data
Fold over basal
1.2
1.1
1
0.9
0.8
0.7
0.6
0
10
20
30
40
50
60
Time (min)

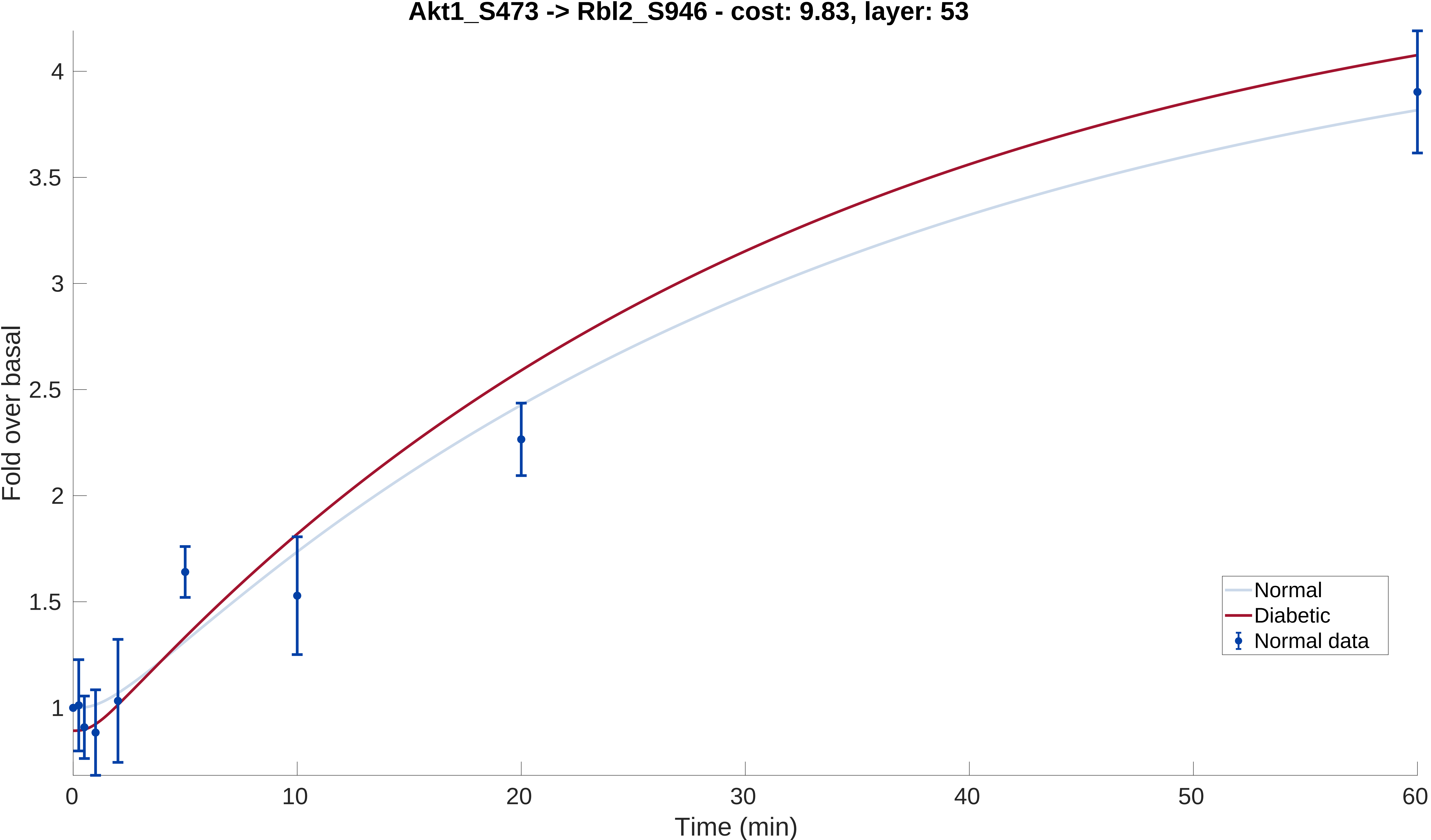
Akt1_S473 -> Rbl2_S946 - cost: 9.83, layer: 53
Fold over basal
4
3.5
3
2.5
2
1.5
1
0
10
20
30
40
50
60
Time (min)
Normal
Diabetic
Normal data

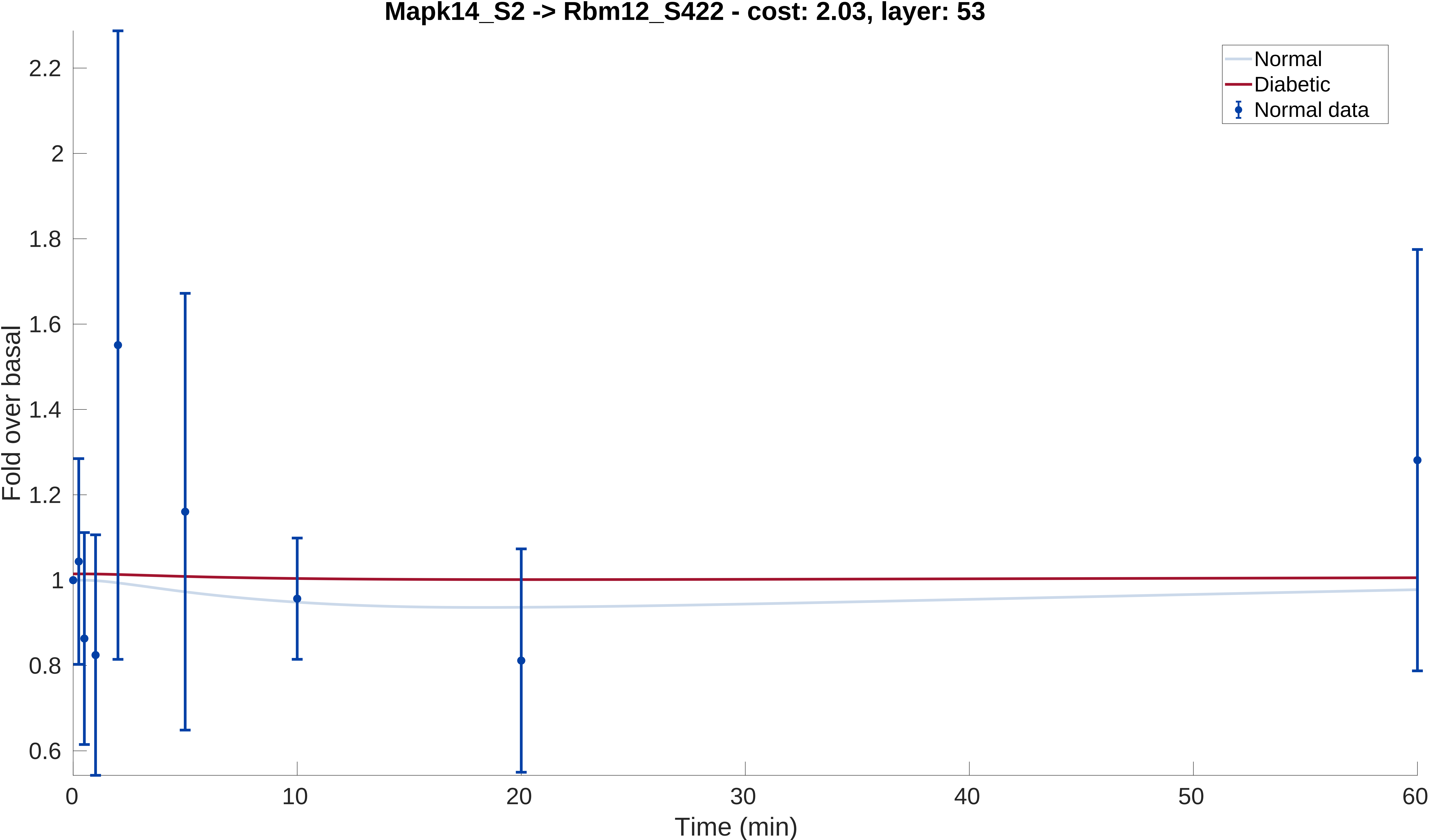

Mapk14_S2 -> Rbm12_S422 - cost: 2.03, layer: 53
Normal
Diabetic
Normal data
Fold over basal
Time (min)
0.6
0.8
1
1.2
1.4
1.6
1.8
2
2.2
0
10
20
30
40
50
60

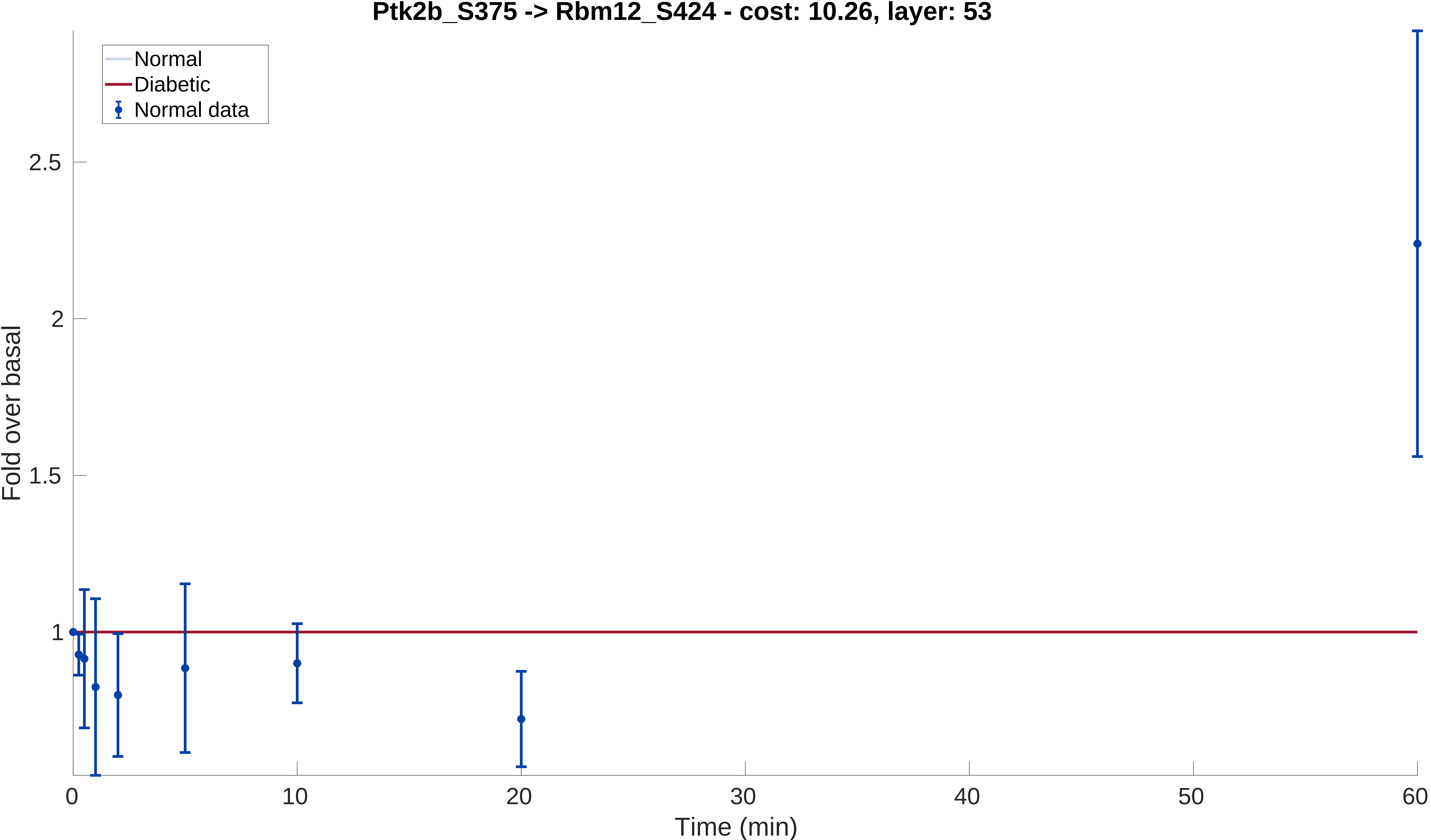
Ptk2b_S375 -> Rbm12_S424 - cost: 10.26, layer: 53
Normal
Diabetic
Normal data
Fold over basal
2.5
2
1.5
1
0
10
20
30
40
50
60
Time (min)

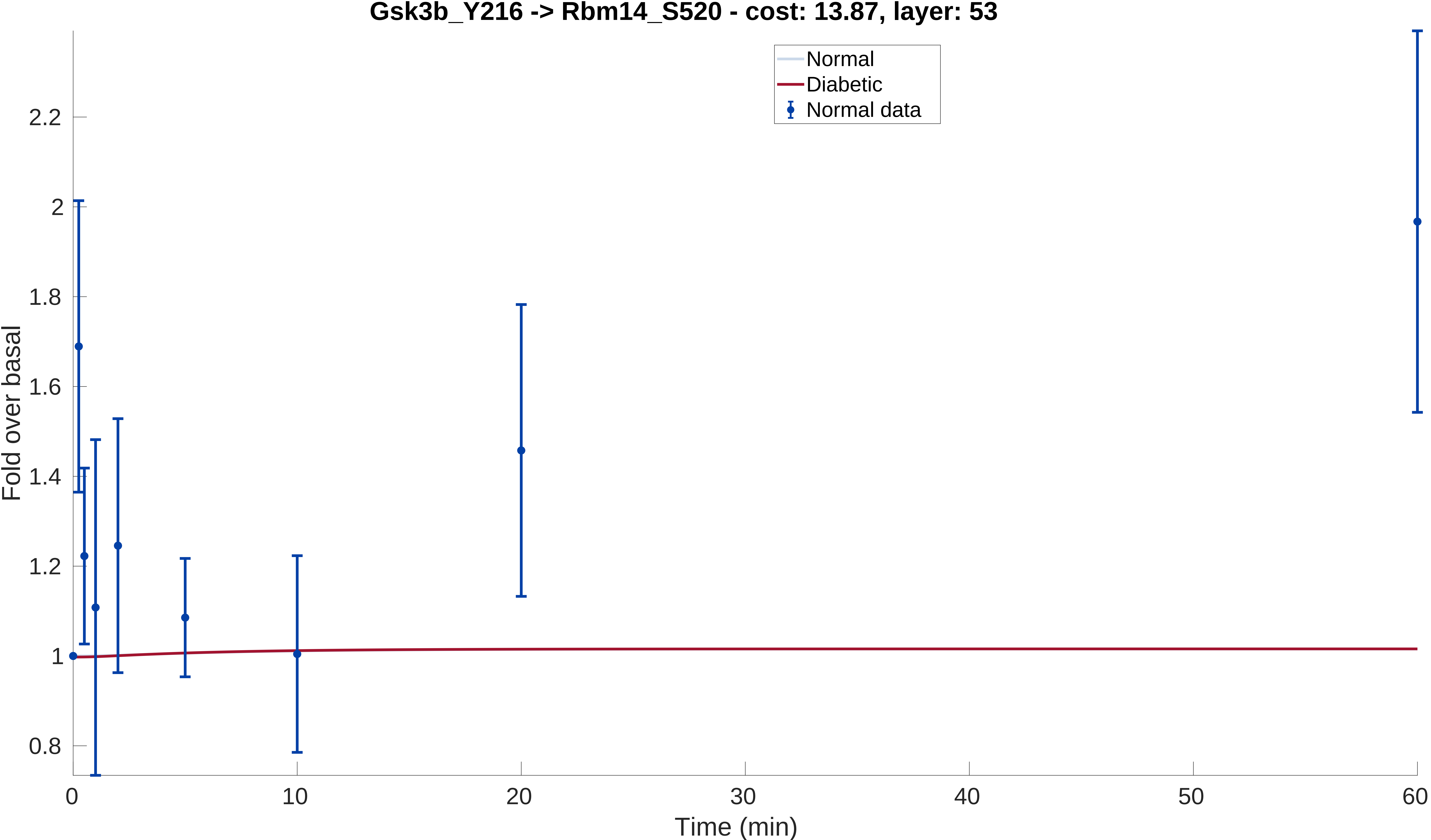

Gsk3b_Y216 -> Rbm14_S520 - cost: 13.87, layer: 53
Normal
Diabetic
Normal data
Fold over basal
0.8
1
1.2
1.4
1.6
1.8
2
2.2
0
10
20
30
40
50
60
Time (min)

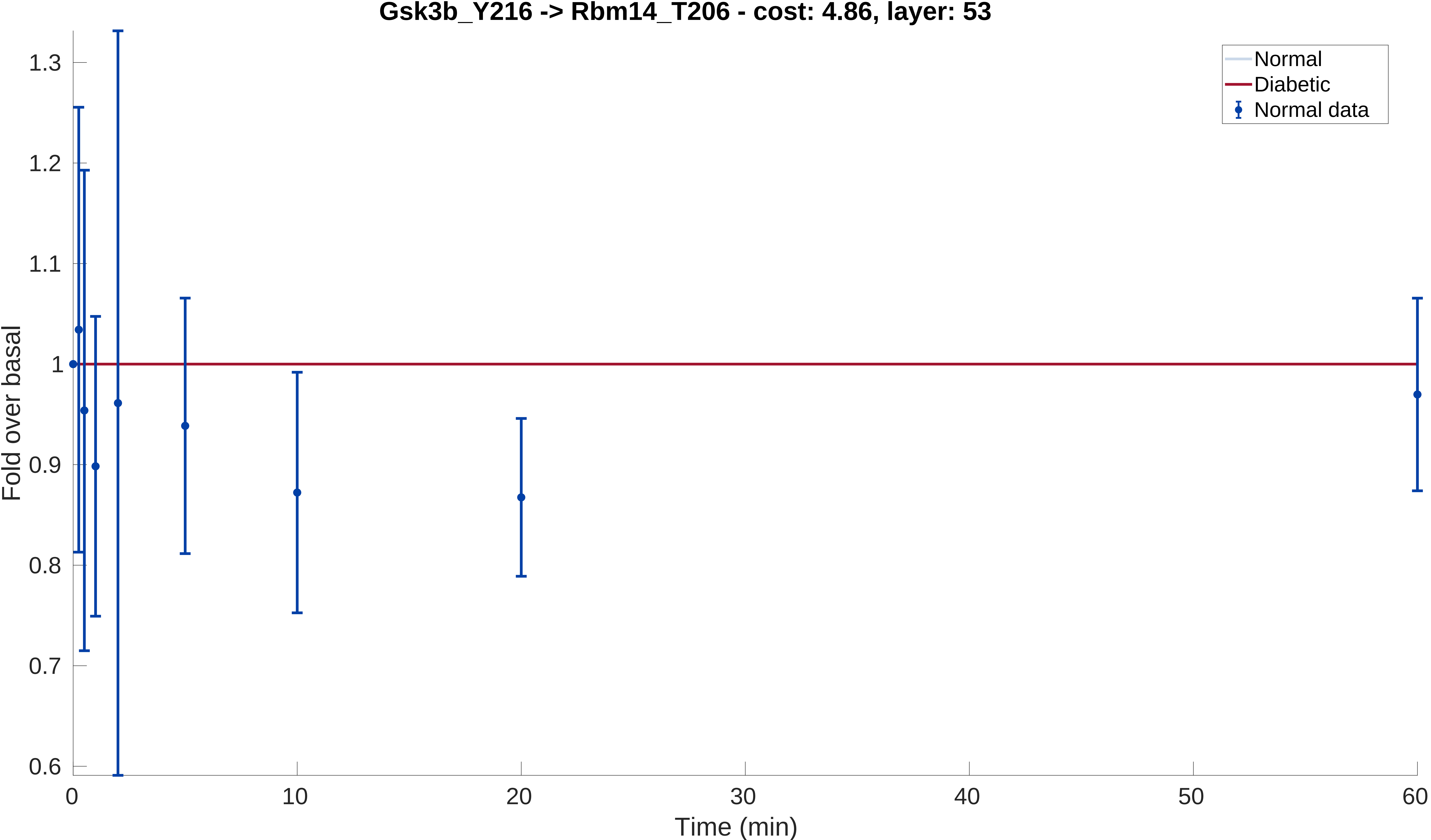

Gsk3b_Y216 -> Rbm14_T206 - cost: 4.86, layer: 53
Normal
Diabetic
Normal data
Fold over basal
Time (min)
1.3
1.2
1.1
1
0.9
0.8
0.7
0.6
0
10
20
30
40
50
60

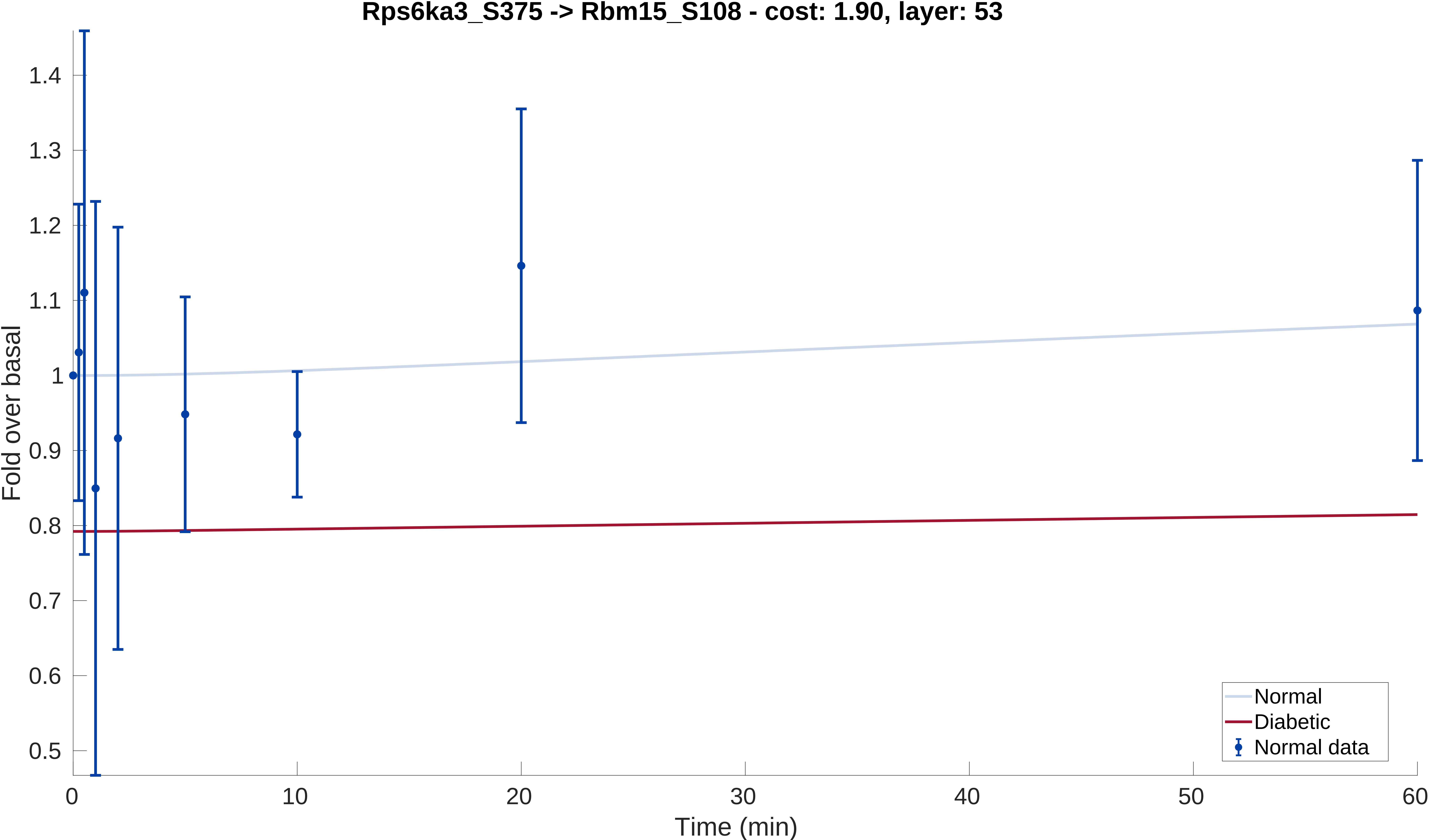

Rps6ka3_S375 -> Rbm15_S108 - cost: 1.90, layer: 53
Fold over basal
1.4
1.3
1.2
1.1
1
0.9
0.8
0.7
0.6
0.5
0
10
20
30
40
50
60
Time (min)
Normal
Diabetic
Normal data

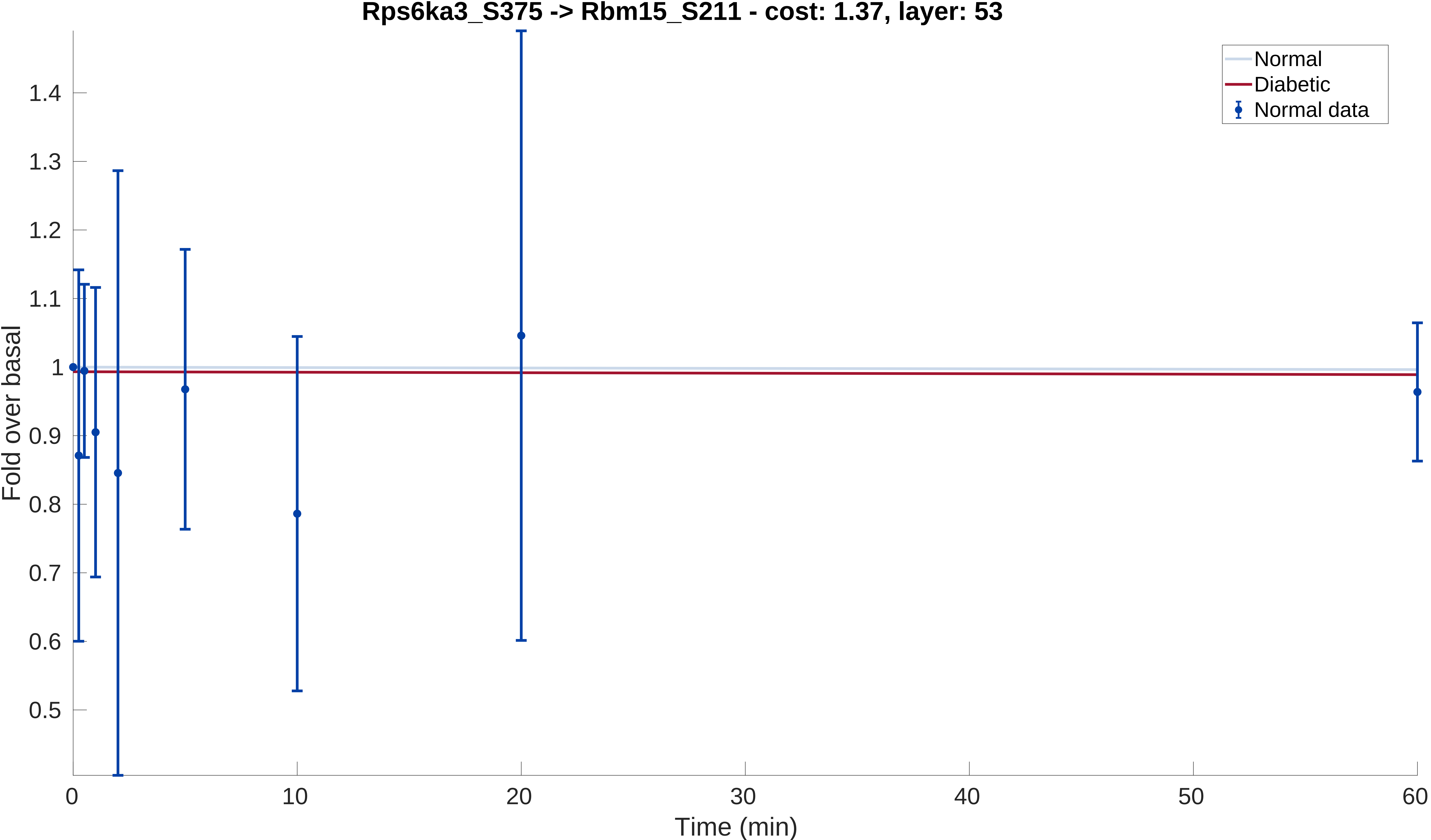
Rps6ka3_S375 -> Rbm15_S211 - cost: 1.37, layer: 53
Normal
Diabetic
Normal data
Fold over basal
1.4
1.3
1.2
1.1
1
0.9
0.8
0.7
0.6
0.5
0
10
20
30
40
50
60
Time (min)

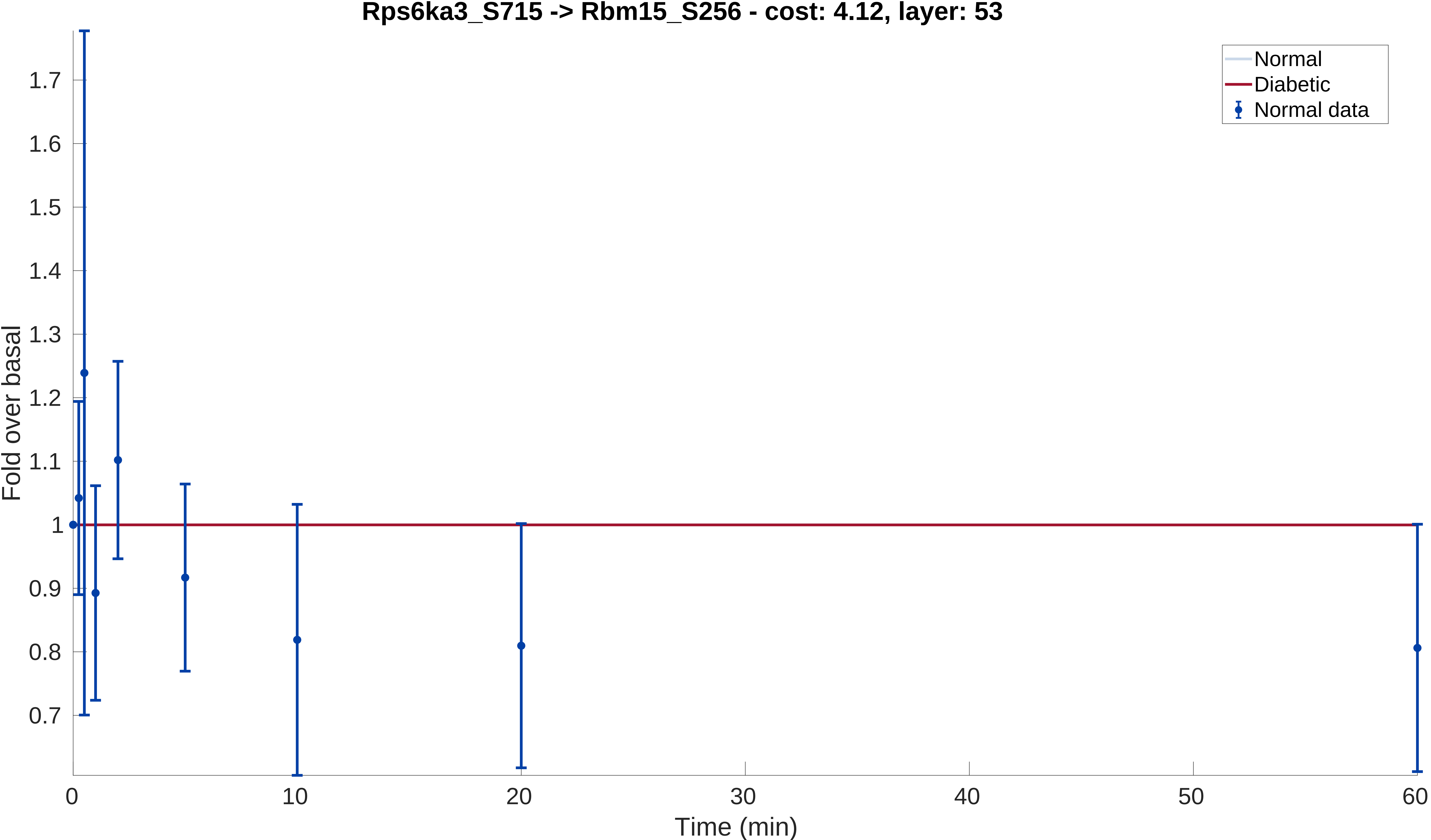
Rps6ka3_S715 -> Rbm15_S256 - cost: 4.12, layer: 53
Normal
Diabetic
Normal data
Fold over basal
1.7
1.6
1.5
1.4
1.3
1.2
1.1
1
0.9
0.8
0.7
0
10
20
30
40
50
60
Time (min)

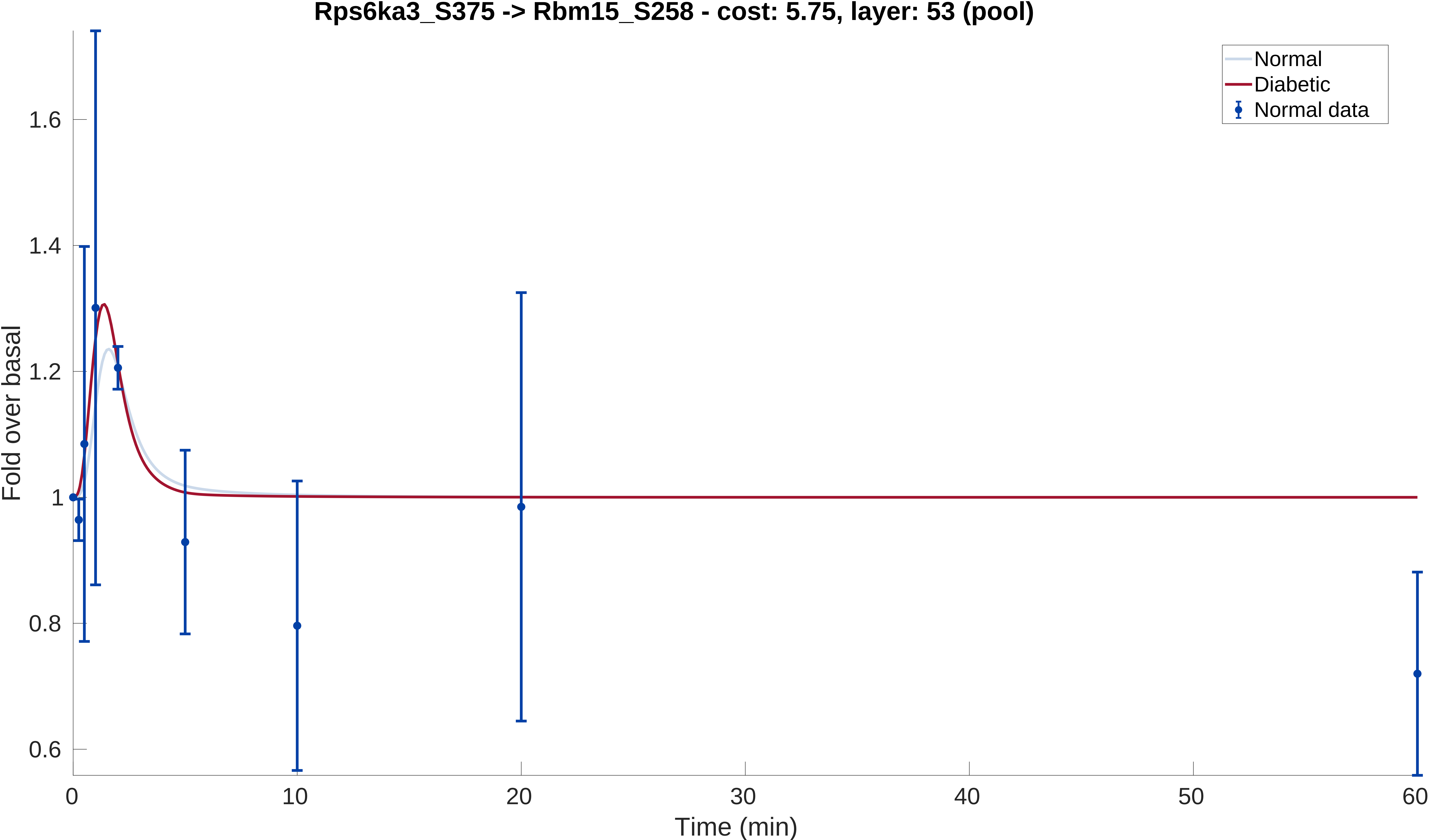
Rps6ka3_S375 -> Rbm15_S258 - cost: 5.75, layer: 53 (pool)
Normal
Diabetic
Normal data
Fold over basal
Time (min)
1.6
1.4
1.2
1
0.8
0.6
0
10
20
30
40
50
60

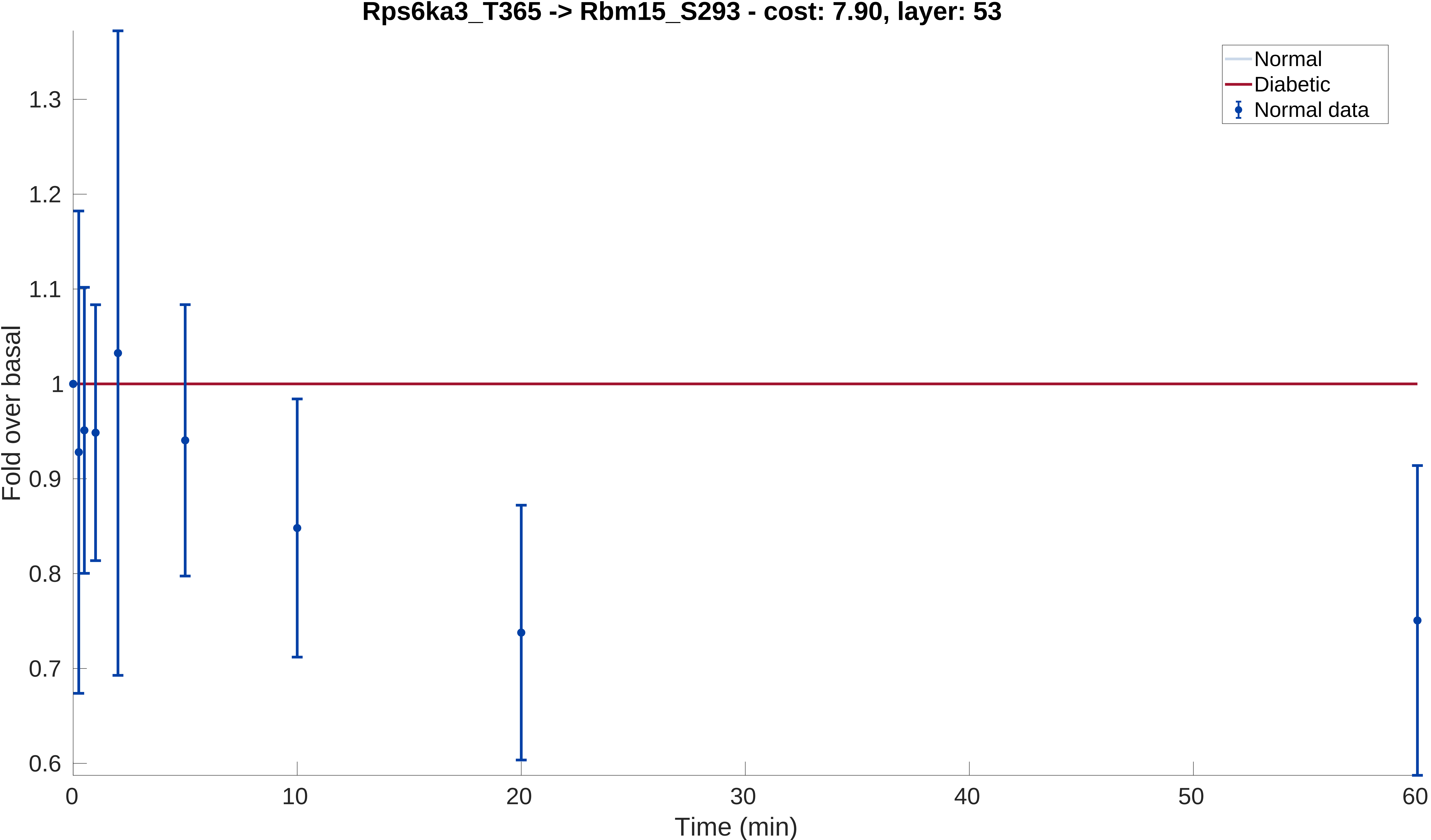

Rps6ka3_T365 -> Rbm15_S293 - cost: 7.90, layer: 53
Normal
Diabetic
Normal data
Fold over basal
Time (min)
1.3
1.2
1.1
1
0.9
0.8
0.7
0.6
0
10
20
30
40
50
60

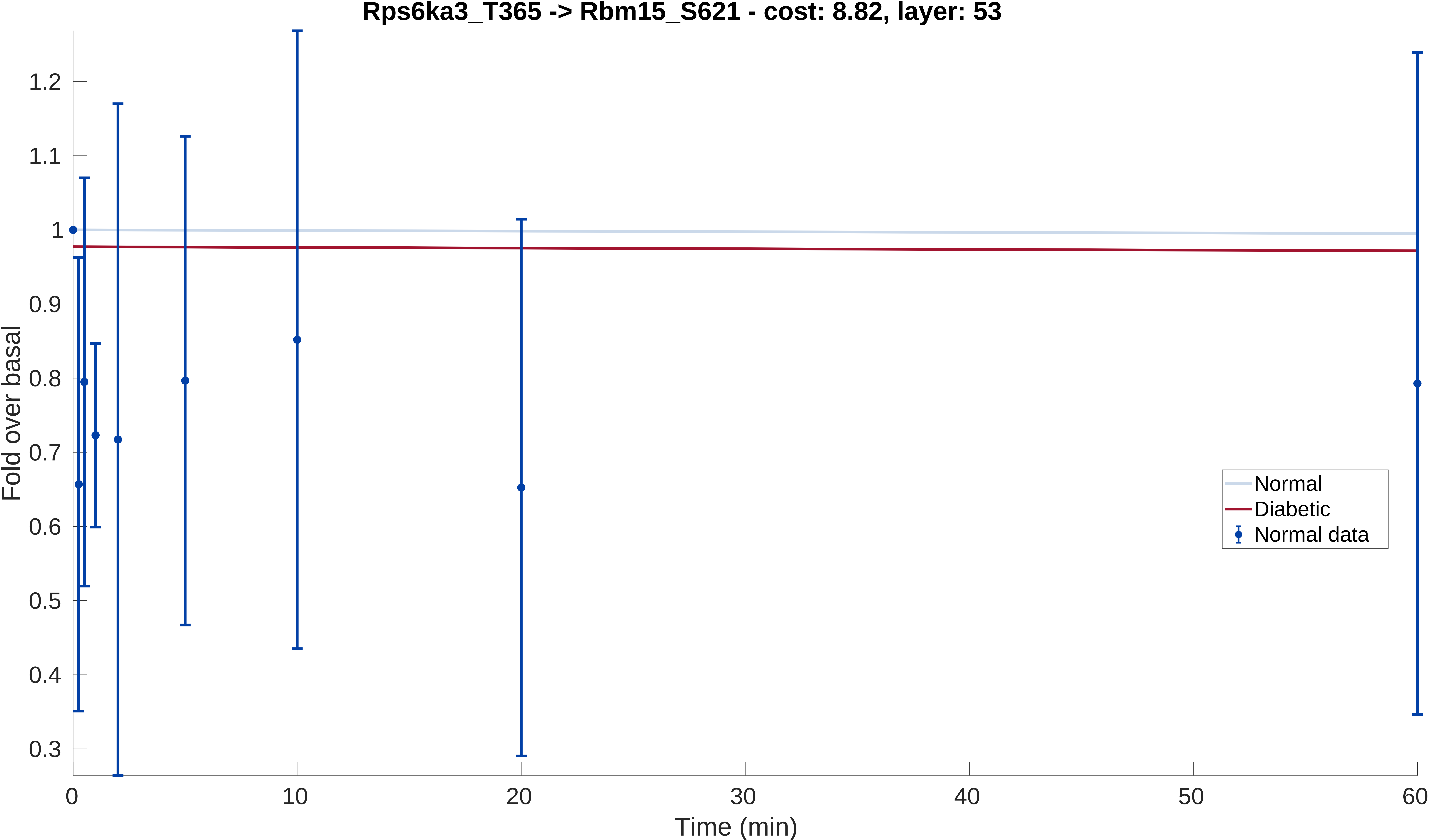

Rps6ka3_T365 -> Rbm15_S621 - cost: 8.82, layer: 53
Fold over basal
1.2
1.1
1
0.9
0.8
0.7
0.6
0.5
0.4
0.3
0
10
20
30
40
50
60
Time (min)
Normal
Diabetic
Normal data

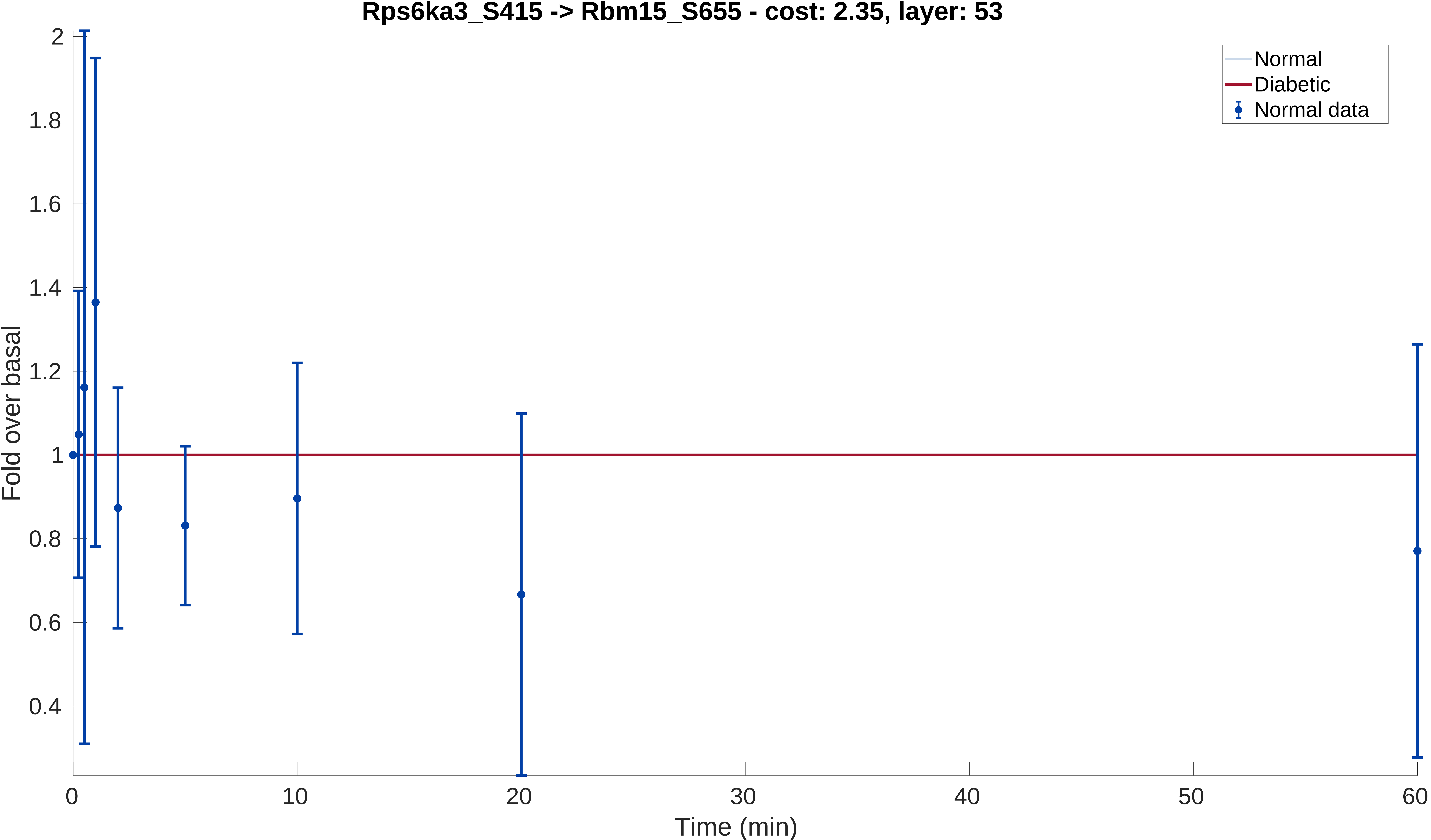

Rps6ka3_S415 -> Rbm15_S655 - cost: 2.35, layer: 53
Normal
Diabetic
Normal data
Fold over basal
Time (min)
2
1.8
1.6
1.4
1.2
1
0.8
0.6
0.4
0
10
20
30
40
50
60

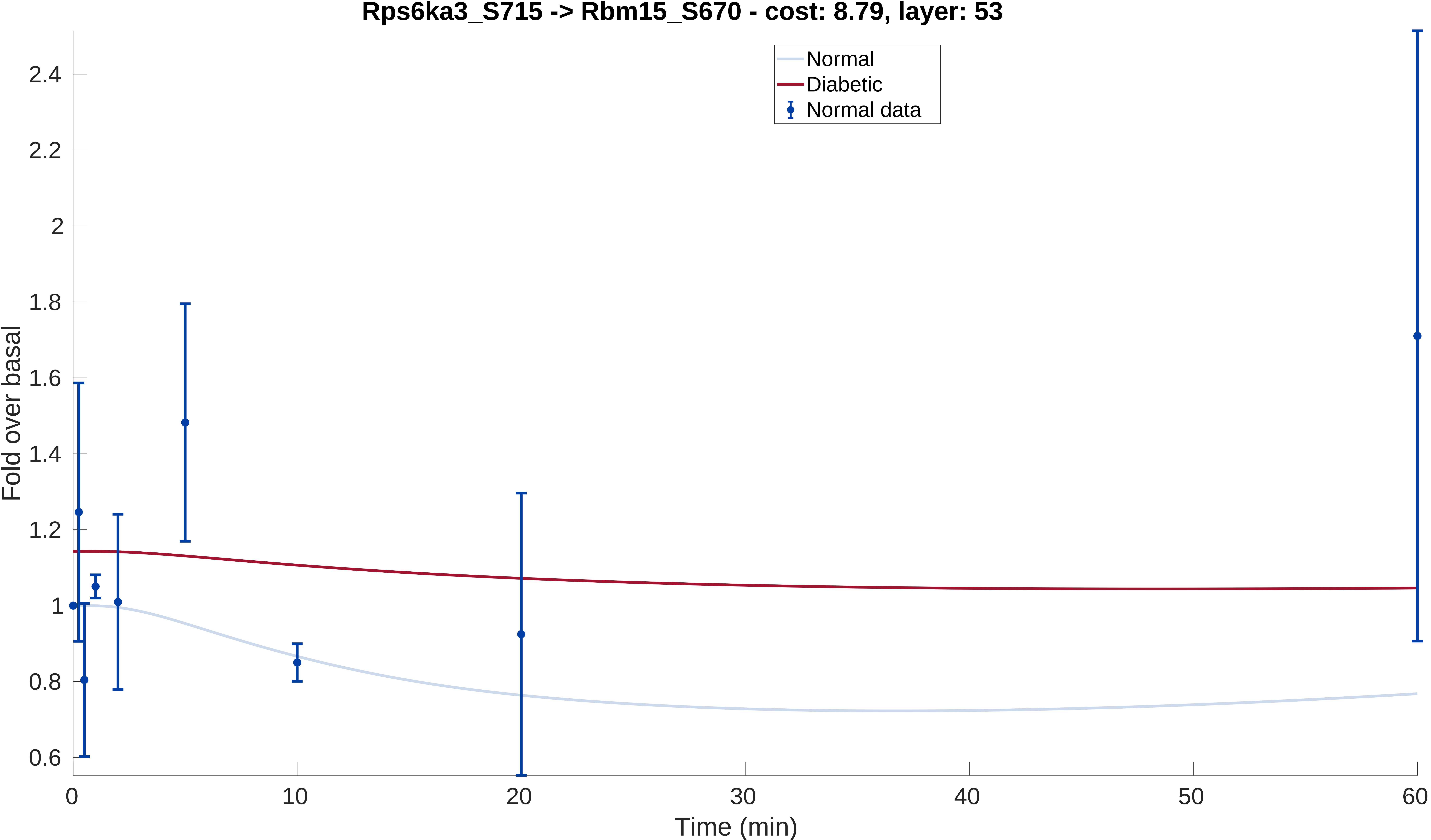

Rps6ka3_S715 -> Rbm15_S670 - cost: 8.79, layer: 53
Normal
Diabetic
Normal data
Fold over basal
2.4
2.2
2
1.8
1.6
1.4
1.2
1
0.8
0.6
0
10
20
30
40
50
60
Time (min)

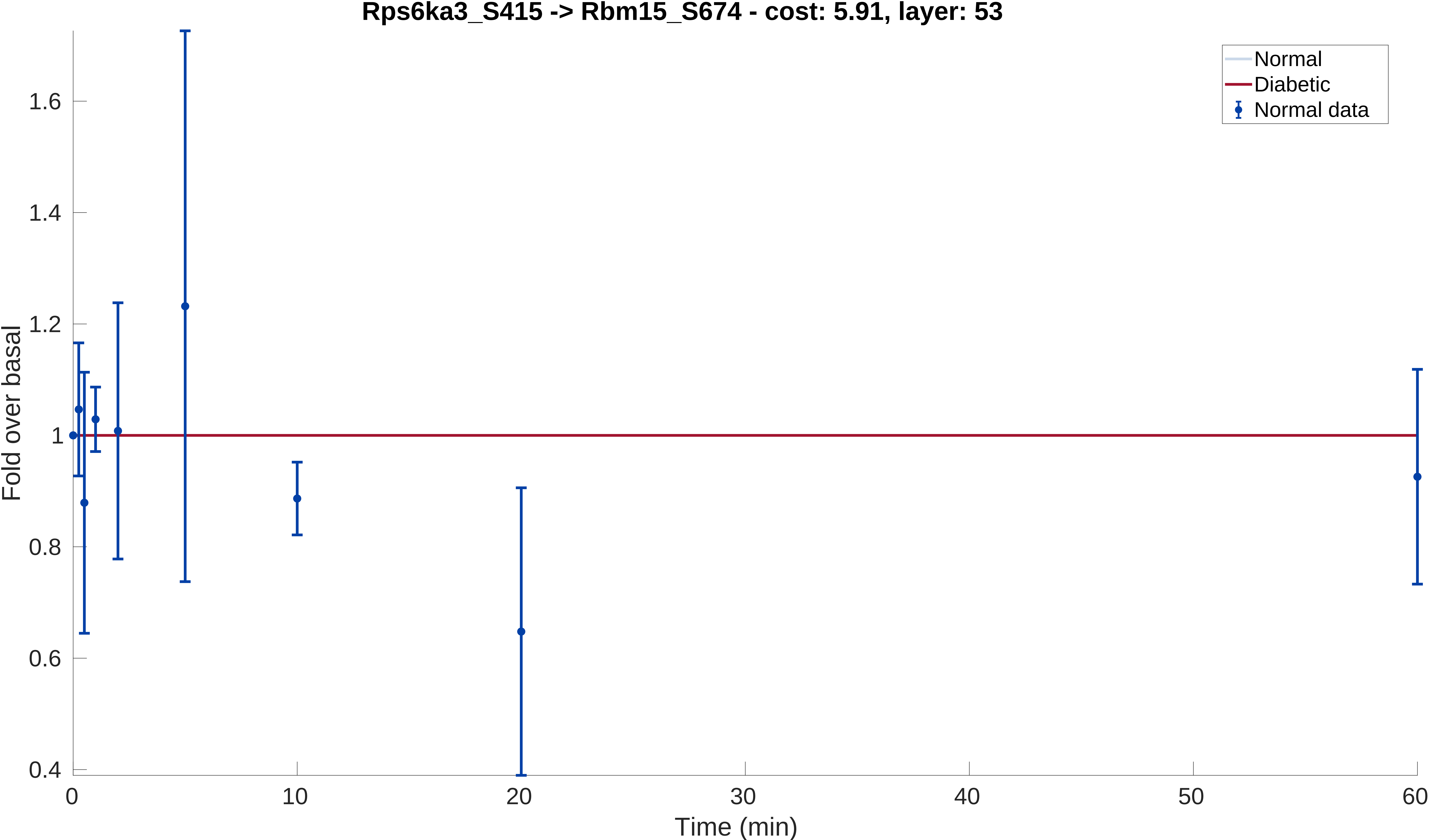

Rps6ka3_S415 -> Rbm15_S674 - cost: 5.91, layer: 53
Normal
Diabetic
Normal data
Fold over basal
1.6
1.4
1.2
1
0.8
0.6
0.4
0
10
20
30
40
50
60
Time (min)

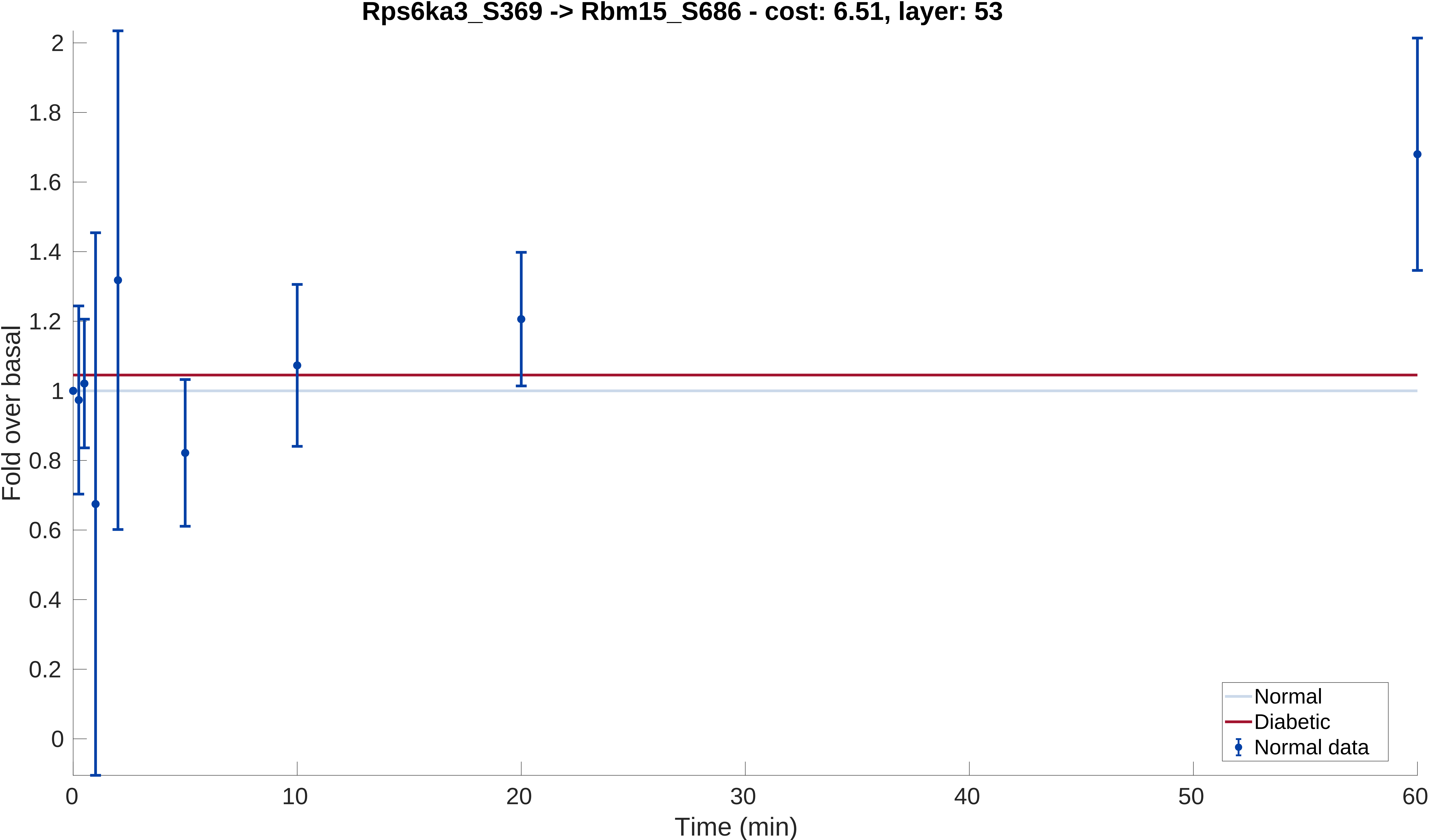

Rps6ka3_S369 -> Rbm15_S686 - cost: 6.51, layer: 53
Fold over basal
Time (min)
0
0.2
0.4
0.6
0.8
1
1.2
1.4
1.6
1.8
2
0
10
20
30
40
50
60
Normal
Diabetic
Normal data

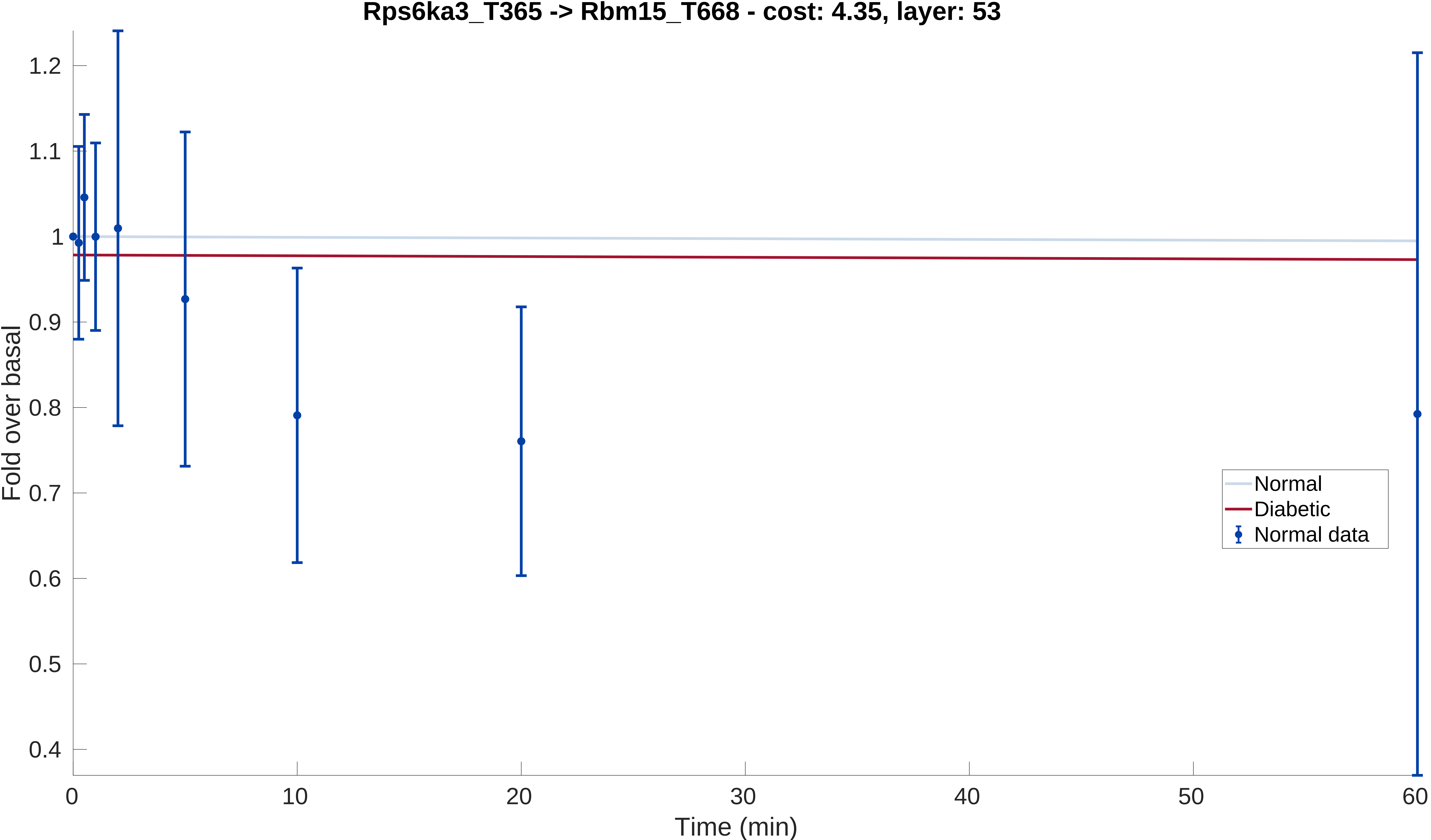

Rps6ka3_T365 -> Rbm15_T668 - cost: 4.35, layer: 53
Fold over basal
1.2
1.1
1
0.9
0.8
0.7
0.6
0.5
0.4
0
10
20
30
40
50
60
Time (min)
Normal
Diabetic
Normal data

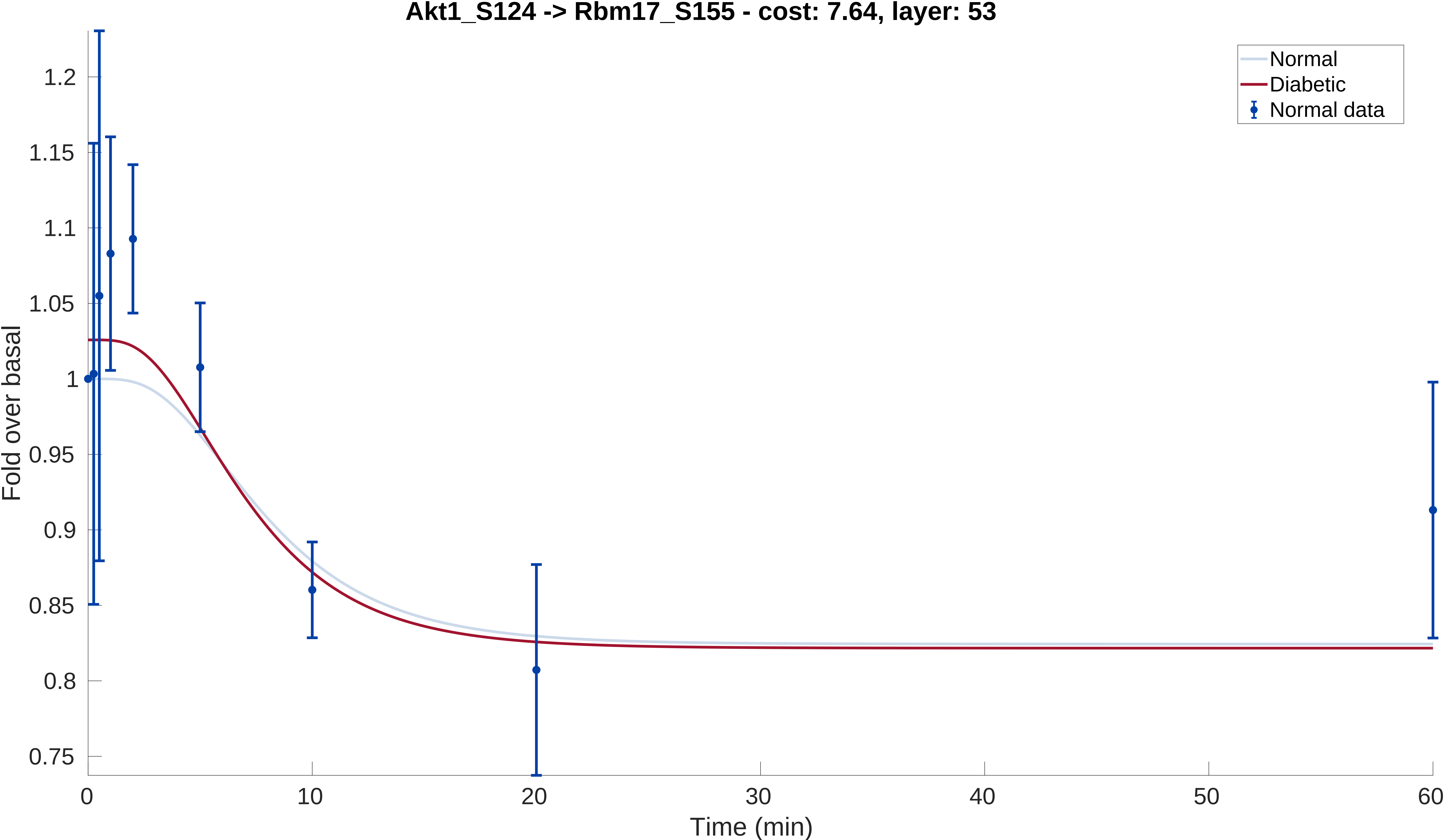

Akt1_S124 -> Rbm17_S155 - cost: 7.64, layer: 53
Normal
Diabetic
Normal data
Fold over basal
Time (min)
1.2
1.15
1.1
1.05
1
0.95
0.9
0.85
0.8
0.75
0
10
20
30
40
50
60

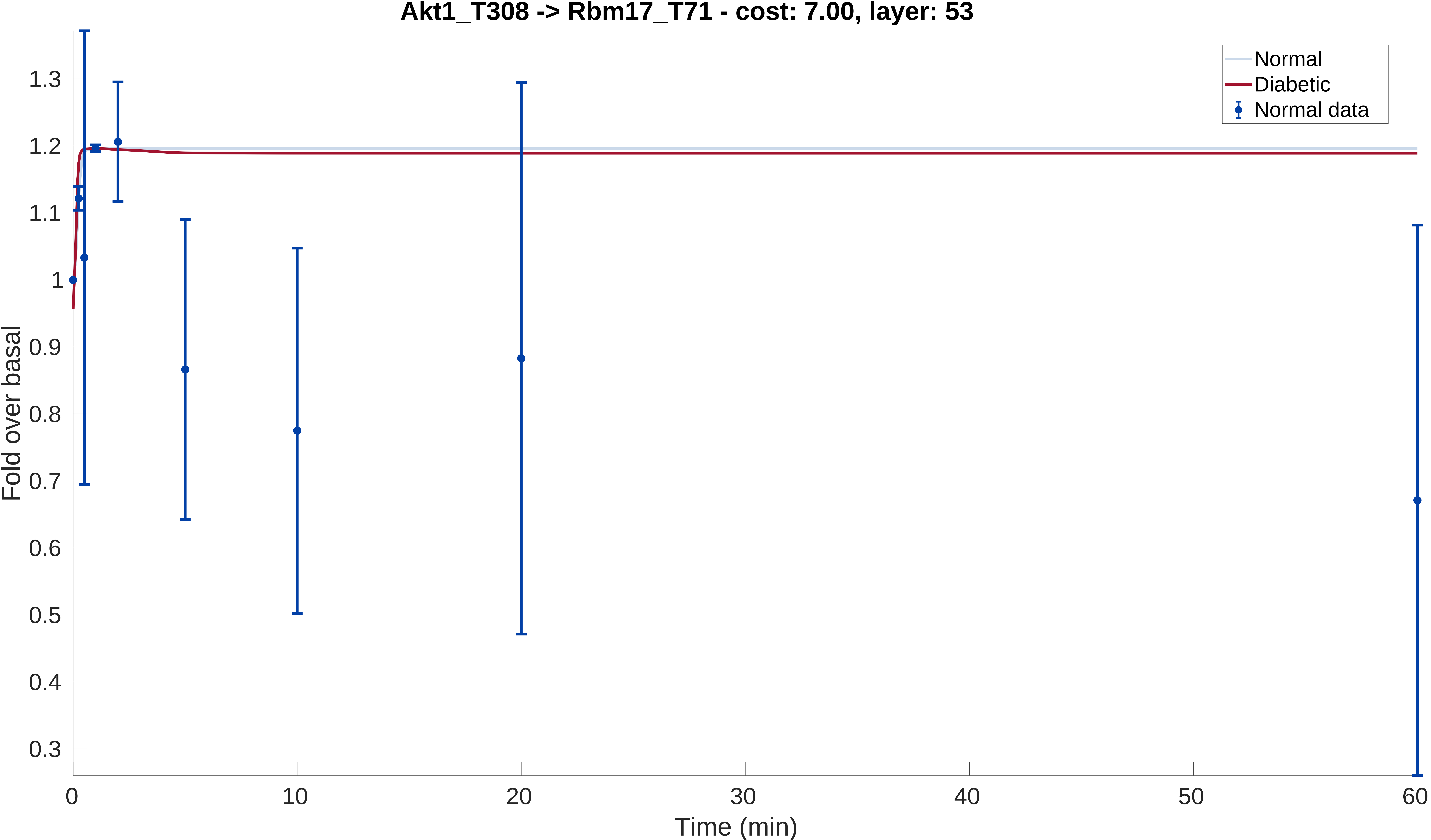

Akt1_T308 -> Rbm17_T71 - cost: 7.00, layer: 53
Normal
Diabetic
Normal data
Fold over basal
1.3
1.2
1.1
1
0.9
0.8
0.7
0.6
0.5
0.4
0.3
0
10
20
30
40
50
60
Time (min)

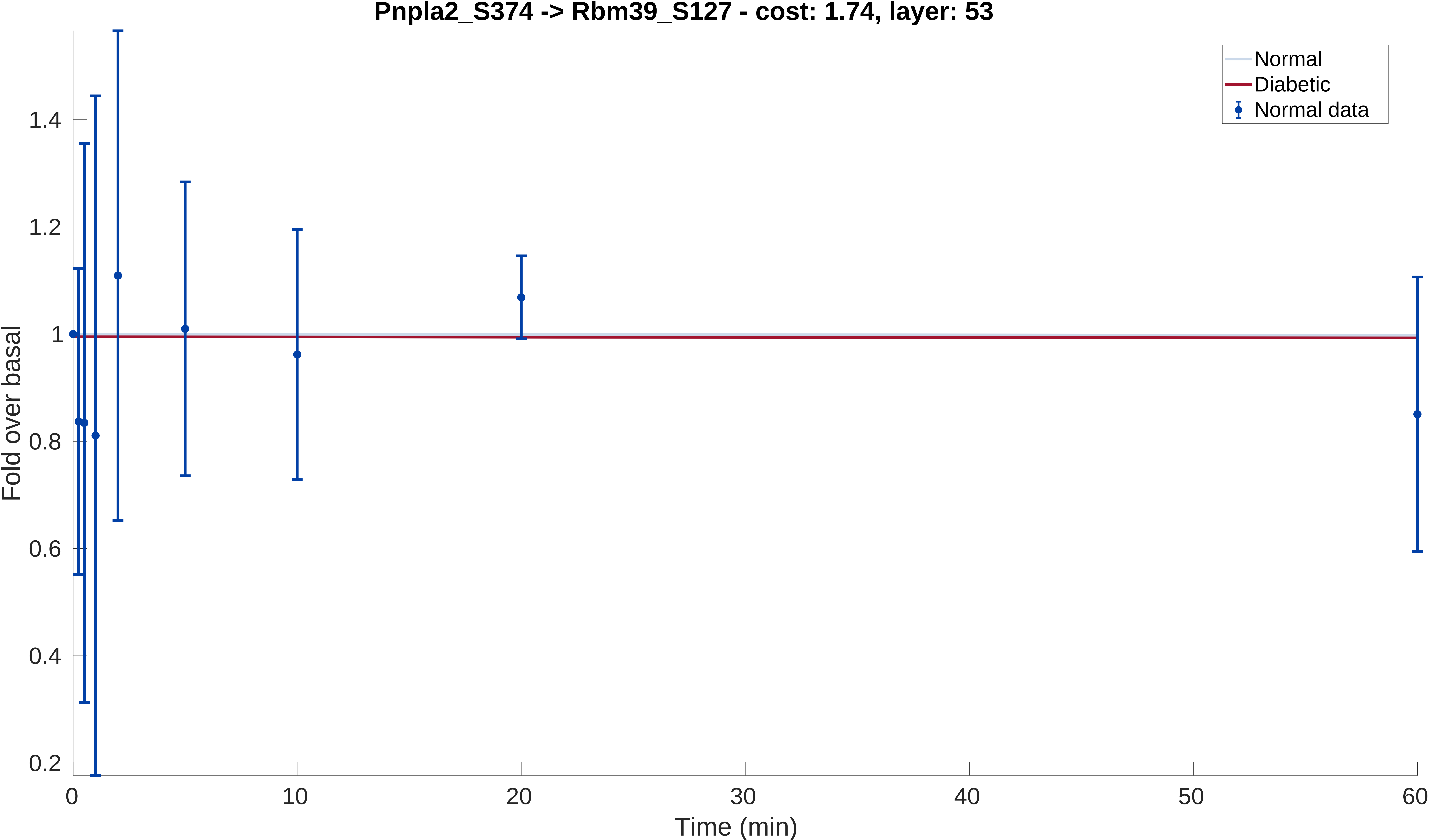

Pnpla2_S374 -> Rbm39_S127 - cost: 1.74, layer: 53
Normal
Diabetic
Normal data
Fold over basal
0.2
0.4
0.6
0.8
1
1.2
1.4
0
10
20
30
40
50
60
Time (min)

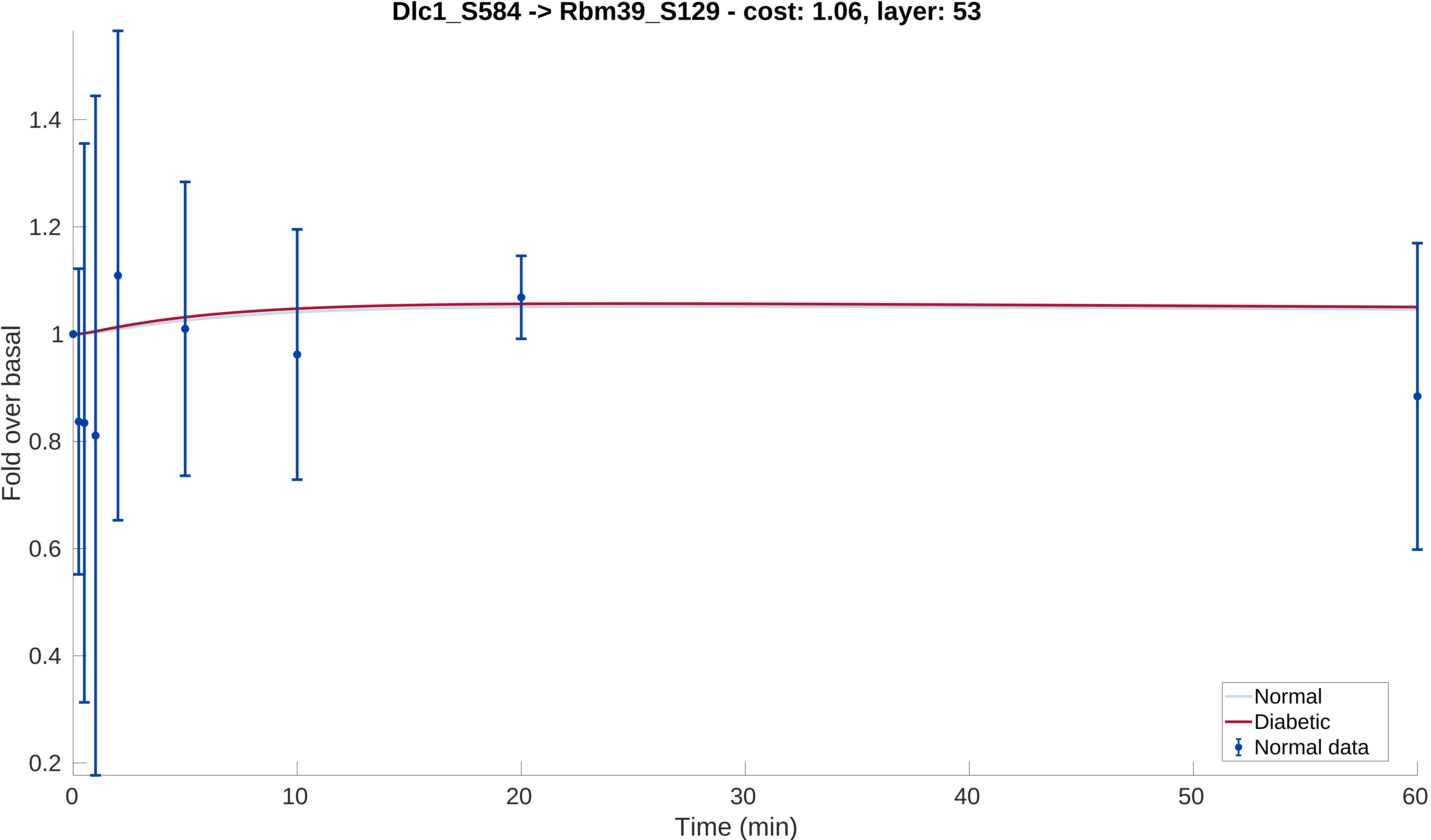
Dlc1_S584 -> Rbm39_S129 - cost: 1.06, layer: 53
Fold over basal
1.4
1.2
1
0.8
0.6
0.4
0.2
0
10
20
30
40
50
60
Time (min)
Normal
Diabetic
Normal data

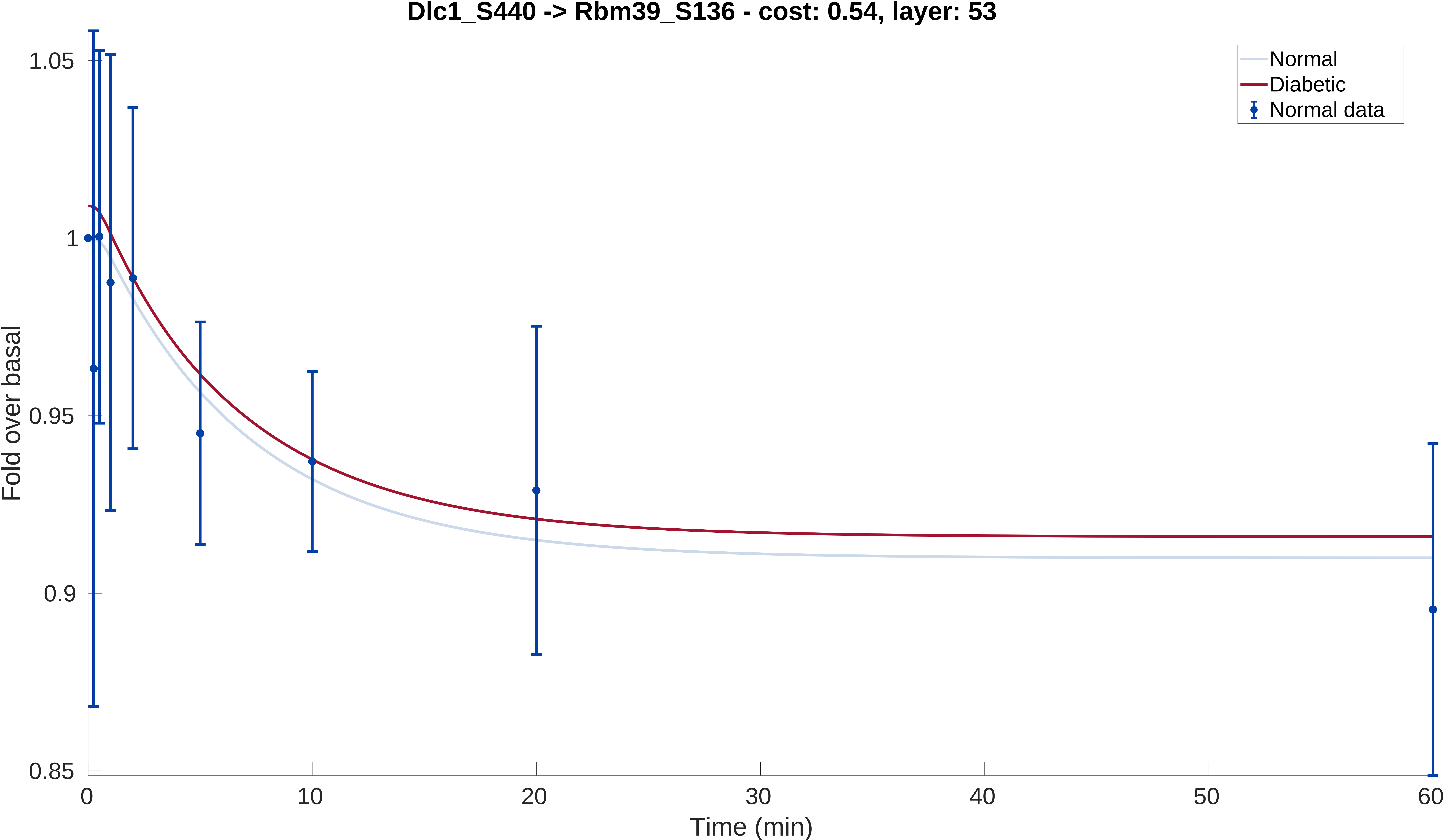

Dlc1_S440 -> Rbm39_S136 - cost: 0.54, layer: 53
Normal
Diabetic
Normal data
Fold over basal
1.05
1
0.95
0.9
0.85
0
10
20
30
40
50
60
Time (min)

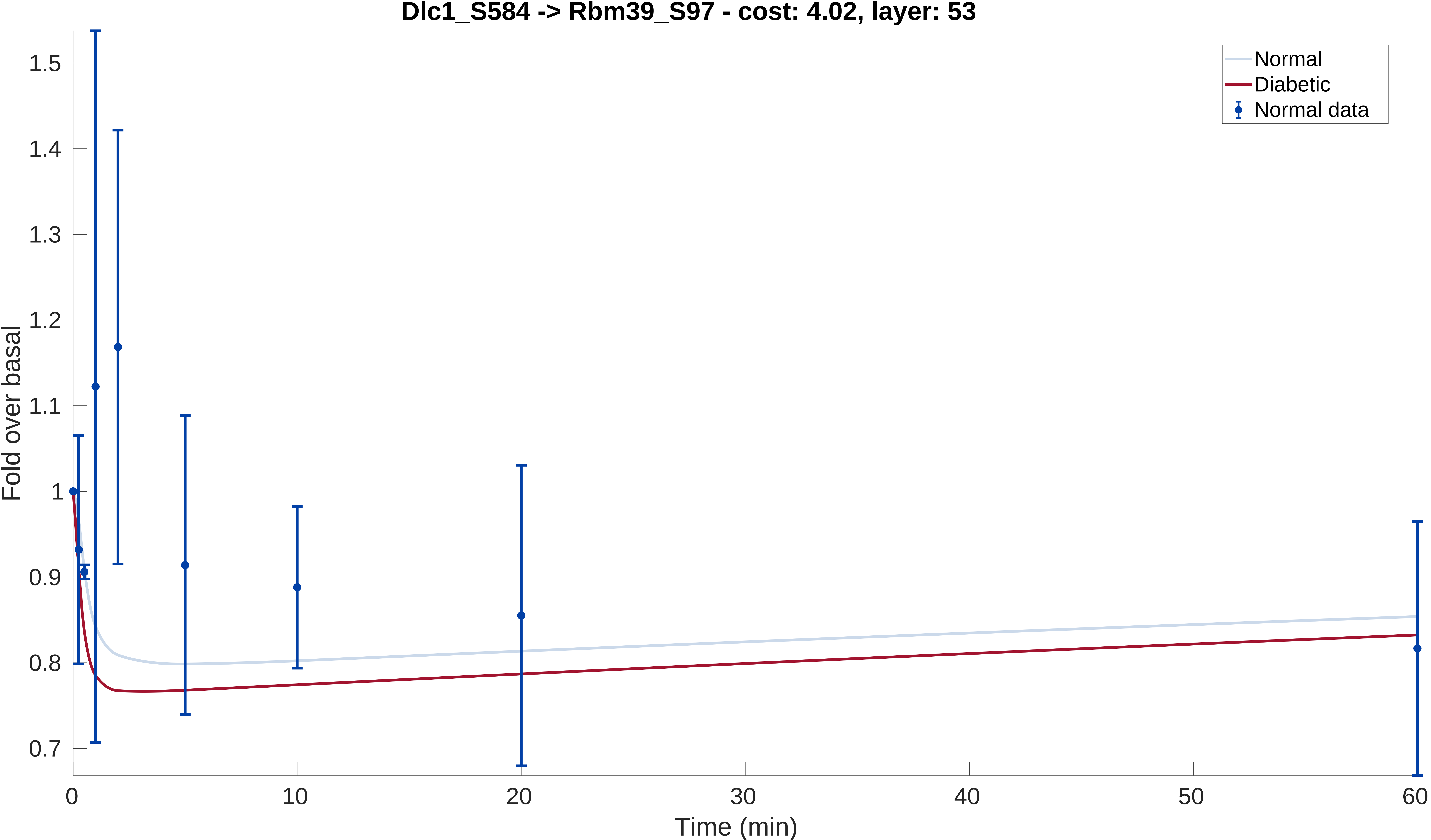

Dlc1_S584 -> Rbm39_S97 - cost: 4.02, layer: 53
Normal
Diabetic
Normal data
Fold over basal
1.5
1.4
1.3
1.2
1.1
1
0.9
0.8
0.7
0
10
20
30
40
50
60
Time (min)

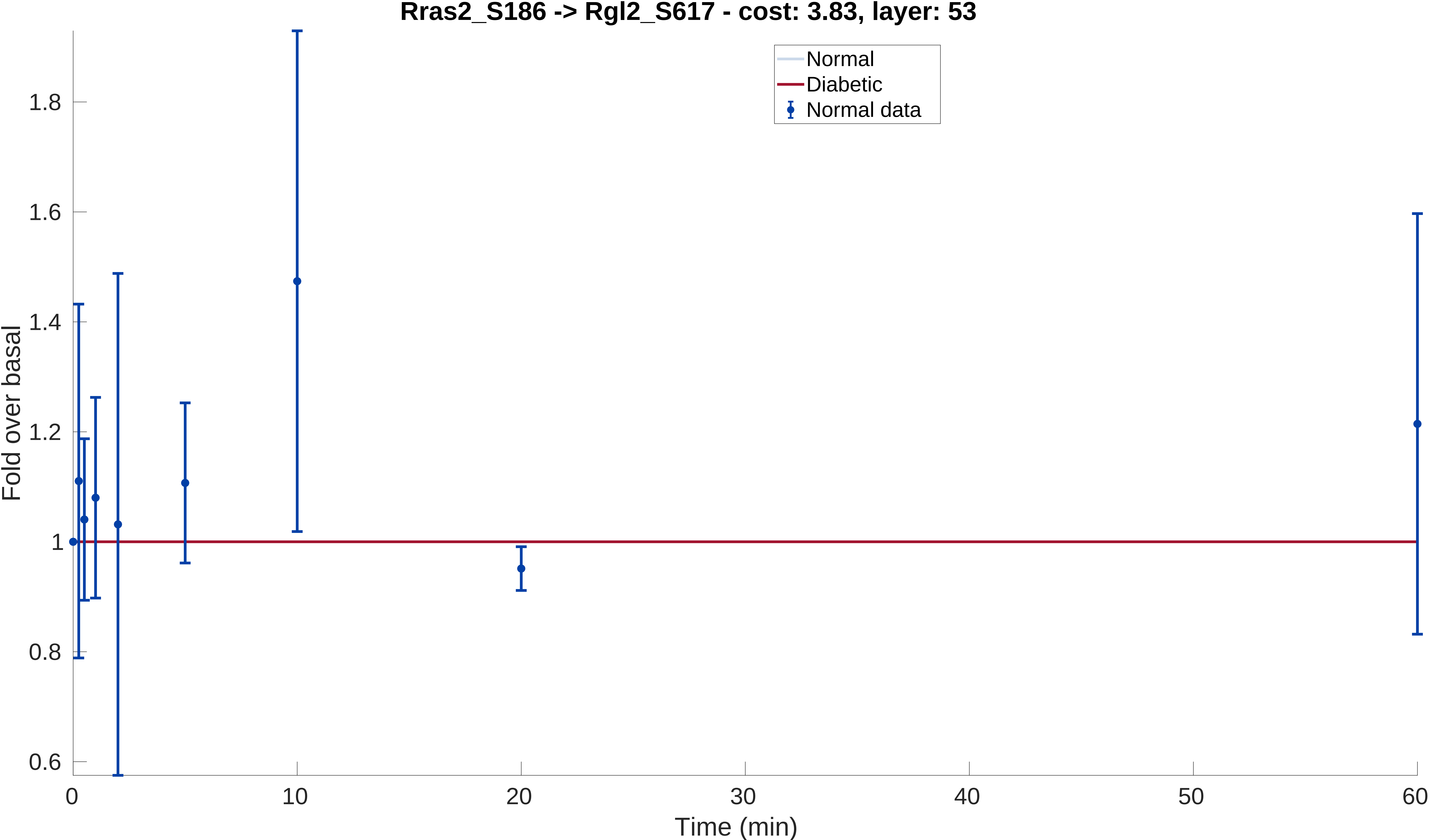

Rras2_S186 -> Rgl2_S617 - cost: 3.83, layer: 53
Normal
Diabetic
Normal data
Fold over basal
Time (min)
0.6
0.8
1
1.2
1.4
1.6
1.8
0
10
20
30
40
50
60

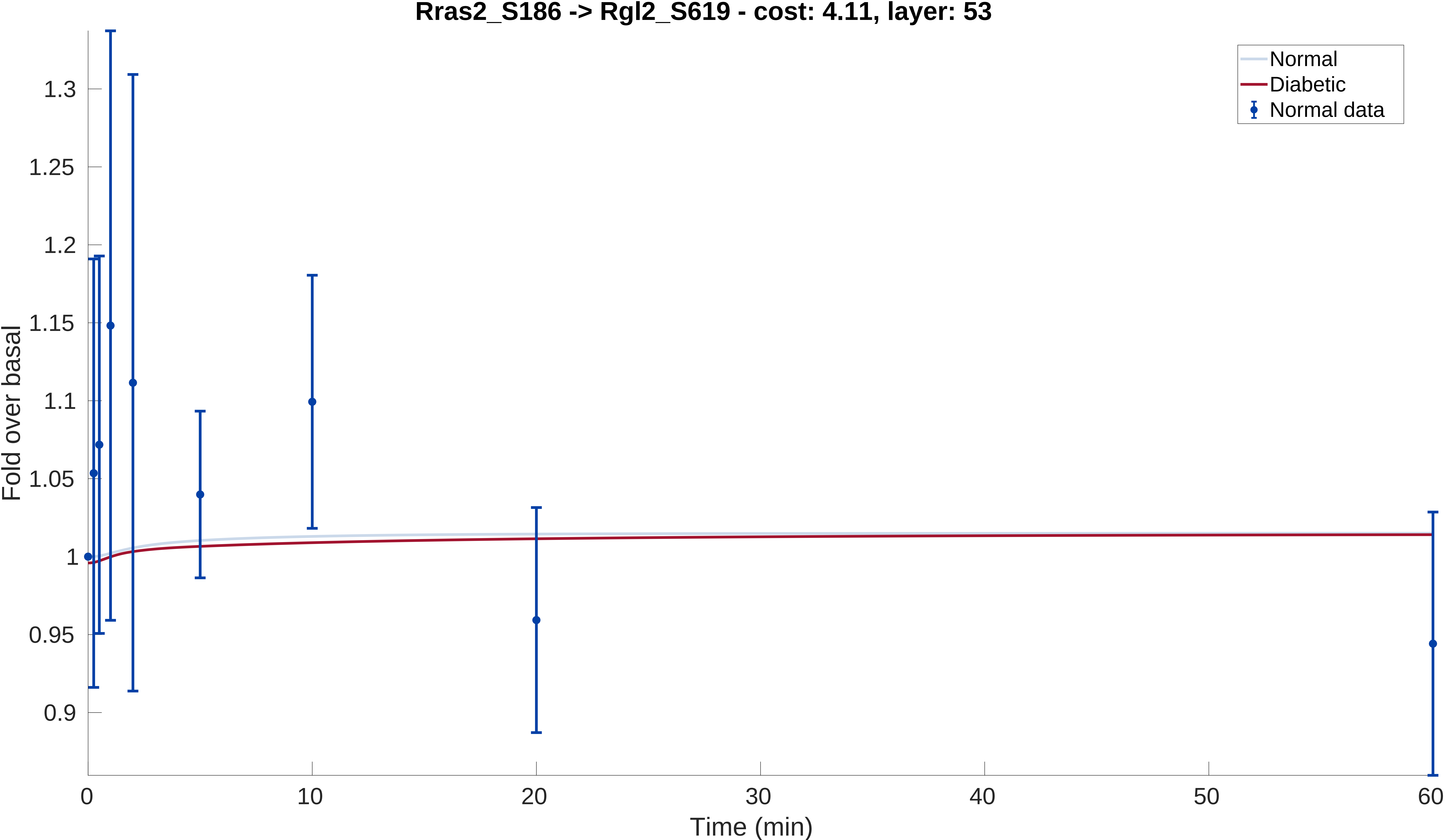

Rras2_S186 -> Rgl2_S619 - cost: 4.11, layer: 53
Fold over basal
1.3
1.25
1.2
1.15
1.1
1.05
1
0.95
0.9
0
10
20
30
40
50
60
Time (min)
Normal
Diabetic
Normal data

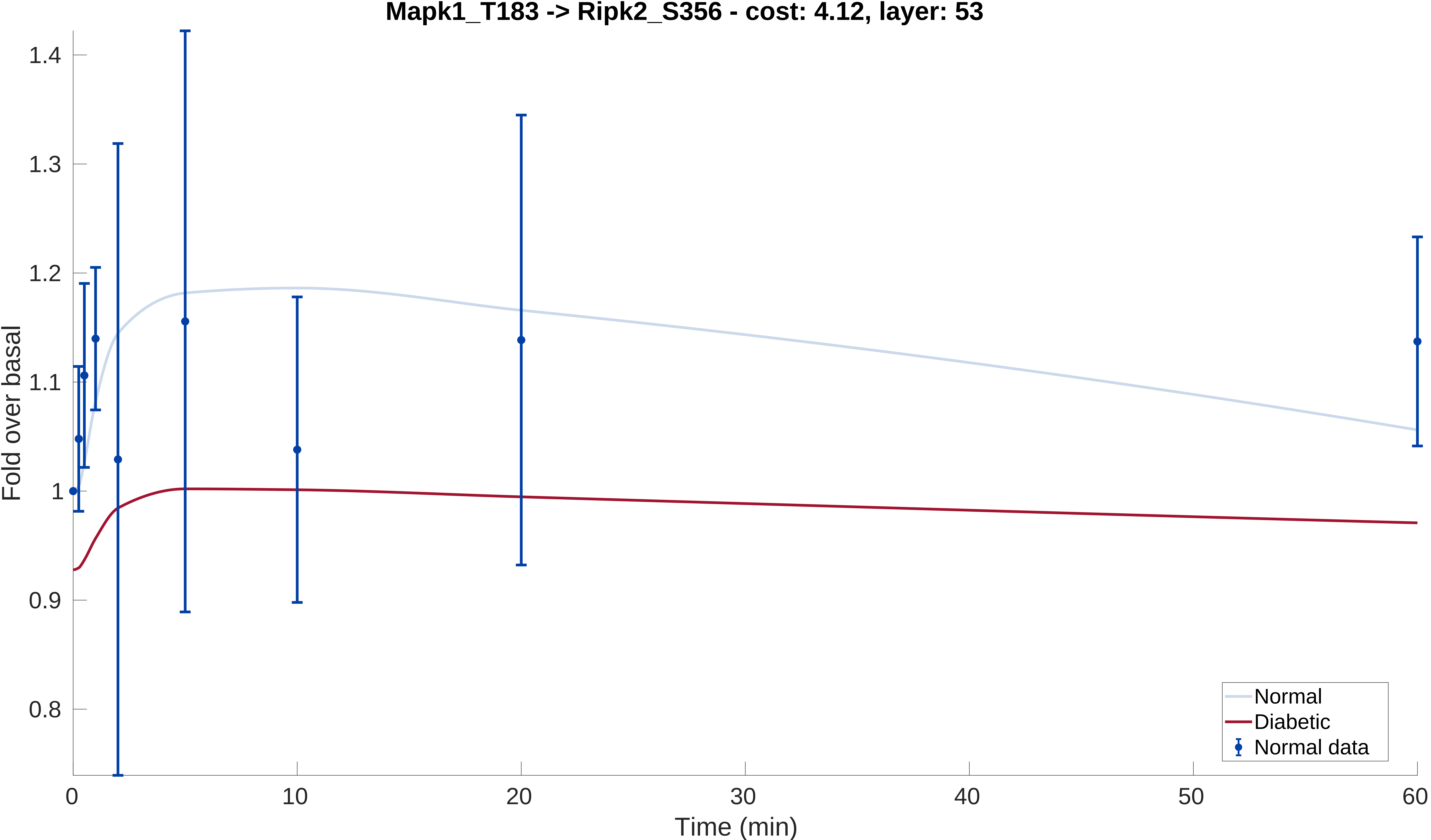
Mapk1_T183 -> Ripk2_S356 - cost: 4.12, layer: 53
Fold over basal
1.4
1.3
1.2
1.1
1
0.9
0.8
0
10
20
30
40
50
60
Time (min)
Normal
Diabetic
Normal data

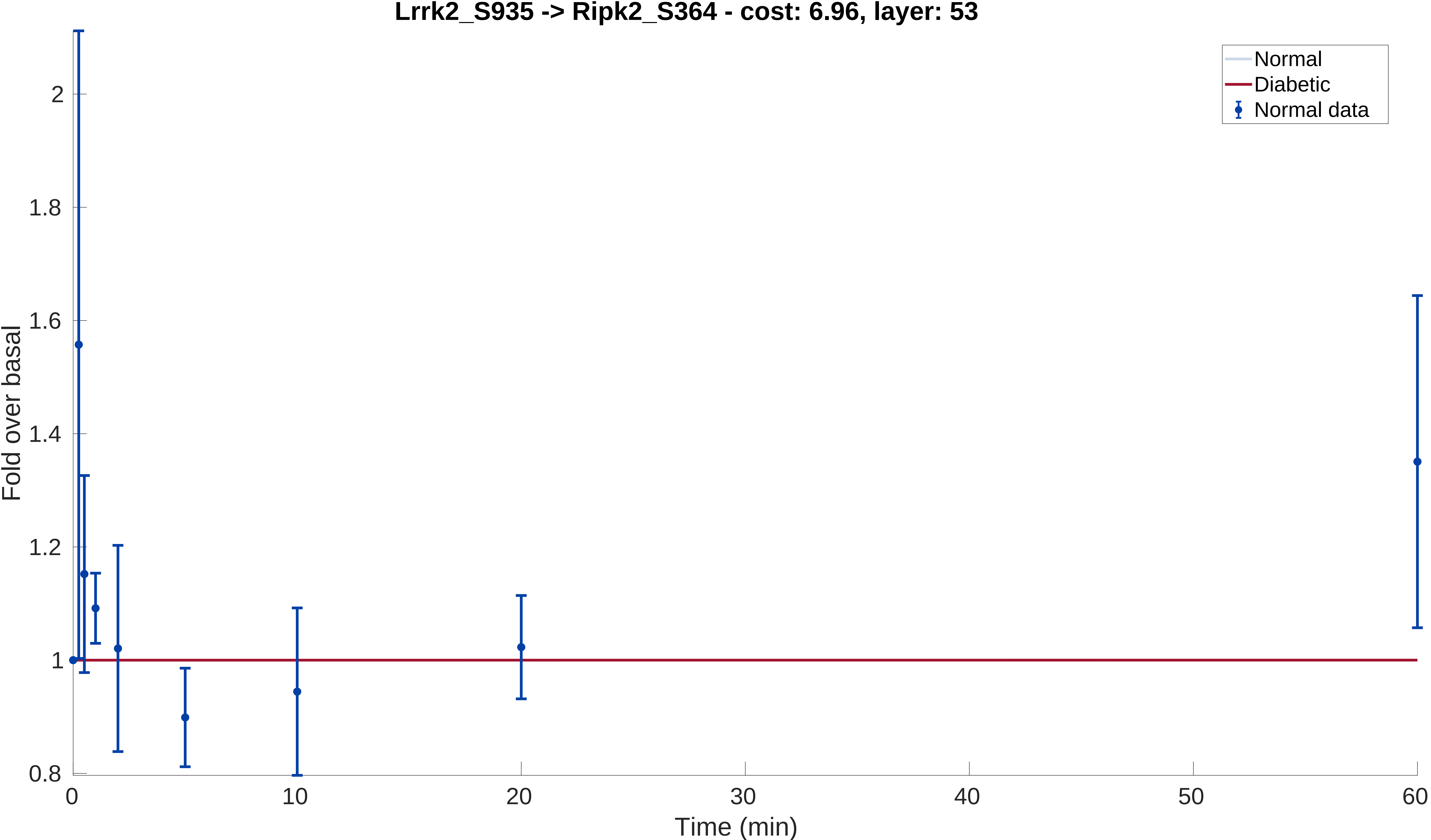

Lrrk2_S935 -> Ripk2_S364 - cost: 6.96, layer: 53
Normal
Diabetic
Normal data
Fold over basal
0.8
1
1.2
1.4
1.6
1.8
2
0
10
20
30
40
50
60
Time (min)

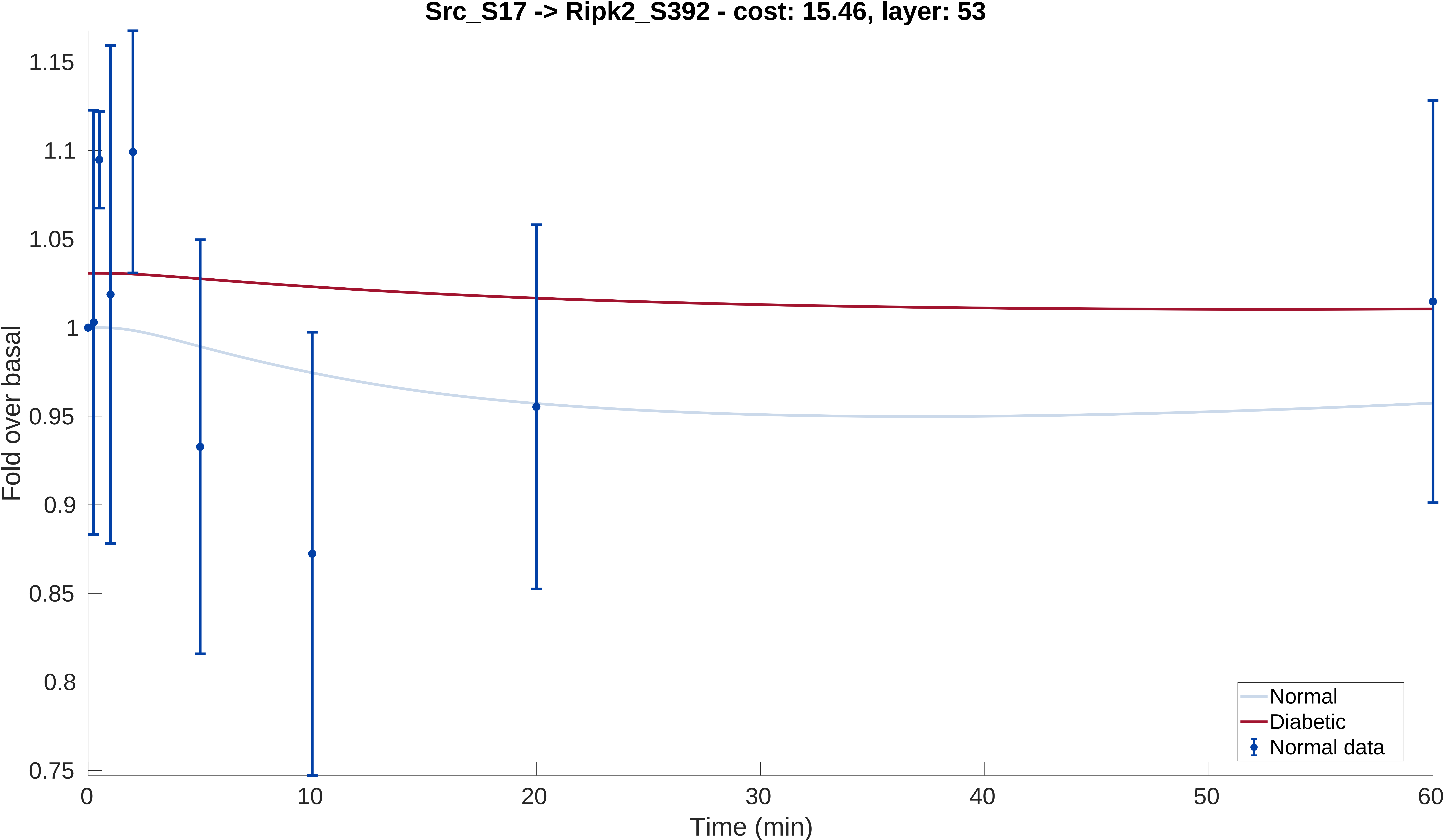
Src_S17 -> Ripk2_S392 - cost: 15.46, layer: 53
Fold over basal
1.15
1.1
1.05
1
0.95
0.9
0.85
0.8
0.75
0
10
20
30
40
50
60
Time (min)
Normal
Diabetic
Normal data

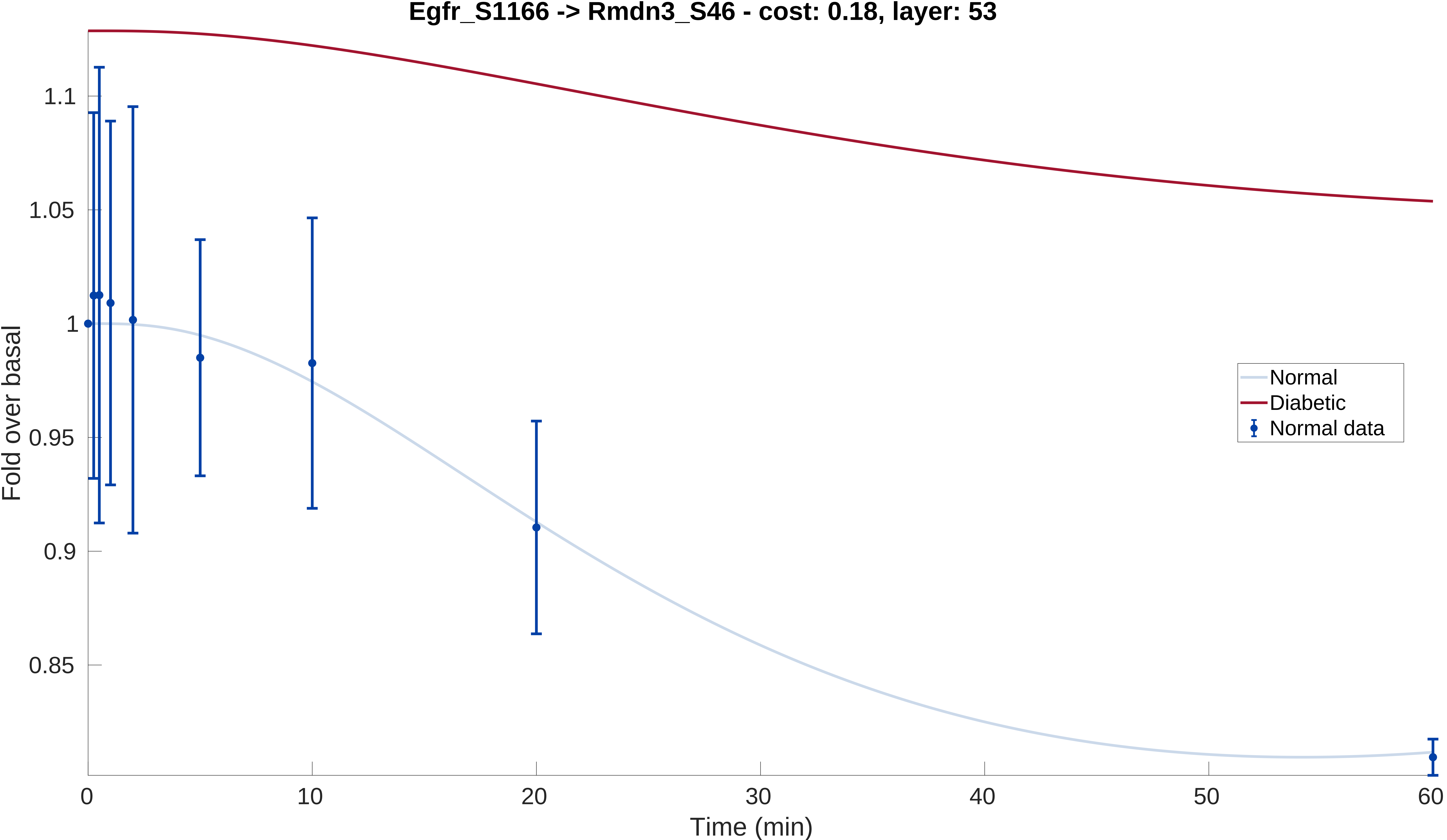

Egfr_S1166 -> Rmdn3_S46 - cost: 0.18, layer: 53
Fold over basal
1.1
1.05
1
0.95
0.9
0.85
0
10
20
30
40
50
60
Time (min)
Normal
Diabetic
Normal data

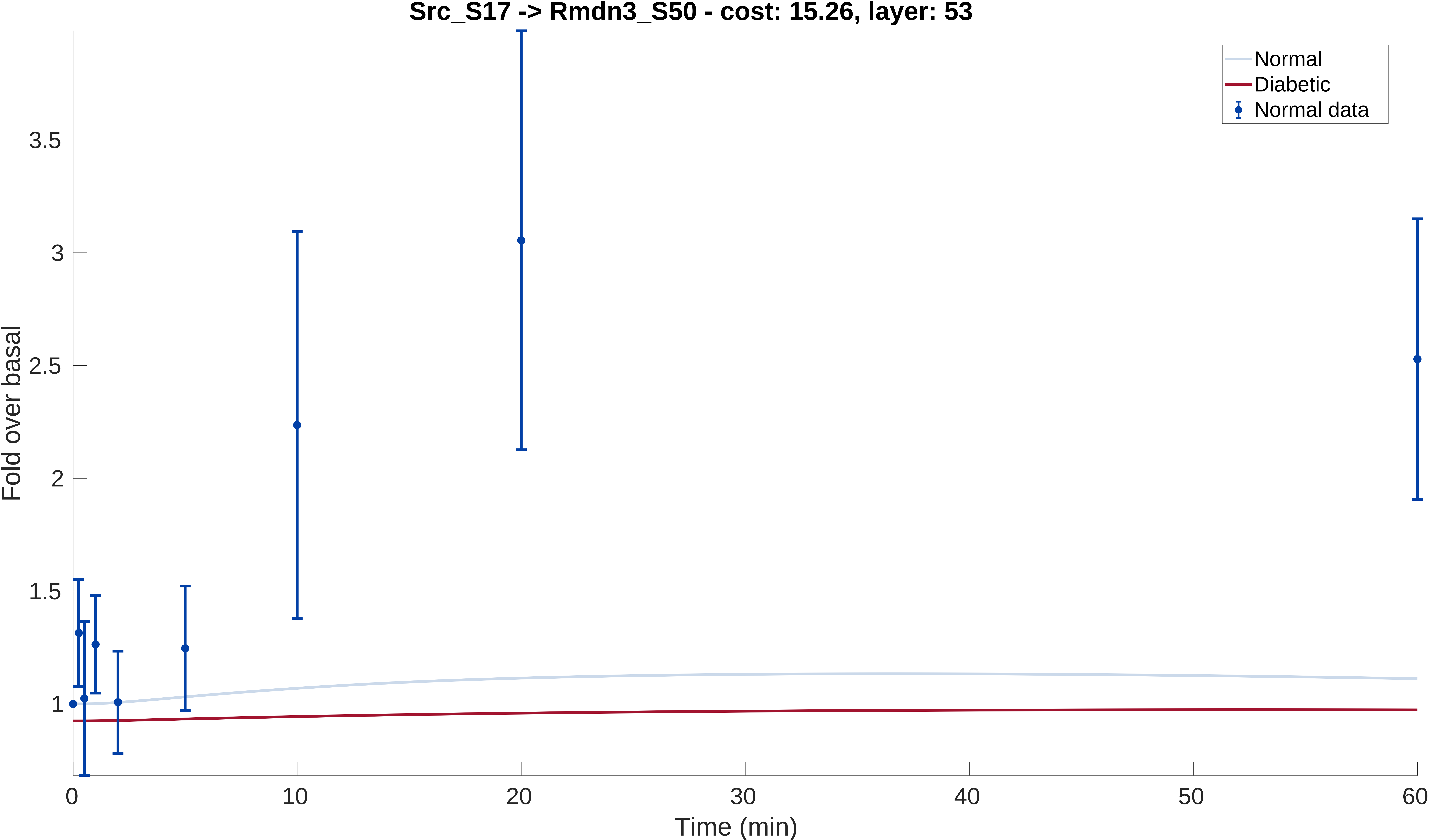
Src_S17 -> Rmdn3_S50 - cost: 15.26, layer: 53
Normal
Diabetic
Normal data
Fold over basal
3.5
3
2.5
2
1.5
1
0
10
20
30
40
50
60
Time (min)

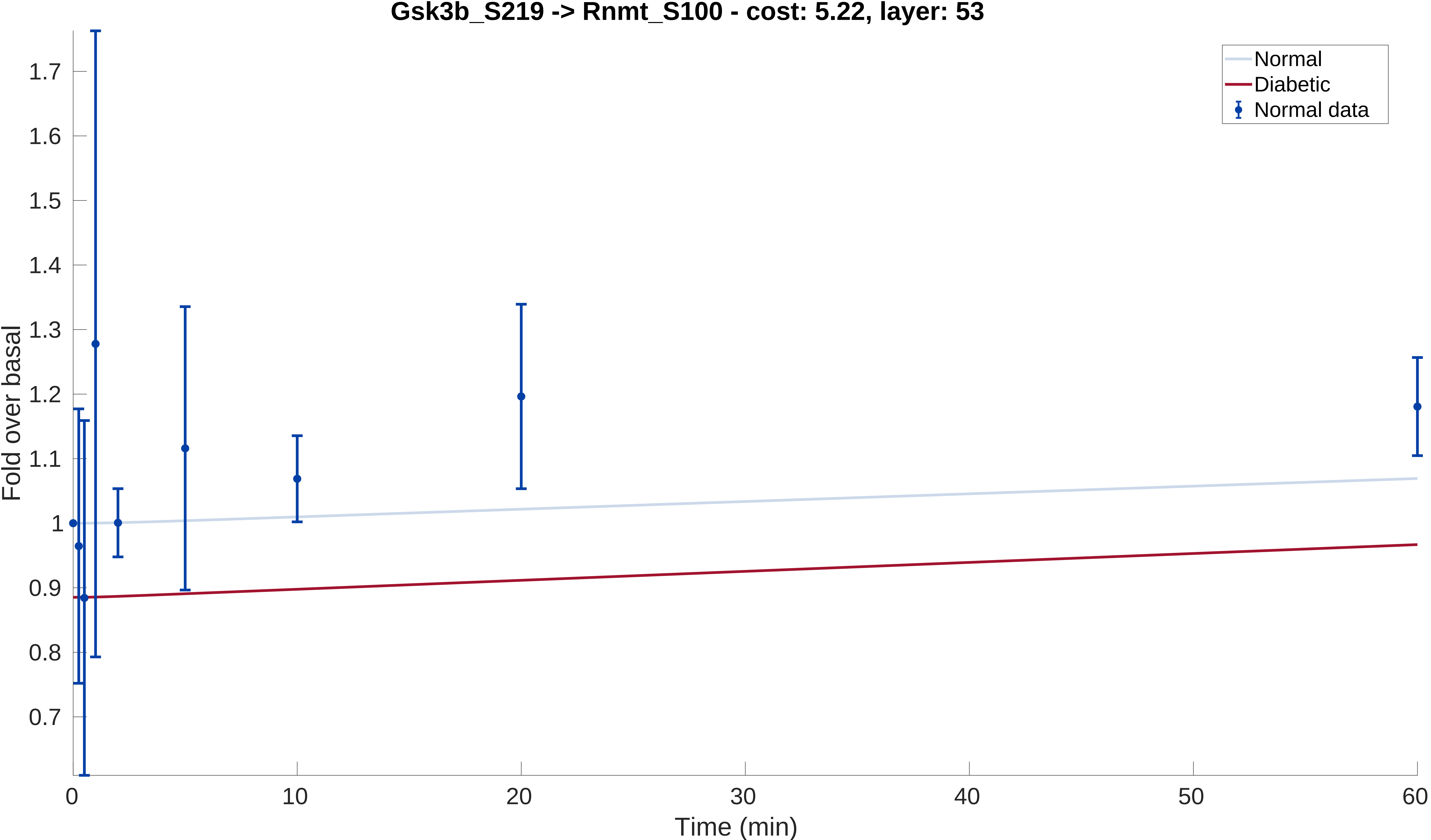

Gsk3b_S219 -> Rnmt_S100 - cost: 5.22, layer: 53
Normal
Diabetic
Normal data
Fold over basal
Time (min)
1.7
1.6
1.5
1.4
1.3
1.2
1.1
1
0.9
0.8
0.7
0
10
20
30
40
50
60

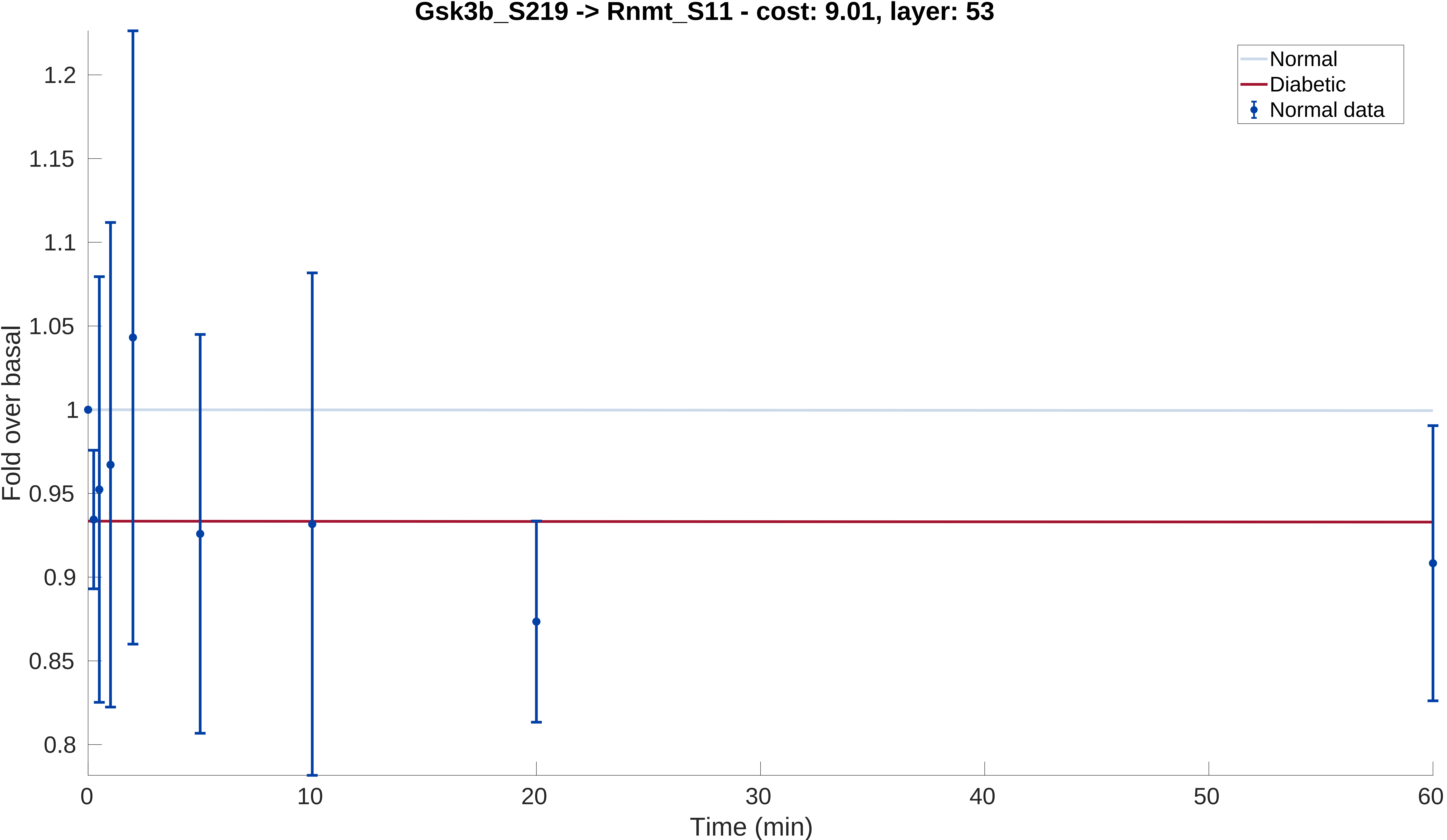

Gsk3b_S219 -> Rnmt_S11 - cost: 9.01, layer: 53
Normal
Diabetic
Normal data
Fold over basal
1.2
1.15
1.1
1.05
1
0.95
0.9
0.85
0.8
0
10
20
30
40
50
60
Time (min)

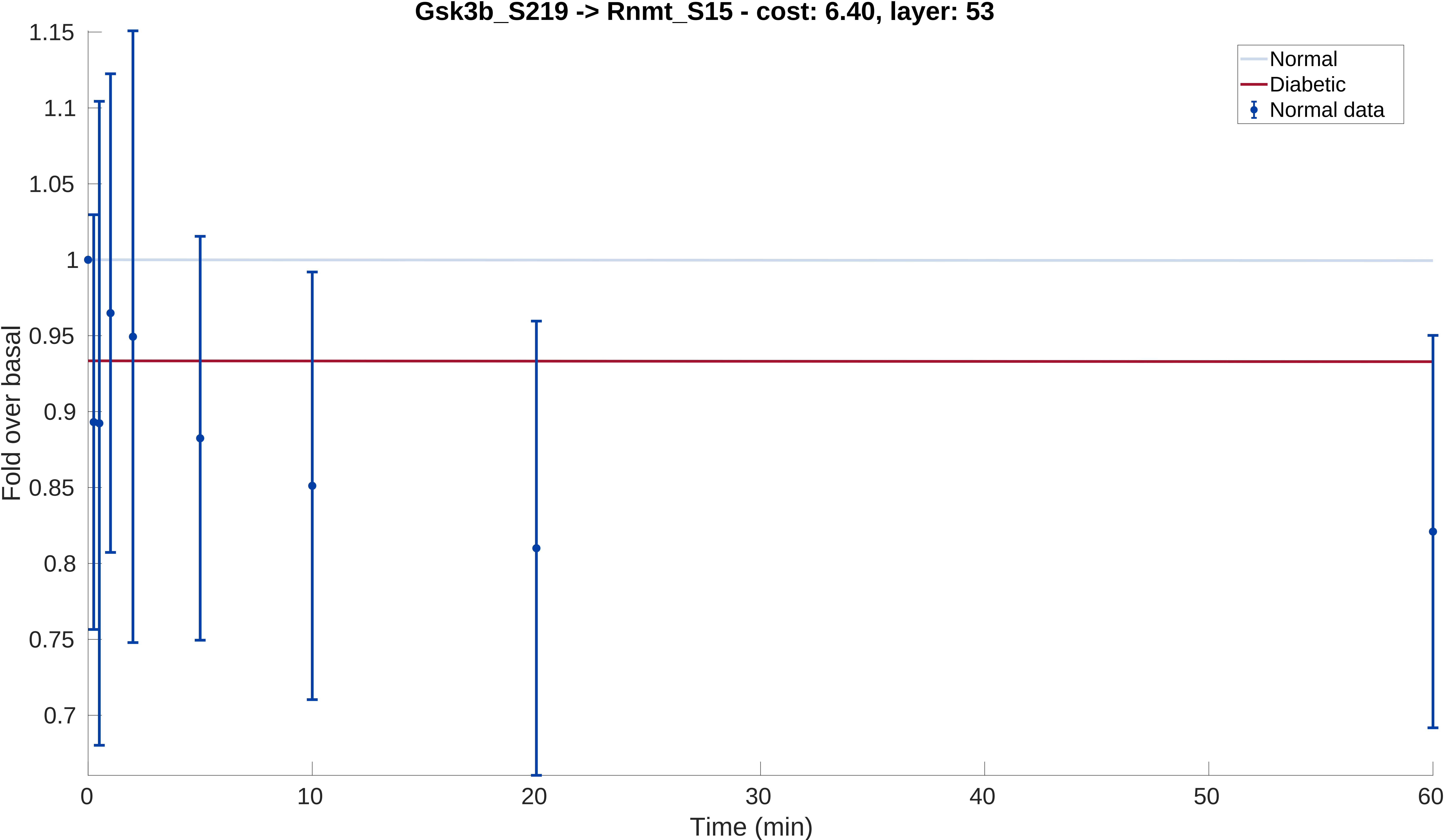

Gsk3b_S219 -> Rnmt_S15 - cost: 6.40, layer: 53
Normal
Diabetic
Normal data
Fold over basal
1.15
1.1
1.05
1
0.95
0.9
0.85
0.8
0.75
0.7
0
10
20
30
40
50
60
Time (min)

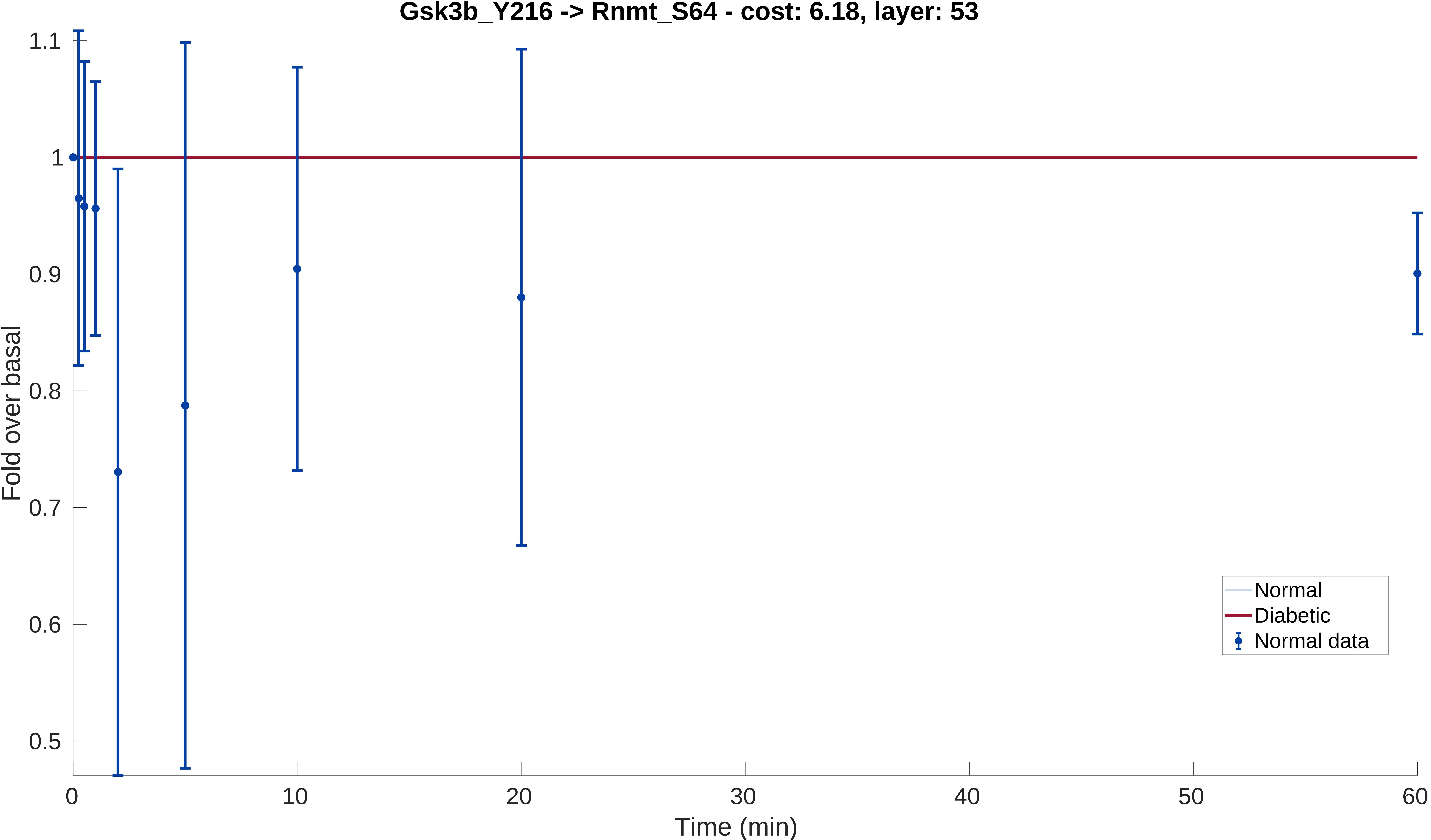

Gsk3b_Y216 -> Rnmt_S64 - cost: 6.18, layer: 53
Fold over basal
Time (min)
1.1
1
0.9
0.8
0.7
0.6
0.5
0
10
20
30
40
50
60
Normal
Diabetic
Normal data

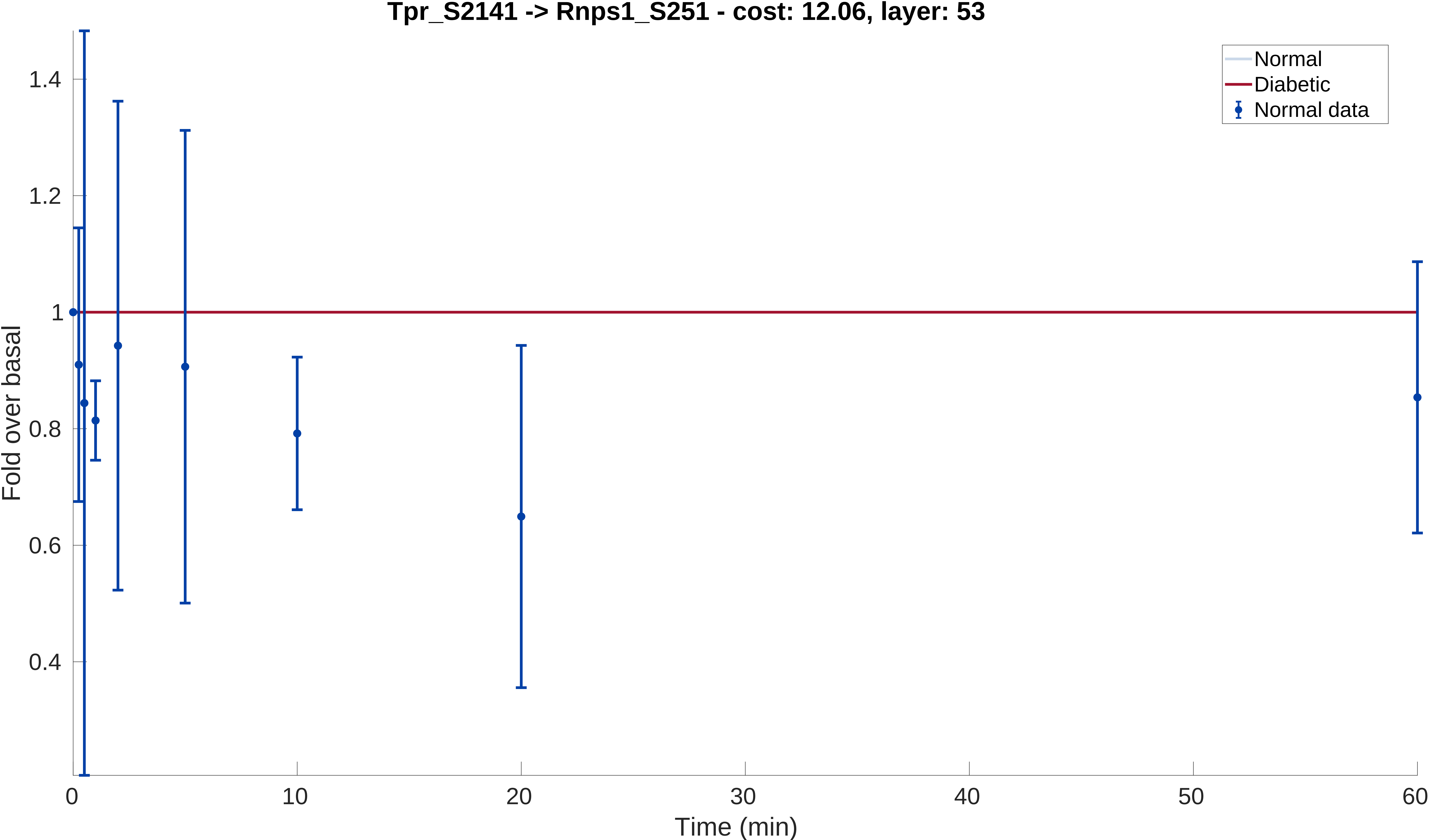
Tpr_S2141 -> Rnps1_S251 - cost: 12.06, layer: 53
Normal
Diabetic
Normal data
Fold over basal
1.4
1.2
1
0.8
0.6
0.4
0
10
20
30
40
50
60
Time (min)

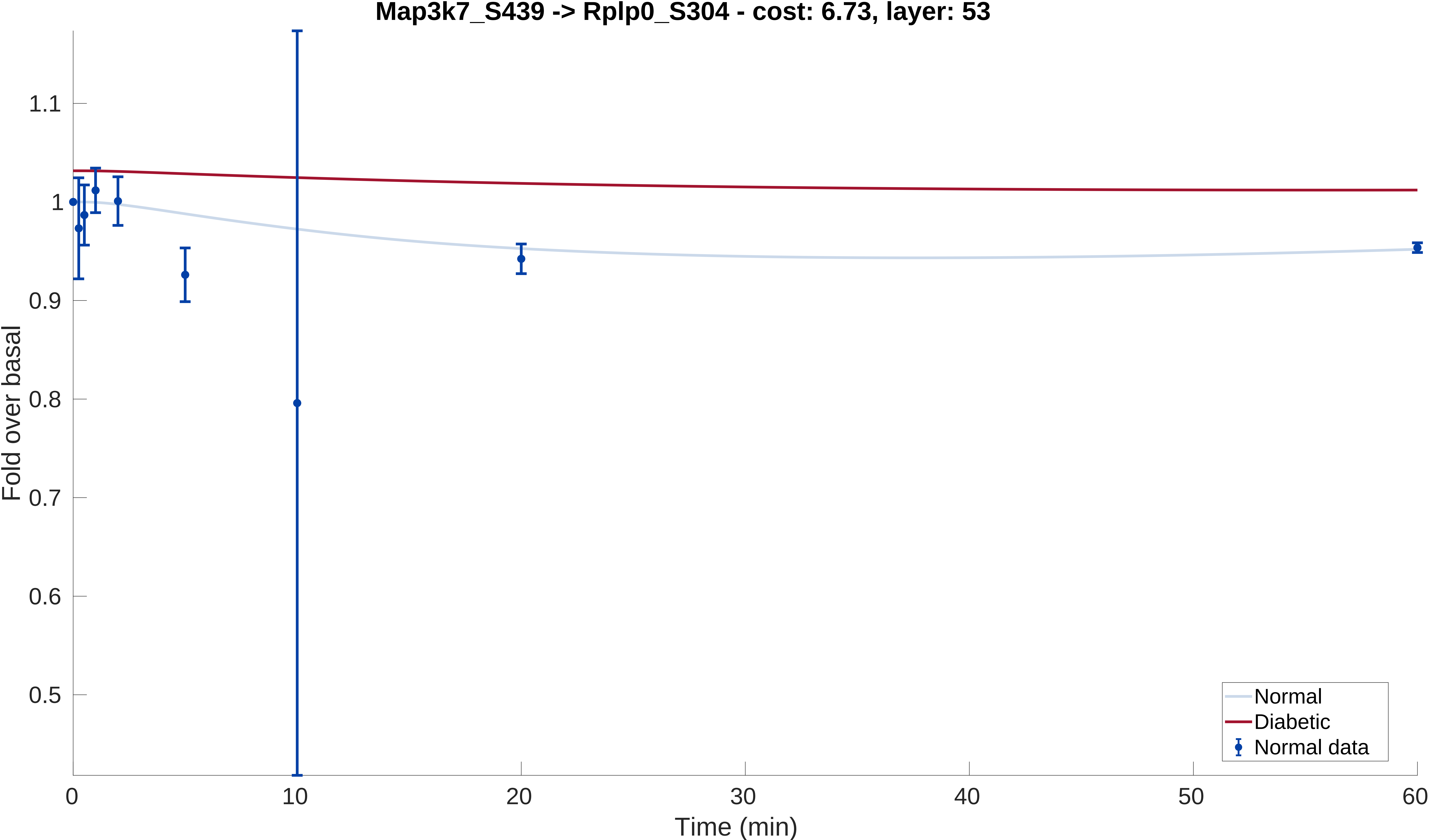

Map3k7_S439 -> Rplp0_S304 - cost: 6.73, layer: 53
1.1
1
0.9
0.8
0.7
0.6
0.5
Fold over basal
0
10
20
30
40
50
60
Time (min)
Normal
Diabetic
Normal data

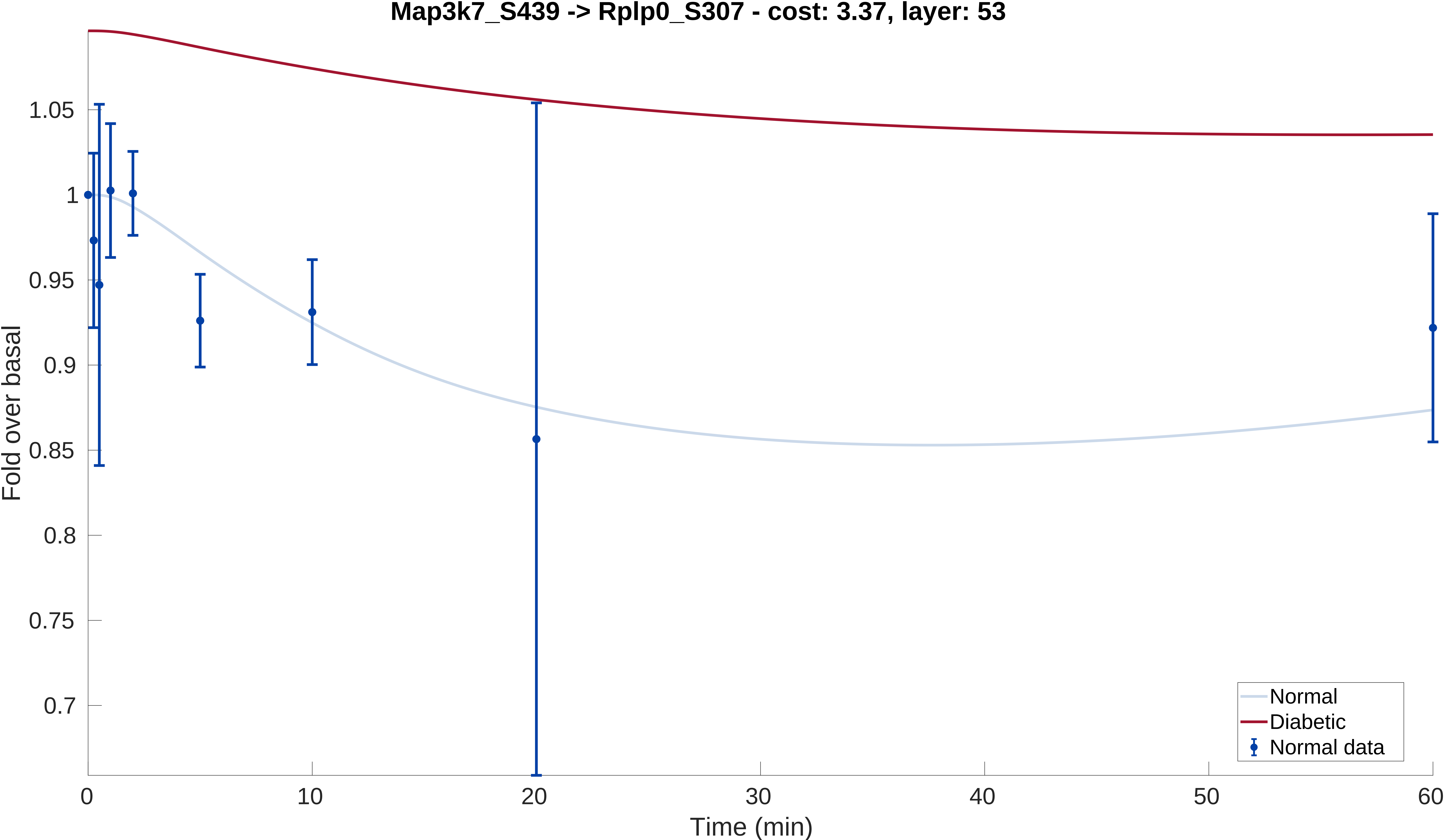

Map3k7_S439 -> Rplp0_S307 - cost: 3.37, layer: 53
Fold over basal
1.05
1
0.95
0.9
0.85
0.8
0.75
0.7
0
10
20
30
40
50
60
Time (min)
Normal
Diabetic
Normal data

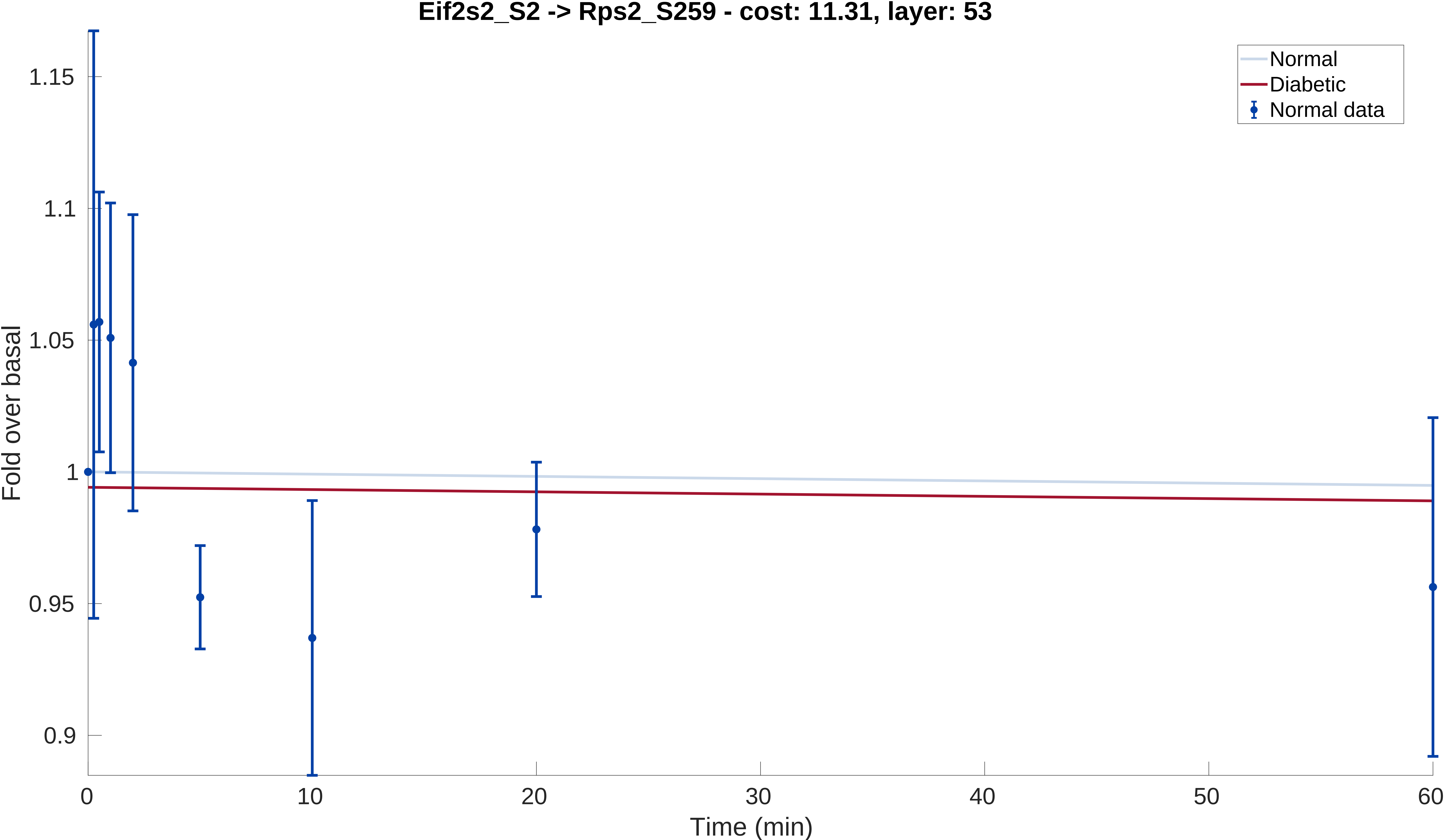

Eif2s2_S2 -> Rps2_S259 - cost: 11.31, layer: 53
Normal
Diabetic
Normal data
1.15
1.1
1.05
1
0.95
0.9
Fold over basal
0
10
20
30
40
50
60
Time (min)

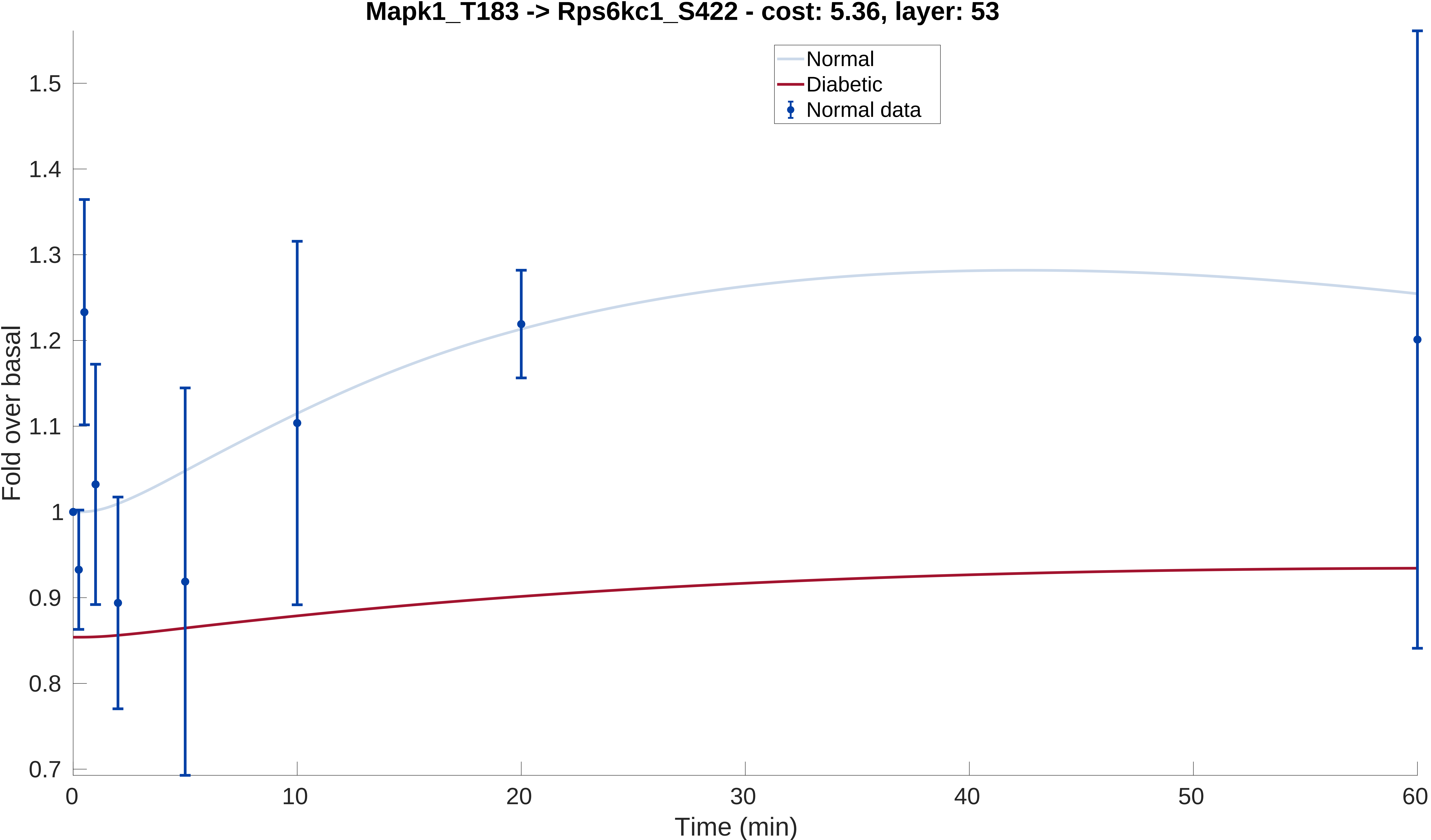

Mapk1_T183 -> Rps6kc1_S422 - cost: 5.36, layer: 53
Normal
Diabetic
Normal data
Fold over basal
1.5
1.4
1.3
1.2
1.1
1
0.9
0.8
0.7
0
10
20
30
40
50
60
Time (min)

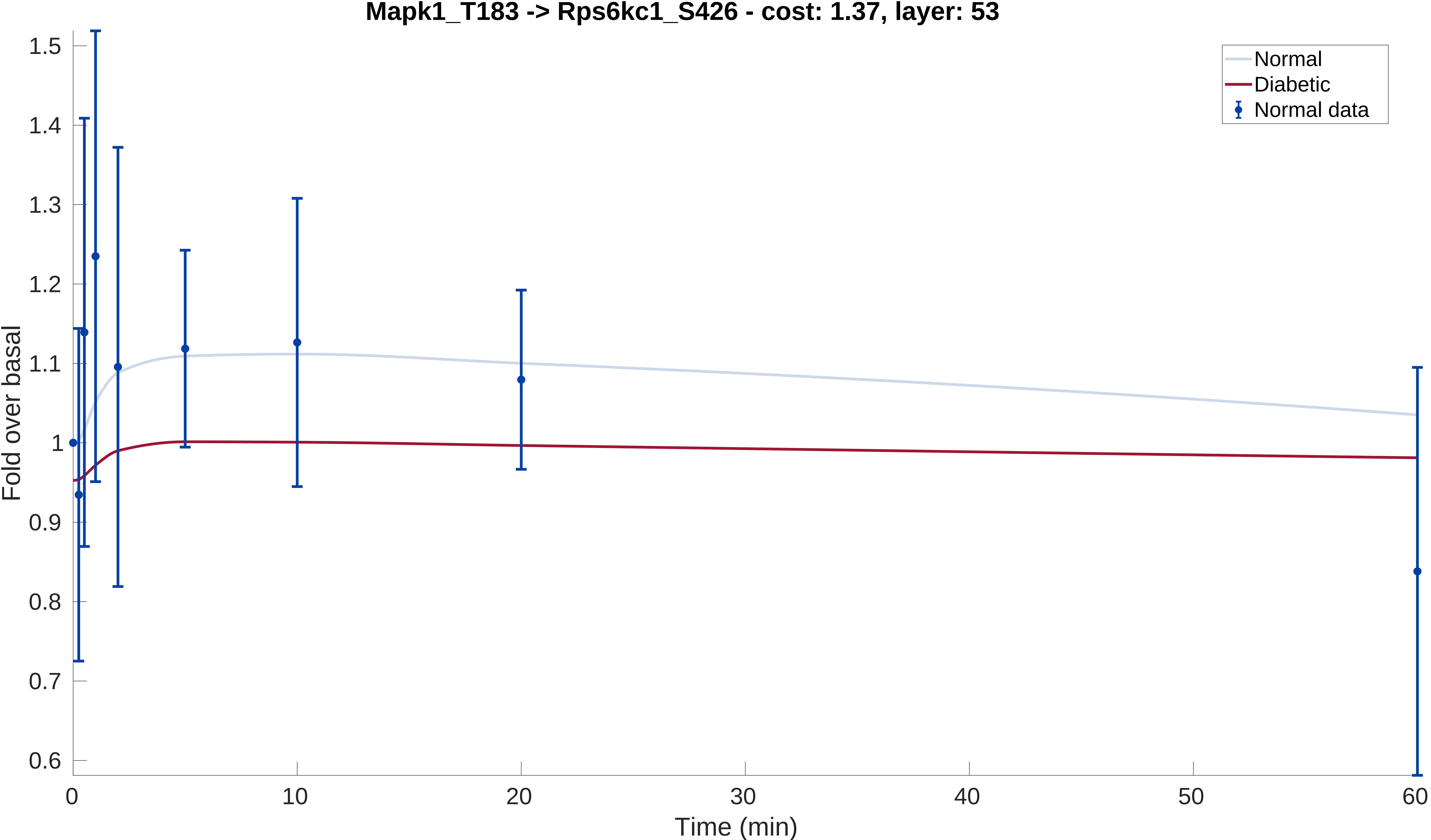

Mapk1_T183 -> Rps6kc1_S426 - cost: 1.37, layer: 53
Normal
Diabetic
Normal data
Fold over basal
1.5
1.4
1.3
1.2
1.1
1
0.9
0.8
0.7
0.6
0
10
20
30
40
50
60
Time (min)

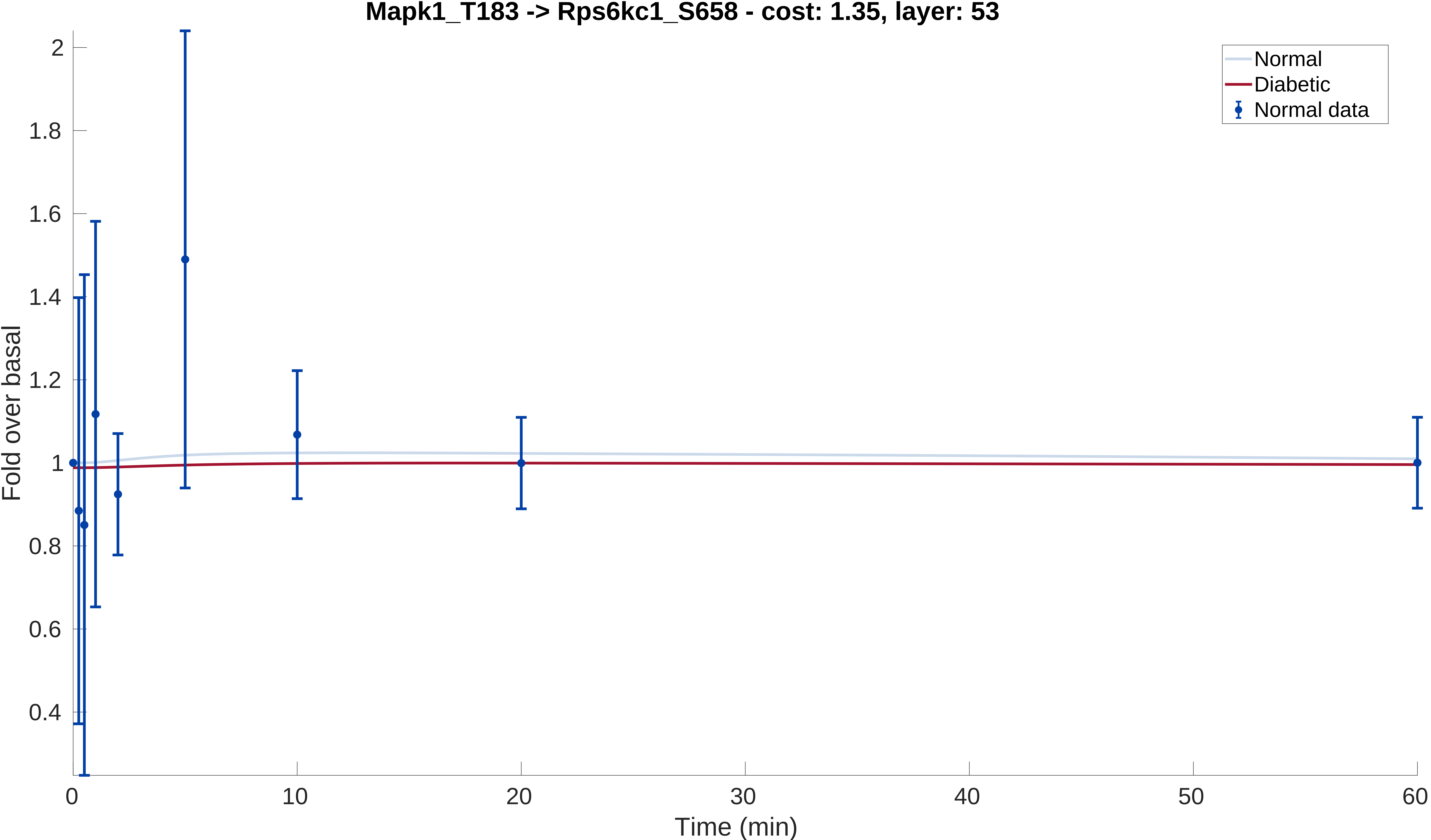

Mapk1_T183 -> Rps6kc1_S658 - cost: 1.35, layer: 53
Normal
Diabetic
Normal data
Fold over basal
2
1.8
1.6
1.4
1.2
1
0.8
0.6
0.4
0
10
20
30
40
50
60
Time (min)

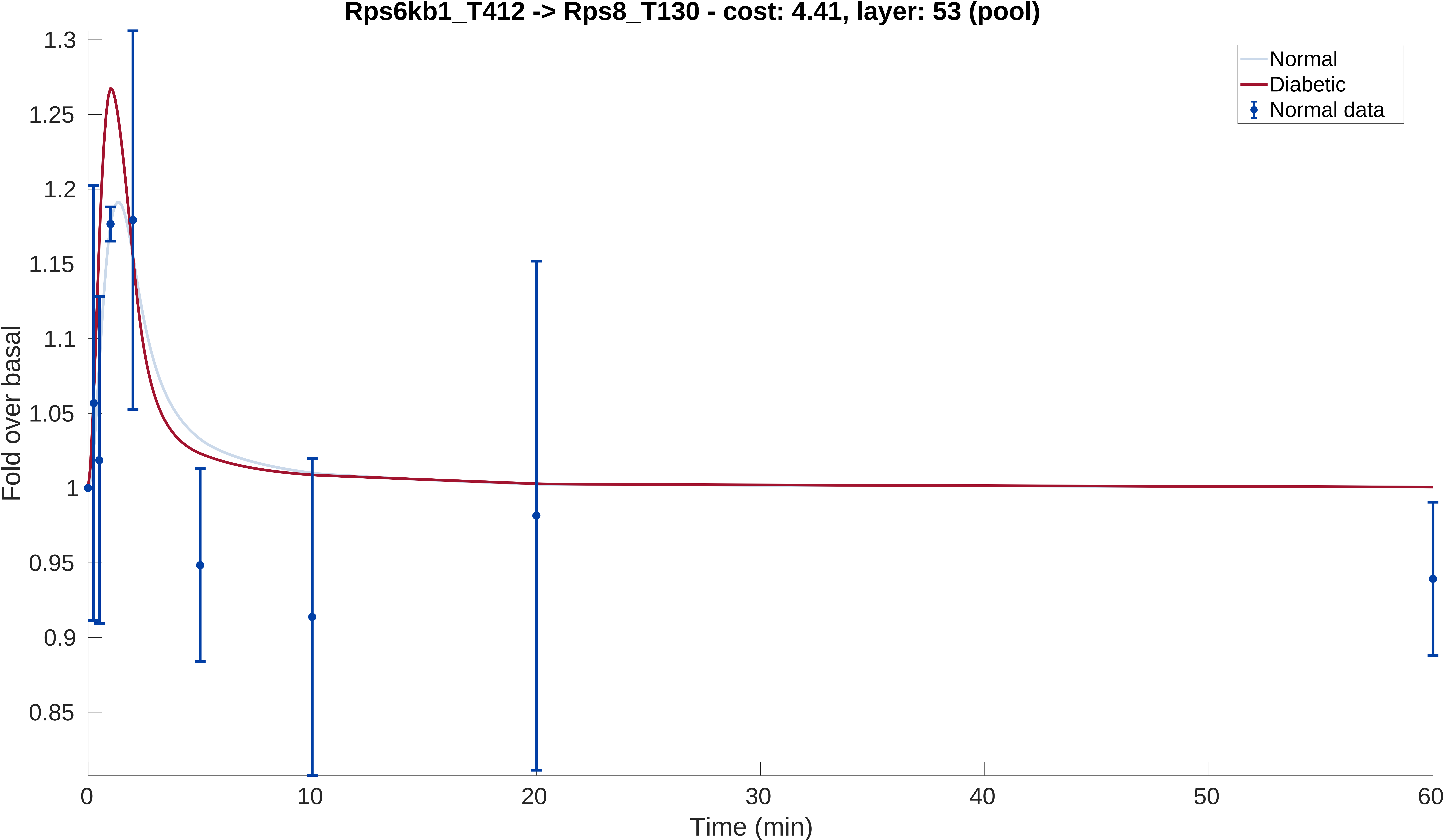

Rps6kb1_T412 -> Rps8_T130 - cost: 4.41, layer: 53 (pool)
Normal
Diabetic
Normal data
Fold over basal
1.3
1.25
1.2
1.15
1.1
1.05
1
0.95
0.9
0.85
0
10
20
30
40
50
60
Time (min)

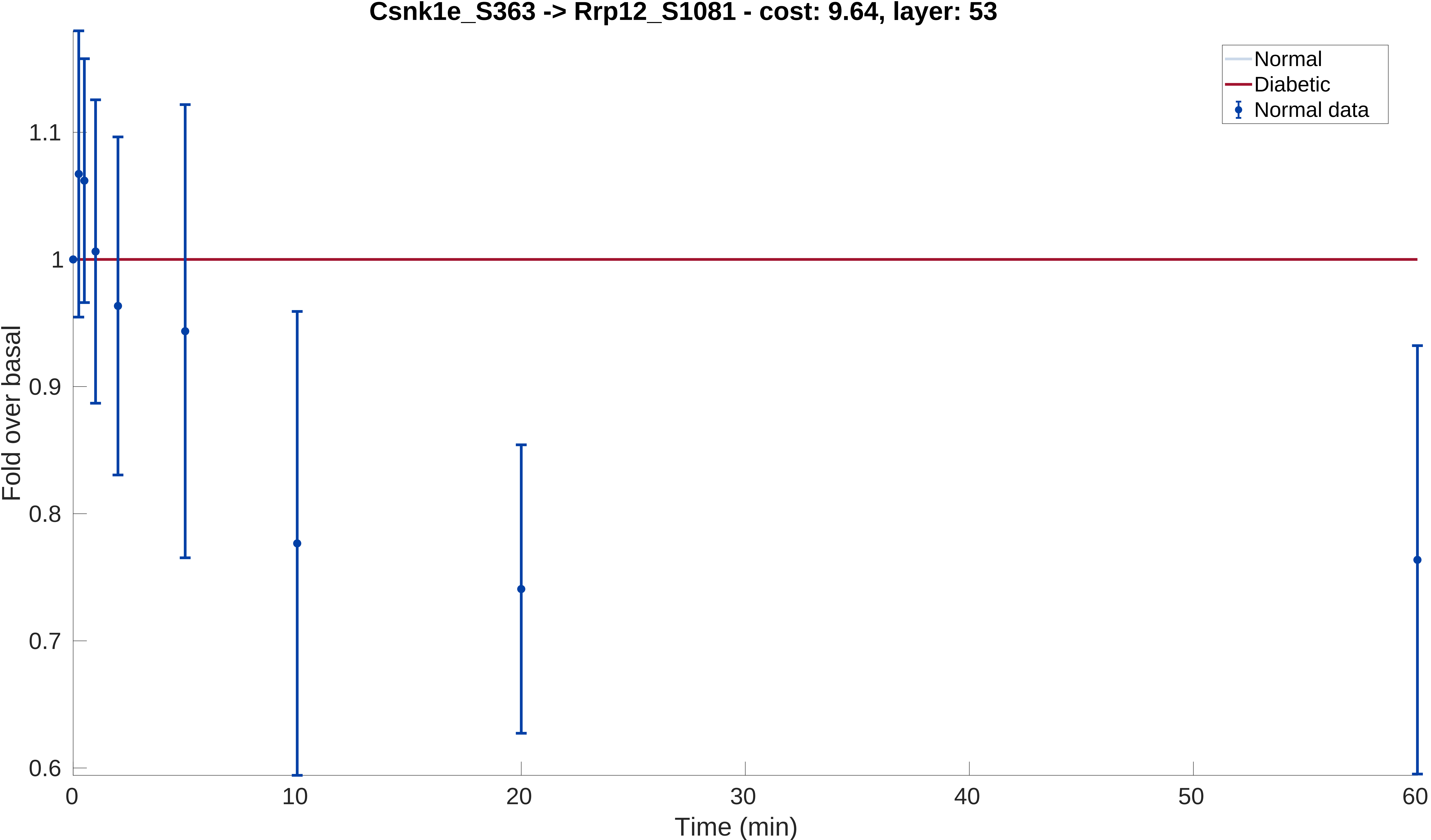
Csnk1e_S363 -> Rrp12_S1081 - cost: 9.64, layer: 53
Normal
Diabetic
Normal data
Fold over basal
1.1
1
0.9
0.8
0.7
0.6
0
10
20
30
40
50
60
Time (min)

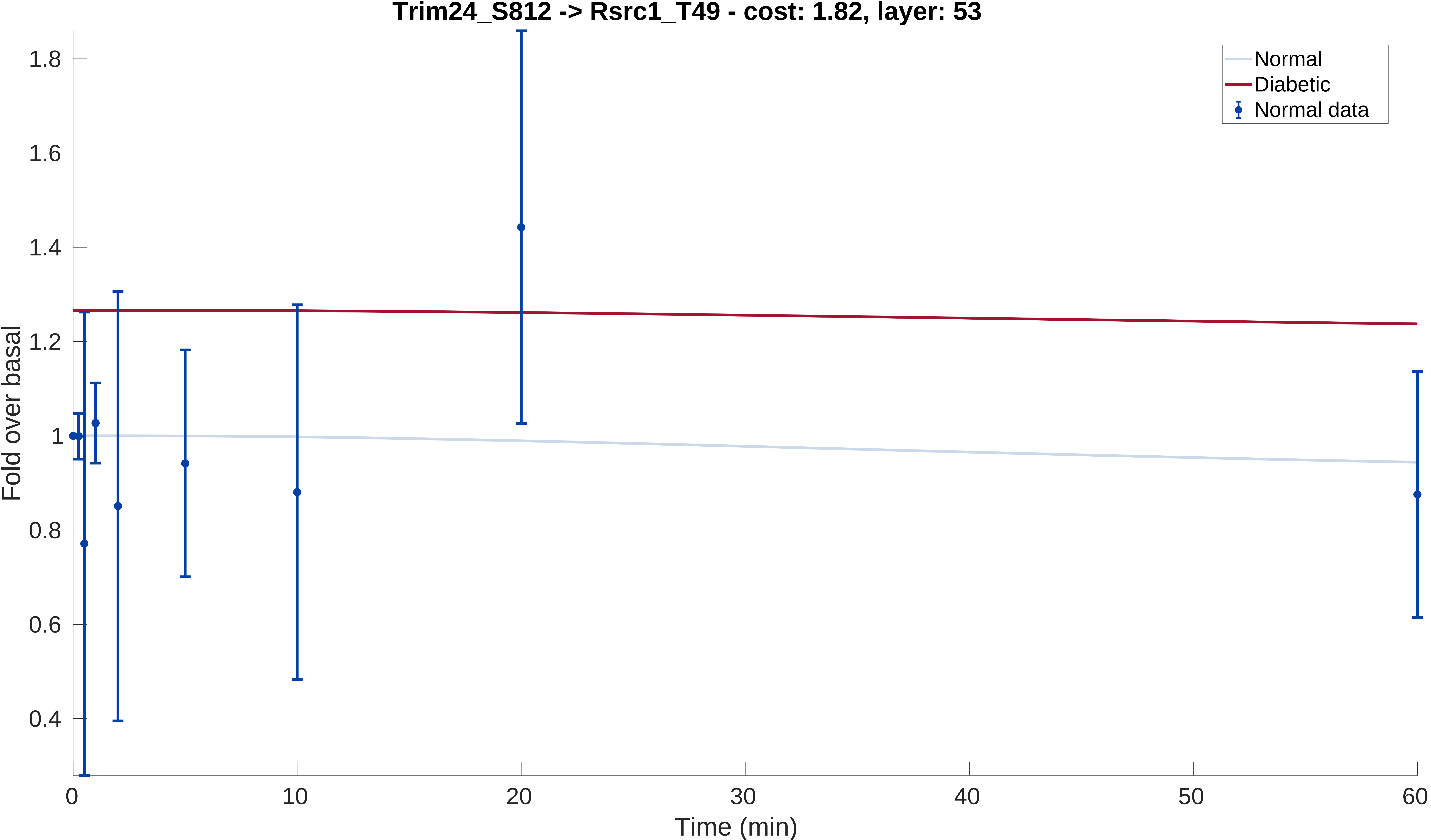

Trim24_S812 -> Rsrc1_T49 - cost: 1.82, layer: 53
Normal
Diabetic
Normal data
Fold over basal
1.8
1.6
1.4
1.2
1
0.8
0.6
0.4
0
10
20
30
40
50
60
Time (min)

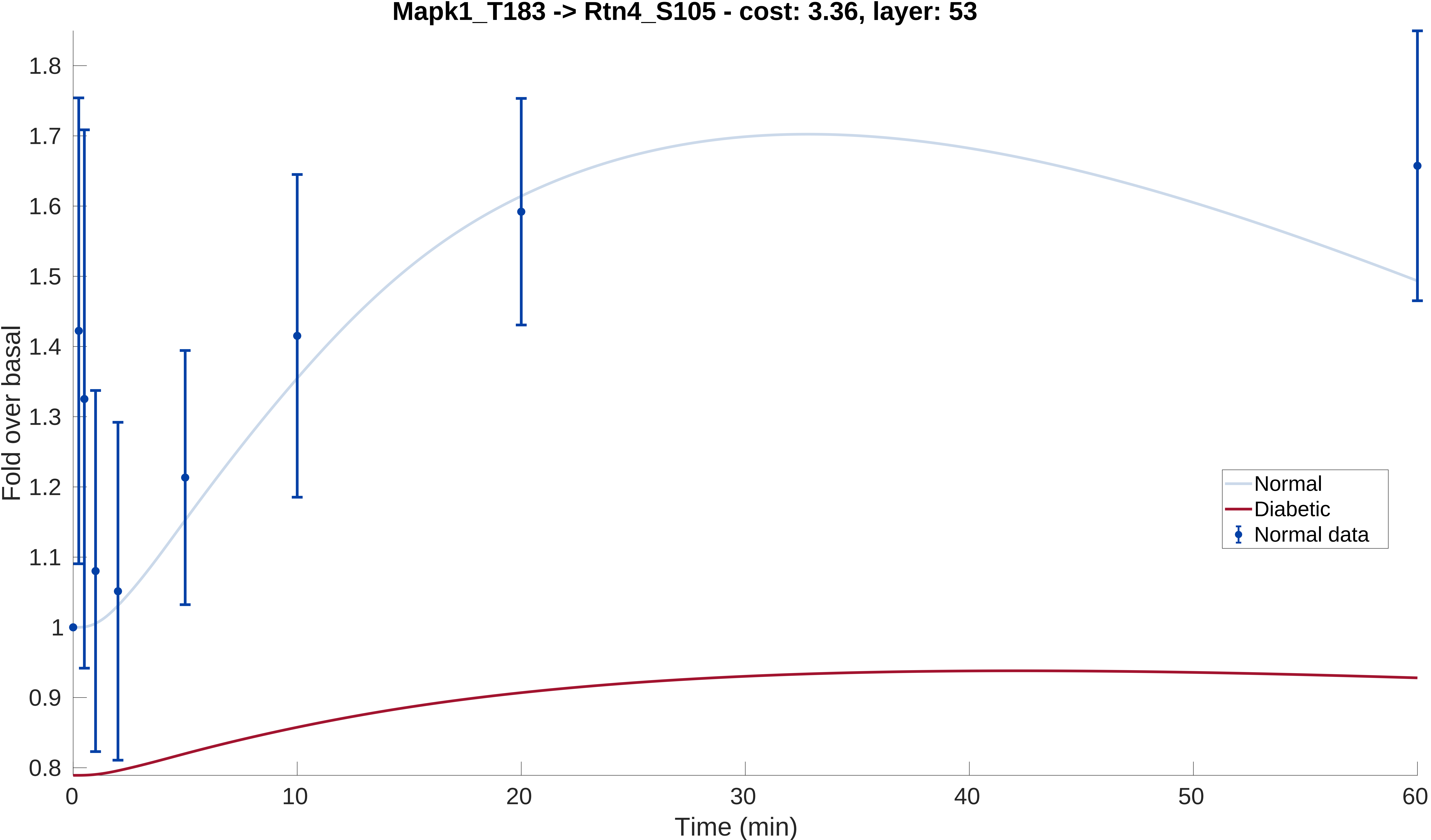

Mapk1_T183 -> Rtn4_S105 - cost: 3.36, layer: 53
Fold over basal
1.8
1.7
1.6
1.5
1.4
1.3
1.2
1.1
1
0.9
0.8
0
10
20
30
40
50
60
Time (min)
Normal
Diabetic
Normal data

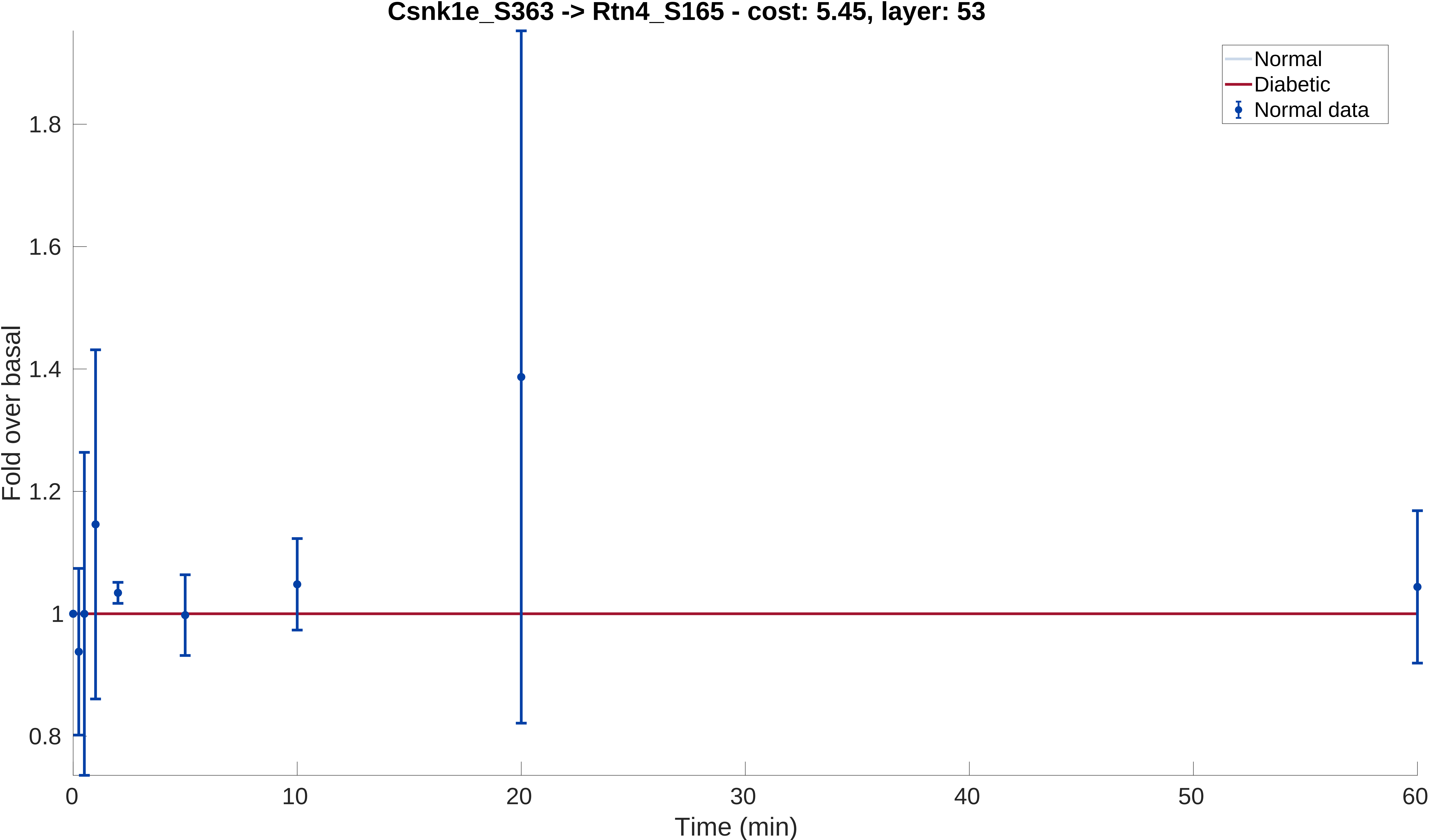

Csnk1e_S363 -> Rtn4_S165 - cost: 5.45, layer: 53
Normal
Diabetic
Normal data
Fold over basal
1.8
1.6
1.4
1.2
1
0.8
0
10
20
30
40
50
60
Time (min)

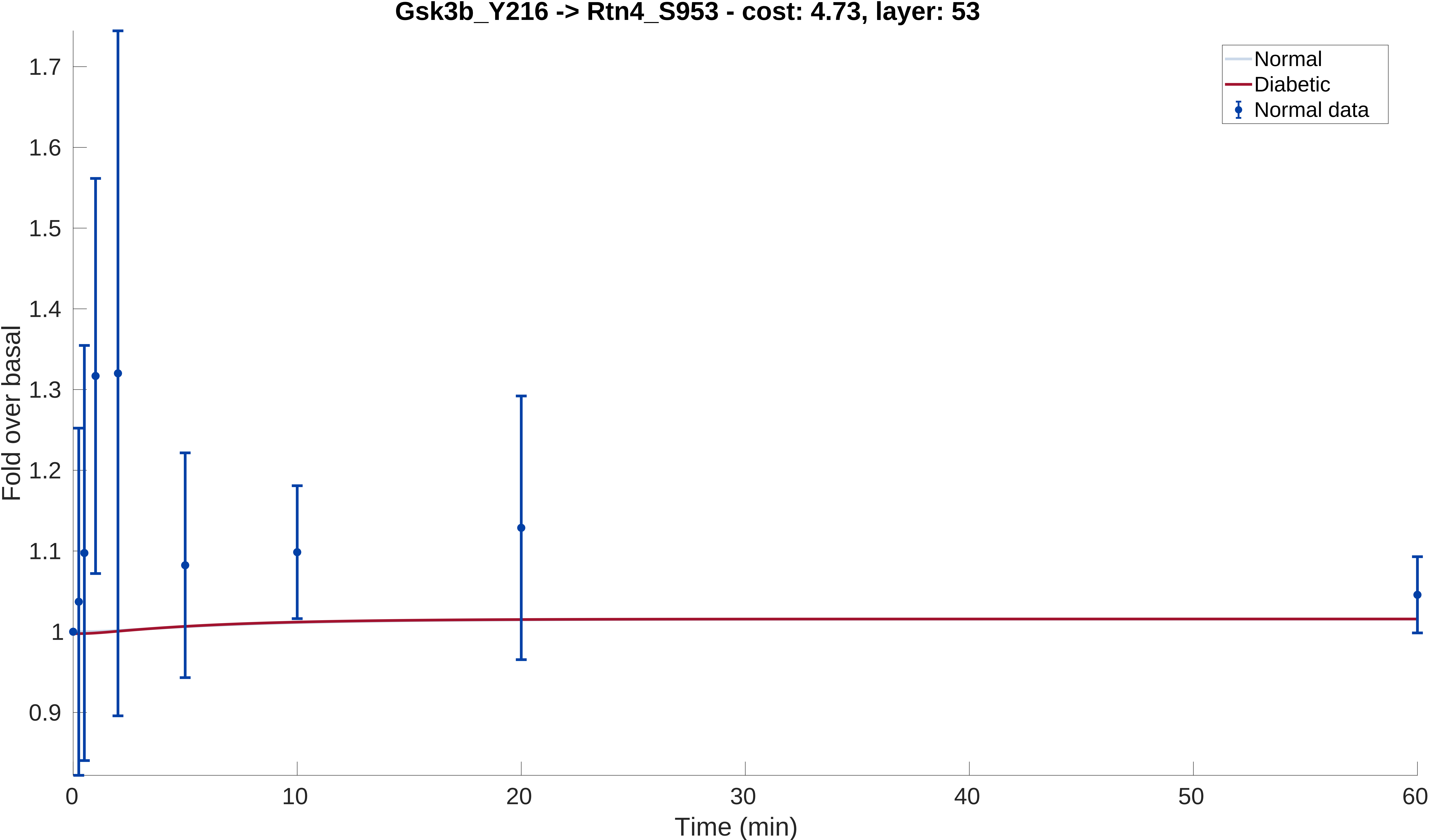

Gsk3b_Y216 -> Rtn4_S953 - cost: 4.73, layer: 53
Normal
Diabetic
Normal data
Fold over basal
1.7
1.6
1.5
1.4
1.3
1.2
1.1
1
0.9
0
10
20
30
40
50
60
Time (min)

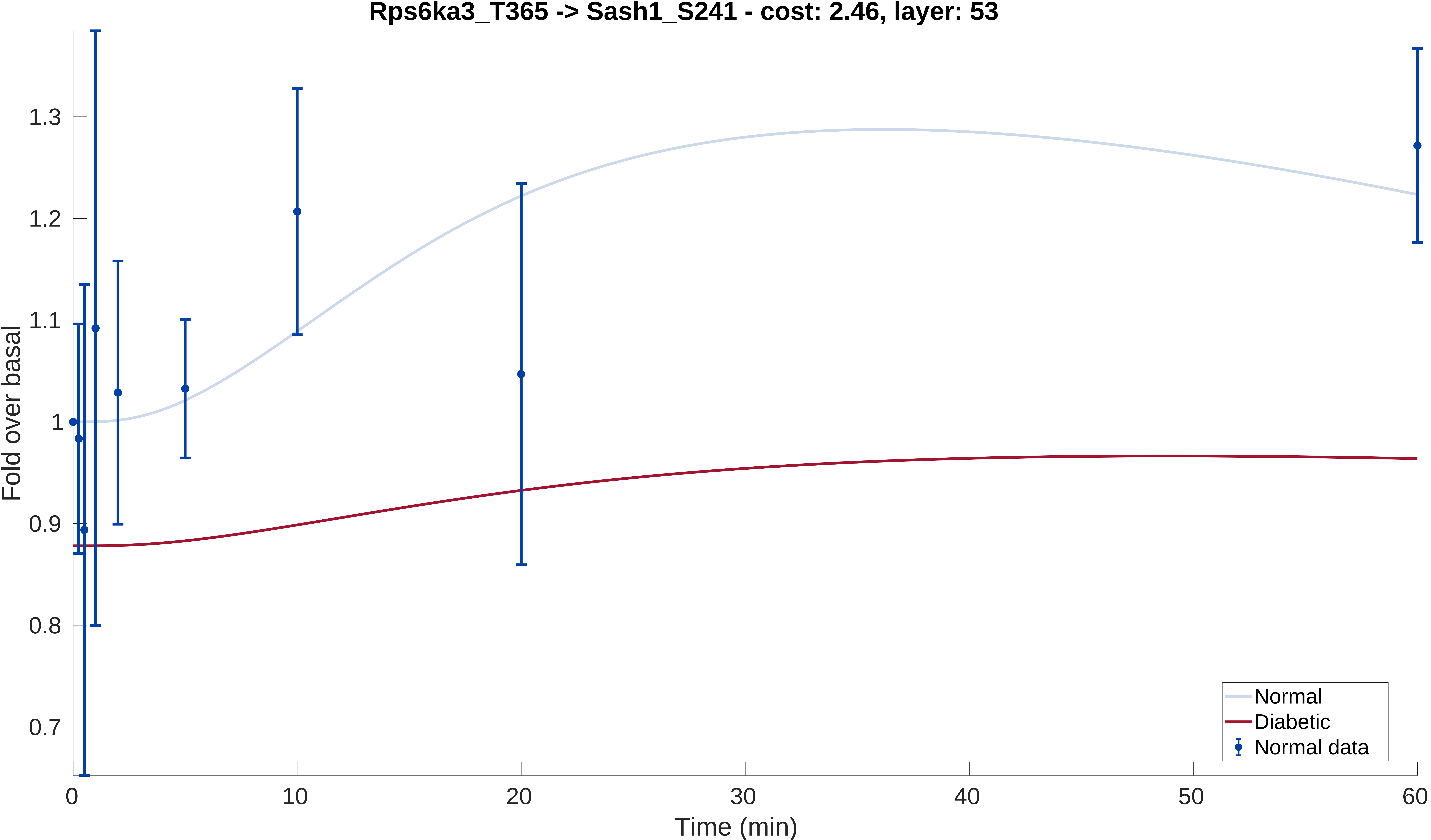
Rps6ka3_T365 -> Sash1_S241 - cost: 2.46, layer: 53
Fold over basal
1.3
1.2
1.1
1
0.9
0.8
0.7
0
10
20
30
40
50
60
Time (min)
Normal
Diabetic
Normal data

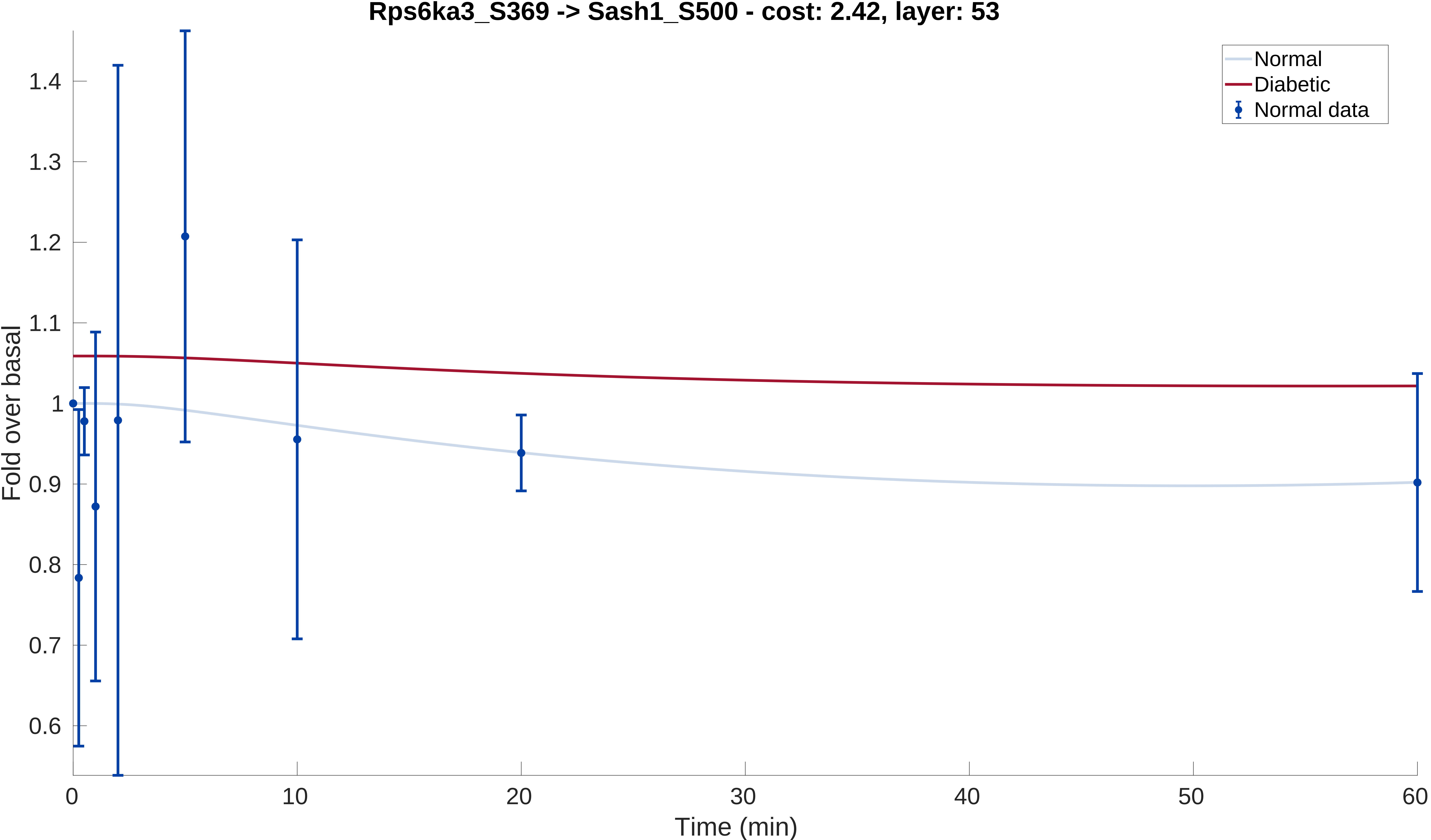

Rps6ka3_S369 -> Sash1_S500 - cost: 2.42, layer: 53
Normal
Diabetic
Normal data
Fold over basal
1.4
1.3
1.2
1.1
1
0.9
0.8
0.7
0.6
0
10
20
30
40
50
60
Time (min)

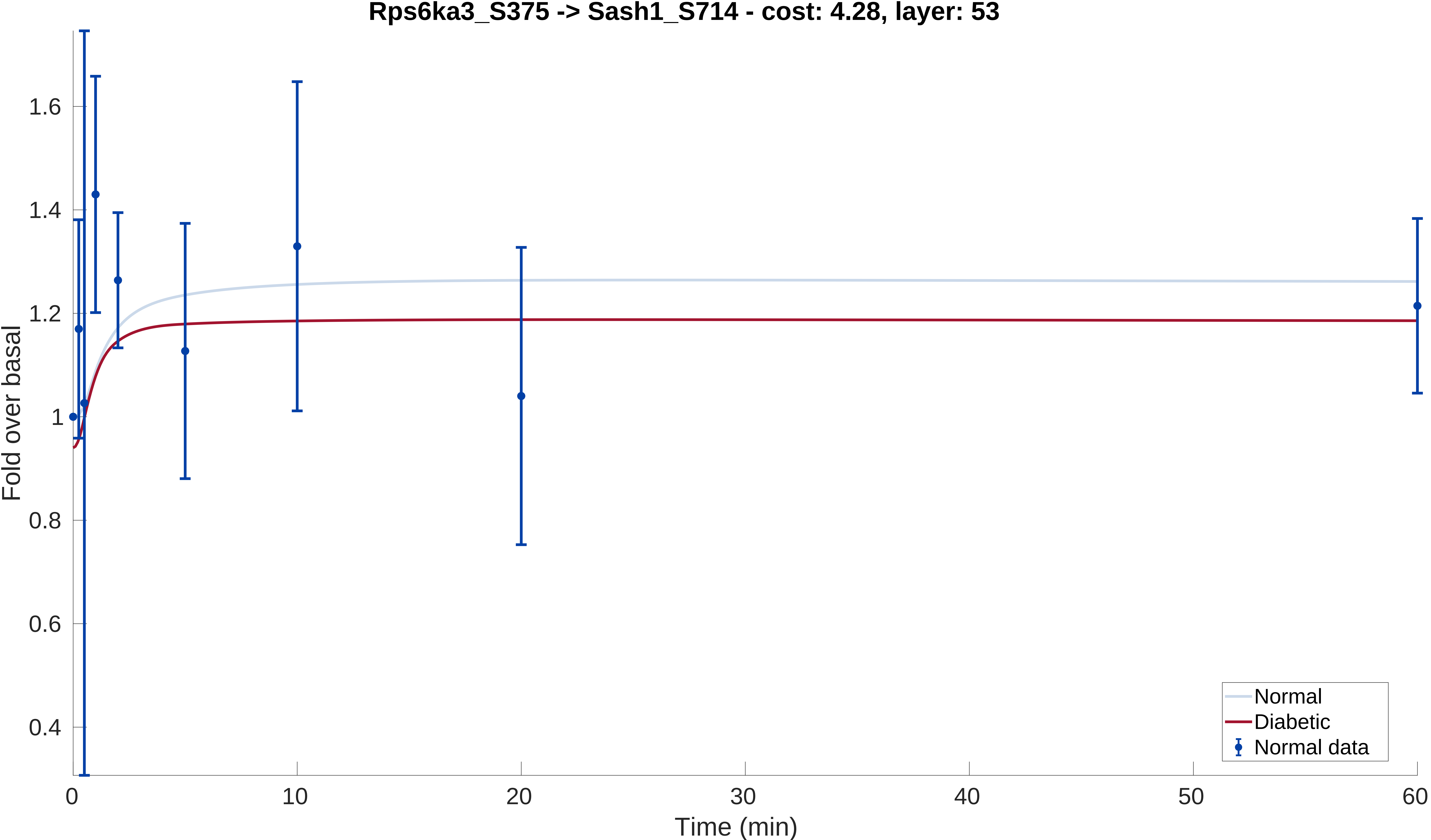

Rps6ka3_S375 -> Sash1_S714 - cost: 4.28, layer: 53
Fold over basal
1.6
1.4
1.2
1
0.8
0.6
0.4
0
10
20
30
40
50
60
Time (min)
Normal
Diabetic
Normal data

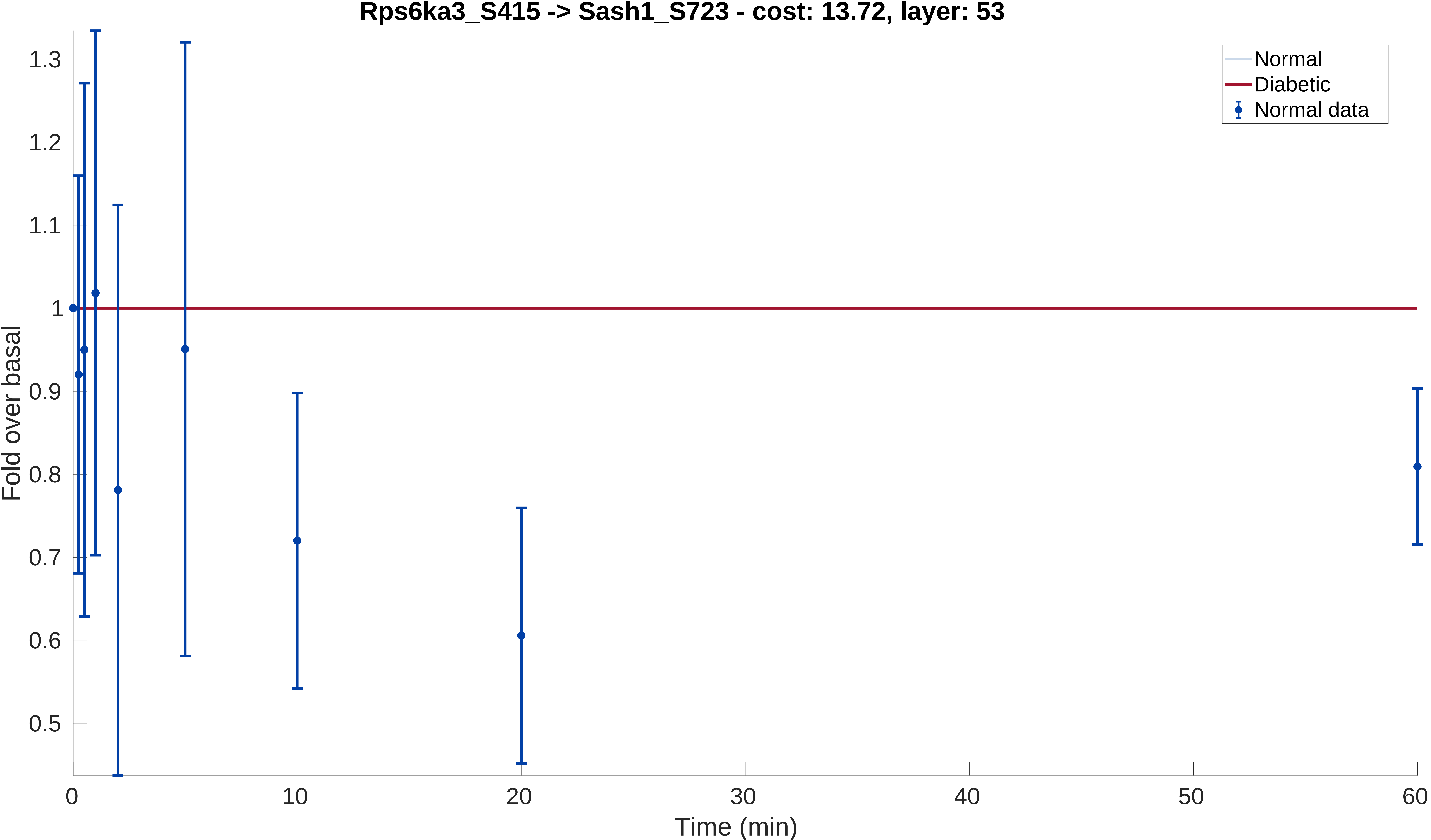

Rps6ka3_S415 -> Sash1_S723 - cost: 13.72, layer: 53
Normal
Diabetic
Normal data
Fold over basal
Time (min)
1.3
1.2
1.1
1
0.9
0.8
0.7
0.6
0.5
0
10
20
30
40
50
60

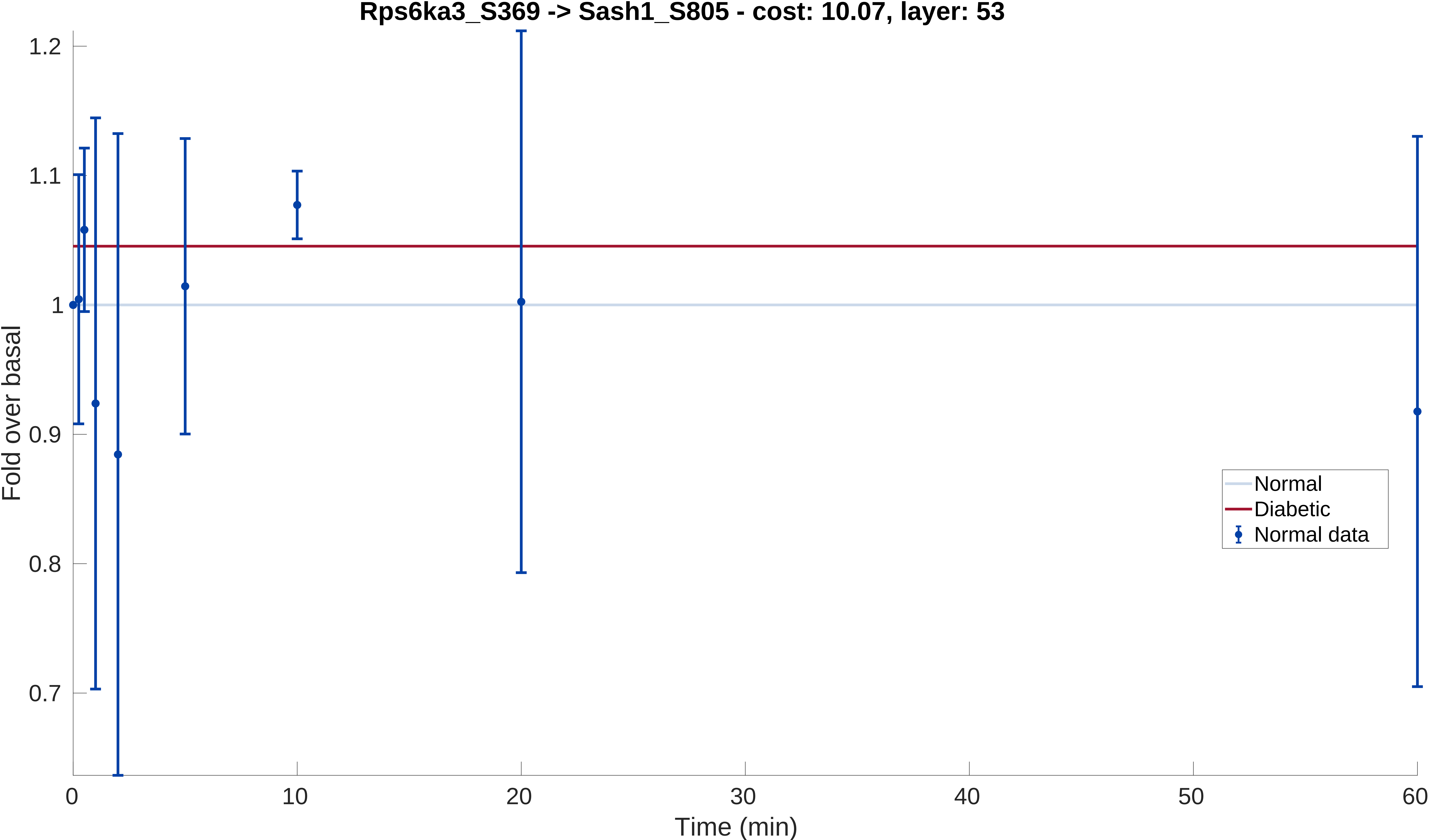

Rps6ka3_S369 -> Sash1_S805 - cost: 10.07, layer: 53
Fold over basal
1.2
1.1
1
0.9
0.8
0.7
0
10
20
30
40
50
60
Time (min)
Normal
Diabetic
Normal data

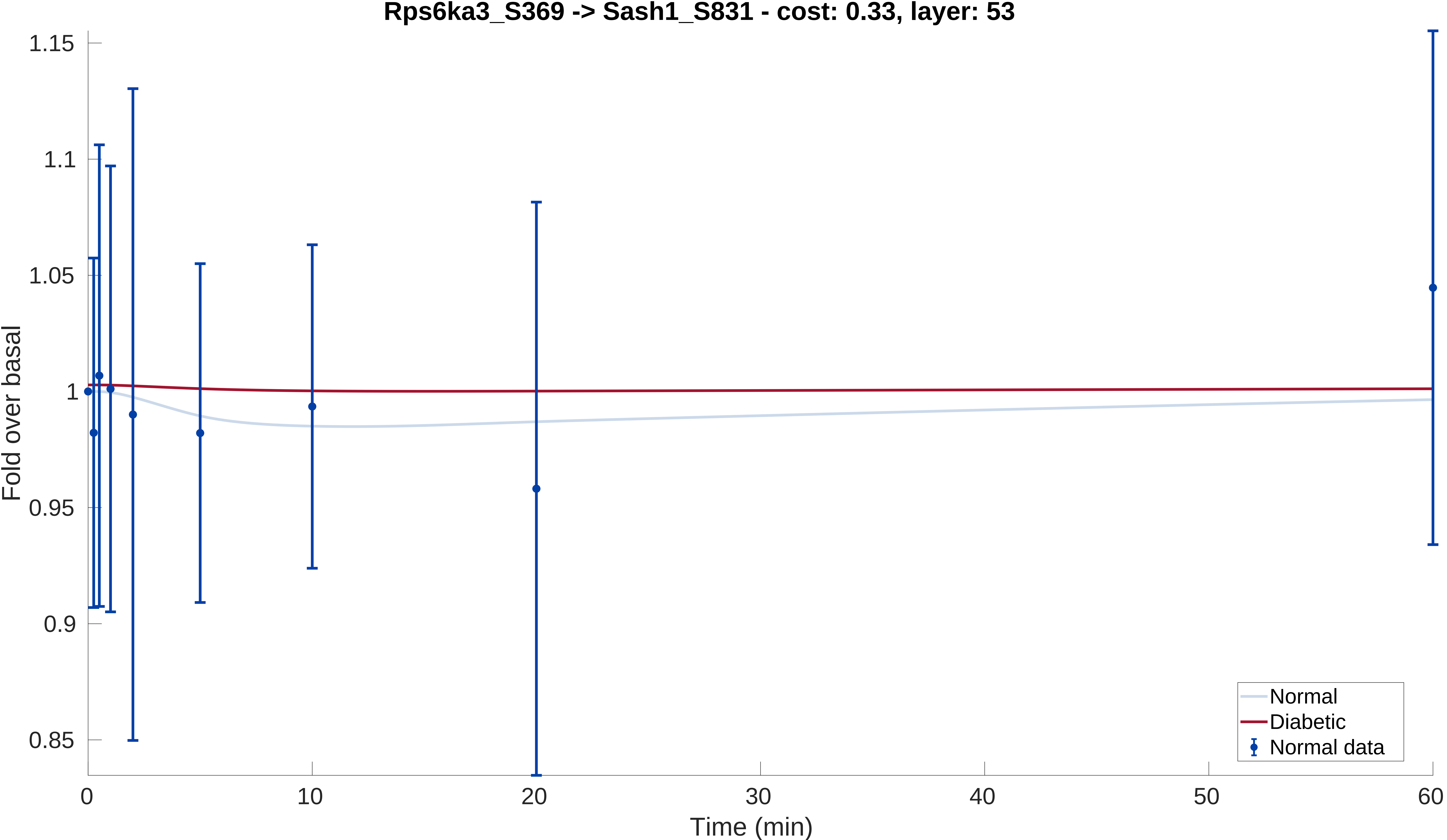

Rps6ka3_S369 -> Sash1_S831 - cost: 0.33, layer: 53
Fold over basal
1.15
1.1
1.05
1
0.95
0.9
0.85
0
10
20
30
40
50
60
Time (min)
Normal
Diabetic
Normal data

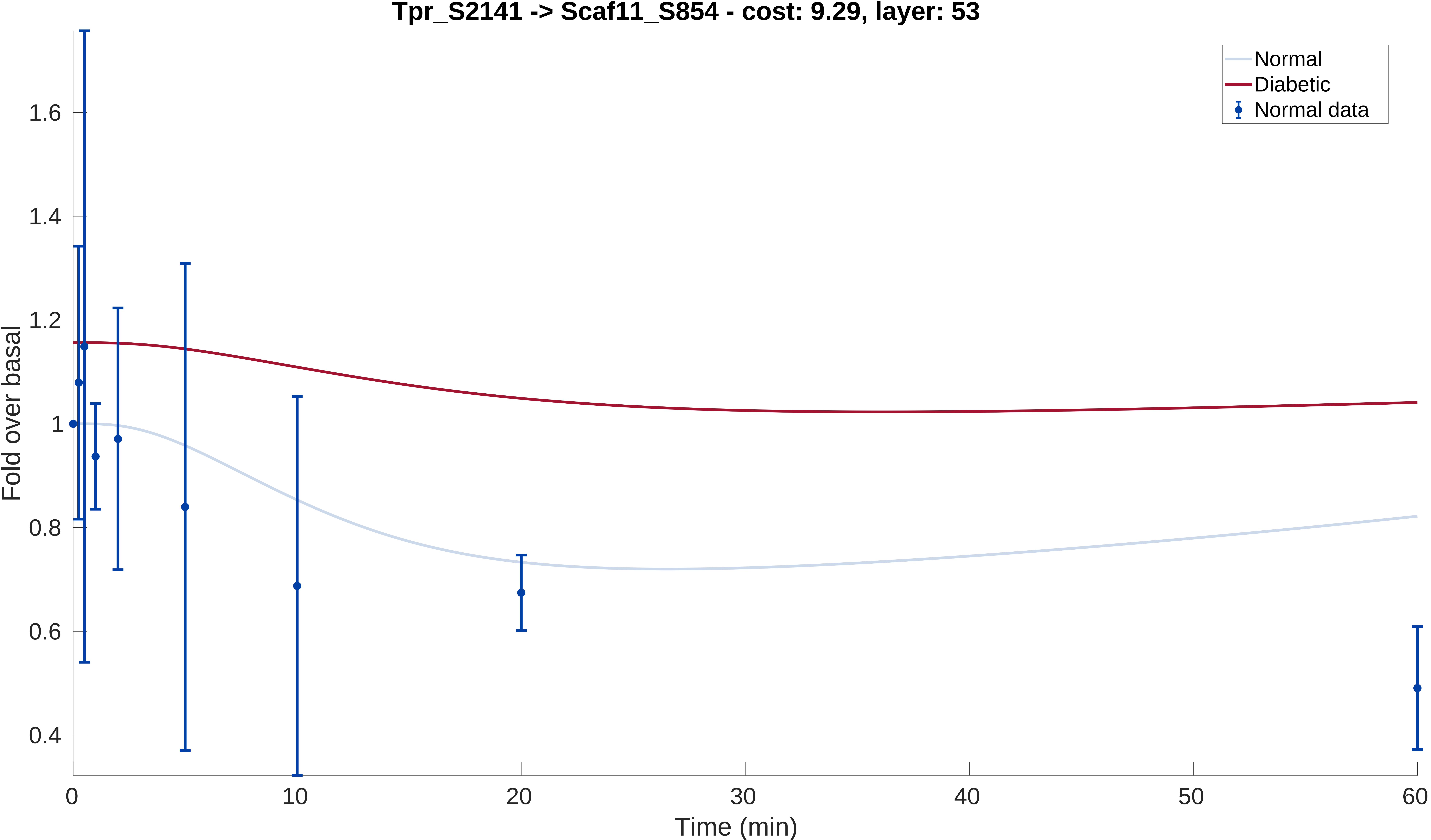

Tpr_S2141 -> Scaf11_S854 - cost: 9.29, layer: 53
Normal
Diabetic
Normal data
1.6
1.4
1.2
1
0.8
0.6
0.4
Fold over basal
0
10
20
30
40
50
60
Time (min)

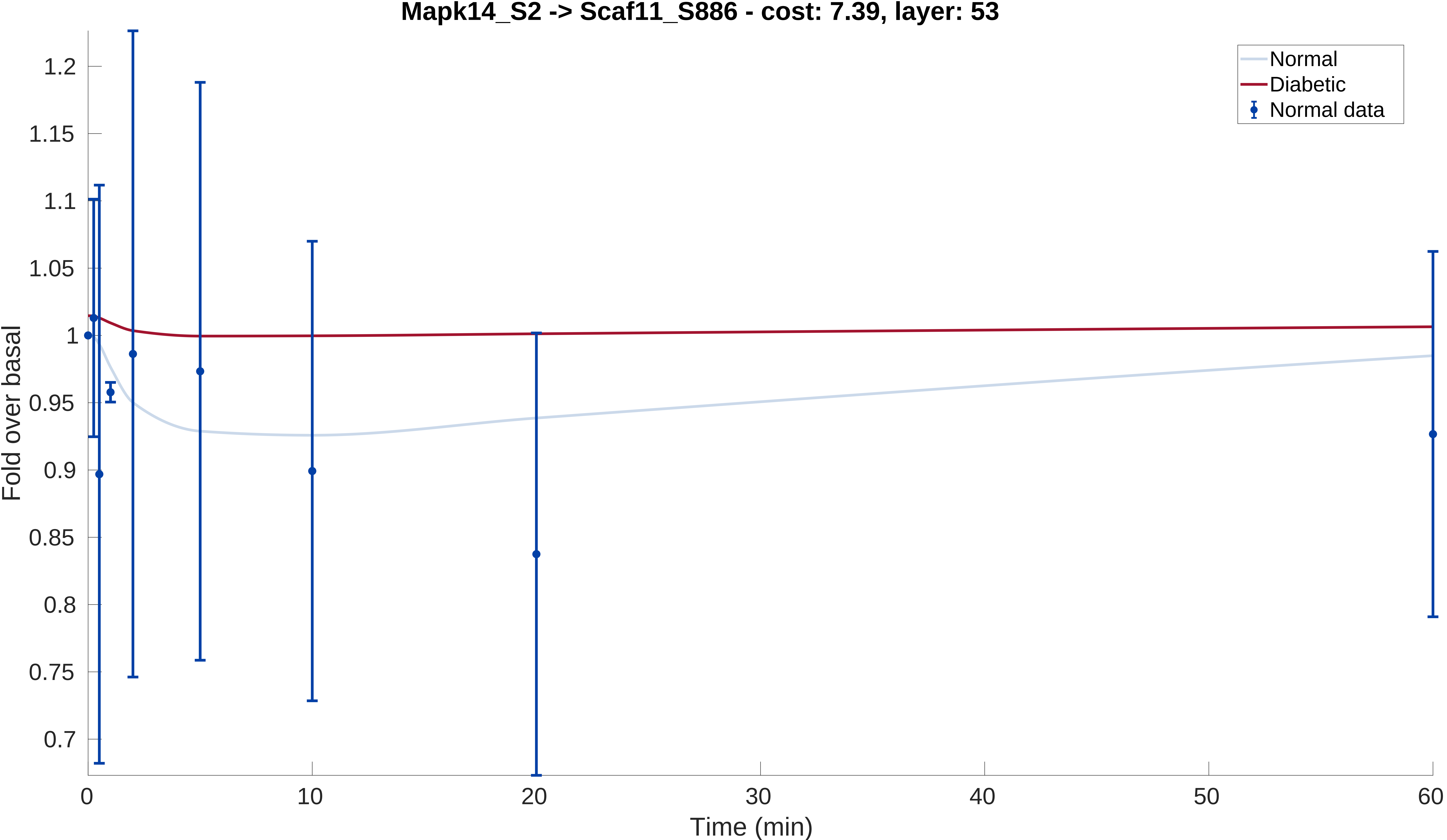

Mapk14_S2 -> Scaf11_S886 - cost: 7.39, layer: 53
Normal
Diabetic
Normal data
Fold over basal
Time (min)
1.2
1.15
1.1
1.05
1
0.95
0.9
0.85
0.8
0.75
0.7
0
10
20
30
40
50
60

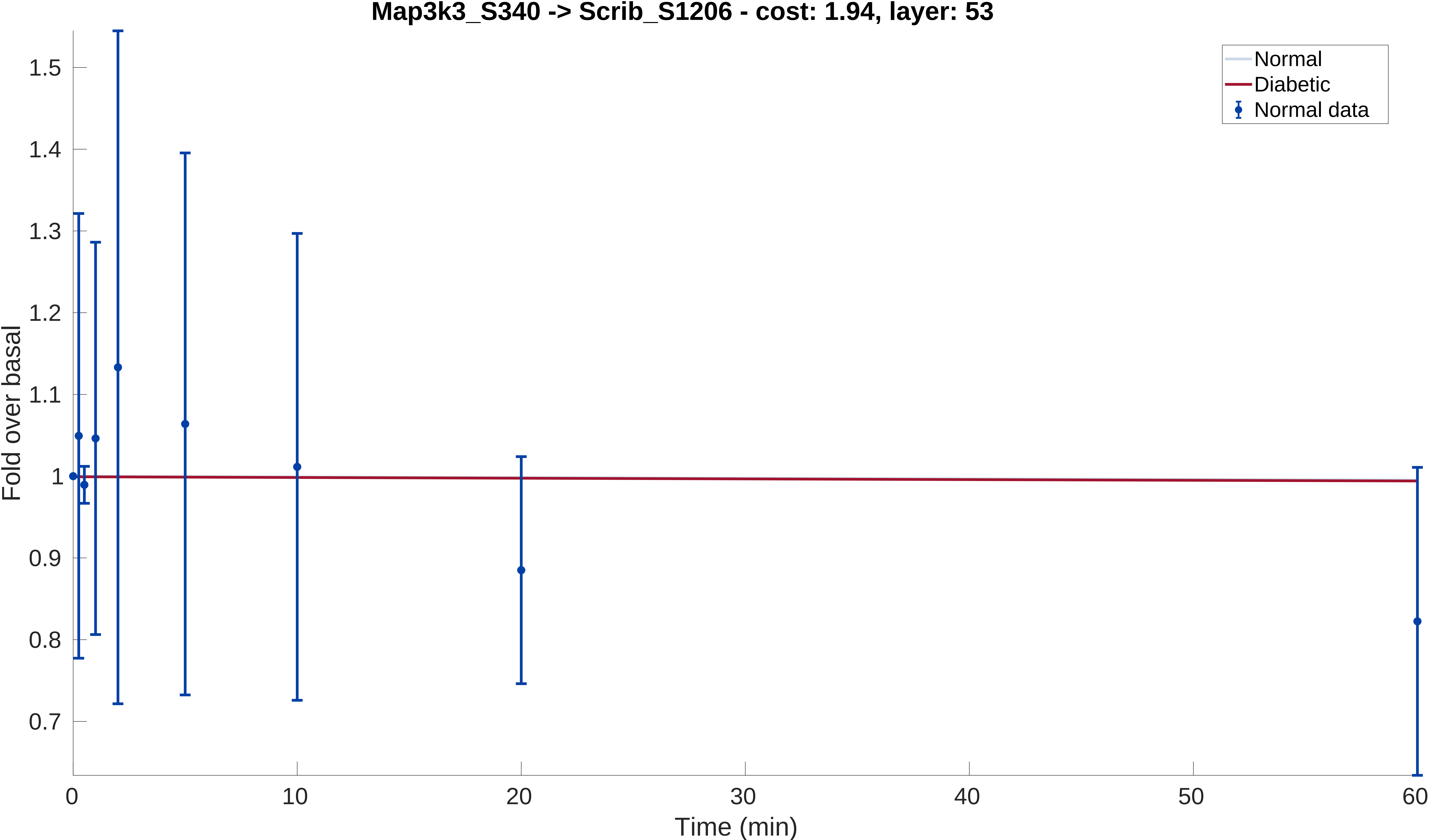

Map3k3_S340 -> Scrib_S1206 - cost: 1.94, layer: 53
Normal
Diabetic
Normal data
Fold over basal
Time (min)
0.7
0.8
0.9
1
1.1
1.2
1.3
1.4
1.5
0
10
20
30
40
50
60

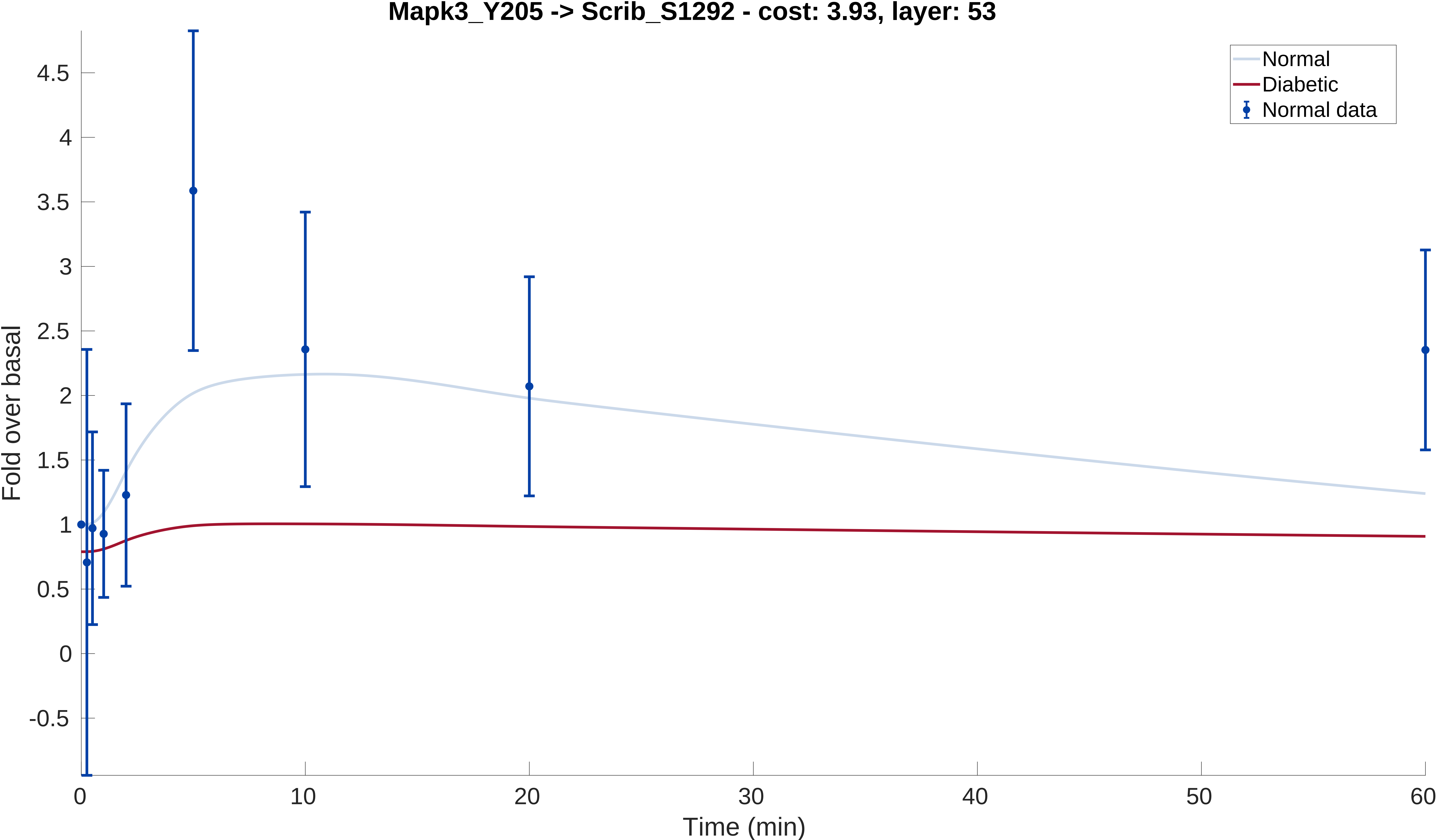

Mapk3_Y205 -> Scrib_S1292 - cost: 3.93, layer: 53
Normal
Diabetic
Normal data
Fold over basal
Time (min)
4.5
4
3.5
3
2.5
2
1.5
1
0.5
0
-0.5
0
10
20
30
40
50
60

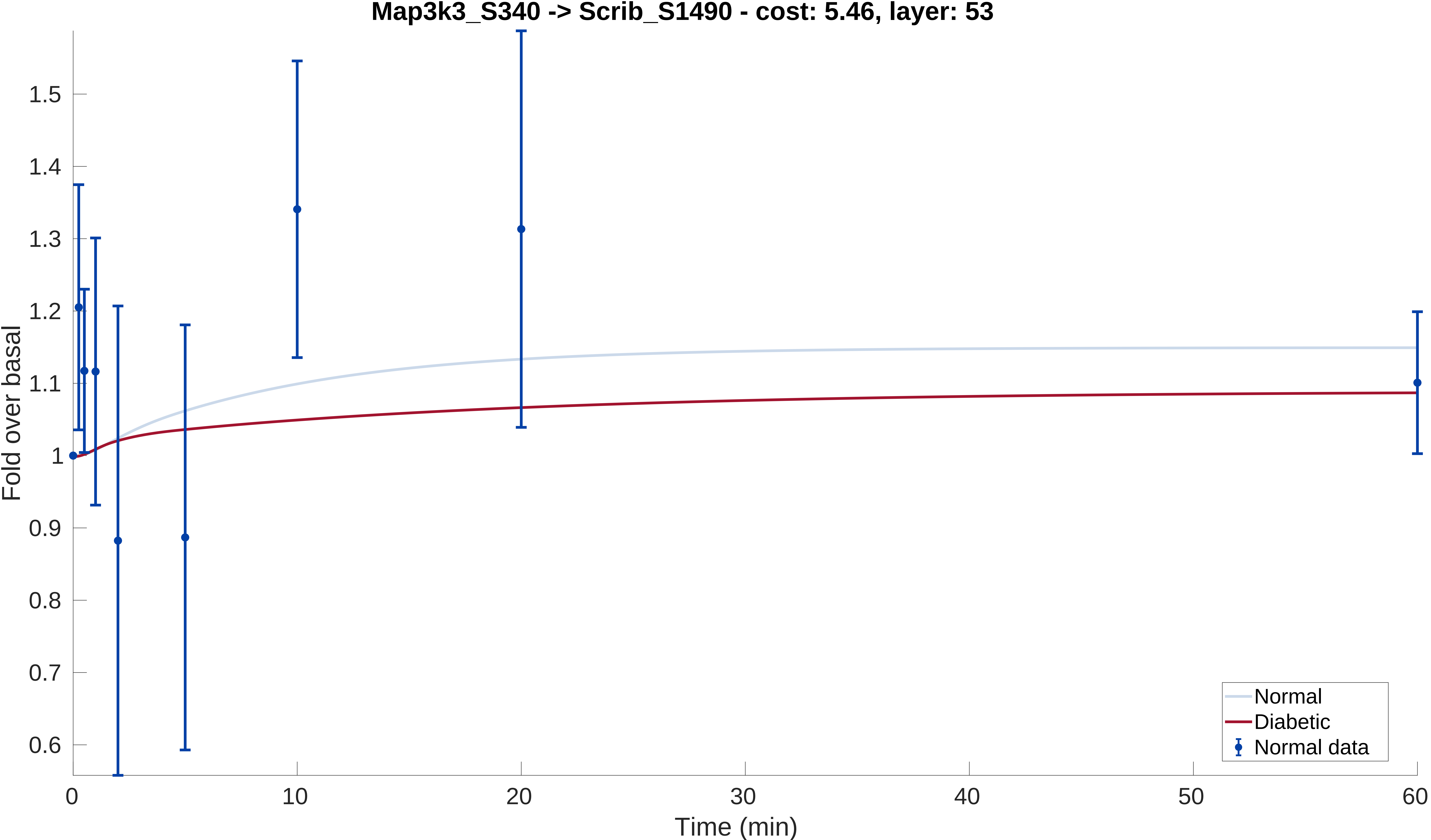
Map3k3_S340 -> Scrib_S1490 - cost: 5.46, layer: 53
Fold over basal
1.5
1.4
1.3
1.2
1.1
1
0.9
0.8
0.7
0.6
0
10
20
30
40
50
60
Time (min)
Normal
Diabetic
Normal data

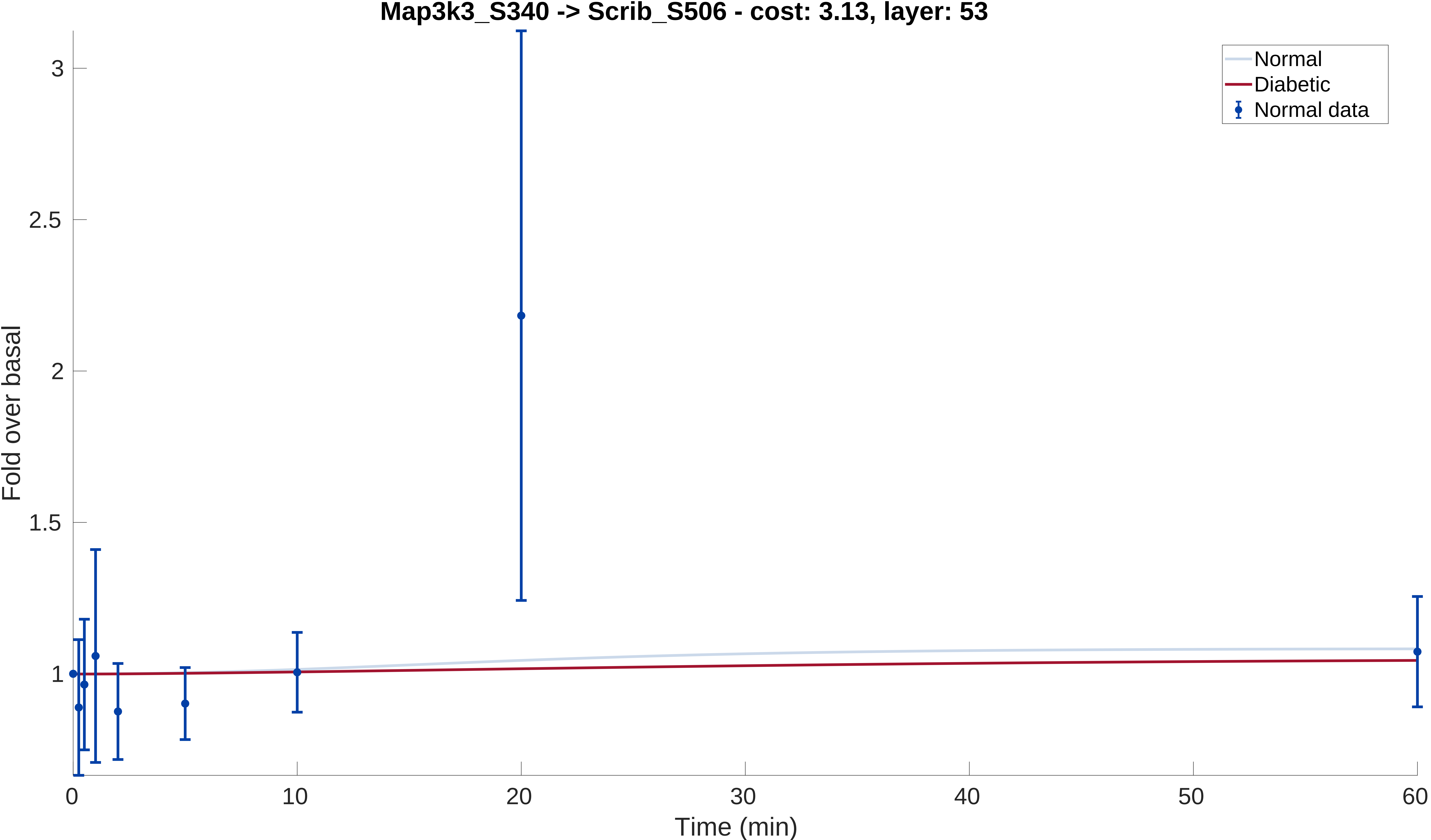

Map3k3_S340 -> Scrib_S506 - cost: 3.13, layer: 53
Normal
Diabetic
Normal data
Fold over basal
3
2.5
2
1.5
1
0
10
20
30
40
50
60
Time (min)

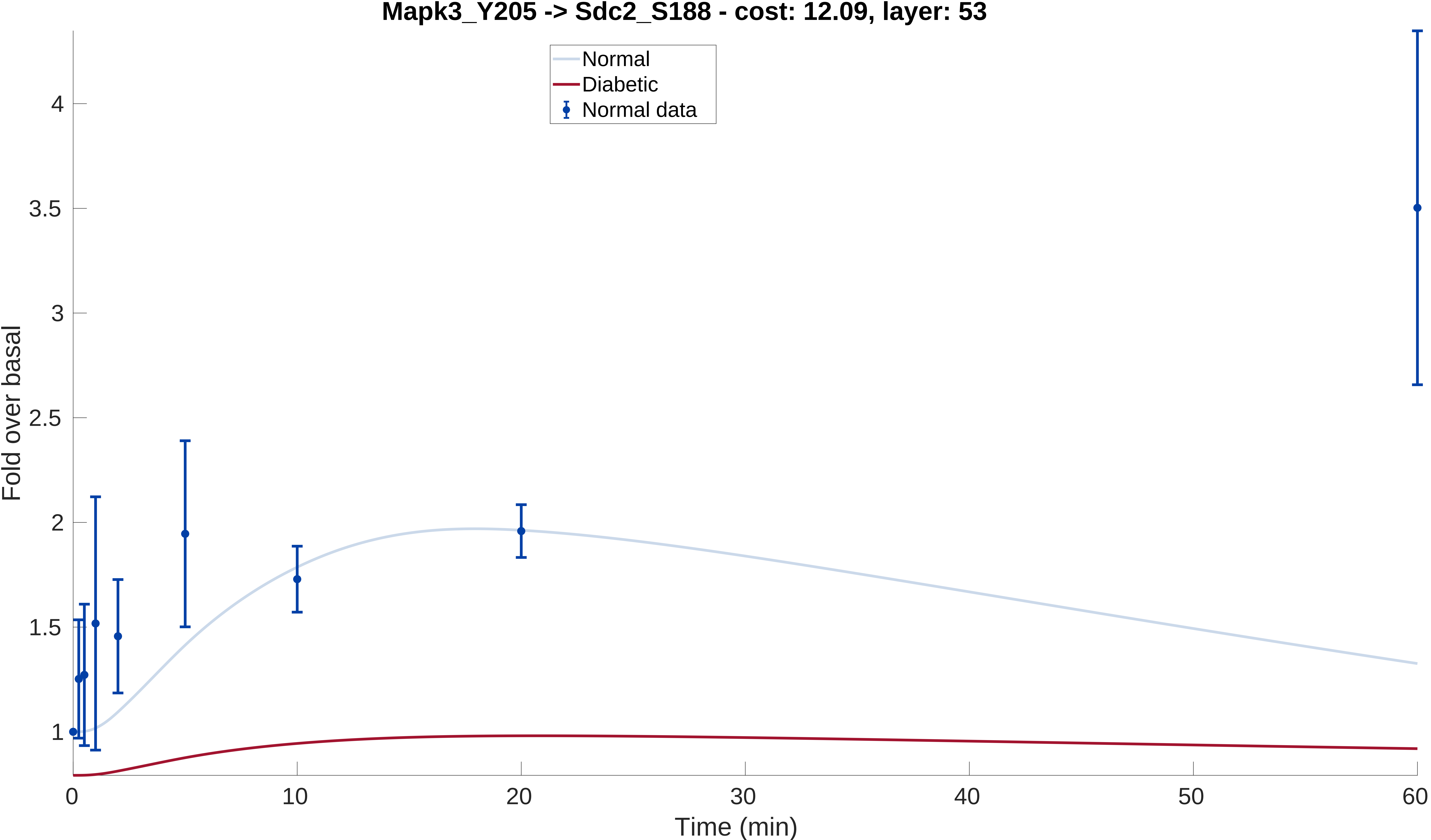

Mapk3_Y205 -> Sdc2_S188 - cost: 12.09, layer: 53
Normal
Diabetic
Normal data
Fold over basal
Time (min)
1
1.5
2
2.5
3
3.5
4
0
10
20
30
40
50
60

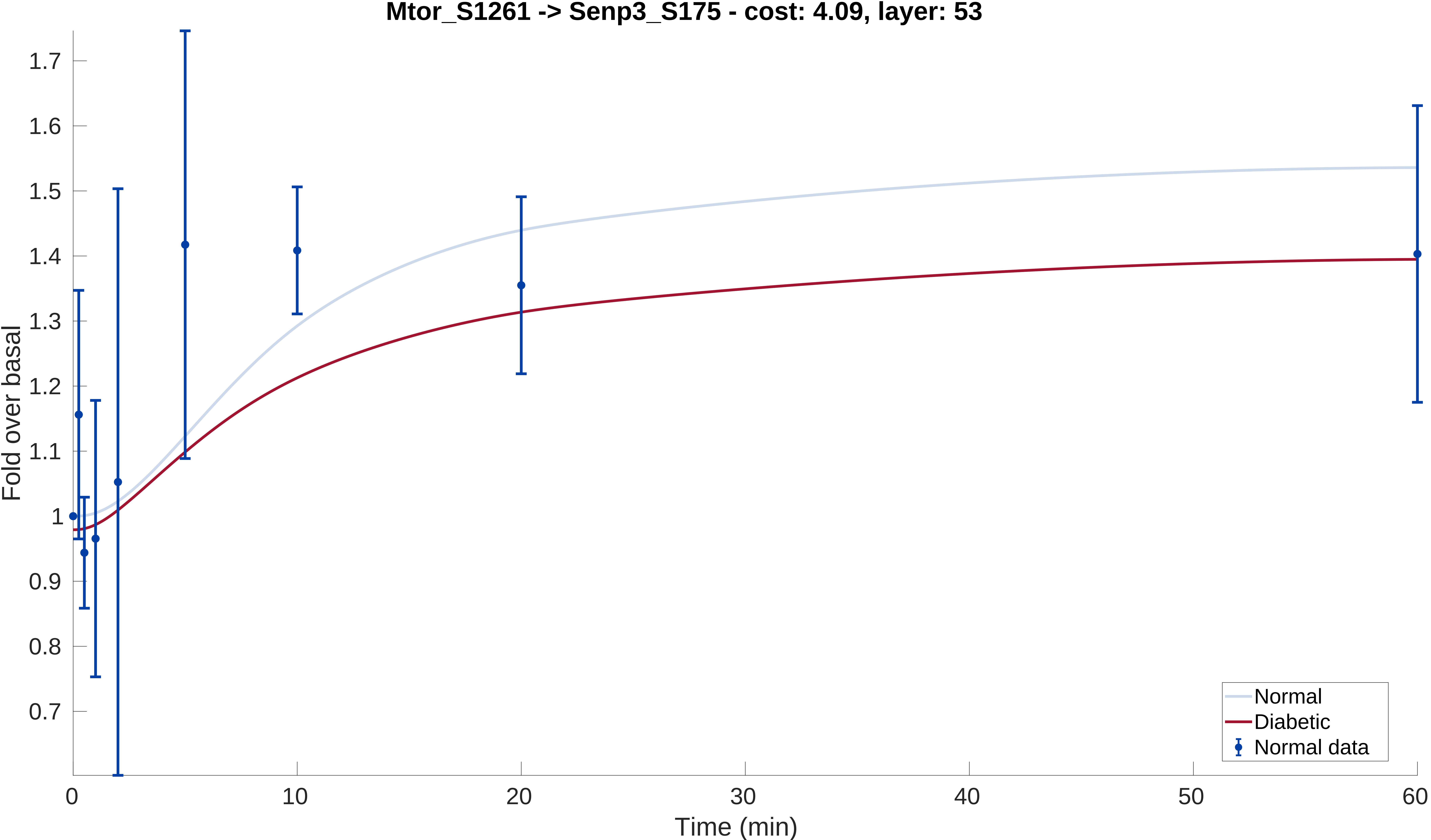
Mtor_S1261 -> Senp3_S175 - cost: 4.09, layer: 53
Fold over basal
Time (min)
1.7
1.6
1.5
1.4
1.3
1.2
1.1
1
0.9
0.8
0.7
0
10
20
30
40
50
60
Normal
Diabetic
Normal data

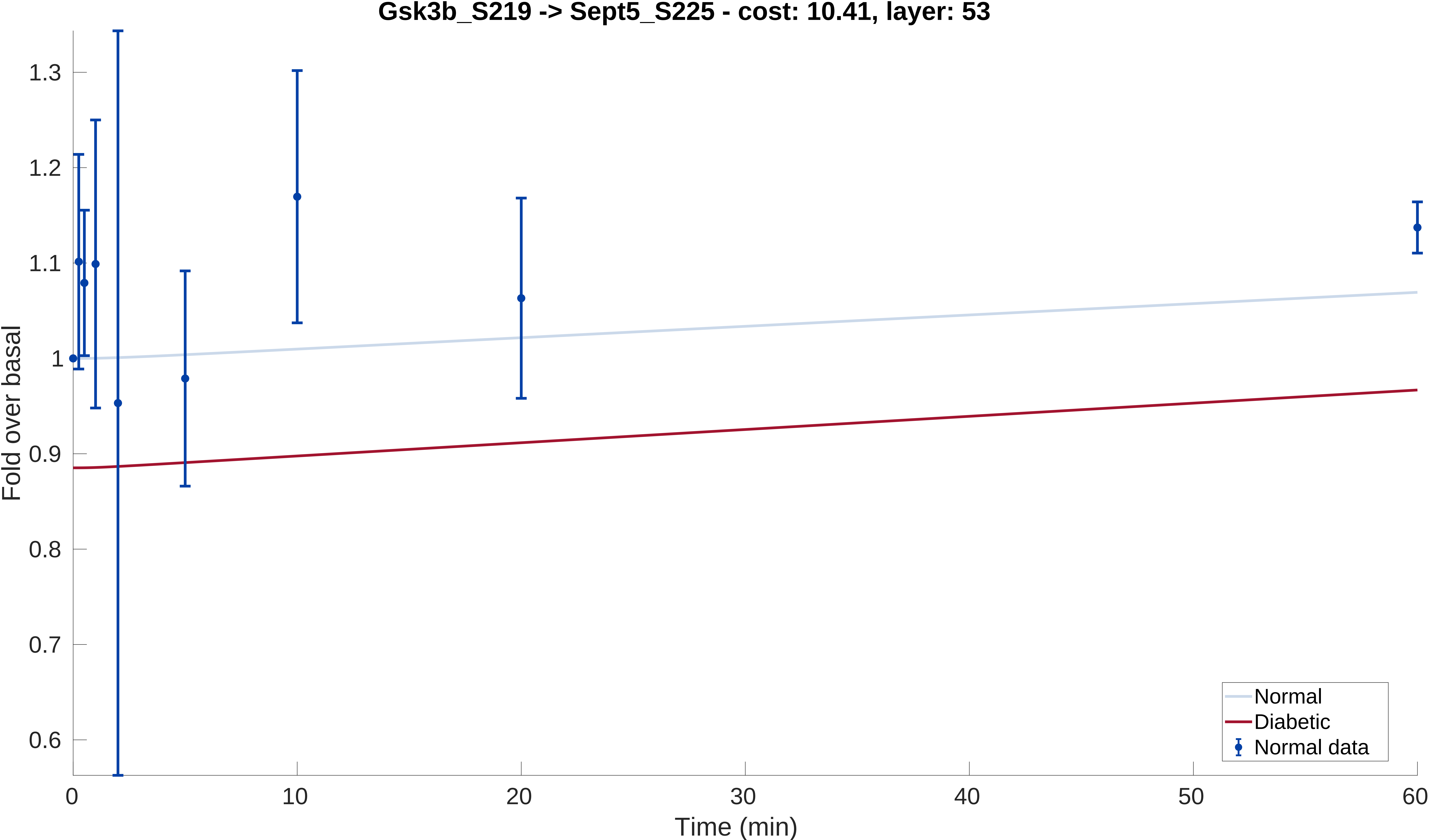

Gsk3b_S219 -> Sept5_S225 - cost: 10.41, layer: 53
Fold over basal
Time (min)
1.3
1.2
1.1
1
0.9
0.8
0.7
0.6
0
10
20
30
40
50
60
Normal
Diabetic
Normal data

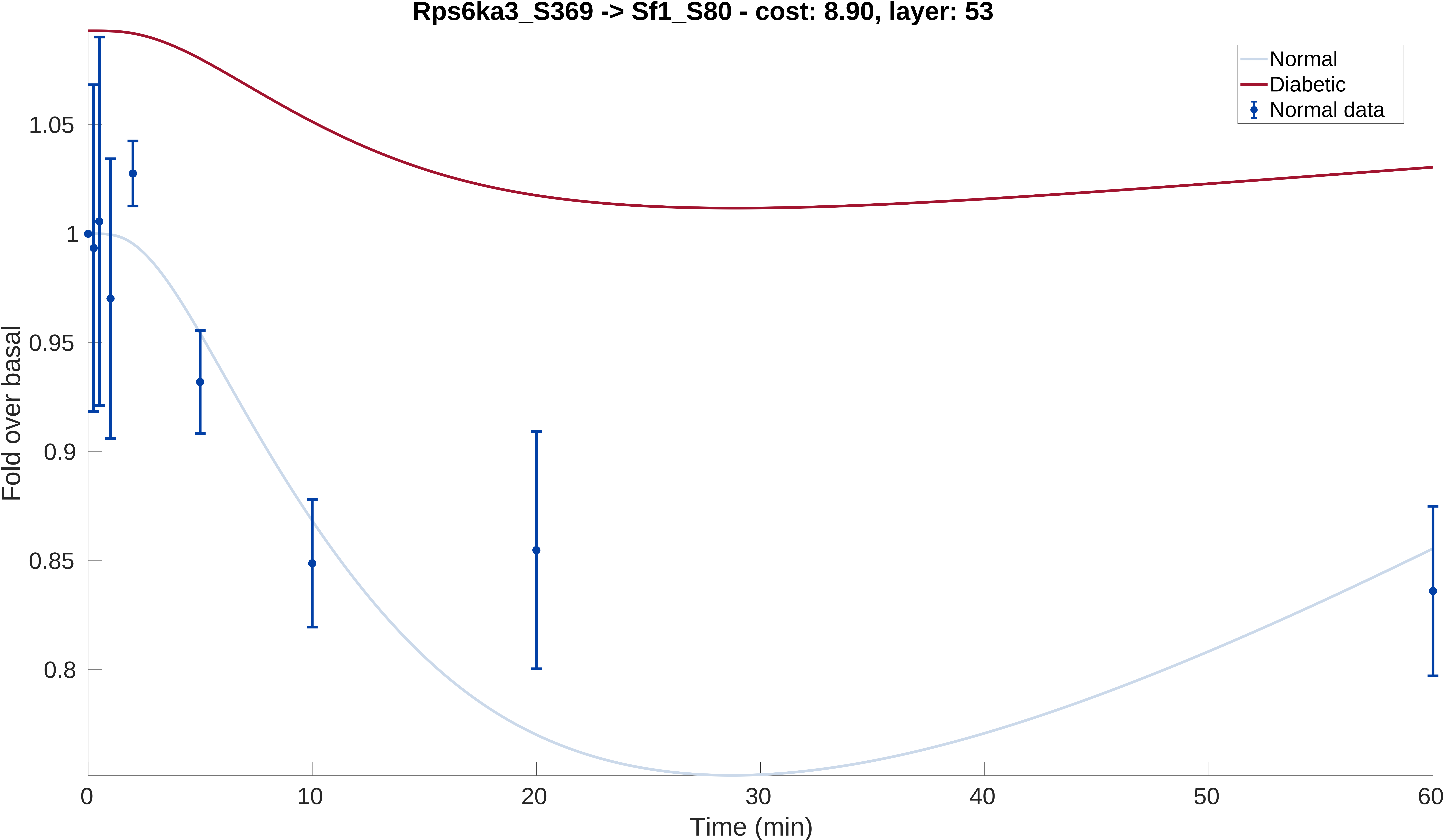
Rps6ka3_S369 -> Sf1_S80 - cost: 8.90, layer: 53
Normal
Diabetic
Normal data
1.05
1
0.95
0.9
0.85
0.8
Fold over basal
0
10
20
30
40
50
60
Time (min)

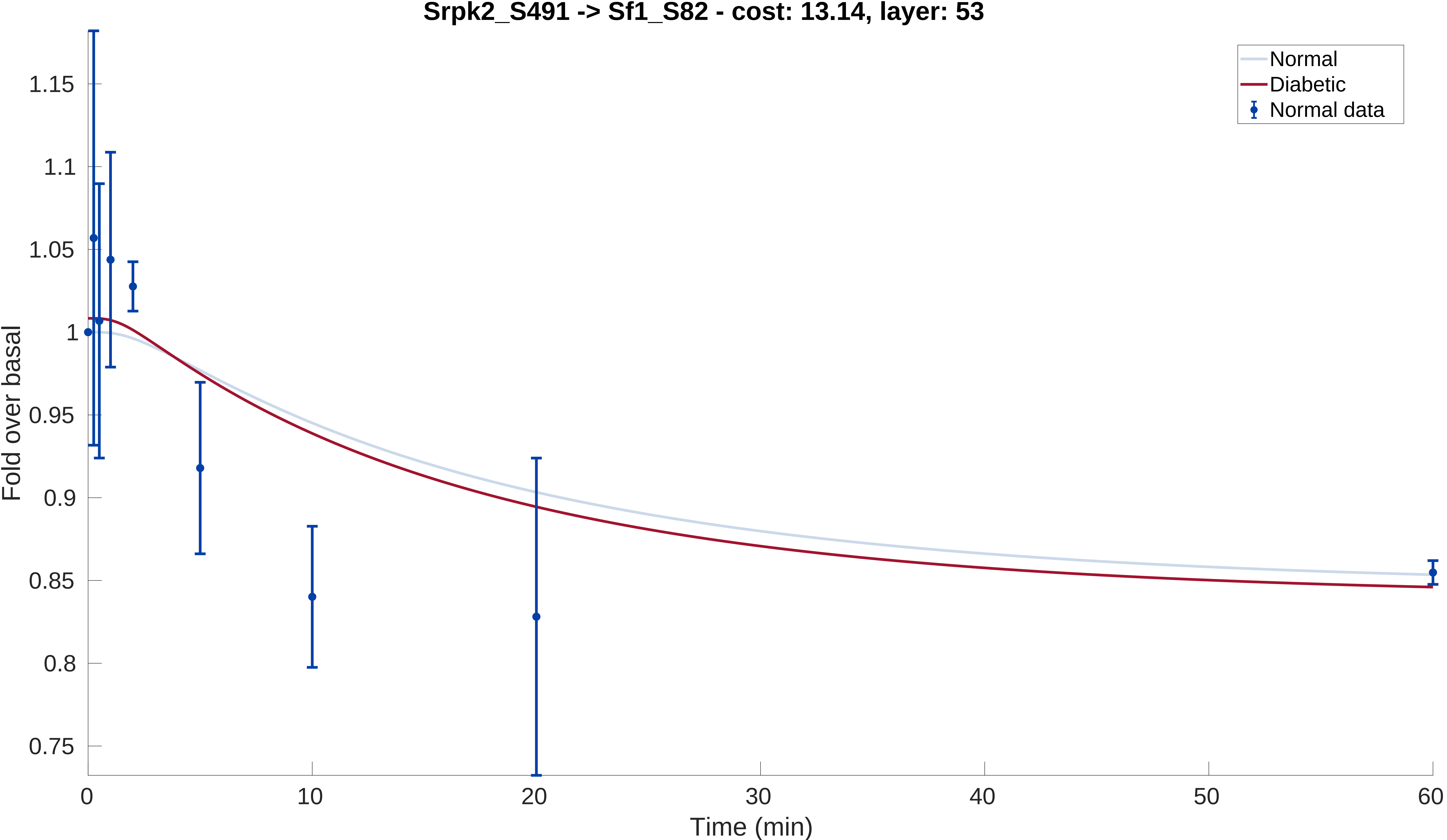
Srpk2_S491 -> Sf1_S82 - cost: 13.14, layer: 53
Normal
Diabetic
Normal data
Fold over basal
1.15
1.1
1.05
1
0.95
0.9
0.85
0.8
0.75
0
10
20
30
40
50
60
Time (min)

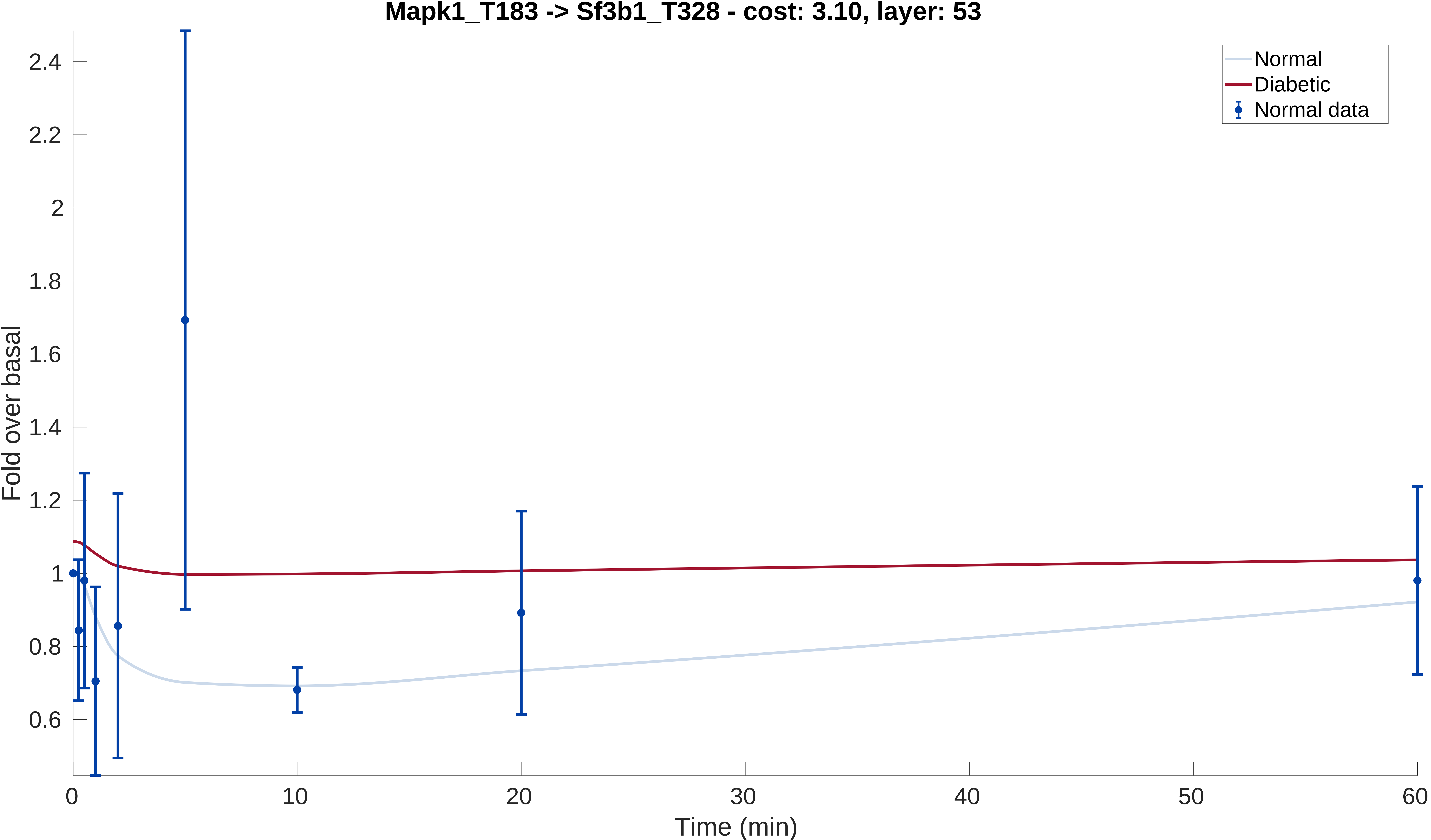
Mapk1_T183 -> Sf3b1_T328 - cost: 3.10, layer: 53
Normal
Diabetic
Normal data
Fold over basal
2.4
2.2
2
1.8
1.6
1.4
1.2
1
0.8
0.6
0
10
20
30
40
50
60
Time (min)

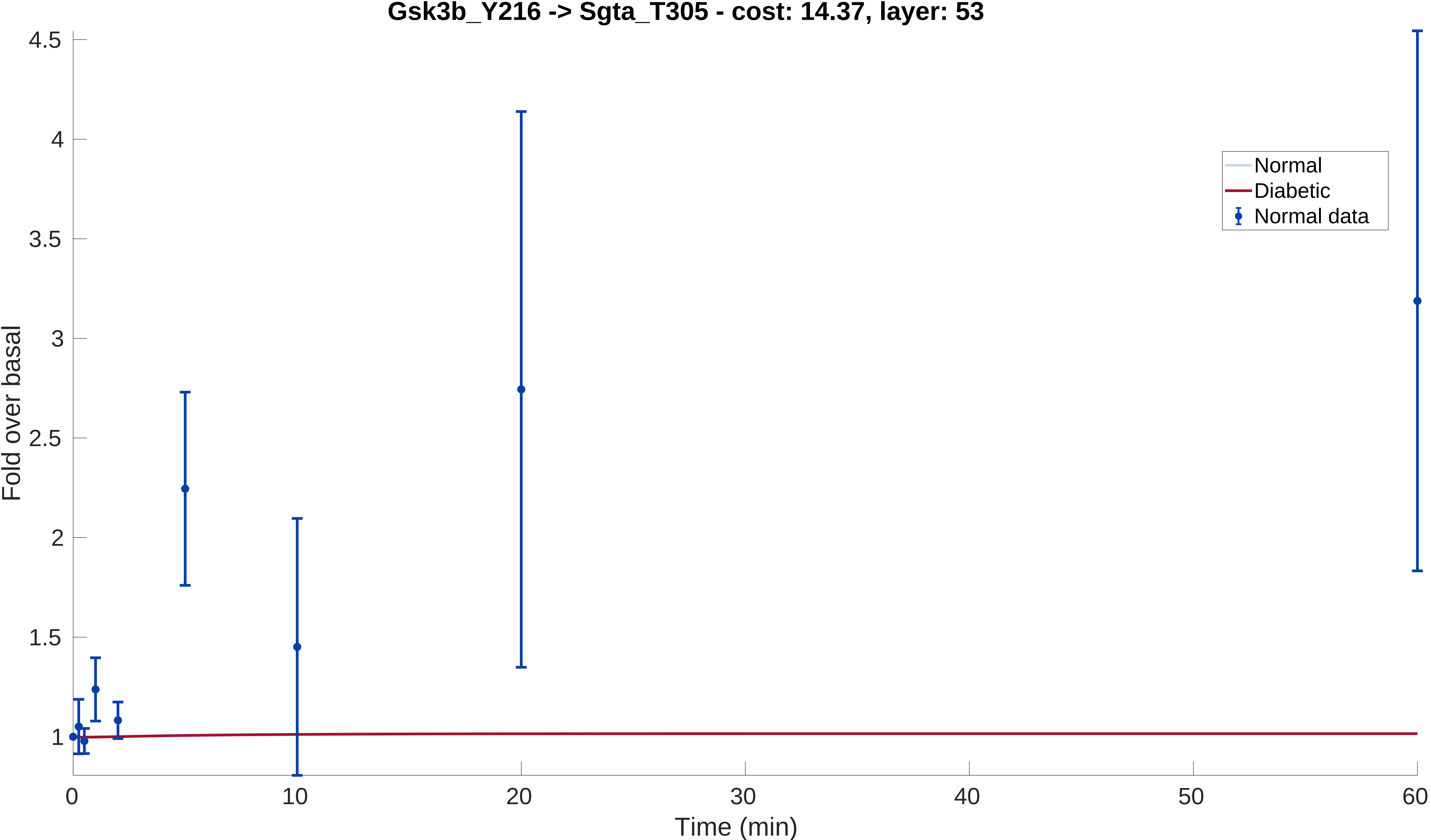
Gsk3b_Y216 -> Sgta_T305 - cost: 14.37, layer: 53
Fold over basal
4.5
4
3.5
3
2.5
2
1.5
1
0
10
20
30
40
50
60
Time (min)
Normal
Diabetic
Normal data

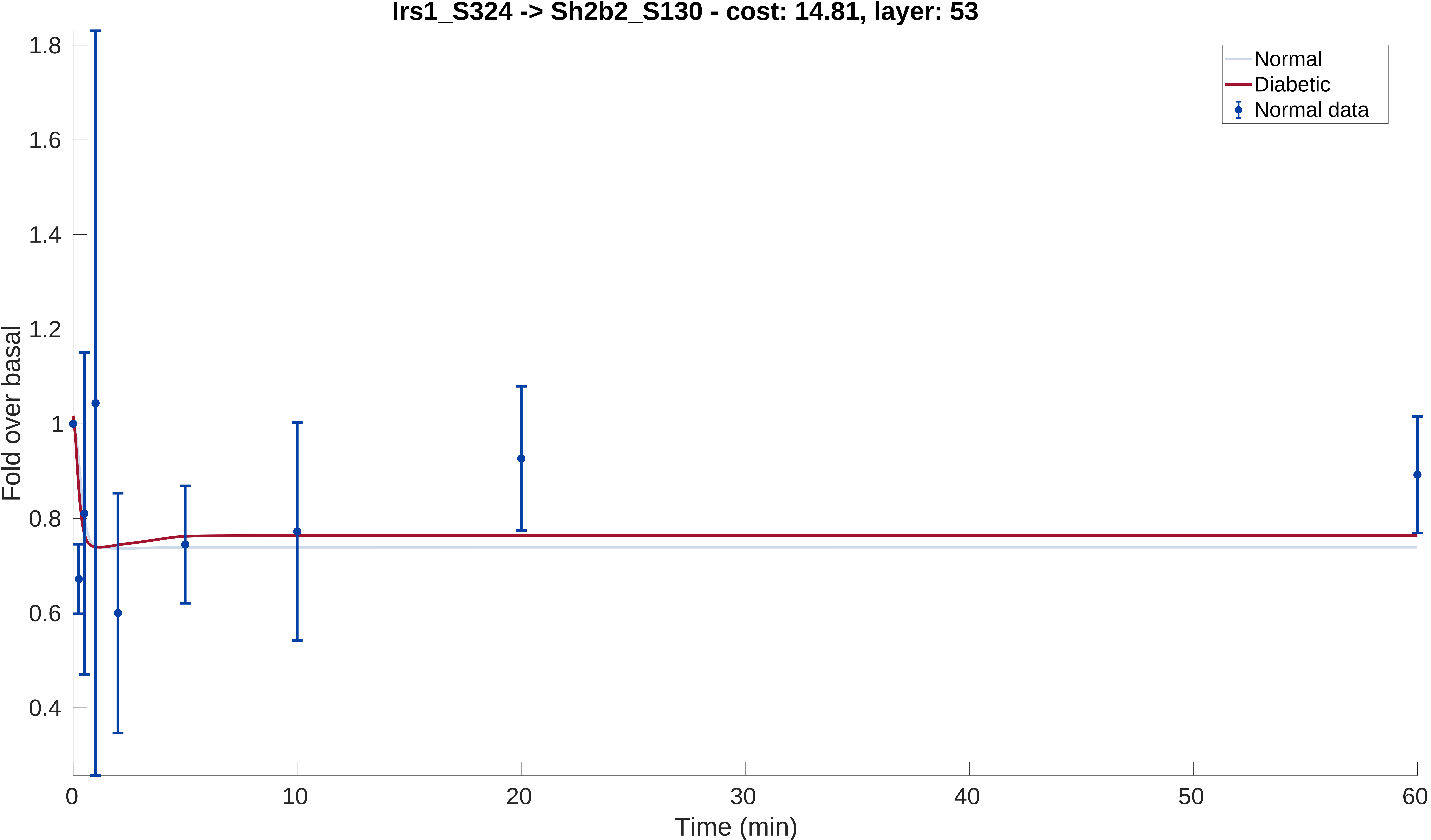
Irs1_S324 -> Sh2b2_S130 - cost: 14.81, layer: 53
Normal
Diabetic
Normal data
Fold over basal
1.8
1.6
1.4
1.2
1
0.8
0.6
0.4
0
10
20
30
40
50
60
Time (min)

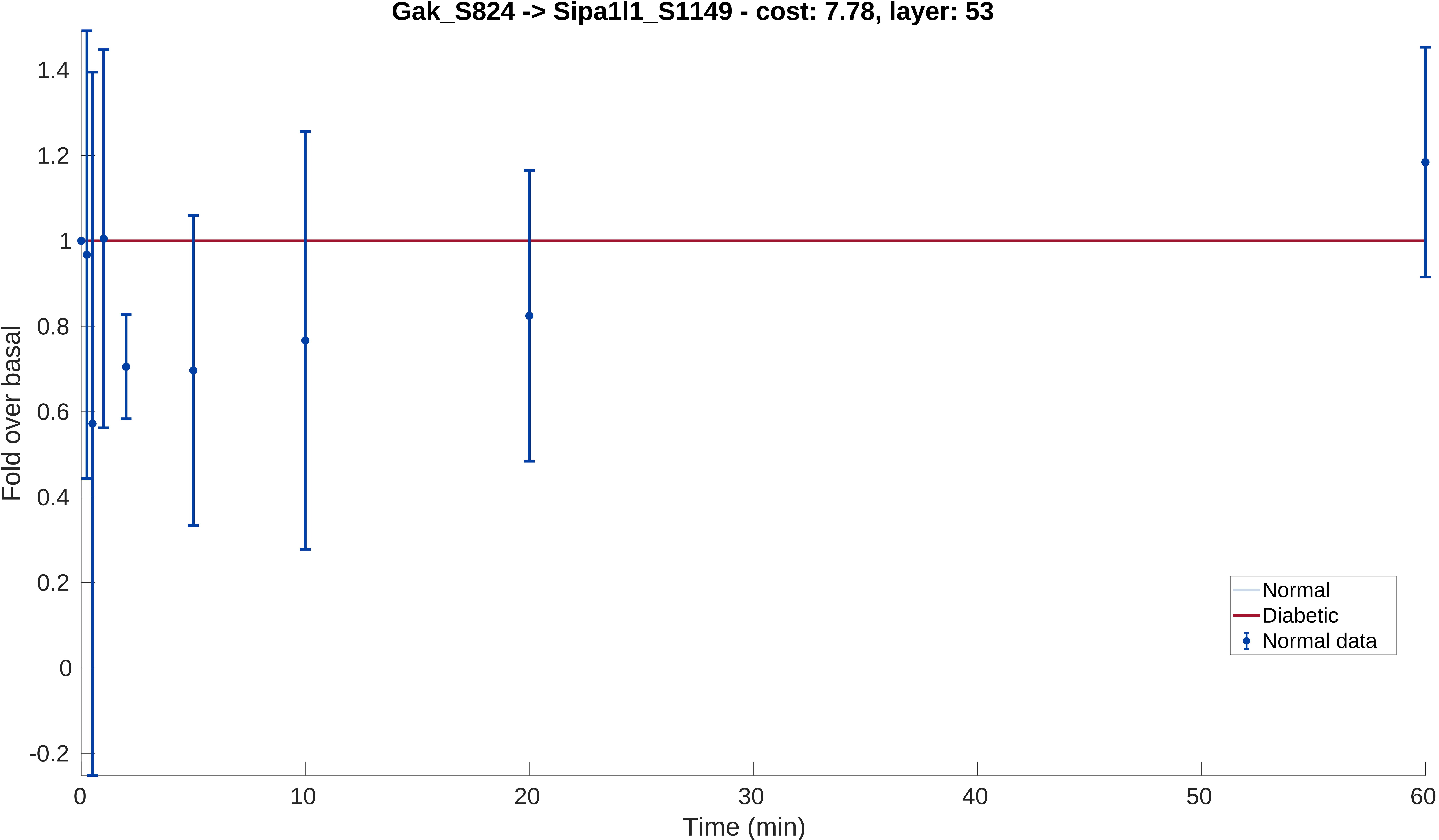

Gak_S824 -> Sipa1l1_S1149 - cost: 7.78, layer: 53
Fold over basal
Time (min)
1.4
1.2
1
0.8
0.6
0.4
0.2
0
-0.2
0
10
20
30
40
50
60
Normal
Diabetic
Normal data

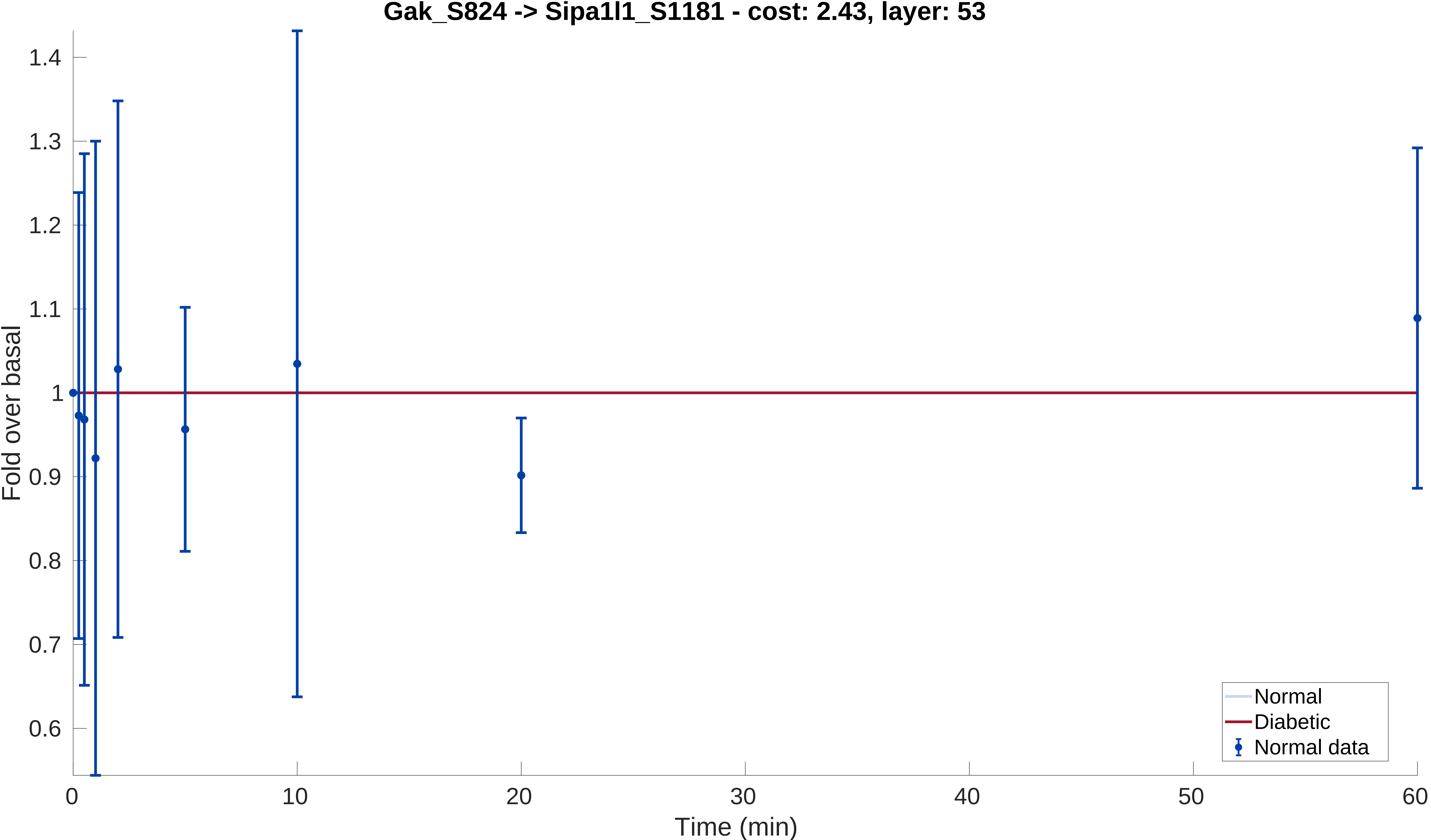

Gak_S824 -> Sipa1l1_S1181 - cost: 2.43, layer: 53
Fold over basal
1.4
1.3
1.2
1.1
1
0.9
0.8
0.7
0.6
0
10
20
30
40
50
60
Time (min)
Normal
Diabetic
Normal data

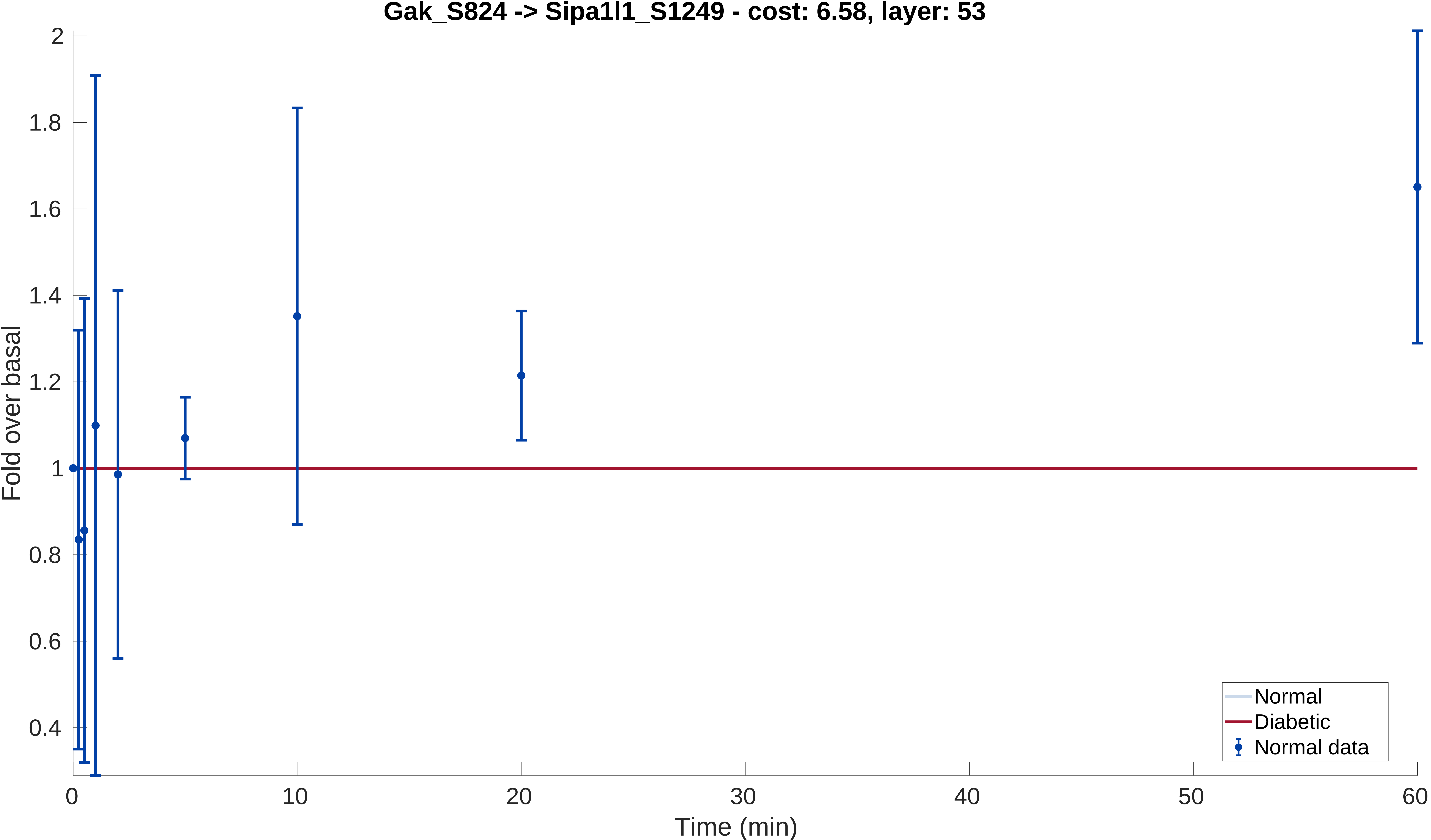

Gak_S824 -> Sipa1l1_S1249 - cost: 6.58, layer: 53
Fold over basal
Time (min)
Normal
Diabetic
Normal data
2
1.8
1.6
1.4
1.2
1
0.8
0.6
0.4
0
10
20
30
40
50
60

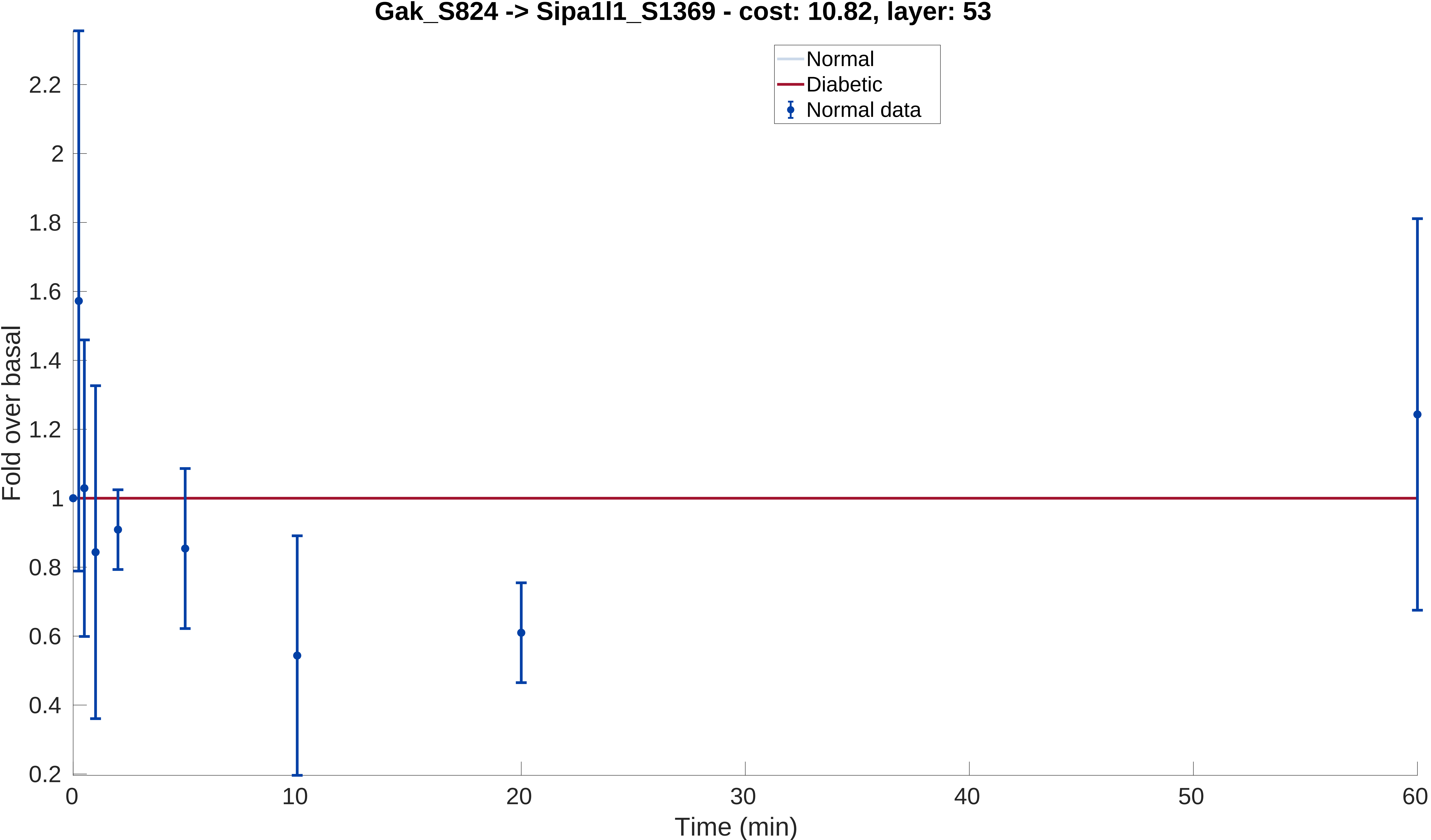

Gak_S824 -> Sipa1l1_S1369 - cost: 10.82, layer: 53
Normal
Diabetic
Normal data
Fold over basal
2.2
2
1.8
1.6
1.4
1.2
1
0.8
0.6
0.4
0.2
0
10
20
30
40
50
60
Time (min)

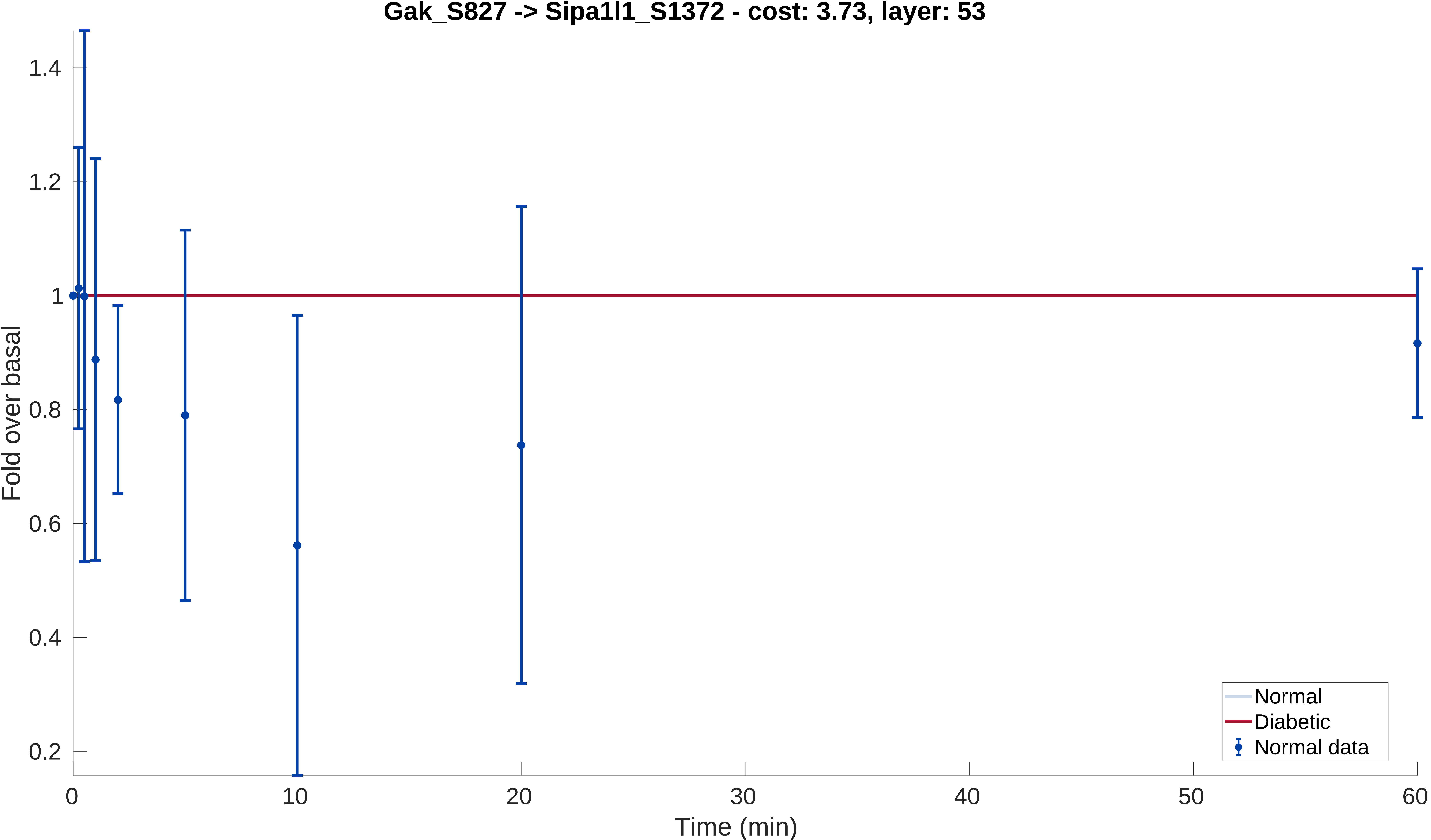

Gak_S827 -> Sipa1l1_S1372 - cost: 3.73, layer: 53
Fold over basal
1.4
1.2
1
0.8
0.6
0.4
0.2
0
10
20
30
40
50
60
Time (min)
Normal
Diabetic
Normal data

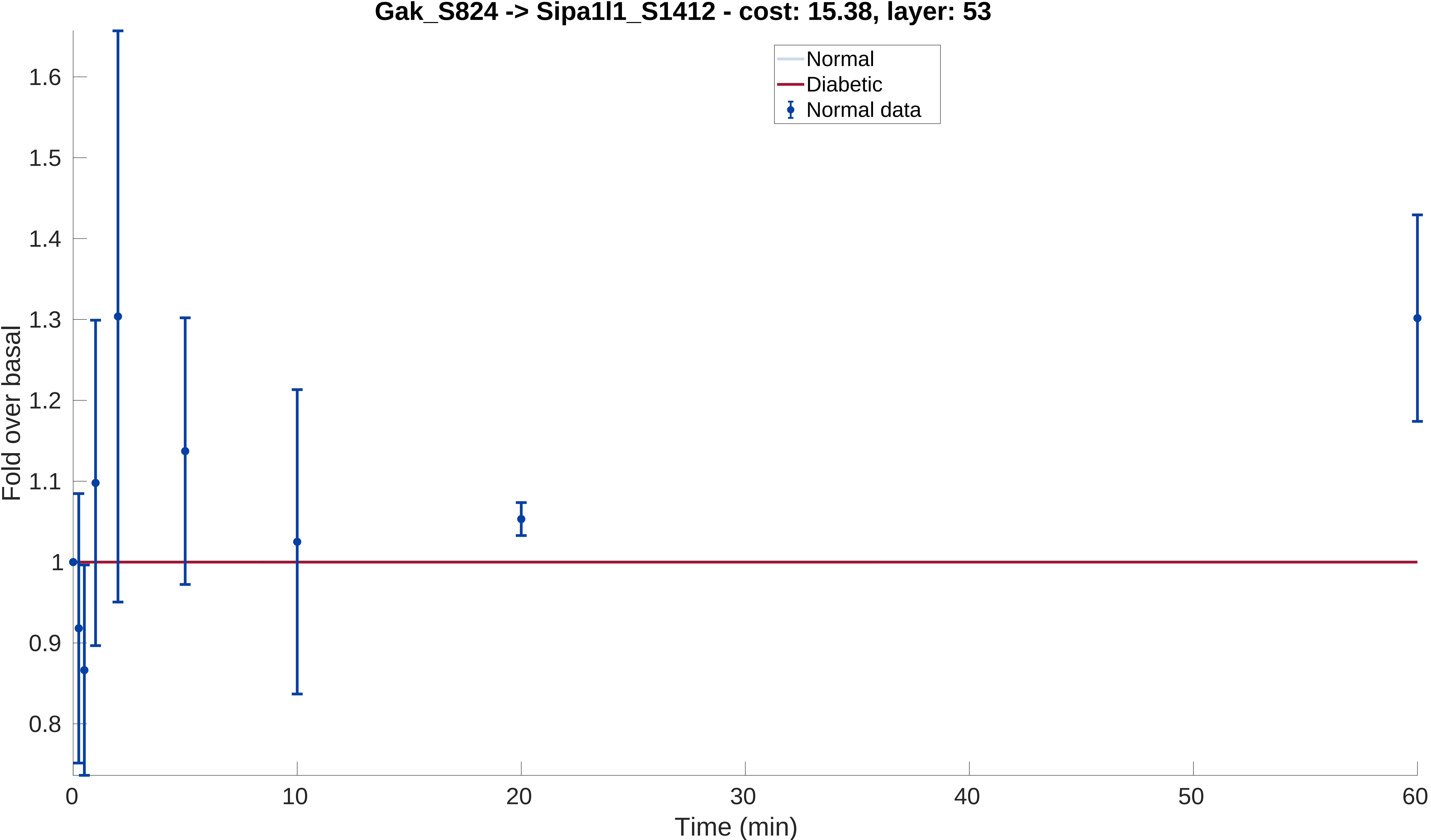

Gak_S824 -> Sipa1l1_S1412 - cost: 15.38, layer: 53
Normal
Diabetic
Normal data
Fold over basal
1.6
1.5
1.4
1.3
1.2
1.1
1
0.9
0.8
0
10
20
30
40
50
60
Time (min)

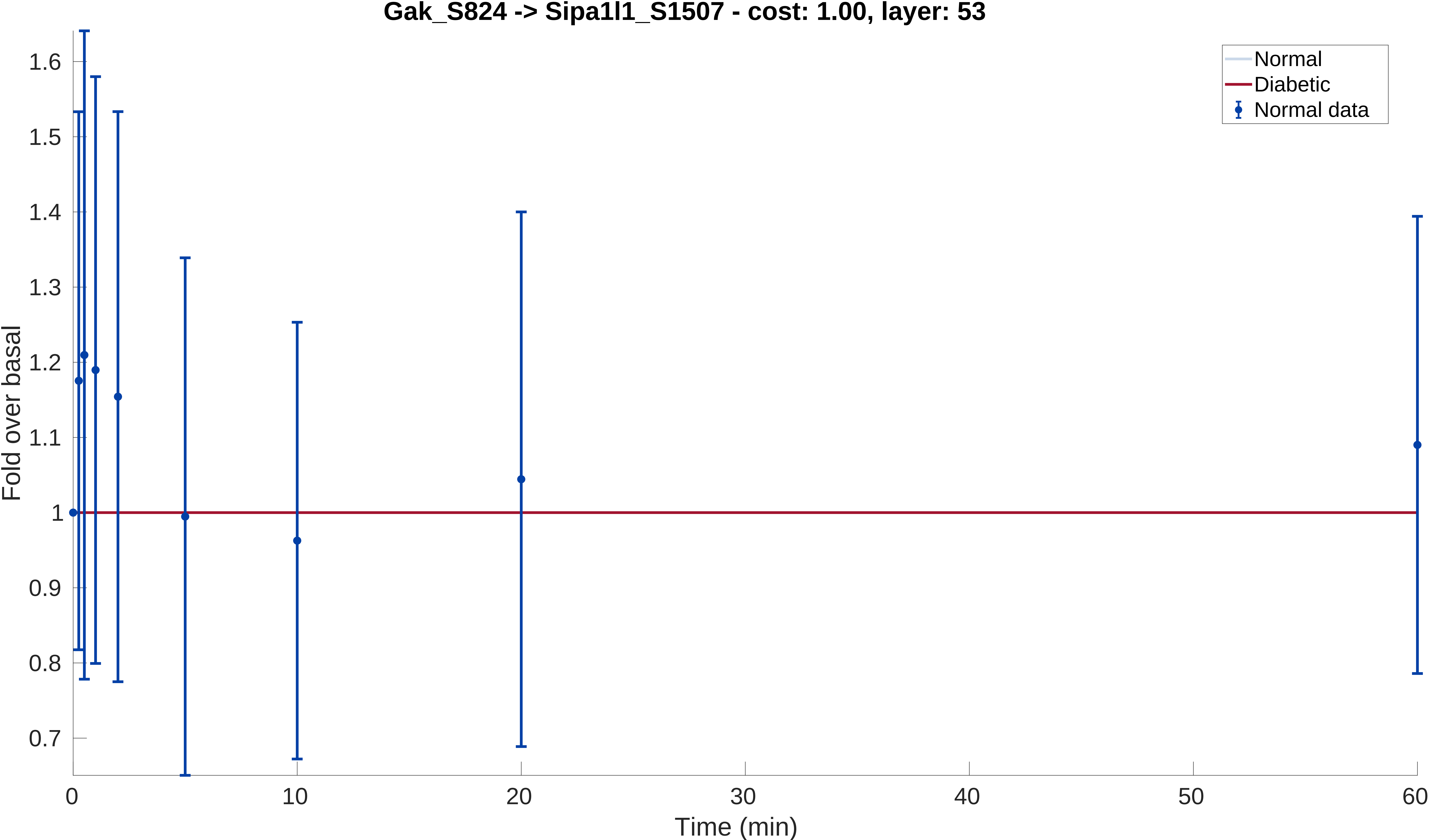
Gak_S824 -> Sipa1l1_S1507 - cost: 1.00, layer: 53
Normal
Diabetic
Normal data
Fold over basal
1.6
1.5
1.4
1.3
1.2
1.1
1
0.9
0.8
0.7
0
10
20
30
40
50
60
Time (min)

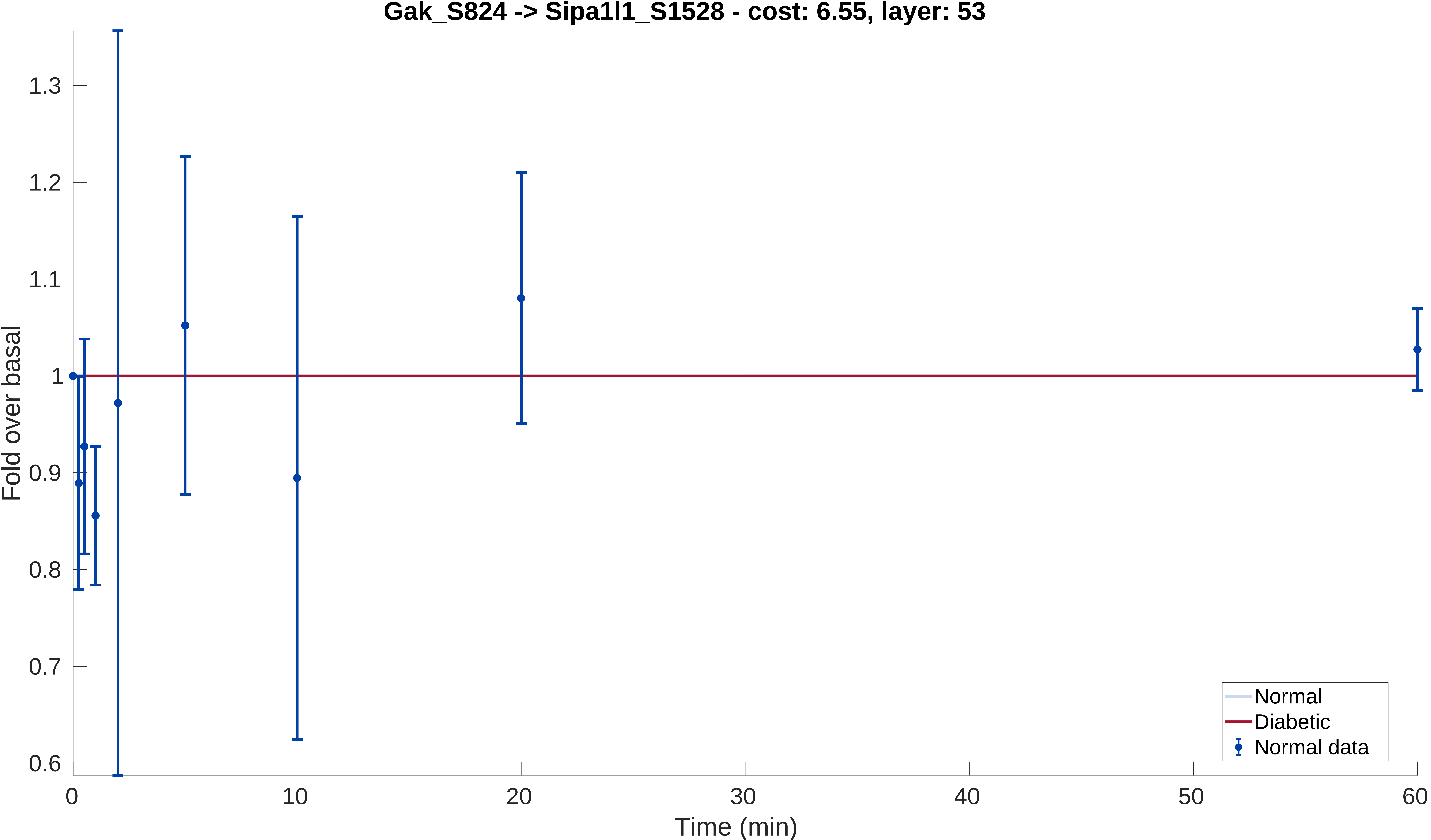
Gak_S824 -> Sipa1l1_S1528 - cost: 6.55, layer: 53
Fold over basal
Time (min)
0.6
0.7
0.8
0.9
1
1.1
1.2
1.3
0
10
20
30
40
50
60
Normal
Diabetic
Normal data

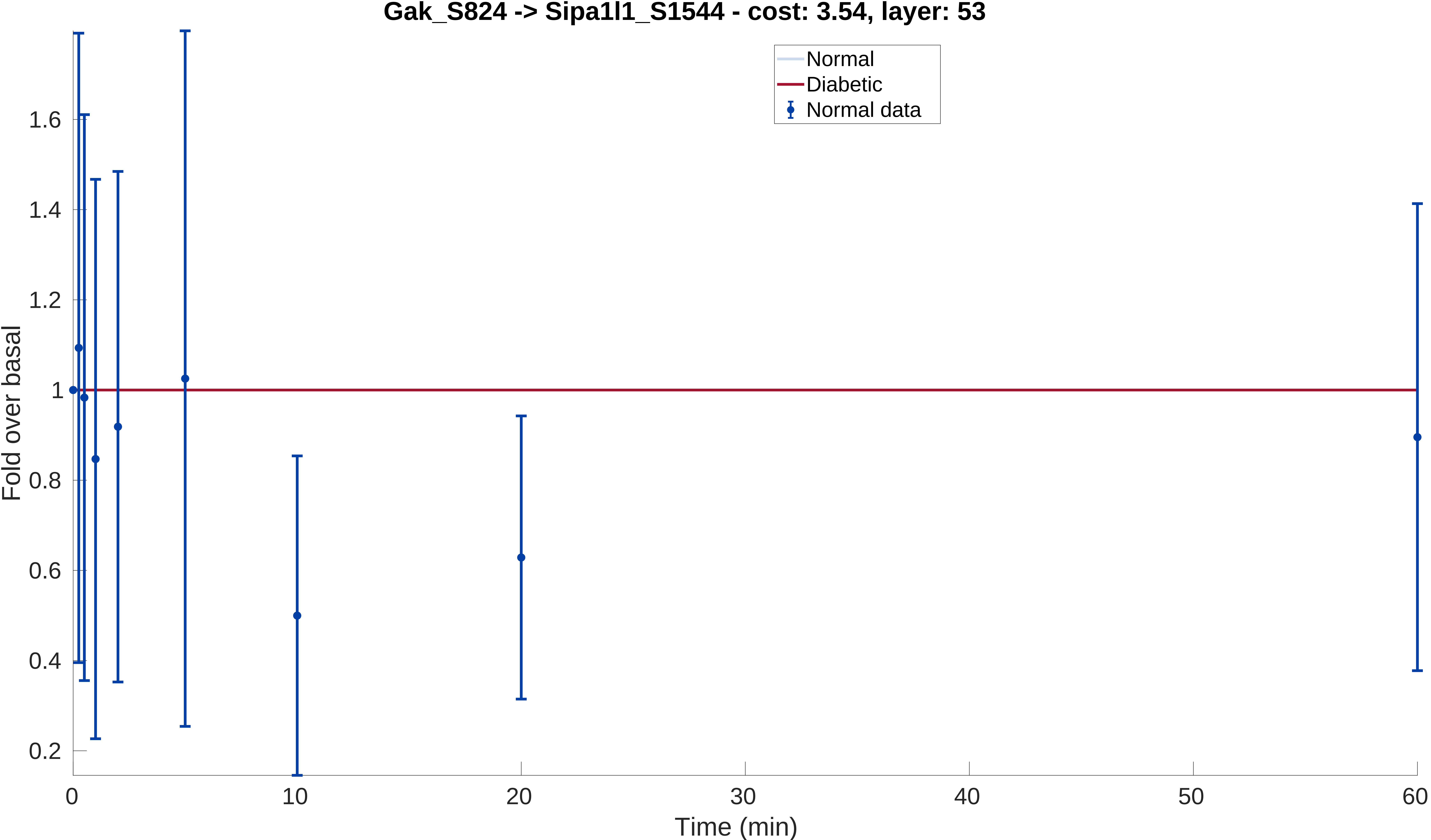

Gak_S824 -> Sipa1l1_S1544 - cost: 3.54, layer: 53
Normal
Diabetic
Normal data
Fold over basal
1.6
1.4
1.2
1
0.8
0.6
0.4
0.2
0
10
20
30
40
50
60
Time (min)

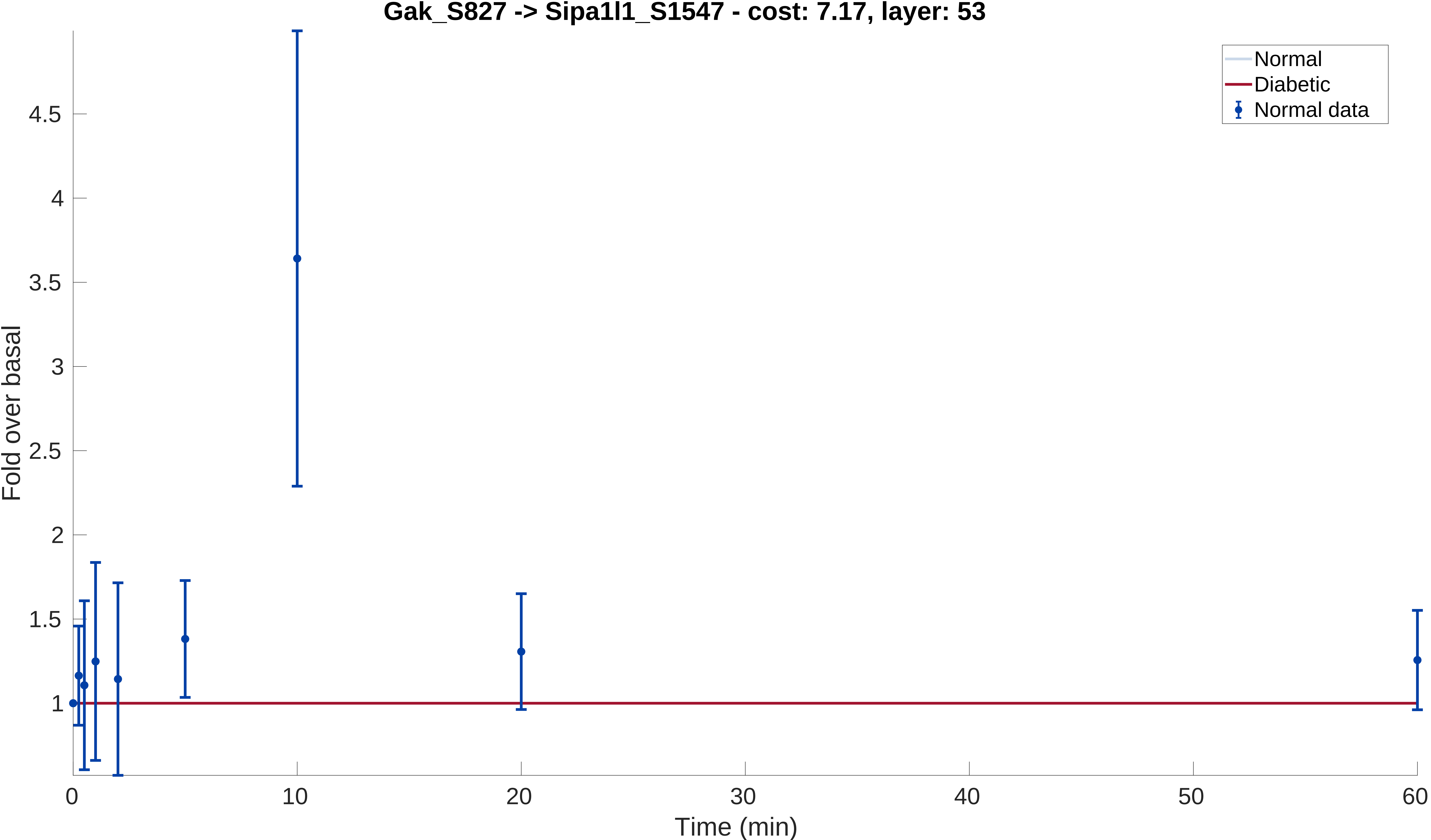

Gak_S827 -> Sipa1l1_S1547 - cost: 7.17, layer: 53
Normal
Diabetic
Normal data
Fold over basal
4.5
4
3.5
3
2.5
2
1.5
1
0
10
20
30
40
50
60
Time (min)

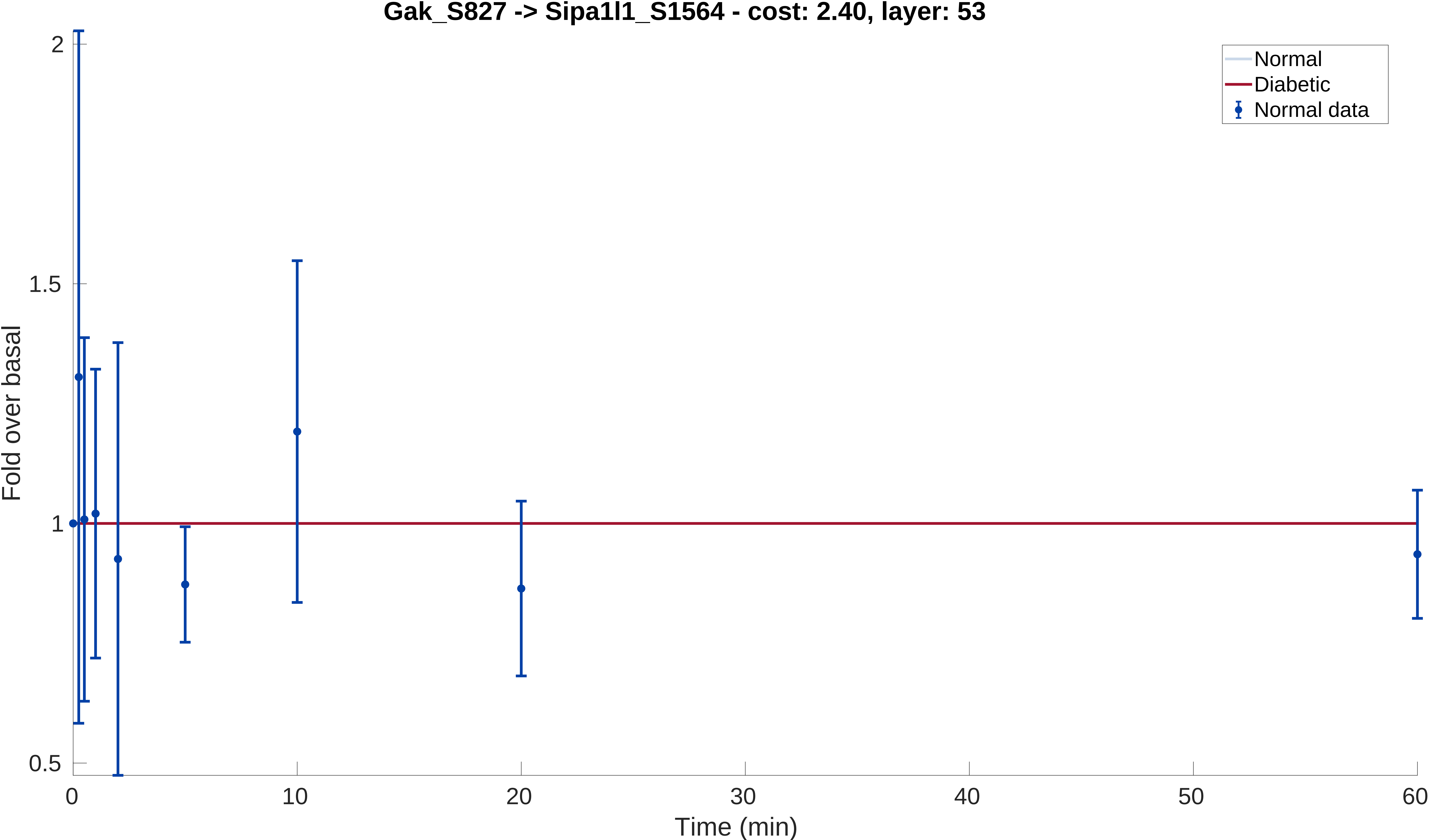

Gak_S827 -> Sipa1l1_S1564 - cost: 2.40, layer: 53
Normal
Diabetic
Normal data
Fold over basal
2
1.5
1
0.5
0
10
20
30
40
50
60
Time (min)

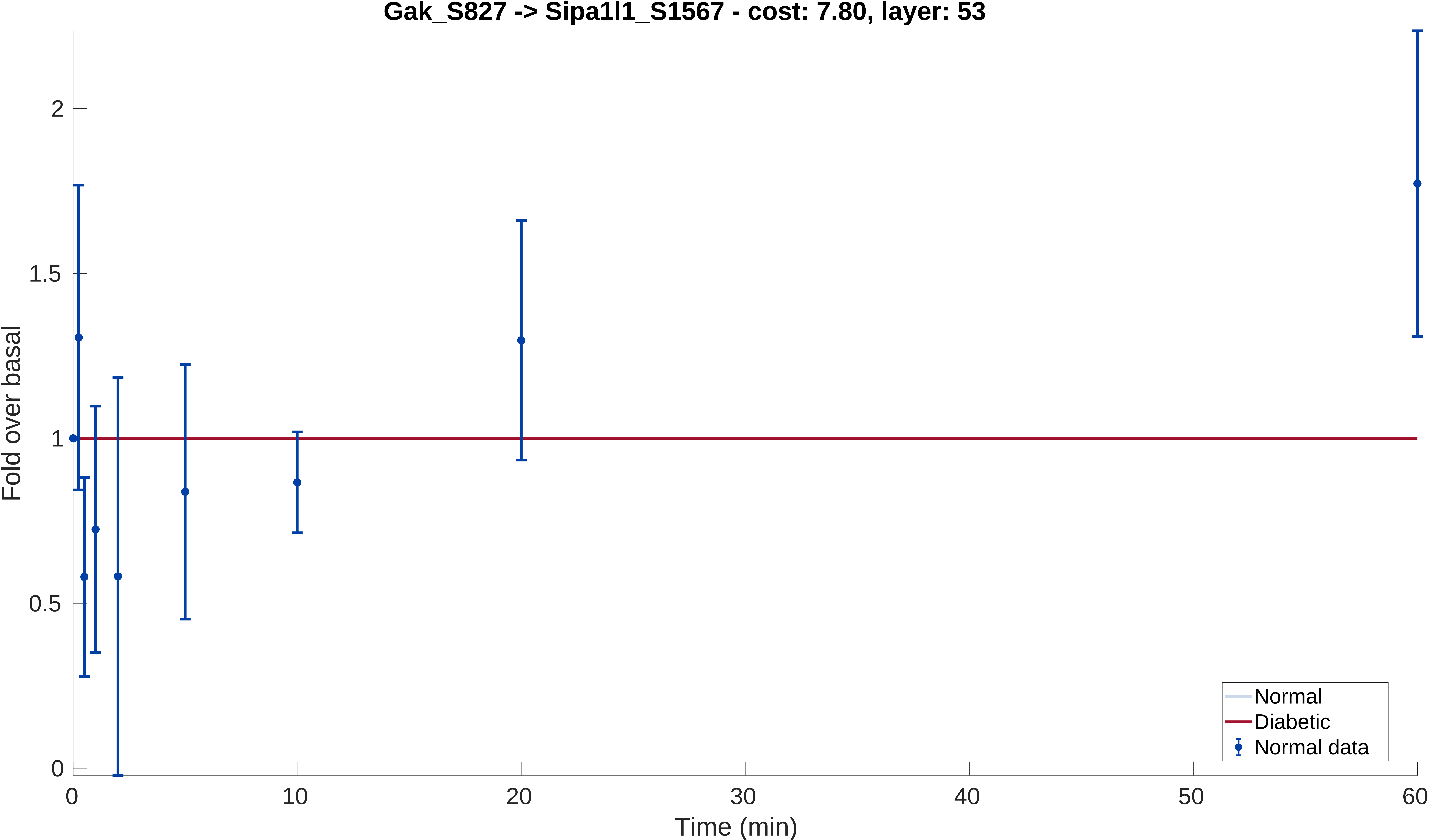

Gak_S827 -> Sipa1l1_S1567 - cost: 7.80, layer: 53
Fold over basal
Time (min)
0
0.5
1
1.5
2
0
10
20
30
40
50
60
Normal
Diabetic
Normal data

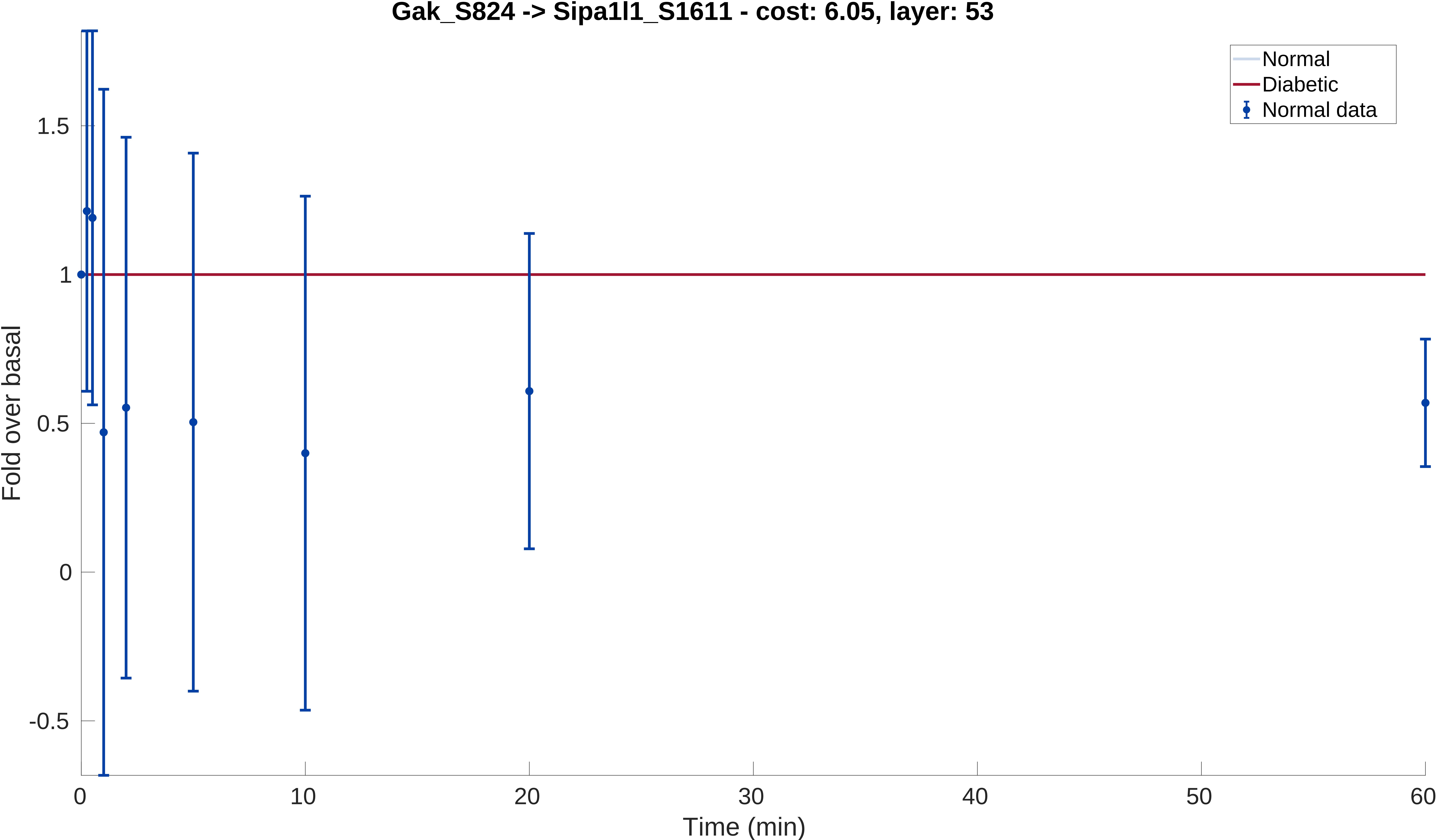

Gak_S824 -> Sipa1l1_S1611 - cost: 6.05, layer: 53
Normal
Diabetic
Normal data
Fold over basal
1.5
1
0.5
0
-0.5
0
10
20
30
40
50
60
Time (min)

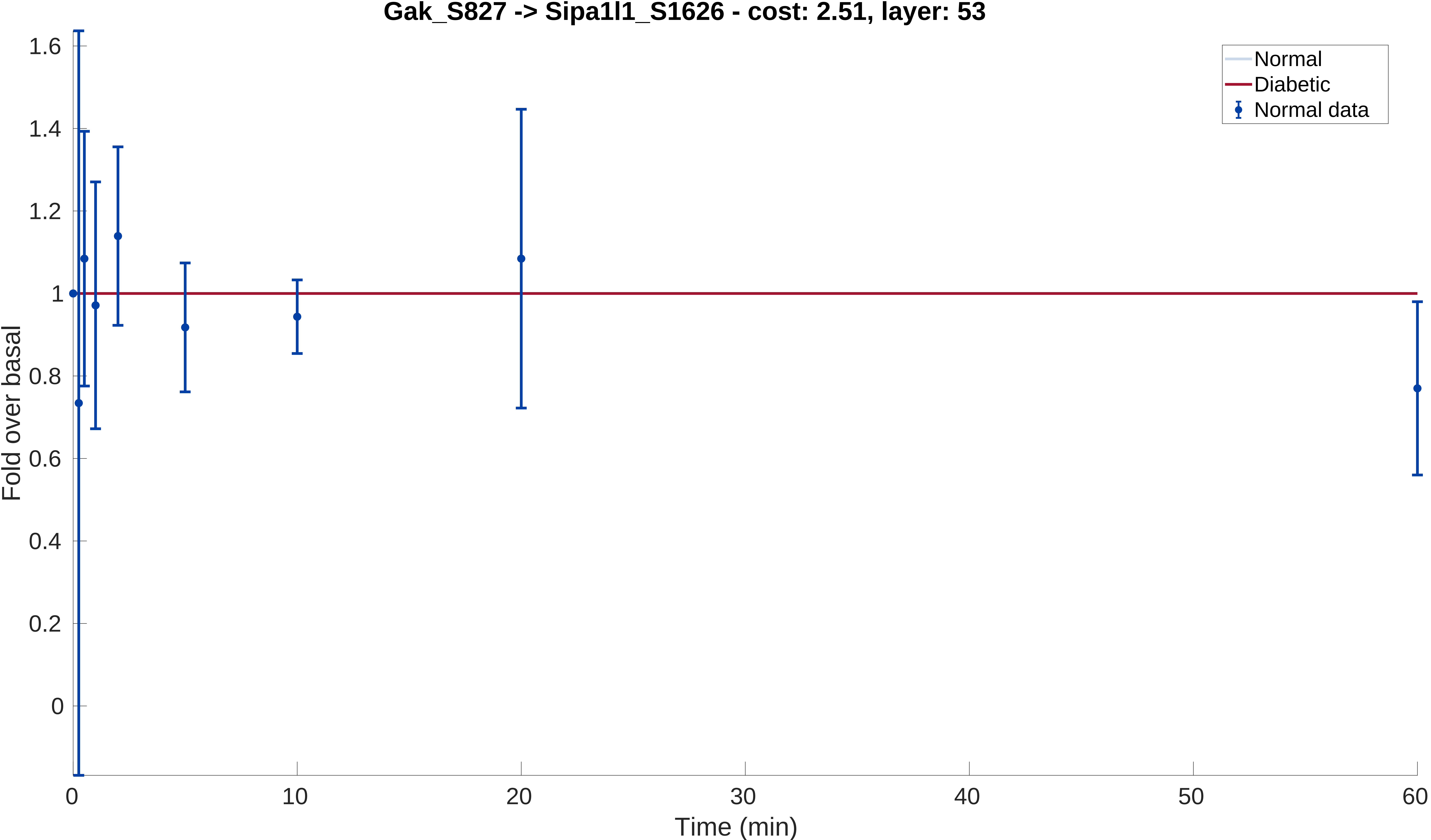

Gak_S827 -> Sipa1l1_S1626 - cost: 2.51, layer: 53
Normal
Diabetic
Normal data
Fold over basal
Time (min)
1.6
1.4
1.2
1
0.8
0.6
0.4
0.2
0
0
10
20
30
40
50
60

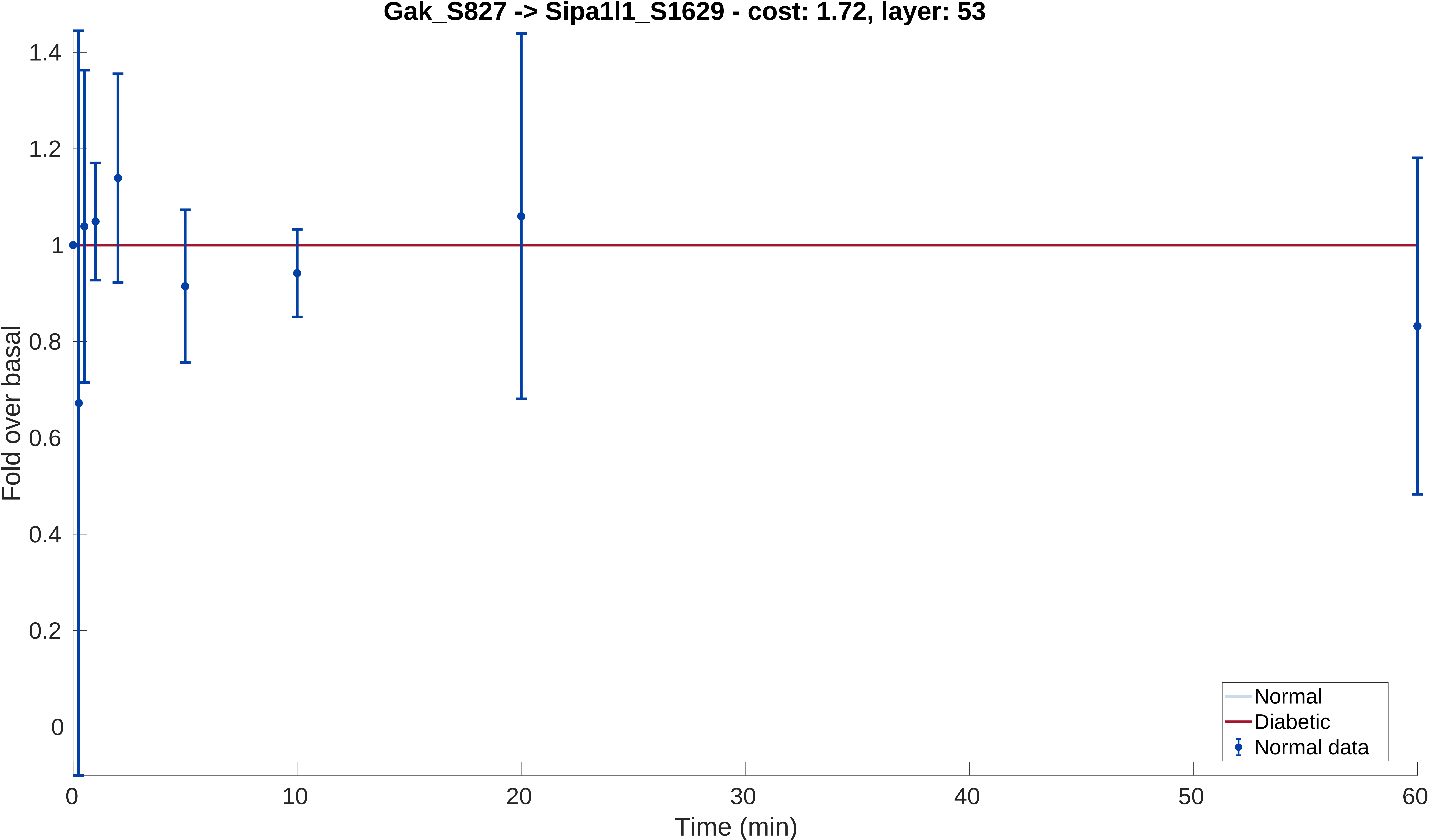

Gak_S827 -> Sipa1l1_S1629 - cost: 1.72, layer: 53
Fold over basal
1.4
1.2
1
0.8
0.6
0.4
0.2
0
0
10
20
30
40
50
60
Time (min)
Normal
Diabetic
Normal data

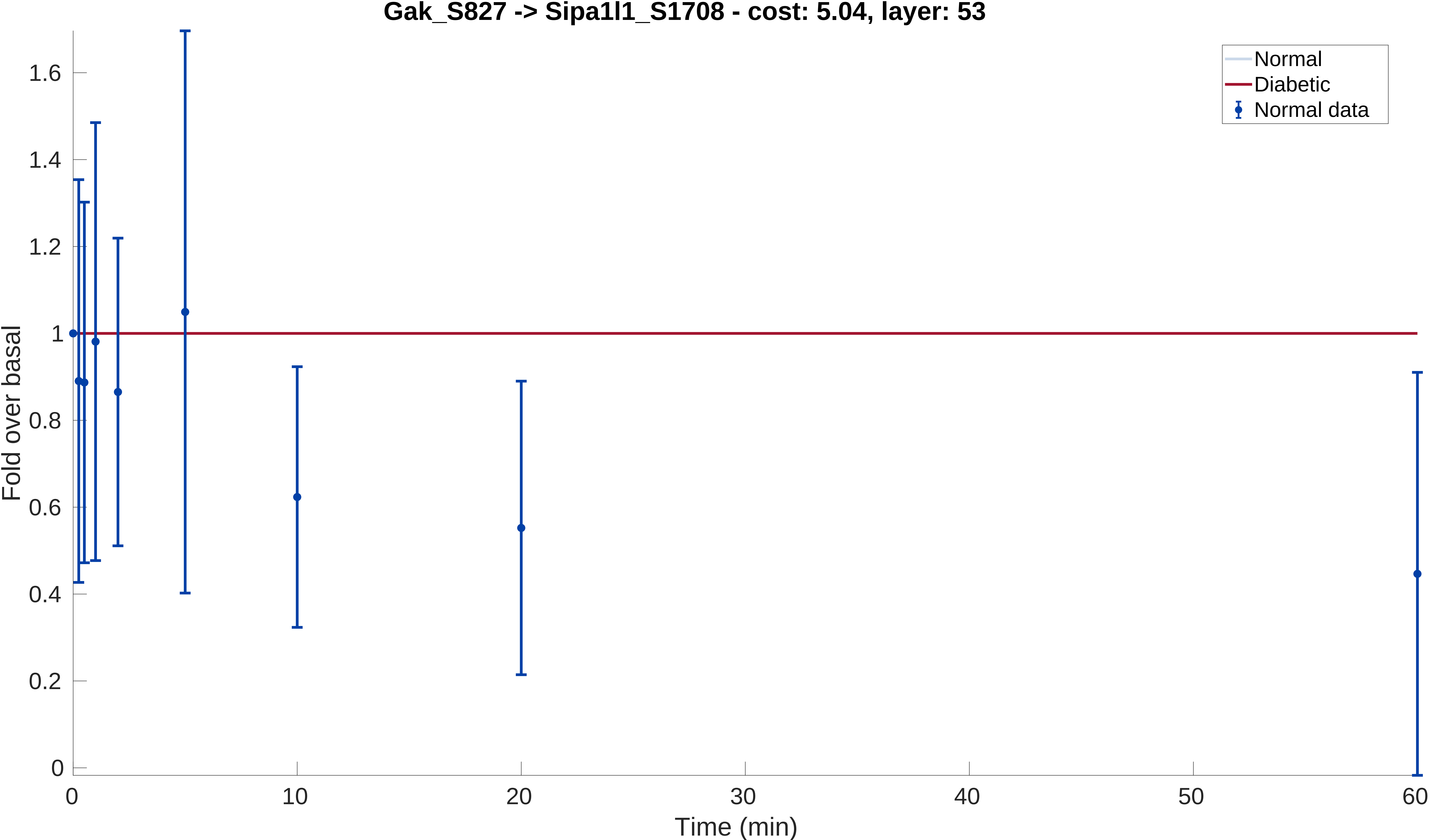

Gak_S827 -> Sipa1l1_S1708 - cost: 5.04, layer: 53
Normal
Diabetic
Normal data
Fold over basal
Time (min)
0
0.2
0.4
0.6
0.8
1
1.2
1.4
1.6
0
10
20
30
40
50
60

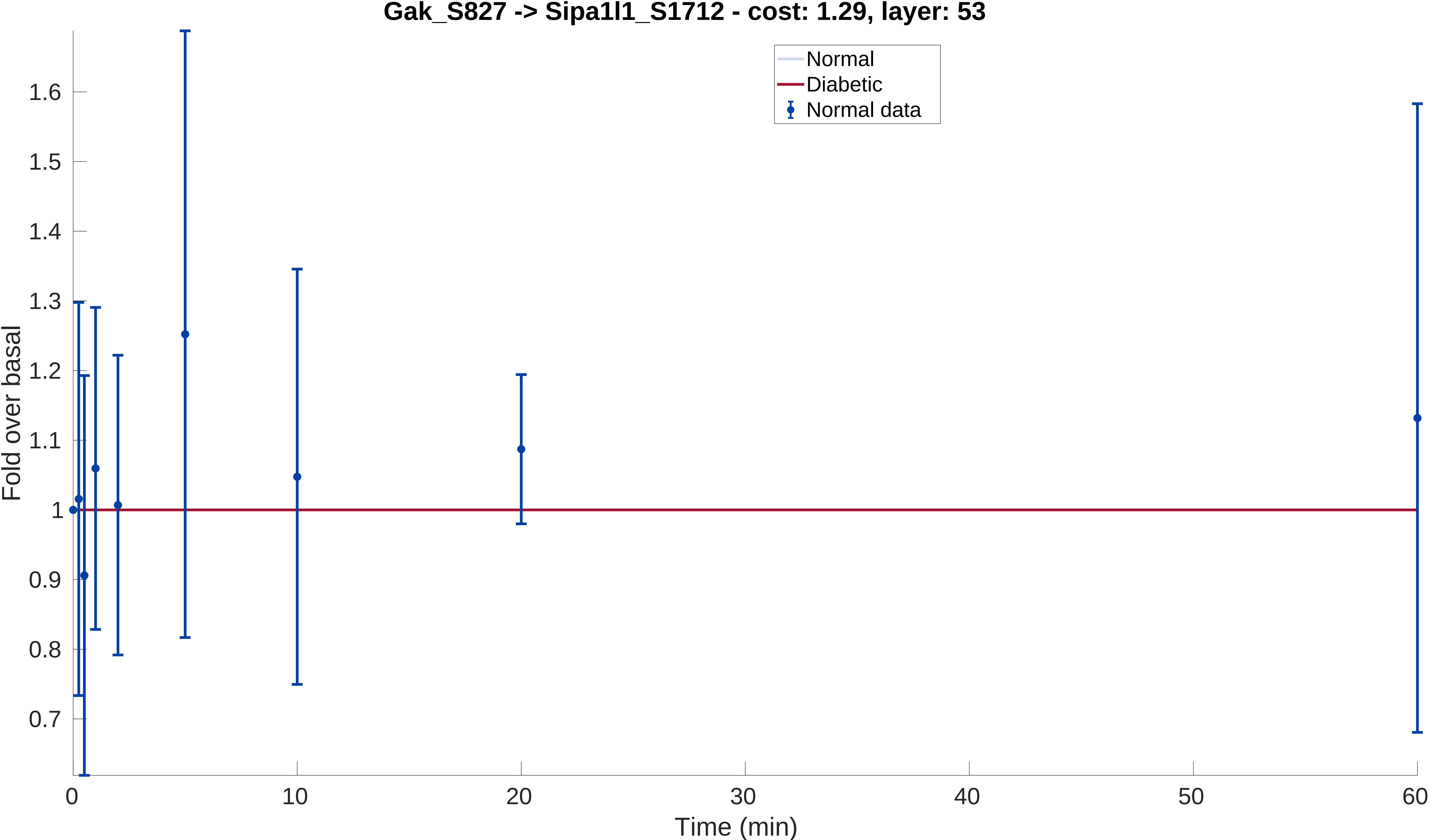

Gak_S827 -> Sipa1l1_S1712 - cost: 1.29, layer: 53
Normal
Diabetic
Normal data
Fold over basal
1.6
1.5
1.4
1.3
1.2
1.1
1
0.9
0.8
0.7
0
10
20
30
40
50
60
Time (min)

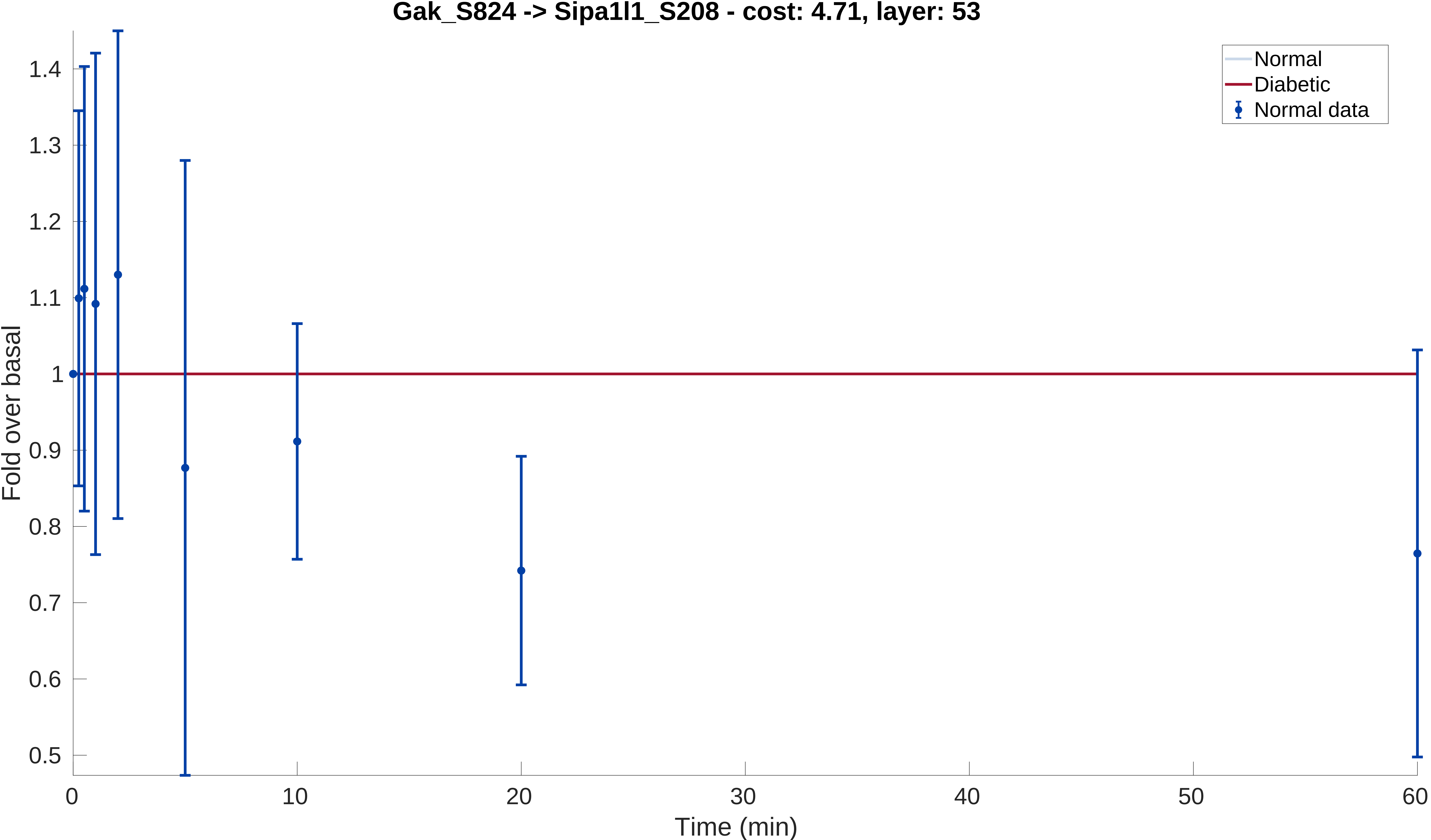

Gak_S824 -> Sipa1l1_S208 - cost: 4.71, layer: 53
Normal
Diabetic
Normal data
Fold over basal
Time (min)
1.4
1.3
1.2
1.1
1
0.9
0.8
0.7
0.6
0.5
0
10
20
30
40
50
60

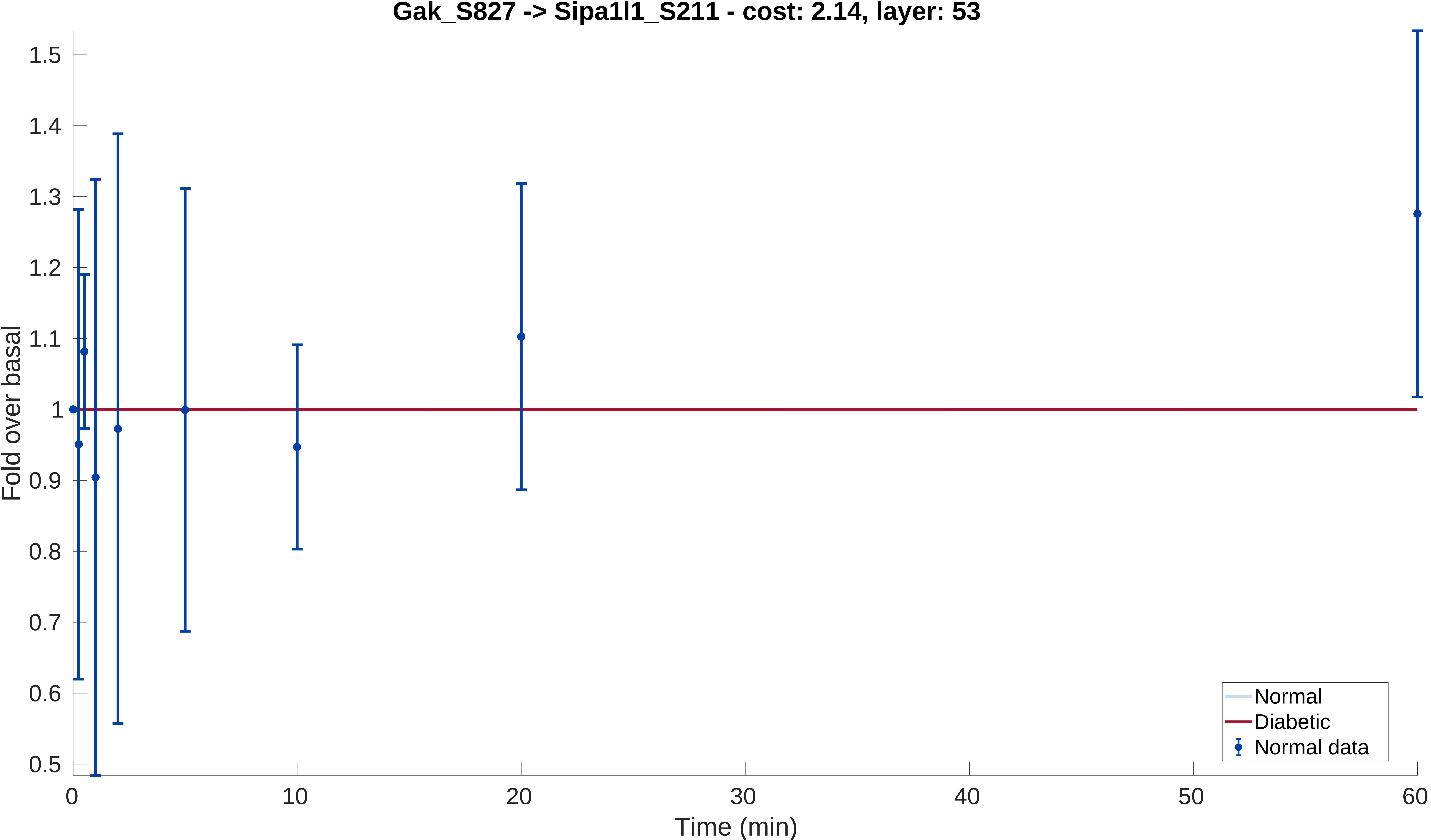
Gak_S827 -> Sipa1l1_S211 - cost: 2.14, layer: 53
Fold over basal
Time (min)
1.5
1.4
1.3
1.2
1.1
1
0.9
0.8
0.7
0.6
0.5
0
10
20
30
40
50
60
Normal
Diabetic
Normal data

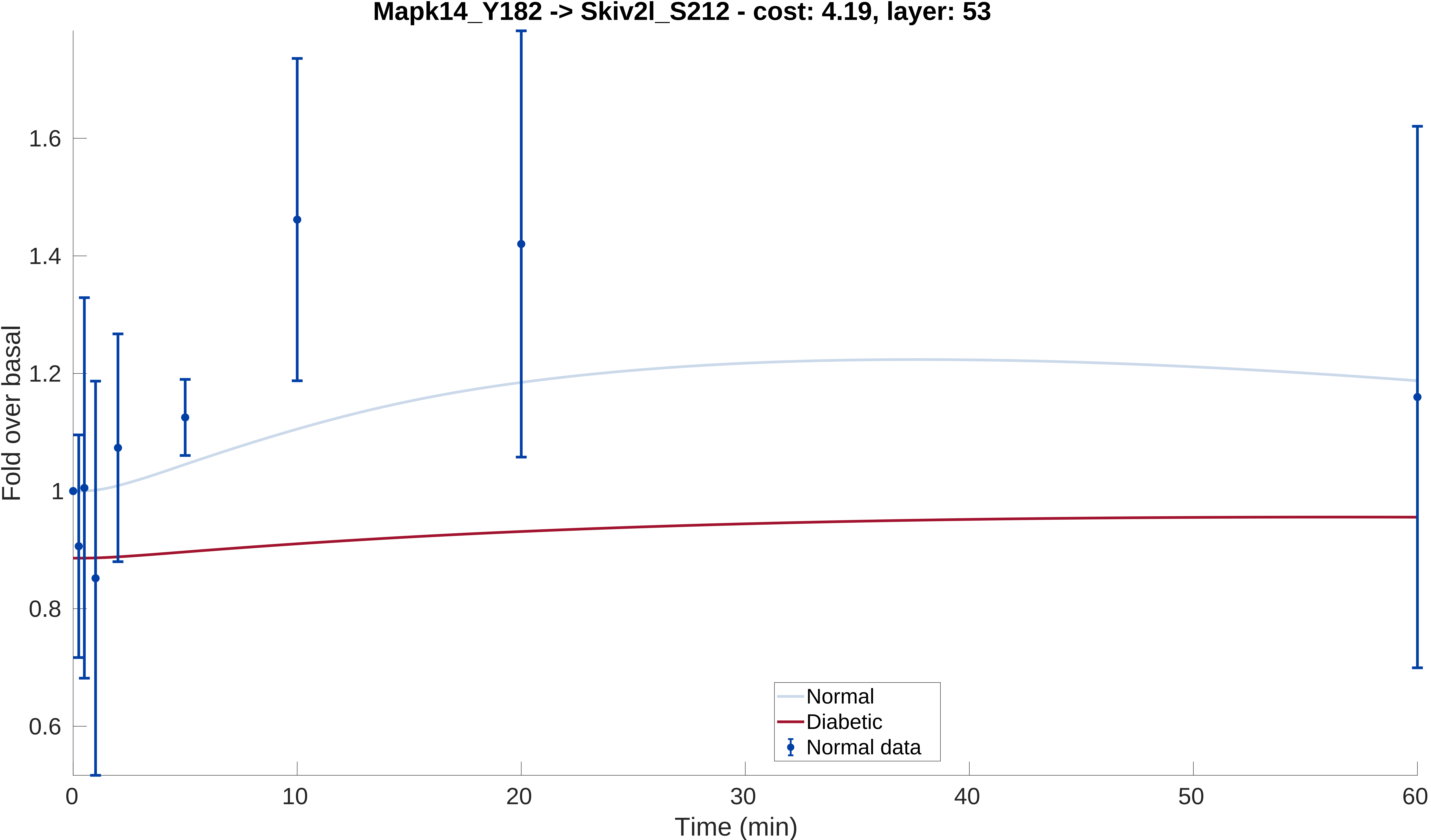

Mapk14_Y182 -> Skiv2l_S212 - cost: 4.19, layer: 53
Fold over basal
1.6
1.4
1.2
1
0.8
0.6
0
10
20
30
40
50
60
Time (min)
Normal
Diabetic
Normal data

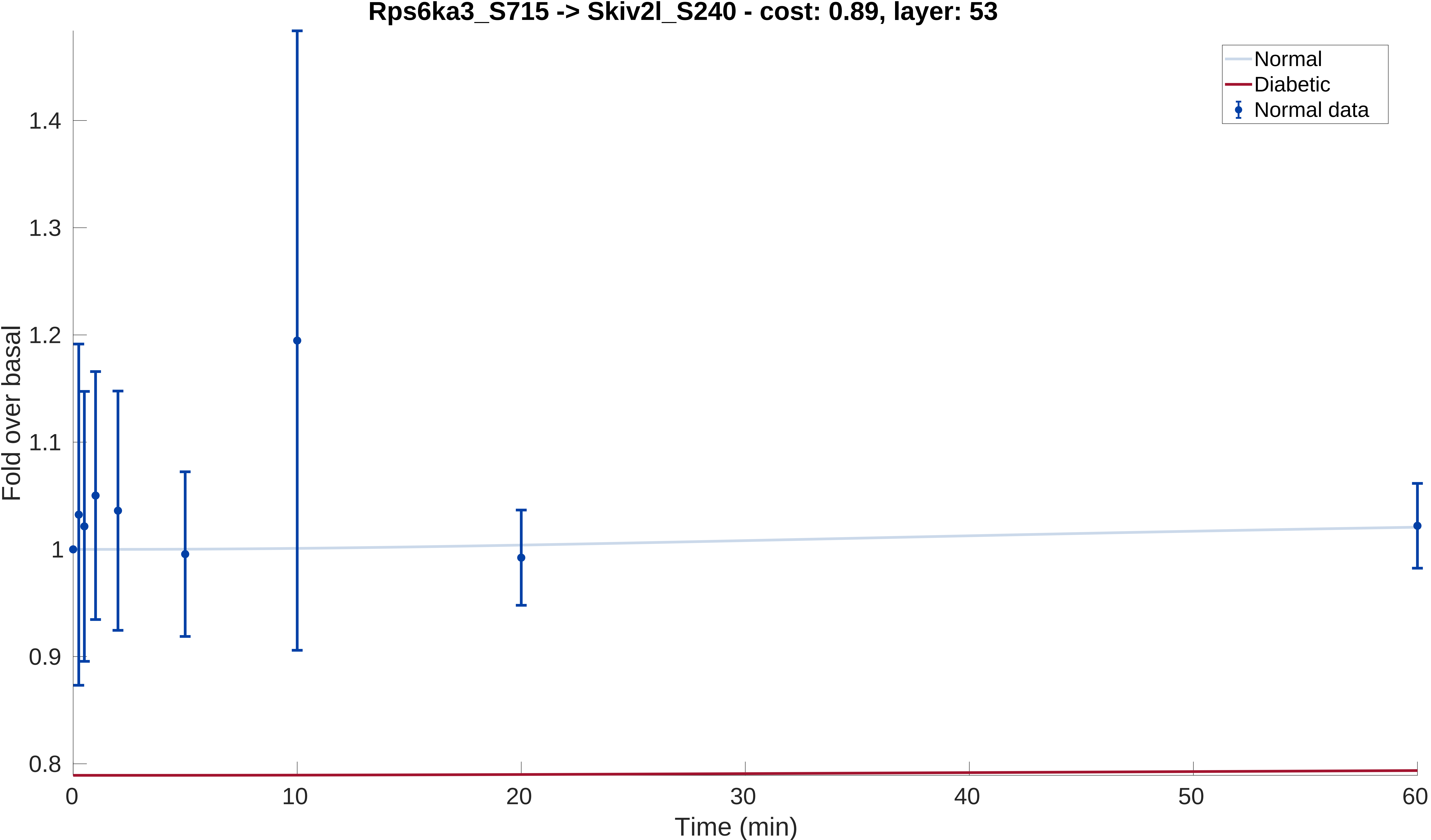

Rps6ka3_S715 -> Skiv2l_S240 - cost: 0.89, layer: 53
Normal
Diabetic
Normal data
Fold over basal
1.4
1.3
1.2
1.1
1
0.9
0.8
0
10
20
30
40
50
60
Time (min)

#### Rps6ka3_S715 -> Skiv2l_S242 - cost: 6.84, layer: 53

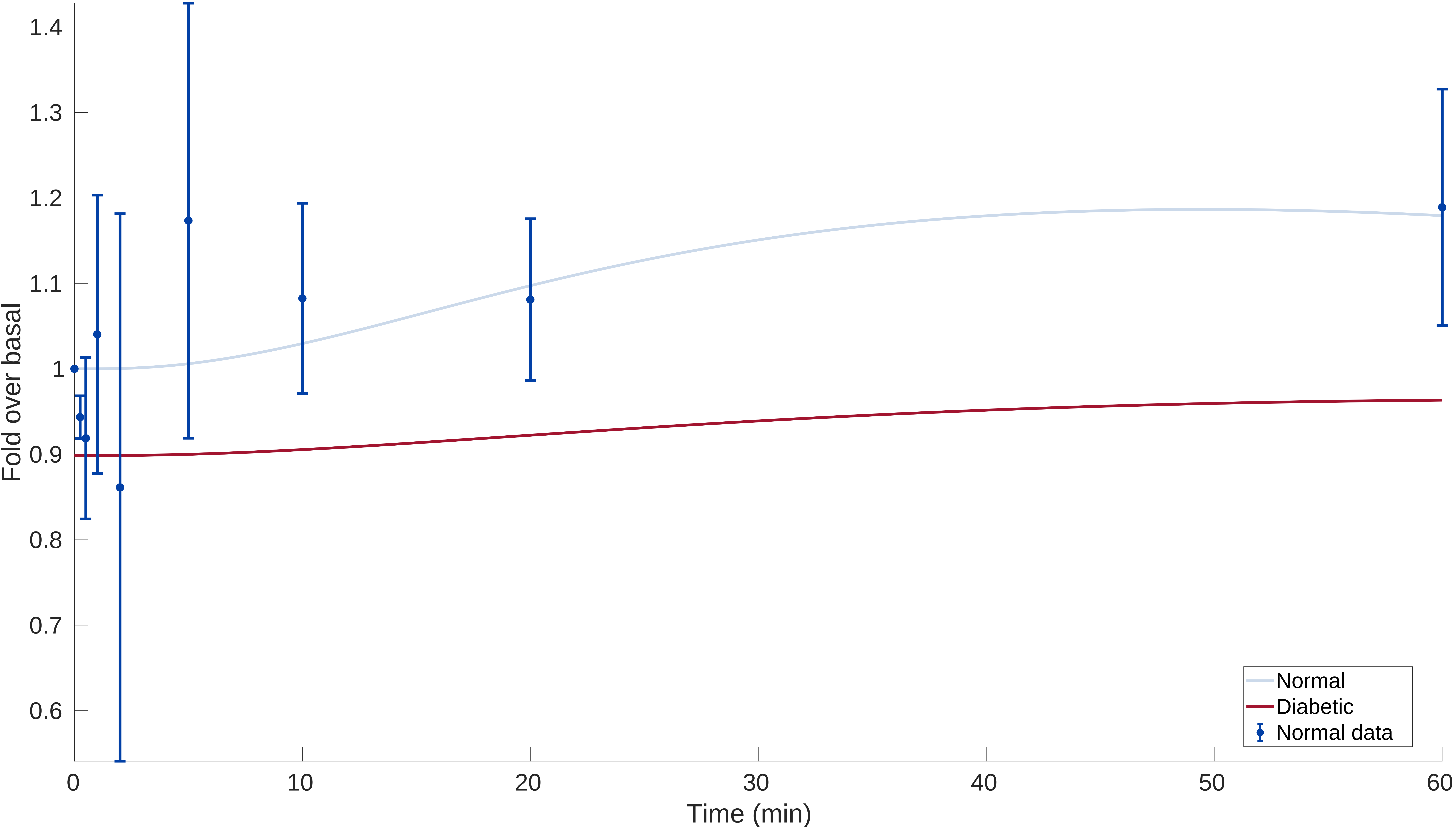

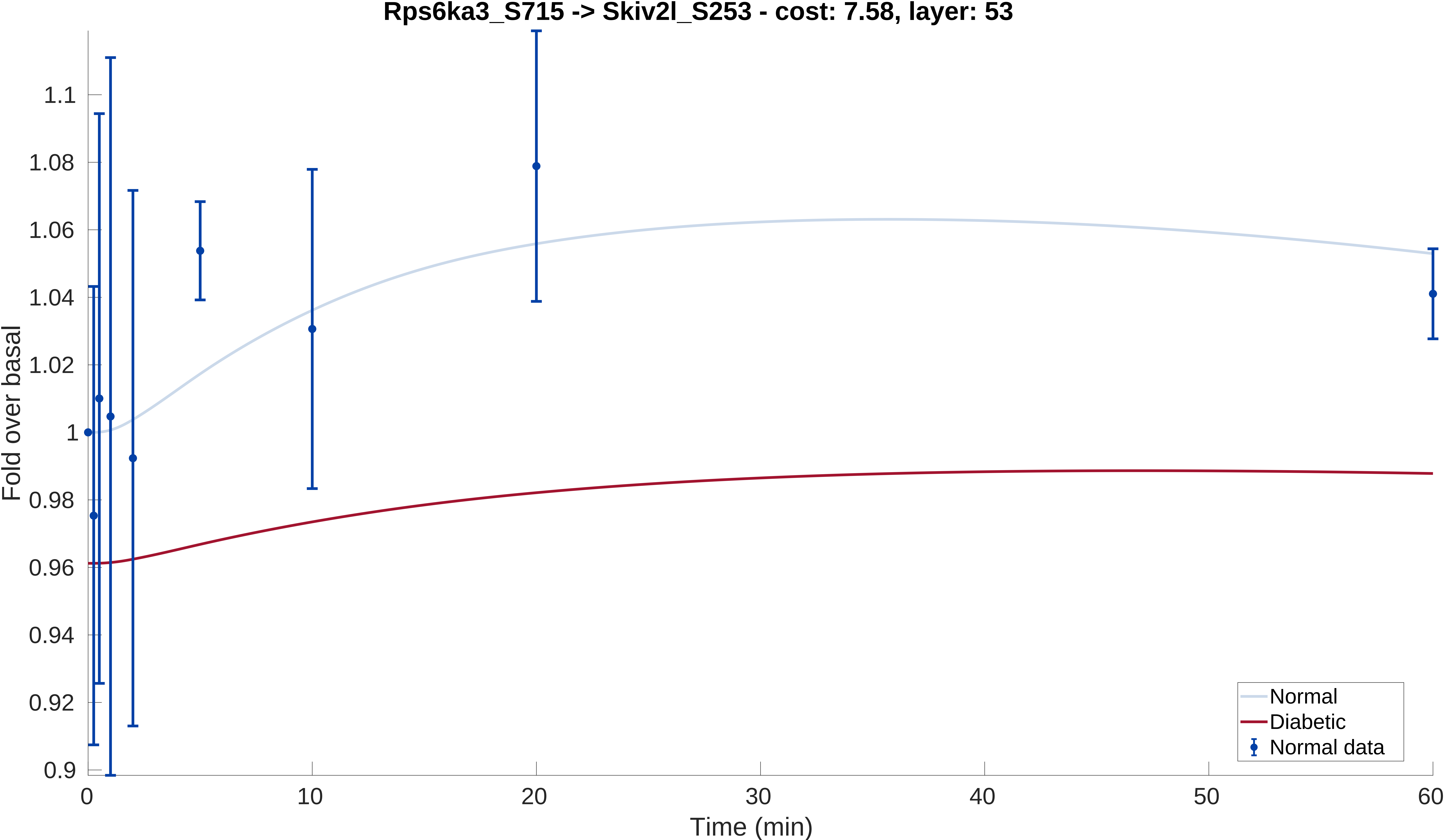

Rps6ka3_S715 -> Skiv2l_S253 - cost: 7.58, layer: 53
Fold over basal
1.1
1.08
1.06
1.04
1.02
1
0.98
0.96
0.94
0.92
0.9
0
10
20
30
40
50
60
Time (min)
Normal
Diabetic
Normal data

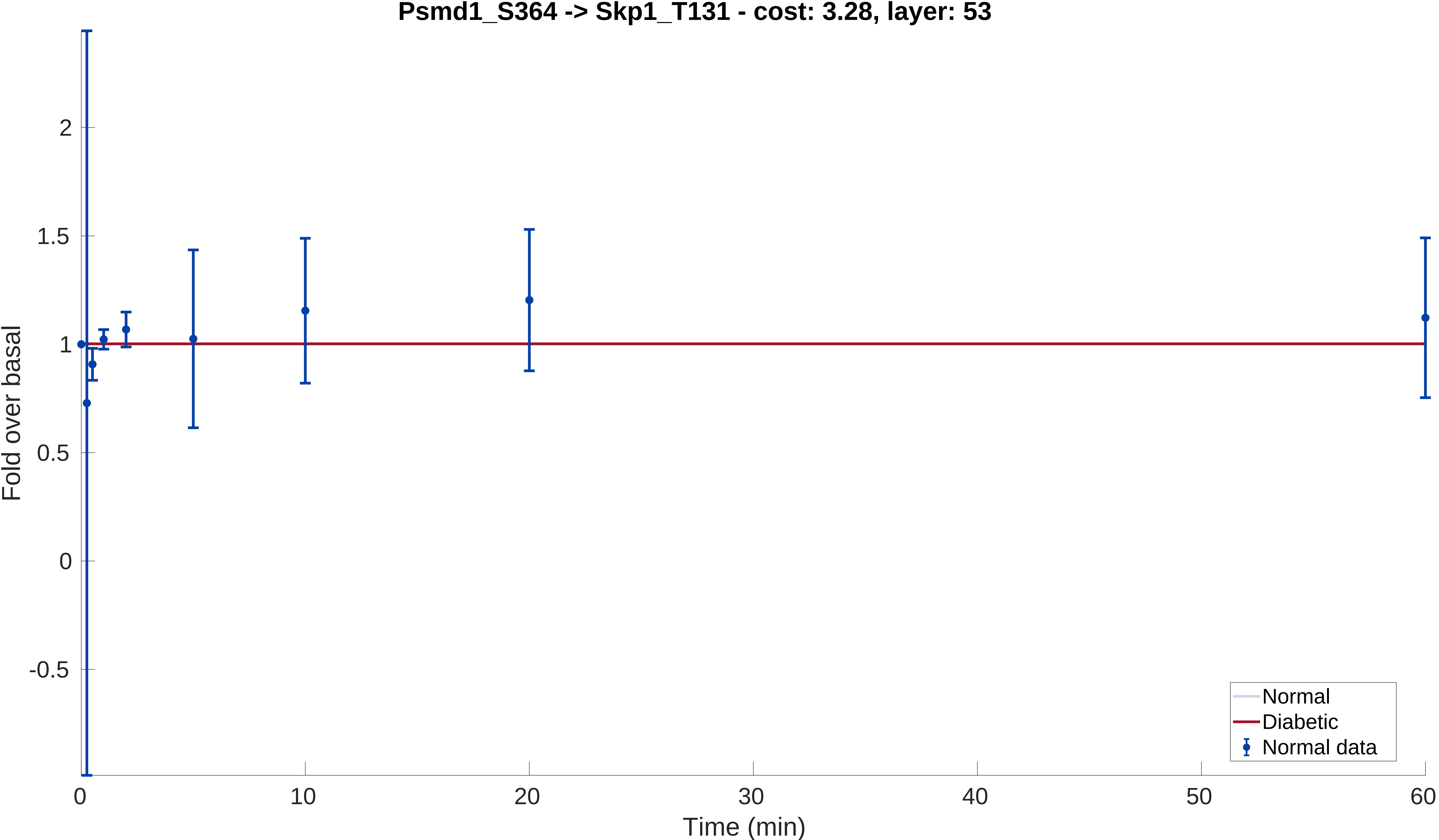
Psmd1_S364 -> Skp1_T131 - cost: 3.28, layer: 53
Fold over basal
2
1.5
1
0.5
0
-0.5
0
10
20
30
40
50
60
Time (min)
Normal
Diabetic
Normal data

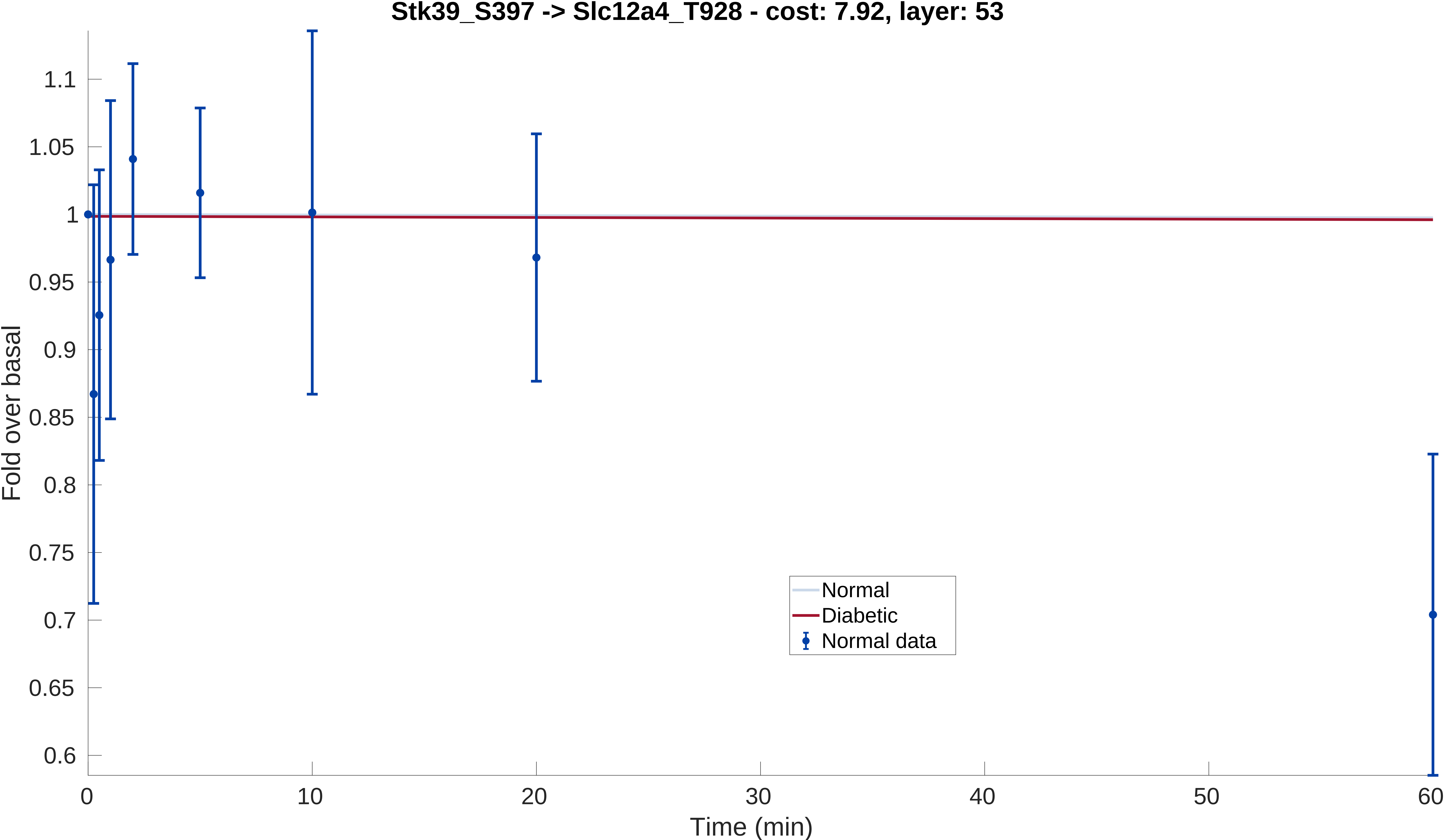

Stk39_S397 -> Slc12a4_T928 - cost: 7.92, layer: 53
Fold over basal
1.1
1.05
1
0.95
0.9
0.85
0.8
0.75
0.7
0.65
0.6
0
10
20
30
40
50
60
Time (min)
Normal
Diabetic
Normal data

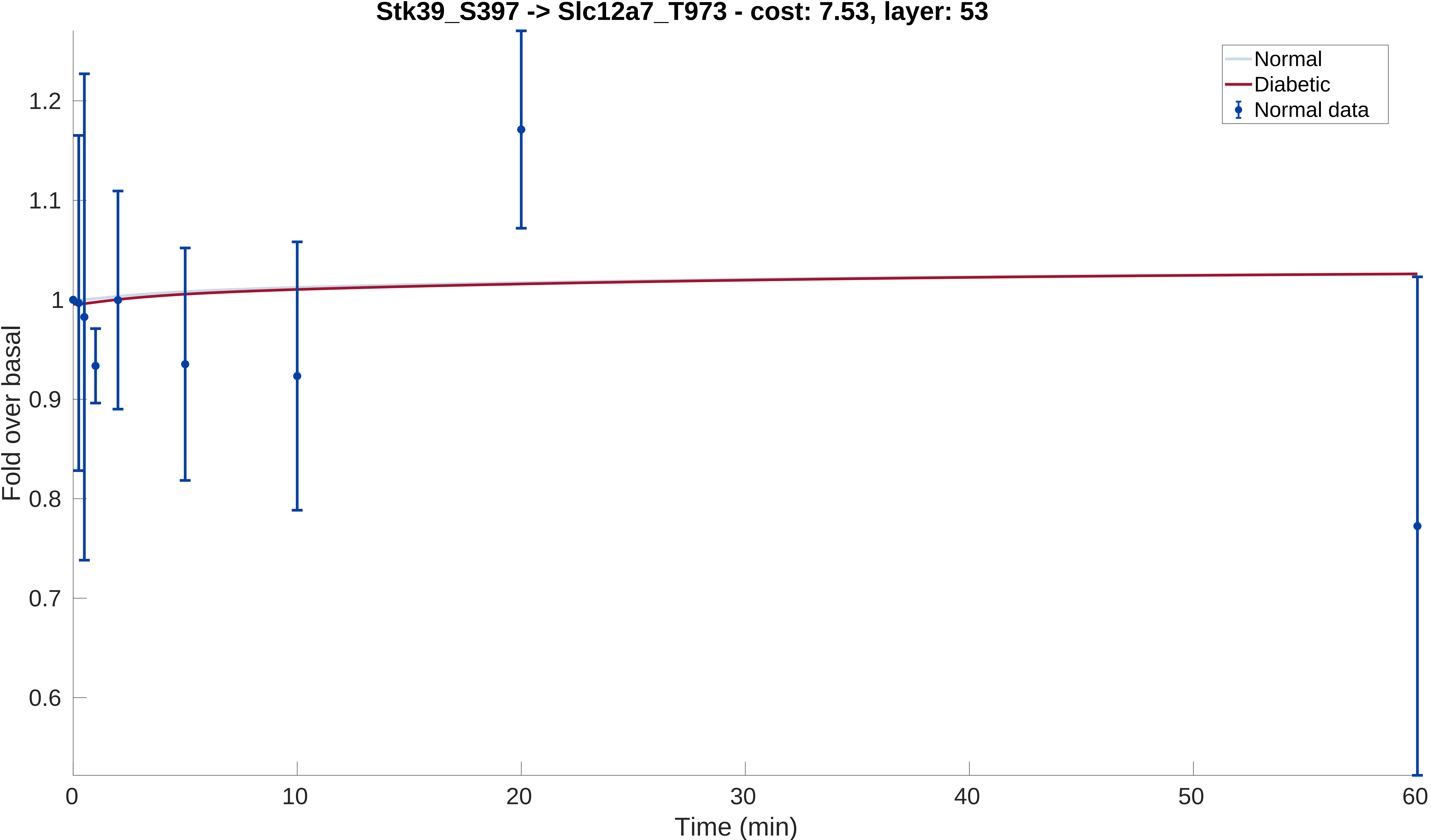
Stk39_S397 -> Slc12a7_T973 - cost: 7.53, layer: 53
Normal
Diabetic
Normal data
Fold over basal
1.2
1.1
1
0.9
0.8
0.7
0.6
0
10
20
30
40
50
60
Time (min)

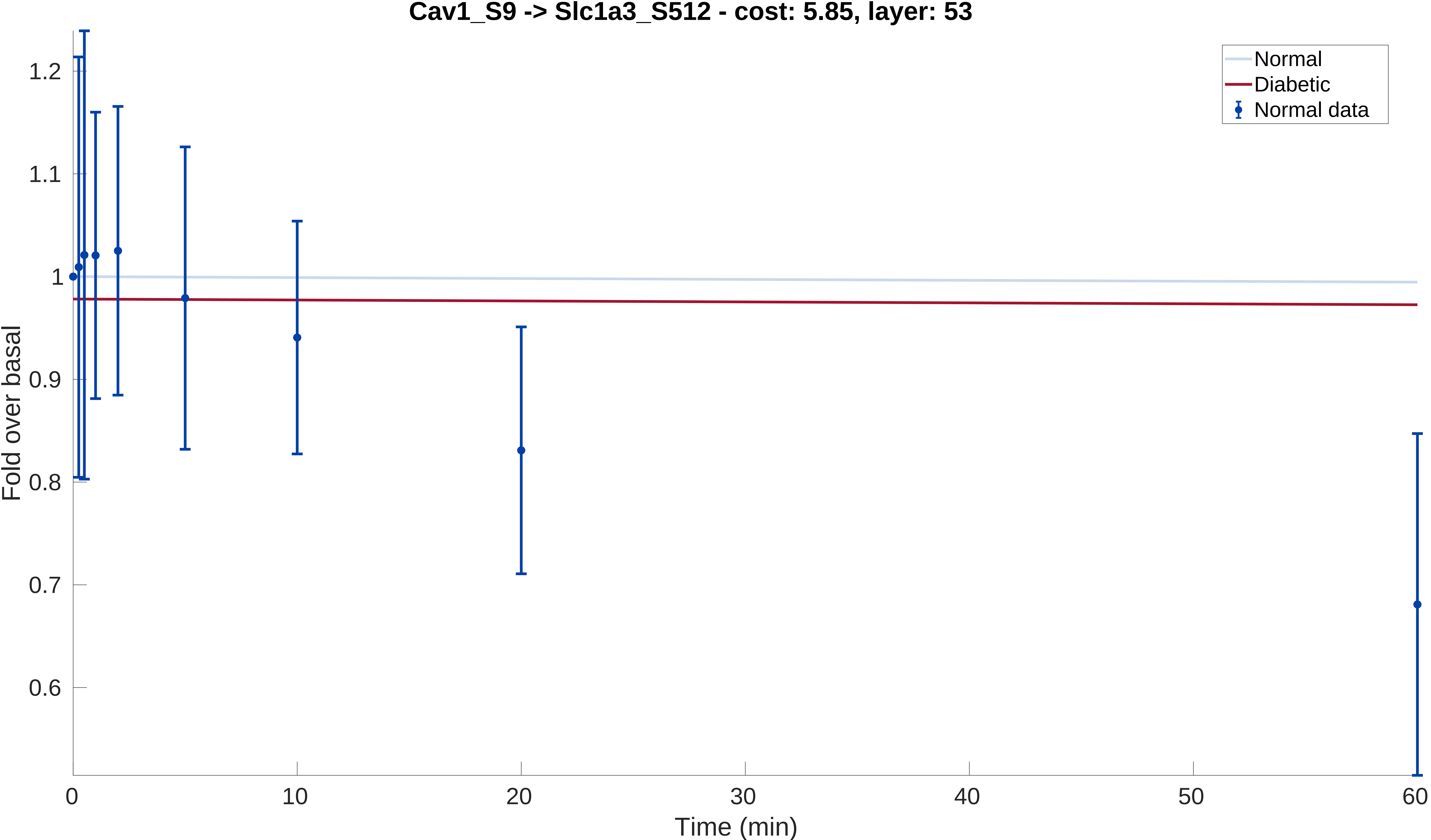

Cav1_S9 -> Slc1a3_S512 - cost: 5.85, layer: 53
Normal
Diabetic
Normal data
1.2
1.1
1
0.9
0.8
0.7
0.6
Fold over basal
0
10
20
30
40
50
60
Time (min)

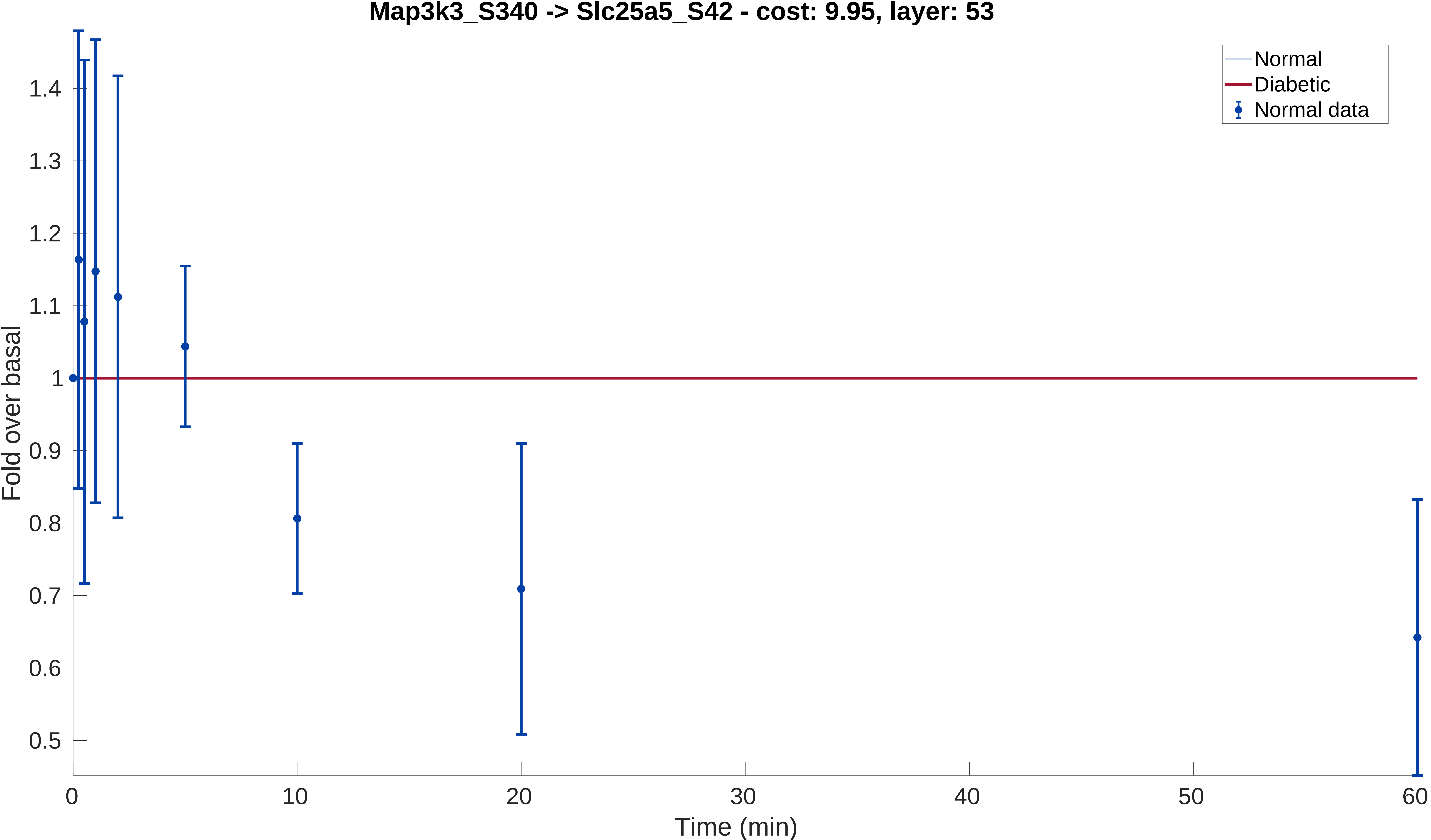

Map3k3_S340 -> Slc25a5_S42 - cost: 9.95, layer: 53
Normal
Diabetic
Normal data
Fold over basal
1.4
1.3
1.2
1.1
1
0.9
0.8
0.7
0.6
0.5
0
10
20
30
40
50
60
Time (min)

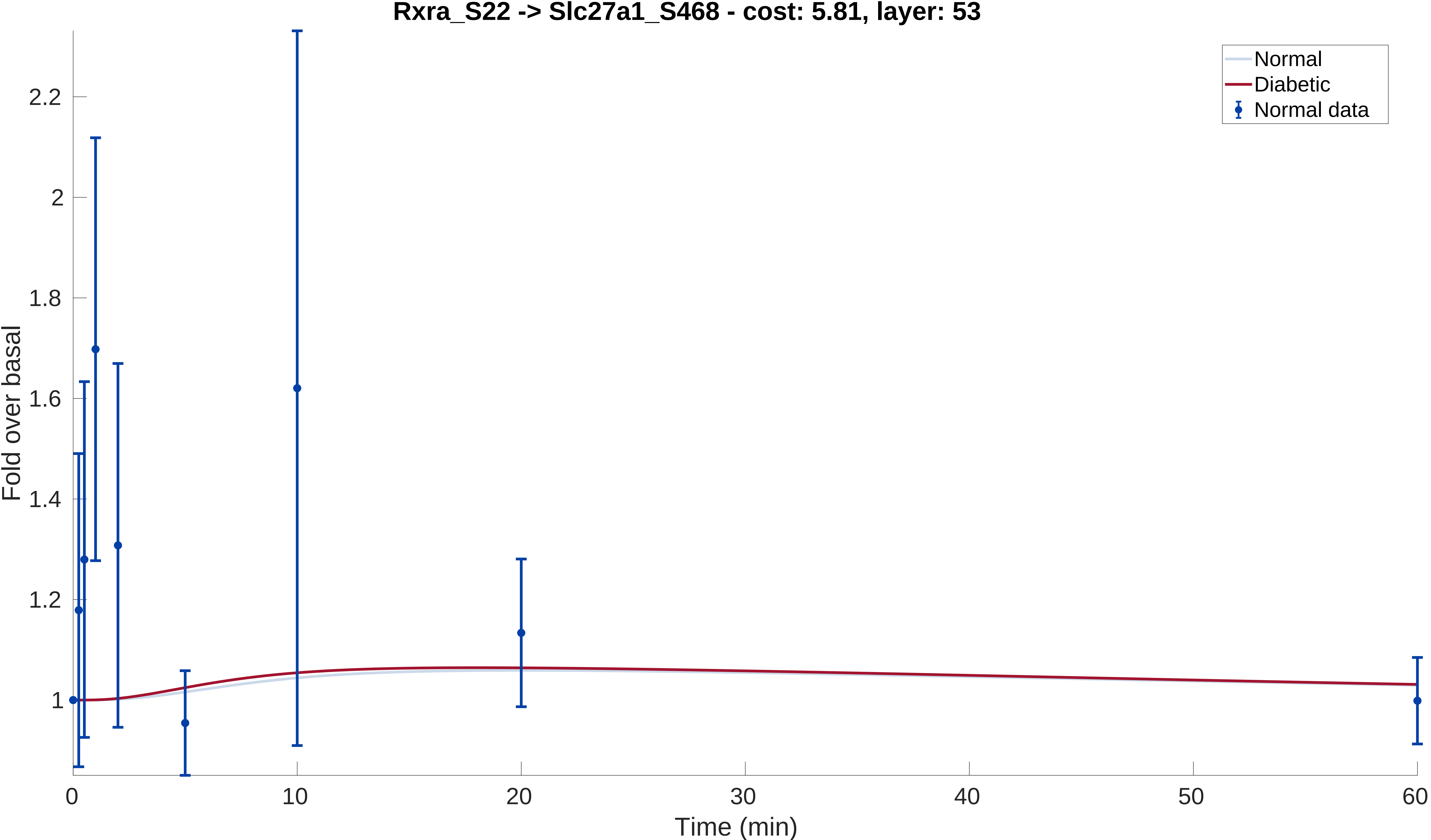

Rxra_S22 -> Slc27a1_S468 - cost: 5.81, layer: 53
Normal
Diabetic
Normal data
Fold over basal
2.2
2
1.8
1.6
1.4
1.2
1
0
10
20
30
40
50
60
Time (min)

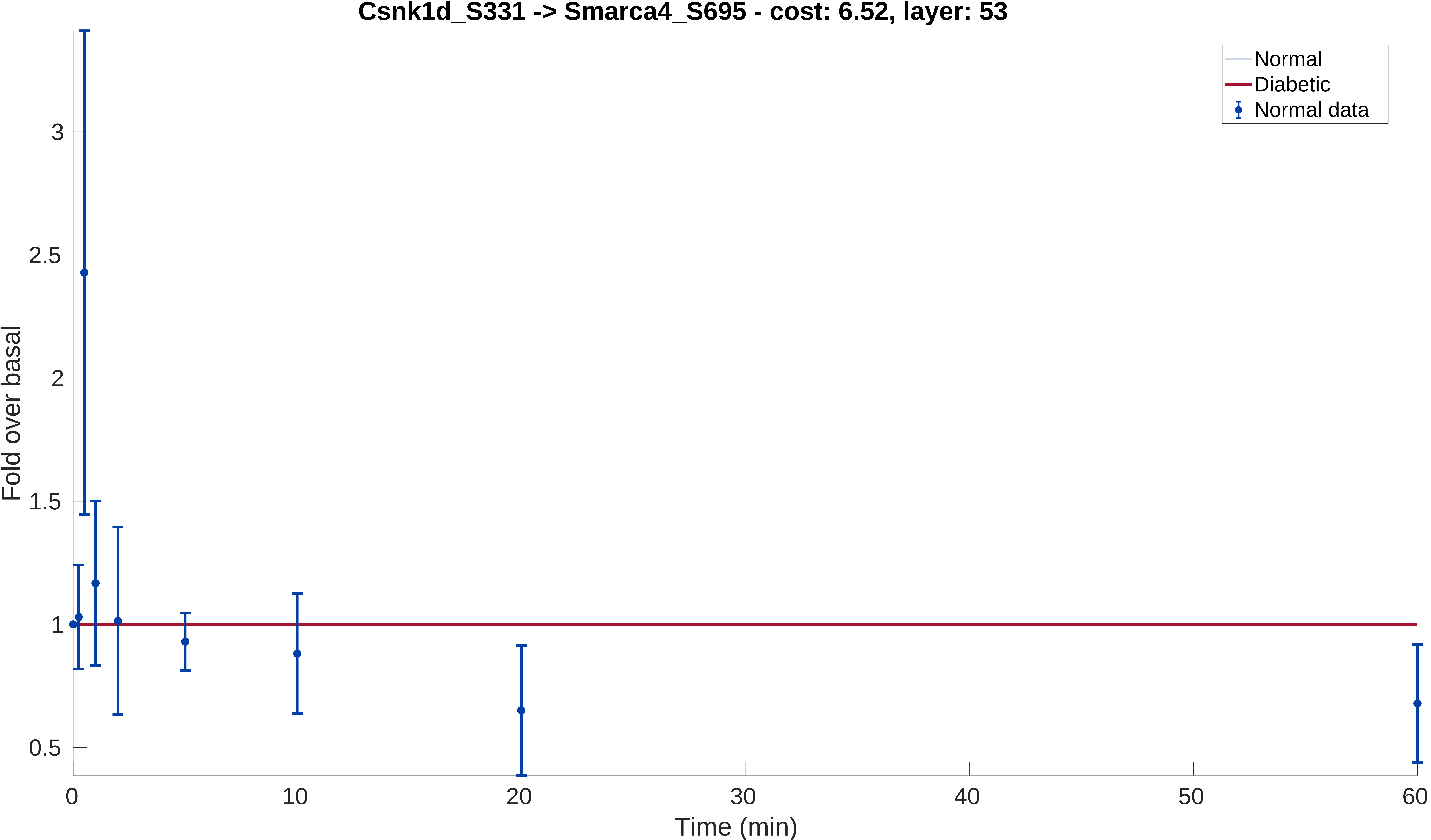
Csnk1d_S331 -> Smarca4_S695 - cost: 6.52, layer: 53
Normal
Diabetic
Normal data
Fold over basal
0.5
1
1.5
2
2.5
3
0
10
20
30
40
50
60
Time (min)

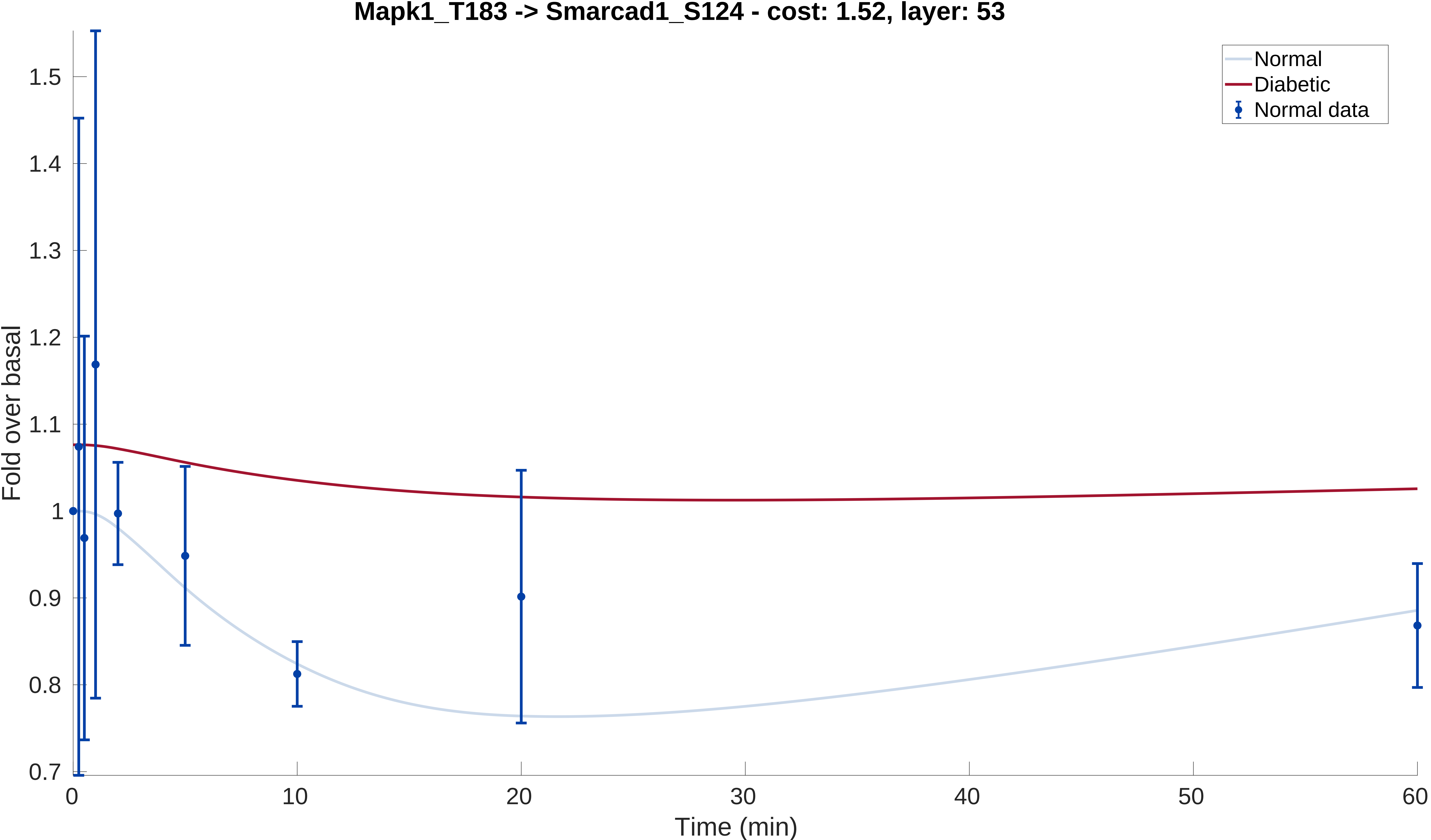
Mapk1_T183 -> Smarcad1_S124 - cost: 1.52, layer: 53
Normal
Diabetic
Normal data
Fold over basal
1.5
1.4
1.3
1.2
1.1
1
0.9
0.8
0.7
0
10
20
30
40
50
60
Time (min)

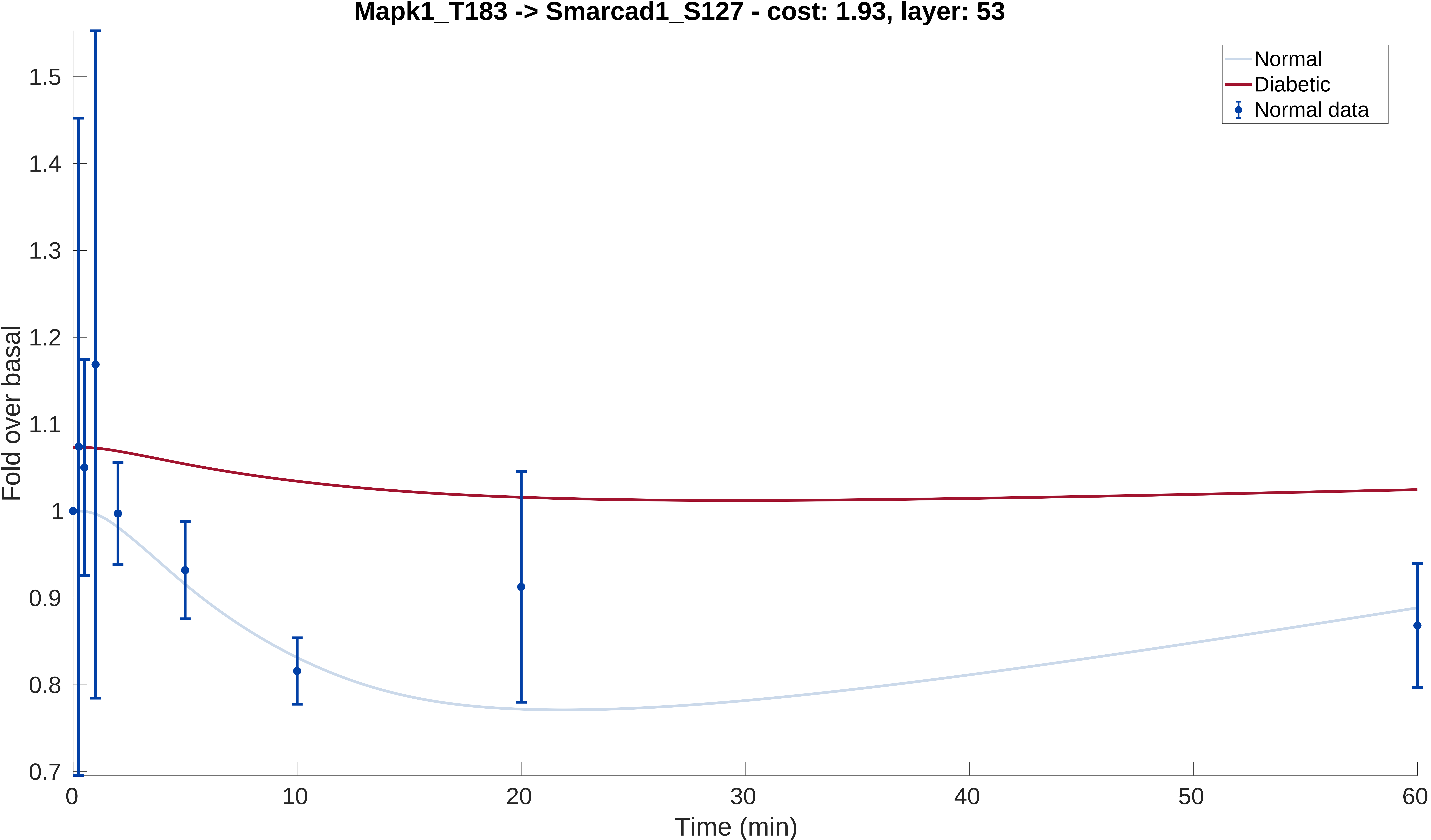

Mapk1_T183 -> Smarcad1_S127 - cost: 1.93, layer: 53
Normal
Diabetic
Normal data
1.5
1.4
1.3
1.2
1.1
1
0.9
0.8
0.7
Fold over basal
0
10
20
30
40
50
60
Time (min)

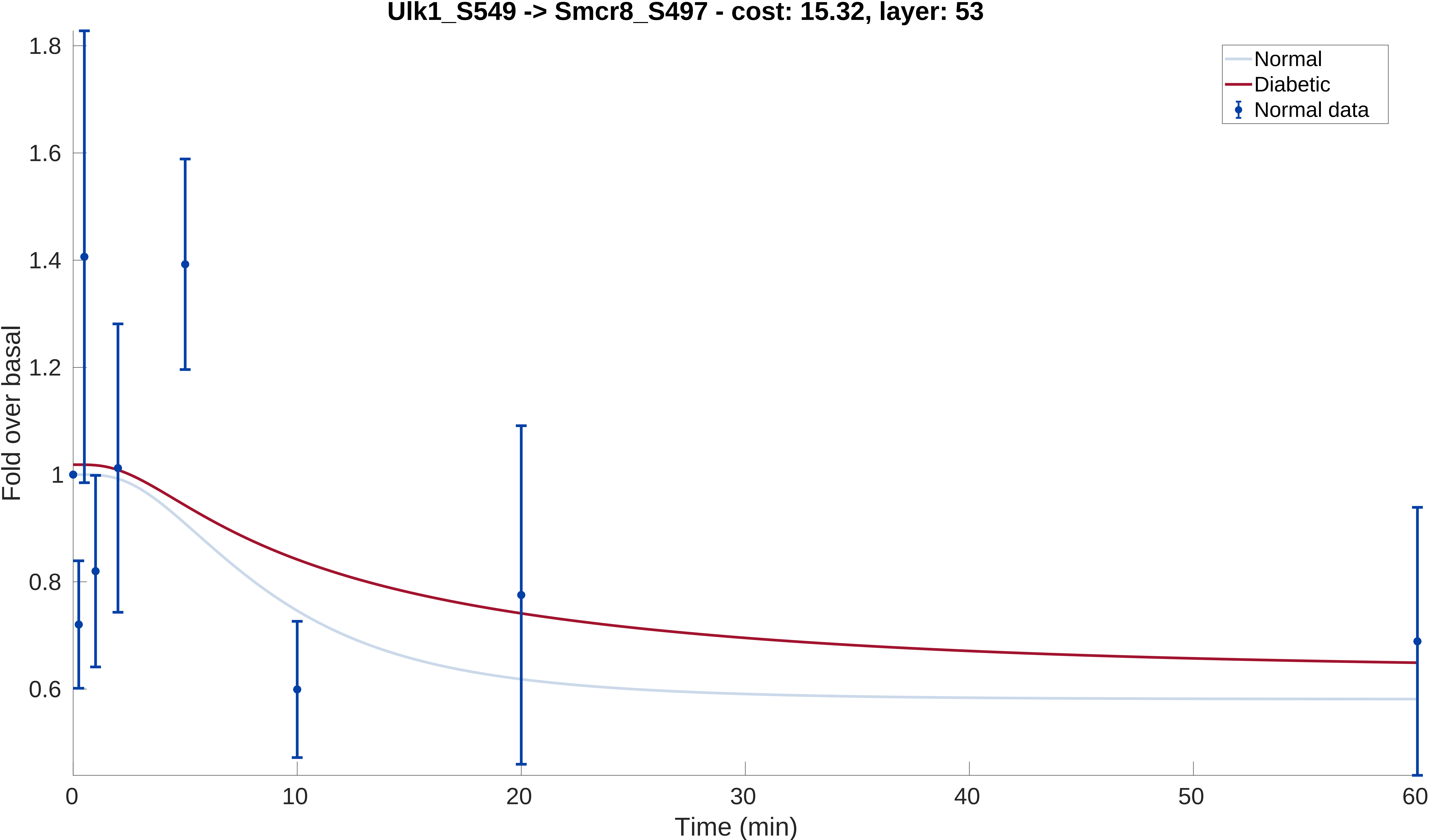
Ulk1_S549 -> Smcr8_S497 - cost: 15.32, layer: 53
Fold over basal
Time (min)
Normal
Diabetic
Normal data
1.8
1.6
1.4
1.2
1
0.8
0.6
0
10
20
30
40
50
60

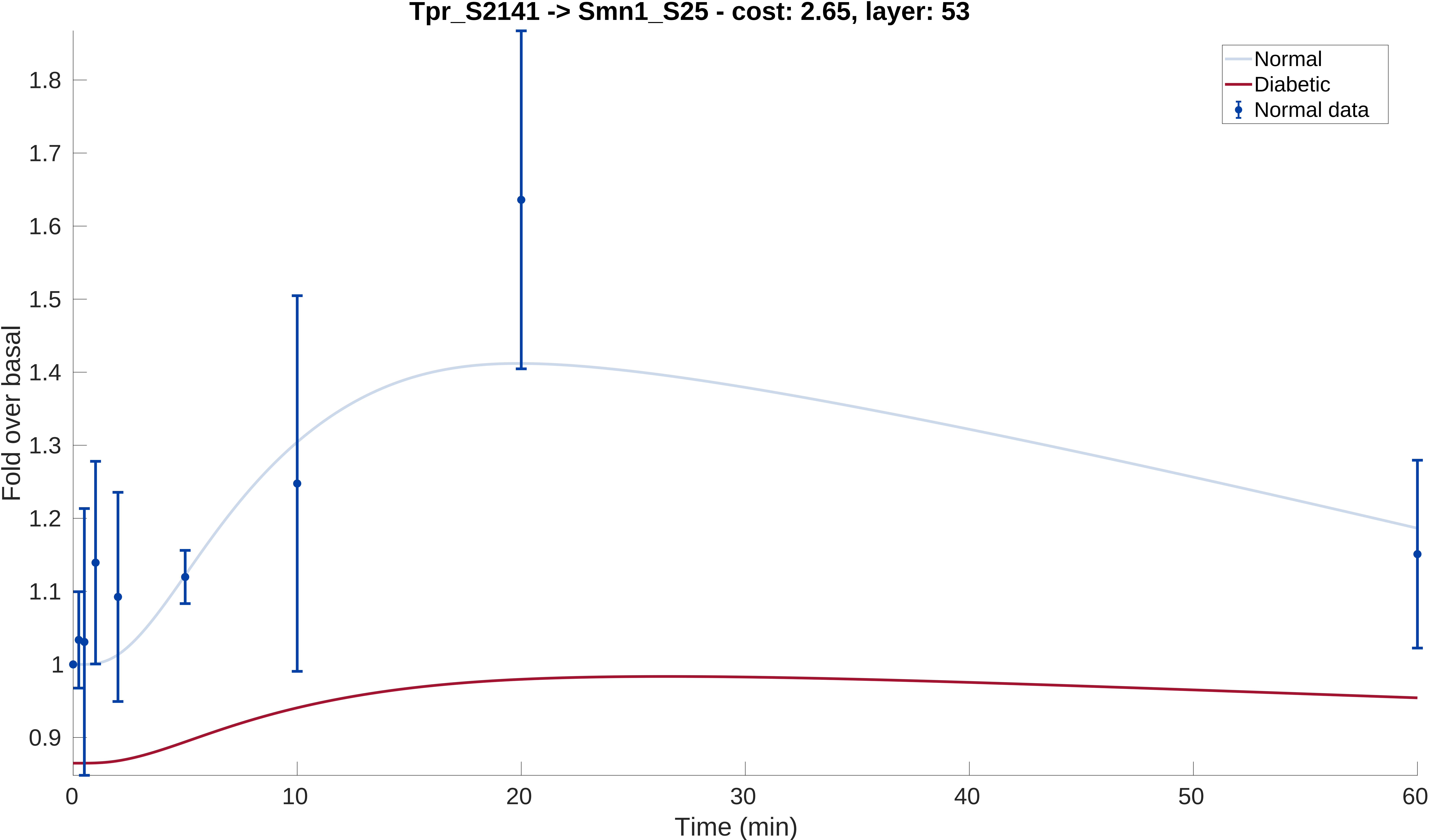

Tpr_S2141 -> Smn1_S25 - cost: 2.65, layer: 53
Normal
Diabetic
Normal data
Fold over basal
1.8
1.7
1.6
1.5
1.4
1.3
1.2
1.1
1
0.9
0
10
20
30
40
50
60
Time (min)

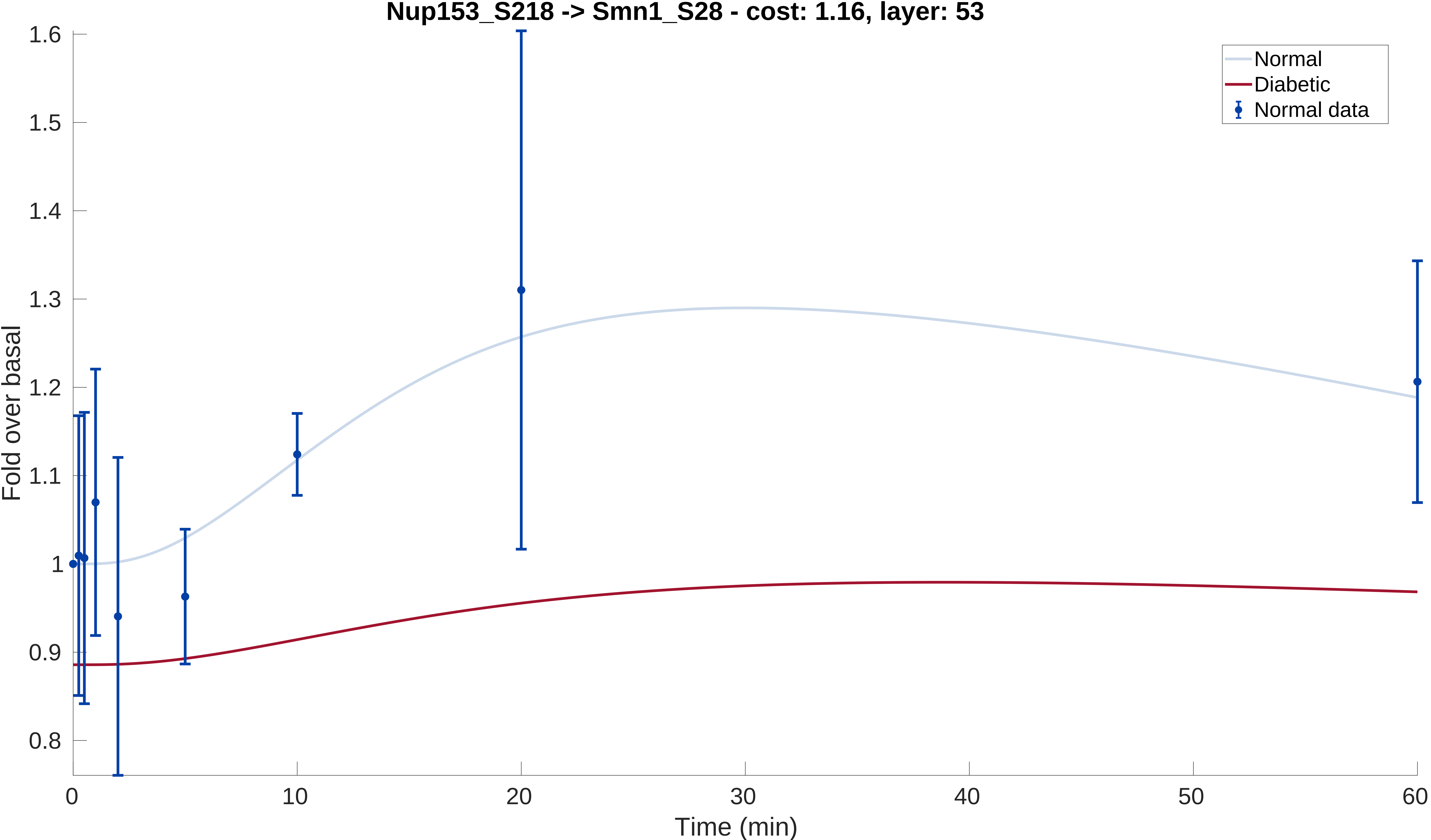

Nup153_S218 -> Smn1_S28 - cost: 1.16, layer: 53
Normal
Diabetic
Normal data
Fold over basal
Time (min)
1.6
1.5
1.4
1.3
1.2
1.1
1
0.9
0.8
0
10
20
30
40
50
60

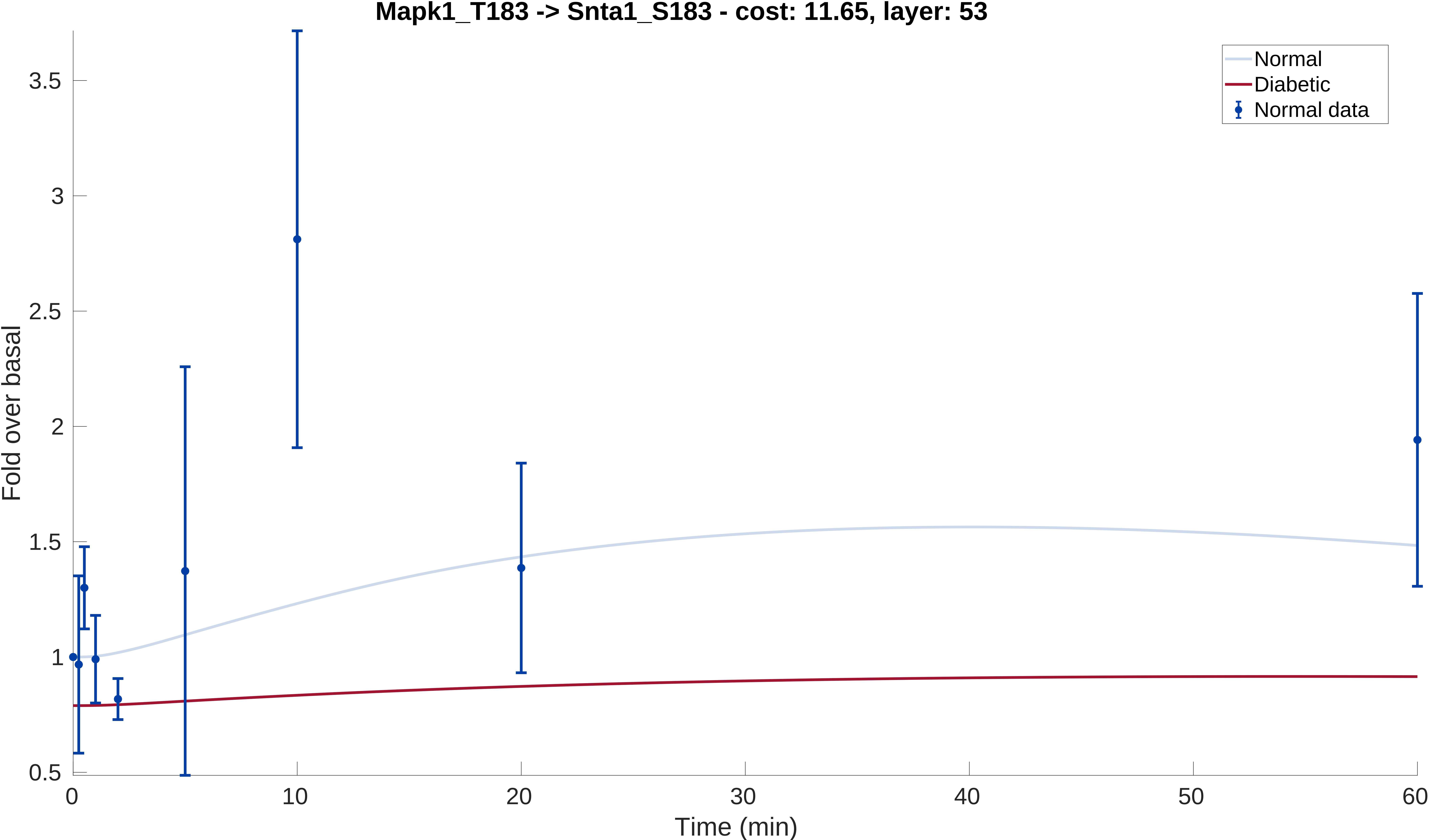

Mapk1_T183 -> Snta1_S183 - cost: 11.65, layer: 53
Normal
Diabetic
Normal data
3.5
3
2.5
2
1.5
1
0.5
0
10
20
30
40
50
60
Fold over basal
Time (min)

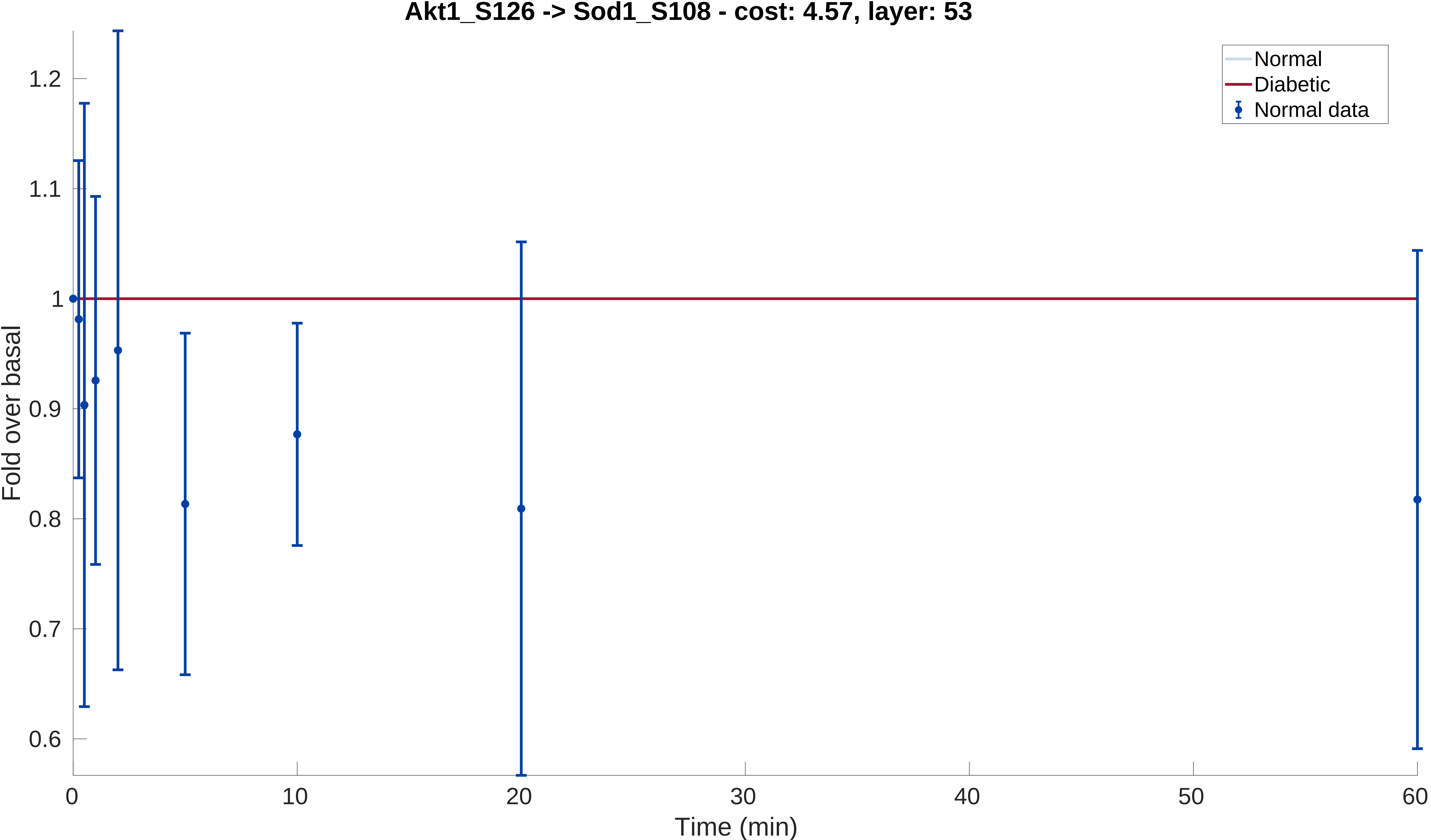
Akt1_S126 -> Sod1_S108 - cost: 4.57, layer: 53
Normal
Diabetic
Normal data
Fold over basal
1.2
1.1
1
0.9
0.8
0.7
0.6
0
10
20
30
40
50
60
Time (min)

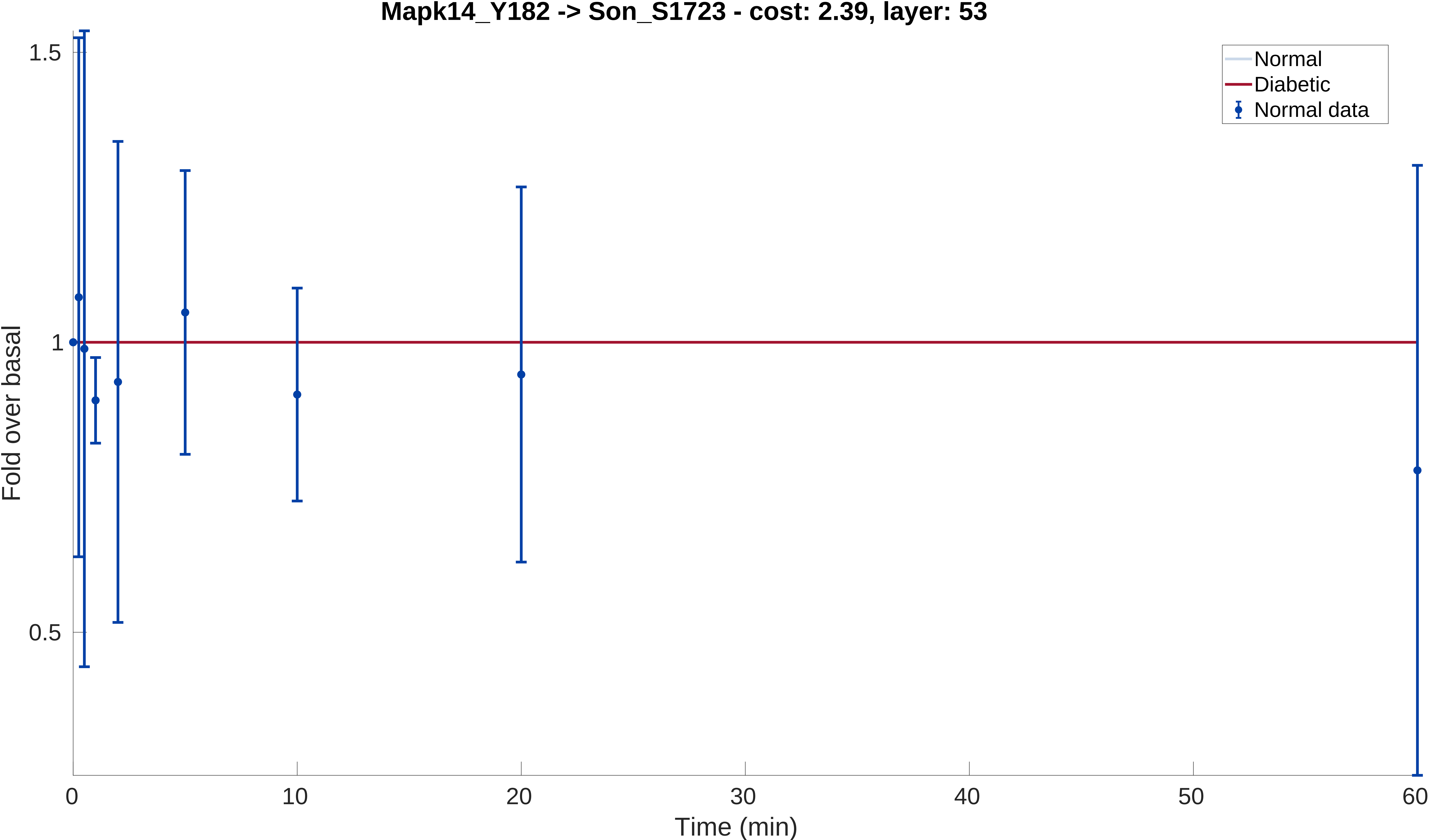

Mapk14_Y182 -> Son_S1723 - cost: 2.39, layer: 53
Normal
Diabetic
Normal data
1.5
1
0.5
Fold over basal
0
10
20
30
40
50
60
Time (min)

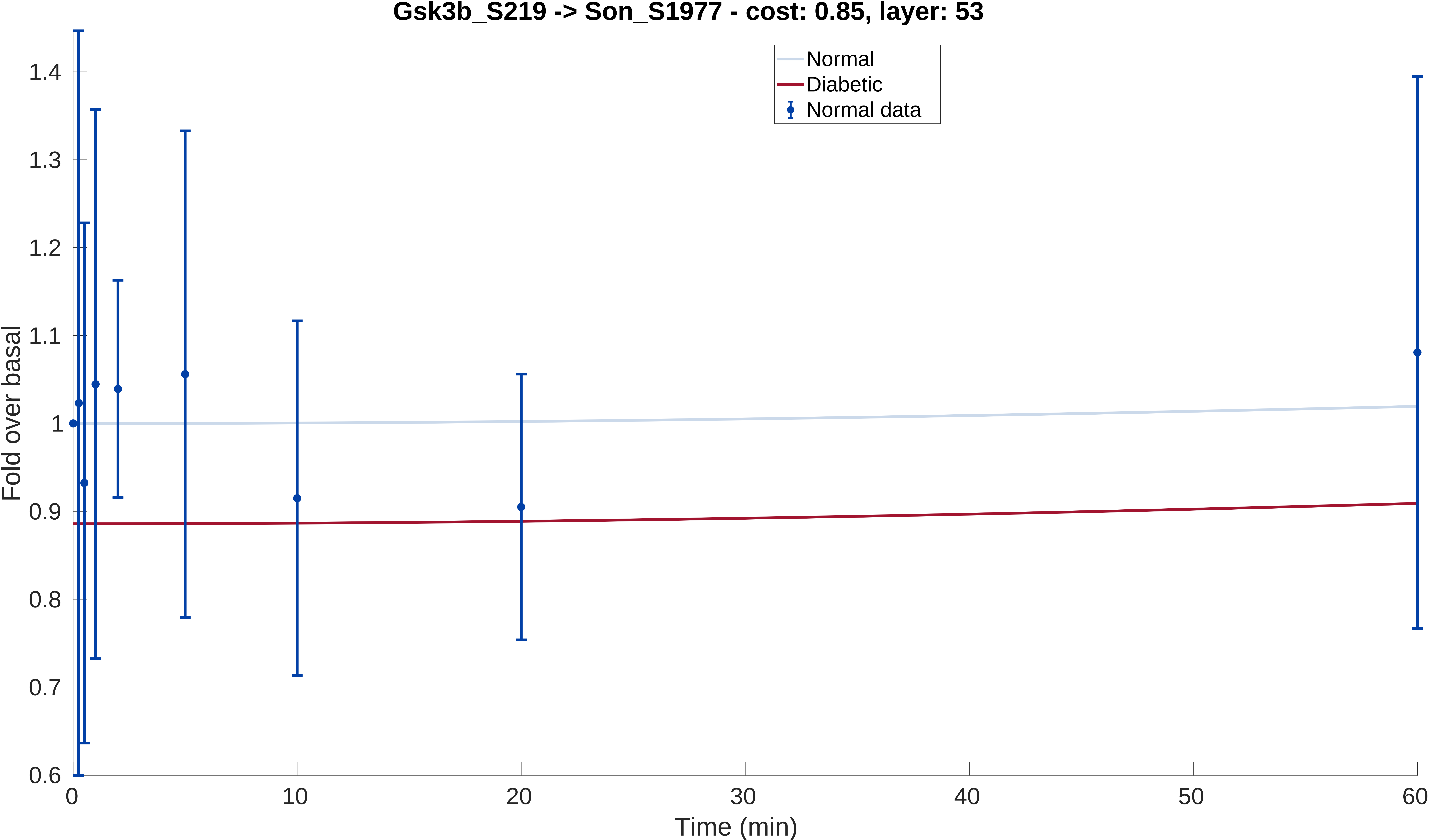

Gsk3b_S219 -> Son_S1977 - cost: 0.85, layer: 53
Normal
Diabetic
Normal data
Fold over basal
1.4
1.3
1.2
1.1
1
0.9
0.8
0.7
0.6
0
10
20
30
40
50
60
Time (min)

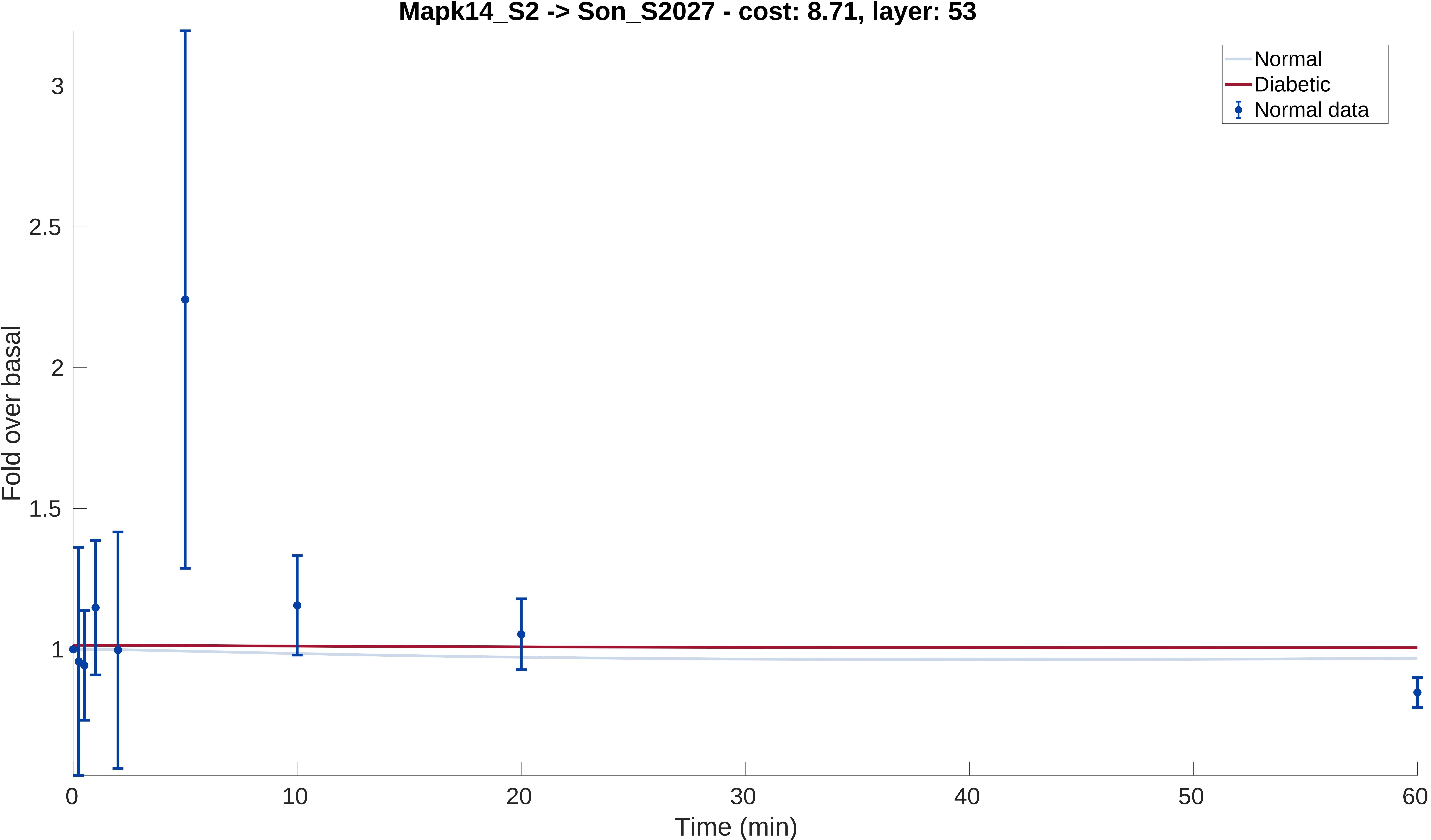

Mapk14_S2 -> Son_S2027 - cost: 8.71, layer: 53
Normal
Diabetic
Normal data
Fold over basal
3
2.5
2
1.5
1
0
10
20
30
40
50
60
Time (min)

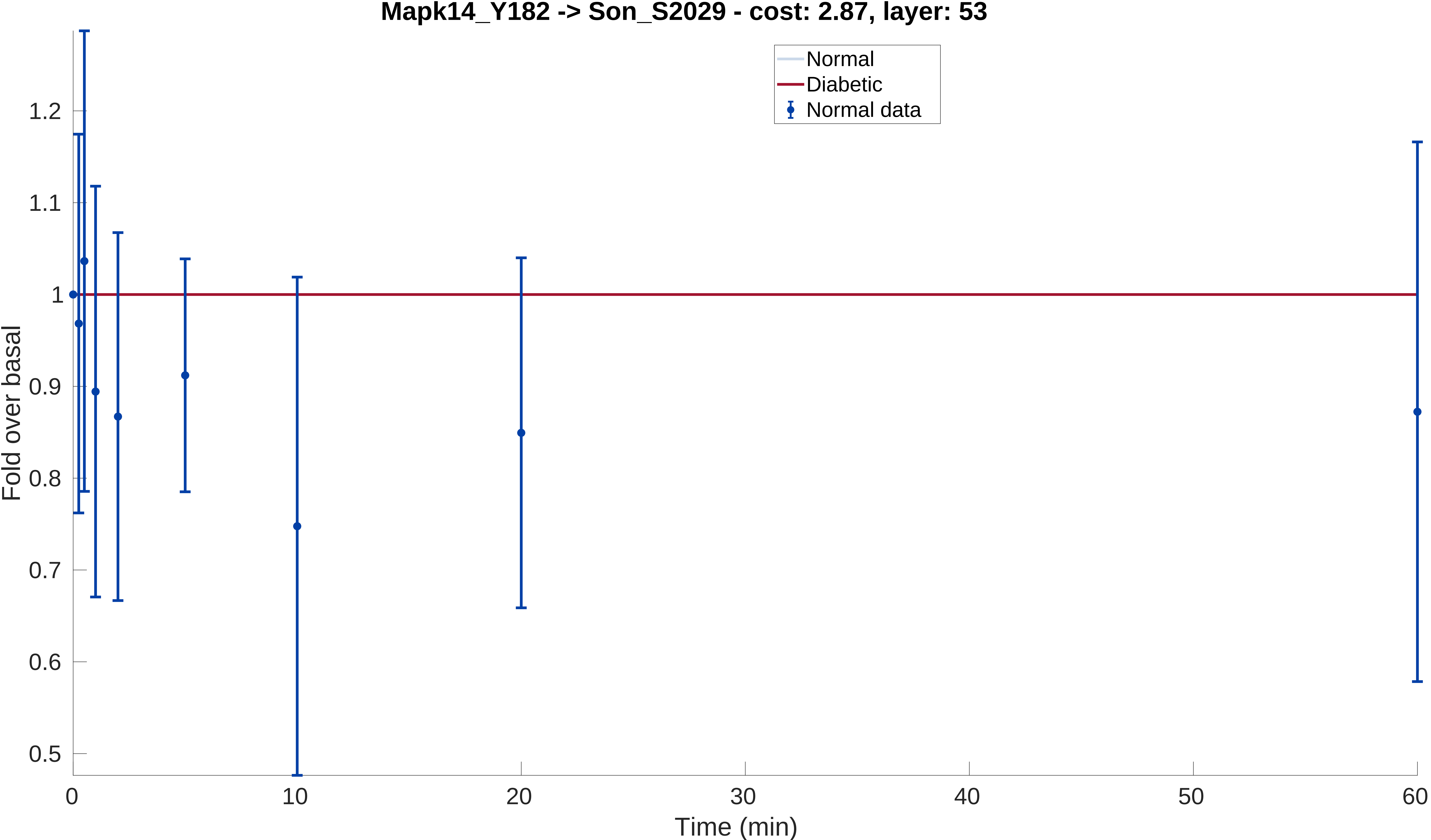
Mapk14_Y182 -> Son_S2029 - cost: 2.87, layer: 53
Normal
Diabetic
Normal data
Fold over basal
1.2
1.1
1
0.9
0.8
0.7
0.6
0.5
0
10
20
30
40
50
60
Time (min)

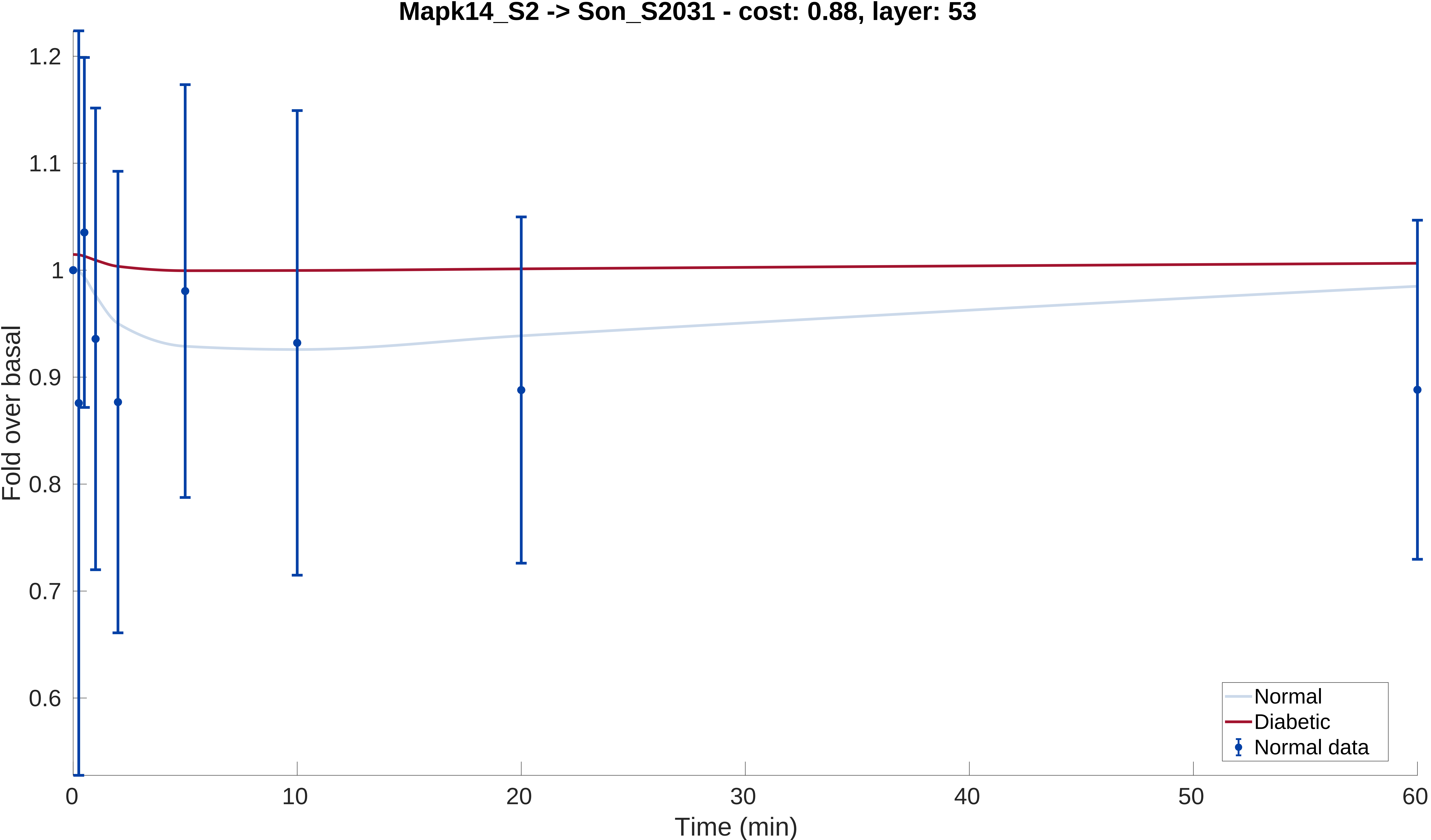

Mapk14_S2 -> Son_S2031 - cost: 0.88, layer: 53
Fold over basal
1.2
1.1
1
0.9
0.8
0.7
0.6
0
10
20
30
40
50
60
Time (min)
Normal
Diabetic
Normal data

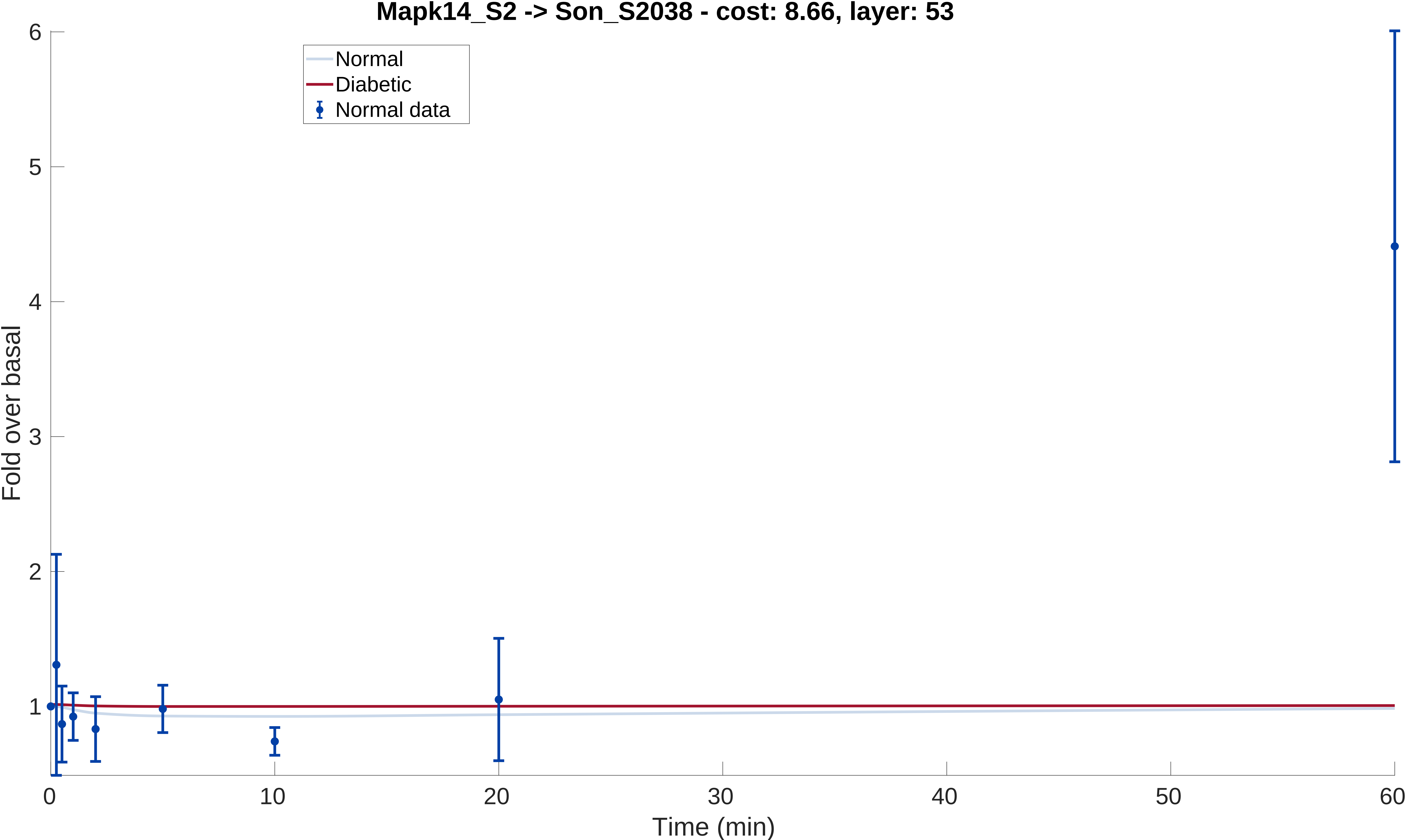

Mapk14_S2 -> Son_S2038 - cost: 8.66, layer: 53
Normal
Diabetic
Normal data
Fold over basal
Time (min)
0
10
20
30
40
50
60
1
2
3
4
5
6

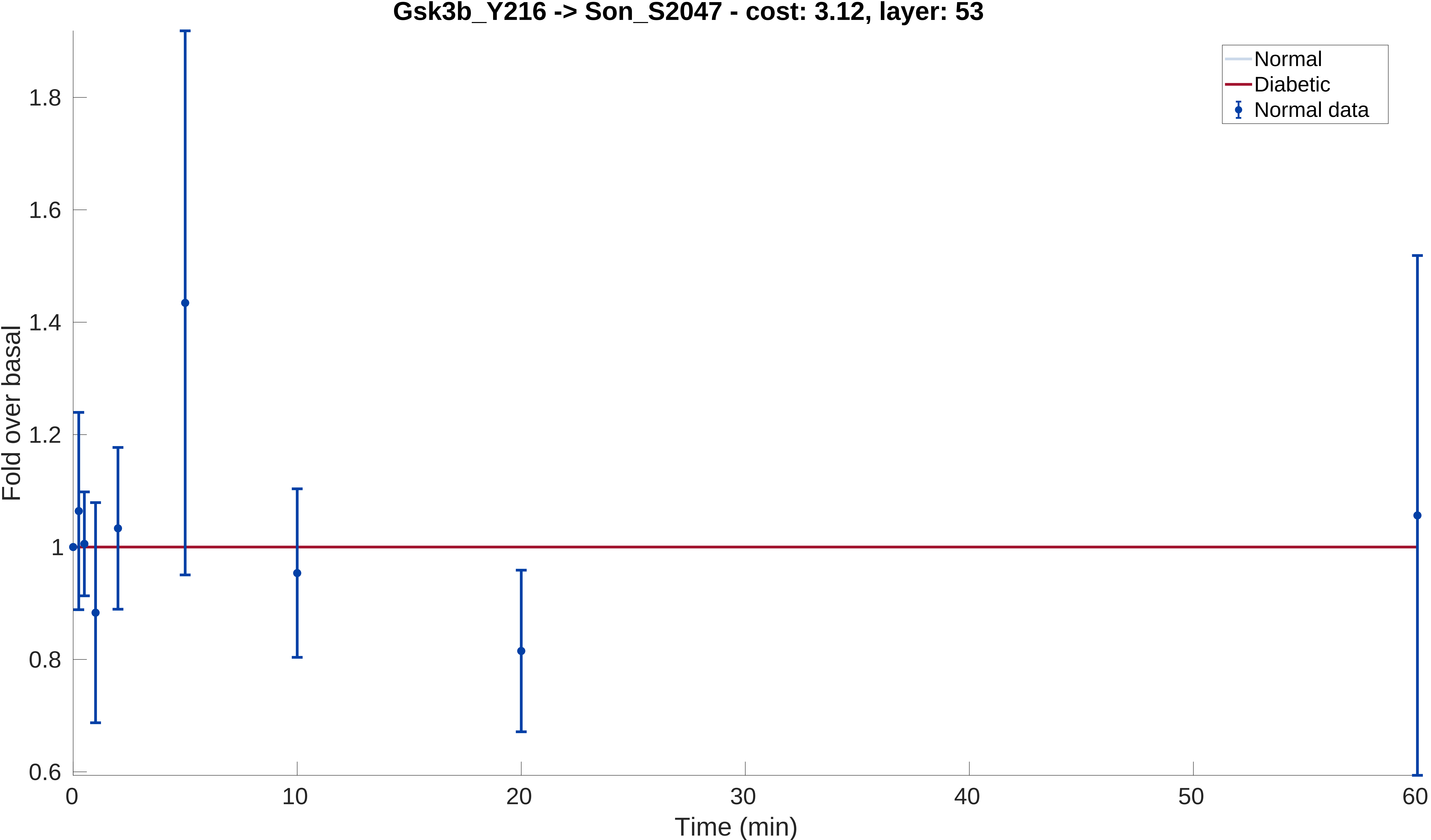

Gsk3b_Y216 -> Son_S2047 - cost: 3.12, layer: 53
Normal
Diabetic
Normal data
Fold over basal
1.8
1.6
1.4
1.2
1
0.8
0.6
0
10
20
30
40
50
60
Time (min)

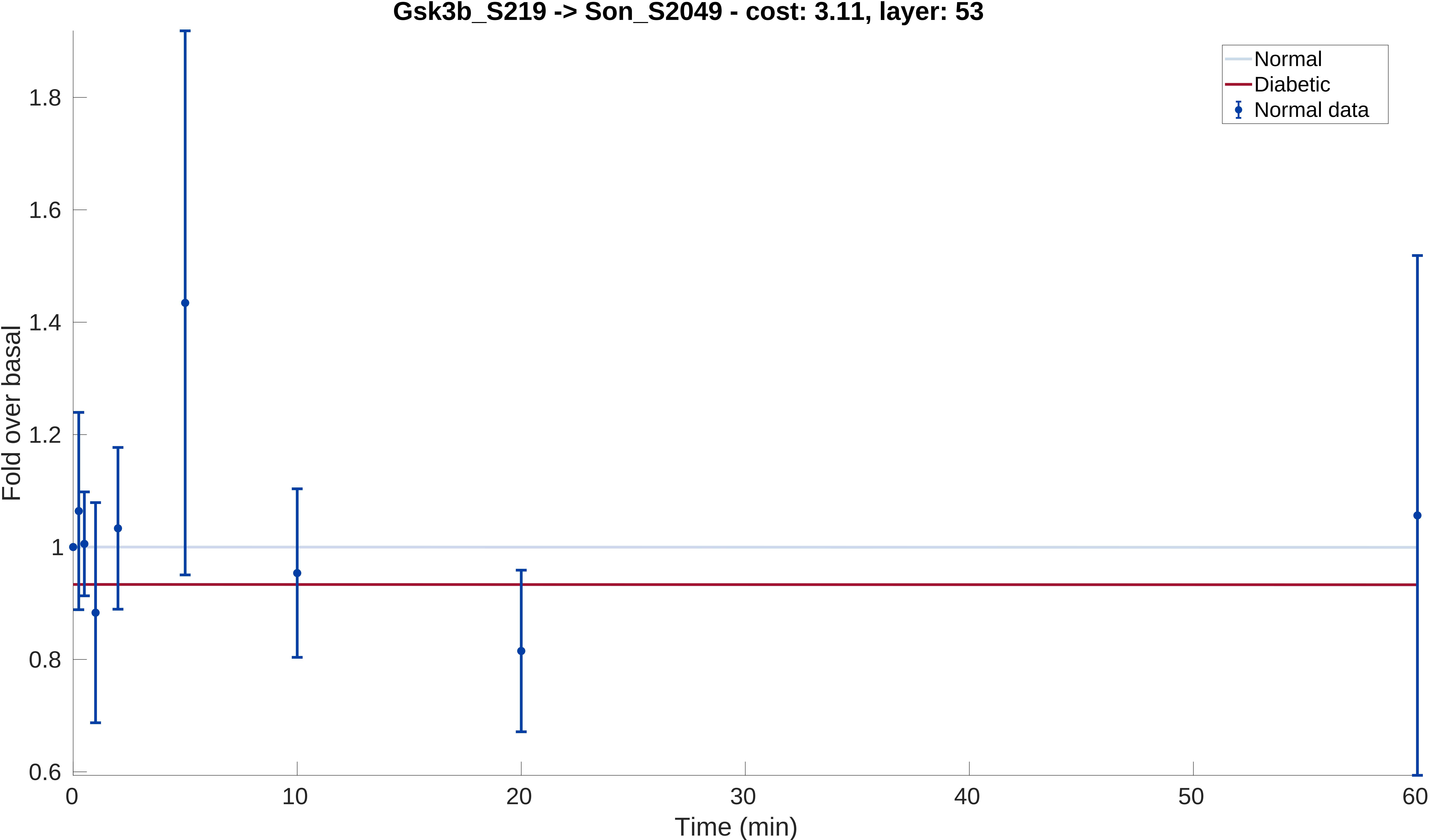

Gsk3b_S219 -> Son_S2049 - cost: 3.11, layer: 53
Normal
Diabetic
Normal data
Fold over basal
1.8
1.6
1.4
1.2
1
0.8
0.6
0
10
20
30
40
50
60
Time (min)

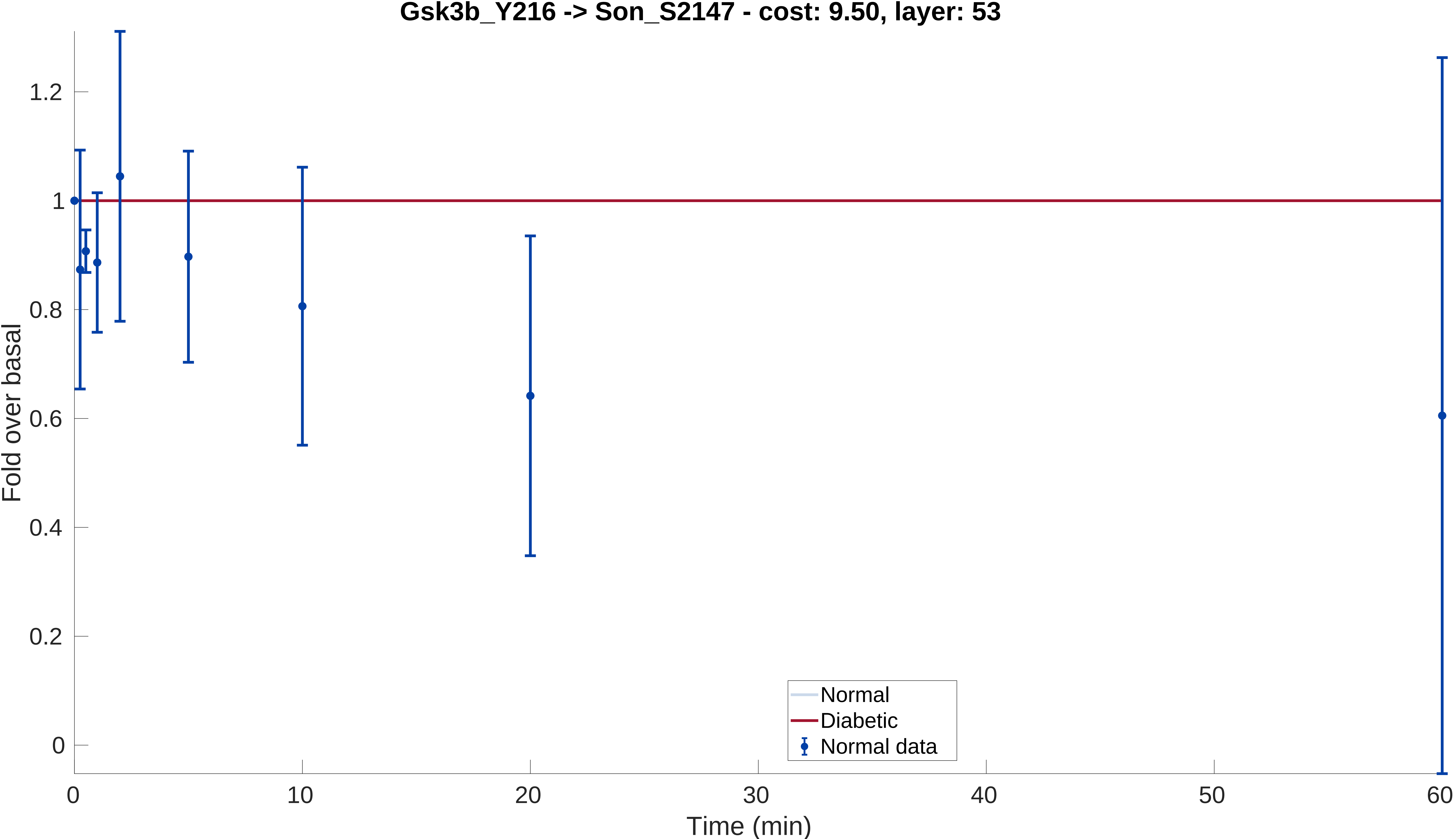
Gsk3b_Y216 -> Son_S2147 - cost: 9.50, layer: 53
Fold over basal
1.2
1
0.8
0.6
0.4
0.2
0
0
10
20
30
40
50
60
Time (min)
Normal
Diabetic
Normal data

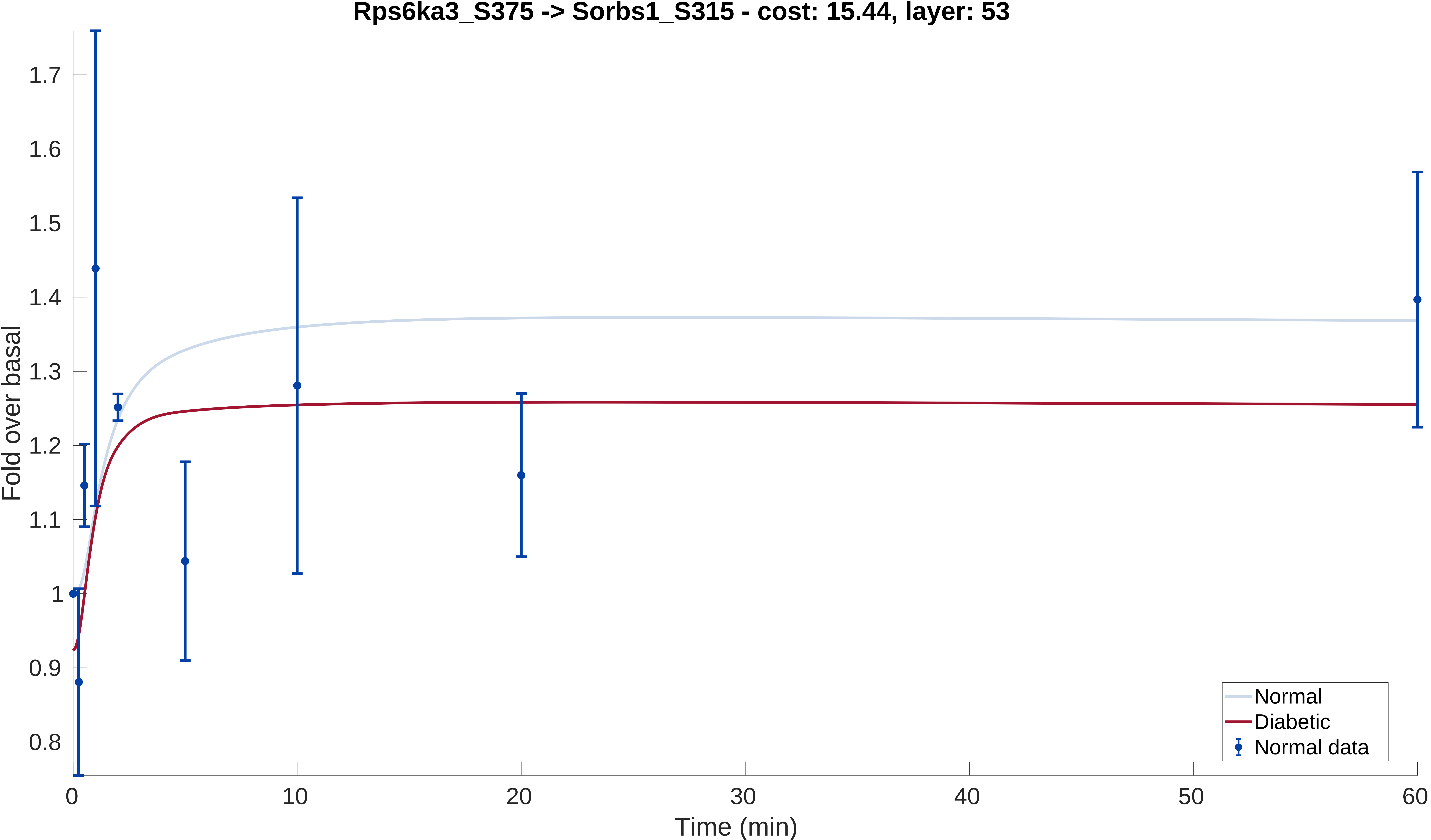

Rps6ka3_S375 -> Sorbs1_S315 - cost: 15.44, layer: 53
Fold over basal
1.7
1.6
1.5
1.4
1.3
1.2
1.1
1
0.9
0.8
0
10
20
30
40
50
60
Time (min)
Normal
Diabetic
Normal data

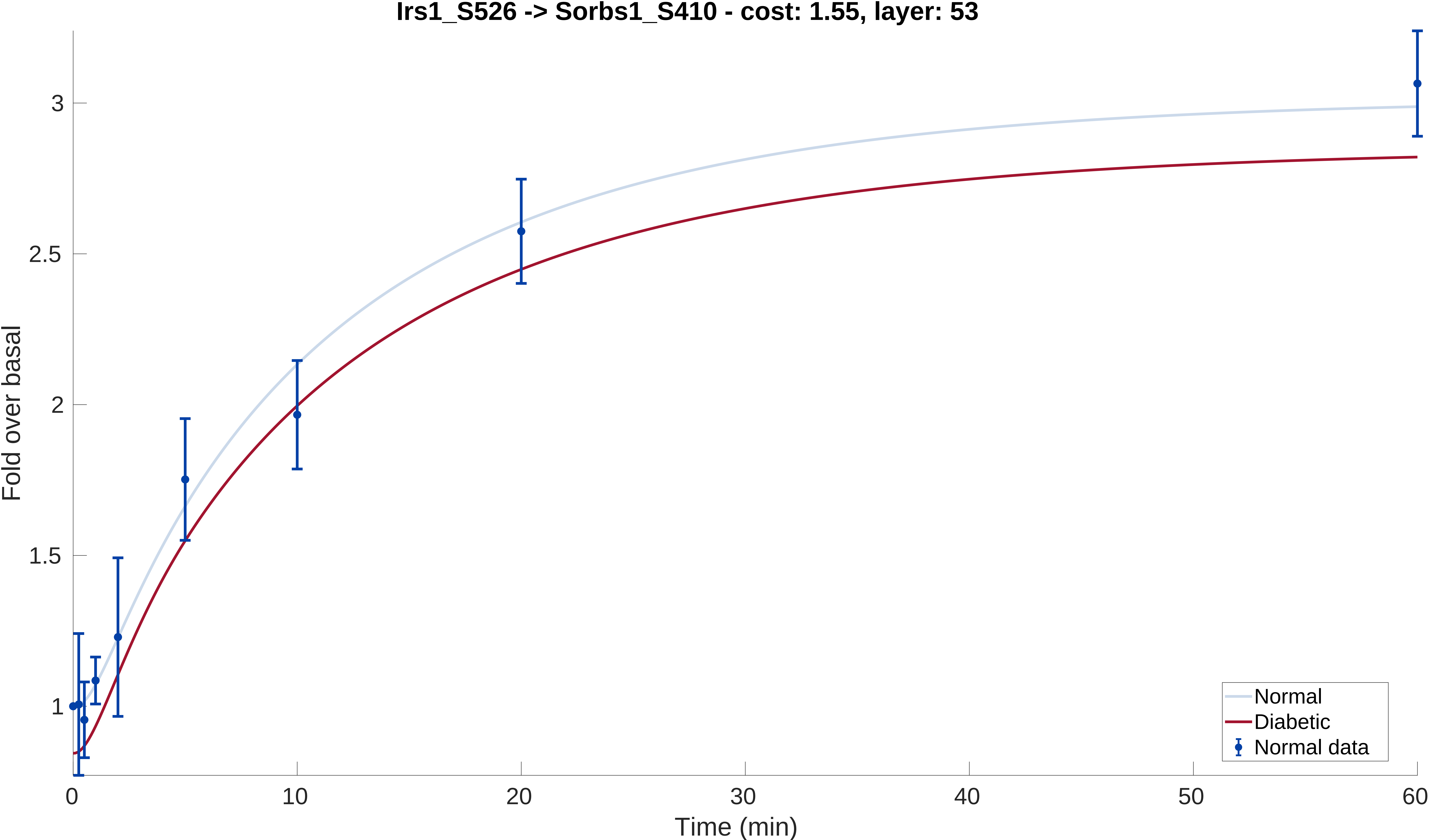

Irs1_S526 -> Sorbs1_S410 - cost: 1.55, layer: 53
Fold over basal
3
2.5
2
1.5
1
0
10
20
30
40
50
60
Time (min)
Normal
Diabetic
Normal data

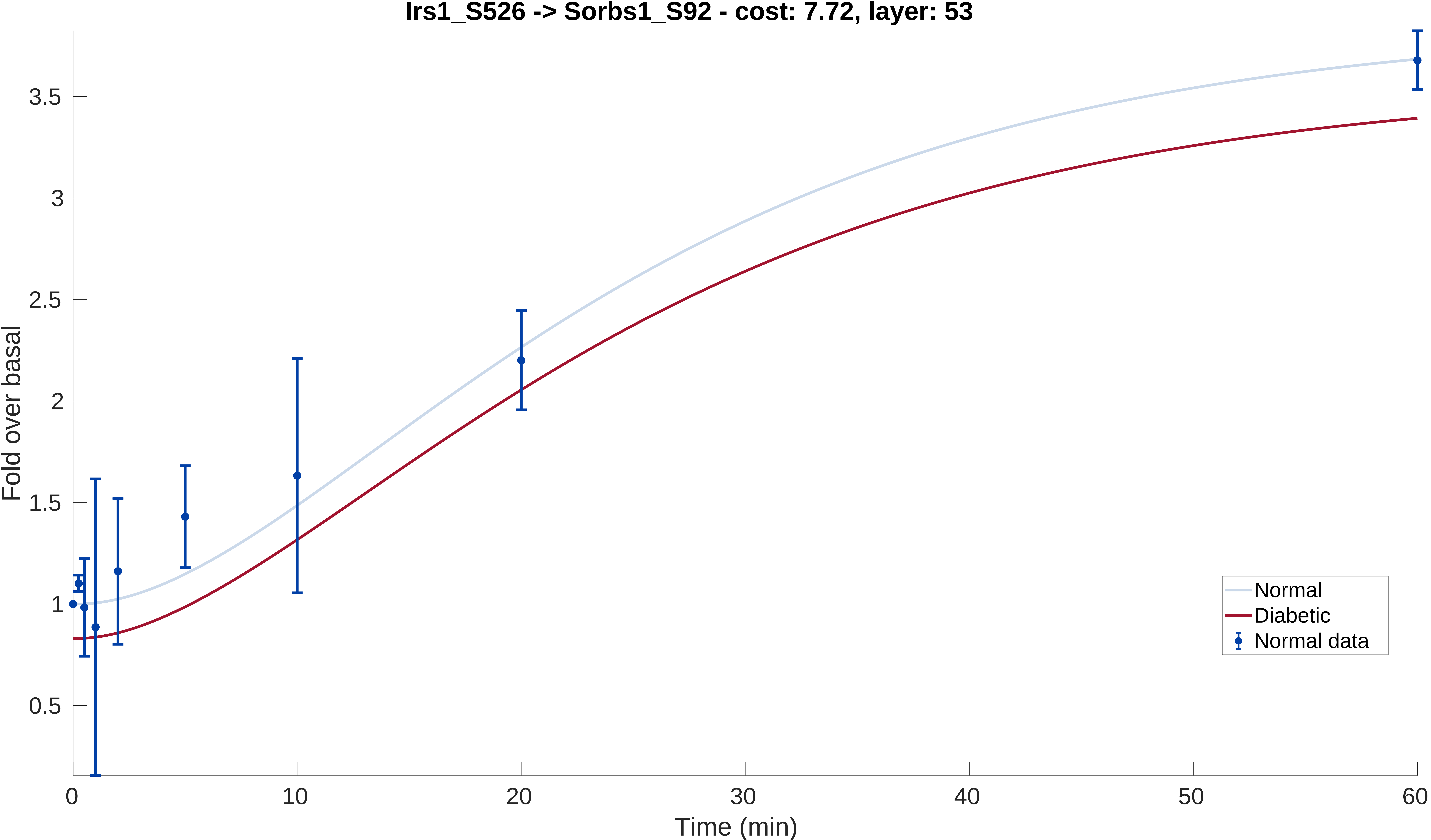
Irs1_S526 -> Sorbs1_S92 - cost: 7.72, layer: 53
Fold over basal
3.5
3
2.5
2
1.5
1
0.5
0
10
20
30
40
50
60
Time (min)
Normal
Diabetic
Normal data

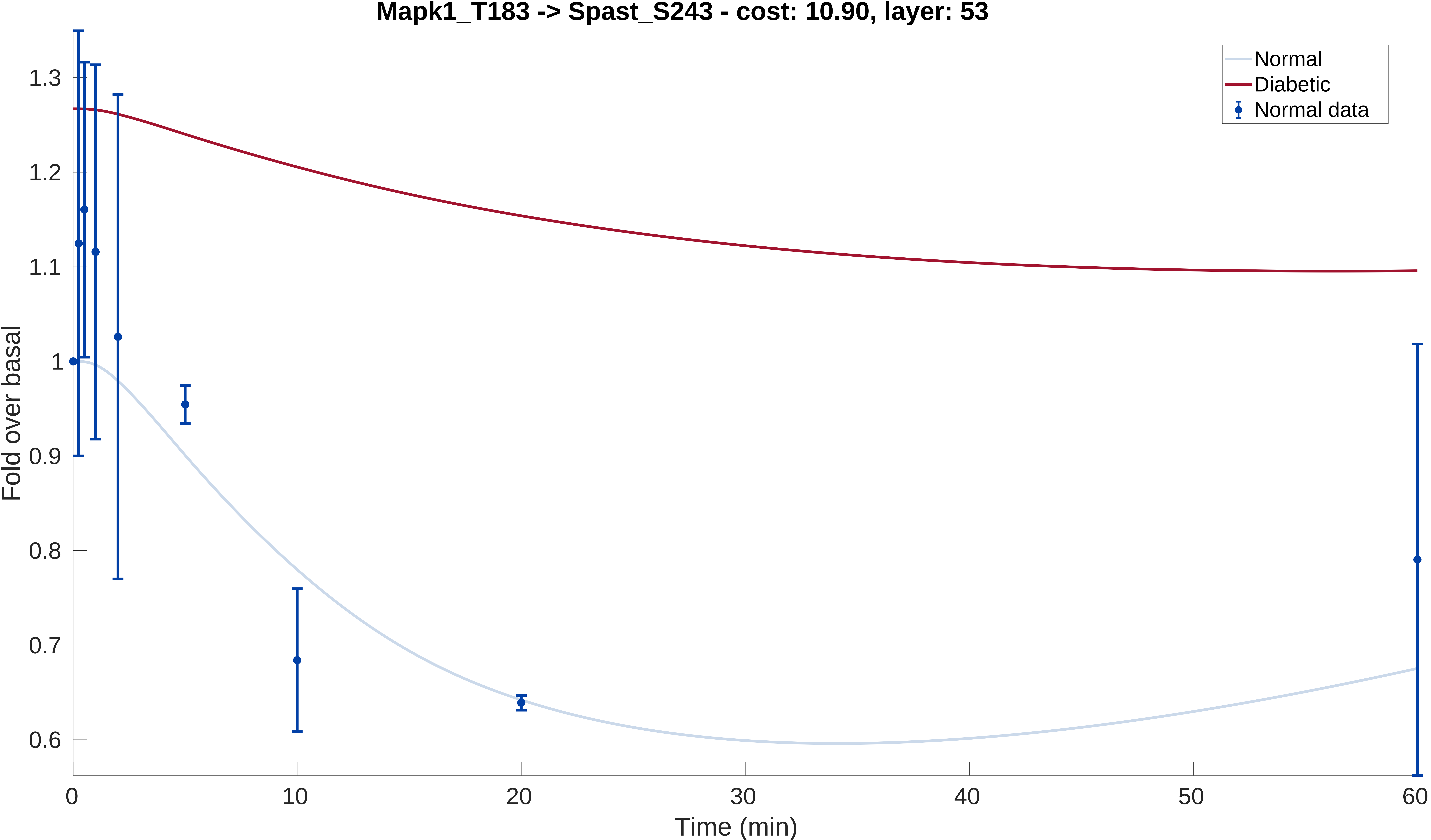

Mapk1_T183 -> Spast_S243 - cost: 10.90, layer: 53
Normal
Diabetic
Normal data
Fold over basal
1.3
1.2
1.1
1
0.9
0.8
0.7
0.6
0
10
20
30
40
50
60
Time (min)

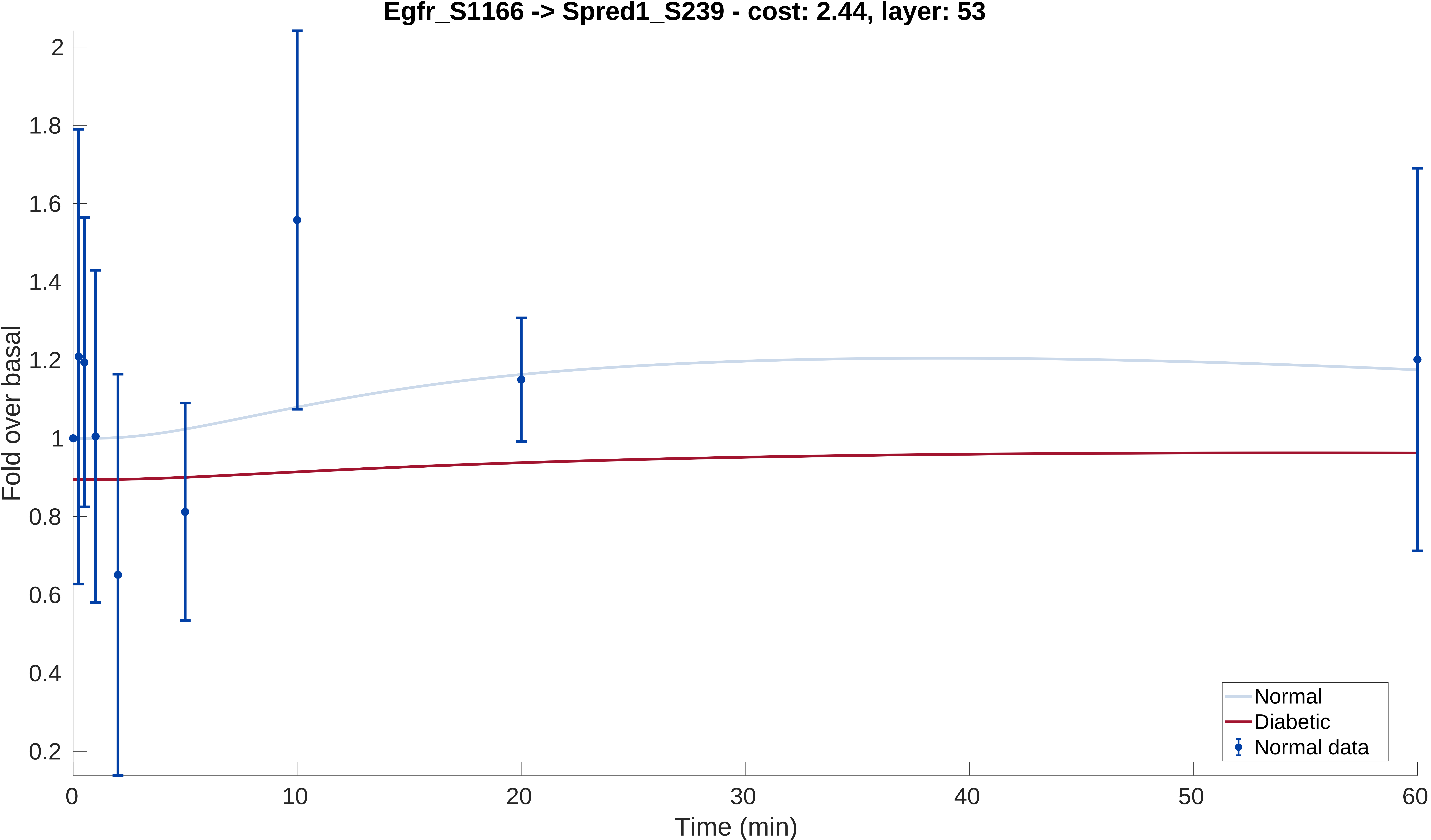

Egfr_S1166 -> Spred1_S239 - cost: 2.44, layer: 53
Fold over basal
Time (min)
0
10
20
30
40
50
60
0.2
0.4
0.6
0.8
1
1.2
1.4
1.6
1.8
2
Normal
Diabetic
Normal data

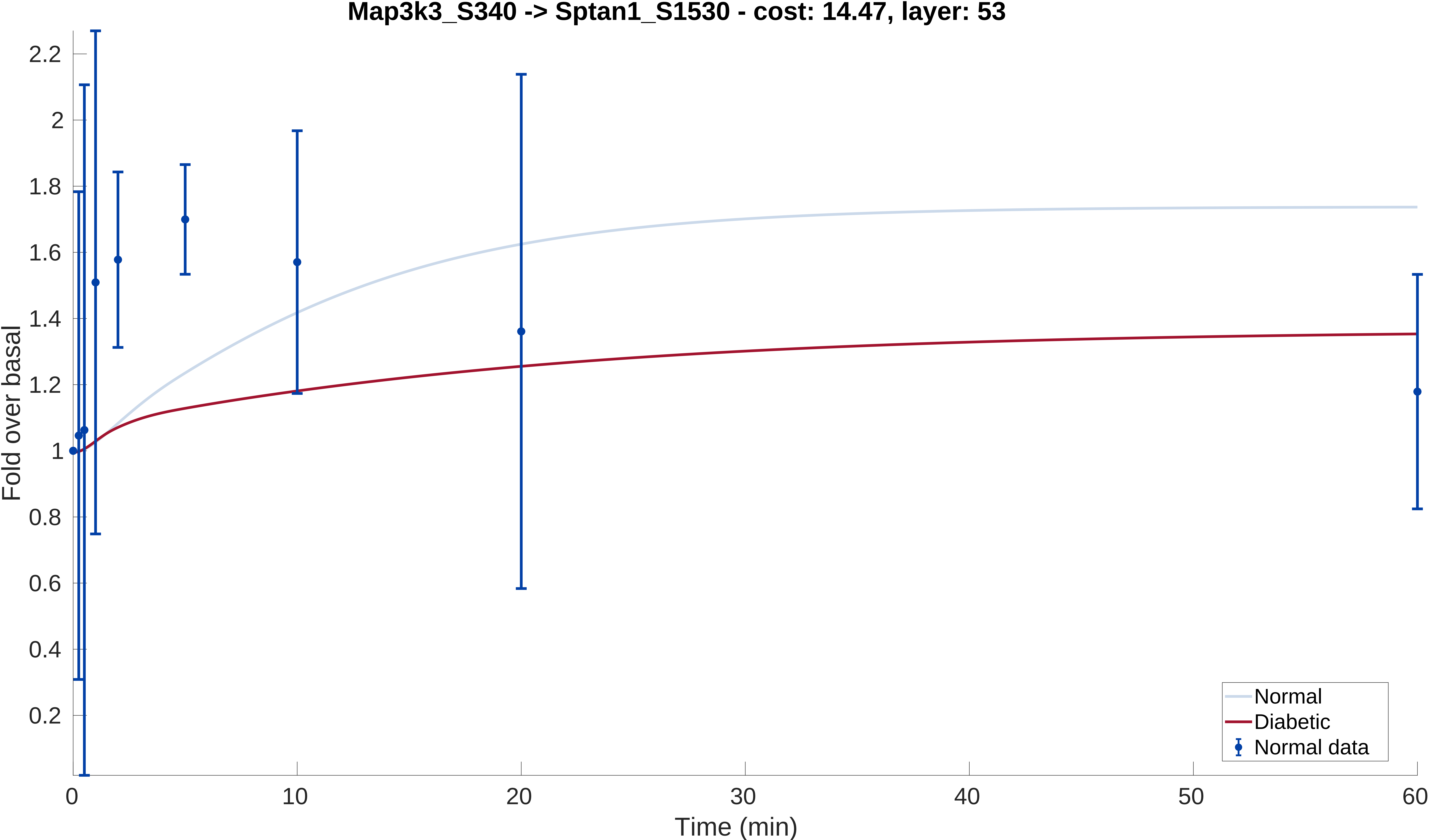

Map3k3_S340 -> Sptan1_S1530 - cost: 14.47, layer: 53
Fold over basal
2.2
2
1.8
1.6
1.4
1.2
1
0.8
0.6
0.4
0.2
0
10
20
30
40
50
60
Time (min)
Normal
Diabetic
Normal data

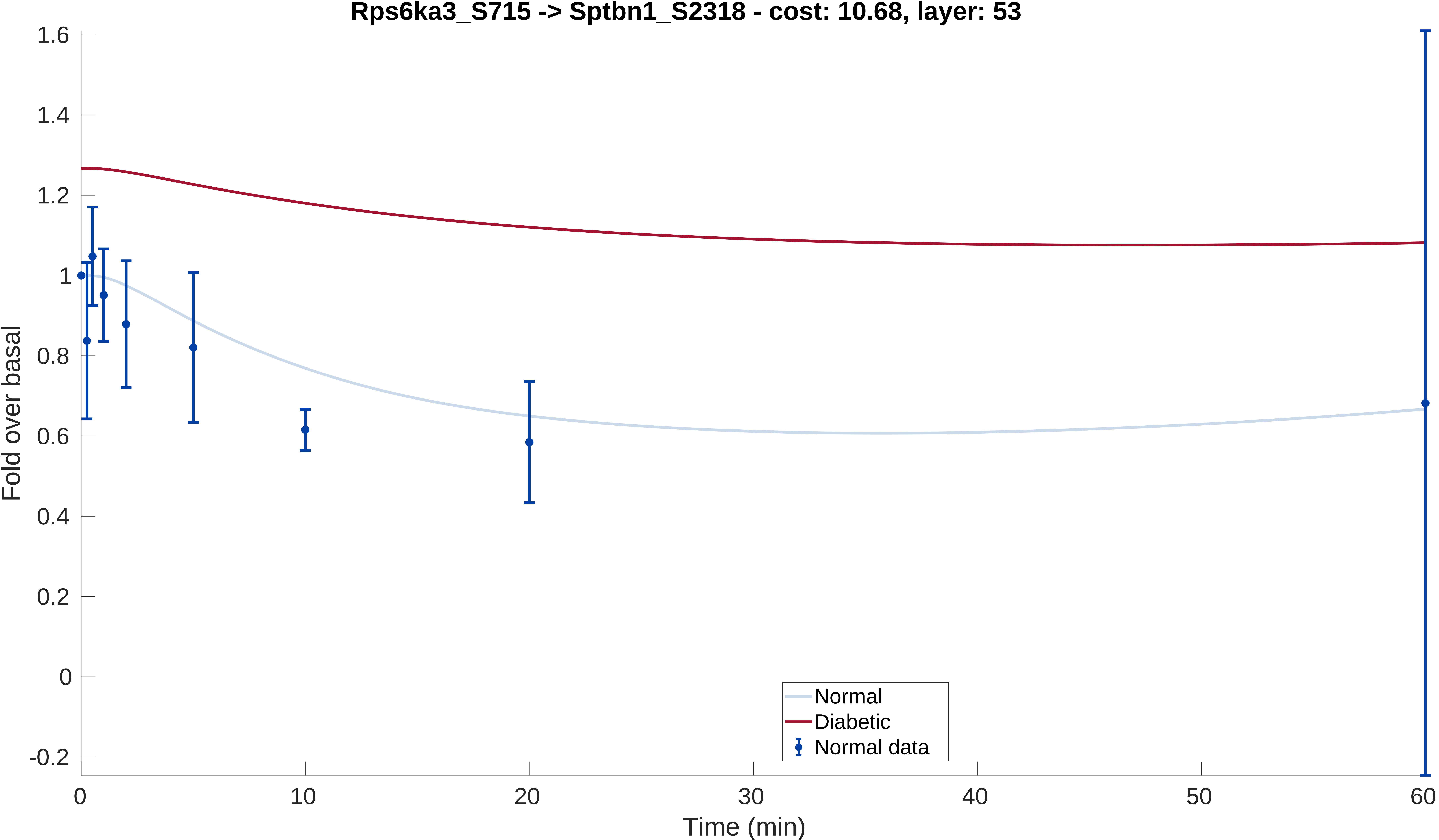

Rps6ka3_S715 -> Sptbn1_S2318 - cost: 10.68, layer: 53
1.6
1.4
1.2
1
0.8
0.6
0.4
0.2
0
-0.2
Fold over basal
0
10
20
30
40
50
60
Time (min)
Normal
Diabetic
Normal data

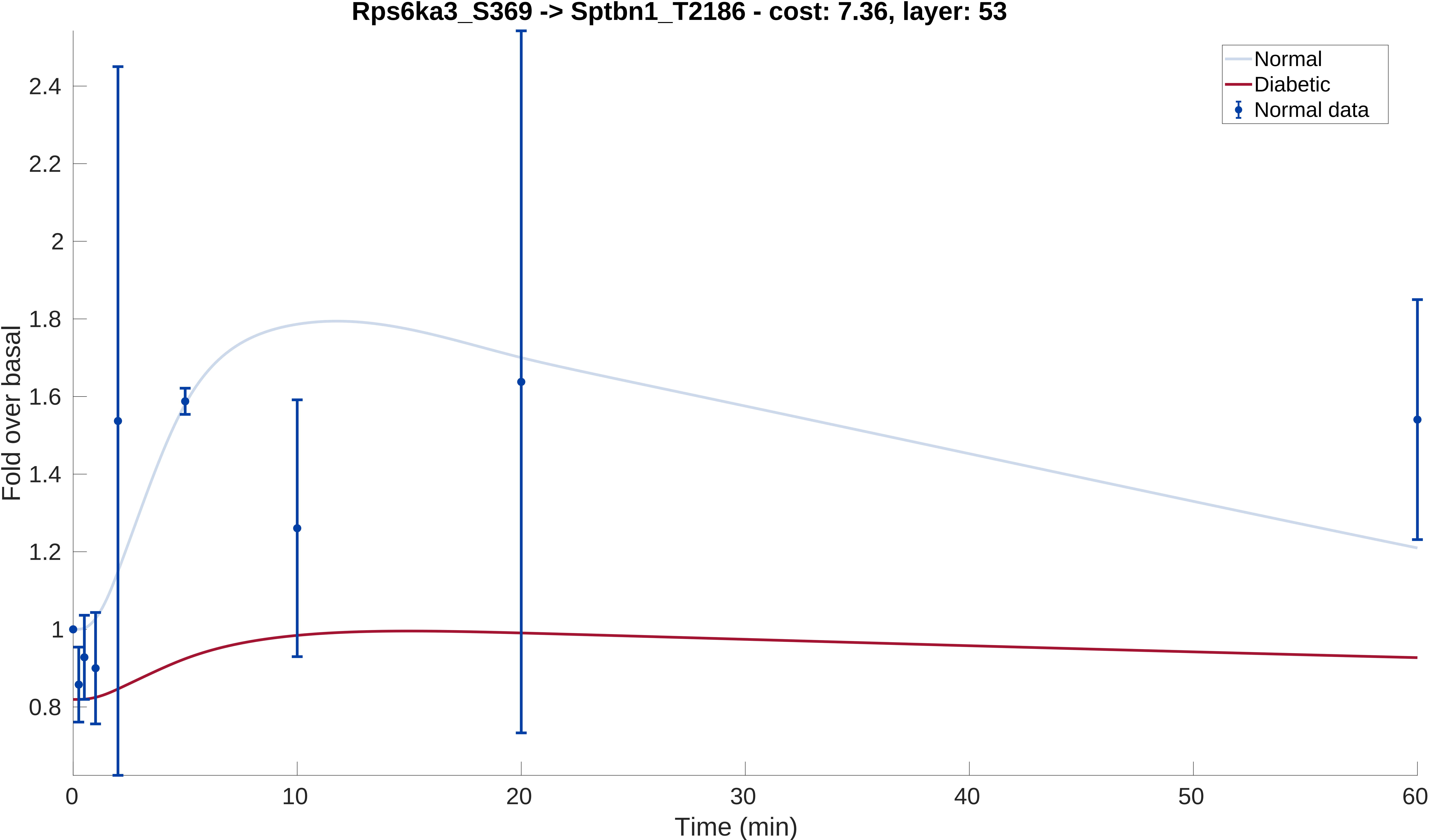
Rps6ka3_S369 -> Sptbn1_T2186 - cost: 7.36, layer: 53
Normal
Diabetic
Normal data
2.4
2.2
2
1.8
1.6
1.4
1.2
1
0.8
Fold over basal
0
10
20
30
40
50
60
Time (min)

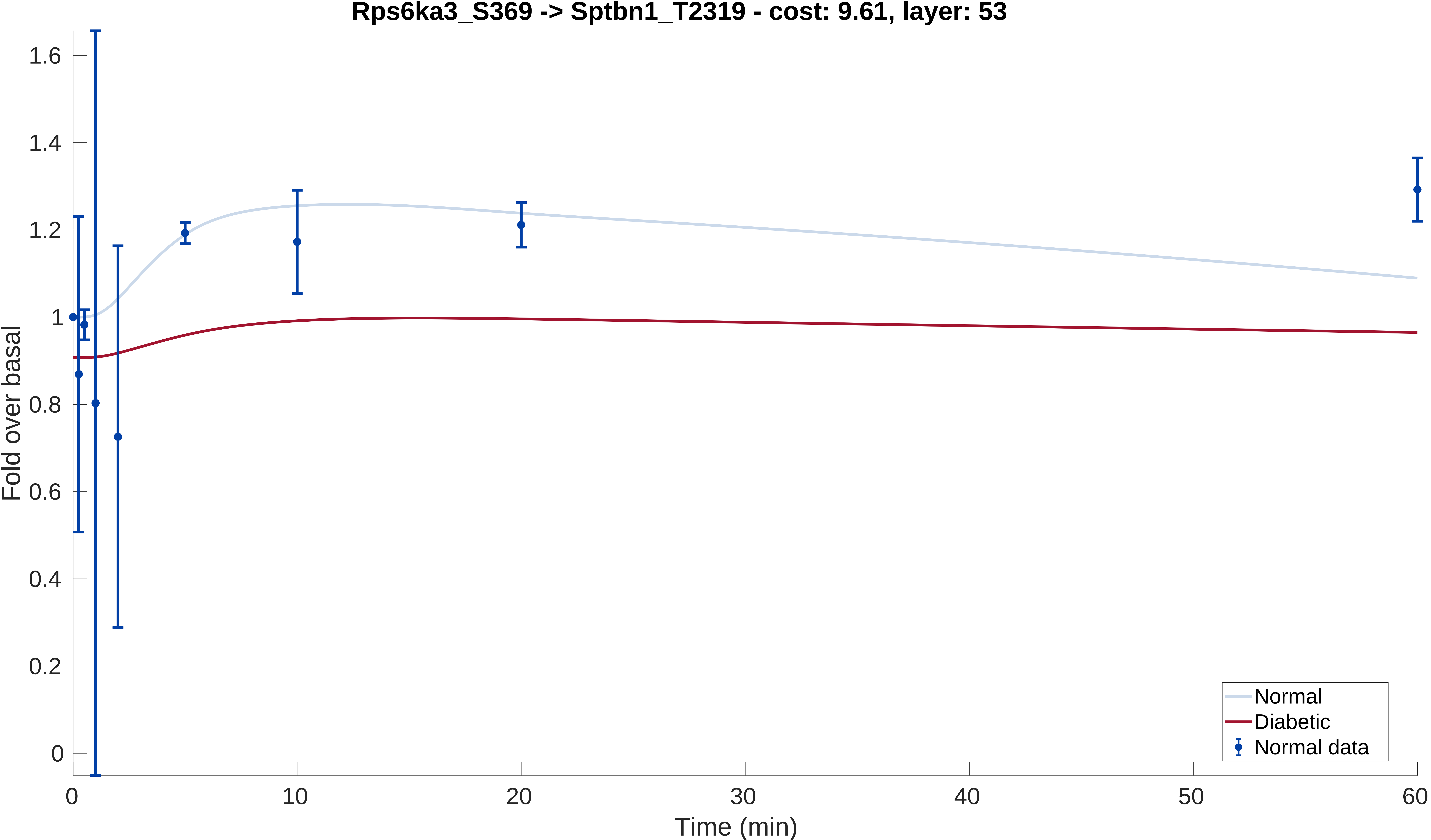

Rps6ka3_S369 -> Sptbn1_T2319 - cost: 9.61, layer: 53
Fold over basal
Time (min)
0
0.2
0.4
0.6
0.8
1
1.2
1.4
1.6
0
10
20
30
40
50
60
Normal
Diabetic
Normal data

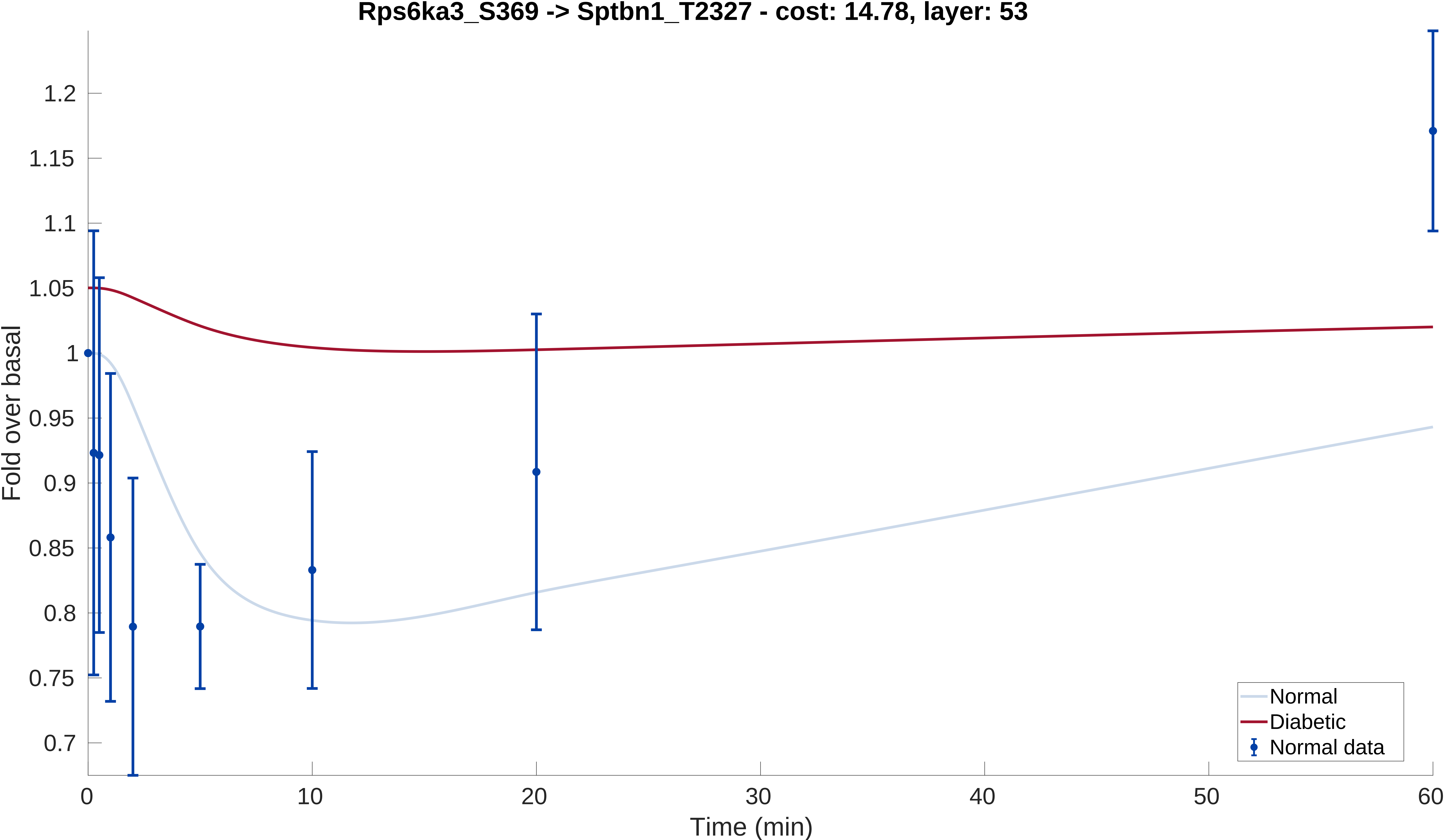

Rps6ka3_S369 -> Sptbn1_T2327 - cost: 14.78, layer: 53
Fold over basal
1.2
1.15
1.1
1.05
1
0.95
0.9
0.85
0.8
0.75
0.7
0
10
20
30
40
50
60
Time (min)
Normal
Diabetic
Normal data

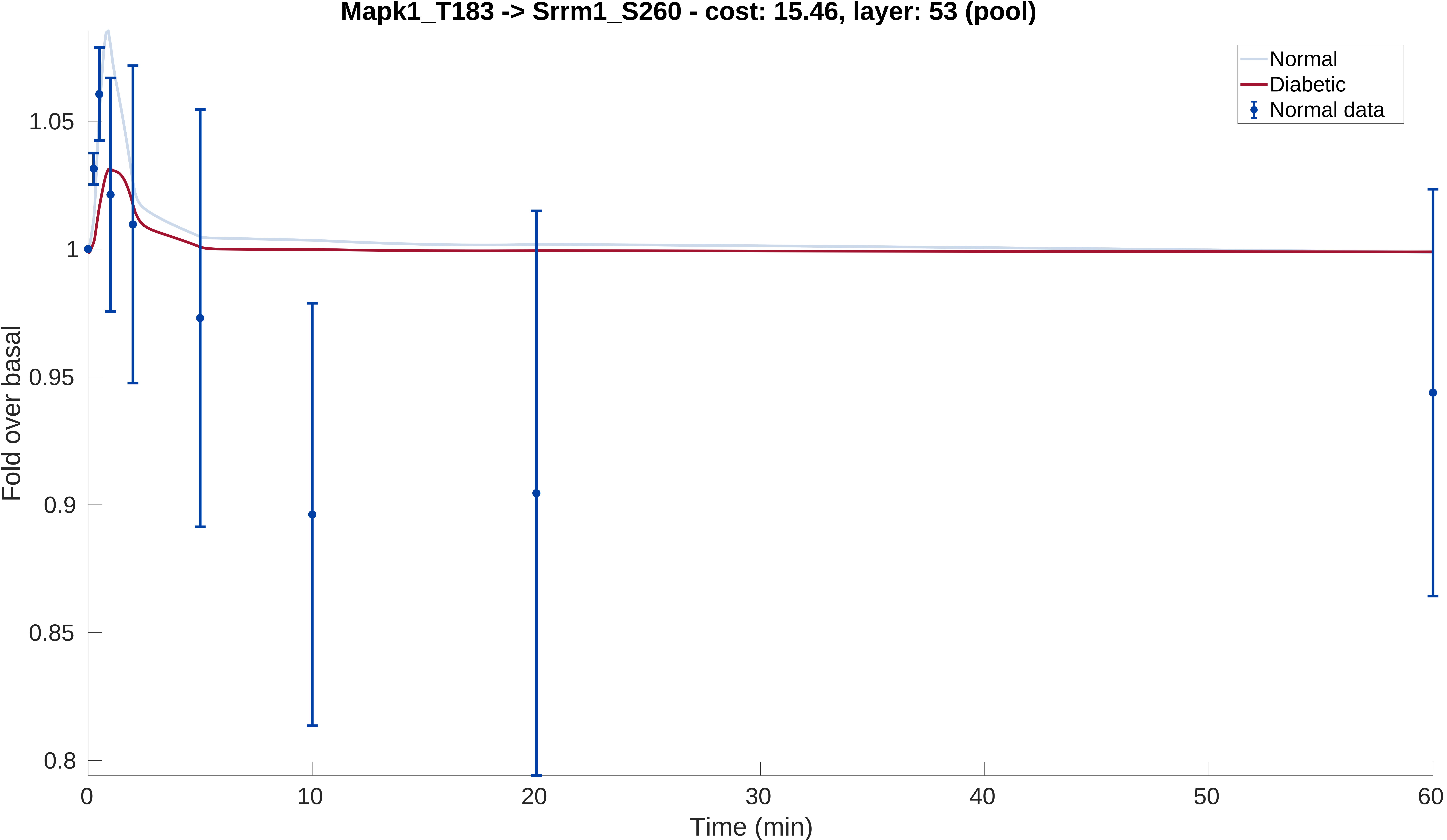
Mapk1_T183 -> Srrm1_S260 - cost: 15.46, layer: 53 (pool)
Normal
Diabetic
Normal data
Fold over basal
1.05
1
0.95
0.9
0.85
0.8
0
10
20
30
40
50
60
Time (min)

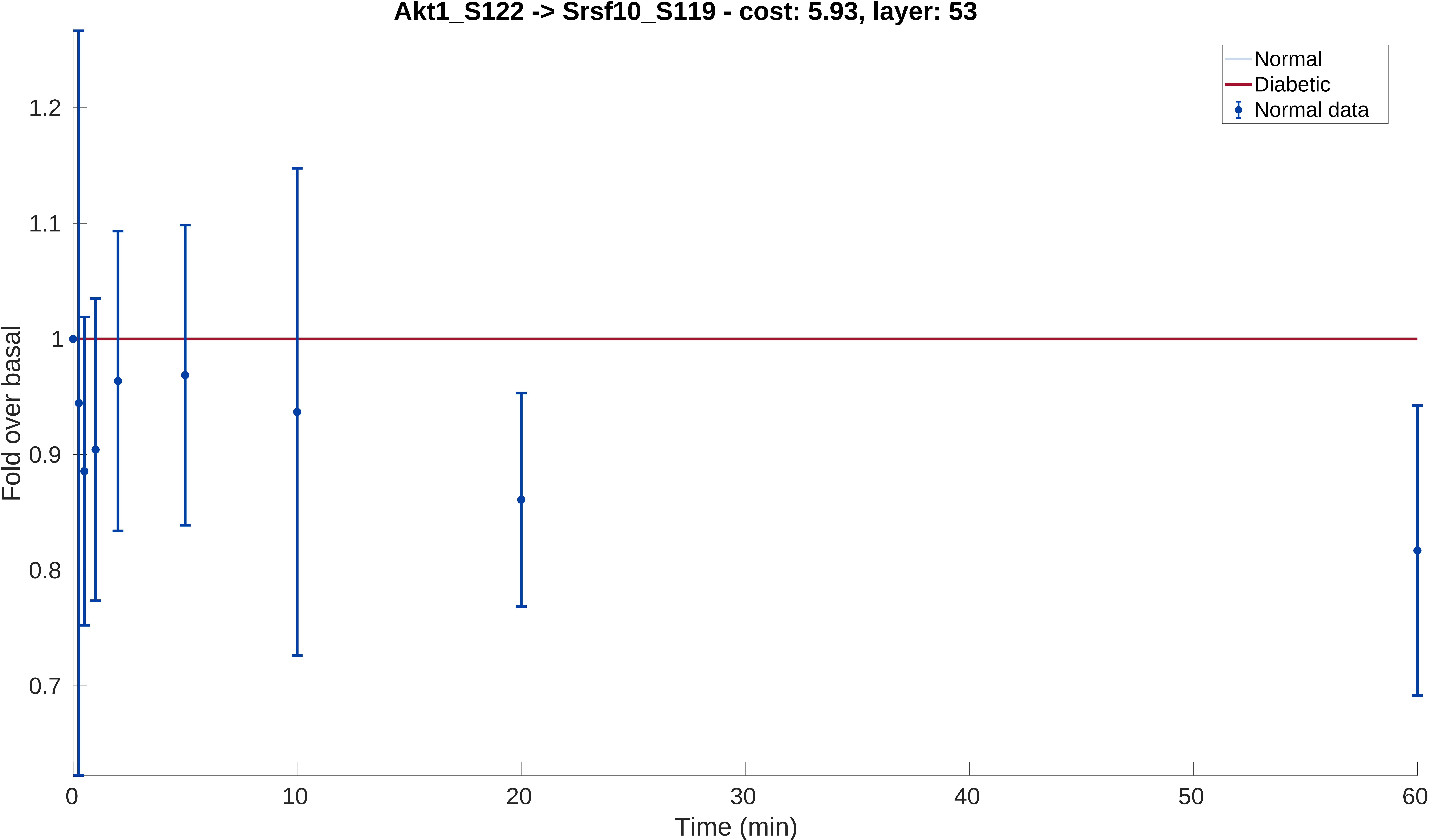

Akt1_S122 -> Srsf10_S119 - cost: 5.93, layer: 53
Normal
Diabetic
Normal data
Fold over basal
1.2
1.1
1
0.9
0.8
0.7
0
10
20
30
40
50
60
Time (min)

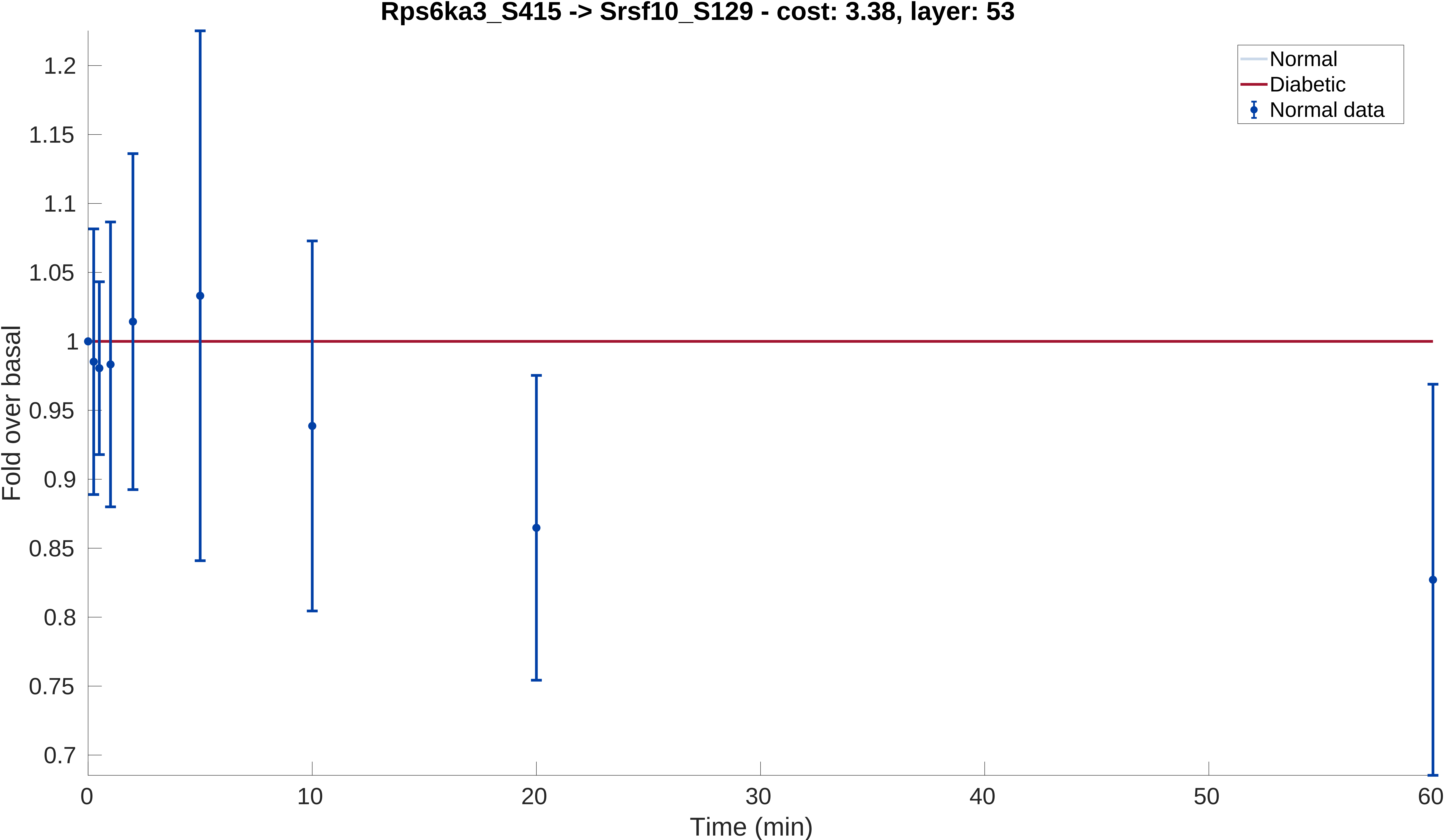

Rps6ka3_S415 -> Srsf10_S129 - cost: 3.38, layer: 53
Normal
Diabetic
Normal data
Fold over basal
1.2
1.15
1.1
1.05
1
0.95
0.9
0.85
0.8
0.75
0.7
0
10
20
30
40
50
60
Time (min)

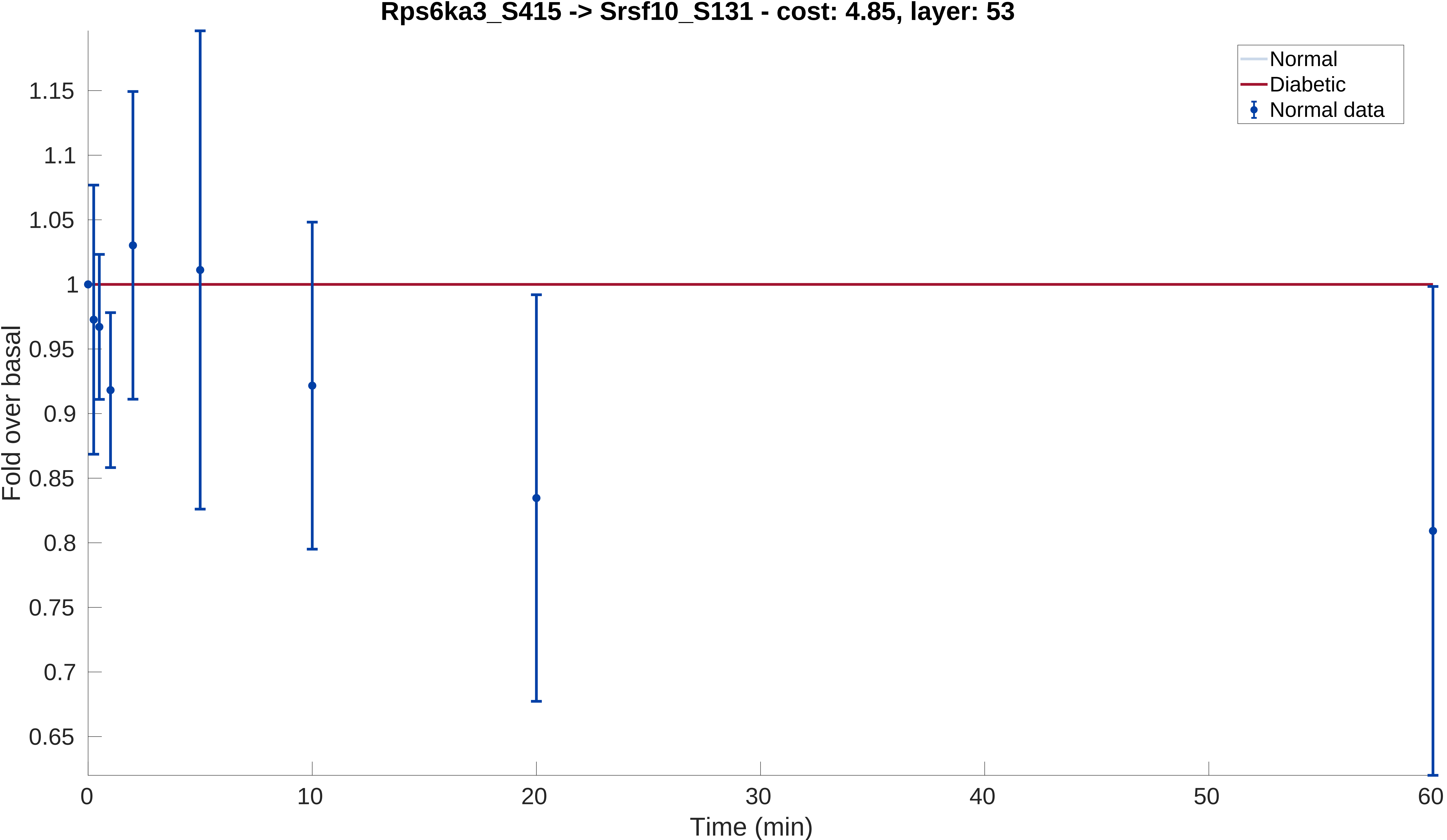
Rps6ka3_S415 -> Srsf10_S131 - cost: 4.85, layer: 53
Normal
Diabetic
Normal data
Fold over basal
1.15
1.1
1.05
1
0.95
0.9
0.85
0.8
0.75
0.7
0.65
0
10
20
30
40
50
60
Time (min)

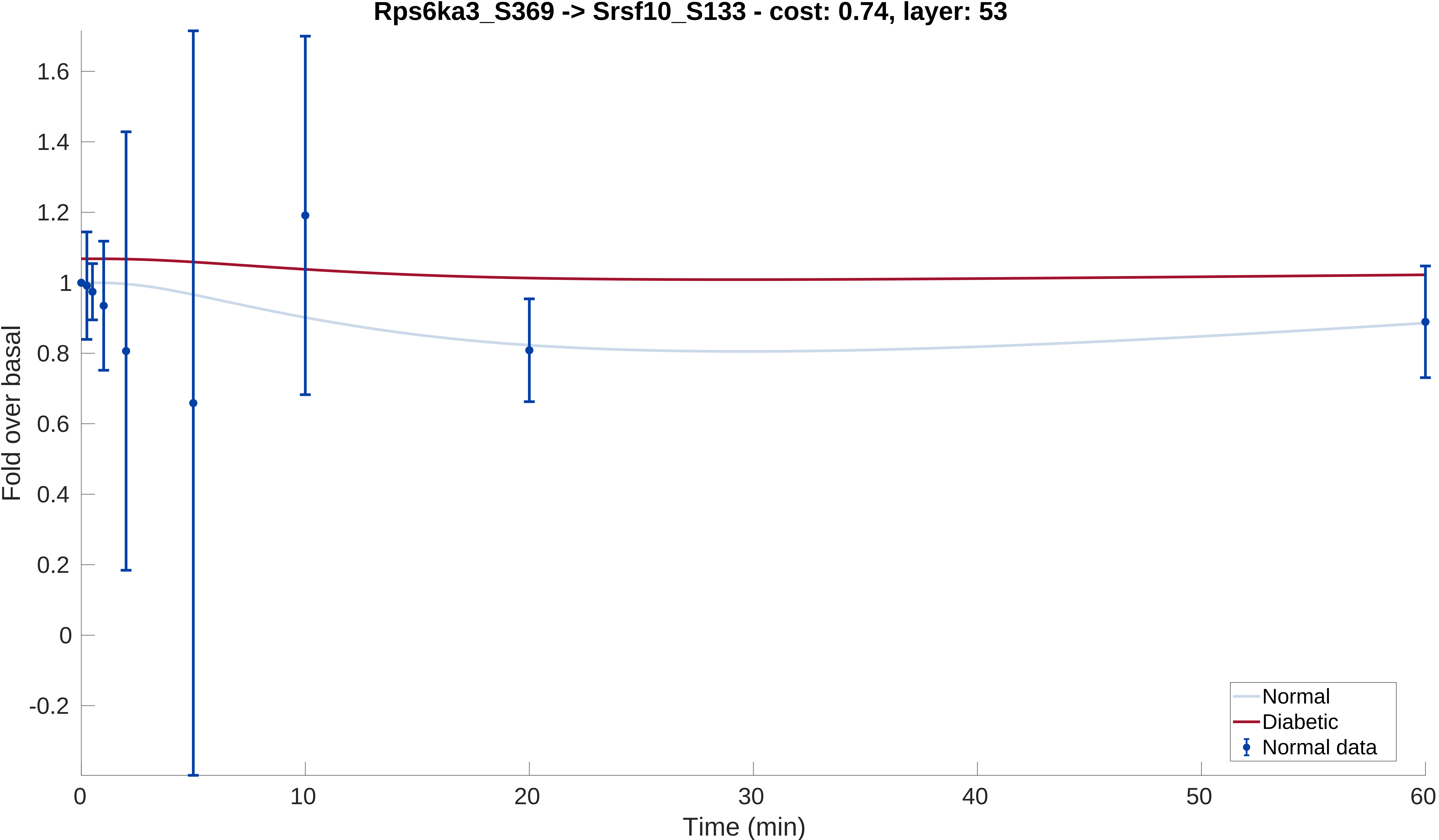

Rps6ka3_S369 -> Srsf10_S133 - cost: 0.74, layer: 53
Fold over basal
1.6
1.4
1.2
1
0.8
0.6
0.4
0.2
0
-0.2
0
10
20
30
40
50
60
Time (min)
Normal
Diabetic
Normal data

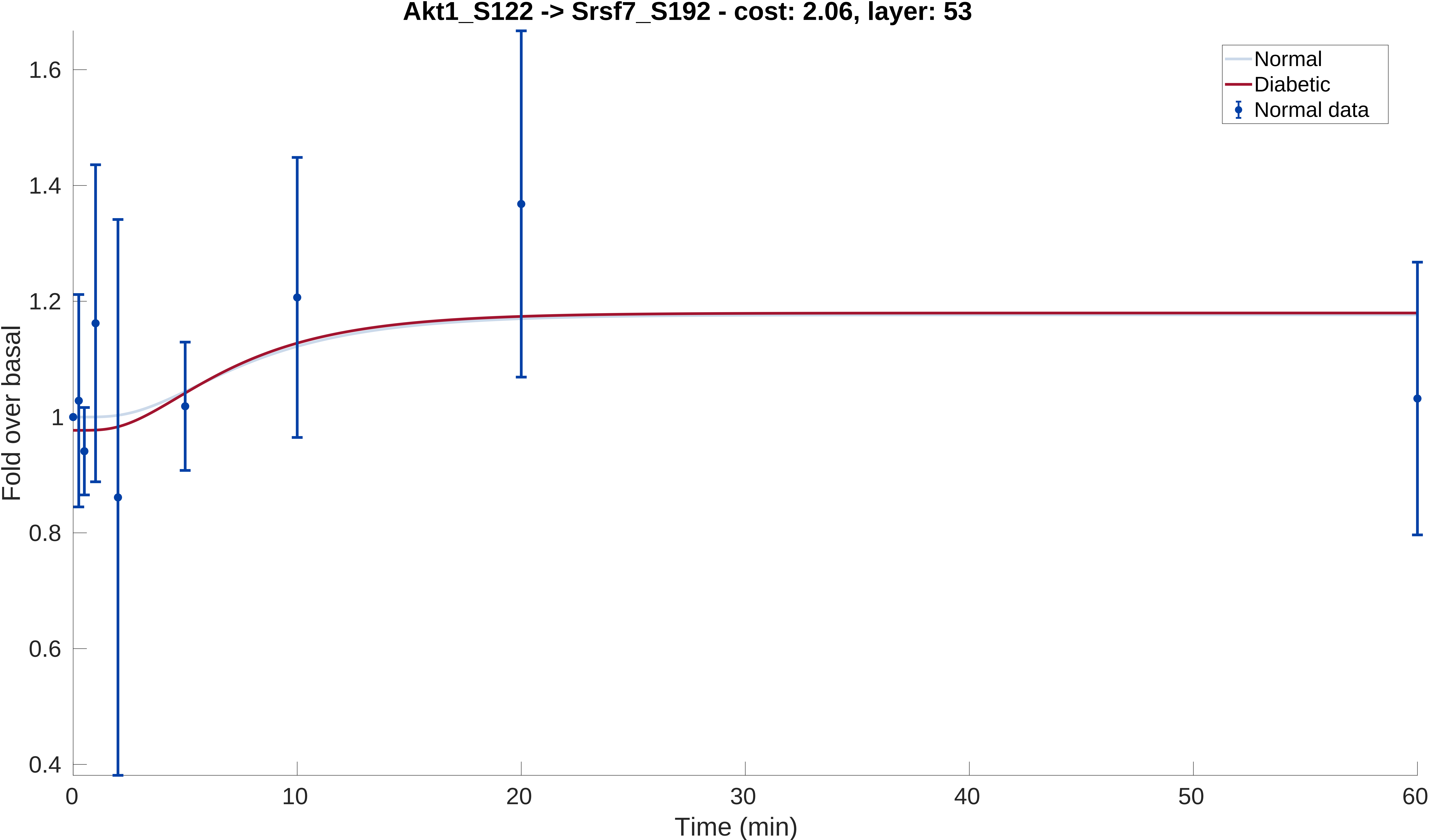
Akt1_S122 -> Srsf7_S192 - cost: 2.06, layer: 53
Normal
Diabetic
Normal data
Fold over basal
Time (min)
0.4
0.6
0.8
1
1.2
1.4
1.6
0
10
20
30
40
50
60

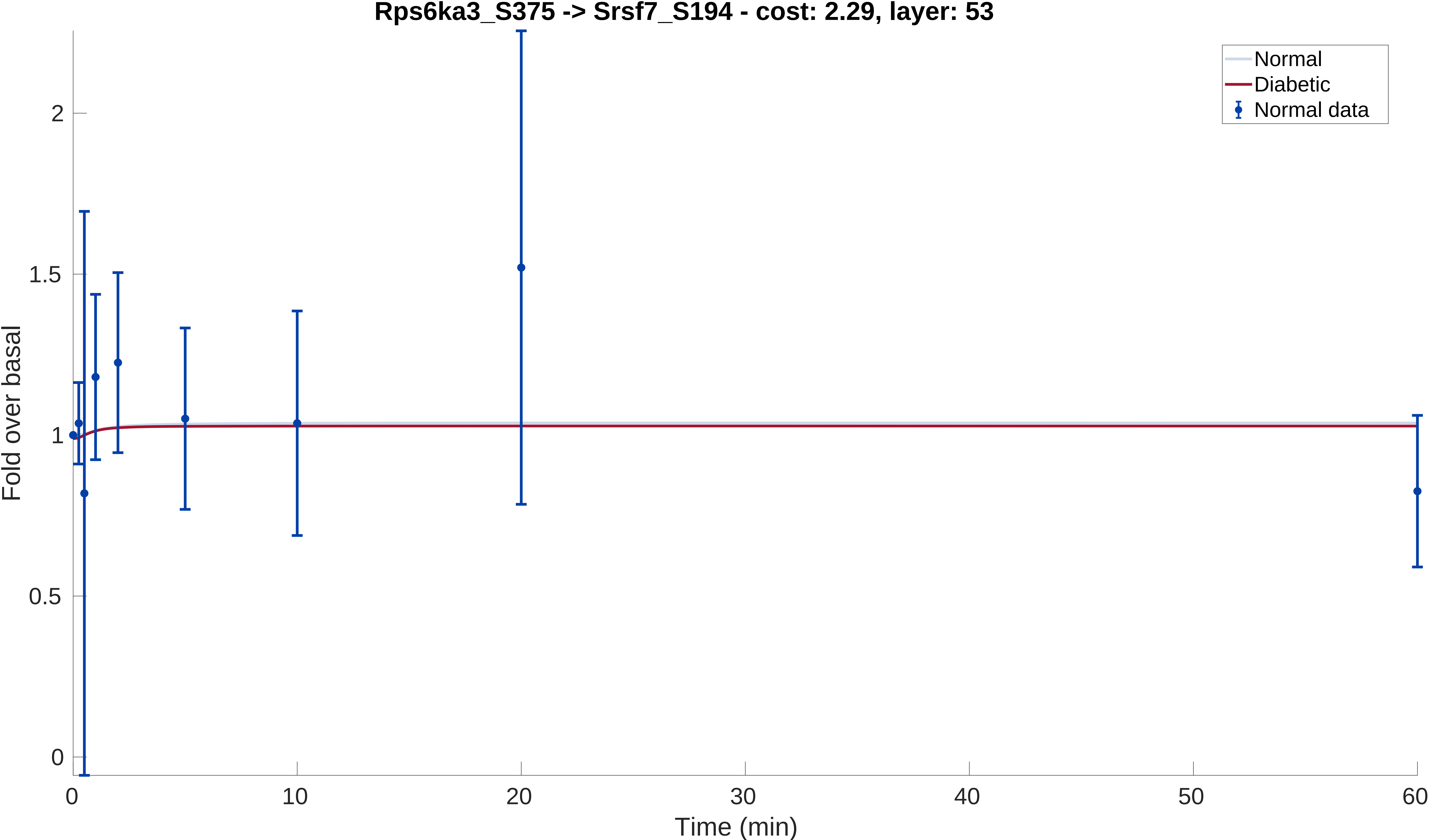

Rps6ka3_S375 -> Srsf7_S194 - cost: 2.29, layer: 53
Normal
Diabetic
Normal data
Fold over basal
0
0.5
1
1.5
2
0
10
20
30
40
50
60
Time (min)

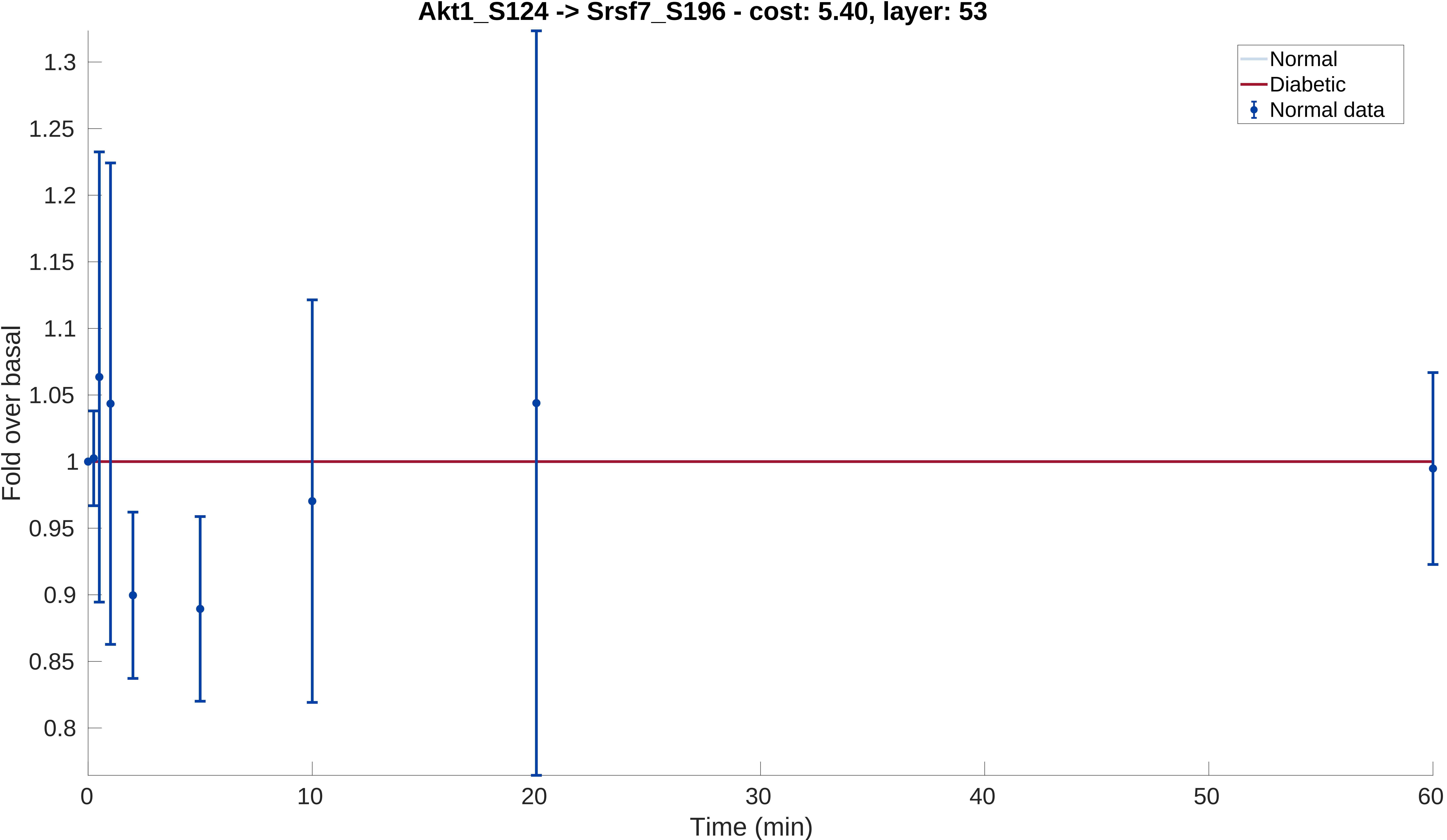

Akt1_S124 -> Srsf7_S196 - cost: 5.40, layer: 53
Normal
Diabetic
Normal data
Fold over basal
1.3
1.25
1.2
1.15
1.1
1.05
1
0.95
0.9
0.85
0.8
0
10
20
30
40
50
60
Time (min)

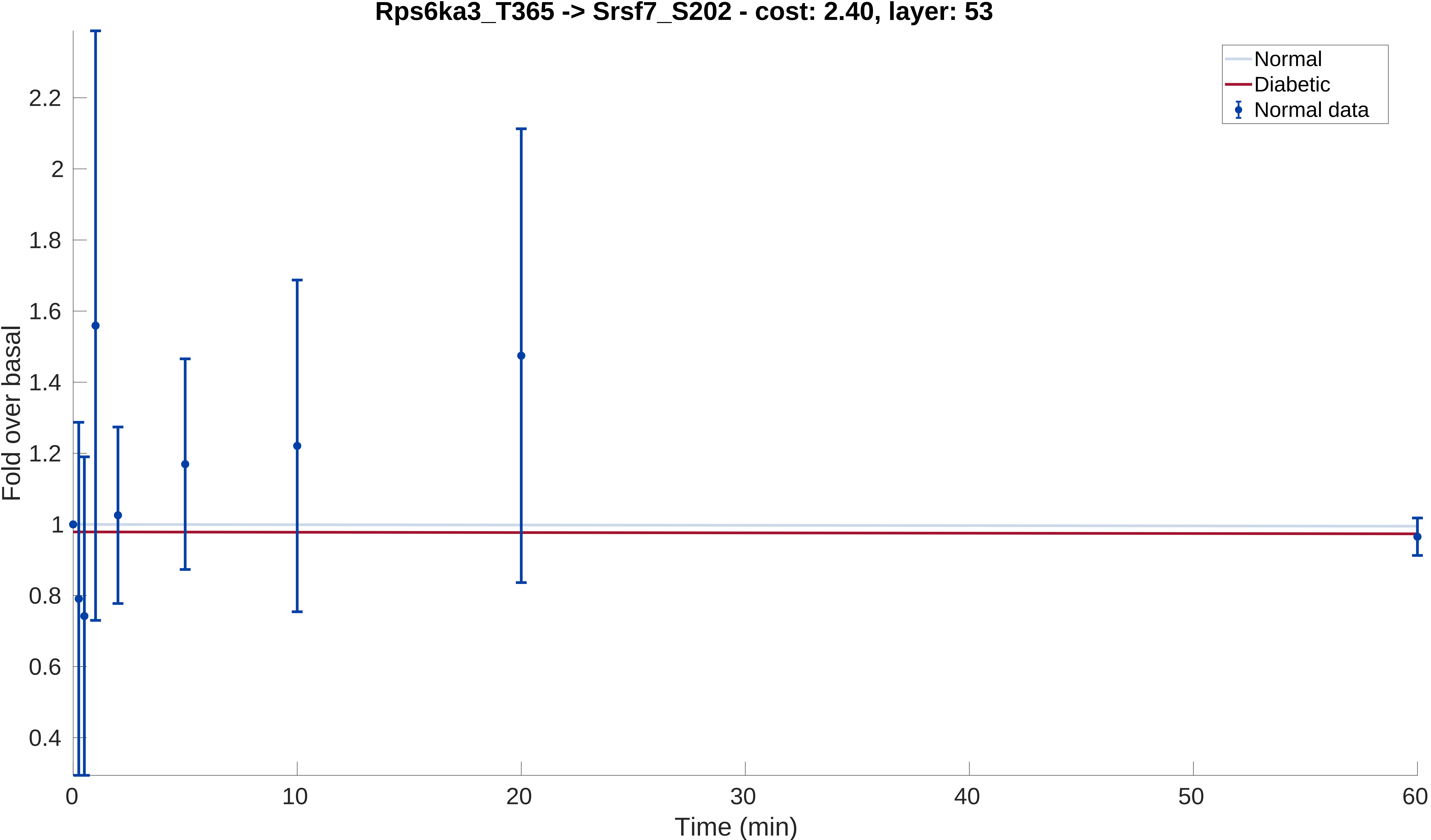

Rps6ka3_T365 -> Srsf7_S202 - cost: 2.40, layer: 53
Normal
Diabetic
Normal data
Fold over basal
Time (min)
0.4
0.6
0.8
1
1.2
1.4
1.6
1.8
2
2.2
0
10
20
30
40
50
60

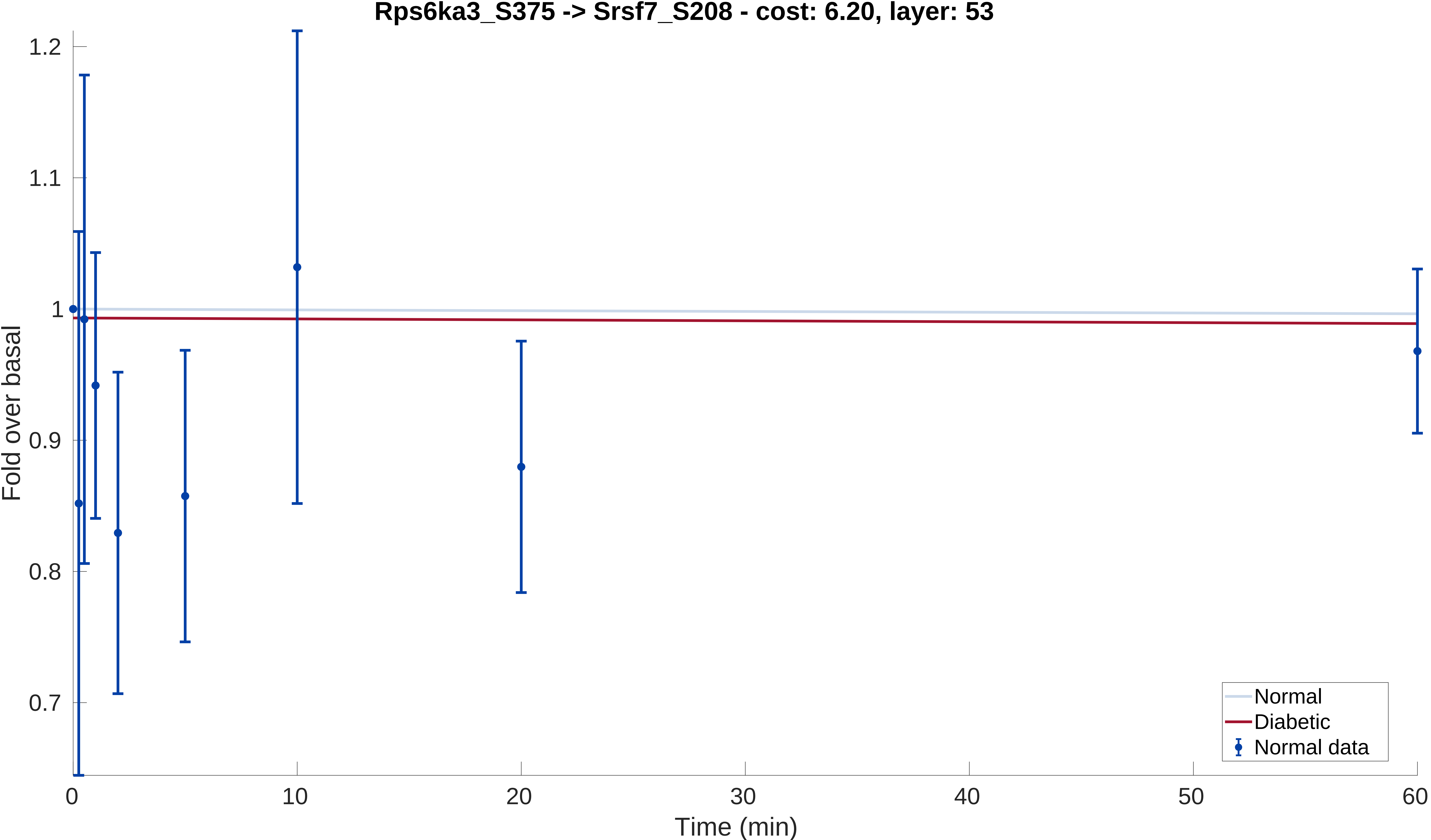

Rps6ka3_S375 -> Srsf7_S208 - cost: 6.20, layer: 53
Fold over basal
1.2
1.1
1
0.9
0.8
0.7
0
10
20
30
40
50
60
Time (min)
Normal
Diabetic
Normal data

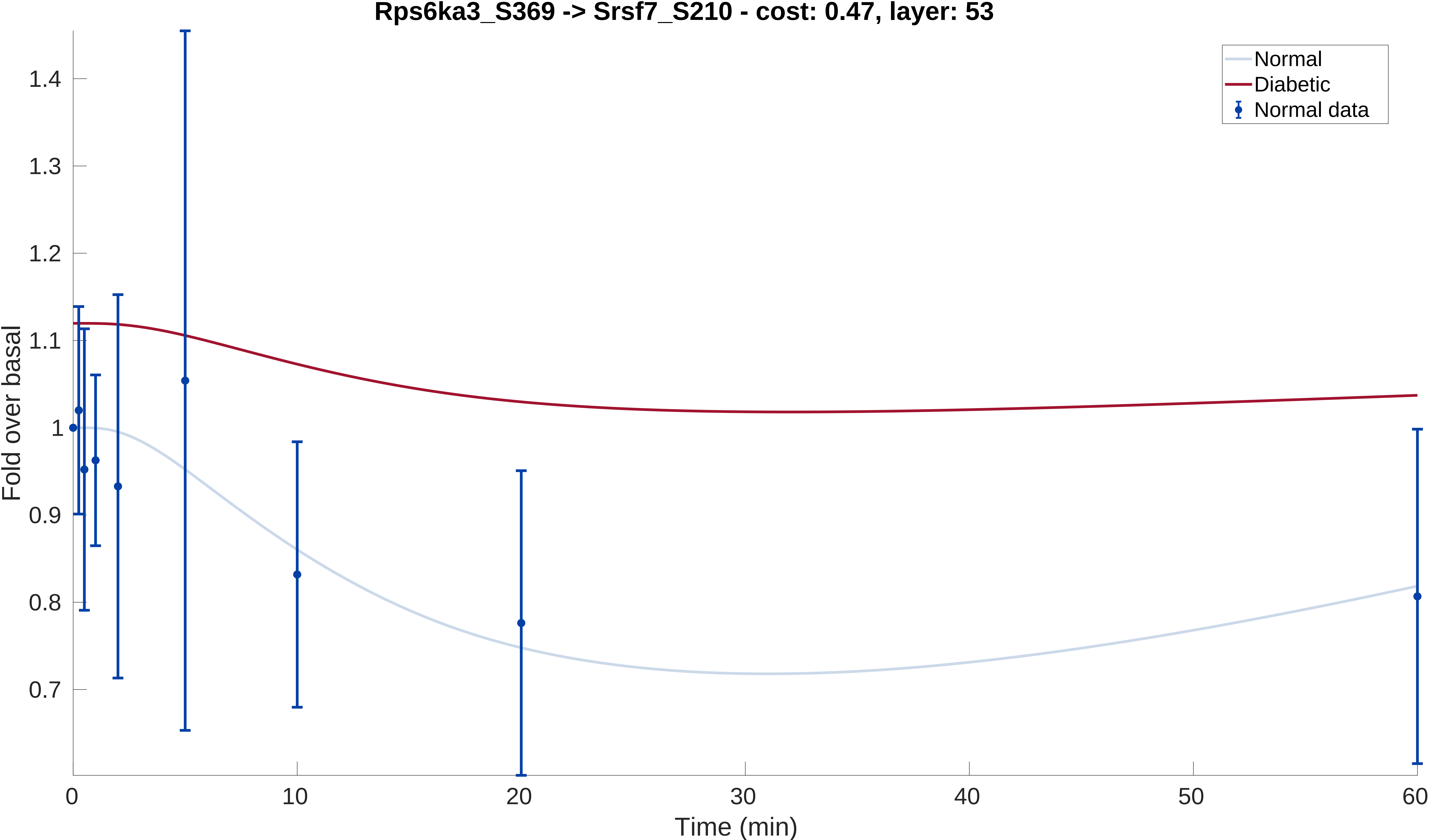
Rps6ka3_S369 -> Srsf7_S210 - cost: 0.47, layer: 53
Normal
Diabetic
Normal data
Fold over basal
Time (min)
1.4
1.3
1.2
1.1
1
0.9
0.8
0.7
0
10
20
30
40
50
60

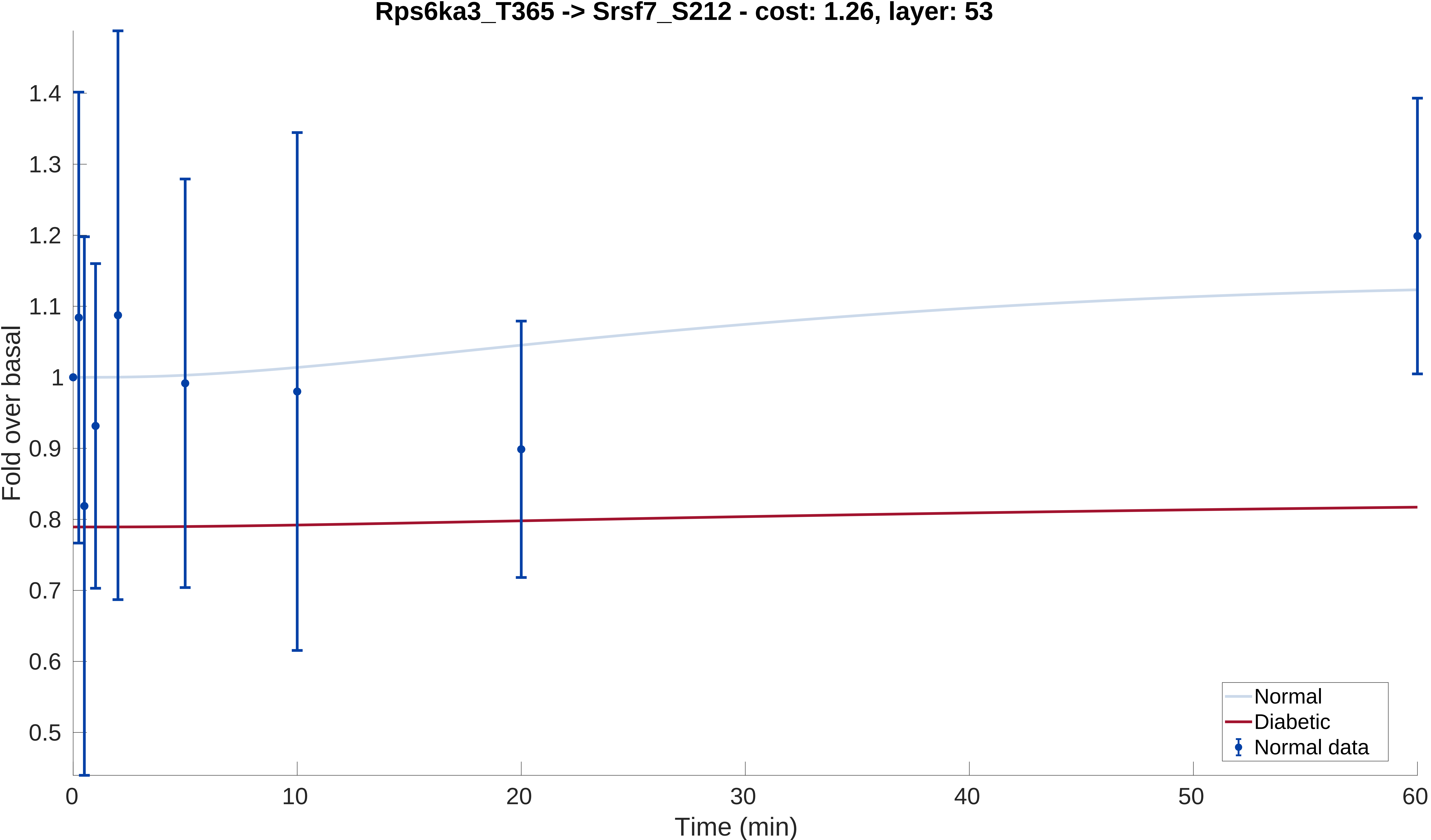

Rps6ka3_T365 -> Srsf7_S212 - cost: 1.26, layer: 53
Fold over basal
Time (min)
1.4
1.3
1.2
1.1
1
0.9
0.8
0.7
0.6
0.5
0
10
20
30
40
50
60
Normal
Diabetic
Normal data

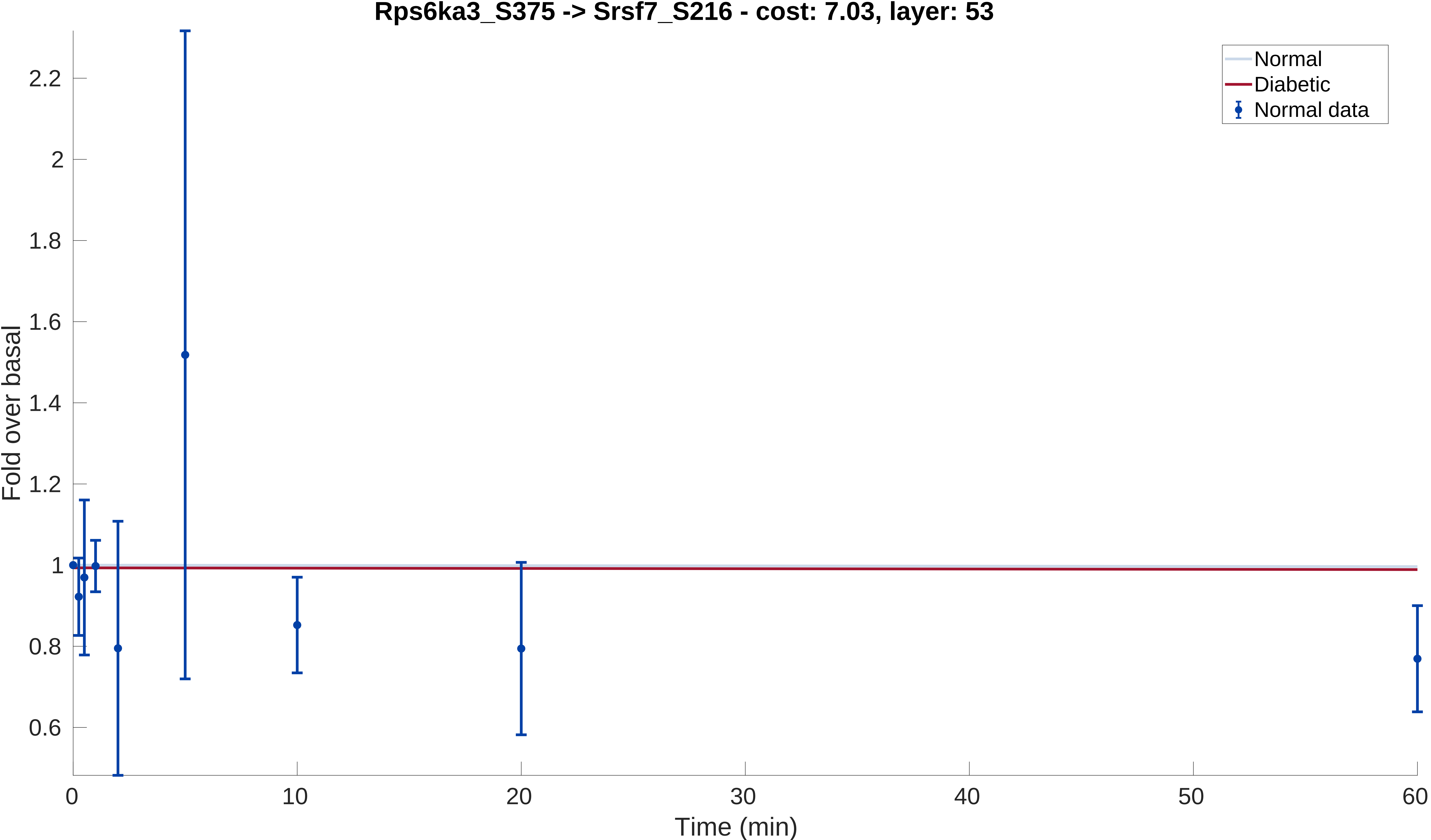

Rps6ka3_S375 -> Srsf7_S216 - cost: 7.03, layer: 53
Normal
Diabetic
Normal data
Fold over basal
Time (min)
0.6
0.8
1
1.2
1.4
1.6
1.8
2
2.2
0
10
20
30
40
50
60

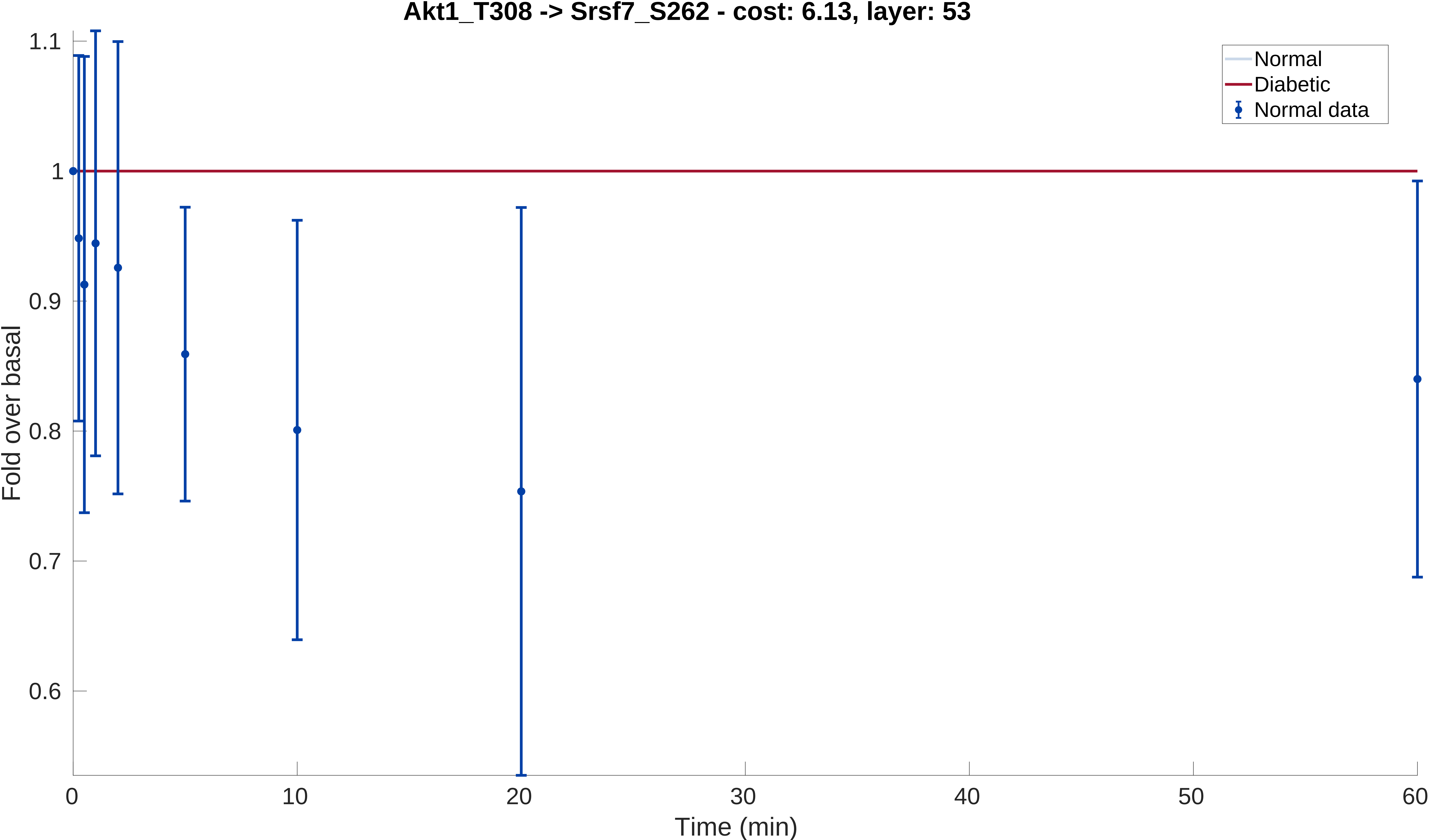

Akt1_T308 -> Srsf7_S262 - cost: 6.13, layer: 53
Normal
Diabetic
Normal data
Fold over basal
1.1
1
0.9
0.8
0.7
0.6
0
10
20
30
40
50
60
Time (min)

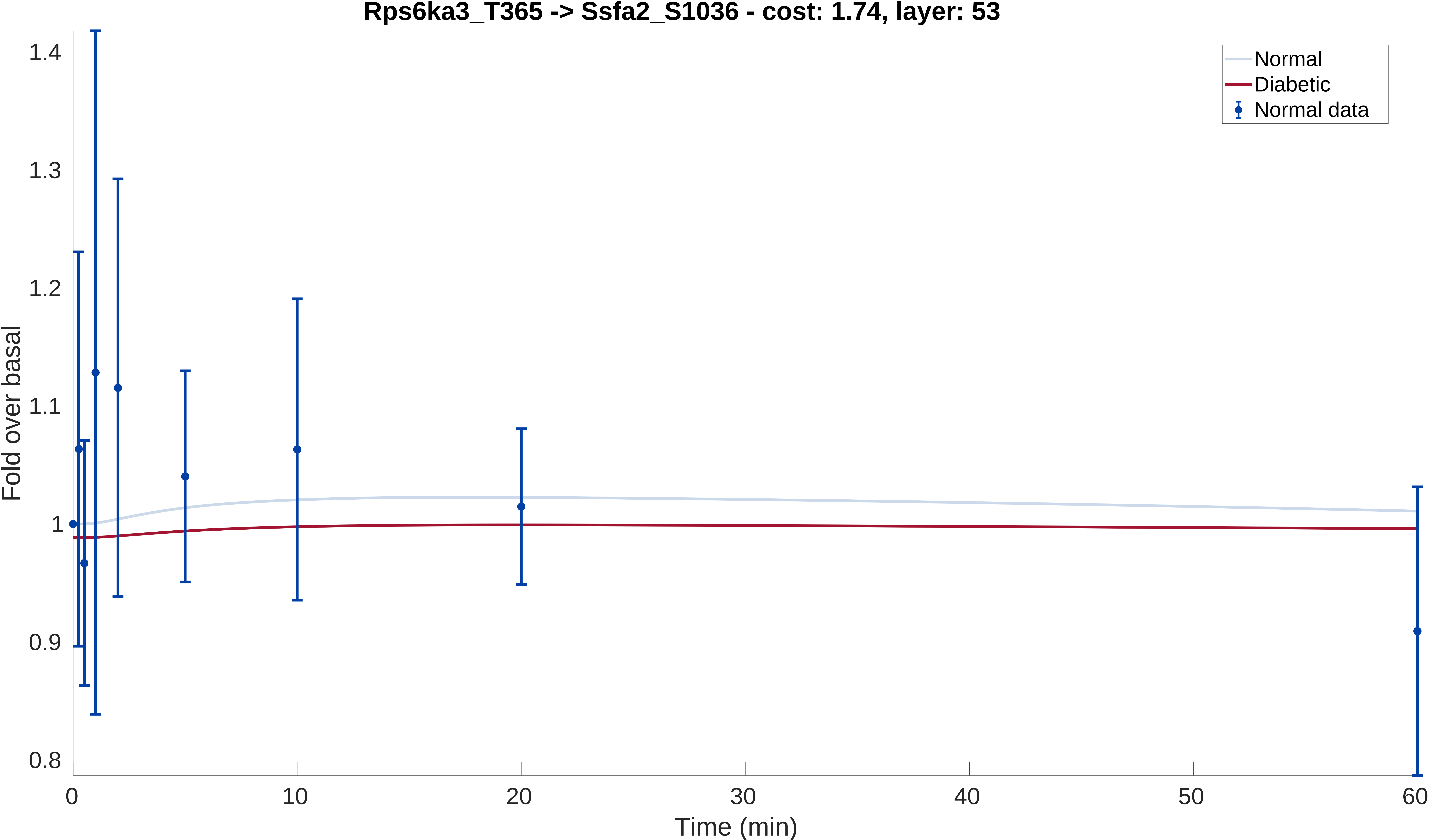

Rps6ka3_T365 -> Ssfa2_S1036 - cost: 1.74, layer: 53
Normal
Diabetic
Normal data
Fold over basal
1.4
1.3
1.2
1.1
1
0.9
0.8
0
10
20
30
40
50
60
Time (min)

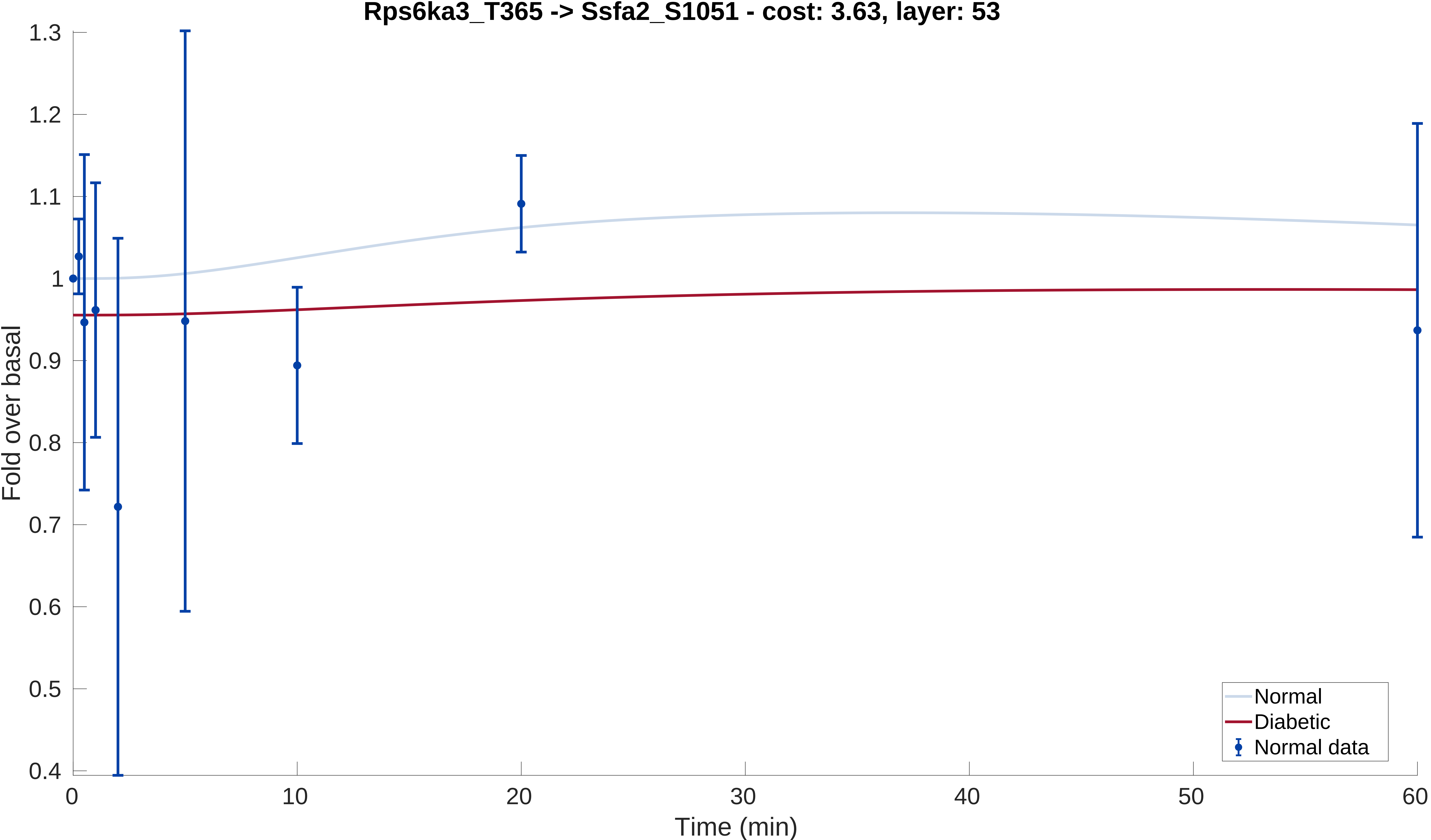
Rps6ka3_T365 -> Ssfa2_S1051 - cost: 3.63, layer: 53
Fold over basal
Time (min)
1.3
1.2
1.1
1
0.9
0.8
0.7
0.6
0.5
0.4
0
10
20
30
40
50
60
Normal
Diabetic
Normal data

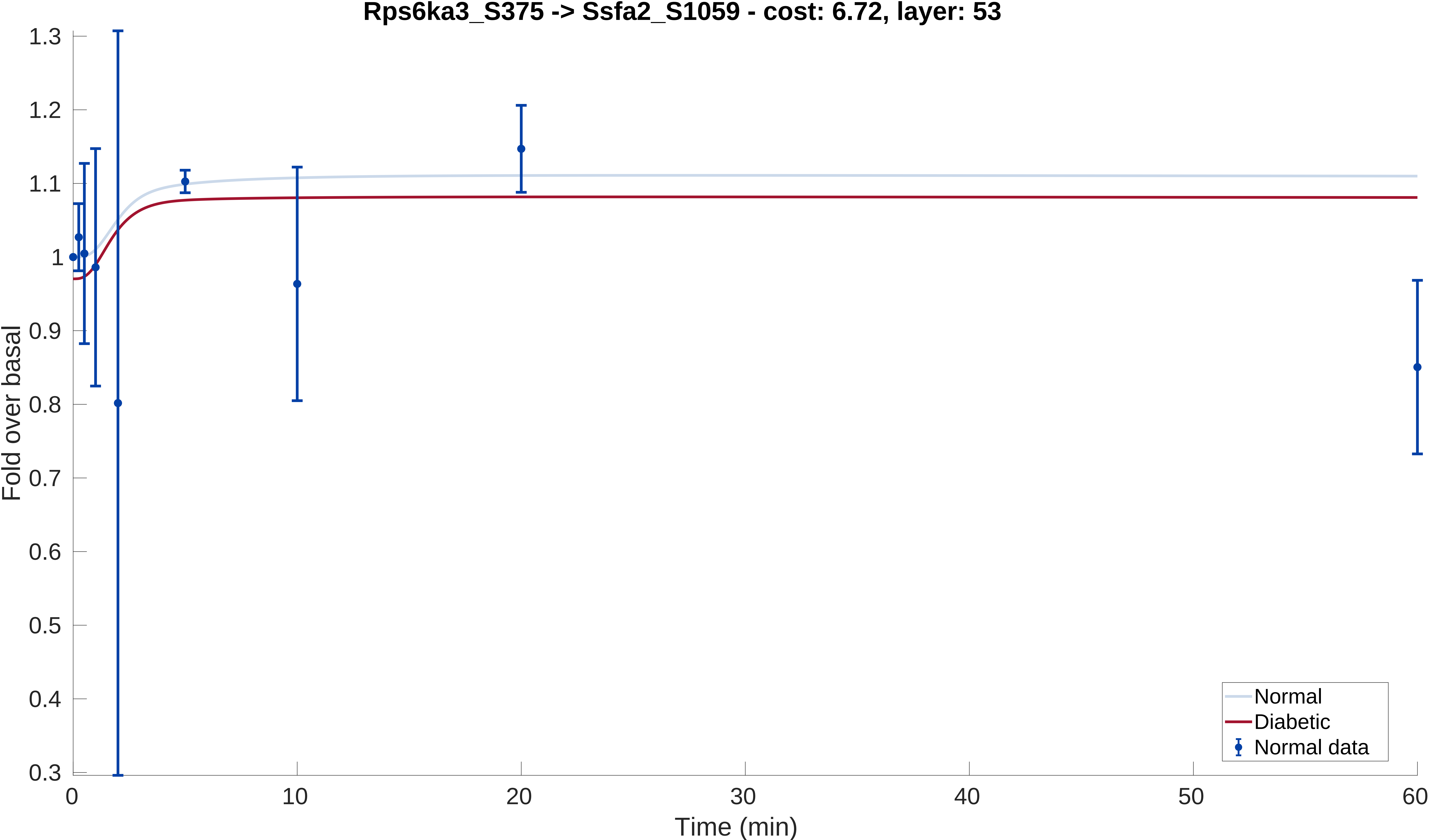
Rps6ka3_S375 -> Ssfa2_S1059 - cost: 6.72, layer: 53
Fold over basal
1.3
1.2
1.1
1
0.9
0.8
0.7
0.6
0.5
0.4
0.3
0
10
20
30
40
50
60
Time (min)
Normal
Diabetic
Normal data

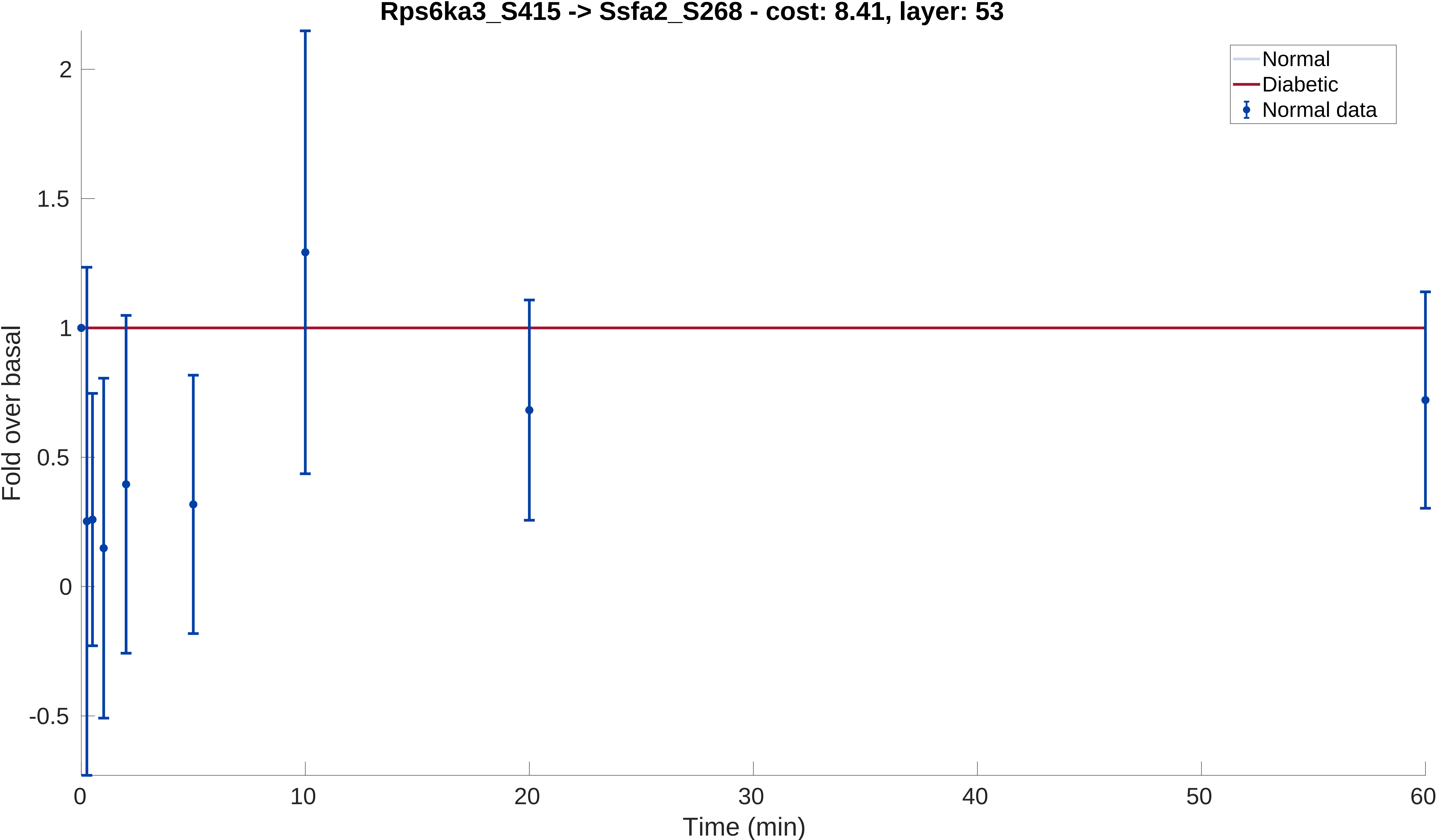

Rps6ka3_S415 -> Ssfa2_S268 - cost: 8.41, layer: 53
Normal
Diabetic
Normal data
Fold over basal
2
1.5
1
0.5
0
-0.5
0
10
20
30
40
50
60
Time (min)

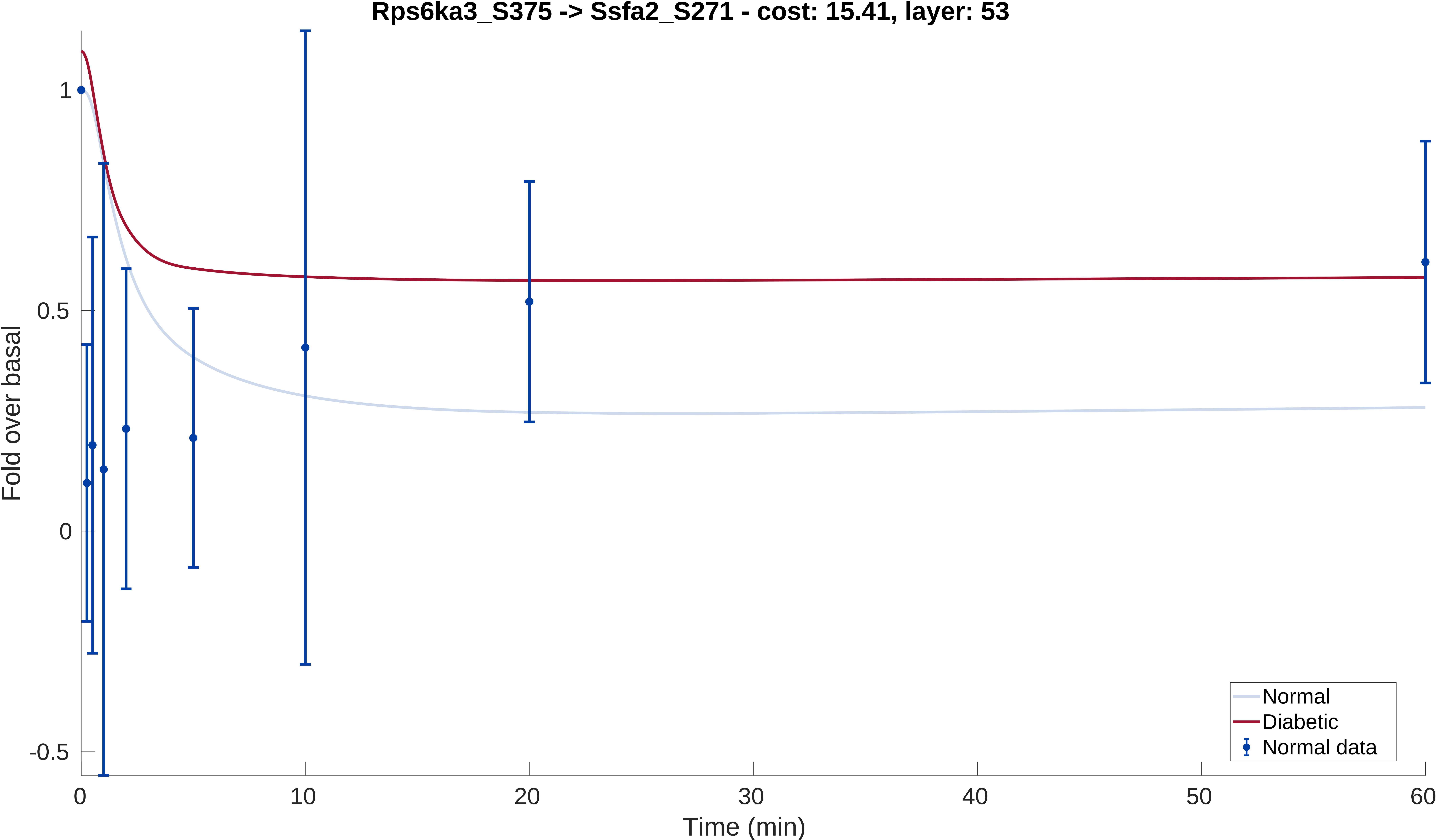
Rps6ka3_S375 -> Ssfa2_S271 - cost: 15.41, layer: 53
Fold over basal
1
0.5
0
-0.5
0
10
20
30
40
50
60
Time (min)
Normal
Diabetic
Normal data

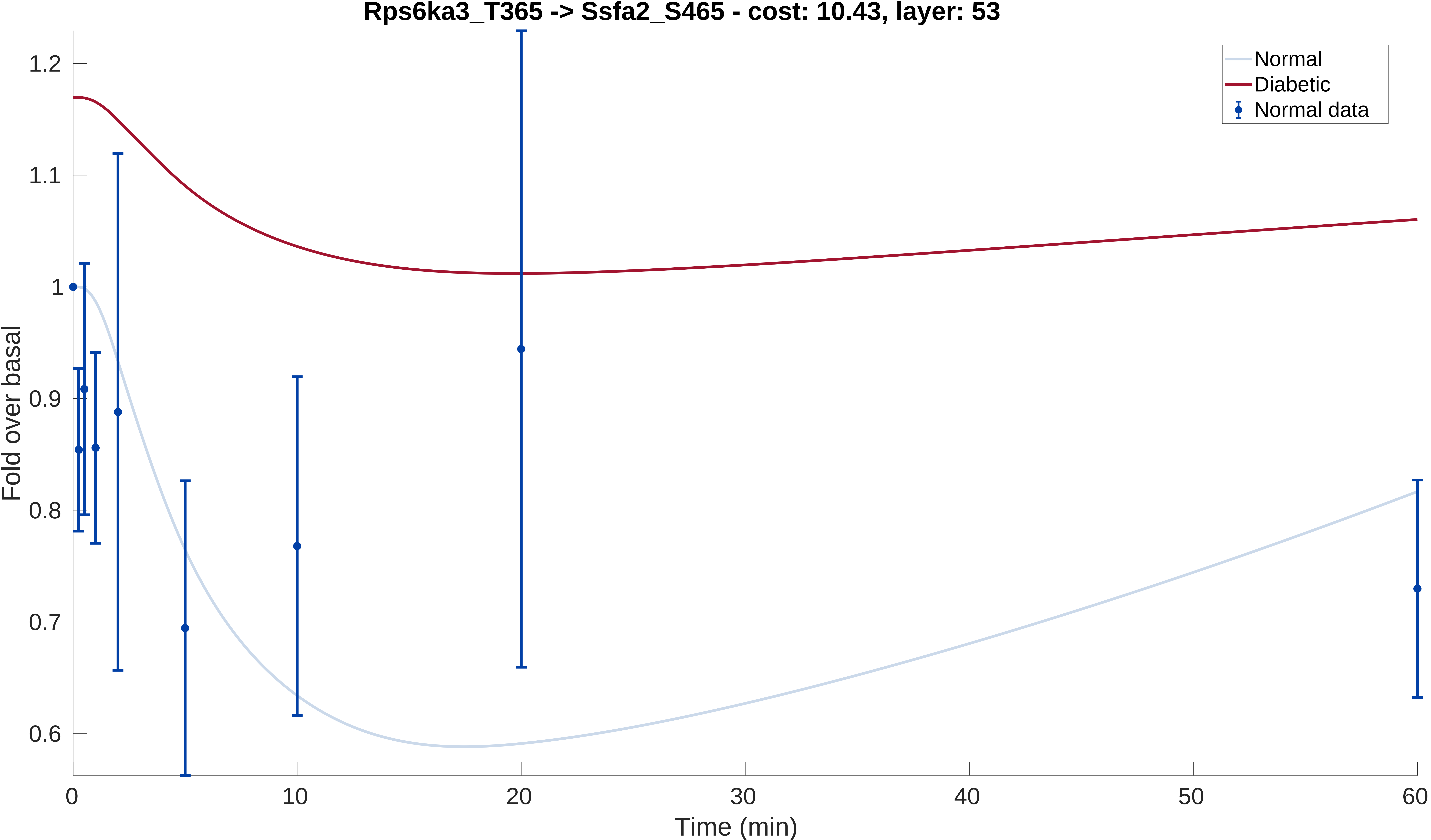
Rps6ka3_T365 -> Ssfa2_S465 - cost: 10.43, layer: 53
Normal
Diabetic
Normal data
Fold over basal
1.2
1.1
1
0.9
0.8
0.7
0.6
0
10
20
30
40
50
60
Time (min)

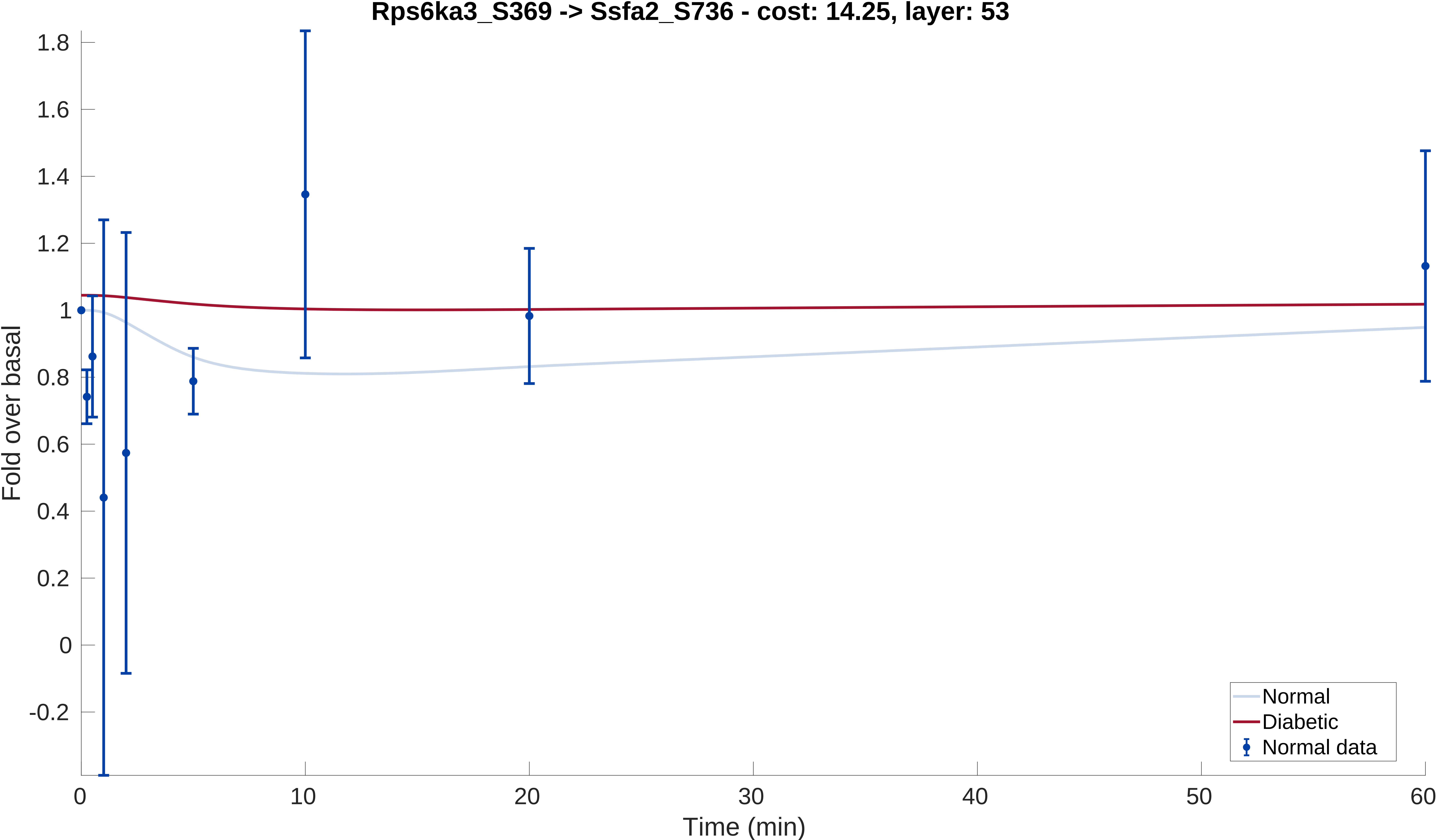
Rps6ka3_S369 -> Ssfa2_S736 - cost: 14.25, layer: 53
Fold over basal
Time (min)
1.8
1.6
1.4
1.2
1
0.8
0.6
0.4
0.2
0
-0.2
0
10
20
30
40
50
60
Normal
Diabetic
Normal data

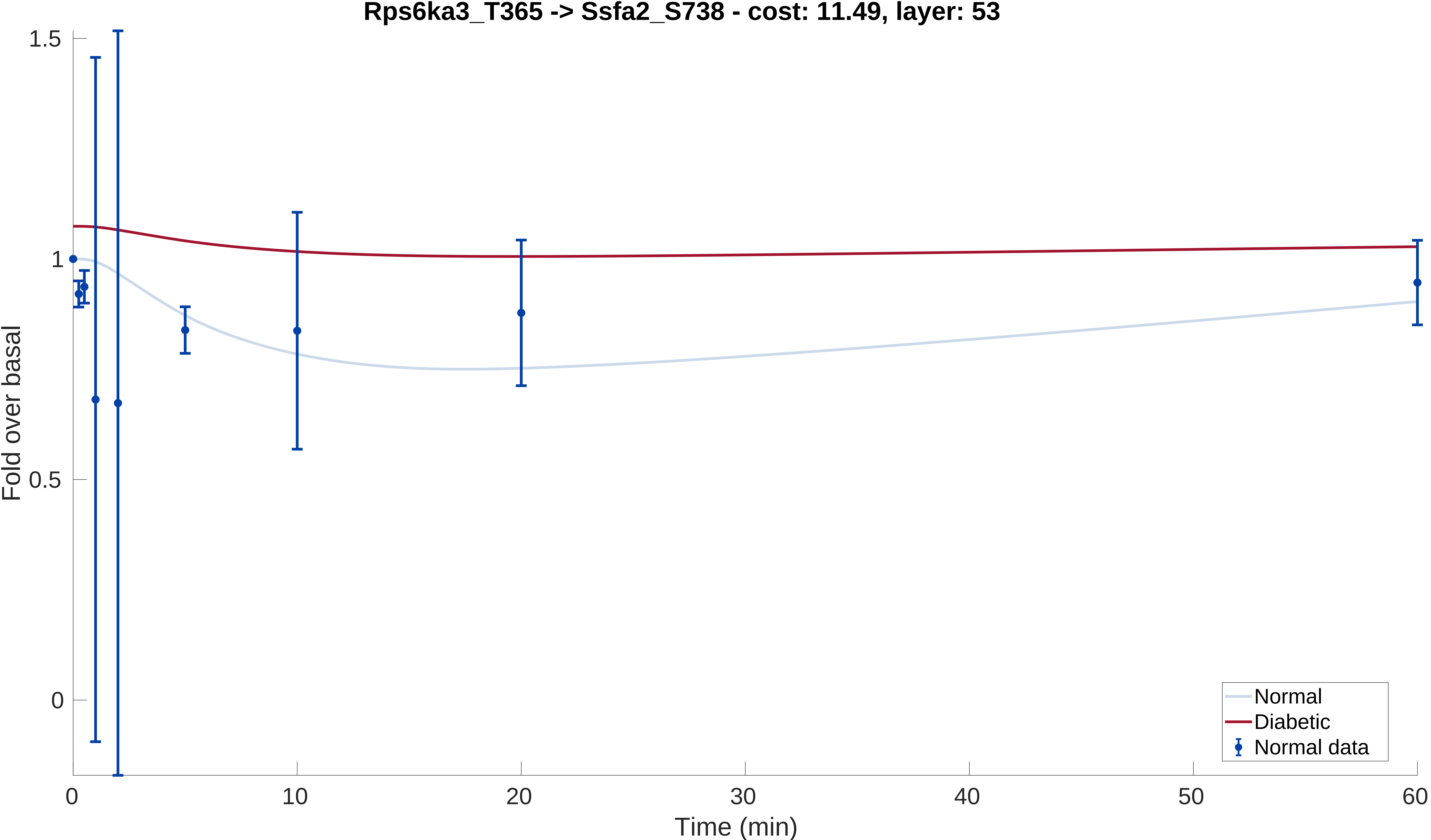

Rps6ka3_T365 -> Ssfa2_S738 - cost: 11.49, layer: 53
1.5
1
0.5
0
Fold over basal
0
10
20
30
40
50
60
Time (min)
Normal
Diabetic
Normal data

#### Rps6ka3_S369 -> Ssfa2_S90 - cost: 3.08, layer: 53

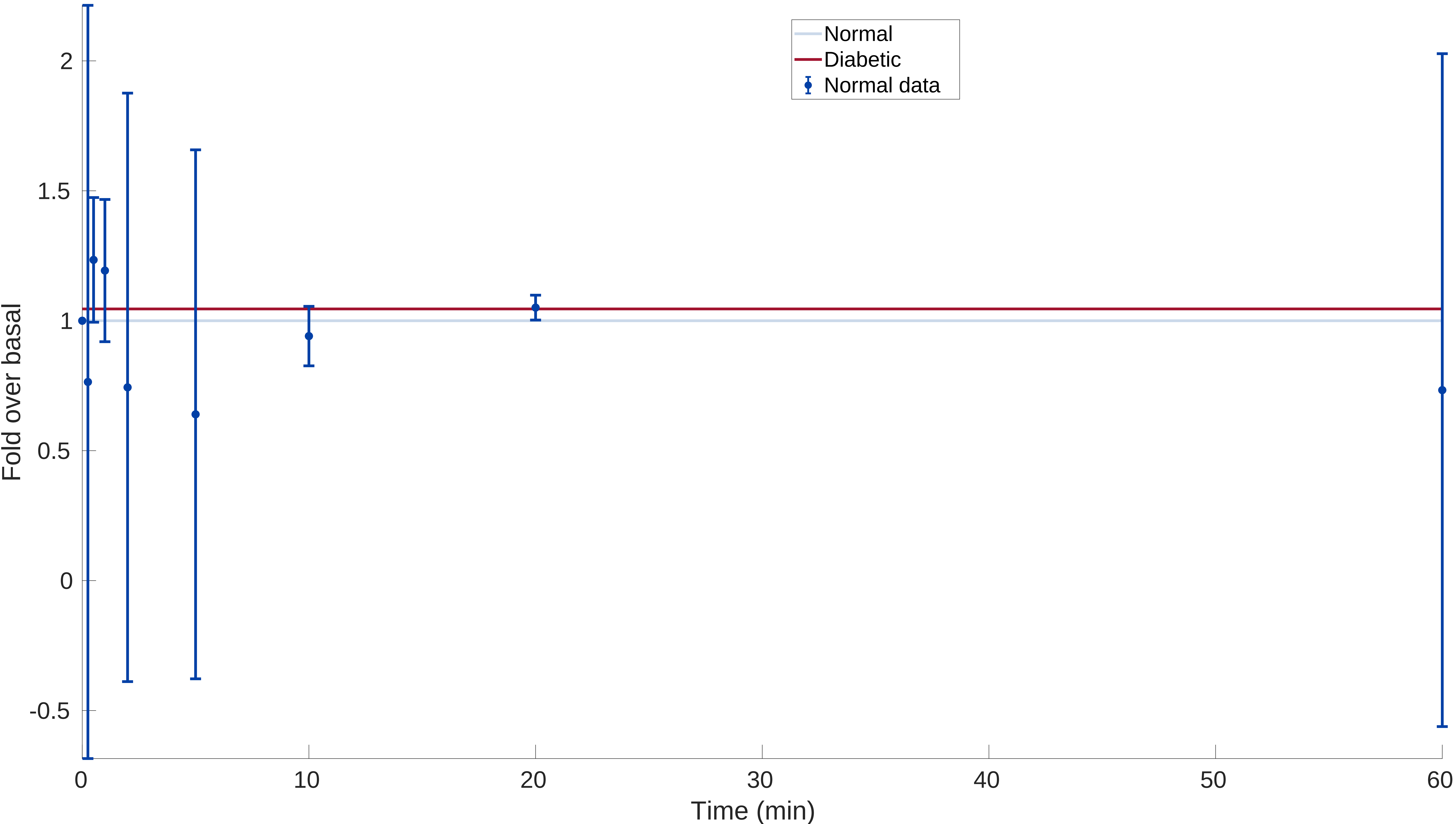

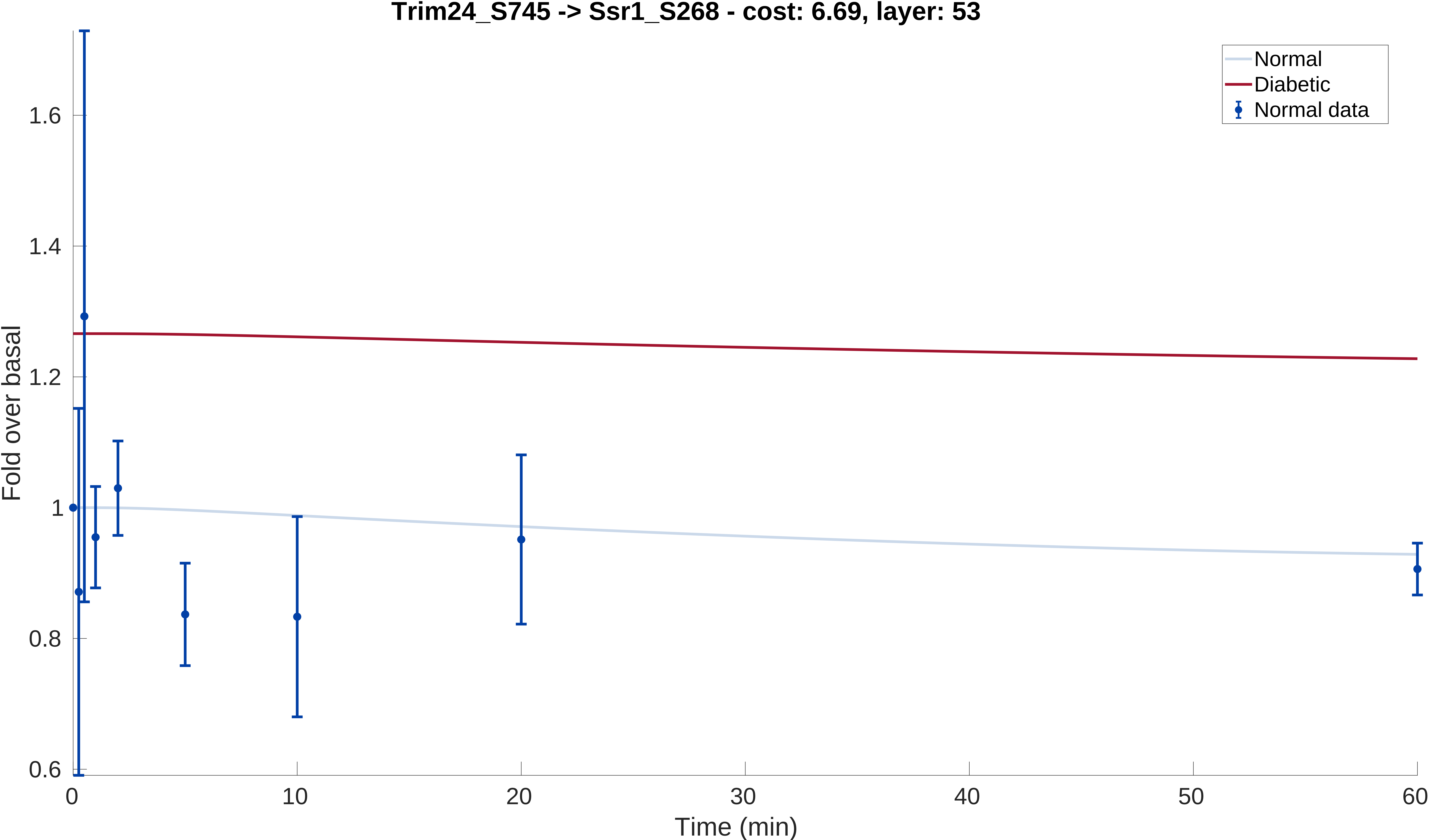

Trim24_S745 -> Ssr1_S268 - cost: 6.69, layer: 53
Normal
Diabetic
Normal data
Fold over basal
1.6
1.4
1.2
1
0.8
0.6
0
10
20
30
40
50
60
Time (min)

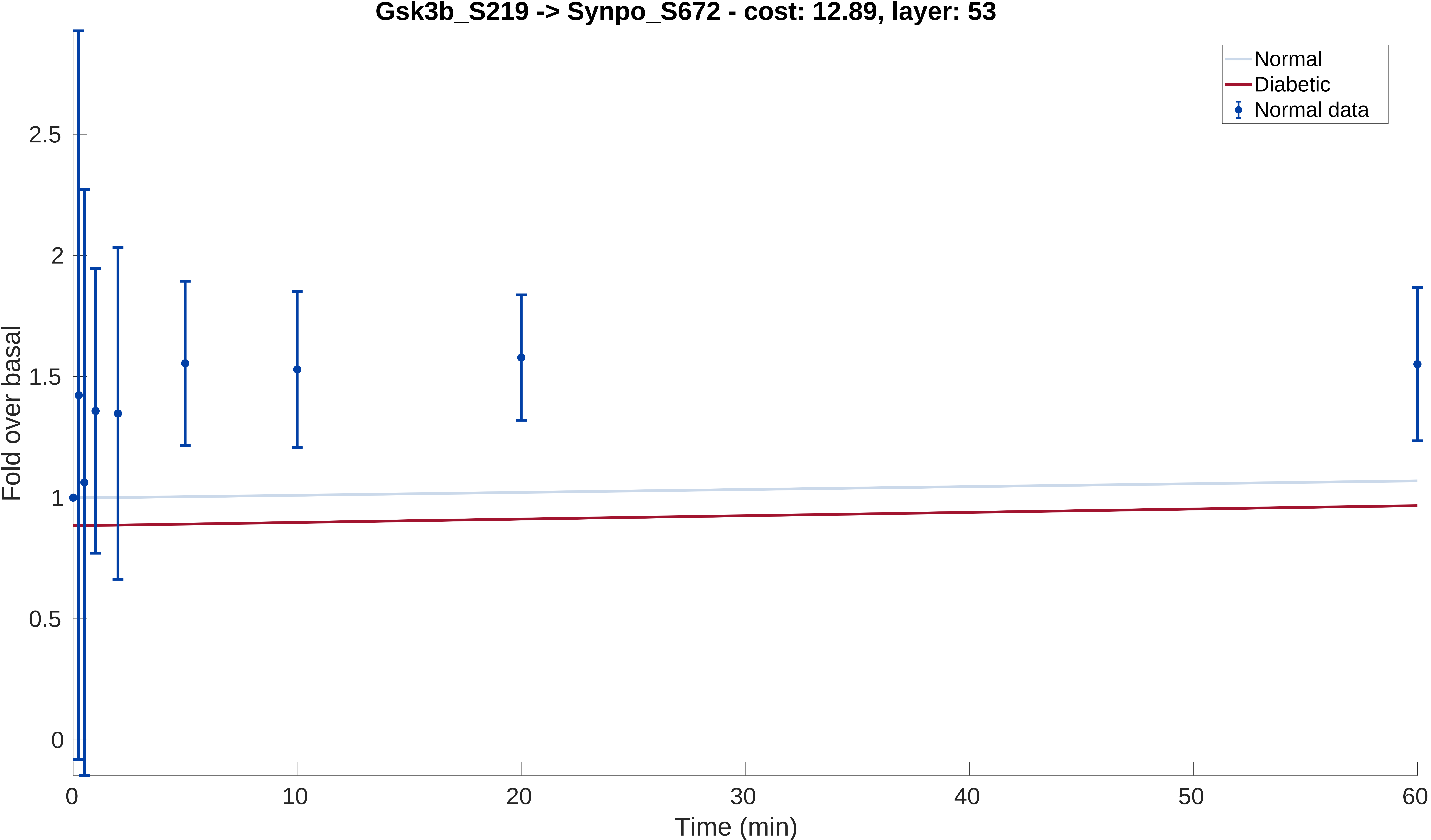

Gsk3b_S219 -> Synpo_S672 - cost: 12.89, layer: 53
Normal
Diabetic
Normal data
Fold over basal
0
0.5
1
1.5
2
2.5
0
10
20
30
40
50
60
Time (min)

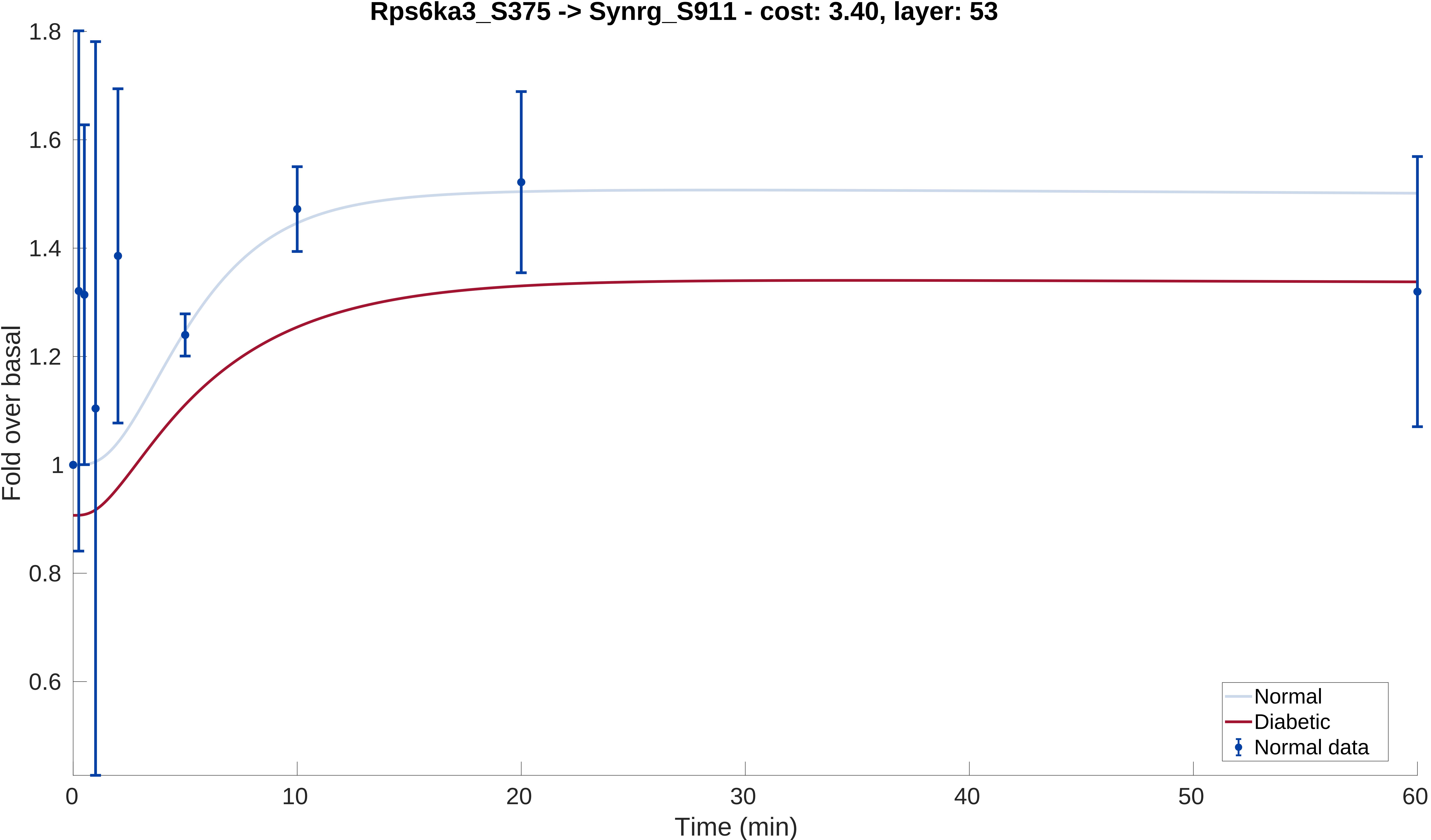
Rps6ka3_S375 -> Synrg_S911 - cost: 3.40, layer: 53
Fold over basal
1.8
1.6
1.4
1.2
1
0.8
0.6
0
10
20
30
40
50
60
Time (min)
Normal
Diabetic
Normal data

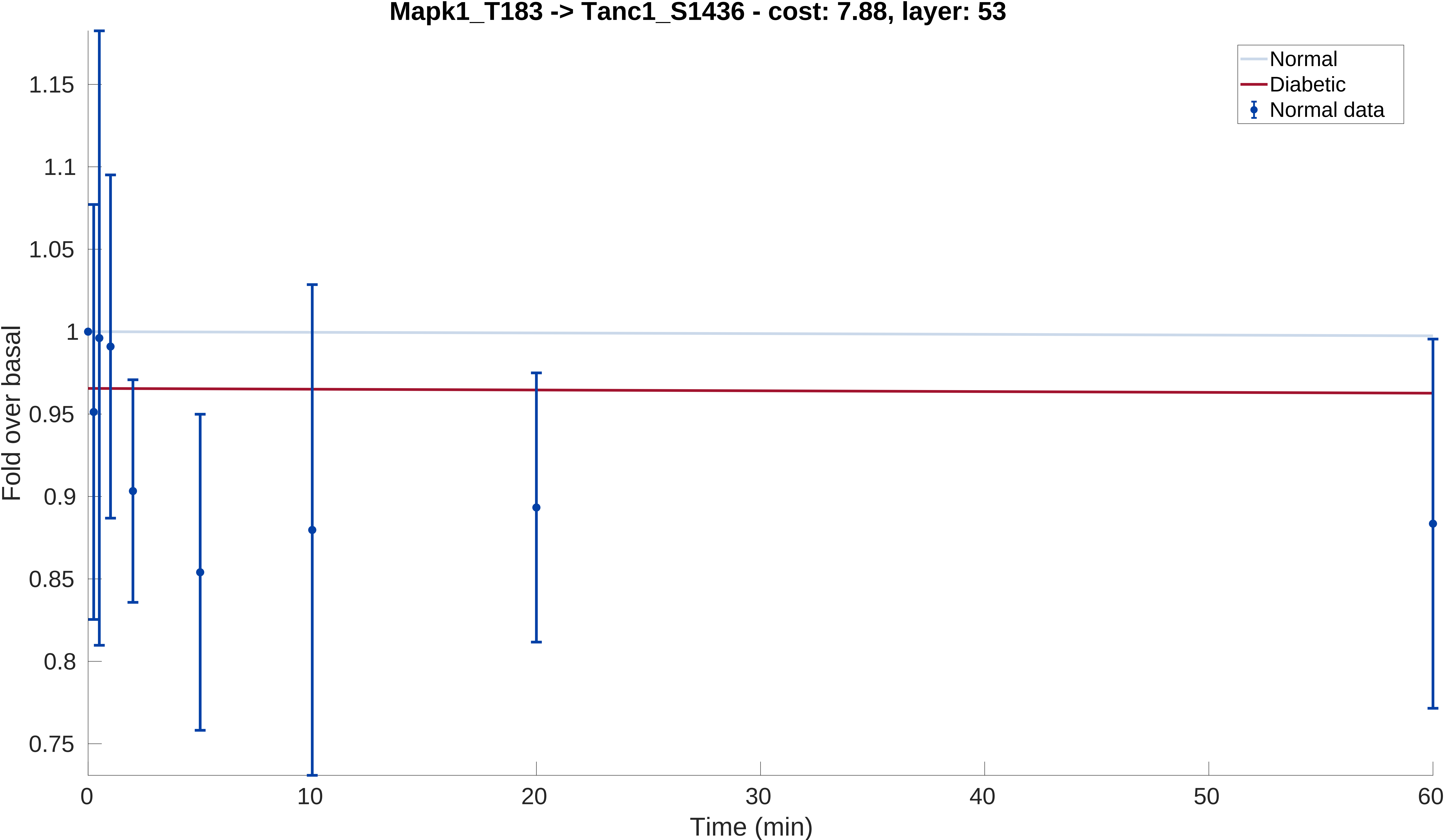

Mapk1_T183 -> Tanc1_S1436 - cost: 7.88, layer: 53
Normal
Diabetic
Normal data
Fold over basal
Time (min)
1.15
1.1
1.05
1
0.95
0.9
0.85
0.8
0.75
0
10
20
30
40
50
60

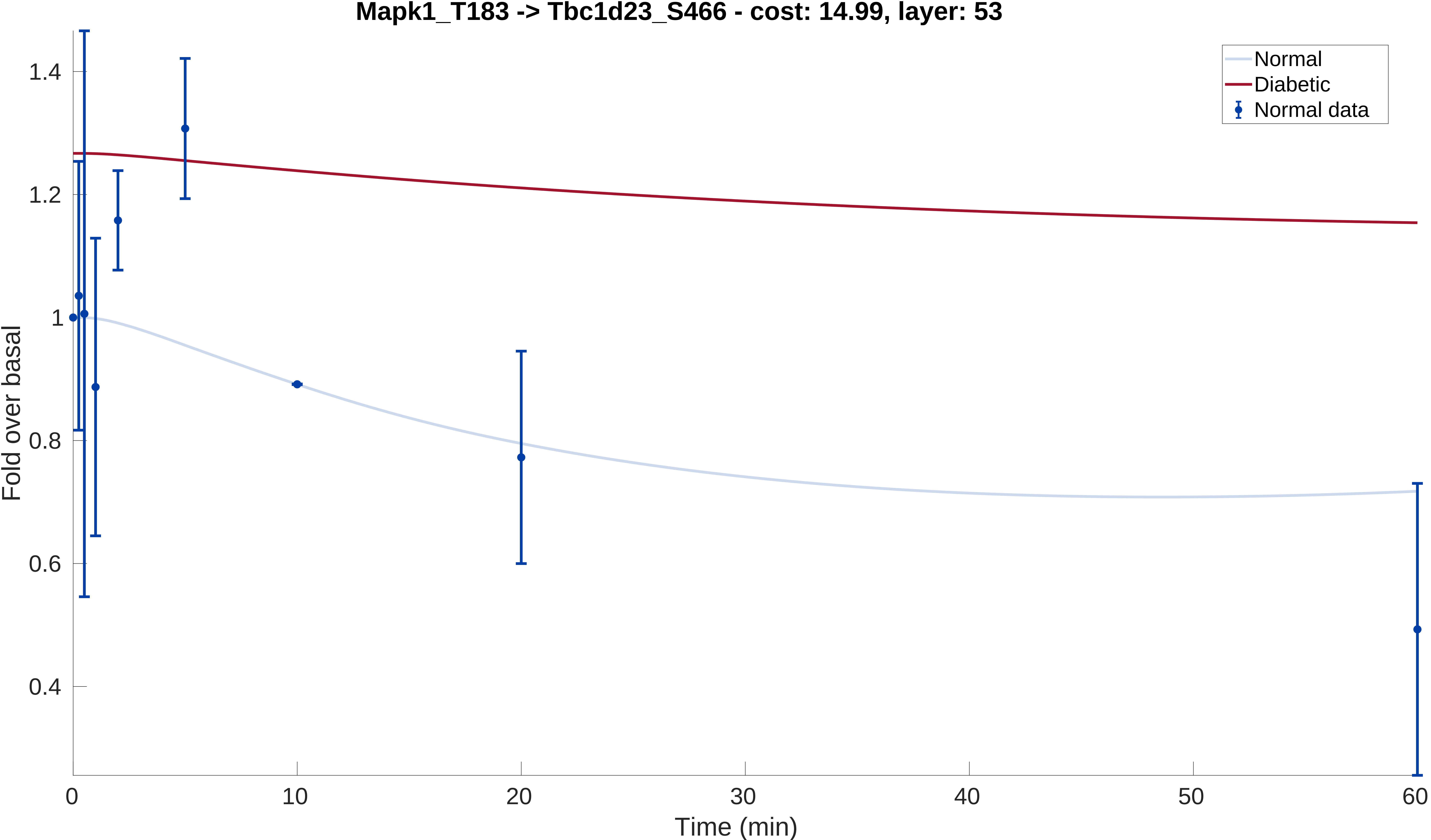
Mapk1_T183 -> Tbc1d23_S466 - cost: 14.99, layer: 53
Normal
Diabetic
Normal data
Fold over basal
1.4
1.2
1
0.8
0.6
0.4
0
10
20
30
40
50
60
Time (min)

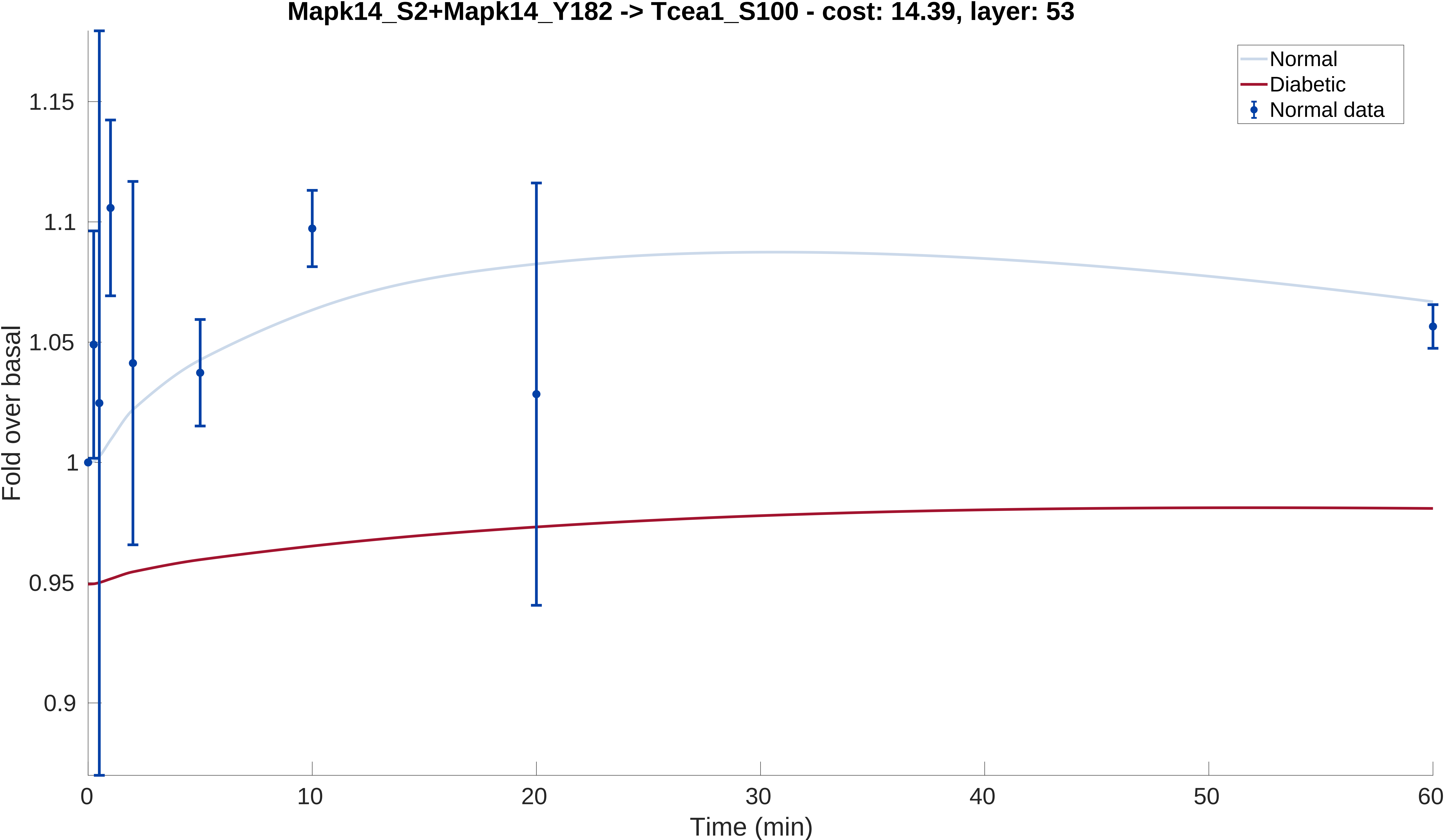

Mapk14_S2+Mapk14_Y182 -> Tcea1_S100 - cost: 14.39, layer: 53
Normal
Diabetic
Normal data
Fold over basal
1.15
1.1
1.05
1
0.95
0.9
0
10
20
30
40
50
60
Time (min)

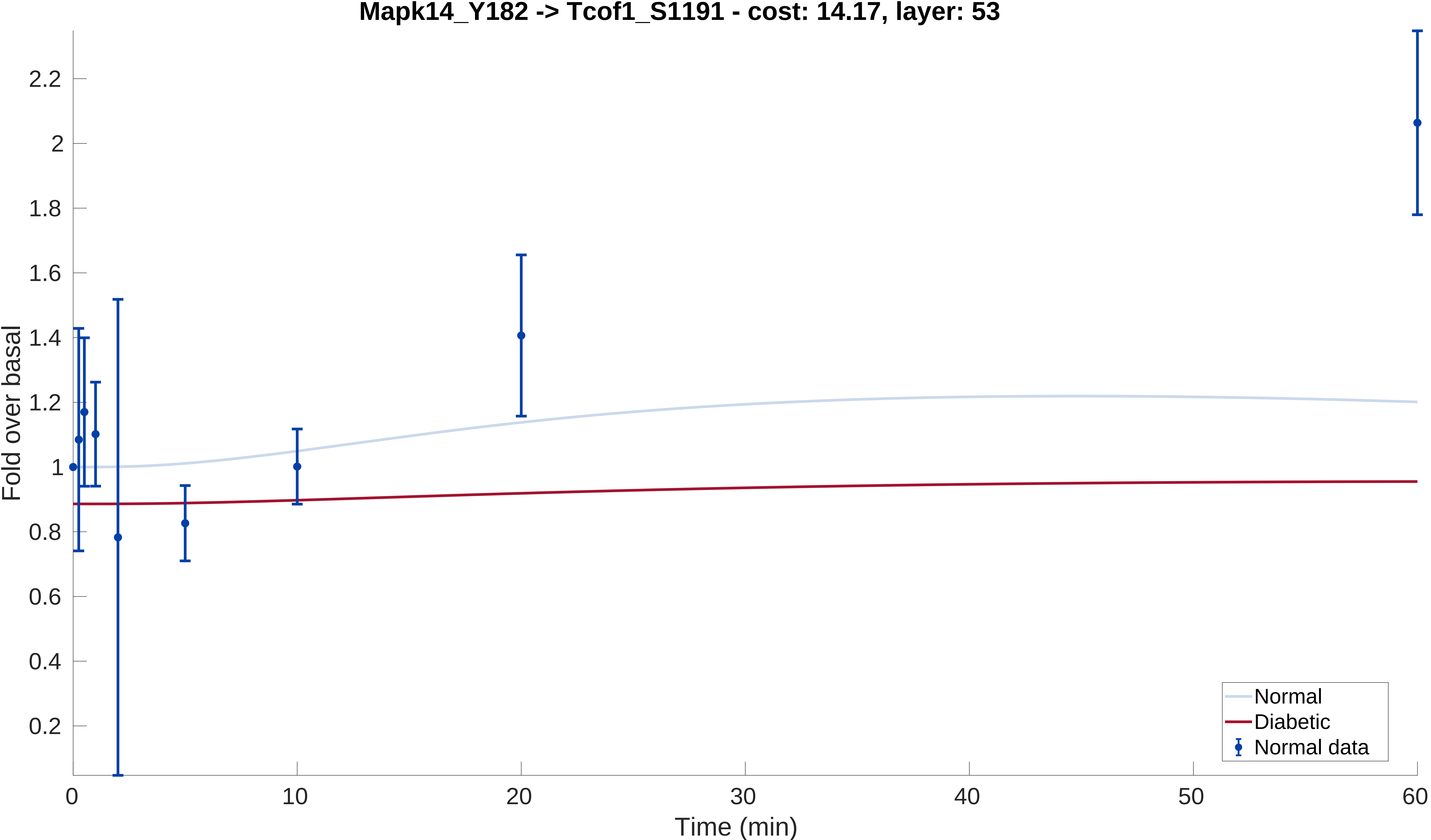

Mapk14_Y182 -> Tcof1_S1191 - cost: 14.17, layer: 53
Fold over basal
2.2
2
1.8
1.6
1.4
1.2
1
0.8
0.6
0.4
0.2
0
10
20
30
40
50
60
Time (min)
Normal
Diabetic
Normal data

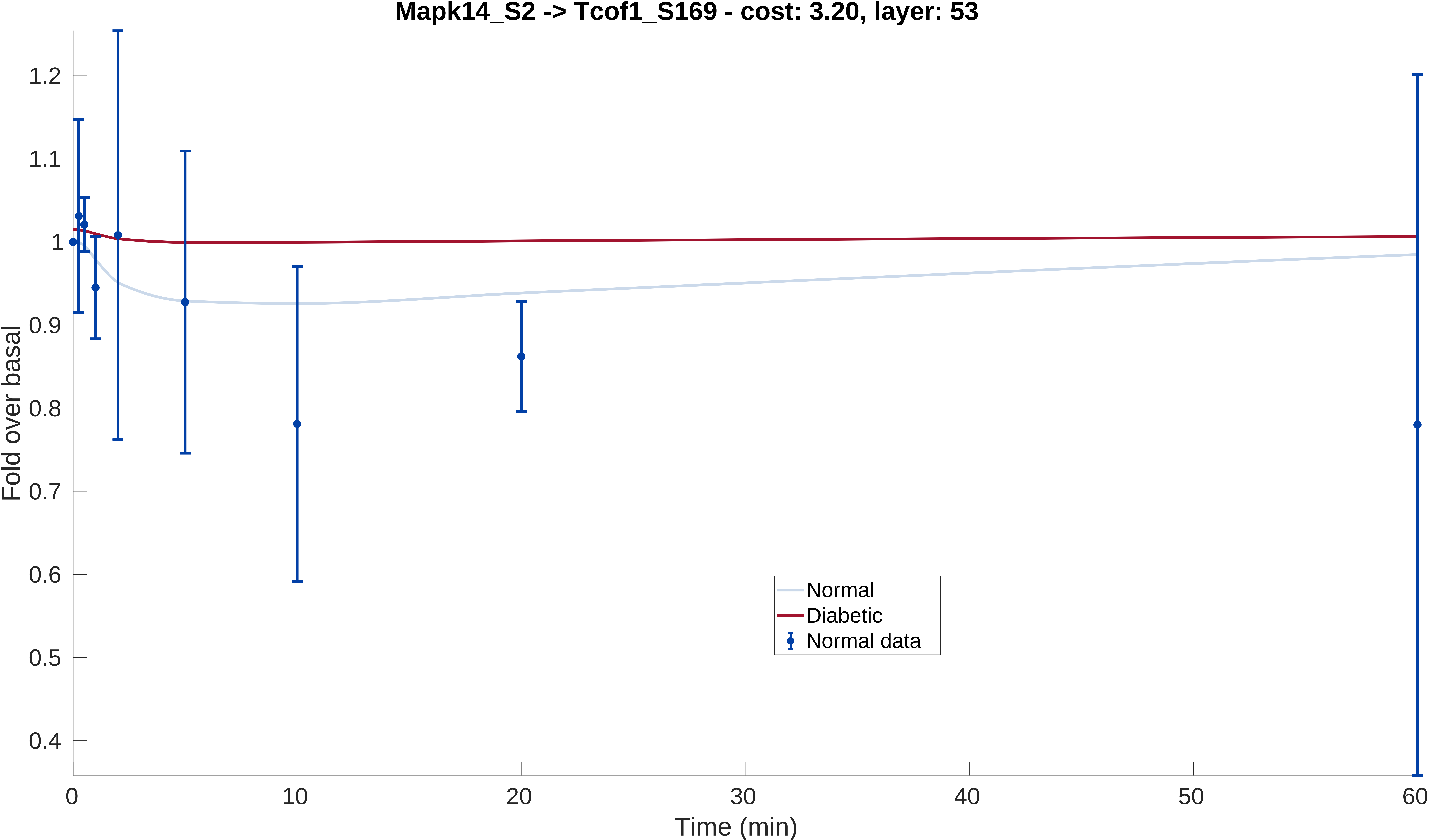
Mapk14_S2 -> Tcof1_S169 - cost: 3.20, layer: 53
Fold over basal
1.2
1.1
1
0.9
0.8
0.7
0.6
0.5
0.4
0
10
20
30
40
50
60
Time (min)
Normal
Diabetic
Normal data

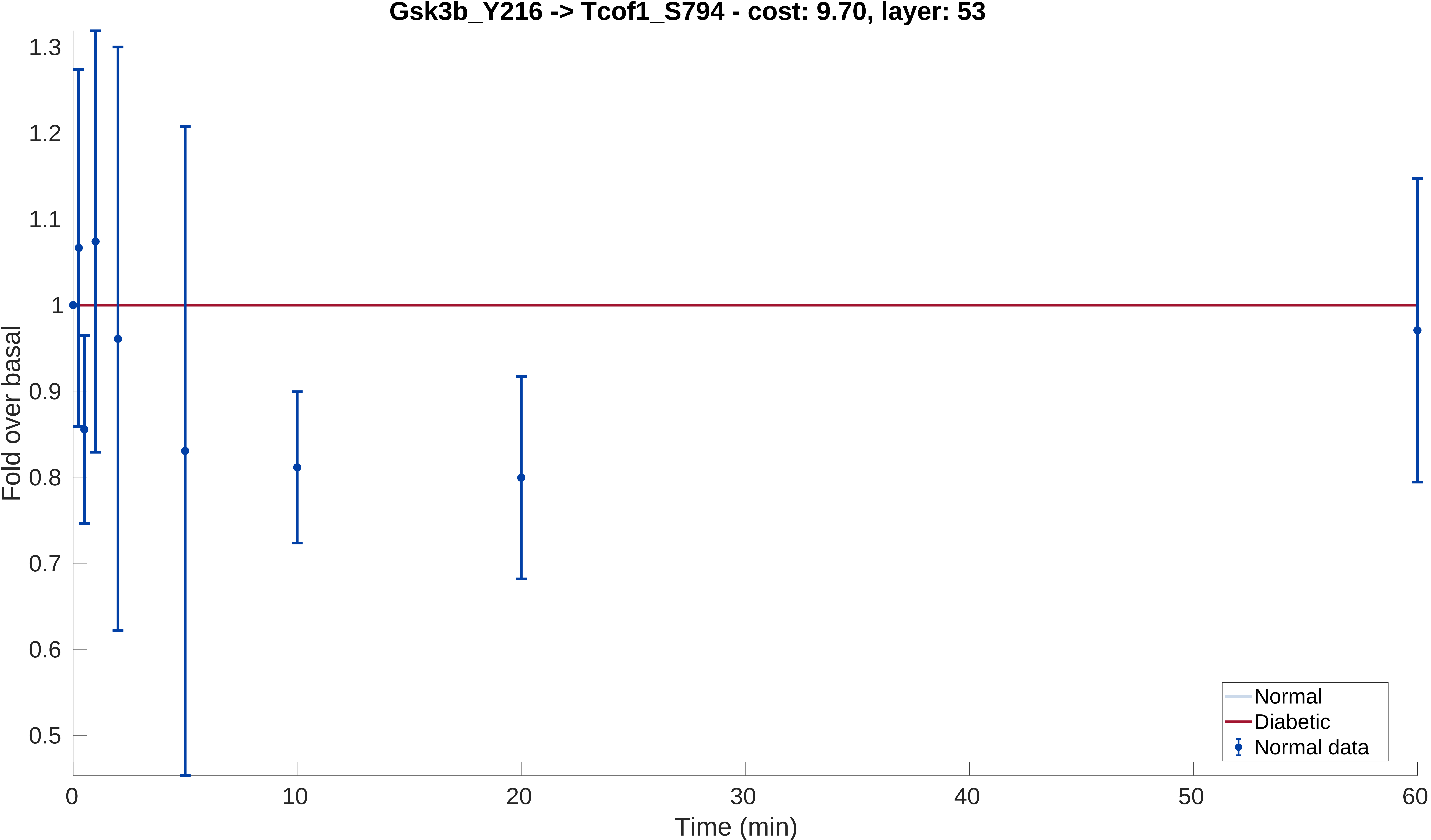

Gsk3b_Y216 -> Tcof1_S794 - cost: 9.70, layer: 53
Fold over basal
Time (min)
1.3
1.2
1.1
1
0.9
0.8
0.7
0.6
0.5
0
10
20
30
40
50
60
Normal
Diabetic
Normal data

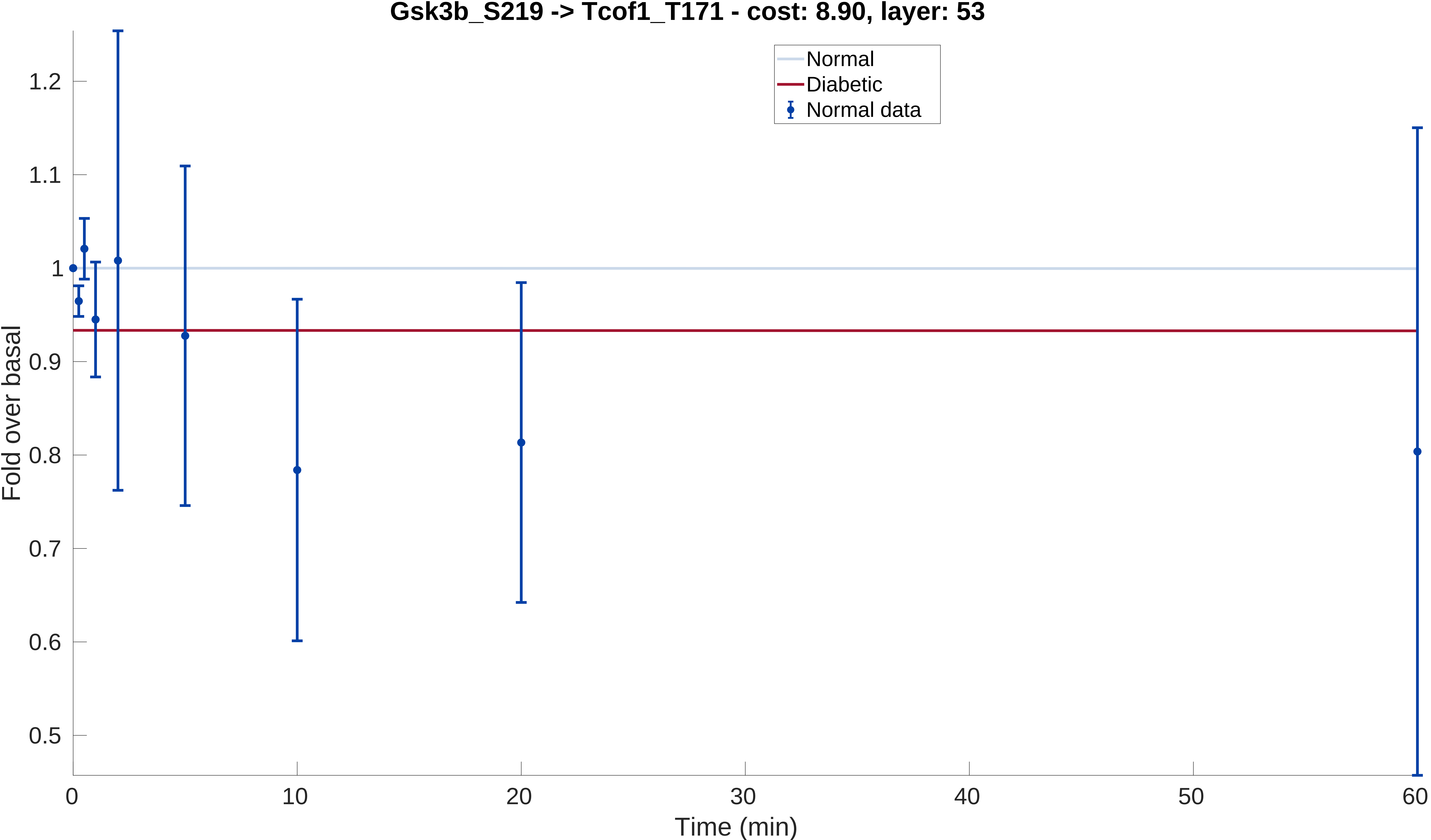

Gsk3b_S219 -> Tcof1_T171 - cost: 8.90, layer: 53
Normal
Diabetic
Normal data
Fold over basal
1.2
1.1
1
0.9
0.8
0.7
0.6
0.5
0
10
20
30
40
50
60
Time (min)

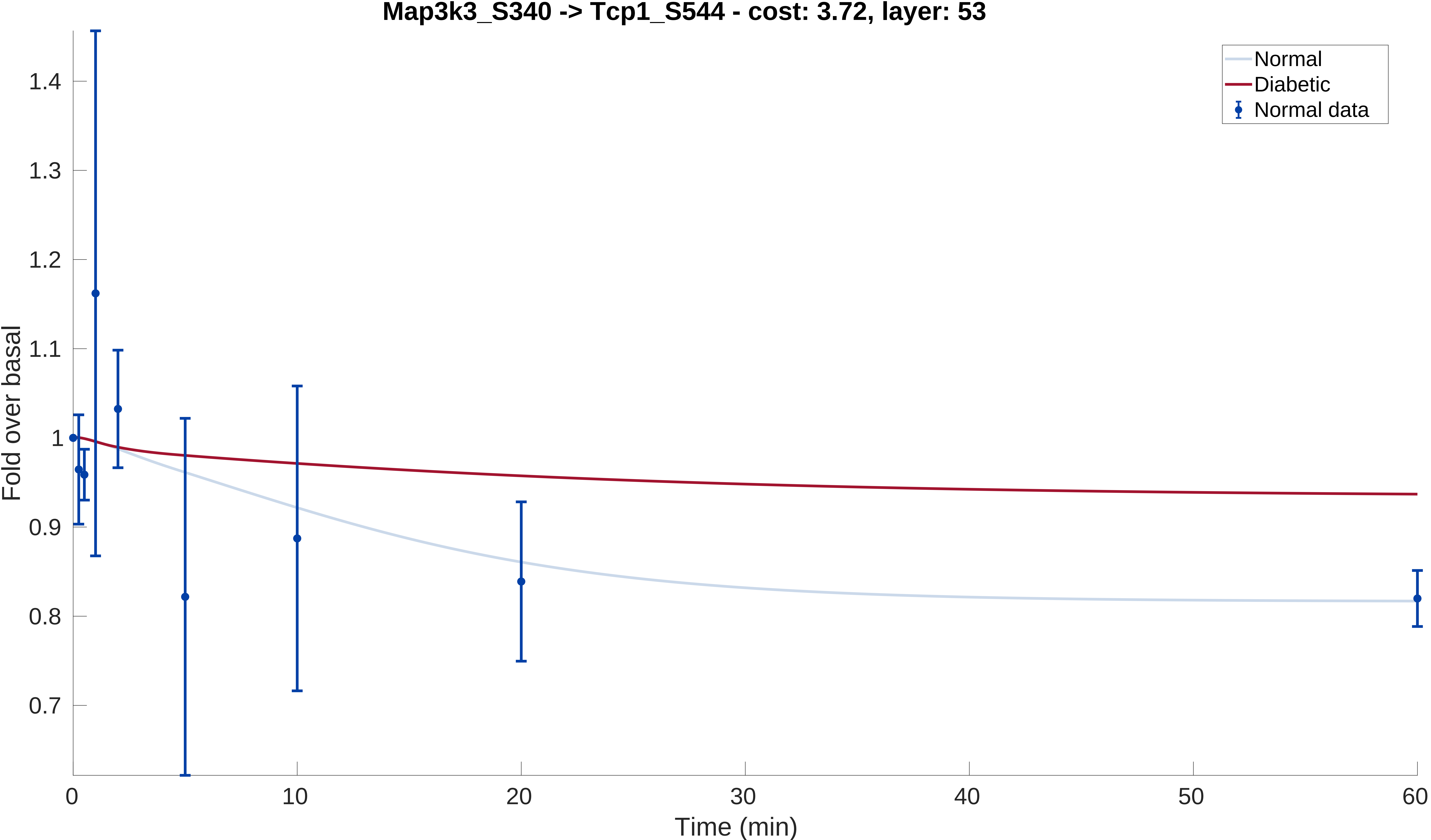

Map3k3_S340 -> Tcp1_S544 - cost: 3.72, layer: 53
Normal
Diabetic
Normal data
Fold over basal
1.4
1.3
1.2
1.1
1
0.9
0.8
0.7
0
10
20
30
40
50
60
Time (min)

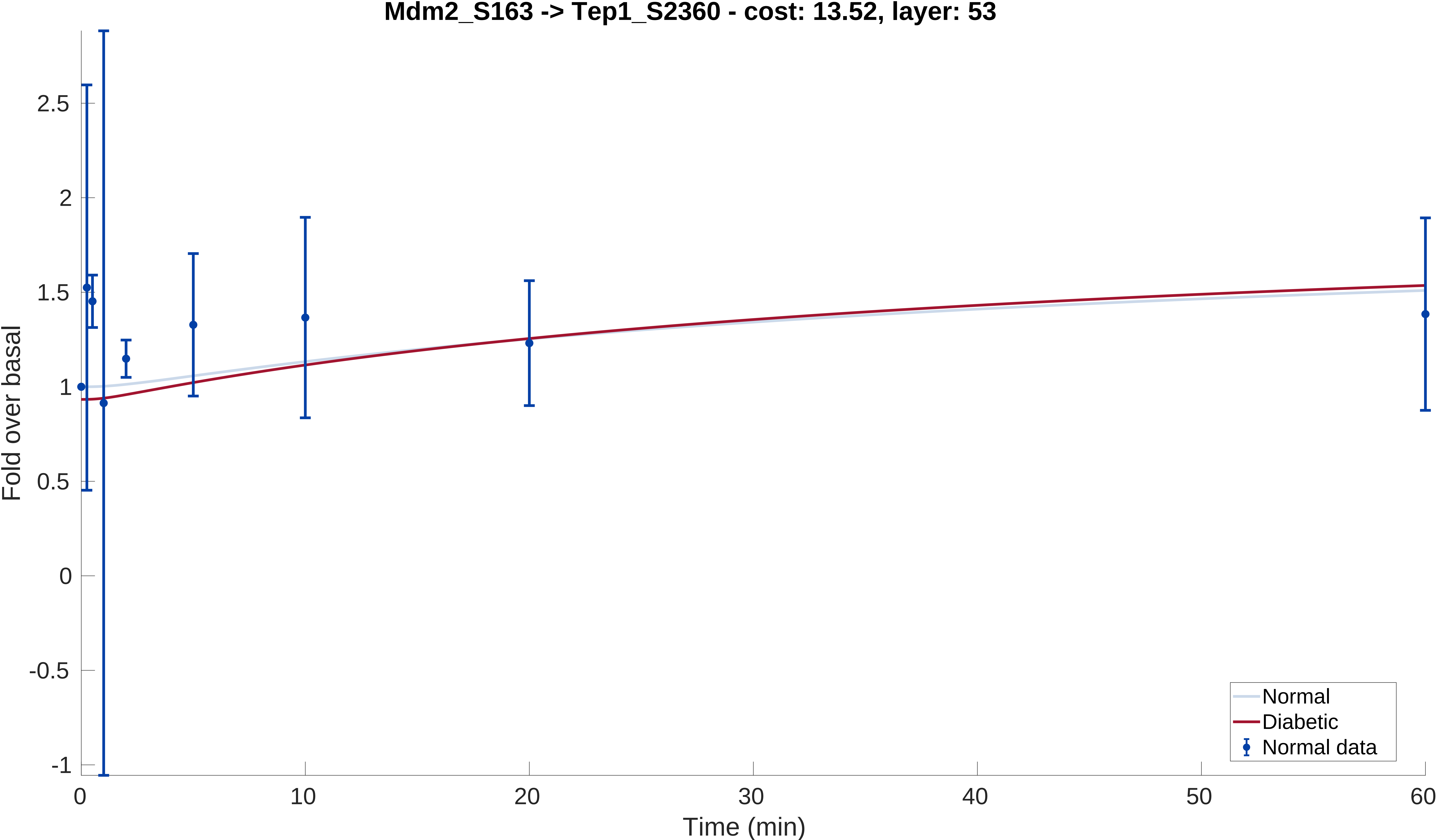

Mdm2_S163 -> Tep1_S2360 - cost: 13.52, layer: 53
Fold over basal
Time (min)
2.5
2
1.5
1
0.5
0
-0.5
-1
0
10
20
30
40
50
60
Normal
Diabetic
Normal data

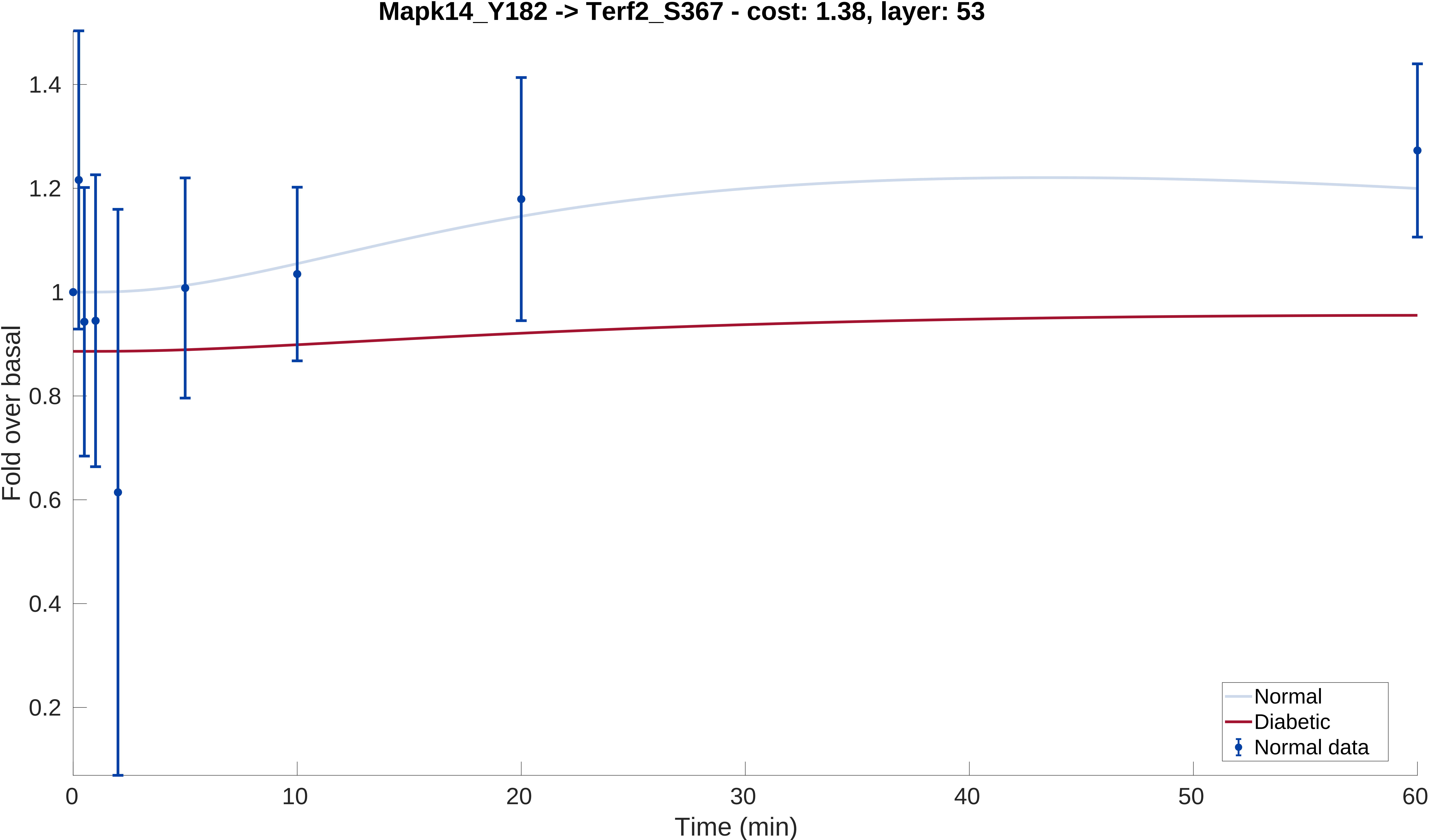

Mapk14_Y182 -> Terf2_S367 - cost: 1.38, layer: 53
Fold over basal
1.4
1.2
1
0.8
0.6
0.4
0.2
0
10
20
30
40
50
60
Time (min)
Normal
Diabetic
Normal data

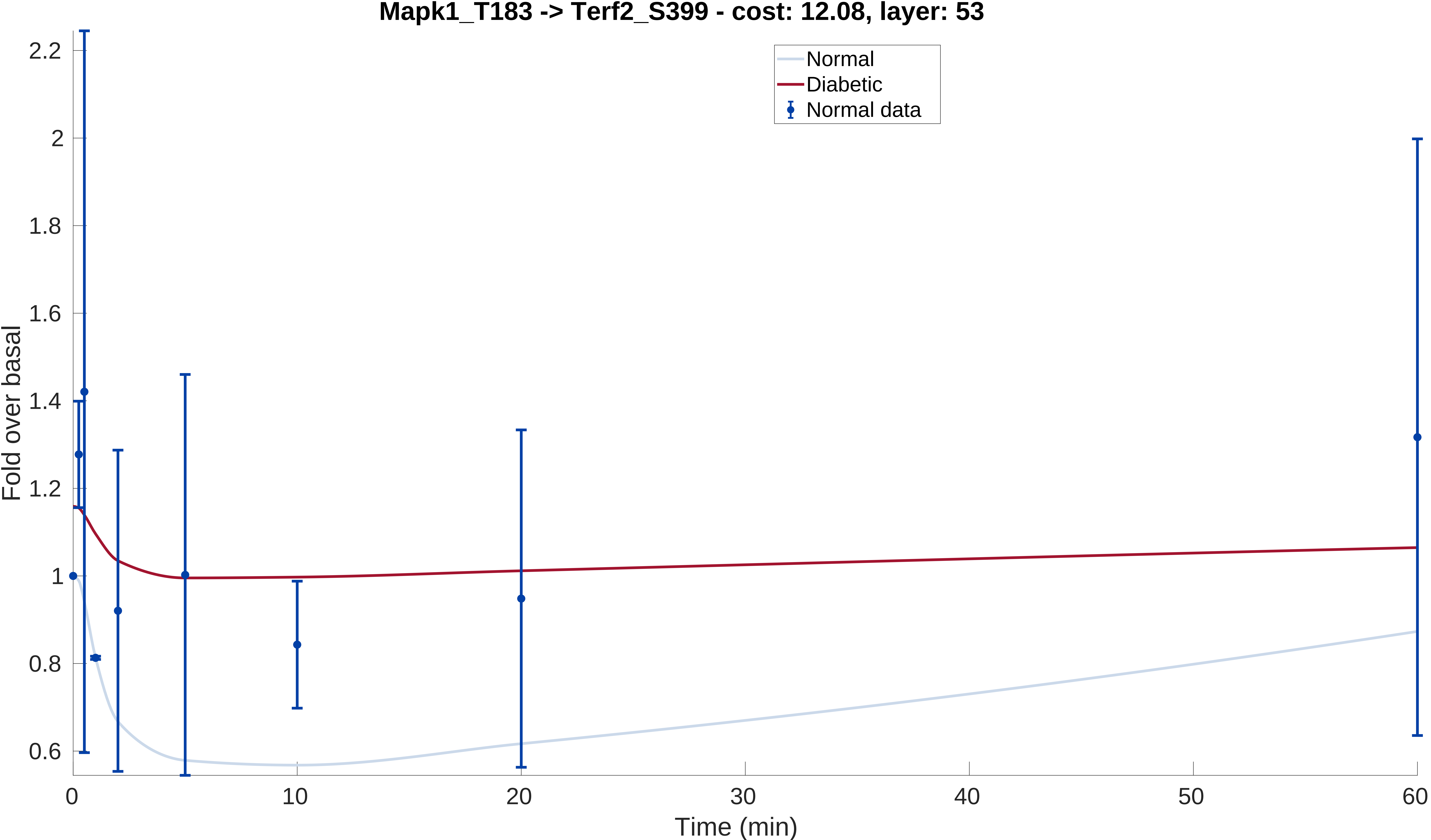

Mapk1_T183 -> Terf2_S399 - cost: 12.08, layer: 53
Normal
Diabetic
Normal data
Fold over basal
2.2
2
1.8
1.6
1.4
1.2
1
0.8
0.6
0
10
20
30
40
50
60
Time (min)

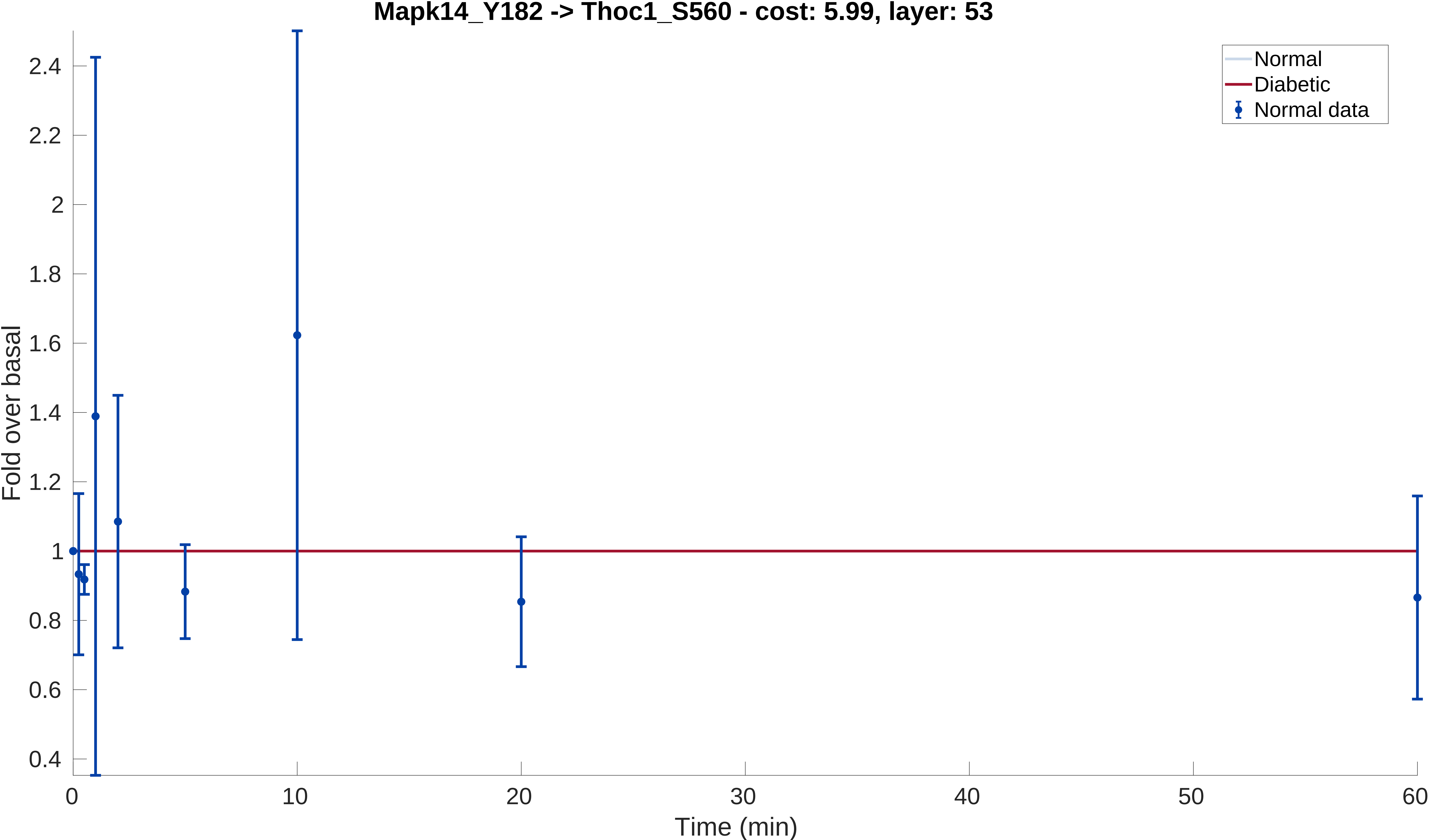

Mapk14_Y182 -> Thoc1_S560 - cost: 5.99, layer: 53
Normal
Diabetic
Normal data
Fold over basal
2.4
2.2
2
1.8
1.6
1.4
1.2
1
0.8
0.6
0.4
0
10
20
30
40
50
60
Time (min)

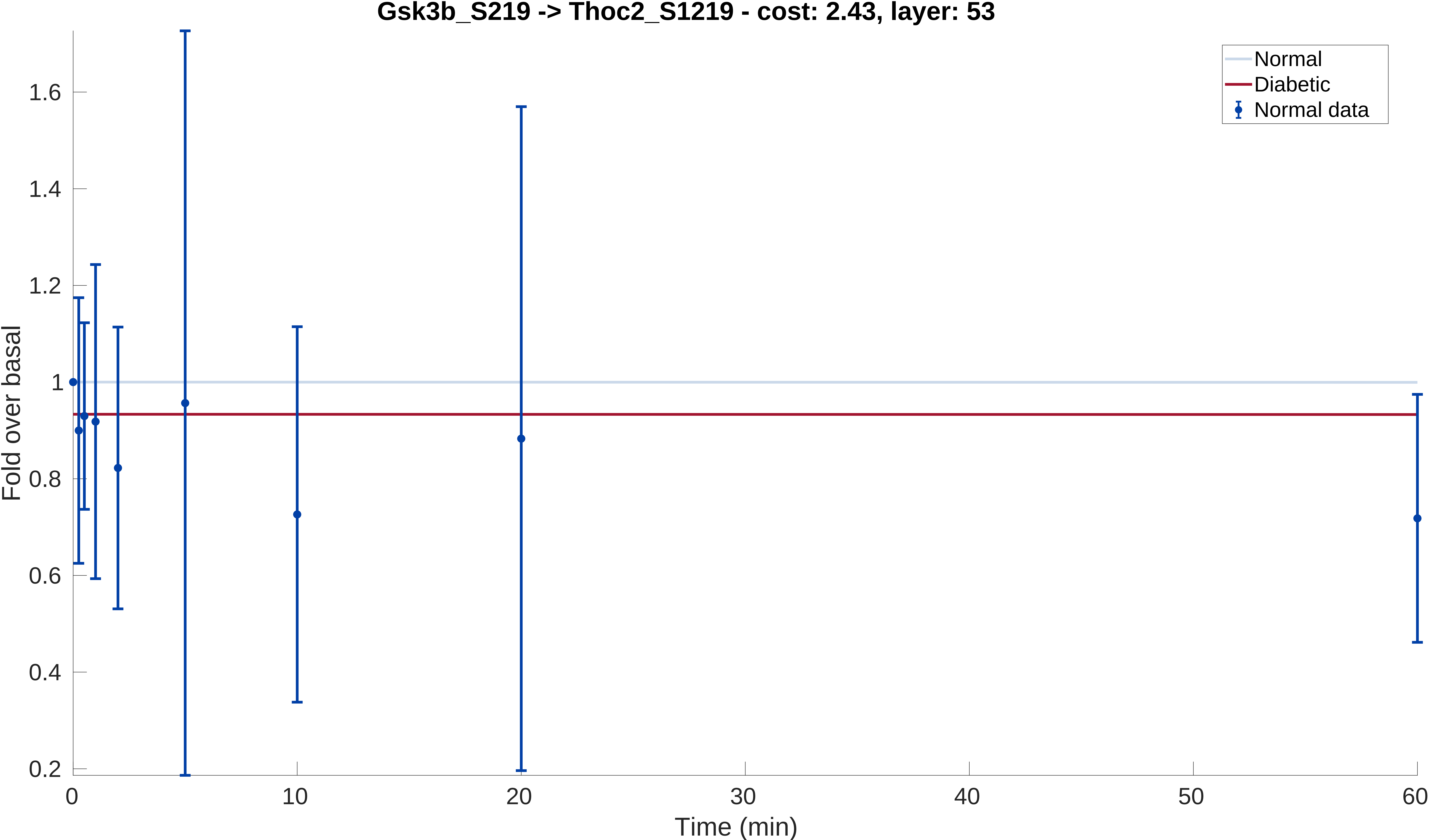

Gsk3b_S219 -> Thoc2_S1219 - cost: 2.43, layer: 53
Normal
Diabetic
Normal data
Fold over basal
1.6
1.4
1.2
1
0.8
0.6
0.4
0.2
0
10
20
30
40
50
60
Time (min)

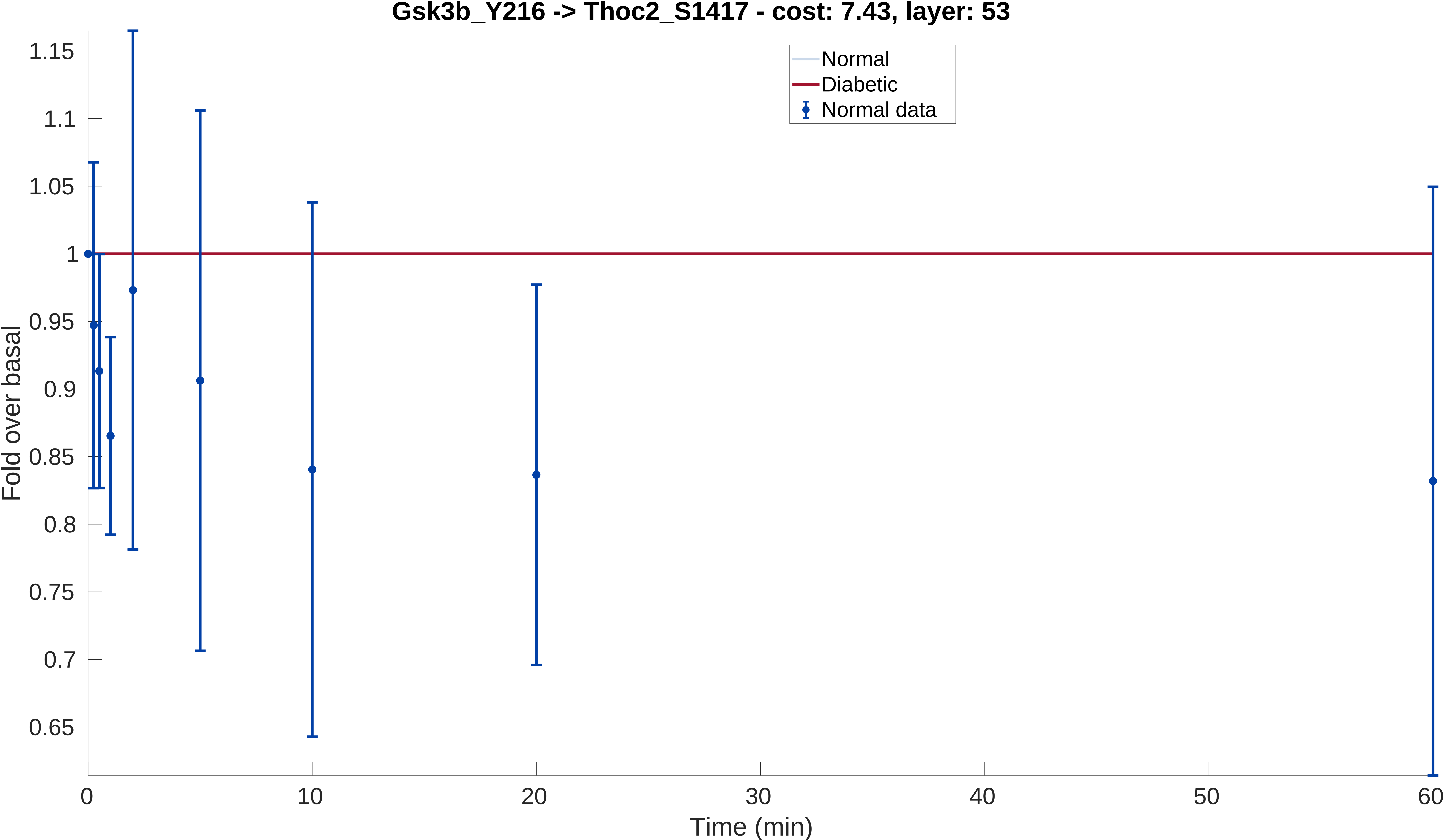

Gsk3b_Y216 -> Thoc2_S1417 - cost: 7.43, layer: 53
Normal
Diabetic
Normal data
Fold over basal
Time (min)
1.15
1.1
1.05
1
0.95
0.9
0.85
0.8
0.75
0.7
0.65
0
10
20
30
40
50
60

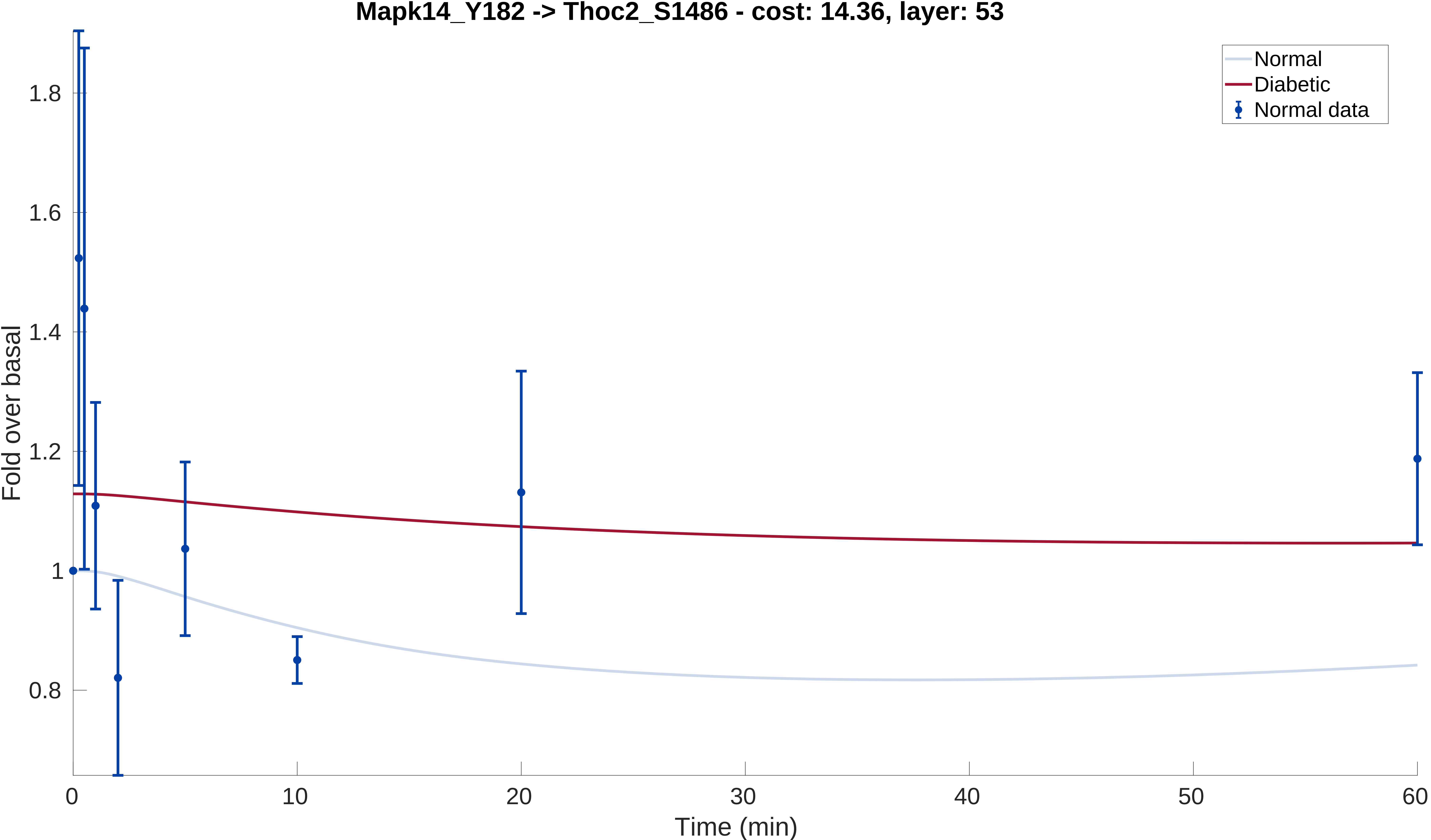

Mapk14_Y182 -> Thoc2_S1486 - cost: 14.36, layer: 53
Normal
Diabetic
Normal data
Fold over basal
1.8
1.6
1.4
1.2
1
0.8
0
10
20
30
40
50
60
Time (min)

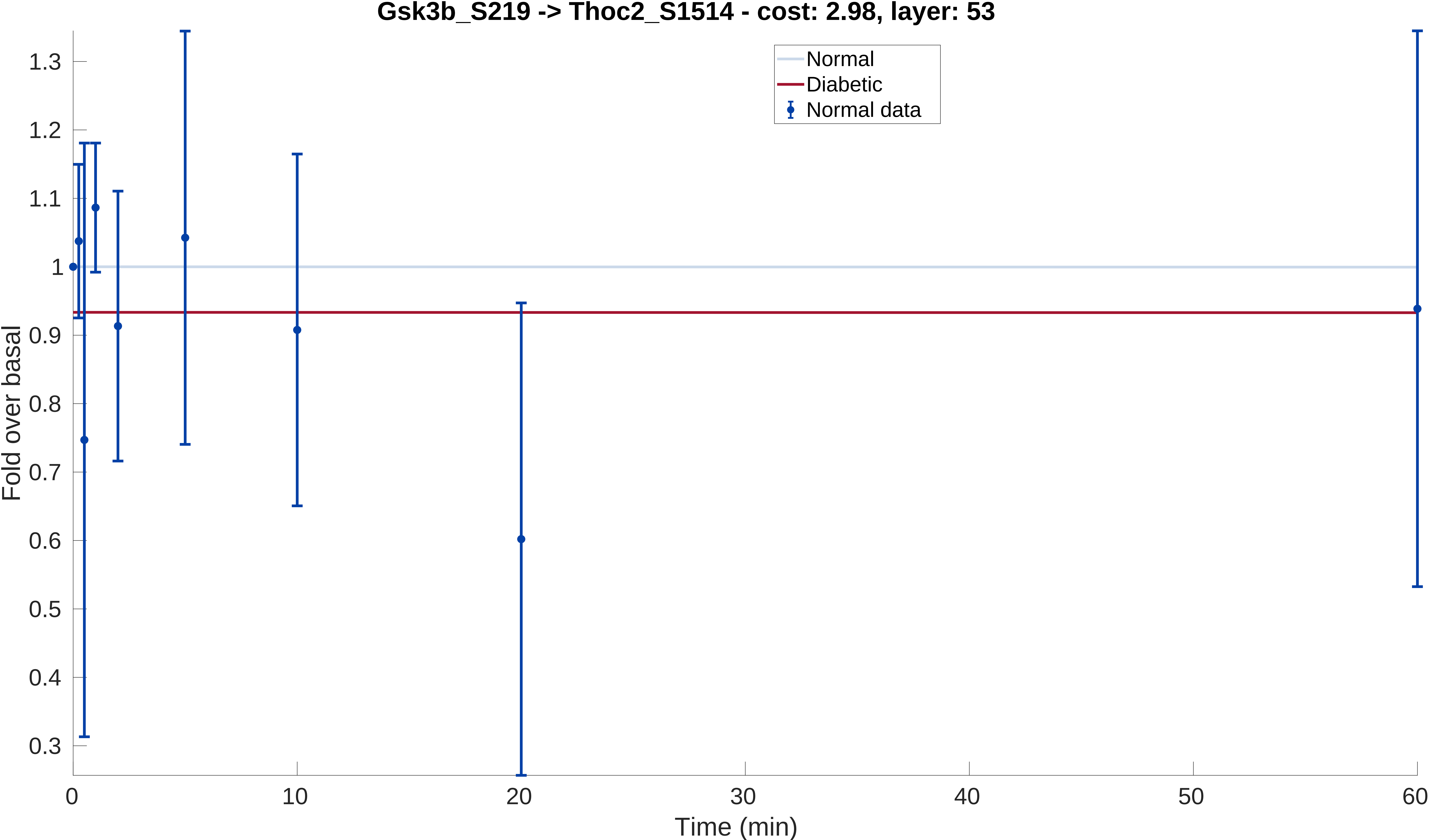

Gsk3b_S219 -> Thoc2_S1514 - cost: 2.98, layer: 53
Normal
Diabetic
Normal data
Fold over basal
1.3
1.2
1.1
1
0.9
0.8
0.7
0.6
0.5
0.4
0.3
0
10
20
30
40
50
60
Time (min)

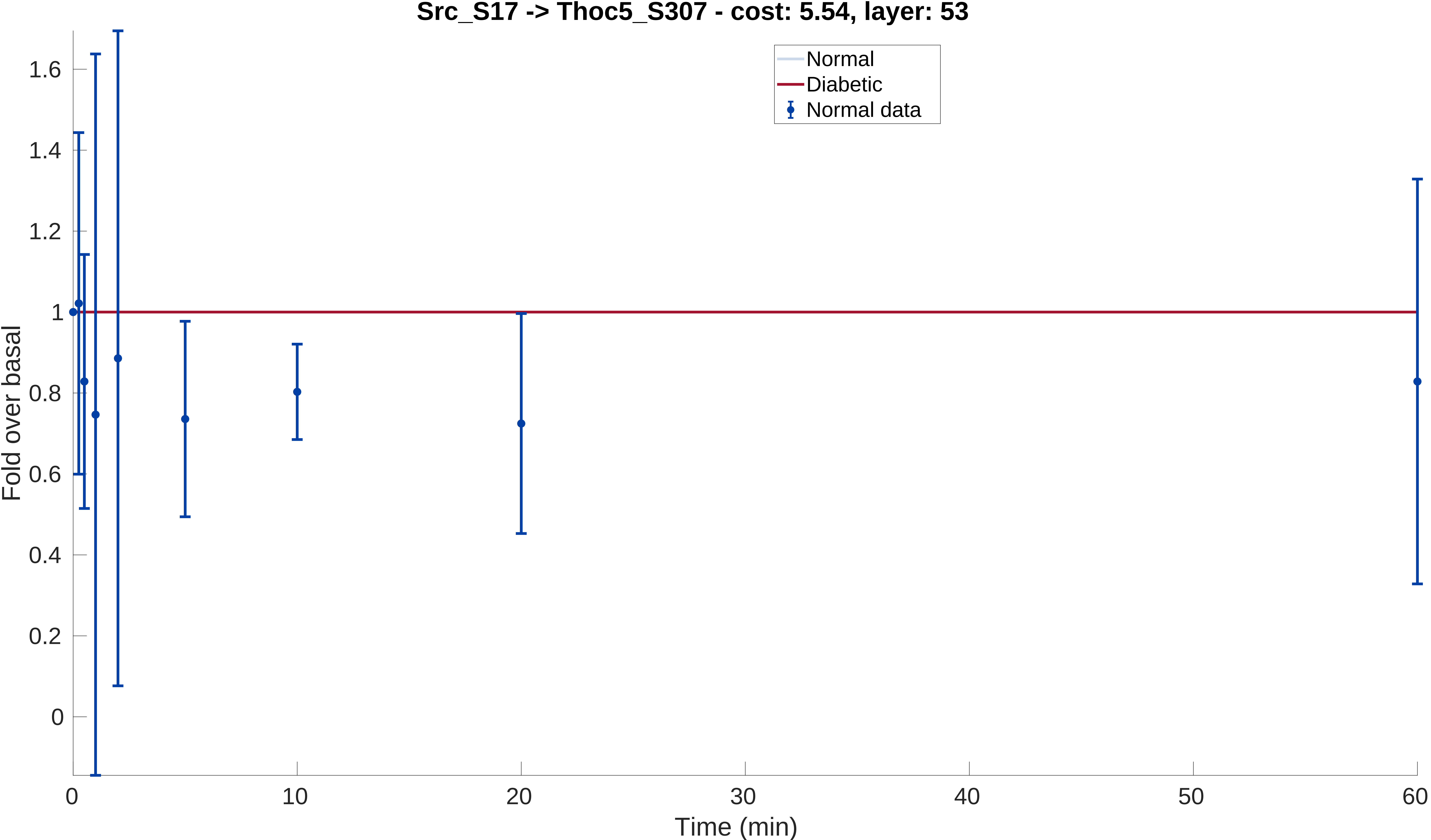

Src_S17 -> Thoc5_S307 - cost: 5.54, layer: 53
Normal
Diabetic
Normal data
Fold over basal
1.6
1.4
1.2
1
0.8
0.6
0.4
0.2
0
0
10
20
30
40
50
60
Time (min)

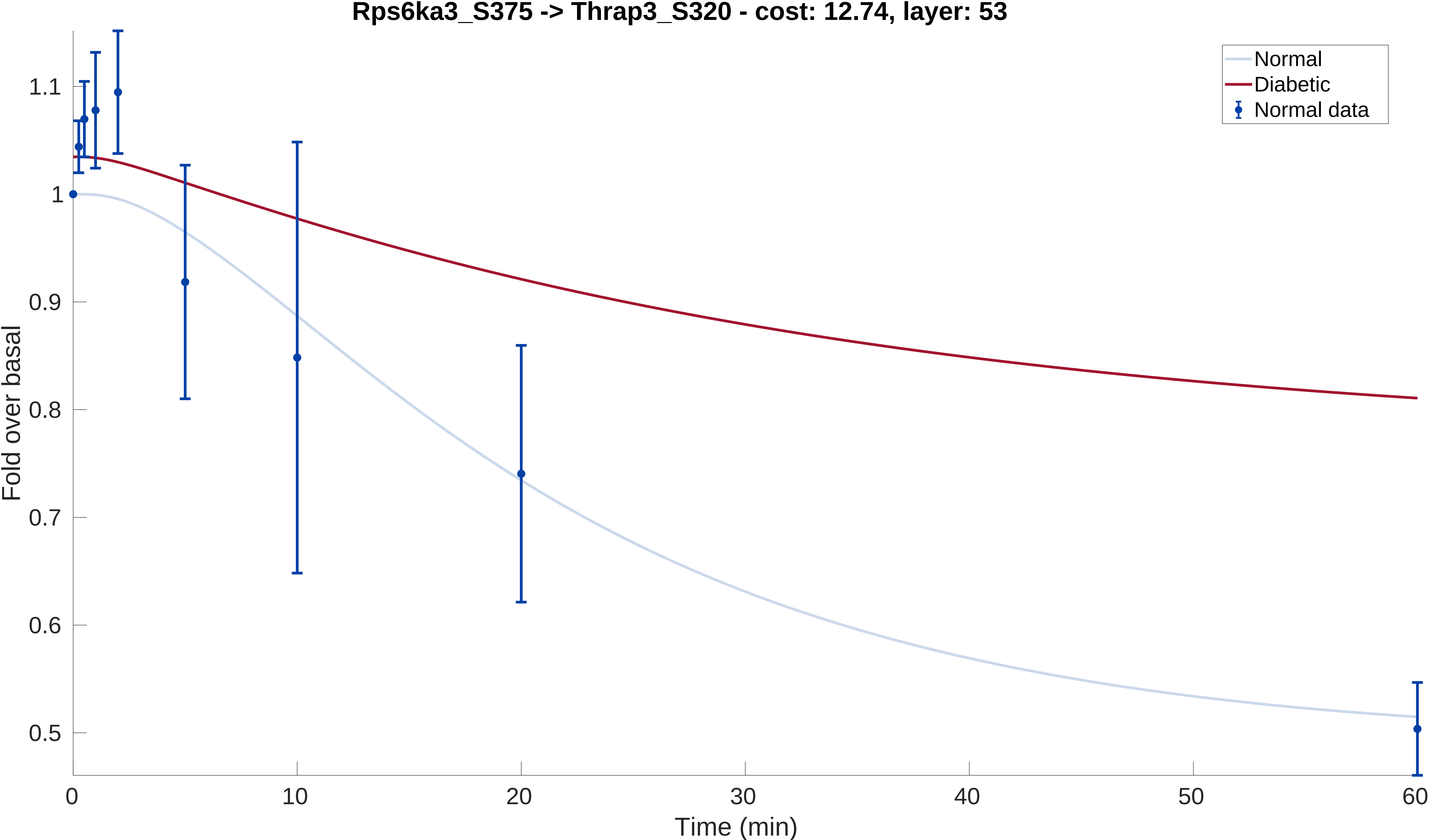

Rps6ka3_S375 -> Thrap3_S320 - cost: 12.74, layer: 53
Normal
Diabetic
Normal data
Fold over basal
1.1
1
0.9
0.8
0.7
0.6
0.5
0
10
20
30
40
50
60
Time (min)

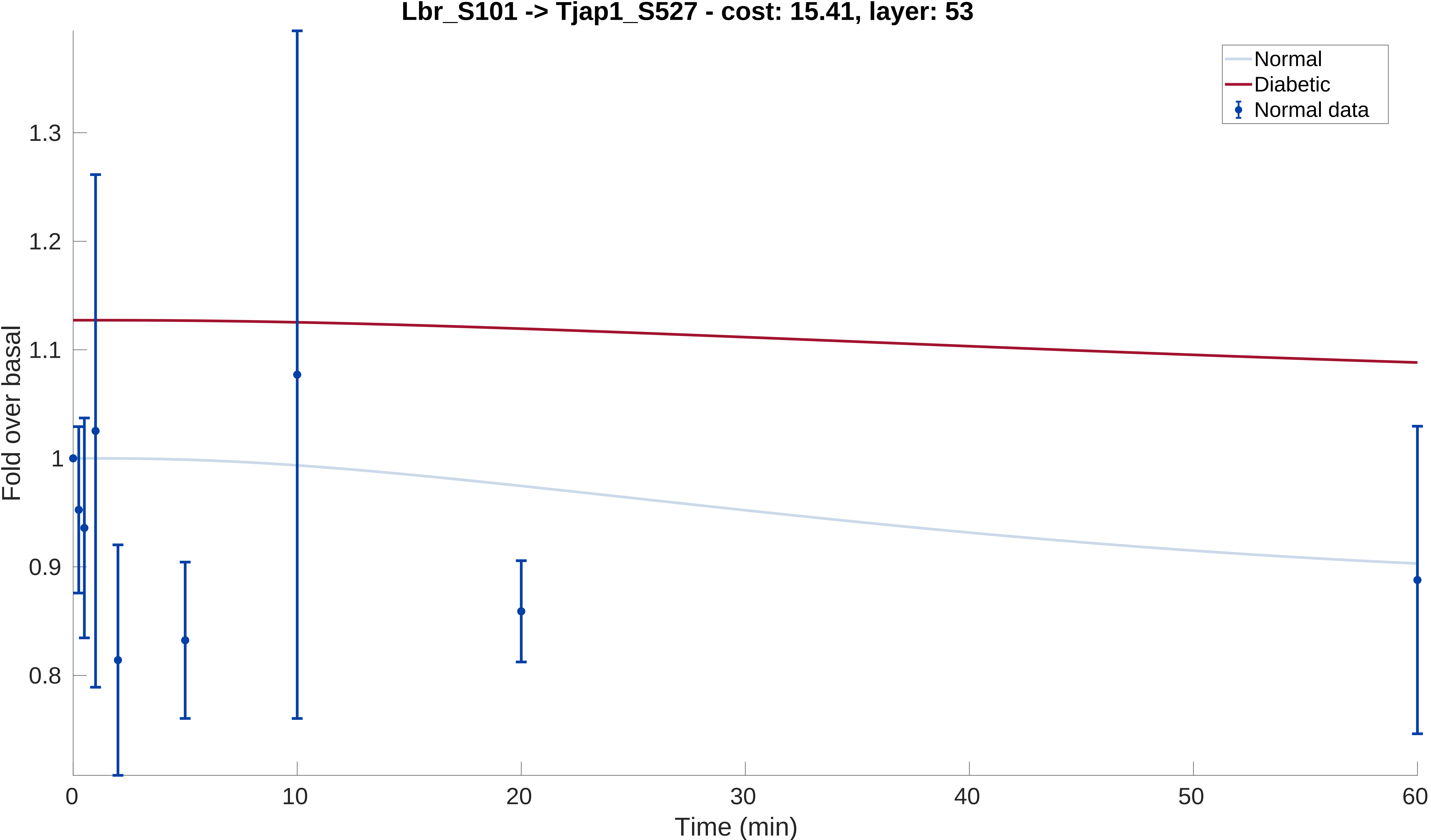

Lbr_S101 -> Tjap1_S527 - cost: 15.41, layer: 53
Normal
Diabetic
Normal data
1.3
1.2
1.1
1
0.9
0.8
Fold over basal
0
10
20
30
40
50
60
Time (min)

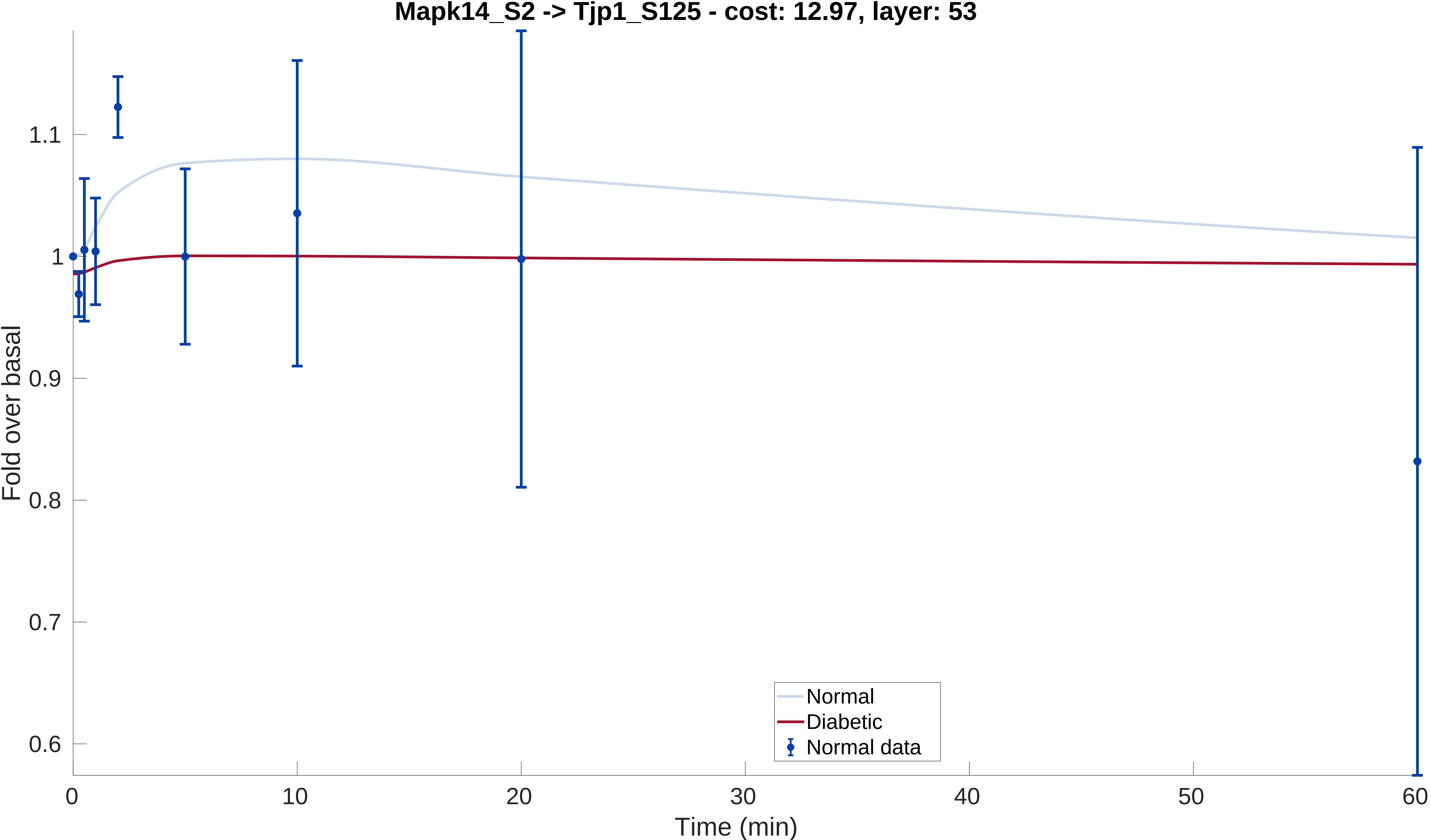

Mapk14_S2 -> Tjp1_S125 - cost: 12.97, layer: 53
Fold over basal
Time (min)
1.1
1
0.9
0.8
0.7
0.6
0
10
20
30
40
50
60
Normal
Diabetic
Normal data

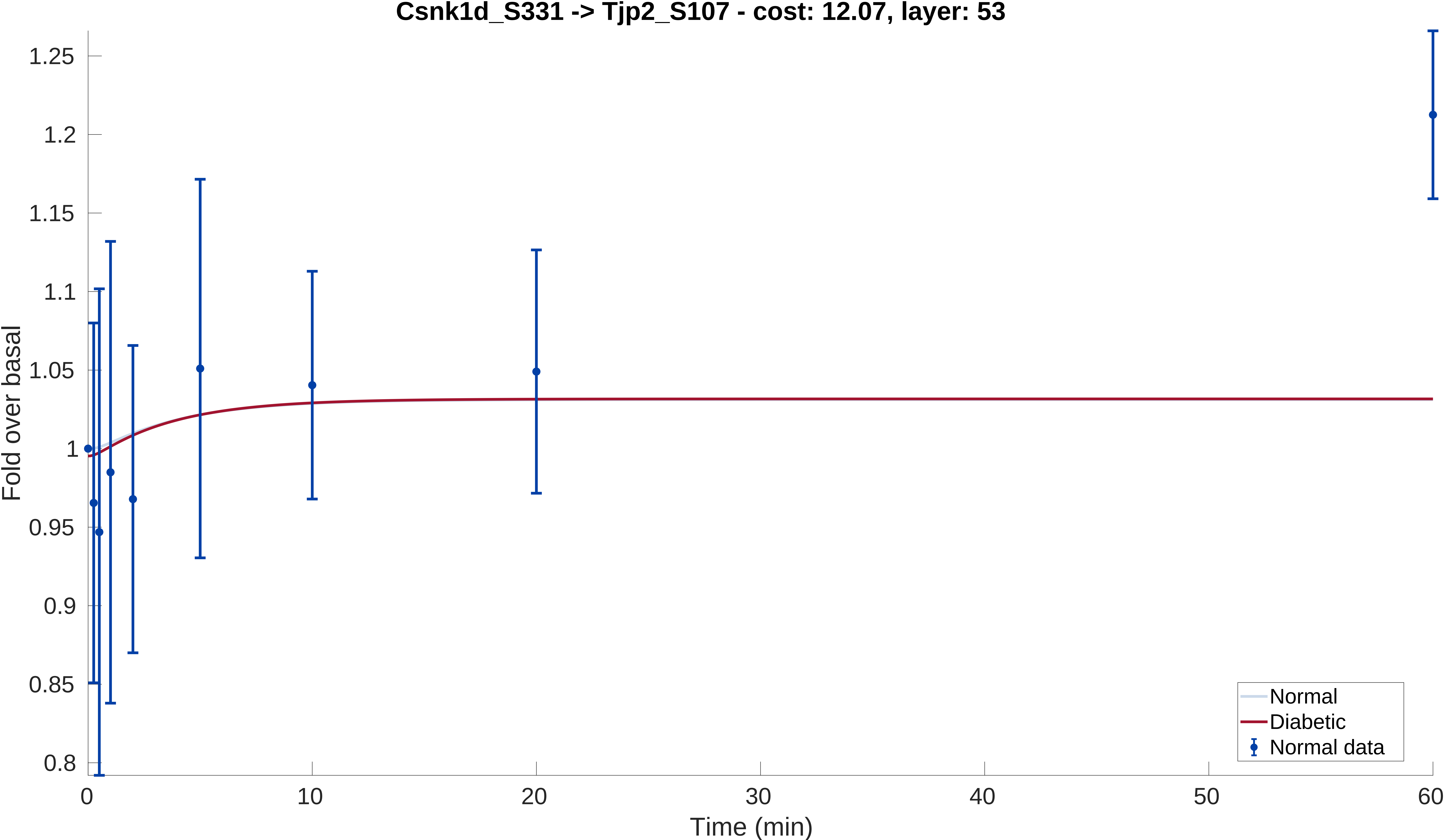

Csnk1d_S331 -> Tjp2_S107 - cost: 12.07, layer: 53
Fold over basal
1.25
1.2
1.15
1.1
1.05
1
0.95
0.9
0.85
0.8
0
10
20
30
40
50
60
Time (min)
Normal
Diabetic
Normal data

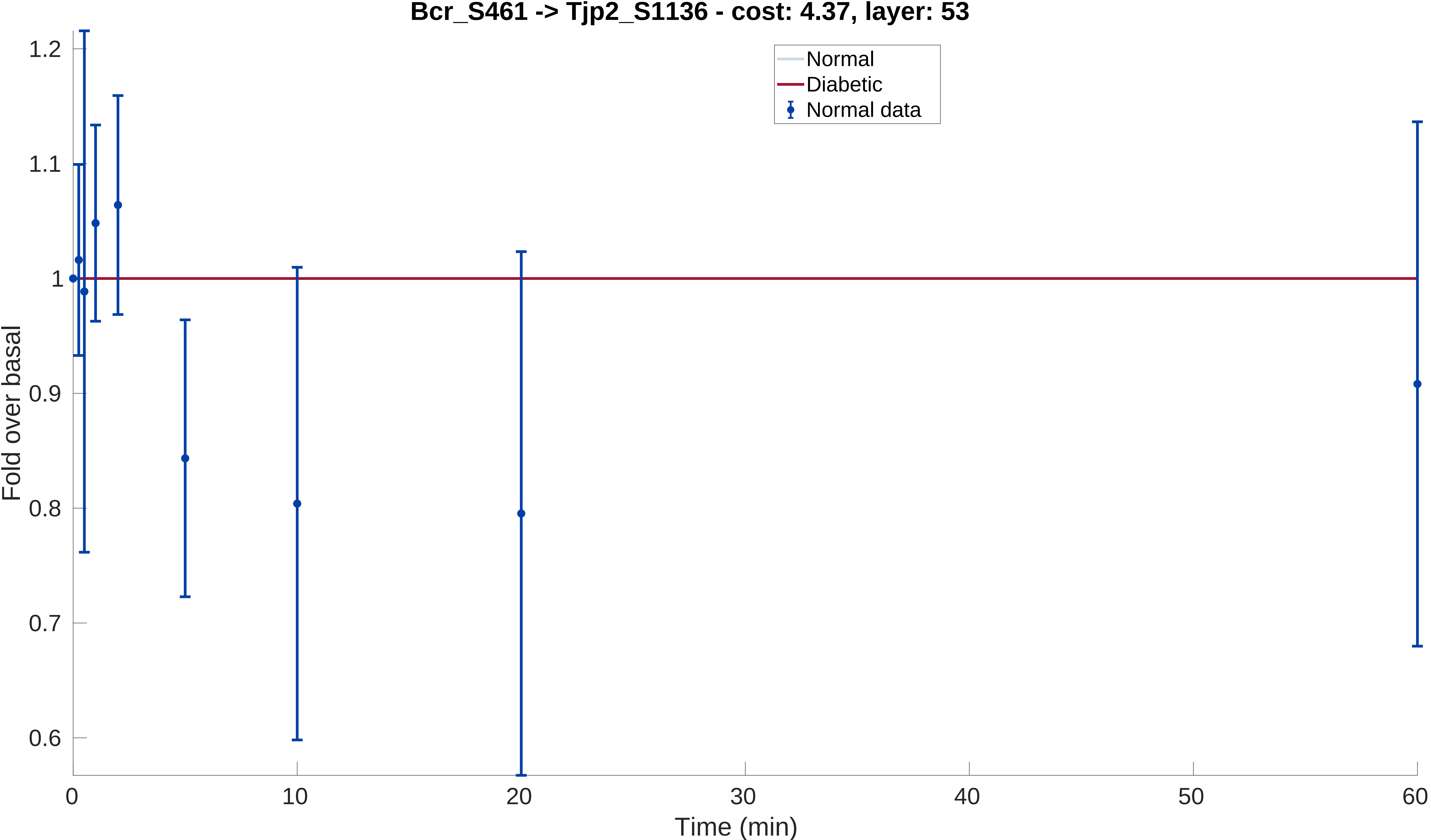

Bcr_S461 -> Tjp2_S1136 - cost: 4.37, layer: 53
Normal
Diabetic
Normal data
Fold over basal
Time (min)
1.2
1.1
1
0.9
0.8
0.7
0.6
0
10
20
30
40
50
60

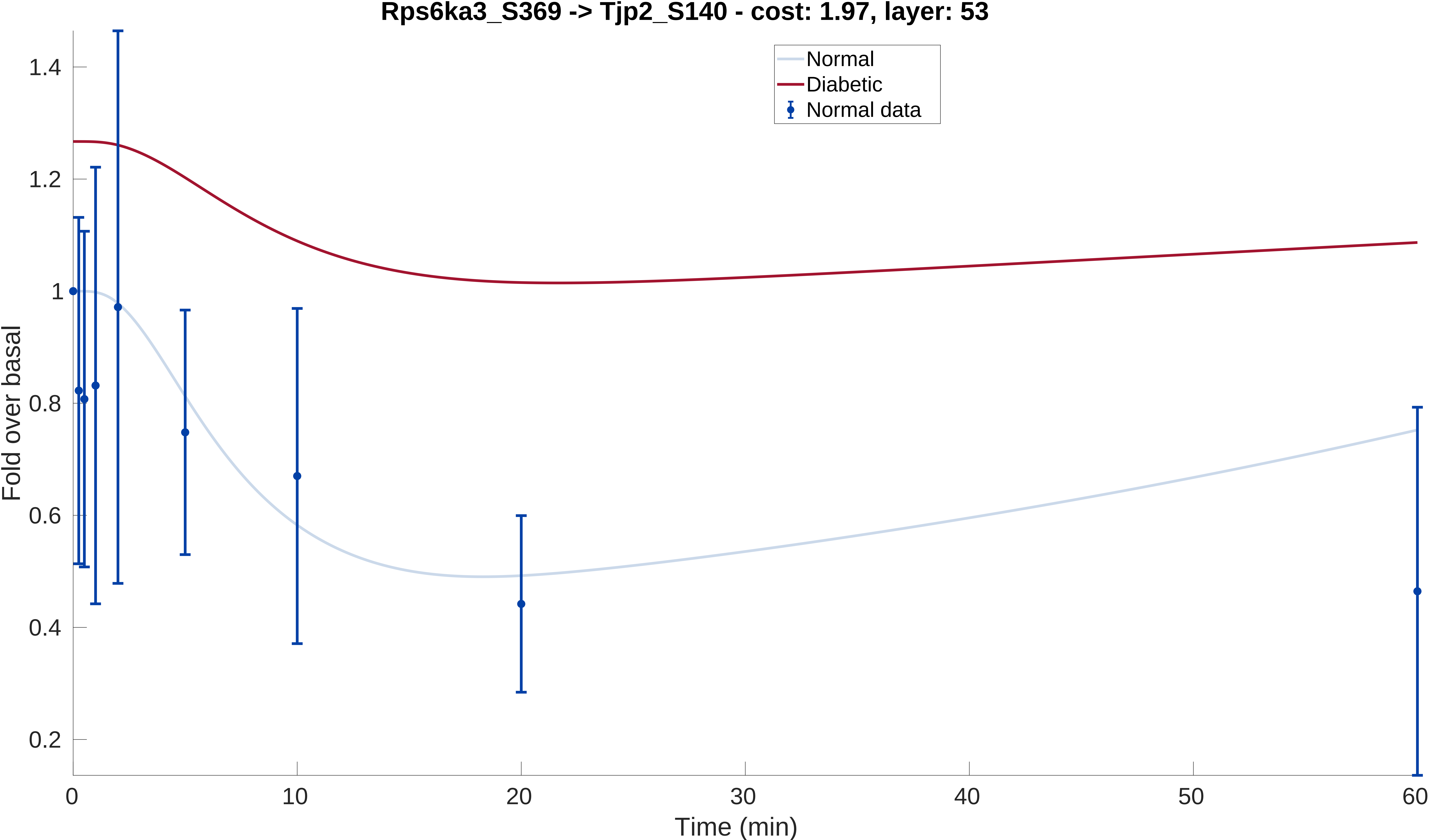

Rps6ka3_S369 -> Tjp2_S140 - cost: 1.97, layer: 53
Normal
Diabetic
Normal data
Fold over basal
Time (min)
1.4
1.2
1
0.8
0.6
0.4
0.2
0
10
20
30
40
50
60

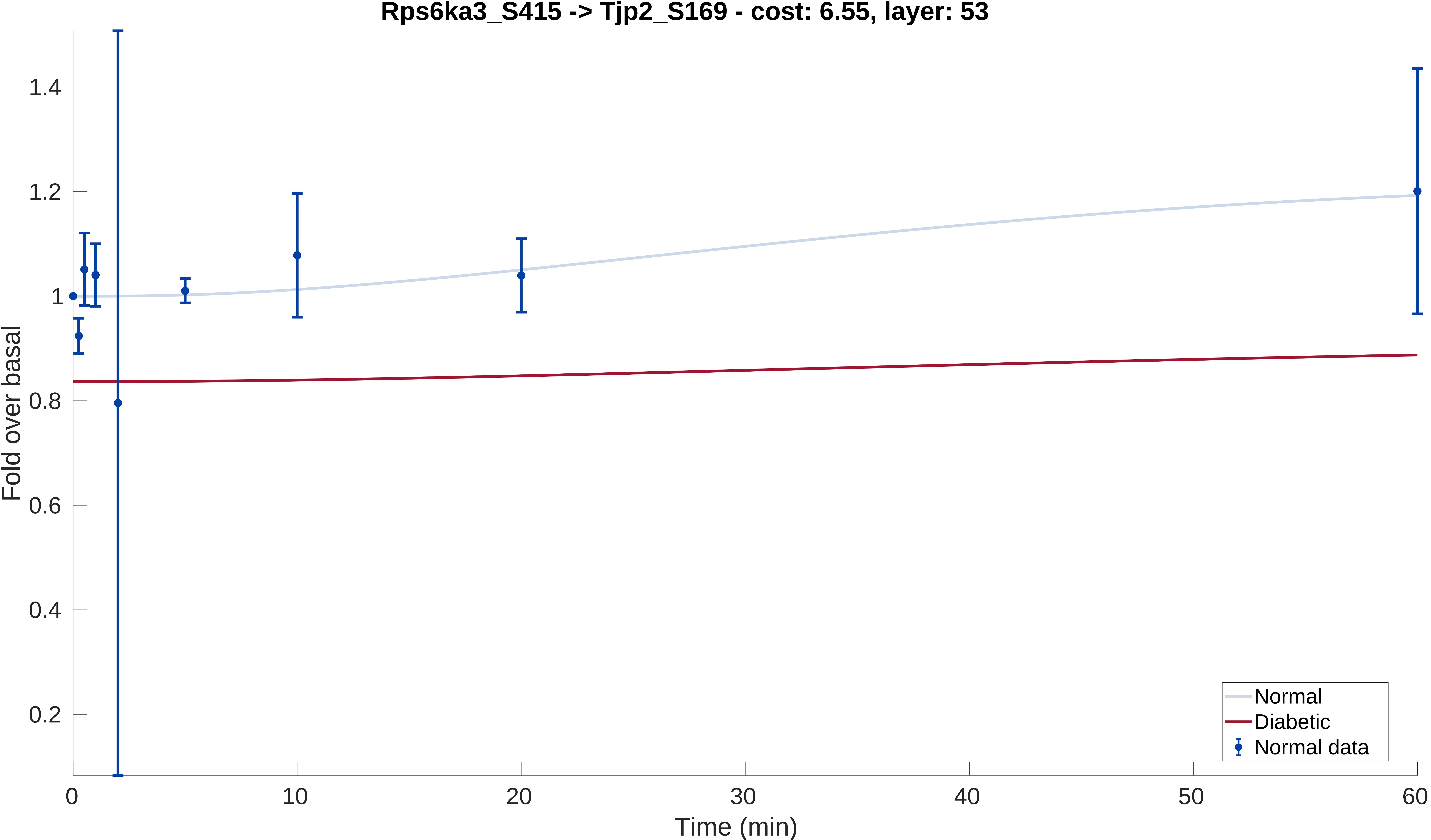
Rps6ka3_S415 -> Tjp2_S169 - cost: 6.55, layer: 53
Fold over basal
1.4
1.2
1
0.8
0.6
0.4
0.2
0
10
20
30
40
50
60
Time (min)
Normal
Diabetic
Normal data

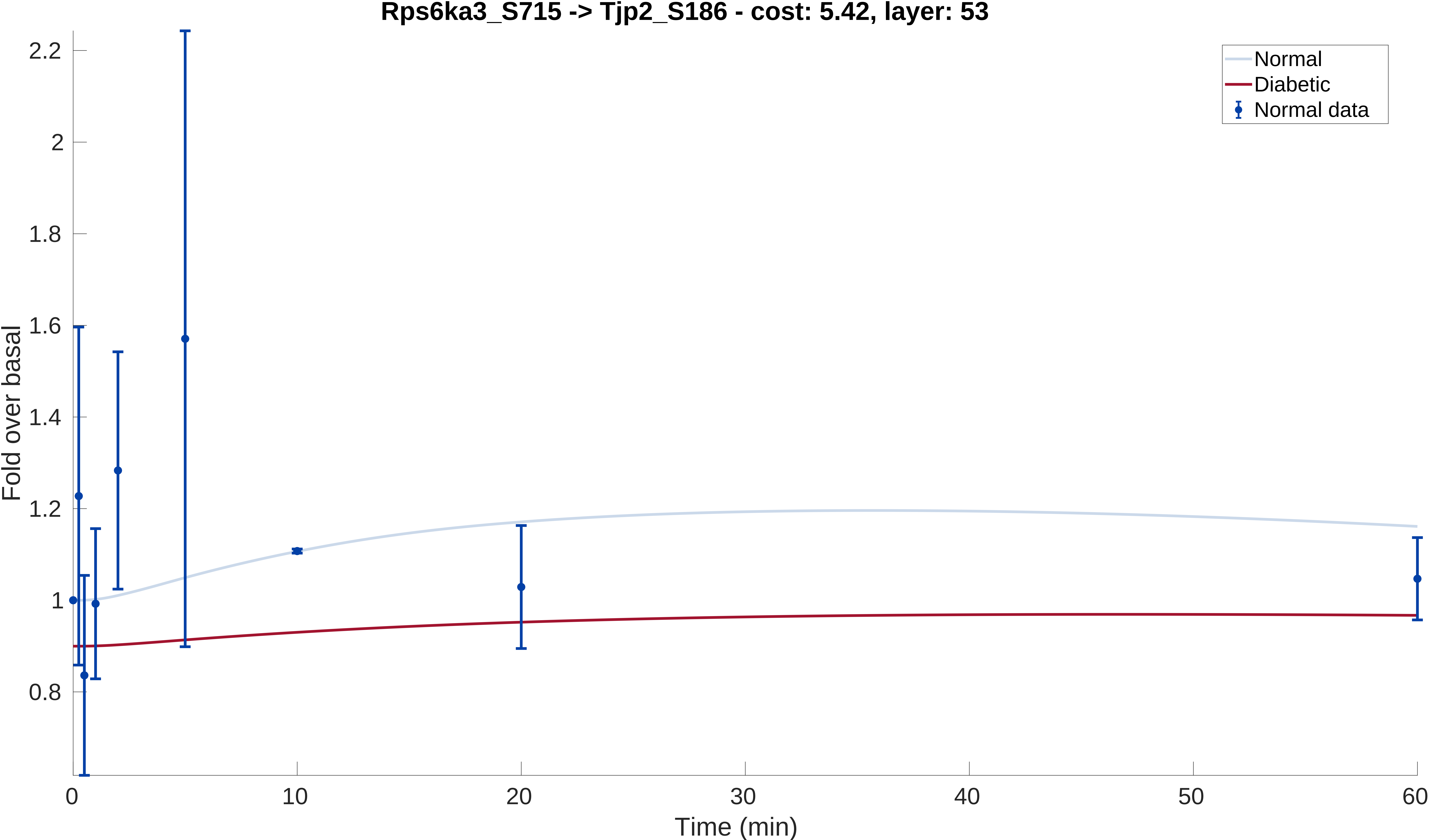

Rps6ka3_S715 -> Tjp2_S186 - cost: 5.42, layer: 53
Normal
Diabetic
Normal data
Fold over basal
2.2
2
1.8
1.6
1.4
1.2
1
0.8
0
10
20
30
40
50
60
Time (min)

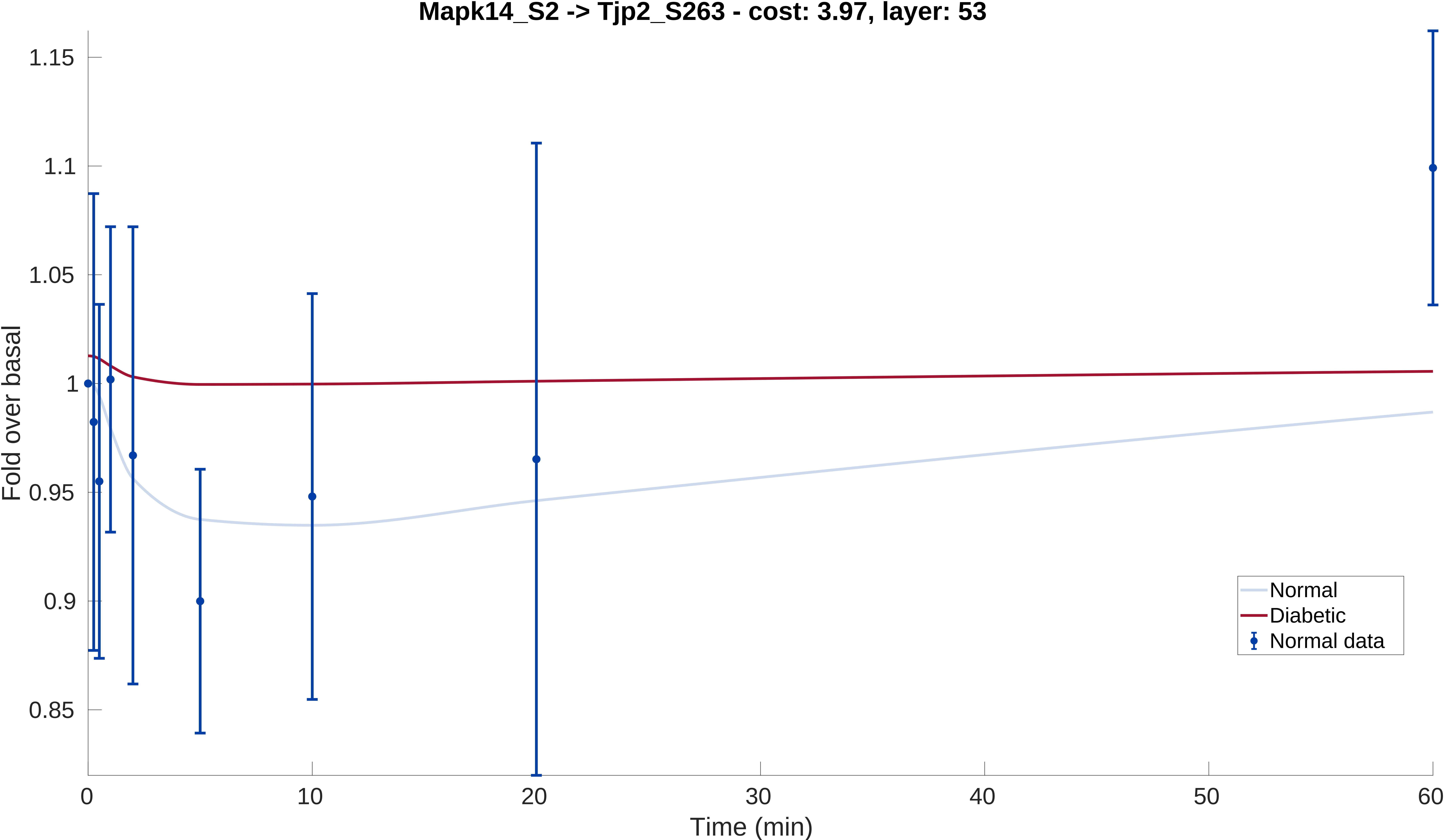
Mapk14_S2 -> Tjp2_S263 - cost: 3.97, layer: 53
1.15
1.1
1.05
1
0.95
0.9
0.85
Fold over basal
0
10
20
30
40
50
60
Time (min)
Normal
Diabetic
Normal data

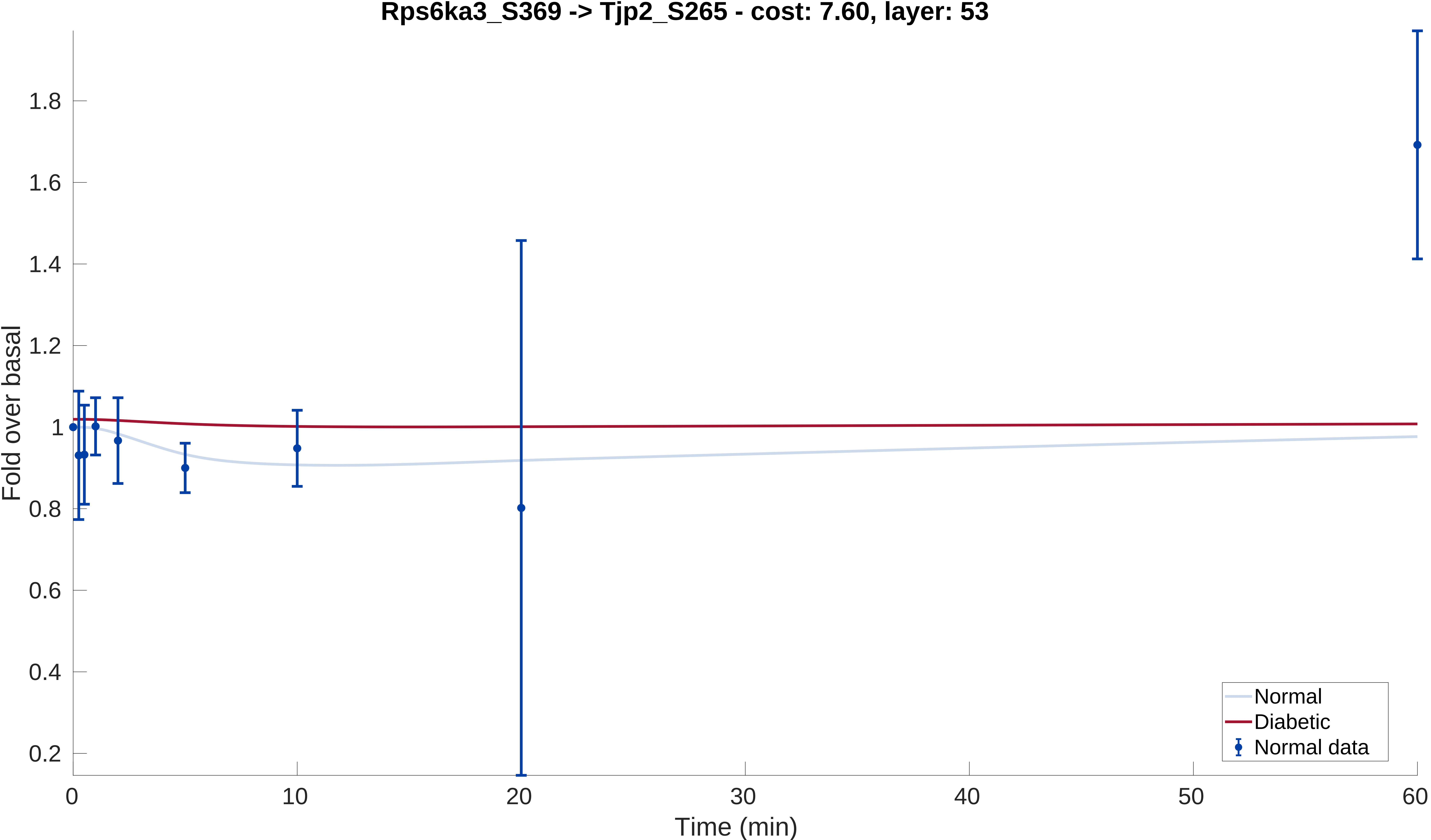

Rps6ka3_S369 -> Tjp2_S265 - cost: 7.60, layer: 53
Fold over basal
1.8
1.6
1.4
1.2
1
0.8
0.6
0.4
0.2
0
10
20
30
40
50
60
Time (min)
Normal
Diabetic
Normal data

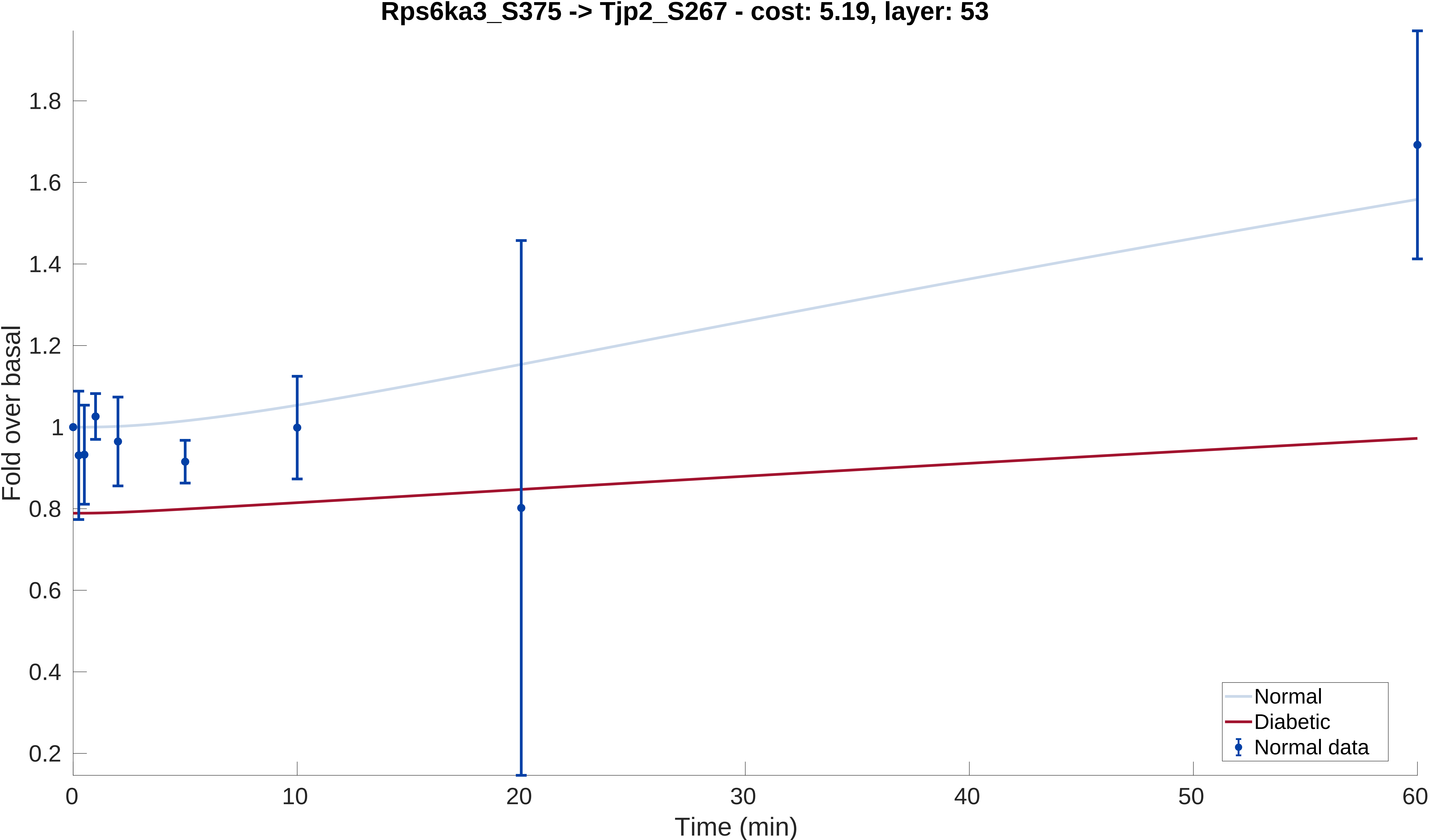
Rps6ka3_S375 -> Tjp2_S267 - cost: 5.19, layer: 53
Fold over basal
1.8
1.6
1.4
1.2
1
0.8
0.6
0.4
0.2
0
10
20
30
40
50
60
Time (min)
Normal
Diabetic
Normal data

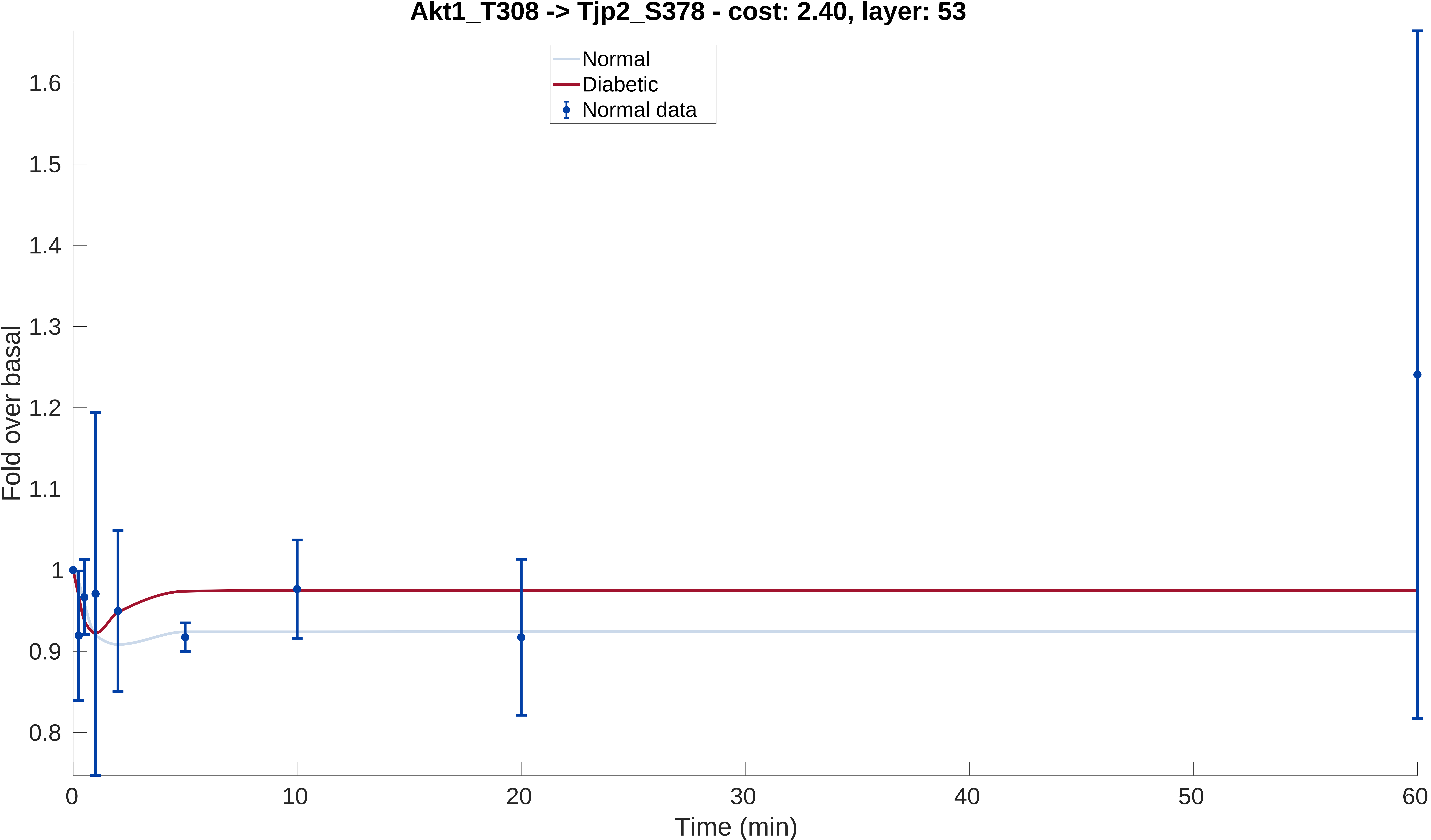

Akt1_T308 -> Tjp2_S378 - cost: 2.40, layer: 53
Normal
Diabetic
Normal data
Fold over basal
1.6
1.5
1.4
1.3
1.2
1.1
1
0.9
0.8
0
10
20
30
40
50
60
Time (min)

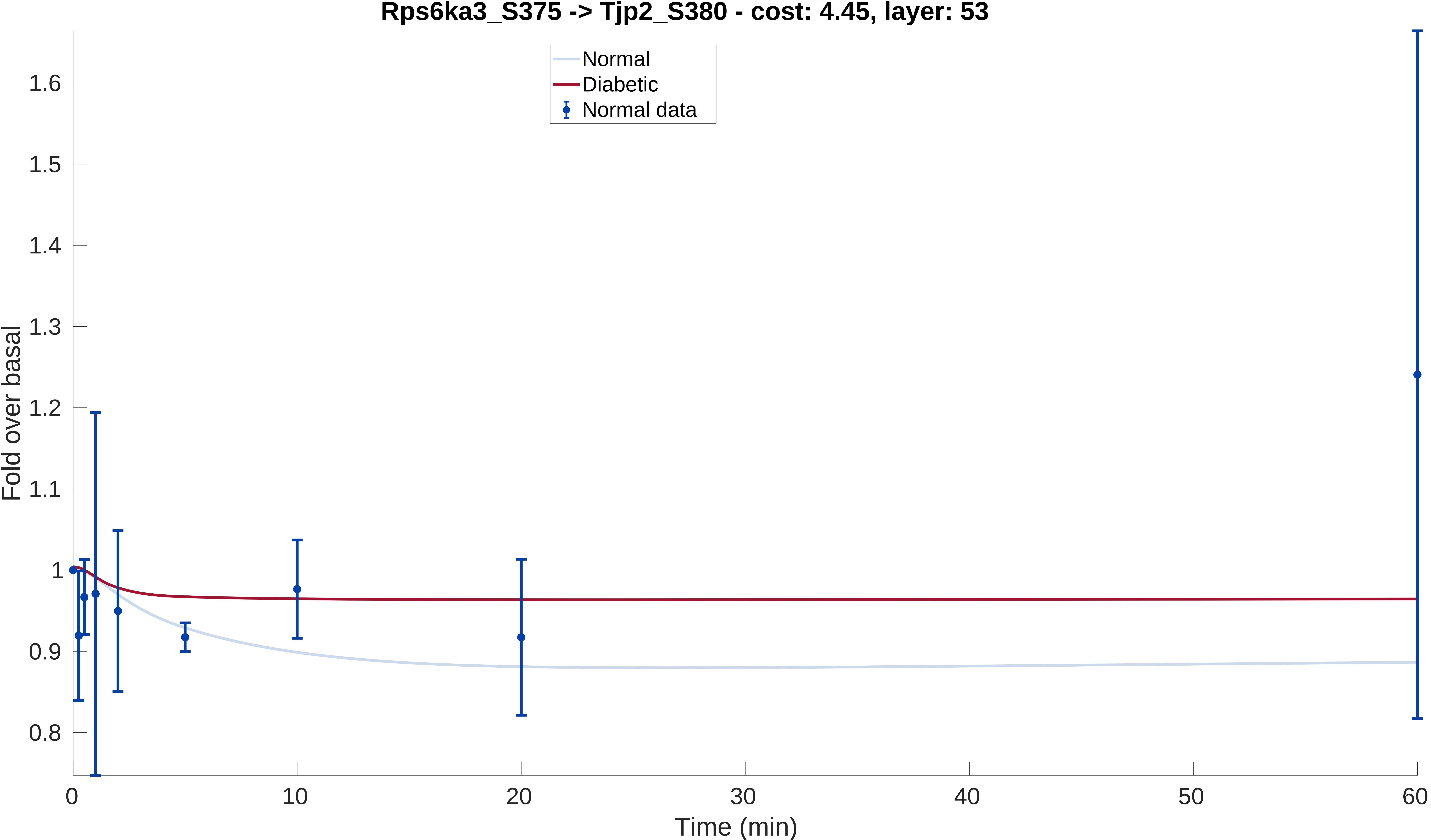
Rps6ka3_S375 -> Tjp2_S380 - cost: 4.45, layer: 53
Normal
Diabetic
Normal data
Fold over basal
1.6
1.5
1.4
1.3
1.2
1.1
1
0.9
0.8
0
10
20
30
40
50
60
Time (min)

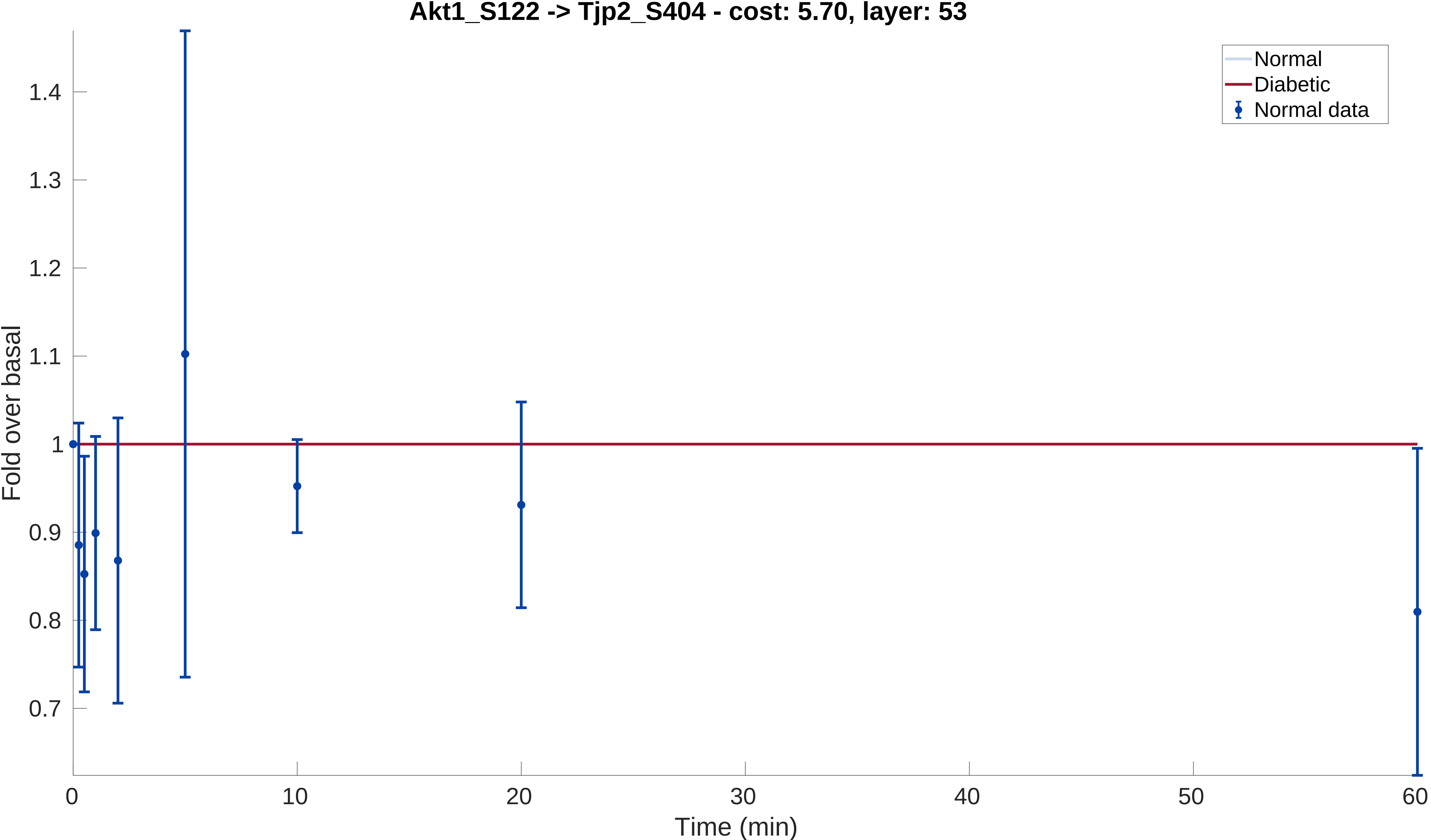

Akt1_S122 -> Tjp2_S404 - cost: 5.70, layer: 53
Normal
Diabetic
Normal data
Fold over basal
1.4
1.3
1.2
1.1
1
0.9
0.8
0.7
0
10
20
30
40
50
60
Time (min)

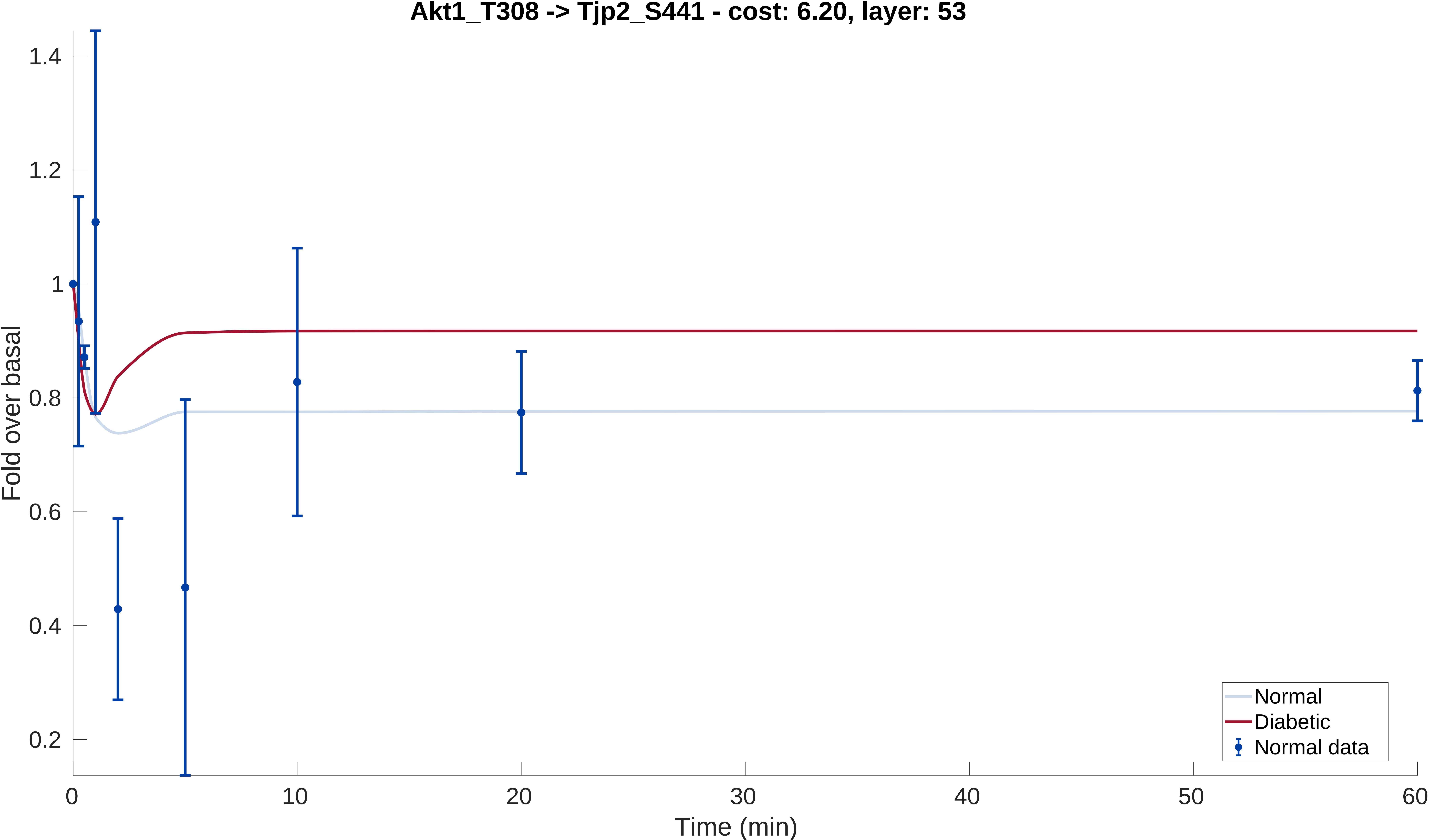

Akt1_T308 -> Tjp2_S441 - cost: 6.20, layer: 53
Fold over basal
1.4
1.2
1
0.8
0.6
0.4
0.2
0
10
20
30
40
50
60
Time (min)
Normal
Diabetic
Normal data

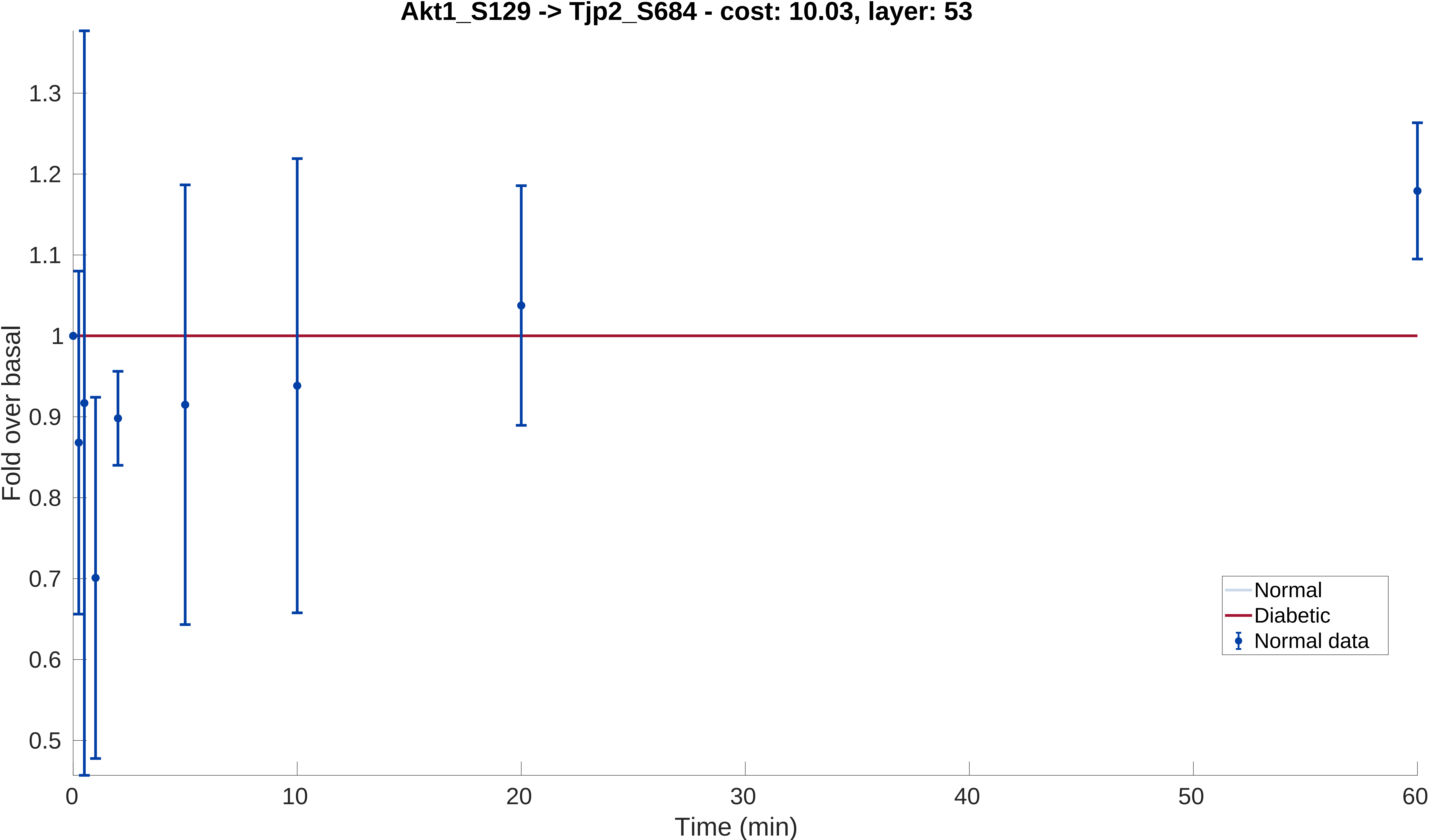
Akt1_S129 -> Tjp2_S684 - cost: 10.03, layer: 53
Fold over basal
Time (min)
0.5
0.6
0.7
0.8
0.9
1
1.1
1.2
1.3
0
10
20
30
40
50
60
Normal
Diabetic
Normal data

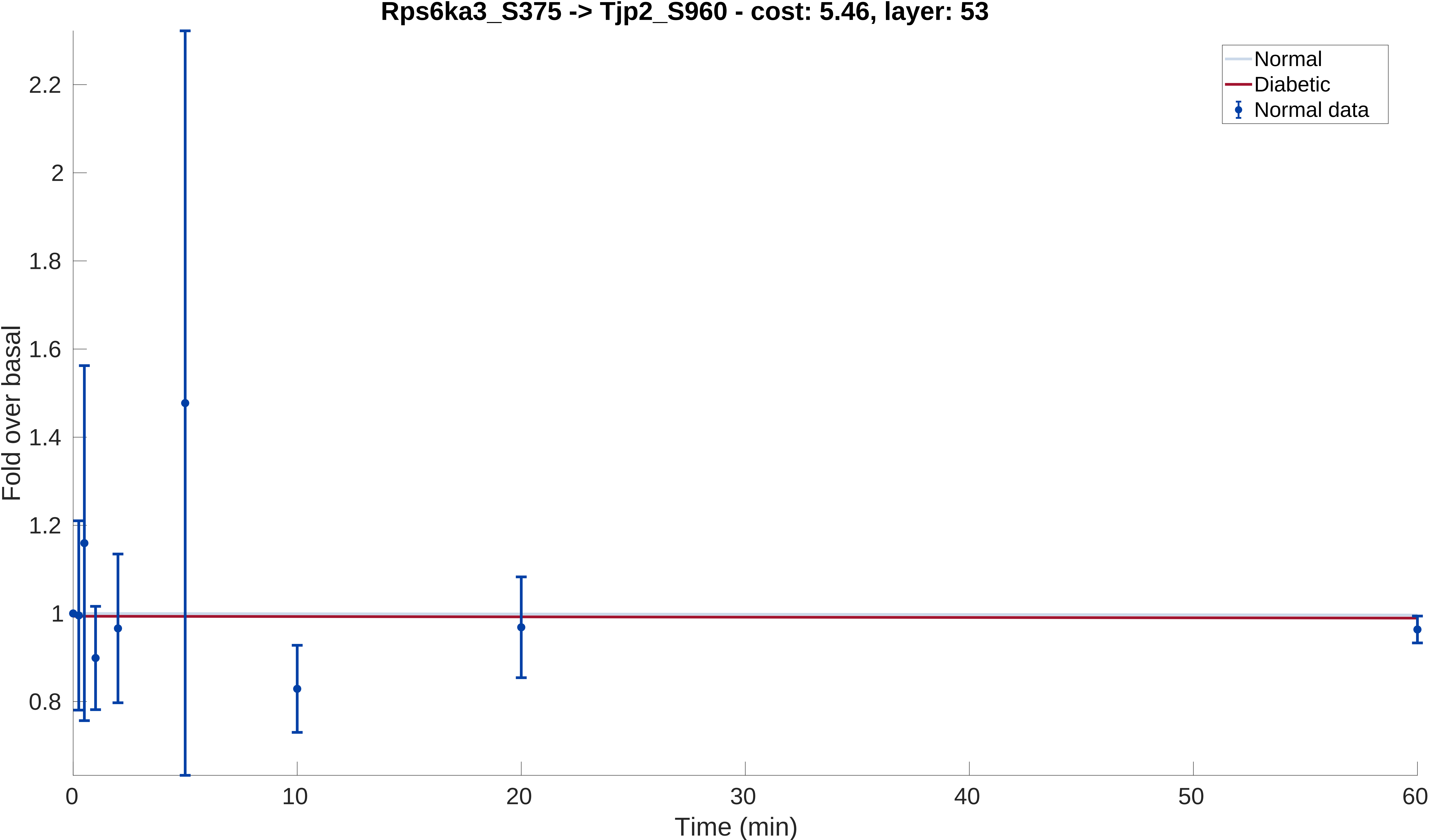
Rps6ka3_S375 -> Tjp2_S960 - cost: 5.46, layer: 53
Normal
Diabetic
Normal data
Fold over basal
2.2
2
1.8
1.6
1.4
1.2
1
0.8
0
10
20
30
40
50
60
Time (min)

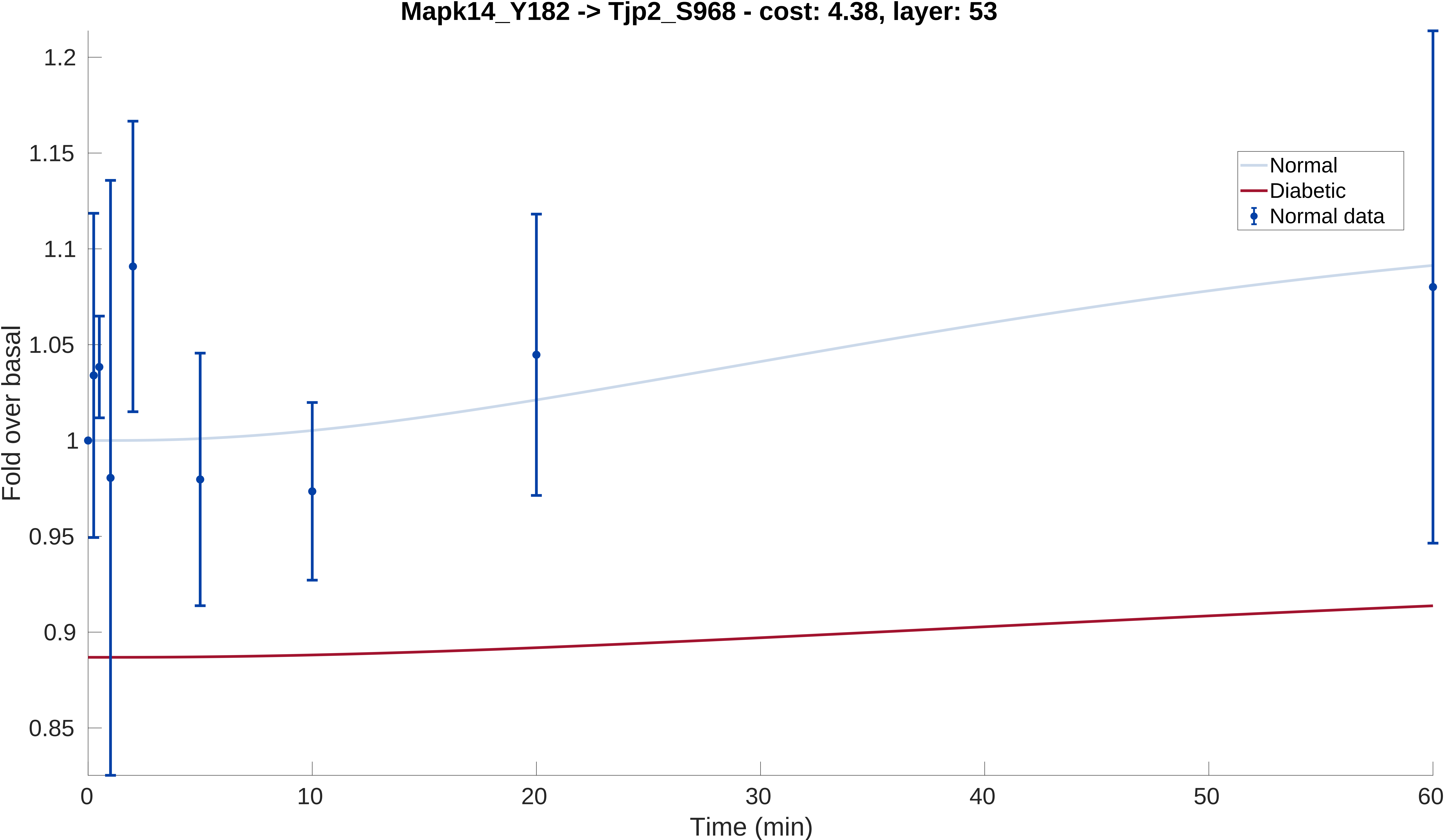
Mapk14_Y182 -> Tjp2_S968 - cost: 4.38, layer: 53
Fold over basal
1.2
1.15
1.1
1.05
1
0.95
0.9
0.85
0
10
20
30
40
50
60
Time (min)
Normal
Diabetic
Normal data

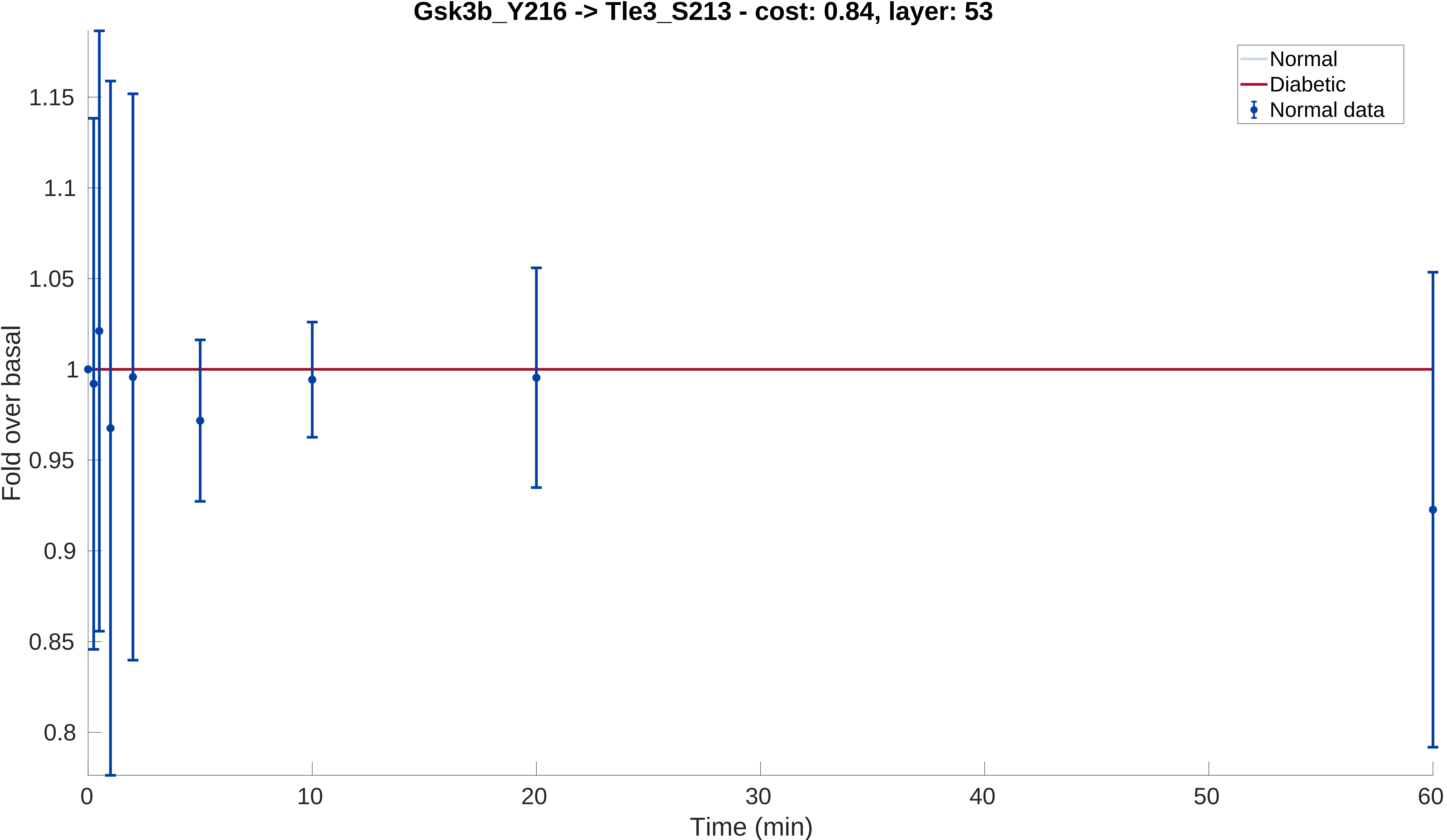

Gsk3b_Y216 -> Tle3_S213 - cost: 0.84, layer: 53
Normal
Diabetic
Normal data
Fold over basal
Time (min)
1.15
1.1
1.05
1
0.95
0.9
0.85
0.8
0
10
20
30
40
50
60

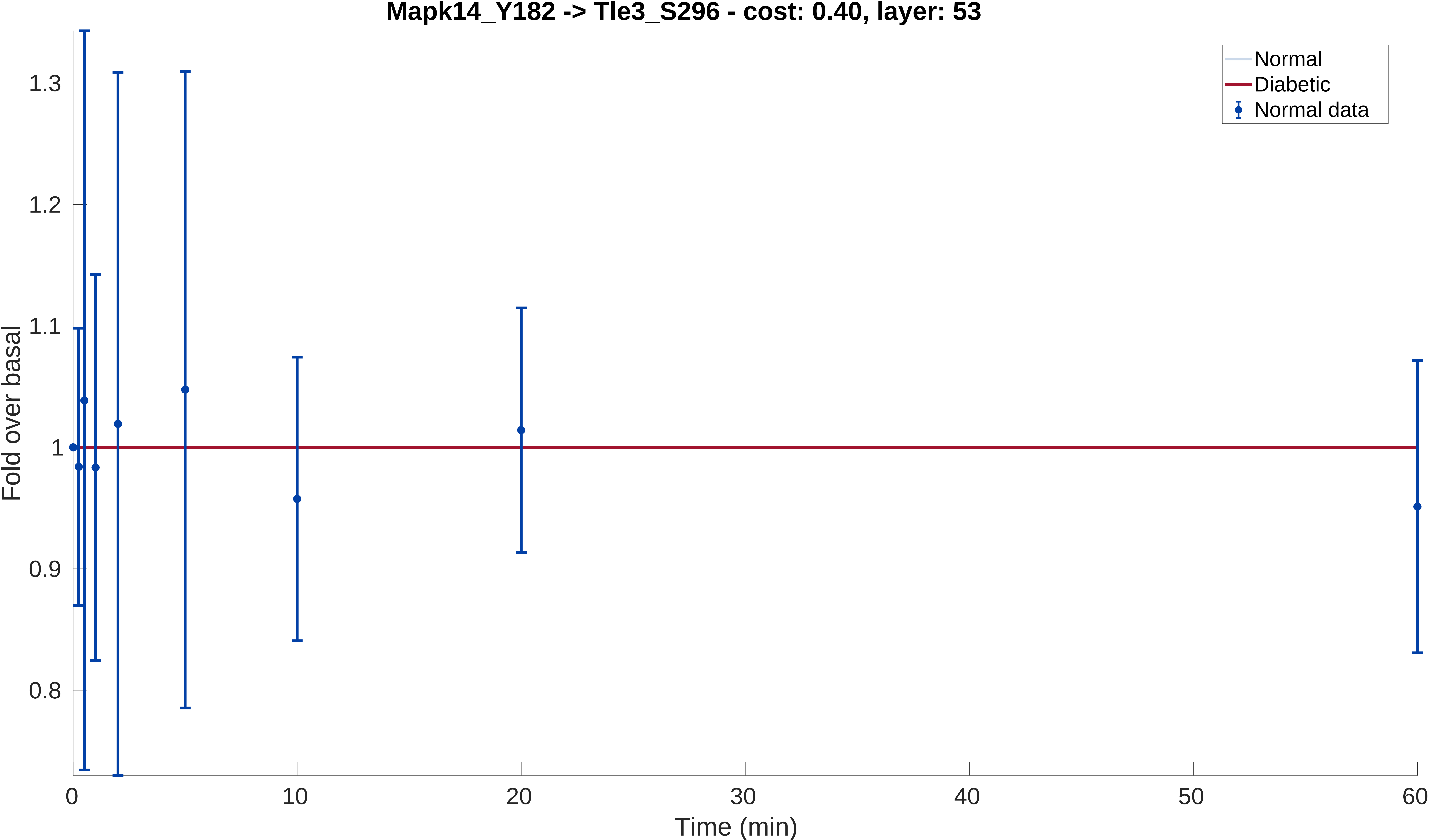

Mapk14_Y182 -> Tle3_S296 - cost: 0.40, layer: 53
Normal
Diabetic
Normal data
Fold over basal
1.3
1.2
1.1
1
0.9
0.8
0
10
20
30
40
50
60
Time (min)

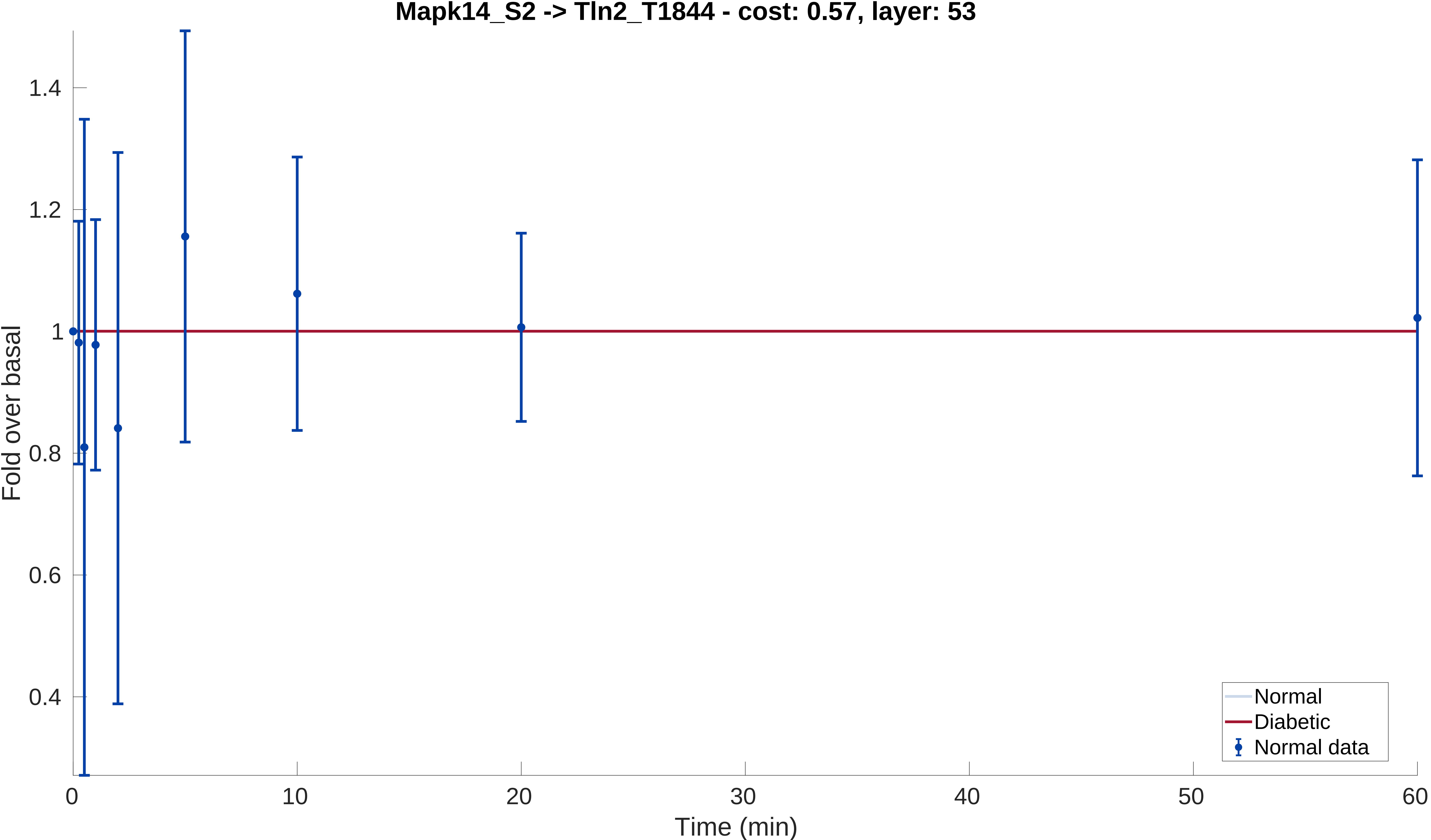

Mapk14_S2 -> Tln2_T1844 - cost: 0.57, layer: 53
Fold over basal
1.4
1.2
1
0.8
0.6
0.4
0
10
20
30
40
50
60
Time (min)
Normal
Diabetic
Normal data

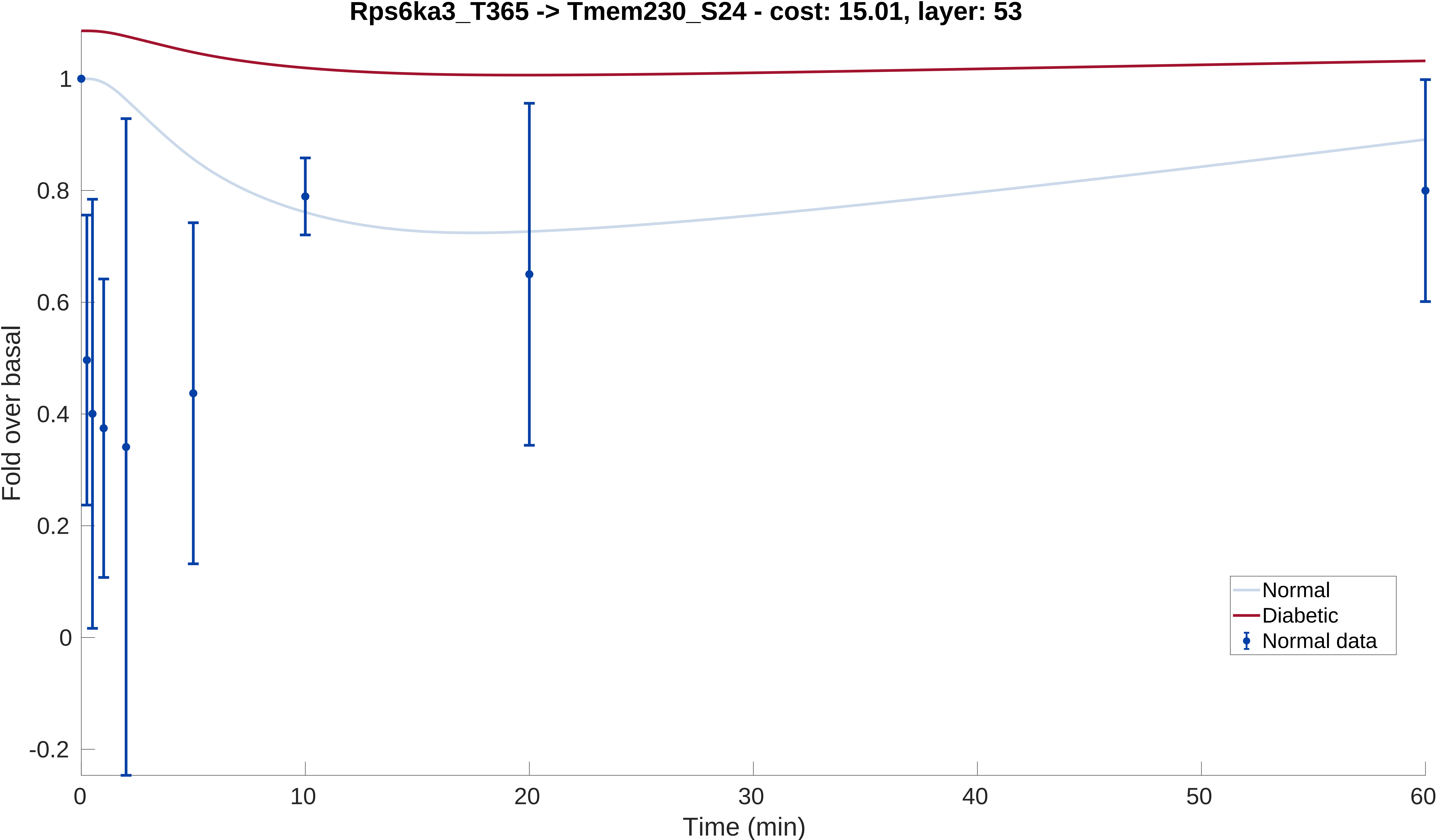

Rps6ka3_T365 -> Tmem230_S24 - cost: 15.01, layer: 53
Fold over basal
1
0.8
0.6
0.4
0.2
0
-0.2
0
10
20
30
40
50
60
Time (min)
Normal
Diabetic
Normal data

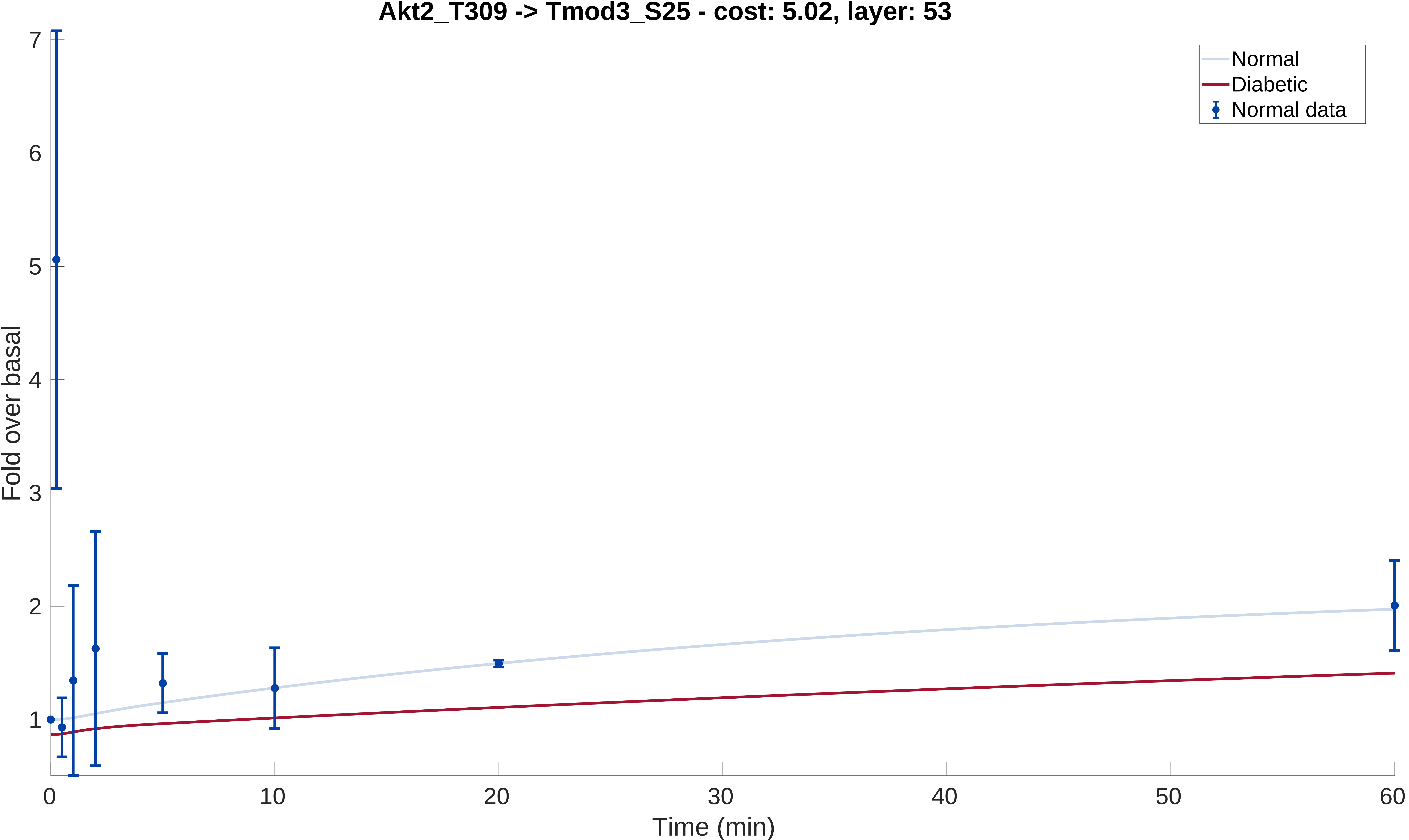
Akt2_T309 -> Tmod3_S25 - cost: 5.02, layer: 53
Fold over basal
Time (min)
Normal
Diabetic
Normal data
0
10
20
30
40
50
60
1
2
3
4
5
6
7

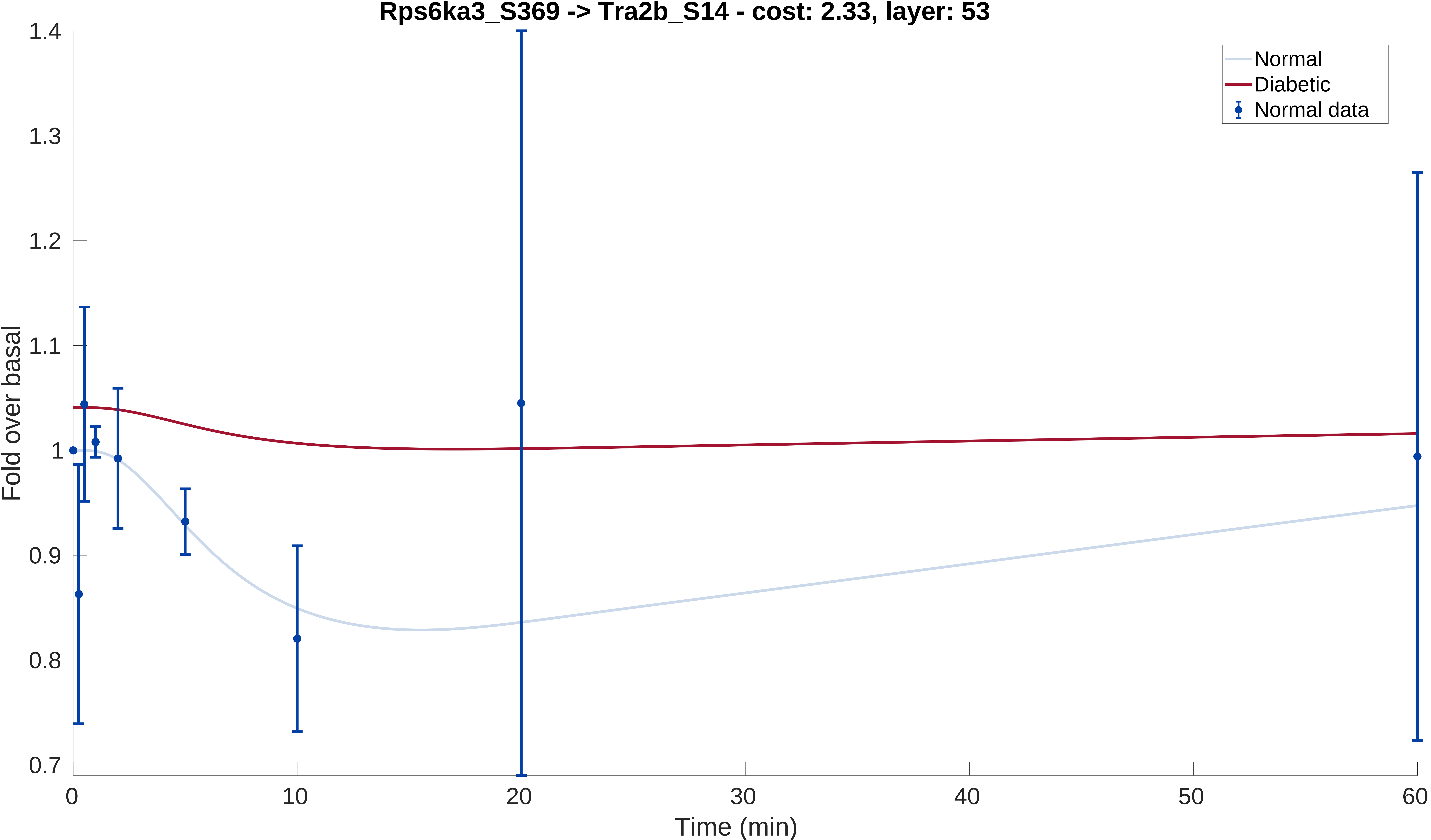

Rps6ka3_S369 -> Tra2b_S14 - cost: 2.33, layer: 53
Normal
Diabetic
Normal data
Fold over basal
1.4
1.3
1.2
1.1
1
0.9
0.8
0.7
0
10
20
30
40
50
60
Time (min)

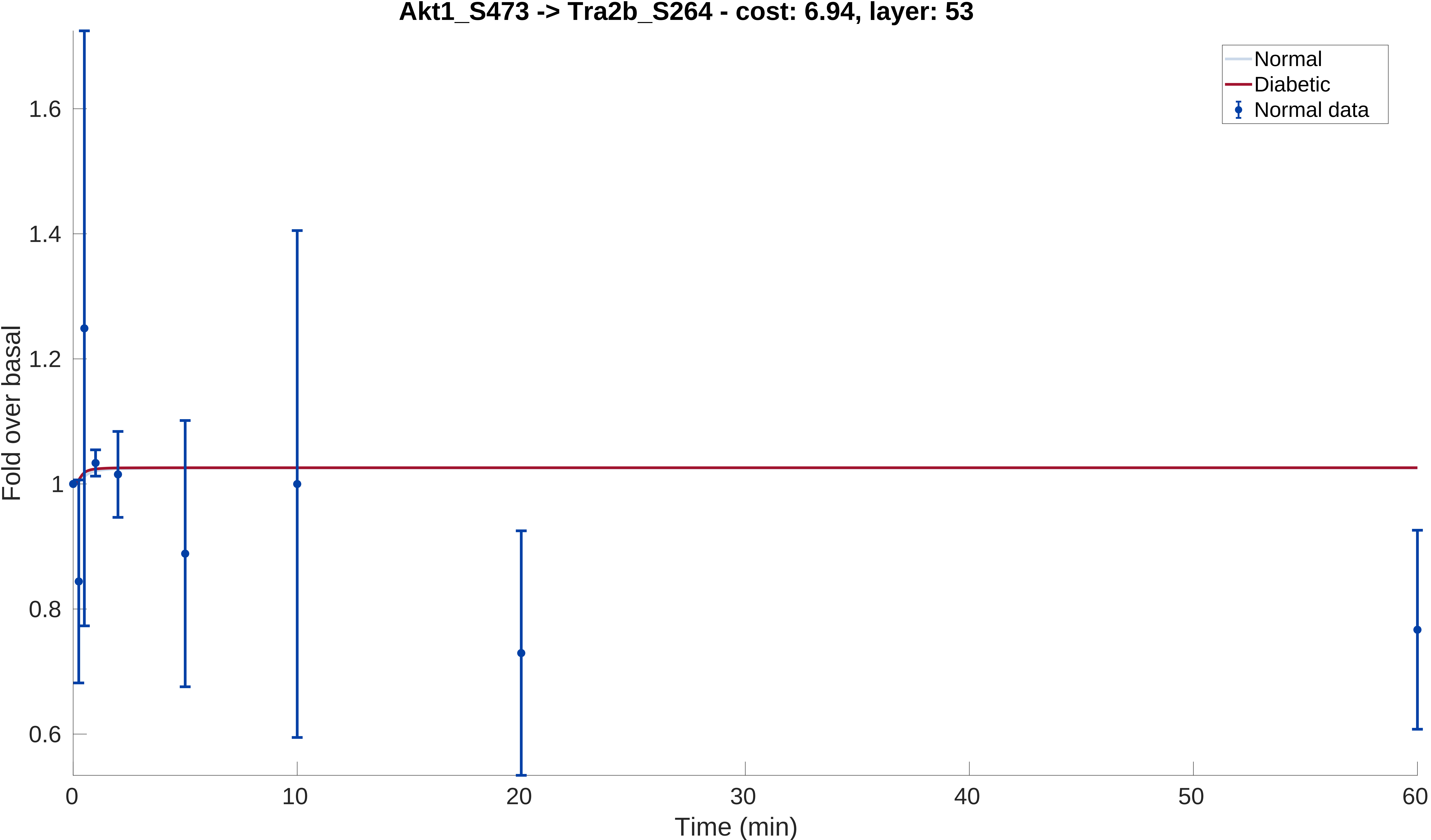

Akt1_S473 -> Tra2b_S264 - cost: 6.94, layer: 53
Normal
Diabetic
Normal data
Fold over basal
1.6
1.4
1.2
1
0.8
0.6
0
10
20
30
40
50
60
Time (min)

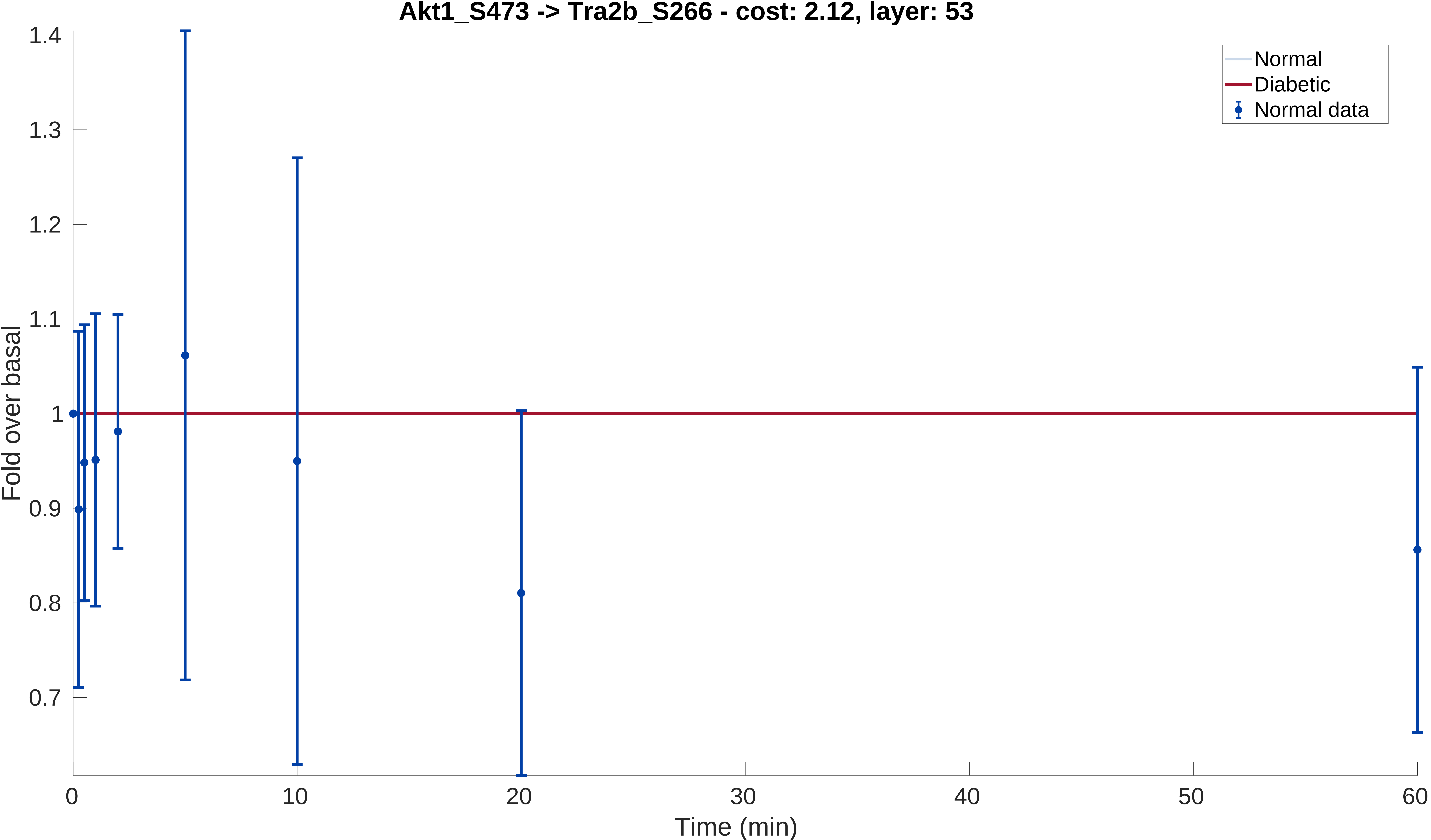

Akt1_S473 -> Tra2b_S266 - cost: 2.12, layer: 53
Normal
Diabetic
Normal data
Fold over basal
1.4
1.3
1.2
1.1
1
0.9
0.8
0.7
0
10
20
30
40
50
60
Time (min)

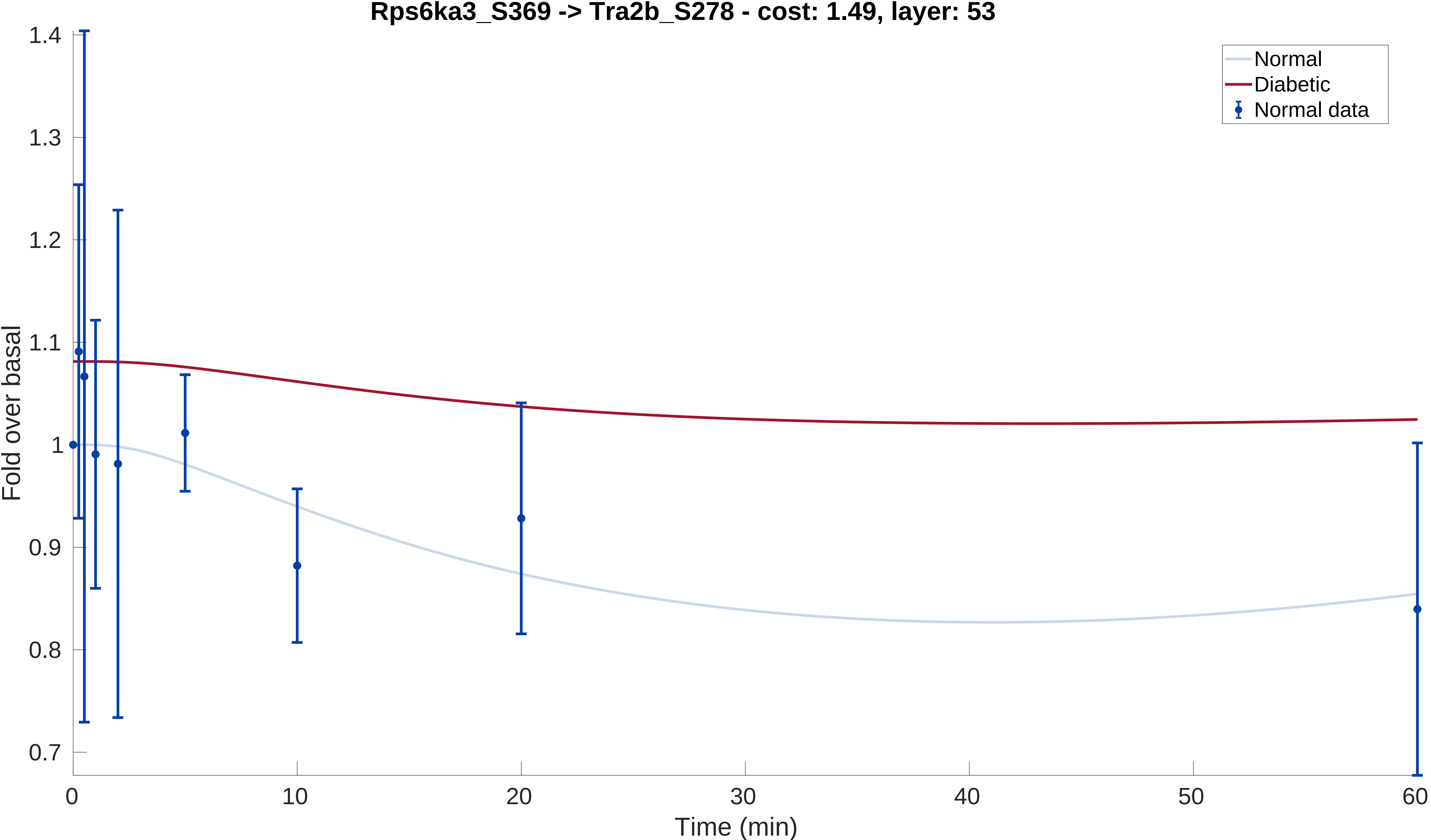

Rps6ka3_S369 -> Tra2b_S278 - cost: 1.49, layer: 53
Normal
Diabetic
Normal data
Fold over basal
Time (min)
1.4
1.3
1.2
1.1
1
0.9
0.8
0.7
0
10
20
30
40
50
60

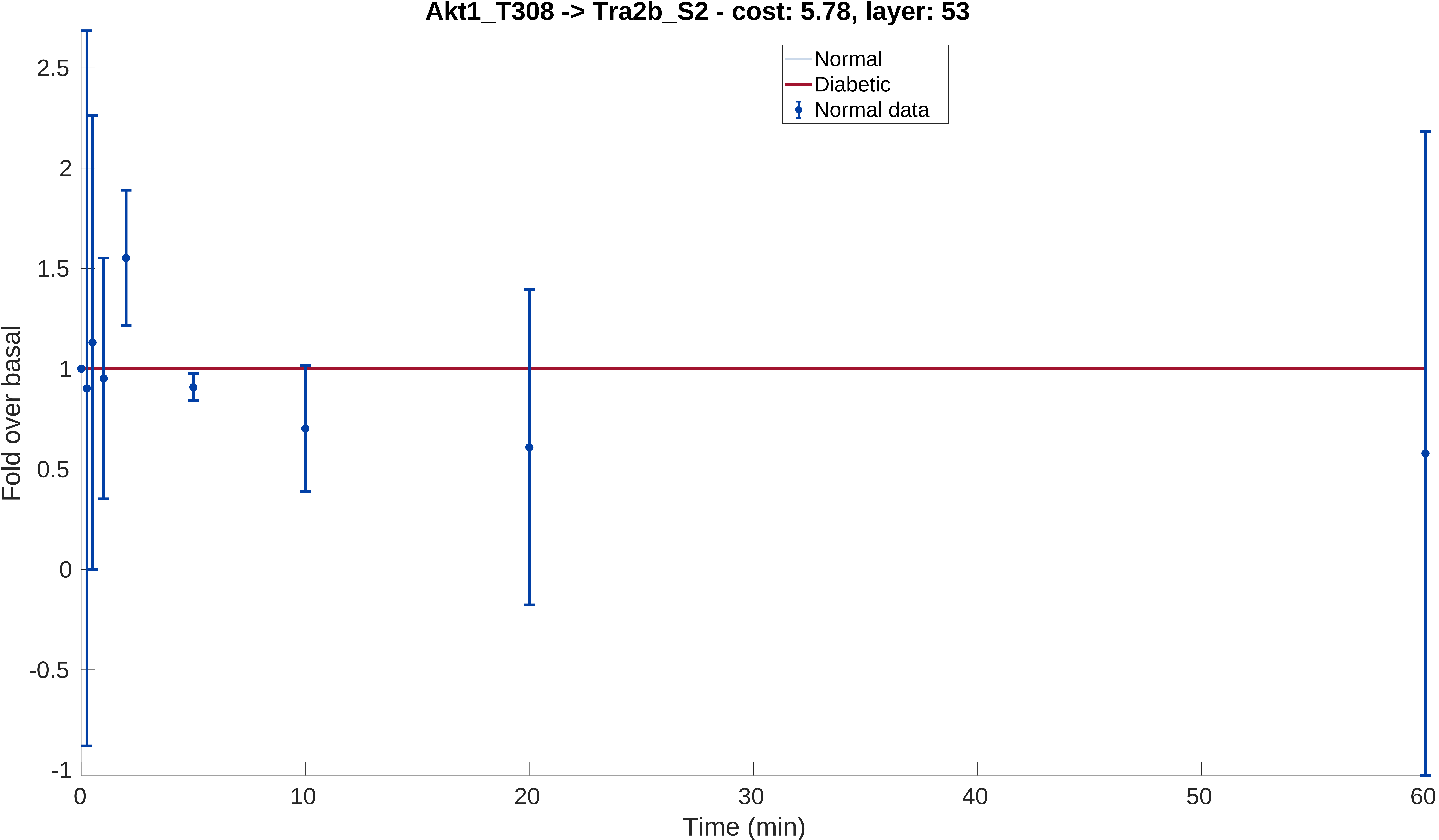

Akt1_T308 -> Tra2b_S2 - cost: 5.78, layer: 53
Normal
Diabetic
Normal data
Fold over basal
Time (min)
2.5
2
1.5
1
0.5
0
-0.5
-1
0
10
20
30
40
50
60

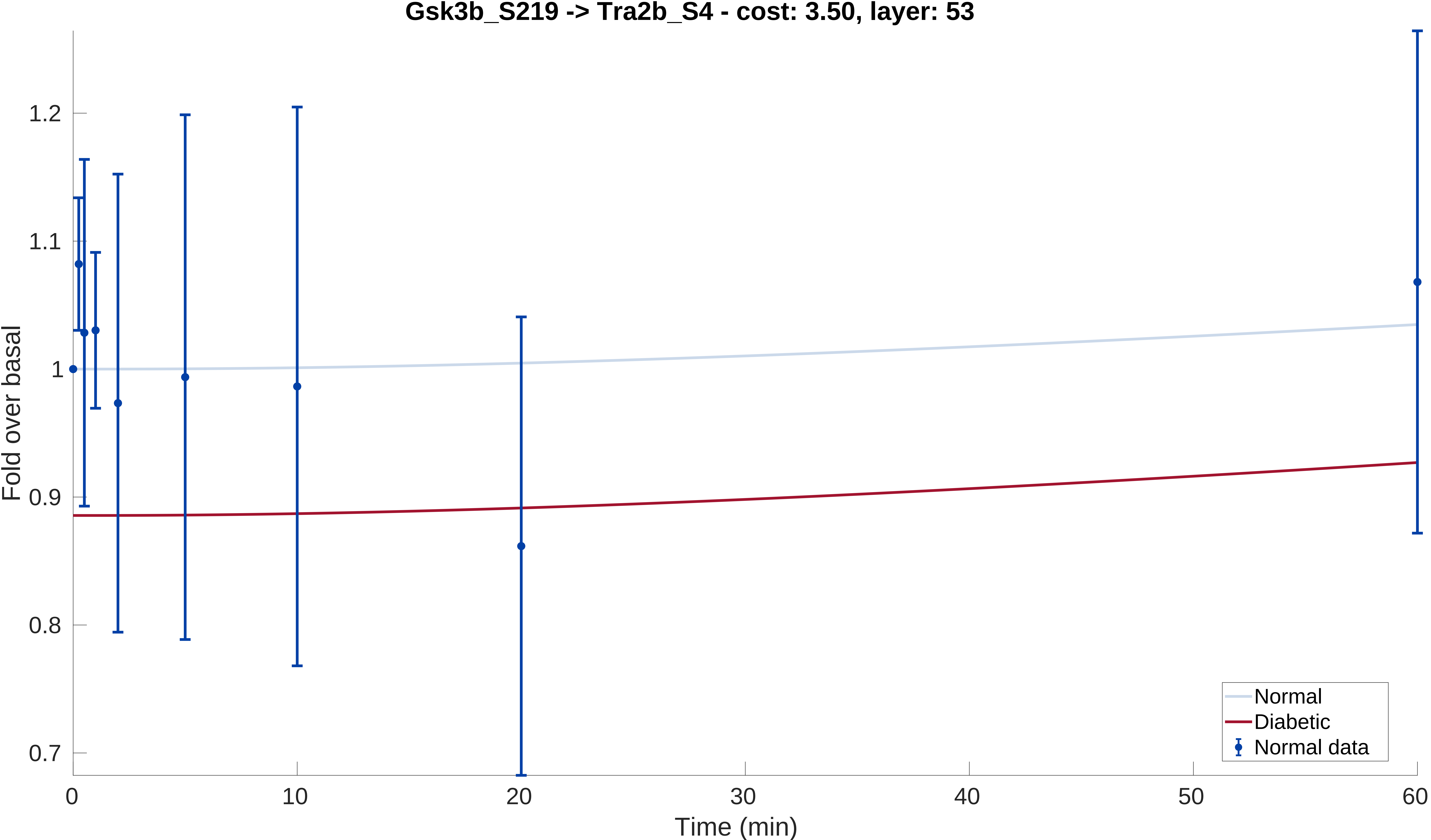

Gsk3b_S219 -> Tra2b_S4 - cost: 3.50, layer: 53
Fold over basal
1.2
1.1
1
0.9
0.8
0.7
0
10
20
30
40
50
60
Time (min)
Normal
Diabetic
Normal data

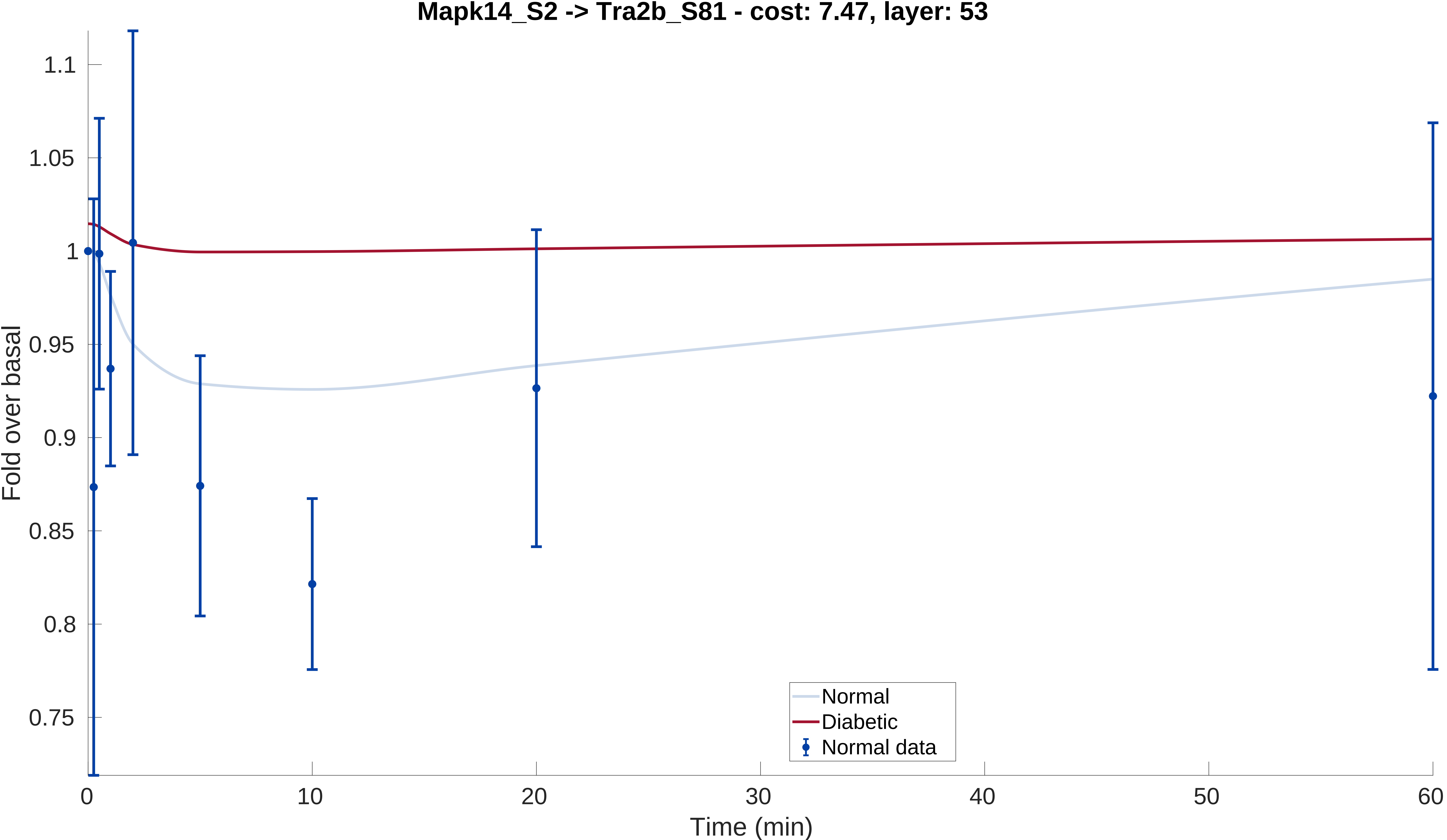
Mapk14_S2 -> Tra2b_S81 - cost: 7.47, layer: 53
Fold over basal
1.1
1.05
1
0.95
0.9
0.85
0.8
0.75
0
10
20
30
40
50
60
Time (min)
Normal
Diabetic
Normal data

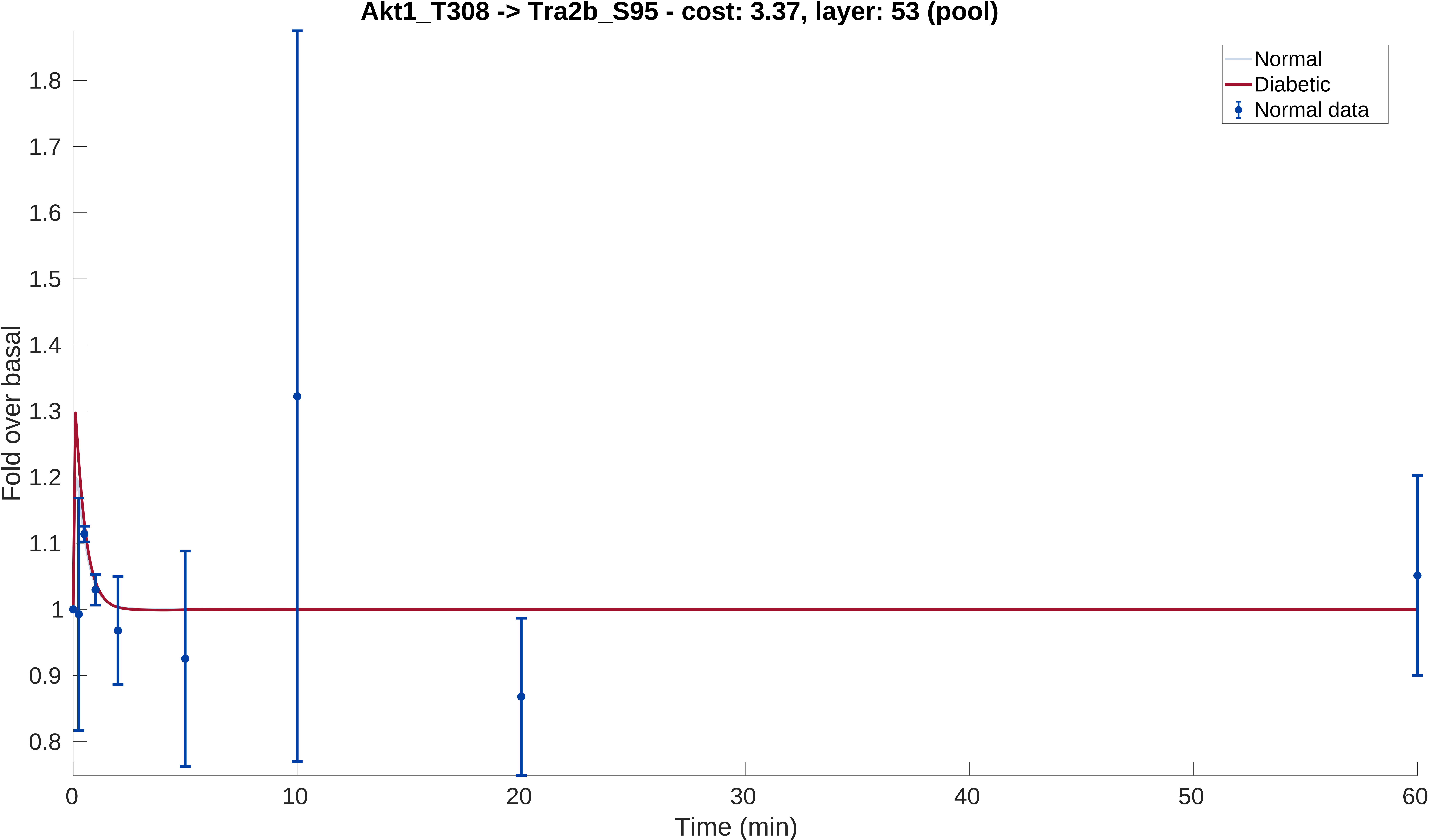
Akt1_T308 -> Tra2b_S95 - cost: 3.37, layer: 53 (pool)
Normal
Diabetic
Normal data
Fold over basal
1.8
1.7
1.6
1.5
1.4
1.3
1.2
1.1
1
0.9
0.8
0
10
20
30
40
50
60
Time (min)

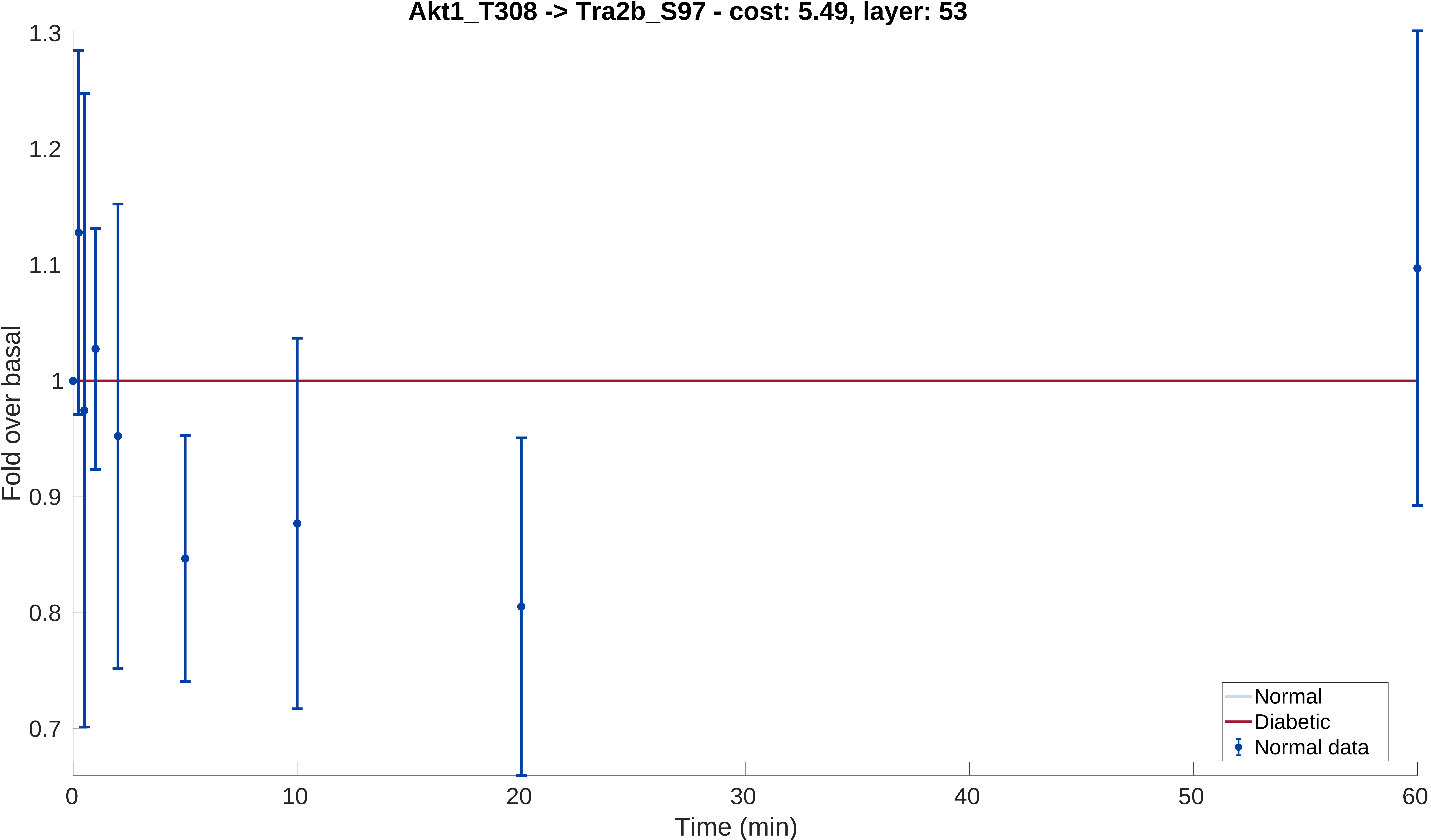

Akt1_T308 -> Tra2b_S97 - cost: 5.49, layer: 53
Fold over basal
1.3
1.2
1.1
1
0.9
0.8
0.7
0
10
20
30
40
50
60
Time (min)
Normal
Diabetic
Normal data

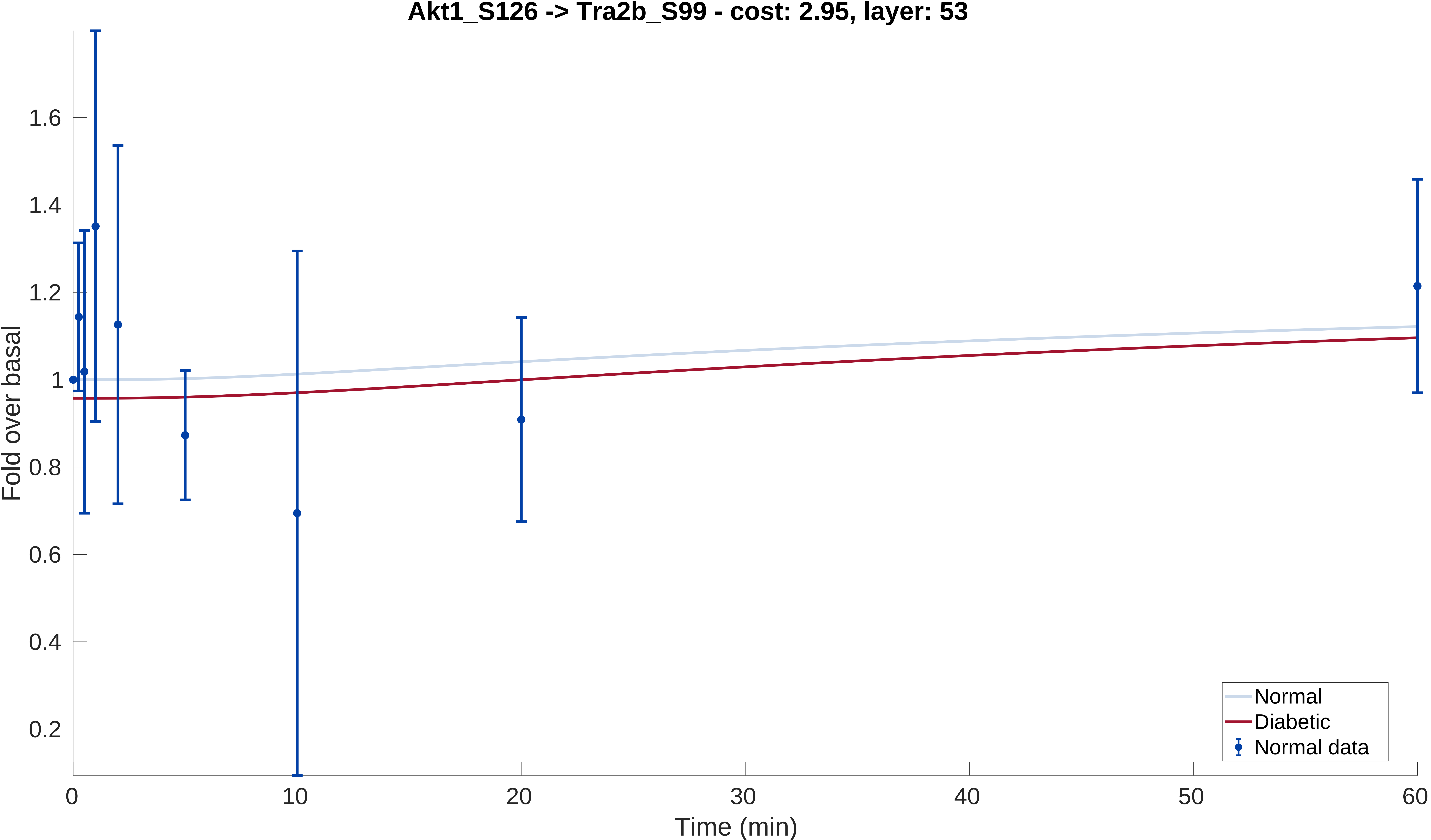
Akt1_S126 -> Tra2b_S99 - cost: 2.95, layer: 53
Fold over basal
1.6
1.4
1.2
1
0.8
0.6
0.4
0.2
0
10
20
30
40
50
60
Time (min)
Normal
Diabetic
Normal data

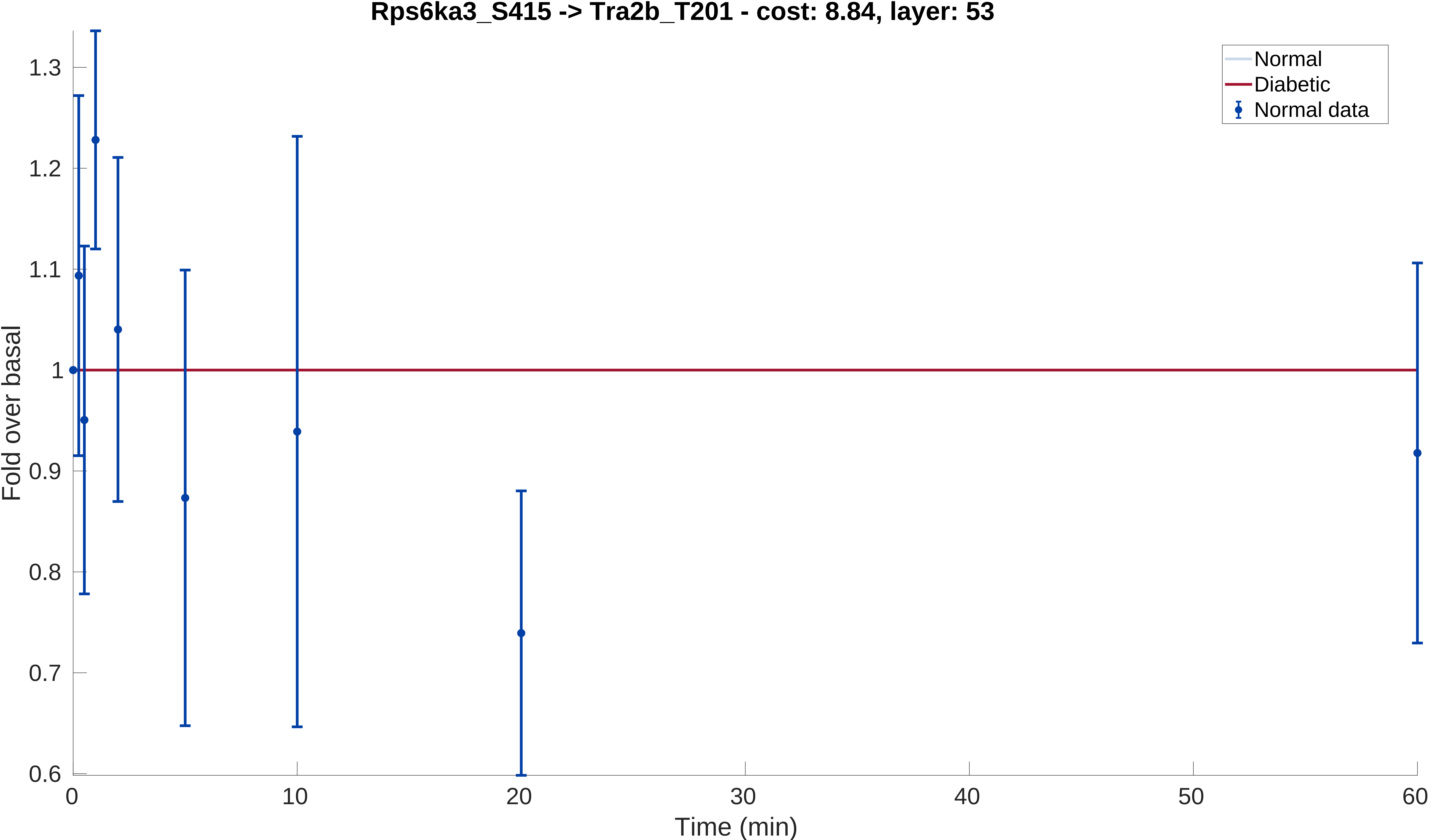

Rps6ka3_S415 -> Tra2b_T201 - cost: 8.84, layer: 53
Normal
Diabetic
Normal data
Fold over basal
1.3
1.2
1.1
1
0.9
0.8
0.7
0.6
0
10
20
30
40
50
60
Time (min)

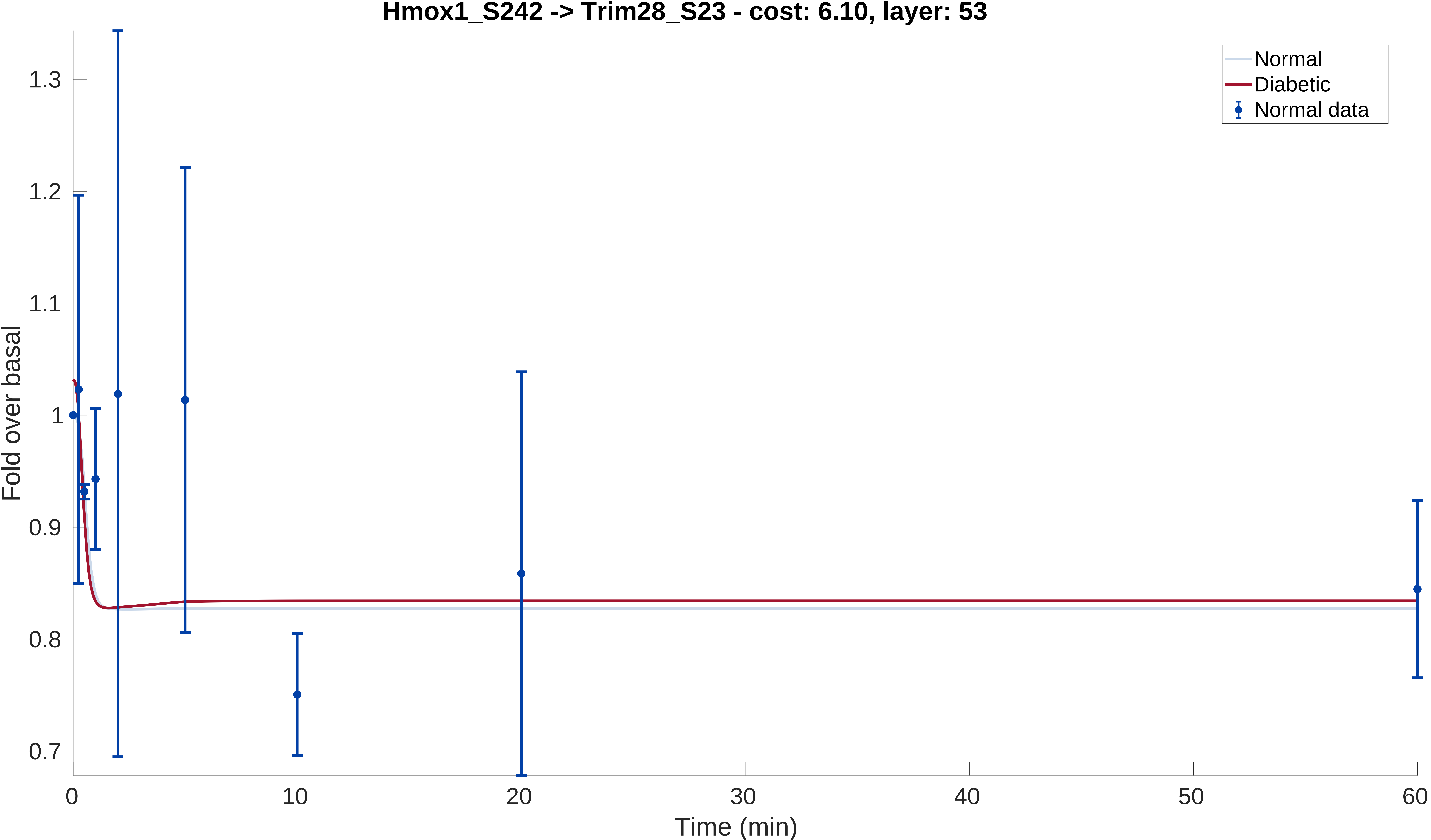
Hmox1_S242 -> Trim28_S23 - cost: 6.10, layer: 53
Normal
Diabetic
Normal data
Fold over basal
1.3
1.2
1.1
1
0.9
0.8
0.7
0
10
20
30
40
50
60
Time (min)

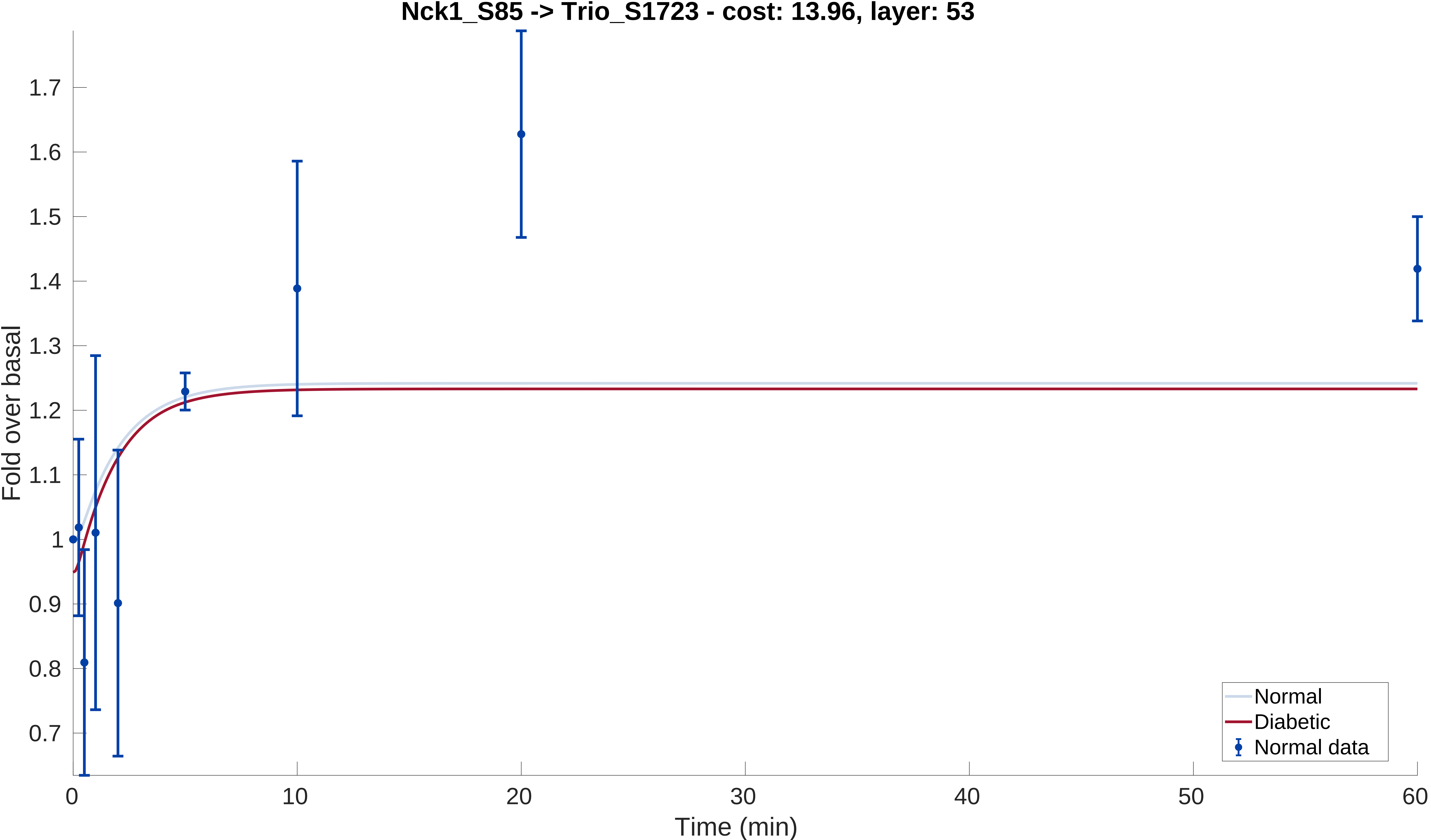

Nck1_S85 -> Trio_S1723 - cost: 13.96, layer: 53
Fold over basal
1.7
1.6
1.5
1.4
1.3
1.2
1.1
1
0.9
0.8
0.7
0
10
20
30
40
50
60
Time (min)
Normal
Diabetic
Normal data

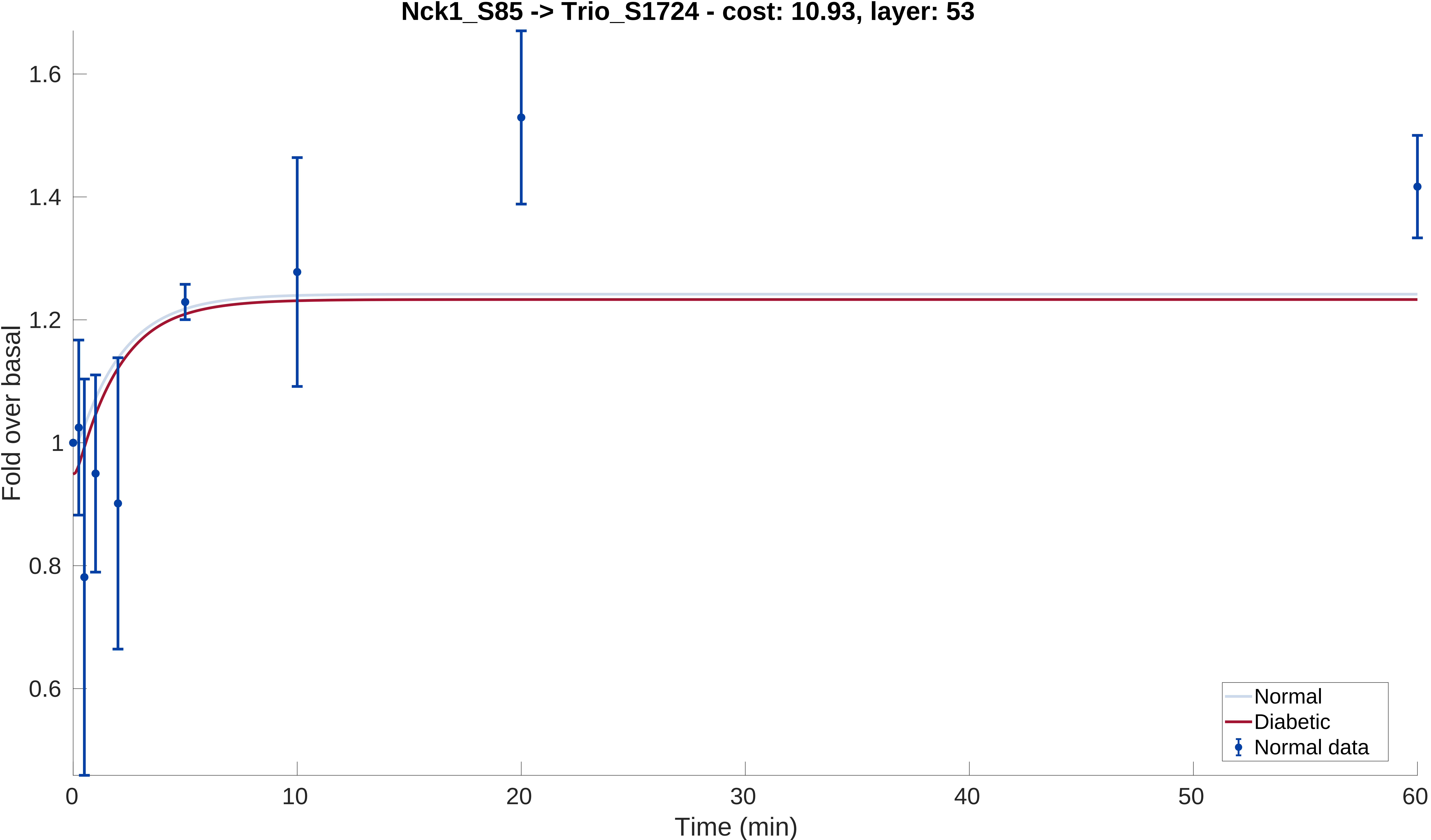

Nck1_S85 -> Trio_S1724 - cost: 10.93, layer: 53
Fold over basal
1.6
1.4
1.2
1
0.8
0.6
0
10
20
30
40
50
60
Time (min)
Normal
Diabetic
Normal data

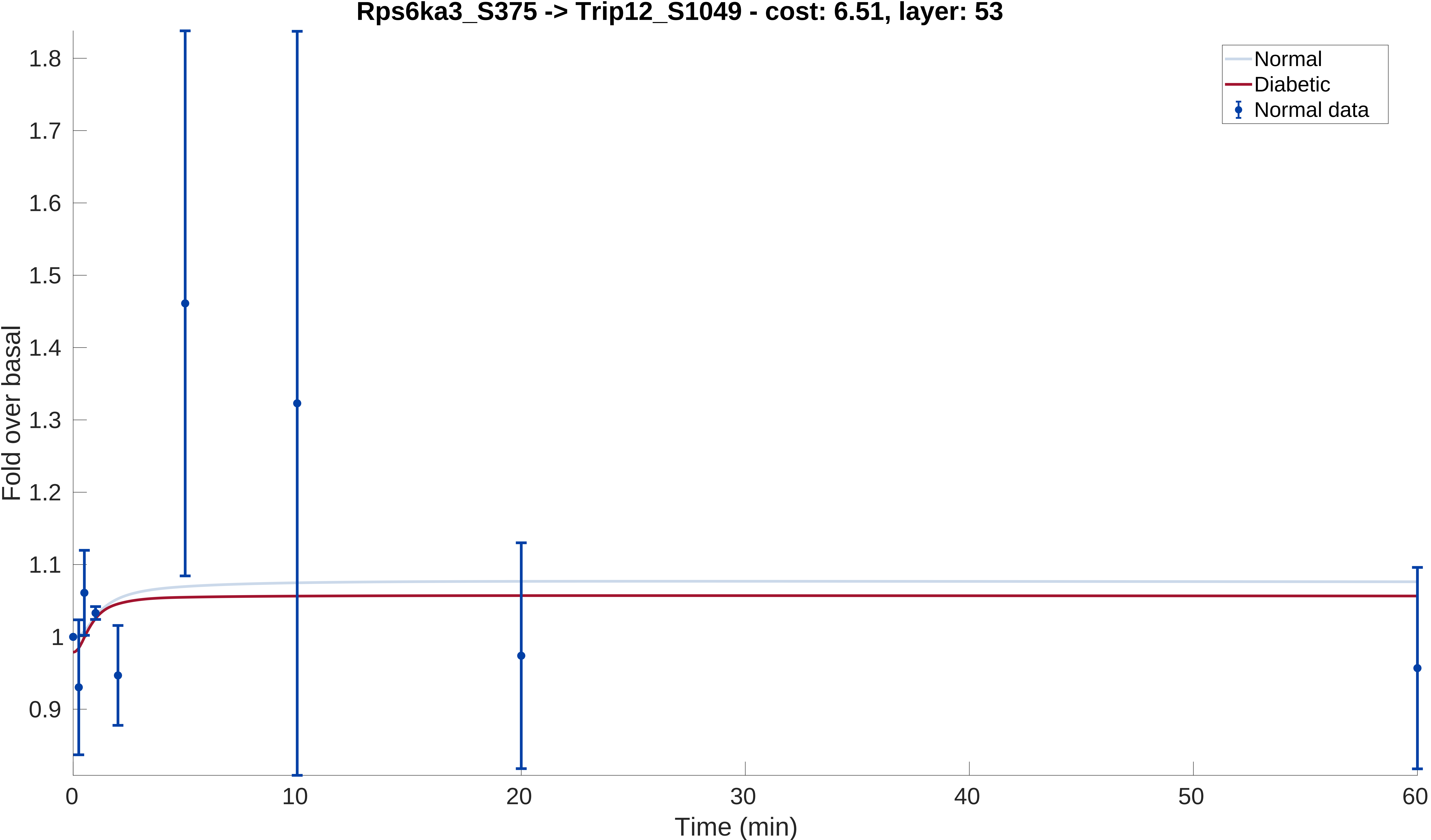

Rps6ka3_S375 -> Trip12_S1049 - cost: 6.51, layer: 53
Normal
Diabetic
Normal data
Fold over basal
1.8
1.7
1.6
1.5
1.4
1.3
1.2
1.1
1
0.9
0
10
20
30
40
50
60
Time (min)

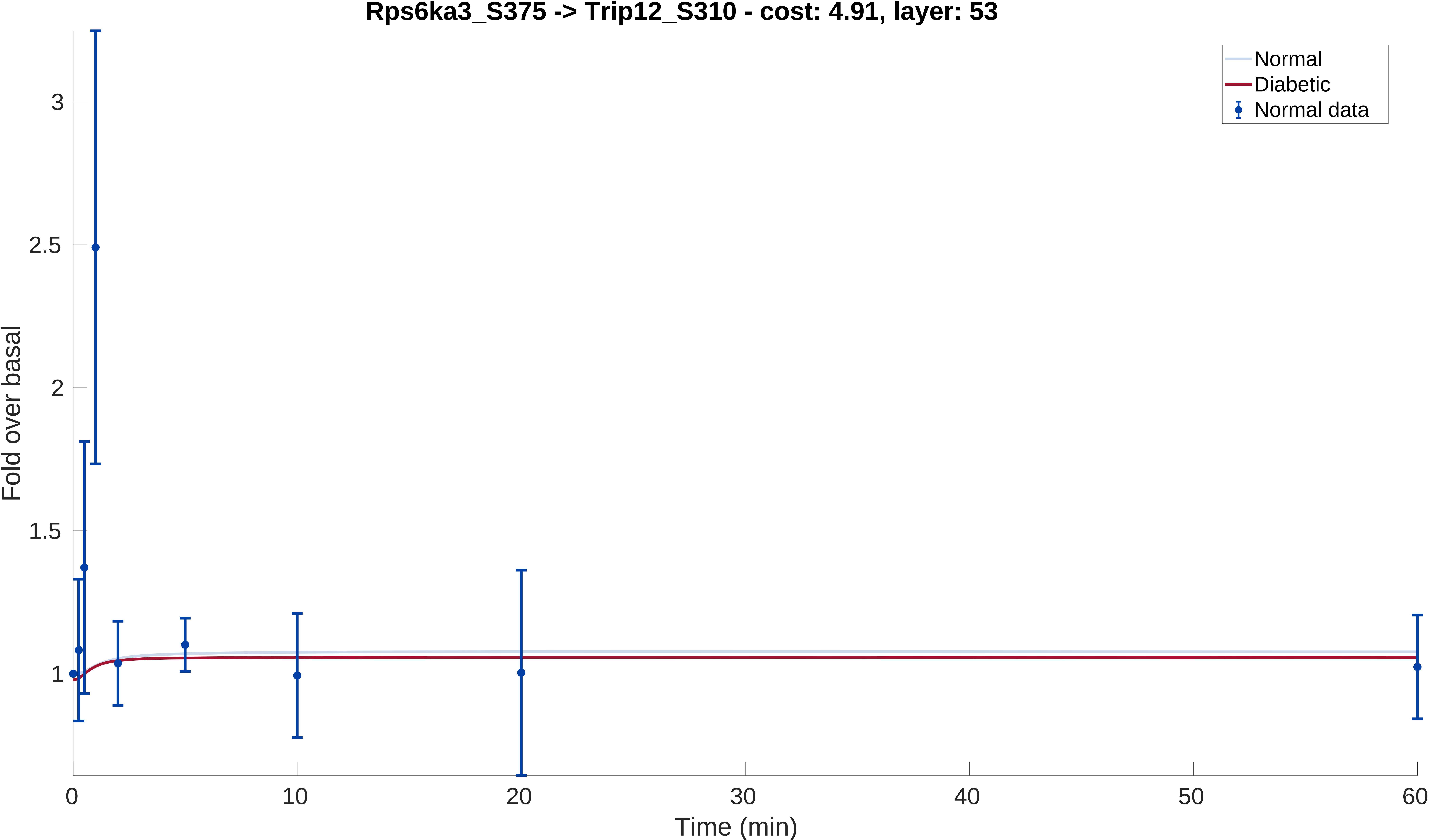

Rps6ka3_S375 -> Trip12_S310 - cost: 4.91, layer: 53
Normal
Diabetic
Normal data
Fold over basal
3
2.5
2
1.5
1
0
10
20
30
40
50
60
Time (min)

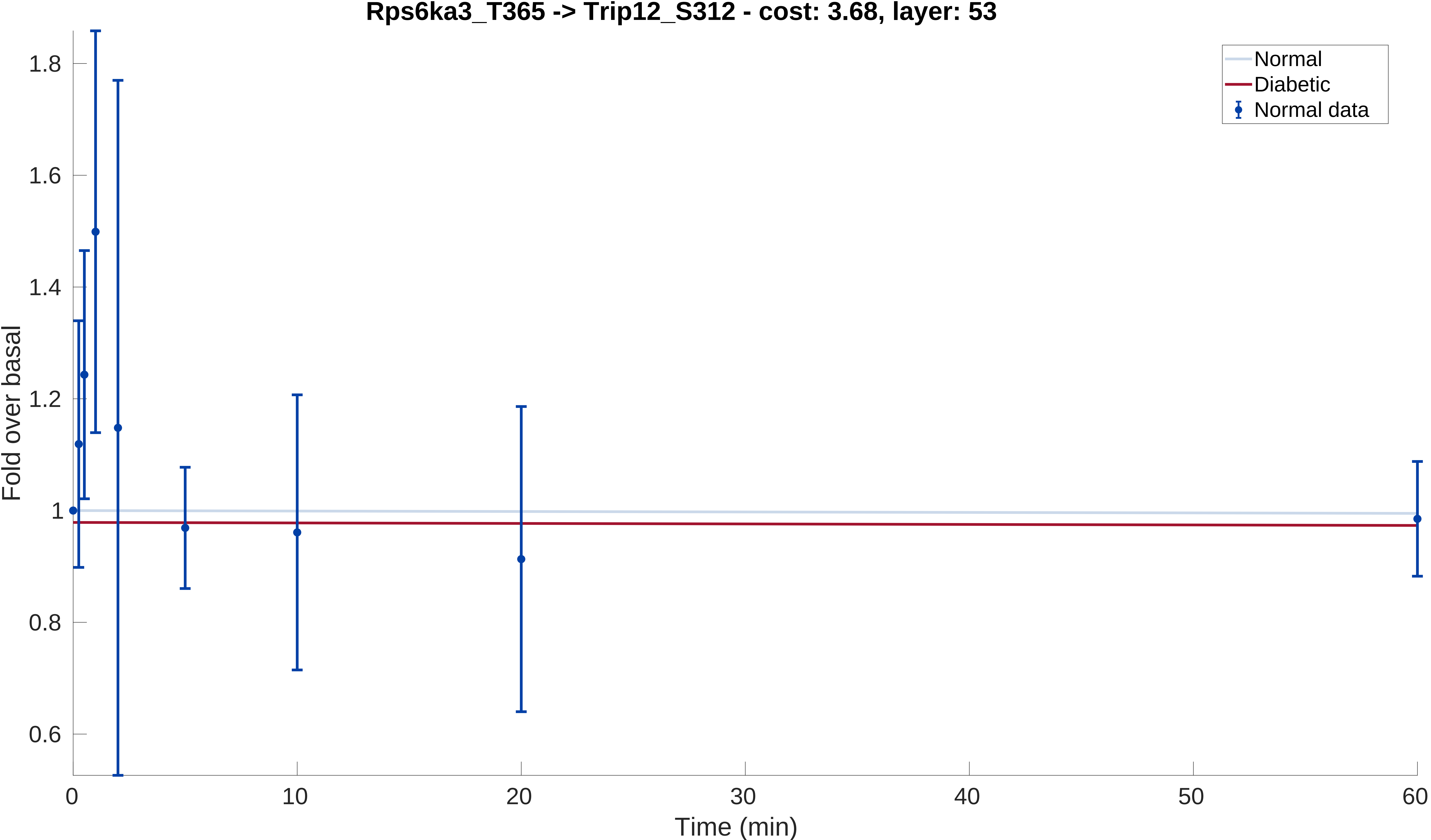

Rps6ka3_T365 -> Trip12_S312 - cost: 3.68, layer: 53
Normal
Diabetic
Normal data
Fold over basal
1.8
1.6
1.4
1.2
1
0.8
0.6
0
10
20
30
40
50
60
Time (min)

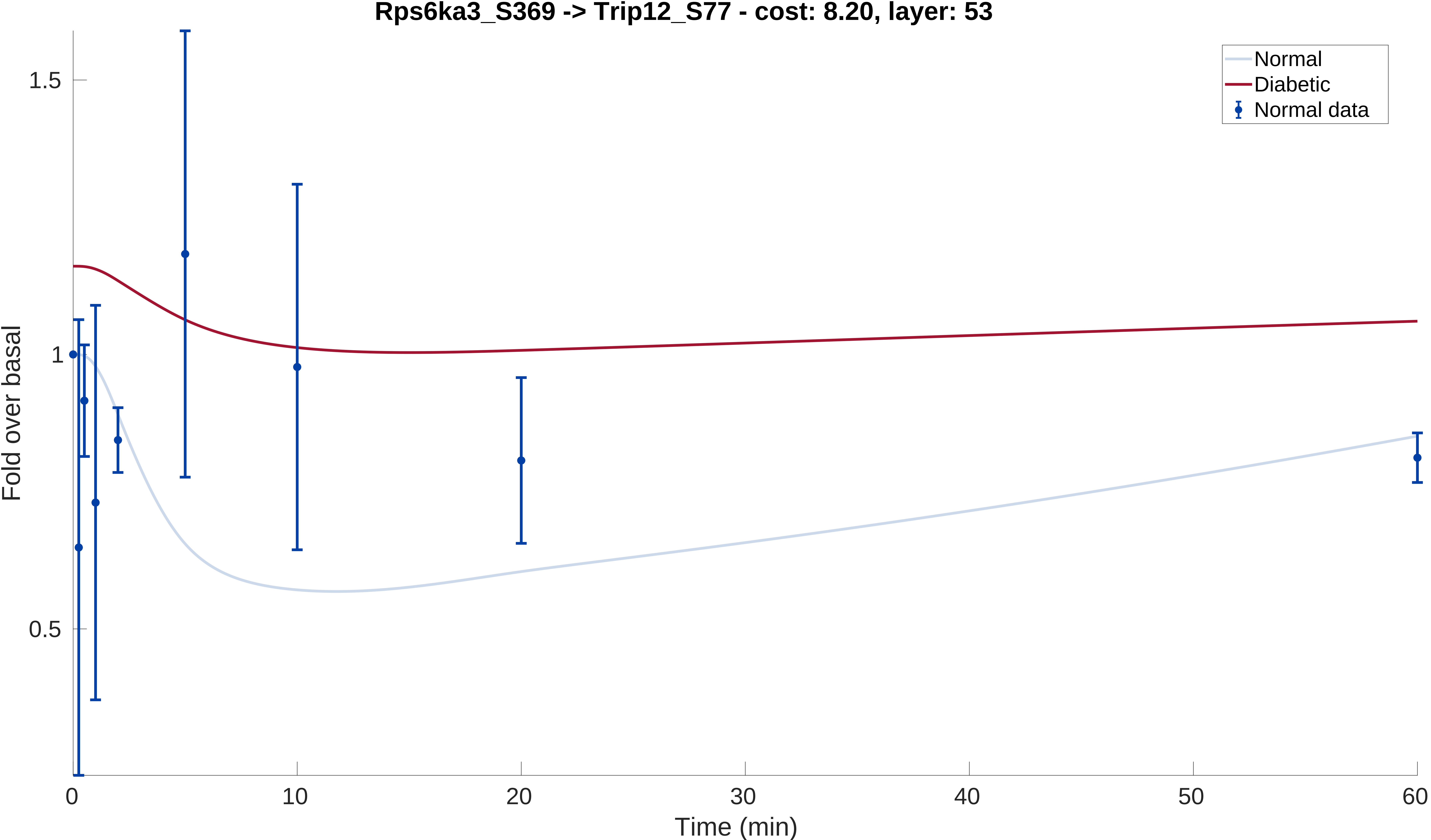

Rps6ka3_S369 -> Trip12_S77 - cost: 8.20, layer: 53
Normal
Diabetic
Normal data
1.5
1
0.5
Fold over basal
0
10
20
30
40
50
60
Time (min)

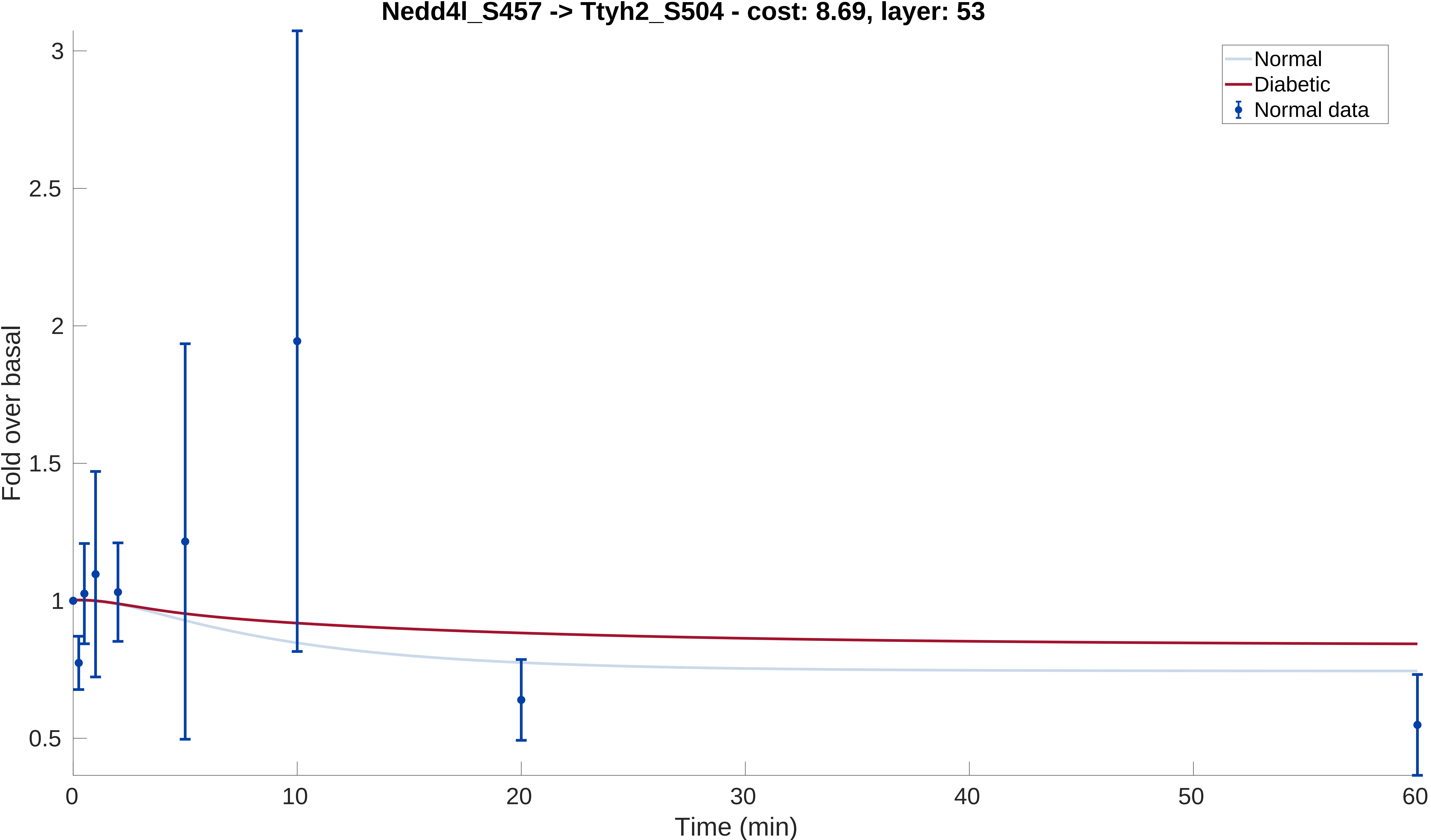

Nedd4l_S457 -> Ttyh2_S504 - cost: 8.69, layer: 53
Normal
Diabetic
Normal data
Fold over basal
Time (min)
0.5
1
1.5
2
2.5
3
0
10
20
30
40
50
60

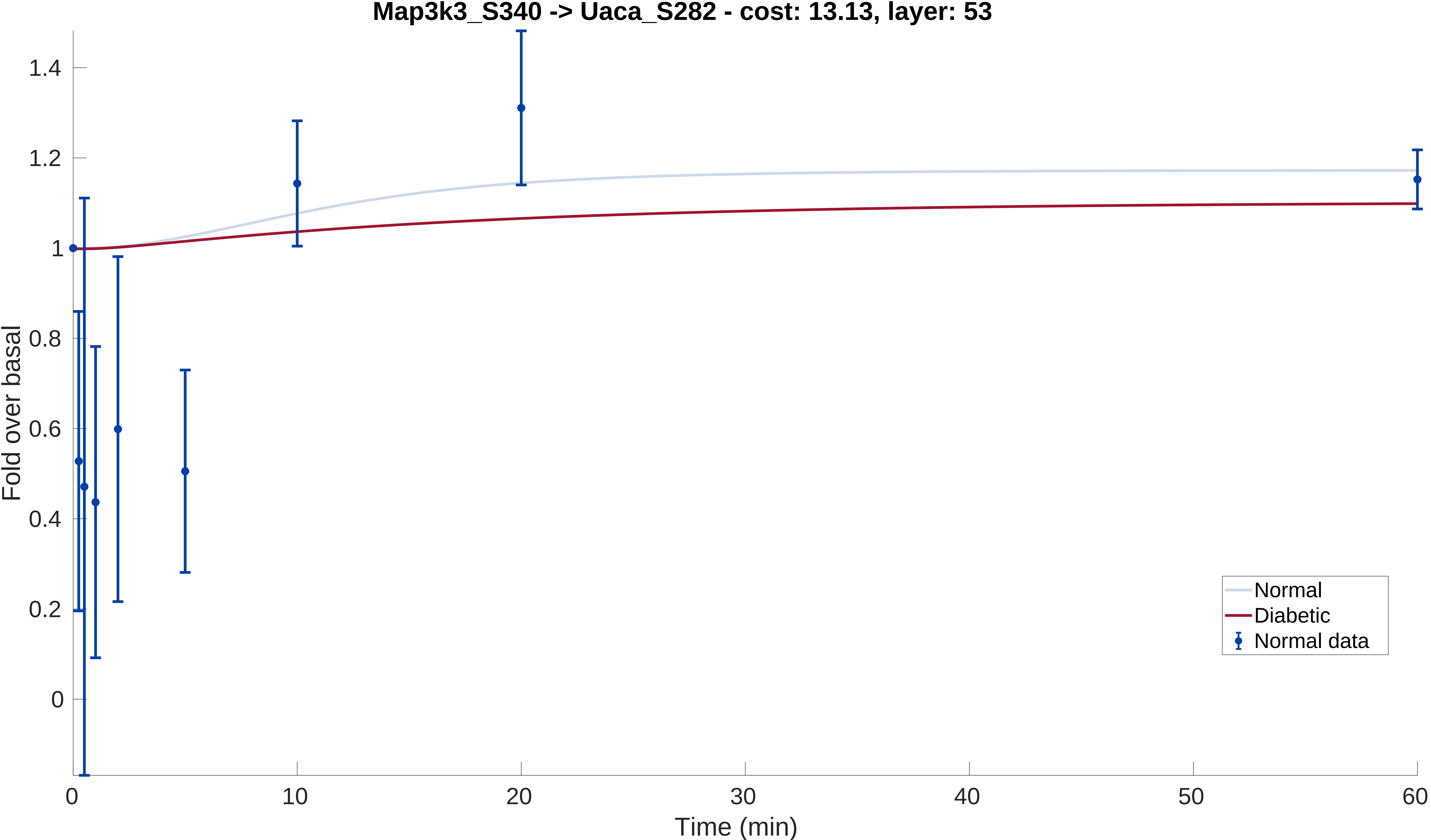

Map3k3_S340 -> Uaca_S282 - cost: 13.13, layer: 53
Fold over basal
1.4
1.2
1
0.8
0.6
0.4
0.2
0
0
10
20
30
40
50
60
Time (min)
Normal
Diabetic
Normal data

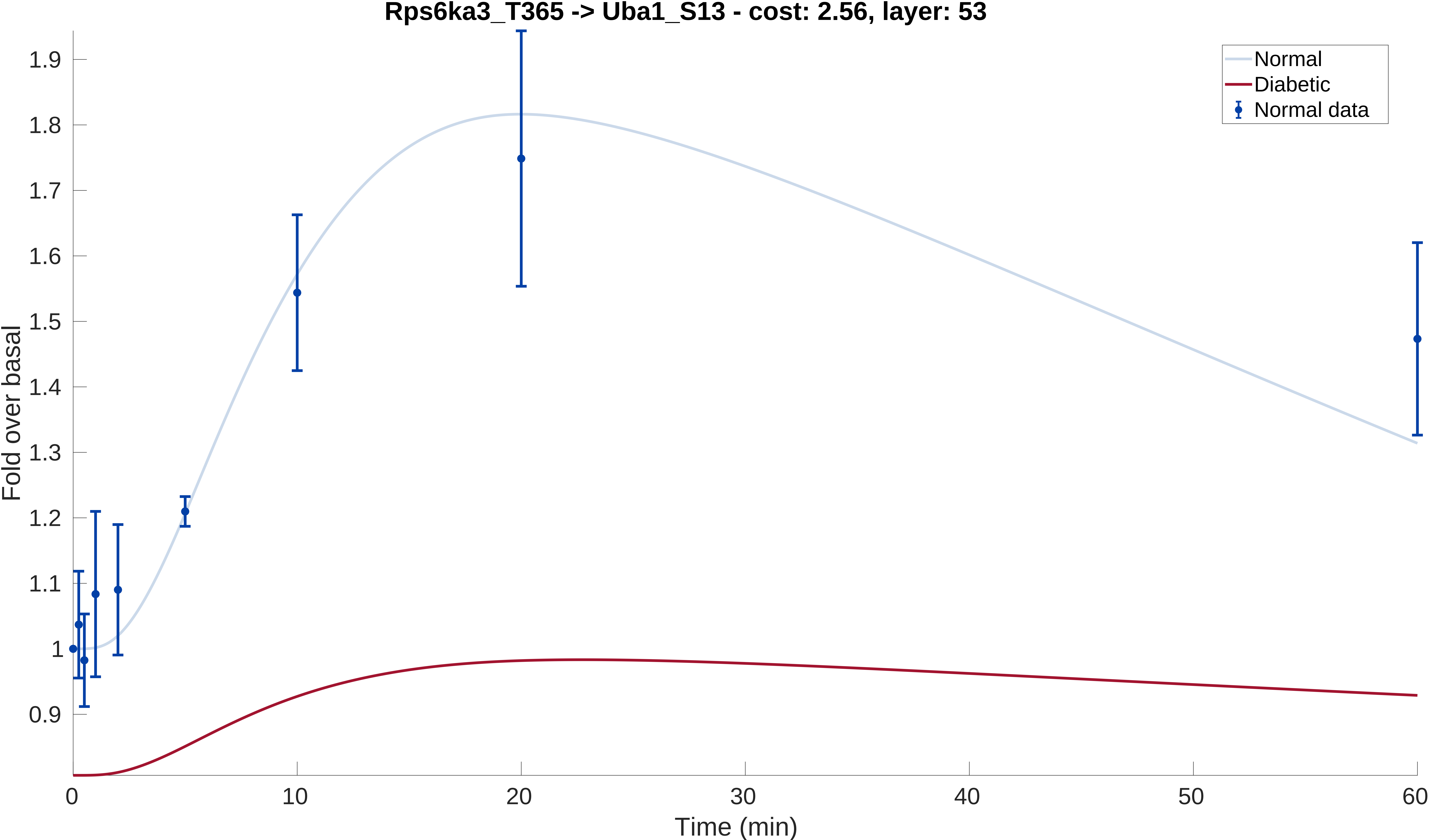
Rps6ka3_T365 -> Uba1_S13 - cost: 2.56, layer: 53
Normal
Diabetic
Normal data
Fold over basal
1.9
1.8
1.7
1.6
1.5
1.4
1.3
1.2
1.1
1
0.9
0
10
20
30
40
50
60
Time (min)

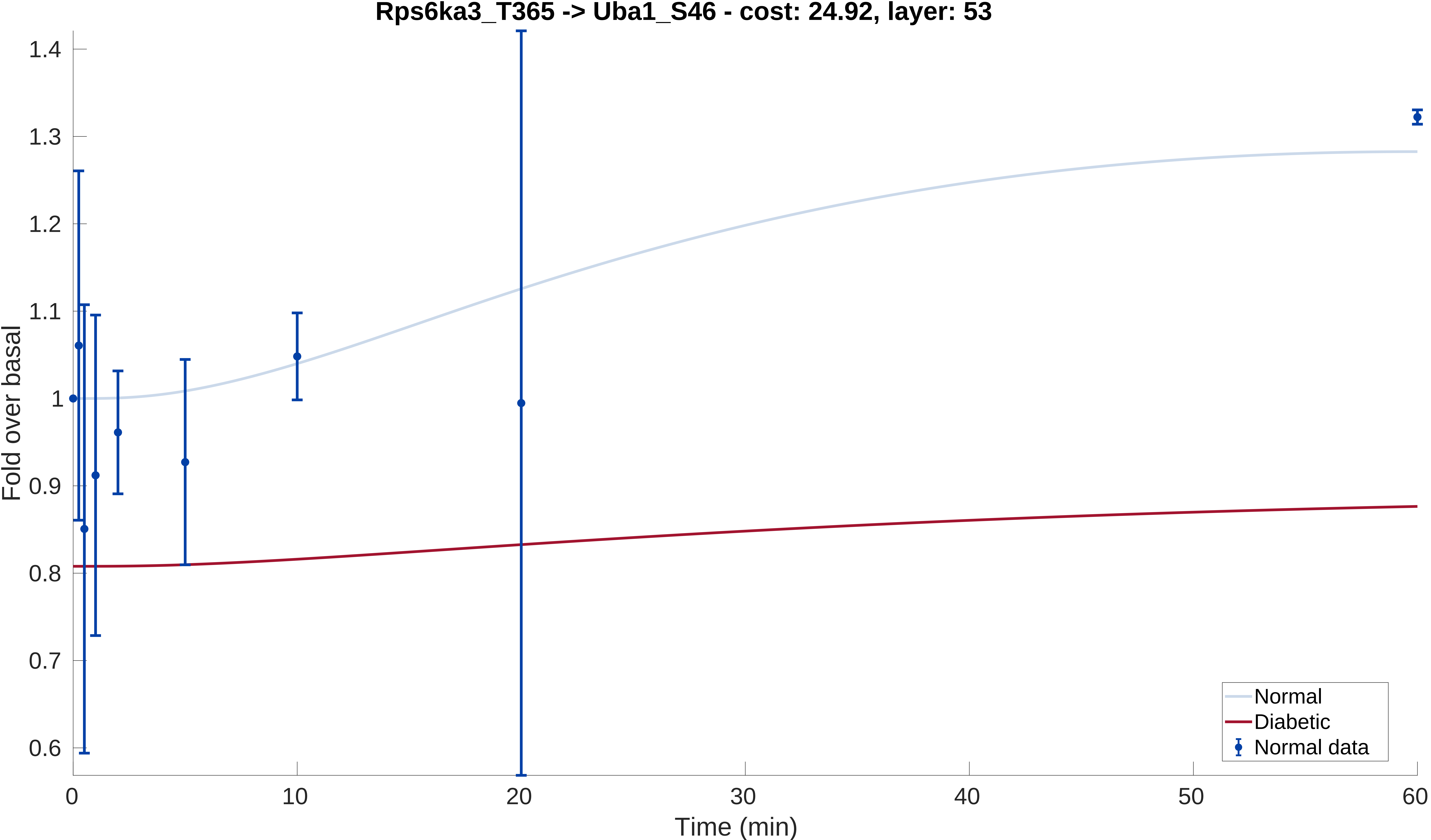

Rps6ka3_T365 -> Uba1_S46 - cost: 24.92, layer: 53
Fold over basal
1.4
1.3
1.2
1.1
1
0.9
0.8
0.7
0.6
0
10
20
30
40
50
60
Time (min)
Normal
Diabetic
Normal data

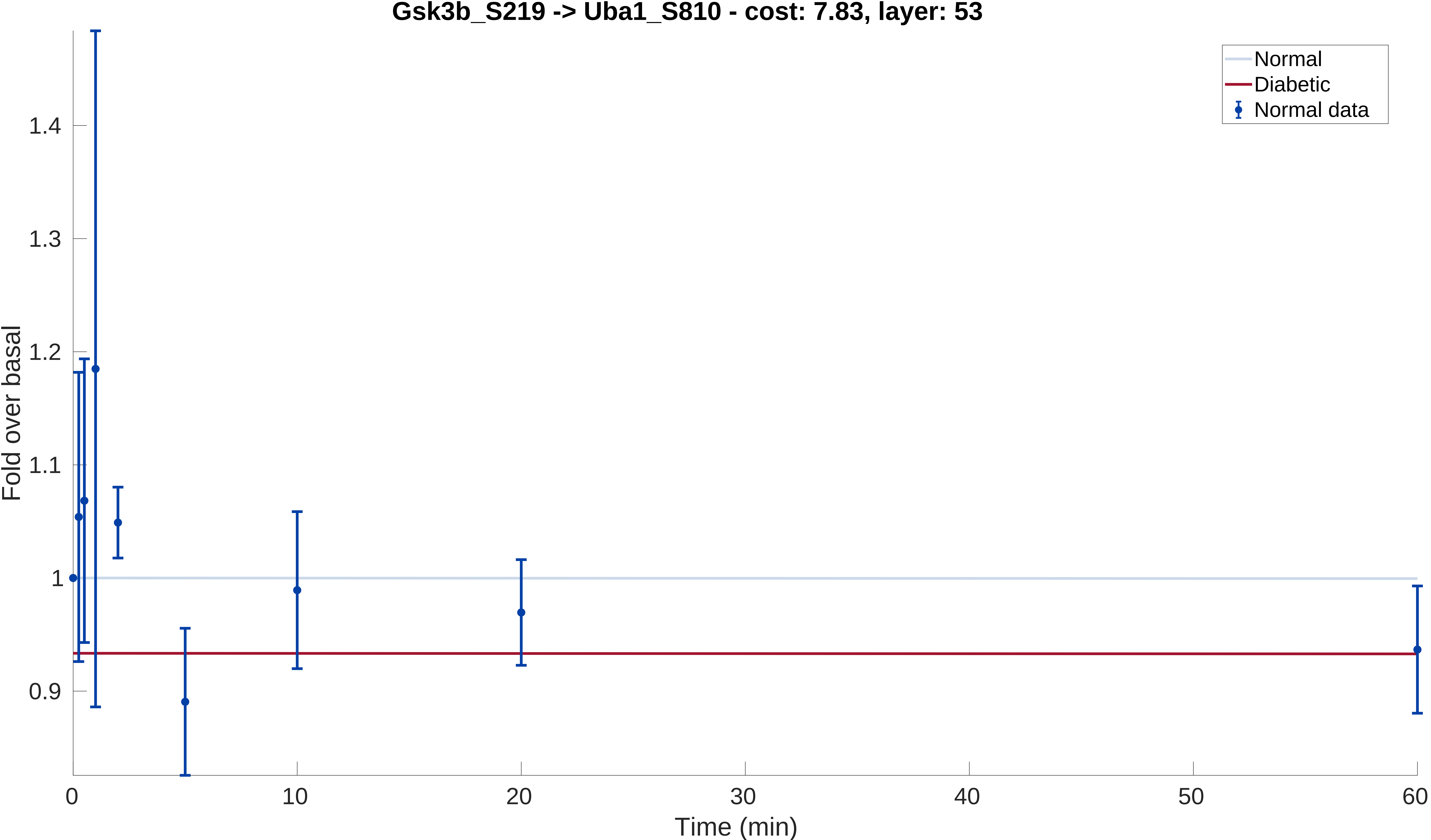

Gsk3b_S219 -> Uba1_S810 - cost: 7.83, layer: 53
Normal
Diabetic
Normal data
Fold over basal
1.4
1.3
1.2
1.1
1
0.9
0
10
20
30
40
50
60
Time (min)

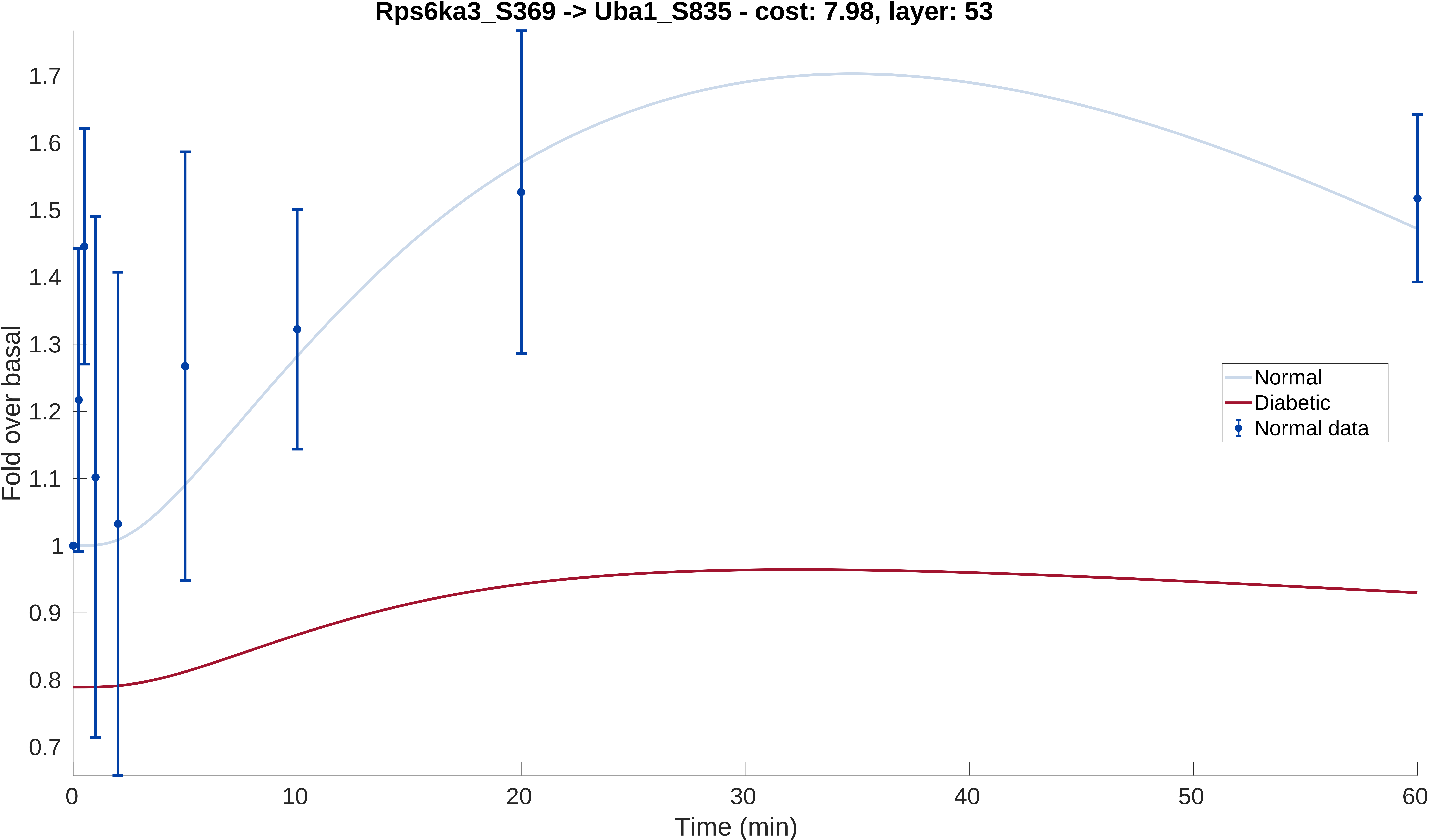
Rps6ka3_S369 -> Uba1_S835 - cost: 7.98, layer: 53
Fold over basal
1.7
1.6
1.5
1.4
1.3
1.2
1.1
1
0.9
0.8
0.7
0
10
20
30
40
50
60
Time (min)
Normal
Diabetic
Normal data

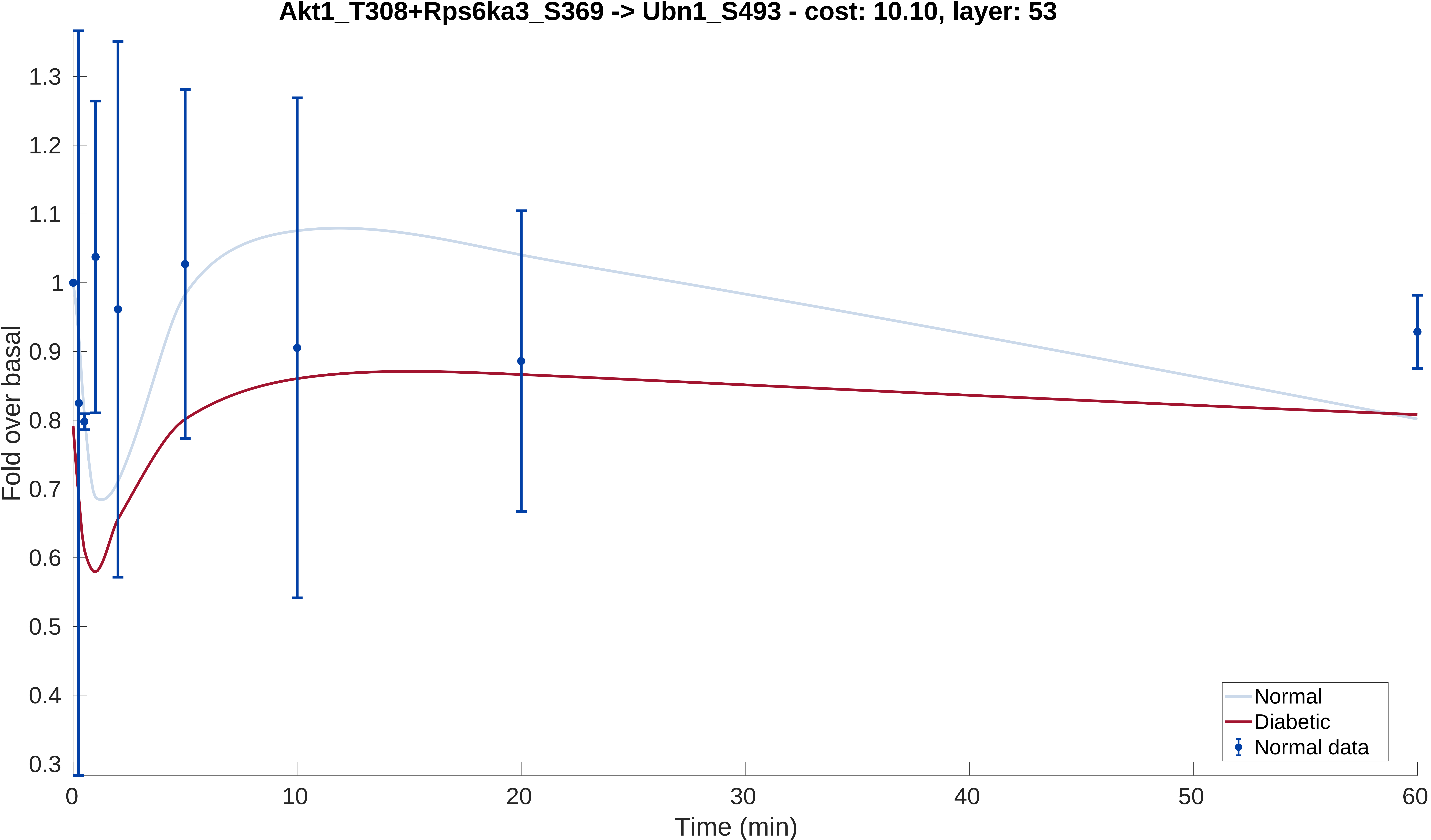

Akt1_T308+Rps6ka3_S369 -> Ubn1_S493 - cost: 10.10, layer: 53
Fold over basal
1.3
1.2
1.1
1
0.9
0.8
0.7
0.6
0.5
0.4
0.3
0
10
20
30
40
50
60
Time (min)
Normal
Diabetic
Normal data

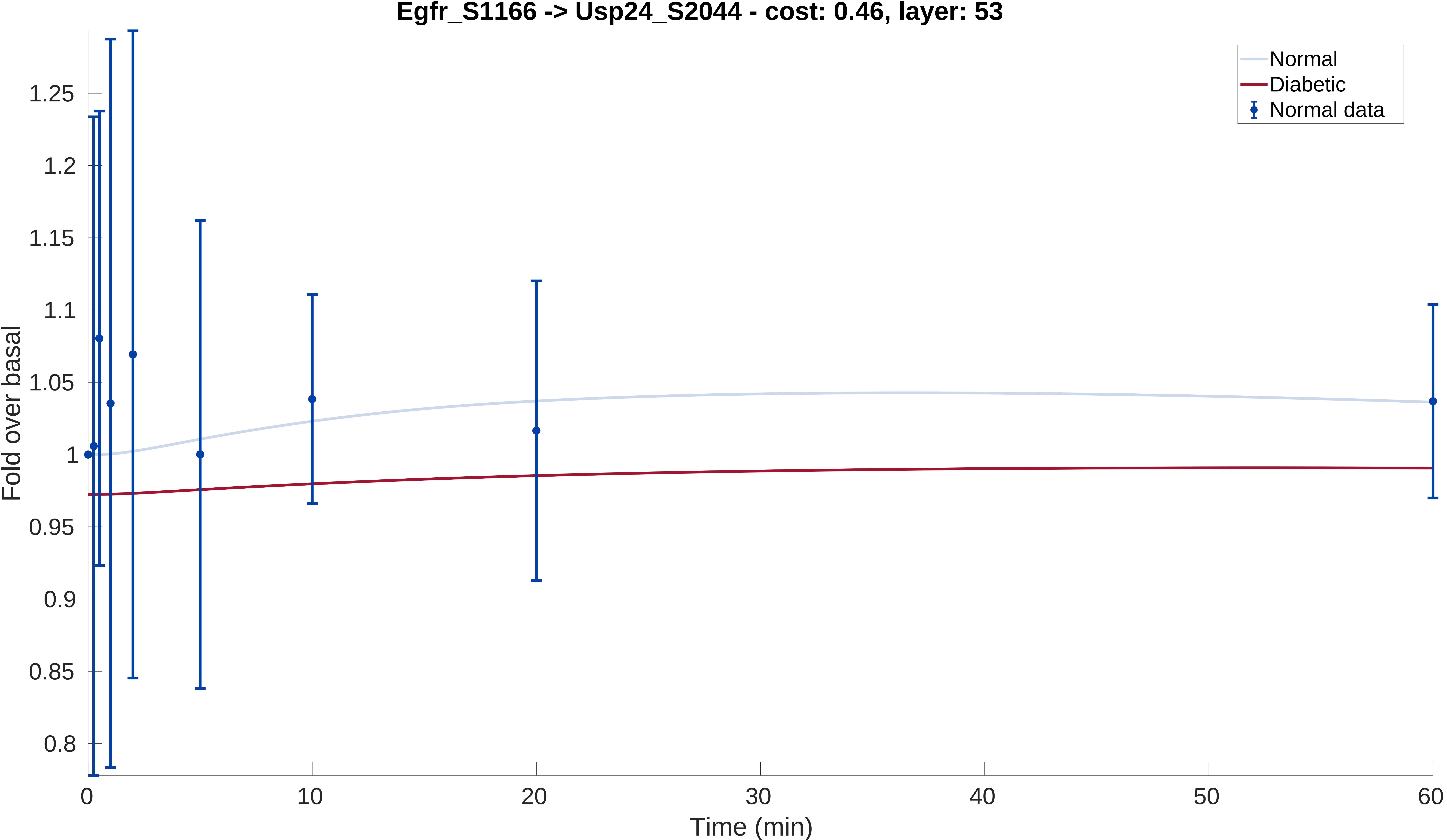

Egfr_S1166 -> Usp24_S2044 - cost: 0.46, layer: 53
Normal
Diabetic
Normal data
Fold over basal
1.25
1.2
1.15
1.1
1.05
1
0.95
0.9
0.85
0.8
0
10
20
30
40
50
60
Time (min)

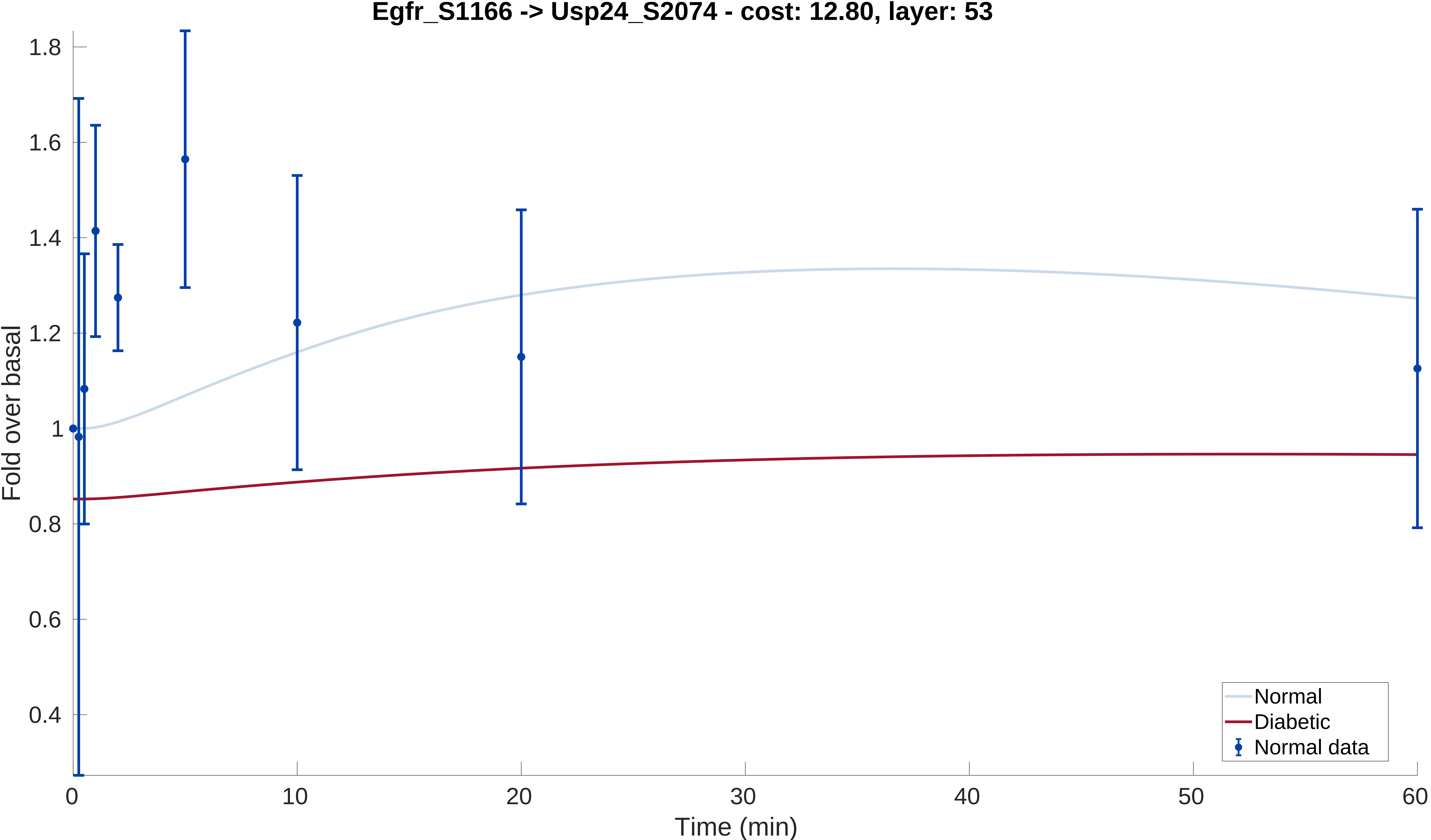
Egfr_S1166 -> Usp24_S2074 - cost: 12.80, layer: 53
Fold over basal
Time (min)
1.8
1.6
1.4
1.2
1
0.8
0.6
0.4
0
10
20
30
40
50
60
Normal
Diabetic
Normal data

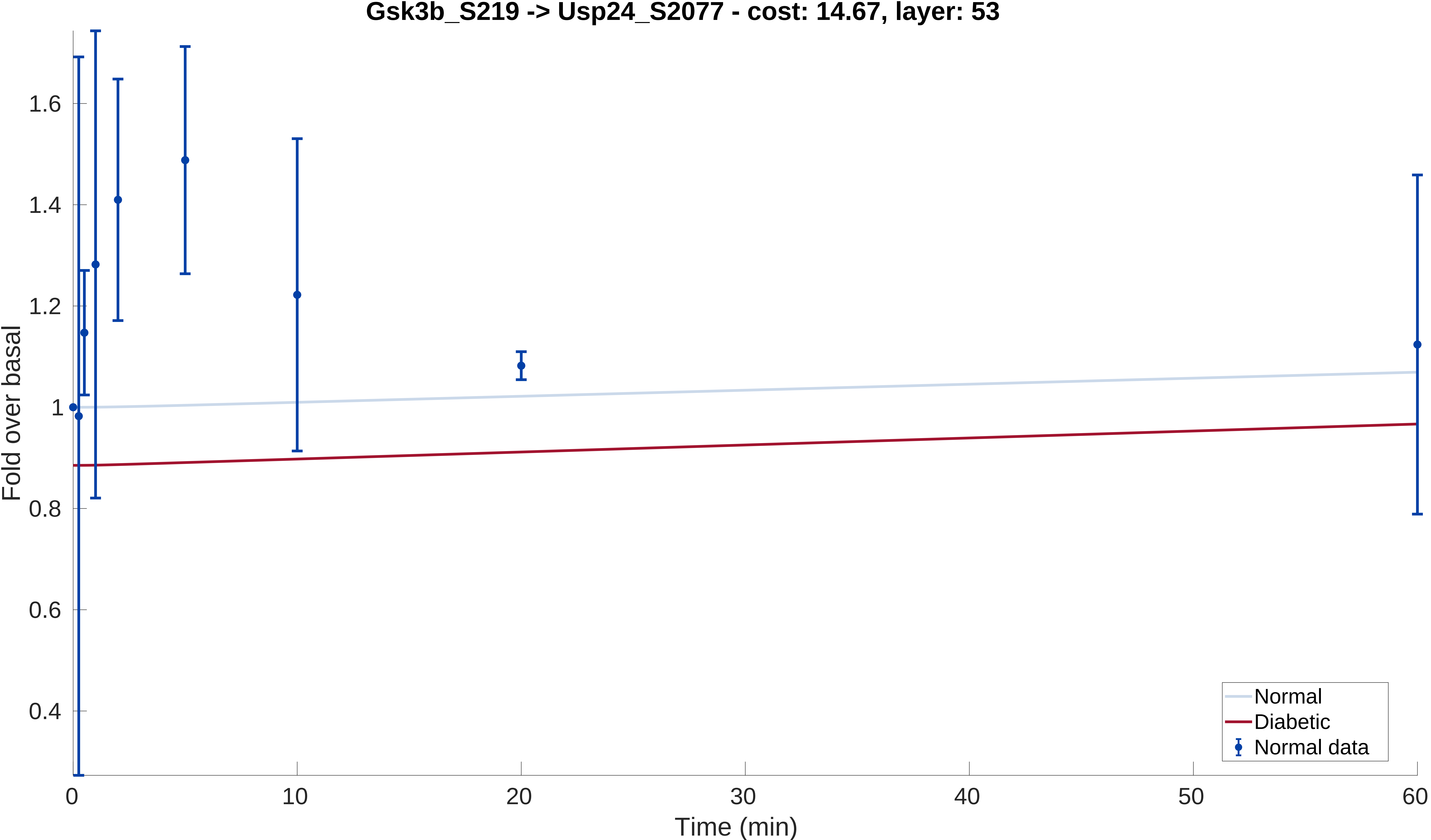
Gsk3b_S219 -> Usp24_S2077 - cost: 14.67, layer: 53
Fold over basal
1.6
1.4
1.2
1
0.8
0.6
0.4
0
10
20
30
40
50
60
Time (min)
Normal
Diabetic
Normal data

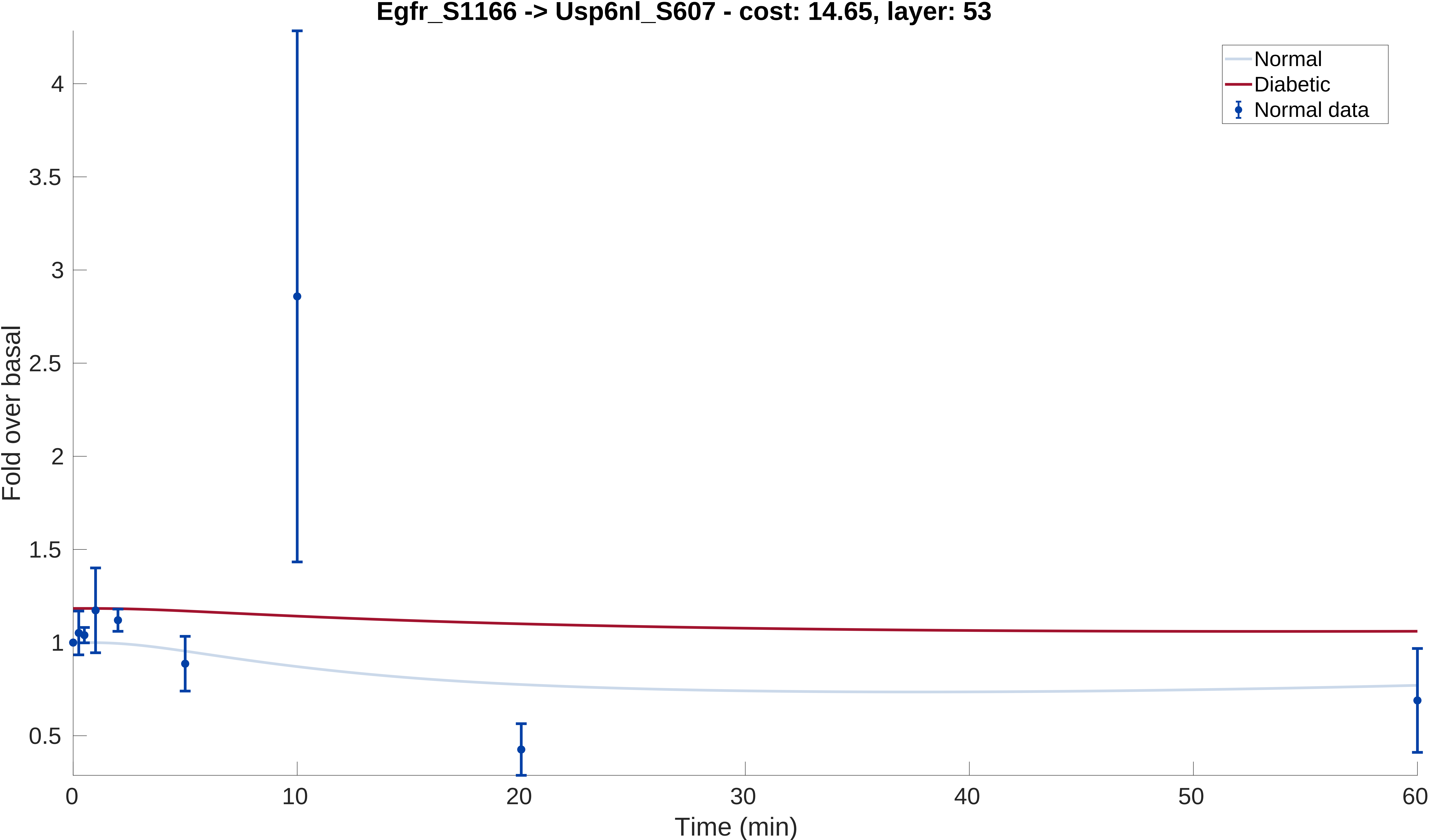

Egfr_S1166 -> Usp6nl_S607 - cost: 14.65, layer: 53
Normal
Diabetic
Normal data
Fold over basal
Time (min)
0.5
1
1.5
2
2.5
3
3.5
4
0
10
20
30
40
50
60

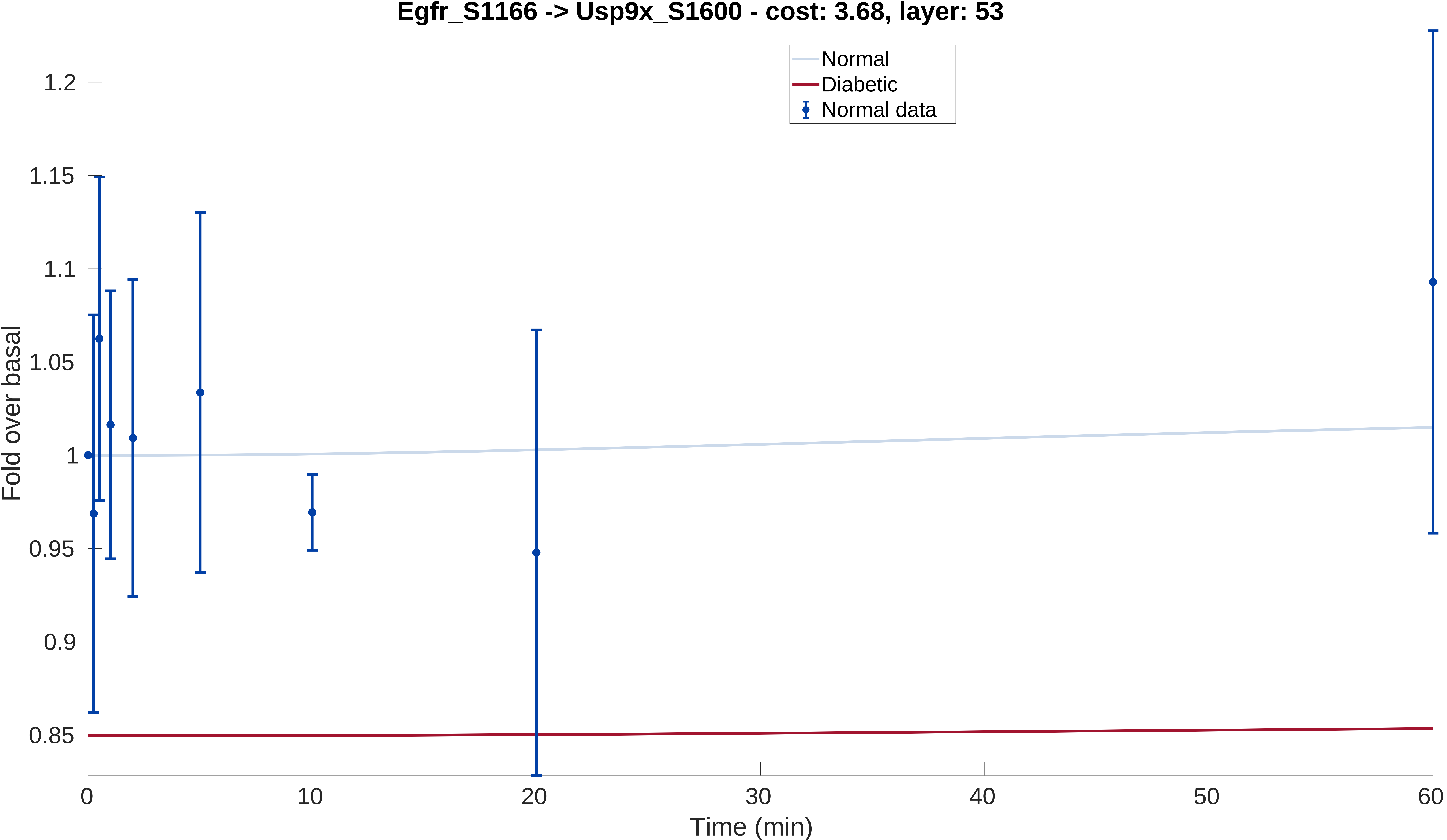
Egfr_S1166 -> Usp9x_S1600 - cost: 3.68, layer: 53
Normal
Diabetic
Normal data
Fold over basal
1.2
1.15
1.1
1.05
1
0.95
0.9
0.85
0
10
20
30
40
50
60
Time (min)

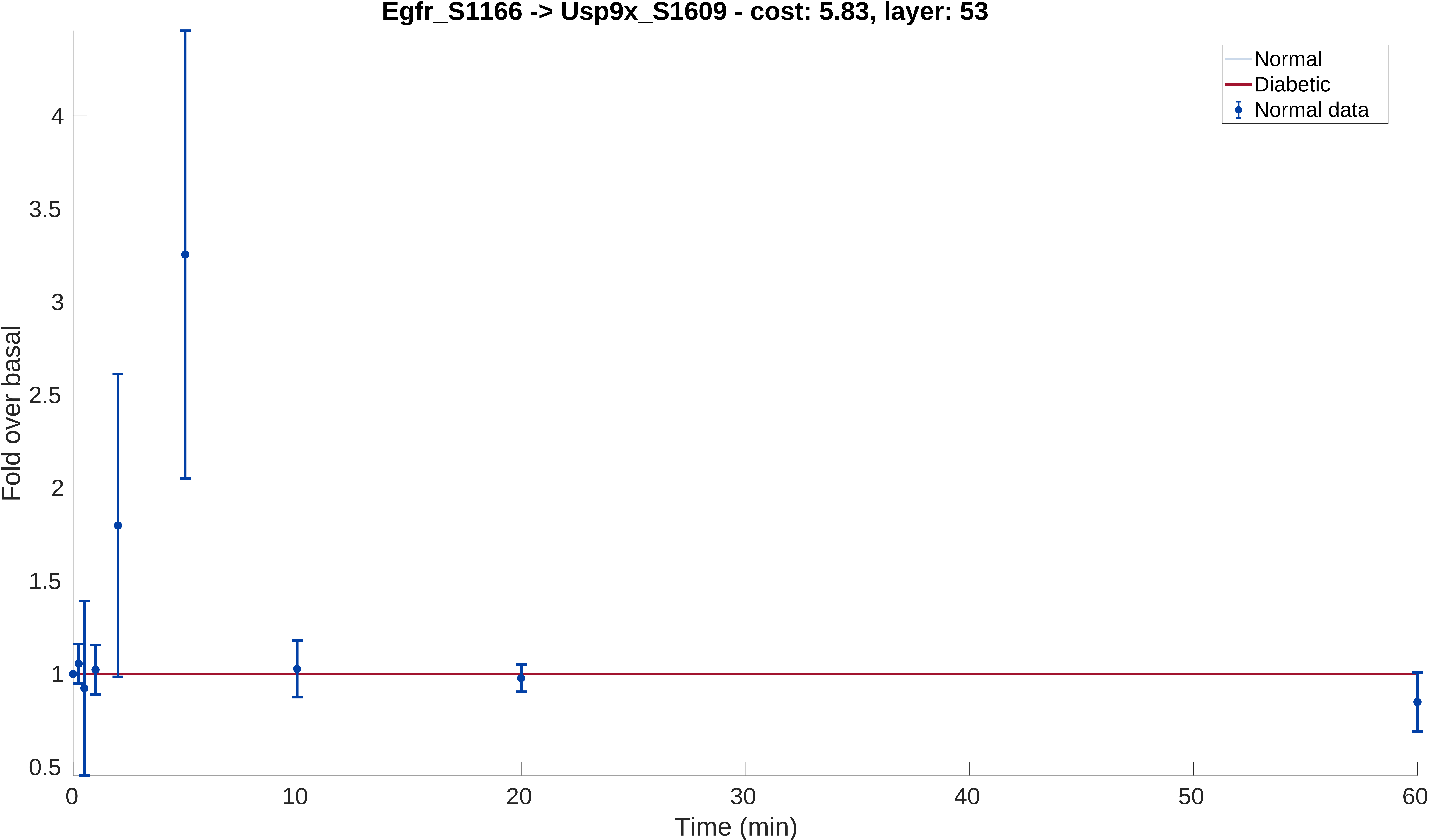

Egfr_S1166 -> Usp9x_S1609 - cost: 5.83, layer: 53
Normal
Diabetic
Normal data
Fold over basal
0.5
1
1.5
2
2.5
3
3.5
4
0
10
20
30
40
50
60
Time (min)

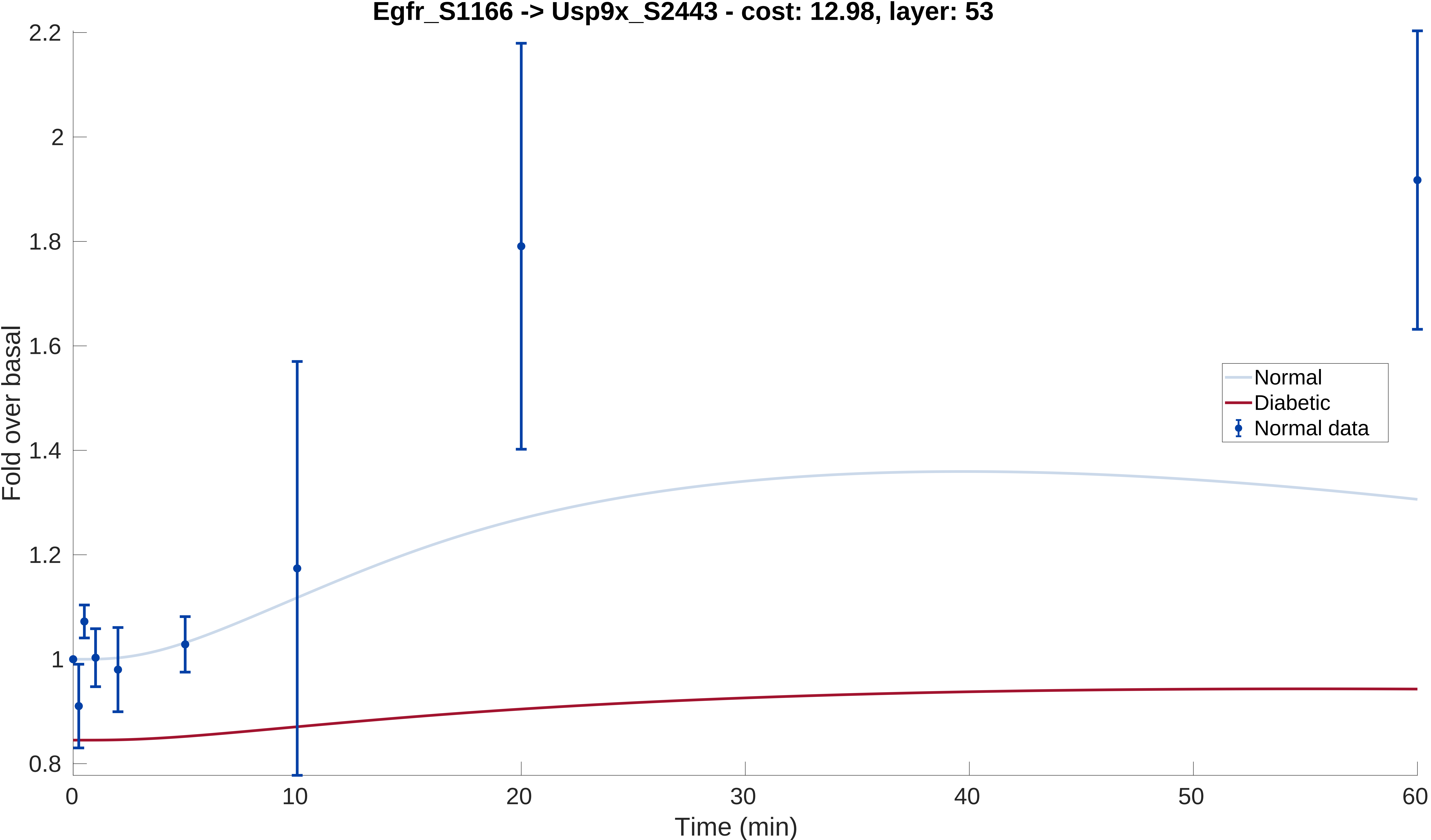

Egfr_S1166 -> Usp9x_S2443 - cost: 12.98, layer: 53
Fold over basal
Time (min)
0.8
1
1.2
1.4
1.6
1.8
2
2.2
0
10
20
30
40
50
60
Normal
Diabetic
Normal data

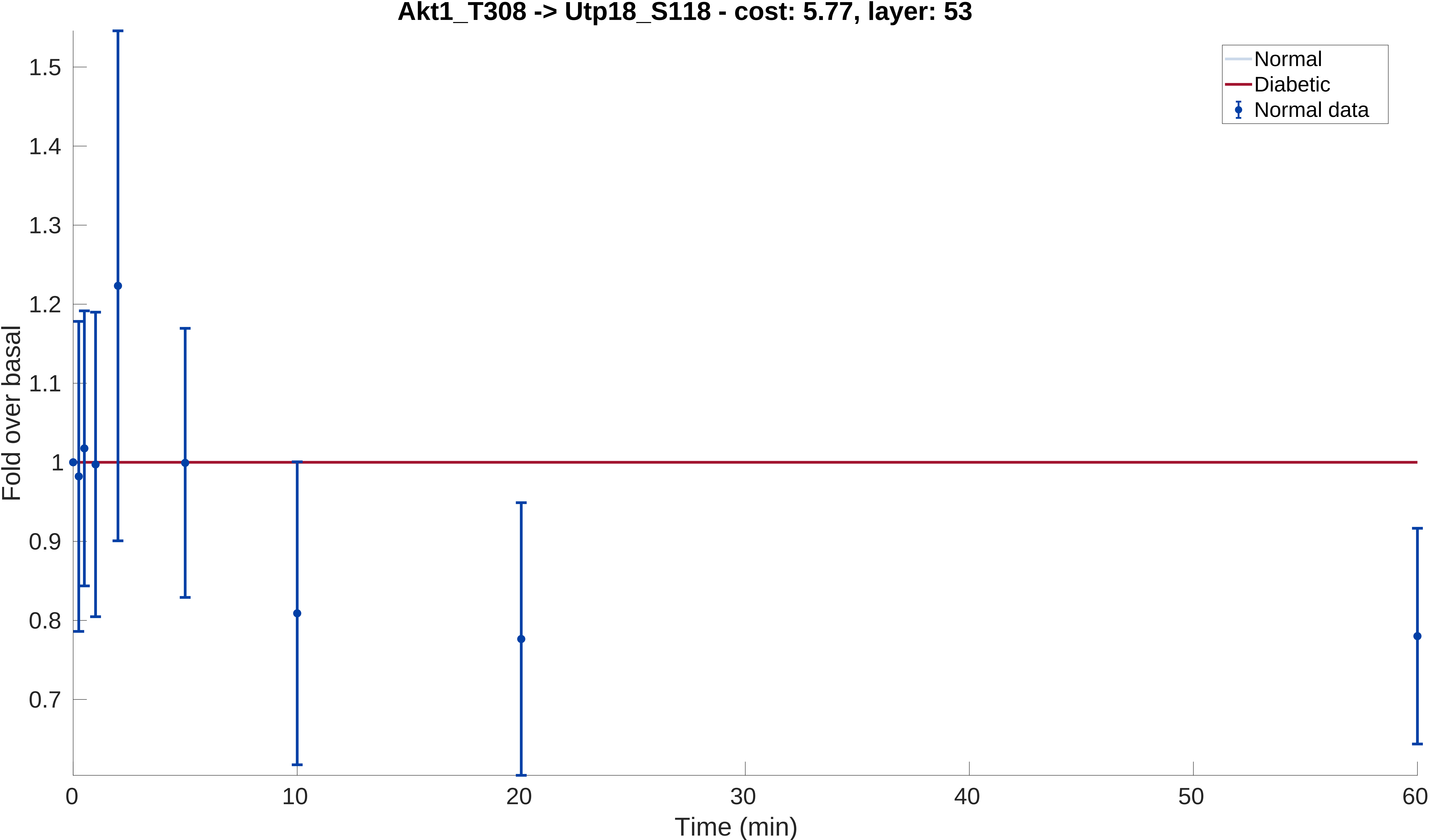

Akt1_T308 -> Utp18_S118 - cost: 5.77, layer: 53
Normal
Diabetic
Normal data
Fold over basal
1.5
1.4
1.3
1.2
1.1
1
0.9
0.8
0.7
0
10
20
30
40
50
60
Time (min)

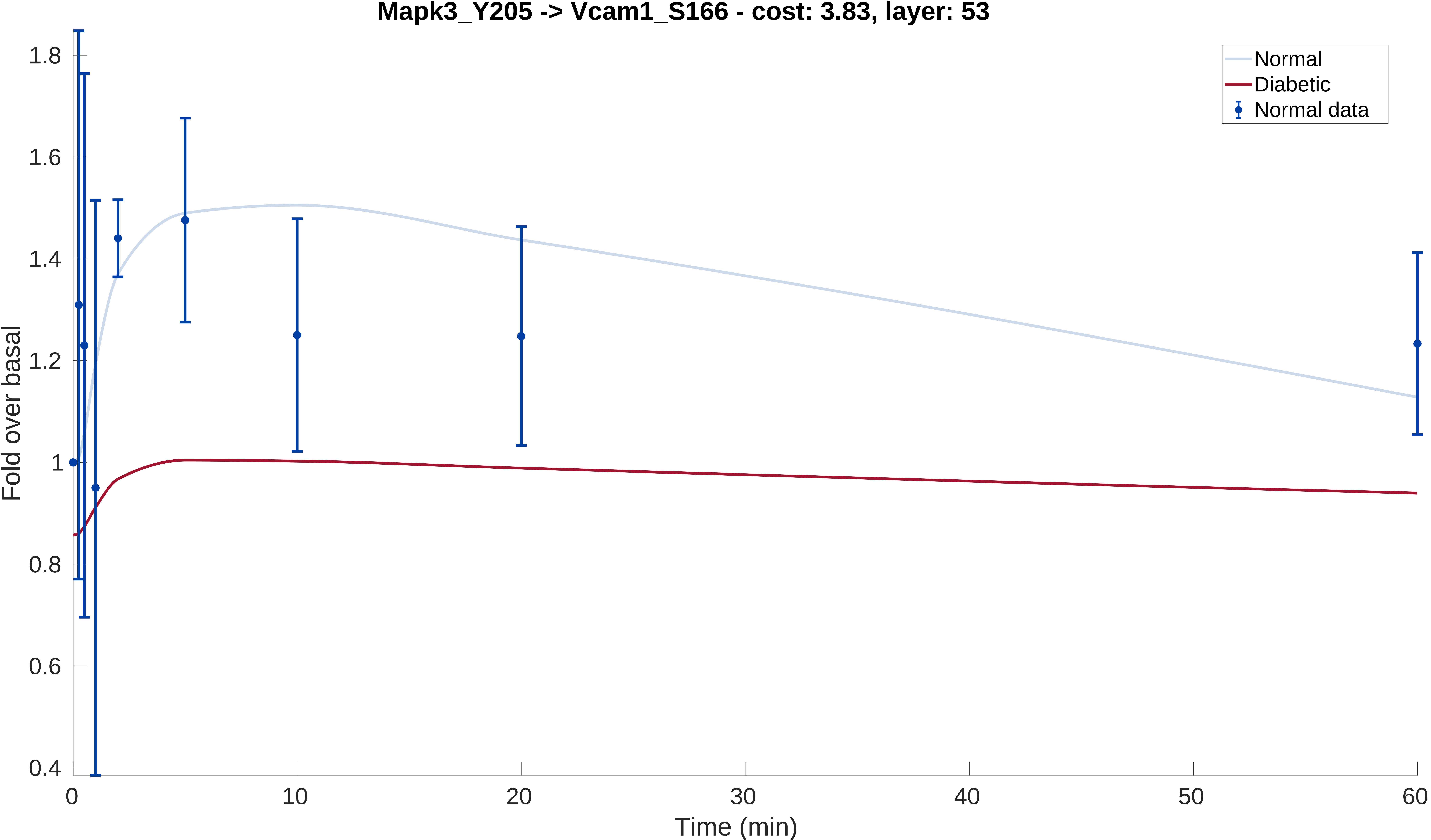

Mapk3_Y205 -> Vcam1_S166 - cost: 3.83, layer: 53
Normal
Diabetic
Normal data
Fold over basal
Time (min)
0.4
0.6
0.8
1
1.2
1.4
1.6
1.8
0
10
20
30
40
50
60

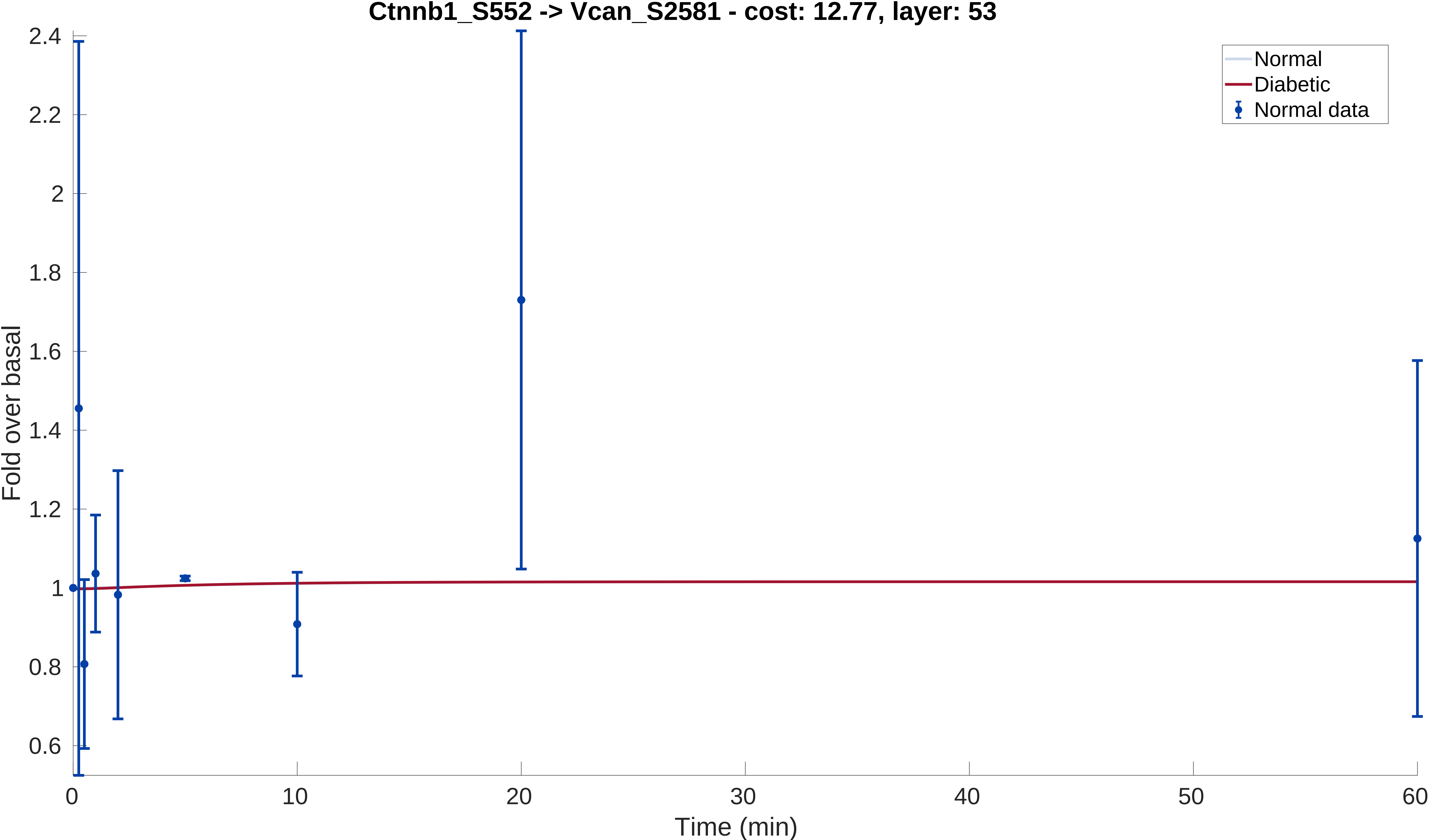

Ctnnb1_S552 -> Vcan_S2581 - cost: 12.77, layer: 53
Normal
Diabetic
Normal data
Fold over basal
2.4
2.2
2
1.8
1.6
1.4
1.2
1
0.8
0.6
0
10
20
30
40
50
60
Time (min)

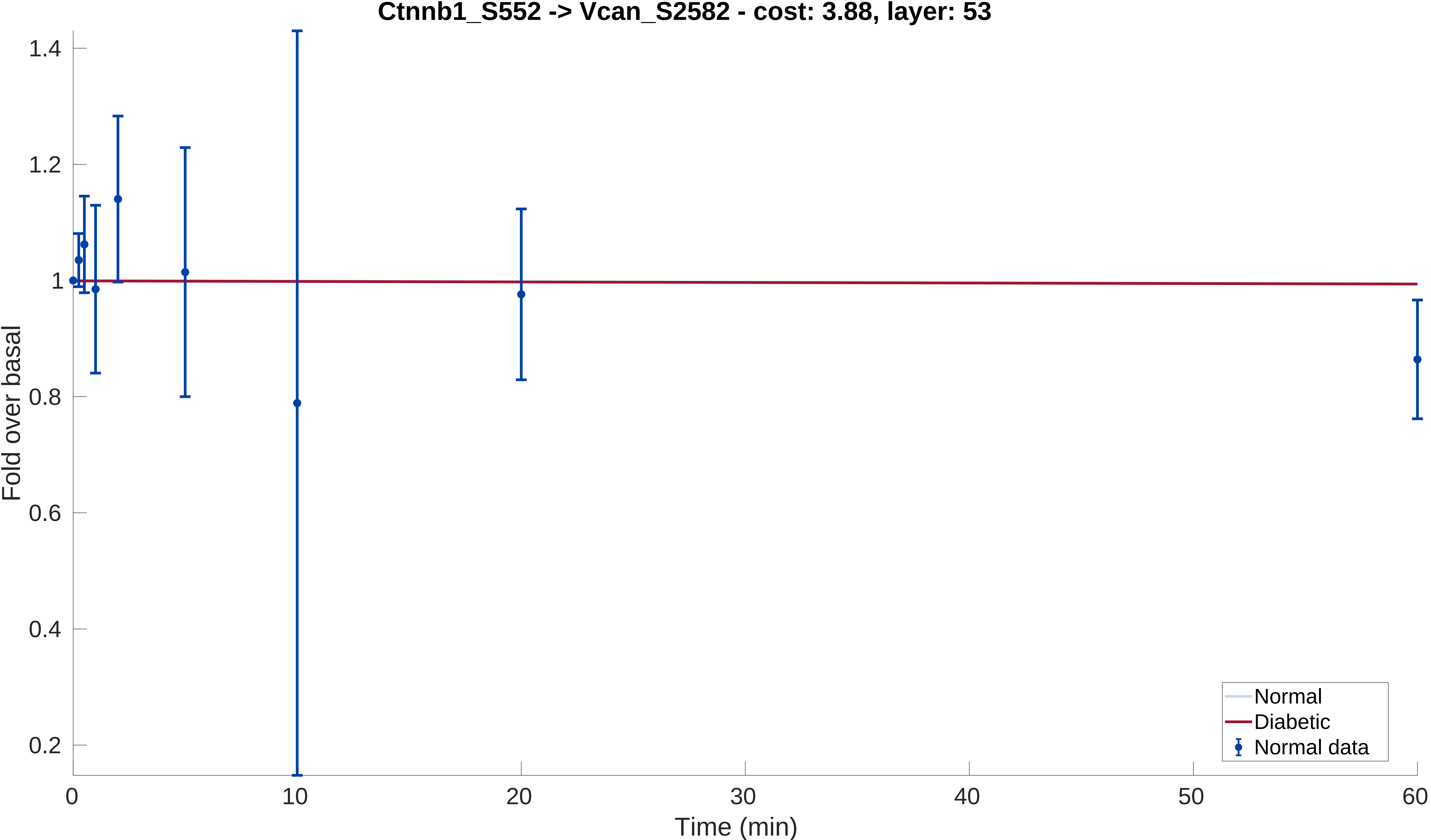

Ctnnb1_S552 -> Vcan_S2582 - cost: 3.88, layer: 53
1.4
1.2
1
0.8
0.6
0.4
0.2
Fold over basal
0
10
20
30
40
50
60
Time (min)
Normal
Diabetic
Normal data

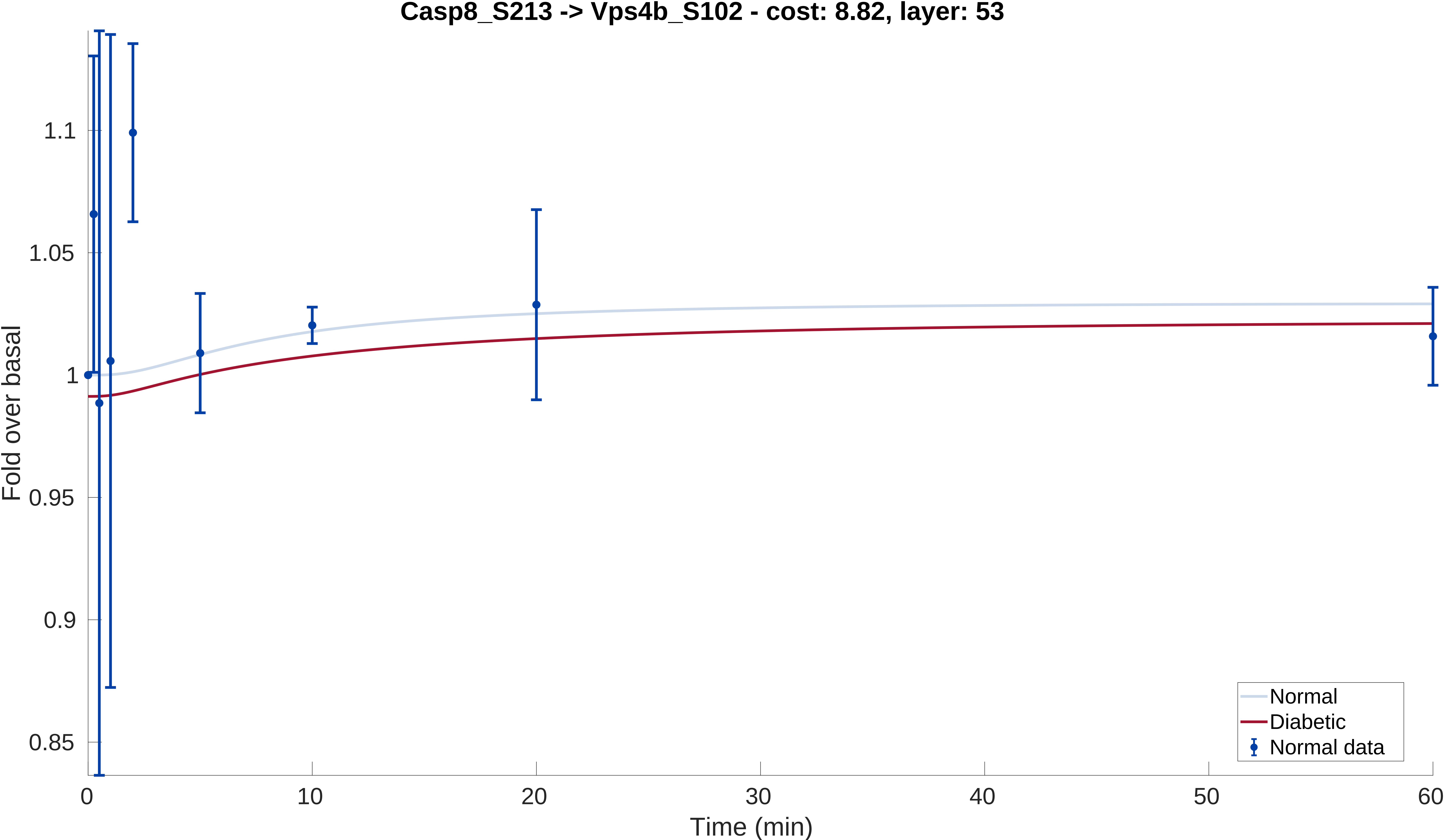

Casp8_S213 -> Vps4b_S102 - cost: 8.82, layer: 53
Fold over basal
1.1
1.05
1
0.95
0.9
0.85
0
10
20
30
40
50
60
Time (min)
Normal
Diabetic
Normal data

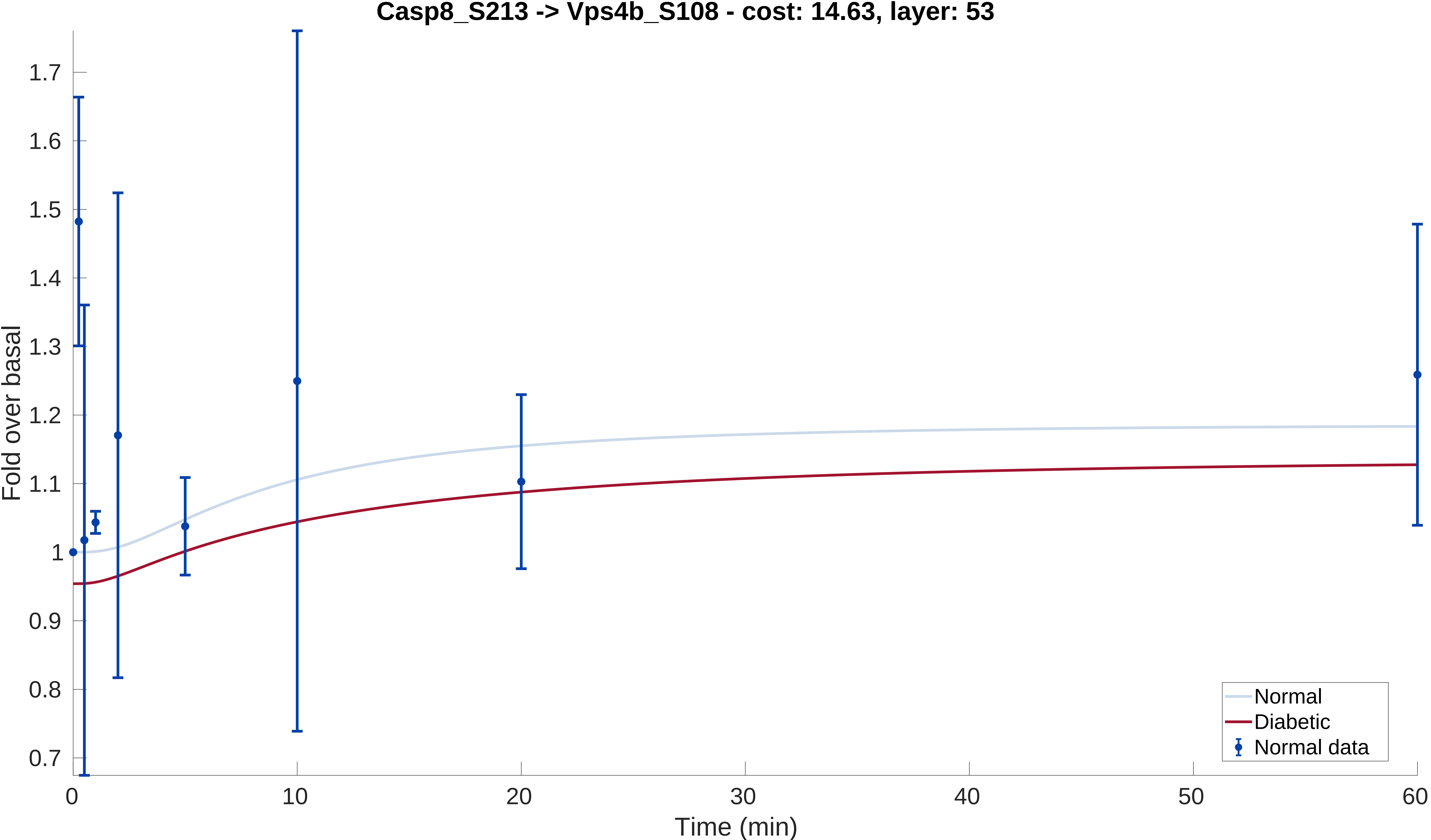
Casp8_S213 -> Vps4b_S108 - cost: 14.63, layer: 53
Fold over basal
1.7
1.6
1.5
1.4
1.3
1.2
1.1
1
0.9
0.8
0.7
0
10
20
30
40
50
60
Time (min)
Normal
Diabetic
Normal data

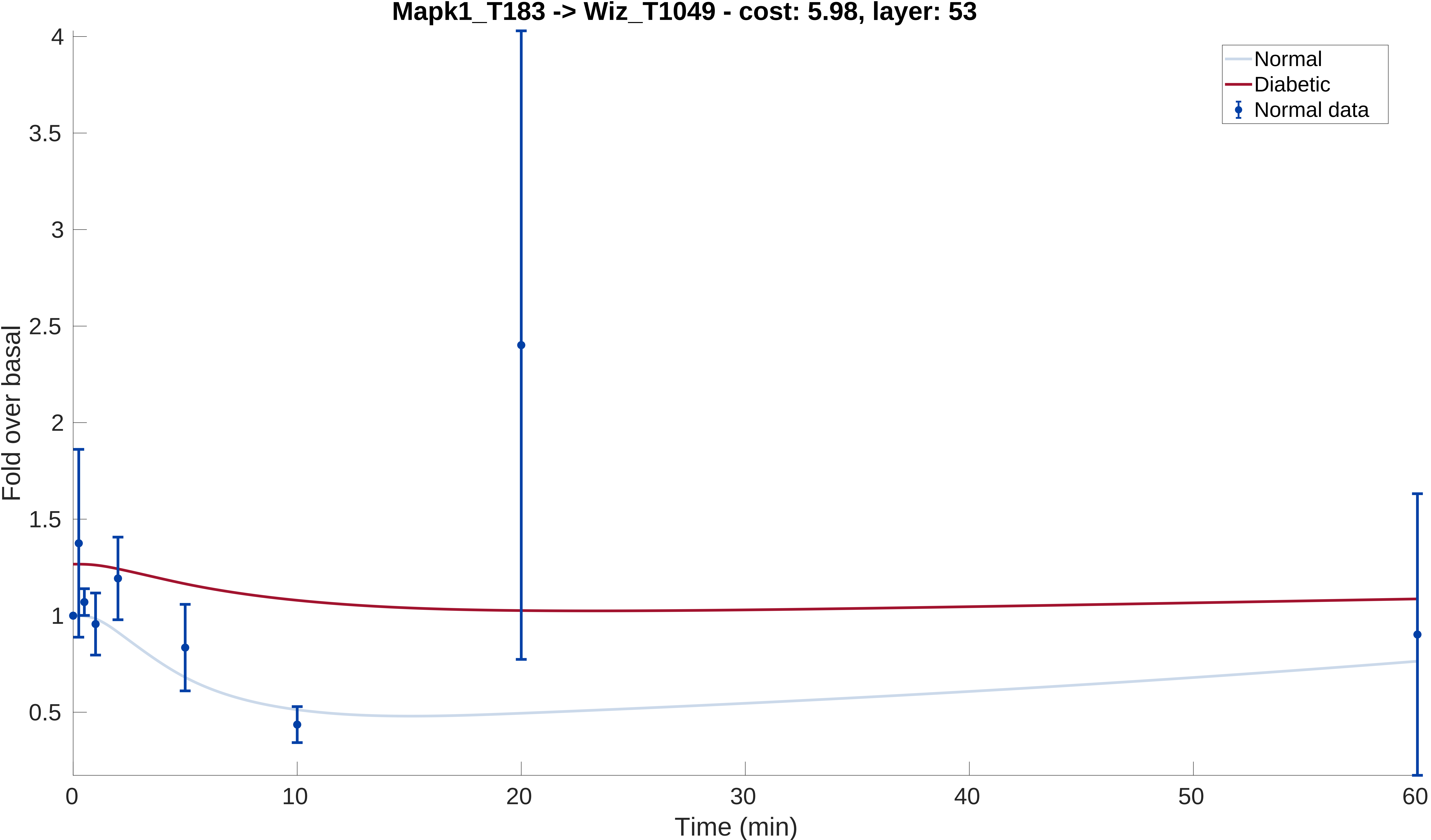

Mapk1_T183 -> Wiz_T1049 - cost: 5.98, layer: 53
Normal
Diabetic
Normal data
Fold over basal
Time (min)
0.5
1
1.5
2
2.5
3
3.5
4
0
10
20
30
40
50
60

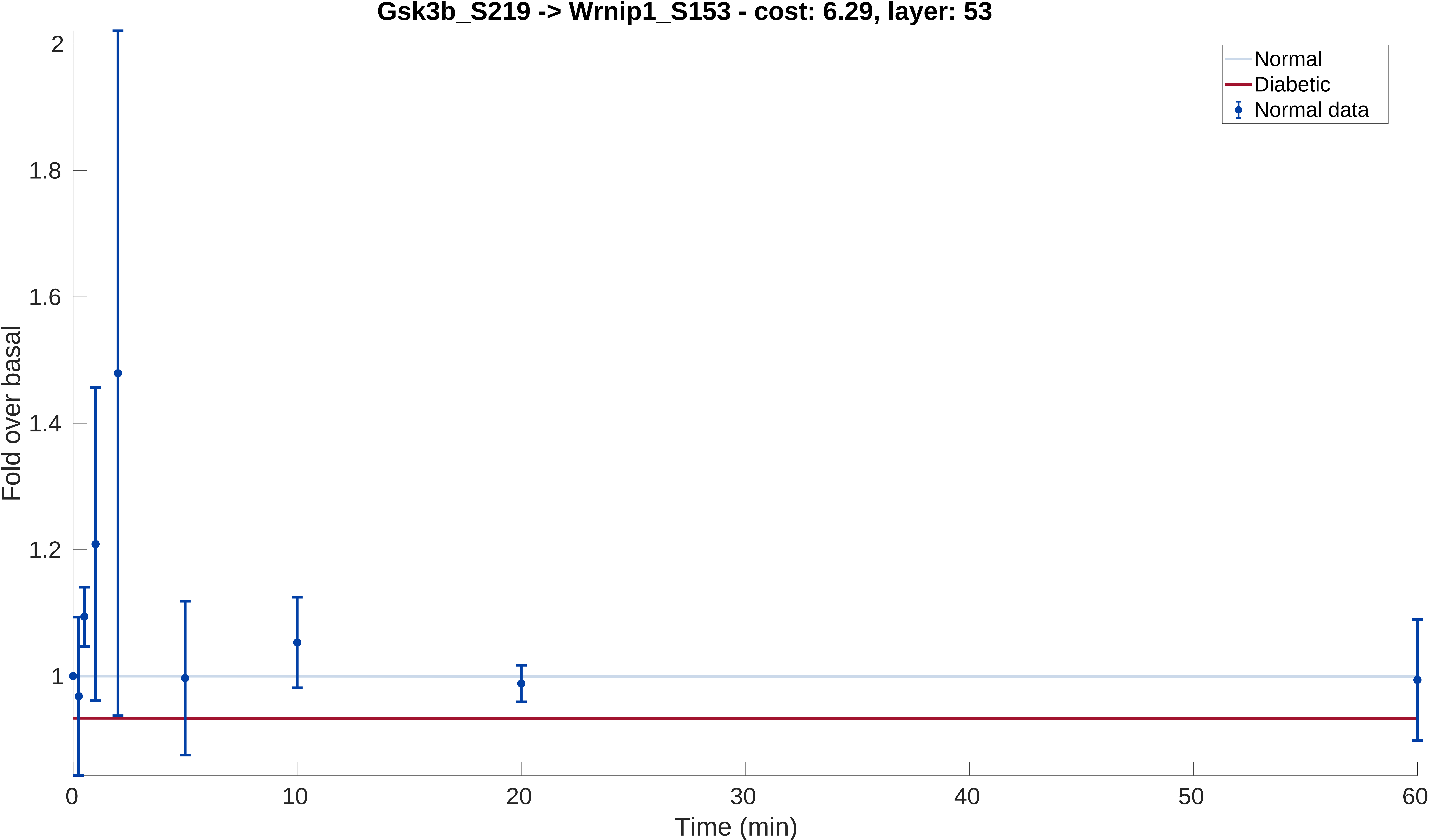

Gsk3b_S219 -> Wrnip1_S153 - cost: 6.29, layer: 53
Normal
Diabetic
Normal data
Fold over basal
2
1.8
1.6
1.4
1.2
1
0
10
20
30
40
50
60
Time (min)

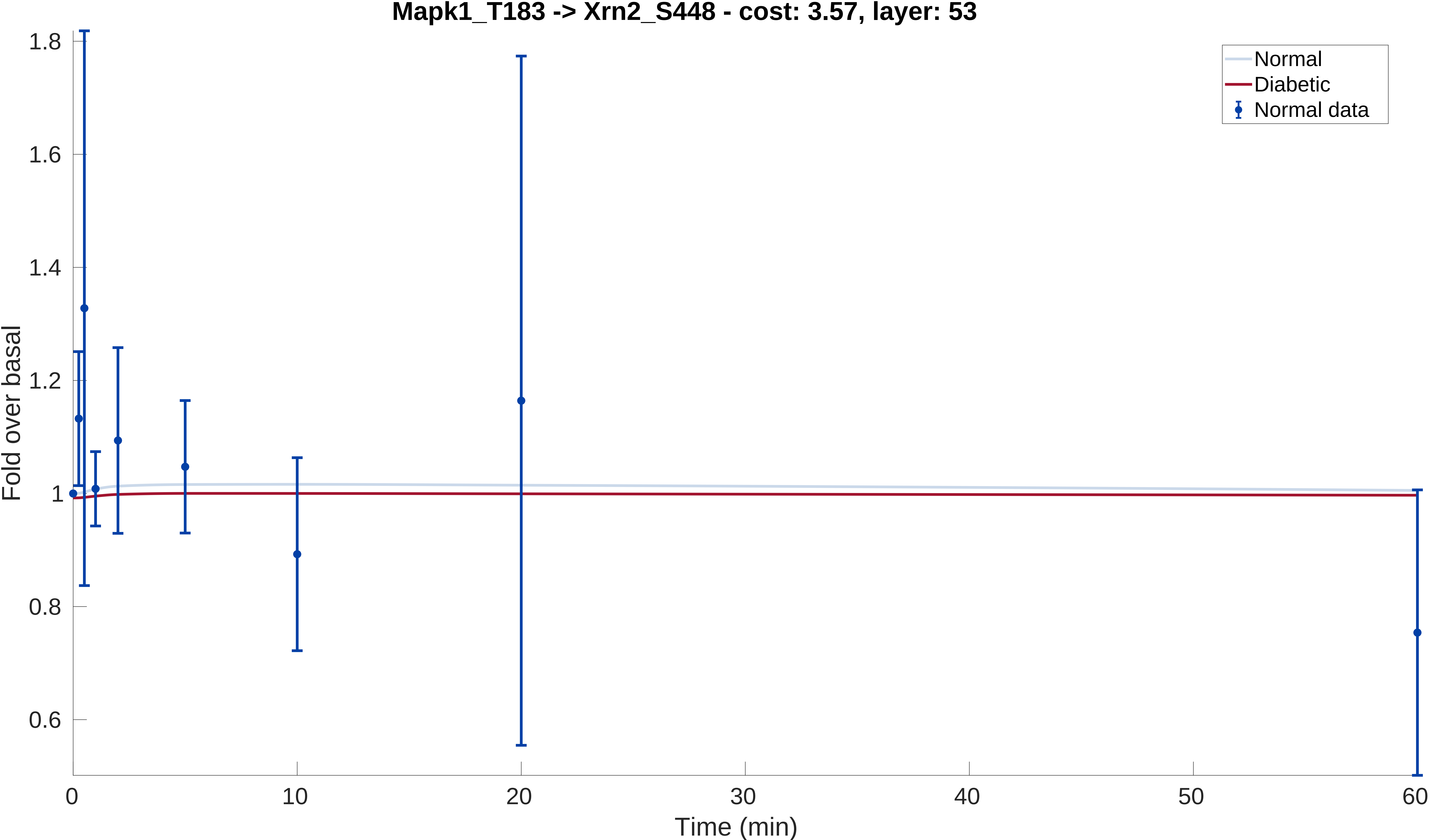

Mapk1_T183 -> Xrn2_S448 - cost: 3.57, layer: 53
Normal
Diabetic
Normal data
Fold over basal
1.8
1.6
1.4
1.2
1
0.8
0.6
0
10
20
30
40
50
60
Time (min)

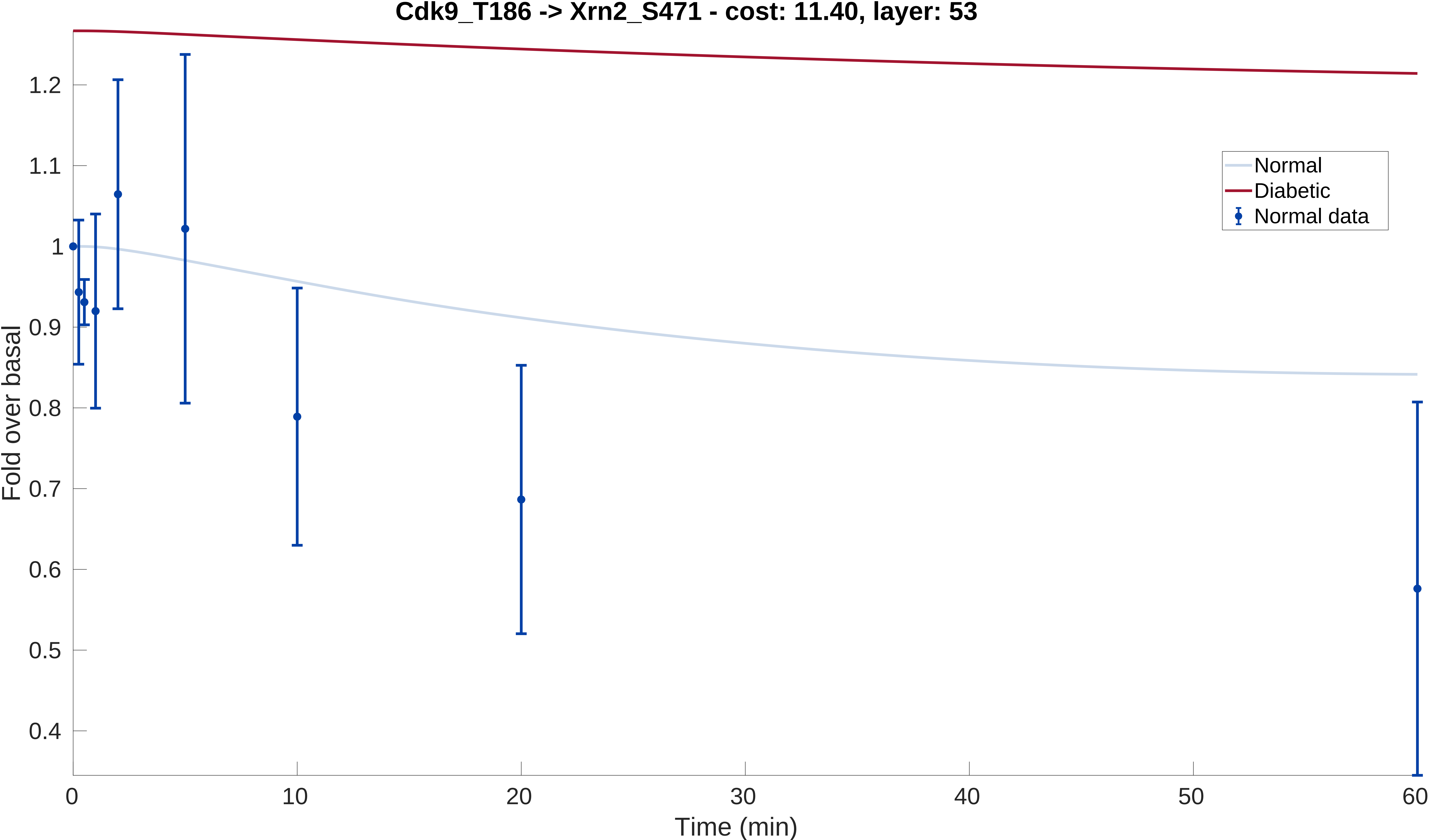

Cdk9_T186 -> Xrn2_S471 - cost: 11.40, layer: 53
Normal
Diabetic
Normal data
Fold over basal
0.4
0.5
0.6
0.7
0.8
0.9
1
1.1
1.2
0
10
20
30
40
50
60
Time (min)

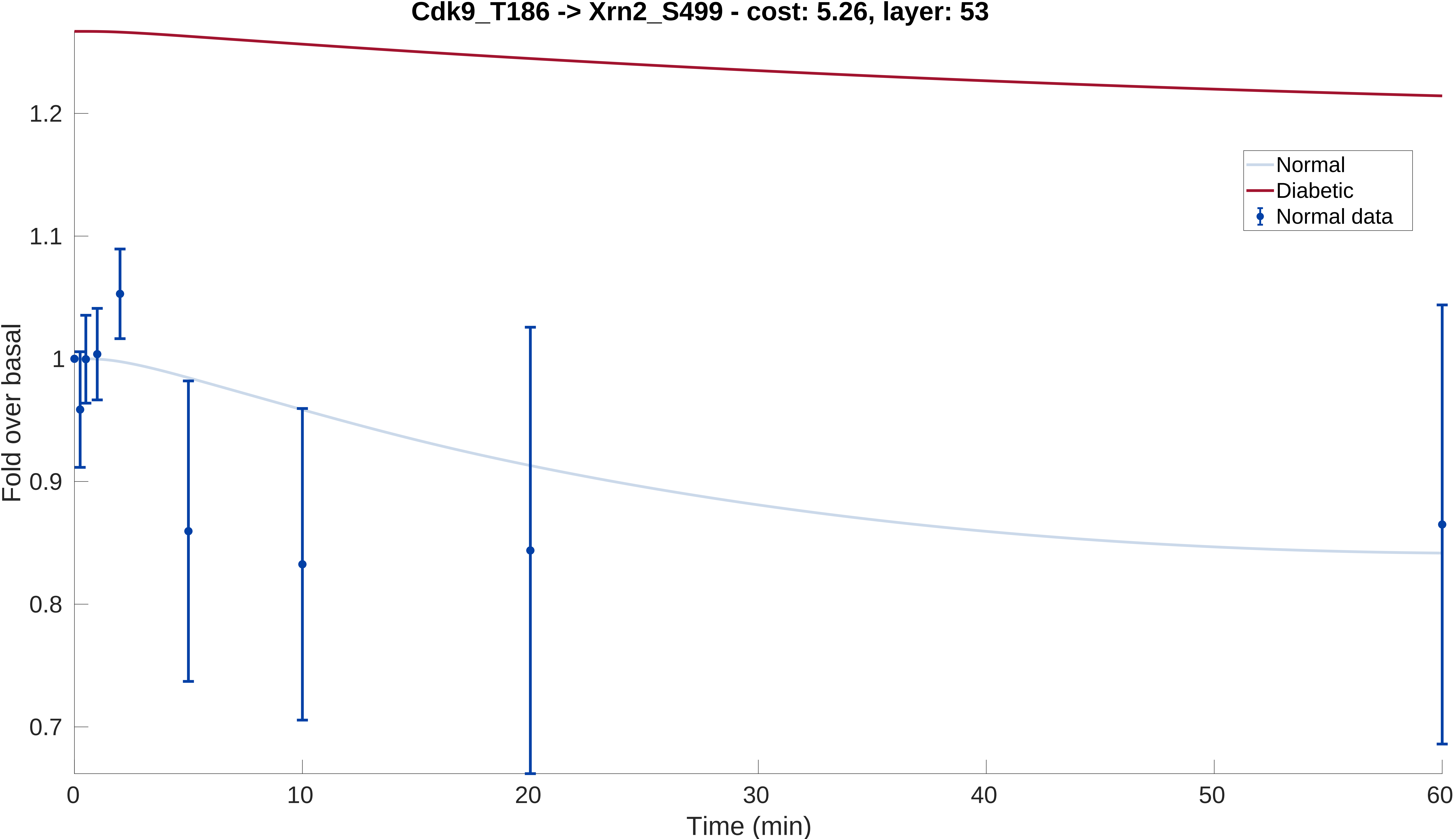
Cdk9_T186 -> Xrn2_S499 - cost: 5.26, layer: 53
Fold over basal
1.2
1.1
1
0.9
0.8
0.7
0
10
20
30
40
50
60
Time (min)
Normal
Diabetic
Normal data

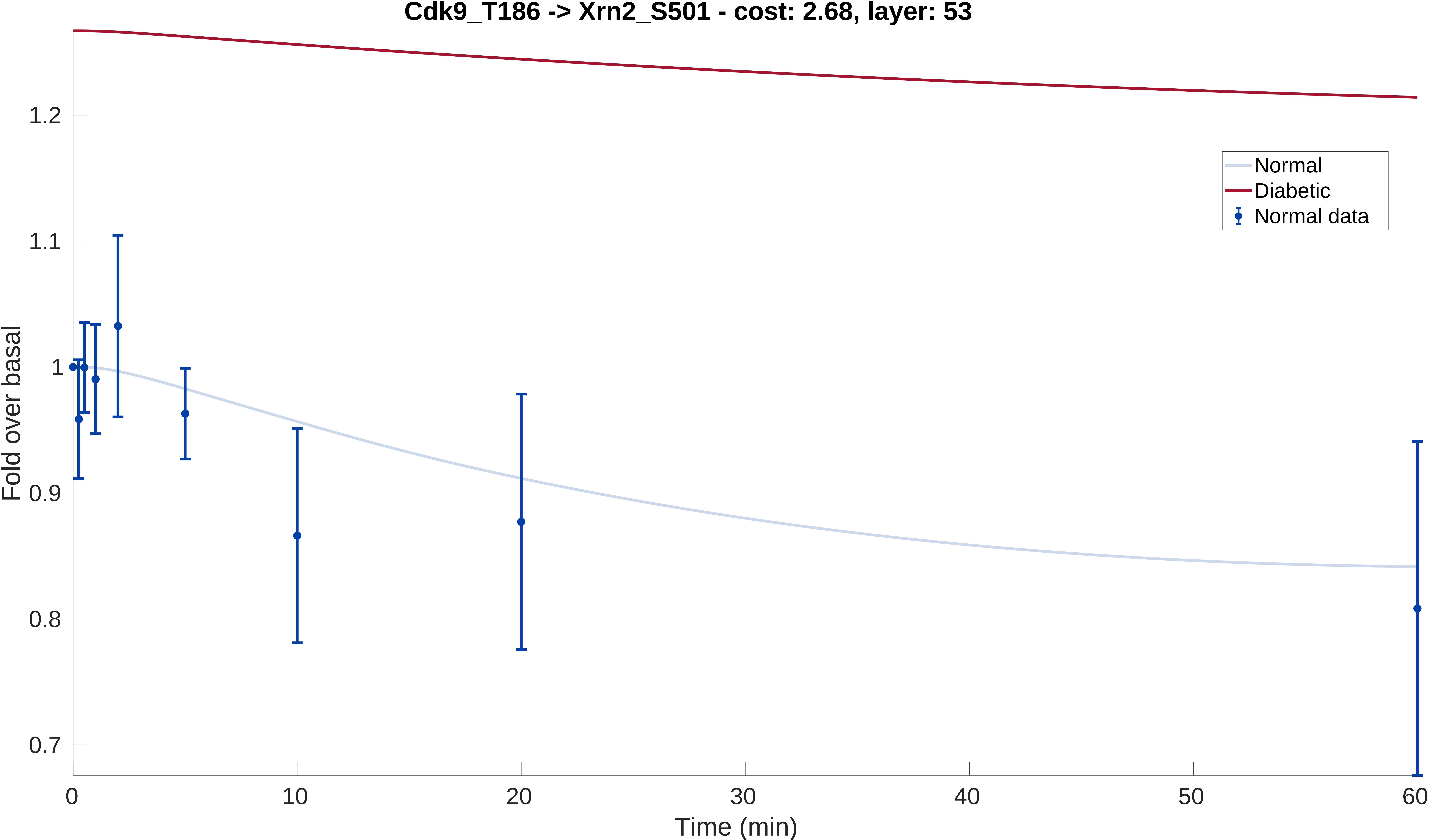

Cdk9_T186 -> Xrn2_S501 - cost: 2.68, layer: 53
Normal
Diabetic
Normal data
Fold over basal
1.2
1.1
1
0.9
0.8
0.7
0
10
20
30
40
50
60
Time (min)

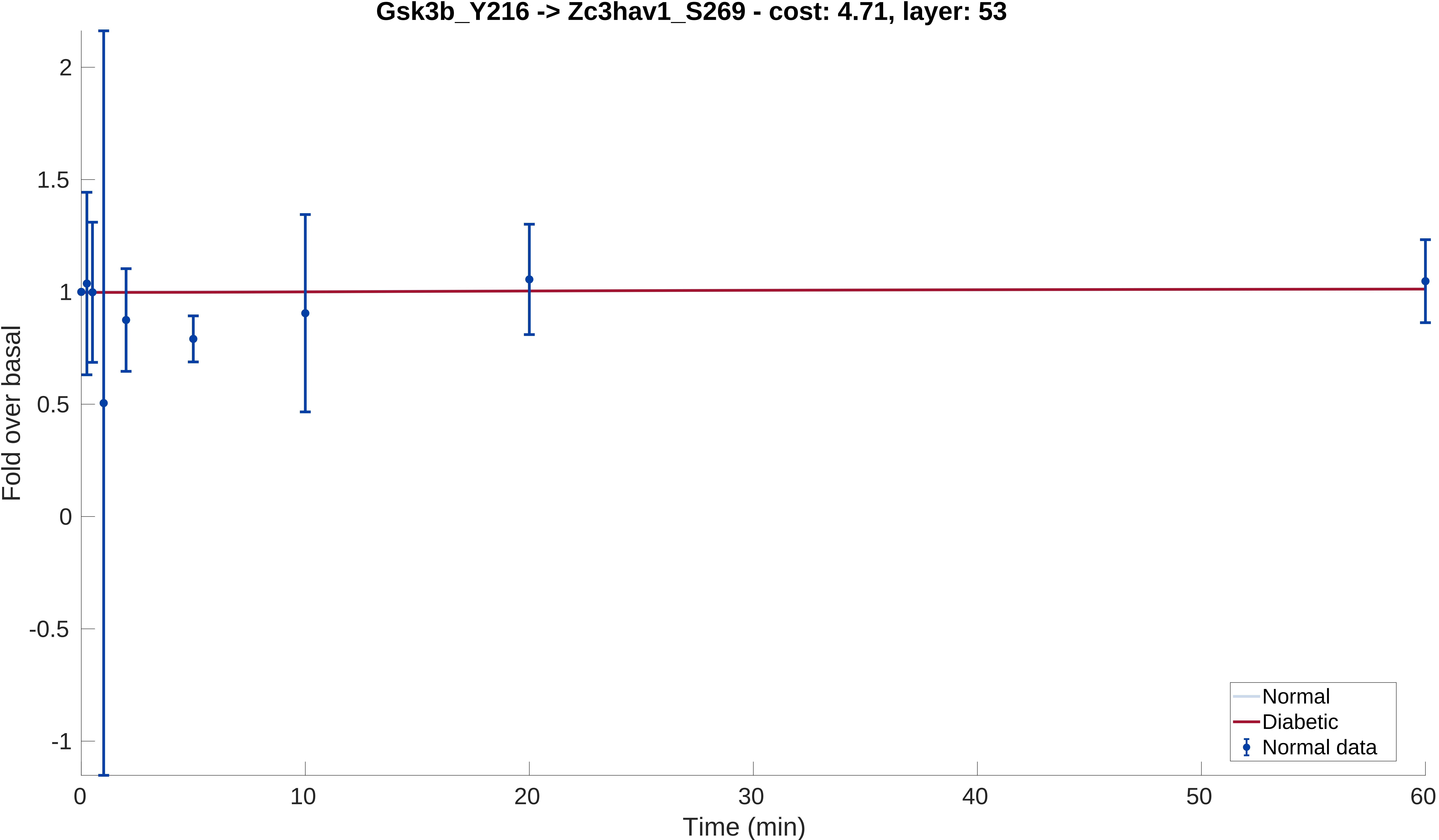

Gsk3b_Y216 -> Zc3hav1_S269 - cost: 4.71, layer: 53
Fold over basal
Time (min)
2
1.5
1
0.5
0
-0.5
-1
0
10
20
30
40
50
60
Normal
Diabetic
Normal data

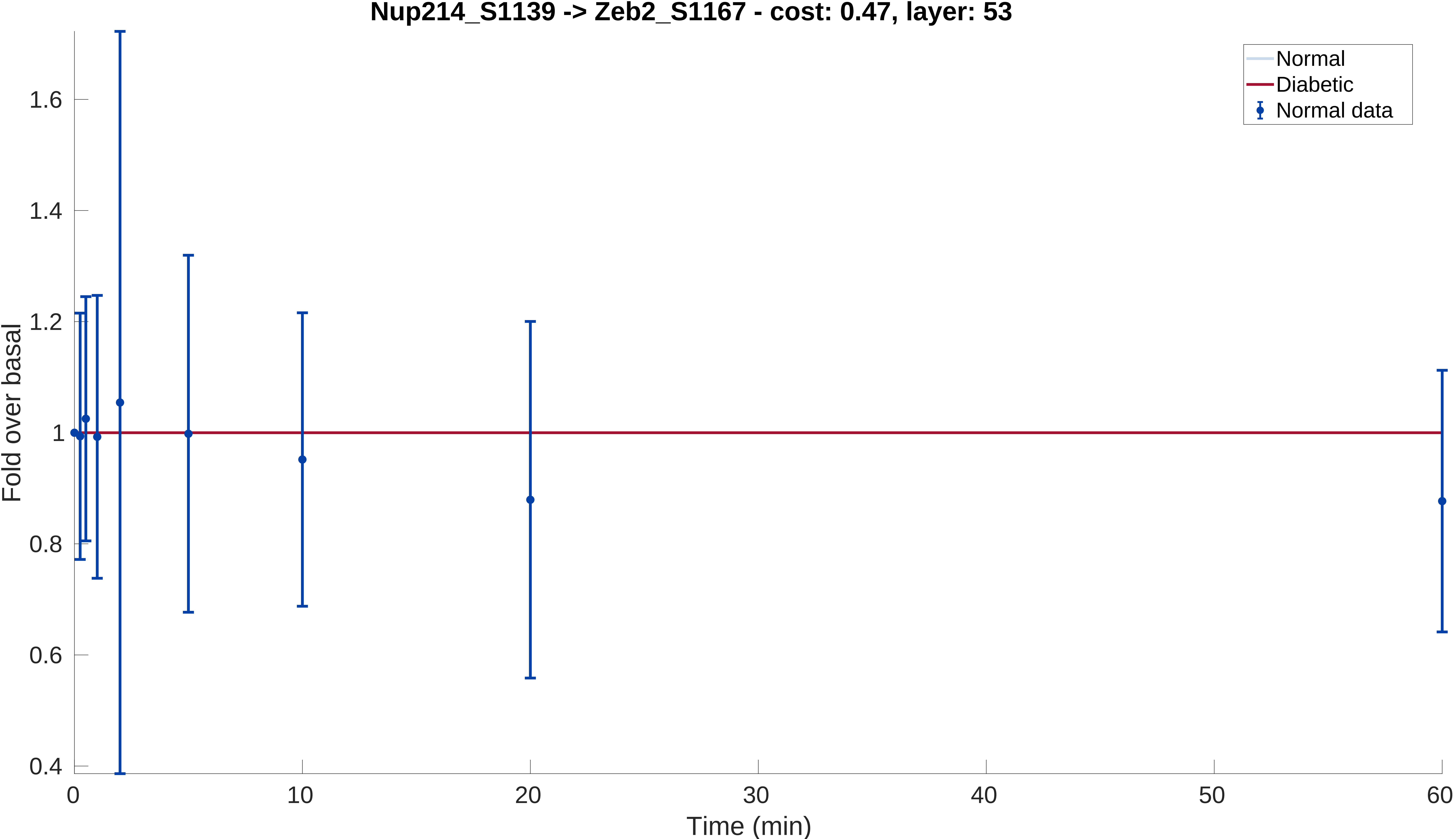

Nup214_S1139 -> Zeb2_S1167 - cost: 0.47, layer: 53
Normal
Diabetic
Normal data
1.6
1.4
1.2
1
0.8
0.6
0.4
Fold over basal
0
10
20
30
40
50
60
Time (min)

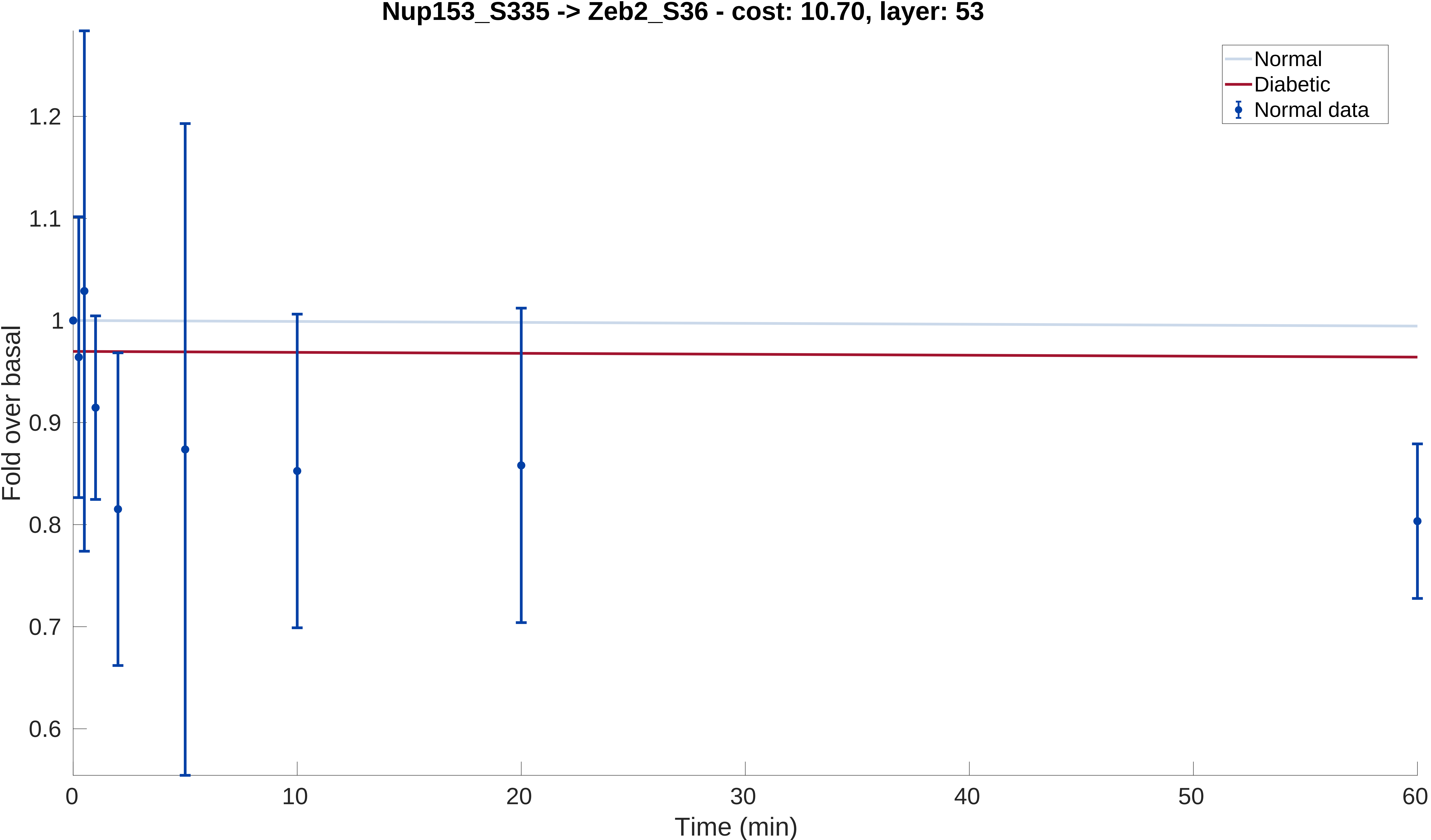
Nup153_S335 -> Zeb2_S36 - cost: 10.70, layer: 53
Normal
Diabetic
Normal data
Fold over basal
Time (min)
0.6
0.7
0.8
0.9
1
1.1
1.2
0
10
20
30
40
50
60

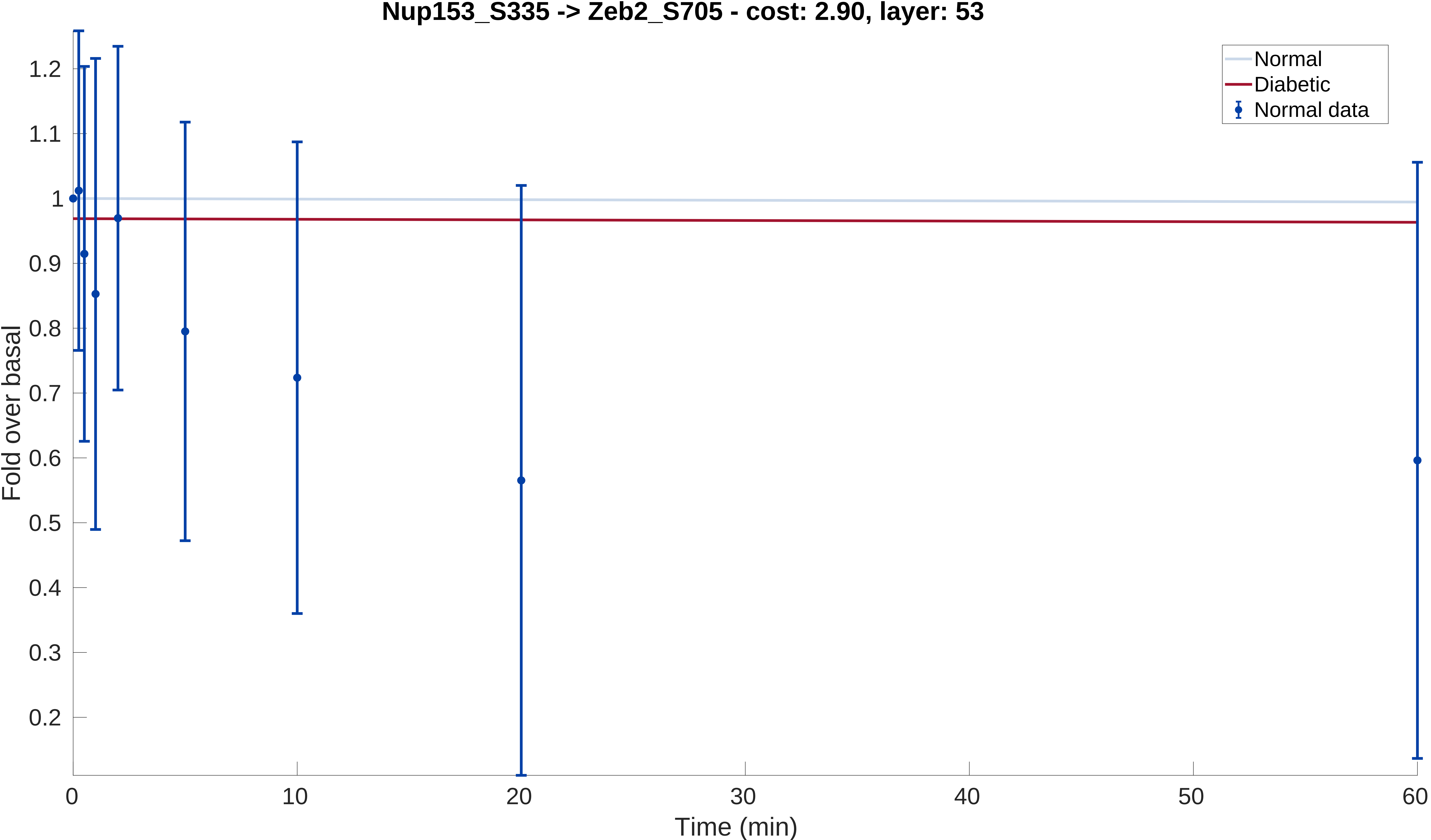

Nup153_S335 -> Zeb2_S705 - cost: 2.90, layer: 53
Normal
Diabetic
Normal data
Fold over basal
Time (min)
0
10
20
30
40
50
60
0.2
0.3
0.4
0.5
0.6
0.7
0.8
0.9
1
1.1
1.2

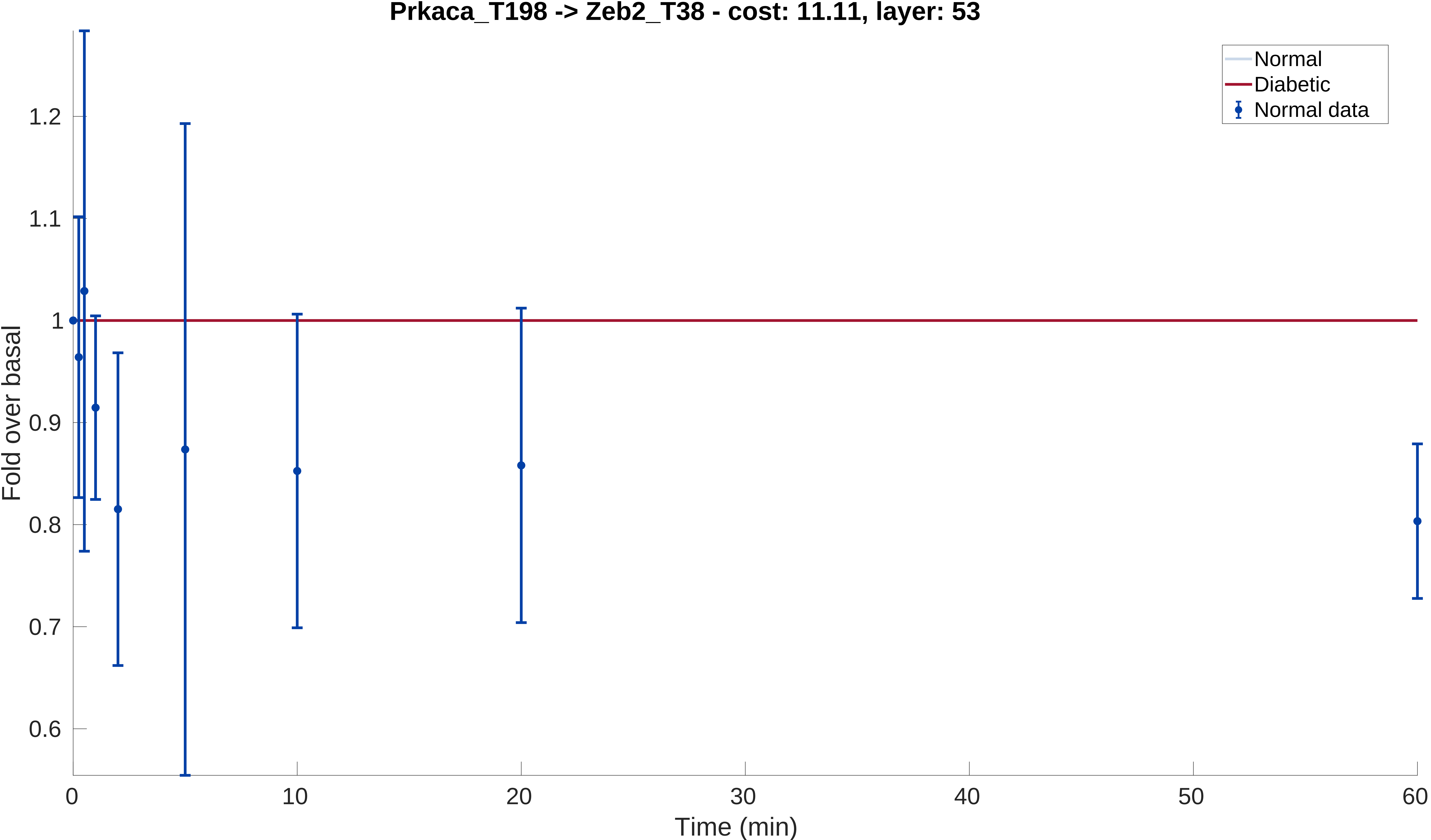

Prkaca_T198 -> Zeb2_T38 - cost: 11.11, layer: 53
Normal
Diabetic
Normal data
Fold over basal
1.2
1.1
1
0.9
0.8
0.7
0.6
0
10
20
30
40
50
60
Time (min)

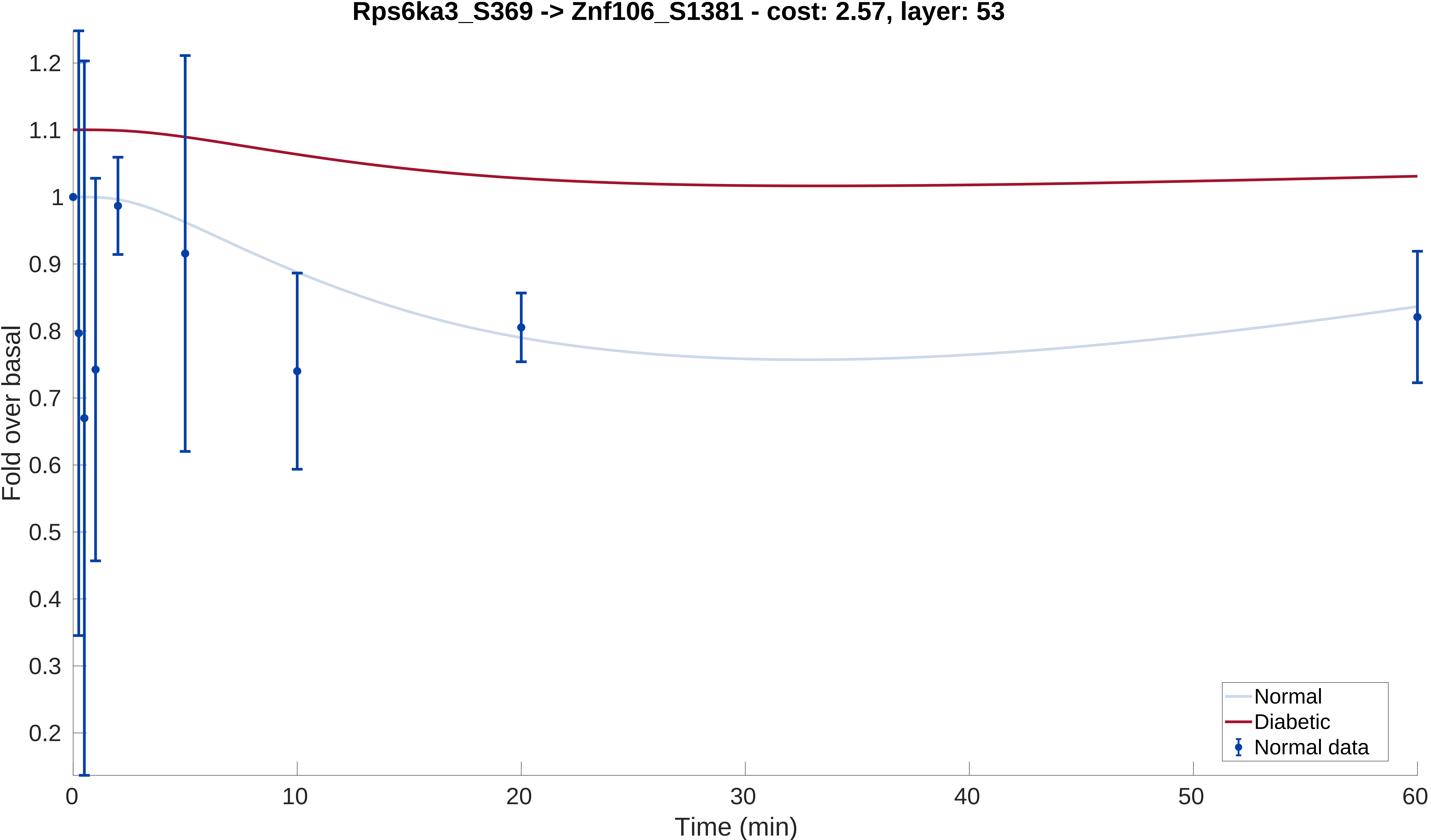

Rps6ka3_S369 -> Znf106_S1381 - cost: 2.57, layer: 53
Fold over basal
Time (min)
1.2
1.1
1
0.9
0.8
0.7
0.6
0.5
0.4
0.3
0.2
0
10
20
30
40
50
60
Normal
Diabetic
Normal data

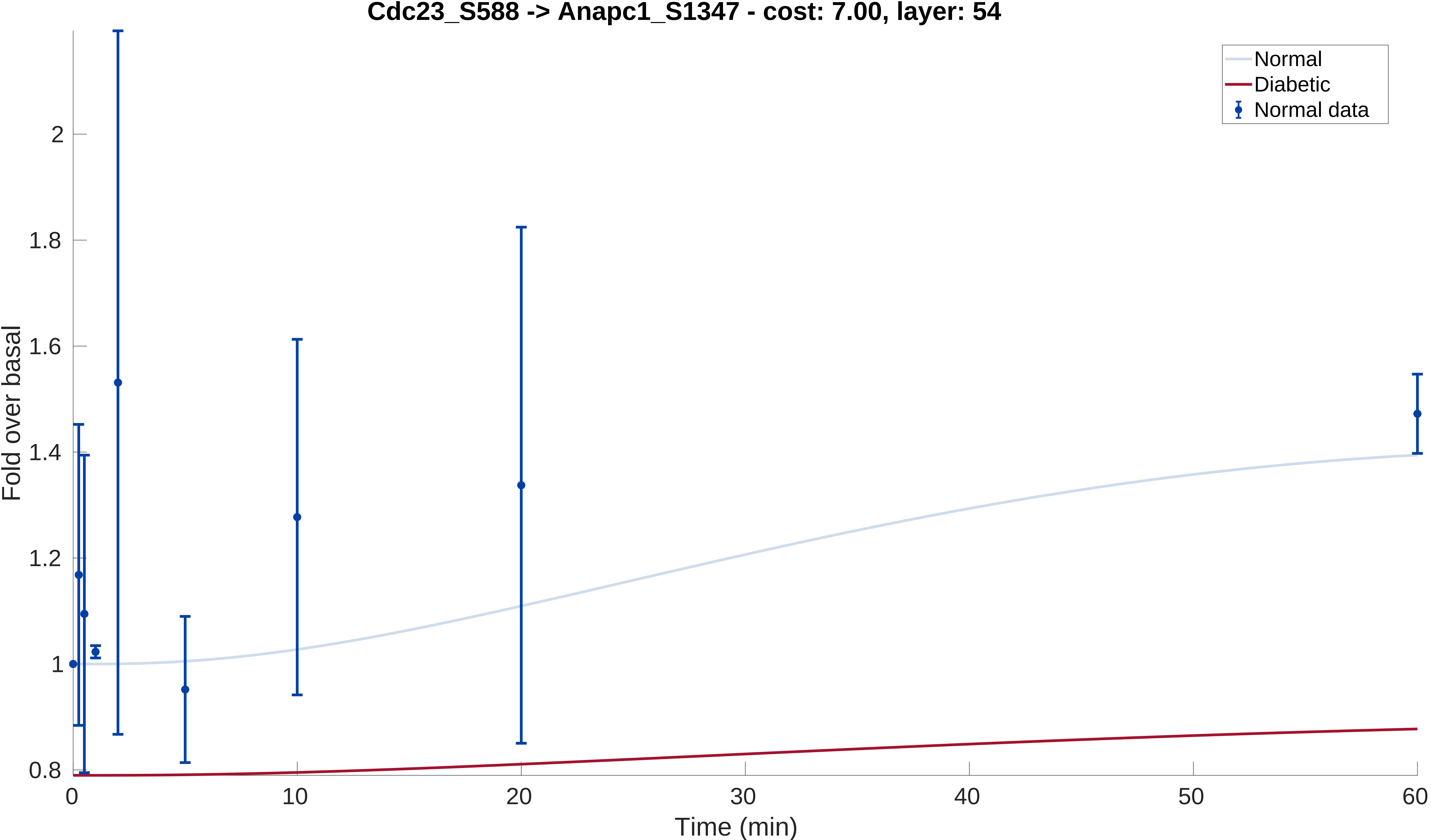

Cdc23_S588 -> Anapc1_S1347 - cost: 7.00, layer: 54
Normal
Diabetic
Normal data
Fold over basal
2
1.8
1.6
1.4
1.2
1
0.8
0
10
20
30
40
50
60
Time (min)

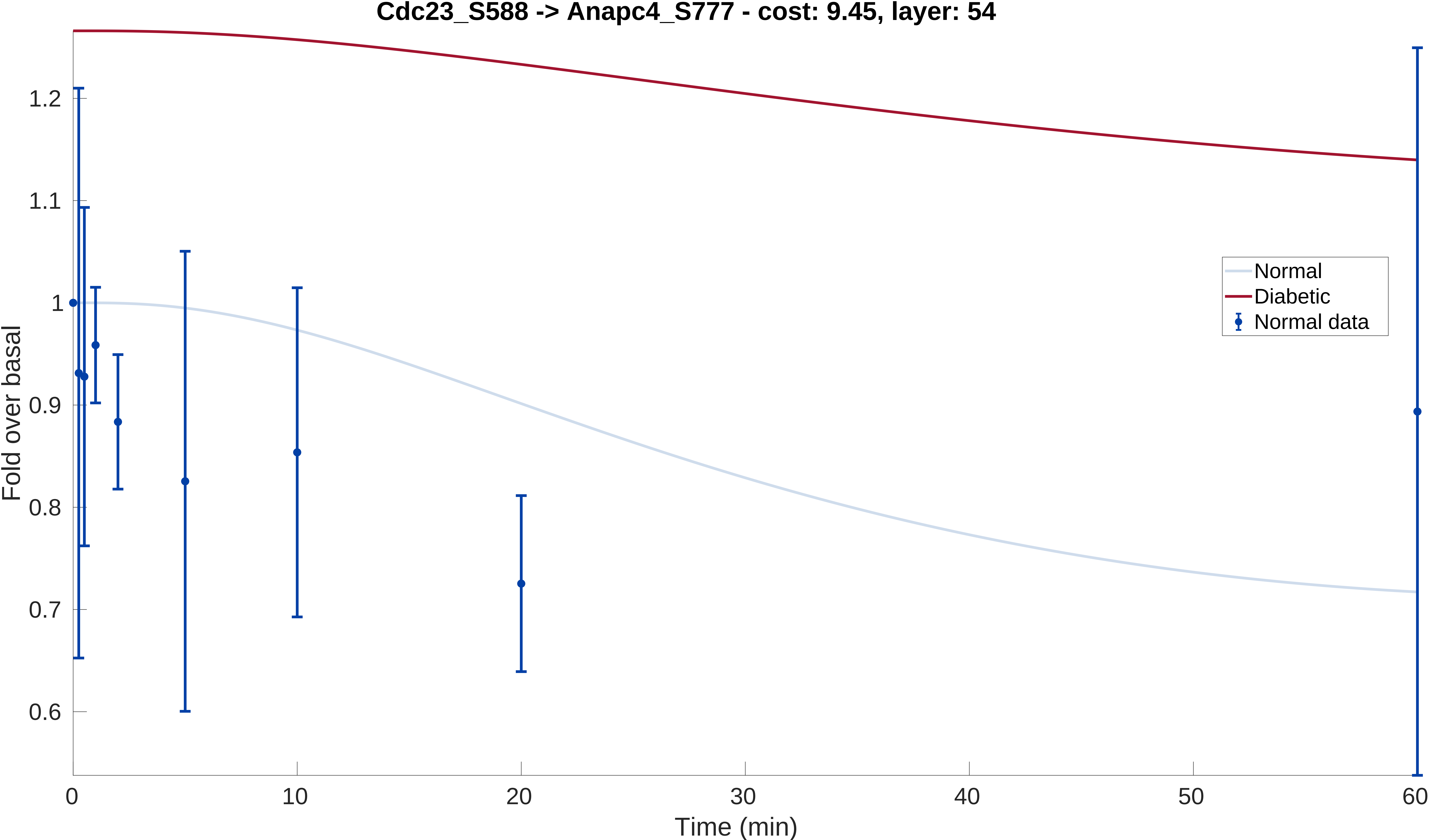

Cdc23_S588 -> Anapc4_S777 - cost: 9.45, layer: 54
Fold over basal
1.2
1.1
1
0.9
0.8
0.7
0.6
0
10
20
30
40
50
60
Time (min)
Normal
Diabetic
Normal data

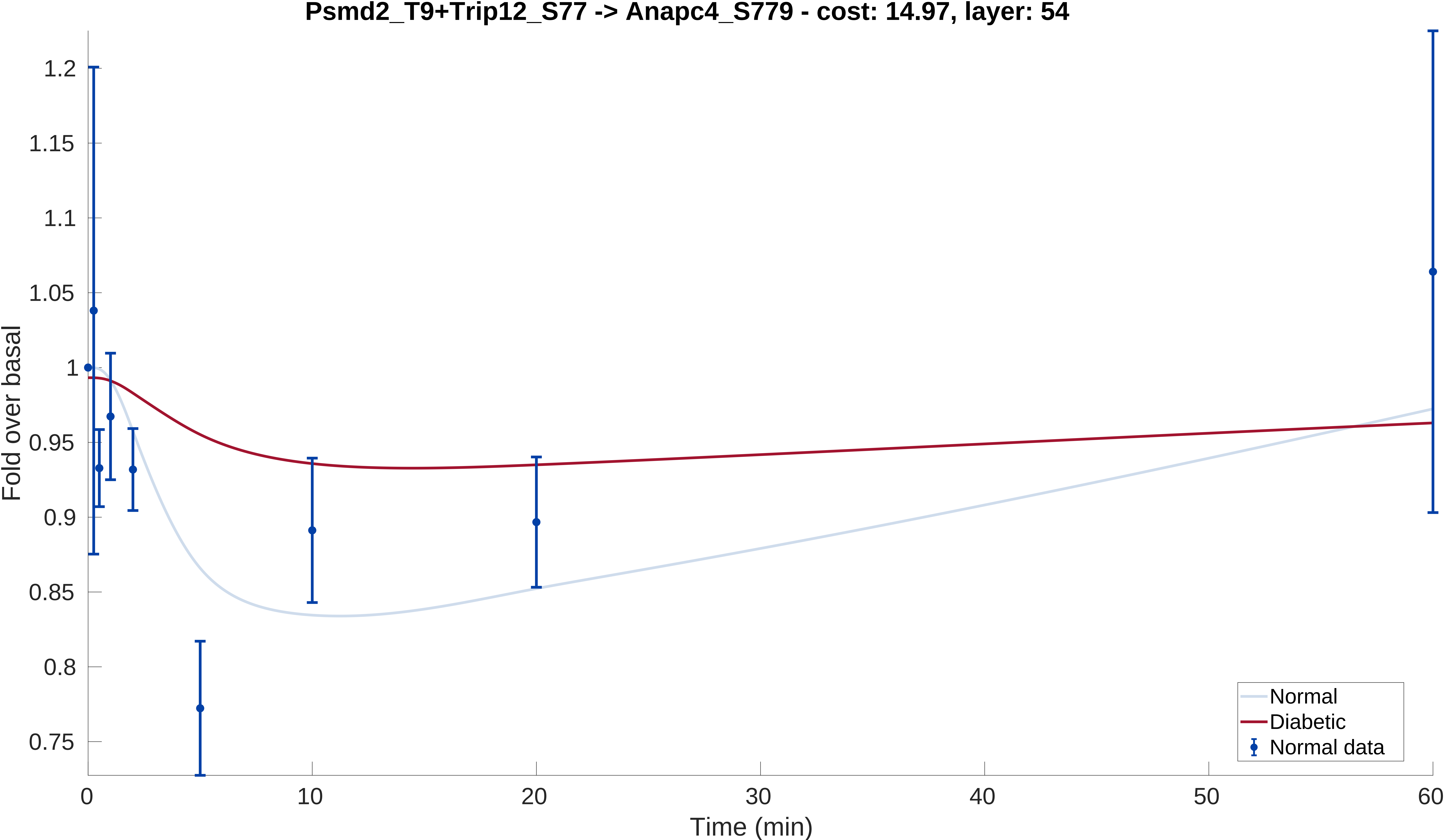
Psmd2_T9+Trip12_S77 -> Anapc4_S779 - cost: 14.97, layer: 54
Fold over basal
1.2
1.15
1.1
1.05
1
0.95
0.9
0.85
0.8
0.75
0
10
20
30
40
50
60
Time (min)
Normal
Diabetic
Normal data

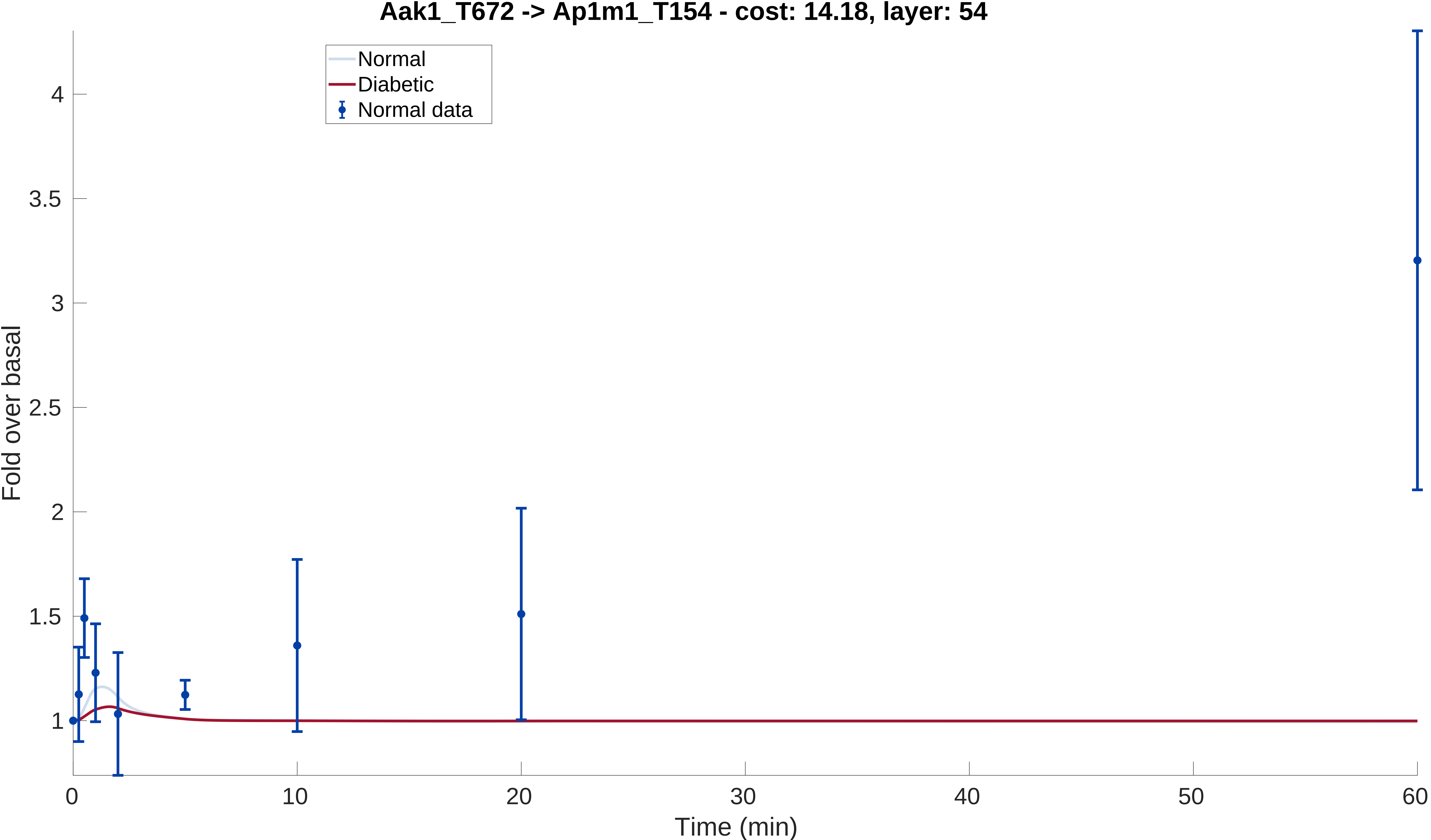

Aak1_T672 -> Ap1m1_T154 - cost: 14.18, layer: 54
Normal
Diabetic
Normal data
Fold over basal
4
3.5
3
2.5
2
1.5
1
0
10
20
30
40
50
60
Time (min)

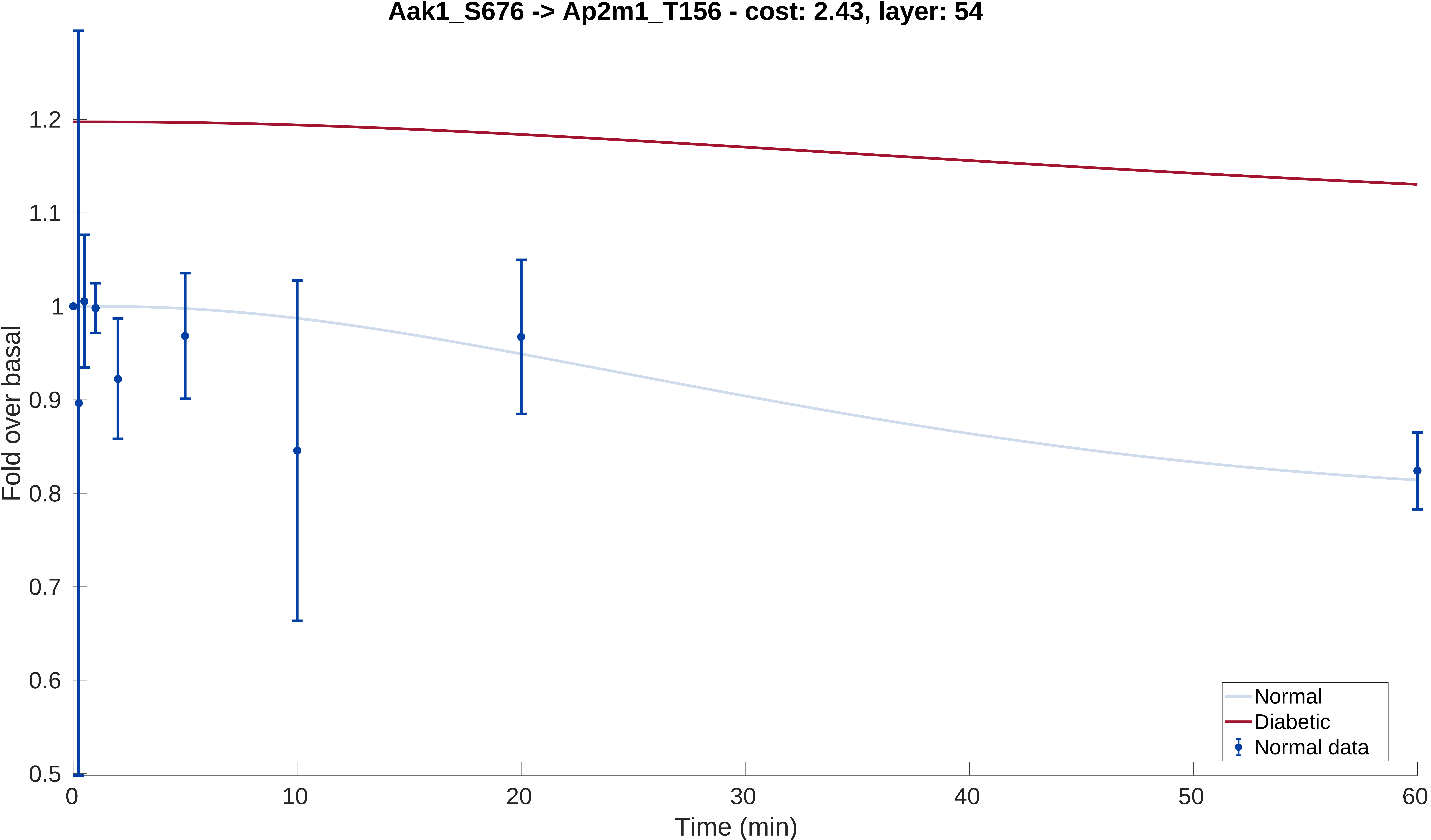

Aak1_S676 -> Ap2m1_T156 - cost: 2.43, layer: 54
Fold over basal
1.2
1.1
1
0.9
0.8
0.7
0.6
0.5
0
10
20
30
40
50
60
Time (min)
Normal
Diabetic
Normal data

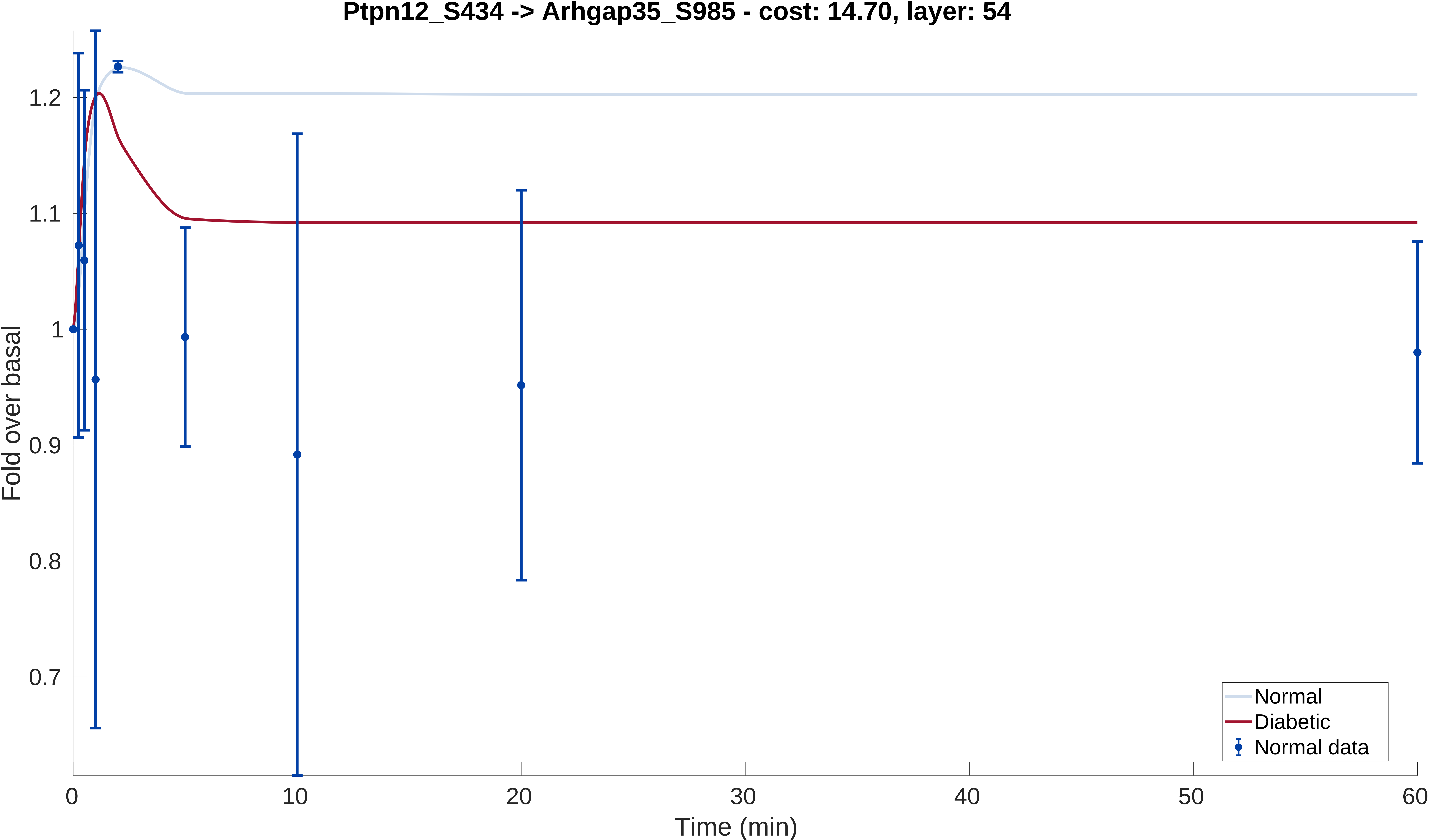
Ptpn12_S434 -> Arhgap35_S985 - cost: 14.70, layer: 54
Fold over basal
1.2
1.1
1
0.9
0.8
0.7
0
10
20
30
40
50
60
Time (min)
Normal
Diabetic
Normal data

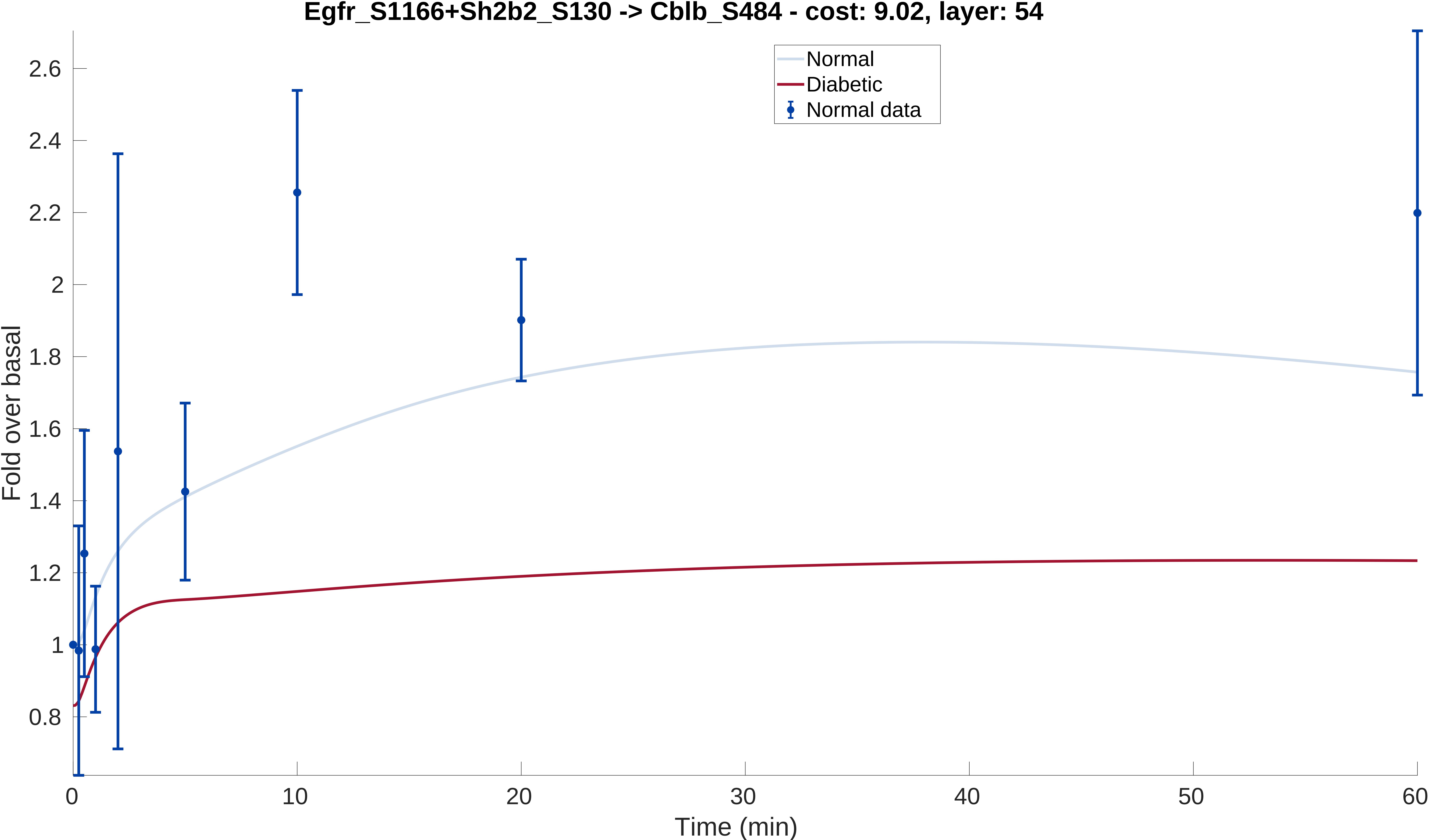
Egfr_S1166+Sh2b2_S130 -> Cblb_S484 - cost: 9.02, layer: 54
Normal
Diabetic
Normal data
Fold over basal
2.6
2.4
2.2
2
1.8
1.6
1.4
1.2
1
0.8
0
10
20
30
40
50
60
Time (min)

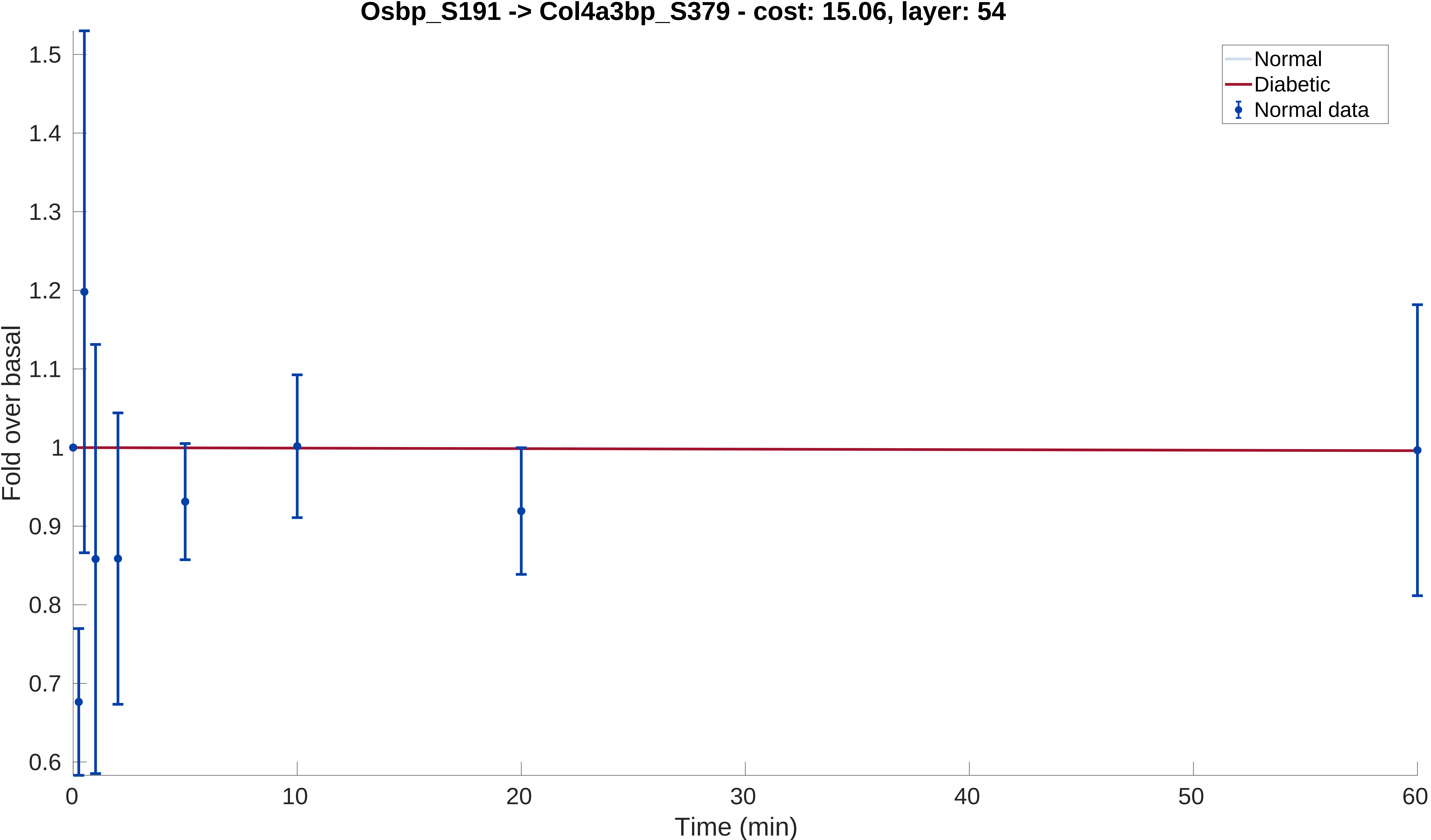
Osbp_S191 -> Col4a3bp_S379 - cost: 15.06, layer: 54
Normal
Diabetic
Normal data
Fold over basal
1.5
1.4
1.3
1.2
1.1
1
0.9
0.8
0.7
0.6
0
10
20
30
40
50
60
Time (min)

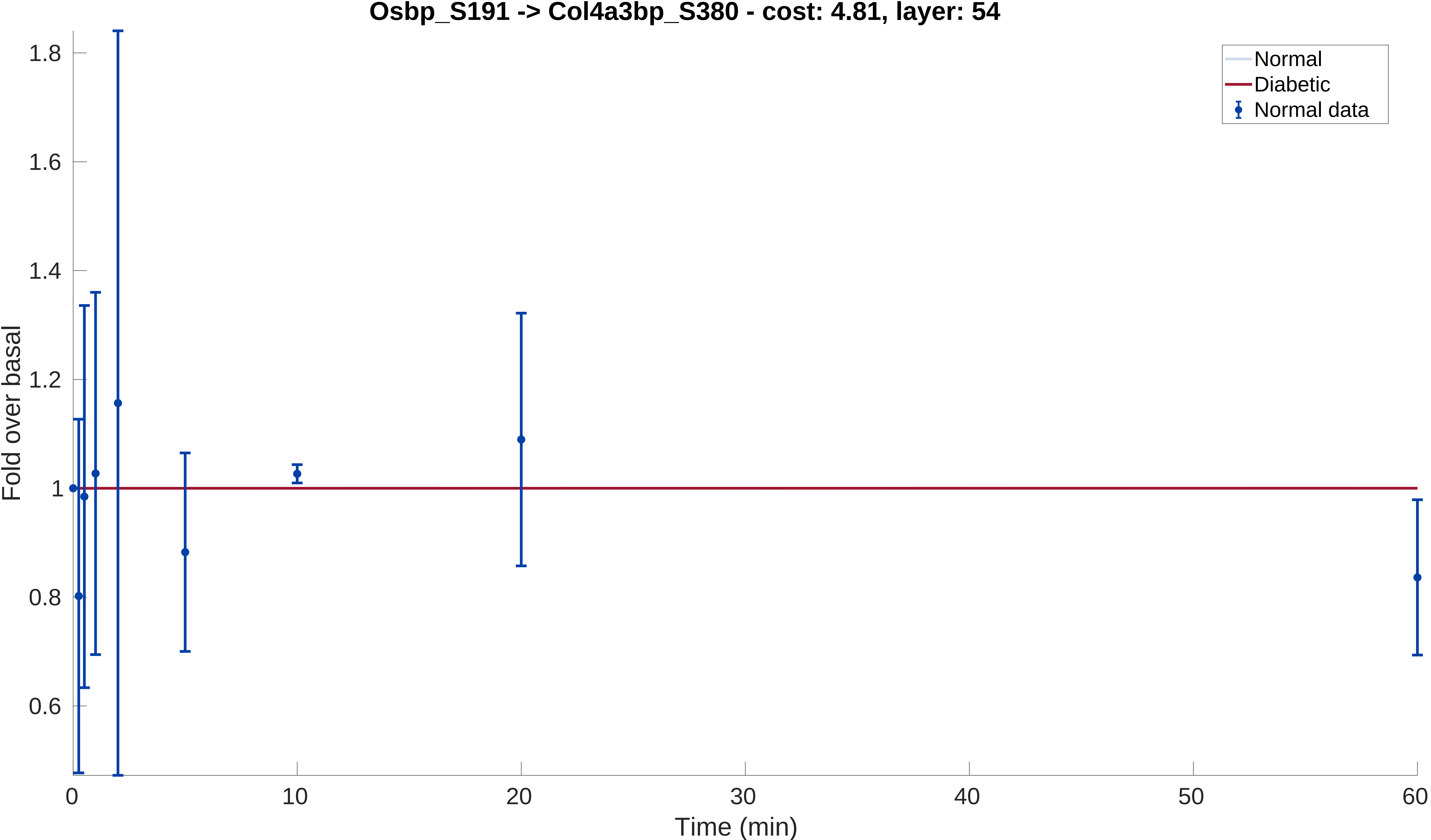

Osbp_S191 -> Col4a3bp_S380 - cost: 4.81, layer: 54
Normal
Diabetic
Normal data
Fold over basal
1.8
1.6
1.4
1.2
1
0.8
0.6
0
10
20
30
40
50
60
Time (min)

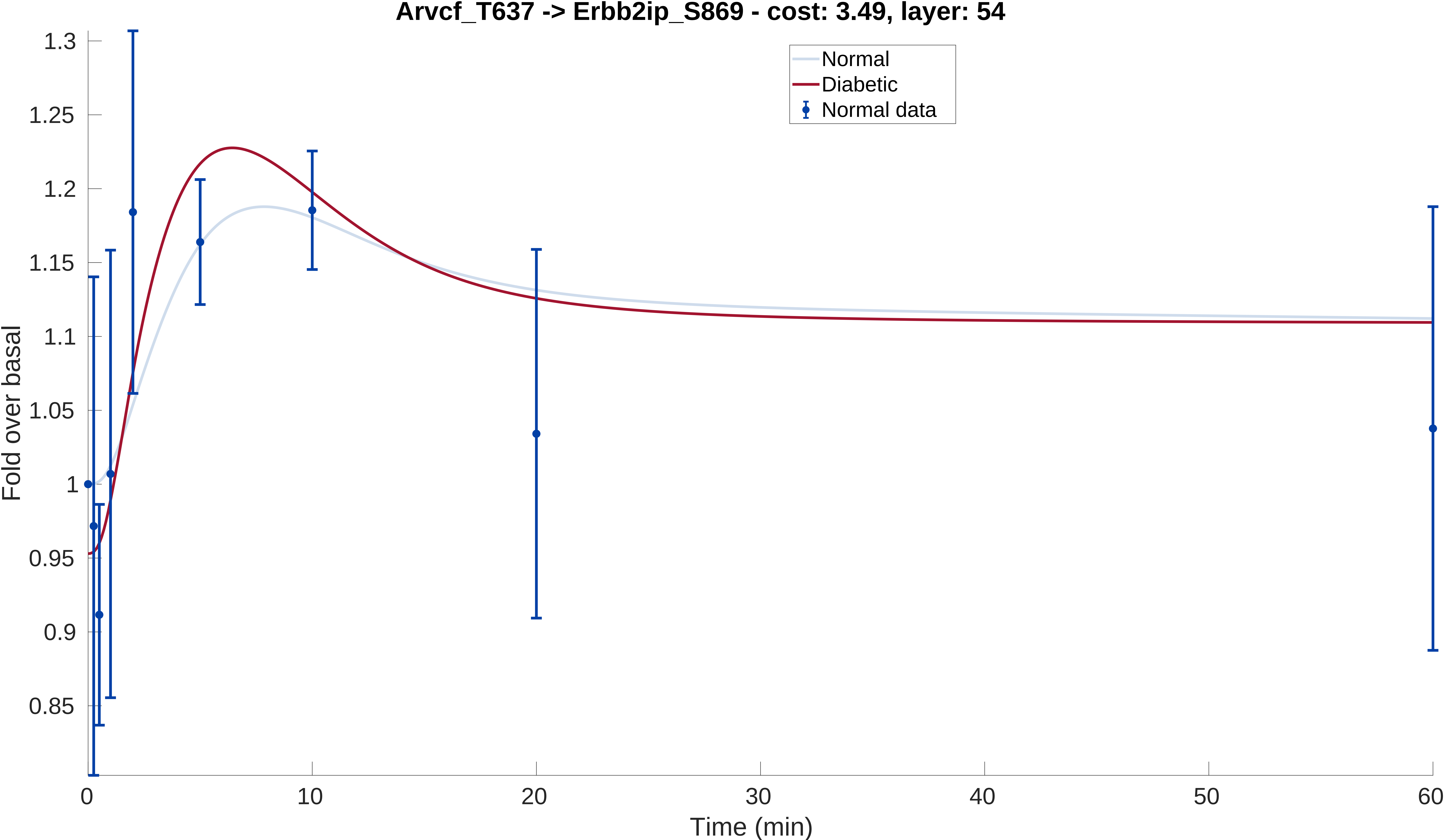

Arvcf_T637 -> Erbb2ip_S869 - cost: 3.49, layer: 54
Normal
Diabetic
Normal data
Fold over basal
1.3
1.25
1.2
1.15
1.1
1.05
1
0.95
0.9
0.85
0
10
20
30
40
50
60
Time (min)

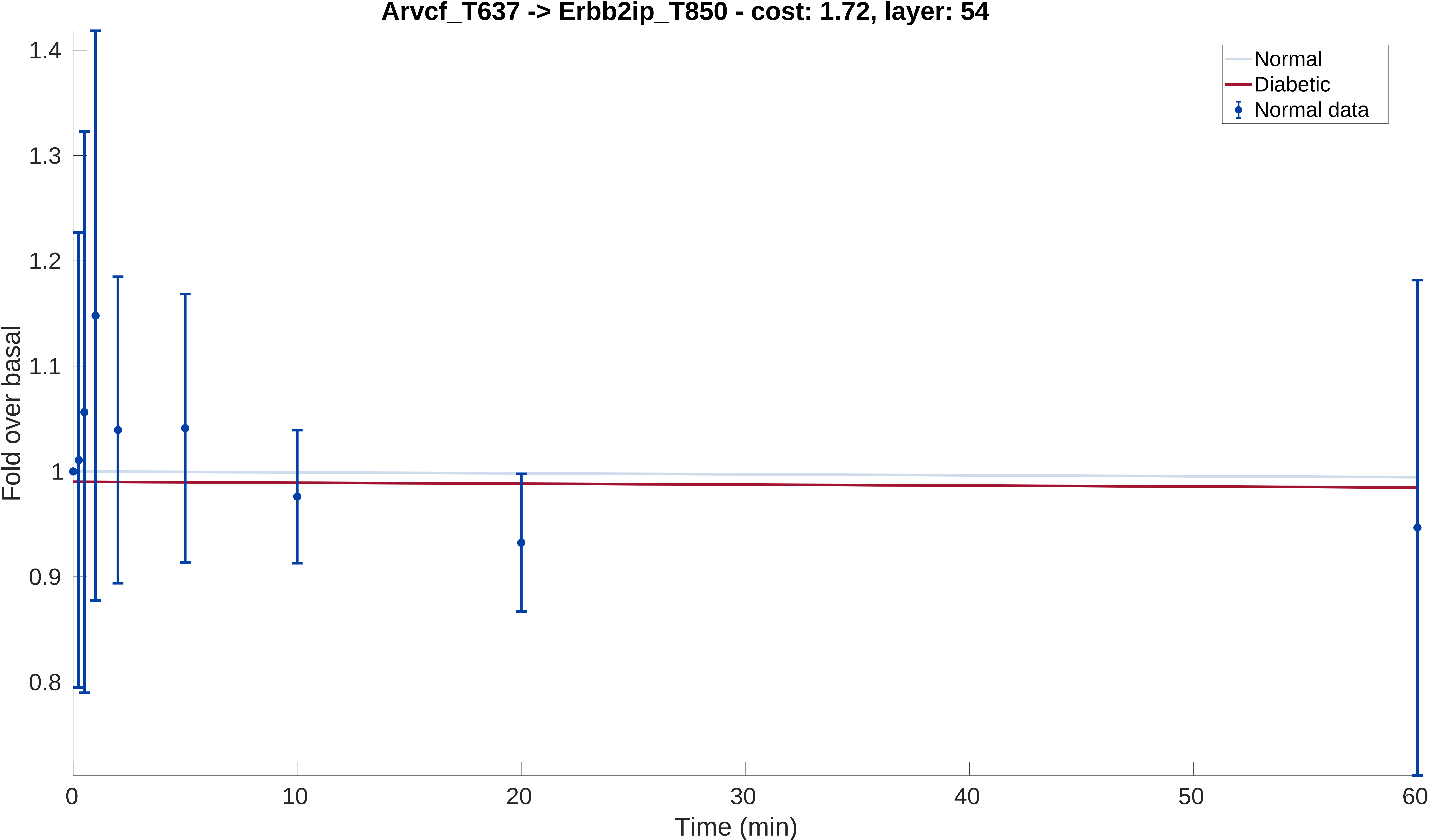

Arvcf_T637 -> Erbb2ip_T850 - cost: 1.72, layer: 54
Normal
Diabetic
Normal data
1.4
1.3
1.2
1.1
1
0.9
0.8
Fold over basal
0
10
20
30
40
50
60
Time (min)

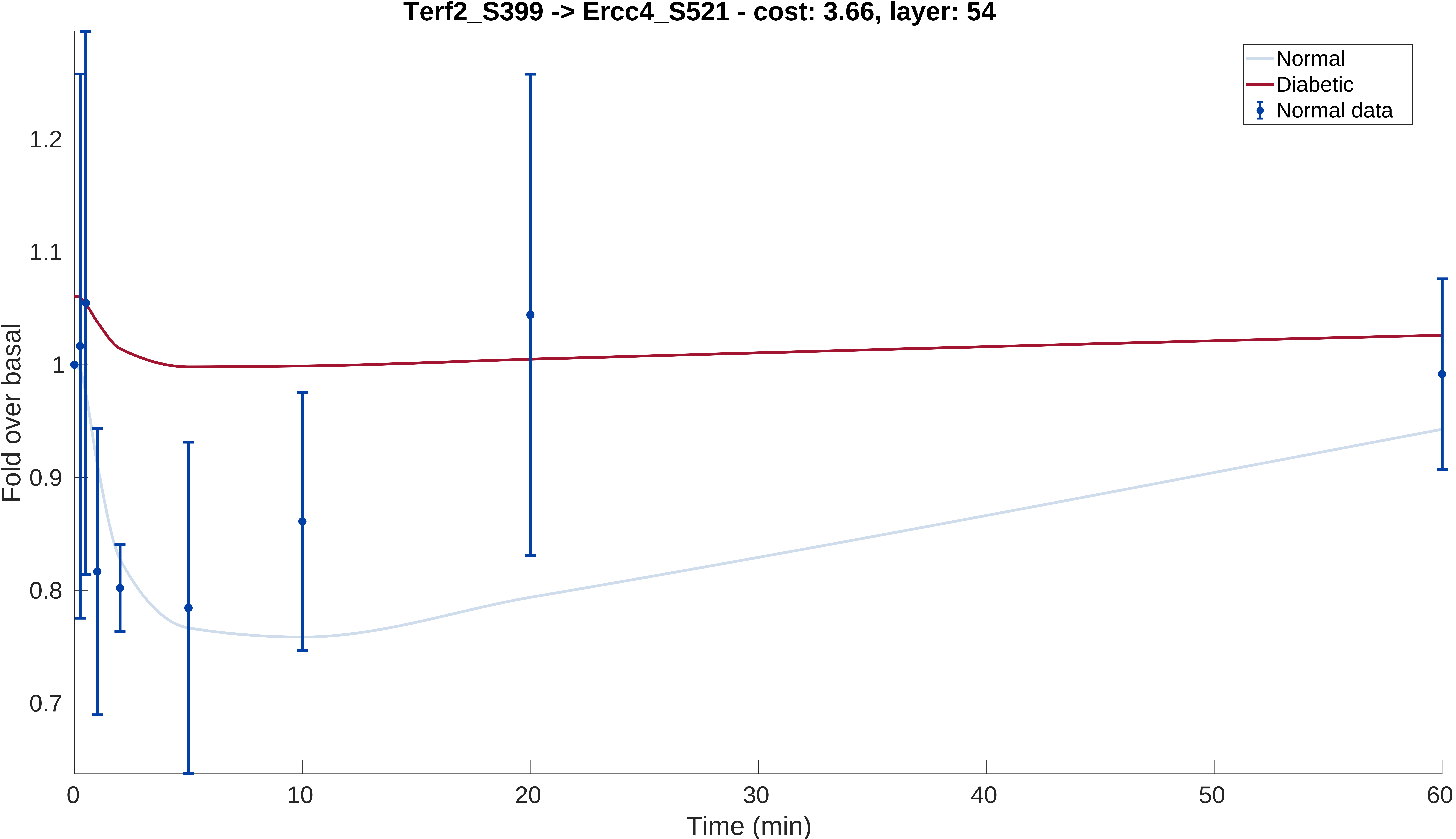

Terf2_S399 -> Ercc4_S521 - cost: 3.66, layer: 54
Normal
Diabetic
Normal data
Fold over basal
1.2
1.1
1
0.9
0.8
0.7
0
10
20
30
40
50
60
Time (min)

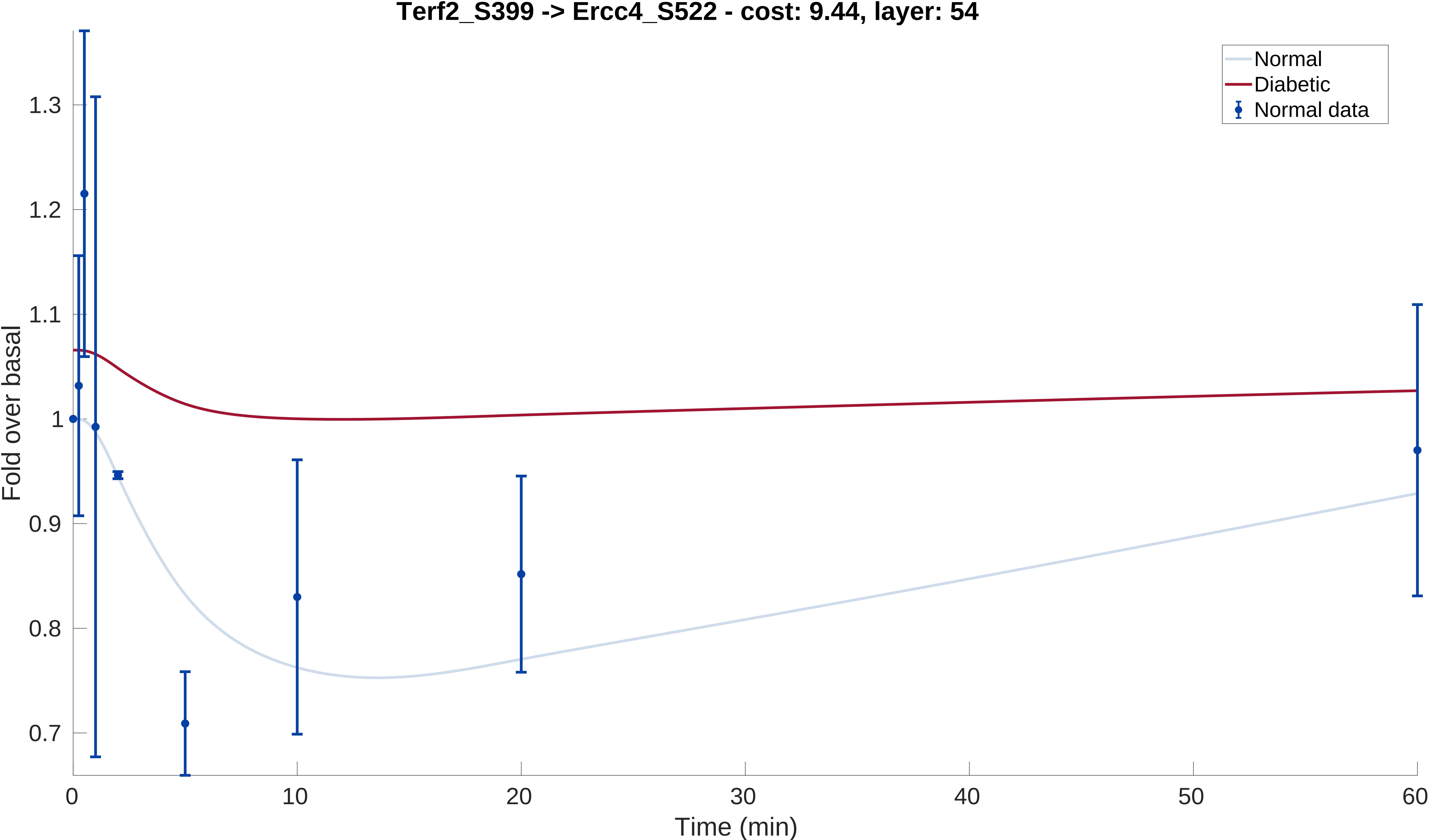

Terf2_S399 -> Ercc4_S522 - cost: 9.44, layer: 54
Normal
Diabetic
Normal data
1.3
1.2
1.1
1
0.9
0.8
0.7
Fold over basal
0
10
20
30
40
50
60
Time (min)

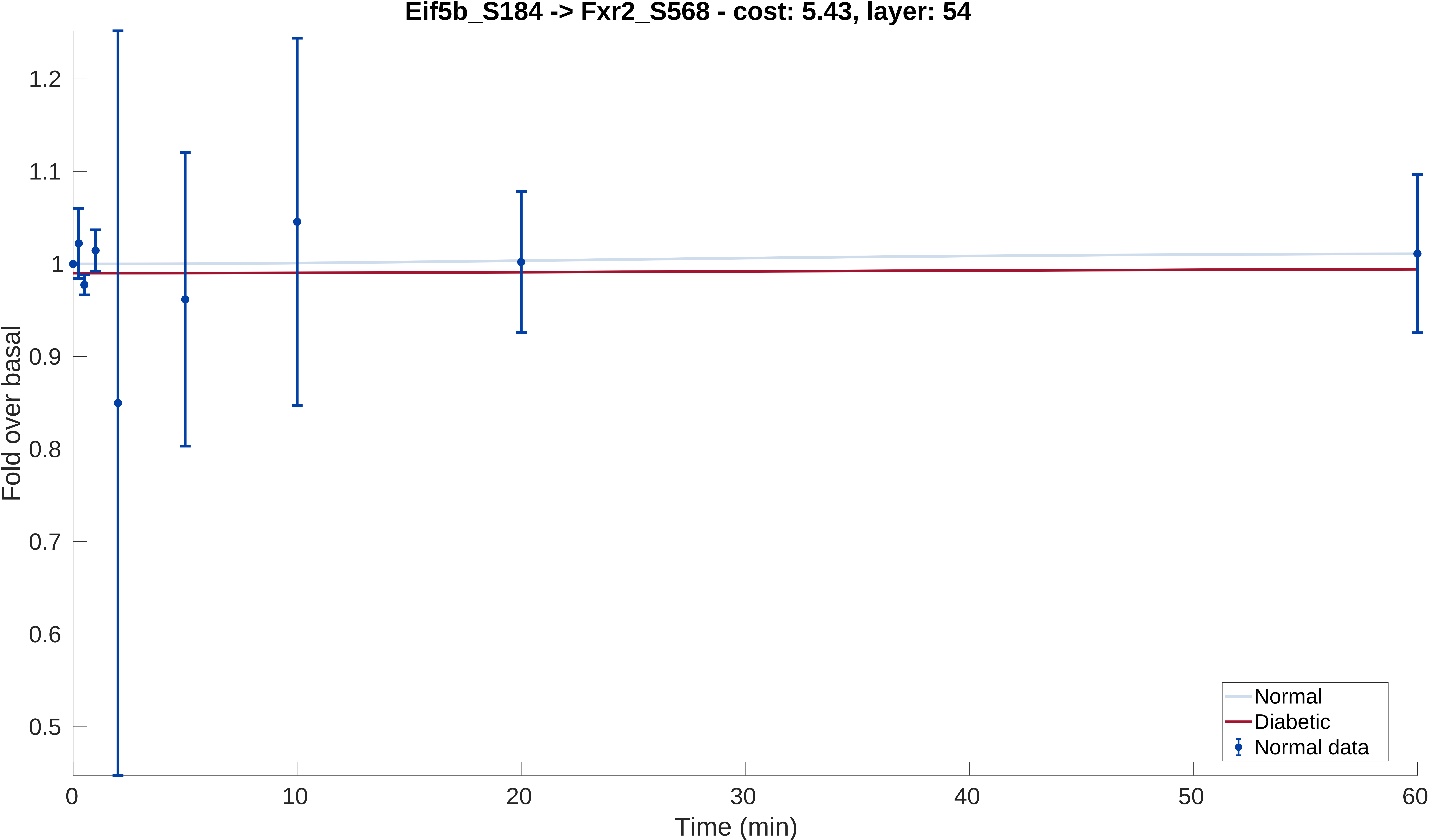

Eif5b_S184 -> Fxr2_S568 - cost: 5.43, layer: 54
Fold over basal
1.2
1.1
1
0.9
0.8
0.7
0.6
0.5
0
10
20
30
40
50
60
Time (min)
Normal
Diabetic
Normal data

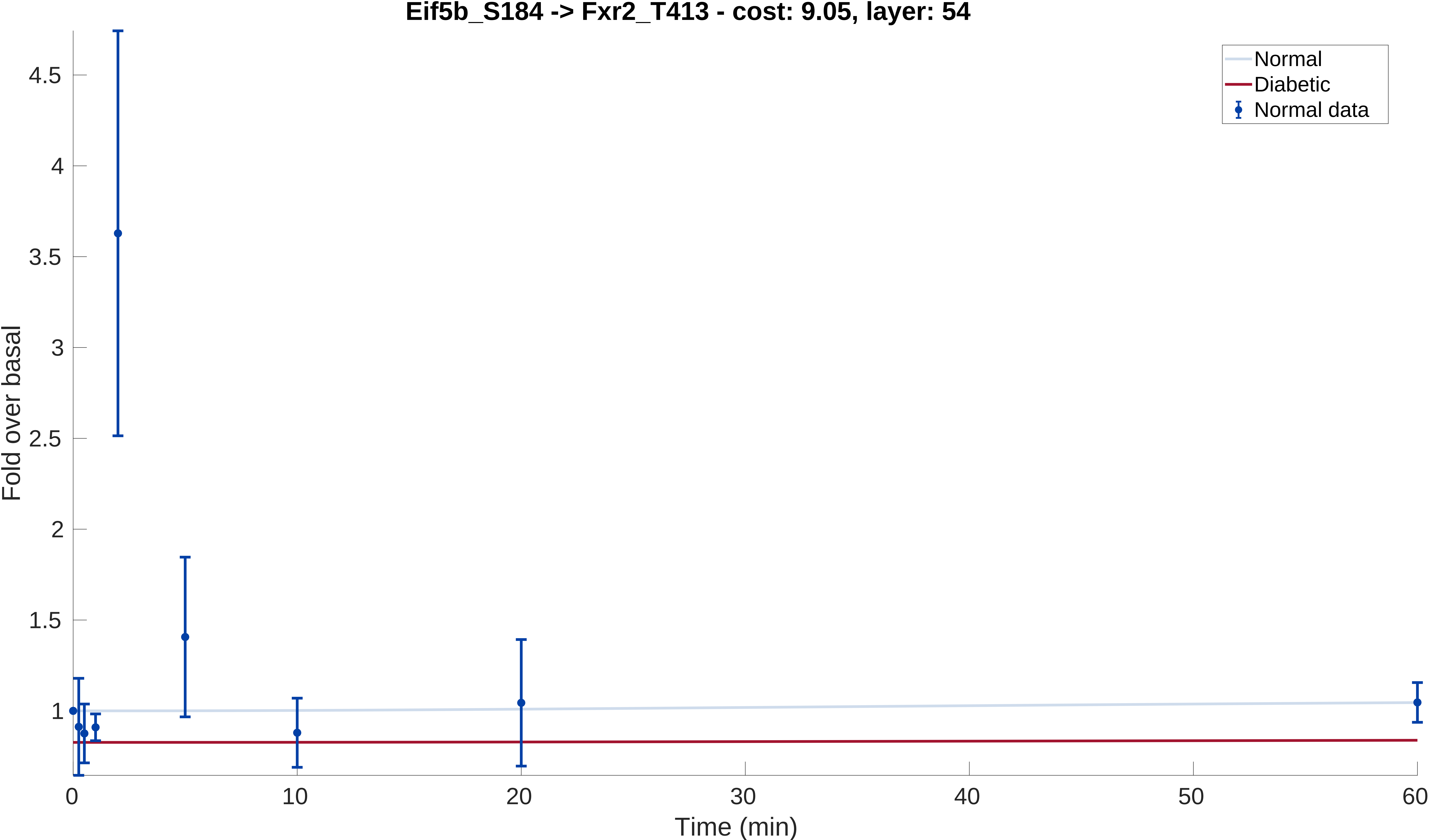

Eif5b_S184 -> Fxr2_T413 - cost: 9.05, layer: 54
Normal
Diabetic
Normal data
Fold over basal
4.5
4
3.5
3
2.5
2
1.5
1
0
10
20
30
40
50
60
Time (min)

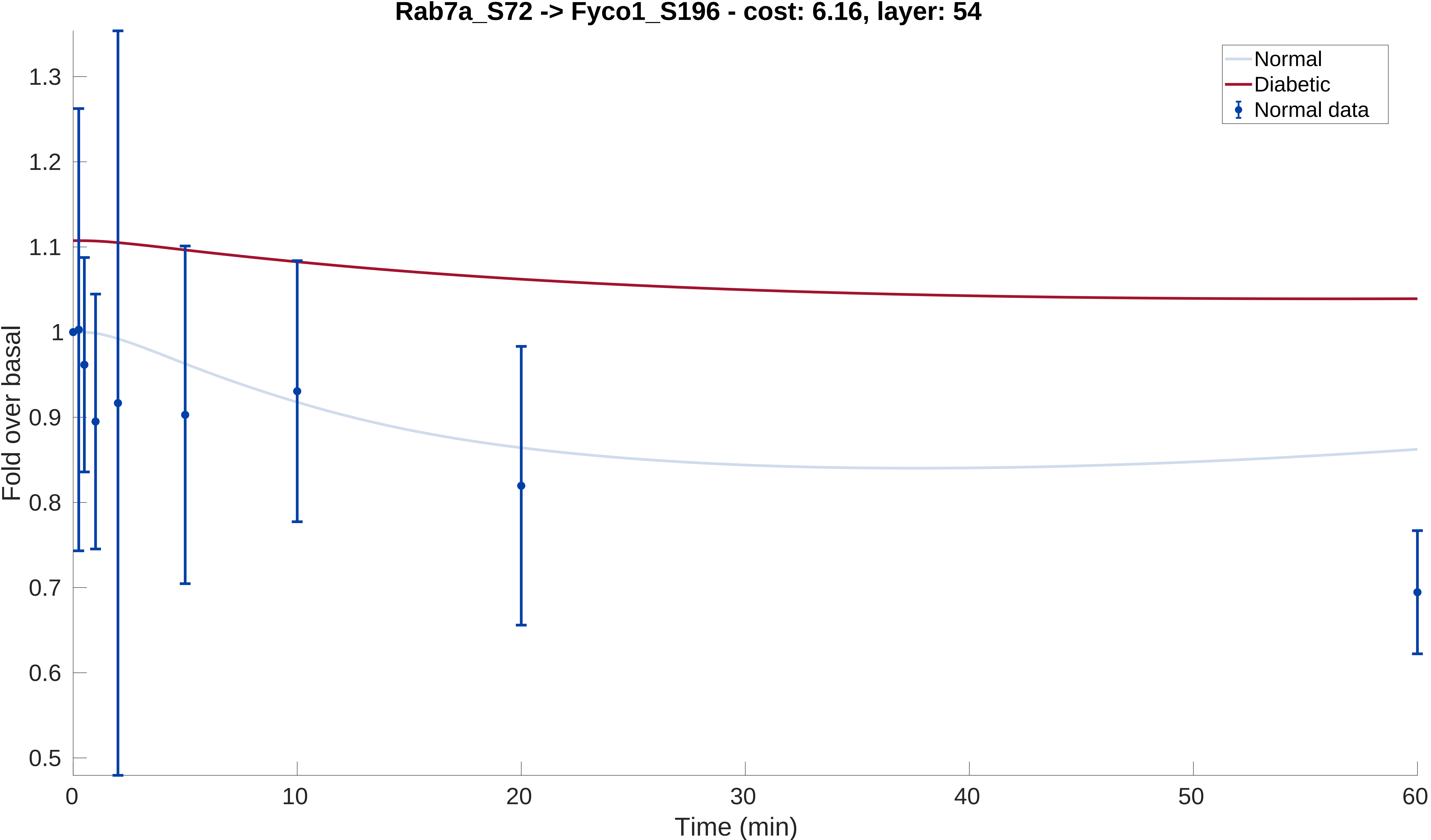
Rab7a_S72 -> Fyco1_S196 - cost: 6.16, layer: 54
Normal
Diabetic
Normal data
Fold over basal
Time (min)
0.5
0.6
0.7
0.8
0.9
1
1.1
1.2
1.3
0
10
20
30
40
50
60

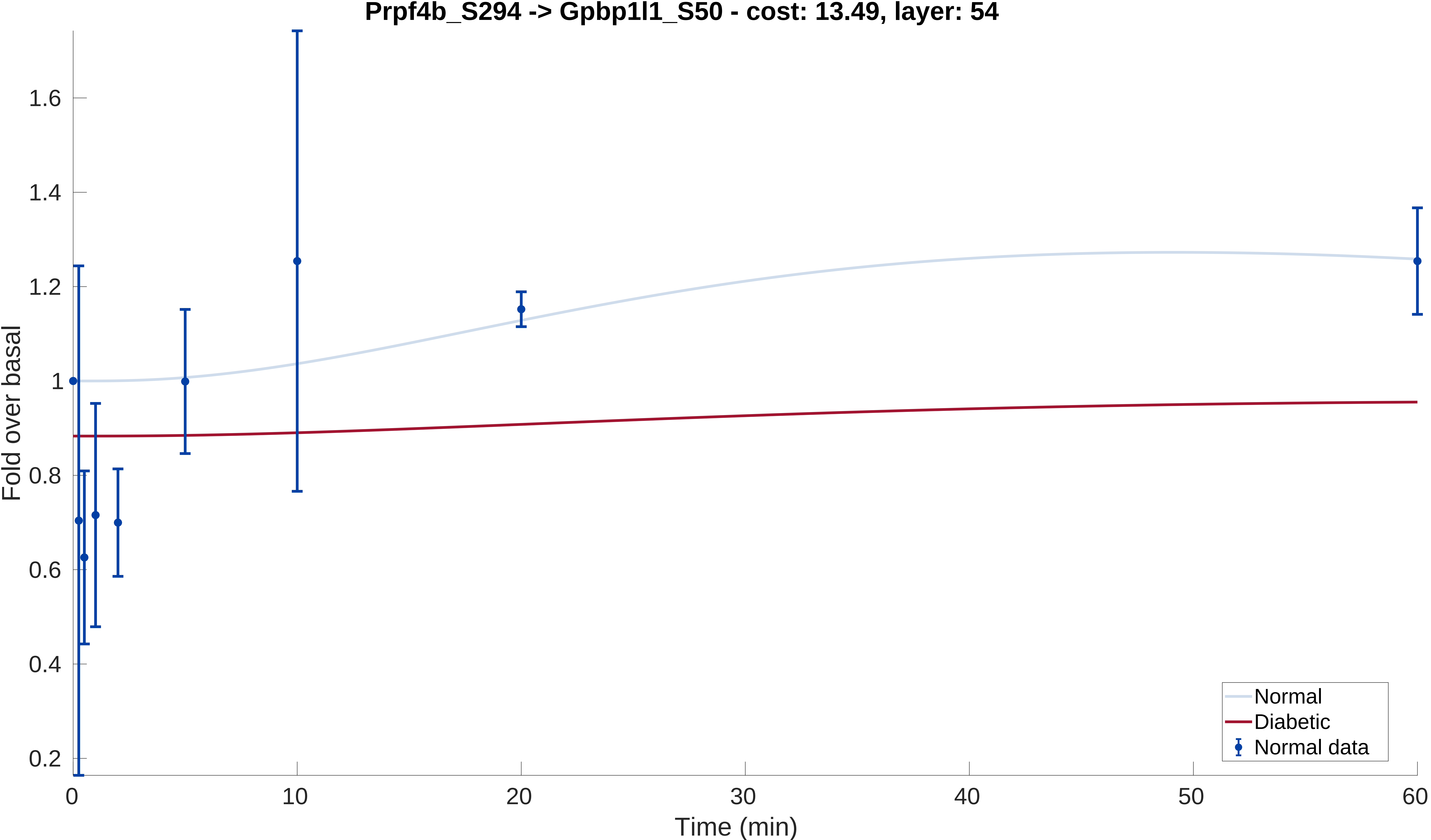

Prpf4b_S294 -> Gpbp1l1_S50 - cost: 13.49, layer: 54
Fold over basal
Time (min)
0.2
0.4
0.6
0.8
1
1.2
1.4
1.6
0
10
20
30
40
50
60
Normal
Diabetic
Normal data

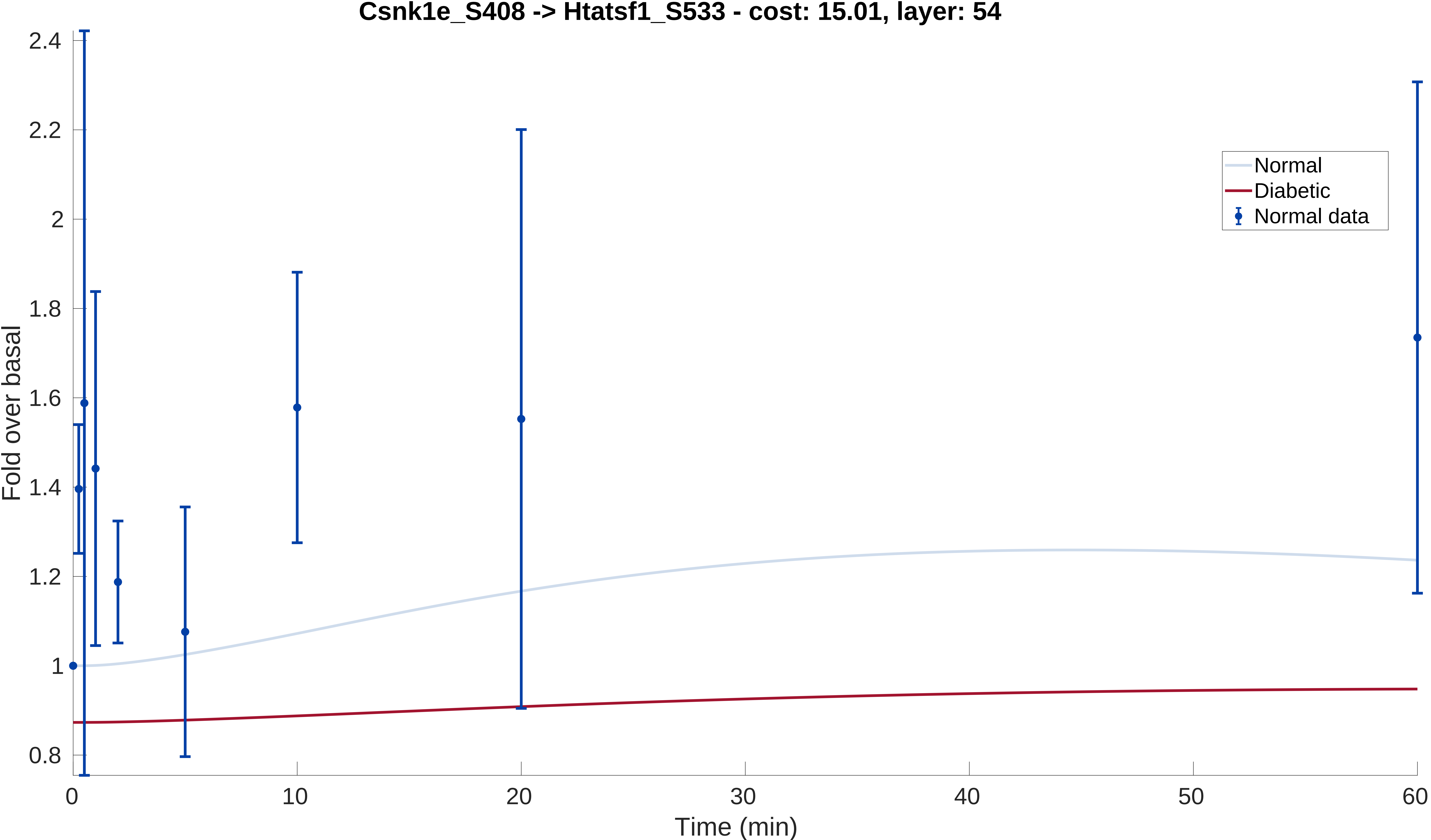

Csnk1e_S408 -> Htatsf1_S533 - cost: 15.01, layer: 54
Fold over basal
Time (min)
Normal
Diabetic
Normal data
2.4
2.2
2
1.8
1.6
1.4
1.2
1
0.8
0
10
20
30
40
50
60

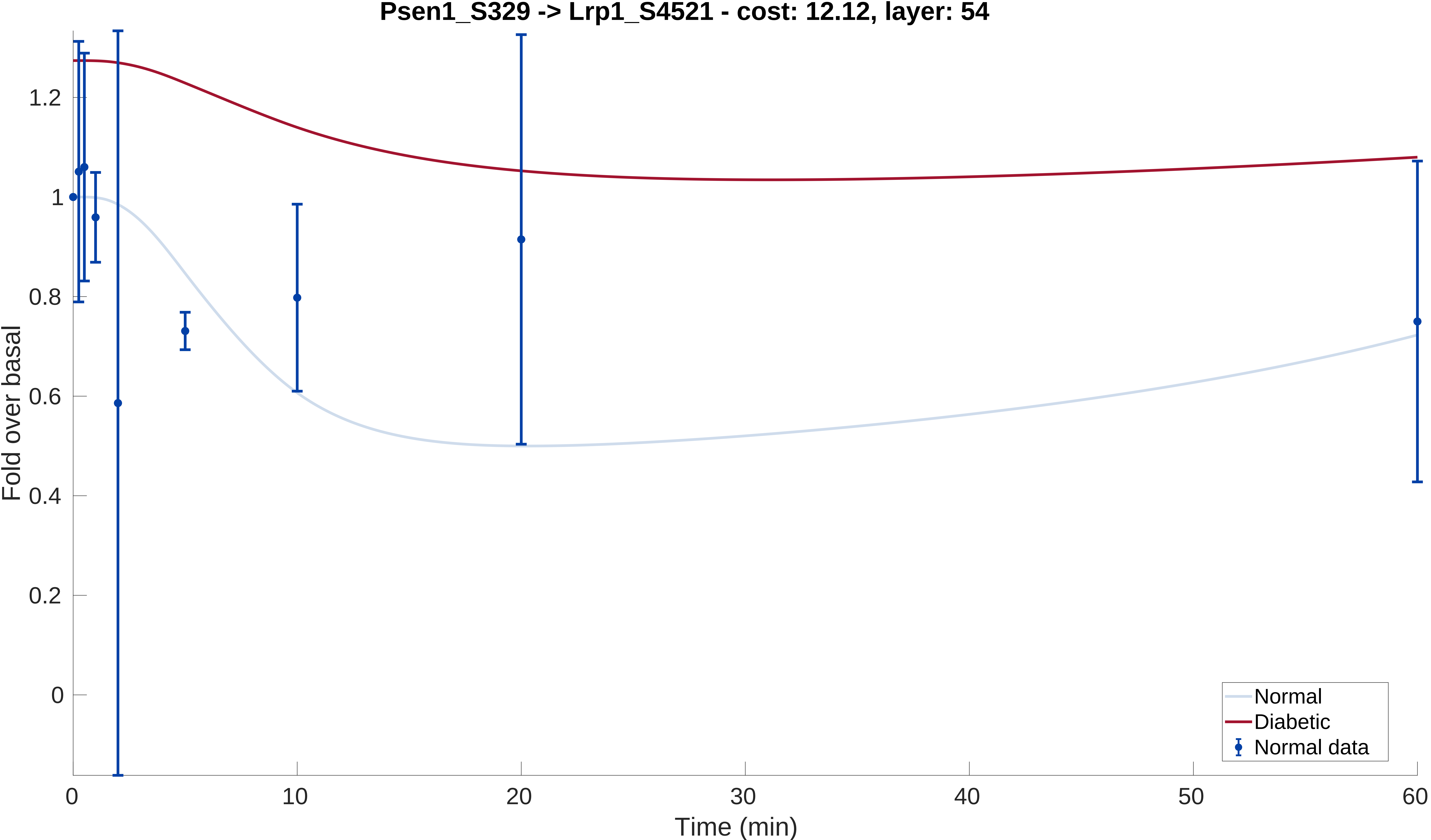

Psen1_S329 -> Lrp1_S4521 - cost: 12.12, layer: 54
Fold over basal
1.2
1
0.8
0.6
0.4
0.2
0
0
10
20
30
40
50
60
Time (min)
Normal
Diabetic
Normal data

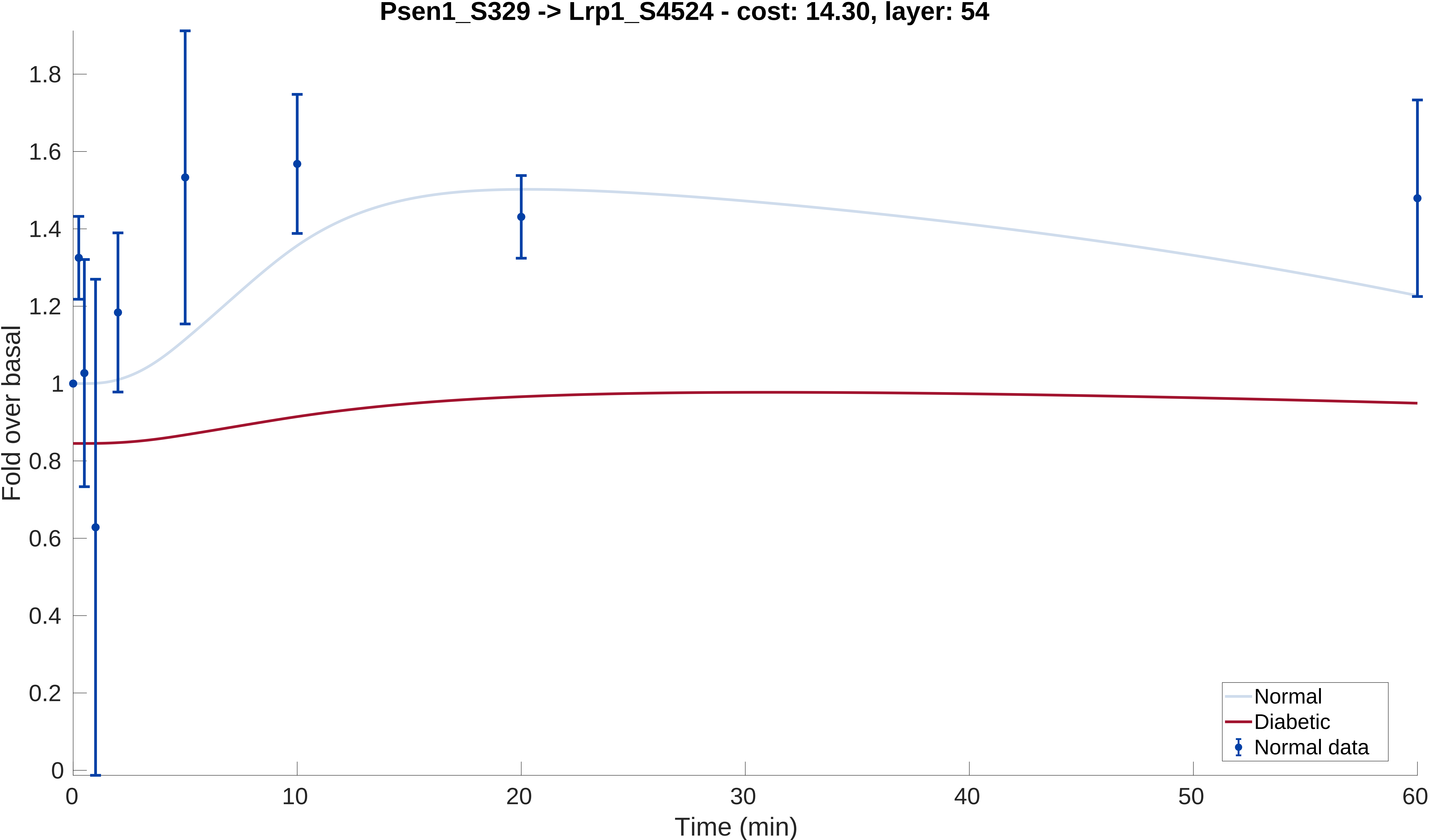

Psen1_S329 -> Lrp1_S4524 - cost: 14.30, layer: 54
Fold over basal
0
0.2
0.4
0.6
0.8
1
1.2
1.4
1.6
1.8
0
10
20
30
40
50
60
Time (min)
Normal
Diabetic
Normal data

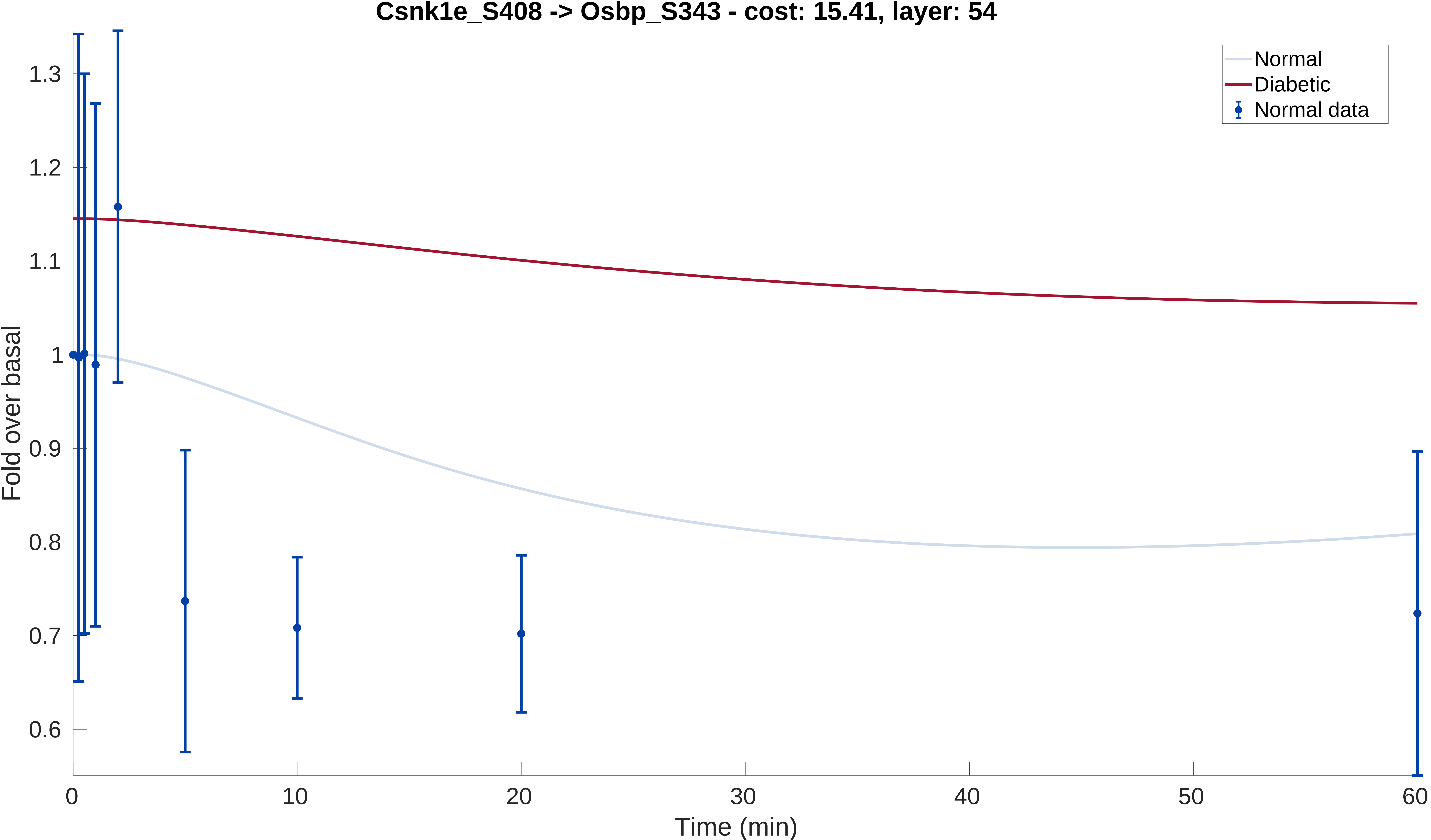
Csnk1e_S408 -> Osbp_S343 - cost: 15.41, layer: 54
Normal
Diabetic
Normal data
Fold over basal
1.3
1.2
1.1
1
0.9
0.8
0.7
0.6
0
10
20
30
40
50
60
Time (min)

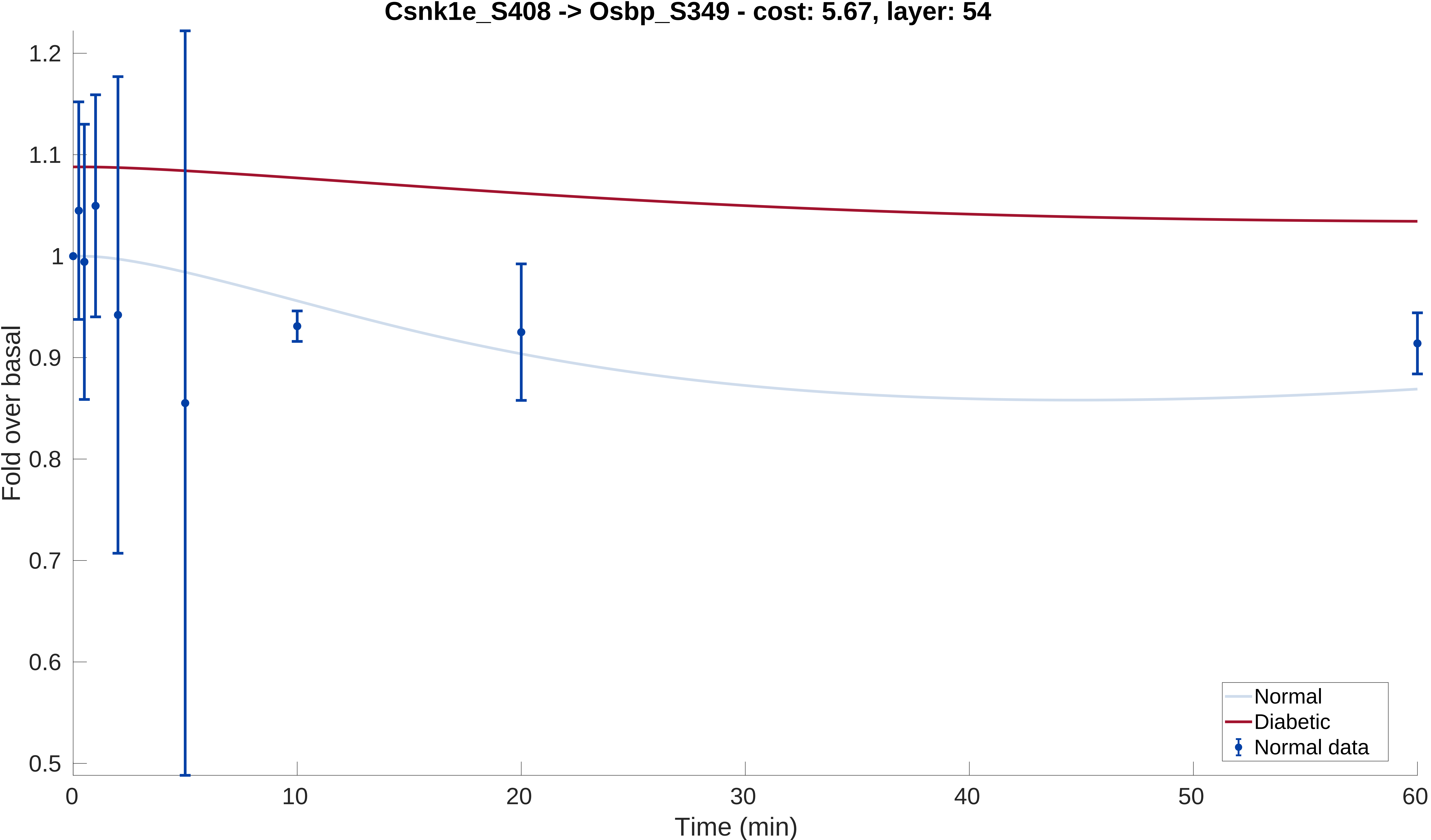

Csnk1e_S408 -> Osbp_S349 - cost: 5.67, layer: 54
Fold over basal
1.2
1.1
1
0.9
0.8
0.7
0.6
0.5
0
10
20
30
40
50
60
Time (min)
Normal
Diabetic
Normal data

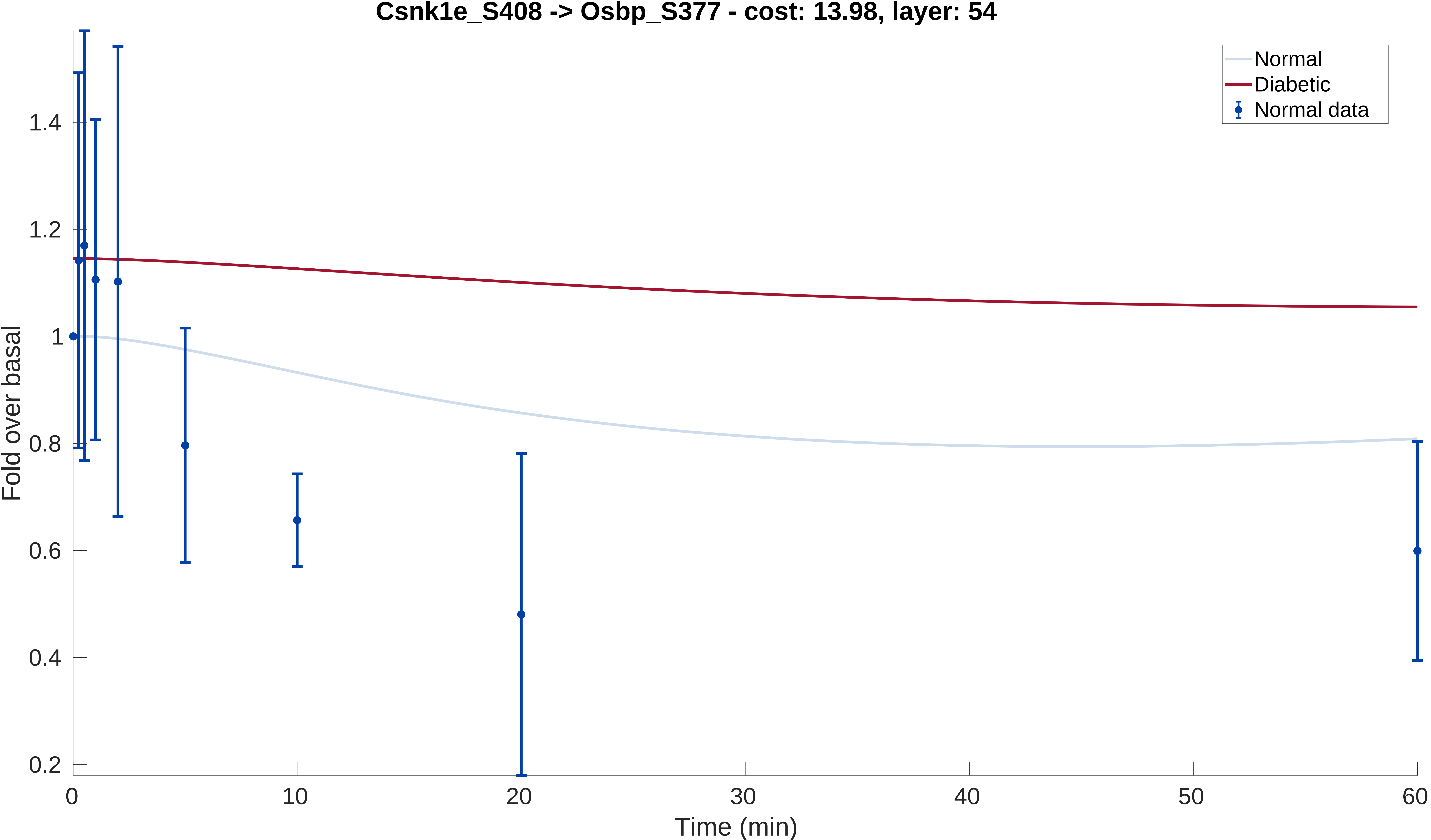

Csnk1e_S408 -> Osbp_S377 - cost: 13.98, layer: 54
Normal
Diabetic
Normal data
Fold over basal
1.4
1.2
1
0.8
0.6
0.4
0.2
0
10
20
30
40
50
60
Time (min)

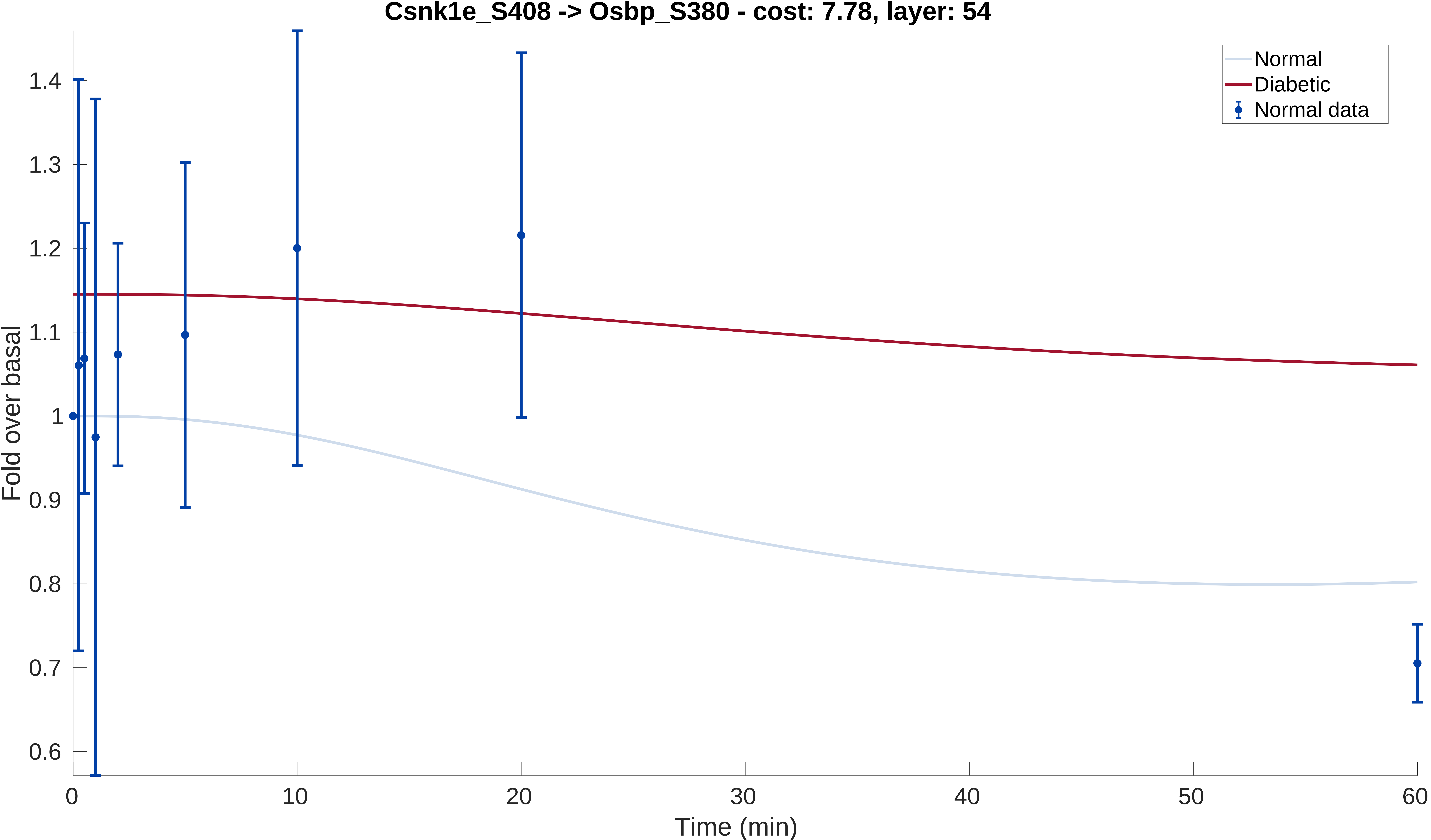

Csnk1e_S408 -> Osbp_S380 - cost: 7.78, layer: 54
Normal
Diabetic
Normal data
Fold over basal
1.4
1.3
1.2
1.1
1
0.9
0.8
0.7
0.6
0
10
20
30
40
50
60
Time (min)

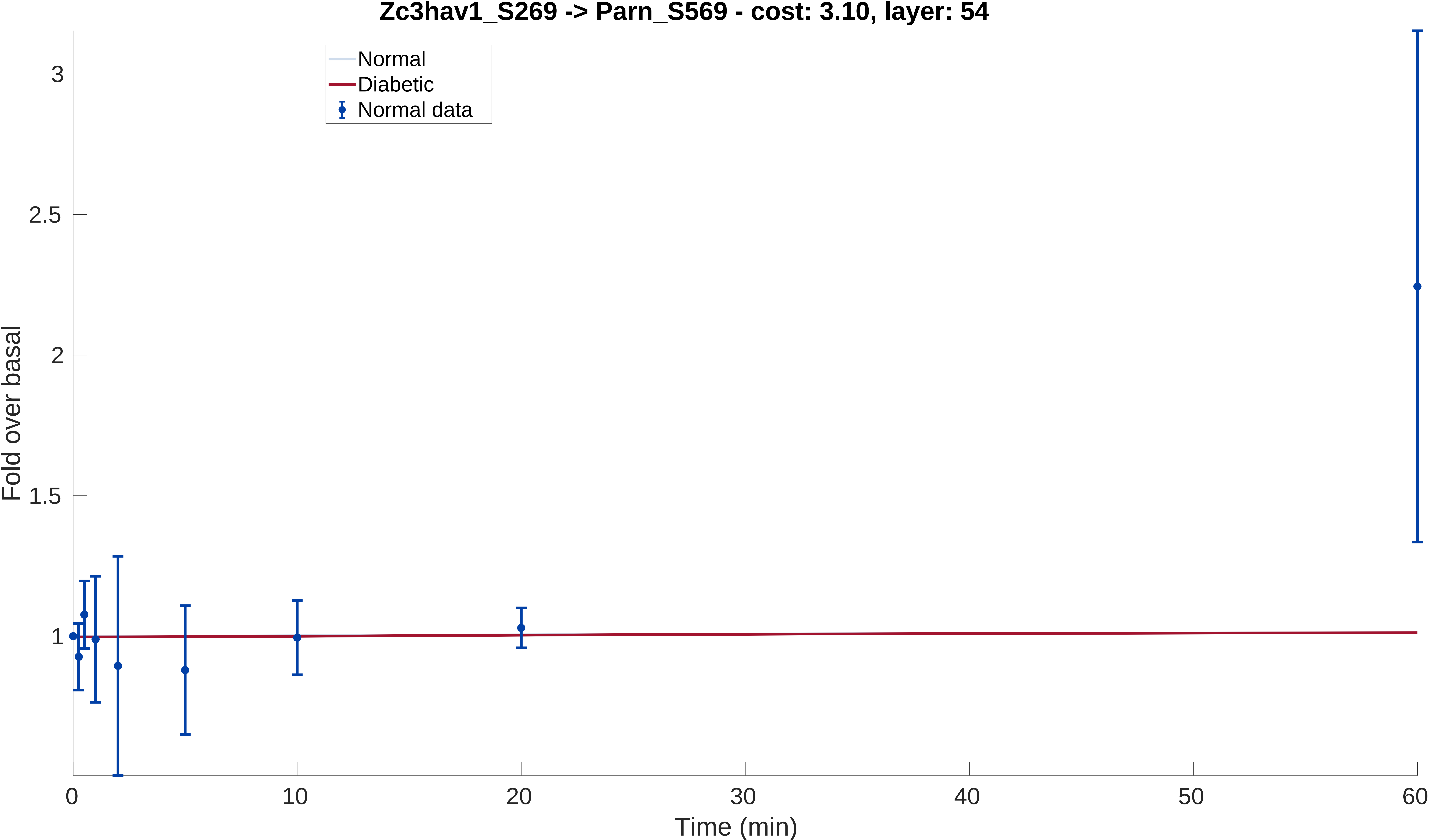
Zc3hav1_S269 -> Parn_S569 - cost: 3.10, layer: 54
Normal
Diabetic
Normal data
Fold over basal
Time (min)
3
2.5
2
1.5
1
0
10
20
30
40
50
60

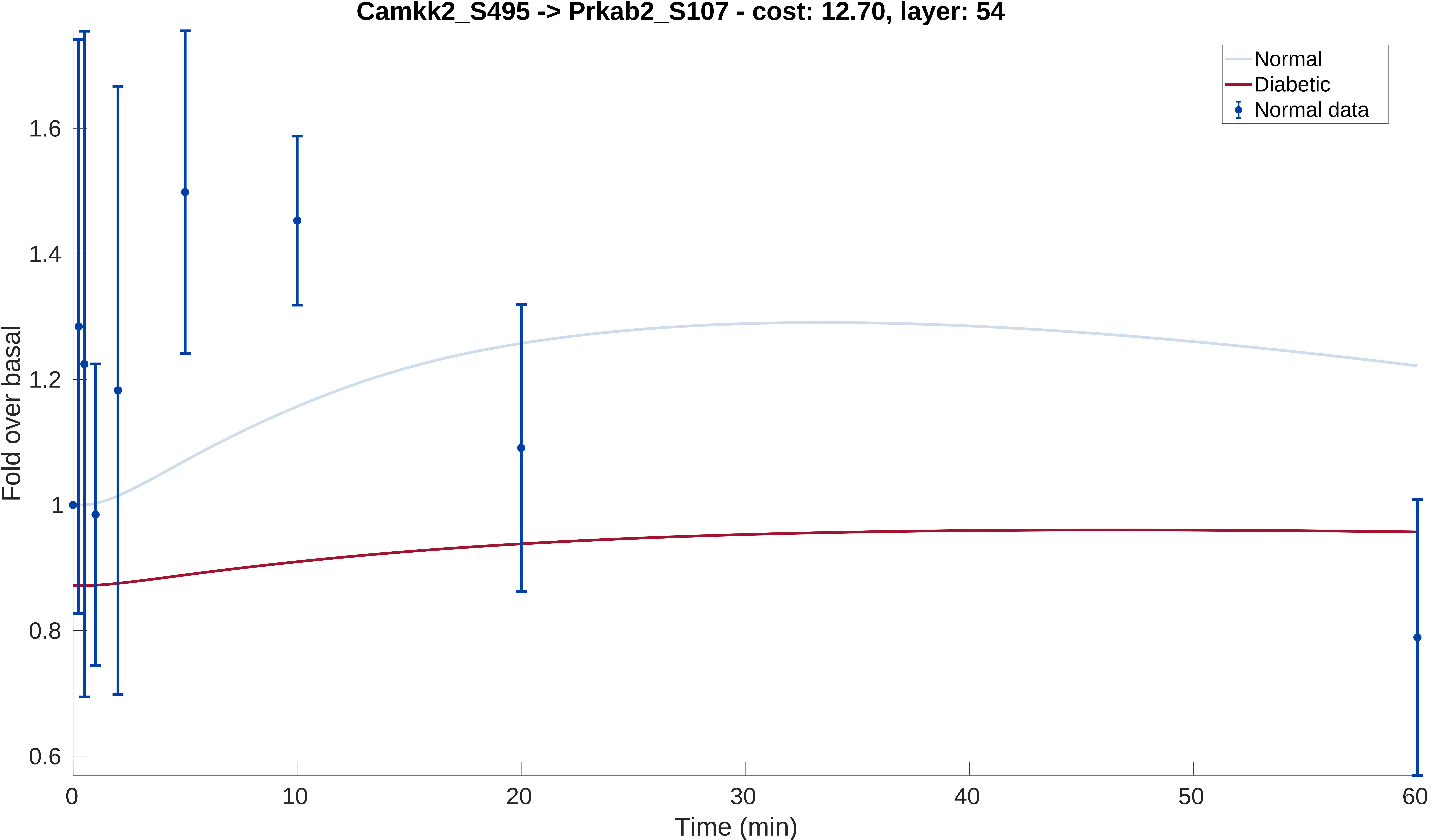
Camkk2_S495 -> Prkab2_S107 - cost: 12.70, layer: 54
Normal
Diabetic
Normal data
Fold over basal
0.6
0.8
1
1.2
1.4
1.6
0
10
20
30
40
50
60
Time (min)

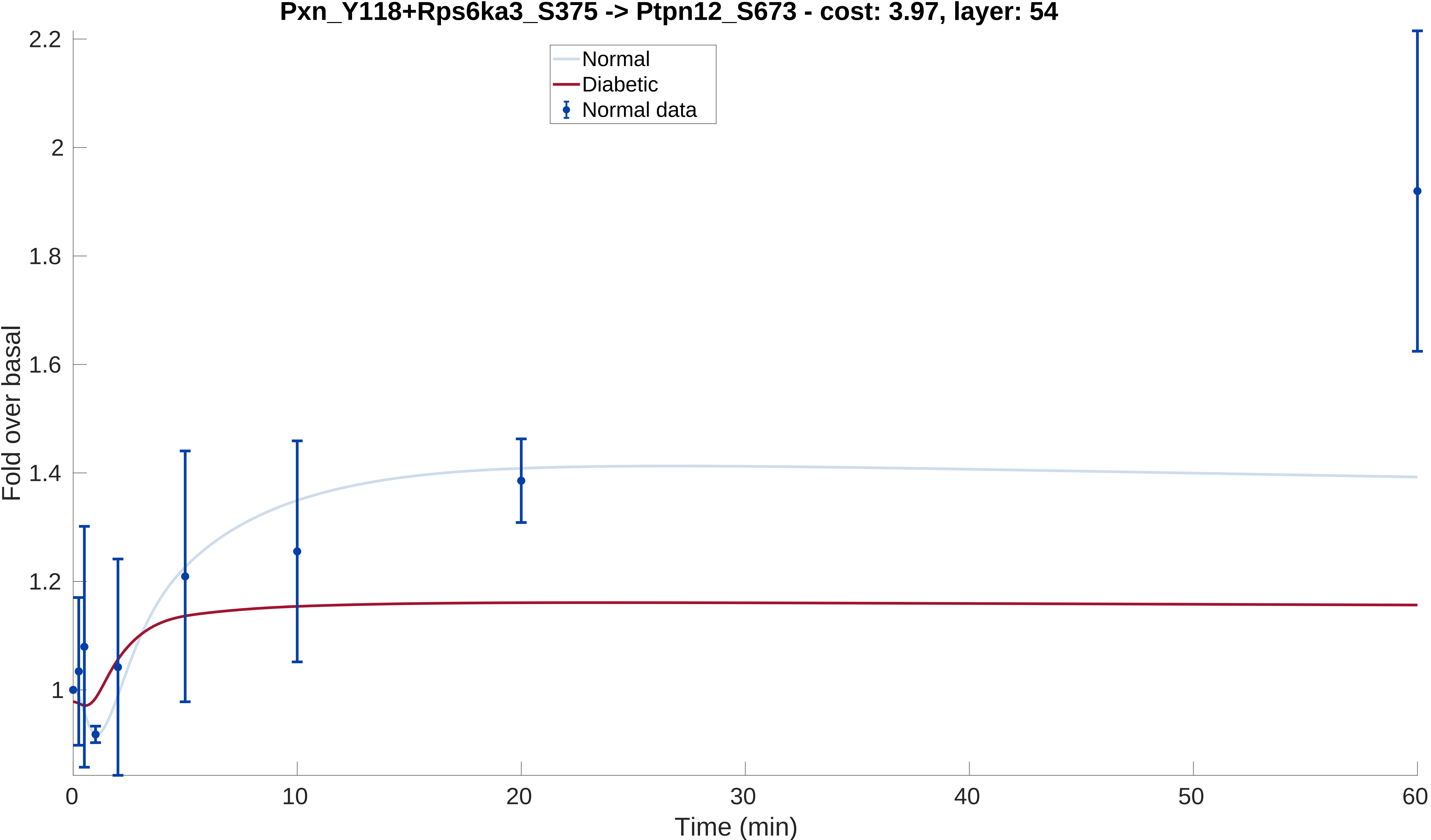

Pxn_Y118+Rps6ka3_S375 -> Ptpn12_S673 - cost: 3.97, layer: 54
Normal
Diabetic
Normal data
Fold over basal
2.2
2
1.8
1.6
1.4
1.2
1
0
10
20
30
40
50
60
Time (min)

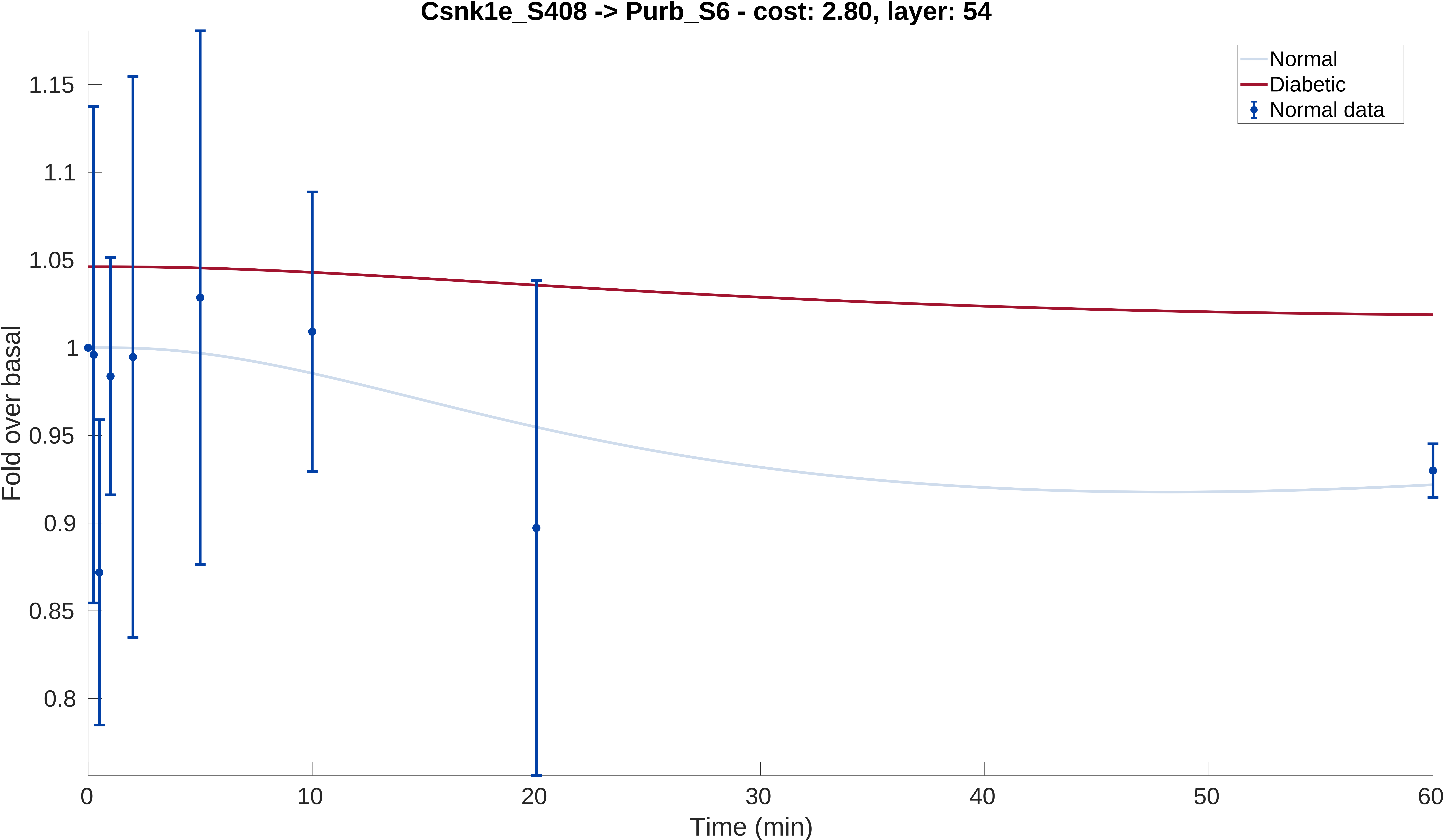
Csnk1e_S408 -> Purb_S6 - cost: 2.80, layer: 54
Normal
Diabetic
Normal data
Fold over basal
1.15
1.1
1.05
1
0.95
0.9
0.85
0.8
0
10
20
30
40
50
60
Time (min)

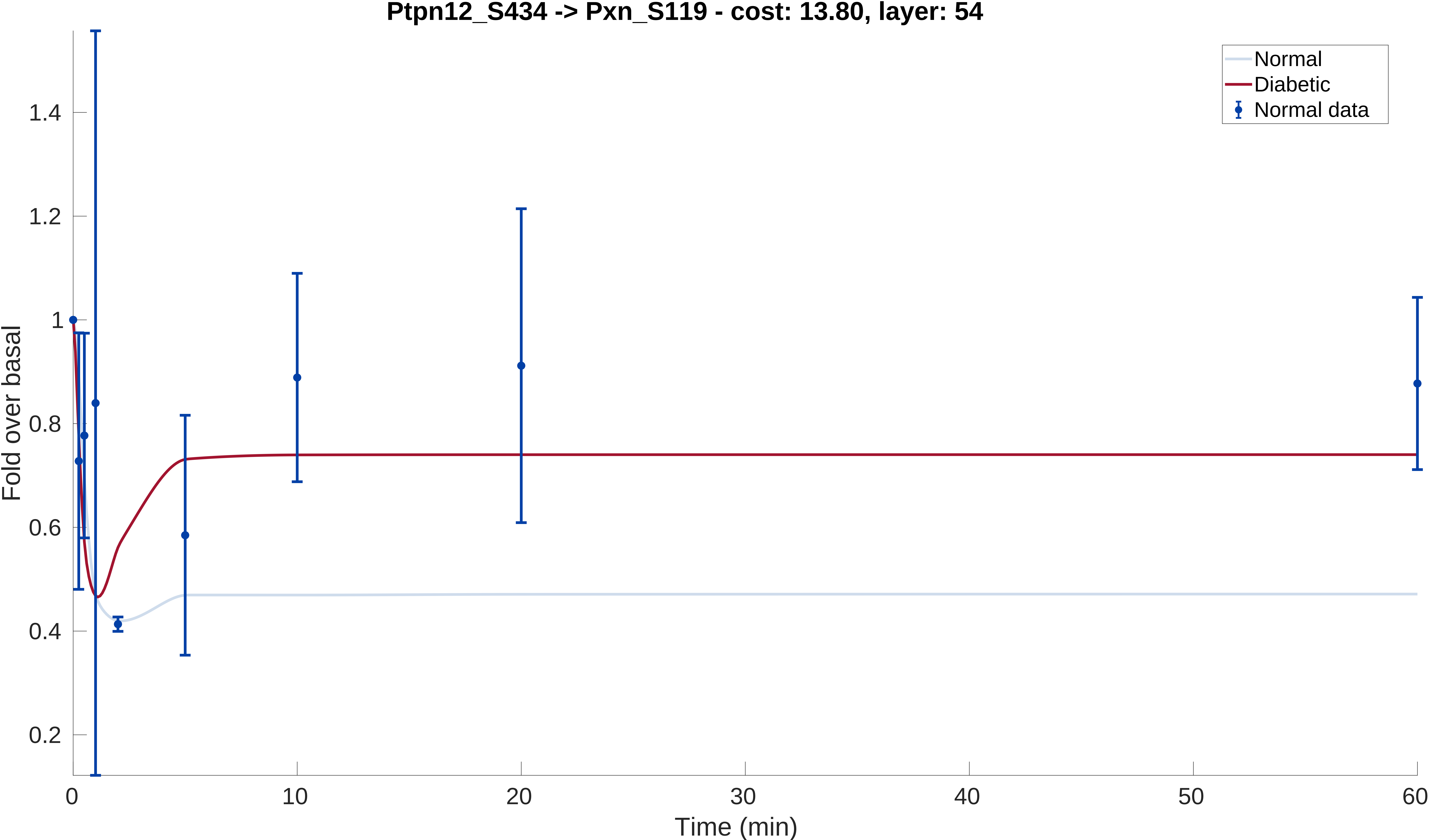

Ptpn12_S434 -> Pxn_S119 - cost: 13.80, layer: 54
Normal
Diabetic
Normal data
Fold over basal
1.4
1.2
1
0.8
0.6
0.4
0.2
0
10
20
30
40
50
60
Time (min)

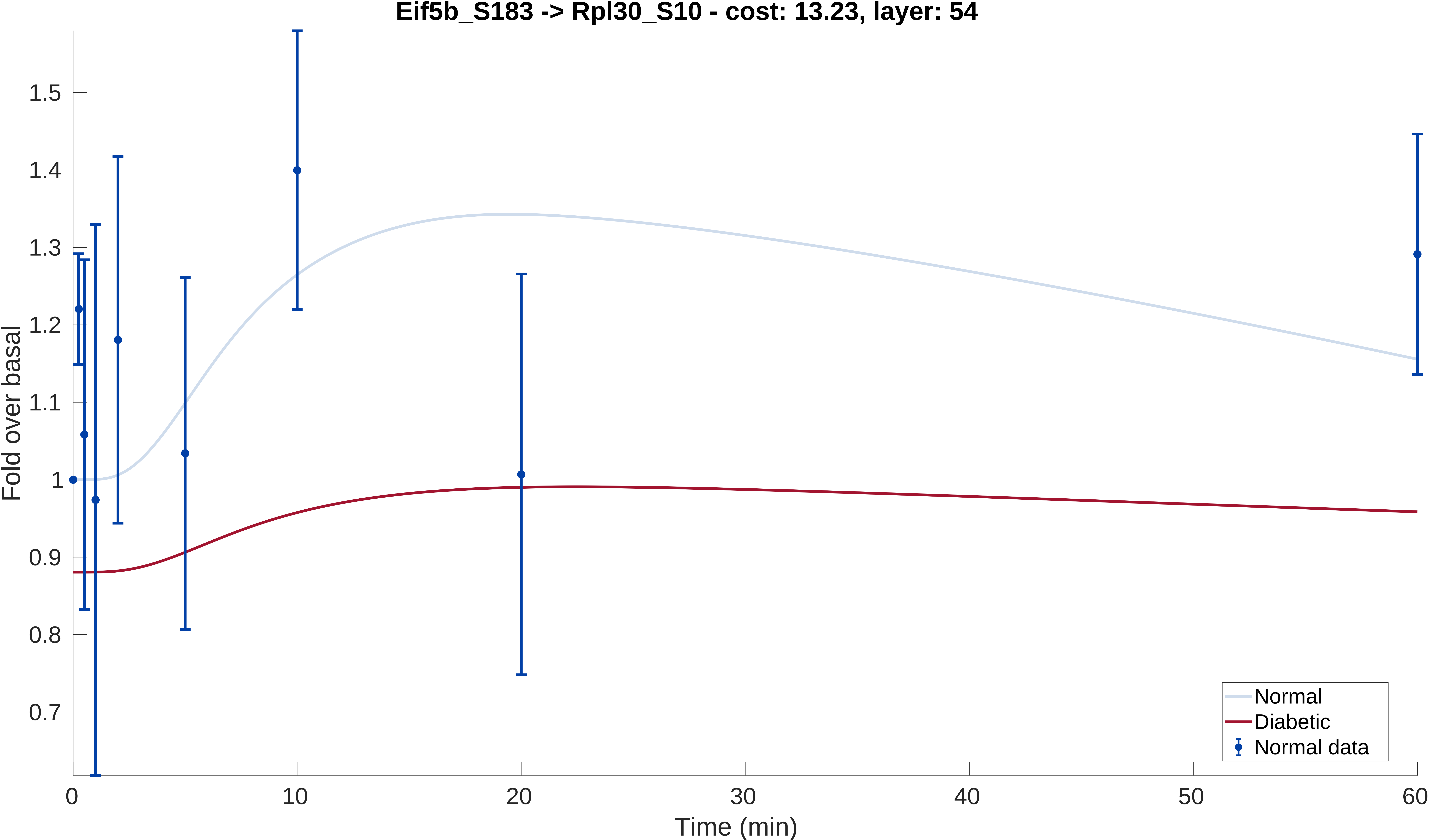

Eif5b_S183 -> Rpl30_S10 - cost: 13.23, layer: 54
Fold over basal
1.5
1.4
1.3
1.2
1.1
1
0.9
0.8
0.7
0
10
20
30
40
50
60
Time (min)
Normal
Diabetic
Normal data

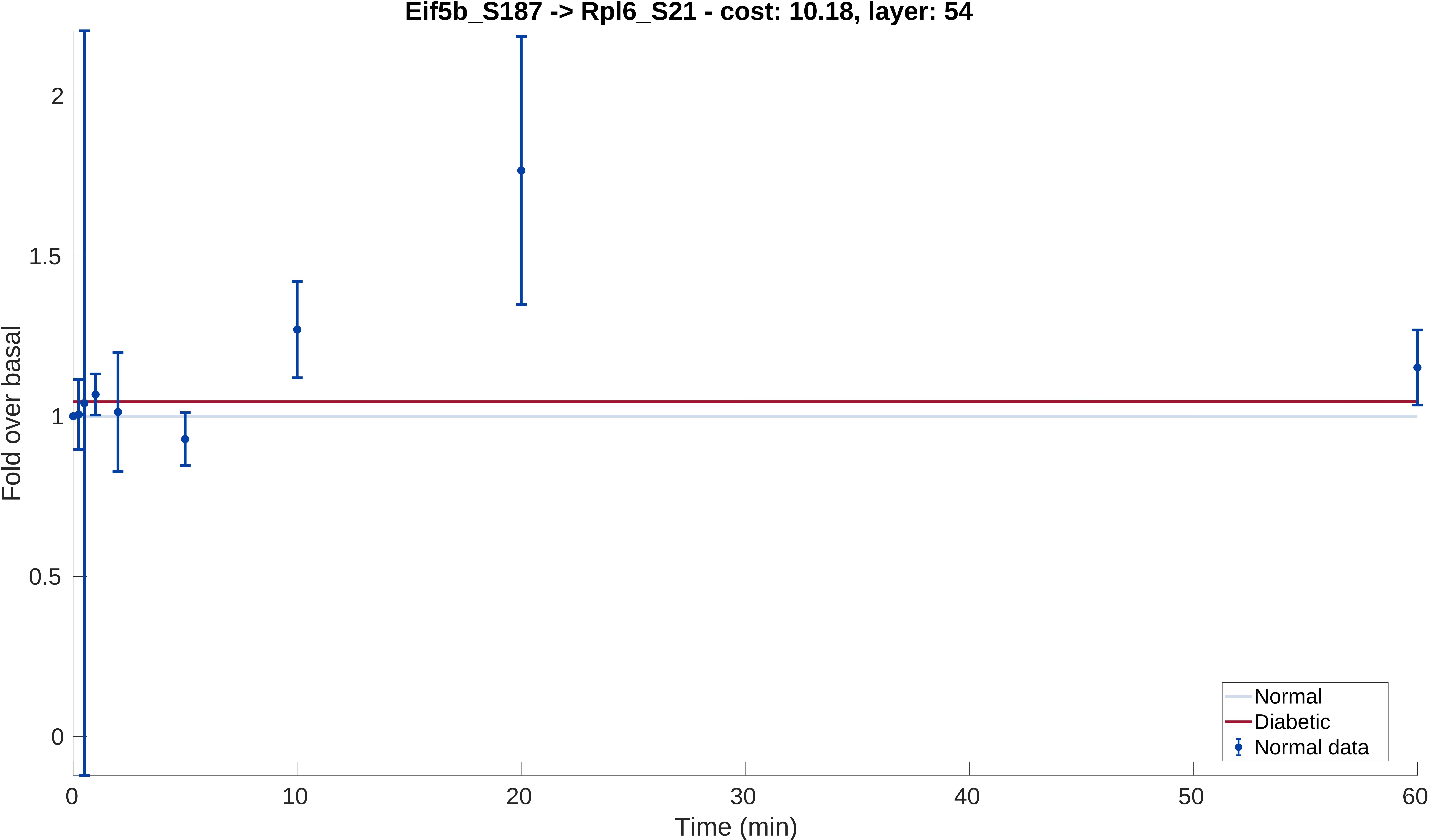
Eif5b_S187 -> Rpl6_S21 - cost: 10.18, layer: 54
Fold over basal
Time (min)
0
0.5
1
1.5
2
0
10
20
30
40
50
60
Normal
Diabetic
Normal data

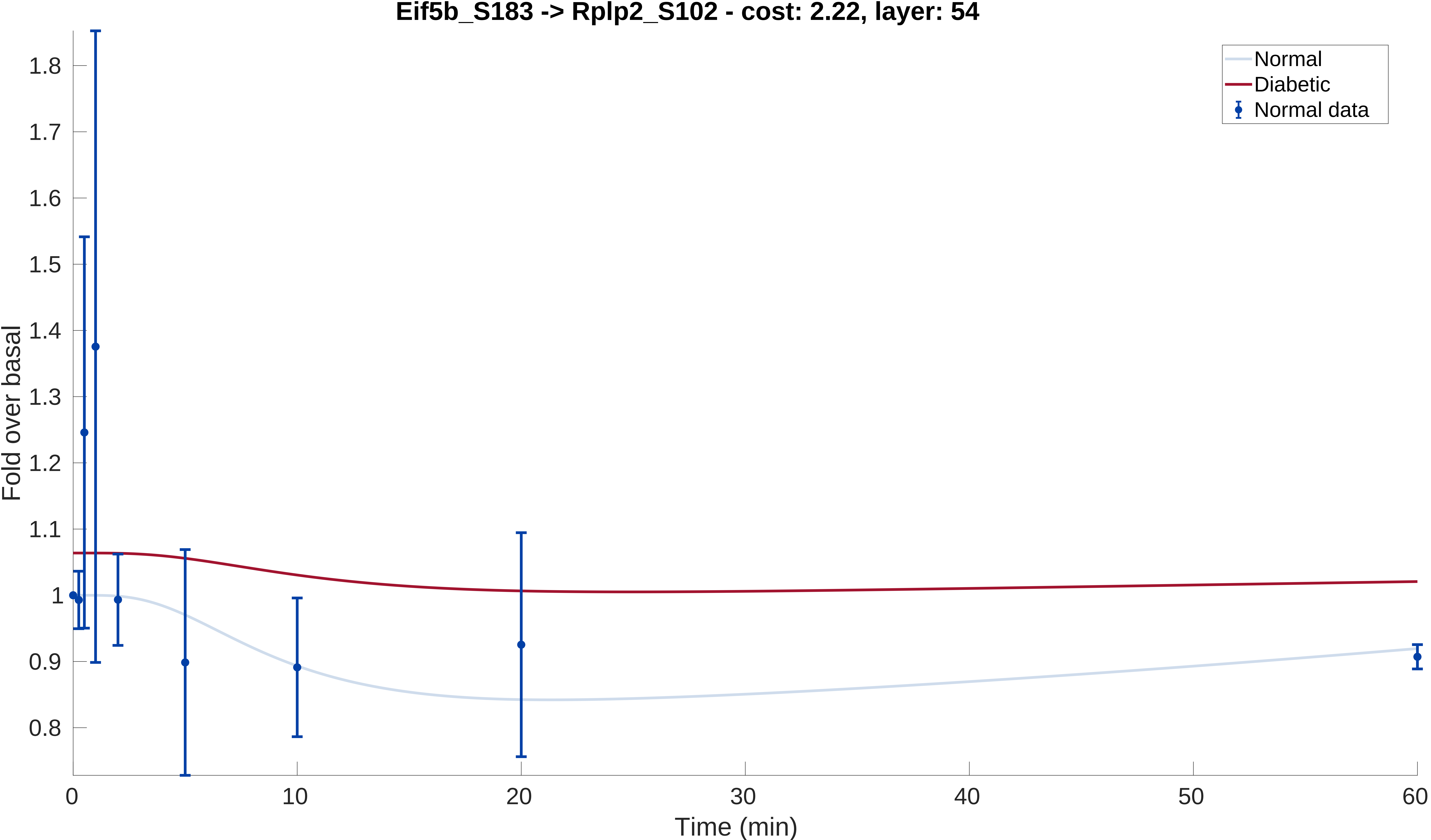

Eif5b_S183 -> Rplp2_S102 - cost: 2.22, layer: 54
Normal
Diabetic
Normal data
Fold over basal
1.8
1.7
1.6
1.5
1.4
1.3
1.2
1.1
1
0.9
0.8
0
10
20
30
40
50
60
Time (min)

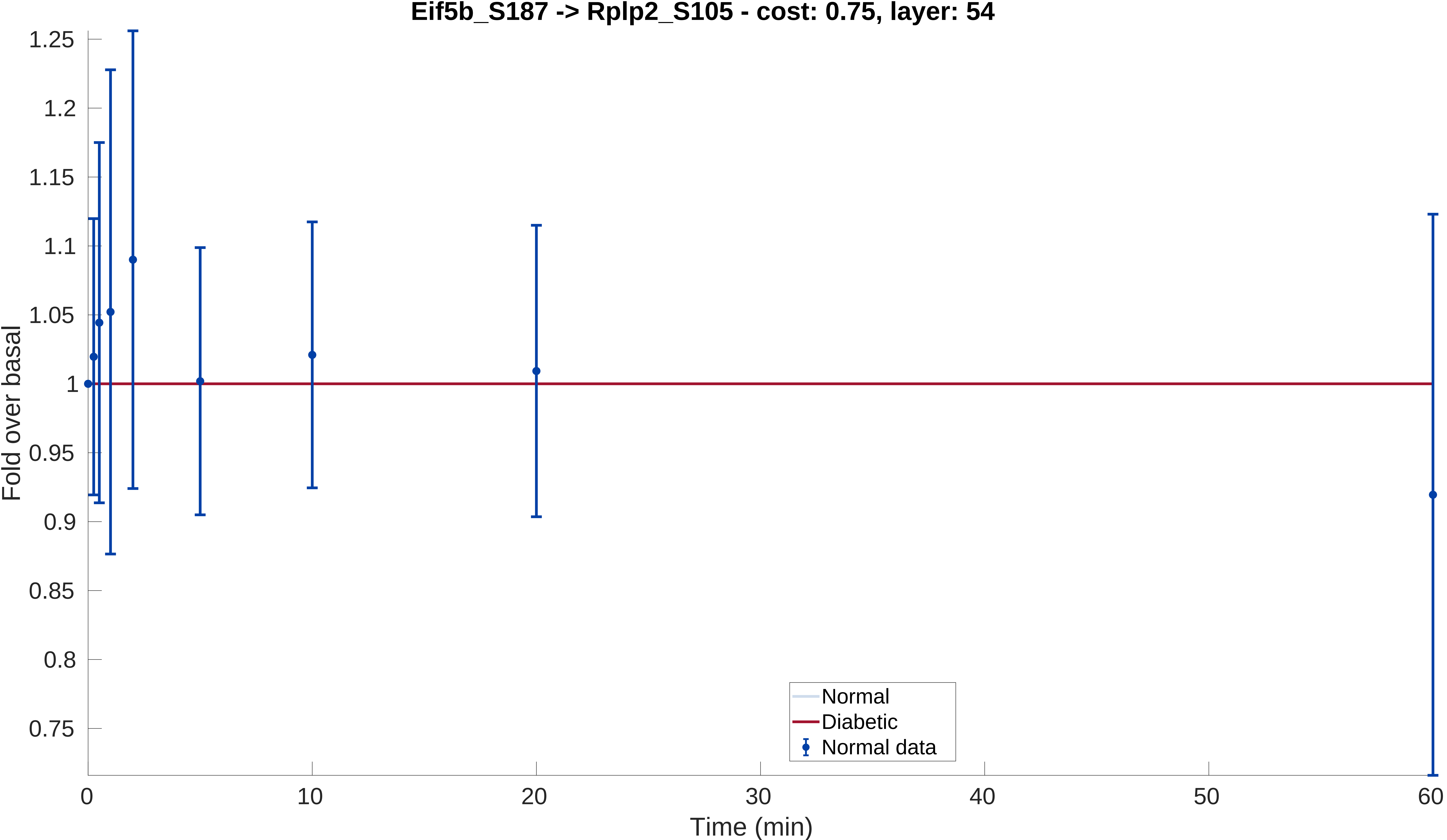

Eif5b_S187 -> Rplp2_S105 - cost: 0.75, layer: 54
Fold over basal
1.25
1.2
1.15
1.1
1.05
1
0.95
0.9
0.85
0.8
0.75
0
10
20
30
40
50
60
Time (min)
Normal
Diabetic
Normal data

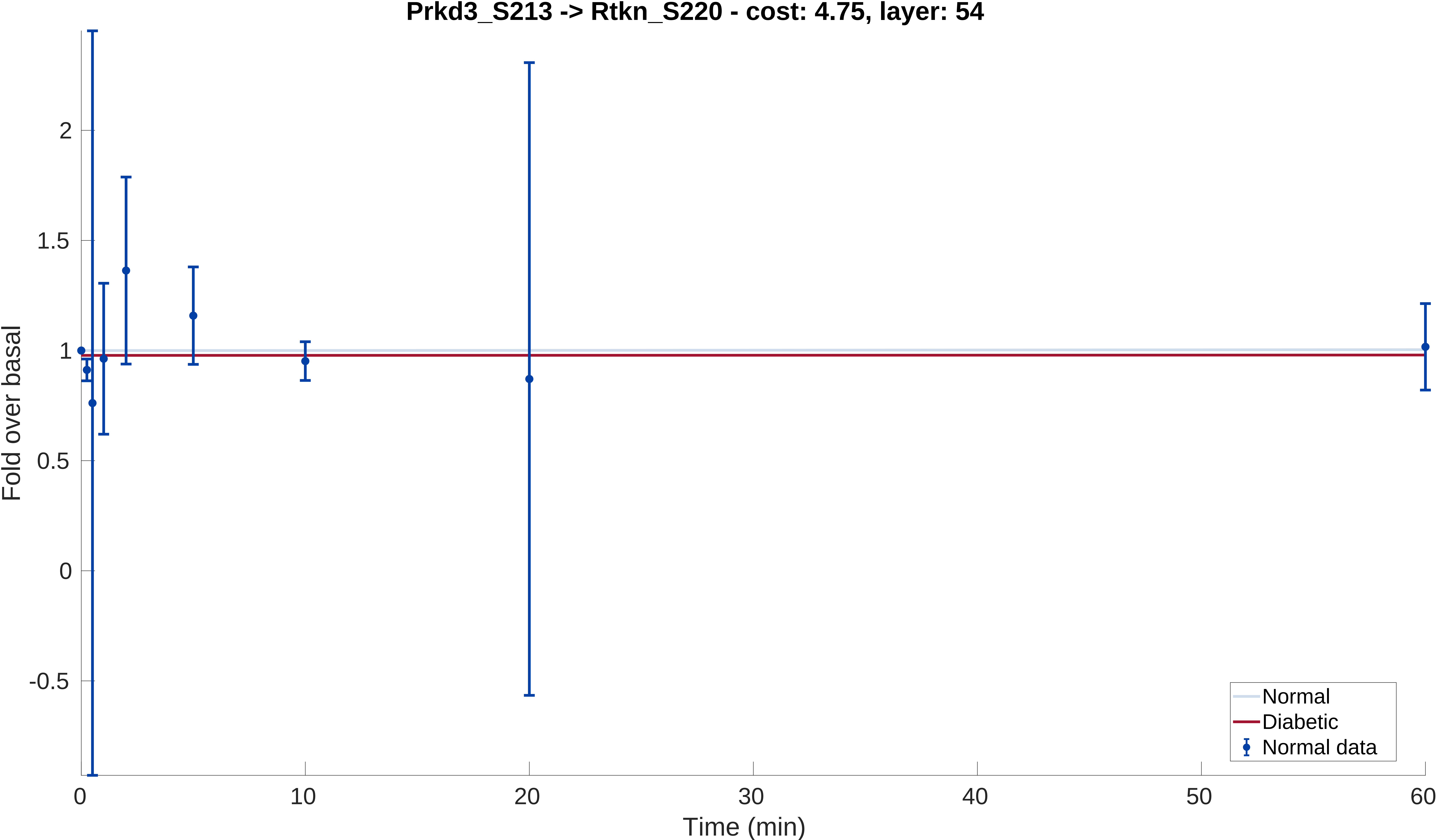

Prkd3_S213 -> Rtkn_S220 - cost: 4.75, layer: 54
Fold over basal
Time (min)
Normal
Diabetic
Normal data
2
1.5
1
0.5
0
-0.5
0
10
20
30
40
50
60

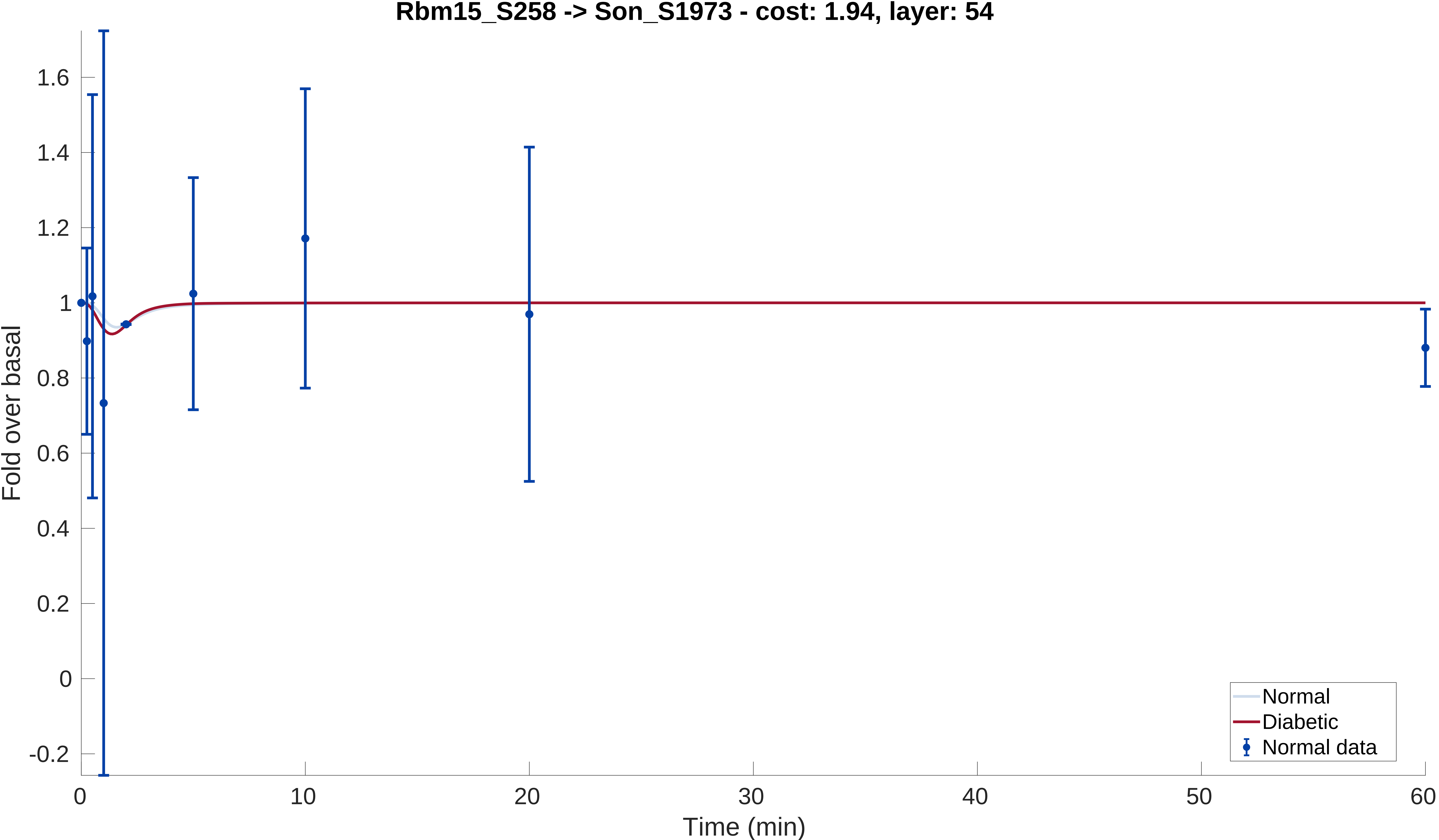

Rbm15_S258 -> Son_S1973 - cost: 1.94, layer: 54
Fold over basal
1.6
1.4
1.2
1
0.8
0.6
0.4
0.2
0
-0.2
0
10
20
30
40
50
60
Time (min)
Normal
Diabetic
Normal data

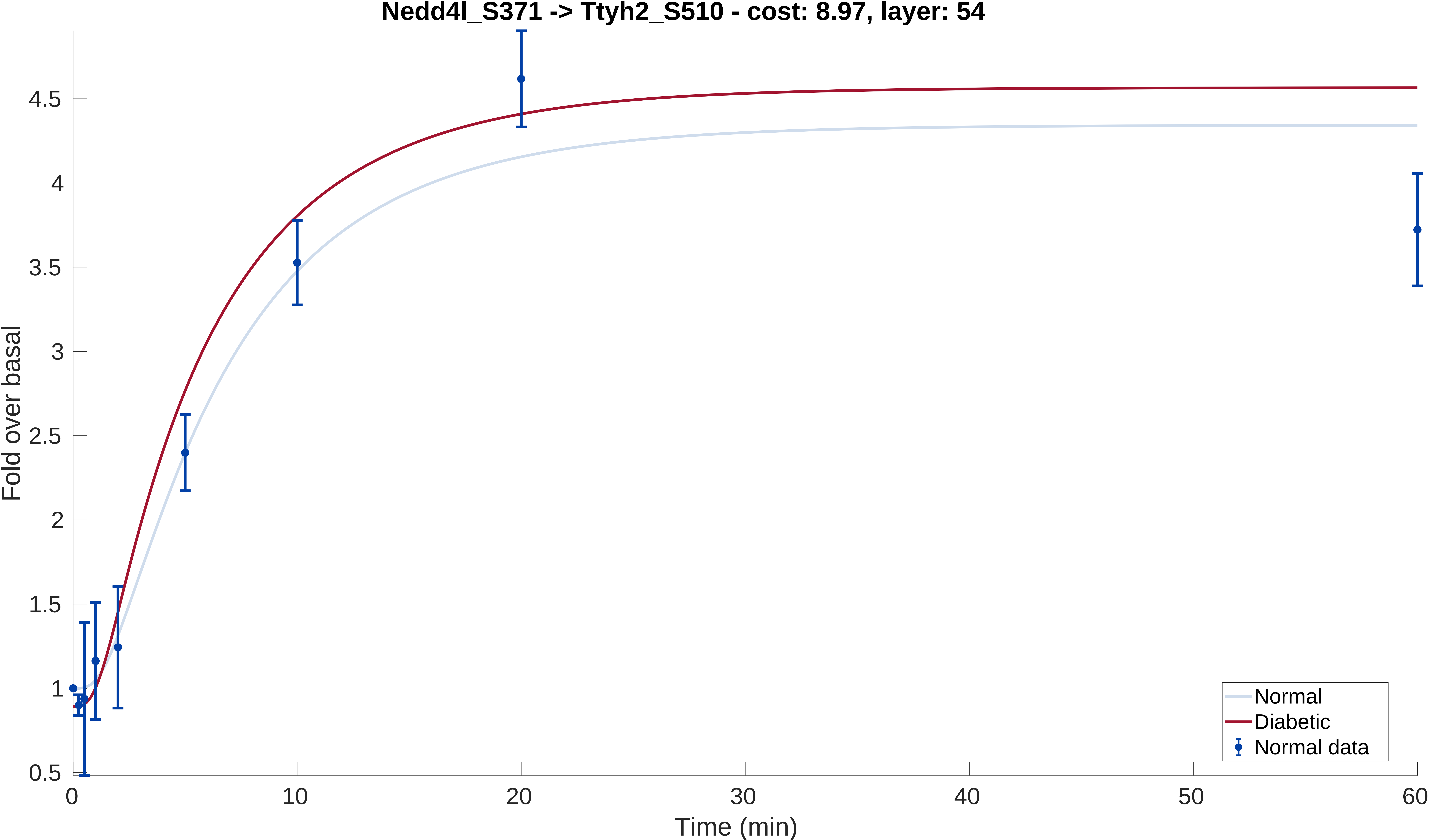
Nedd4l_S371 -> Ttyh2_S510 - cost: 8.97, layer: 54
Fold over basal
Time (min)
0.5
1
1.5
2
2.5
3
3.5
4
4.5
0
10
20
30
40
50
60
Normal
Diabetic
Normal data

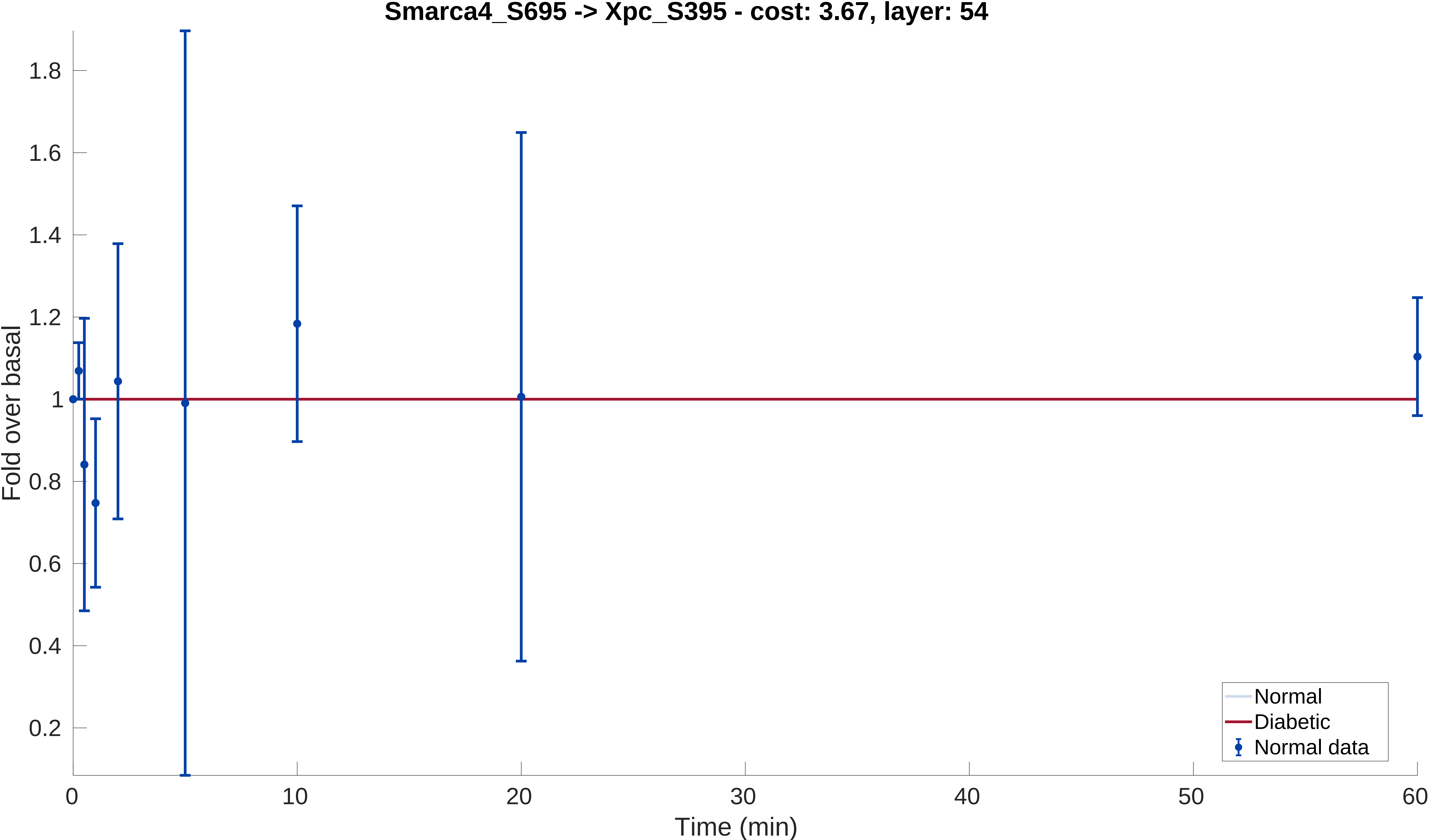

Smarca4_S695 -> Xpc_S395 - cost: 3.67, layer: 54
Fold over basal
1.8
1.6
1.4
1.2
1
0.8
0.6
0.4
0.2
0
10
20
30
40
50
60
Time (min)
Normal
Diabetic
Normal data

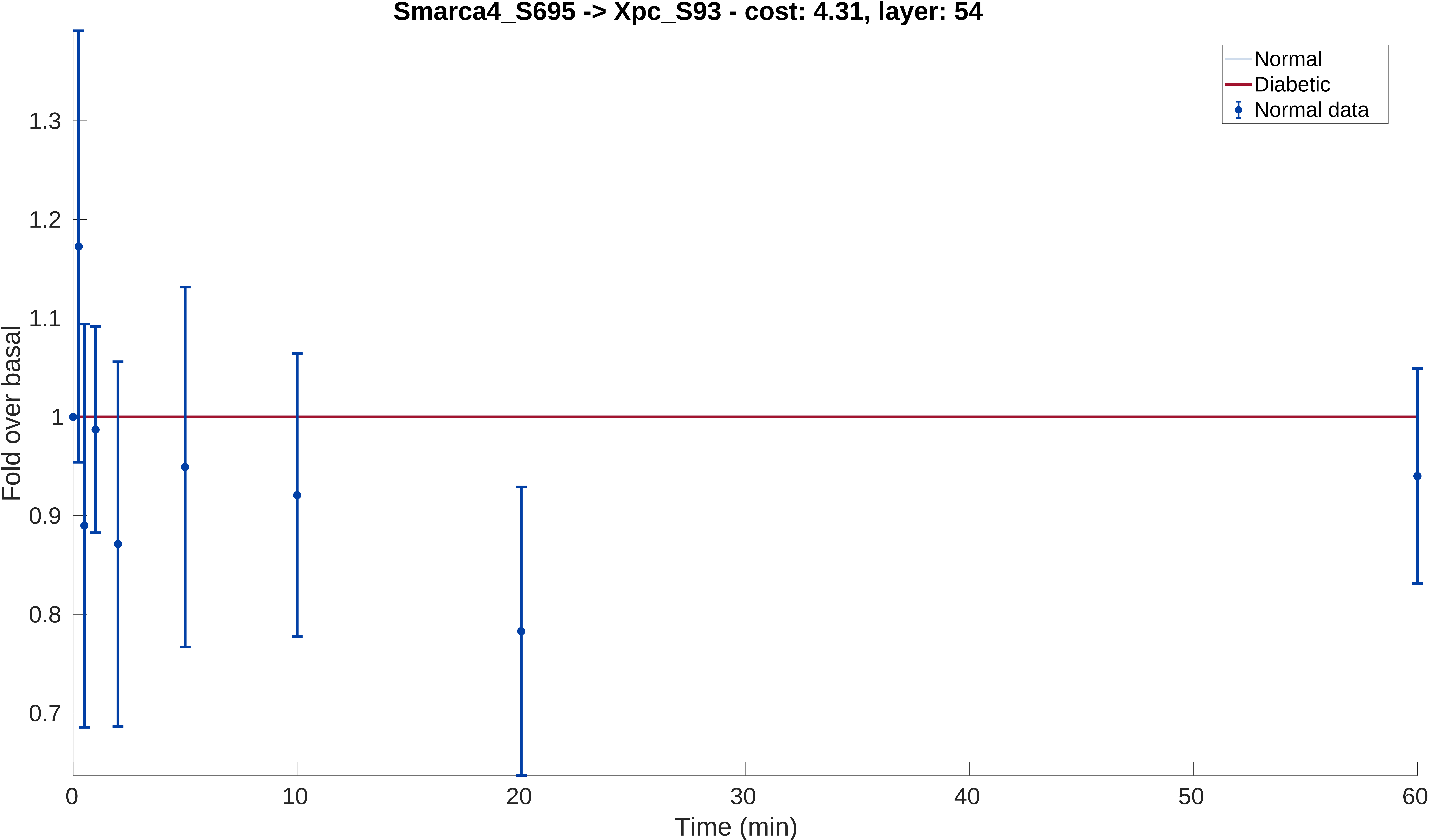

Smarca4_S695 -> Xpc_S93 - cost: 4.31, layer: 54
Normal
Diabetic
Normal data
Fold over basal
1.3
1.2
1.1
1
0.9
0.8
0.7
0
10
20
30
40
50
60
Time (min)

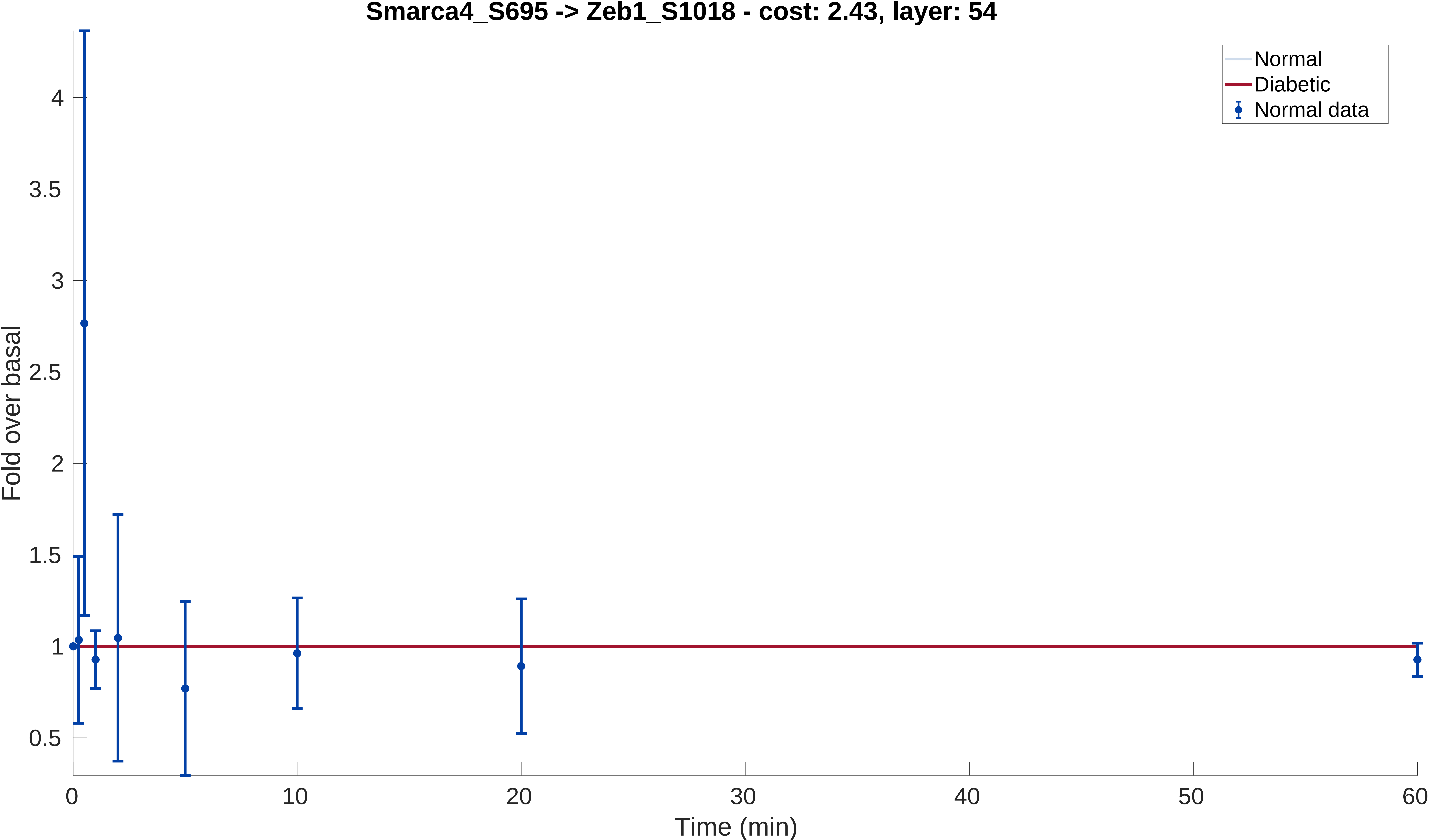

Smarca4_S695 -> Zeb1_S1018 - cost: 2.43, layer: 54
Normal
Diabetic
Normal data
Fold over basal
4
3.5
3
2.5
2
1.5
1
0.5
0
10
20
30
40
50
60
Time (min)

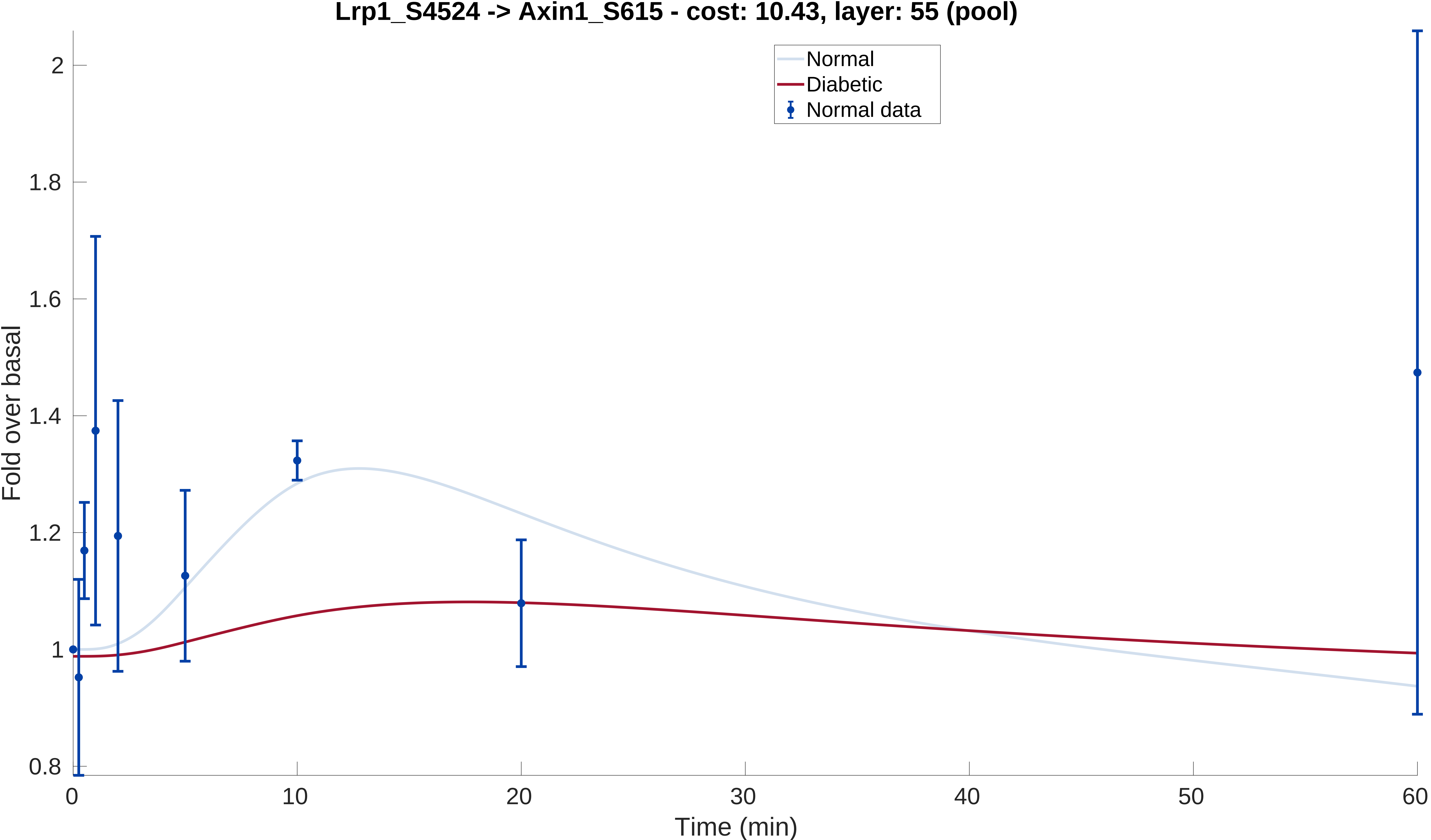
Lrp1_S4524 -> Axin1_S615 - cost: 10.43, layer: 55 (pool)
Normal
Diabetic
Normal data
Fold over basal
Time (min)
0.8
1
1.2
1.4
1.6
1.8
2
0
10
20
30
40
50
60

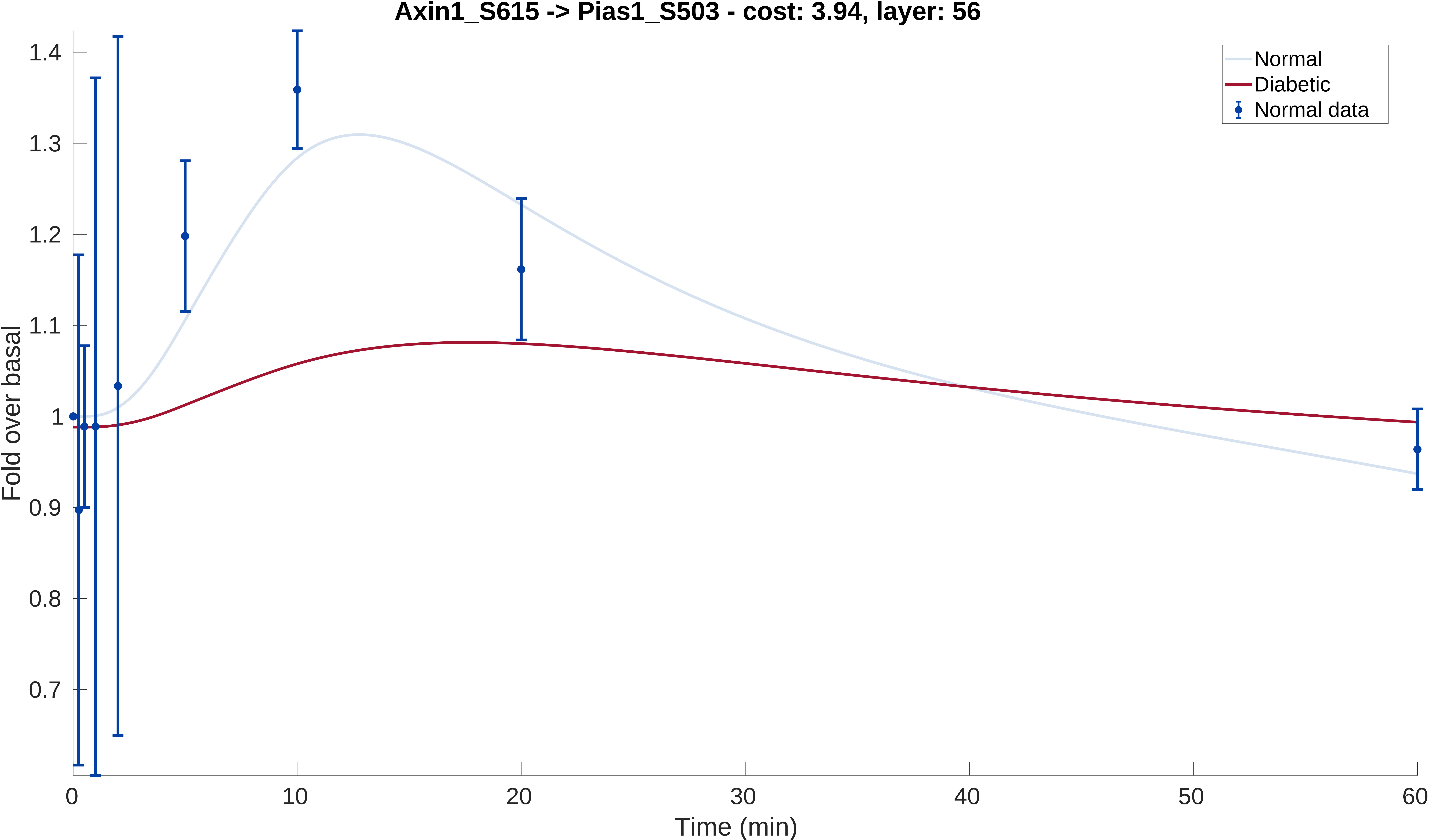

Axin1_S615 -> Pias1_S503 - cost: 3.94, layer: 56
Normal
Diabetic
Normal data
Fold over basal
1.4
1.3
1.2
1.1
1
0.9
0.8
0.7
0
10
20
30
40
50
60
Time (min)

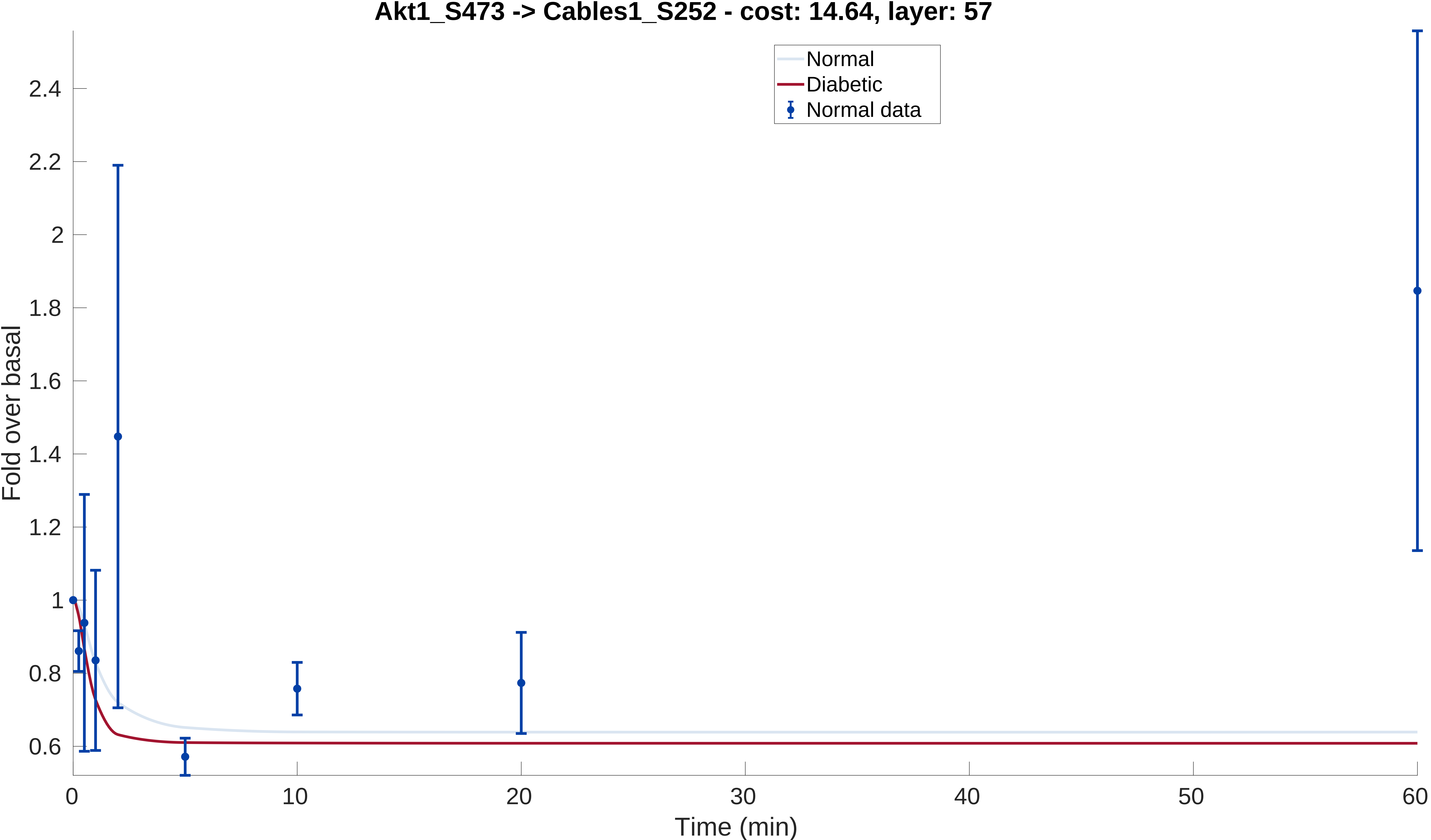

Akt1_S473 -> Cables1_S252 - cost: 14.64, layer: 57
Normal
Diabetic
Normal data
Fold over basal
Time (min)
0.6
0.8
1
1.2
1.4
1.6
1.8
2
2.2
2.4
0
10
20
30
40
50
60

**Akt1_S126 -> Rpl13_S106 - cost: 7.84, layer: 57**

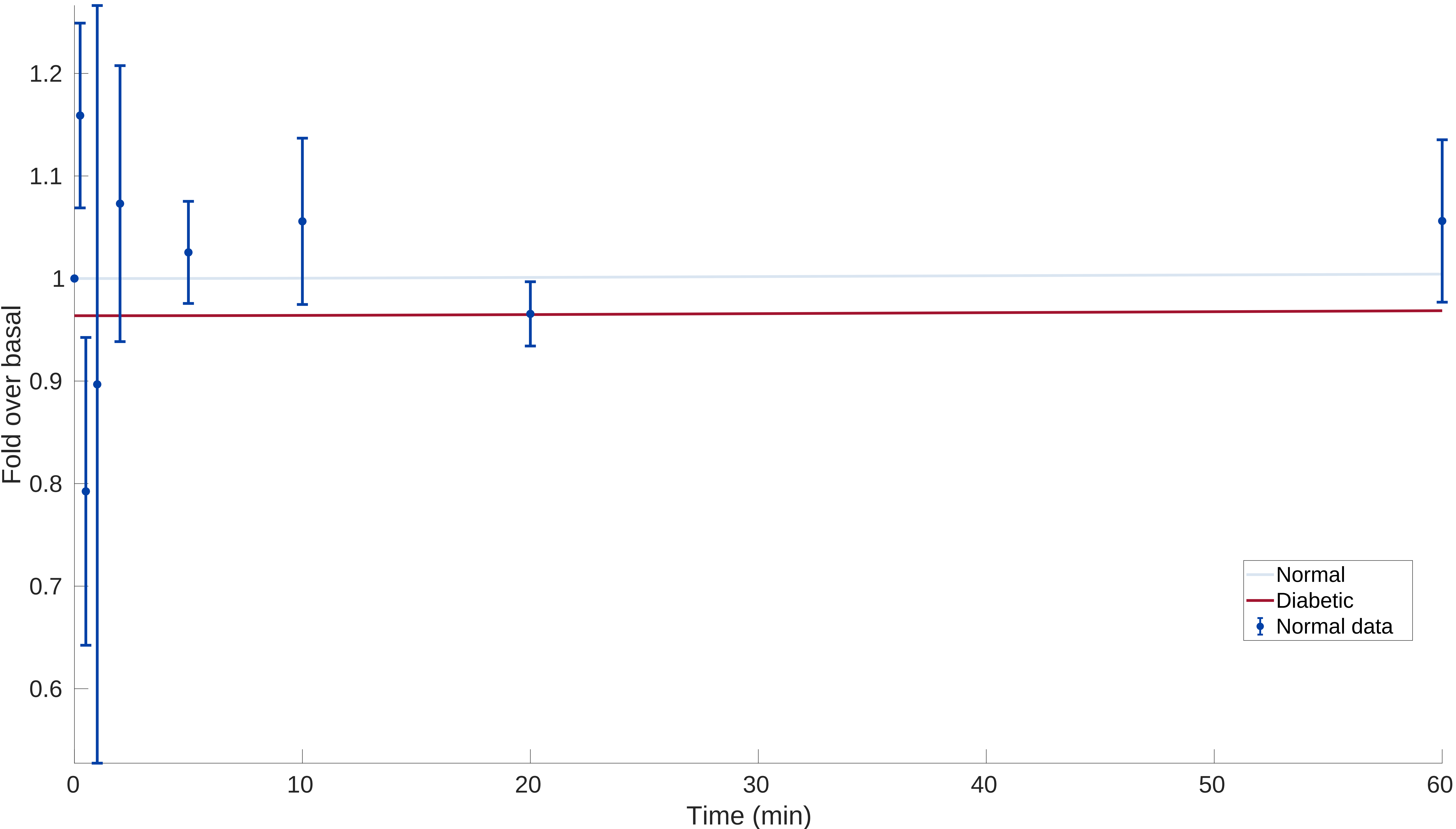

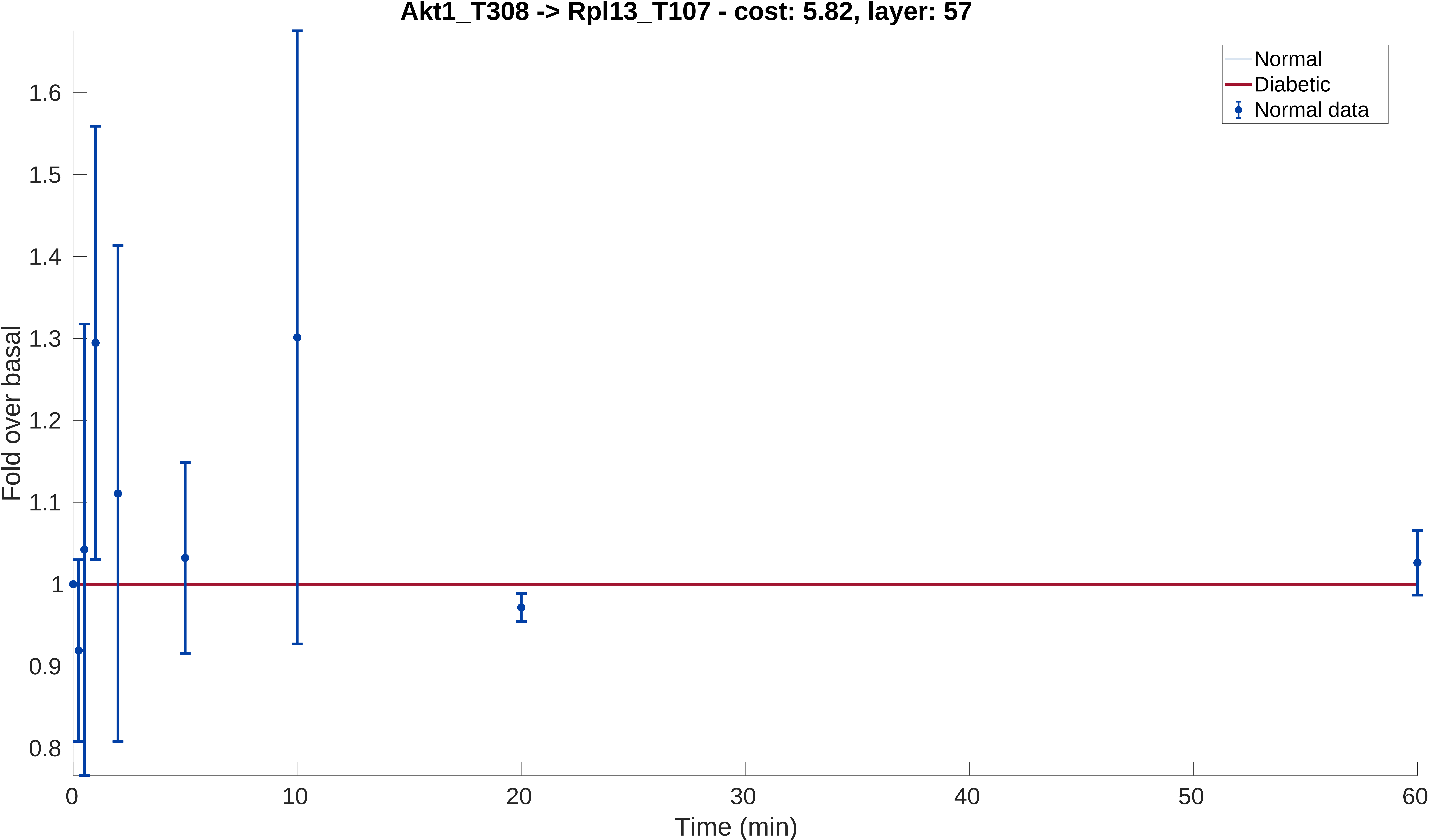
Akt1_T308 -> Rpl13_T107 - cost: 5.82, layer: 57
Normal
Diabetic
Normal data
Fold over basal
Time (min)
0
10
20
30
40
50
60
0.8
0.9
1
1.1
1.2
1.3
1.4
1.5
1.6

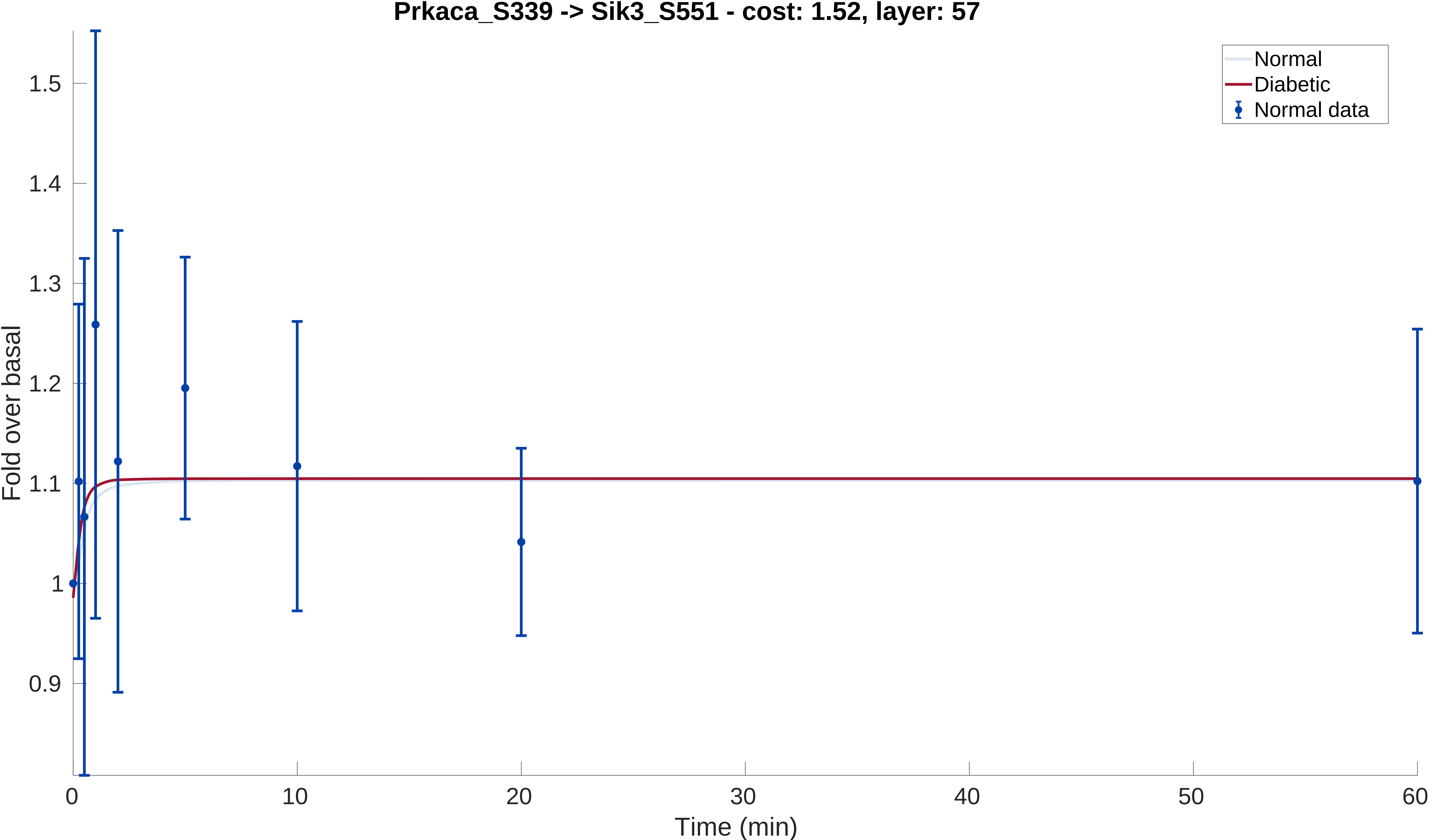

Prkaca_S339 -> Sik3_S551 - cost: 1.52, layer: 57
Normal
Diabetic
Normal data
Fold over basal
1.5
1.4
1.3
1.2
1.1
1
0.9
0
10
20
30
40
50
60
Time (min)

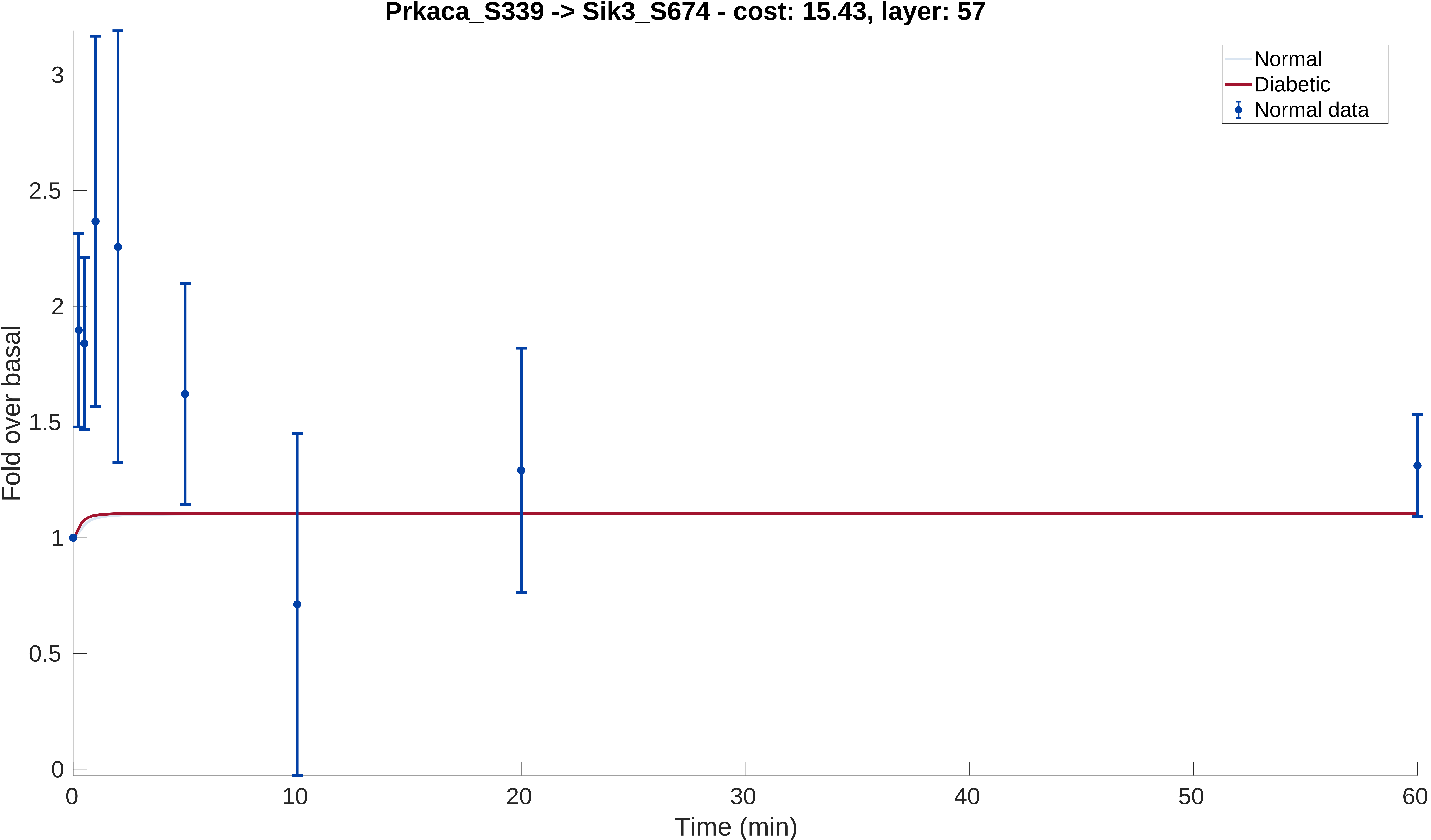

Prkaca_S339 -> Sik3_S674 - cost: 15.43, layer: 57
Normal
Diabetic
Normal data
Fold over basal
Time (min)
0
0.5
1
1.5
2
2.5
3
0
10
20
30
40
50
60

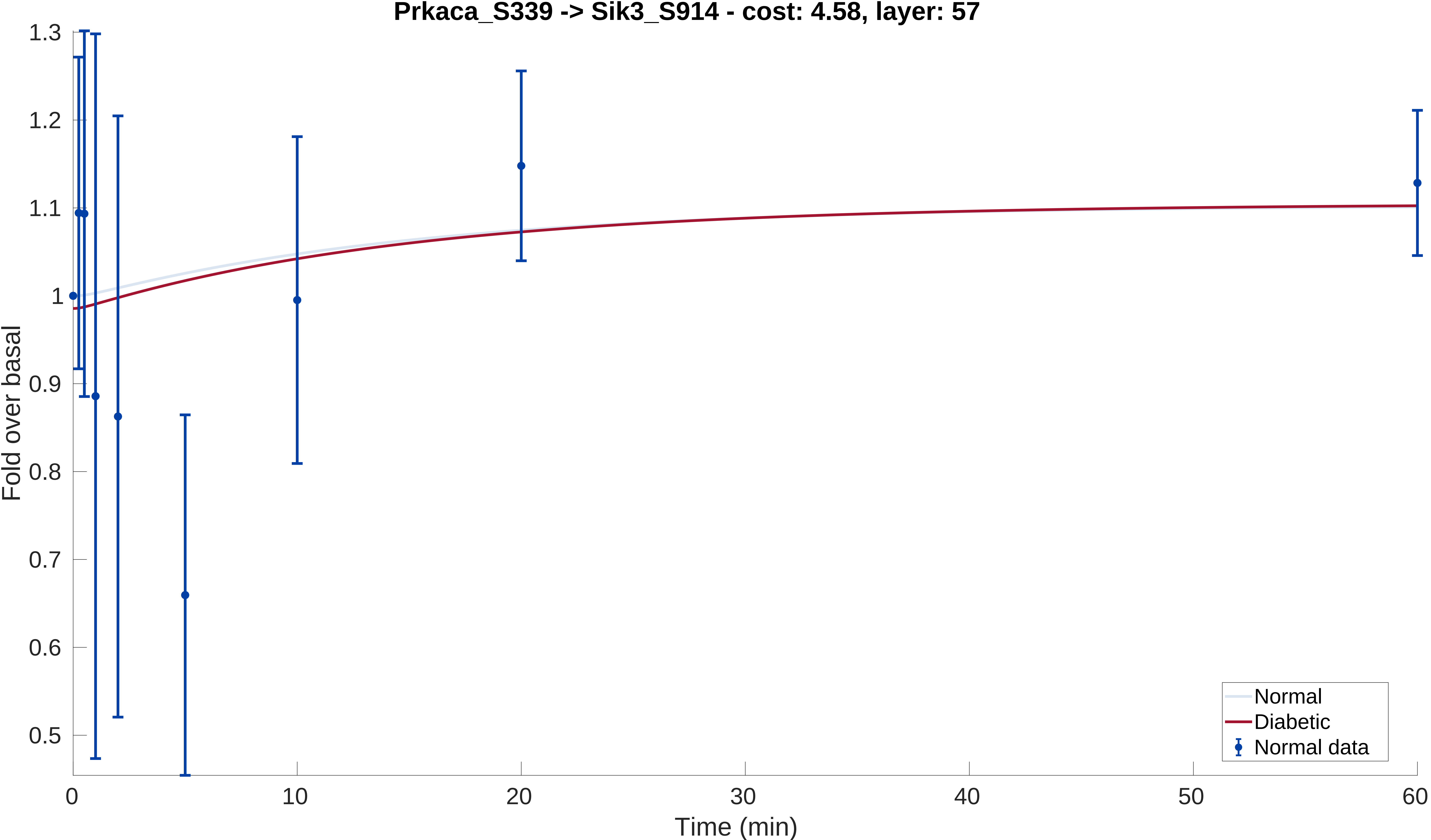
Prkaca_S339 -> Sik3_S914 - cost: 4.58, layer: 57
1.3
1.2
1.1
1
0.9
0.8
0.7
0.6
0.5
Fold over basal
0
10
20
30
40
50
60
Time (min)
Normal
Diabetic
Normal data

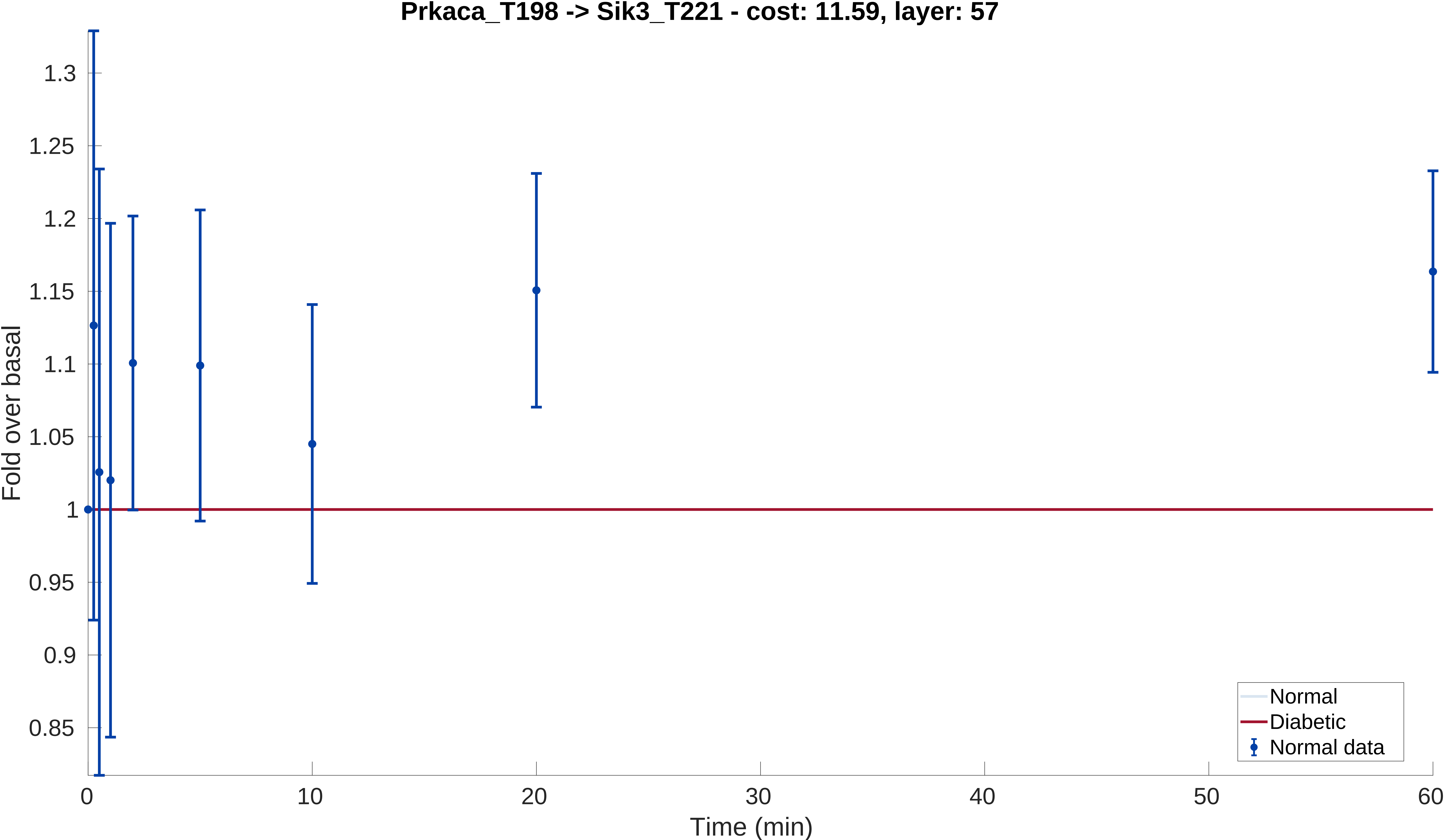

Prrkaca_T198 -> Sik3_T221 - cost: 11.59, layer: 57
Fold over basal
1.3
1.25
1.2
1.15
1.1
1.05
1
0.95
0.9
0.85
0
10
20
30
40
50
60
Time (min)
Normal
Diabetic
Normal data

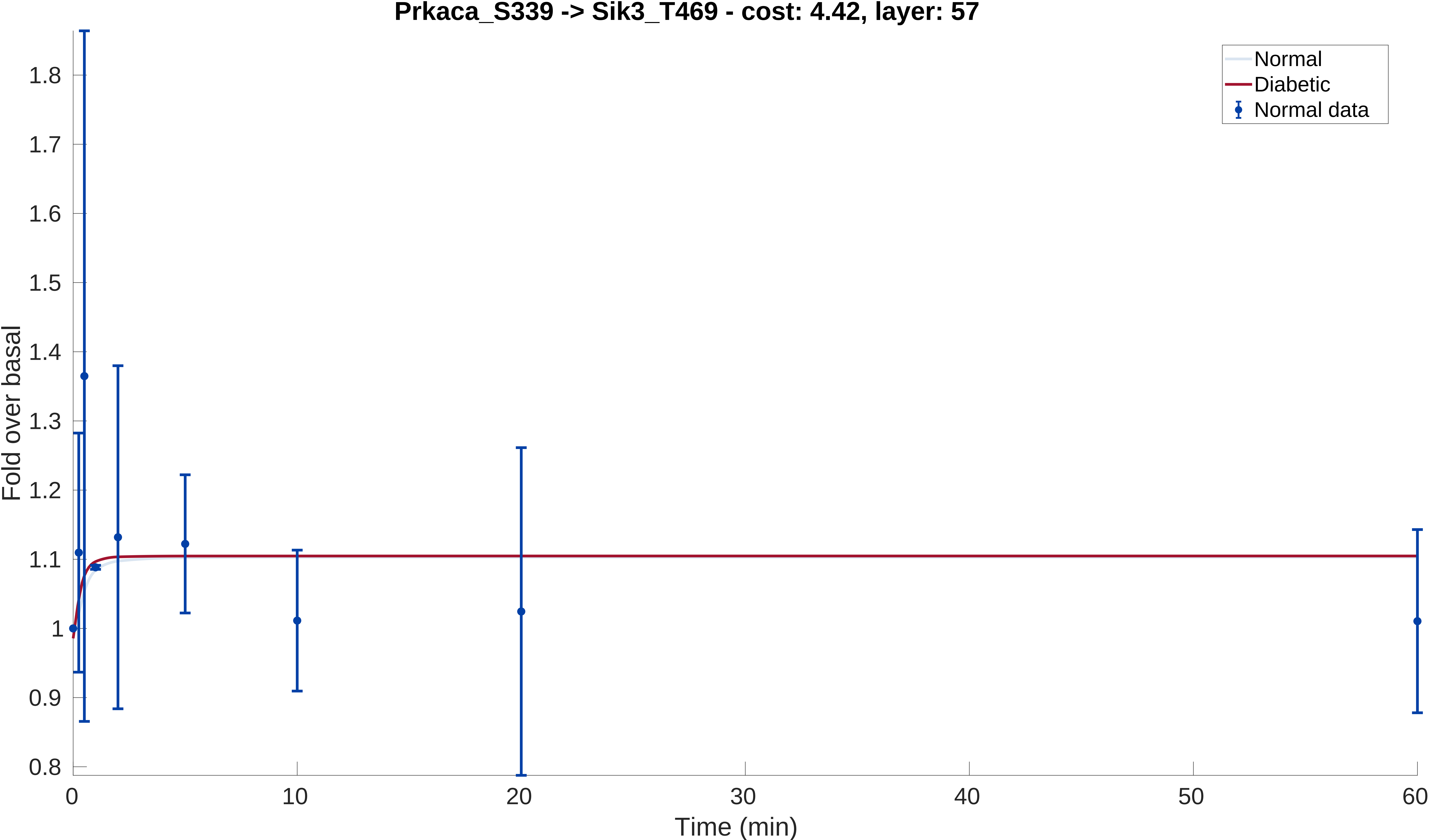
Prkaca_S339 -> Sik3_T469 - cost: 4.42, layer: 57
Normal
Diabetic
Normal data
Fold over basal
0.8
0.9
1
1.1
1.2
1.3
1.4
1.5
1.6
1.7
1.8
0
10
20
30
40
50
60
Time (min)

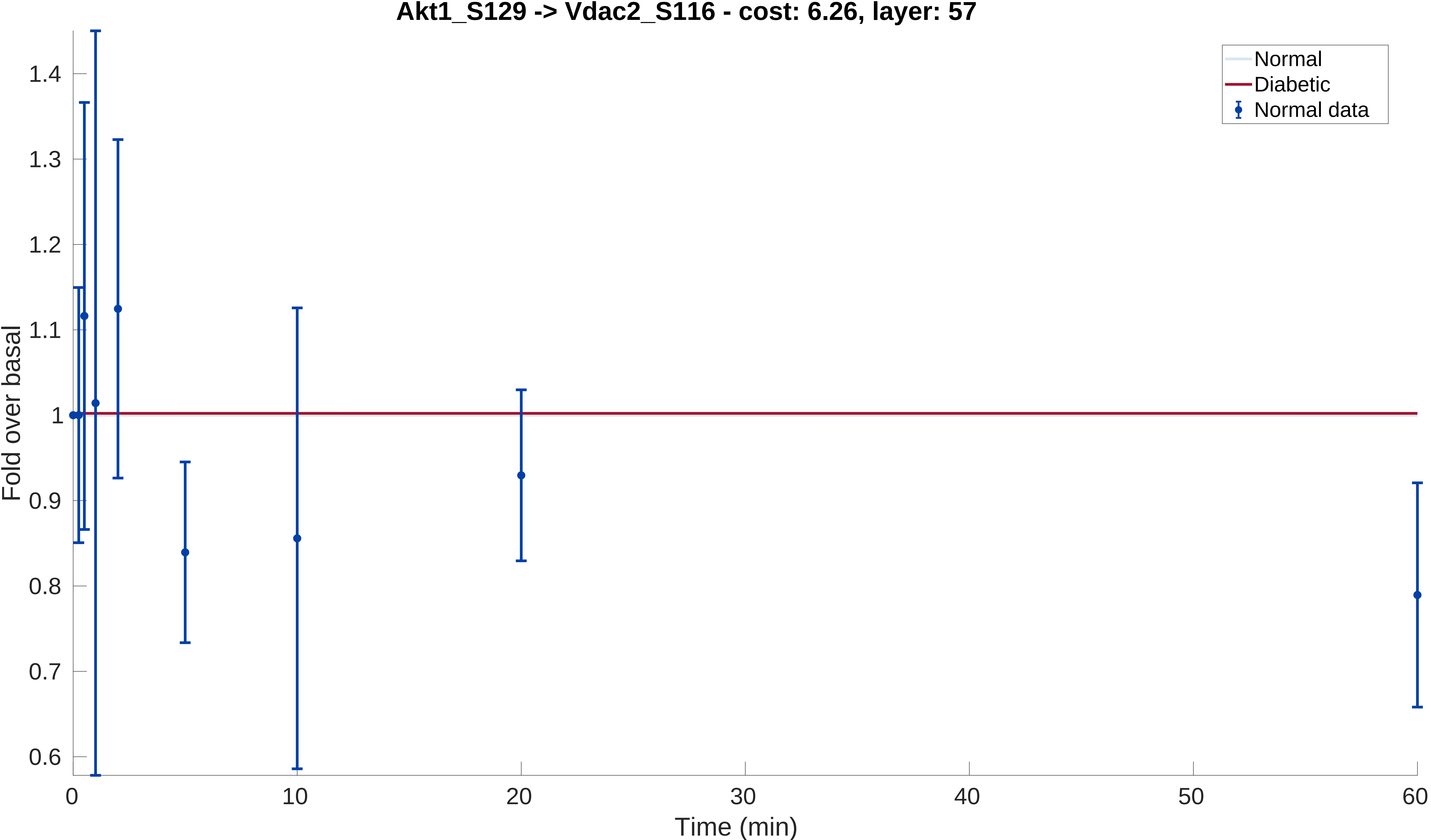

Akt1_S129 -> Vdac2_S116 - cost: 6.26, layer: 57
Normal
Diabetic
Normal data
Fold over basal
1.4
1.3
1.2
1.1
1
0.9
0.8
0.7
0.6
0
10
20
30
40
50
60
Time (min)

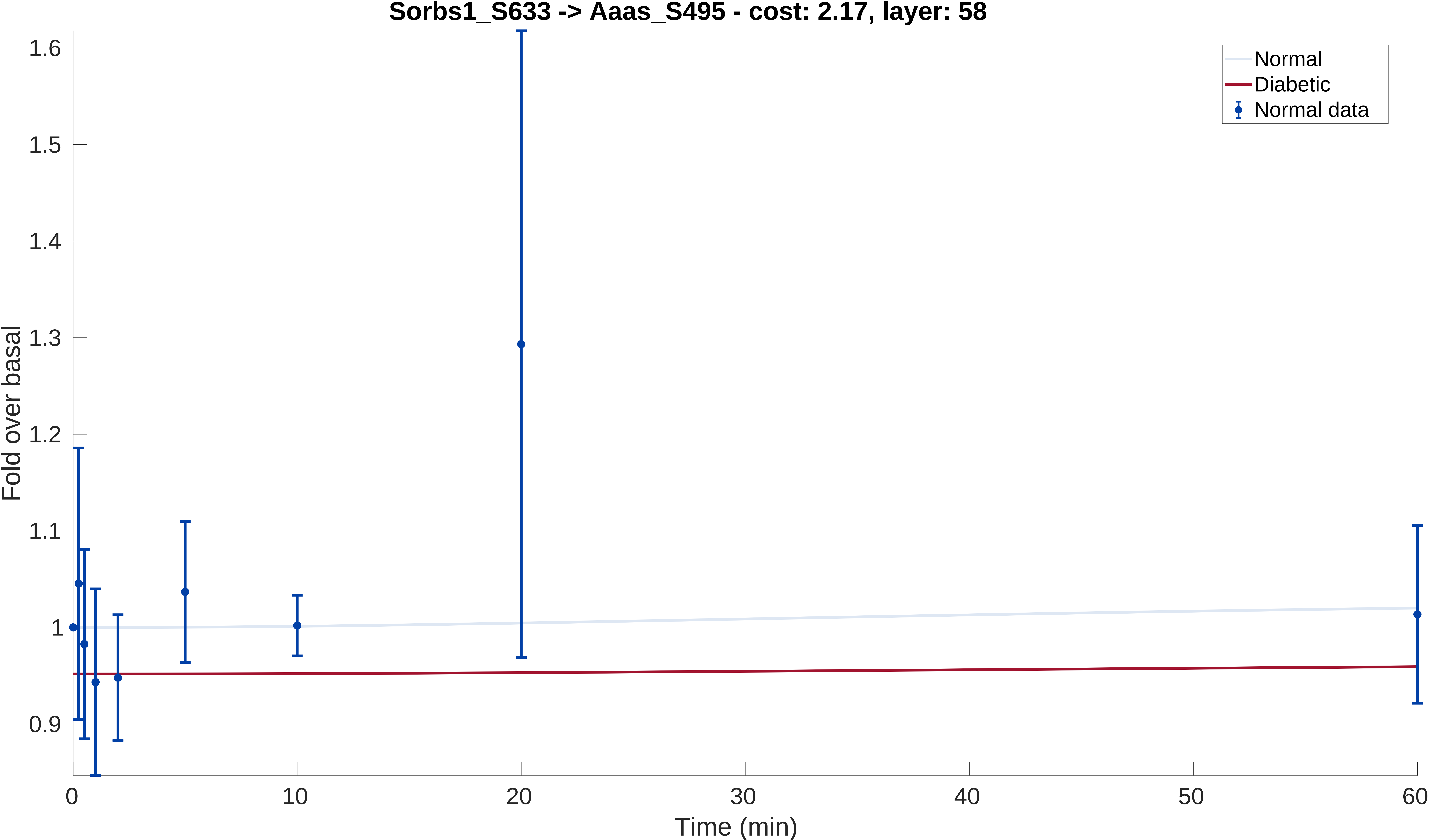
Sorbs1_S633 -> Aaas_S495 - cost: 2.17, layer: 58
Fold over basal
Time (min)
Normal
Diabetic
Normal data
1.6
1.5
1.4
1.3
1.2
1.1
1
0.9
0
10
20
30
40
50
60

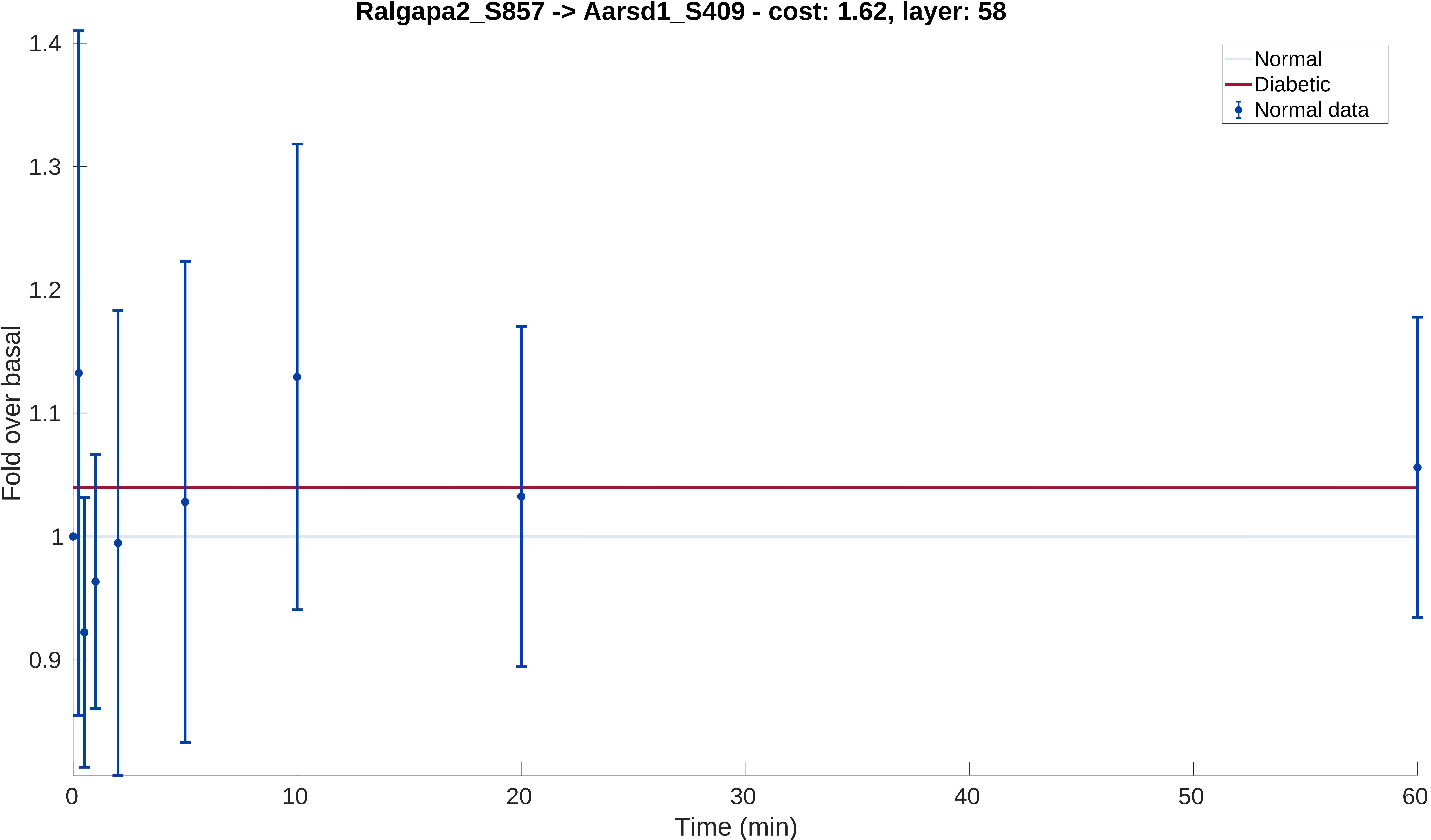

Ralgapa2_S857 -> Aarsd1_S409 - cost: 1.62, layer: 58
Fold over basal
Time (min)
1.4
1.3
1.2
1.1
1
0.9
0
10
20
30
40
50
60
Normal
Diabetic
Normal data

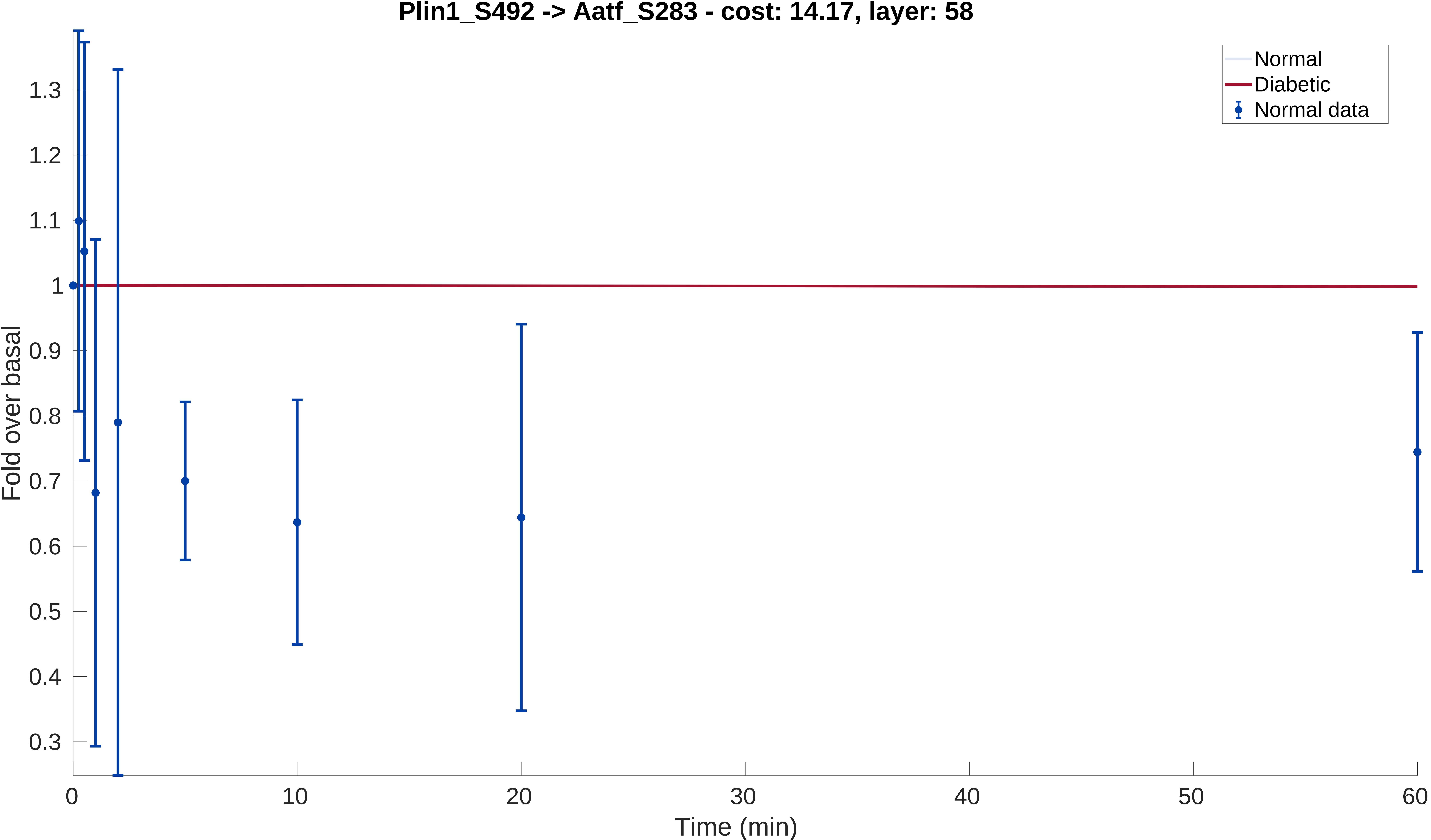

Plin1_S492 -> Aatf_S283 - cost: 14.17, layer: 58
Normal
Diabetic
Normal data
Fold over basal
1.3
1.2
1.1
1
0.9
0.8
0.7
0.6
0.5
0.4
0.3
0
10
20
30
40
50
60
Time (min)

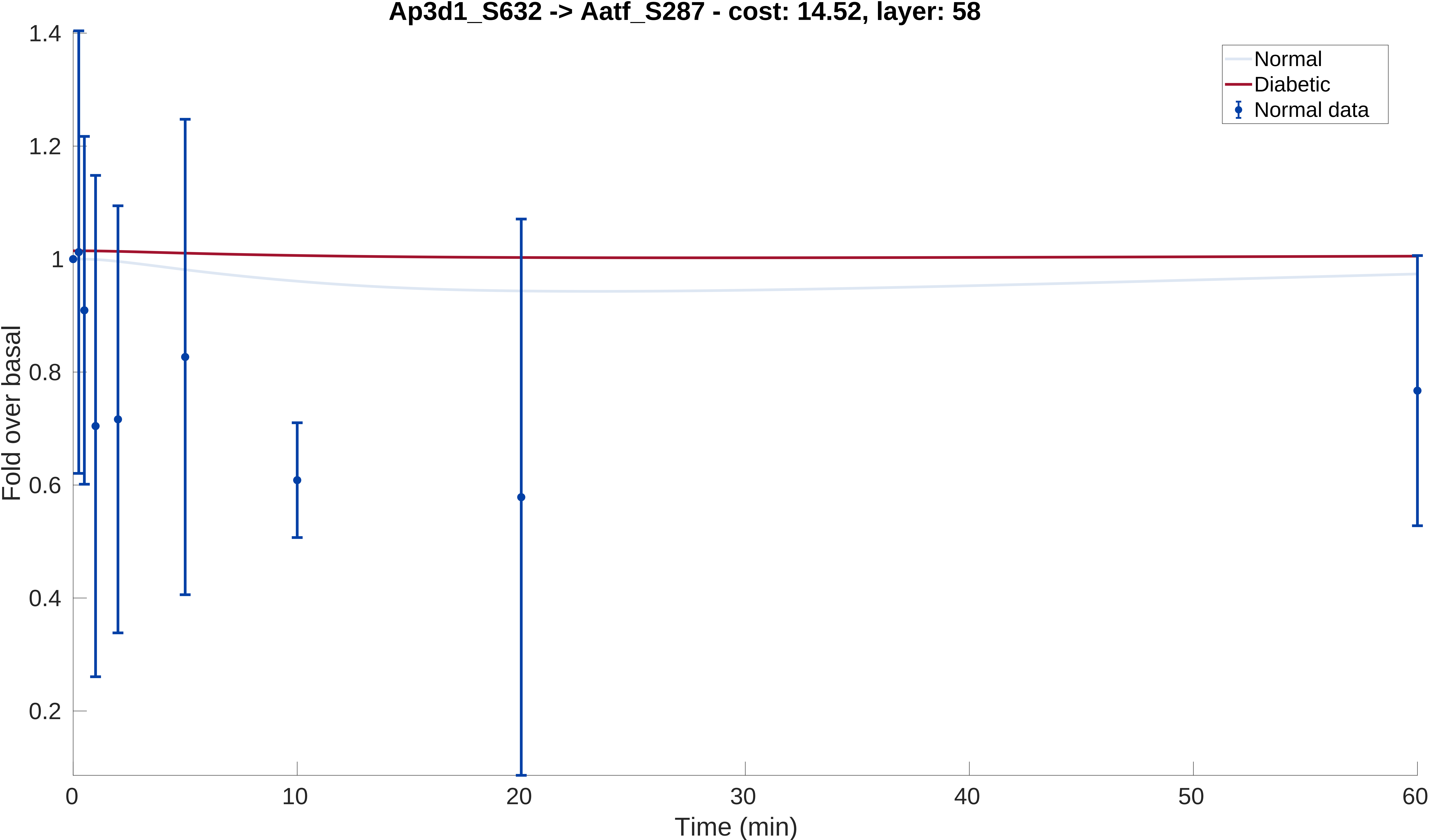

Ap3d1_S632 -> Aatf_S287 - cost: 14.52, layer: 58
Normal
Diabetic
Normal data
Fold over basal
1.4
1.2
1
0.8
0.6
0.4
0.2
0
10
20
30
40
50
60
Time (min)

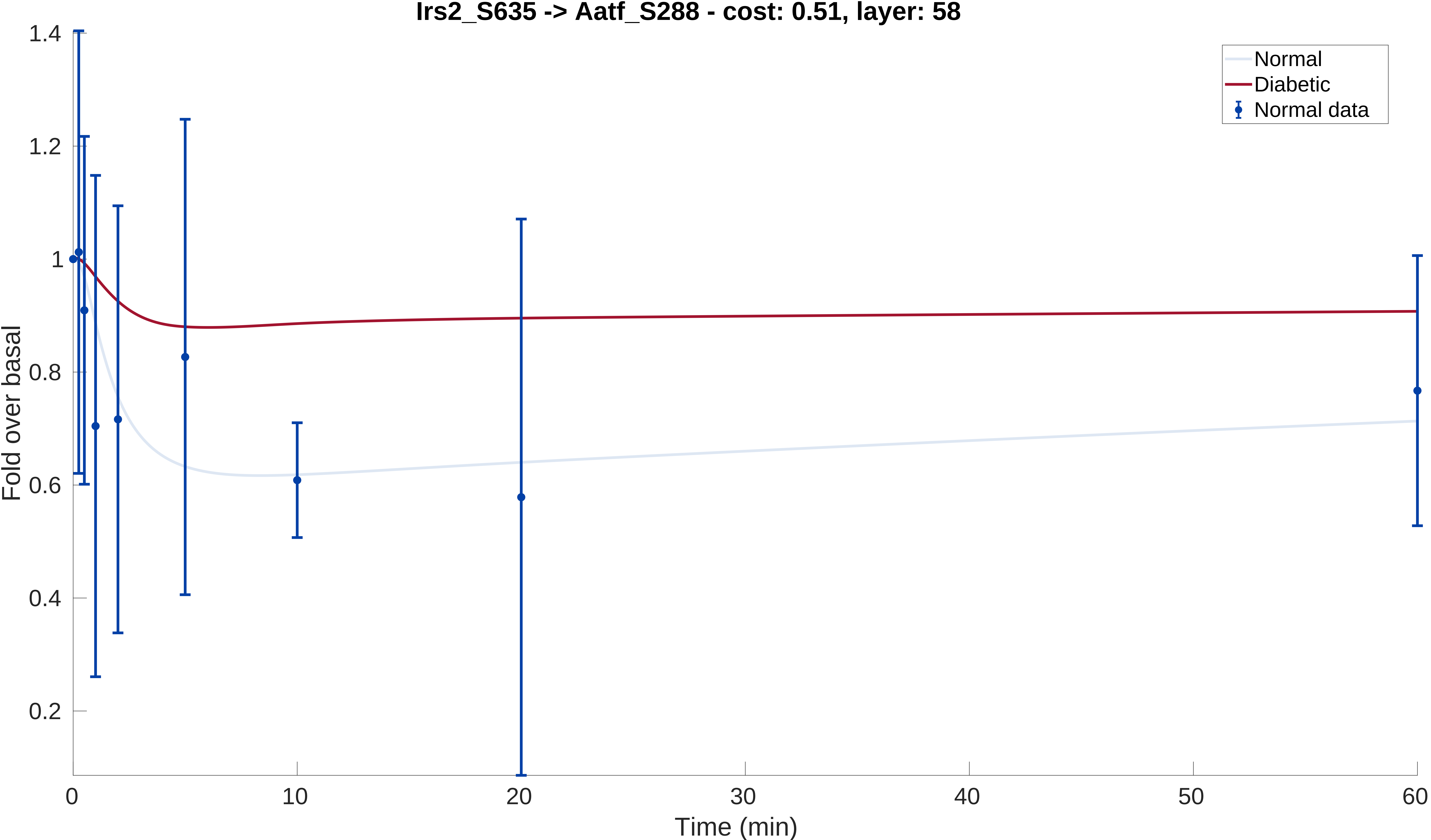

Irs2_S635 -> Aatf_S288 - cost: 0.51, layer: 58
1.4
1.2
1
0.8
0.6
0.4
0.2
Fold over basal
0
10
20
30
40
50
60
Time (min)
Normal
Diabetic
Normal data

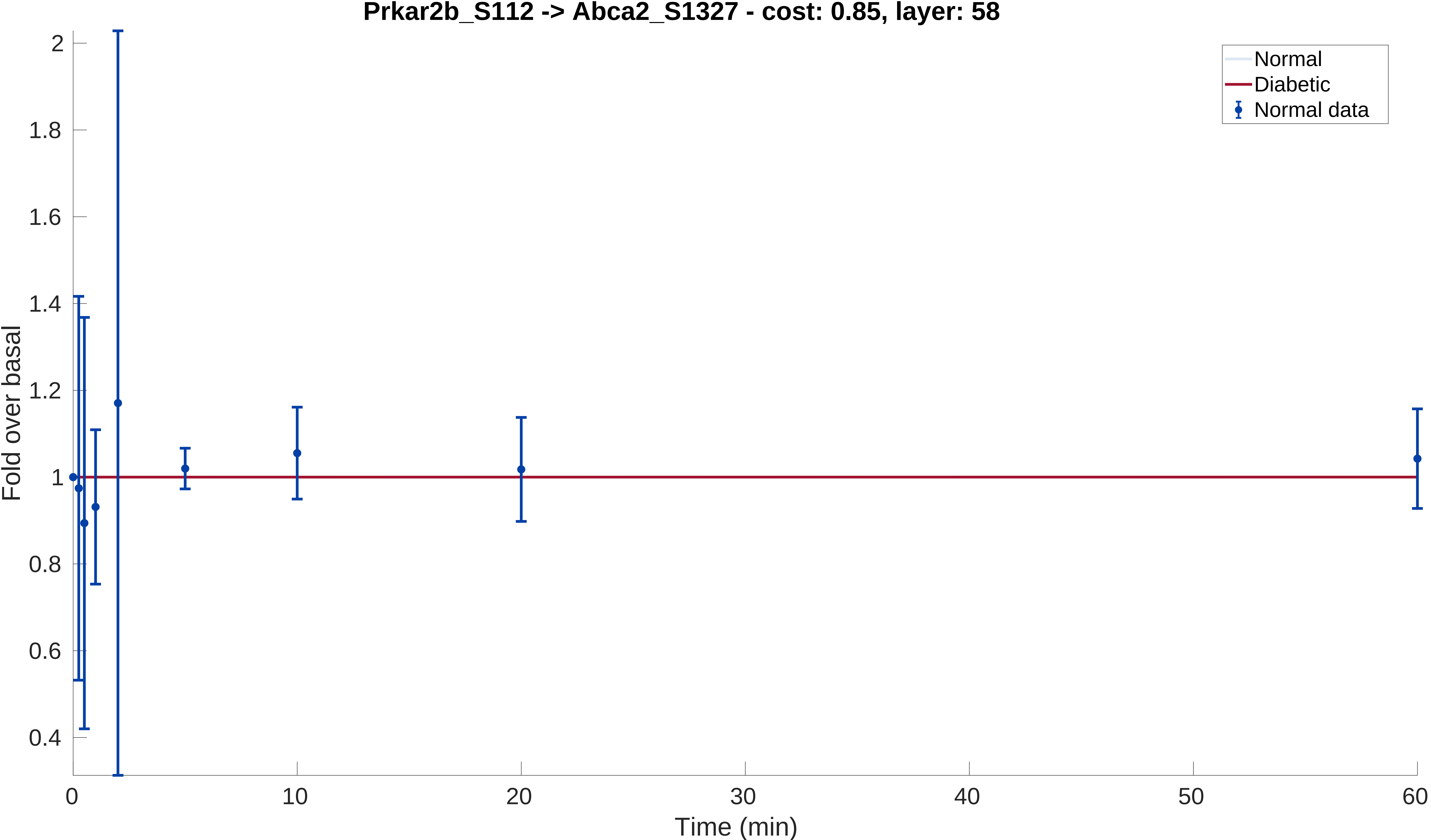

Prkar2b_S112 -> Abca2_S1327 - cost: 0.85, layer: 58
Normal
Diabetic
Normal data
Fold over basal
Time (min)
0.4
0.6
0.8
1
1.2
1.4
1.6
1.8
2
0
10
20
30
40
50
60

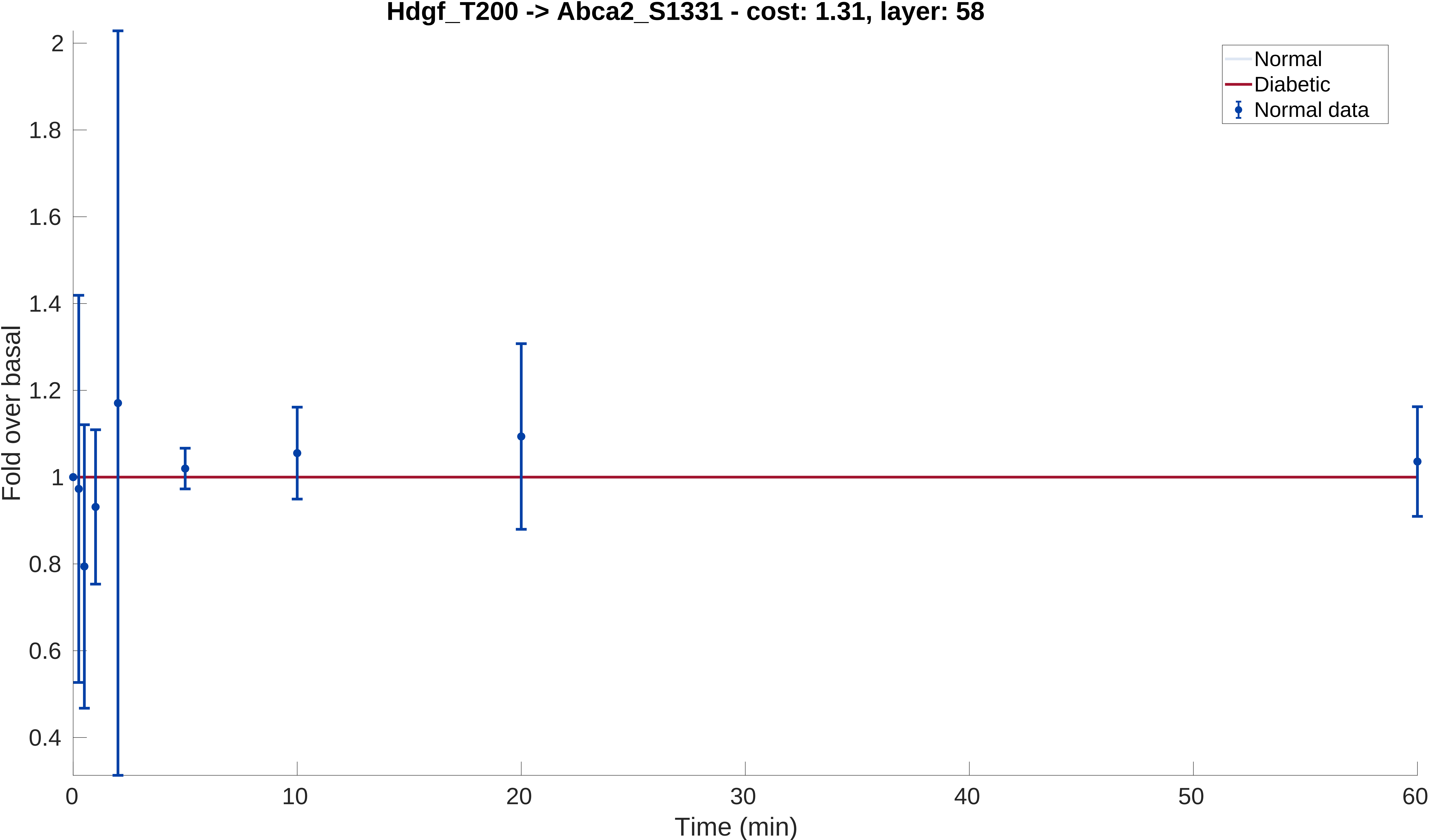

Hdgf_T200 -> Abca2_S1331 - cost: 1.31, layer: 58
Normal
Diabetic
Normal data
Fold over basal
Time (min)
0.4
0.6
0.8
1
1.2
1.4
1.6
1.8
2
0
10
20
30
40
50
60

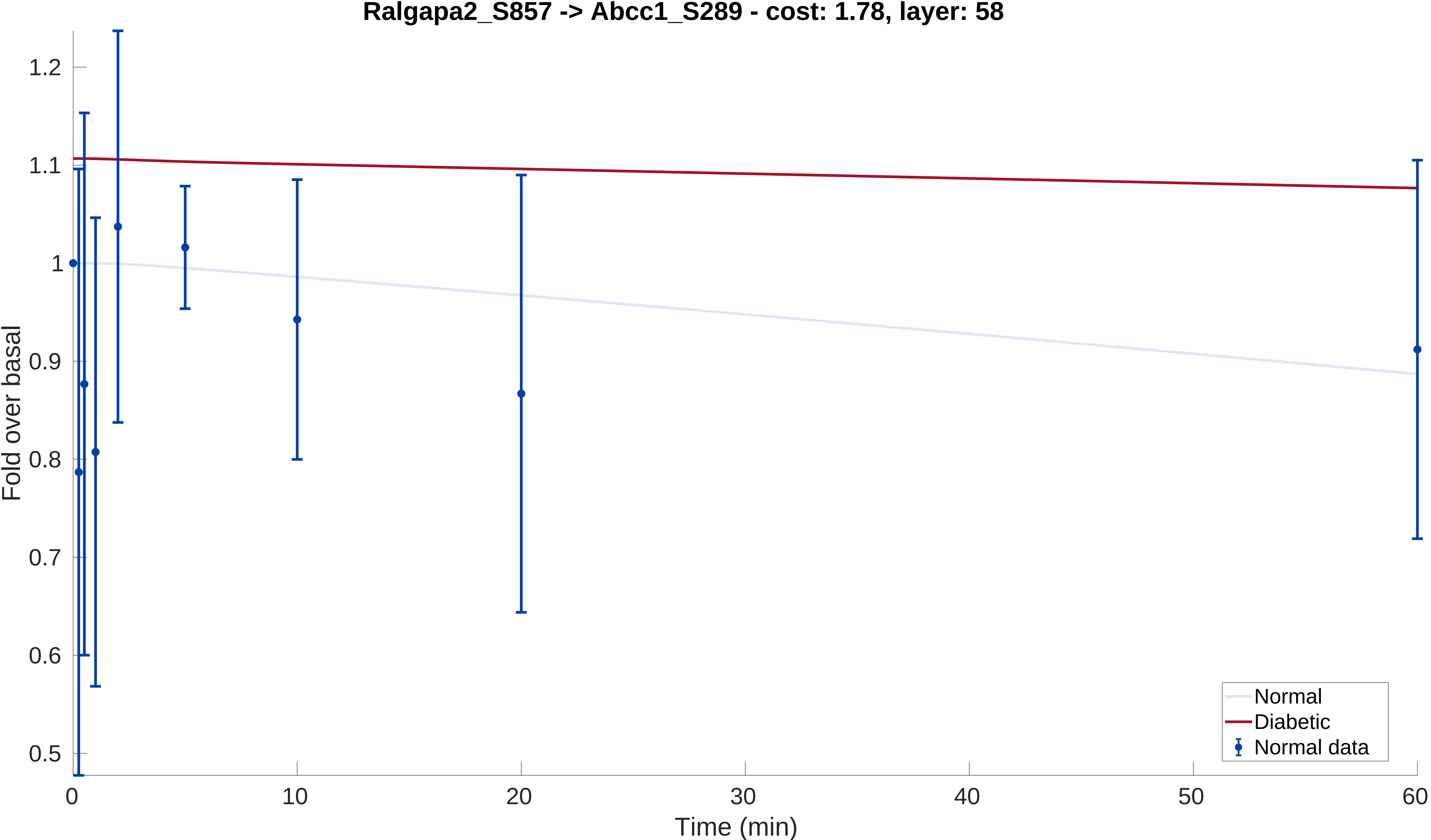

Ralgapa2_S857 -> Abcc1_S289 - cost: 1.78, layer: 58
Fold over basal
1.2
1.1
1
0.9
0.8
0.7
0.6
0.5
0
10
20
30
40
50
60
Time (min)
Normal
Diabetic
Normal data

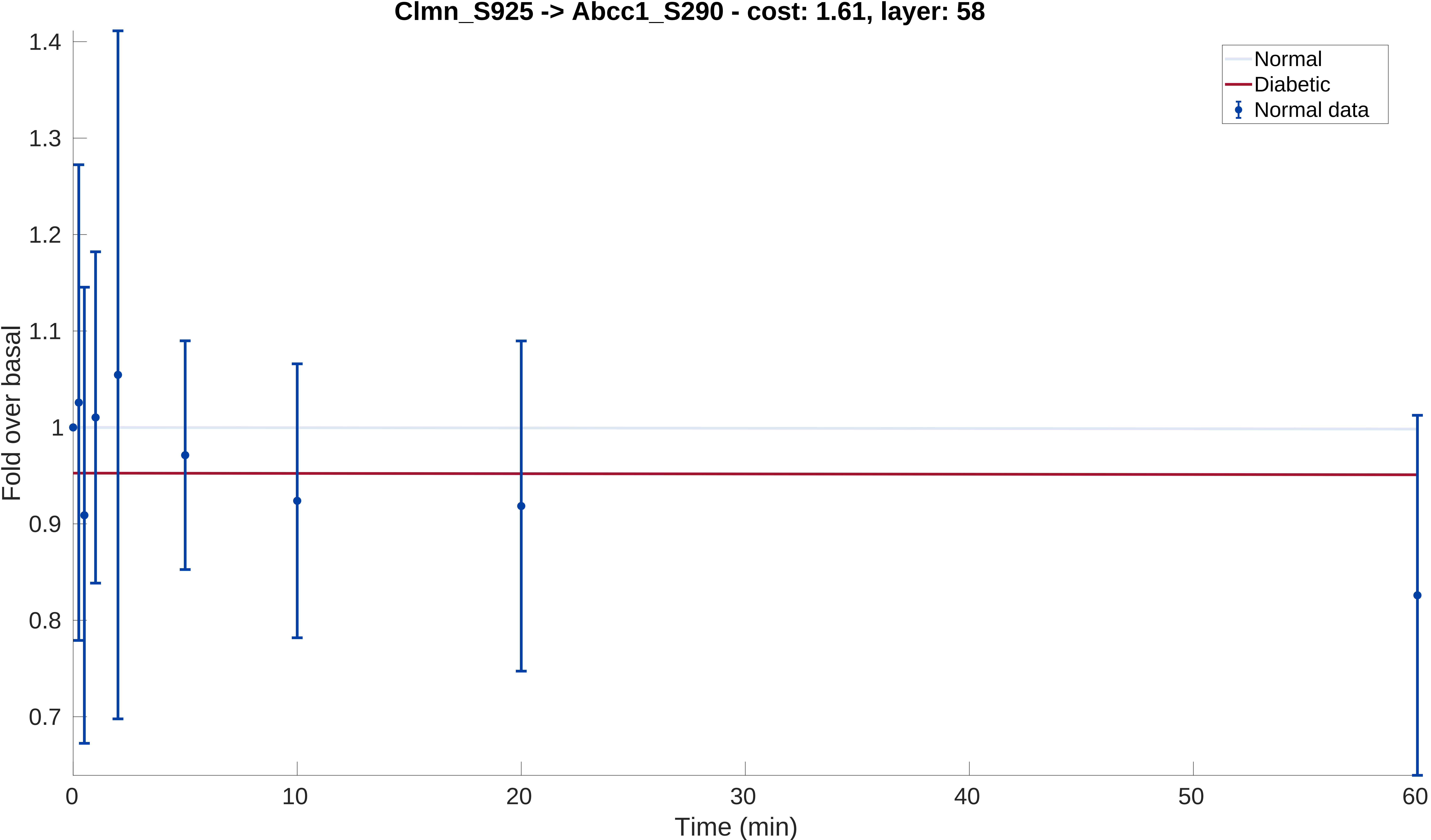

Clmn_S925 -> Abcc1_S290 - cost: 1.61, layer: 58
Normal
Diabetic
Normal data
Fold over basal
1.4
1.3
1.2
1.1
1
0.9
0.8
0.7
0
10
20
30
40
50
60
Time (min)

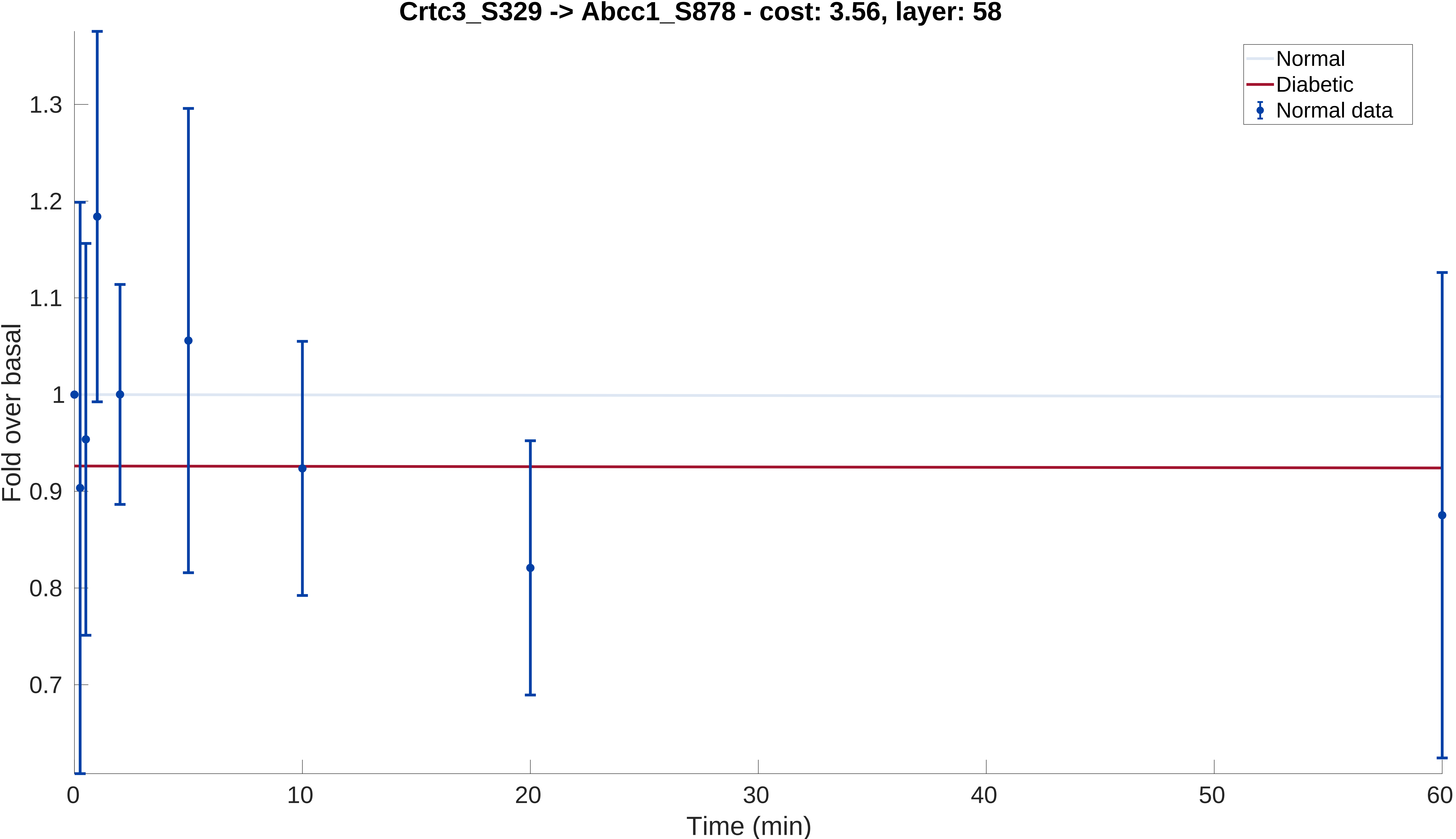

Crtc3_S329 -> Abcc1_S878 - cost: 3.56, layer: 58
Normal
Diabetic
Normal data
Fold over basal
Time (min)
1.3
1.2
1.1
1
0.9
0.8
0.7
0
10
20
30
40
50
60

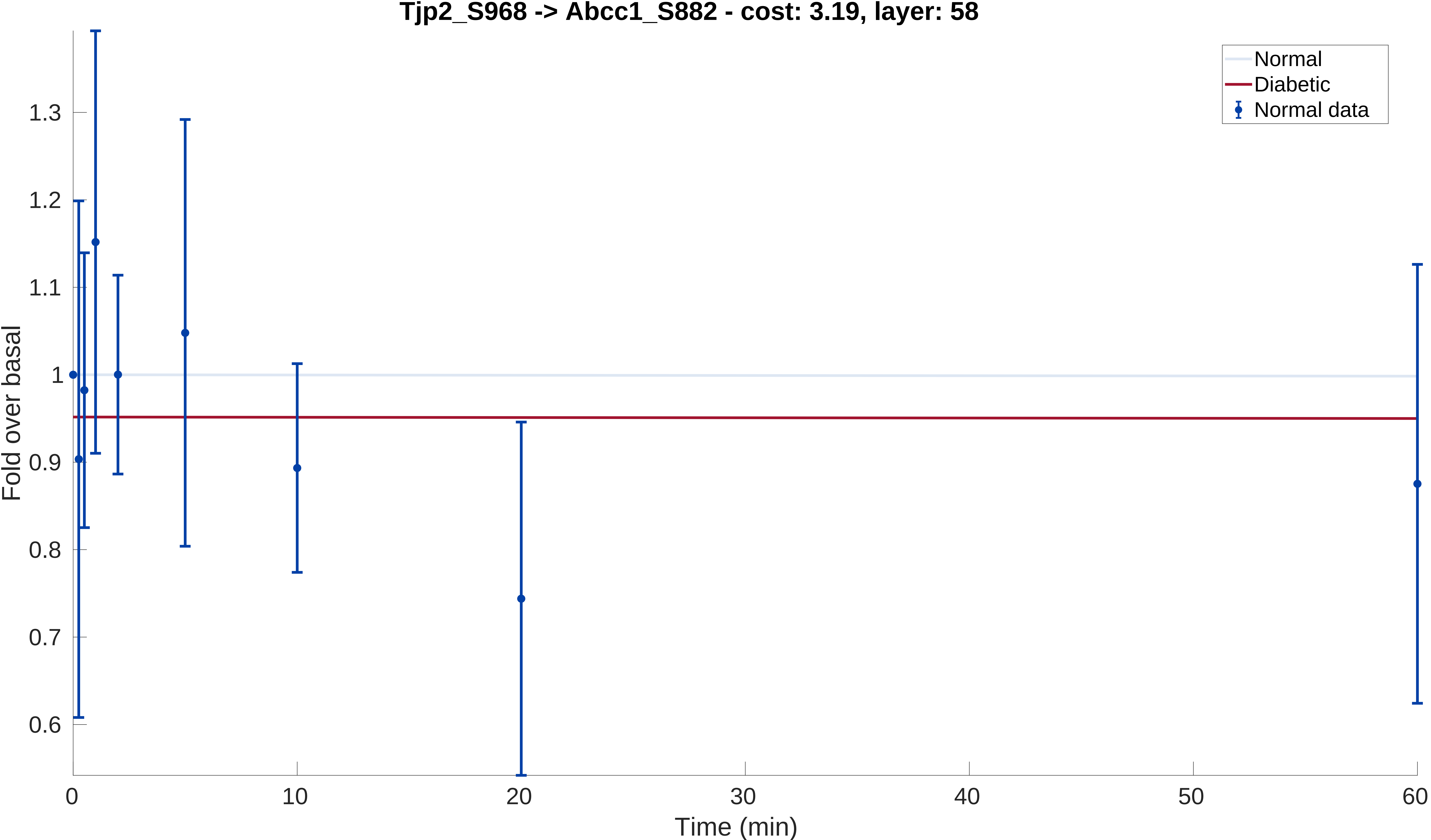

Tjp2_S968 -> Abcc1_S882 - cost: 3.19, layer: 58
Normal
Diabetic
Normal data
Fold over basal
1.3
1.2
1.1
1
0.9
0.8
0.7
0.6
0
10
20
30
40
50
60
Time (min)

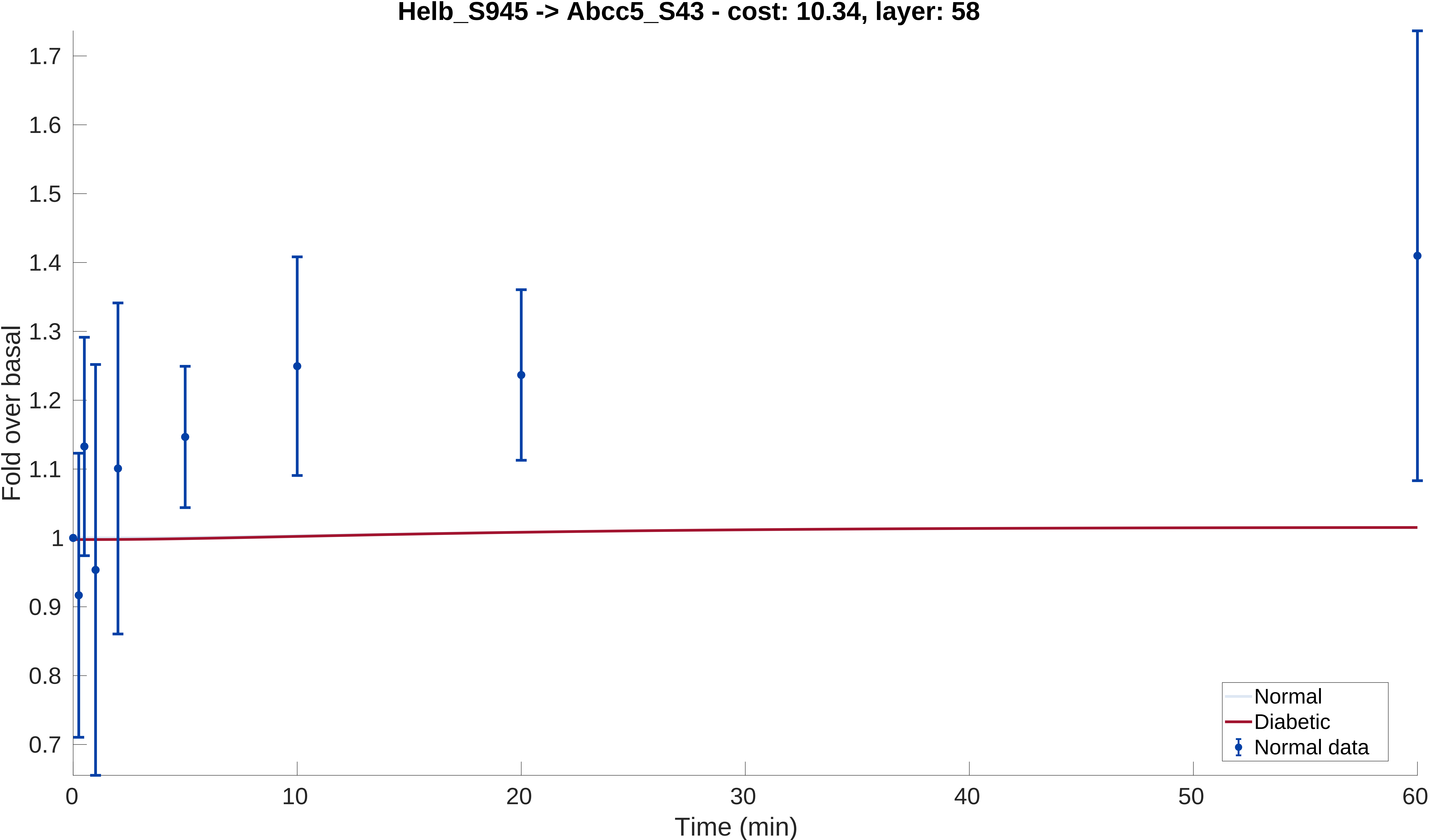
Helb_S945 -> Abcc5_S43 - cost: 10.34, layer: 58
Fold over basal
1.7
1.6
1.5
1.4
1.3
1.2
1.1
1
0.9
0.8
0.7
0
10
20
30
40
50
60
Time (min)
Normal
Diabetic
Normal data

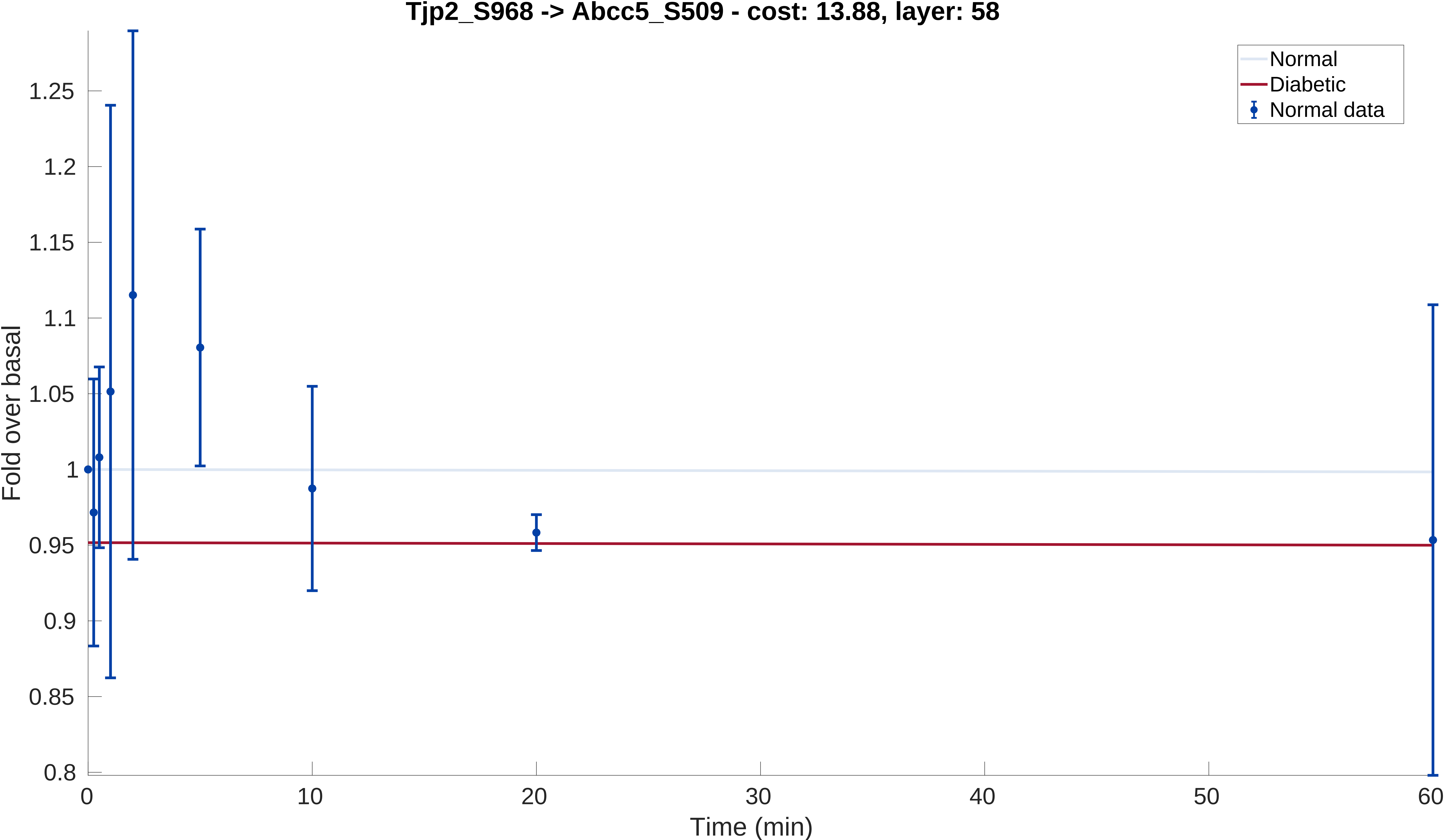

Tjp2_S968 -> Abcc5_S509 - cost: 13.88, layer: 58
Normal
Diabetic
Normal data
Fold over basal
1.25
1.2
1.15
1.1
1.05
1
0.95
0.9
0.85
0.8
0
10
20
30
40
50
60
Time (min)

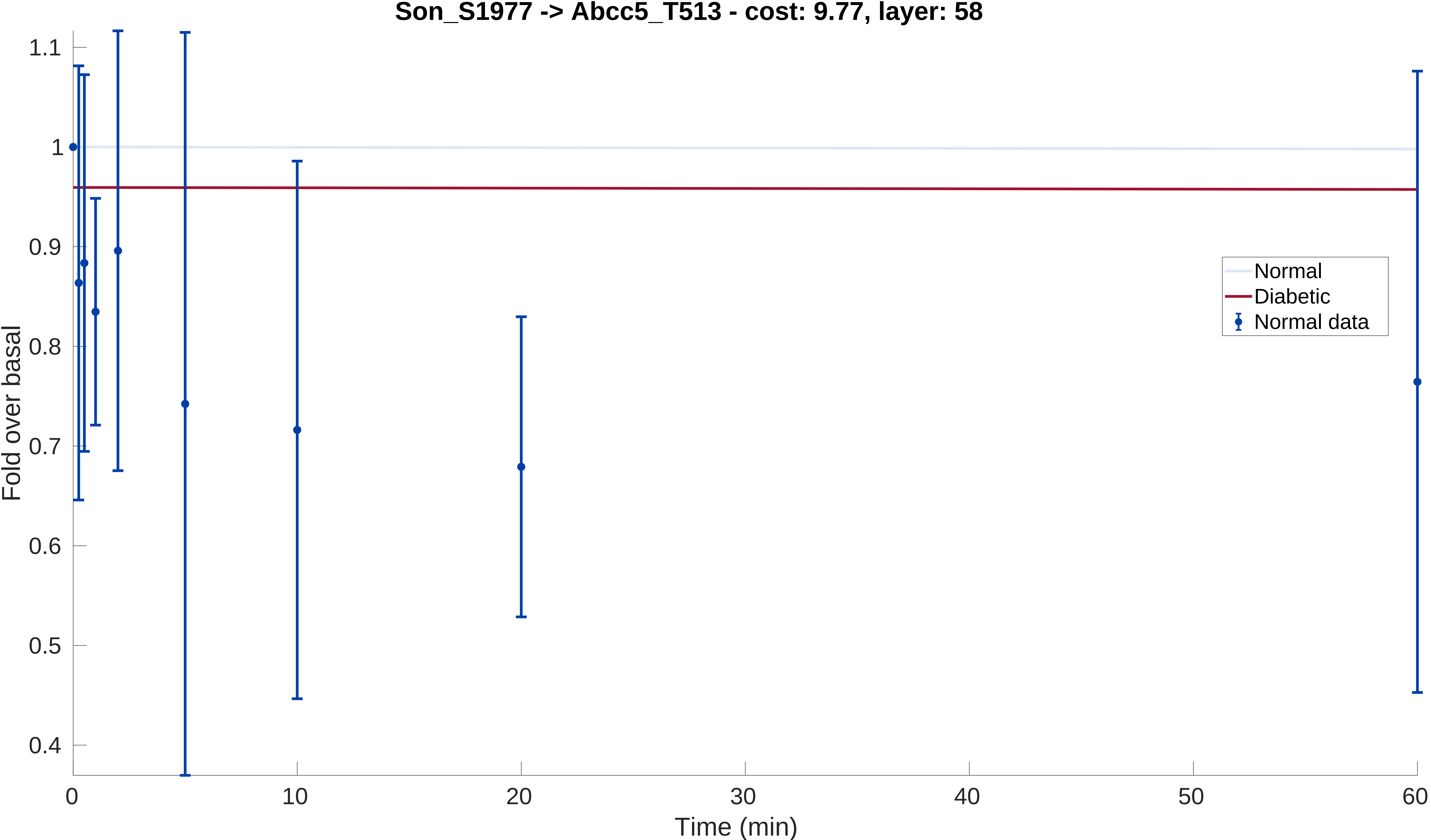

Son_S1977 -> Abcc5_T513 - cost: 9.77, layer: 58
1.1
1
0.9
0.8
0.7
0.6
0.5
0.4
Fold over basal
0
10
20
30
40
50
60
Time (min)
Normal
Diabetic
Normal data

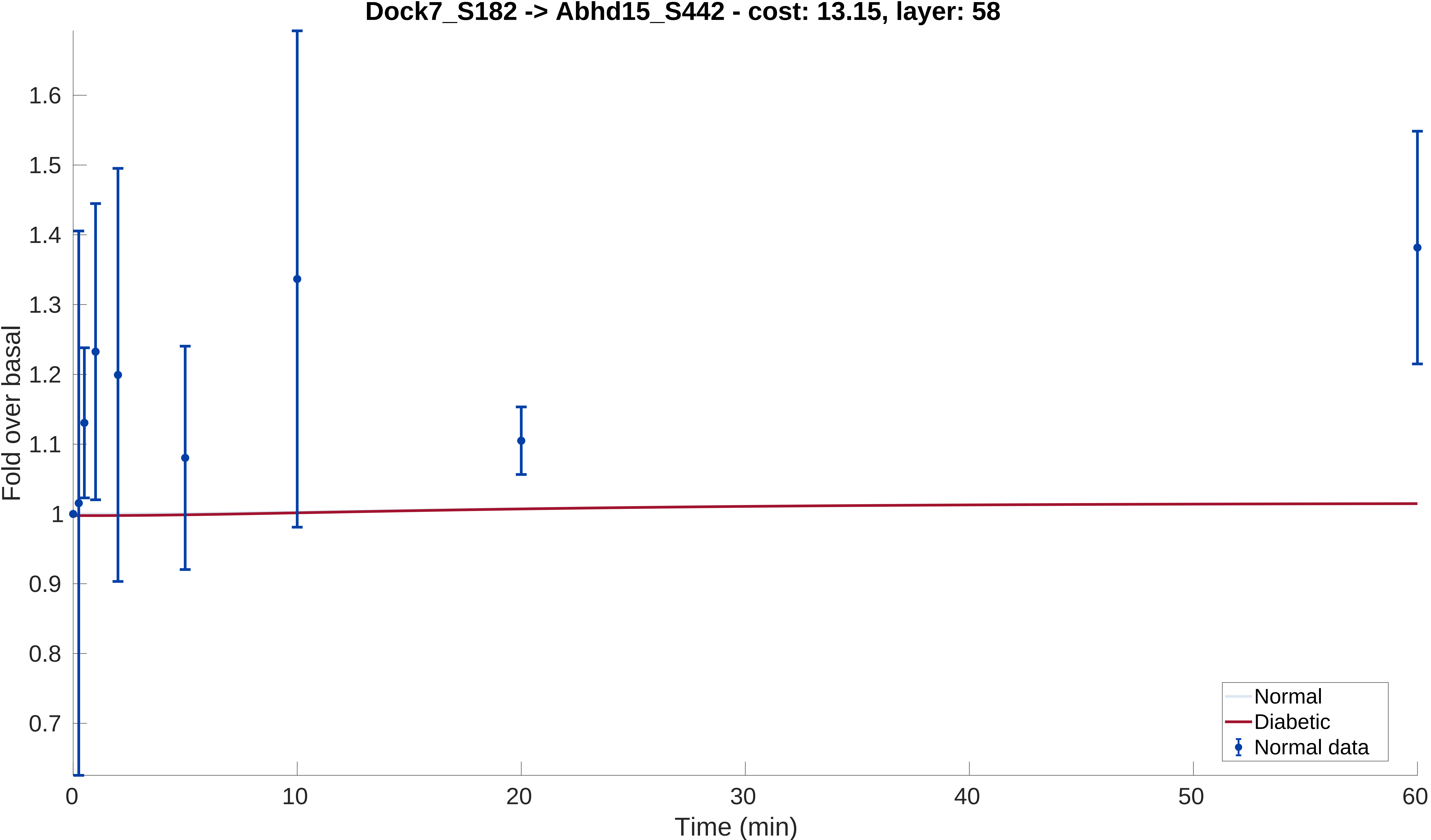

Dock7_S182 -> Abhd15_S442 - cost: 13.15, layer: 58
Fold over basal
Time (min)
1.6
1.5
1.4
1.3
1.2
1.1
1
0.9
0.8
0.7
0
10
20
30
40
50
60
Normal
Diabetic
Normal data

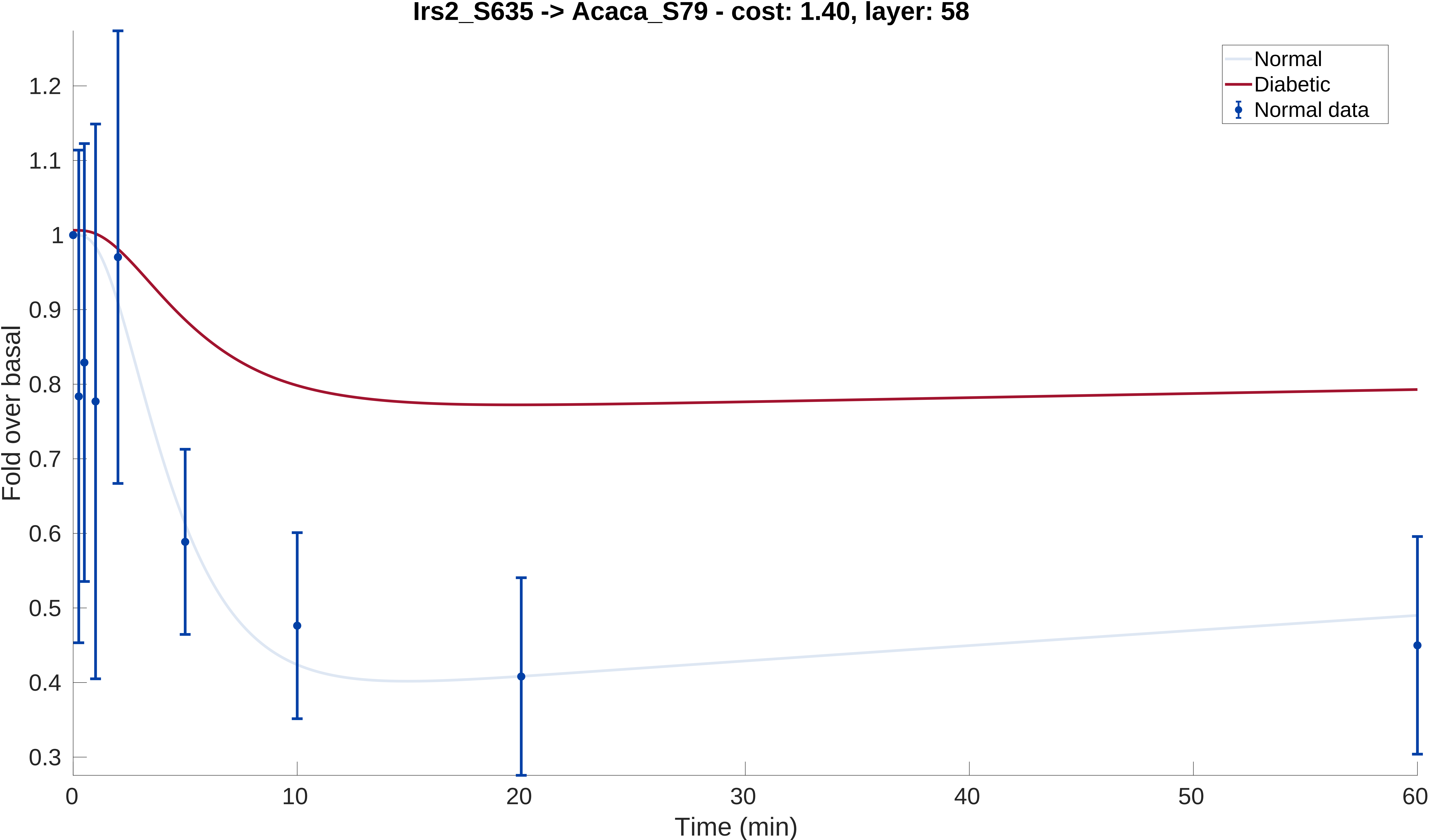

Irs2_S635 -> Acaca_S79 - cost: 1.40, layer: 58
Normal
Diabetic
Normal data
Fold over basal
1.2
1.1
1
0.9
0.8
0.7
0.6
0.5
0.4
0.3
0
10
20
30
40
50
60
Time (min)

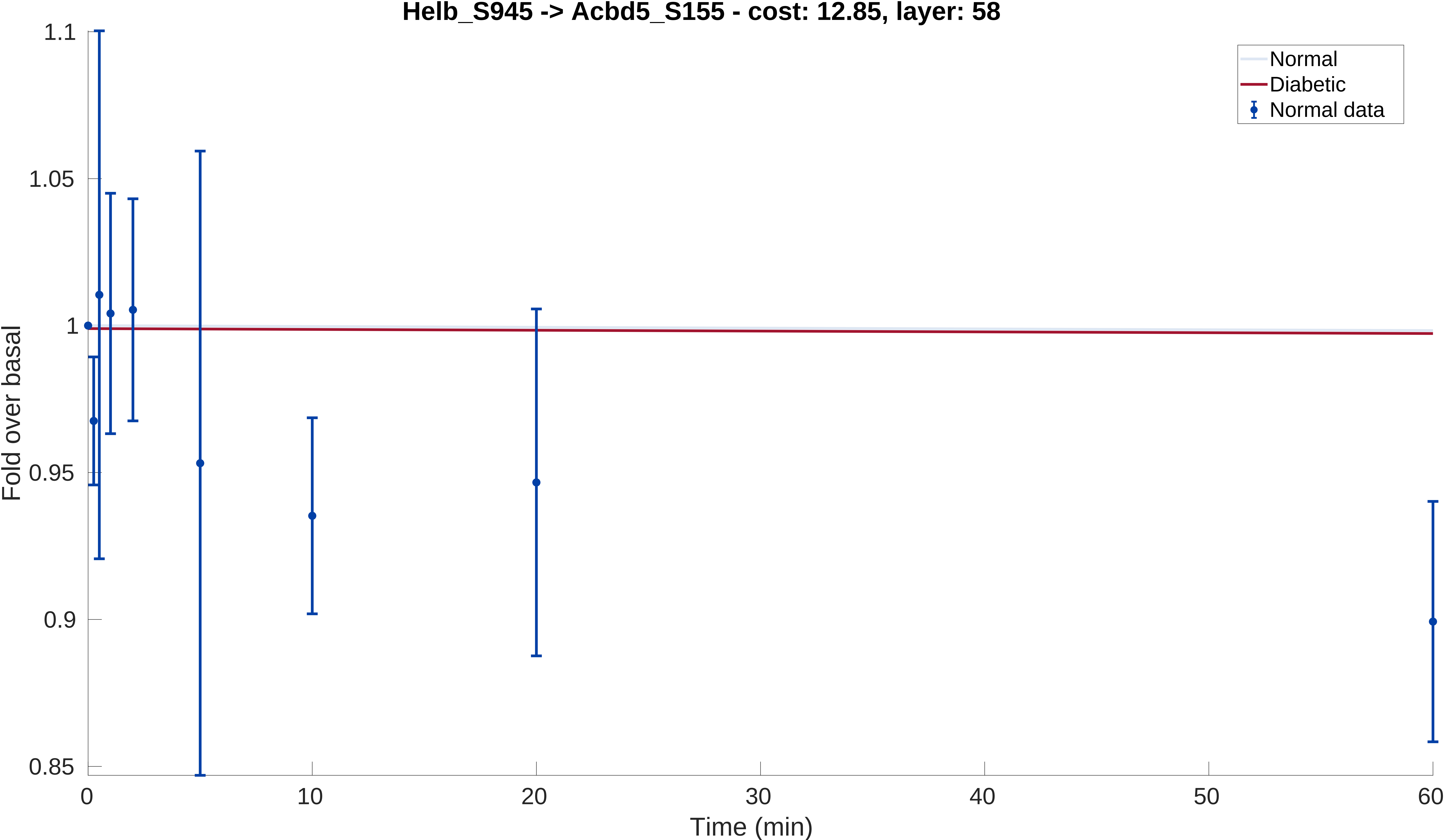

Helb_S945 -> Acbd5_S155 - cost: 12.85, layer: 58
Normal
Diabetic
Normal data
Fold over basal
Time (min)
1.1
1.05
1
0.95
0.9
0.85
0
10
20
30
40
50
60

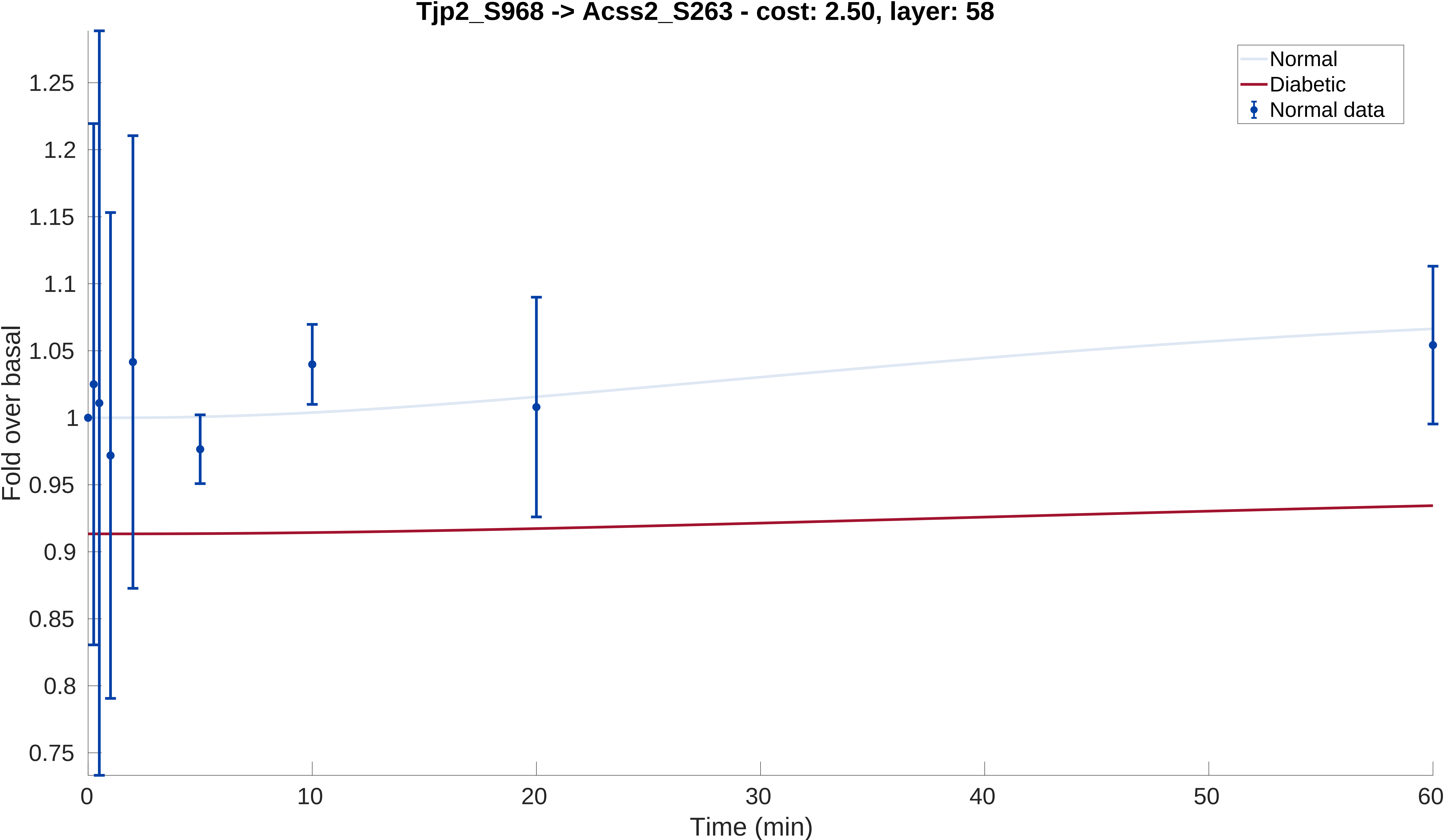

Tjp2_S968 -> Acss2_S263 - cost: 2.50, layer: 58
Normal
Diabetic
Normal data
1.25
1.2
1.15
1.1
1.05
1
0.95
0.9
0.85
0.8
0.75
Fold over basal
0
10
20
30
40
50
60
Time (min)

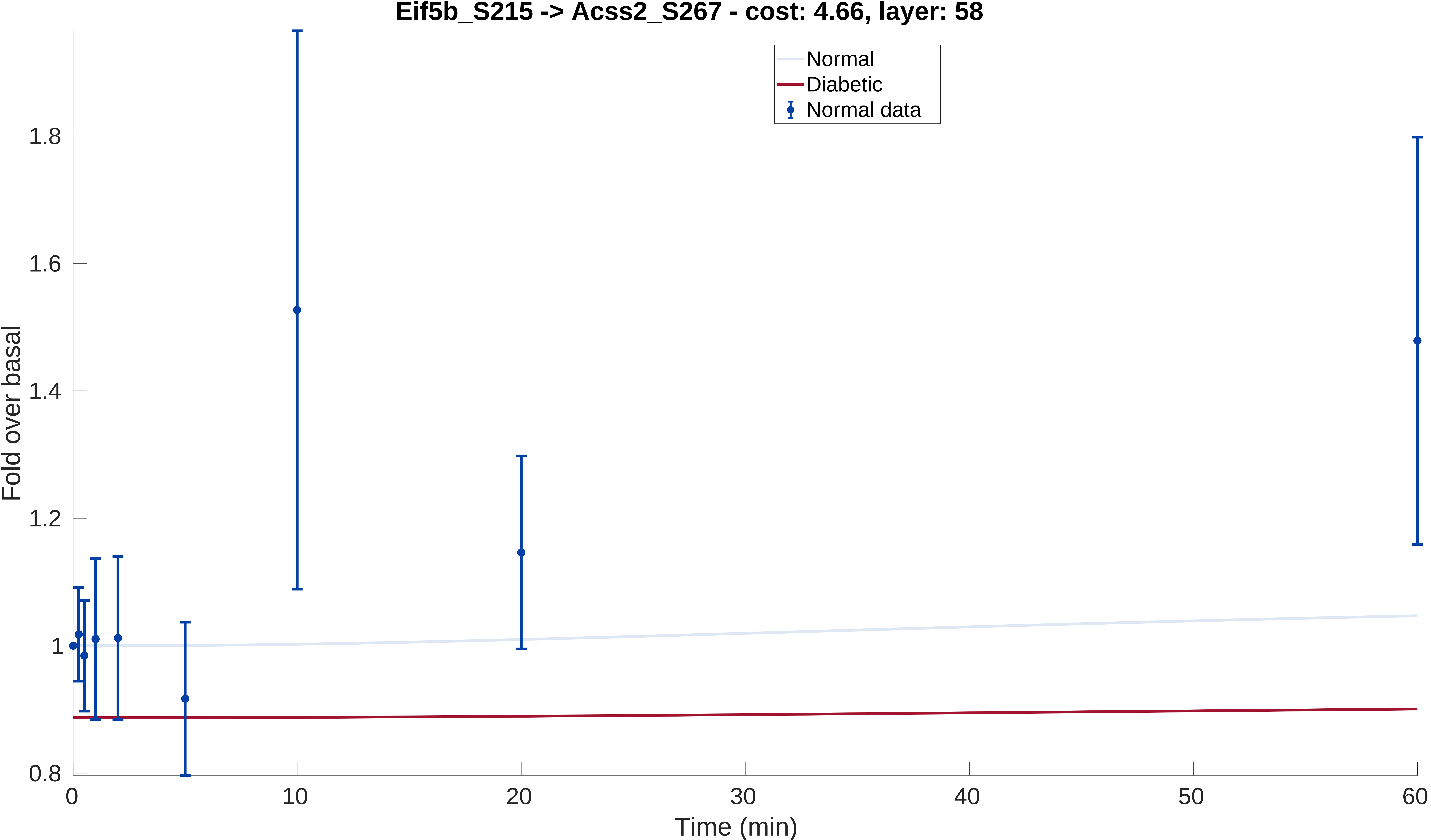
Eif5b_S215 -> Acss2_S267 - cost: 4.66, layer: 58
Normal
Diabetic
Normal data
Fold over basal
1.8
1.6
1.4
1.2
1
0.8
0
10
20
30
40
50
60
Time (min)

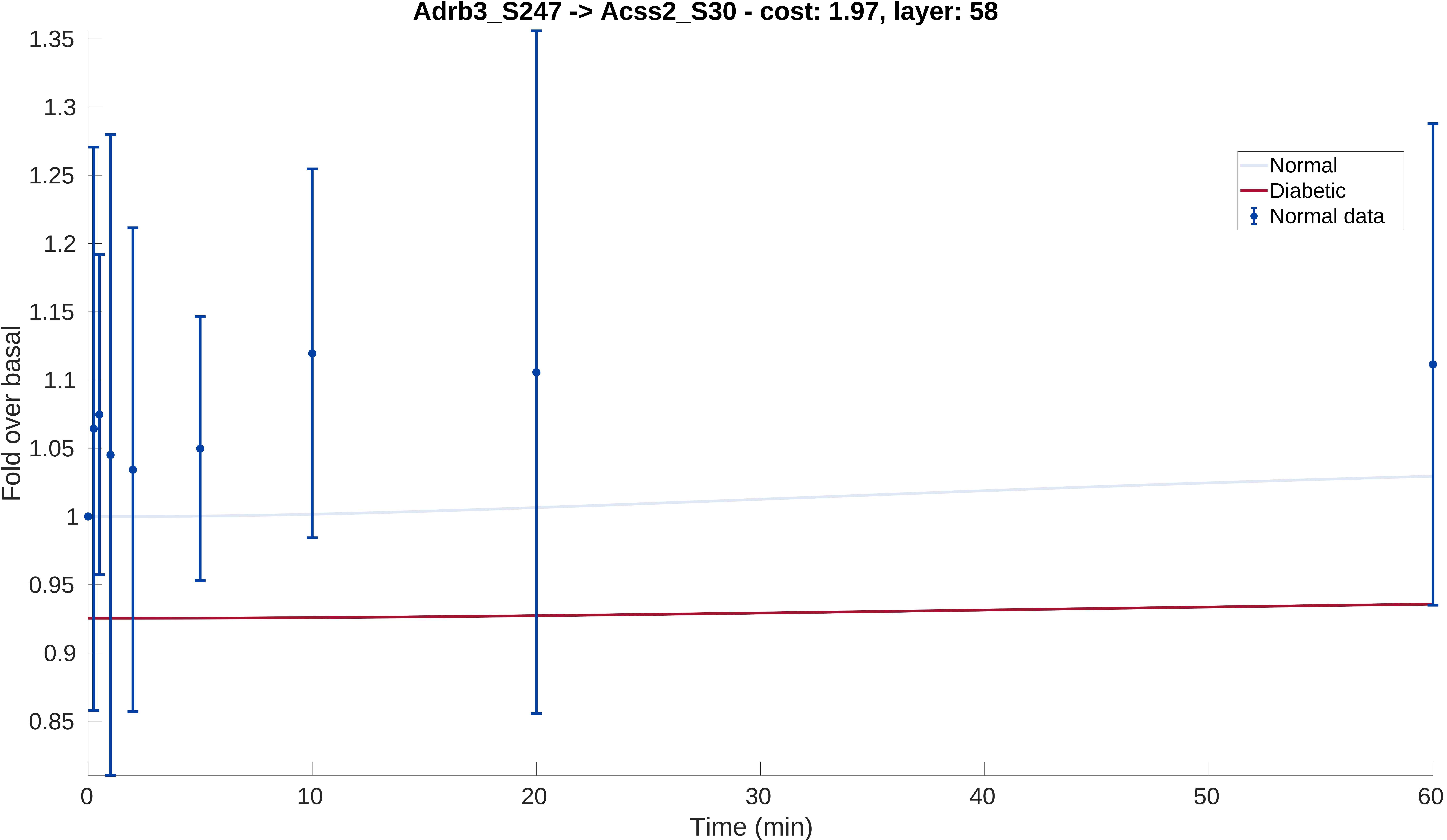

Adrb3_S247 -> Acss2_S30 - cost: 1.97, layer: 58
Fold over basal
1.35
1.3
1.25
1.2
1.15
1.1
1.05
1
0.95
0.9
0.85
Normal
Diabetic
Normal data
0
10
20
30
40
50
60
Time (min)

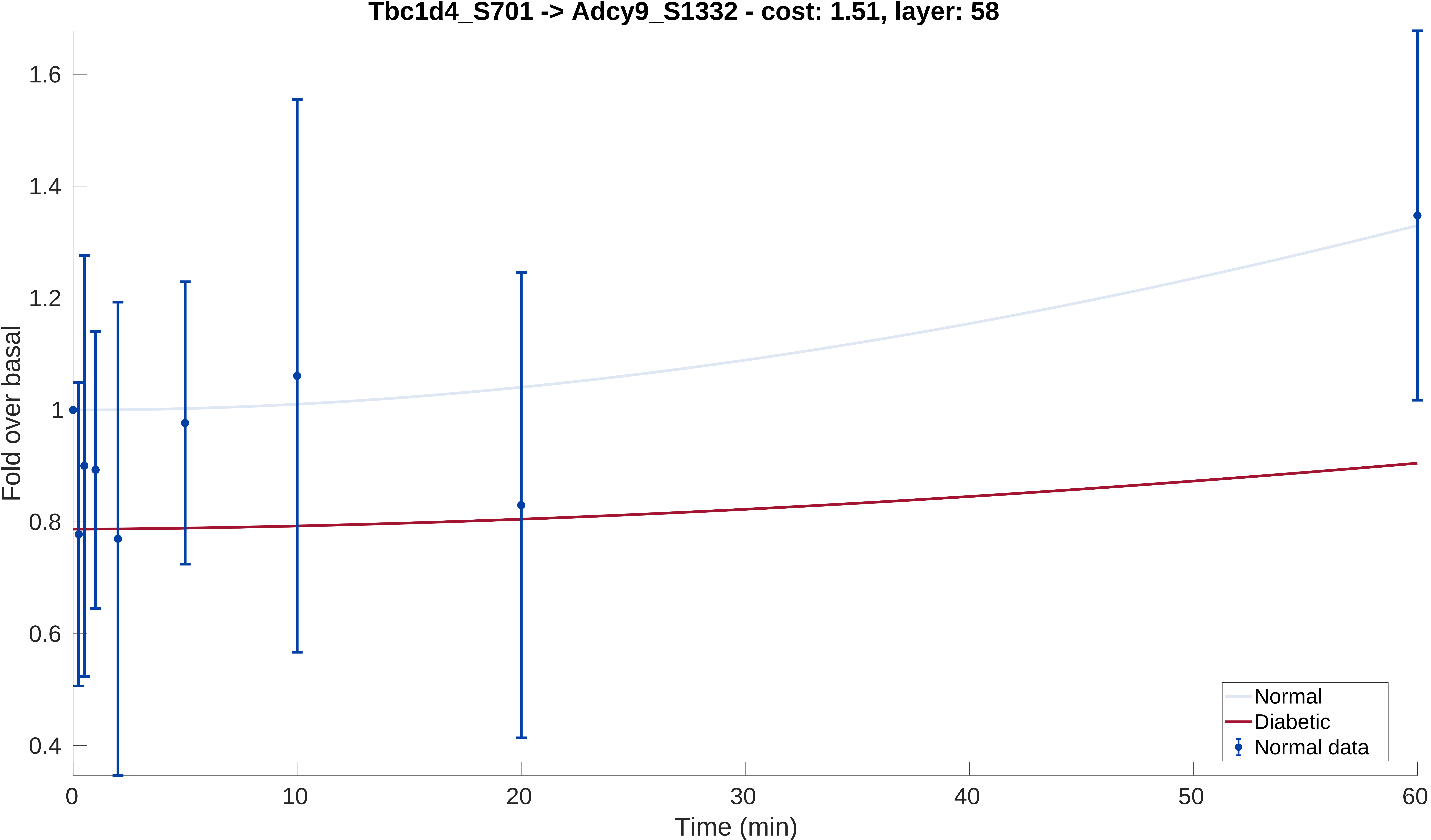

Tbc1d4_S701 -> Adcy9_S1332 - cost: 1.51, layer: 58
Fold over basal
1.6
1.4
1.2
1
0.8
0.6
0.4
0
10
20
30
40
50
60
Time (min)
Normal
Diabetic
Normal data

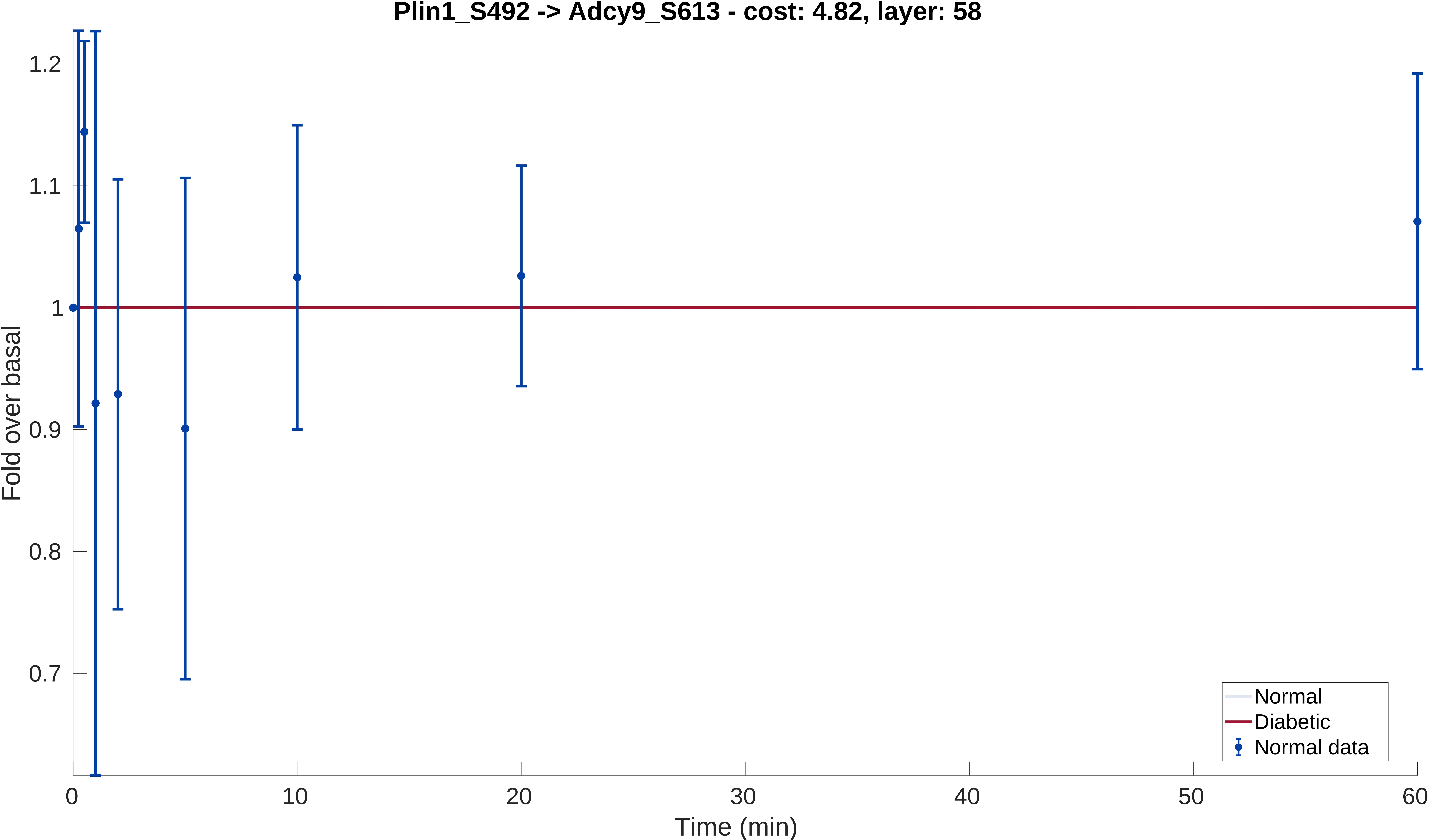

Plin1_S492 -> Adcy9_S613 - cost: 4.82, layer: 58
Fold over basal
1.2
1.1
1
0.9
0.8
0.7
0
10
20
30
40
50
60
Time (min)
Normal
Diabetic
Normal data

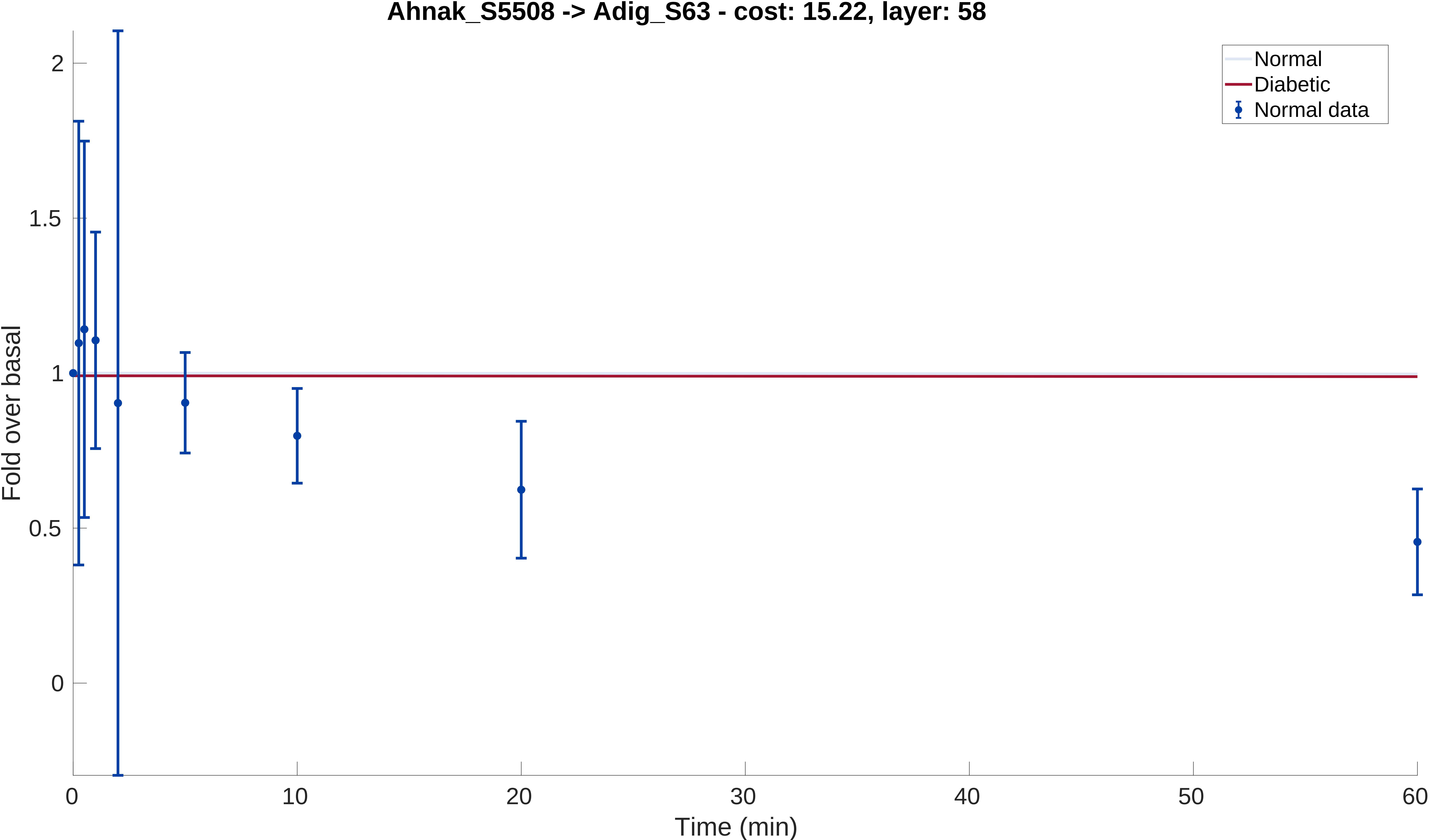

Ahnak_S5508 -> Adig_S63 - cost: 15.22, layer: 58
Normal
Diabetic
Normal data
Fold over basal
Time (min)
0
0.5
1
1.5
2
0
10
20
30
40
50
60

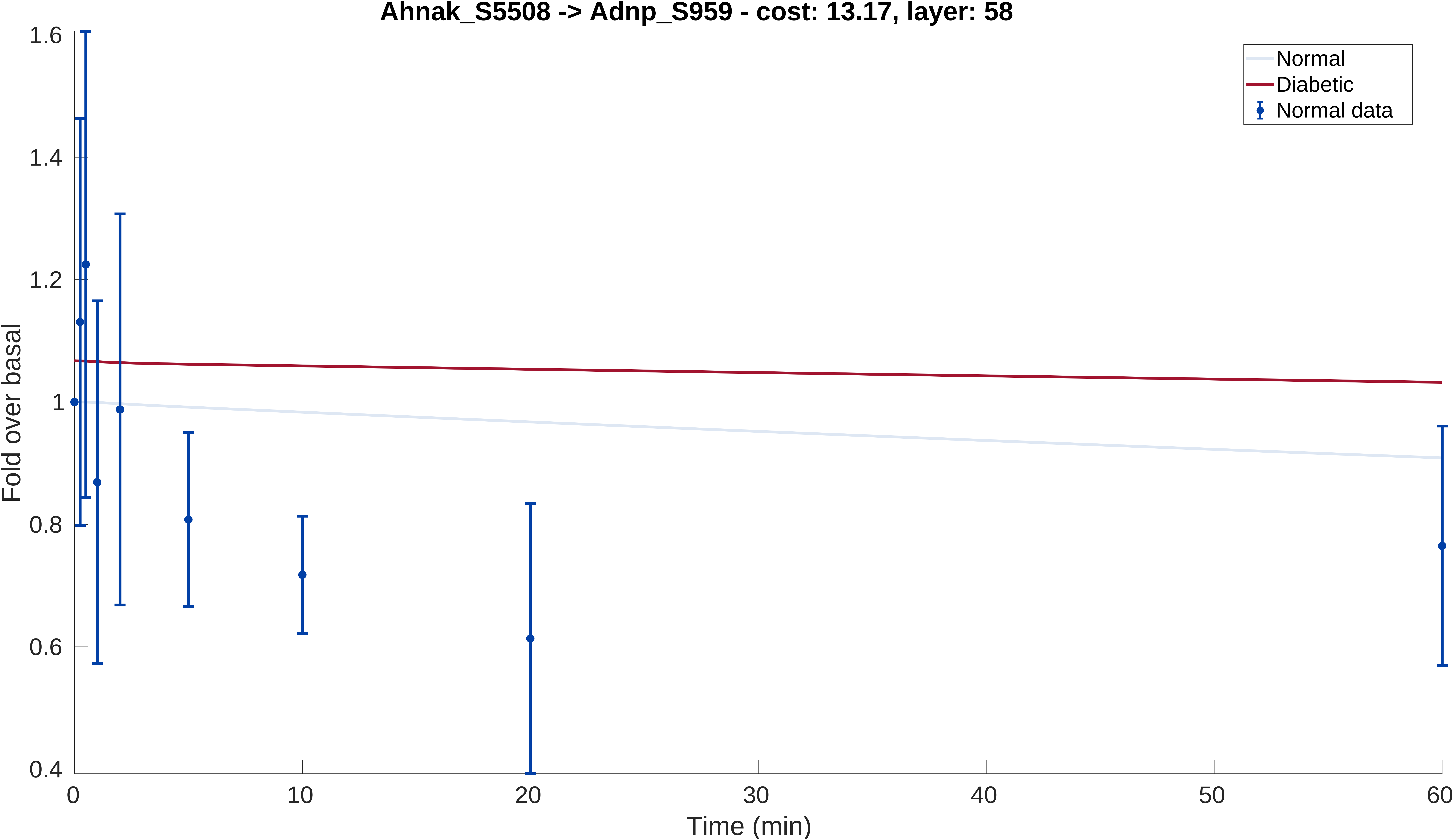

Ahnak_S5508 -> Adnp_S959 - cost: 13.17, layer: 58
Normal
Diabetic
Normal data
Fold over basal
Time (min)
1.6
1.4
1.2
1
0.8
0.6
0.4
0
10
20
30
40
50
60

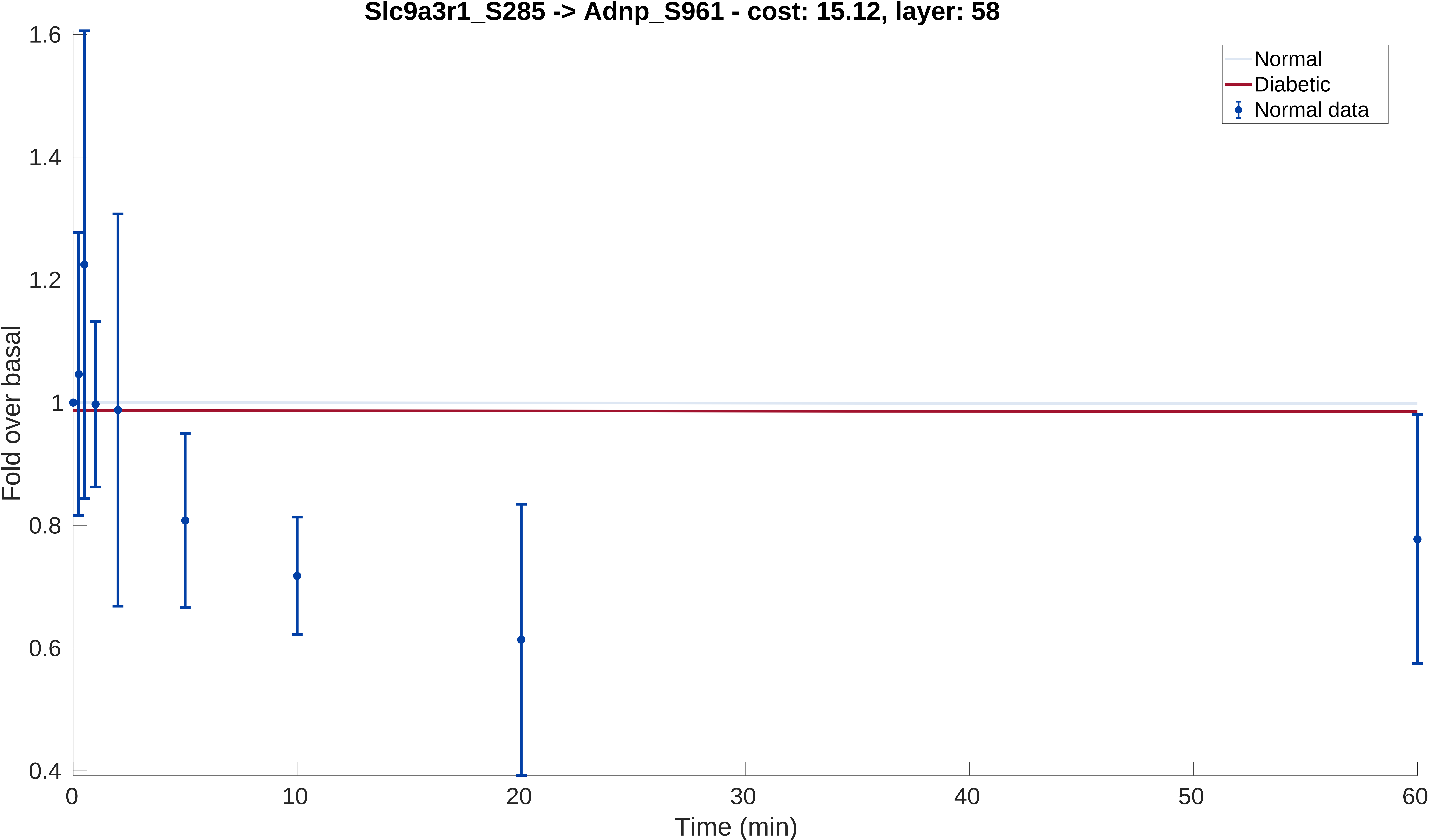
Slc9a3r1_S285 -> Adnp_S961 - cost: 15.12, layer: 58
Normal
Diabetic
Normal data
Fold over basal
1.6
1.4
1.2
1
0.8
0.6
0.4
0
10
20
30
40
50
60
Time (min)

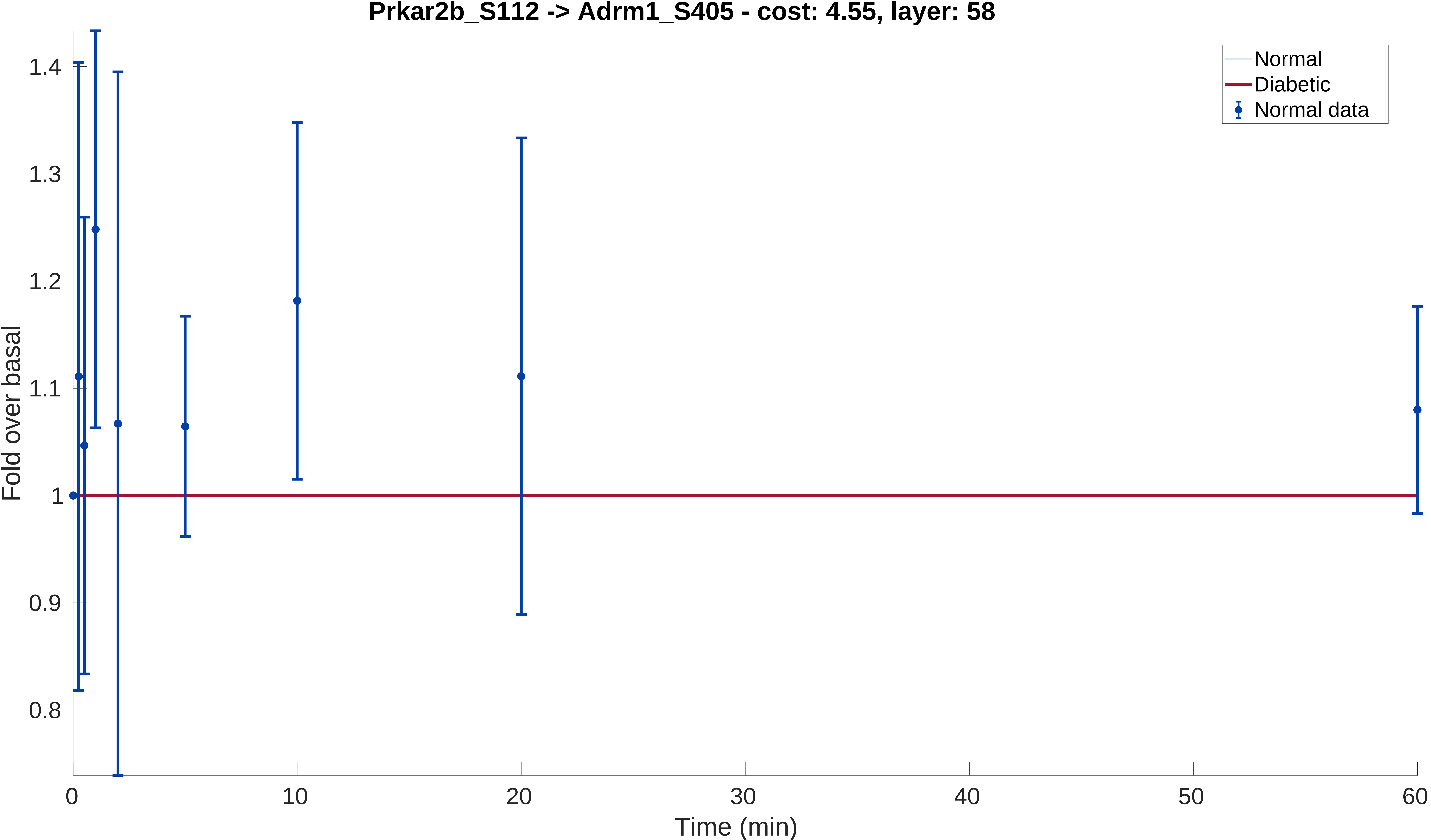

Prkar2b_S112 -> Adrm1_S405 - cost: 4.55, layer: 58
Normal
Diabetic
Normal data
Fold over basal
1.4
1.3
1.2
1.1
1
0.9
0.8
0
10
20
30
40
50
60
Time (min)

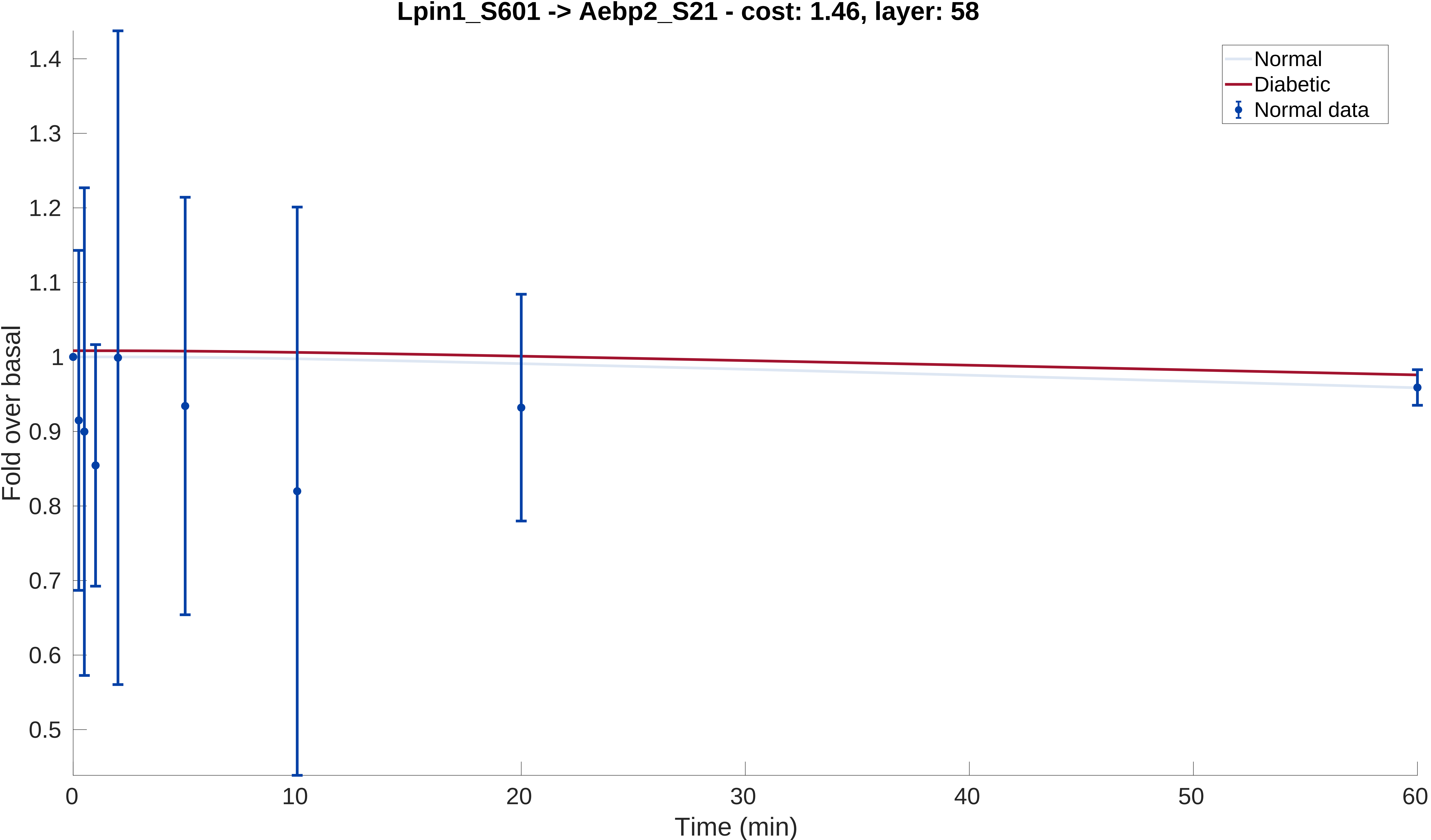

Lpin1_S601 -> Aebp2_S21 - cost: 1.46, layer: 58
Normal
Diabetic
Normal data
Fold over basal
Time (min)
1.4
1.3
1.2
1.1
1
0.9
0.8
0.7
0.6
0.5
0
10
20
30
40
50
60

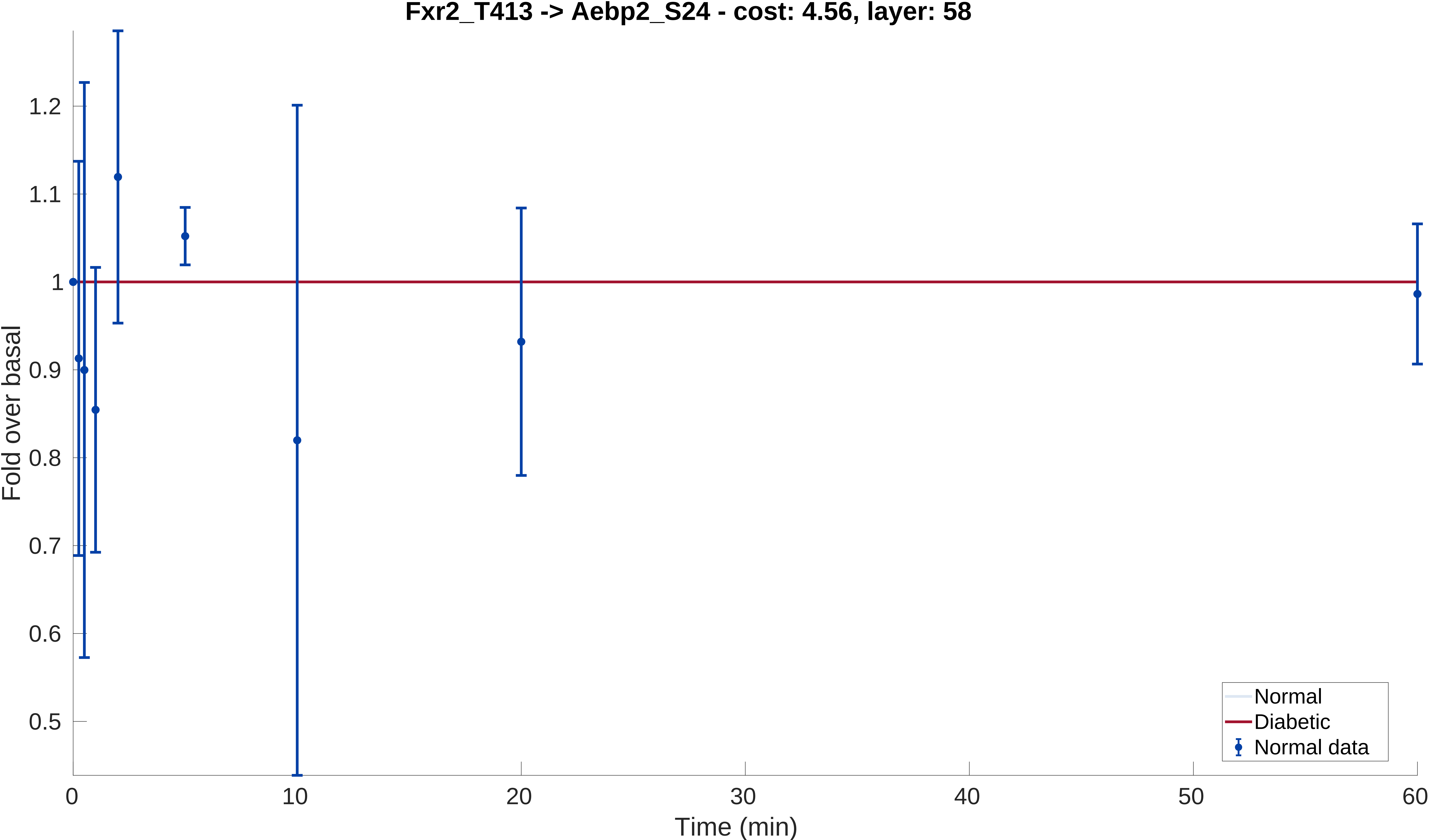

Fxr2_T413 -> Aebp2_S24 - cost: 4.56, layer: 58
Fold over basal
1.2
1.1
1
0.9
0.8
0.7
0.6
0.5
0
10
20
30
40
50
60
Time (min)
Normal
Diabetic
Normal data

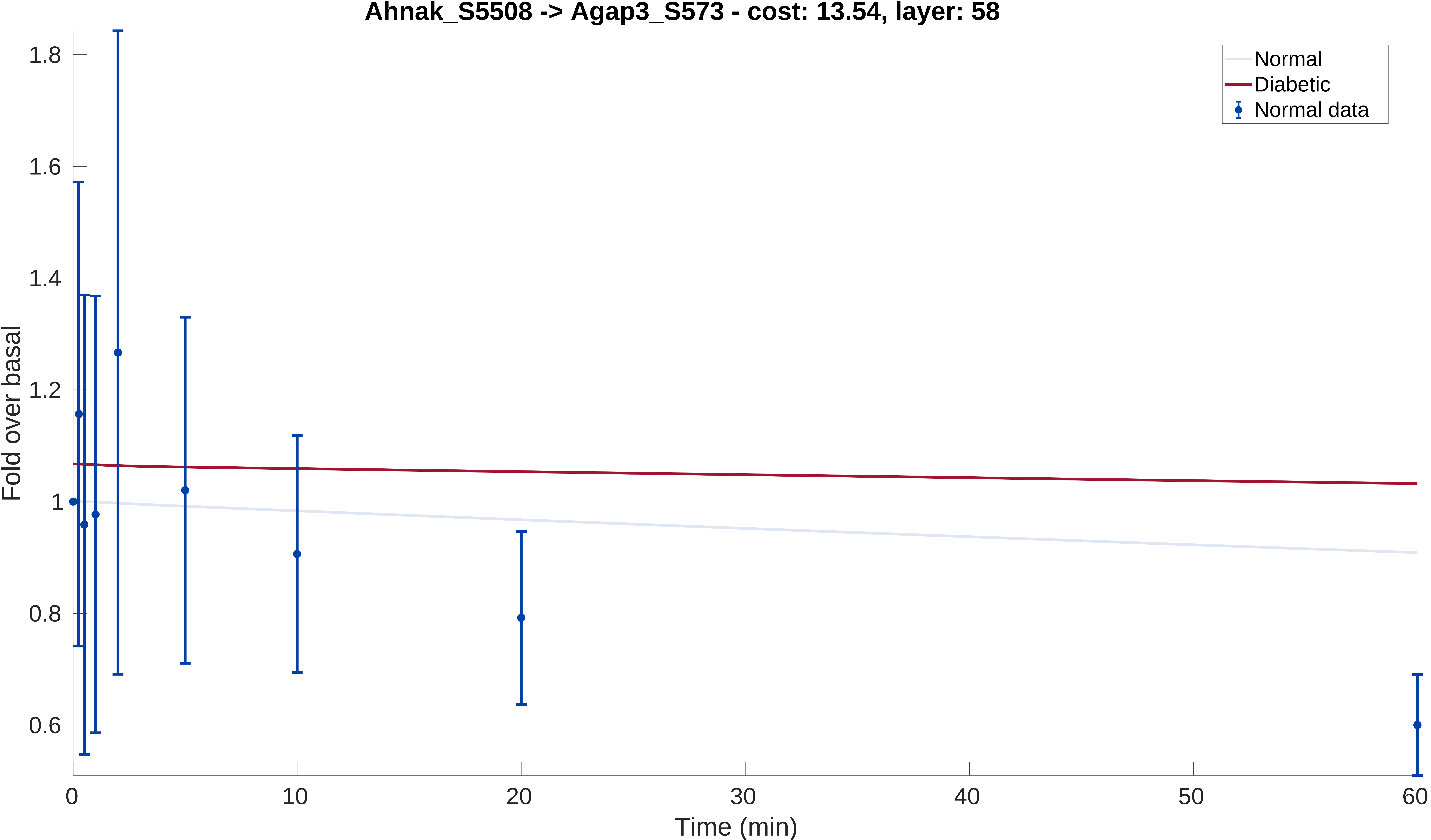

Ahnak_S5508 -> Agap3_S573 - cost: 13.54, layer: 58
Normal
Diabetic
Normal data
Fold over basal
Time (min)
1.8
1.6
1.4
1.2
1
0.8
0.6
0
10
20
30
40
50
60

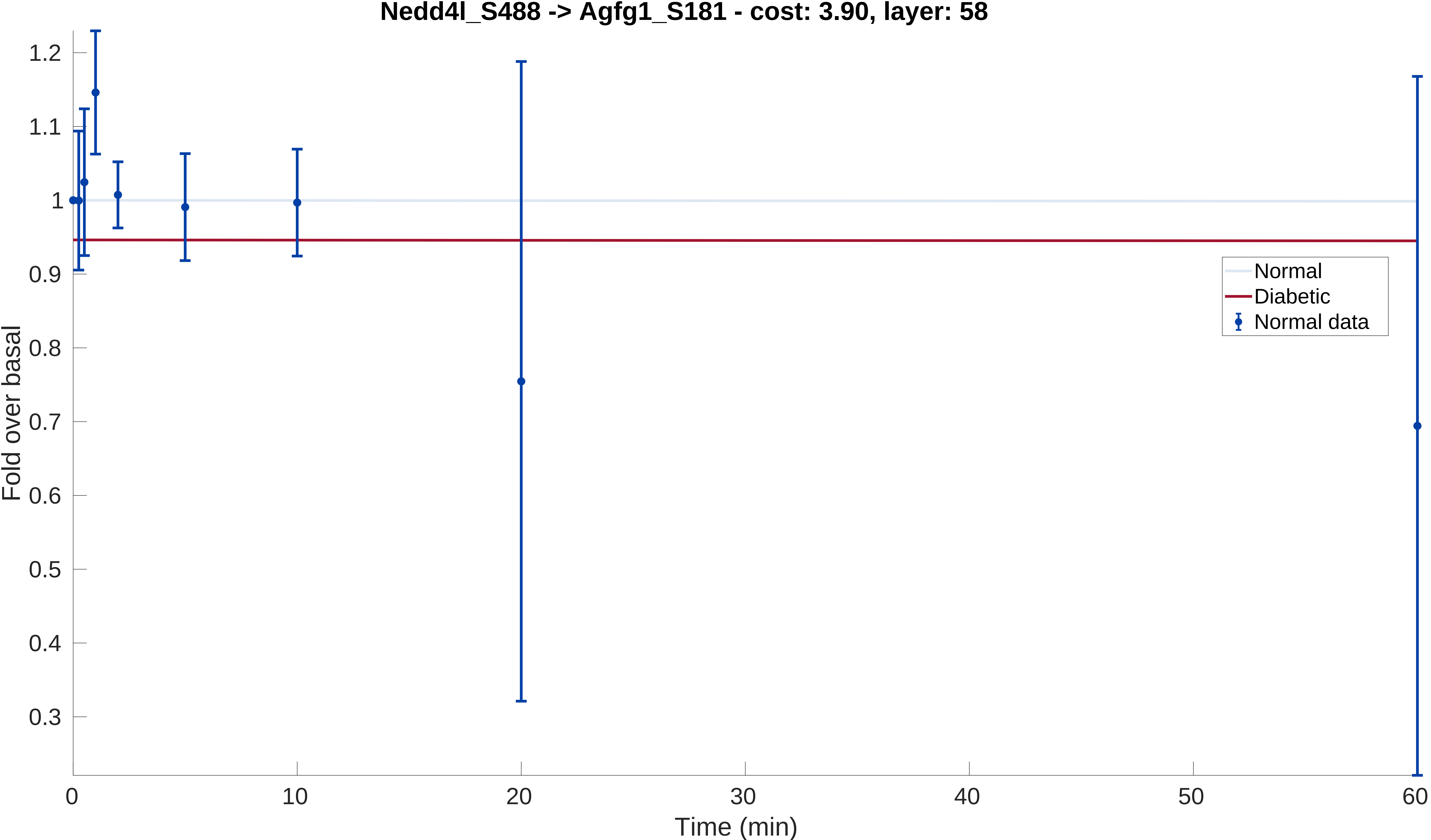

Nedd4l_S488 -> Agfg1_S181 - cost: 3.90, layer: 58
Fold over basal
1.2
1.1
1
0.9
0.8
0.7
0.6
0.5
0.4
0.3
0
10
20
30
40
50
60
Time (min)
Normal
Diabetic
Normal data

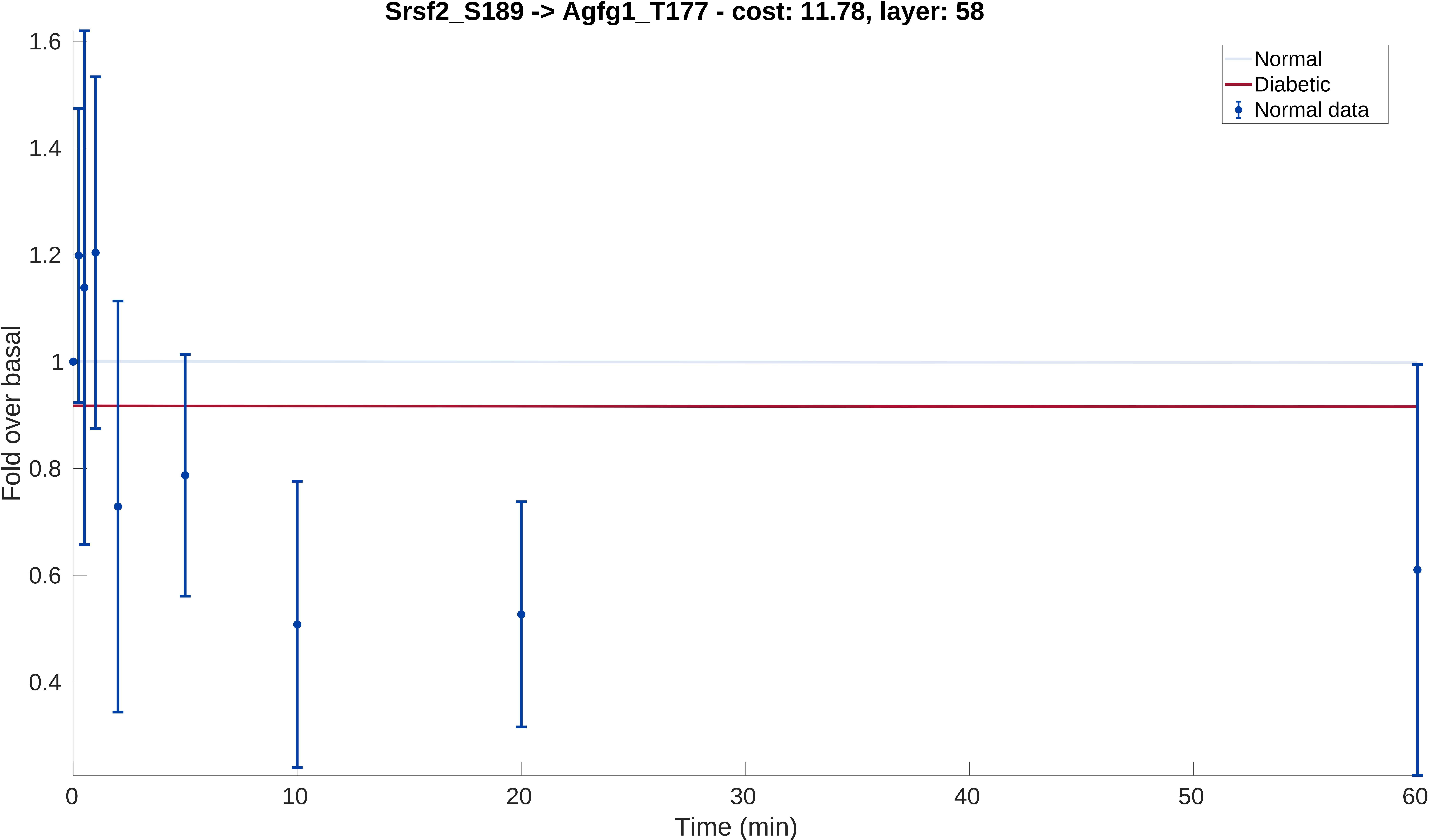

Srsf2_S189 -> Agfg1_T177 - cost: 11.78, layer: 58
Normal
Diabetic
Normal data
Fold over basal
1.6
1.4
1.2
1
0.8
0.6
0.4
0
10
20
30
40
50
60
Time (min)

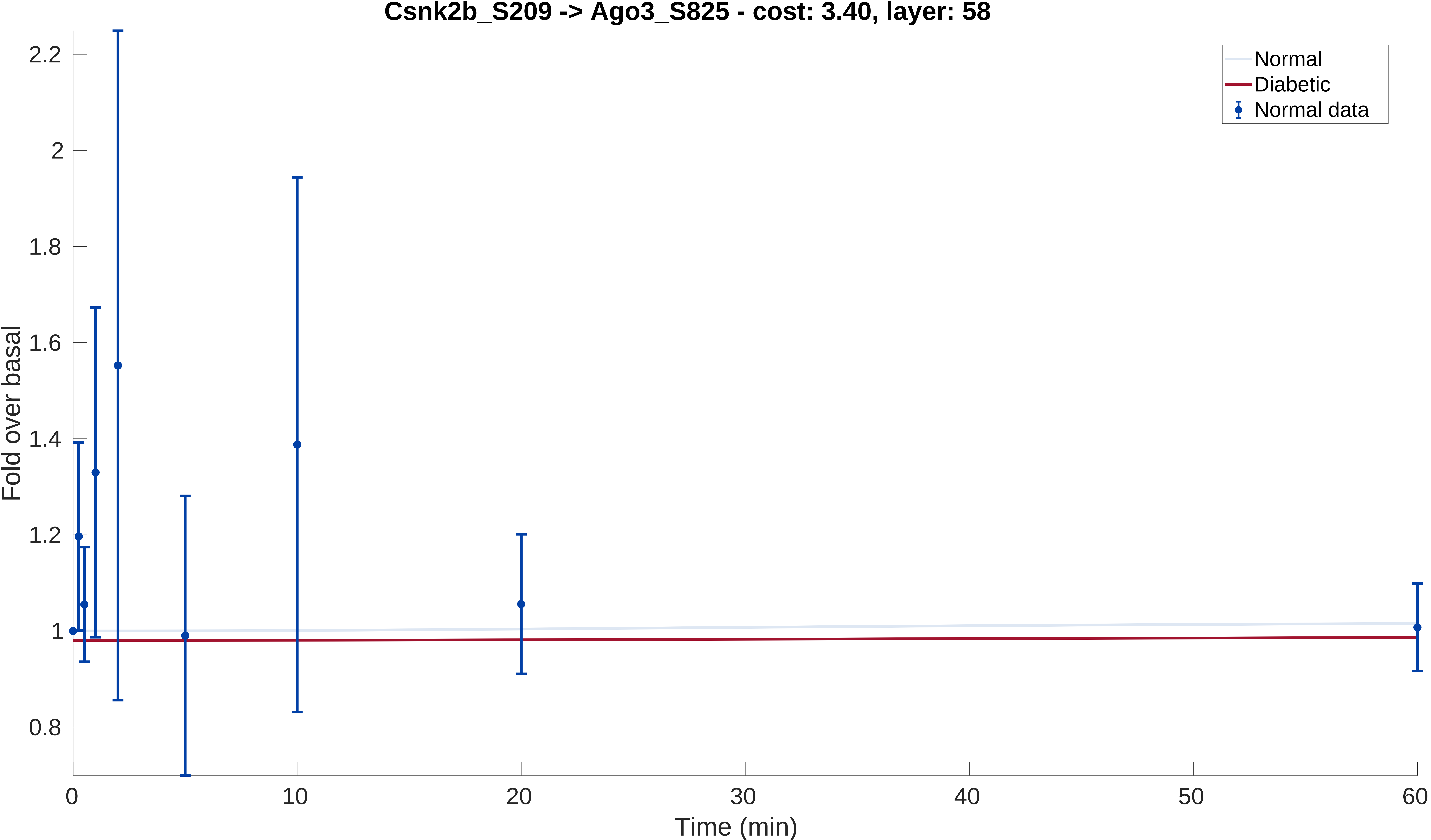

Csnk2b_S209 -> Ago3_S825 - cost: 3.40, layer: 58
Normal
Diabetic
Normal data
Fold over basal
Time (min)
0.8
1
1.2
1.4
1.6
1.8
2
2.2
0
10
20
30
40
50
60

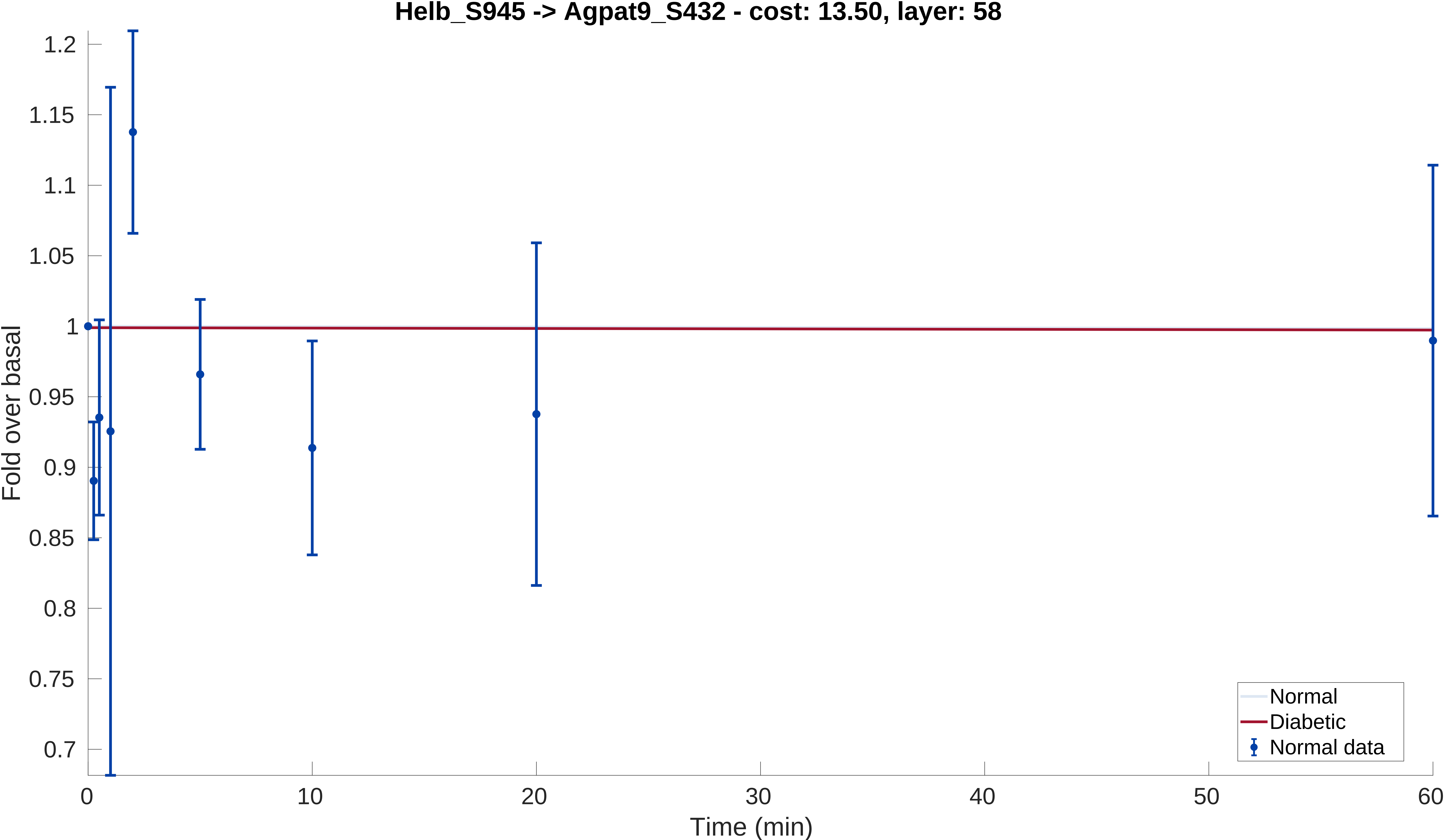

Helb_S945 -> Agpat9_S432 - cost: 13.50, layer: 58
1.2
1.15
1.1
1.05
1
0.95
0.9
0.85
0.8
0.75
0.7
Fold over basal
0
10
20
30
40
50
60
Time (min)
Normal
Diabetic
Normal data

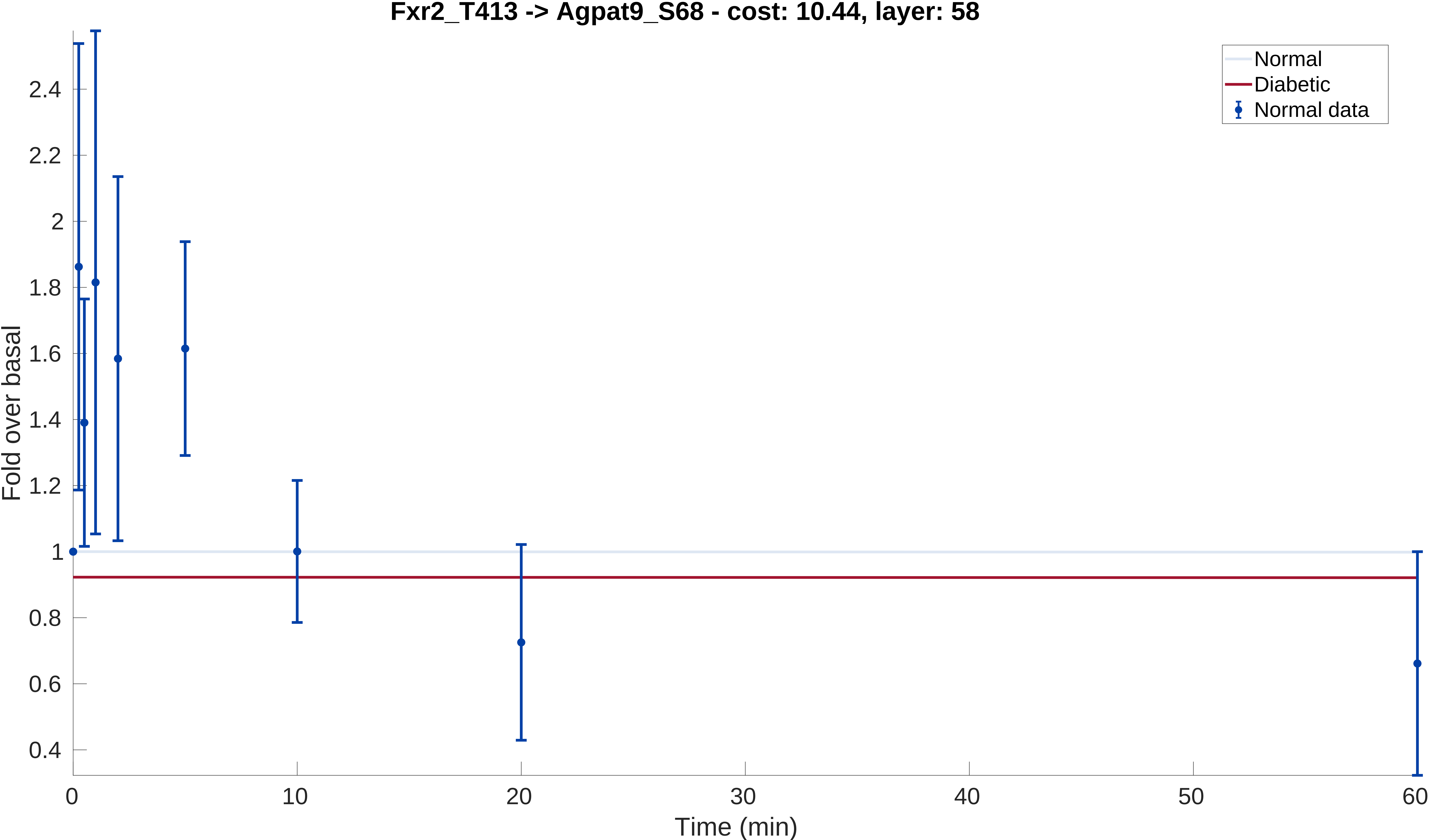

Fxr2_T413 -> Agpat9_S68 - cost: 10.44, layer: 58
Normal
Diabetic
Normal data
Fold over basal
2.4
2.2
2
1.8
1.6
1.4
1.2
1
0.8
0.6
0.4
0
10
20
30
40
50
60
Time (min)

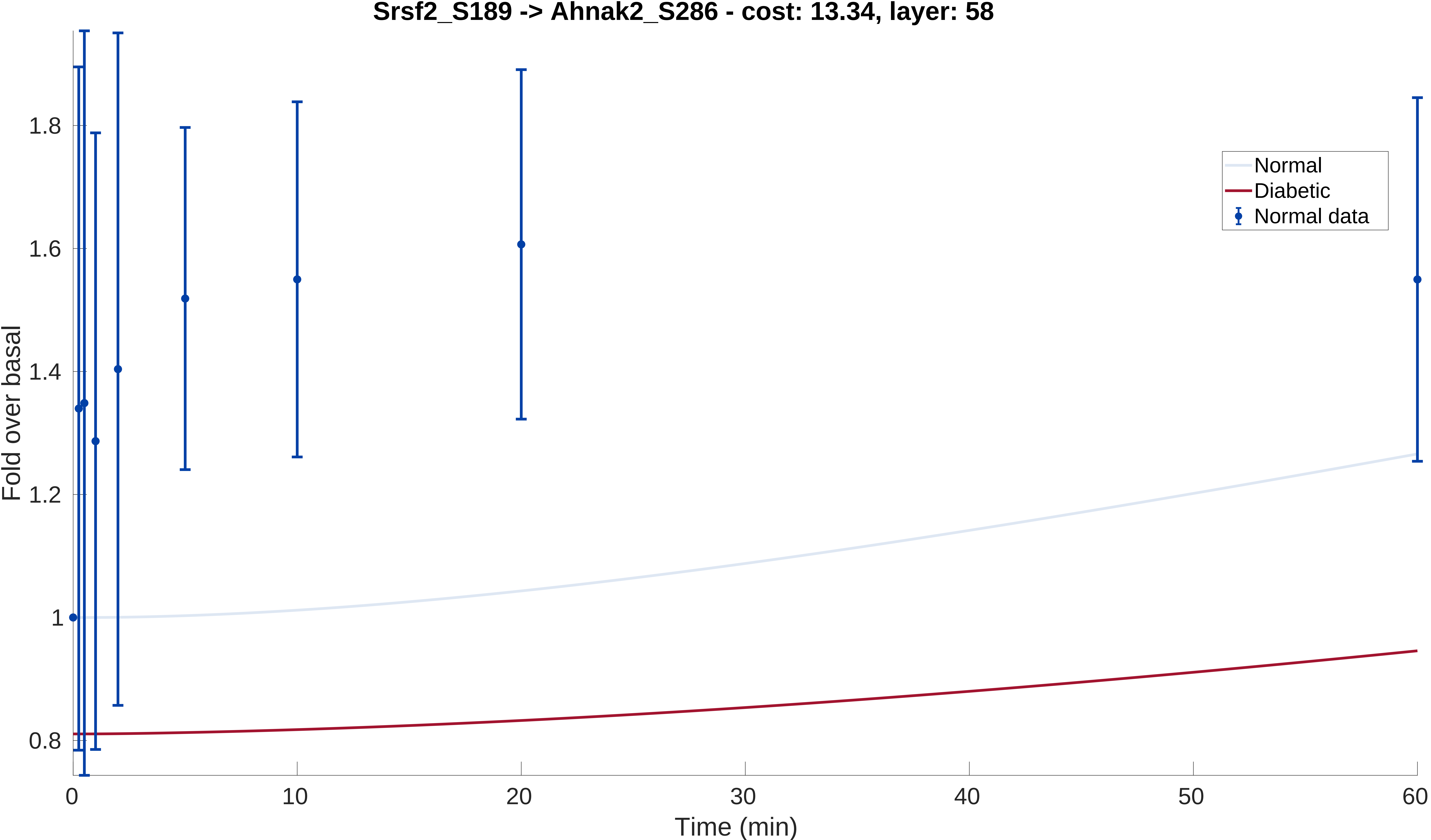
Srsf2_S189 -> Ahnak2_S286 - cost: 13.34, layer: 58
Fold over basal
1.8
1.6
1.4
1.2
1
0.8
0
10
20
30
40
50
60
Time (min)
Normal
Diabetic
Normal data

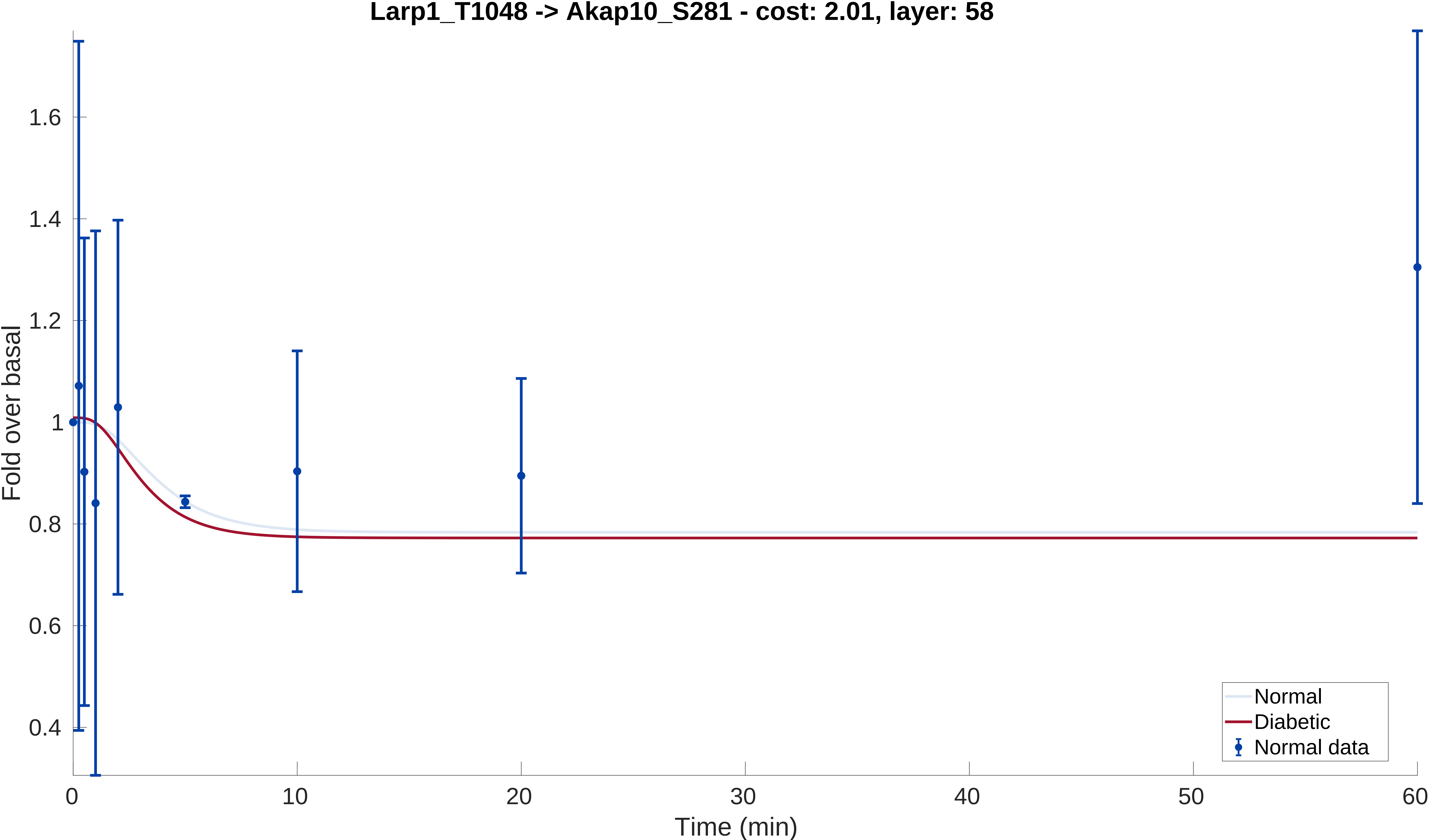

Larp1_T1048 -> Akap10_S281 - cost: 2.01, layer: 58
Fold over basal
1.6
1.4
1.2
1
0.8
0.6
0.4
0
10
20
30
40
50
60
Time (min)
Normal
Diabetic
Normal data

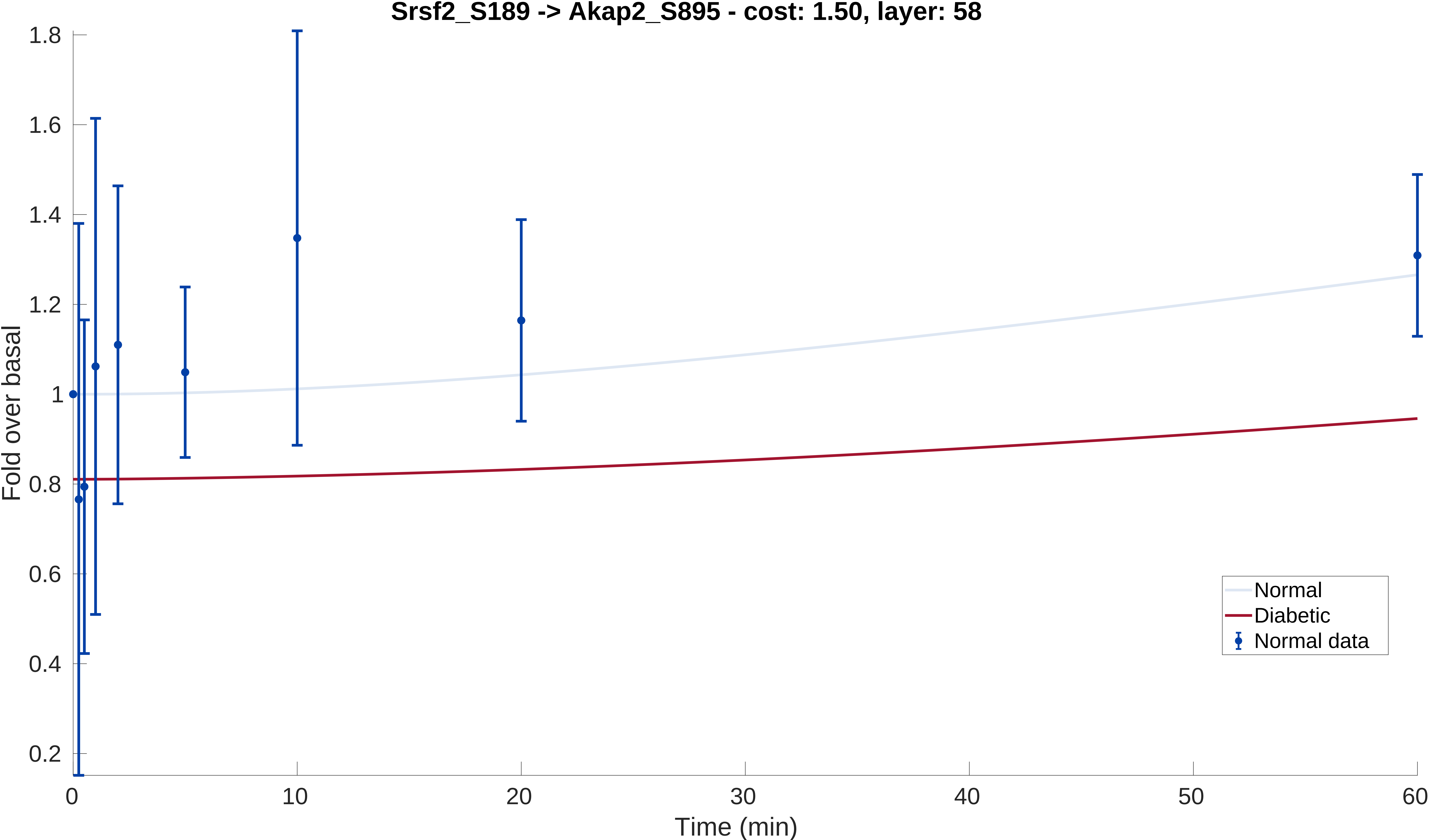

Srsf2_S189 -> Akap2_S895 - cost: 1.50, layer: 58
Fold over basal
1.8
1.6
1.4
1.2
1
0.8
0.6
0.4
0.2
0
10
20
30
40
50
60
Time (min)
Normal
Diabetic
Normal data

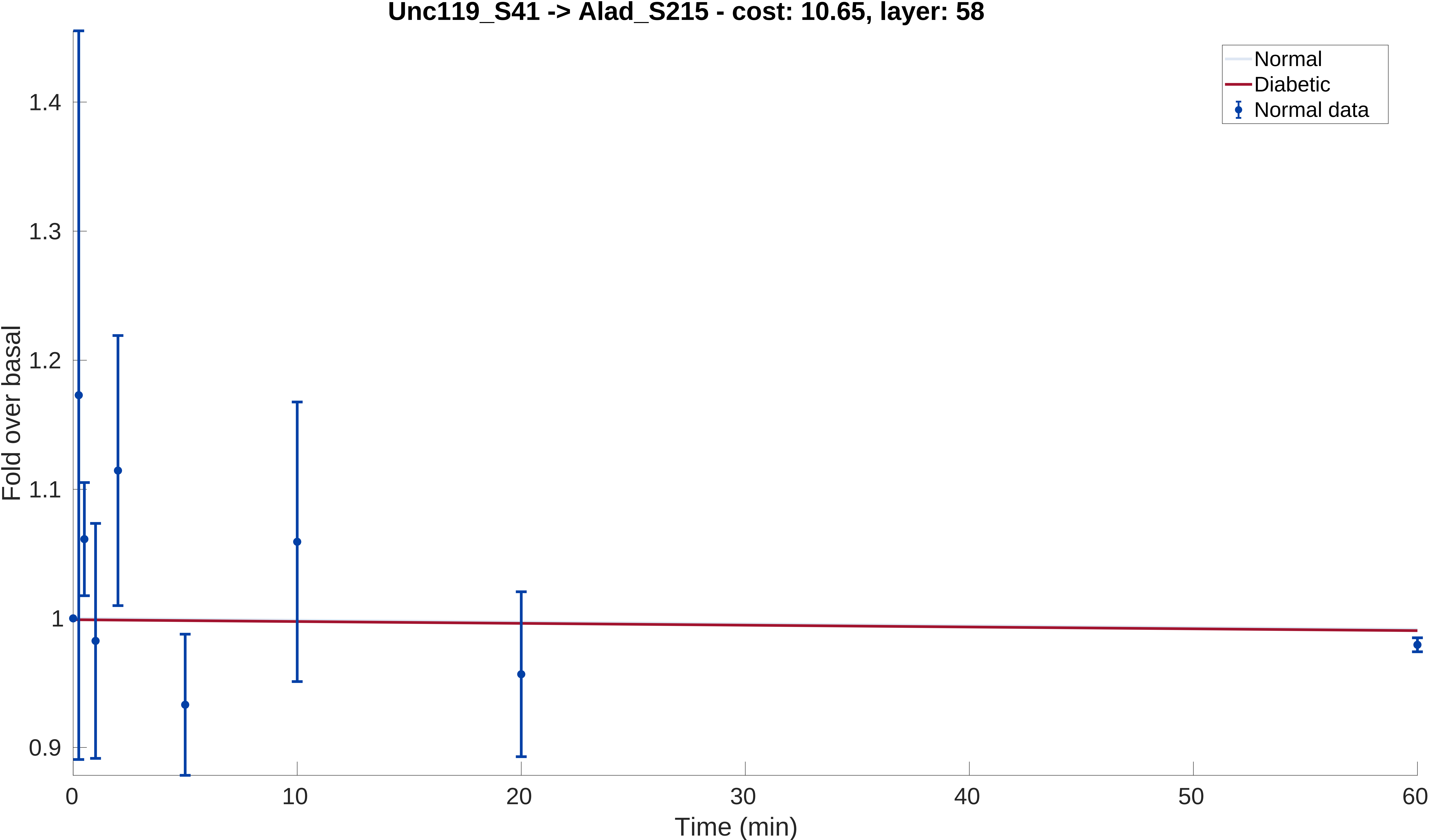

Unc119_S41 -> Alad_S215 - cost: 10.65, layer: 58
Normal
Diabetic
Normal data
1.4
1.3
1.2
1.1
1
0.9
Fold over basal
0
10
20
30
40
50
60
Time (min)

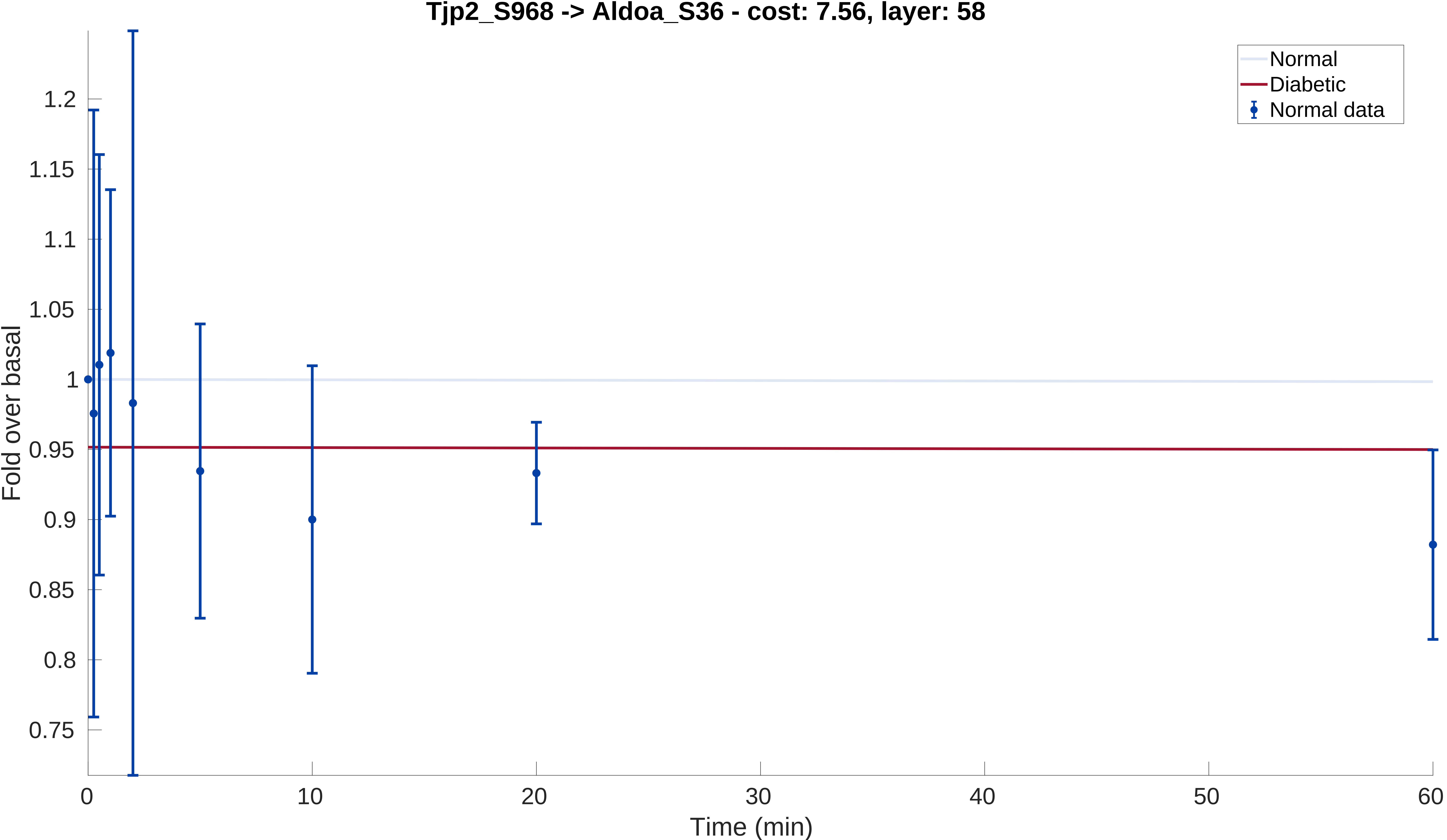

Tjp2_S968 -> Aldoa_S36 - cost: 7.56, layer: 58
Normal
Diabetic
Normal data
Fold over basal
1.2
1.15
1.1
1.05
1
0.95
0.9
0.85
0.8
0.75
0
10
20
30
40
50
60
Time (min)

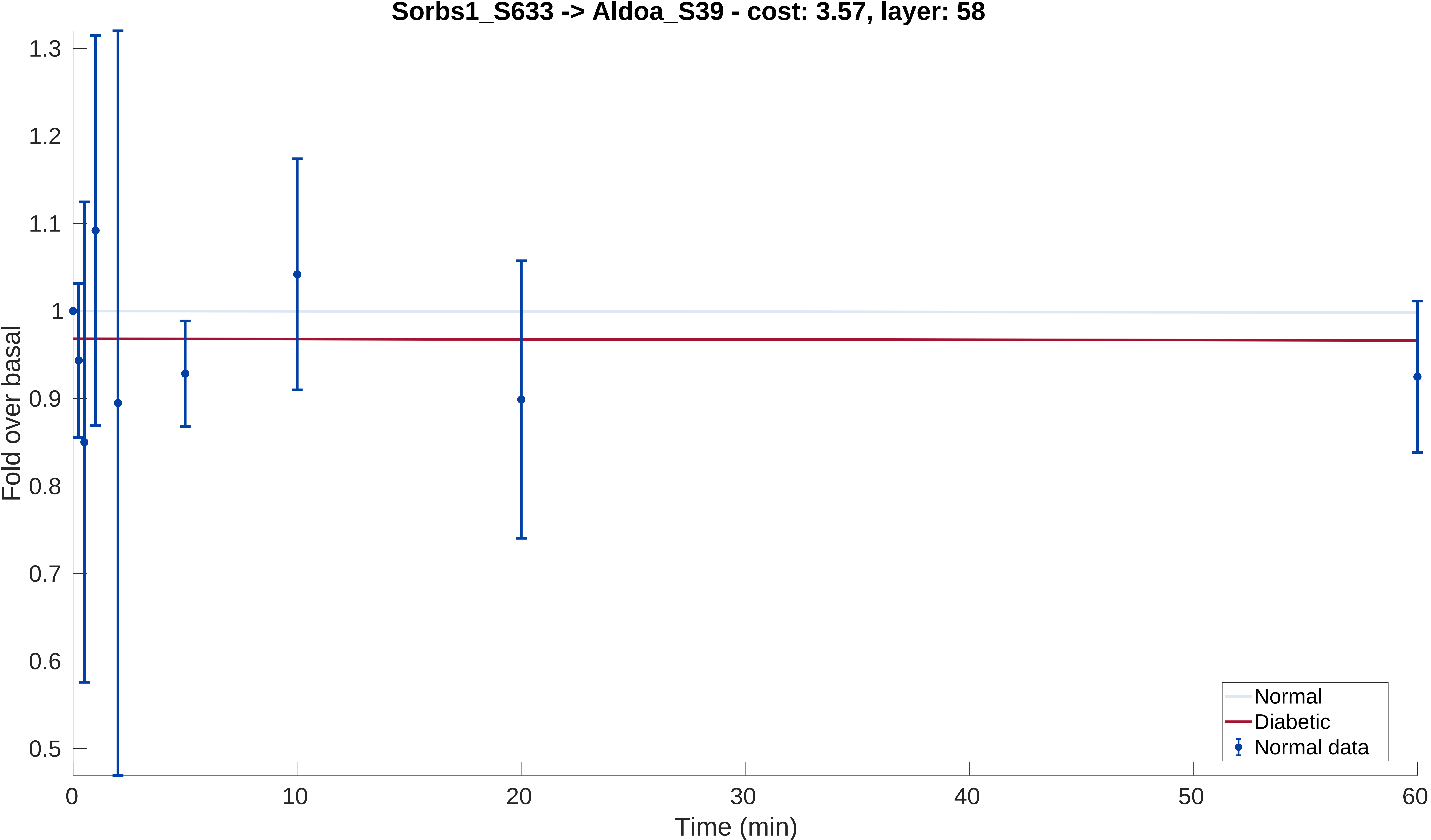

Sorbs1_S633 -> Aldoa_S39 - cost: 3.57, layer: 58
Fold over basal
Time (min)
1.3
1.2
1.1
1
0.9
0.8
0.7
0.6
0.5
0
10
20
30
40
50
60
Normal
Diabetic
Normal data

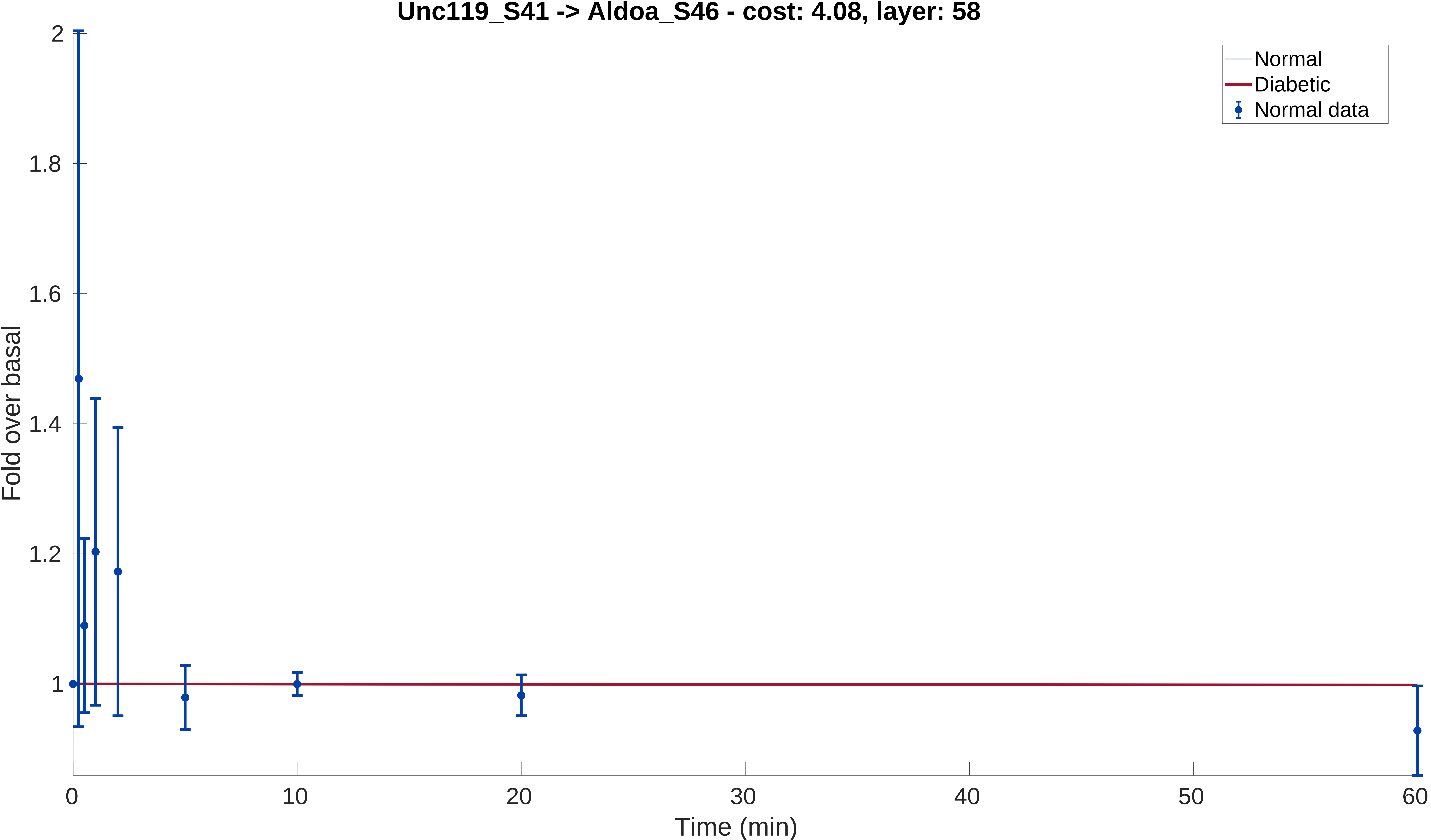
Unc119_S41 -> Aldoa_S46 - cost: 4.08, layer: 58
Normal
Diabetic
Normal data
Fold over basal
2
1.8
1.6
1.4
1.2
1
0
10
20
30
40
50
60
Time (min)

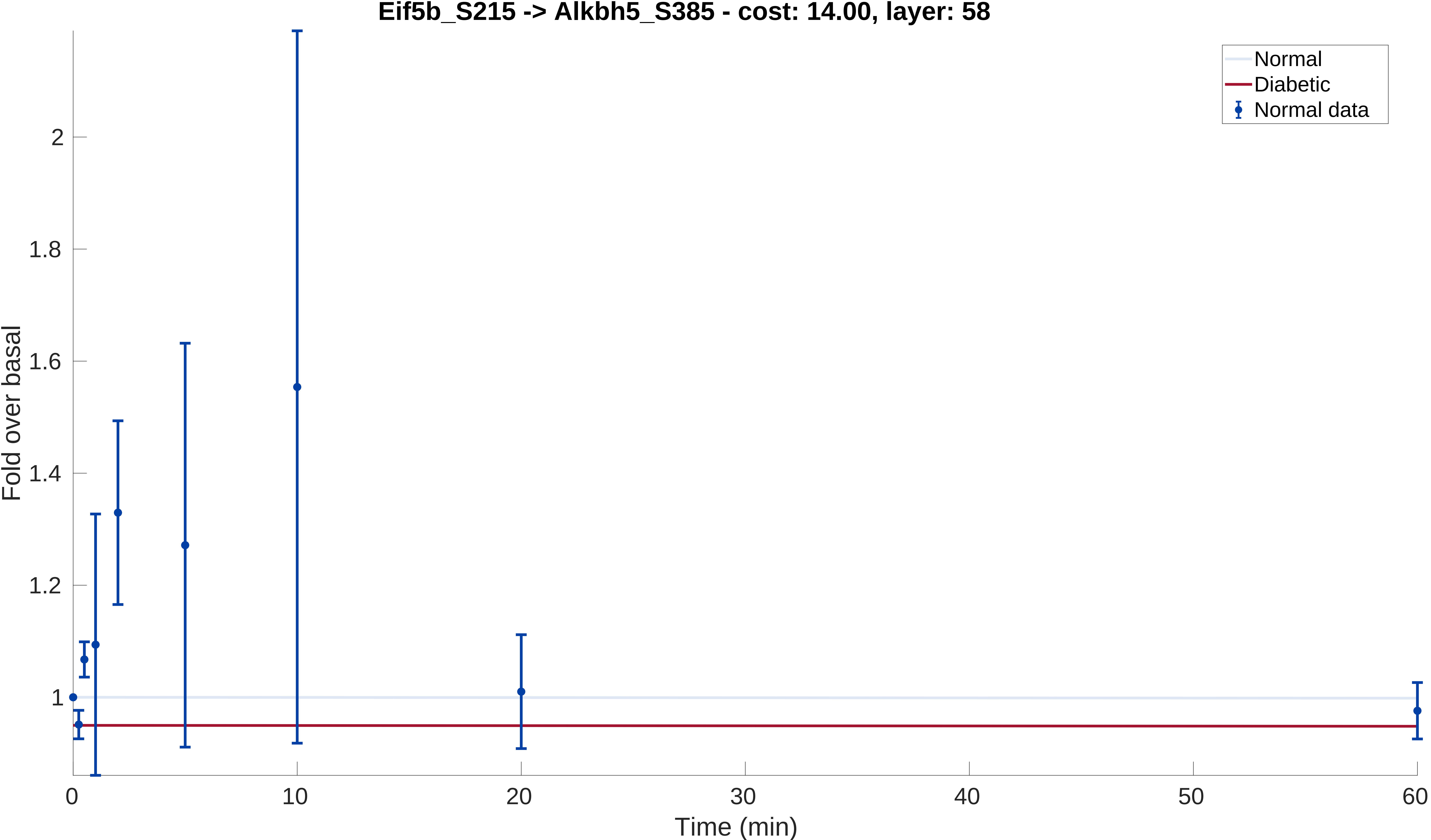

Eif5b_S215 -> Alkbh5_S385 - cost: 14.00, layer: 58
Normal
Diabetic
Normal data
Fold over basal
2
1.8
1.6
1.4
1.2
1
0
10
20
30
40
50
60
Time (min)

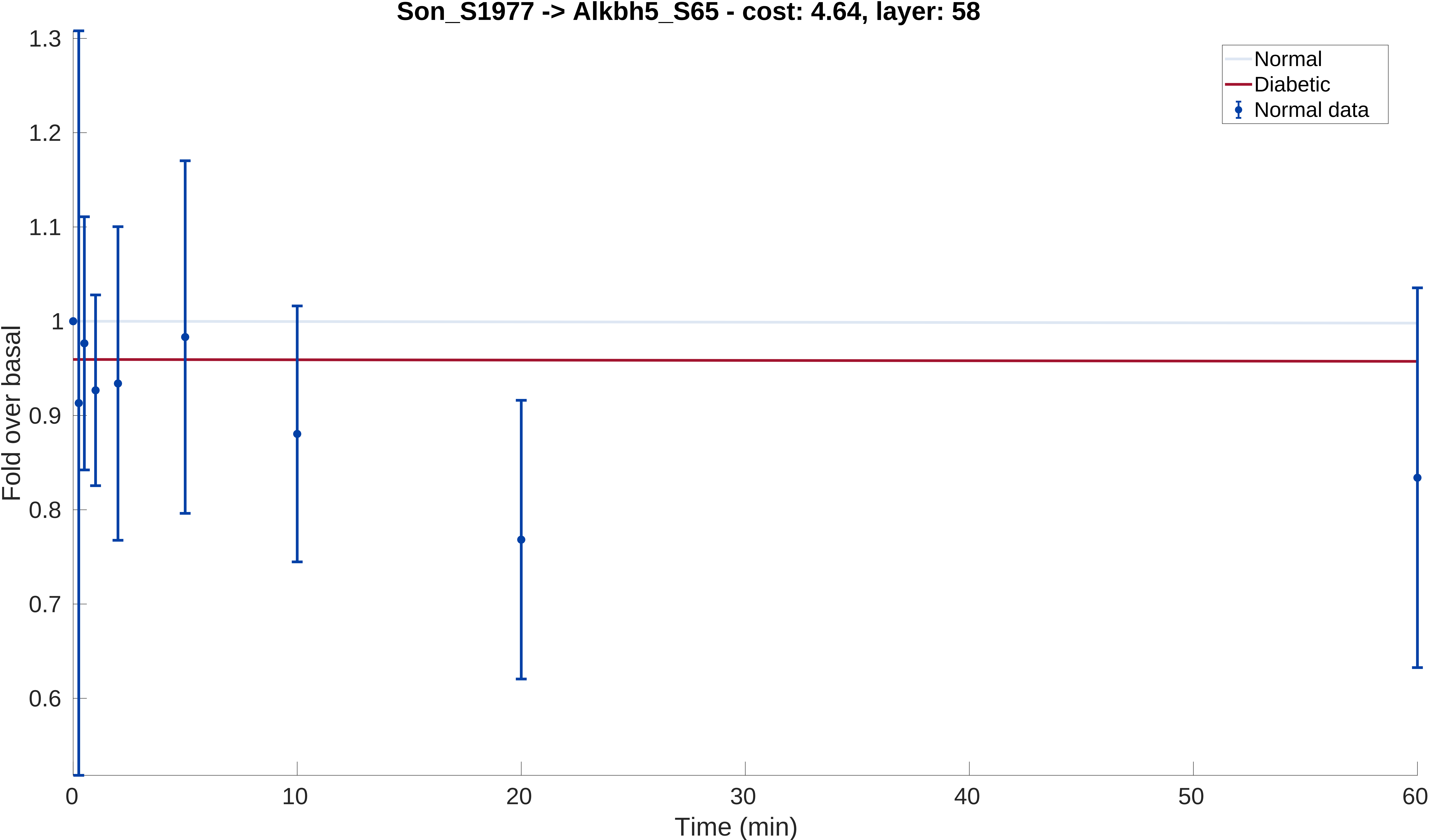

Son_S1977 -> Alkbh5_S65 - cost: 4.64, layer: 58
Normal
Diabetic
Normal data
Fold over basal
1.3
1.2
1.1
1
0.9
0.8
0.7
0.6
0
10
20
30
40
50
60
Time (min)

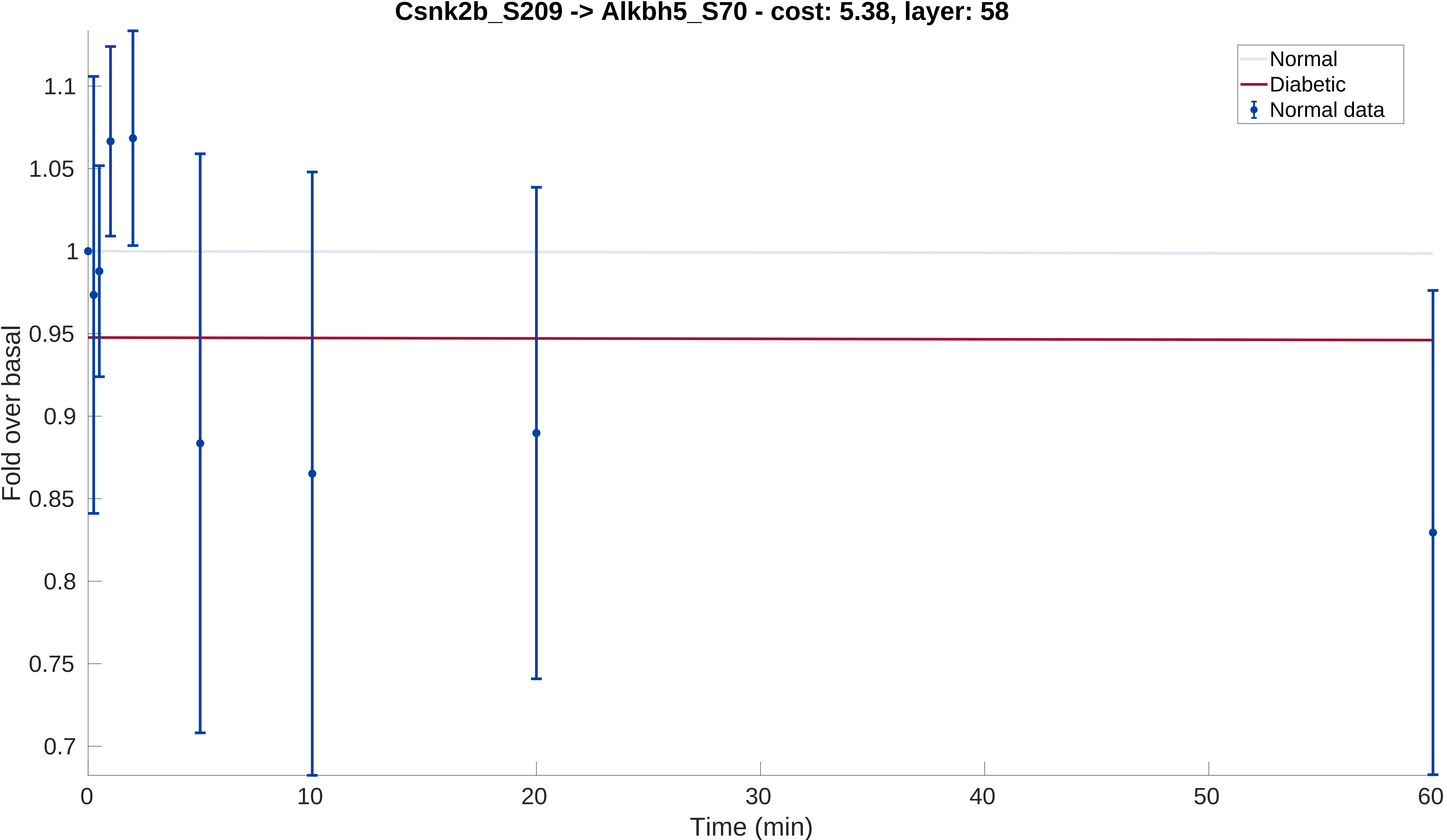

Csnk2b_S209 -> Alkbh5_S70 - cost: 5.38, layer: 58
Normal
Diabetic
Normal data
Fold over basal
Time (min)
1.1
1.05
1
0.95
0.9
0.85
0.8
0.75
0.7
0
10
20
30
40
50
60

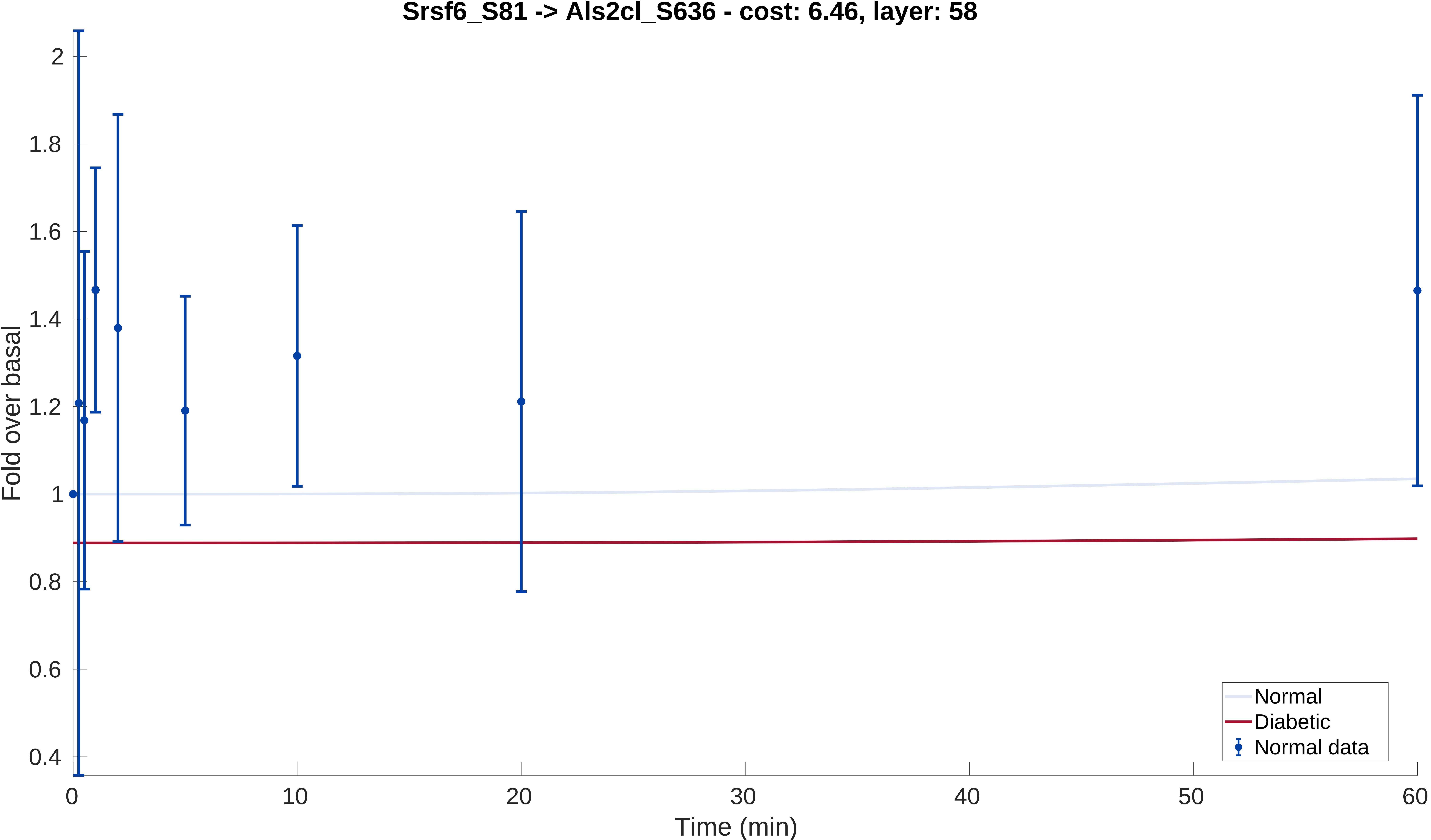

Srsf6_S81 -> Als2cl_S636 - cost: 6.46, layer: 58
Fold over basal
Time (min)
0.4
0.6
0.8
1
1.2
1.4
1.6
1.8
2
0
10
20
30
40
50
60
Normal
Diabetic
Normal data

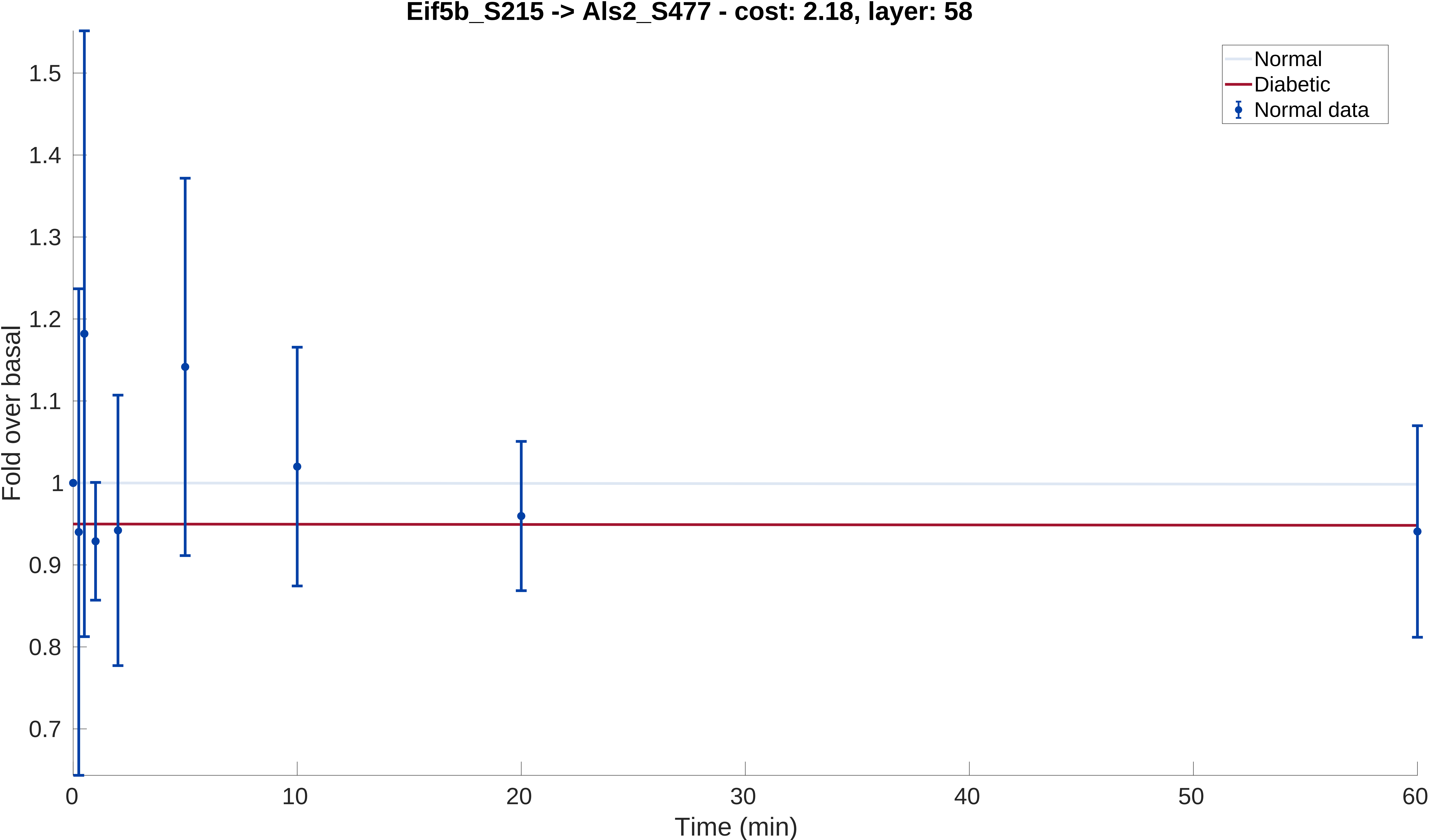

Eif5b_S215 -> Als2_S477 - cost: 2.18, layer: 58
Normal
Diabetic
Normal data
Fold over basal
1.5
1.4
1.3
1.2
1.1
1
0.9
0.8
0.7
0
10
20
30
40
50
60
Time (min)

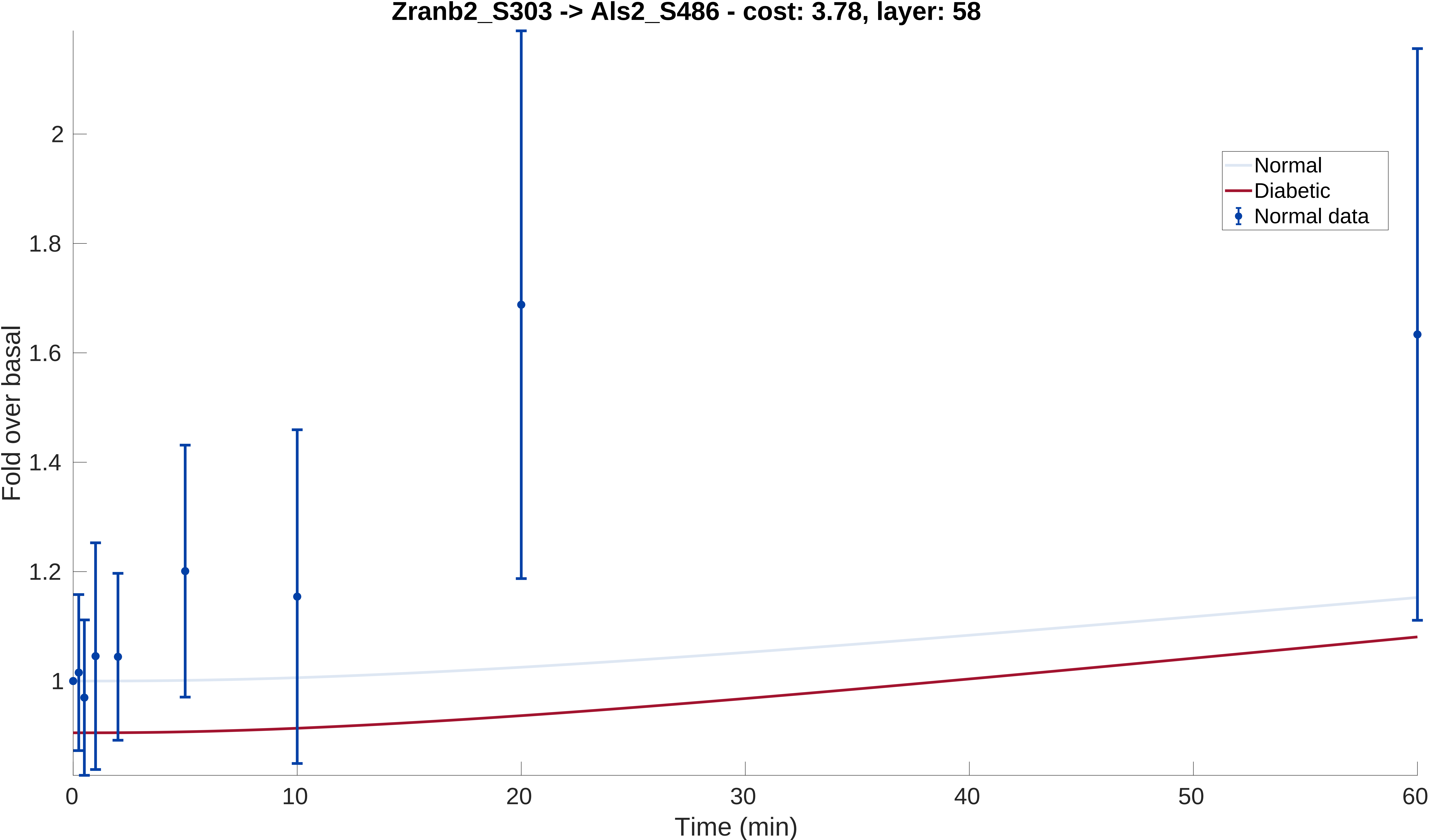
Zranb2_S303 -> Als2_S486 - cost: 3.78, layer: 58
Fold over basal
Time (min)
Normal
Diabetic
Normal data
1
1.2
1.4
1.6
1.8
2
0
10
20
30
40
50
60

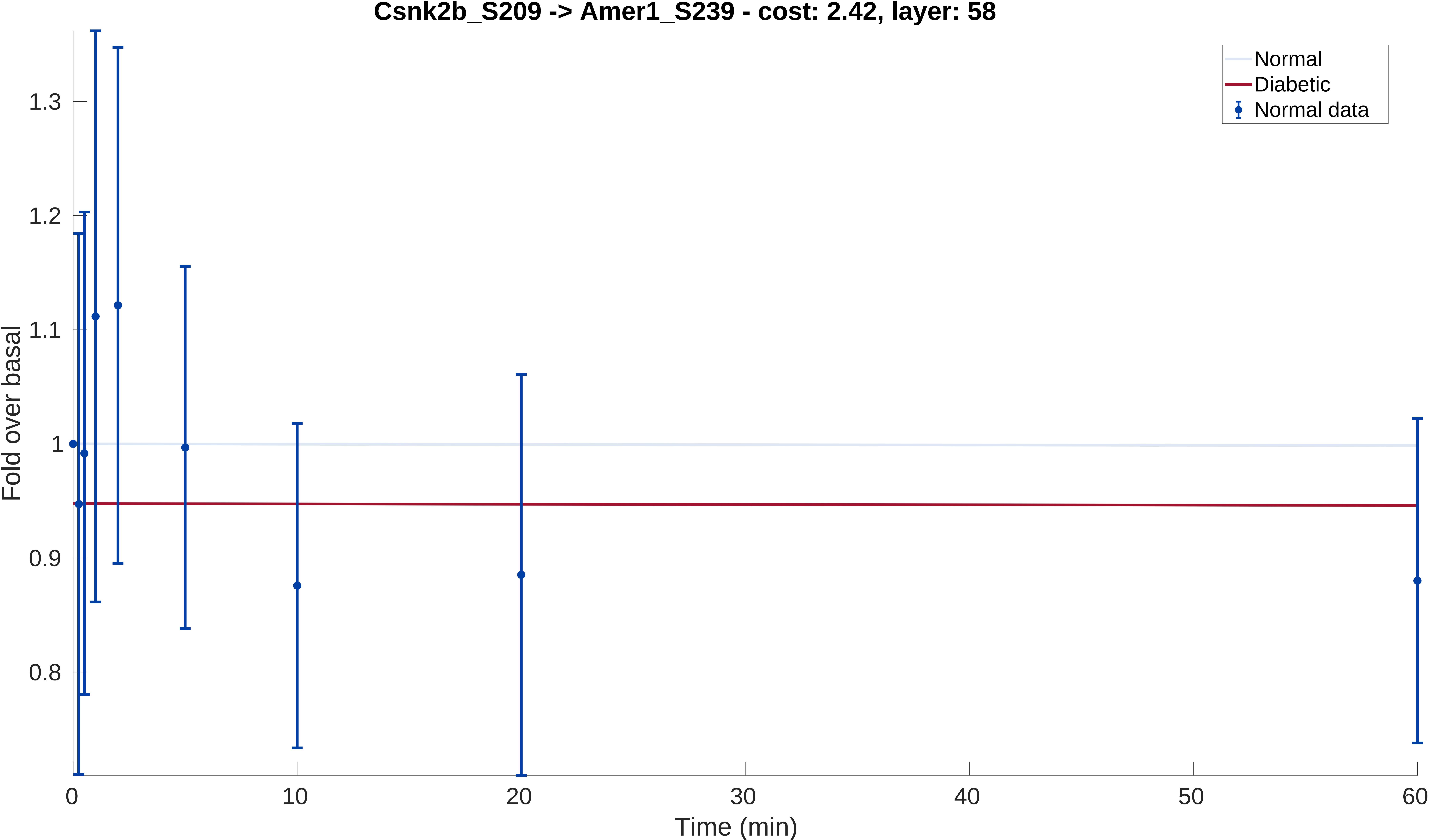
Csnk2b_S209 -> Amer1_S239 - cost: 2.42, layer: 58
Normal
Diabetic
Normal data
Fold over basal
1.3
1.2
1.1
1
0.9
0.8
0
10
20
30
40
50
60
Time (min)

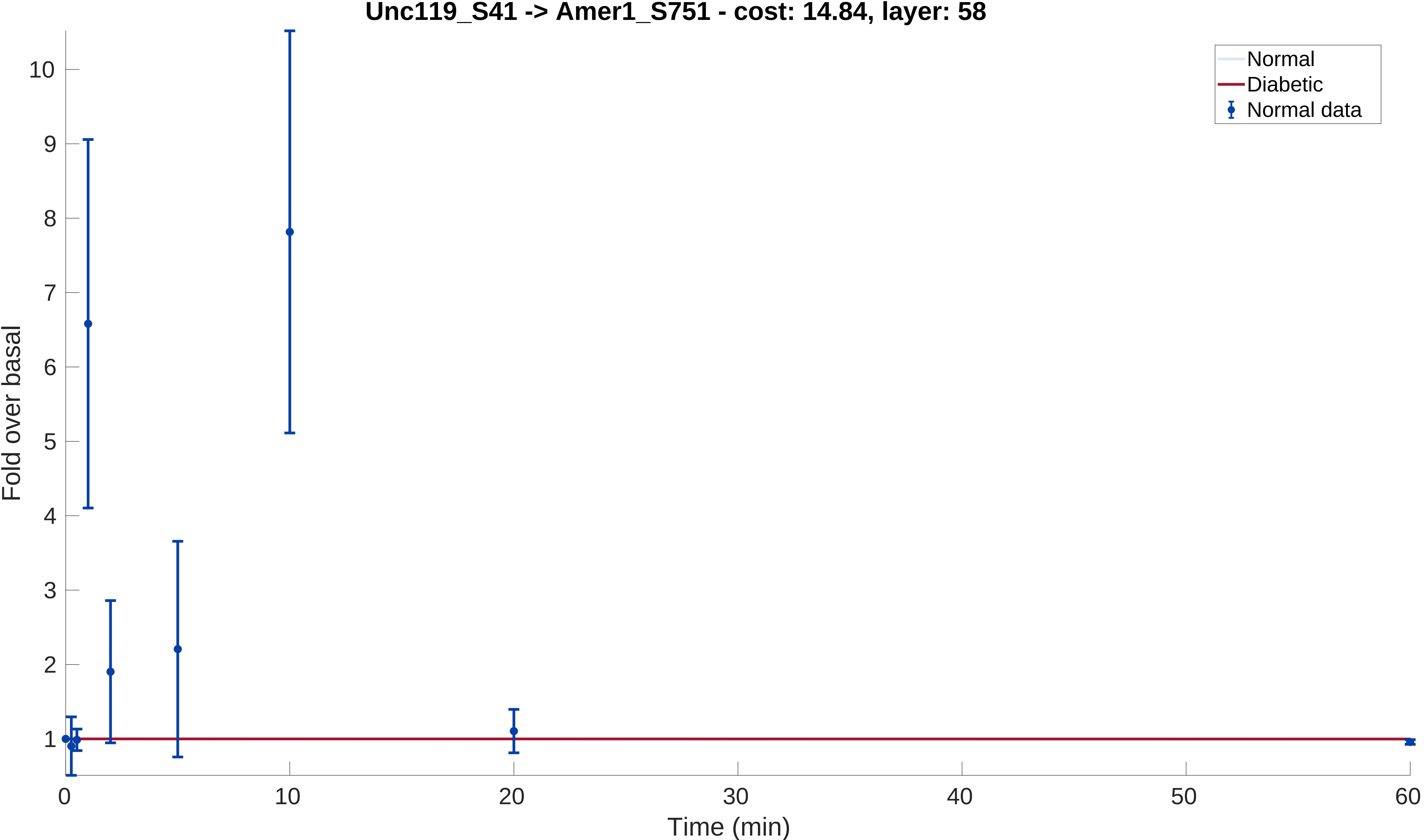

Unc119_S41 -> Amer1_S751 - cost: 14.84, layer: 58
Normal
Diabetic
Normal data
Fold over basal
Time (min)
0
10
20
30
40
50
60
1
2
3
4
5
6
7
8
9
10

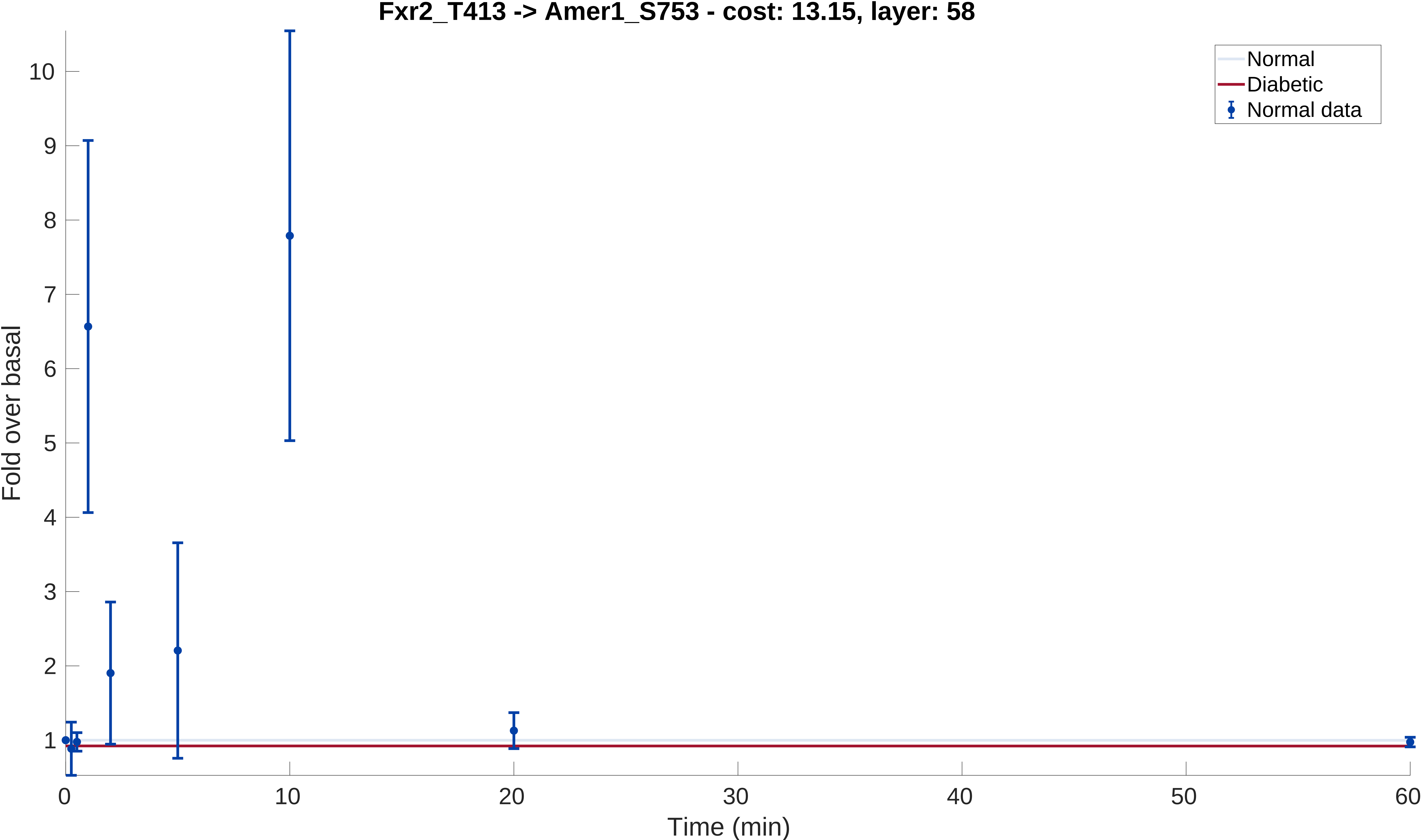

Fxr2_T413 -> Amer1_S753 - cost: 13.15, layer: 58
Normal
Diabetic
Normal data
Fold over basal
Time (min)
0
10
20
30
40
50
60
1
2
3
4
5
6
7
8
9
10

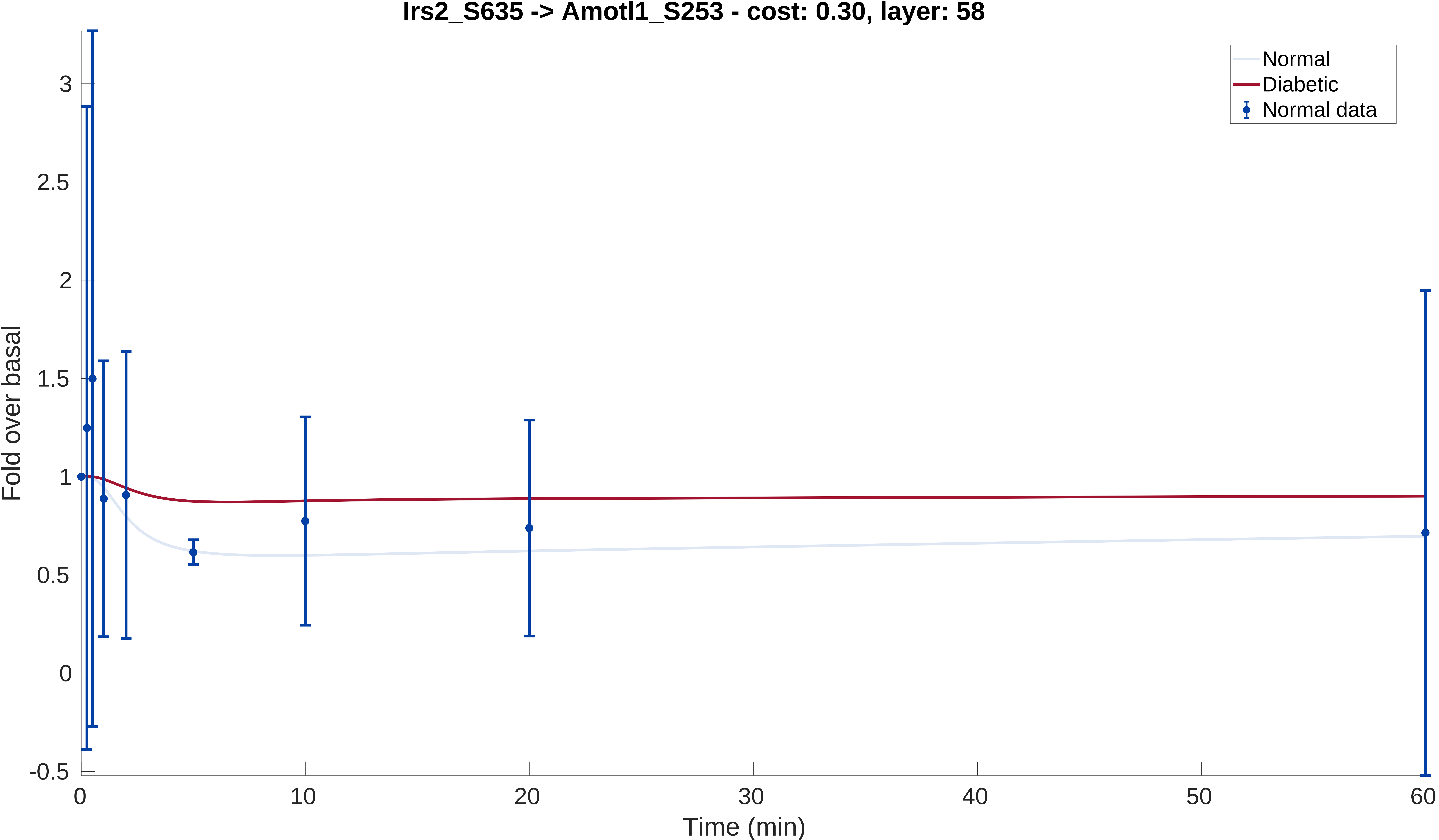

Irs2_S635 -> Amotl1_S253 - cost: 0.30, layer: 58
Normal
Diabetic
Normal data
Fold over basal
Time (min)
3
2.5
2
1.5
1
0.5
0
-0.5
0
10
20
30
40
50
60

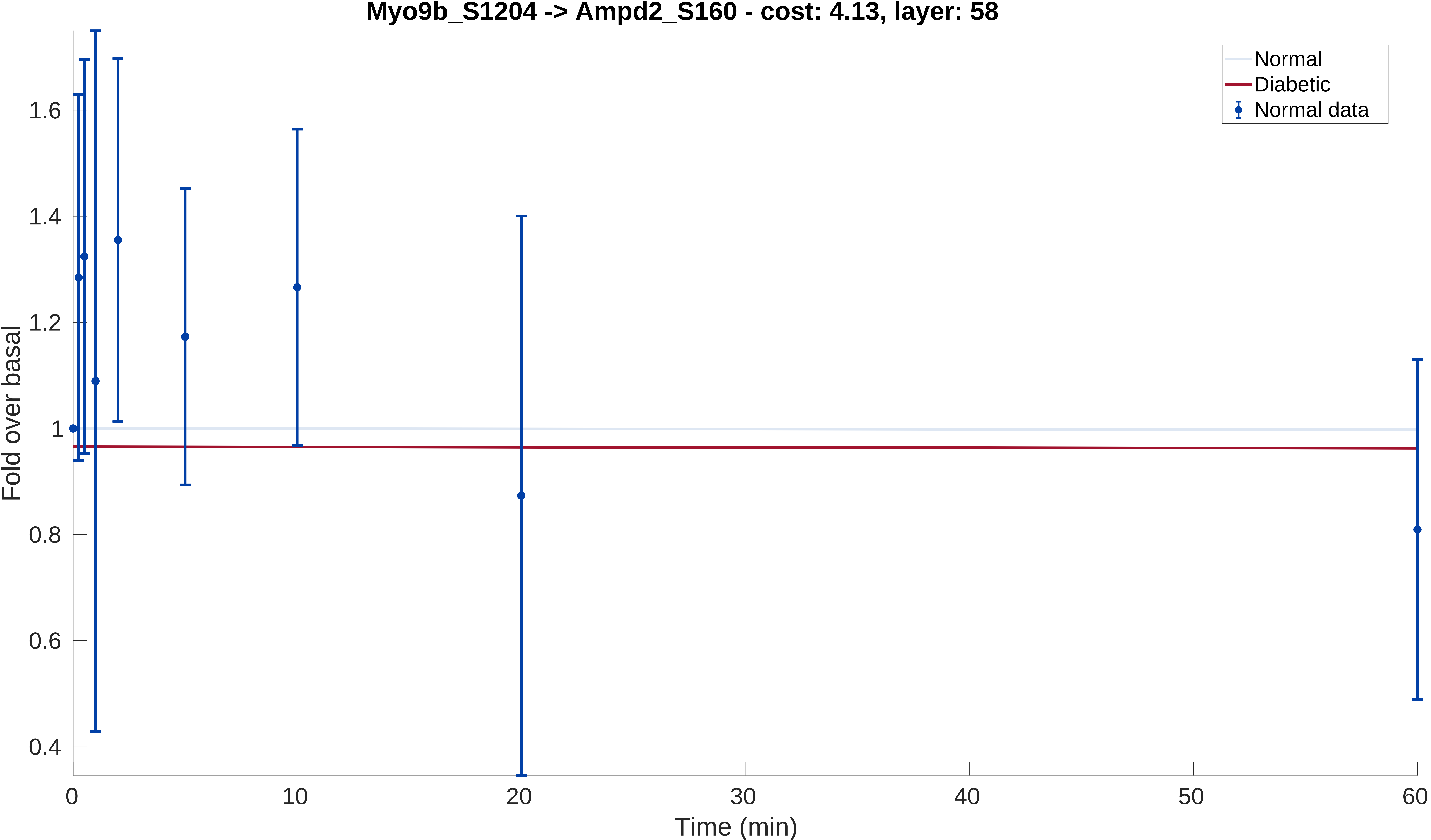

Myo9b_S1204 -> Ampd2_S160 - cost: 4.13, layer: 58
Normal
Diabetic
Normal data
Fold over basal
Time (min)
0.4
0.6
0.8
1
1.2
1.4
1.6
0
10
20
30
40
50
60

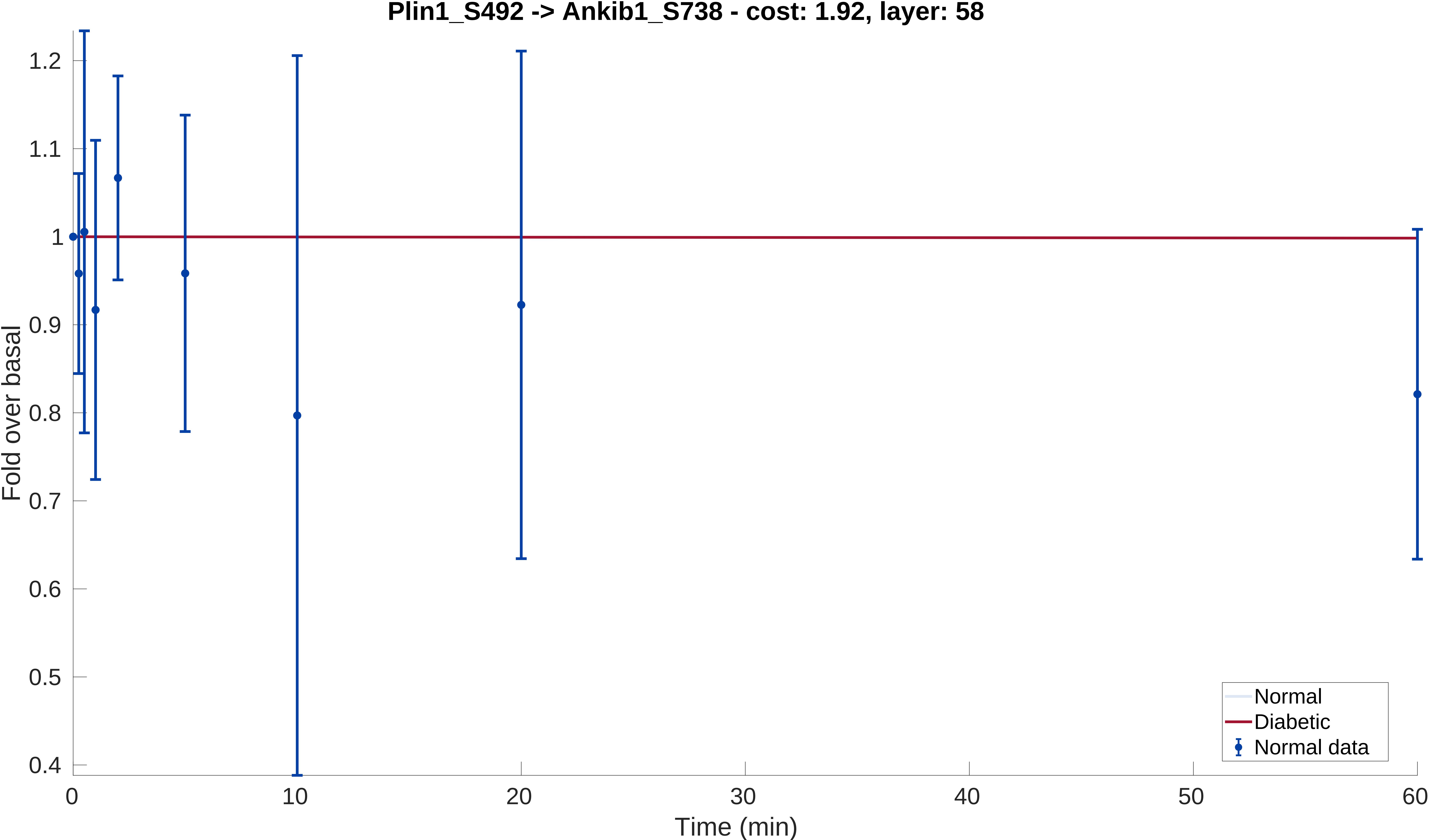

Plin1_S492 -> Ankib1_S738 - cost: 1.92, layer: 58
Fold over basal
1.2
1.1
1
0.9
0.8
0.7
0.6
0.5
0.4
0
10
20
30
40
50
60
Time (min)
Normal
Diabetic
Normal data

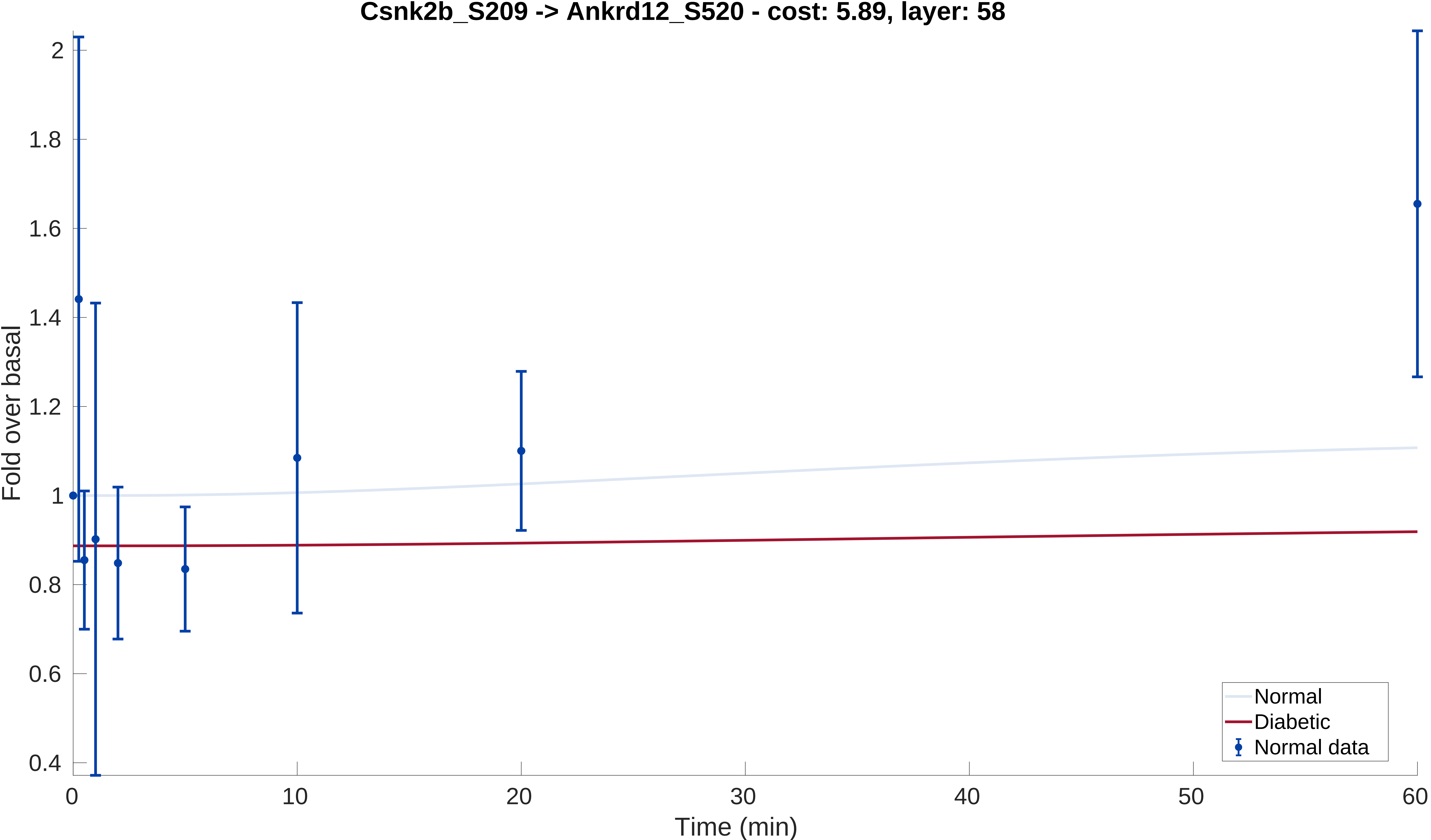

Csnk2b_S209 -> Ankrd12_S520 - cost: 5.89, layer: 58
Fold over basal
Time (min)
2
1.8
1.6
1.4
1.2
1
0.8
0.6
0.4
0
10
20
30
40
50
60
Normal
Diabetic
Normal data

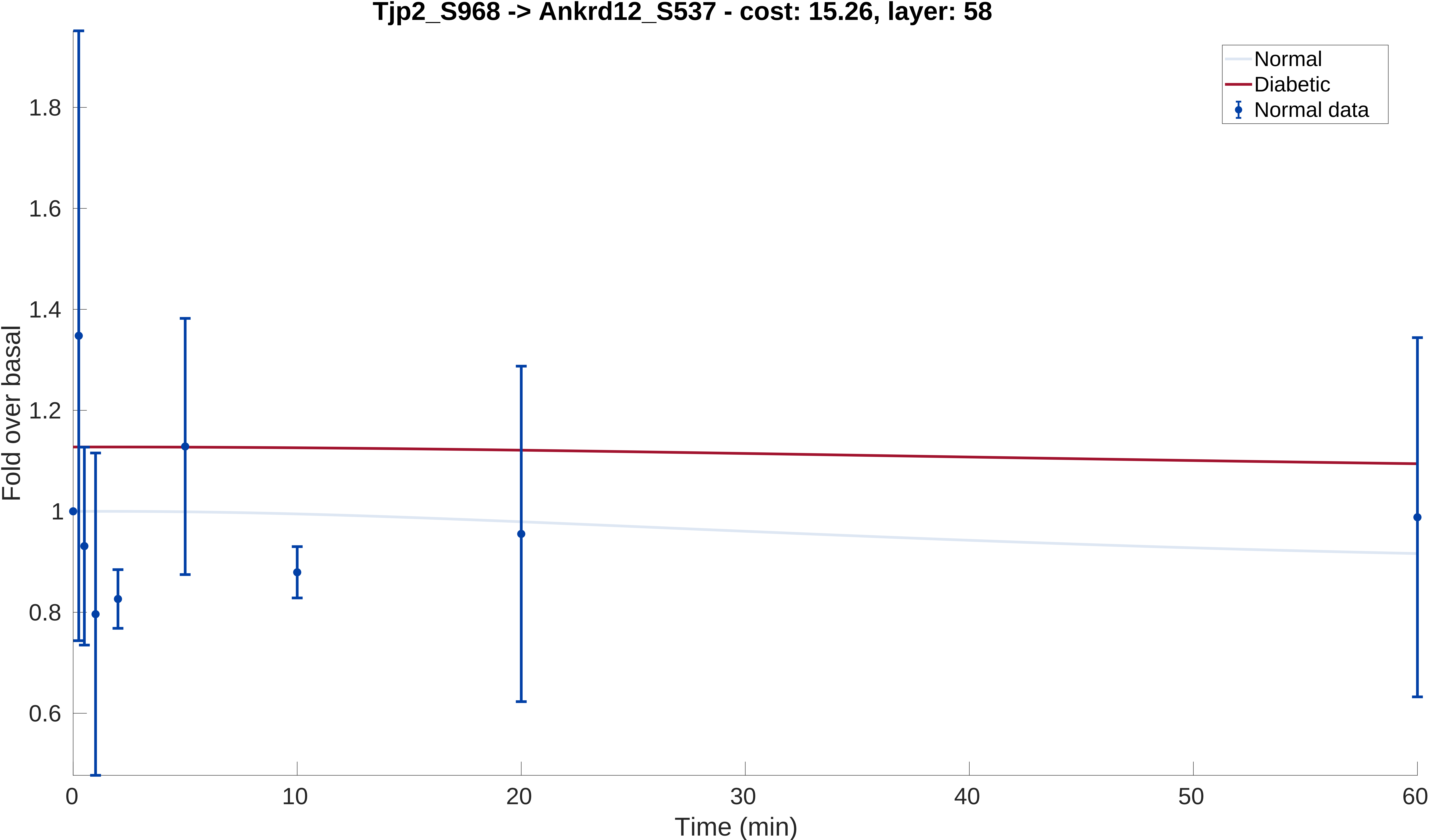

Tjp2_S968 -> Ankrd12_S537 - cost: 15.26, layer: 58
Normal
Diabetic
Normal data
Fold over basal
1.8
1.6
1.4
1.2
1
0.8
0.6
0
10
20
30
40
50
60
Time (min)

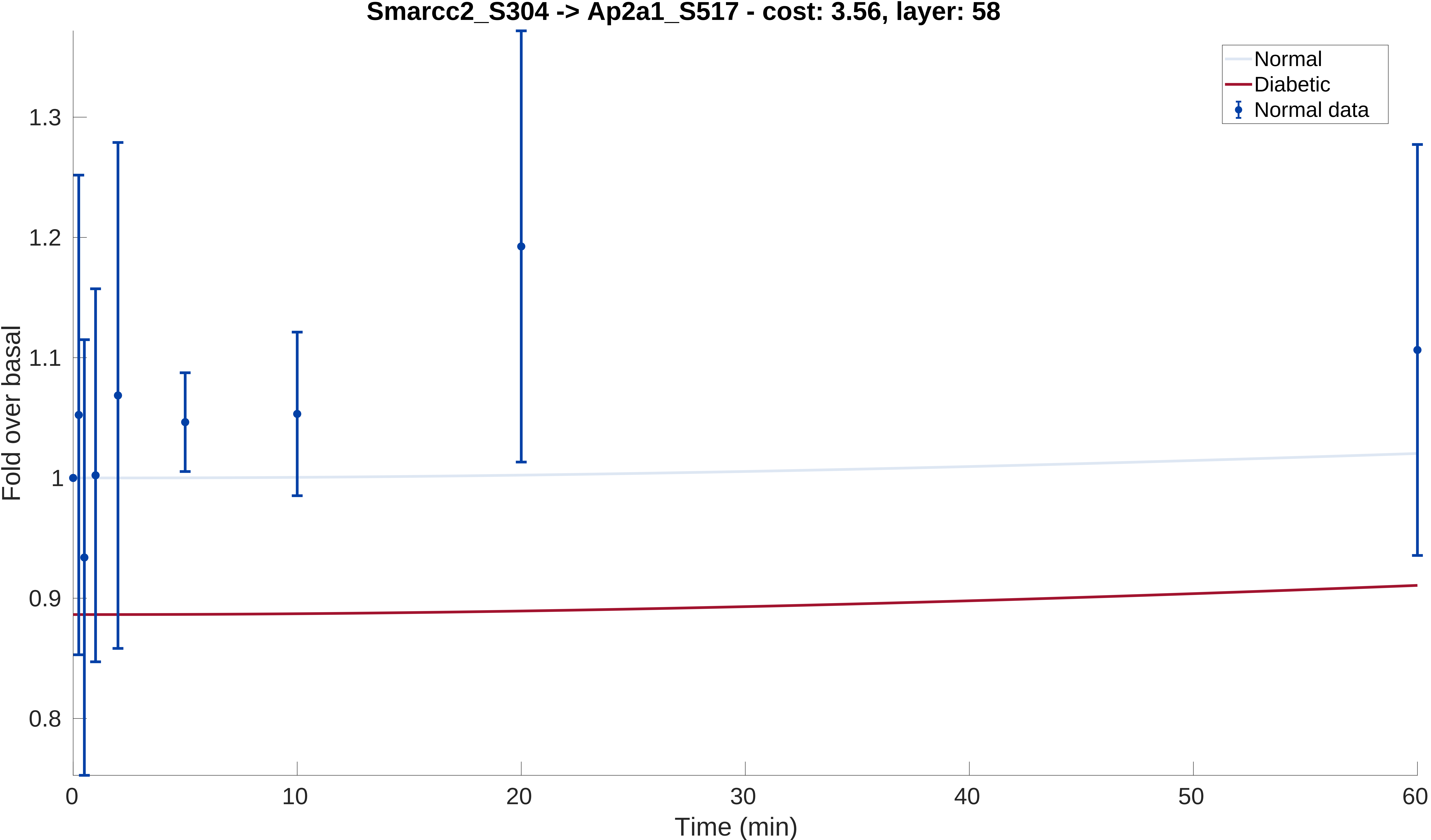
Smarcc2_S304 -> Ap2a1_S517 - cost: 3.56, layer: 58
Normal
Diabetic
Normal data
Fold over basal
Time (min)
1.3
1.2
1.1
1
0.9
0.8
0
10
20
30
40
50
60

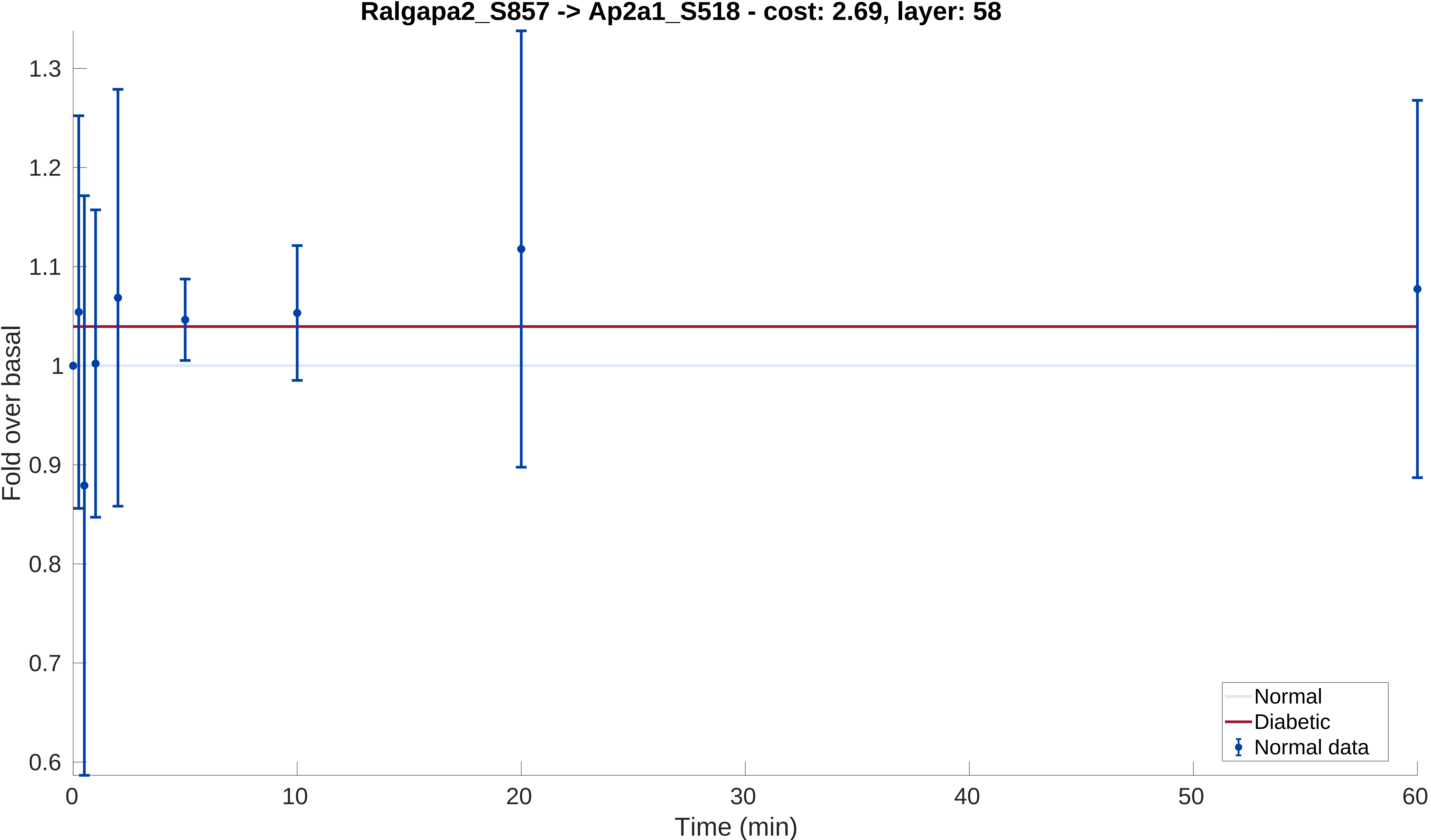

Ralgapa2_S857 -> Ap2a1_S518 - cost: 2.69, layer: 58
Fold over basal
1.3
1.2
1.1
1
0.9
0.8
0.7
0.6
0
10
20
30
40
50
60
Time (min)
Normal
Diabetic
Normal data

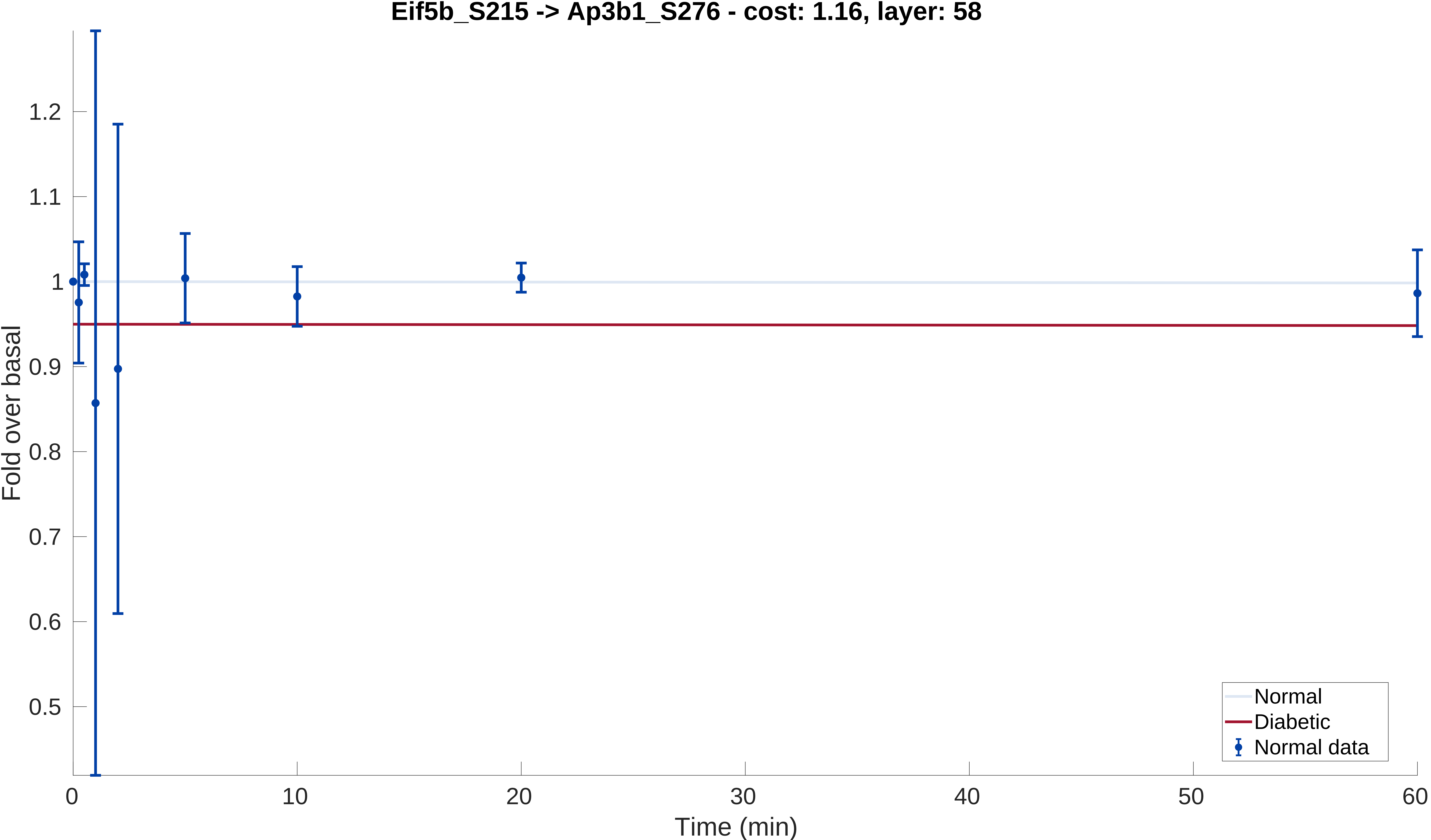

Eif5b_S215 -> Ap3b1_S276 - cost: 1.16, layer: 58
Fold over basal
1.2
1.1
1
0.9
0.8
0.7
0.6
0.5
0
10
20
30
40
50
60
Time (min)
Normal
Diabetic
Normal data

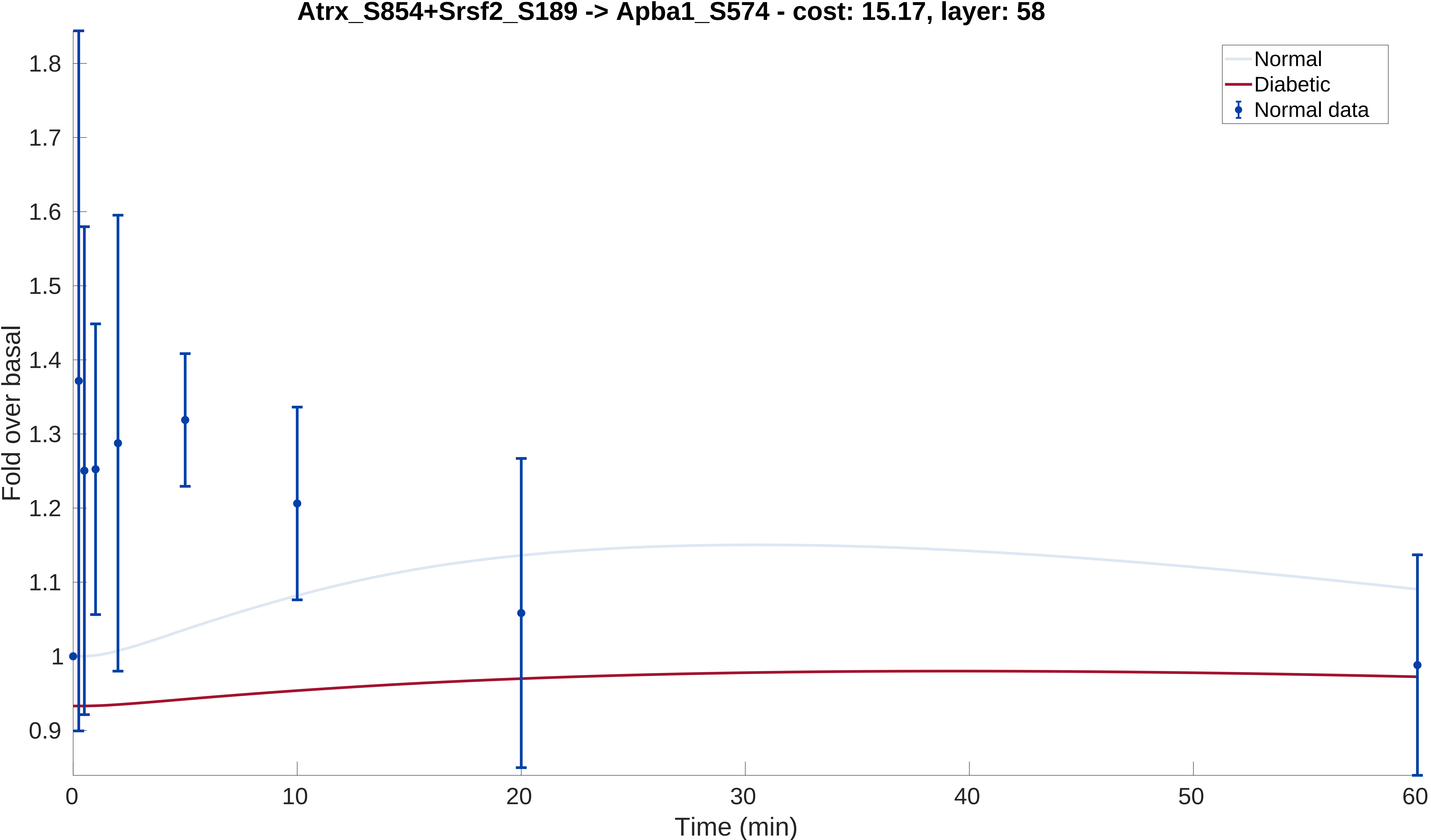

Atrx_S854+Srsf2_S189 -> Apba1_S574 - cost: 15.17, layer: 58
Normal
Diabetic
Normal data
Fold over basal
1.8
1.7
1.6
1.5
1.4
1.3
1.2
1.1
1
0.9
0
10
20
30
40
50
60
Time (min)

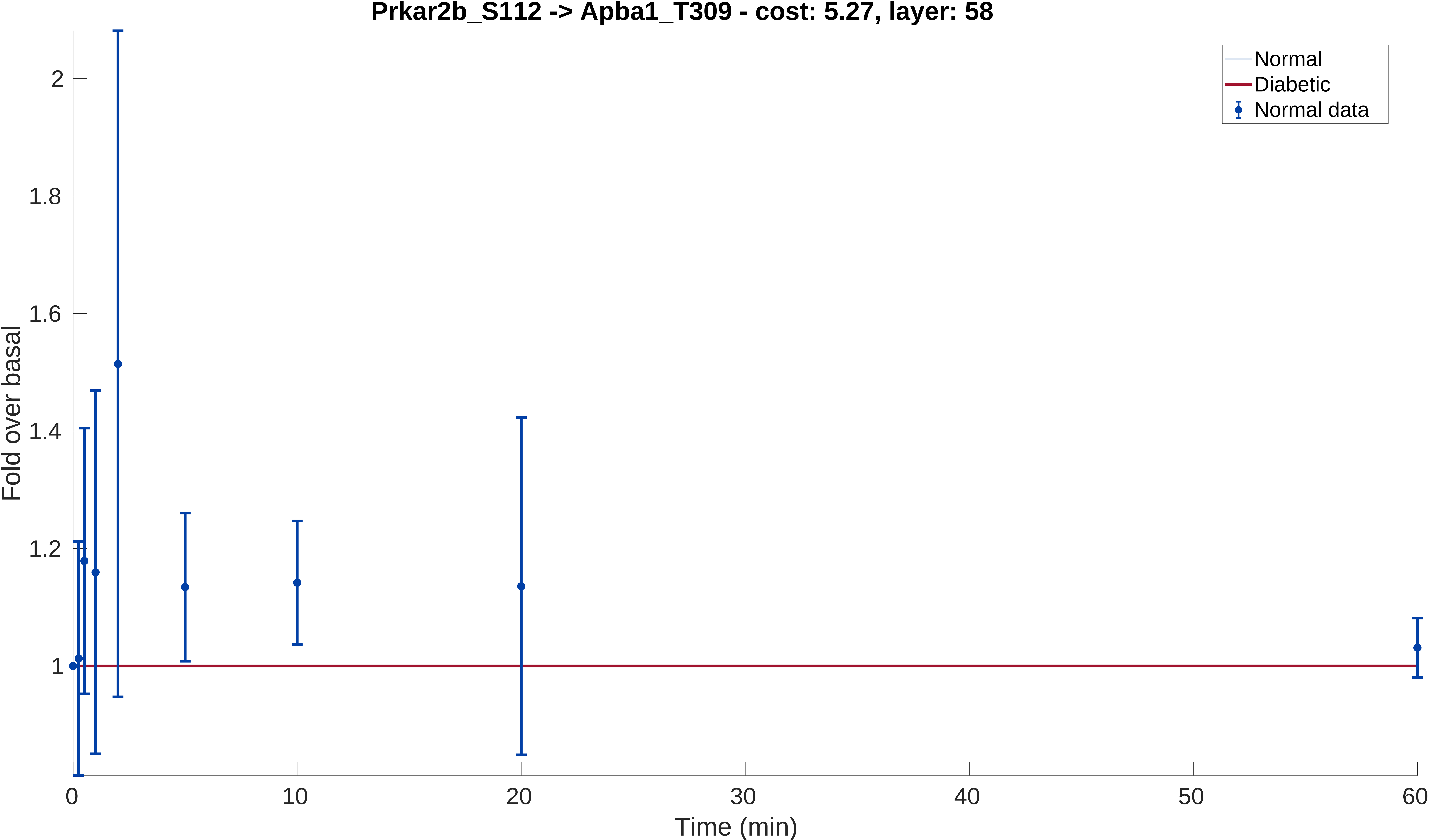
Prkar2b_S112 -> Apba1_T309 - cost: 5.27, layer: 58
Normal
Diabetic
Normal data
Fold over basal
2
1.8
1.6
1.4
1.2
1
0
10
20
30
40
50
60
Time (min)

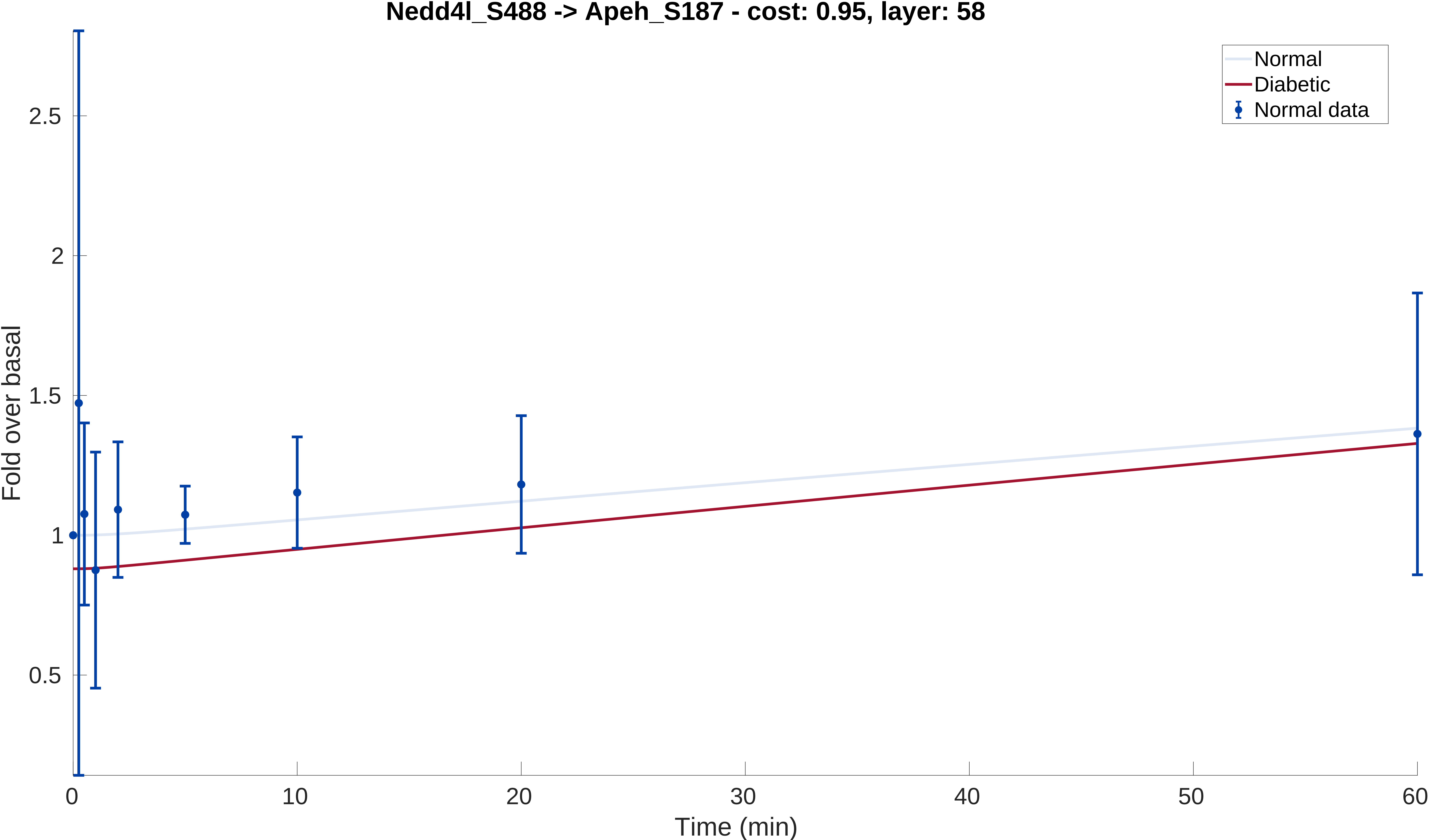

Nedd4l_S488 -> Apeh_S187 - cost: 0.95, layer: 58
Normal
Diabetic
Normal data
Fold over basal
2.5
2
1.5
1
0.5
0
10
20
30
40
50
60
Time (min)

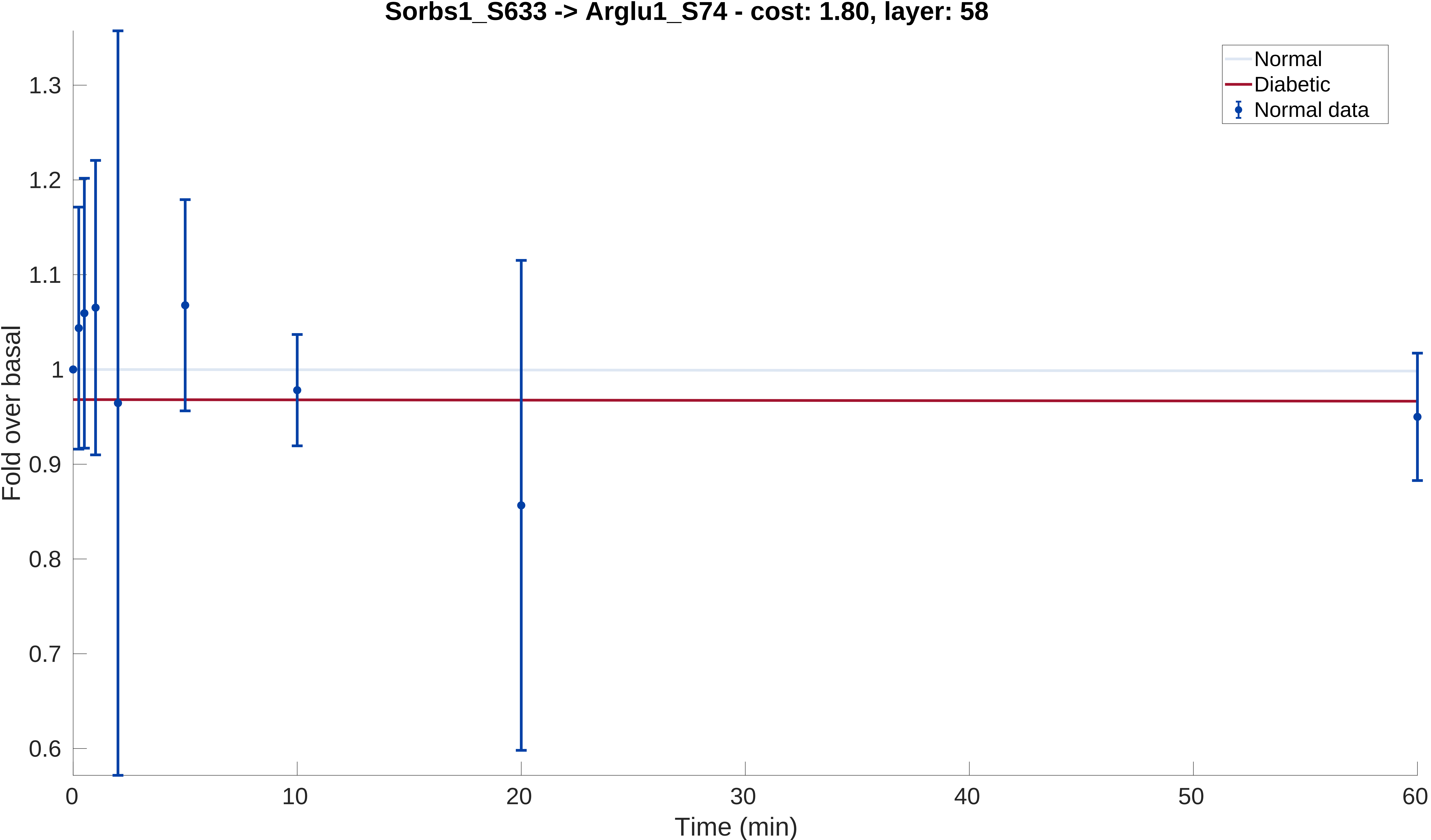

Sorbs1_S633 -> Arglu1_S74 - cost: 1.80, layer: 58
Normal
Diabetic
Normal data
Fold over basal
1.3
1.2
1.1
1
0.9
0.8
0.7
0.6
0
10
20
30
40
50
60
Time (min)

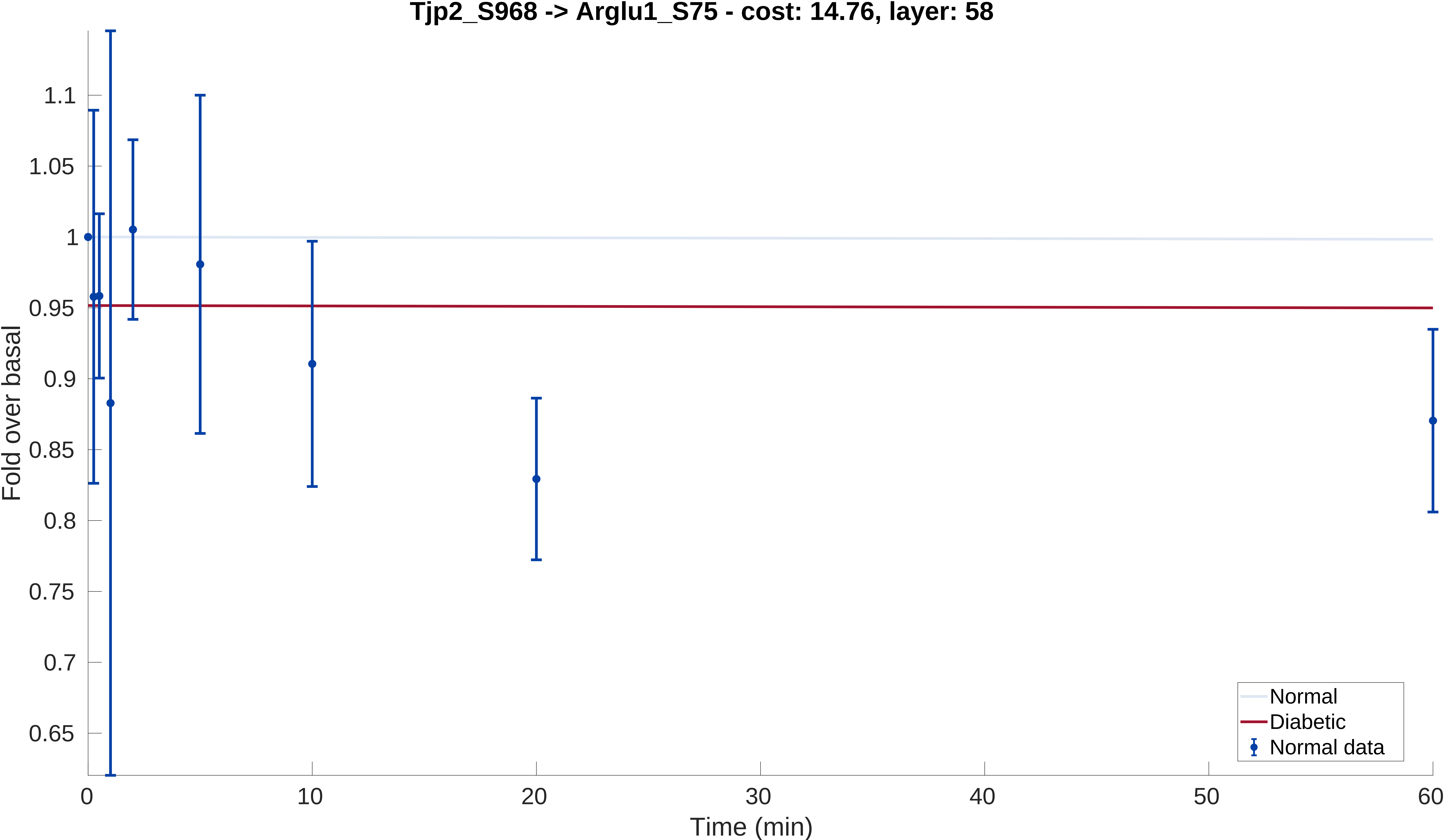
Tjp2_S968 -> Arglu1_S75 - cost: 14.76, layer: 58
Fold over basal
1.1
1.05
1
0.95
0.9
0.85
0.8
0.75
0.7
0.65
0
10
20
30
40
50
60
Time (min)
Normal
Diabetic
Normal data

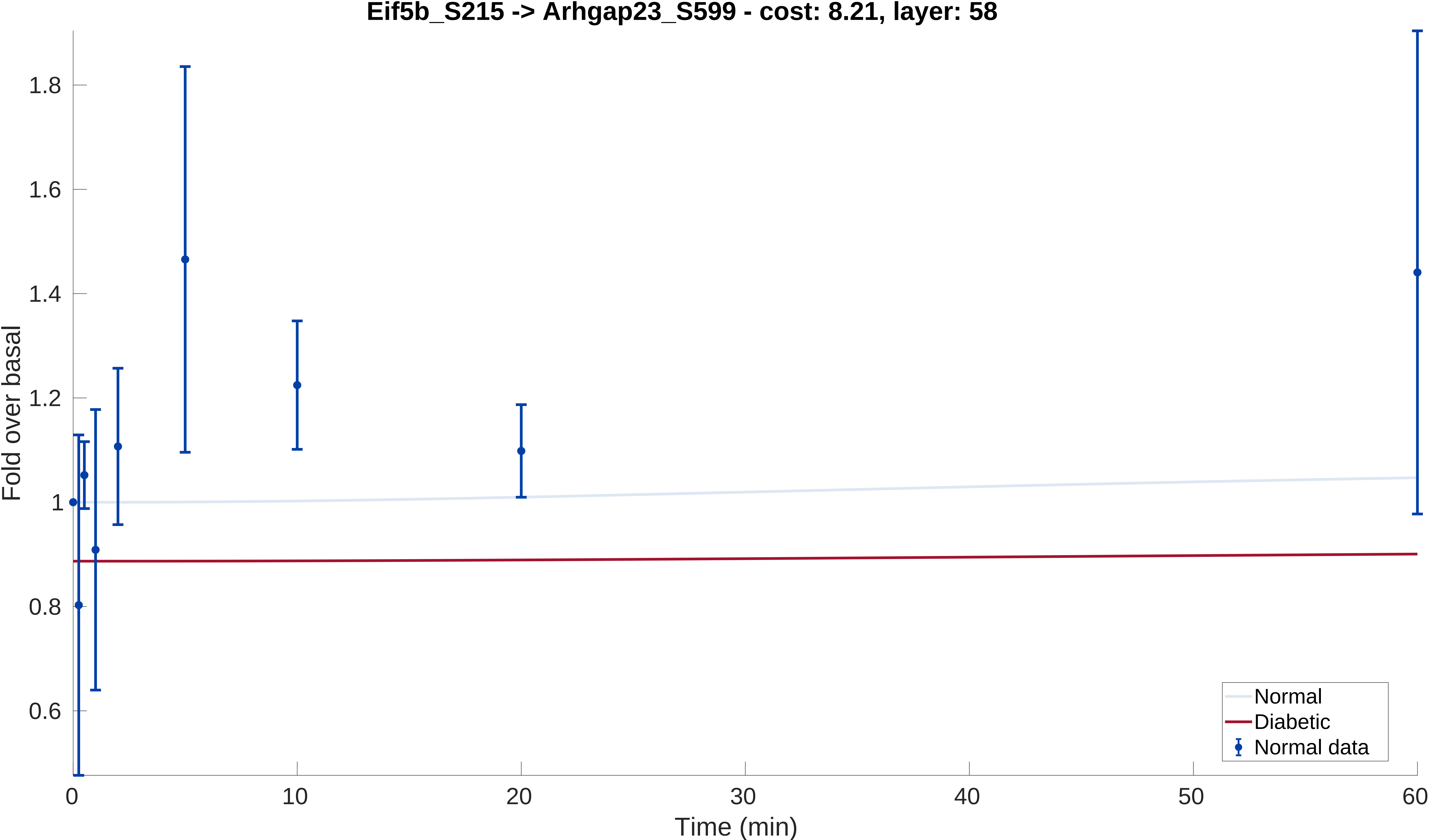
Eif5b_S215 -> Arhgap23_S599 - cost: 8.21, layer: 58
Fold over basal
1.8
1.6
1.4
1.2
1
0.8
0.6
0
10
20
30
40
50
60
Time (min)
Normal
Diabetic
Normal data

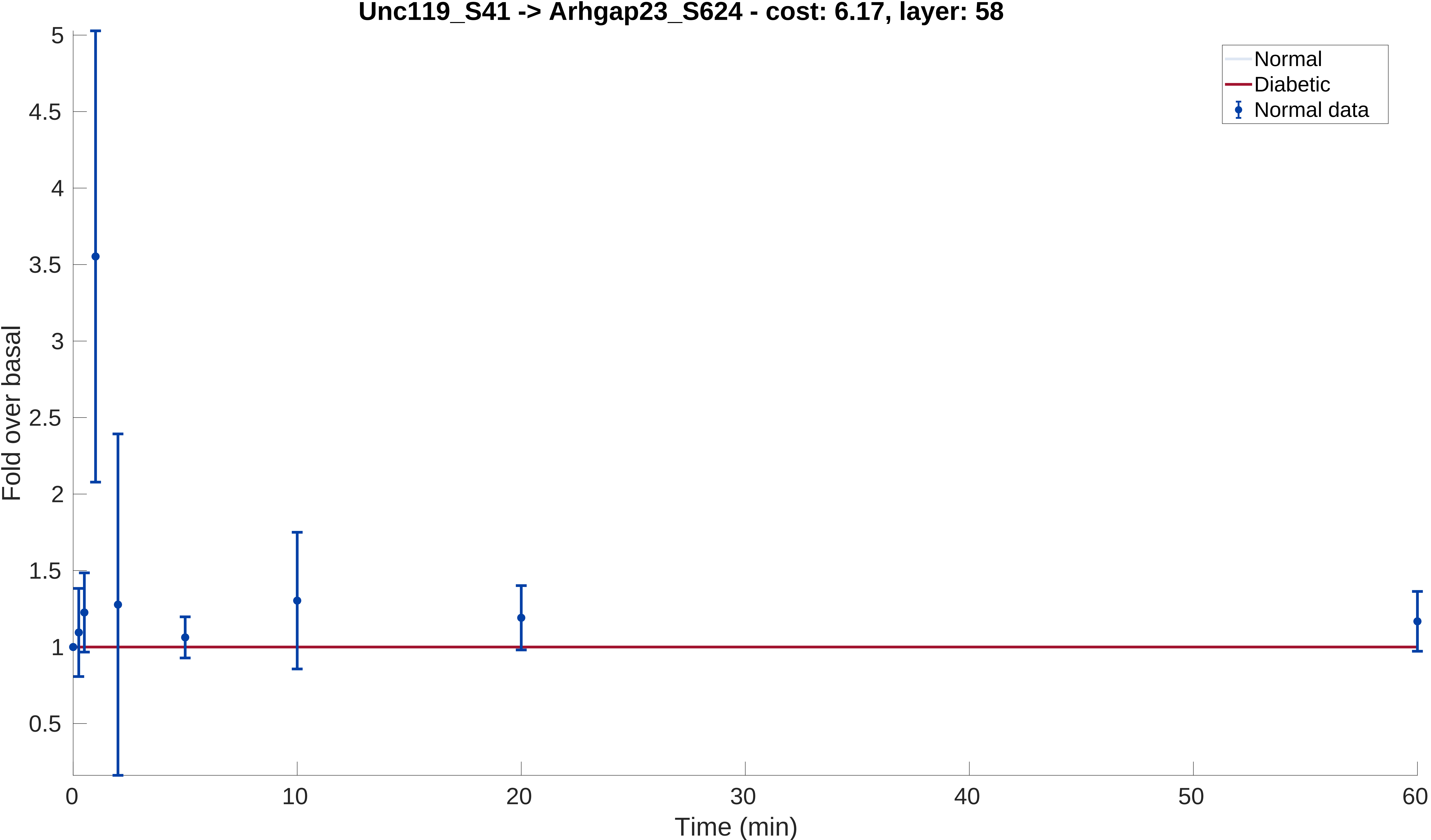
Unc119_S41 -> Arhgap23_S624 - cost: 6.17, layer: 58
Normal
Diabetic
Normal data
Fold over basal
5
4.5
4
3.5
3
2.5
2
1.5
1
0.5
0
10
20
30
40
50
60
Time (min)

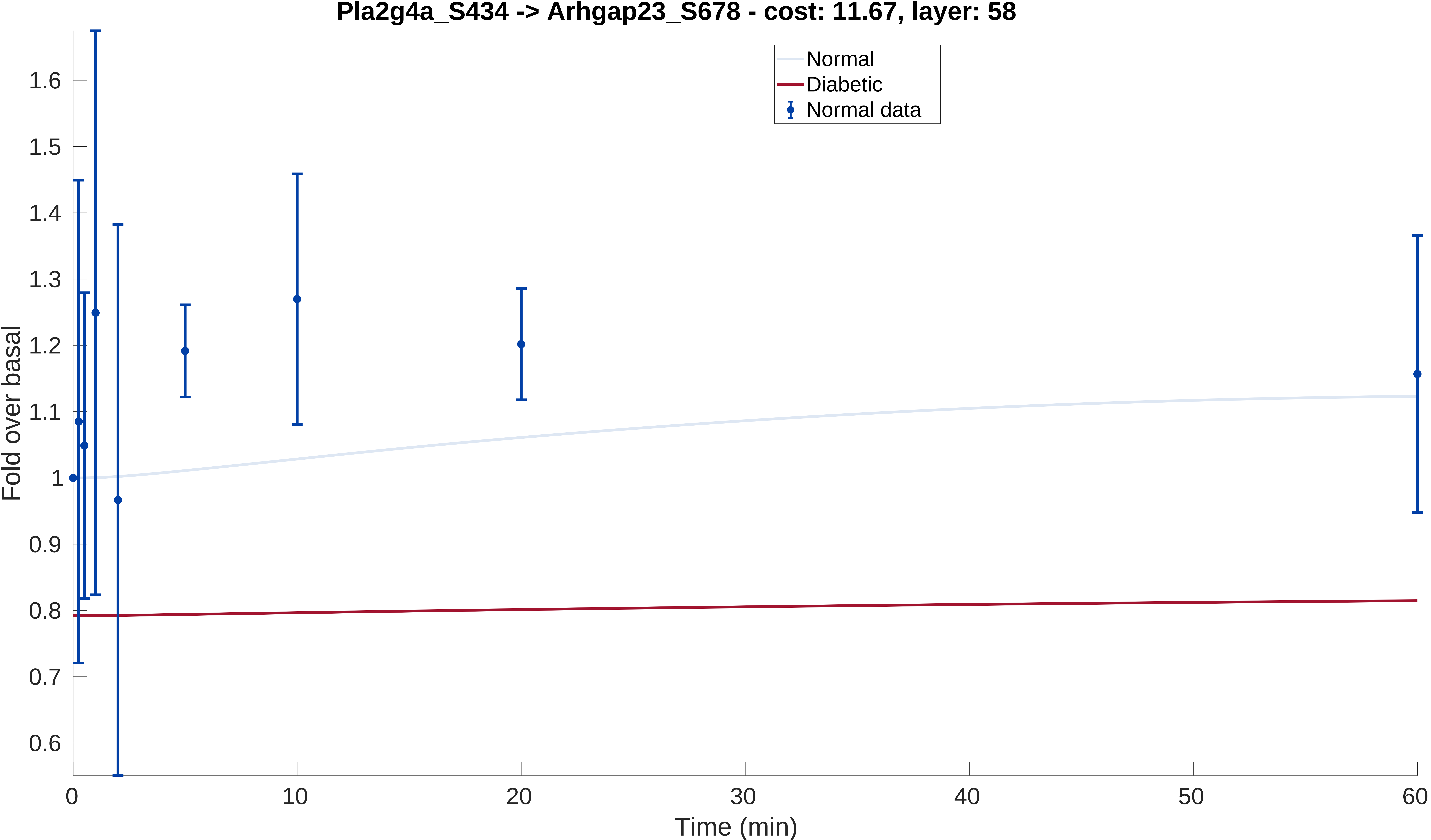

Pla2g4a_S434 -> Arhgap23_S678 - cost: 11.67, layer: 58
Normal
Diabetic
Normal data
Fold over basal
Time (min)
0
10
20
30
40
50
60
0.6
0.7
0.8
0.9
1
1.1
1.2
1.3
1.4
1.5
1.6

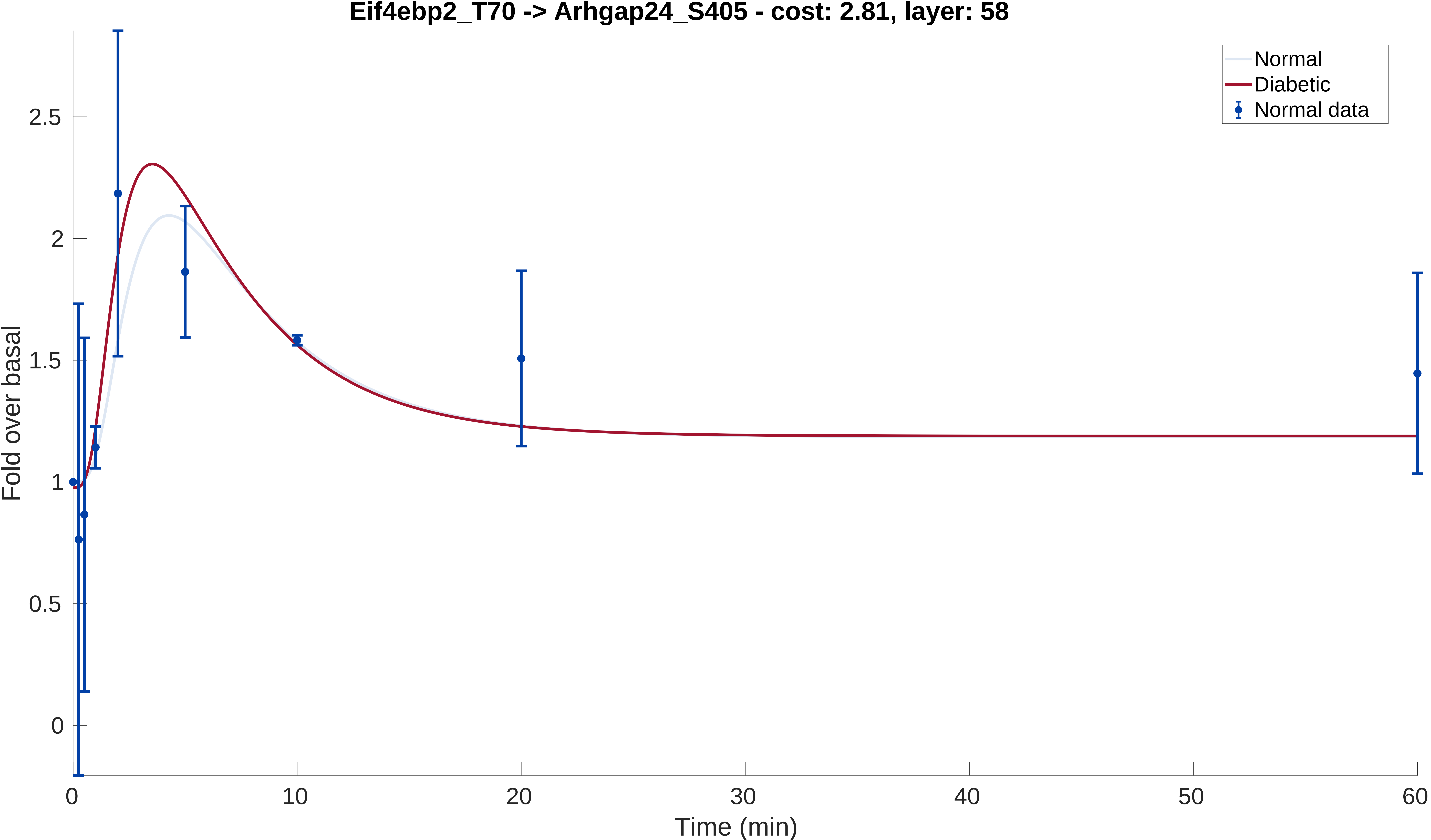

Eif4ebp2_T70 -> Arhgap24_S405 - cost: 2.81, layer: 58
Normal
Diabetic
Normal data
Fold over basal
0
0.5
1
1.5
2
2.5
0
10
20
30
40
50
60
Time (min)

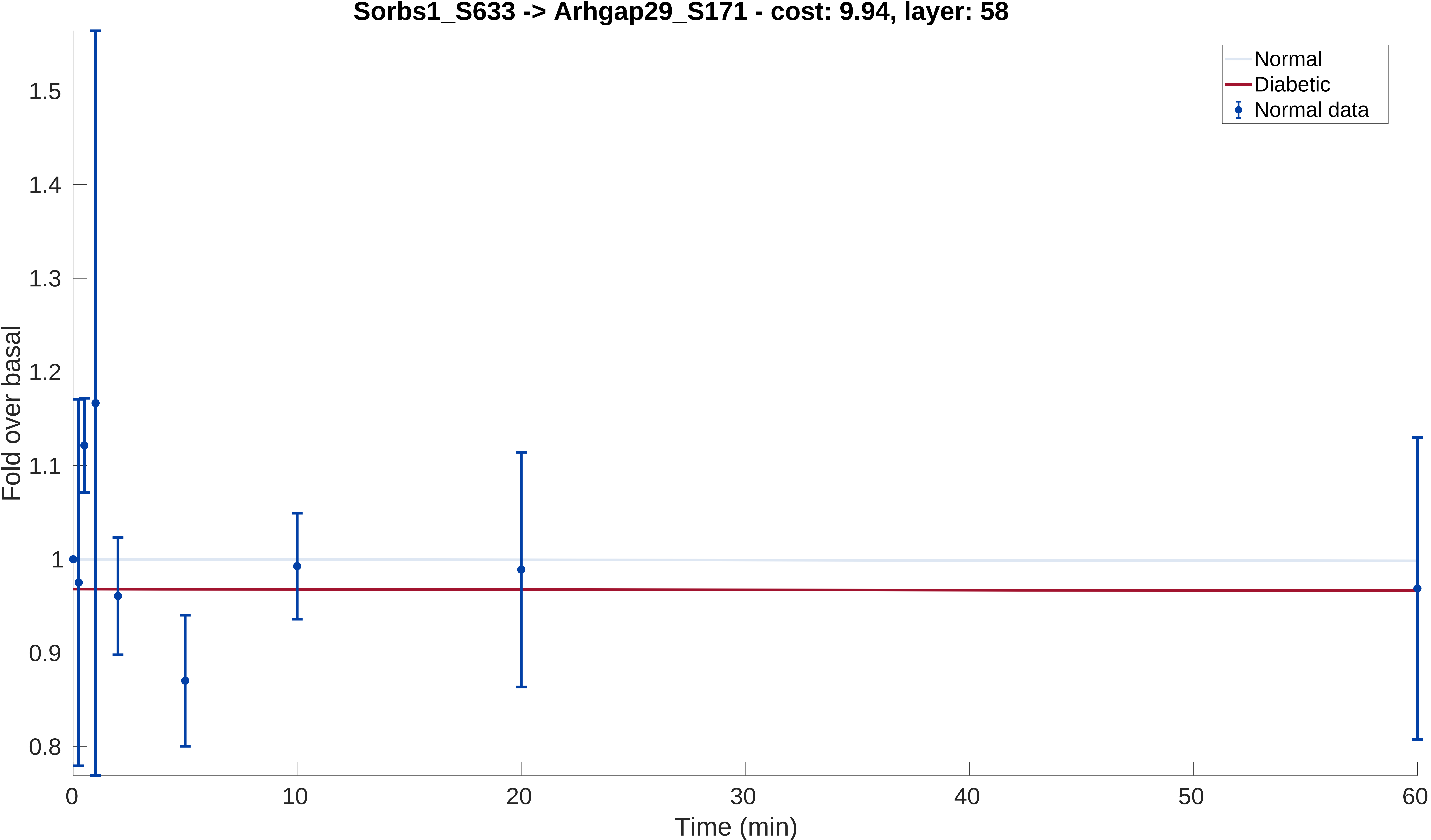

Sorbs1_S633 -> Arhgap29_S171 - cost: 9.94, layer: 58
Normal
Diabetic
Normal data
Fold over basal
1.5
1.4
1.3
1.2
1.1
1
0.9
0.8
0
10
20
30
40
50
60
Time (min)

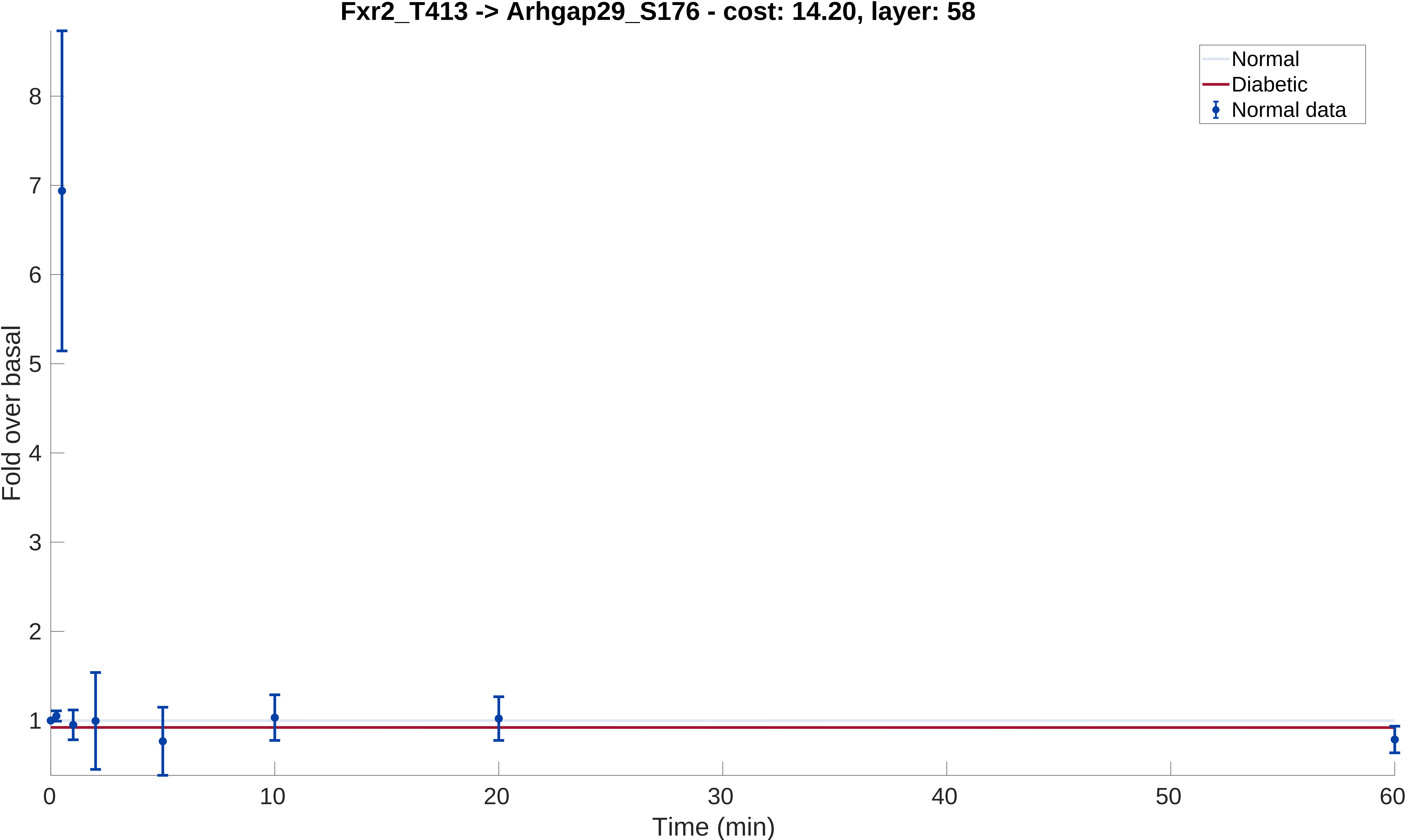

Fxr2_T413 -> Arhgap29_S176 - cost: 14.20, layer: 58
Normal
Diabetic
Normal data
Fold over basal
Time (min)
1
2
3
4
5
6
7
8
0
10
20
30
40
50
60

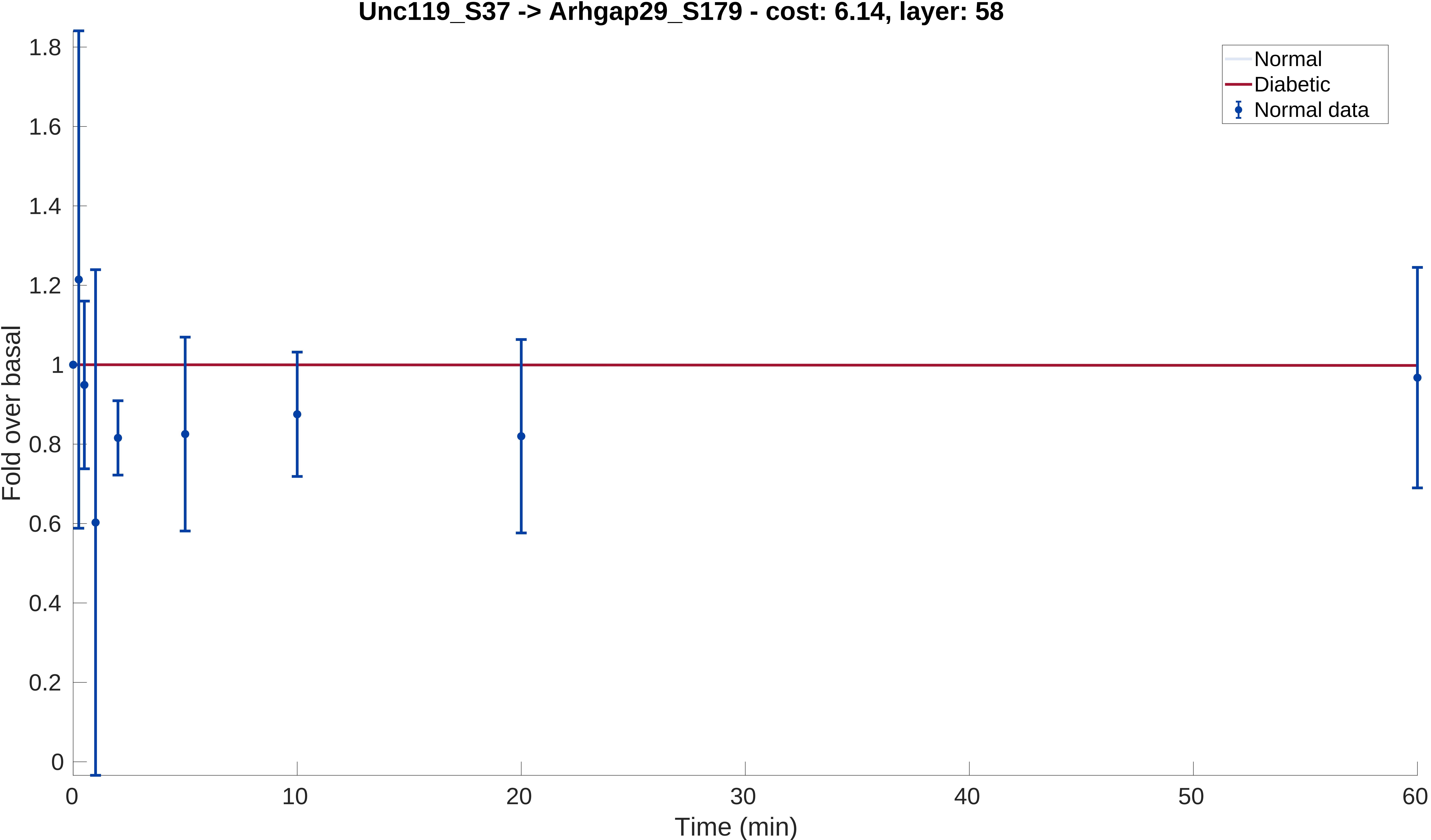

Unc119_S37 -> Arhgap29_S179 - cost: 6.14, layer: 58
Normal
Diabetic
Normal data
Fold over basal
Time (min)
0
0.2
0.4
0.6
0.8
1
1.2
1.4
1.6
1.8
0
10
20
30
40
50
60

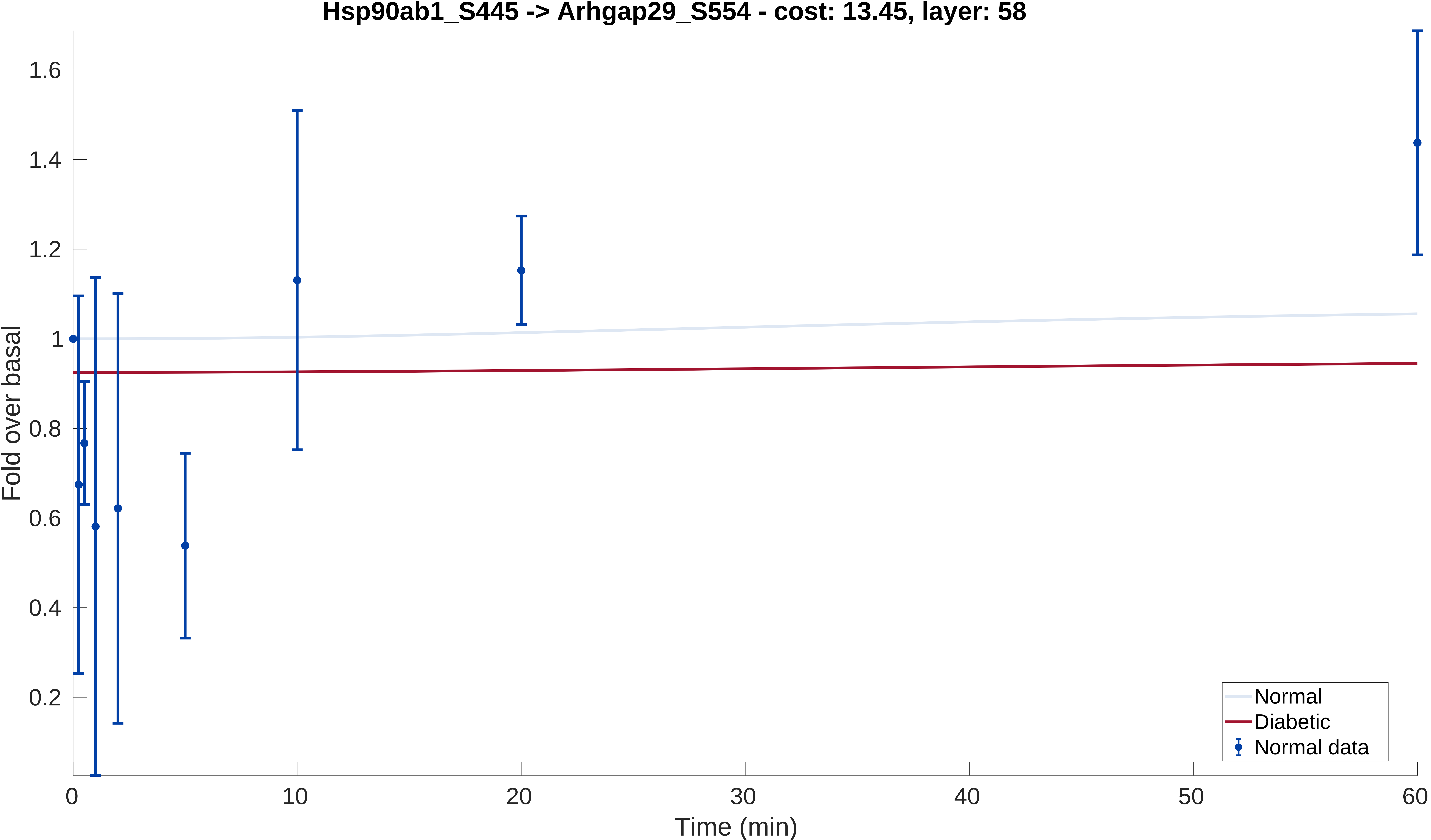

Hsp90ab1_S445 -> Arhgap29_S554 - cost: 13.45, layer: 58
Fold over basal
Time (min)
1.6
1.4
1.2
1
0.8
0.6
0.4
0.2
0
10
20
30
40
50
60
Normal
Diabetic
Normal data

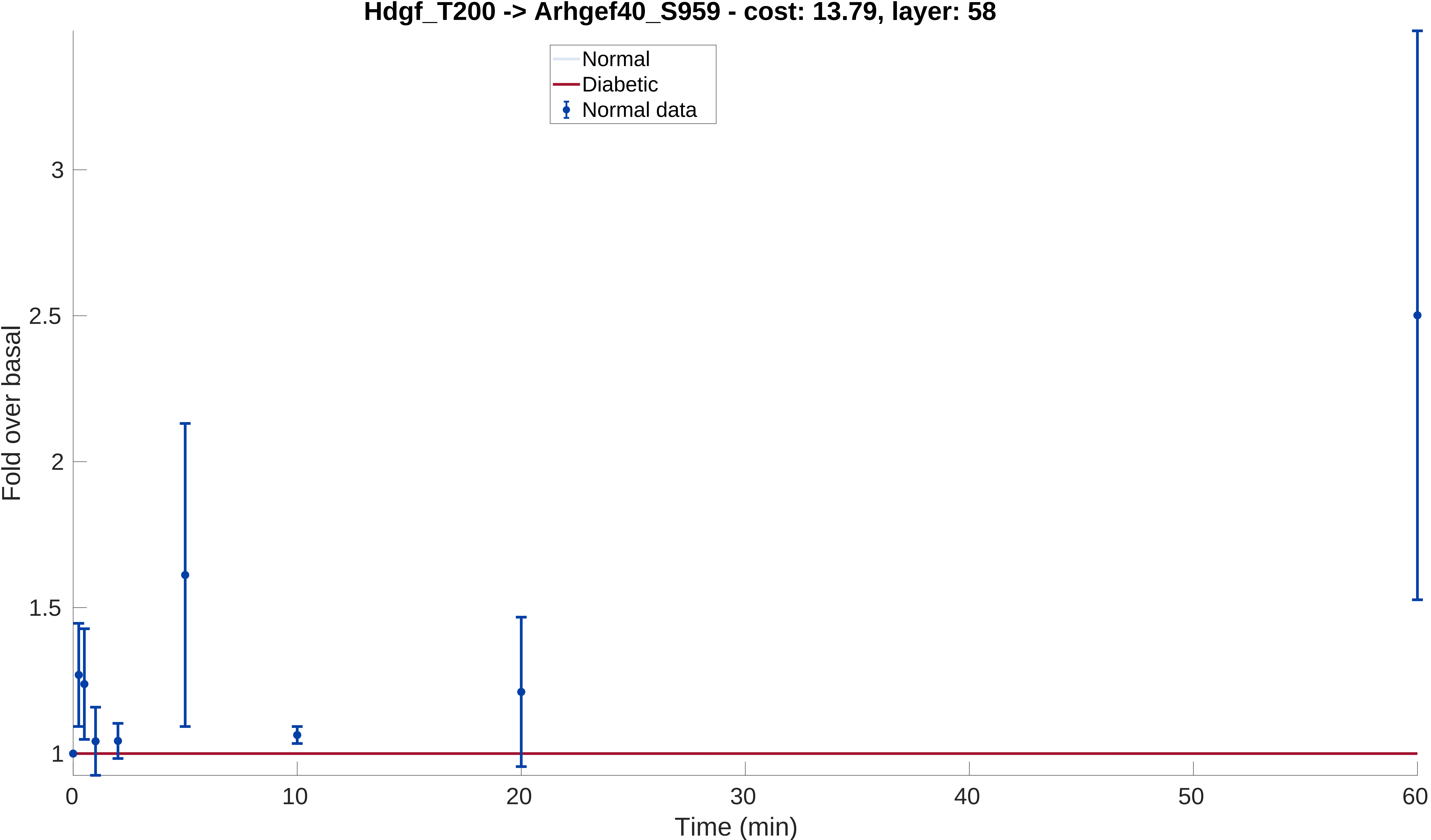

Hdgf_T200 -> Arhgef40_S959 - cost: 13.79, layer: 58
Normal
Diabetic
Normal data
Fold over basal
3
2.5
2
1.5
1
0
10
20
30
40
50
60
Time (min)

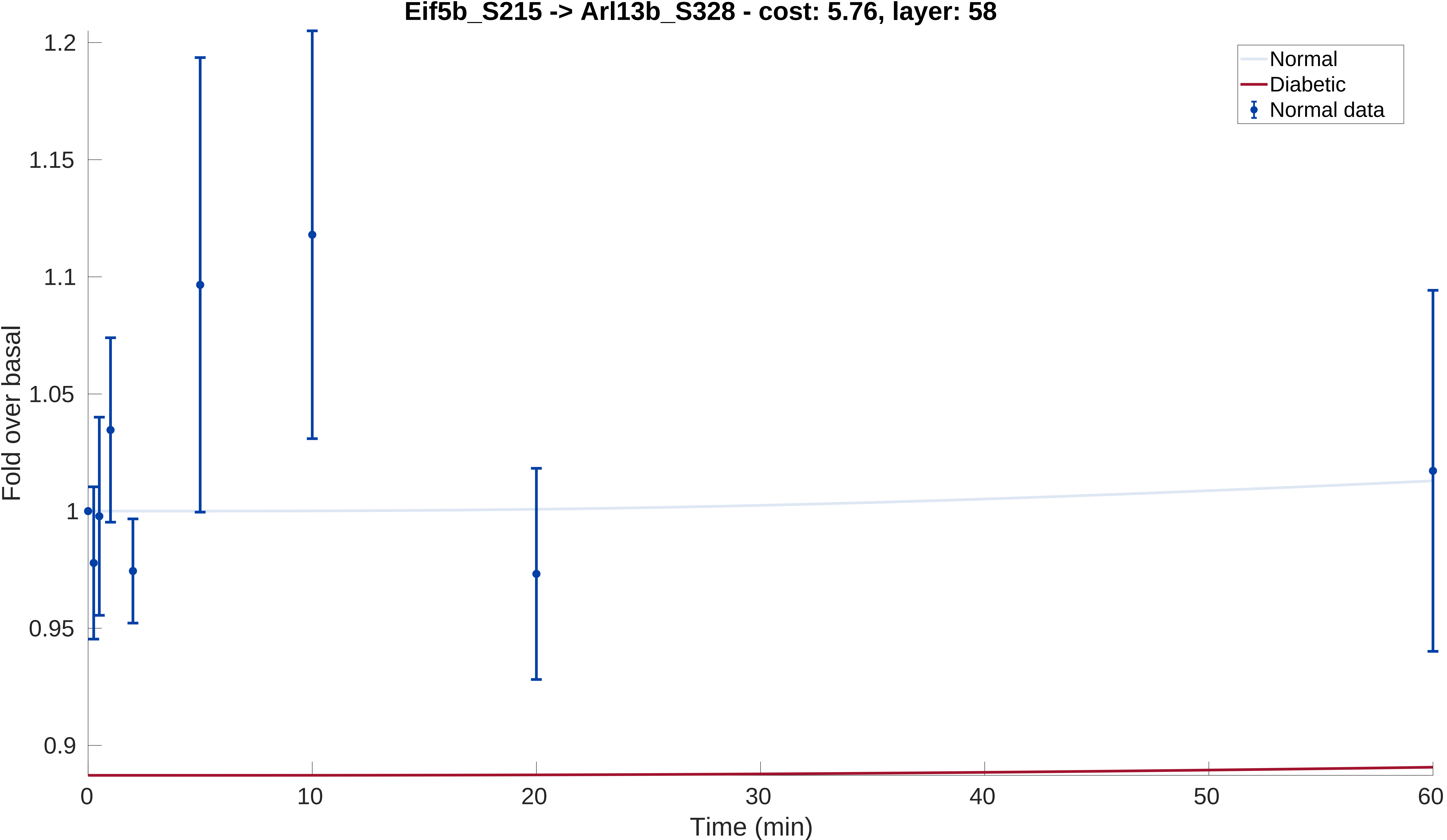

Eif5b_S215 -> Arl13b_S328 - cost: 5.76, layer: 58
Normal
Diabetic
Normal data
Fold over basal
1.2
1.15
1.1
1.05
1
0.95
0.9
0
10
20
30
40
50
60
Time (min)

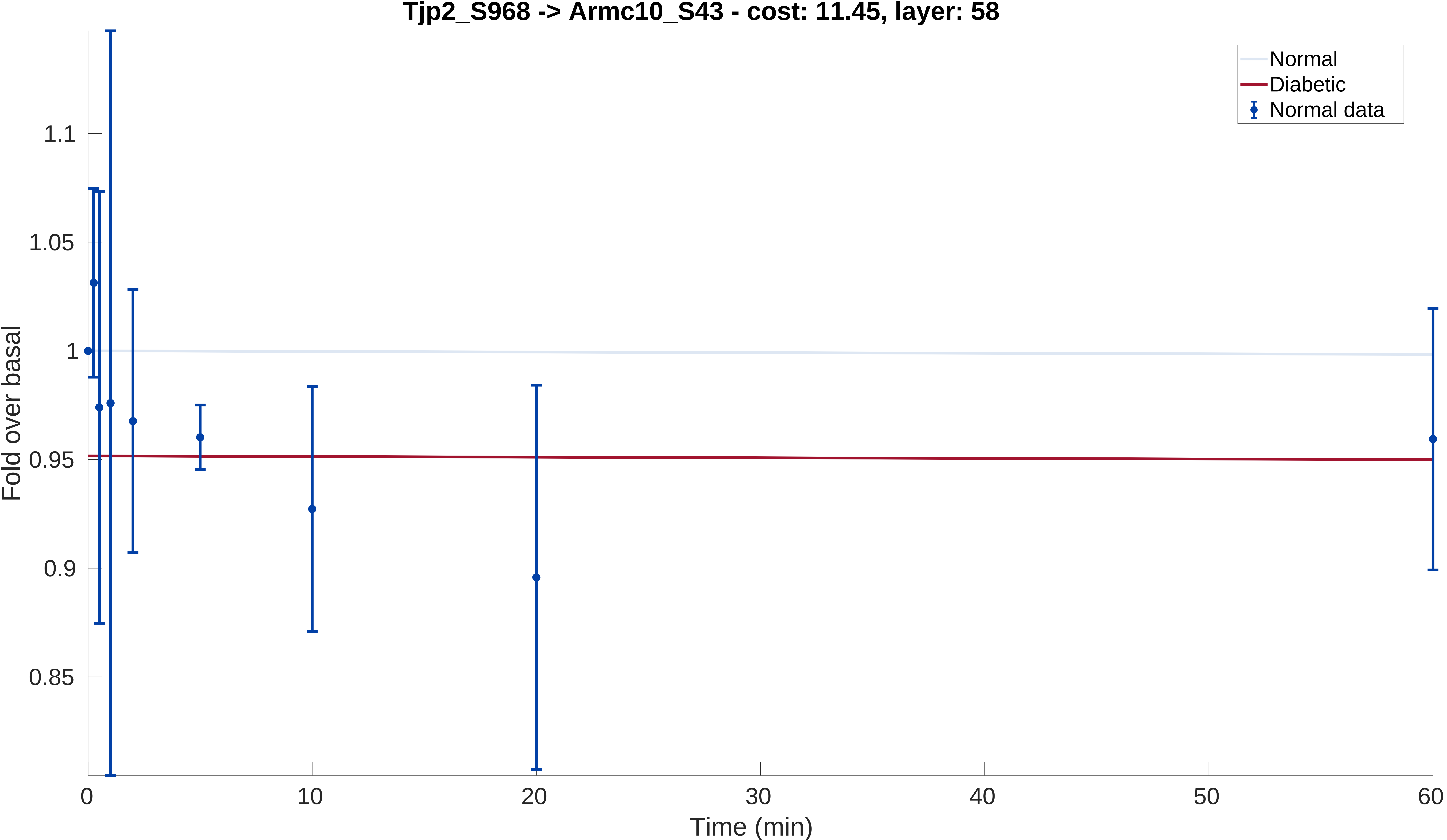

Tjp2_S968 -> Armc10_S43 - cost: 11.45, layer: 58
Normal
Diabetic
Normal data
1.1
1.05
1
0.95
0.9
0.85
Fold over basal
0
10
20
30
40
50
60
Time (min)

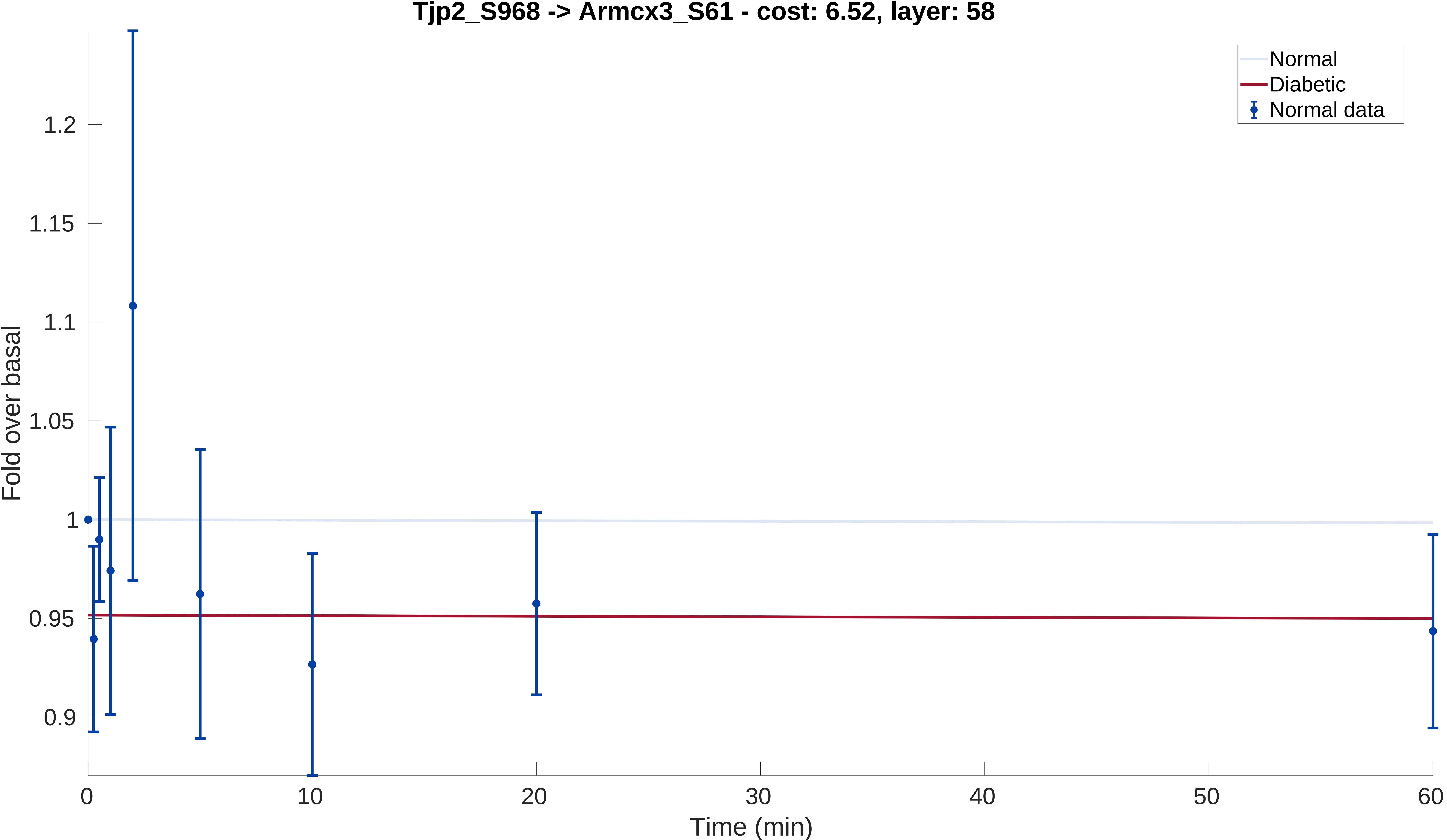

Tjp2_S968 -> Armcx3_S61 - cost: 6.52, layer: 58
Normal
Diabetic
Normal data
Fold over basal
1.2
1.15
1.1
1.05
1
0.95
0.9
0
10
20
30
40
50
60
Time (min)

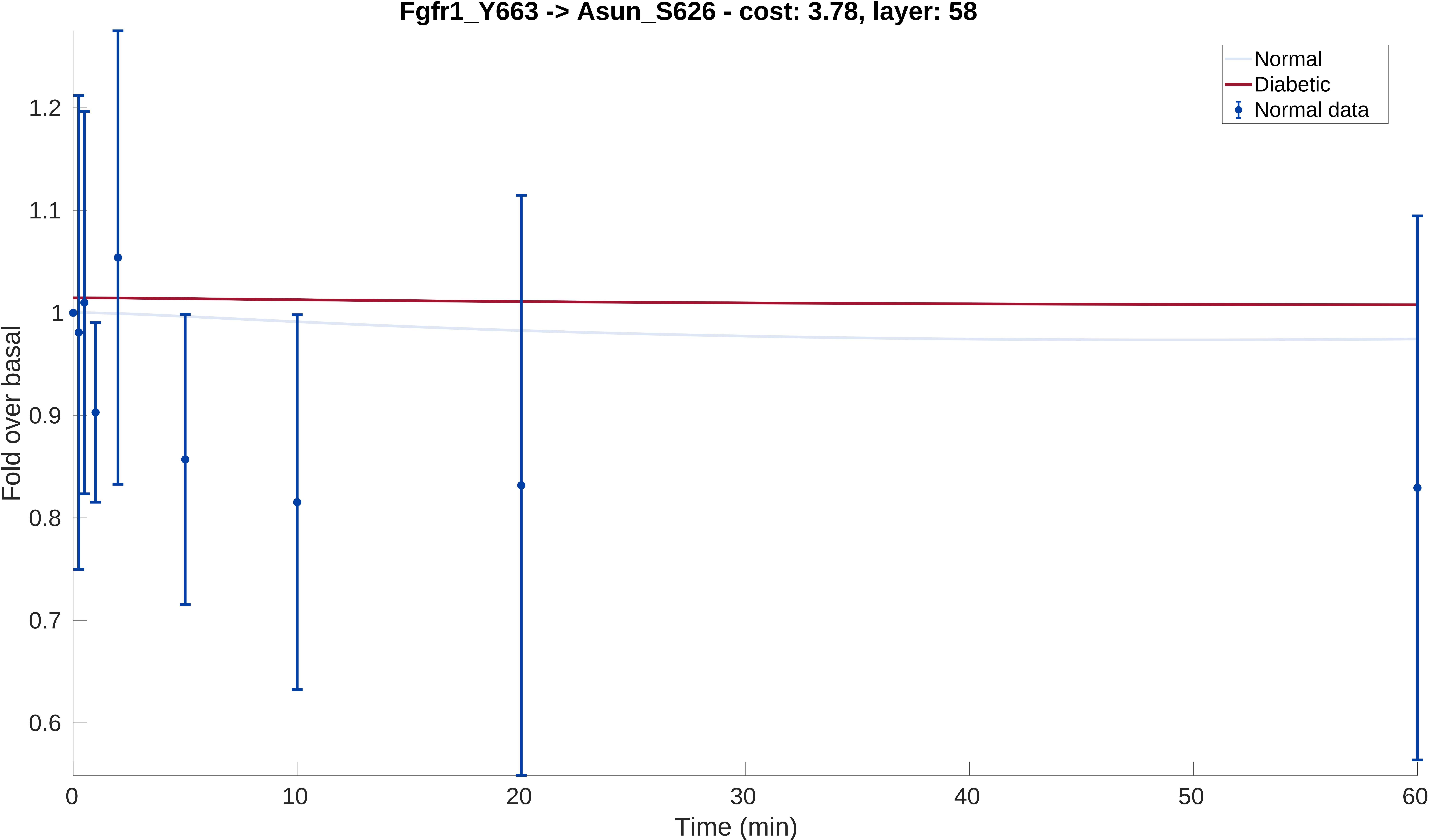

Fgfr1_Y663 -> Asun_S626 - cost: 3.78, layer: 58
Normal
Diabetic
Normal data
Fold over basal
1.2
1.1
1
0.9
0.8
0.7
0.6
0
10
20
30
40
50
60
Time (min)

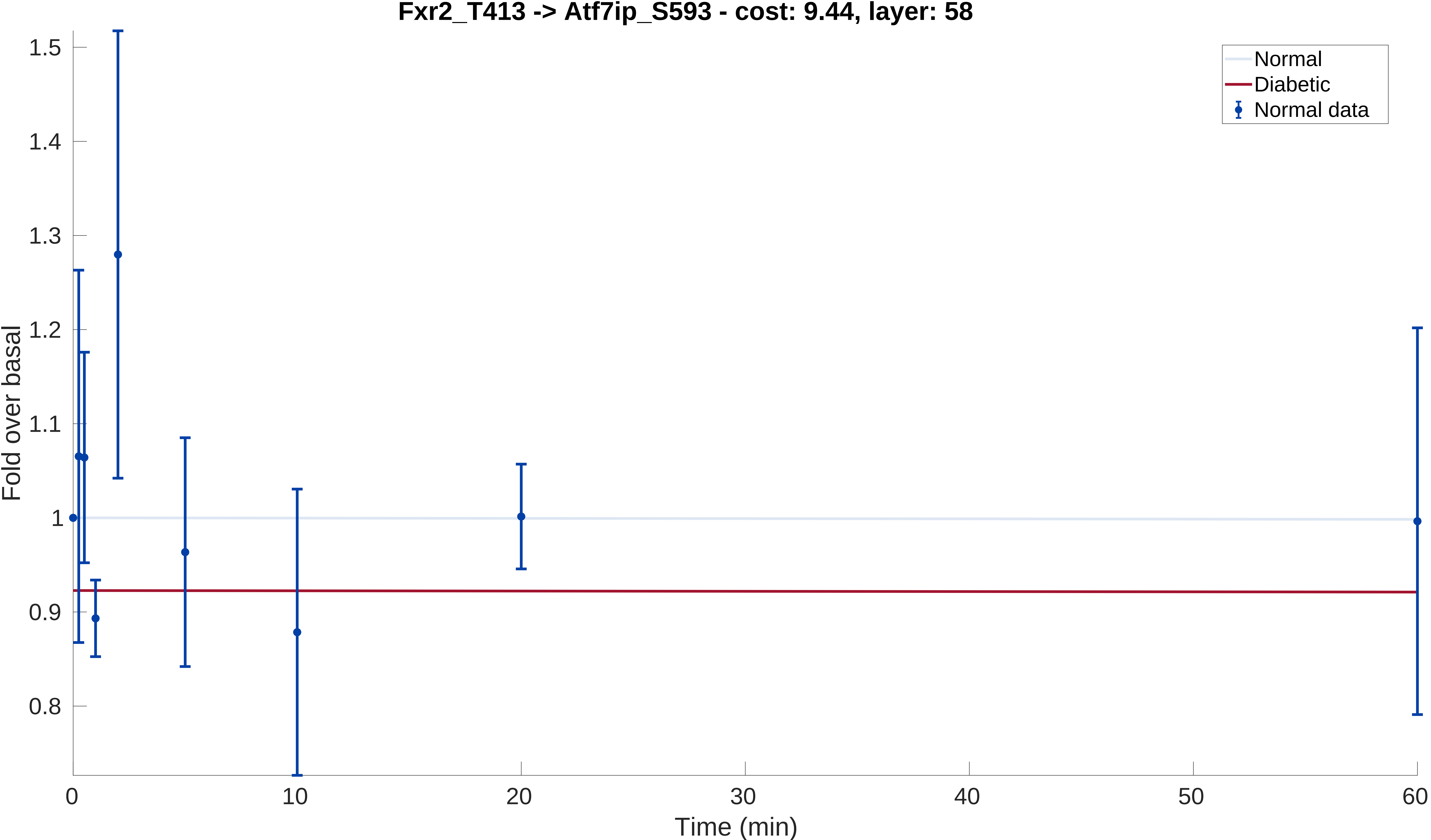
Fxr2_T413 -> Atf7ip_S593 - cost: 9.44, layer: 58
Normal
Diabetic
Normal data
Fold over basal
1.5
1.4
1.3
1.2
1.1
1
0.9
0.8
0
10
20
30
40
50
60
Time (min)

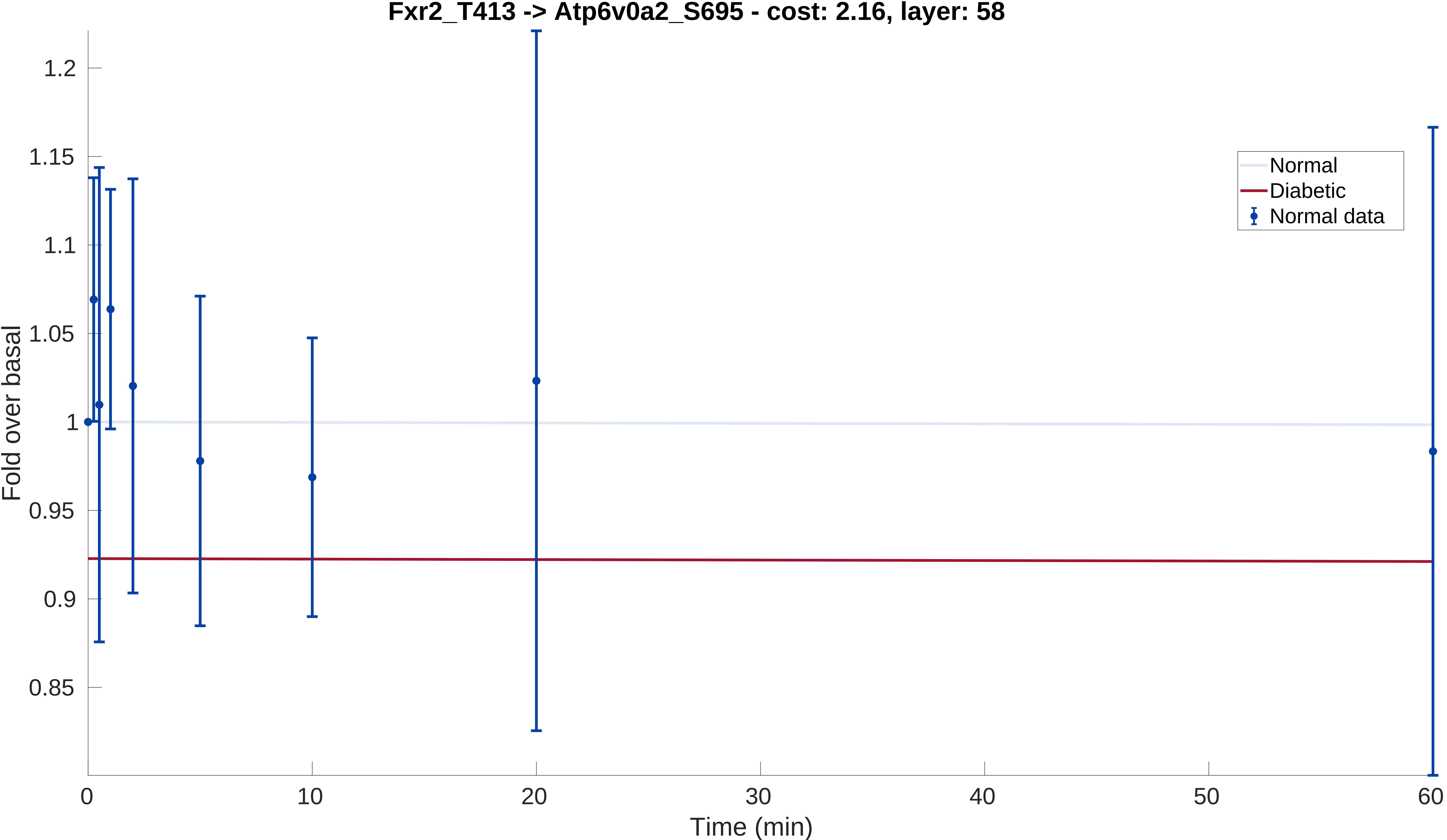
Fxr2_T413 -> Atp6v0a2_S695 - cost: 2.16, layer: 58
Fold over basal
1.2
1.15
1.1
1.05
1
0.95
0.9
0.85
0
10
20
30
40
50
60
Time (min)
Normal
Diabetic
Normal data

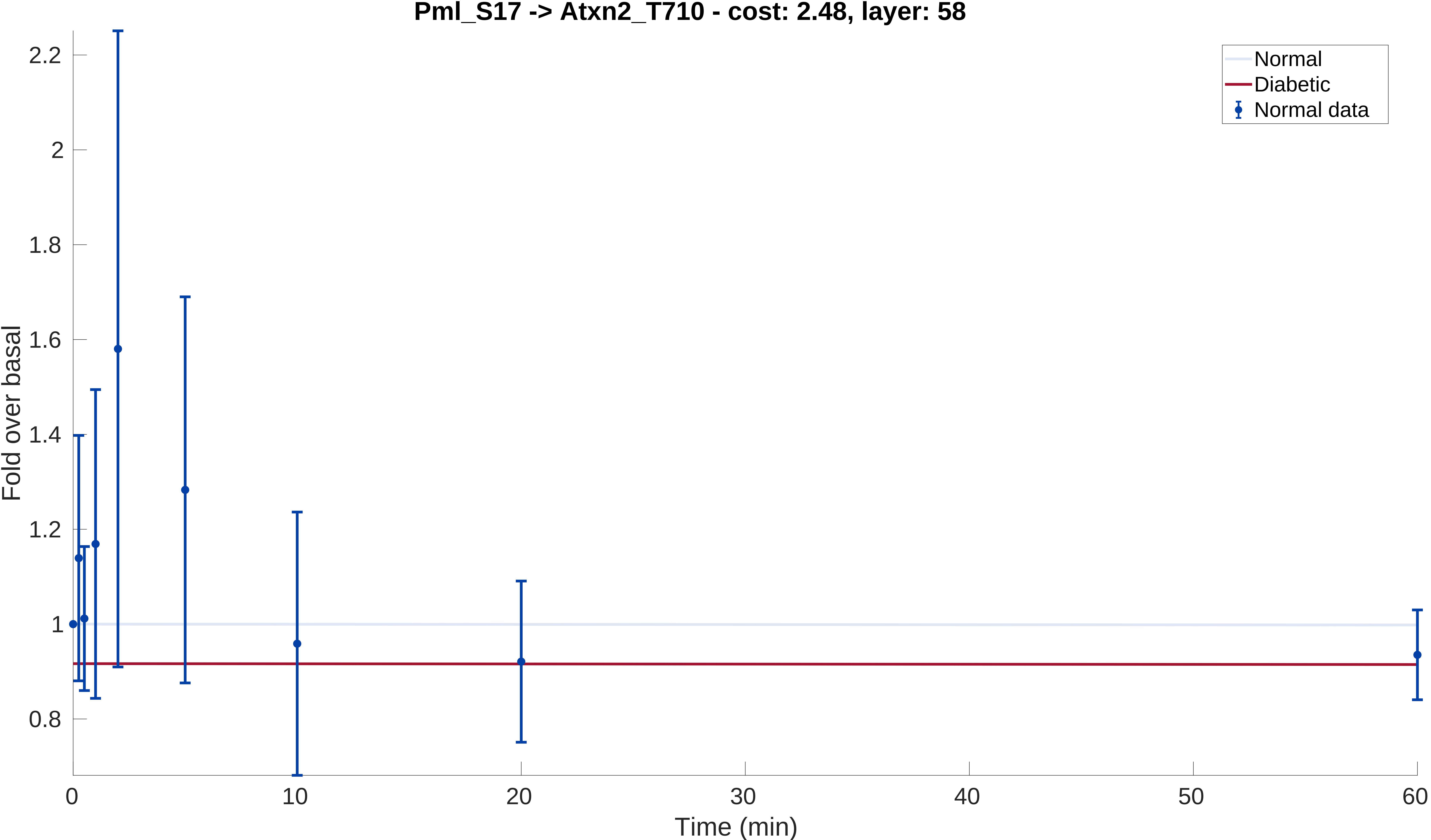

Pml_S17 -> Atxn2_T710 - cost: 2.48, layer: 58
Fold over basal
Time (min)
Normal
Diabetic
Normal data
2.2
2
1.8
1.6
1.4
1.2
1
0.8
0
10
20
30
40
50
60

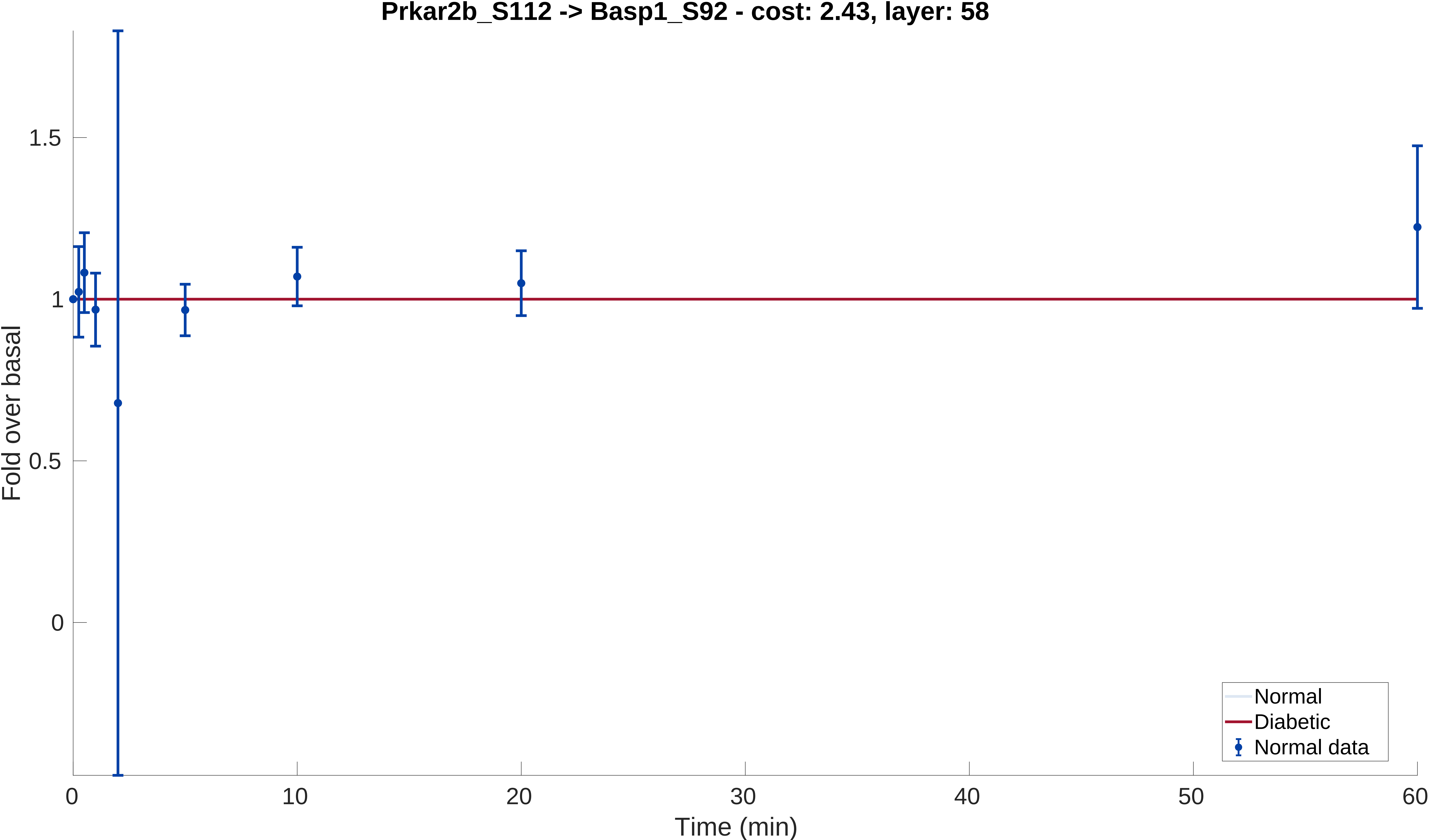
Prrkar2b_S112 -> Basp1_S92 - cost: 2.43, layer: 58
Fold over basal
1.5
1
0.5
0
0
10
20
30
40
50
60
Time (min)
Normal
Diabetic
Normal data

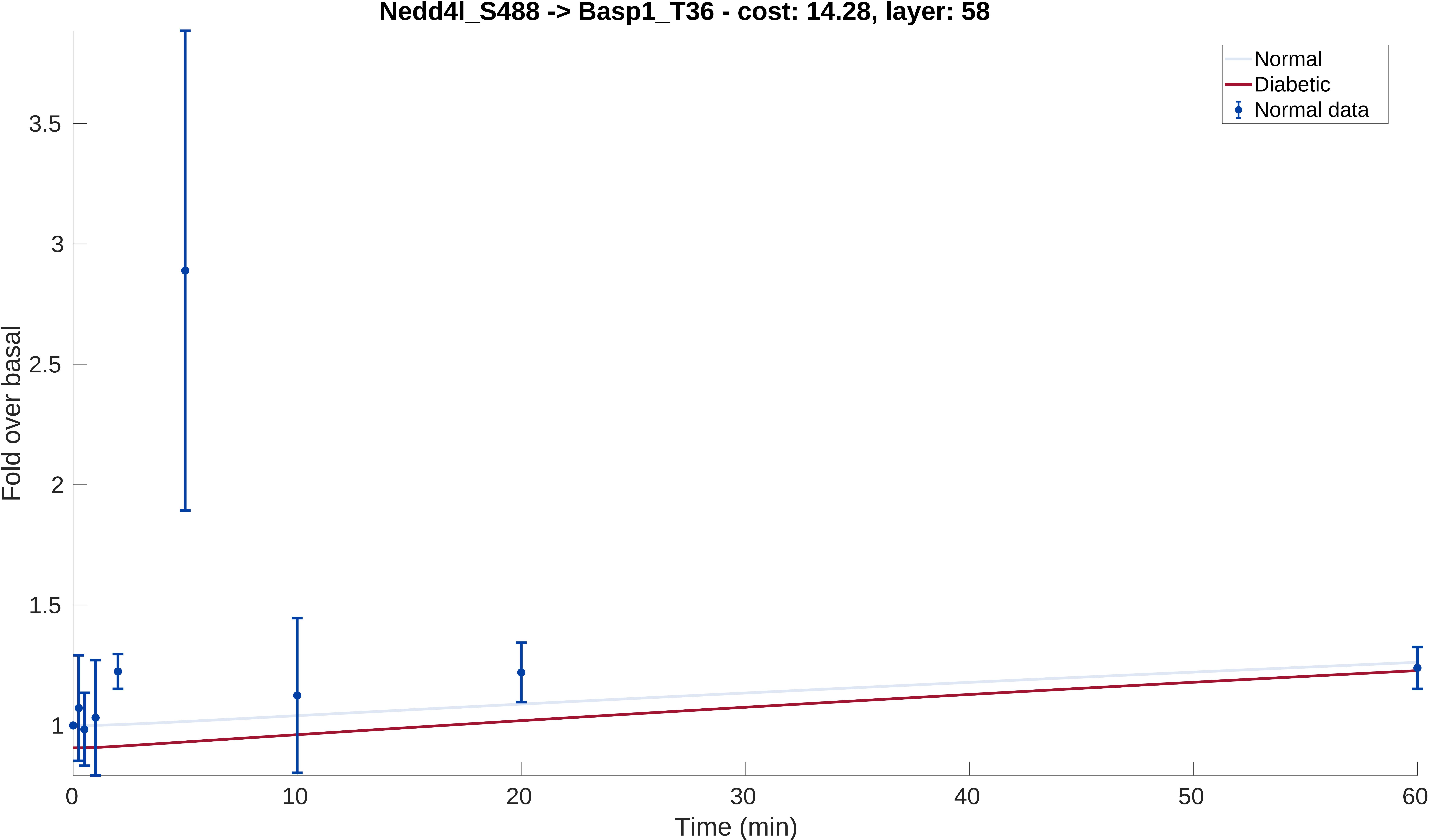
Nedd4l_S488 -> Basp1_T36 - cost: 14.28, layer: 58
Normal
Diabetic
Normal data
3.5
3
2.5
2
1.5
1
Fold over basal
0
10
20
30
40
50
60
Time (min)

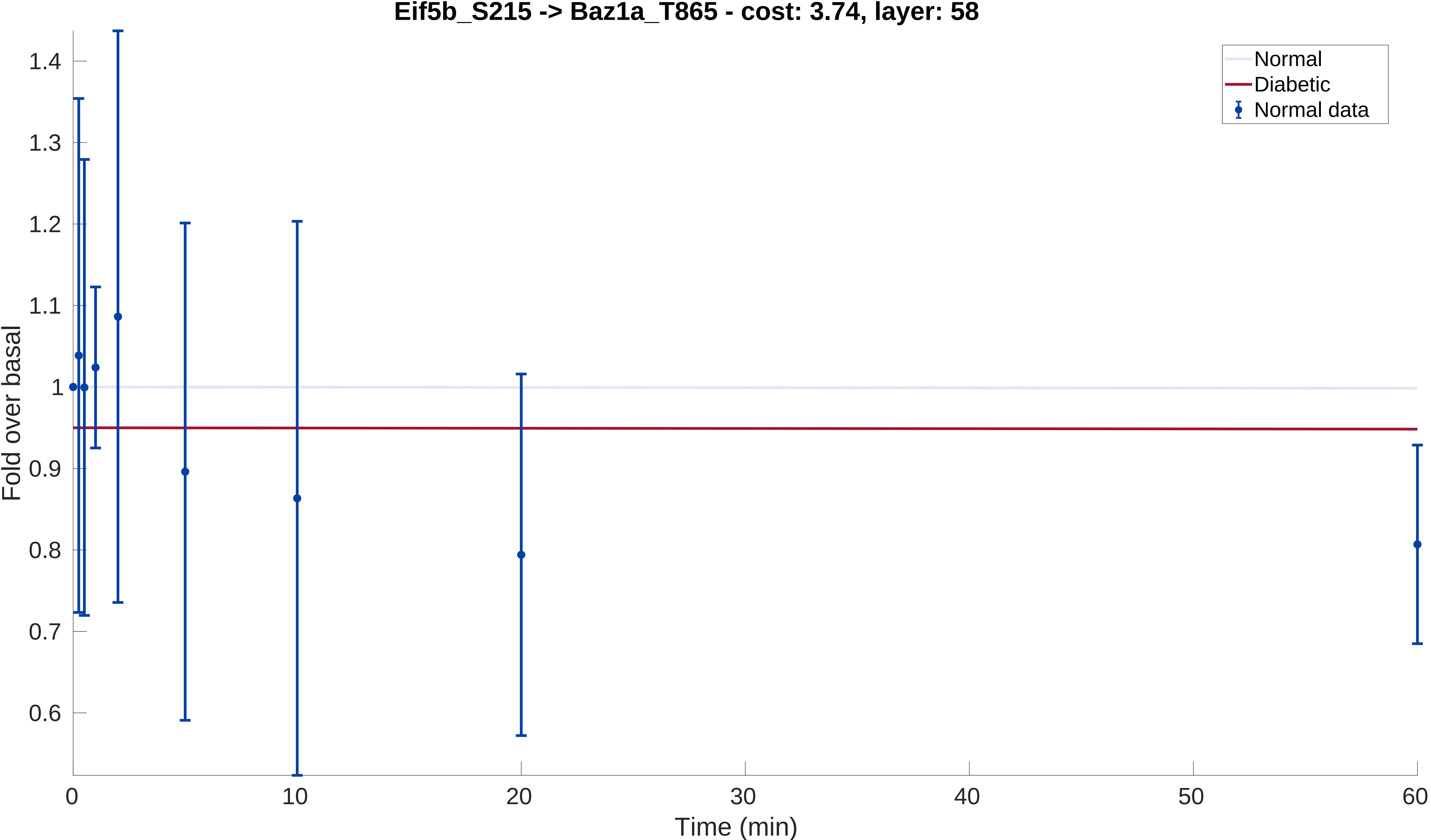

Eif5b_S215 -> Baz1a_T865 - cost: 3.74, layer: 58
Normal
Diabetic
Normal data
Fold over basal
1.4
1.3
1.2
1.1
1
0.9
0.8
0.7
0.6
0
10
20
30
40
50
60
Time (min)

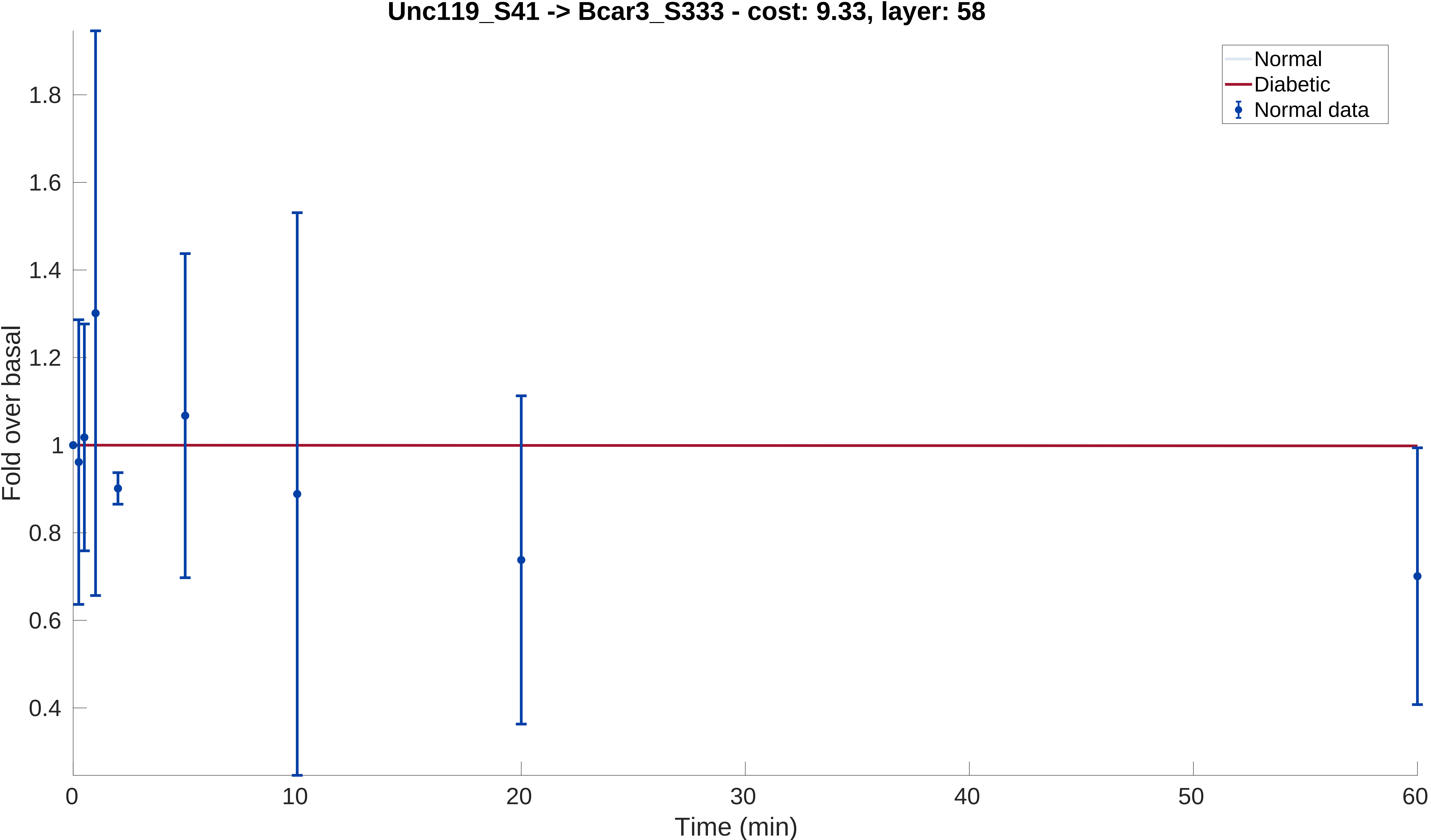

Unc119_S41 -> Bcar3_S333 - cost: 9.33, layer: 58
Normal
Diabetic
Normal data
Fold over basal
1.8
1.6
1.4
1.2
1
0.8
0.6
0.4
0
10
20
30
40
50
60
Time (min)

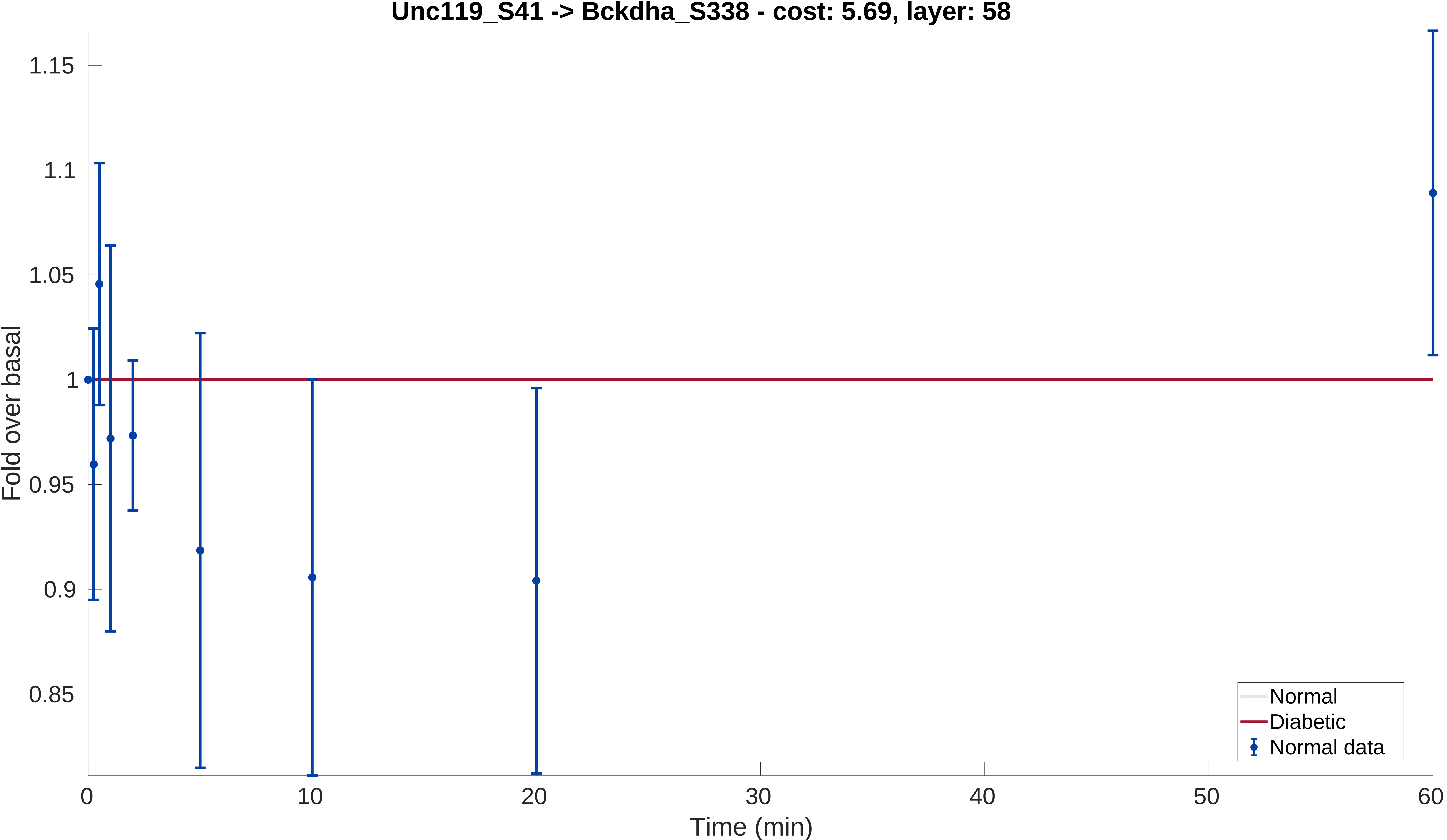

Unc119_S41 -> Bckdha_S338 - cost: 5.69, layer: 58
Fold over basal
1.15
1.1
1.05
1
0.95
0.9
0.85
0
10
20
30
40
50
60
Time (min)
Normal
Diabetic
Normal data

#### Eif4ebp2_T70 -> Bckdha_S343 - cost: 6.68, layer: 58

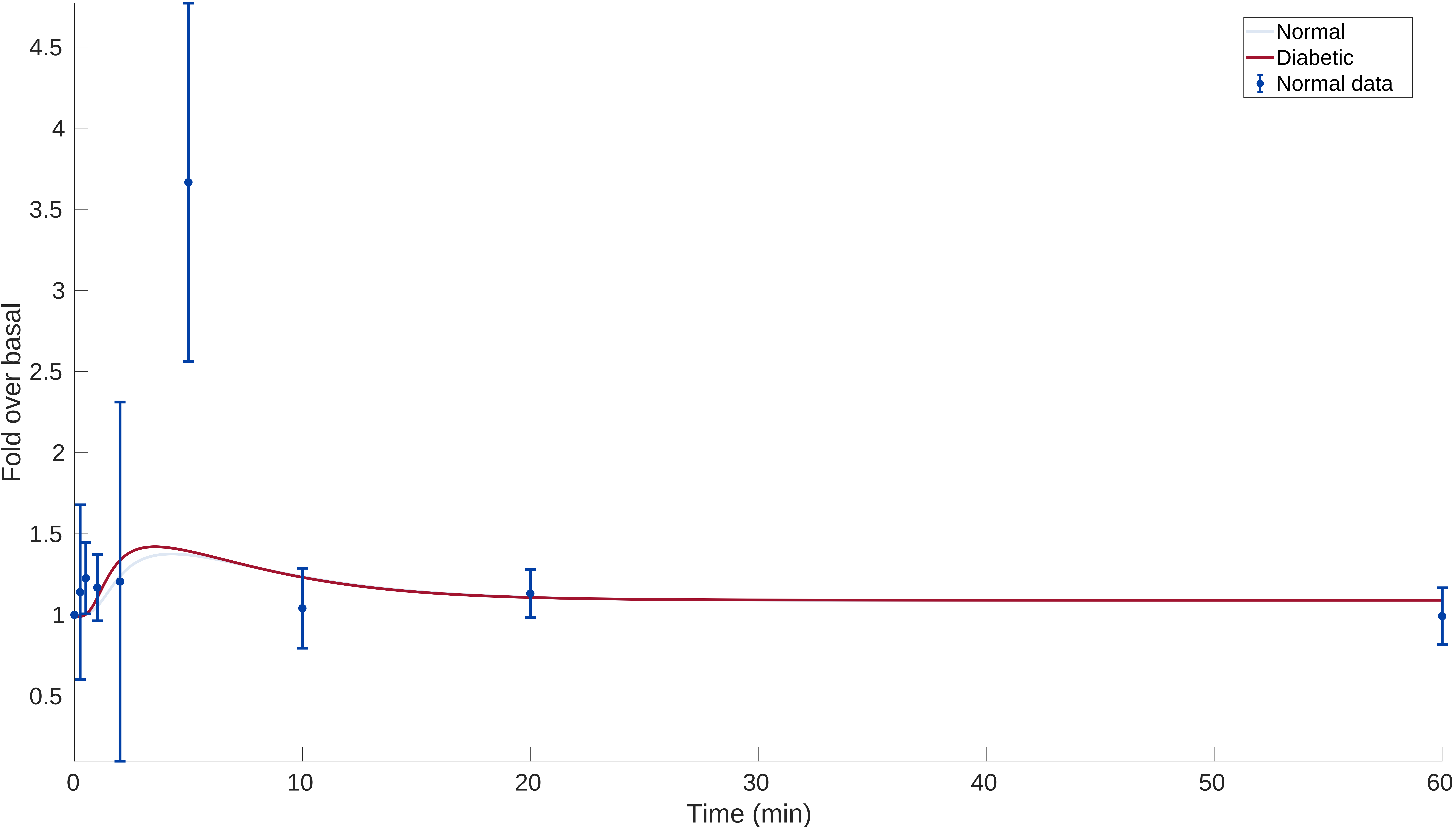

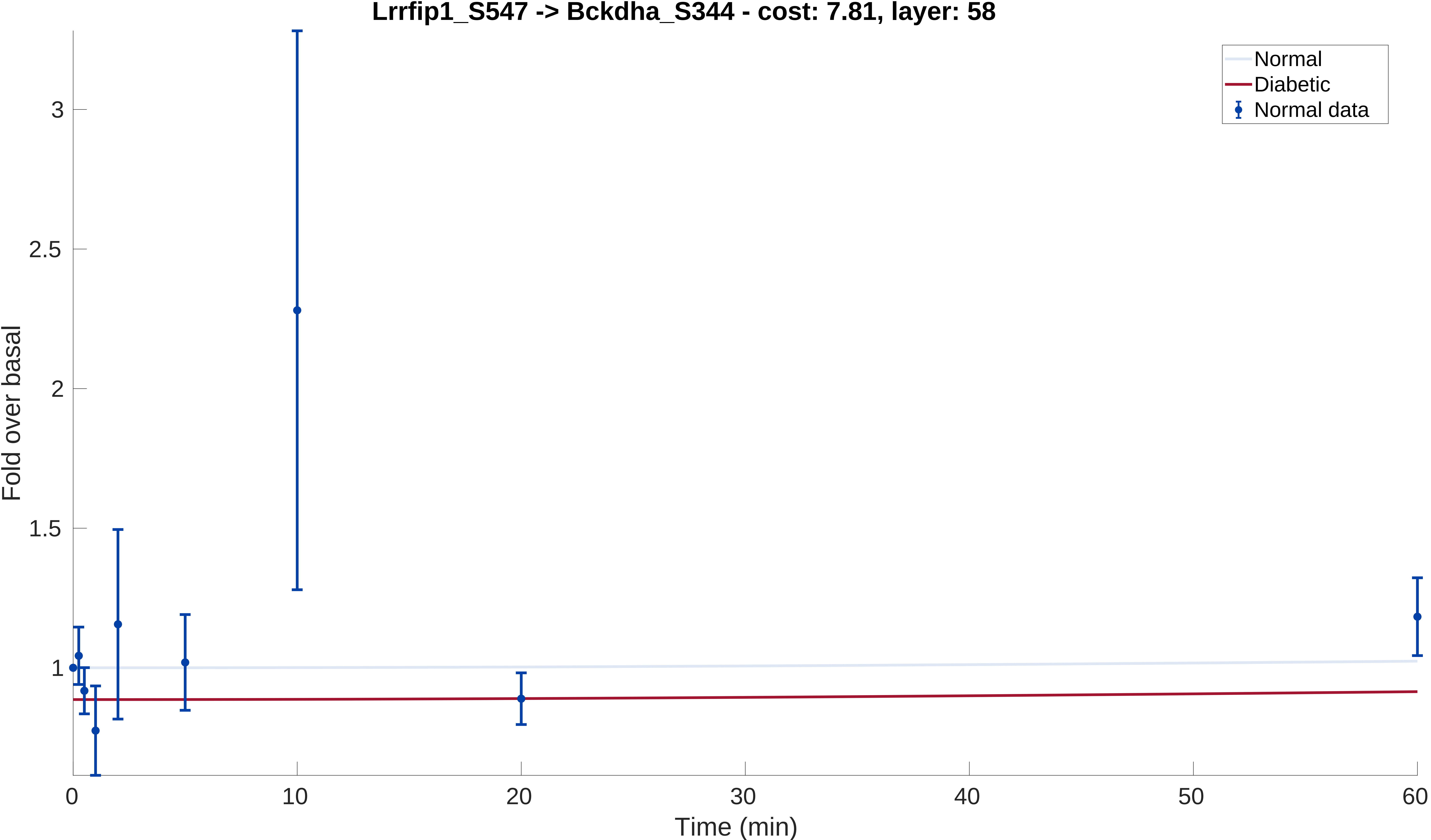
Lrrfip1_S547 -> Bckdha_S344 - cost: 7.81, layer: 58
Normal
Diabetic
Normal data
Fold over basal
3
2.5
2
1.5
1
0
10
20
30
40
50
60
Time (min)

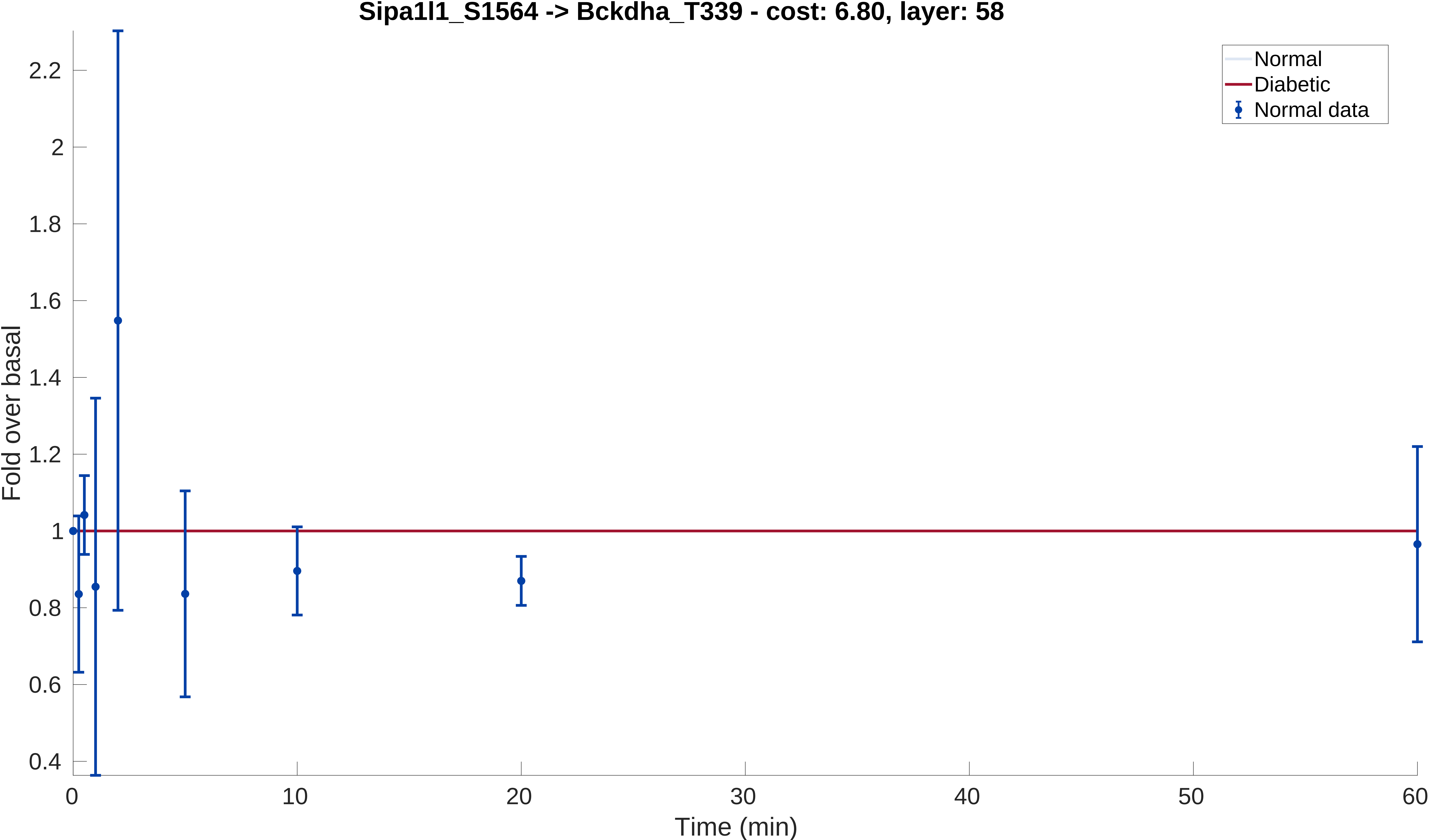

Sipa1l1_S1564 -> Bckdha_T339 - cost: 6.80, layer: 58
Normal
Diabetic
Normal data
Fold over basal
2.2
2
1.8
1.6
1.4
1.2
1
0.8
0.6
0.4
0
10
20
30
40
50
60
Time (min)

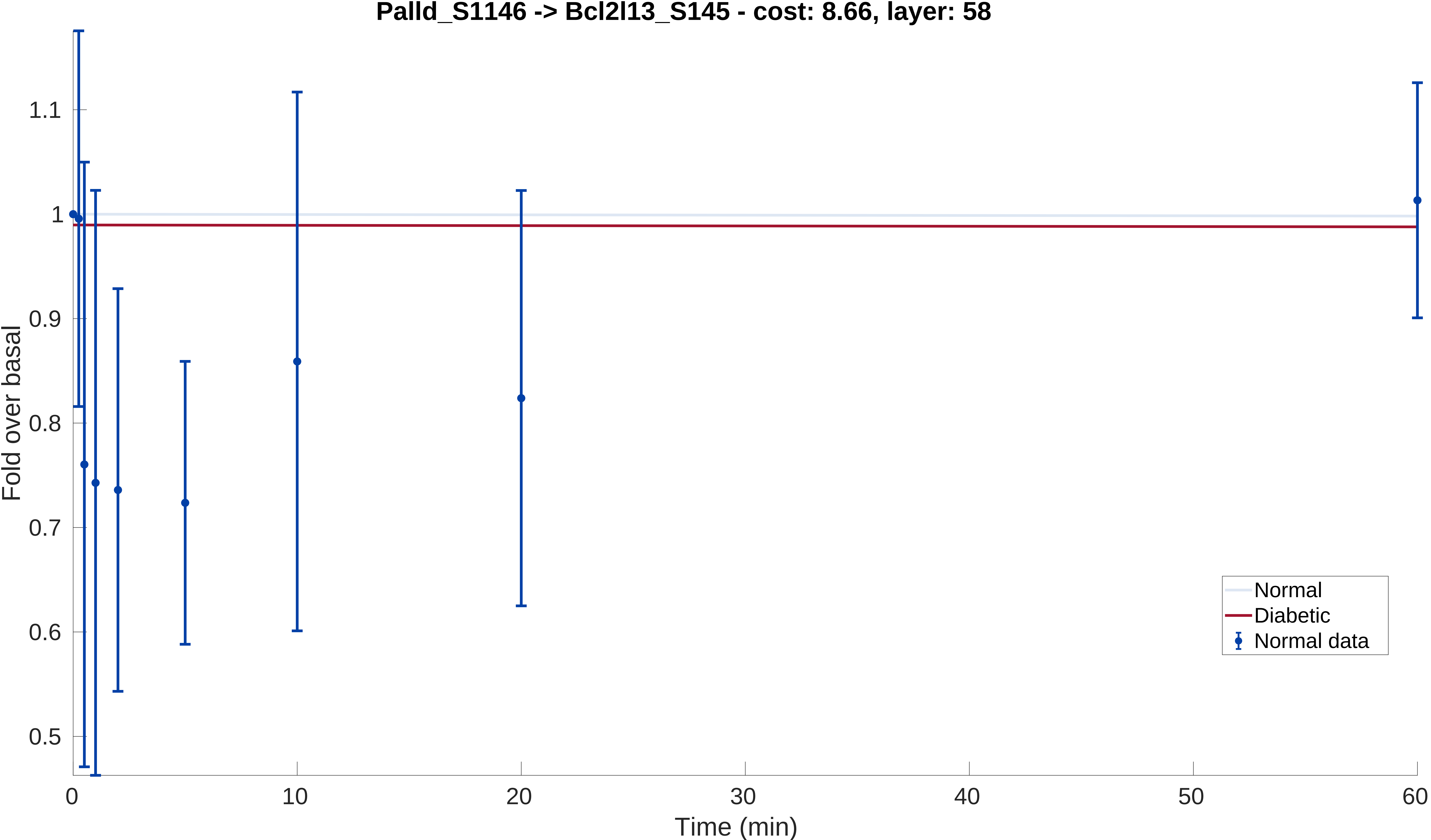
Palld_S1146 -> Bcl2l13_S145 - cost: 8.66, layer: 58
Fold over basal
1.1
1
0.9
0.8
0.7
0.6
0.5
0
10
20
30
40
50
60
Time (min)
Normal
Diabetic
Normal data

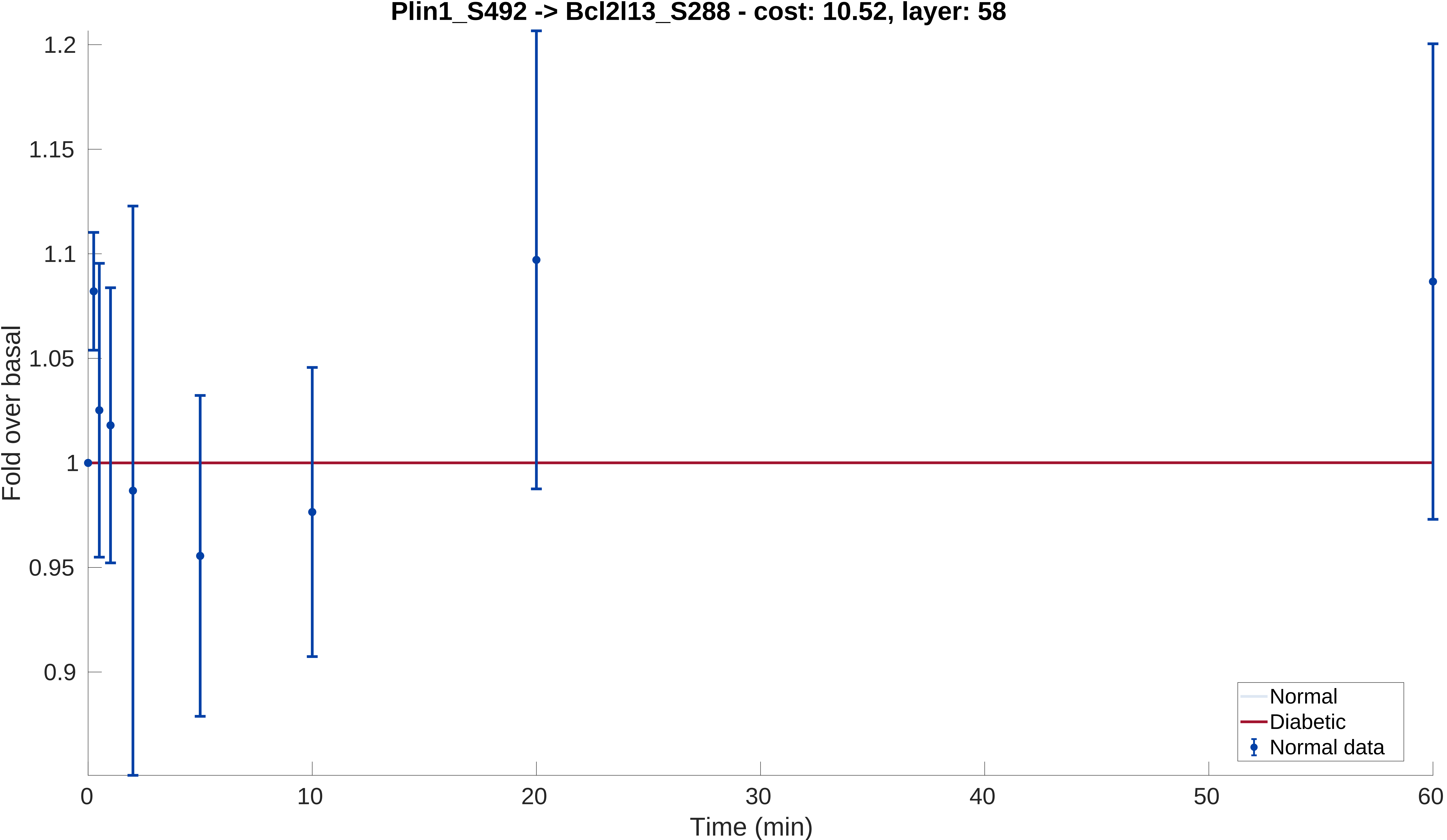

Plin1_S492 -> Bcl2l13_S288 - cost: 10.52, layer: 58
Fold over basal
1.2
1.15
1.1
1.05
1
0.95
0.9
0
10
20
30
40
50
60
Time (min)
Normal
Diabetic
Normal data

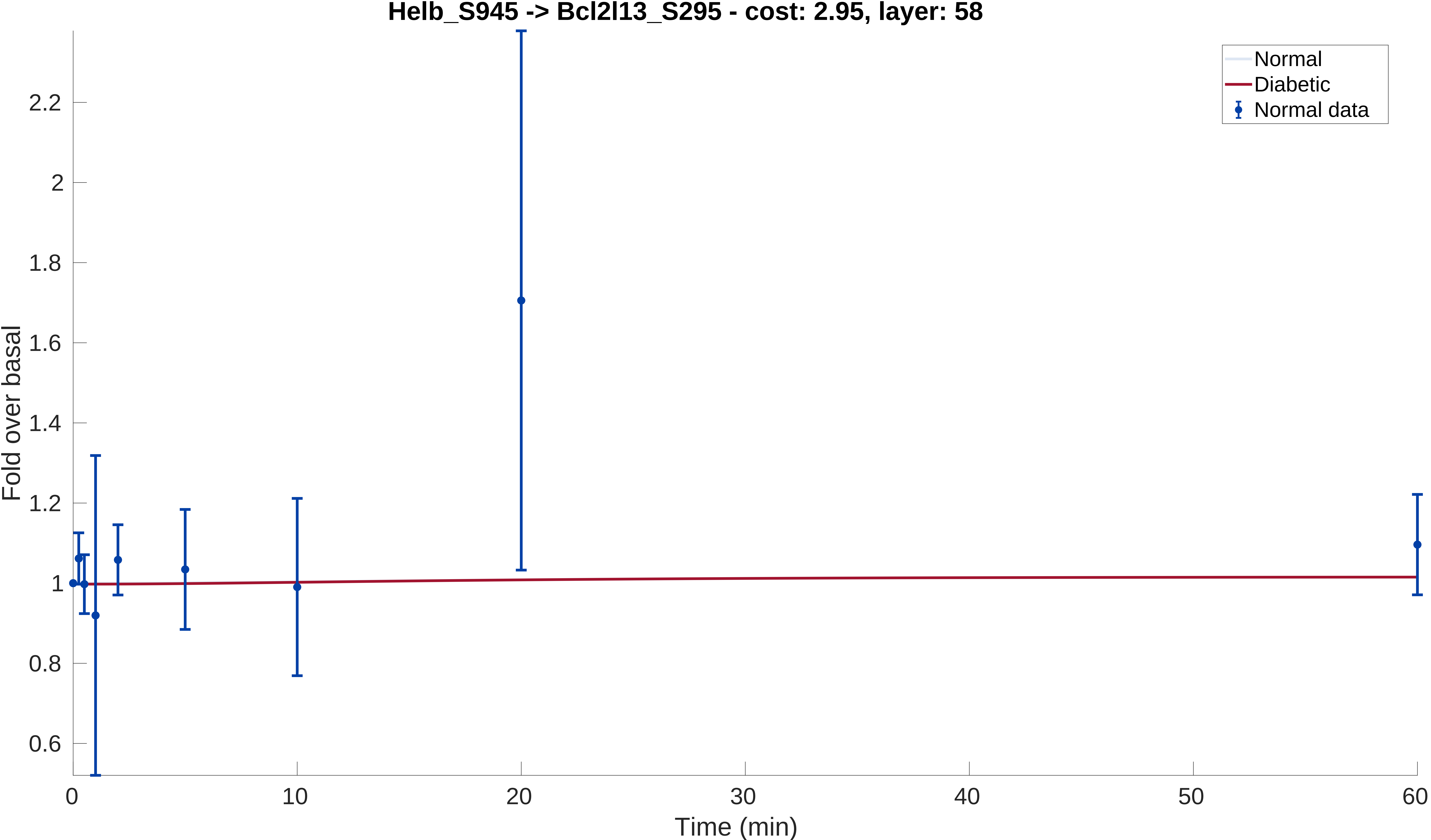

Helb_S945 -> Bcl2l13_S295 - cost: 2.95, layer: 58
Normal
Diabetic
Normal data
Fold over basal
2.2
2
1.8
1.6
1.4
1.2
1
0.8
0.6
0
10
20
30
40
50
60
Time (min)

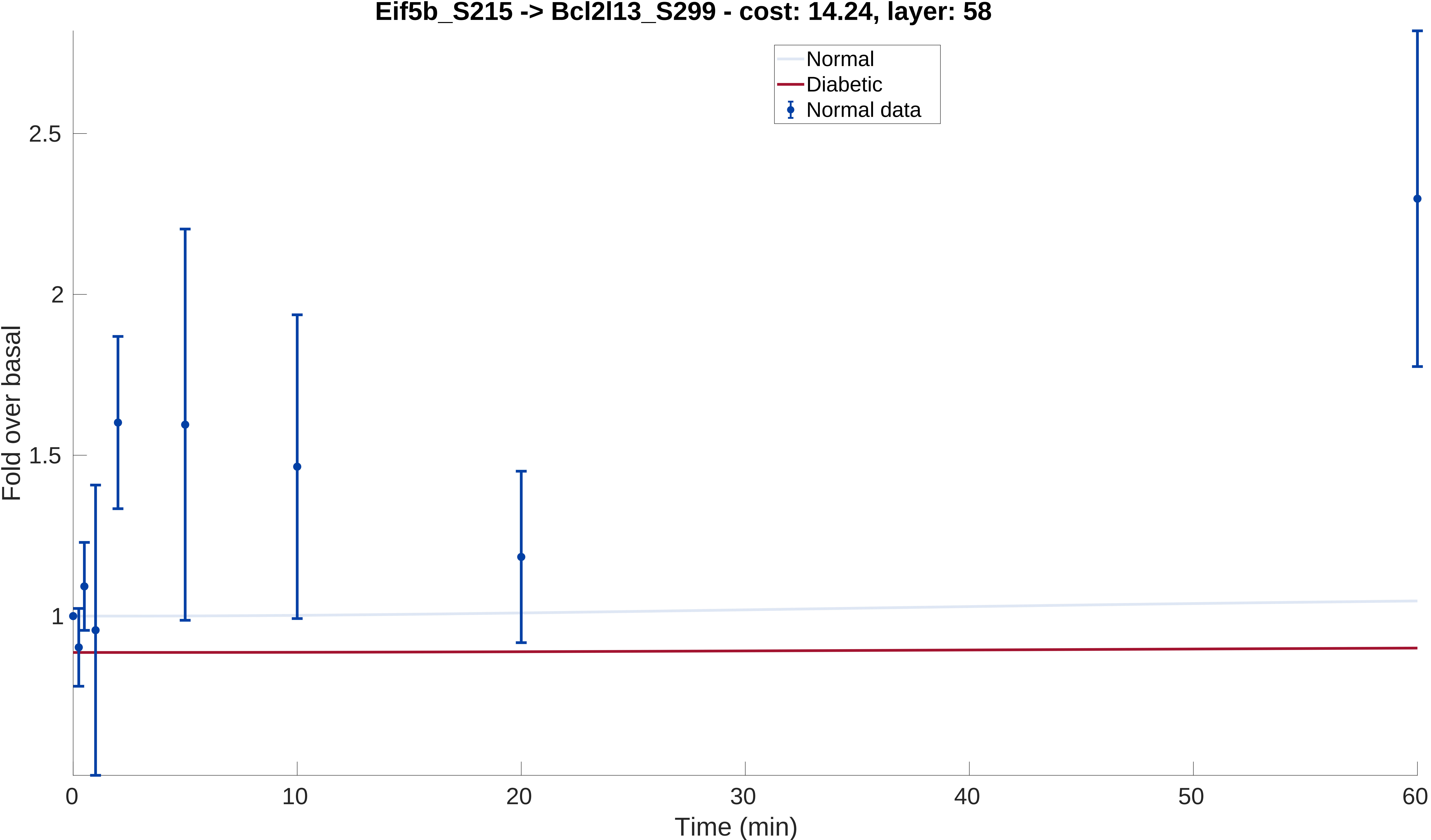
Eif5b_S215 -> Bcl2l13_S299 - cost: 14.24, layer: 58
Normal
Diabetic
Normal data
Fold over basal
2.5
2
1.5
1
0
10
20
30
40
50
60
Time (min)

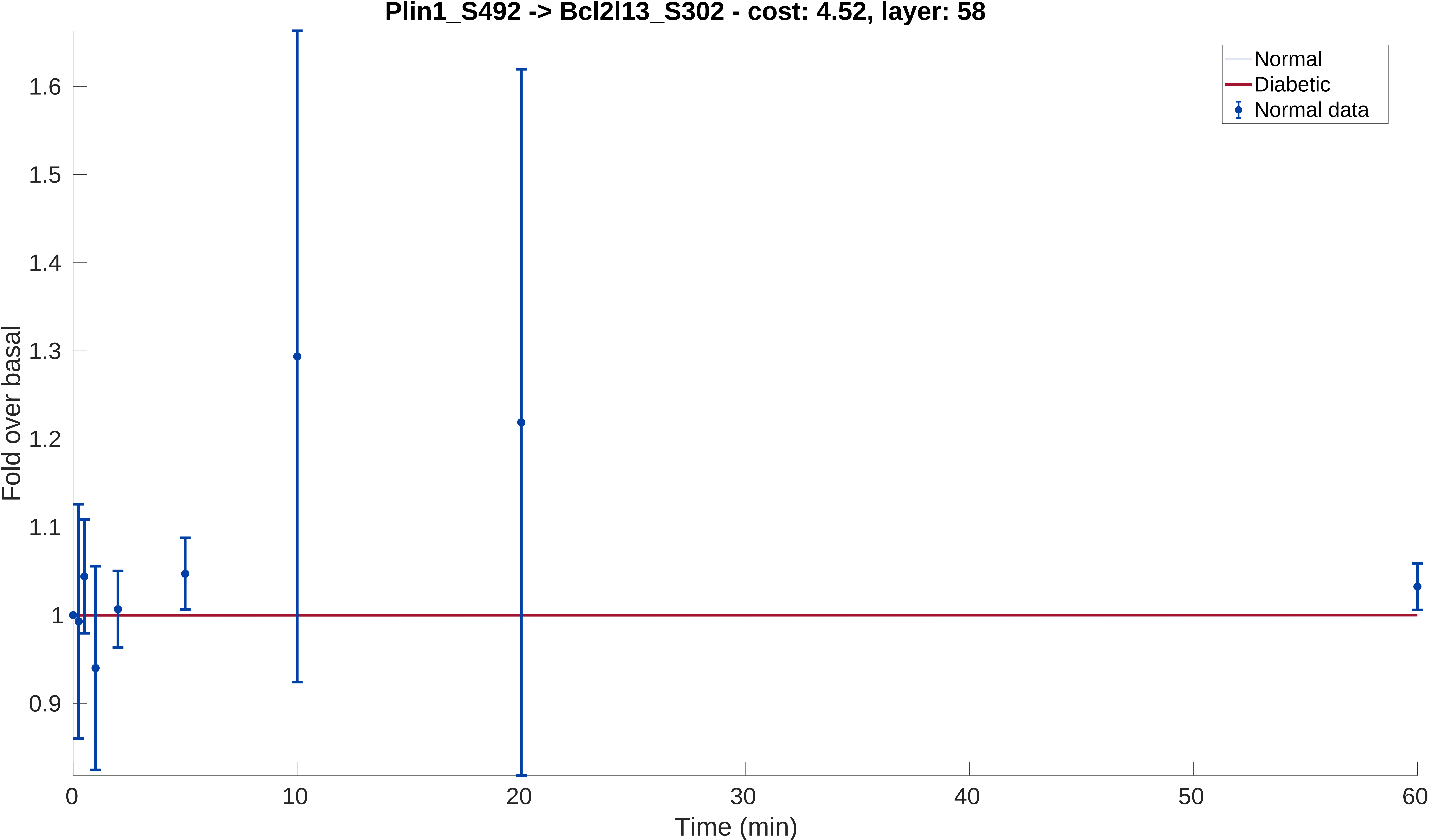
Plin1_S492 -> Bcl2l13_S302 - cost: 4.52, layer: 58
Normal
Diabetic
Normal data
Fold over basal
1.6
1.5
1.4
1.3
1.2
1.1
1
0.9
0
10
20
30
40
50
60
Time (min)

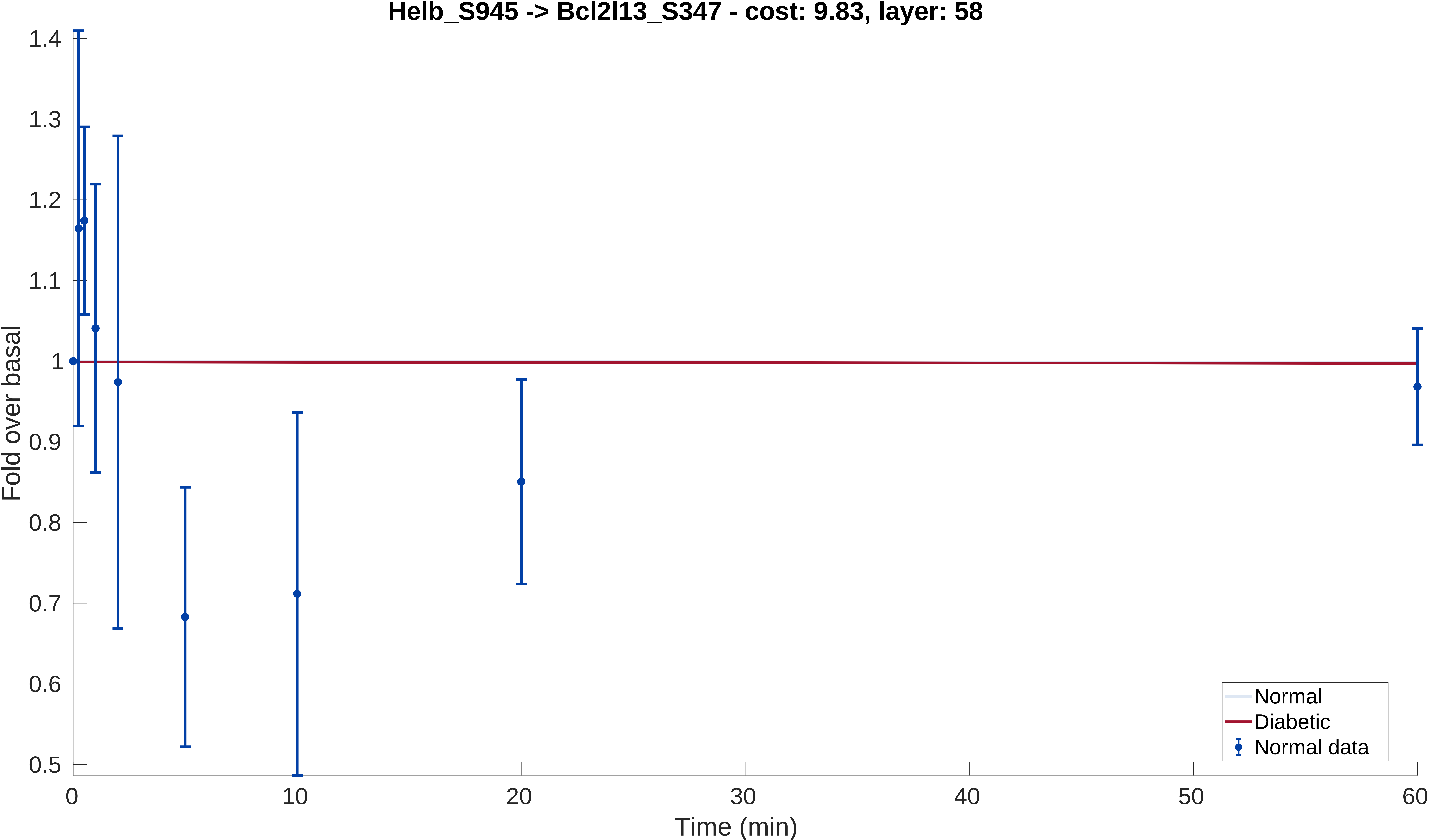

Helb_S945 -> Bcl2l13_S347 - cost: 9.83, layer: 58
Fold over basal
1.4
1.3
1.2
1.1
1
0.9
0.8
0.7
0.6
0.5
0
10
20
30
40
50
60
Time (min)
Normal
Diabetic
Normal data

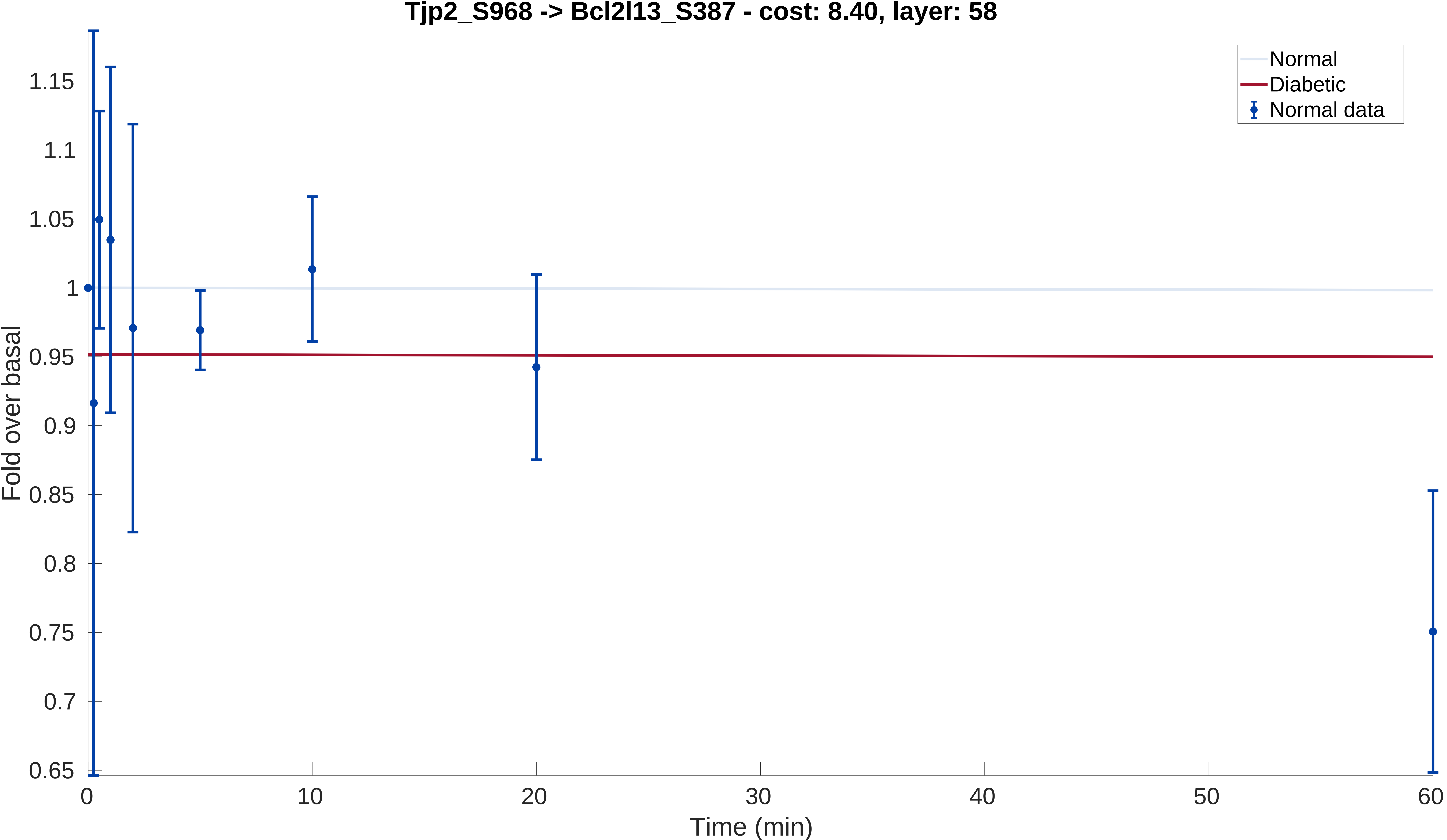

Tjp2_S968 -> Bcl2l13_S387 - cost: 8.40, layer: 58
Normal
Diabetic
Normal data
Fold over basal
1.15
1.1
1.05
1
0.95
0.9
0.85
0.8
0.75
0.7
0.65
0
10
20
30
40
50
60
Time (min)

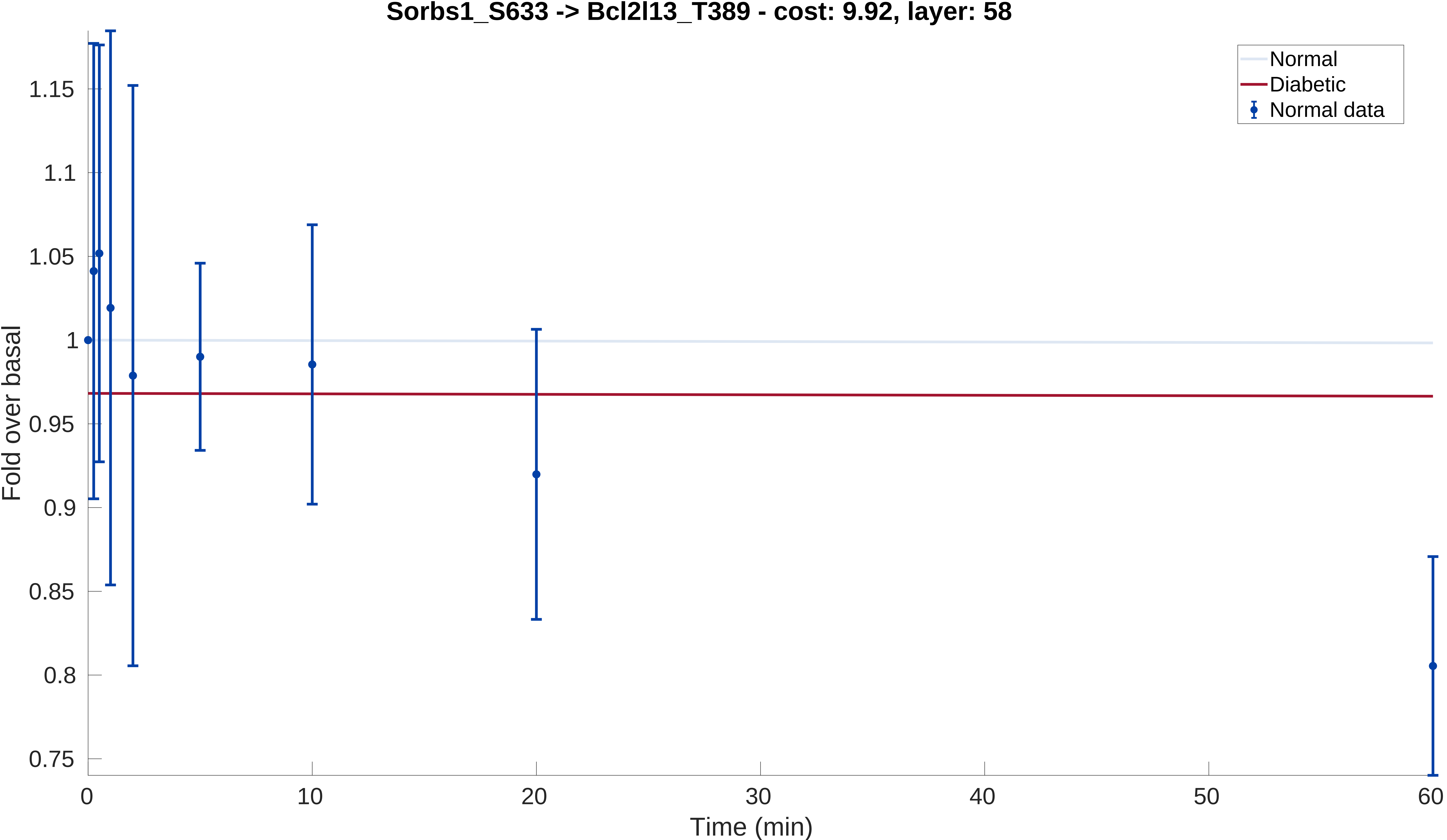

Sorbs1_S633 -> Bcl2l13_T389 - cost: 9.92, layer: 58
Normal
Diabetic
Normal data
Fold over basal
1.15
1.1
1.05
1
0.95
0.9
0.85
0.8
0.75
0
10
20
30
40
50
60
Time (min)

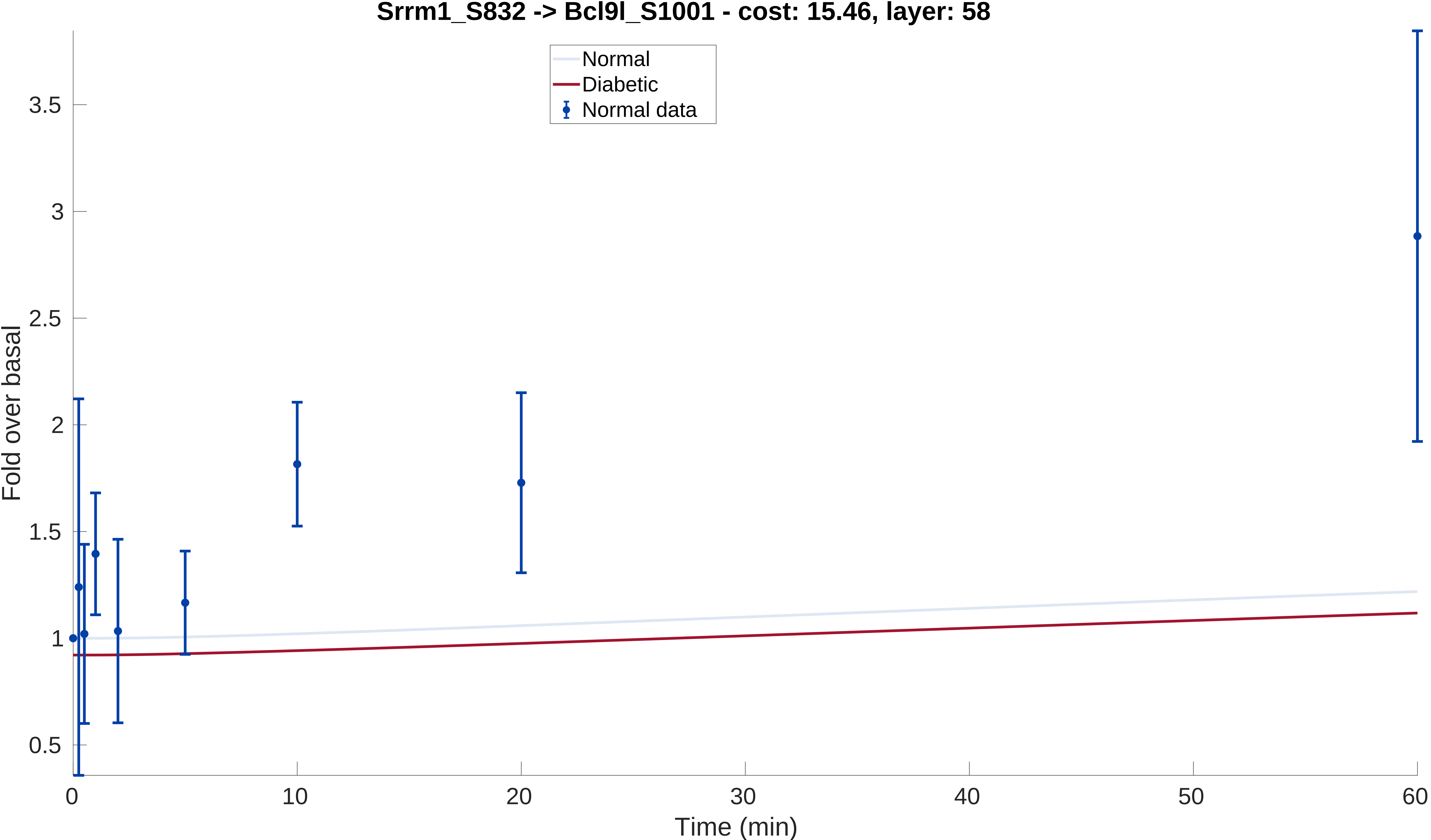

Srrm1_S832 -> Bcl9l_S1001 - cost: 15.46, layer: 58
Normal
Diabetic
Normal data
Fold over basal
Time (min)
0.5
1
1.5
2
2.5
3
3.5
0
10
20
30
40
50
60

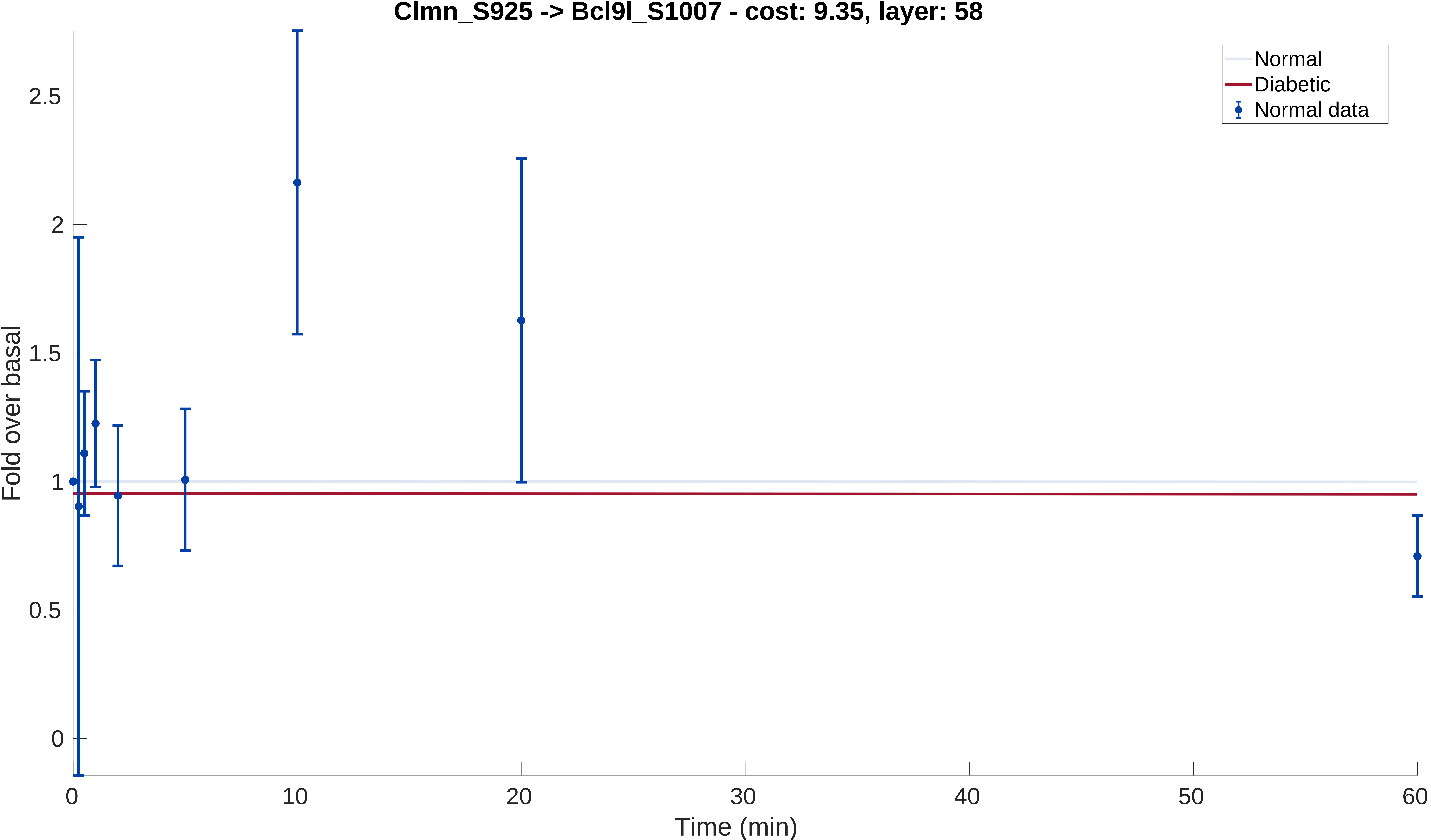

Clmn_S925 -> Bcl9l_S1007 - cost: 9.35, layer: 58
Normal
Diabetic
Normal data
Fold over basal
Time (min)
0
0.5
1
1.5
2
2.5
0
10
20
30
40
50
60

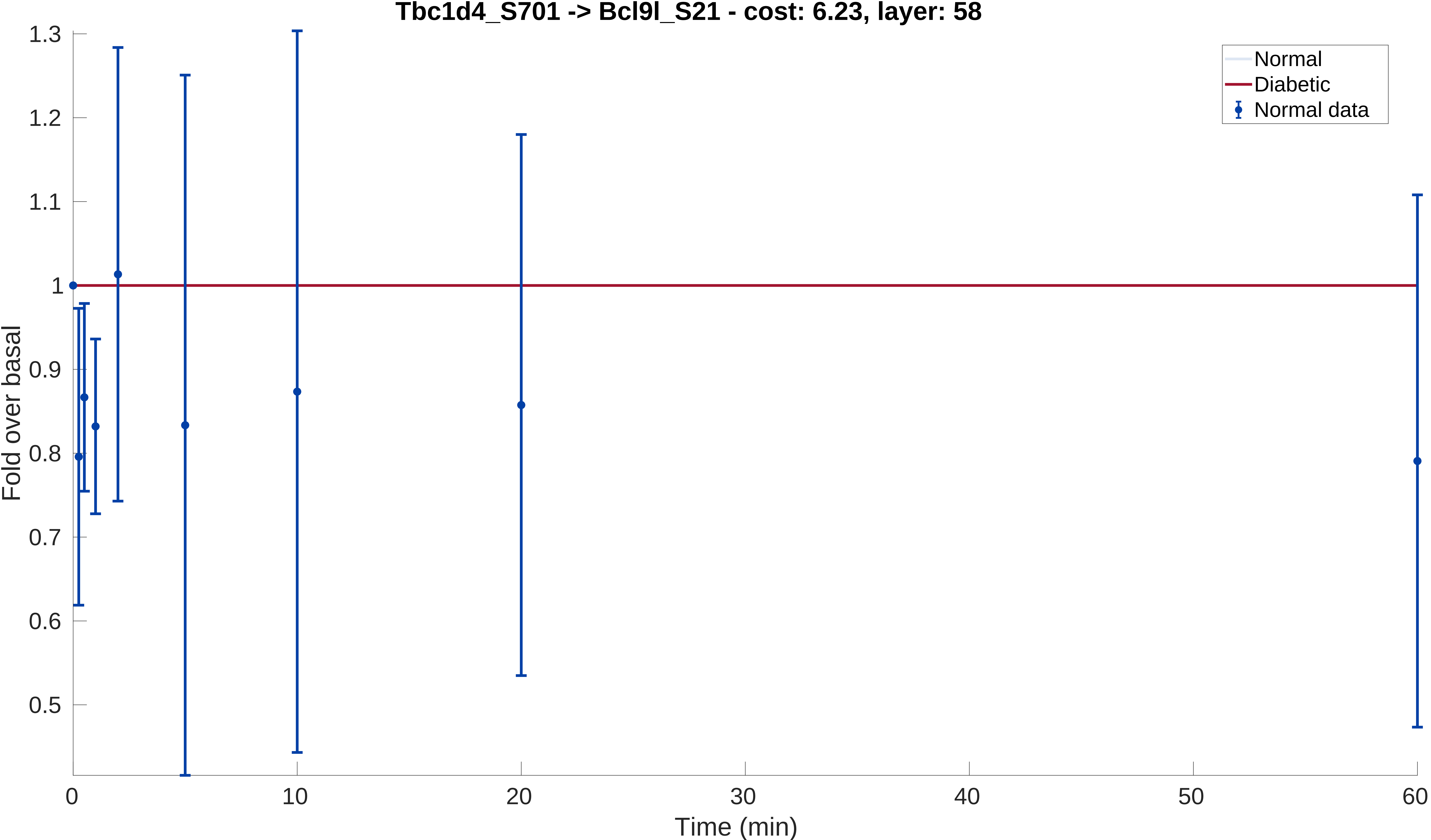
Tbc1d4_S701 -> Bcl9l_S21 - cost: 6.23, layer: 58
Normal
Diabetic
Normal data
Fold over basal
Time (min)
1.3
1.2
1.1
1
0.9
0.8
0.7
0.6
0.5
0
10
20
30
40
50
60

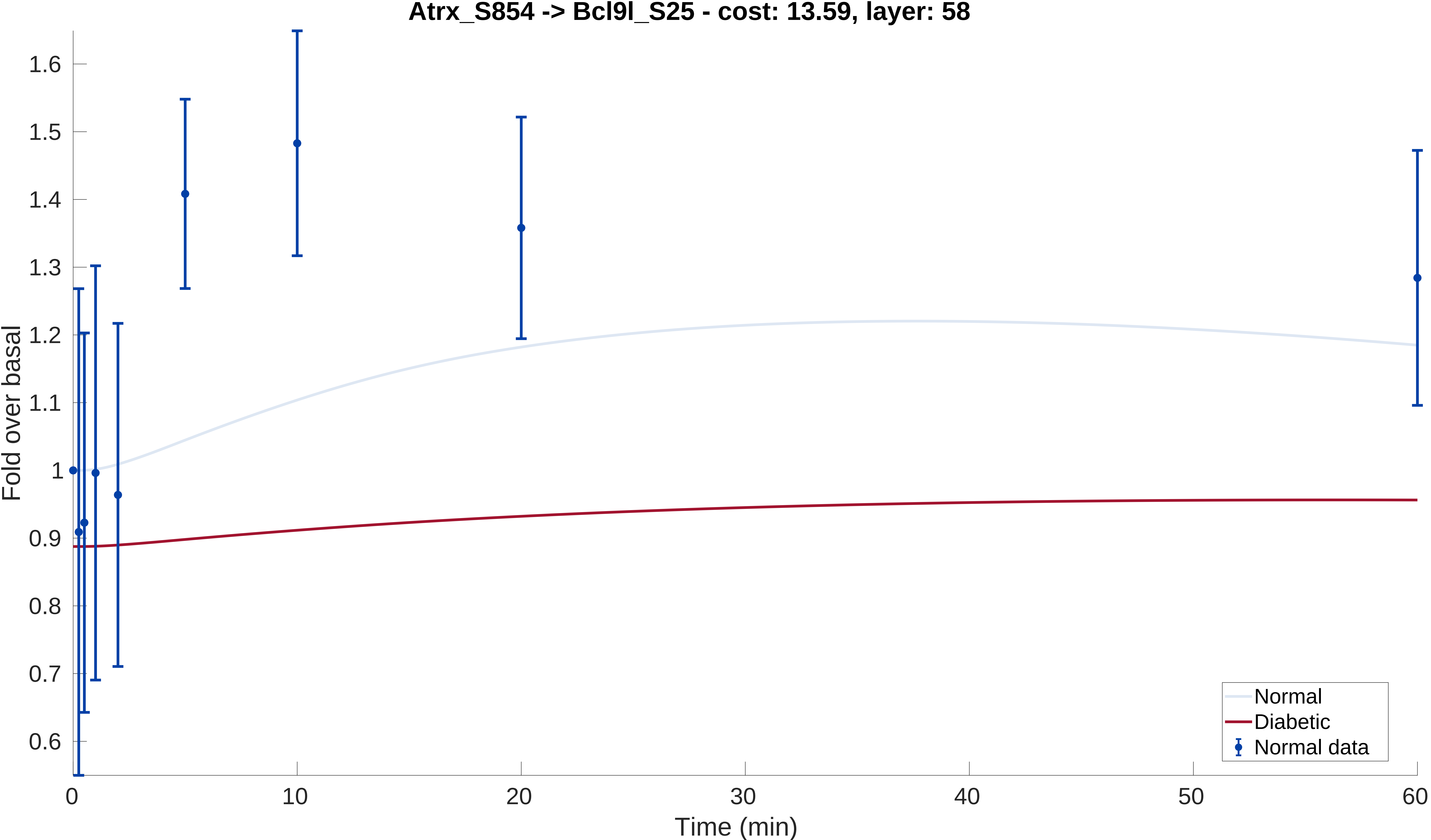
Atrx_S854 -> Bcl9l_S25 - cost: 13.59, layer: 58
Fold over basal
Time (min)
1.6
1.5
1.4
1.3
1.2
1.1
1
0.9
0.8
0.7
0.6
0
10
20
30
40
50
60
Normal
Diabetic
Normal data

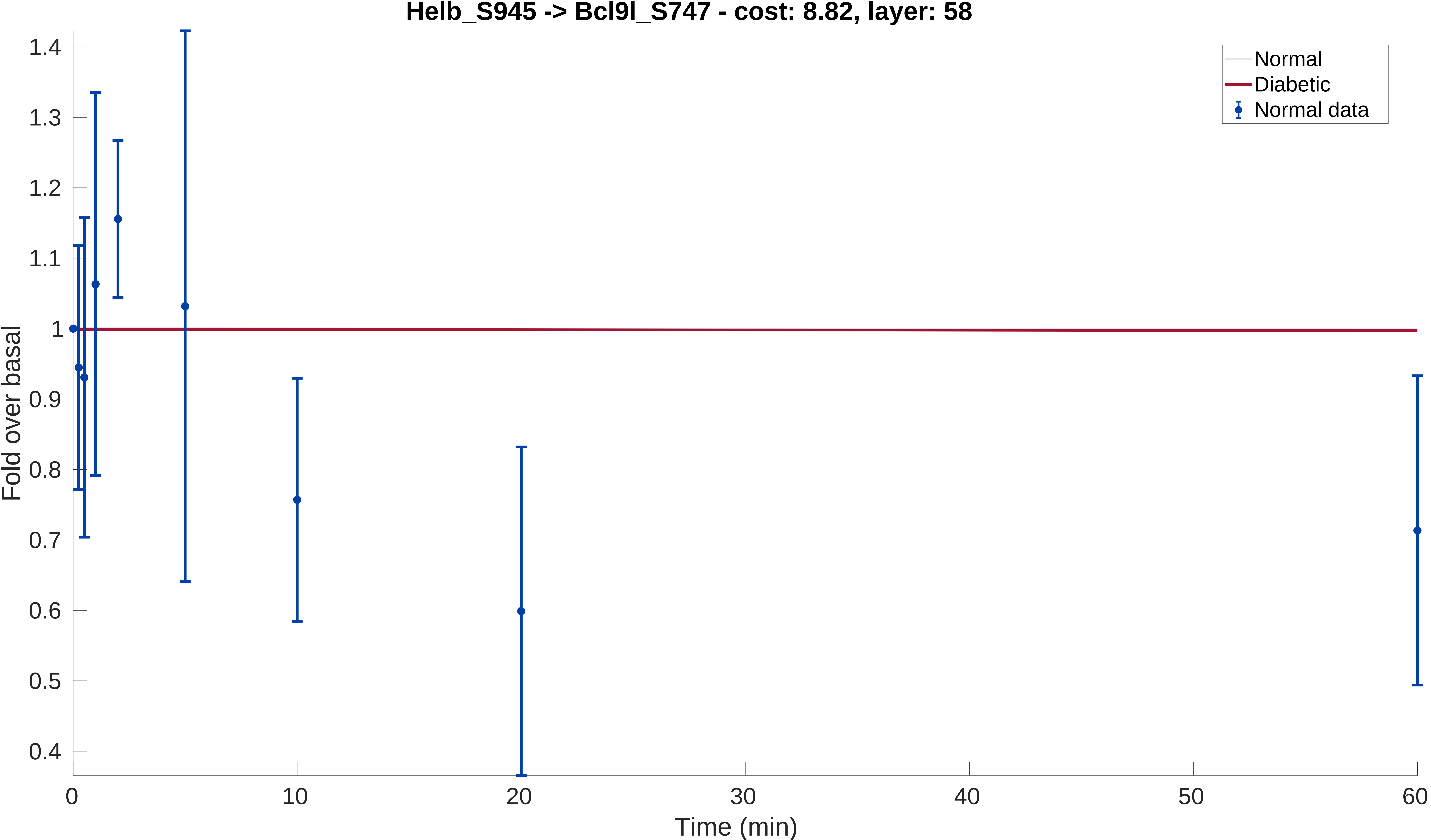

Helb_S945 -> Bcl9l_S747 - cost: 8.82, layer: 58
Normal
Diabetic
Normal data
Fold over basal
1.4
1.3
1.2
1.1
1
0.9
0.8
0.7
0.6
0.5
0.4
0
10
20
30
40
50
60
Time (min)

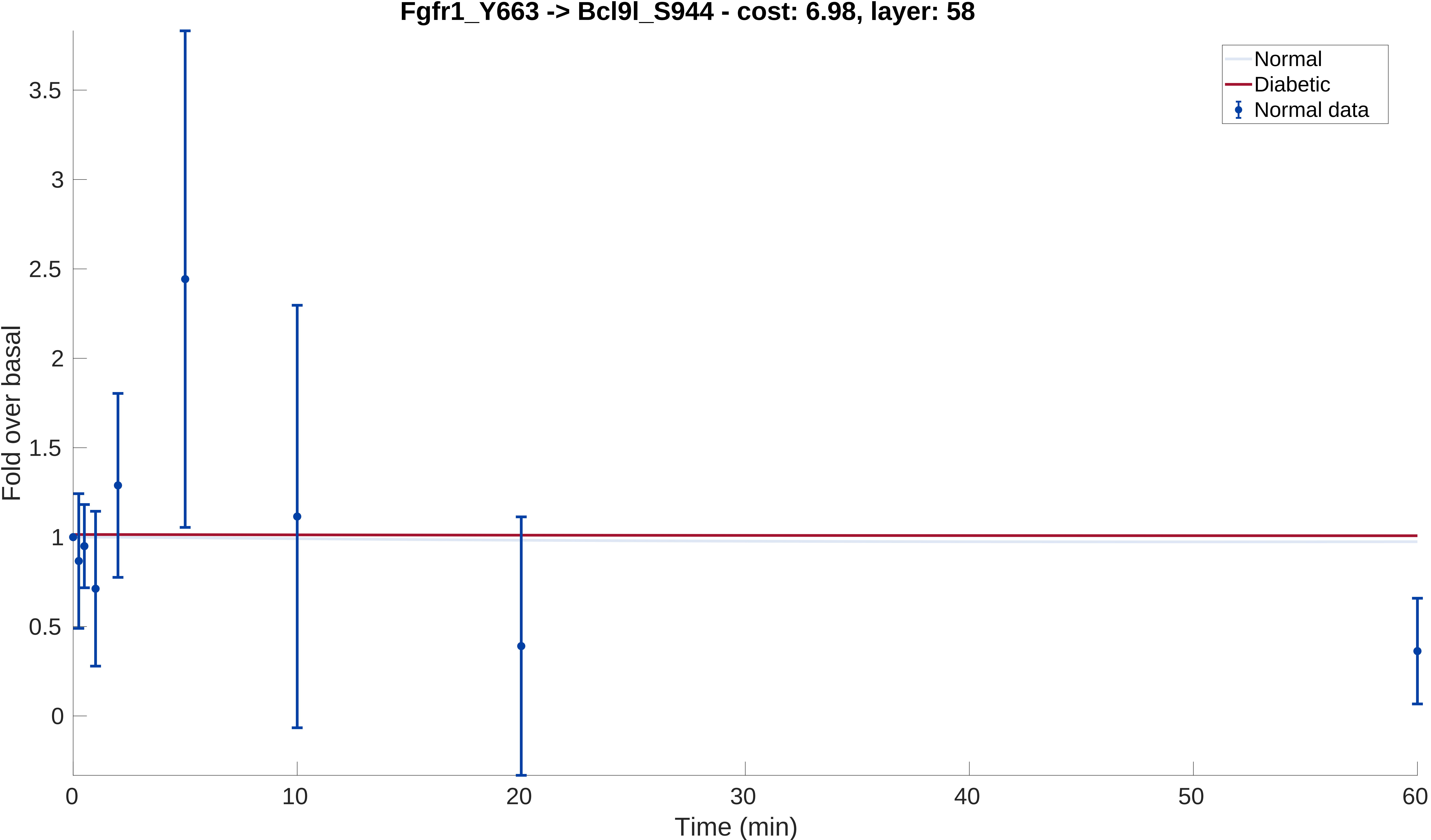

Fgfr1_Y663 -> Bcl9l_S944 - cost: 6.98, layer: 58
Normal
Diabetic
Normal data
Fold over basal
Time (min)
0
0.5
1
1.5
2
2.5
3
3.5
0
10
20
30
40
50
60

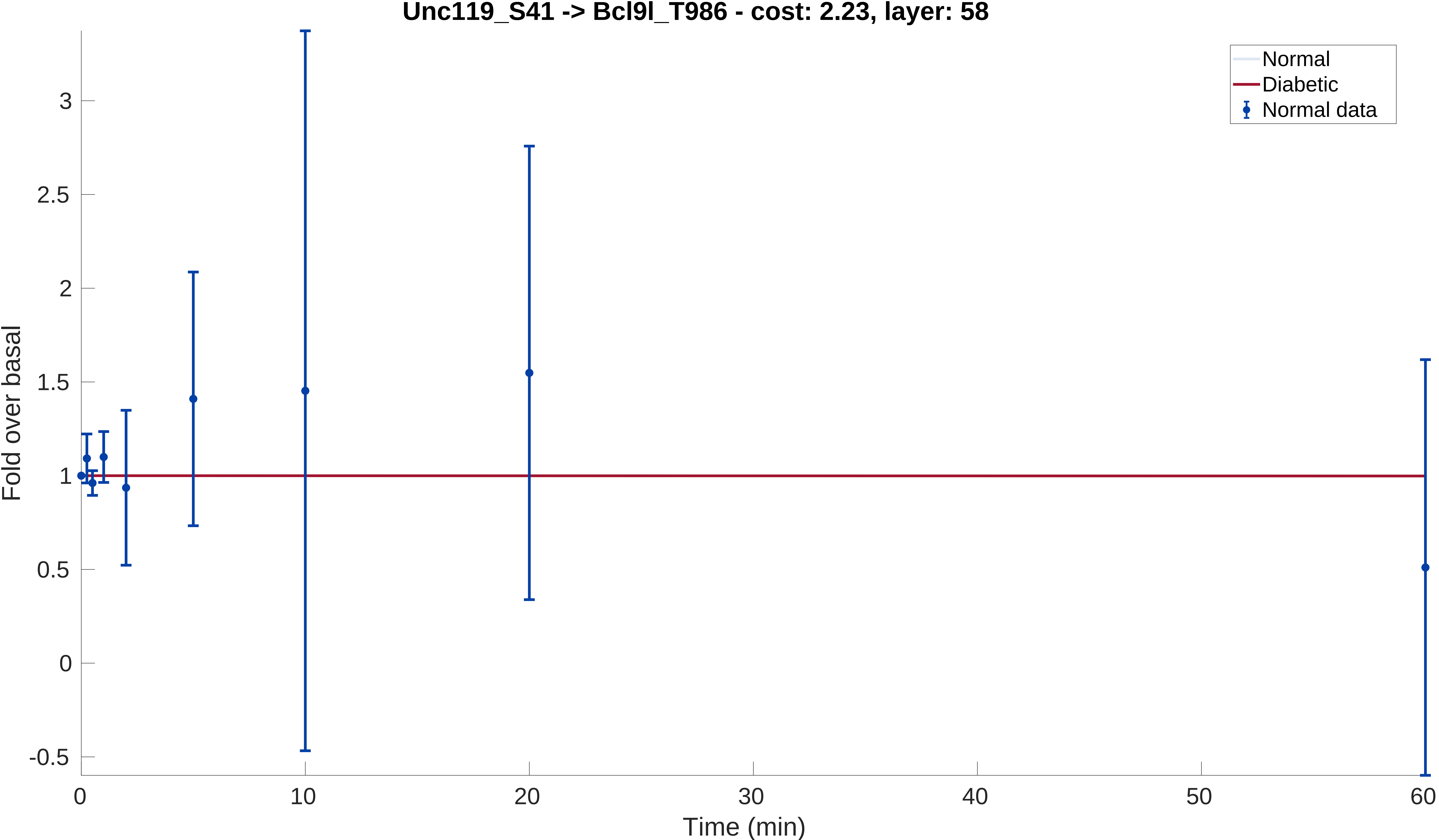
Unc119_S41 -> Bcl9l_T986 - cost: 2.23, layer: 58
Normal
Diabetic
Normal data
Fold over basal
Time (min)
3
2.5
2
1.5
1
0.5
0
-0.5
0
10
20
30
40
50
60

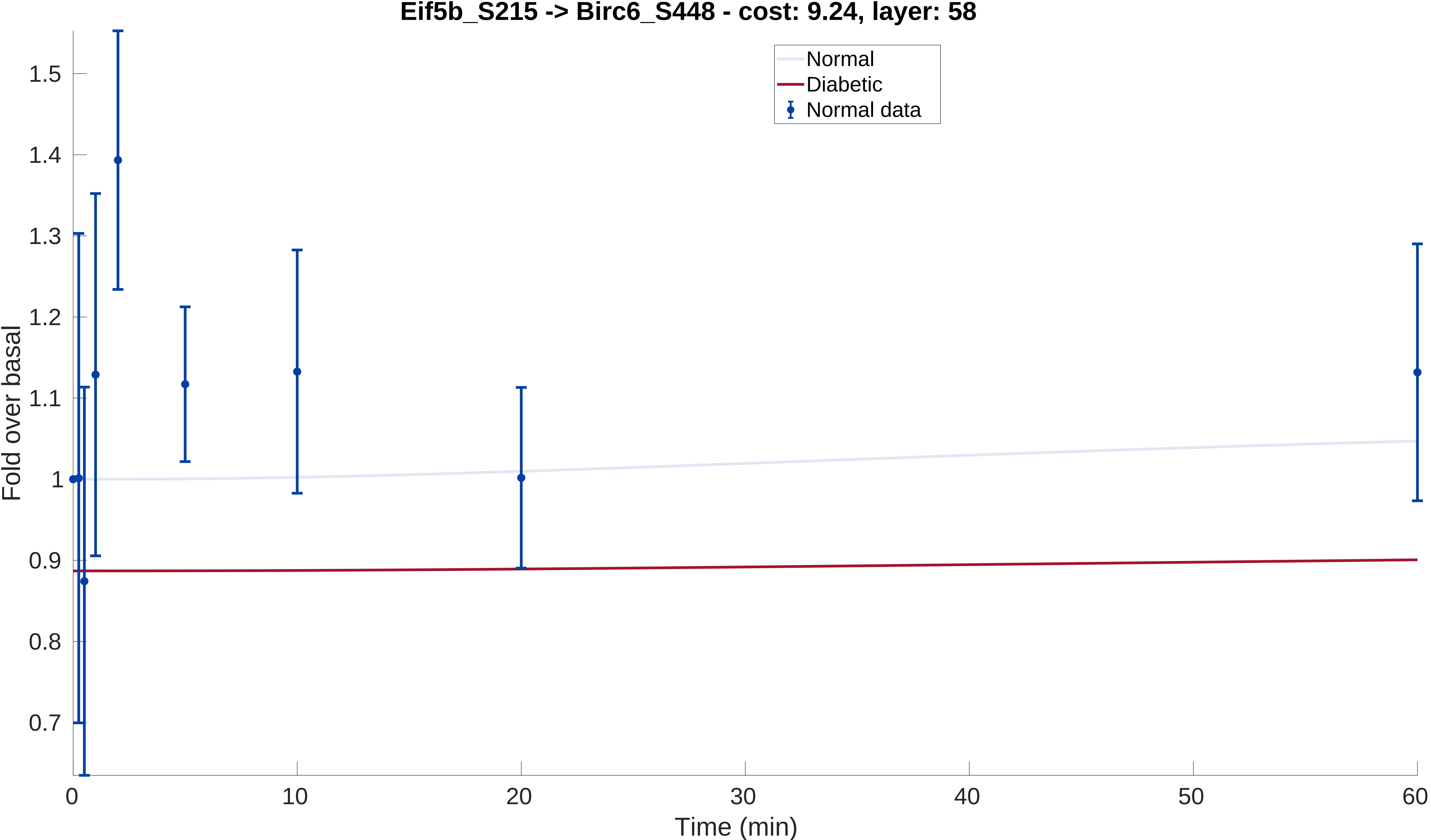
Eif5b_S215 -> Birc6_S448 - cost: 9.24, layer: 58
Normal
Diabetic
Normal data
Fold over basal
1.5
1.4
1.3
1.2
1.1
1
0.9
0.8
0.7
0
10
20
30
40
50
60
Time (min)

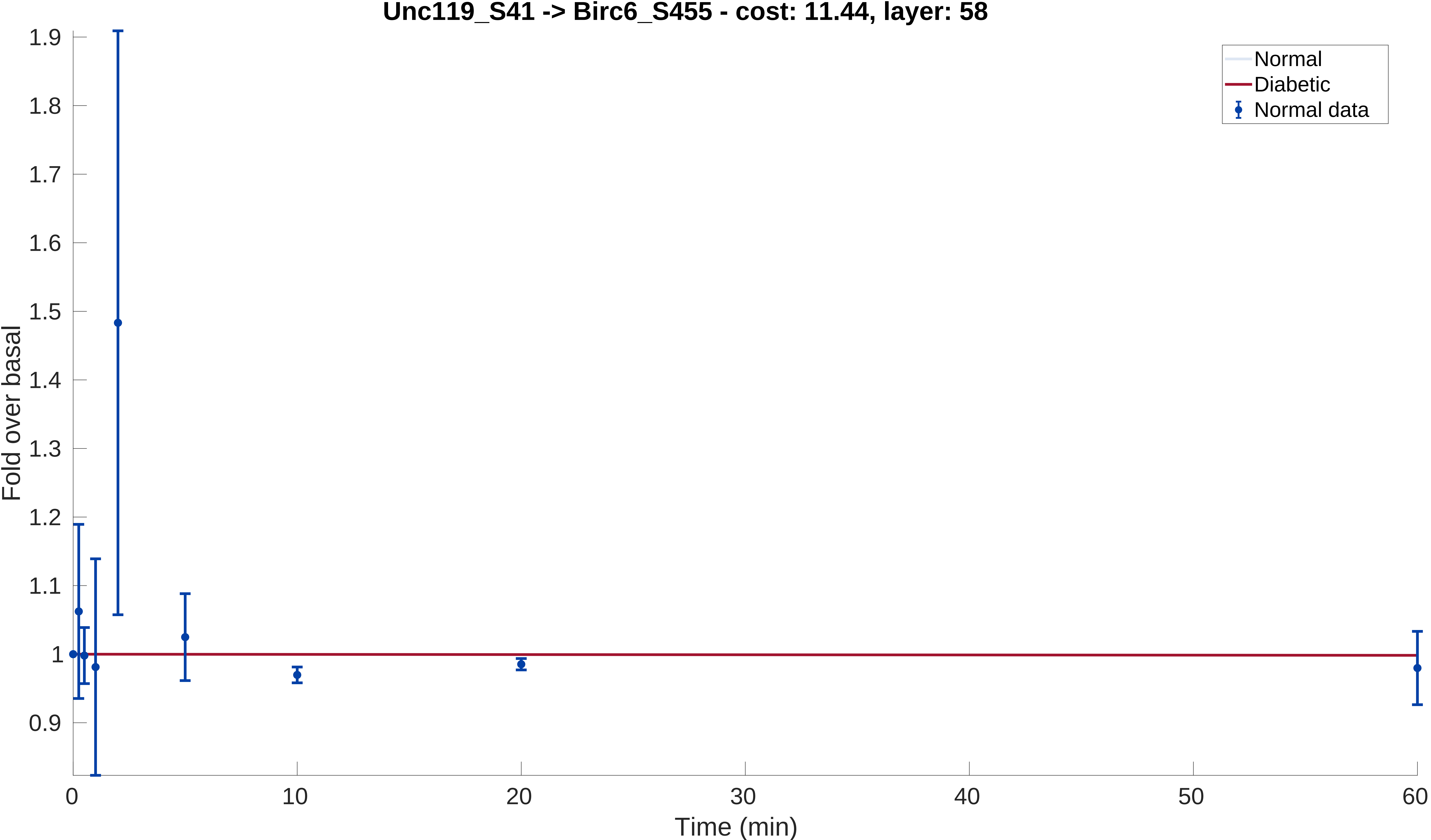

Unc119_S41 -> Birc6_S455 - cost: 11.44, layer: 58
Normal
Diabetic
Normal data
Fold over basal
1.9
1.8
1.7
1.6
1.5
1.4
1.3
1.2
1.1
1
0.9
0
10
20
30
40
50
60
Time (min)

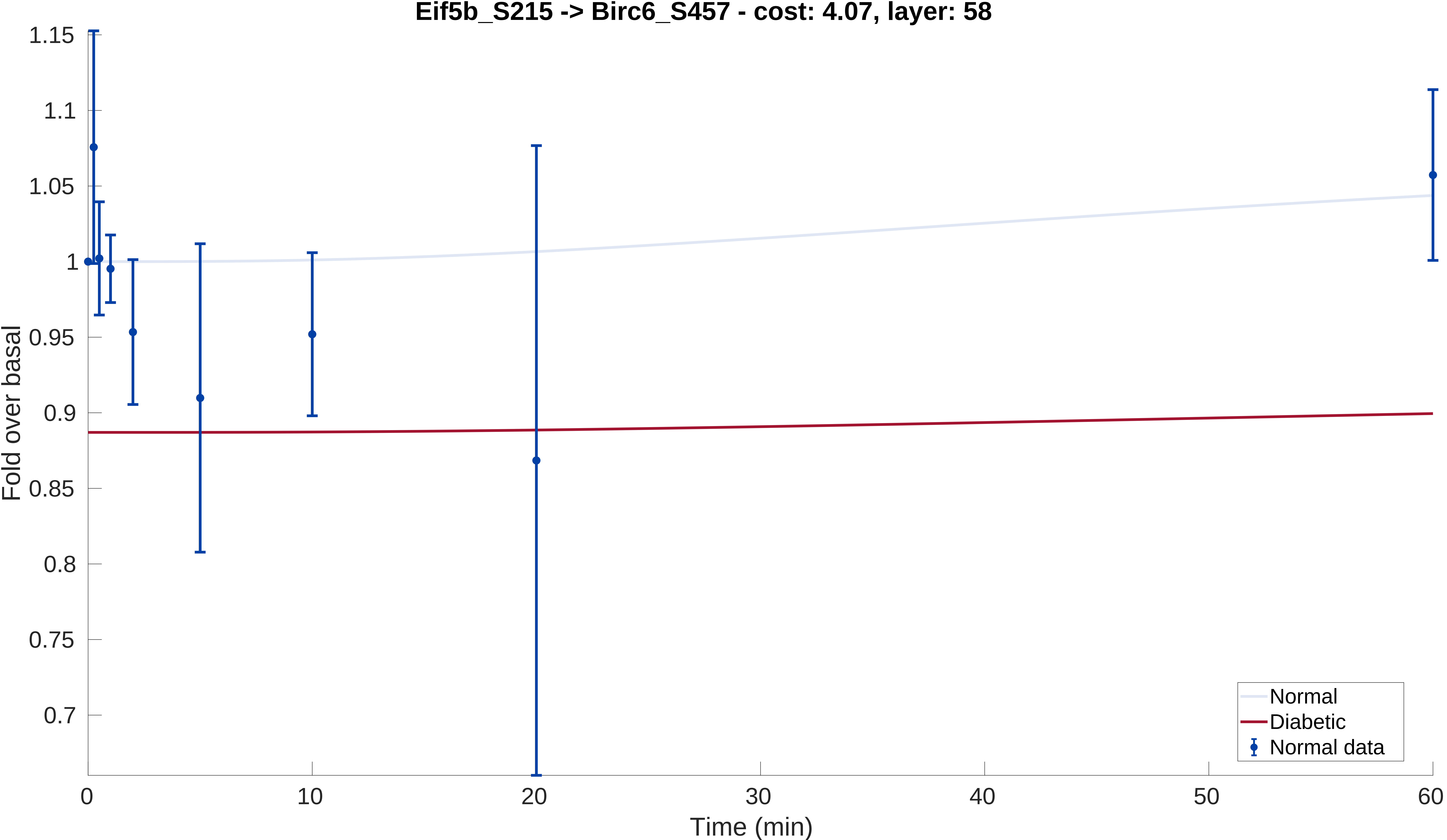

Eif5b_S215 -> Birc6_S457 - cost: 4.07, layer: 58
1.15
1.1
1.05
1
0.95
0.9
0.85
0.8
0.75
0.7
Fold over basal
0
10
20
30
40
50
60
Time (min)
Normal
Diabetic
Normal data

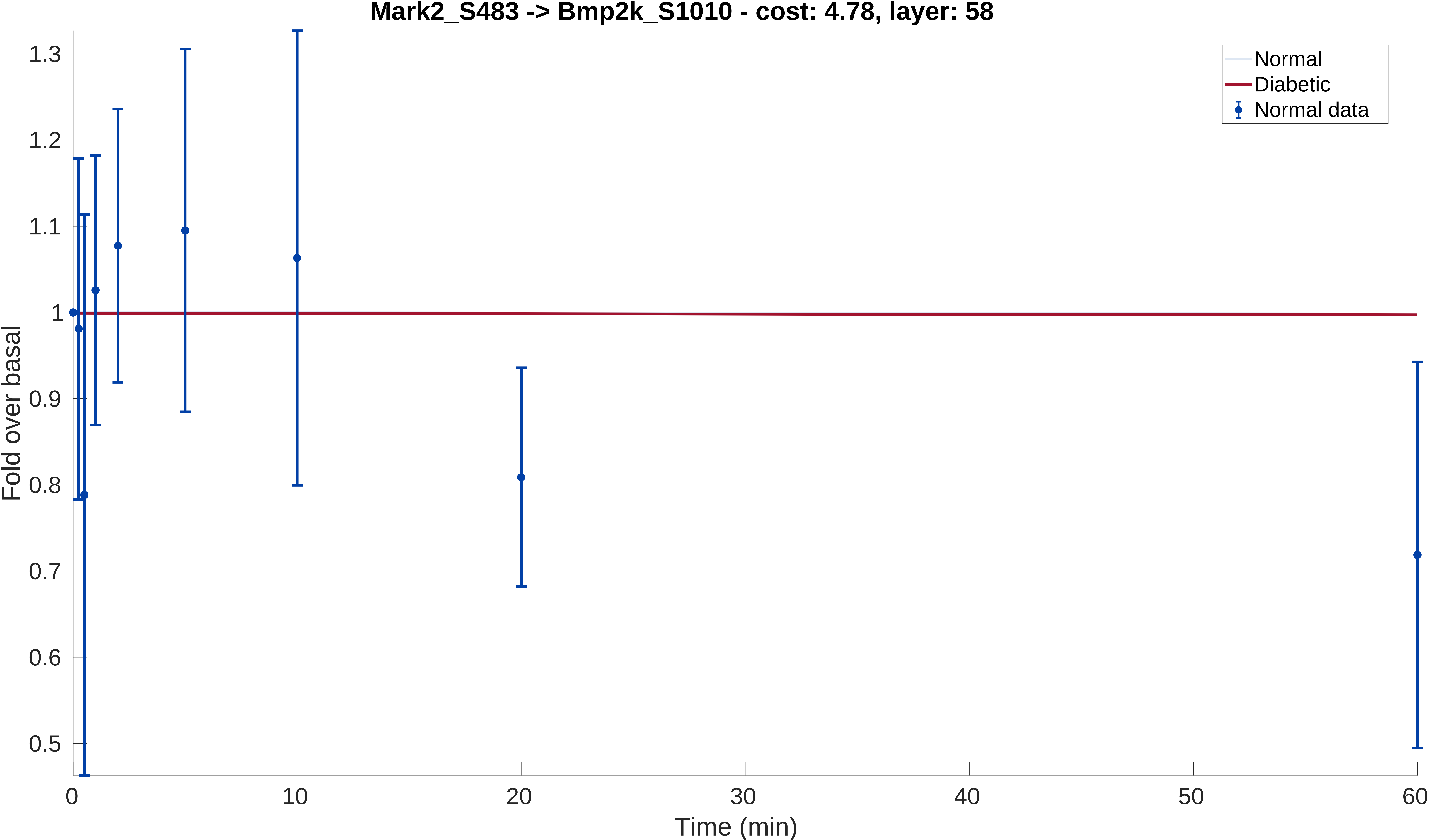

Mark2_S483 -> Bmp2k_S1010 - cost: 4.78, layer: 58
Normal
Diabetic
Normal data
Fold over basal
1.3
1.2
1.1
1
0.9
0.8
0.7
0.6
0.5
0
10
20
30
40
50
60
Time (min)

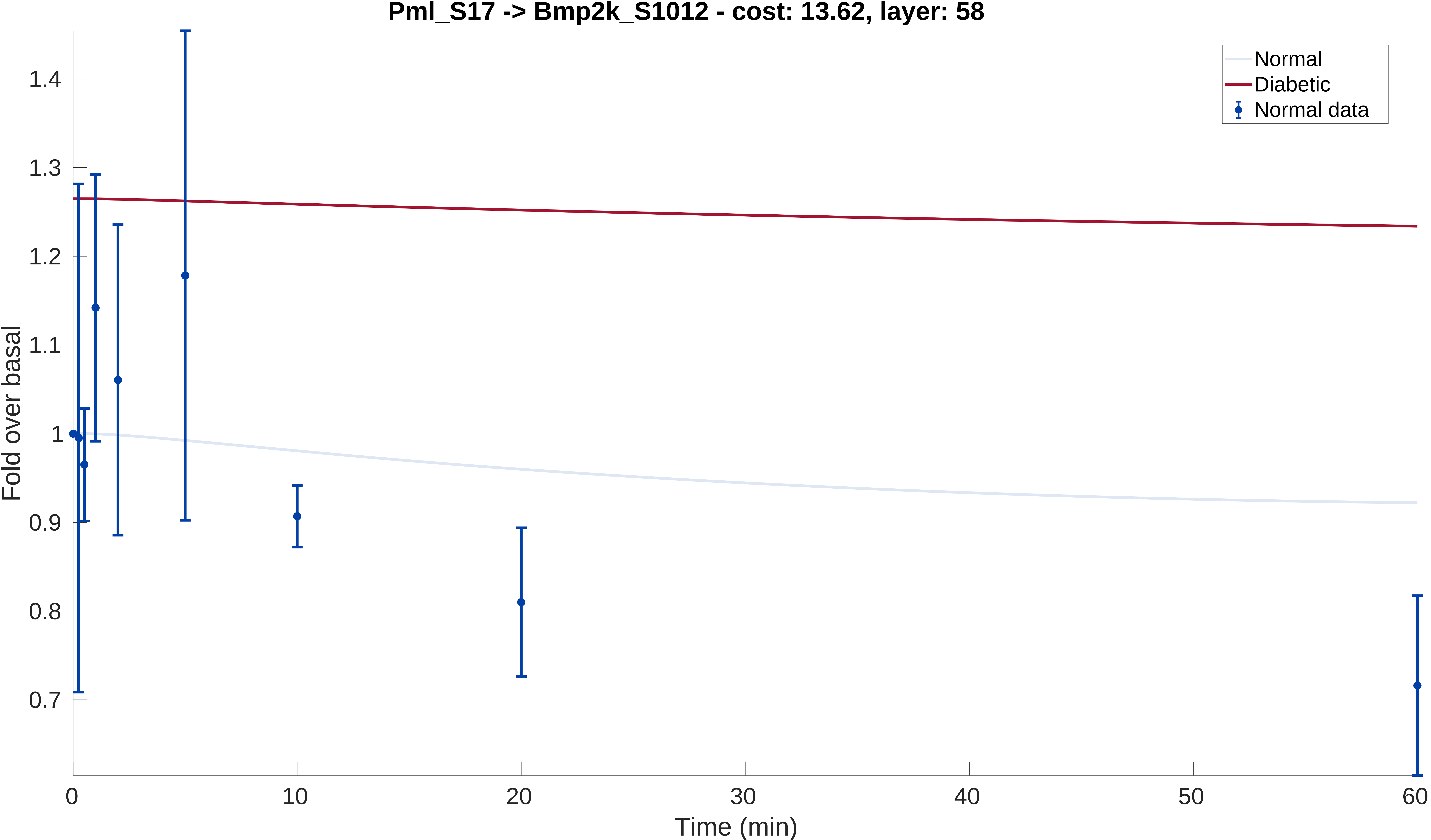

Pml_S17 -> Bmp2k_S1012 - cost: 13.62, layer: 58
Normal
Diabetic
Normal data
Fold over basal
1.4
1.3
1.2
1.1
1
0.9
0.8
0.7
0
10
20
30
40
50
60
Time (min)

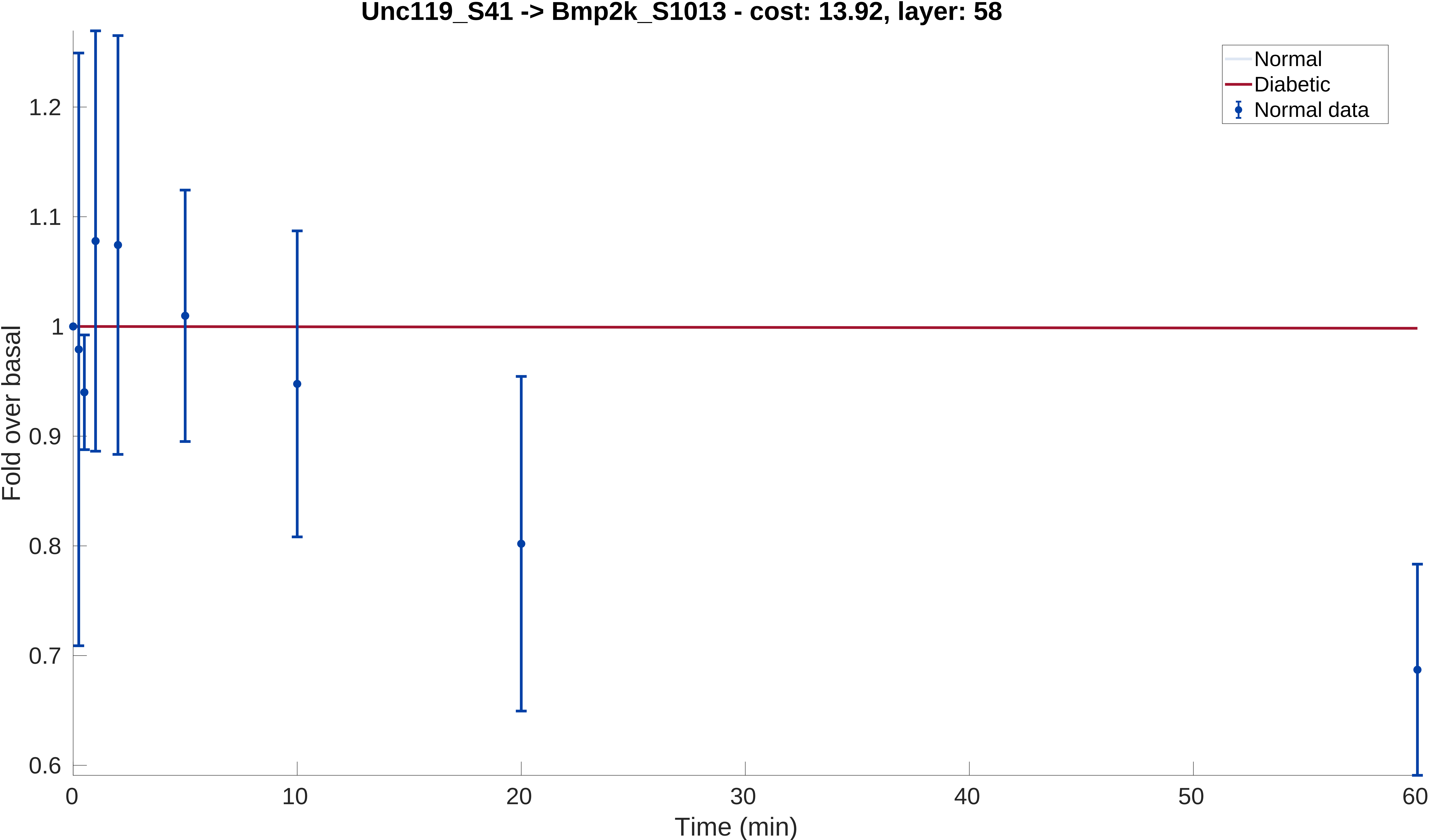
Unc119_S41 -> Bmp2k_S1013 - cost: 13.92, layer: 58
Normal
Diabetic
Normal data
Fold over basal
1.2
1.1
1
0.9
0.8
0.7
0.6
0
10
20
30
40
50
60
Time (min)

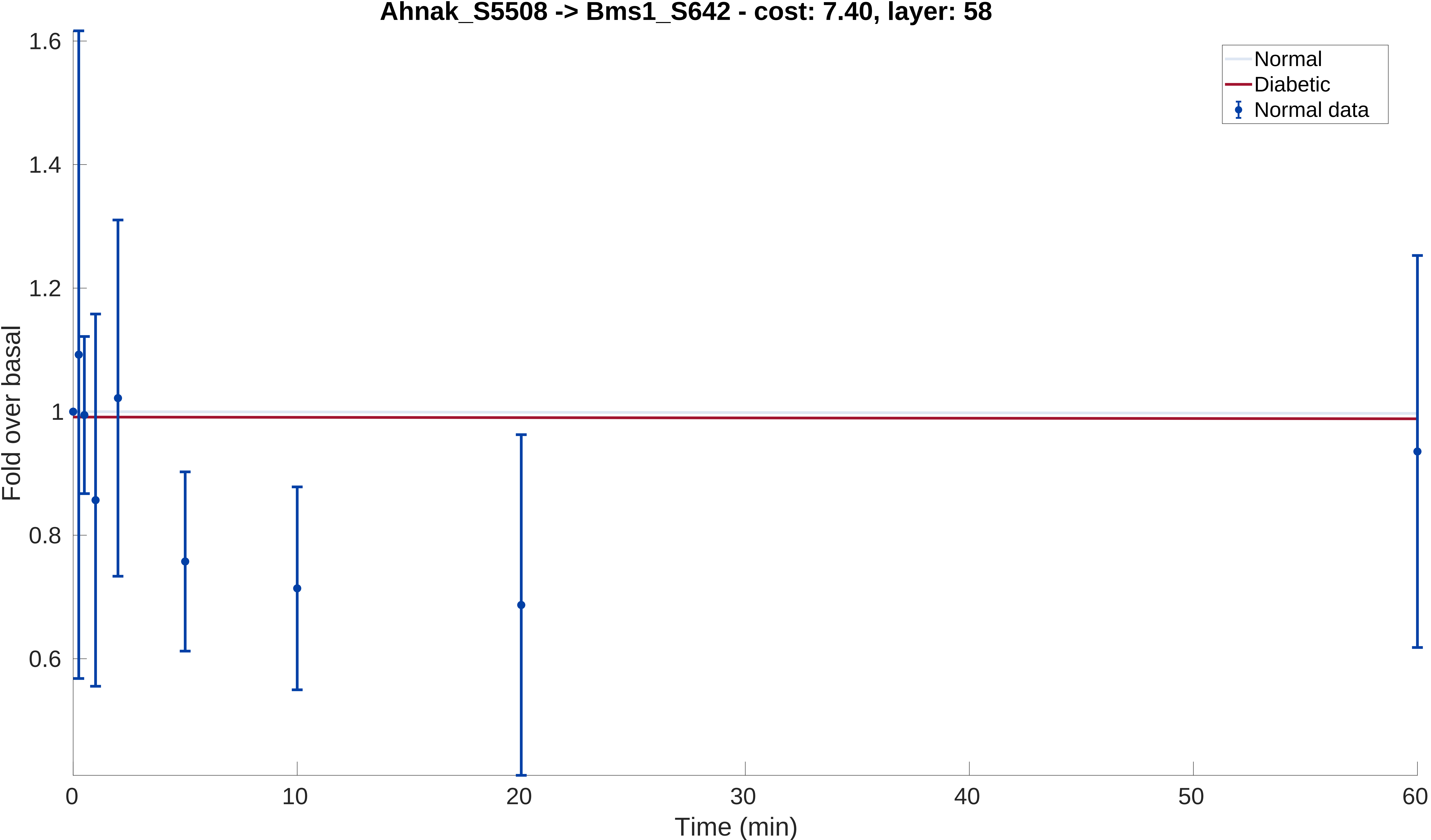

Ahnak_S5508 -> Bms1_S642 - cost: 7.40, layer: 58
Normal
Diabetic
Normal data
1.6
1.4
1.2
1
0.8
0.6
Fold over basal
0
10
20
30
40
50
60
Time (min)

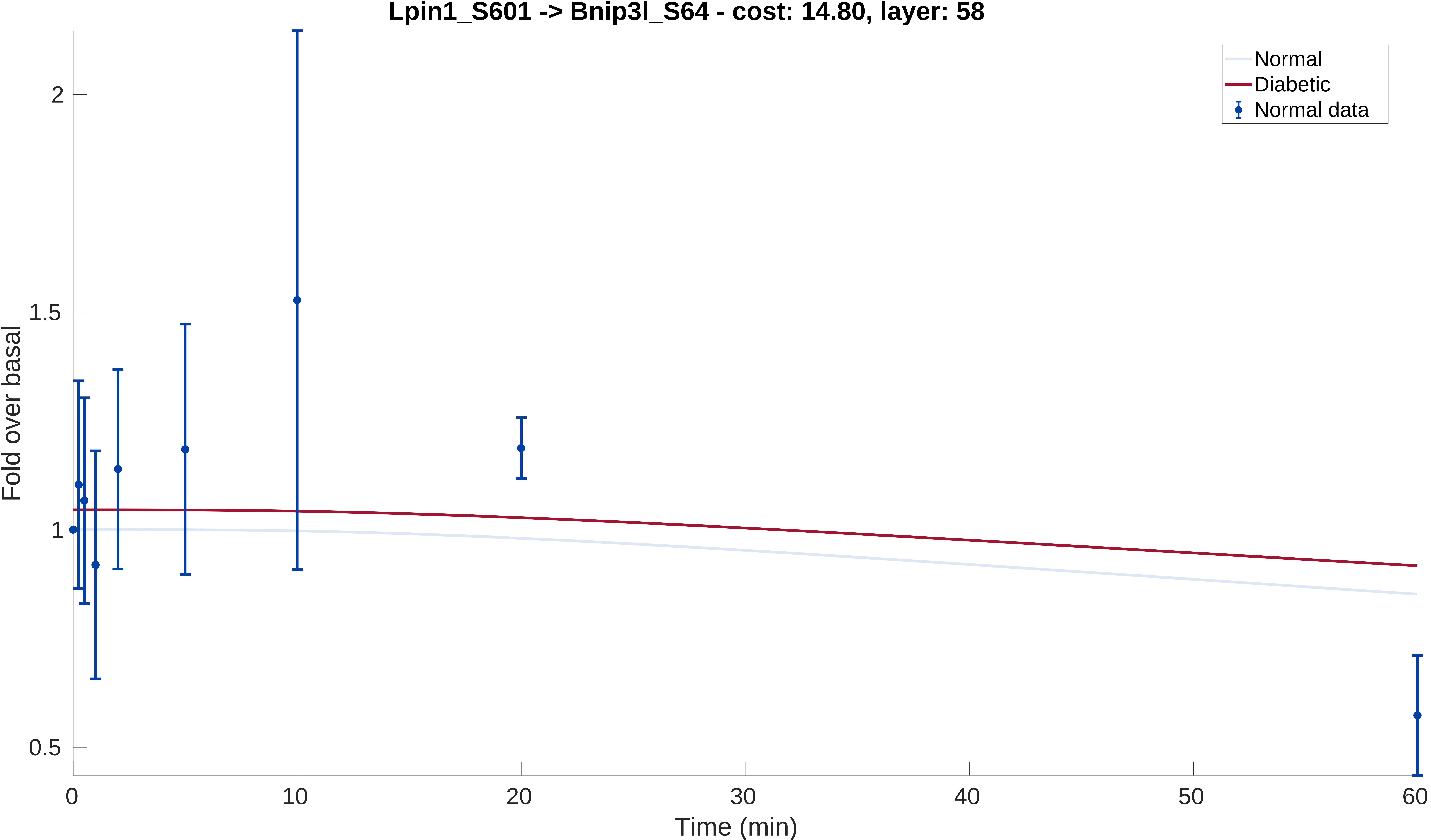

Lpin1_S601 -> Bnip3l_S64 - cost: 14.80, layer: 58
Normal
Diabetic
Normal data
Fold over basal
Time (min)
0.5
1
1.5
2
0
10
20
30
40
50
60

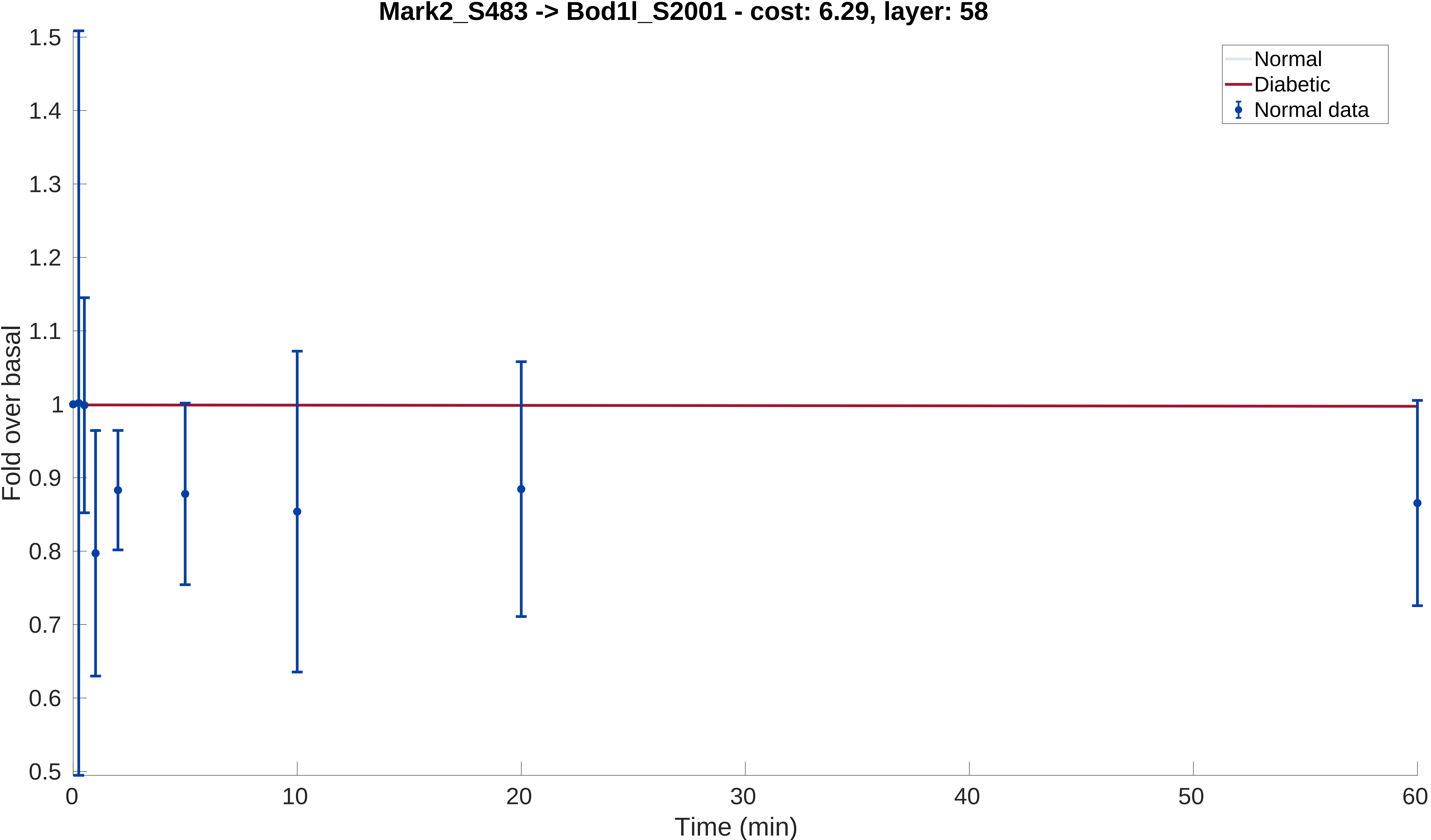
Mark2_S483 -> Bod1l_S2001 - cost: 6.29, layer: 58
Normal
Diabetic
Normal data
Fold over basal
Time (min)
1.5
1.4
1.3
1.2
1.1
1
0.9
0.8
0.7
0.6
0.5
0
10
20
30
40
50
60

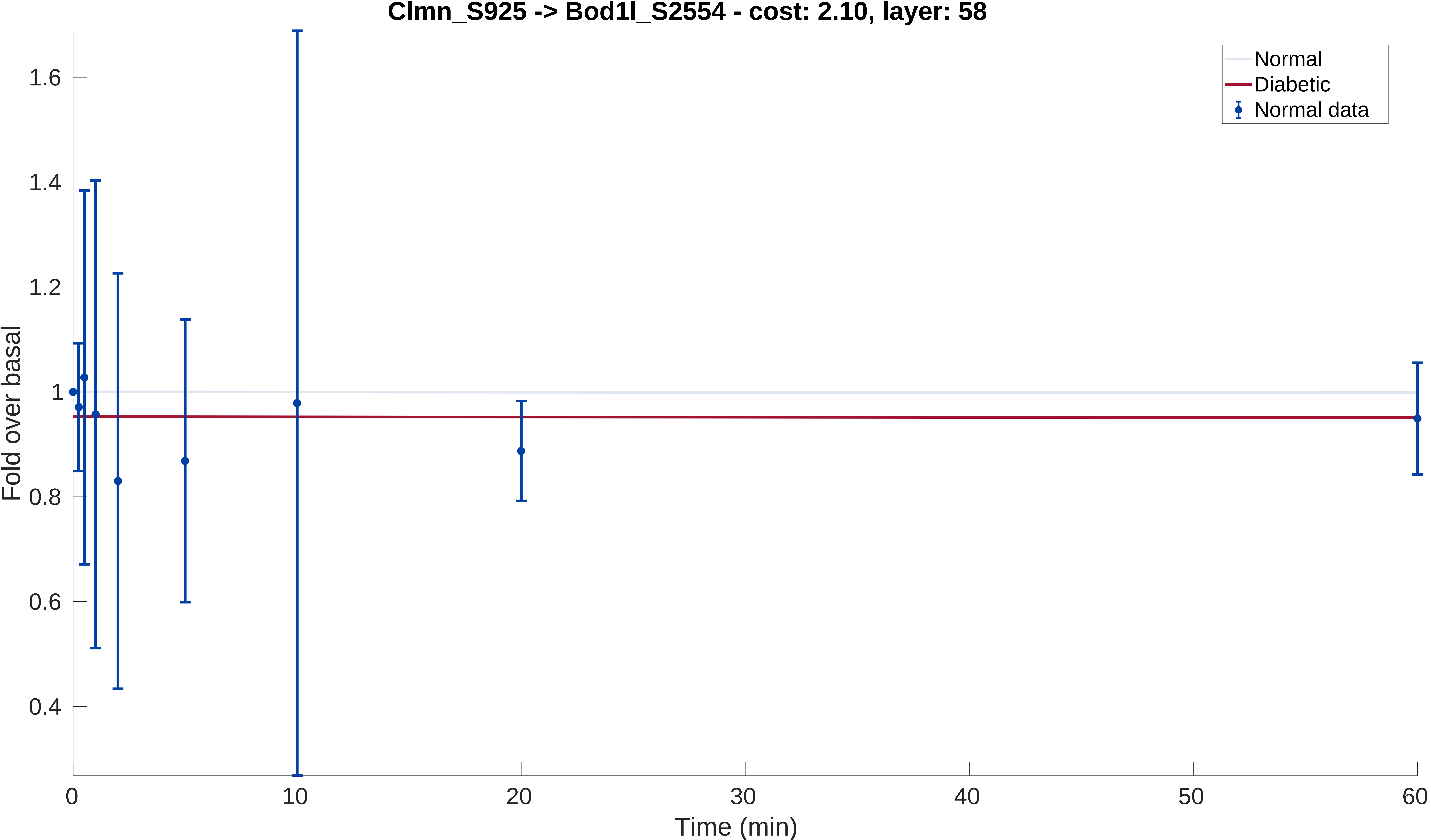

Clmn_S925 -> Bod1l_S2554 - cost: 2.10, layer: 58
Normal
Diabetic
Normal data
Fold over basal
1.6
1.4
1.2
1
0.8
0.6
0.4
0
10
20
30
40
50
60
Time (min)

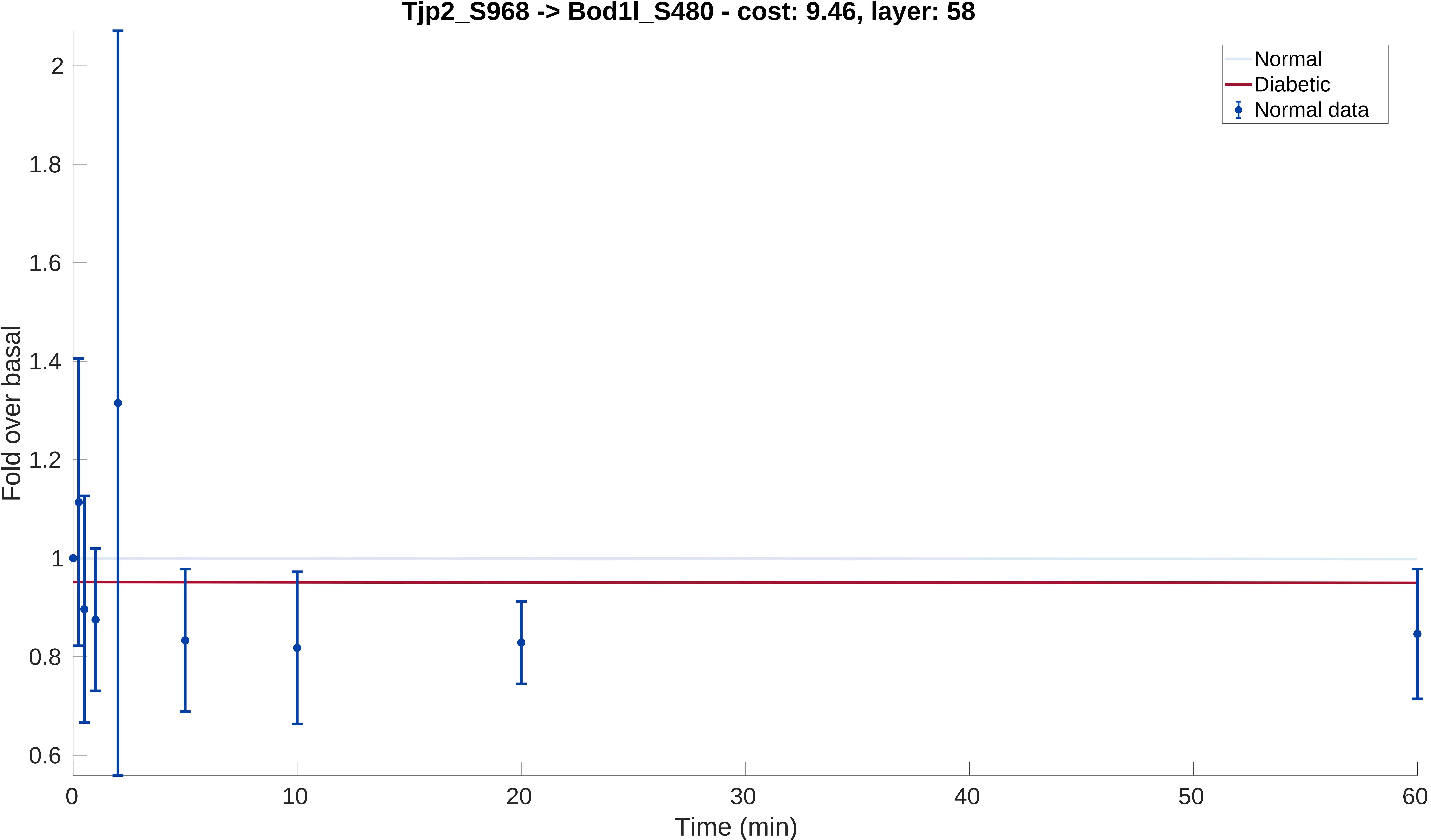
Tjp2_S968 -> Bod1l_S480 - cost: 9.46, layer: 58
Normal
Diabetic
Normal data
Fold over basal
0.6
0.8
1
1.2
1.4
1.6
1.8
2
0
10
20
30
40
50
60
Time (min)

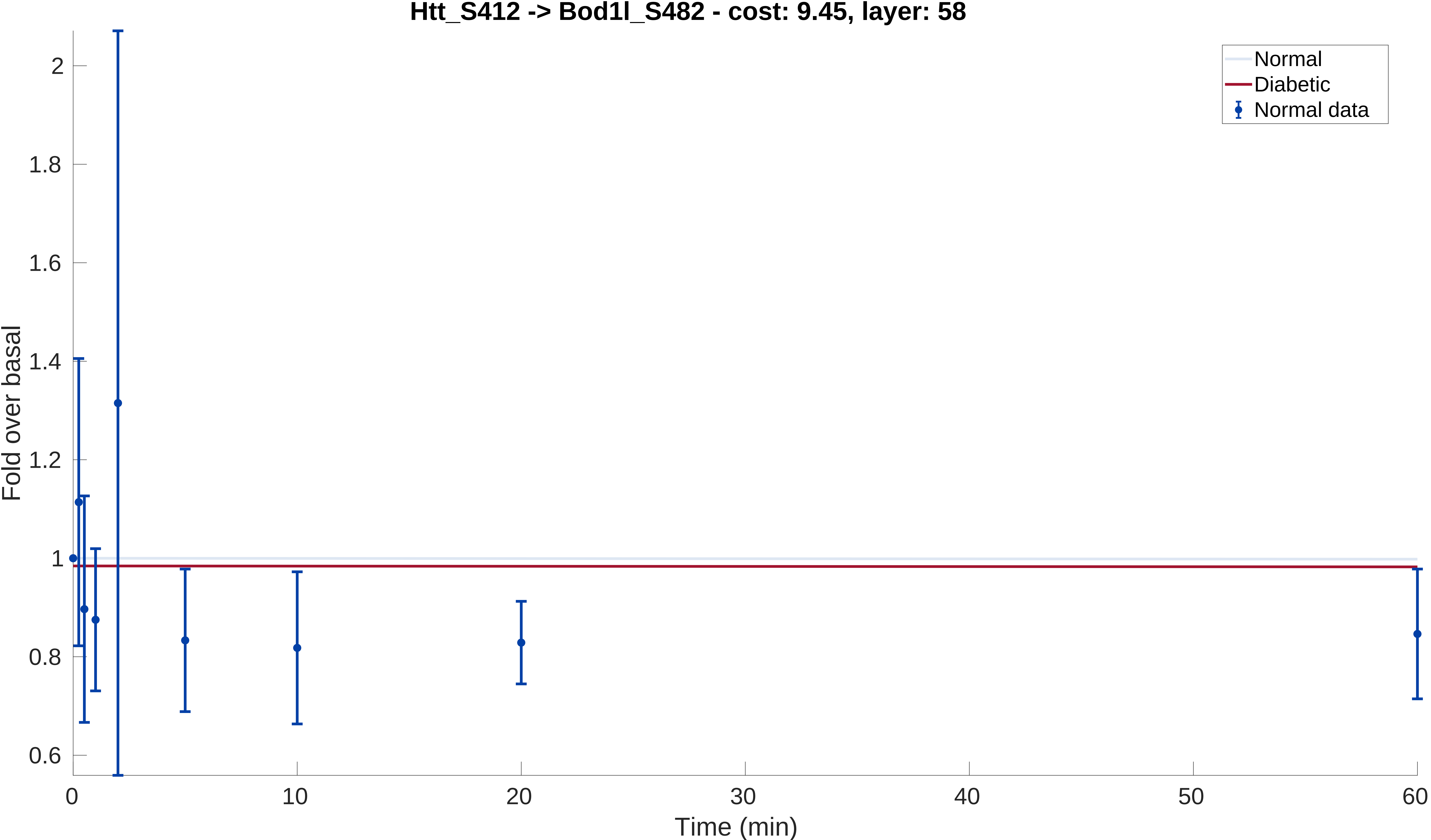
Htt_S412 -> Bod1l_S482 - cost: 9.45, layer: 58
Normal
Diabetic
Normal data
Fold over basal
2
1.8
1.6
1.4
1.2
1
0.8
0.6
0
10
20
30
40
50
60
Time (min)

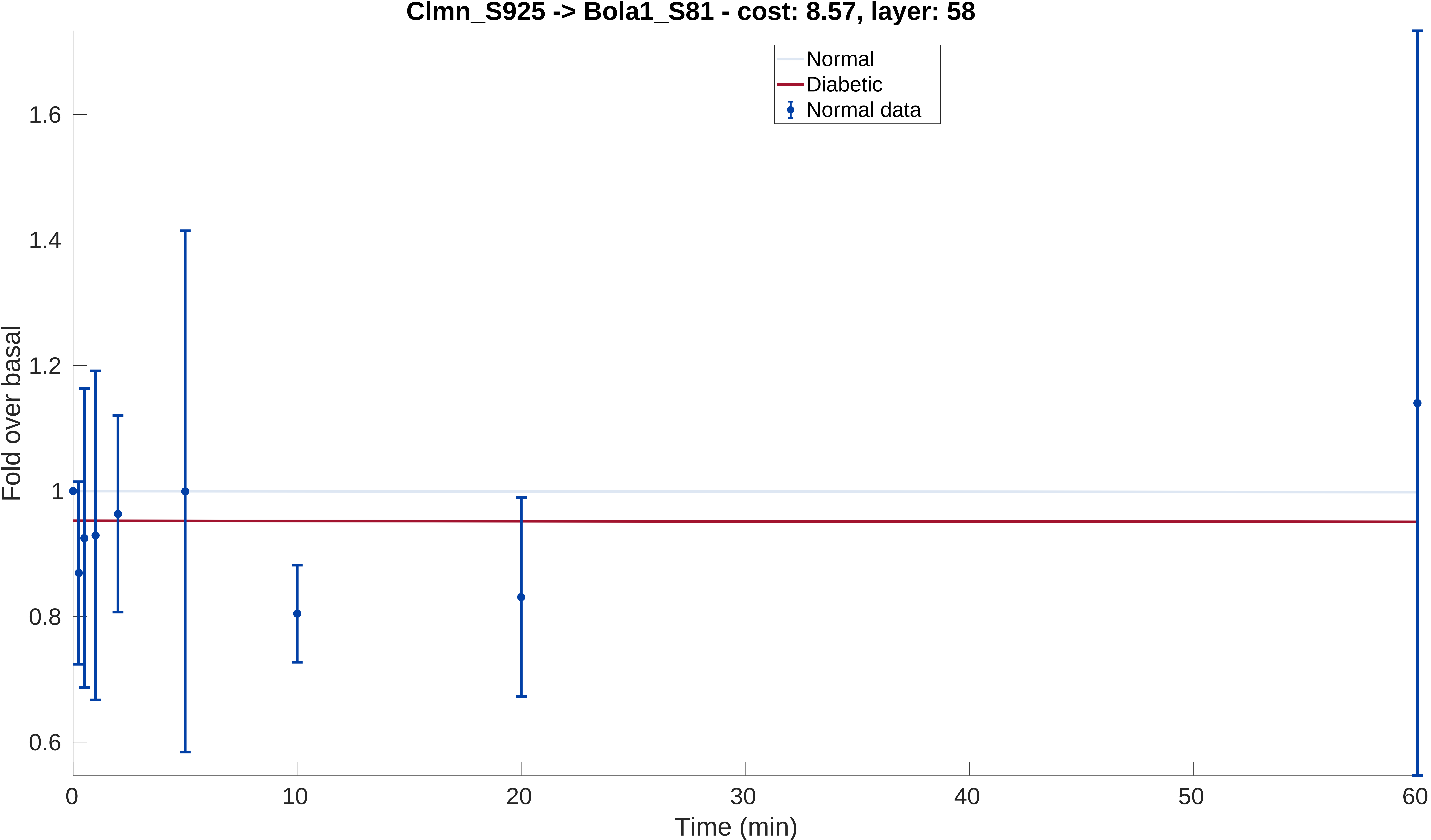

Clmn_S925 -> Bola1_S81 - cost: 8.57, layer: 58
Normal
Diabetic
Normal data
Fold over basal
1.6
1.4
1.2
1
0.8
0.6
0
10
20
30
40
50
60
Time (min)

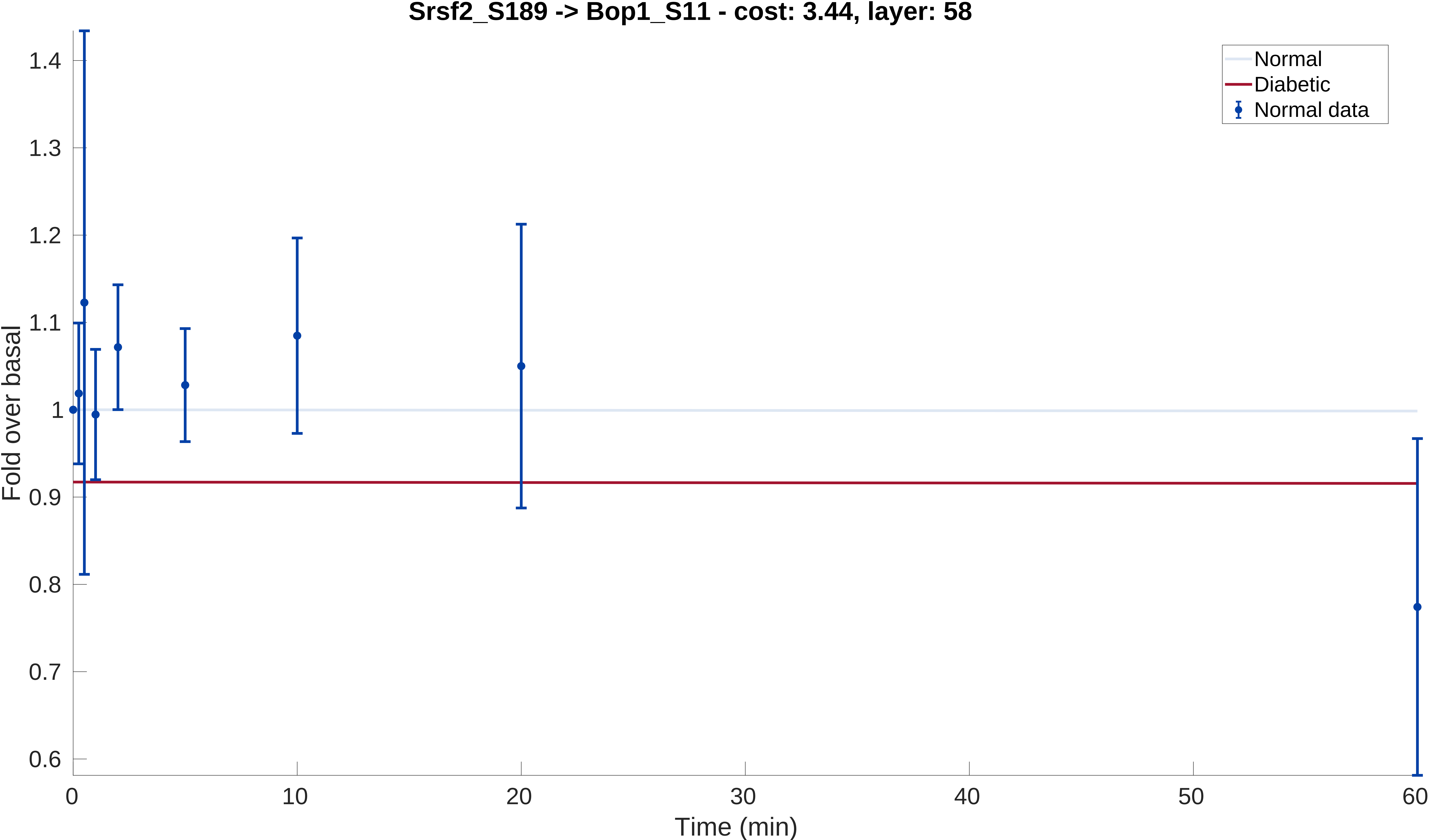
Srsf2_S189 -> Bop1_S11 - cost: 3.44, layer: 58
Fold over basal
1.4
1.3
1.2
1.1
1
0.9
0.8
0.7
0.6
0
10
20
30
40
50
60
Time (min)
Normal
Diabetic
Normal data

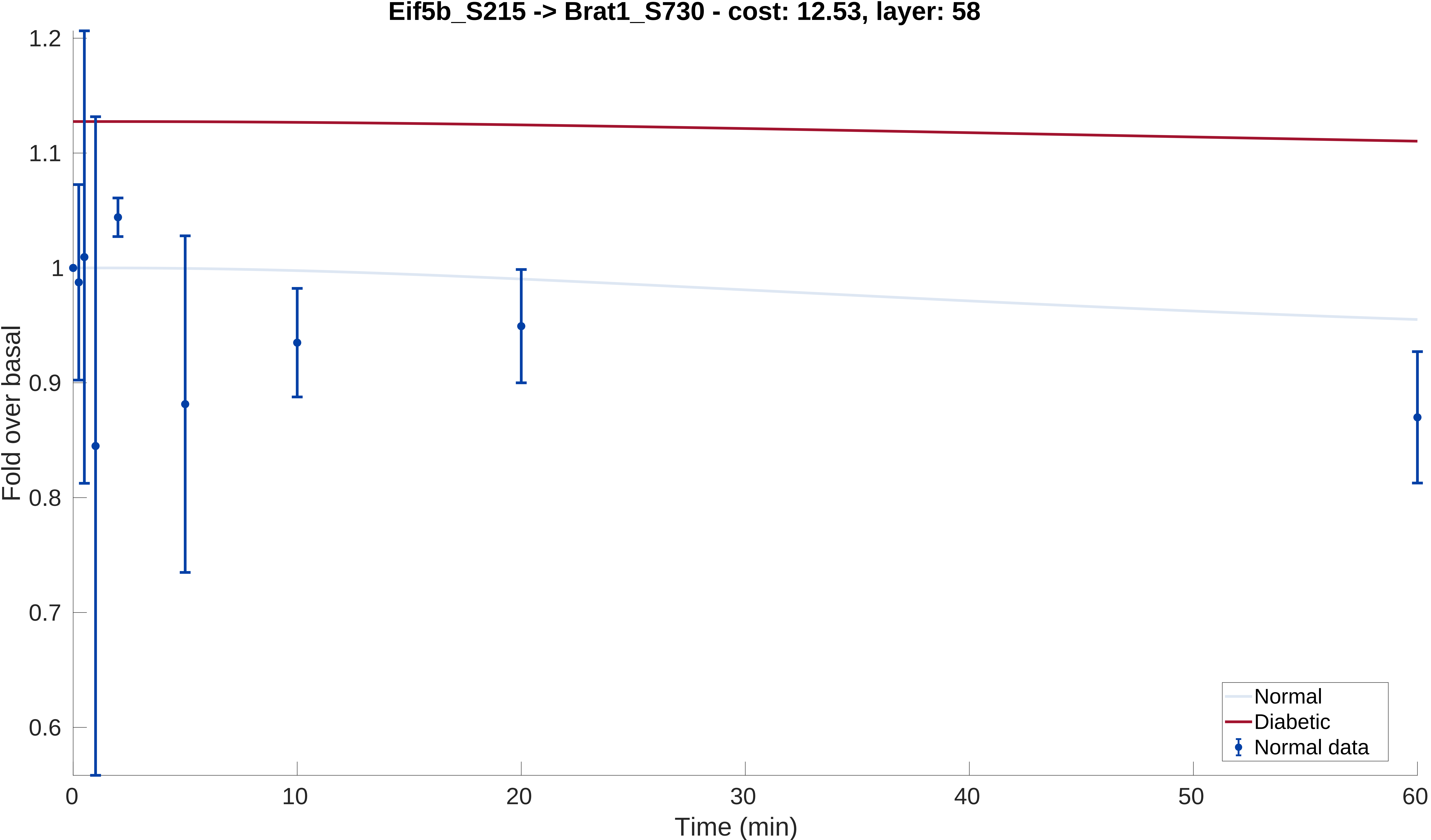

Eif5b_S215 -> Brat1_S730 - cost: 12.53, layer: 58
Fold over basal
1.2
1.1
1
0.9
0.8
0.7
0.6
0
10
20
30
40
50
60
Time (min)
Normal
Diabetic
Normal data

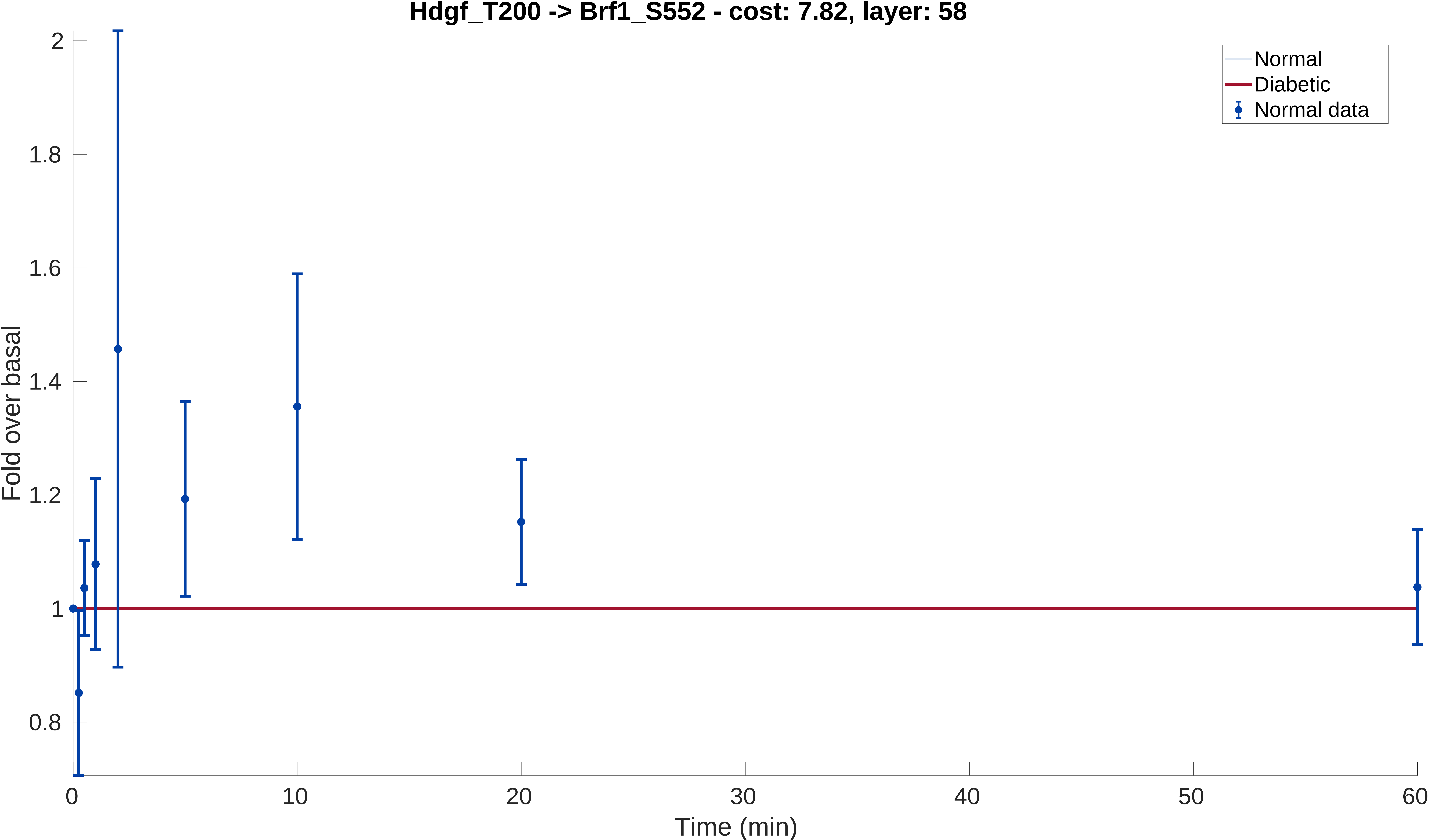

Hdgf_T200 -> Brf1_S552 - cost: 7.82, layer: 58
Normal
Diabetic
Normal data
Fold over basal
2
1.8
1.6
1.4
1.2
1
0.8
0
10
20
30
40
50
60
Time (min)

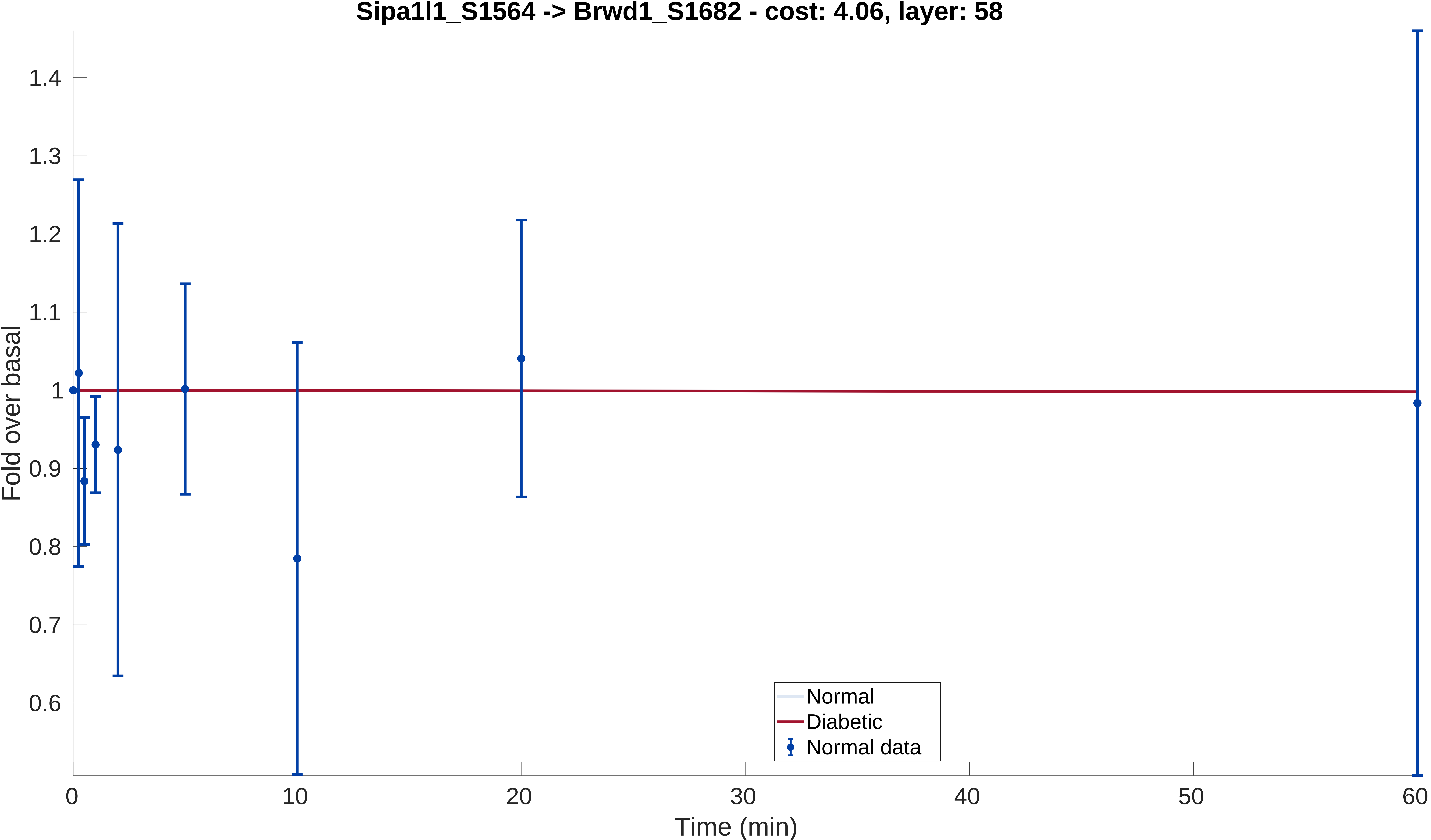

Sipa1l1_S1564 -> Brwd1_S1682 - cost: 4.06, layer: 58
Fold over basal
1.4
1.3
1.2
1.1
1
0.9
0.8
0.7
0.6
0
10
20
30
40
50
60
Time (min)
Normal
Diabetic
Normal data

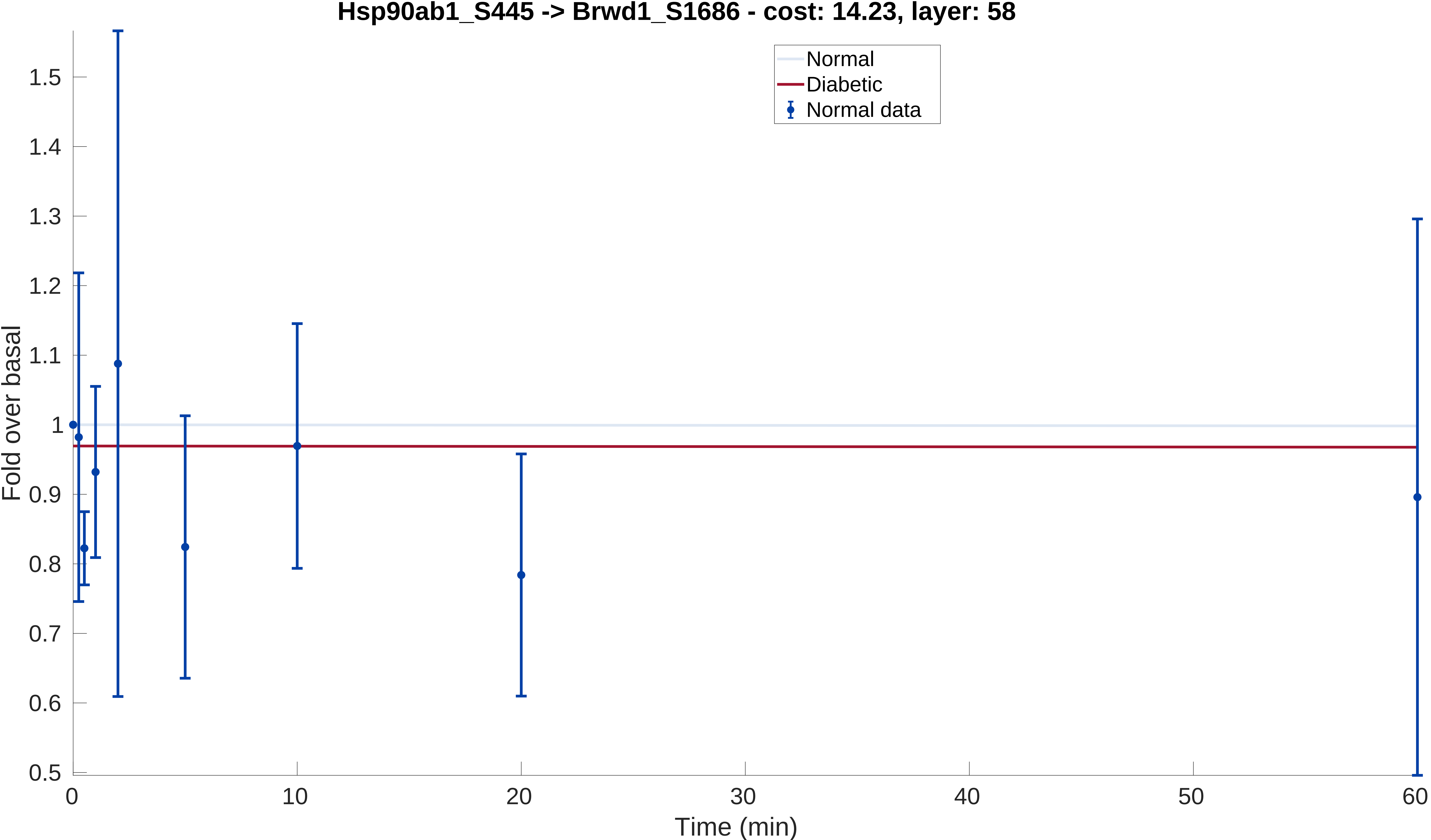

Hsp90ab1_S445 -> Brwd1_S1686 - cost: 14.23, layer: 58
Normal
Diabetic
Normal data
Fold over basal
1.5
1.4
1.3
1.2
1.1
1
0.9
0.8
0.7
0.6
0.5
0
10
20
30
40
50
60
Time (min)

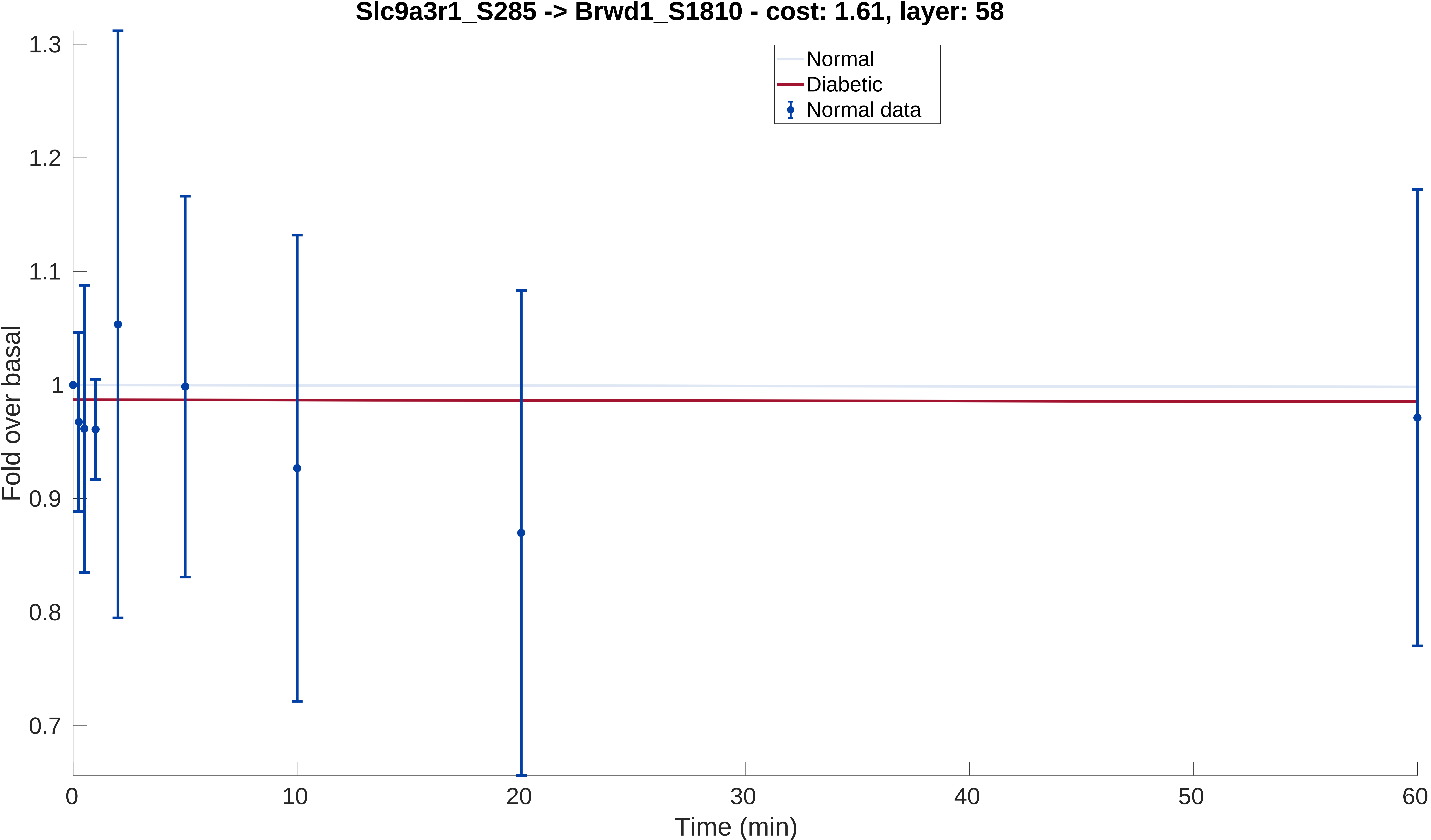

Slc9a3r1_S285 -> Brwd1_S1810 - cost: 1.61, layer: 58
Normal
Diabetic
Normal data
Fold over basal
Time (min)
0.7
0.8
0.9
1
1.1
1.2
1.3
0
10
20
30
40
50
60

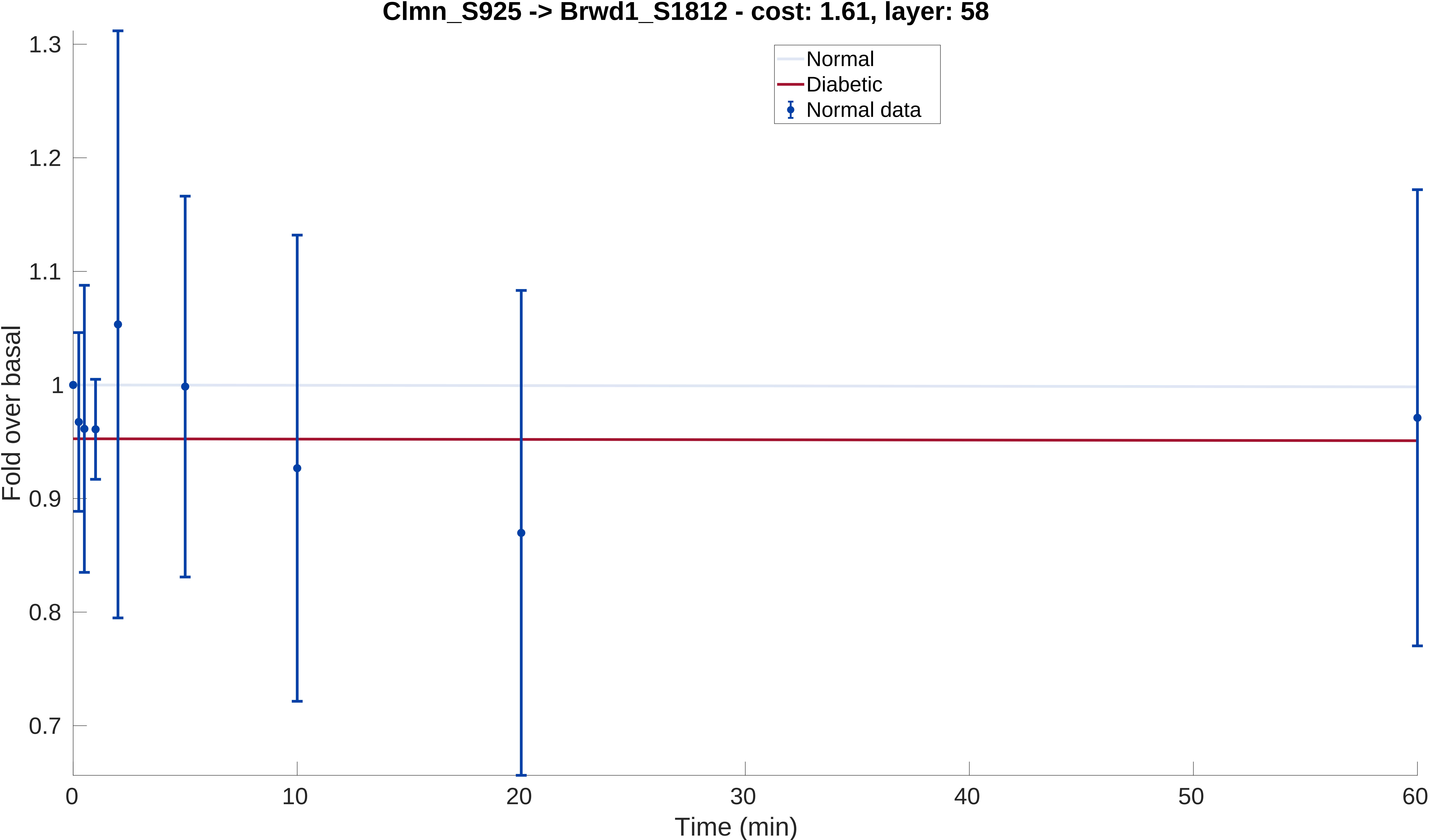

Clmn_S925 -> Brwd1_S1812 - cost: 1.61, layer: 58
Normal
Diabetic
Normal data
1.3
1.2
1.1
1
0.9
0.8
0.7
Fold over basal
0
10
20
30
40
50
60
Time (min)

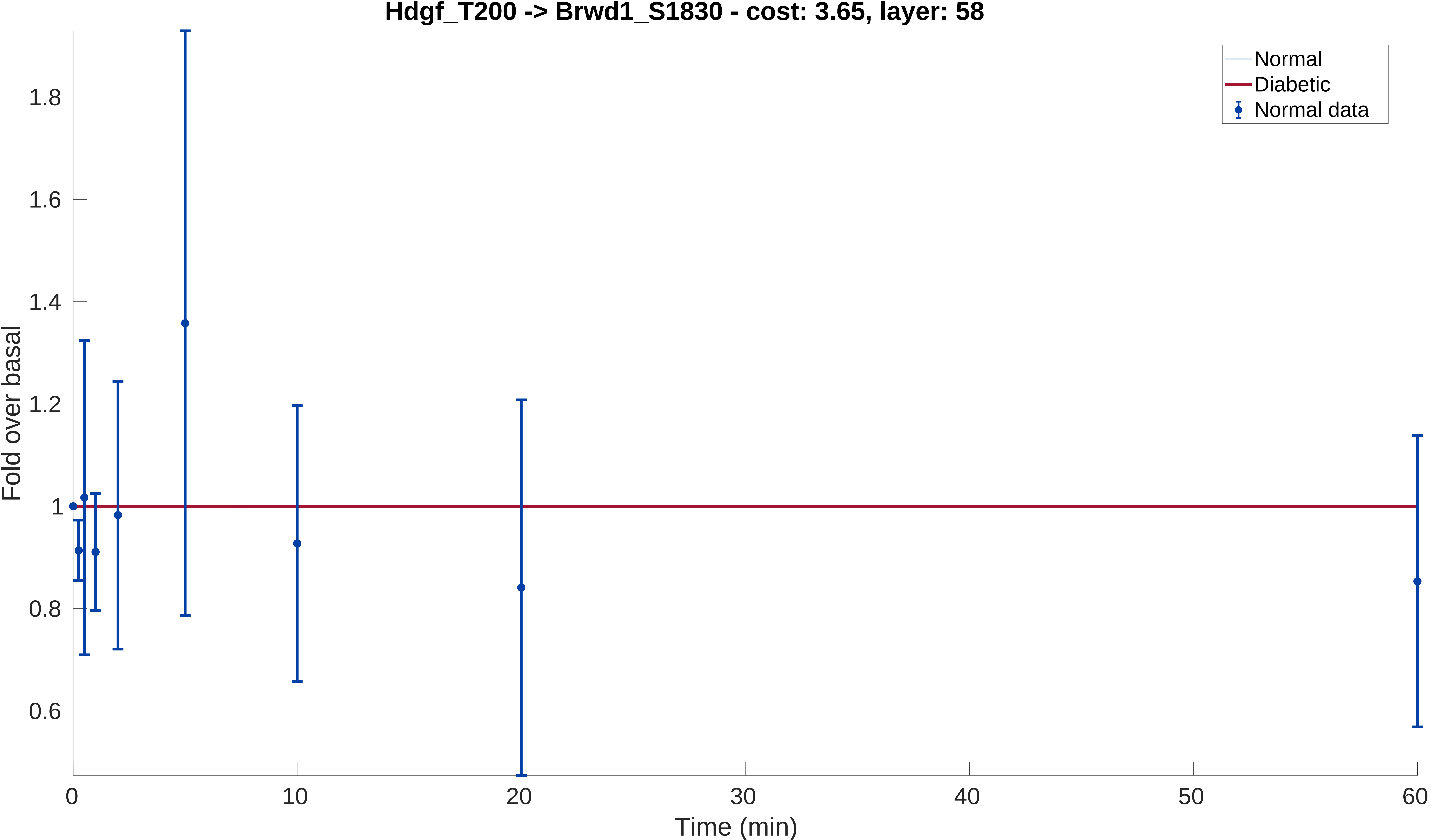
Hdgf_T200 -> Brwd1_S1830 - cost: 3.65, layer: 58
Normal
Diabetic
Normal data
Fold over basal
1.8
1.6
1.4
1.2
1
0.8
0.6
0
10
20
30
40
50
60
Time (min)

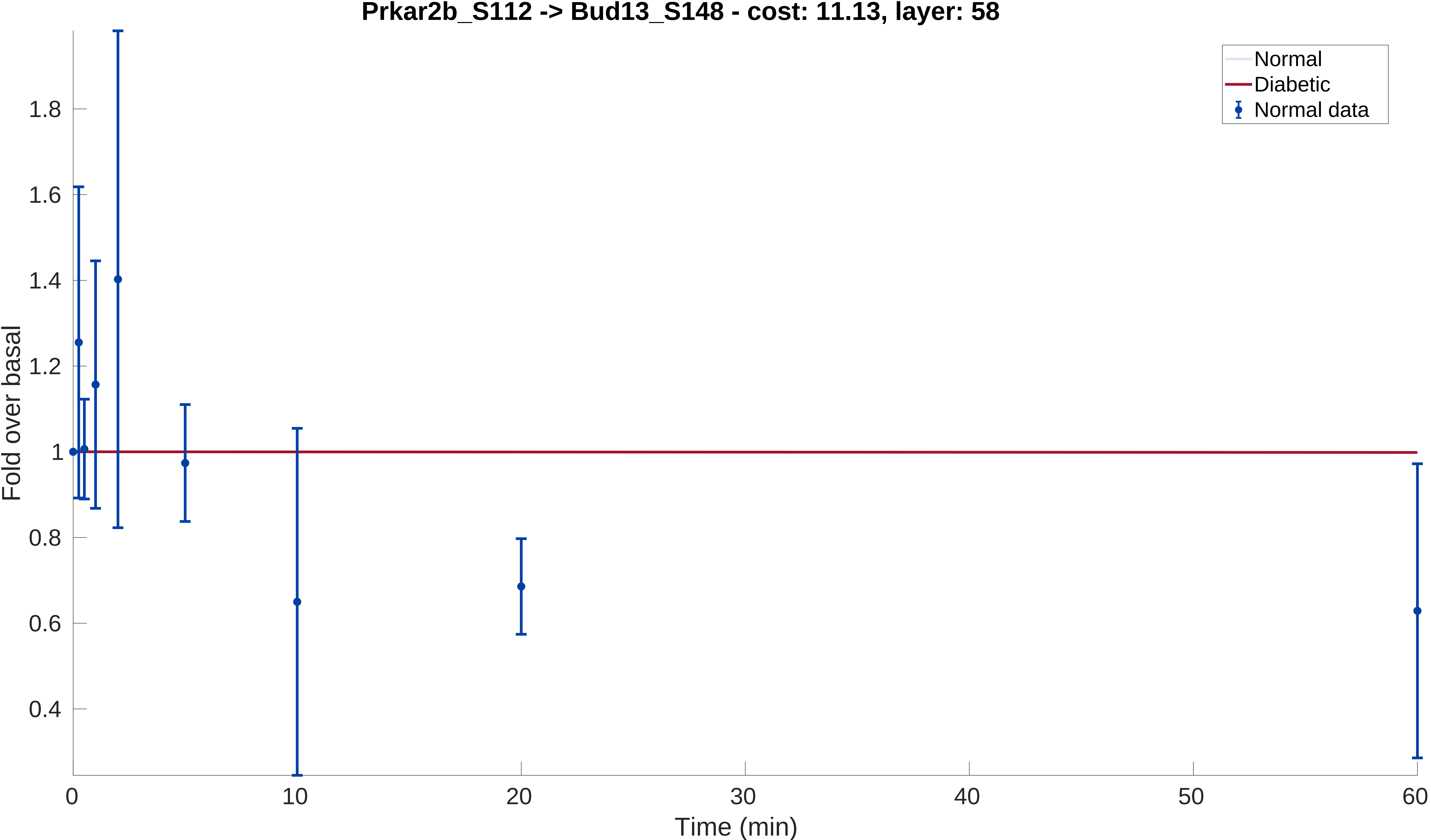

Prkar2b_S112 -> Bud13_S148 - cost: 11.13, layer: 58
Normal
Diabetic
Normal data
1.8
1.6
1.4
1.2
1
0.8
0.6
0.4
Fold over basal
0
10
20
30
40
50
60
Time (min)

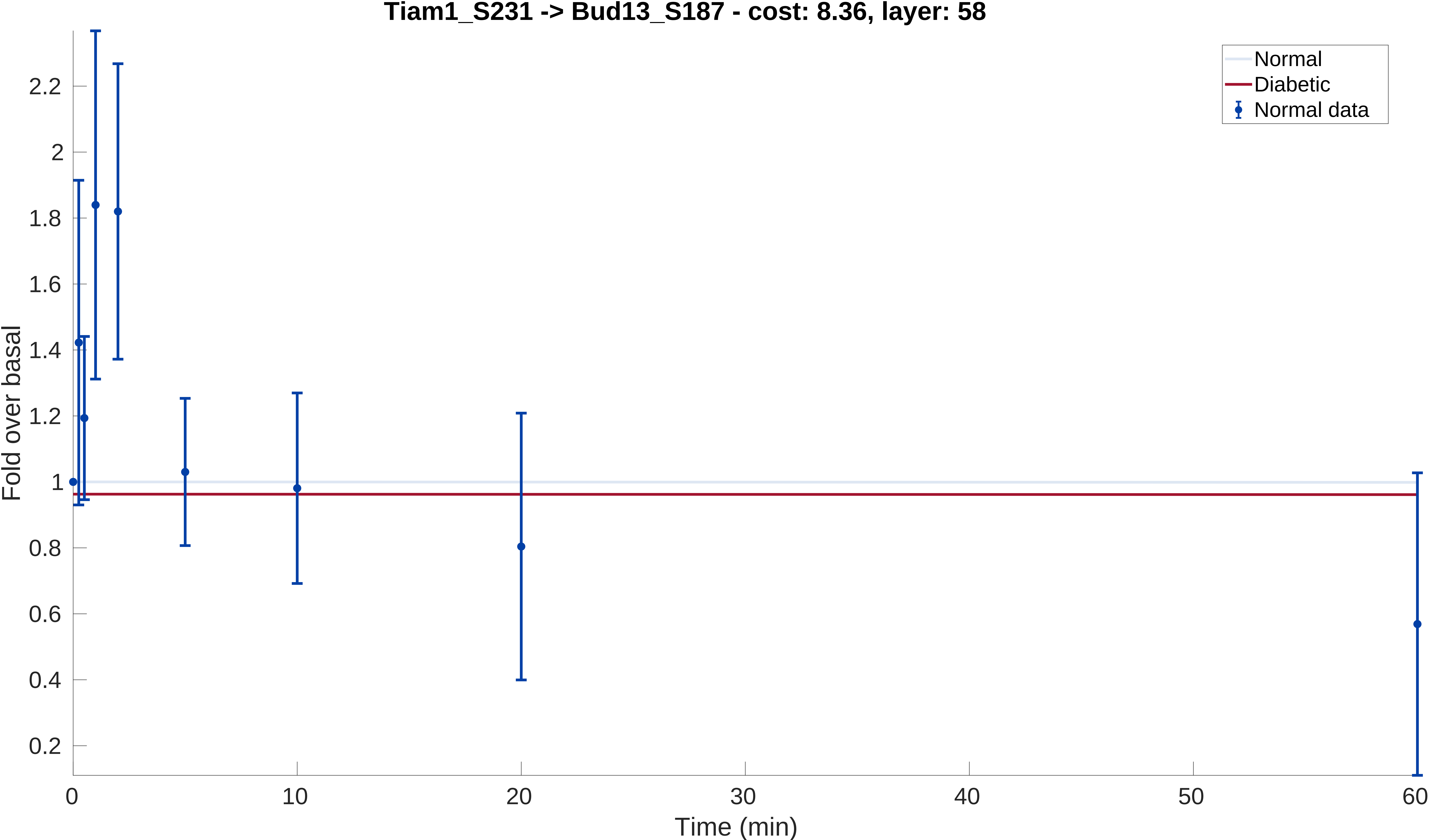

Tiam1_S231 -> Bud13_S187 - cost: 8.36, layer: 58
Normal
Diabetic
Normal data
Fold over basal
2.2
2
1.8
1.6
1.4
1.2
1
0.8
0.6
0.4
0.2
0
10
20
30
40
50
60
Time (min)

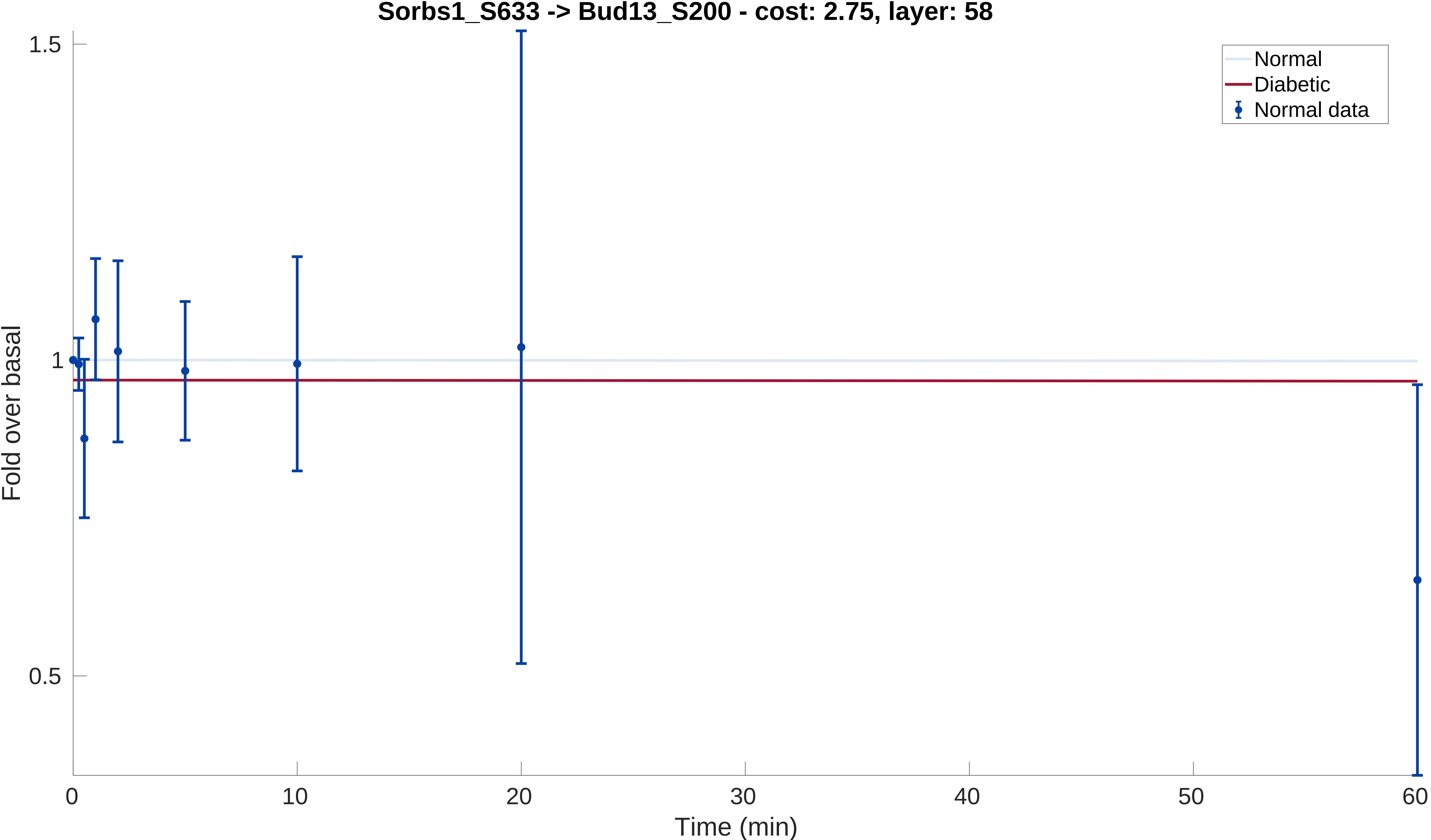

Sorbs1_S633 -> Bud13_S200 - cost: 2.75, layer: 58
Normal
Diabetic
Normal data
Fold over basal
1.5
1
0.5
0
10
20
30
40
50
60
Time (min)

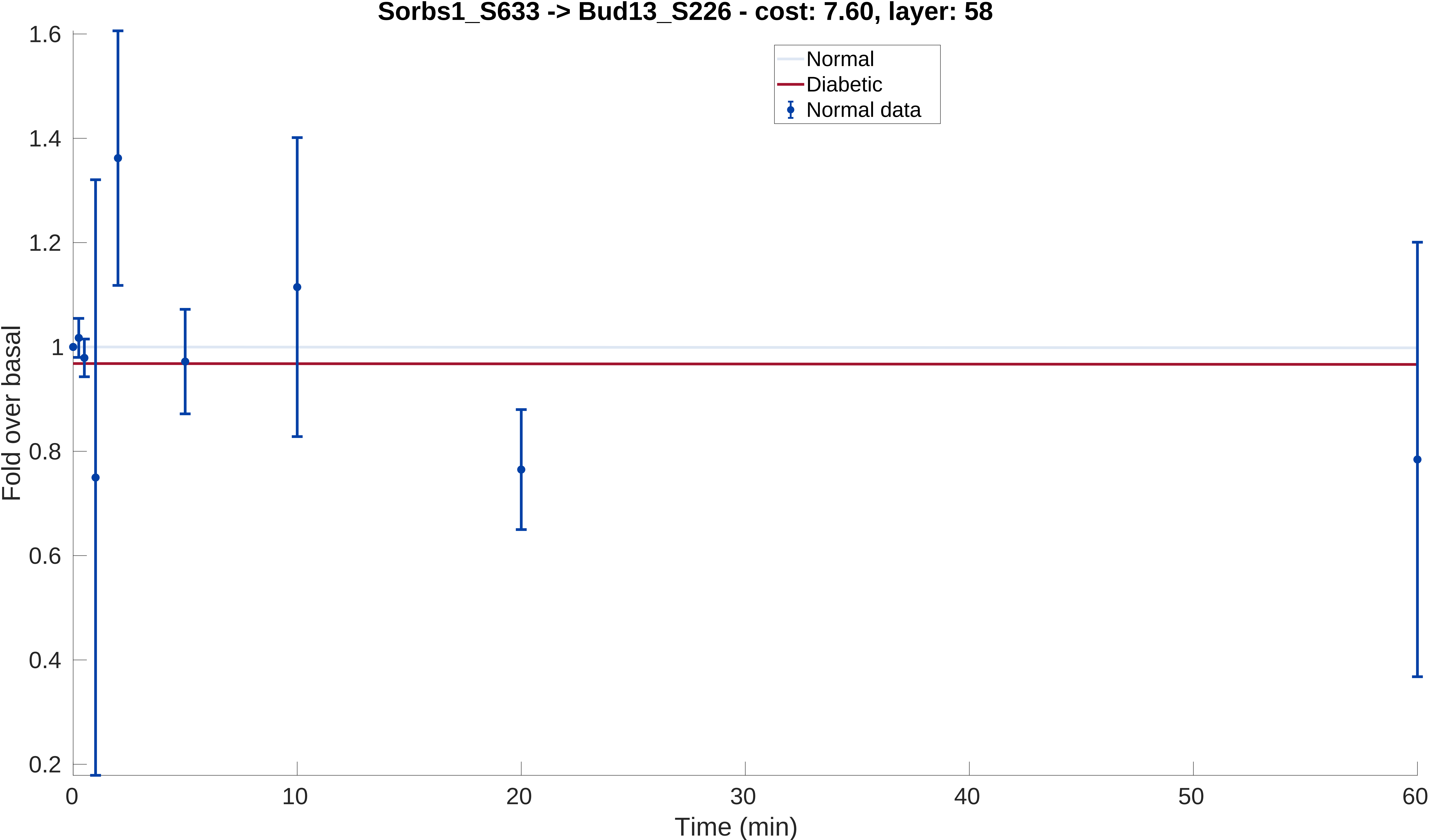

Sorbs1_S633 -> Bud13_S226 - cost: 7.60, layer: 58
Normal
Diabetic
Normal data
Fold over basal
1.6
1.4
1.2
1
0.8
0.6
0.4
0.2
0
10
20
30
40
50
60
Time (min)

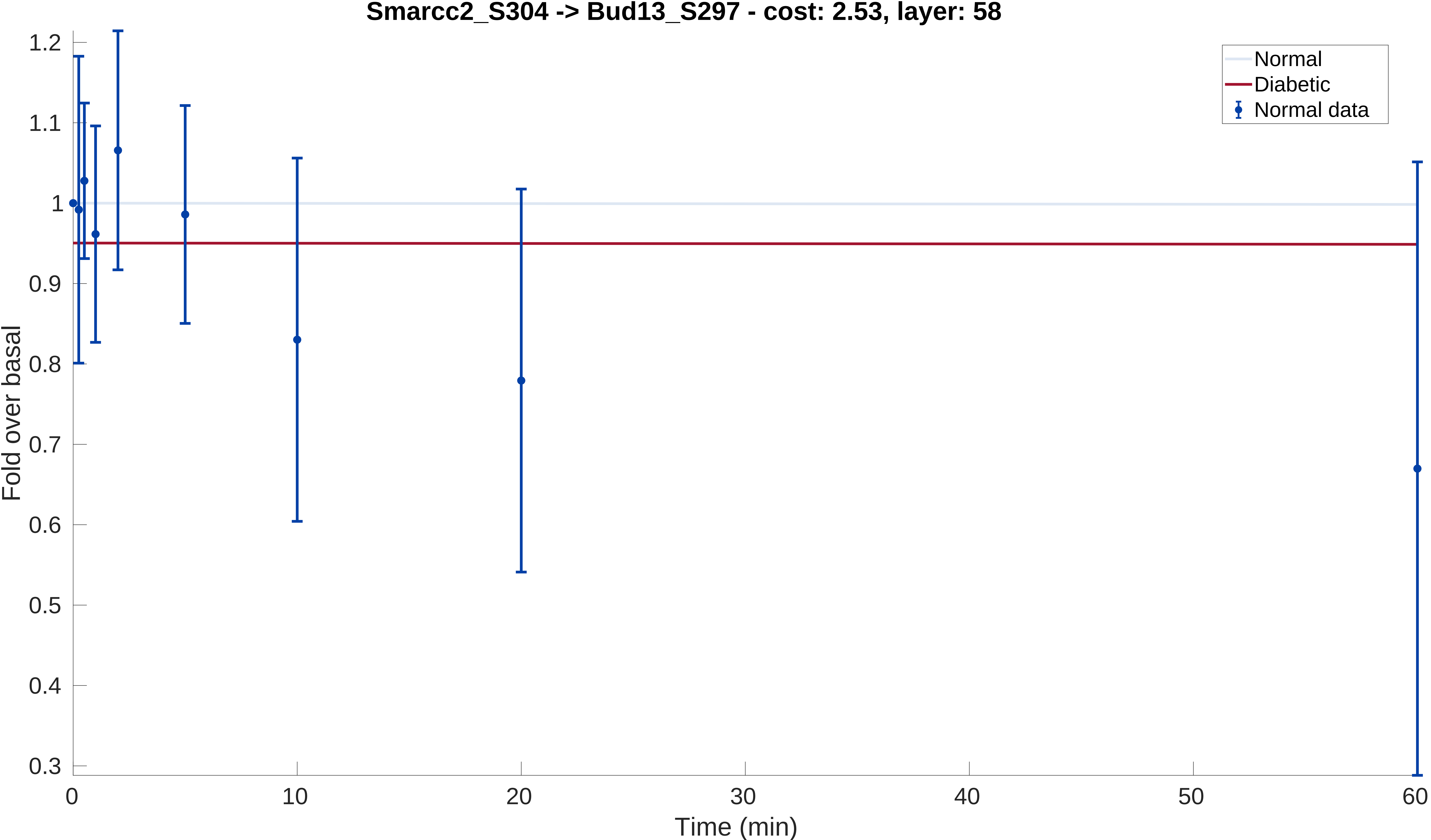

Smarcc2_S304 -> Bud13_S297 - cost: 2.53, layer: 58
Normal
Diabetic
Normal data
Fold over basal
Time (min)
1.2
1.1
1
0.9
0.8
0.7
0.6
0.5
0.4
0.3
0
10
20
30
40
50
60

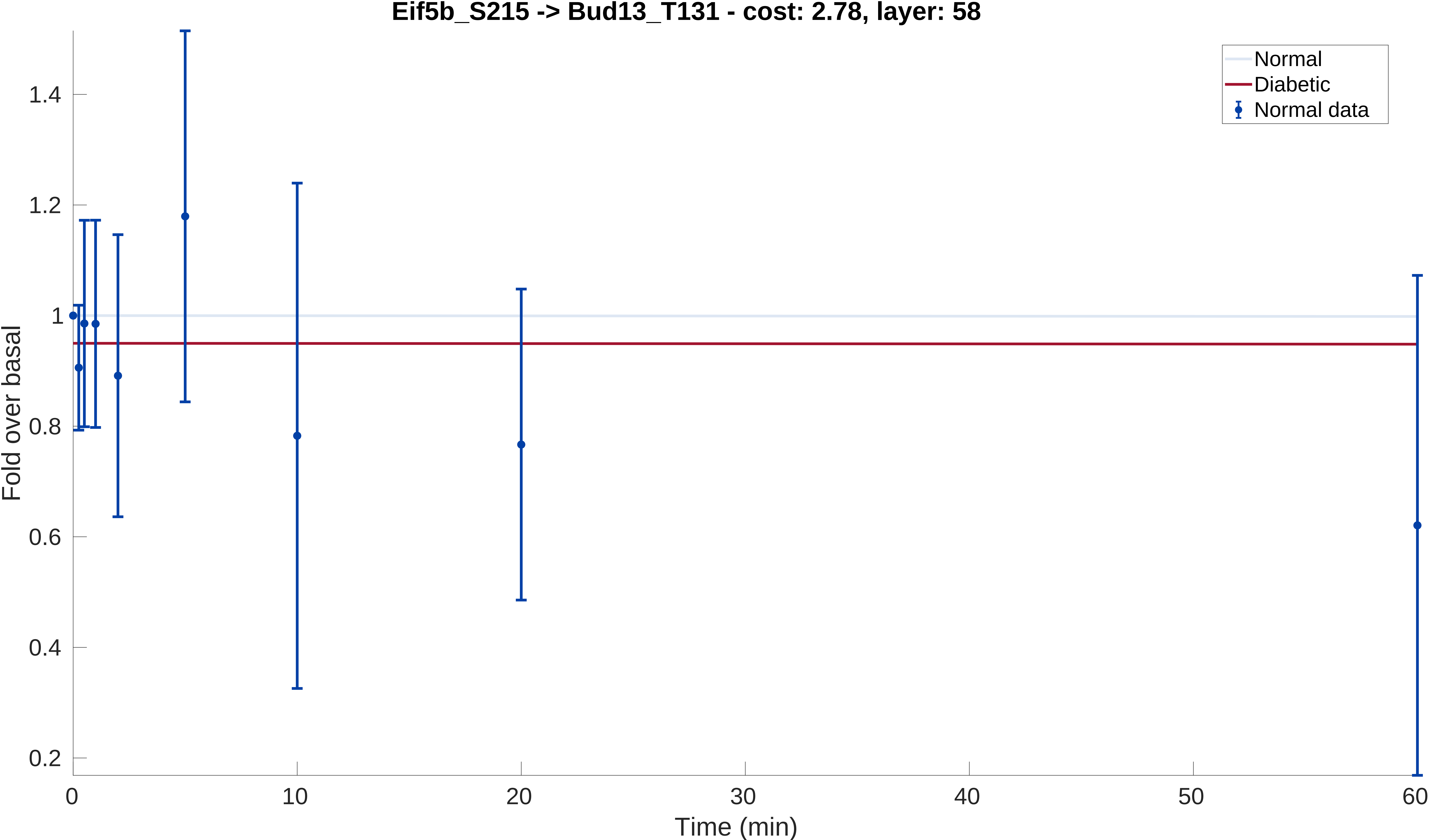

Eif5b_S215 -> Bud13_T131 - cost: 2.78, layer: 58
Normal
Diabetic
Normal data
Fold over basal
1.4
1.2
1
0.8
0.6
0.4
0.2
0
10
20
30
40
50
60
Time (min)

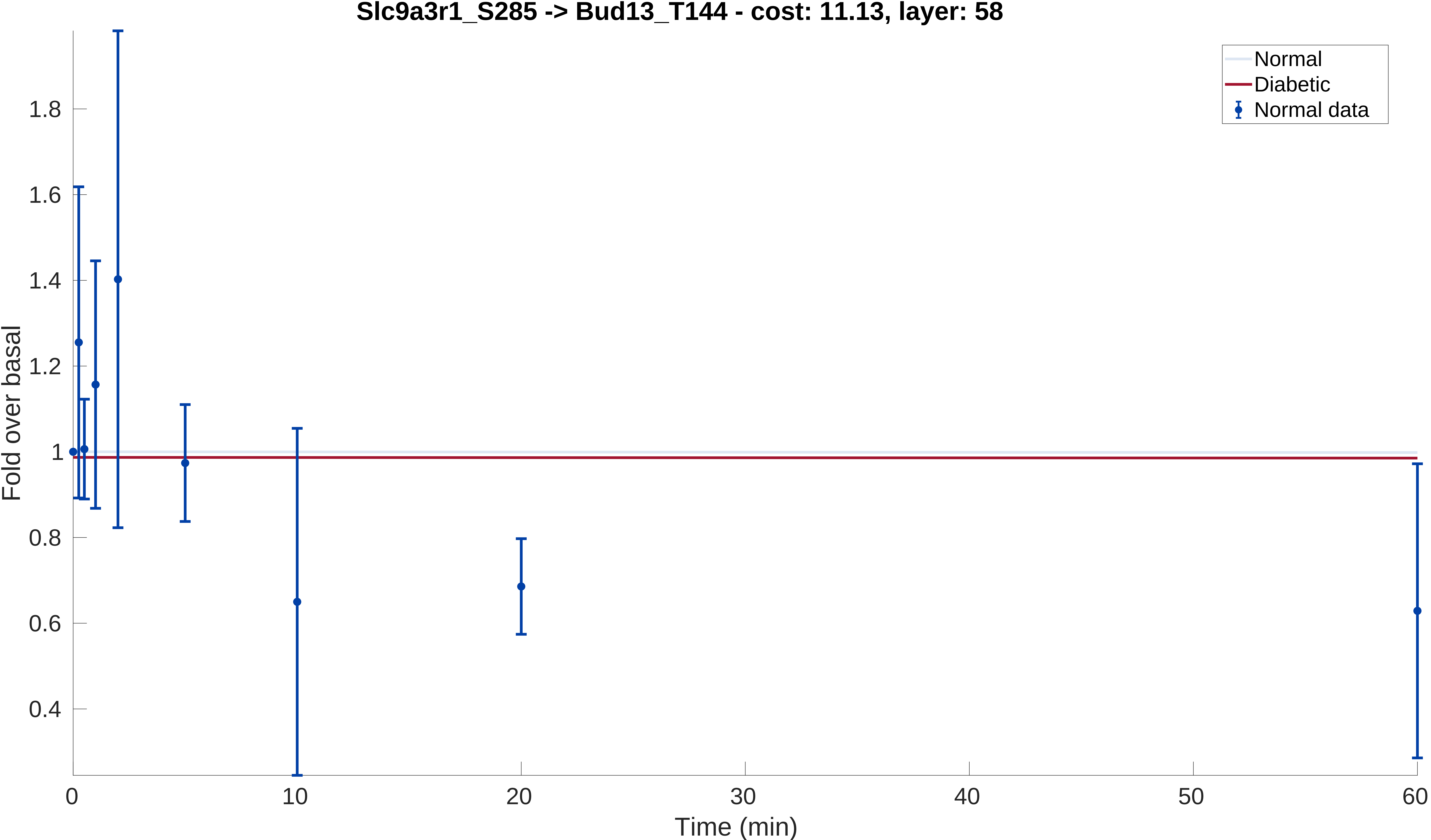

Slc9a3r1_S285 -> Bud13_T144 - cost: 11.13, layer: 58
Normal
Diabetic
Normal data
1.8
1.6
1.4
1.2
1
0.8
0.6
0.4
Fold over basal
0
10
20
30
40
50
60
Time (min)

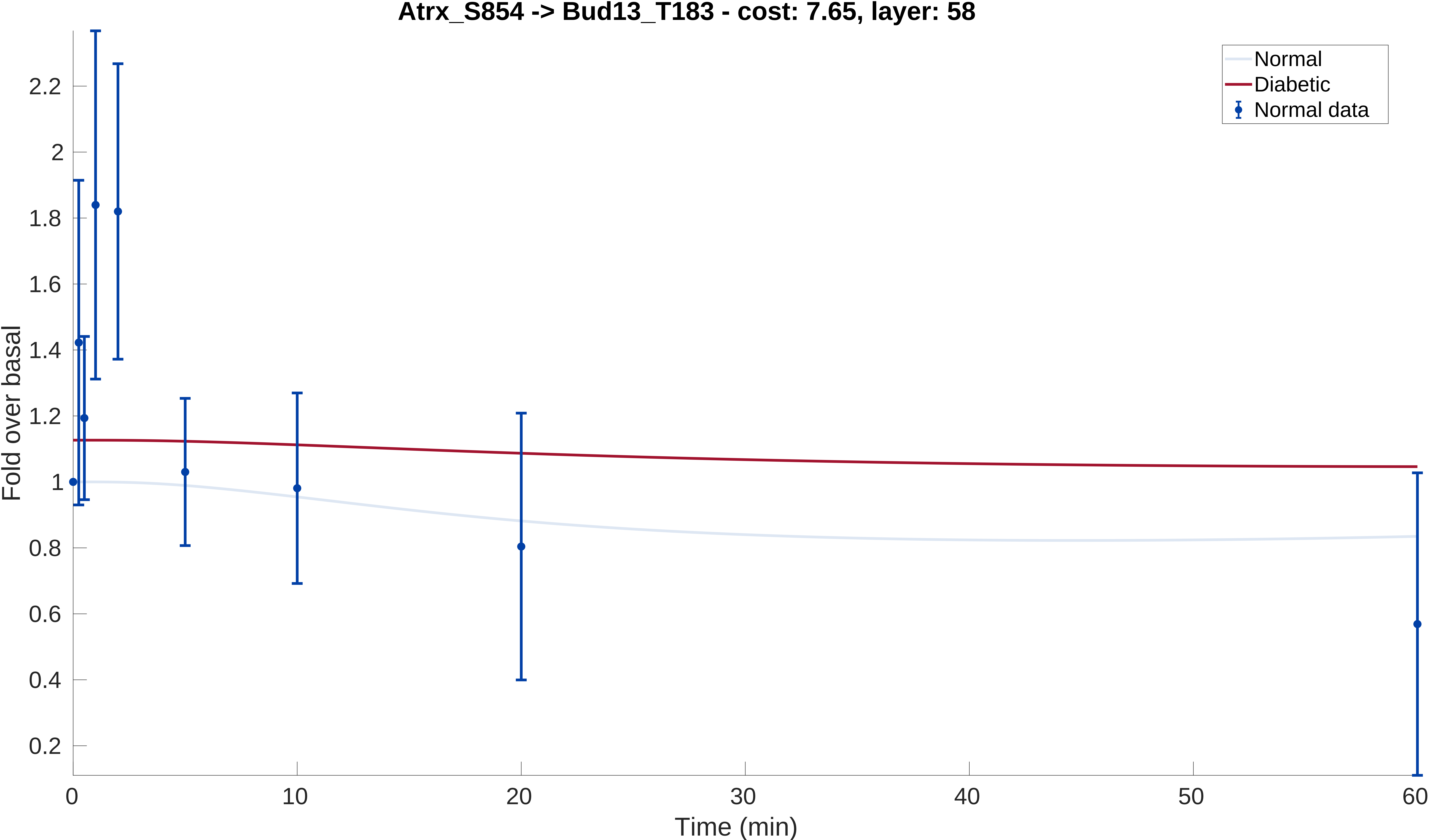

Atrx_S854 -> Bud13_T183 - cost: 7.65, layer: 58
Normal
Diabetic
Normal data
Fold over basal
0.2
0.4
0.6
0.8
1
1.2
1.4
1.6
1.8
2
2.2
0
10
20
30
40
50
60
Time (min)

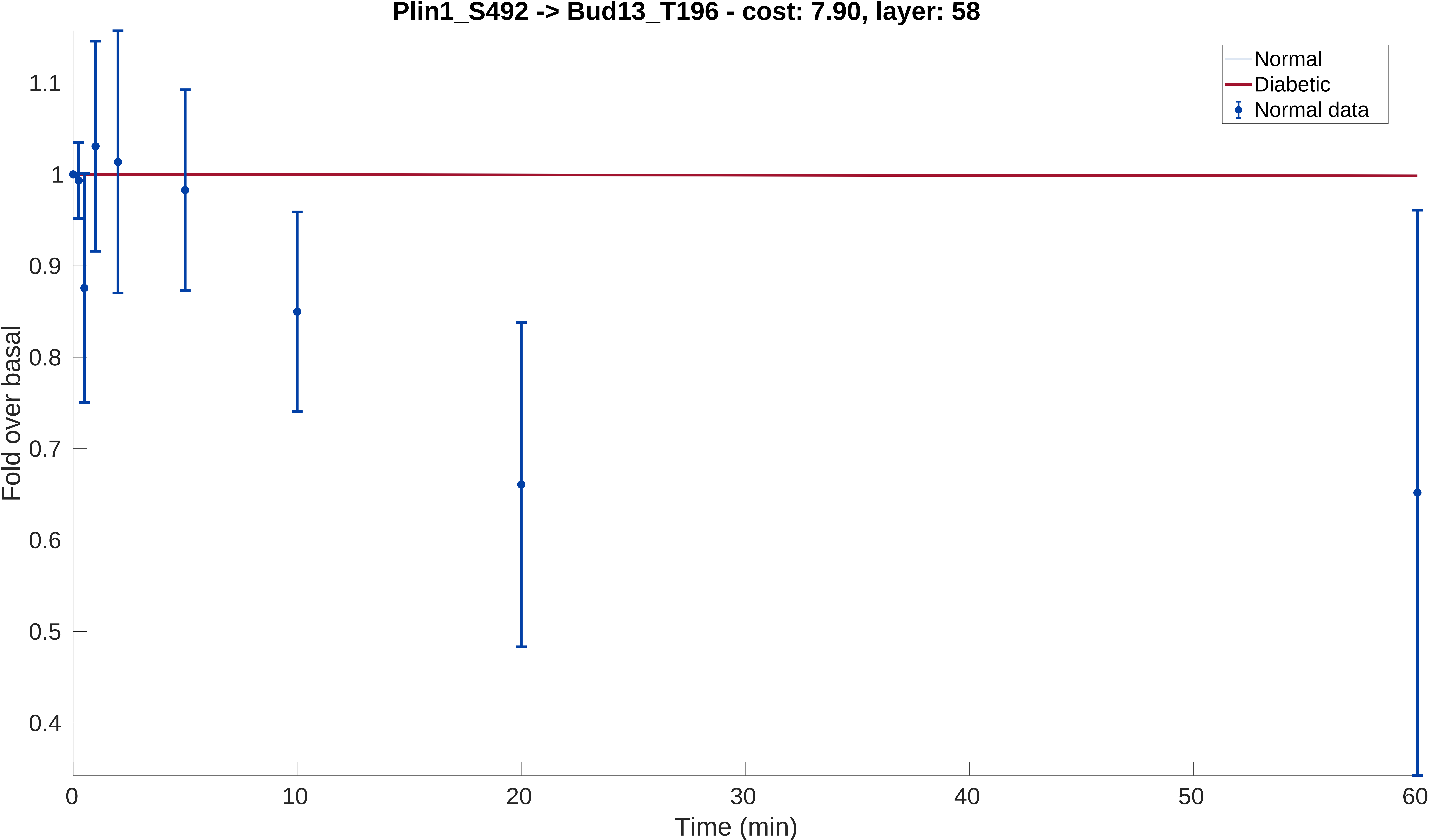

Plin1_S492 -> Bud13_T196 - cost: 7.90, layer: 58
Normal
Diabetic
Normal data
Fold over basal
1.1
1
0.9
0.8
0.7
0.6
0.5
0.4
0
10
20
30
40
50
60
Time (min)

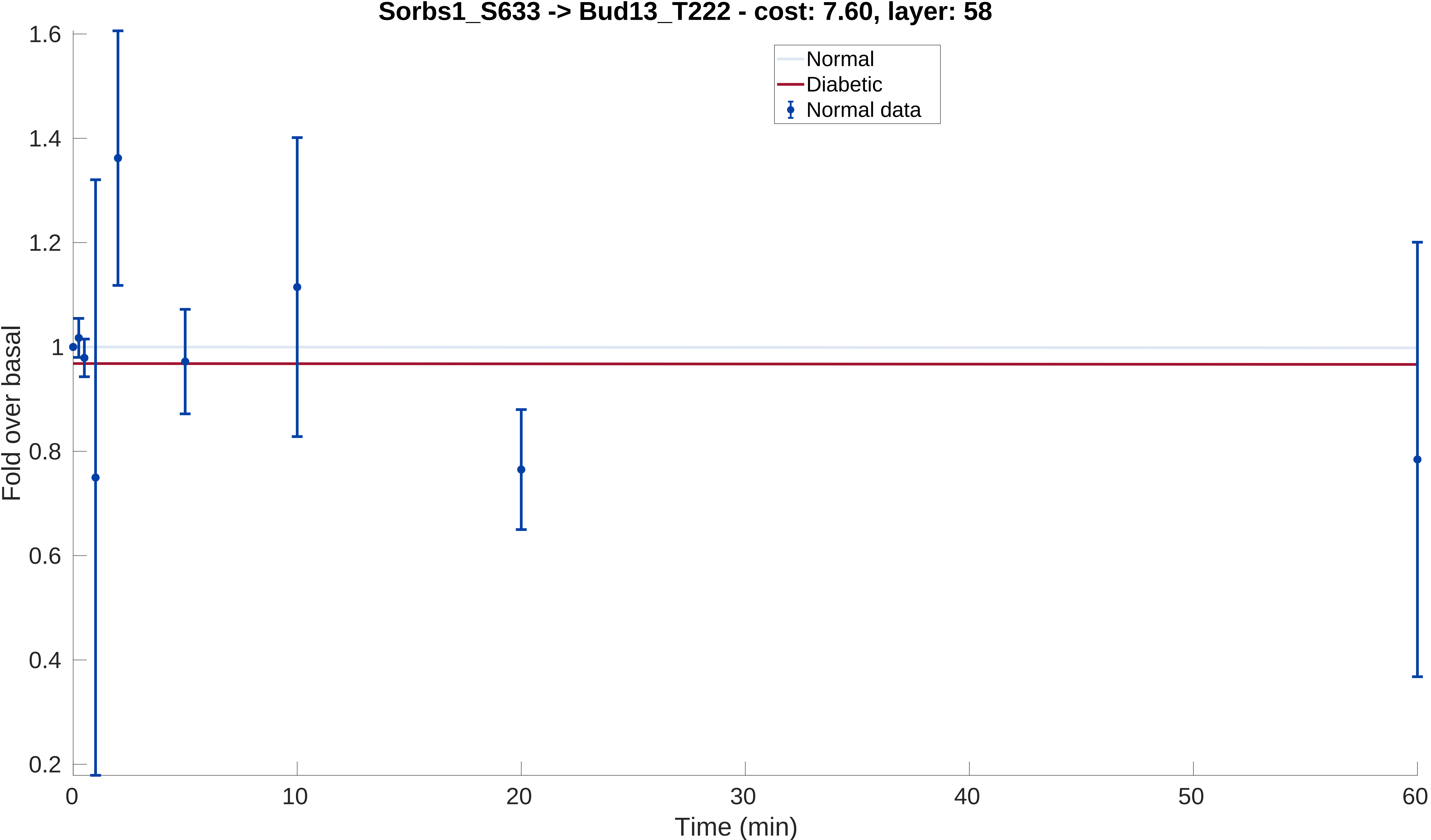
Sorbs1_S633 -> Bud13_T222 - cost: 7.60, layer: 58
Normal
Diabetic
Normal data
Fold over basal
Time (min)
1.6
1.4
1.2
1
0.8
0.6
0.4
0.2
0
10
20
30
40
50
60

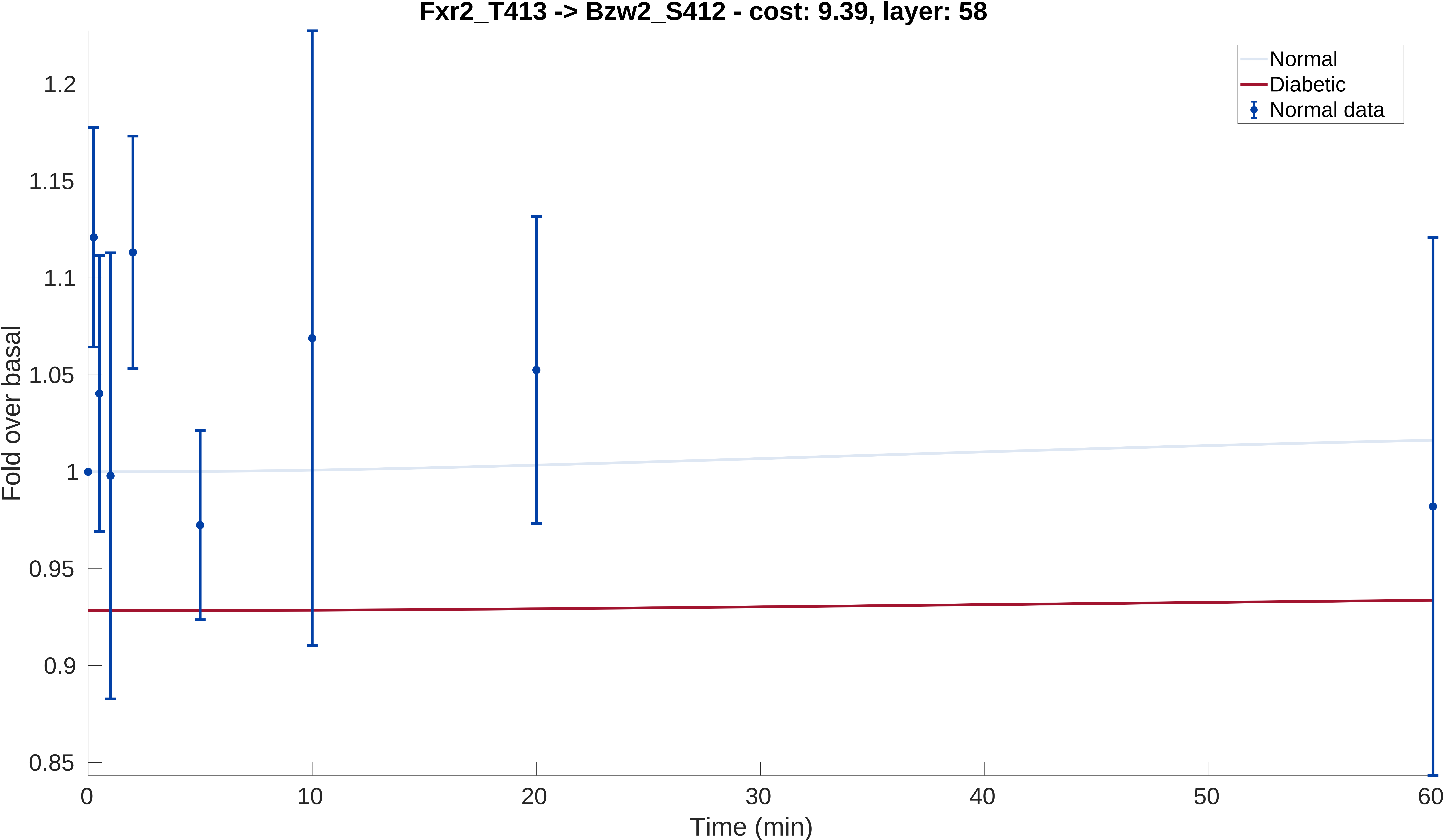
Fxr2_T413 -> Bzw2_S412 - cost: 9.39, layer: 58
Normal
Diabetic
Normal data
Fold over basal
1.2
1.15
1.1
1.05
1
0.95
0.9
0.85
0
10
20
30
40
50
60
Time (min)

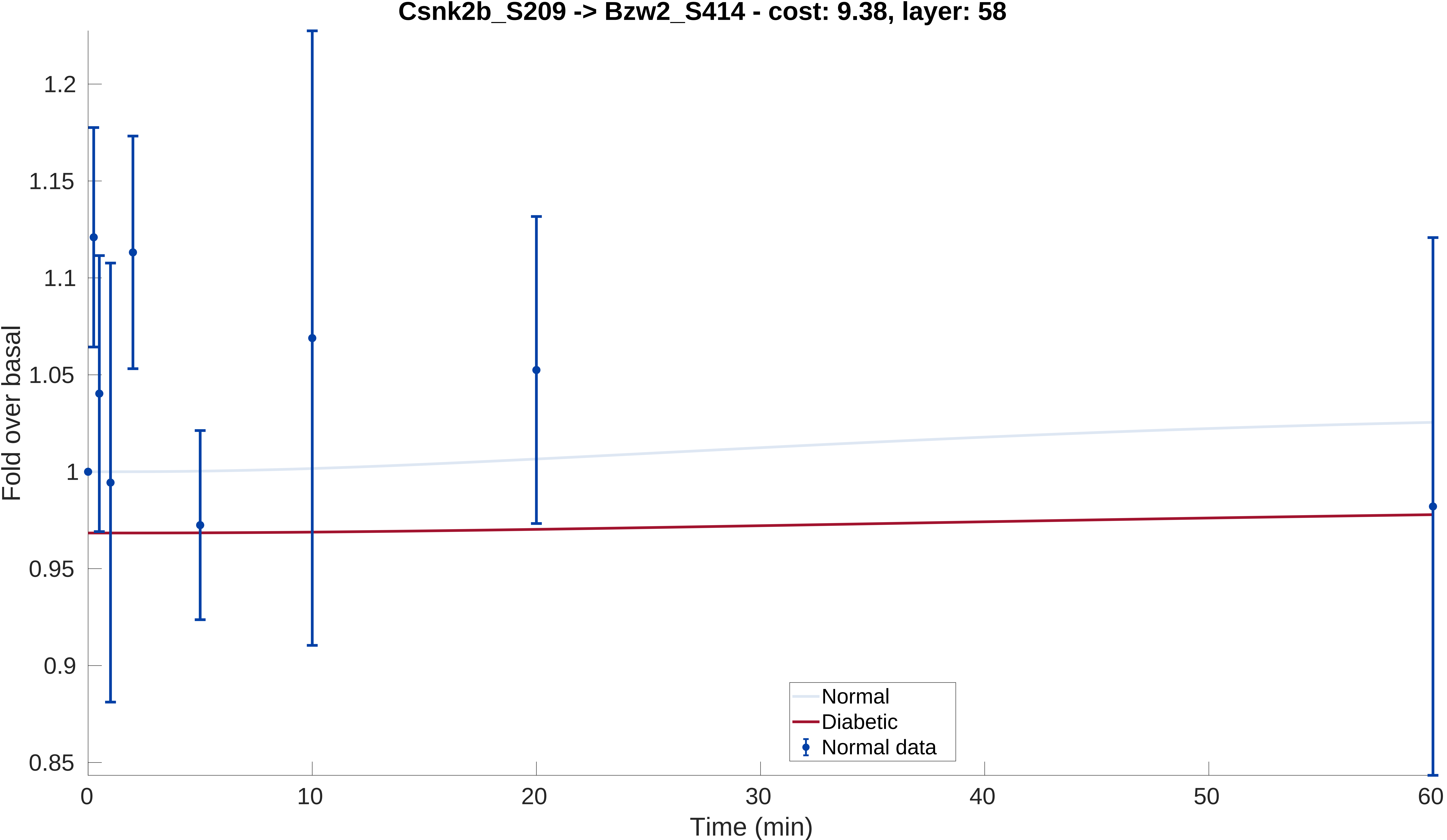

Csnk2b_S209 -> Bzw2_S414 - cost: 9.38, layer: 58
Fold over basal
1.2
1.15
1.1
1.05
1
0.95
0.9
0.85
0
10
20
30
40
50
60
Time (min)
Normal
Diabetic
Normal data

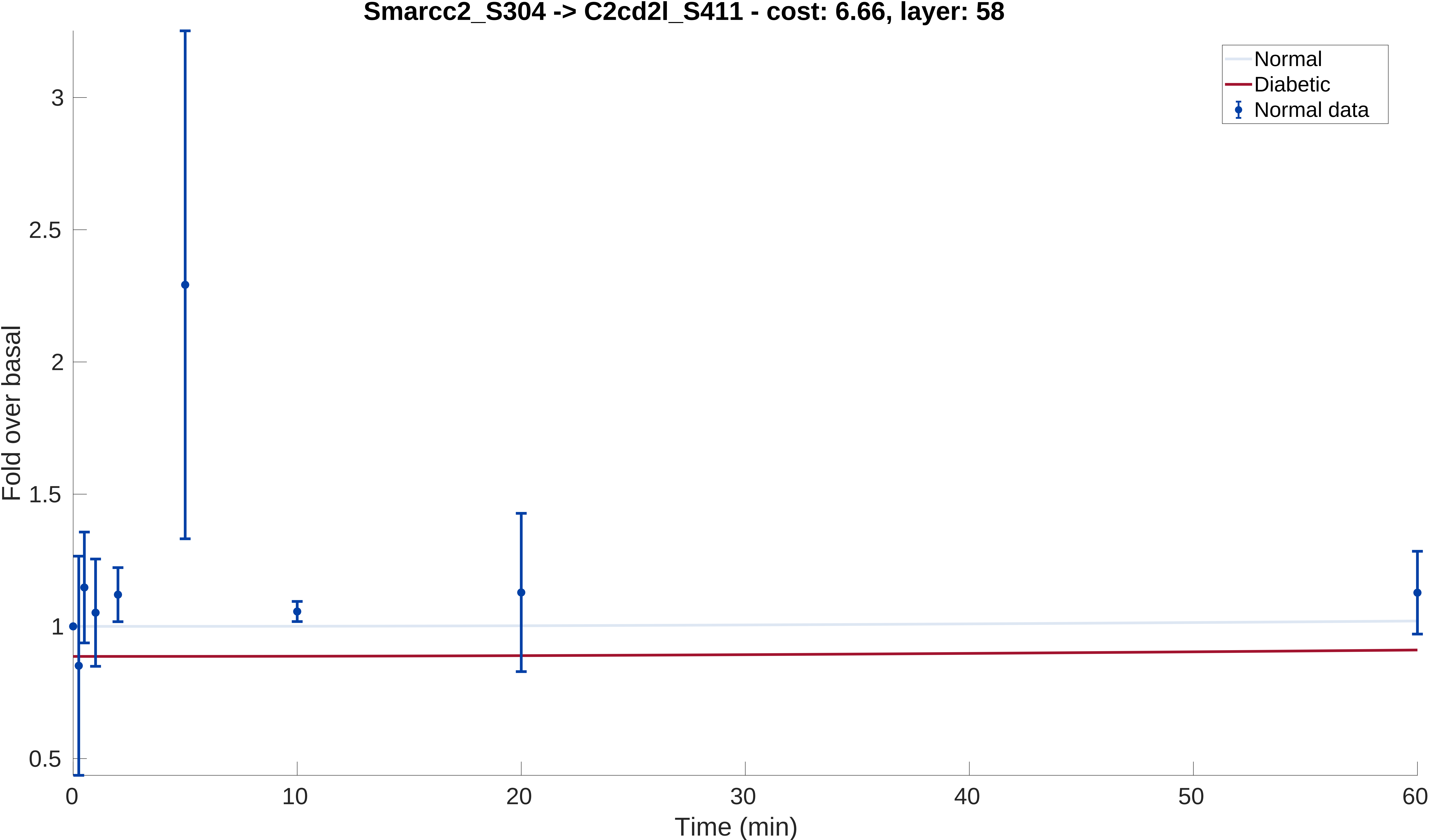
Smarcc2_S304 -> C2cd2l_S411 - cost: 6.66, layer: 58
Fold over basal
Time (min)
Normal
Diabetic
Normal data
0.5
1
1.5
2
2.5
3
0
10
20
30
40
50
60

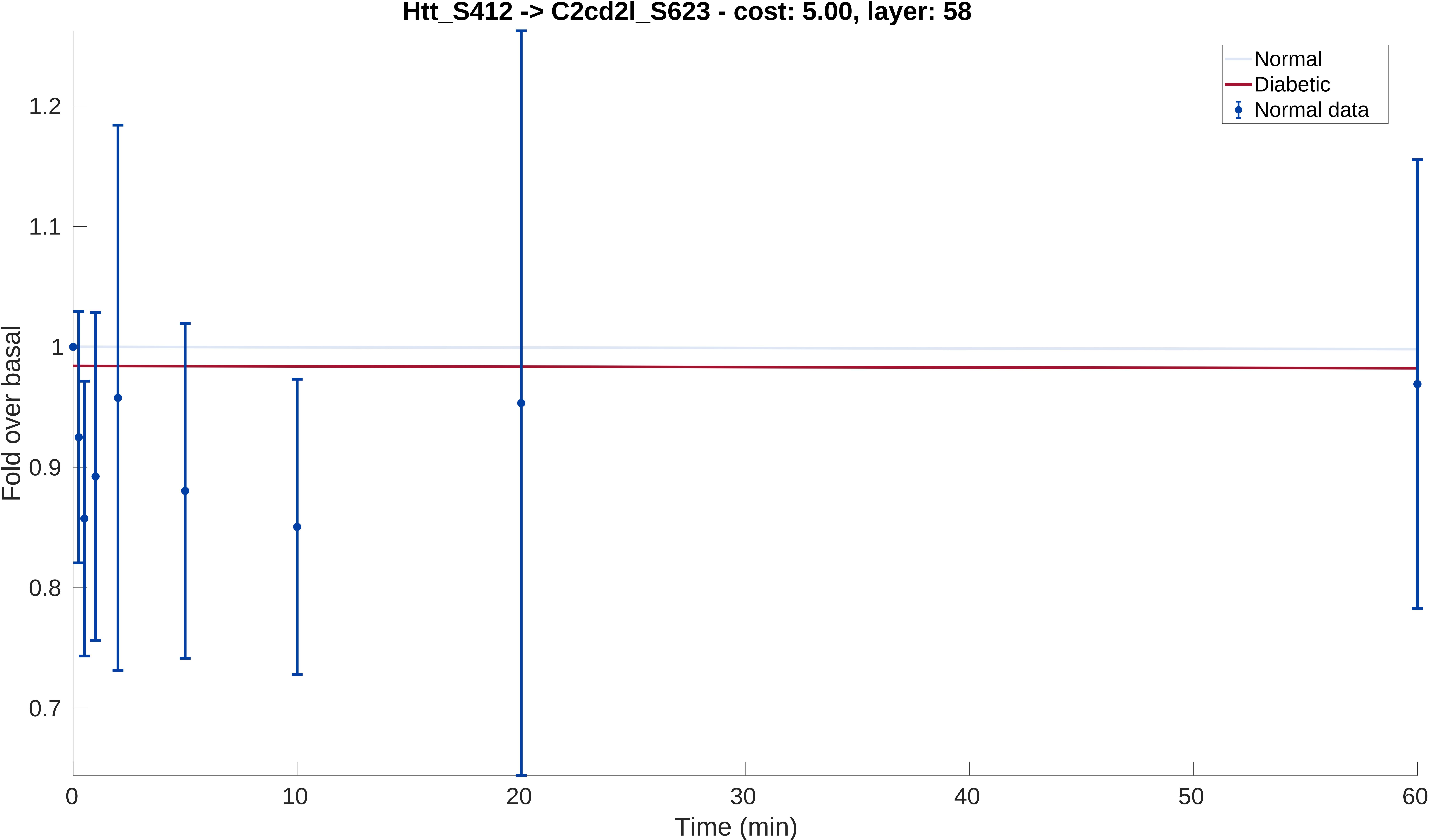

Htt_S412 -> C2cd2l_S623 - cost: 5.00, layer: 58
Normal
Diabetic
Normal data
Fold over basal
1.2
1.1
1
0.9
0.8
0.7
0
10
20
30
40
50
60
Time (min)

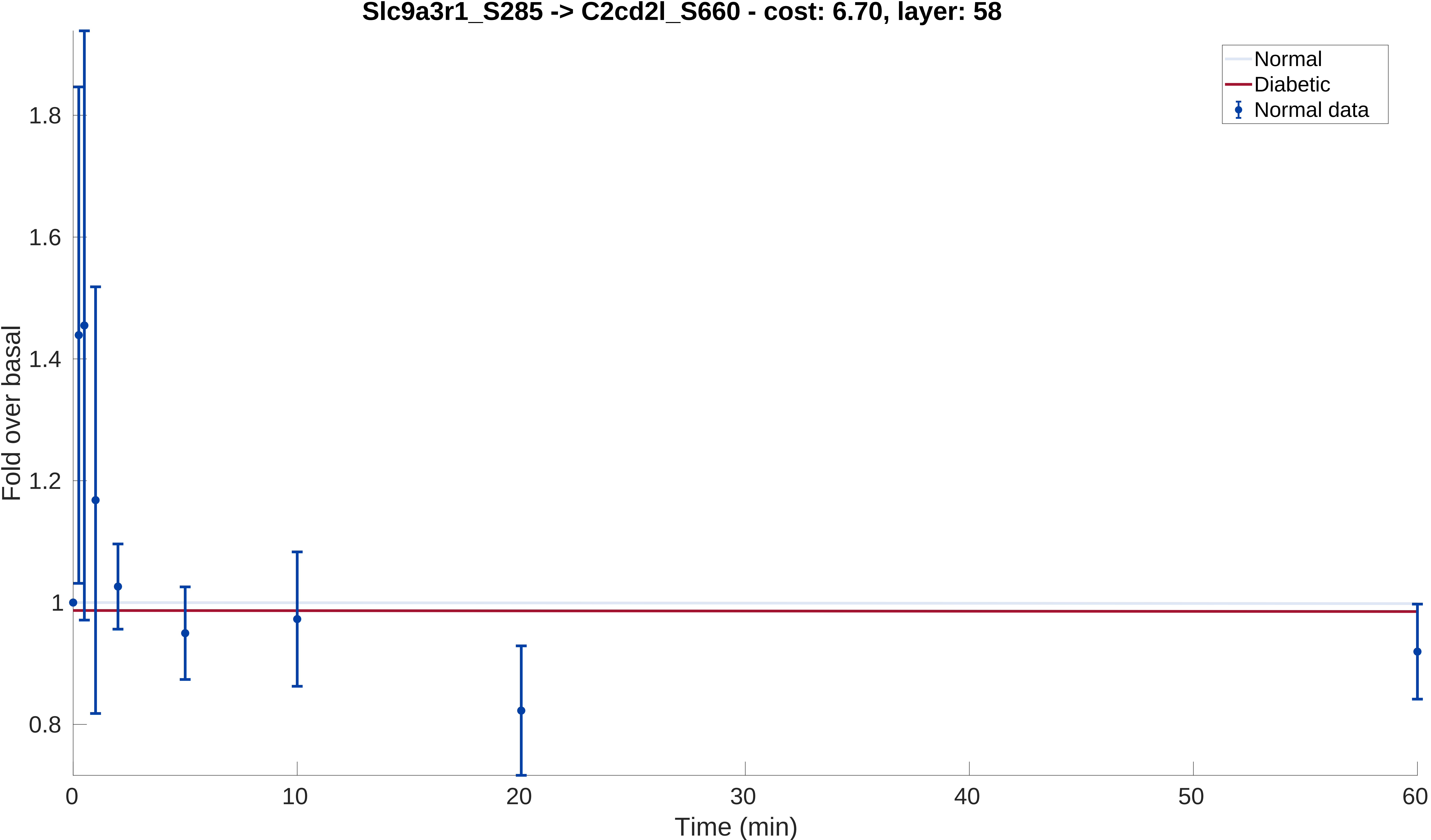

Slc9a3r1_S285 -> C2cd2l_S660 - cost: 6.70, layer: 58
Normal
Diabetic
Normal data
Fold over basal
1.8
1.6
1.4
1.2
1
0.8
0
10
20
30
40
50
60
Time (min)

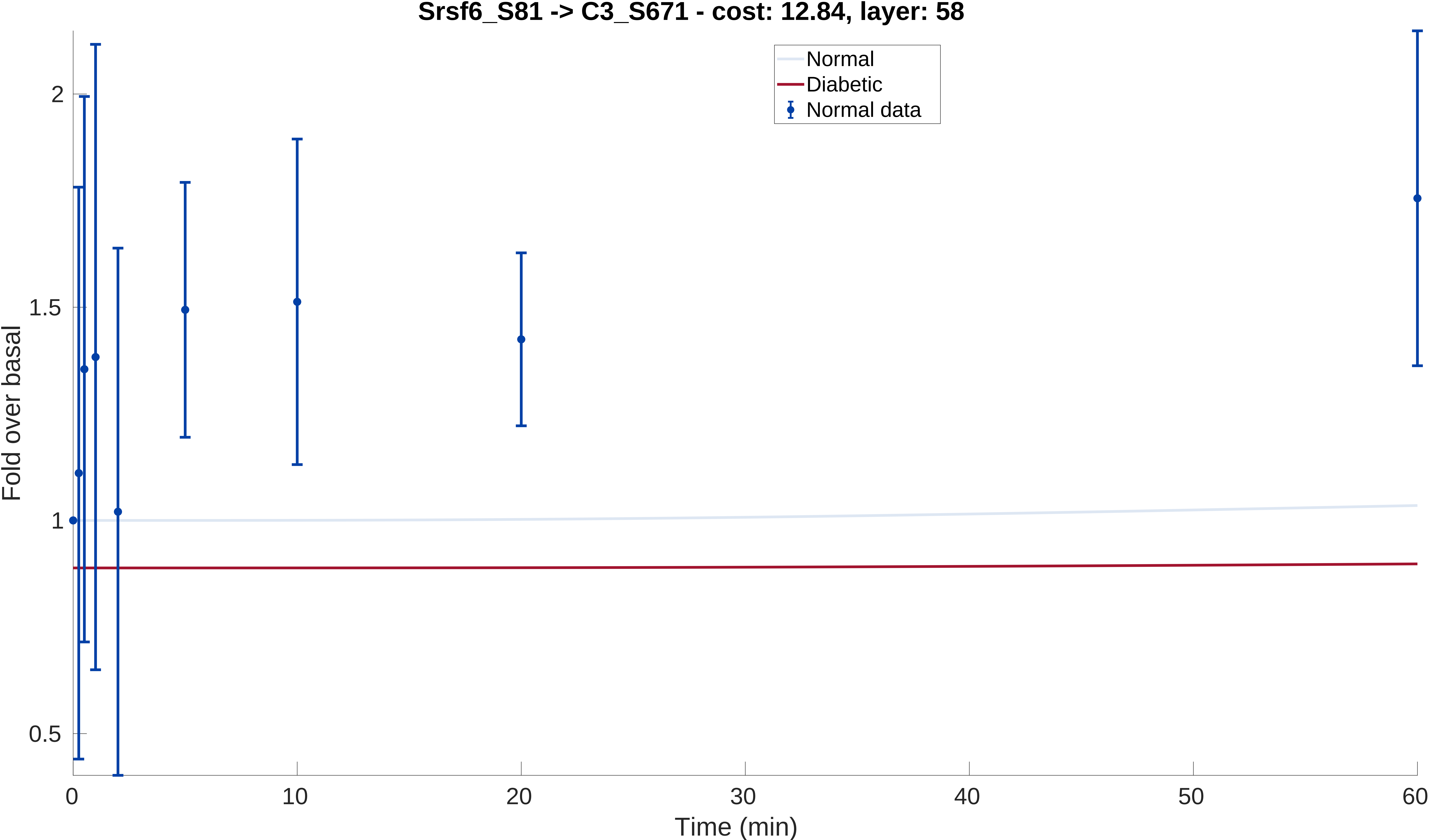

Srsf6_S81 -> C3_S671 - cost: 12.84, layer: 58
Normal
Diabetic
Normal data
Fold over basal
2
1.5
1
0.5
0
10
20
30
40
50
60
Time (min)

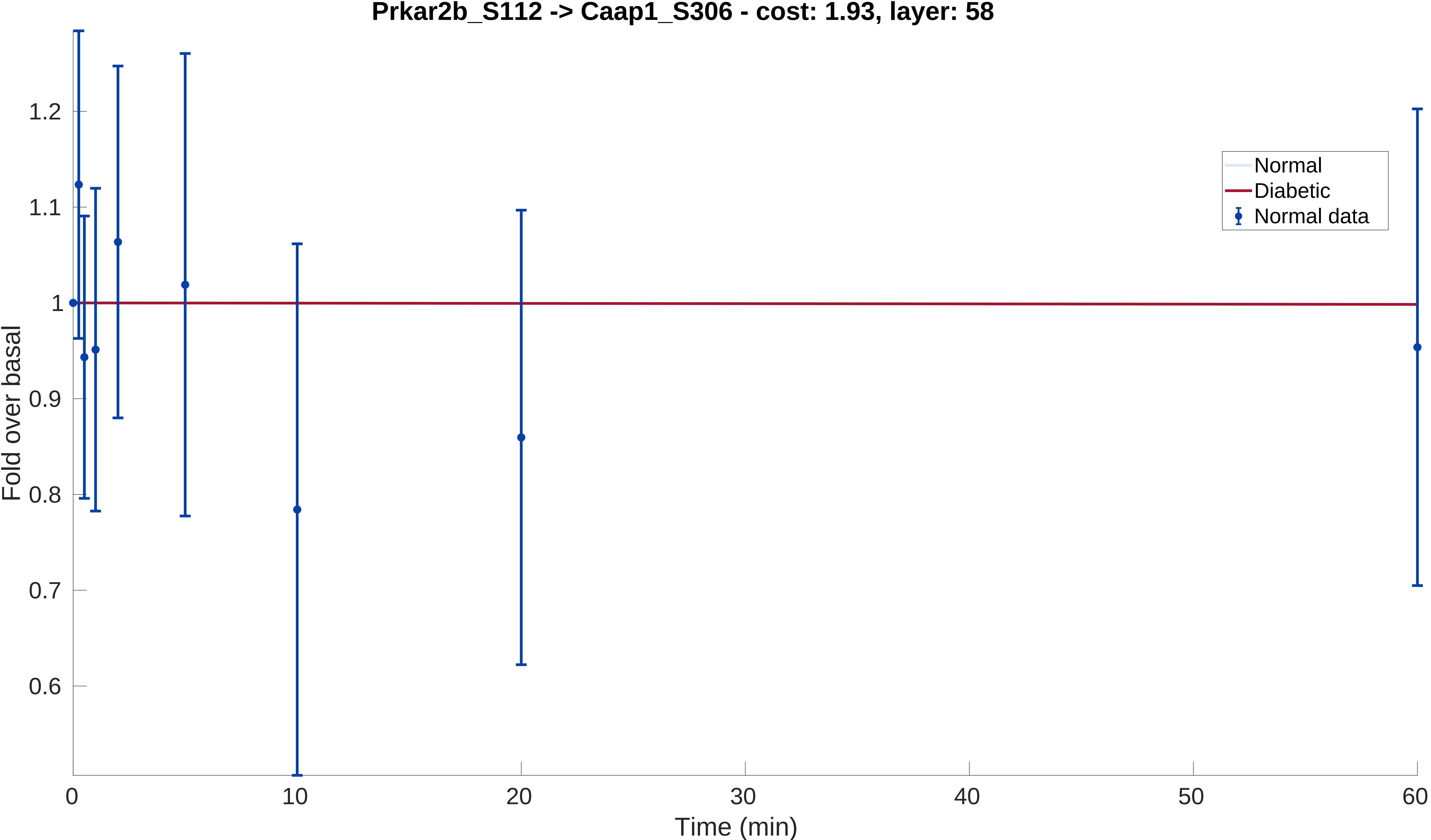

Prkar2b_S112 -> Caap1_S306 - cost: 1.93, layer: 58
Fold over basal
Time (min)
Normal
Diabetic
Normal data
1.2
1.1
1
0.9
0.8
0.7
0.6
0
10
20
30
40
50
60

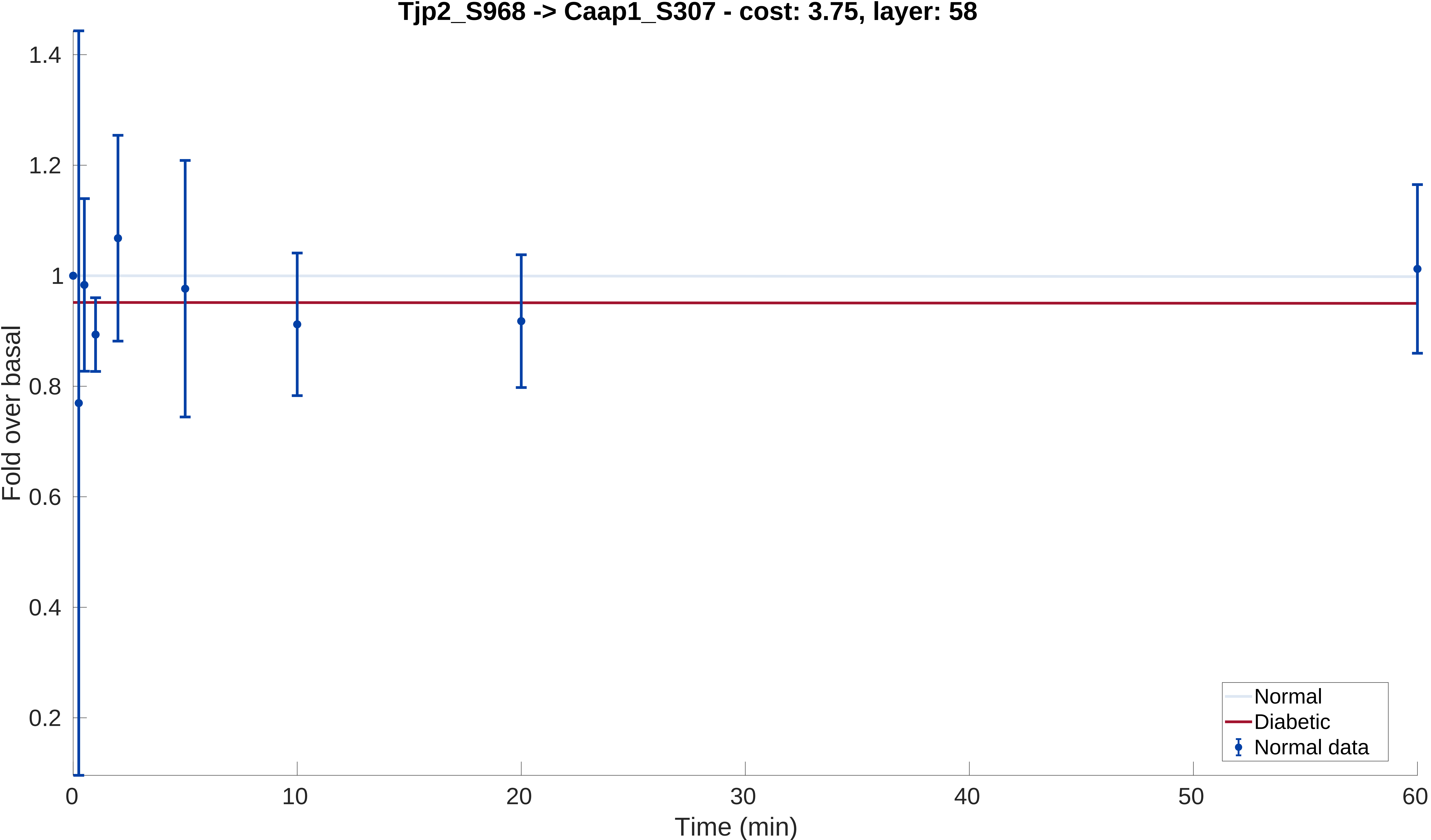

Tjp2_S968 -> Caap1_S307 - cost: 3.75, layer: 58
Fold over basal
1.4
1.2
1
0.8
0.6
0.4
0.2
0
10
20
30
40
50
60
Time (min)
Normal
Diabetic
Normal data

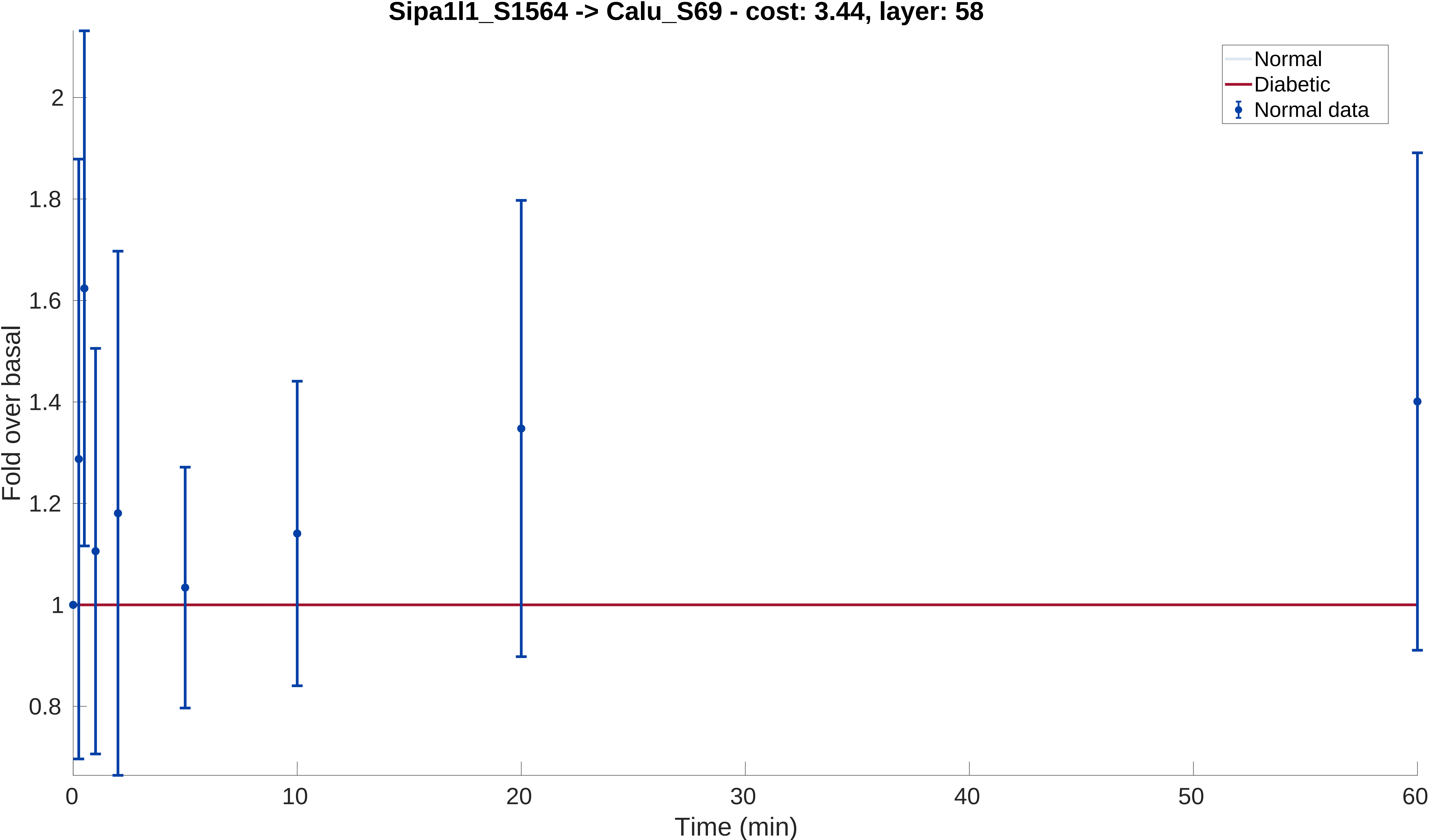

Sipa1l1_S1564 -> Calu_S69 - cost: 3.44, layer: 58
Normal
Diabetic
Normal data
Fold over basal
Time (min)
0.8
1
1.2
1.4
1.6
1.8
2
0
10
20
30
40
50
60

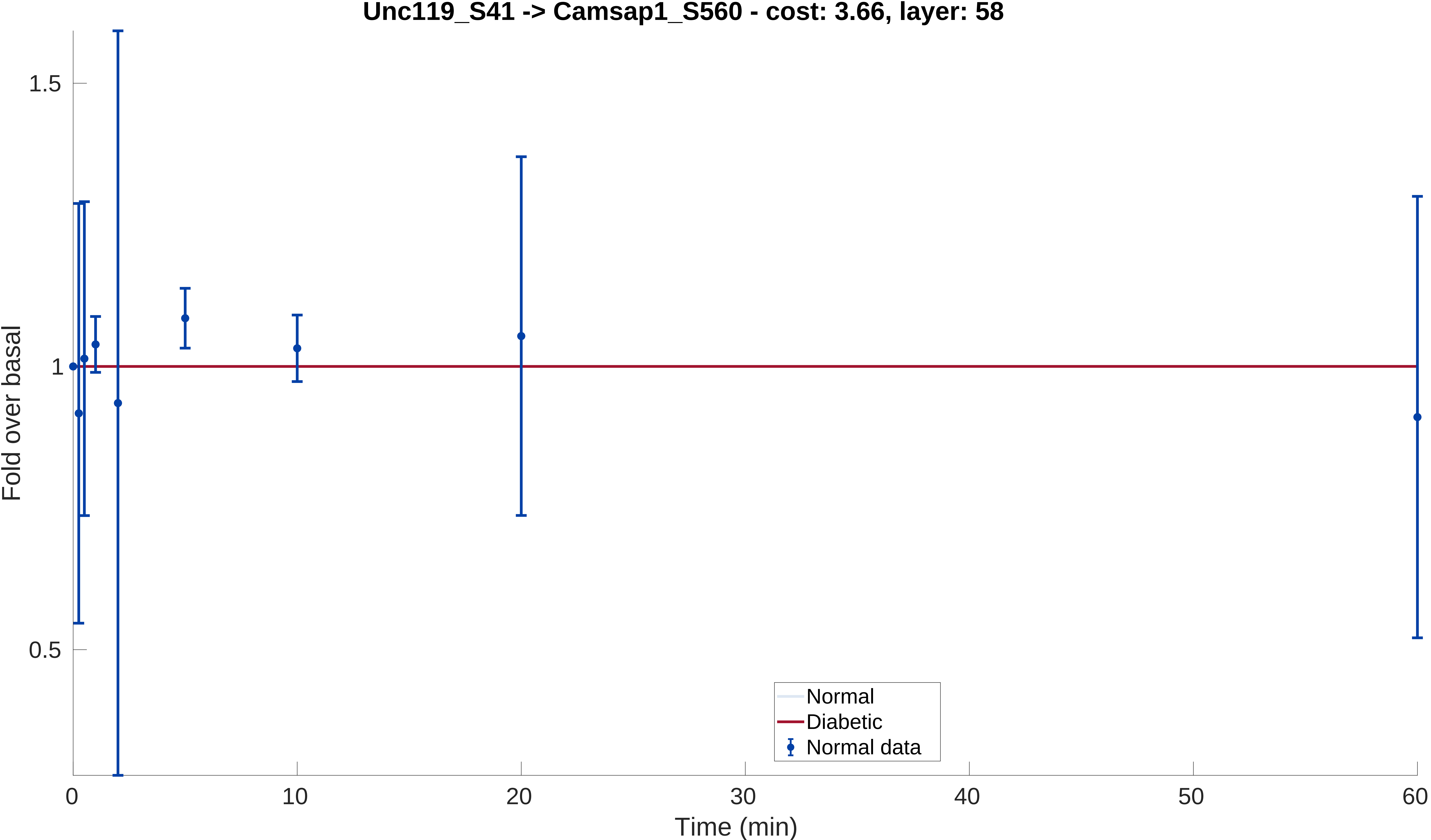

Unc119_S41 -> Camsap1_S560 - cost: 3.66, layer: 58
Fold over basal
1.5
1
0.5
0
10
20
30
40
50
60
Time (min)
Normal
Diabetic
Normal data

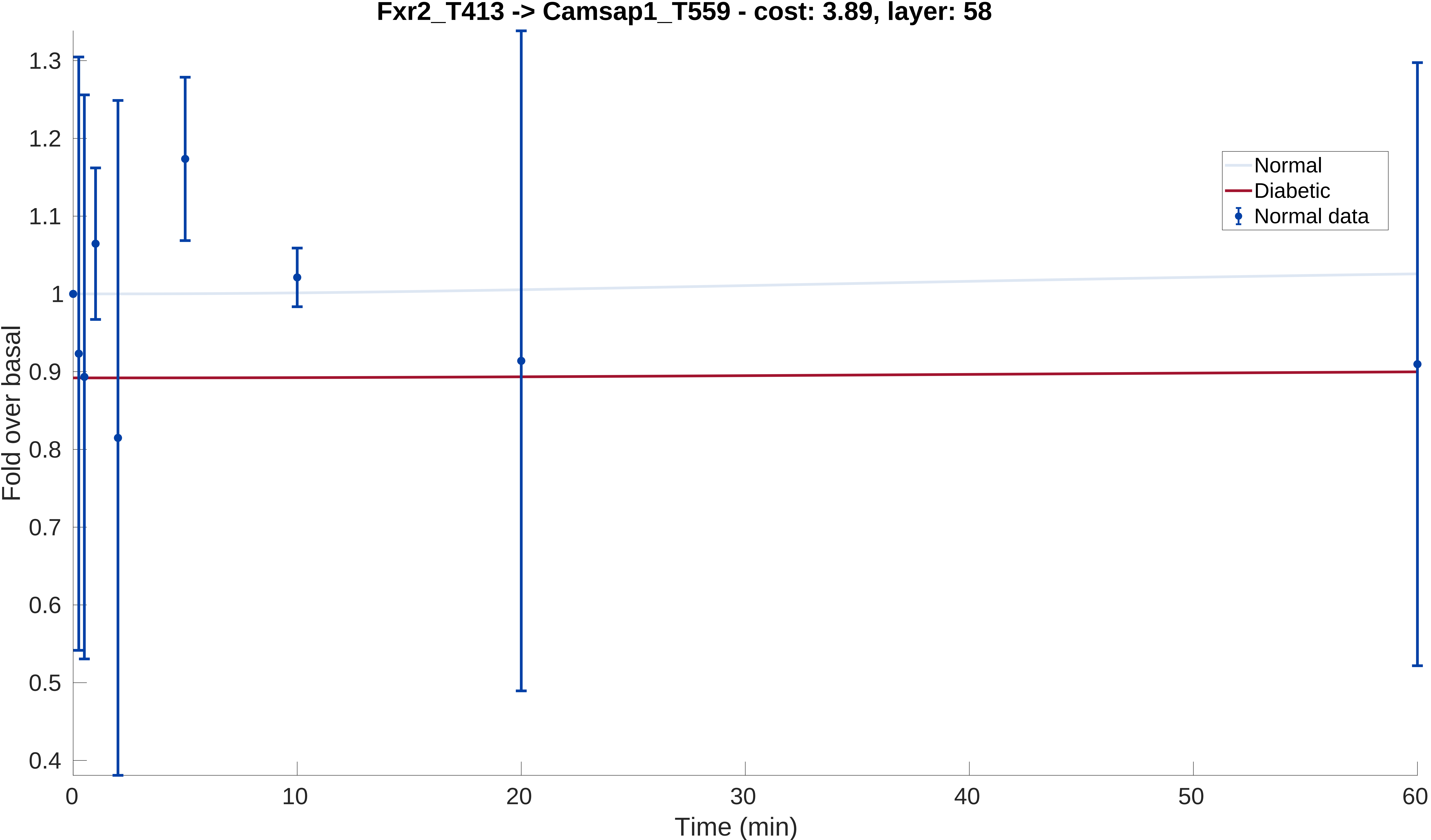

Fxr2_T413 -> Camsap1_T559 - cost: 3.89, layer: 58
Fold over basal
1.3
1.2
1.1
1
0.9
0.8
0.7
0.6
0.5
0.4
0
10
20
30
40
50
60
Time (min)
Normal
Diabetic
Normal data

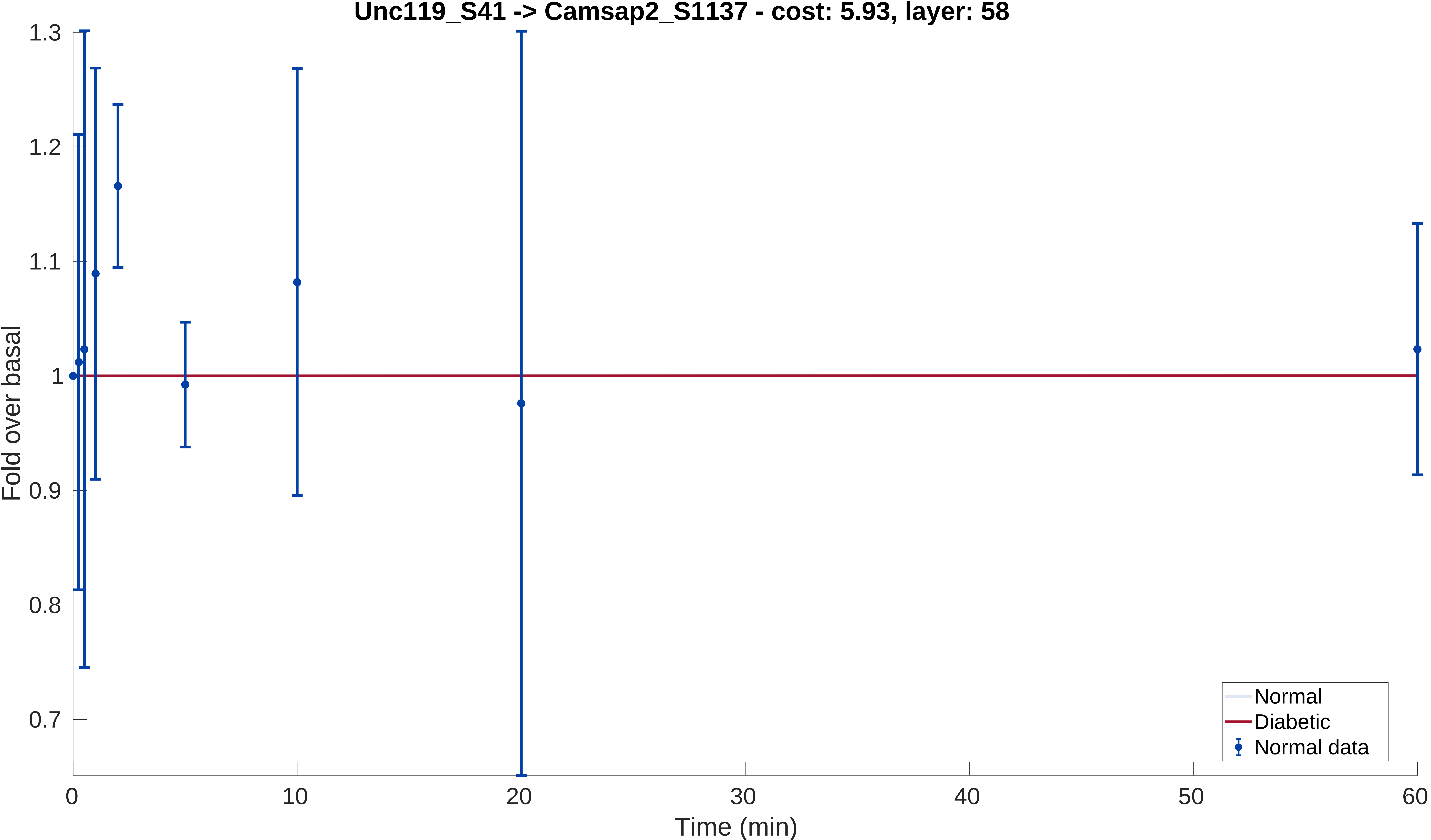

Unc119_S41 -> Camsap2_S1137 - cost: 5.93, layer: 58
Fold over basal
Time (min)
1.3
1.2
1.1
1
0.9
0.8
0.7
0
10
20
30
40
50
60
Normal
Diabetic
Normal data

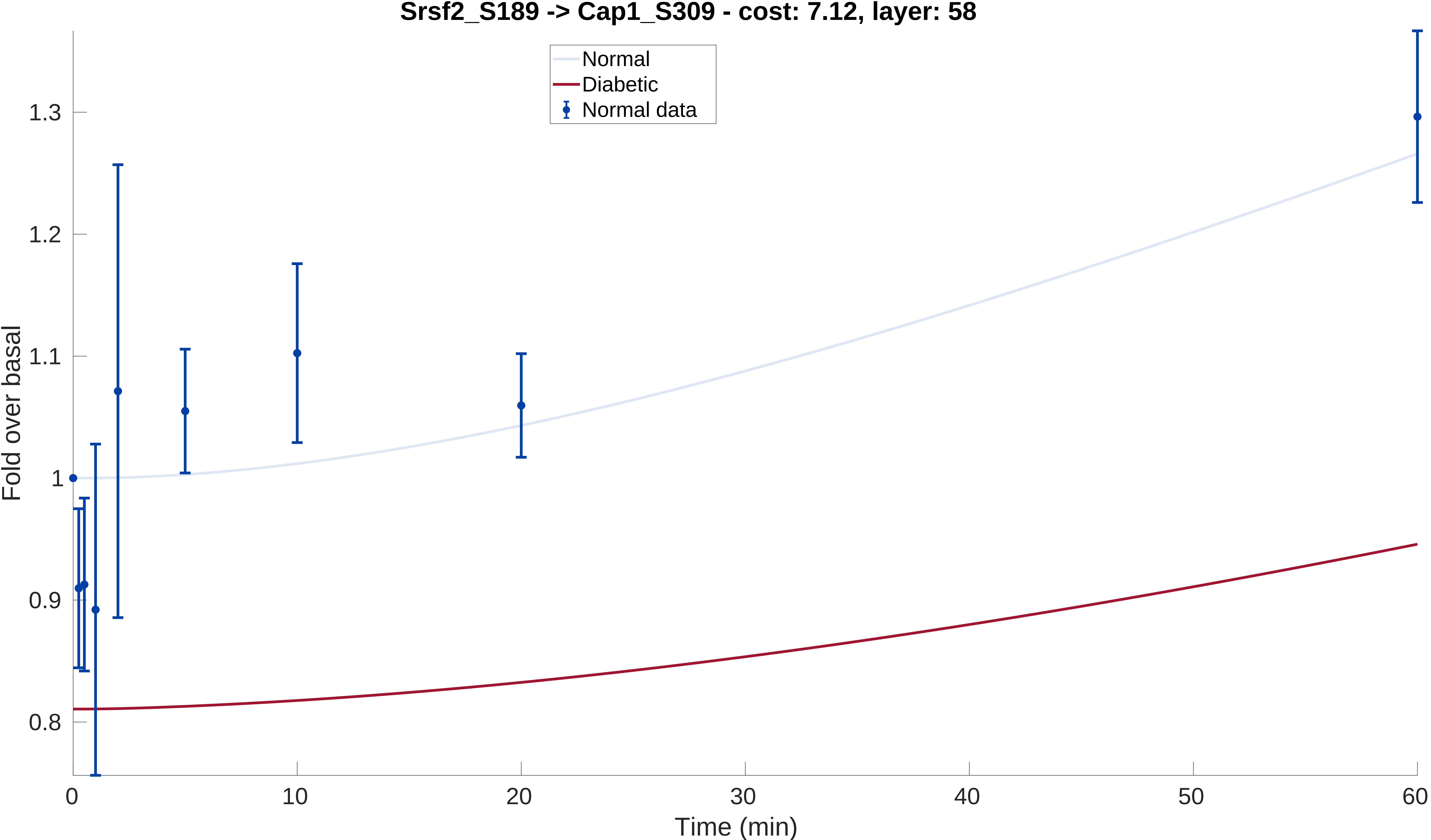
Srsf2_S189 -> Cap1_S309 - cost: 7.12, layer: 58
Normal
Diabetic
Normal data
Fold over basal
1.3
1.2
1.1
1
0.9
0.8
0
10
20
30
40
50
60
Time (min)

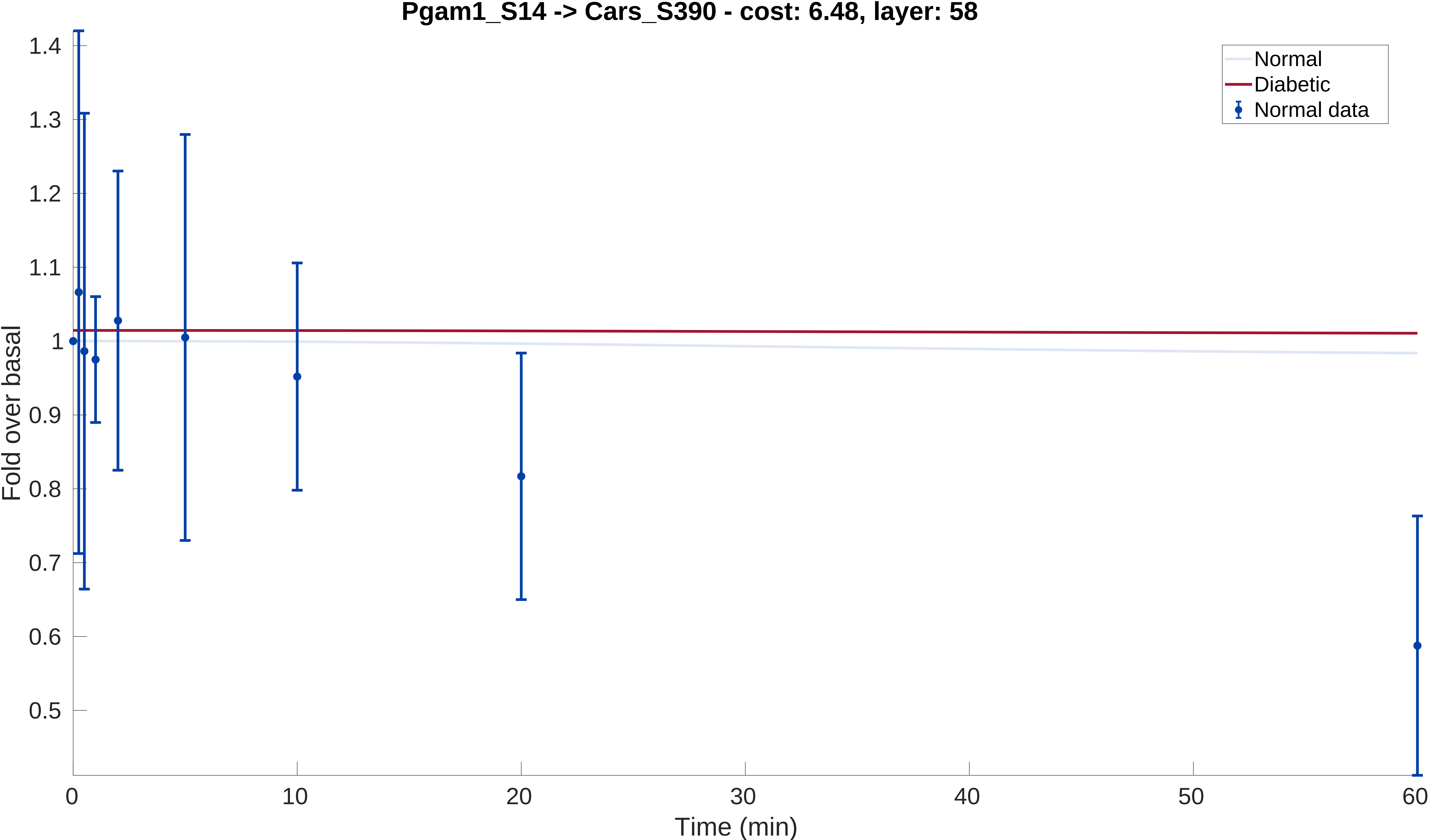

Pgam1_S14 -> Cars_S390 - cost: 6.48, layer: 58
Normal
Diabetic
Normal data
Fold over basal
Time (min)
1.4
1.3
1.2
1.1
1
0.9
0.8
0.7
0.6
0.5
0
10
20
30
40
50
60

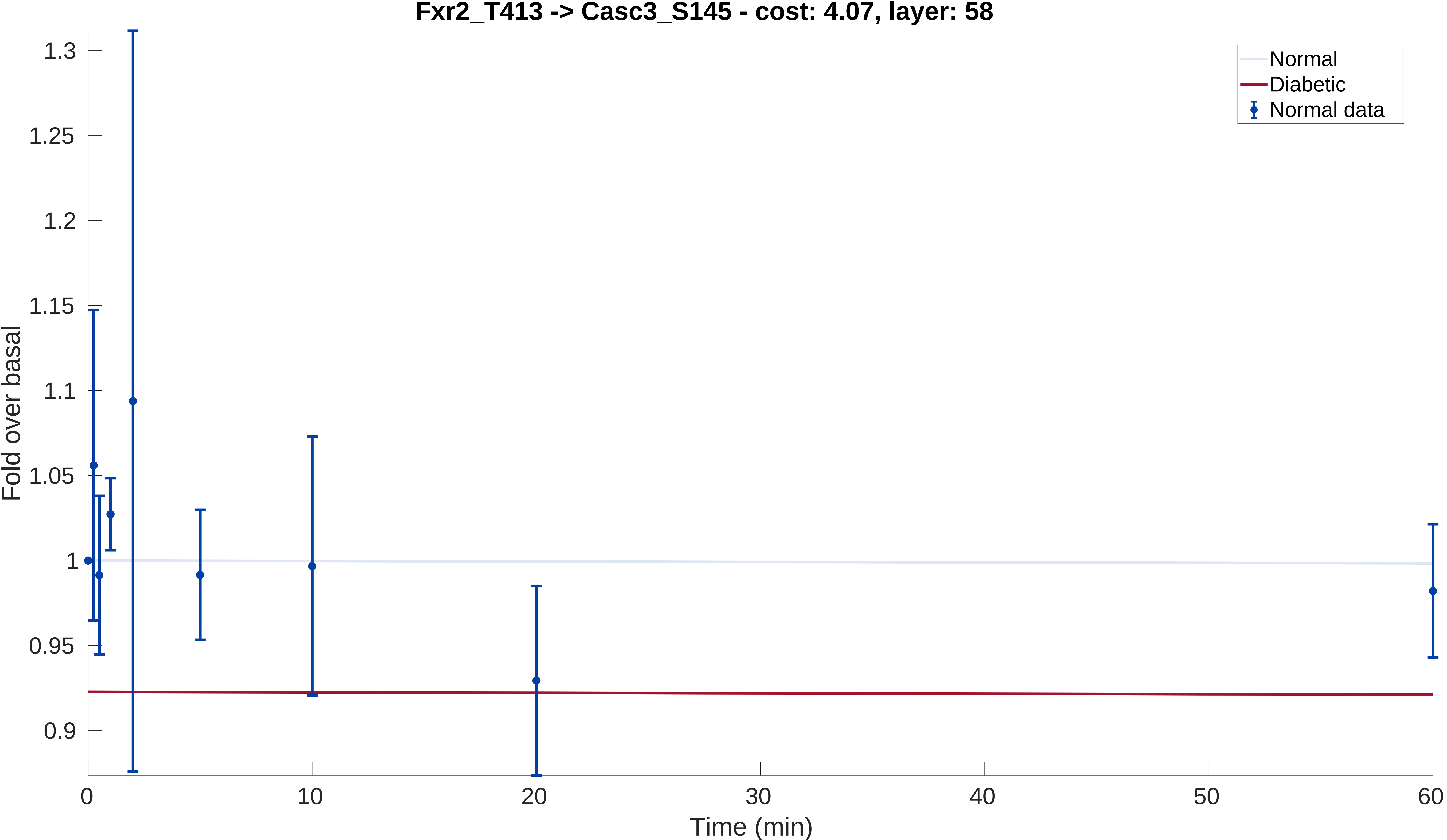

Fxr2_T413 -> Casc3_S145 - cost: 4.07, layer: 58
Normal
Diabetic
Normal data
Fold over basal
1.3
1.25
1.2
1.15
1.1
1.05
1
0.95
0.9
0
10
20
30
40
50
60
Time (min)

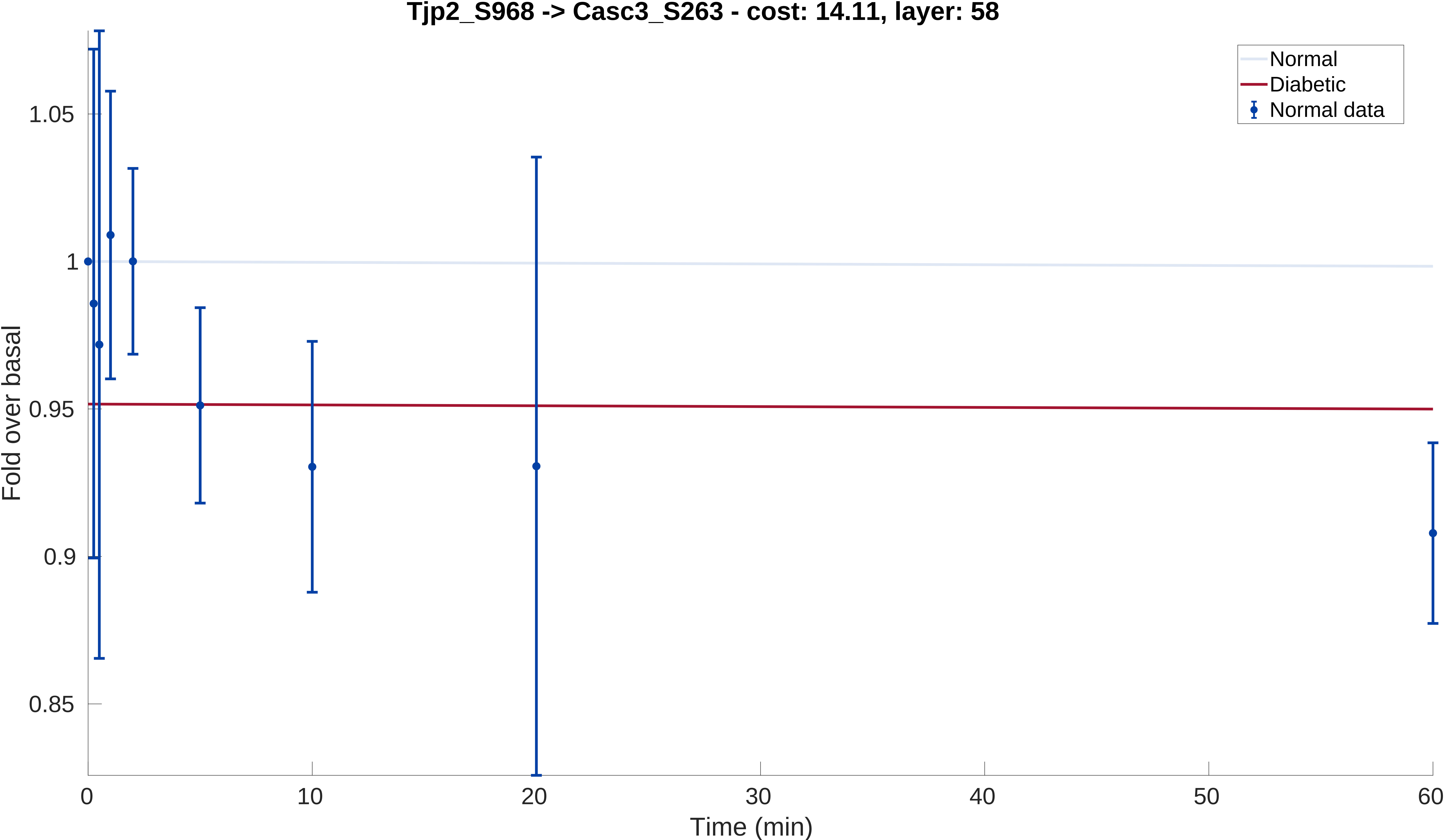

Tjp2_S968 -> Casc3_S263 - cost: 14.11, layer: 58
Normal
Diabetic
Normal data
Fold over basal
1.05
1
0.95
0.9
0.85
0
10
20
30
40
50
60
Time (min)

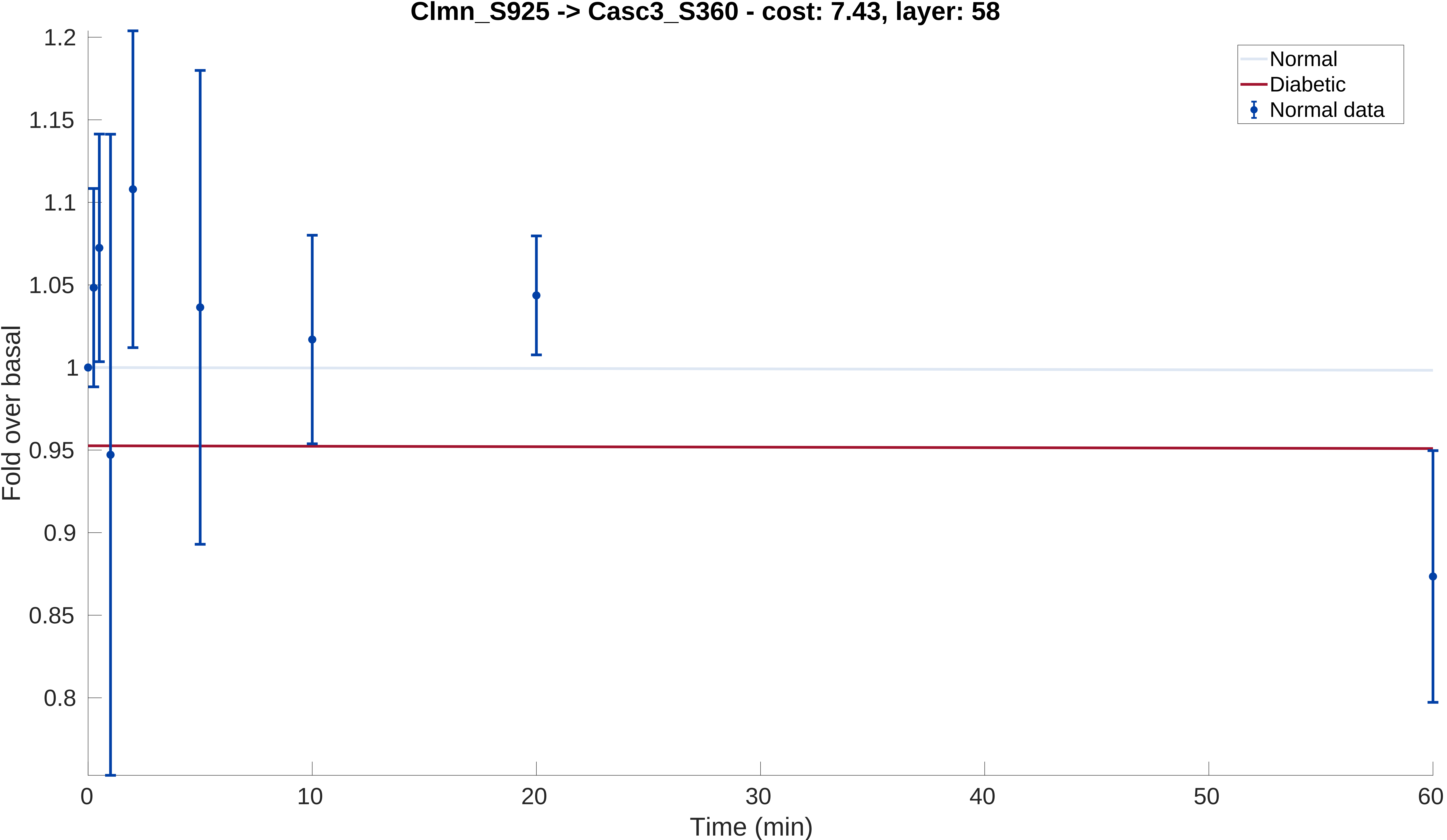

Clmn_S925 -> Casc3_S360 - cost: 7.43, layer: 58
Normal
Diabetic
Normal data
Fold over basal
Time (min)
1.2
1.15
1.1
1.05
1
0.95
0.9
0.85
0.8
0
10
20
30
40
50
60

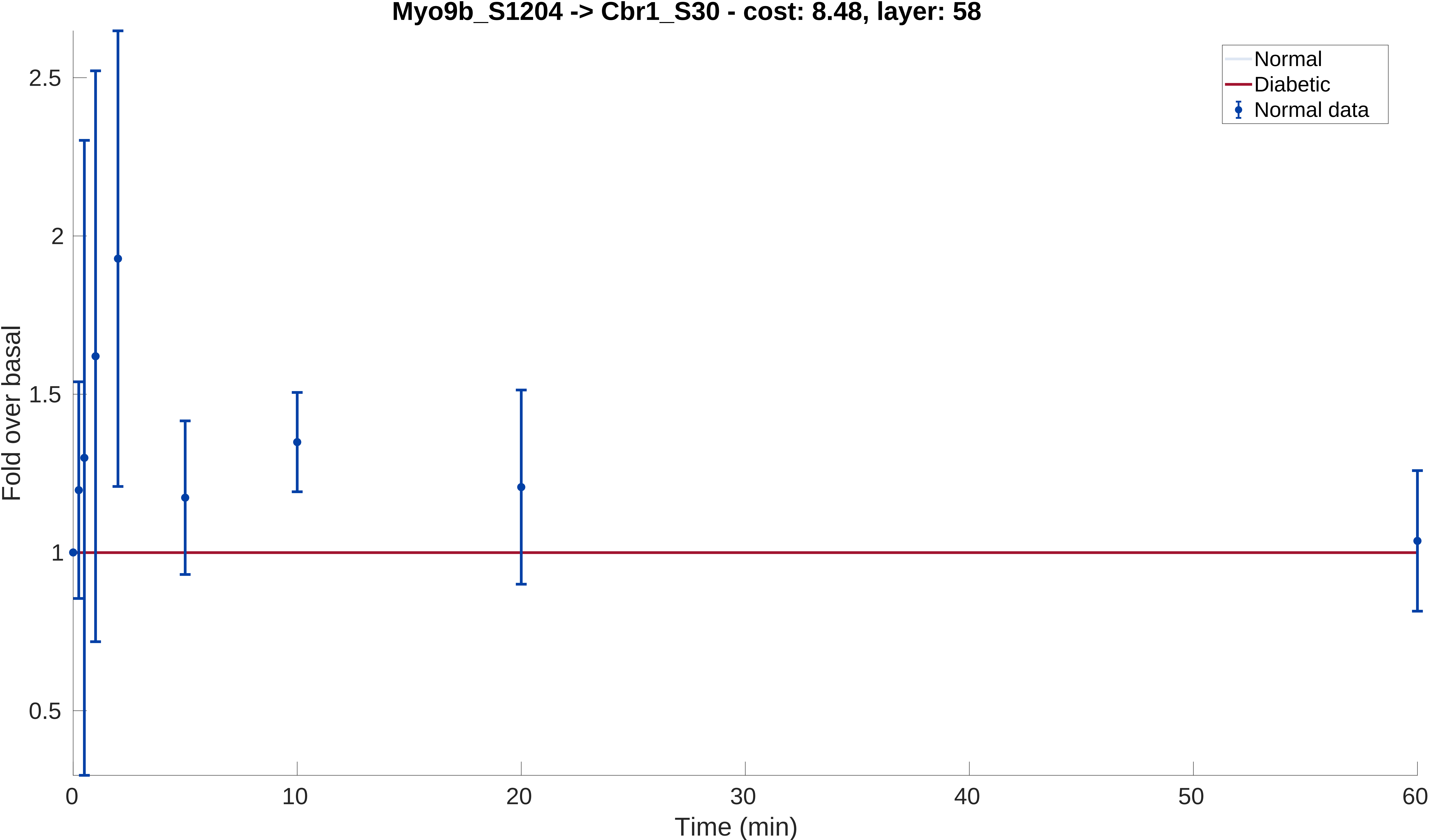
Myo9b_S1204 -> Cbr1_S30 - cost: 8.48, layer: 58
Normal
Diabetic
Normal data
Fold over basal
Time (min)
0.5
1
1.5
2
2.5
0
10
20
30
40
50
60

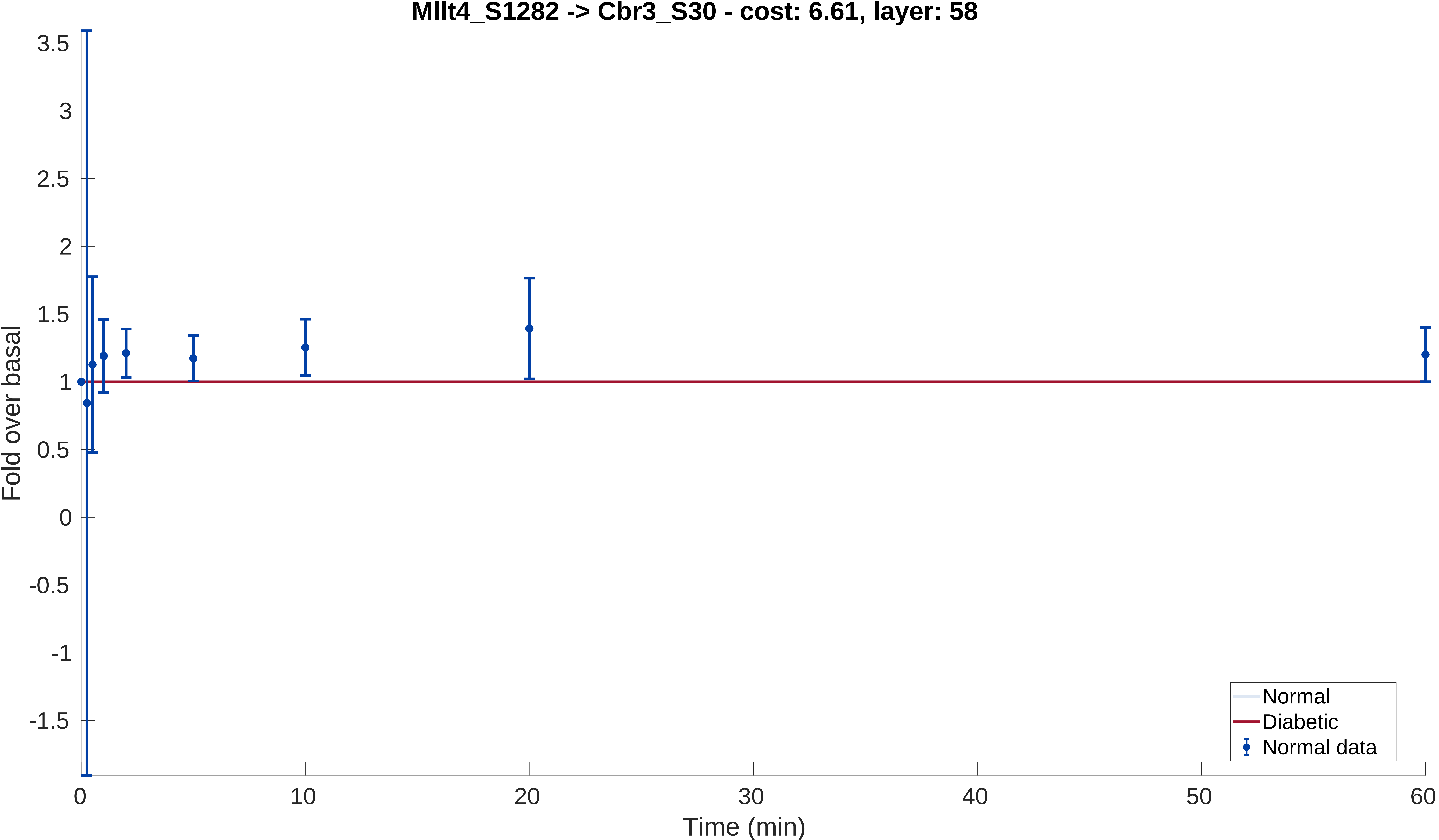

Mllt4_S1282 -> Cbr3_S30 - cost: 6.61, layer: 58
Fold over basal
3.5
3
2.5
2
1.5
1
0.5
0
-0.5
-1
-1.5
0
10
20
30
40
50
60
Time (min)
Normal
Diabetic
Normal data

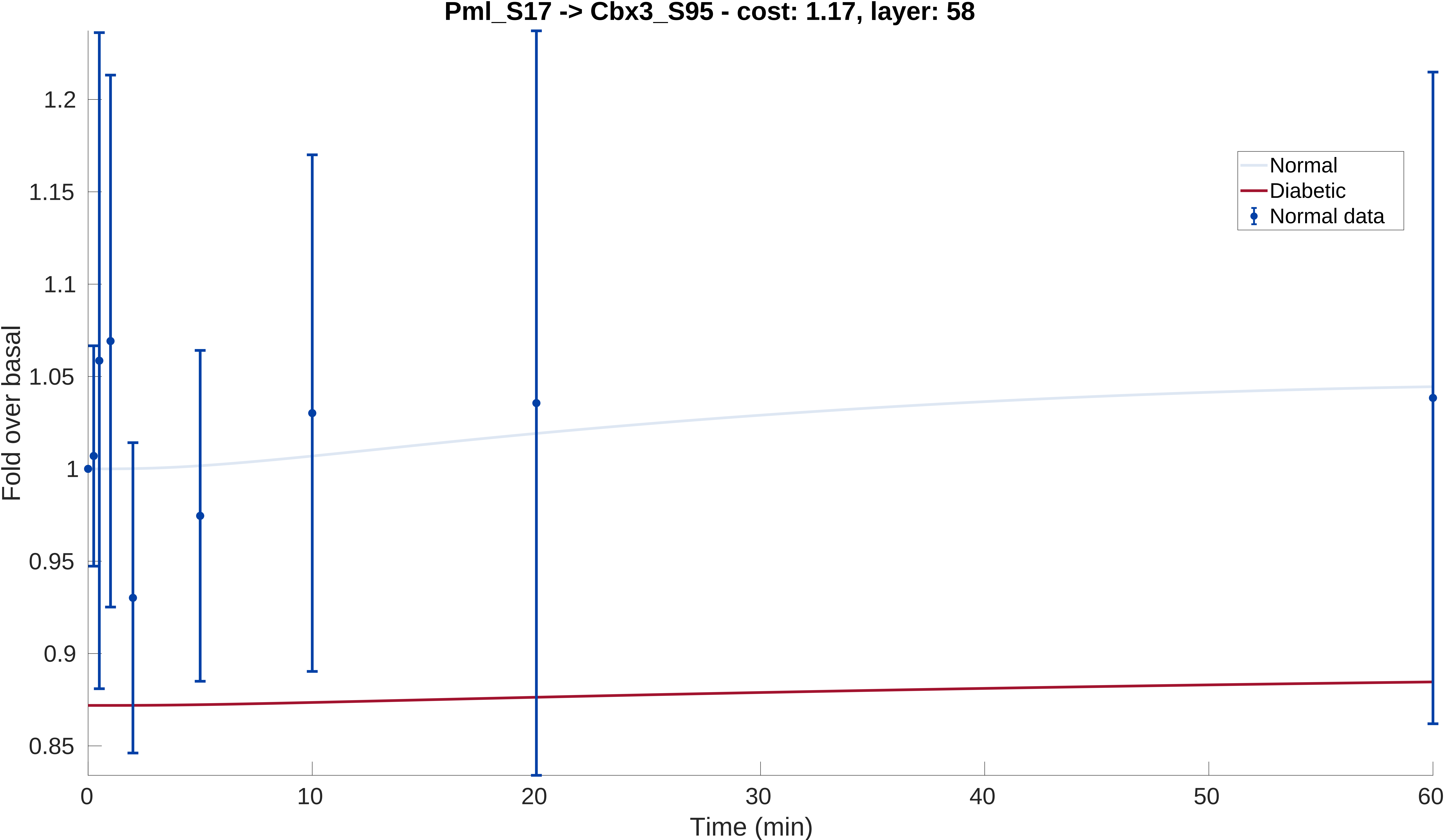

Pml_S17 -> Cbx3_S95 - cost: 1.17, layer: 58
Fold over basal
1.2
1.15
1.1
1.05
1
0.95
0.9
0.85
0
10
20
30
40
50
60
Time (min)
Normal
Diabetic
Normal data

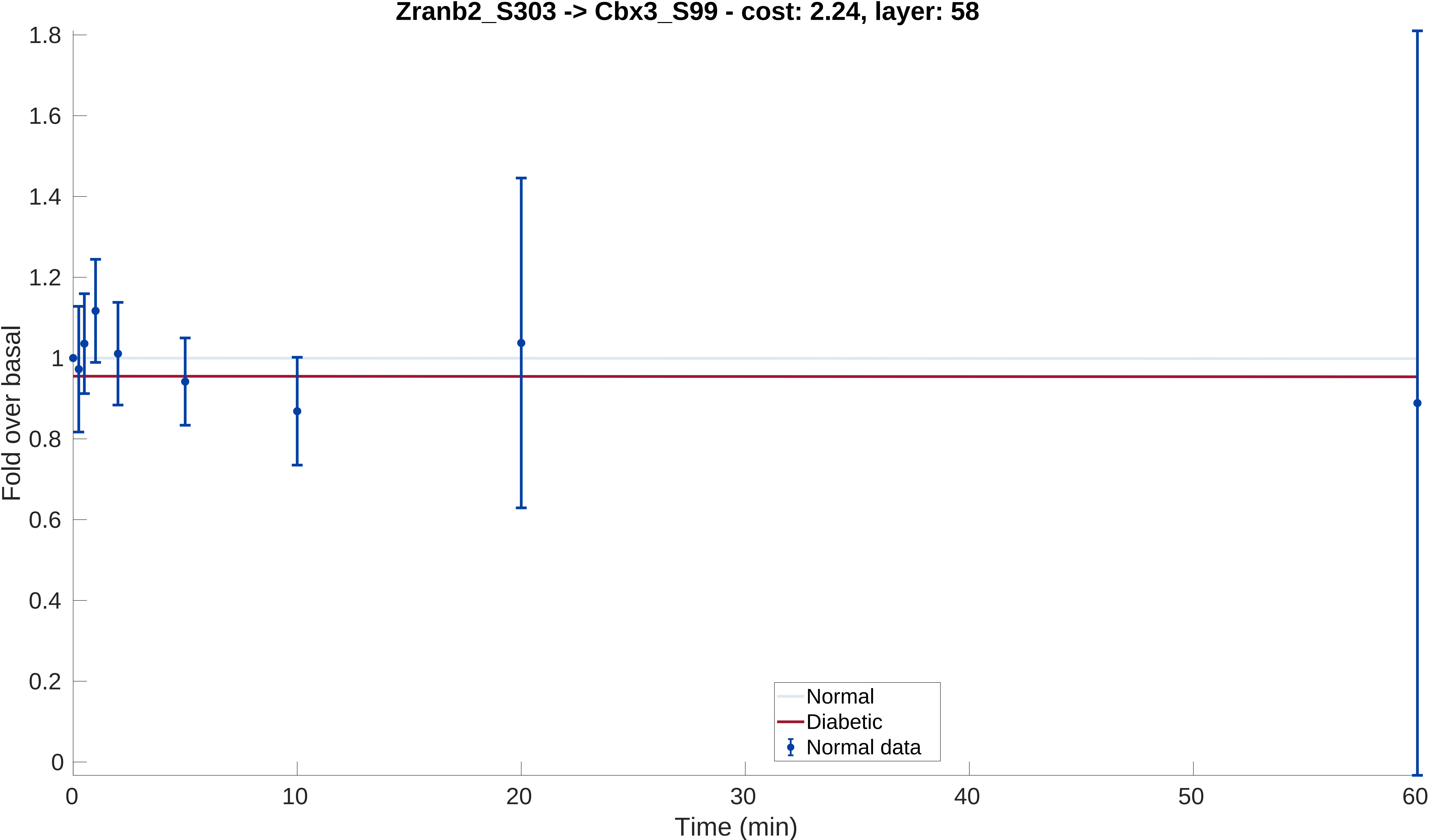
Zranb2_S303 -> Cbx3_S99 - cost: 2.24, layer: 58
Fold over basal
Time (min)
0
0.2
0.4
0.6
0.8
1
1.2
1.4
1.6
1.8
0
10
20
30
40
50
60
Normal
Diabetic
Normal data

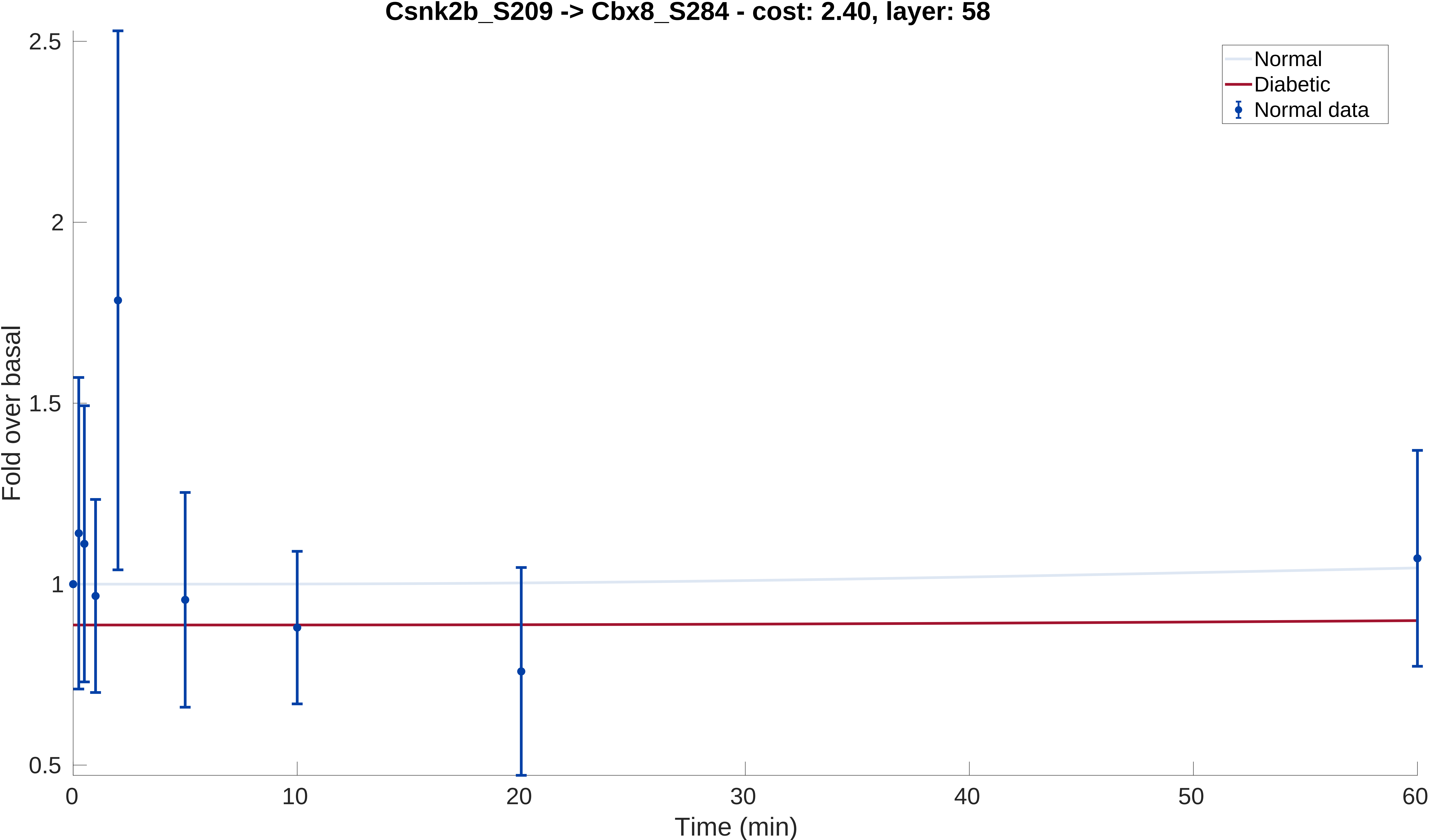

Csnk2b_S209 -> Cbx8_S284 - cost: 2.40, layer: 58
Normal
Diabetic
Normal data
Fold over basal
2.5
2
1.5
1
0.5
0
10
20
30
40
50
60
Time (min)

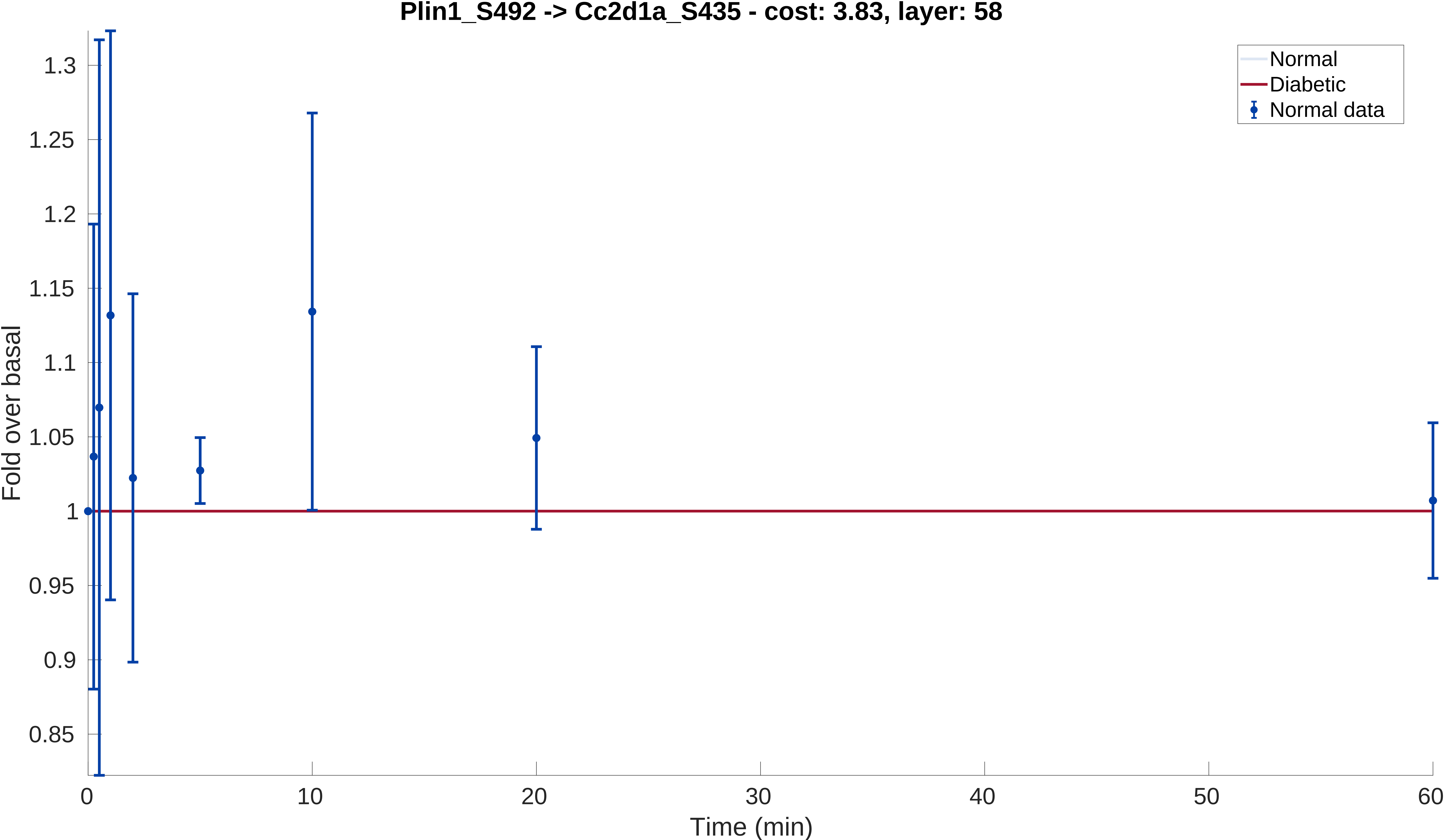
Plin1_S492 -> Cc2d1a_S435 - cost: 3.83, layer: 58
Normal
Diabetic
Normal data
Fold over basal
1.3
1.25
1.2
1.15
1.1
1.05
1
0.95
0.9
0.85
0
10
20
30
40
50
60
Time (min)

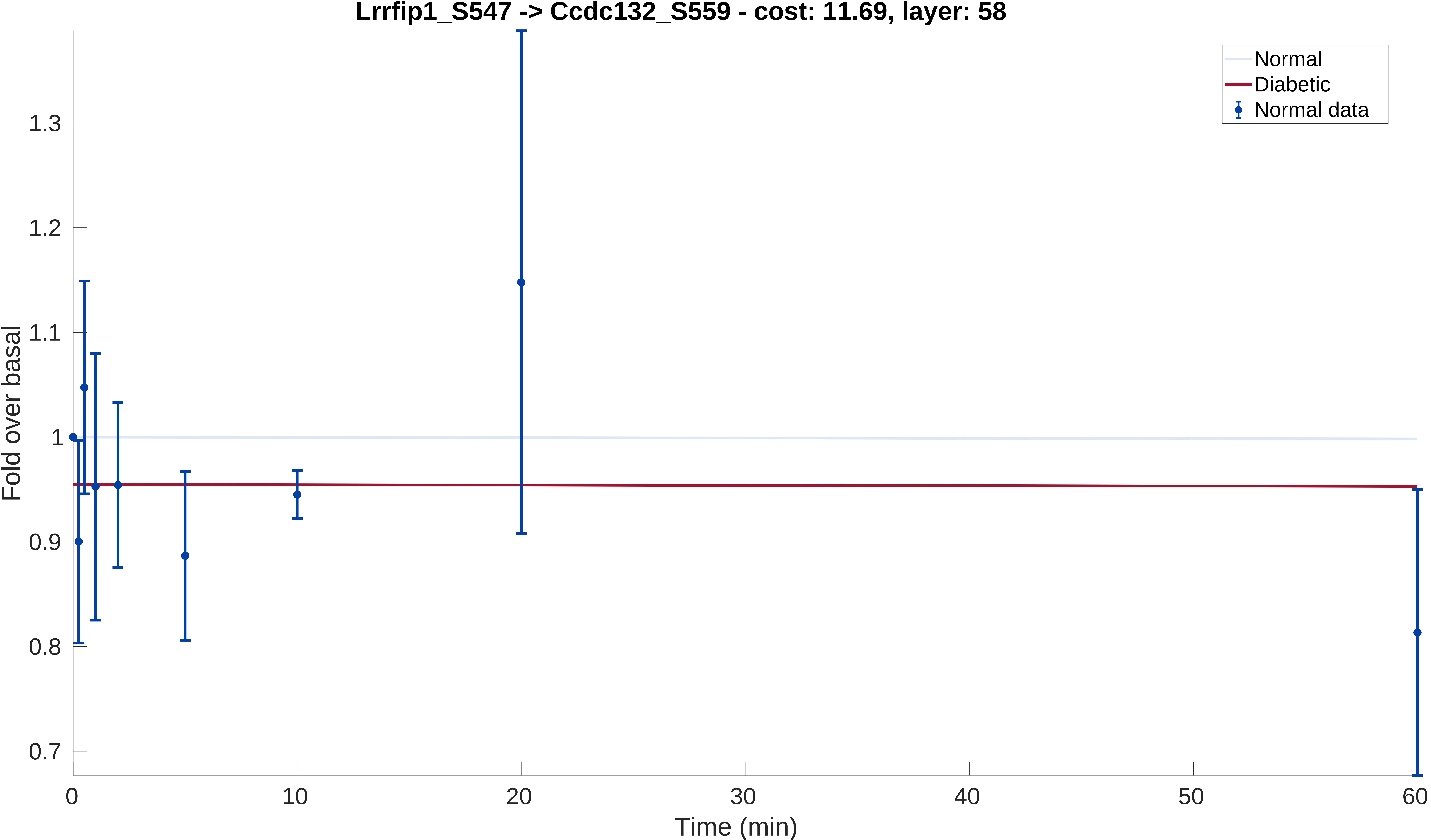

Lrrfip1_S547 -> Ccdc132_S559 - cost: 11.69, layer: 58
Normal
Diabetic
Normal data
Fold over basal
Time (min)
0.7
0.8
0.9
1
1.1
1.2
1.3
0
10
20
30
40
50
60

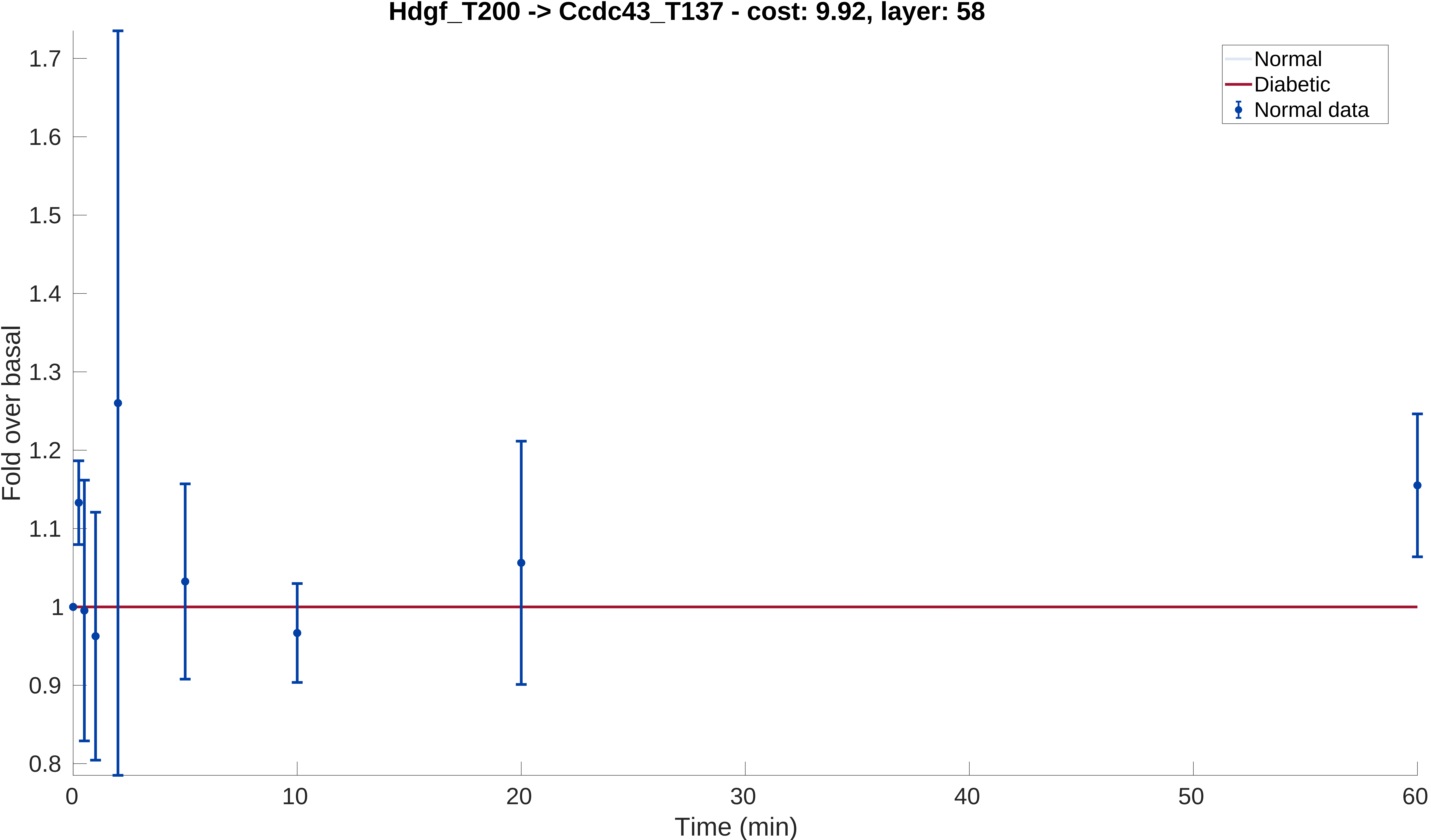

Hdgf_T200 -> Ccdc43_T137 - cost: 9.92, layer: 58
Normal
Diabetic
Normal data
Fold over basal
1.7
1.6
1.5
1.4
1.3
1.2
1.1
1
0.9
0.8
0
10
20
30
40
50
60
Time (min)

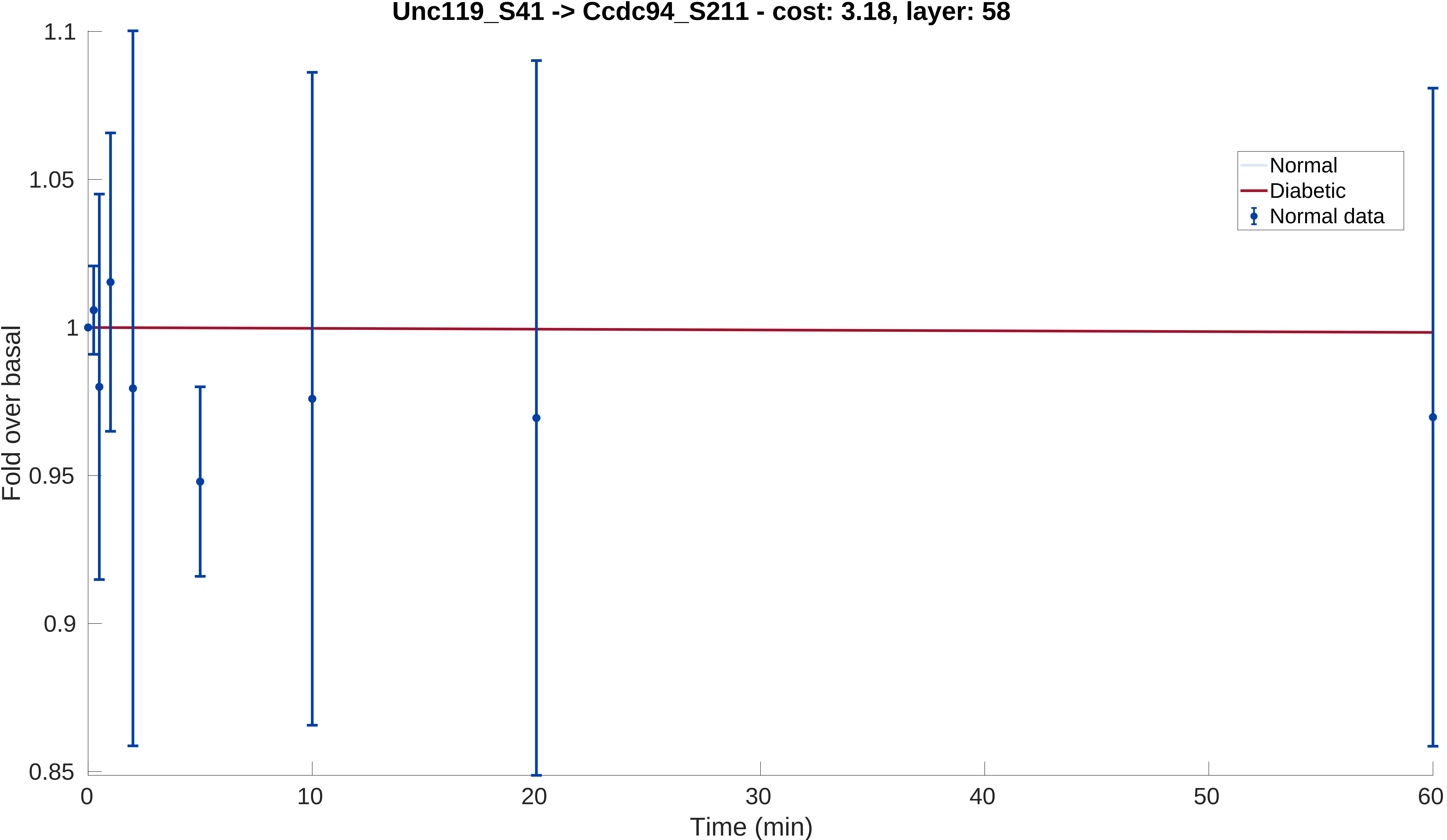

Unc119_S41 -> Ccdc94_S211 - cost: 3.18, layer: 58
Fold over basal
1.1
1.05
1
0.95
0.9
0.85
0
10
20
30
40
50
60
Time (min)
Normal
Diabetic
Normal data

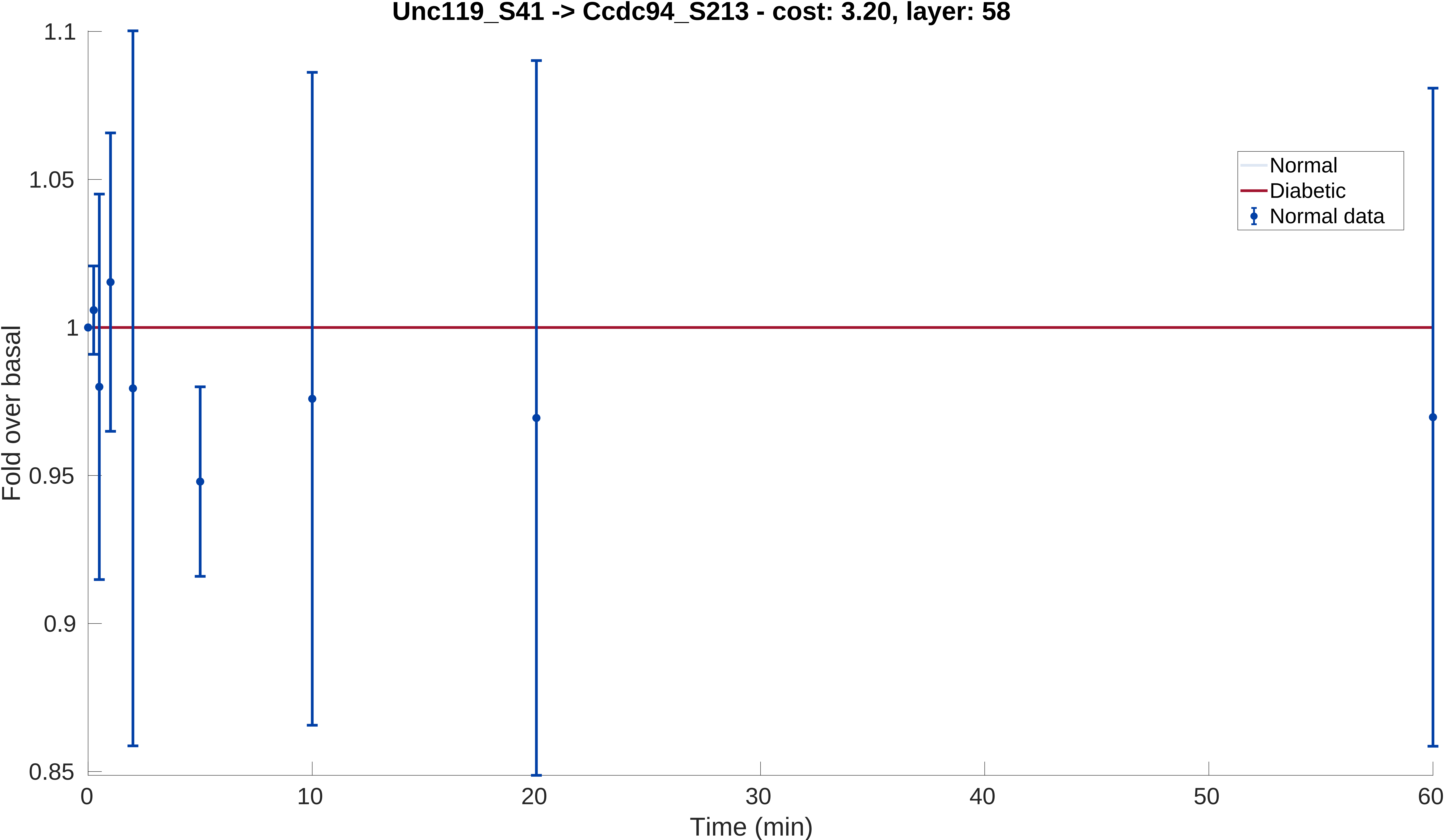

Unc119_S41 -> Ccdc94_S213 - cost: 3.20, layer: 58
Fold over basal
1.1
1.05
1
0.95
0.9
0.85
0
10
20
30
40
50
60
Time (min)
Normal
Diabetic
Normal data

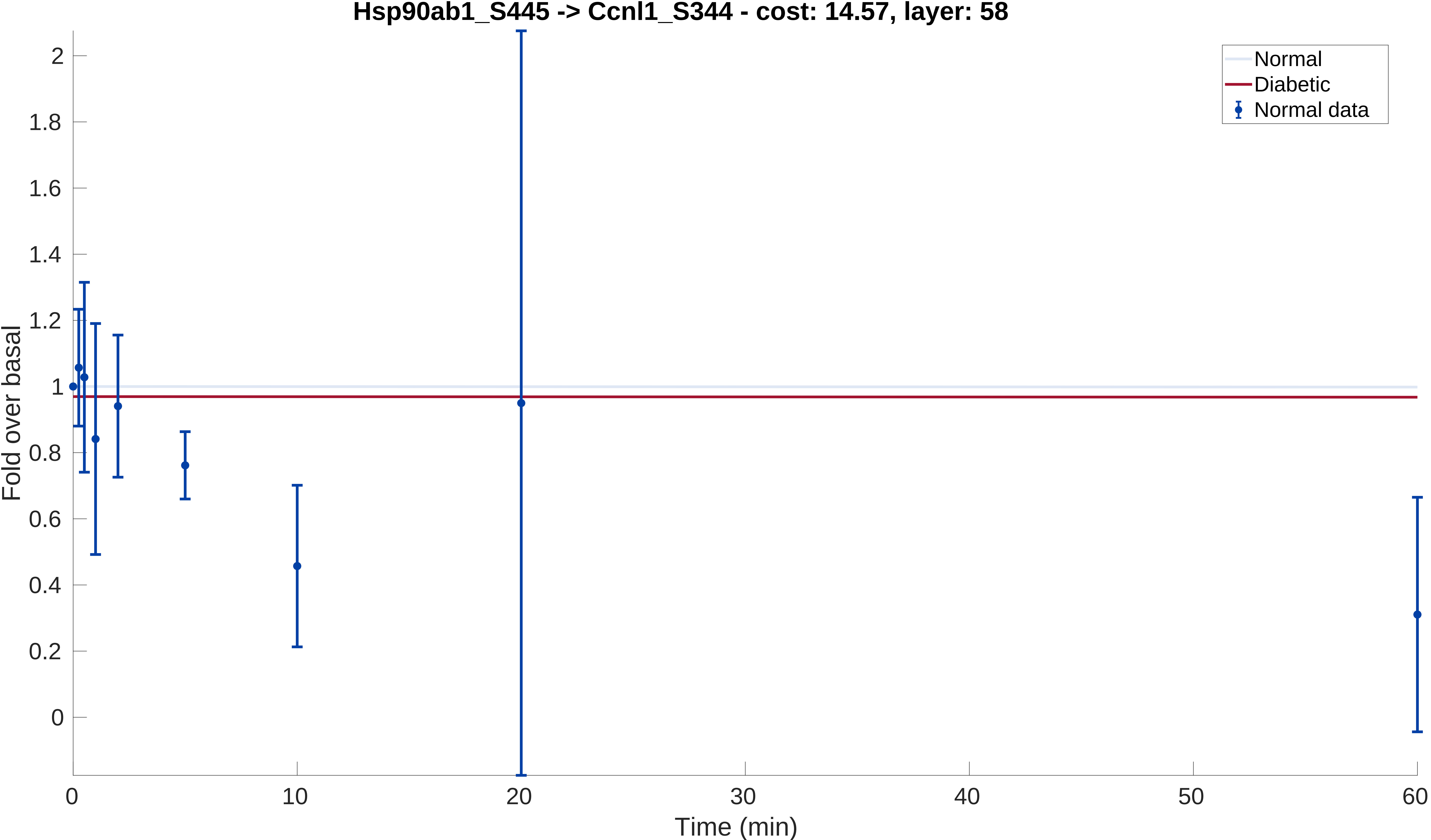
Hsp90ab1_S445 -> Ccnl1_S344 - cost: 14.57, layer: 58
Normal
Diabetic
Normal data
Fold over basal
Time (min)
2
1.8
1.6
1.4
1.2
1
0.8
0.6
0.4
0.2
0
0
10
20
30
40
50
60

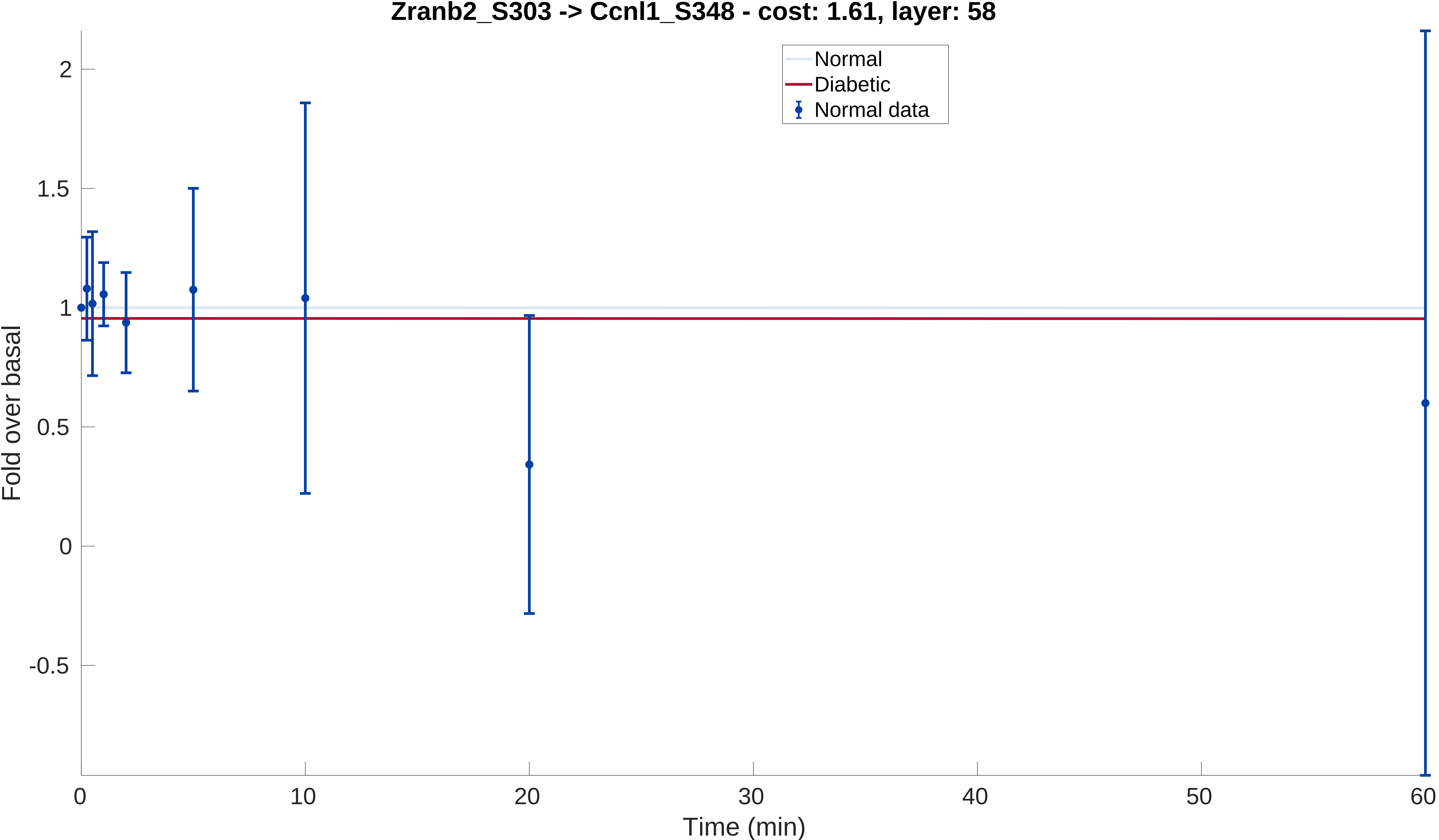

Zranb2_S303 -> Ccnl1_S348 - cost: 1.61, layer: 58
Normal
Diabetic
Normal data
Fold over basal
Time (min)
2
1.5
1
0.5
0
-0.5
0
10
20
30
40
50
60

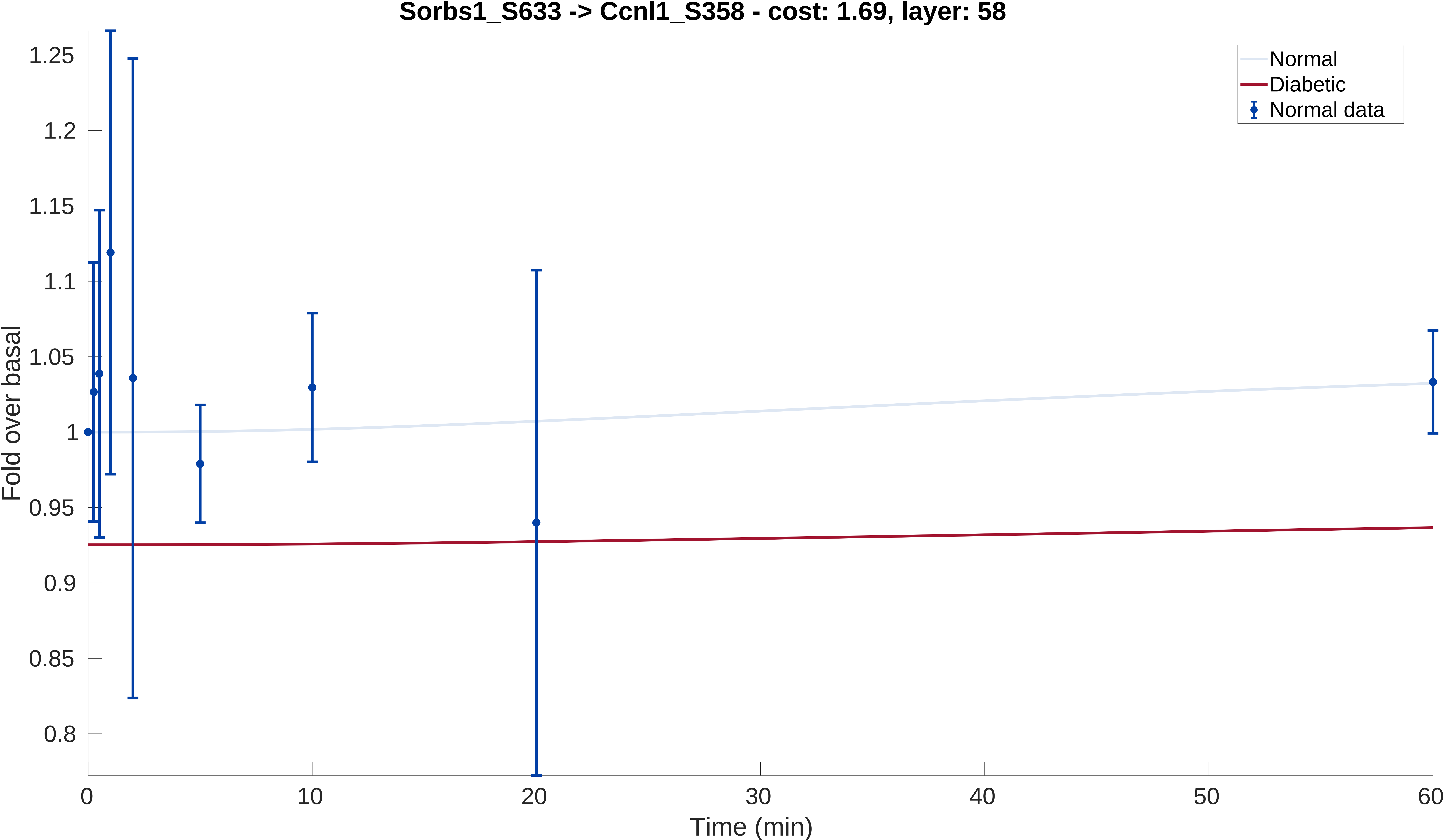

Sorbs1_S633 -> Ccnl1_S358 - cost: 1.69, layer: 58
Normal
Diabetic
Normal data
Fold over basal
1.25
1.2
1.15
1.1
1.05
1
0.95
0.9
0.85
0.8
0
10
20
30
40
50
60
Time (min)

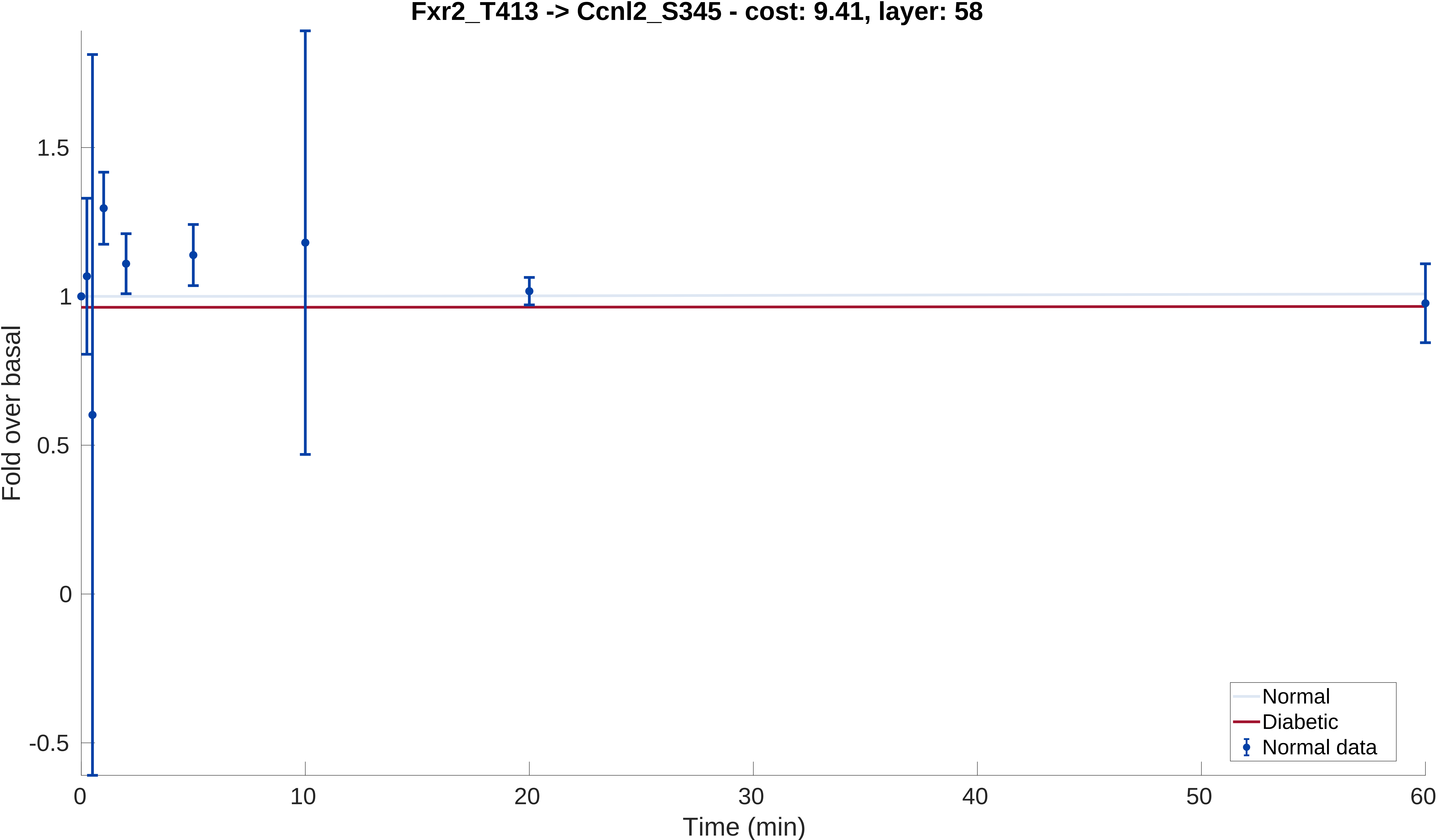

Fxr2_T413 -> Ccnl2_S345 - cost: 9.41, layer: 58
Fold over basal
1.5
1
0.5
0
-0.5
0
10
20
30
40
50
60
Time (min)
Normal
Diabetic
Normal data

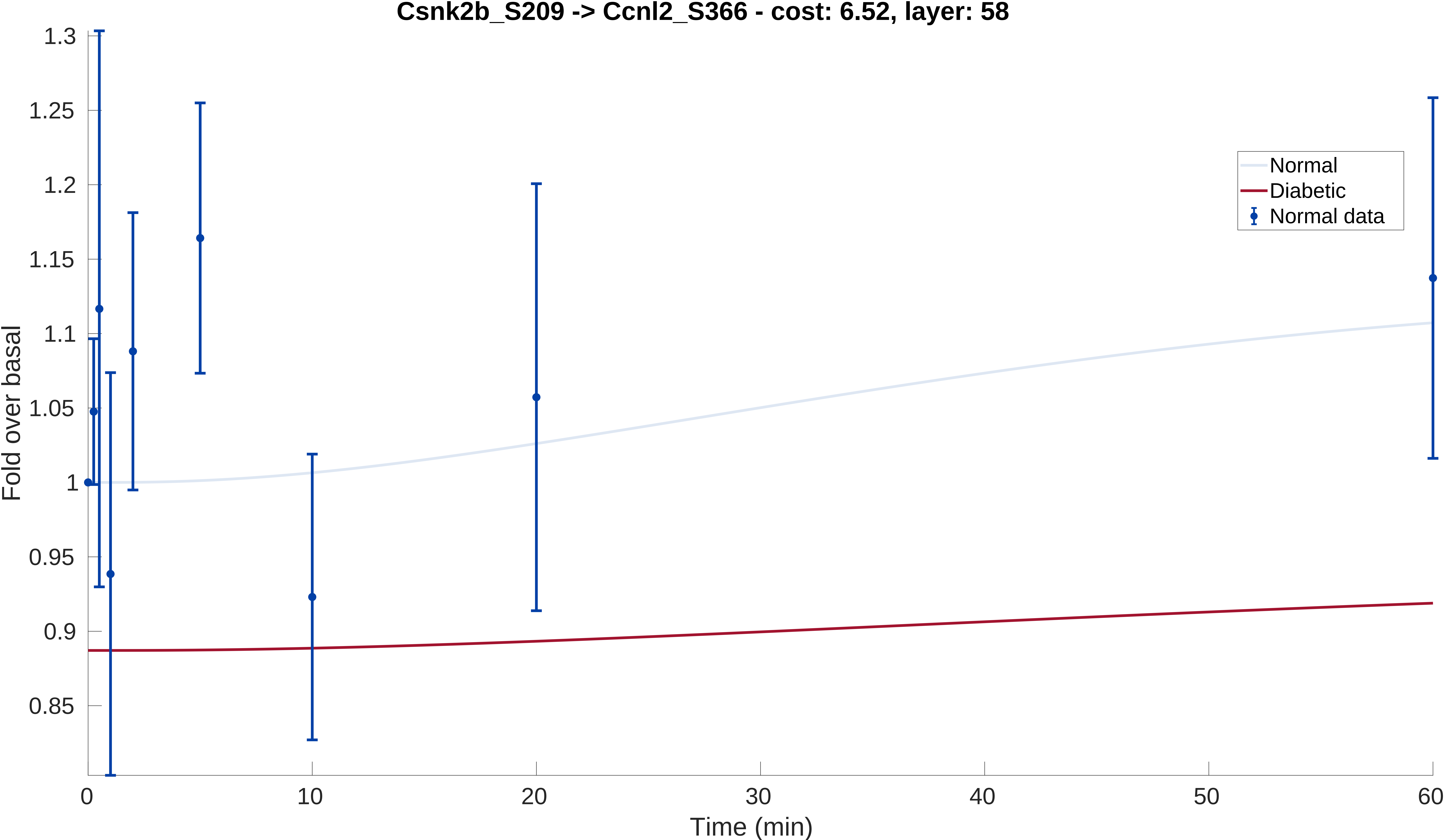

Csnk2b_S209 -> Ccnl2_S366 - cost: 6.52, layer: 58
Normal
Diabetic
Normal data
Fold over basal
1.3
1.25
1.2
1.15
1.1
1.05
1
0.95
0.9
0.85
0
10
20
30
40
50
60
Time (min)

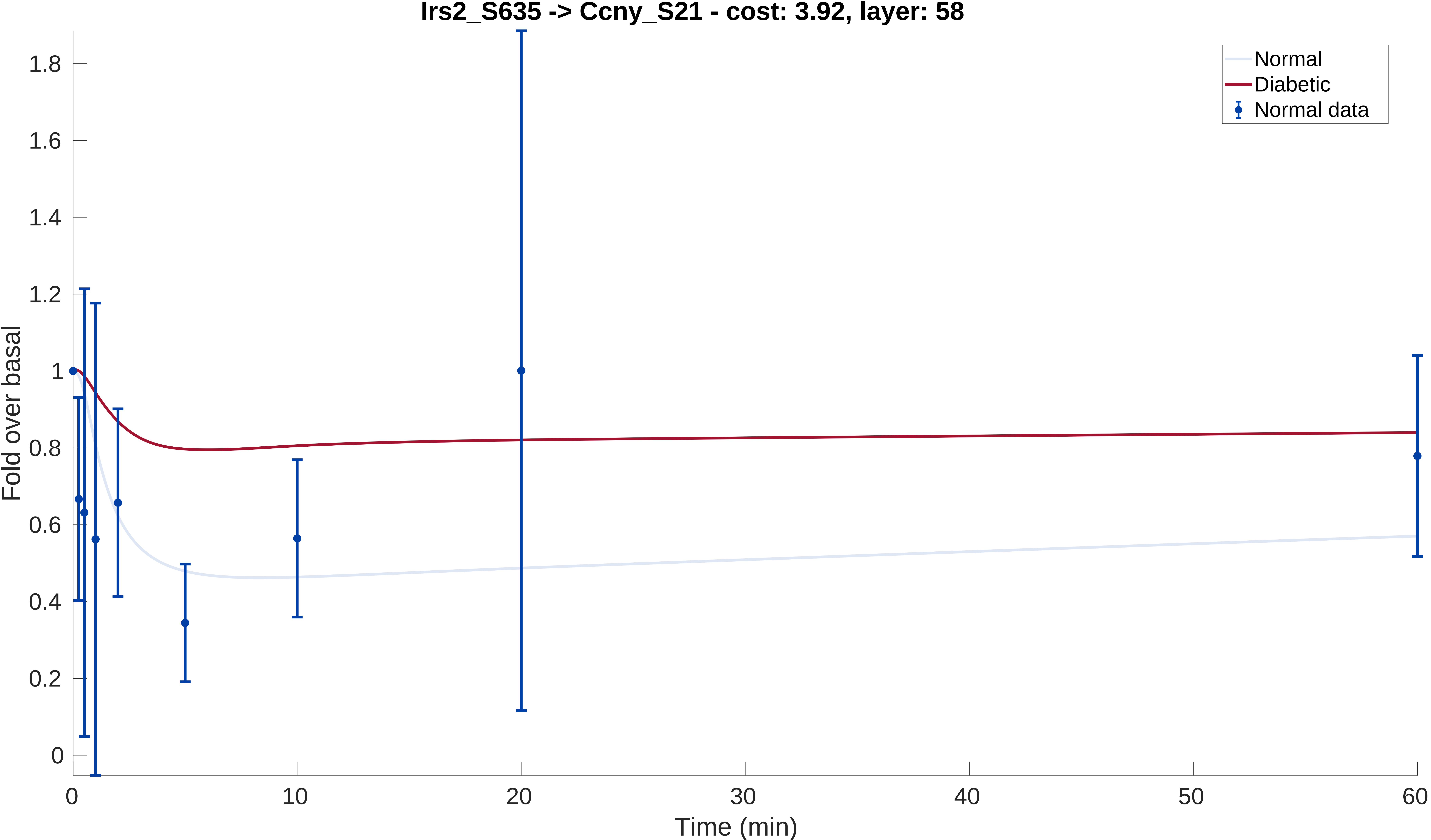

Irs2_S635 -> Ccny_S21 - cost: 3.92, layer: 58
Normal
Diabetic
Normal data
Fold over basal
1.8
1.6
1.4
1.2
1
0.8
0.6
0.4
0.2
0
0
10
20
30
40
50
60
Time (min)

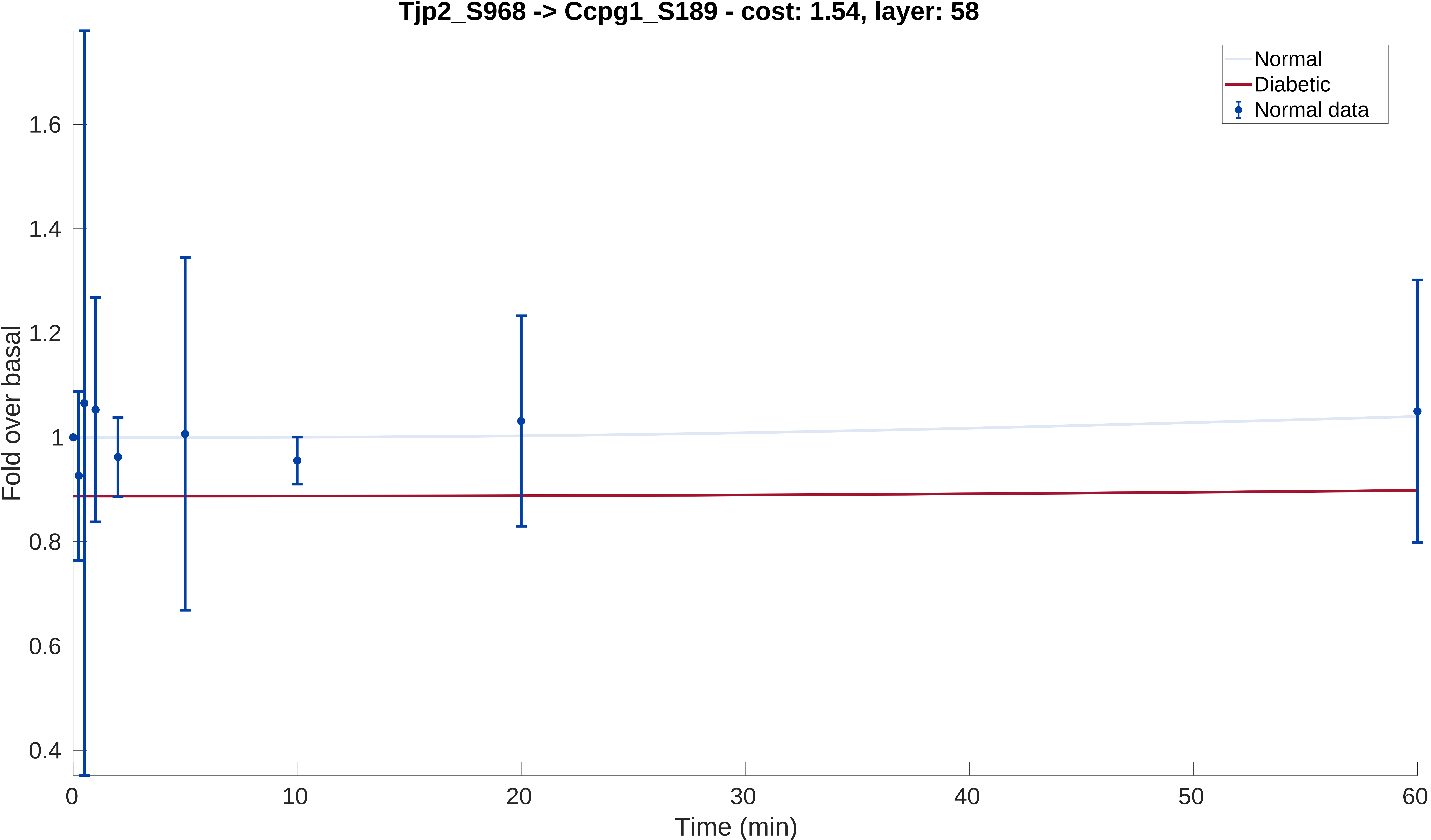

Tjp2_S968 -> Ccpg1_S189 - cost: 1.54, layer: 58
Normal
Diabetic
Normal data
Fold over basal
1.6
1.4
1.2
1
0.8
0.6
0.4
0
10
20
30
40
50
60
Time (min)

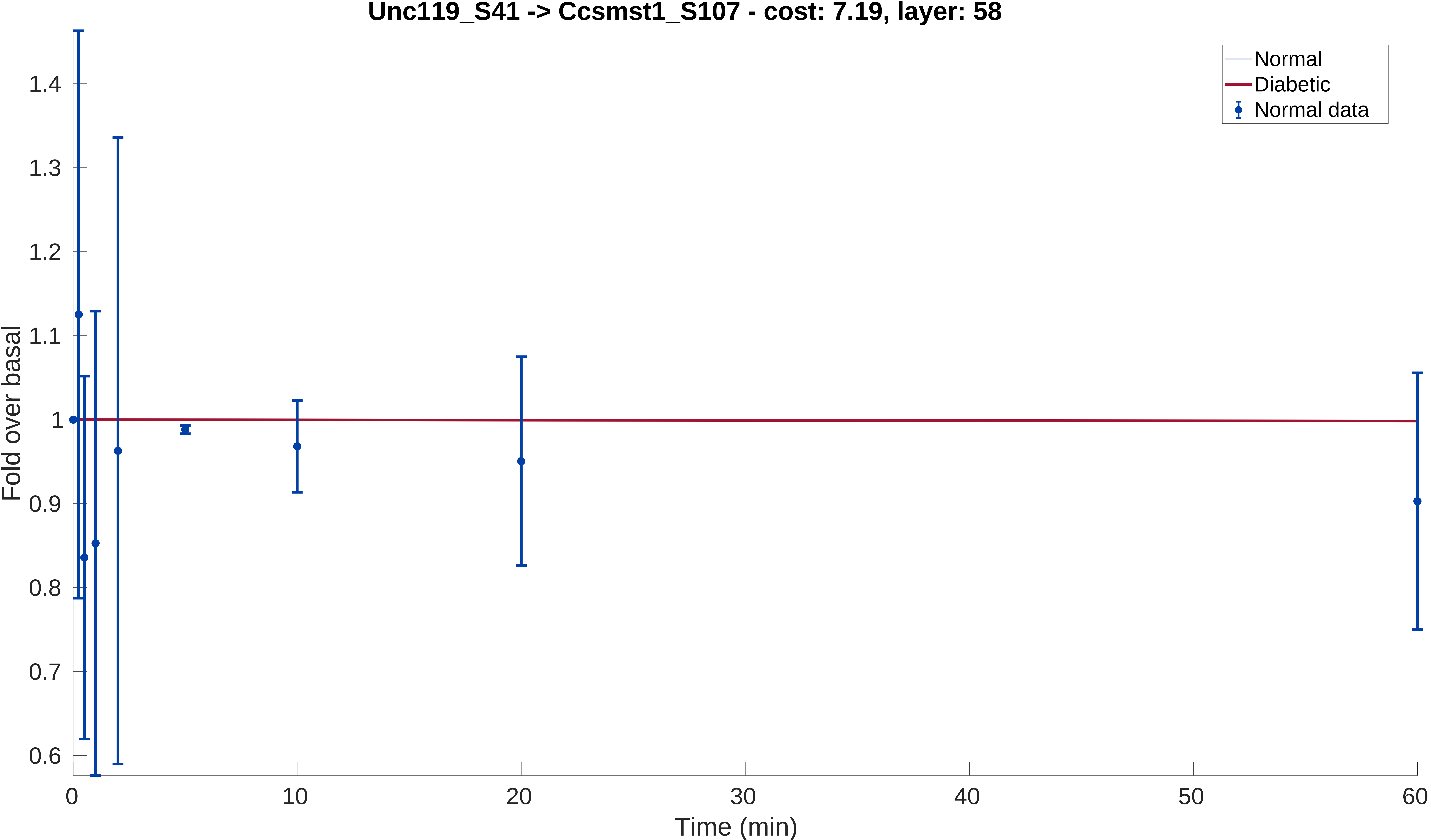

Unc119_S41 -> Ccsmst1_S107 - cost: 7.19, layer: 58
Normal
Diabetic
Normal data
Fold over basal
1.4
1.3
1.2
1.1
1
0.9
0.8
0.7
0.6
0
10
20
30
40
50
60
Time (min)

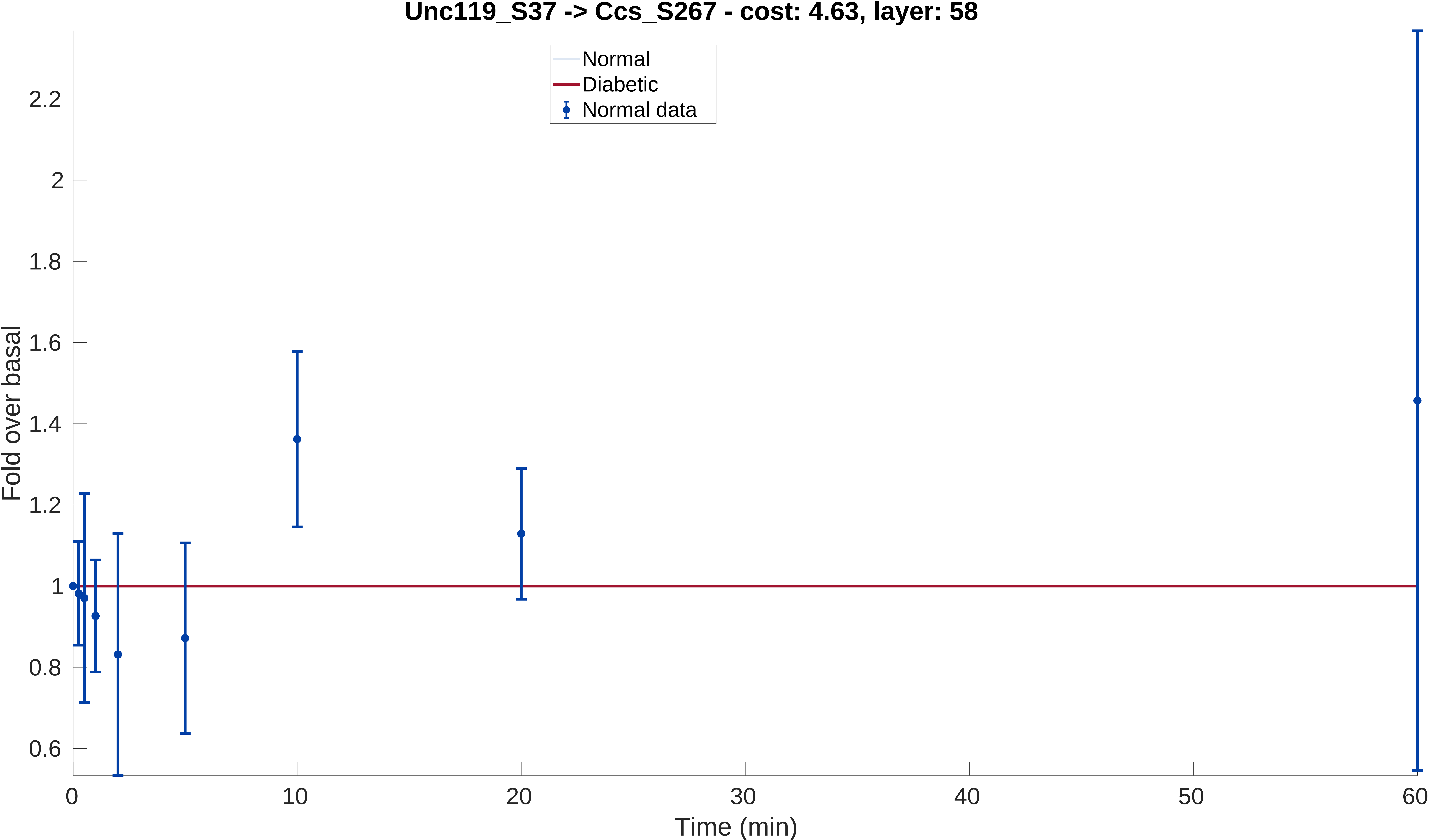

Unc119_S37 -> Ccs_S267 - cost: 4.63, layer: 58
Normal
Diabetic
Normal data
Fold over basal
2.2
2
1.8
1.6
1.4
1.2
1
0.8
0.6
0
10
20
30
40
50
60
Time (min)

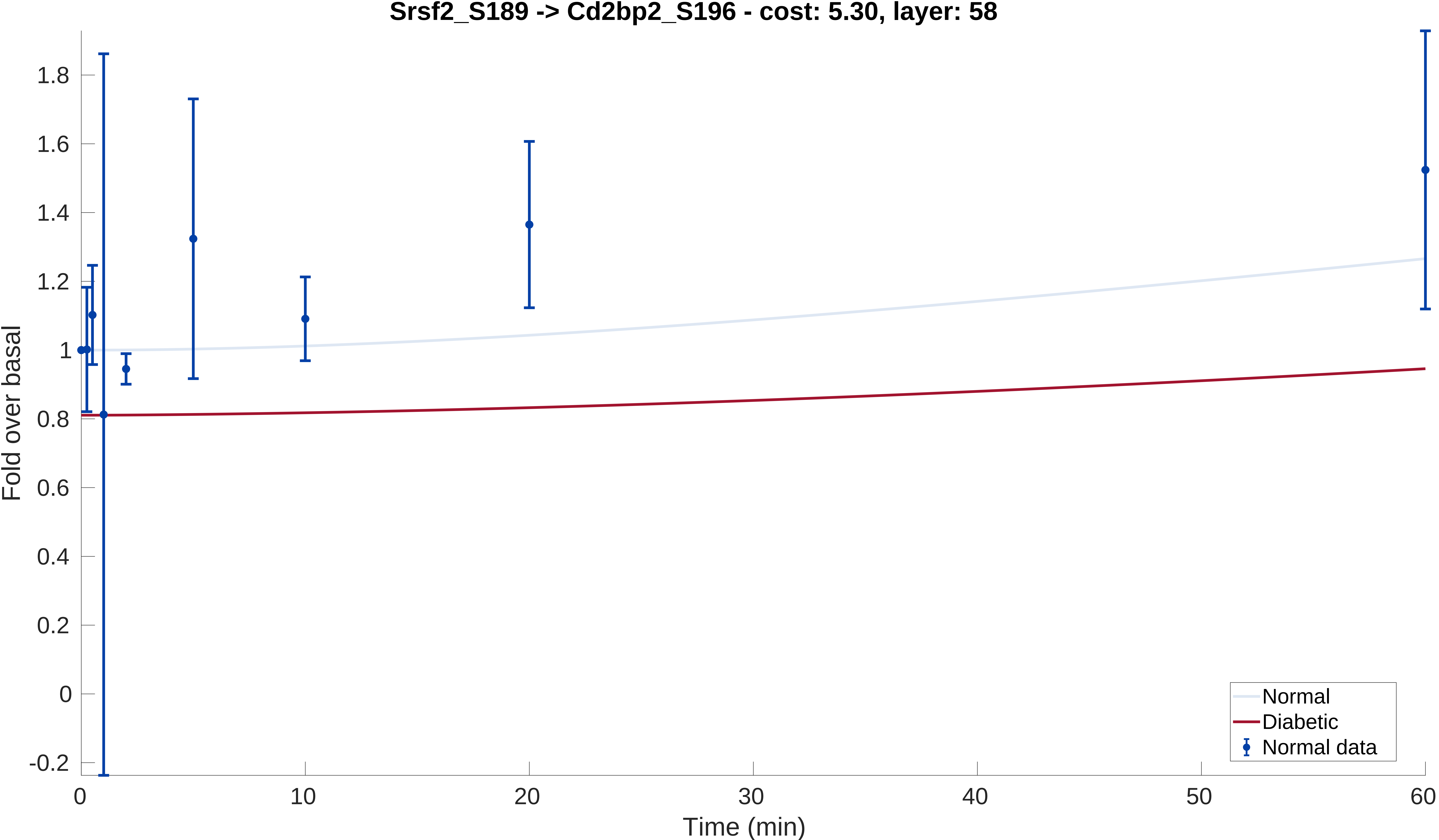

Srsf2_S189 -> Cd2bp2_S196 - cost: 5.30, layer: 58
Fold over basal
1.8
1.6
1.4
1.2
1
0.8
0.6
0.4
0.2
0
-0.2
0
10
20
30
40
50
60
Time (min)
Normal
Diabetic
Normal data

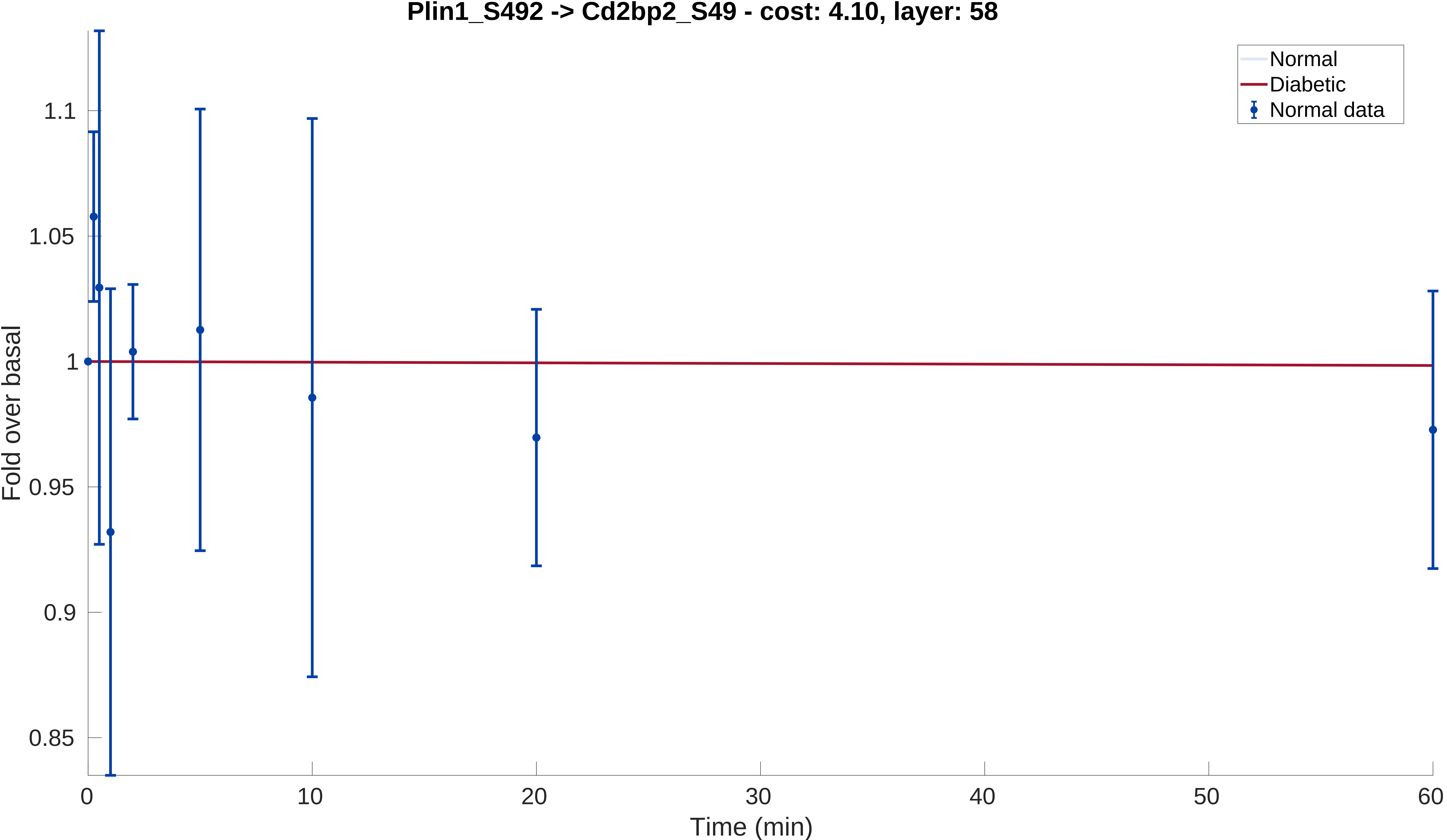
Plin1_S492 -> Cd2bp2_S49 - cost: 4.10, layer: 58
Normal
Diabetic
Normal data
Fold over basal
1.1
1.05
1
0.95
0.9
0.85
0
10
20
30
40
50
60
Time (min)

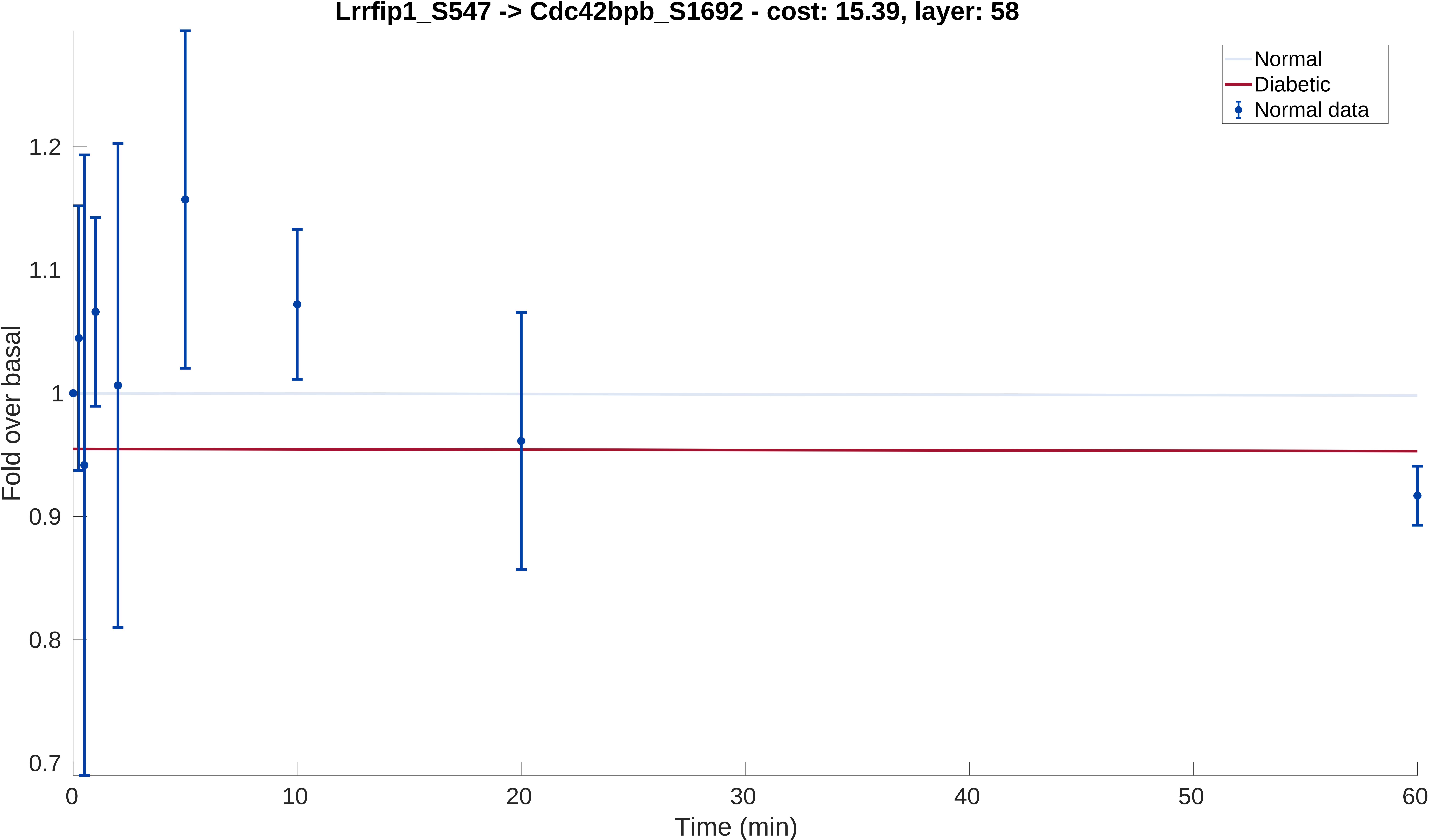

Lrrfip1_S547 -> Cdc42bpb_S1692 - cost: 15.39, layer: 58
Normal
Diabetic
Normal data
1.2
1.1
1
0.9
0.8
0.7
Fold over basal
0
10
20
30
40
50
60
Time (min)

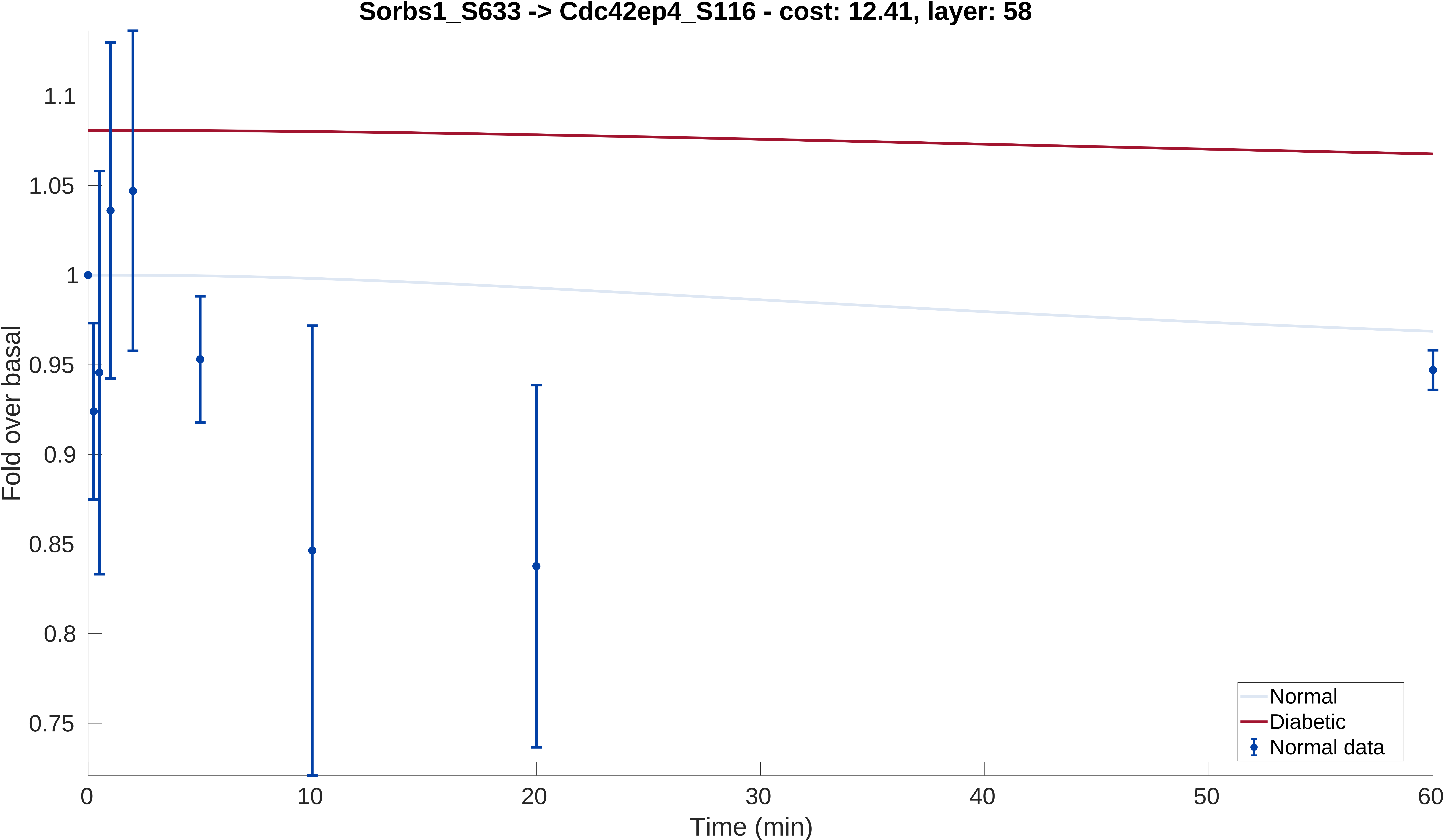

Sorbs1_S633 -> Cdc42ep4_S116 - cost: 12.41, layer: 58
Fold over basal
1.1
1.05
1
0.95
0.9
0.85
0.8
0.75
0
10
20
30
40
50
60
Time (min)
Normal
Diabetic
Normal data

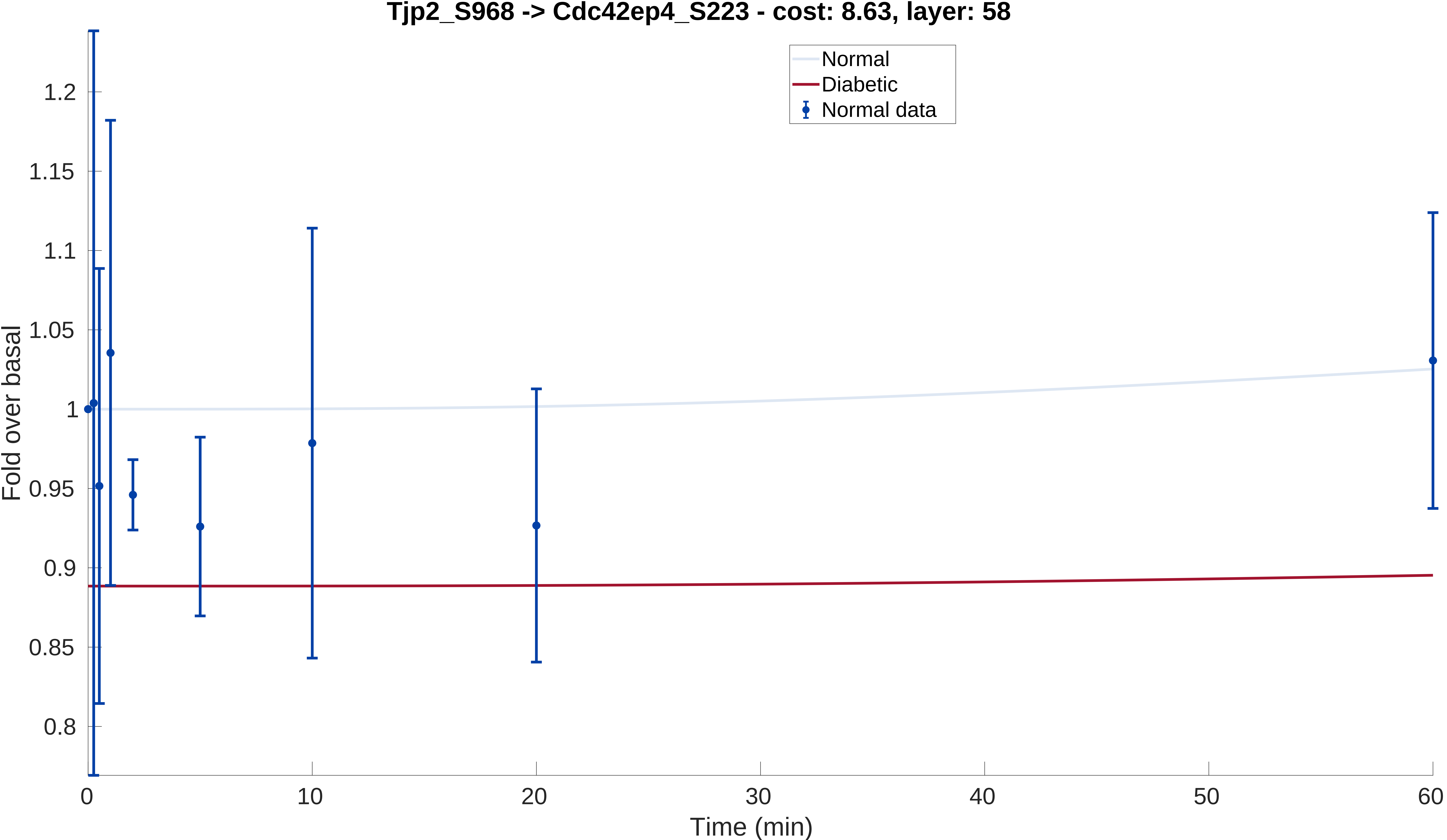

Tjp2_S968 -> Cdc42ep4_S223 - cost: 8.63, layer: 58
Normal
Diabetic
Normal data
Fold over basal
1.2
1.15
1.1
1.05
1
0.95
0.9
0.85
0.8
0
10
20
30
40
50
60
Time (min)

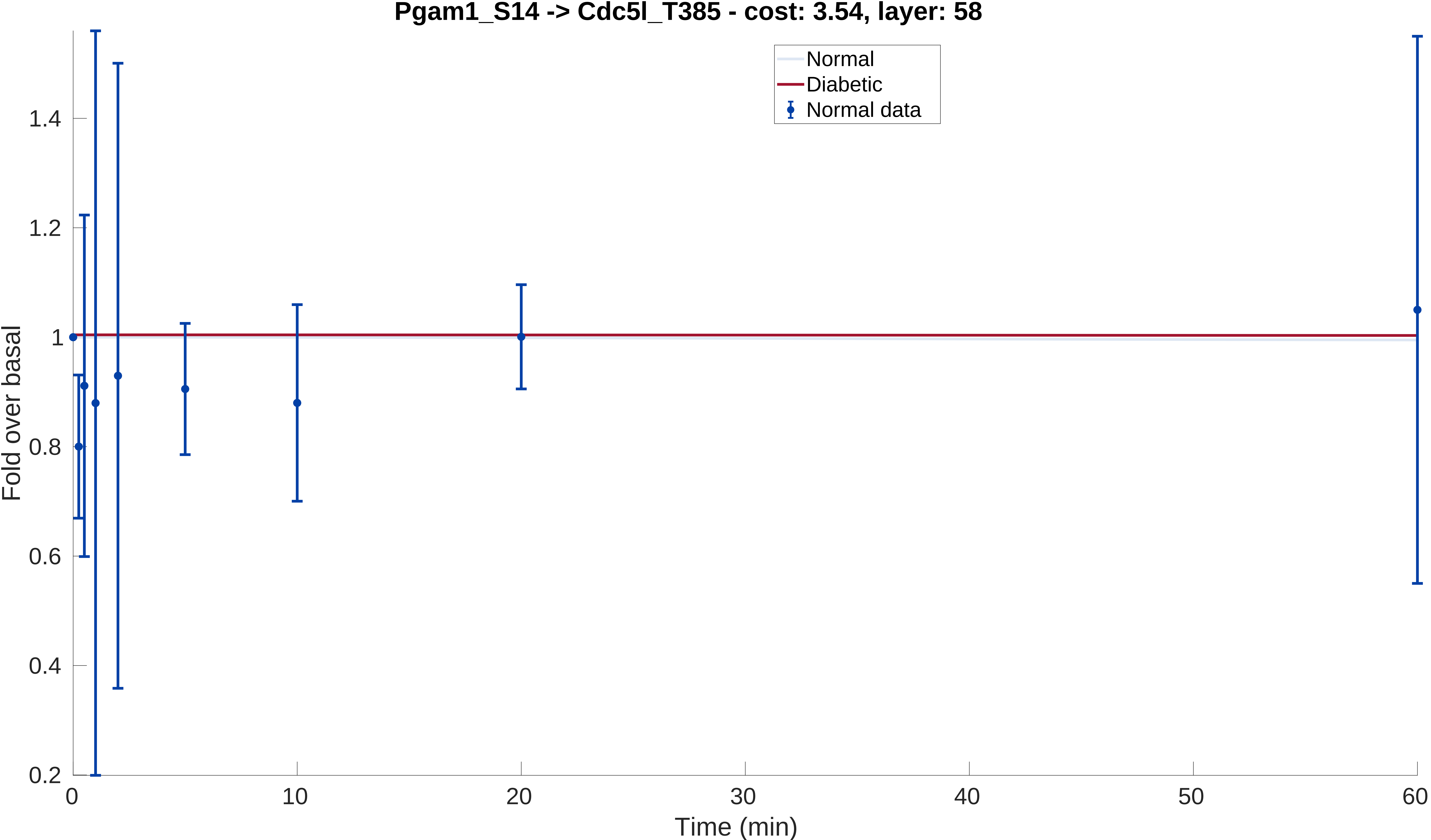

Pgam1_S14 -> Cdc5l_T385 - cost: 3.54, layer: 58
Normal
Diabetic
Normal data
Fold over basal
1.4
1.2
1
0.8
0.6
0.4
0.2
0
10
20
30
40
50
60
Time (min)

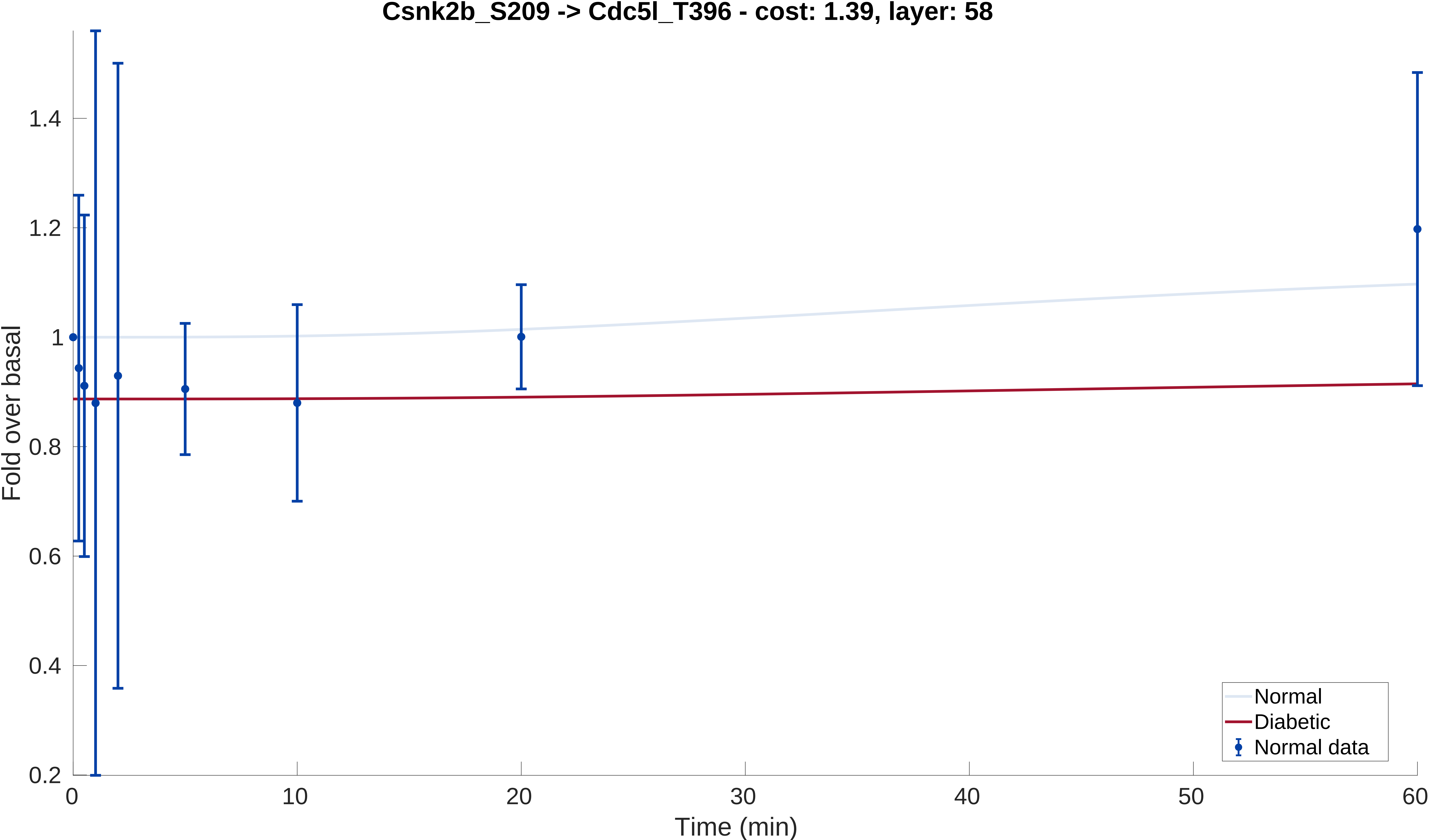
Csnk2b_S209 -> Cdc5l_T396 - cost: 1.39, layer: 58
Fold over basal
Time (min)
Normal
Diabetic
Normal data
0.2
0.4
0.6
0.8
1
1.2
1.4
0
10
20
30
40
50
60

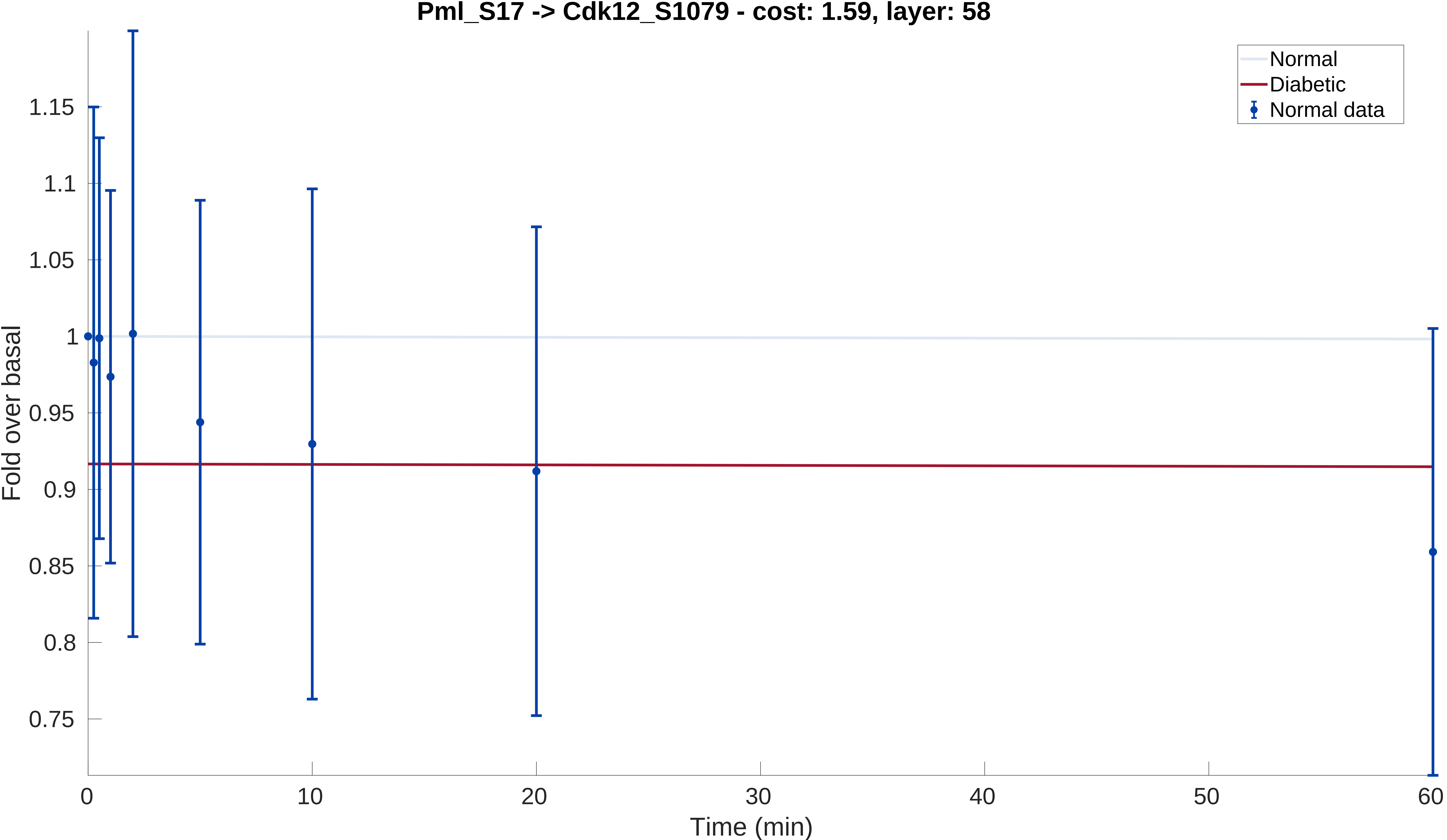

Pml_S17 -> Cdk12_S1079 - cost: 1.59, layer: 58
Normal
Diabetic
Normal data
Fold over basal
1.15
1.1
1.05
1
0.95
0.9
0.85
0.8
0.75
0
10
20
30
40
50
60
Time (min)

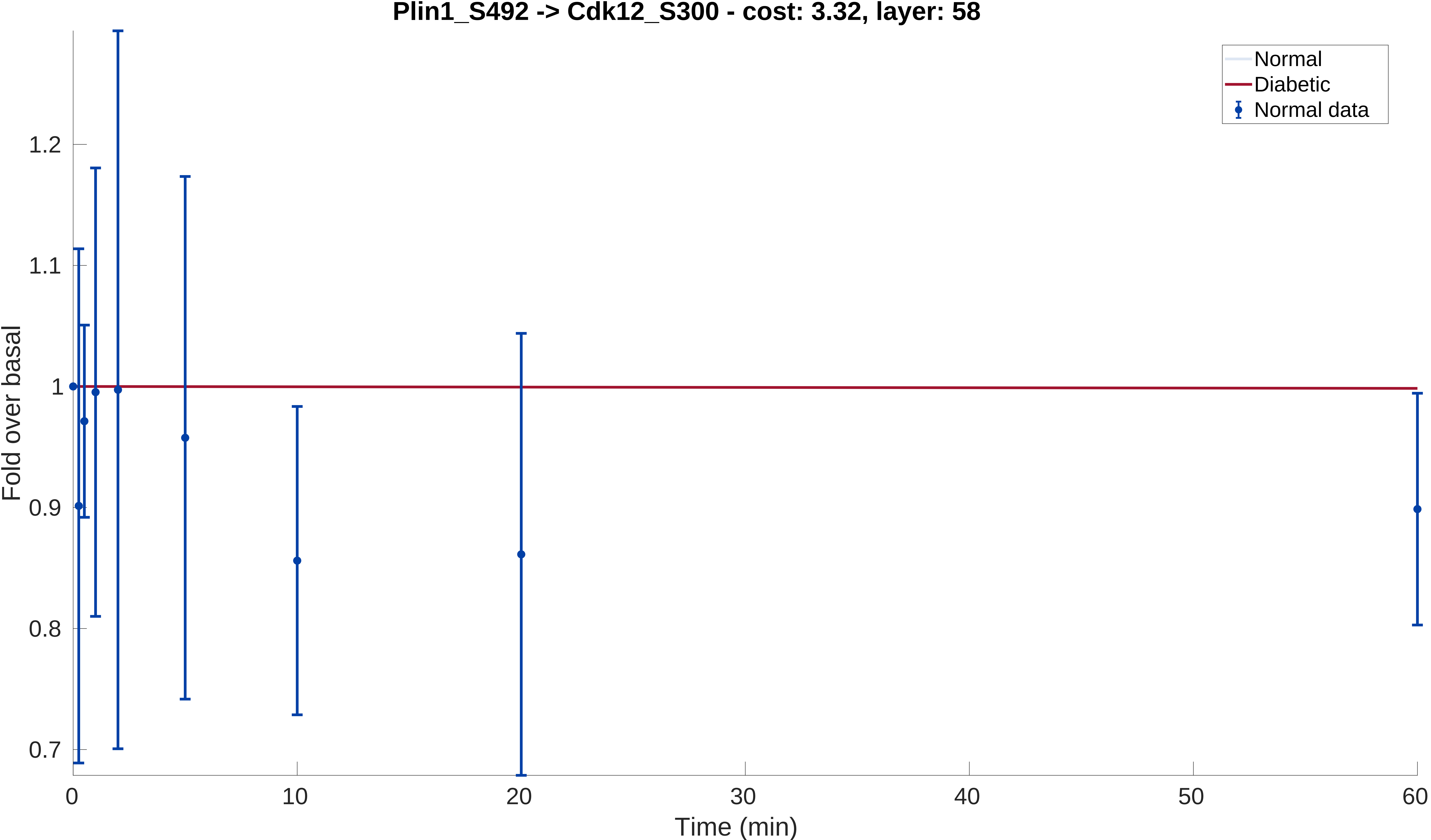

Plin1_S492 -> Cdk12_S300 - cost: 3.32, layer: 58
Normal
Diabetic
Normal data
Fold over basal
1.2
1.1
1
0.9
0.8
0.7
0
10
20
30
40
50
60
Time (min)

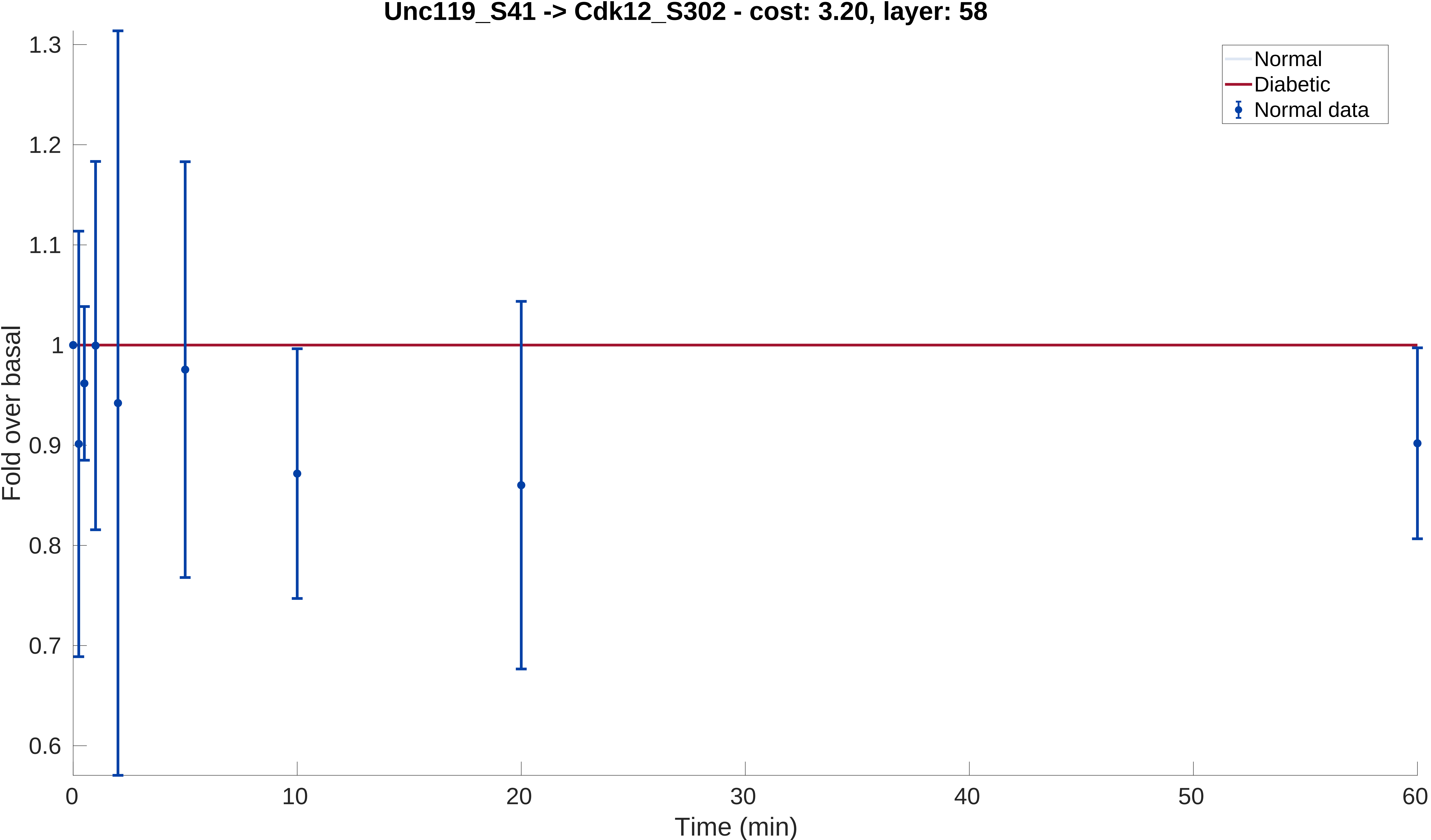

Unc119_S41 -> Cdk12_S302 - cost: 3.20, layer: 58
Normal
Diabetic
Normal data
Fold over basal
1.3
1.2
1.1
1
0.9
0.8
0.7
0.6
0
10
20
30
40
50
60
Time (min)

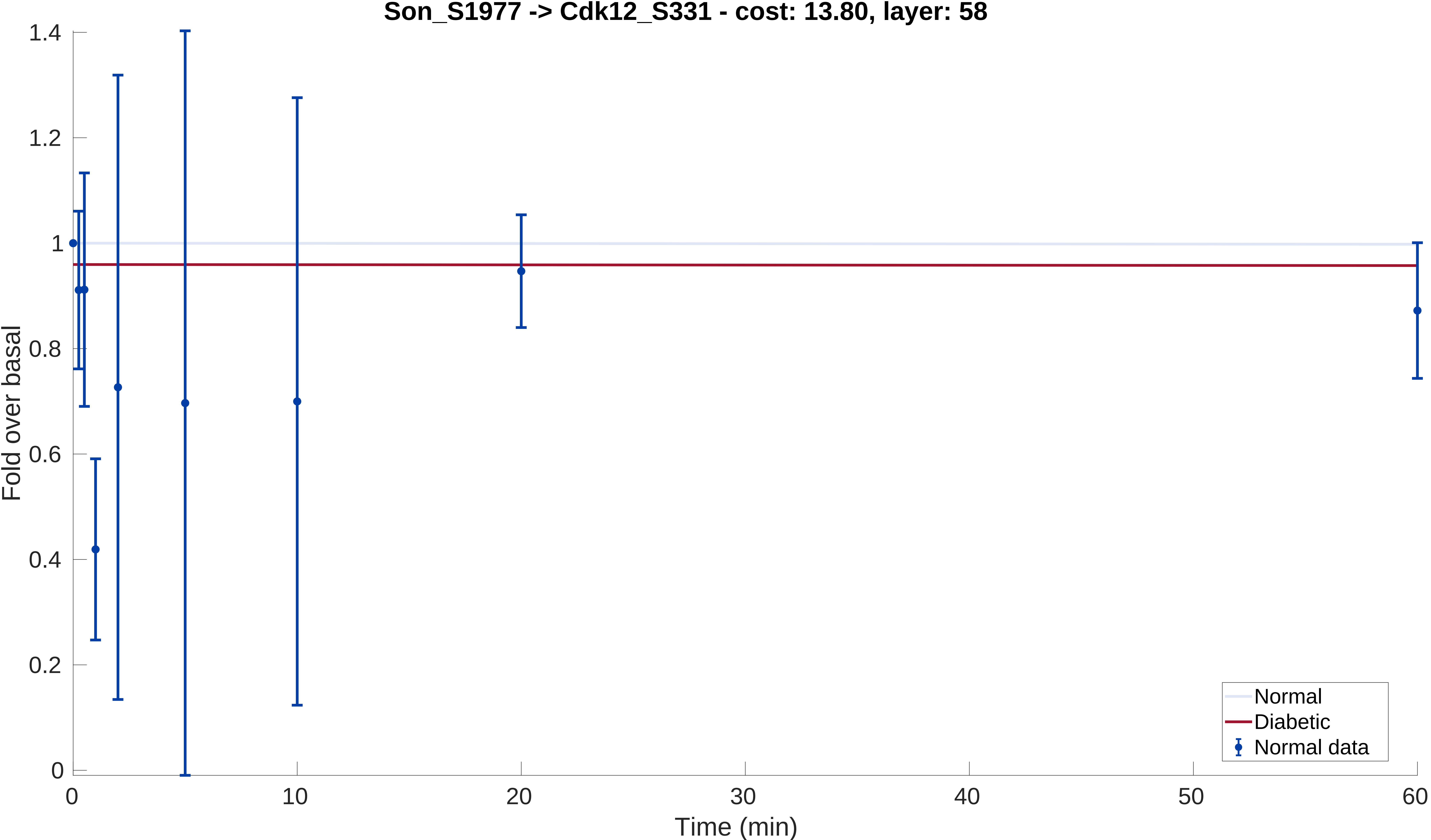
Son_S1977 -> Cdk12_S331 - cost: 13.80, layer: 58
Fold over basal
Time (min)
0
0.2
0.4
0.6
0.8
1
1.2
1.4
0
10
20
30
40
50
60
Normal
Diabetic
Normal data

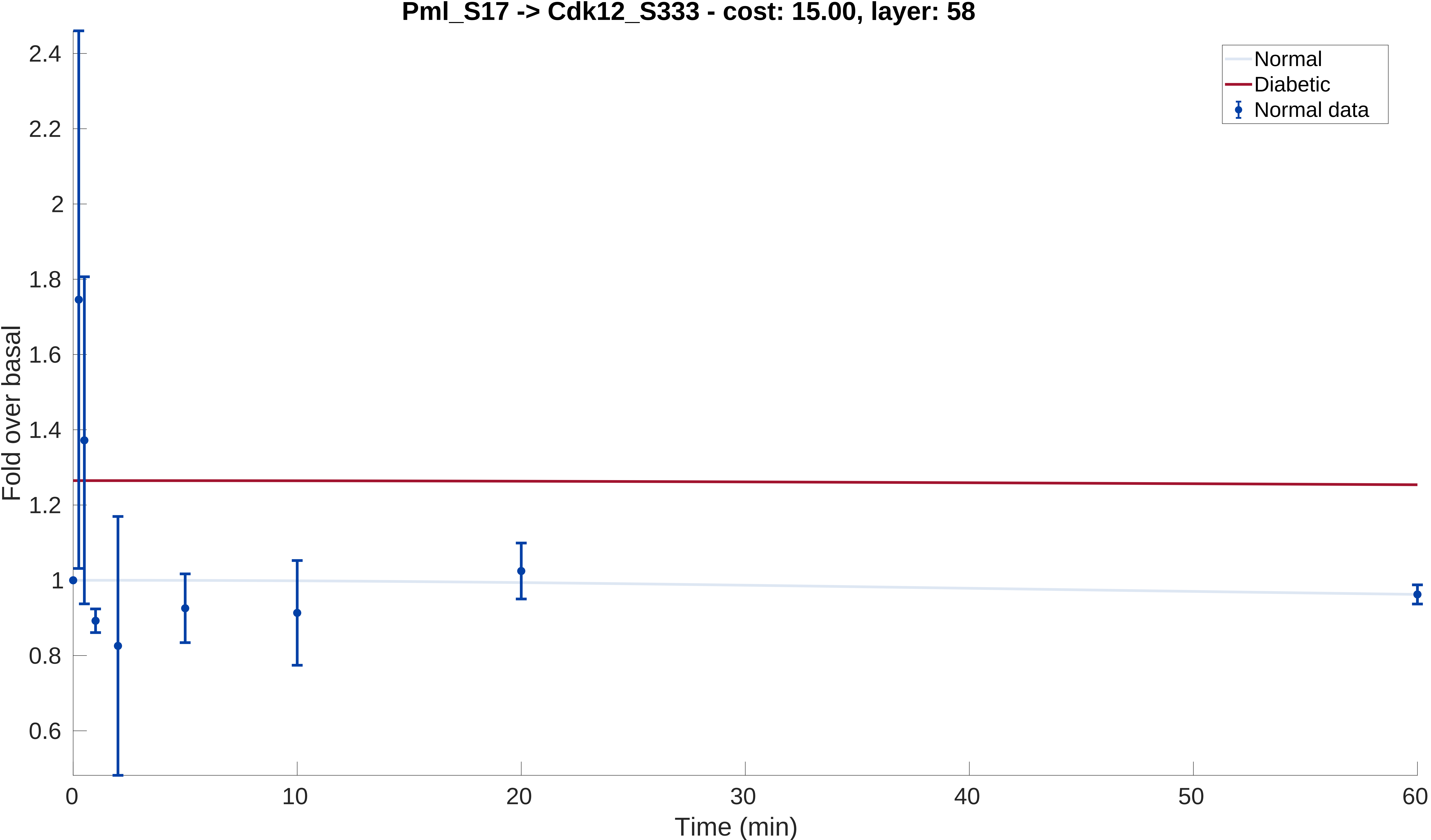

Pml_S17 -> Cdk12_S333 - cost: 15.00, layer: 58
Normal
Diabetic
Normal data
Fold over basal
Time (min)
2.4
2.2
2
1.8
1.6
1.4
1.2
1
0.8
0.6
0
10
20
30
40
50
60

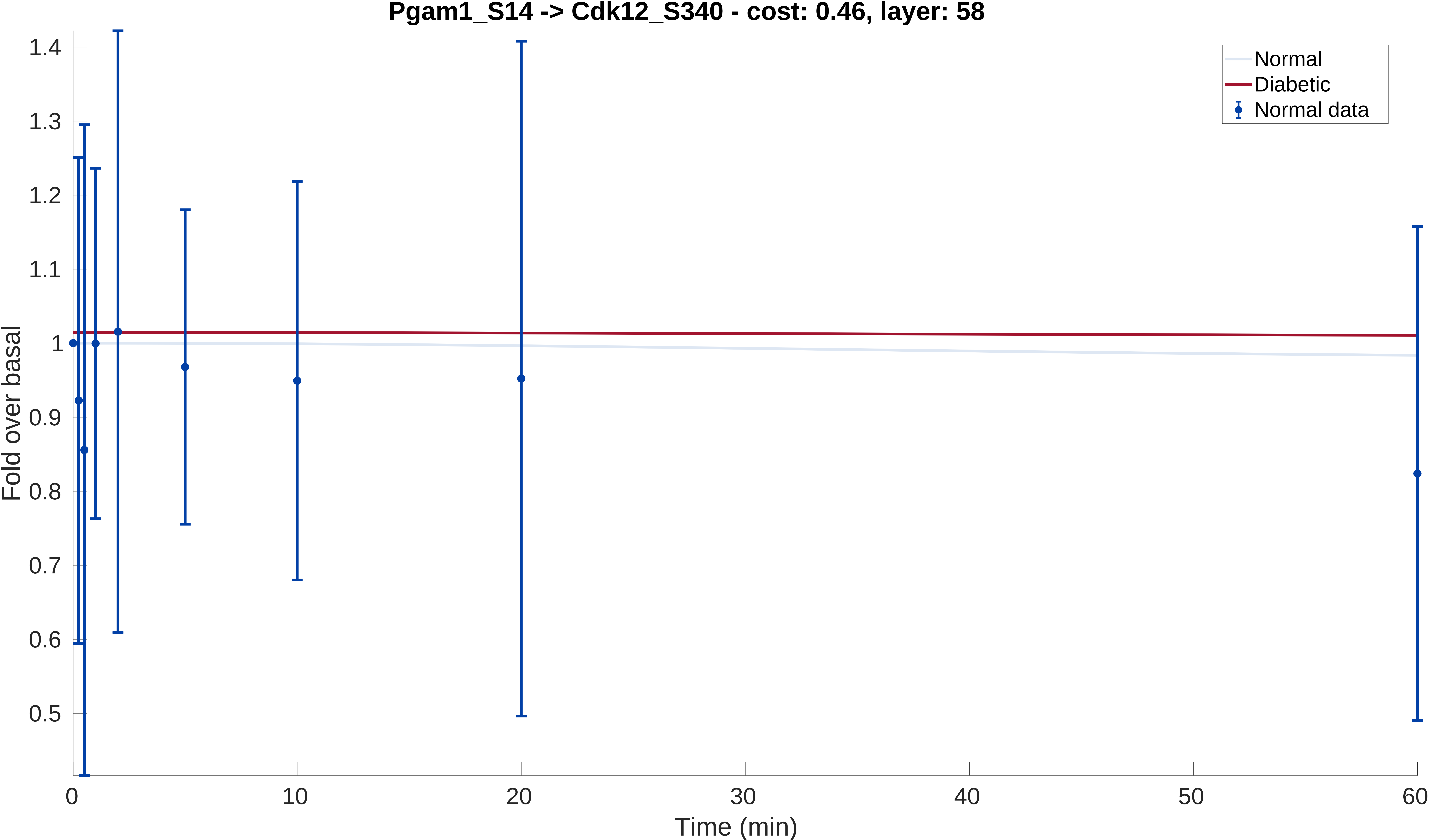

Pgam1_S14 -> Cdk12_S340 - cost: 0.46, layer: 58
Normal
Diabetic
Normal data
Fold over basal
Time (min)
1.4
1.3
1.2
1.1
1
0.9
0.8
0.7
0.6
0.5
0
10
20
30
40
50
60

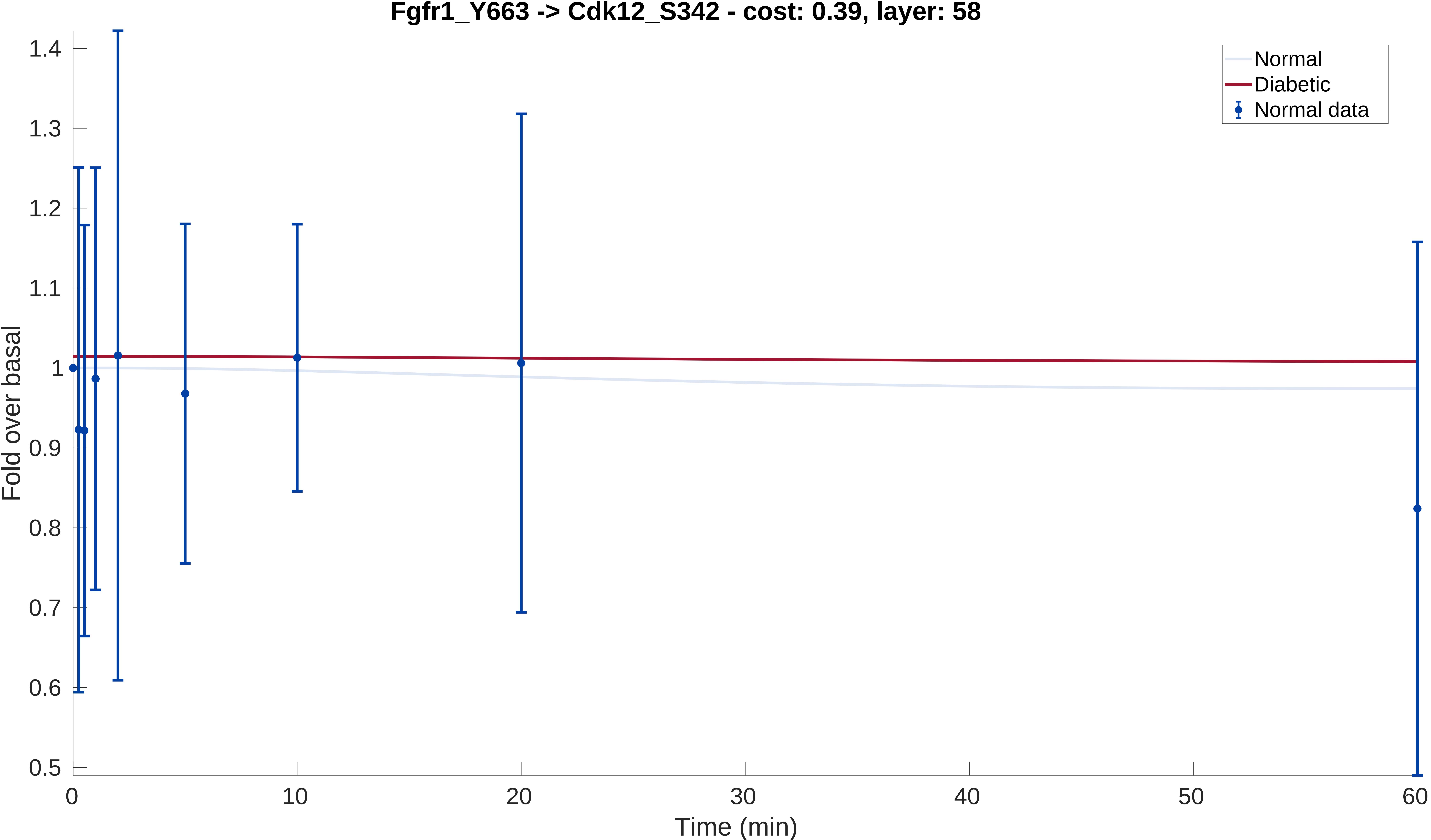

Fgfr1_Y663 -> Cdk12_S342 - cost: 0.39, layer: 58
Normal
Diabetic
Normal data
Fold over basal
0.5
0.6
0.7
0.8
0.9
1
1.1
1.2
1.3
1.4
0
10
20
30
40
50
60
Time (min)

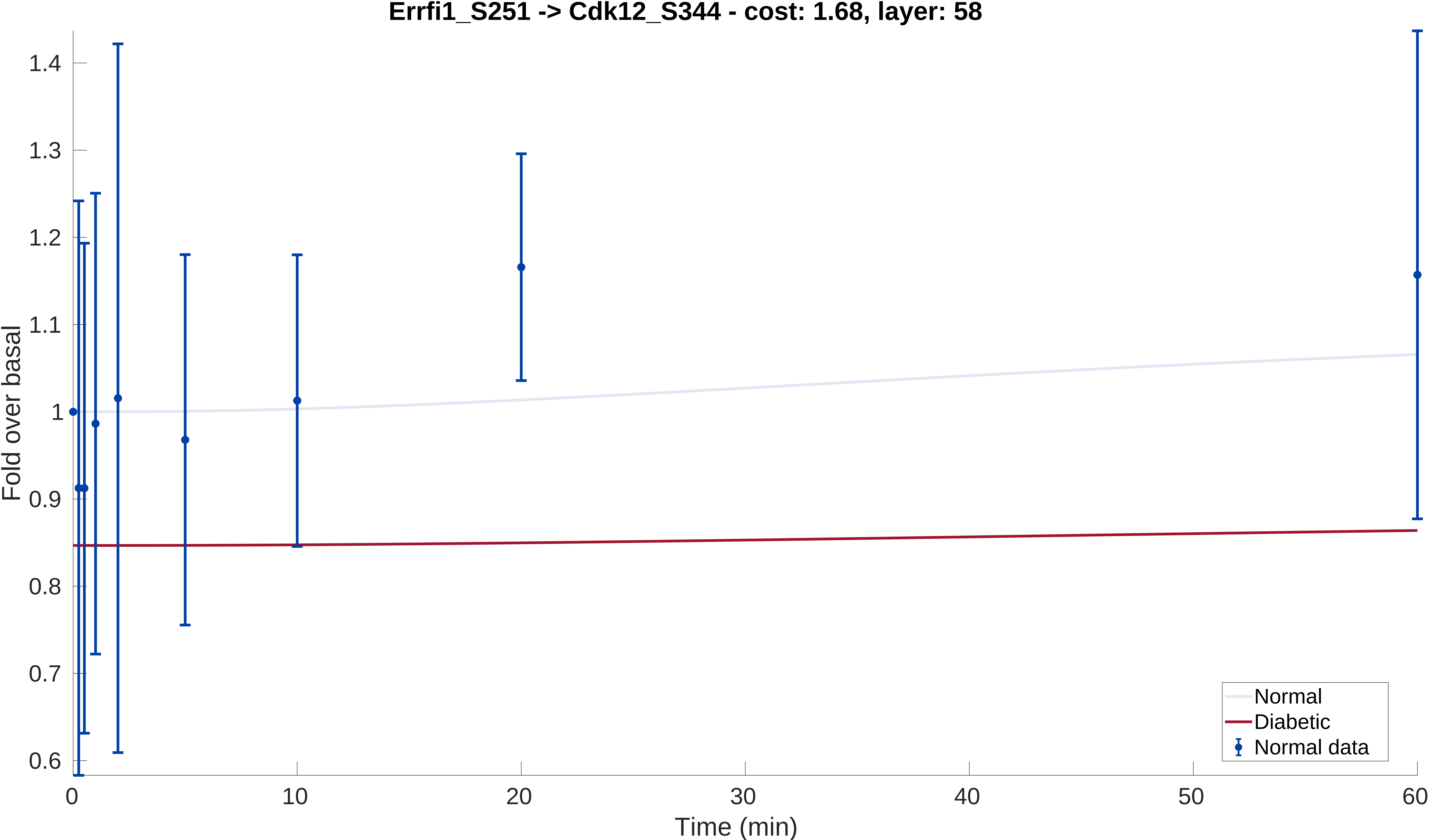

Errfi1_S251 -> Cdk12_S344 - cost: 1.68, layer: 58
Fold over basal
1.4
1.3
1.2
1.1
1
0.9
0.8
0.7
0.6
0
10
20
30
40
50
60
Time (min)
Normal
Diabetic
Normal data

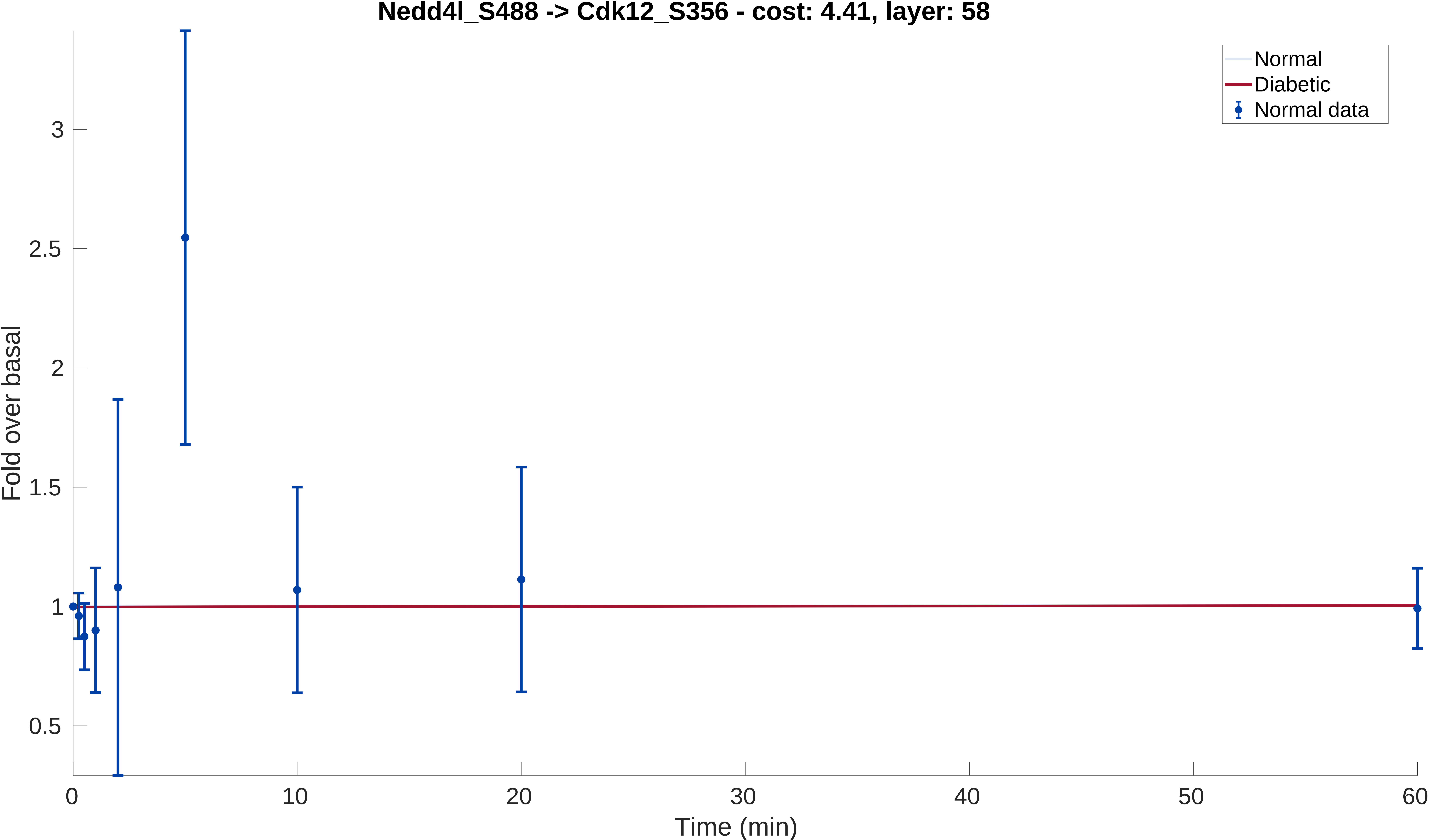

Nedd4l_S488 -> Cdk12_S356 - cost: 4.41, layer: 58
Normal
Diabetic
Normal data
Fold over basal
3
2.5
2
1.5
1
0.5
0
10
20
30
40
50
60
Time (min)

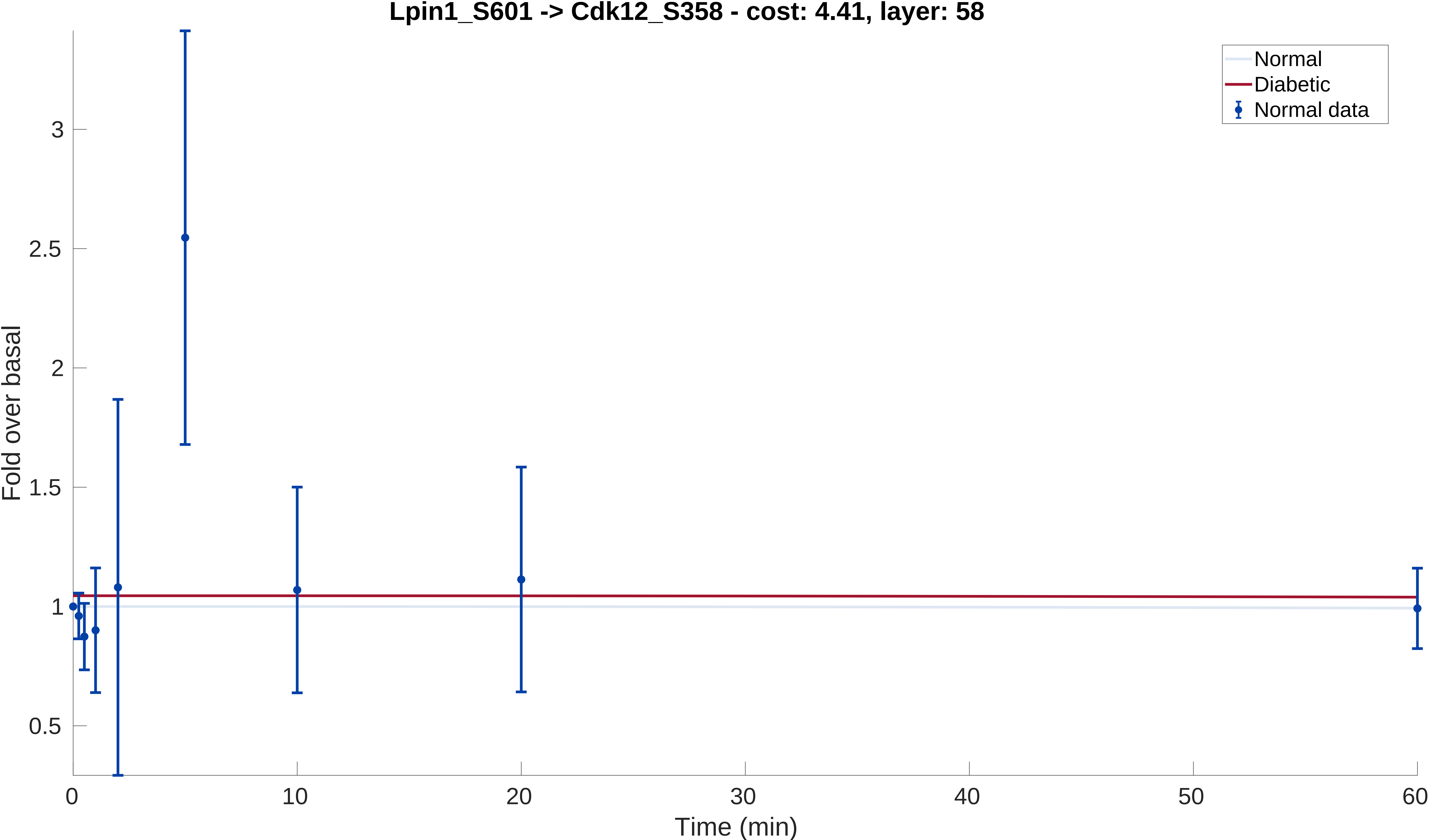

Lpin1_S601 -> Cdk12_S358 - cost: 4.41, layer: 58
Normal
Diabetic
Normal data
Fold over basal
3
2.5
2
1.5
1
0.5
0
10
20
30
40
50
60
Time (min)

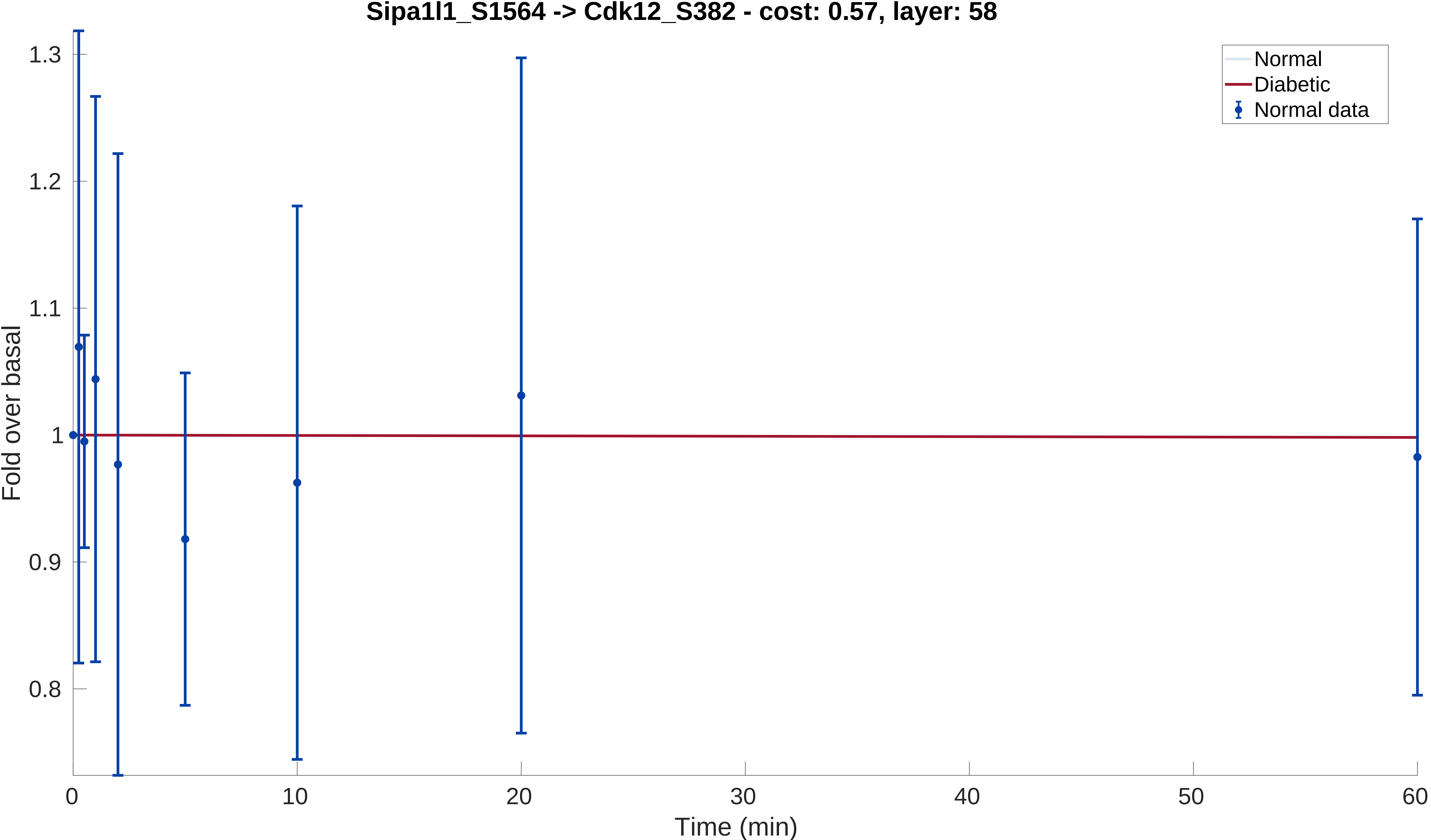

Sipa1l1_S1564 -> Cdk12_S382 - cost: 0.57, layer: 58
Fold over basal
1.3
1.2
1.1
1
0.9
0.8
0
10
20
30
40
50
60
Time (min)
Normal
Diabetic
Normal data

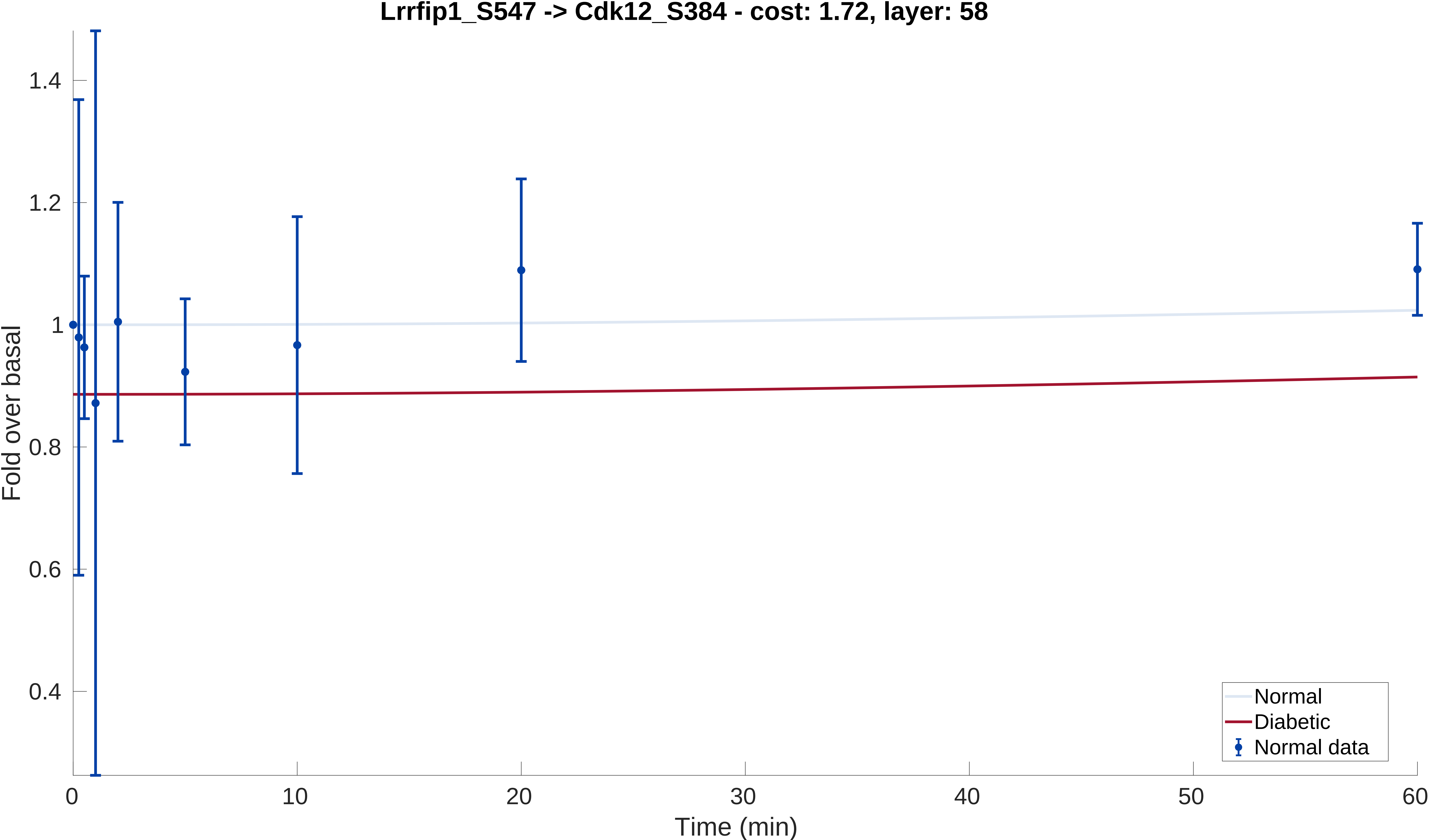

Lrrfip1_S547 -> Cdk12_S384 - cost: 1.72, layer: 58
1.4
1.2
1
0.8
0.6
0.4
Fold over basal
0
10
20
30
40
50
60
Time (min)
Normal
Diabetic
Normal data

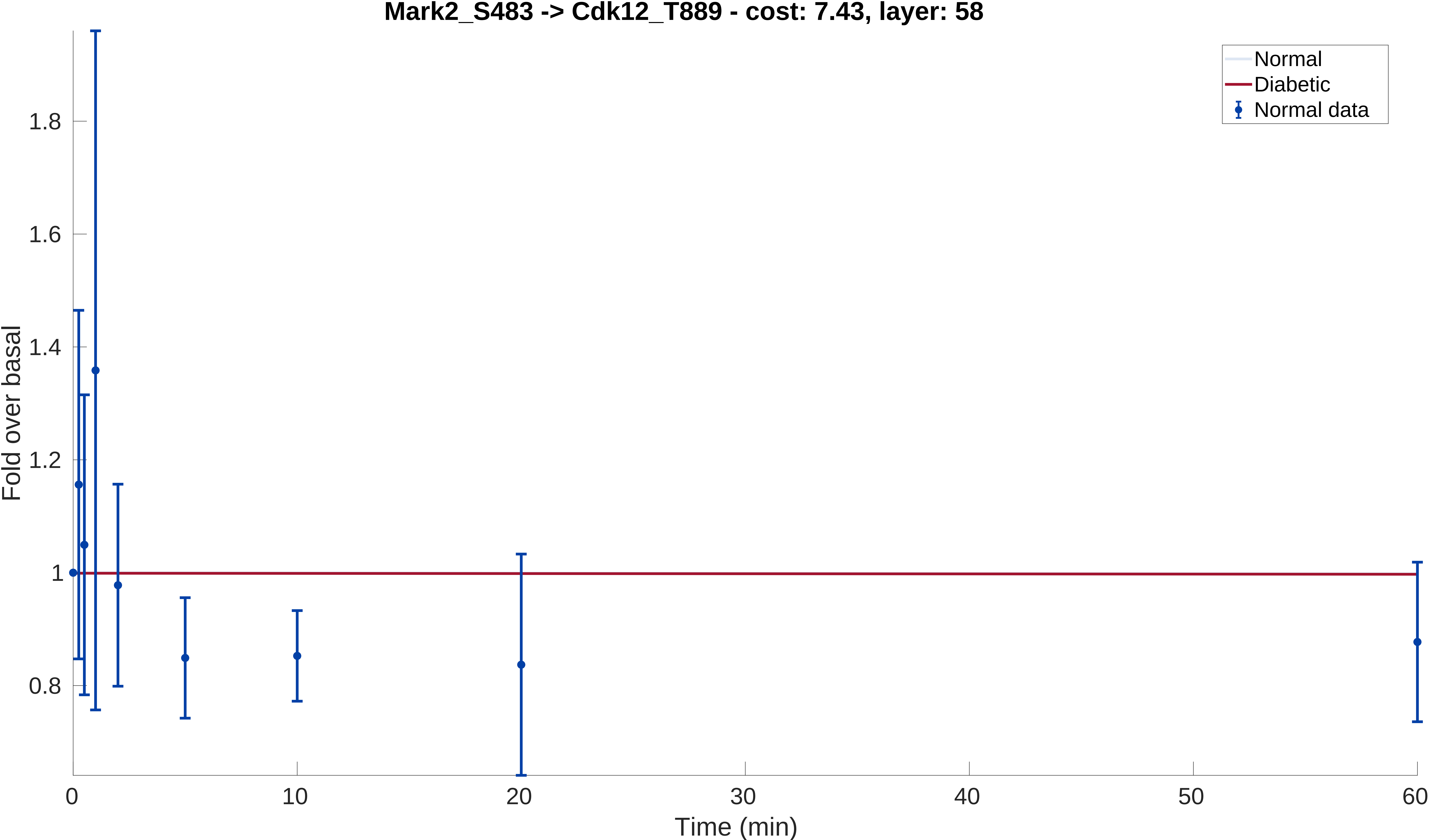

Mark2_S483 -> Cdk12_T889 - cost: 7.43, layer: 58
Normal
Diabetic
Normal data
Fold over basal
1.8
1.6
1.4
1.2
1
0.8
0
10
20
30
40
50
60
Time (min)

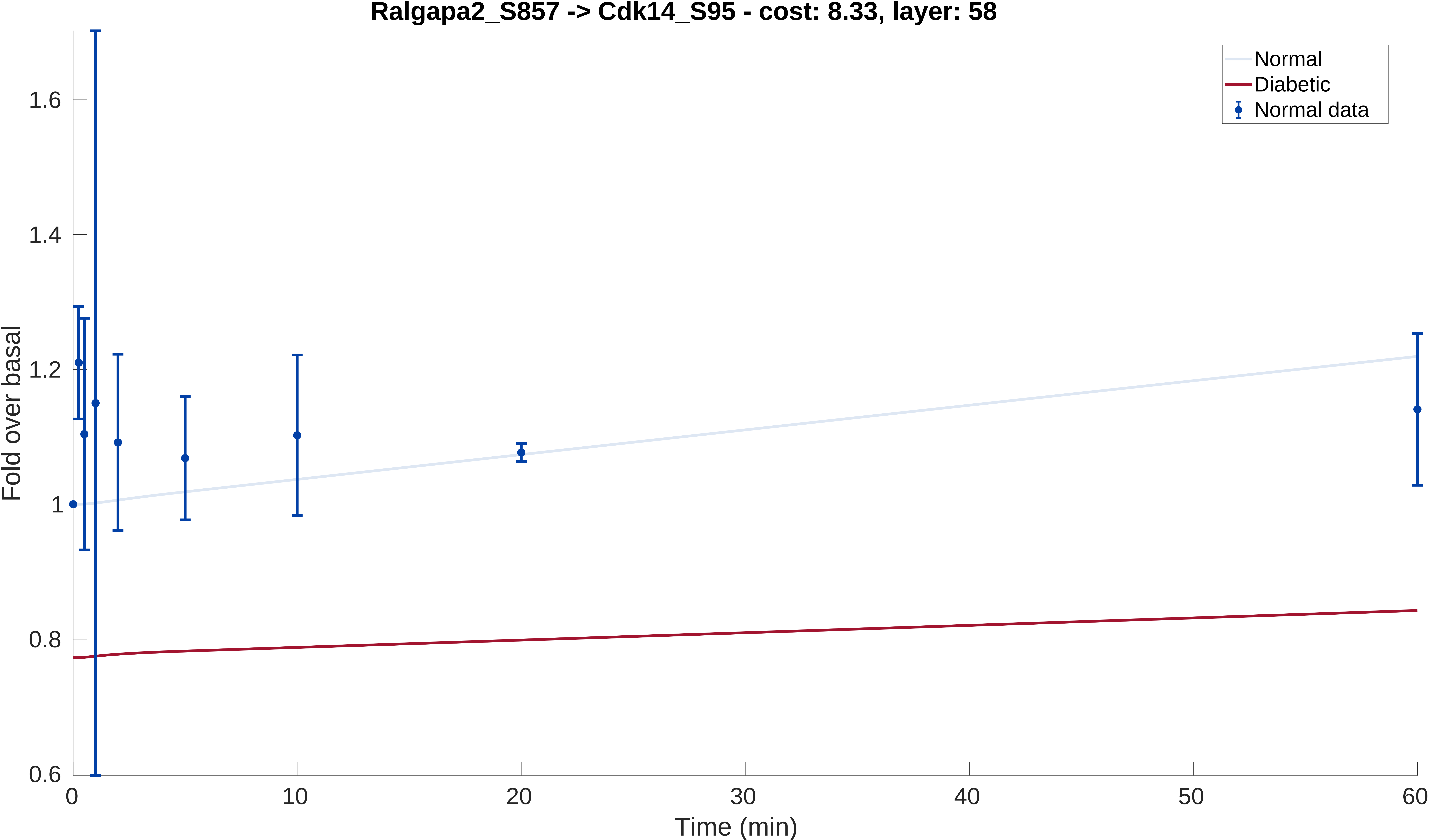

Ralgapa2_S857 -> Cdk14_S95 - cost: 8.33, layer: 58
Normal
Diabetic
Normal data
Fold over basal
1.6
1.4
1.2
1
0.8
0.6
0
10
20
30
40
50
60
Time (min)

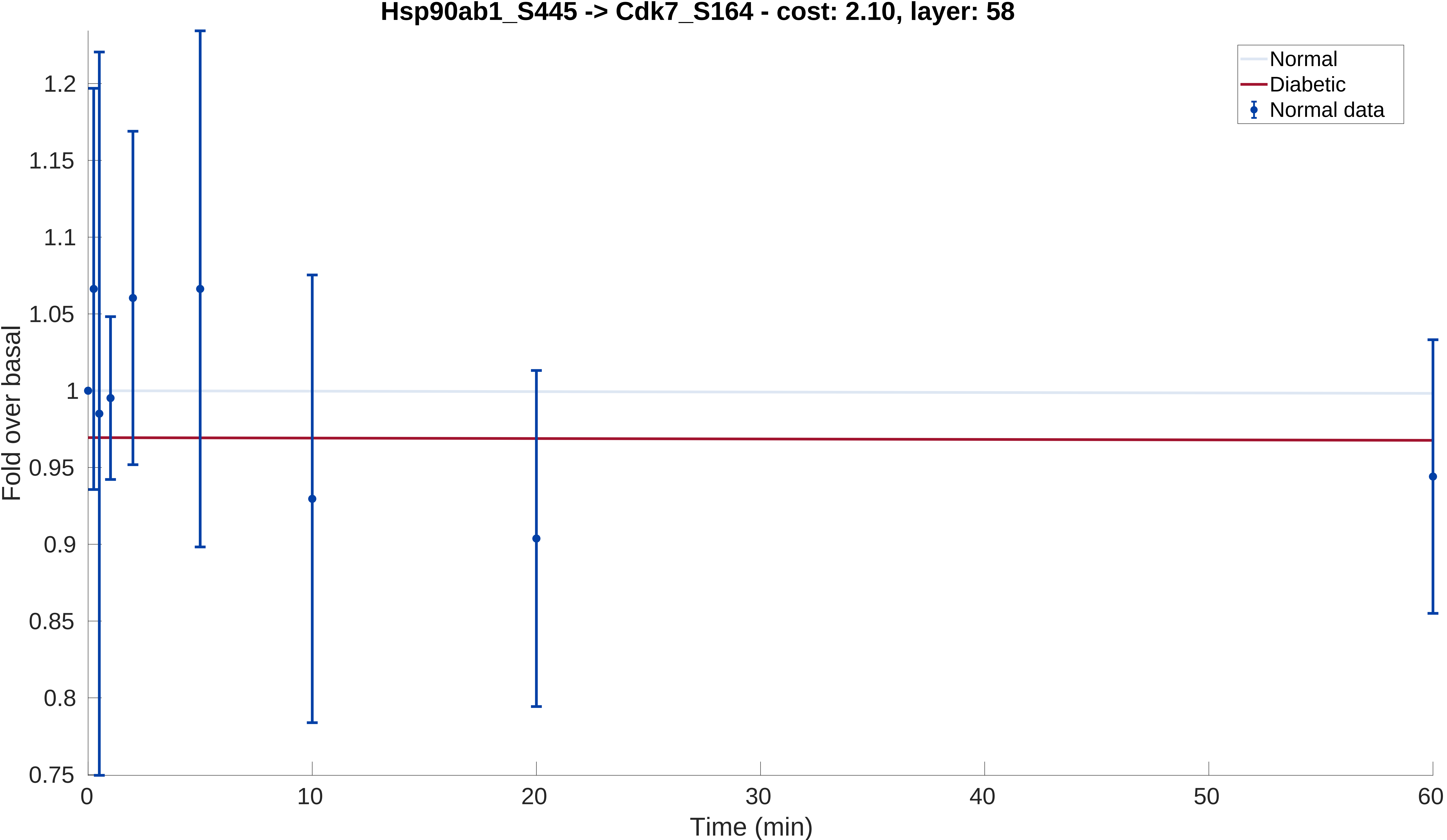
Hsp90ab1_S445 -> Cdk7_S164 - cost: 2.10, layer: 58
Normal
Diabetic
Normal data
Fold over basal
1.2
1.15
1.1
1.05
1
0.95
0.9
0.85
0.8
0.75
0
10
20
30
40
50
60
Time (min)

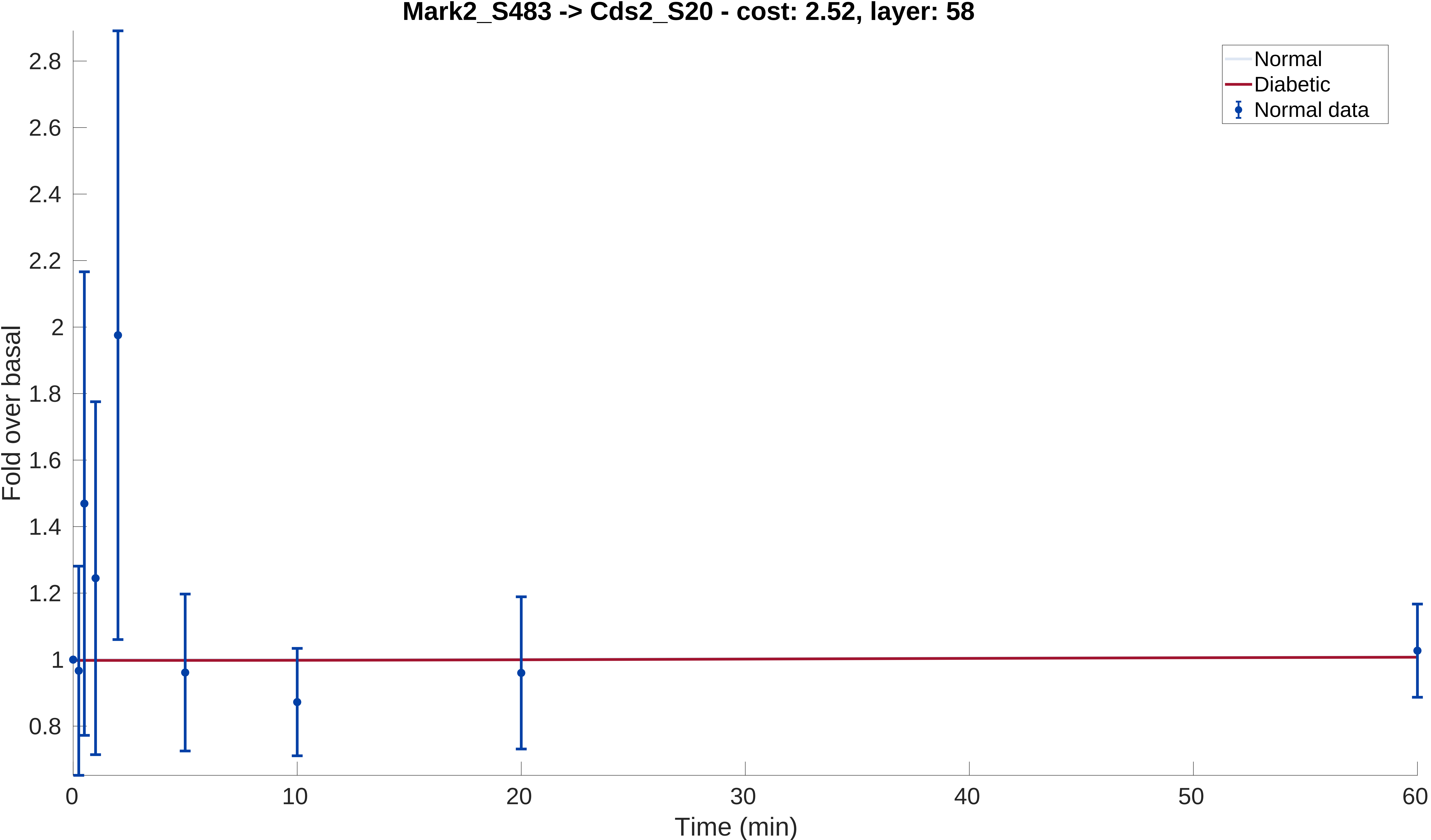

Mark2_S483 -> Cds2_S20 - cost: 2.52, layer: 58
Normal
Diabetic
Normal data
Fold over basal
Time (min)
0.8
1
1.2
1.4
1.6
1.8
2
2.2
2.4
2.6
2.8
0
10
20
30
40
50
60

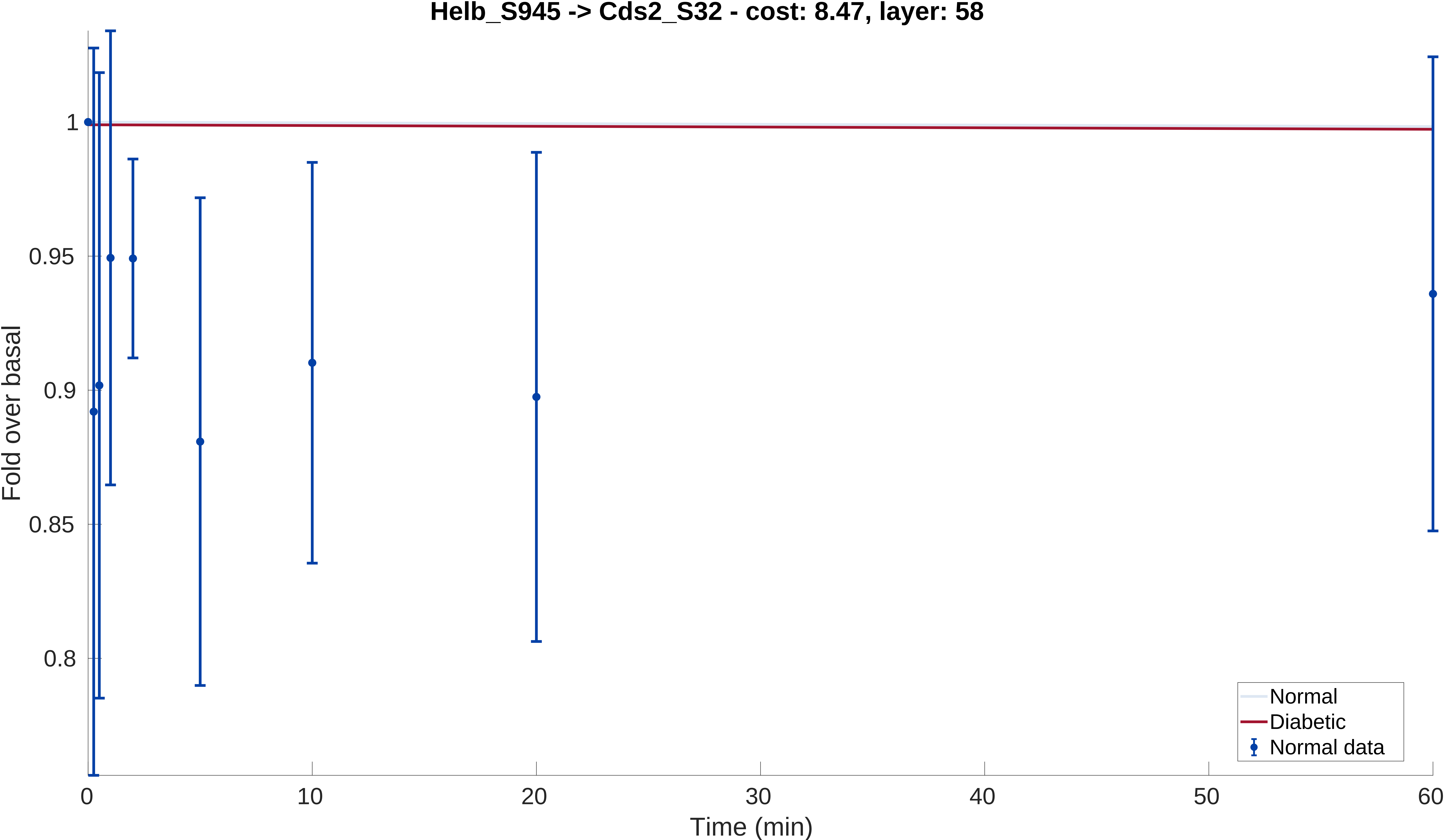

Helb_S945 -> Cds2_S32 - cost: 8.47, layer: 58
Fold over basal
1
0.95
0.9
0.85
0.8
0
10
20
30
40
50
60
Time (min)
Normal
Diabetic
Normal data

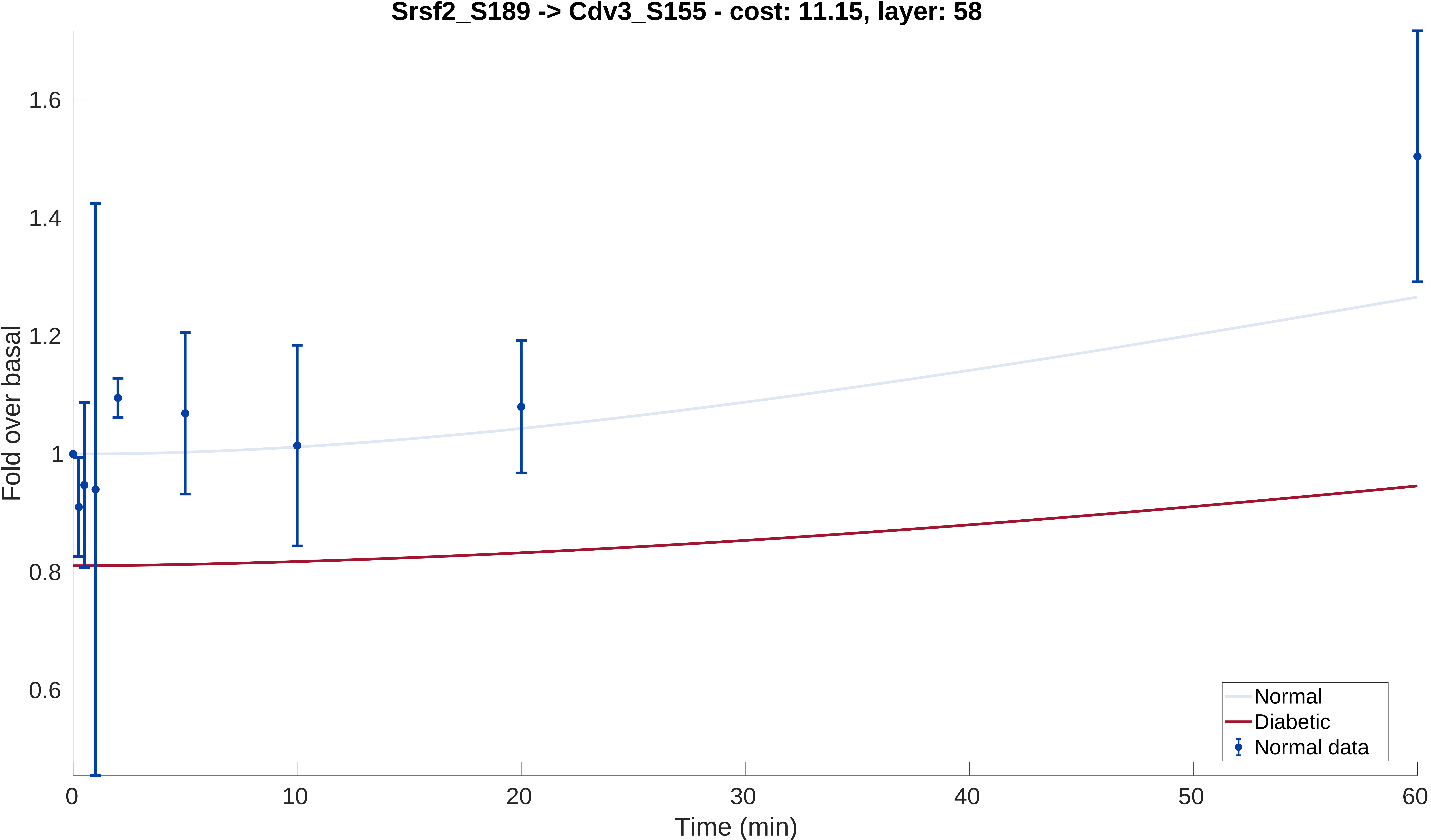

Srsf2_S189 -> Cdv3_S155 - cost: 11.15, layer: 58
1.6
1.4
1.2
1
0.8
0.6
Fold over basal
0
10
20
30
40
50
60
Time (min)
Normal
Diabetic
Normal data

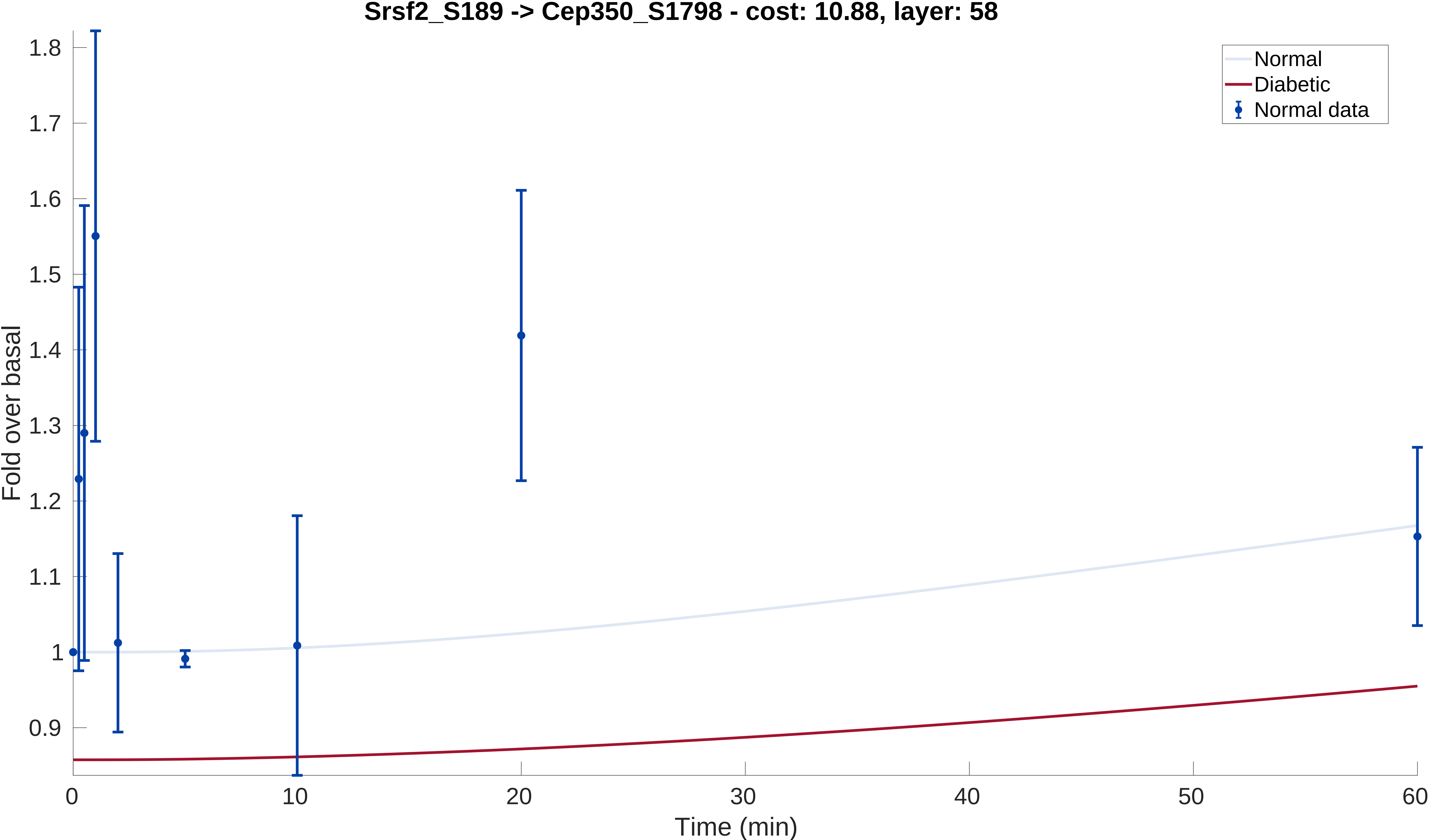

Srsf2_S189 -> Cep350_S1798 - cost: 10.88, layer: 58
Normal
Diabetic
Normal data
Fold over basal
1.8
1.7
1.6
1.5
1.4
1.3
1.2
1.1
1
0.9
0
10
20
30
40
50
60
Time (min)

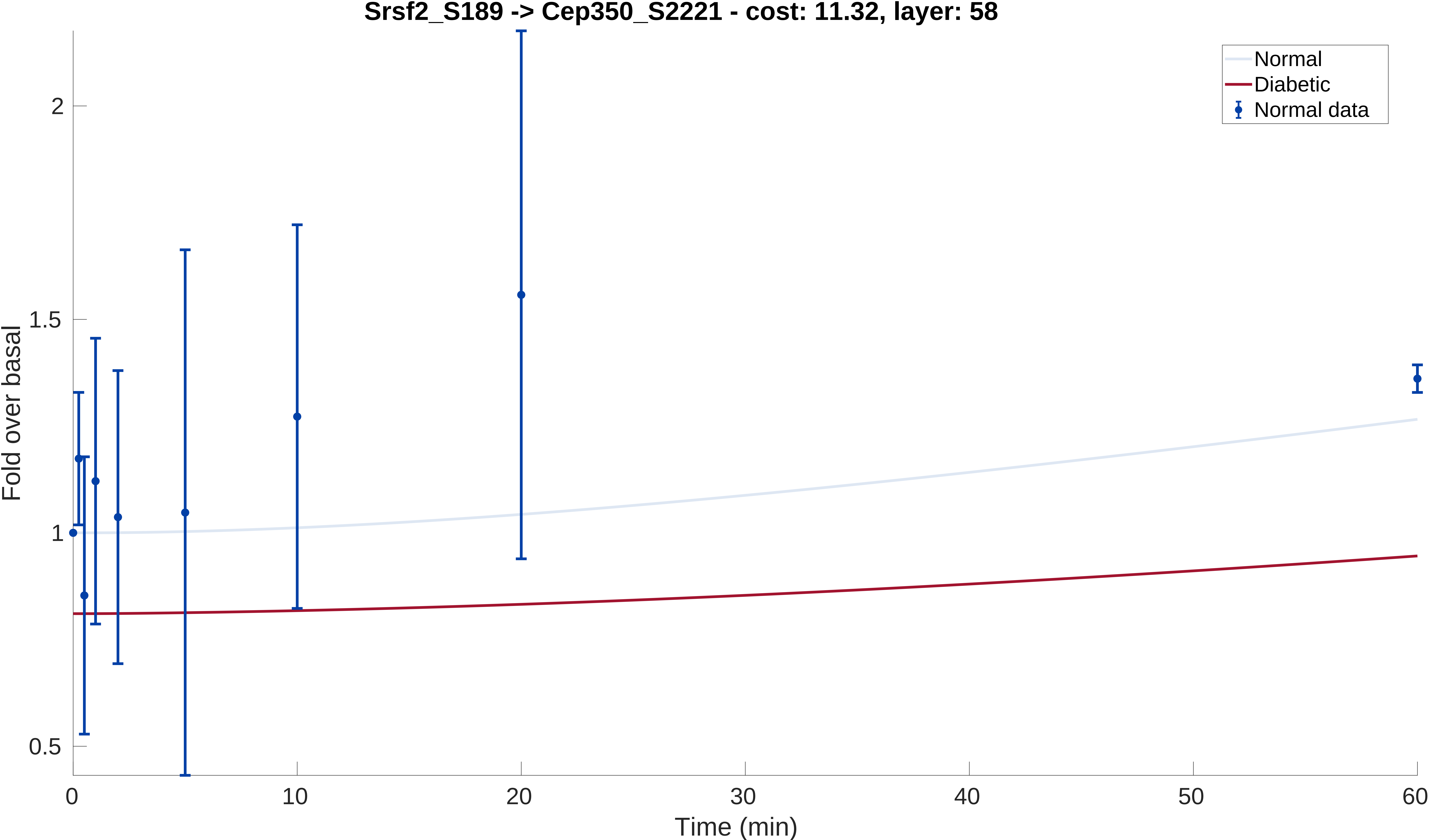

Srsf2_S189 -> Cep350_S2221 - cost: 11.32, layer: 58
Normal
Diabetic
Normal data
Fold over basal
0.5
1
1.5
2
0
10
20
30
40
50
60
Time (min)

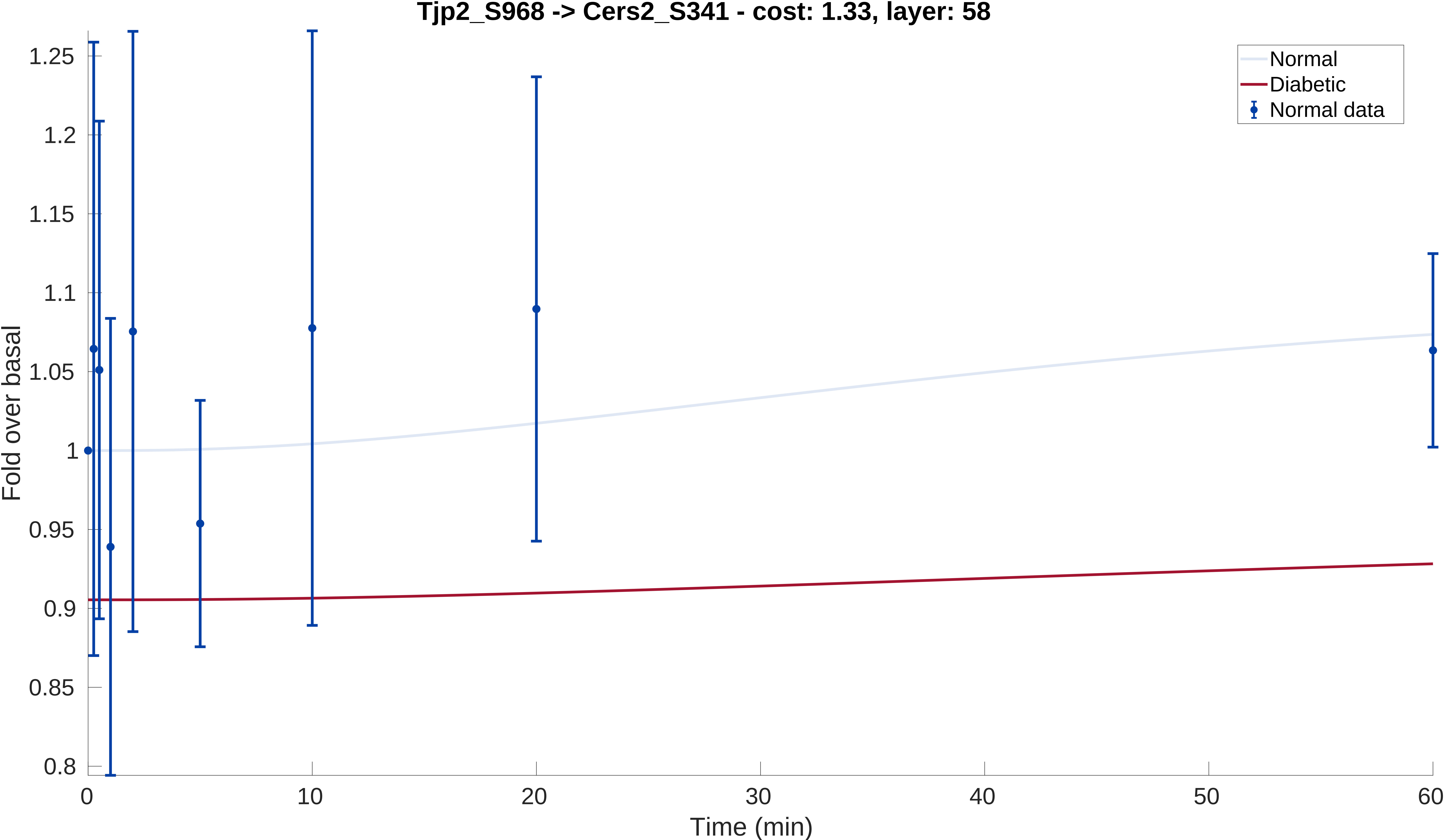

Tjp2_S968 -> Cers2_S341 - cost: 1.33, layer: 58
Normal
Diabetic
Normal data
Fold over basal
1.25
1.2
1.15
1.1
1.05
1
0.95
0.9
0.85
0.8
0
10
20
30
40
50
60
Time (min)

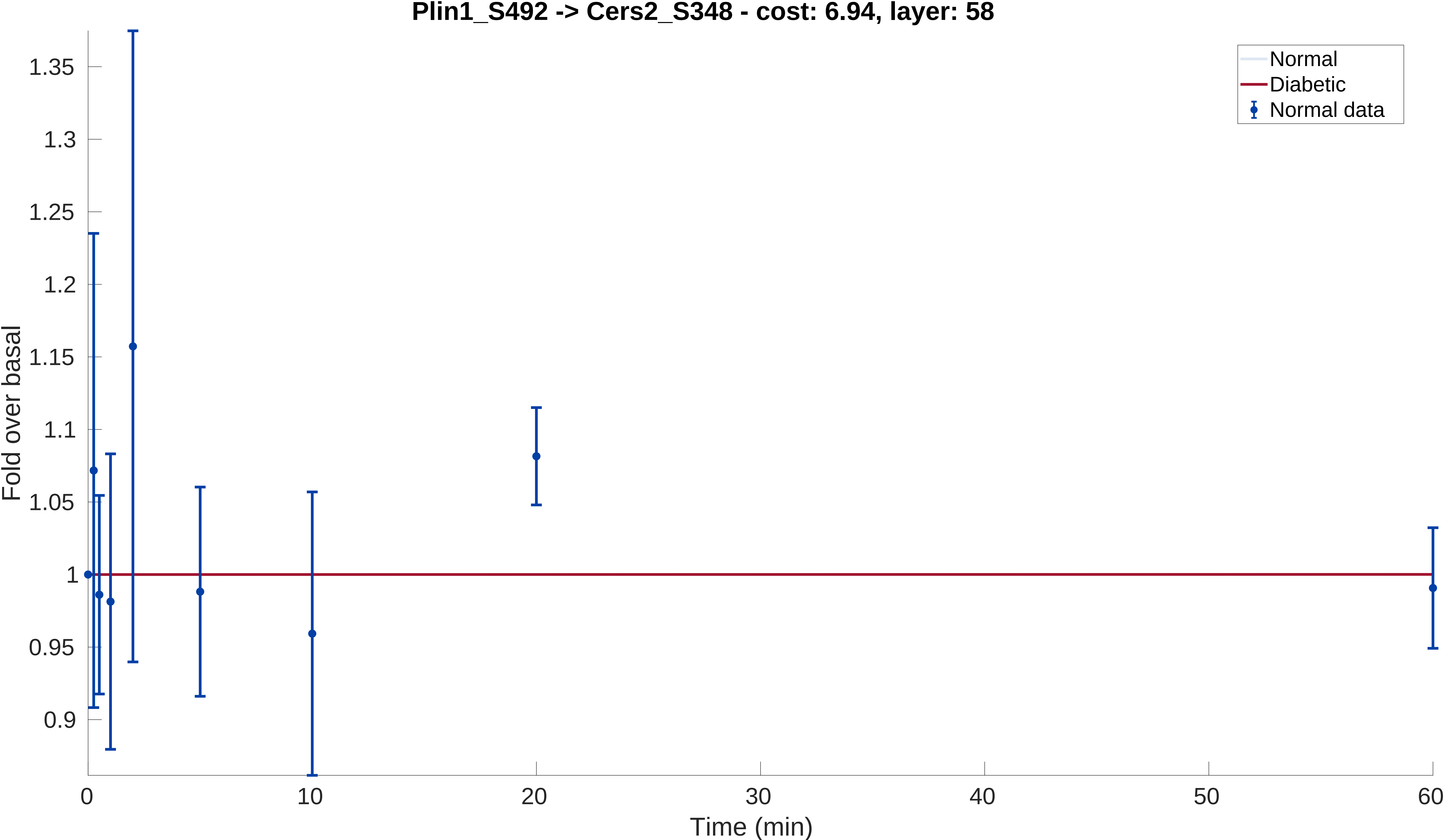

Plin1_S492 -> Cers2_S348 - cost: 6.94, layer: 58
Normal
Diabetic
Normal data
Fold over basal
1.35
1.3
1.25
1.2
1.15
1.1
1.05
1
0.95
0.9
0
10
20
30
40
50
60
Time (min)

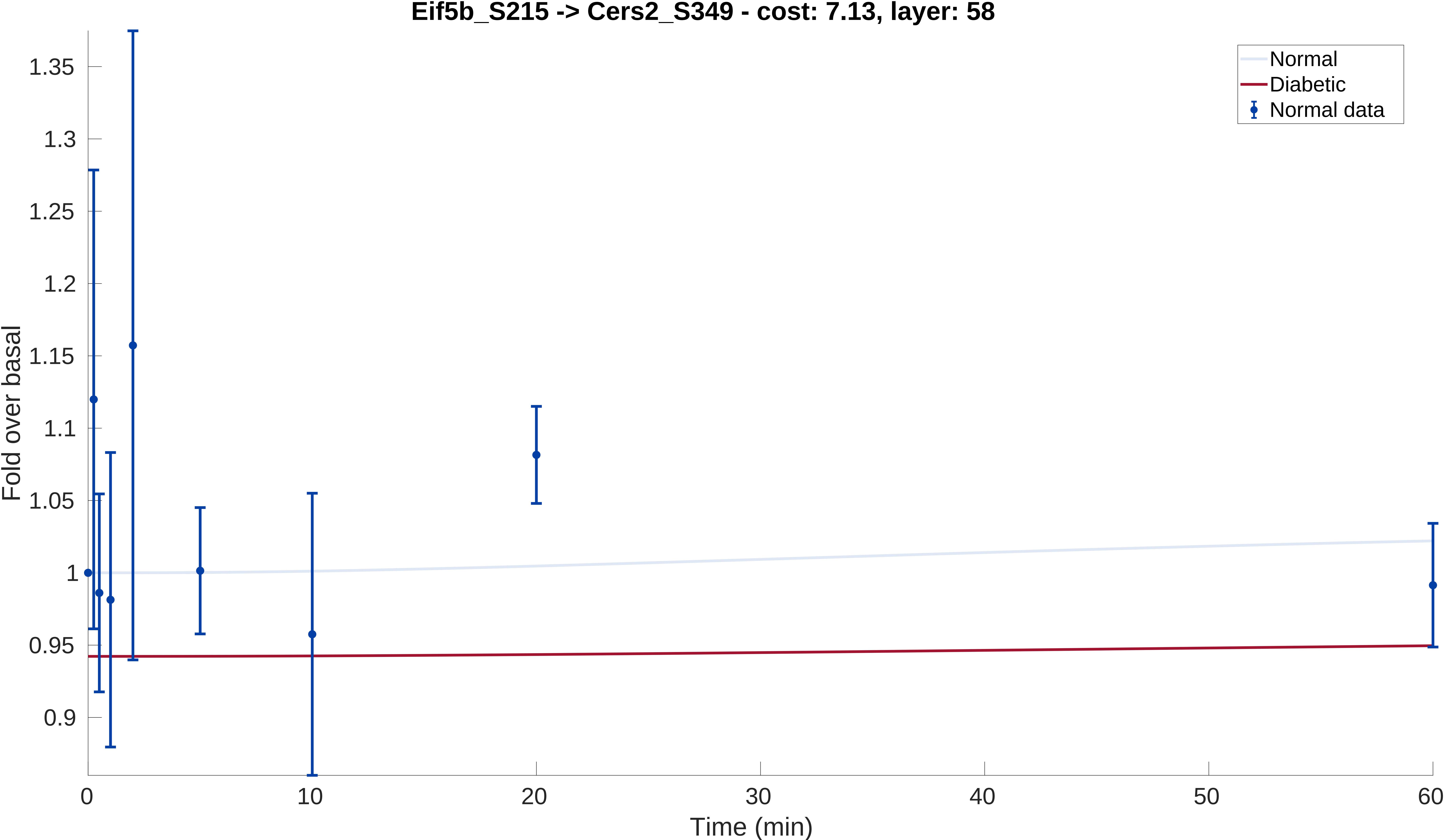

Eif5b_S215 -> Cers2_S349 - cost: 7.13, layer: 58
Normal
Diabetic
Normal data
Fold over basal
Time (min)
1.35
1.3
1.25
1.2
1.15
1.1
1.05
1
0.95
0.9
0
10
20
30
40
50
60

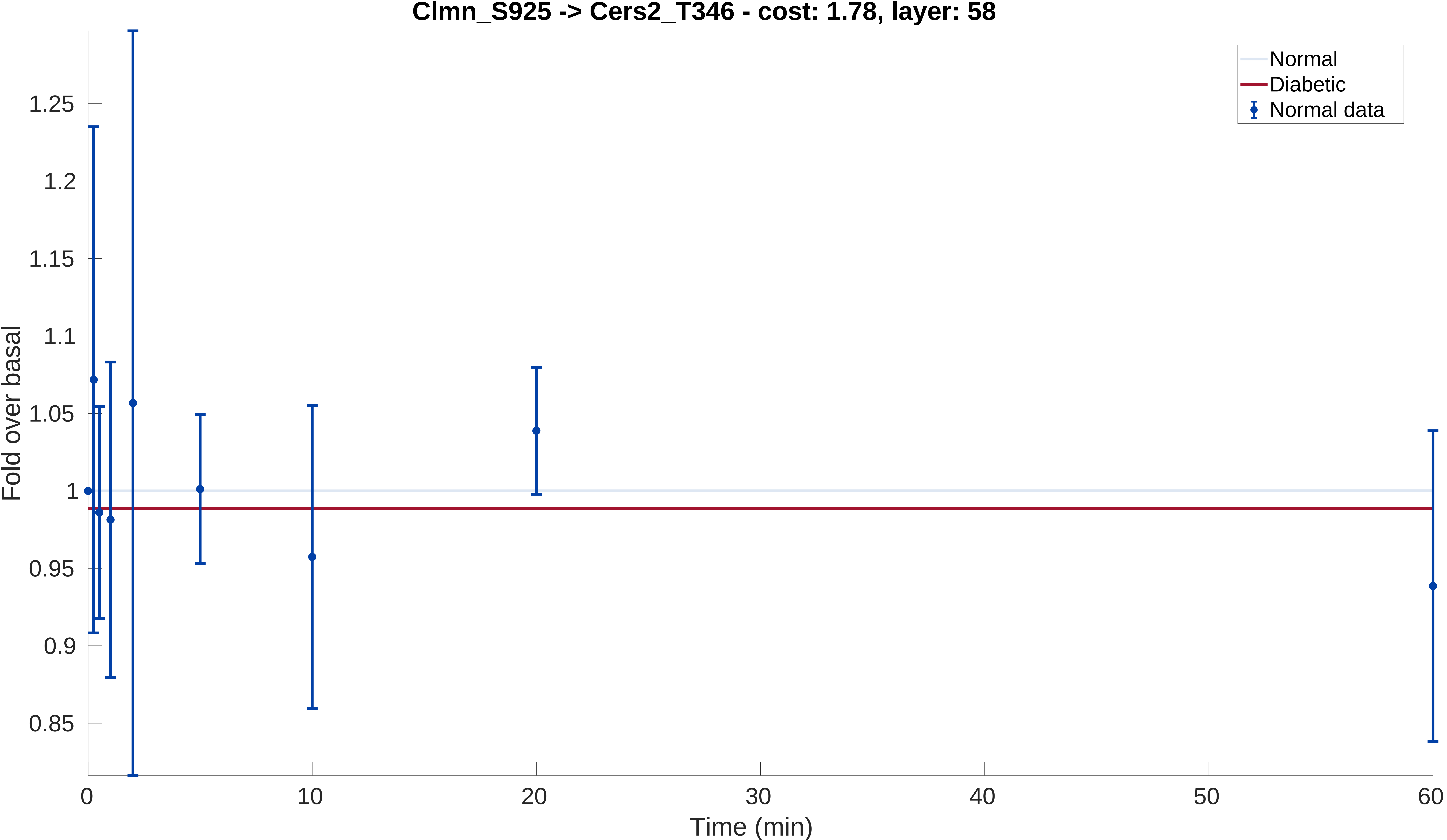

Clmn_S925 -> Cers2_T346 - cost: 1.78, layer: 58
Normal
Diabetic
Normal data
Fold over basal
1.25
1.2
1.15
1.1
1.05
1
0.95
0.9
0.85
0
10
20
30
40
50
60
Time (min)

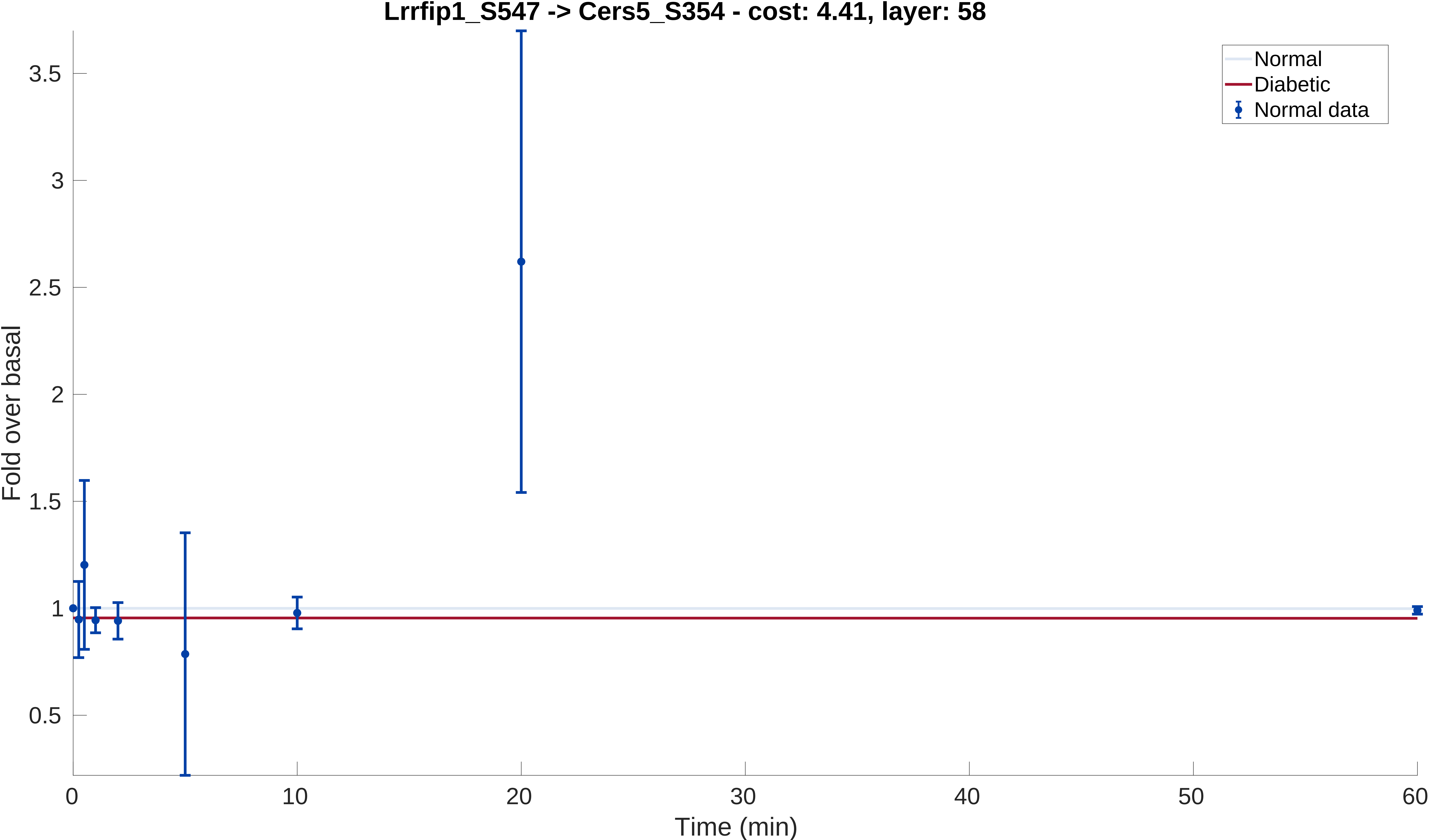

Lrrfip1_S547 -> Cers5_S354 - cost: 4.41, layer: 58
Normal
Diabetic
Normal data
Fold over basal
3.5
3
2.5
2
1.5
1
0.5
0
10
20
30
40
50
60
Time (min)

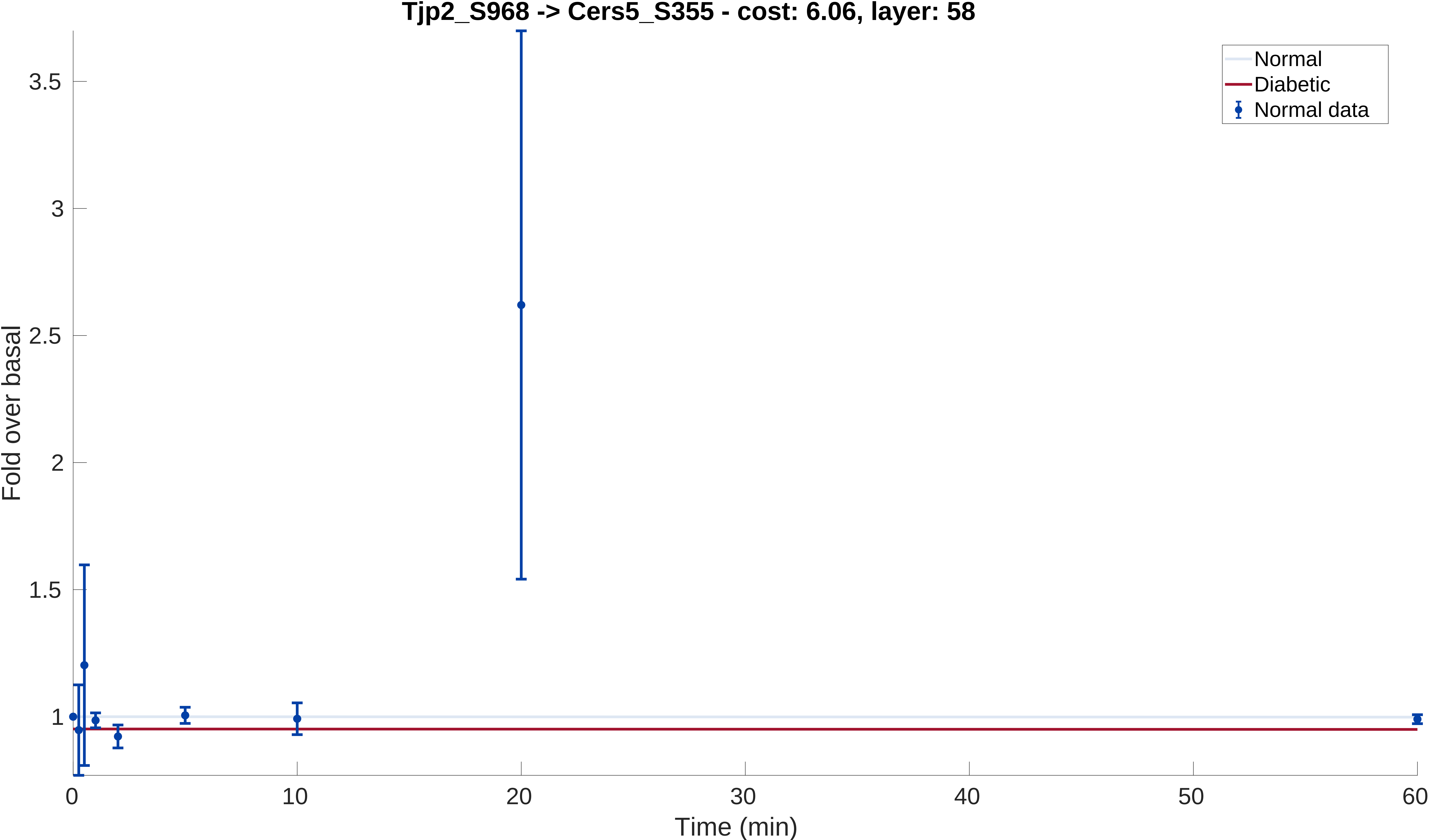

Tjp2_S968 -> Cers5_S355 - cost: 6.06, layer: 58
Normal
Diabetic
Normal data
Fold over basal
Time (min)
3.5
3
2.5
2
1.5
1
0
10
20
30
40
50
60

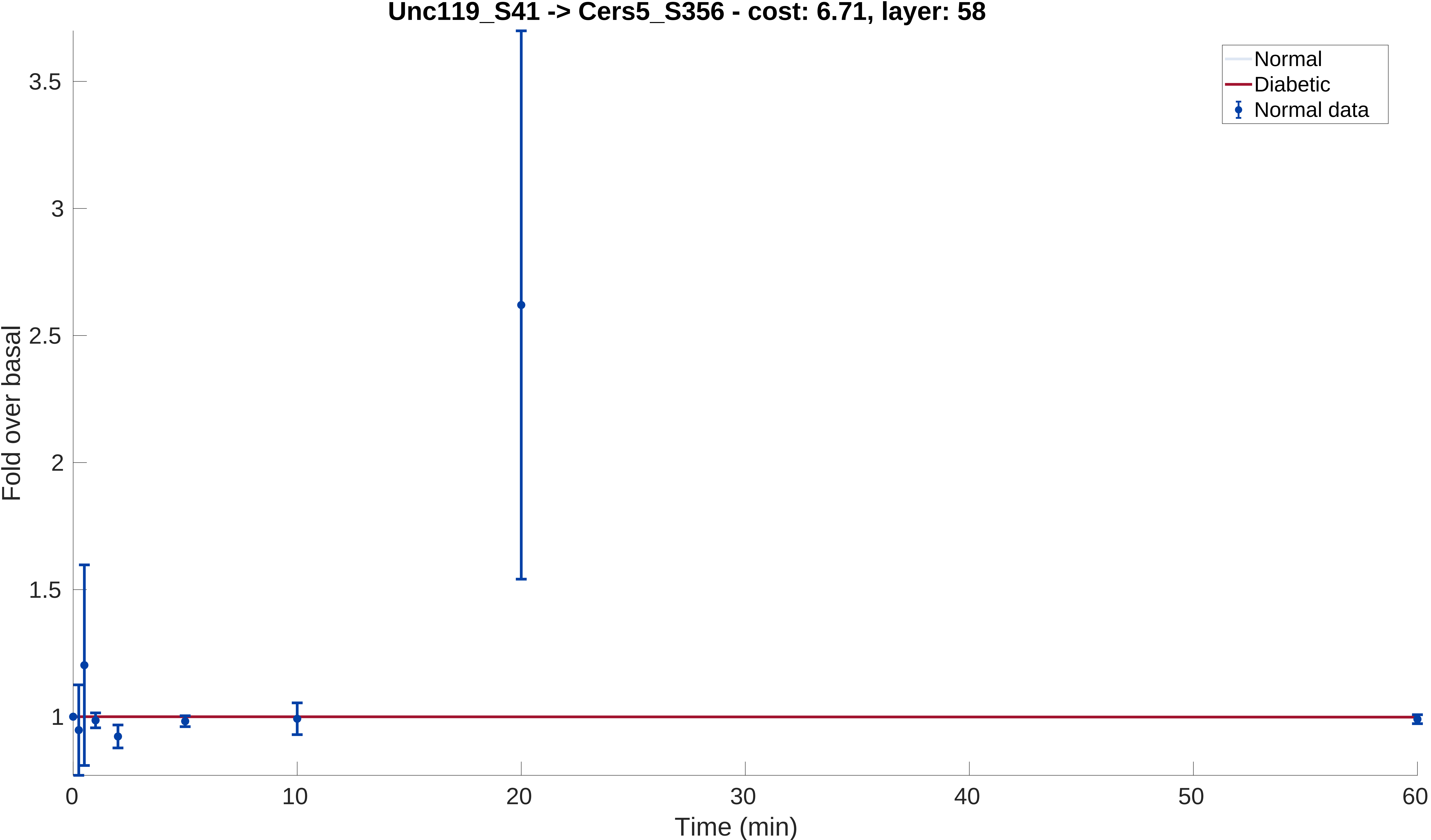

Unc119_S41 -> Cers5_S356 - cost: 6.71, layer: 58
Normal
Diabetic
Normal data
Fold over basal
3.5
3
2.5
2
1.5
1
0
10
20
30
40
50
60
Time (min)

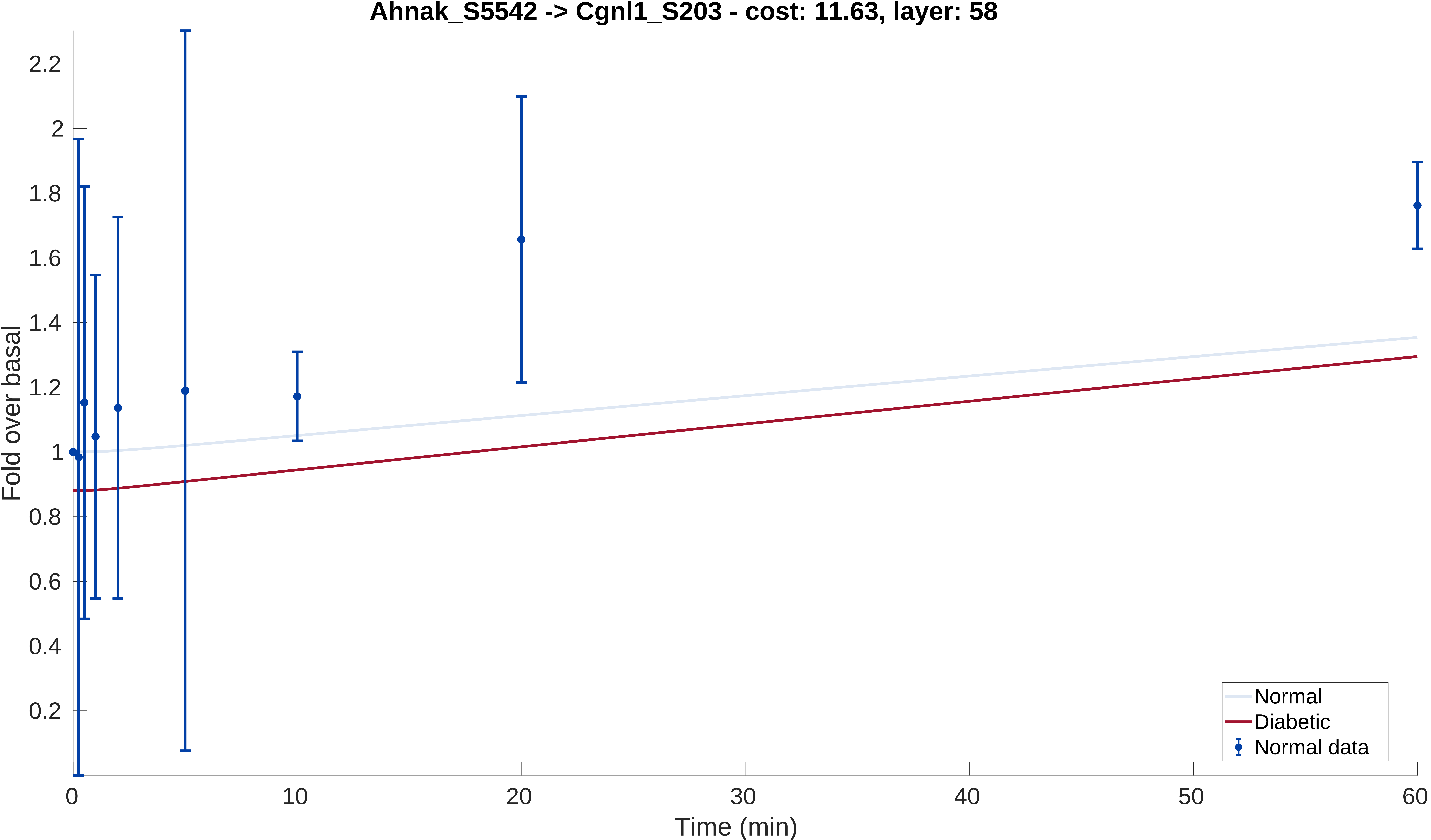

Ahnak_S5542 -> Cgnl1_S203 - cost: 11.63, layer: 58
Fold over basal
Time (min)
0.2
0.4
0.6
0.8
1
1.2
1.4
1.6
1.8
2
2.2
0
10
20
30
40
50
60
Normal
Diabetic
Normal data

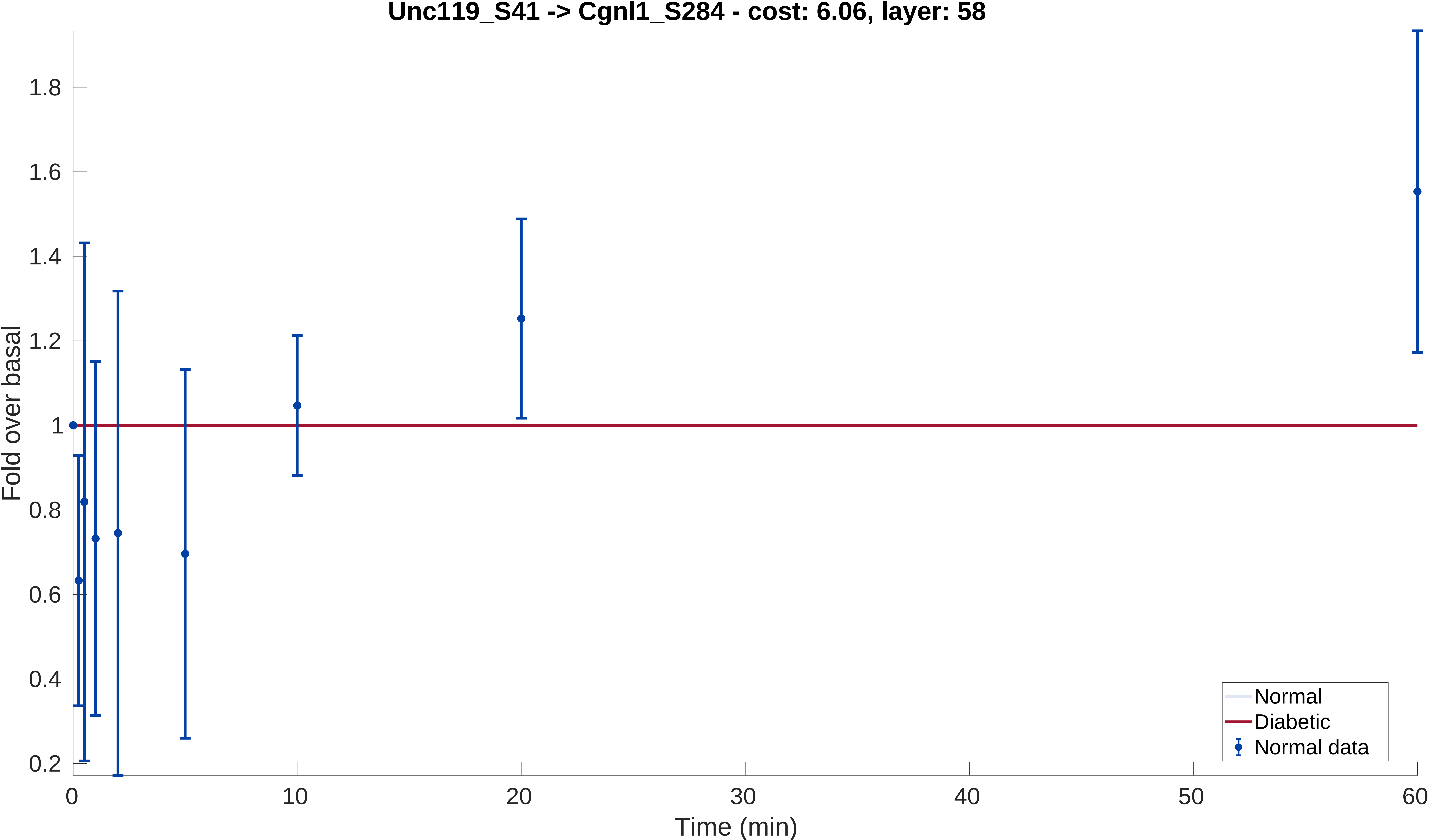

Unc119_S41 -> Cgnl1_S284 - cost: 6.06, layer: 58
Fold over basal
1.8
1.6
1.4
1.2
1
0.8
0.6
0.4
0.2
0
10
20
30
40
50
60
Time (min)
Normal
Diabetic
Normal data

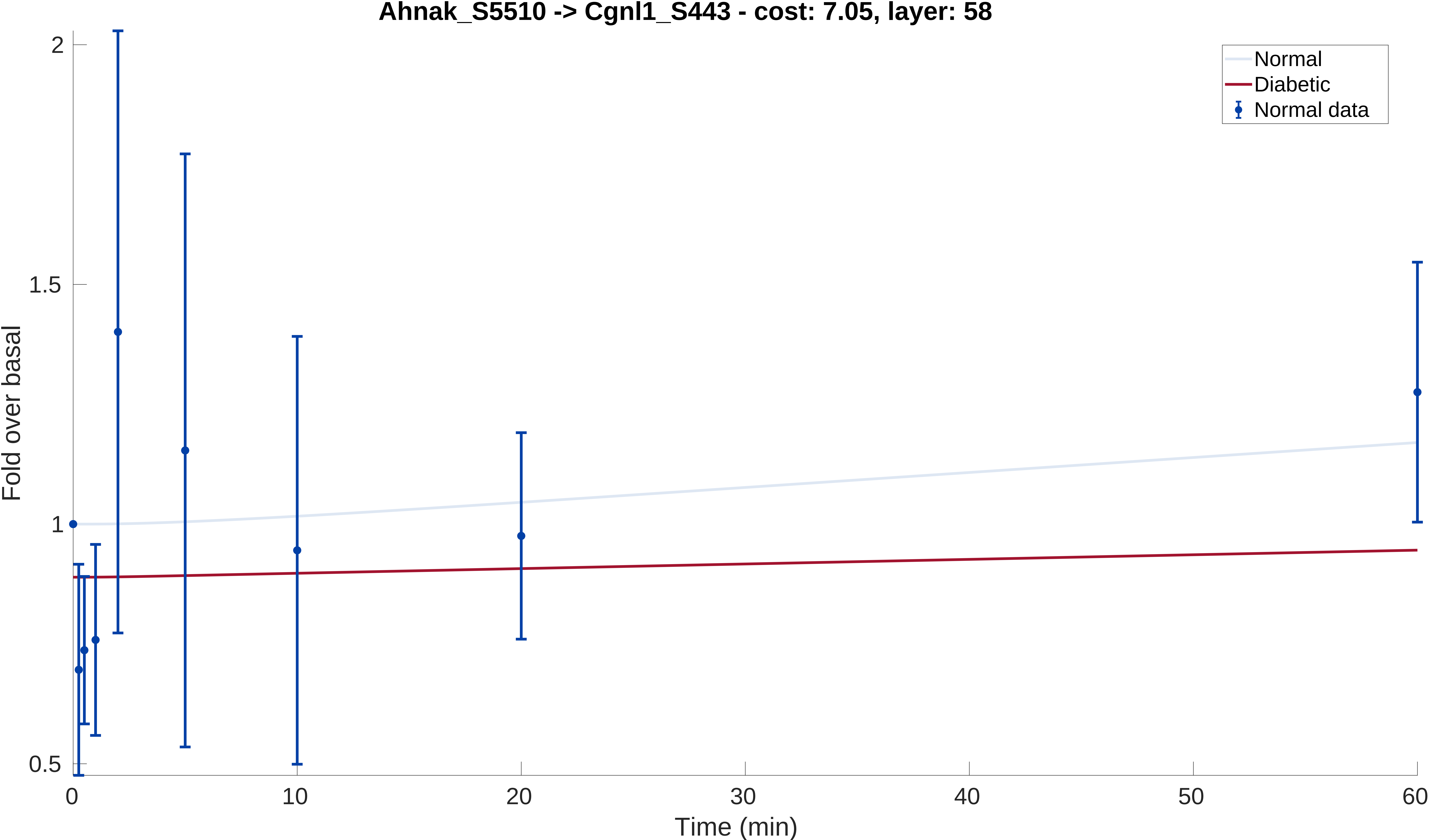
Ahnak_S5510 -> Cgnl1_S443 - cost: 7.05, layer: 58
Normal
Diabetic
Normal data
Fold over basal
Time (min)
0.5
1
1.5
2
0
10
20
30
40
50
60

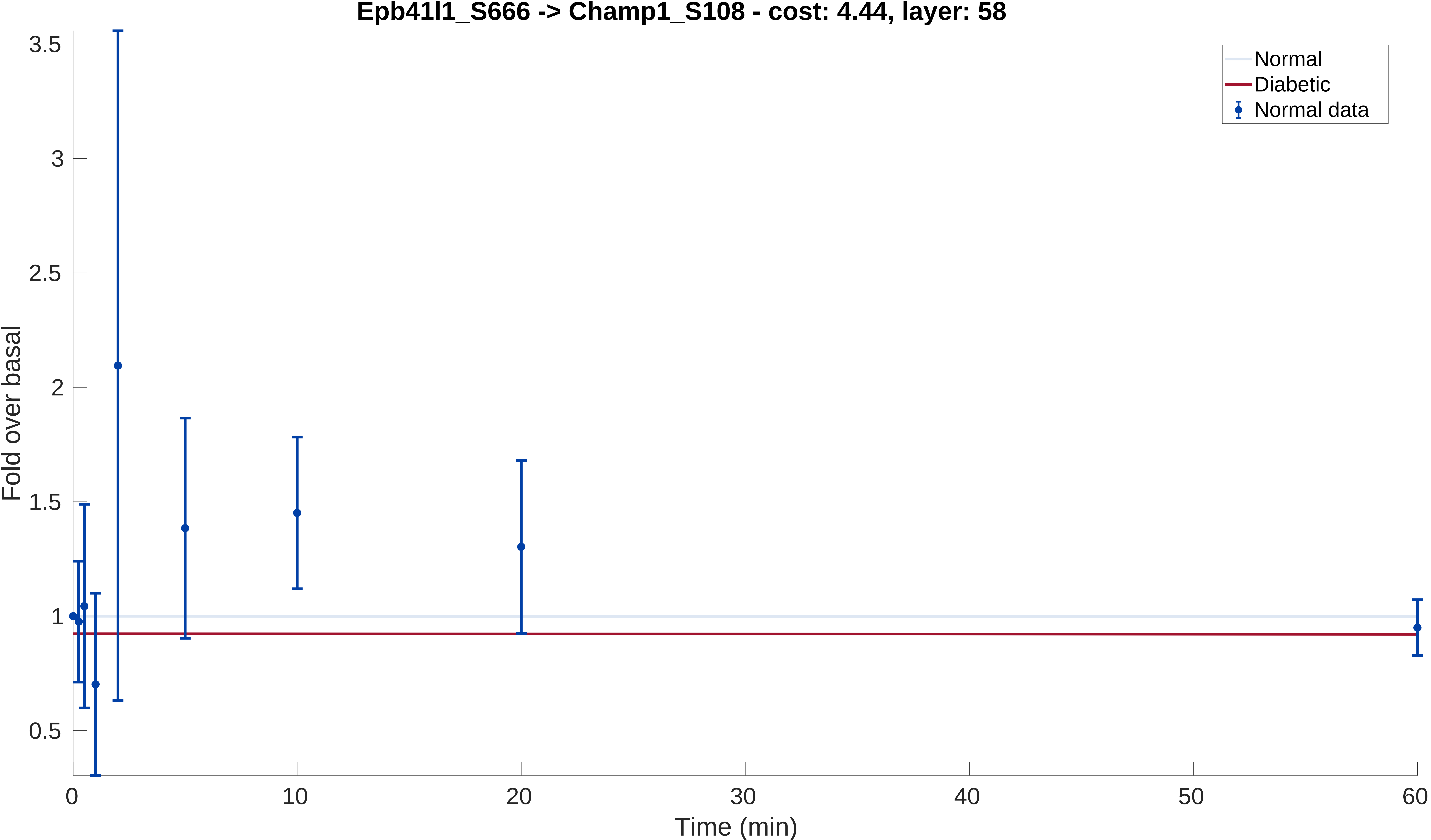

Epb41l1_S666 -> Champ1_S108 - cost: 4.44, layer: 58
Normal
Diabetic
Normal data
Fold over basal
Time (min)
0.5
1
1.5
2
2.5
3
3.5
0
10
20
30
40
50
60

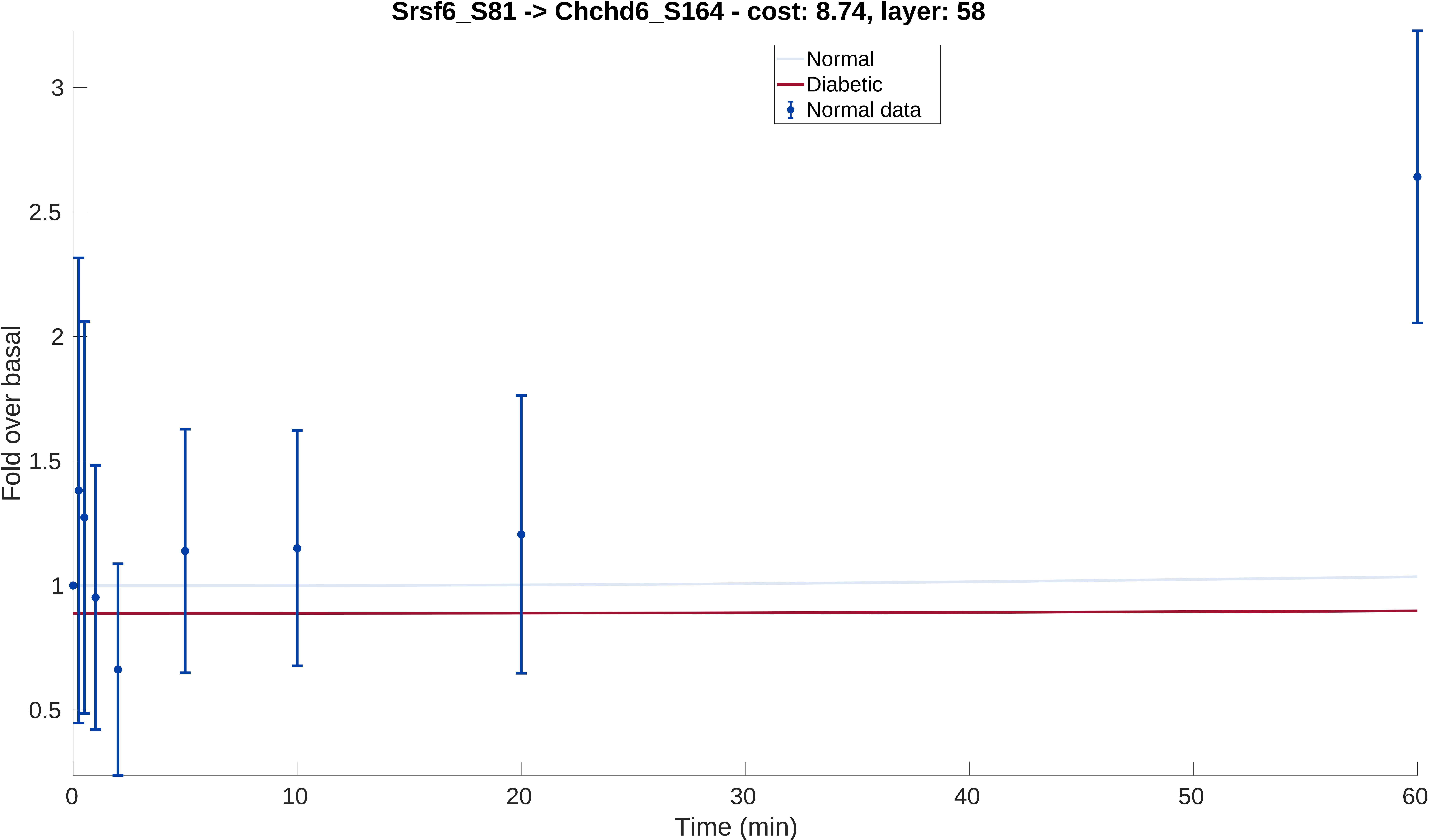

Srsf6_S81 -> Chchd6_S164 - cost: 8.74, layer: 58
Normal
Diabetic
Normal data
Fold over basal
3
2.5
2
1.5
1
0.5
0
10
20
30
40
50
60
Time (min)

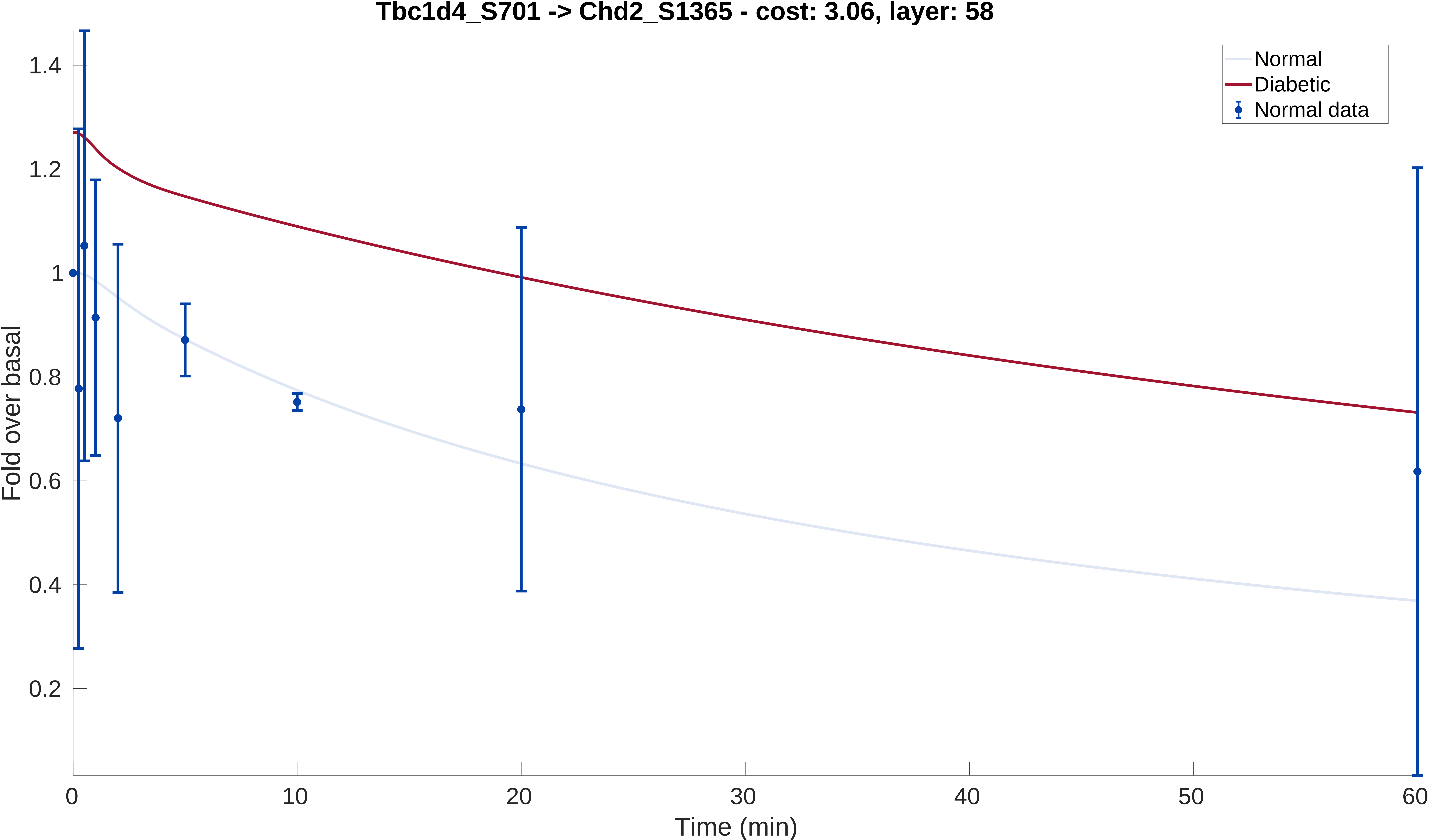

Tbc1d4_S701 -> Chd2_S1365 - cost: 3.06, layer: 58
Normal
Diabetic
Normal data
Fold over basal
1.4
1.2
1
0.8
0.6
0.4
0.2
0
10
20
30
40
50
60
Time (min)

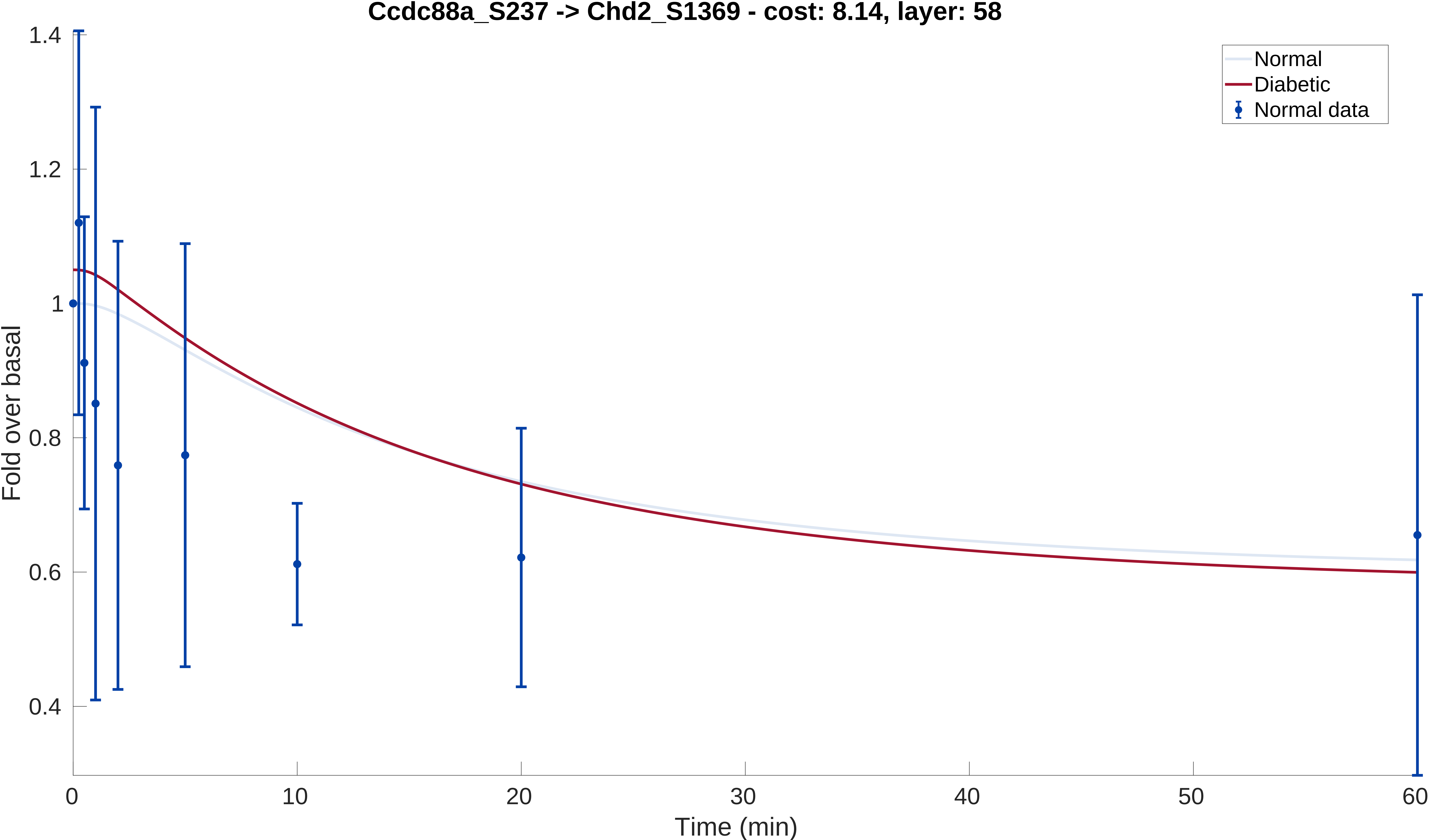
Ccdc88a_S237 -> Chd2_S1369 - cost: 8.14, layer: 58
Normal
Diabetic
Normal data
Fold over basal
Time (min)
1.4
1.2
1
0.8
0.6
0.4
0
10
20
30
40
50
60

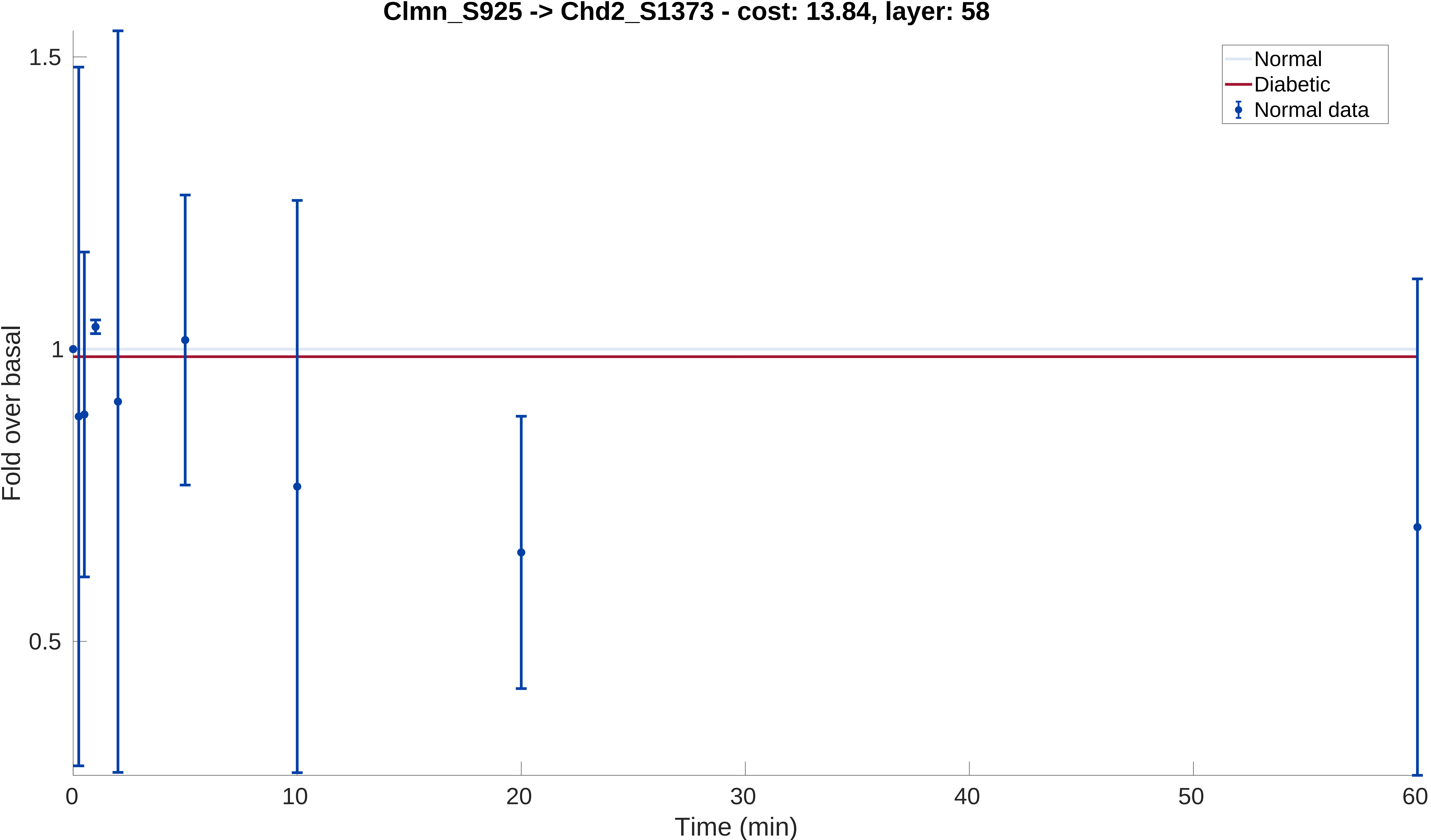

Clmn_S925 -> Chd2_S1373 - cost: 13.84, layer: 58
Normal
Diabetic
Normal data
Fold over basal
Time (min)
1.5
1
0.5
0
10
20
30
40
50
60

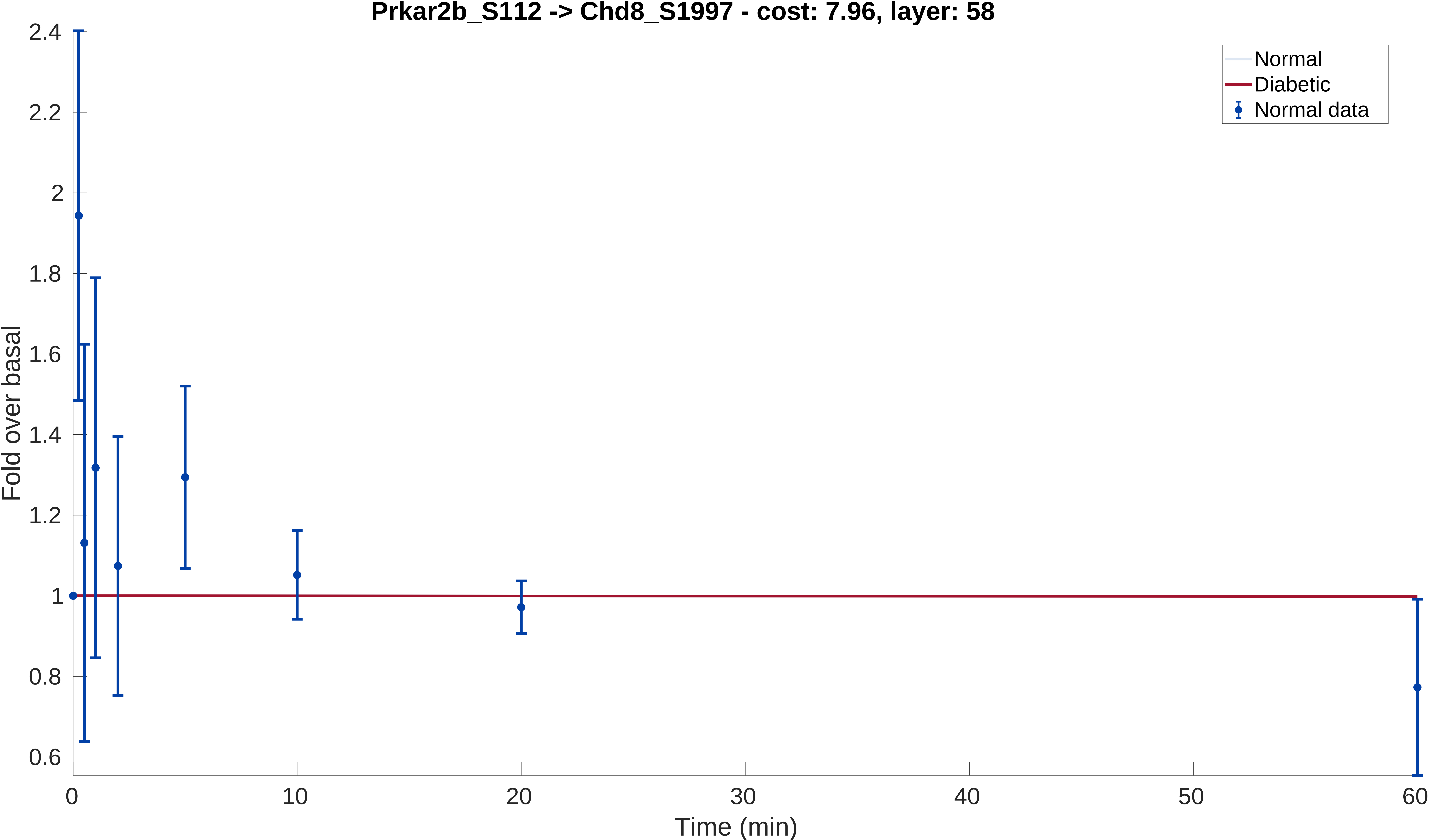

Prkar2b_S112 -> Chd8_S1997 - cost: 7.96, layer: 58
Normal
Diabetic
Normal data
Fold over basal
Time (min)
2.4
2.2
2
1.8
1.6
1.4
1.2
1
0.8
0.6
0
10
20
30
40
50
60

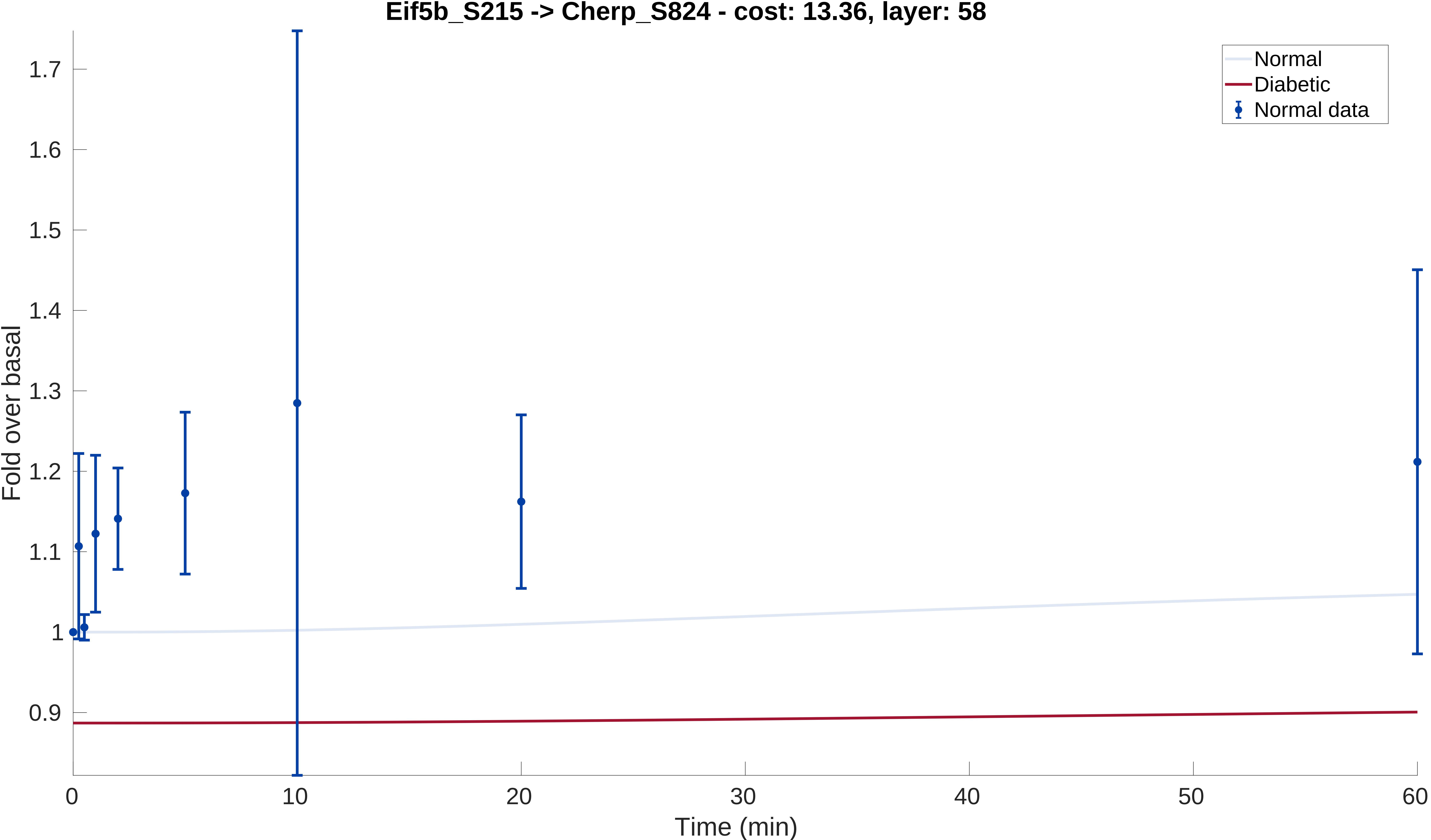

Eif5b_S215 -> Cherp_S824 - cost: 13.36, layer: 58
Normal
Diabetic
Normal data
Fold over basal
1.7
1.6
1.5
1.4
1.3
1.2
1.1
1
0.9
0
10
20
30
40
50
60
Time (min)

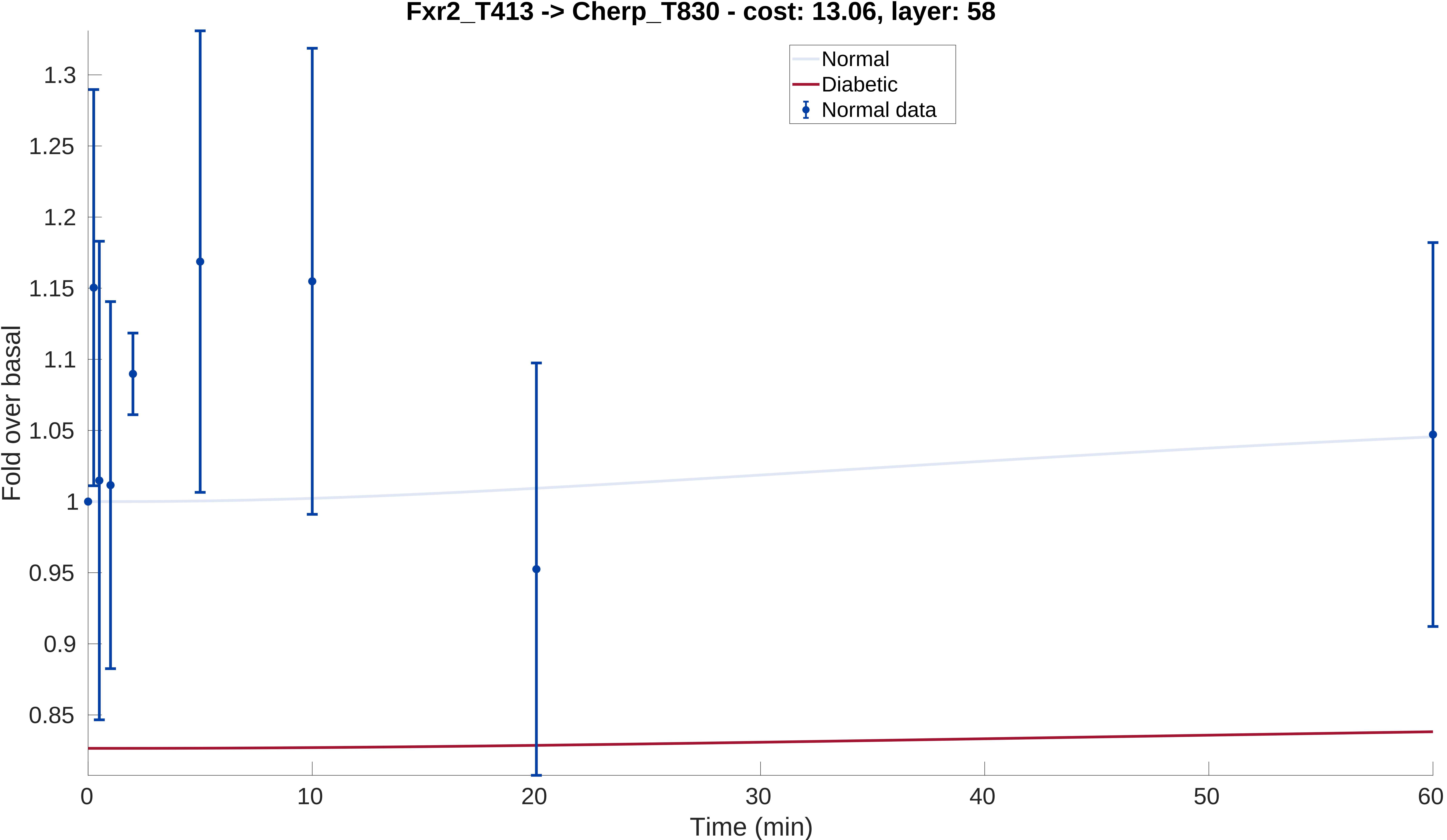
Fxr2_T413 -> Cherp_T830 - cost: 13.06, layer: 58
Normal
Diabetic
Normal data
Fold over basal
1.3
1.25
1.2
1.15
1.1
1.05
1
0.95
0.9
0.85
0
10
20
30
40
50
60
Time (min)

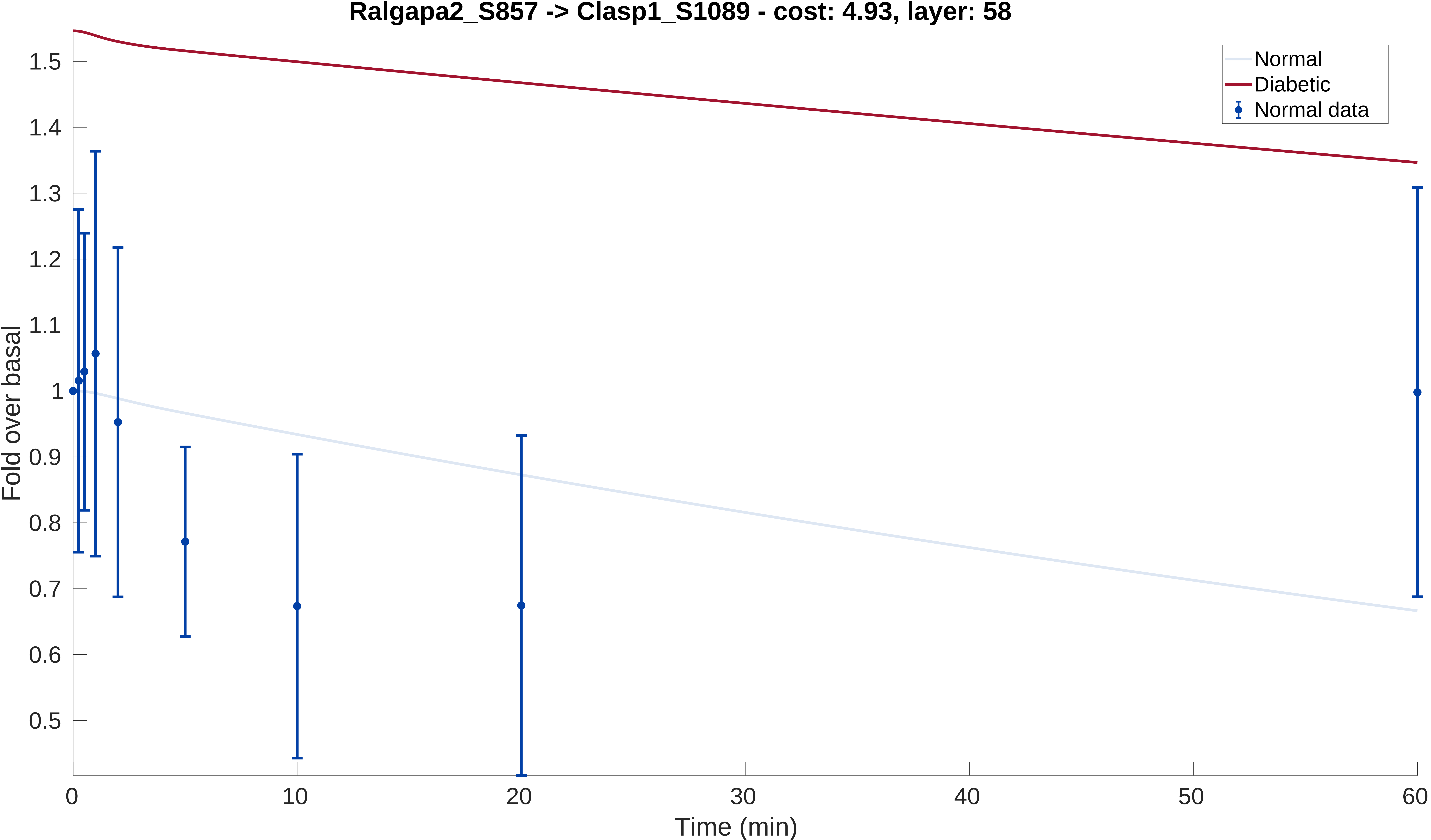

Ralgapa2_S857 -> Clasp1_S1089 - cost: 4.93, layer: 58
Normal
Diabetic
Normal data
Fold over basal
Time (min)
1.5
1.4
1.3
1.2
1.1
1
0.9
0.8
0.7
0.6
0.5
0
10
20
30
40
50
60

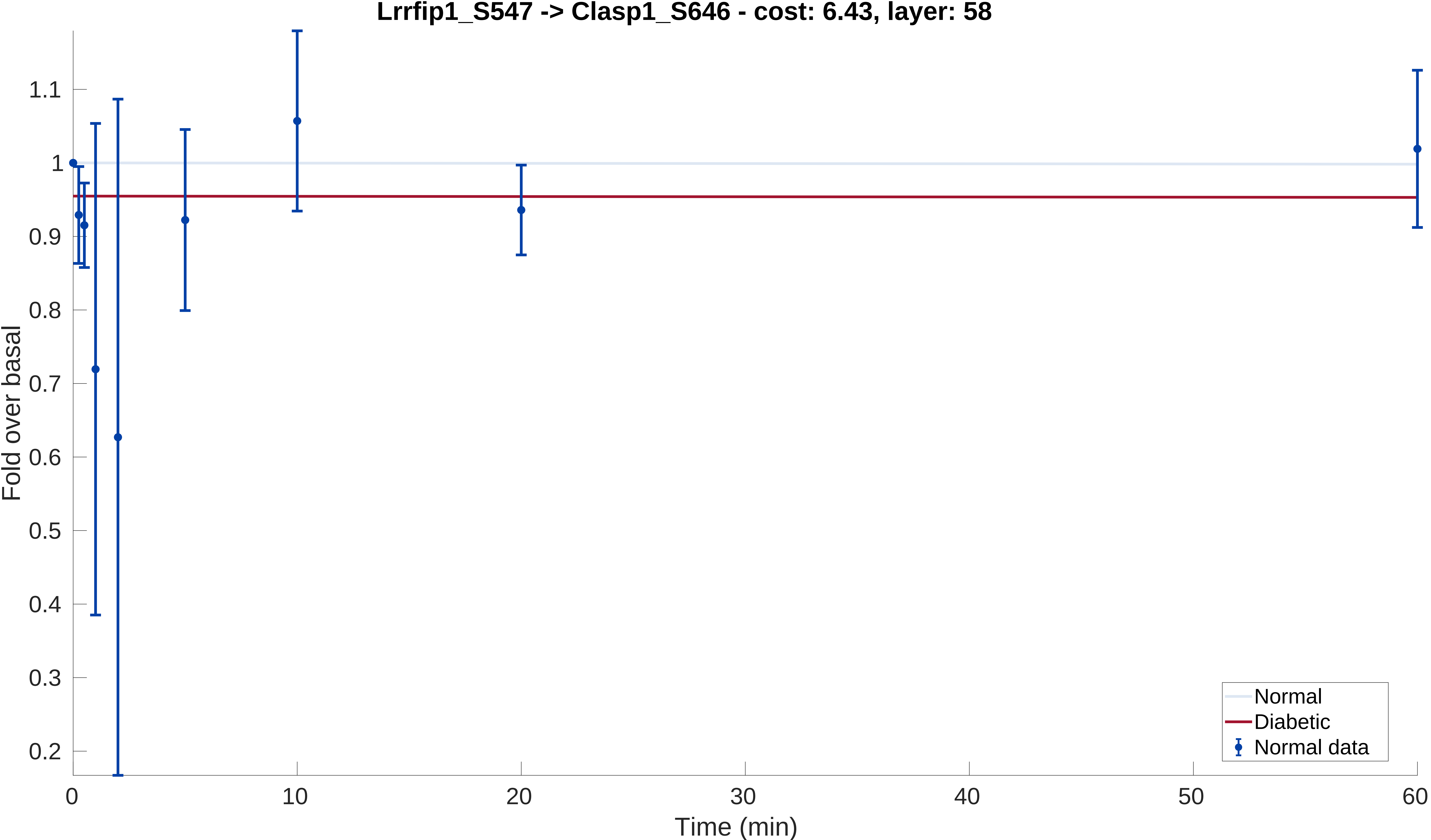

Lrrfip1_S547 -> Clasp1_S646 - cost: 6.43, layer: 58
Fold over basal
1.1
1
0.9
0.8
0.7
0.6
0.5
0.4
0.3
0.2
0
10
20
30
40
50
60
Time (min)
Normal
Diabetic
Normal data

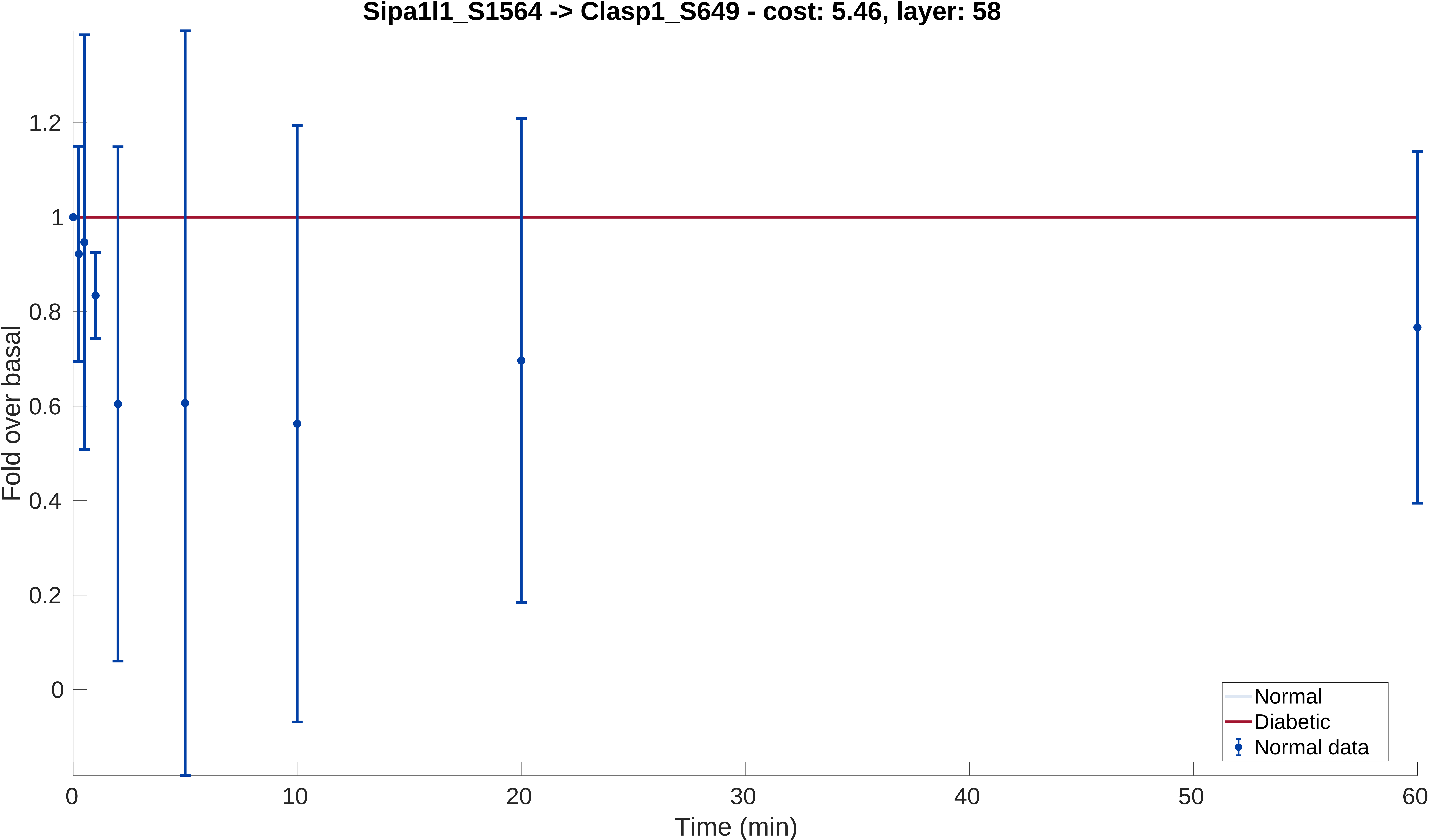

Sipa1l1_S1564 -> Clasp1_S649 - cost: 5.46, layer: 58
Fold over basal
1.2
1
0.8
0.6
0.4
0.2
0
0
10
20
30
40
50
60
Time (min)
Normal
Diabetic
Normal data

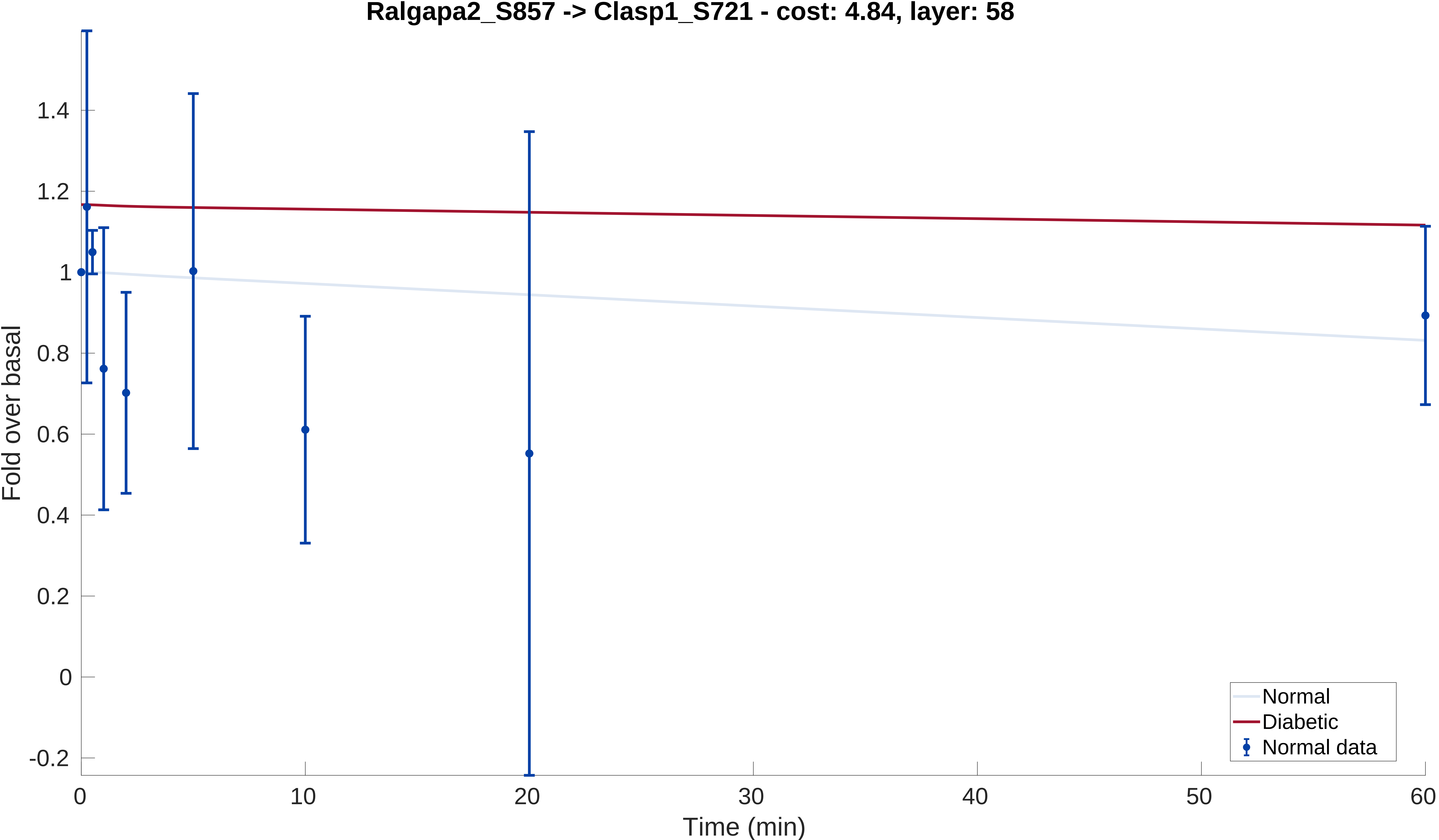

Ralgapa2_S857 -> Clasp1_S721 - cost: 4.84, layer: 58
Fold over basal
Time (min)
1.4
1.2
1
0.8
0.6
0.4
0.2
0
-0.2
0
10
20
30
40
50
60
Normal
Diabetic
Normal data

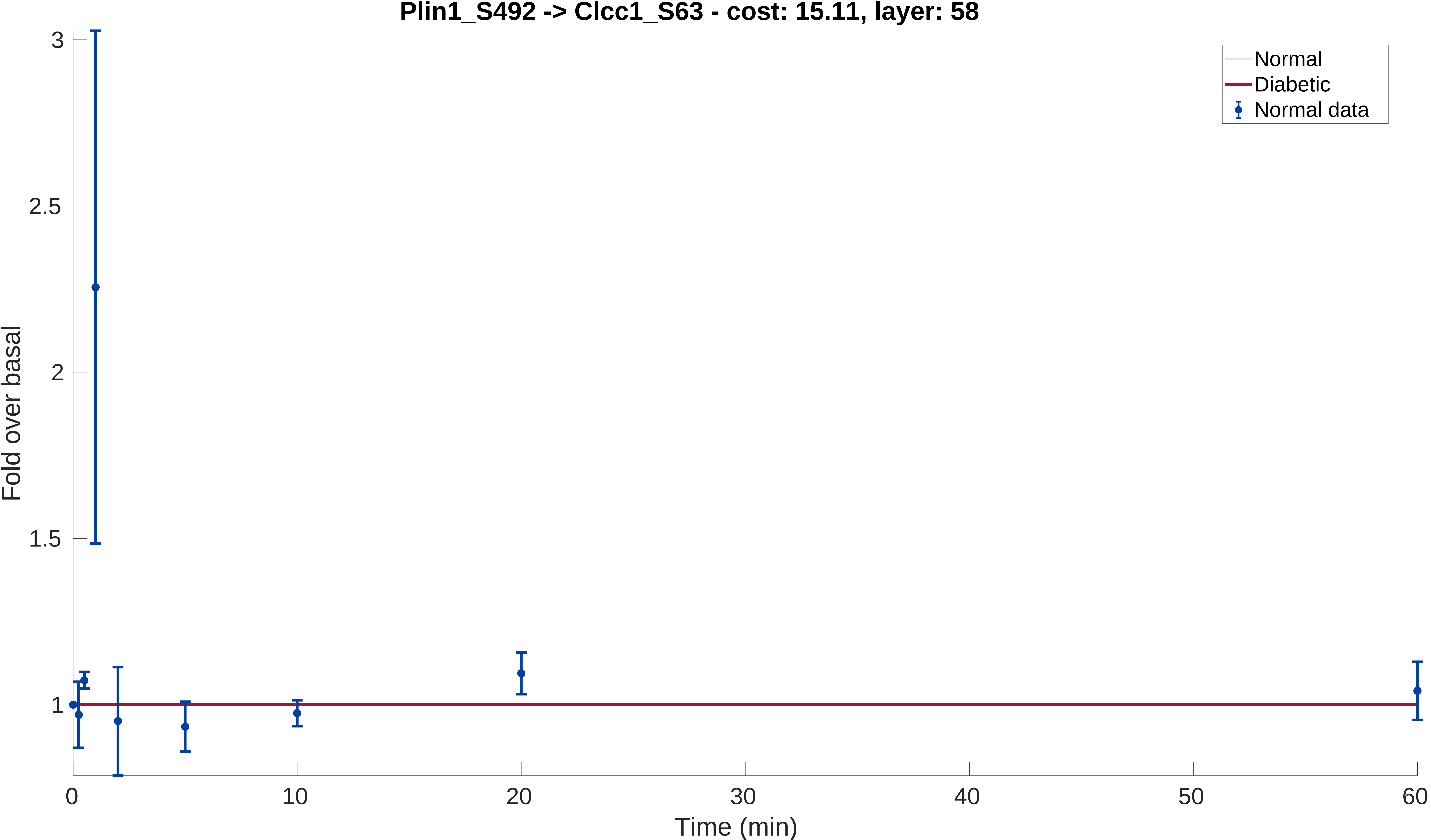

Plin1_S492 -> Clcc1_S63 - cost: 15.11, layer: 58
Normal
Diabetic
Normal data
Fold over basal
Time (min)
3
2.5
2
1.5
1
0
10
20
30
40
50
60

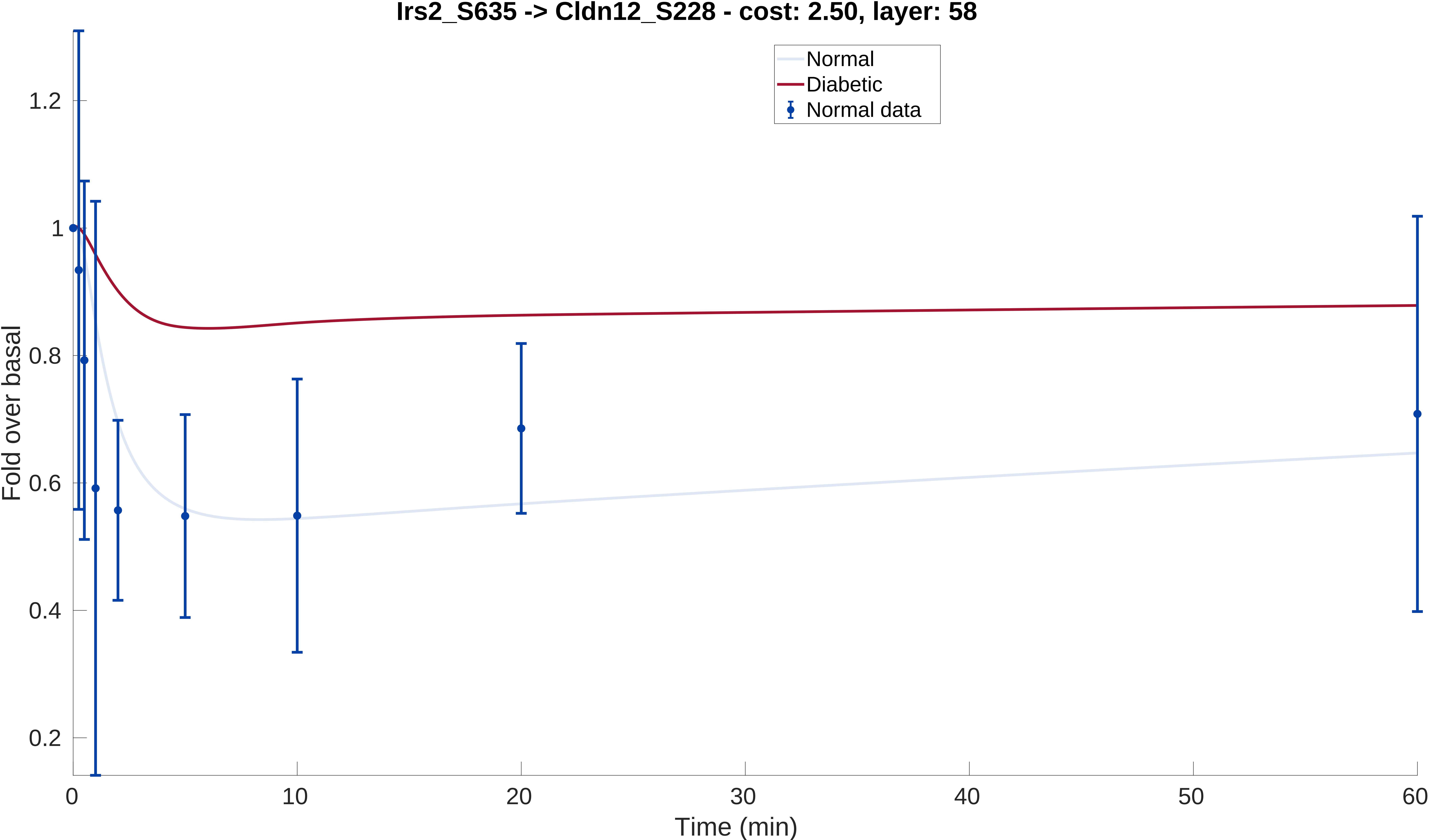
Irs2_S635 -> Cldn12_S228 - cost: 2.50, layer: 58
Normal
Diabetic
Normal data
Fold over basal
0.2
0.4
0.6
0.8
1
1.2
0
10
20
30
40
50
60
Time (min)

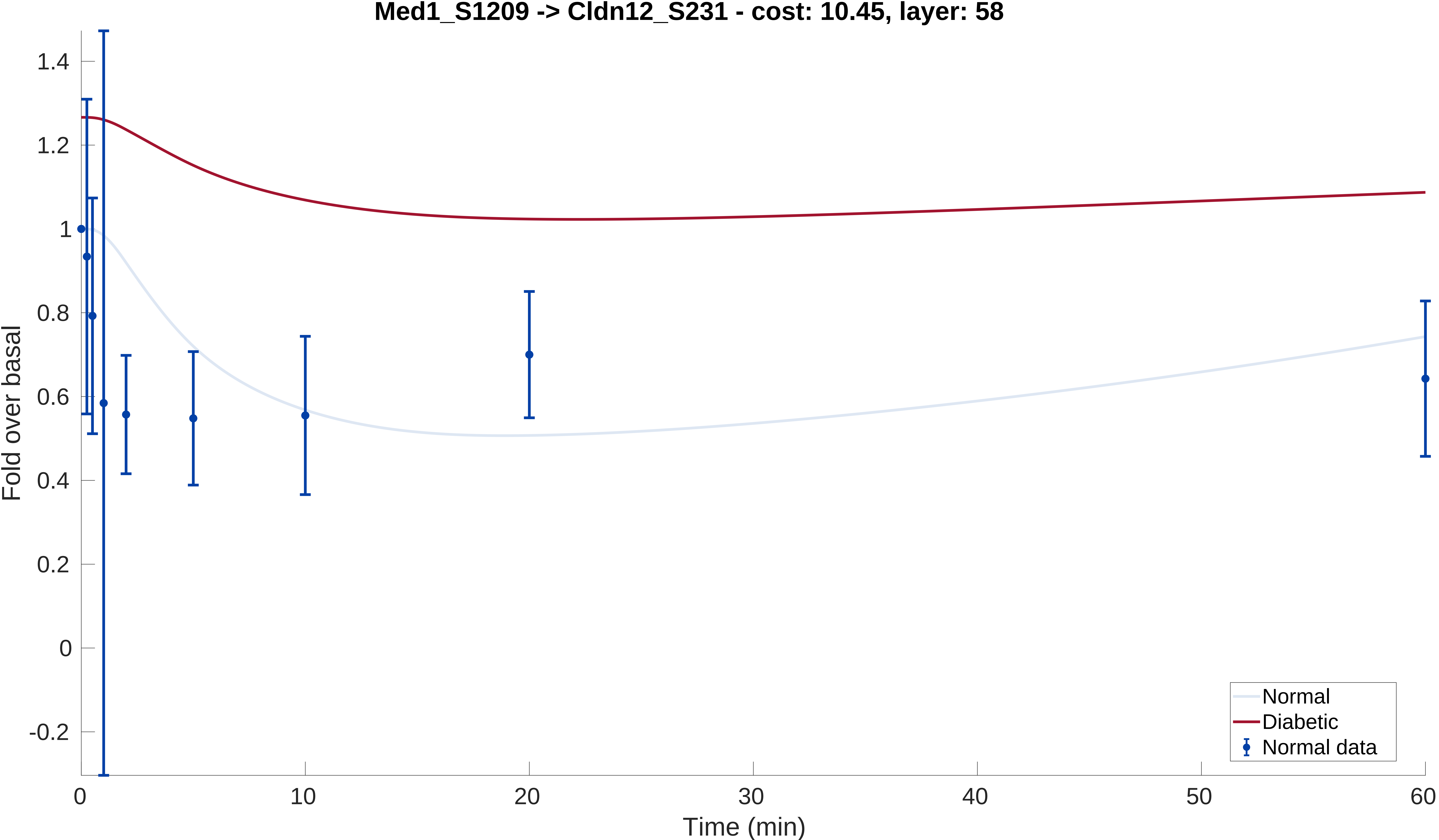
Med1_S1209 -> Cldn12_S231 - cost: 10.45, layer: 58
Fold over basal
1.4
1.2
1
0.8
0.6
0.4
0.2
0
-0.2
0
10
20
30
40
50
60
Time (min)
Normal
Diabetic
Normal data

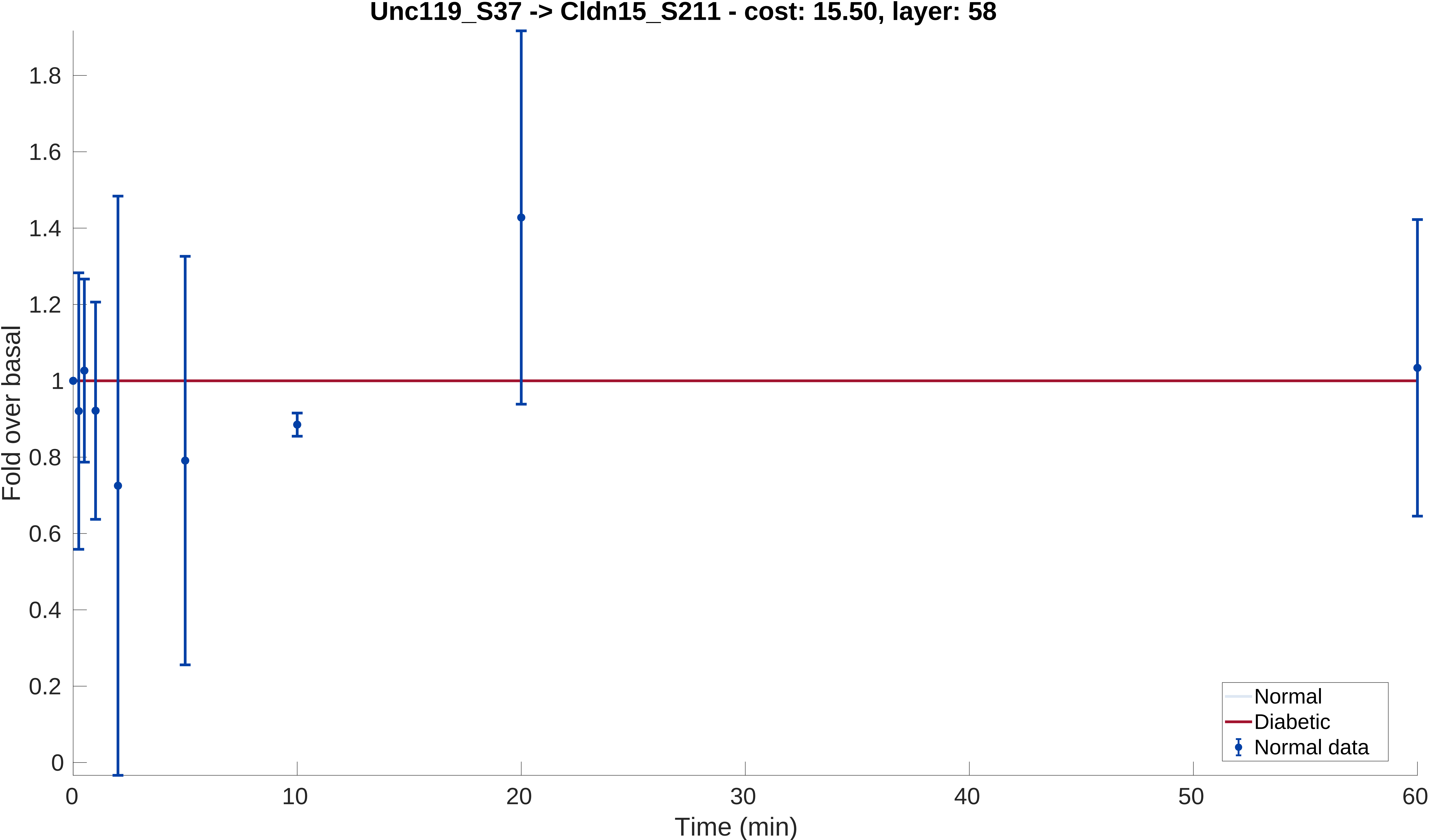

Unc119_S37 -> Cldn15_S211 - cost: 15.50, layer: 58
Fold over basal
Time (min)
0
0.2
0.4
0.6
0.8
1
1.2
1.4
1.6
1.8
0
10
20
30
40
50
60
Normal
Diabetic
Normal data

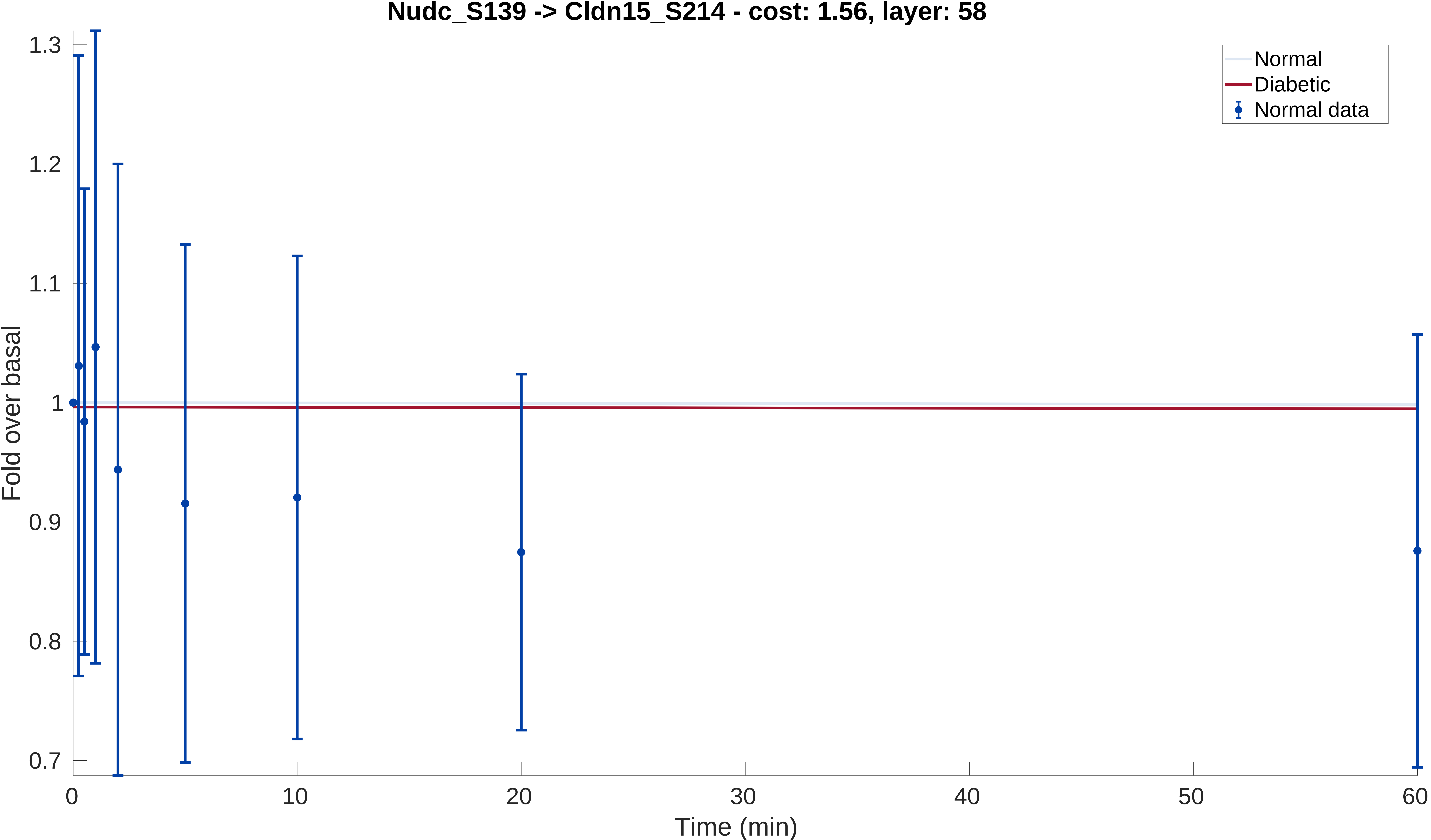
Nudc_S139 -> Cldn15_S214 - cost: 1.56, layer: 58
Normal
Diabetic
Normal data
Fold over basal
Time (min)
1.3
1.2
1.1
1
0.9
0.8
0.7
0
10
20
30
40
50
60

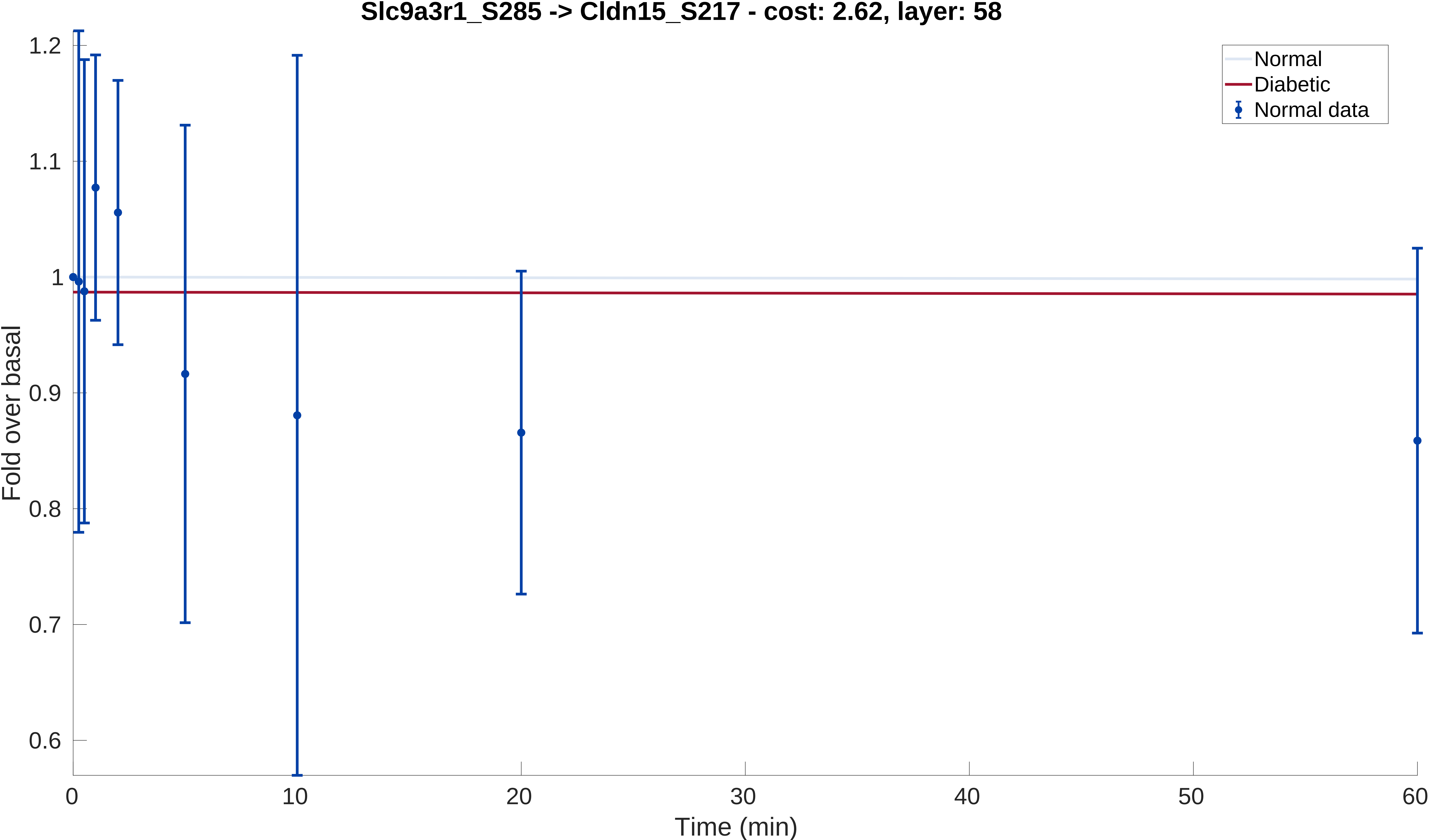

Slc9a3r1_S285 -> Cldn15_S217 - cost: 2.62, layer: 58
Normal
Diabetic
Normal data
Fold over basal
Time (min)
1.2
1.1
1
0.9
0.8
0.7
0.6
0
10
20
30
40
50
60

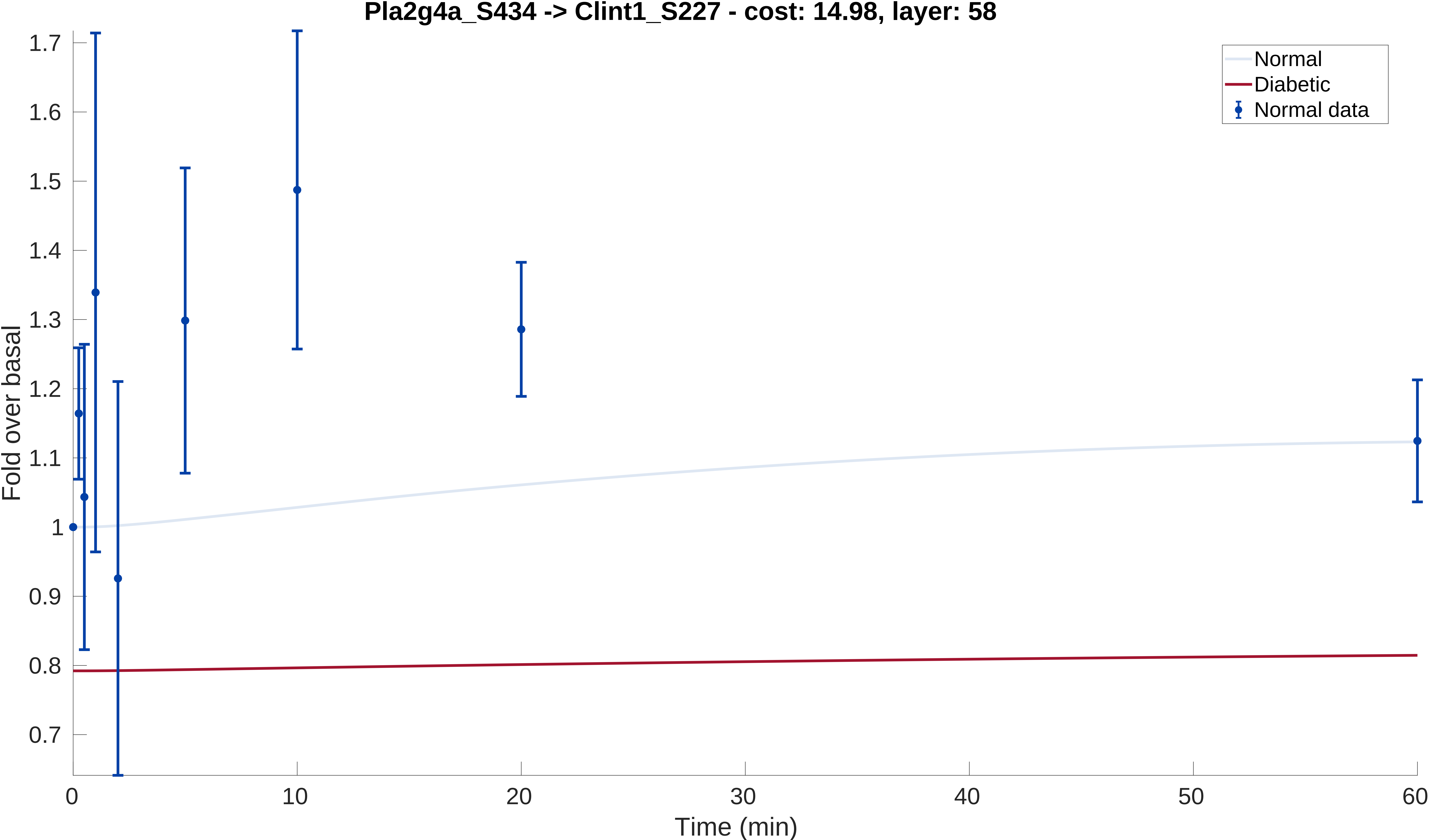

Pla2g4a_S434 -> Clint1_S227 - cost: 14.98, layer: 58
Normal
Diabetic
Normal data
Fold over basal
1.7
1.6
1.5
1.4
1.3
1.2
1.1
1
0.9
0.8
0.7
0
10
20
30
40
50
60
Time (min)

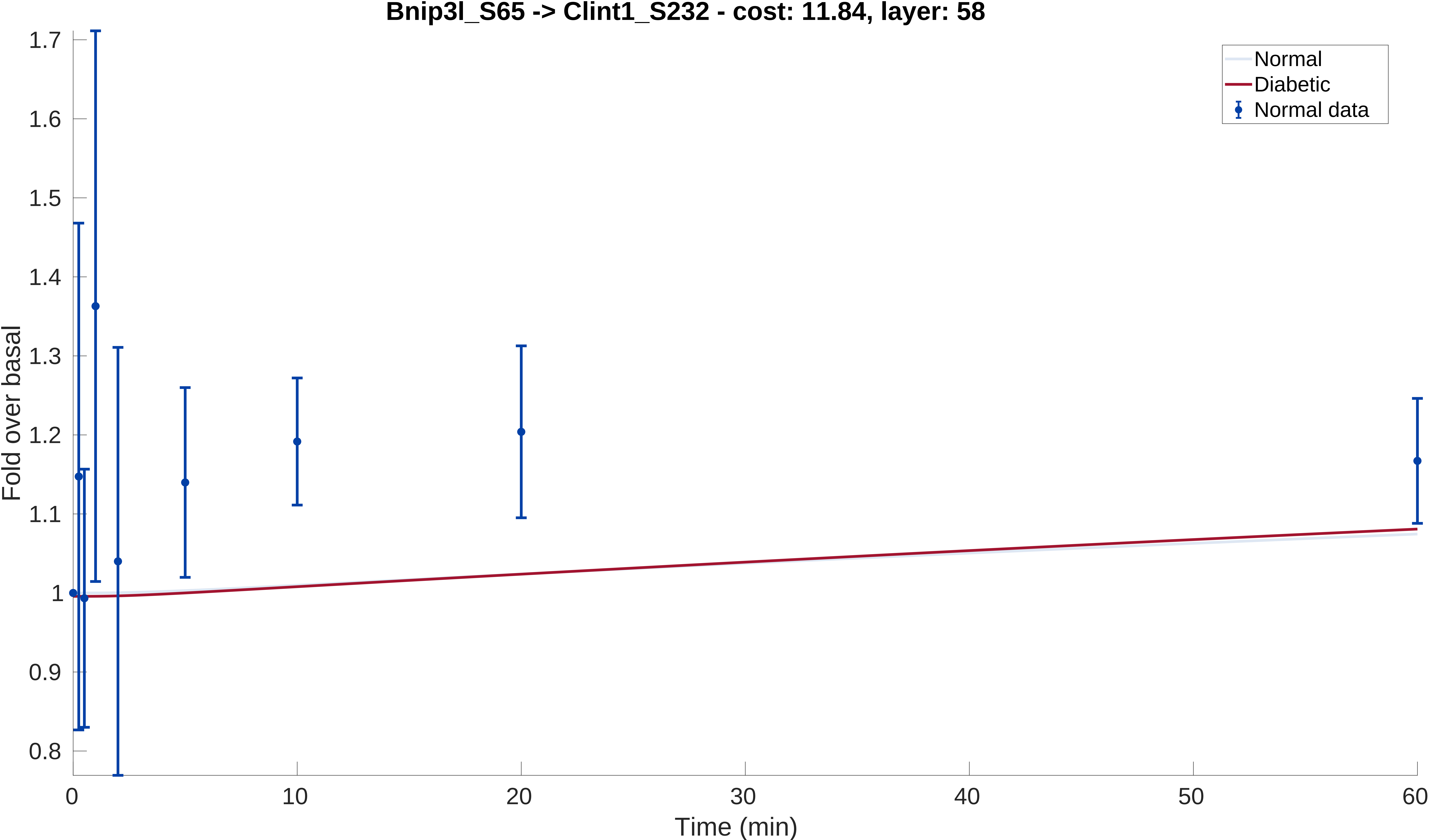
Bnip3l_S65 -> Clint1_S232 - cost: 11.84, layer: 58
Normal
Diabetic
Normal data
Fold over basal
1.7
1.6
1.5
1.4
1.3
1.2
1.1
1
0.9
0.8
0
10
20
30
40
50
60
Time (min)

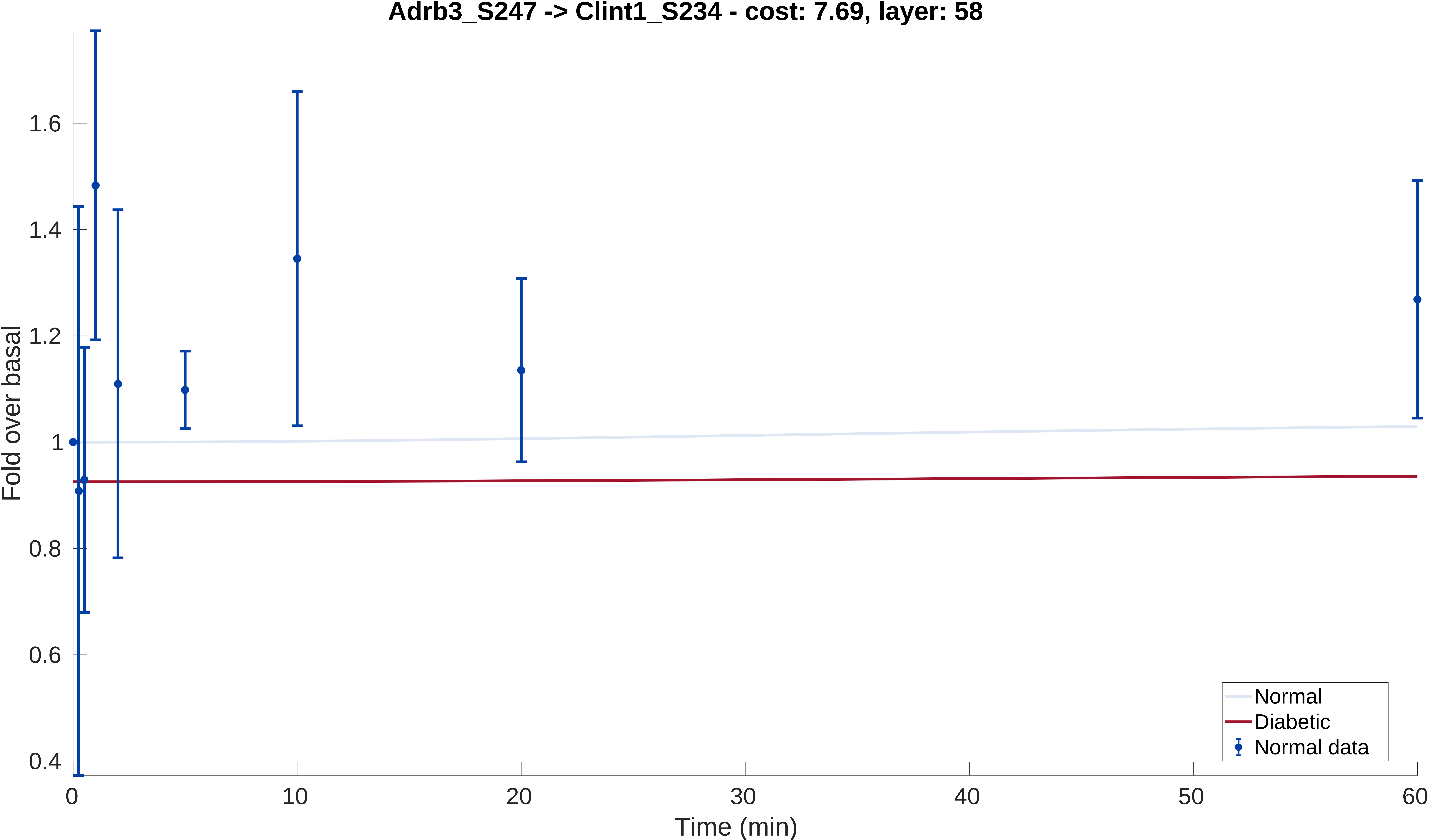

Adrb3_S247 -> Clint1_S234 - cost: 7.69, layer: 58
Fold over basal
1.6
1.4
1.2
1
0.8
0.6
0.4
0
10
20
30
40
50
60
Time (min)
Normal
Diabetic
Normal data

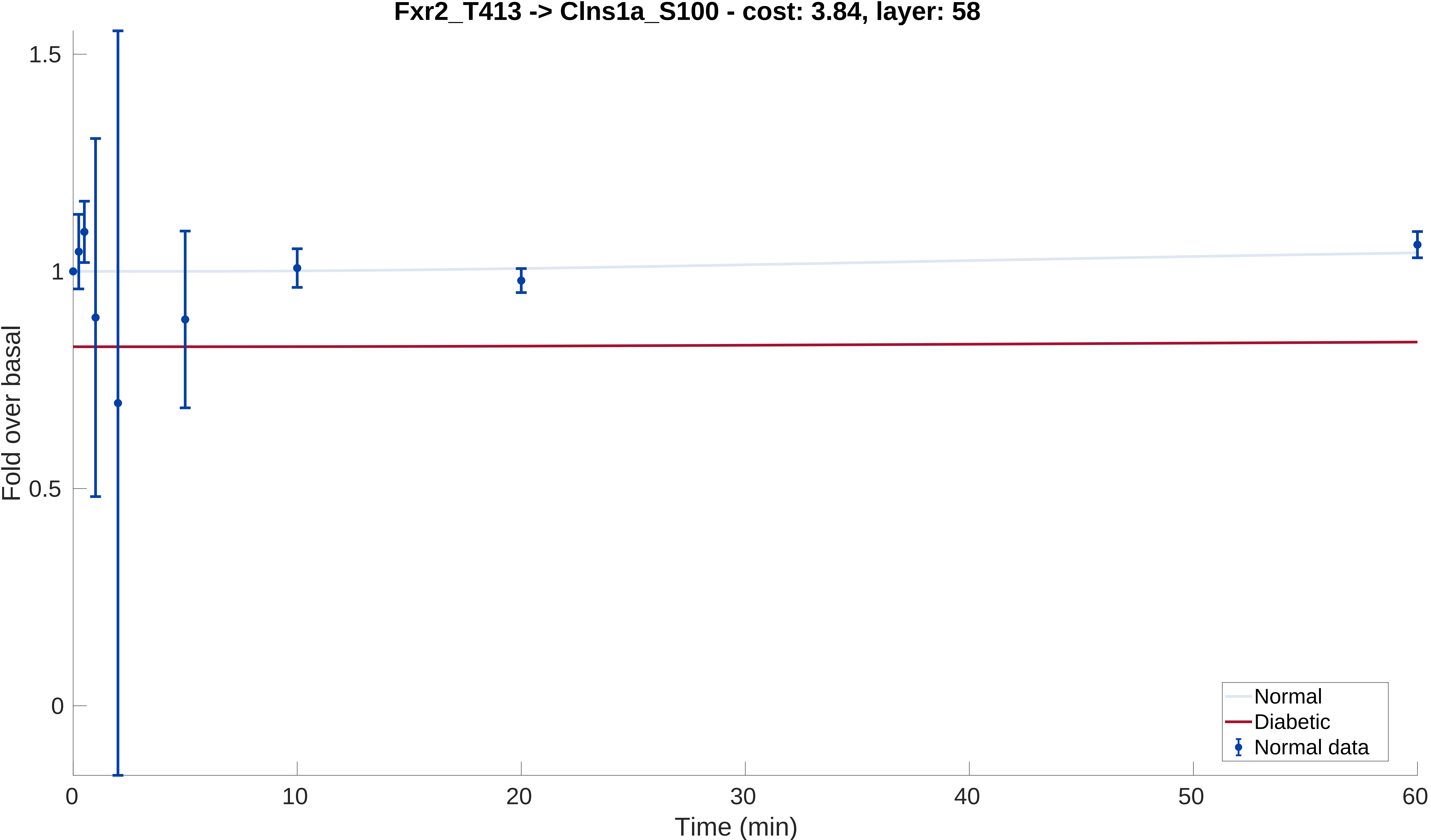

Fxr2_T413 -> Clns1a_S100 - cost: 3.84, layer: 58
1.5
1
0.5
0
Fold over basal
0
10
20
30
40
50
60
Time (min)
Normal
Diabetic
Normal data

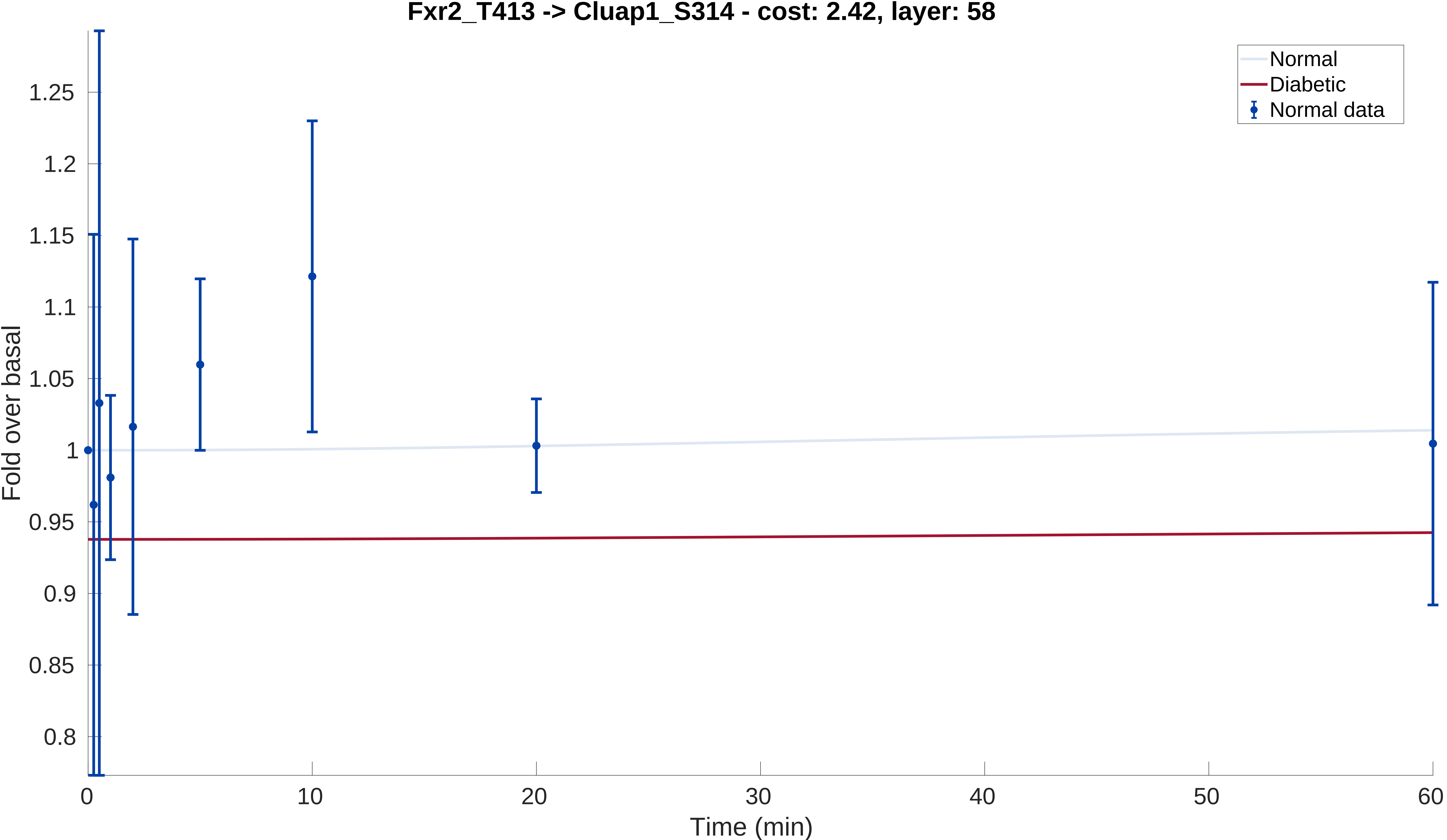
Fxr2_T413 -> Cluap1_S314 - cost: 2.42, layer: 58
Normal
Diabetic
Normal data
Fold over basal
1.25
1.2
1.15
1.1
1.05
1
0.95
0.9
0.85
0.8
0
10
20
30
40
50
60
Time (min)

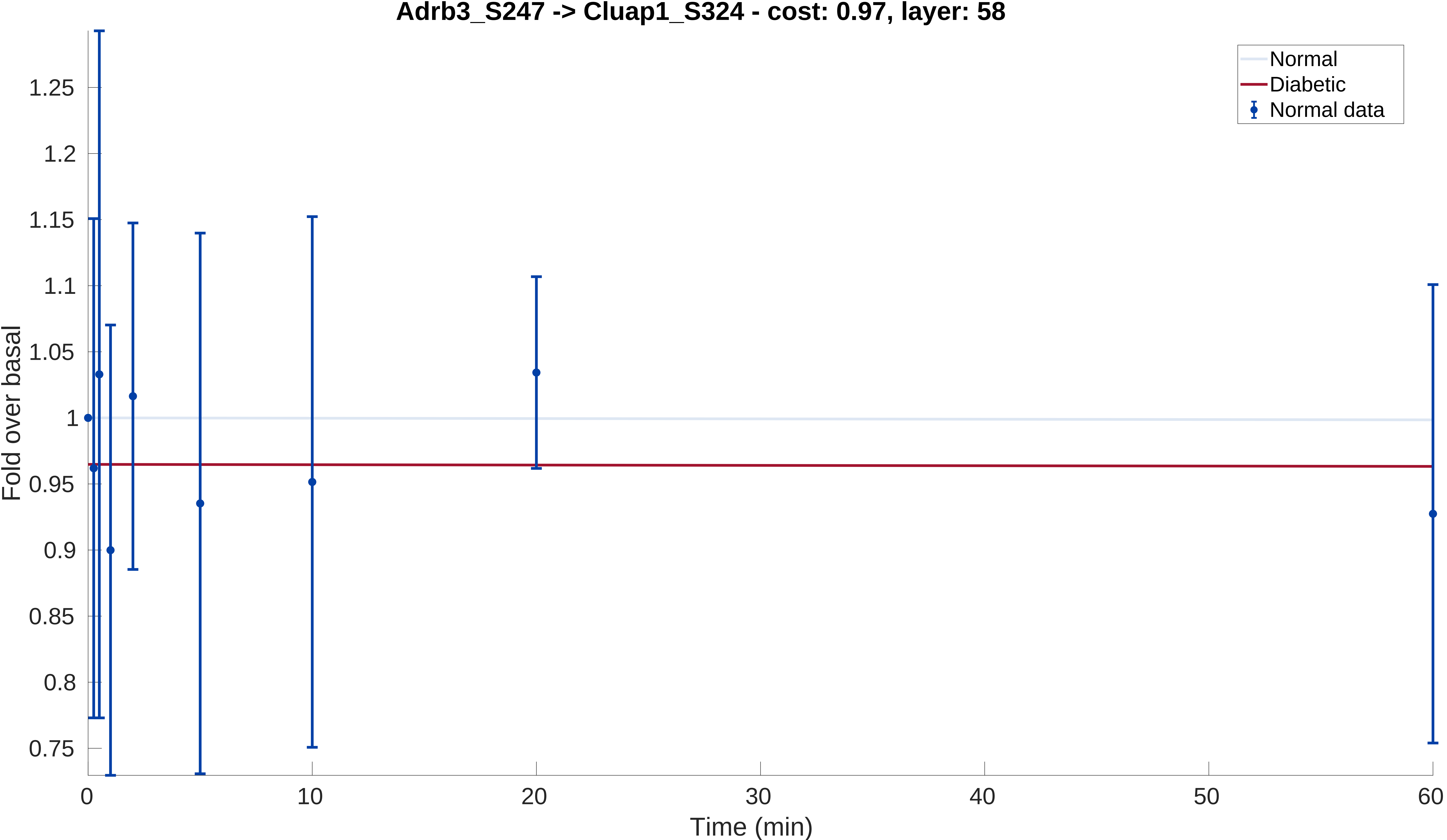
Adrb3_S247 -> Cluap1_S324 - cost: 0.97, layer: 58
Normal
Diabetic
Normal data
Fold over basal
1.25
1.2
1.15
1.1
1.05
1
0.95
0.9
0.85
0.8
0.75
0
10
20
30
40
50
60
Time (min)

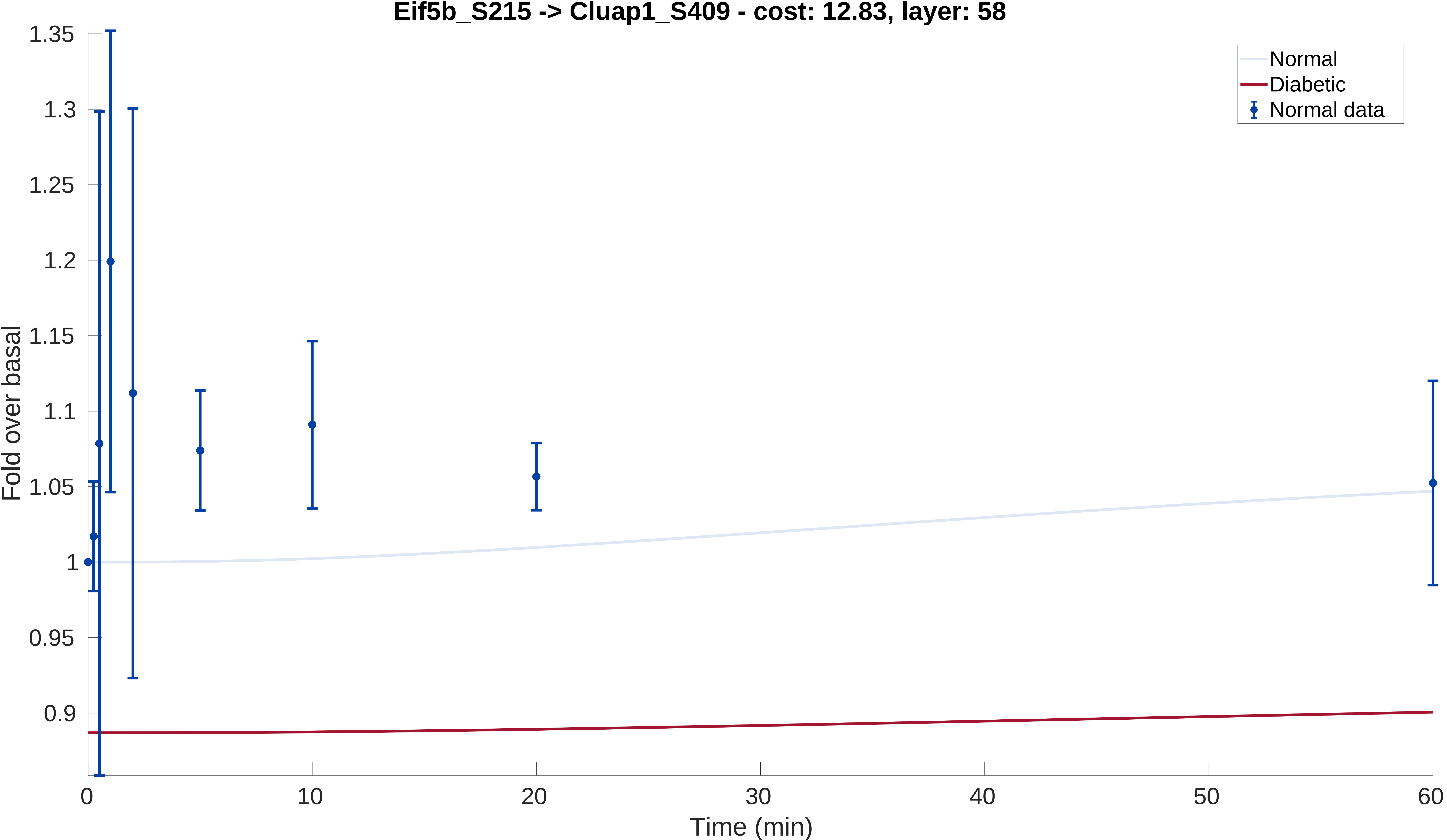
Eif5b_S215 -> Cluap1_S409 - cost: 12.83, layer: 58
Fold over basal
Time (min)
1.35
1.3
1.25
1.2
1.15
1.1
1.05
1
0.95
0.9
0
10
20
30
40
50
60
Normal
Diabetic
Normal data

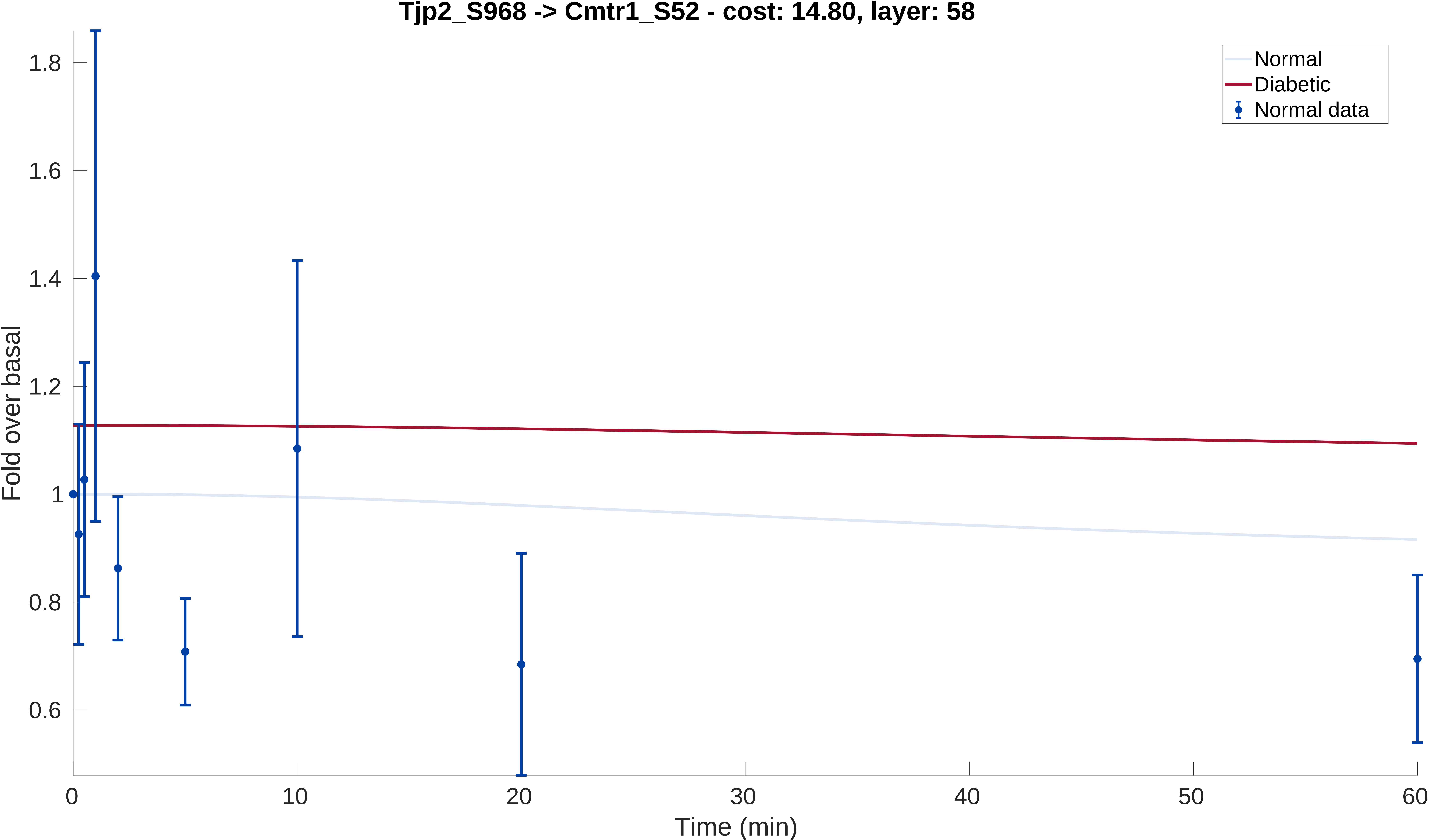

Tjp2_S968 -> Cmtr1_S52 - cost: 14.80, layer: 58
Normal
Diabetic
Normal data
1.8
1.6
1.4
1.2
1
0.8
0.6
Fold over basal
0
10
20
30
40
50
60
Time (min)

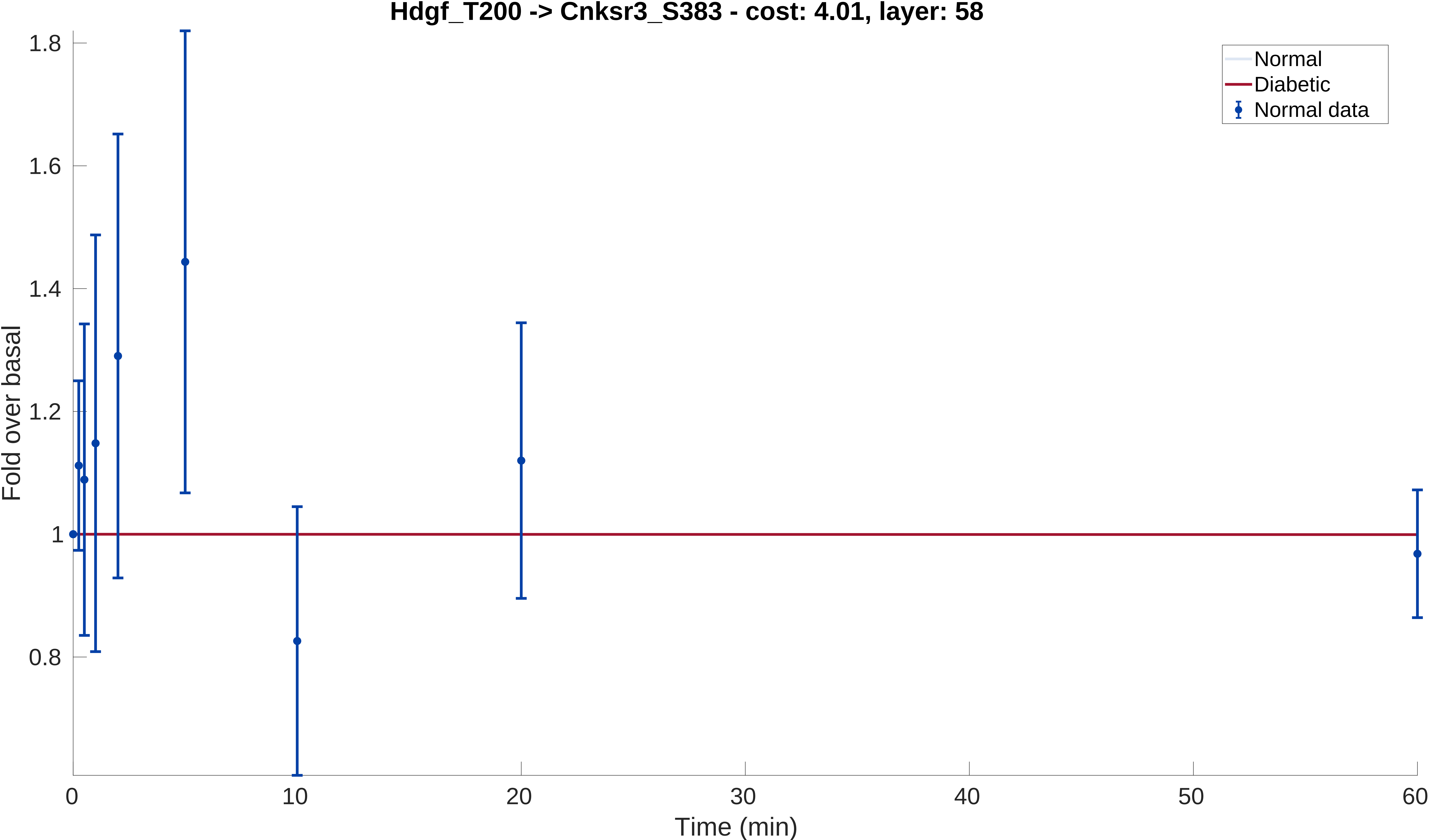
Hdgf_T200 -> Cnksr3_S383 - cost: 4.01, layer: 58
Normal
Diabetic
Normal data
Fold over basal
1.8
1.6
1.4
1.2
1
0.8
0
10
20
30
40
50
60
Time (min)

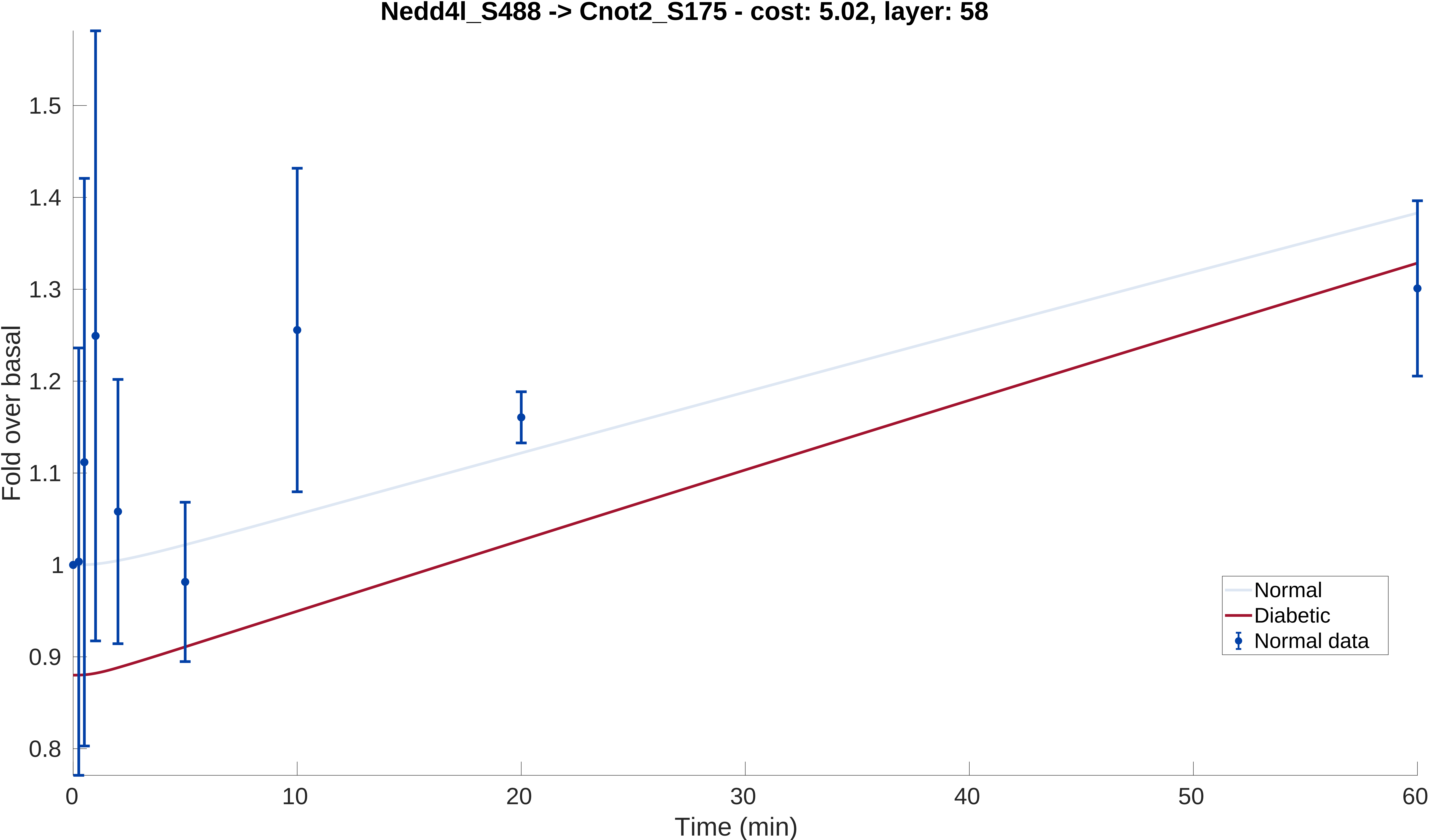
Nedd4l_S488 -> Cnot2_S175 - cost: 5.02, layer: 58
Fold over basal
1.5
1.4
1.3
1.2
1.1
1
0.9
0.8
0
10
20
30
40
50
60
Time (min)
Normal
Diabetic
Normal data

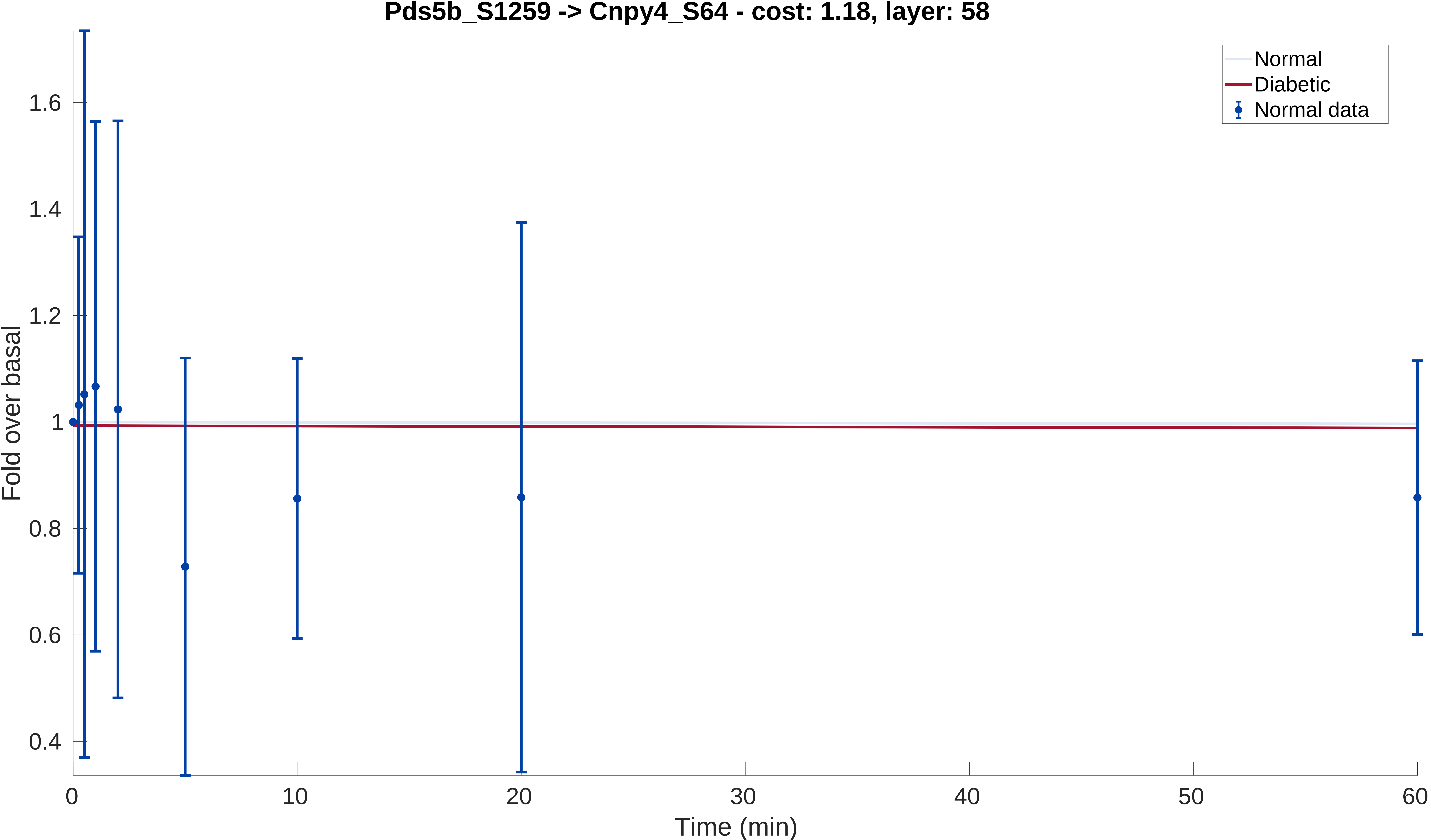
Pds5b_S1259 -> Cnpy4_S64 - cost: 1.18, layer: 58
Normal
Diabetic
Normal data
Fold over basal
Time (min)
0.4
0.6
0.8
1
1.2
1.4
1.6
0
10
20
30
40
50
60

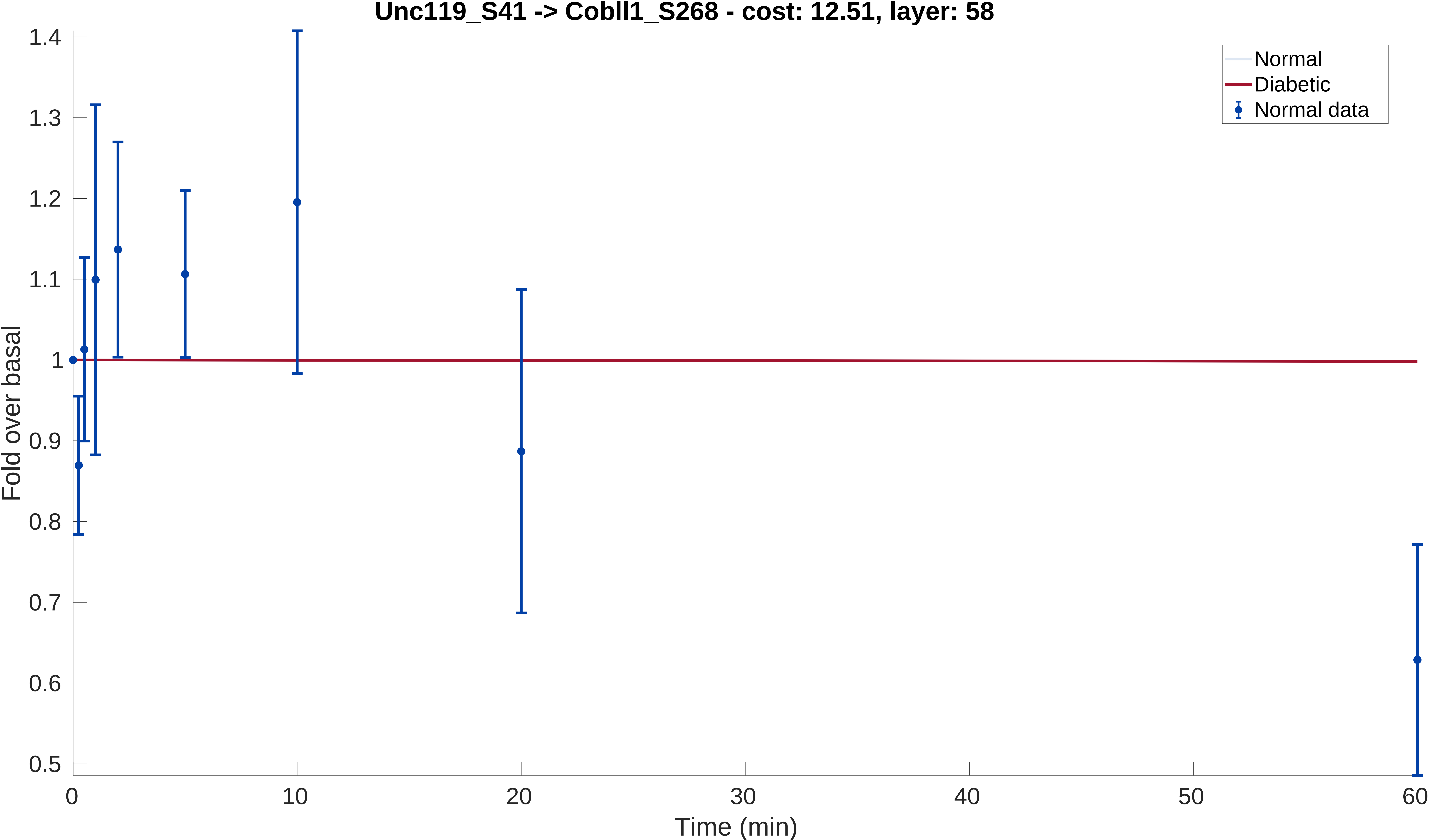

Unc119_S41 -> Cobll1_S268 - cost: 12.51, layer: 58
Fold over basal
Time (min)
Normal
Diabetic
Normal data
1.4
1.3
1.2
1.1
1
0.9
0.8
0.7
0.6
0.5
0
10
20
30
40
50
60

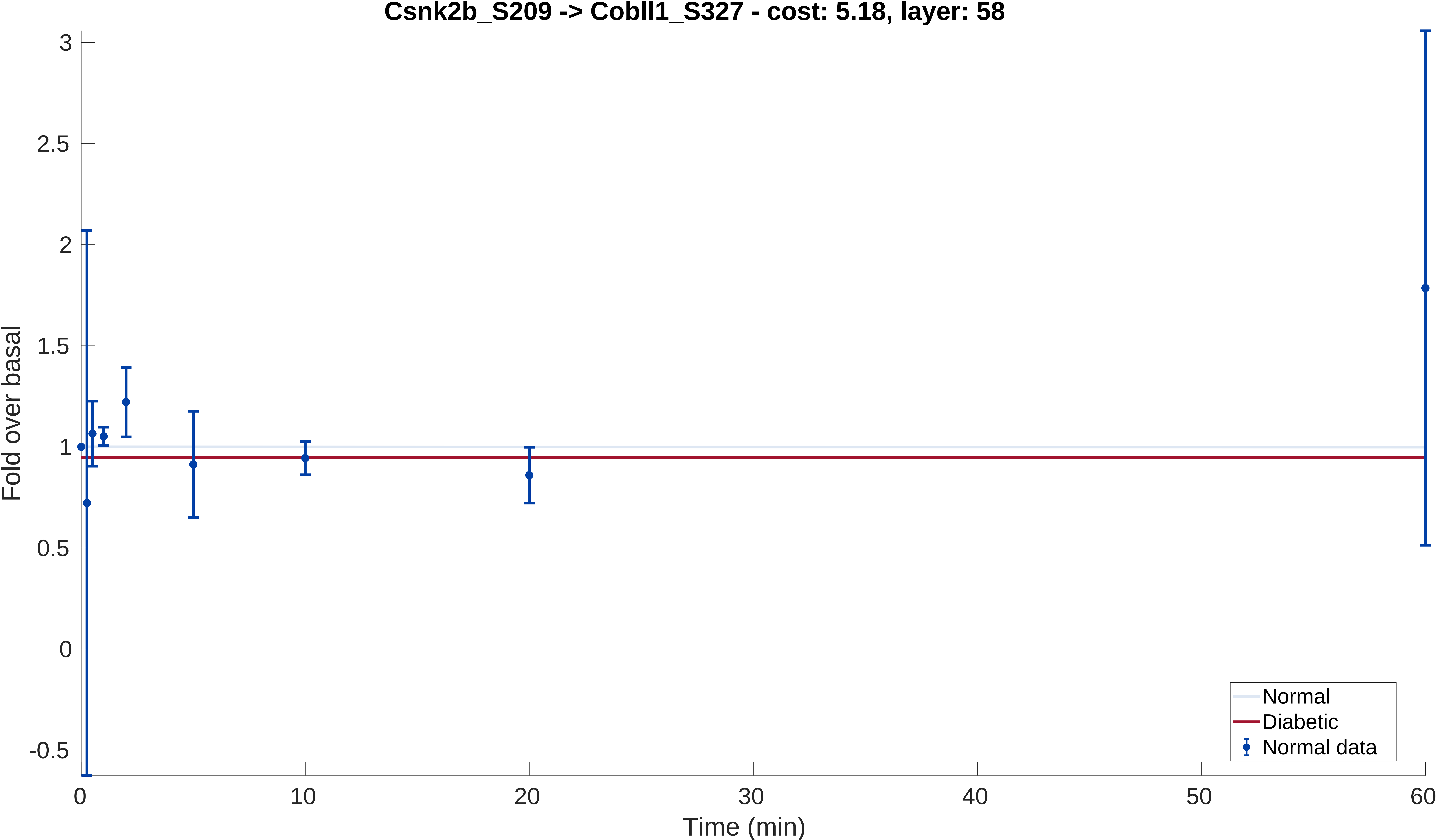

Csnk2b_S209 -> Cobll1_S327 - cost: 5.18, layer: 58
Fold over basal
3
2.5
2
1.5
1
0.5
0
-0.5
0
10
20
30
40
50
60
Time (min)
Normal
Diabetic
Normal data

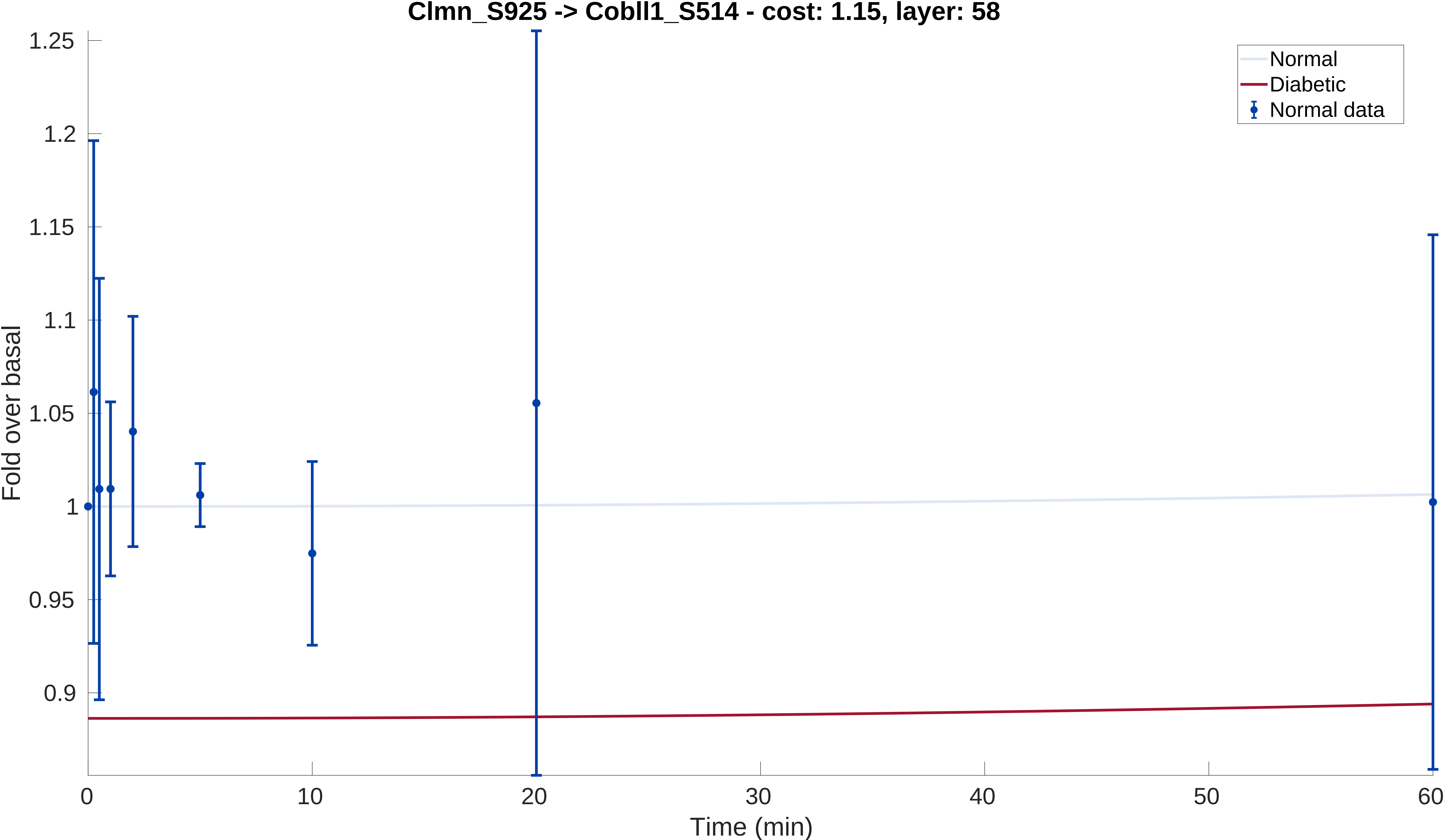

Clmn_S925 -> Cobll1_S514 - cost: 1.15, layer: 58
Normal
Diabetic
Normal data
Fold over basal
Time (min)
1.25
1.2
1.15
1.1
1.05
1
0.95
0.9
0
10
20
30
40
50
60

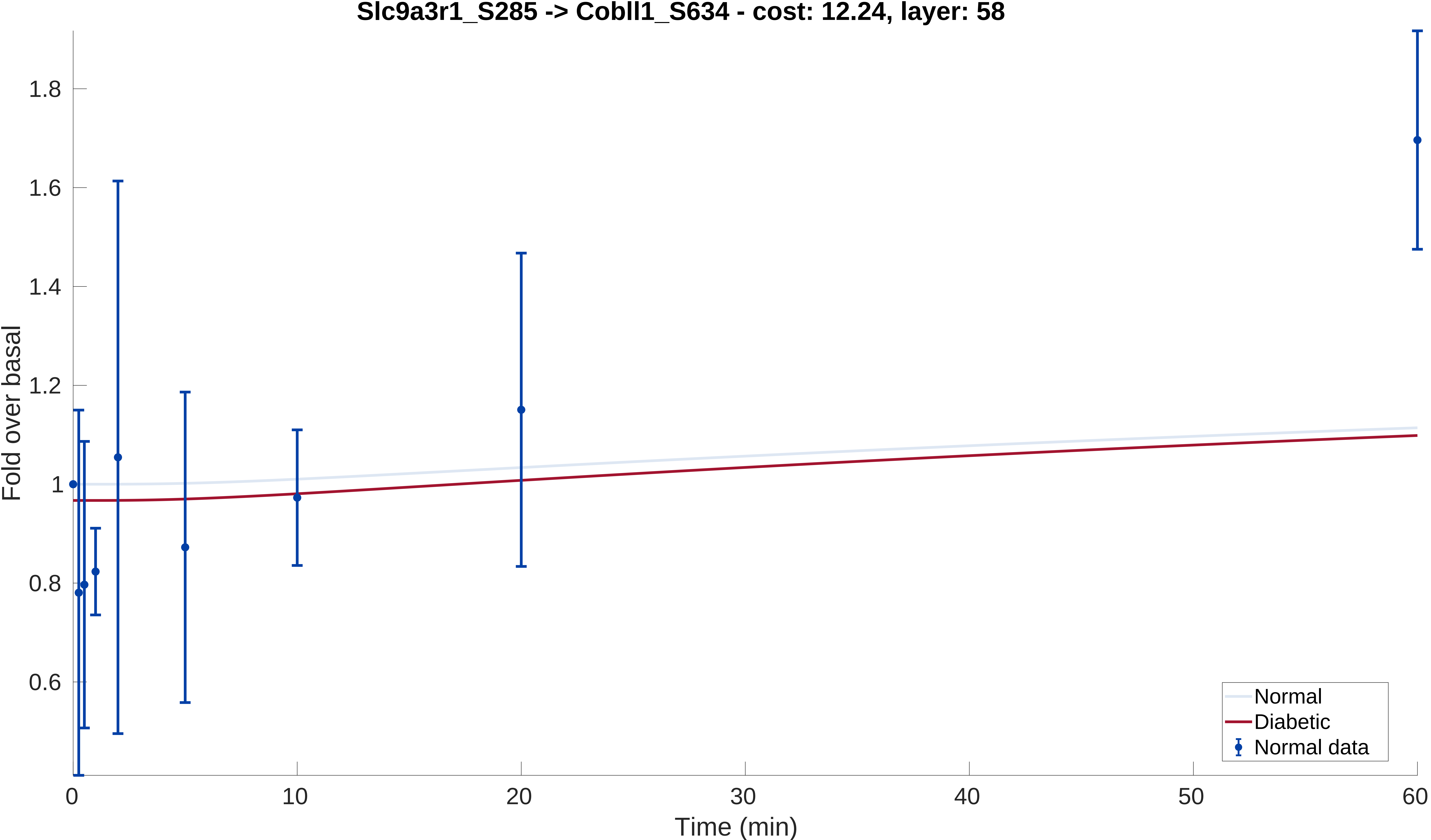

Slc9a3r1_S285 -> Cobll1_S634 - cost: 12.24, layer: 58
Fold over basal
1.8
1.6
1.4
1.2
1
0.8
0.6
0
10
20
30
40
50
60
Time (min)
Normal
Diabetic
Normal data

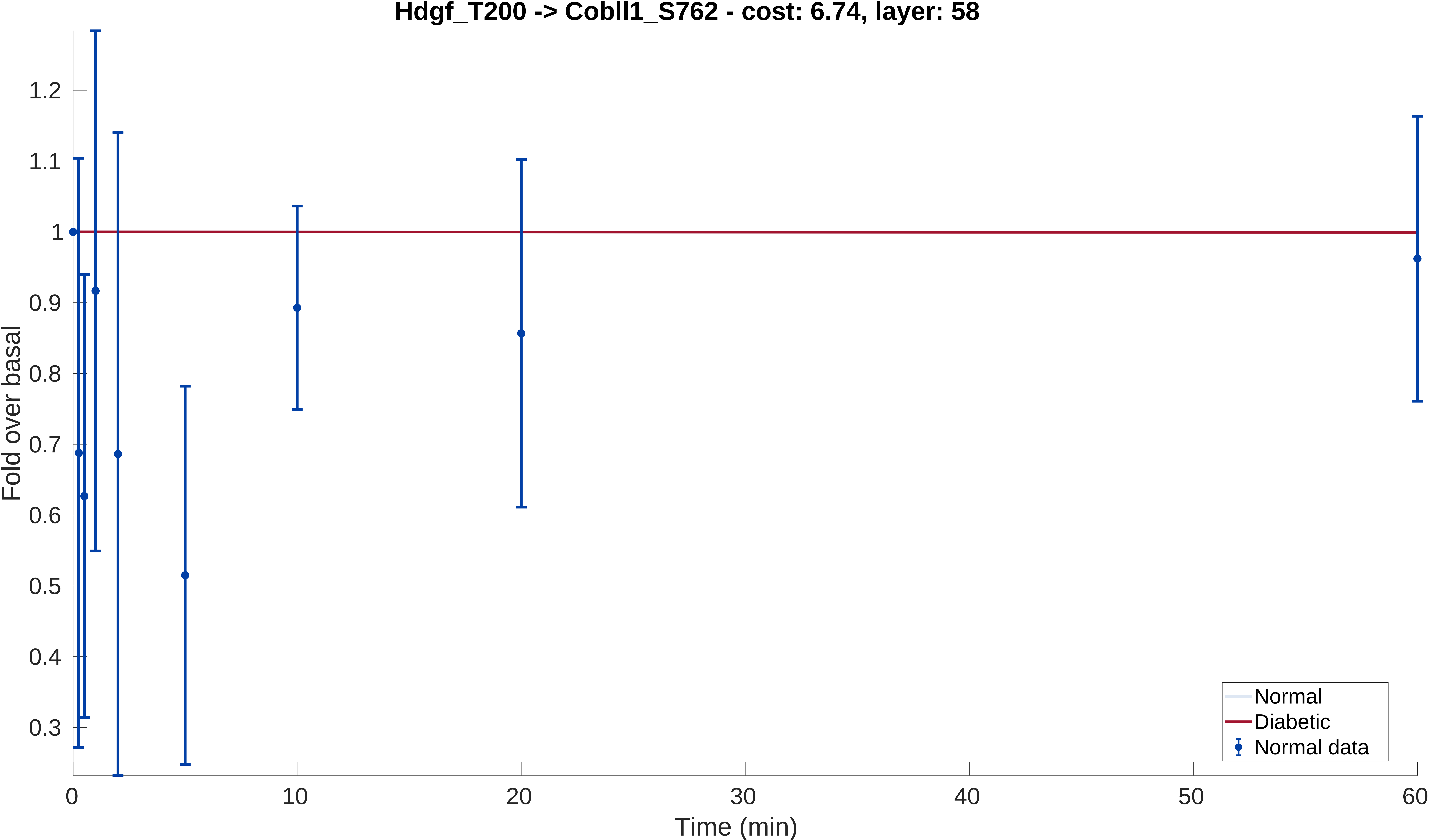

Hdgf_T200 -> Cobll1_S762 - cost: 6.74, layer: 58
Fold over basal
1.2
1.1
1
0.9
0.8
0.7
0.6
0.5
0.4
0.3
0
10
20
30
40
50
60
Time (min)
Normal
Diabetic
Normal data

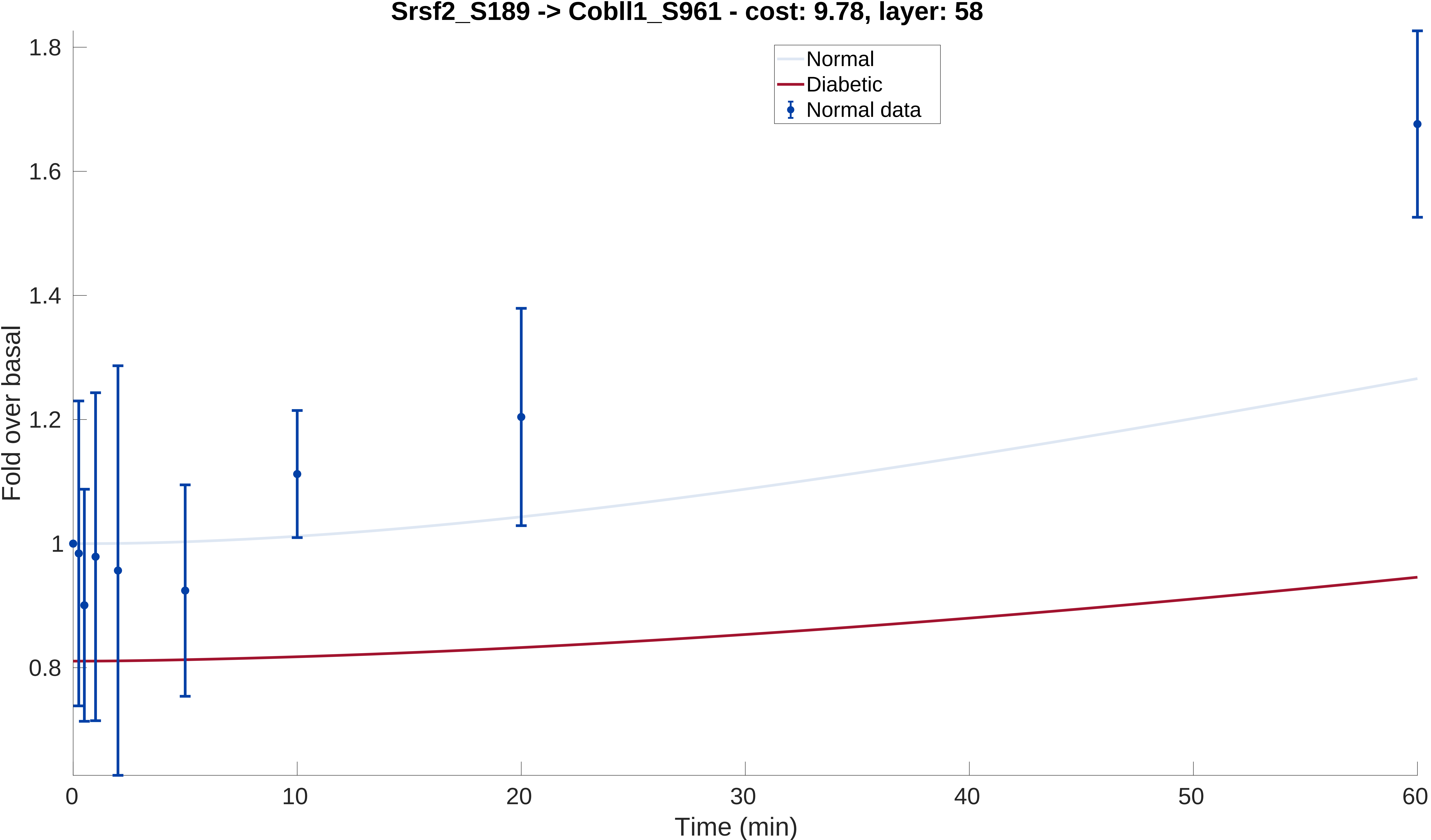

Srsf2_S189 -> Cobll1_S961 - cost: 9.78, layer: 58
Normal
Diabetic
Normal data
Fold over basal
1.8
1.6
1.4
1.2
1
0.8
0
10
20
30
40
50
60
Time (min)

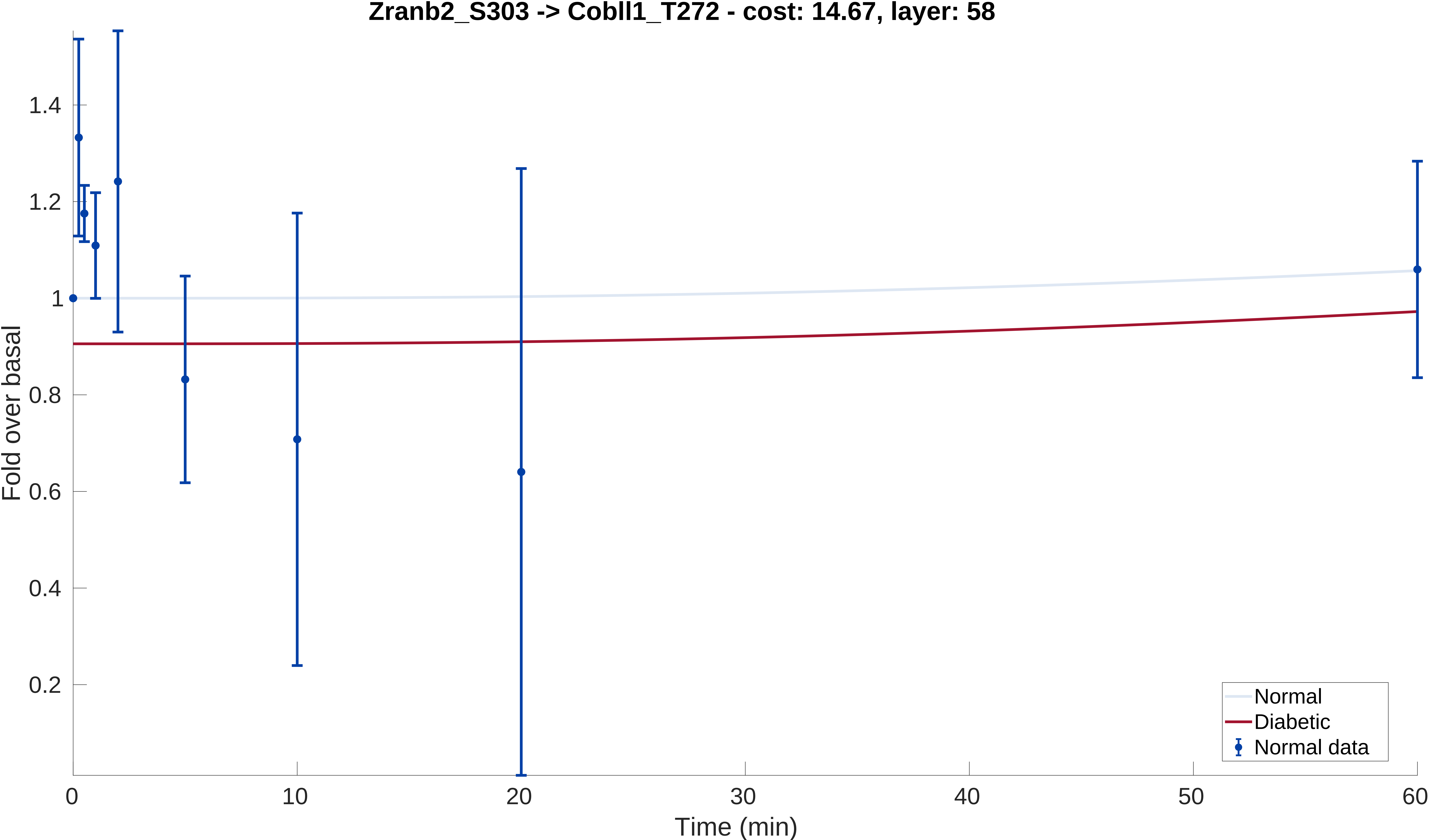

Zranb2_S303 -> Cobll1_T272 - cost: 14.67, layer: 58
Fold over basal
1.4
1.2
1
0.8
0.6
0.4
0.2
0
10
20
30
40
50
60
Time (min)
Normal
Diabetic
Normal data

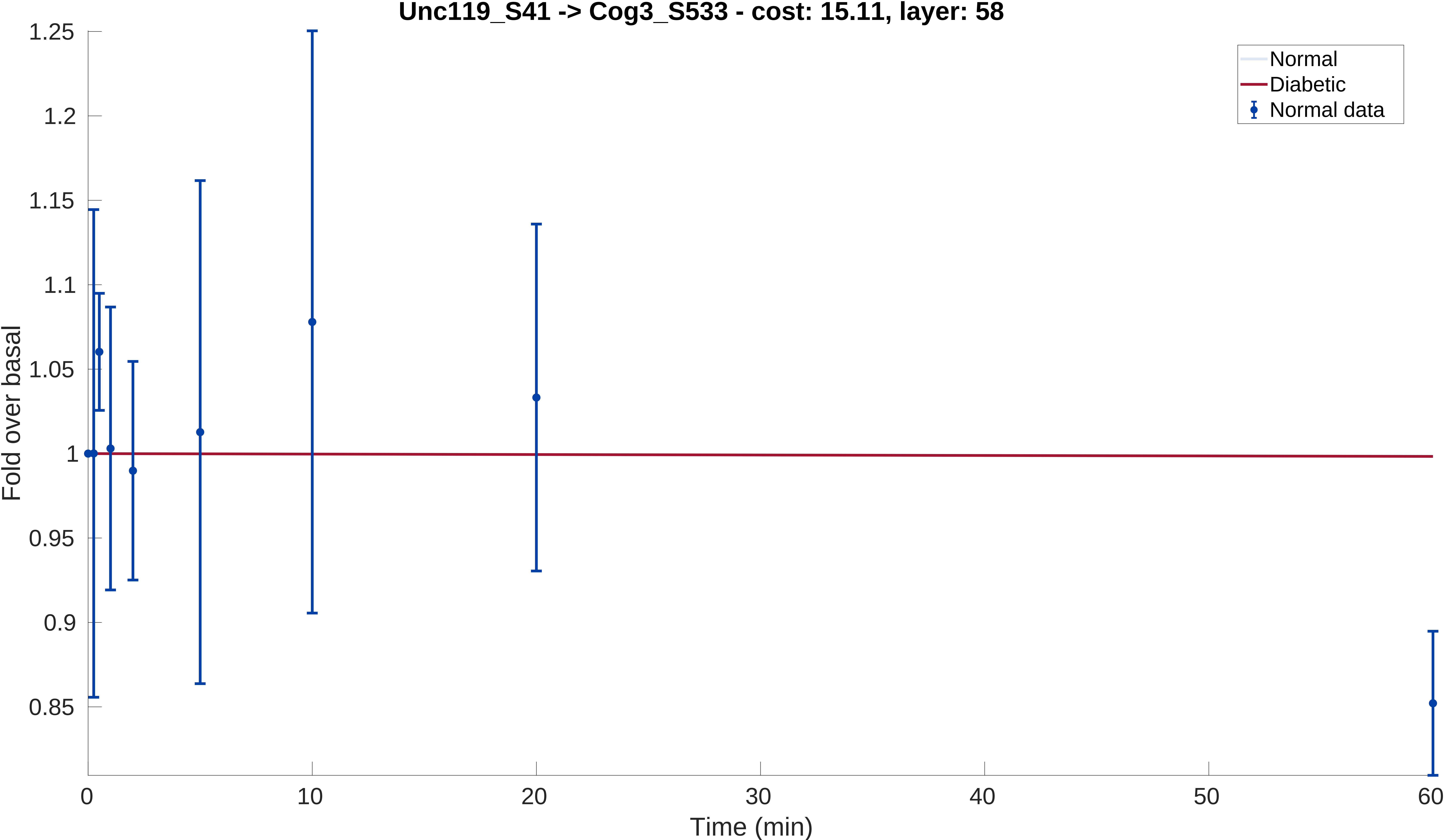
Unc119_S41 -> Cog3_S533 - cost: 15.11, layer: 58
1.25
1.2
1.15
1.1
1.05
1
0.95
0.9
0.85
Fold over basal
0
10
20
30
40
50
60
Time (min)
Normal
Diabetic
Normal data

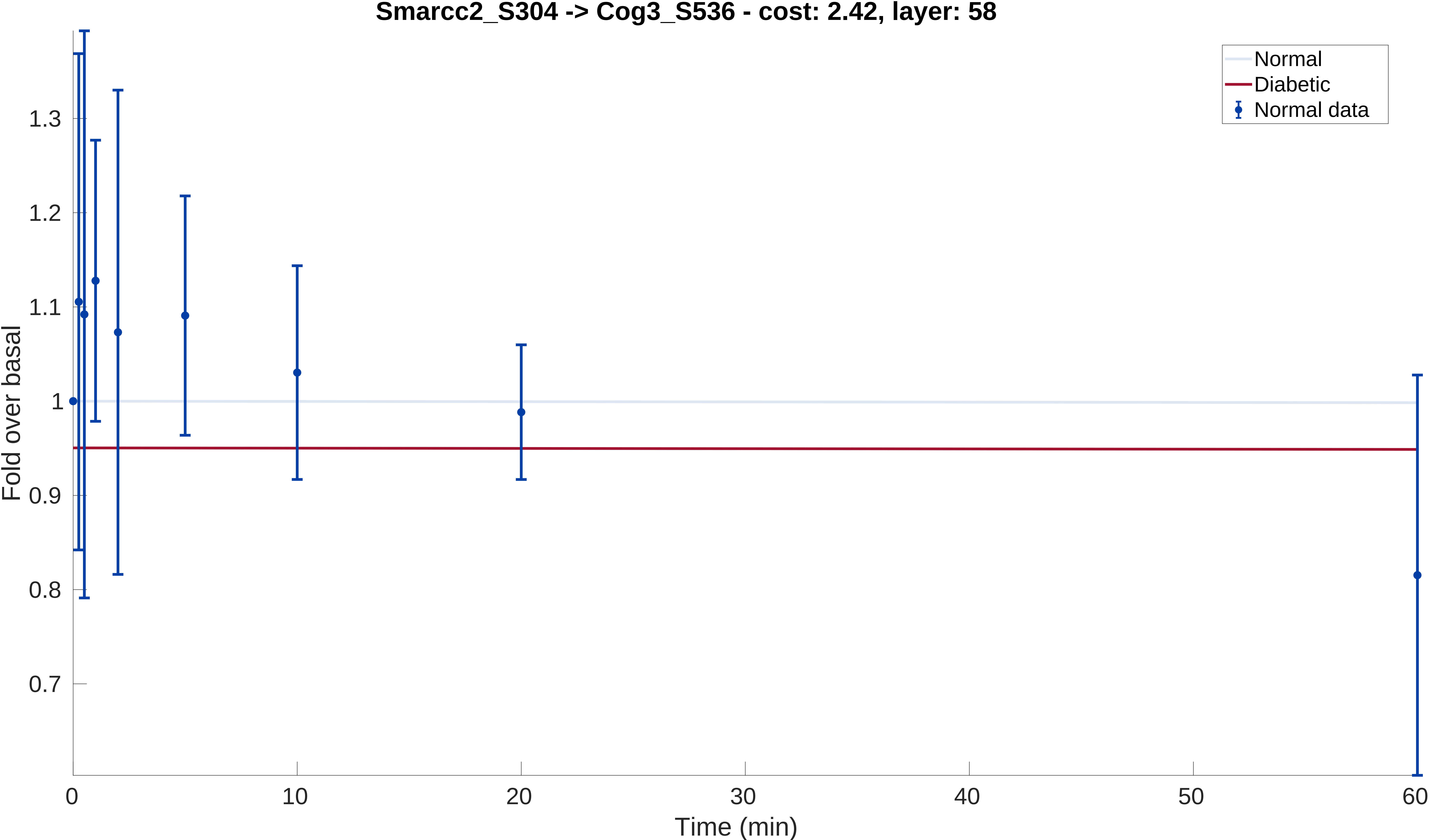

Smarcc2_S304 -> Cog3_S536 - cost: 2.42, layer: 58
Normal
Diabetic
Normal data
Fold over basal
1.3
1.2
1.1
1
0.9
0.8
0.7
0
10
20
30
40
50
60
Time (min)

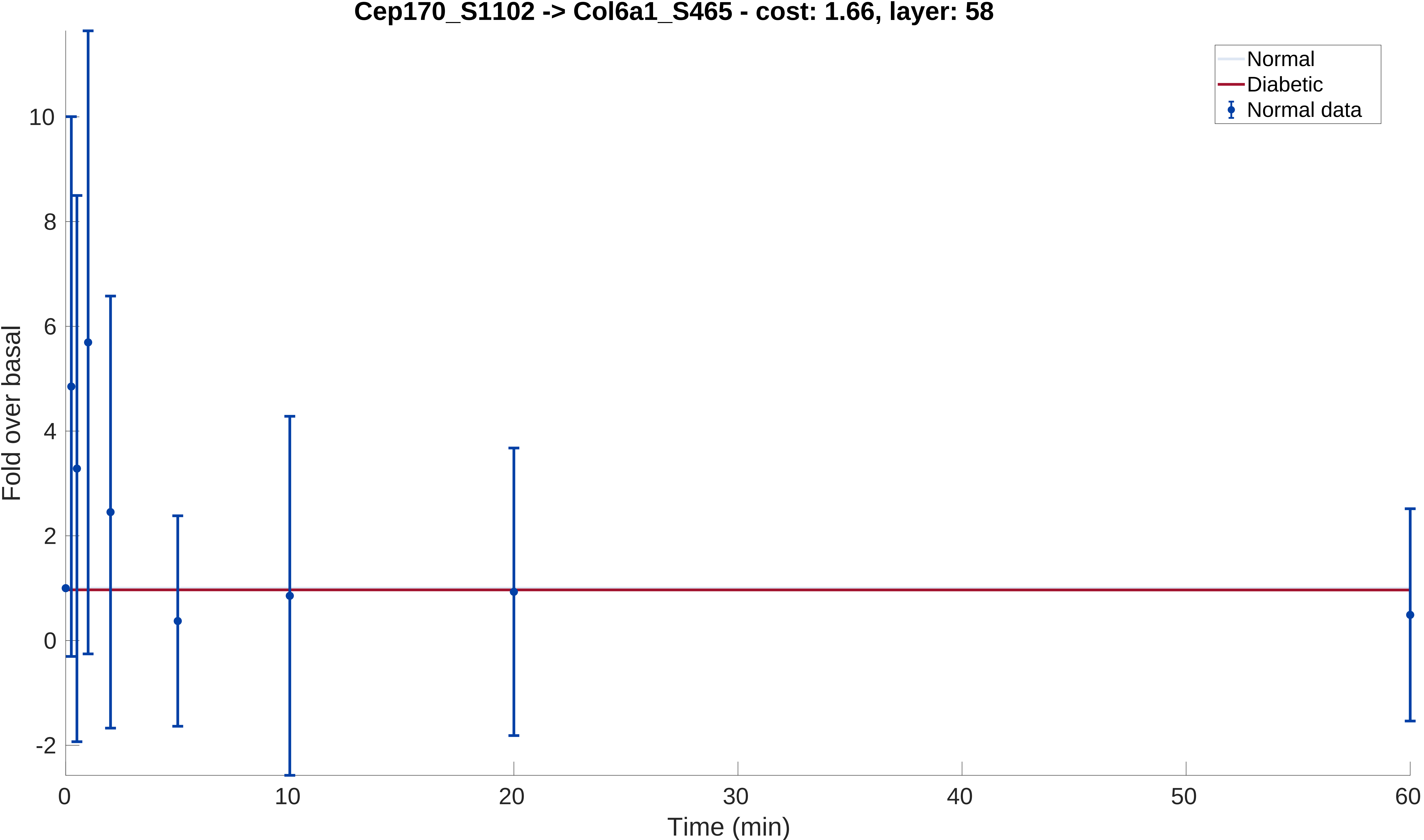

Cep170_S1102 -> Col6a1_S465 - cost: 1.66, layer: 58
Normal
Diabetic
Normal data
Fold over basal
Time (min)
10
8
6
4
2
0
-2
0
10
20
30
40
50
60

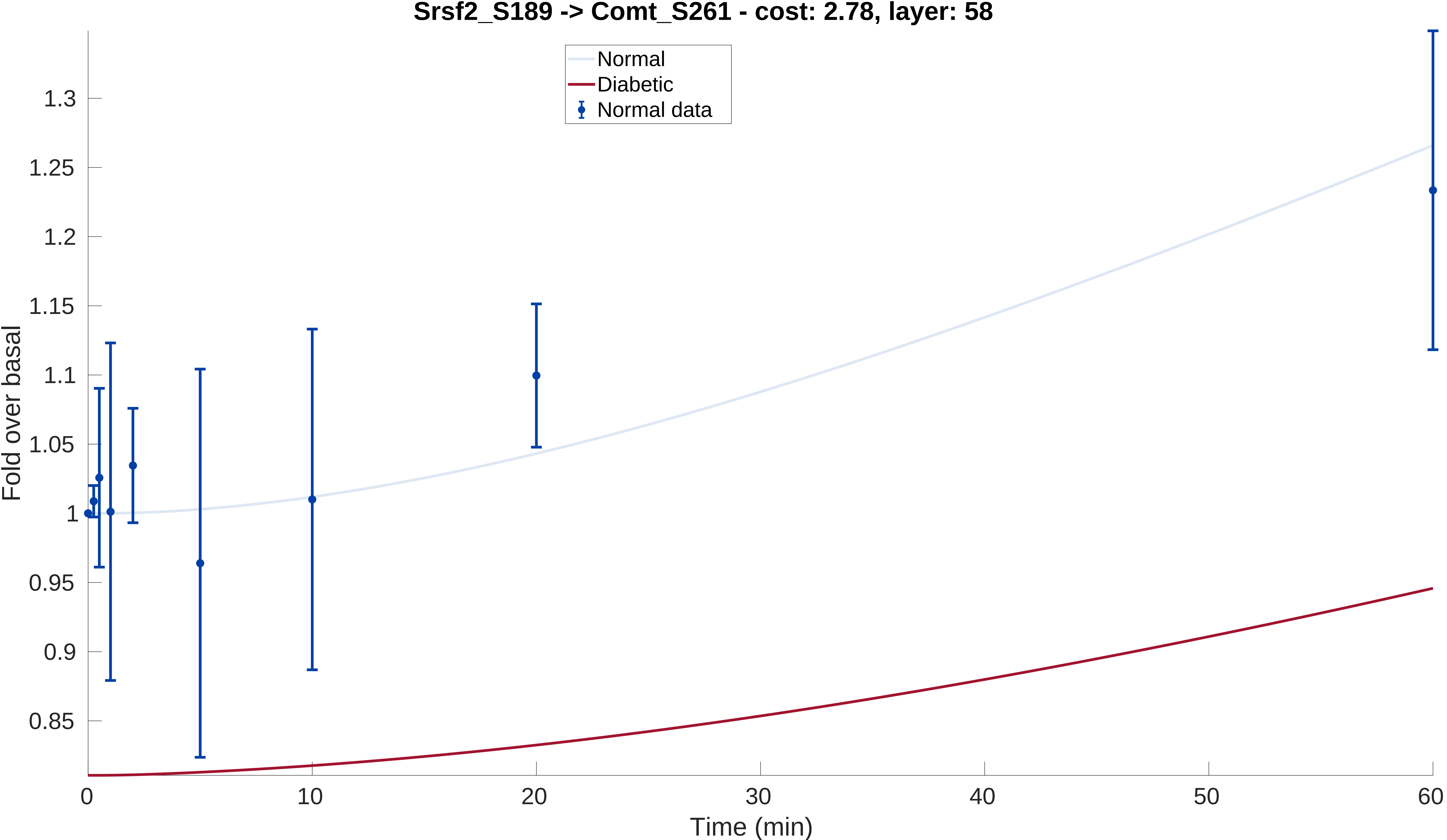
Srsf2_S189 -> Comt_S261 - cost: 2.78, layer: 58
Normal
Diabetic
Normal data
Fold over basal
1.3
1.25
1.2
1.15
1.1
1.05
1
0.95
0.9
0.85
0
10
20
30
40
50
60
Time (min)

#### Prkar2b_S112 -> Coq9_S81 - cost: 0.77, layer: 58

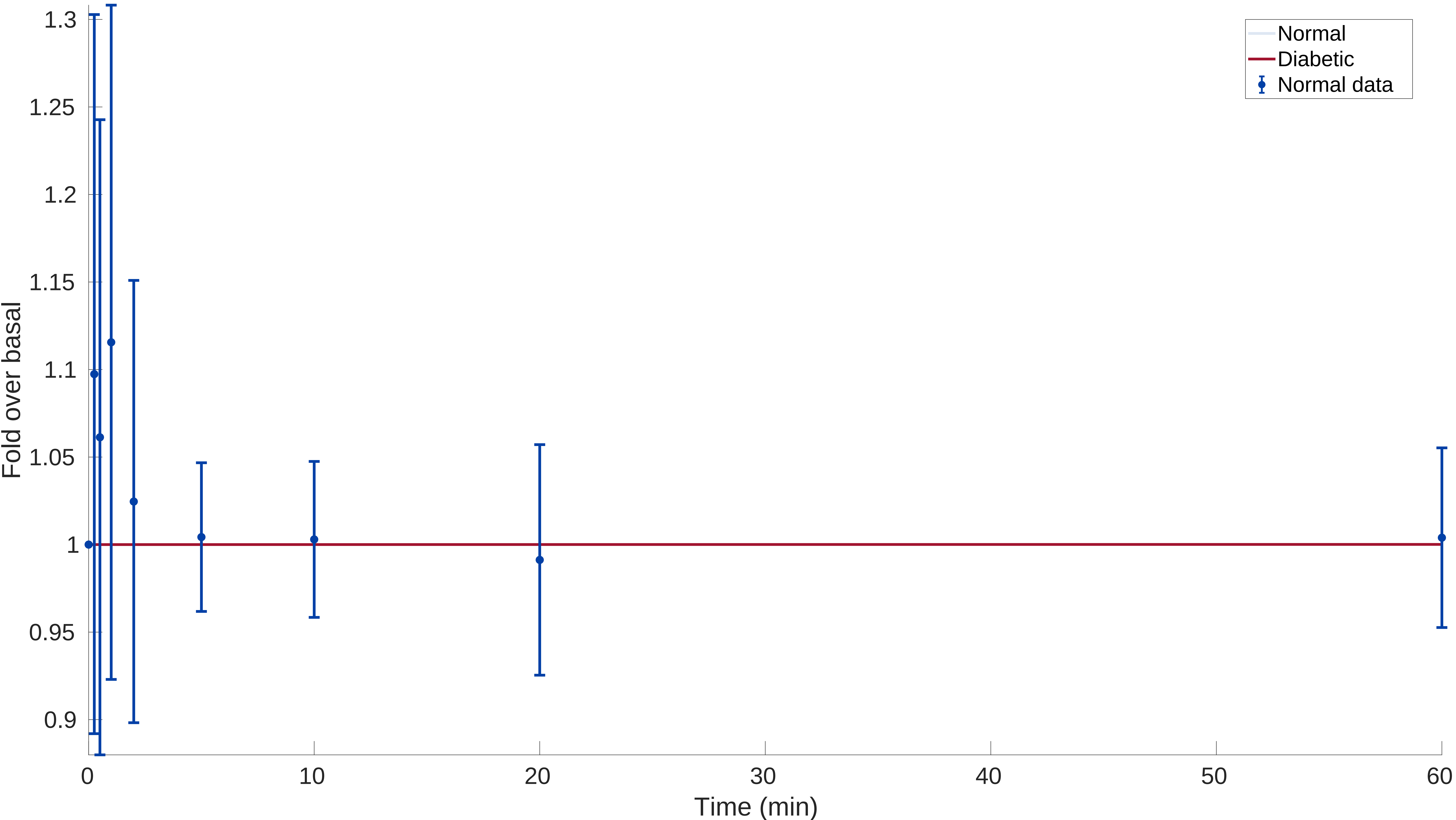

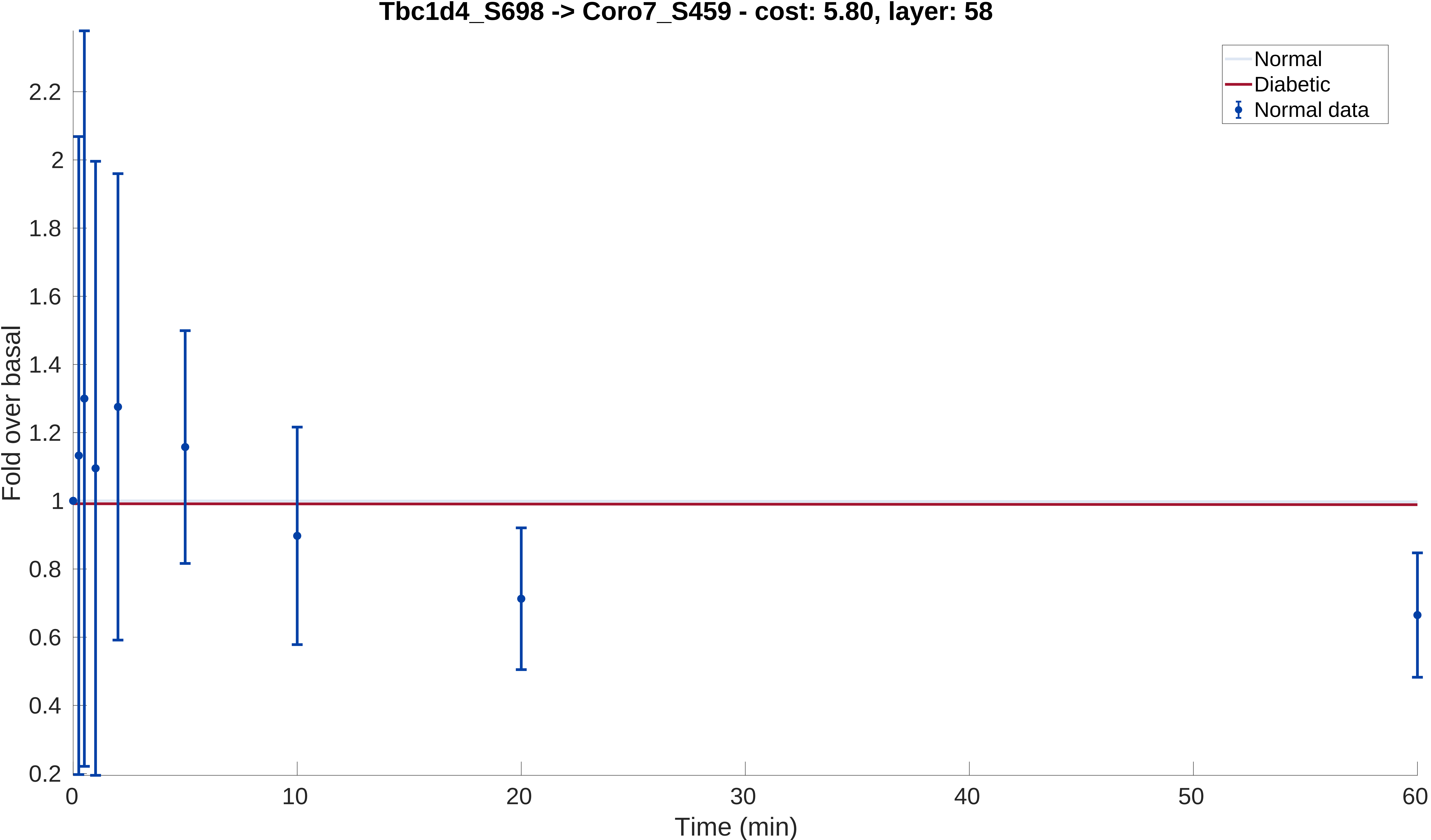

Tbc1d4_S698 -> Coro7_S459 - cost: 5.80, layer: 58
Normal
Diabetic
Normal data
Fold over basal
2.2
2
1.8
1.6
1.4
1.2
1
0.8
0.6
0.4
0.2
0
10
20
30
40
50
60
Time (min)

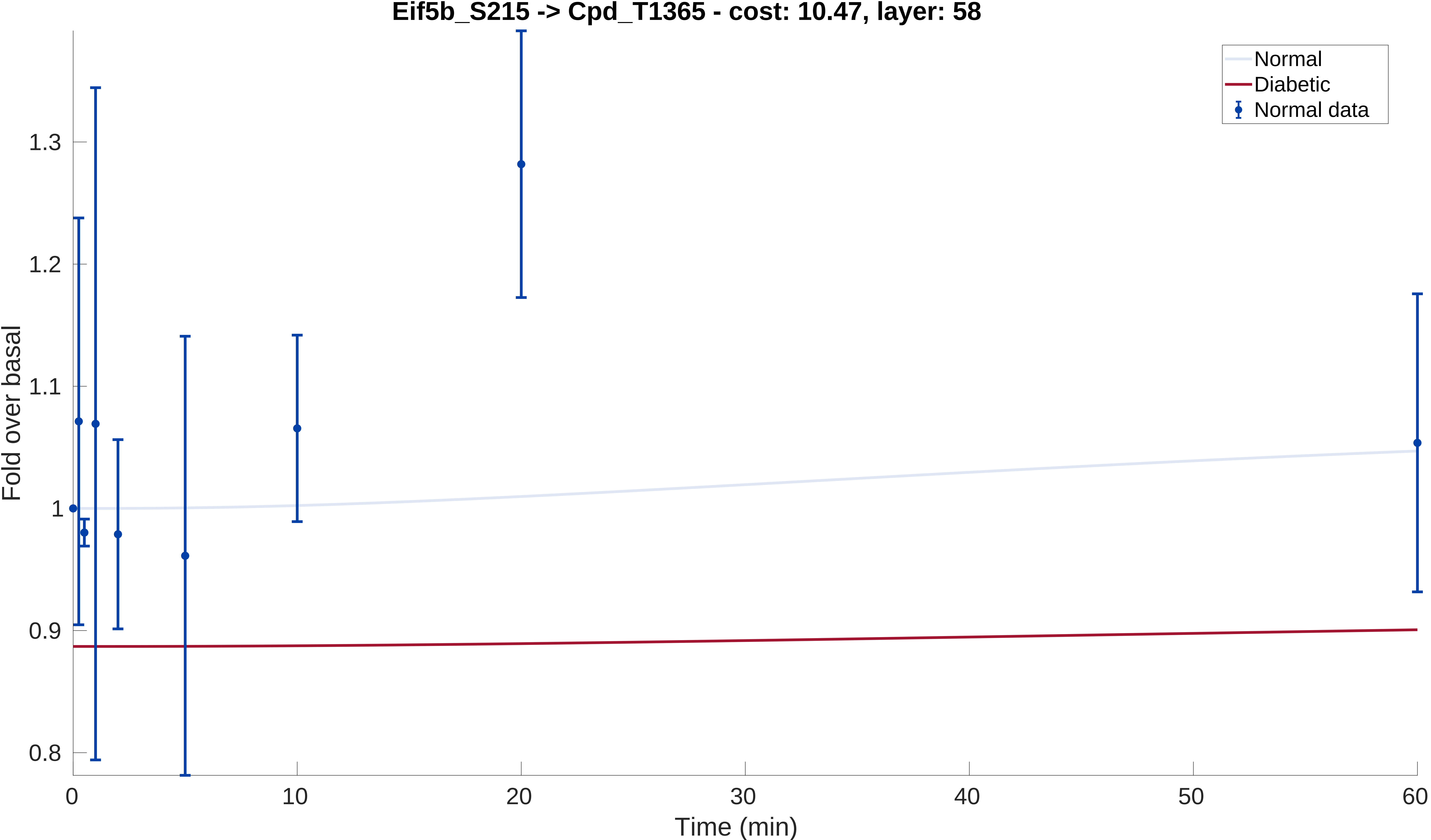

Eif5b_S215 -> Cpd_T1365 - cost: 10.47, layer: 58
Normal
Diabetic
Normal data
1.3
1.2
1.1
1
0.9
0.8
Fold over basal
0
10
20
30
40
50
60
Time (min)

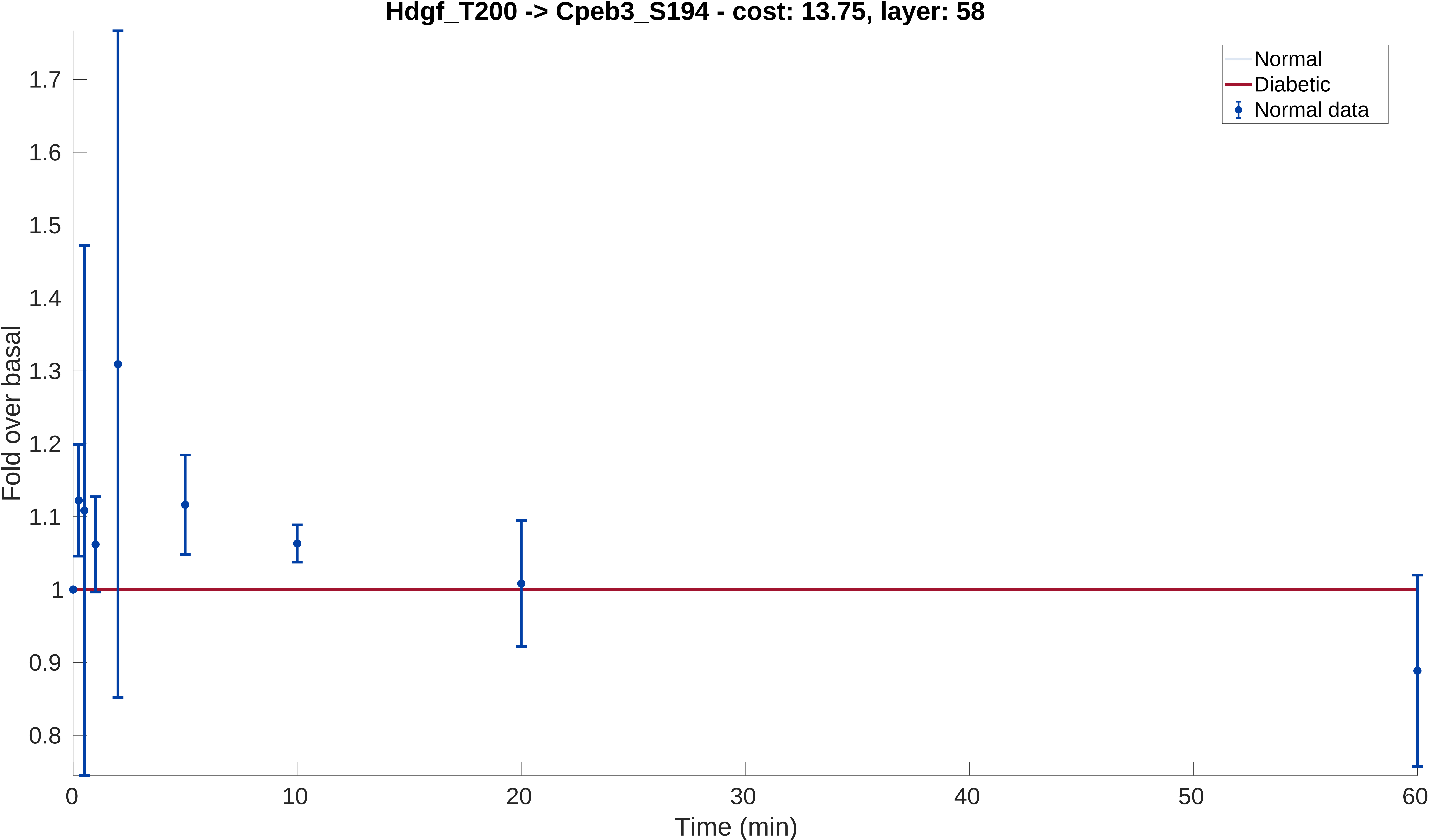

Hdgf_T200 -> Cpeb3_S194 - cost: 13.75, layer: 58
Normal
Diabetic
Normal data
Fold over basal
1.7
1.6
1.5
1.4
1.3
1.2
1.1
1
0.9
0.8
0
10
20
30
40
50
60
Time (min)

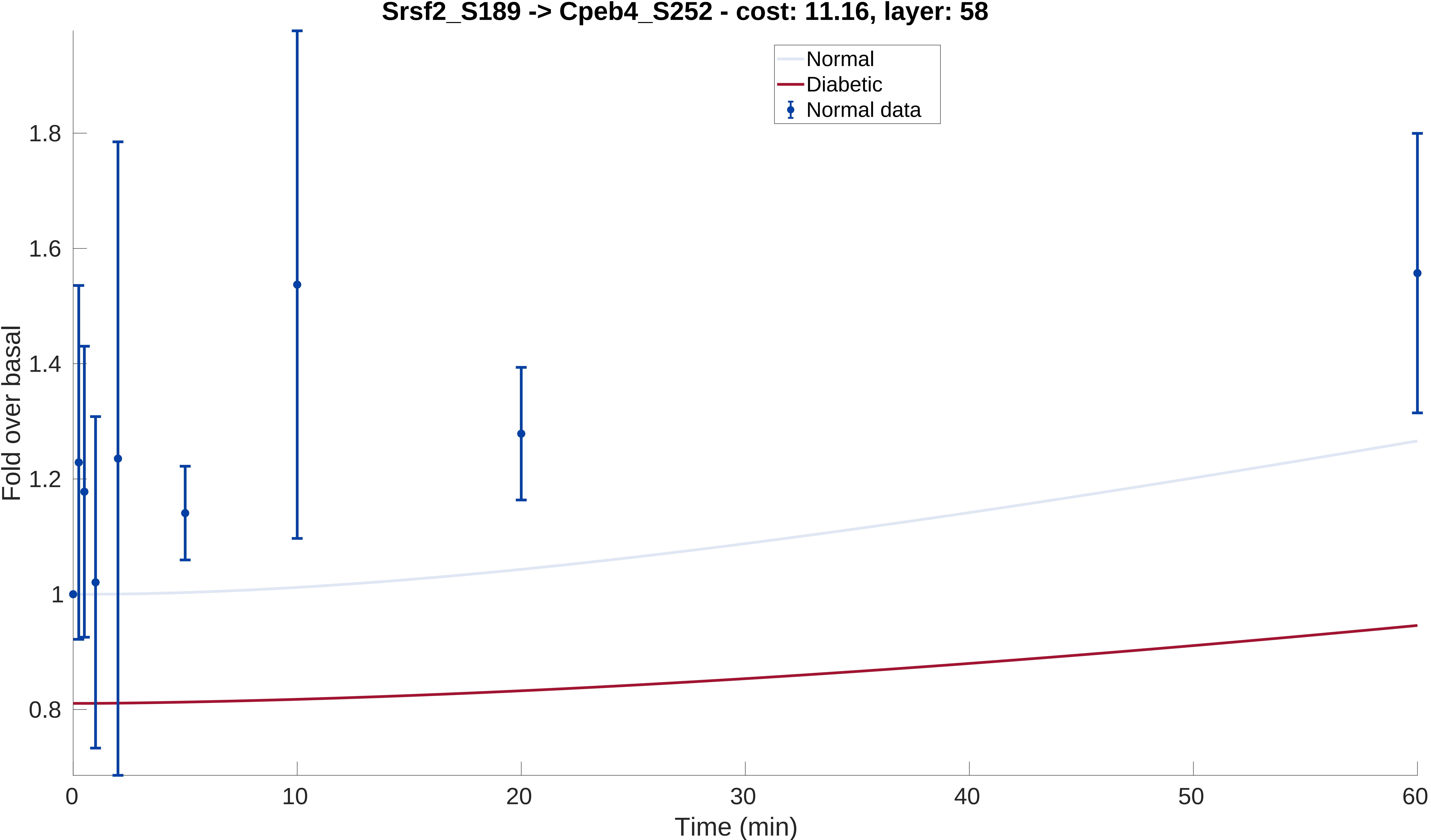

Srsf2_S189 -> Cpeb4_S252 - cost: 11.16, layer: 58
Normal
Diabetic
Normal data
Fold over basal
1.8
1.6
1.4
1.2
1
0.8
0
10
20
30
40
50
60
Time (min)

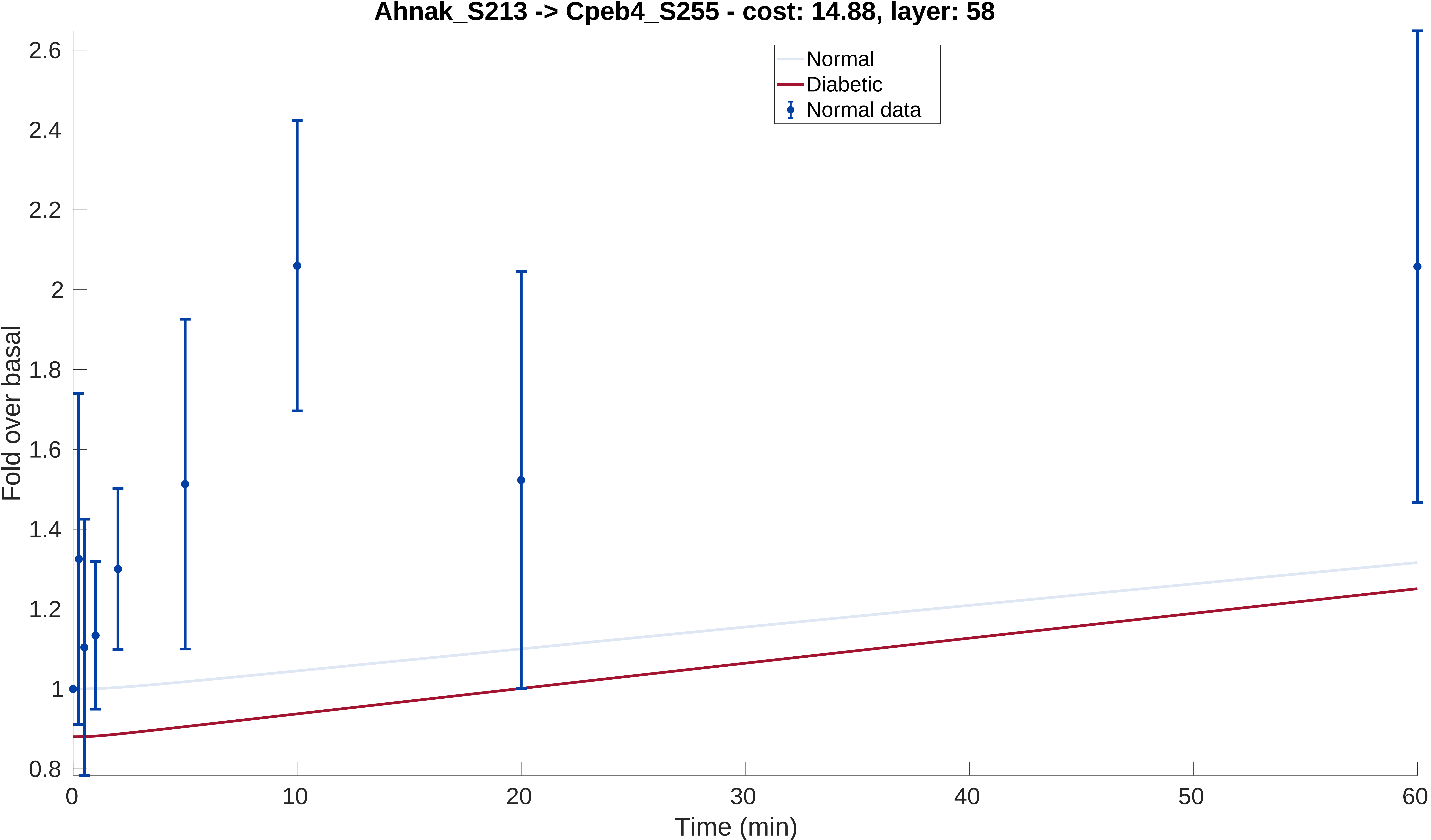

Ahnak_S213 -> Cpeb4_S255 - cost: 14.88, layer: 58
Normal
Diabetic
Normal data
Fold over basal
Time (min)
0.8
1
1.2
1.4
1.6
1.8
2
2.2
2.4
2.6
0
10
20
30
40
50
60

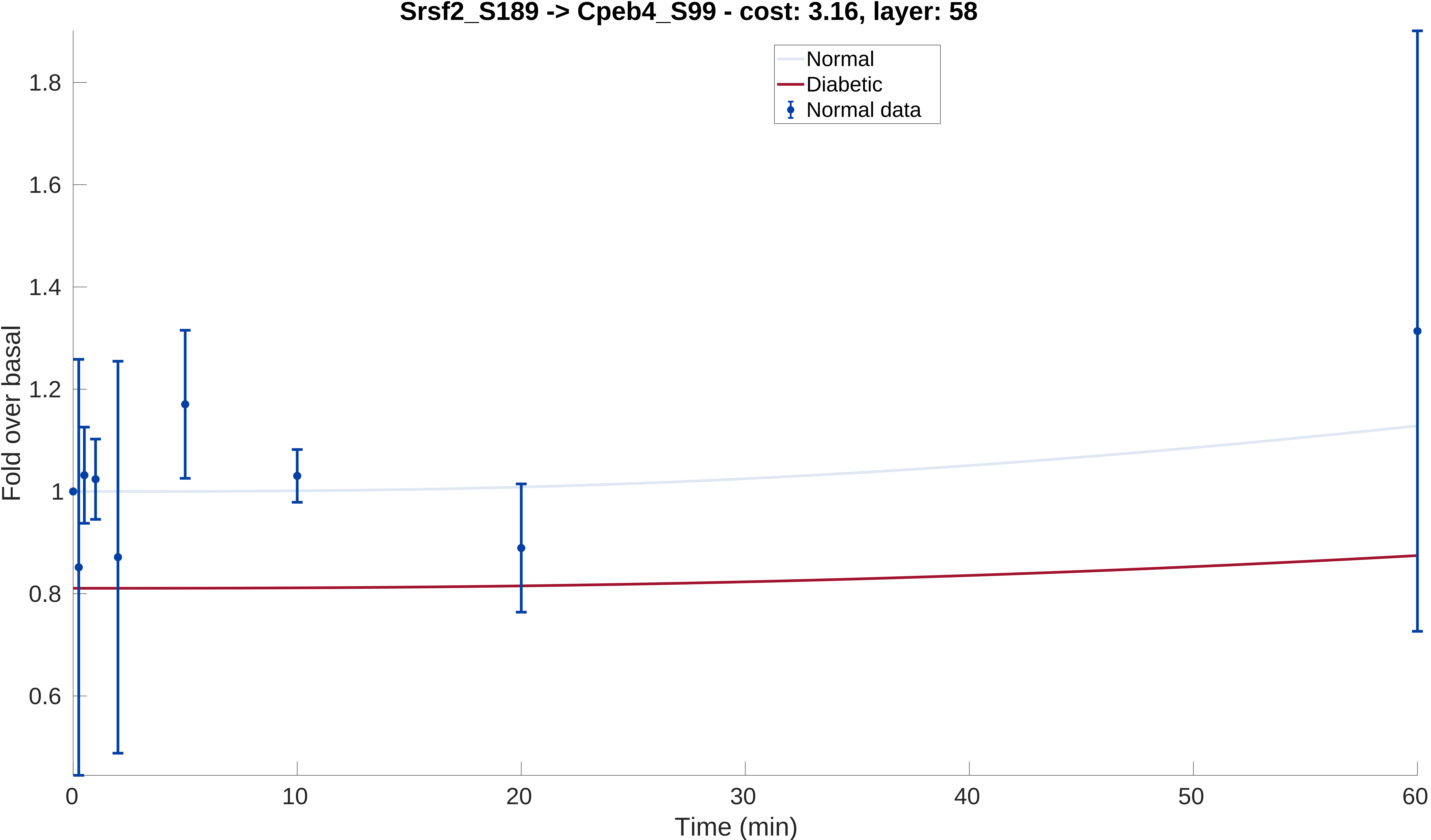
Srsf2_S189 -> Cpeb4_S99 - cost: 3.16, layer: 58
Normal
Diabetic
Normal data
Fold over basal
1.8
1.6
1.4
1.2
1
0.8
0.6
0
10
20
30
40
50
60
Time (min)

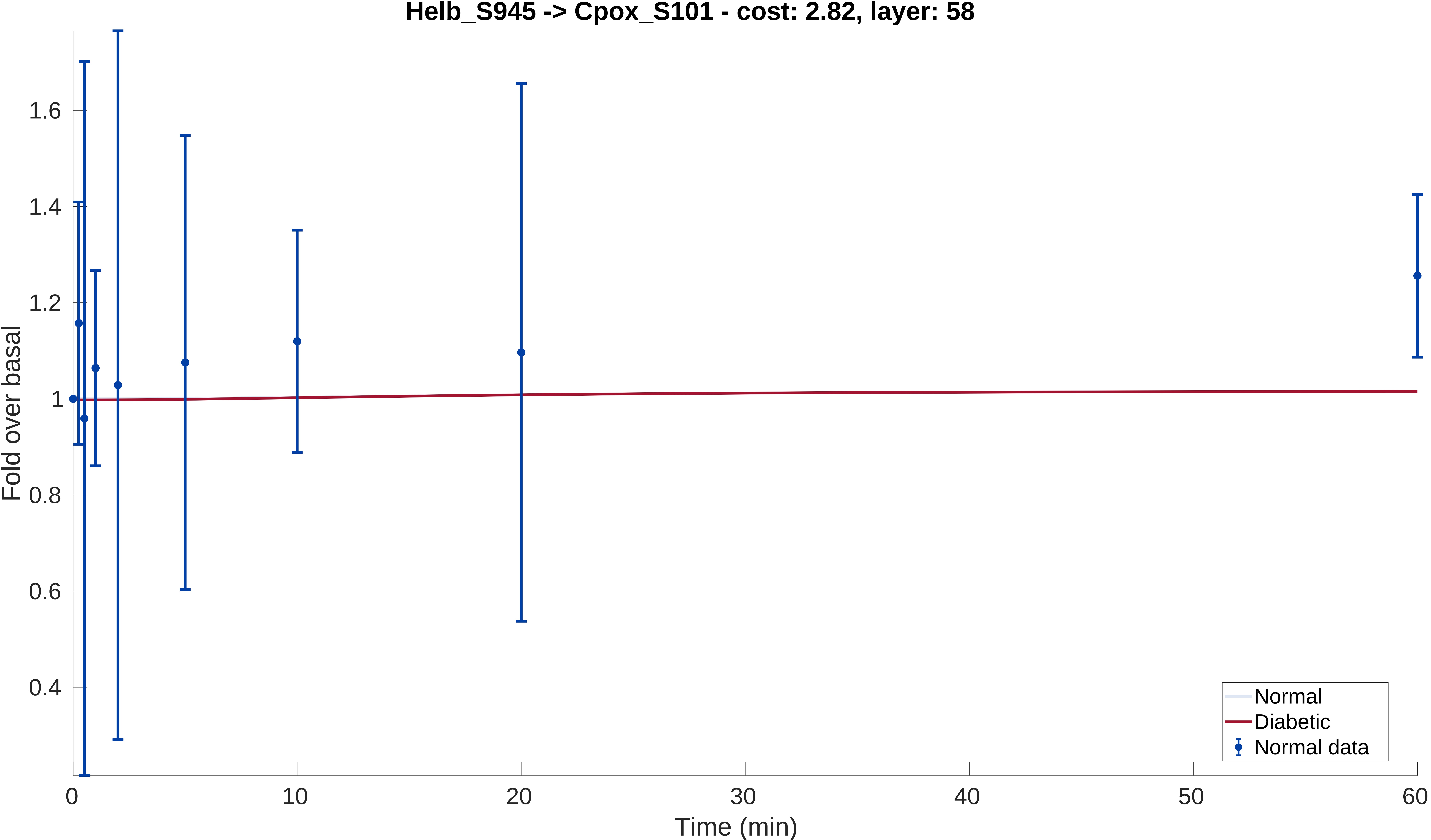

Helb_S945 -> Cpox_S101 - cost: 2.82, layer: 58
Fold over basal
1.6
1.4
1.2
1
0.8
0.6
0.4
0
10
20
30
40
50
60
Time (min)
Normal
Diabetic
Normal data

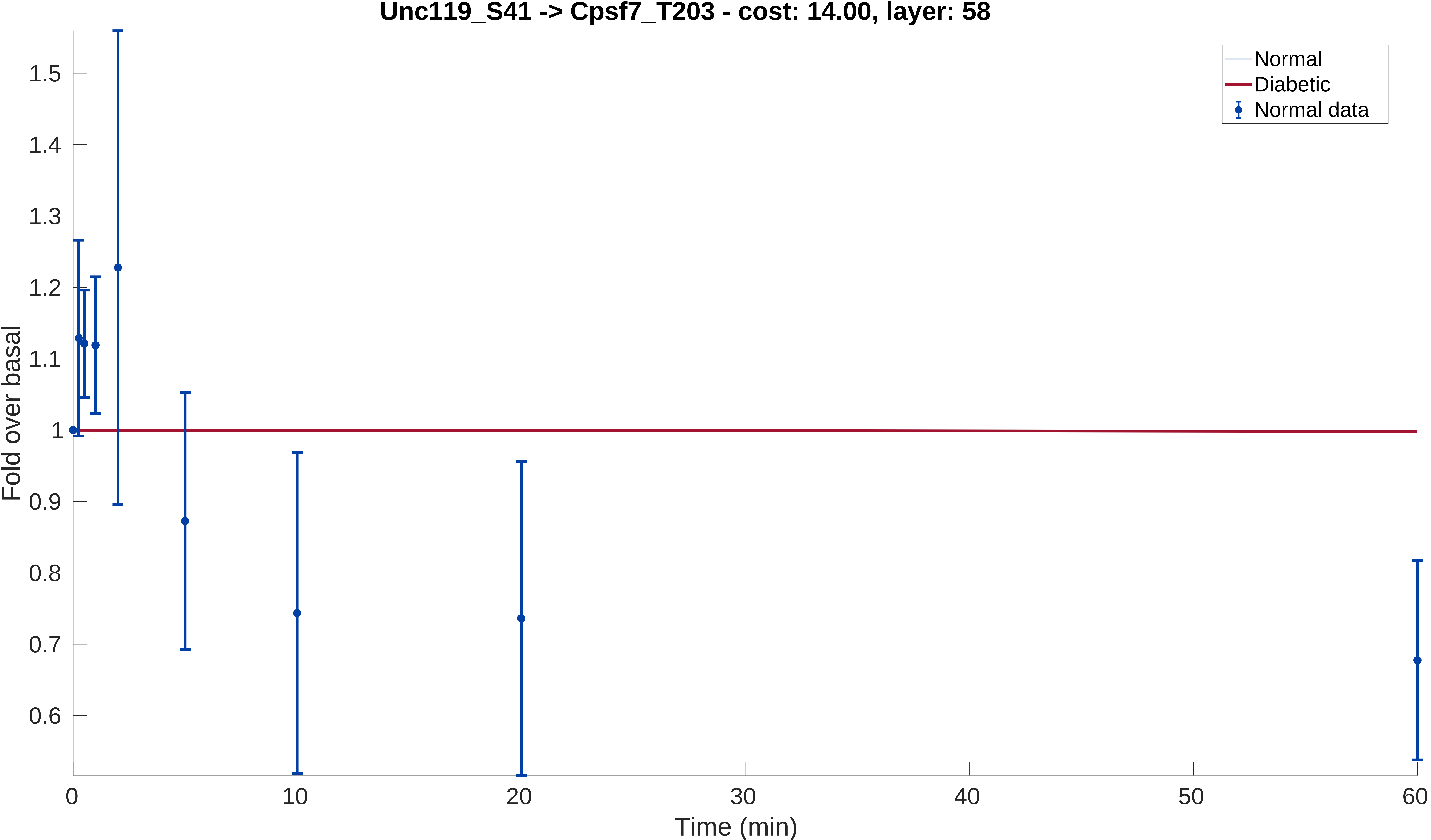

Unc119_S41 -> Cpsf7_T203 - cost: 14.00, layer: 58
Normal
Diabetic
Normal data
Fold over basal
1.5
1.4
1.3
1.2
1.1
1
0.9
0.8
0.7
0.6
0
10
20
30
40
50
60
Time (min)

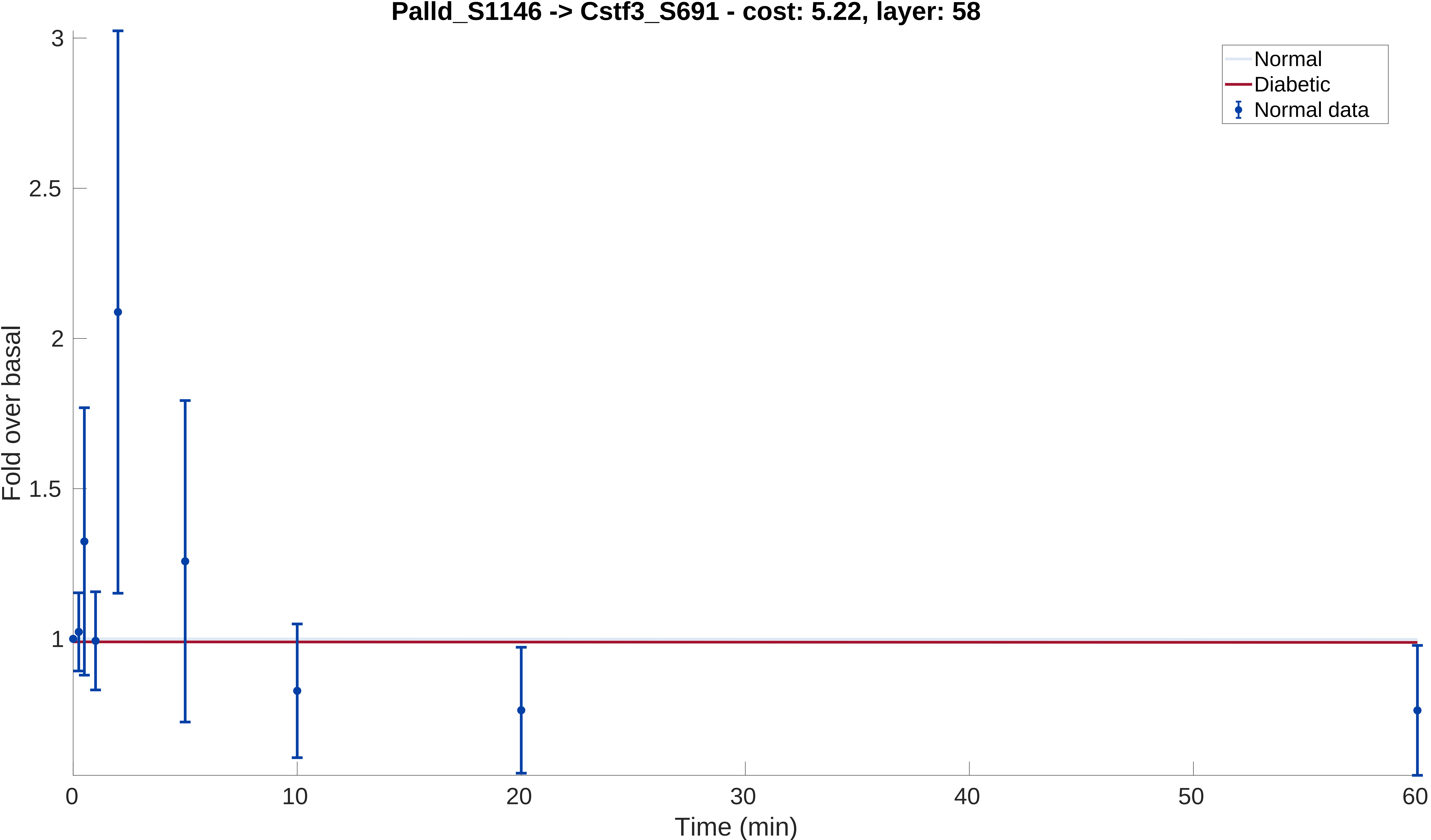
Palld_S1146 -> Cstf3_S691 - cost: 5.22, layer: 58
Fold over basal
Time (min)
Normal
Diabetic
Normal data
0
10
20
30
40
50
60
1
1.5
2
2.5
3

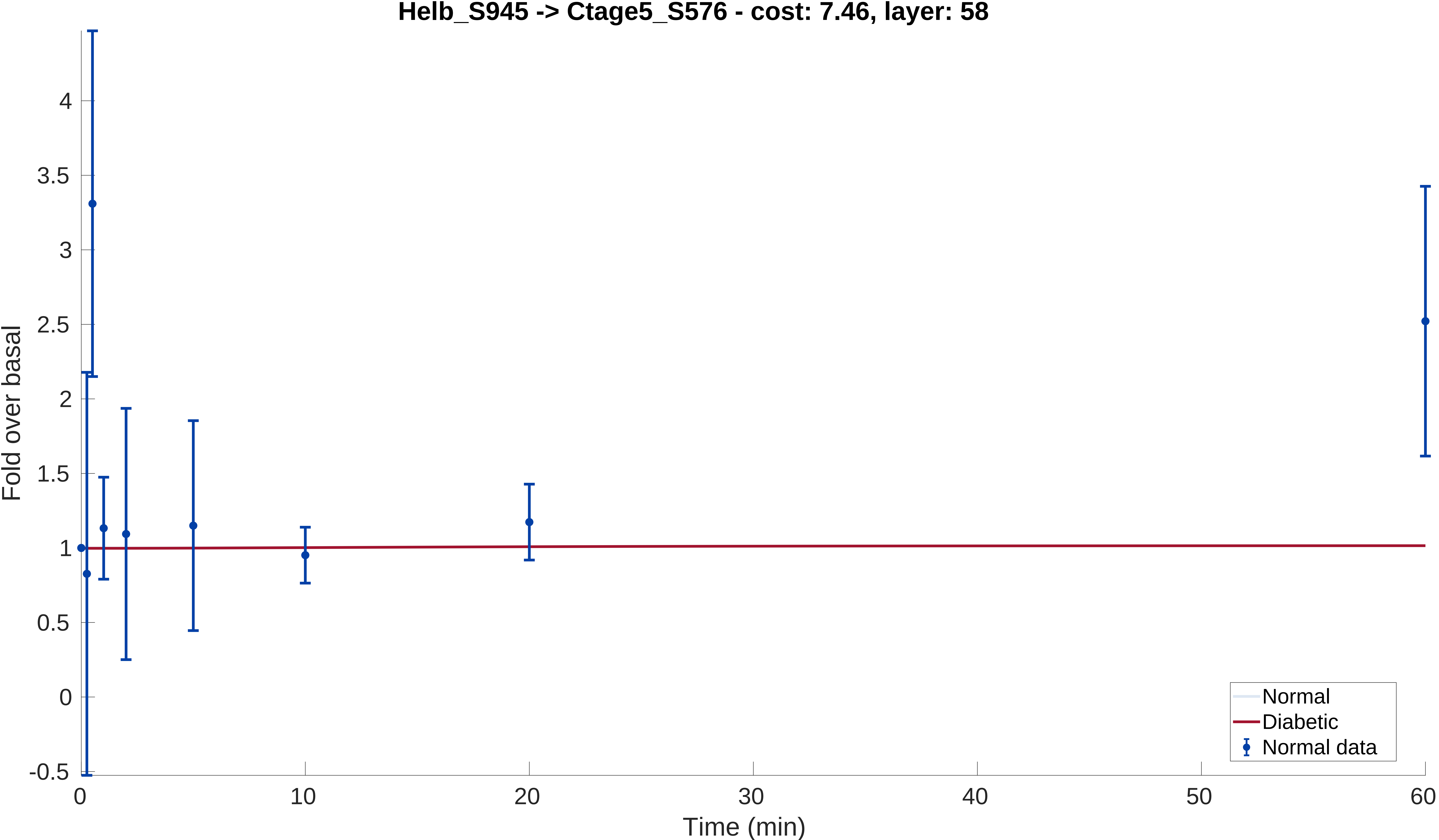
Helb_S945 -> Ctage5_S576 - cost: 7.46, layer: 58
Fold over basal
4
3.5
3
2.5
2
1.5
1
0.5
0
-0.5
0
10
20
30
40
50
60
Time (min)
Normal
Diabetic
Normal data

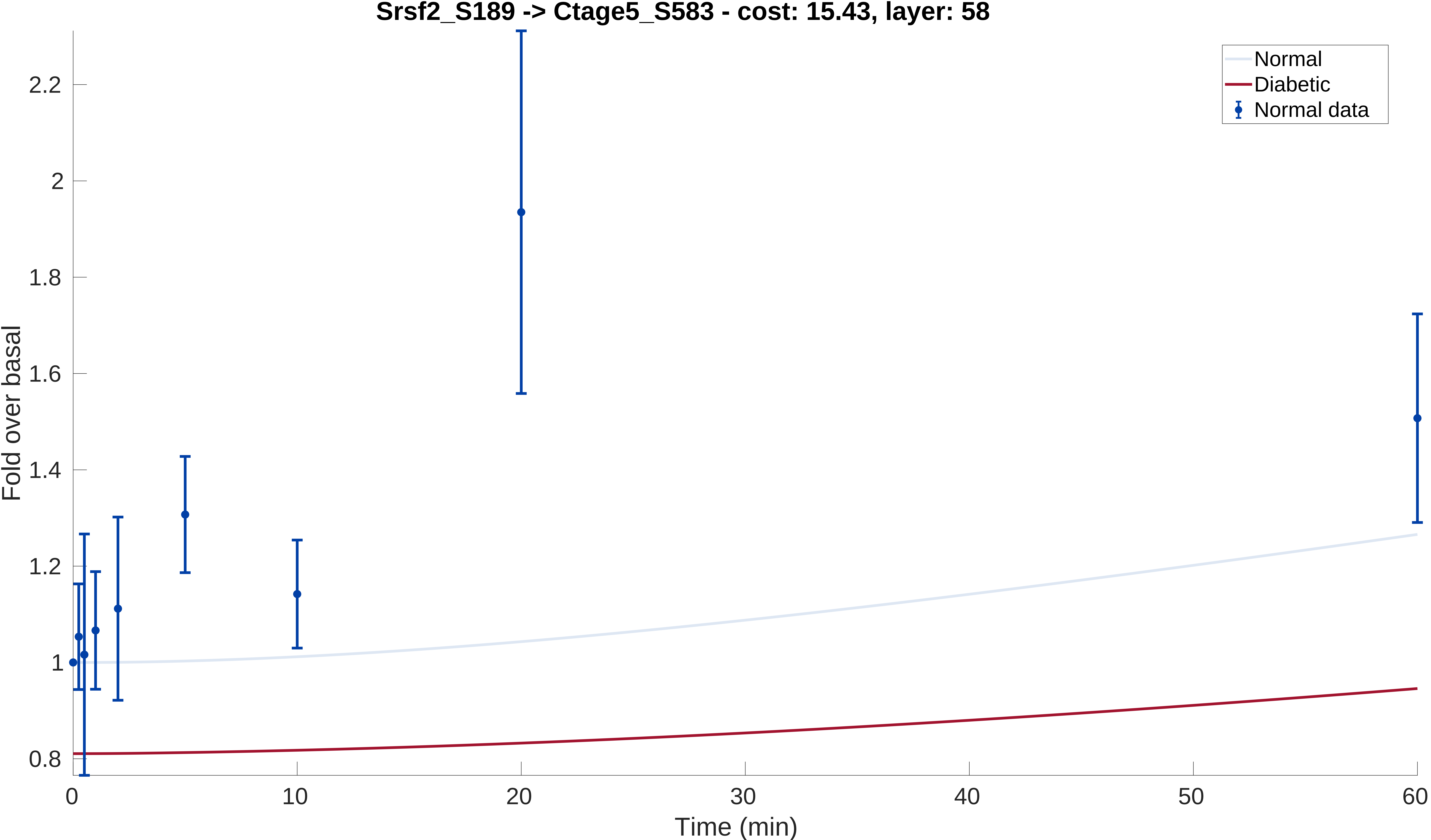

Srsf2_S189 -> Ctage5_S583 - cost: 15.43, layer: 58
Normal
Diabetic
Normal data
Fold over basal
Time (min)
0.8
1
1.2
1.4
1.6
1.8
2
2.2
0
10
20
30
40
50
60

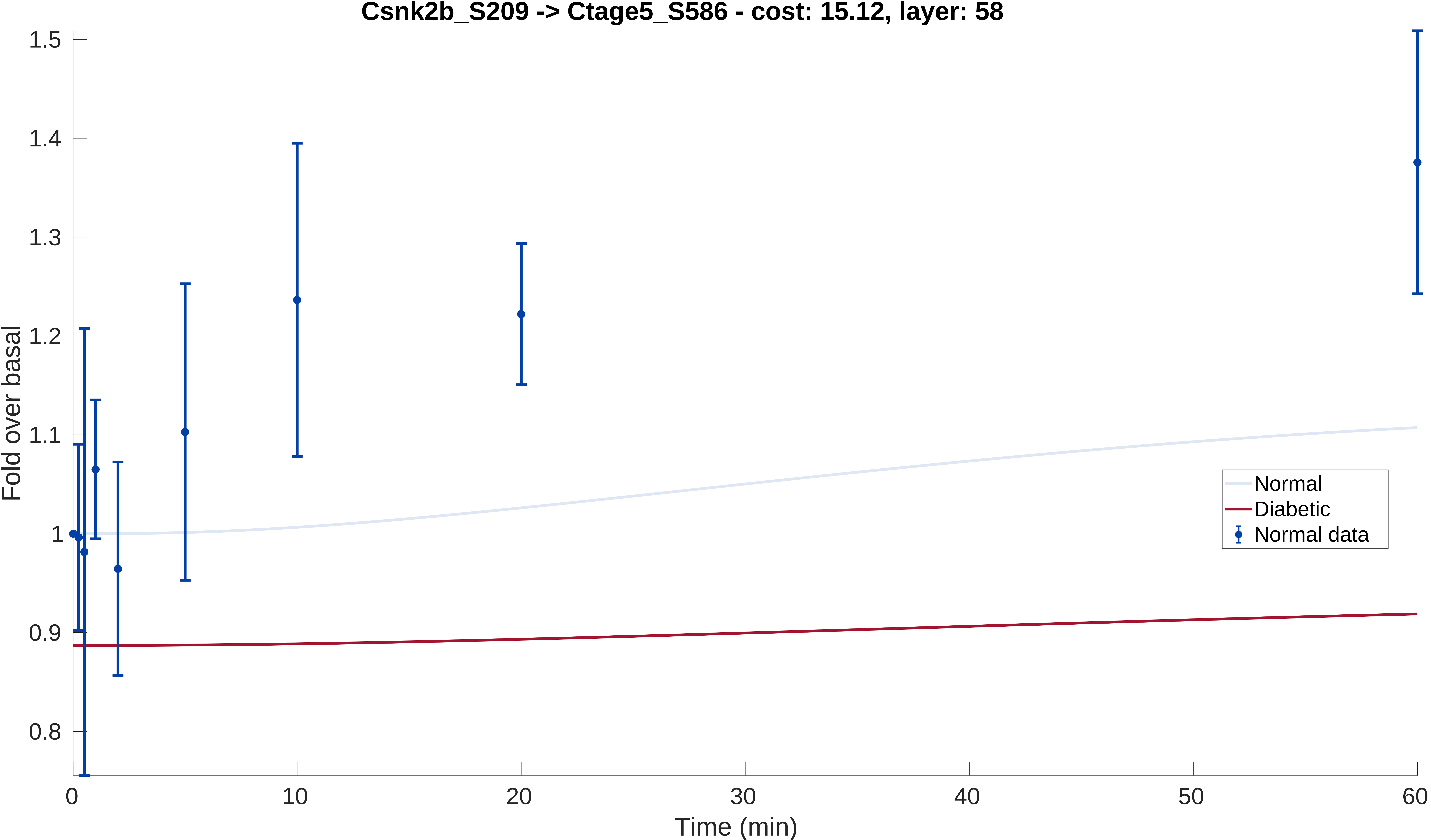

Csnk2b_S209 -> Ctage5_S586 - cost: 15.12, layer: 58
Fold over basal
1.5
1.4
1.3
1.2
1.1
1
0.9
0.8
0
10
20
30
40
50
60
Time (min)
Normal
Diabetic
Normal data

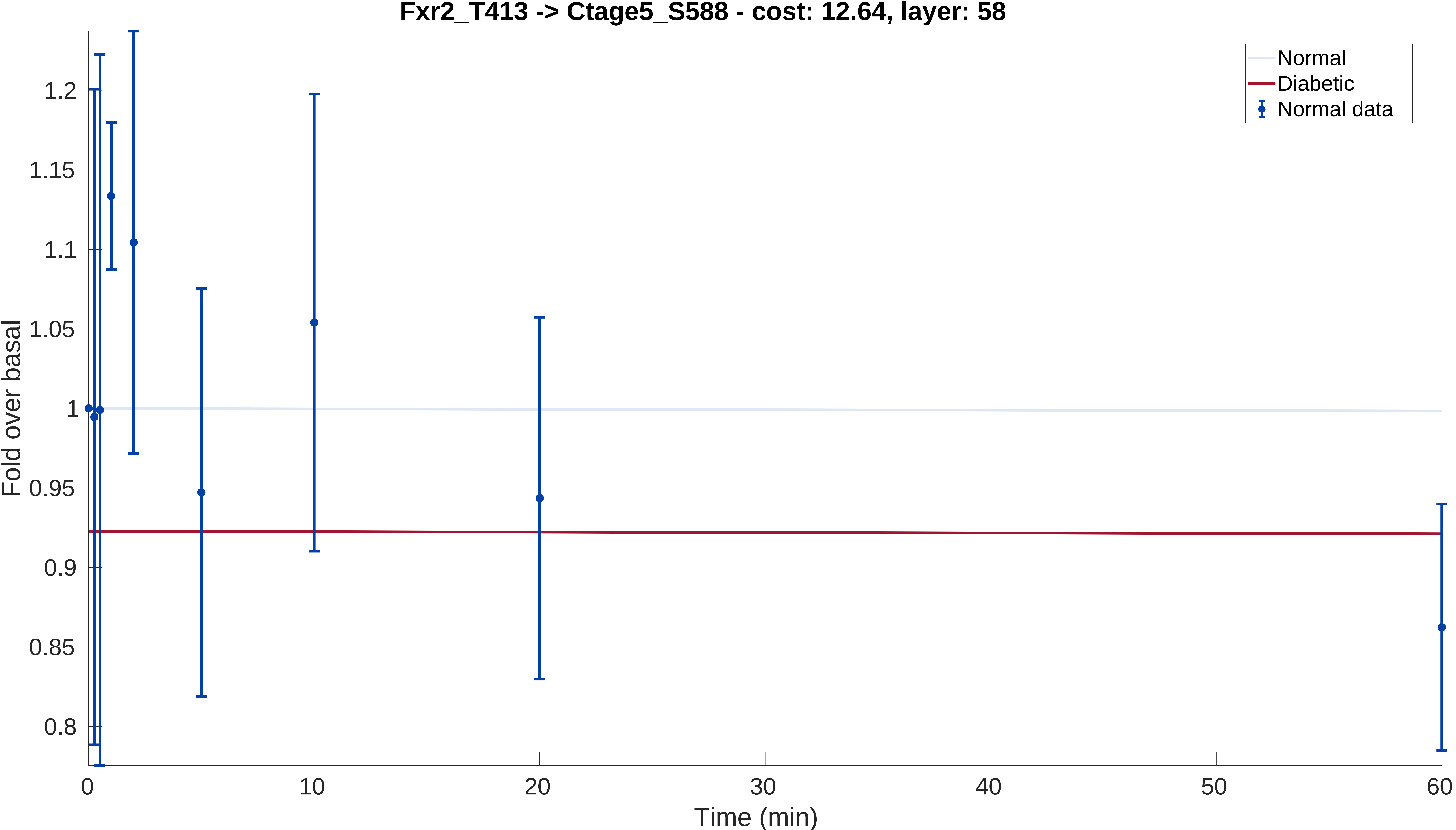

Fxr2_T413 -> Ctage5_S588 - cost: 12.64, layer: 58
Normal
Diabetic
Normal data
Fold over basal
1.2
1.15
1.1
1.05
1
0.95
0.9
0.85
0.8
0
10
20
30
40
50
60
Time (min)

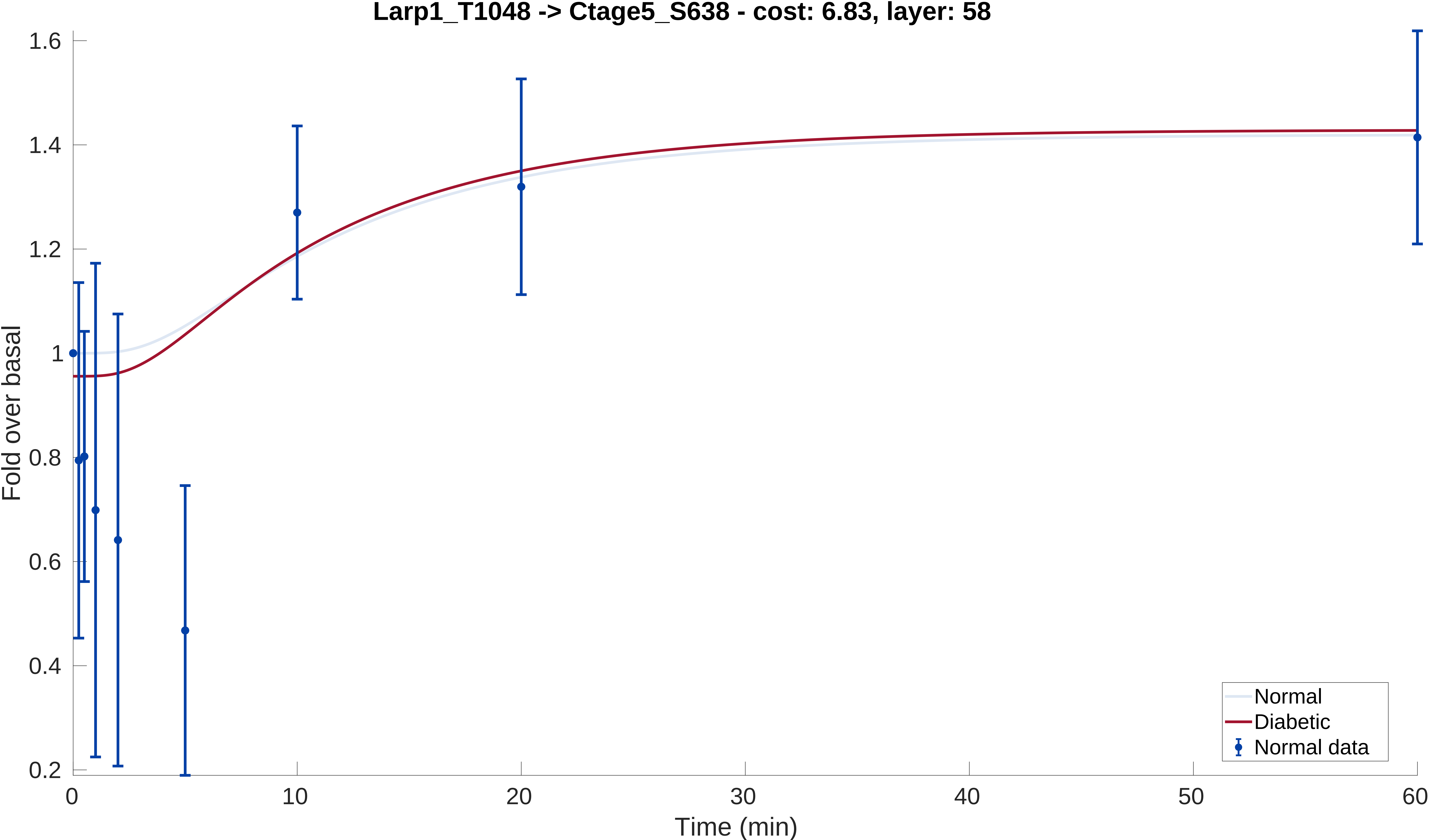

Larp1_T1048 -> Ctage5_S638 - cost: 6.83, layer: 58
1.6
1.4
1.2
1
0.8
0.6
0.4
0.2
Fold over basal
0
10
20
30
40
50
60
Time (min)
Normal
Diabetic
Normal data

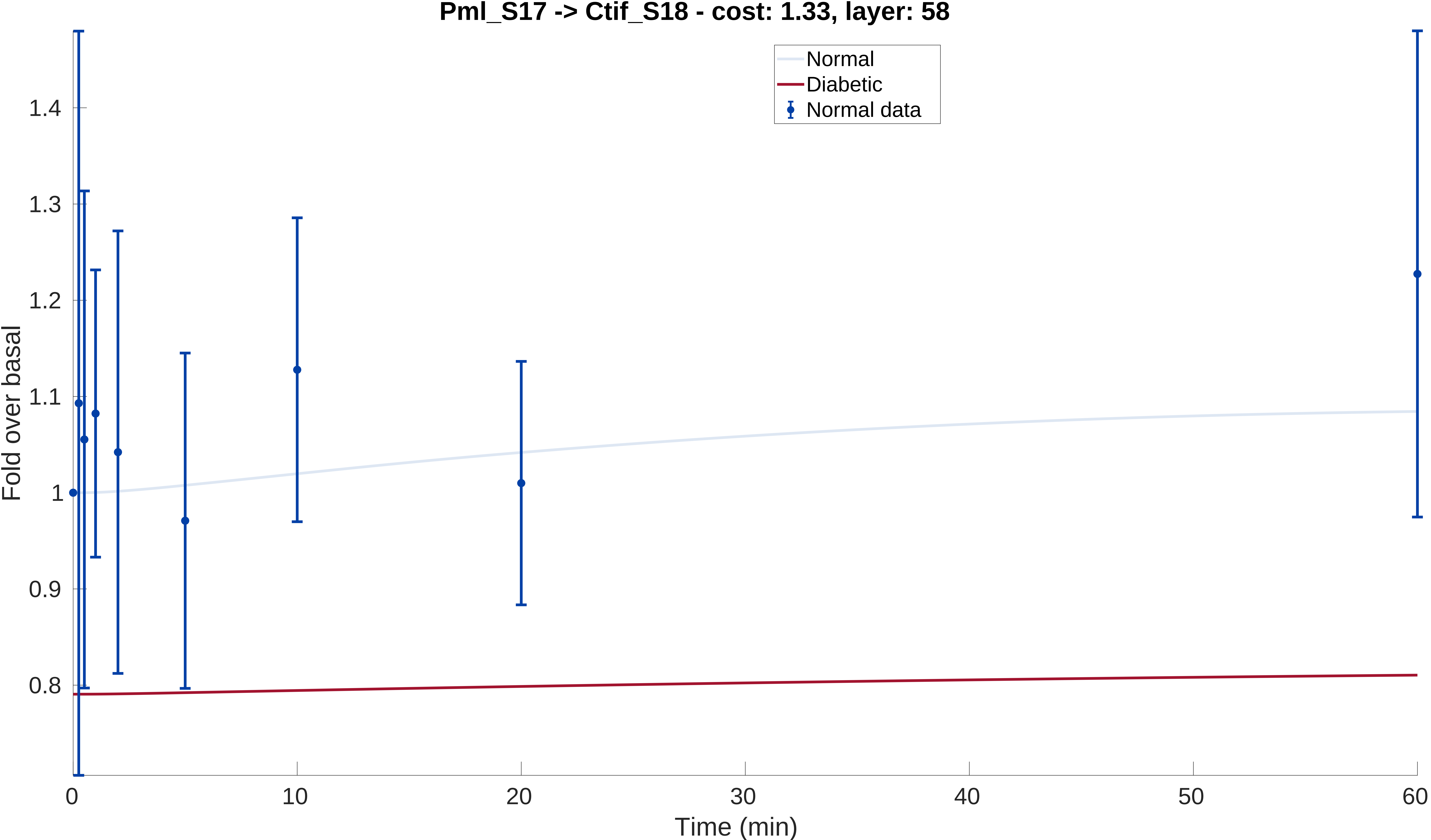
Pml_S17 -> Ctif_S18 - cost: 1.33, layer: 58
Normal
Diabetic
Normal data
Fold over basal
1.4
1.3
1.2
1.1
1
0.9
0.8
0
10
20
30
40
50
60
Time (min)

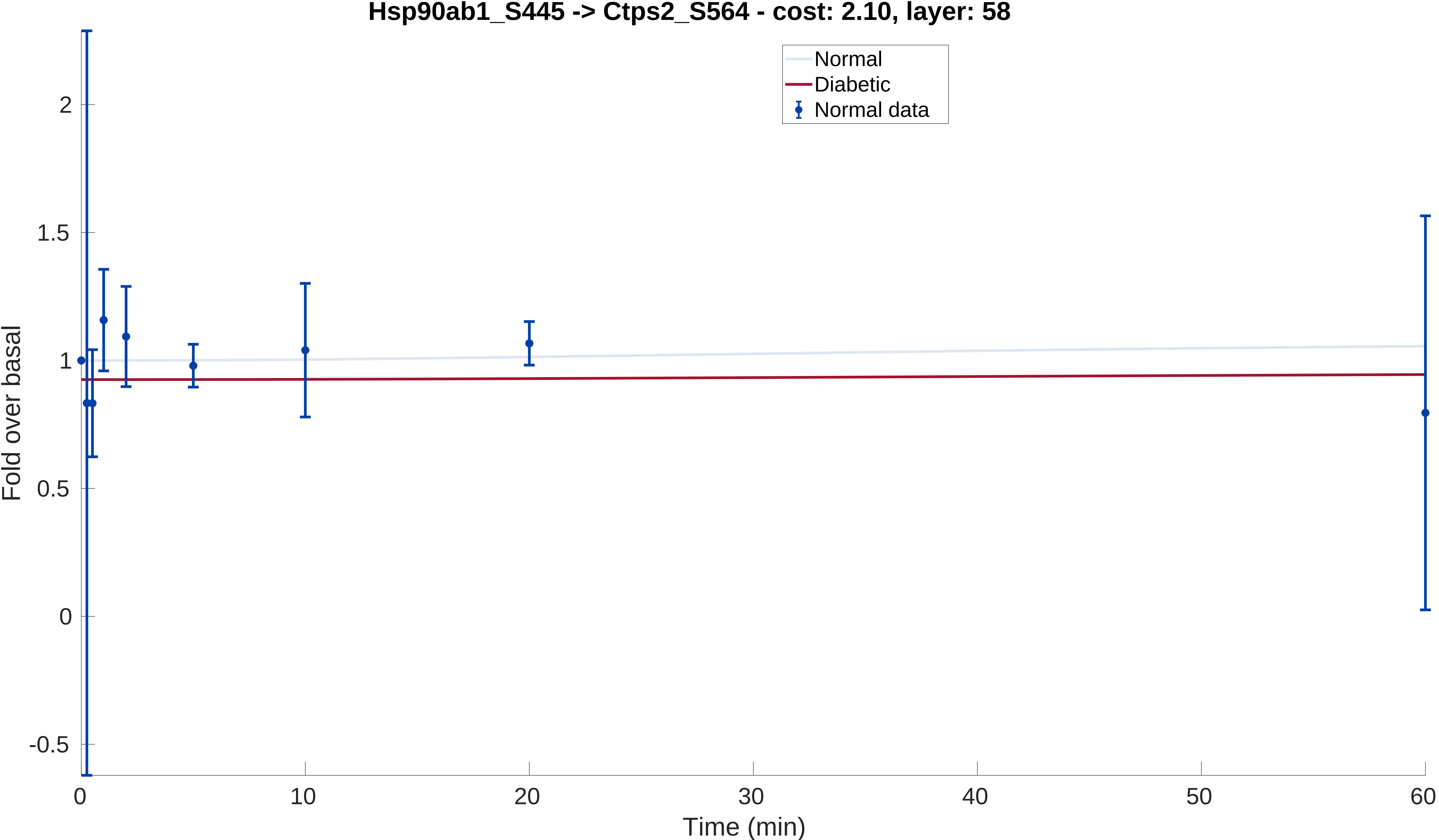
Hsp90ab1_S445 -> Ctps2_S564 - cost: 2.10, layer: 58
Normal
Diabetic
Normal data
Fold over basal
2
1.5
1
0.5
0
-0.5
0
10
20
30
40
50
60
Time (min)

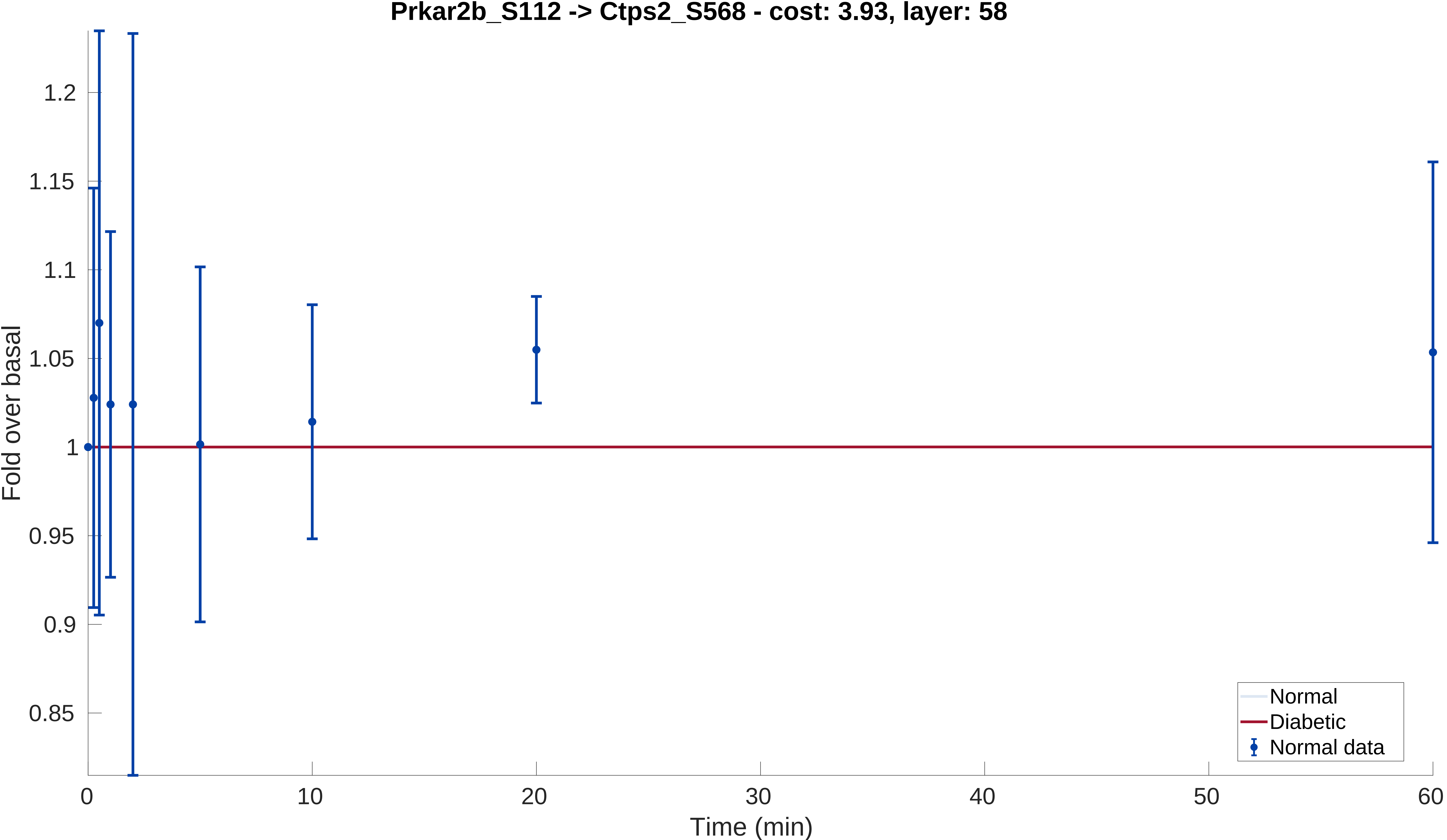
Prkar2b_S112 -> Ctps2_S568 - cost: 3.93, layer: 58
Fold over basal
1.2
1.15
1.1
1.05
1
0.95
0.9
0.85
0
10
20
30
40
50
60
Time (min)
Normal
Diabetic
Normal data

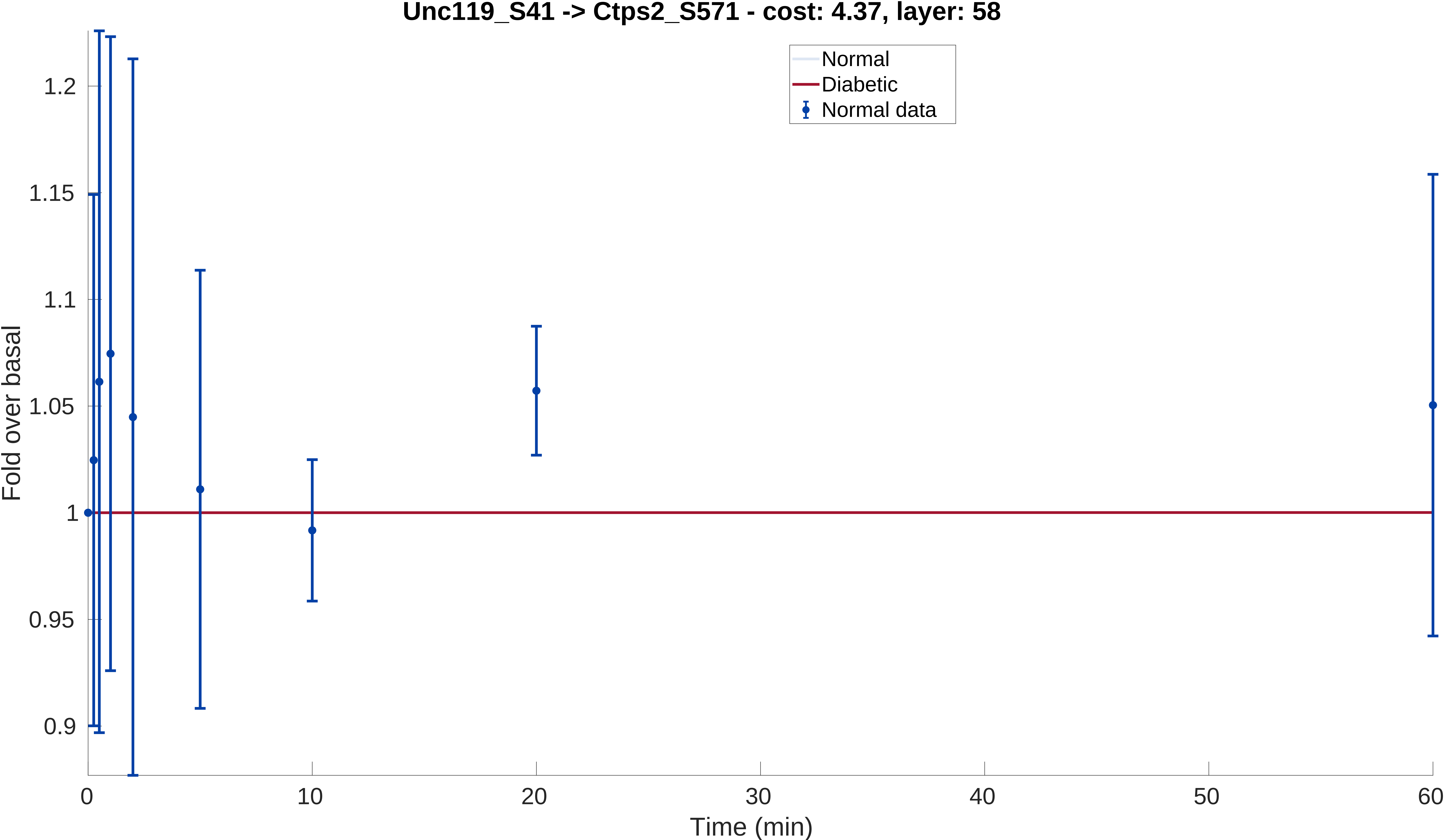

Unc119_S41 -> Ctps2_S571 - cost: 4.37, layer: 58
Normal
Diabetic
Normal data
Fold over basal
1.2
1.15
1.1
1.05
1
0.95
0.9
0
10
20
30
40
50
60
Time (min)

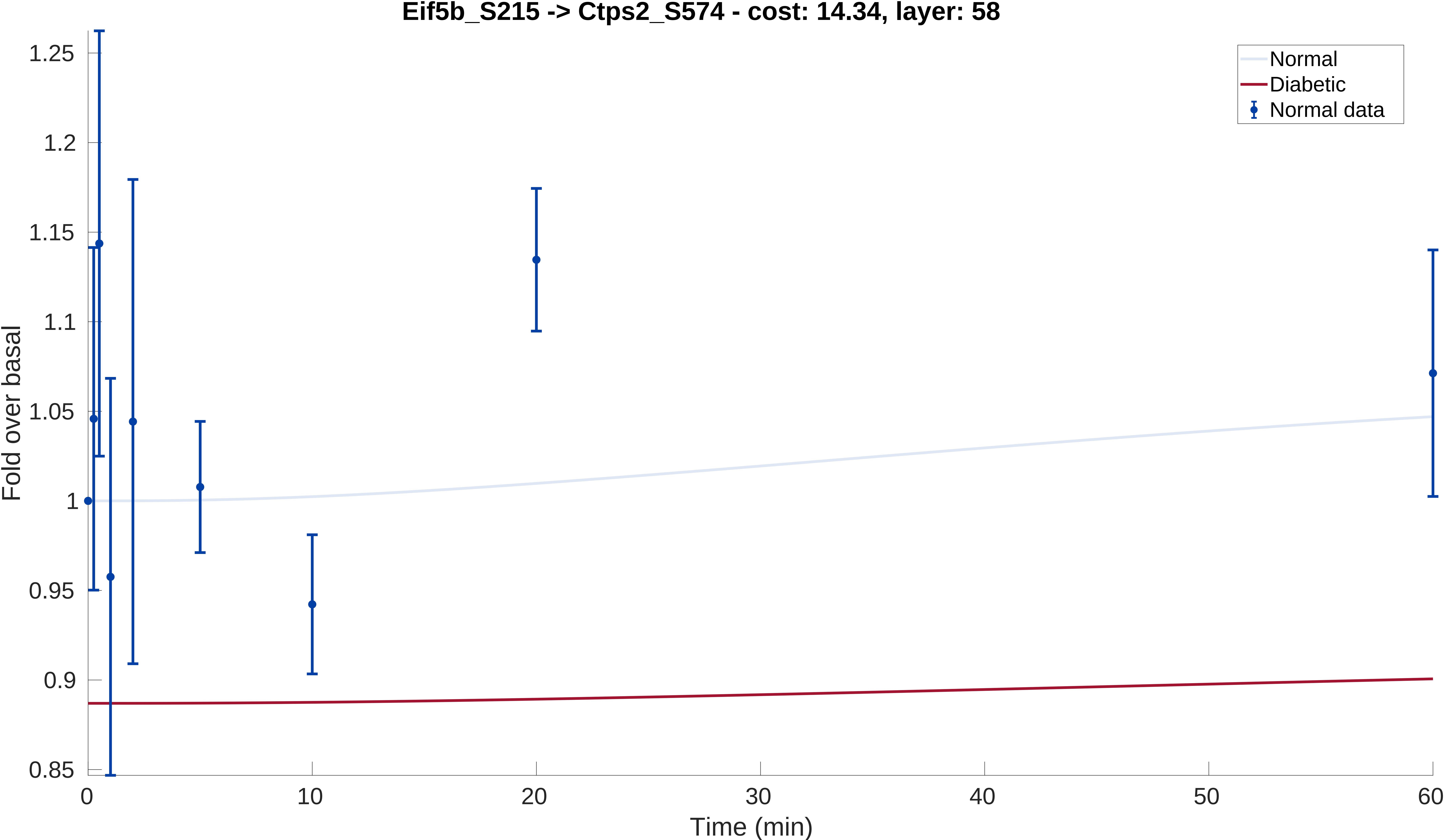

Eif5b_S215 -> Ctps2_S574 - cost: 14.34, layer: 58
Normal
Diabetic
Normal data
Fold over basal
1.25
1.2
1.15
1.1
1.05
1
0.95
0.9
0.85
0
10
20
30
40
50
60
Time (min)

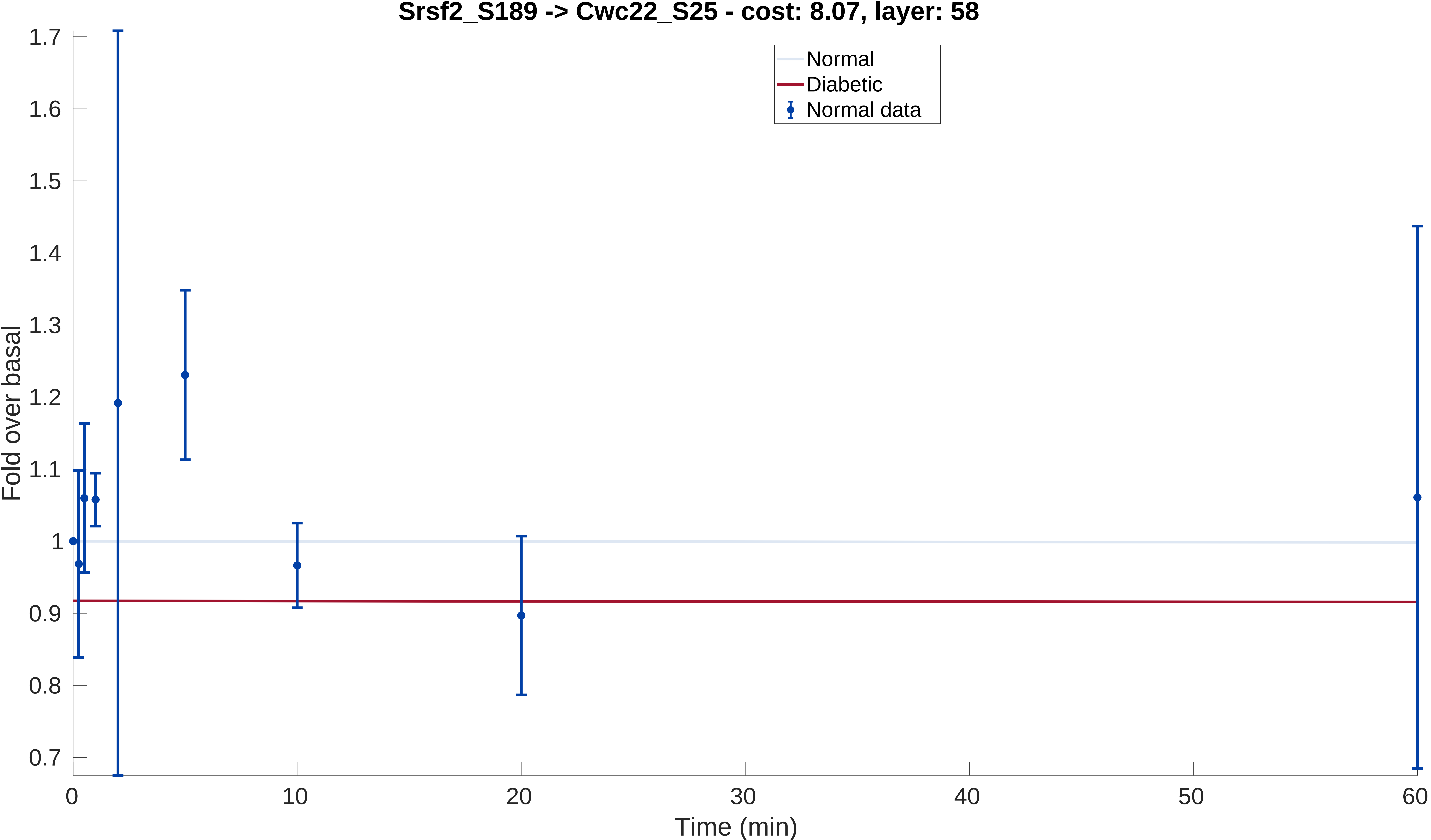

Srsf2_S189 -> Cwc22_S25 - cost: 8.07, layer: 58
Normal
Diabetic
Normal data
Fold over basal
Time (min)
0.7
0.8
0.9
1
1.1
1.2
1.3
1.4
1.5
1.6
1.7
0
10
20
30
40
50
60

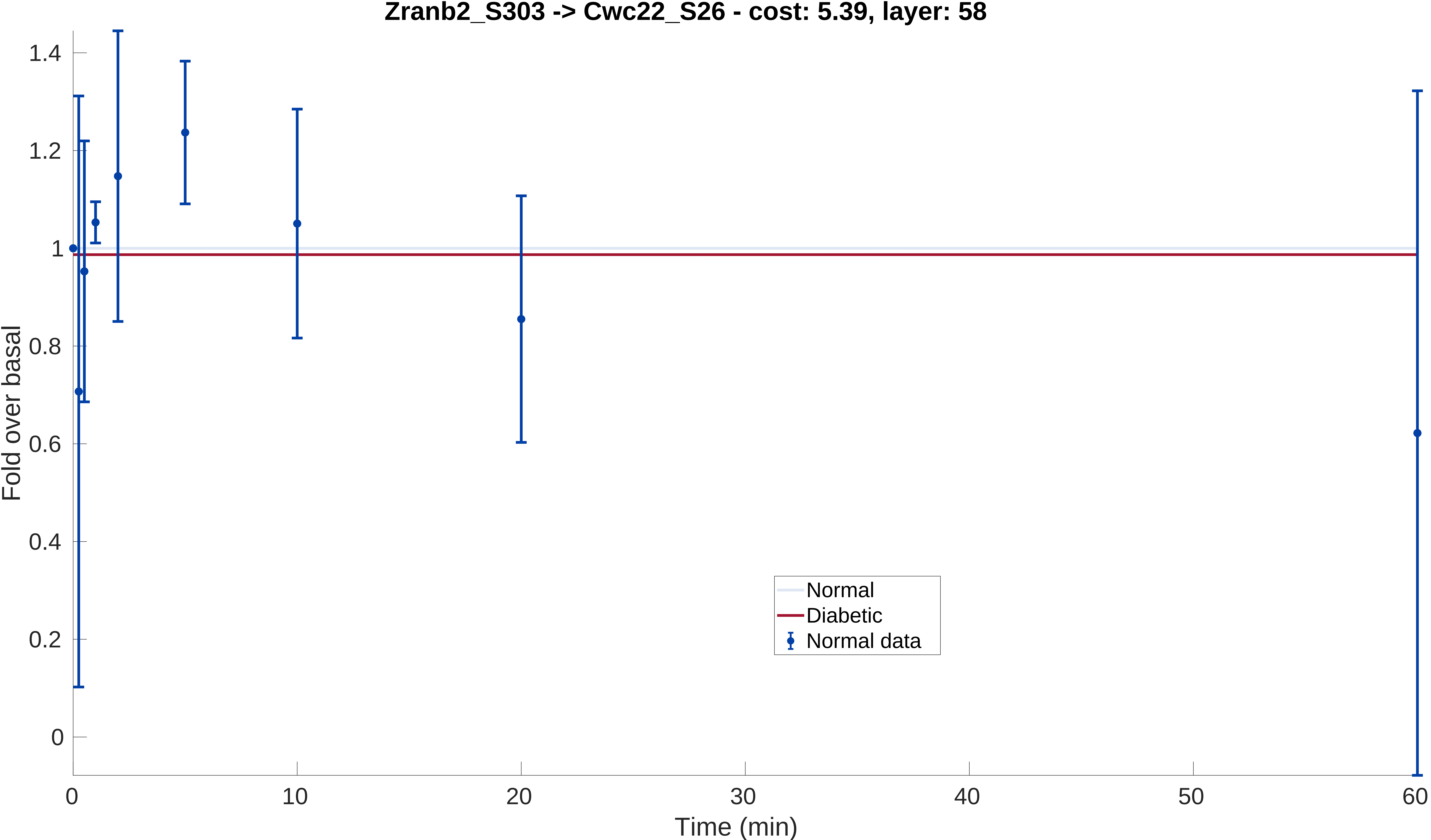

Zranb2_S303 -> Cwc22_S26 - cost: 5.39, layer: 58
Fold over basal
1.4
1.2
1
0.8
0.6
0.4
0.2
0
0
10
20
30
40
50
60
Time (min)
Normal
Diabetic
Normal data

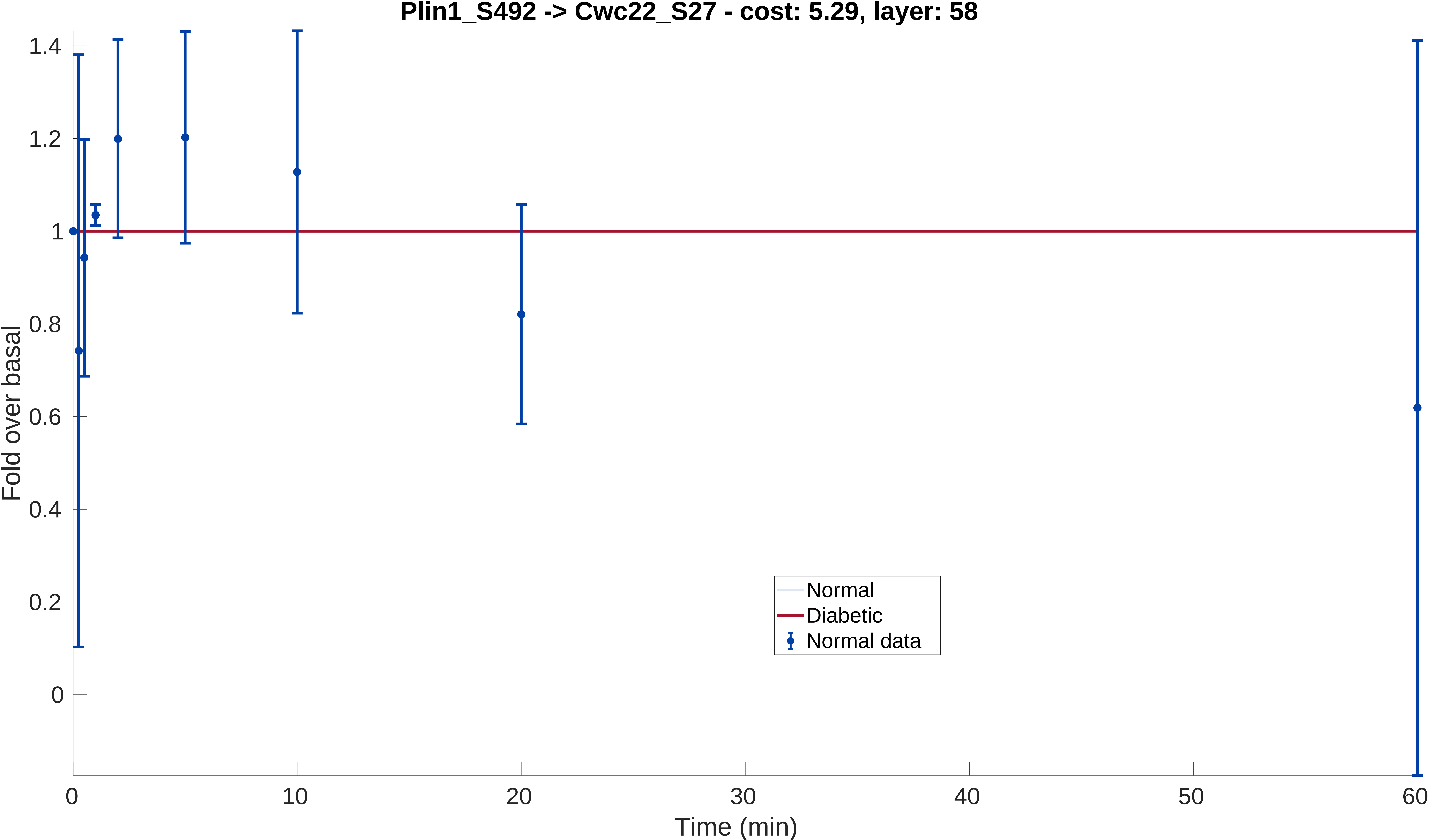
Plin1_S492 -> Cwc22_S27 - cost: 5.29, layer: 58
Fold over basal
1.4
1.2
1
0.8
0.6
0.4
0.2
0
0
10
20
30
40
50
60
Time (min)
Normal
Diabetic
Normal data

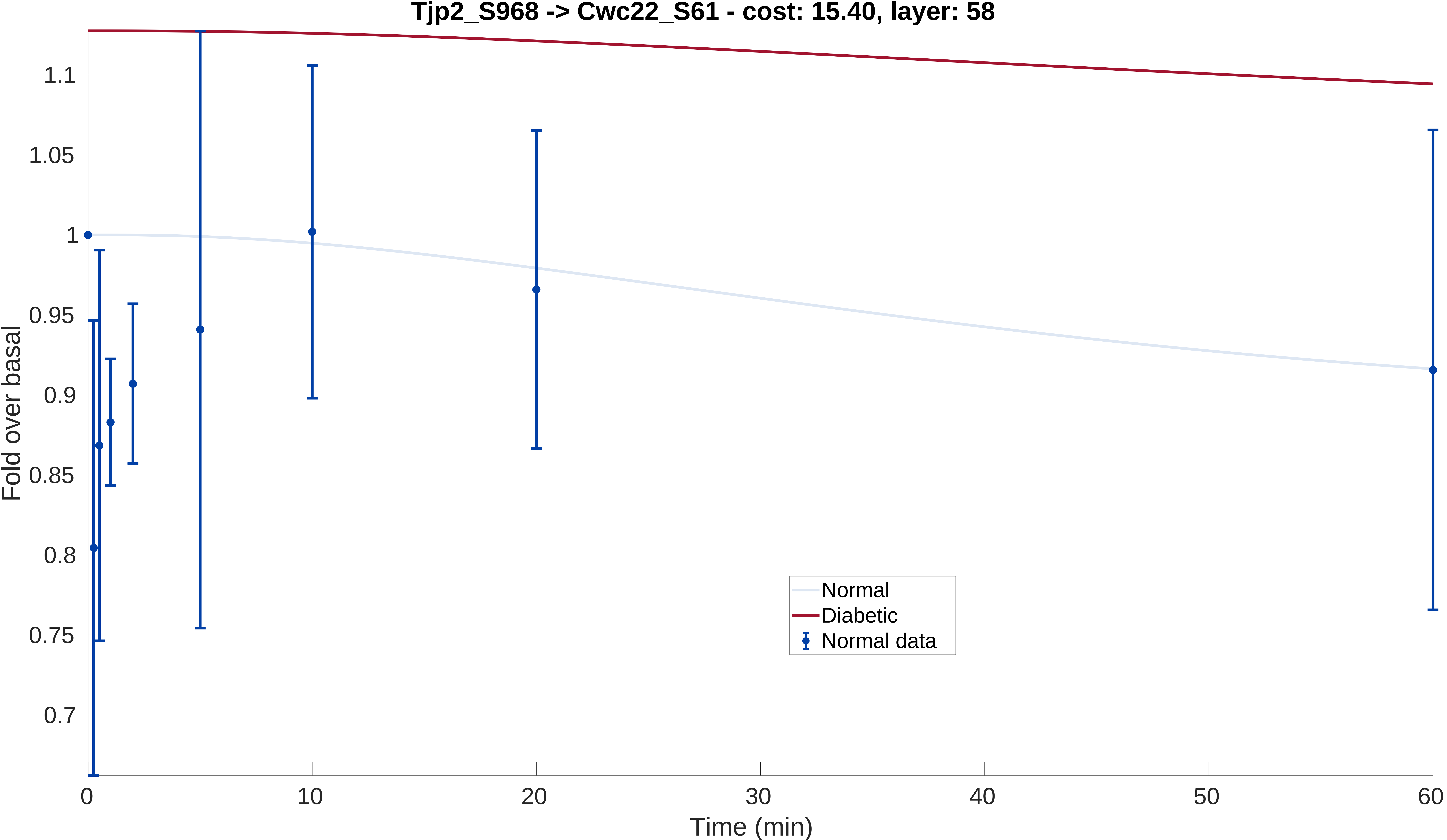

Tjp2_S968 -> Cwc22_S61 - cost: 15.40, layer: 58
Fold over basal
1.1
1.05
1
0.95
0.9
0.85
0.8
0.75
0.7
0
10
20
30
40
50
60
Time (min)
Normal
Diabetic
Normal data

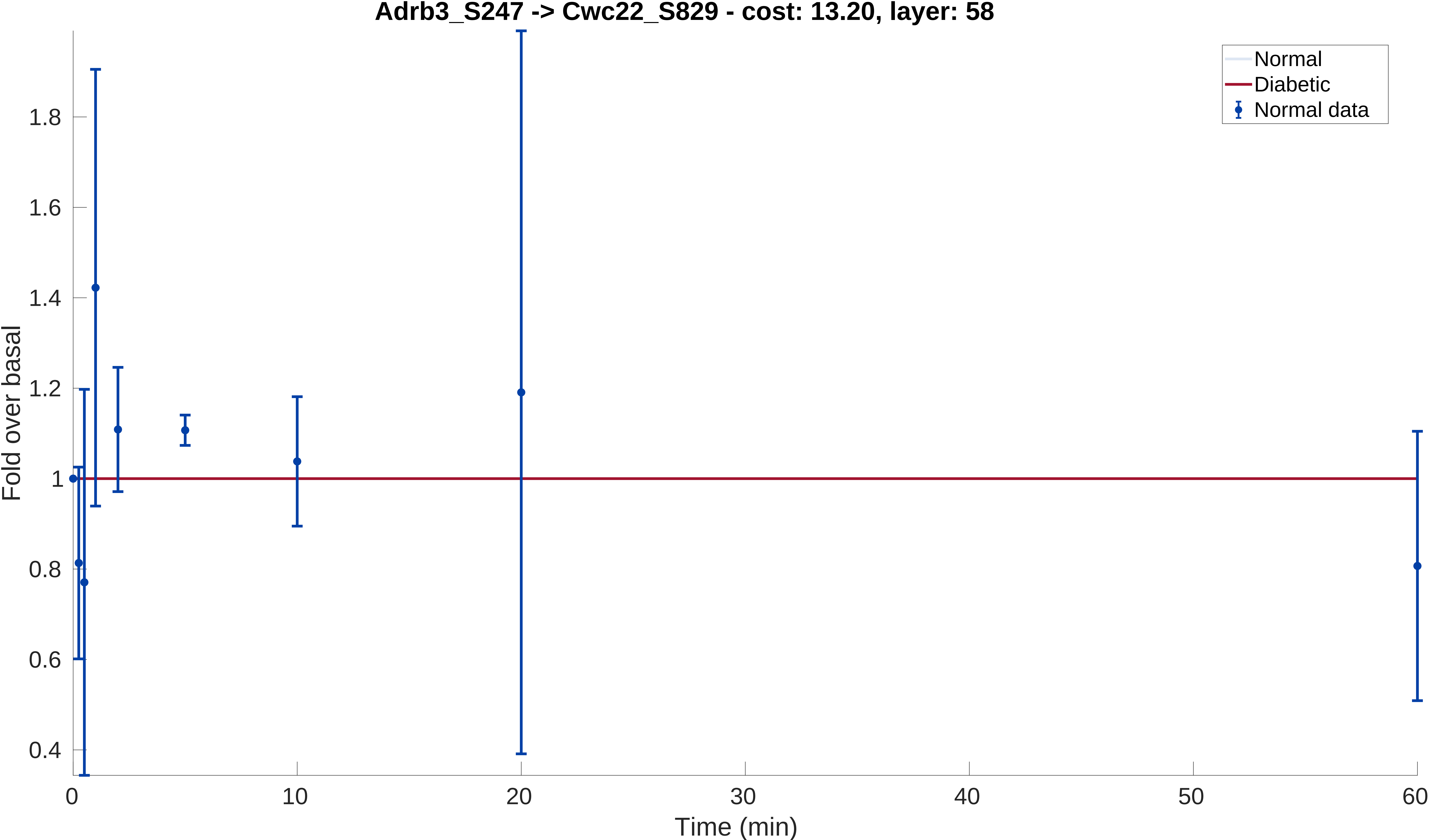

Adrb3_S247 -> Cwc22_S829 - cost: 13.20, layer: 58
Normal
Diabetic
Normal data
Fold over basal
1.8
1.6
1.4
1.2
1
0.8
0.6
0.4
0
10
20
30
40
50
60
Time (min)

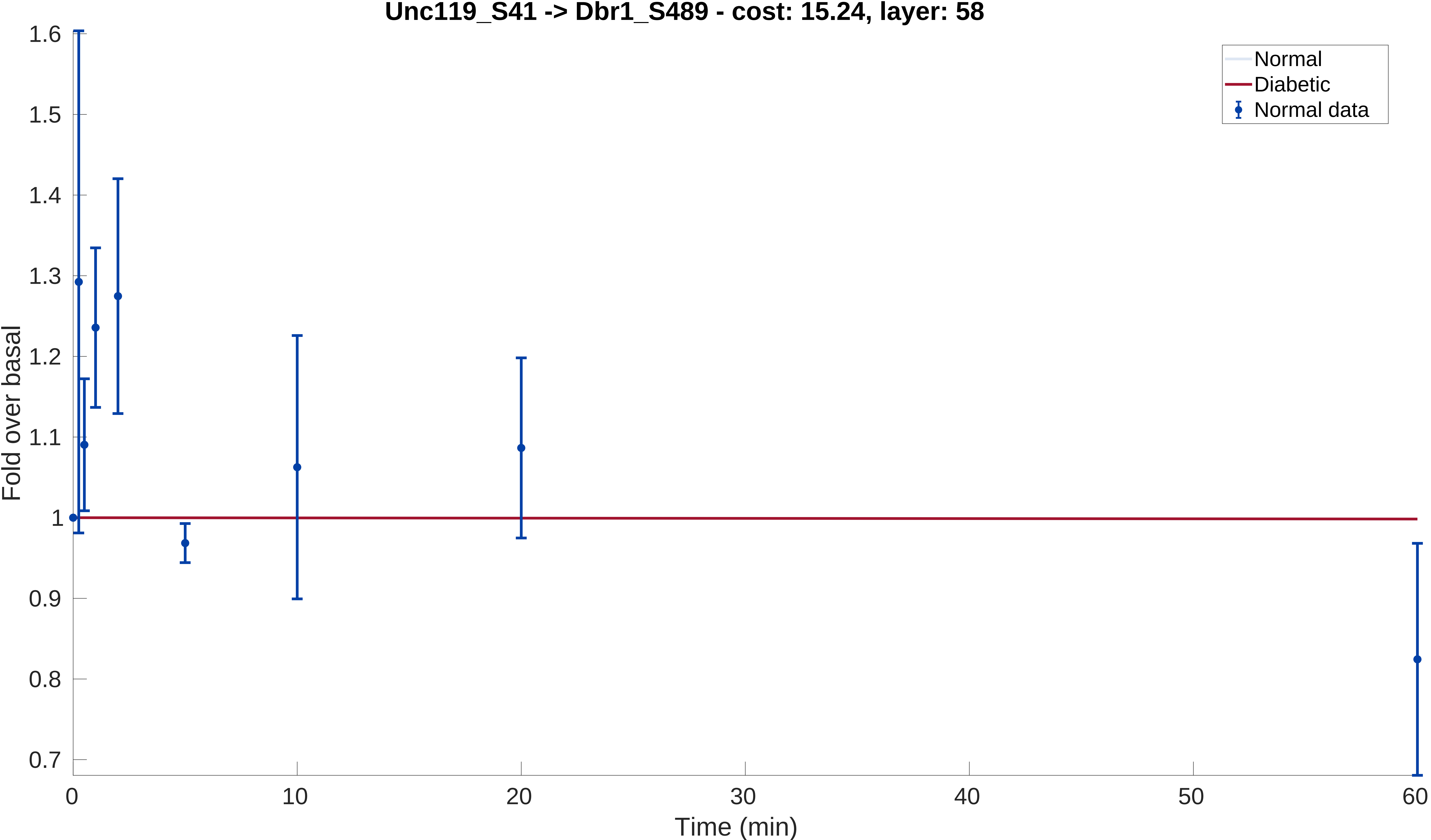
Unc119_S41 -> Dbr1_S489 - cost: 15.24, layer: 58
Fold over basal
Time (min)
Normal
Diabetic
Normal data
0.7
0.8
0.9
1
1.1
1.2
1.3
1.4
1.5
1.6
0
10
20
30
40
50
60

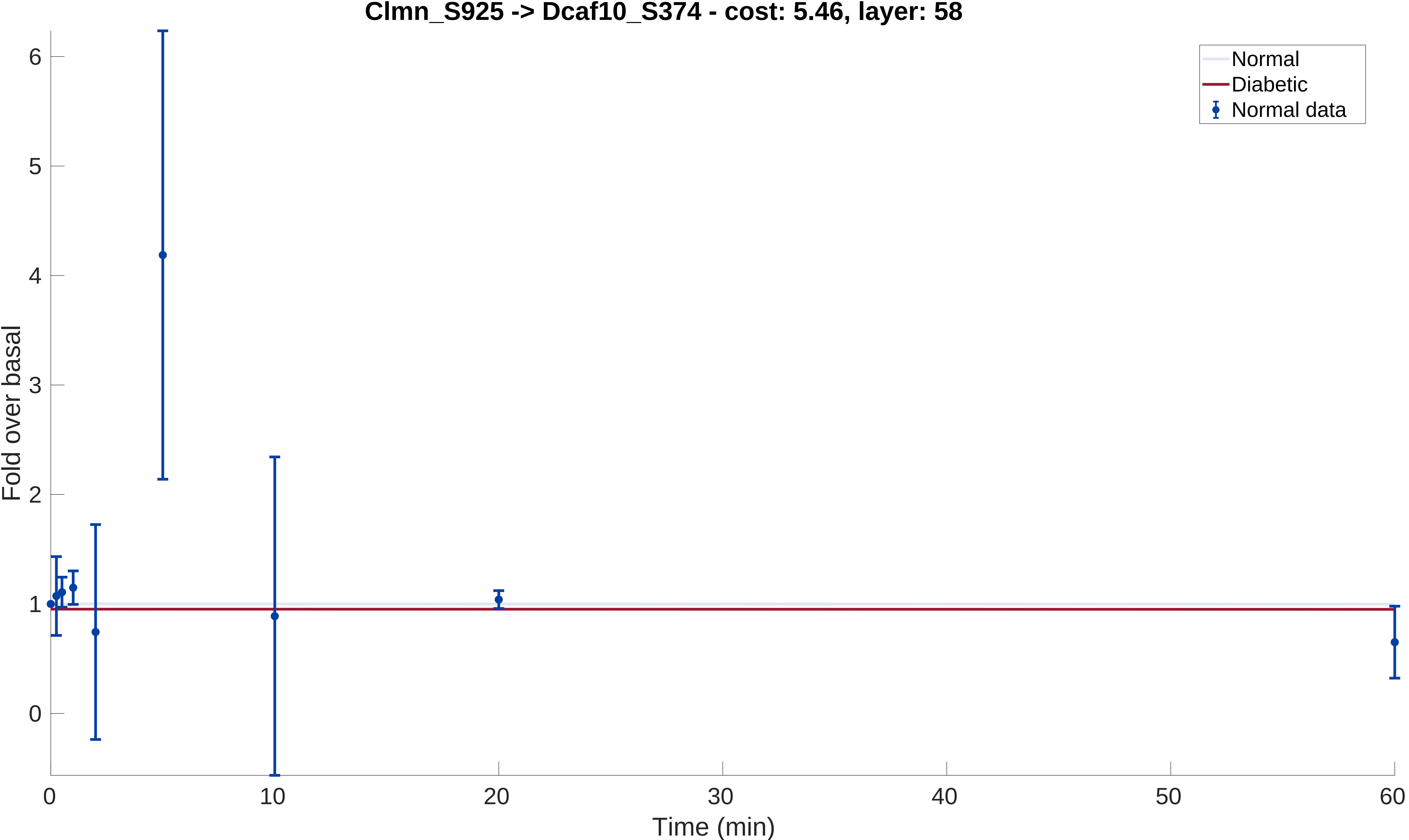

Clmn_S925 -> Dcaf10_S374 - cost: 5.46, layer: 58
Normal
Diabetic
Normal data
Fold over basal
Time (min)
0
1
2
3
4
5
6
0
10
20
30
40
50
60

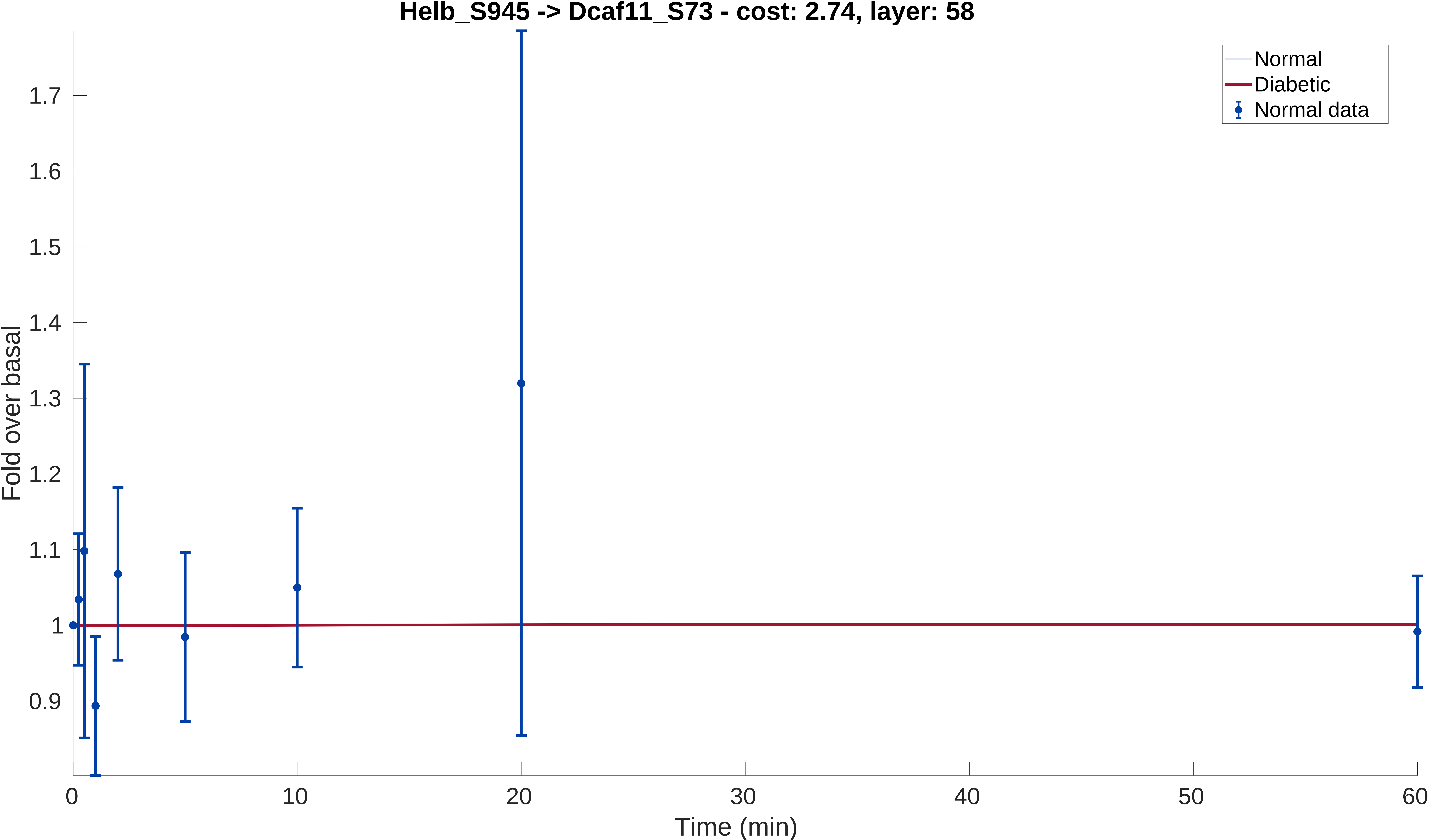

Helb_S945 -> Dcaf11_S73 - cost: 2.74, layer: 58
Normal
Diabetic
Normal data
Fold over basal
1.7
1.6
1.5
1.4
1.3
1.2
1.1
1
0.9
0
10
20
30
40
50
60
Time (min)

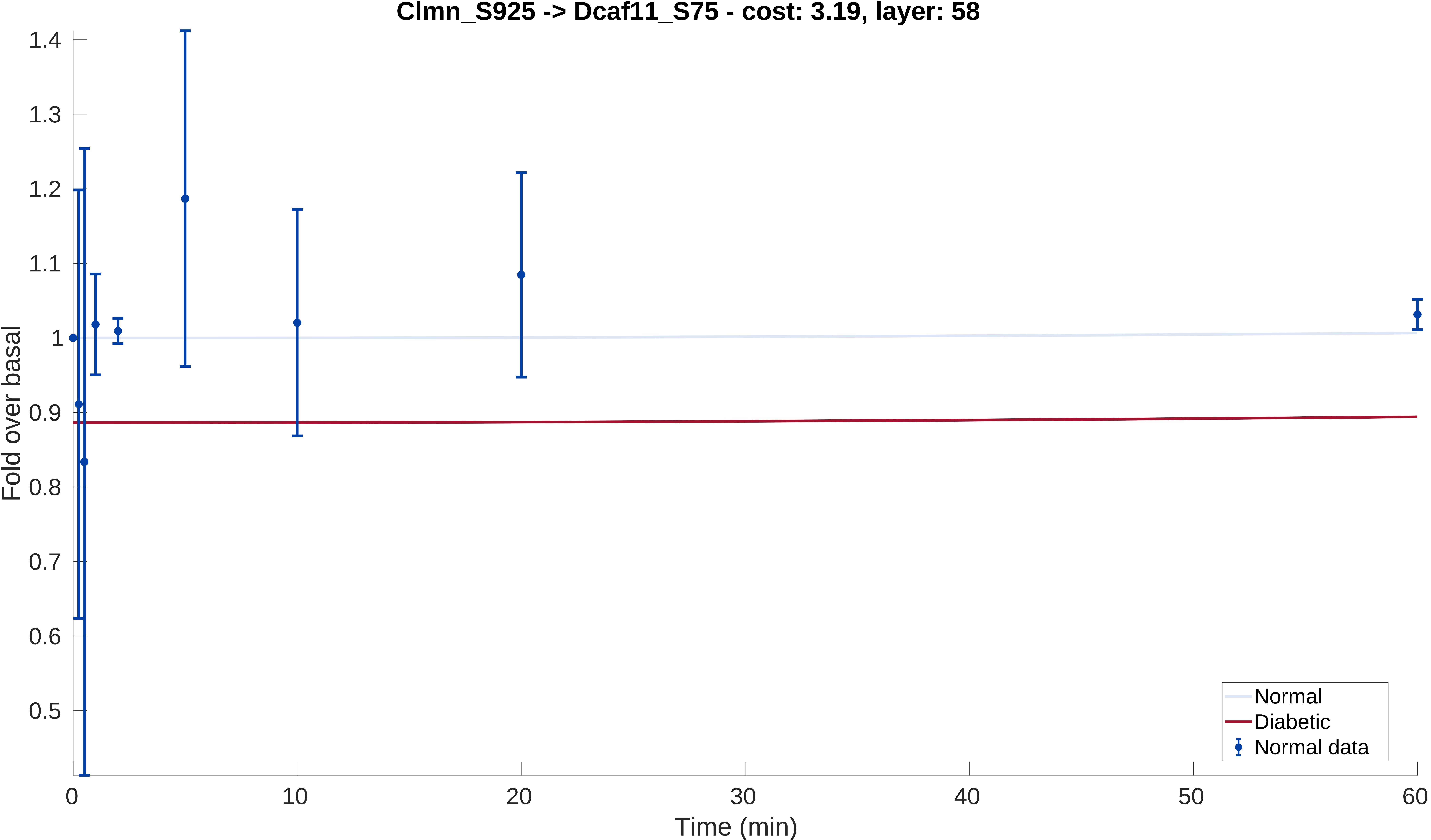

Clmn_S925 -> Dcaf11_S75 - cost: 3.19, layer: 58
Fold over basal
1.4
1.3
1.2
1.1
1
0.9
0.8
0.7
0.6
0.5
0
10
20
30
40
50
60
Time (min)
Normal
Diabetic
Normal data

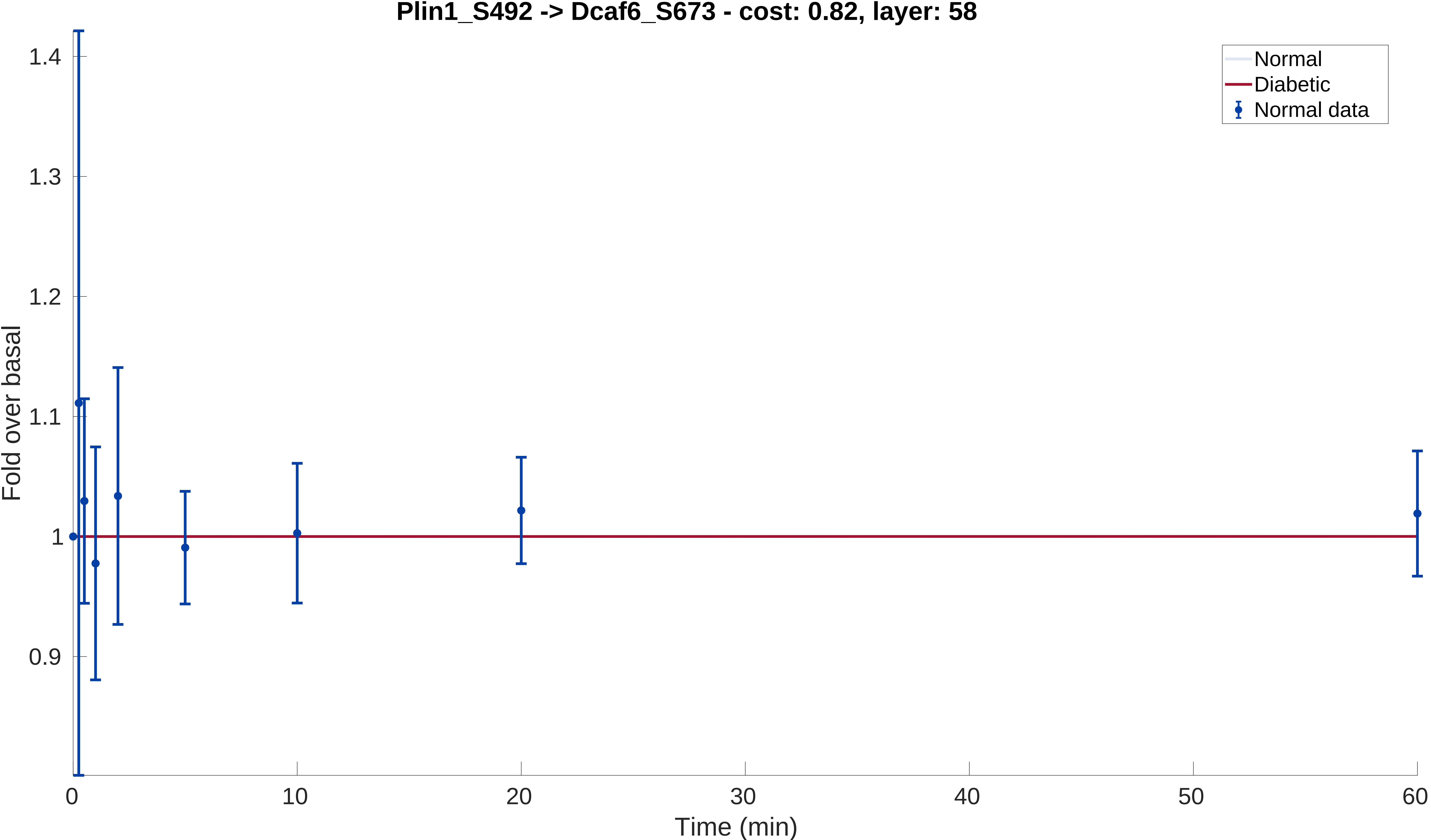

Plin1_S492 -> Dcaf6_S673 - cost: 0.82, layer: 58
Normal
Diabetic
Normal data
Fold over basal
1.4
1.3
1.2
1.1
1
0.9
0
10
20
30
40
50
60
Time (min)

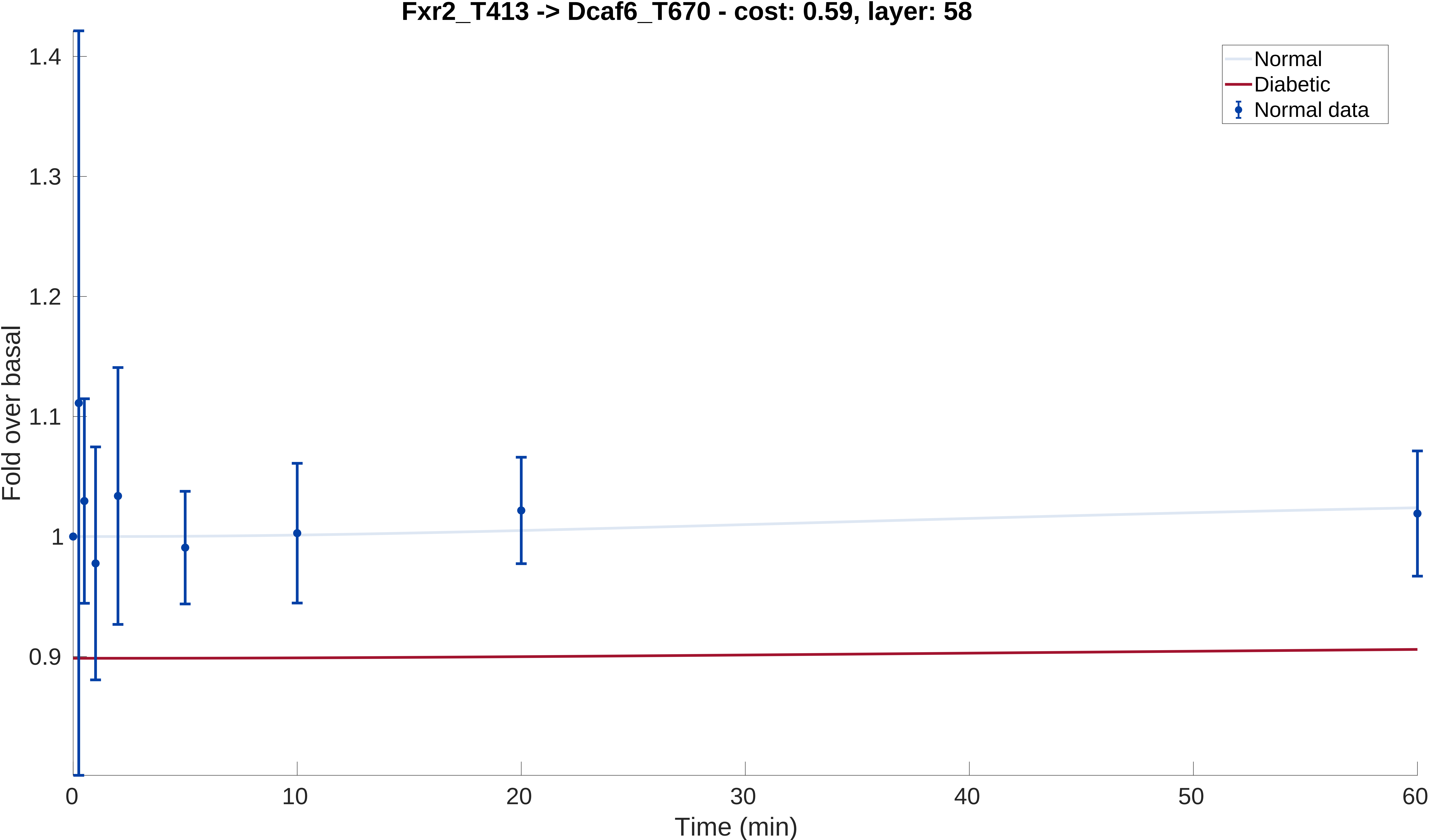

Fxr2_T413 -> Dcaf6_T670 - cost: 0.59, layer: 58
Normal
Diabetic
Normal data
Fold over basal
1.4
1.3
1.2
1.1
1
0.9
0
10
20
30
40
50
60
Time (min)

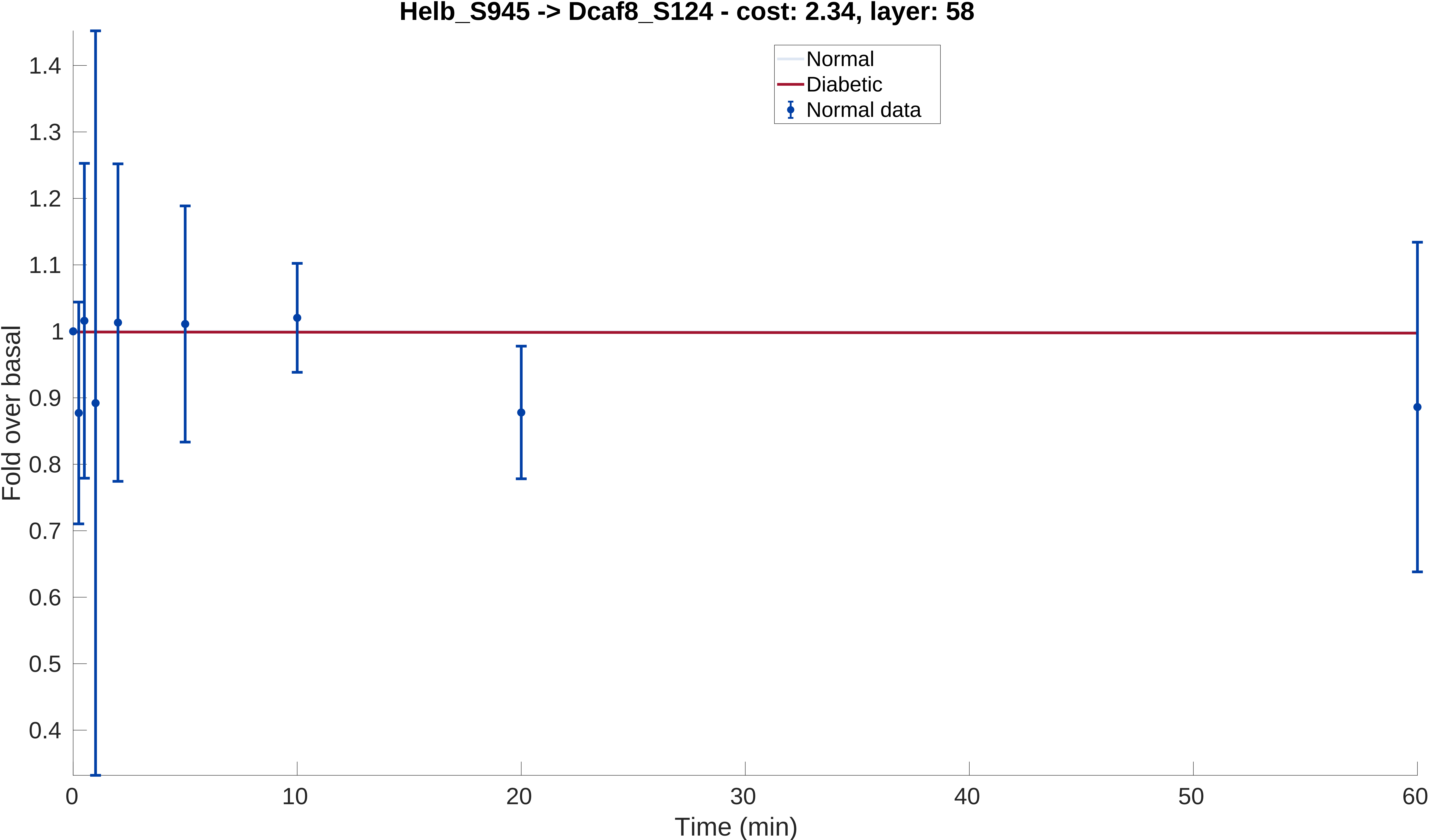
Helb_S945 -> Dcaf8_S124 - cost: 2.34, layer: 58
Normal
Diabetic
Normal data
Fold over basal
1.4
1.3
1.2
1.1
1
0.9
0.8
0.7
0.6
0.5
0.4
0
10
20
30
40
50
60
Time (min)

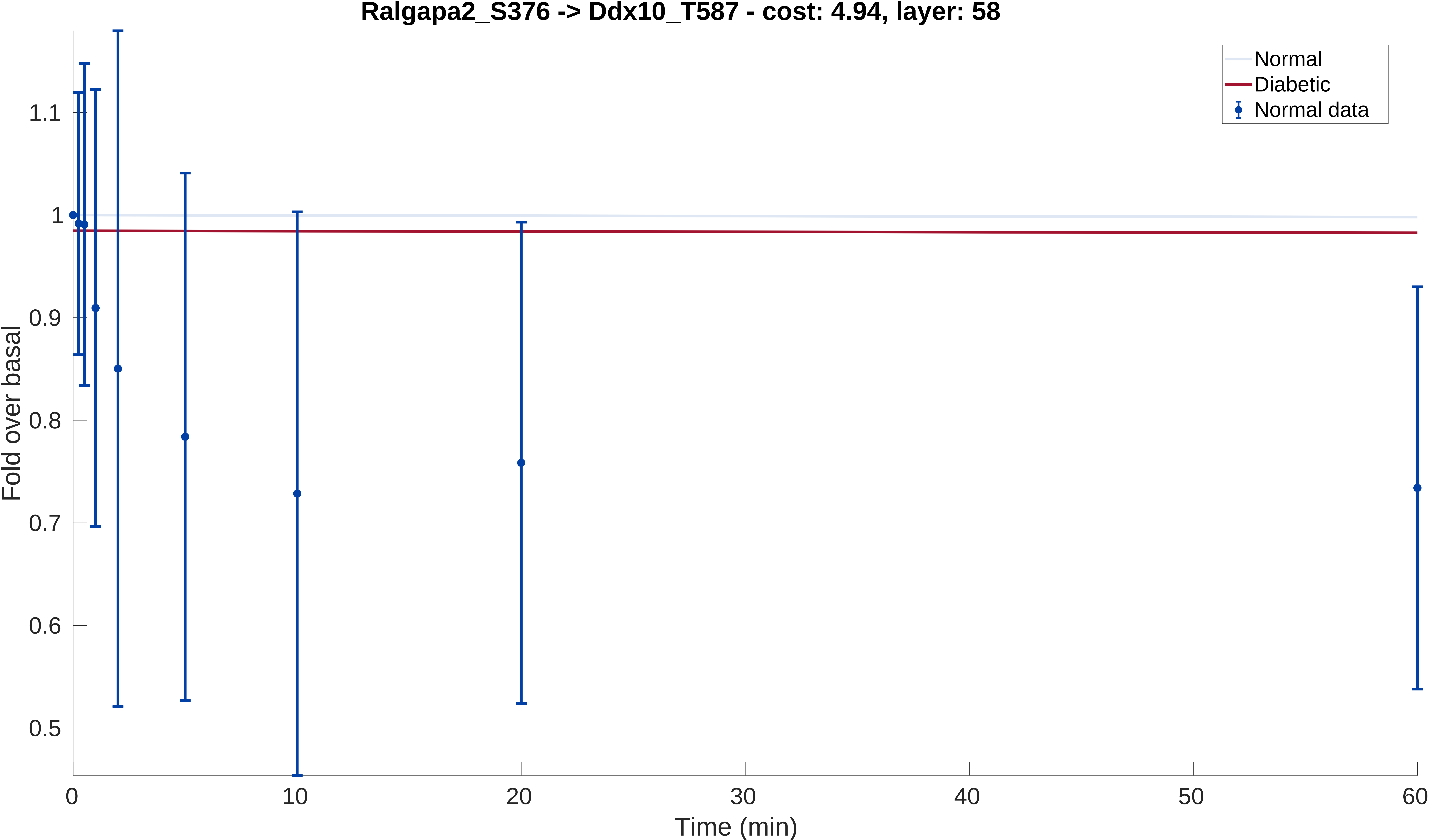

Ralgapa2_S376 -> Ddx10_T587 - cost: 4.94, layer: 58
Normal
Diabetic
Normal data
Fold over basal
1.1
1
0.9
0.8
0.7
0.6
0.5
0
10
20
30
40
50
60
Time (min)

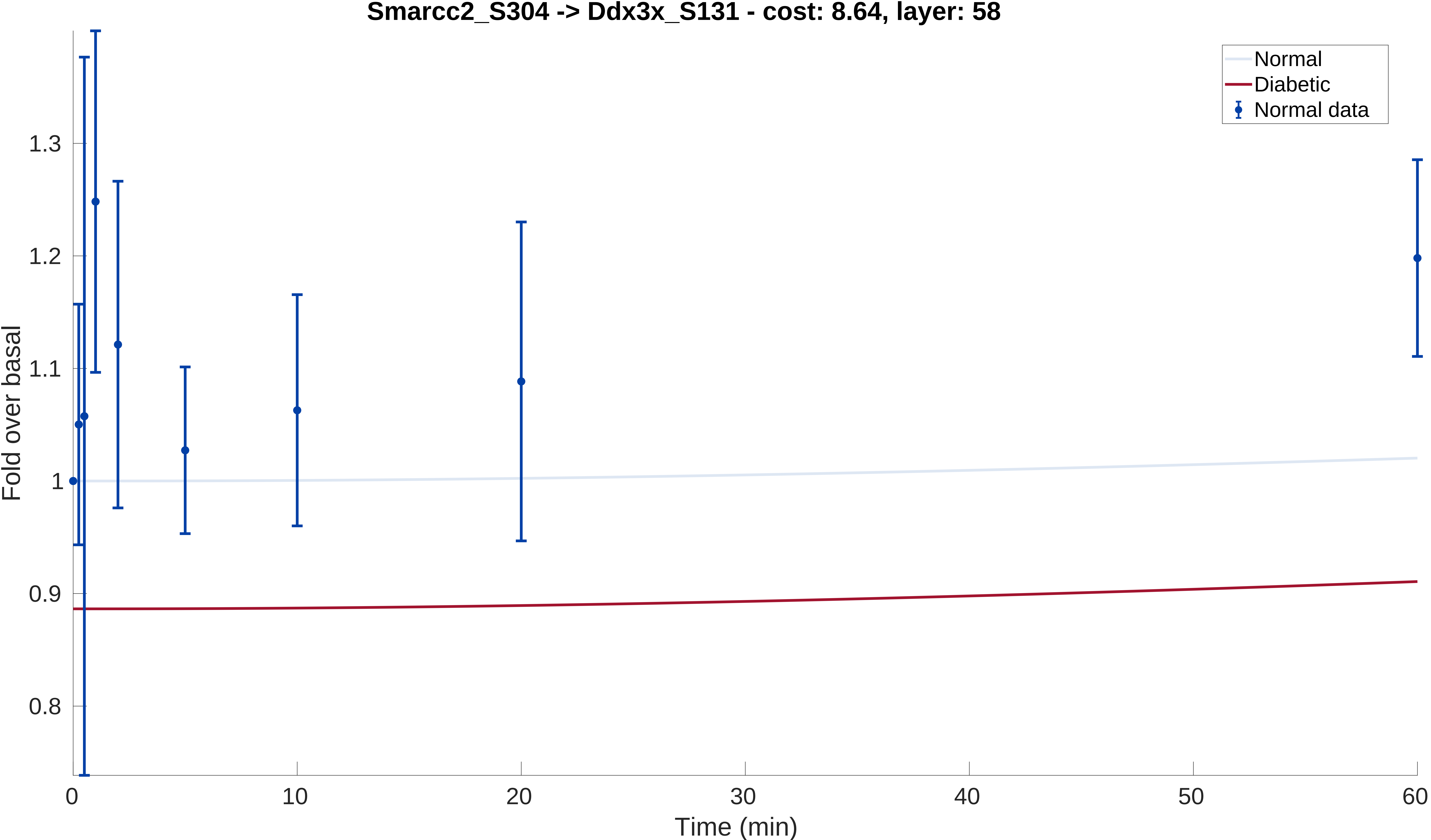

Smarcc2_S304 -> Ddx3x_S131 - cost: 8.64, layer: 58
Normal
Diabetic
Normal data
Fold over basal
1.3
1.2
1.1
1
0.9
0.8
0
10
20
30
40
50
60
Time (min)

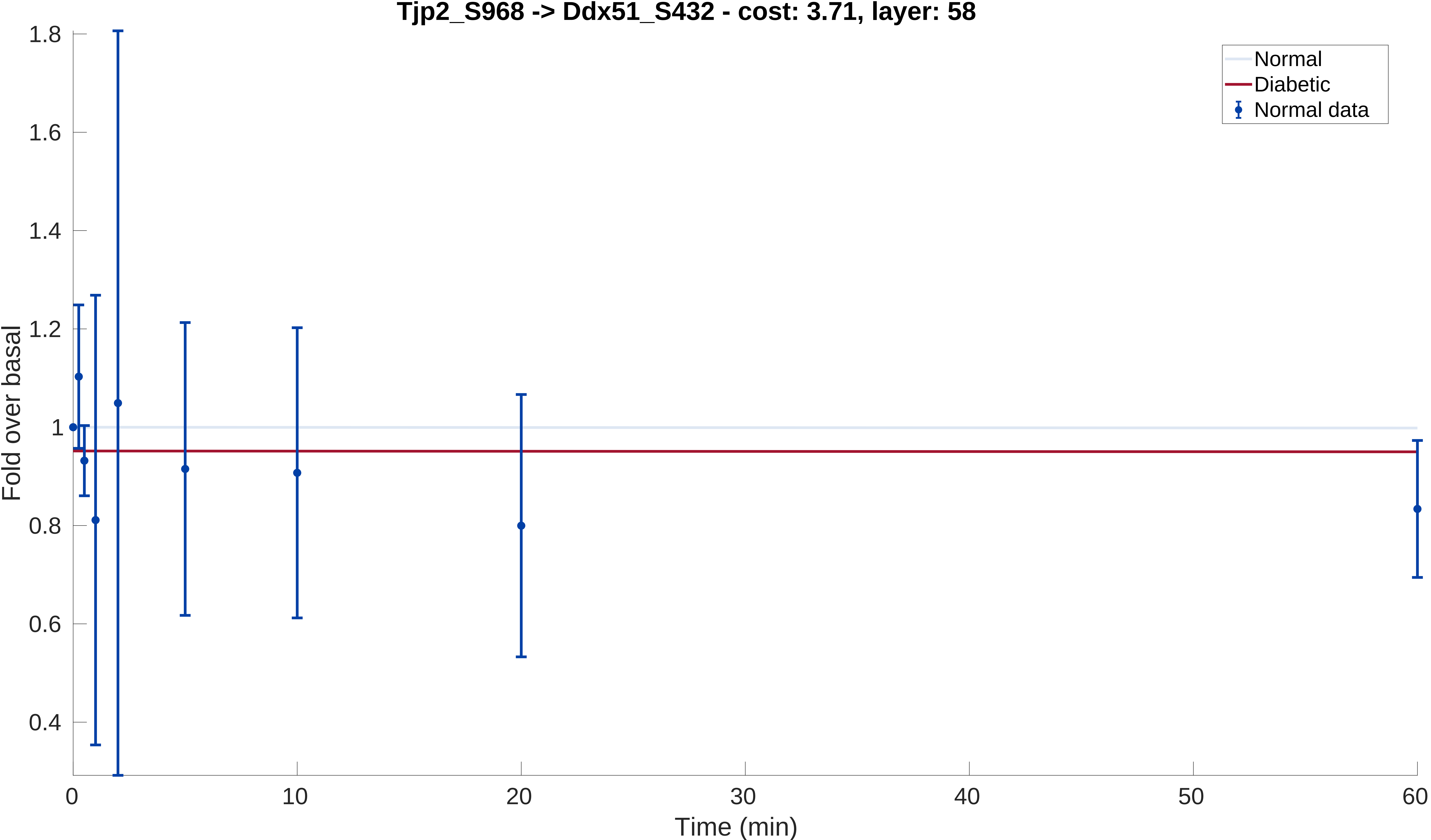

Tjp2_S968 -> Ddx51_S432 - cost: 3.71, layer: 58
Normal
Diabetic
Normal data
Fold over basal
1.8
1.6
1.4
1.2
1
0.8
0.6
0.4
0
10
20
30
40
50
60
Time (min)

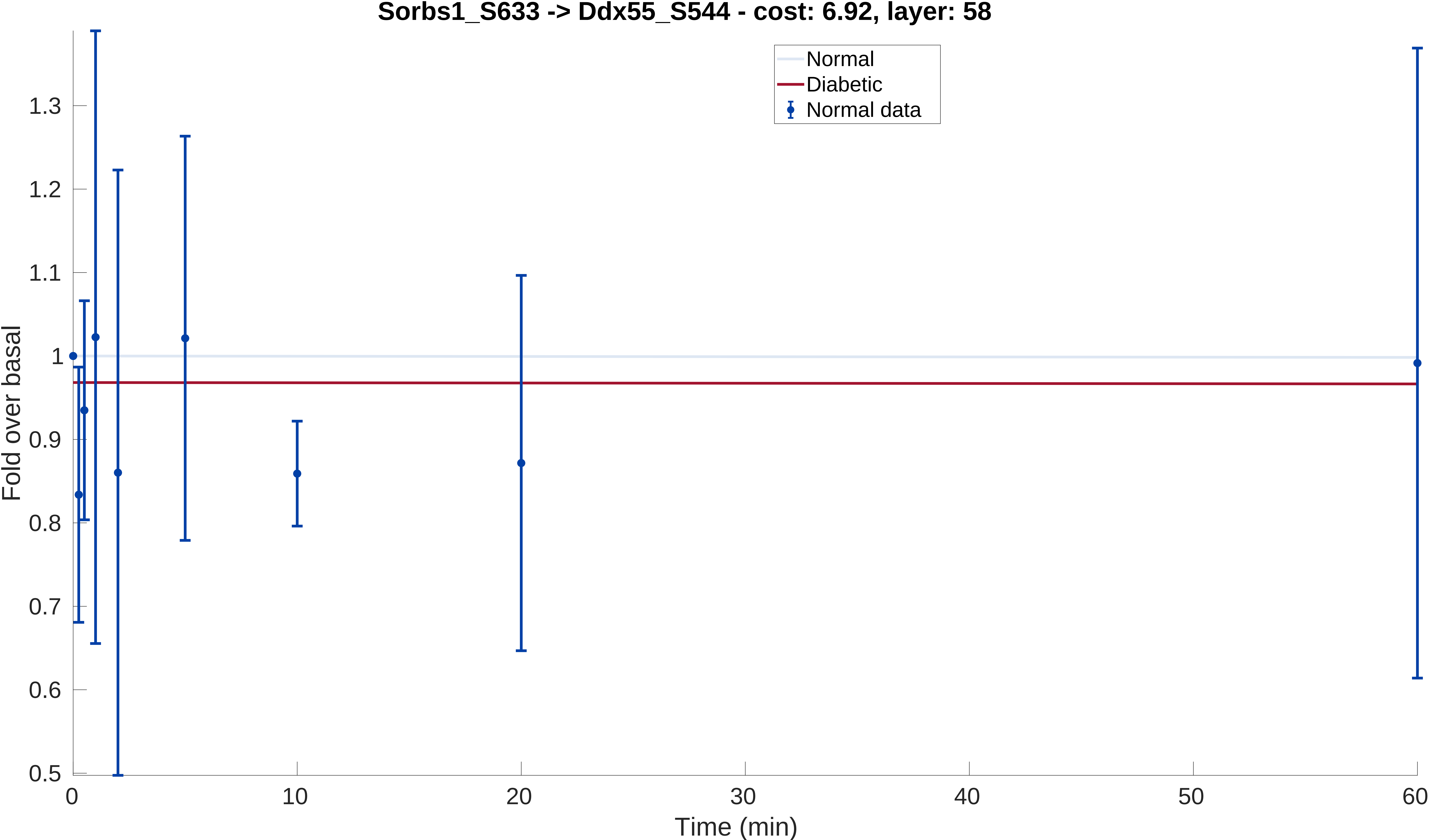

Sorbs1_S633 -> Ddx55_S544 - cost: 6.92, layer: 58
Normal
Diabetic
Normal data
1.3
1.2
1.1
1
0.9
0.8
0.7
0.6
0.5
Fold over basal
0
10
20
30
40
50
60
Time (min)

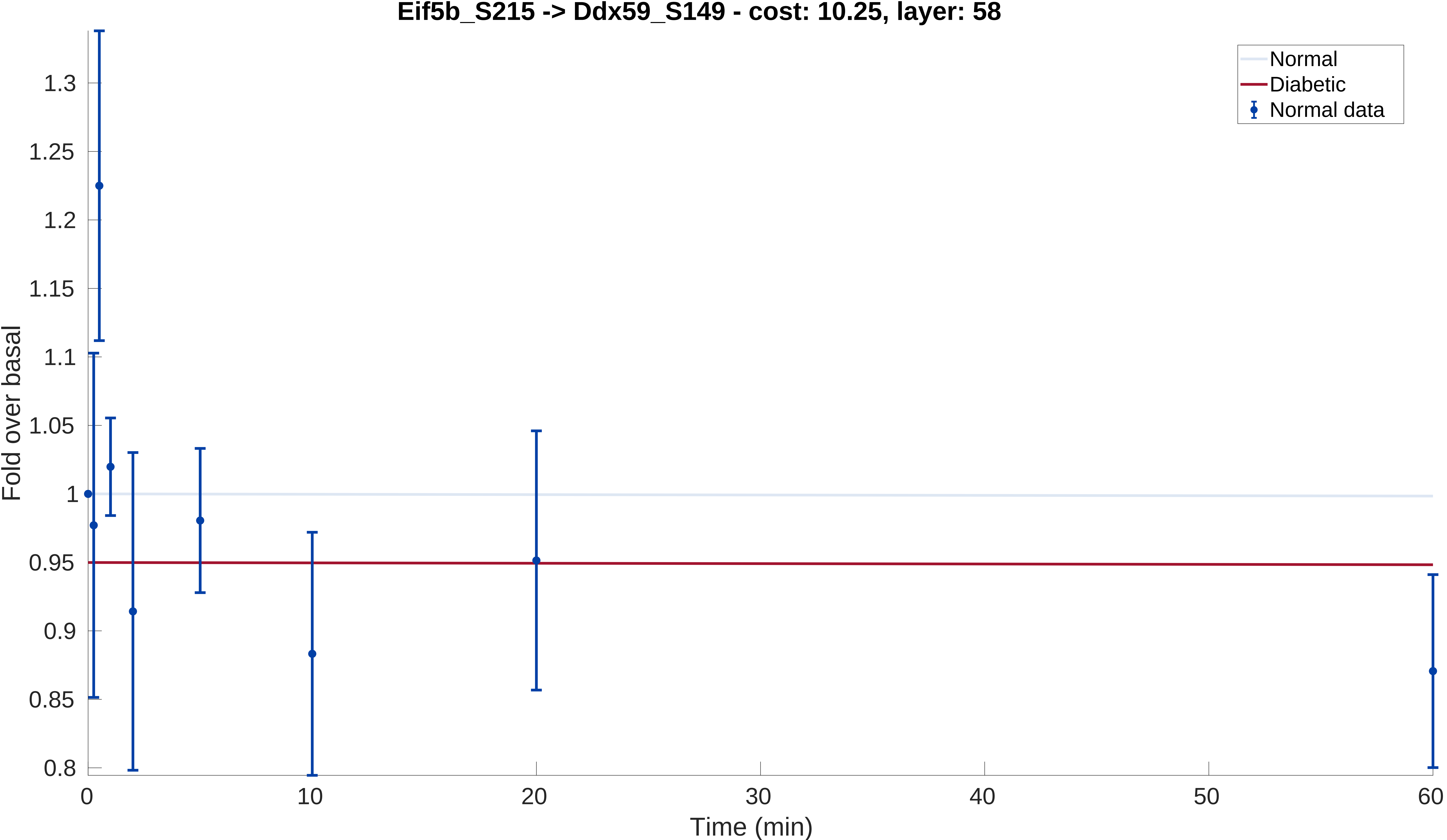

Eif5b_S215 -> Ddx59_S149 - cost: 10.25, layer: 58
Normal
Diabetic
Normal data
Fold over basal
Time (min)
1.3
1.25
1.2
1.15
1.1
1.05
1
0.95
0.9
0.85
0.8
0
10
20
30
40
50
60

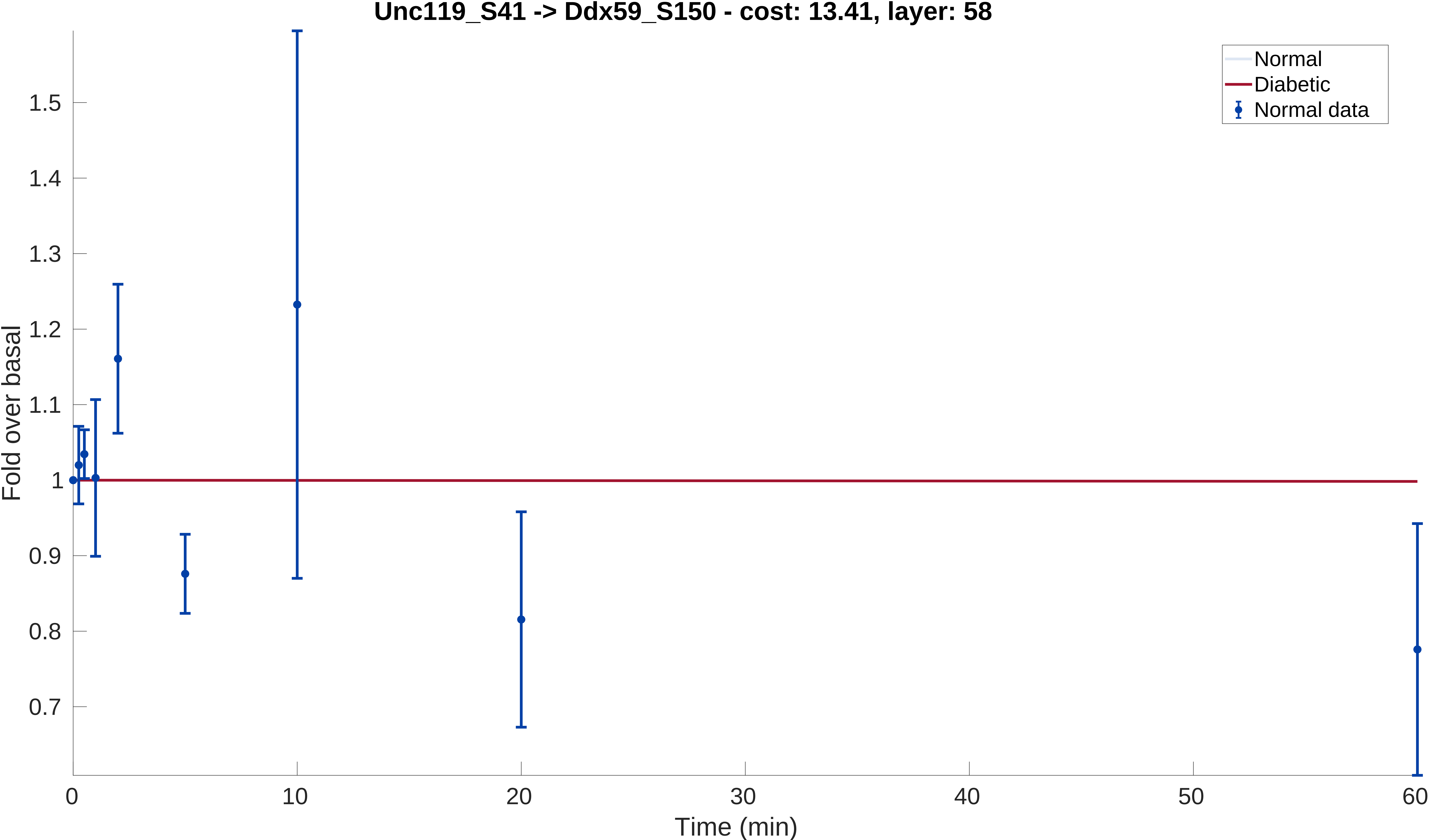

Unc119_S41 -> Ddx59_S150 - cost: 13.41, layer: 58
Normal
Diabetic
Normal data
Fold over basal
1.5
1.4
1.3
1.2
1.1
1
0.9
0.8
0.7
0
10
20
30
40
50
60
Time (min)

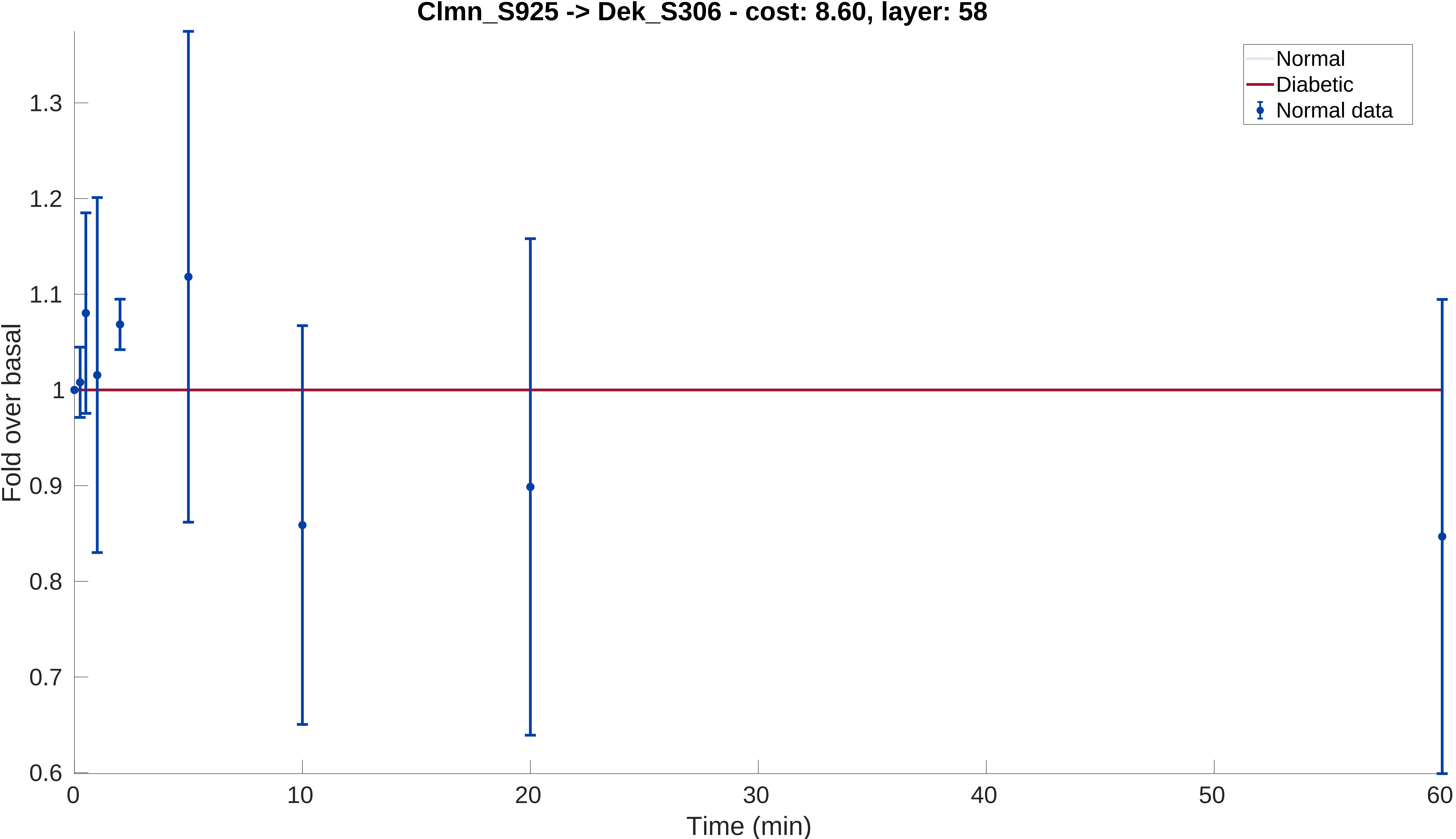

Clmn_S925 -> Dek_S306 - cost: 8.60, layer: 58
Normal
Diabetic
Normal data
Fold over basal
1.3
1.2
1.1
1
0.9
0.8
0.7
0.6
0
10
20
30
40
50
60
Time (min)

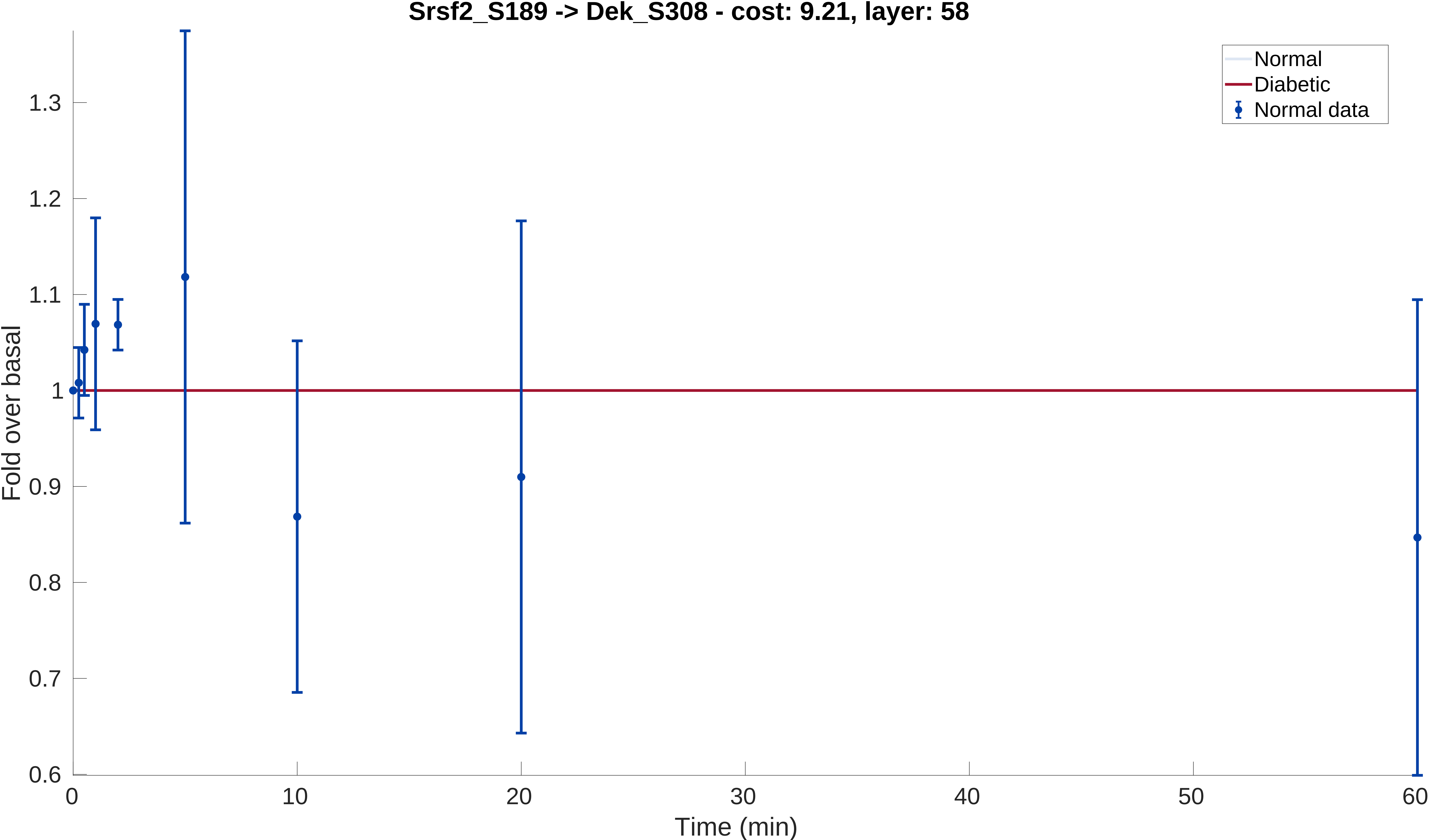

Srsf2_S189 -> Dek_S308 - cost: 9.21, layer: 58
Normal
Diabetic
Normal data
1.3
1.2
1.1
1
0.9
0.8
0.7
0.6
Fold over basal
0
10
20
30
40
50
60
Time (min)

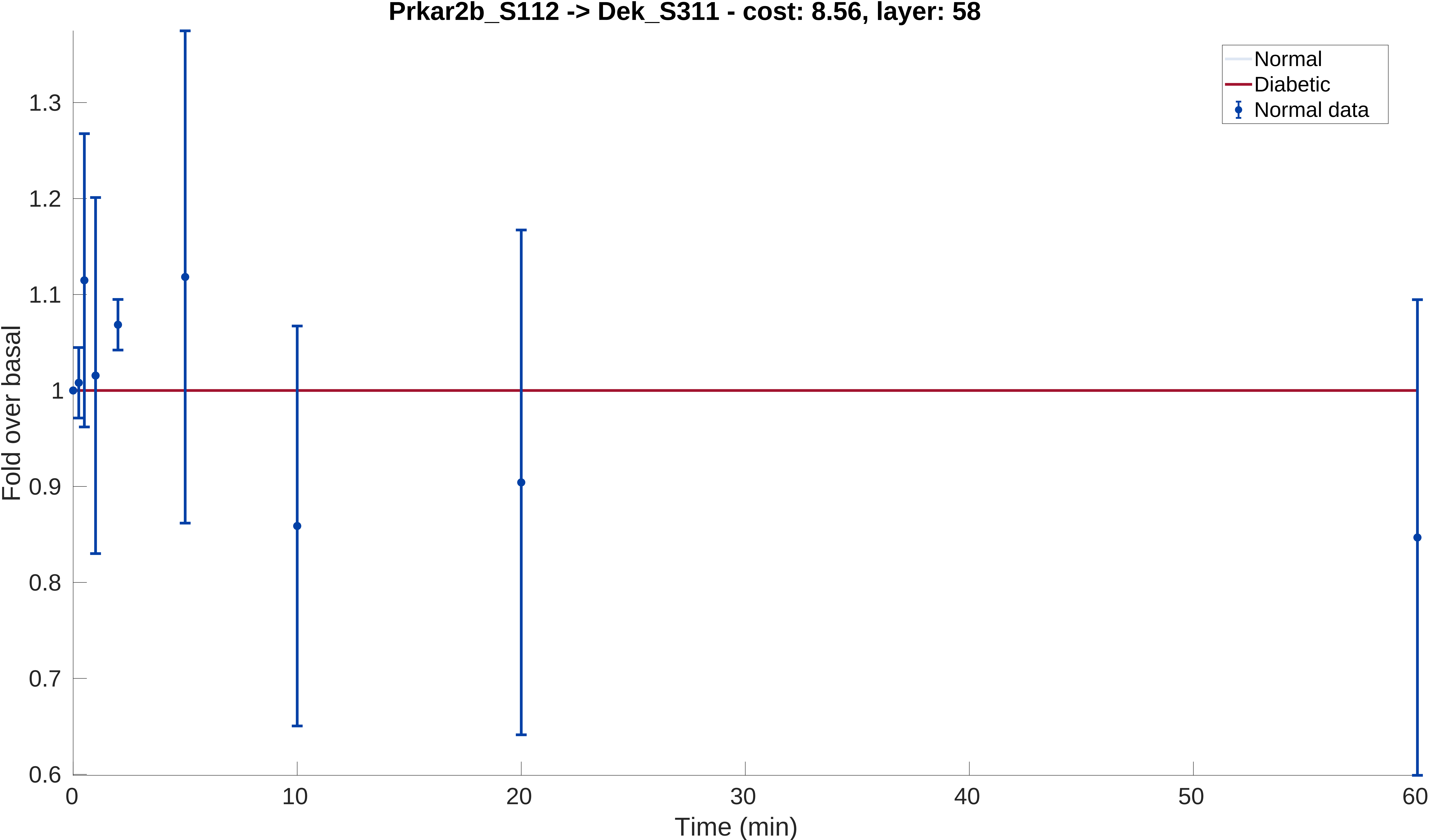

Prkar2b_S112 -> Dek_S311 - cost: 8.56, layer: 58
Normal
Diabetic
Normal data
Fold over basal
1.3
1.2
1.1
1
0.9
0.8
0.7
0.6
0
10
20
30
40
50
60
Time (min)

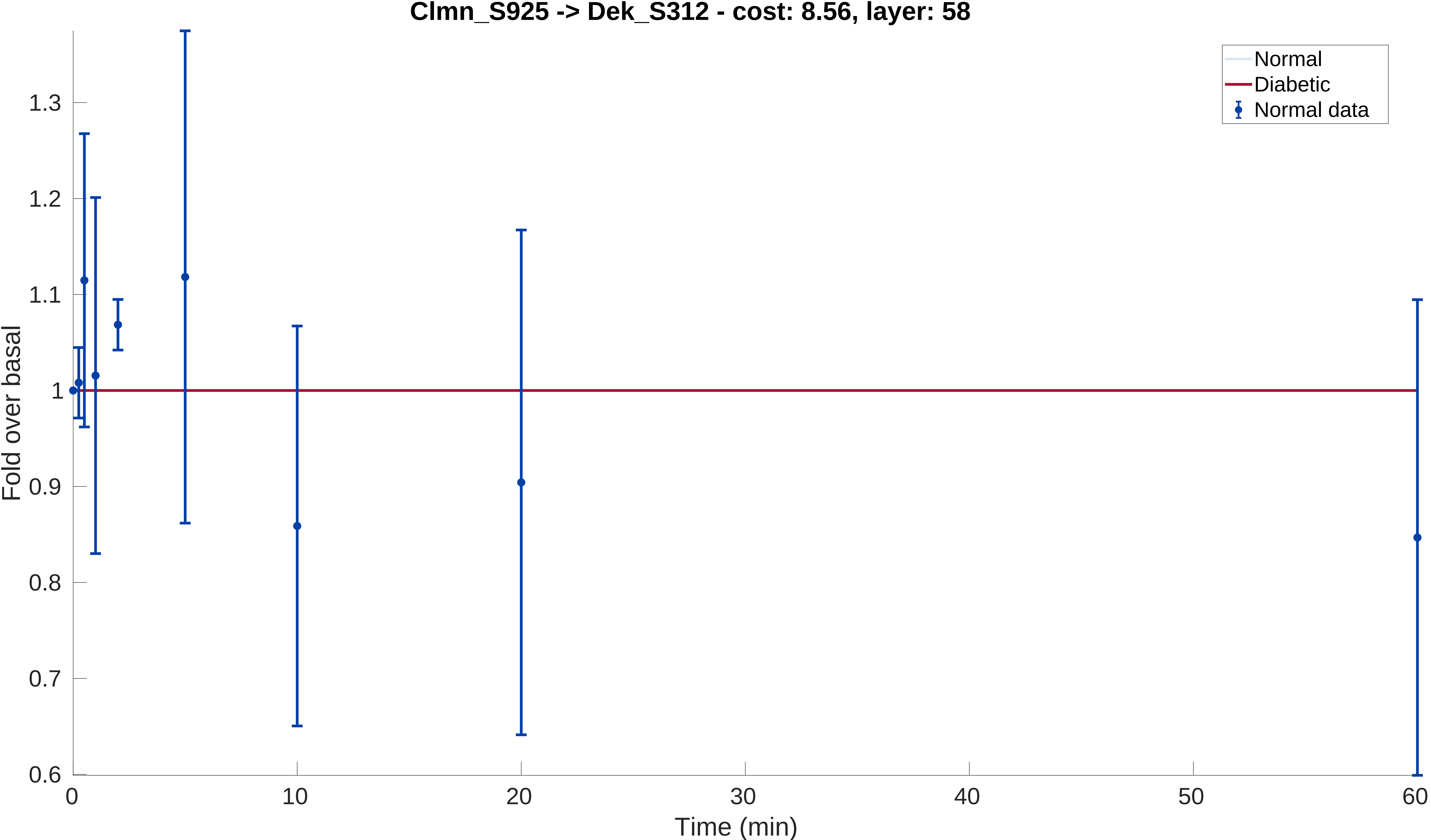

Clmn_S925 -> Dek_S312 - cost: 8.56, layer: 58
Normal
Diabetic
Normal data
Fold over basal
1.3
1.2
1.1
1
0.9
0.8
0.7
0.6
0
10
20
30
40
50
60
Time (min)

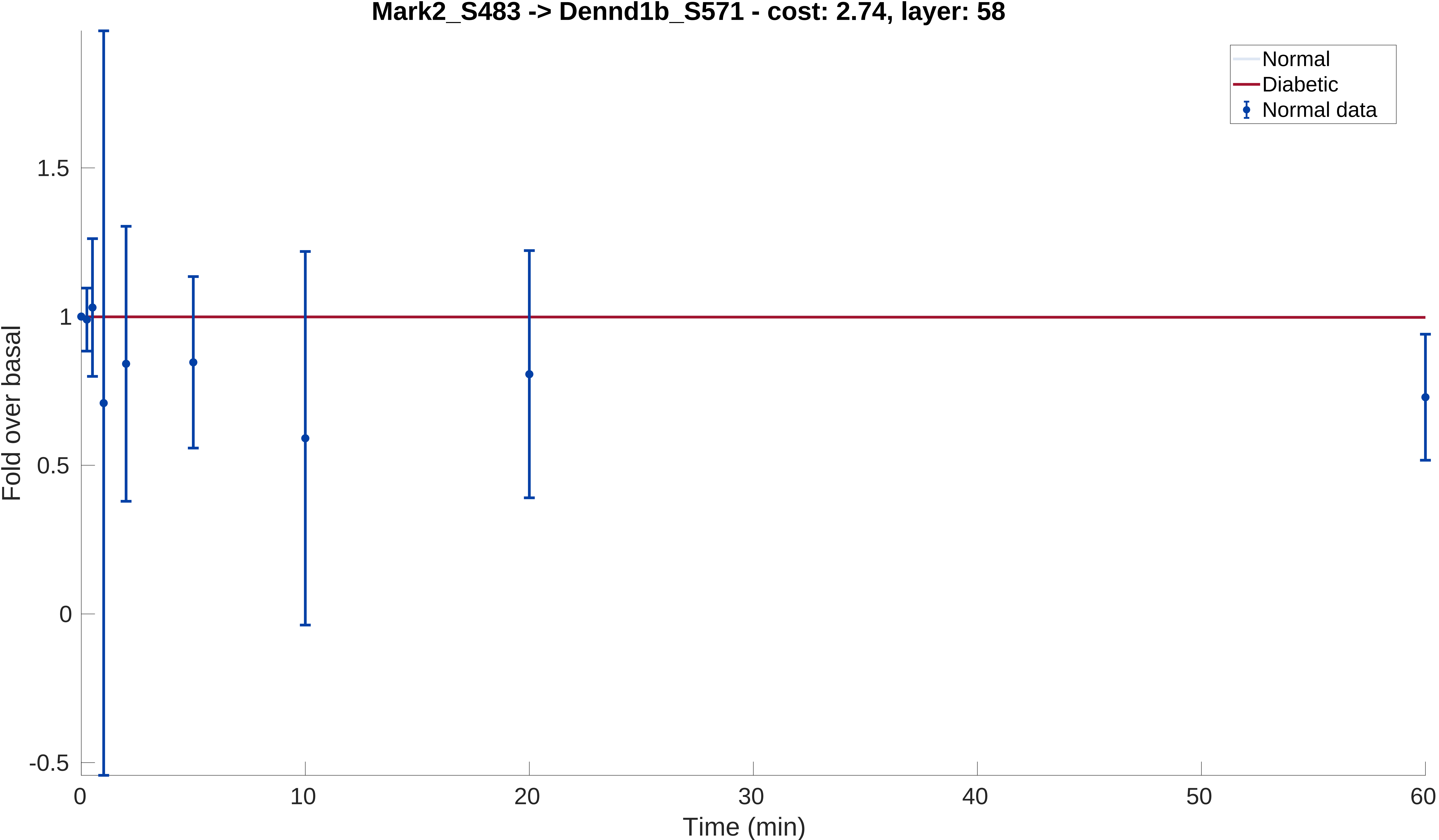

Mark2_S483 -> Dennd1b_S571 - cost: 2.74, layer: 58
Normal
Diabetic
Normal data
Fold over basal
1.5
1
0.5
0
-0.5
0
10
20
30
40
50
60
Time (min)

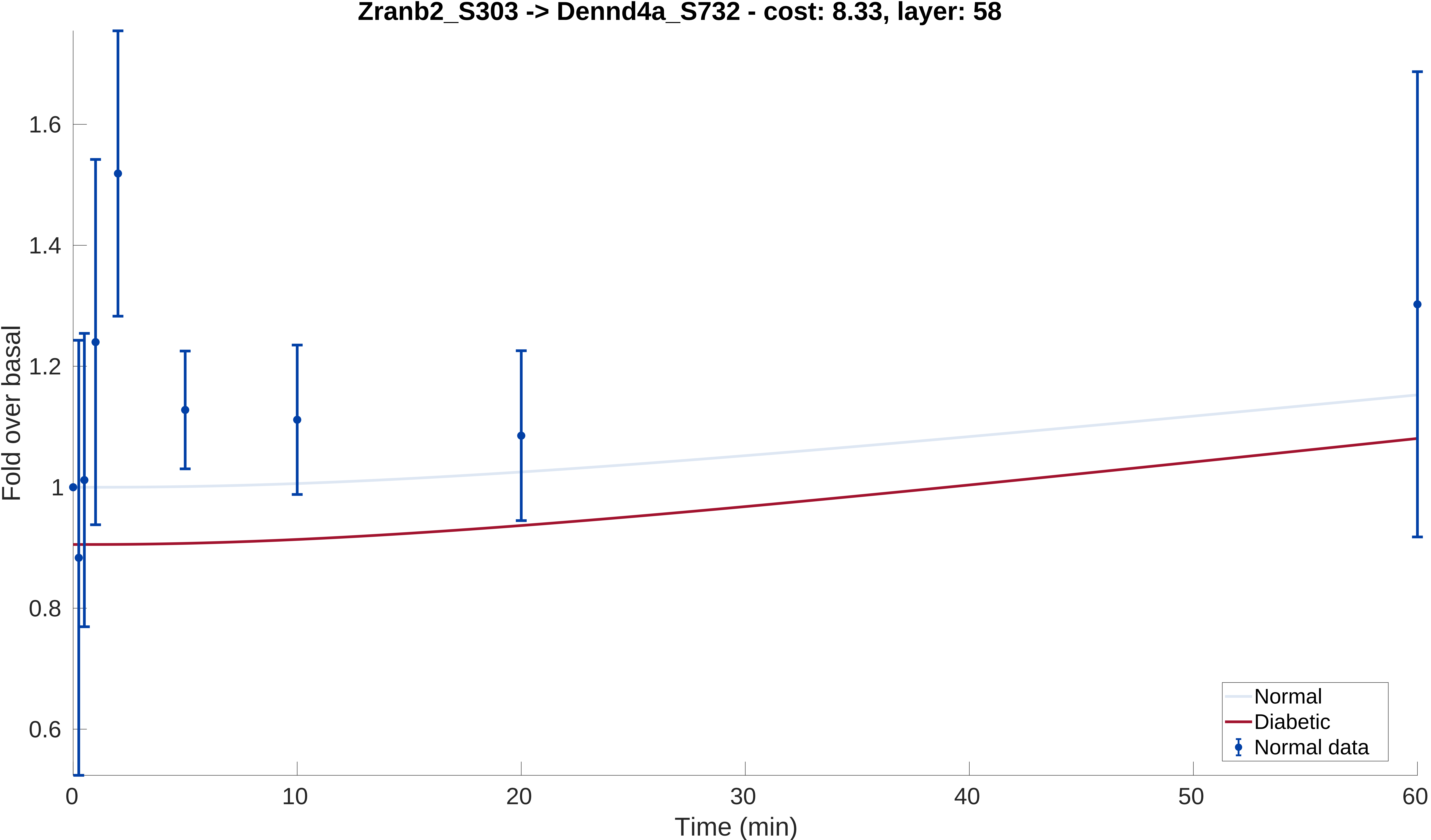

Zranb2_S303 -> Dennd4a_S732 - cost: 8.33, layer: 58
Fold over basal
1.6
1.4
1.2
1
0.8
0.6
0
10
20
30
40
50
60
Time (min)
Normal
Diabetic
Normal data

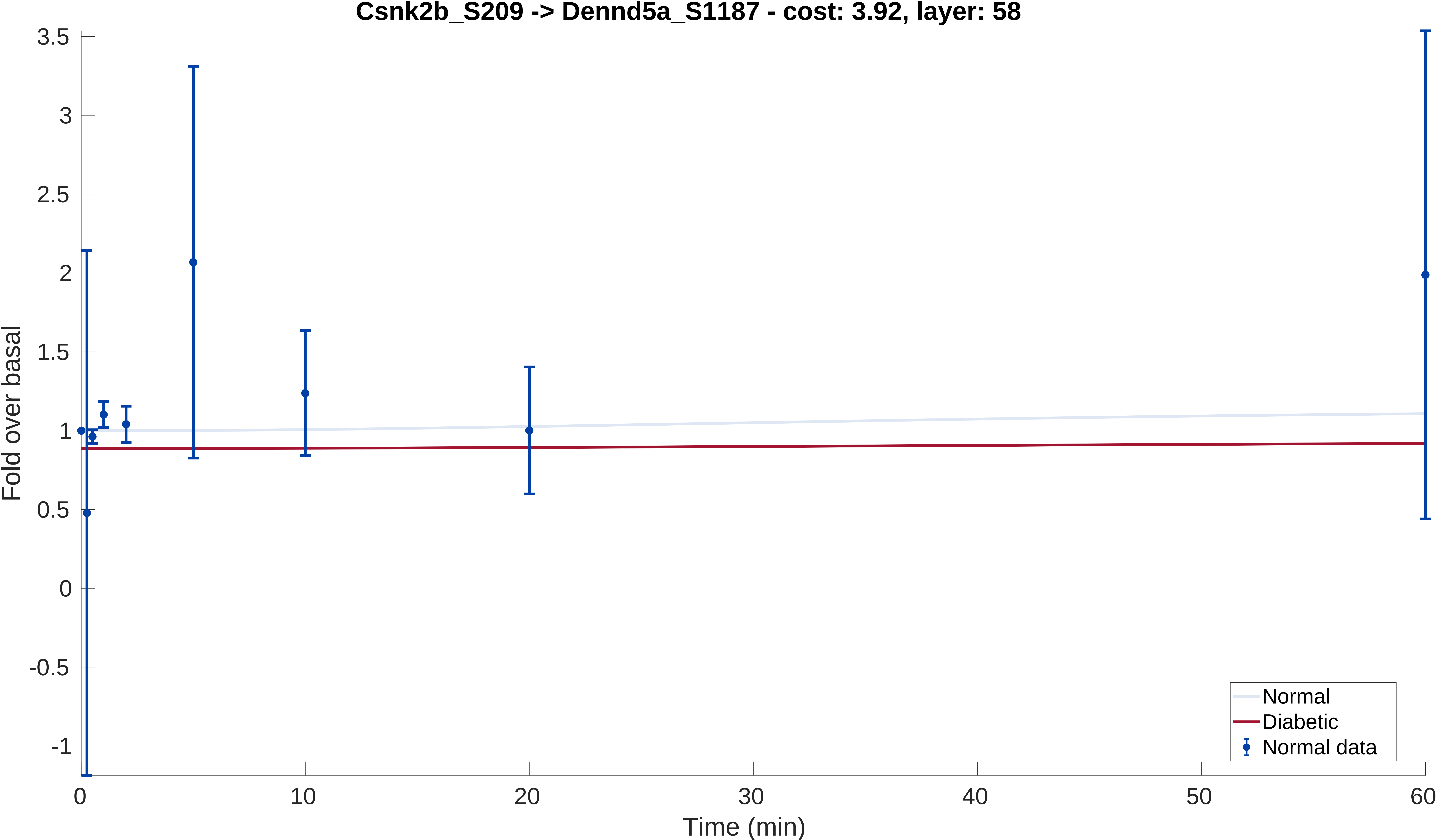

Csnk2b_S209 -> Dennd5a_S1187 - cost: 3.92, layer: 58
Fold over basal
3.5
3
2.5
2
1.5
1
0.5
0
-0.5
-1
0
10
20
30
40
50
60
Time (min)
Normal
Diabetic
Normal data

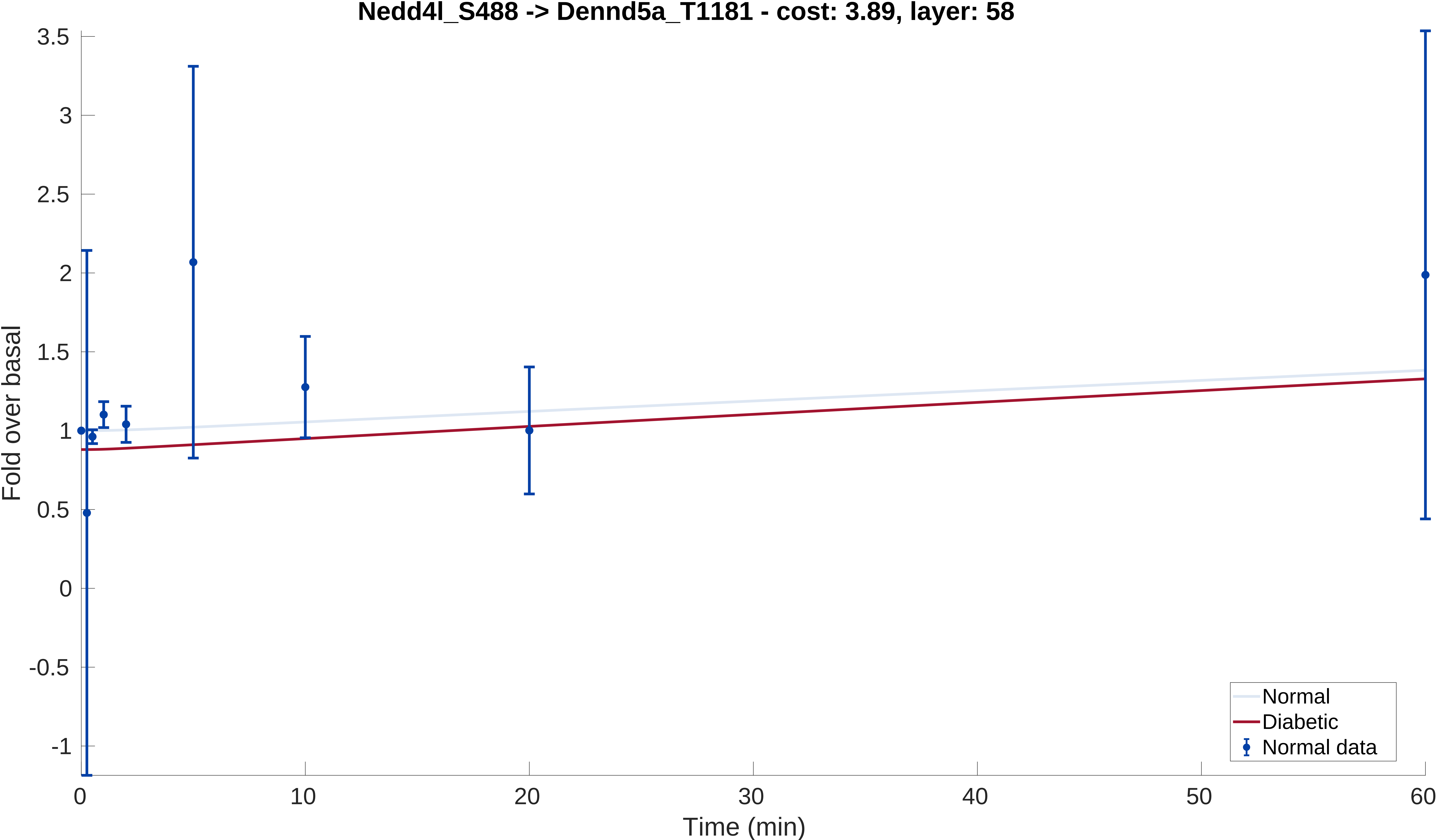
Nedd4l_S488 -> Dennd5a_T1181 - cost: 3.89, layer: 58
Fold over basal
3.5
3
2.5
2
1.5
1
0.5
0
-0.5
-1
0
10
20
30
40
50
60
Time (min)
Normal
Diabetic
Normal data

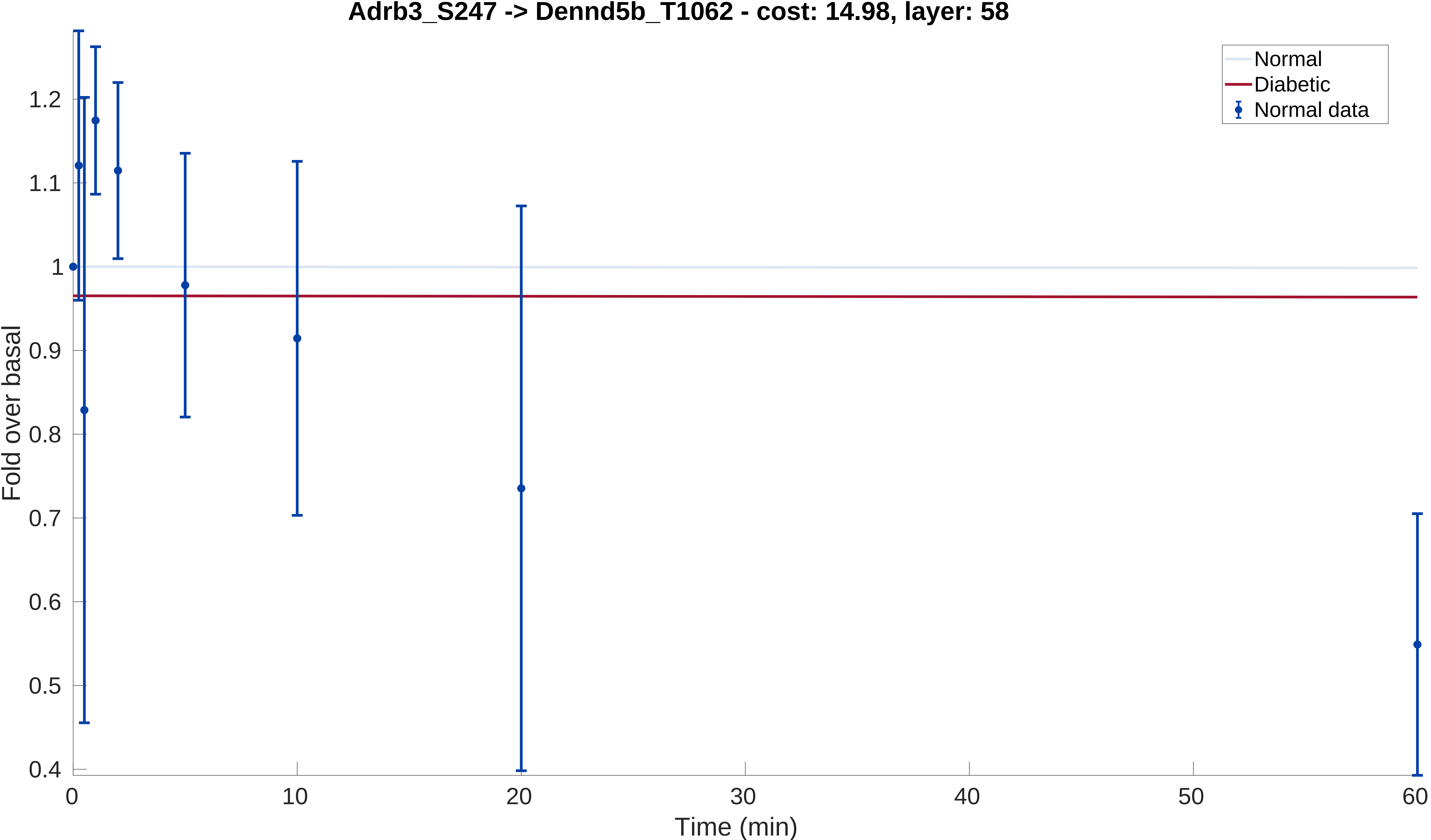
Adrb3_S247 -> Dennd5b_T1062 - cost: 14.98, layer: 58
Normal
Diabetic
Normal data
1.2
1.1
1
0.9
0.8
0.7
0.6
0.5
0.4
Fold over basal
0
10
20
30
40
50
60
Time (min)

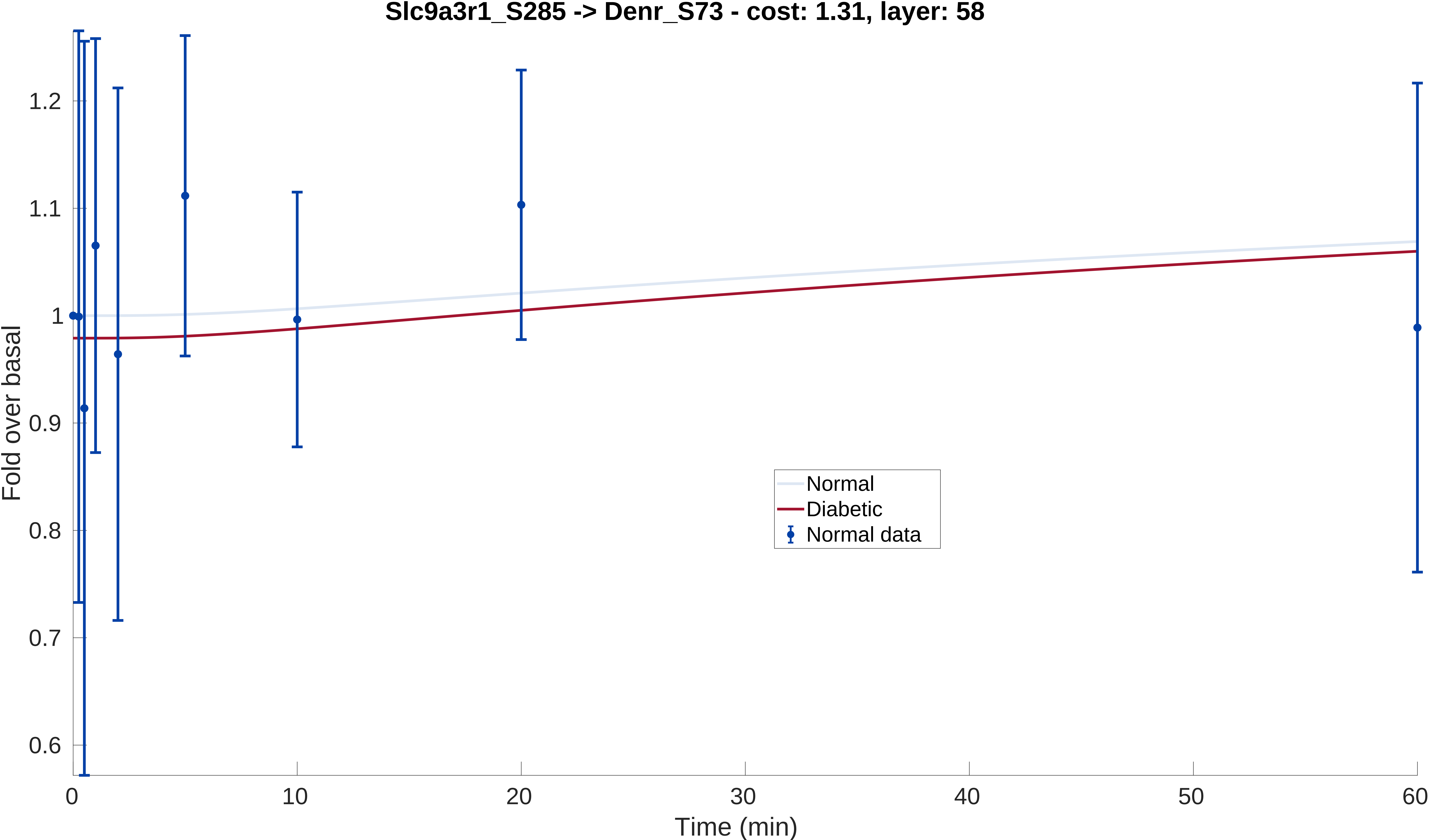

Slc9a3r1_S285 -> Denr_S73 - cost: 1.31, layer: 58
Fold over basal
1.2
1.1
1
0.9
0.8
0.7
0.6
0
10
20
30
40
50
60
Time (min)
Normal
Diabetic
Normal data

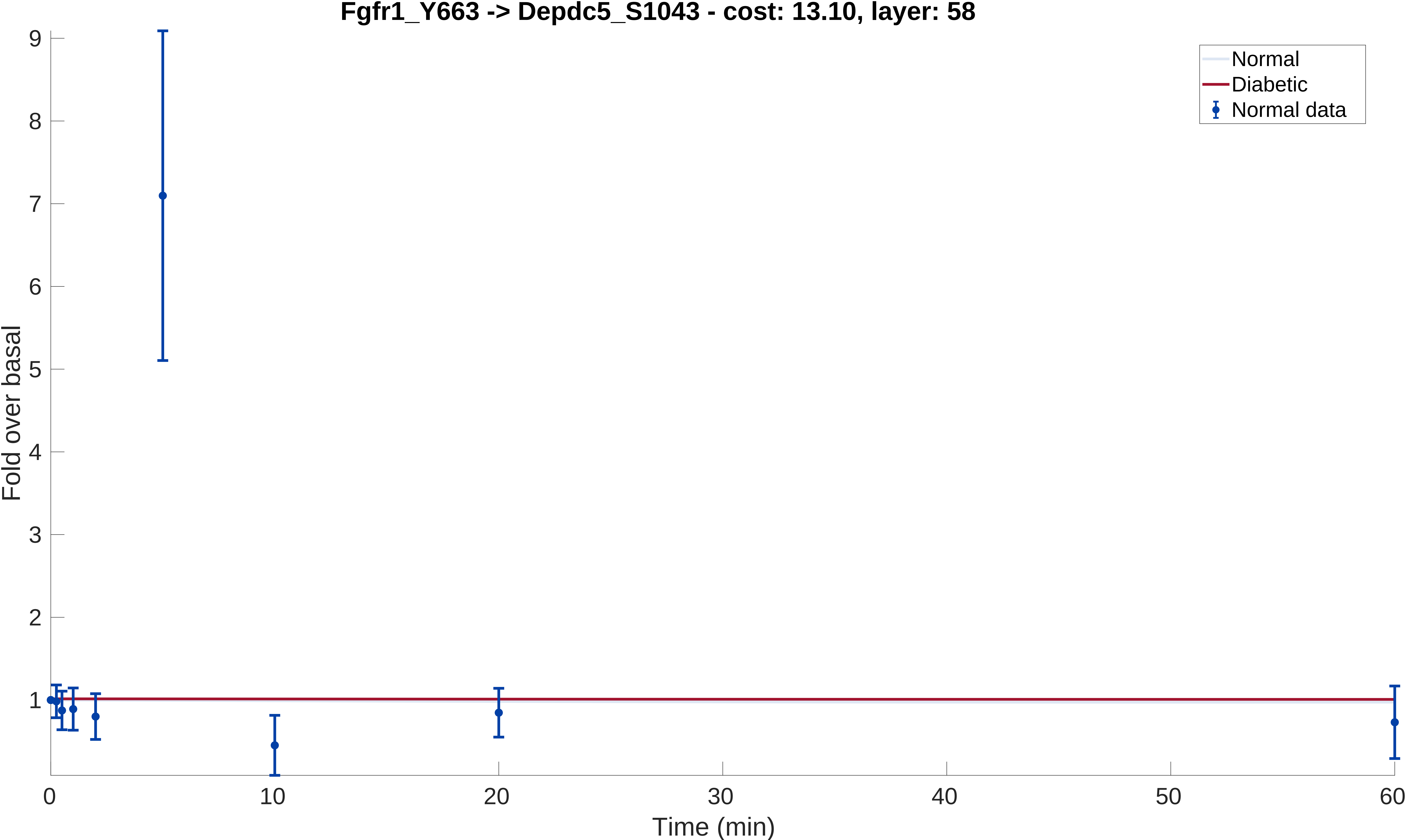

Fgfr1_Y663 -> Depdc5_S1043 - cost: 13.10, layer: 58
Normal
Diabetic
Normal data
Fold over basal
Time (min)
0
10
20
30
40
50
60
1
2
3
4
5
6
7
8
9

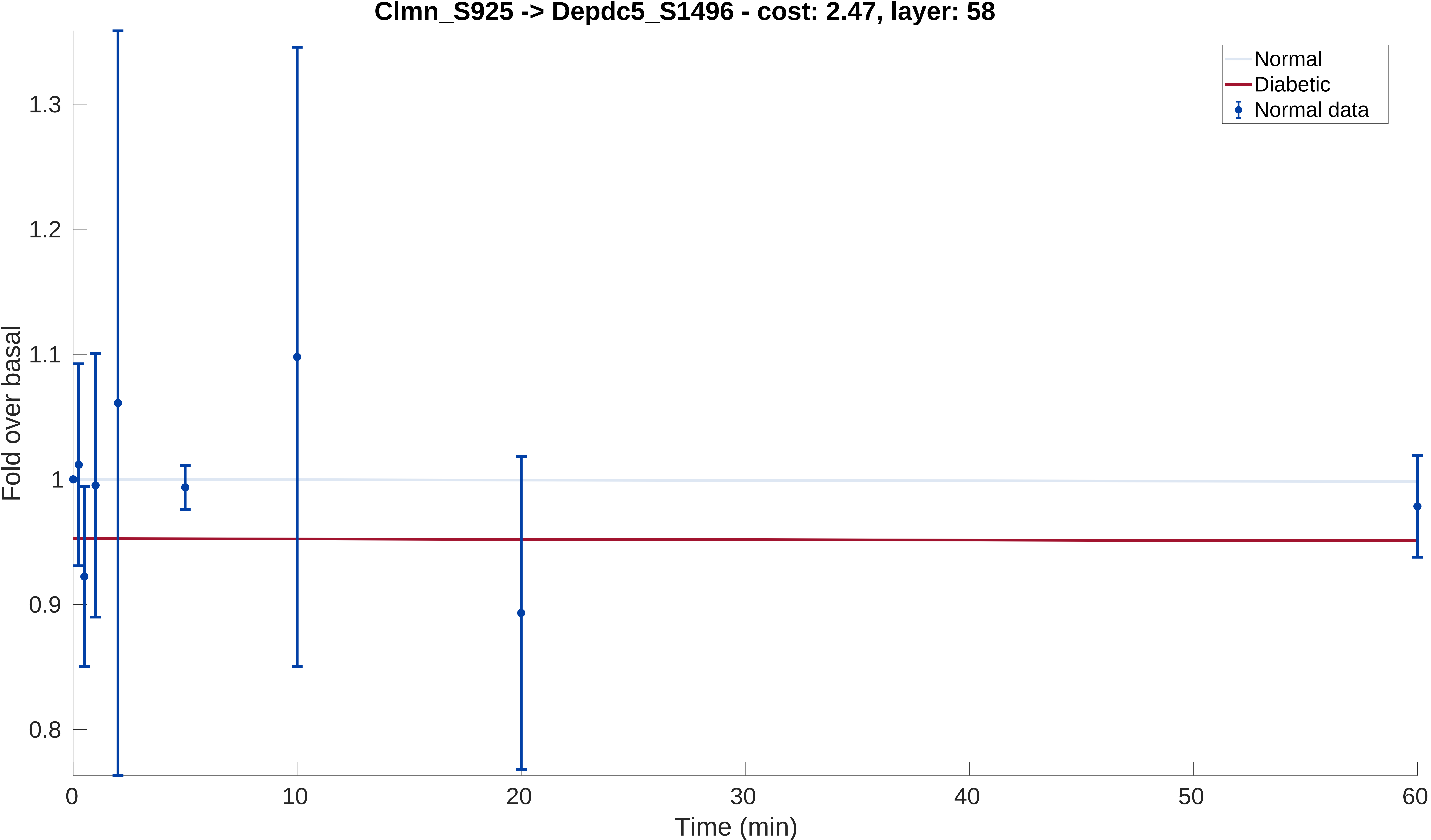

Clmn_S925 -> Depdc5_S1496 - cost: 2.47, layer: 58
Normal
Diabetic
Normal data
Fold over basal
1.3
1.2
1.1
1
0.9
0.8
0
10
20
30
40
50
60
Time (min)

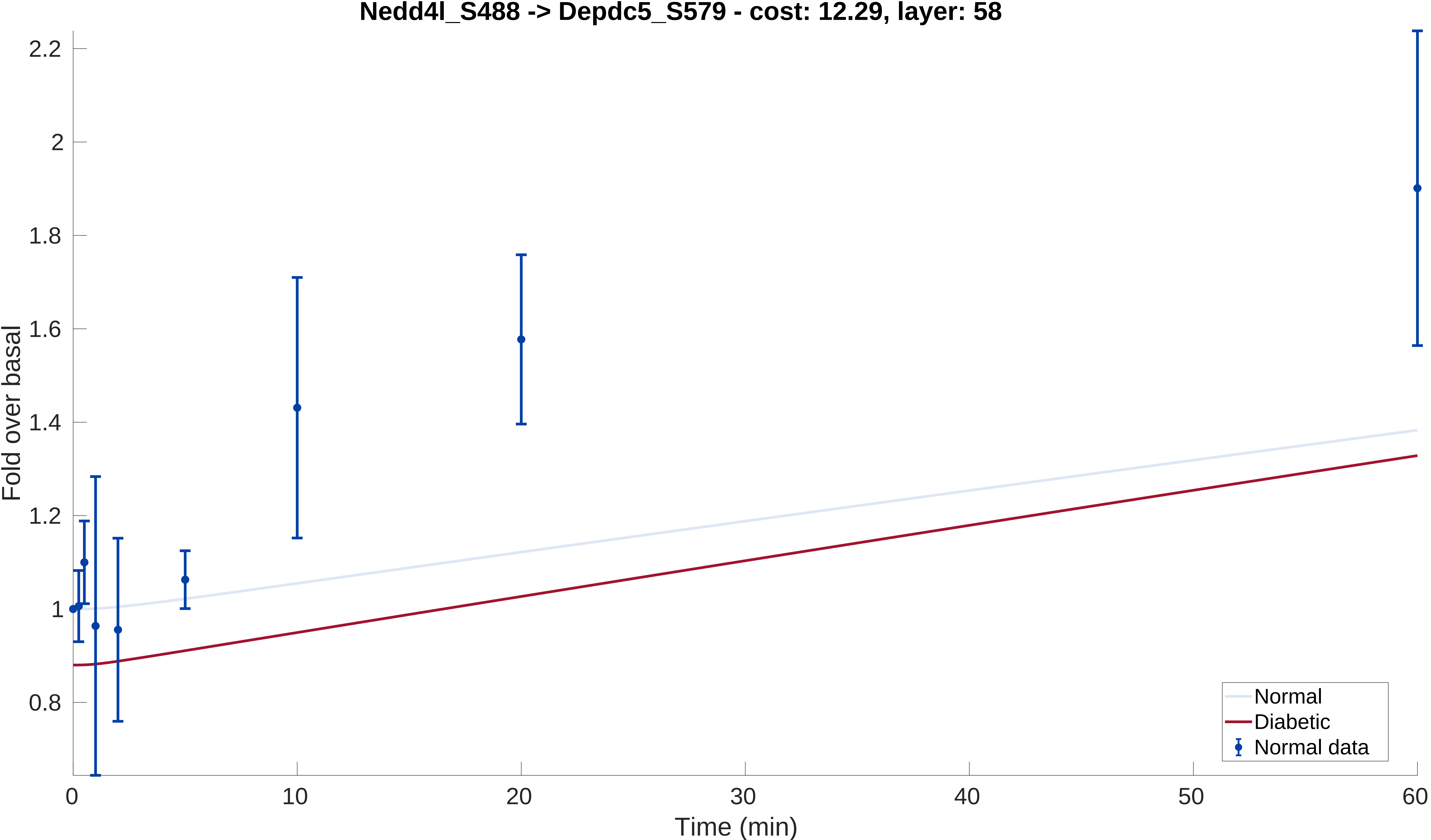

Nedd4l_S488 -> Depdc5_S579 - cost: 12.29, layer: 58
Fold over basal
2.2
2
1.8
1.6
1.4
1.2
1
0.8
0
10
20
30
40
50
60
Time (min)
Normal
Diabetic
Normal data

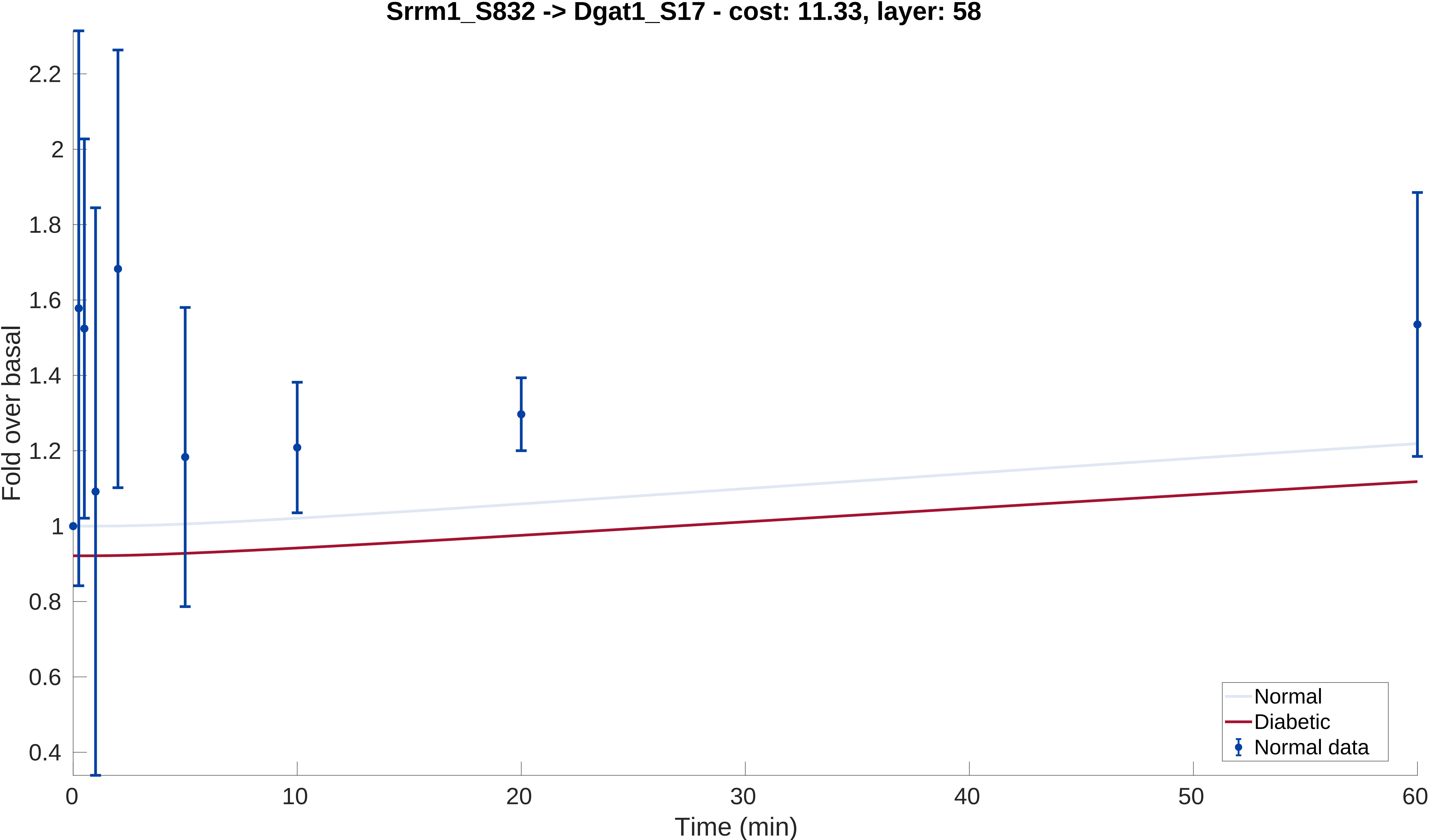
Srrm1_S832 -> Dgat1_S17 - cost: 11.33, layer: 58
Fold over basal
Time (min)
0.4
0.6
0.8
1
1.2
1.4
1.6
1.8
2
2.2
0
10
20
30
40
50
60
Normal
Diabetic
Normal data

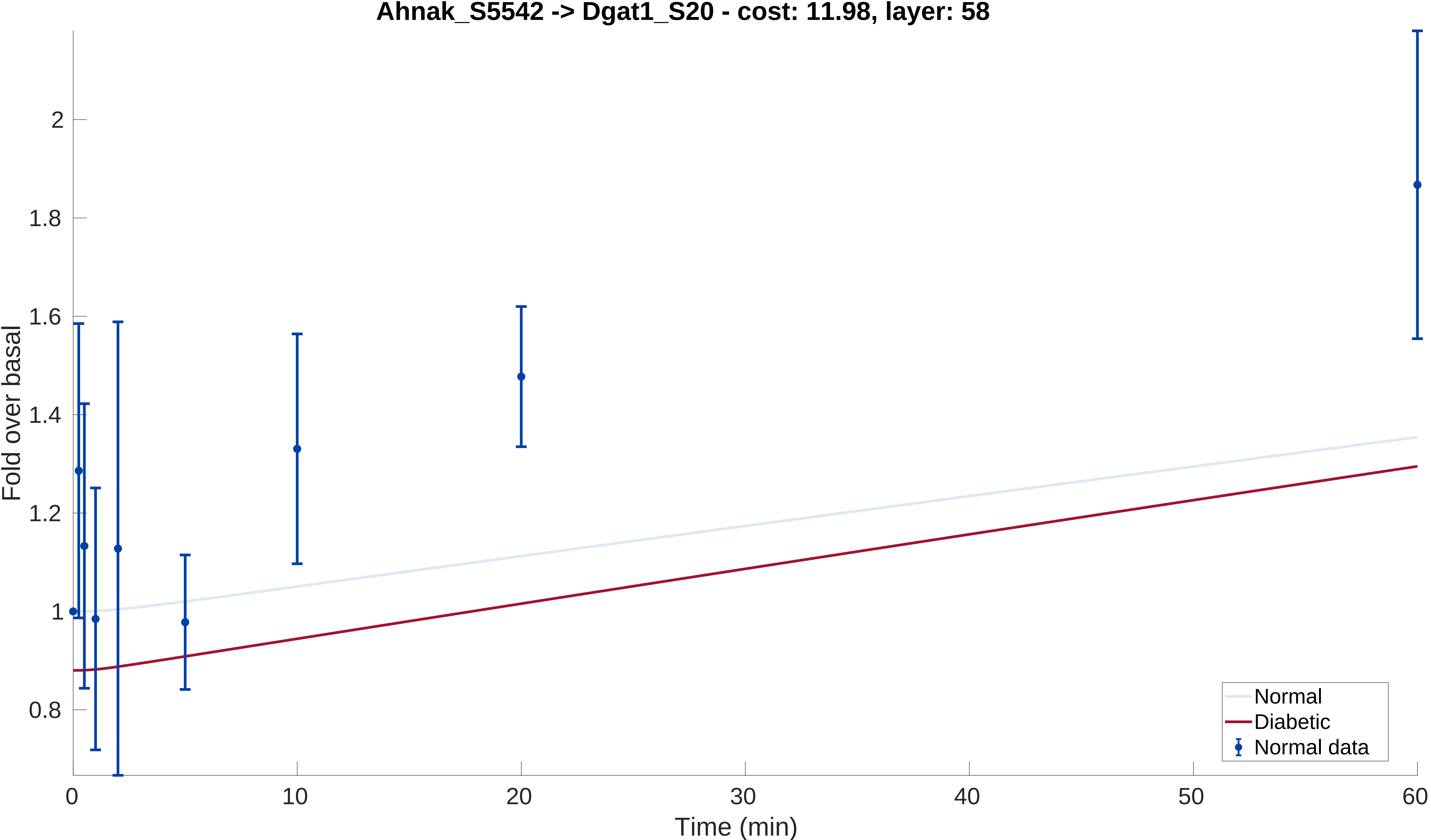
Ahnak_S5542 -> Dgat1_S20 - cost: 11.98, layer: 58
Fold over basal
2
1.8
1.6
1.4
1.2
1
0.8
0
10
20
30
40
50
60
Time (min)
Normal
Diabetic
Normal data

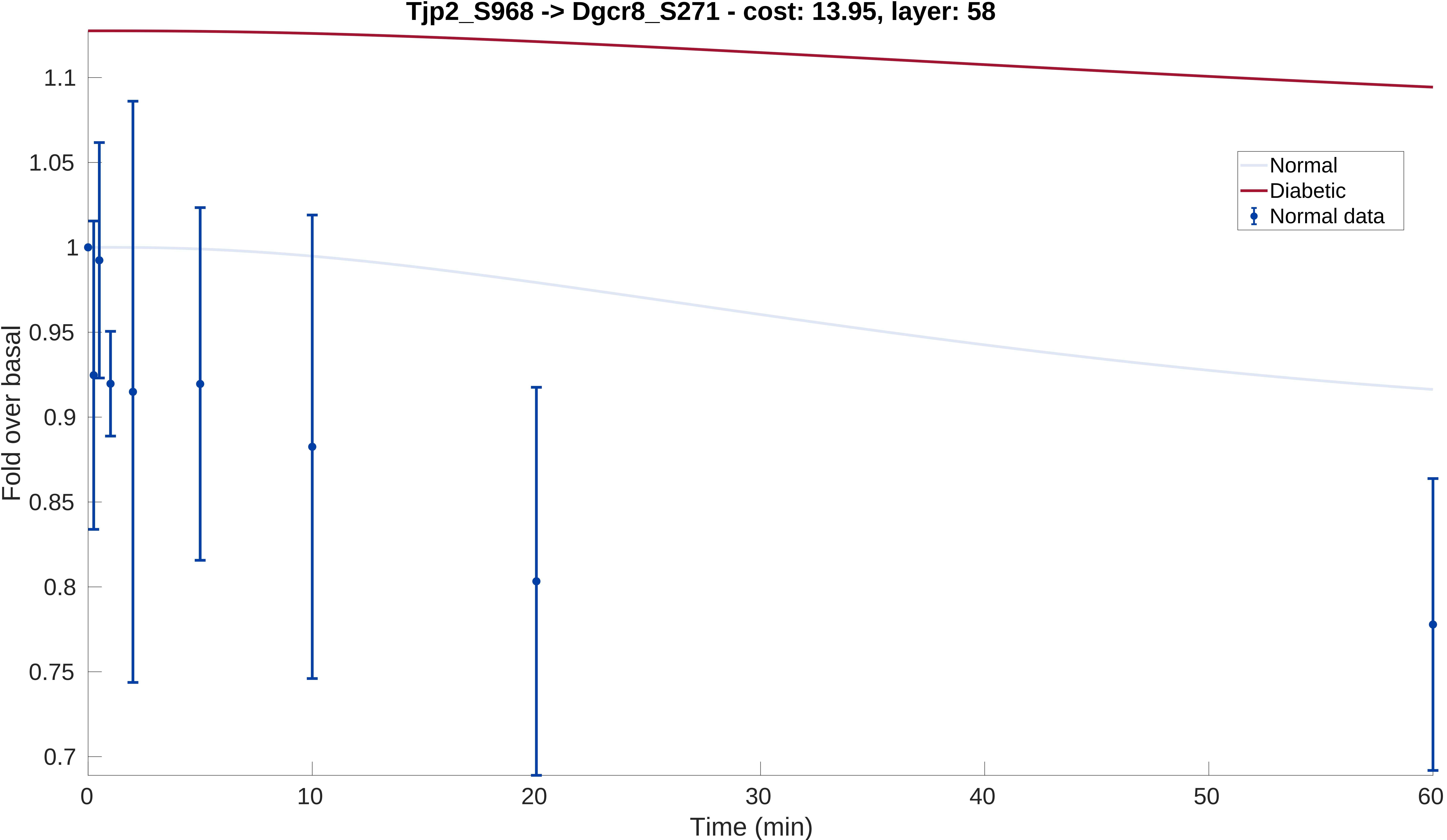

Tjp2_S968 -> Dgcr8_S271 - cost: 13.95, layer: 58
Normal
Diabetic
Normal data
Fold over basal
1.1
1.05
1
0.95
0.9
0.85
0.8
0.75
0.7
0
10
20
30
40
50
60
Time (min)

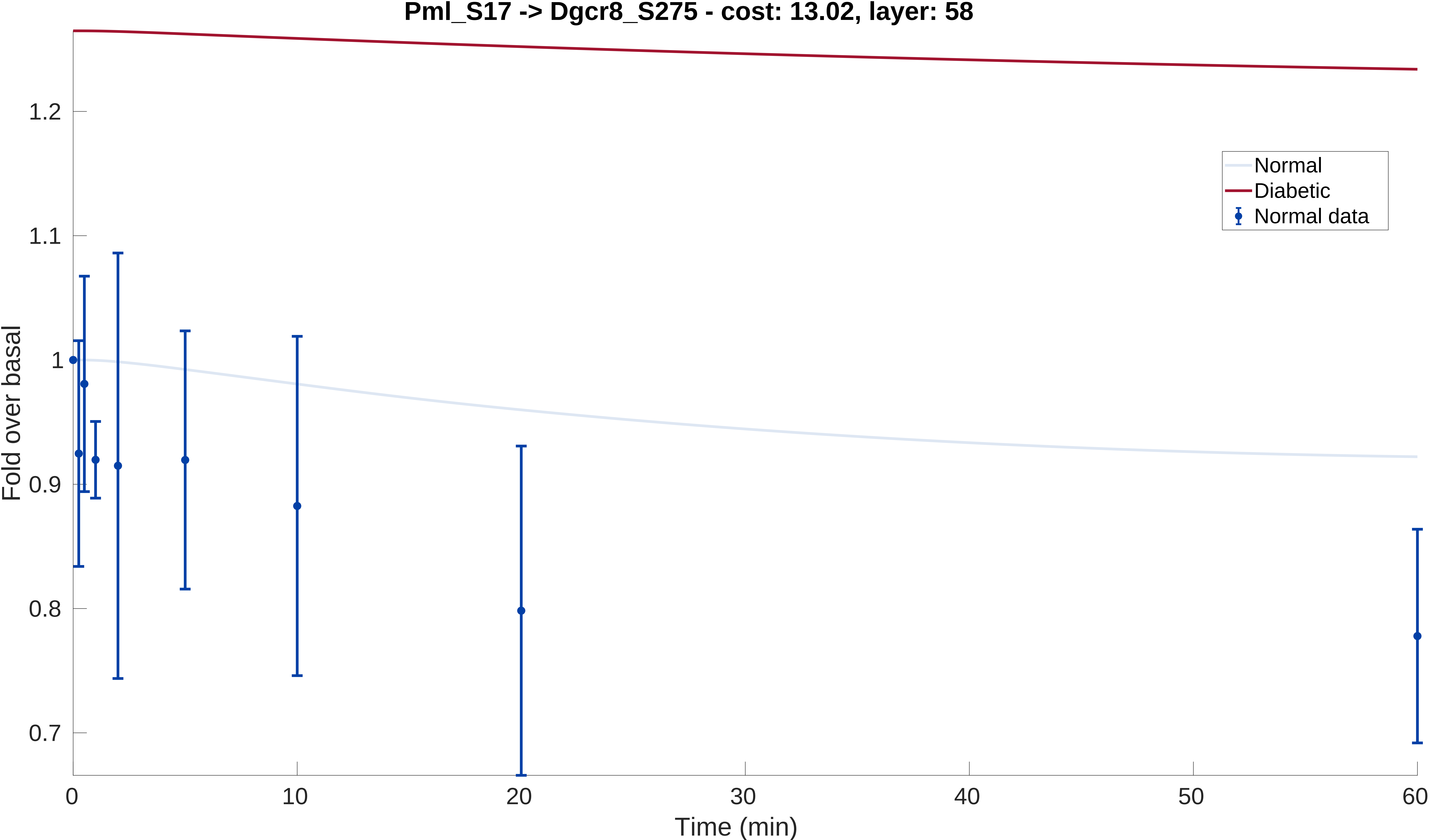

Pml_S17 -> Dgcr8_S275 - cost: 13.02, layer: 58
Normal
Diabetic
Normal data
Fold over basal
1.2
1.1
1
0.9
0.8
0.7
0
10
20
30
40
50
60
Time (min)

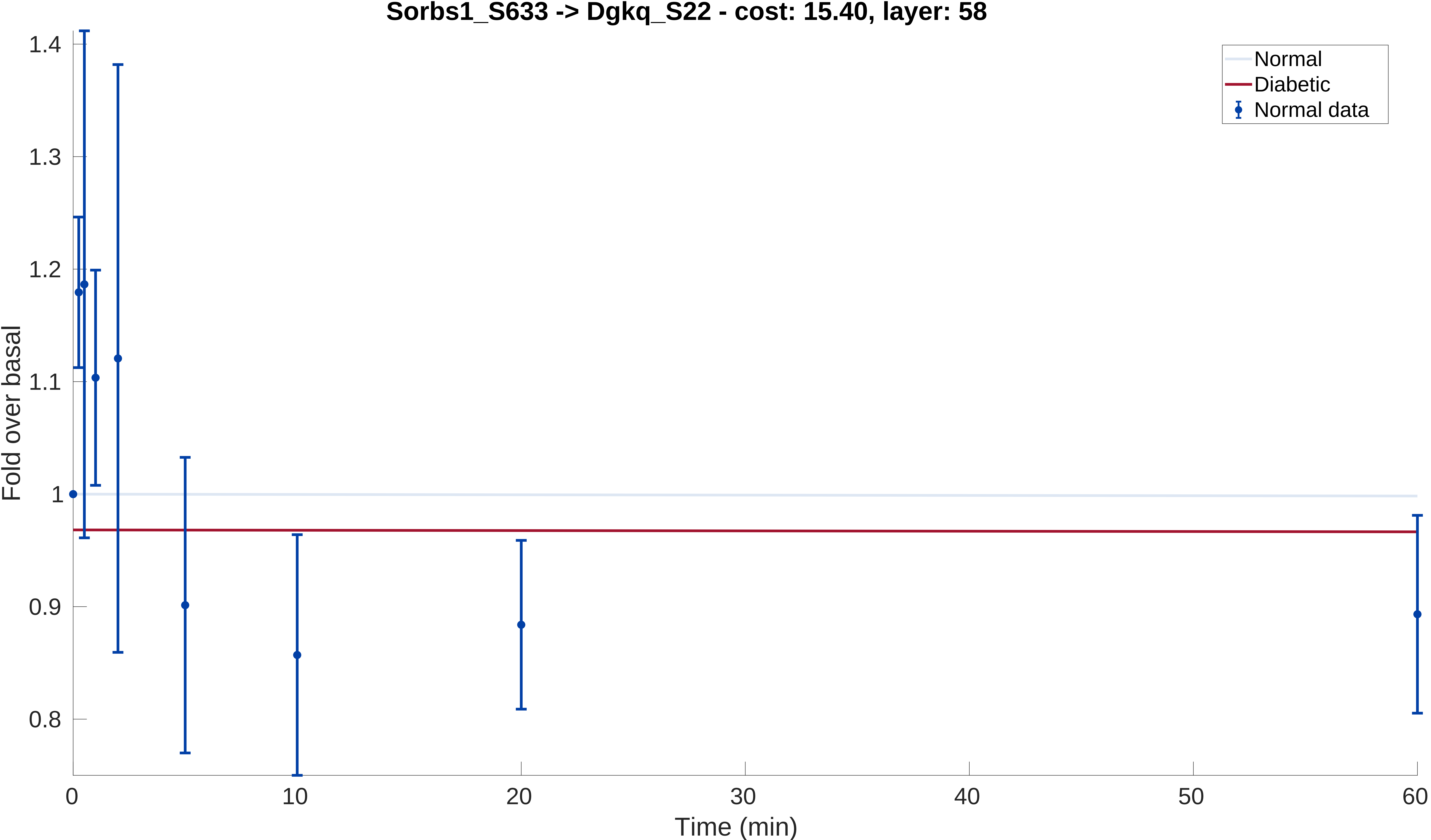
Sorbs1_S633 -> Dgkq_S22 - cost: 15.40, layer: 58
Normal
Diabetic
Normal data
Fold over basal
1.4
1.3
1.2
1.1
1
0.9
0.8
0
10
20
30
40
50
60
Time (min)

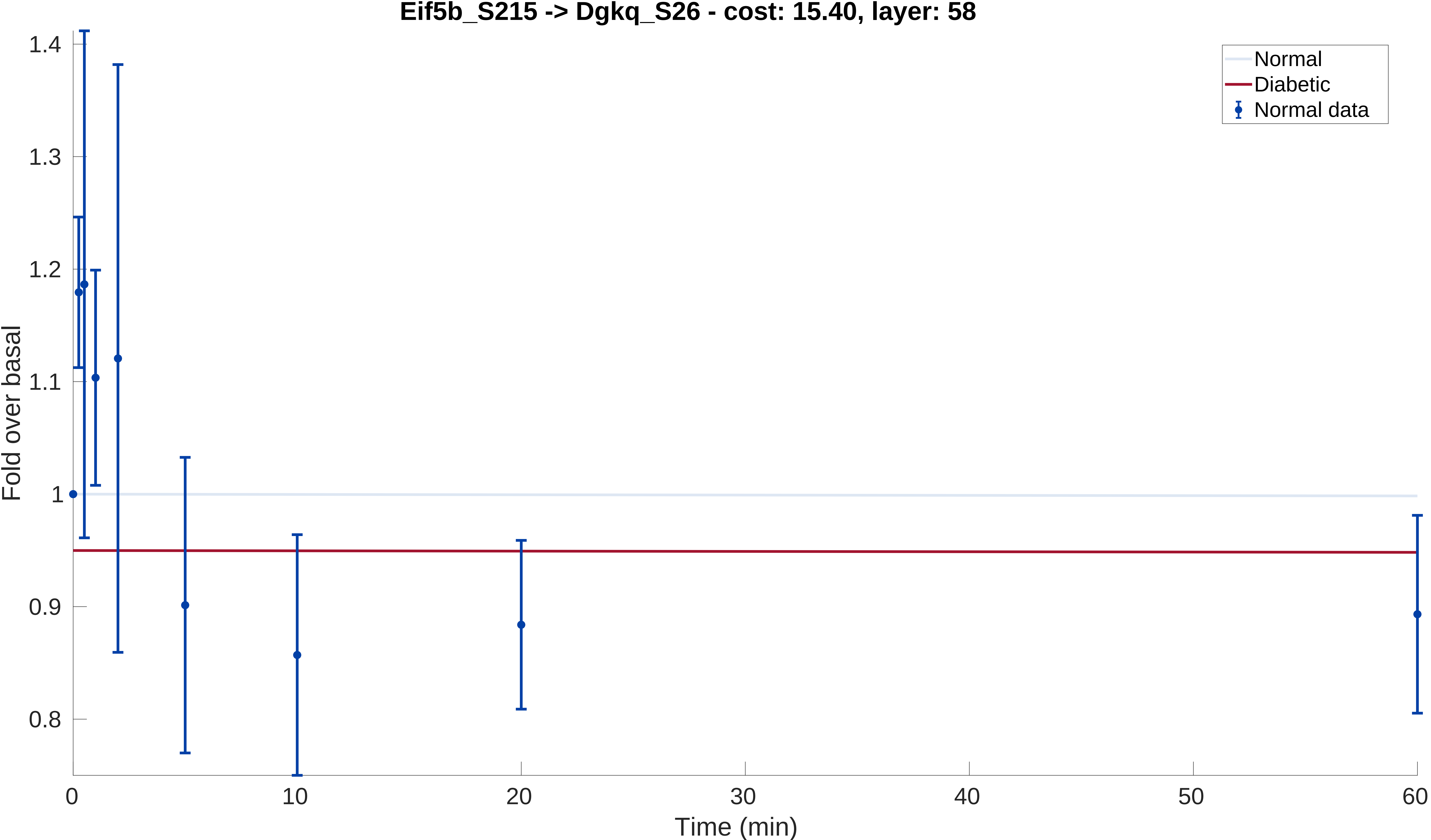

Eif5b_S215 -> Dgkq_S26 - cost: 15.40, layer: 58
Normal
Diabetic
Normal data
Fold over basal
1.4
1.3
1.2
1.1
1
0.9
0.8
0
10
20
30
40
50
60
Time (min)

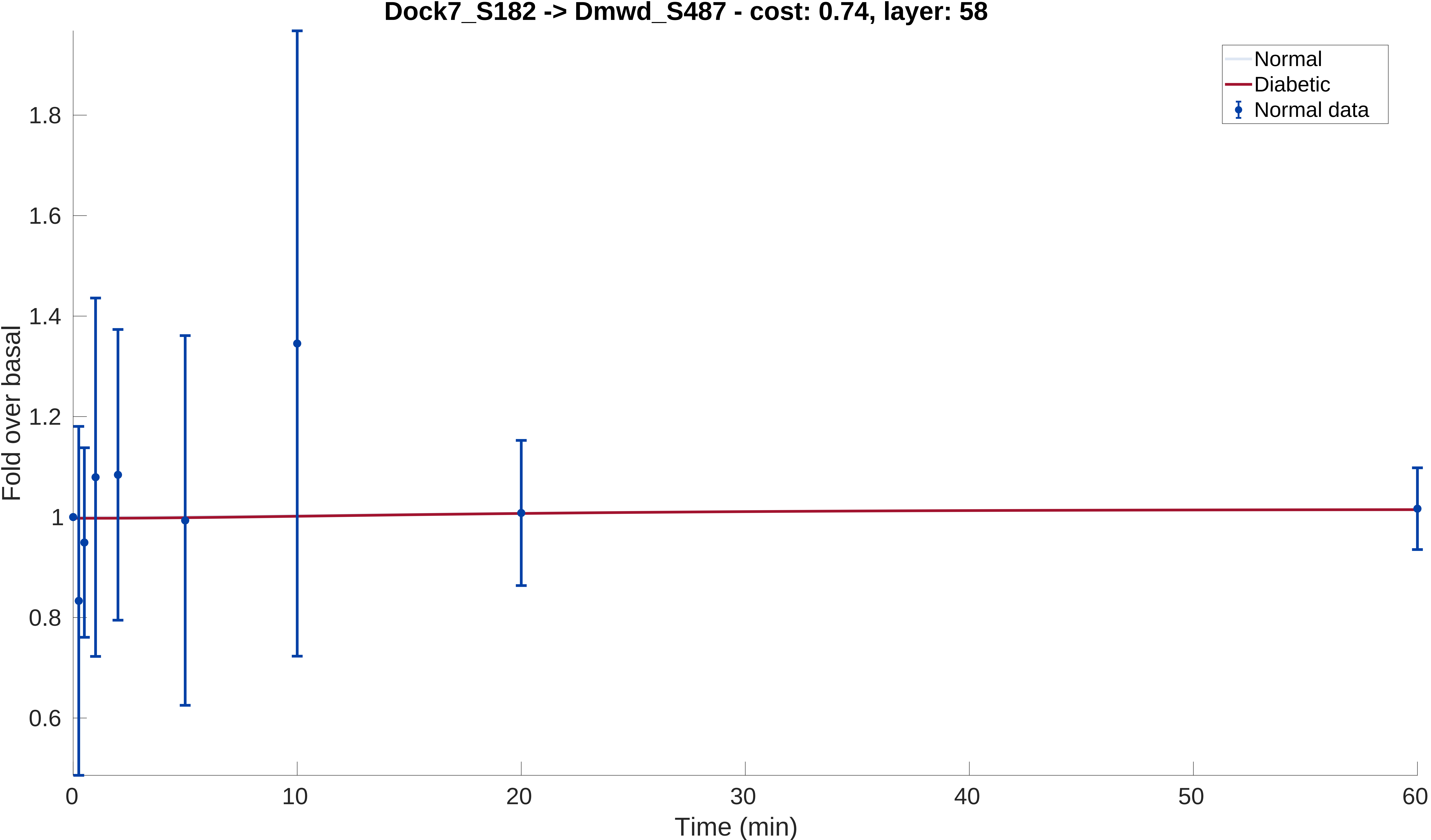

Dock7_S182 -> Dmwd_S487 - cost: 0.74, layer: 58
Normal
Diabetic
Normal data
Fold over basal
Time (min)
1.8
1.6
1.4
1.2
1
0.8
0.6
0
10
20
30
40
50
60

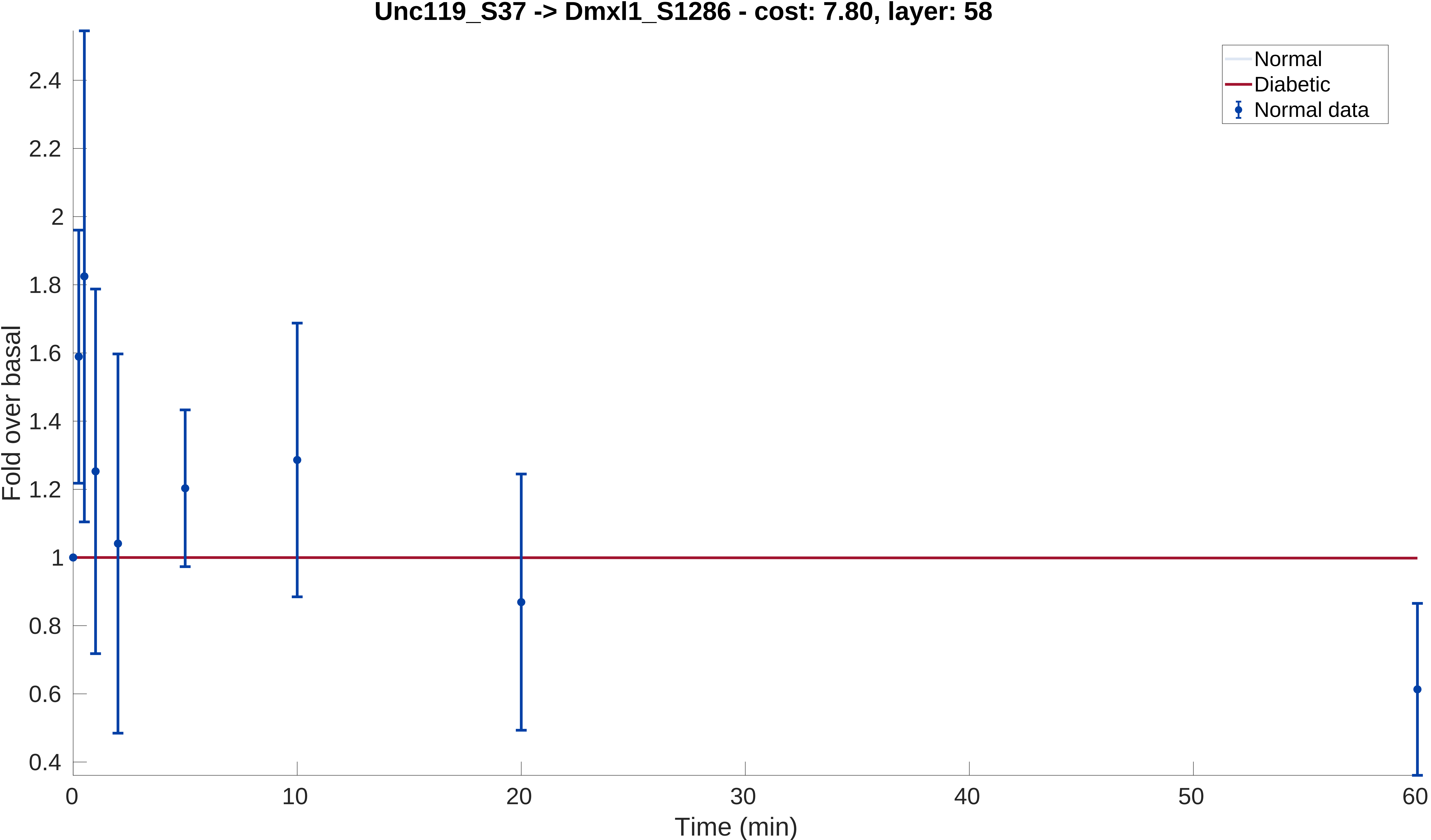

Unc119_S37 -> Dmxl1_S1286 - cost: 7.80, layer: 58
Normal
Diabetic
Normal data
Fold over basal
Time (min)
2.4
2.2
2
1.8
1.6
1.4
1.2
1
0.8
0.6
0.4
0
10
20
30
40
50
60

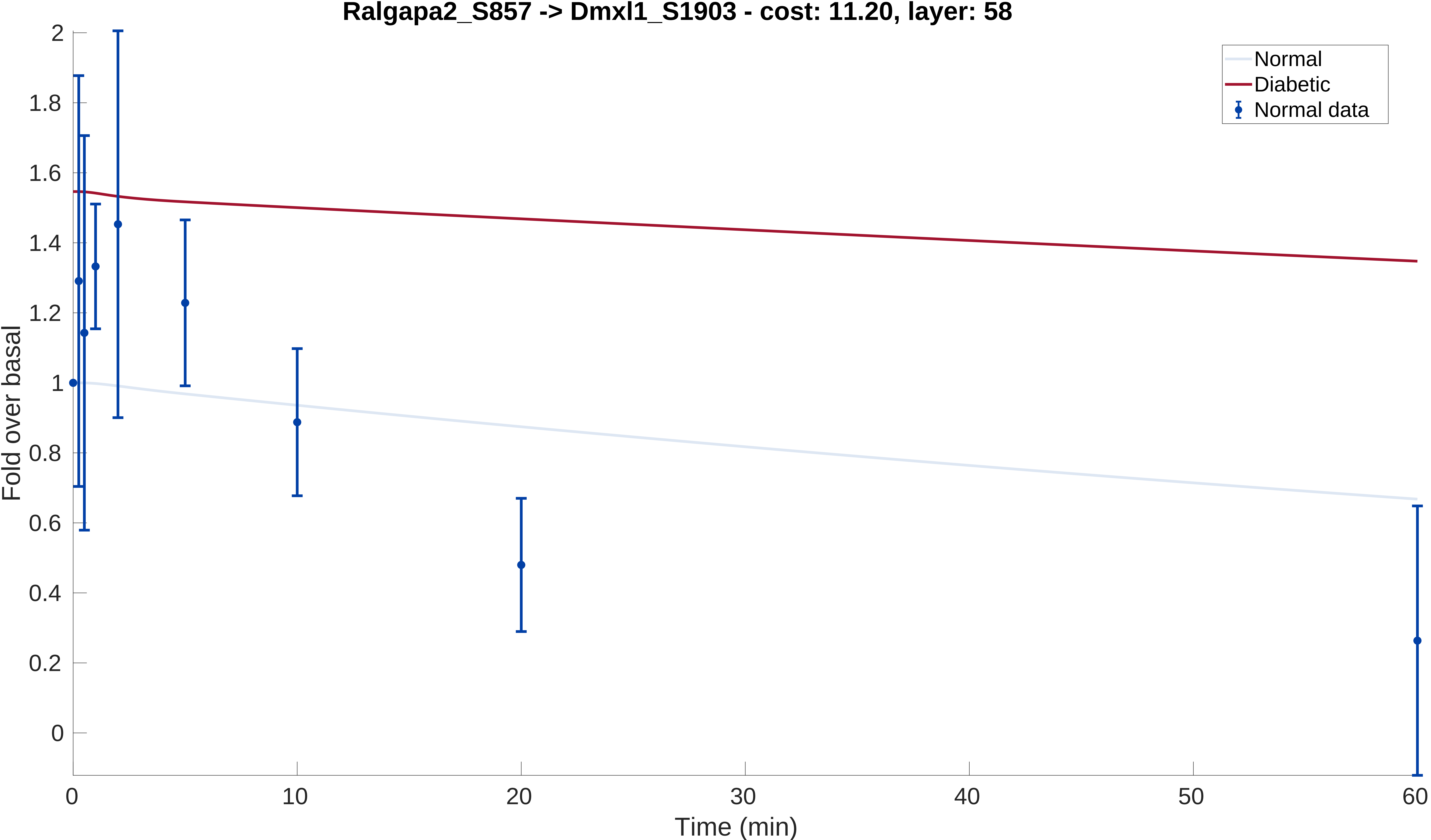
Ralgapa2_S857 -> Dmxl1_S1903 - cost: 11.20, layer: 58
Normal
Diabetic
Normal data
Fold over basal
Time (min)
2
1.8
1.6
1.4
1.2
1
0.8
0.6
0.4
0.2
0
10
20
30
40
50
60

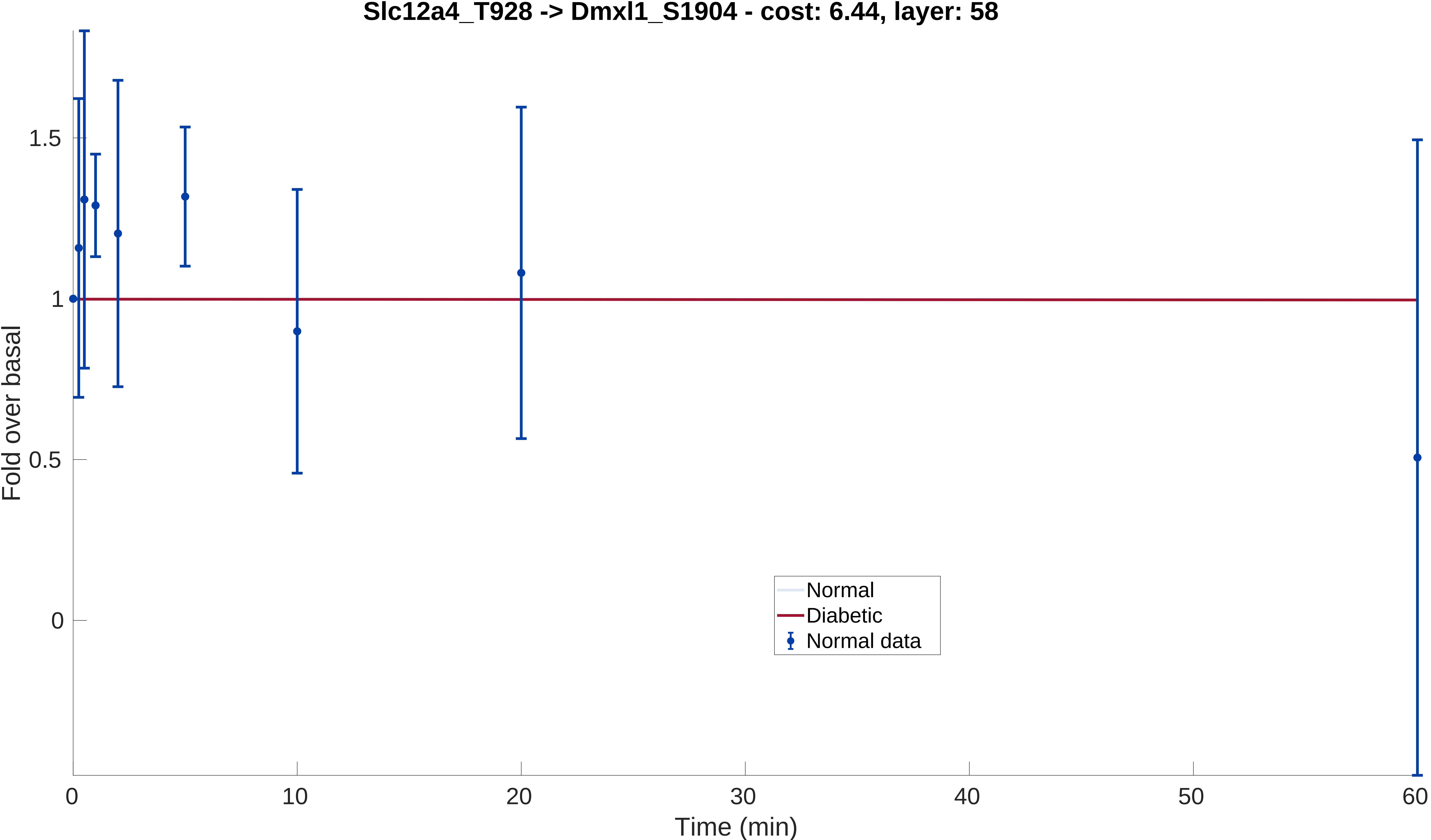
Slc12a4_T928 -> Dmxl1_S1904 - cost: 6.44, layer: 58
Fold over basal
1.5
1
0.5
0
0
10
20
30
40
50
60
Time (min)
Normal
Diabetic
Normal data

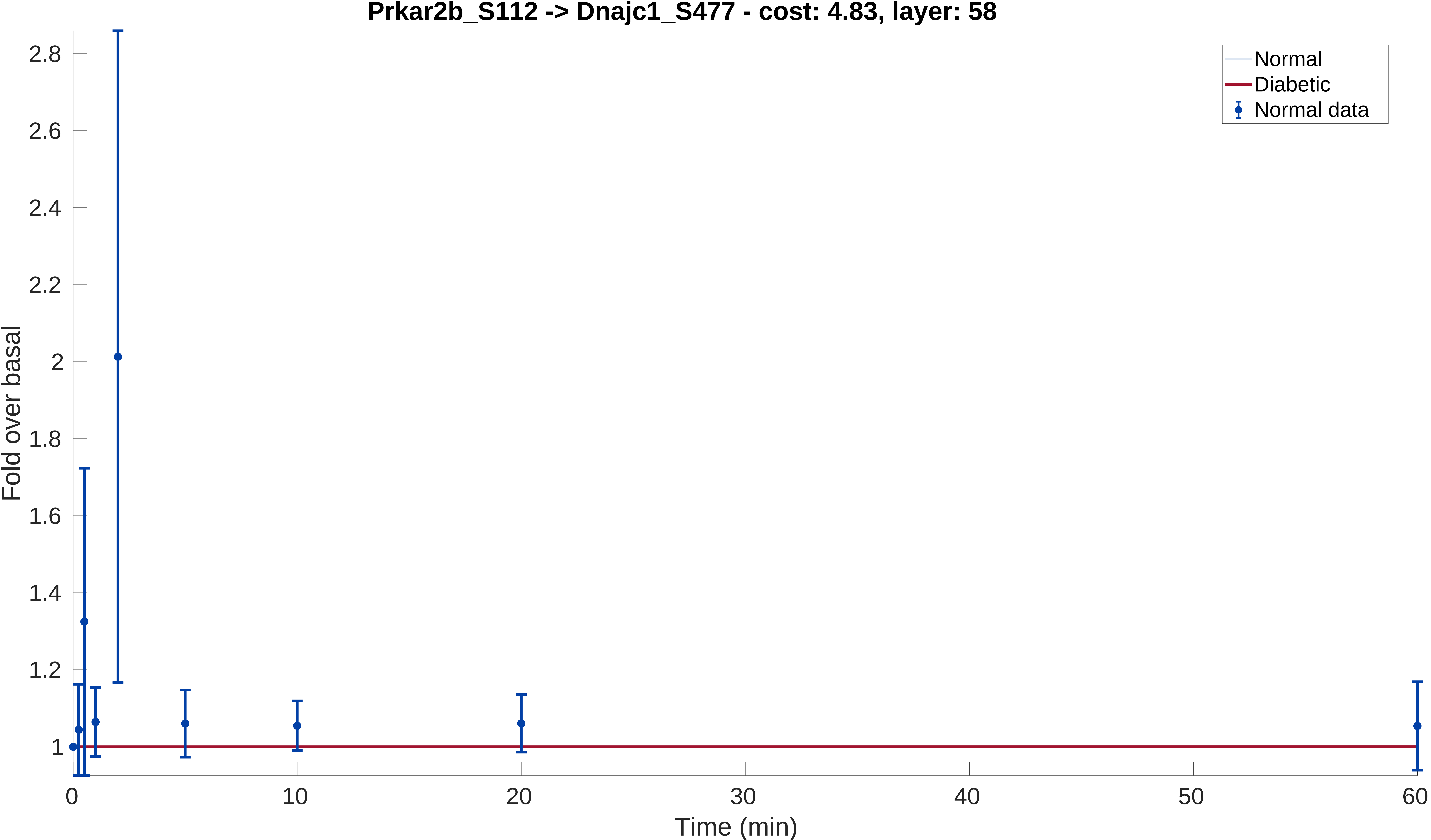

Prkar2b_S112 -> Dnajc1_S477 - cost: 4.83, layer: 58
Normal
Diabetic
Normal data
Fold over basal
2.8
2.6
2.4
2.2
2
1.8
1.6
1.4
1.2
1
0
10
20
30
40
50
60
Time (min)

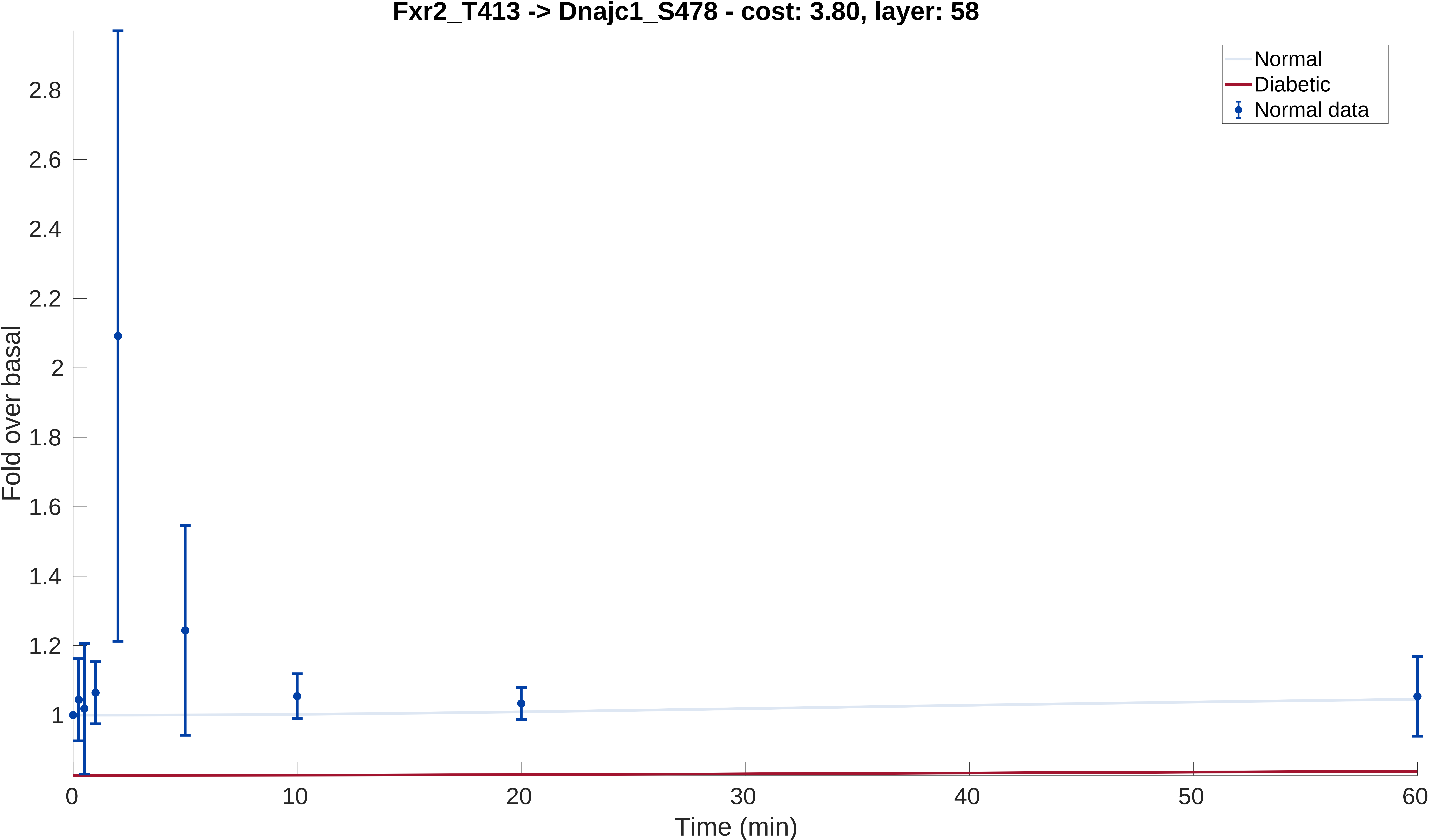

Fxr2_T413 -> Dnajc1_S478 - cost: 3.80, layer: 58
Normal
Diabetic
Normal data
Fold over basal
2.8
2.6
2.4
2.2
2
1.8
1.6
1.4
1.2
1
0
10
20
30
40
50
60
Time (min)

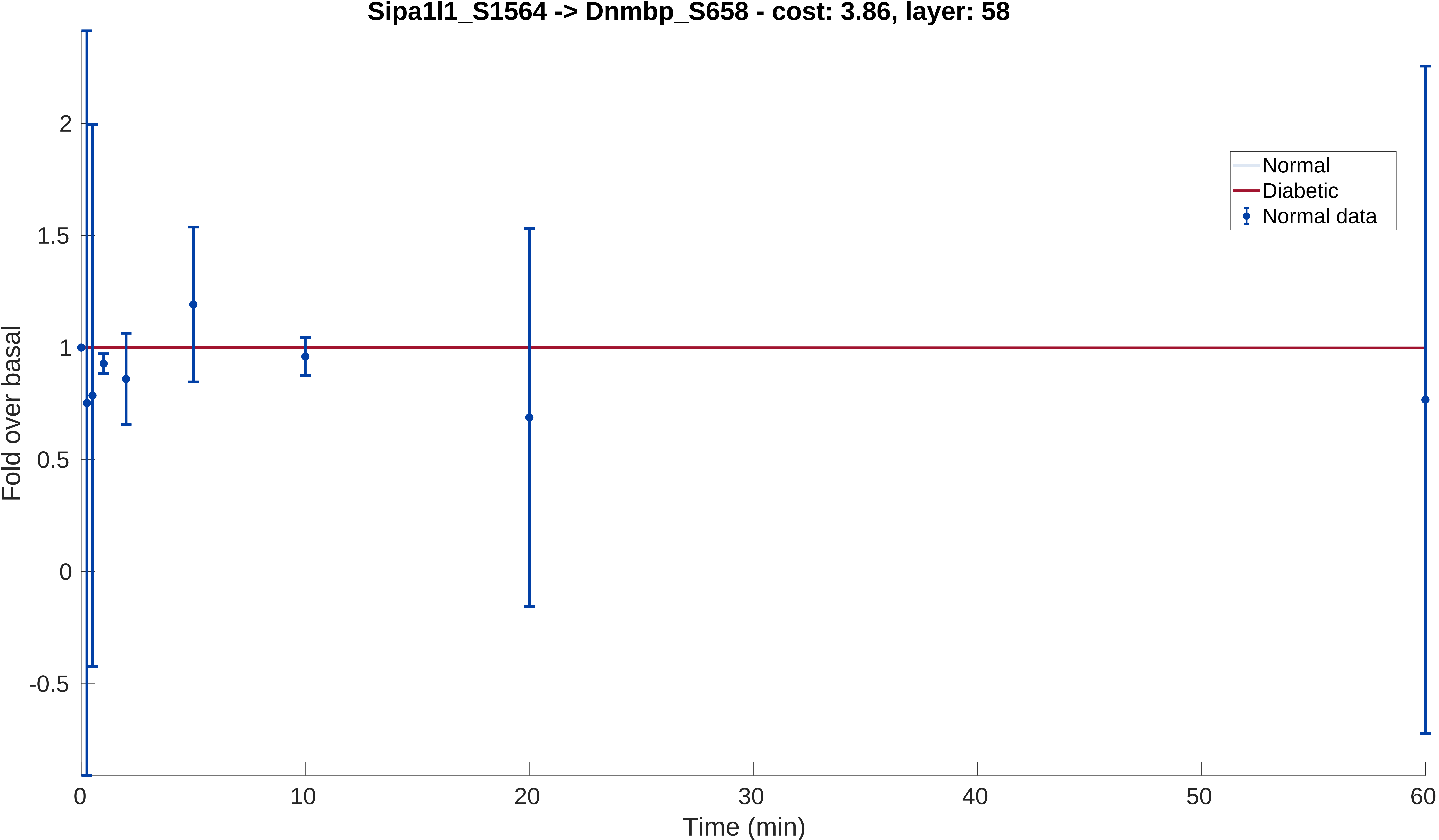

Sipa1l1_S1564 -> Dnmbp_S658 - cost: 3.86, layer: 58
Fold over basal
2
1.5
1
0.5
0
-0.5
0
10
20
30
40
50
60
Time (min)
Normal
Diabetic
Normal data

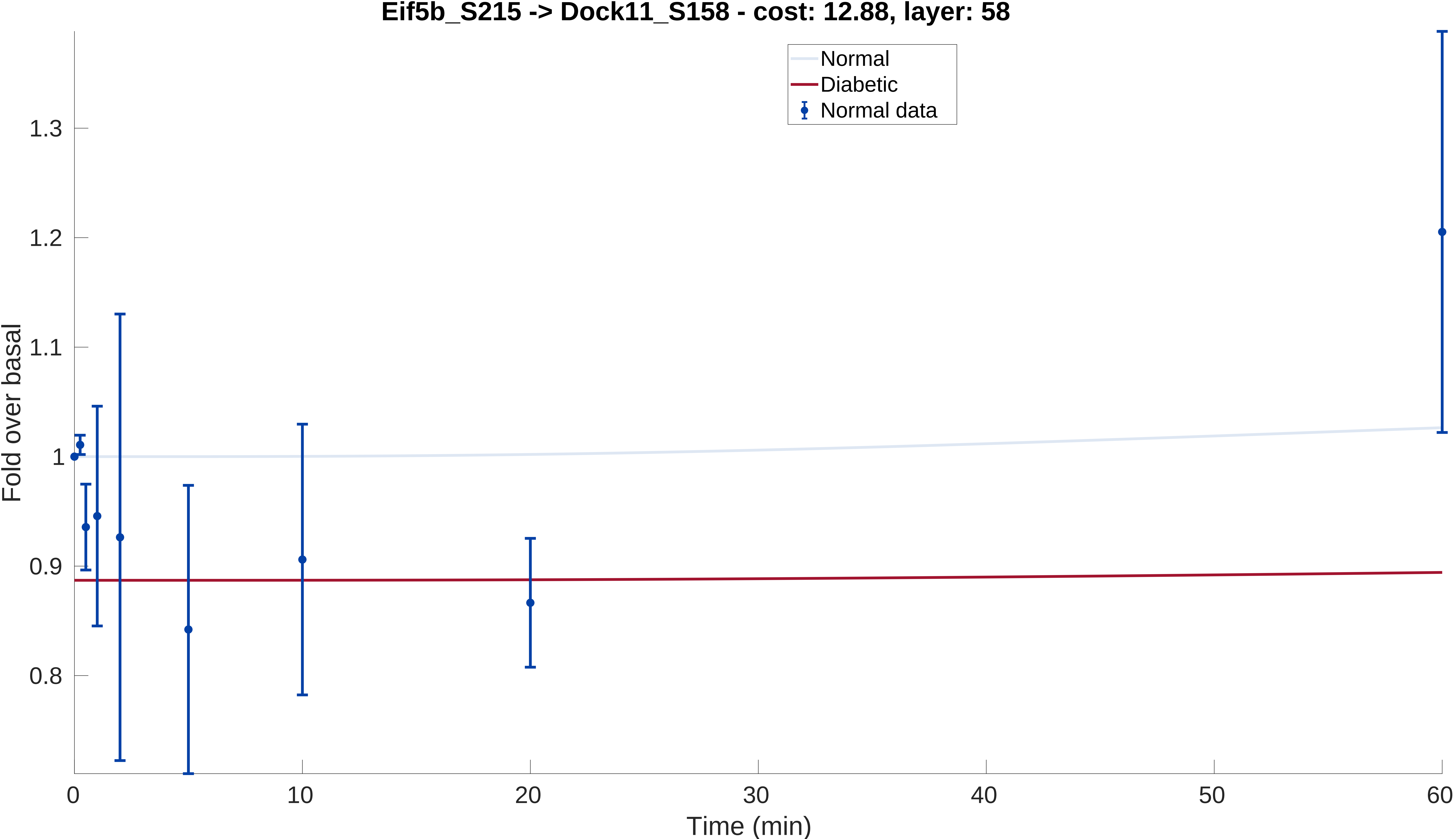
Eif5b_S215 -> Dock11_S158 - cost: 12.88, layer: 58
Normal
Diabetic
Normal data
Fold over basal
1.3
1.2
1.1
1
0.9
0.8
0
10
20
30
40
50
60
Time (min)

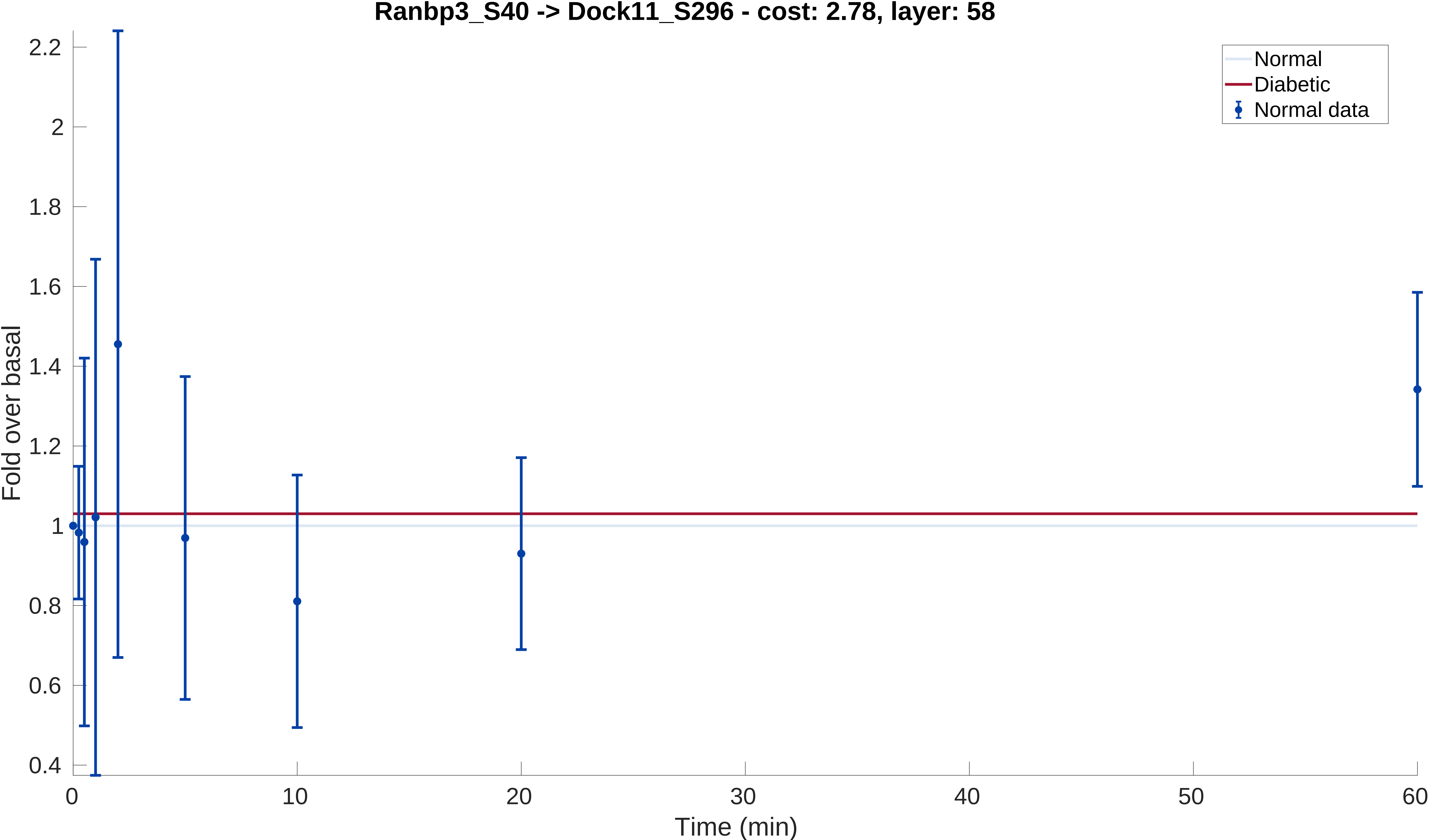
Ranbp3_S40 -> Dock11_S296 - cost: 2.78, layer: 58
Normal
Diabetic
Normal data
Fold over basal
Time (min)
2.2
2
1.8
1.6
1.4
1.2
1
0.8
0.6
0.4
0
10
20
30
40
50
60

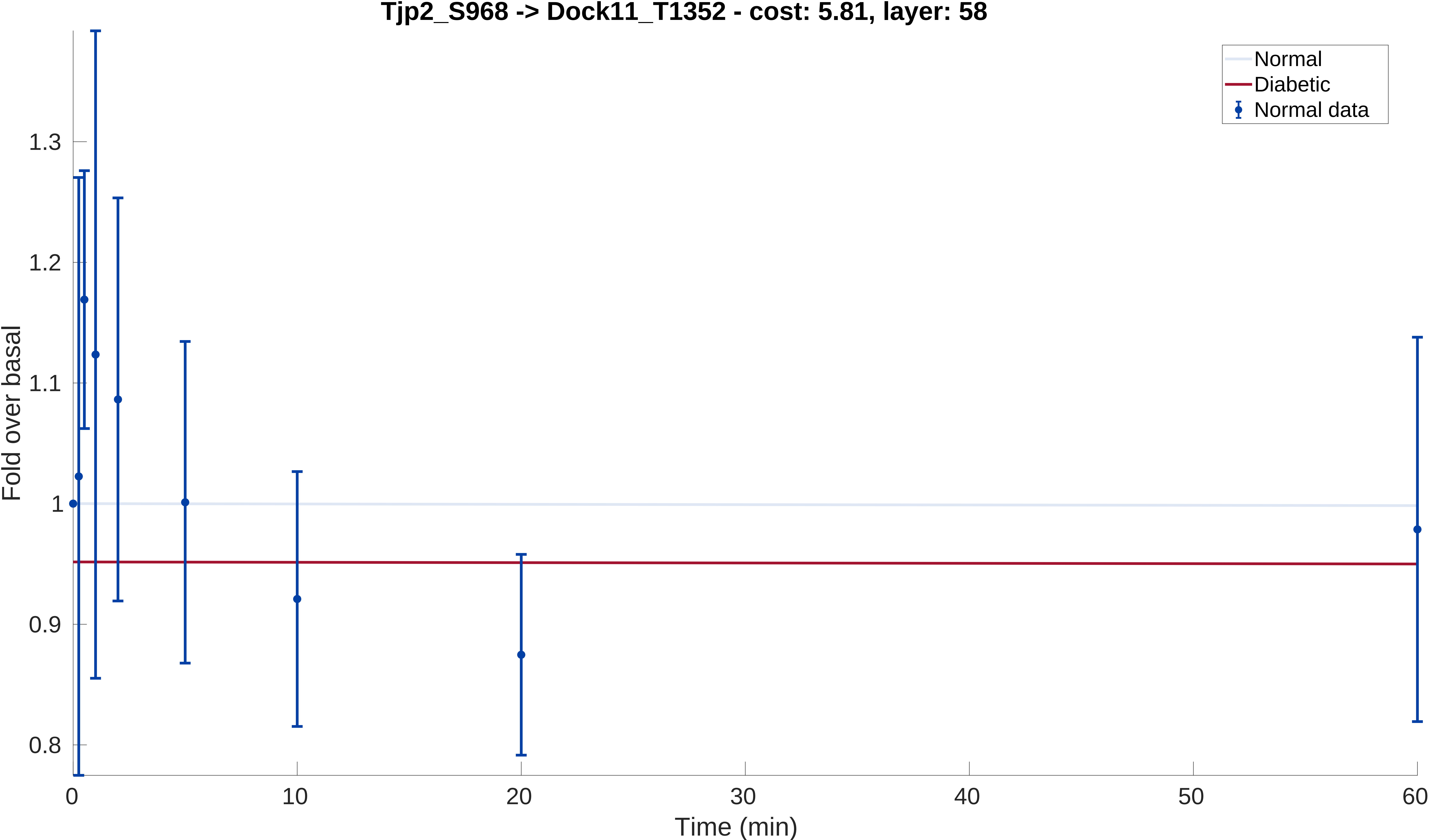
Tjp2_S968 -> Dock11_T1352 - cost: 5.81, layer: 58
Normal
Diabetic
Normal data
Fold over basal
1.3
1.2
1.1
1
0.9
0.8
0
10
20
30
40
50
60
Time (min)

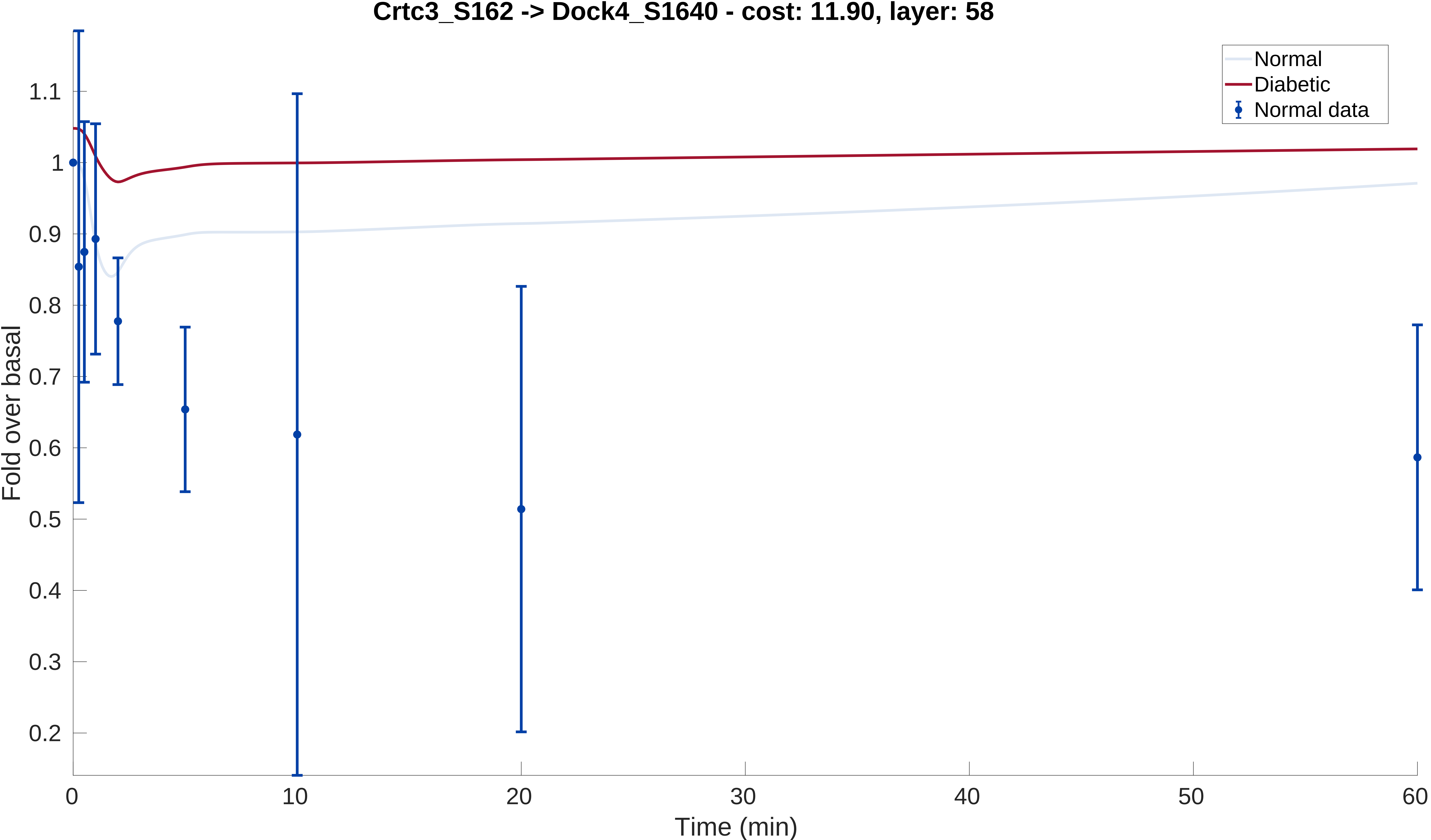
Crtc3_S162 -> Dock4_S1640 - cost: 11.90, layer: 58
Normal
Diabetic
Normal data
Fold over basal
1.1
1
0.9
0.8
0.7
0.6
0.5
0.4
0.3
0.2
0
10
20
30
40
50
60
Time (min)

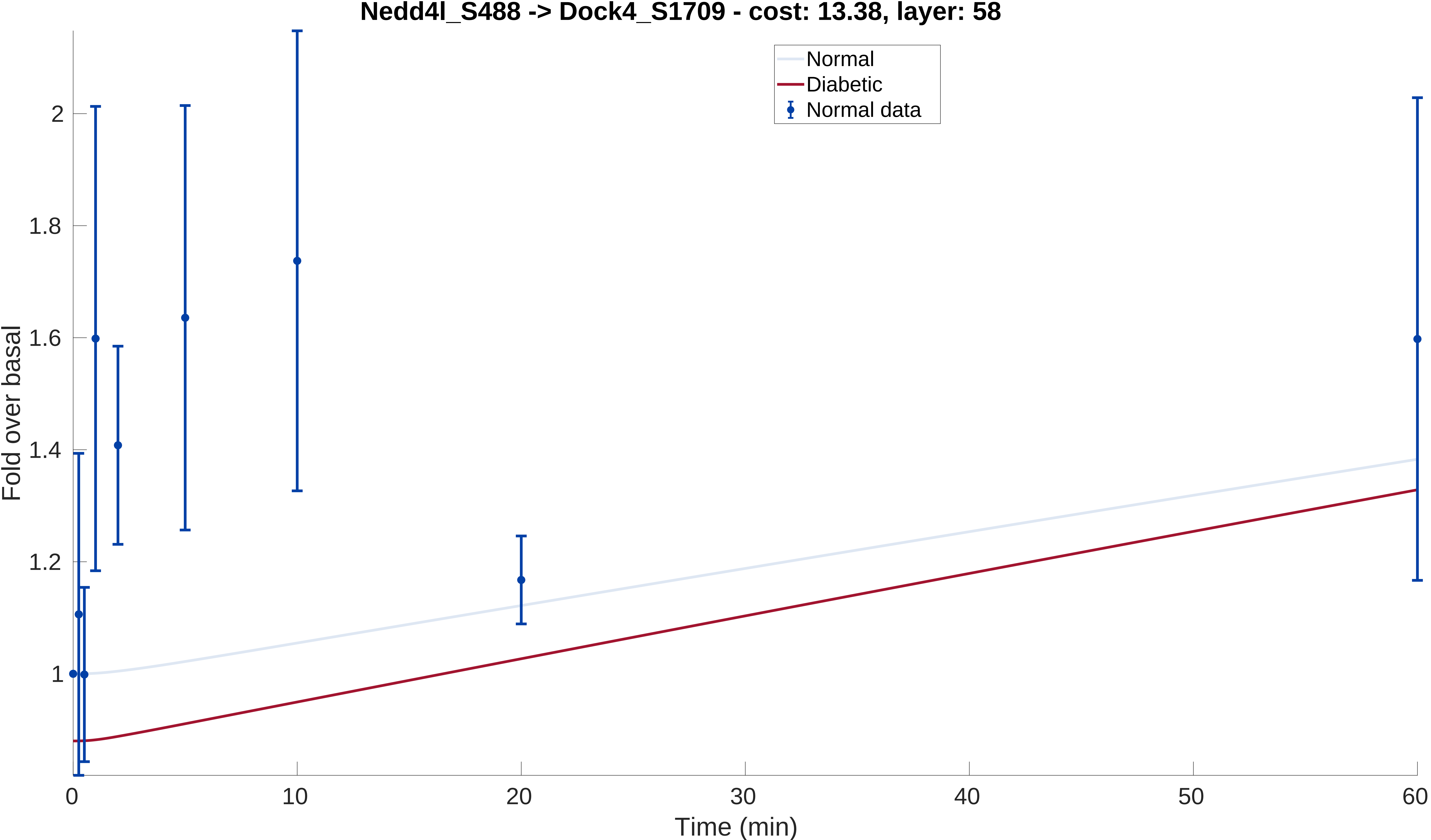

Nedd4l_S488 -> Dock4_S1709 - cost: 13.38, layer: 58
Normal
Diabetic
Normal data
Fold over basal
2
1.8
1.6
1.4
1.2
1
0
10
20
30
40
50
60
Time (min)

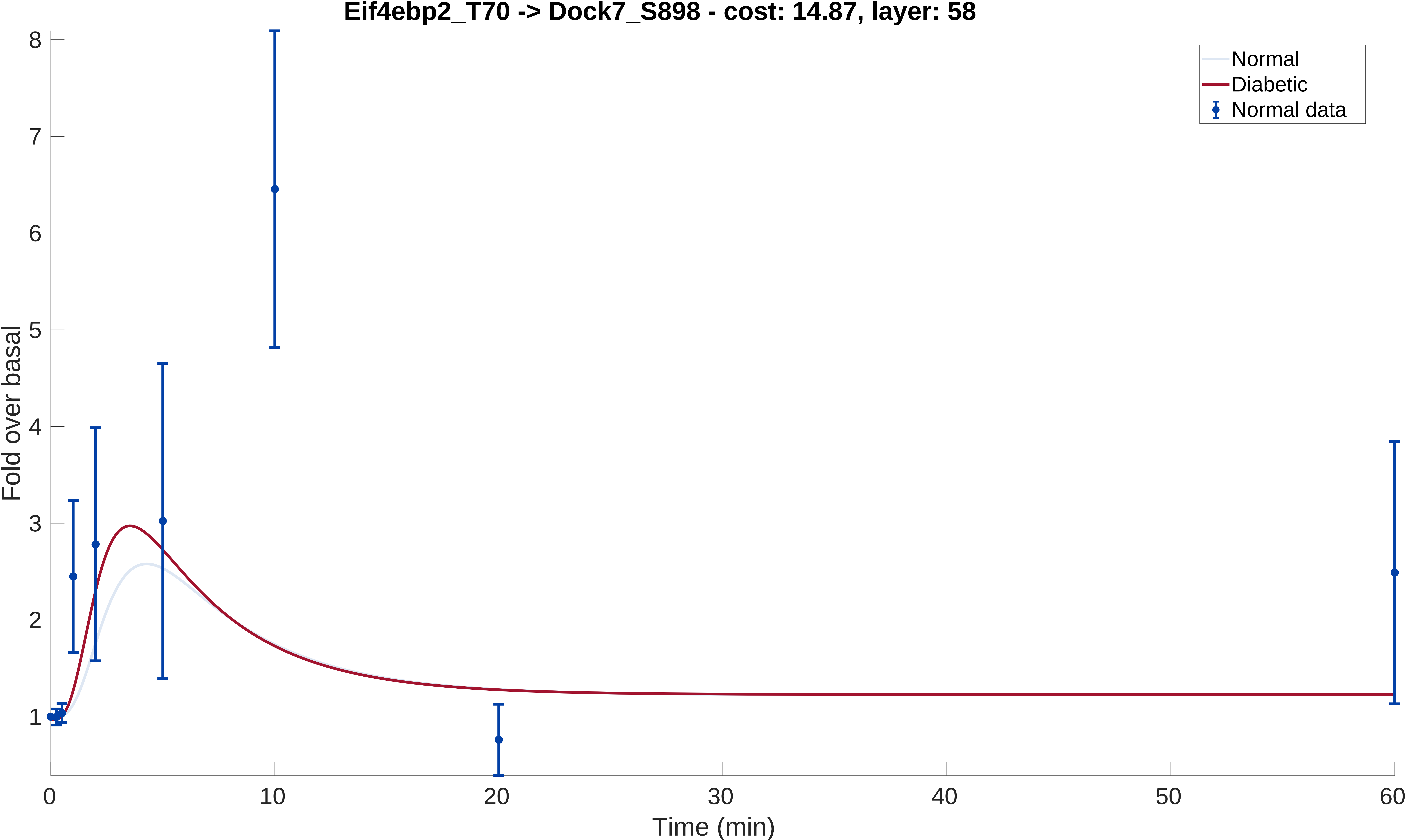
Eif4ebp2_T70 -> Dock7_S898 - cost: 14.87, layer: 58
Normal
Diabetic
Normal data
Fold over basal
Time (min)
0
10
20
30
40
50
60
1
2
3
4
5
6
7
8

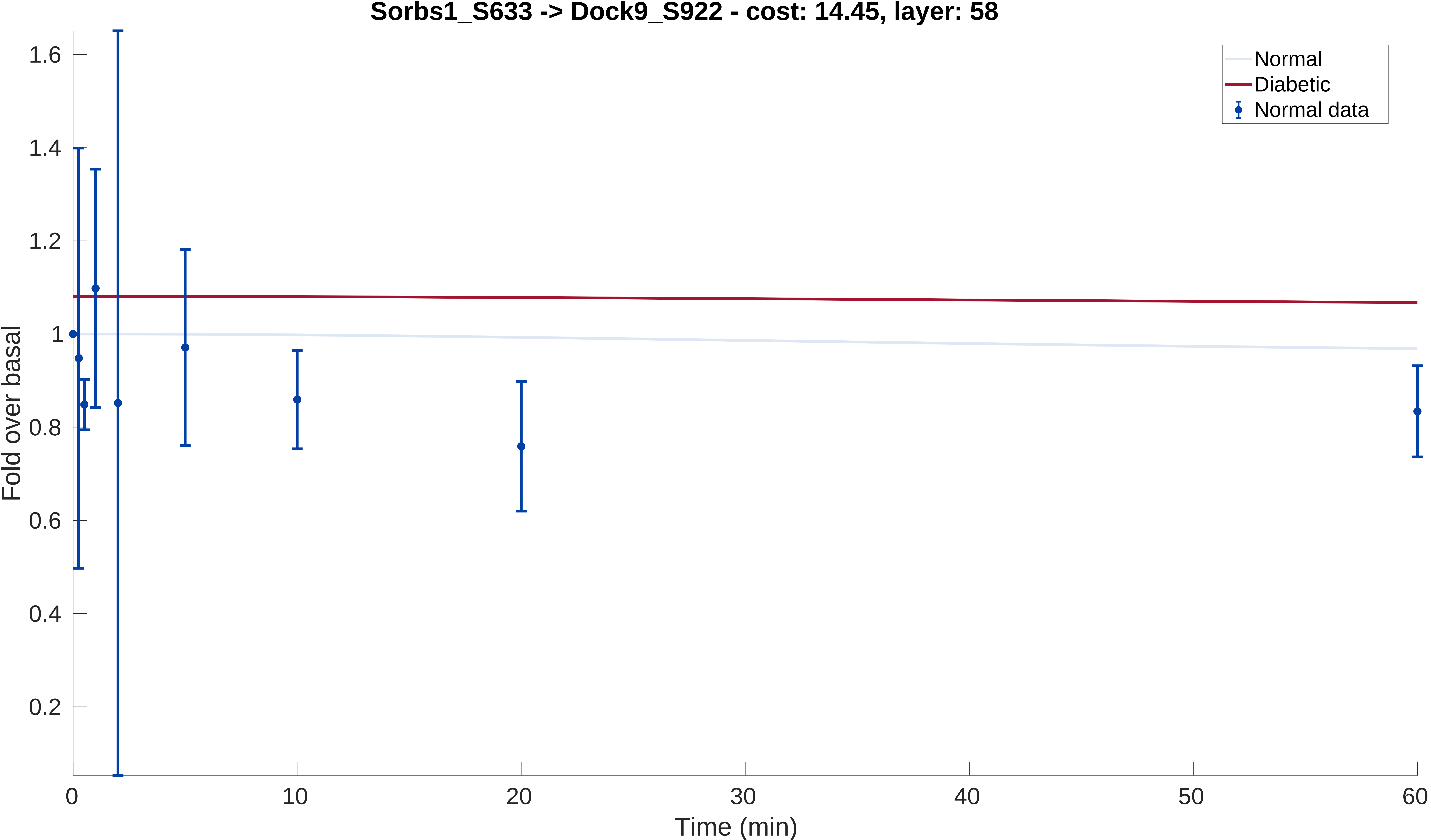
Sorbs1_S633 -> Dock9_S922 - cost: 14.45, layer: 58
Normal
Diabetic
Normal data
Fold over basal
Time (min)
0
10
20
30
40
50
60
0.2
0.4
0.6
0.8
1
1.2
1.4
1.6

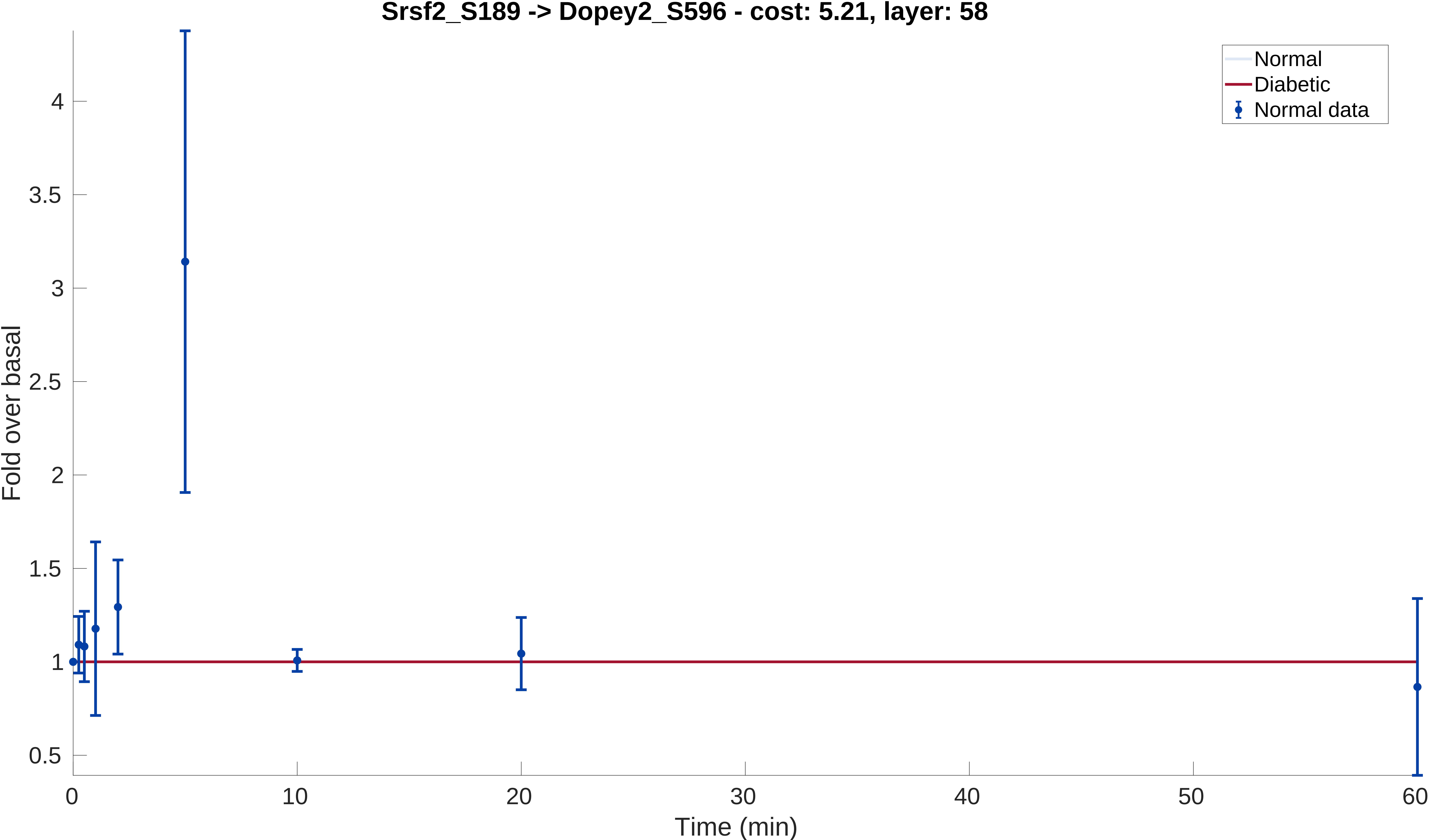

Srsf2_S189 -> Dopey2_S596 - cost: 5.21, layer: 58
Normal
Diabetic
Normal data
Fold over basal
Time (min)
0.5
1
1.5
2
2.5
3
3.5
4
0
10
20
30
40
50
60

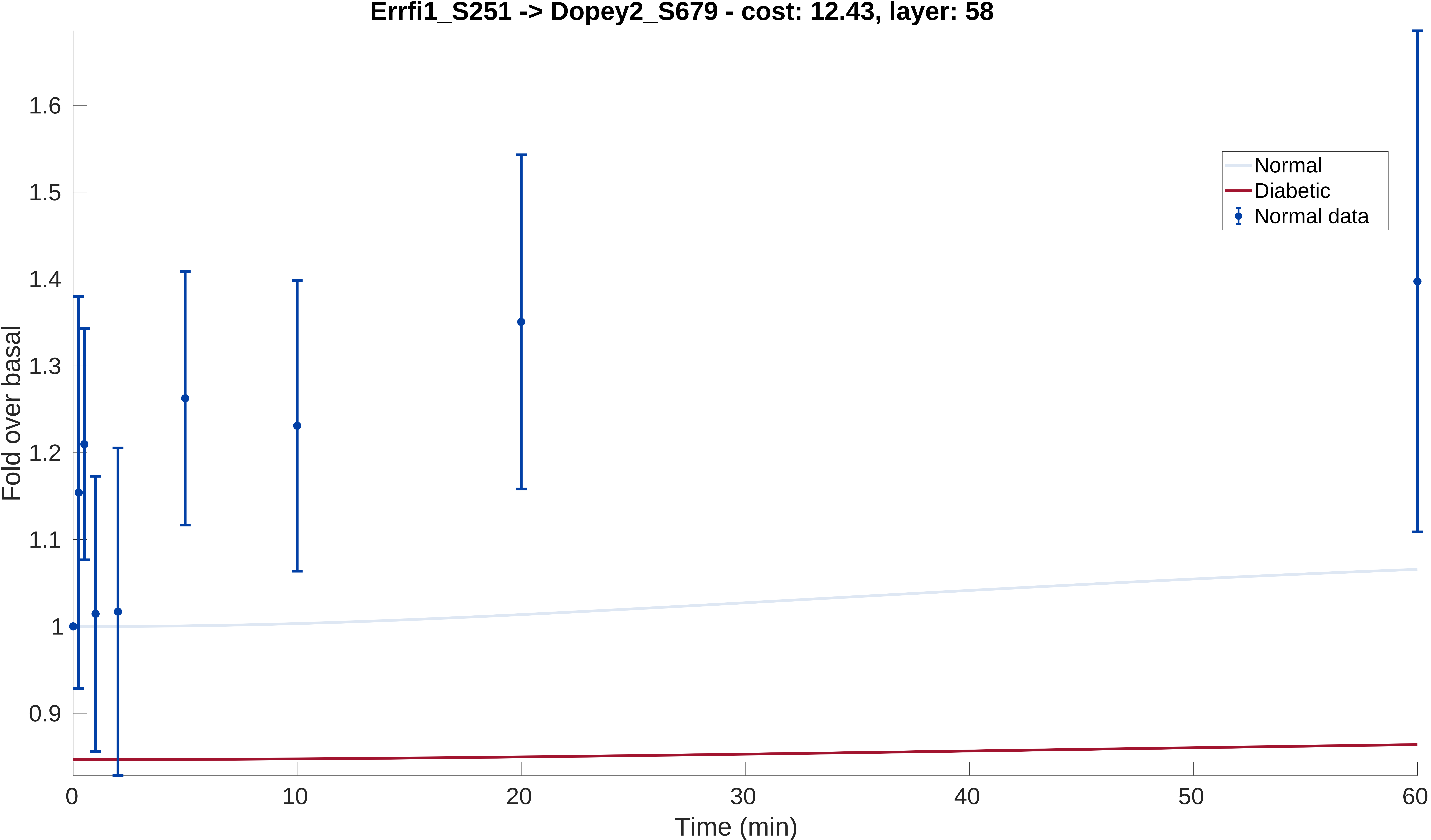
Errfi1_S251 -> Dopey2_S679 - cost: 12.43, layer: 58
Fold over basal
1.6
1.5
1.4
1.3
1.2
1.1
1
0.9
0
10
20
30
40
50
60
Time (min)
Normal
Diabetic
Normal data

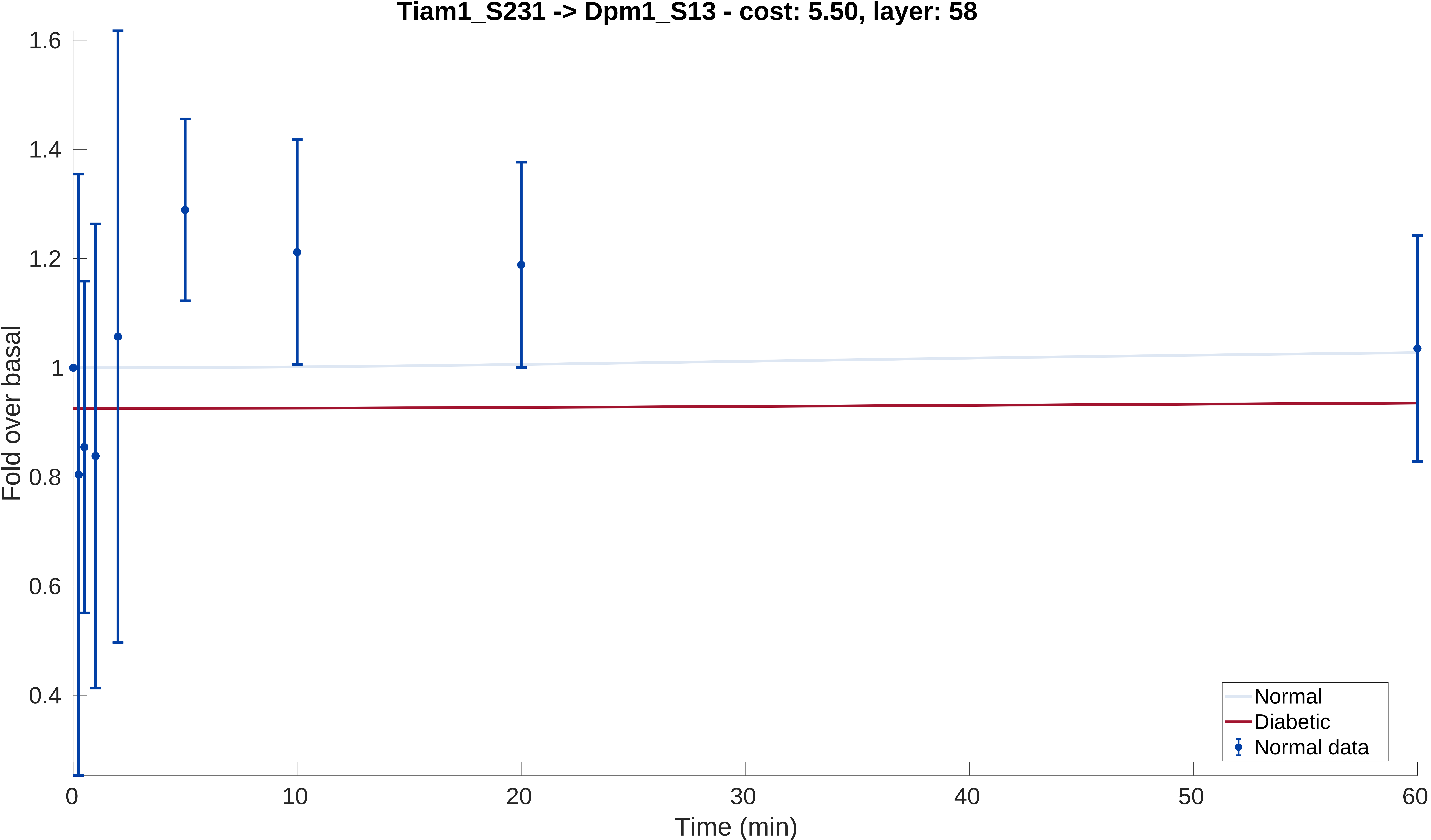

Tiam1_S231 -> Dpm1_S13 - cost: 5.50, layer: 58
Fold over basal
1.6
1.4
1.2
1
0.8
0.6
0.4
0
10
20
30
40
50
60
Time (min)
Normal
Diabetic
Normal data

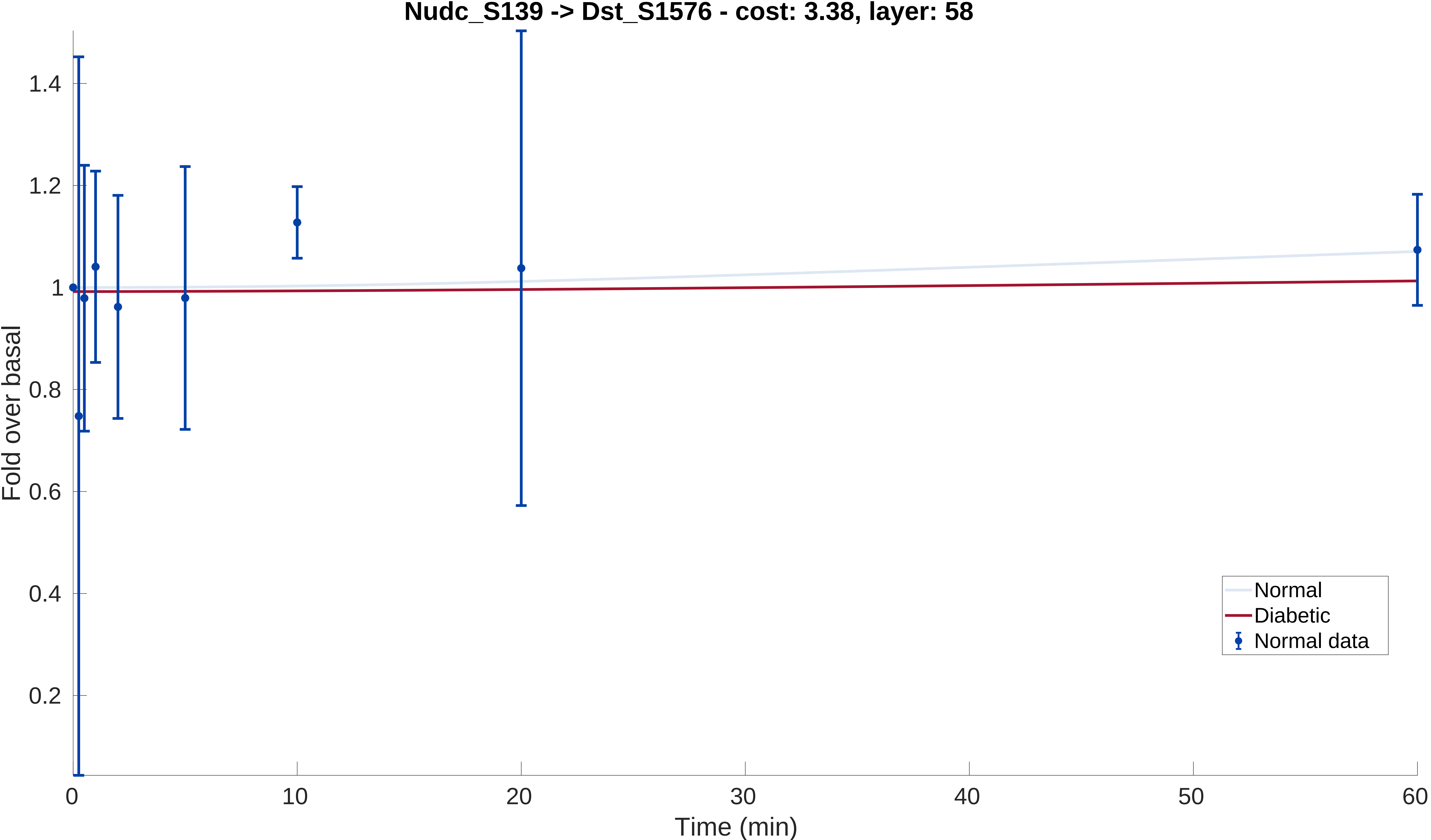
Nudc_S139 -> Dst_S1576 - cost: 3.38, layer: 58
Fold over basal
1.4
1.2
1
0.8
0.6
0.4
0.2
0
10
20
30
40
50
60
Time (min)
Normal
Diabetic
Normal data

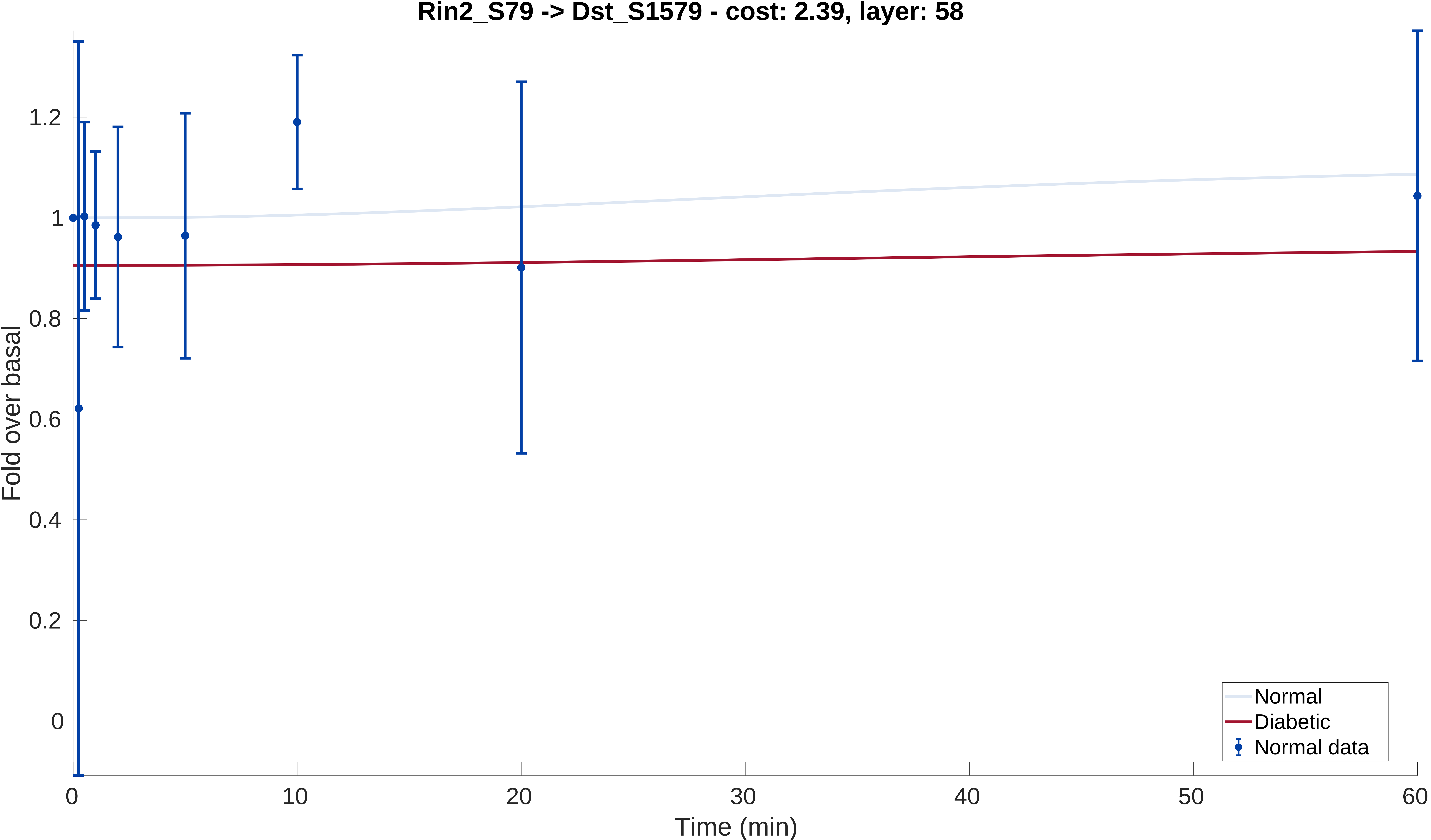

Rin2_S79 -> Dst_S1579 - cost: 2.39, layer: 58
Fold over basal
1.2
1
0.8
0.6
0.4
0.2
0
0
10
20
30
40
50
60
Time (min)
Normal
Diabetic
Normal data

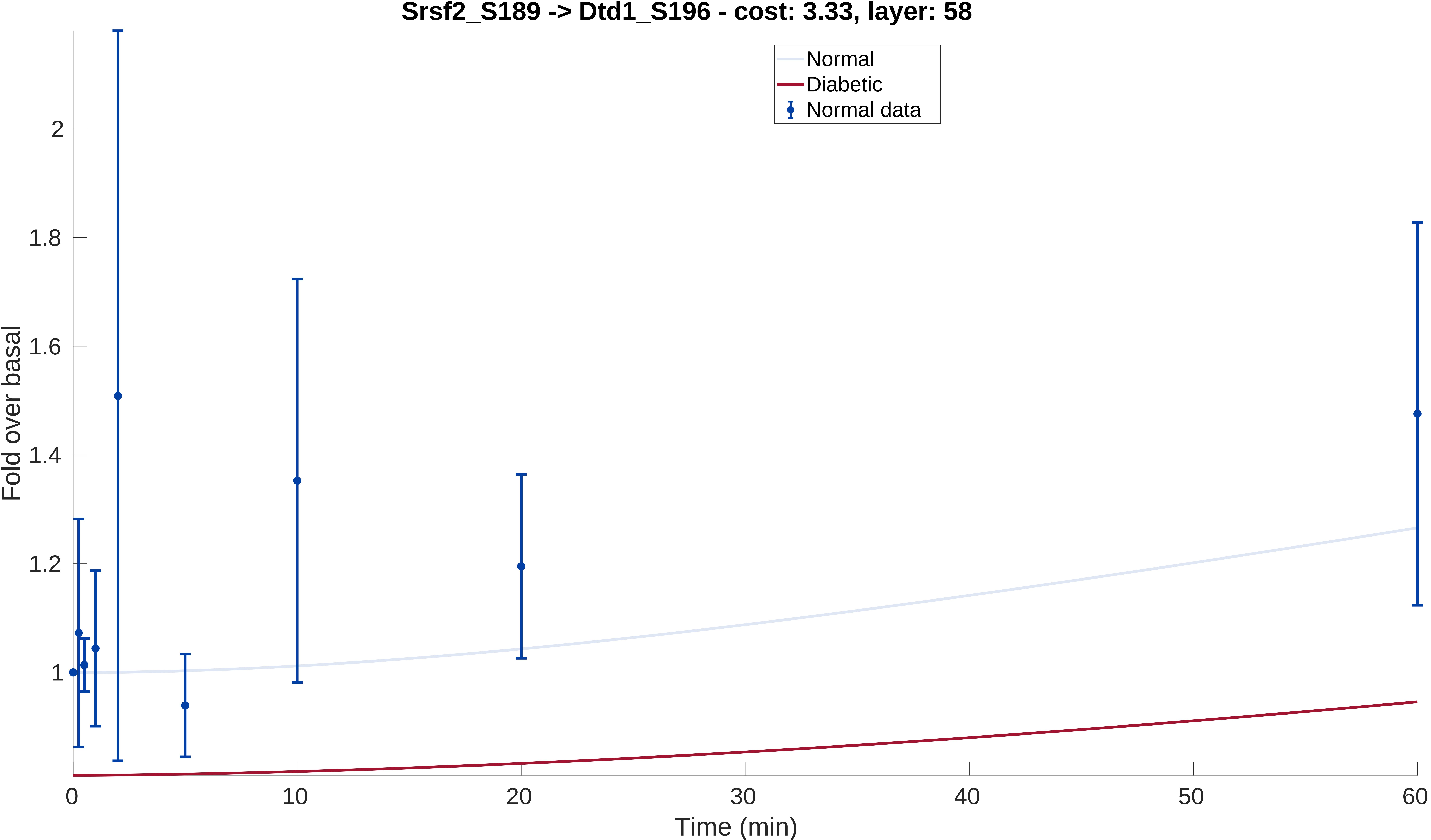

Srsf2_S189 -> Dtd1_S196 - cost: 3.33, layer: 58
Normal
Diabetic
Normal data
Fold over basal
2
1.8
1.6
1.4
1.2
1
0
10
20
30
40
50
60
Time (min)

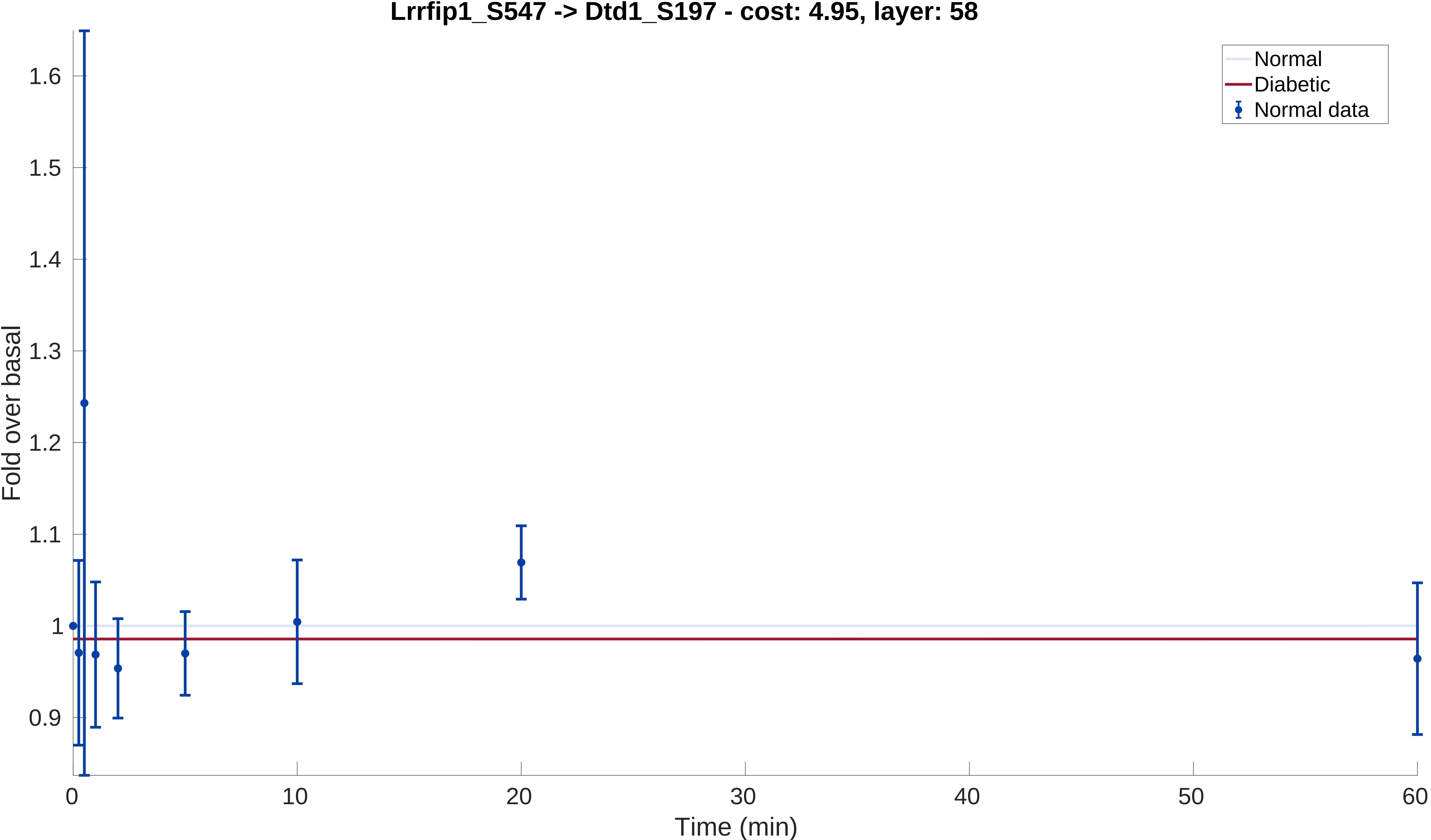

Lrrfip1_S547 -> Dtd1_S197 - cost: 4.95, layer: 58
Normal
Diabetic
Normal data
Fold over basal
1.6
1.5
1.4
1.3
1.2
1.1
1
0.9
0
10
20
30
40
50
60
Time (min)

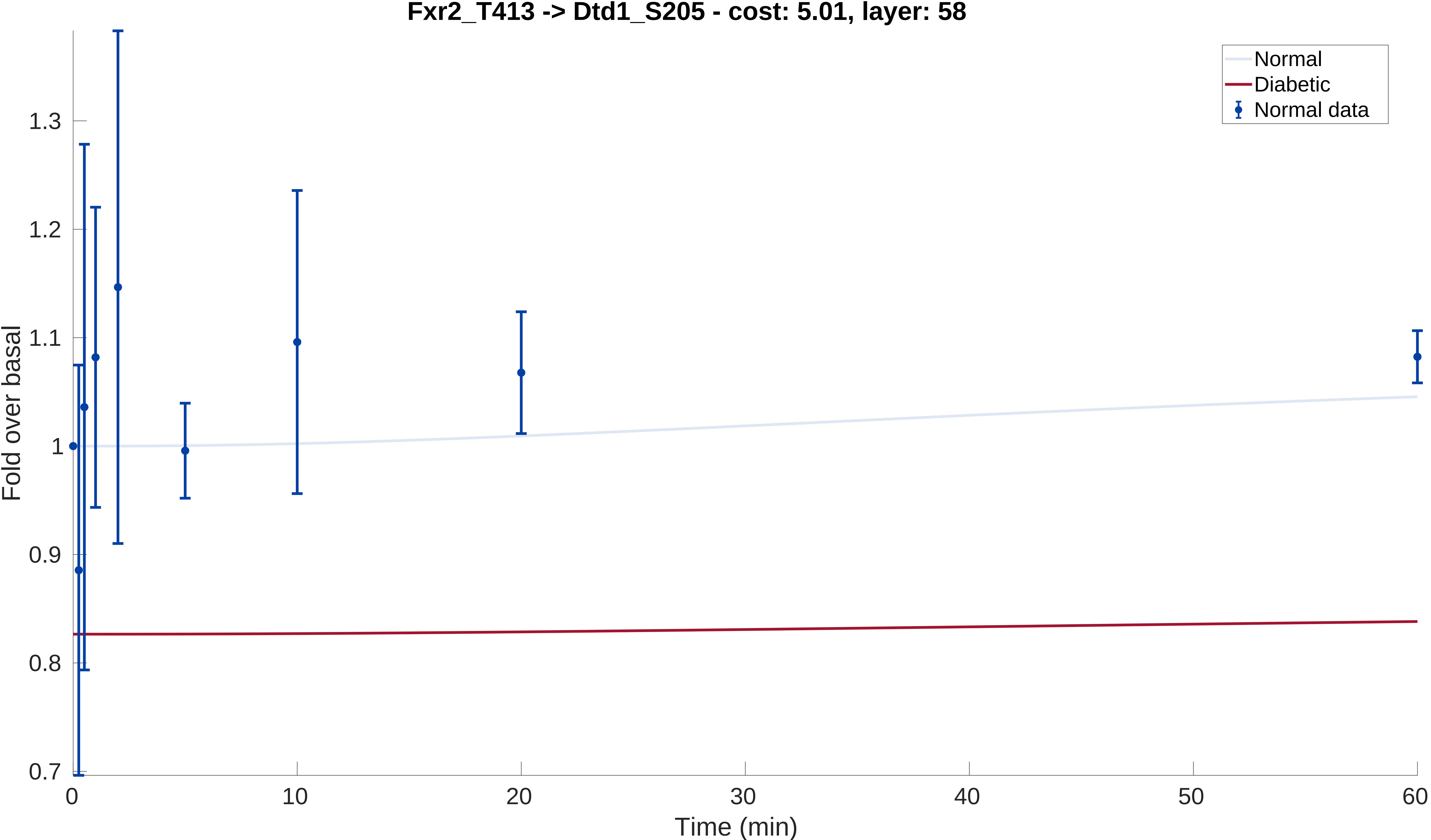

Fxr2_T413 -> Dtd1_S205 - cost: 5.01, layer: 58
Normal
Diabetic
Normal data
Fold over basal
1.3
1.2
1.1
1
0.9
0.8
0.7
0
10
20
30
40
50
60
Time (min)

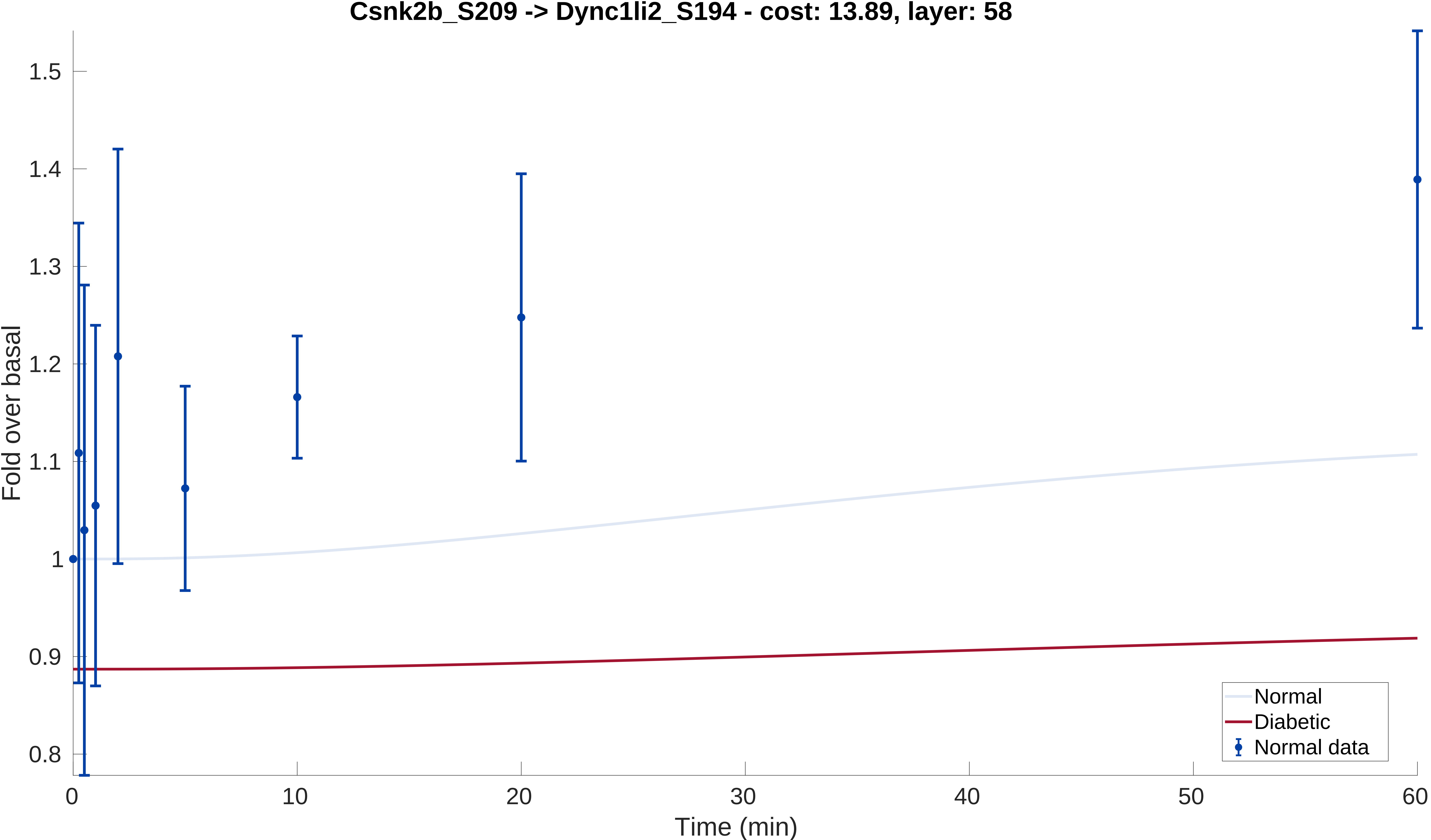

Csnk2b_S209 -> Dync1li2_S194 - cost: 13.89, layer: 58
Fold over basal
1.5
1.4
1.3
1.2
1.1
1
0.9
0.8
0
10
20
30
40
50
60
Time (min)
Normal
Diabetic
Normal data

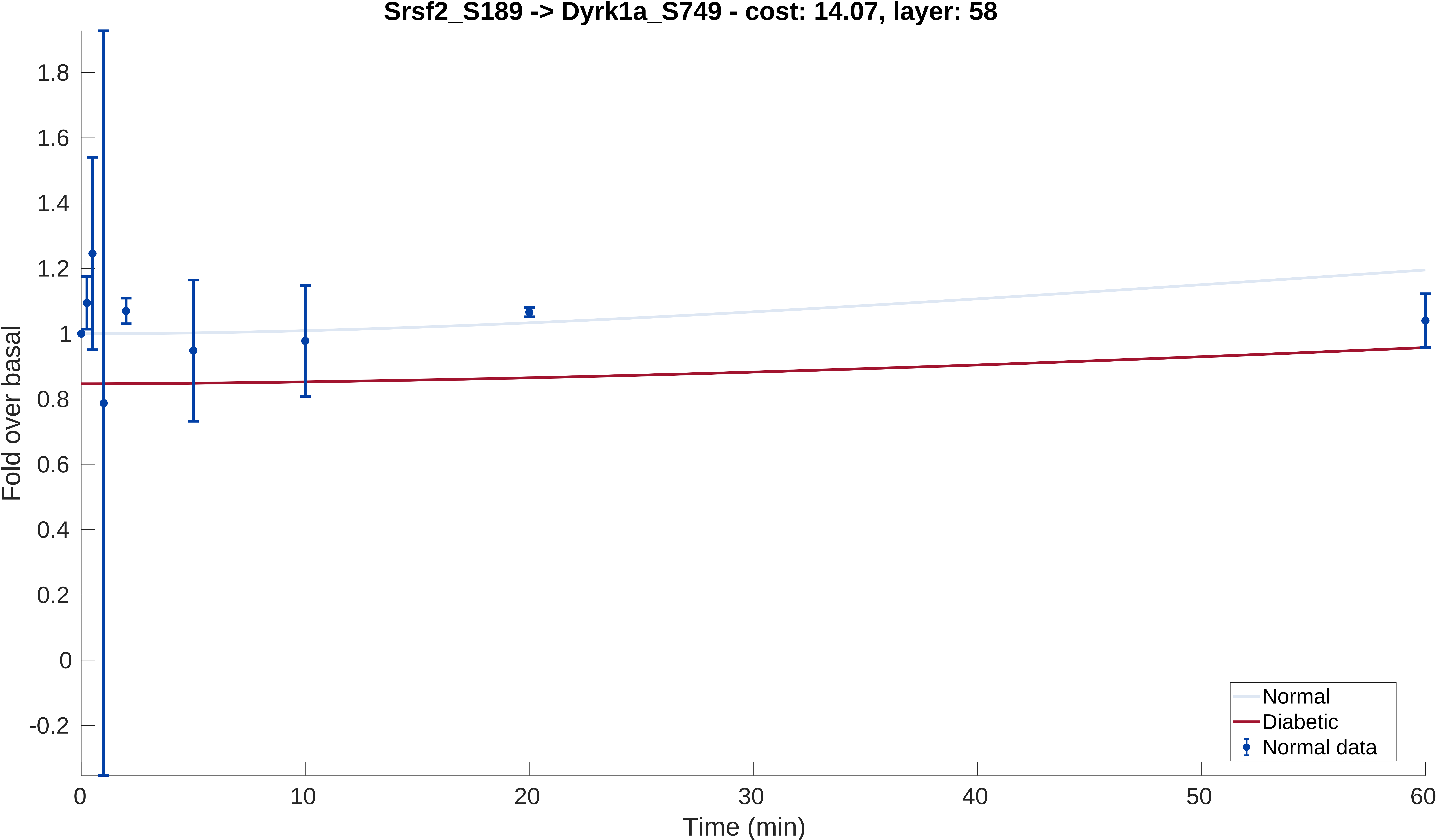

Srsf2_S189 -> Dyrk1a_S749 - cost: 14.07, layer: 58
Fold over basal
1.8
1.6
1.4
1.2
1
0.8
0.6
0.4
0.2
0
-0.2
0
10
20
30
40
50
60
Time (min)
Normal
Diabetic
Normal data

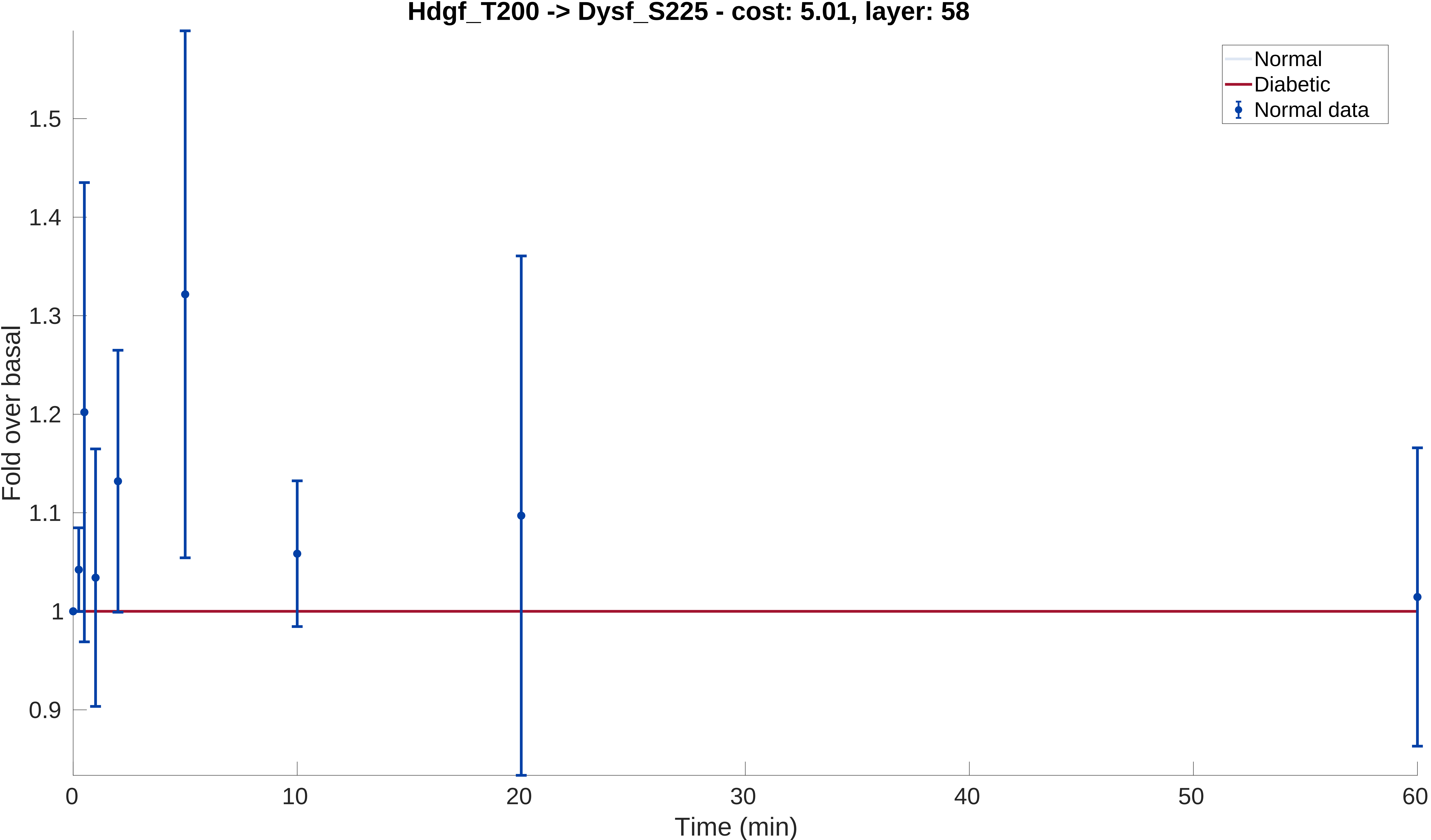

Hdgf_T200 -> Dysf_S225 - cost: 5.01, layer: 58
Normal
Diabetic
Normal data
Fold over basal
Time (min)
1.5
1.4
1.3
1.2
1.1
1
0.9
0
10
20
30
40
50
60

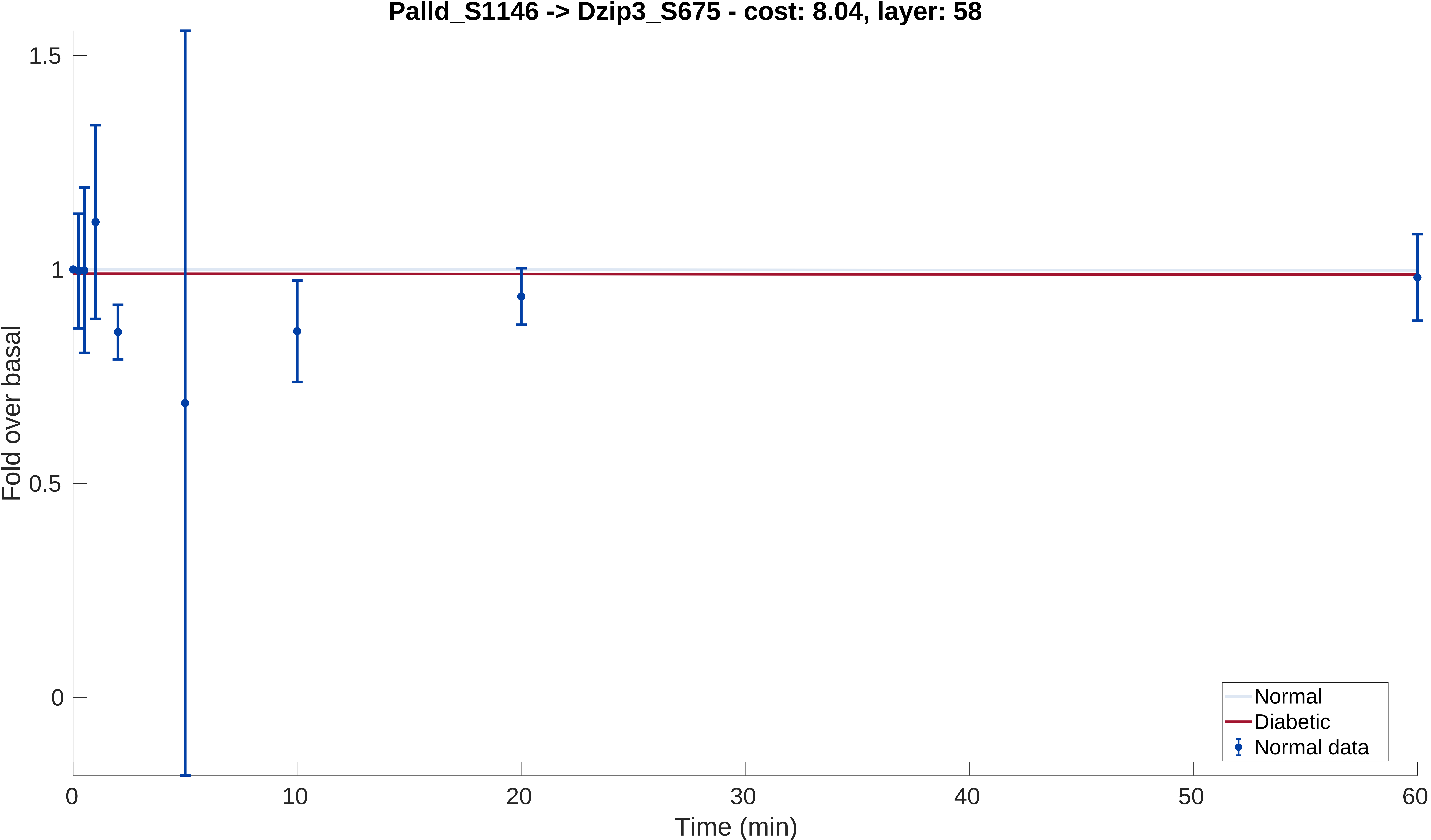

Palld_S1146 -> Dzip3_S675 - cost: 8.04, layer: 58
1.5
1
0.5
0
Fold over basal
0
10
20
30
40
50
60
Time (min)
Normal
Diabetic
Normal data

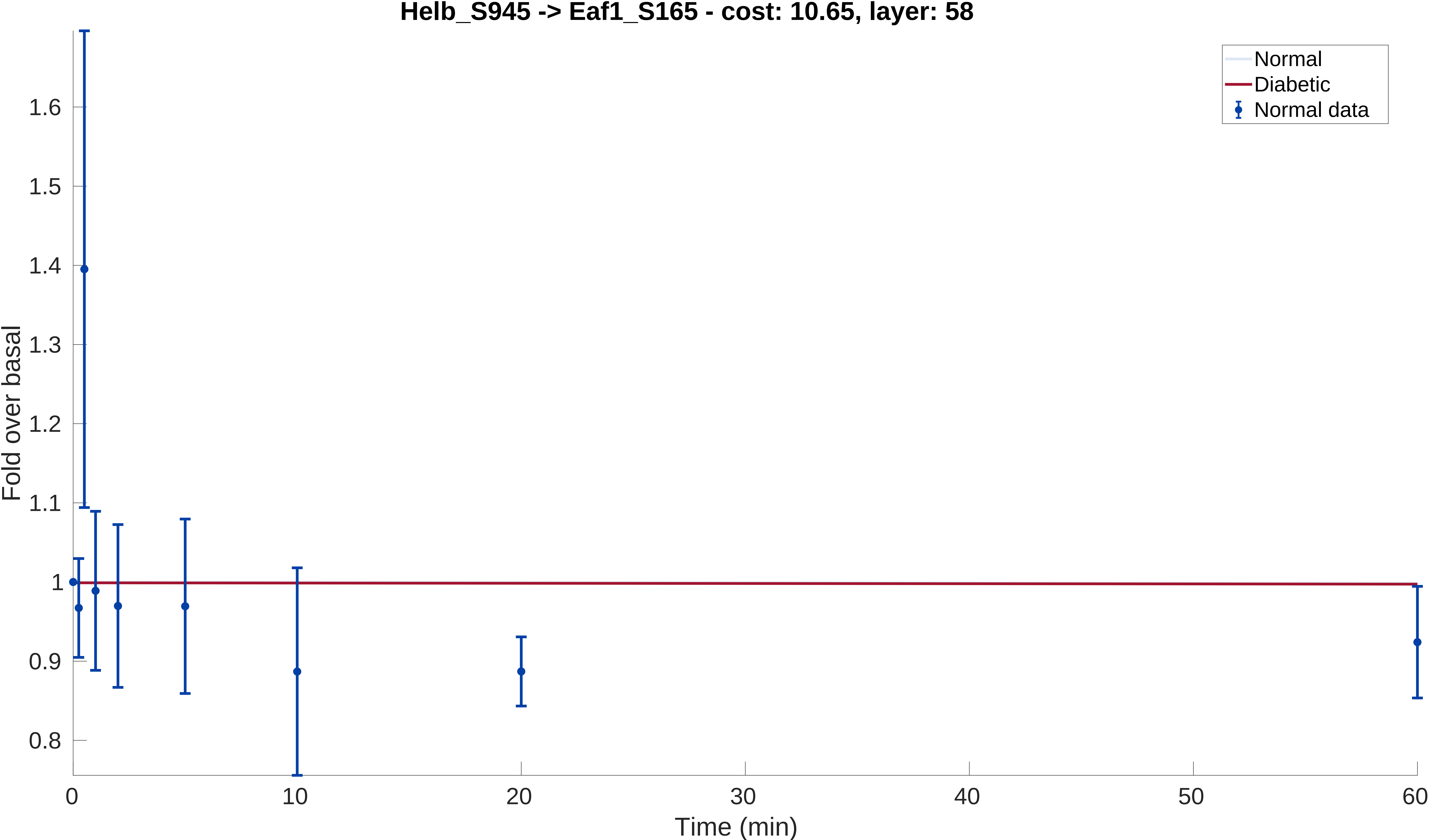

Helb_S945 -> Eaf1_S165 - cost: 10.65, layer: 58
Normal
Diabetic
Normal data
Fold over basal
1.6
1.5
1.4
1.3
1.2
1.1
1
0.9
0.8
0
10
20
30
40
50
60
Time (min)

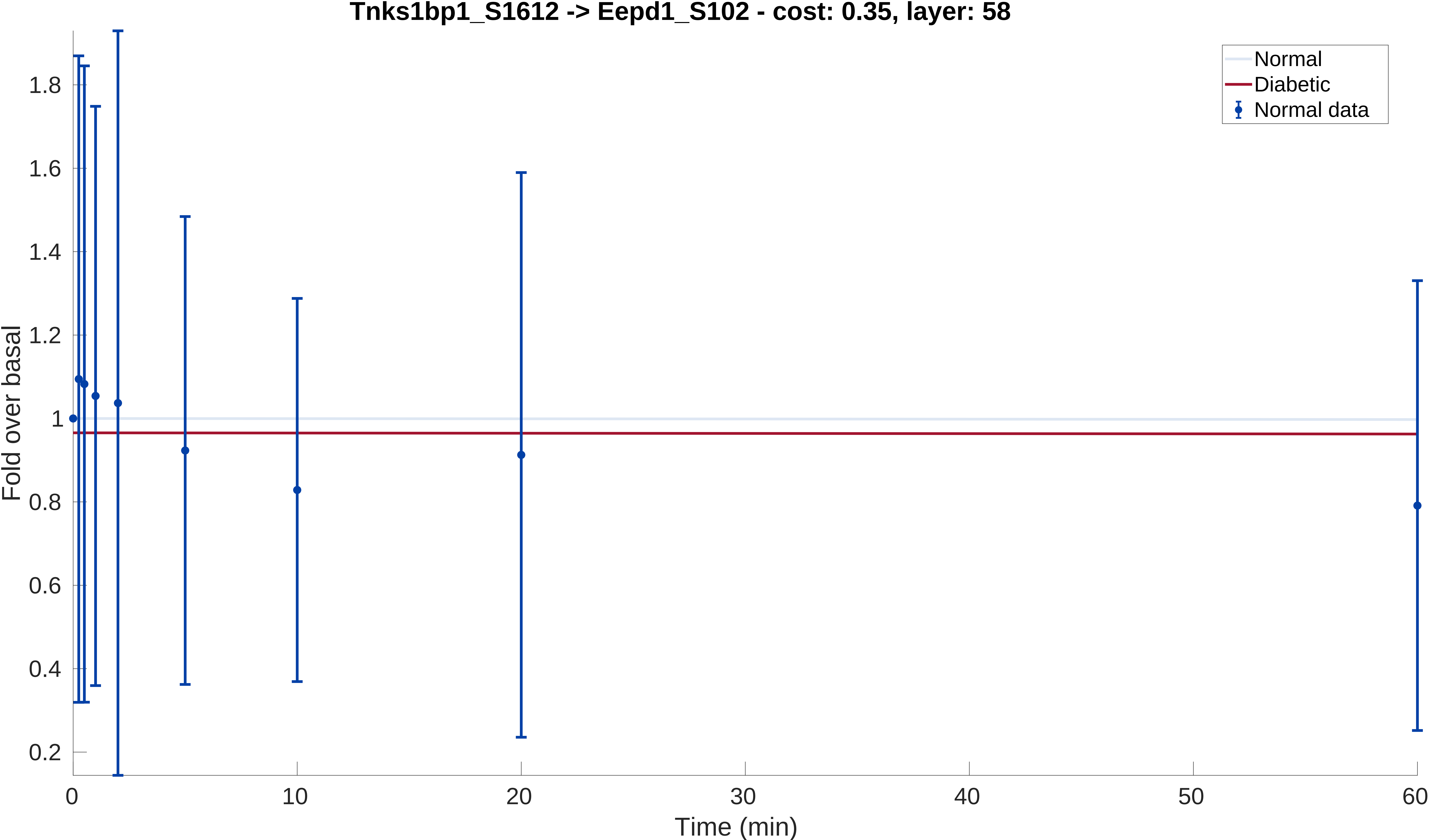
Tnks1bp1_S1612 -> Eepd1_S102 - cost: 0.35, layer: 58
Fold over basal
Time (min)
Normal
Diabetic
Normal data
0.2
0.4
0.6
0.8
1
1.2
1.4
1.6
1.8
0
10
20
30
40
50
60

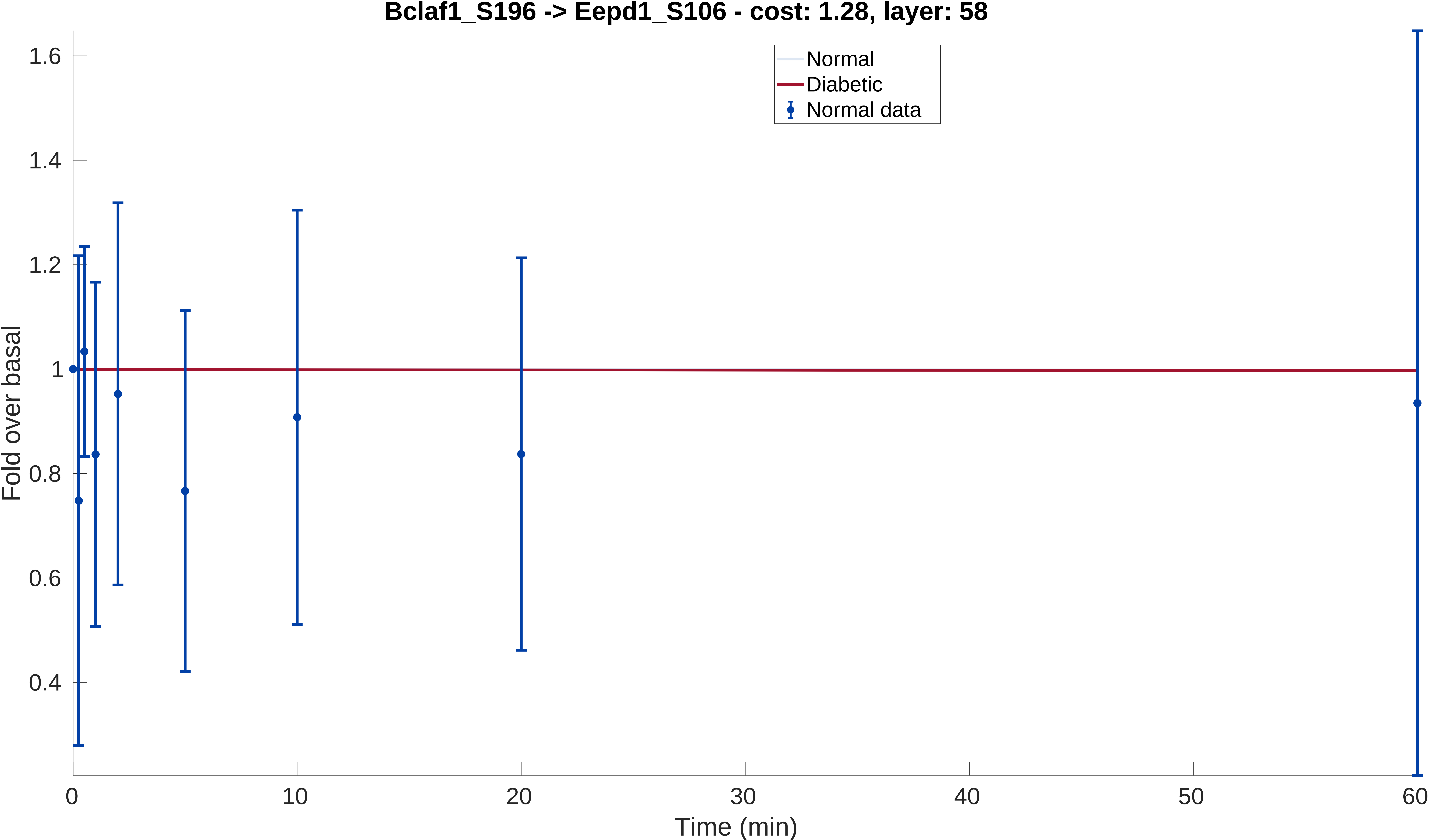
Bclaf1_S196 -> Eepd1_S106 - cost: 1.28, layer: 58
Normal
Diabetic
Normal data
Fold over basal
Time (min)
1.6
1.4
1.2
1
0.8
0.6
0.4
0
10
20
30
40
50
60

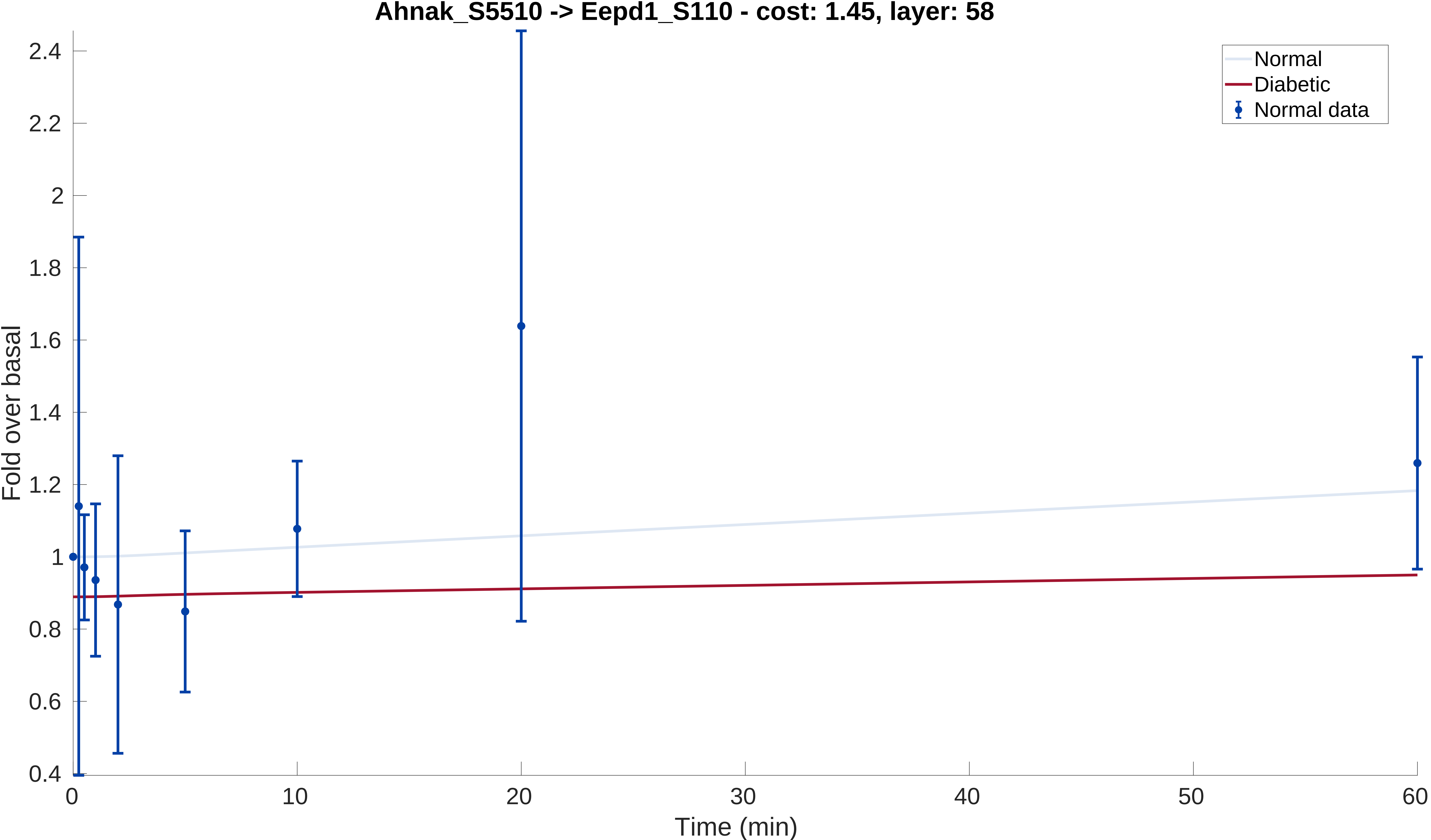

Ahnak_S5510 -> Eepd1_S110 - cost: 1.45, layer: 58
Normal
Diabetic
Normal data
Fold over basal
Time (min)
2.4
2.2
2
1.8
1.6
1.4
1.2
1
0.8
0.6
0.4
0
10
20
30
40
50
60

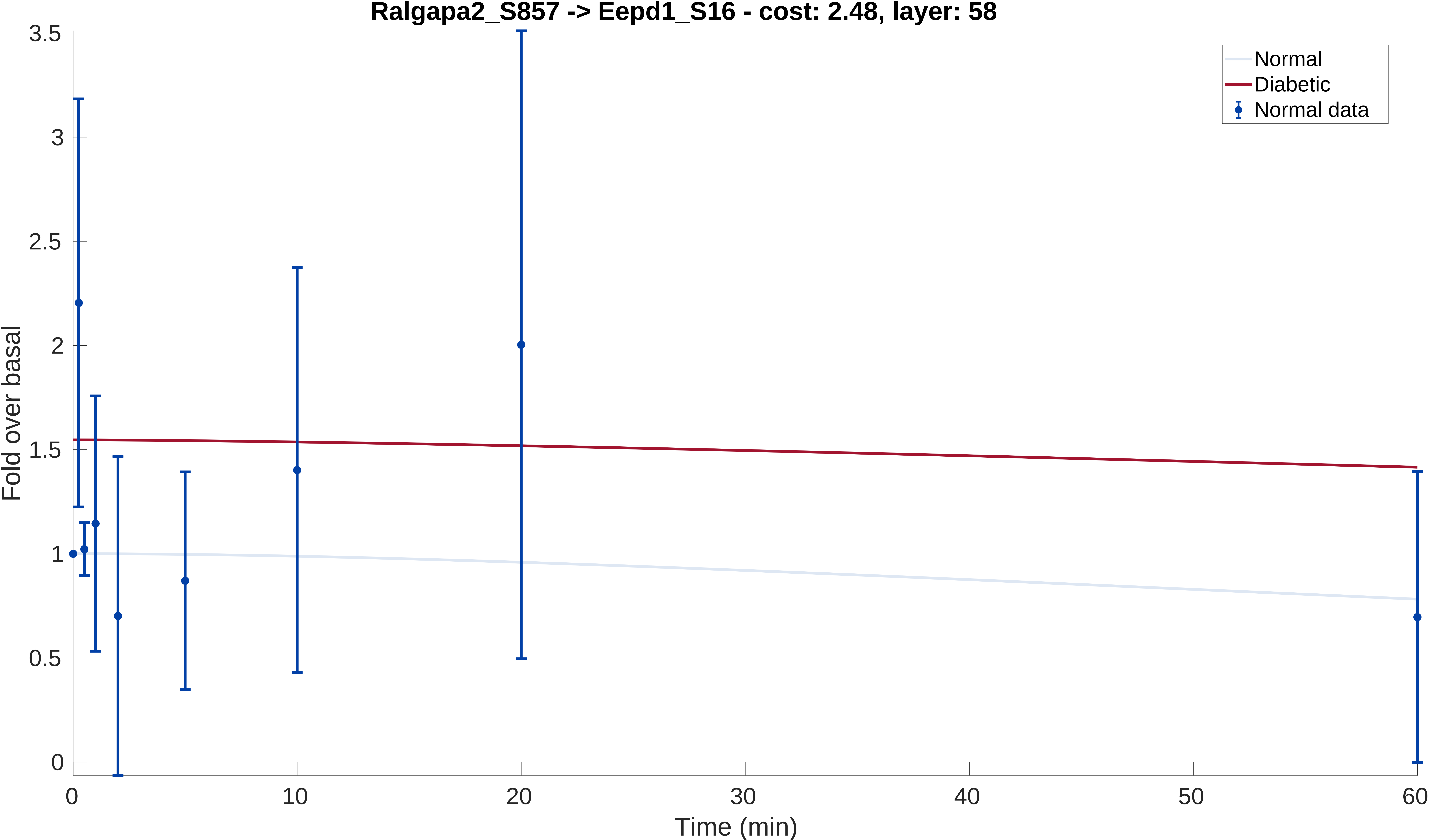

Ralgapa2_S857 -> Eepd1_S16 - cost: 2.48, layer: 58
Normal
Diabetic
Normal data
Fold over basal
Time (min)
0
0.5
1
1.5
2
2.5
3
3.5
0
10
20
30
40
50
60

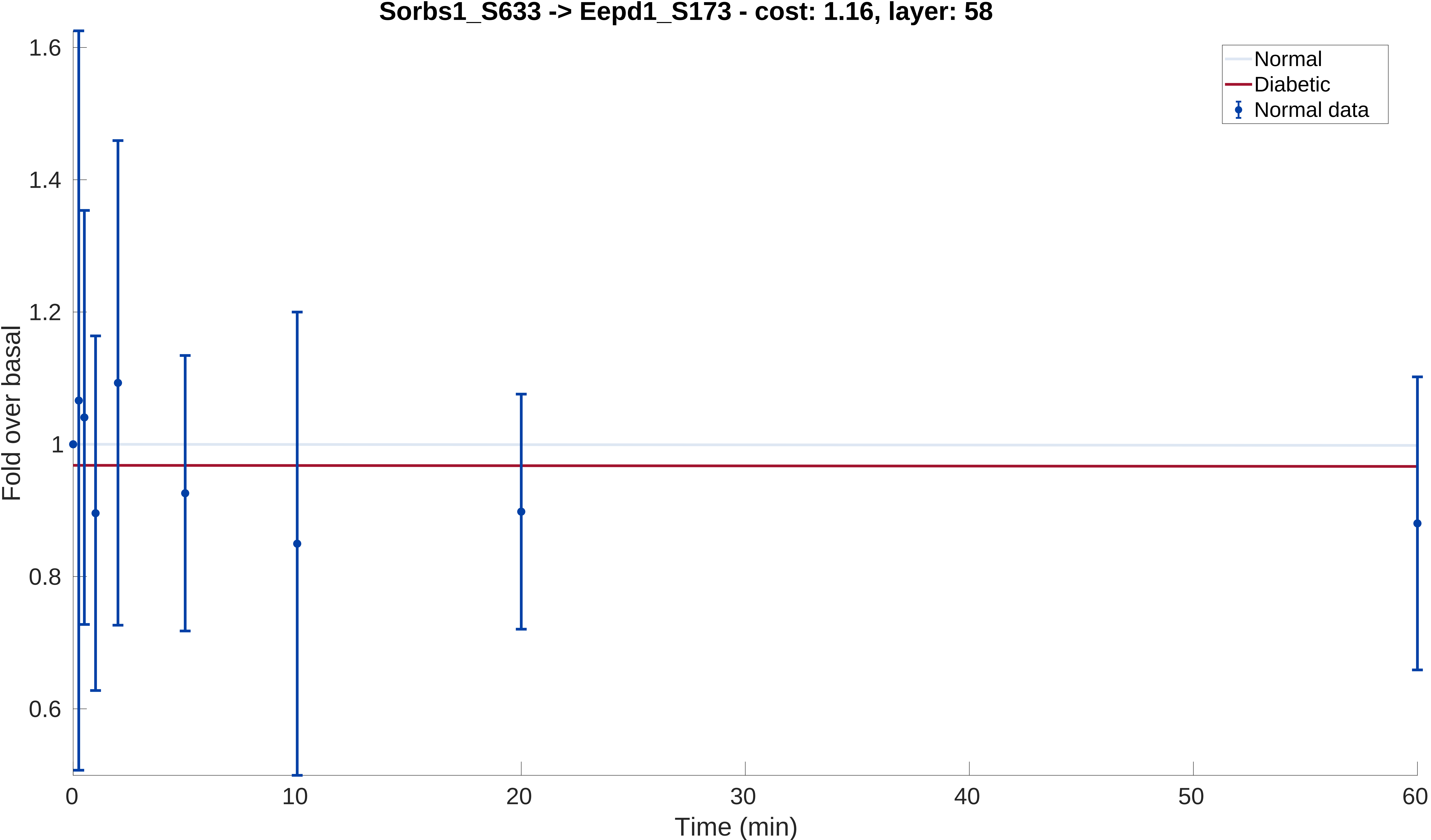

Sorbs1_S633 -> Eepd1_S173 - cost: 1.16, layer: 58
Normal
Diabetic
Normal data
Fold over basal
1.6
1.4
1.2
1
0.8
0.6
0
10
20
30
40
50
60
Time (min)

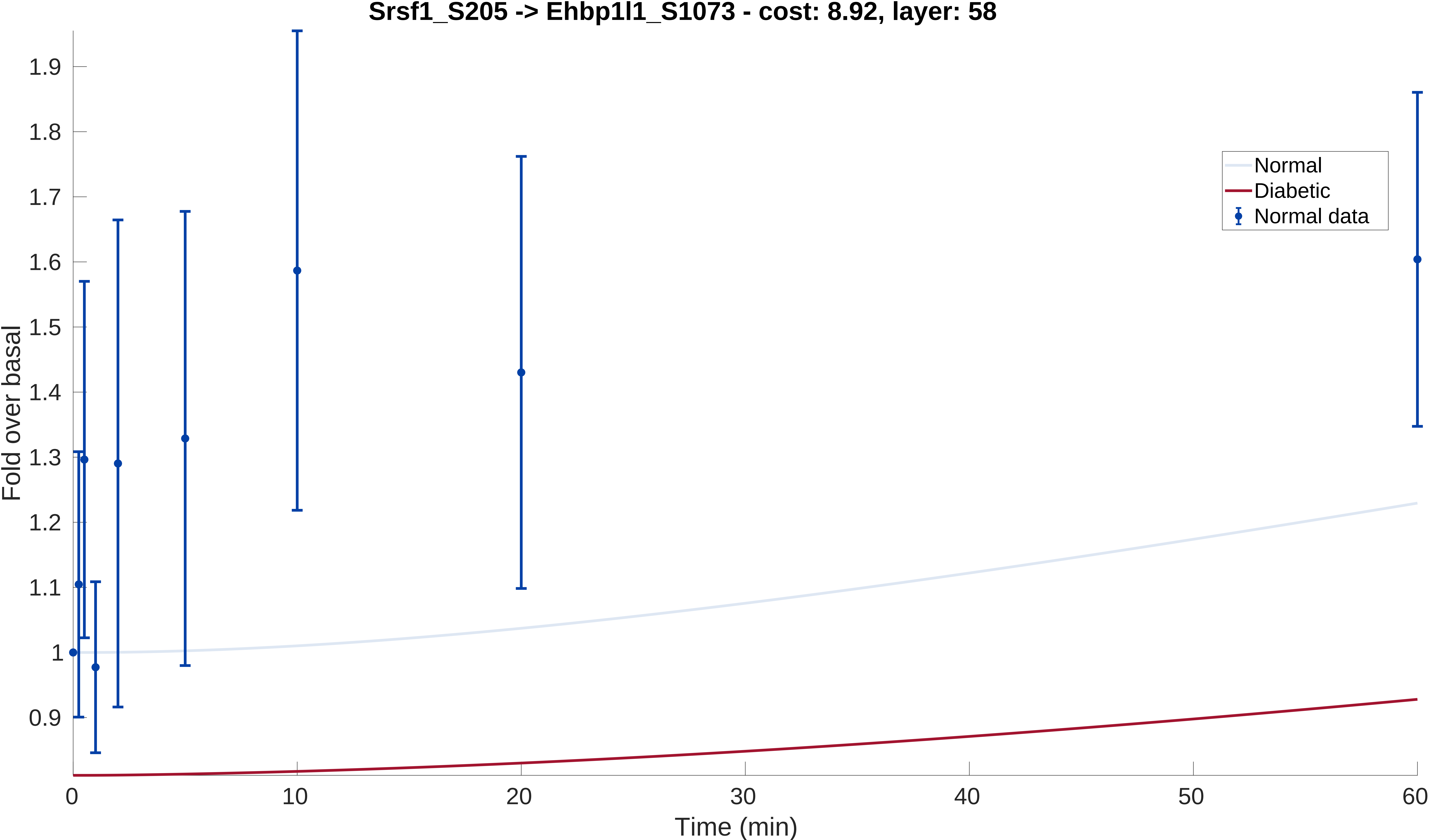

Srsf1_S205 -> Ehbp1l1_S1073 - cost: 8.92, layer: 58
Fold over basal
1.9
1.8
1.7
1.6
1.5
1.4
1.3
1.2
1.1
1
0.9
0
10
20
30
40
50
60
Time (min)
Normal
Diabetic
Normal data

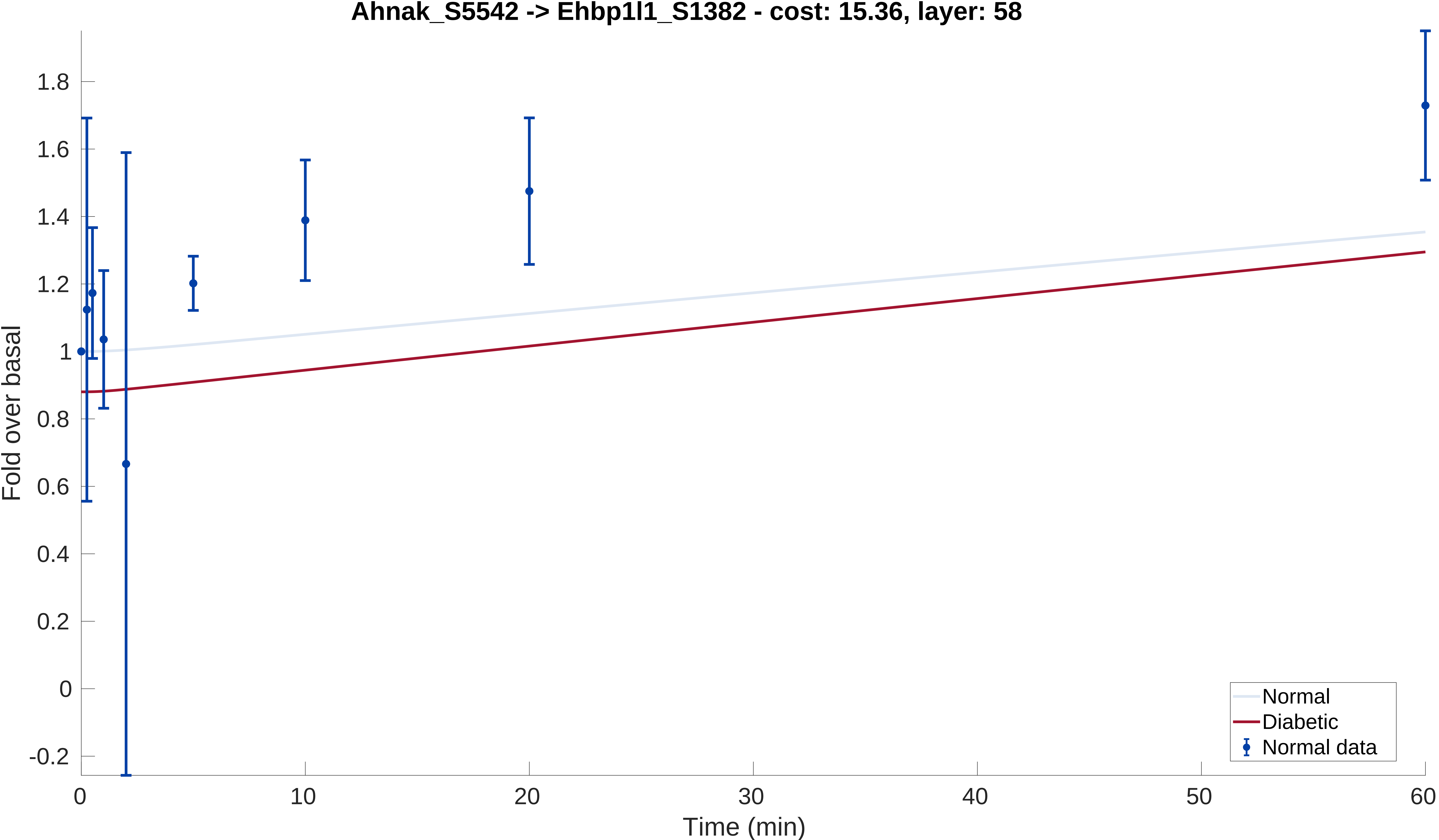

Ahnak_S5542 -> Ehbp1l1_S1382 - cost: 15.36, layer: 58
Fold over basal
1.8
1.6
1.4
1.2
1
0.8
0.6
0.4
0.2
0
-0.2
0
10
20
30
40
50
60
Time (min)
Normal
Diabetic
Normal data

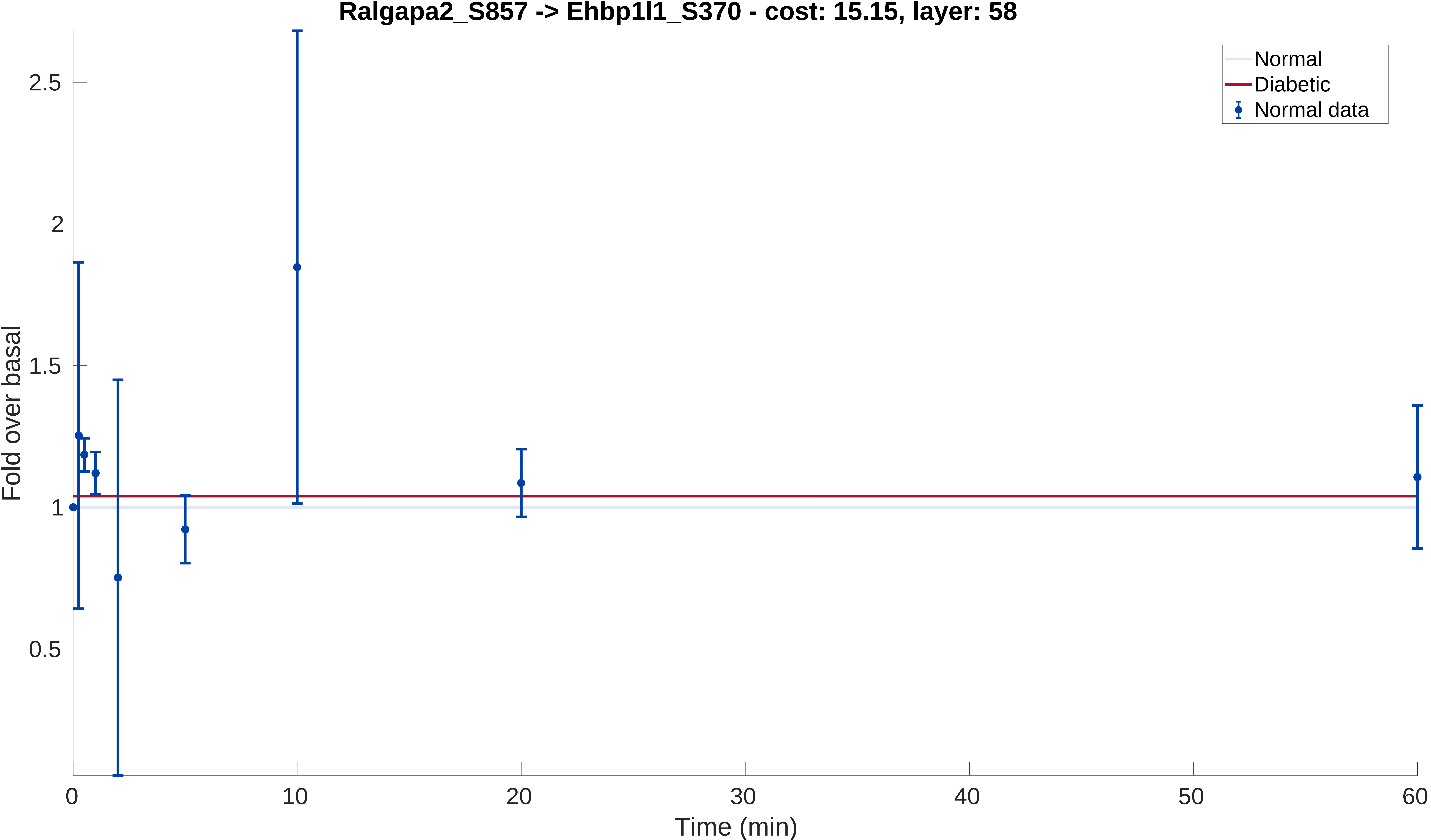

Ralgapa2_S857 -> Ehbp1l1_S370 - cost: 15.15, layer: 58
Normal
Diabetic
Normal data
Fold over basal
2.5
2
1.5
1
0.5
0
10
20
30
40
50
60
Time (min)

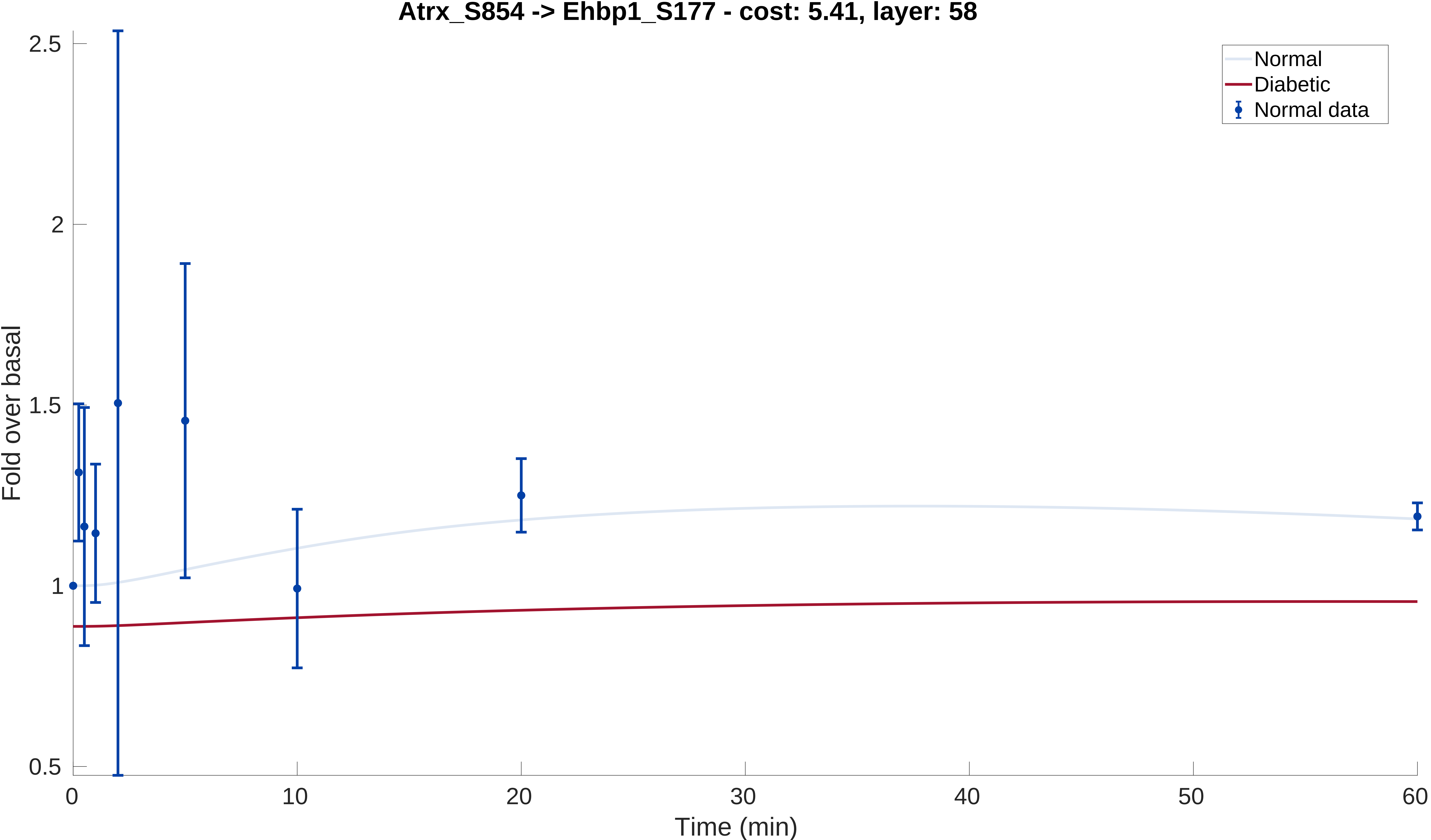

Atrx_S854 -> Ehbp1_S177 - cost: 5.41, layer: 58
Normal
Diabetic
Normal data
Fold over basal
Time (min)
0.5
1
1.5
2
2.5
0
10
20
30
40
50
60

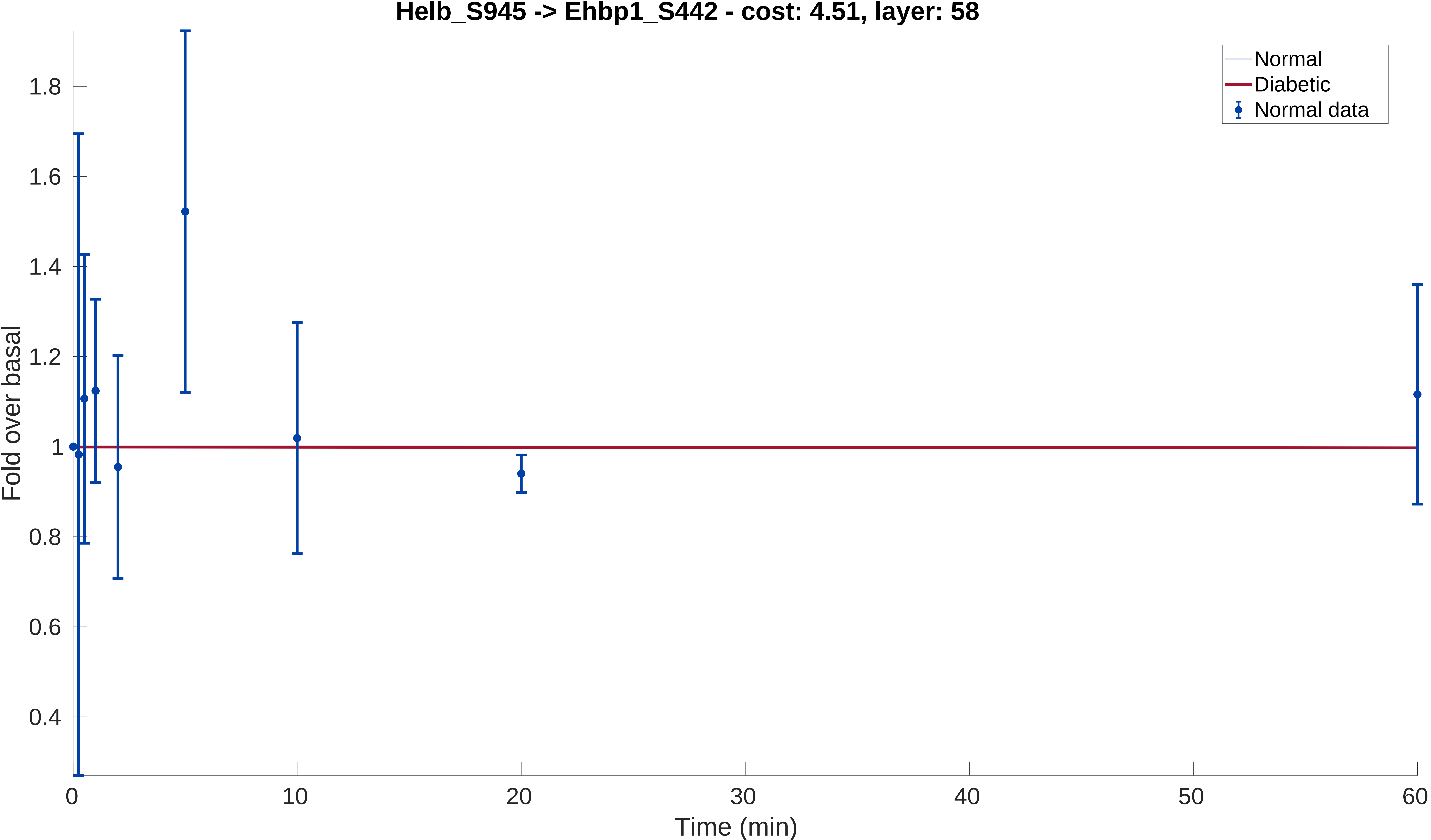

Helb_S945 -> Ehbp1_S442 - cost: 4.51, layer: 58
Normal
Diabetic
Normal data
Fold over basal
1.8
1.6
1.4
1.2
1
0.8
0.6
0.4
0
10
20
30
40
50
60
Time (min)

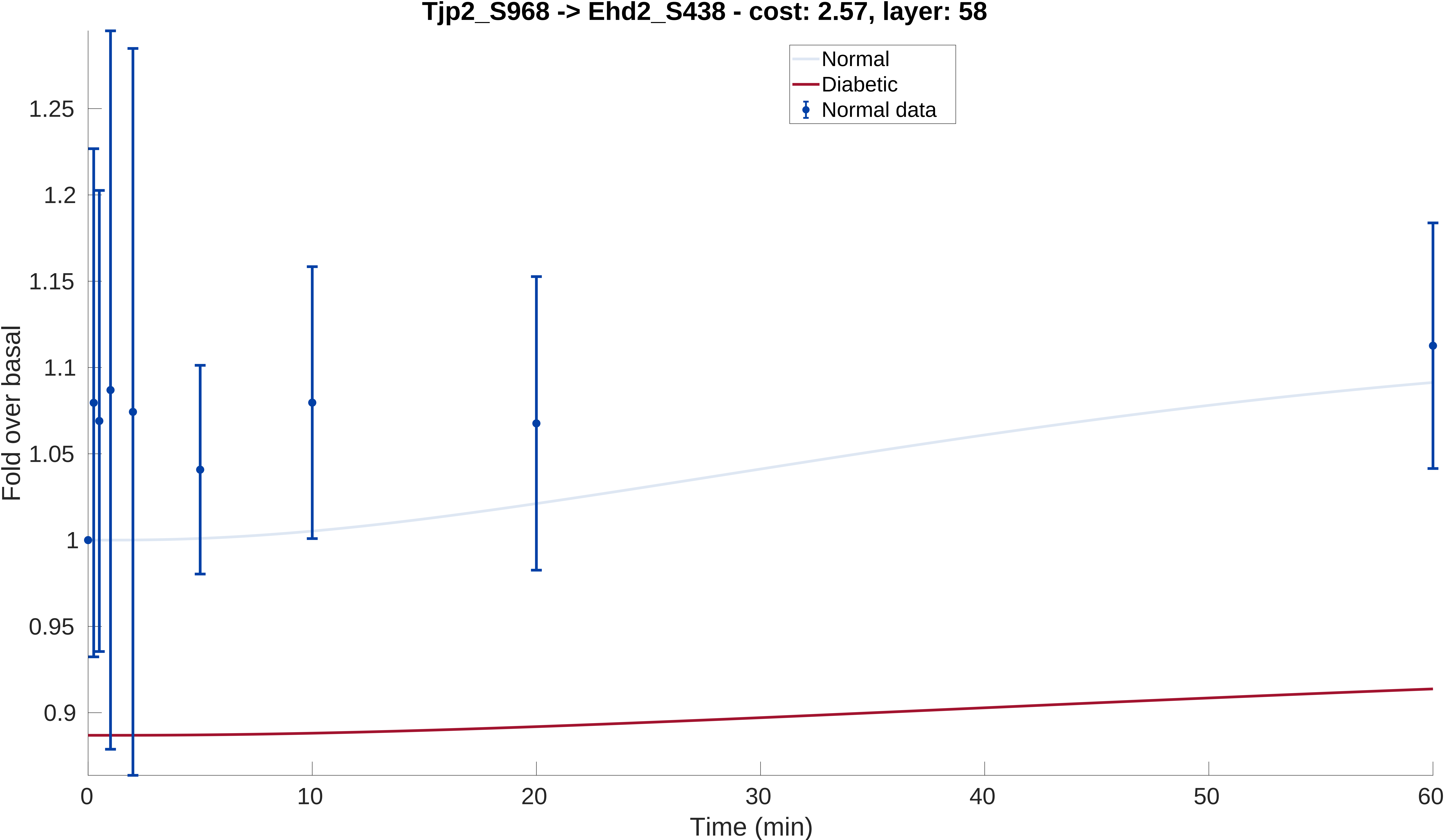

Tjp2_S968 -> Ehd2_S438 - cost: 2.57, layer: 58
Normal
Diabetic
Normal data
Fold over basal
1.25
1.2
1.15
1.1
1.05
1
0.95
0.9
0
10
20
30
40
50
60
Time (min)

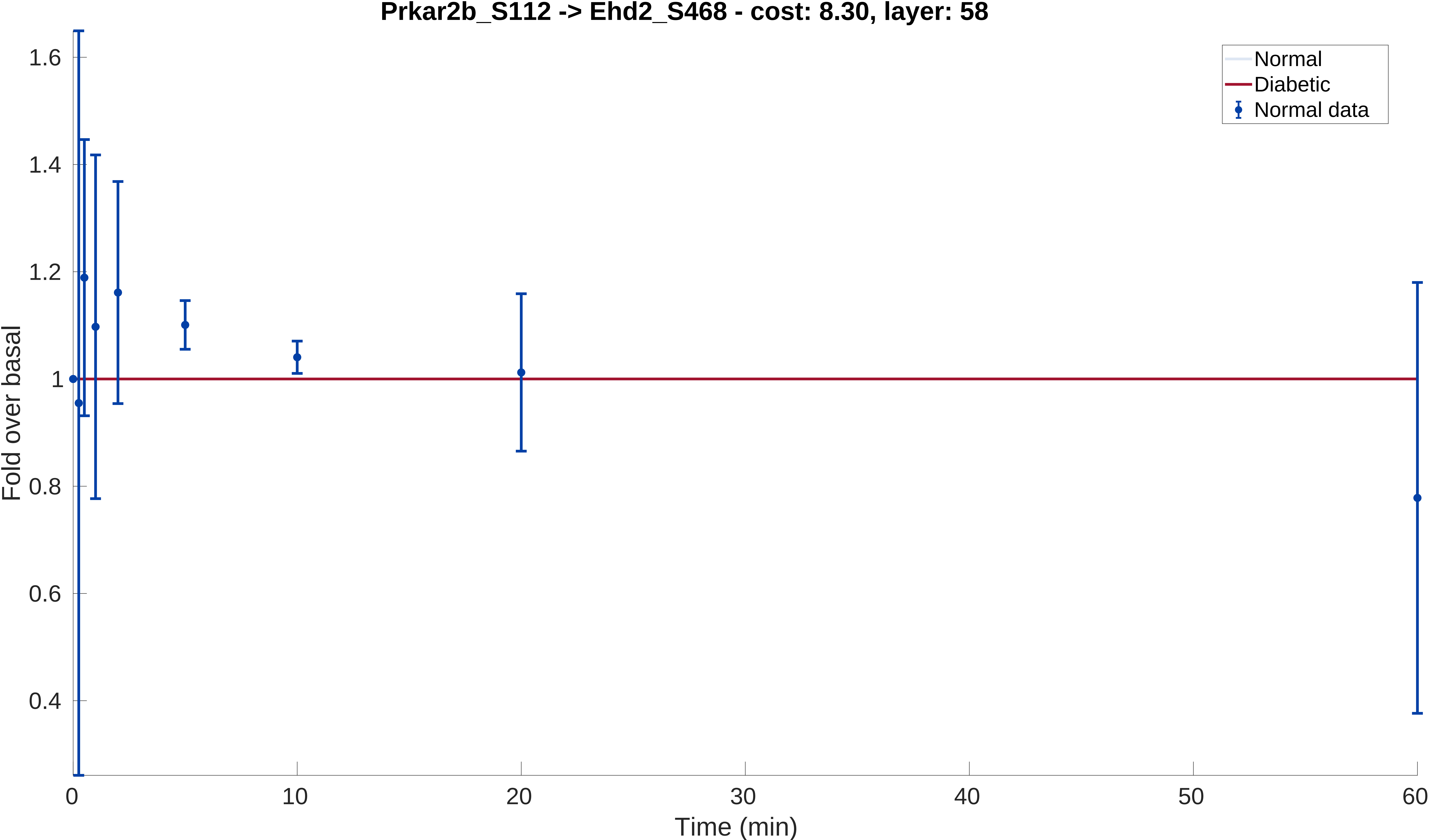
Prkar2b_S112 -> Ehd2_S468 - cost: 8.30, layer: 58
Normal
Diabetic
Normal data
Fold over basal
1.6
1.4
1.2
1
0.8
0.6
0.4
0
10
20
30
40
50
60
Time (min)

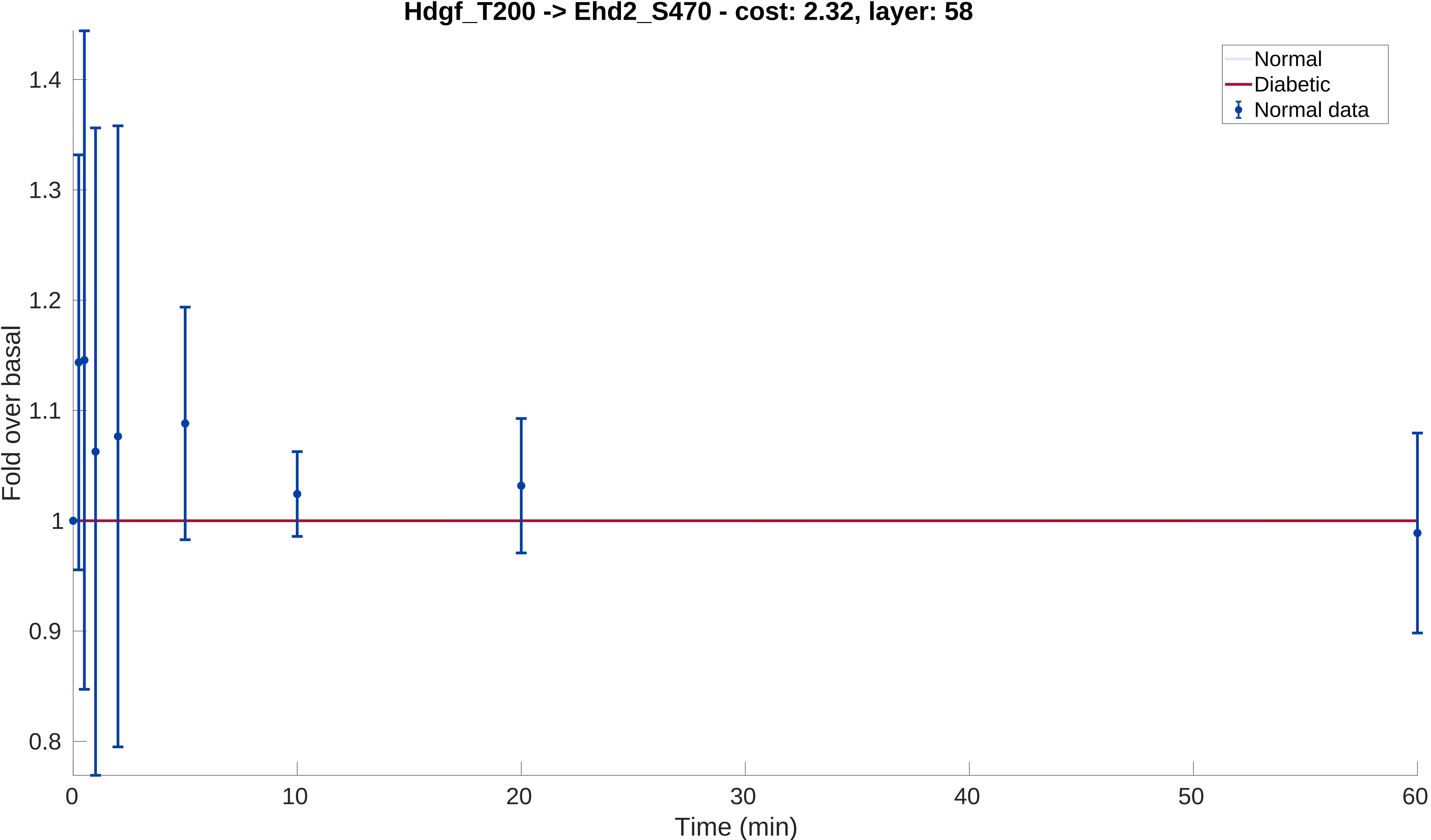
Hdgf_T200 -> Ehd2_S470 - cost: 2.32, layer: 58
Normal
Diabetic
Normal data
Fold over basal
1.4
1.3
1.2
1.1
1
0.9
0.8
0
10
20
30
40
50
60
Time (min)

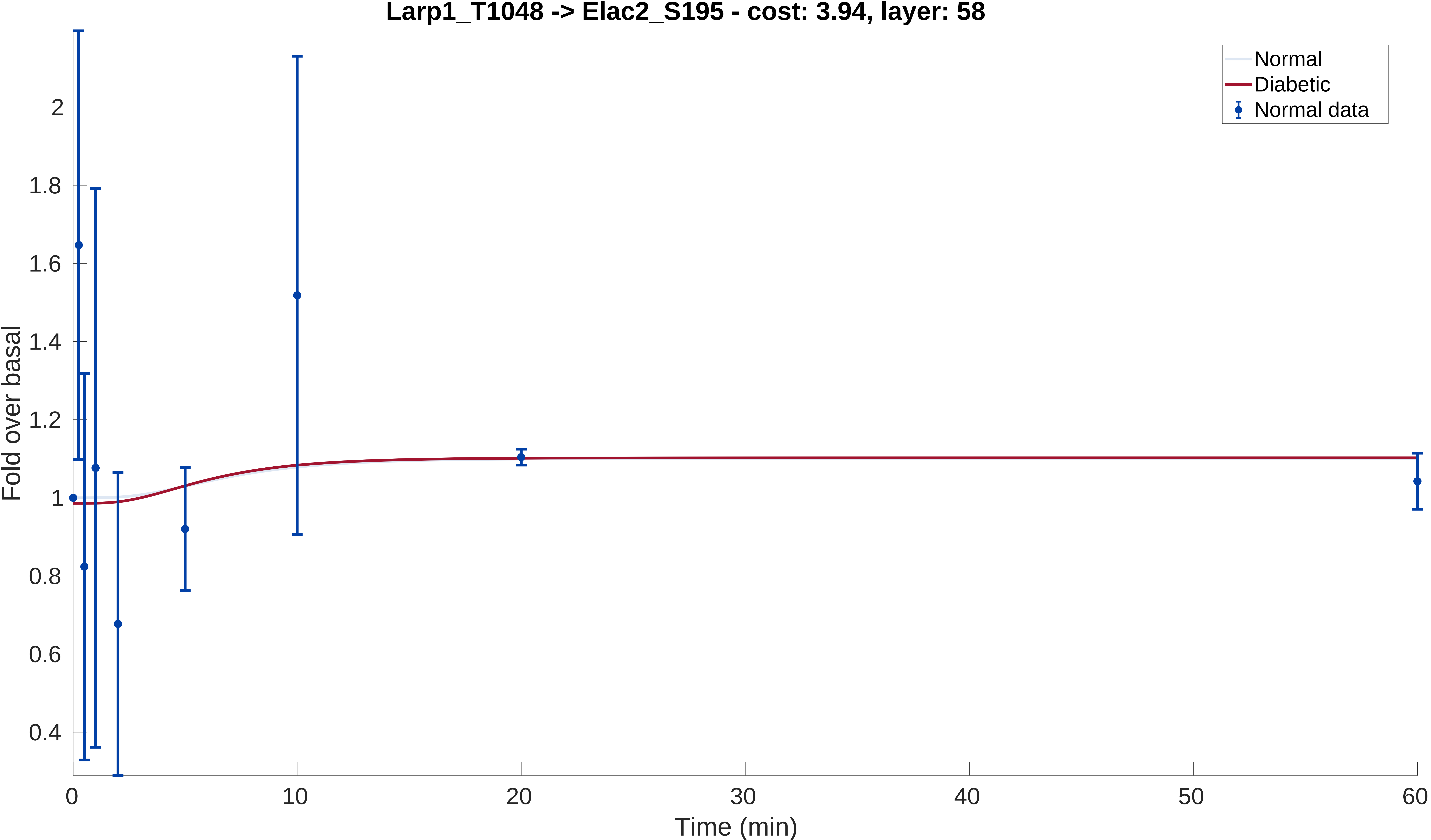

Larp1_T1048 -> Elac2_S195 - cost: 3.94, layer: 58
Normal
Diabetic
Normal data
Fold over basal
2
1.8
1.6
1.4
1.2
1
0.8
0.6
0.4
0
10
20
30
40
50
60
Time (min)

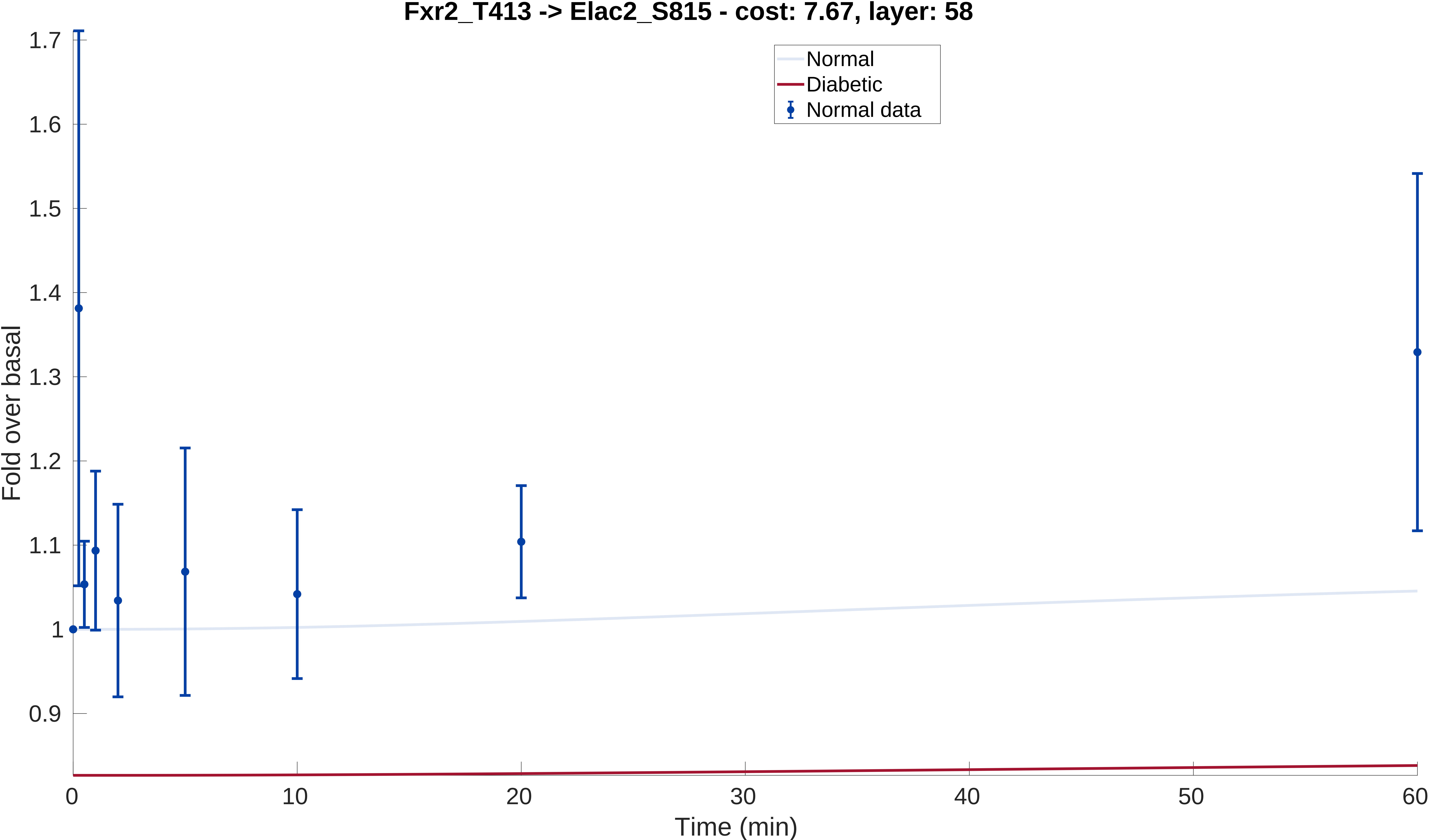

Fxr2_T413 -> Elac2_S815 - cost: 7.67, layer: 58
Normal
Diabetic
Normal data
1.7
1.6
1.5
1.4
1.3
1.2
1.1
1
0.9
Fold over basal
0
10
20
30
40
50
60
Time (min)

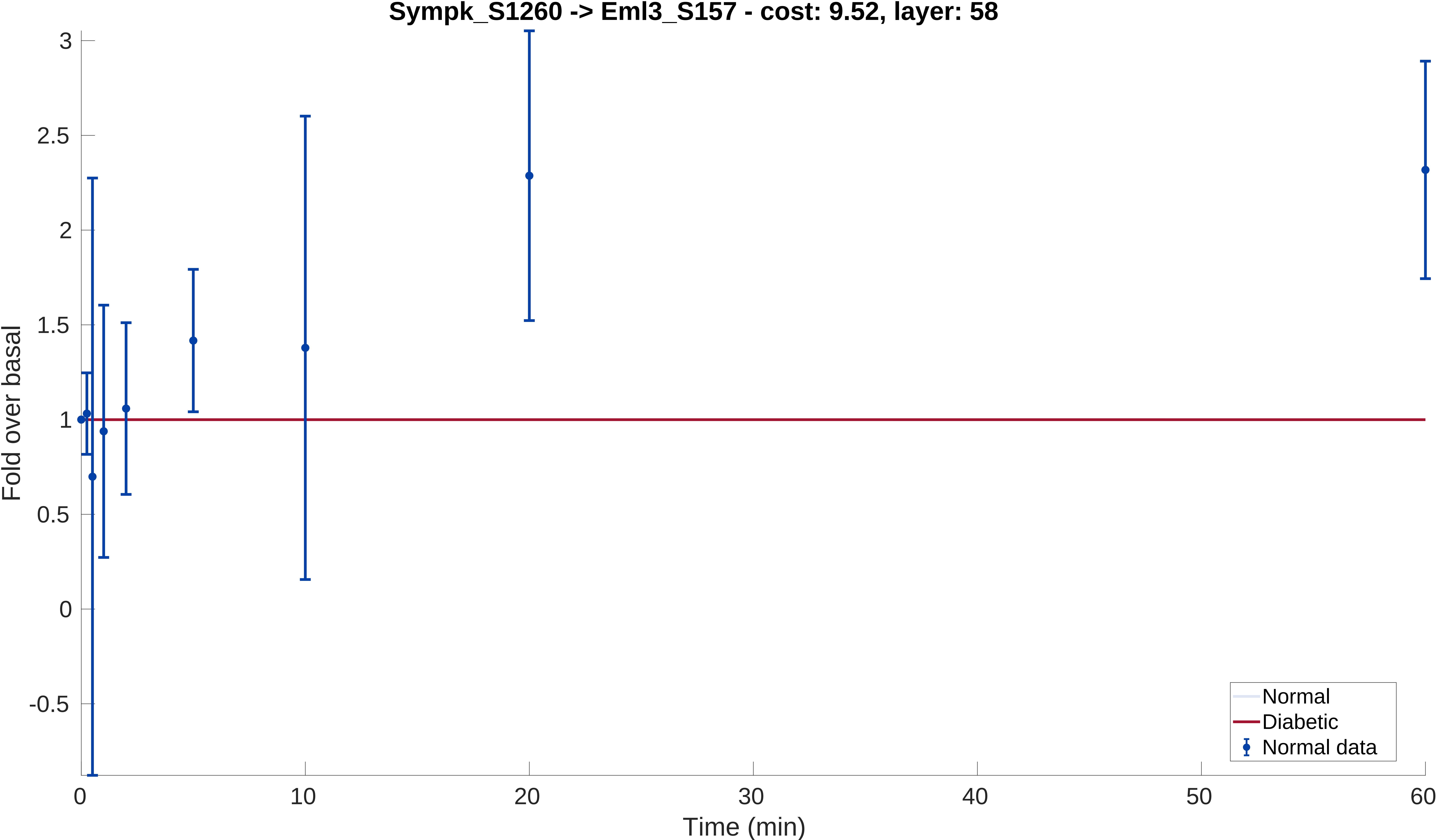
Sympk_S1260 -> Eml3_S157 - cost: 9.52, layer: 58
Fold over basal
Time (min)
Normal
Diabetic
Normal data
3
2.5
2
1.5
1
0.5
0
-0.5
0
10
20
30
40
50
60

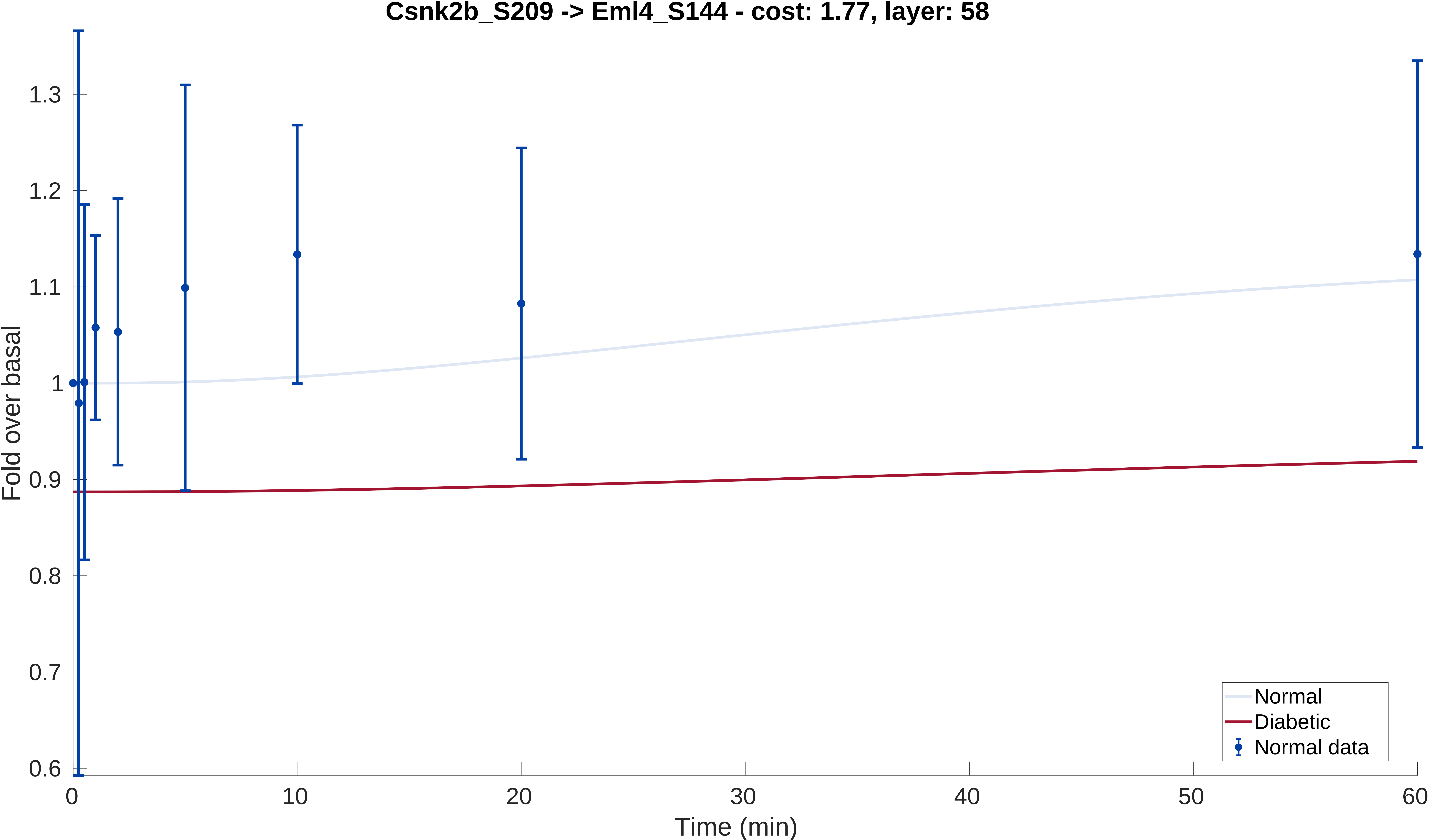

Csnk2b_S209 -> Eml4_S144 - cost: 1.77, layer: 58
Fold over basal
1.3
1.2
1.1
1
0.9
0.8
0.7
0.6
0
10
20
30
40
50
60
Time (min)
Normal
Diabetic
Normal data

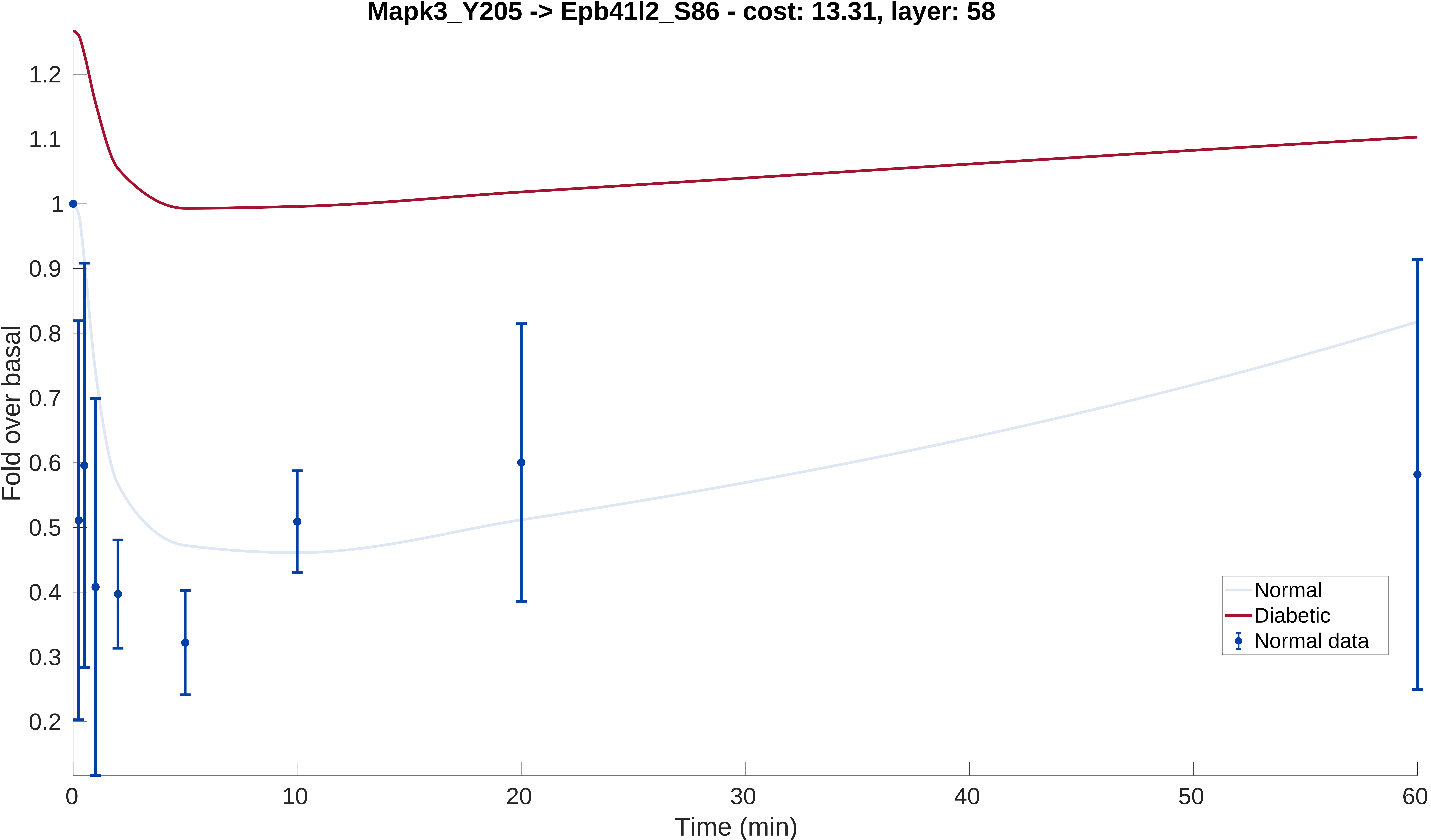
Mapk3_Y205 -> Epb41l2_S86 - cost: 13.31, layer: 58
Fold over basal
1.2
1.1
1
0.9
0.8
0.7
0.6
0.5
0.4
0.3
0.2
0
10
20
30
40
50
60
Time (min)
Normal
Diabetic
Normal data

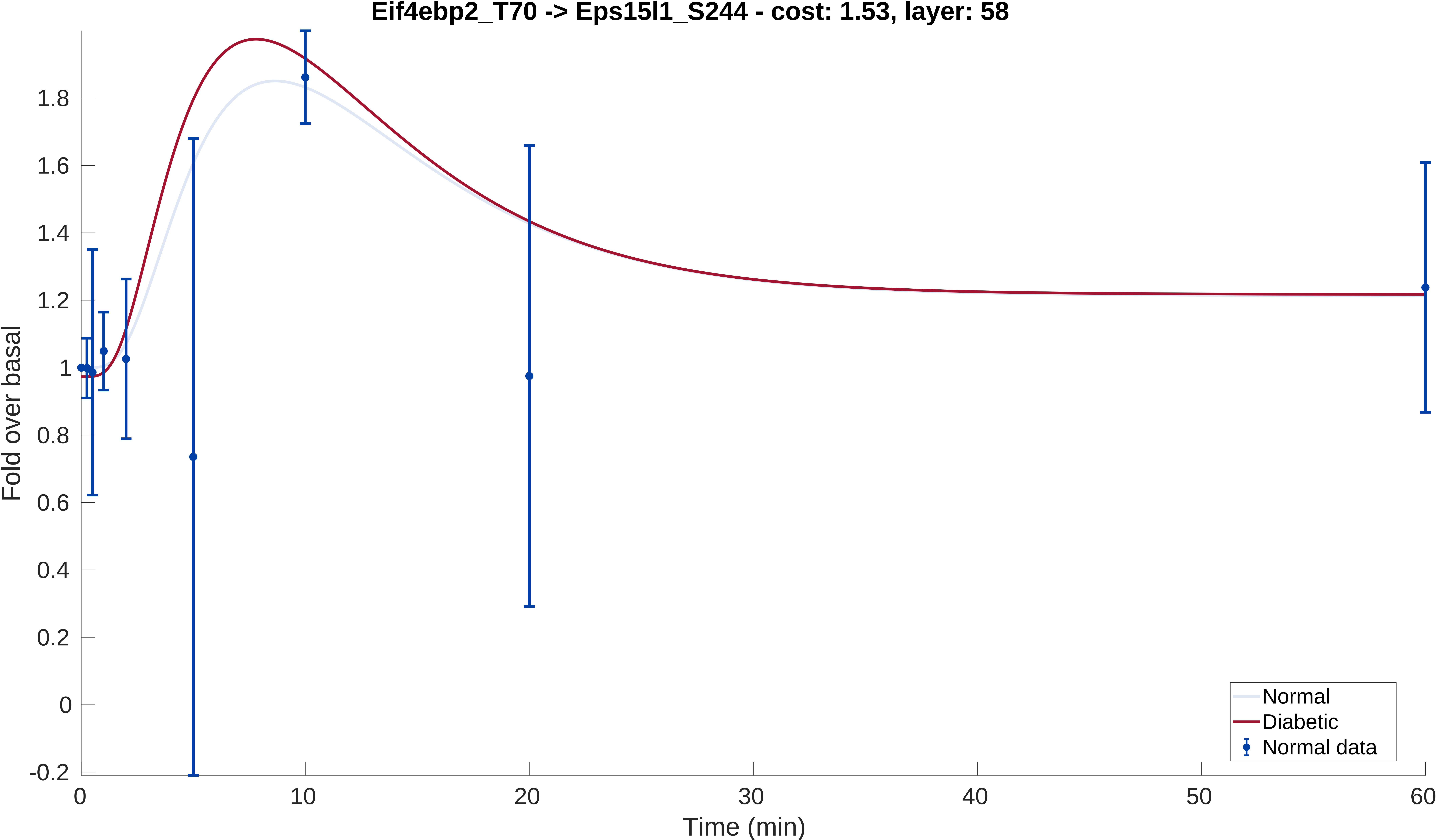
Eif4ebp2_T70 -> Eps15l1_S244 - cost: 1.53, layer: 58
Fold over basal
Time (min)
-0.2
0
0.2
0.4
0.6
0.8
1
1.2
1.4
1.6
1.8
0
10
20
30
40
50
60
Normal
Diabetic
Normal data

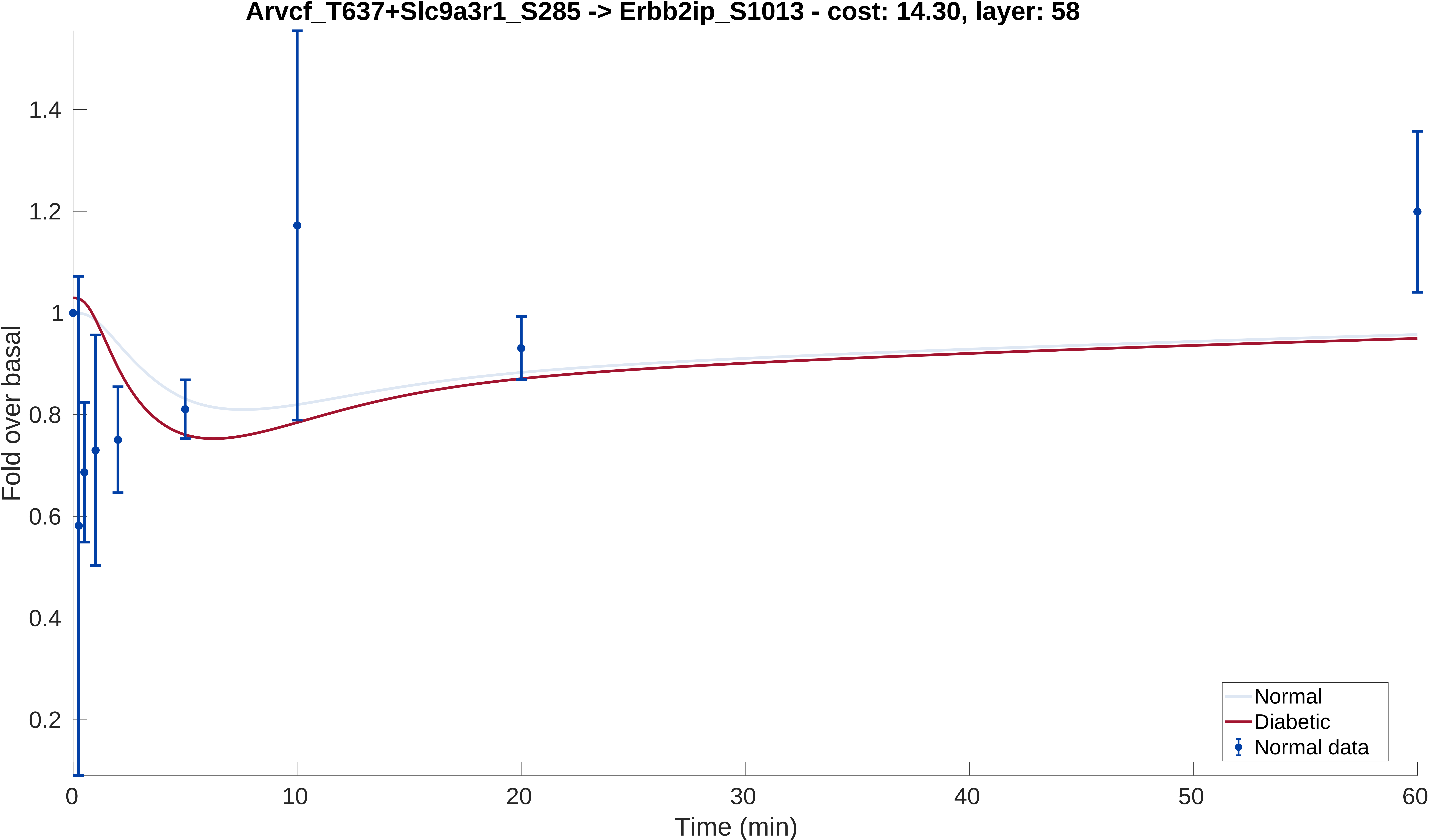

Arvcf_T637+Slc9a3r1_S285 -> Erbb2ip_S1013 - cost: 14.30, layer: 58
Fold over basal
1.4
1.2
1
0.8
0.6
0.4
0.2
0
10
20
30
40
50
60
Time (min)
Normal
Diabetic
Normal data

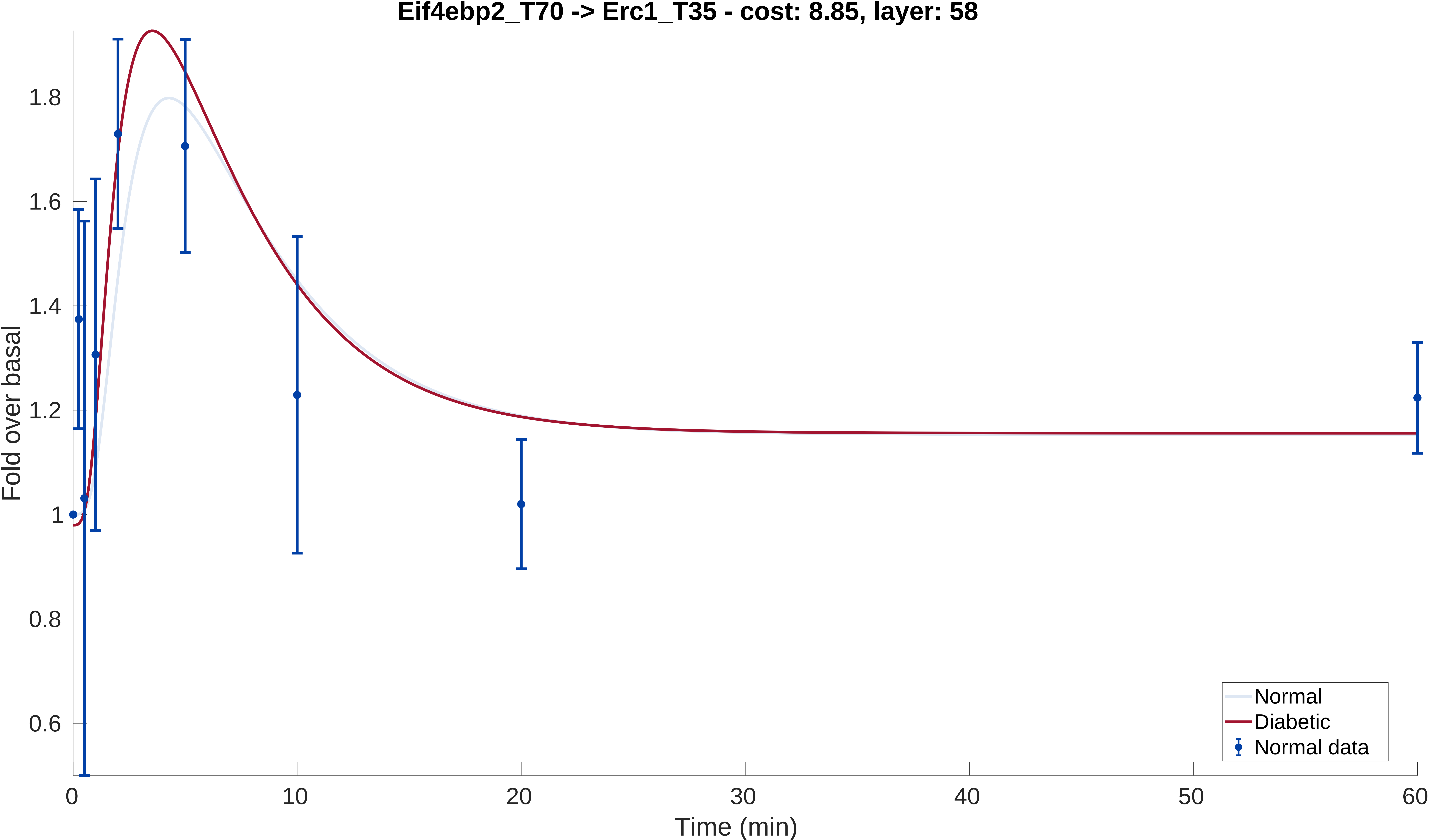
Eif4ebp2_T70 -> Erc1_T35 - cost: 8.85, layer: 58
Fold over basal
1.8
1.6
1.4
1.2
1
0.8
0.6
0
10
20
30
40
50
60
Time (min)
Normal
Diabetic
Normal data

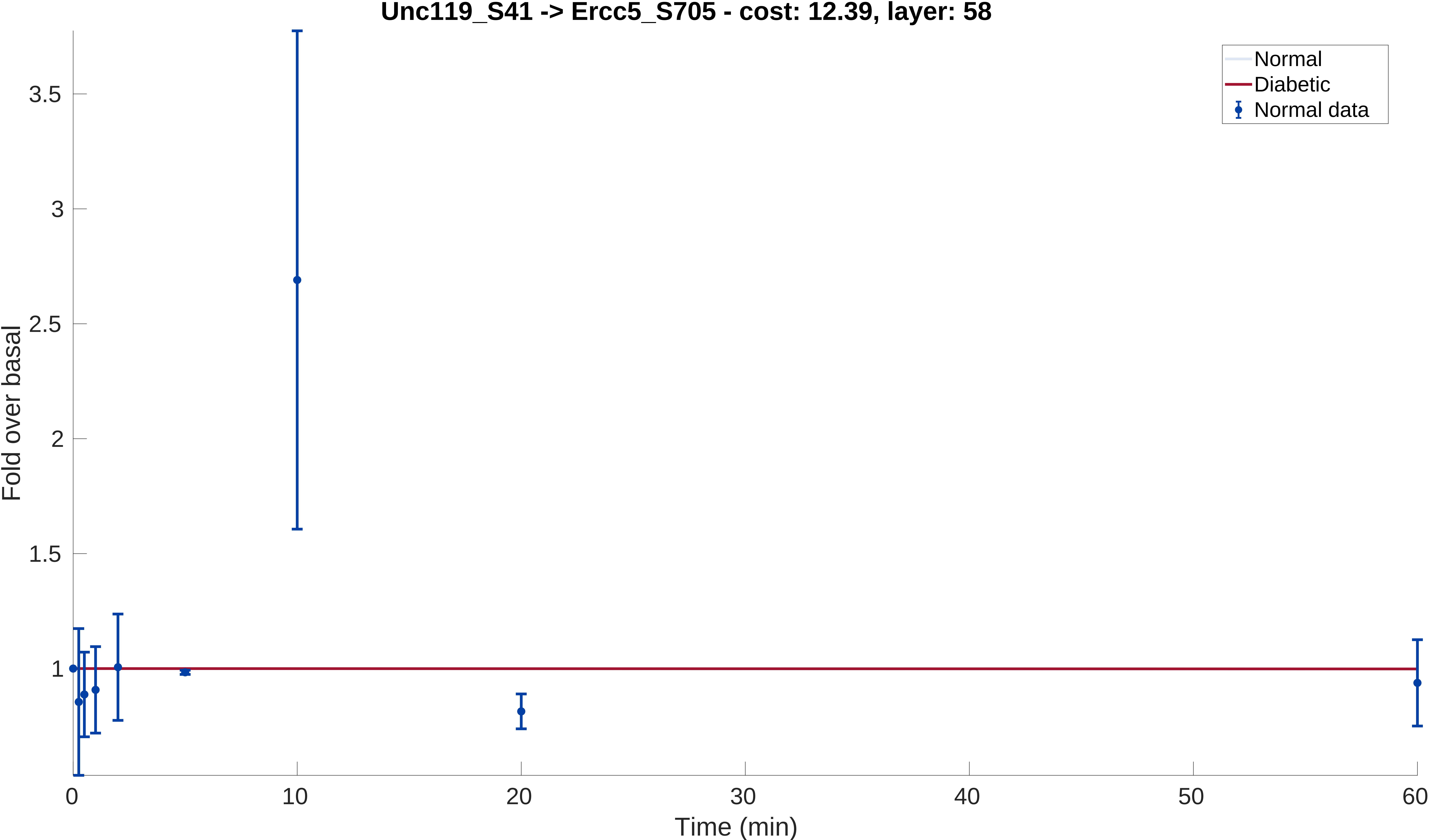

Unc119_S41 -> Ercc5_S705 - cost: 12.39, layer: 58
Normal
Diabetic
Normal data
Fold over basal
3.5
3
2.5
2
1.5
1
0
10
20
30
40
50
60
Time (min)

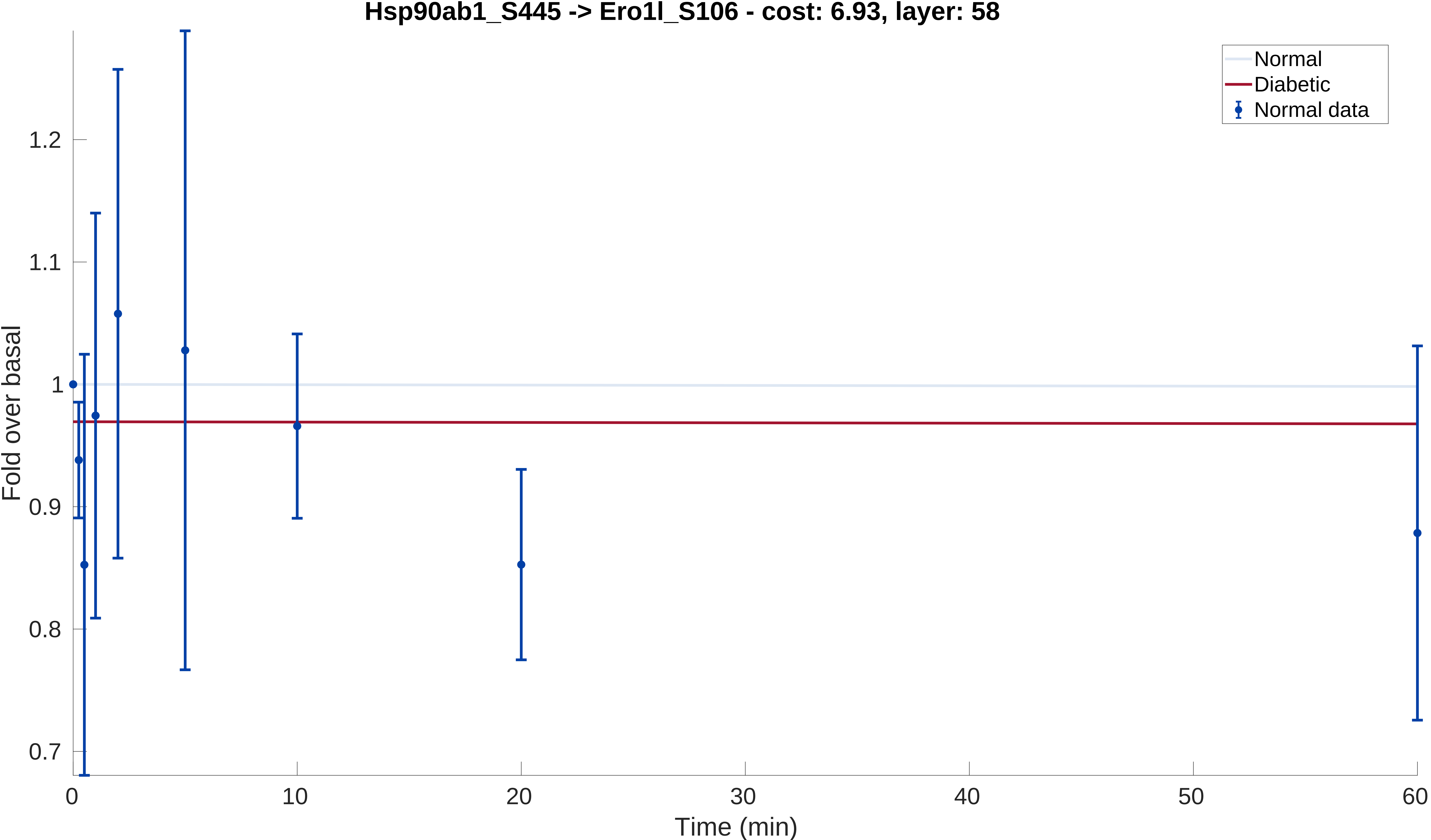

Hsp90ab1_S445 -> Ero1l_S106 - cost: 6.93, layer: 58
Normal
Diabetic
Normal data
Fold over basal
1.2
1.1
1
0.9
0.8
0.7
0
10
20
30
40
50
60
Time (min)

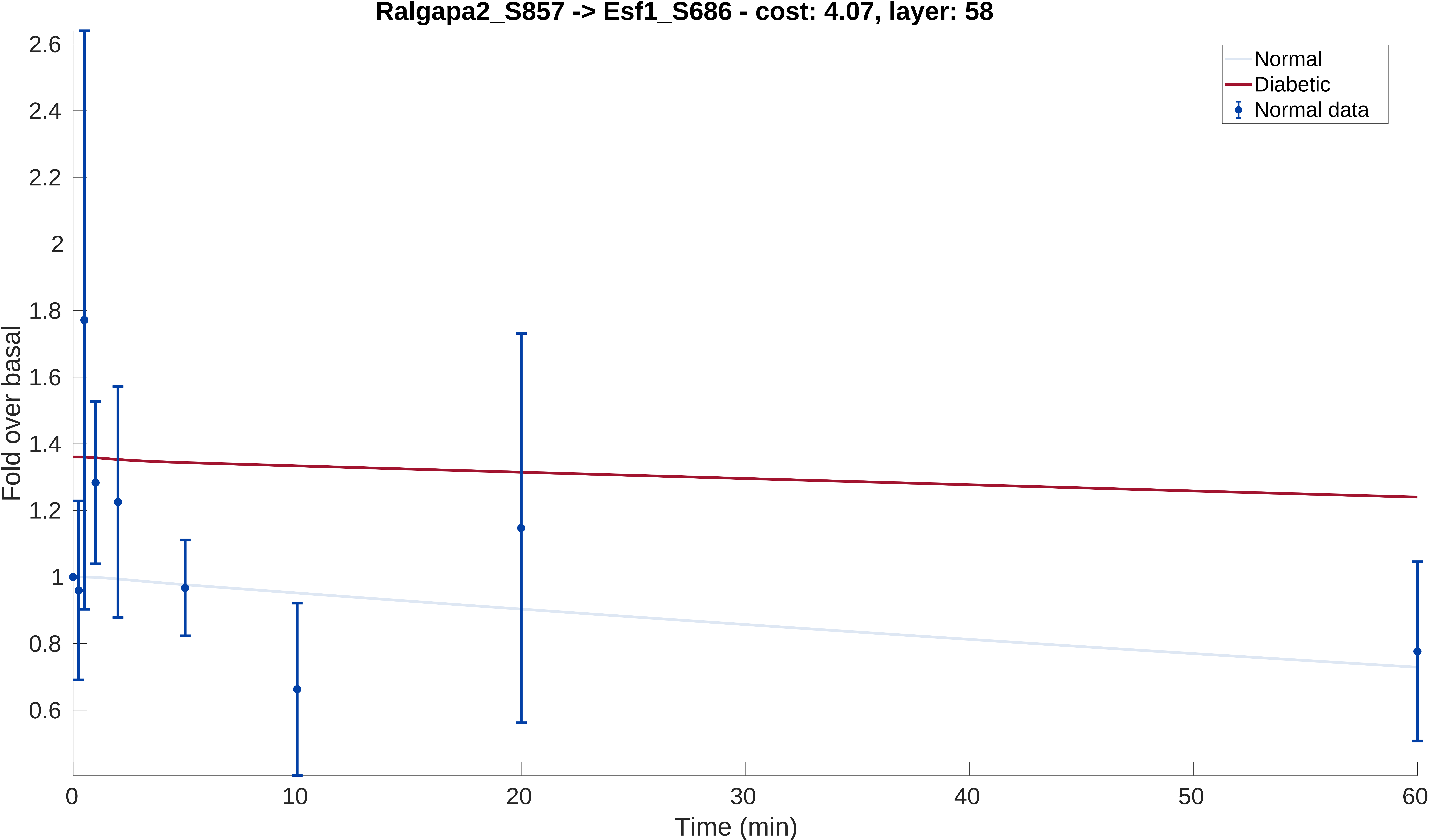
Ralgapa2_S857 -> Esf1_S686 - cost: 4.07, layer: 58
Normal
Diabetic
Normal data
Fold over basal
2.6
2.4
2.2
2
1.8
1.6
1.4
1.2
1
0.8
0.6
0
10
20
30
40
50
60
Time (min)

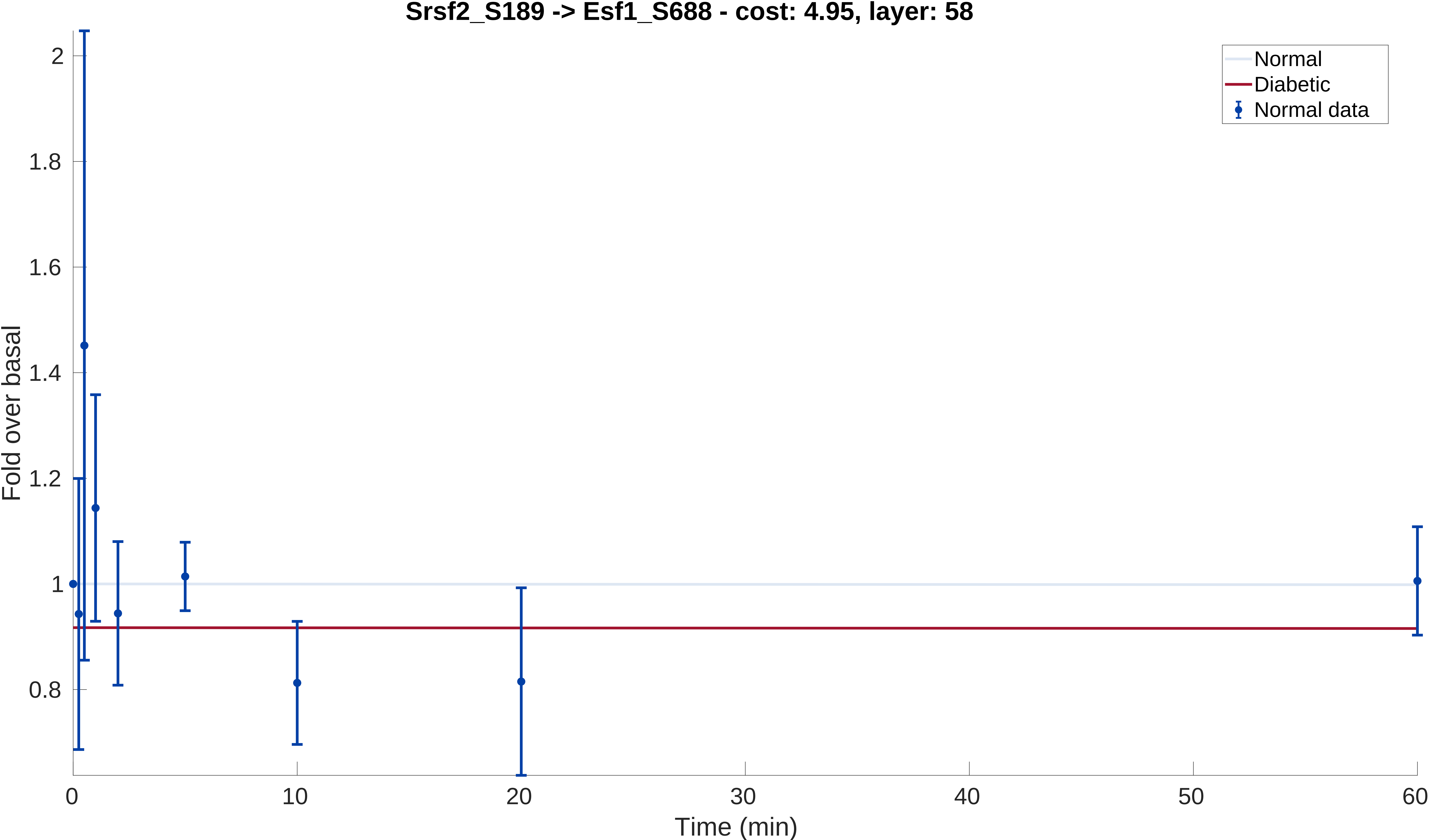
Srsf2_S189 -> Esf1_S688 - cost: 4.95, layer: 58
Fold over basal
Time (min)
Normal
Diabetic
Normal data
2
1.8
1.6
1.4
1.2
1
0.8
0
10
20
30
40
50
60

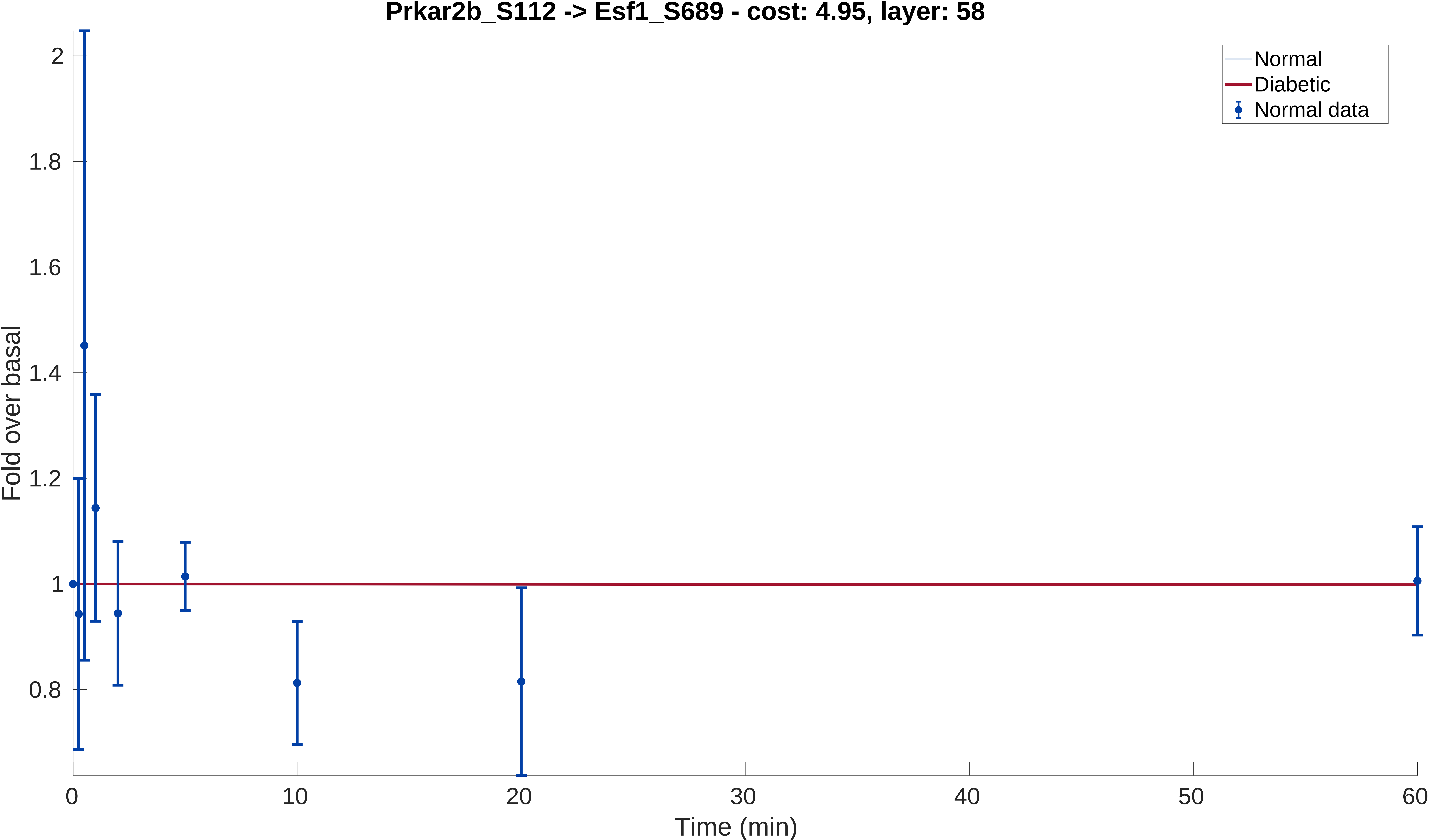

Prkar2b_S112 -> Esf1_S689 - cost: 4.95, layer: 58
Normal
Diabetic
Normal data
Fold over basal
2
1.8
1.6
1.4
1.2
1
0.8
0
10
20
30
40
50
60
Time (min)

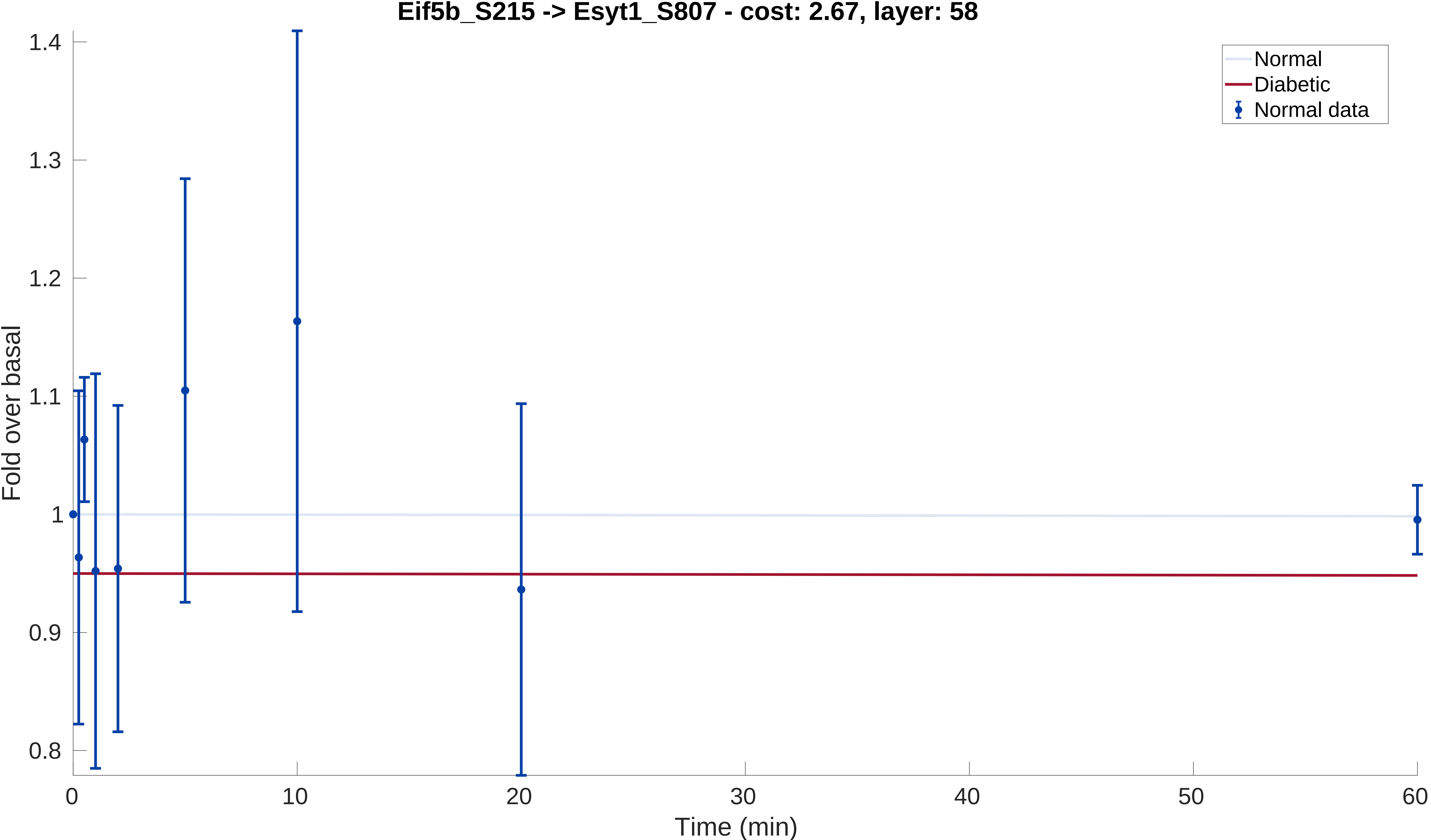
Eif5b_S215 -> Esyt1_S807 - cost: 2.67, layer: 58
Fold over basal
Time (min)
Normal
Diabetic
Normal data
1.4
1.3
1.2
1.1
1
0.9
0.8
0
10
20
30
40
50
60

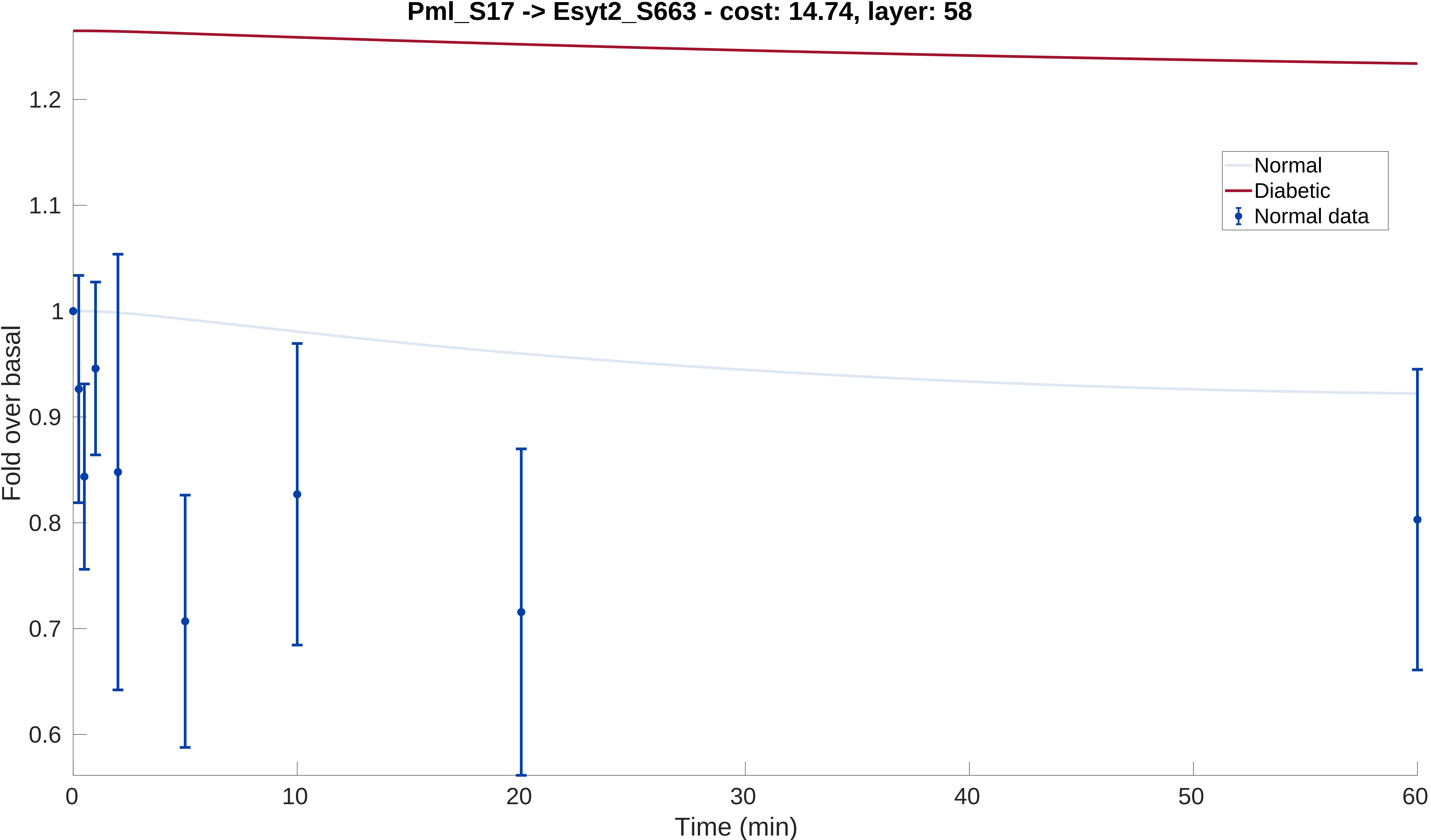

Pml_S17 -> Esyt2_S663 - cost: 14.74, layer: 58
Normal
Diabetic
Normal data
Fold over basal
1.2
1.1
1
0.9
0.8
0.7
0.6
0
10
20
30
40
50
60
Time (min)

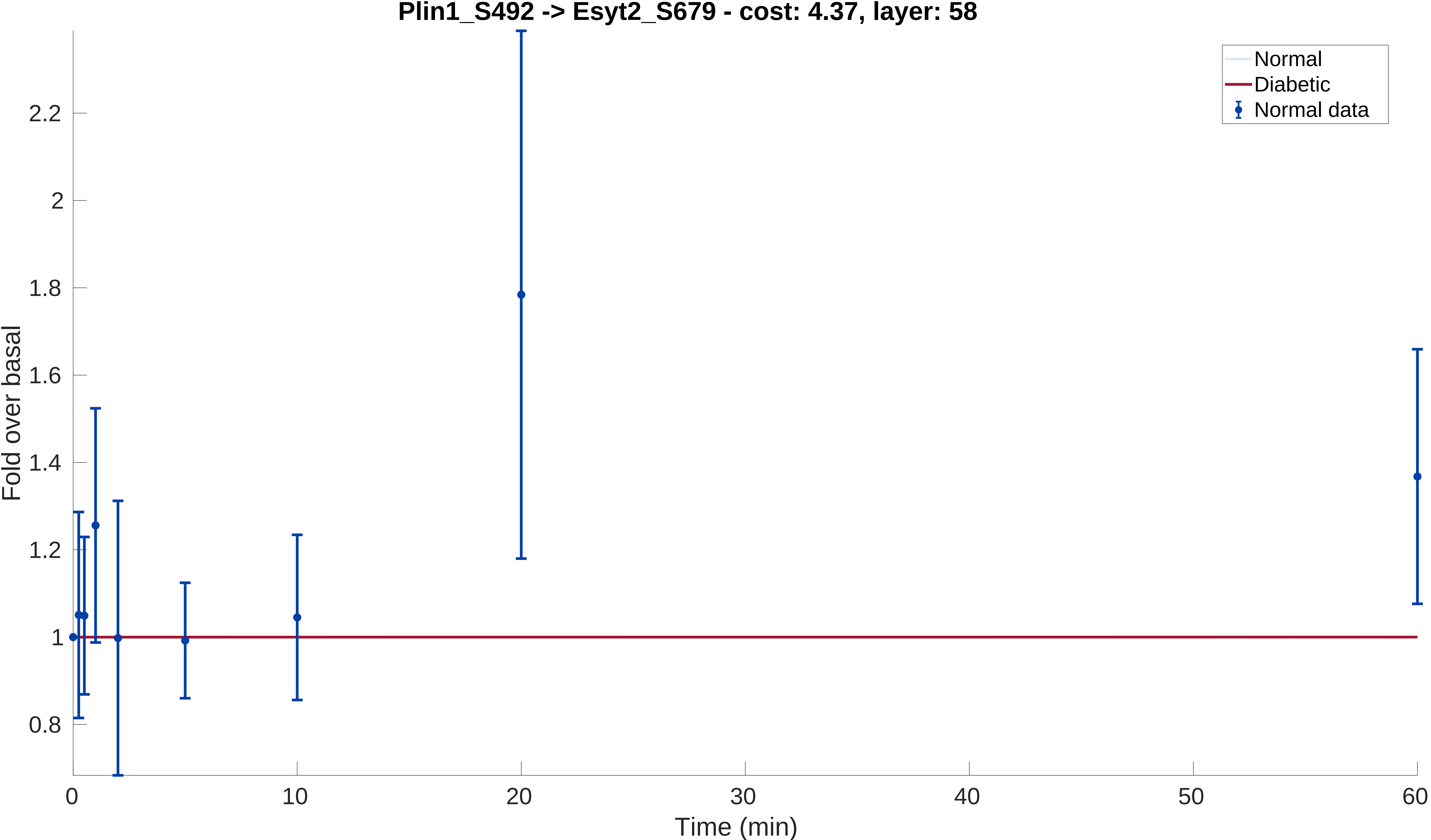

Plin1_S492 -> Esyt2_S679 - cost: 4.37, layer: 58
Normal
Diabetic
Normal data
Fold over basal
Time (min)
0.8
1
1.2
1.4
1.6
1.8
2
2.2
0
10
20
30
40
50
60

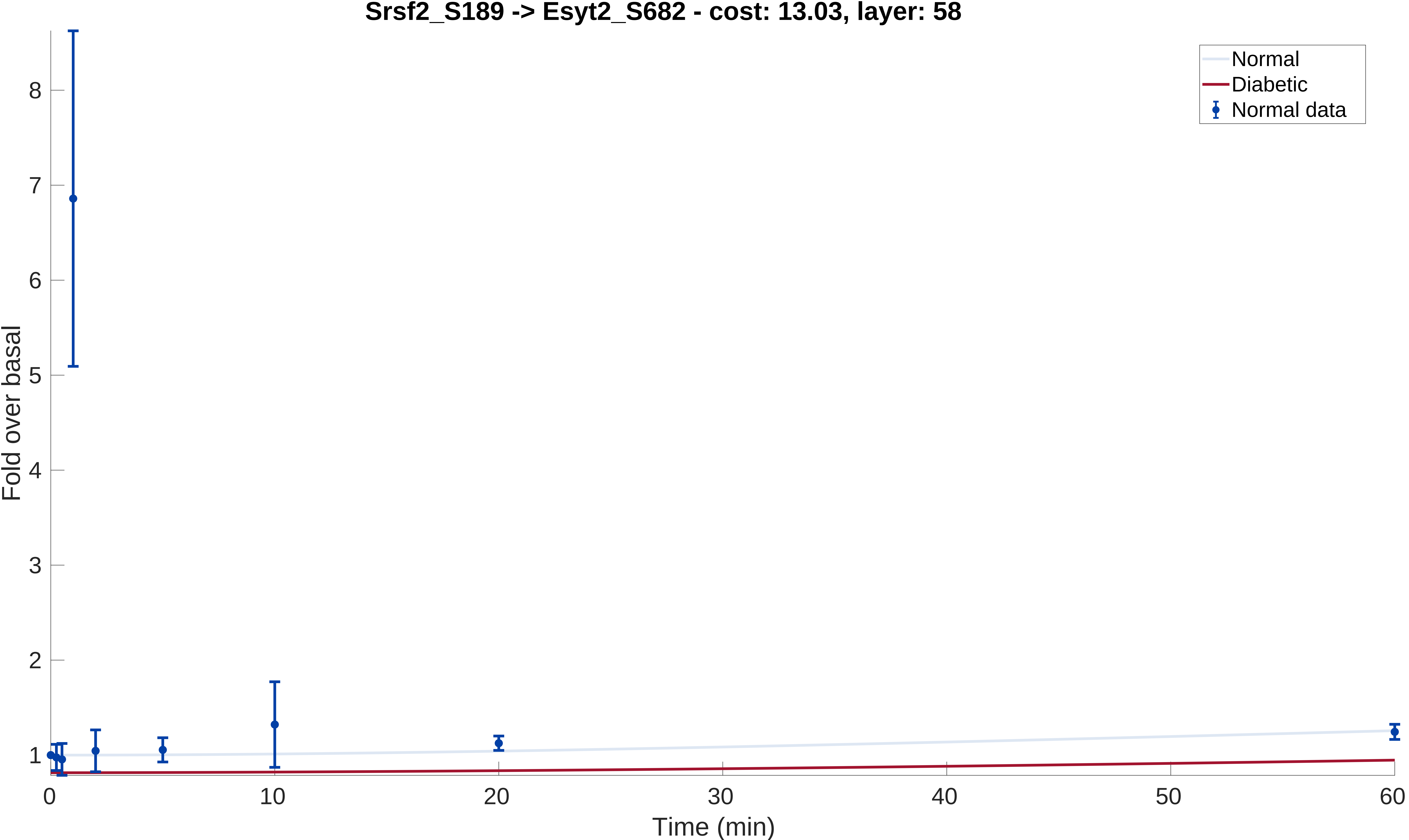
Srsf2_S189 -> Esyt2_S682 - cost: 13.03, layer: 58
Normal
Diabetic
Normal data
Fold over basal
1
2
3
4
5
6
7
8
0
10
20
30
40
50
60
Time (min)

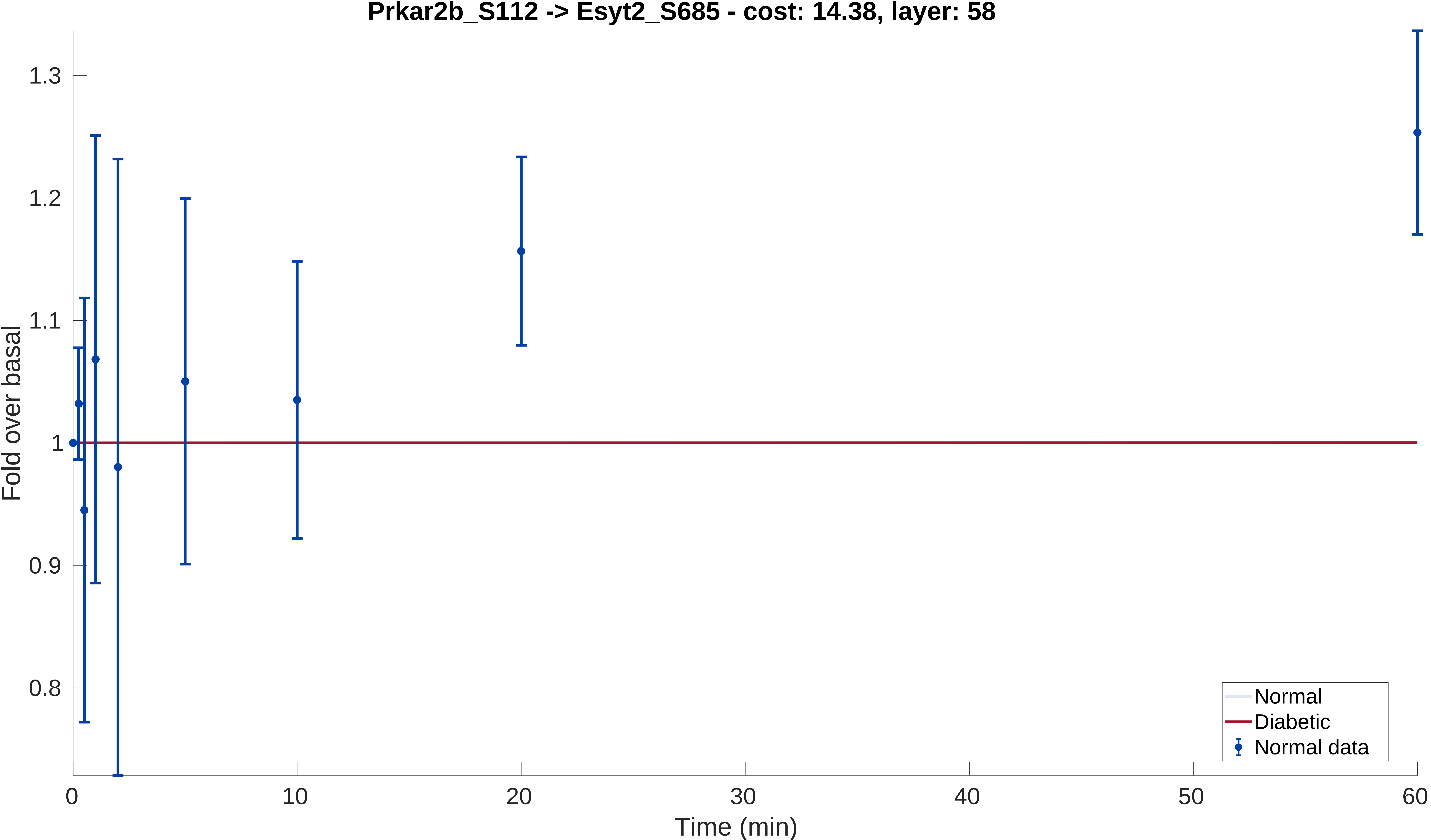

Prkar2b_S112 -> Esyt2_S685 - cost: 14.38, layer: 58
1.3
1.2
1.1
1
0.9
0.8
Fold over basal
0
10
20
30
40
50
60
Time (min)
Normal
Diabetic
Normal data

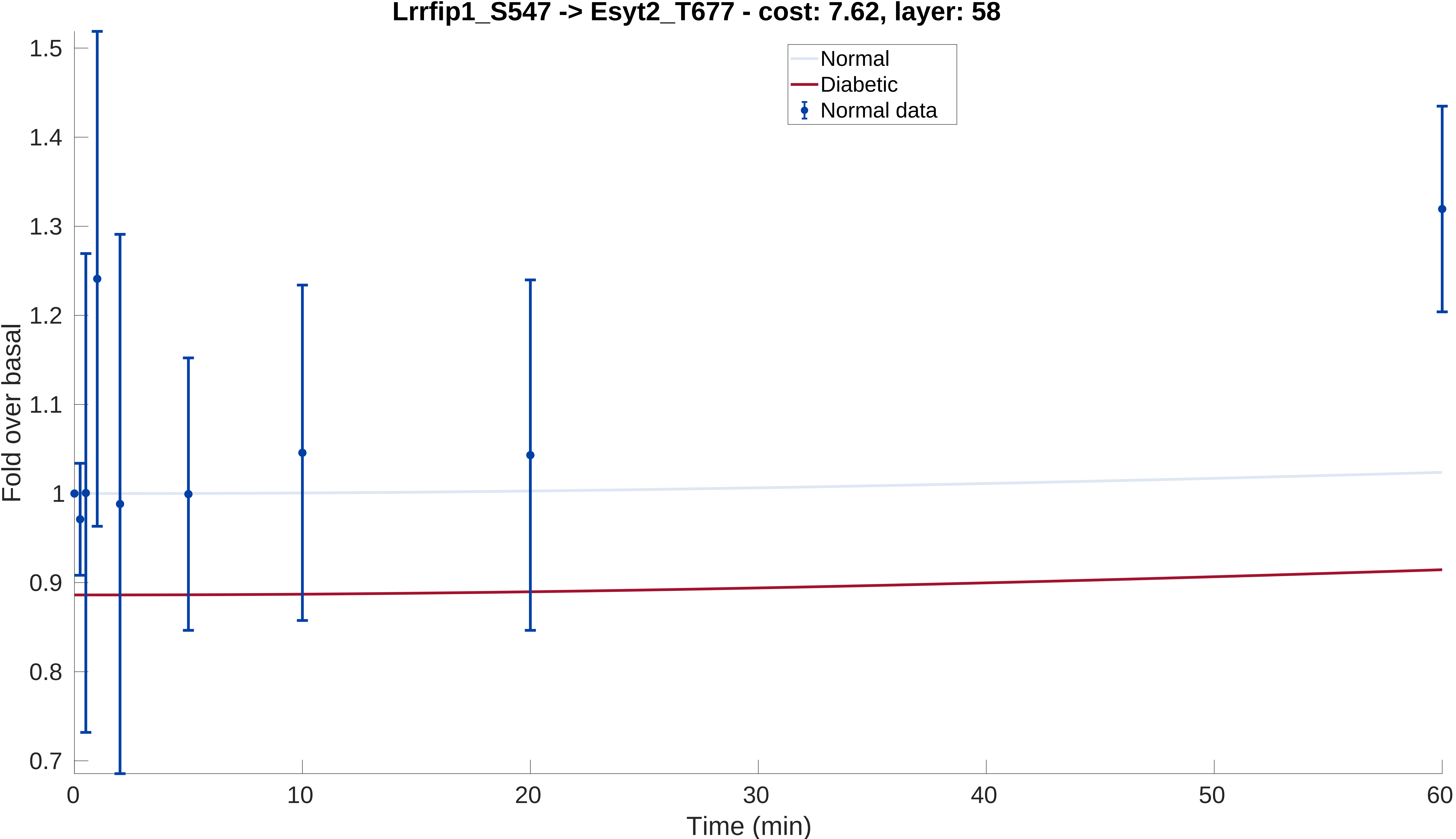
Lrrfip1_S547 -> Esyt2_T677 - cost: 7.62, layer: 58
Normal
Diabetic
Normal data
Fold over basal
Time (min)
1.5
1.4
1.3
1.2
1.1
1
0.9
0.8
0.7
0
10
20
30
40
50
60

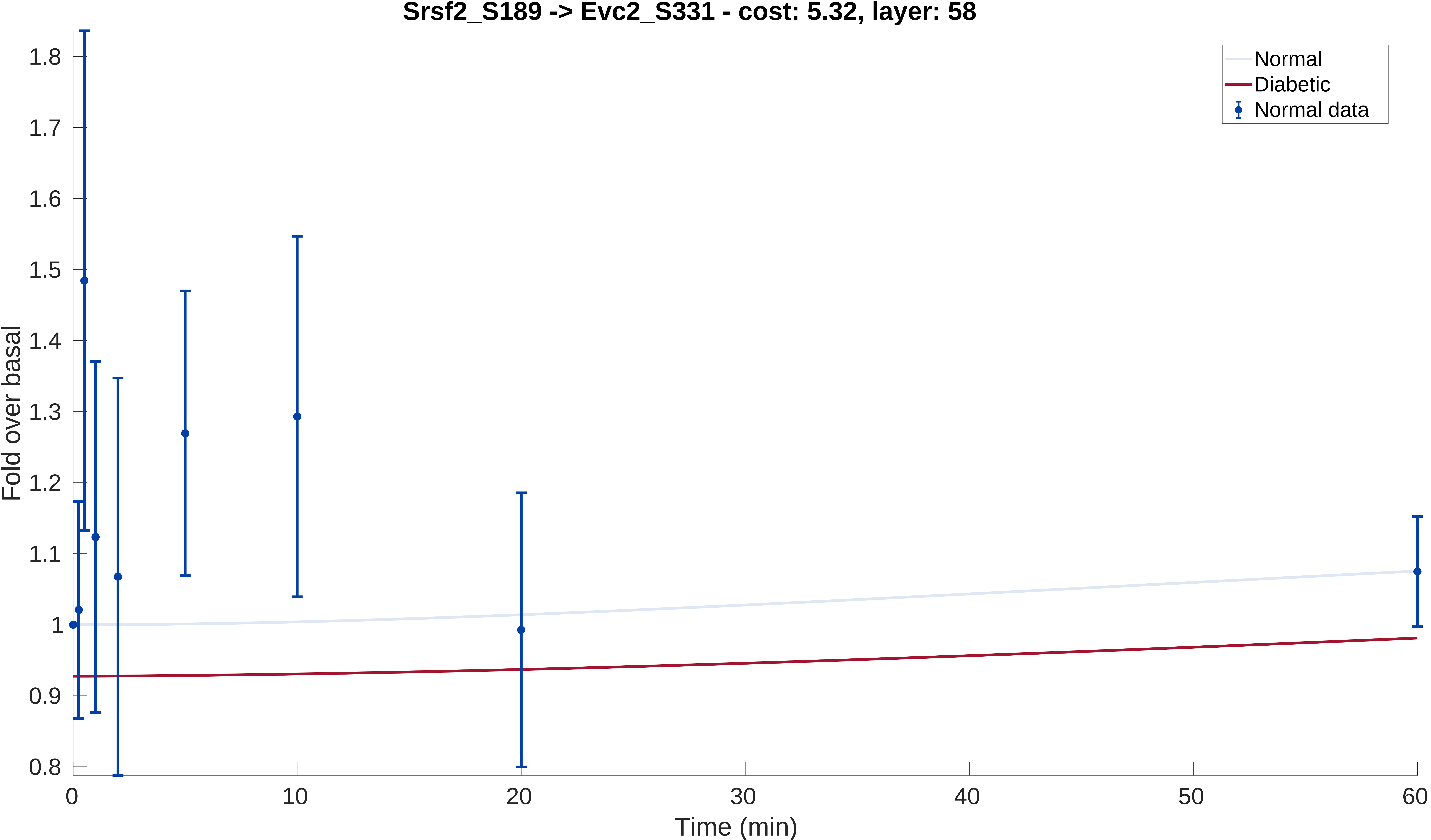

Srsf2_S189 -> Evc2_S331 - cost: 5.32, layer: 58
Normal
Diabetic
Normal data
Fold over basal
Time (min)
0.8
0.9
1
1.1
1.2
1.3
1.4
1.5
1.6
1.7
1.8
0
10
20
30
40
50
60

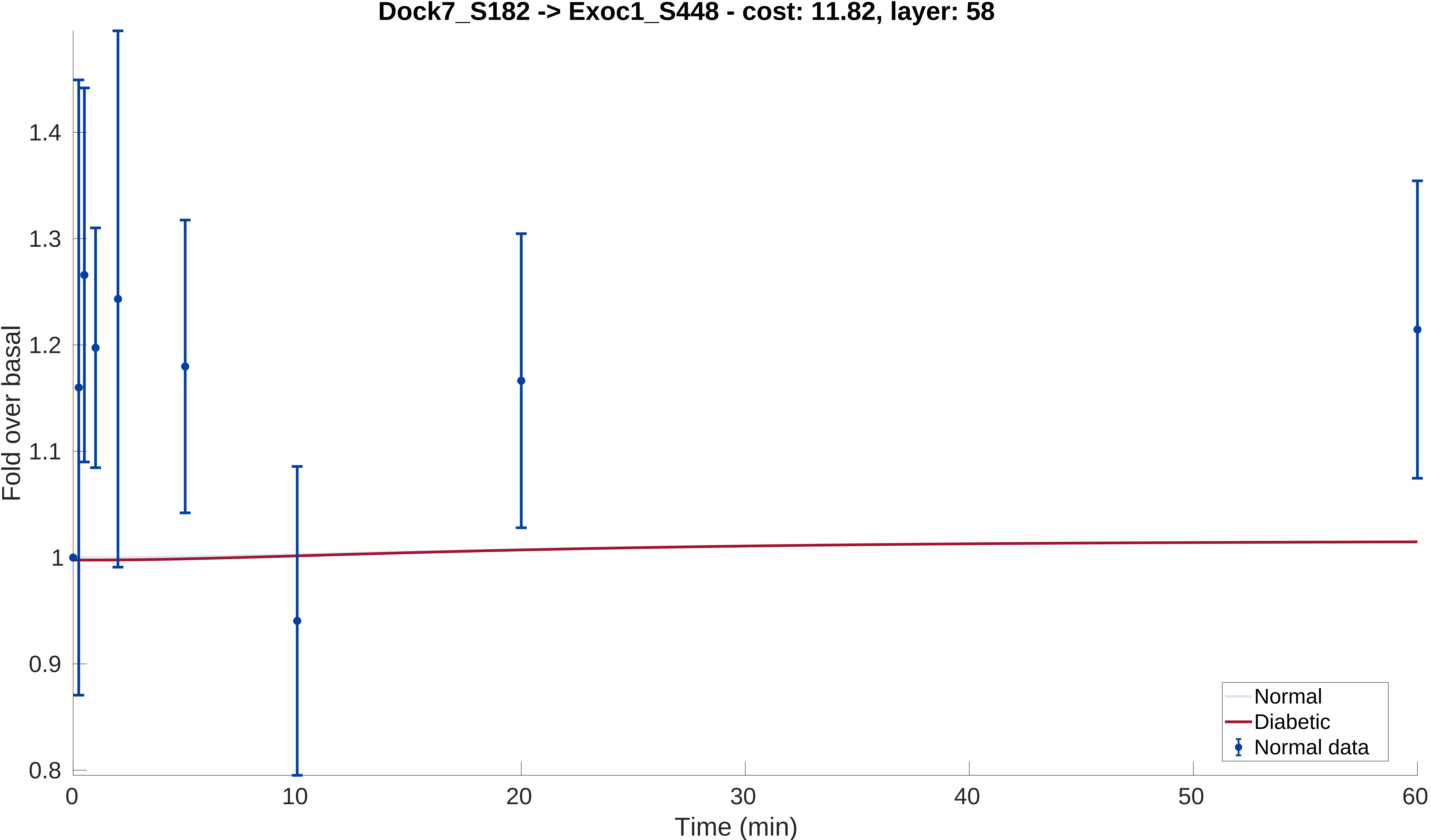

Dock7_S182 -> Exoc1_S448 - cost: 11.82, layer: 58
Fold over basal
1.4
1.3
1.2
1.1
1
0.9
0.8
0
10
20
30
40
50
60
Time (min)
Normal
Diabetic
Normal data

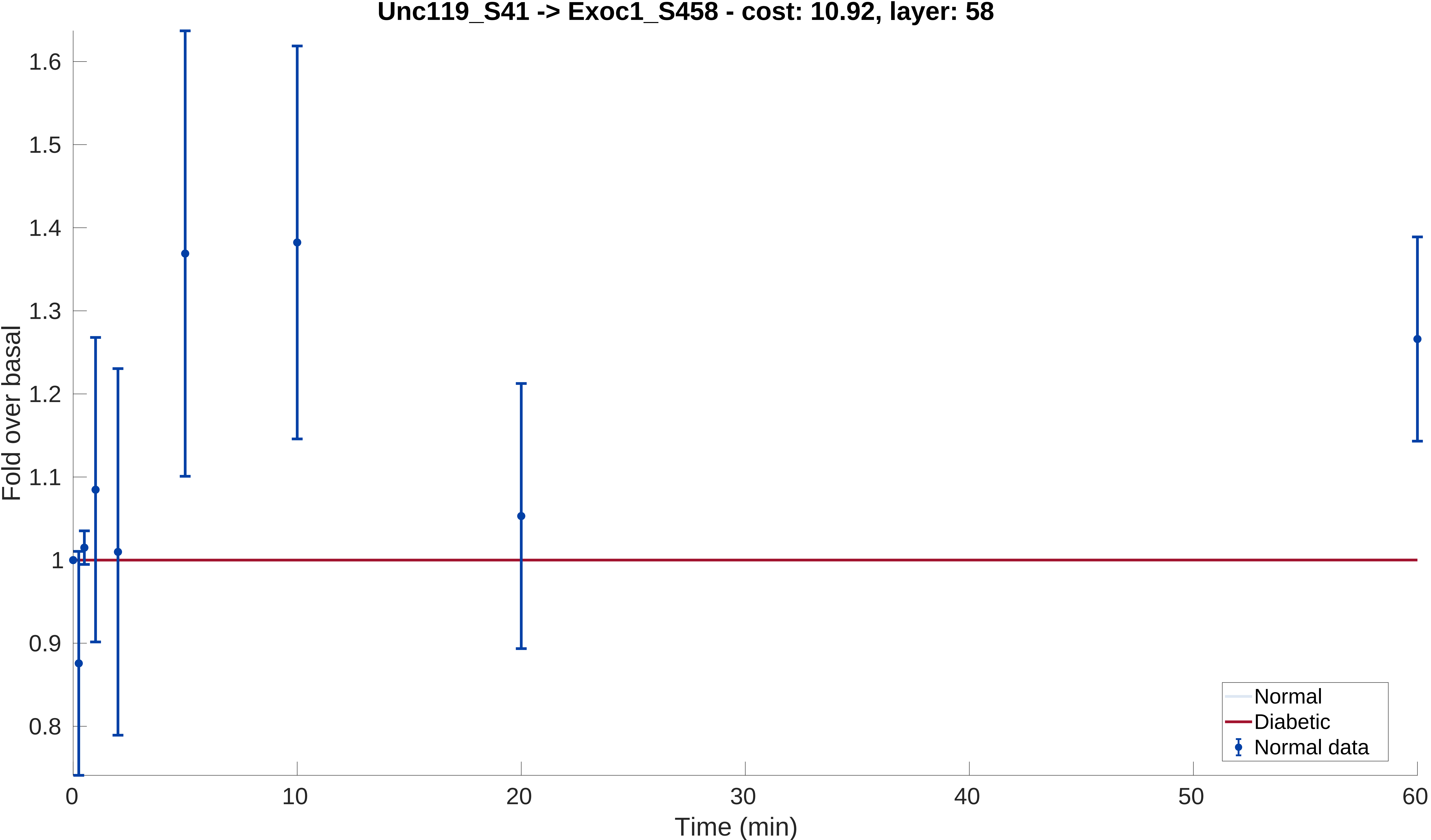

Unc119_S41 -> Exoc1_S458 - cost: 10.92, layer: 58
Fold over basal
1.6
1.5
1.4
1.3
1.2
1.1
1
0.9
0.8
0
10
20
30
40
50
60
Time (min)
Normal
Diabetic
Normal data

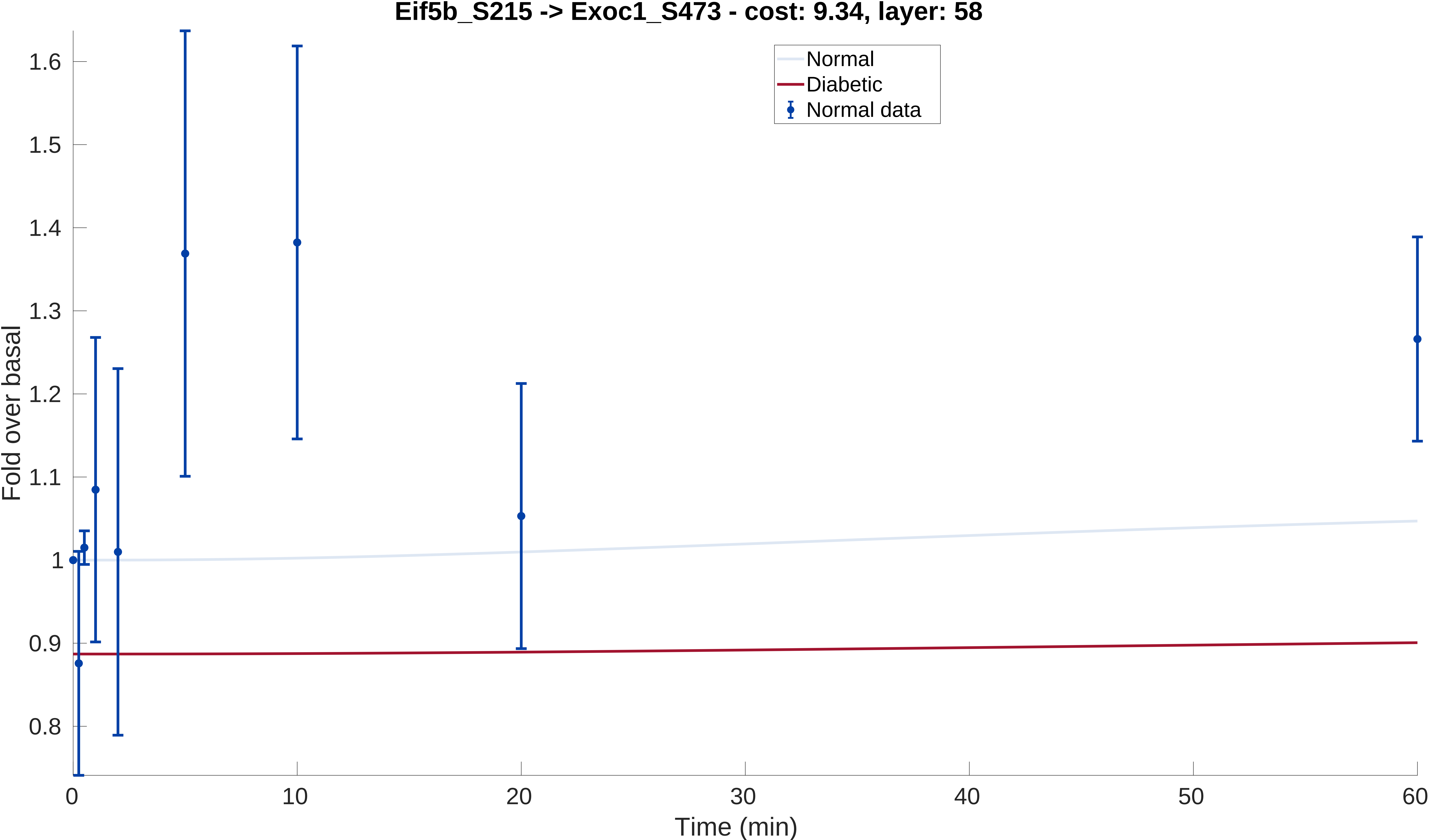
Eif5b_S215 -> Exoc1_S473 - cost: 9.34, layer: 58
Normal
Diabetic
Normal data
Fold over basal
1.6
1.5
1.4
1.3
1.2
1.1
1
0.9
0.8
0
10
20
30
40
50
60
Time (min)

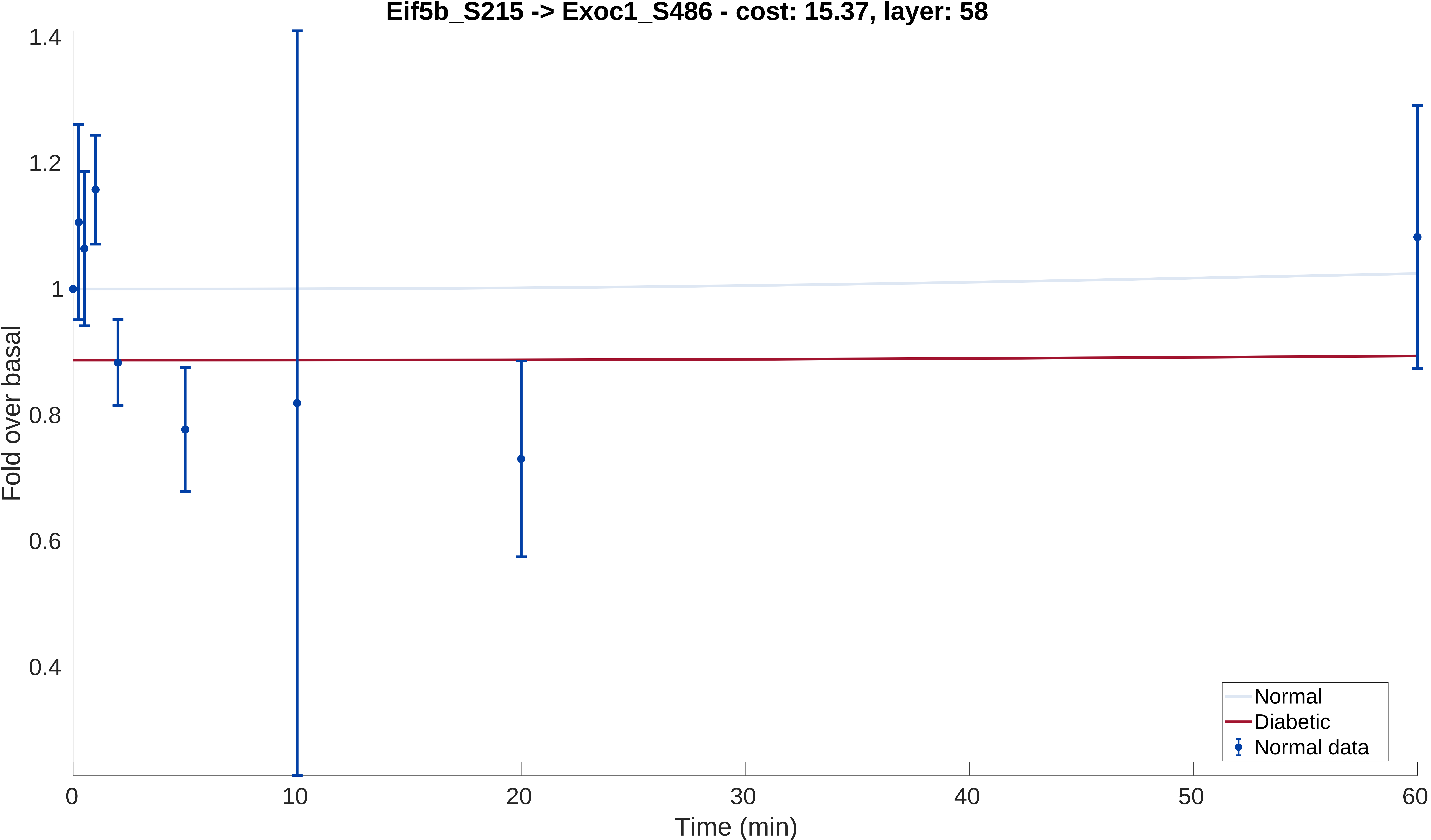

Eif5b_S215 -> Exoc1_S486 - cost: 15.37, layer: 58
Fold over basal
1.4
1.2
1
0.8
0.6
0.4
0
10
20
30
40
50
60
Time (min)
Normal
Diabetic
Normal data

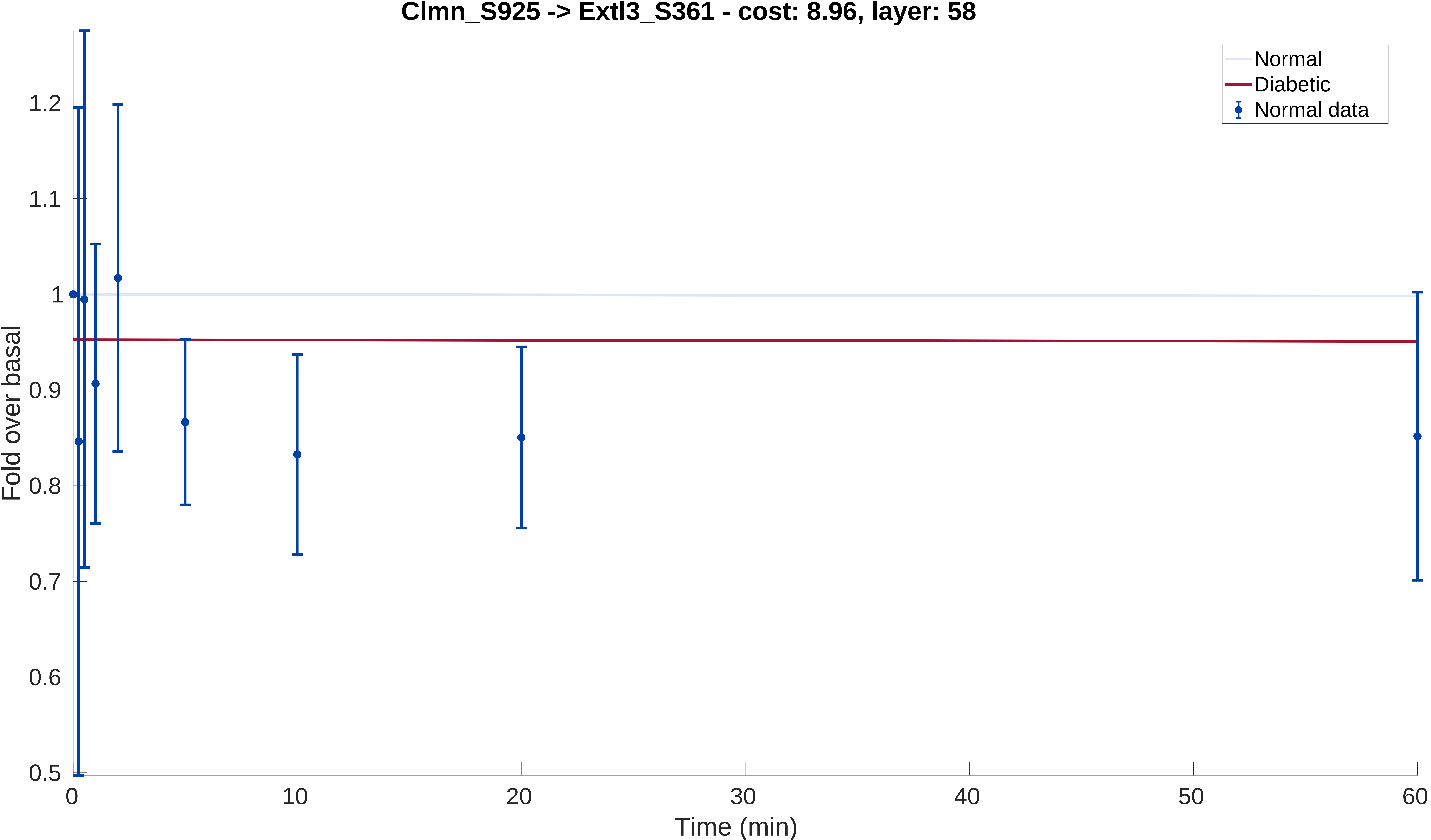

Clmn_S925 -> Extl3_S361 - cost: 8.96, layer: 58
Normal
Diabetic
Normal data
Fold over basal
1.2
1.1
1
0.9
0.8
0.7
0.6
0.5
0
10
20
30
40
50
60
Time (min)

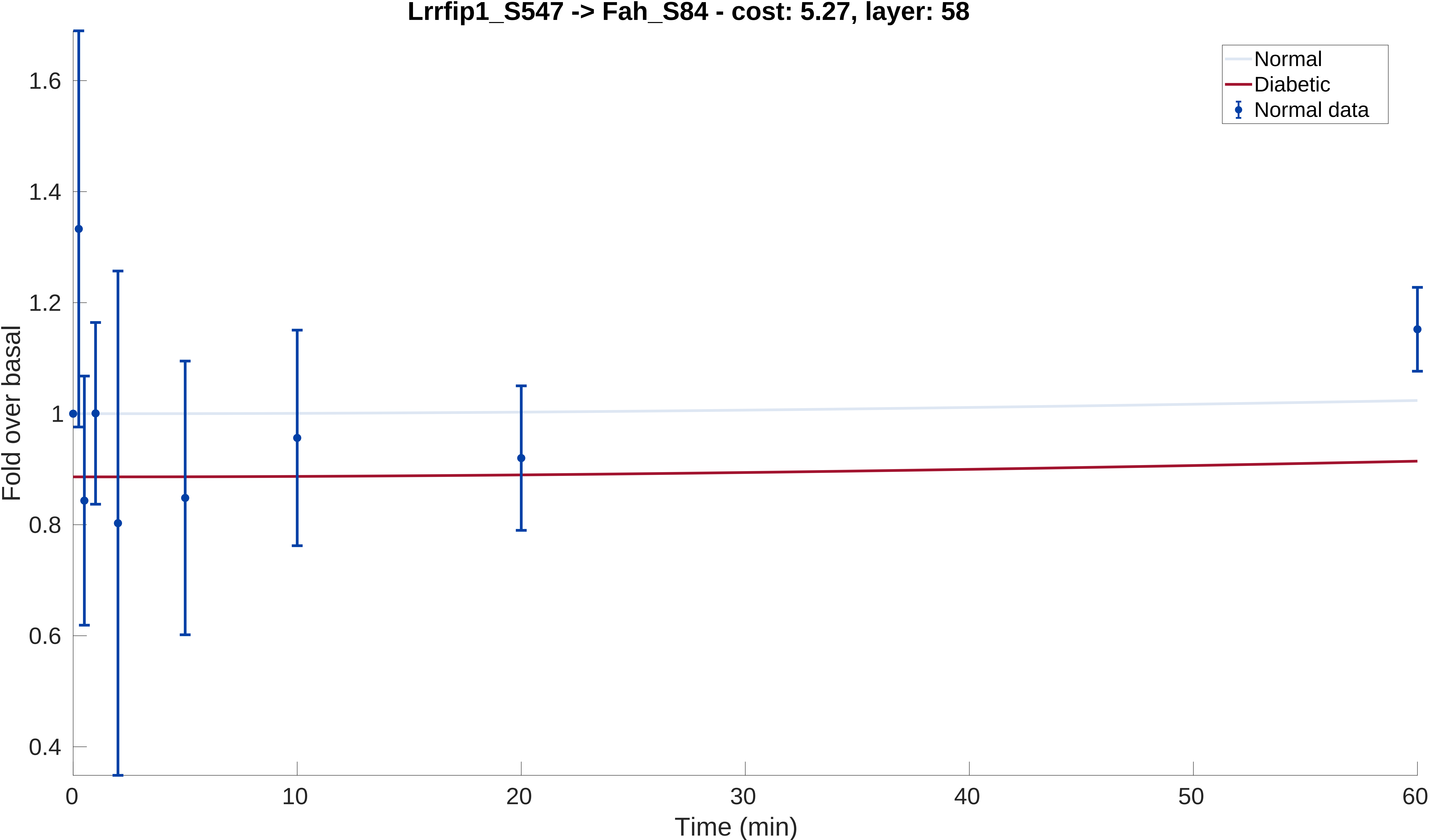

Lrrfip1_S547 -> Fah_S84 - cost: 5.27, layer: 58
Normal
Diabetic
Normal data
Fold over basal
1.6
1.4
1.2
1
0.8
0.6
0.4
0
10
20
30
40
50
60
Time (min)

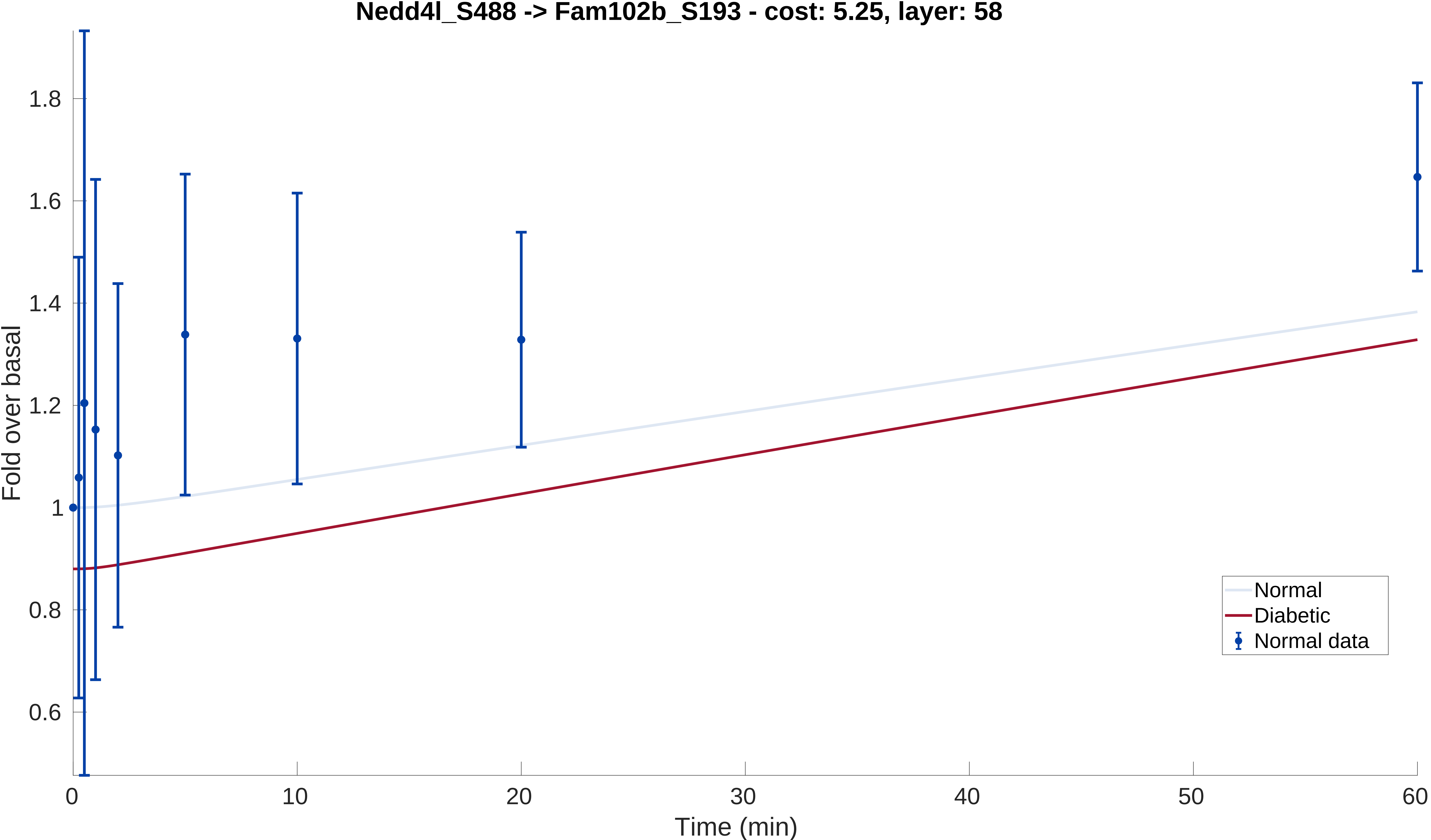

Nedd4l_S488 -> Fam102b_S193 - cost: 5.25, layer: 58
Fold over basal
1.8
1.6
1.4
1.2
1
0.8
0.6
0
10
20
30
40
50
60
Time (min)
Normal
Diabetic
Normal data

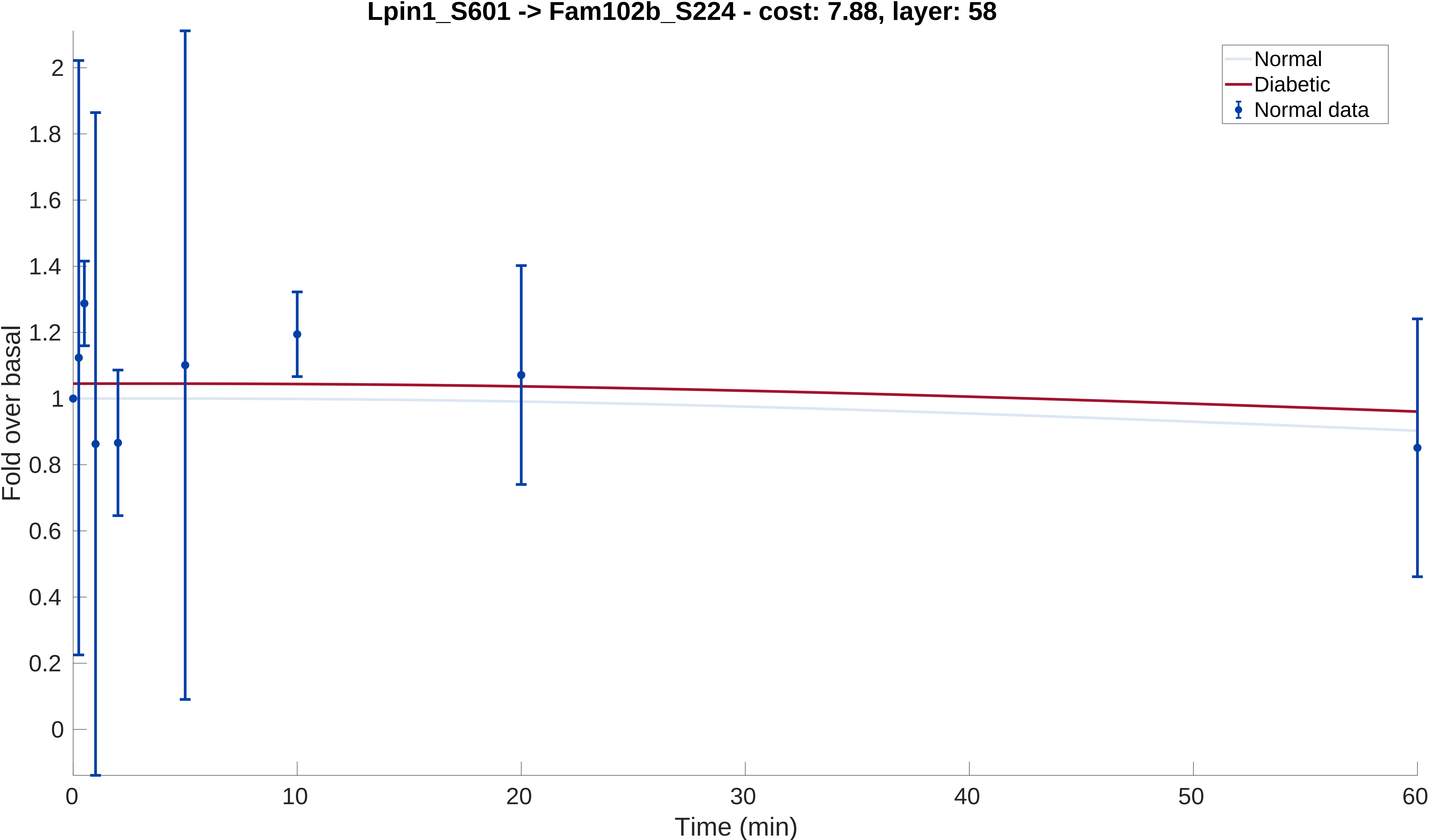

Lpin1_S601 -> Fam102b_S224 - cost: 7.88, layer: 58
Normal
Diabetic
Normal data
Fold over basal
2
1.8
1.6
1.4
1.2
1
0.8
0.6
0.4
0.2
0
0
10
20
30
40
50
60
Time (min)

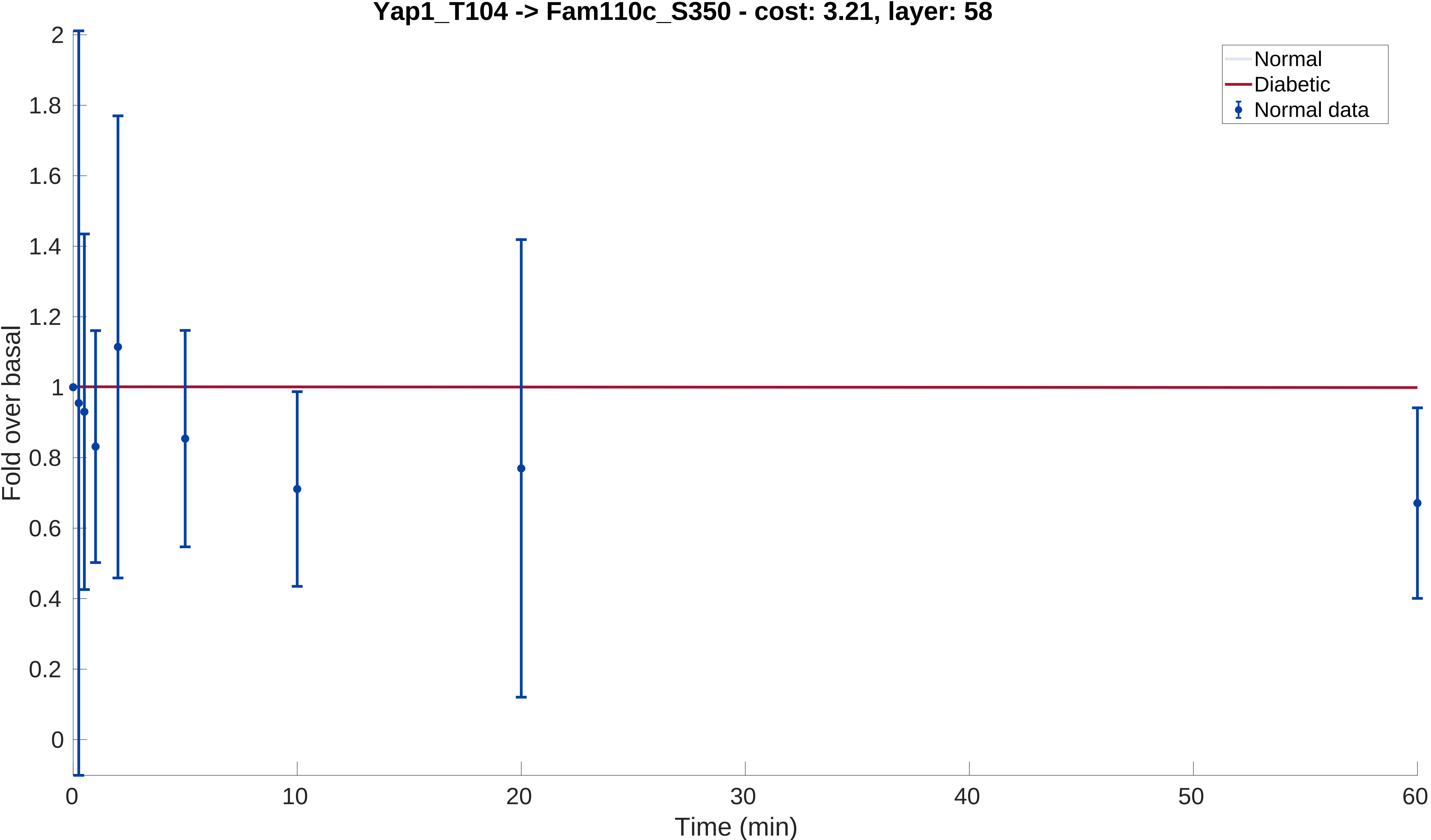

Yap1_T104 -> Fam110c_S350 - cost: 3.21, layer: 58
Normal
Diabetic
Normal data
Fold over basal
2
1.8
1.6
1.4
1.2
1
0.8
0.6
0.4
0.2
0
0
10
20
30
40
50
60
Time (min)

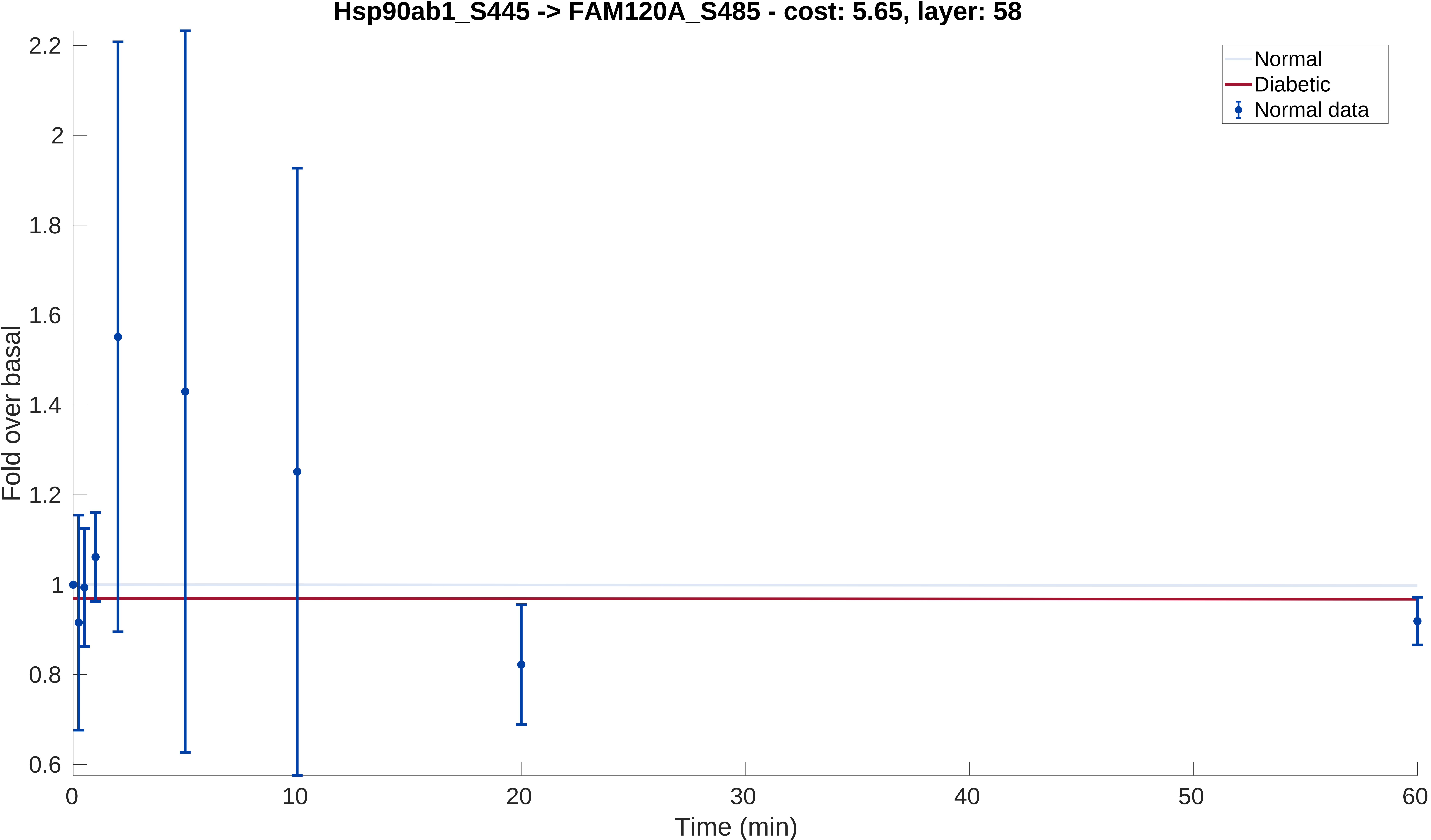
Hsp90ab1_S445 -> FAM120A_S485 - cost: 5.65, layer: 58
Normal
Diabetic
Normal data
Fold over basal
Time (min)
0.6
0.8
1
1.2
1.4
1.6
1.8
2
2.2
0
10
20
30
40
50
60

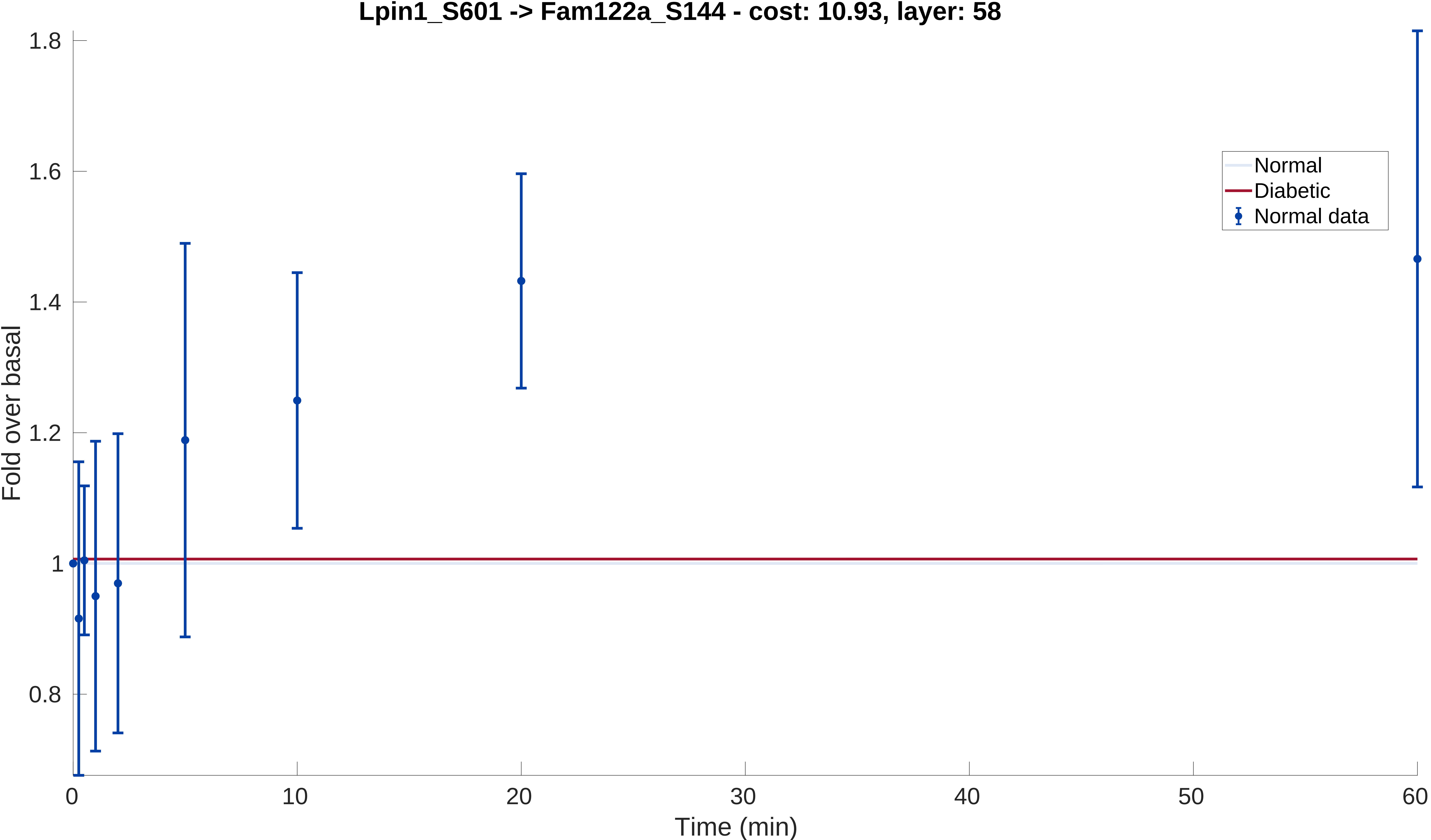

Lpin1_S601 -> Fam122a_S144 - cost: 10.93, layer: 58
Fold over basal
Time (min)
Normal
Diabetic
Normal data
0.8
1
1.2
1.4
1.6
1.8
0
10
20
30
40
50
60

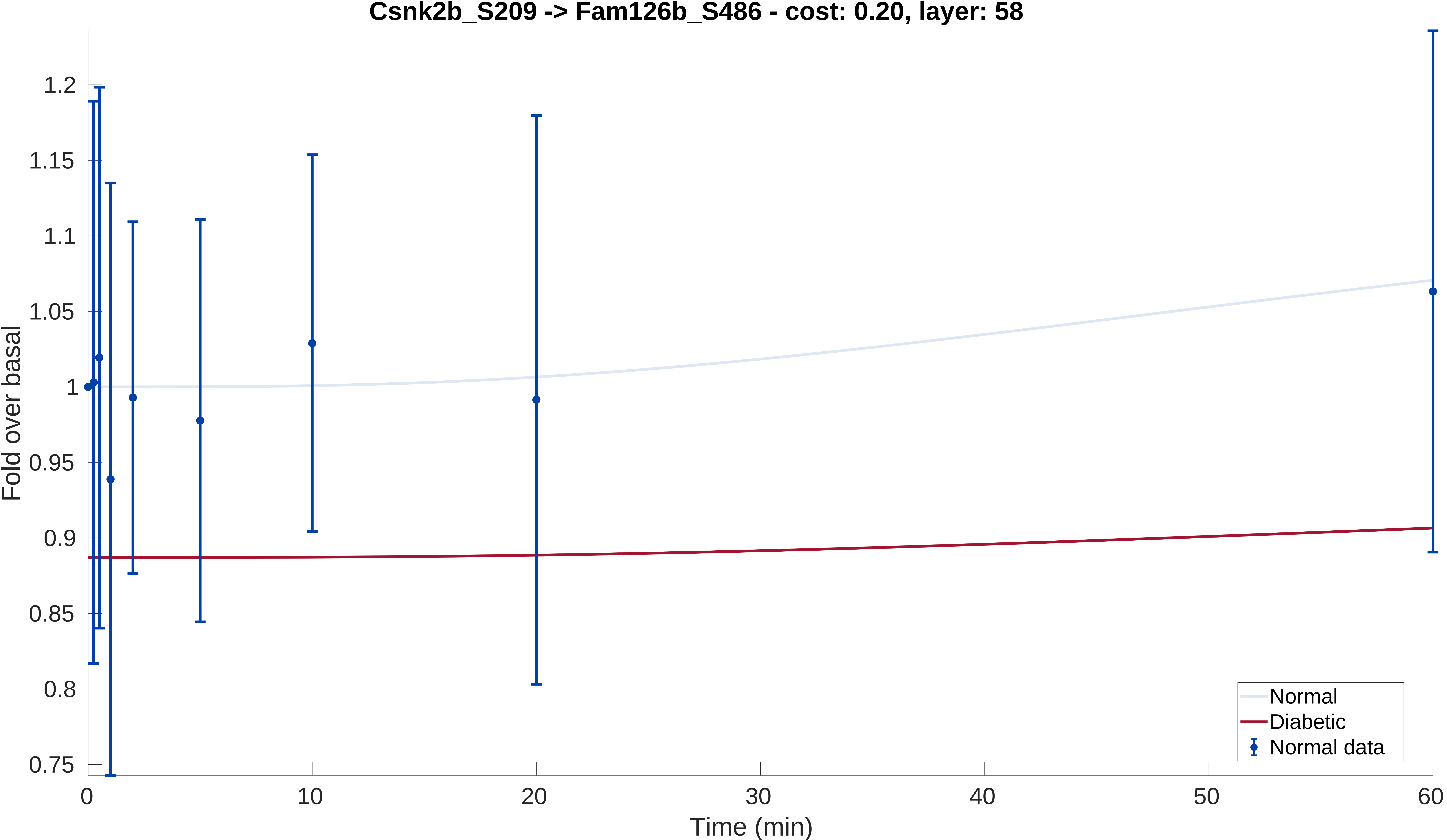

Csnk2b_S209 -> Fam126b_S486 - cost: 0.20, layer: 58
Fold over basal
1.2
1.15
1.1
1.05
1
0.95
0.9
0.85
0.8
0.75
0
10
20
30
40
50
60
Time (min)
Normal
Diabetic
Normal data

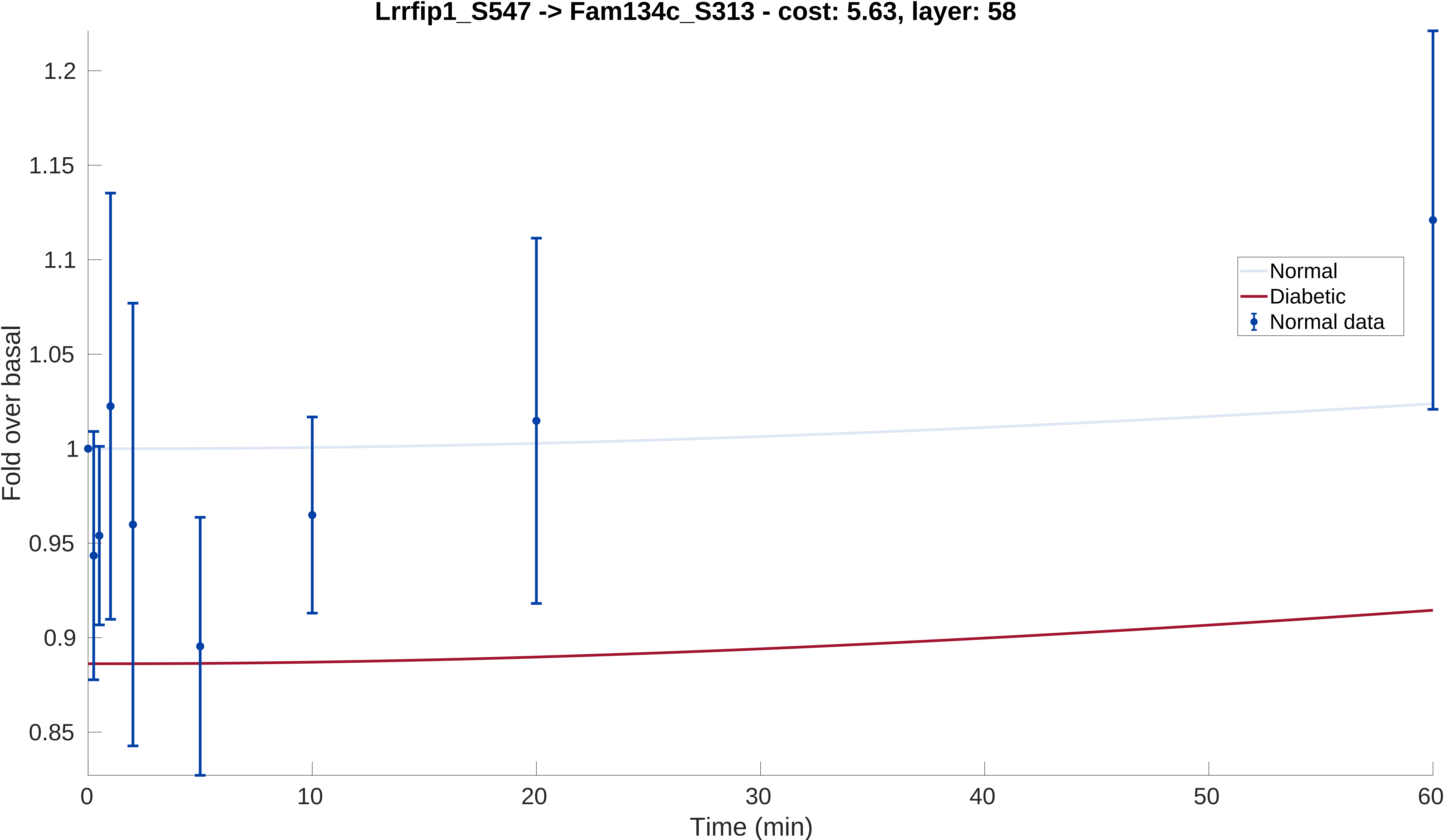
Lrrfip1_S547 -> Fam134c_S313 - cost: 5.63, layer: 58
Fold over basal
1.2
1.15
1.1
1.05
1
0.95
0.9
0.85
0
10
20
30
40
50
60
Time (min)
Normal
Diabetic
Normal data

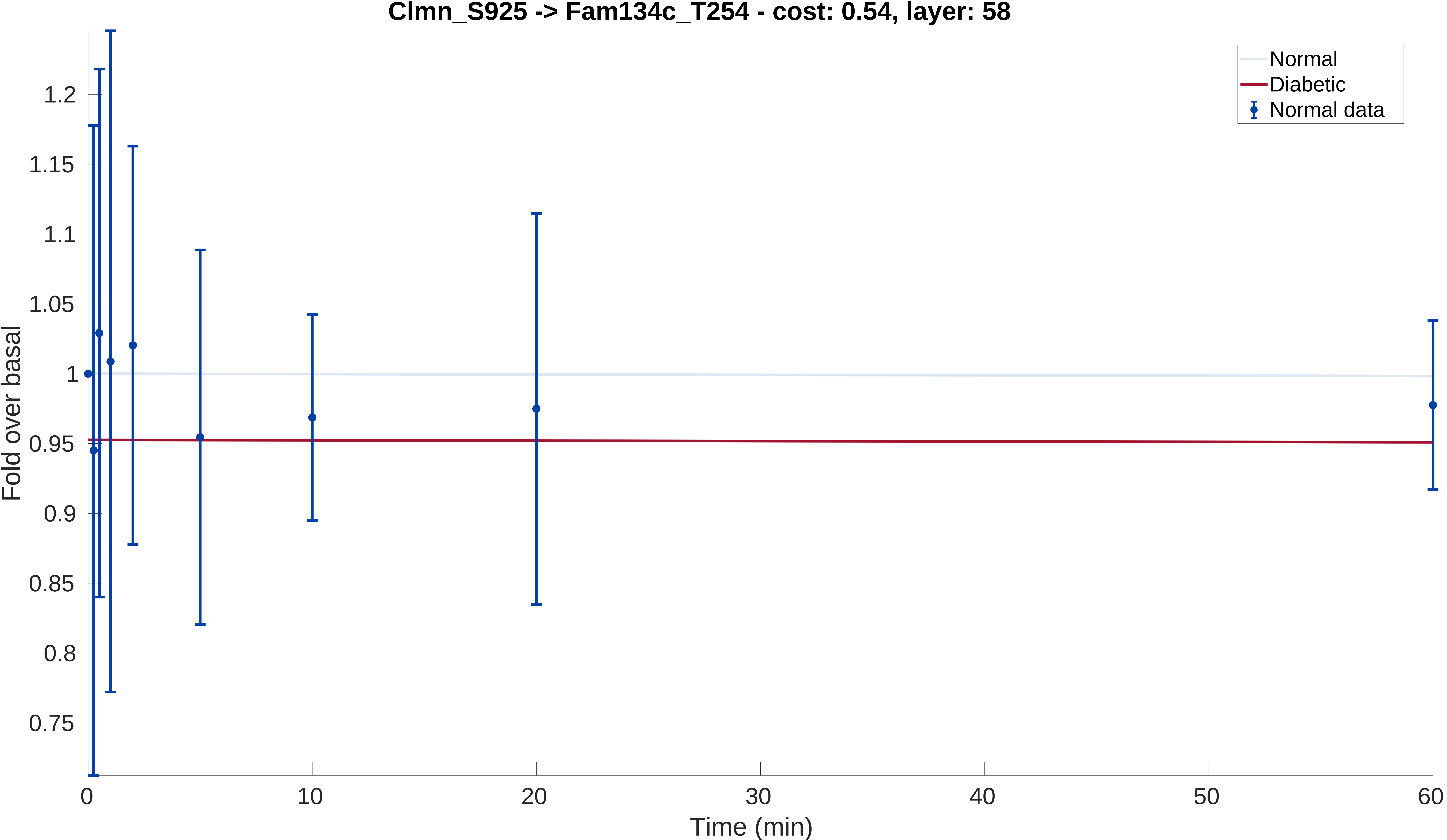

Clmn_S925 -> Fam134c_T254 - cost: 0.54, layer: 58
Normal
Diabetic
Normal data
Fold over basal
1.2
1.15
1.1
1.05
1
0.95
0.9
0.85
0.8
0.75
0
10
20
30
40
50
60
Time (min)

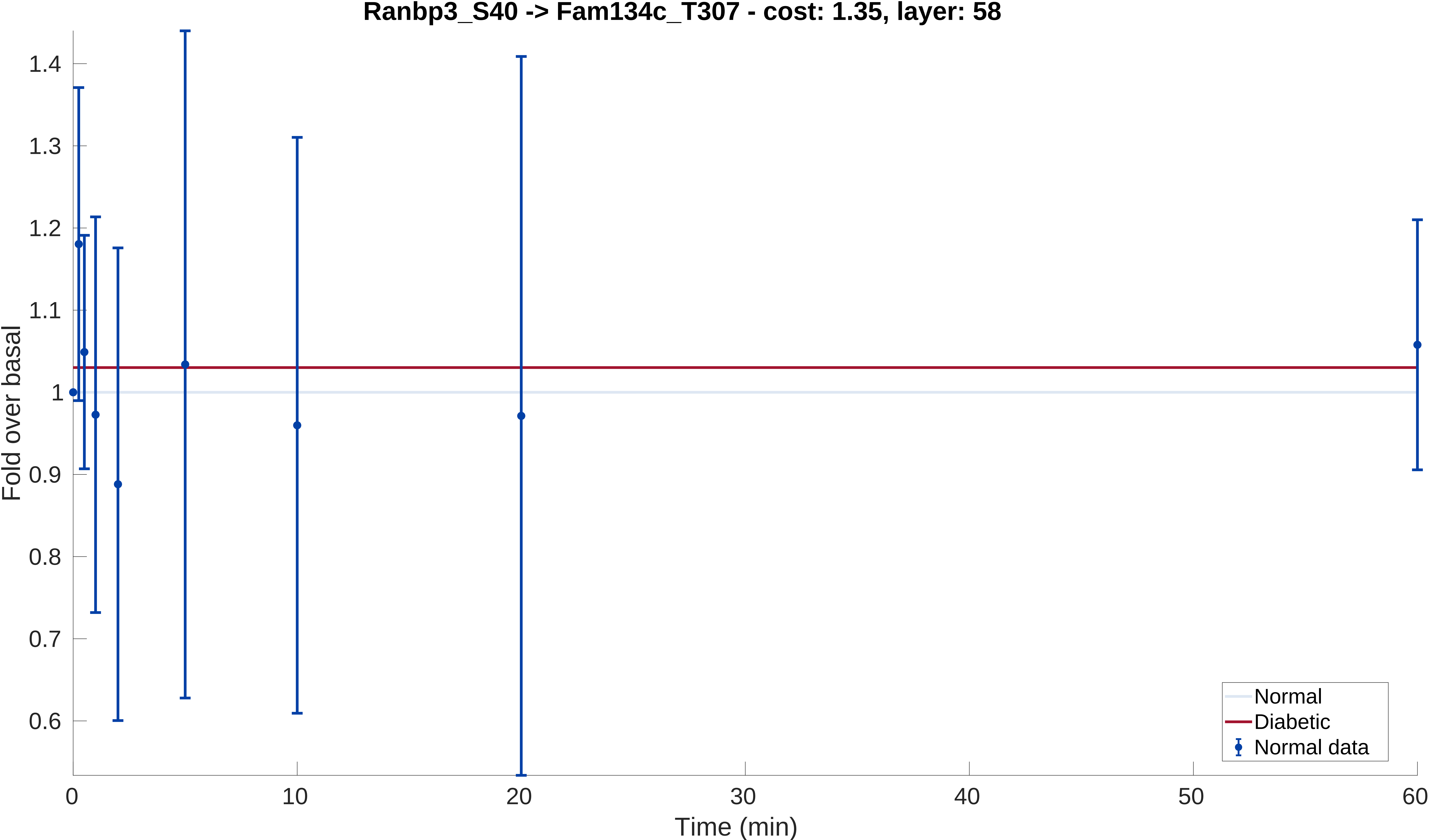

Ranbp3_S40 -> Fam134c_T307 - cost: 1.35, layer: 58
Fold over basal
Time (min)
1.4
1.3
1.2
1.1
1
0.9
0.8
0.7
0.6
0
10
20
30
40
50
60
Normal
Diabetic
Normal data

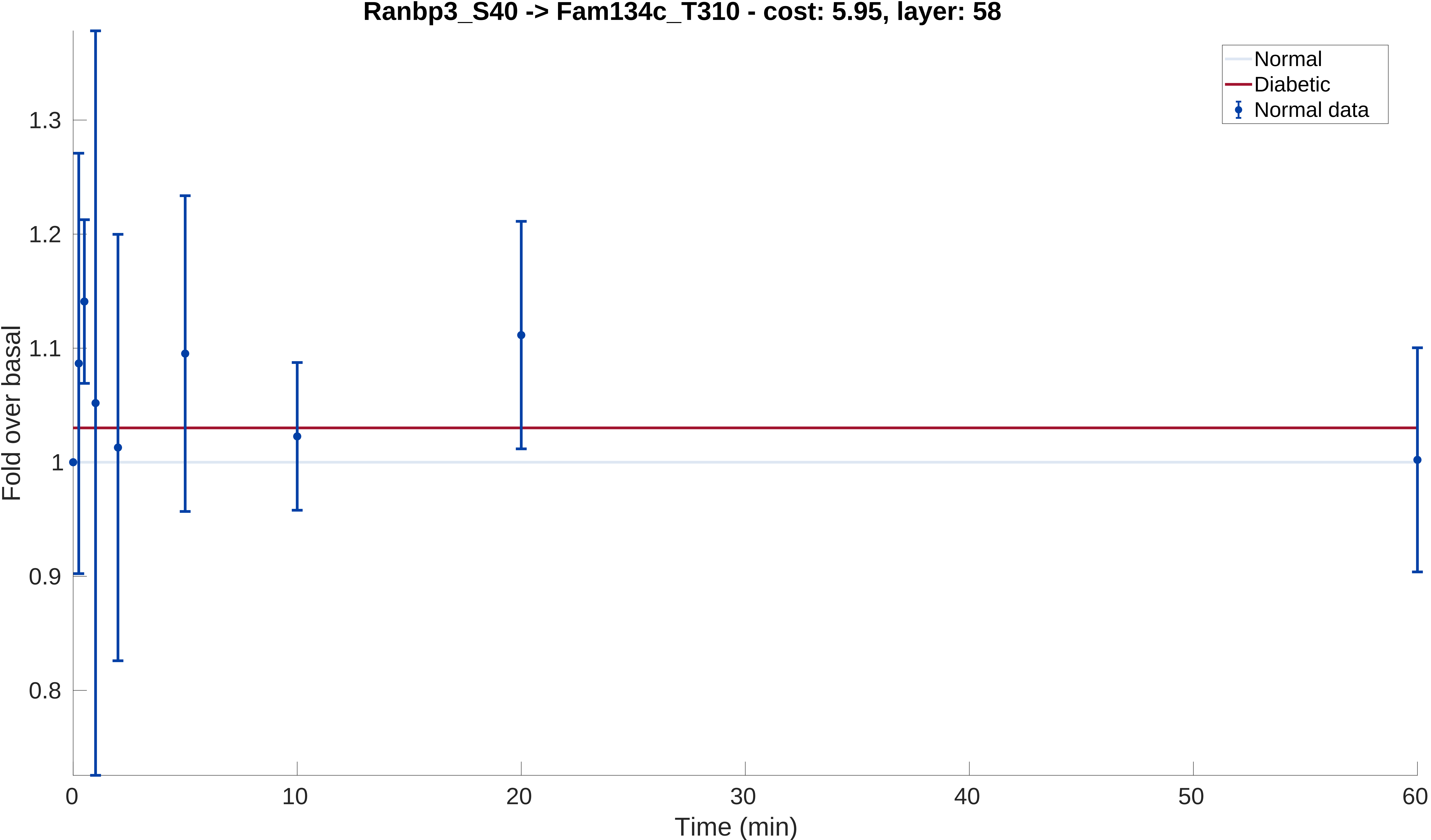
Ranbp3_S40 -> Fam134c_T310 - cost: 5.95, layer: 58
Normal
Diabetic
Normal data
Fold over basal
Time (min)
1.3
1.2
1.1
1
0.9
0.8
0
10
20
30
40
50
60

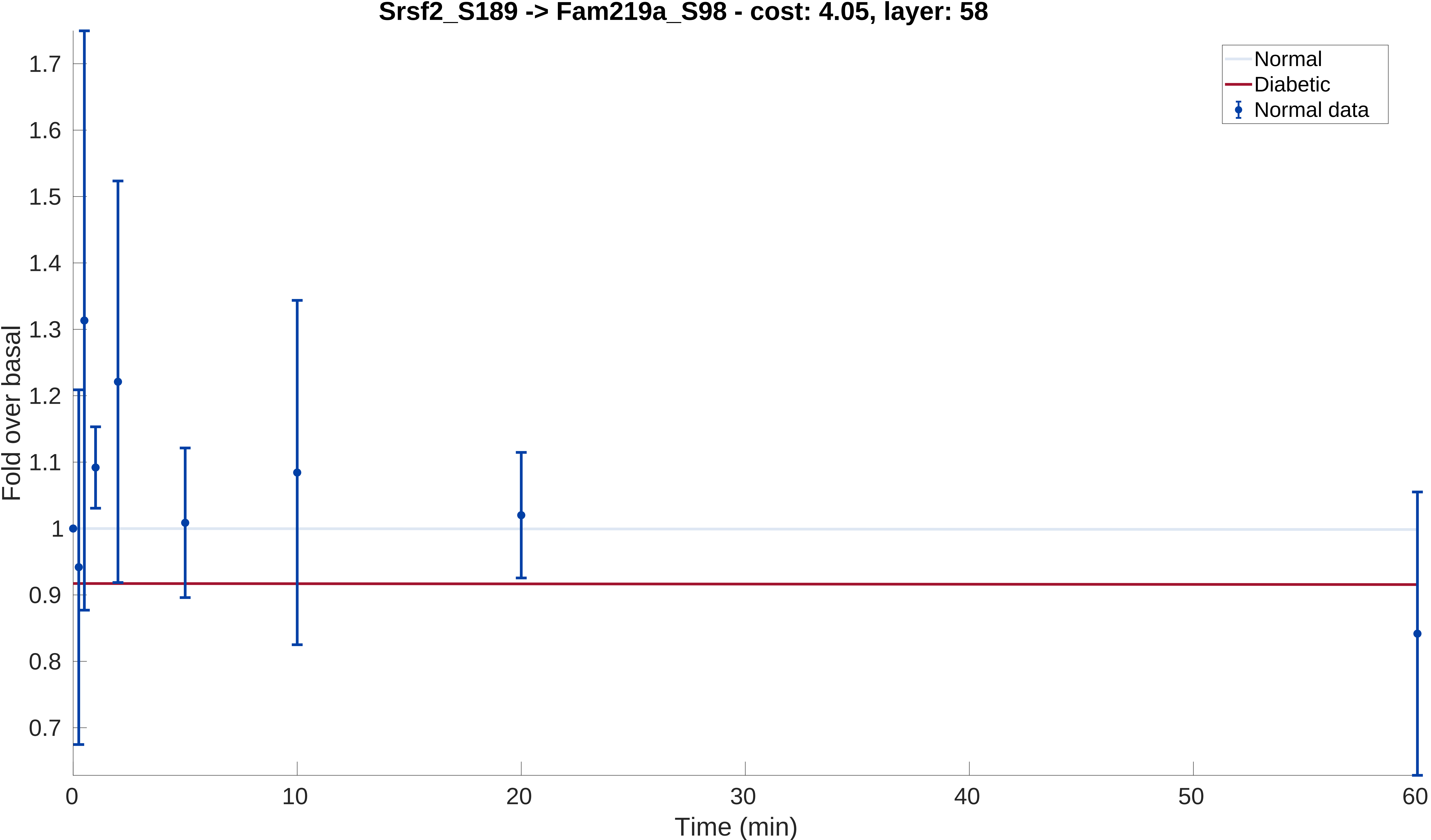

Srsf2_S189 -> Fam219a_S98 - cost: 4.05, layer: 58
Normal
Diabetic
Normal data
Fold over basal
1.7
1.6
1.5
1.4
1.3
1.2
1.1
1
0.9
0.8
0.7
0
10
20
30
40
50
60
Time (min)

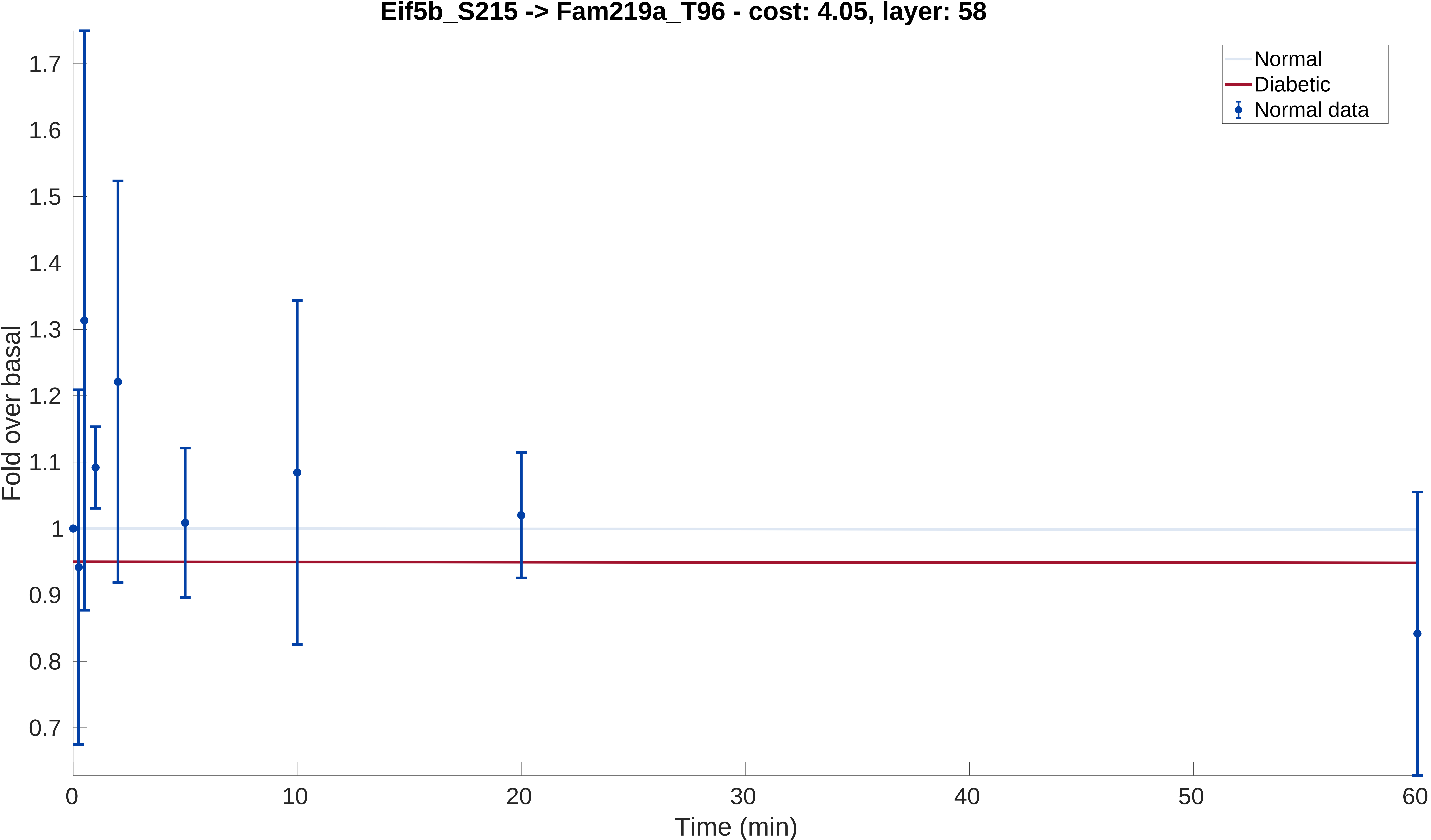

Eif5b_S215 -> Fam219a_T96 - cost: 4.05, layer: 58
Normal
Diabetic
Normal data
Fold over basal
1.7
1.6
1.5
1.4
1.3
1.2
1.1
1
0.9
0.8
0.7
0
10
20
30
40
50
60
Time (min)

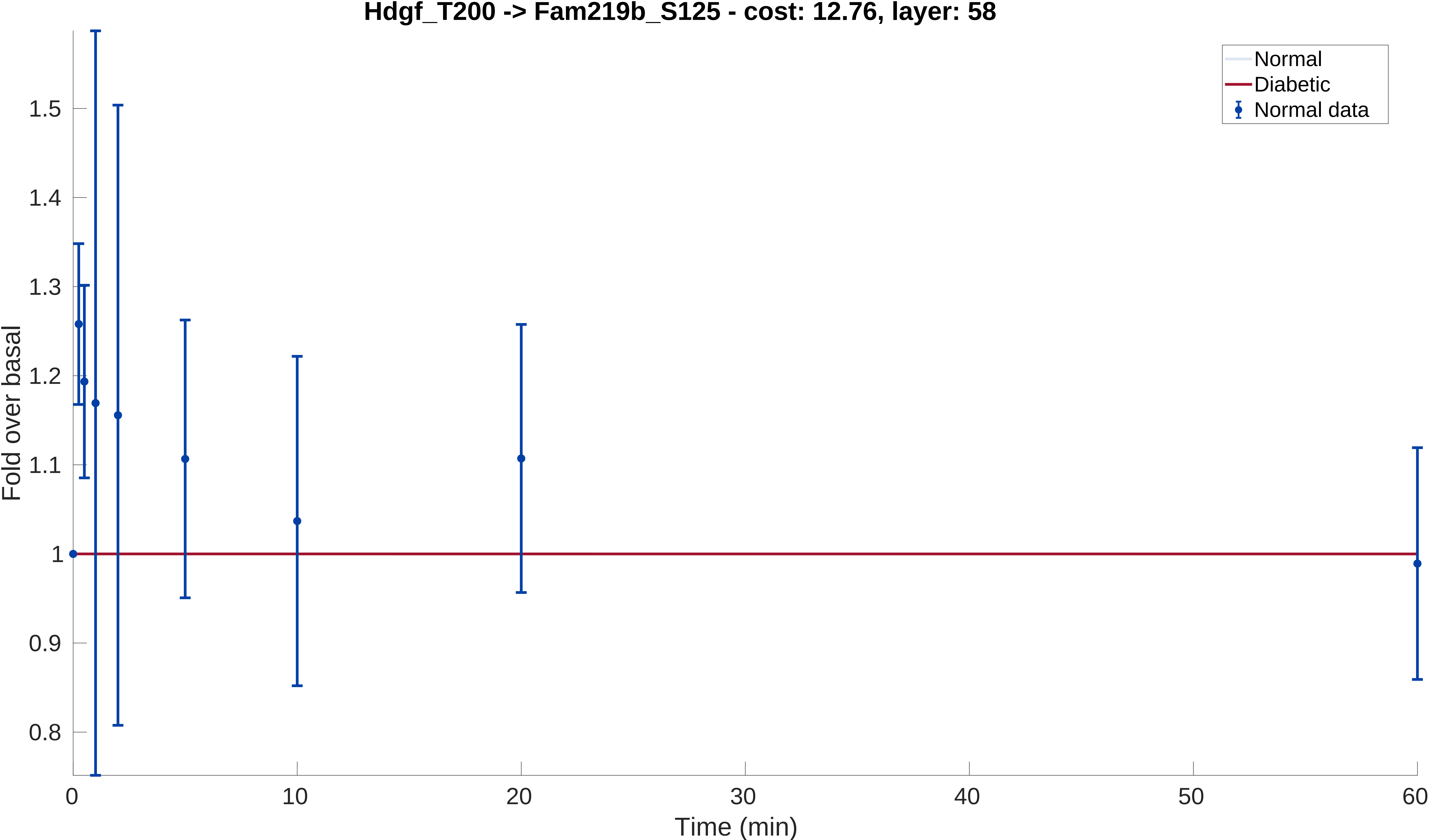
Hdgf_T200 -> Fam219b_S125 - cost: 12.76, layer: 58
Normal
Diabetic
Normal data
Fold over basal
1.5
1.4
1.3
1.2
1.1
1
0.9
0.8
0
10
20
30
40
50
60
Time (min)

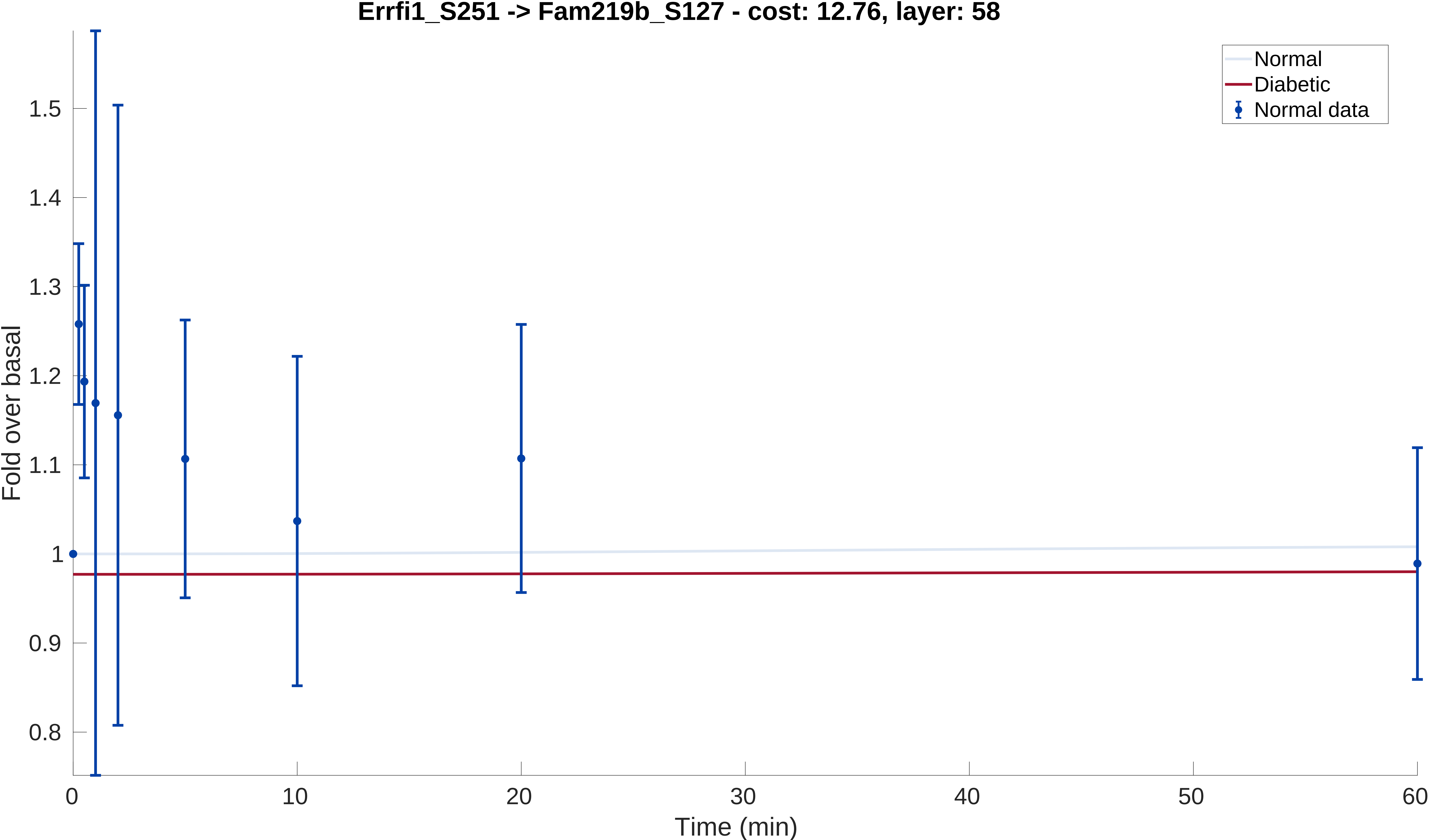

Errfi1_S251 -> Fam219b_S127 - cost: 12.76, layer: 58
Normal
Diabetic
Normal data
Fold over basal
1.5
1.4
1.3
1.2
1.1
1
0.9
0.8
0
10
20
30
40
50
60
Time (min)

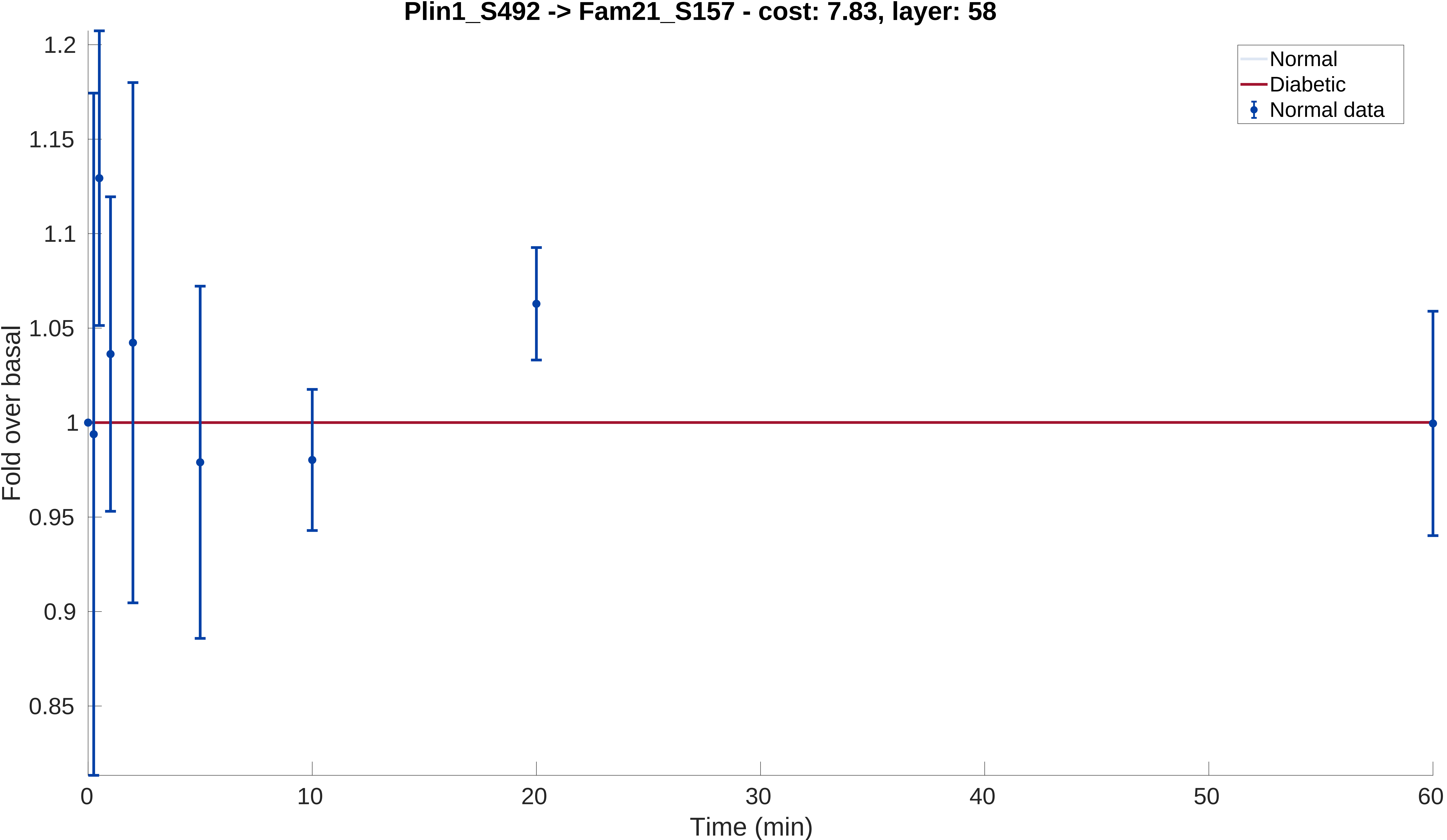
Plin1_S492 -> Fam21_S157 - cost: 7.83, layer: 58
Normal
Diabetic
Normal data
Fold over basal
Time (min)
1.2
1.15
1.1
1.05
1
0.95
0.9
0.85
0
10
20
30
40
50
60

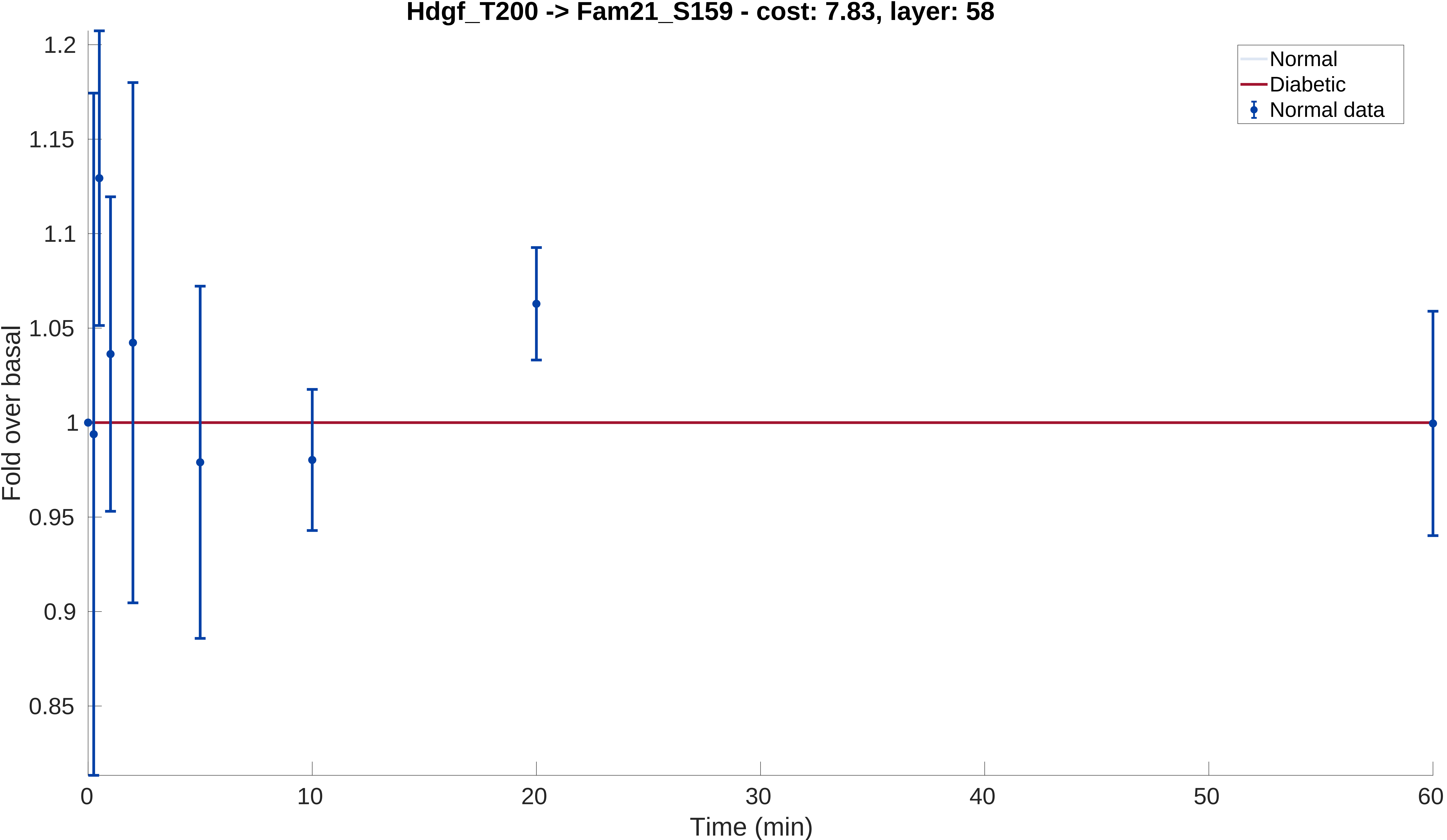
Hdgf_T200 -> Fam21_S159 - cost: 7.83, layer: 58
Normal
Diabetic
Normal data
Fold over basal
1.2
1.15
1.1
1.05
1
0.95
0.9
0.85
Time (min)
0
10
20
30
40
50
60

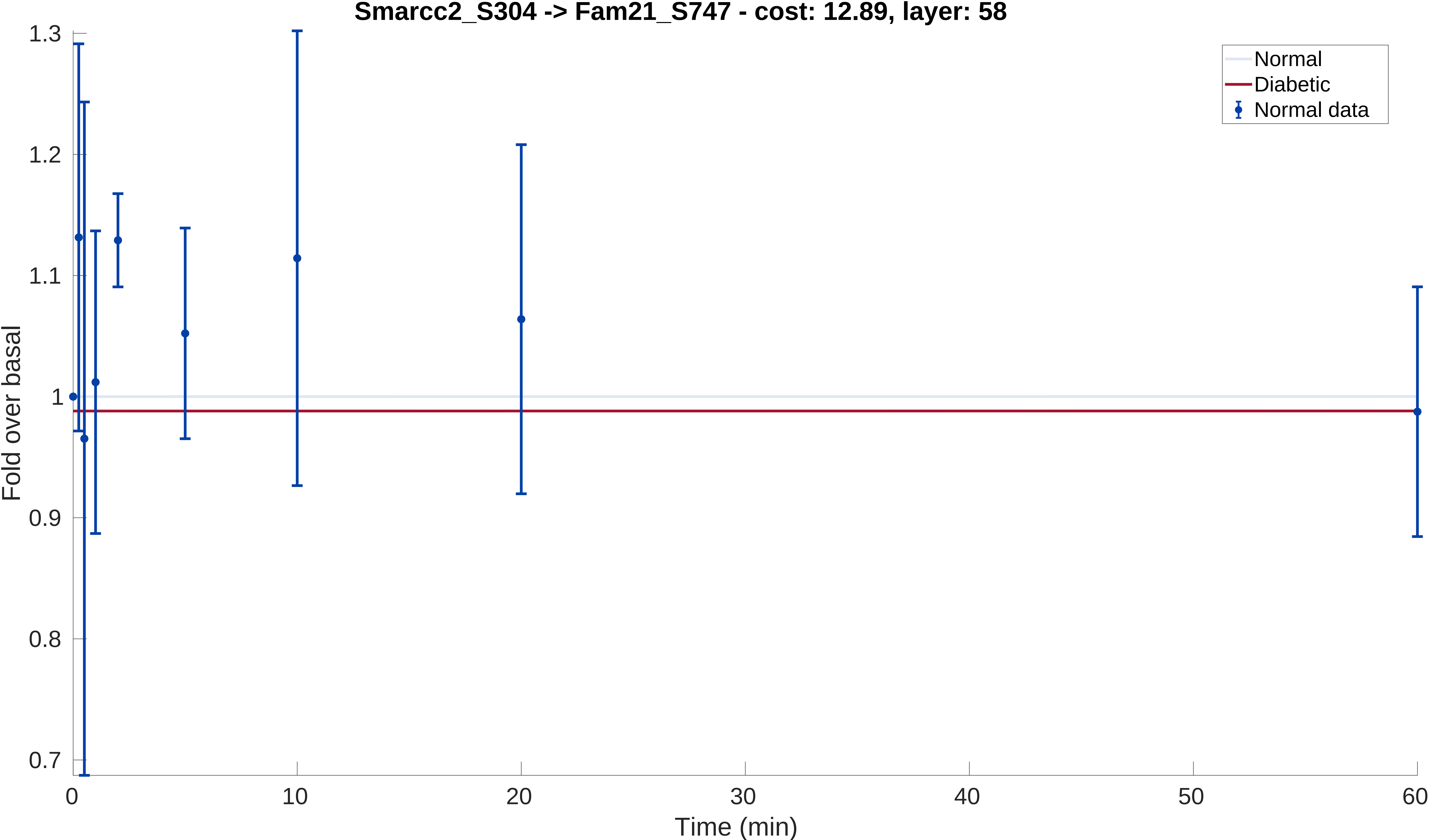

Smarcc2_S304 -> Fam21_S747 - cost: 12.89, layer: 58
Normal
Diabetic
Normal data
Fold over basal
Time (min)
1.3
1.2
1.1
1
0.9
0.8
0.7
0
10
20
30
40
50
60

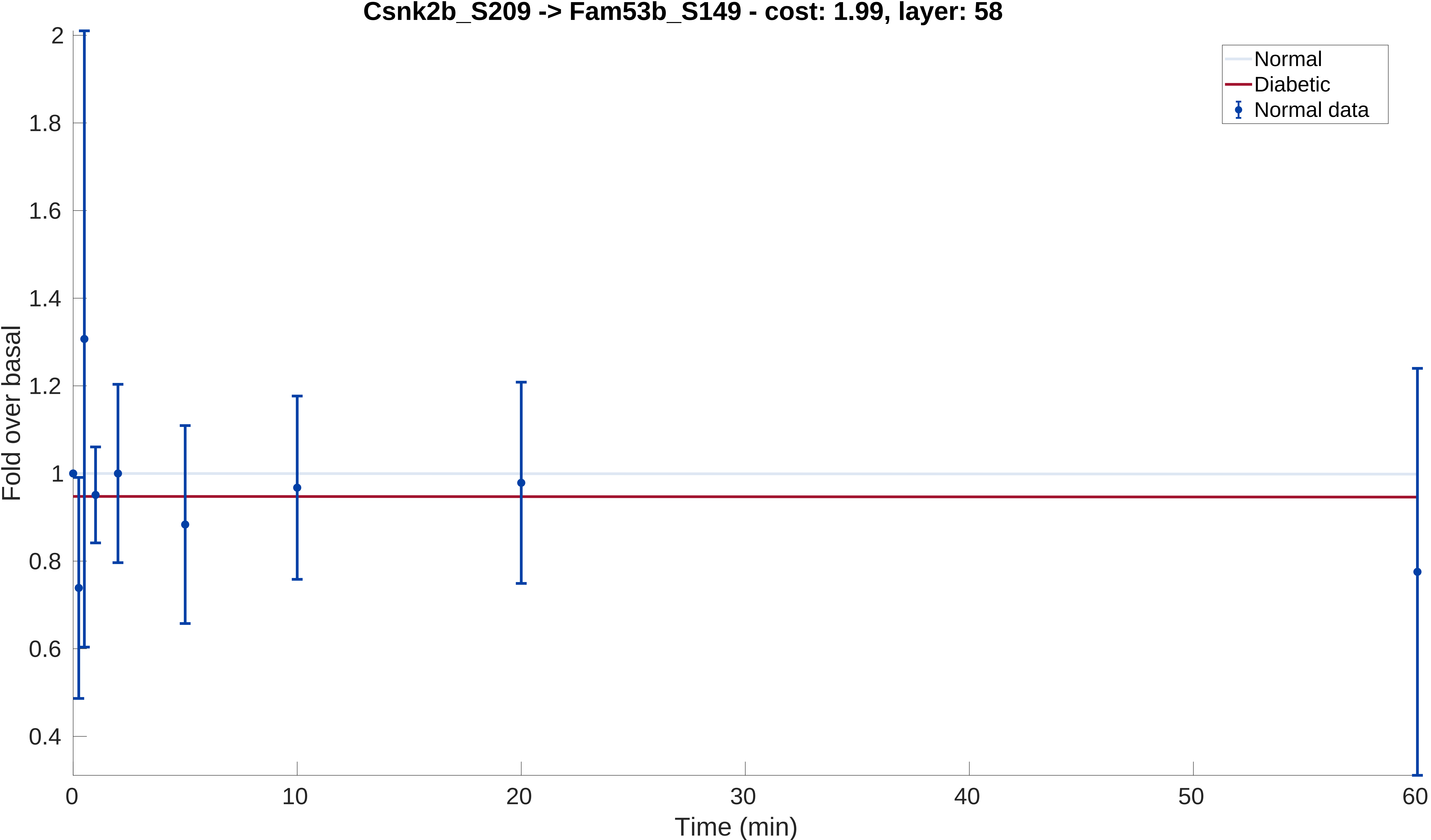

Csnk2b_S209 -> Fam53b_S149 - cost: 1.99, layer: 58
Normal
Diabetic
Normal data
Fold over basal
Time (min)
2
1.8
1.6
1.4
1.2
1
0.8
0.6
0.4
0
10
20
30
40
50
60

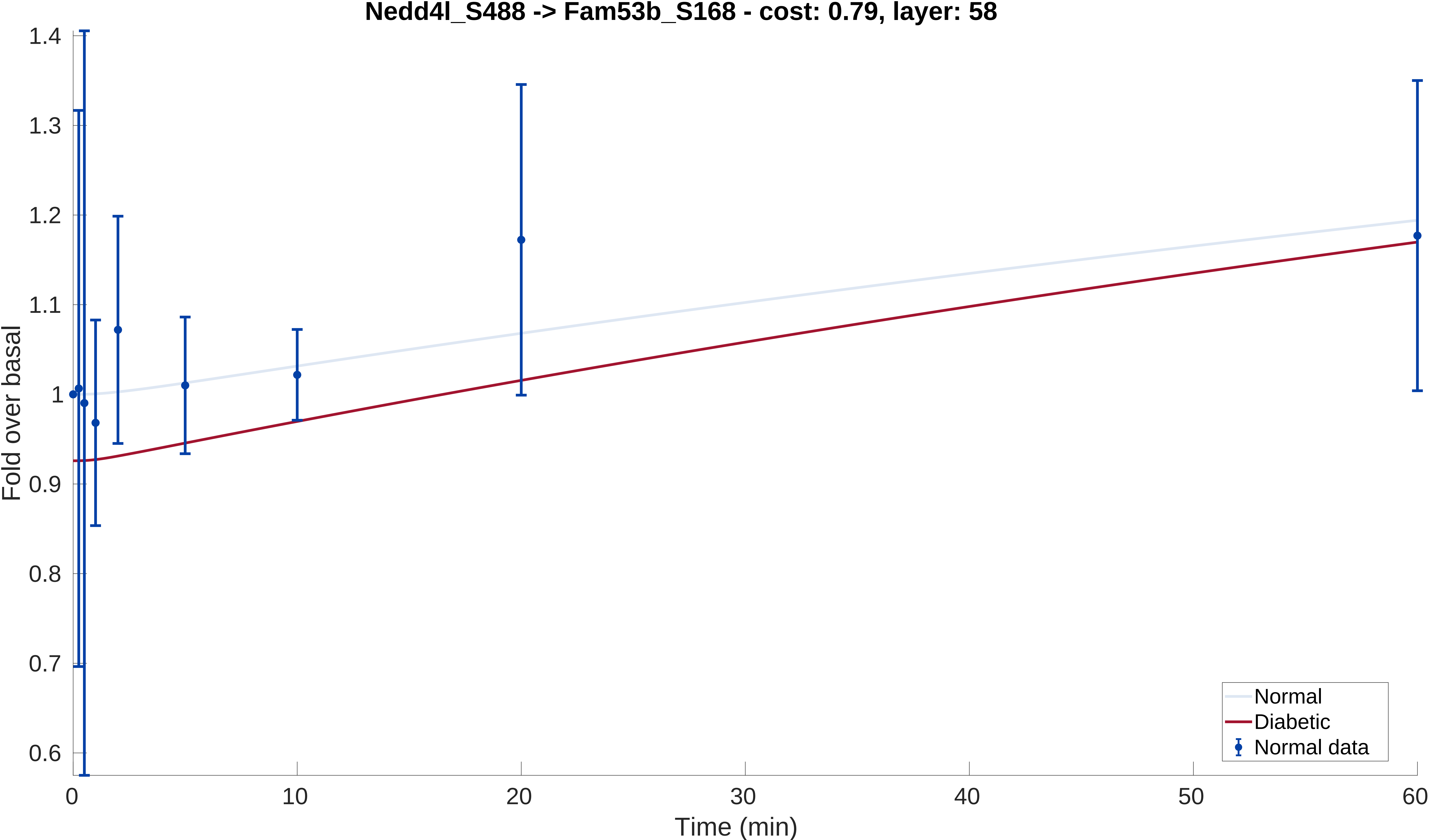
Nedd4l_S488 -> Fam53b_S168 - cost: 0.79, layer: 58
Fold over basal
1.4
1.3
1.2
1.1
1
0.9
0.8
0.7
0.6
0
10
20
30
40
50
60
Time (min)
Normal
Diabetic
Normal data

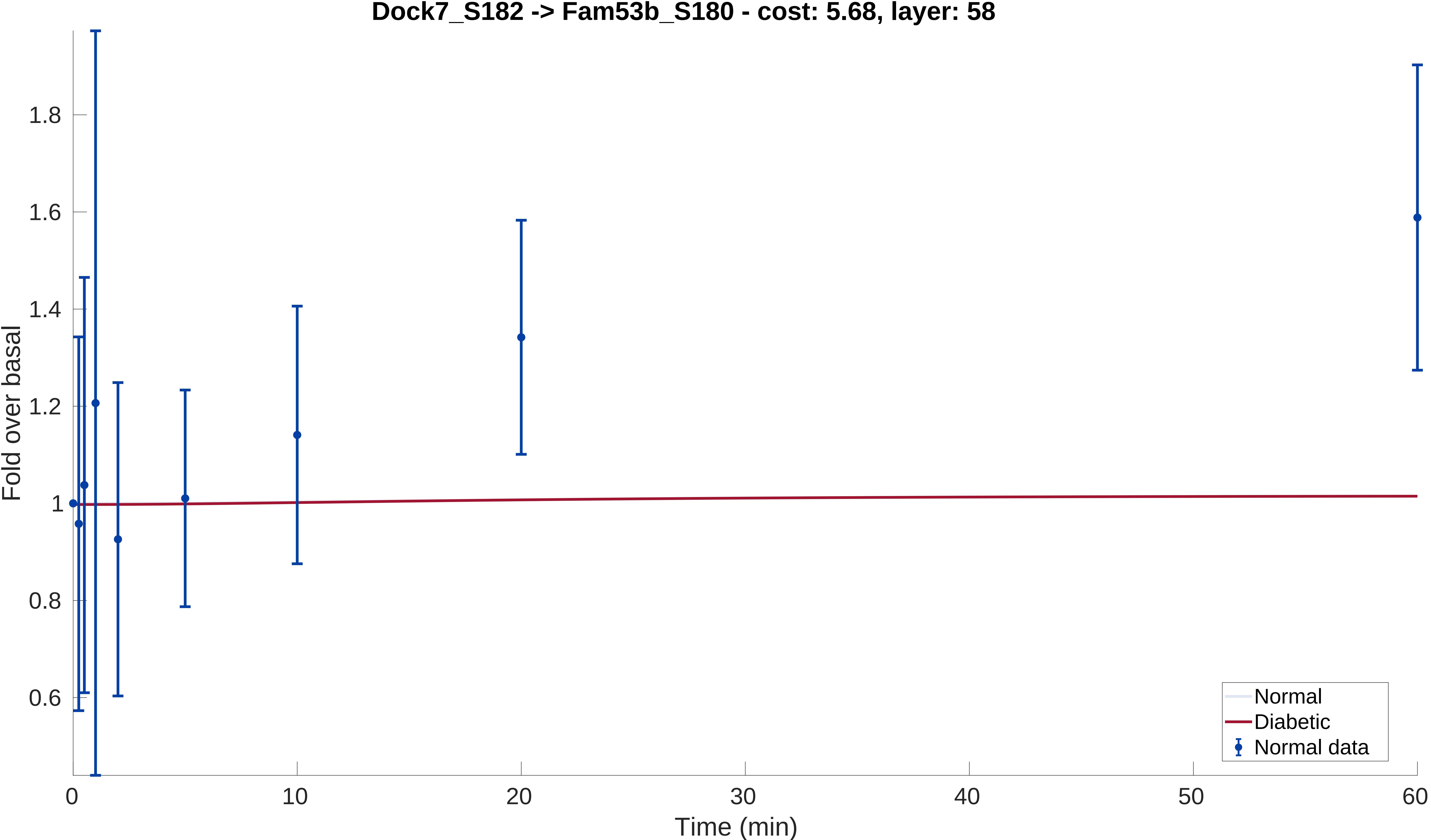
Dock7_S182 -> Fam53b_S180 - cost: 5.68, layer: 58
Fold over basal
1.8
1.6
1.4
1.2
1
0.8
0.6
0
10
20
30
40
50
60
Time (min)
Normal
Diabetic
Normal data

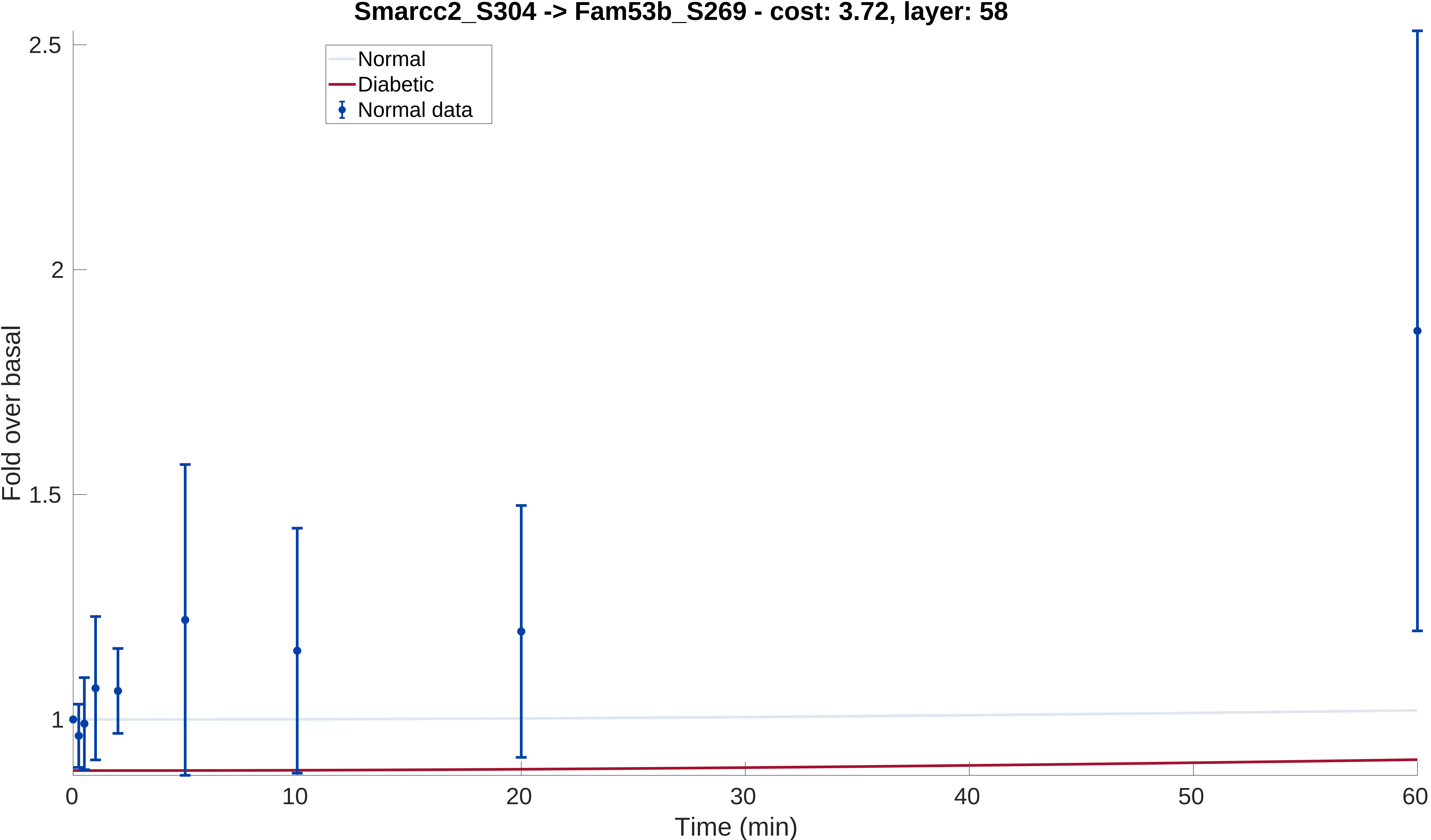

Smarcc2_S304 -> Fam53b_S269 - cost: 3.72, layer: 58
Normal
Diabetic
Normal data
Fold over basal
2.5
2
1.5
1
0
10
20
30
40
50
60
Time (min)

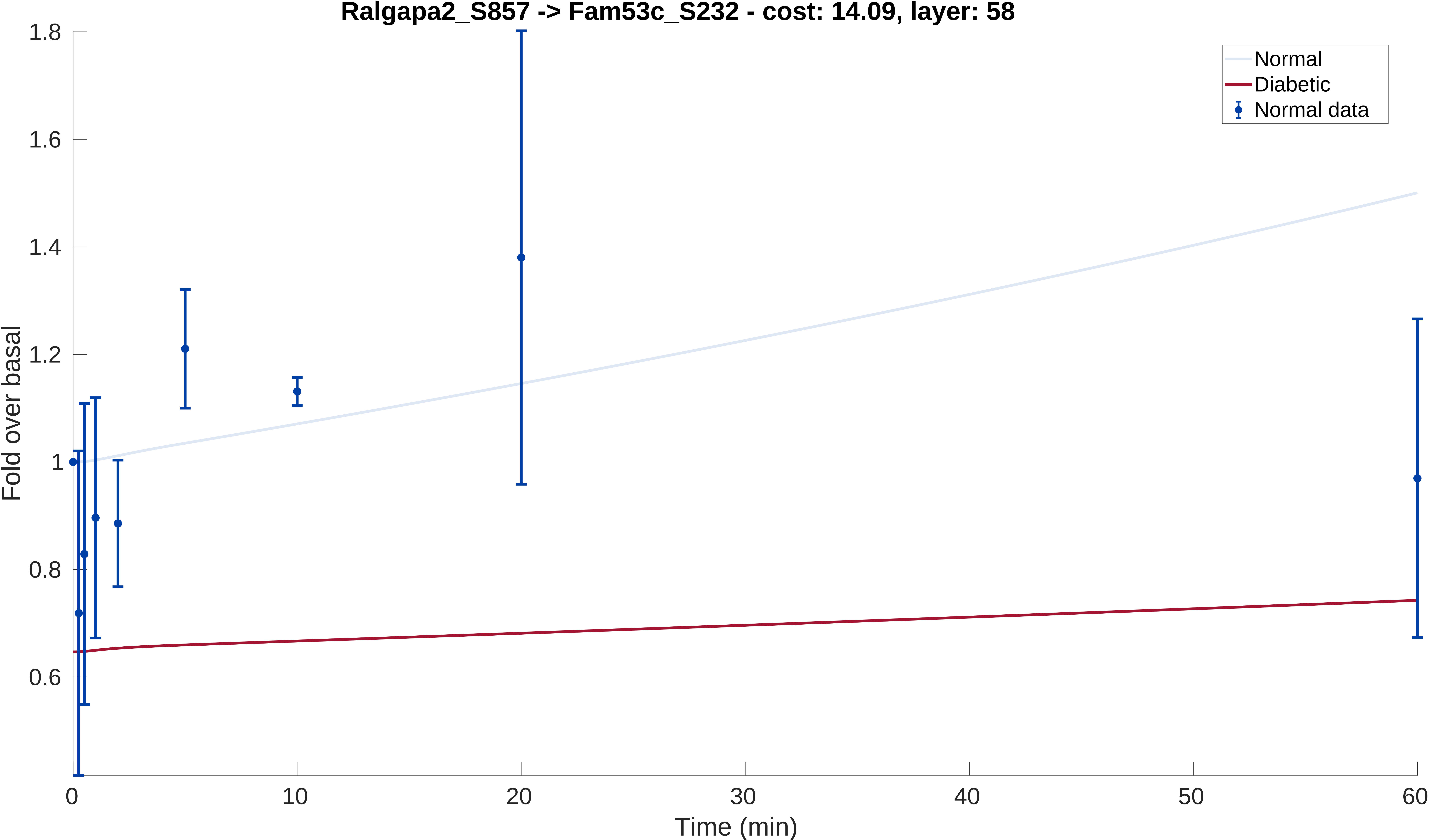

Ralgapa2_S857 -> Fam53c_S232 - cost: 14.09, layer: 58
Normal
Diabetic
Normal data
Fold over basal
1.8
1.6
1.4
1.2
1
0.8
0.6
0
10
20
30
40
50
60
Time (min)

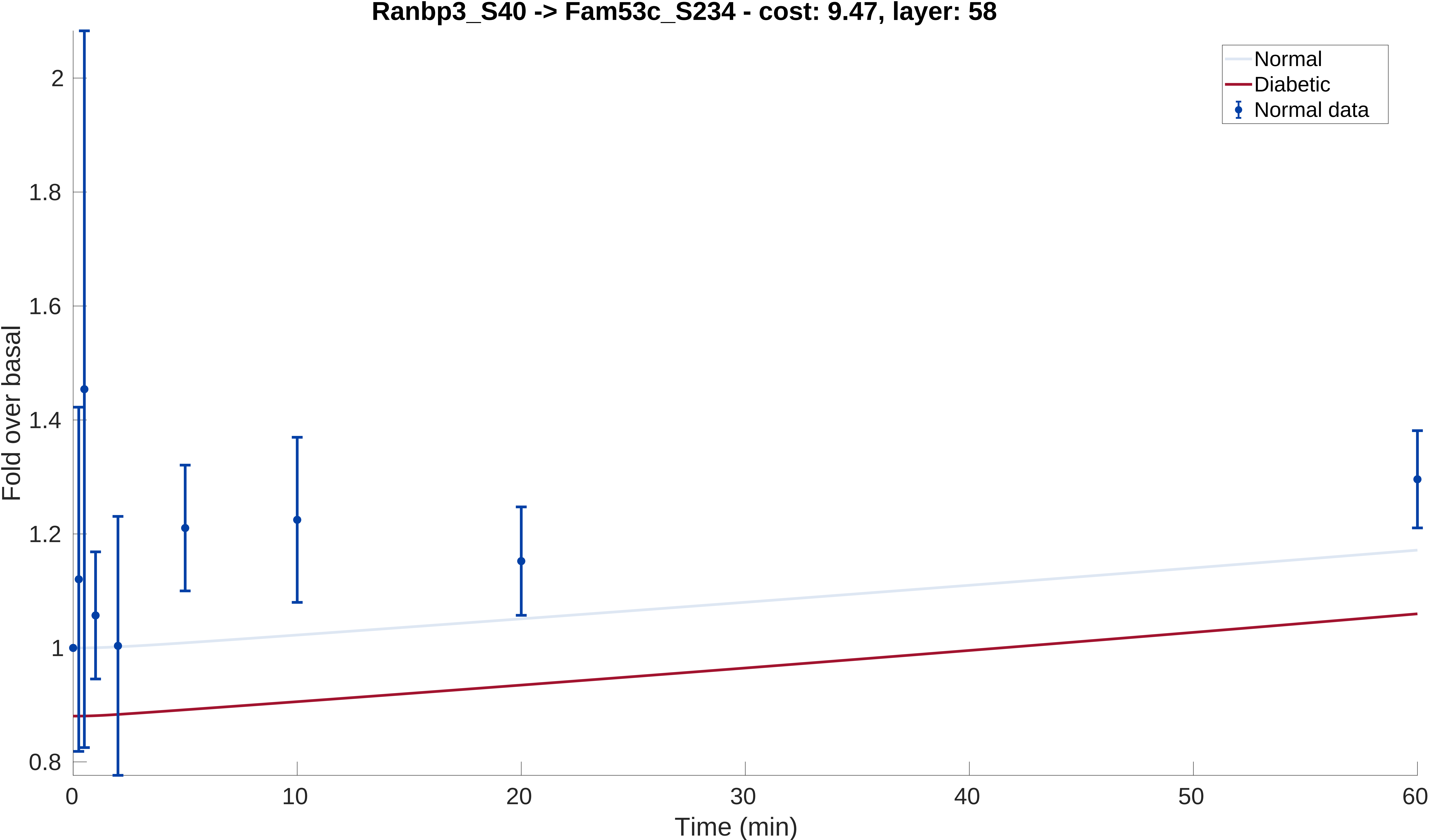

Ranbp3_S40 -> Fam53c_S234 - cost: 9.47, layer: 58
Normal
Diabetic
Normal data
Fold over basal
0.8
1
1.2
1.4
1.6
1.8
2
0
10
20
30
40
50
60
Time (min)

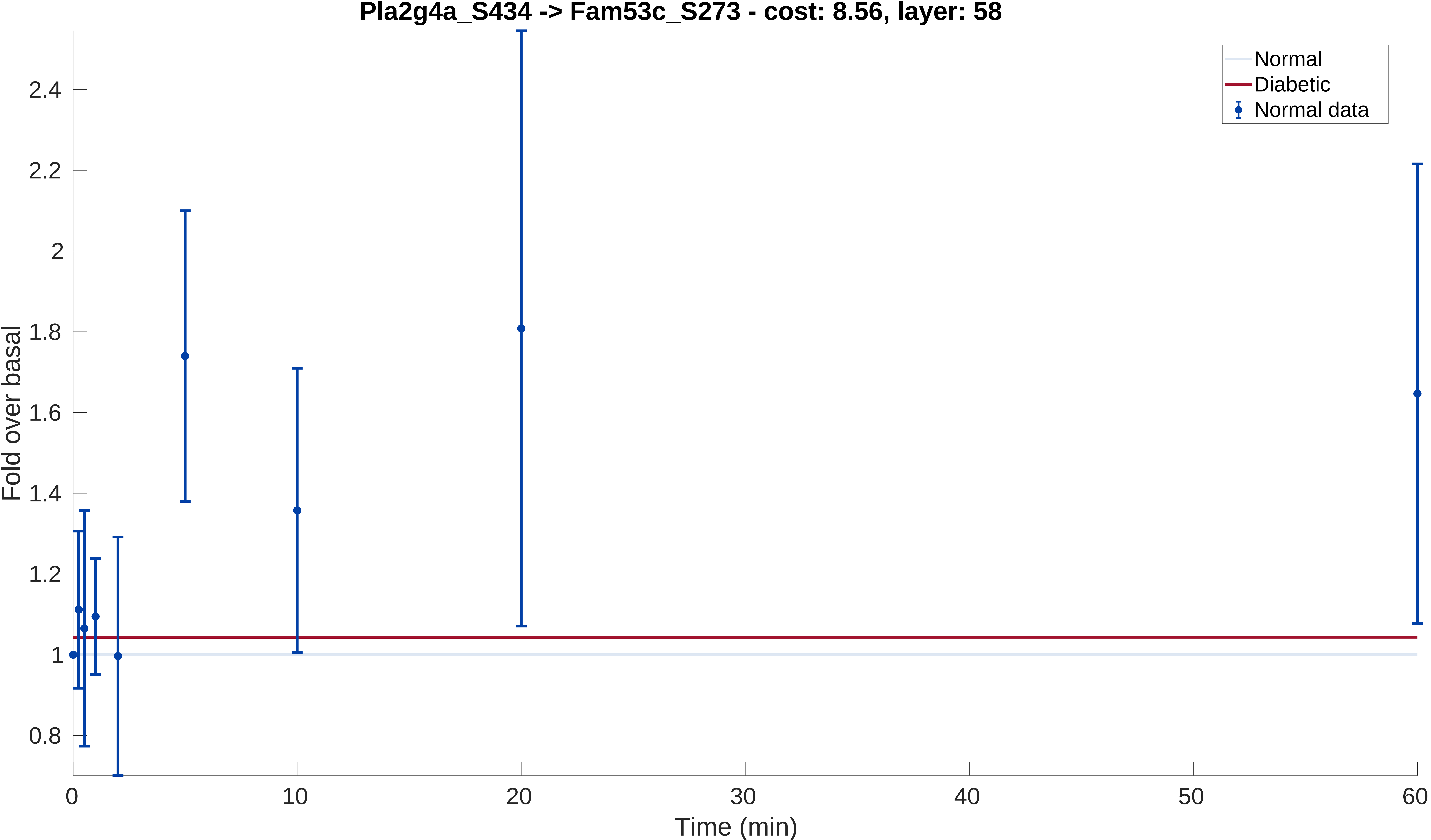

Pla2g4a_S434 -> Fam53c_S273 - cost: 8.56, layer: 58
Normal
Diabetic
Normal data
Fold over basal
2.4
2.2
2
1.8
1.6
1.4
1.2
1
0.8
0
10
20
30
40
50
60
Time (min)

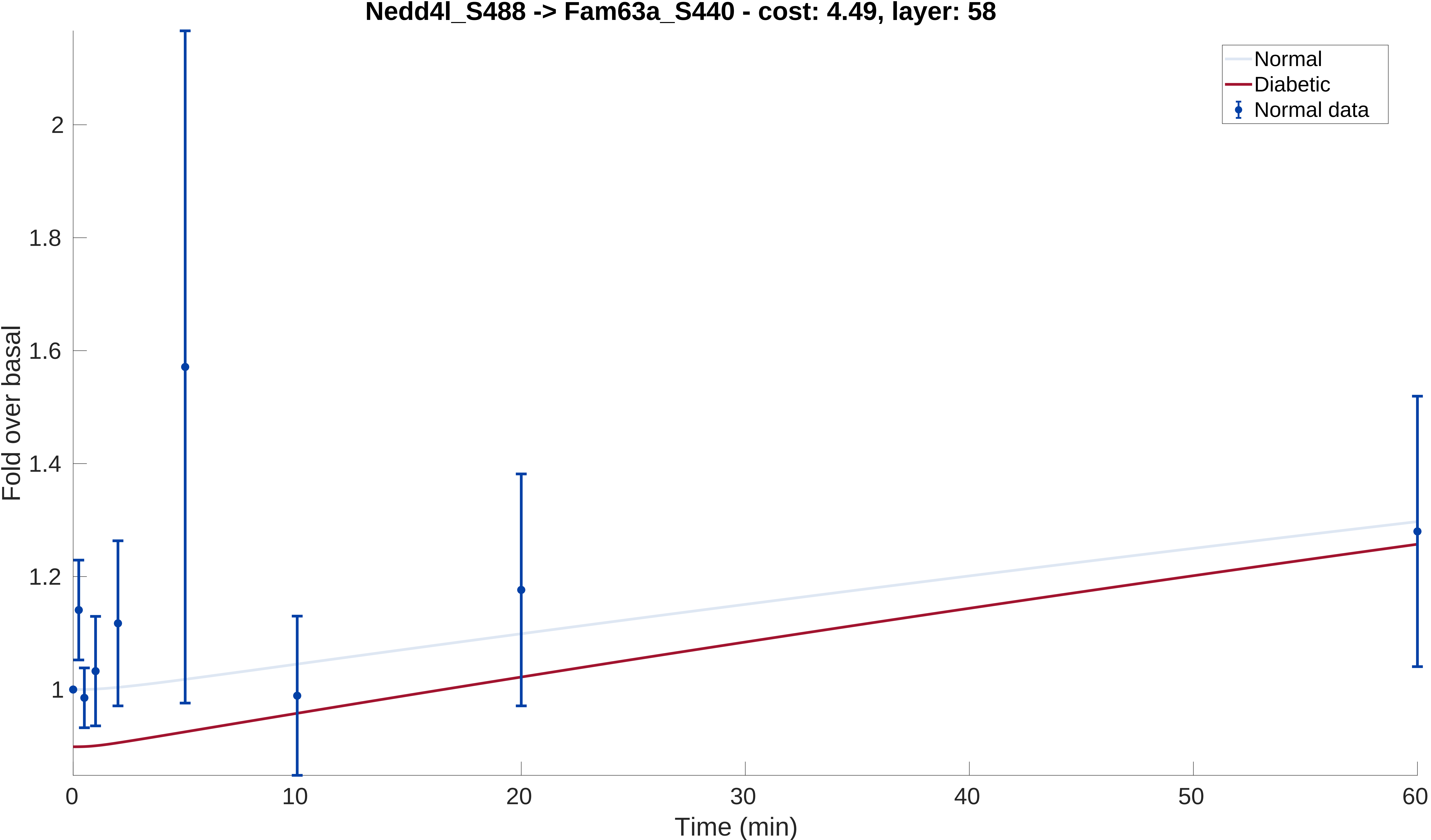
Nedd4l_S488 -> Fam63a_S440 - cost: 4.49, layer: 58
Normal
Diabetic
Normal data
Fold over basal
2
1.8
1.6
1.4
1.2
1
0
10
20
30
40
50
60
Time (min)

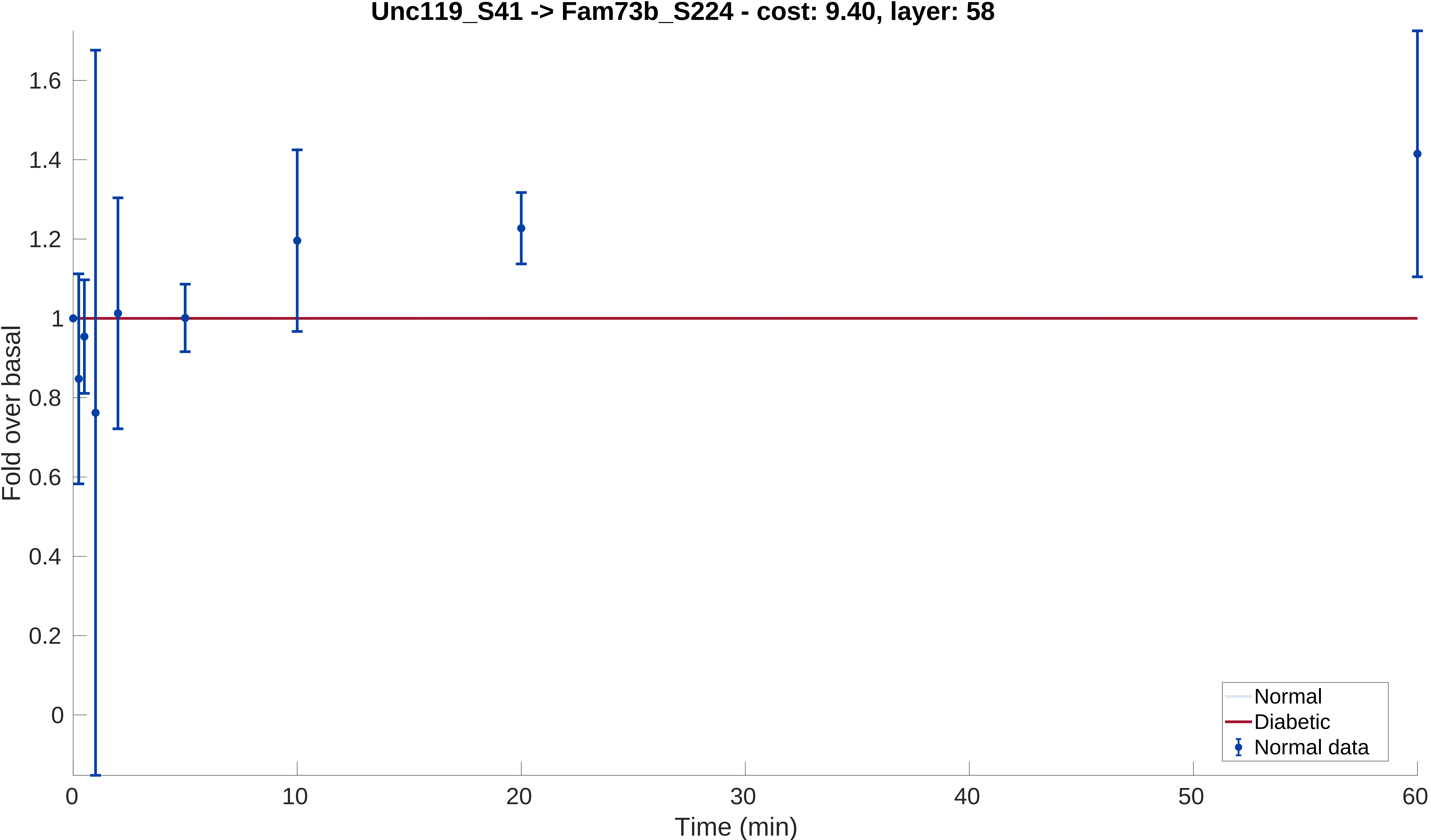
Unc119_S41 -> Fam73b_S224 - cost: 9.40, layer: 58
Fold over basal
1.6
1.4
1.2
1
0.8
0.6
0.4
0.2
0
0
10
20
30
40
50
60
Time (min)
Normal
Diabetic
Normal data

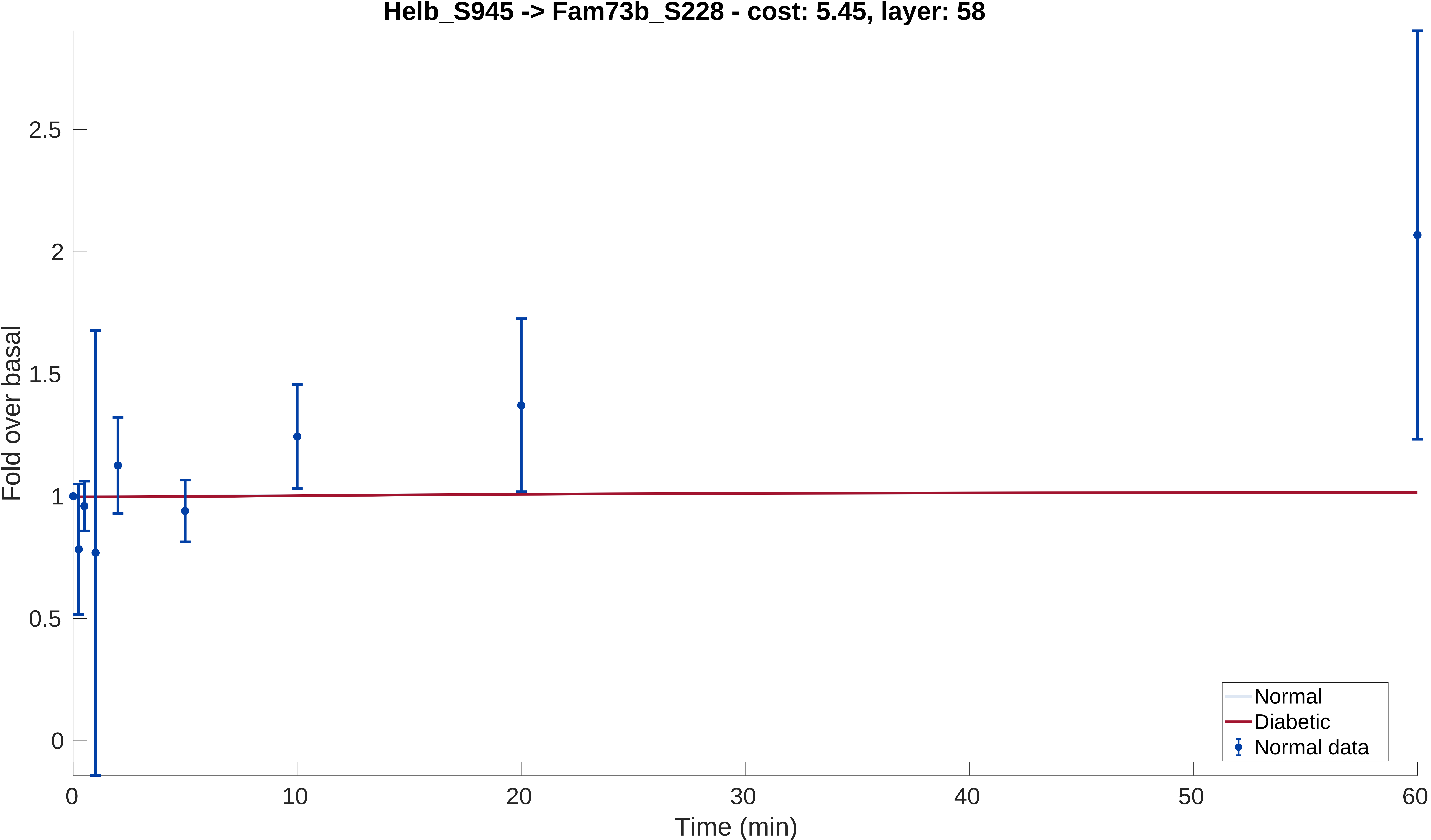

Helb_S945 -> Fam73b_S228 - cost: 5.45, layer: 58
Fold over basal
2.5
2
1.5
1
0.5
0
0
10
20
30
40
50
60
Time (min)
Normal
Diabetic
Normal data

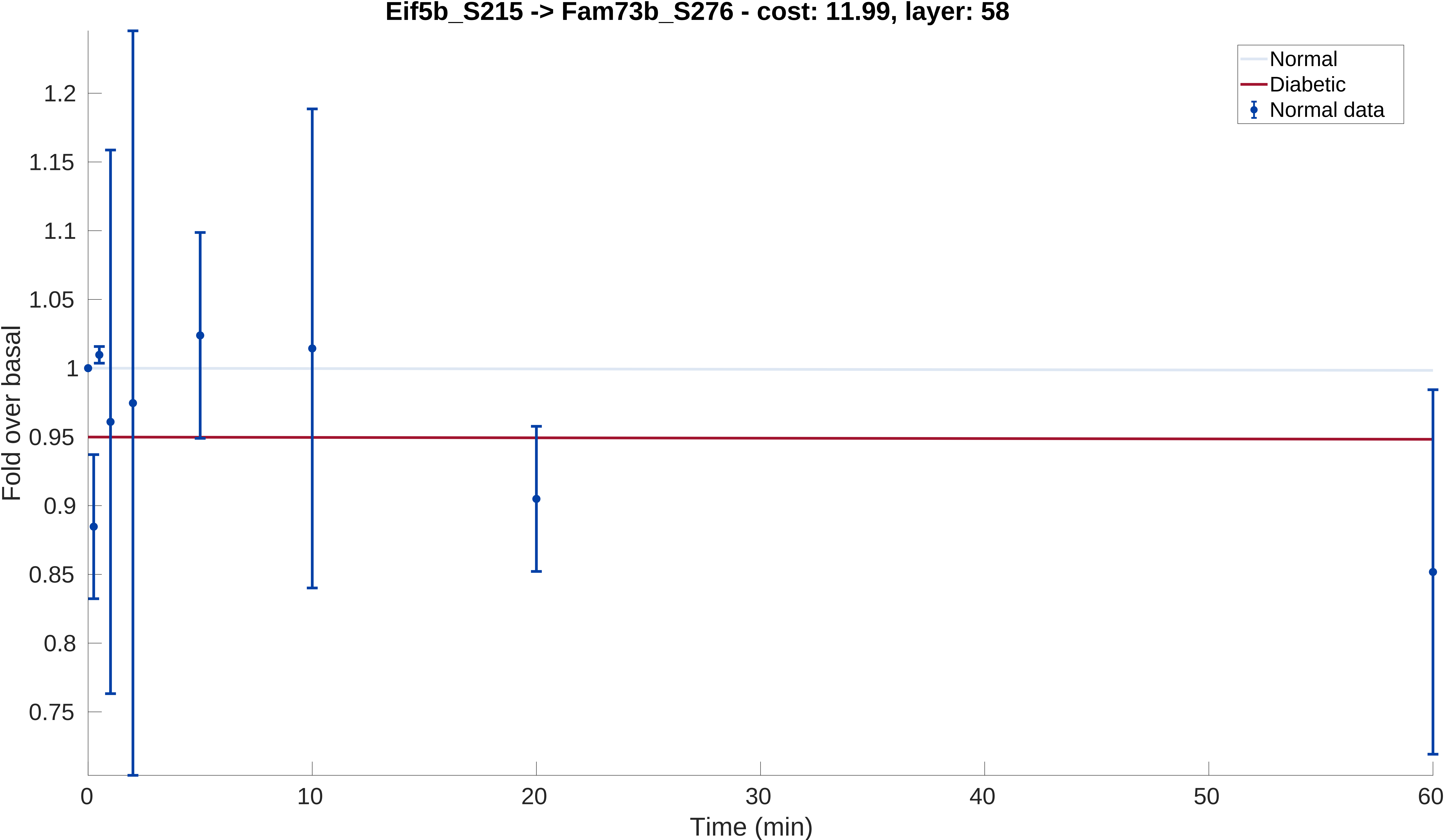

Eif5b_S215 -> Fam73b_S276 - cost: 11.99, layer: 58
Normal
Diabetic
Normal data
Fold over basal
Time (min)
1.2
1.15
1.1
1.05
1
0.95
0.9
0.85
0.8
0.75
0
10
20
30
40
50
60

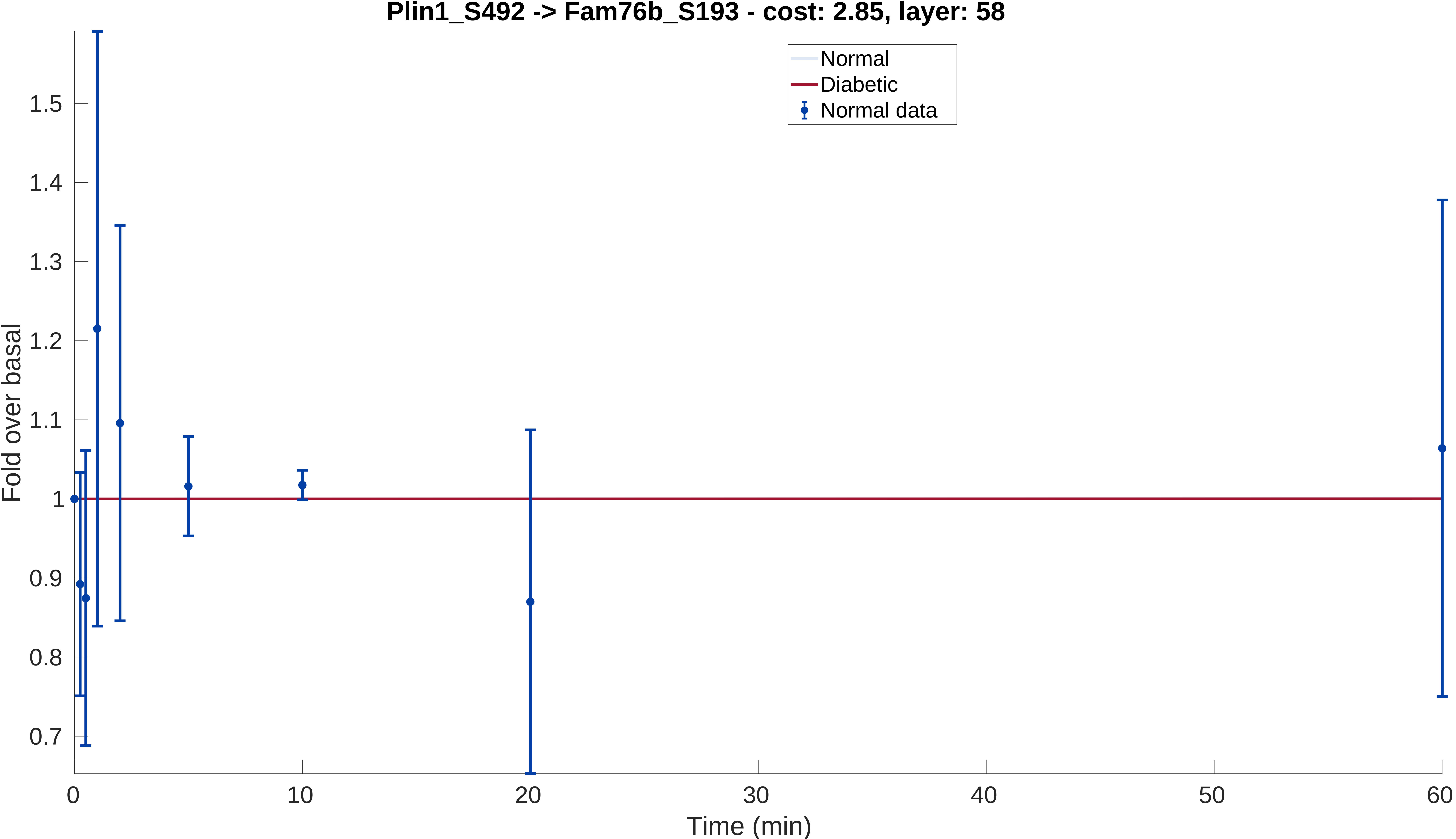

Plin1_S492 -> Fam76b_S193 - cost: 2.85, layer: 58
Normal
Diabetic
Normal data
Fold over basal
1.5
1.4
1.3
1.2
1.1
1
0.9
0.8
0.7
0
10
20
30
40
50
60
Time (min)

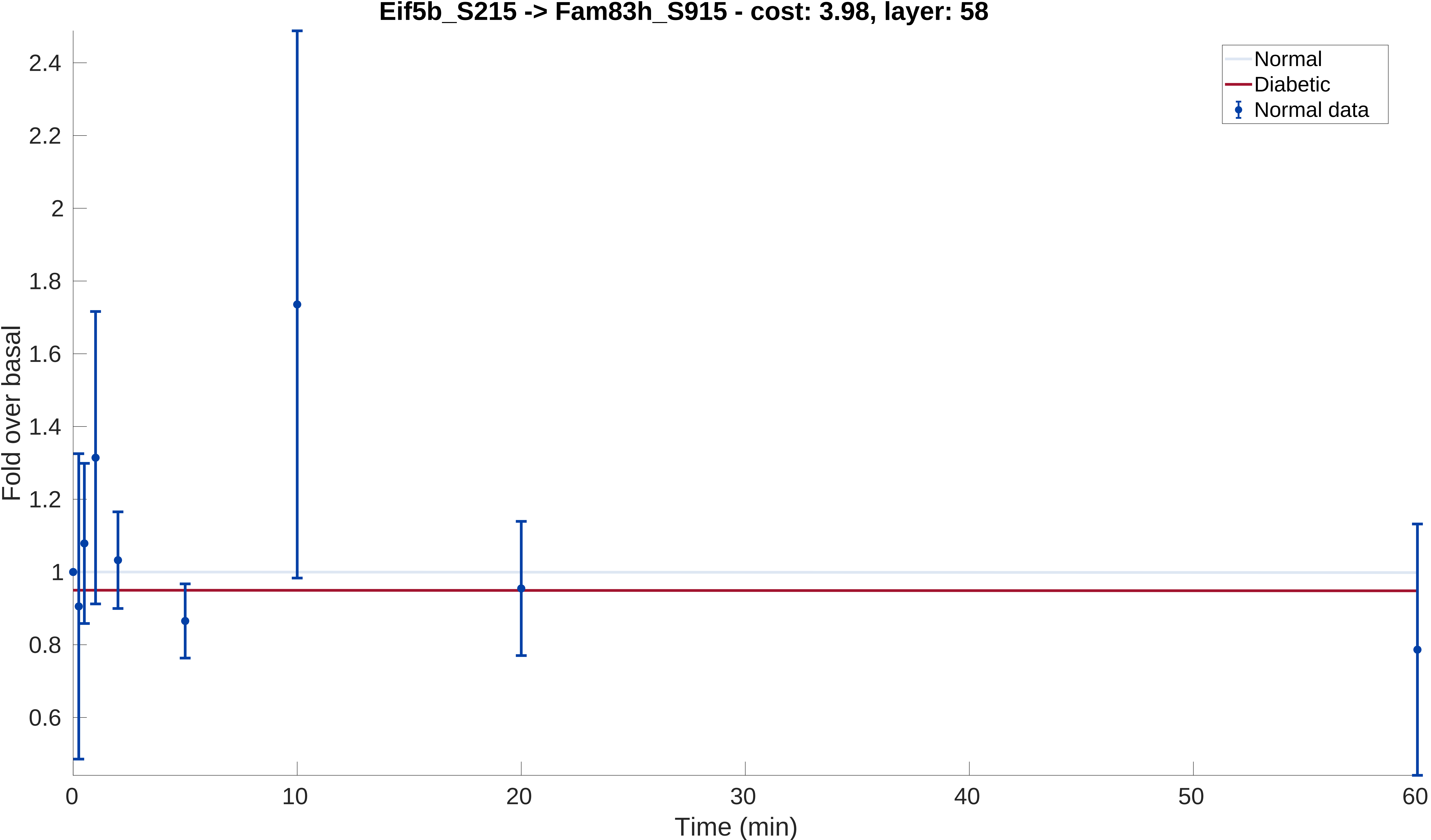

Eif5b_S215 -> Fam83h_S915 - cost: 3.98, layer: 58
Fold over basal
Time (min)
Normal
Diabetic
Normal data
2.4
2.2
2
1.8
1.6
1.4
1.2
1
0.8
0.6
0
10
20
30
40
50
60

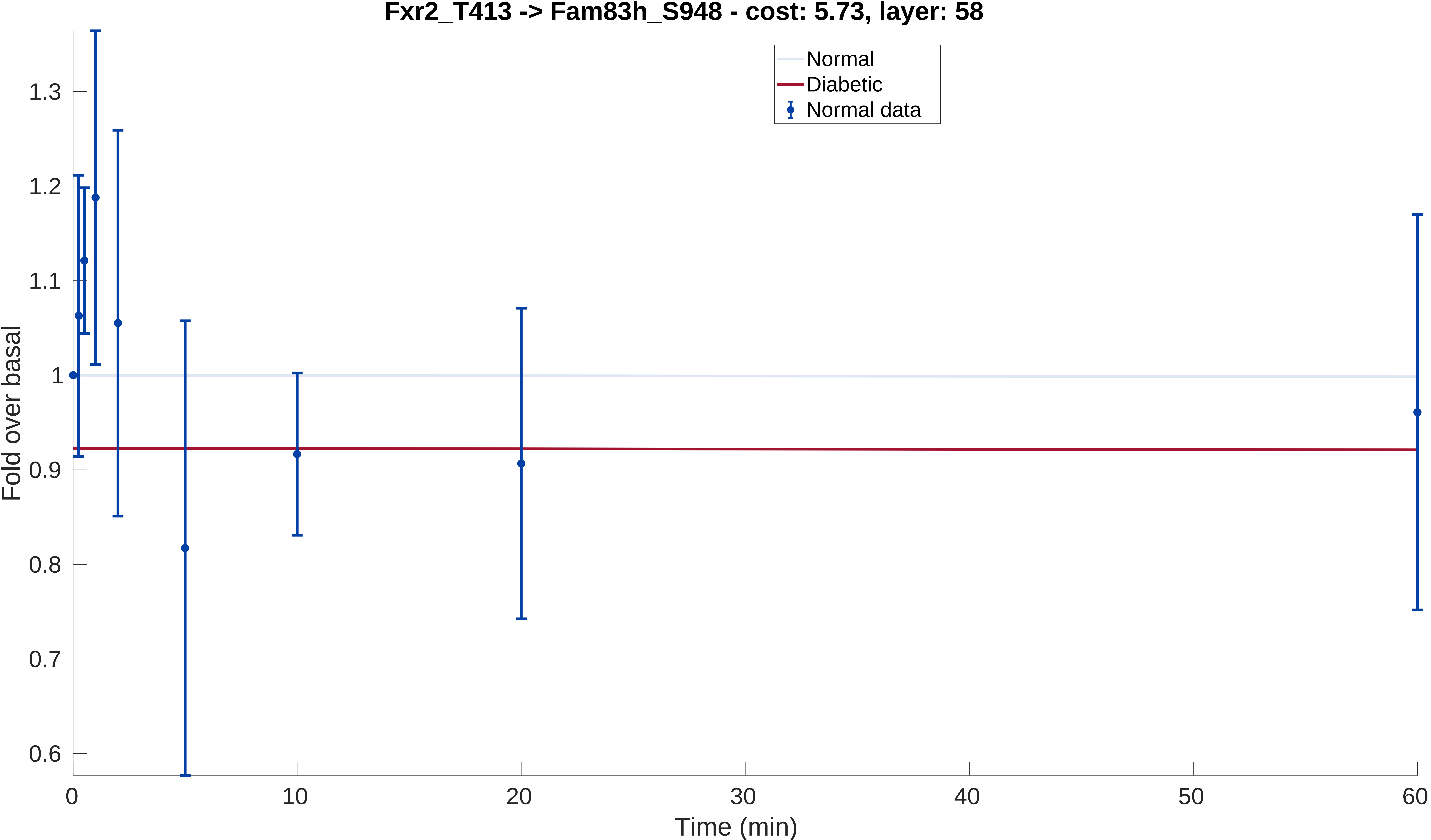

Fxr2_T413 -> Fam83h_S948 - cost: 5.73, layer: 58
Normal
Diabetic
Normal data
1.3
1.2
1.1
1
0.9
0.8
0.7
0.6
Fold over basal
0
10
20
30
40
50
60
Time (min)

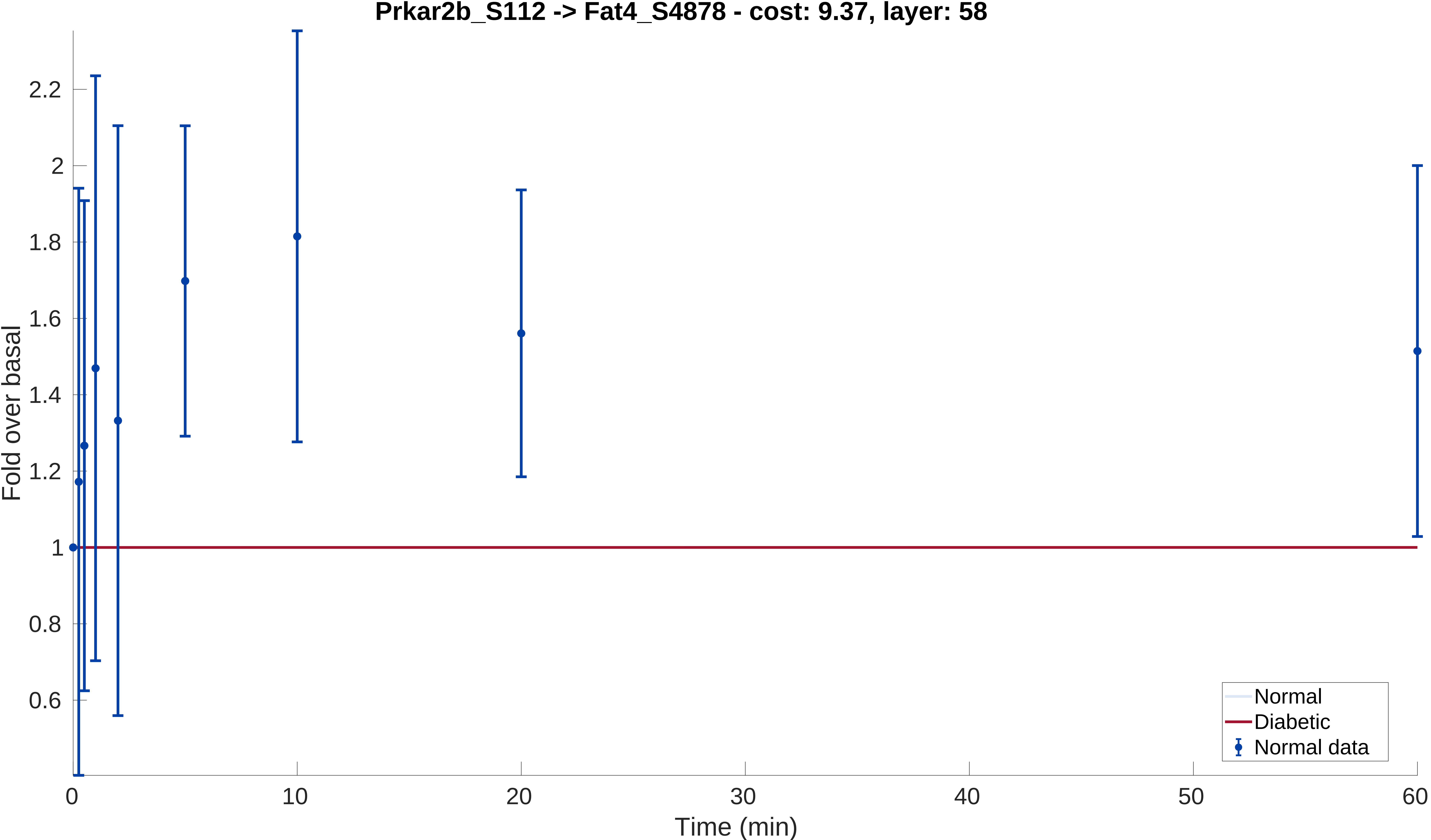

Prkar2b_S112 -> Fat4_S4878 - cost: 9.37, layer: 58
Fold over basal
2.2
2
1.8
1.6
1.4
1.2
1
0.8
0.6
0
10
20
30
40
50
60
Time (min)
Normal
Diabetic
Normal data

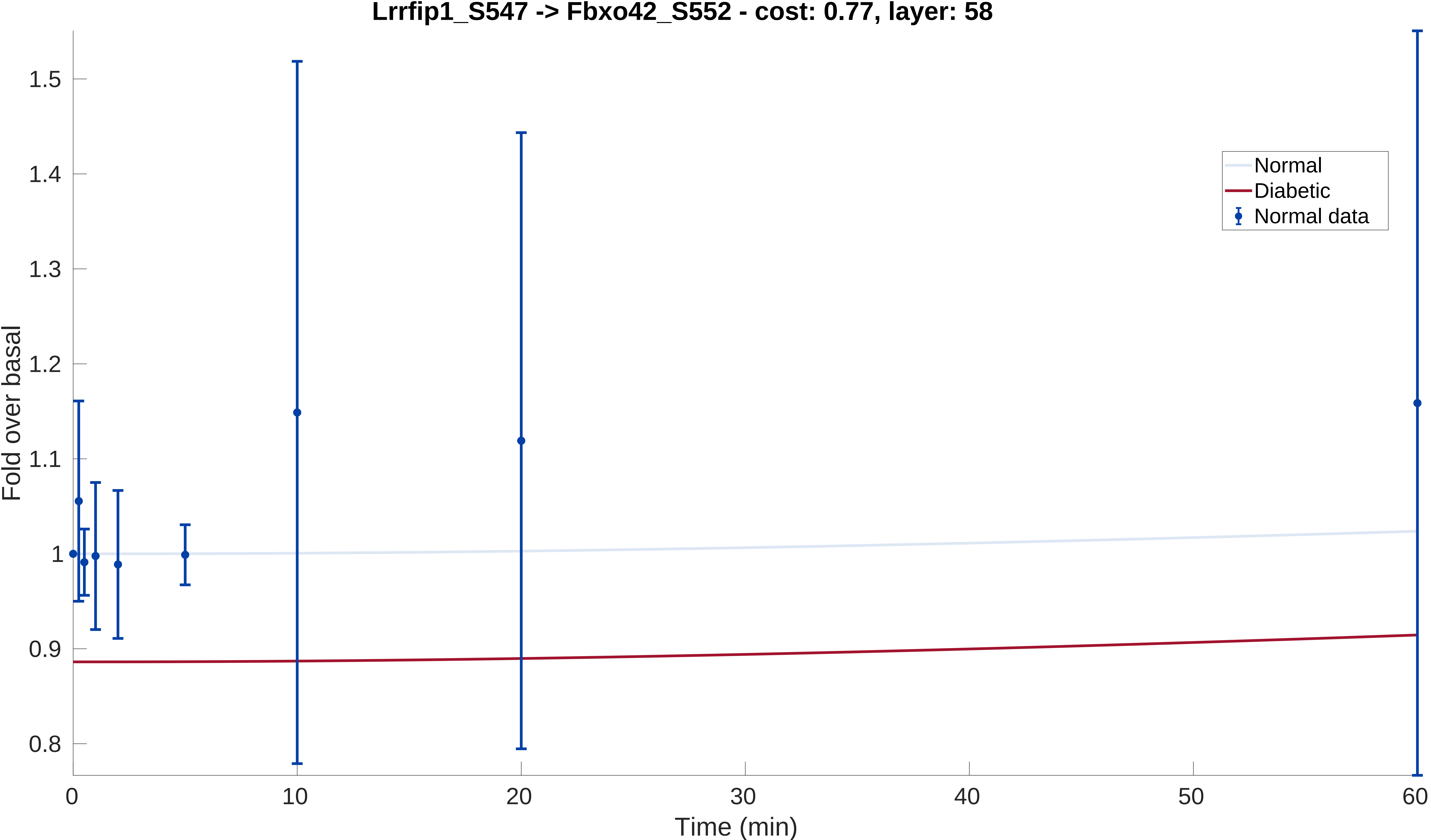
Lrrfip1_S547 -> Fbxo42_S552 - cost: 0.77, layer: 58
Fold over basal
Time (min)
Normal
Diabetic
Normal data
0.8
0.9
1
1.1
1.2
1.3
1.4
1.5
0
10
20
30
40
50
60

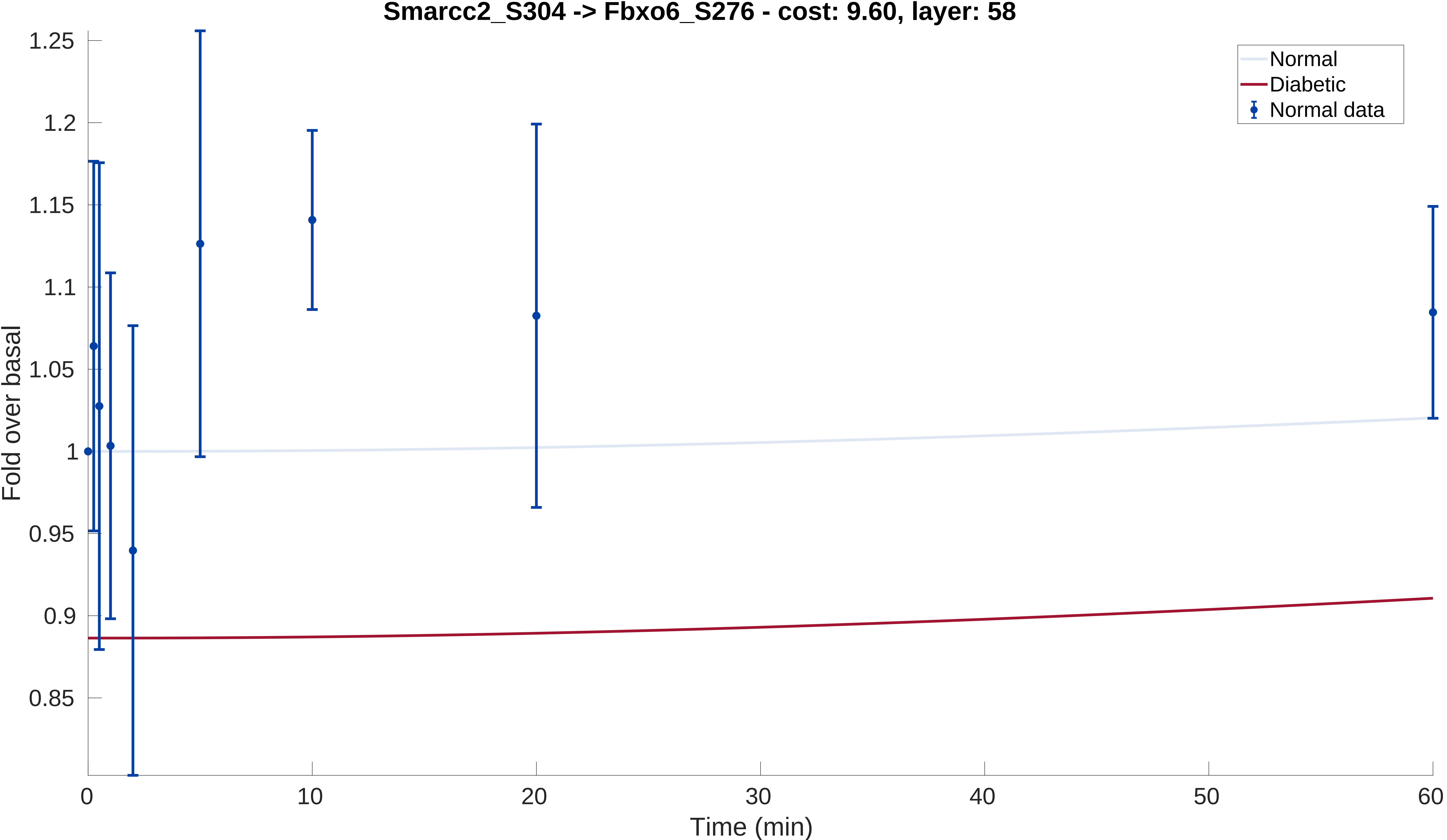

Smarcc2_S304 -> Fbxo6_S276 - cost: 9.60, layer: 58
Normal
Diabetic
Normal data
1.25
1.2
1.15
1.1
1.05
1
0.95
0.9
0.85
Fold over basal
0
10
20
30
40
50
60
Time (min)

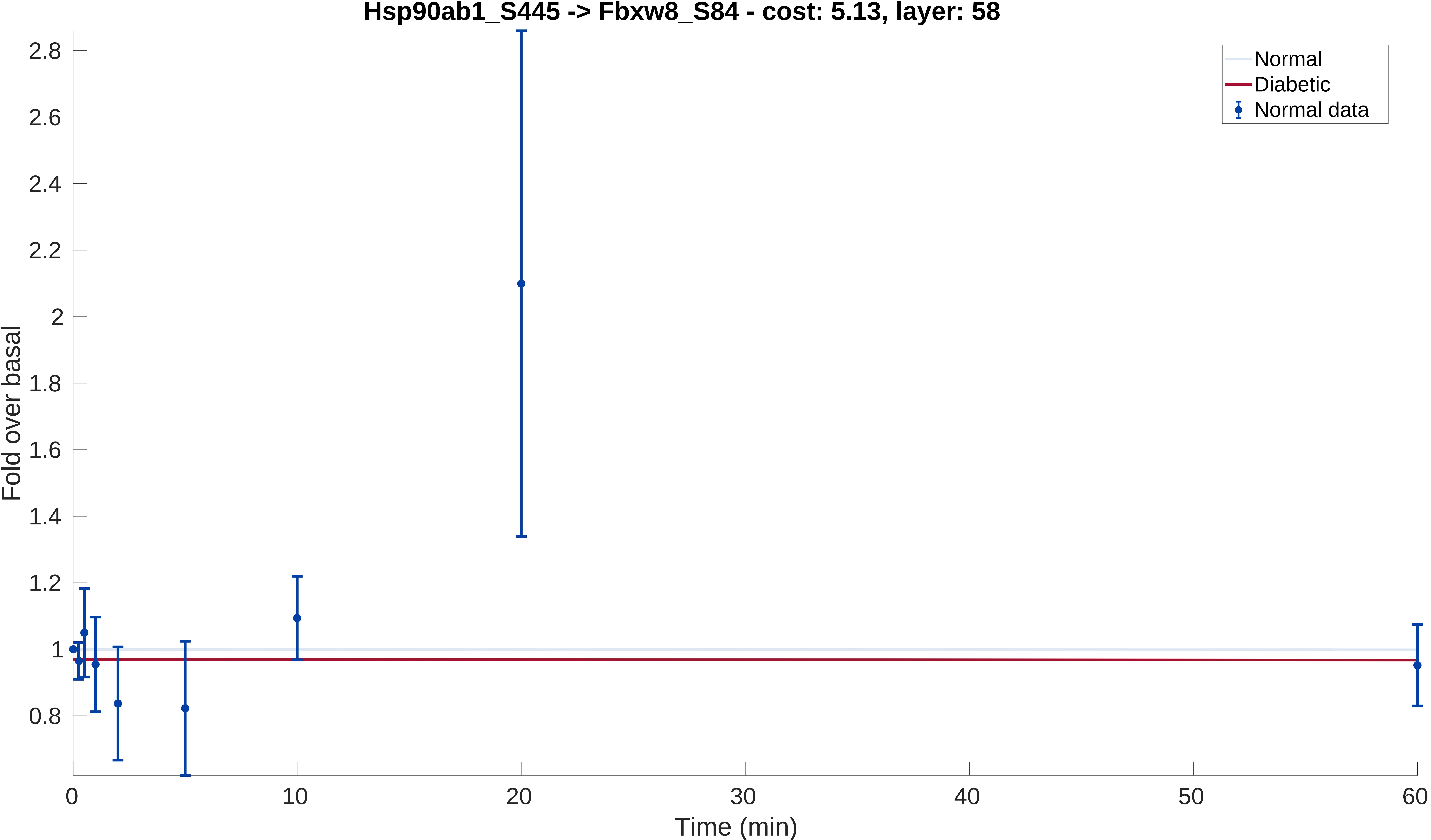

Hsp90ab1_S445 -> Fbxw8_S84 - cost: 5.13, layer: 58
Normal
Diabetic
Normal data
Fold over basal
2.8
2.6
2.4
2.2
2
1.8
1.6
1.4
1.2
1
0.8
0
10
20
30
40
50
60
Time (min)

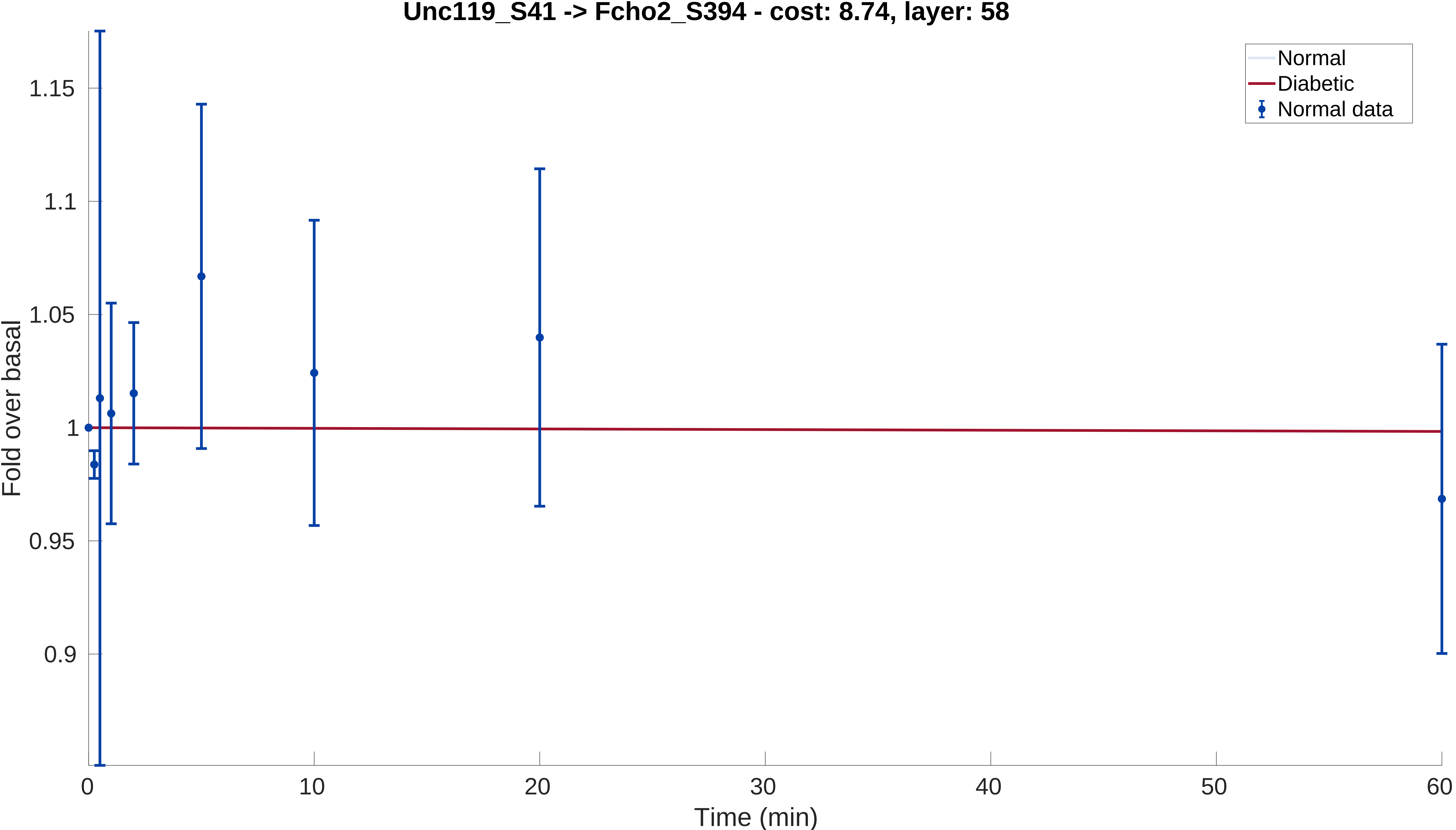
Unc119_S41 -> Fcho2_S394 - cost: 8.74, layer: 58
Normal
Diabetic
Normal data
Fold over basal
1.15
1.1
1.05
1
0.95
0.9
Time (min)
0
10
20
30
40
50
60

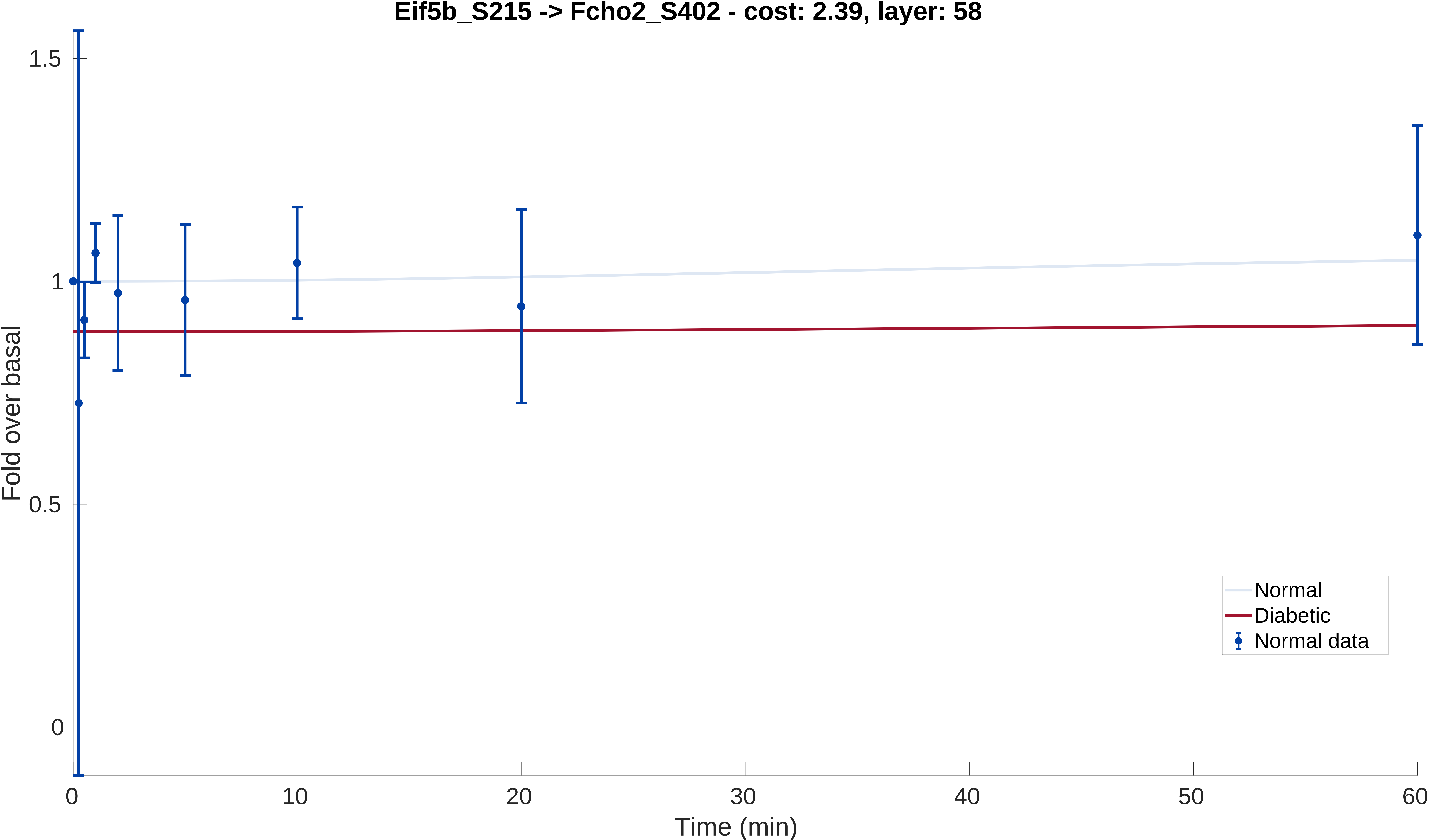
Eif5b_S215 -> Fcho2_S402 - cost: 2.39, layer: 58
Fold over basal
1.5
1
0.5
0
0
10
20
30
40
50
60
Time (min)
Normal
Diabetic
Normal data

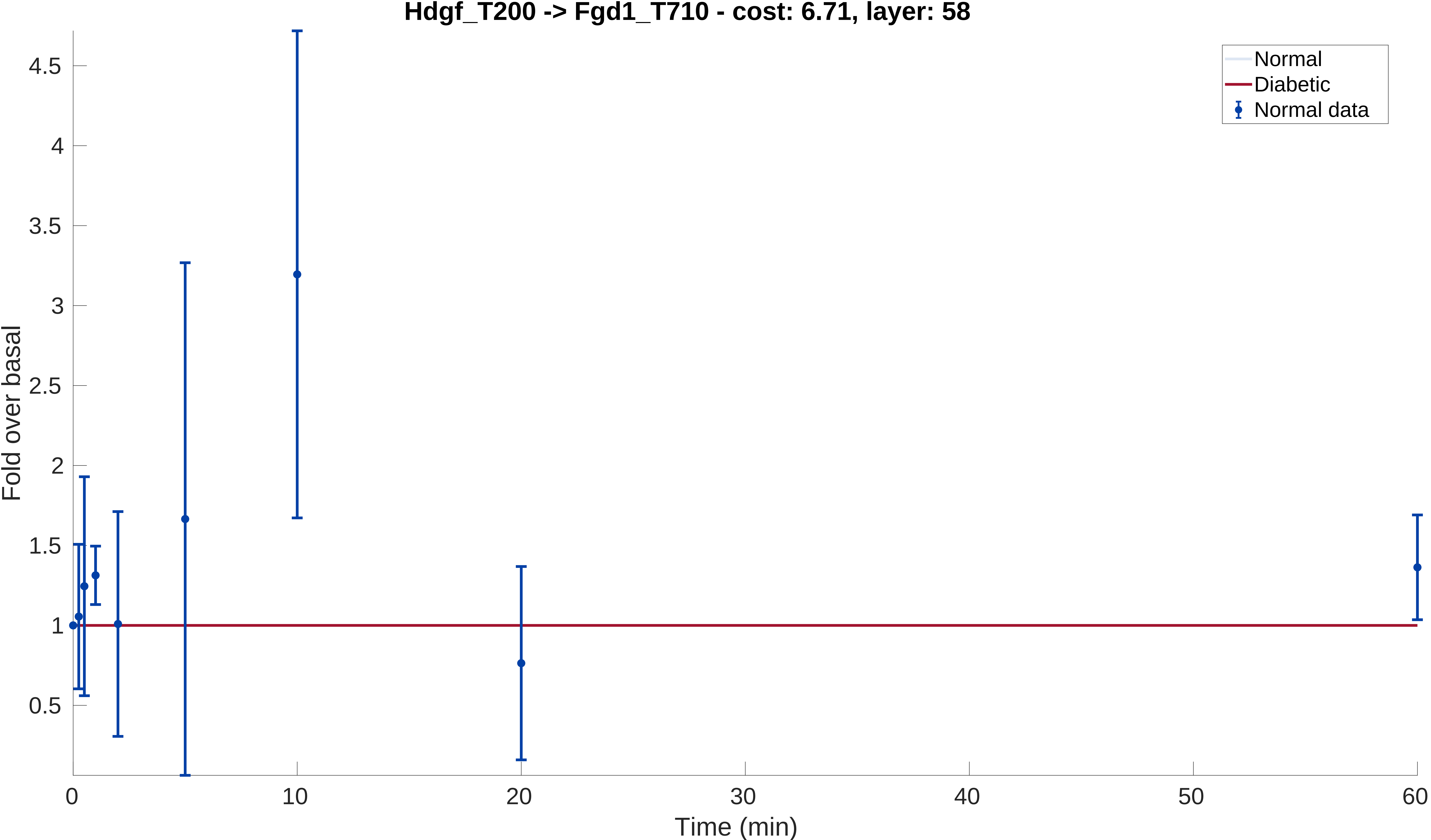
Hdgf_T200 -> Fgd1_T710 - cost: 6.71, layer: 58
Normal
Diabetic
Normal data
Fold over basal
Time (min)
0.5
1
1.5
2
2.5
3
3.5
4
4.5
0
10
20
30
40
50
60

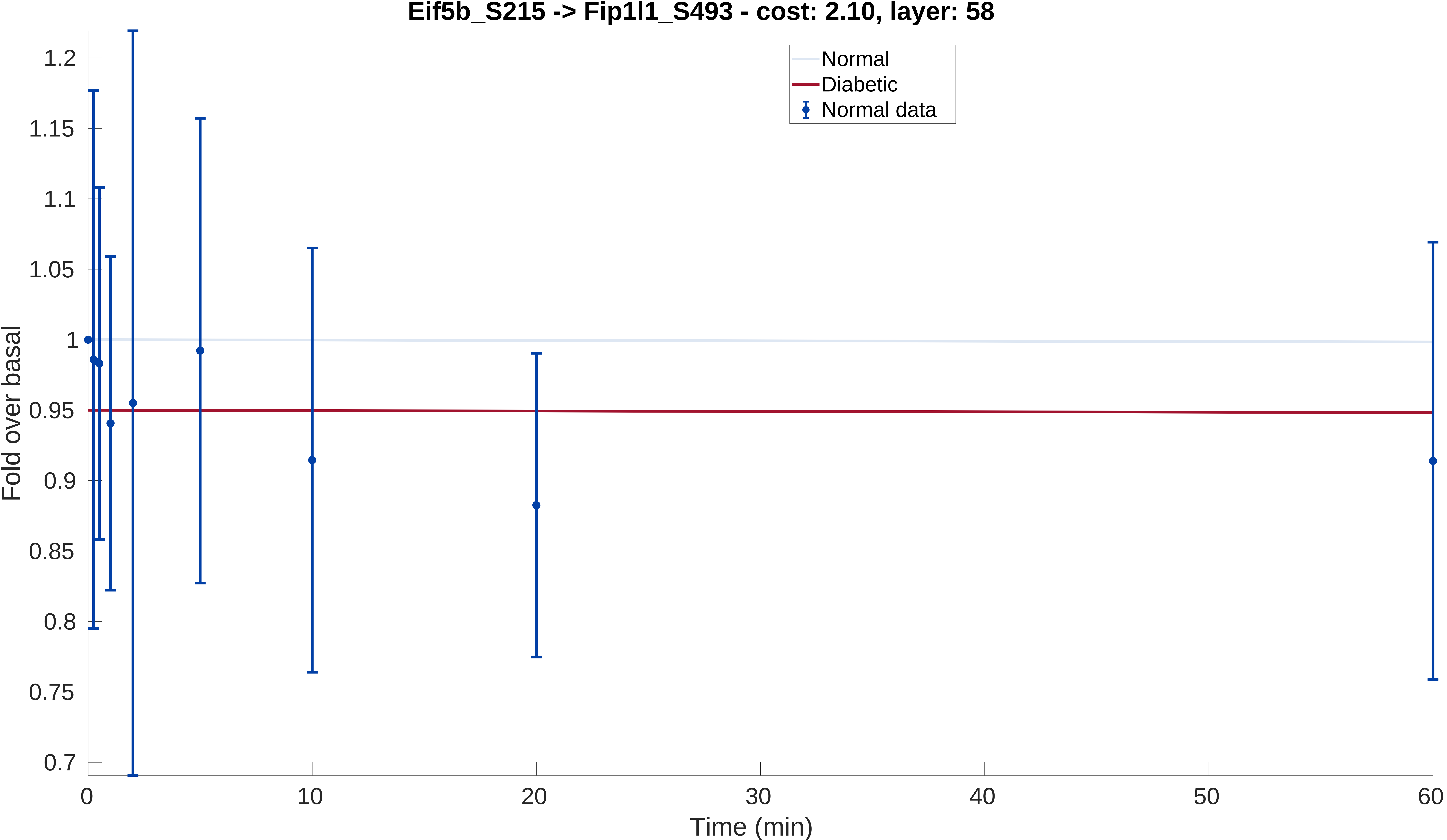
Eif5b_S215 -> Fip1l1_S493 - cost: 2.10, layer: 58
Normal
Diabetic
Normal data
Fold over basal
1.2
1.15
1.1
1.05
1
0.95
0.9
0.85
0.8
0.75
0.7
0
10
20
30
40
50
60
Time (min)

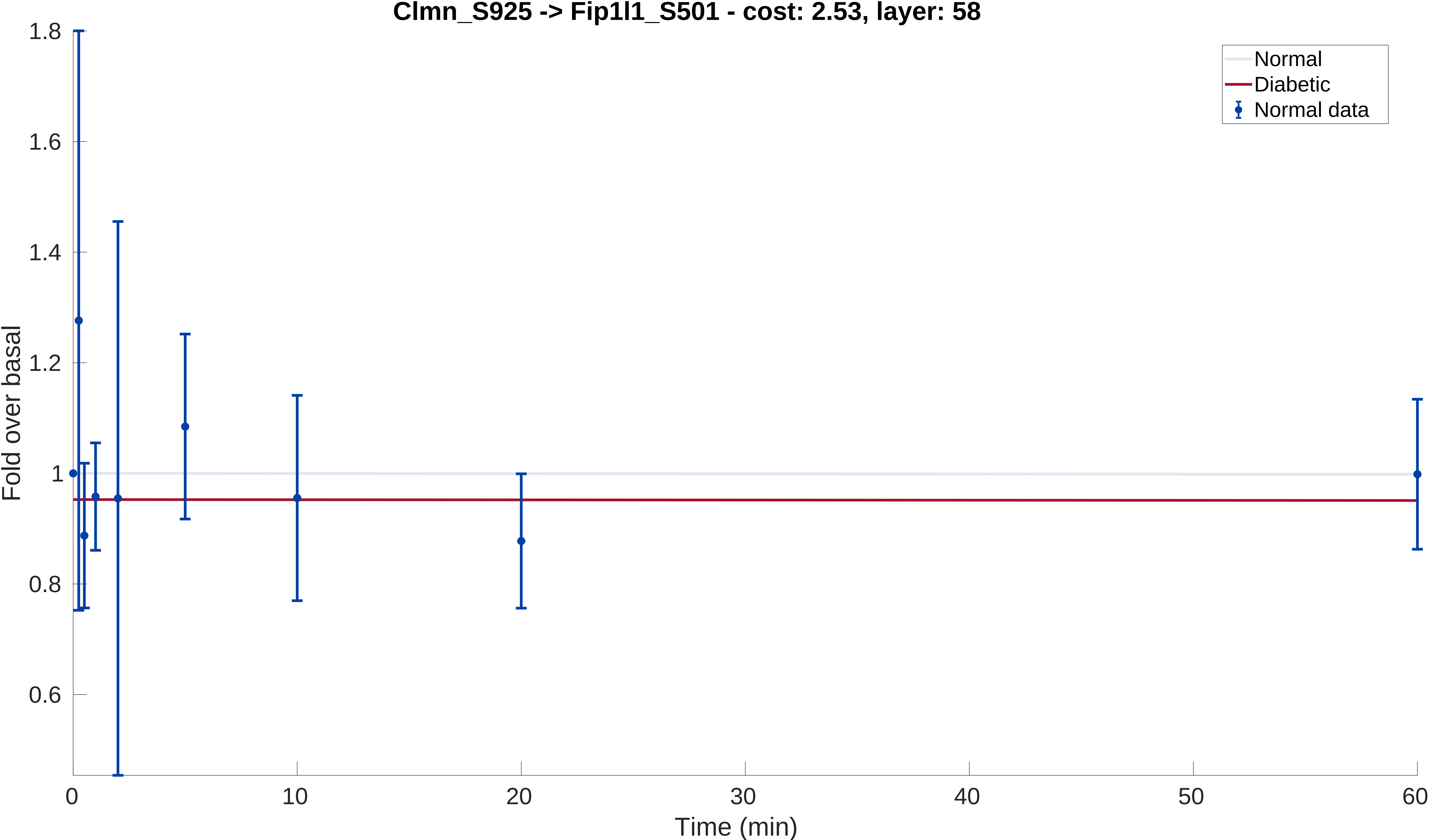

Clmn_S925 -> Fip1l1_S501 - cost: 2.53, layer: 58
Normal
Diabetic
Normal data
Fold over basal
1.8
1.6
1.4
1.2
1
0.8
0.6
0
10
20
30
40
50
60
Time (min)

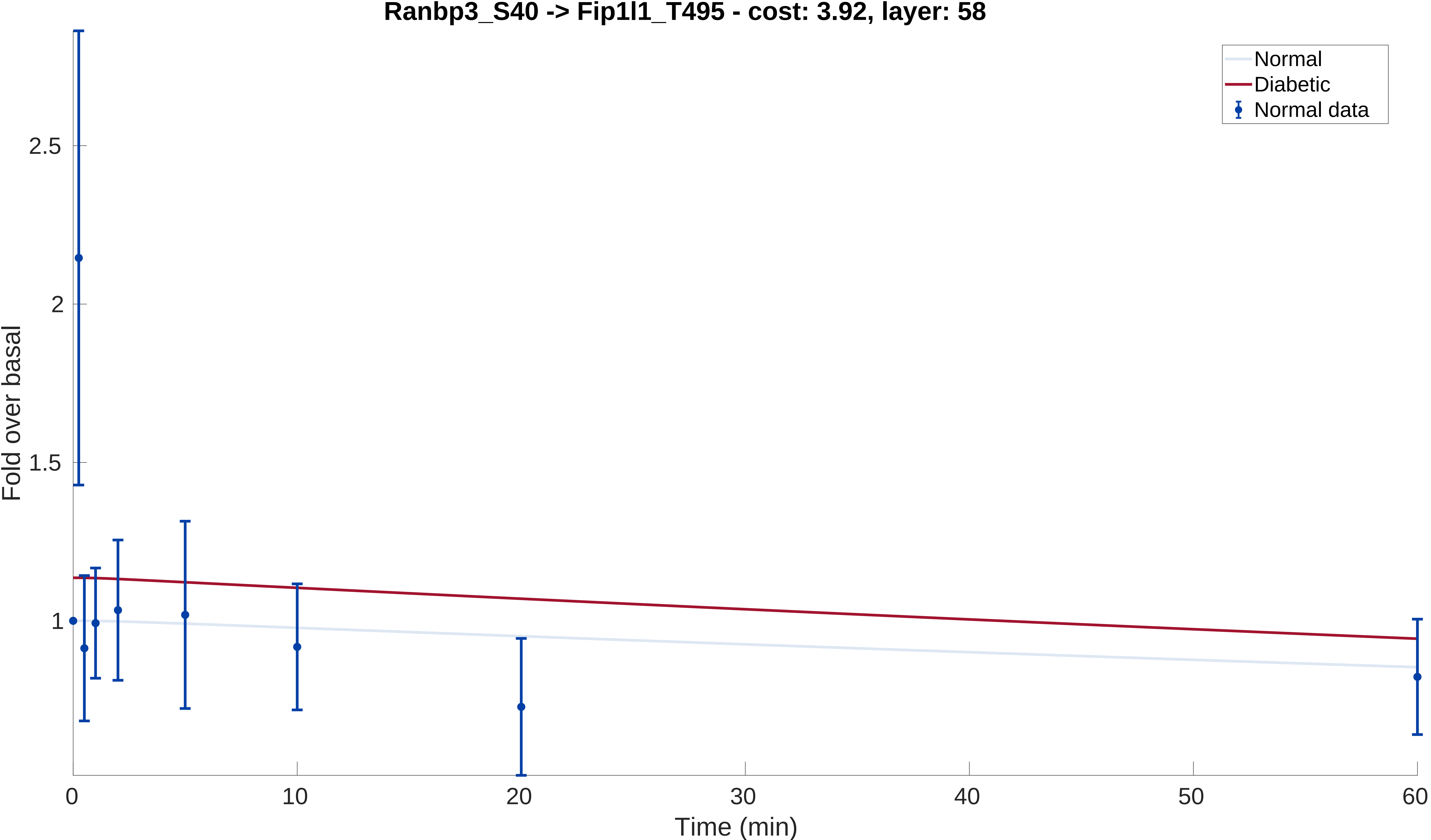

Ranbp3_S40 -> Fip1l1_T495 - cost: 3.92, layer: 58
Normal
Diabetic
Normal data
Fold over basal
2.5
2
1.5
1
0
10
20
30
40
50
60
Time (min)

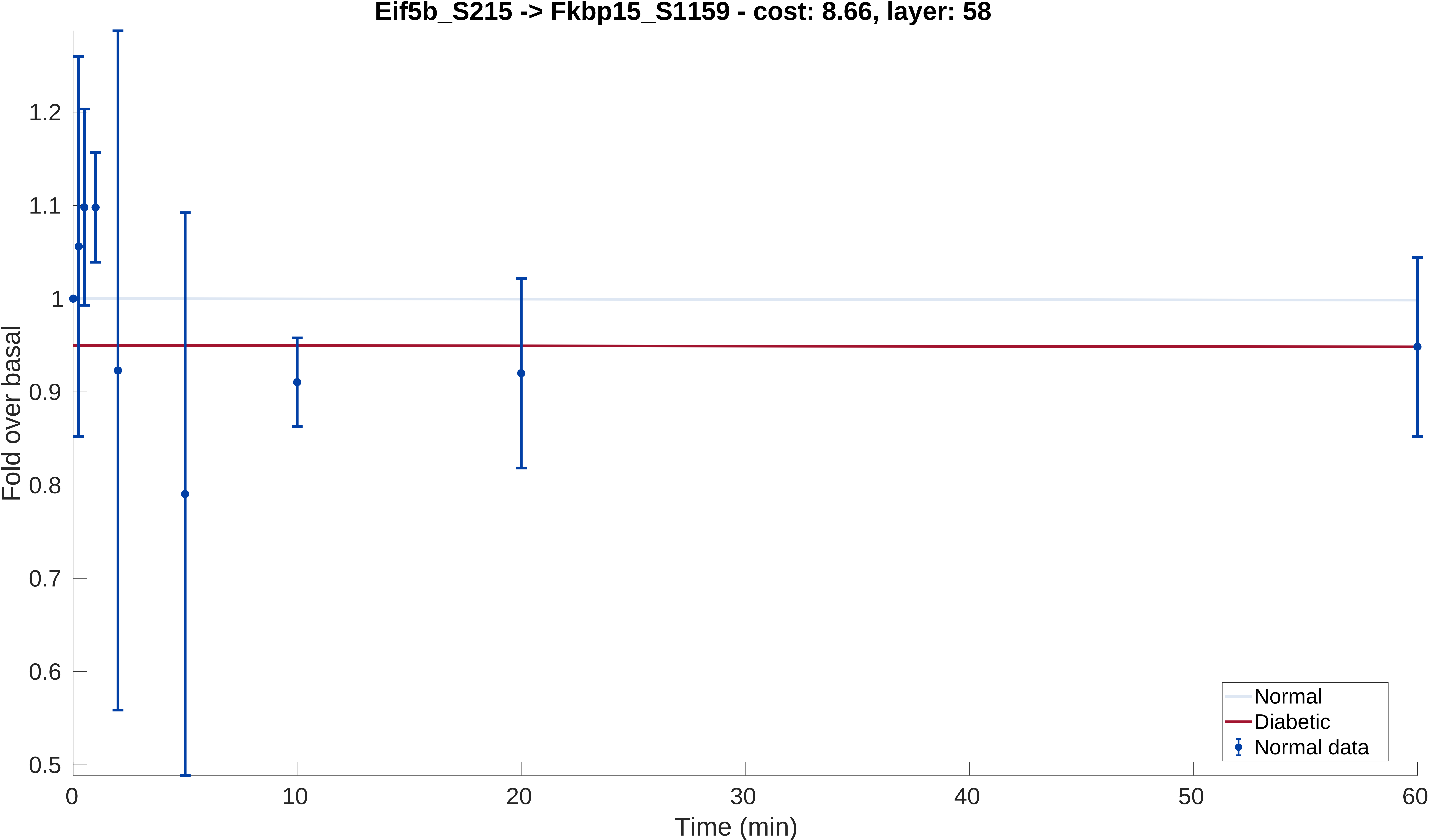

Eif5b_S215 -> Fkbp15_S1159 - cost: 8.66, layer: 58
Fold over basal
0.5
0.6
0.7
0.8
0.9
1
1.1
1.2
0
10
20
30
40
50
60
Time (min)
Normal
Diabetic
Normal data

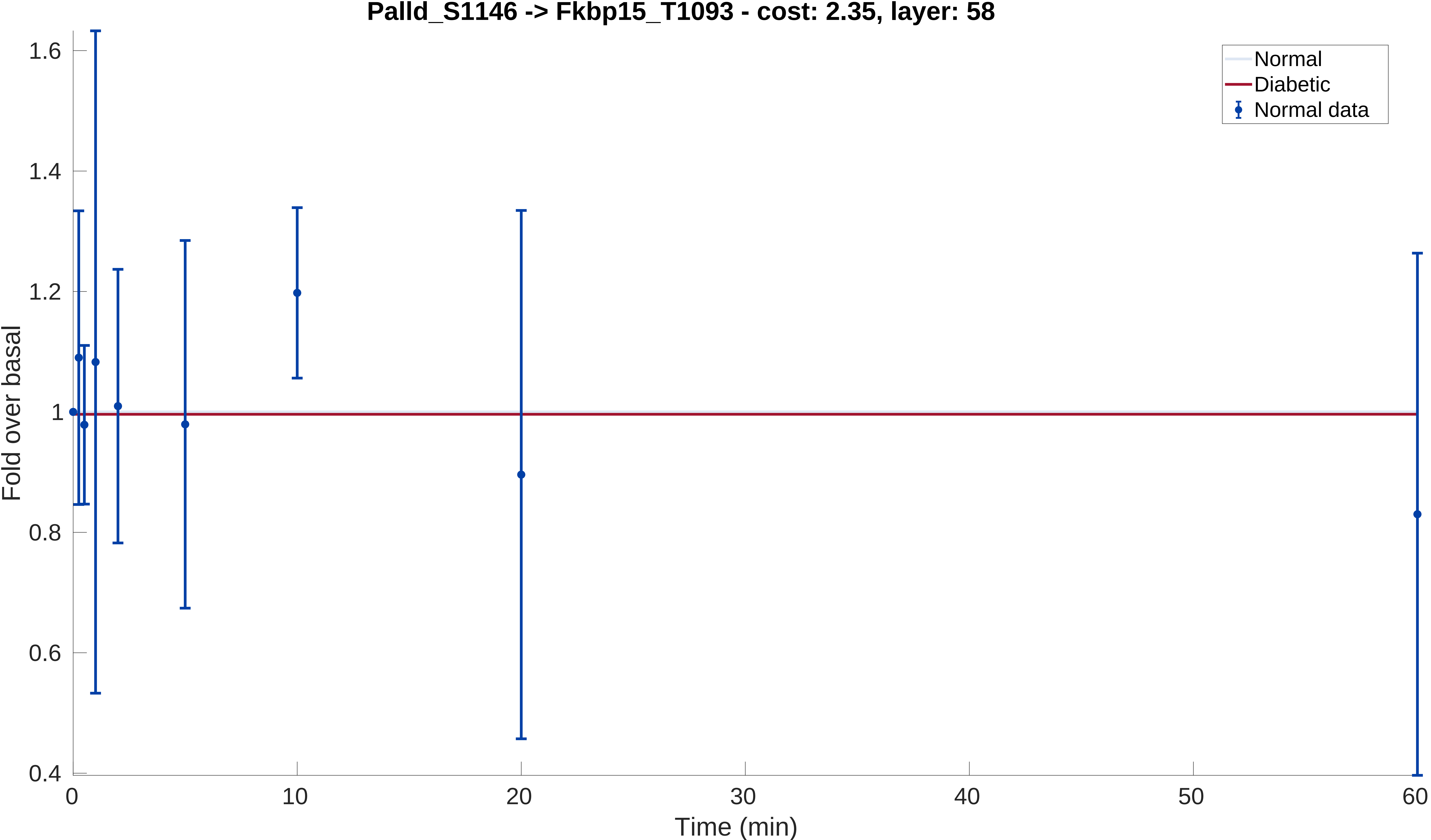
Palld_S1146 -> Fkbp15_T1093 - cost: 2.35, layer: 58
Normal
Diabetic
Normal data
Fold over basal
1.6
1.4
1.2
1
0.8
0.6
0.4
0
10
20
30
40
50
60
Time (min)

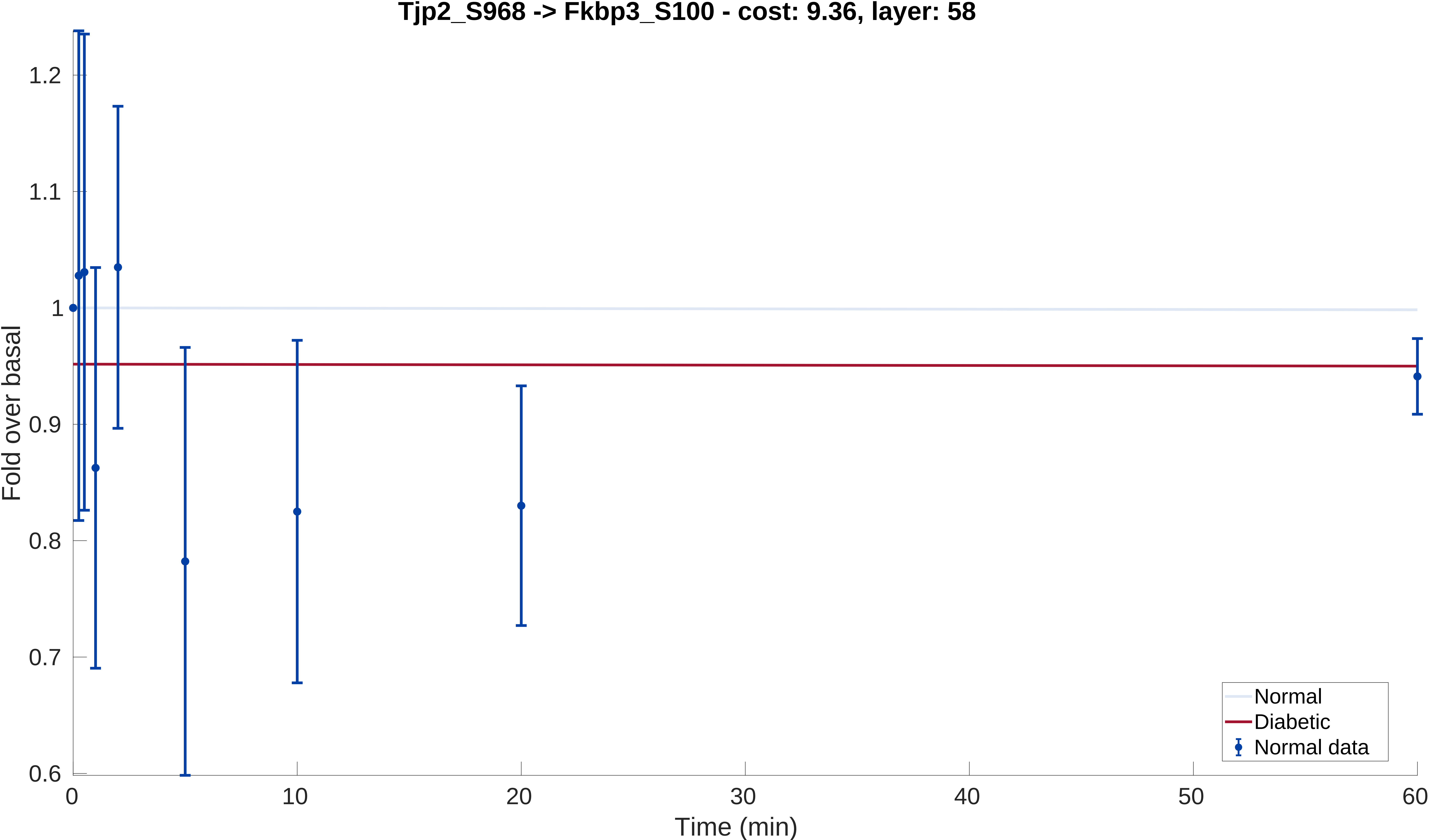

Tjp2_S968 -> Fkbp3_S100 - cost: 9.36, layer: 58
Fold over basal
1.2
1.1
1
0.9
0.8
0.7
0.6
0
10
20
30
40
50
60
Time (min)
Normal
Diabetic
Normal data

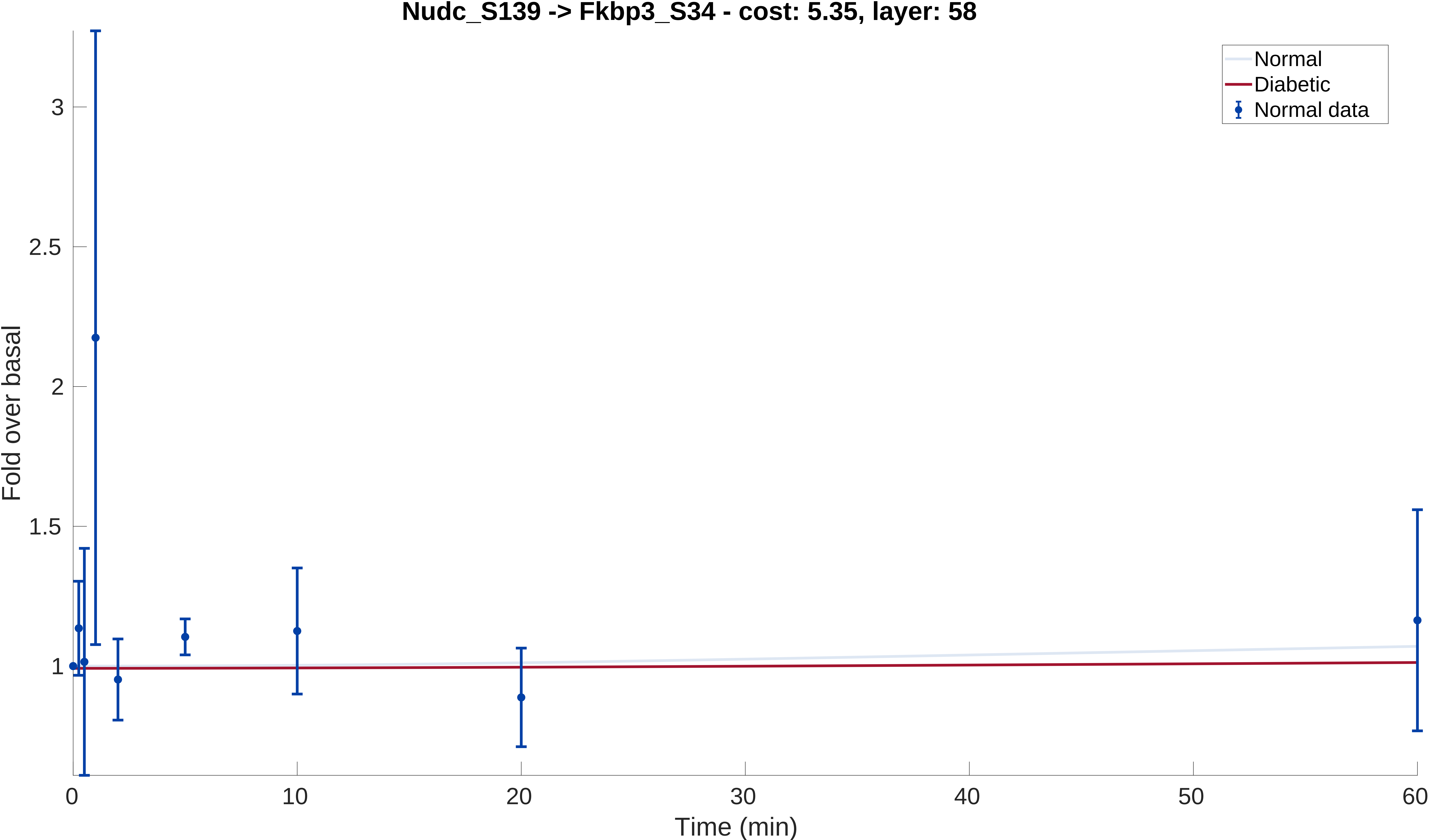

Nudc_S139 -> Fkbp3_S34 - cost: 5.35, layer: 58
Normal
Diabetic
Normal data
Fold over basal
3
2.5
2
1.5
1
0
10
20
30
40
50
60
Time (min)

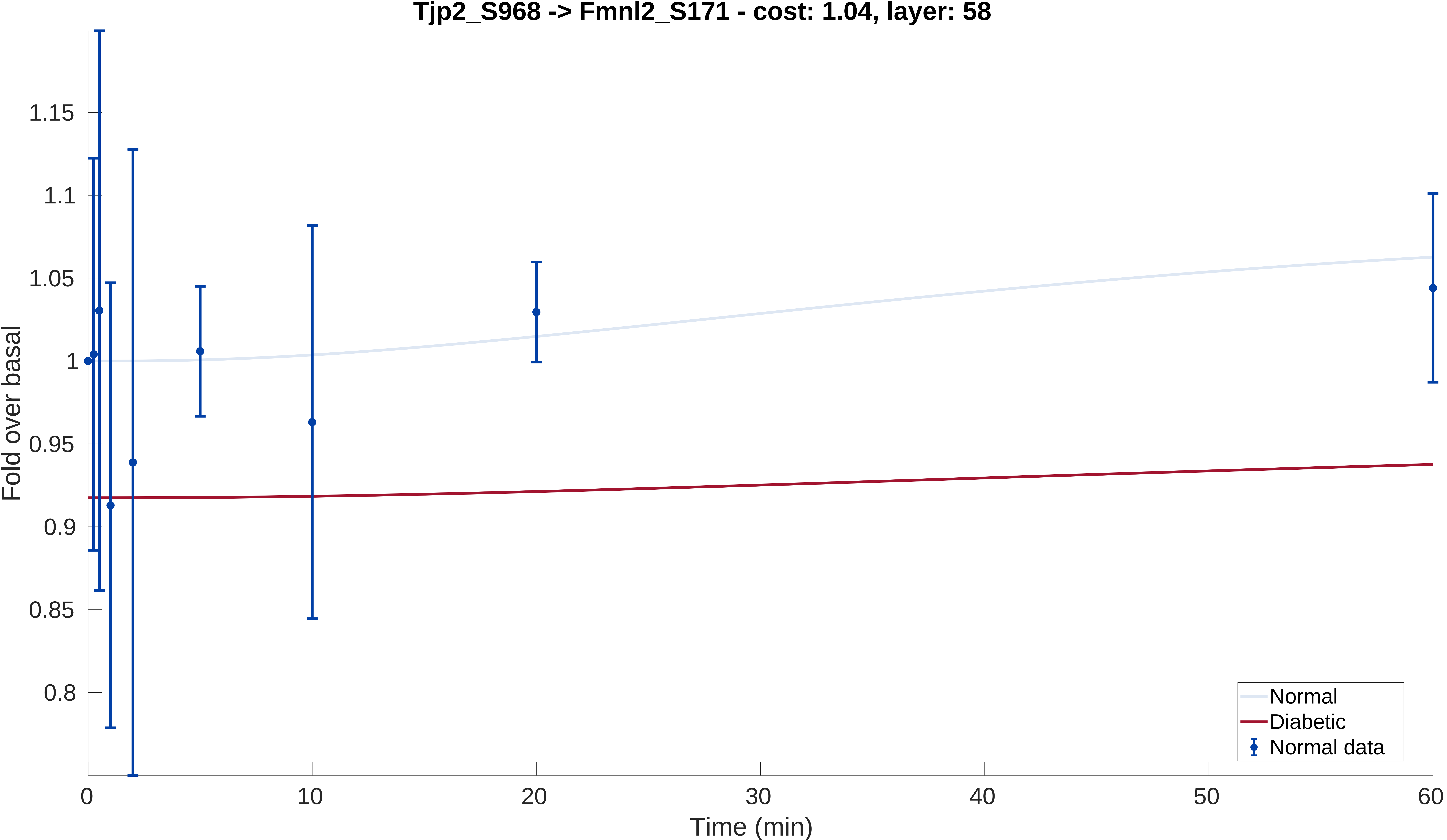

Tjp2_S968 -> Fmnl2_S171 - cost: 1.04, layer: 58
Fold over basal
1.15
1.1
1.05
1
0.95
0.9
0.85
0.8
0
10
20
30
40
50
60
Time (min)
Normal
Diabetic
Normal data

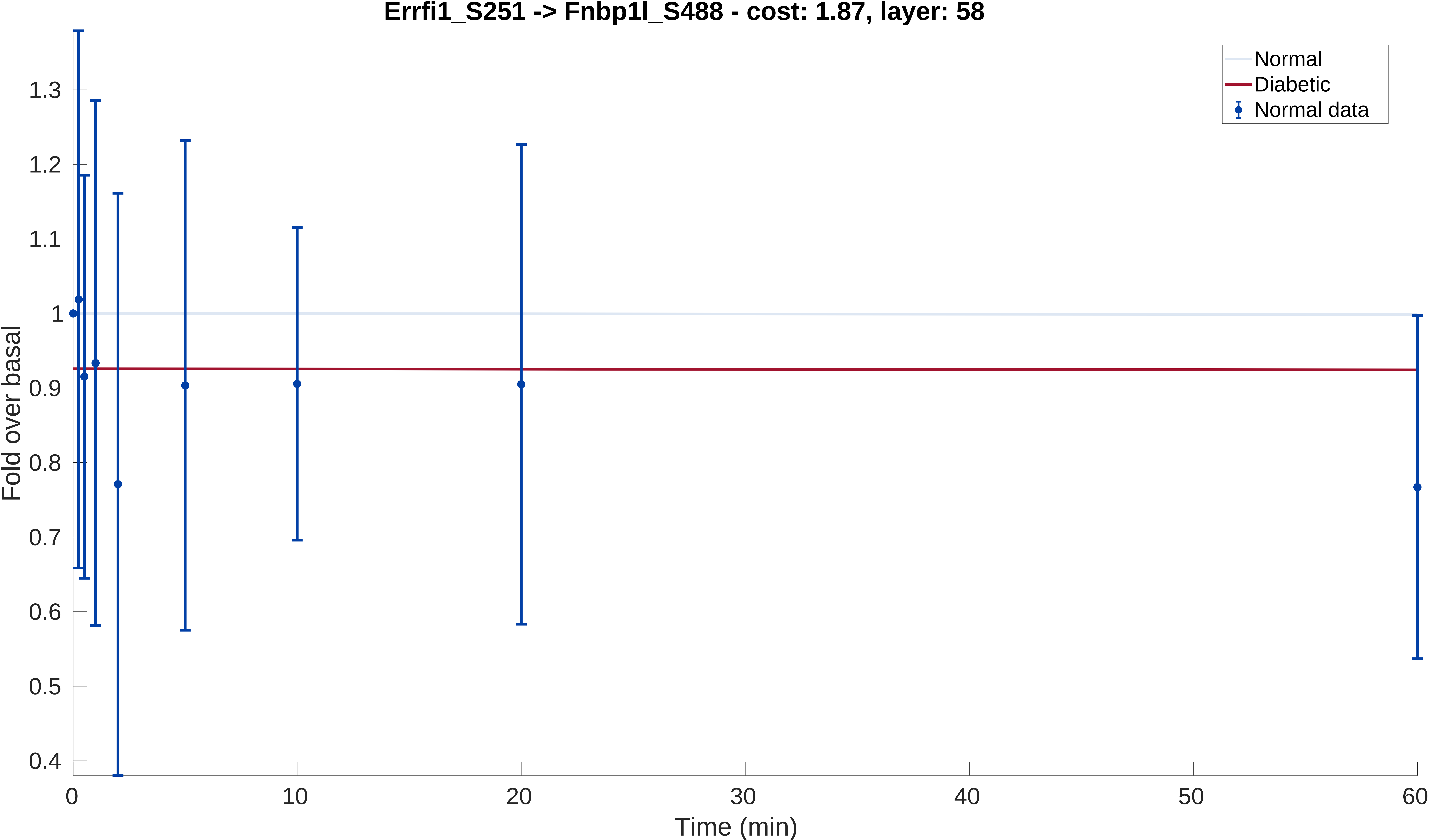

Errfi1_S251 -> Fnbp1l_S488 - cost: 1.87, layer: 58
Normal
Diabetic
Normal data
Fold over basal
1.3
1.2
1.1
1
0.9
0.8
0.7
0.6
0.5
0.4
0
10
20
30
40
50
60
Time (min)

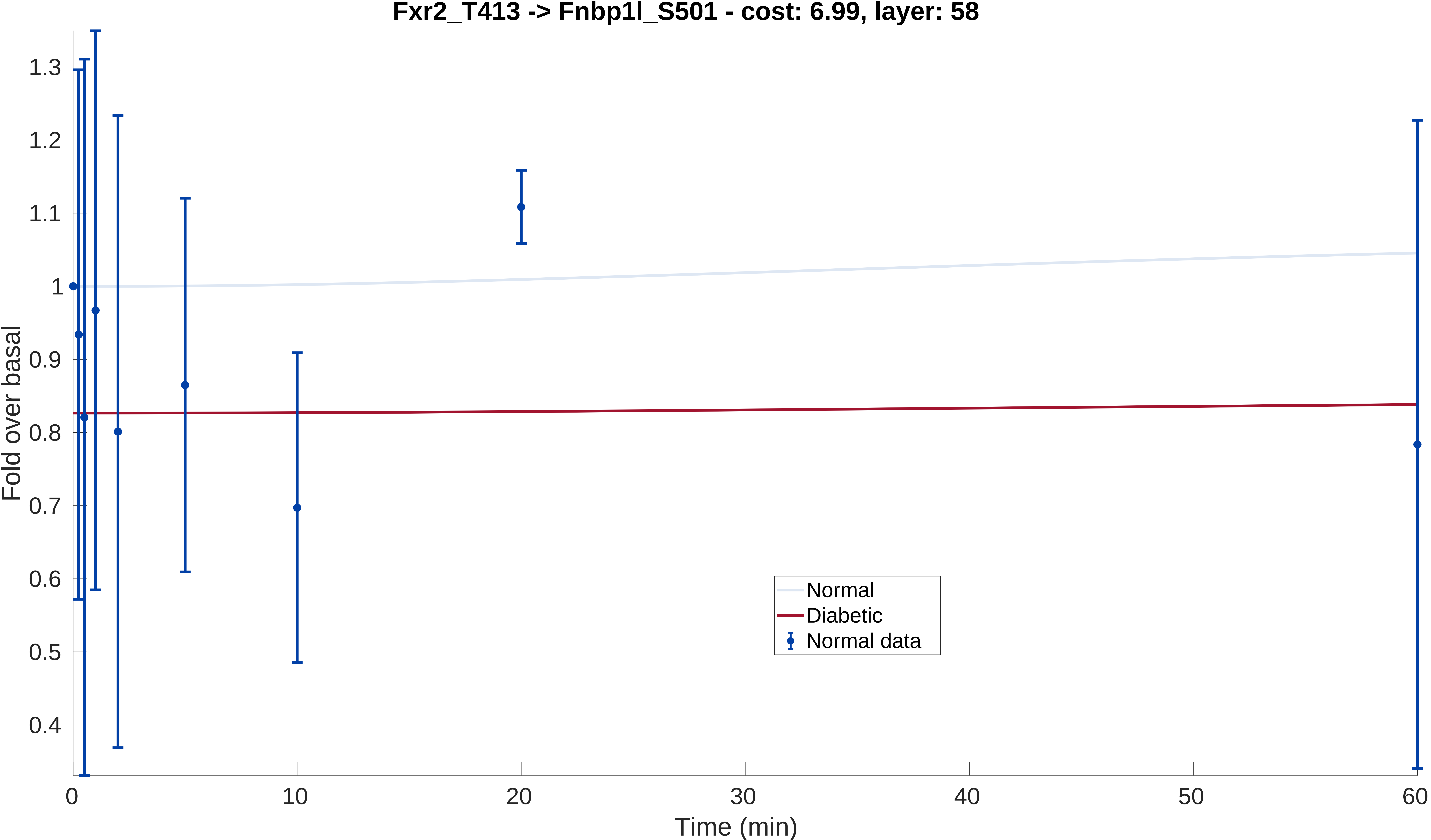

Fxr2_T413 -> Fnbp1l_S501 - cost: 6.99, layer: 58
Fold over basal
1.3
1.2
1.1
1
0.9
0.8
0.7
0.6
0.5
0.4
0
10
20
30
40
50
60
Time (min)
Normal
Diabetic
Normal data

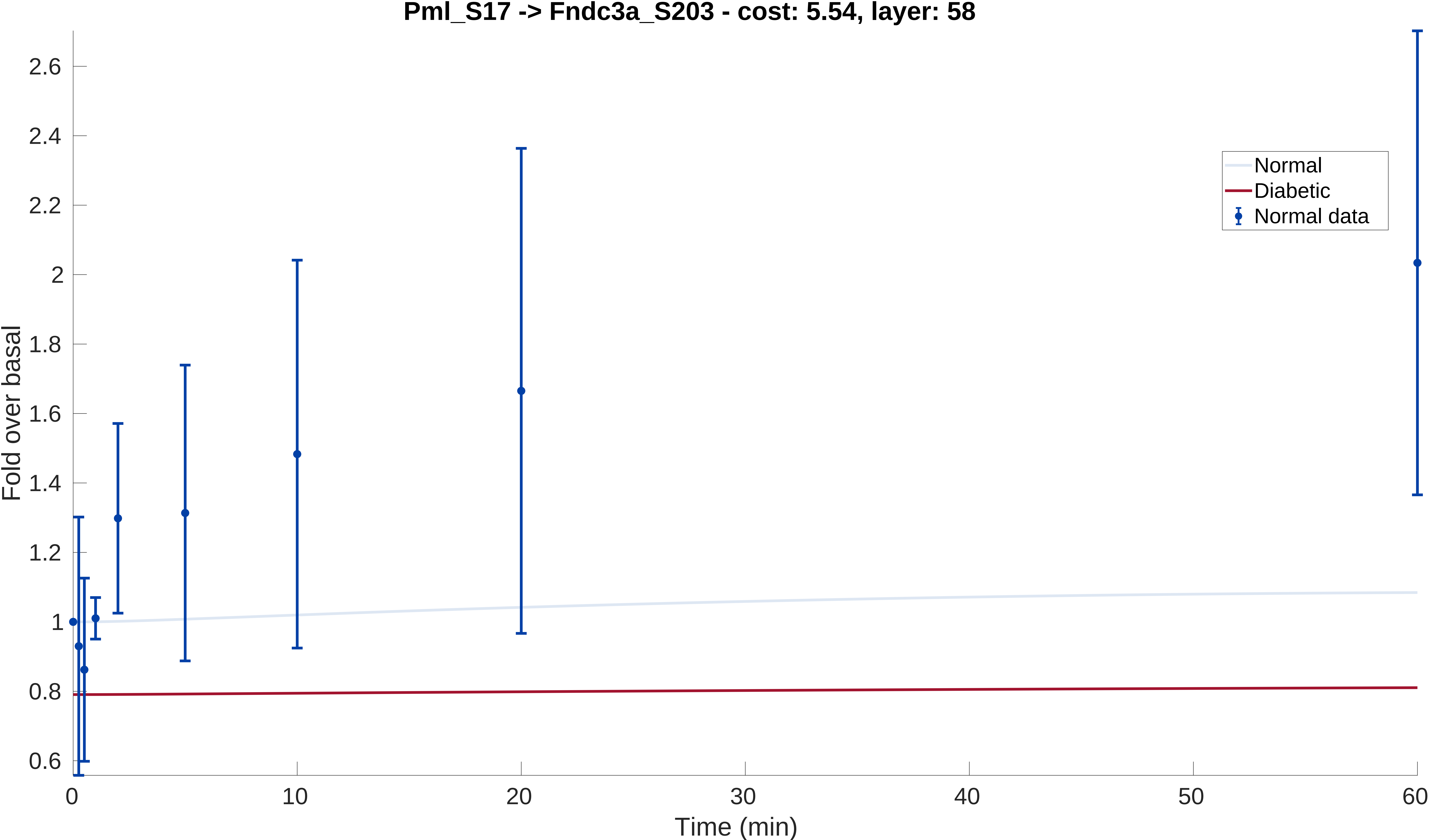

Pml_S17 -> Fndc3a_S203 - cost: 5.54, layer: 58
Fold over basal
Time (min)
Normal
Diabetic
Normal data
2.6
2.4
2.2
2
1.8
1.6
1.4
1.2
1
0.8
0.6
0
10
20
30
40
50
60

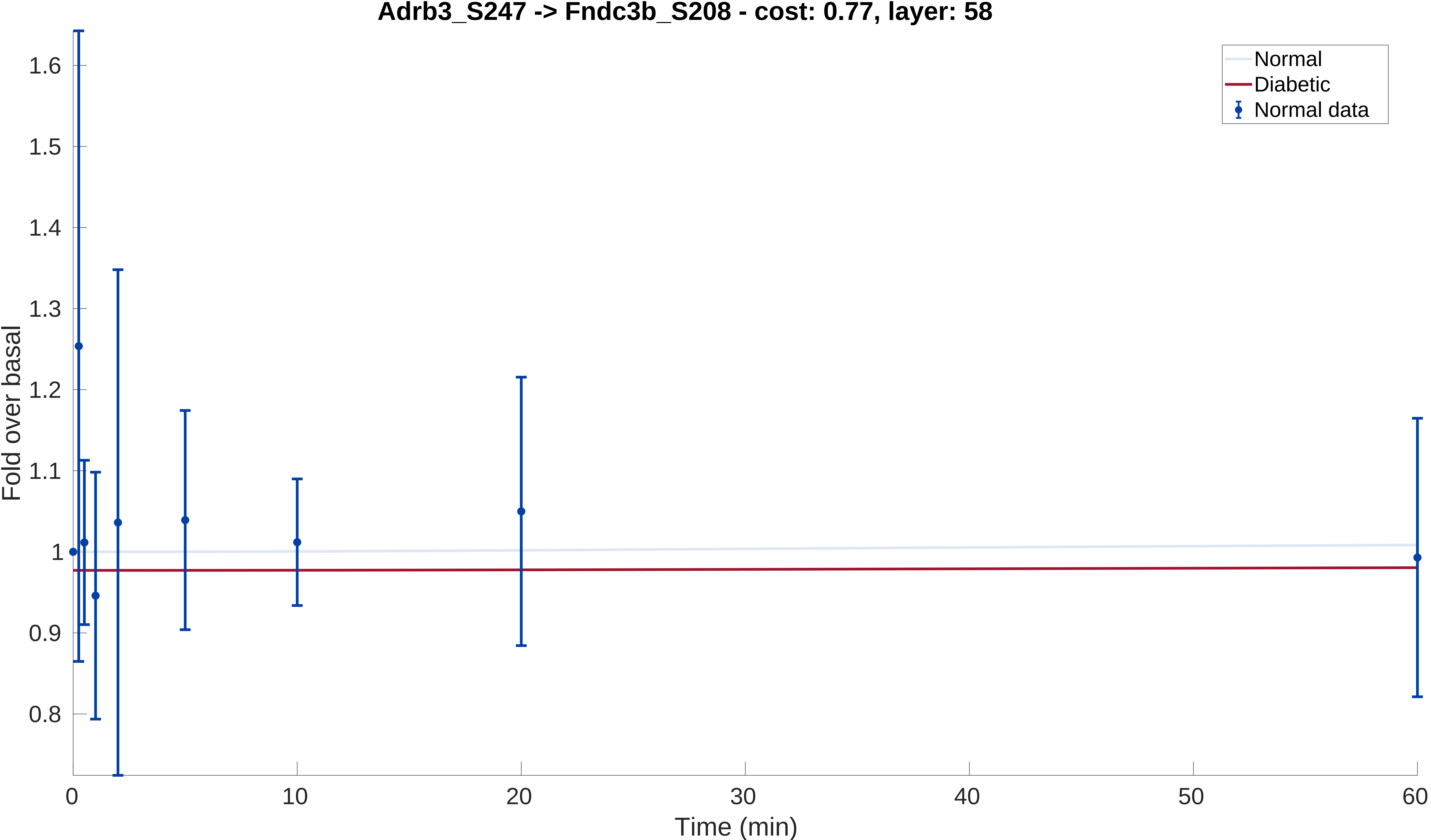

Adrb3_S247 -> Fndc3b_S208 - cost: 0.77, layer: 58
Normal
Diabetic
Normal data
Fold over basal
1.6
1.5
1.4
1.3
1.2
1.1
1
0.9
0.8
0
10
20
30
40
50
60
Time (min)

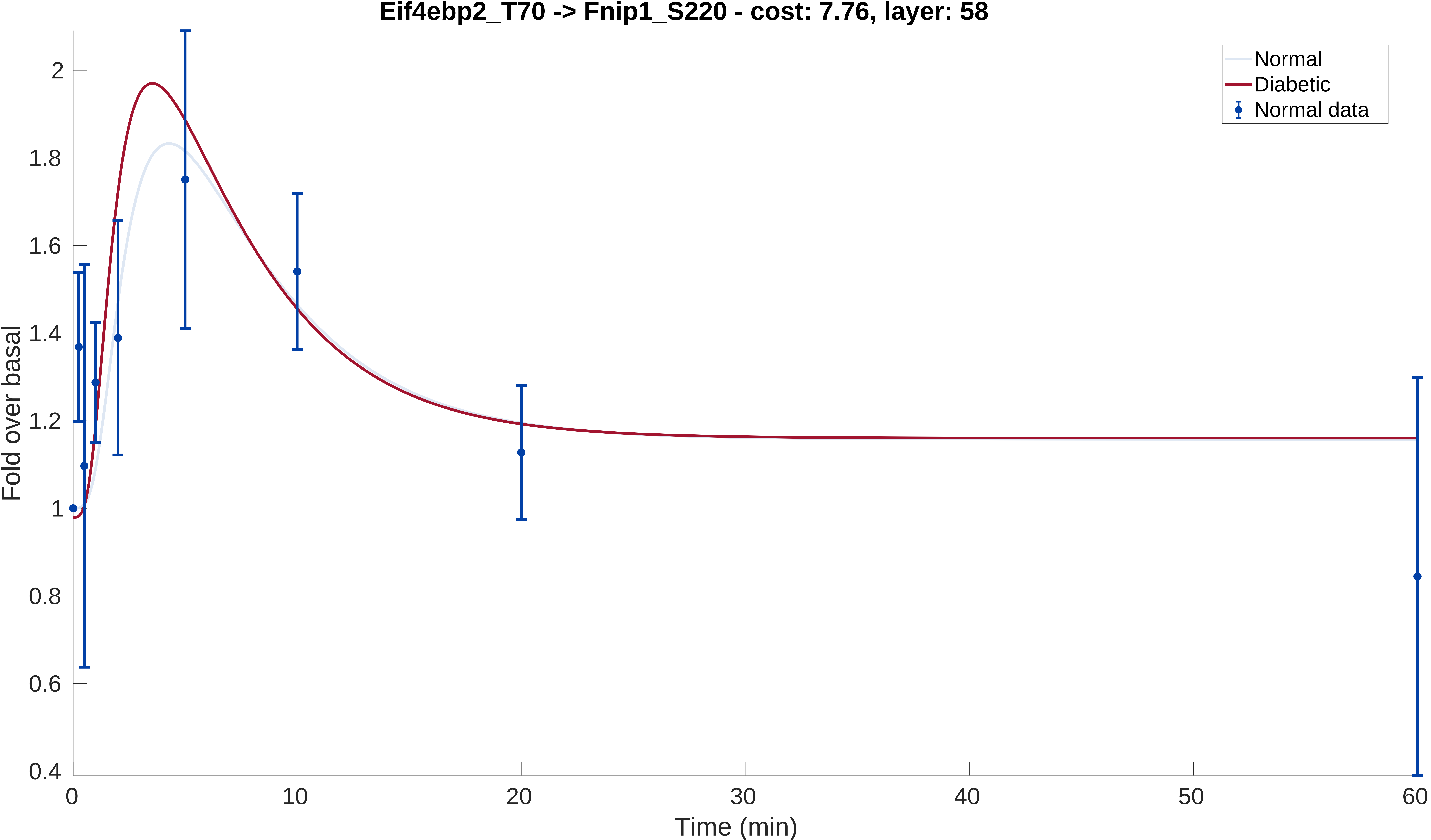

Eif4ebp2_T70 -> Fnip1_S220 - cost: 7.76, layer: 58
Normal
Diabetic
Normal data
Fold over basal
2
1.8
1.6
1.4
1.2
1
0.8
0.6
0.4
0
10
20
30
40
50
60
Time (min)

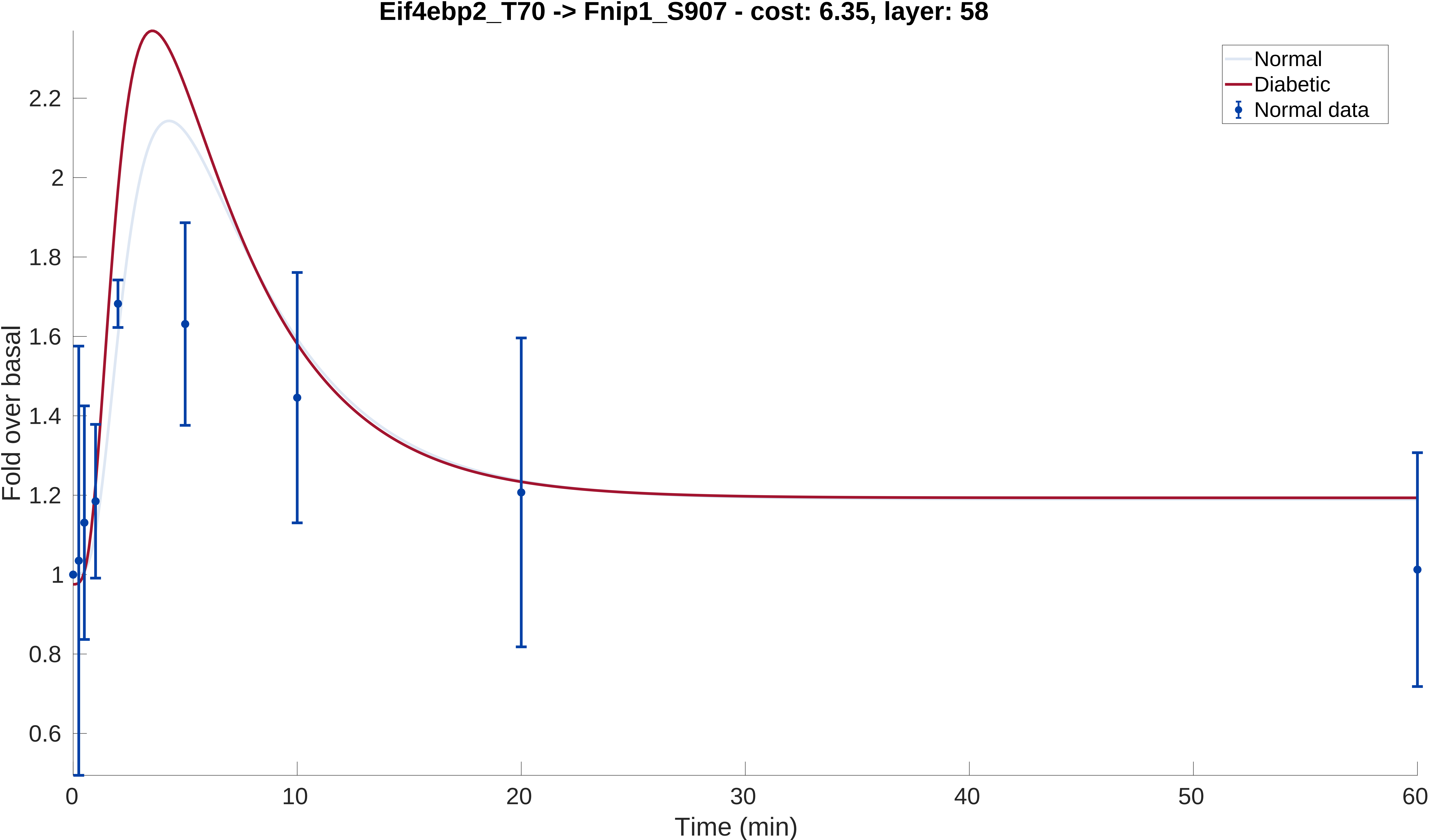
Eif4ebp2_T70 -> Fnip1_S907 - cost: 6.35, layer: 58
Normal
Diabetic
Normal data
Fold over basal
0.6
0.8
1
1.2
1.4
1.6
1.8
2
2.2
0
10
20
30
40
50
60
Time (min)

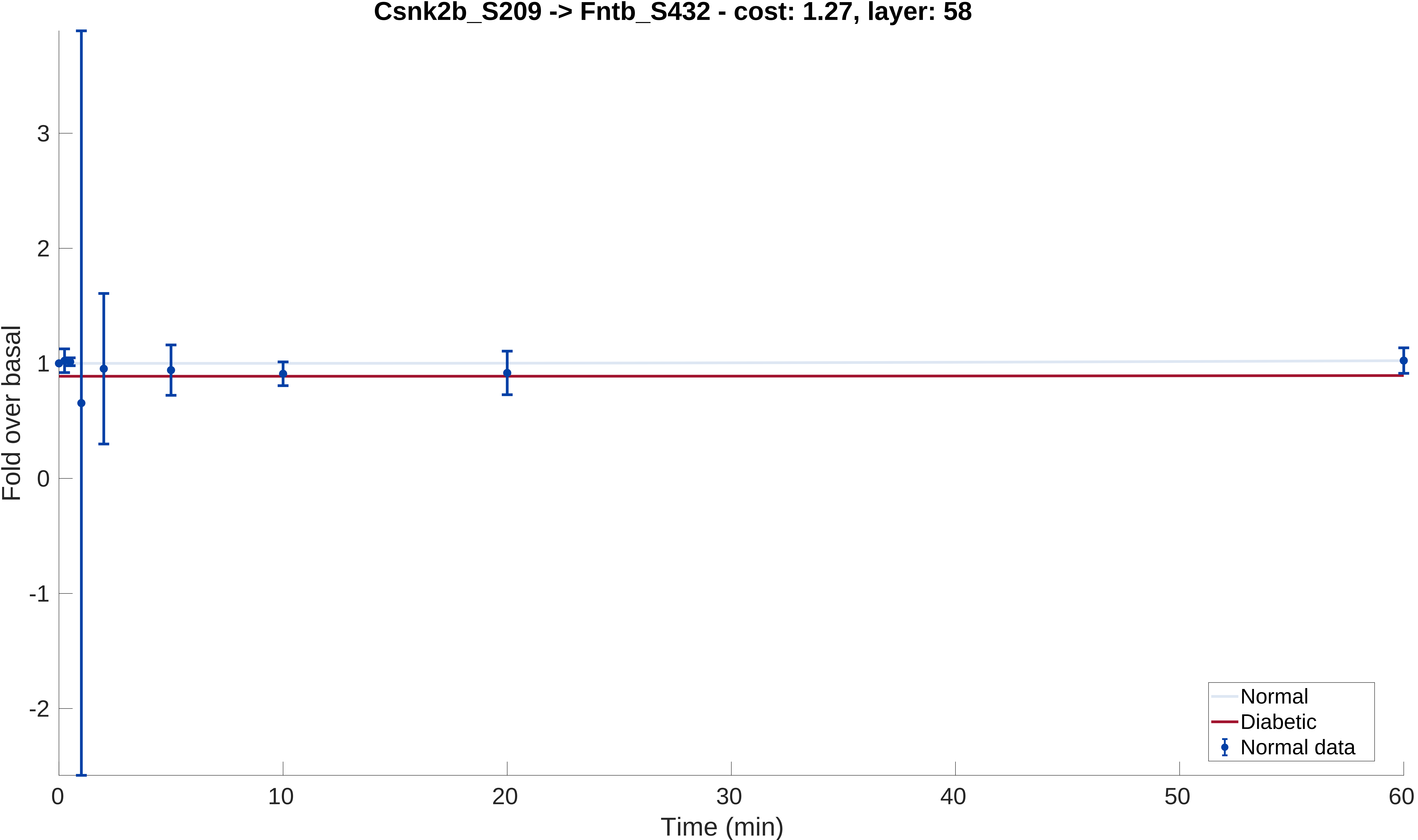

Csnk2b_S209 -> Fntb_S432 - cost: 1.27, layer: 58
Fold over basal
3
2
1
0
-1
-2
0
10
20
30
40
50
60
Time (min)
Normal
Diabetic
Normal data

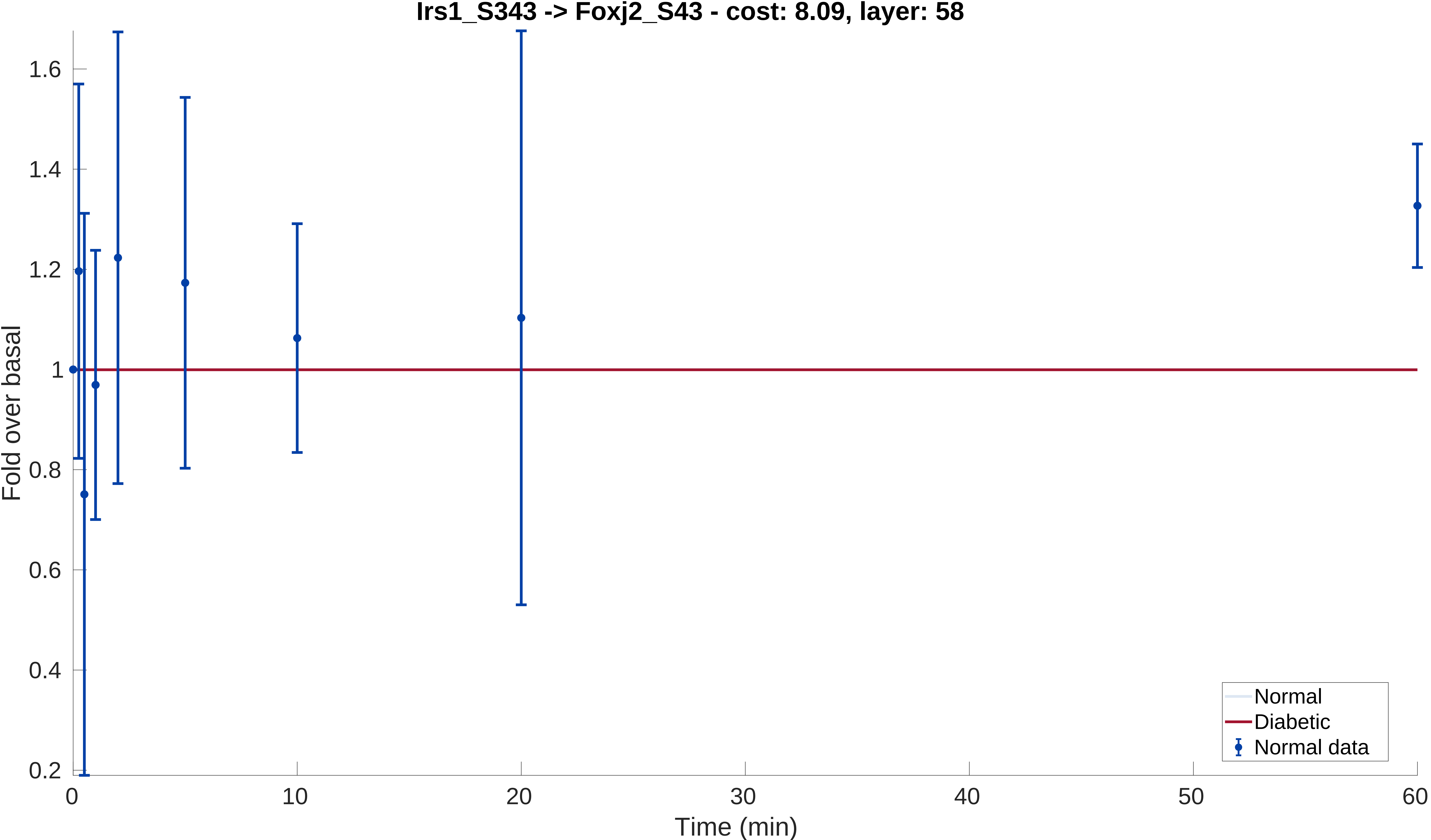

Irs1_S343 -> Foxj2_S43 - cost: 8.09, layer: 58
Fold over basal
1.6
1.4
1.2
1
0.8
0.6
0.4
0.2
0
10
20
30
40
50
60
Time (min)
Normal
Diabetic
Normal data

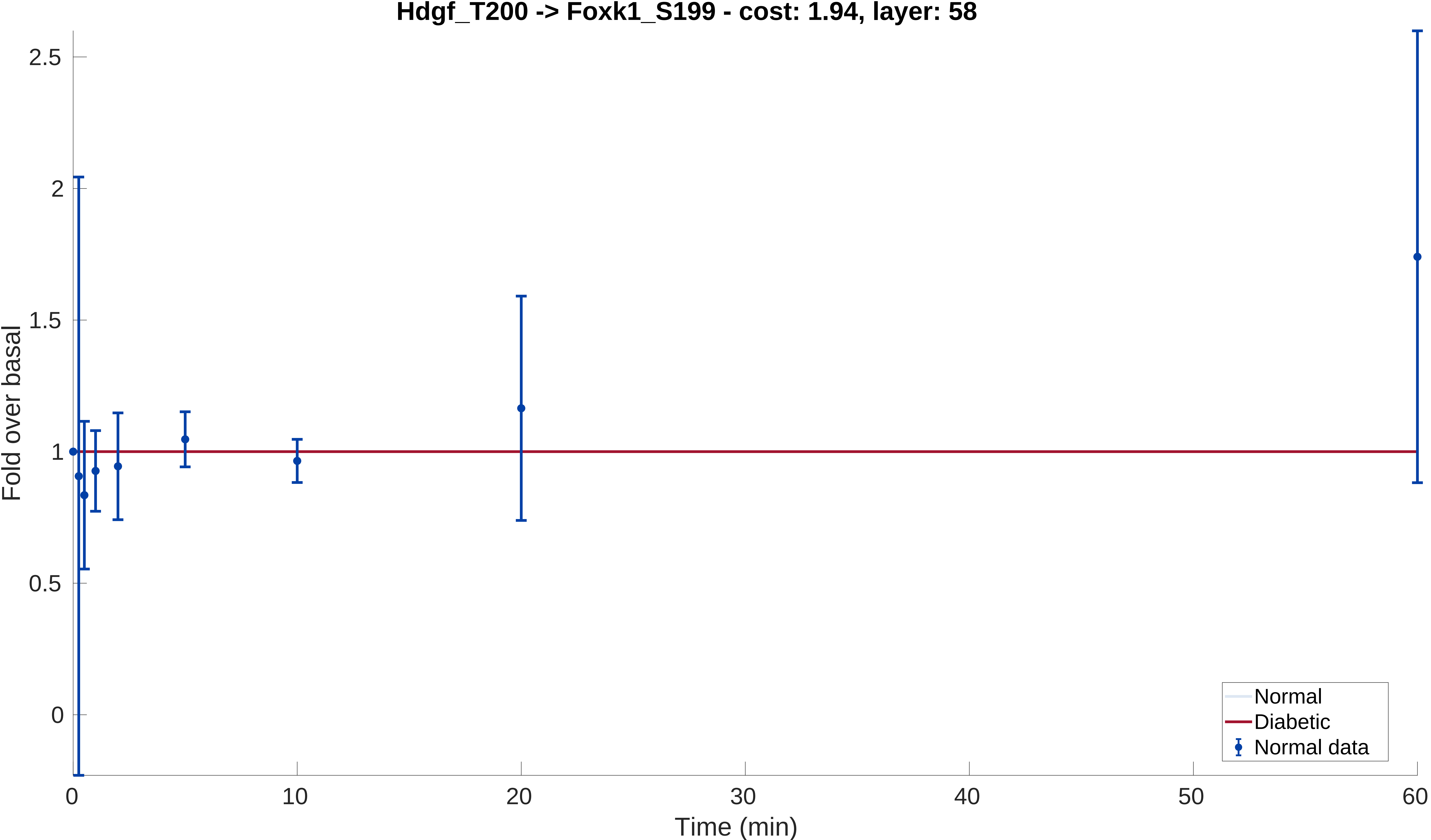

Hdgf_T200 -> Foxk1_S199 - cost: 1.94, layer: 58
Fold over basal
Time (min)
0
0.5
1
1.5
2
2.5
0
10
20
30
40
50
60
Normal
Diabetic
Normal data

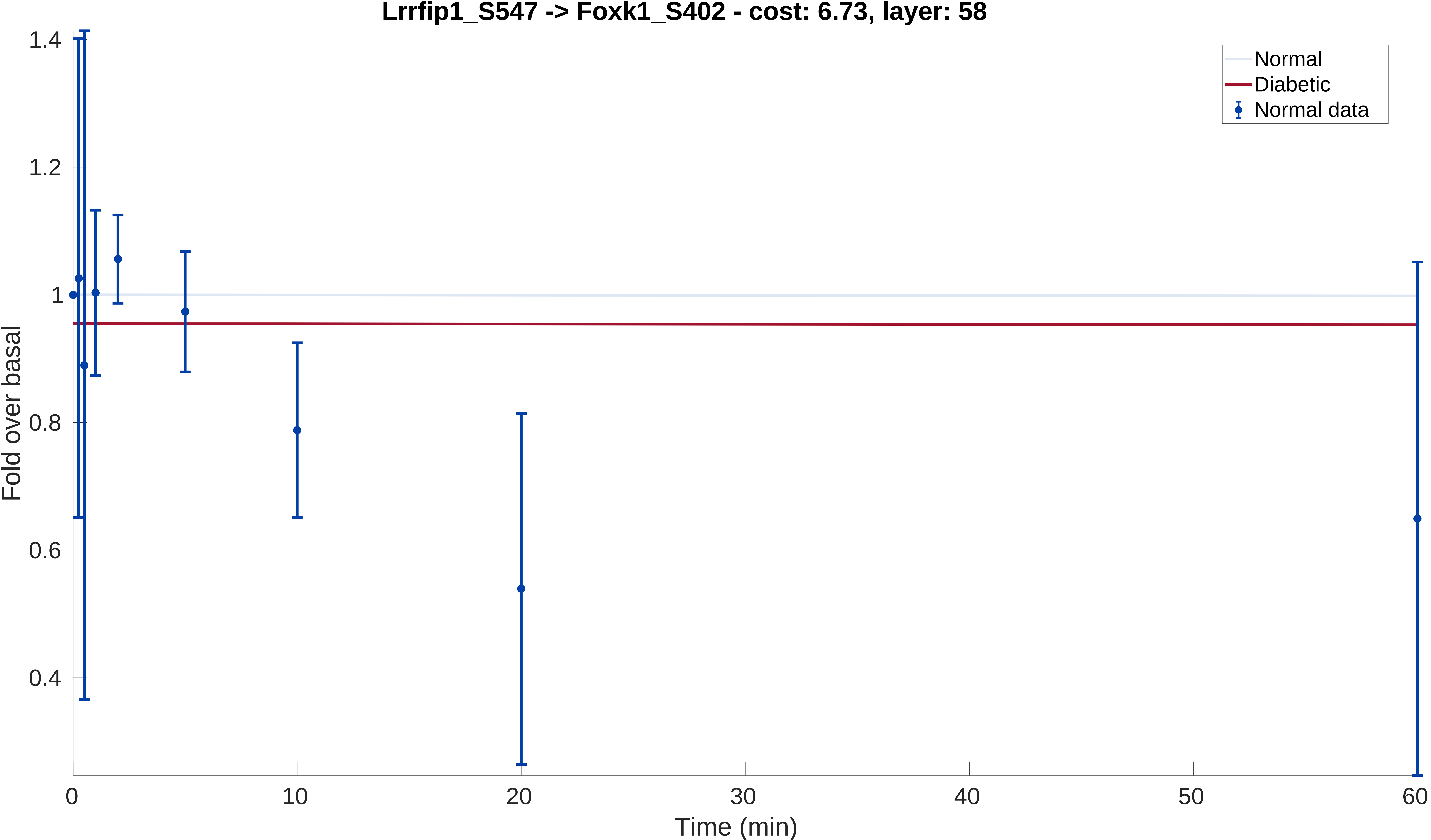

Lrrfip1_S547 -> Foxk1_S402 - cost: 6.73, layer: 58
Normal
Diabetic
Normal data
Fold over basal
1.4
1.2
1
0.8
0.6
0.4
0
10
20
30
40
50
60
Time (min)

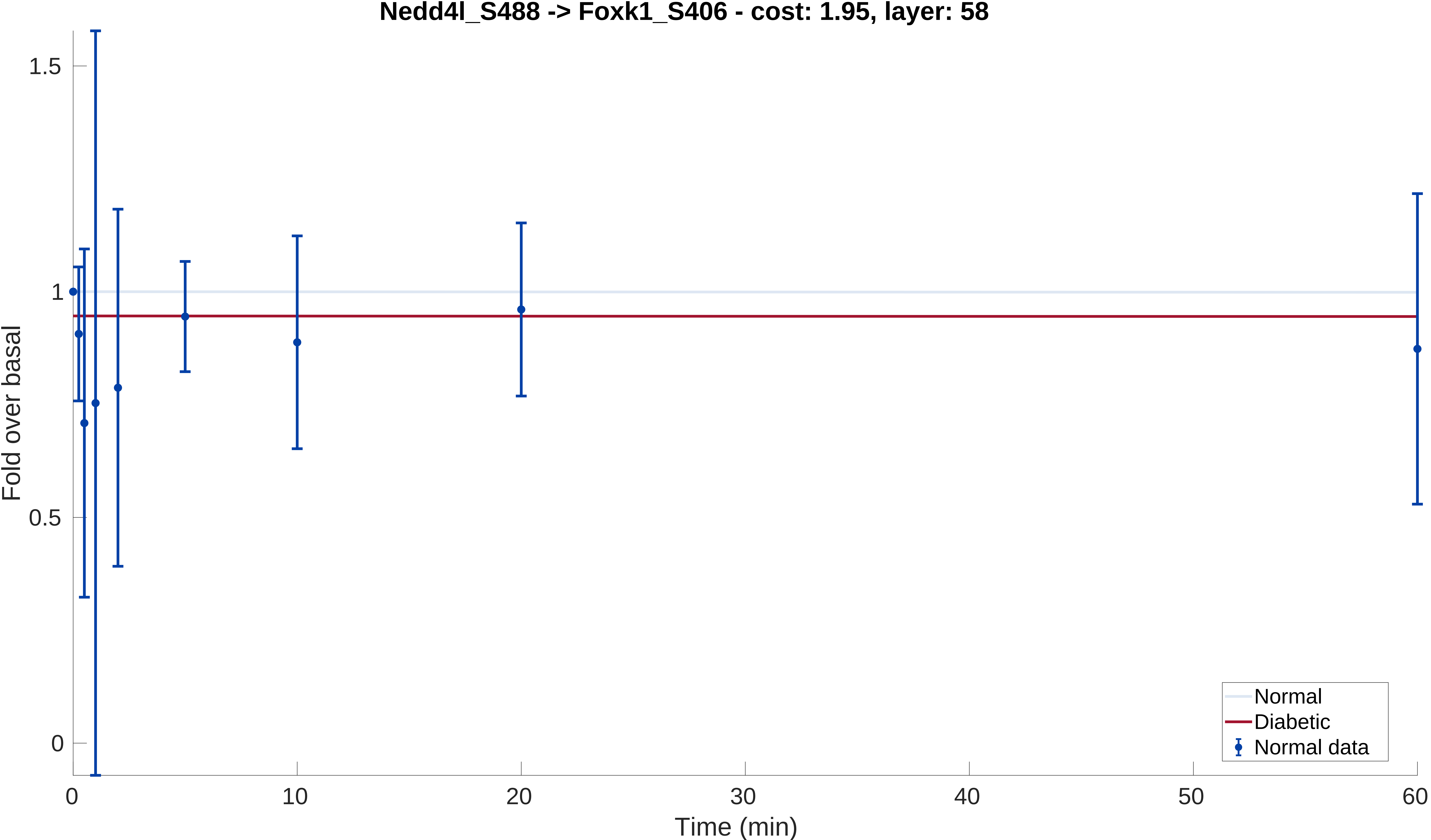
Nedd4l_S488 -> Foxk1_S406 - cost: 1.95, layer: 58
Fold over basal
Time (min)
1.5
1
0.5
0
0
10
20
30
40
50
60
Normal
Diabetic
Normal data

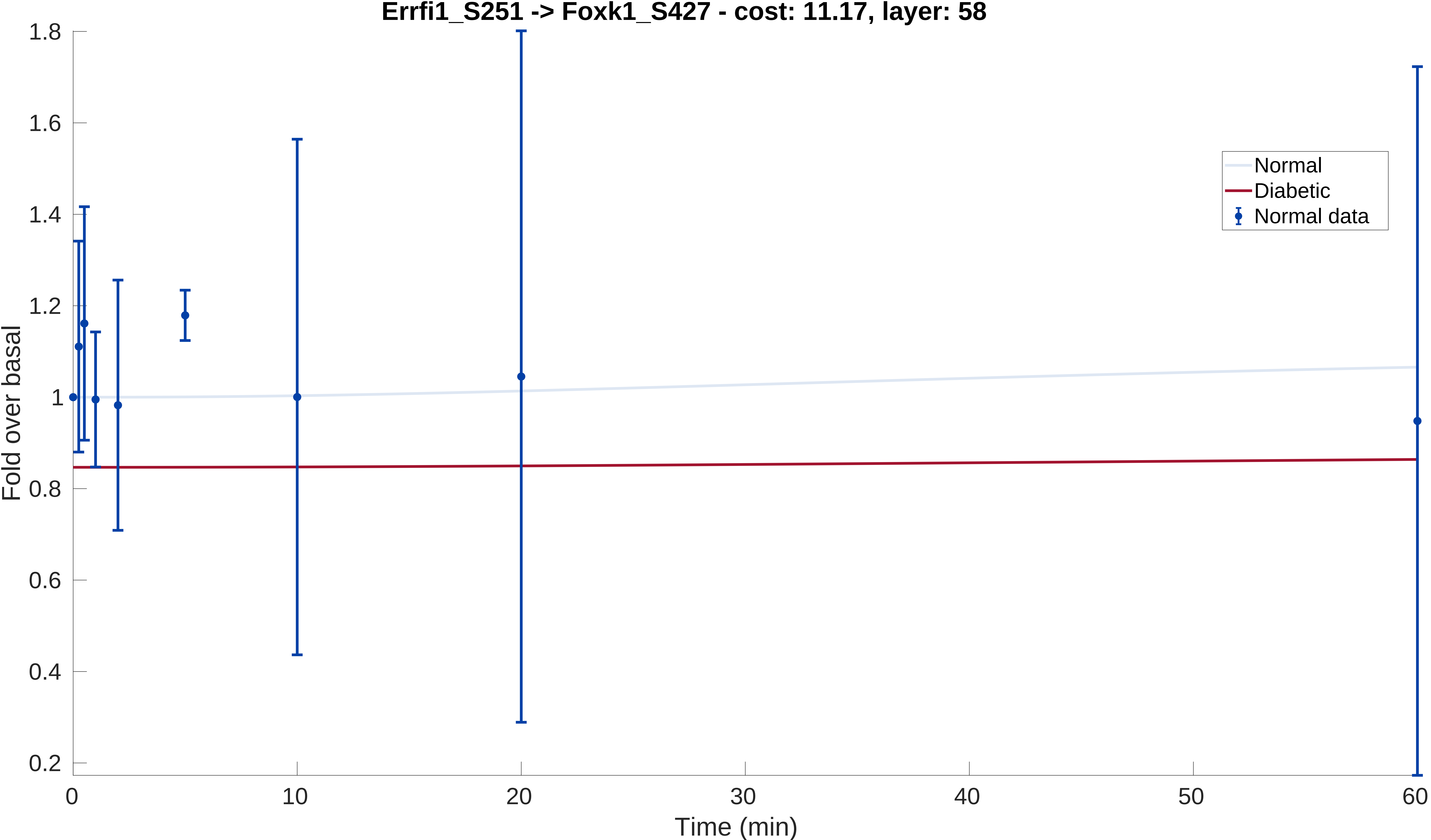

Errfi1_S251 -> Foxk1_S427 - cost: 11.17, layer: 58
Fold over basal
Time (min)
Normal
Diabetic
Normal data
1.8
1.6
1.4
1.2
1
0.8
0.6
0.4
0.2
0
10
20
30
40
50
60

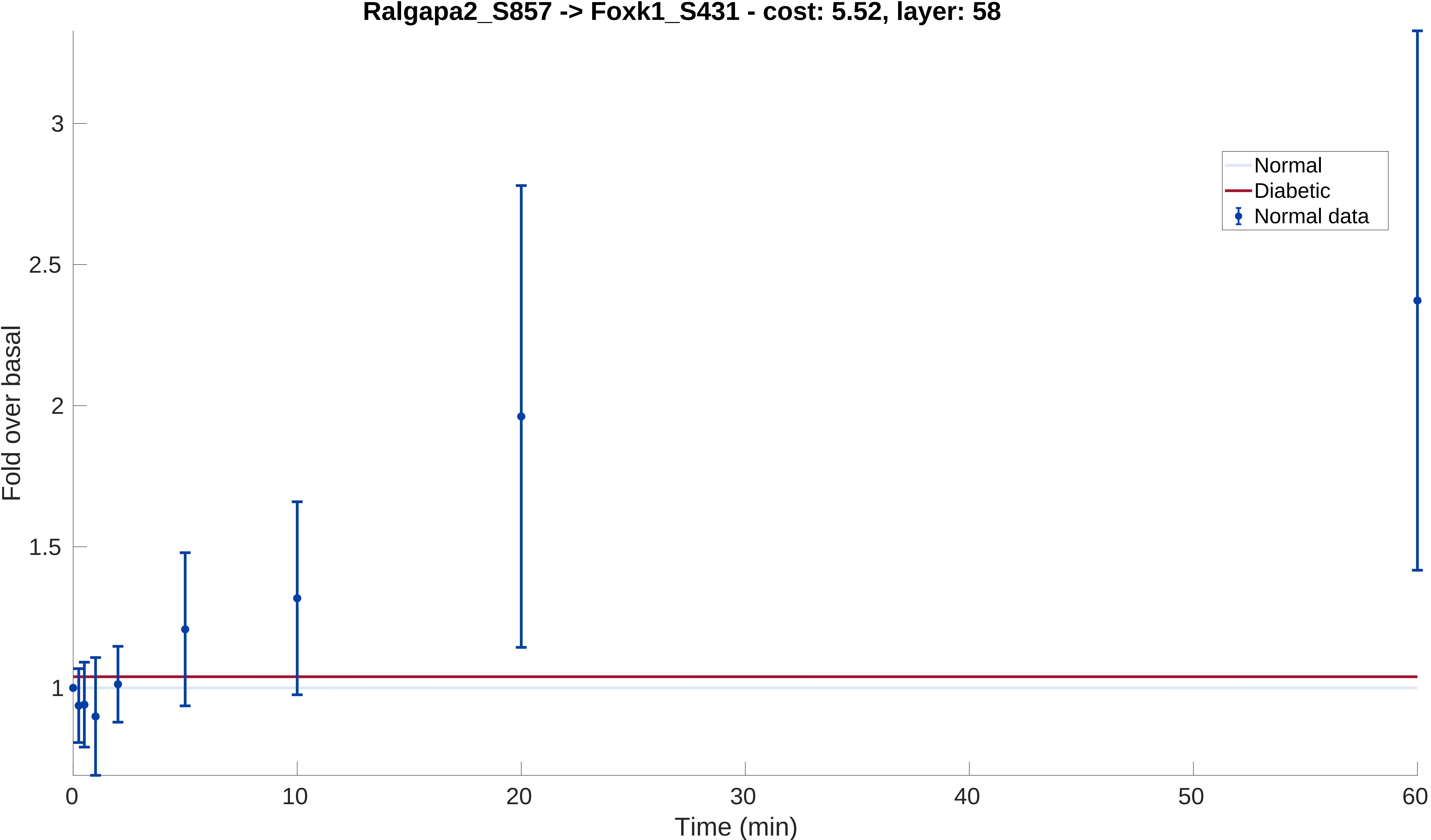
Ralgapa2_S857 -> Foxk1_S431 - cost: 5.52, layer: 58
Fold over basal
Time (min)
Normal
Diabetic
Normal data
3
2.5
2
1.5
1
0
10
20
30
40
50
60

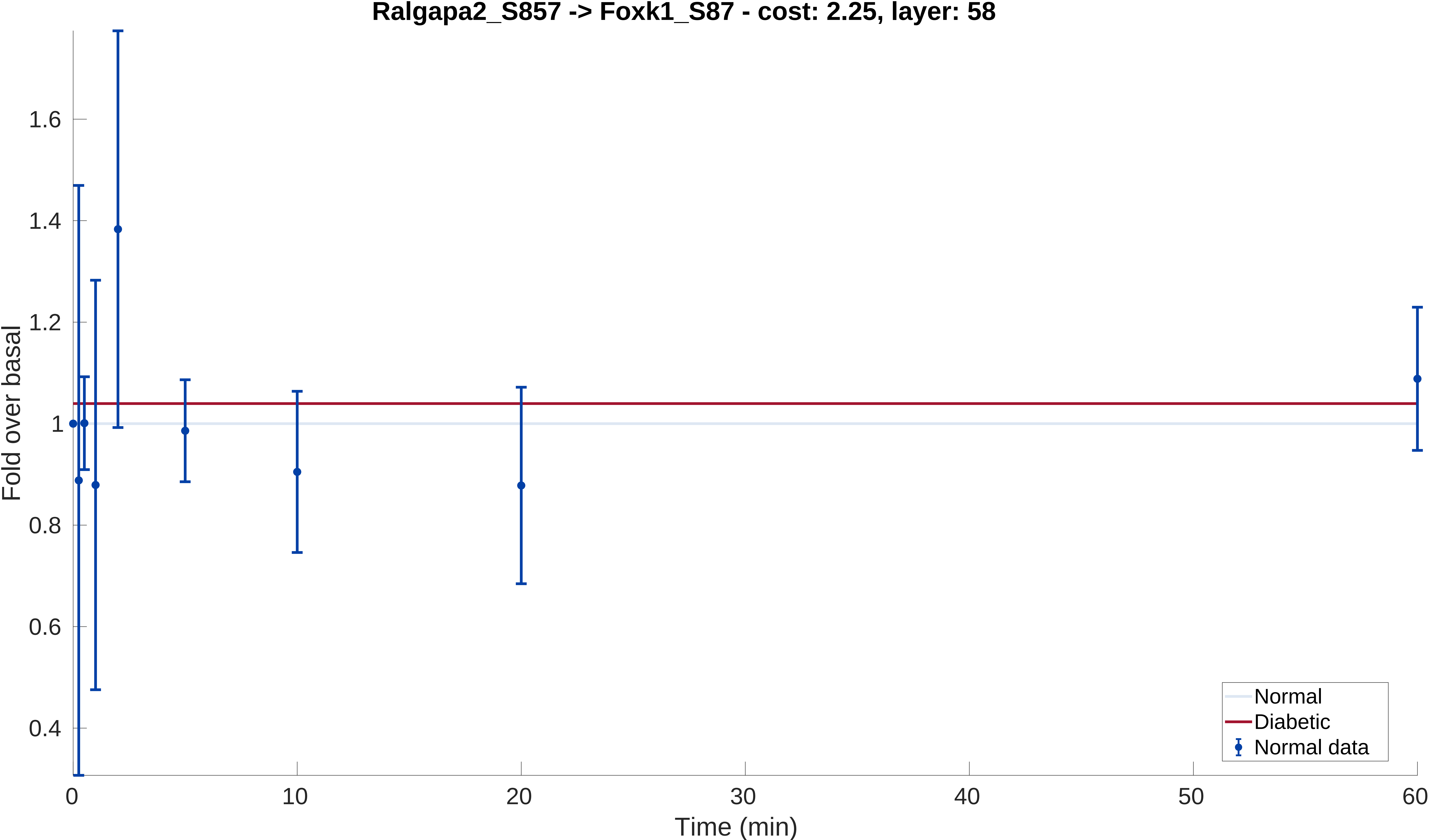

Ralgapa2_S857 -> Foxk1_S87 - cost: 2.25, layer: 58
Fold over basal
Time (min)
0
10
20
30
40
50
60
0.4
0.6
0.8
1
1.2
1.4
1.6
Normal
Diabetic
Normal data

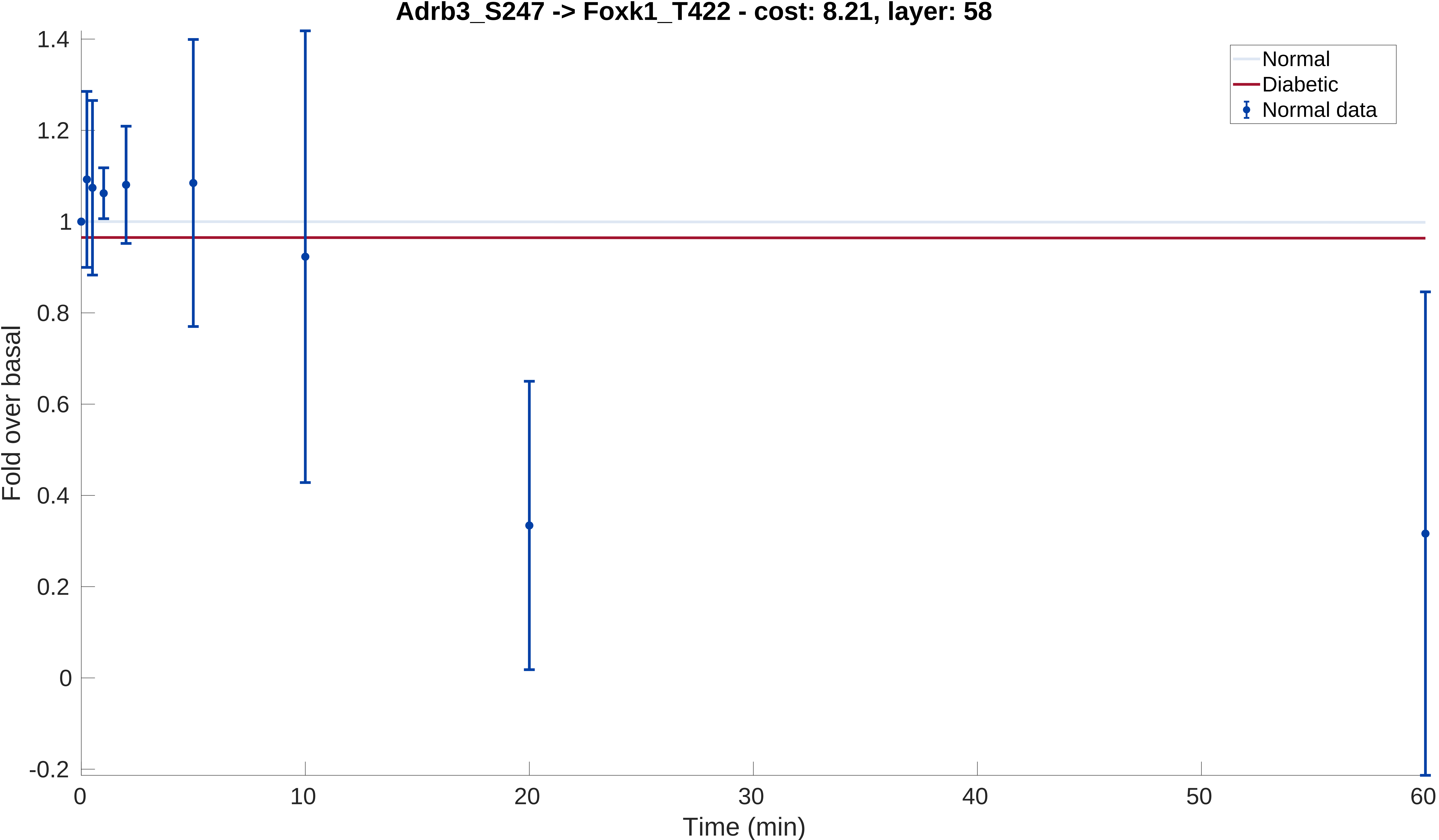

Adrb3_S247 -> Foxk1_T422 - cost: 8.21, layer: 58
Normal
Diabetic
Normal data
Fold over basal
Time (min)
1.4
1.2
1
0.8
0.6
0.4
0.2
0
-0.2
0
10
20
30
40
50
60

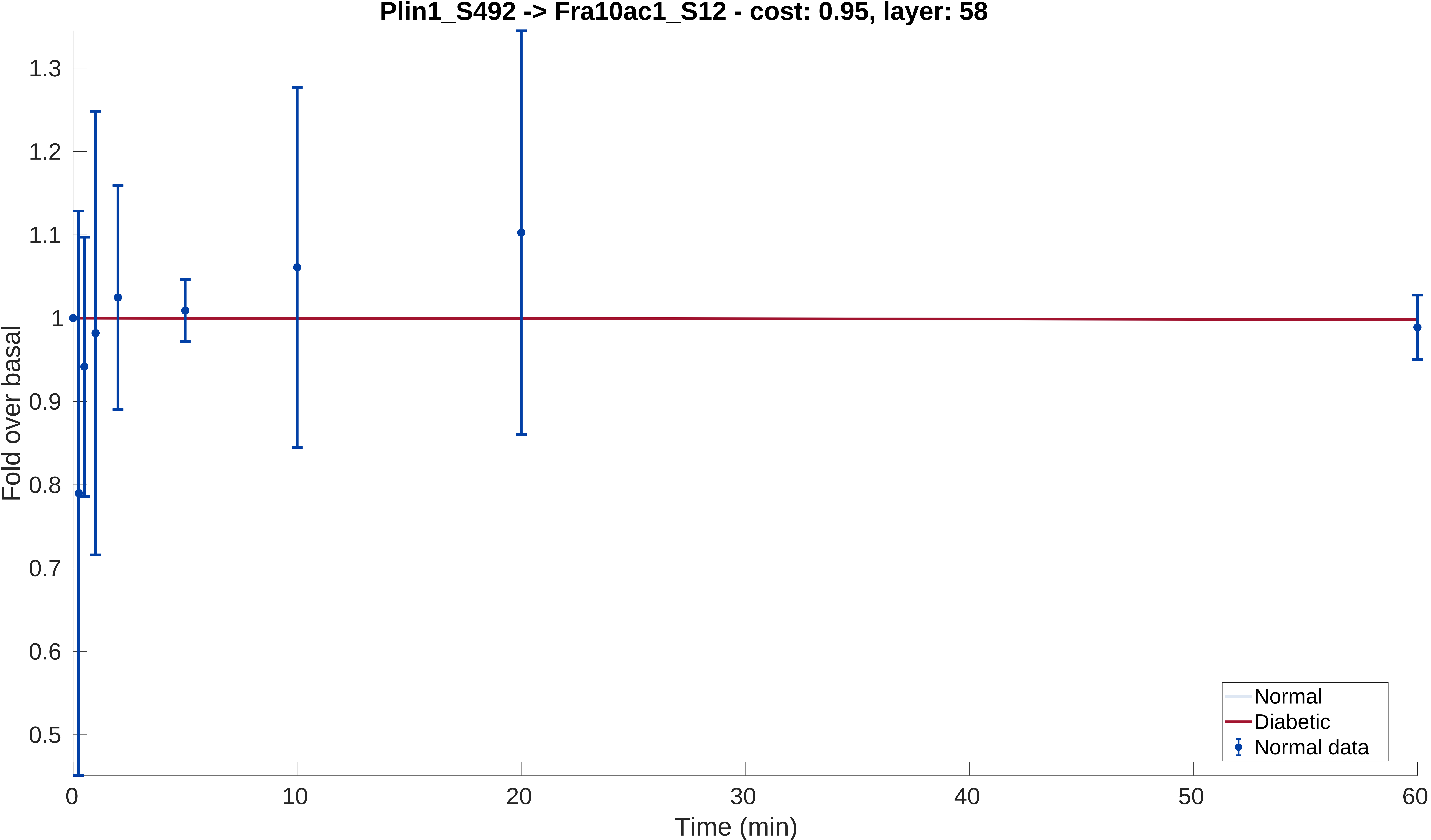

Plin1_S492 -> Fra10ac1_S12 - cost: 0.95, layer: 58
Fold over basal
1.3
1.2
1.1
1
0.9
0.8
0.7
0.6
0.5
0
10
20
30
40
50
60
Time (min)
Normal
Diabetic
Normal data

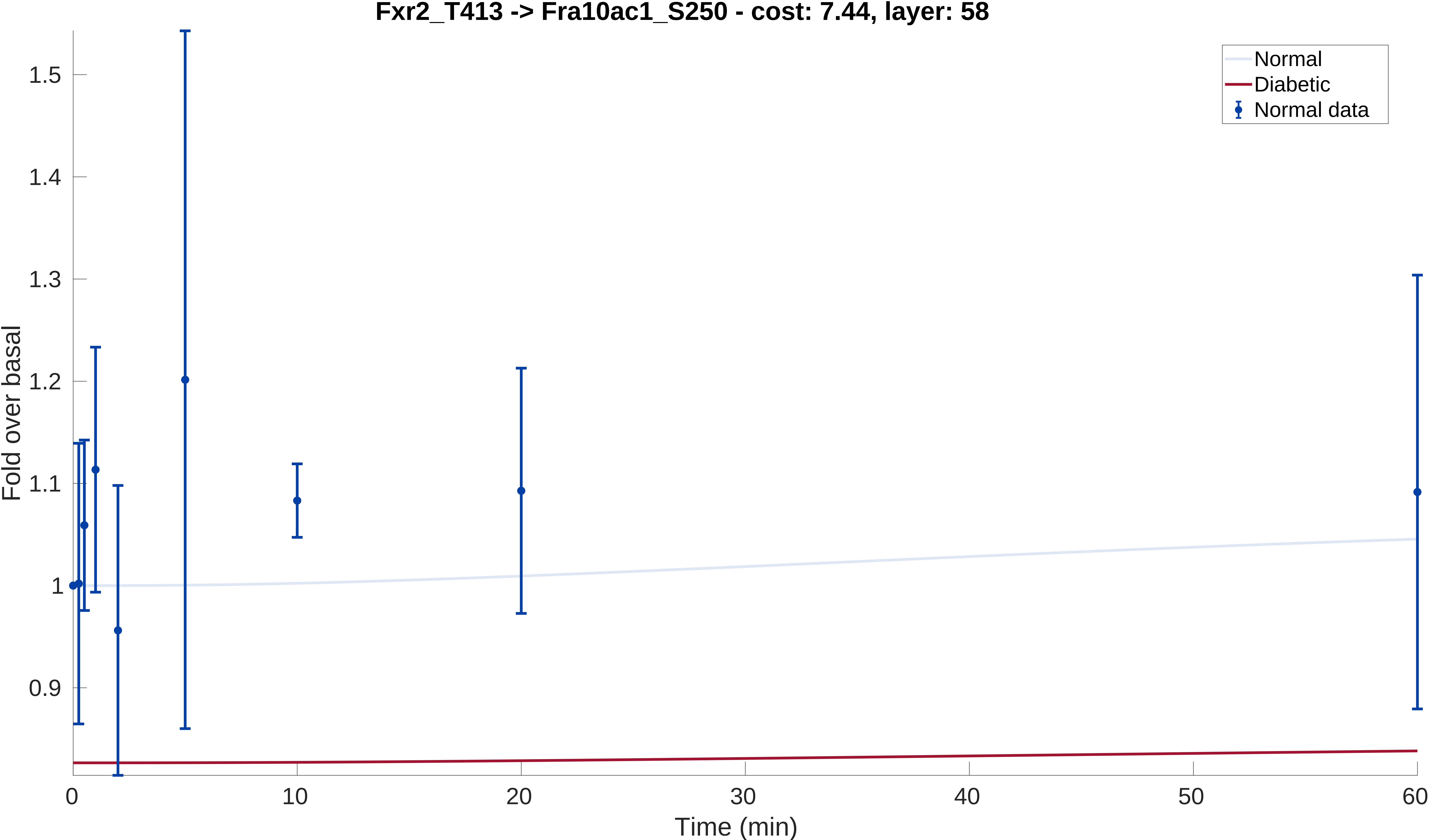
Fxr2_T413 -> Fra10ac1_S250 - cost: 7.44, layer: 58
Normal
Diabetic
Normal data
Fold over basal
1.5
1.4
1.3
1.2
1.1
1
0.9
0
10
20
30
40
50
60
Time (min)

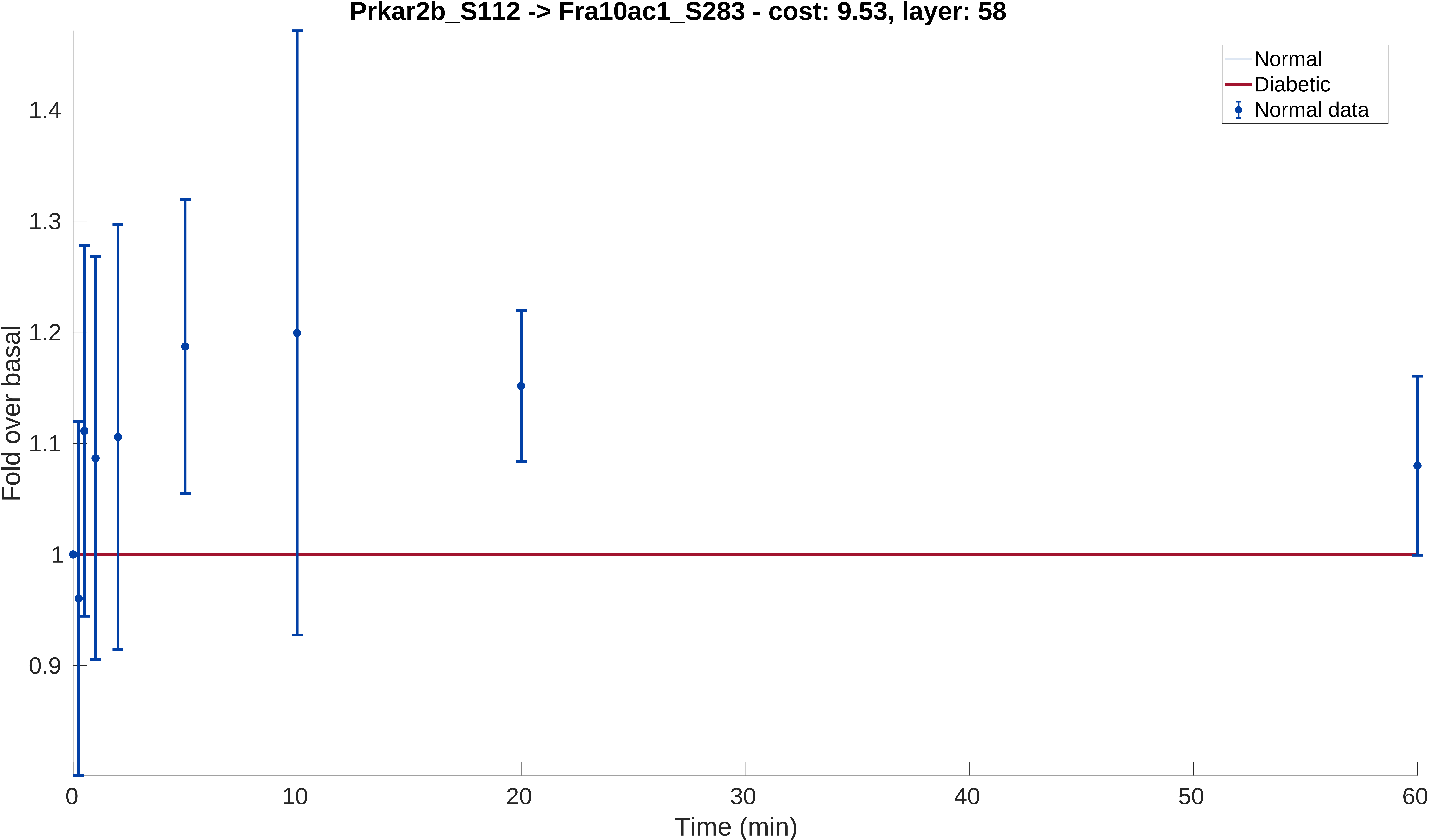

Prkar2b_S112 -> Fra10ac1_S283 - cost: 9.53, layer: 58
Normal
Diabetic
Normal data
Fold over basal
1.4
1.3
1.2
1.1
1
0.9
0
10
20
30
40
50
60
Time (min)

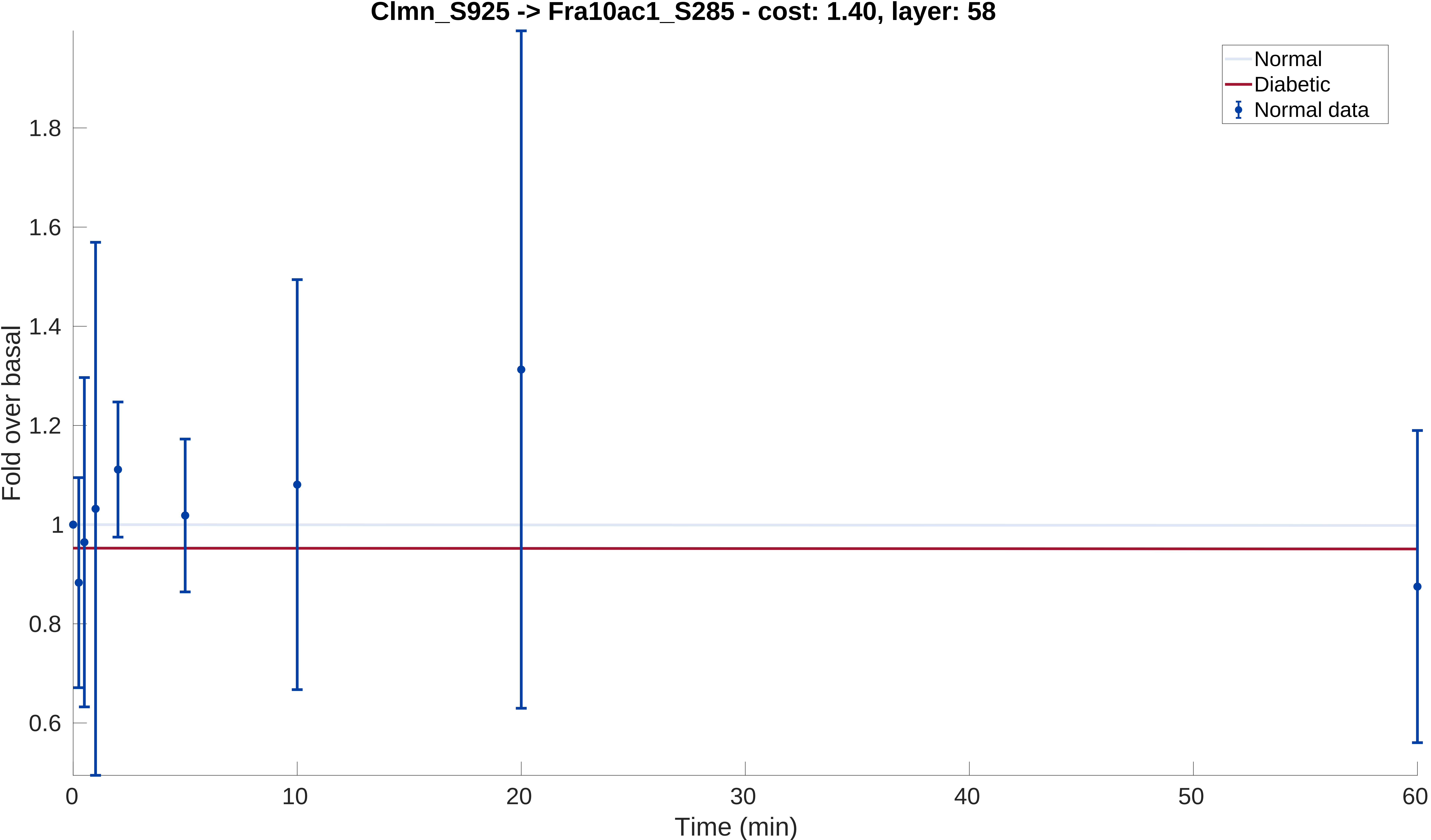

Clmn_S925 -> Fra10ac1_S285 - cost: 1.40, layer: 58
Normal
Diabetic
Normal data
1.8
1.6
1.4
1.2
1
0.8
0.6
Fold over basal
0
10
20
30
40
50
60
Time (min)

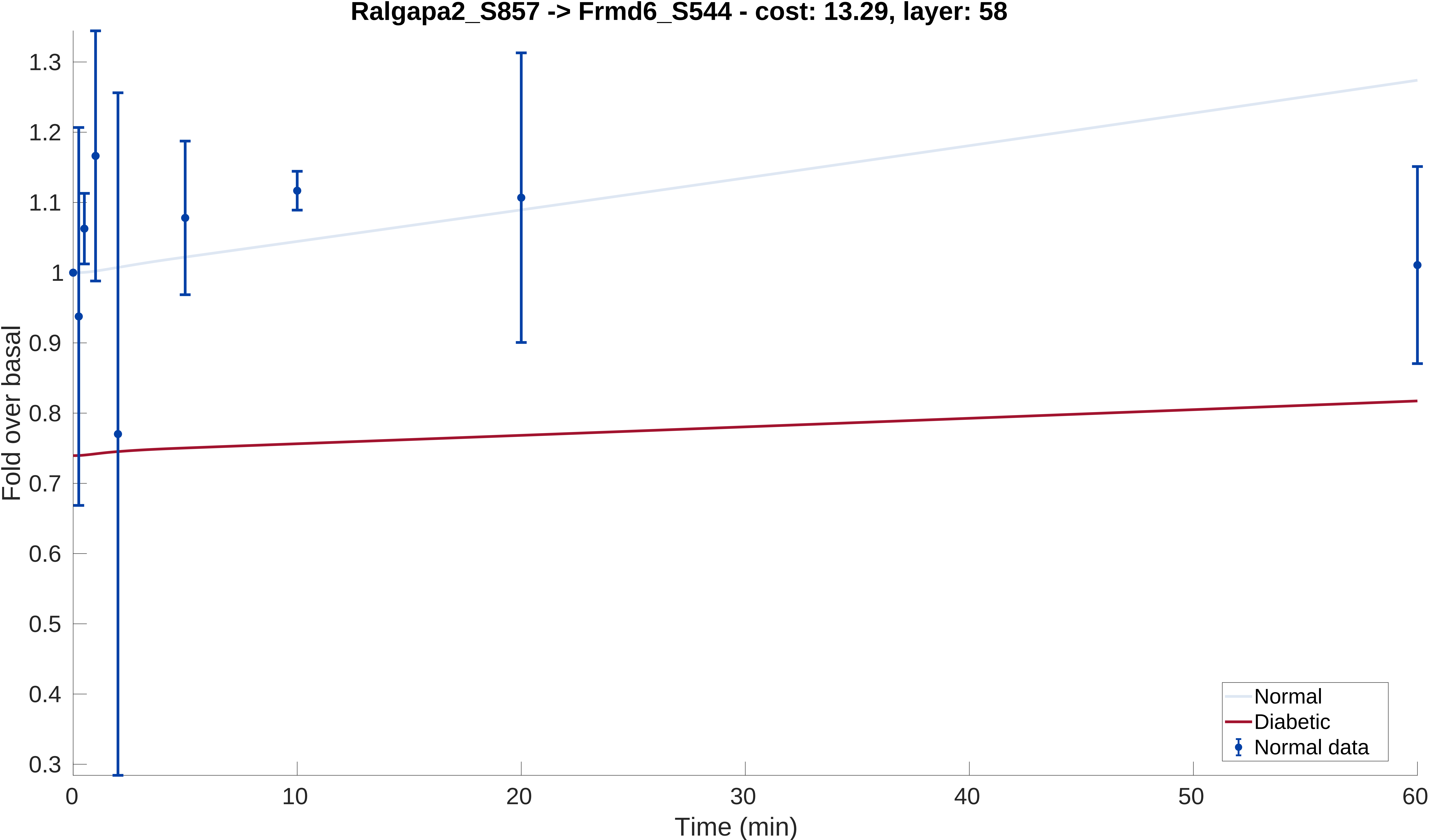

Ralgapa2_S857 -> Frmd6_S544 - cost: 13.29, layer: 58
Fold over basal
1.3
1.2
1.1
1
0.9
0.8
0.7
0.6
0.5
0.4
0.3
0
10
20
30
40
50
60
Time (min)
Normal
Diabetic
Normal data

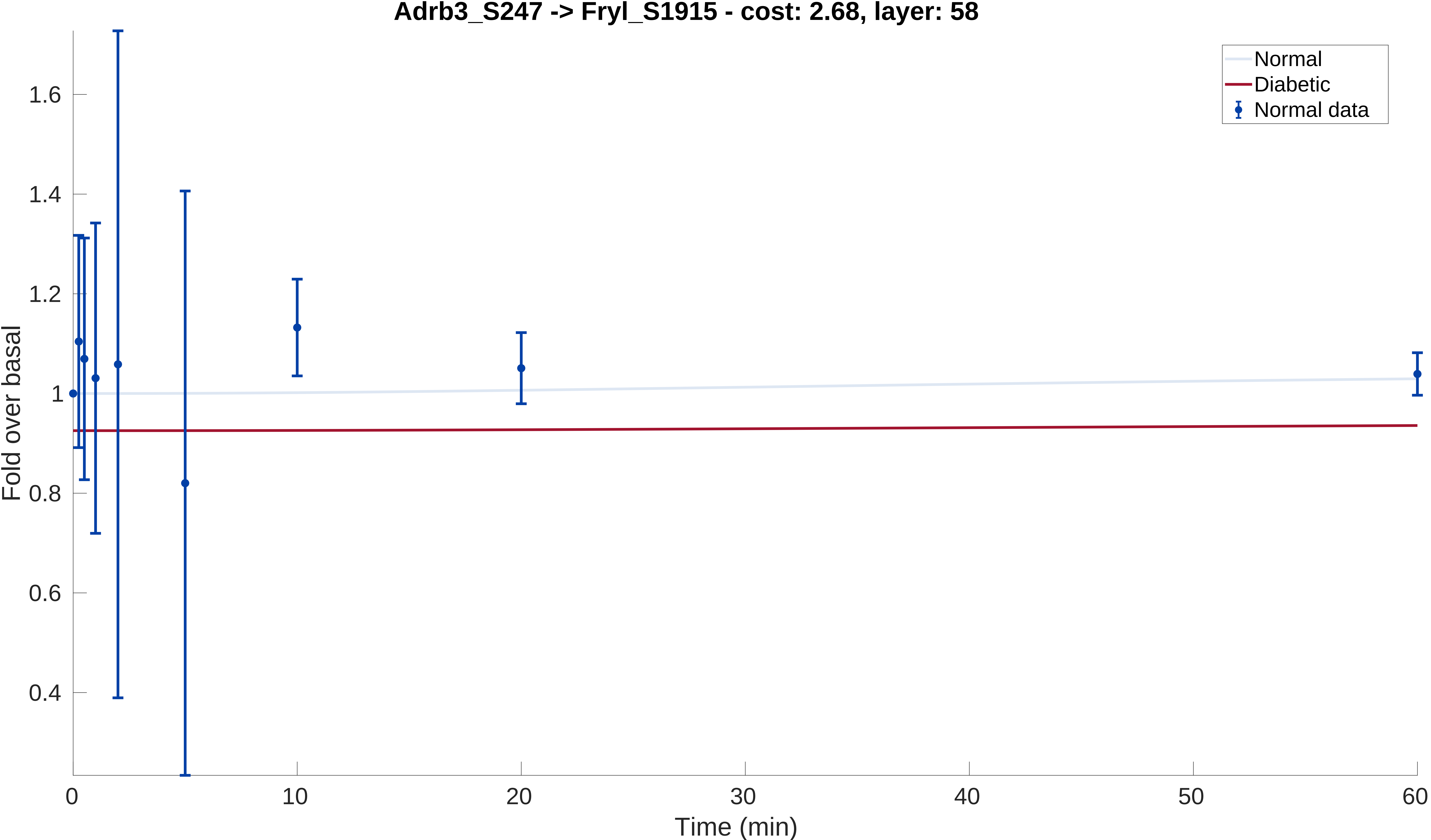

Adrb3_S247 -> Fryl_S1915 - cost: 2.68, layer: 58
Normal
Diabetic
Normal data
Fold over basal
1.6
1.4
1.2
1
0.8
0.6
0.4
0
10
20
30
40
50
60
Time (min)

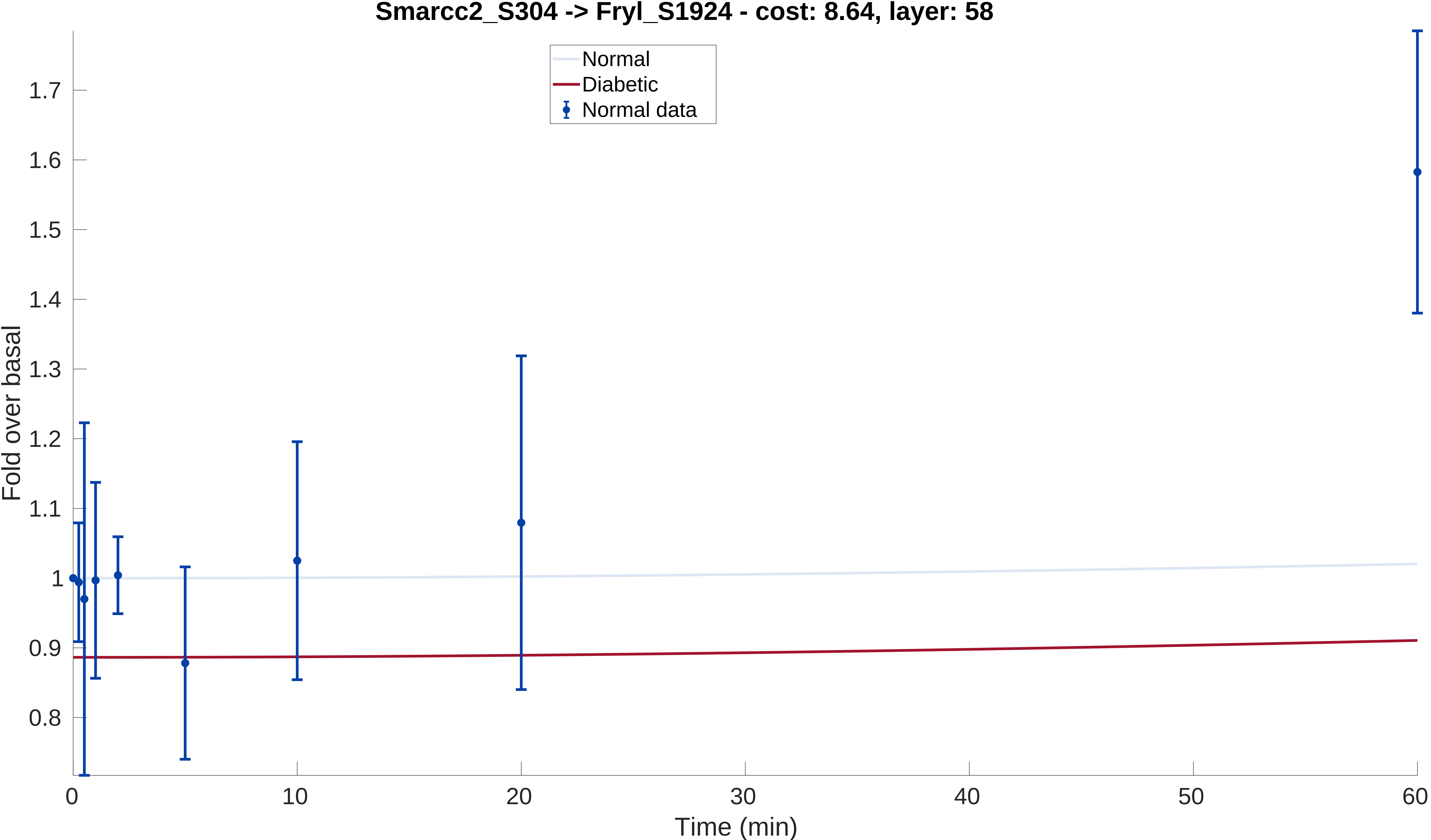

Smarcc2_S304 -> Fryl_S1924 - cost: 8.64, layer: 58
Normal
Diabetic
Normal data
Fold over basal
1.7
1.6
1.5
1.4
1.3
1.2
1.1
1
0.9
0.8
0
10
20
30
40
50
60
Time (min)

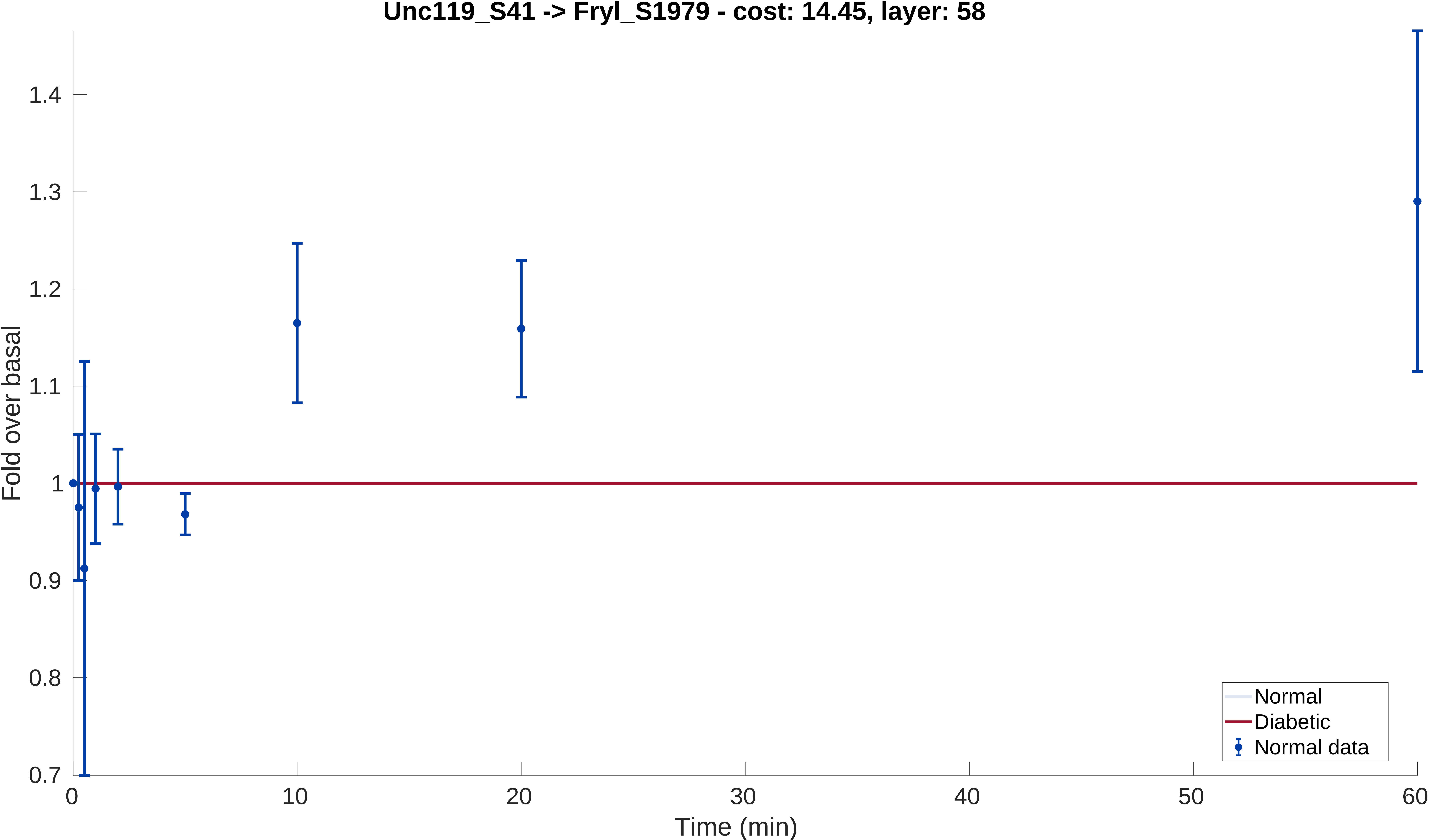

Unc119_S41 -> Fryl_S1979 - cost: 14.45, layer: 58
Fold over basal
1.4
1.3
1.2
1.1
1
0.9
0.8
0.7
0
10
20
30
40
50
60
Time (min)
Normal
Diabetic
Normal data

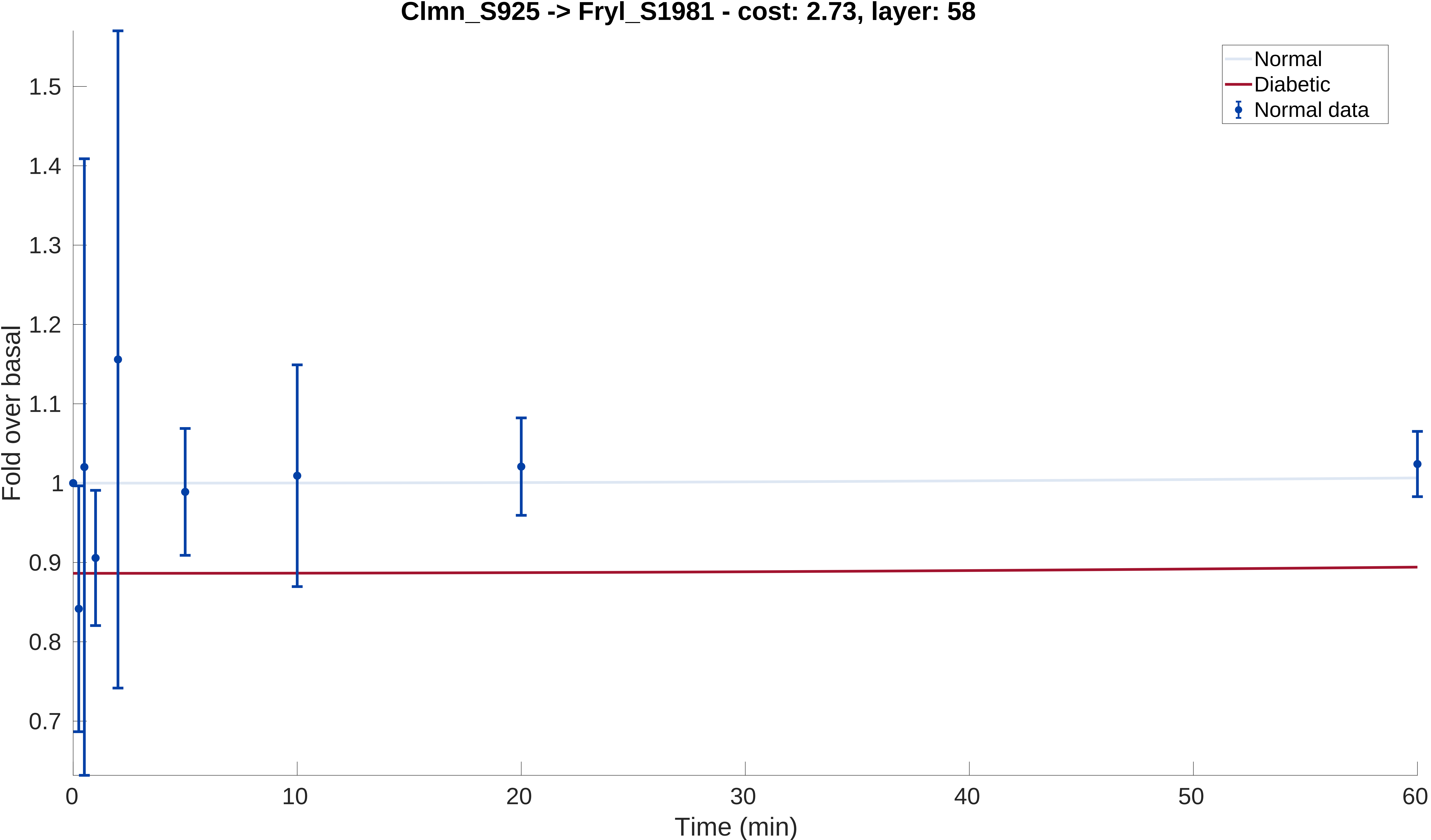
Clmn_S925 -> Fryl_S1981 - cost: 2.73, layer: 58
Normal
Diabetic
Normal data
Fold over basal
1.5
1.4
1.3
1.2
1.1
1
0.9
0.8
0.7
0
10
20
30
40
50
60
Time (min)

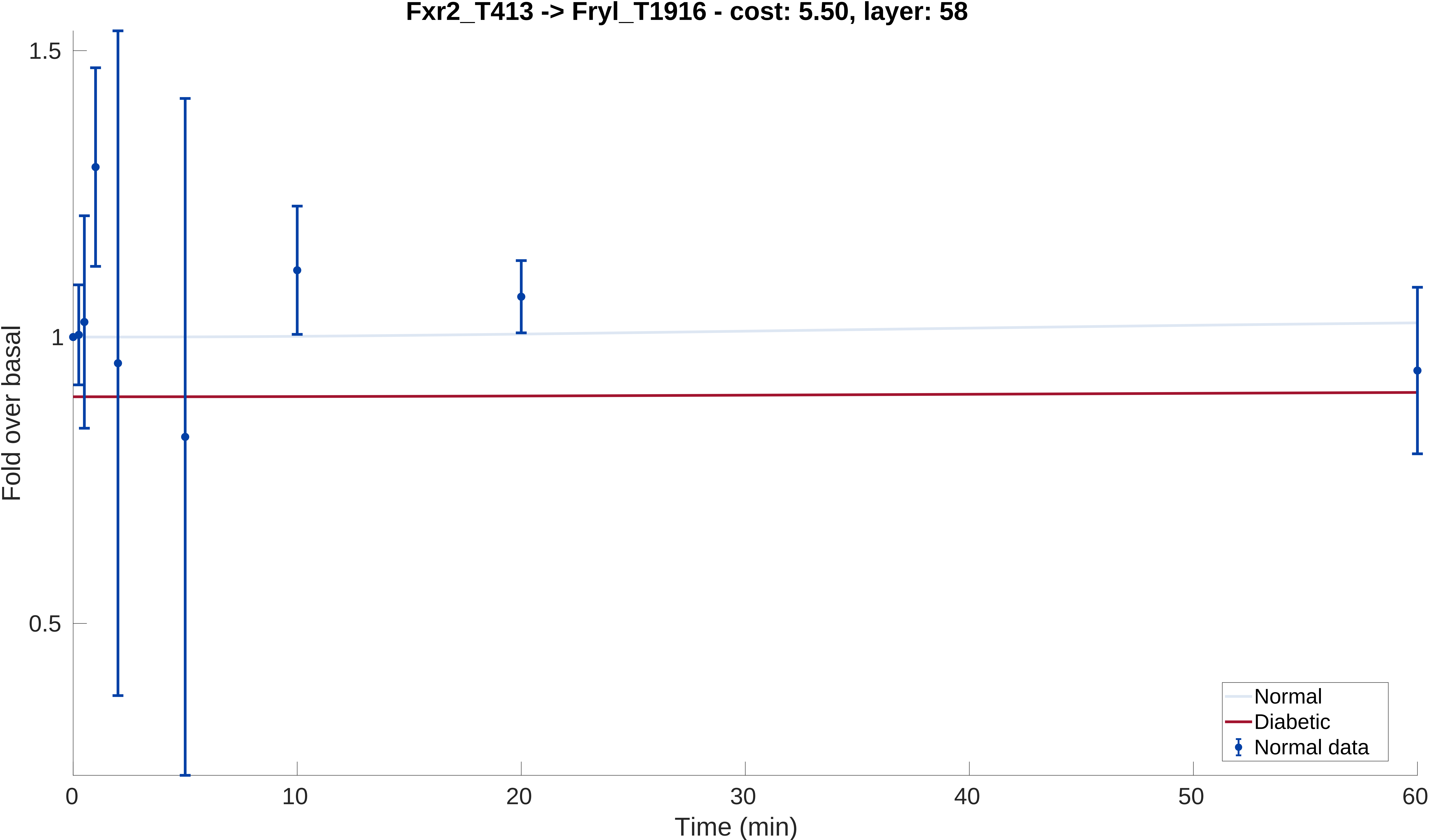

Fxr2_T413 -> Fryl_T1916 - cost: 5.50, layer: 58
1.5
1
0.5
Fold over basal
0
10
20
30
40
50
60
Time (min)
Normal
Diabetic
Normal data

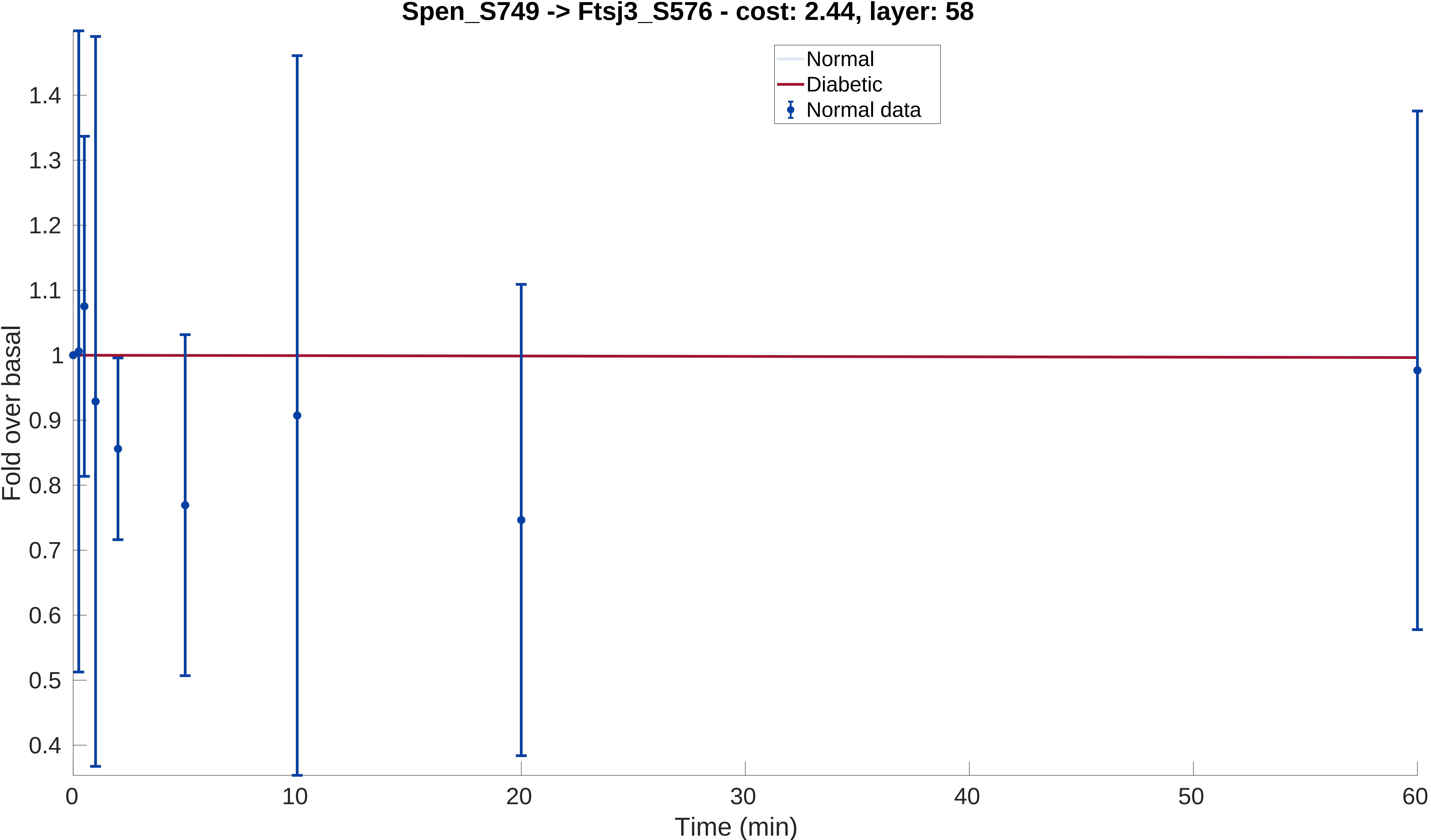
Spen_S749 -> Ftsj3_S576 - cost: 2.44, layer: 58
Normal
Diabetic
Normal data
Fold over basal
Time (min)
0.4
0.5
0.6
0.7
0.8
0.9
1
1.1
1.2
1.3
1.4
0
10
20
30
40
50
60

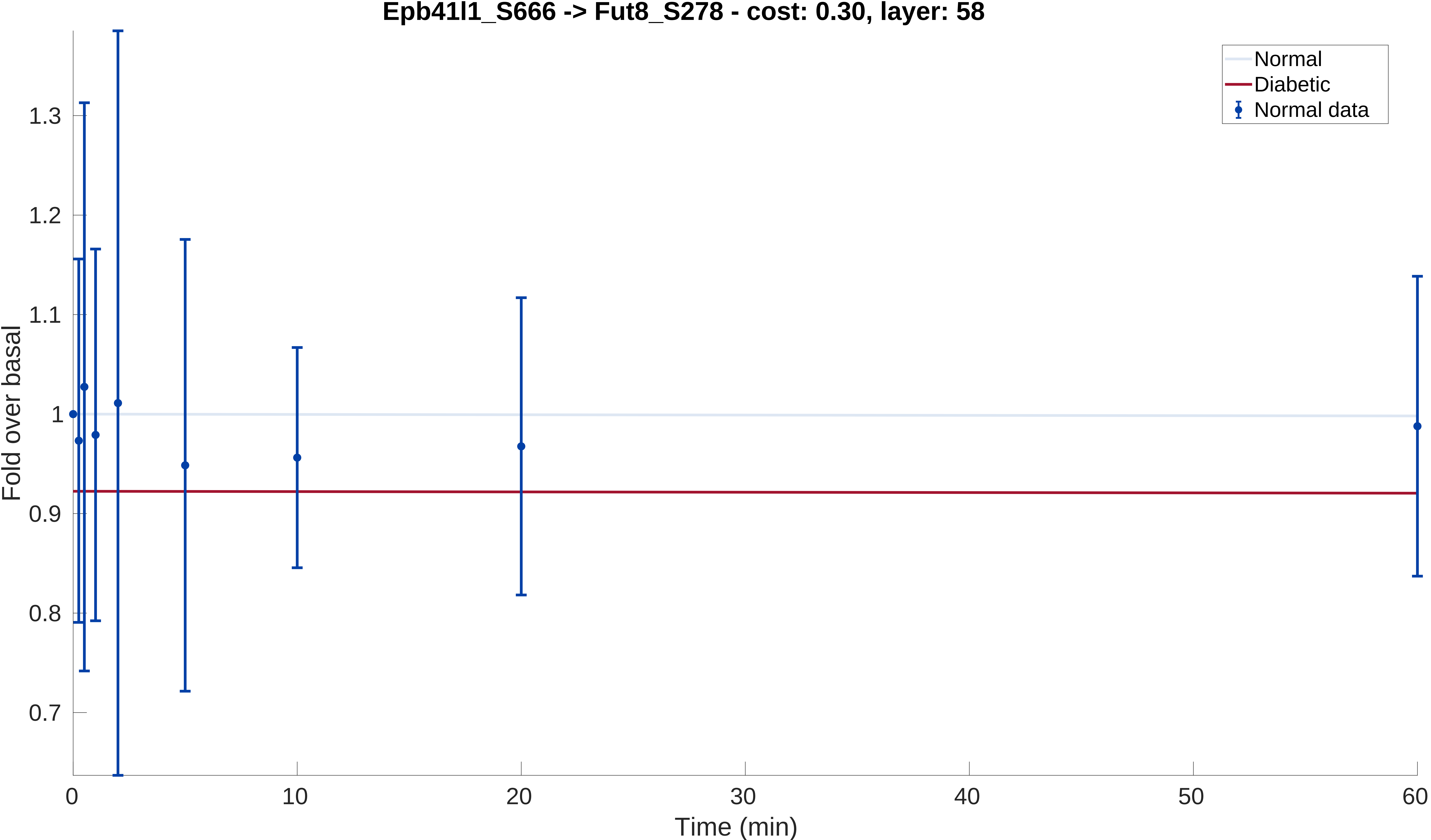

Epb41l1_S666 -> Fut8_S278 - cost: 0.30, layer: 58
Normal
Diabetic
Normal data
Fold over basal
1.3
1.2
1.1
1
0.9
0.8
0.7
0
10
20
30
40
50
60
Time (min)

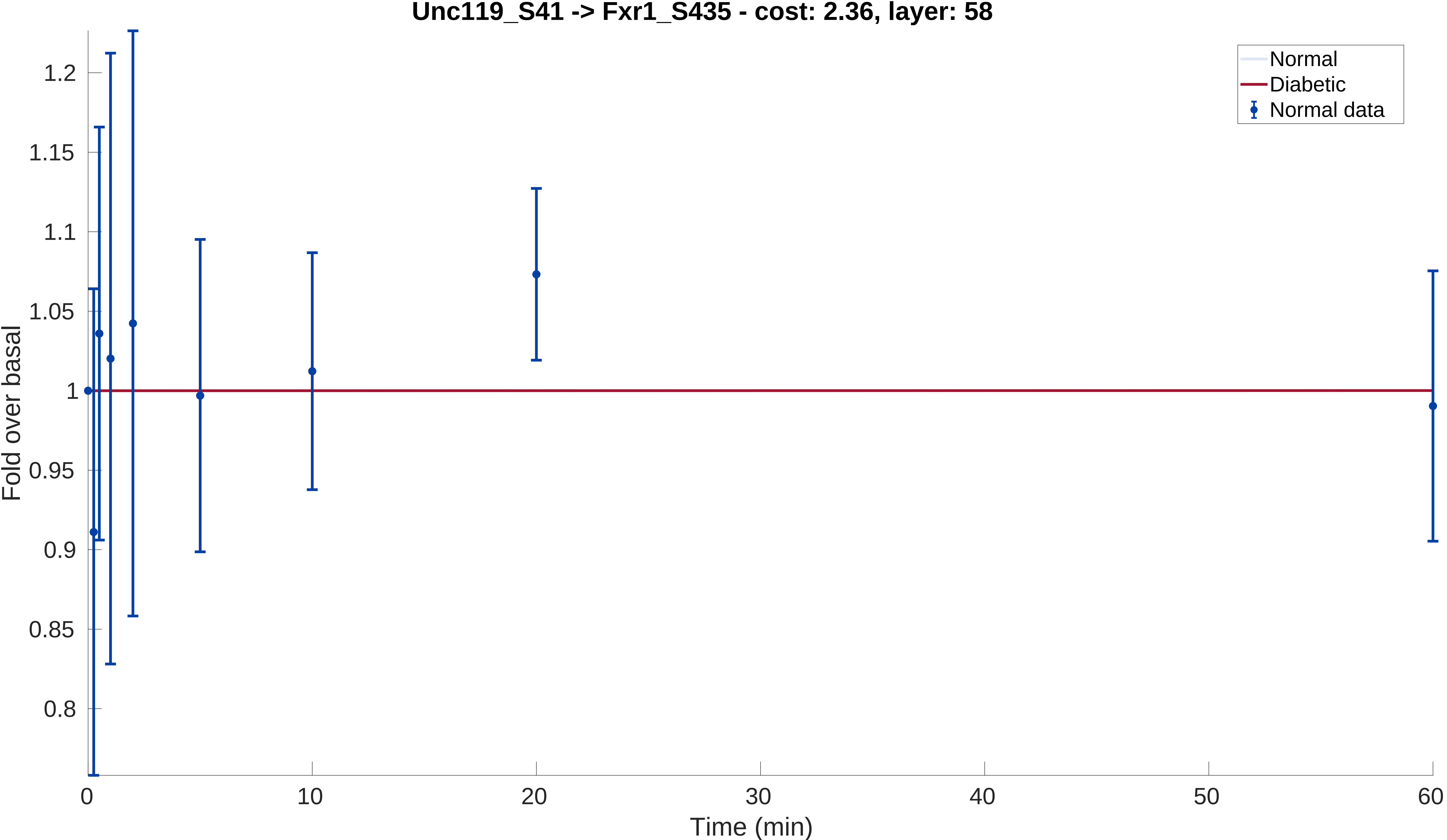

Unc119_S41 -> Fxr1_S435 - cost: 2.36, layer: 58
Normal
Diabetic
Normal data
Fold over basal
Time (min)
1.2
1.15
1.1
1.05
1
0.95
0.9
0.85
0.8
0
10
20
30
40
50
60

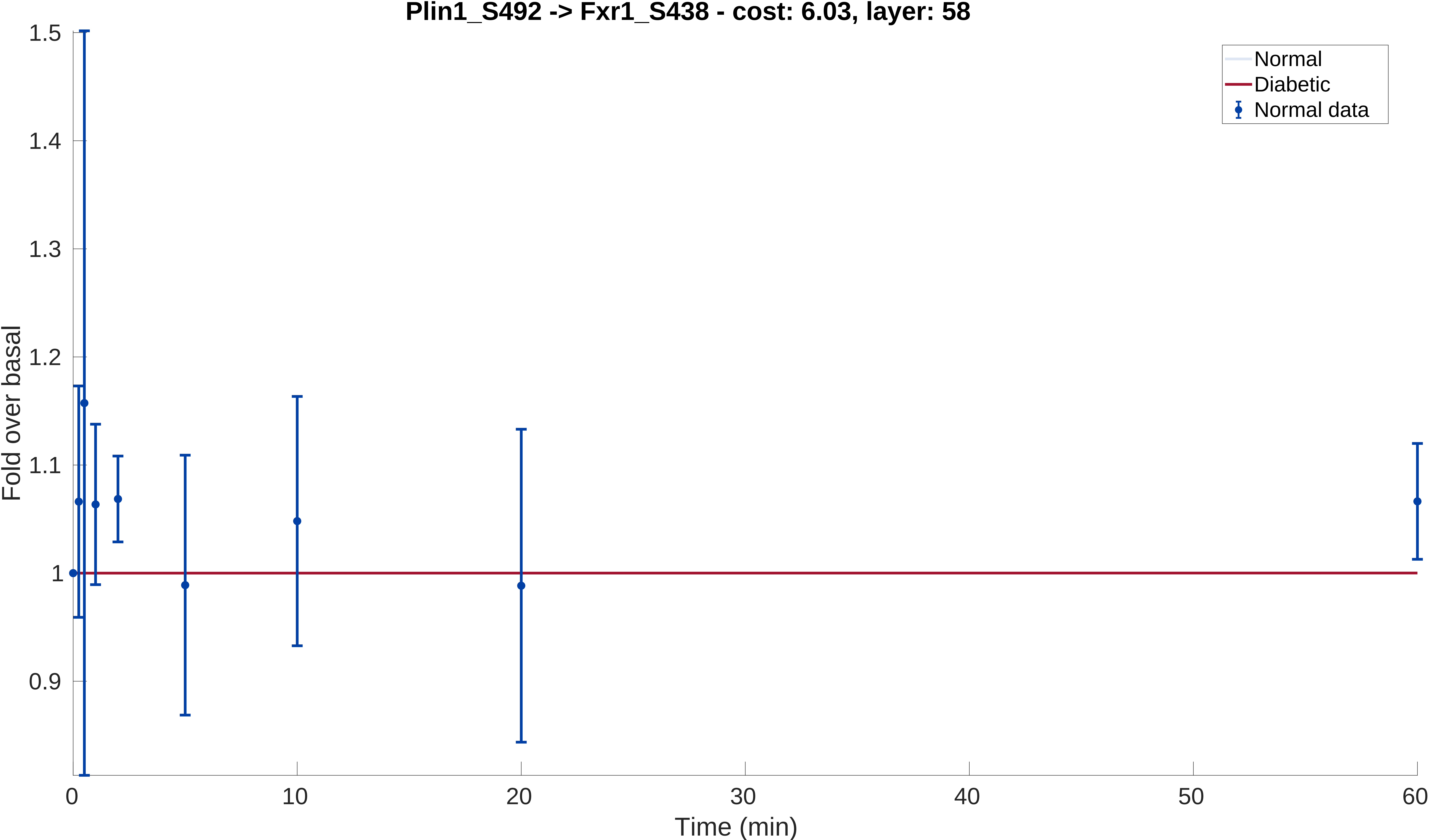

Plin1_S492 -> Fxr1_S438 - cost: 6.03, layer: 58
Normal
Diabetic
Normal data
1.5
1.4
1.3
1.2
1.1
1
0.9
Fold over basal
0
10
20
30
40
50
60
Time (min)

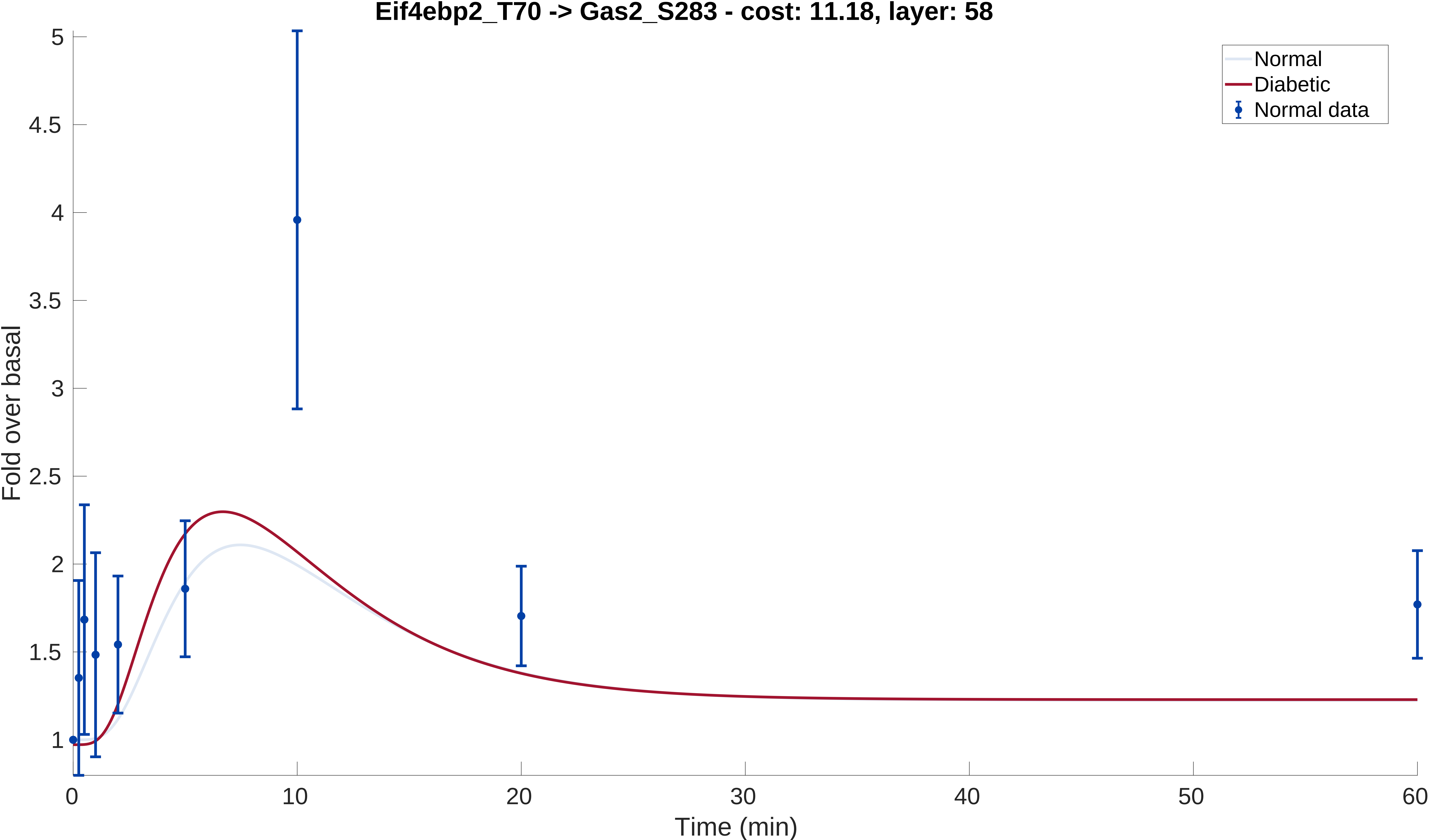

Eif4ebp2_T70 -> Gas2_S283 - cost: 11.18, layer: 58
Normal
Diabetic
Normal data
Fold over basal
Time (min)
5
4.5
4
3.5
3
2.5
2
1.5
1
0
10
20
30
40
50
60

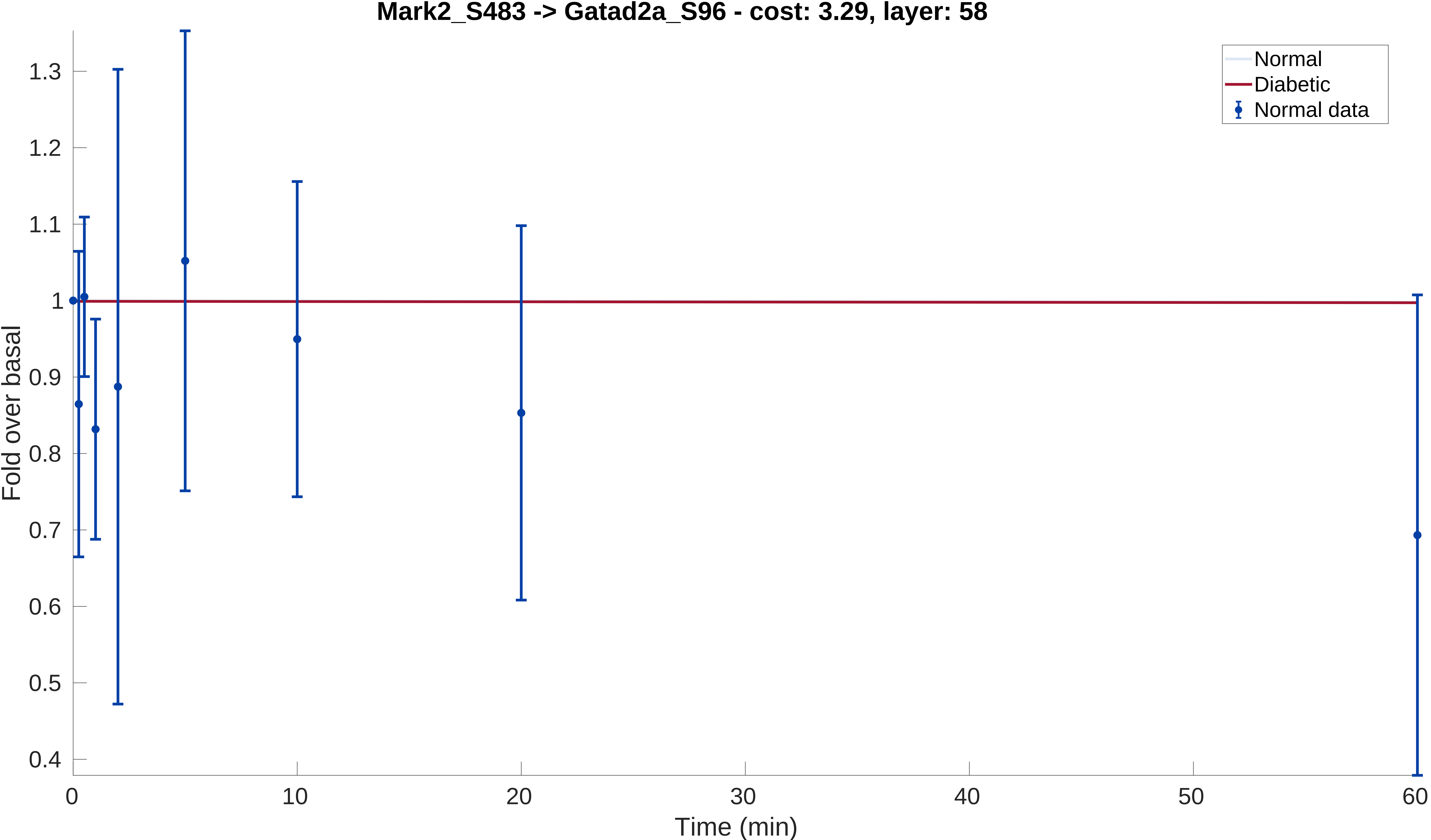
Mark2_S483 -> Gatad2a_S96 - cost: 3.29, layer: 58
Normal
Diabetic
Normal data
Fold over basal
1.3
1.2
1.1
1
0.9
0.8
0.7
0.6
0.5
0.4
0
10
20
30
40
50
60
Time (min)

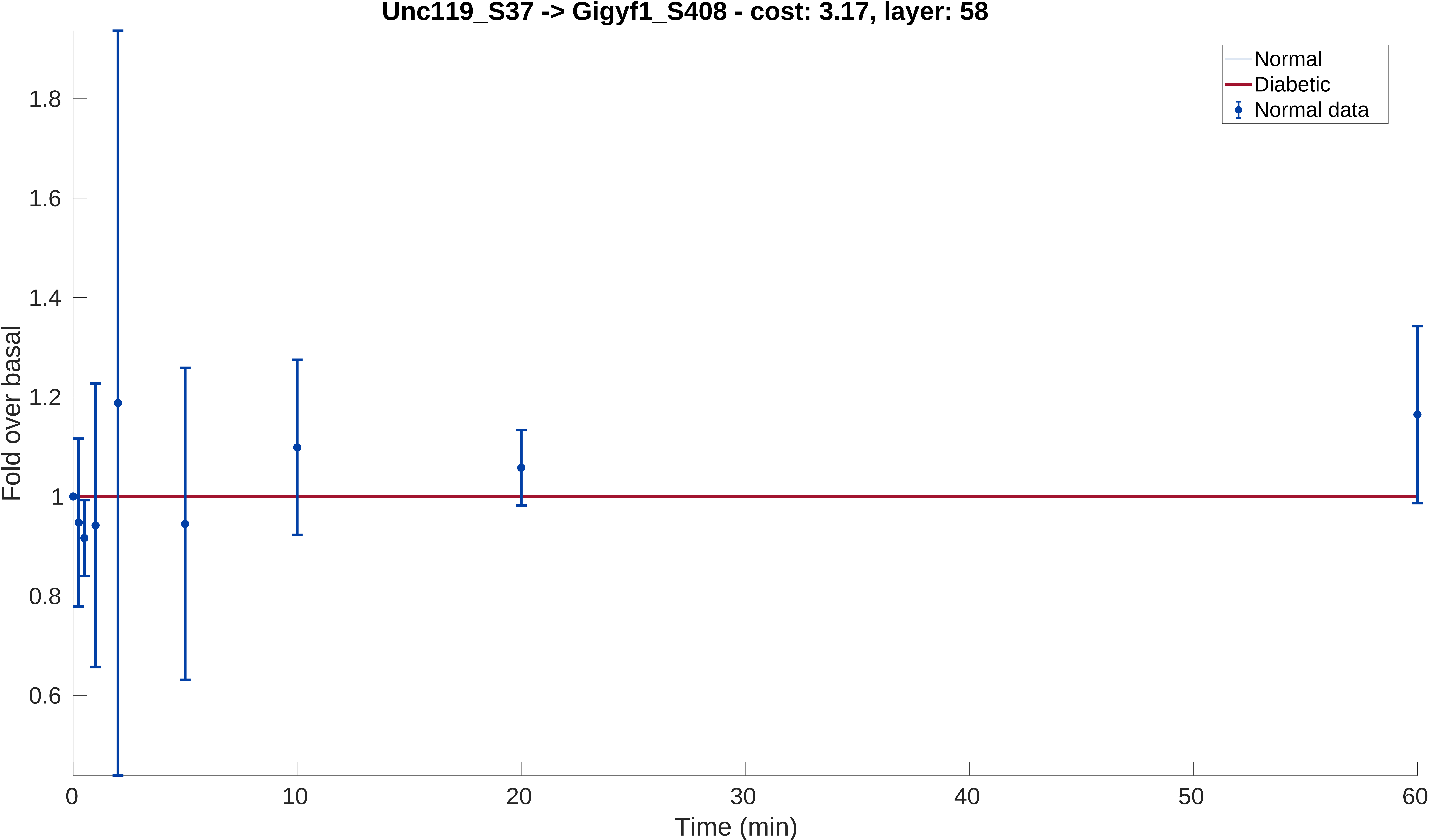

Unc119_S37 -> Gigyf1_S408 - cost: 3.17, layer: 58
Normal
Diabetic
Normal data
Fold over basal
1.8
1.6
1.4
1.2
1
0.8
0.6
0
10
20
30
40
50
60
Time (min)

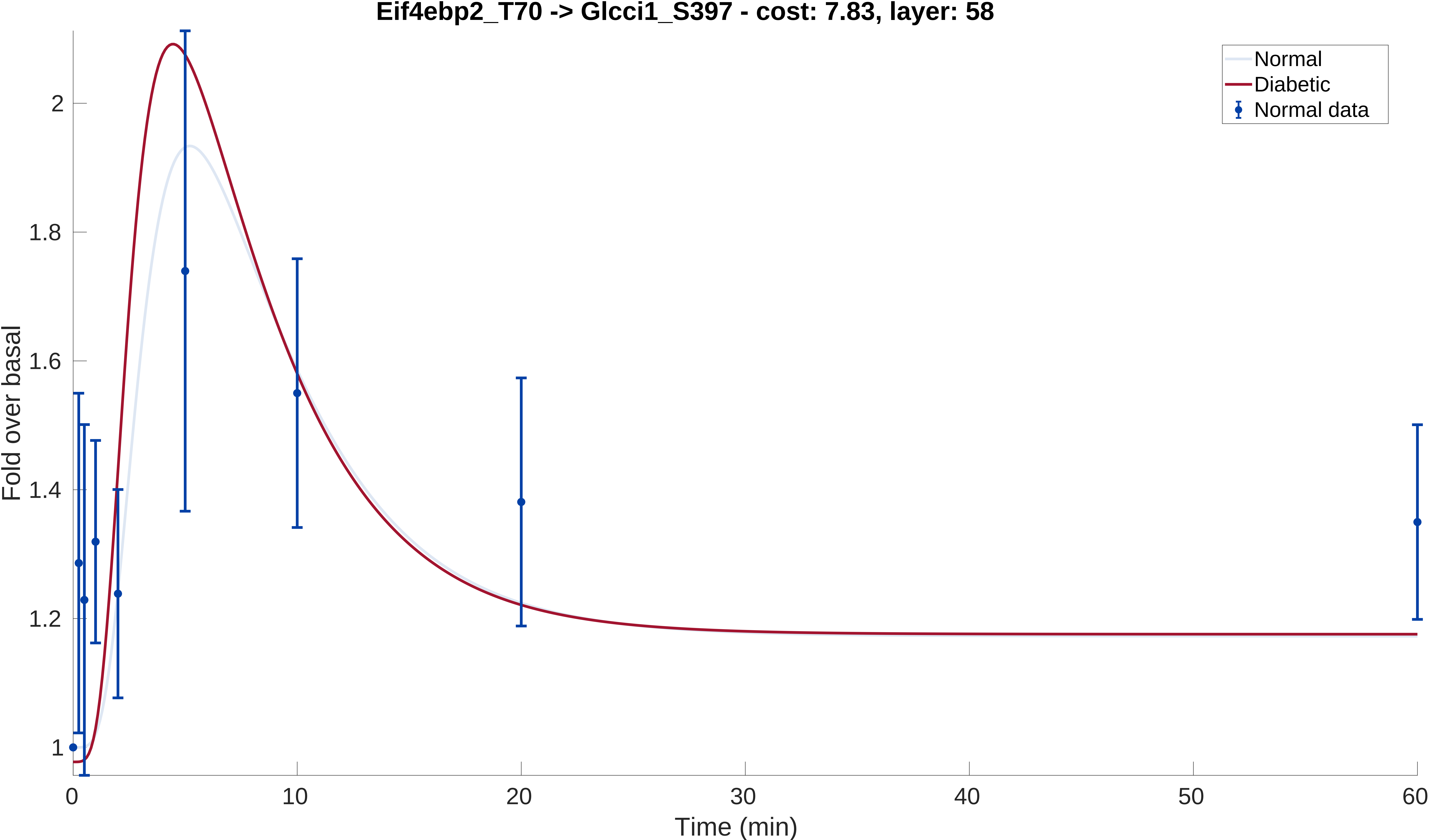

Eif4ebp2_T70 -> Glcci1_S397 - cost: 7.83, layer: 58
Normal
Diabetic
Normal data
Fold over basal
2
1.8
1.6
1.4
1.2
1
0
10
20
30
40
50
60
Time (min)

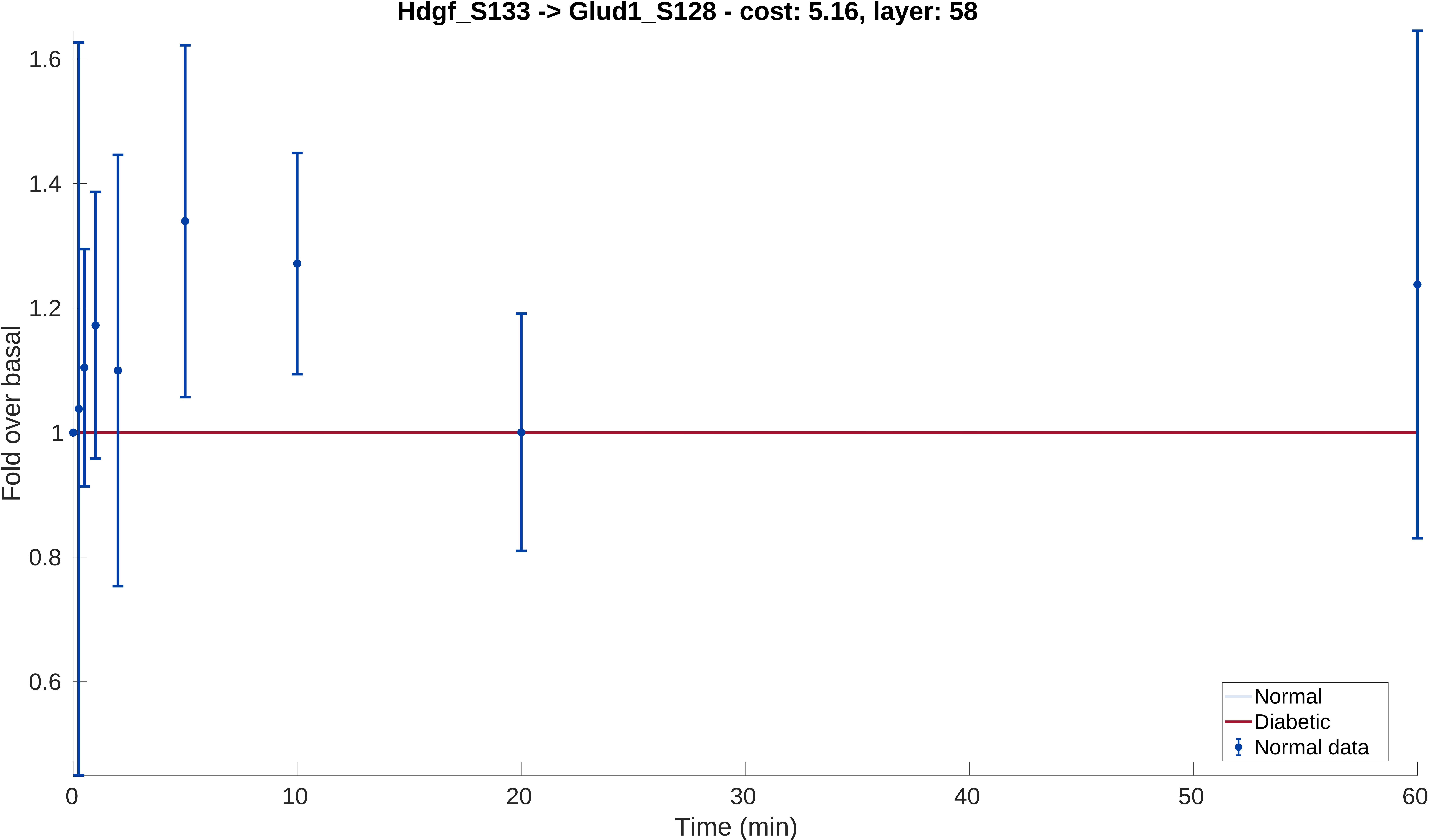

Hdgf_S133 -> Glud1_S128 - cost: 5.16, layer: 58
Fold over basal
1.6
1.4
1.2
1
0.8
0.6
0
10
20
30
40
50
60
Time (min)
Normal
Diabetic
Normal data

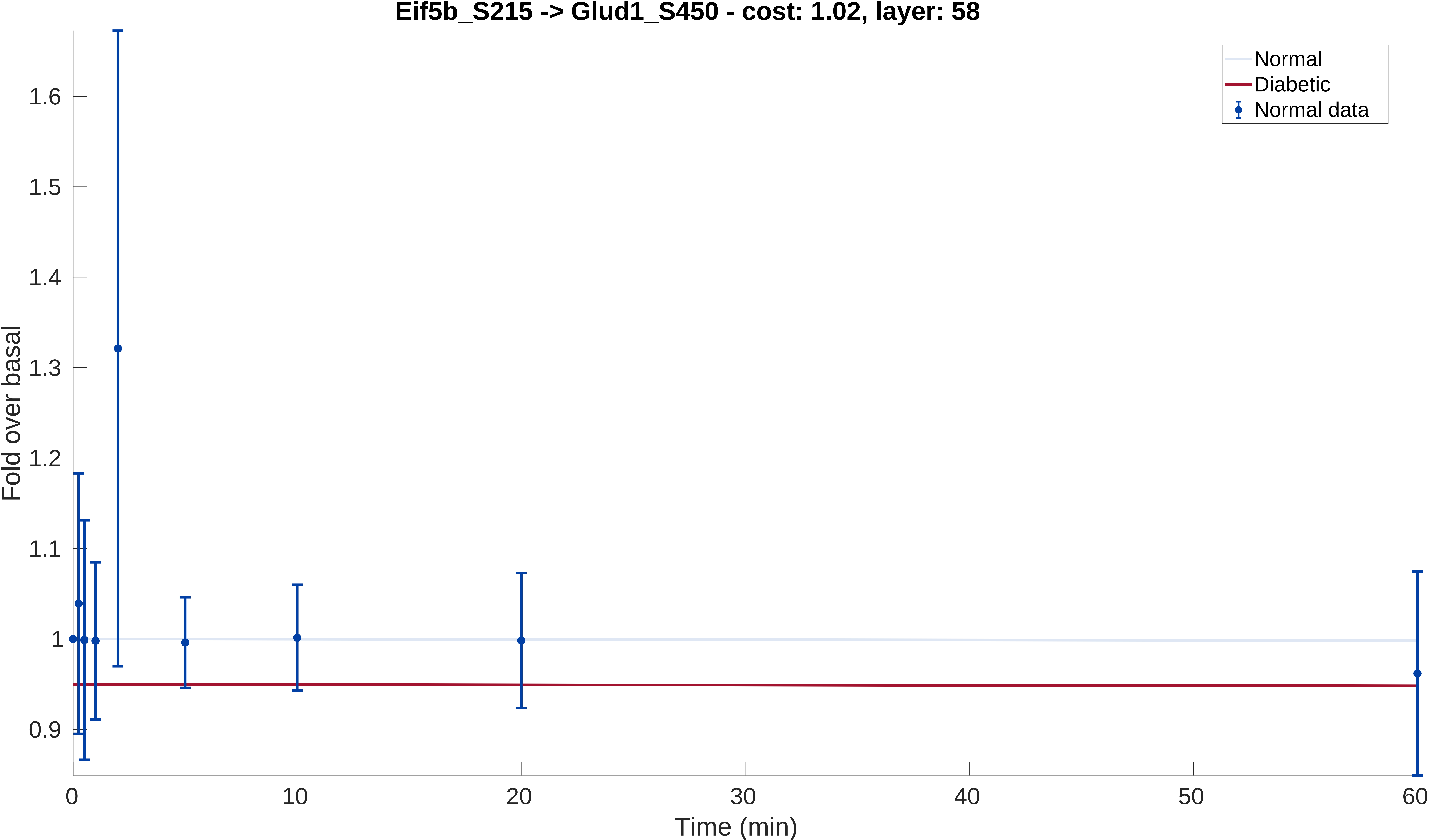

Eif5b_S215 -> Glud1_S450 - cost: 1.02, layer: 58
Normal
Diabetic
Normal data
Fold over basal
1.6
1.5
1.4
1.3
1.2
1.1
1
0.9
0
10
20
30
40
50
60
Time (min)

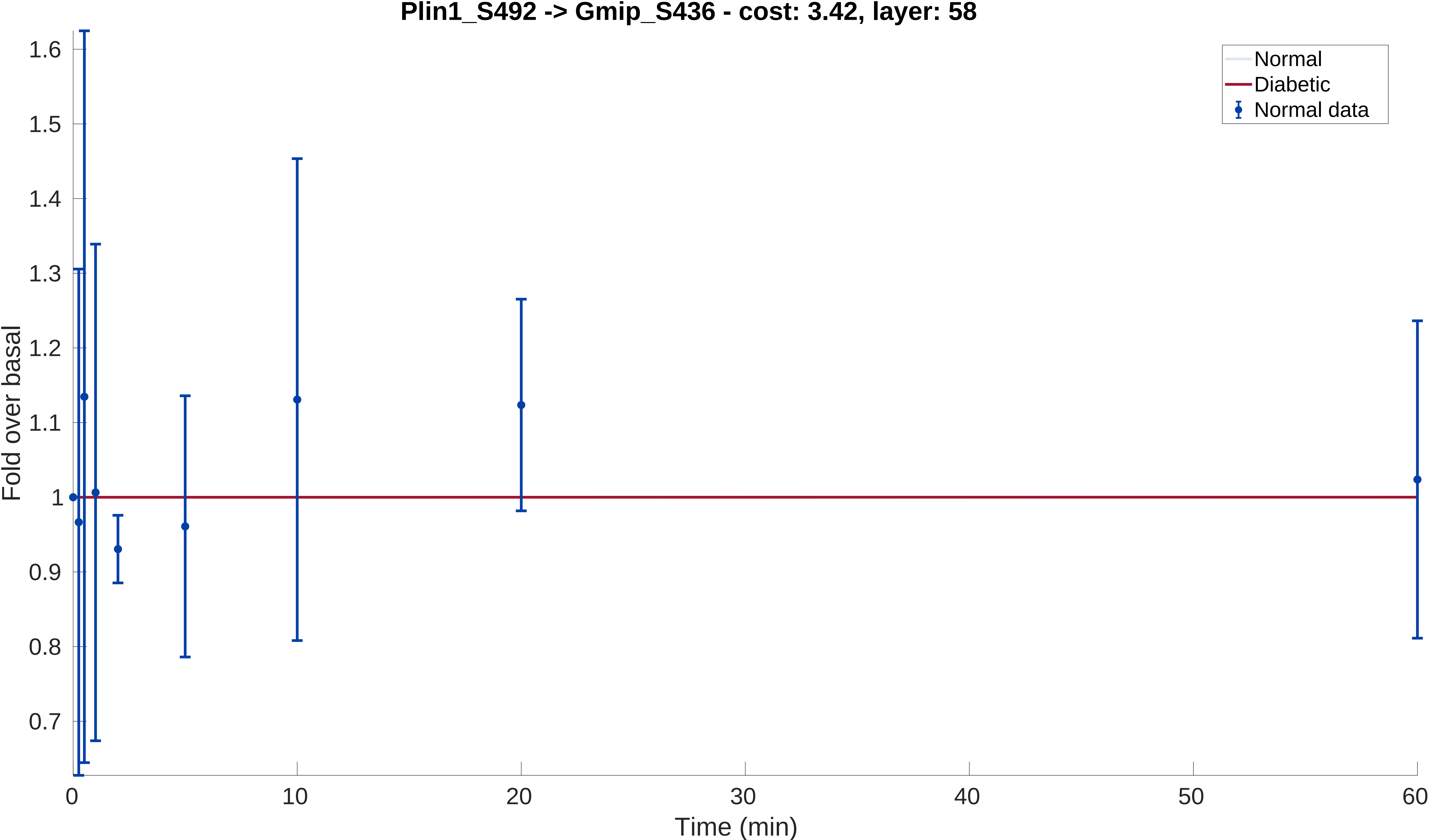
Plin1_S492 -> Gmip_S436 - cost: 3.42, layer: 58
Normal
Diabetic
Normal data
Fold over basal
1.6
1.5
1.4
1.3
1.2
1.1
1
0.9
0.8
0.7
0
10
20
30
40
50
60
Time (min)

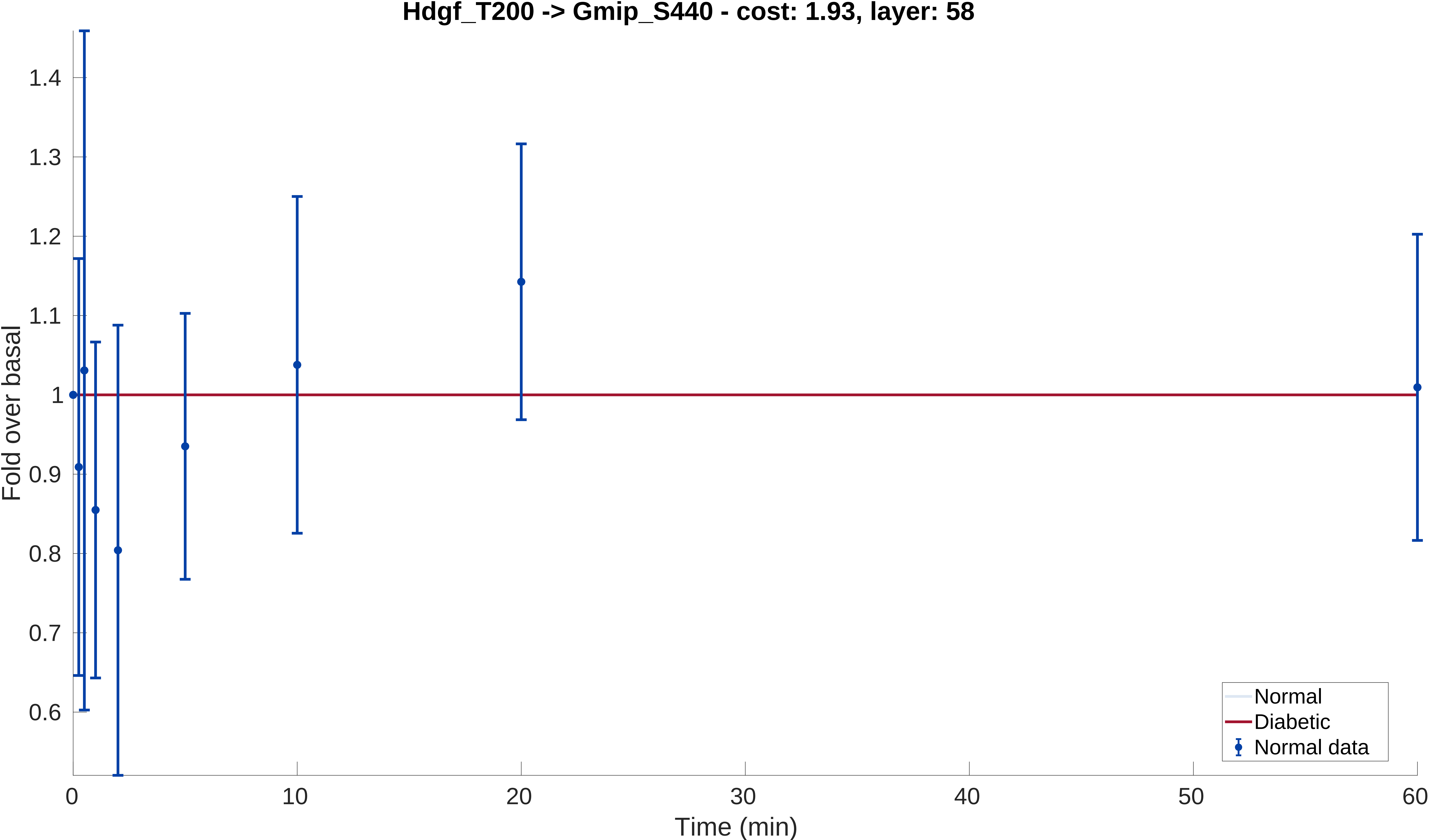
Hdgf_T200 -> Gmip_S440 - cost: 1.93, layer: 58
Fold over basal
1.4
1.3
1.2
1.1
1
0.9
0.8
0.7
0.6
0
10
20
30
40
50
60
Time (min)
Normal
Diabetic
Normal data

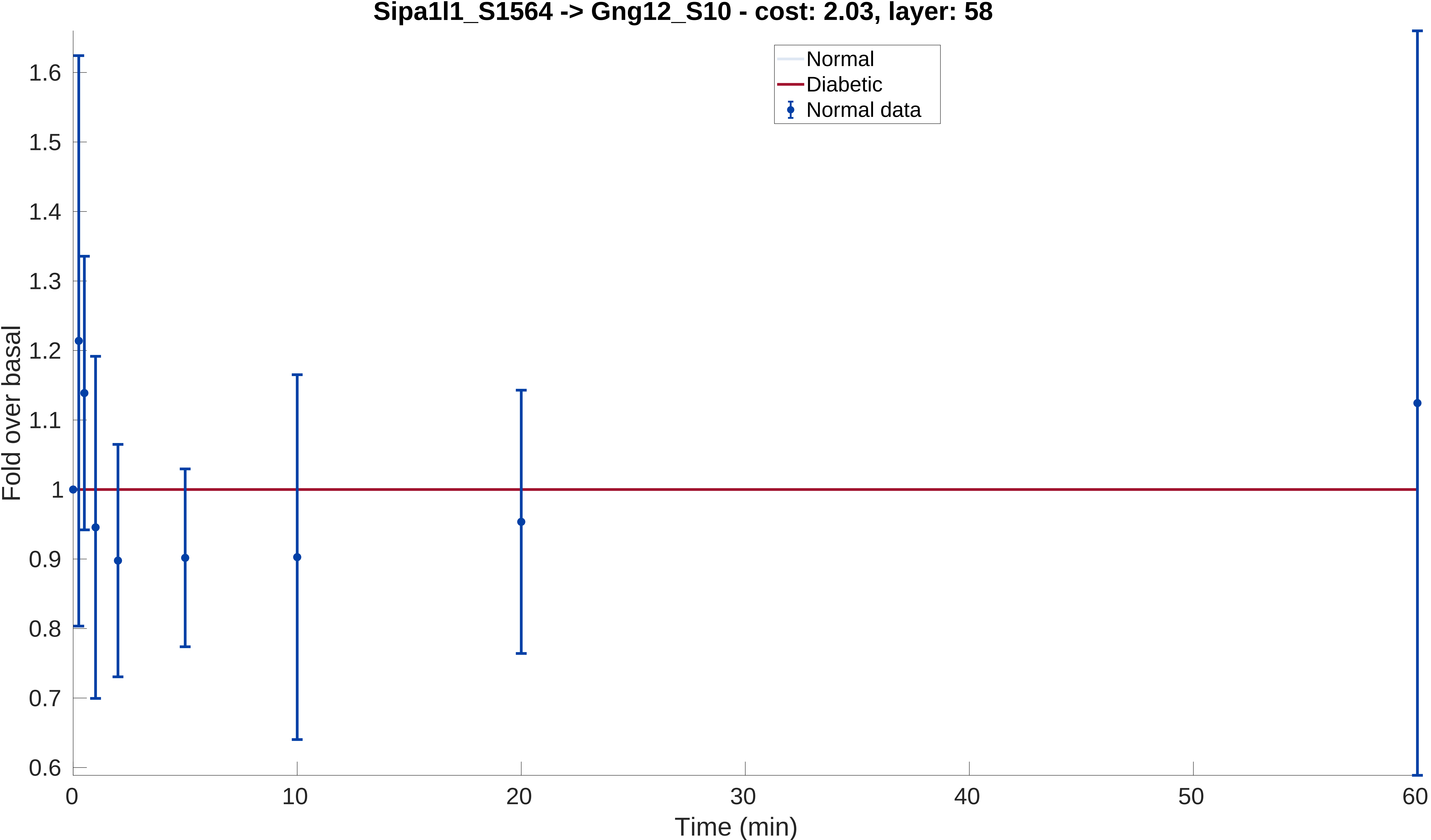
Sipa1l1_S1564 -> Gng12_S10 - cost: 2.03, layer: 58
Normal
Diabetic
Normal data
Fold over basal
1.6
1.5
1.4
1.3
1.2
1.1
1
0.9
0.8
0.7
0.6
0
10
20
30
40
50
60
Time (min)

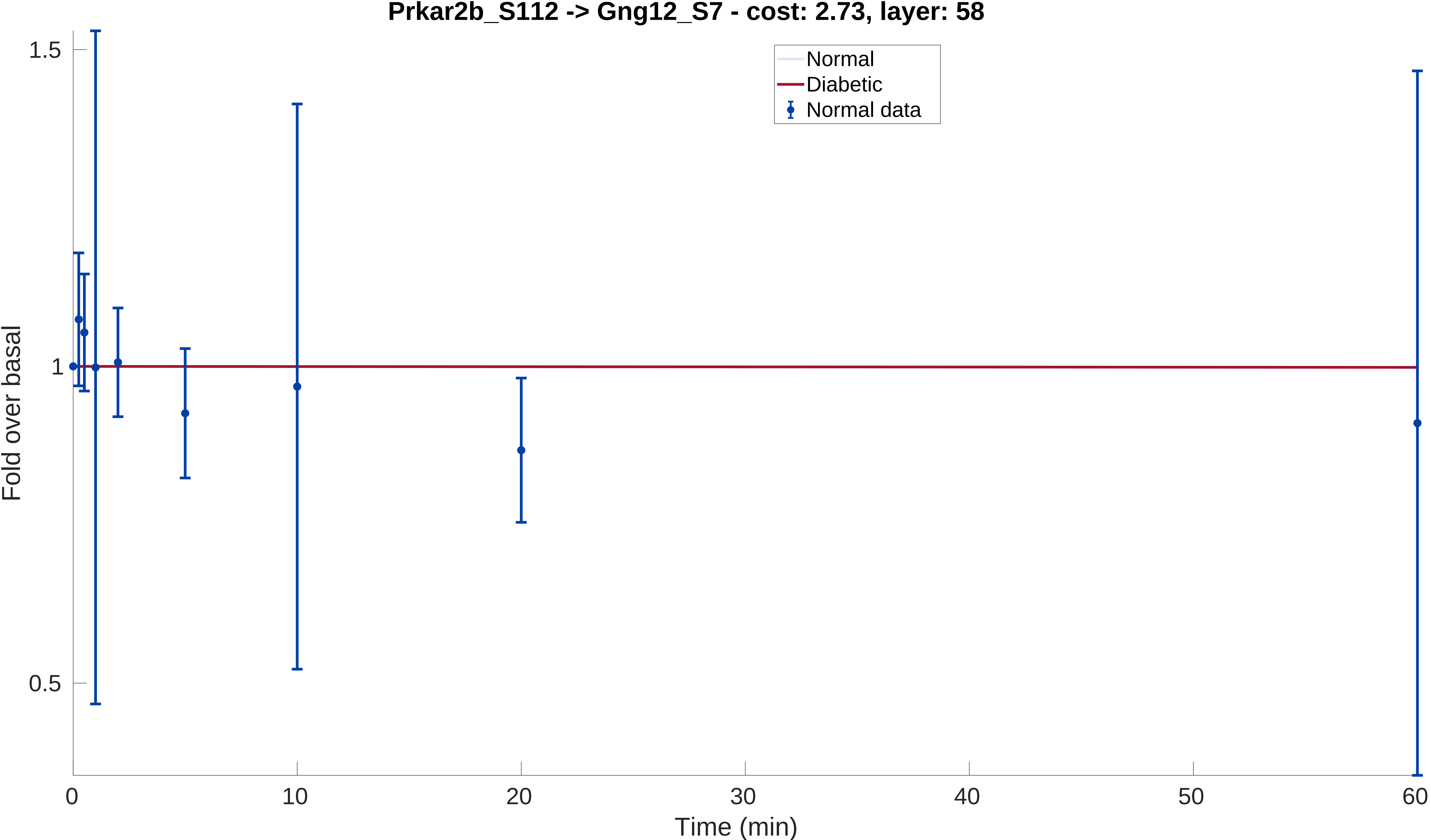

Prkar2b_S112 -> Gng12_S7 - cost: 2.73, layer: 58
Normal
Diabetic
Normal data
1.5
1
0.5
Fold over basal
0
10
20
30
40
50
60
Time (min)

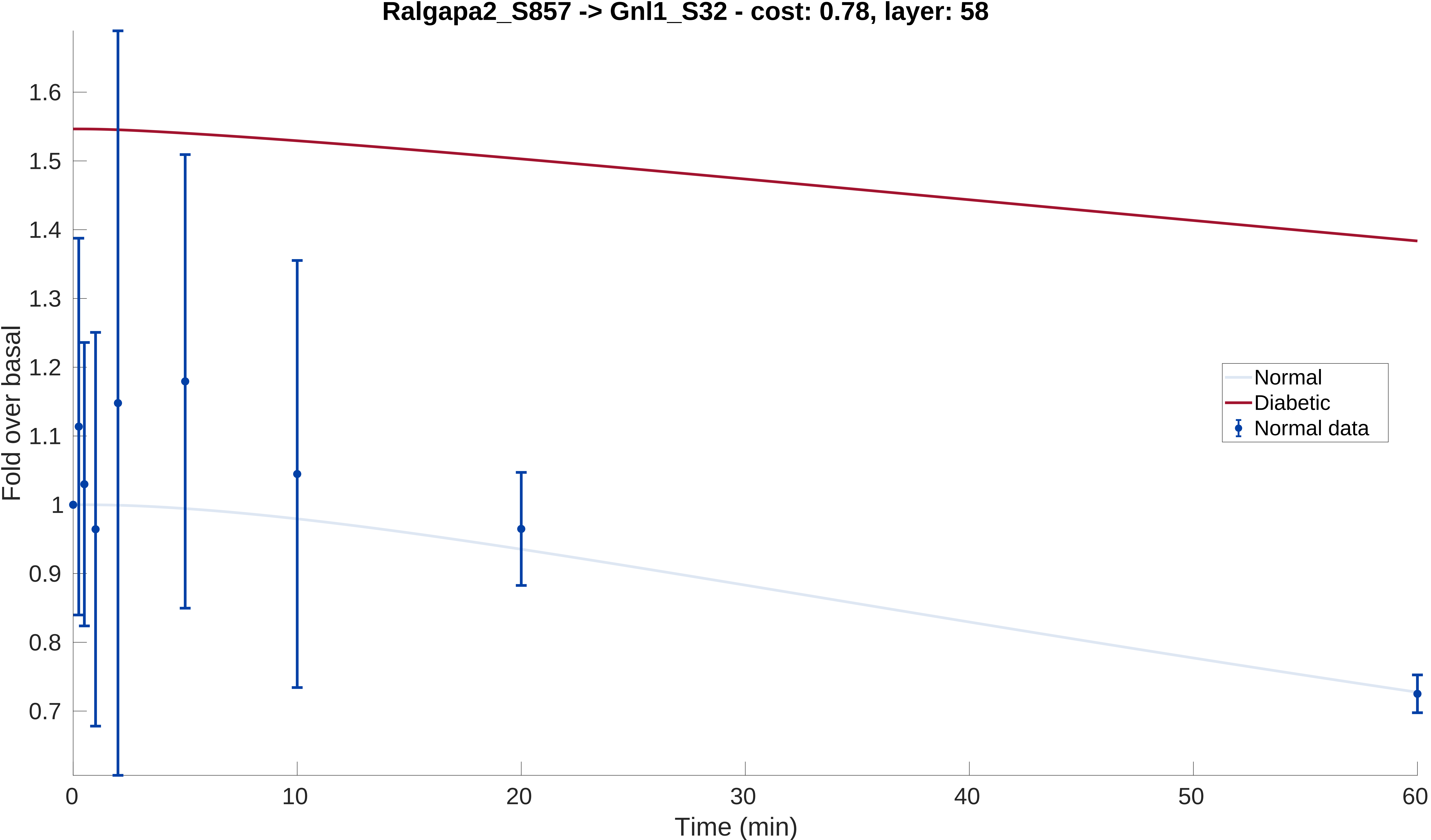

Ralgapa2_S857 -> Gnl1_S32 - cost: 0.78, layer: 58
Fold over basal
1.6
1.5
1.4
1.3
1.2
1.1
1
0.9
0.8
0.7
0
10
20
30
40
50
60
Time (min)
Normal
Diabetic
Normal data

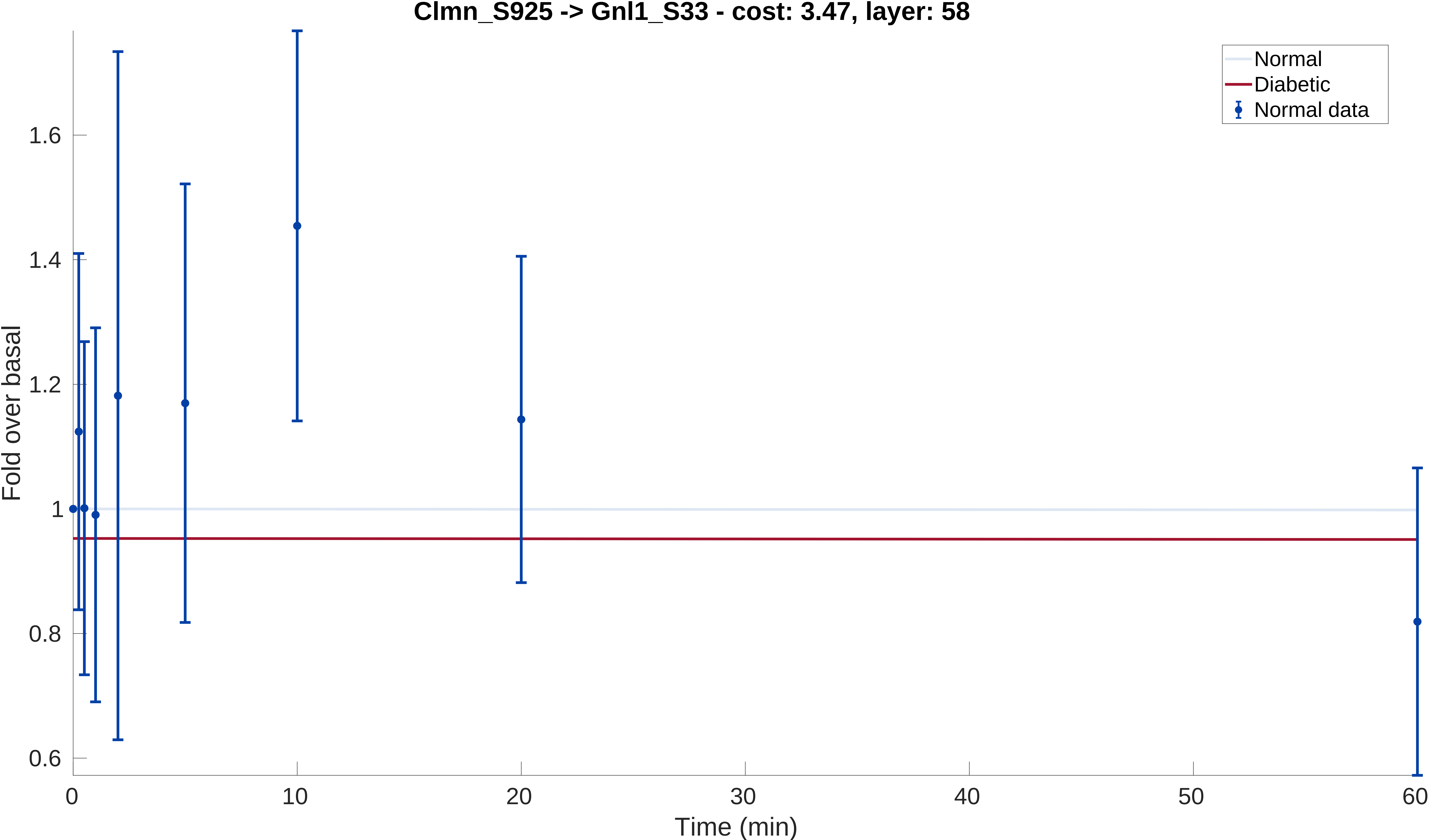
Clmn_S925 -> Gnl1_S33 - cost: 3.47, layer: 58
Normal
Diabetic
Normal data
Fold over basal
1.6
1.4
1.2
1
0.8
0.6
0
10
20
30
40
50
60
Time (min)

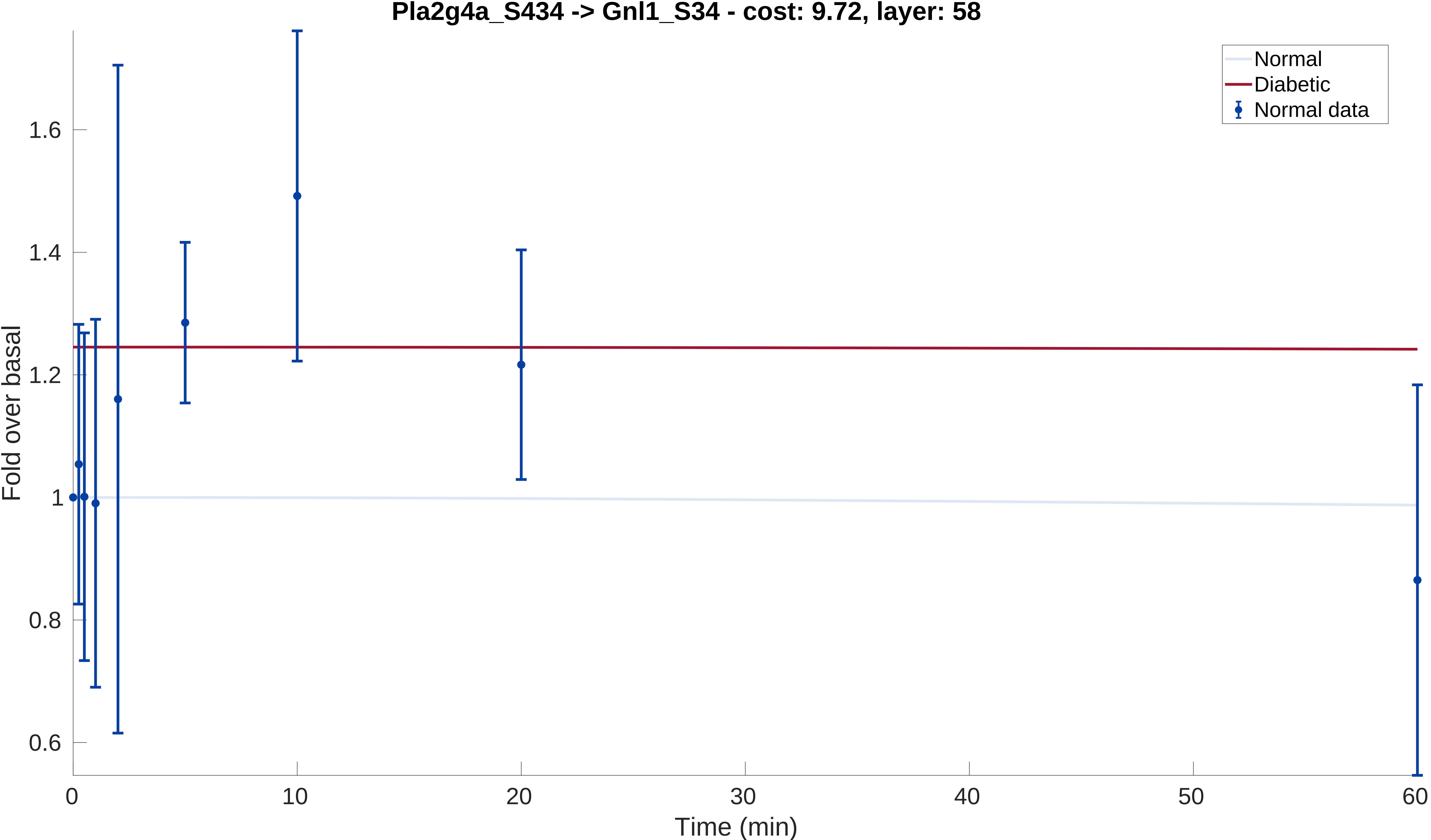

Pla2g4a_S434 -> Gnl1_S34 - cost: 9.72, layer: 58
Normal
Diabetic
Normal data
Fold over basal
1.6
1.4
1.2
1
0.8
0.6
0
10
20
30
40
50
60
Time (min)

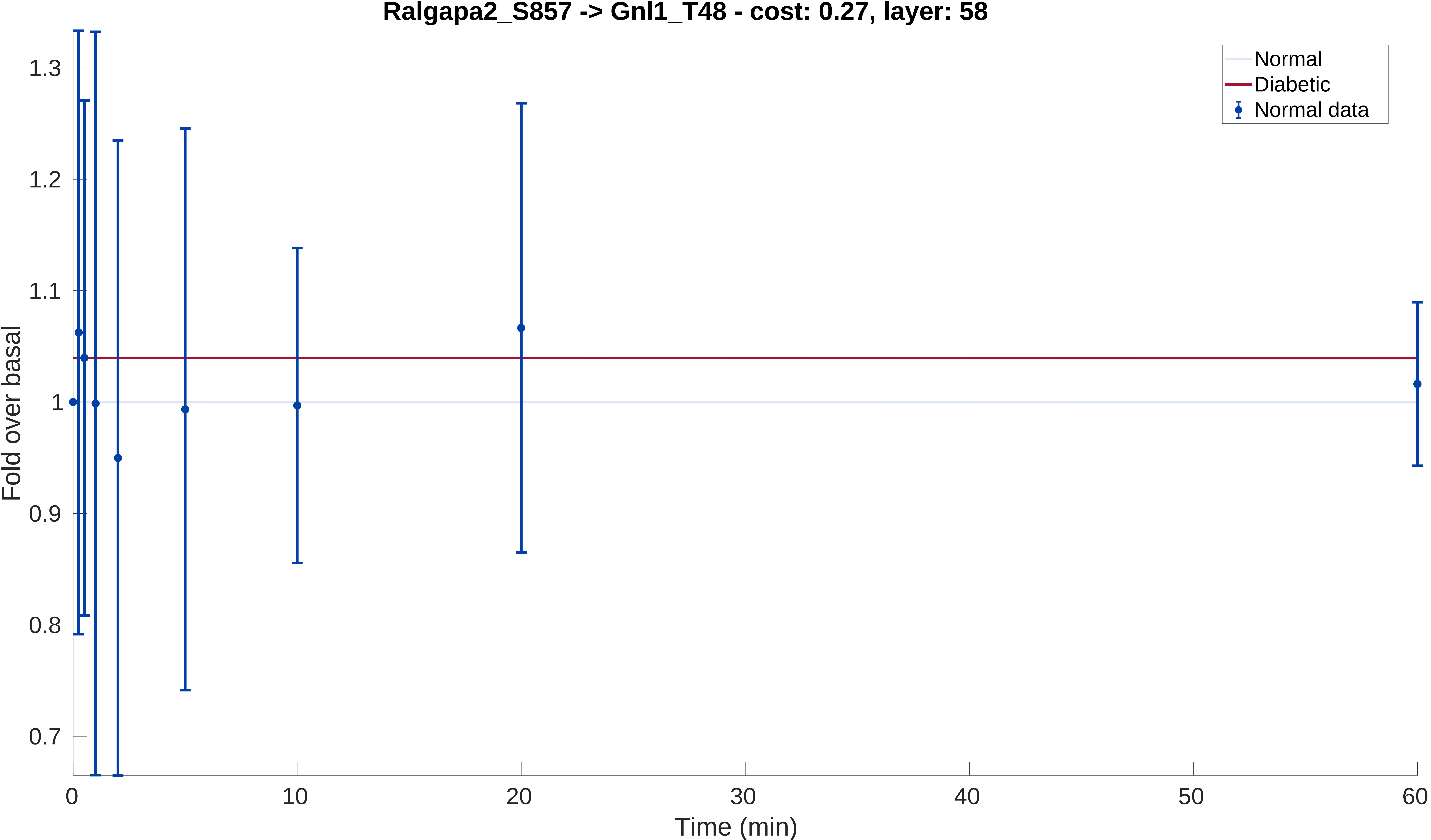
Ralgapa2_S857 -> Gnl1_T48 - cost: 0.27, layer: 58
Normal
Diabetic
Normal data
Fold over basal
1.3
1.2
1.1
1
0.9
0.8
0.7
0
10
20
30
40
50
60
Time (min)

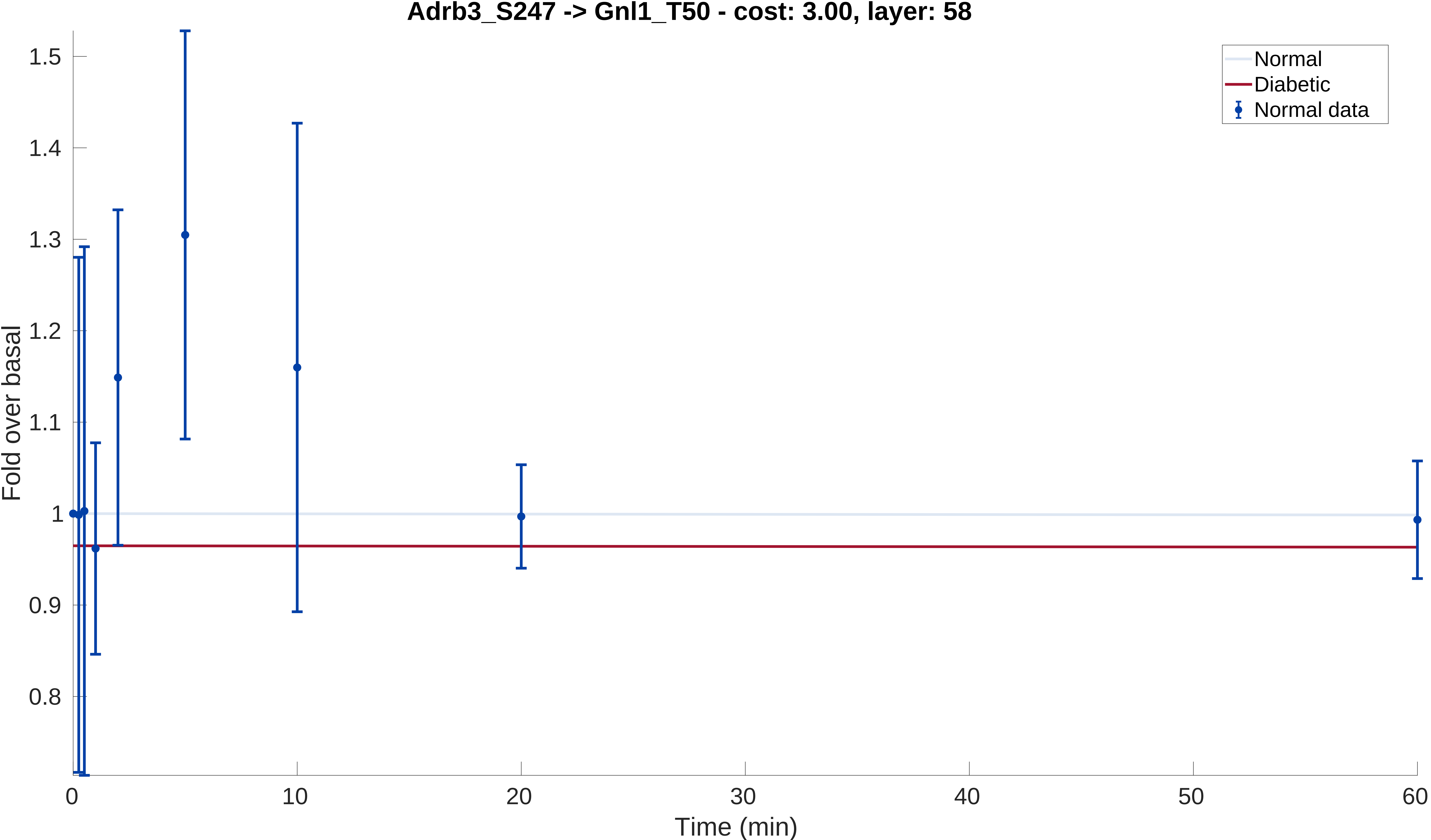
Adrb3_S247 -> Gnl1_T50 - cost: 3.00, layer: 58
Normal
Diabetic
Normal data
1.5
1.4
1.3
1.2
1.1
1
0.9
0.8
Fold over basal
0
10
20
30
40
50
60
Time (min)

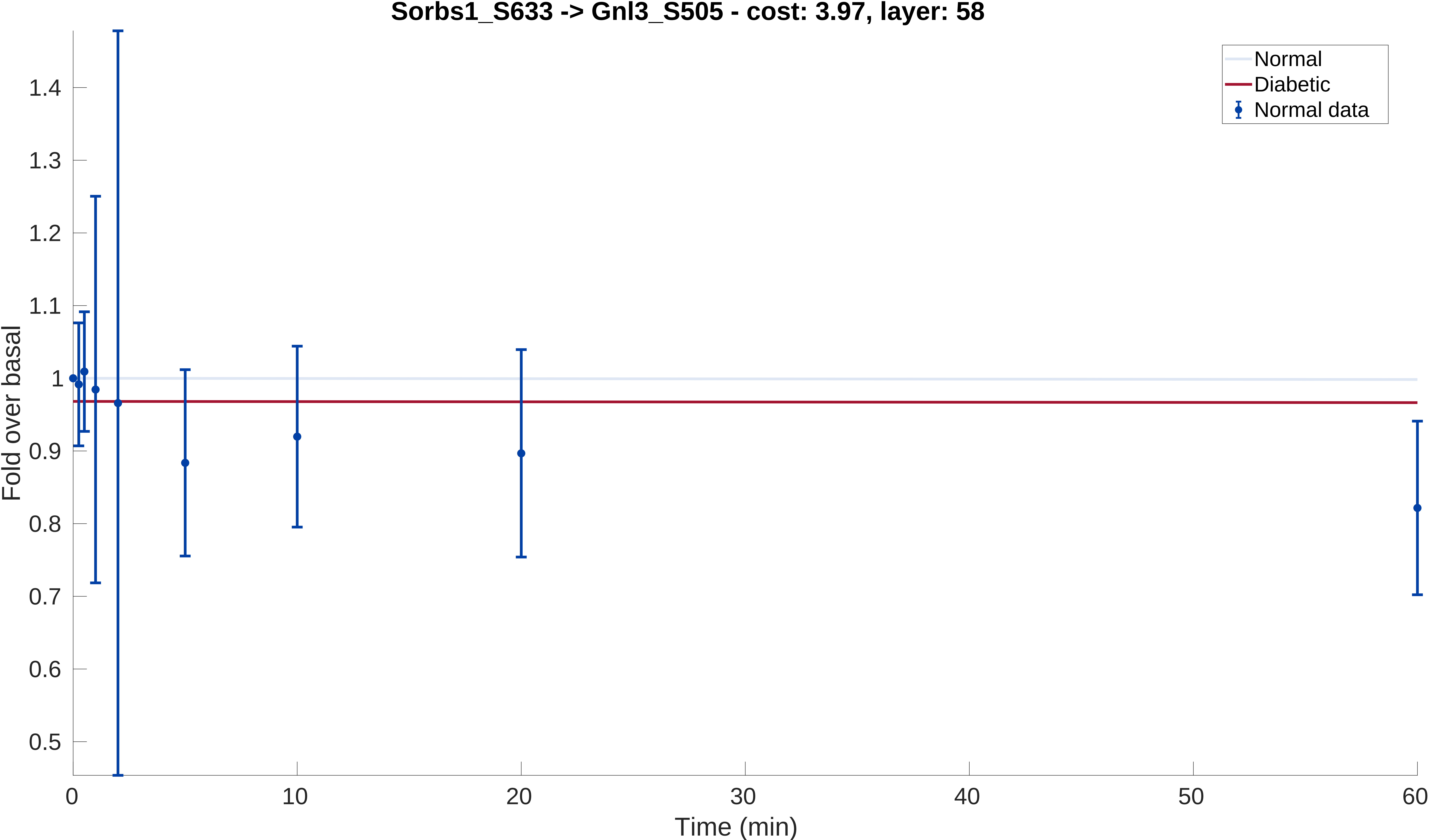
Sorbs1_S633 -> Gnl3_S505 - cost: 3.97, layer: 58
Normal
Diabetic
Normal data
Fold over basal
1.4
1.3
1.2
1.1
1
0.9
0.8
0.7
0.6
0.5
0
10
20
30
40
50
60
Time (min)

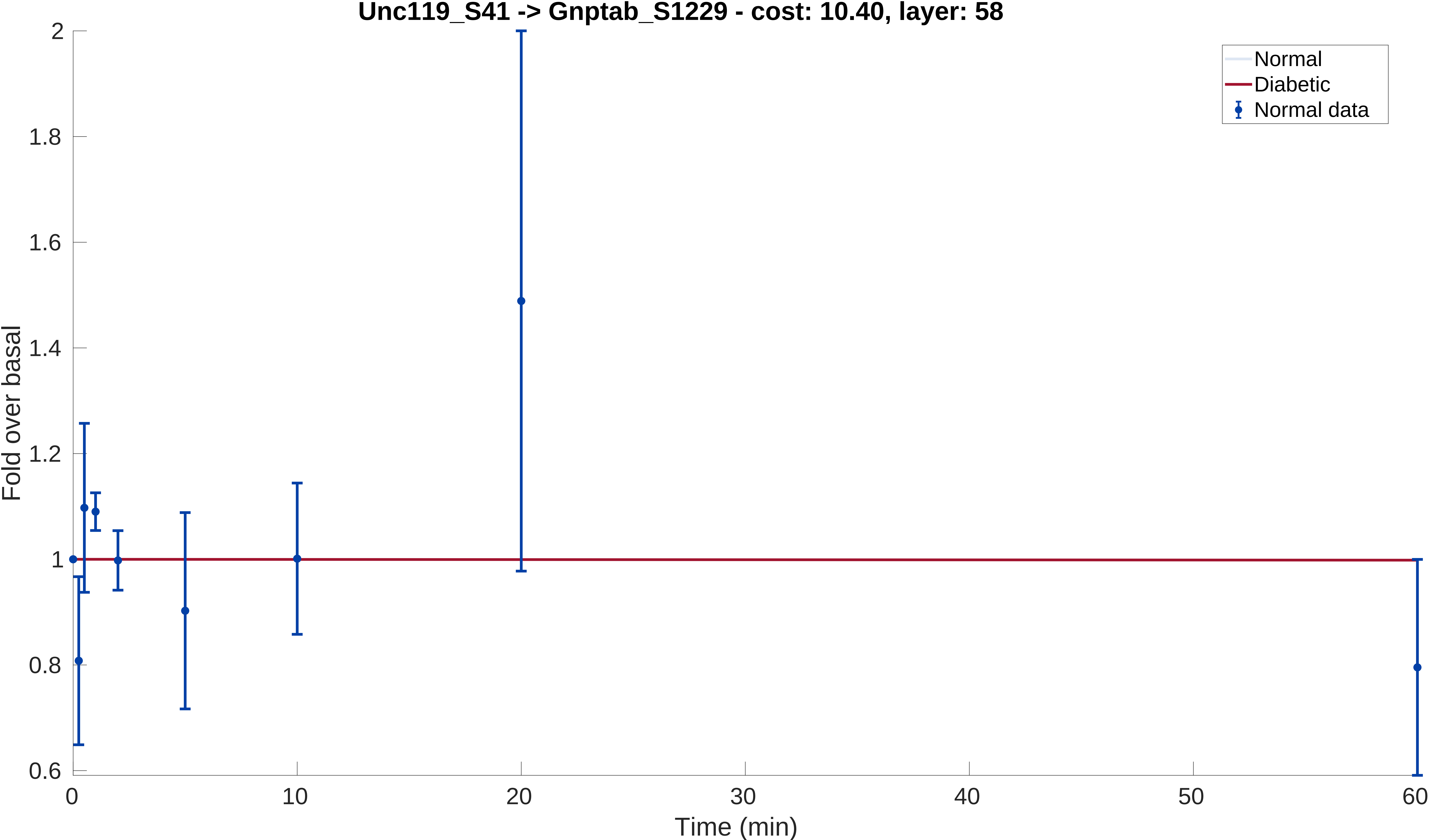

Unc119_S41 -> Gnptab_S1229 - cost: 10.40, layer: 58
Normal
Diabetic
Normal data
Fold over basal
Time (min)
2
1.8
1.6
1.4
1.2
1
0.8
0.6
0
10
20
30
40
50
60

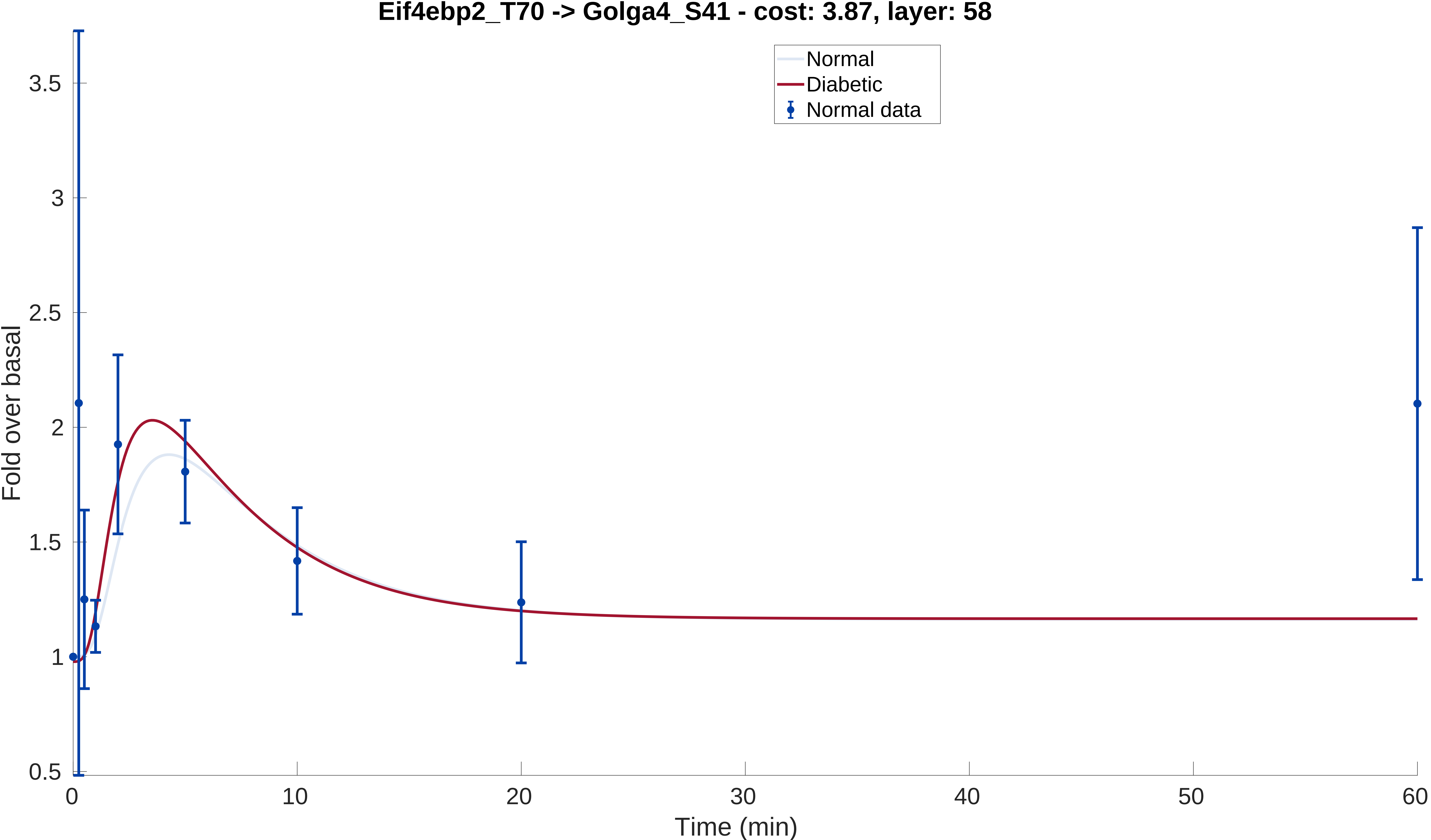

Eif4ebp2_T70 -> Golga4_S41 - cost: 3.87, layer: 58
Normal
Diabetic
Normal data
Fold over basal
Time (min)
0.5
1
1.5
2
2.5
3
3.5
0
10
20
30
40
50
60

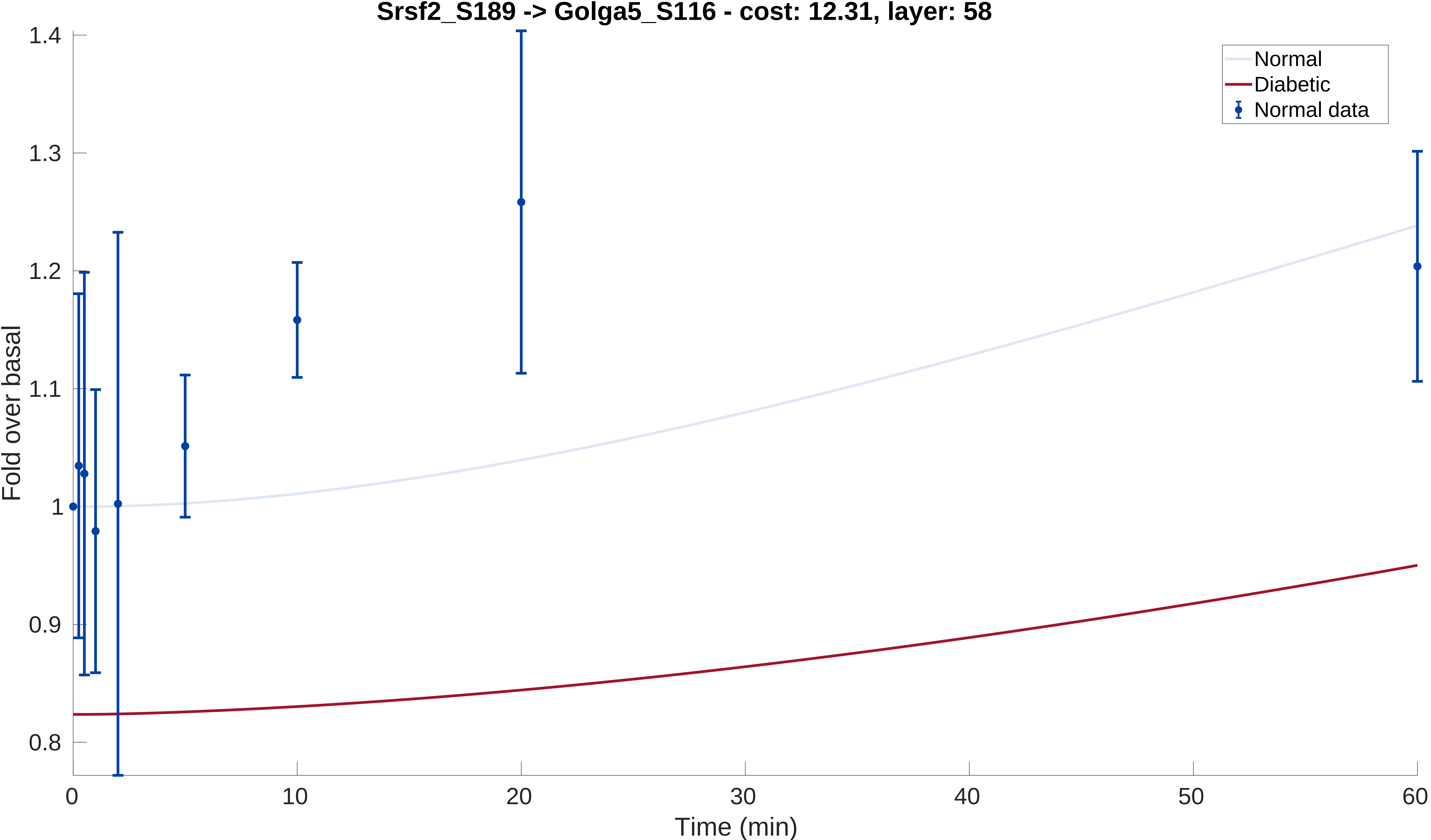

Srsf2_S189 -> Golga5_S116 - cost: 12.31, layer: 58
Normal
Diabetic
Normal data
Fold over basal
Time (min)
1.4
1.3
1.2
1.1
1
0.9
0.8
0
10
20
30
40
50
60

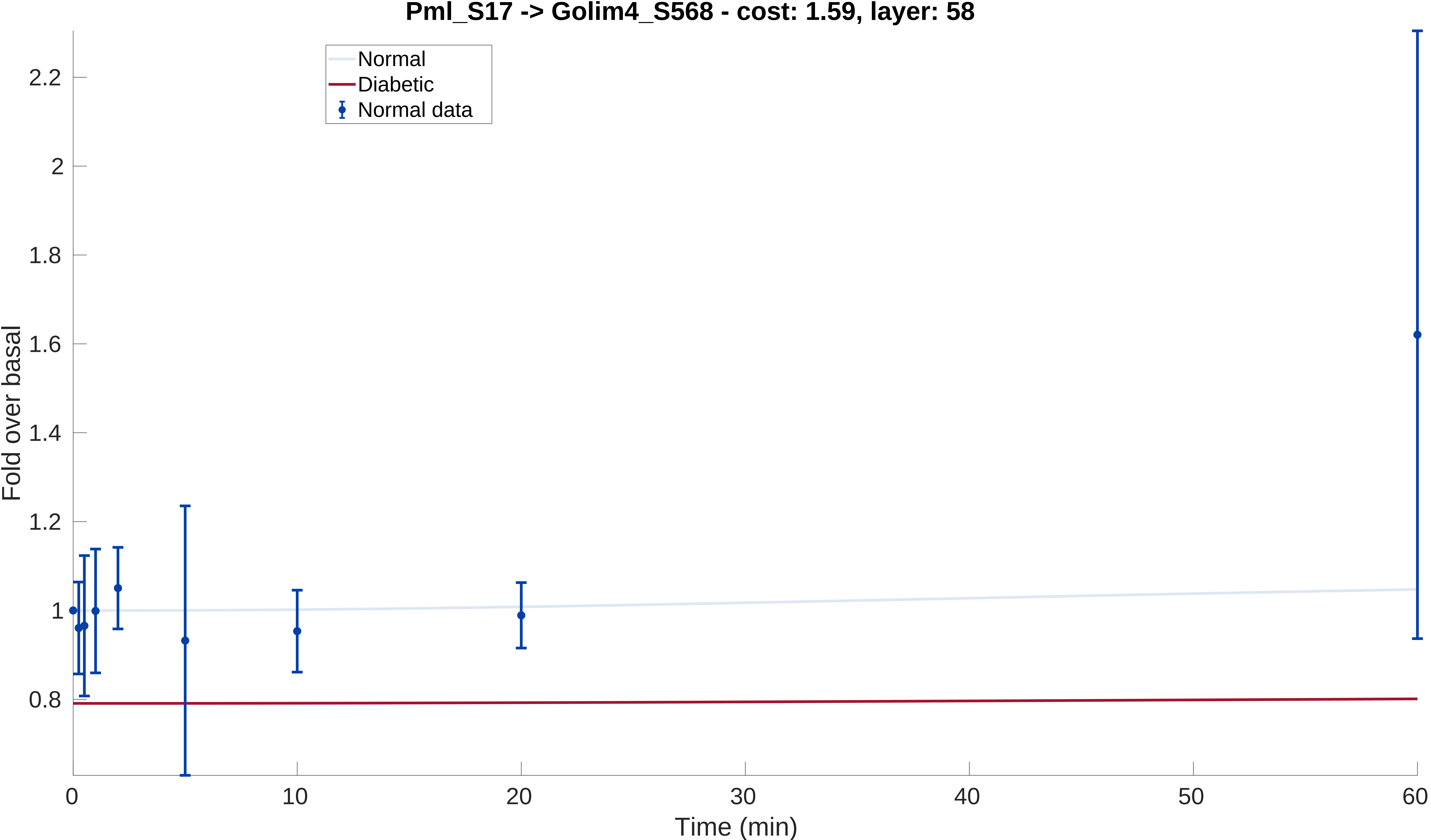
Pml_S17 -> Golim4_S568 - cost: 1.59, layer: 58
Normal
Diabetic
Normal data
Fold over basal
2.2
2
1.8
1.6
1.4
1.2
1
0.8
0
10
20
30
40
50
60
Time (min)

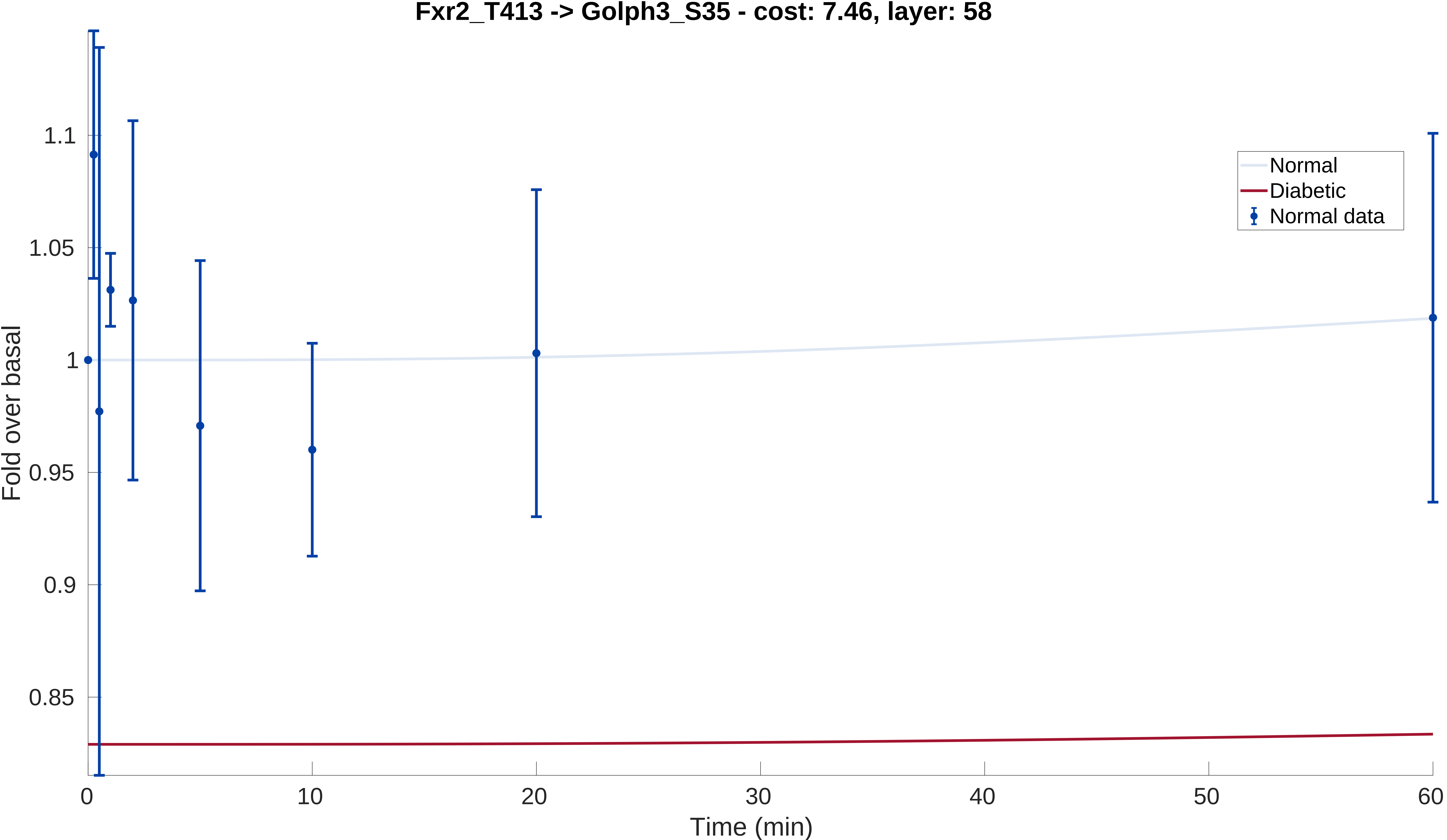

Fxr2_T413 -> Golph3_S35 - cost: 7.46, layer: 58
Fold over basal
1.1
1.05
1
0.95
0.9
0.85
0
10
20
30
40
50
60
Time (min)
Normal
Diabetic
Normal data

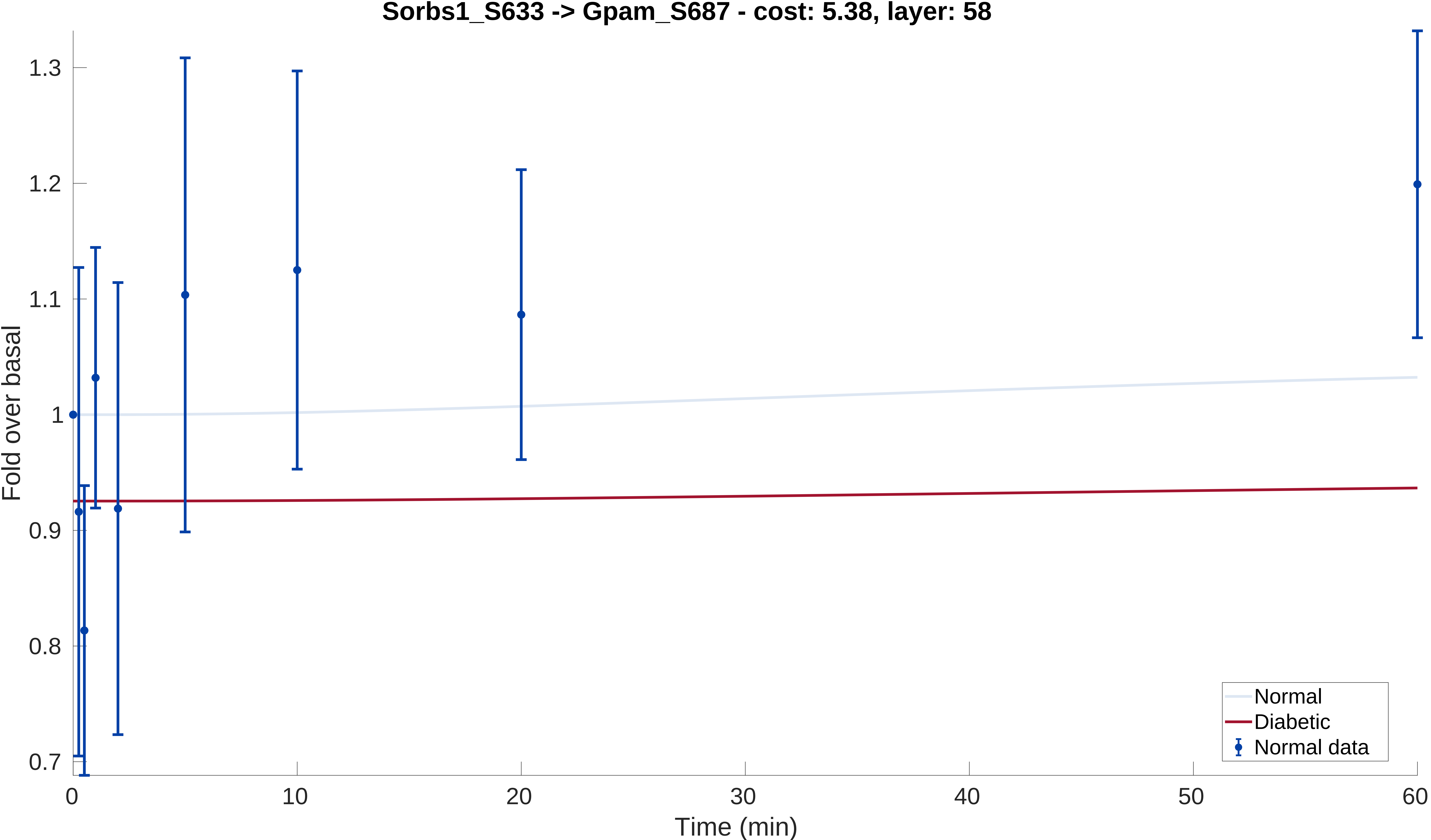

Sorbs1_S633 -> Gpam_S687 - cost: 5.38, layer: 58
Fold over basal
1.3
1.2
1.1
1
0.9
0.8
0.7
0
10
20
30
40
50
60
Time (min)
Normal
Diabetic
Normal data

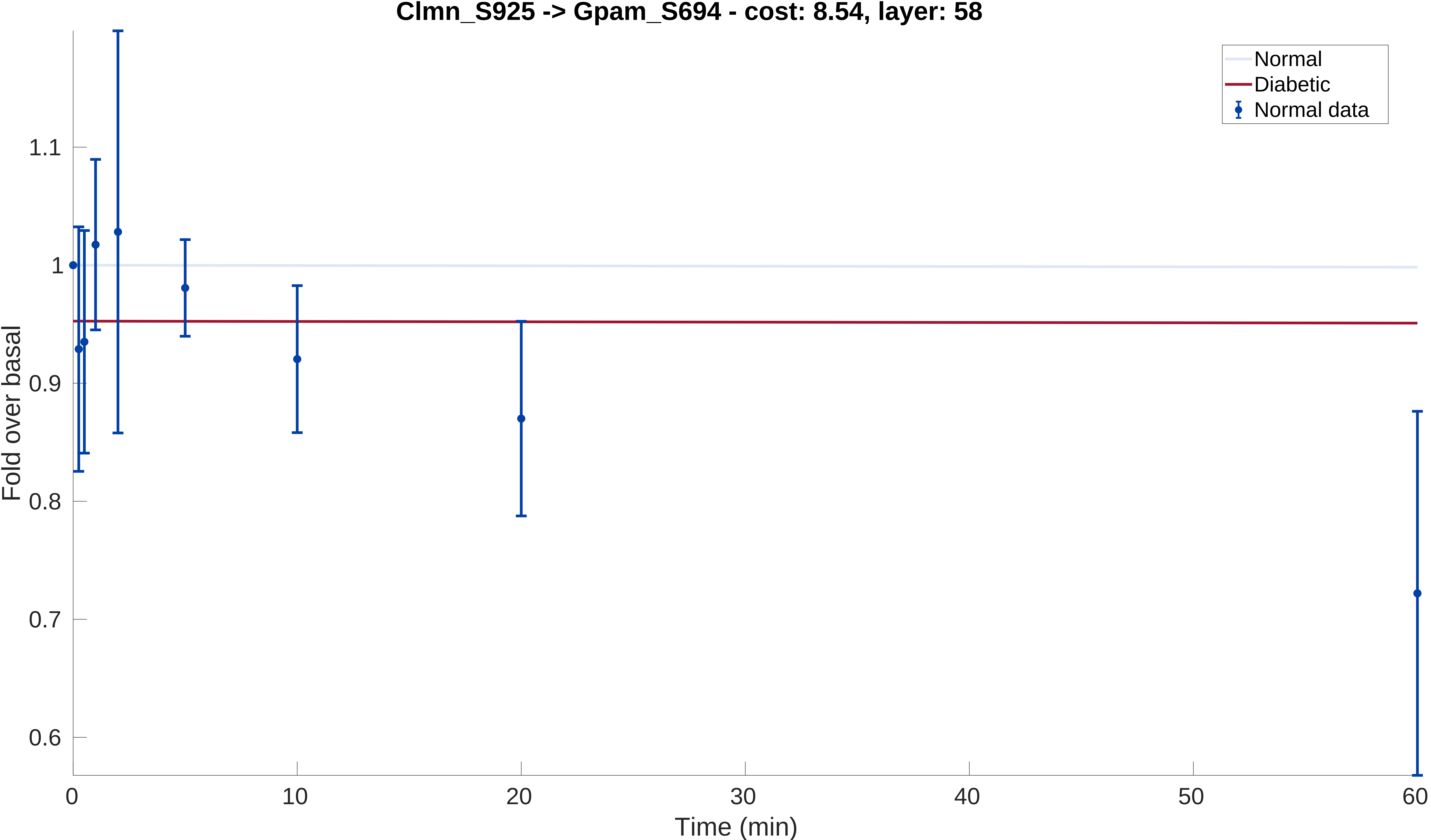

Clmn_S925 -> Gpam_S694 - cost: 8.54, layer: 58
Normal
Diabetic
Normal data
Fold over basal
1.1
1
0.9
0.8
0.7
0.6
0
10
20
30
40
50
60
Time (min)

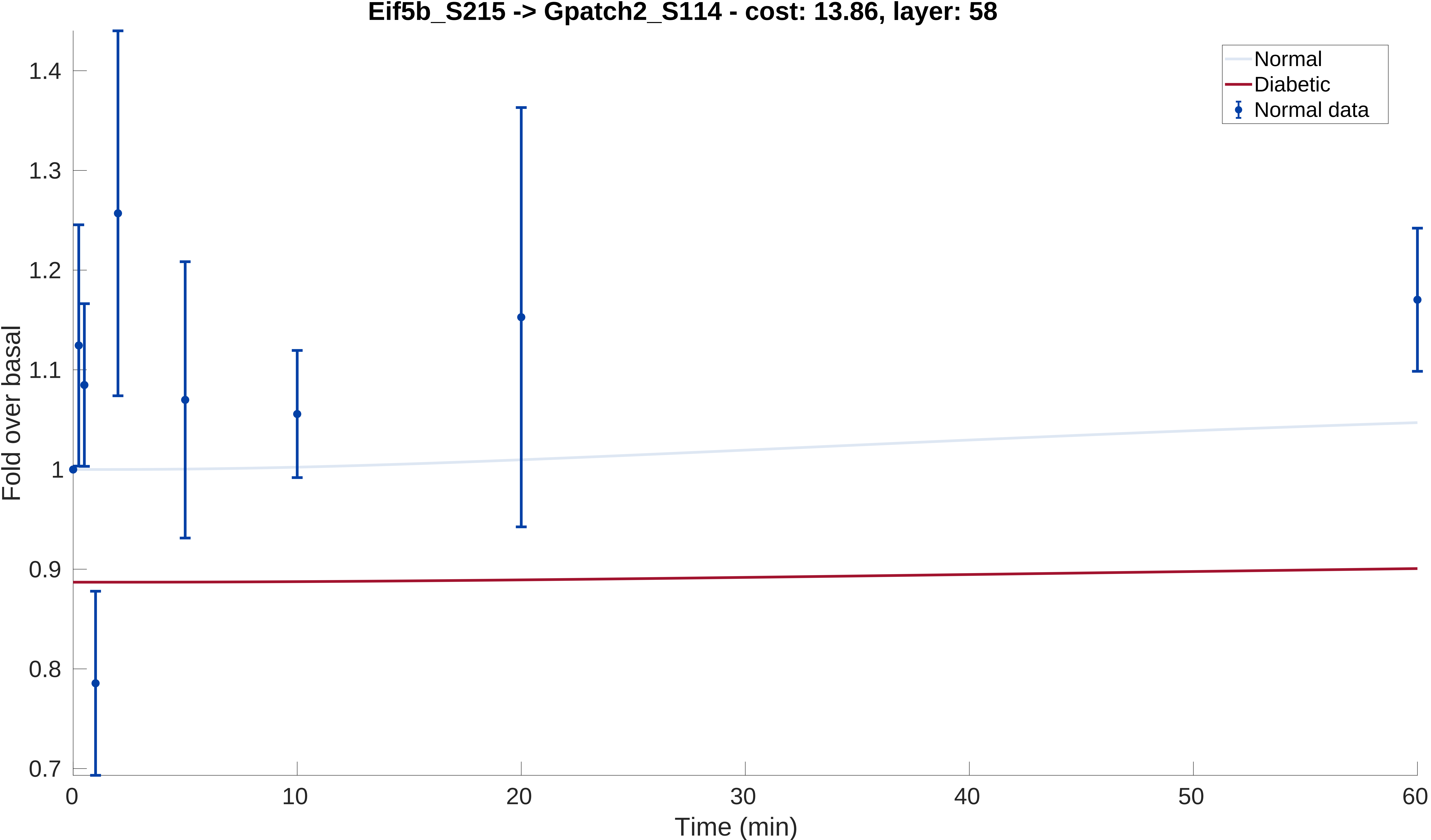

Eif5b_S215 -> Gpatch2_S114 - cost: 13.86, layer: 58
Normal
Diabetic
Normal data
1.4
1.3
1.2
1.1
1
0.9
0.8
0.7
Fold over basal
0
10
20
30
40
50
60
Time (min)

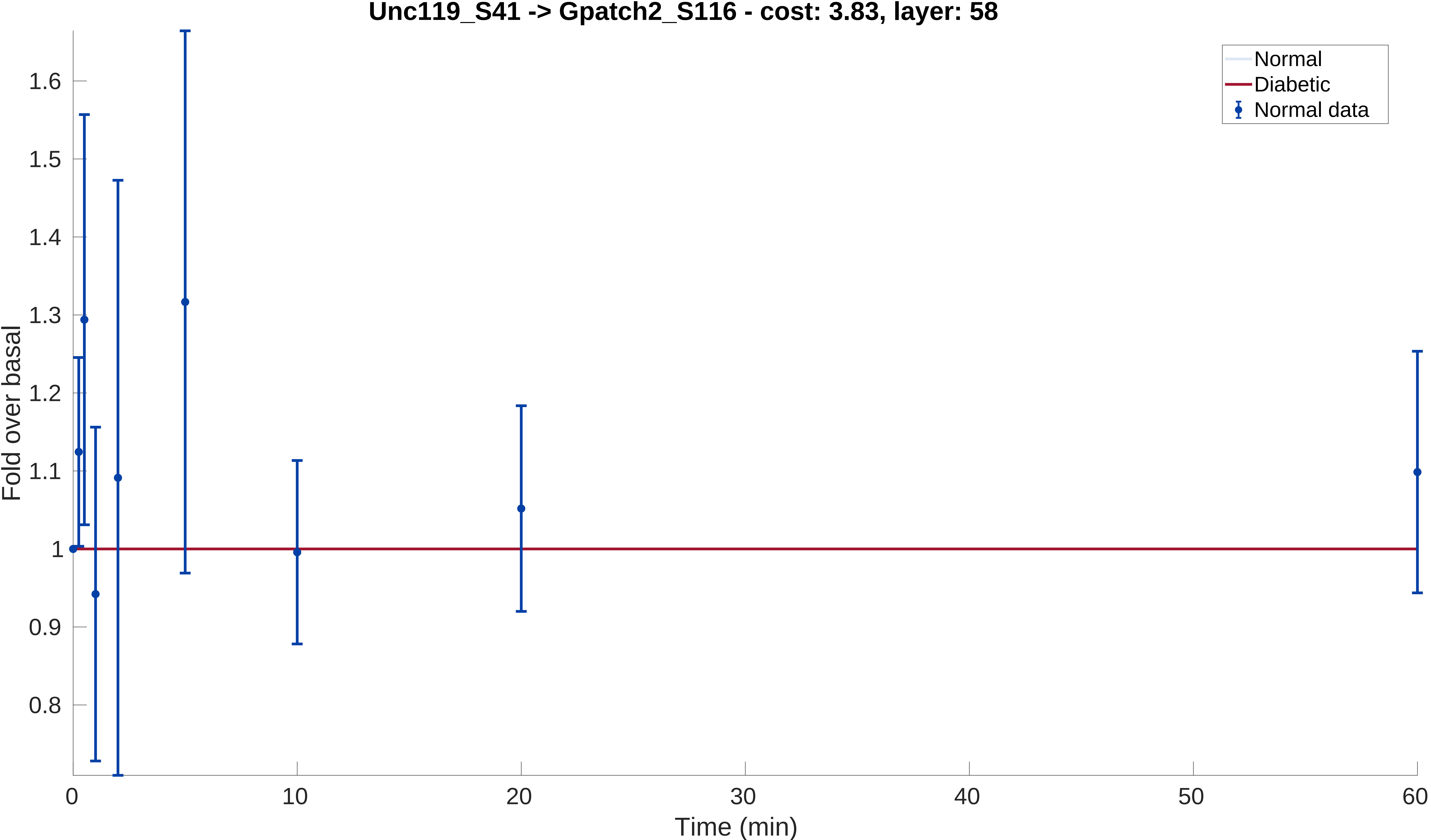
Unc119_S41 -> Gpatch2_S116 - cost: 3.83, layer: 58
Normal
Diabetic
Normal data
Fold over basal
1.6
1.5
1.4
1.3
1.2
1.1
1
0.9
0.8
0
10
20
30
40
50
60
Time (min)

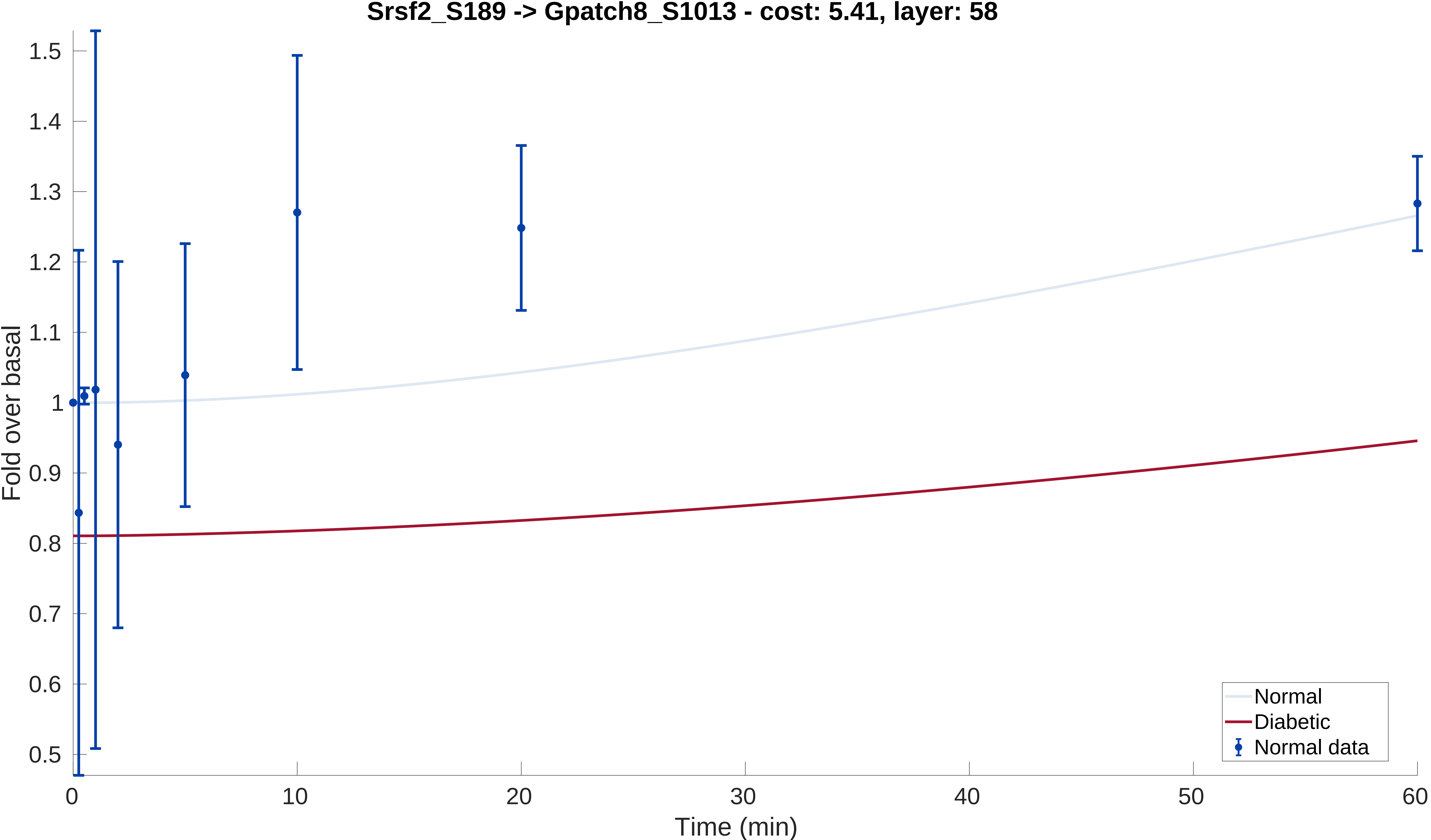

Srsf2_S189 -> Gpatch8_S1013 - cost: 5.41, layer: 58
Fold over basal
1.5
1.4
1.3
1.2
1.1
1
0.9
0.8
0.7
0.6
0.5
0
10
20
30
40
50
60
Time (min)
Normal
Diabetic
Normal data

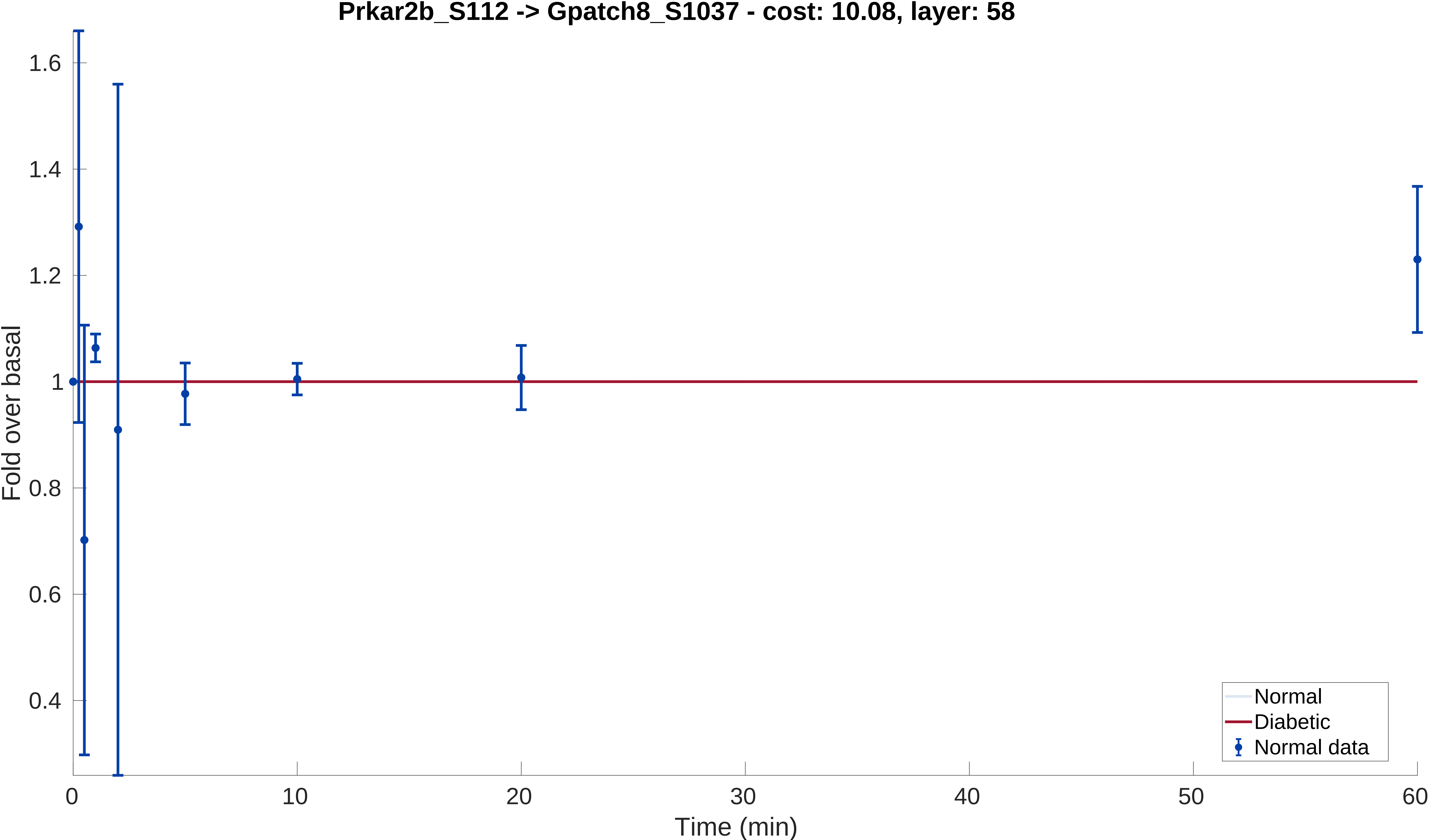
Prrkar2b_S112 -> Gpatch8_S1037 - cost: 10.08, layer: 58
Fold over basal
1.6
1.4
1.2
1
0.8
0.6
0.4
0
10
20
30
40
50
60
Time (min)
Normal
Diabetic
Normal data

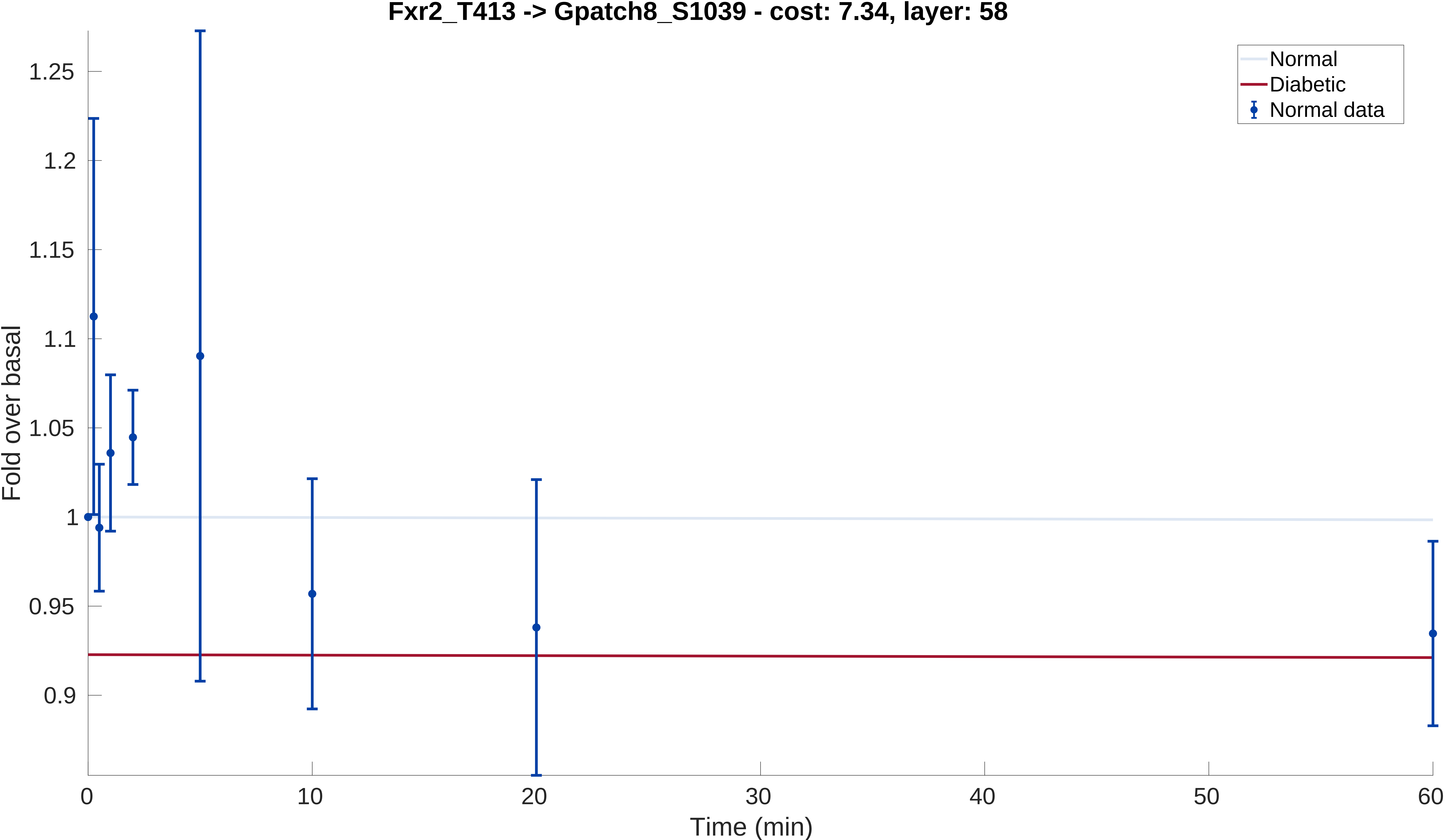

Fxr2_T413 -> Gpatch8_S1039 - cost: 7.34, layer: 58
Normal
Diabetic
Normal data
Fold over basal
1.25
1.2
1.15
1.1
1.05
1
0.95
0.9
0
10
20
30
40
50
60
Time (min)

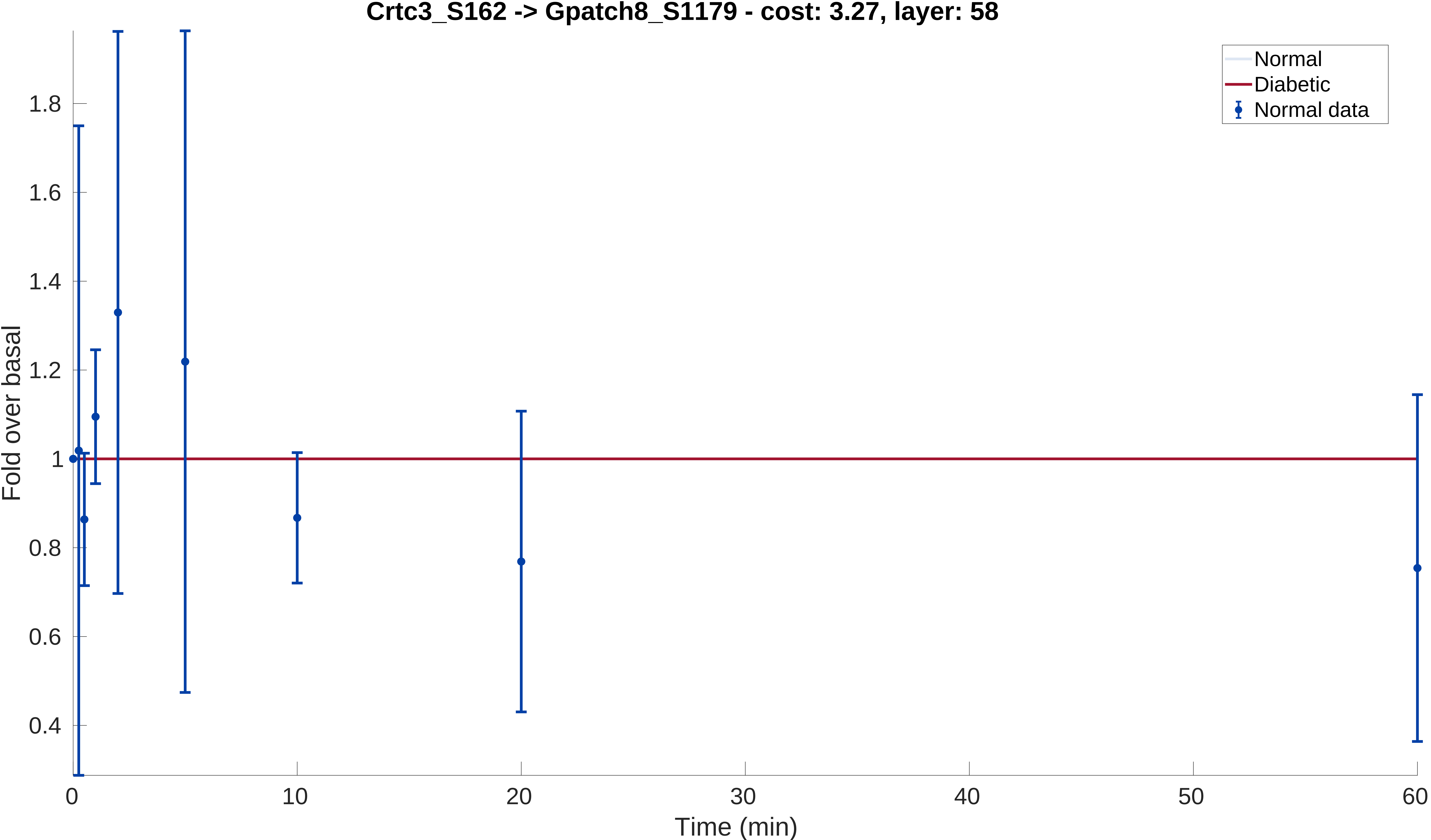

Crtc3_S162 -> Gpatch8_S1179 - cost: 3.27, layer: 58
Normal
Diabetic
Normal data
Fold over basal
1.8
1.6
1.4
1.2
1
0.8
0.6
0.4
0
10
20
30
40
50
60
Time (min)

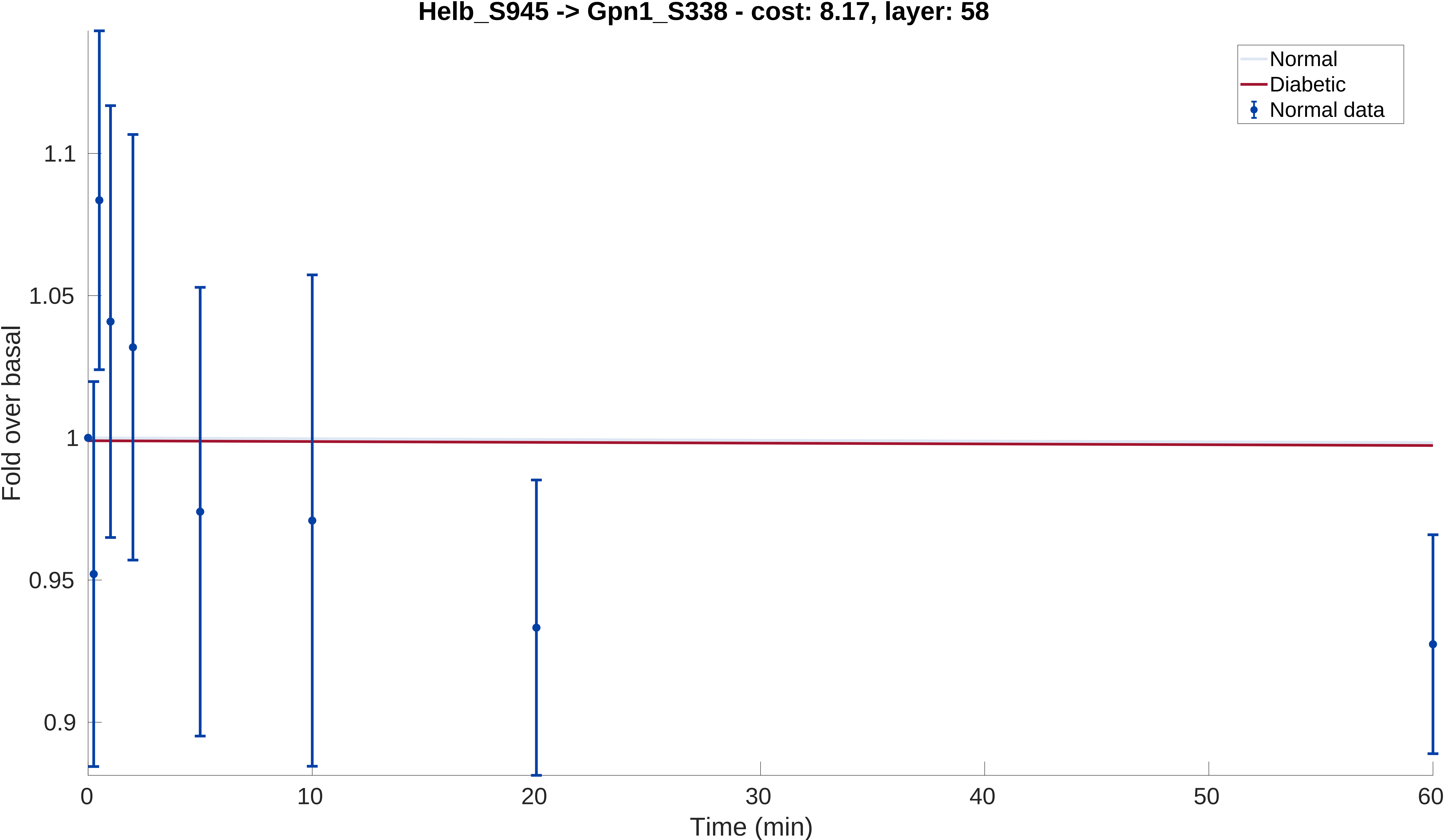

Helb_S945 -> Gpn1_S338 - cost: 8.17, layer: 58
Normal
Diabetic
Normal data
Fold over basal
1.1
1.05
1
0.95
0.9
0
10
20
30
40
50
60
Time (min)

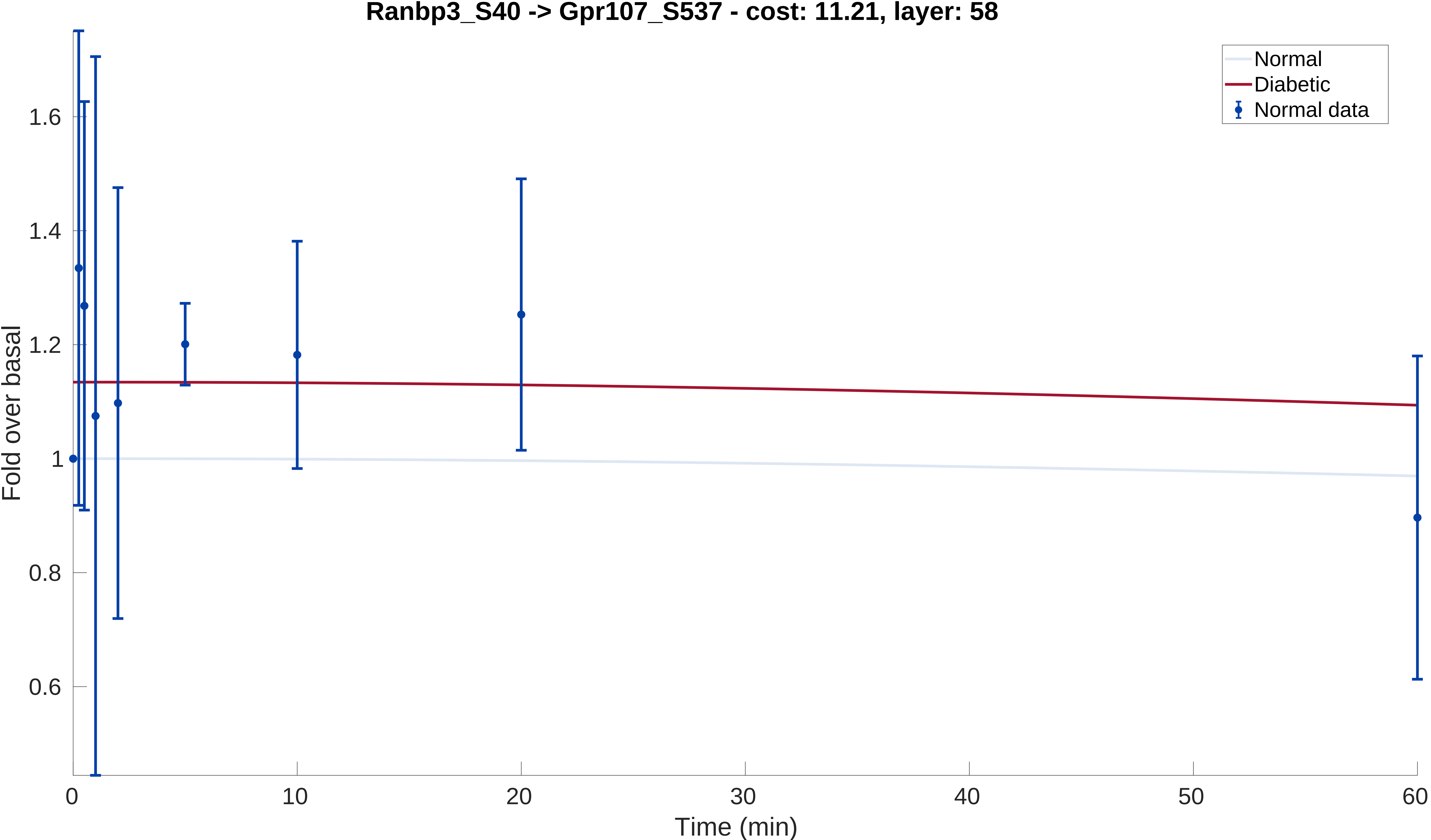
Ranbp3_S40 -> Gpr107_S537 - cost: 11.21, layer: 58
Normal
Diabetic
Normal data
Fold over basal
1.6
1.4
1.2
1
0.8
0.6
0
10
20
30
40
50
60
Time (min)

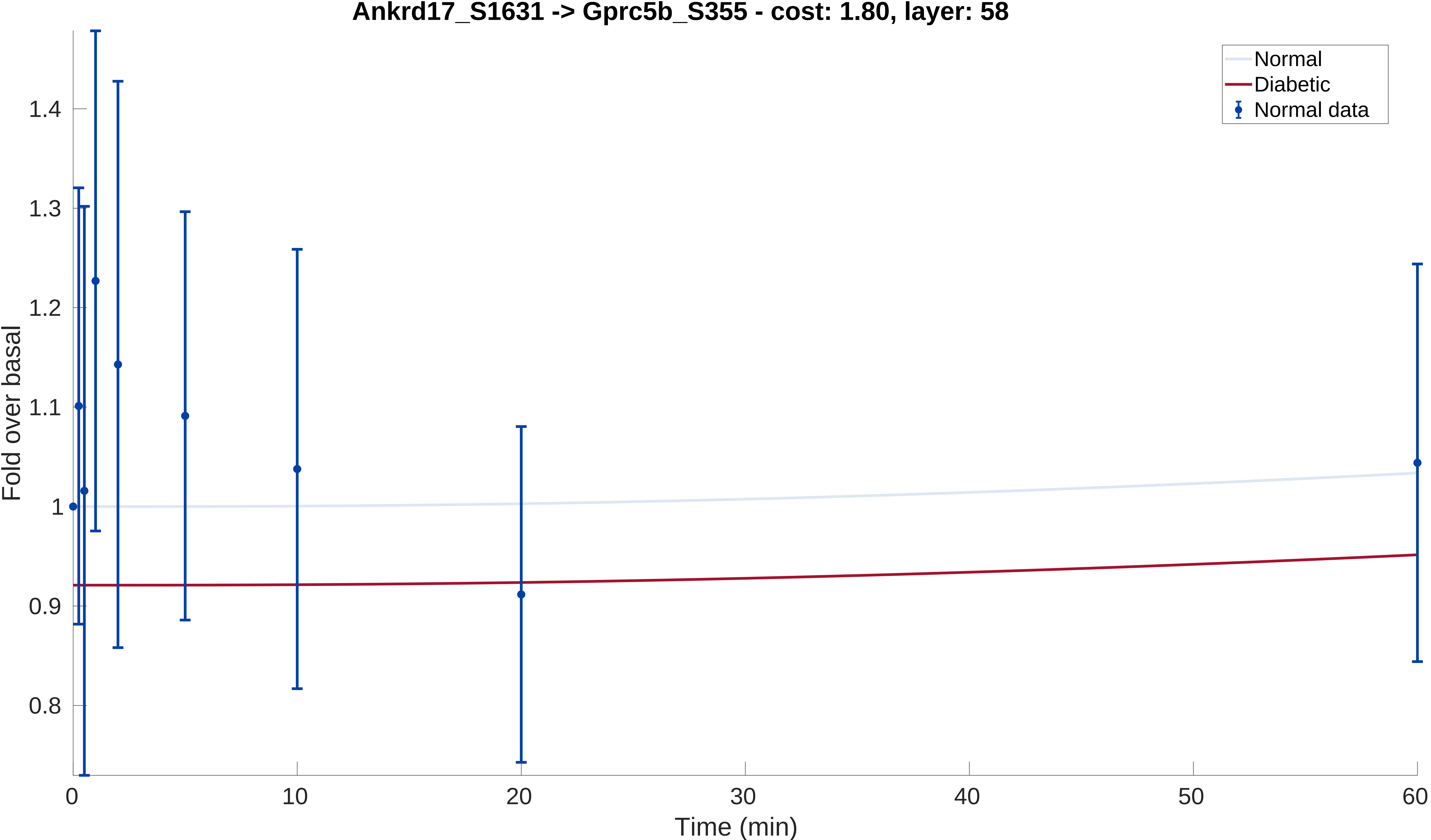

Ankrd17_S1631 -> Gprc5b_S355 - cost: 1.80, layer: 58
Normal
Diabetic
Normal data
Fold over basal
1.4
1.3
1.2
1.1
1
0.9
0.8
0
10
20
30
40
50
60
Time (min)

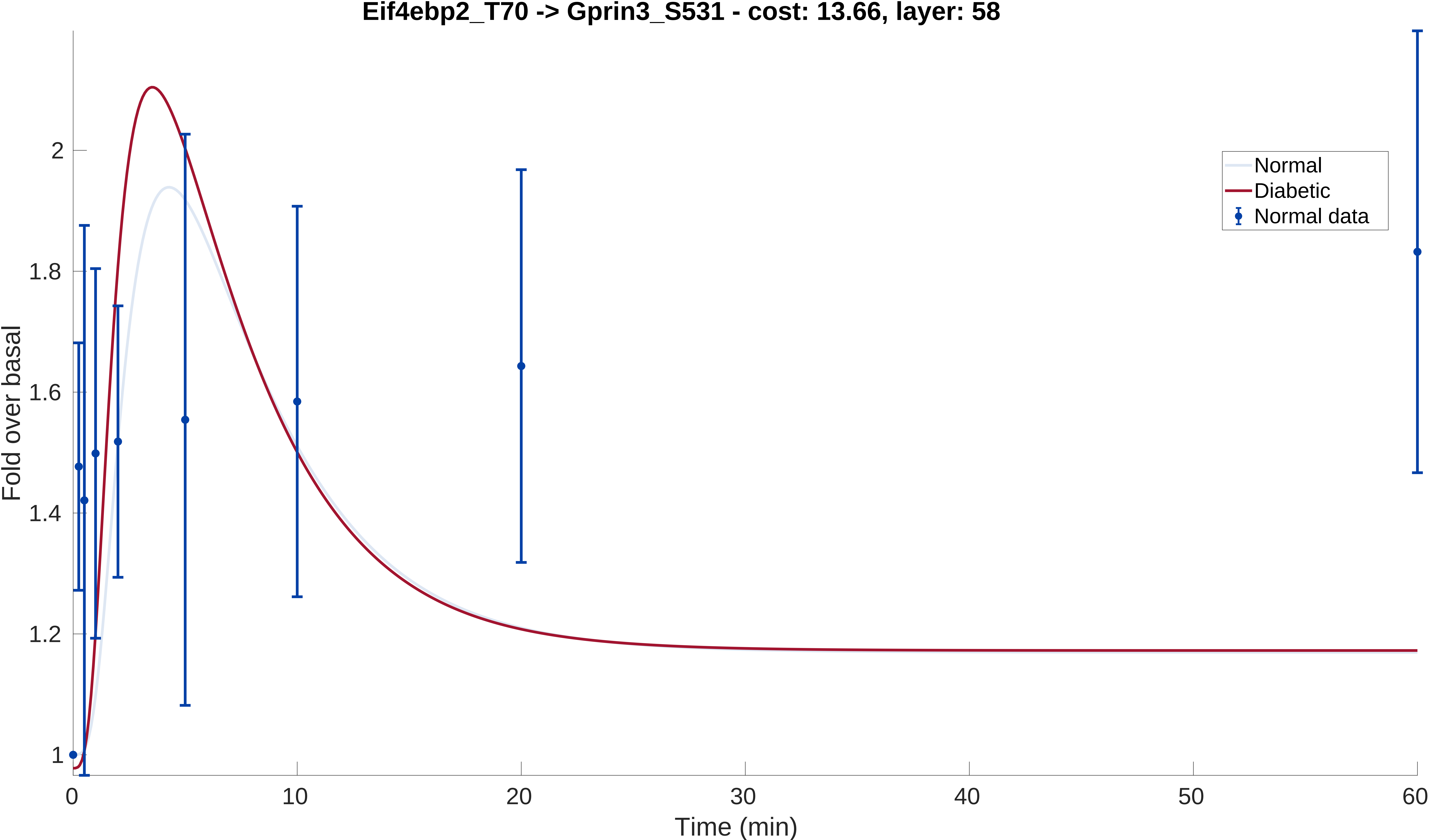
Eif4ebp2_T70 -> Gprin3_S531 - cost: 13.66, layer: 58
Fold over basal
Time (min)
Normal
Diabetic
Normal data
2
1.8
1.6
1.4
1.2
1
0
10
20
30
40
50
60

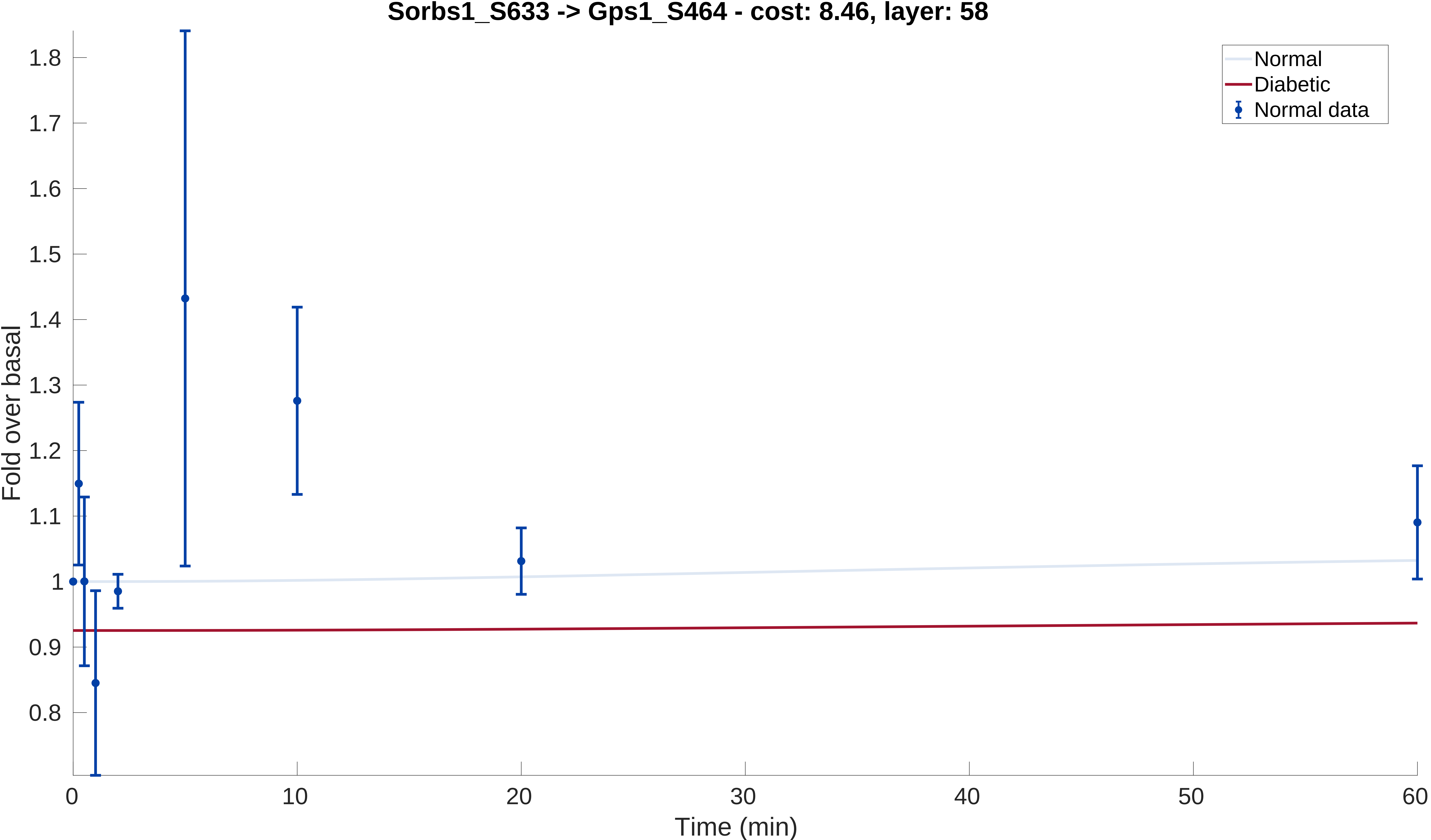

Sorbs1_S633 -> Gps1_S464 - cost: 8.46, layer: 58
Normal
Diabetic
Normal data
Fold over basal
1.8
1.7
1.6
1.5
1.4
1.3
1.2
1.1
1
0.9
0.8
0
10
20
30
40
50
60
Time (min)

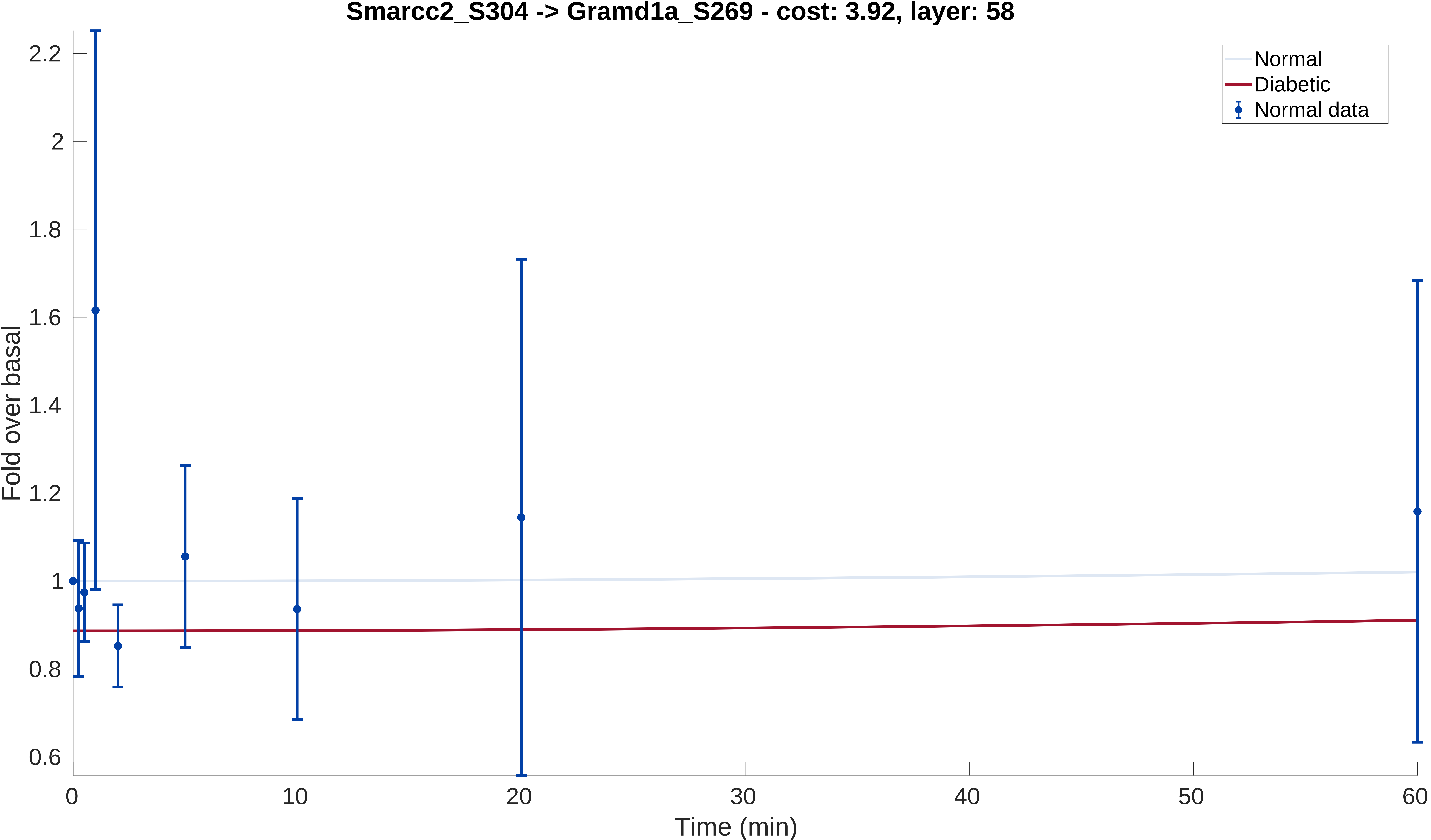
Smarcc2_S304 -> Gramd1a_S269 - cost: 3.92, layer: 58
Fold over basal
Time (min)
Normal
Diabetic
Normal data
0.6
0.8
1
1.2
1.4
1.6
1.8
2
2.2
0
10
20
30
40
50
60

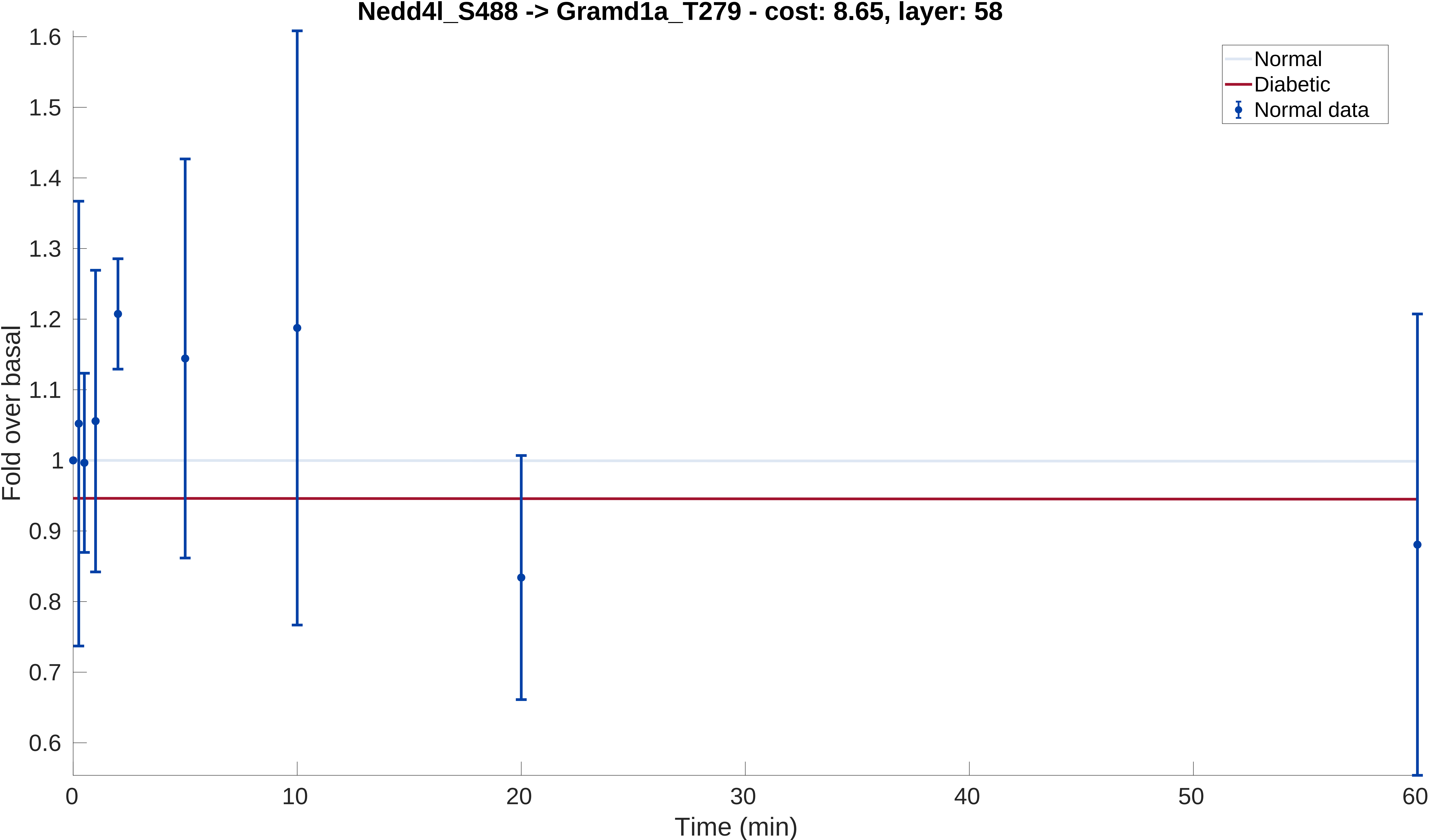
Nedd4l_S488 -> Gramd1a_T279 - cost: 8.65, layer: 58
Fold over basal
Time (min)
Normal
Diabetic
Normal data
0.6
0.7
0.8
0.9
1
1.1
1.2
1.3
1.4
1.5
1.6
0
10
20
30
40
50
60

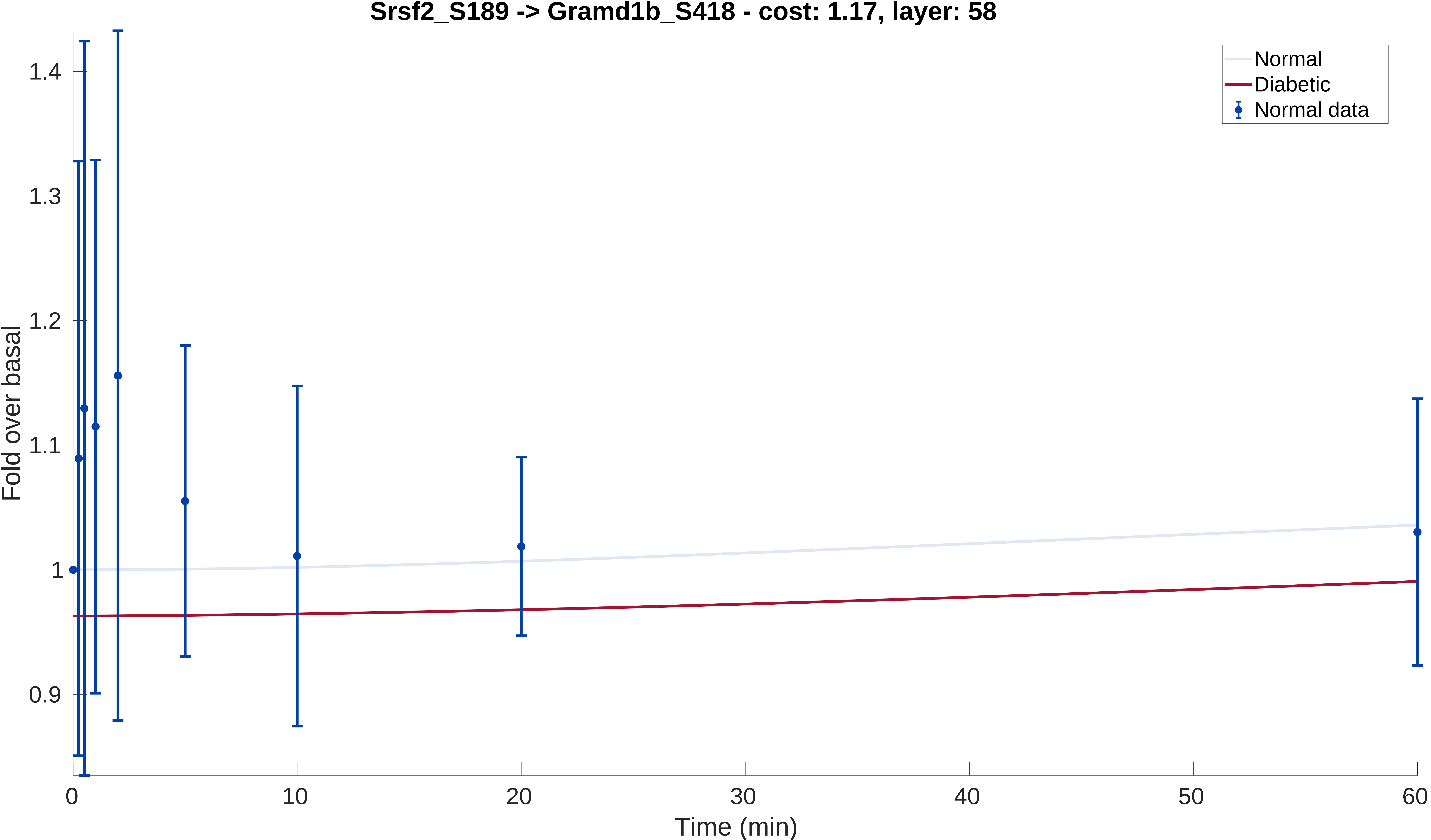

Srsf2_S189 -> Gramd1b_S418 - cost: 1.17, layer: 58
Normal
Diabetic
Normal data
Fold over basal
1.4
1.3
1.2
1.1
1
0.9
0
10
20
30
40
50
60
Time (min)

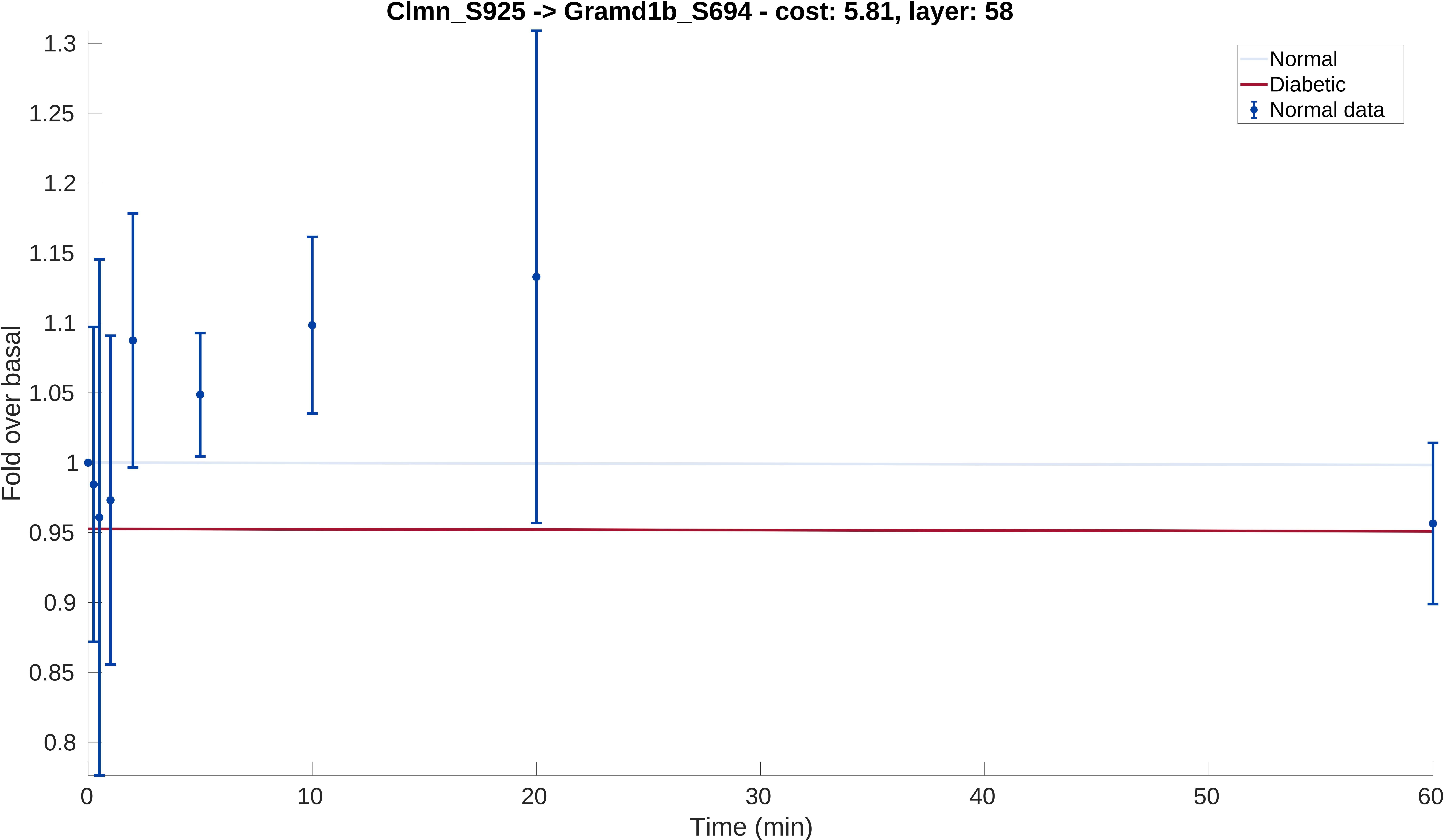
Clmn_S925 -> Gramd1b_S694 - cost: 5.81, layer: 58
Normal
Diabetic
Normal data
Fold over basal
Time (min)
1.3
1.25
1.2
1.15
1.1
1.05
1
0.95
0.9
0.85
0.8
0
10
20
30
40
50
60

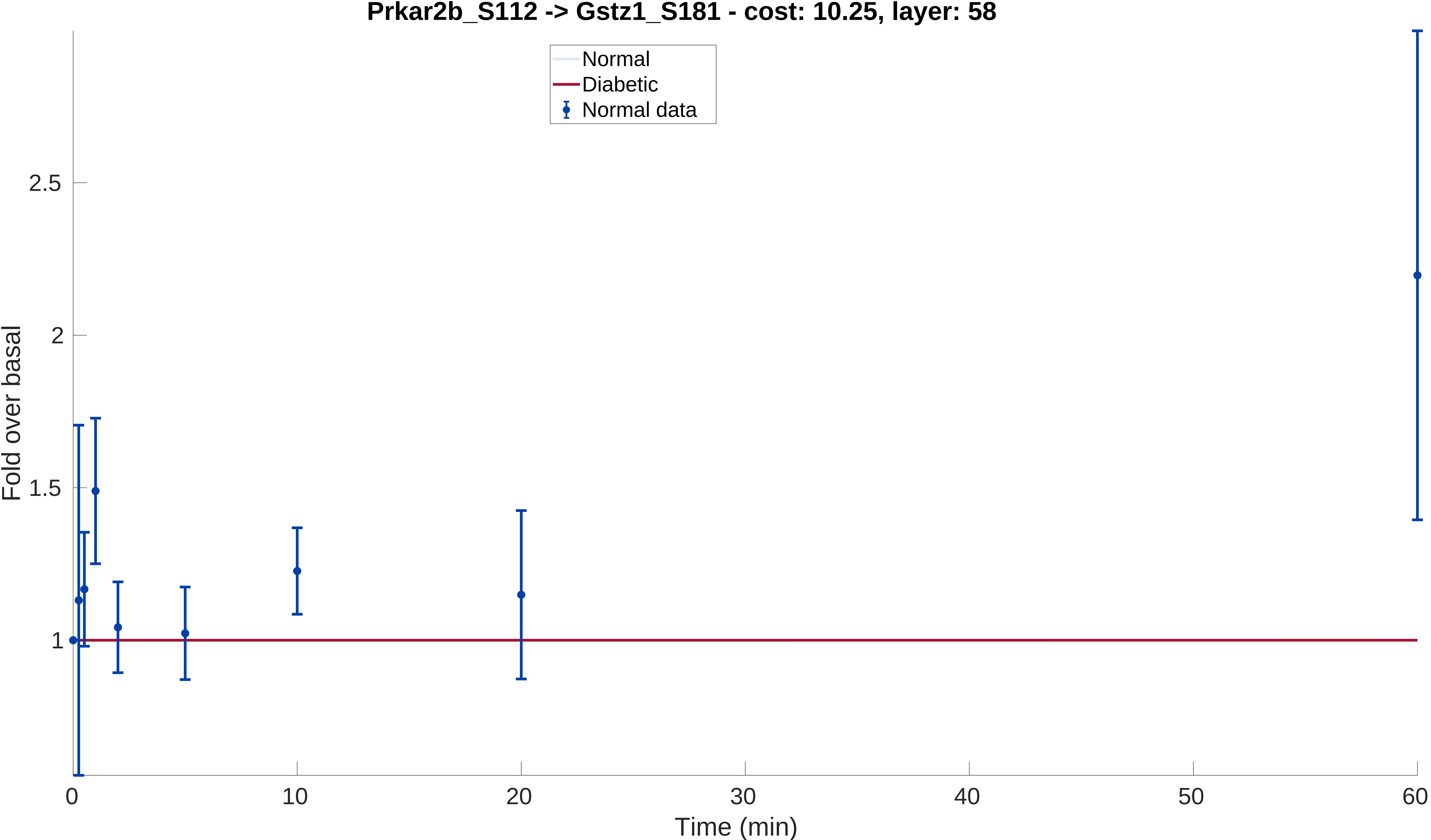

Prkar2b_S112 -> Gstz1_S181 - cost: 10.25, layer: 58
Normal
Diabetic
Normal data
Fold over basal
2.5
2
1.5
1
0
10
20
30
40
50
60
Time (min)

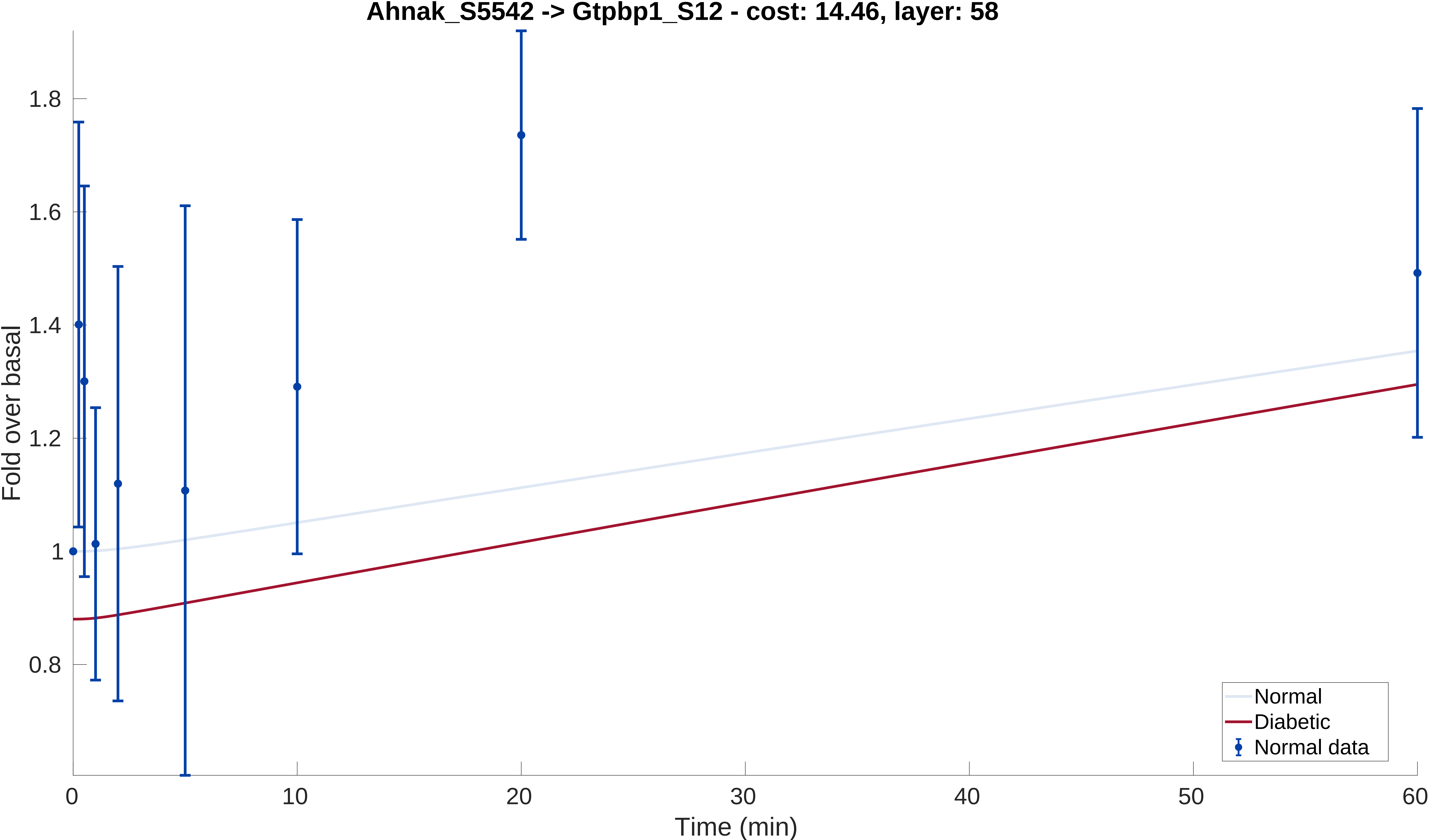
Ahnak_S5542 -> Gtpbp1_S12 - cost: 14.46, layer: 58
Fold over basal
1.8
1.6
1.4
1.2
1
0.8
0
10
20
30
40
50
60
Time (min)
Normal
Diabetic
Normal data

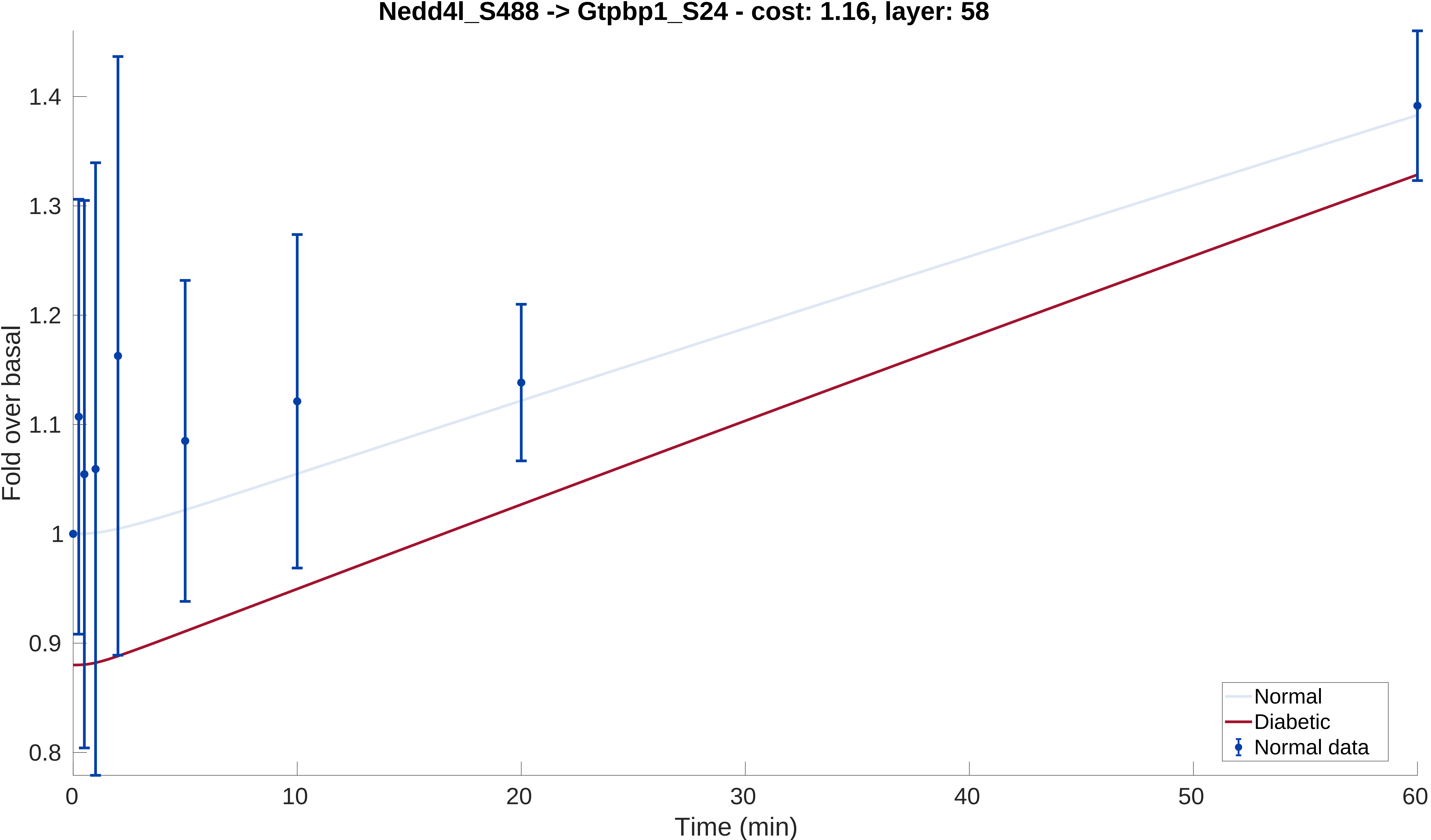

Nedd4l_S488 -> Gtpbp1_S24 - cost: 1.16, layer: 58
Fold over basal
Time (min)
1.4
1.3
1.2
1.1
1
0.9
0.8
0
10
20
30
40
50
60
Normal
Diabetic
Normal data

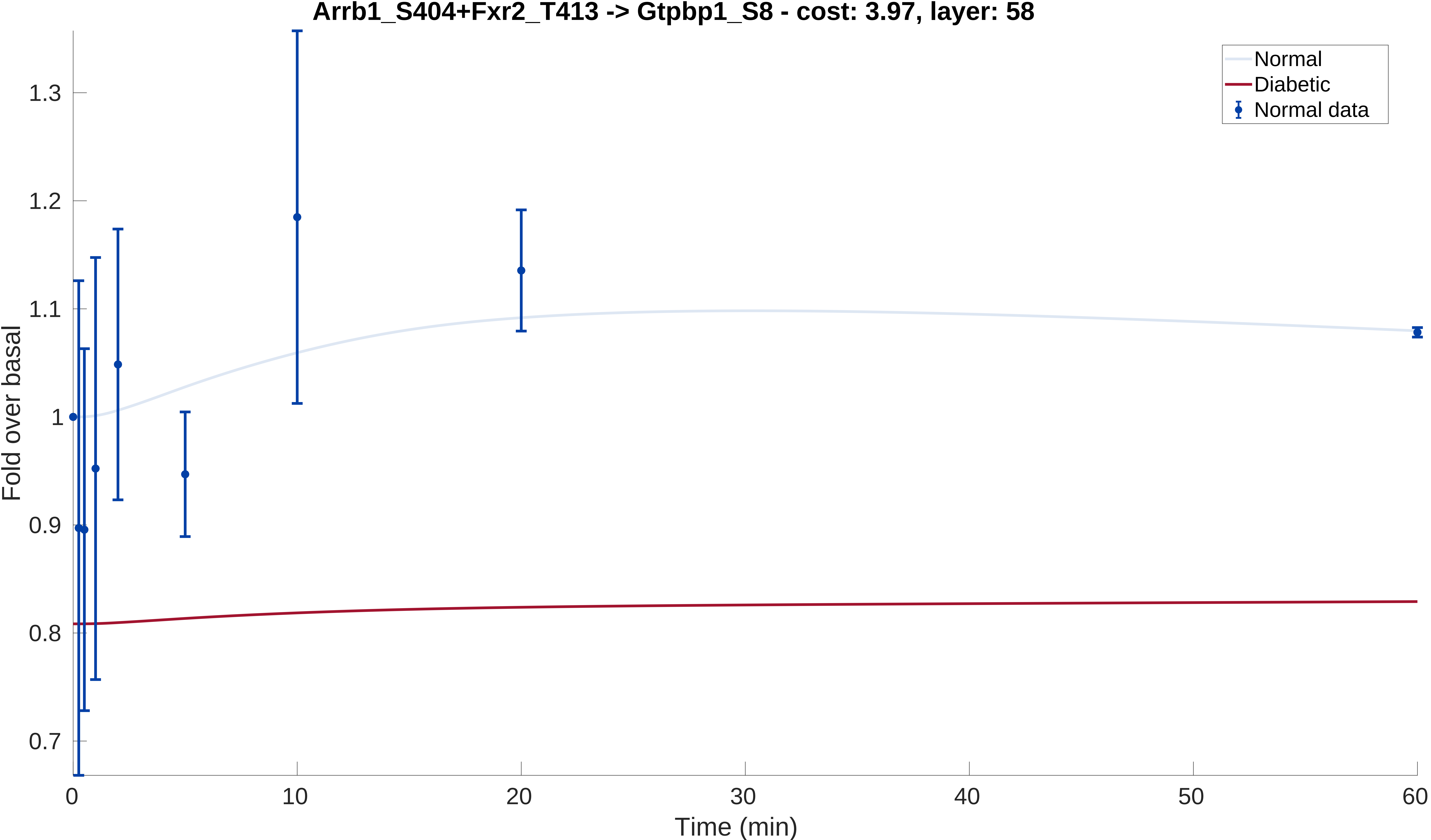

Arrb1_S404+Fxr2_T413 -> Gtpbp1_S8 - cost: 3.97, layer: 58
Normal
Diabetic
Normal data
Fold over basal
1.3
1.2
1.1
1
0.9
0.8
0.7
0
10
20
30
40
50
60
Time (min)

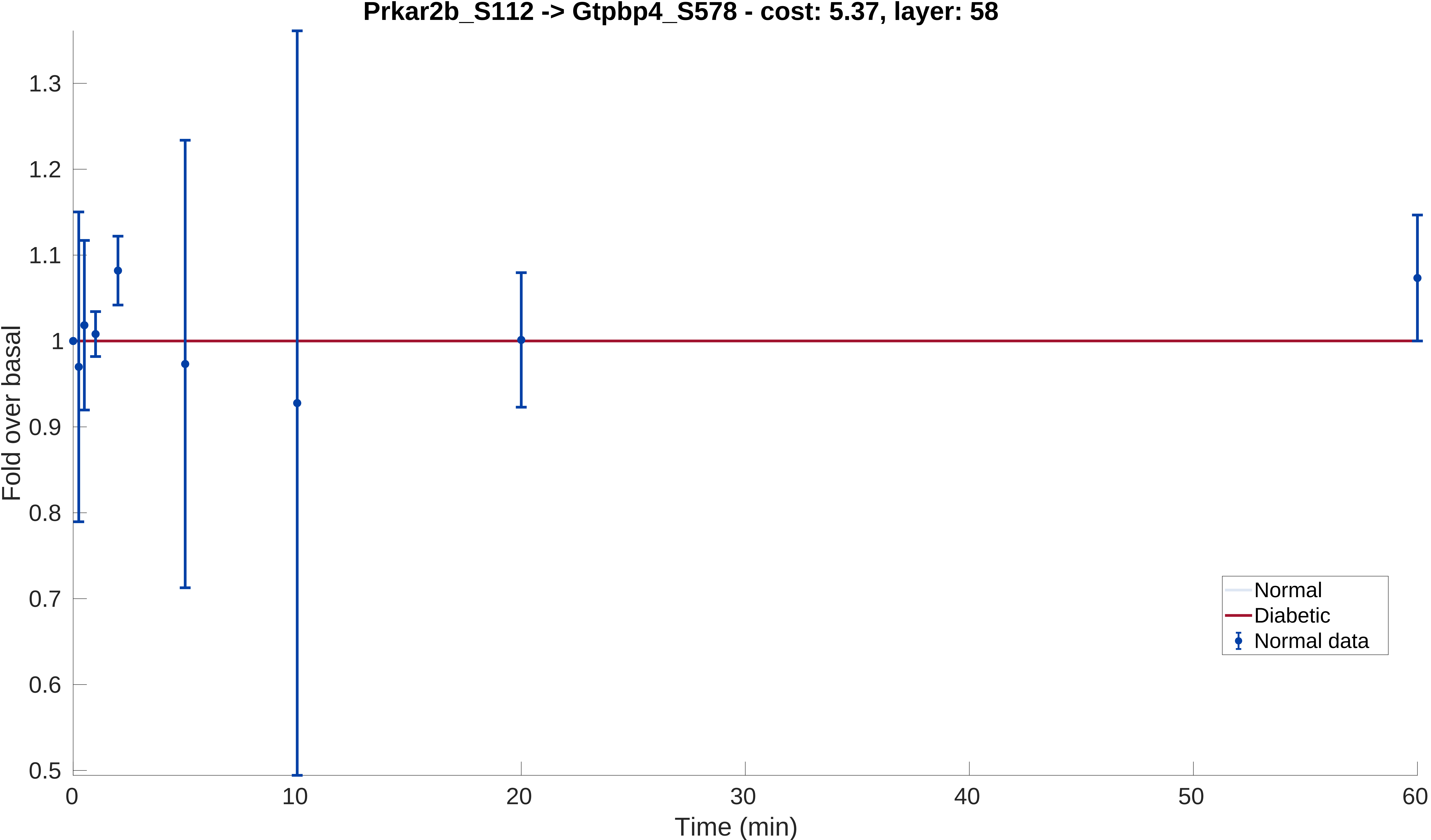

Prkar2b_S112 -> Gtpbp4_S578 - cost: 5.37, layer: 58
Fold over basal
1.3
1.2
1.1
1
0.9
0.8
0.7
0.6
0.5
0
10
20
30
40
50
60
Time (min)
Normal
Diabetic
Normal data

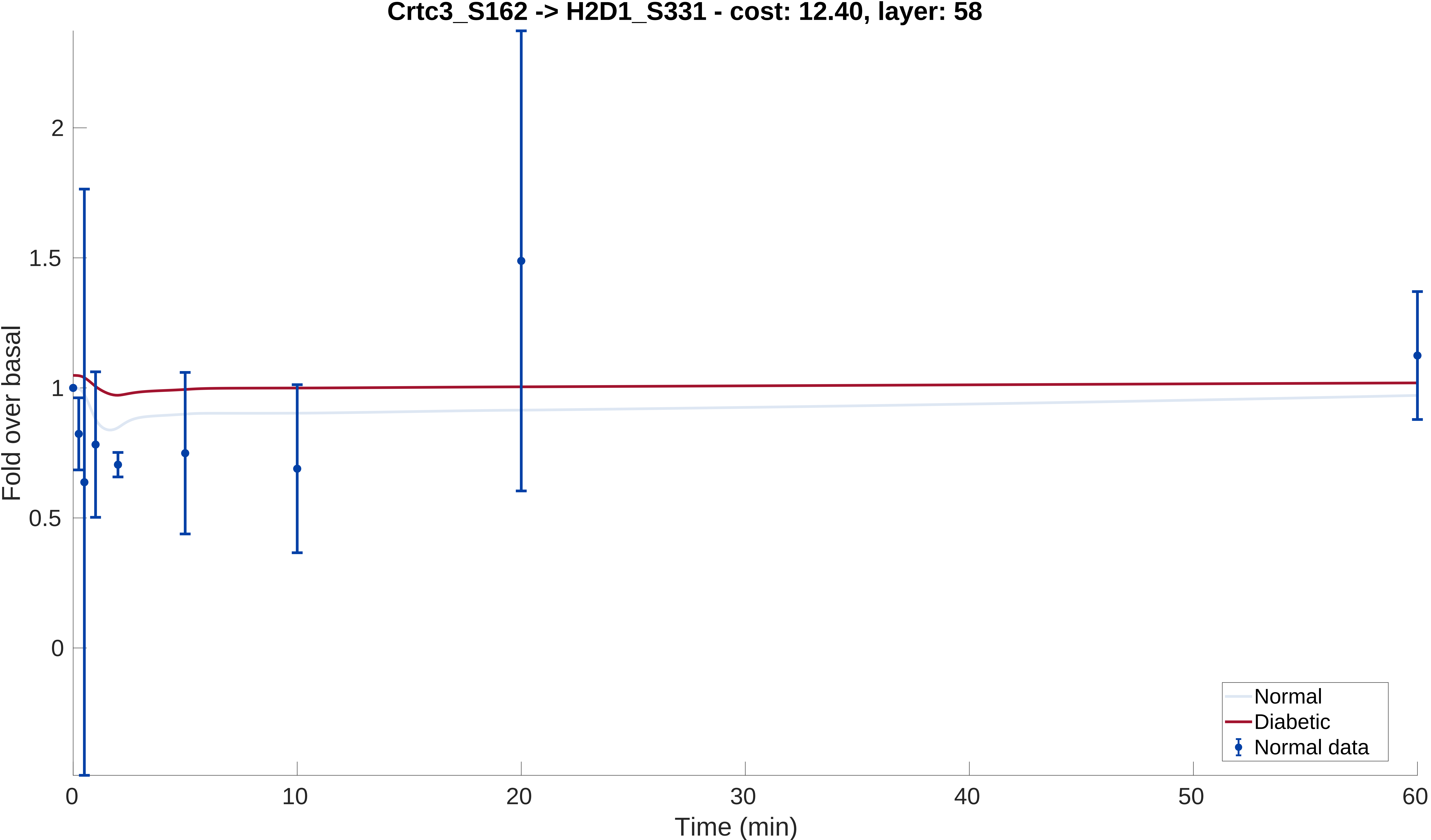

Crtc3_S162 -> H2D1_S331 - cost: 12.40, layer: 58
Fold over basal
2
1.5
1
0.5
0
0
10
20
30
40
50
60
Time (min)
Normal
Diabetic
Normal data

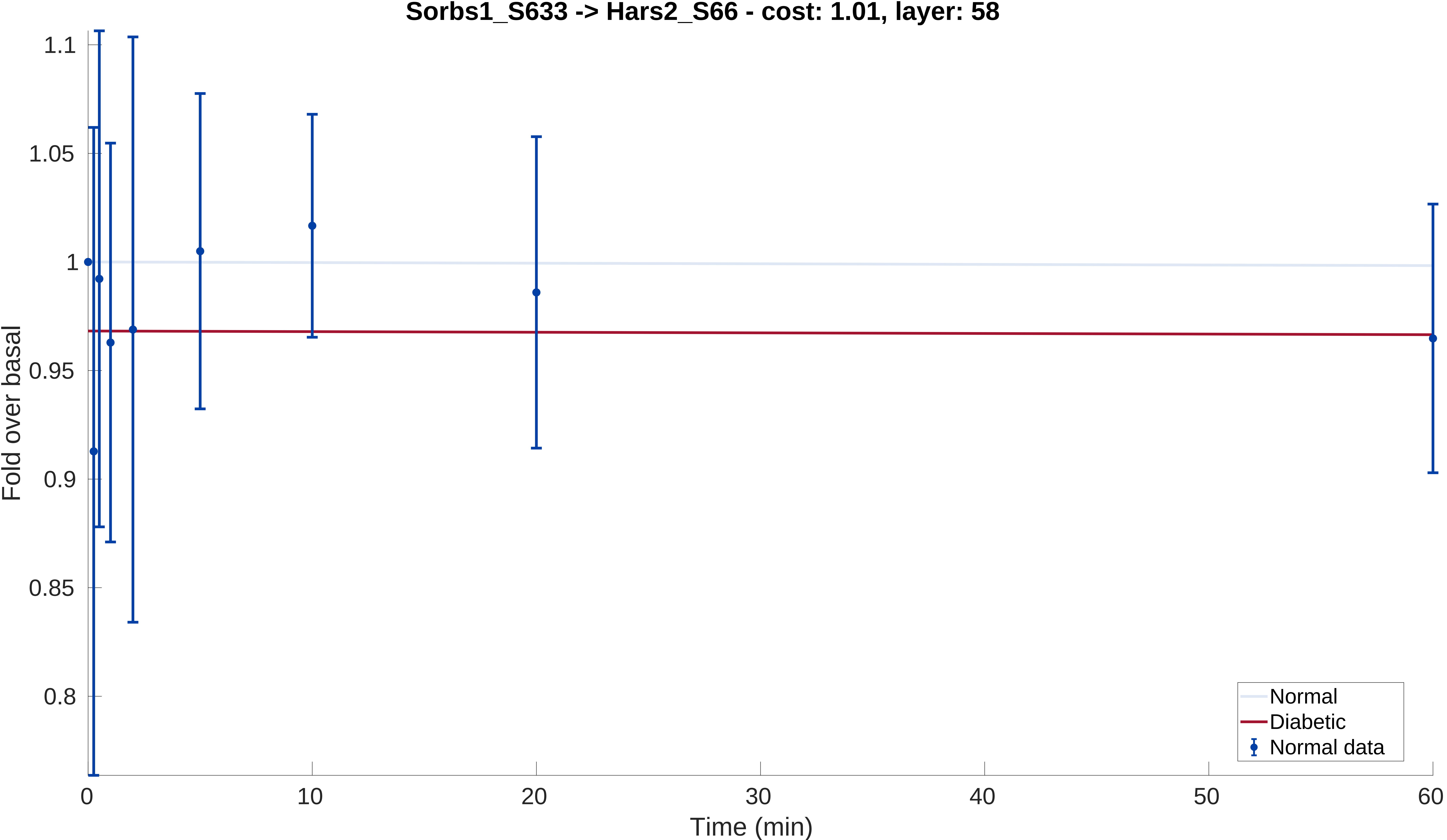

Sorbs1_S633 -> Hars2_S66 - cost: 1.01, layer: 58
Fold over basal
1.1
1.05
1
0.95
0.9
0.85
0.8
0
10
20
30
40
50
60
Time (min)
Normal
Diabetic
Normal data

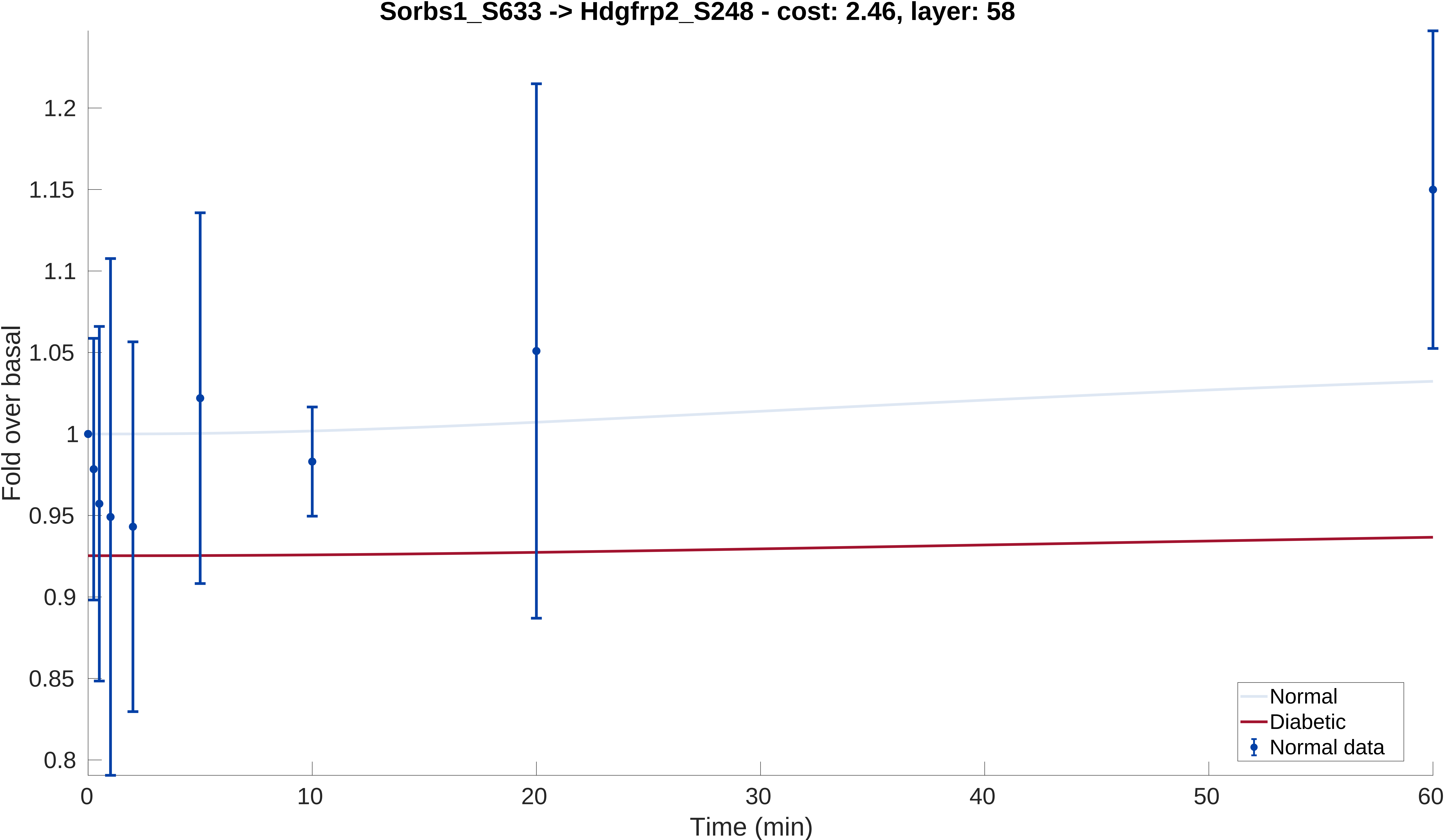
Sorbs1_S633 -> Hdgfrp2_S248 - cost: 2.46, layer: 58
Fold over basal
1.2
1.15
1.1
1.05
1
0.95
0.9
0.85
0.8
0
10
20
30
40
50
60
Time (min)
Normal
Diabetic
Normal data

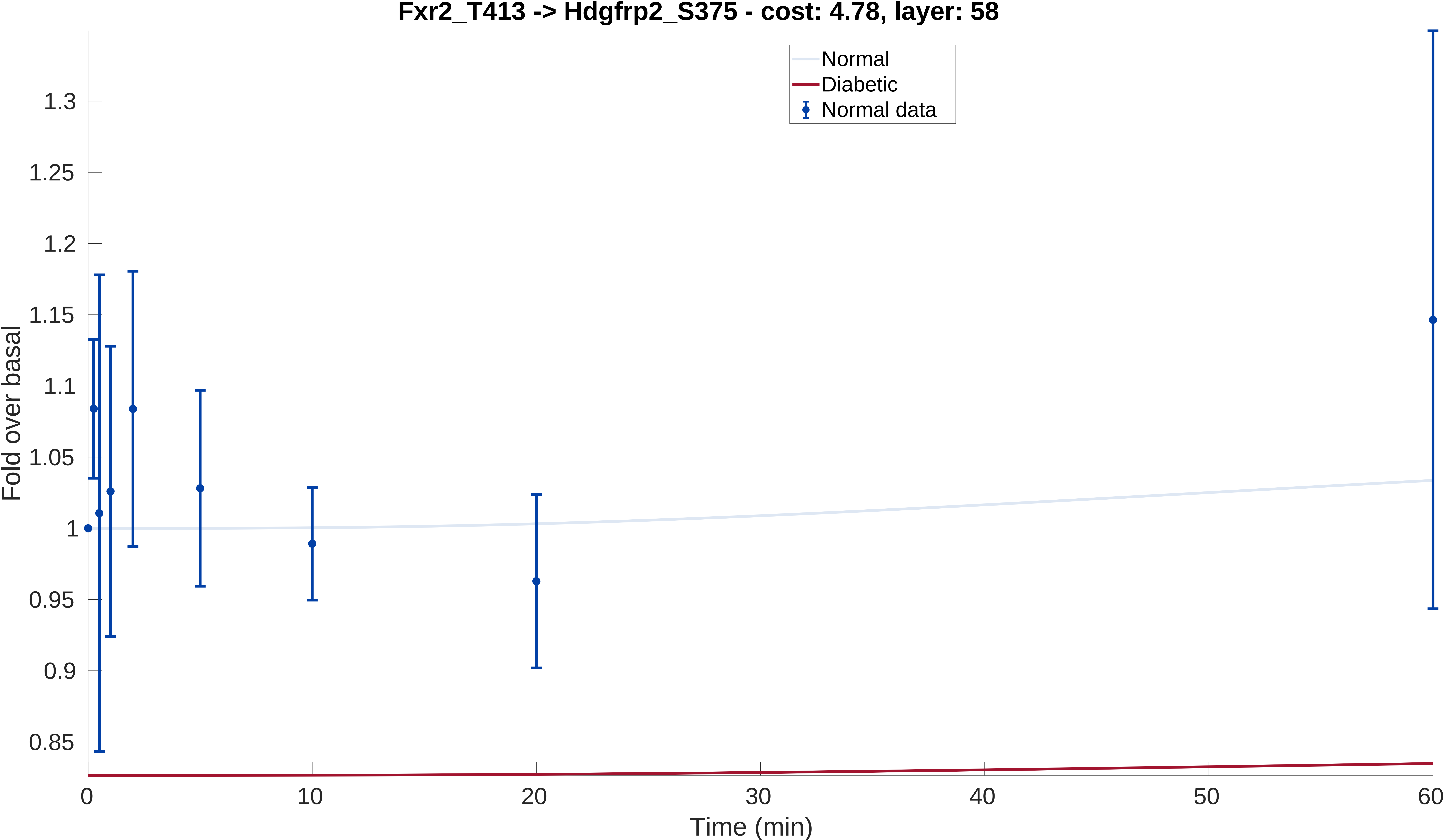

Fxr2_T413 -> Hdgfrp2_S375 - cost: 4.78, layer: 58
Normal
Diabetic
Normal data
Fold over basal
Time (min)
0.85
0.9
0.95
1
1.05
1.1
1.15
1.2
1.25
1.3
0
10
20
30
40
50
60

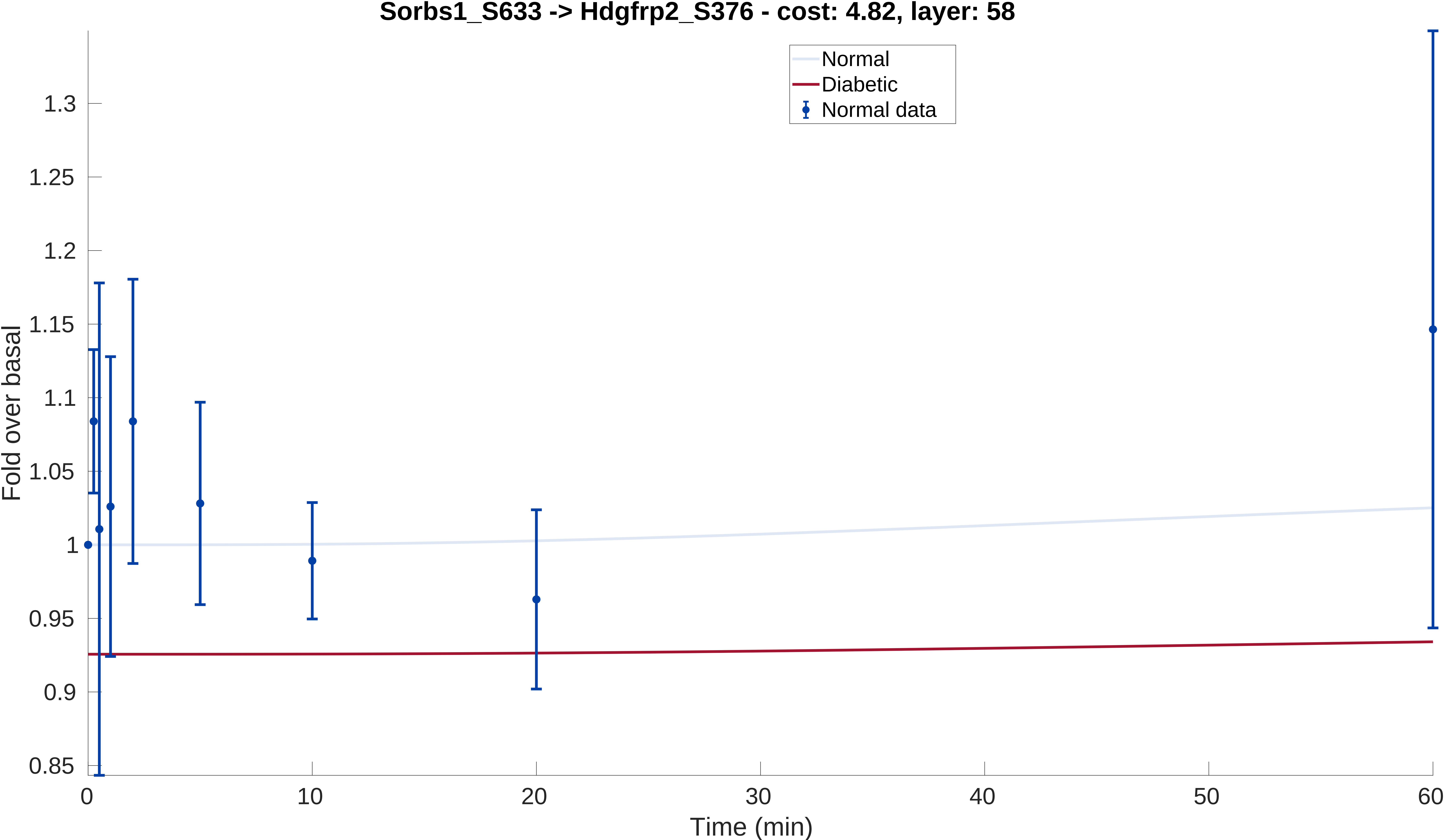

Sorbs1_S633 -> Hdgfrp2_S376 - cost: 4.82, layer: 58
Normal
Diabetic
Normal data
Fold over basal
1.3
1.25
1.2
1.15
1.1
1.05
1
0.95
0.9
0.85
0
10
20
30
40
50
60
Time (min)

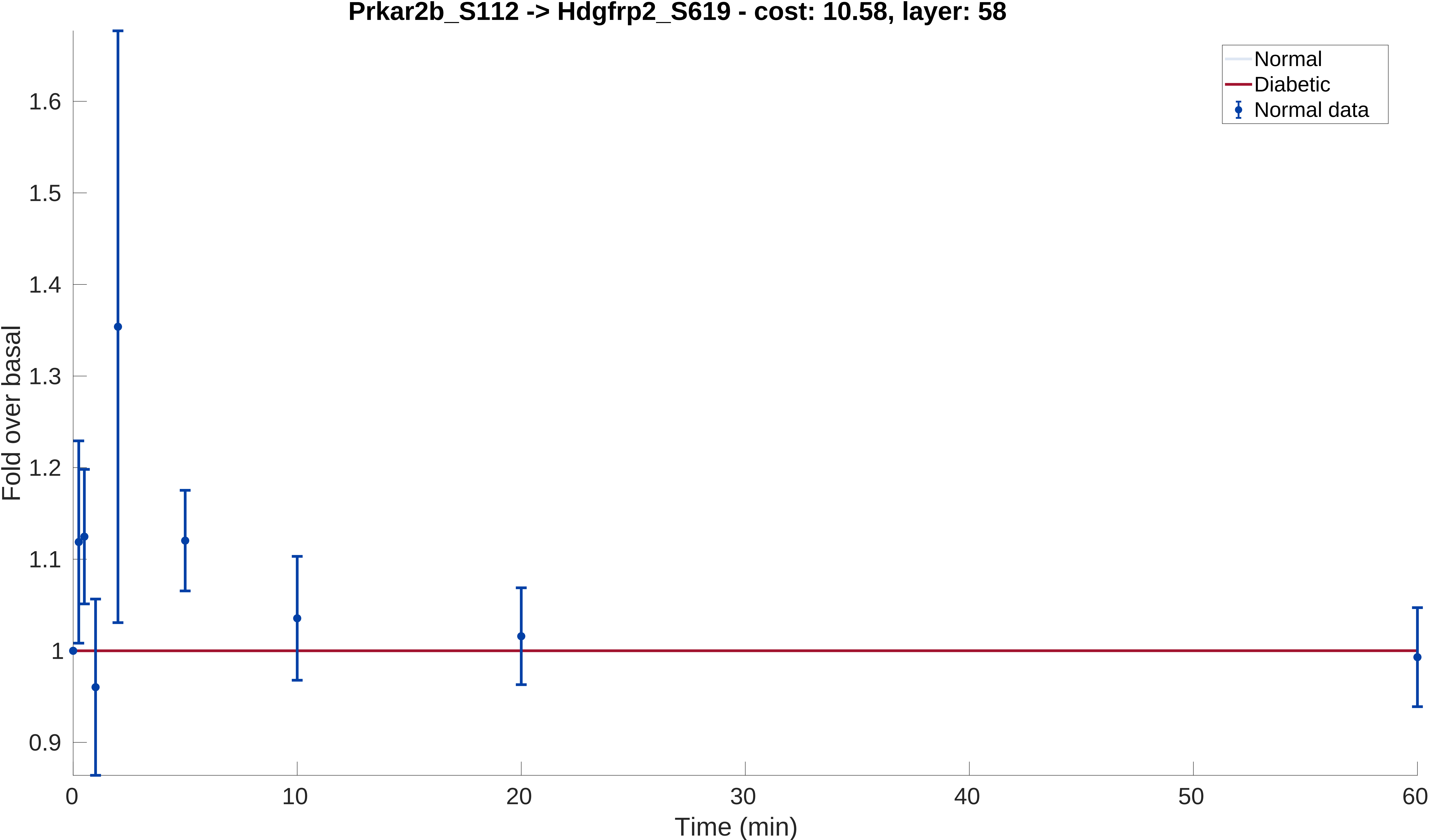

Prkar2b_S112 -> Hdgfrp2_S619 - cost: 10.58, layer: 58
Normal
Diabetic
Normal data
Fold over basal
1.6
1.5
1.4
1.3
1.2
1.1
1
0.9
0
10
20
30
40
50
60
Time (min)

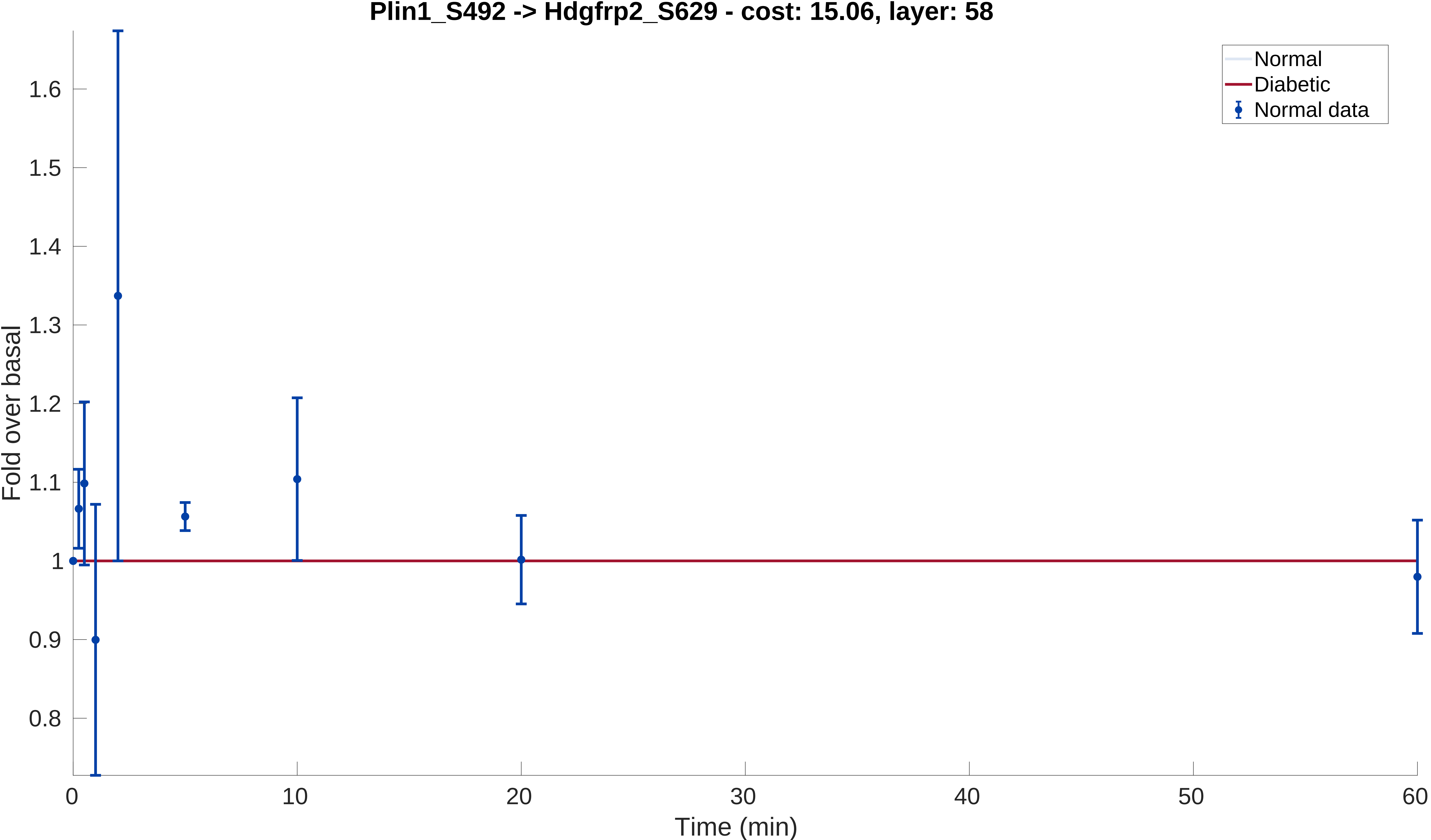

Plin1_S492 -> Hdgfrp2_S629 - cost: 15.06, layer: 58
Normal
Diabetic
Normal data
Fold over basal
Time (min)
0.8
0.9
1
1.1
1.2
1.3
1.4
1.5
1.6
0
10
20
30
40
50
60

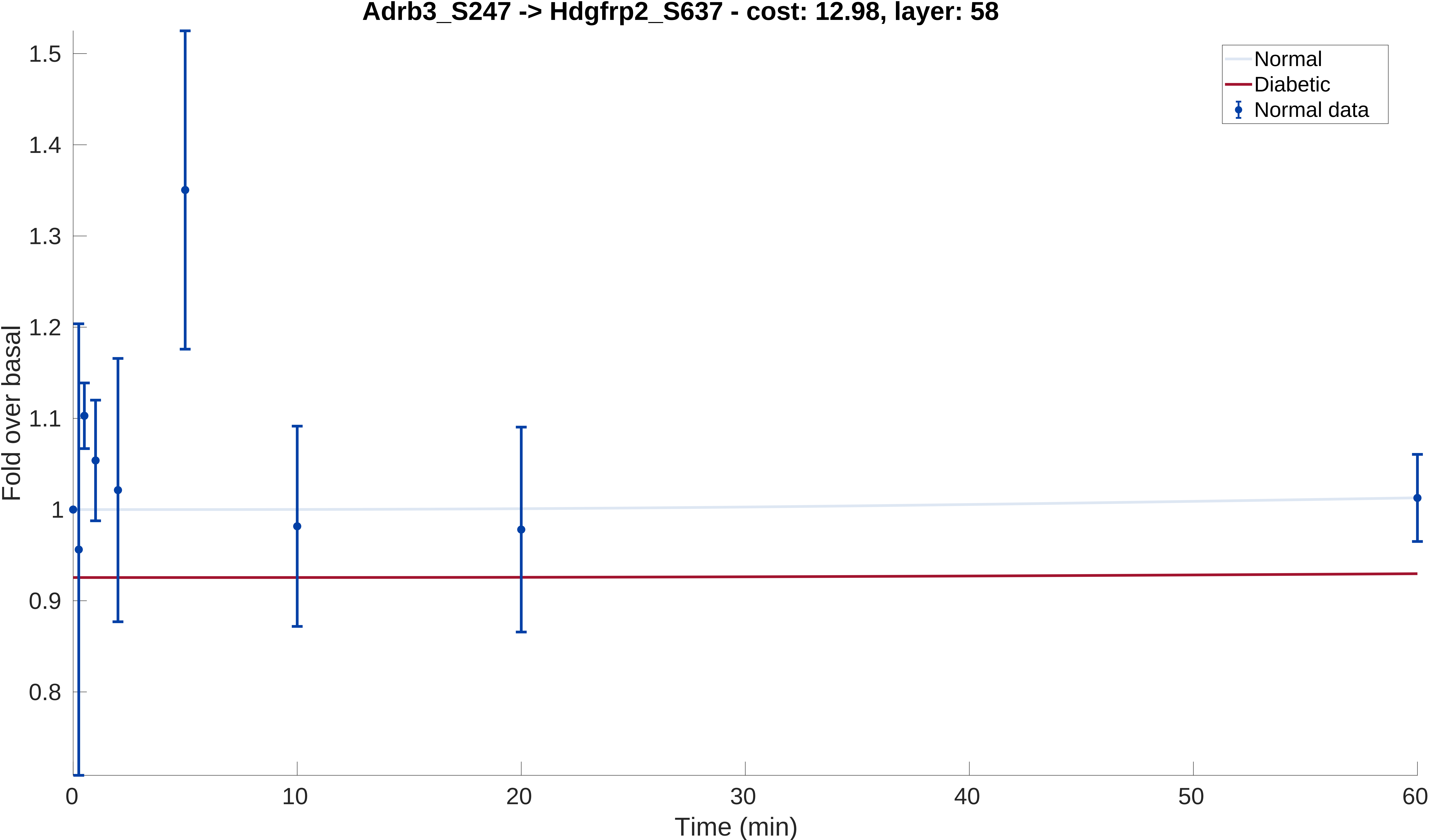

Adrb3_S247 -> Hdgfrp2_S637 - cost: 12.98, layer: 58
Normal
Diabetic
Normal data
Fold over basal
1.5
1.4
1.3
1.2
1.1
1
0.9
0.8
0
10
20
30
40
50
60
Time (min)

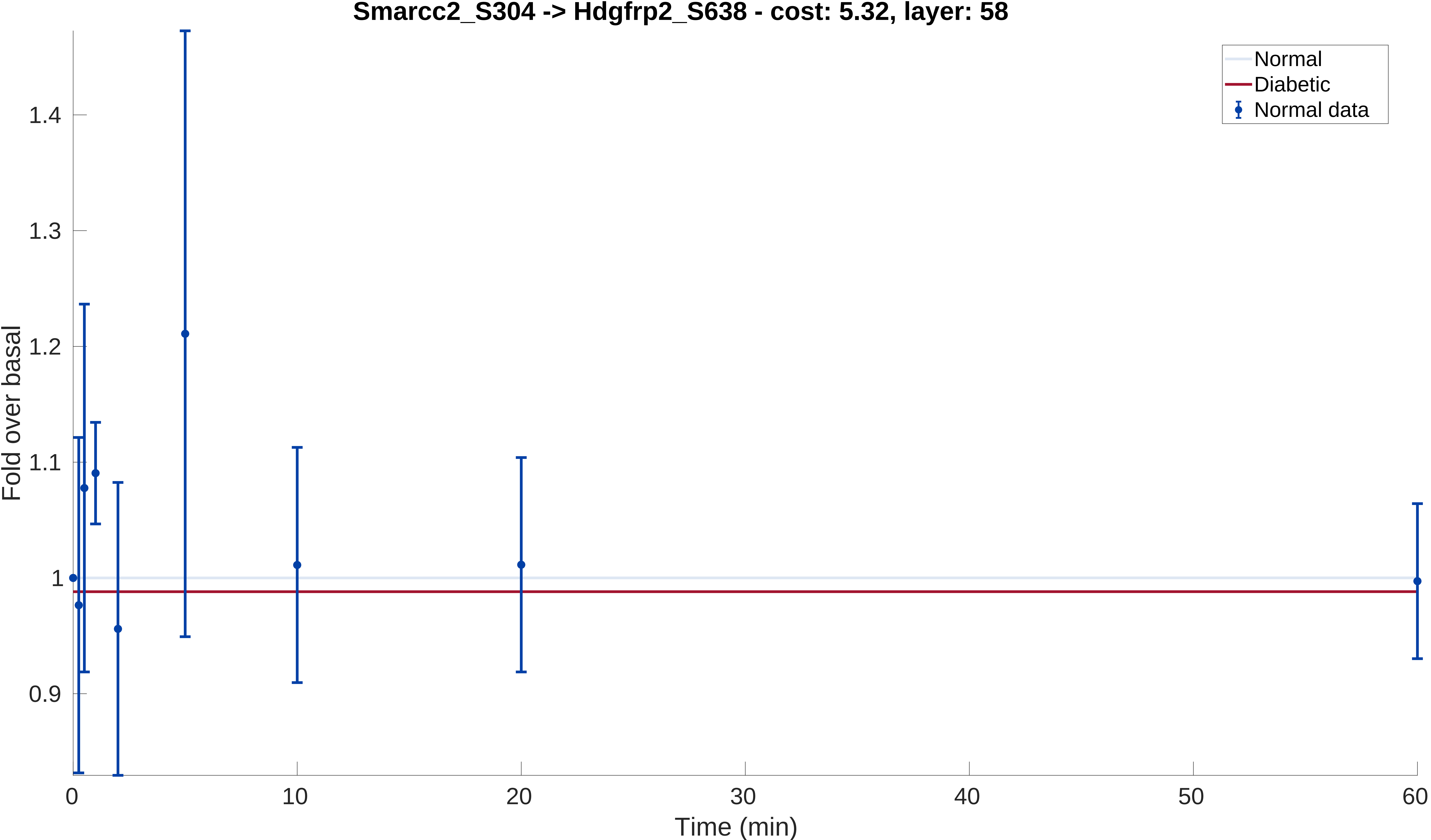

Smarcc2_S304 -> Hdgfrp2_S638 - cost: 5.32, layer: 58
Normal
Diabetic
Normal data
Fold over basal
1.4
1.3
1.2
1.1
1
0.9
0
10
20
30
40
50
60
Time (min)

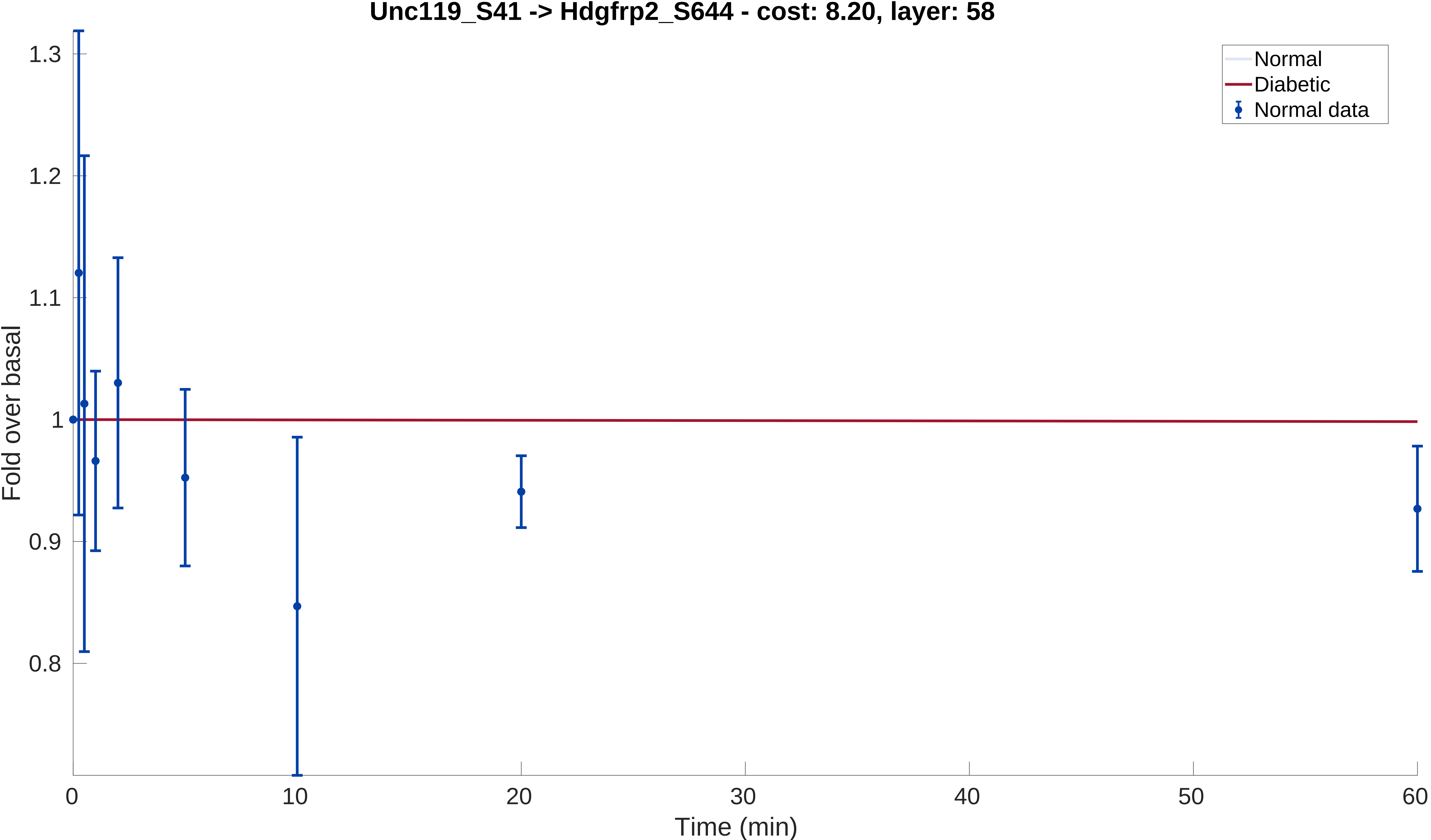
Unc119_S41 -> Hdgfrp2_S644 - cost: 8.20, layer: 58
Normal
Diabetic
Normal data
Fold over basal
1.3
1.2
1.1
1
0.9
0.8
0
10
20
30
40
50
60
Time (min)

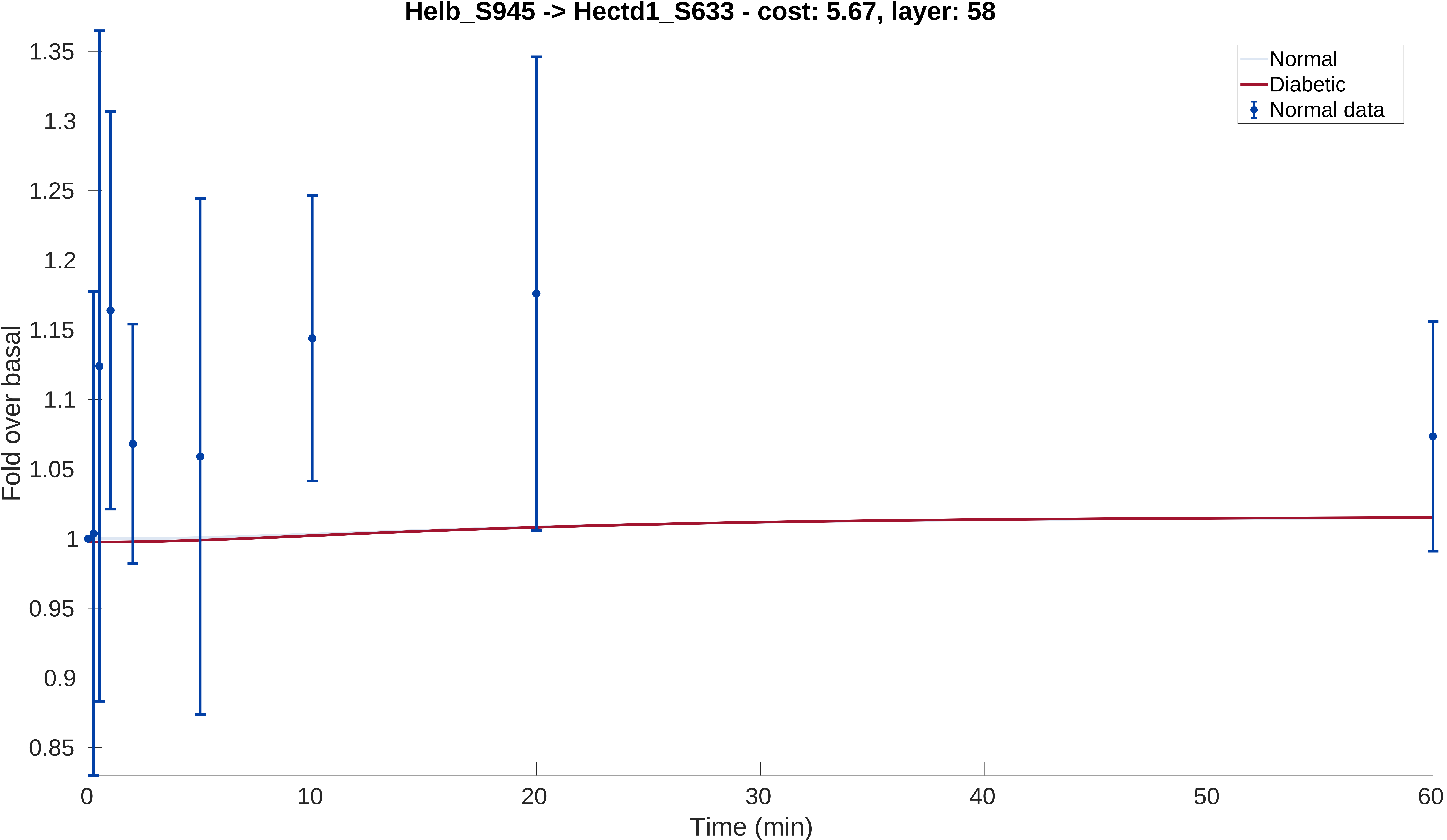
Helb_S945 -> Hectd1_S633 - cost: 5.67, layer: 58
Normal
Diabetic
Normal data
Fold over basal
Time (min)
1.35
1.3
1.25
1.2
1.15
1.1
1.05
1
0.95
0.9
0.85
0
10
20
30
40
50
60

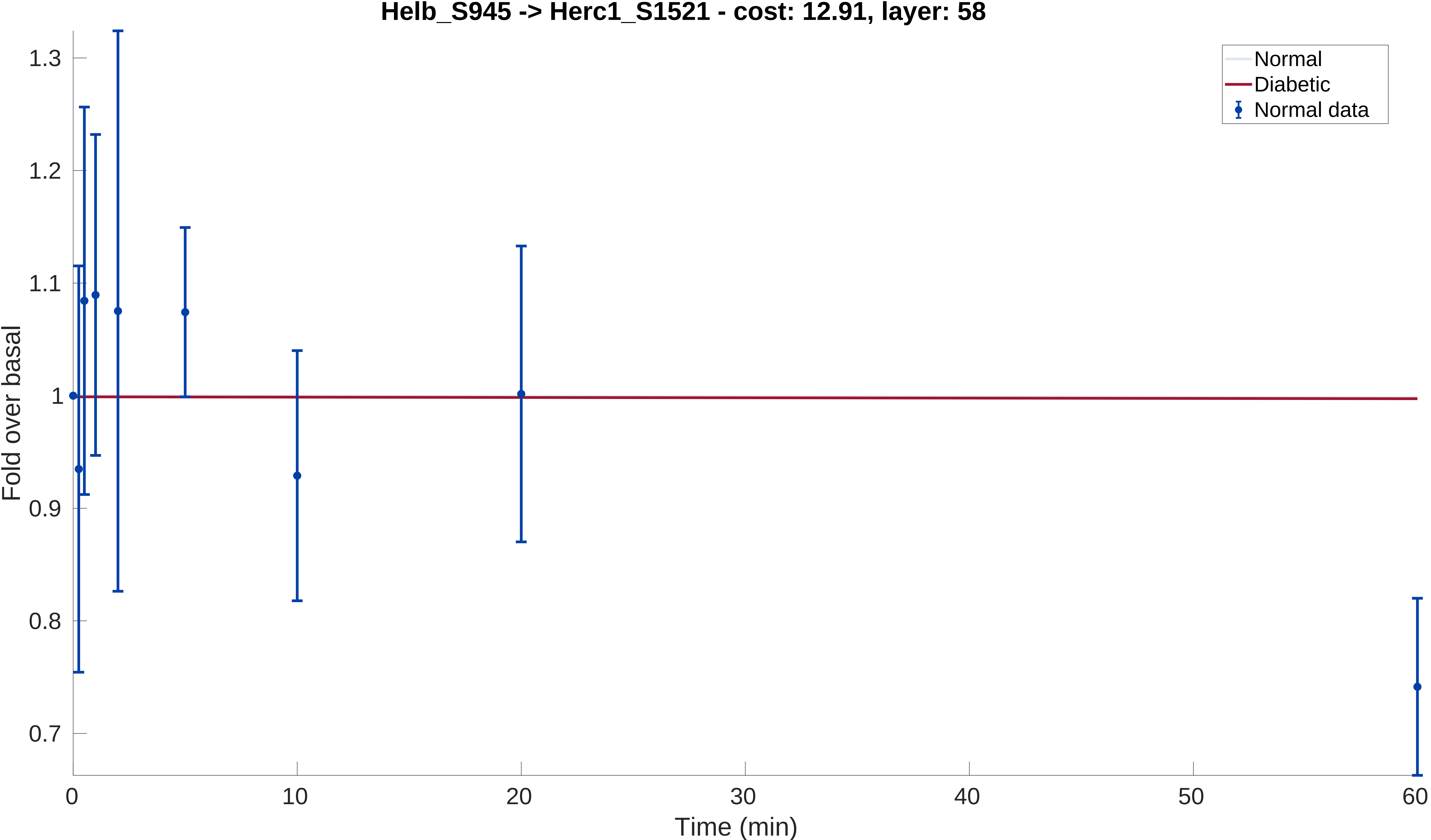

Helb_S945 -> Herc1_S1521 - cost: 12.91, layer: 58
Normal
Diabetic
Normal data
Fold over basal
1.3
1.2
1.1
1
0.9
0.8
0.7
0
10
20
30
40
50
60
Time (min)

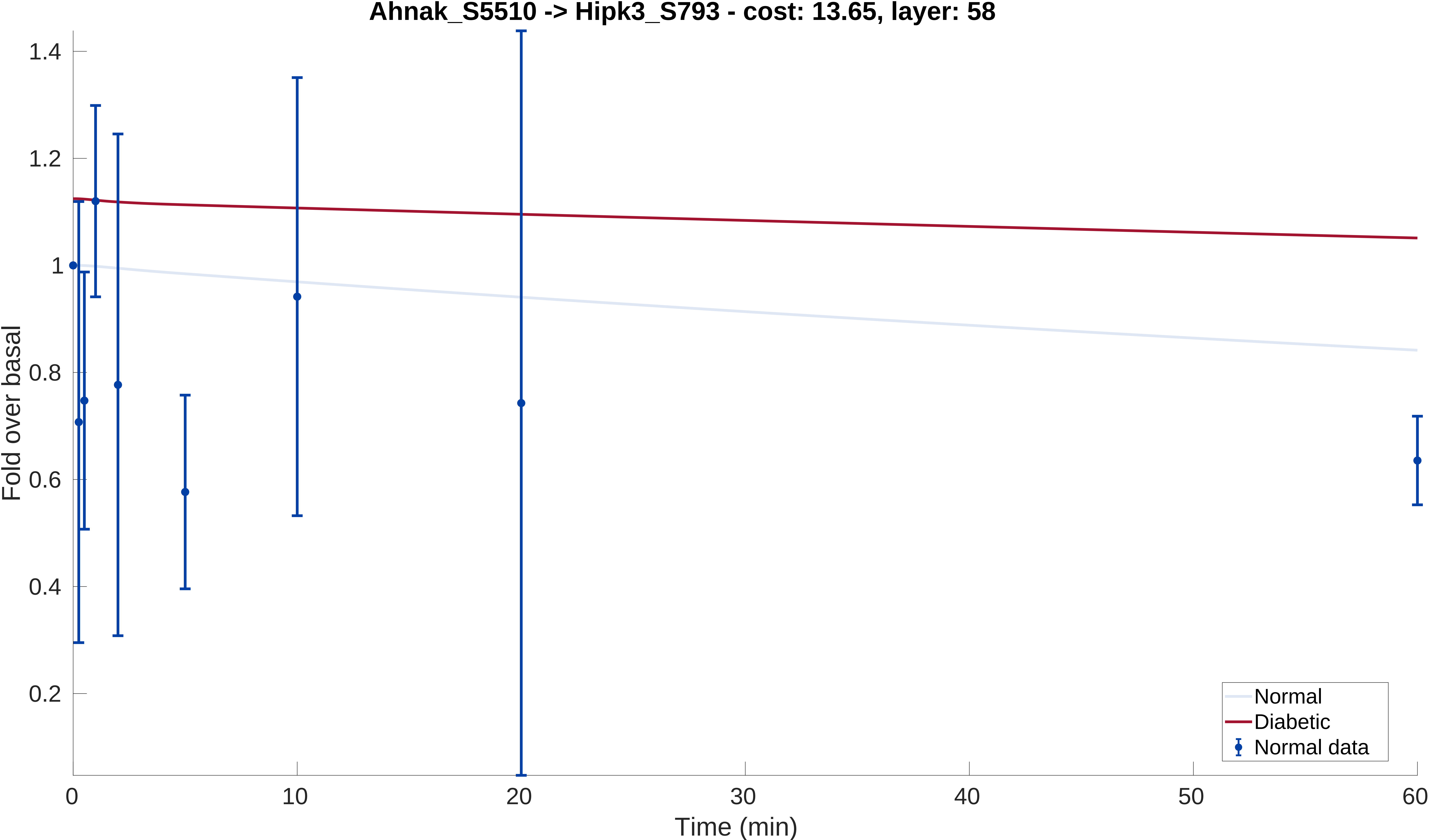

Ahnak_S5510 -> Hipk3_S793 - cost: 13.65, layer: 58
Fold over basal
1.4
1.2
1
0.8
0.6
0.4
0.2
0
10
20
30
40
50
60
Time (min)
Normal
Diabetic
Normal data

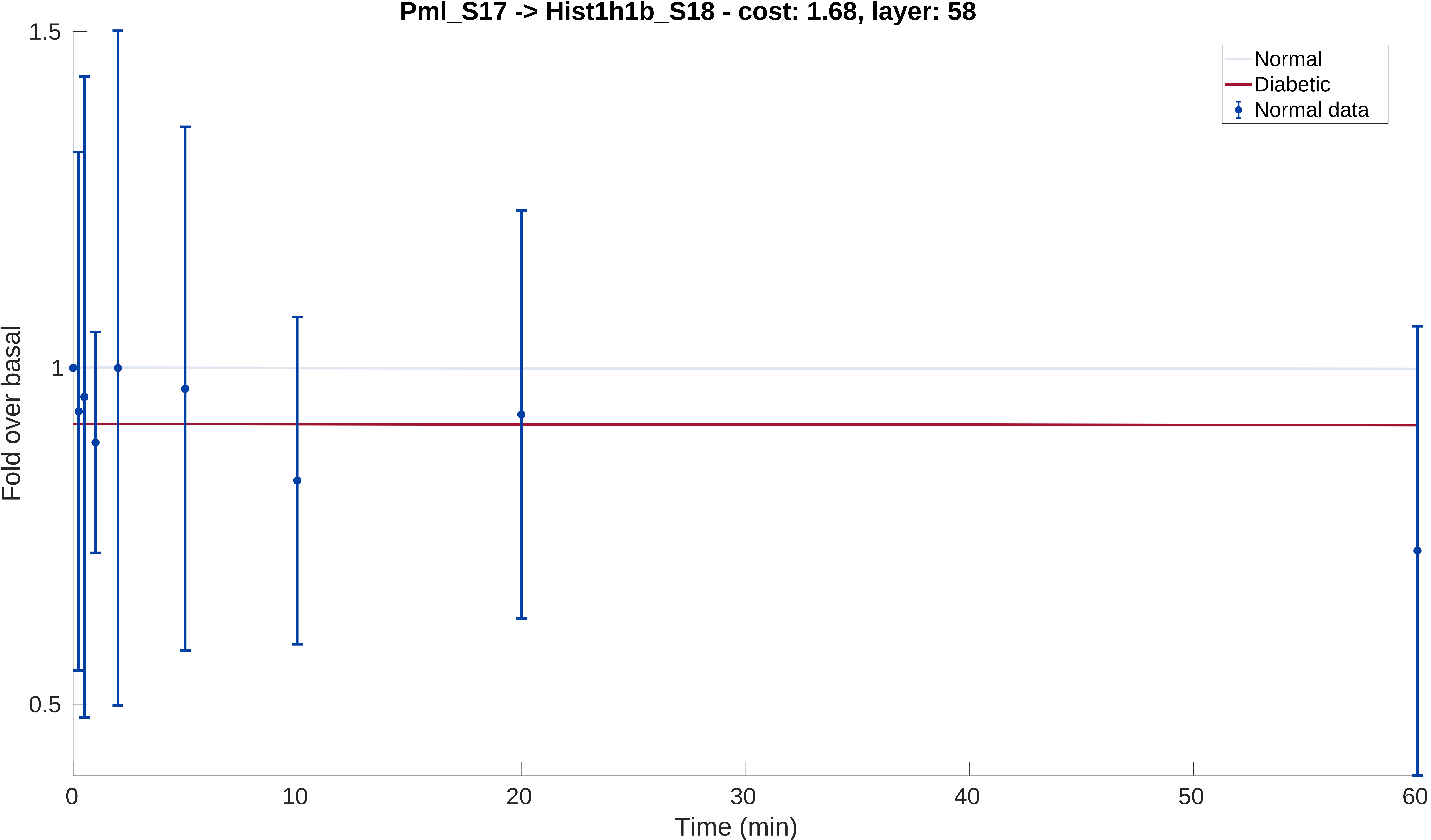

Pml_S17 -> Hist1h1b_S18 - cost: 1.68, layer: 58
Fold over basal
Time (min)
Normal
Diabetic
Normal data
1.5
1
0.5
0
10
20
30
40
50
60

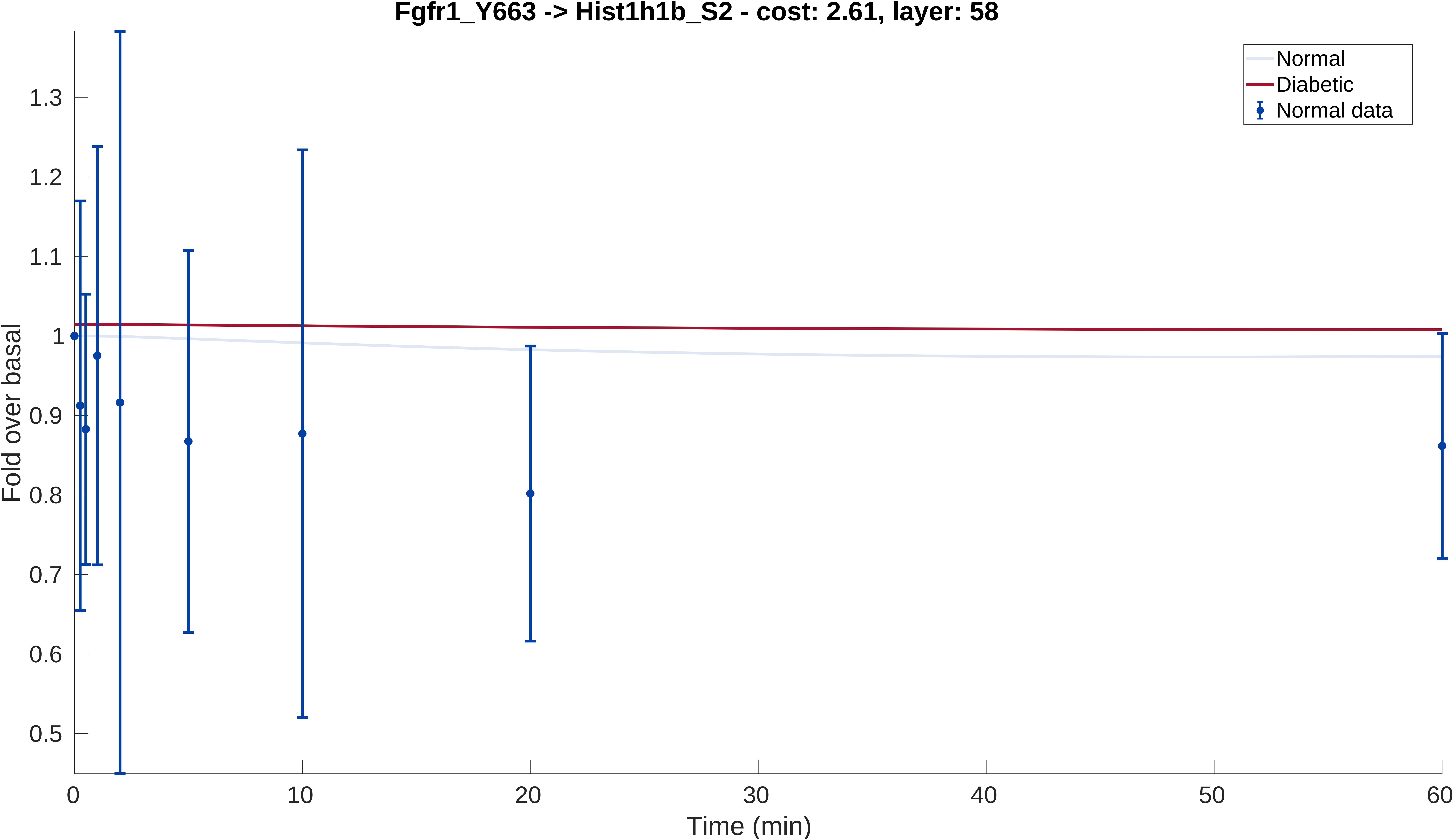
Fgfr1_Y663 -> Hist1h1b_S2 - cost: 2.61, layer: 58
Normal
Diabetic
Normal data
Fold over basal
1.3
1.2
1.1
1
0.9
0.8
0.7
0.6
0.5
0
10
20
30
40
50
60
Time (min)

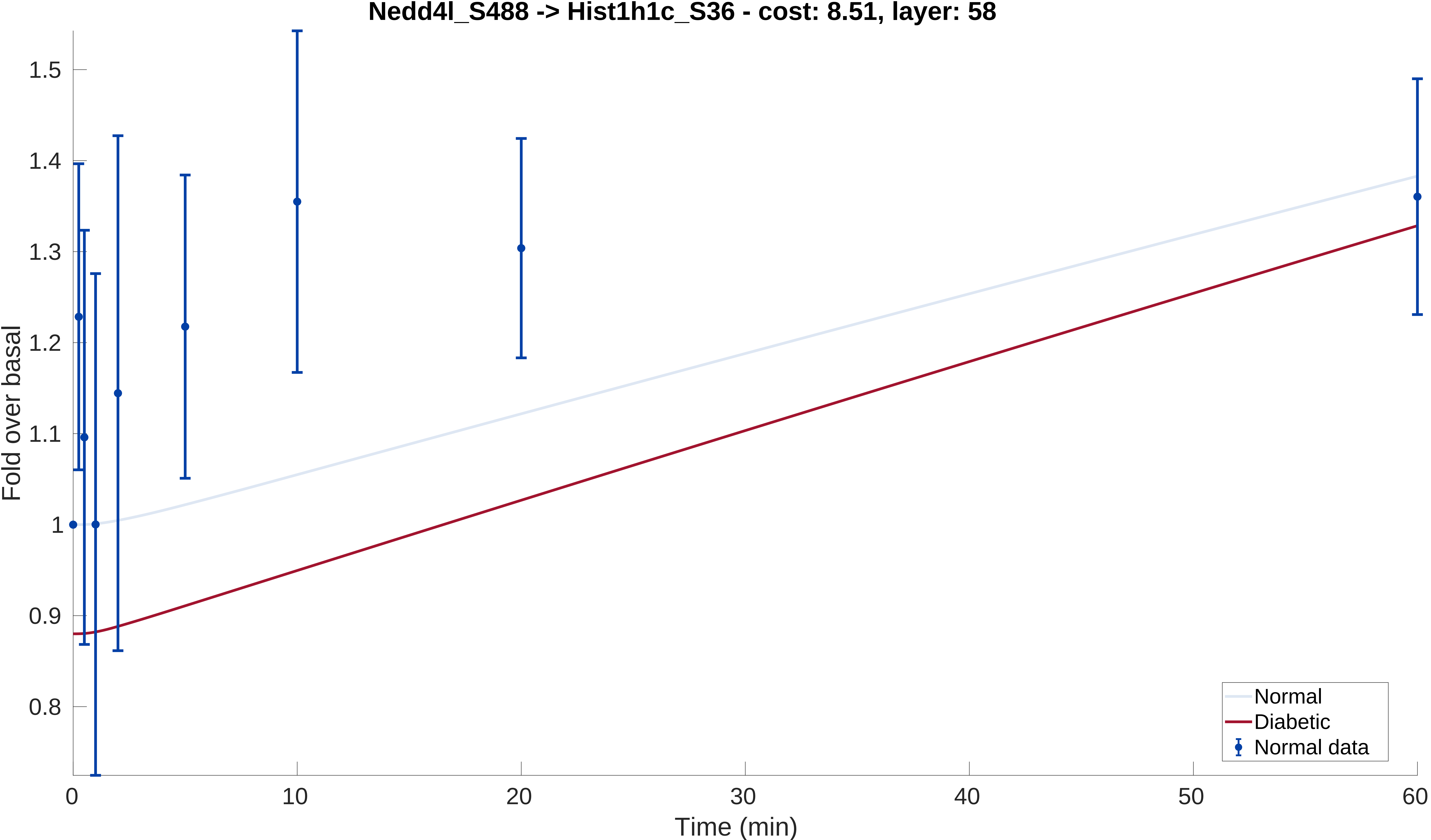

Nedd4l_S488 -> Hist1h1c_S36 - cost: 8.51, layer: 58
Fold over basal
Time (min)
0.8
0.9
1
1.1
1.2
1.3
1.4
1.5
0
10
20
30
40
50
60
Normal
Diabetic
Normal data

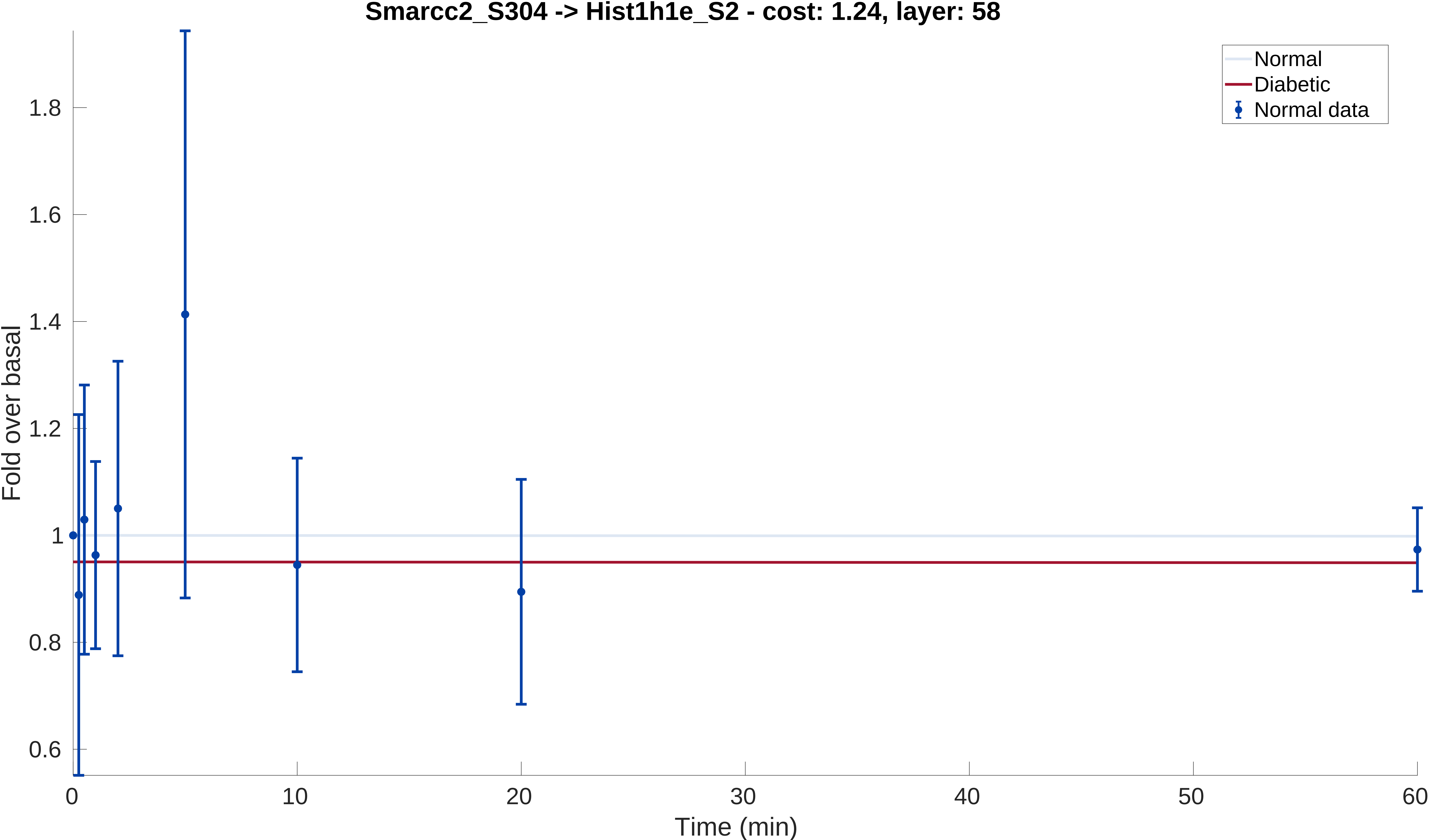

Smarcc2_S304 -> Hist1h1e_S2 - cost: 1.24, layer: 58
Normal
Diabetic
Normal data
1.8
1.6
1.4
1.2
1
0.8
0.6
Fold over basal
0
10
20
30
40
50
60
Time (min)

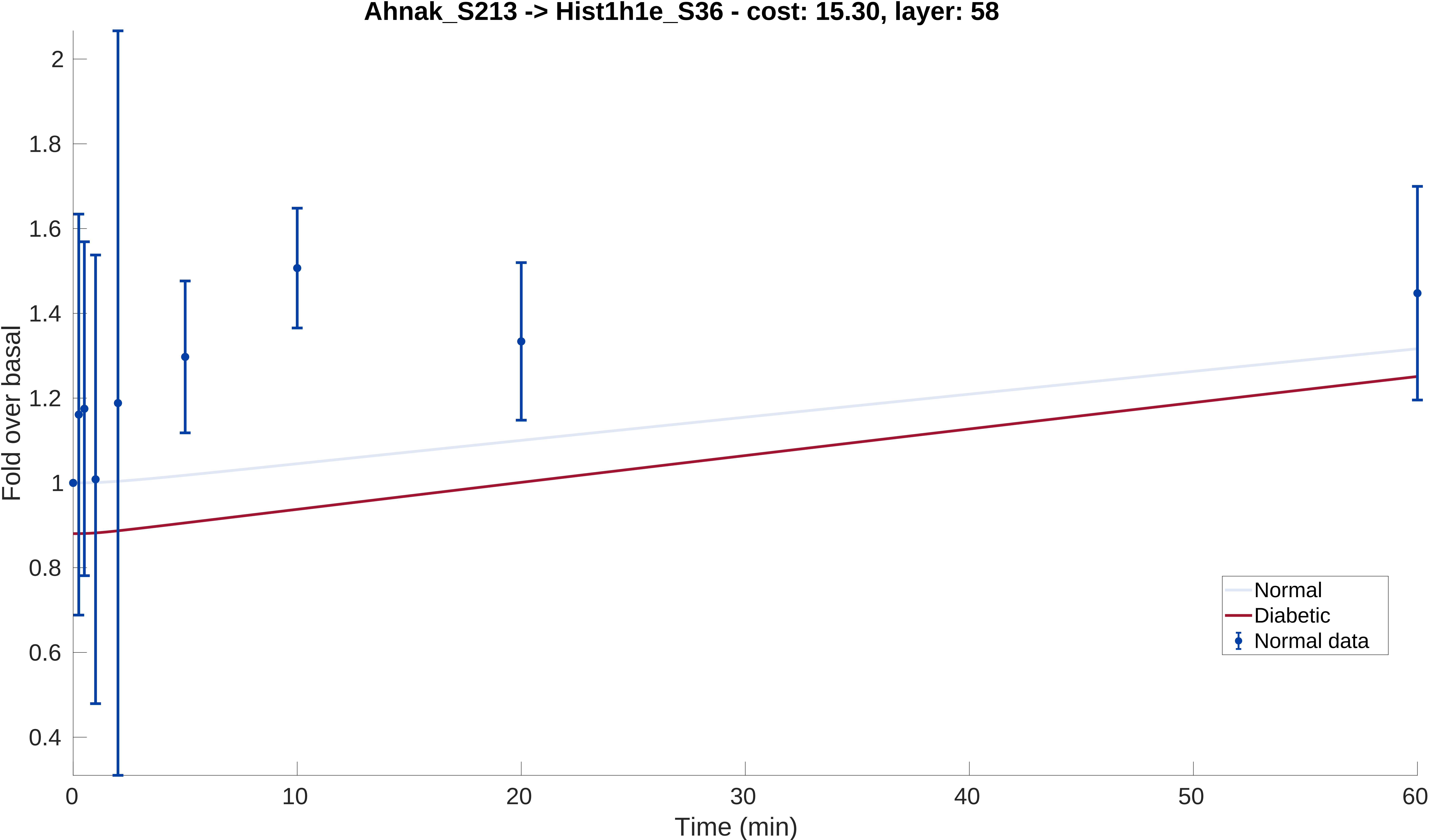
Ahnak_S213 -> Hist1h1e_S36 - cost: 15.30, layer: 58
Fold over basal
2
1.8
1.6
1.4
1.2
1
0.8
0.6
0.4
0
10
20
30
40
50
60
Time (min)
Normal
Diabetic
Normal data

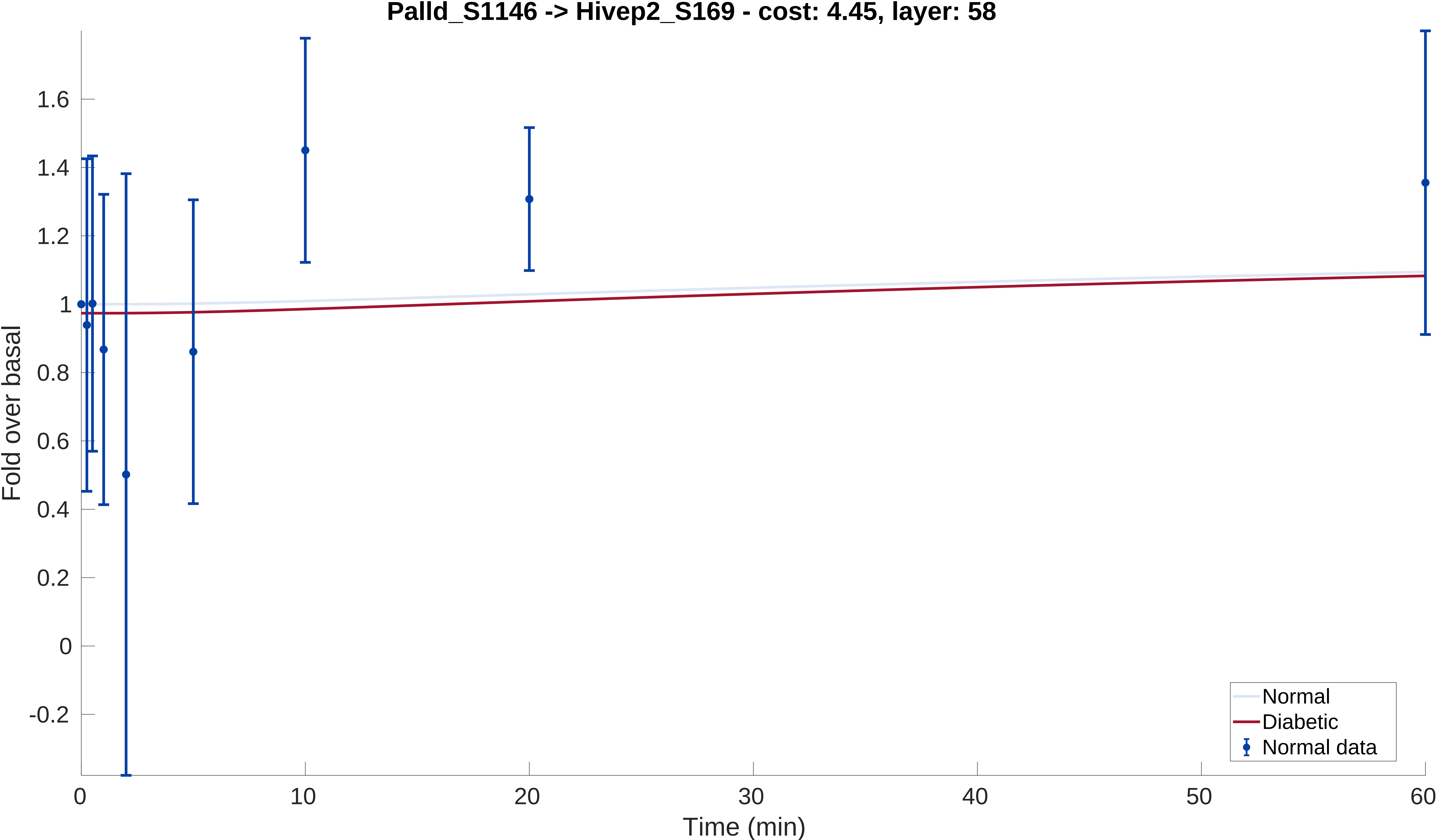

Palld_S1146 -> Hivep2_S169 - cost: 4.45, layer: 58
Fold over basal
1.6
1.4
1.2
1
0.8
0.6
0.4
0.2
0
-0.2
0
10
20
30
40
50
60
Time (min)
Normal
Diabetic
Normal data

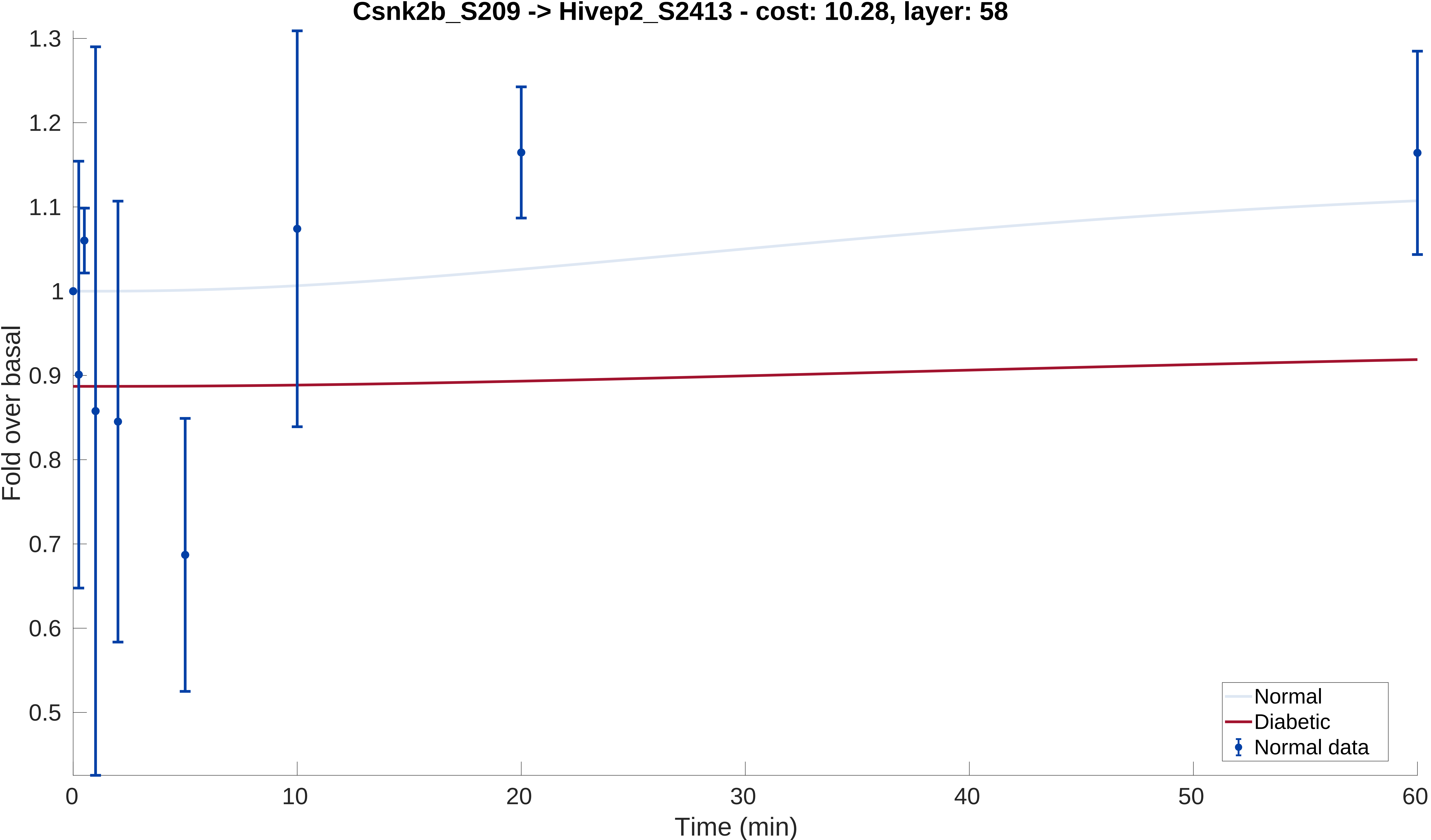

Csnk2b_S209 -> Hivep2_S2413 - cost: 10.28, layer: 58
1.3
1.2
1.1
1
0.9
0.8
0.7
0.6
0.5
Fold over basal
0
10
20
30
40
50
60
Time (min)
Normal
Diabetic
Normal data

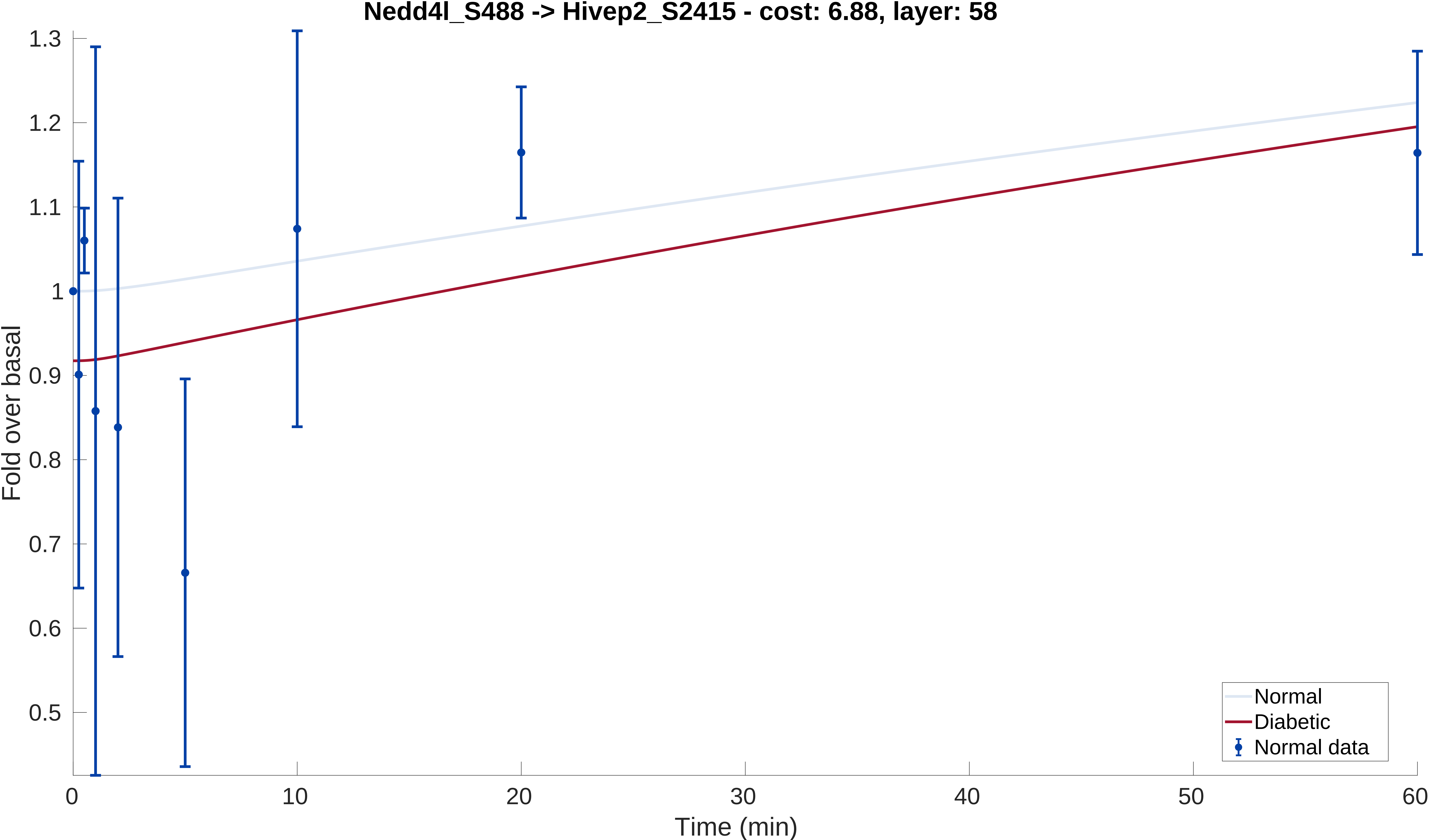

Nedd4l_S488 -> Hivep2_S2415 - cost: 6.88, layer: 58
Fold over basal
Time (min)
1.3
1.2
1.1
1
0.9
0.8
0.7
0.6
0.5
0
10
20
30
40
50
60
Normal
Diabetic
Normal data

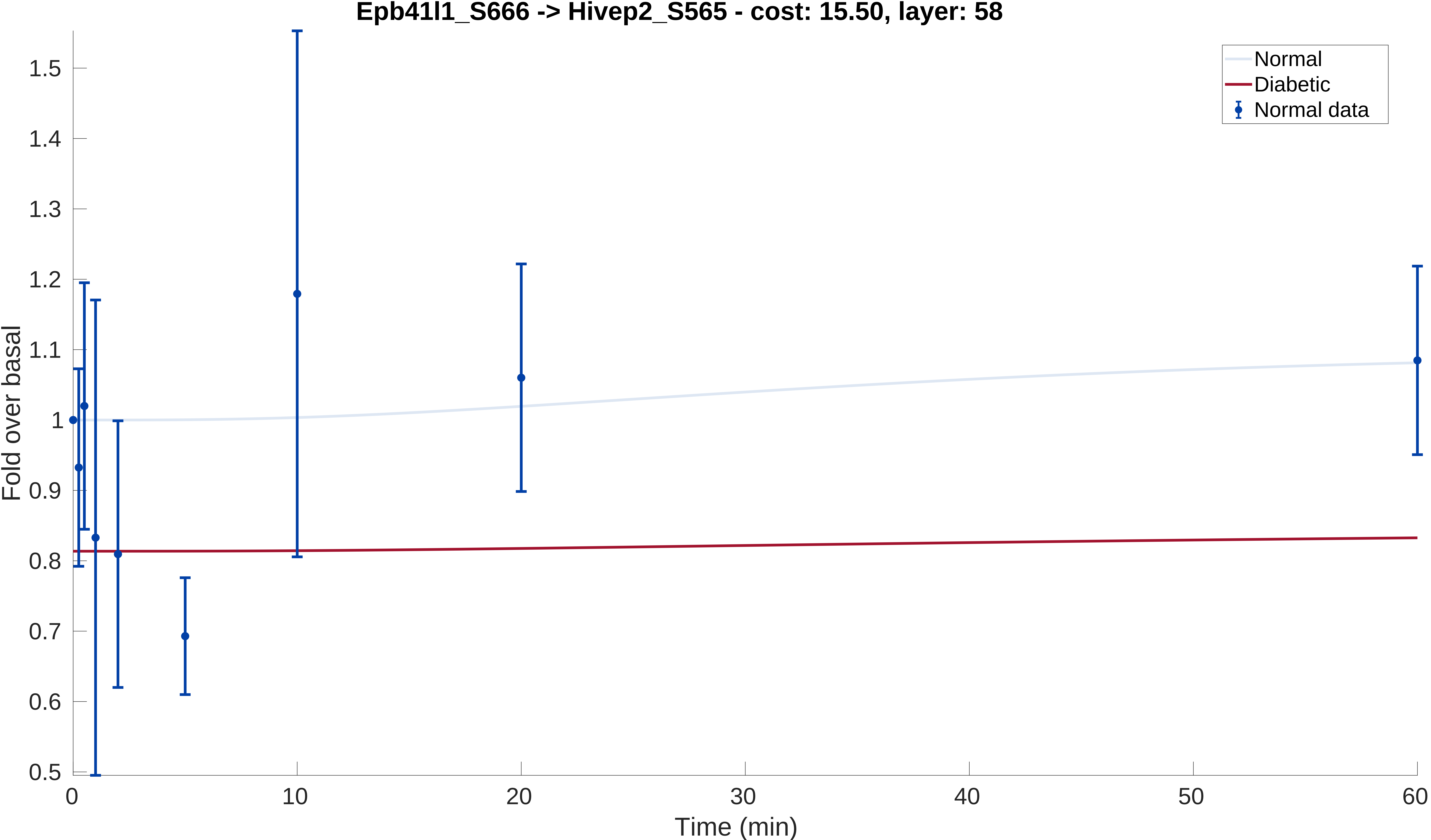

Epb41l1_S666 -> Hivep2_S565 - cost: 15.50, layer: 58
Normal
Diabetic
Normal data
Fold over basal
1.5
1.4
1.3
1.2
1.1
1
0.9
0.8
0.7
0.6
0.5
0
10
20
30
40
50
60
Time (min)

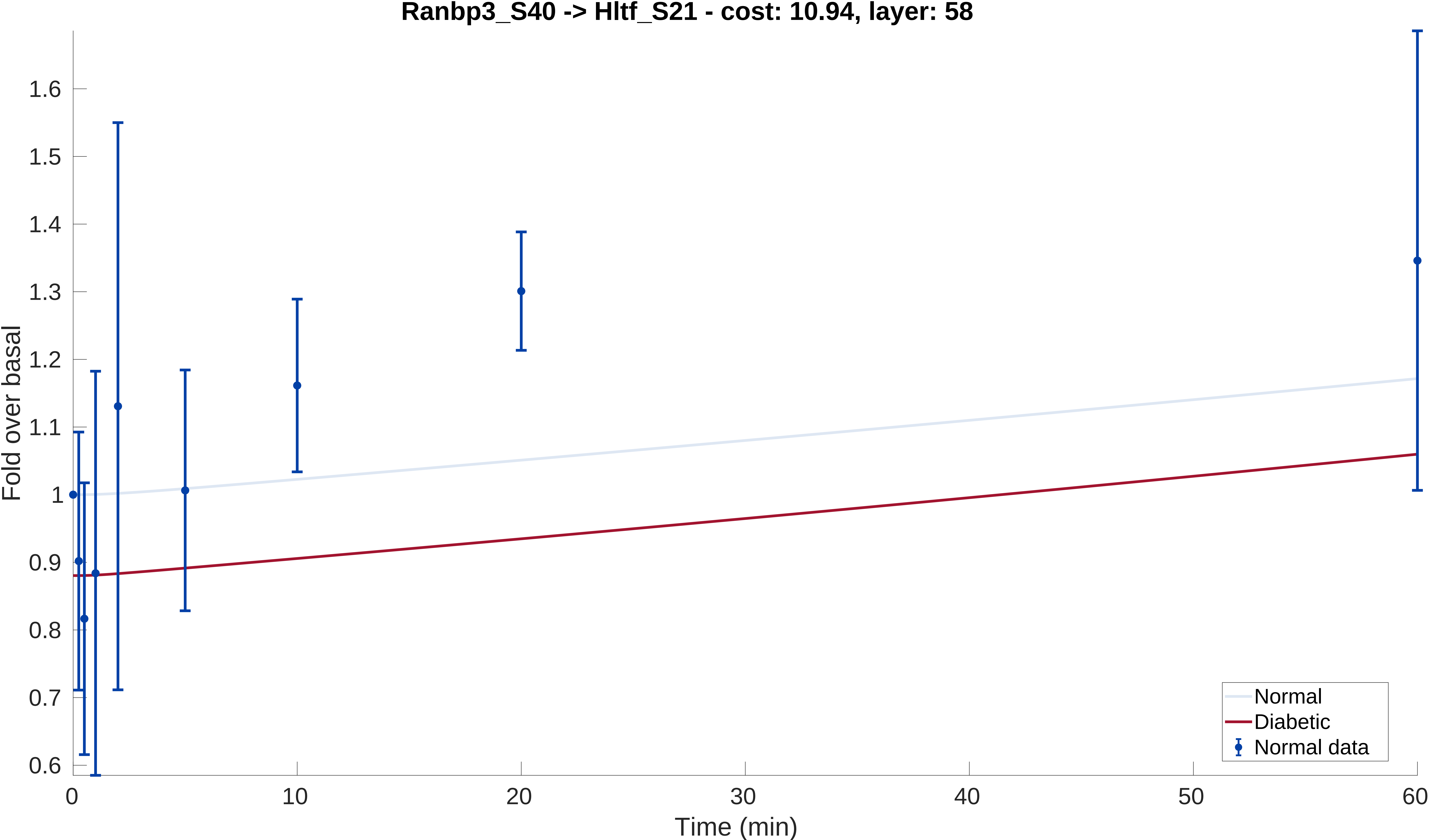

Ranbp3_S40 -> Hltf_S21 - cost: 10.94, layer: 58
Fold over basal
1.6
1.5
1.4
1.3
1.2
1.1
1
0.9
0.8
0.7
0.6
0
10
20
30
40
50
60
Time (min)
Normal
Diabetic
Normal data

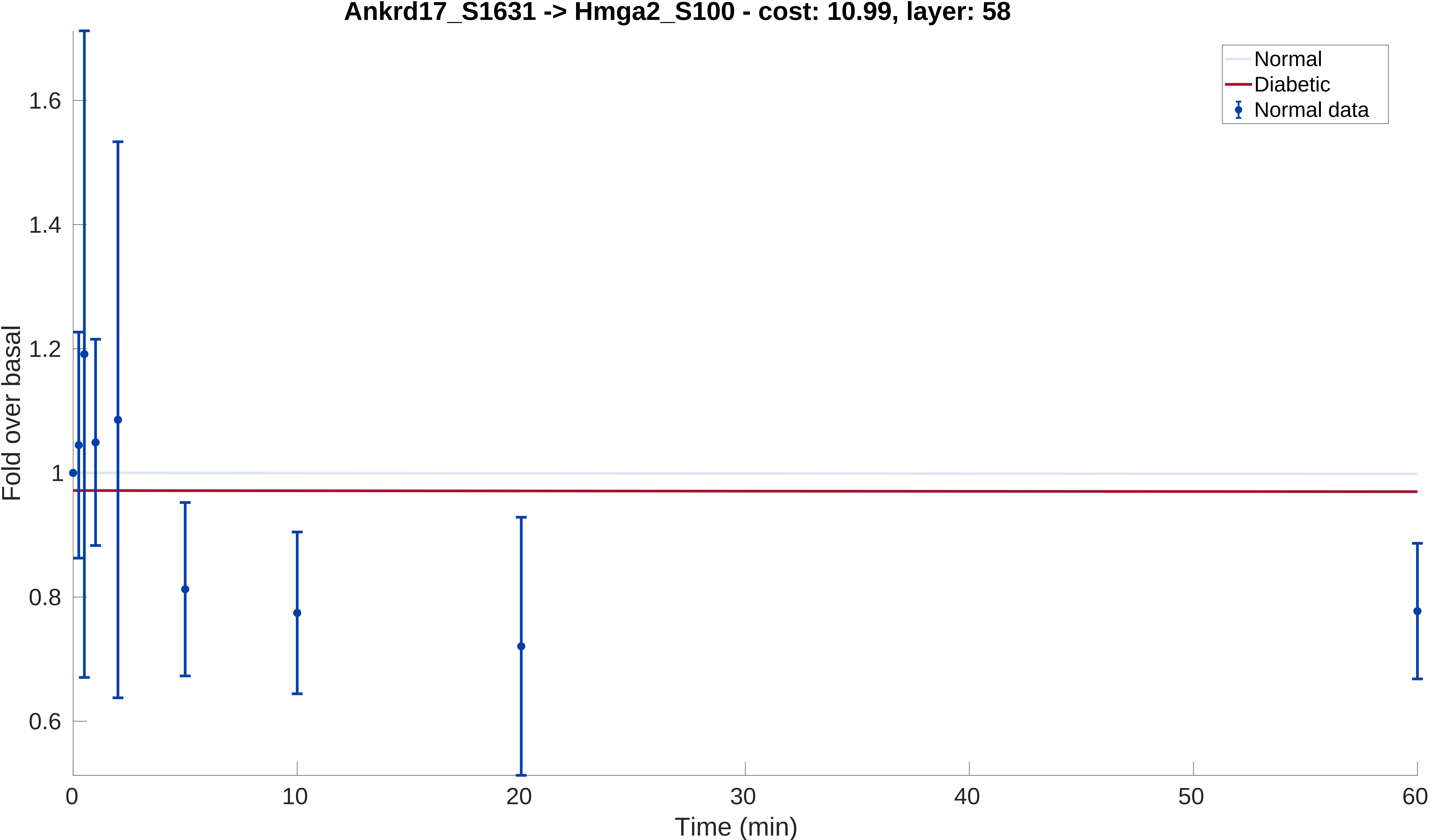

Ankrd17_S1631 -> Hmga2_S100 - cost: 10.99, layer: 58
Normal
Diabetic
Normal data
Fold over basal
1.6
1.4
1.2
1
0.8
0.6
0
10
20
30
40
50
60
Time (min)

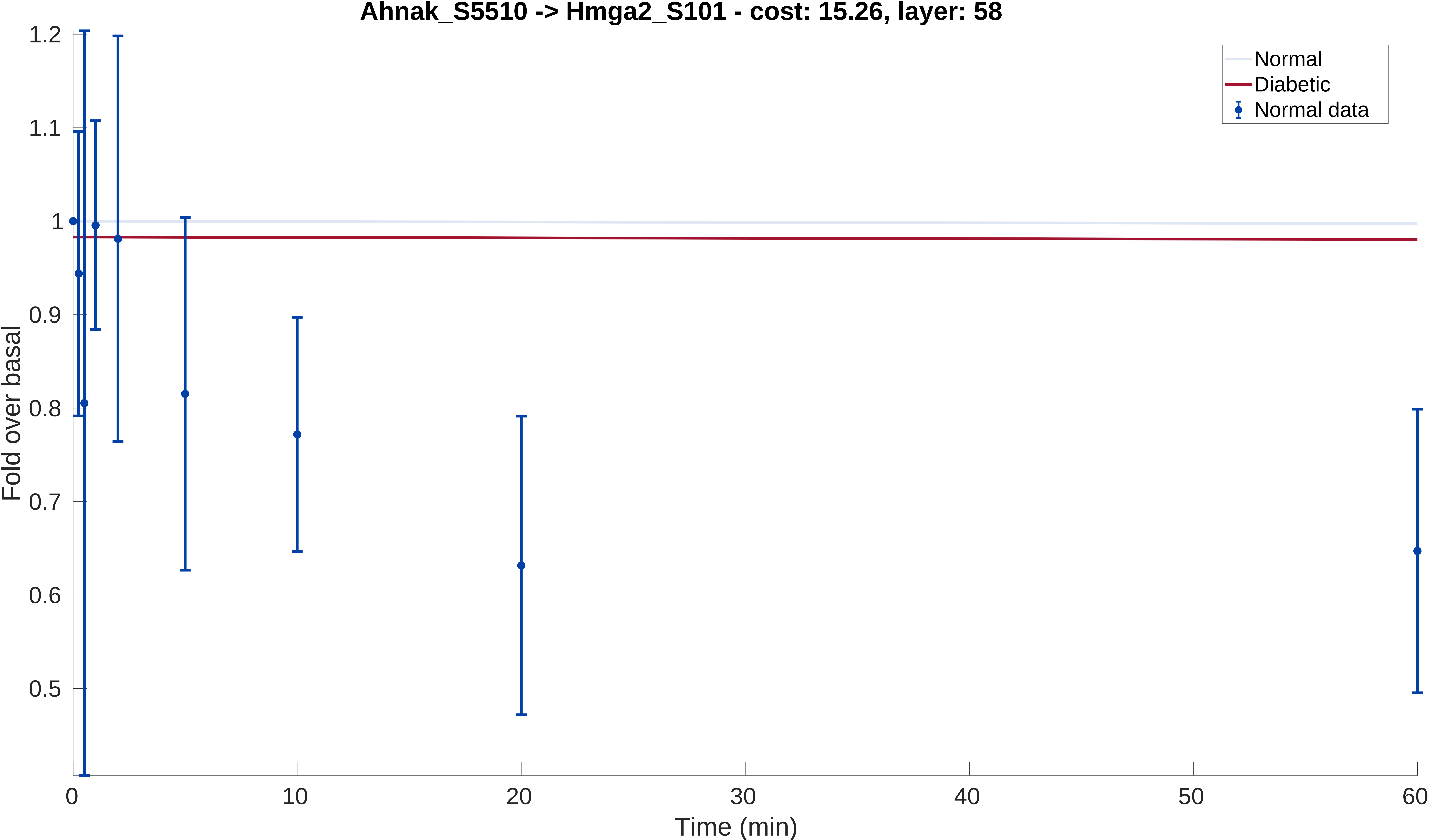

Ahnak_S5510 -> Hmga2_S101 - cost: 15.26, layer: 58
Normal
Diabetic
Normal data
Fold over basal
1.2
1.1
1
0.9
0.8
0.7
0.6
0.5
0
10
20
30
40
50
60
Time (min)

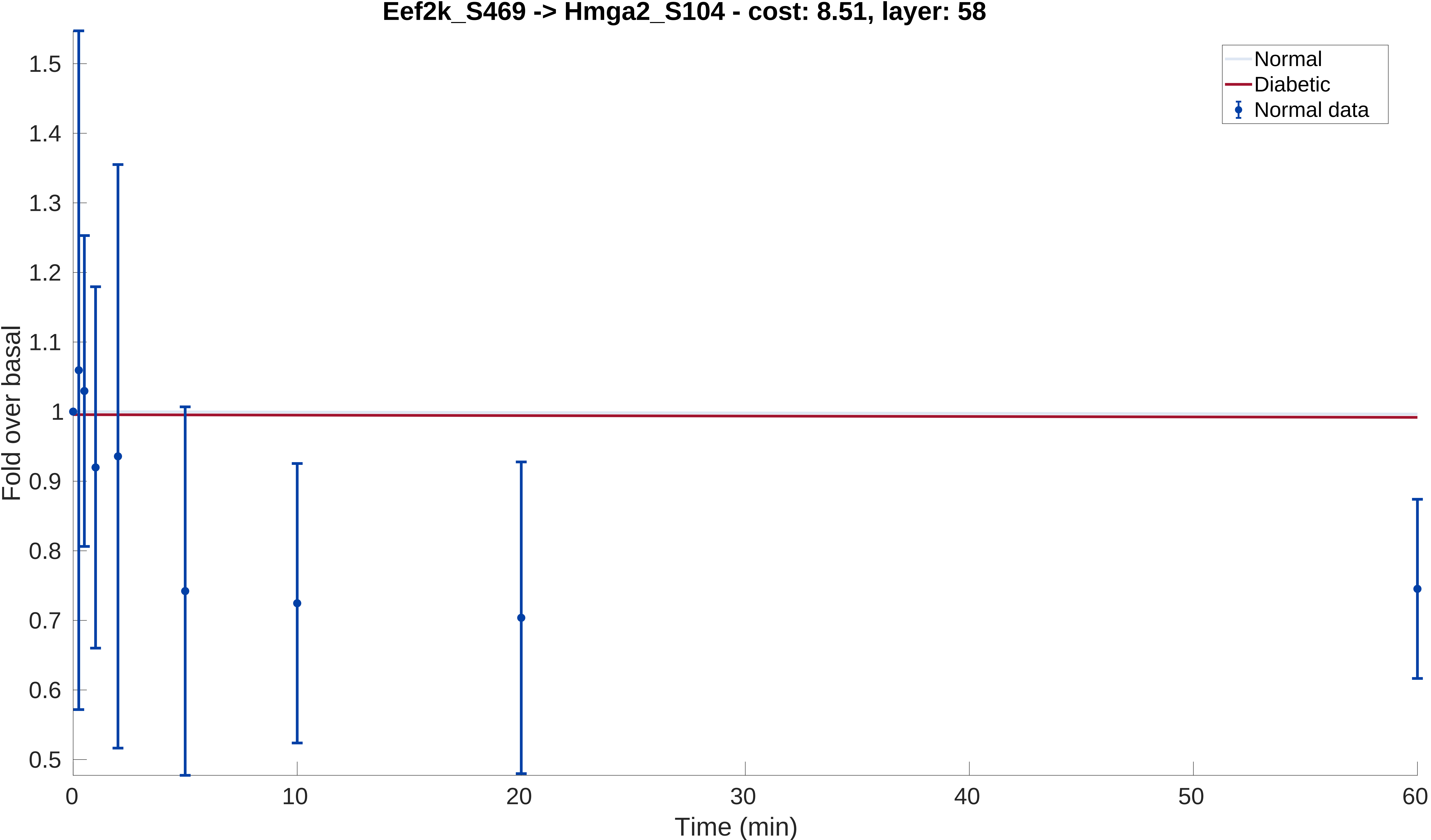

Eef2k_S469 -> Hmga2_S104 - cost: 8.51, layer: 58
Normal
Diabetic
Normal data
Fold over basal
Time (min)
1.5
1.4
1.3
1.2
1.1
1
0.9
0.8
0.7
0.6
0.5
0
10
20
30
40
50
60

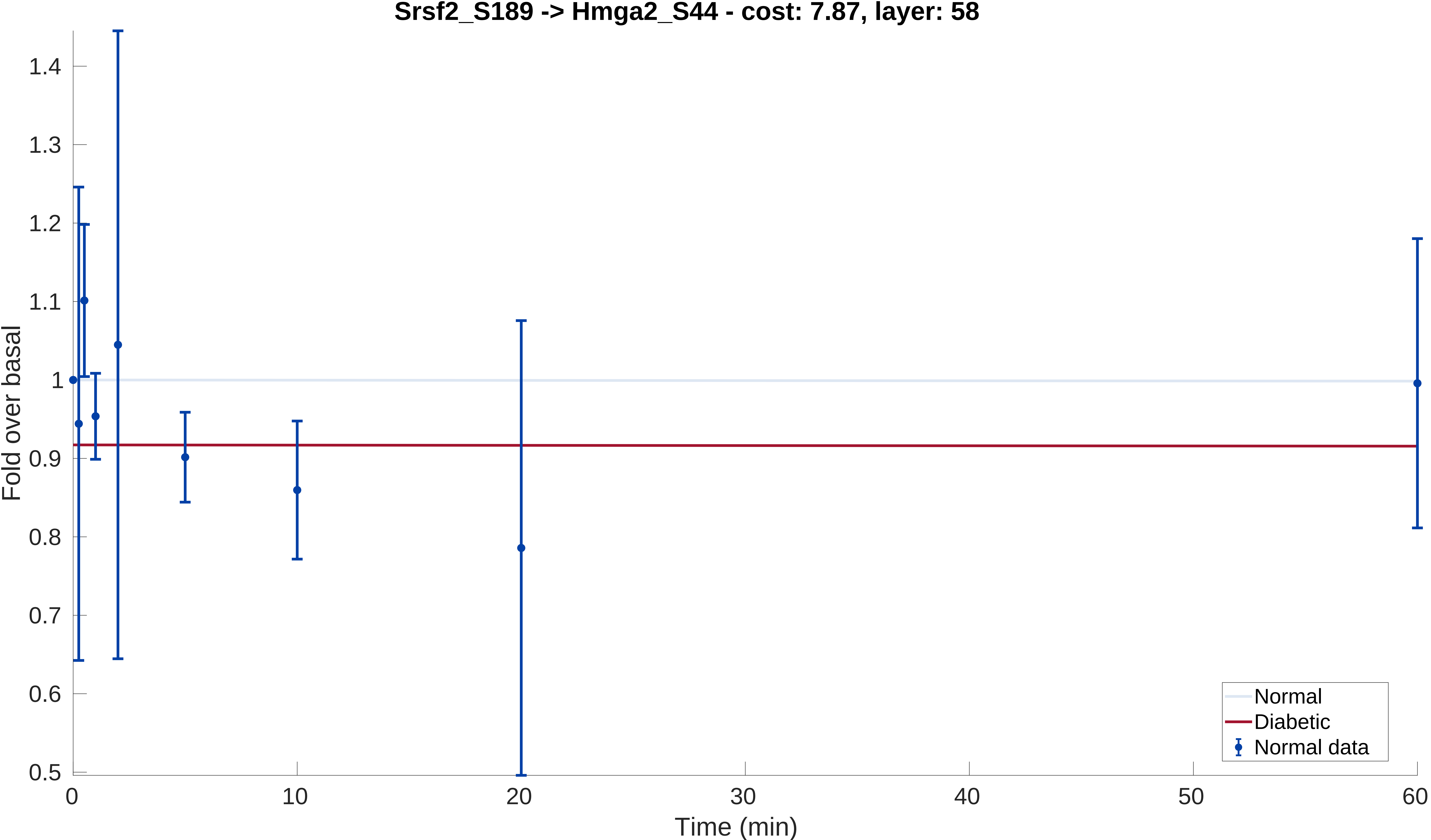

Srsf2_S189 -> Hmga2_S44 - cost: 7.87, layer: 58
Fold over basal
1.4
1.3
1.2
1.1
1
0.9
0.8
0.7
0.6
0.5
0
10
20
30
40
50
60
Time (min)
Normal
Diabetic
Normal data

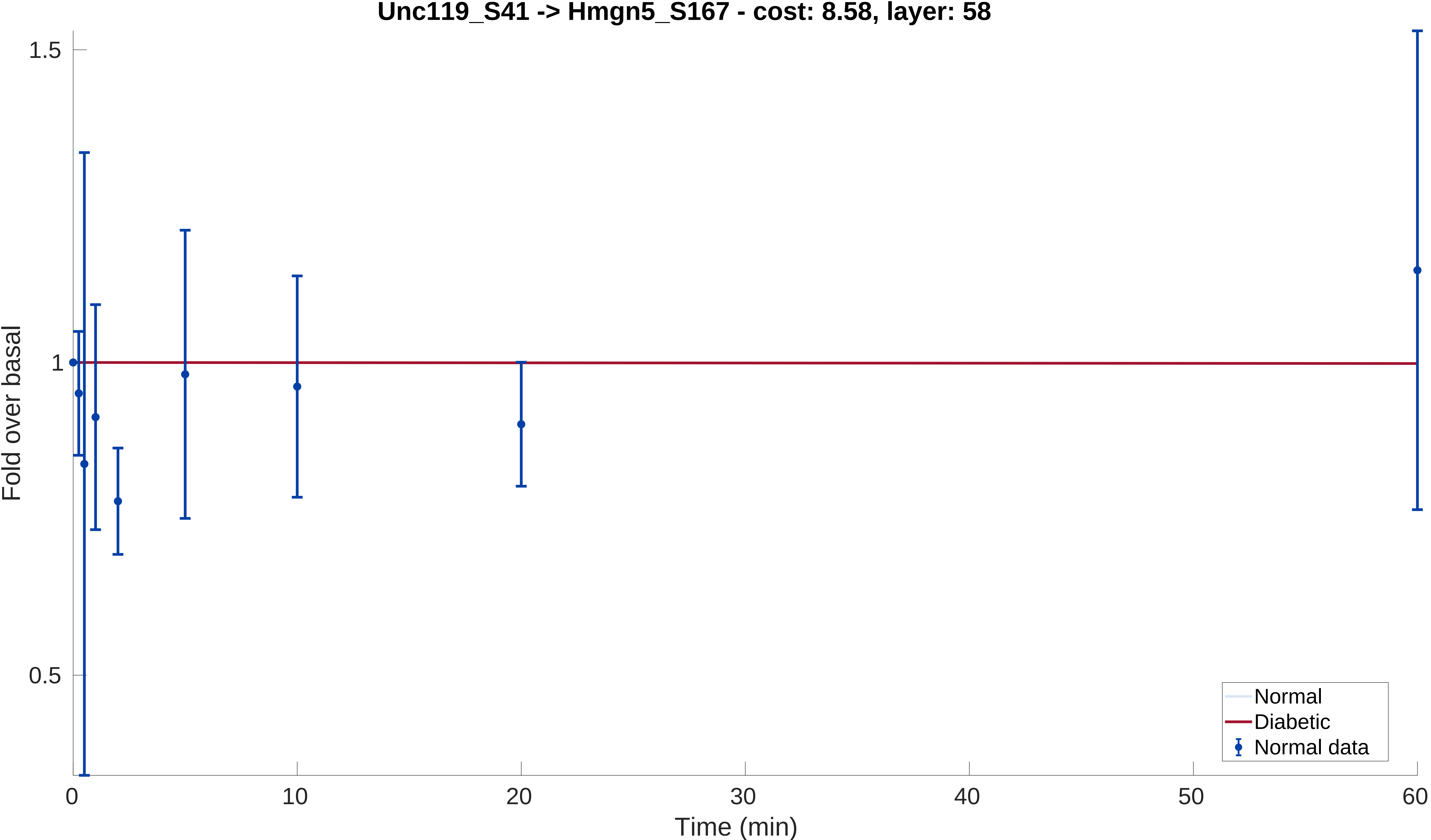

Unc119_S41 -> Hmgn5_S167 - cost: 8.58, layer: 58
1.5
1
0.5
Fold over basal
0
10
20
30
40
50
60
Time (min)
Normal
Diabetic
Normal data

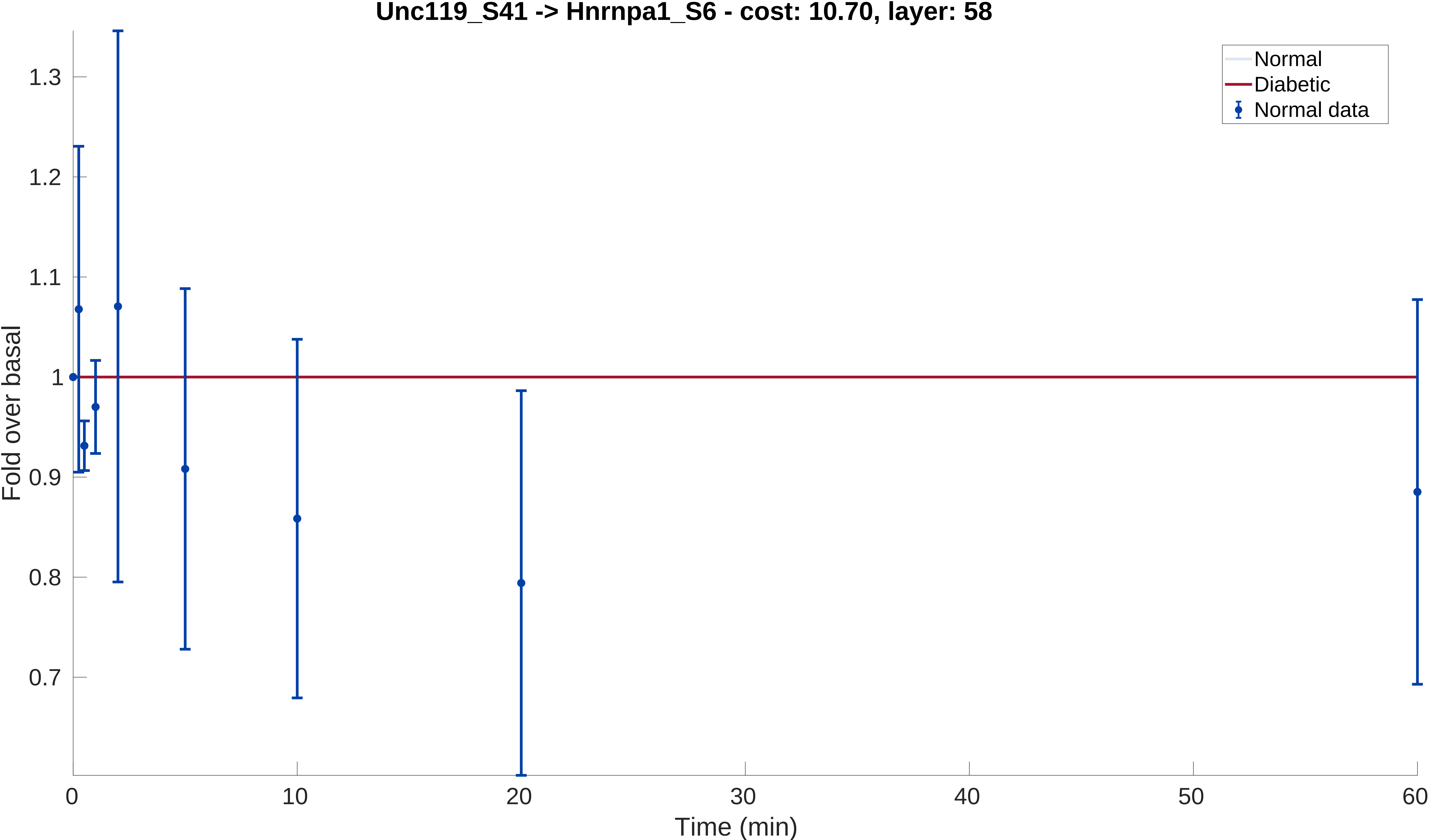

Unc119_S41 -> Hnrnpa1_S6 - cost: 10.70, layer: 58
Normal
Diabetic
Normal data
Fold over basal
1.3
1.2
1.1
1
0.9
0.8
0.7
0
10
20
30
40
50
60
Time (min)

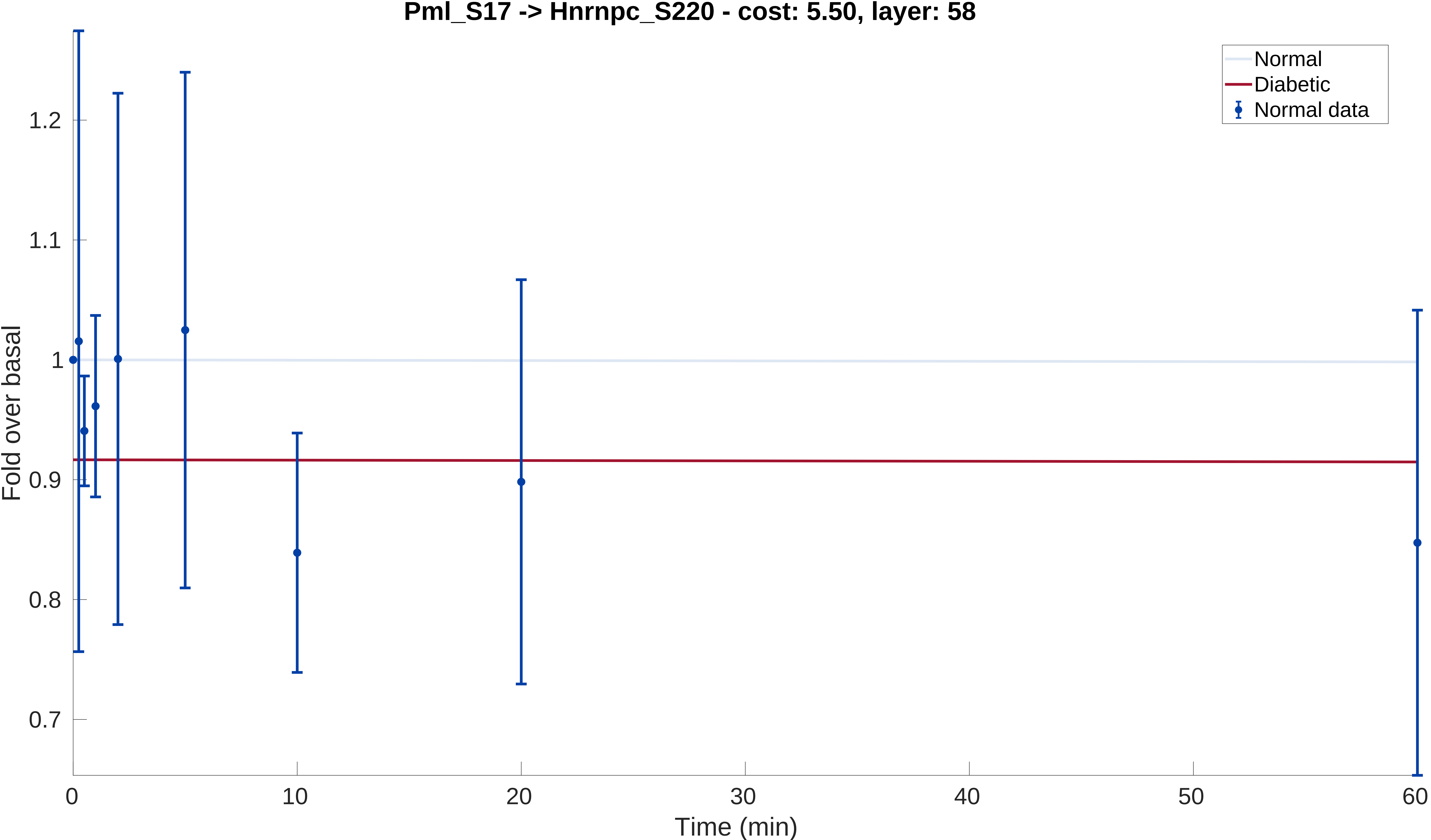
Pml_S17 -> Hnrnpc_S220 - cost: 5.50, layer: 58
Normal
Diabetic
Normal data
Fold over basal
1.2
1.1
1
0.9
0.8
0.7
0
10
20
30
40
50
60
Time (min)

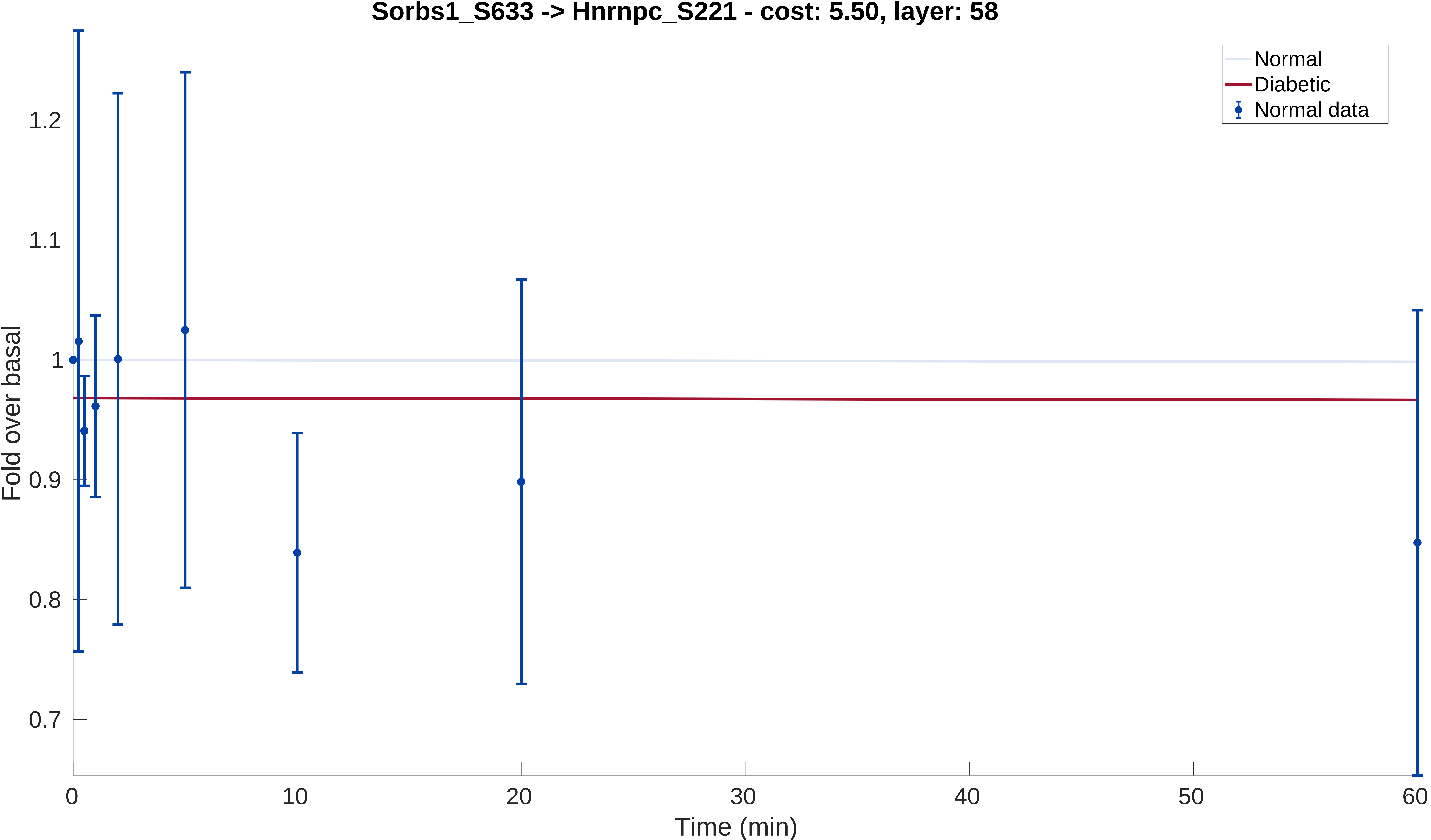
Sorbs1_S633 -> Hnrnpc_S221 - cost: 5.50, layer: 58
Normal
Diabetic
Normal data
Fold over basal
1.2
1.1
1
0.9
0.8
0.7
0
10
20
30
40
50
60
Time (min)

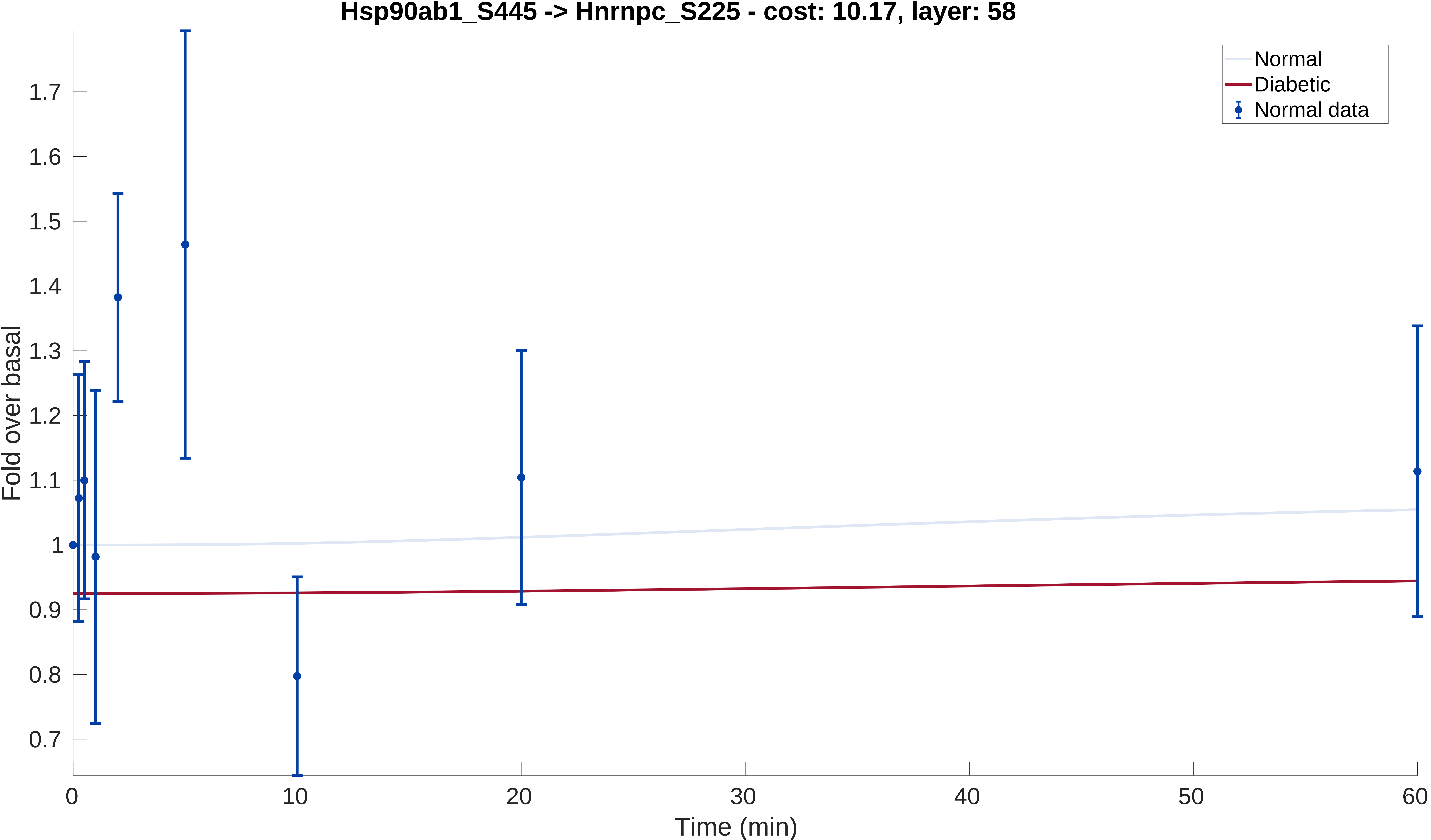
Hsp90ab1_S445 -> Hnrnpc_S225 - cost: 10.17, layer: 58
Normal
Diabetic
Normal data
Fold over basal
1.7
1.6
1.5
1.4
1.3
1.2
1.1
1
0.9
0.8
0.7
0
10
20
30
40
50
60
Time (min)

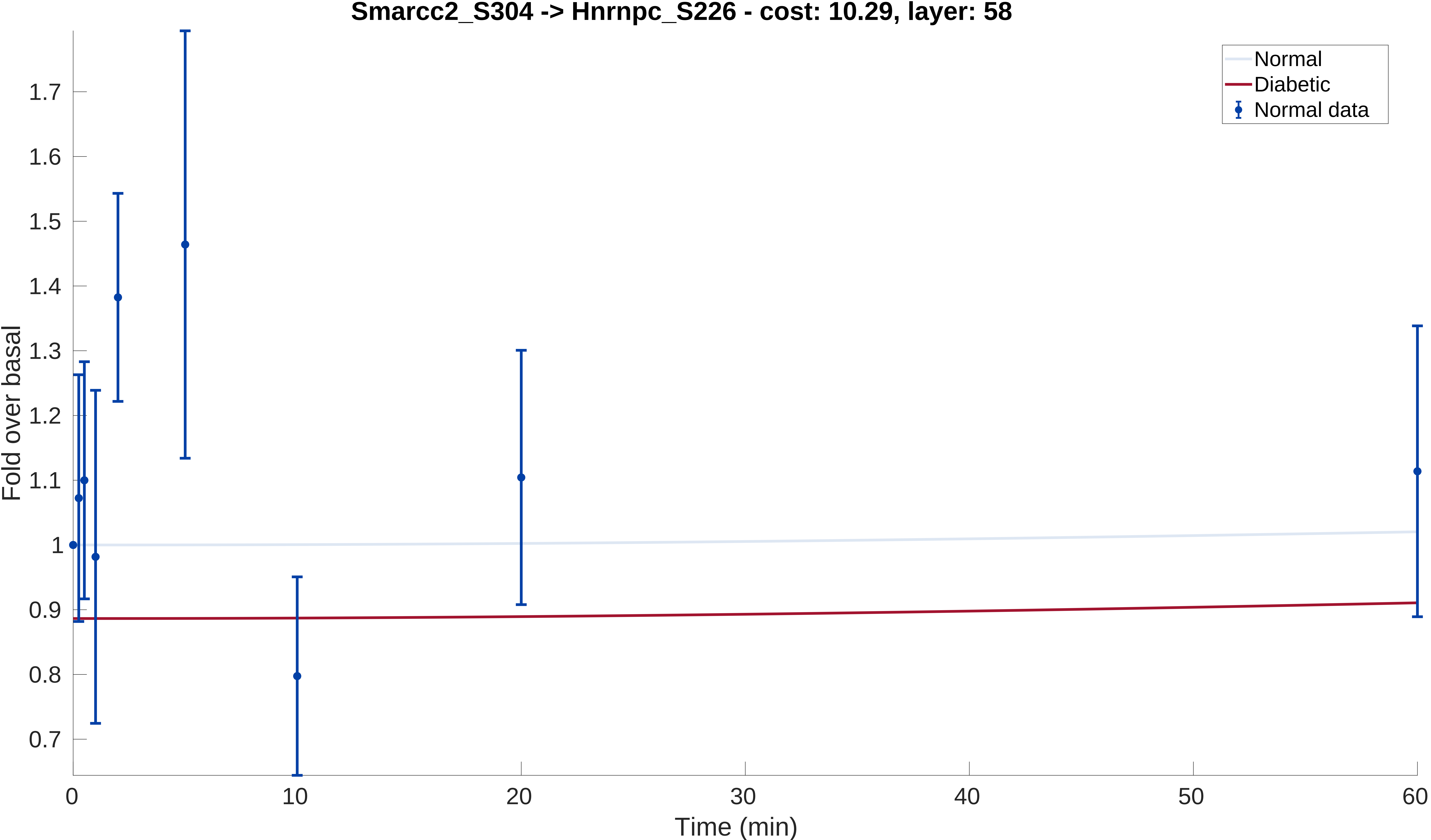

Smarcc2_S304 -> Hnrnpc_S226 - cost: 10.29, layer: 58
Normal
Diabetic
Normal data
Fold over basal
Time (min)
0.7
0.8
0.9
1
1.1
1.2
1.3
1.4
1.5
1.6
1.7
0
10
20
30
40
50
60

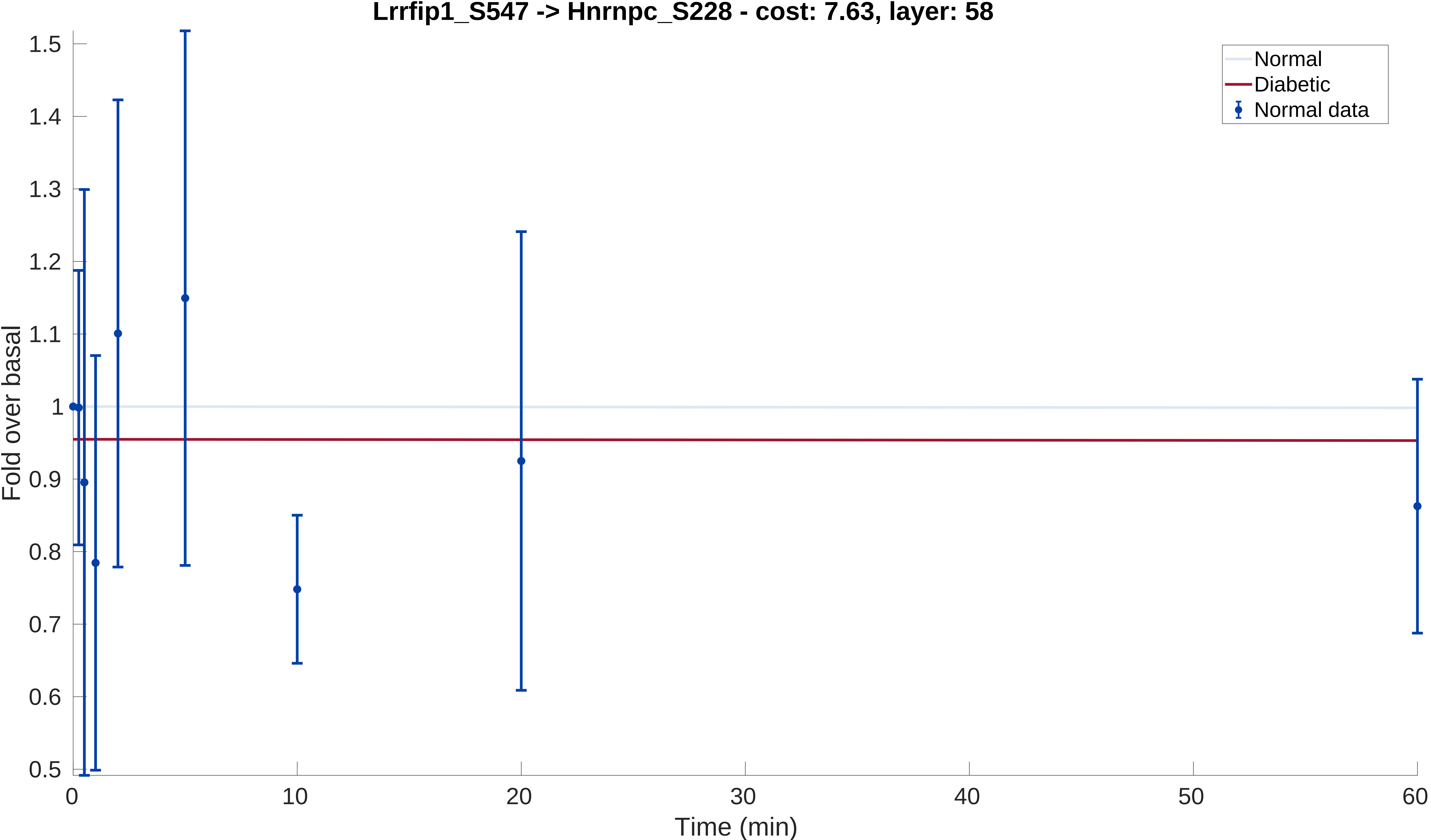
Lrrfip1_S547 -> Hnrnpc_S228 - cost: 7.63, layer: 58
Normal
Diabetic
Normal data
Fold over basal
1.5
1.4
1.3
1.2
1.1
1
0.9
0.8
0.7
0.6
0.5
0
10
20
30
40
50
60
Time (min)

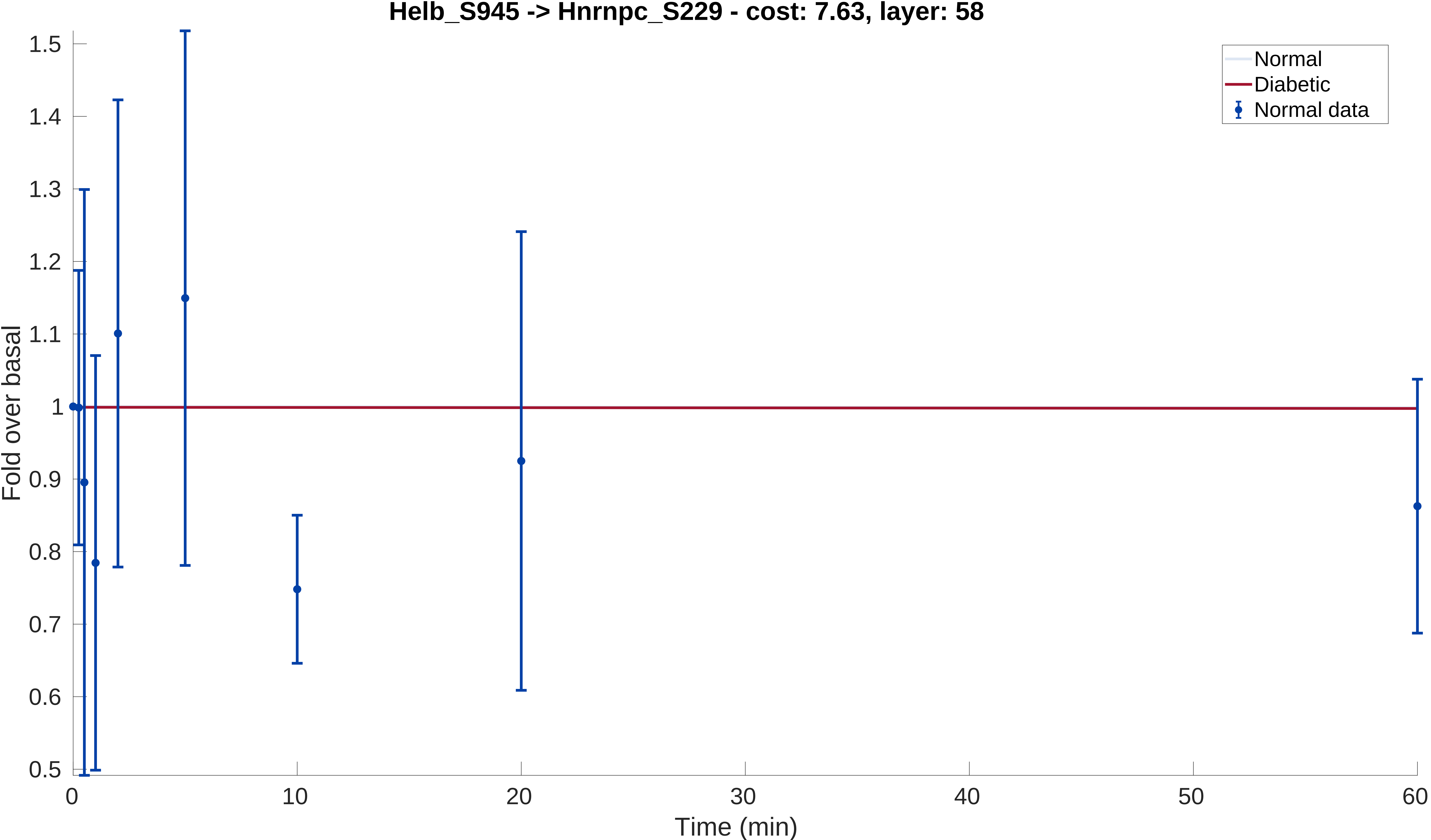

Helb_S945 -> Hnrnpc_S229 - cost: 7.63, layer: 58
Normal
Diabetic
Normal data
Fold over basal
Time (min)
1.5
1.4
1.3
1.2
1.1
1
0.9
0.8
0.7
0.6
0.5
0
10
20
30
40
50
60

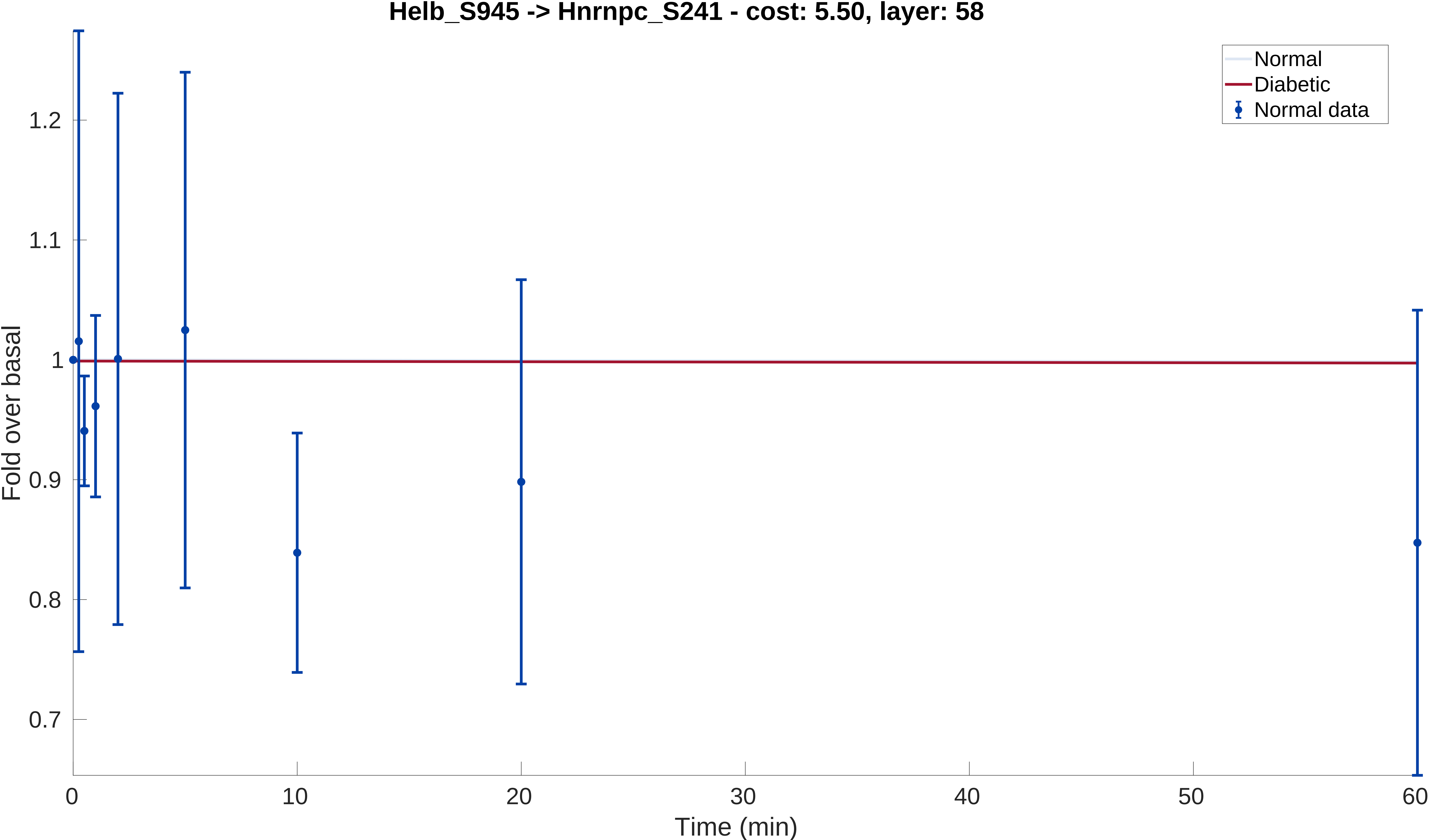

Helb_S945 -> Hnrnpc_S241 - cost: 5.50, layer: 58
Normal
Diabetic
Normal data
Fold over basal
1.2
1.1
1
0.9
0.8
0.7
0
10
20
30
40
50
60
Time (min)

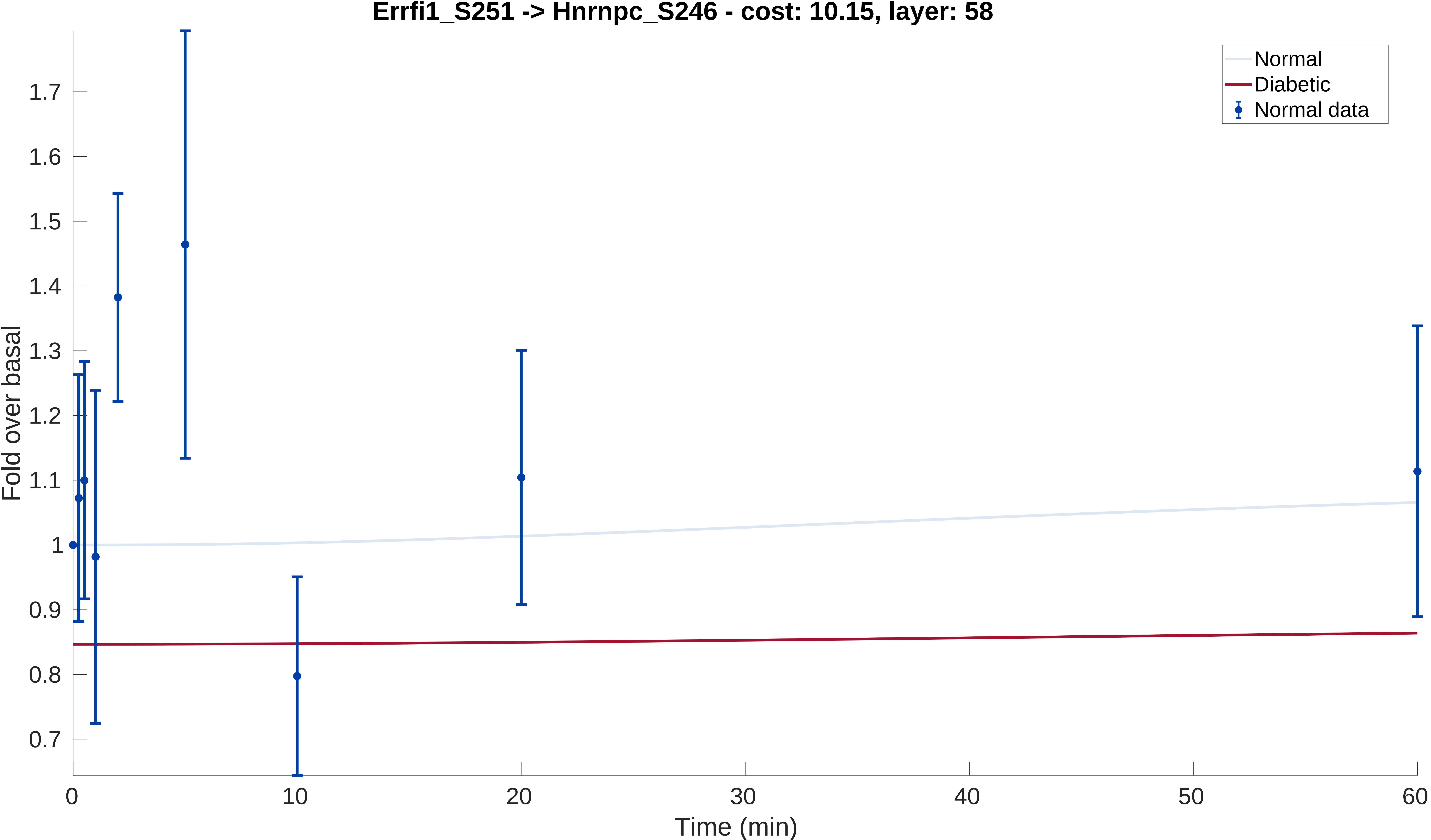

Errfi1_S251 -> Hnrnpc_S246 - cost: 10.15, layer: 58
Normal
Diabetic
Normal data
Fold over basal
1.7
1.6
1.5
1.4
1.3
1.2
1.1
1
0.9
0.8
0.7
0
10
20
30
40
50
60
Time (min)

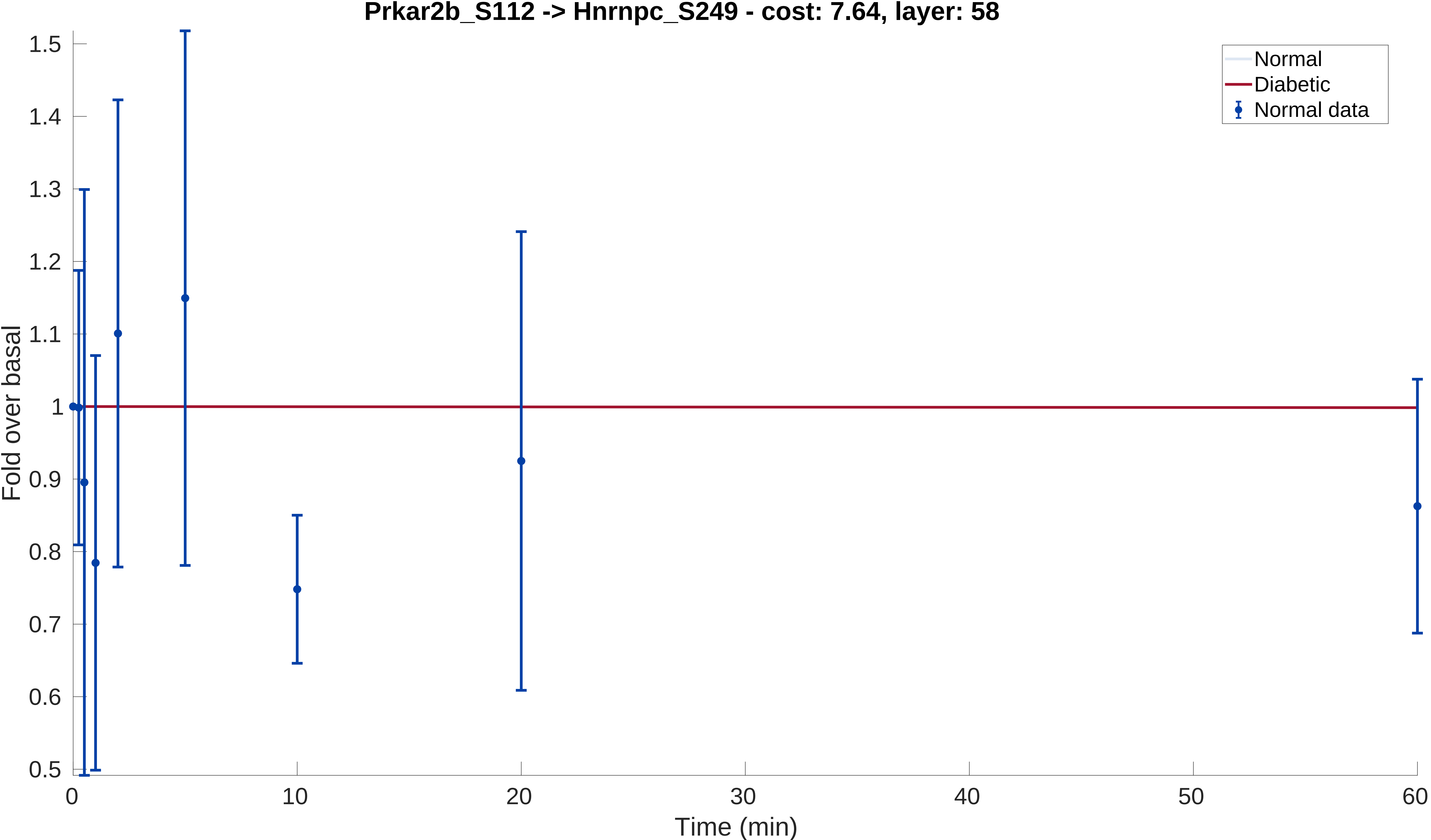

Prkar2b_S112 -> Hnrnpc_S249 - cost: 7.64, layer: 58
Fold over basal
Time (min)
Normal
Diabetic
Normal data
1.5
1.4
1.3
1.2
1.1
1
0.9
0.8
0.7
0.6
0.5
0
10
20
30
40
50
60

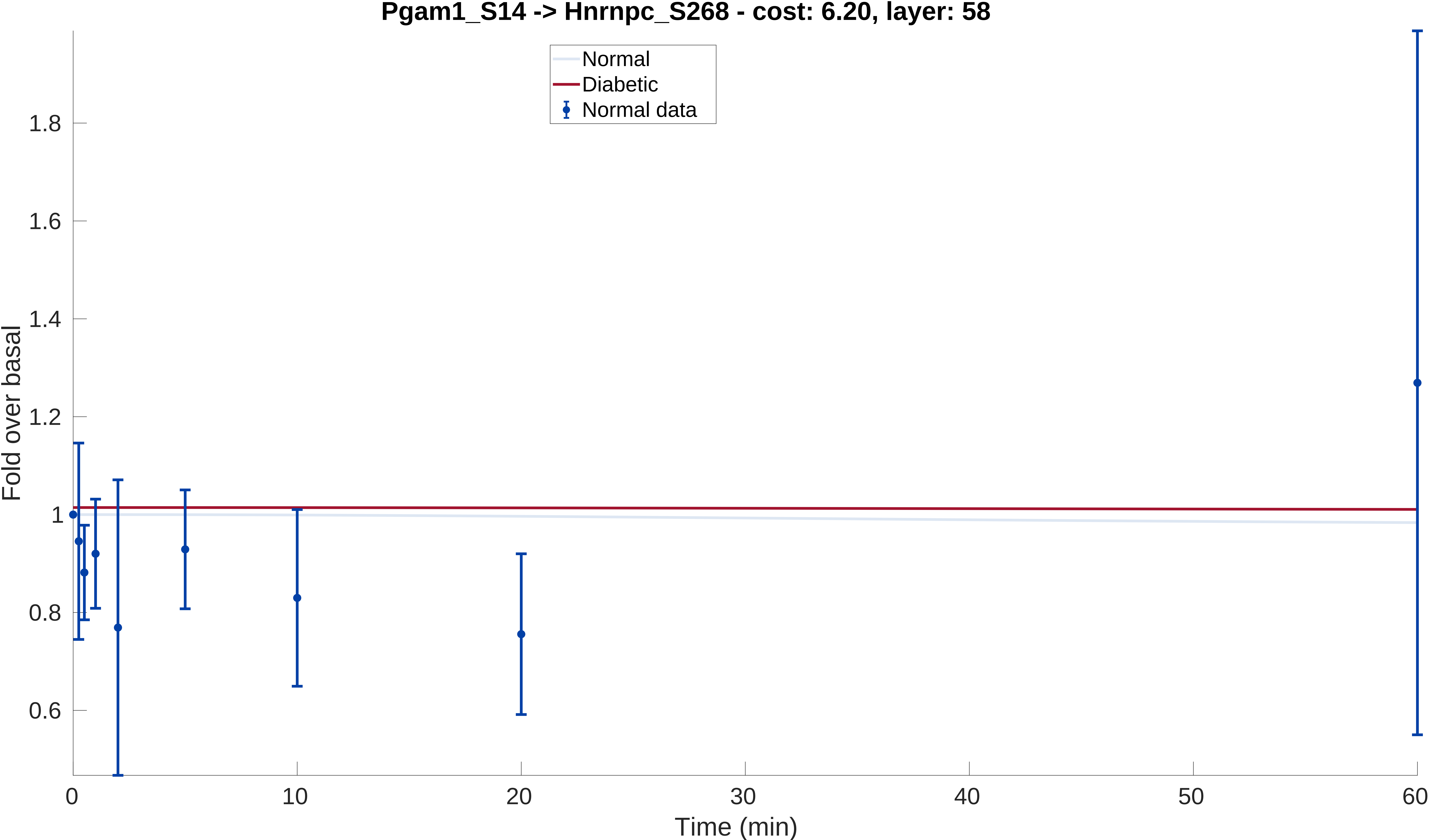

Pgam1_S14 -> Hnrnpc_S268 - cost: 6.20, layer: 58
Normal
Diabetic
Normal data
Fold over basal
1.8
1.6
1.4
1.2
1
0.8
0.6
0
10
20
30
40
50
60
Time (min)

#### Tbc1d4_S701 -> Hnrnpll_S37 - cost: 10.30, layer: 58

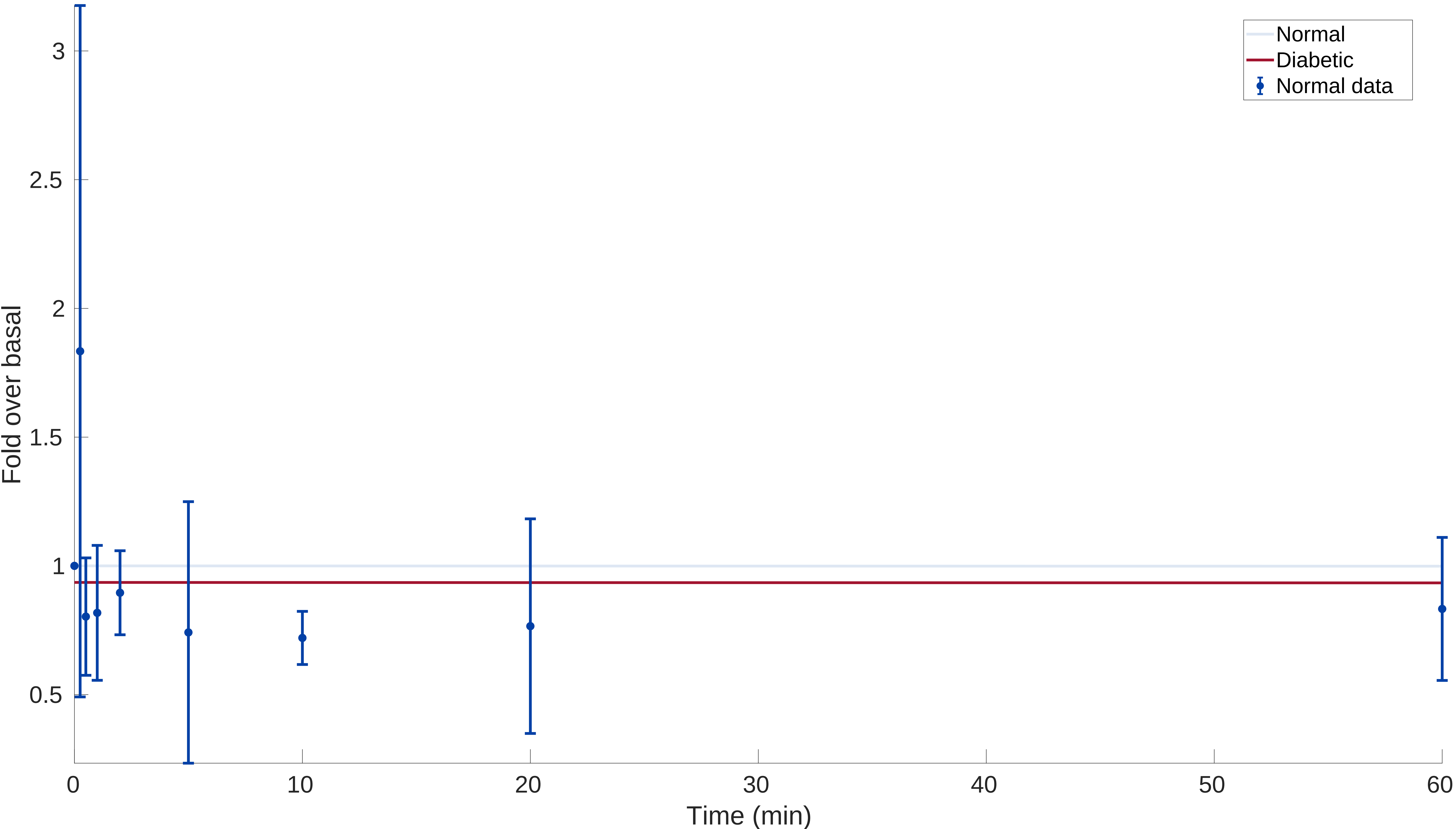

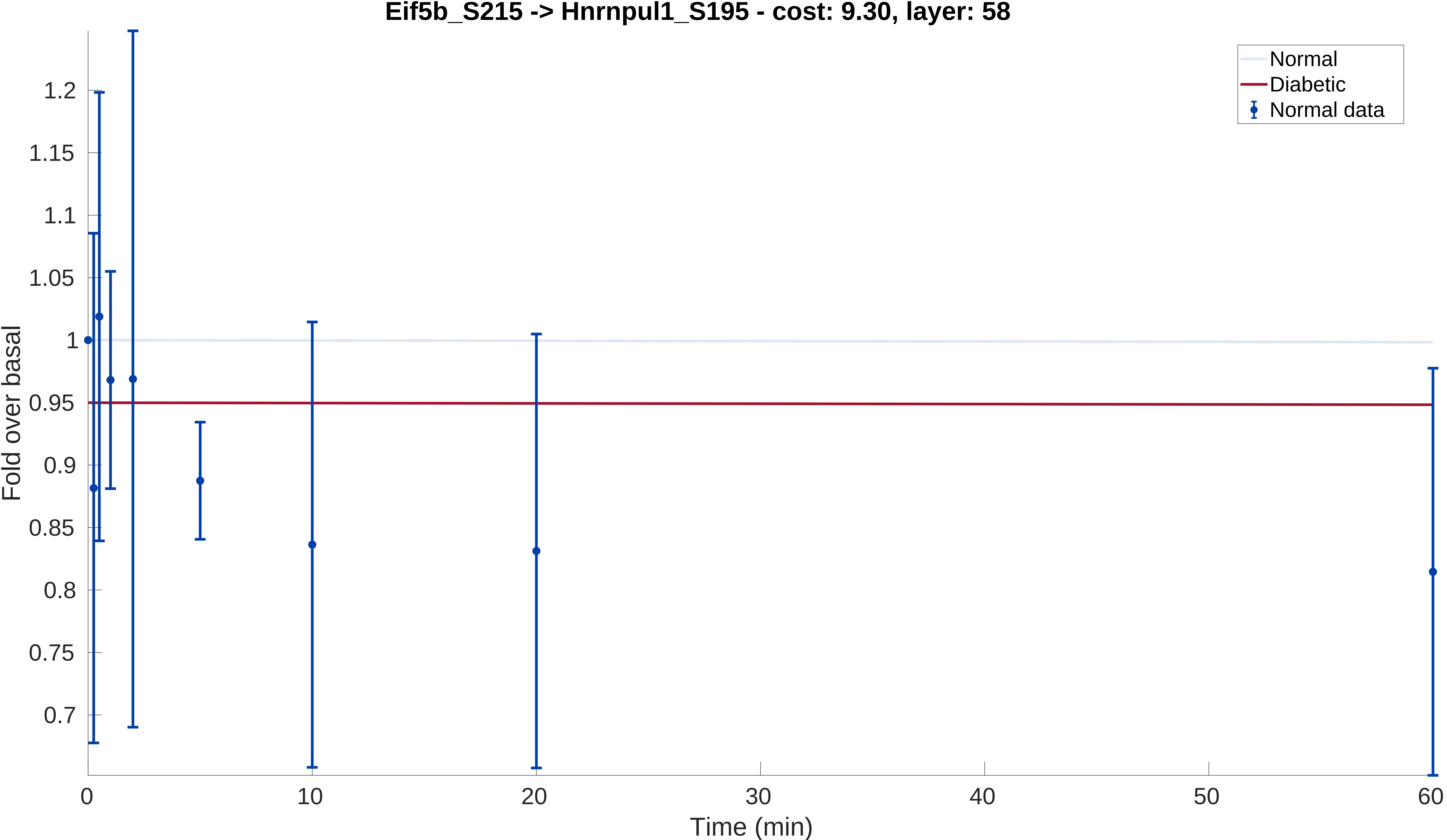

Eif5b_S215 -> Hnrnpul1_S195 - cost: 9.30, layer: 58
Normal
Diabetic
Normal data
Fold over basal
1.2
1.15
1.1
1.05
1
0.95
0.9
0.85
0.8
0.75
0.7
0
10
20
30
40
50
60
Time (min)

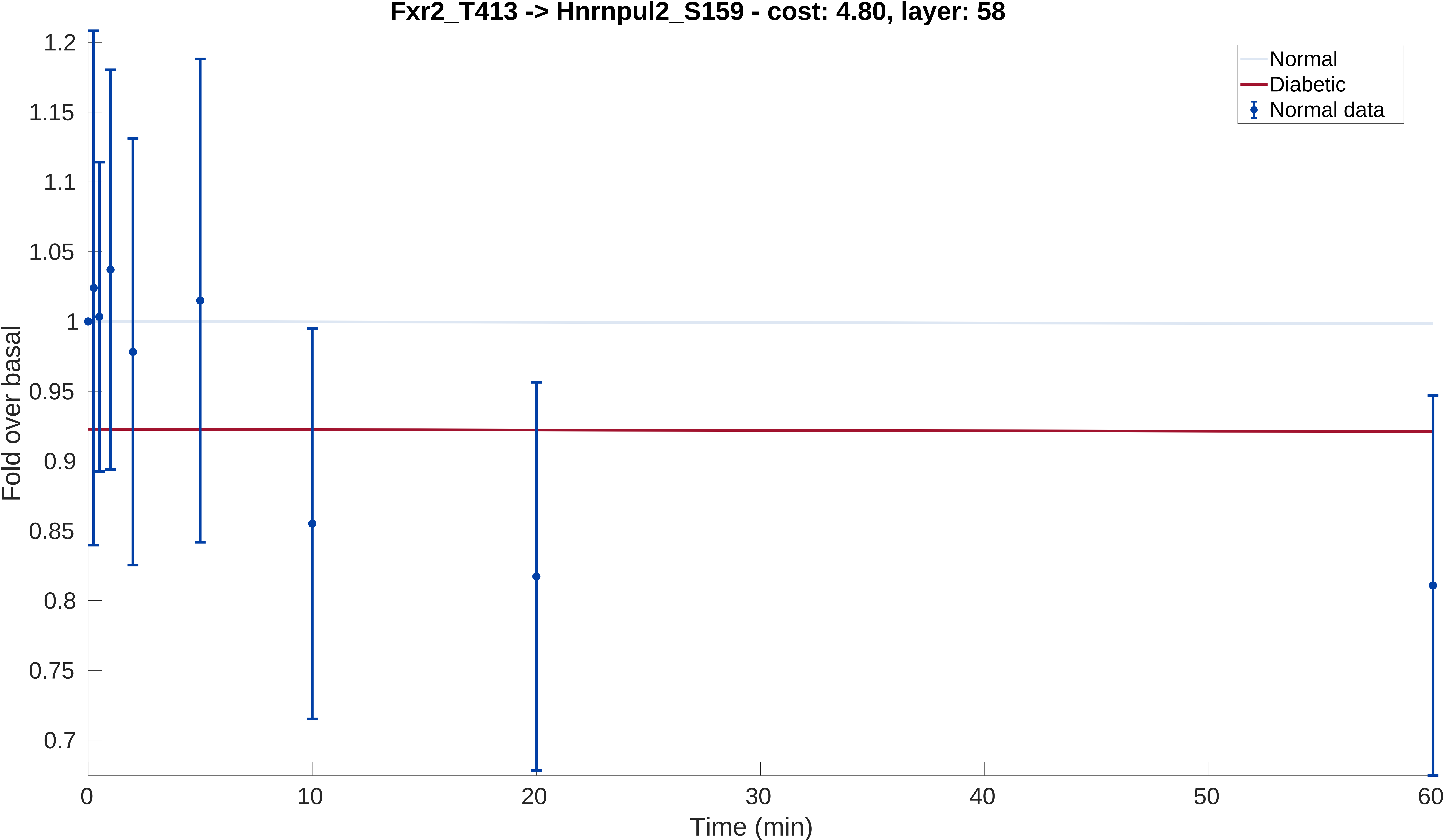

Fxr2_T413 -> Hnrnpul2_S159 - cost: 4.80, layer: 58
Normal
Diabetic
Normal data
Fold over basal
1.2
1.15
1.1
1.05
1
0.95
0.9
0.85
0.8
0.75
0.7
0
10
20
30
40
50
60
Time (min)

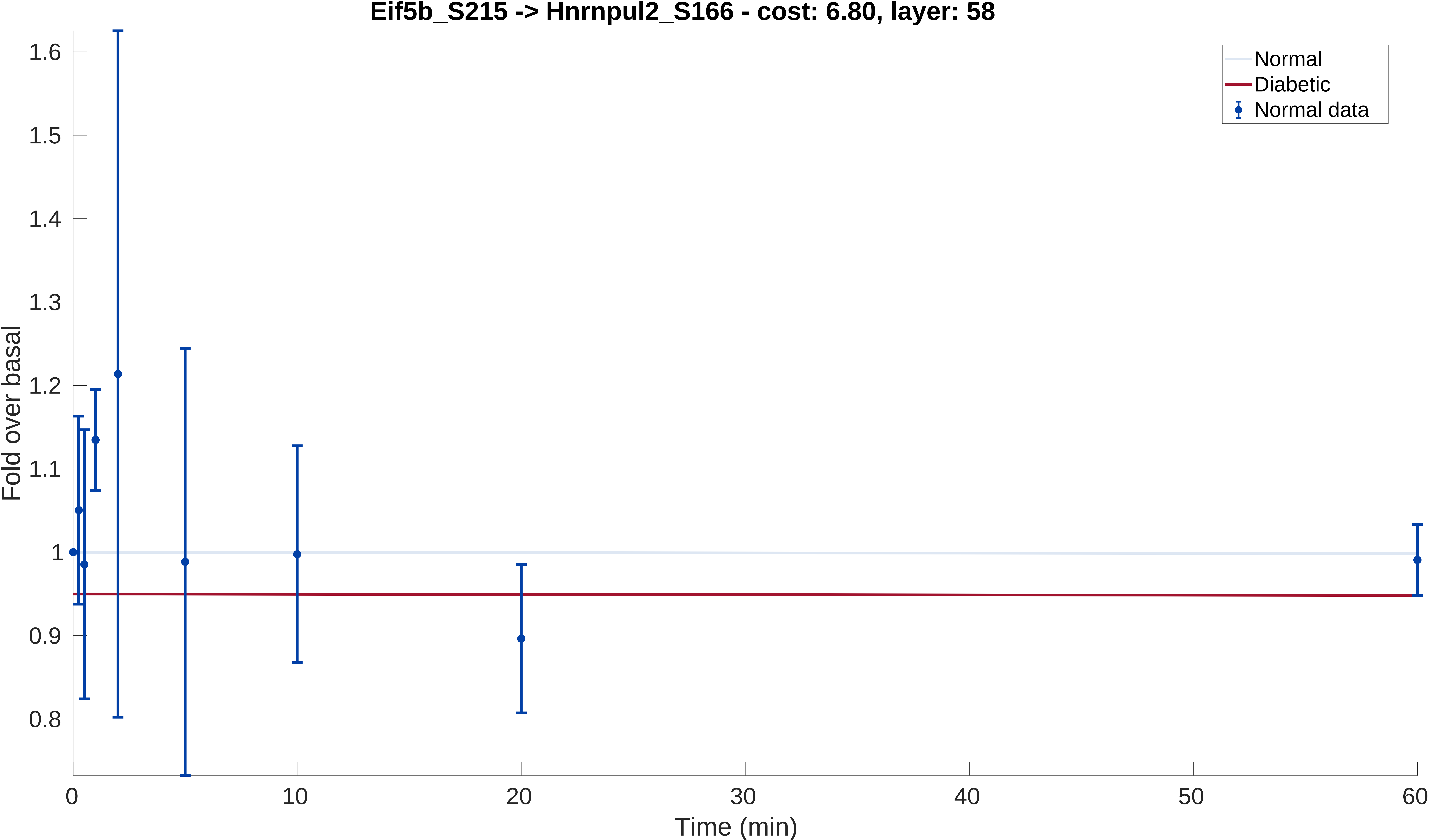

Eif5b_S215 -> Hnrnpul2_S166 - cost: 6.80, layer: 58
Normal
Diabetic
Normal data
Fold over basal
1.6
1.5
1.4
1.3
1.2
1.1
1
0.9
0.8
0
10
20
30
40
50
60
Time (min)

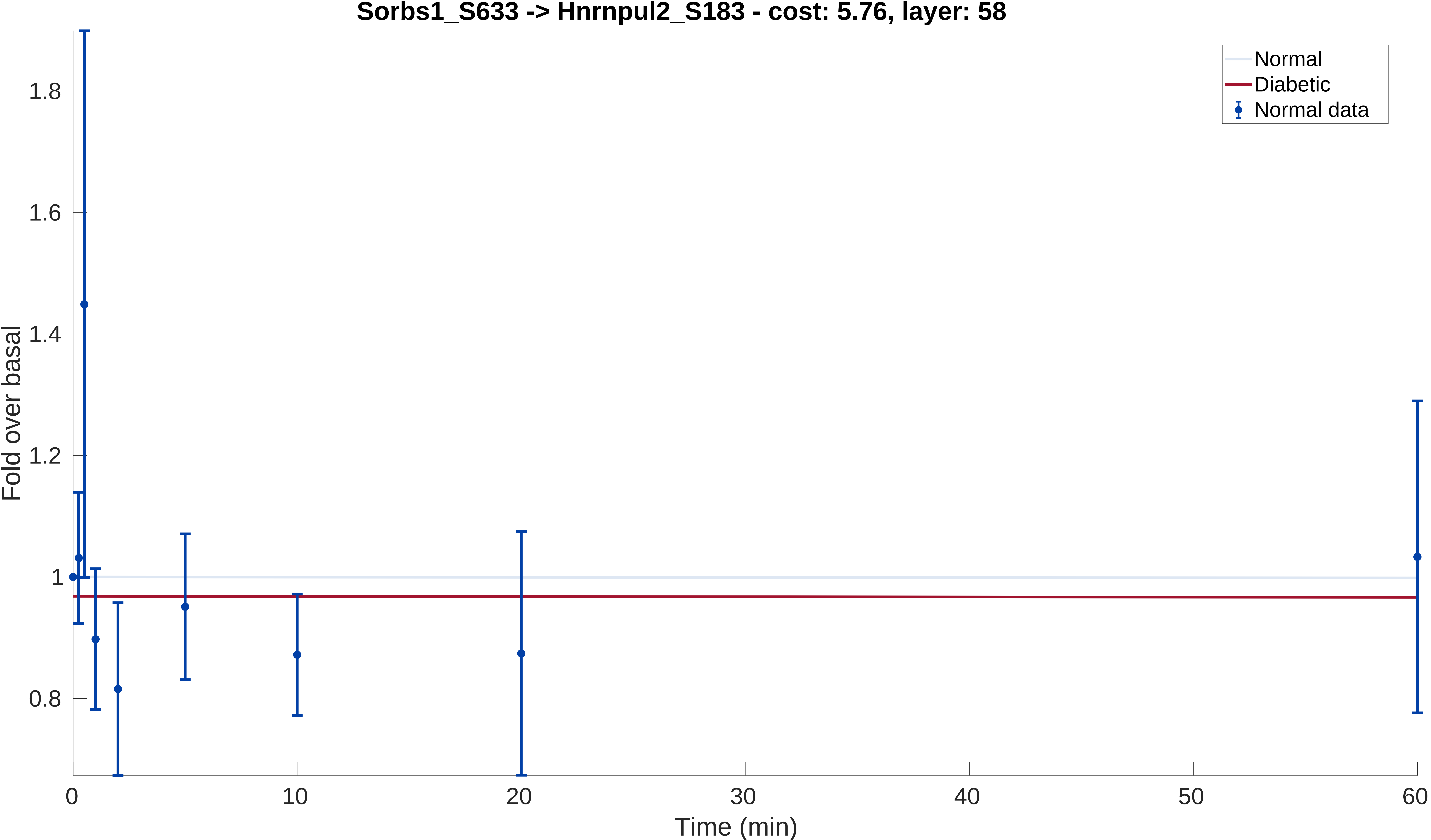

Sorbs1_S633 -> Hnrnpul2_S183 - cost: 5.76, layer: 58
Normal
Diabetic
Normal data
Fold over basal
1.8
1.6
1.4
1.2
1
0.8
0
10
20
30
40
50
60
Time (min)

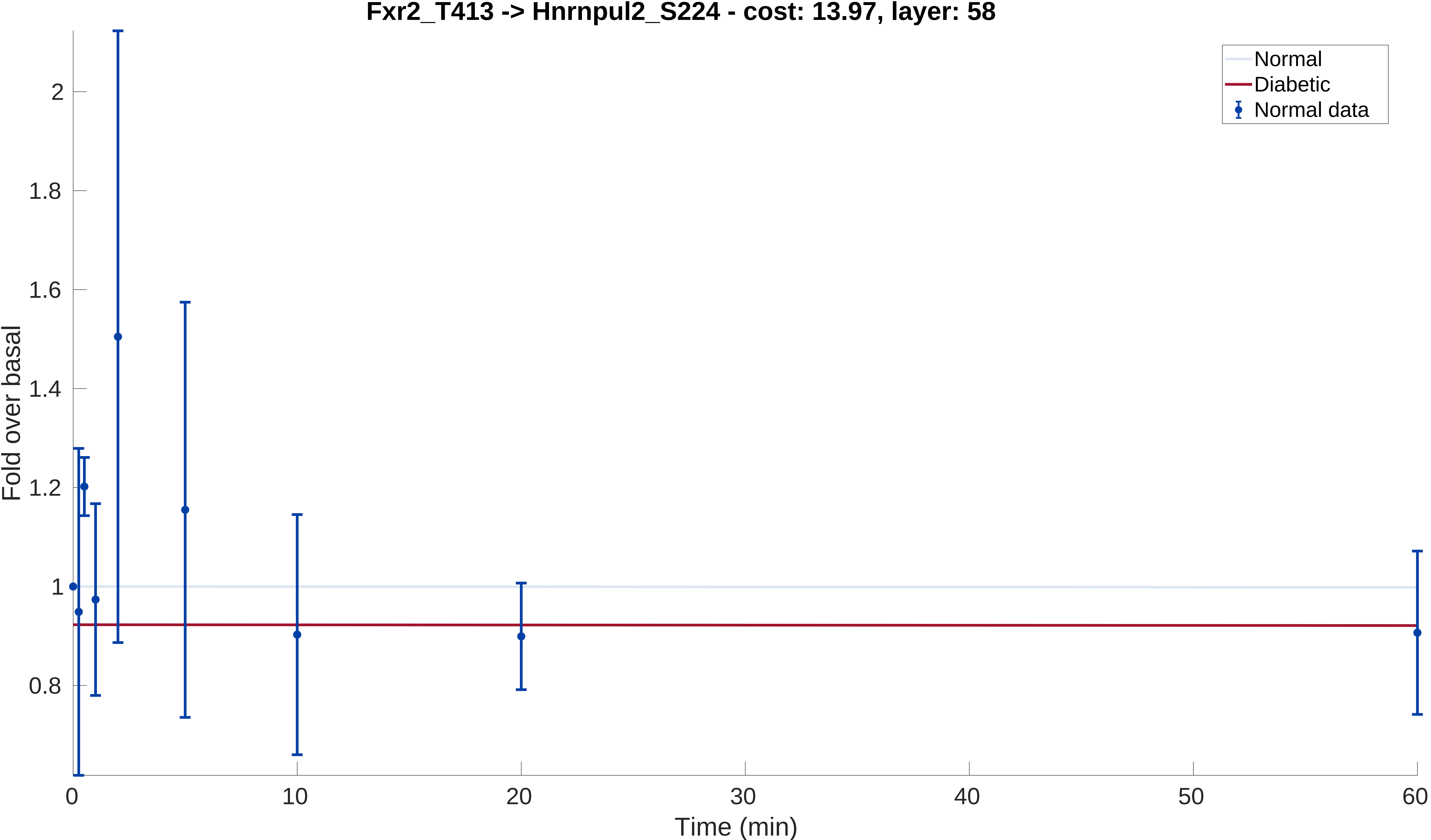

Fxr2_T413 -> Hnrnpul2_S224 - cost: 13.97, layer: 58
Normal
Diabetic
Normal data
Fold over basal
2
1.8
1.6
1.4
1.2
1
0.8
0
10
20
30
40
50
60
Time (min)

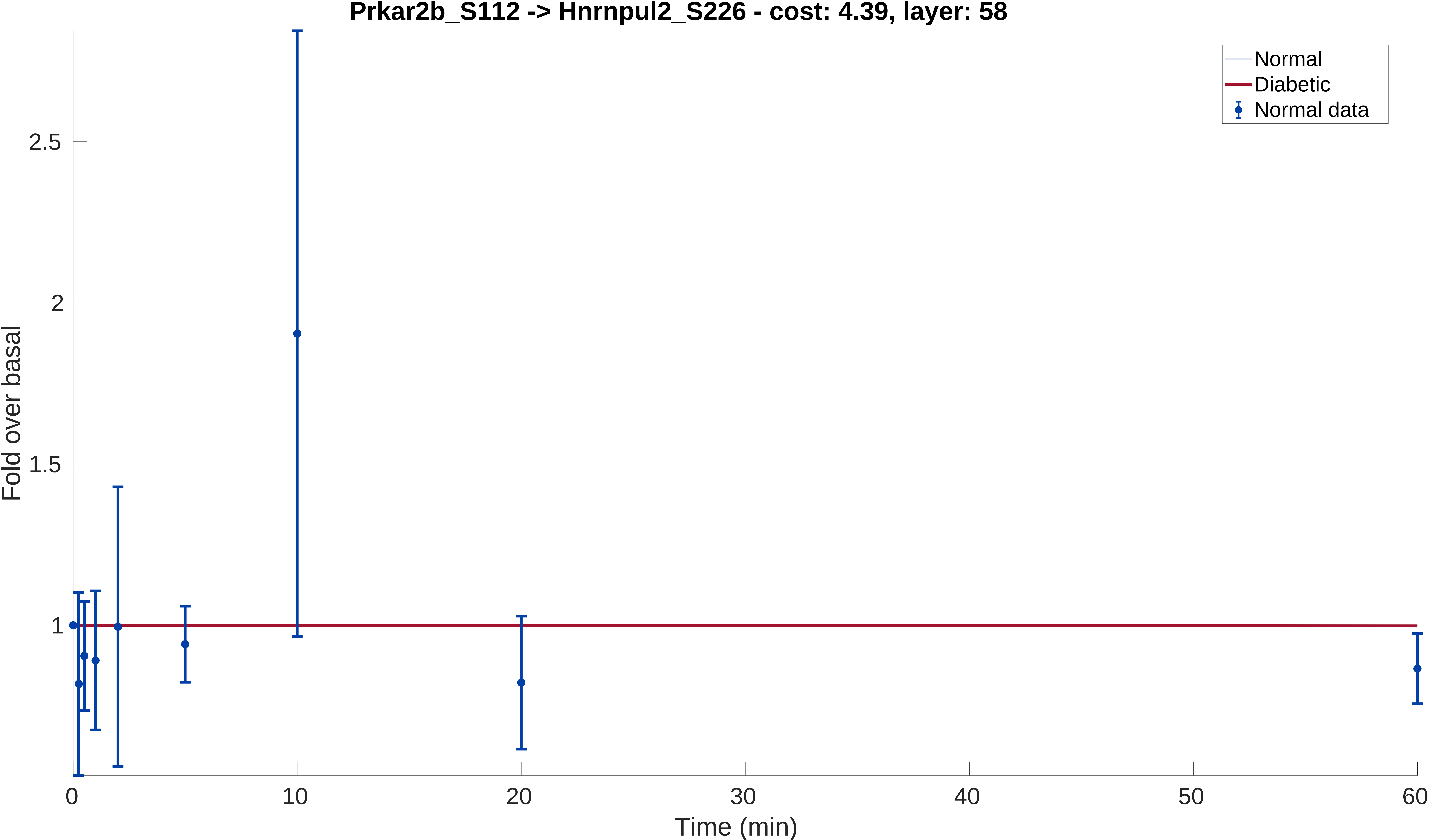

Prkar2b_S112 -> Hnrnpul2_S226 - cost: 4.39, layer: 58
Normal
Diabetic
Normal data
2.5
2
1.5
1
Fold over basal
0
10
20
30
40
50
60
Time (min)

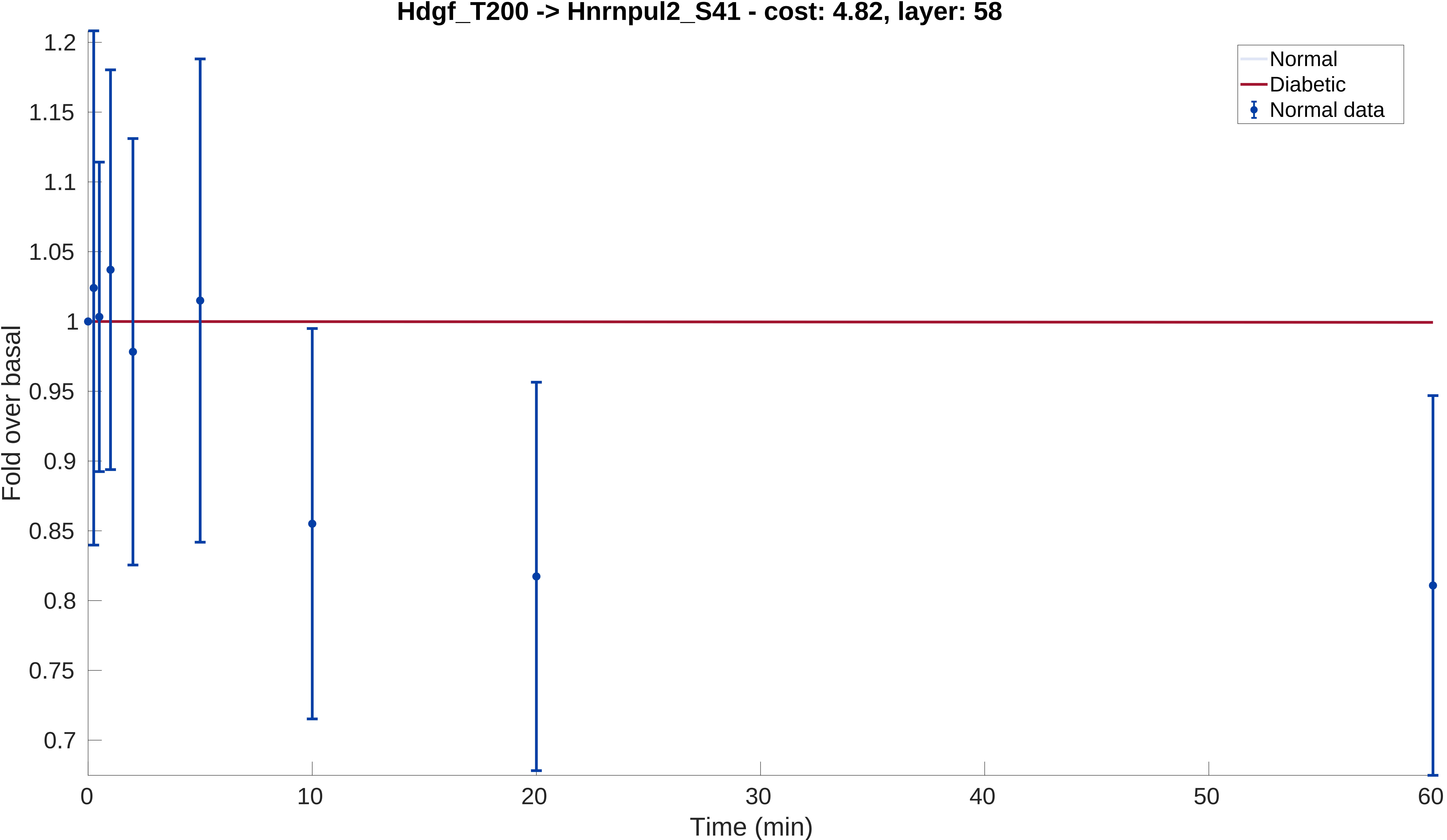

Hdgf_T200 -> Hnrnpul2_S41 - cost: 4.82, layer: 58
Normal
Diabetic
Normal data
1.2
1.15
1.1
1.05
1
0.95
0.9
0.85
0.8
0.75
0.7
Fold over basal
0
10
20
30
40
50
60
Time (min)

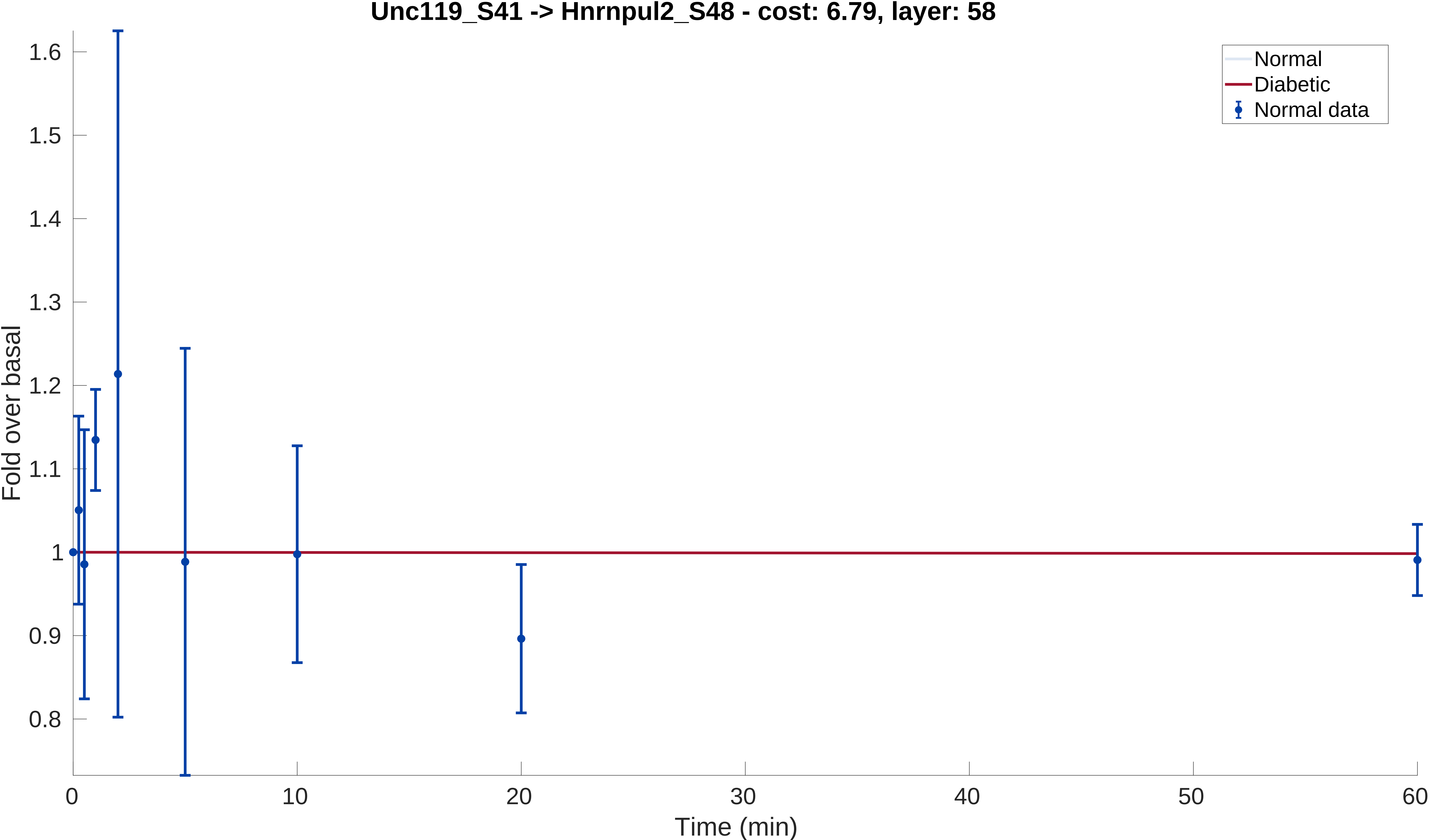

Unc119_S41 -> Hnrnpul2_S48 - cost: 6.79, layer: 58
Normal
Diabetic
Normal data
Fold over basal
1.6
1.5
1.4
1.3
1.2
1.1
1
0.9
0.8
0
10
20
30
40
50
60
Time (min)

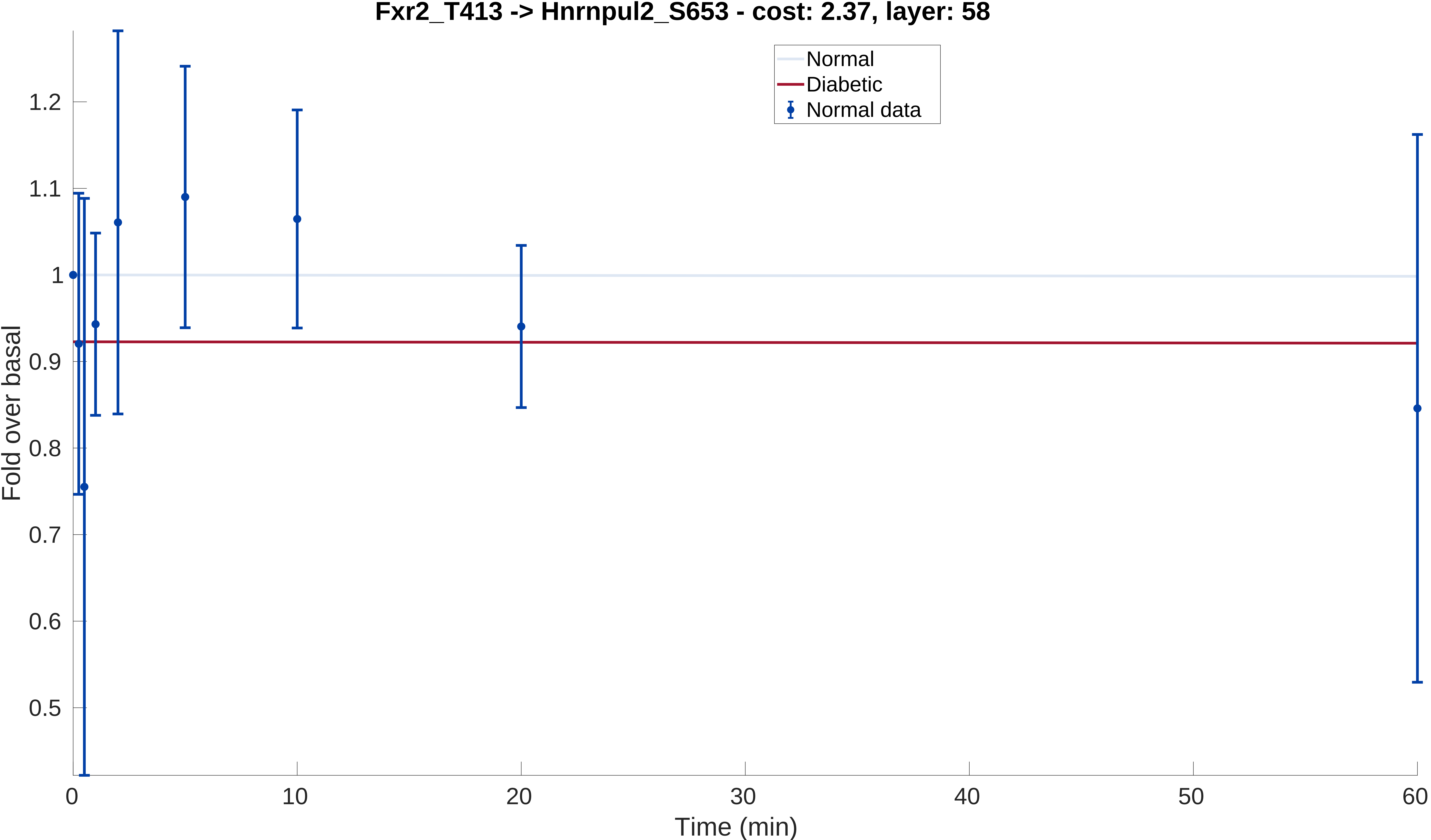

Fxr2_T413 -> Hnrnpul2_S653 - cost: 2.37, layer: 58
Normal
Diabetic
Normal data
Fold over basal
1.2
1.1
1
0.9
0.8
0.7
0.6
0.5
0
10
20
30
40
50
60
Time (min)

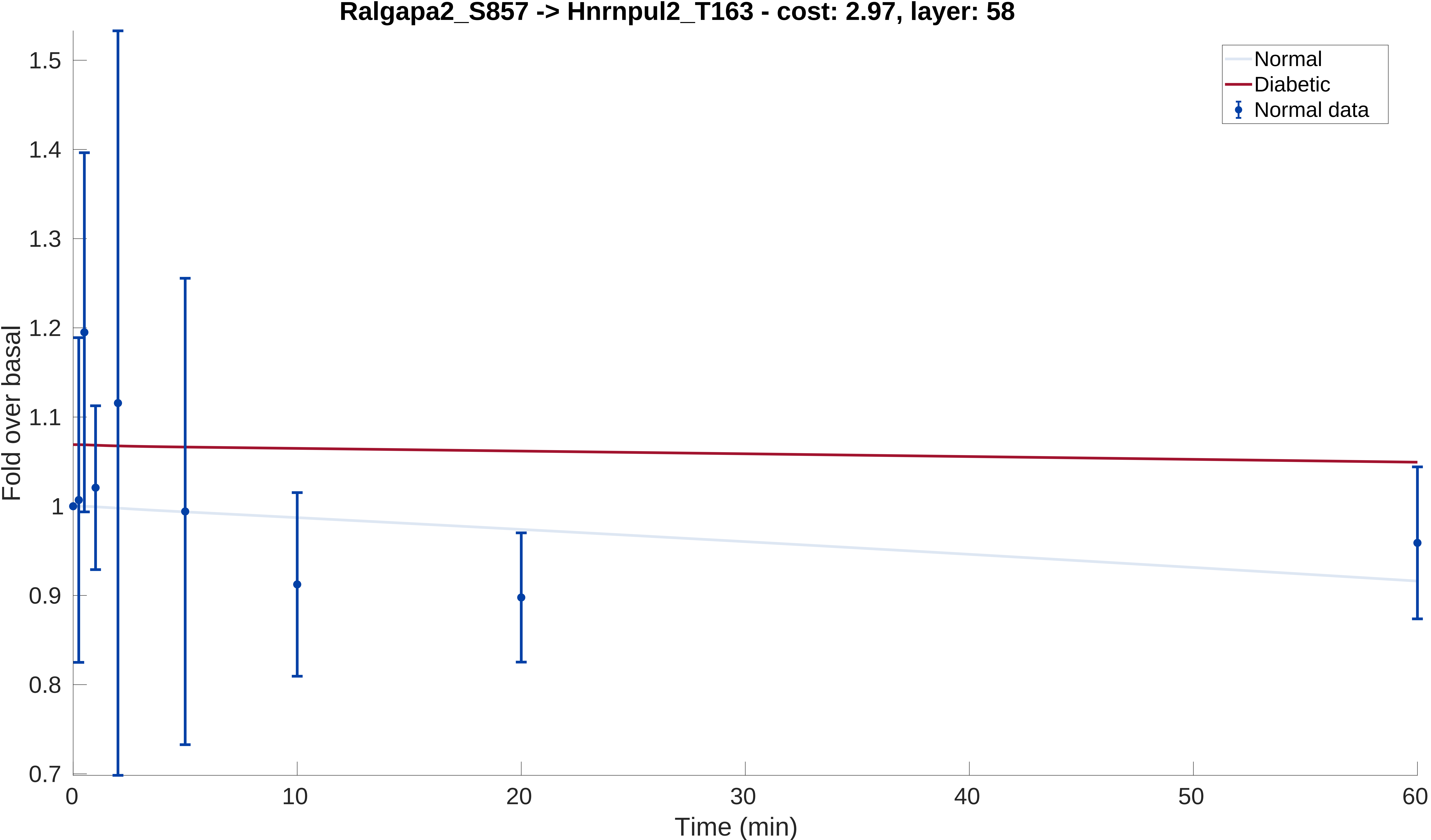

Ralgapa2_S857 -> Hnrnpul2_T163 - cost: 2.97, layer: 58
Normal
Diabetic
Normal data
Fold over basal
1.5
1.4
1.3
1.2
1.1
1
0.9
0.8
0.7
0
10
20
30
40
50
60
Time (min)

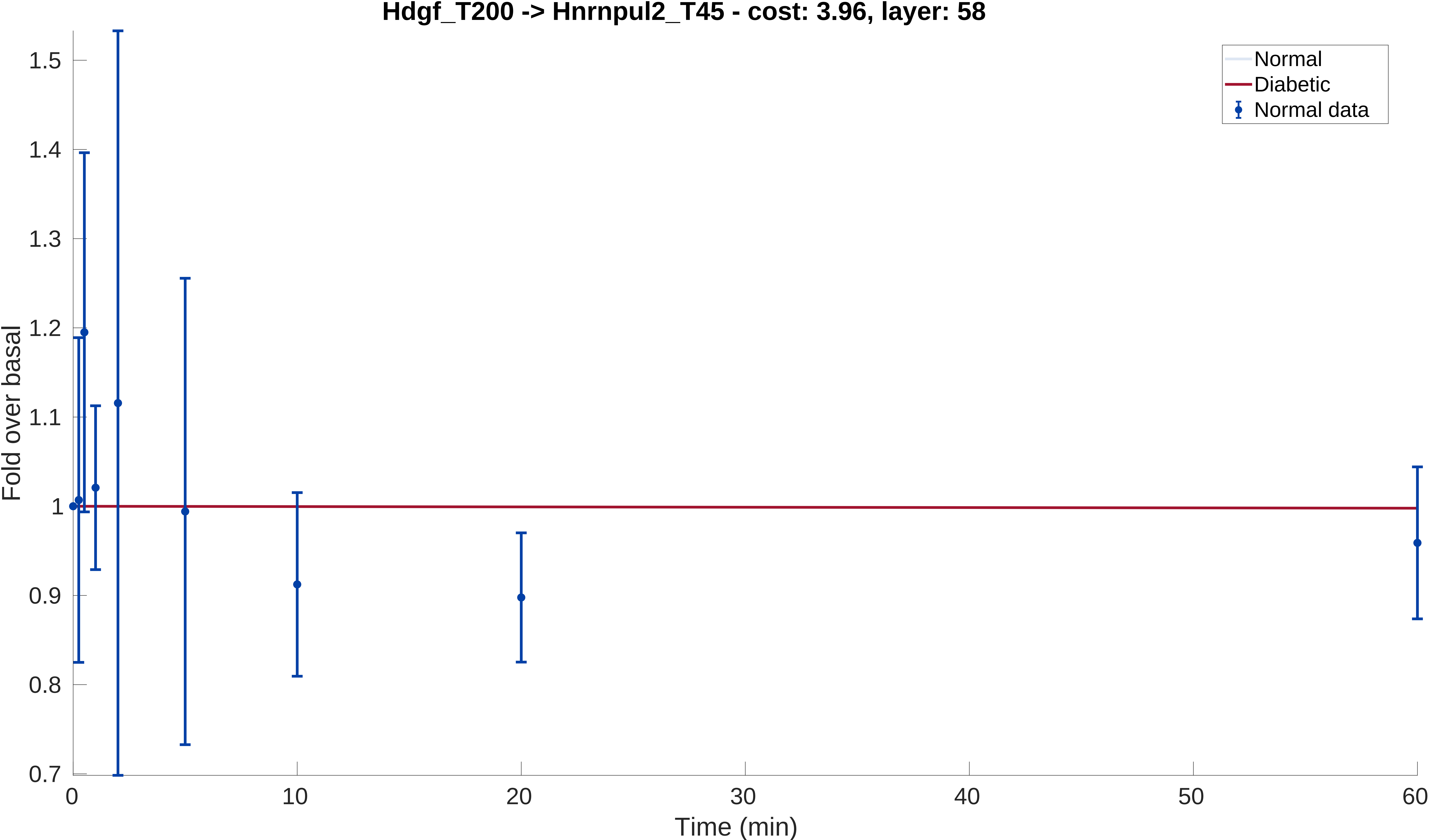

Hdgf_T200 -> Hnrnpul2_T45 - cost: 3.96, layer: 58
Normal
Diabetic
Normal data
Fold over basal
1.5
1.4
1.3
1.2
1.1
1
0.9
0.8
0.7
0
10
20
30
40
50
60
Time (min)

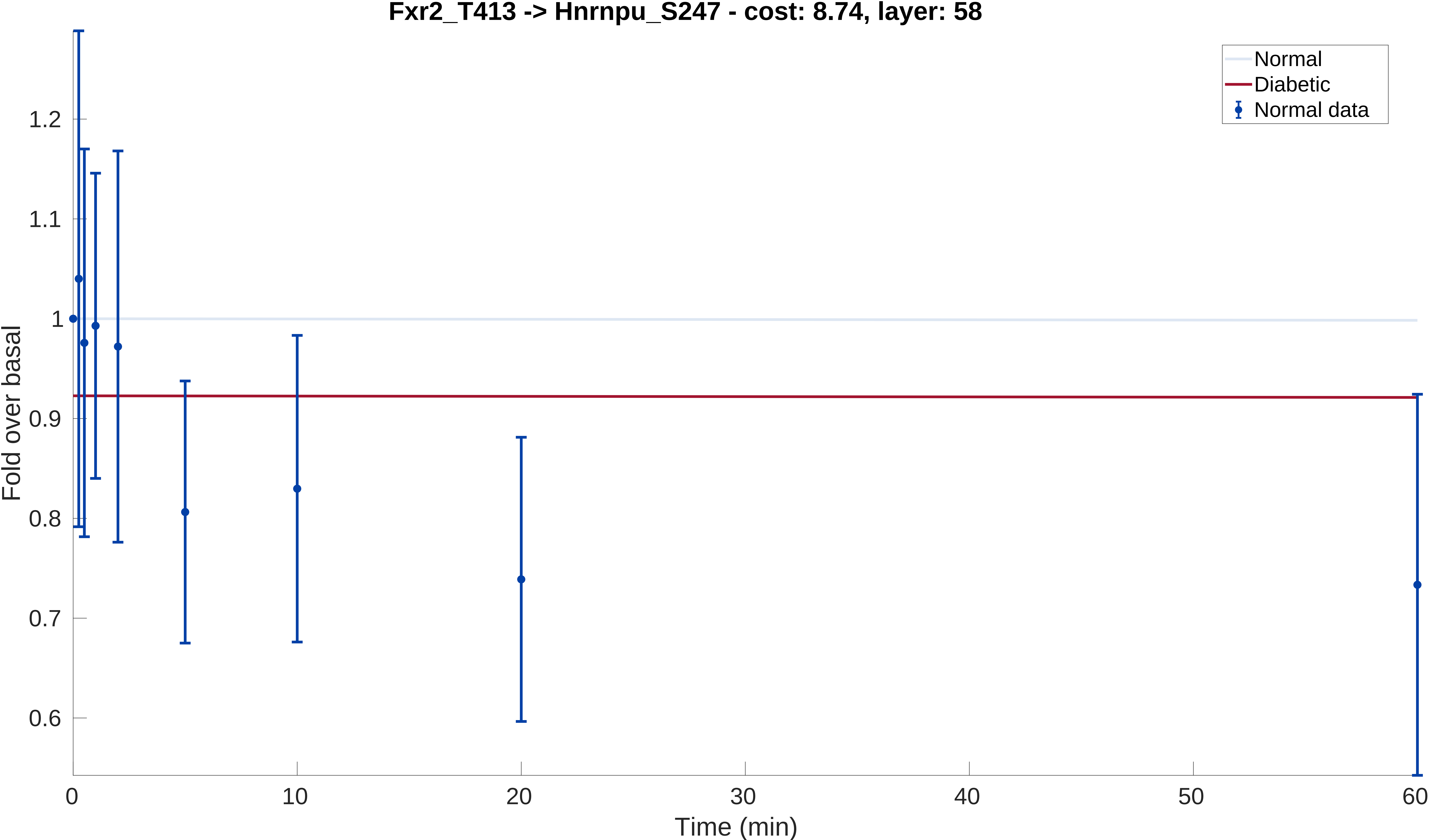

Fxr2_T413 -> Hnrnpu_S247 - cost: 8.74, layer: 58
Normal
Diabetic
Normal data
Fold over basal
1.2
1.1
1
0.9
0.8
0.7
0.6
0
10
20
30
40
50
60
Time (min)

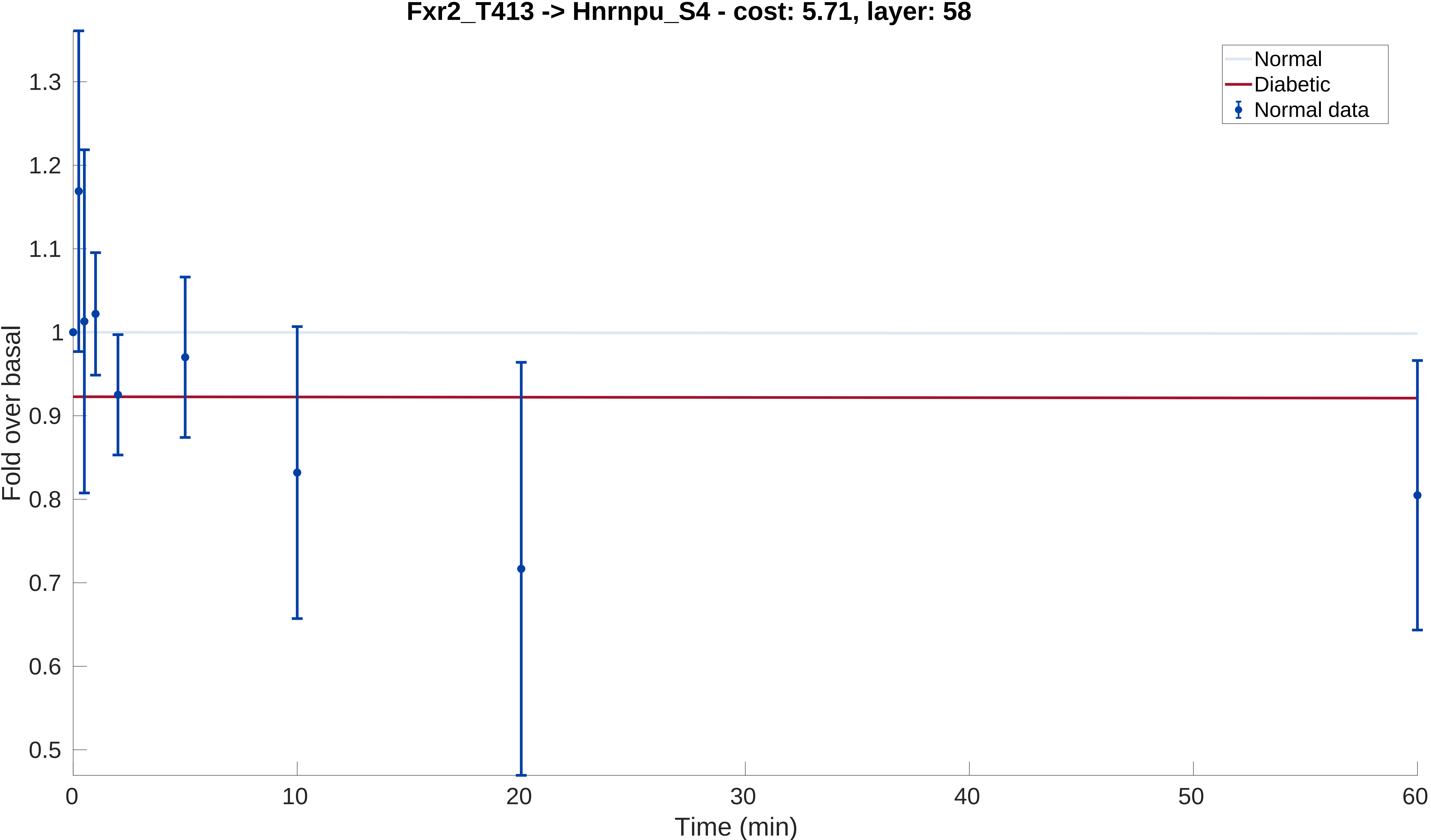

Fxr2_T413 -> Hnrnpu_S4 - cost: 5.71, layer: 58
Normal
Diabetic
Normal data
1.3
1.2
1.1
1
0.9
0.8
0.7
0.6
0.5
Fold over basal
0
10
20
30
40
50
60
Time (min)

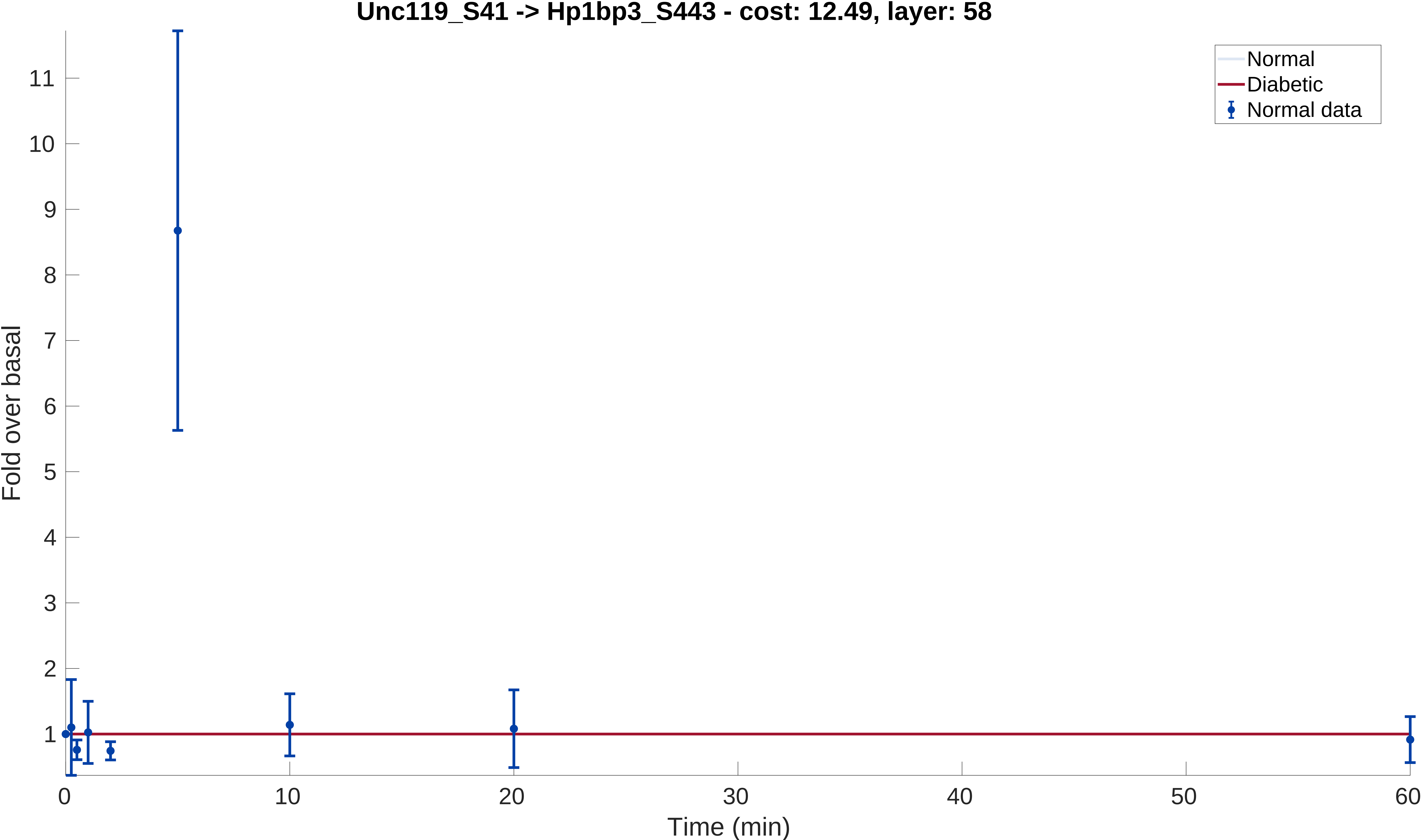
Unc119_S41 -> Hp1bp3_S443 - cost: 12.49, layer: 58
Normal
Diabetic
Normal data
Fold over basal
11
10
9
8
7
6
5
4
3
2
1
0
10
20
30
40
50
60
Time (min)

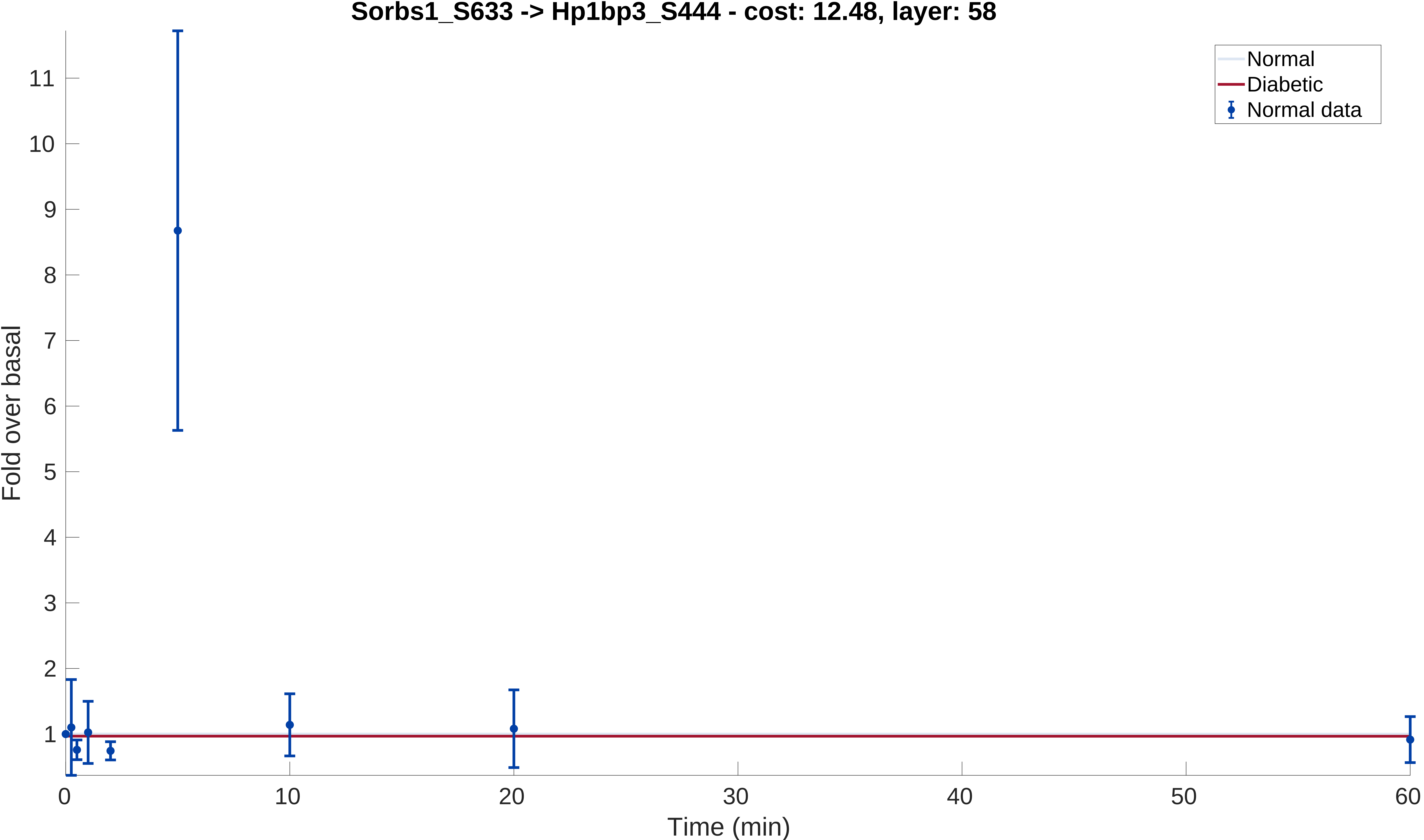

Sorbs1_S633 -> Hp1bp3_S444 - cost: 12.48, layer: 58
Normal
Diabetic
Normal data
Fold over basal
Time (min)
0
10
20
30
40
50
60
1
2
3
4
5
6
7
8
9
10
11

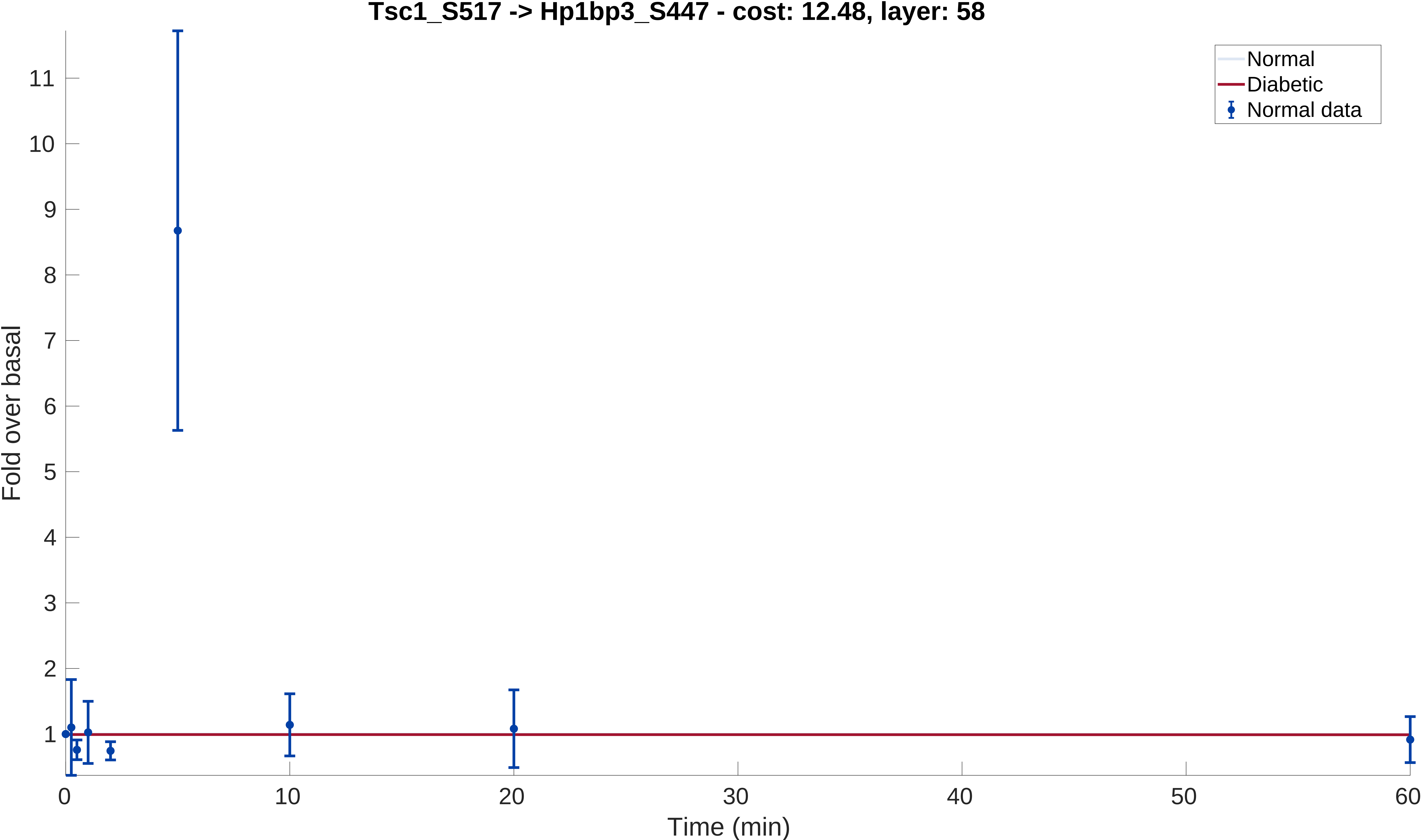
Tsc1_S517 -> Hp1bp3_S447 - cost: 12.48, layer: 58
Normal
Diabetic
Normal data
Fold over basal
Time (min)
1
2
3
4
5
6
7
8
9
10
11
0
10
20
30
40
50
60

Plin1_S492 -> Hp1bp3_T51 - cost: 1.95, layer: 58

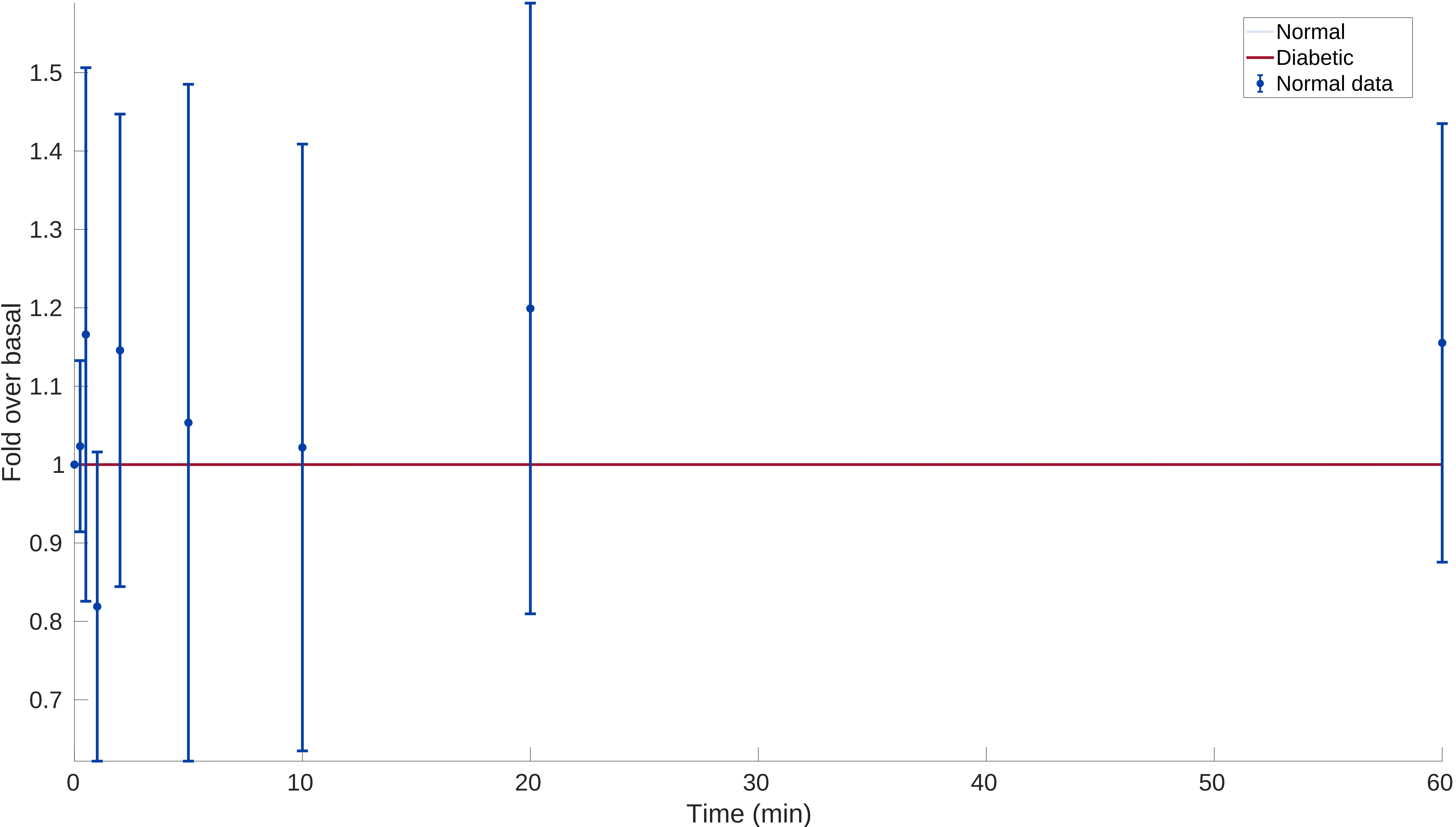

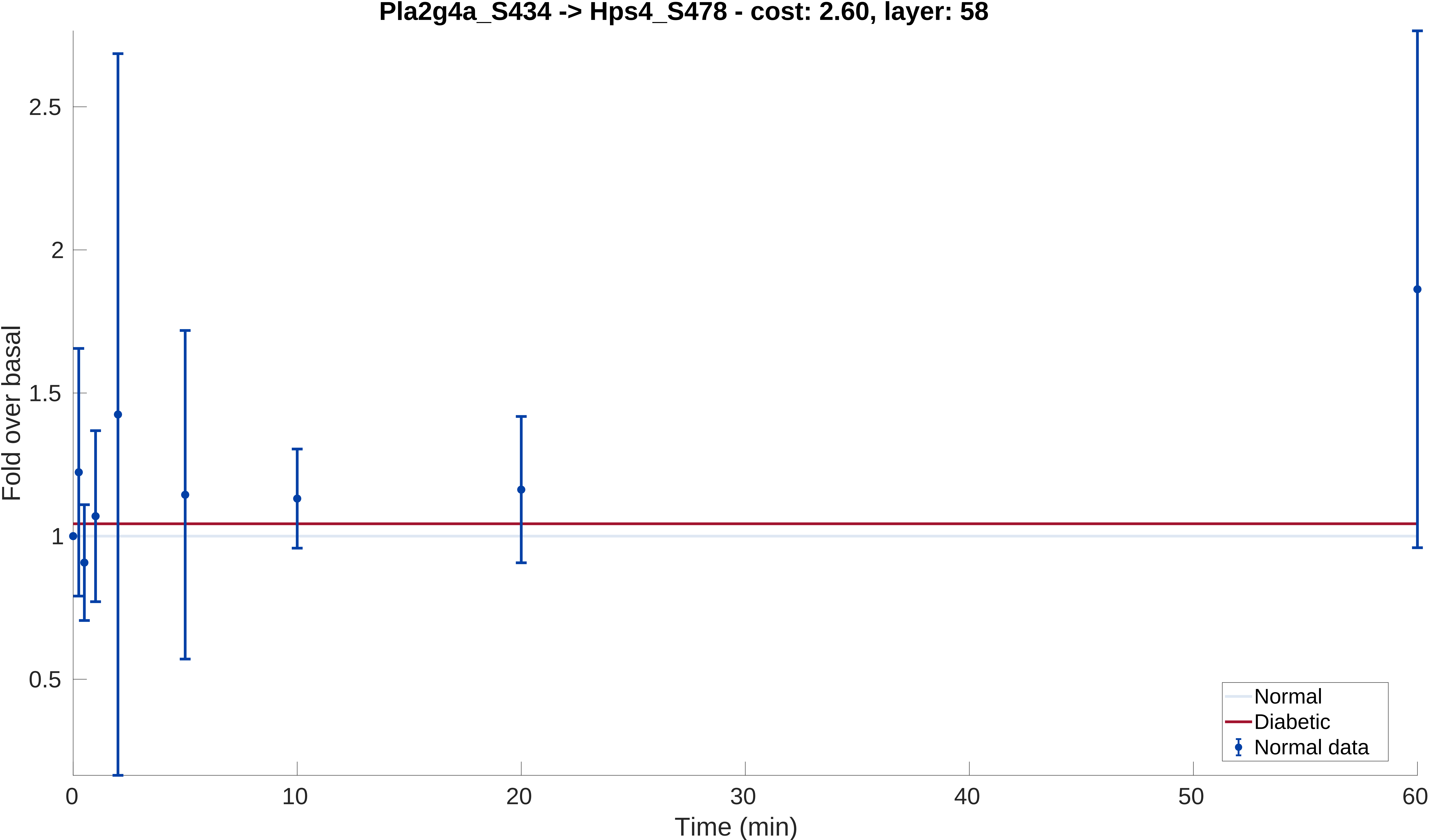

Pla2g4a_S434 -> Hps4_S478 - cost: 2.60, layer: 58
Fold over basal
2.5
2
1.5
1
0.5
0
10
20
30
40
50
60
Time (min)
Normal
Diabetic
Normal data

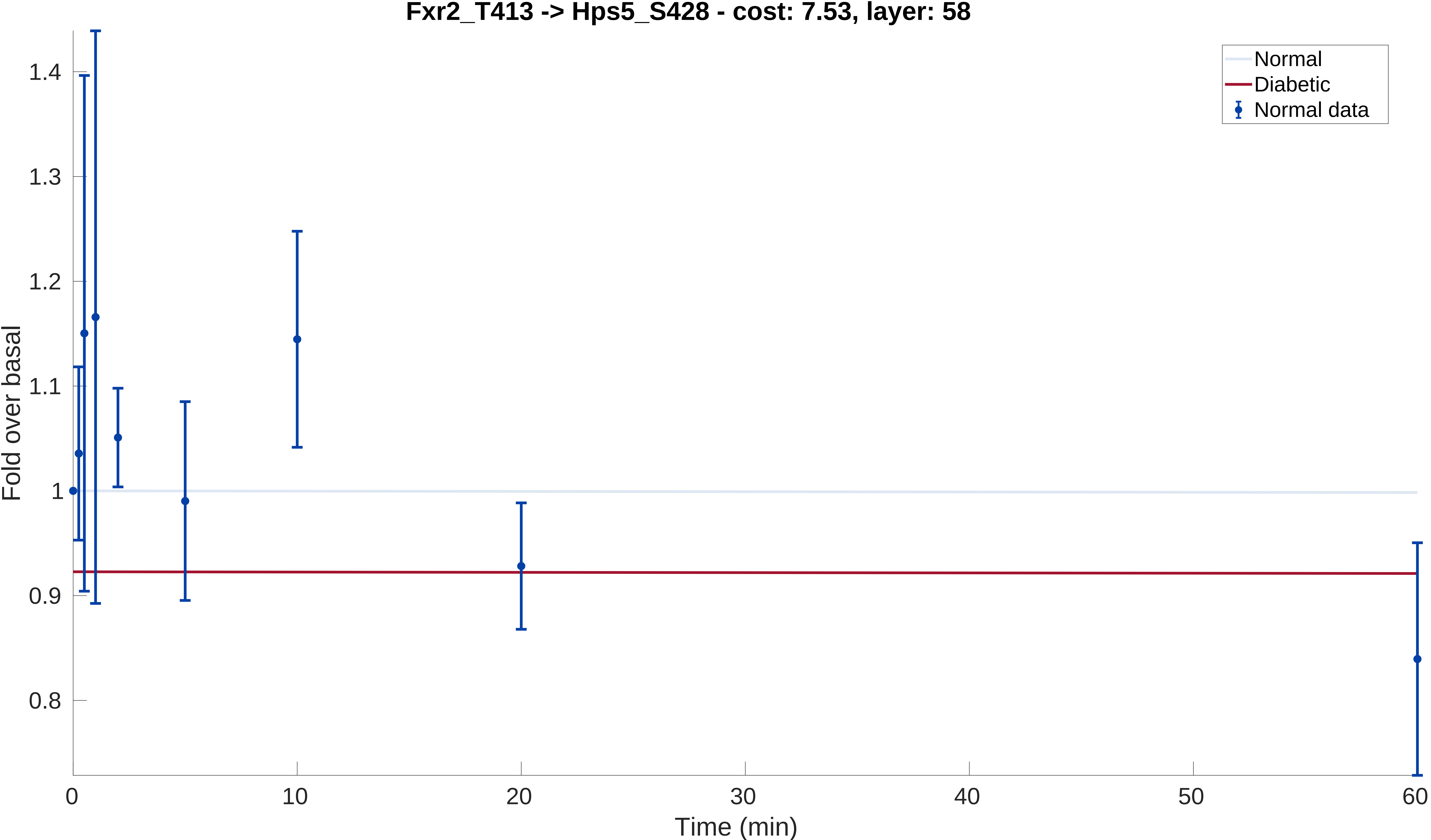

Fxr2_T413 -> Hps5_S428 - cost: 7.53, layer: 58
Normal
Diabetic
Normal data
Fold over basal
1.4
1.3
1.2
1.1
1
0.9
0.8
0
10
20
30
40
50
60
Time (min)

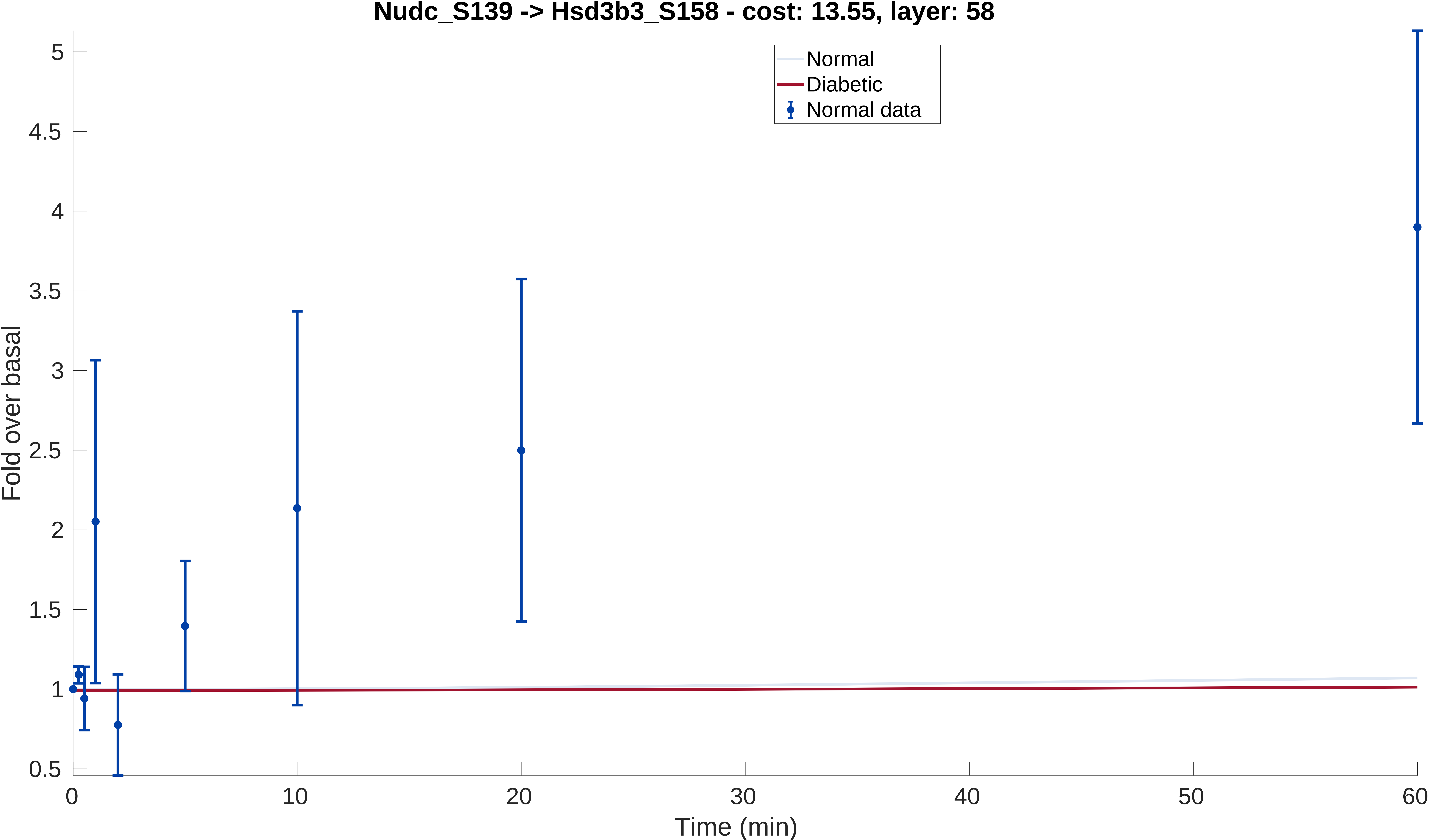

Nudc_S139 -> Hsd3b3_S158 - cost: 13.55, layer: 58
Normal
Diabetic
Normal data
Fold over basal
Time (min)
0.5
1
1.5
2
2.5
3
3.5
4
4.5
5
0
10
20
30
40
50
60

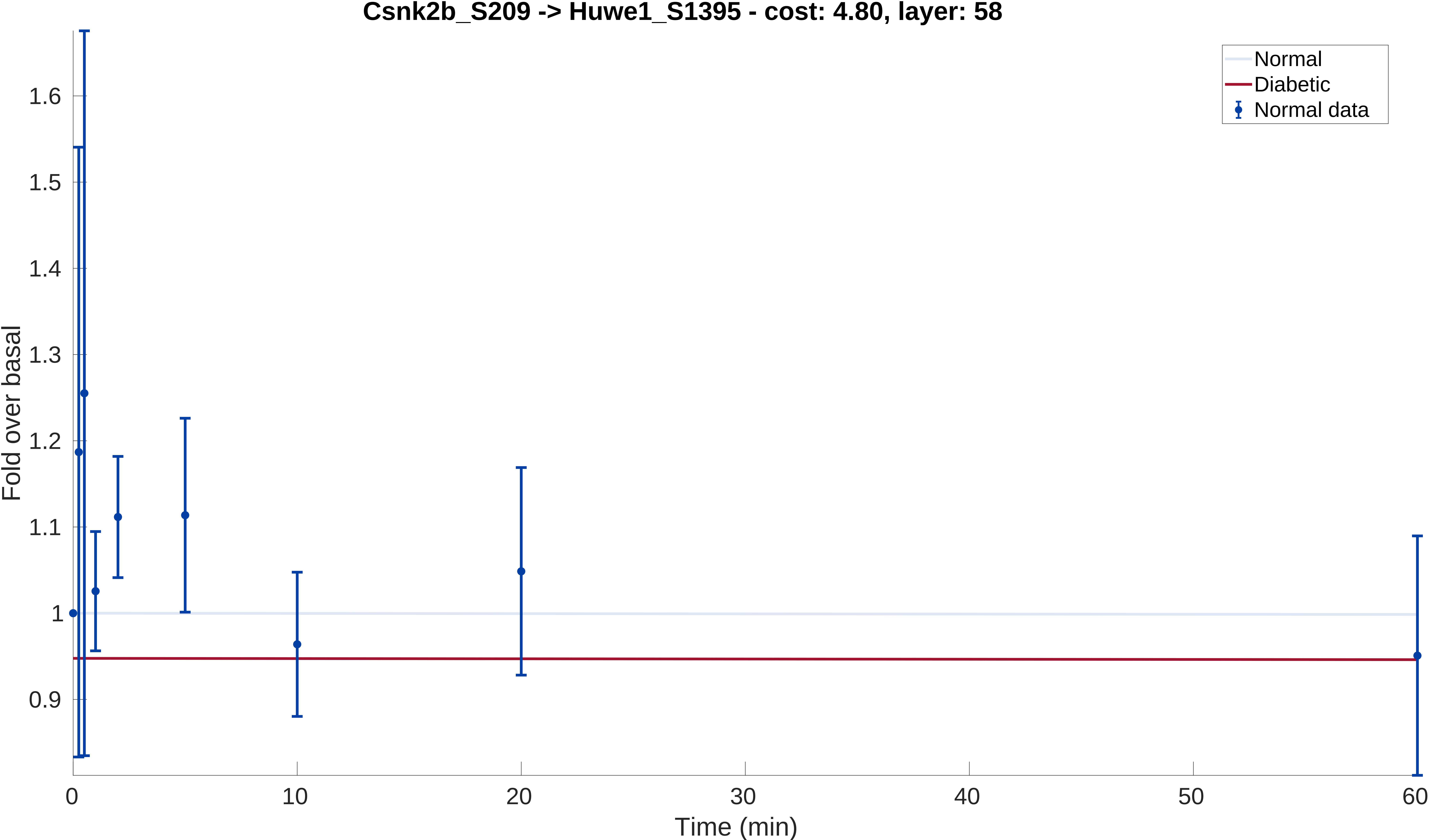

Csnk2b_S209 -> Huwe1_S1395 - cost: 4.80, layer: 58
Normal
Diabetic
Normal data
Fold over basal
1.6
1.5
1.4
1.3
1.2
1.1
1
0.9
0
10
20
30
40
50
60
Time (min)

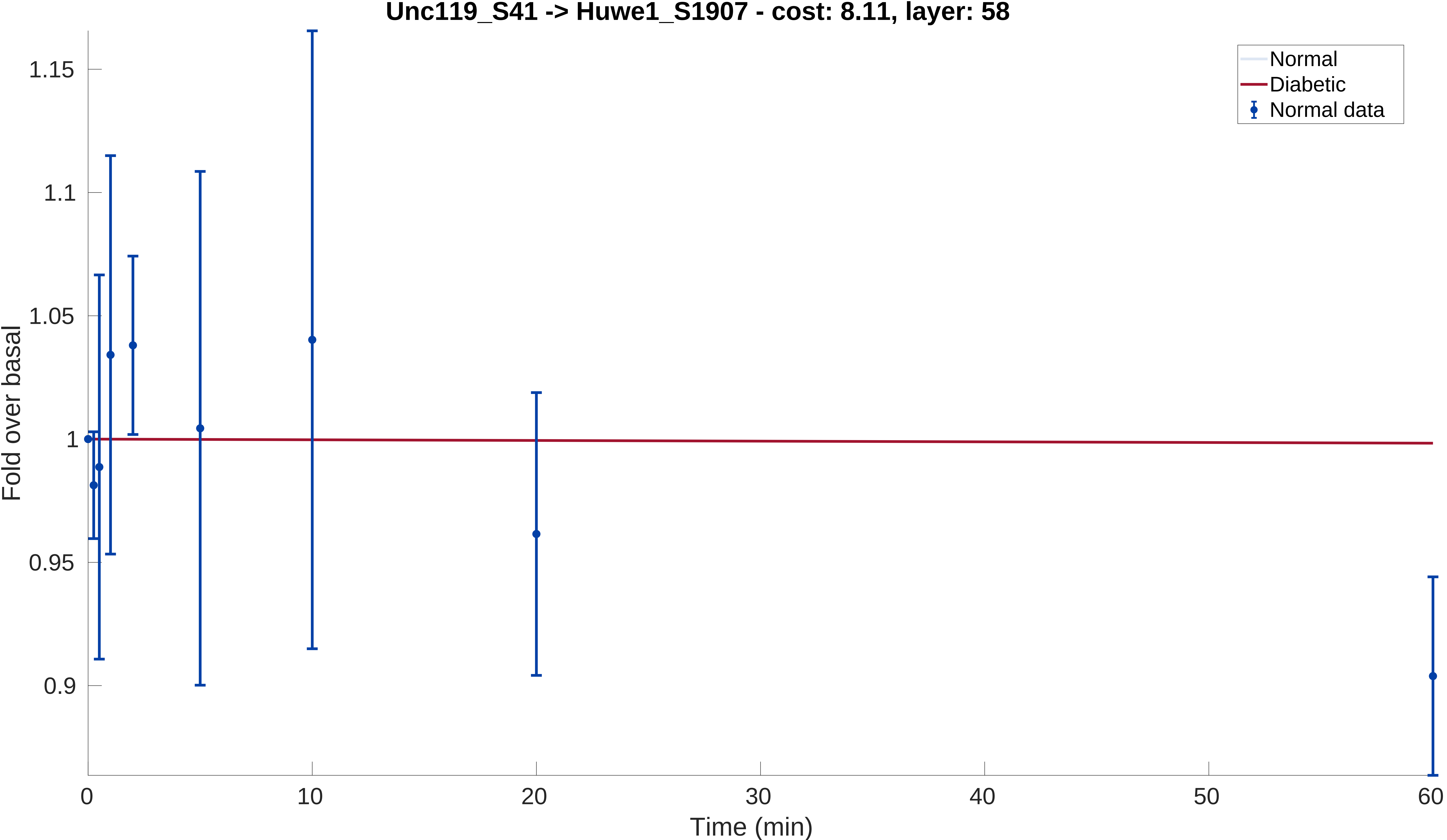

Unc119_S41 -> Huwe1_S1907 - cost: 8.11, layer: 58
Normal
Diabetic
Normal data
1.15
1.1
1.05
1
0.95
0.9
Fold over basal
0
10
20
30
40
50
60
Time (min)

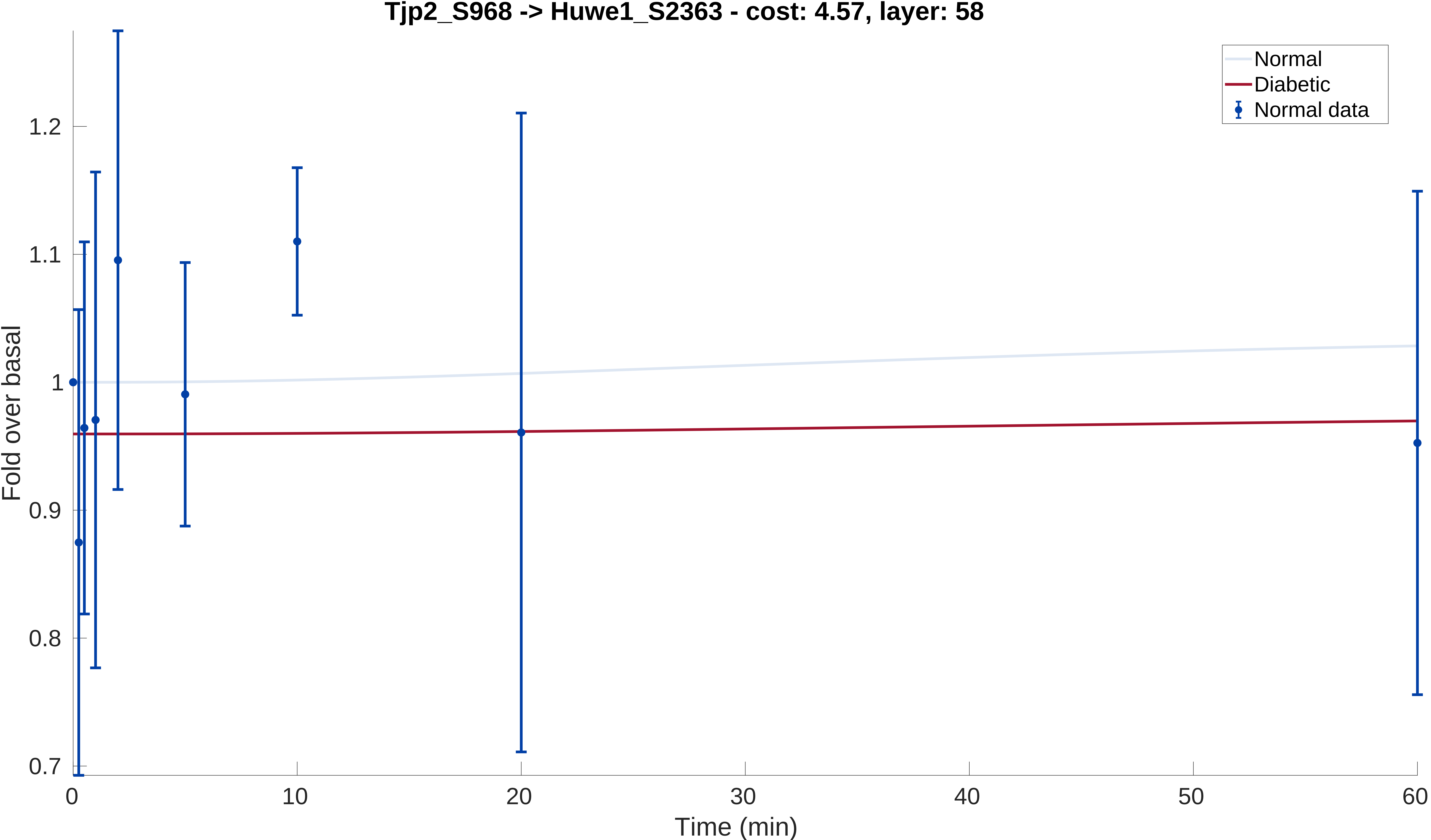

Tjp2_S968 -> Huwe1_S2363 - cost: 4.57, layer: 58
Normal
Diabetic
Normal data
Fold over basal
1.2
1.1
1
0.9
0.8
0.7
0
10
20
30
40
50
60
Time (min)

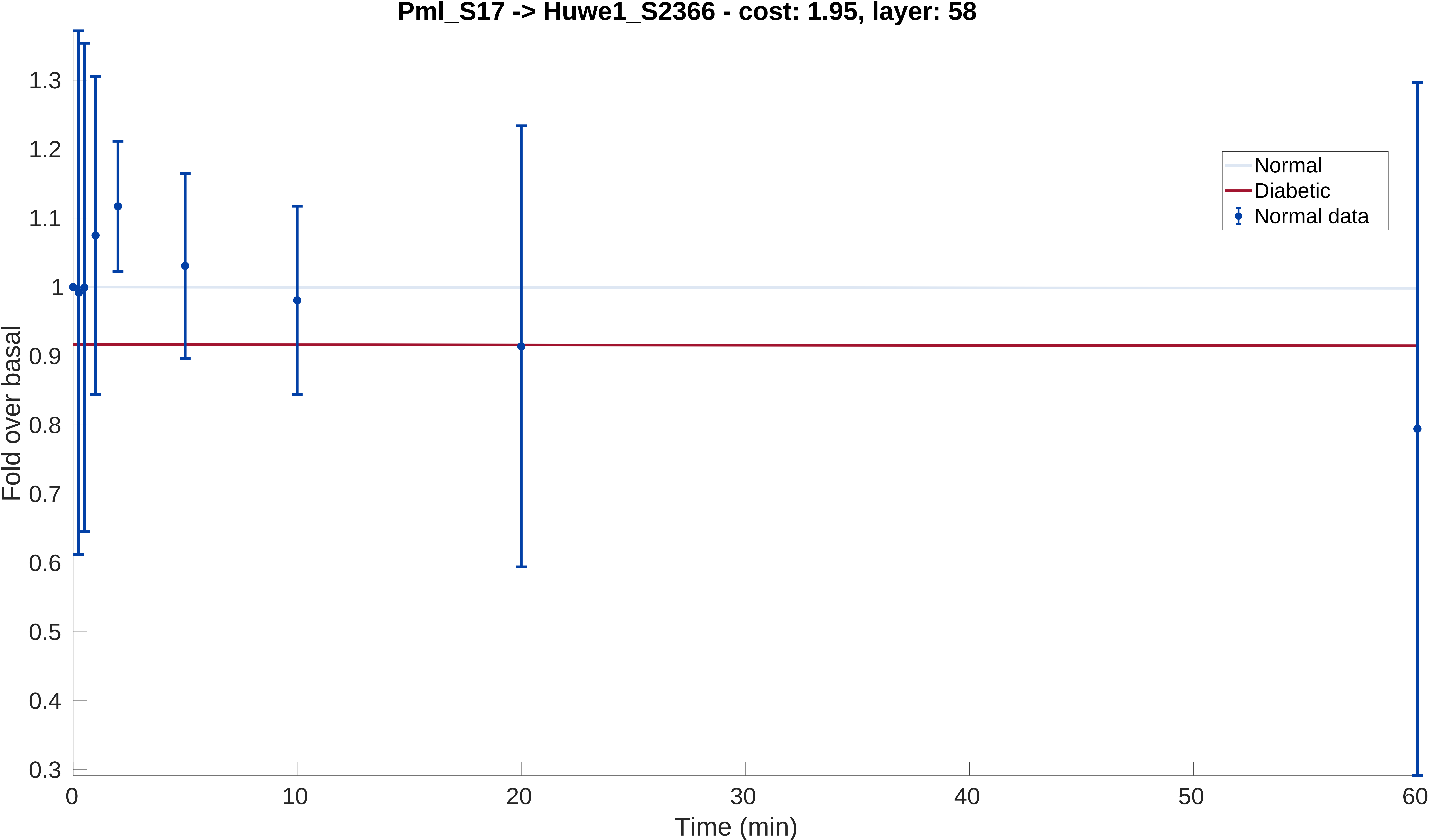

Pml_S17 -> Huwe1_S2366 - cost: 1.95, layer: 58
Fold over basal
1.3
1.2
1.1
1
0.9
0.8
0.7
0.6
0.5
0.4
0.3
0
10
20
30
40
50
60
Time (min)
Normal
Diabetic
Normal data

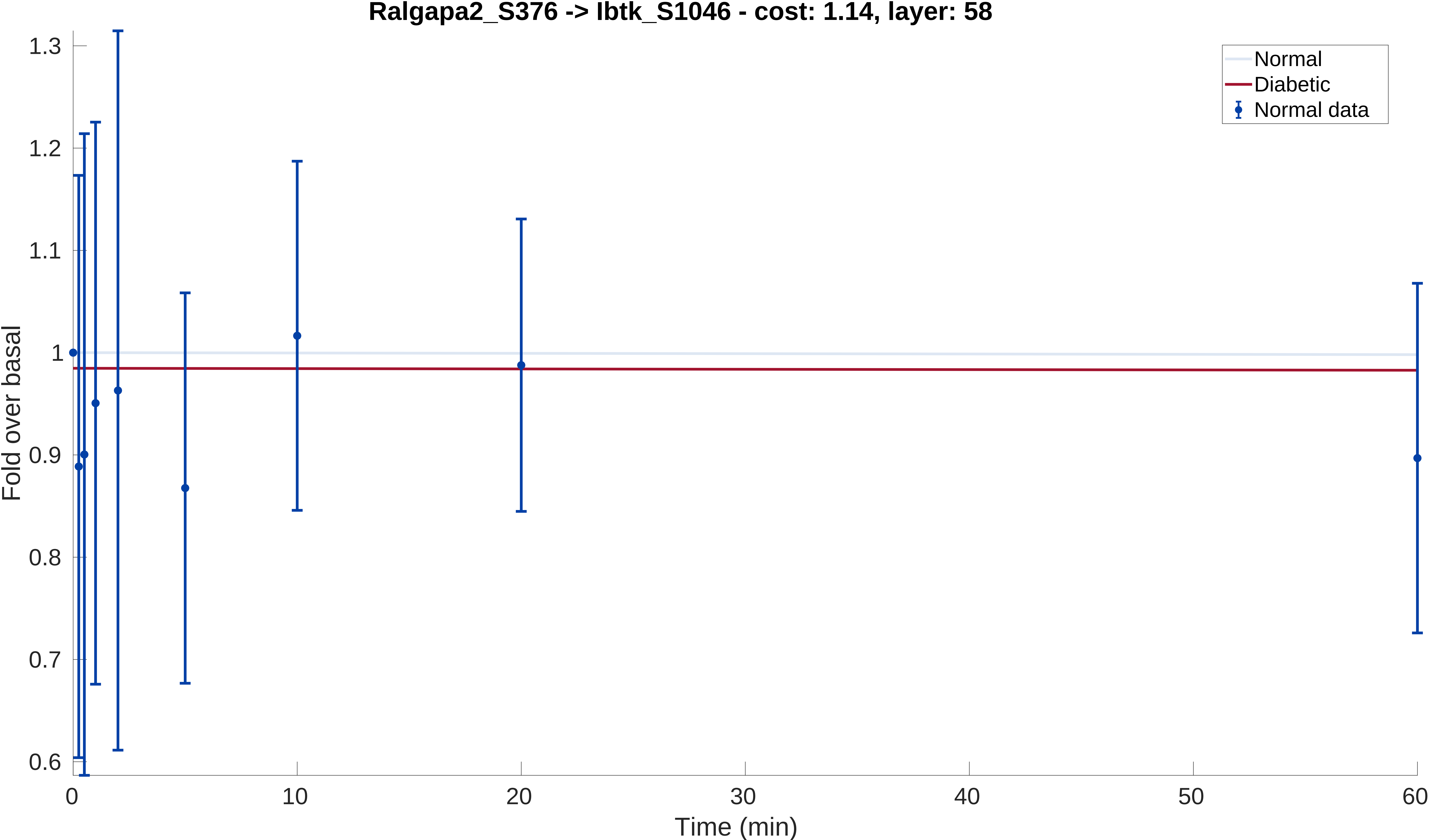

Ralgapa2_S376 -> Ibtk_S1046 - cost: 1.14, layer: 58
Normal
Diabetic
Normal data
Fold over basal
Time (min)
1.3
1.2
1.1
1
0.9
0.8
0.7
0.6
0
10
20
30
40
50
60

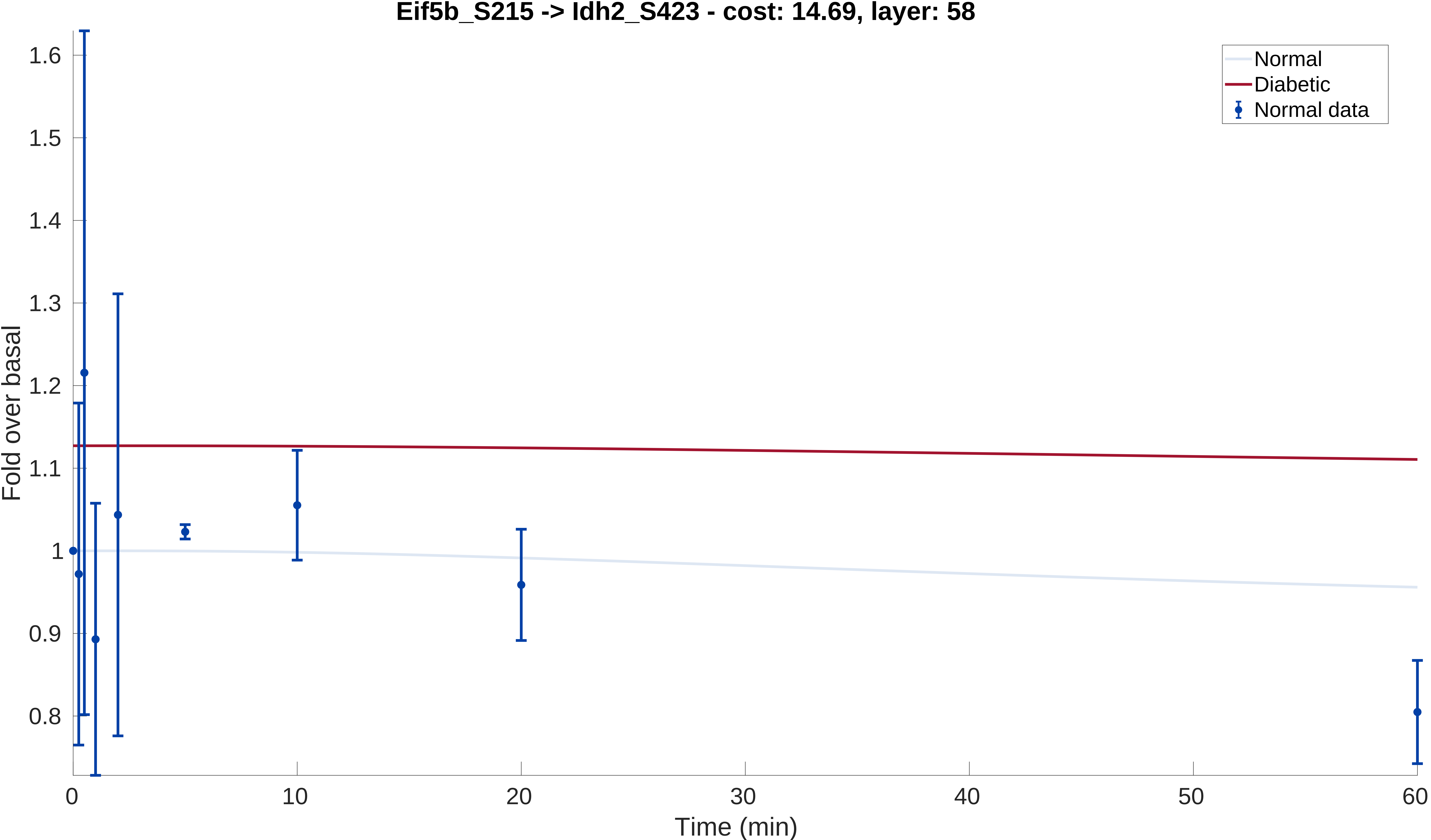
Eif5b_S215 -> Idh2_S423 - cost: 14.69, layer: 58
Fold over basal
1.6
1.5
1.4
1.3
1.2
1.1
1
0.9
0.8
Normal
Diabetic
Normal data
0
10
20
30
40
50
60
Time (min)

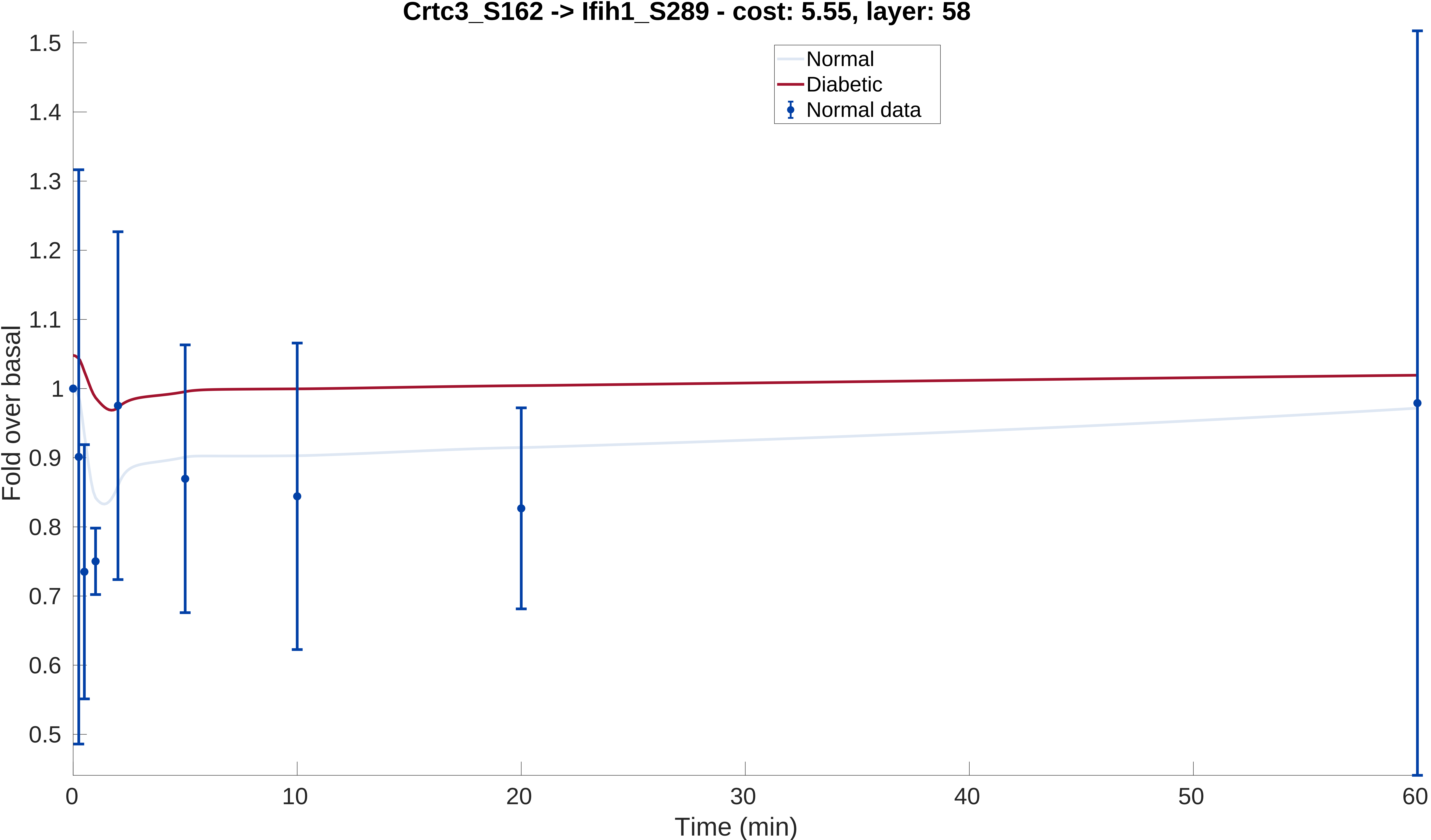

Crtc3_S162 -> Ifih1_S289 - cost: 5.55, layer: 58
Normal
Diabetic
Normal data
Fold over basal
Time (min)
1.5
1.4
1.3
1.2
1.1
1
0.9
0.8
0.7
0.6
0.5
0
10
20
30
40
50
60

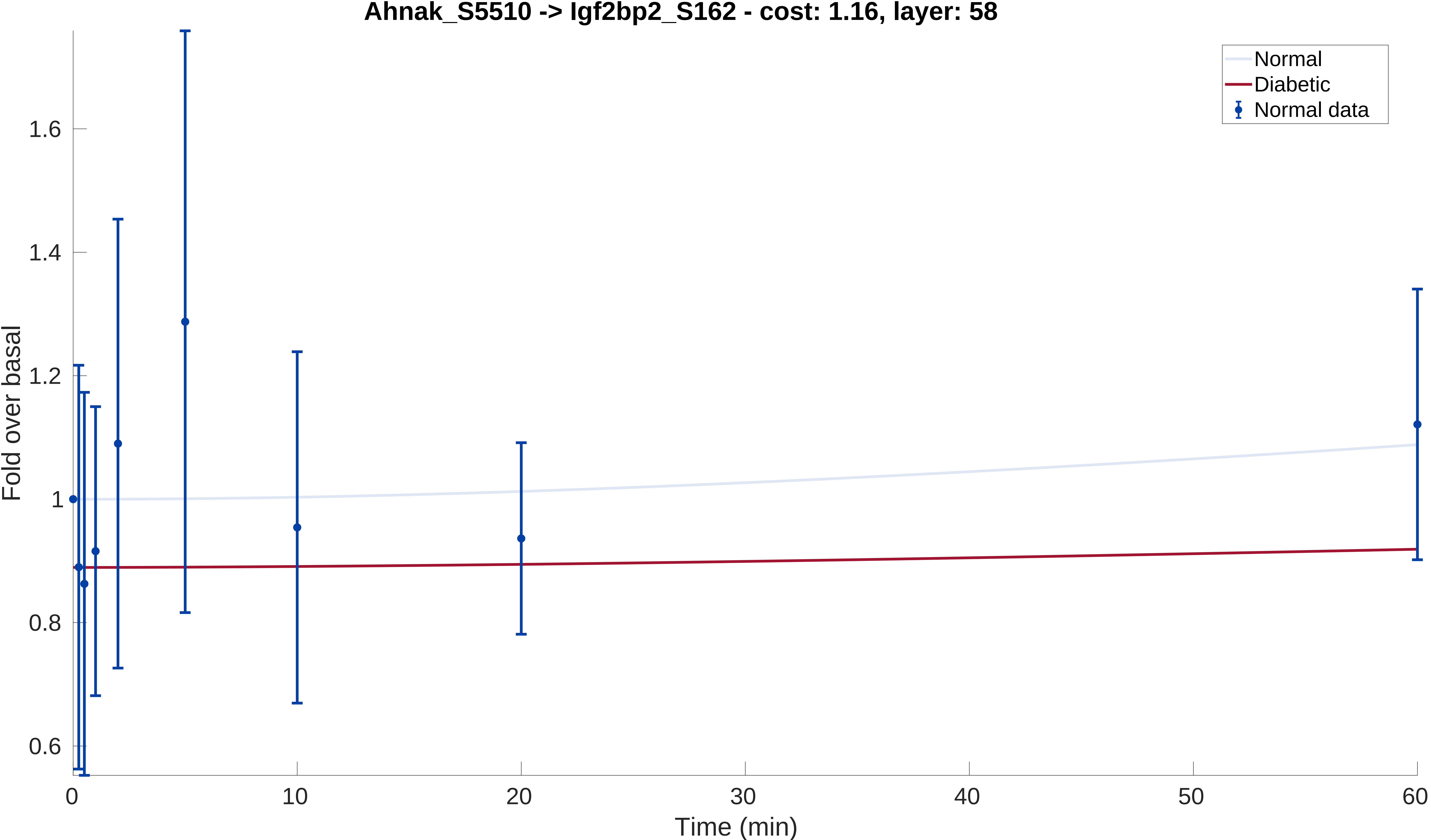

Ahnak_S5510 -> Igf2bp2_S162 - cost: 1.16, layer: 58
Normal
Diabetic
Normal data
Fold over basal
1.6
1.4
1.2
1
0.8
0.6
0
10
20
30
40
50
60
Time (min)

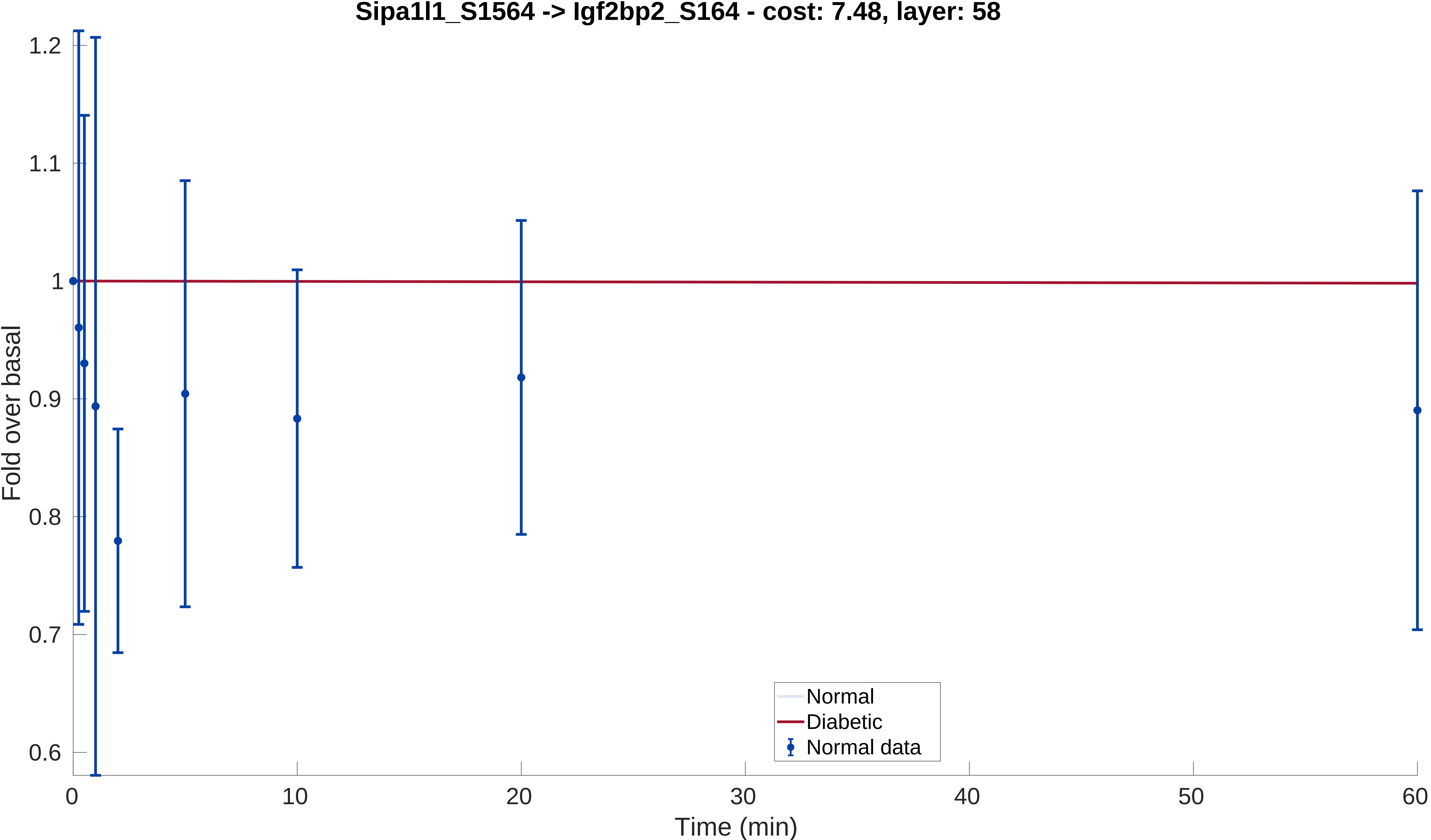

Sipa1l1_S1564 -> Igf2bp2_S164 - cost: 7.48, layer: 58
Fold over basal
1.2
1.1
1
0.9
0.8
0.7
0.6
0
10
20
30
40
50
60
Time (min)
Normal
Diabetic
Normal data

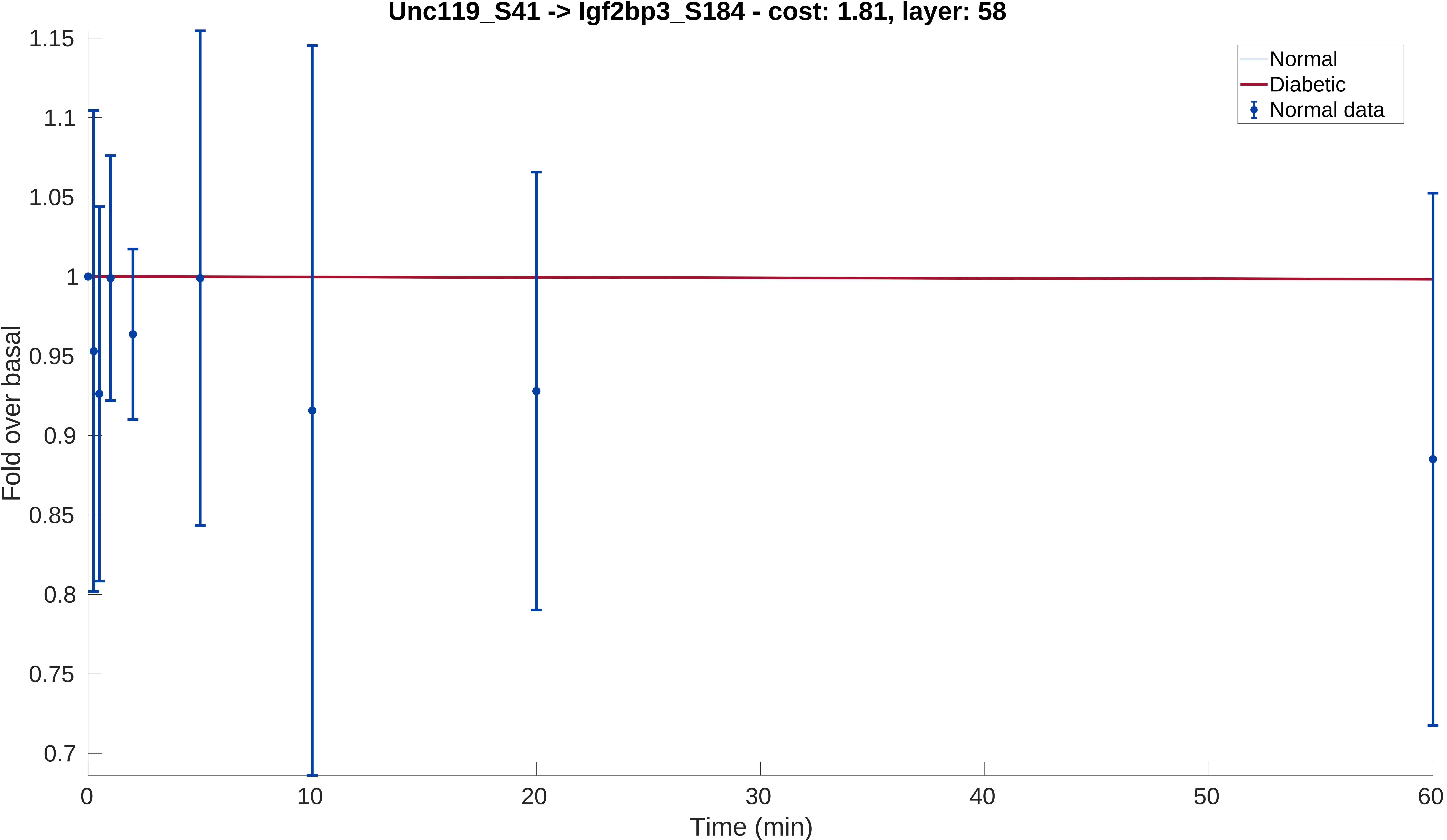

Unc119_S41 -> Igf2bp3_S184 - cost: 1.81, layer: 58
Normal
Diabetic
Normal data
Fold over basal
Time (min)
1.15
1.1
1.05
1
0.95
0.9
0.85
0.8
0.75
0.7
0
10
20
30
40
50
60

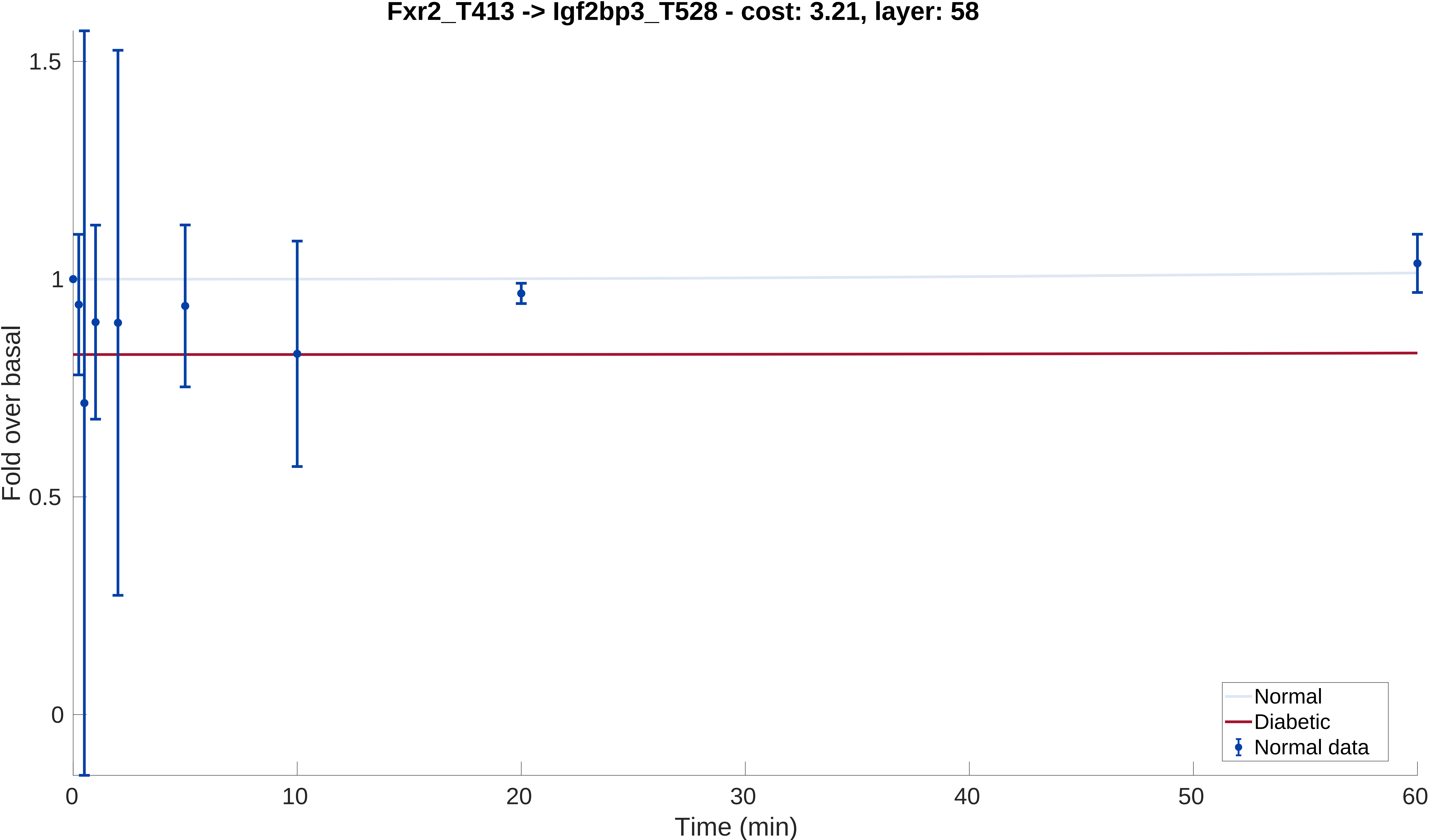

Fxr2_T413 -> Igf2bp3_T528 - cost: 3.21, layer: 58
Fold over basal
Time (min)
1.5
1
0.5
0
0
10
20
30
40
50
60
Normal
Diabetic
Normal data

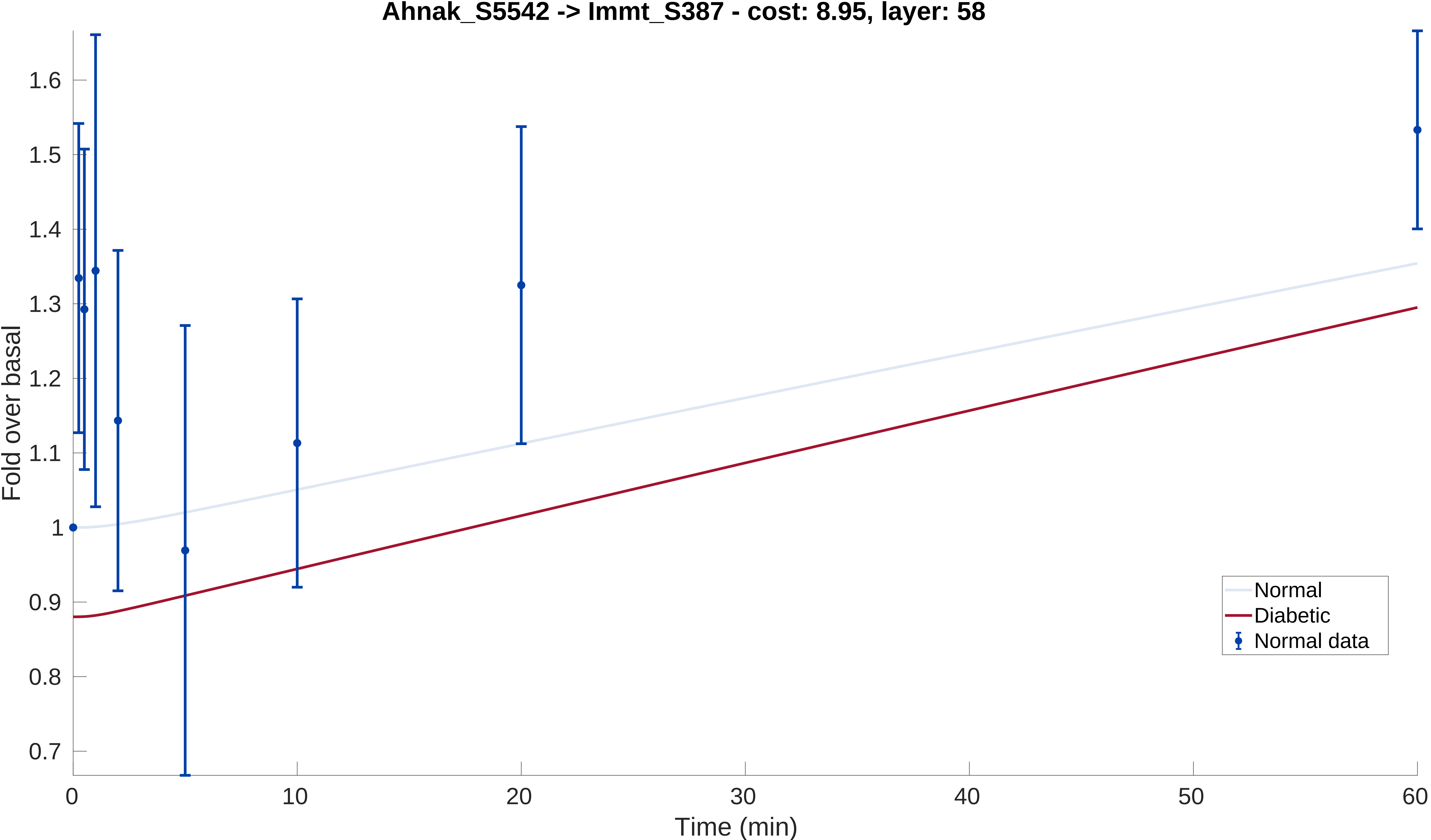

Ahnak_S5542 -> Immt_S387 - cost: 8.95, layer: 58
1.6
1.5
1.4
1.3
1.2
1.1
1
0.9
0.8
0.7
Fold over basal
0
10
20
30
40
50
60
Time (min)
Normal
Diabetic
Normal data

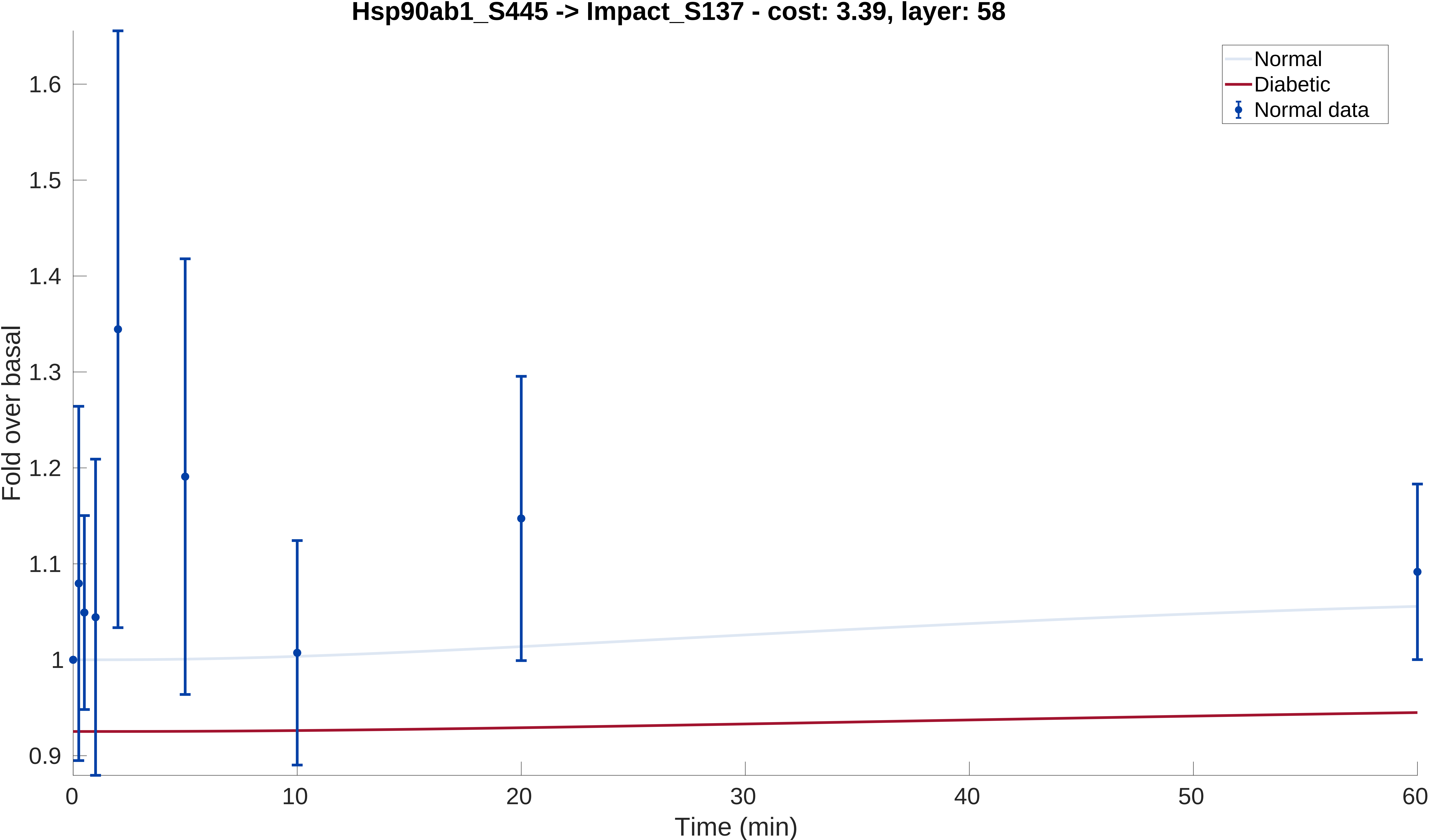
Hsp90ab1_S445 -> Impact_S137 - cost: 3.39, layer: 58
Normal
Diabetic
Normal data
Fold over basal
1.6
1.5
1.4
1.3
1.2
1.1
1
0.9
0
10
20
30
40
50
60
Time (min)

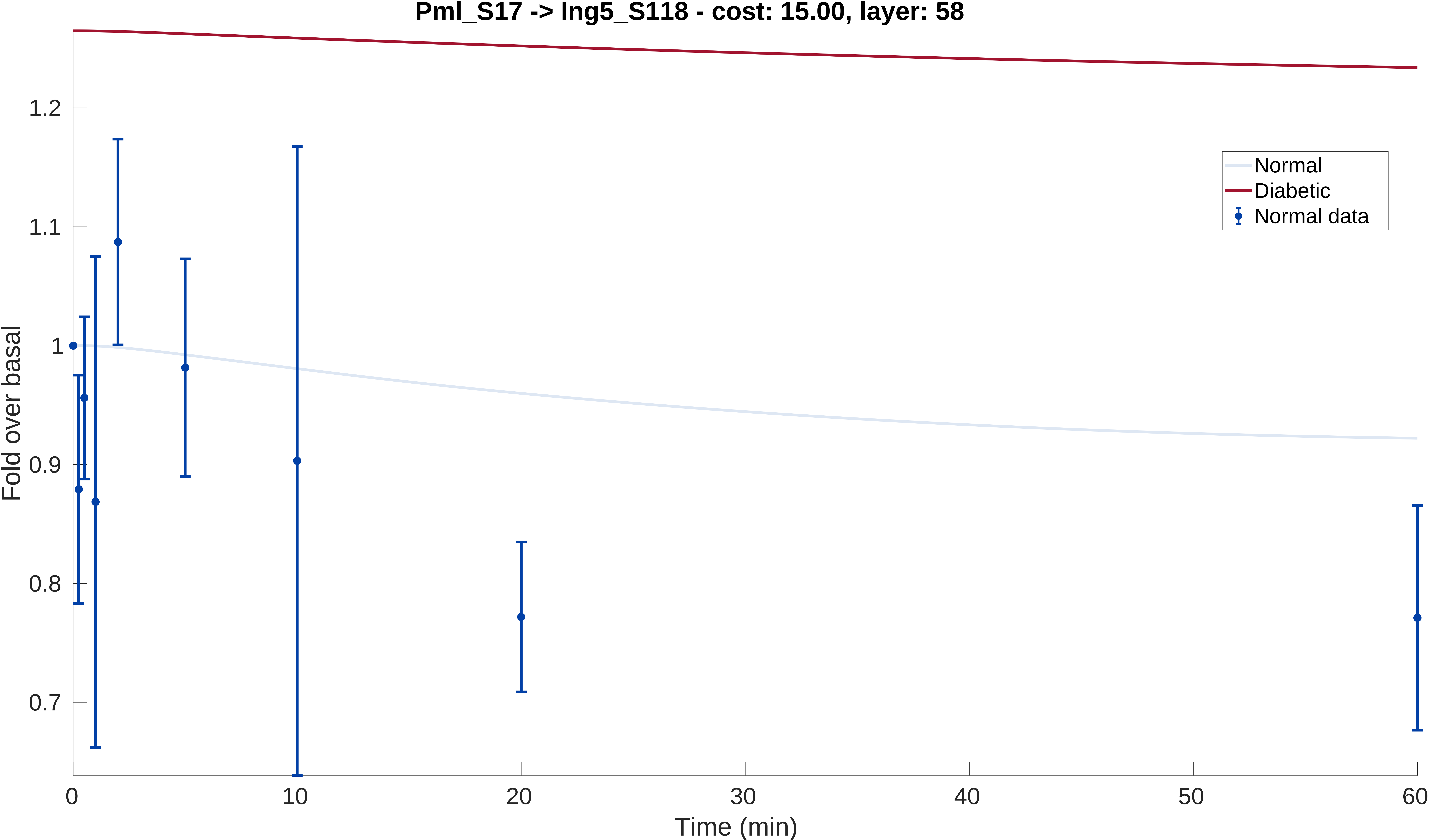
Pml_S17 -> Ing5_S118 - cost: 15.00, layer: 58
Normal
Diabetic
Normal data
Fold over basal
1.2
1.1
1
0.9
0.8
0.7
0
10
20
30
40
50
60
Time (min)

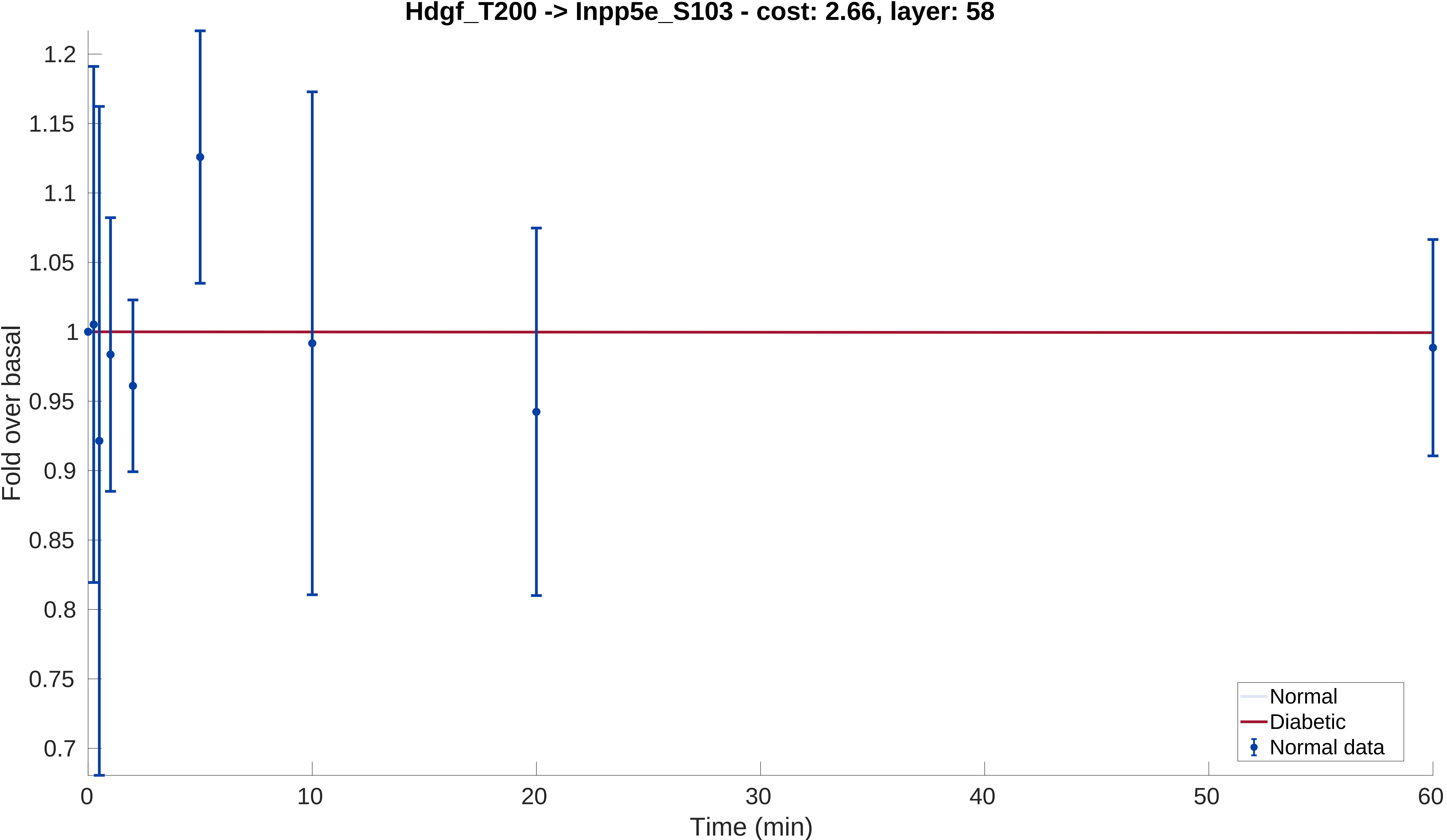

Hdgf_T200 -> Inpp5e_S103 - cost: 2.66, layer: 58
Fold over basal
1.2
1.15
1.1
1.05
1
0.95
0.9
0.85
0.8
0.75
0.7
0
10
20
30
40
50
60
Time (min)
Normal
Diabetic
Normal data

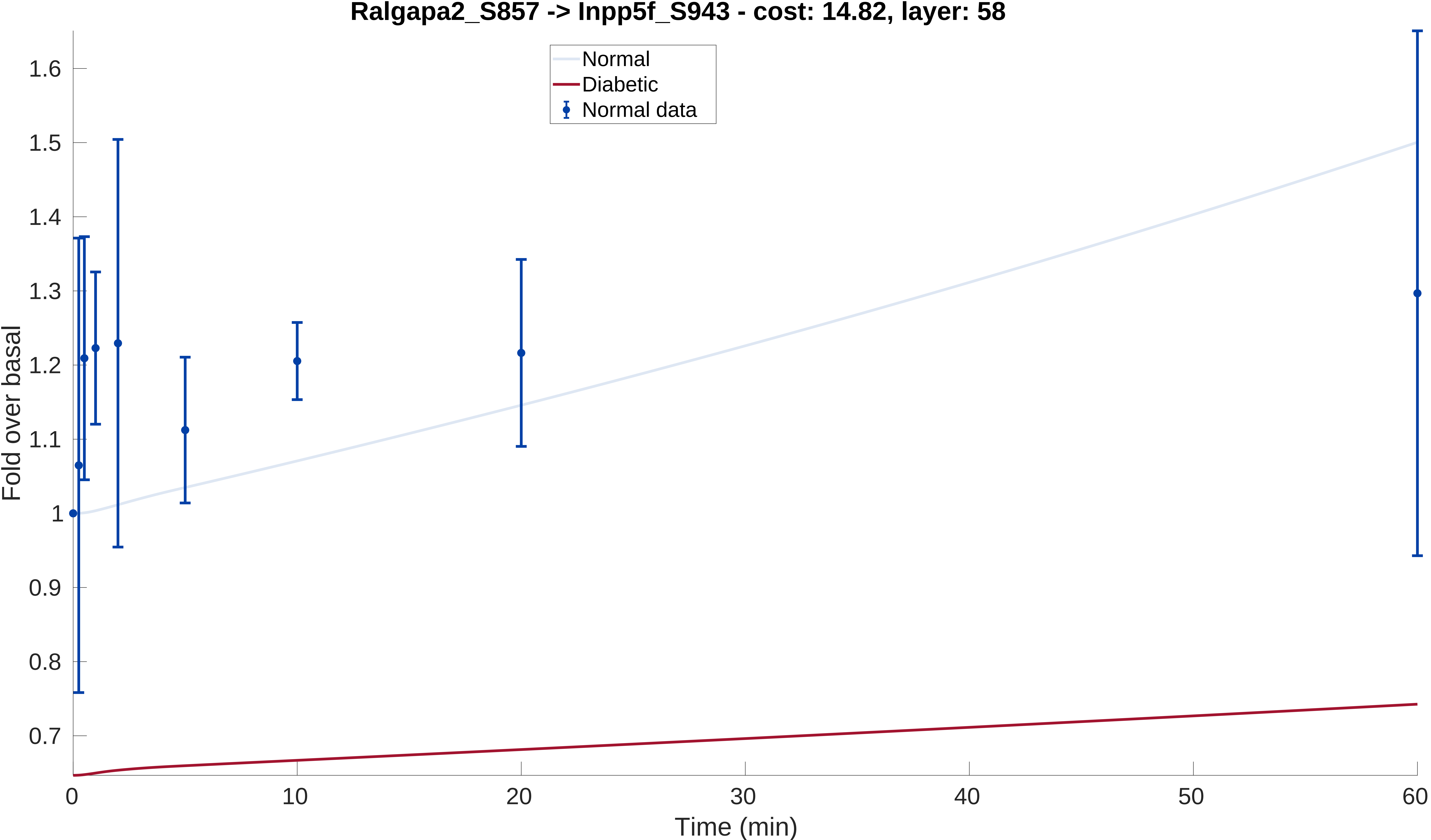

Ralgapa2_S857 -> Inpp5f_S943 - cost: 14.82, layer: 58
Normal
Diabetic
Normal data
Fold over basal
1.6
1.5
1.4
1.3
1.2
1.1
1
0.9
0.8
0.7
0
10
20
30
40
50
60
Time (min)

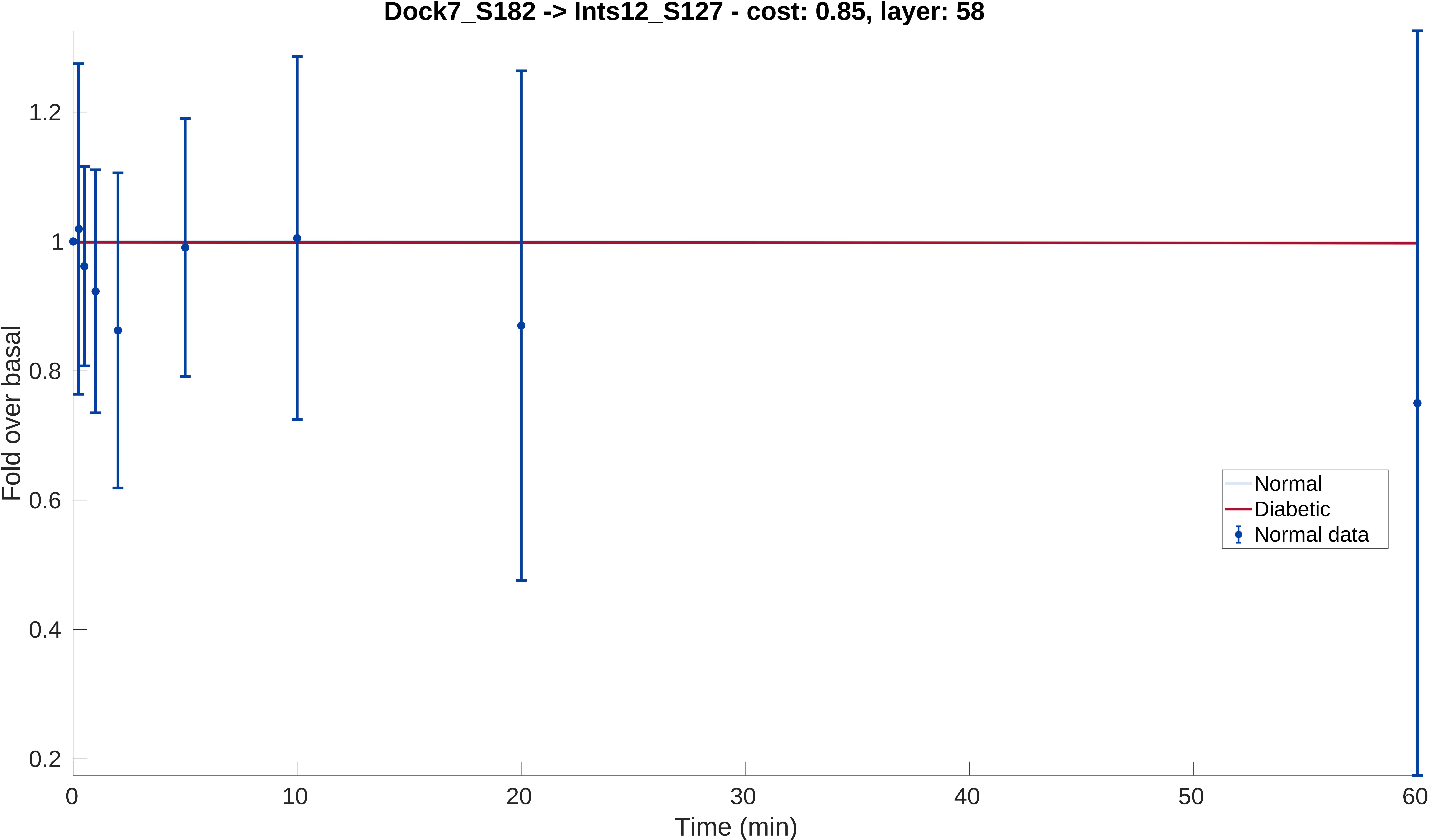

Dock7_S182 -> Ints12_S127 - cost: 0.85, layer: 58
Fold over basal
Time (min)
0.2
0.4
0.6
0.8
1
1.2
0
10
20
30
40
50
60
Normal
Diabetic
Normal data

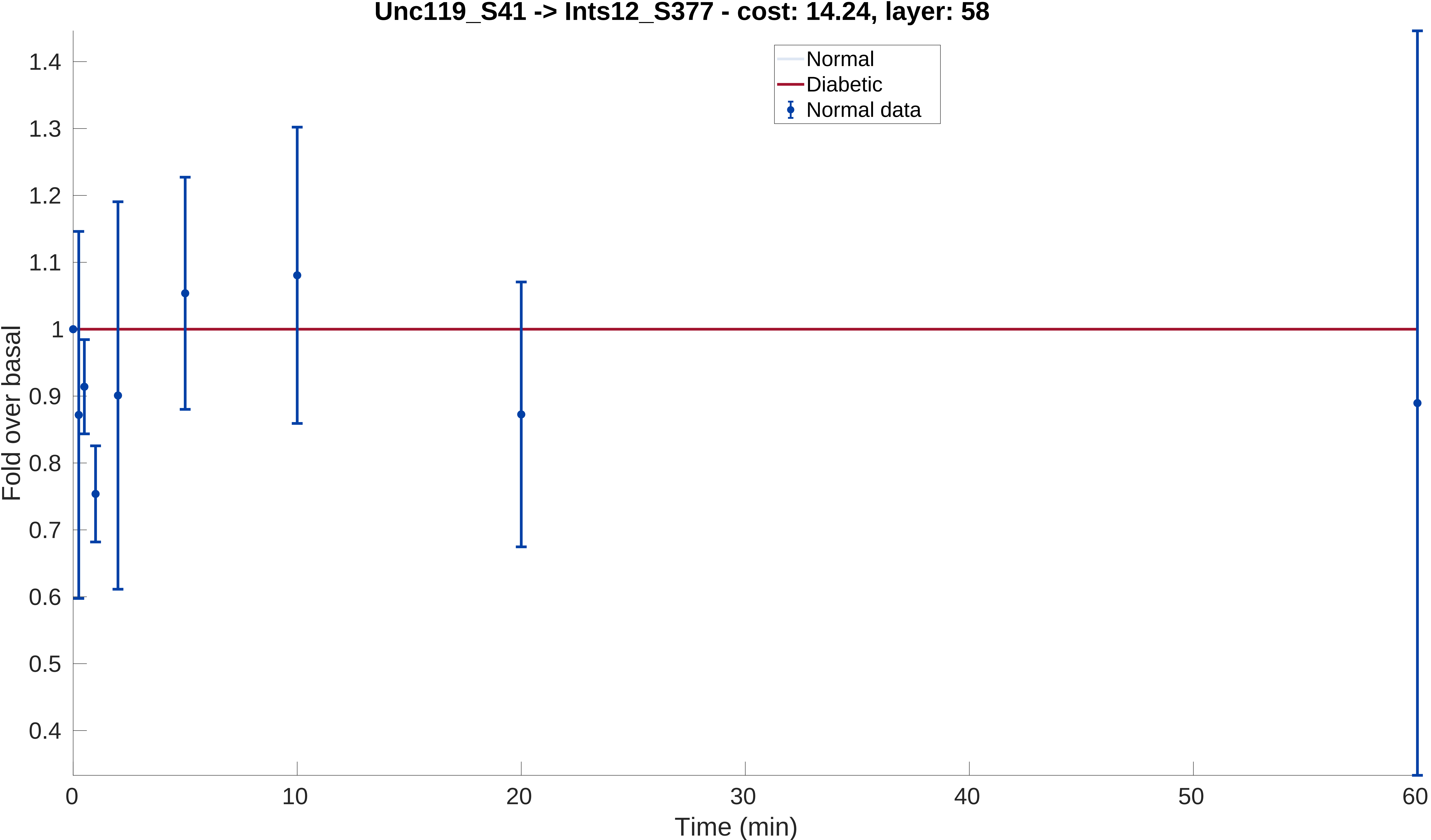

Unc119_S41 -> Ints12_S377 - cost: 14.24, layer: 58
Normal
Diabetic
Normal data
Fold over basal
1.4
1.3
1.2
1.1
1
0.9
0.8
0.7
0.6
0.5
0.4
0
10
20
30
40
50
60
Time (min)

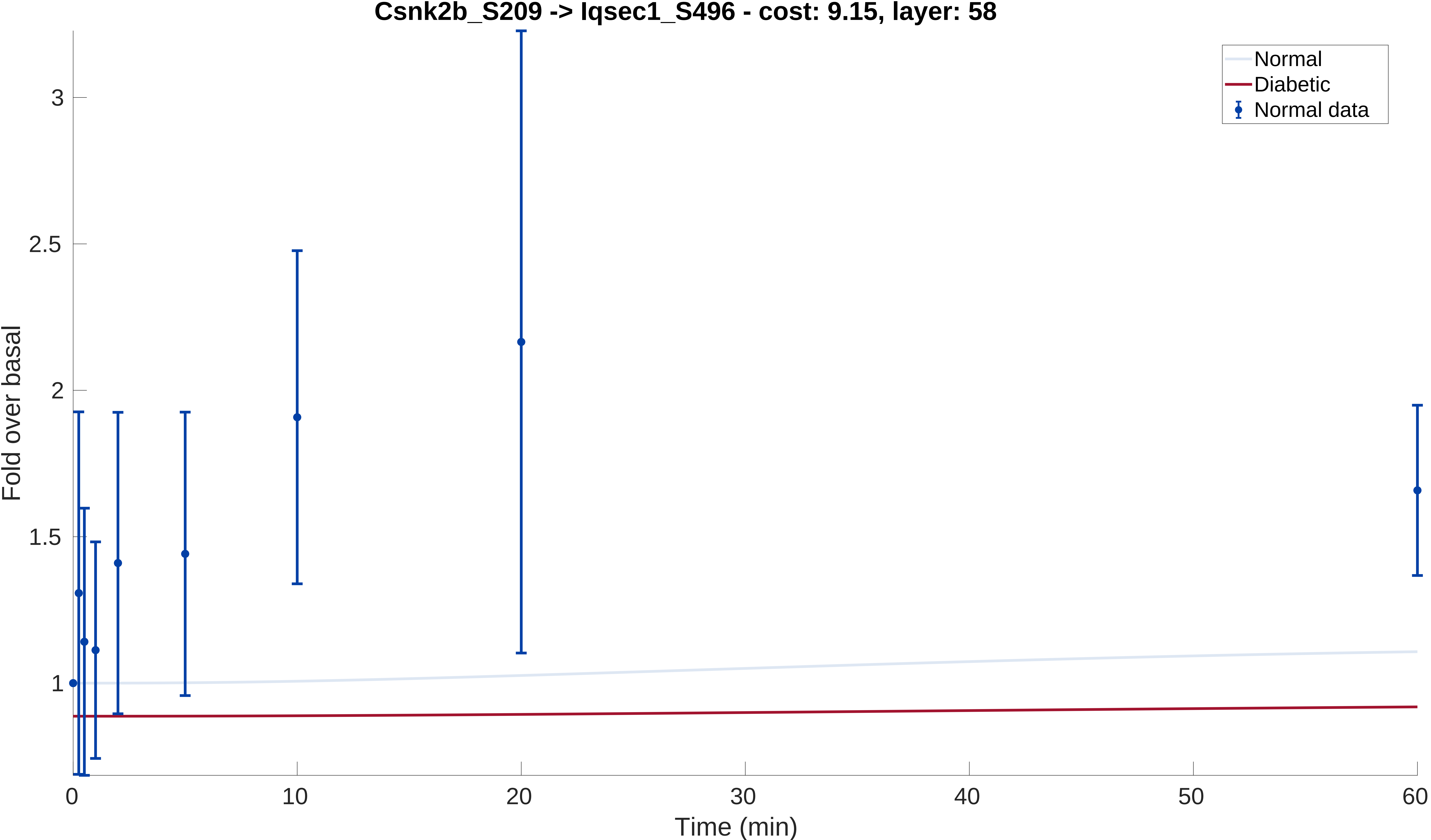
Csnk2b_S209 -> Iqsec1_S496 - cost: 9.15, layer: 58
Normal
Diabetic
Normal data
Fold over basal
3
2.5
2
1.5
1
0
10
20
30
40
50
60
Time (min)

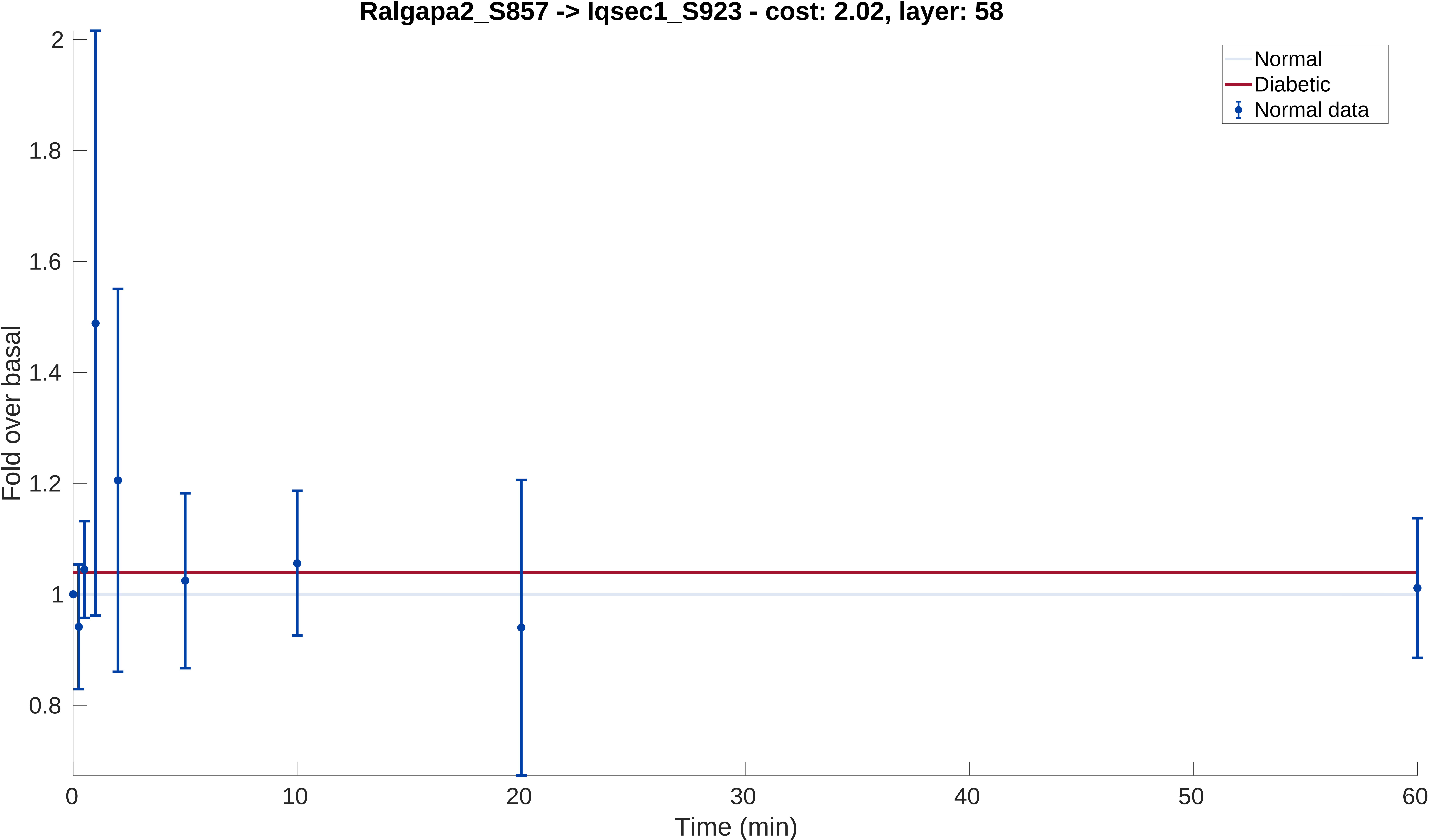
Ralgapa2_S857 -> Iqsec1_S923 - cost: 2.02, layer: 58
Normal
Diabetic
Normal data
Fold over basal
2
1.8
1.6
1.4
1.2
1
0.8
0
10
20
30
40
50
60
Time (min)

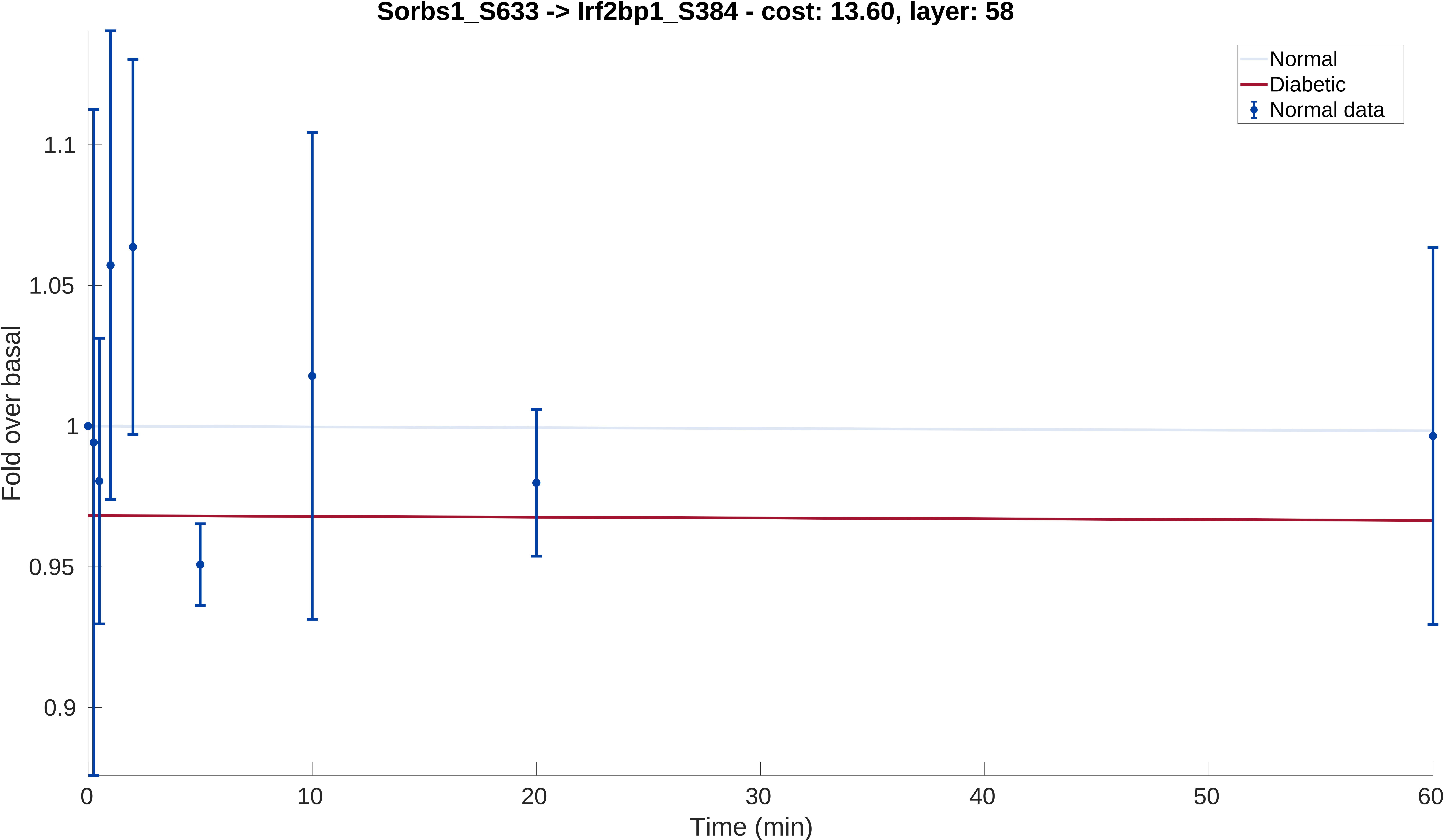

Sorbs1_S633 -> Irf2bp1_S384 - cost: 13.60, layer: 58
Normal
Diabetic
Normal data
1.1
1.05
1
0.95
0.9
Fold over basal
0
10
20
30
40
50
60
Time (min)

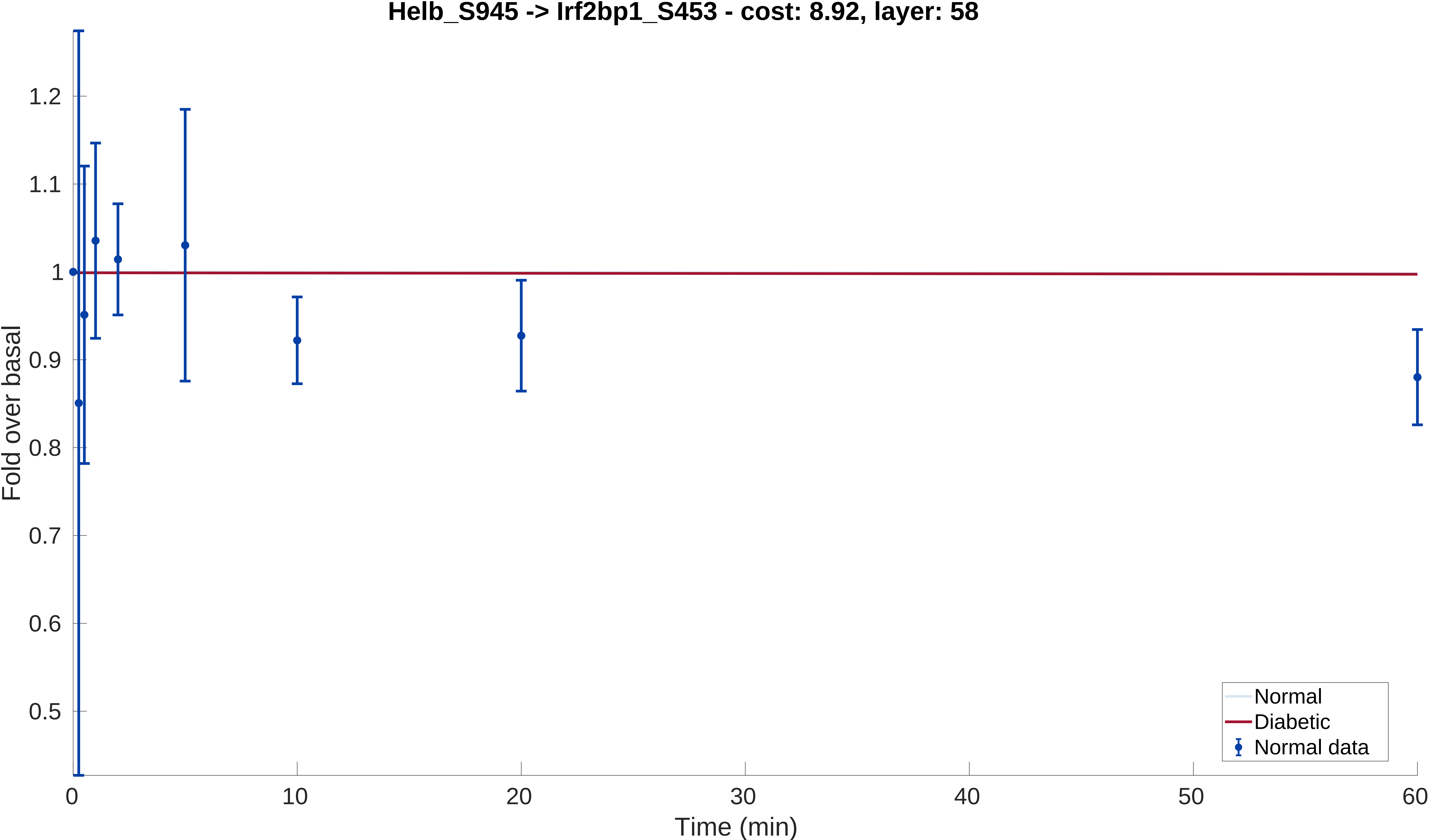

Helb_S945 -> Irf2bp1_S453 - cost: 8.92, layer: 58
Fold over basal
1.2
1.1
1
0.9
0.8
0.7
0.6
0.5
0
10
20
30
40
50
60
Time (min)
Normal
Diabetic
Normal data

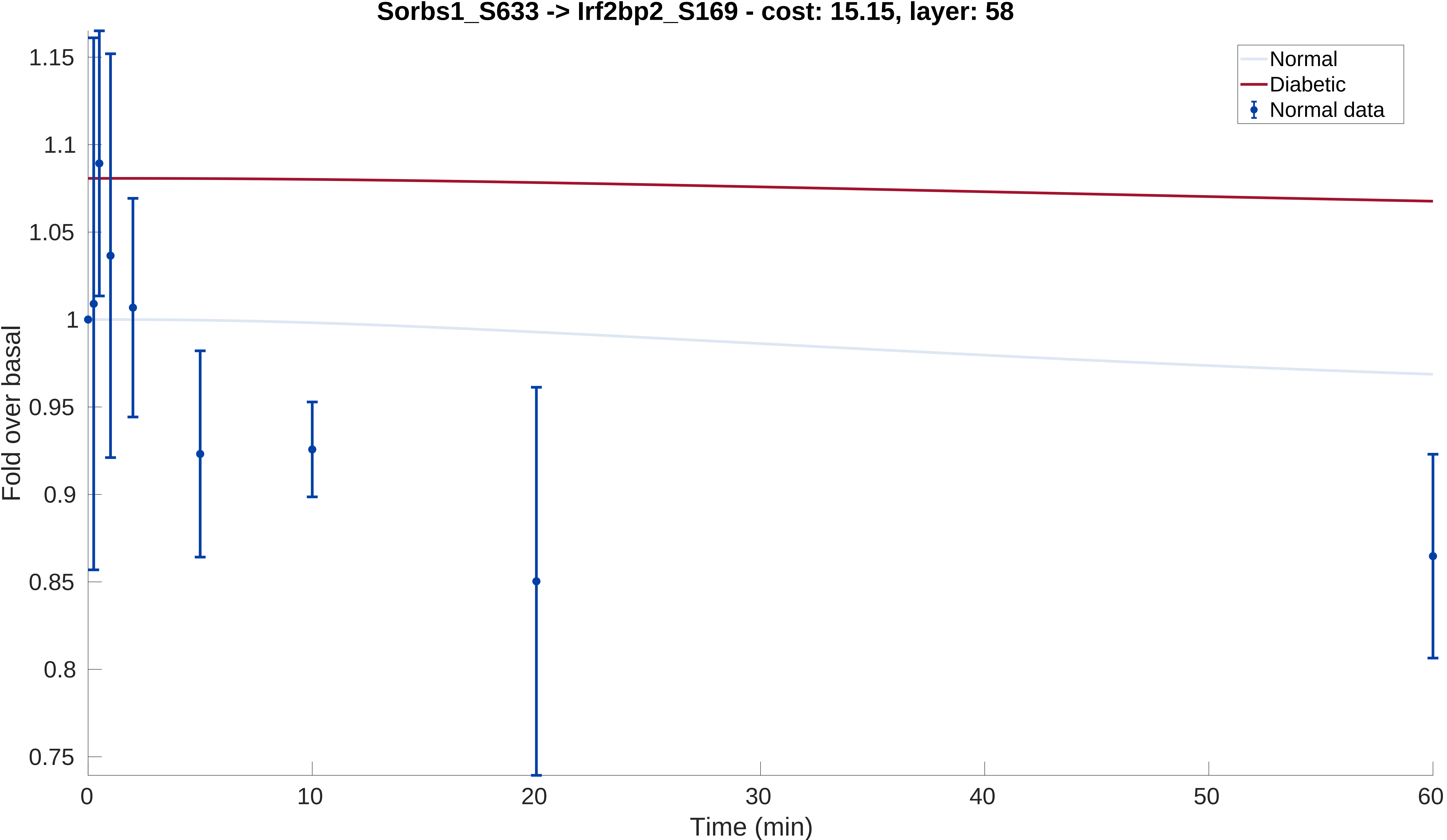
Sorbs1_S633 -> Irf2bp2_S169 - cost: 15.15, layer: 58
Normal
Diabetic
Normal data
Fold over basal
Time (min)
1.15
1.1
1.05
1
0.95
0.9
0.85
0.8
0.75
0
10
20
30
40
50
60

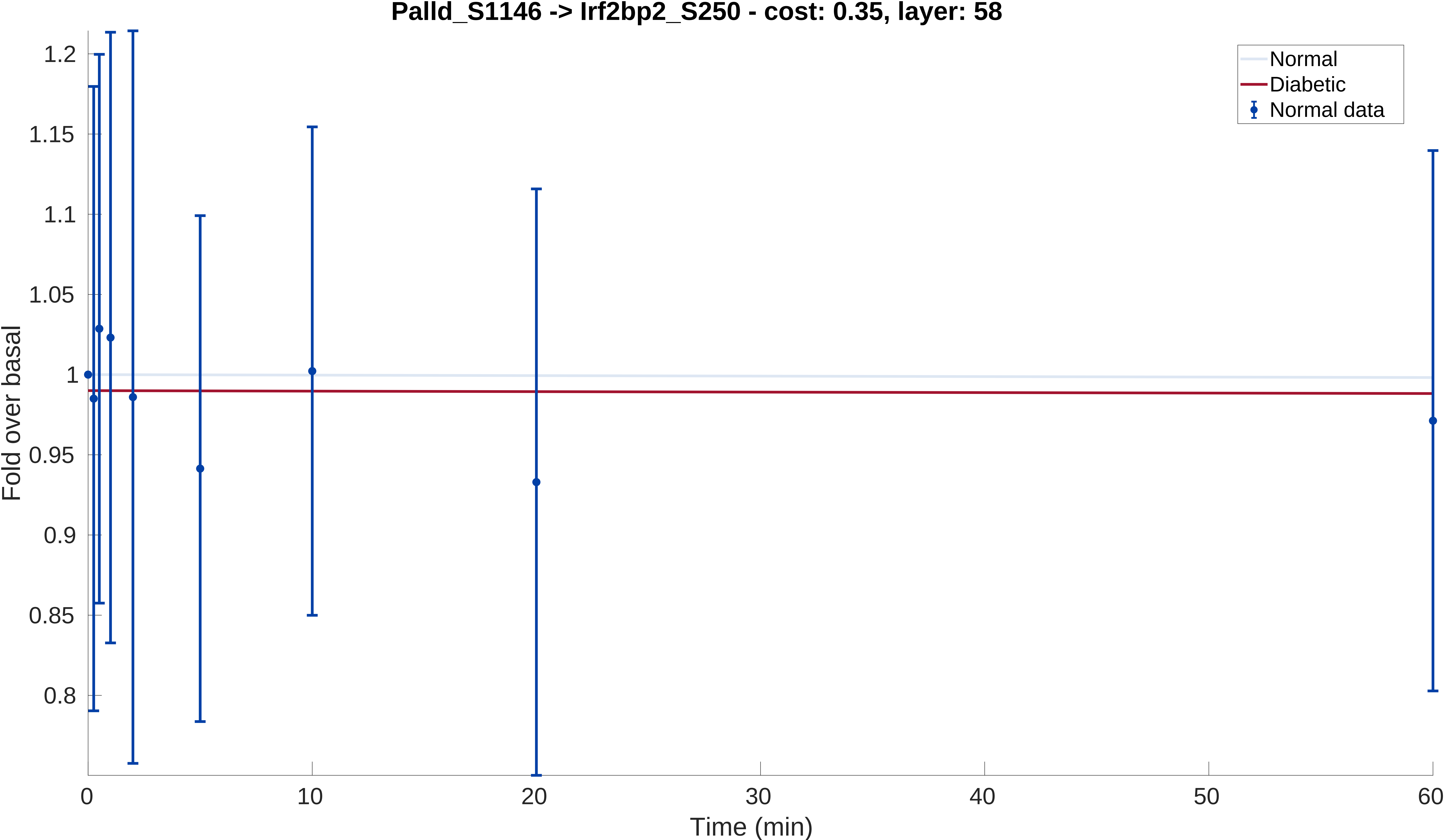
Palld_S1146 -> Irf2bp2_S250 - cost: 0.35, layer: 58
Normal
Diabetic
Normal data
Fold over basal
1.2
1.15
1.1
1.05
1
0.95
0.9
0.85
0.8
0
10
20
30
40
50
60
Time (min)

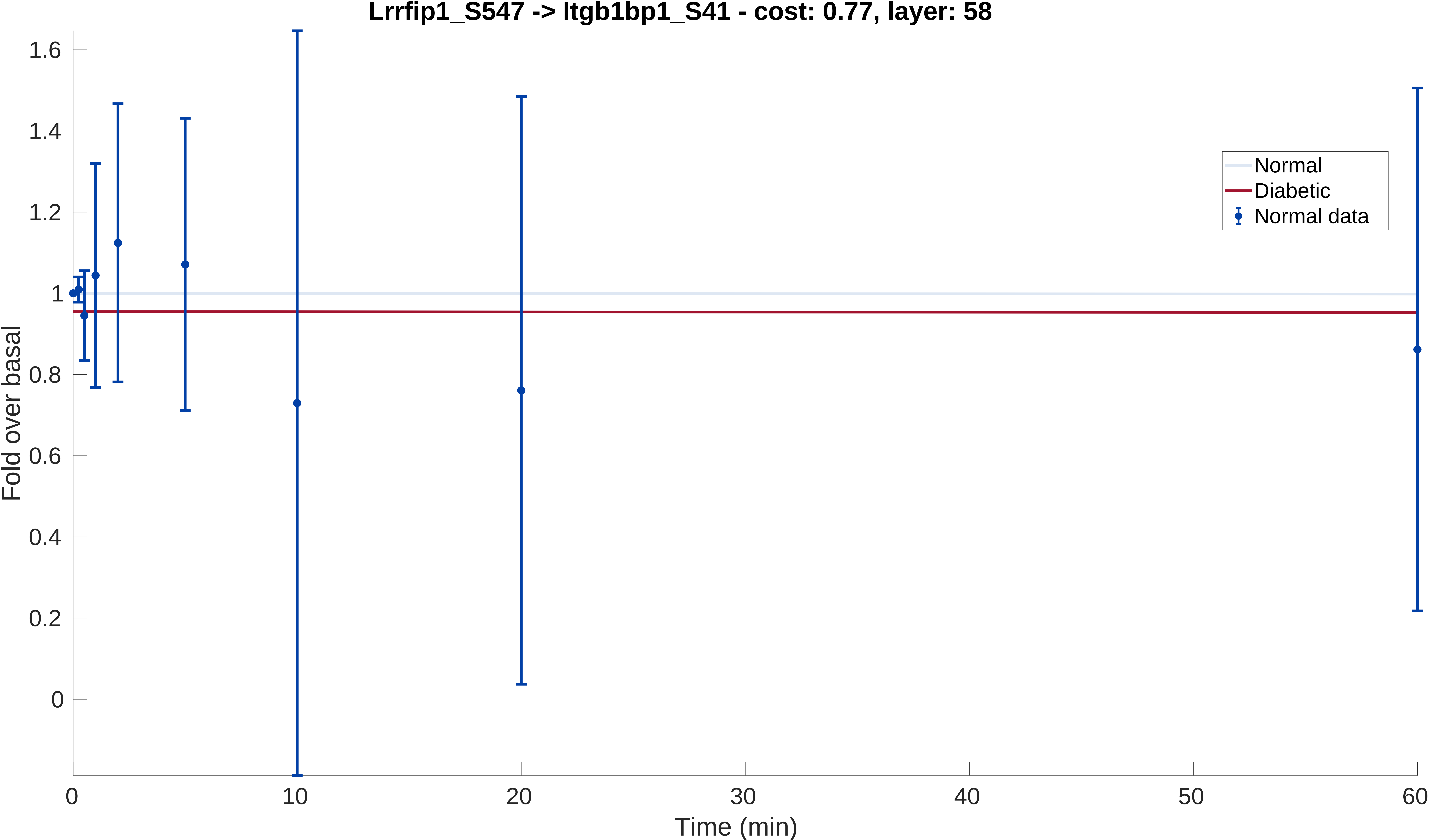

Lrrfip1_S547 -> Itgb1bp1_S41 - cost: 0.77, layer: 58
Fold over basal
Time (min)
Normal
Diabetic
Normal data
1.6
1.4
1.2
1
0.8
0.6
0.4
0.2
0
0
10
20
30
40
50
60

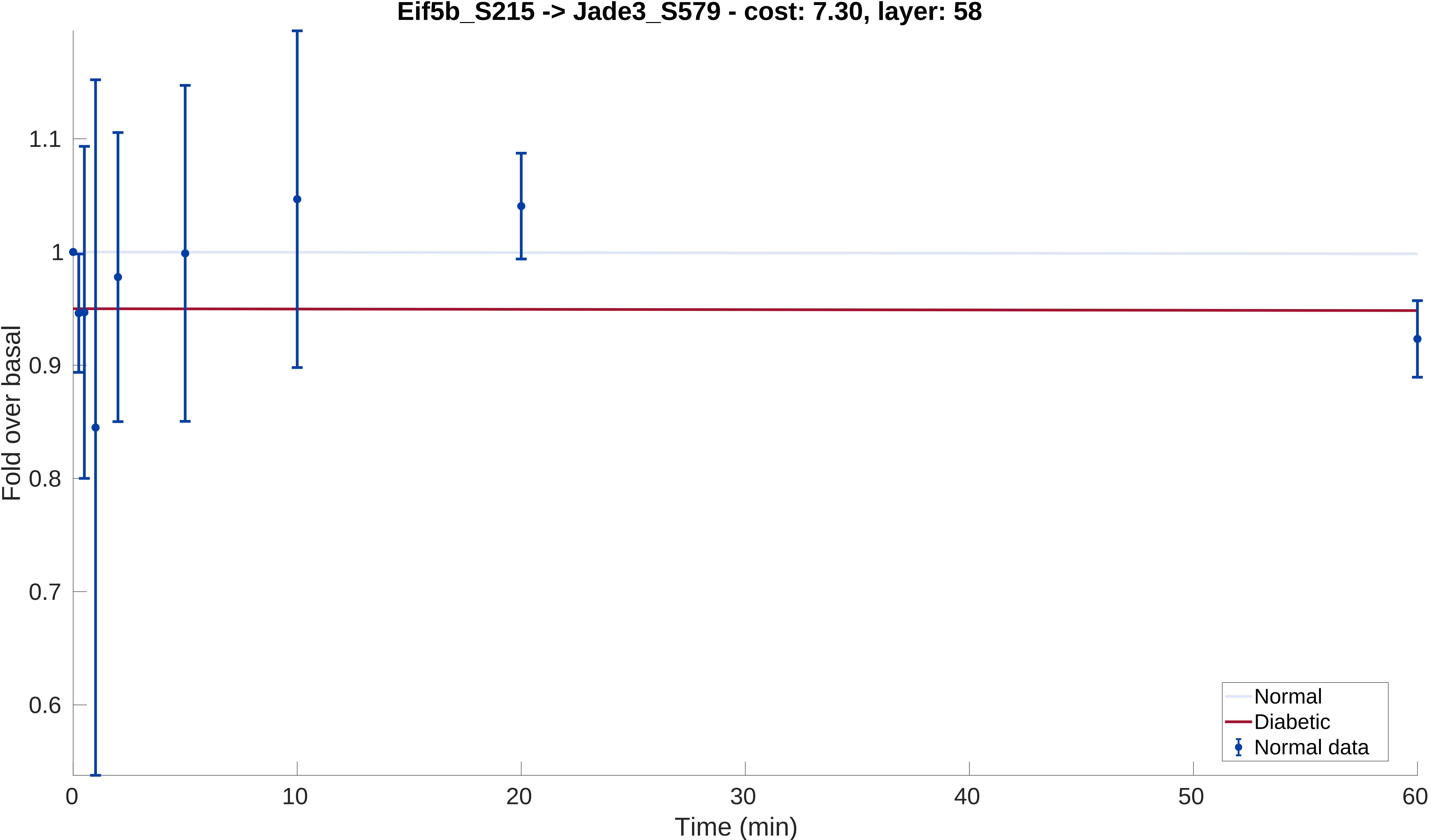

Eif5b_S215 -> Jade3_S579 - cost: 7.30, layer: 58
Fold over basal
1.1
1
0.9
0.8
0.7
0.6
0
10
20
30
40
50
60
Time (min)
Normal
Diabetic
Normal data

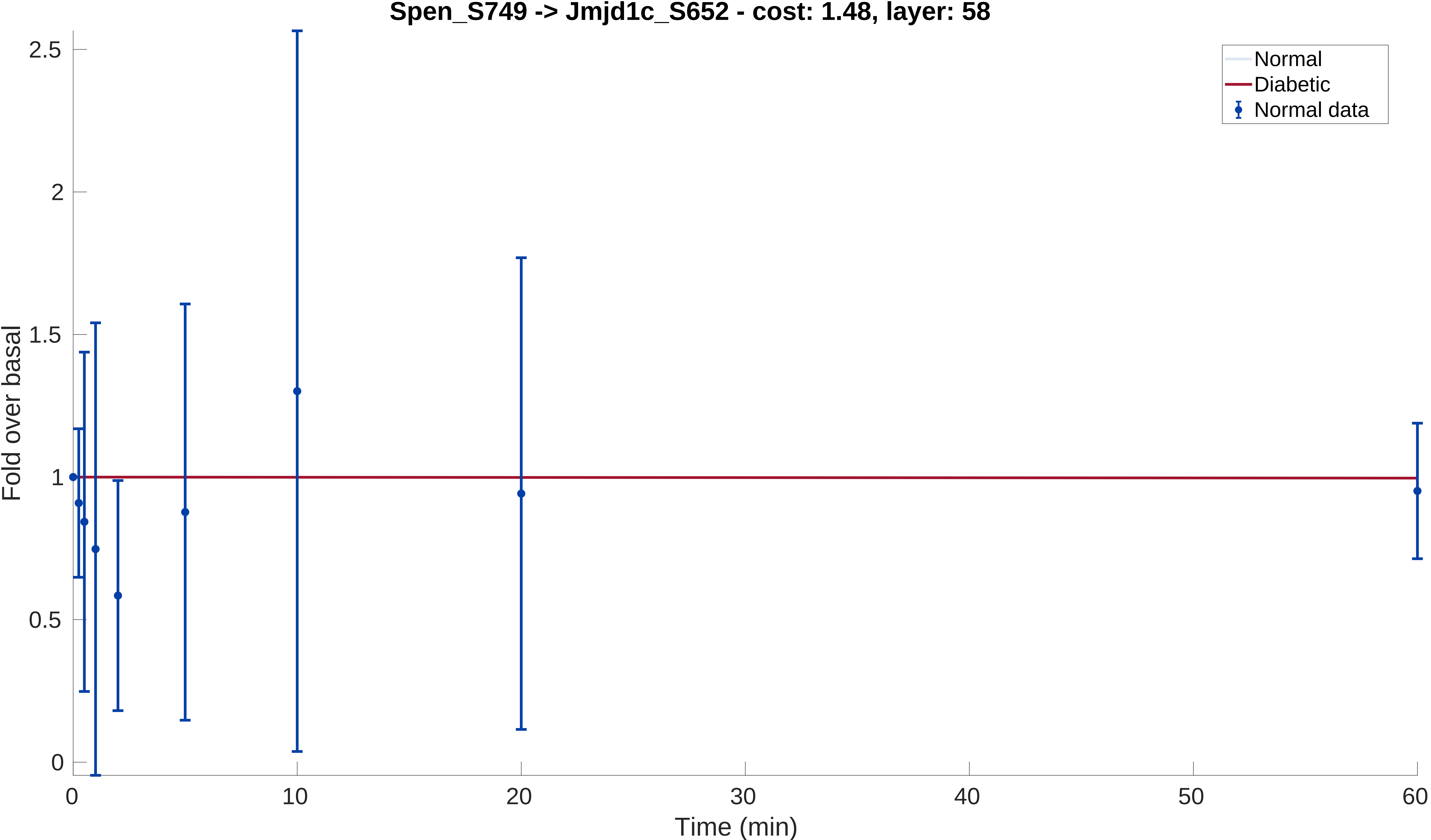

Spen_S749 -> Jmjd1c_S652 - cost: 1.48, layer: 58
Normal
Diabetic
Normal data
Fold over basal
Time (min)
0
0.5
1
1.5
2
2.5
0
10
20
30
40
50
60

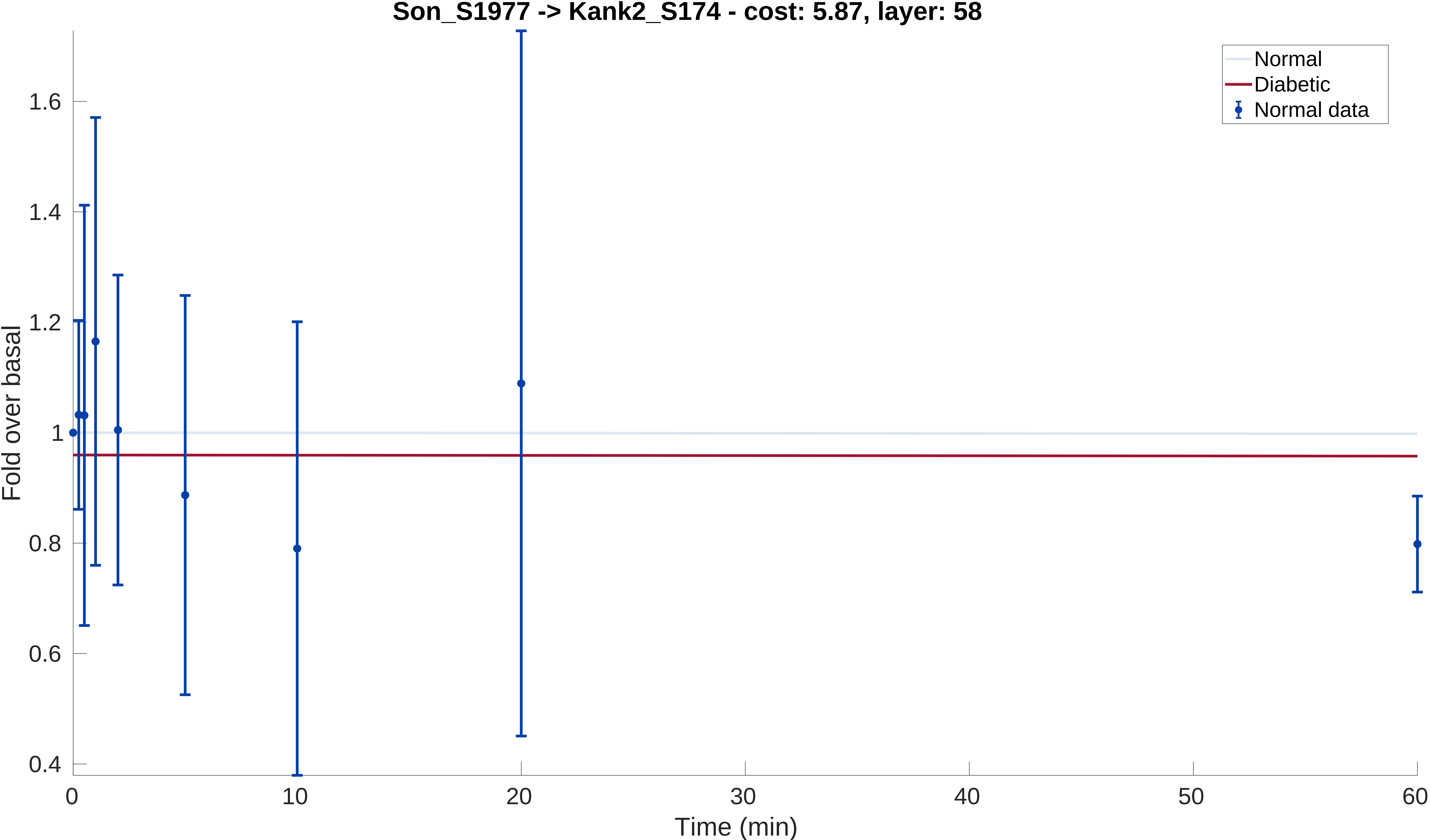

Son_S1977 -> Kank2_S174 - cost: 5.87, layer: 58
Normal
Diabetic
Normal data
Fold over basal
1.6
1.4
1.2
1
0.8
0.6
0.4
0
10
20
30
40
50
60
Time (min)

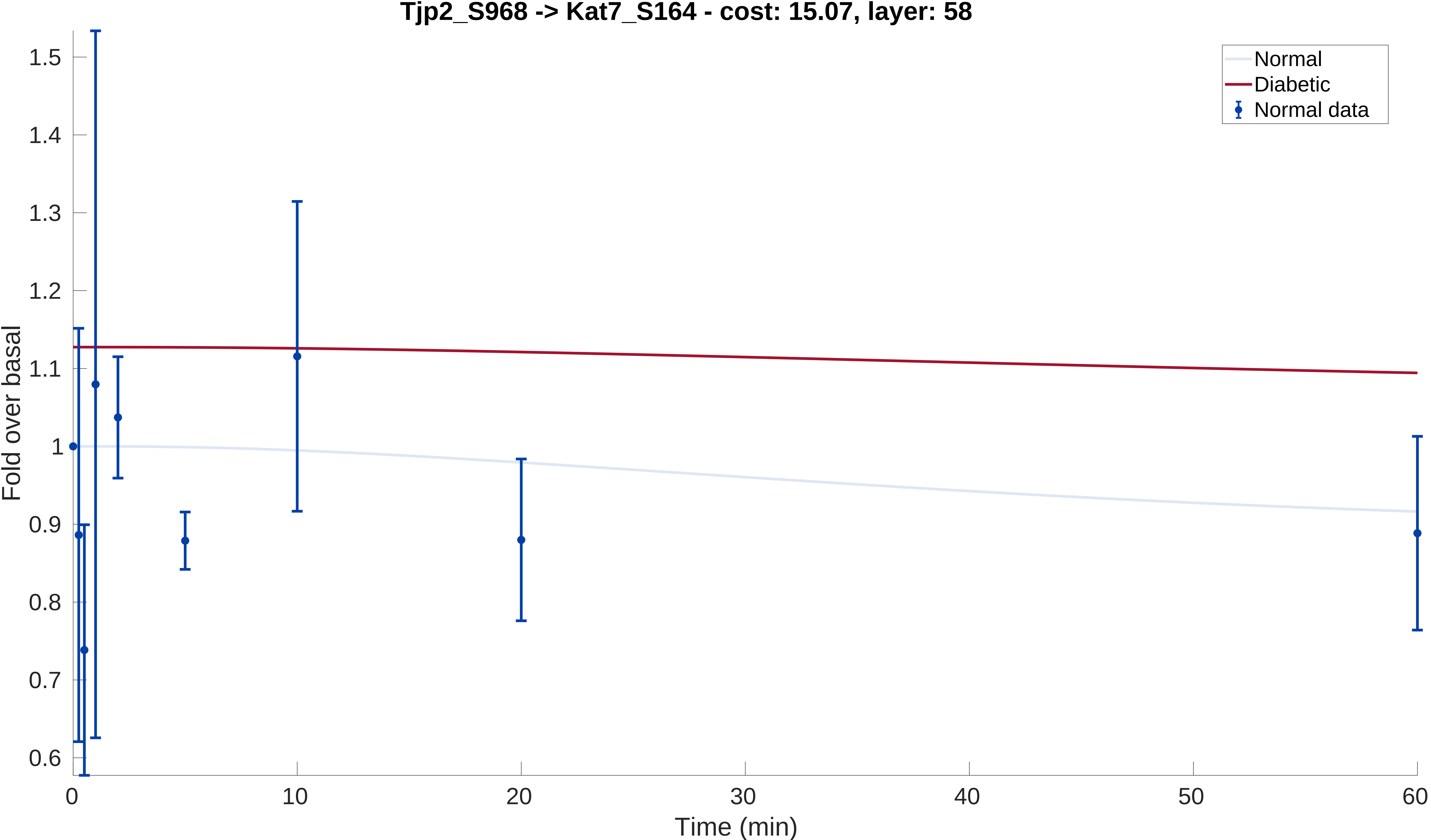
Tjp2_S968 -> Kat7_S164 - cost: 15.07, layer: 58
Fold over basal
Time (min)
Normal
Diabetic
Normal data
0.6
0.7
0.8
0.9
1
1.1
1.2
1.3
1.4
1.5
0
10
20
30
40
50
60

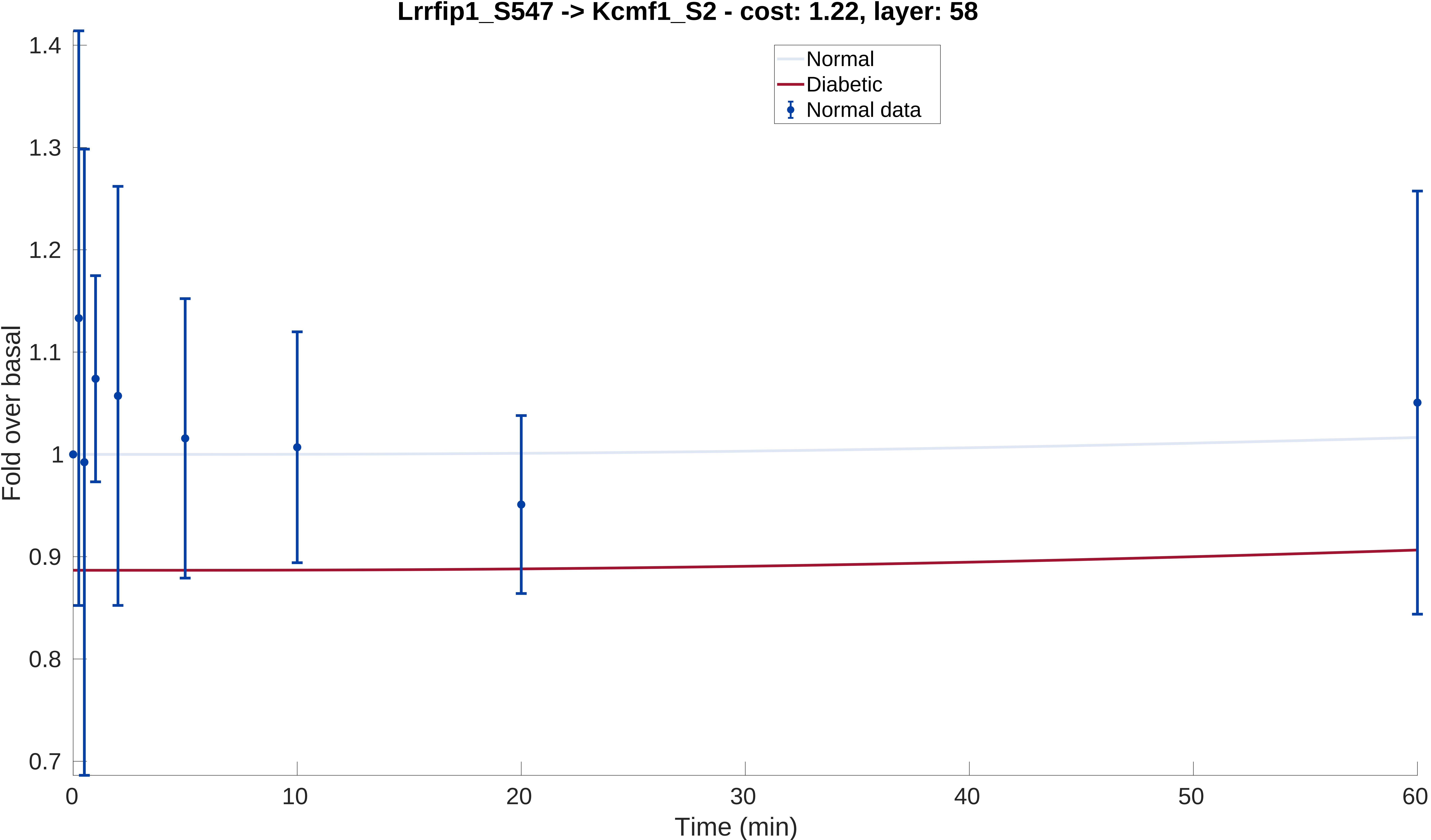

Lrrfip1_S547 -> Kcmf1_S2 - cost: 1.22, layer: 58
Normal
Diabetic
Normal data
Fold over basal
Time (min)
0.7
0.8
0.9
1
1.1
1.2
1.3
1.4
0
10
20
30
40
50
60

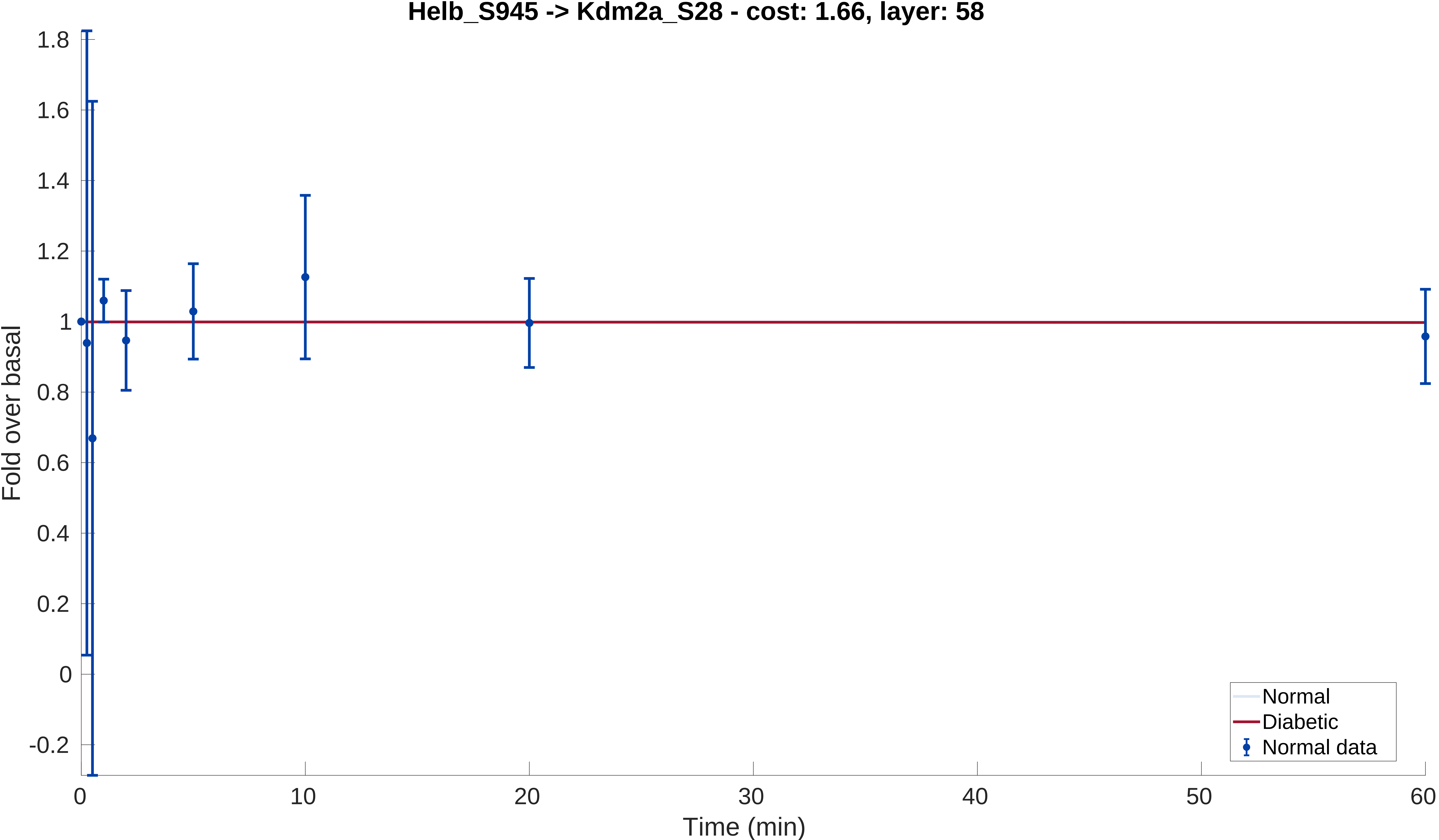

Helb_S945 -> Kdm2a_S28 - cost: 1.66, layer: 58
Fold over basal
1.8
1.6
1.4
1.2
1
0.8
0.6
0.4
0.2
0
-0.2
0
10
20
30
40
50
60
Time (min)
Normal
Diabetic
Normal data

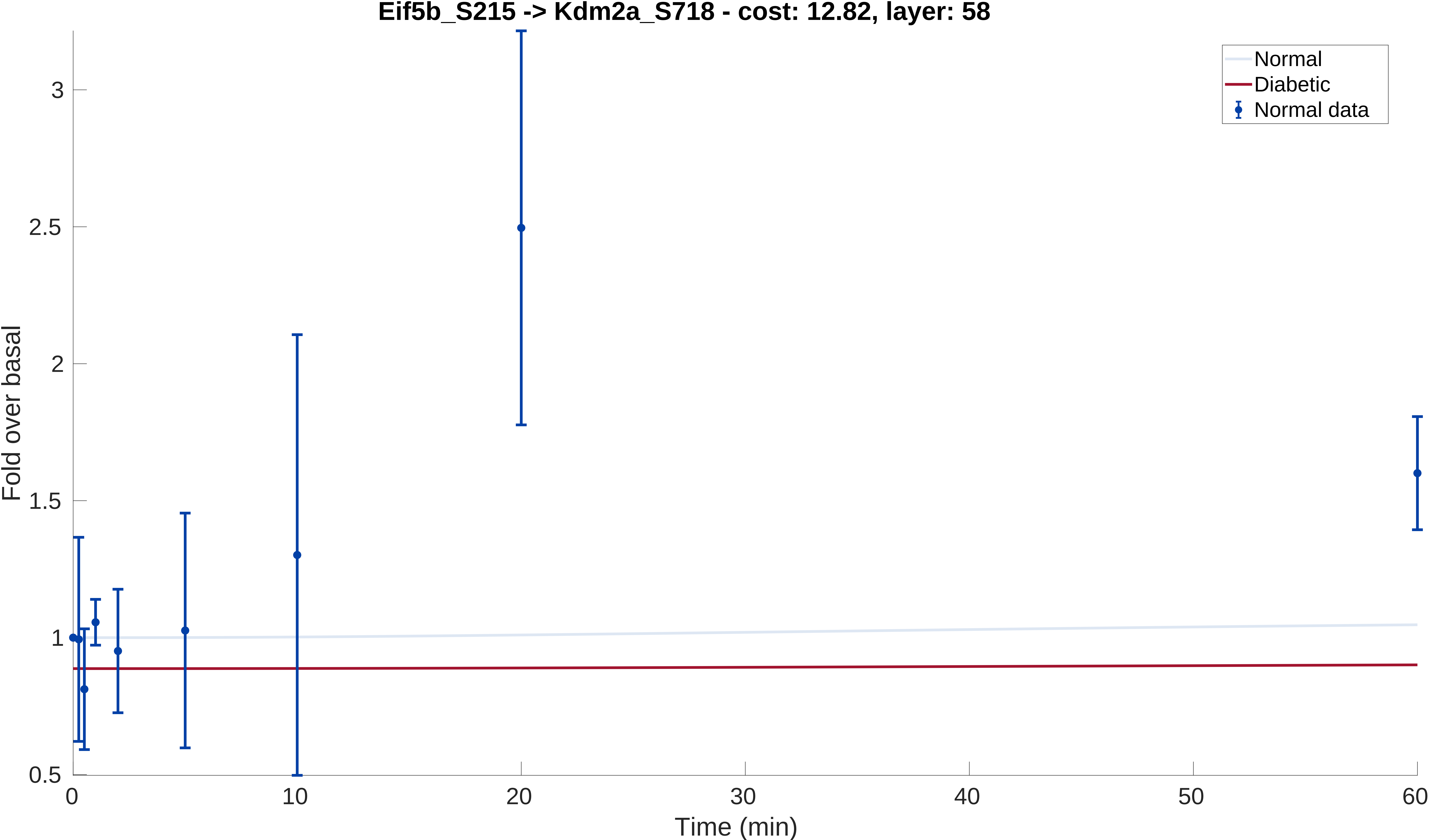

Eif5b_S215 -> Kdm2a_S718 - cost: 12.82, layer: 58
Normal
Diabetic
Normal data
Fold over basal
3
2.5
2
1.5
1
0.5
0
10
20
30
40
50
60
Time (min)

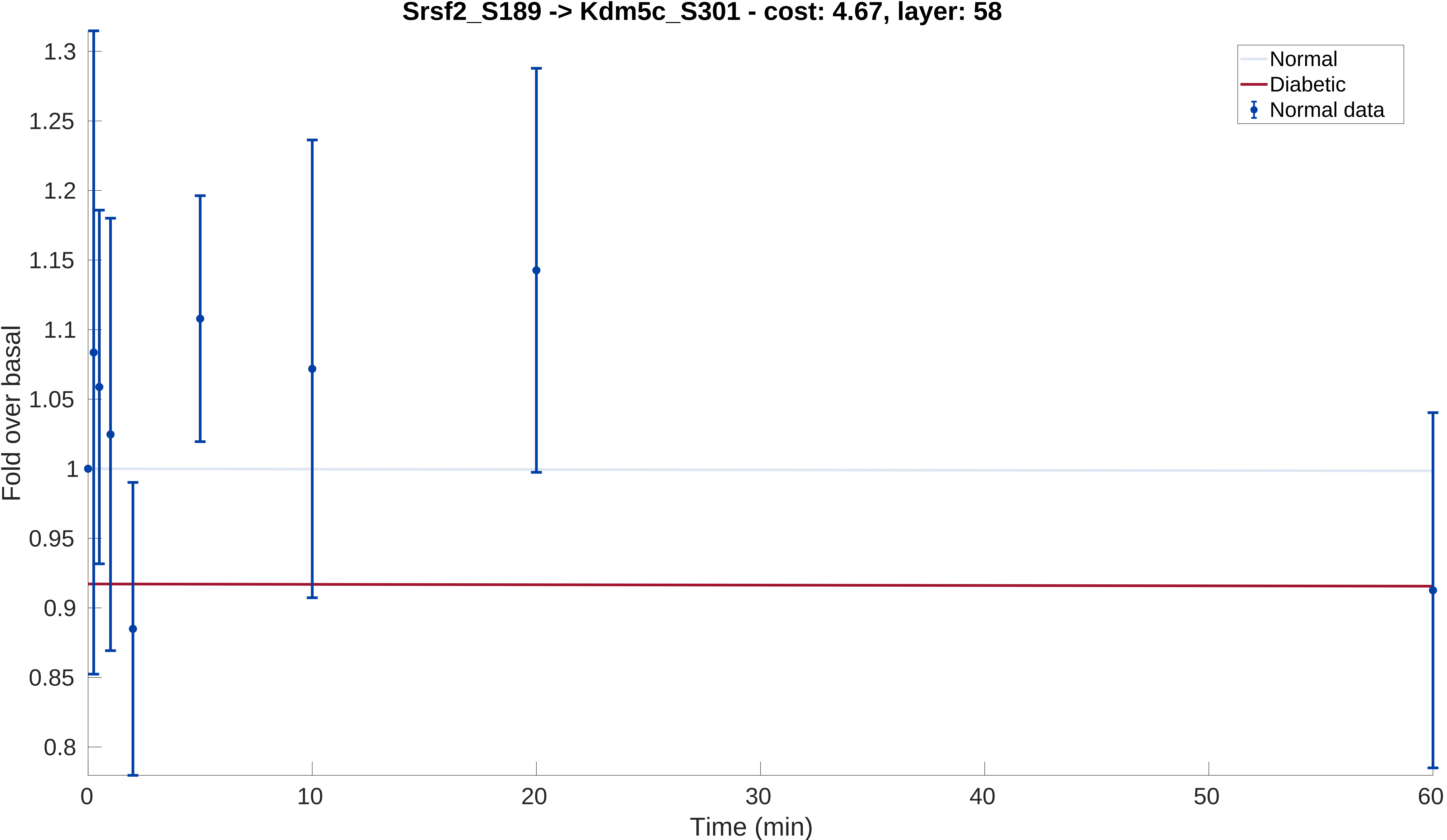

Srsf2_S189 -> Kdm5c_S301 - cost: 4.67, layer: 58
Normal
Diabetic
Normal data
Fold over basal
1.3
1.25
1.2
1.15
1.1
1.05
1
0.95
0.9
0.85
0.8
0
10
20
30
40
50
60
Time (min)

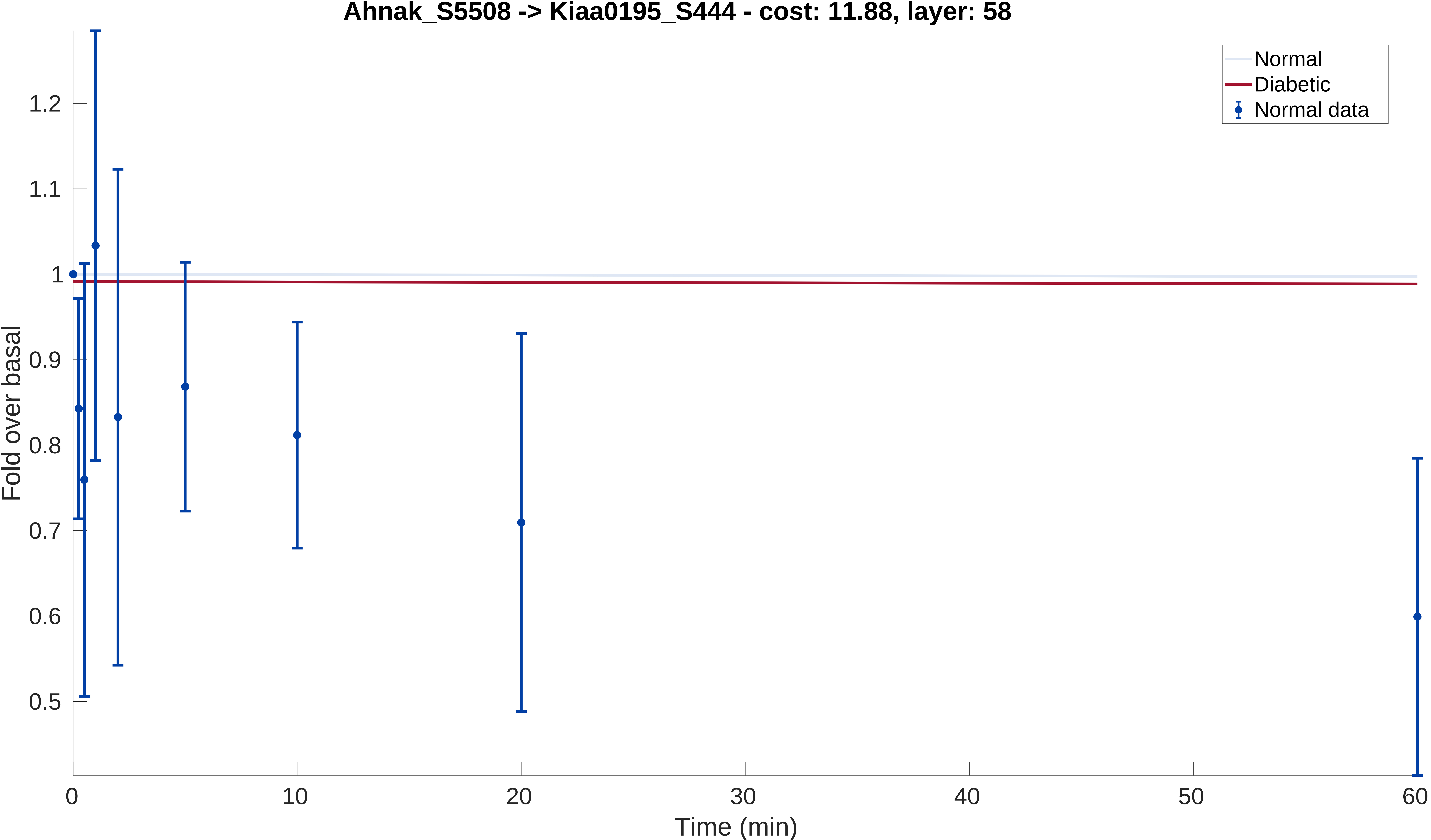

Ahnak_S5508 -> Kiaa0195_S444 - cost: 11.88, layer: 58
Normal
Diabetic
Normal data
Fold over basal
1.2
1.1
1
0.9
0.8
0.7
0.6
0.5
0
10
20
30
40
50
60
Time (min)

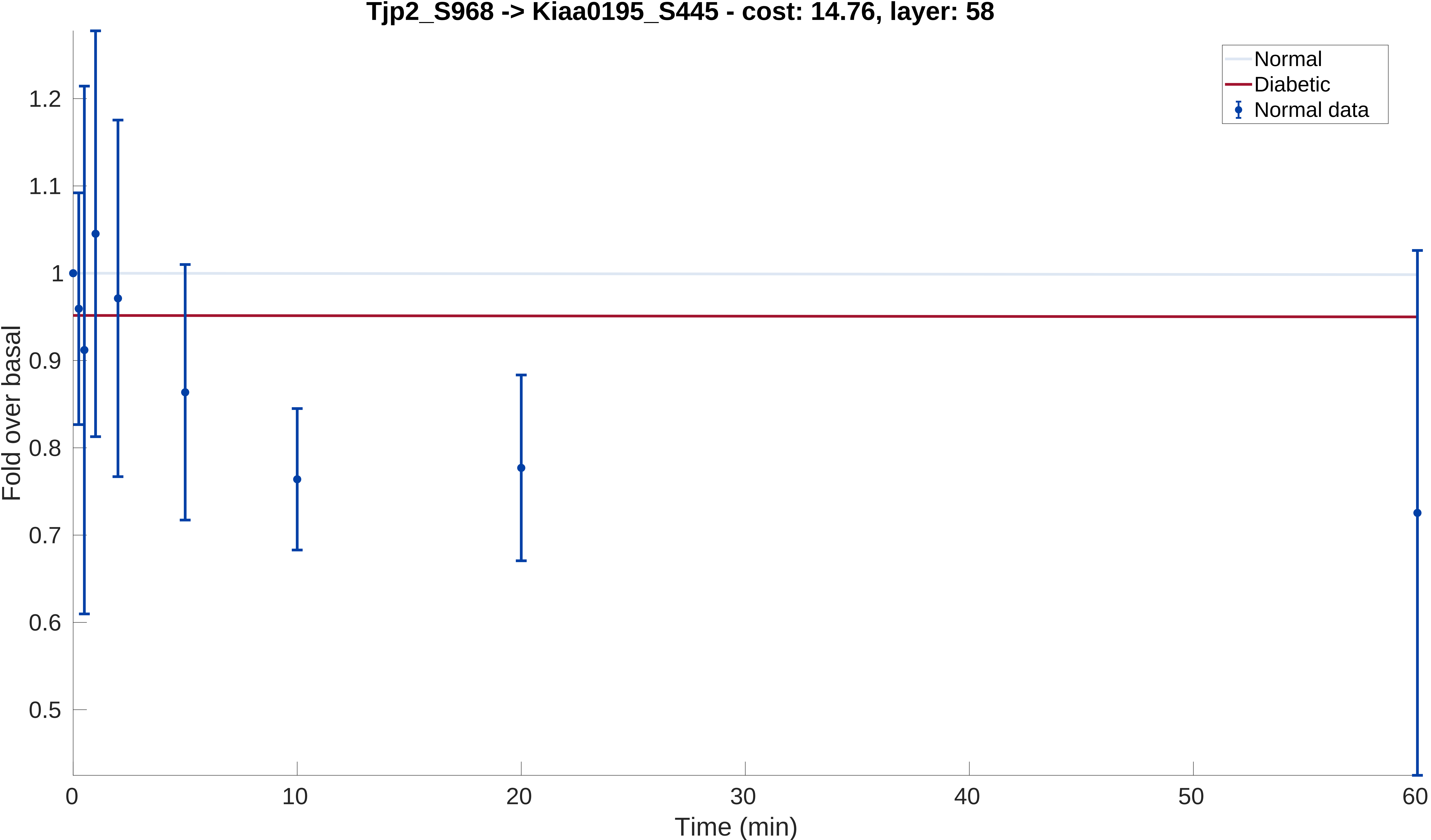

Tjp2_S968 -> Kiaa0195_S445 - cost: 14.76, layer: 58
Normal
Diabetic
Normal data
Fold over basal
Time (min)
0.5
0.6
0.7
0.8
0.9
1
1.1
1.2
0
10
20
30
40
50
60

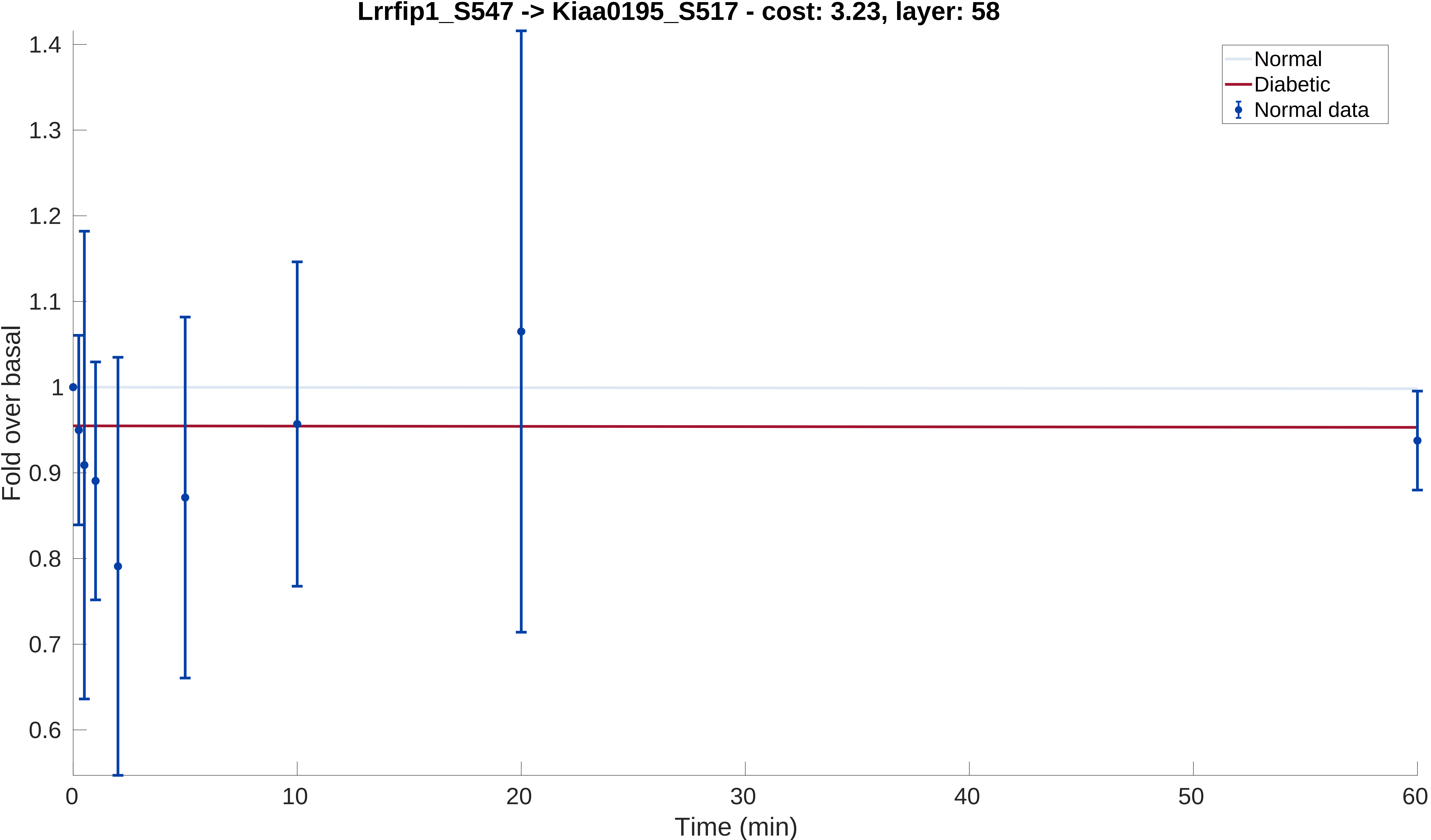

Lrrfip1_S547 -> Kiaa0195_S517 - cost: 3.23, layer: 58
Normal
Diabetic
Normal data
Fold over basal
1.4
1.3
1.2
1.1
1
0.9
0.8
0.7
0.6
0
10
20
30
40
50
60
Time (min)

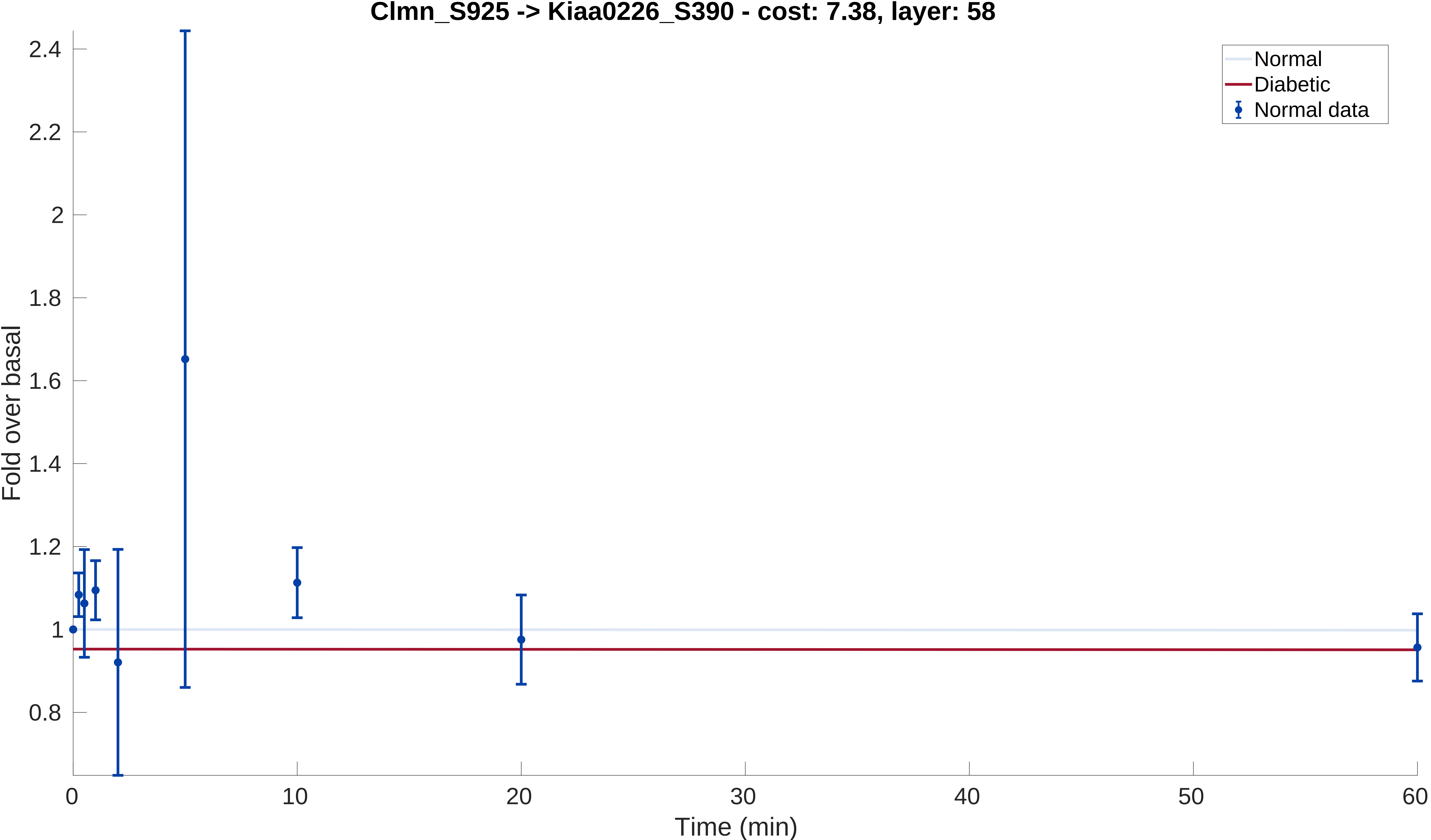

Clmn_S925 -> Kiaa0226_S390 - cost: 7.38, layer: 58
Normal
Diabetic
Normal data
Fold over basal
Time (min)
2.4
2.2
2
1.8
1.6
1.4
1.2
1
0.8
0
10
20
30
40
50
60

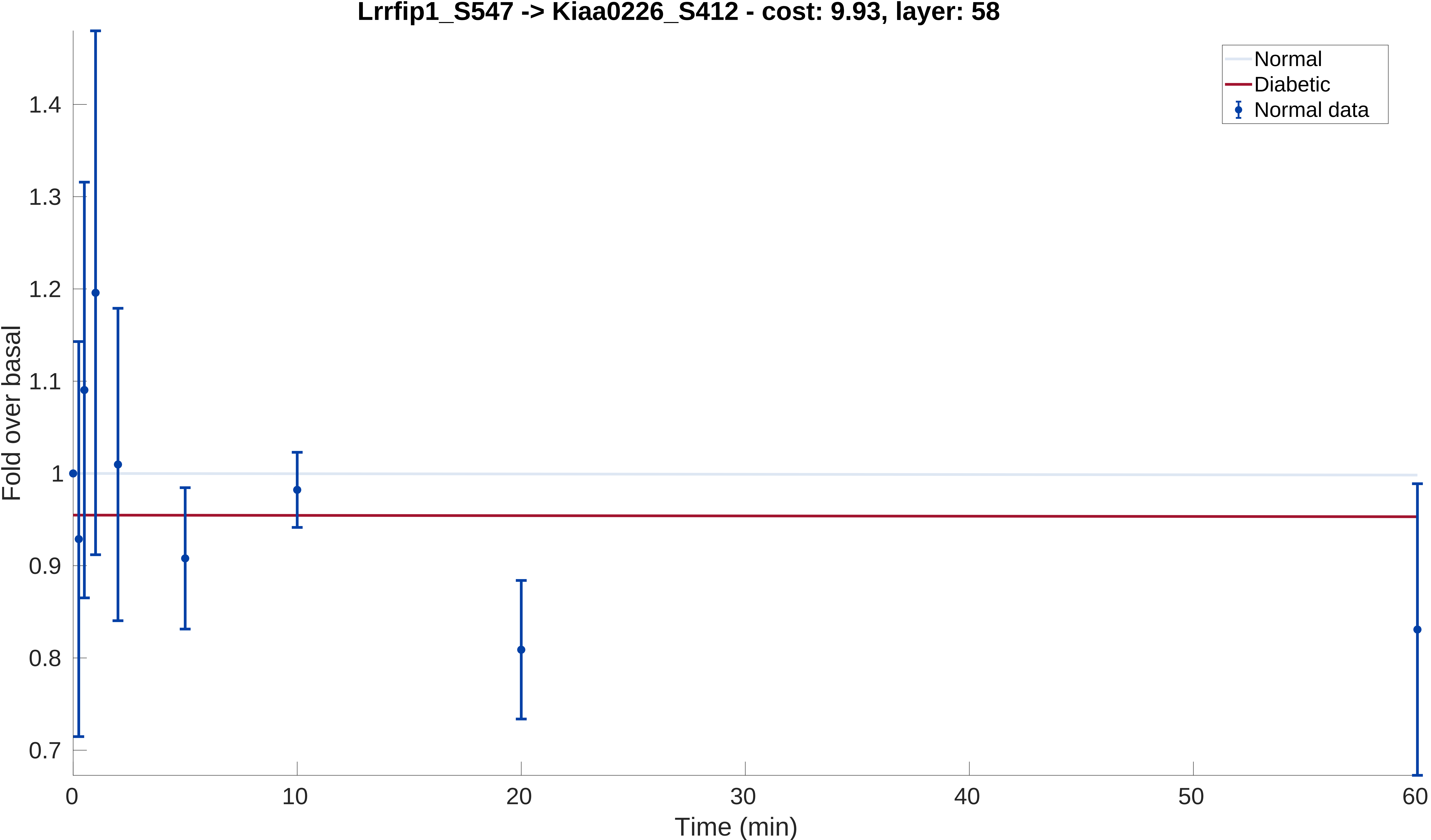

Lrrfip1_S547 -> Kiaa0226_S412 - cost: 9.93, layer: 58
Normal
Diabetic
Normal data
Fold over basal
1.4
1.3
1.2
1.1
1
0.9
0.8
0.7
0
10
20
30
40
50
60
Time (min)

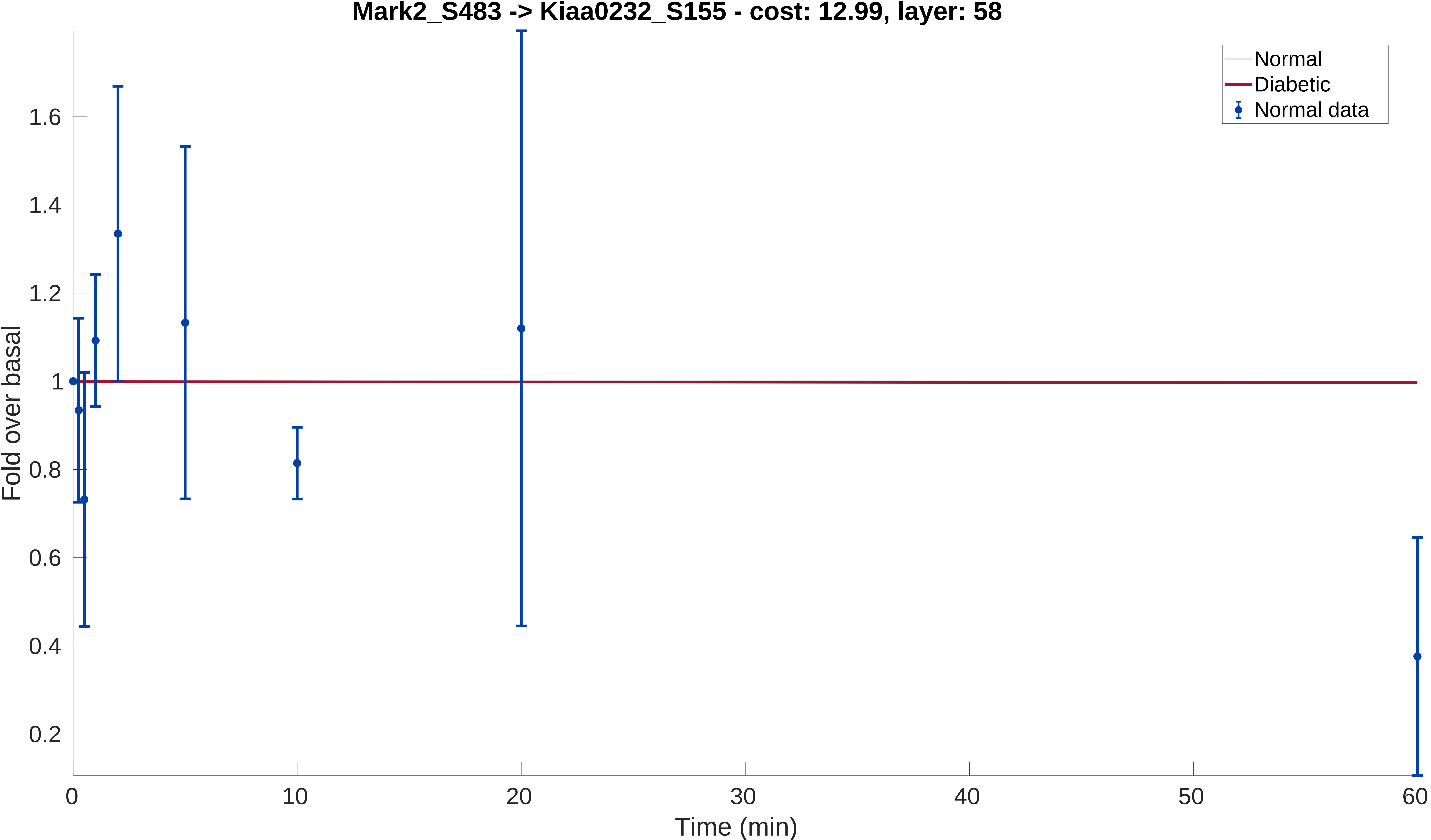

Mark2_S483 -> Kiaa0232_S155 - cost: 12.99, layer: 58
Normal
Diabetic
Normal data
Fold over basal
0.2
0.4
0.6
0.8
1
1.2
1.4
1.6
0
10
20
30
40
50
60
Time (min)

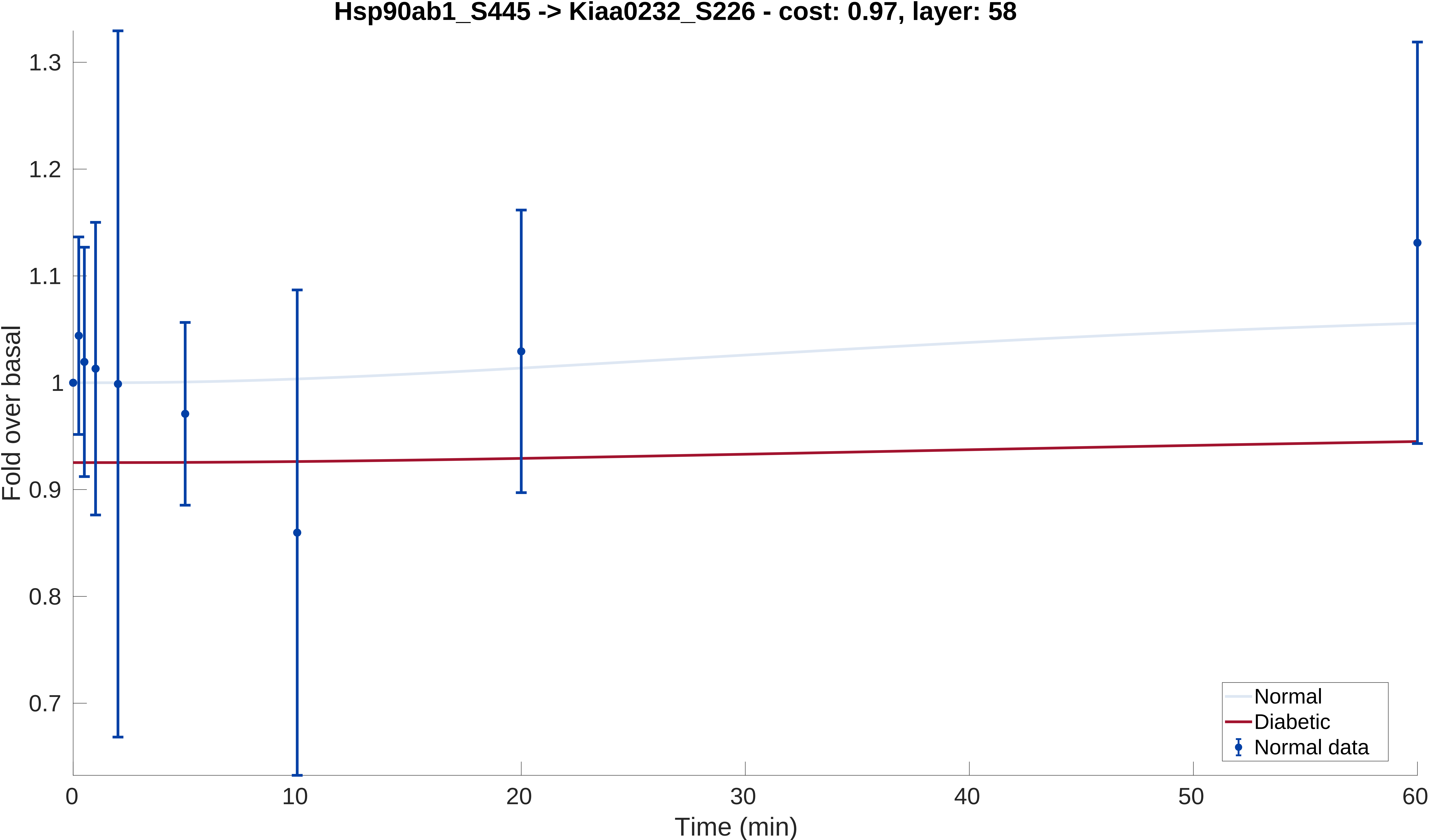
Hsp90ab1_S445 -> Kiaa0232_S226 - cost: 0.97, layer: 58
Fold over basal
Time (min)
1.3
1.2
1.1
1
0.9
0.8
0.7
0
10
20
30
40
50
60
Normal
Diabetic
Normal data

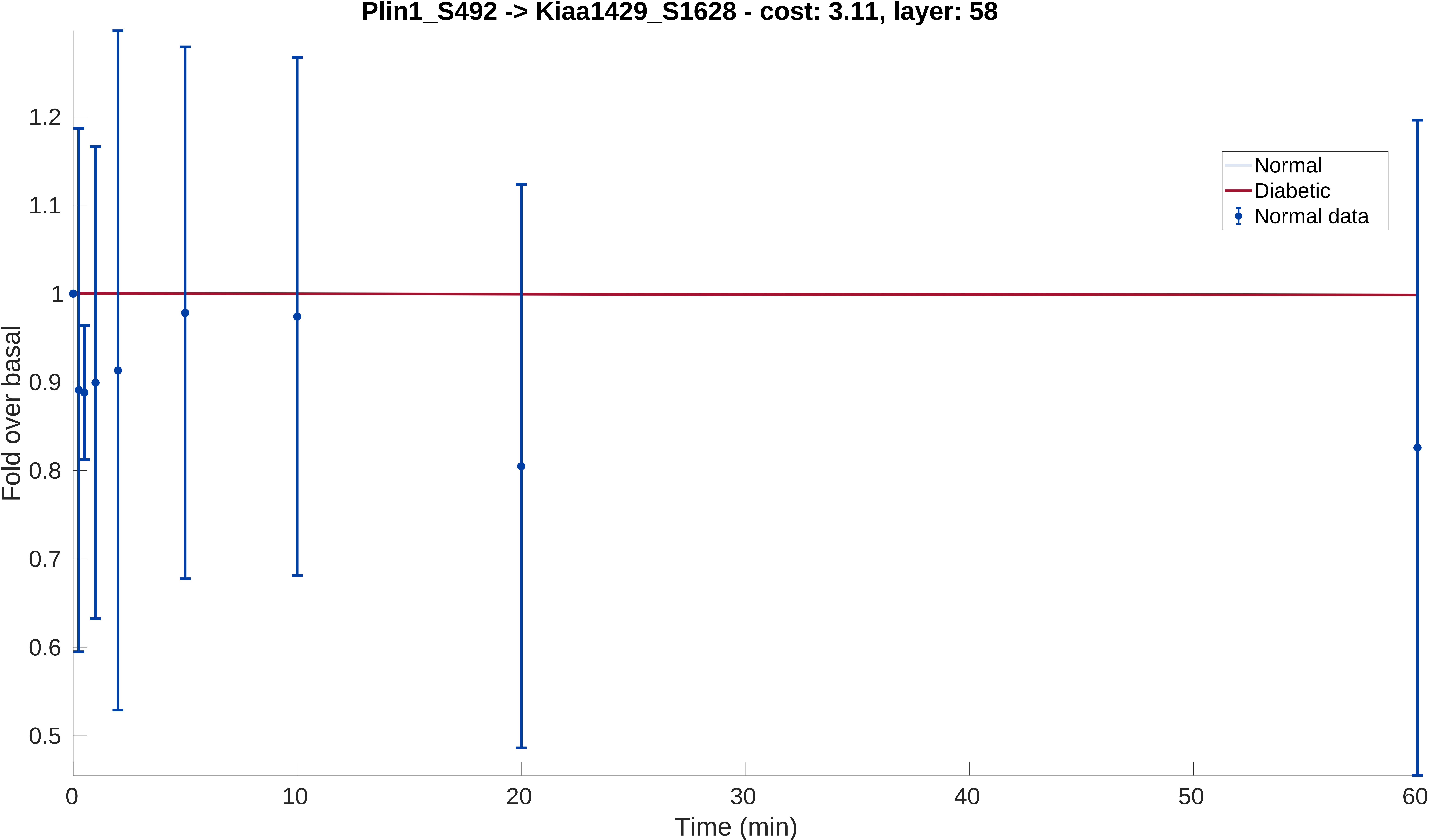

Plin1_S492 -> Kiaa1429_S1628 - cost: 3.11, layer: 58
Fold over basal
1.2
1.1
1
0.9
0.8
0.7
0.6
0.5
0
10
20
30
40
50
60
Time (min)
Normal
Diabetic
Normal data

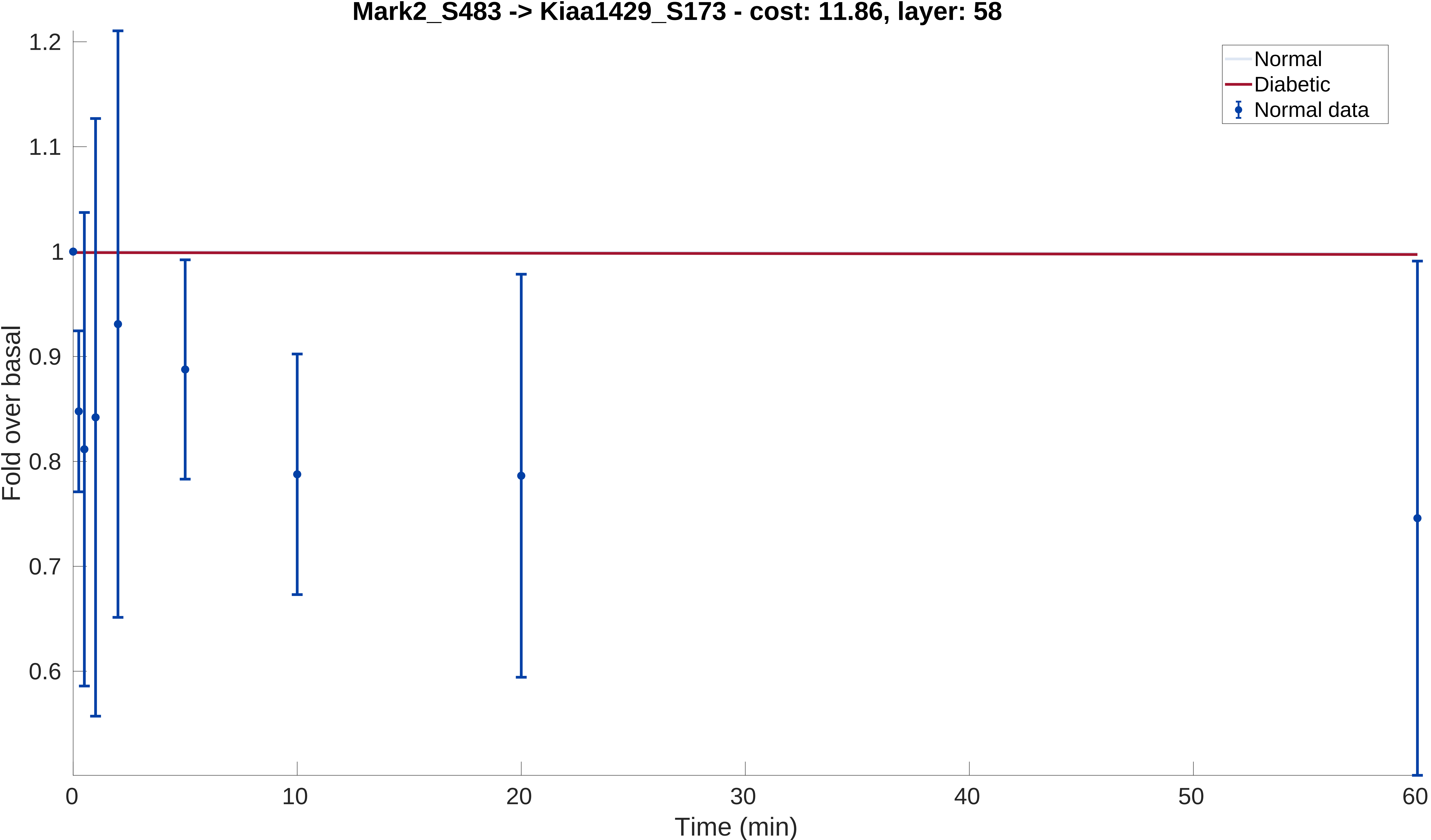

Mark2_S483 -> Kiaa1429_S173 - cost: 11.86, layer: 58
Normal
Diabetic
Normal data
Fold over basal
1.2
1.1
1
0.9
0.8
0.7
0.6
0
10
20
30
40
50
60
Time (min)

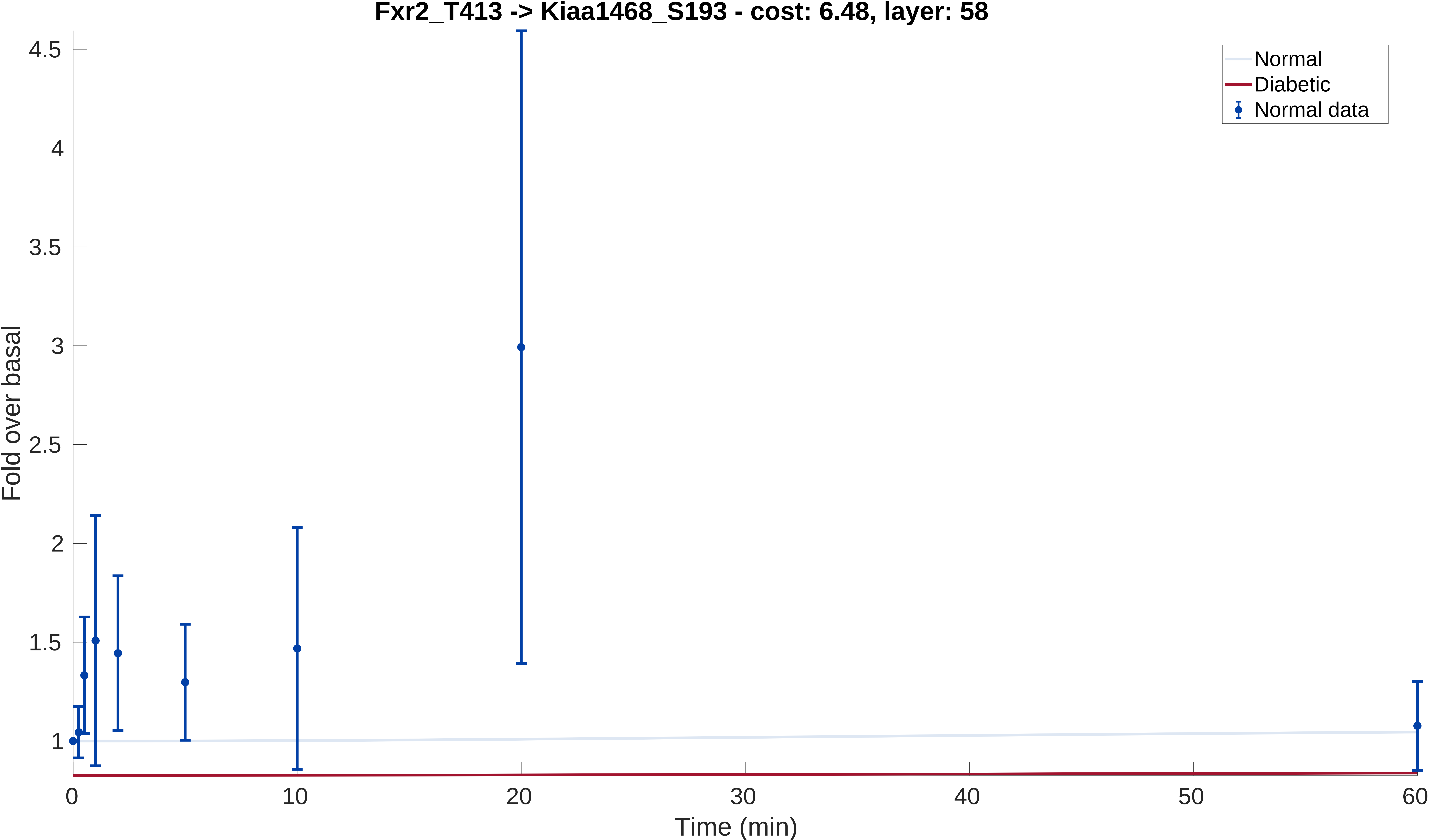
Fxr2_T413 -> Kiaa1468_S193 - cost: 6.48, layer: 58
Normal
Diabetic
Normal data
Fold over basal
Time (min)
4.5
4
3.5
3
2.5
2
1.5
1
0
10
20
30
40
50
60

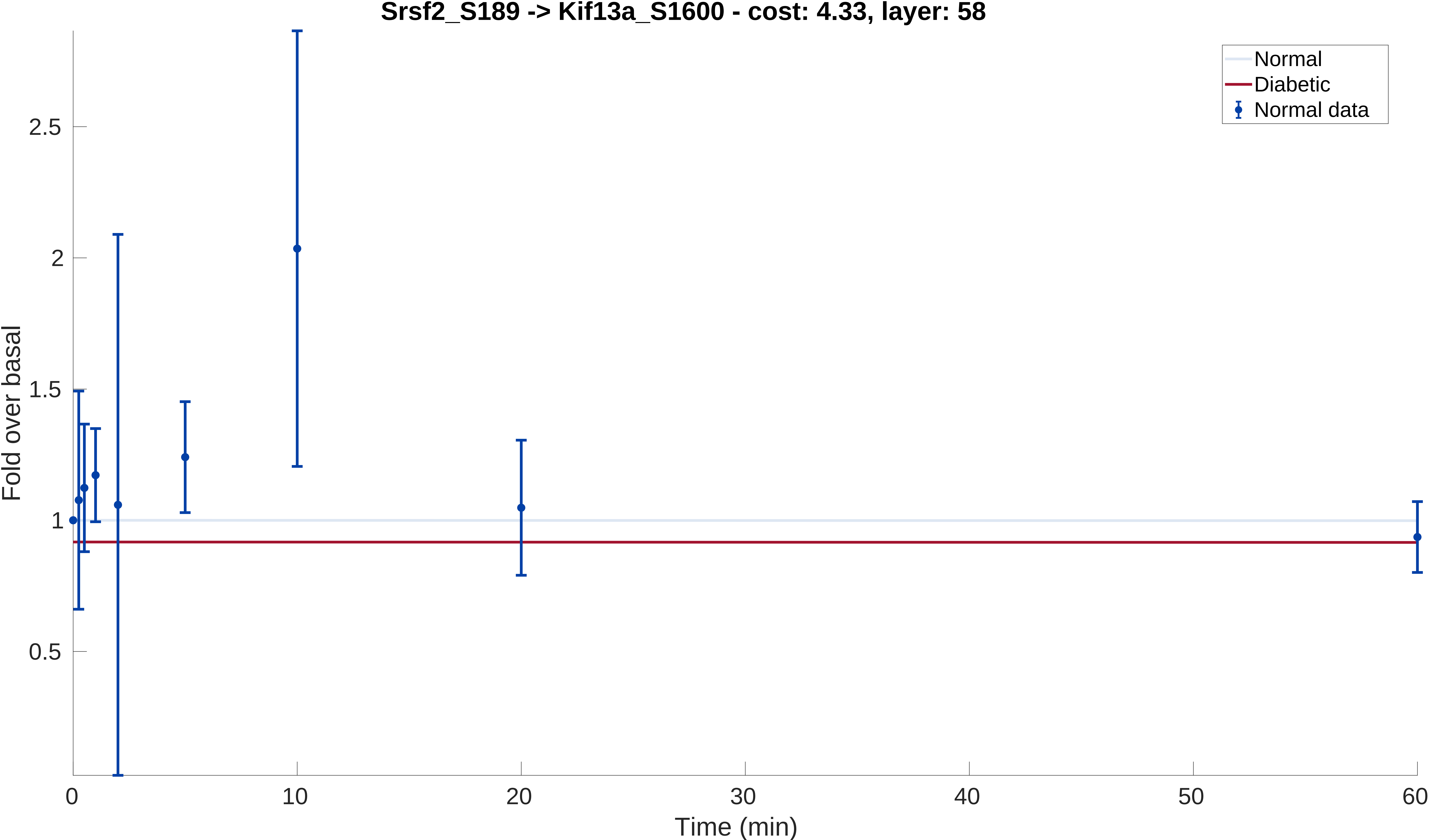
Srsf2_S189 -> Kif13a_S1600 - cost: 4.33, layer: 58
Normal
Diabetic
Normal data
Fold over basal
2.5
2
1.5
1
0.5
0
10
20
30
40
50
60
Time (min)

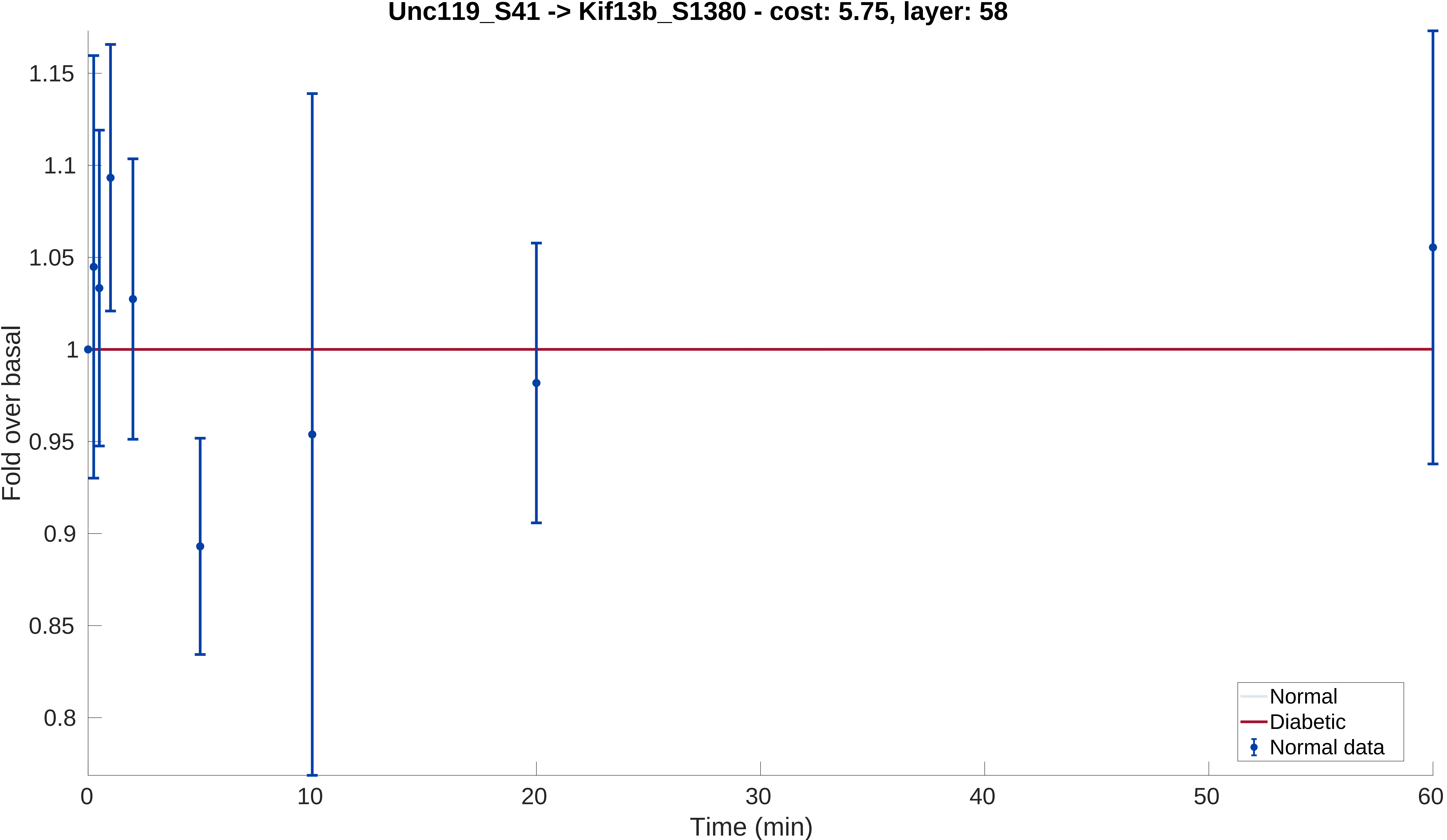

Unc119_S41 -> Kif13b_S1380 - cost: 5.75, layer: 58
Fold over basal
1.15
1.1
1.05
1
0.95
0.9
0.85
0.8
0
10
20
30
40
50
60
Time (min)
Normal
Diabetic
Normal data

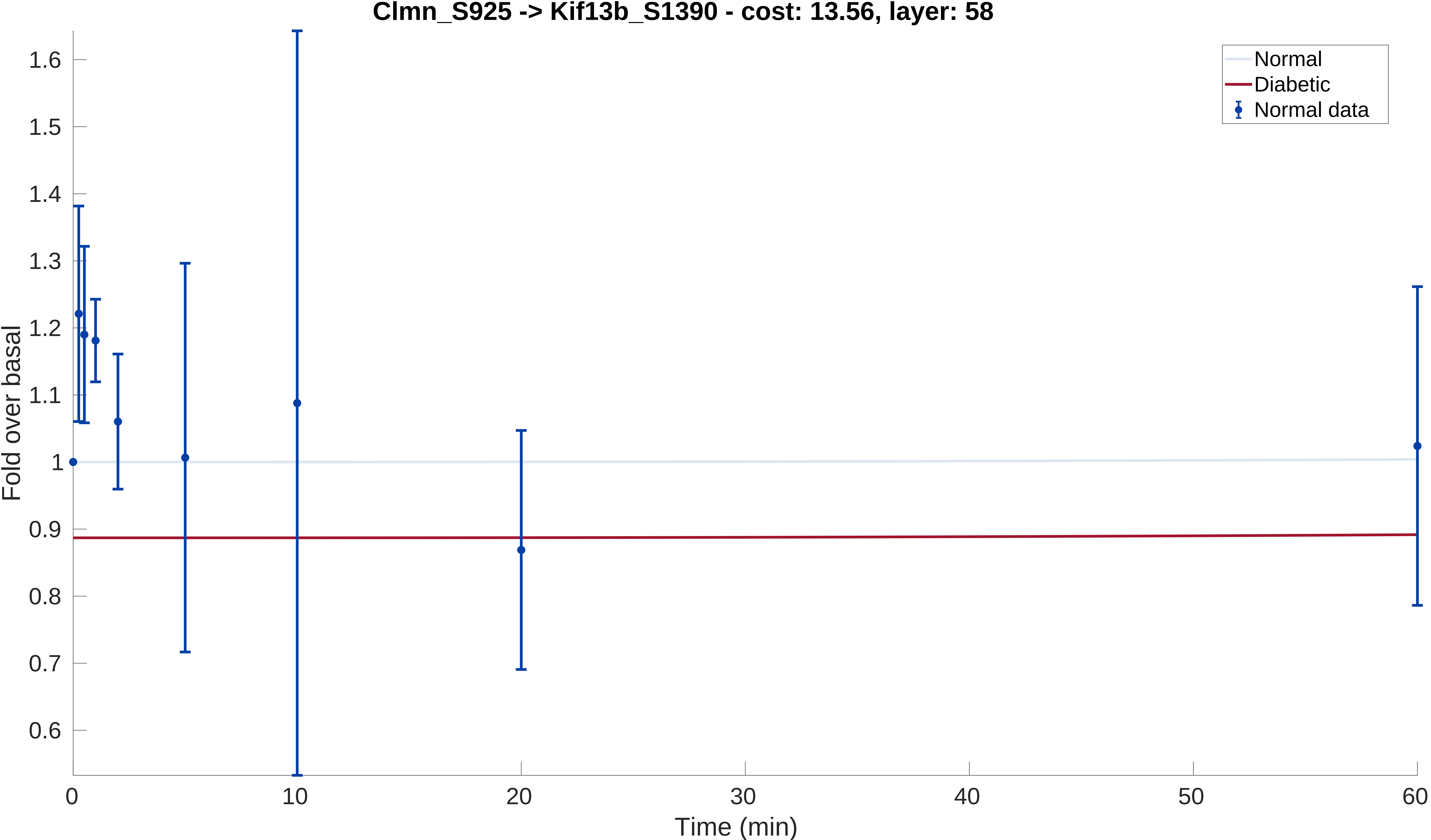
Clmn_S925 -> Kif13b_S1390 - cost: 13.56, layer: 58
Normal
Diabetic
Normal data
Fold over basal
1.6
1.5
1.4
1.3
1.2
1.1
1
0.9
0.8
0.7
0.6
0
10
20
30
40
50
60
Time (min)

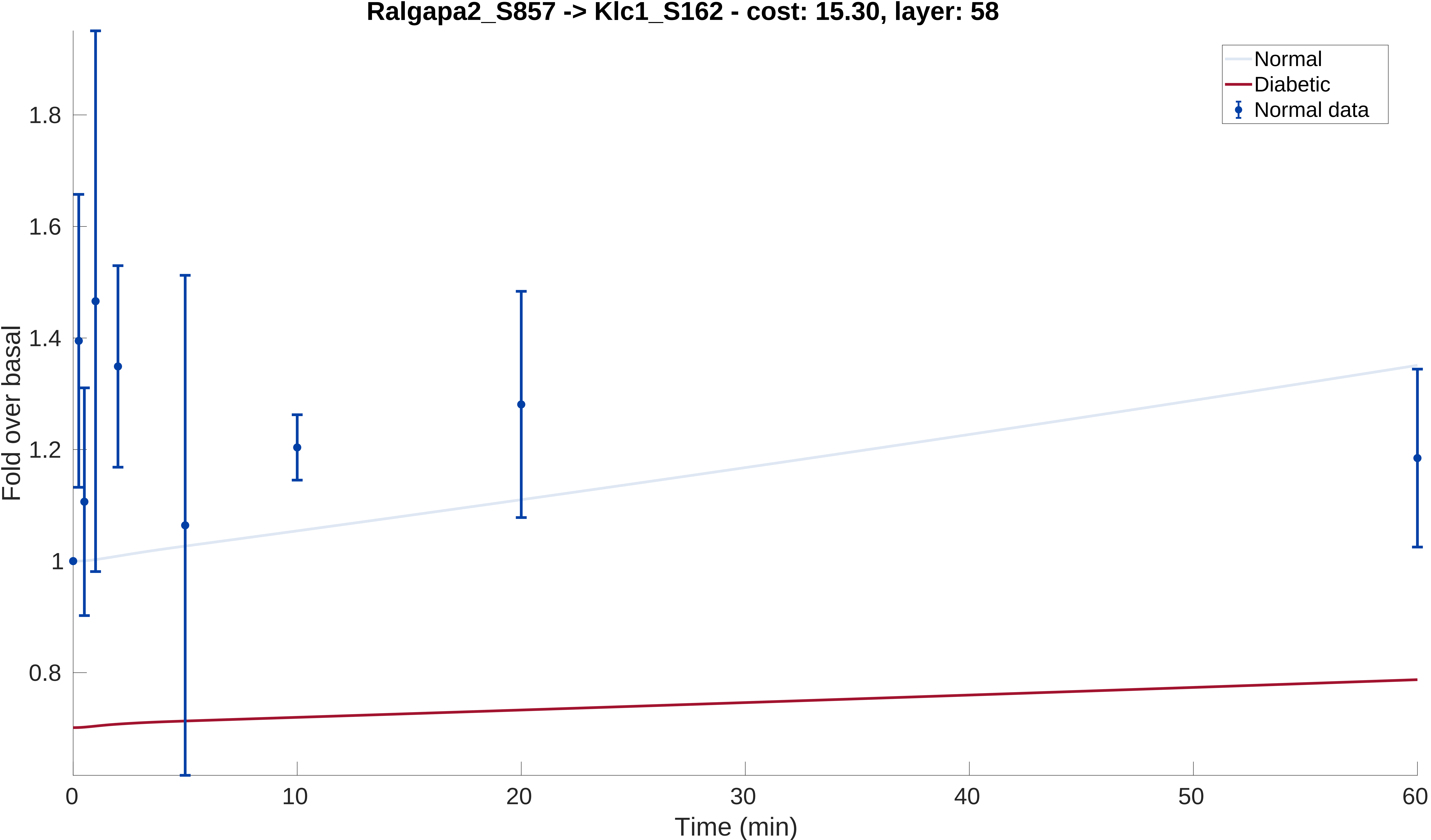

Ralgapa2_S857 -> Klc1_S162 - cost: 15.30, layer: 58
Normal
Diabetic
Normal data
Fold over basal
1.8
1.6
1.4
1.2
1
0.8
0
10
20
30
40
50
60
Time (min)

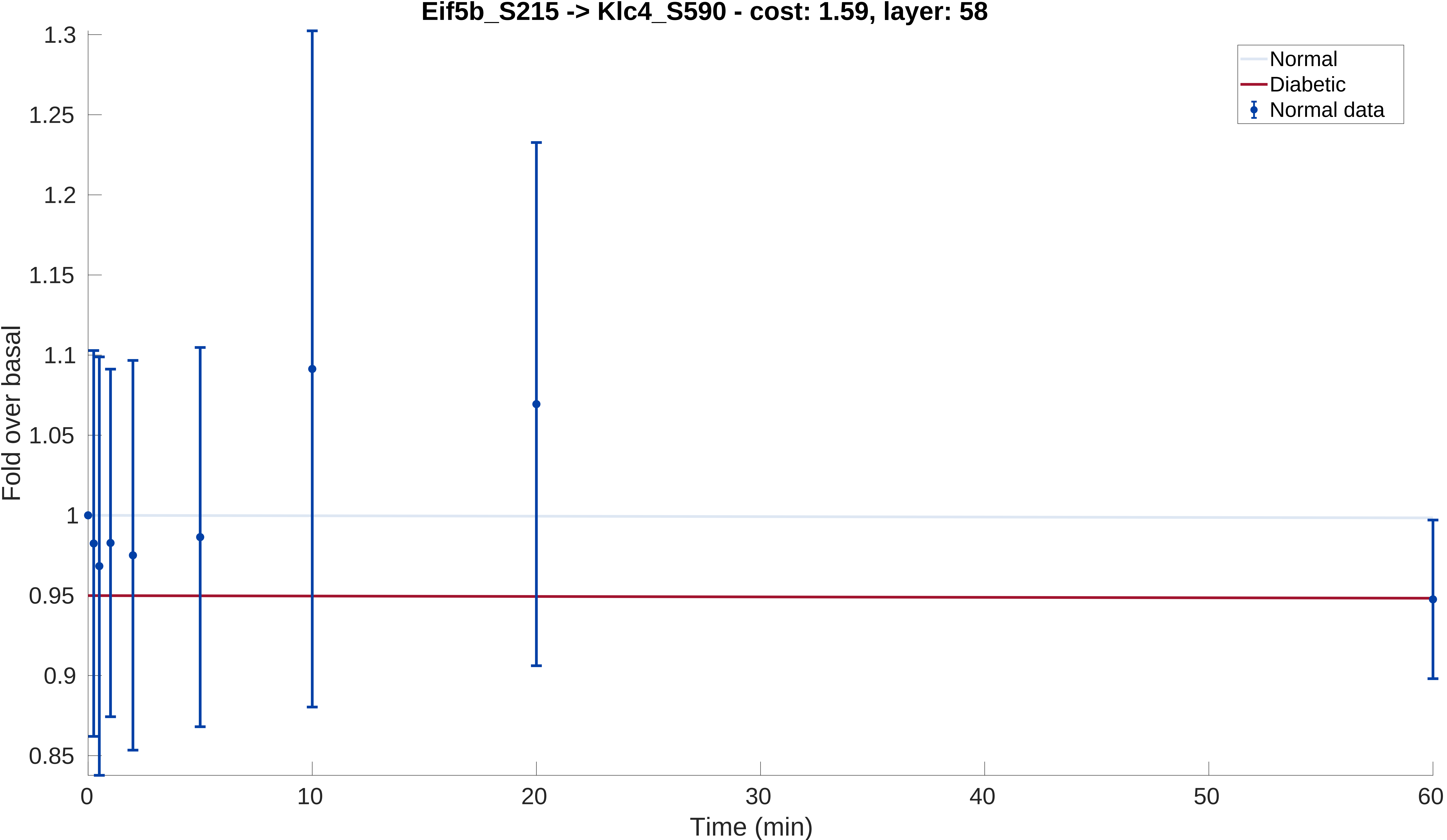
Eif5b_S215 -> Klc4_S590 - cost: 1.59, layer: 58
Normal
Diabetic
Normal data
Fold over basal
Time (min)
1.3
1.25
1.2
1.15
1.1
1.05
1
0.95
0.9
0.85
0
10
20
30
40
50
60

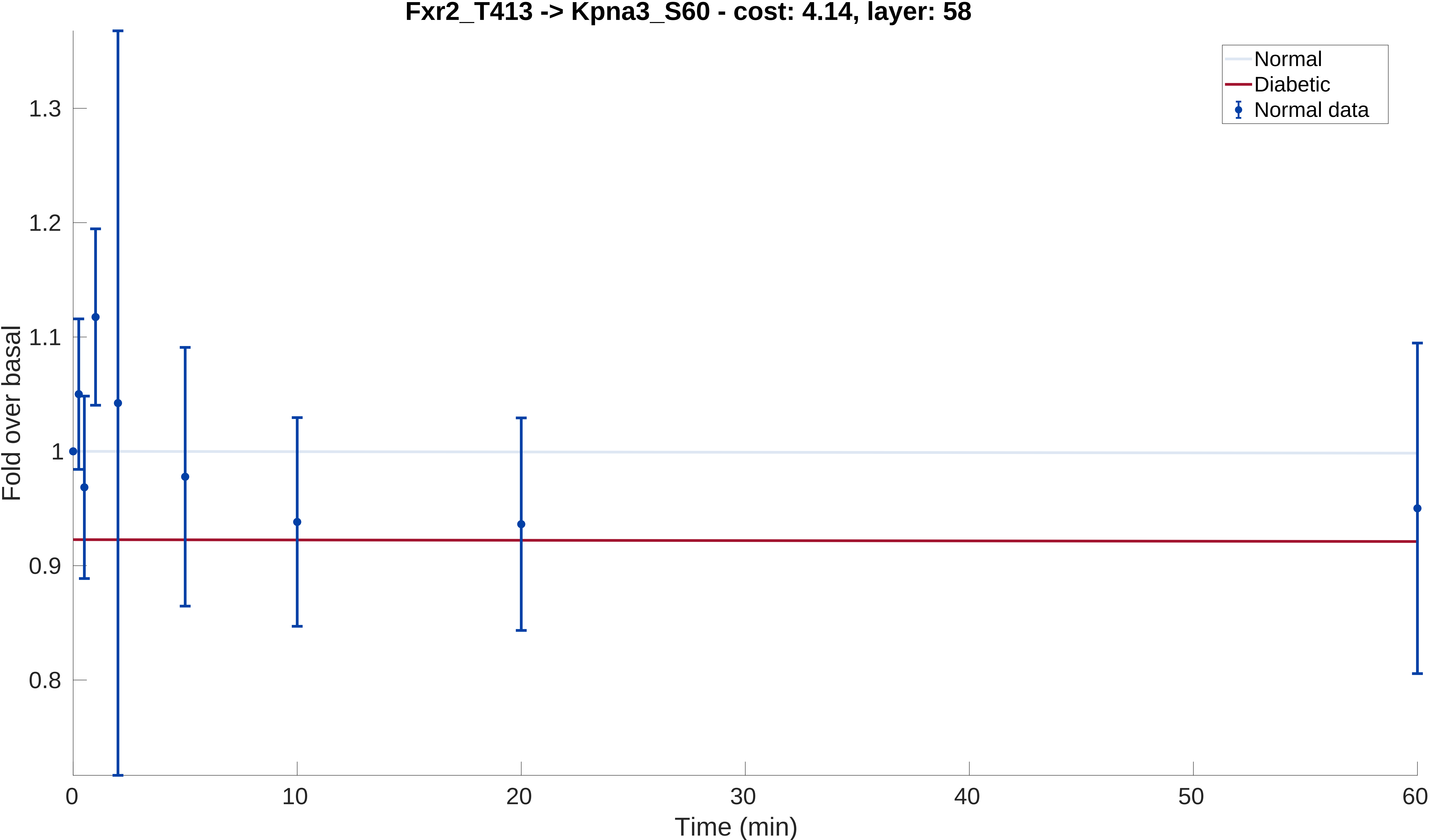

Fxr2_T413 -> Kpna3_S60 - cost: 4.14, layer: 58
Normal
Diabetic
Normal data
Fold over basal
1.3
1.2
1.1
1
0.9
0.8
0
10
20
30
40
50
60
Time (min)

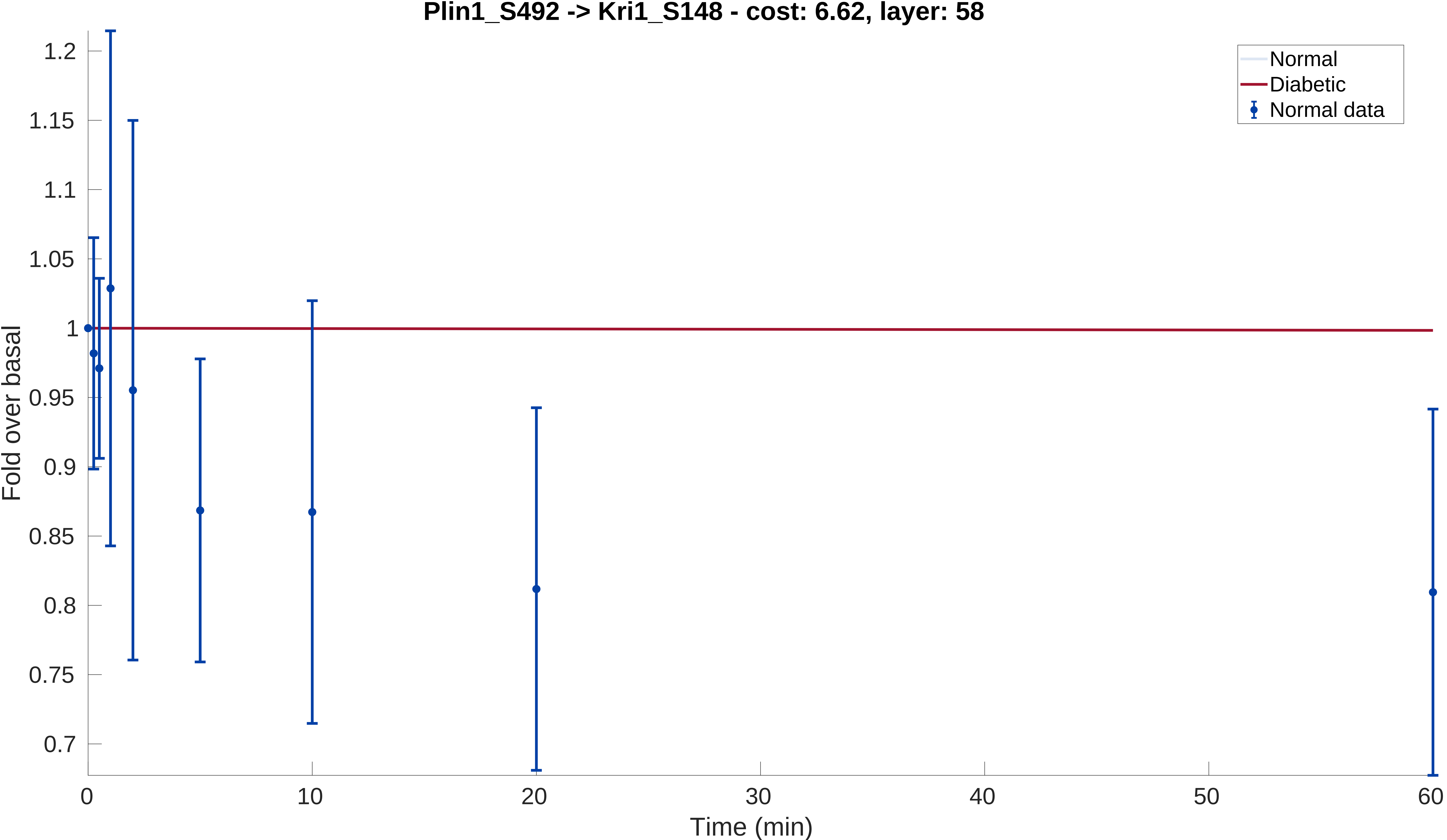
Plin1_S492 -> Kri1_S148 - cost: 6.62, layer: 58
Fold over basal
Time (min)
1.2
1.15
1.1
1.05
1
0.95
0.9
0.85
0.8
0.75
0.7
0
10
20
30
40
50
60
Normal
Diabetic
Normal data

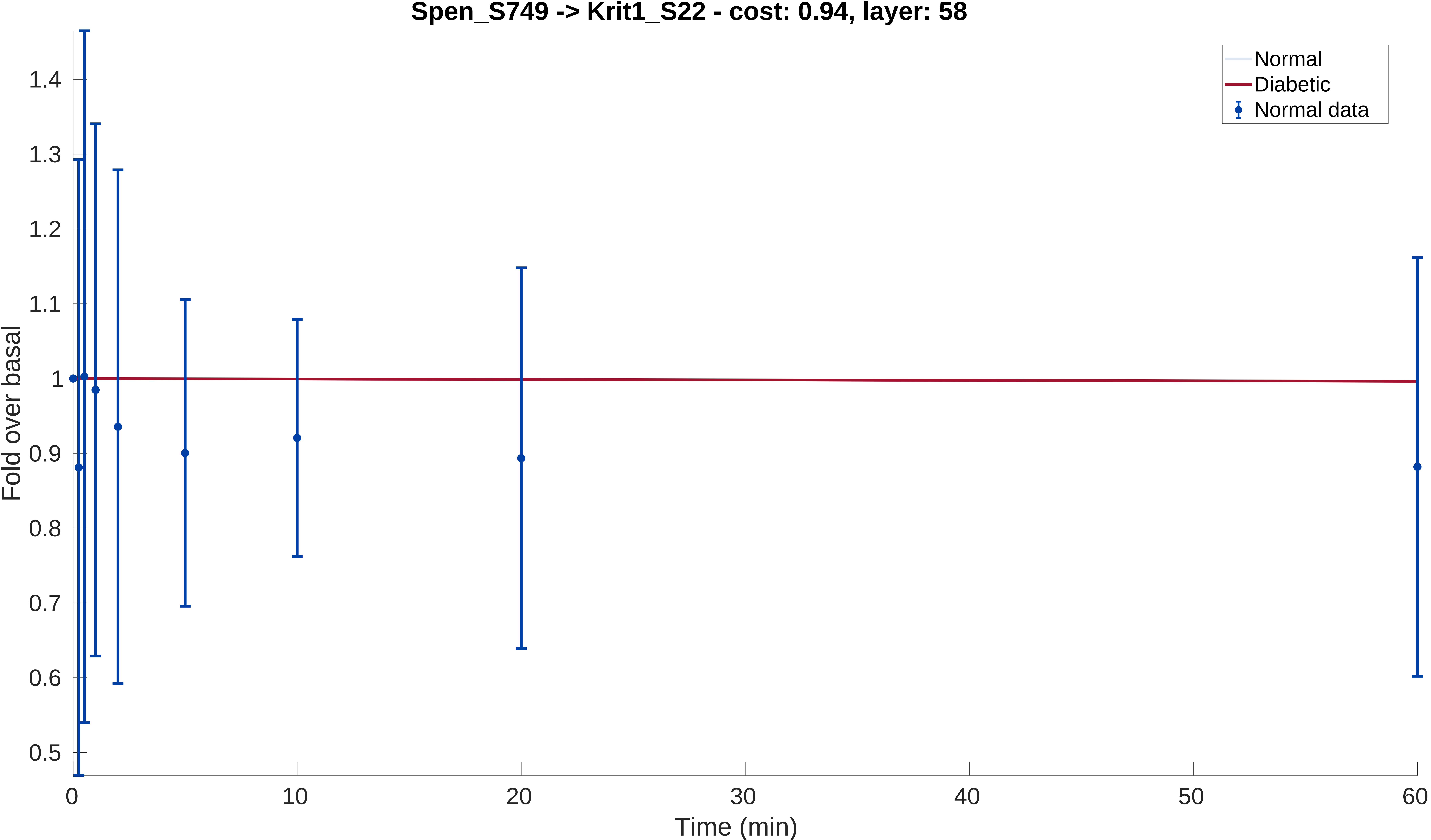
Spen_S749 -> Krit1_S22 - cost: 0.94, layer: 58
Normal
Diabetic
Normal data
Fold over basal
1.4
1.3
1.2
1.1
1
0.9
0.8
0.7
0.6
0.5
0
10
20
30
40
50
60
Time (min)

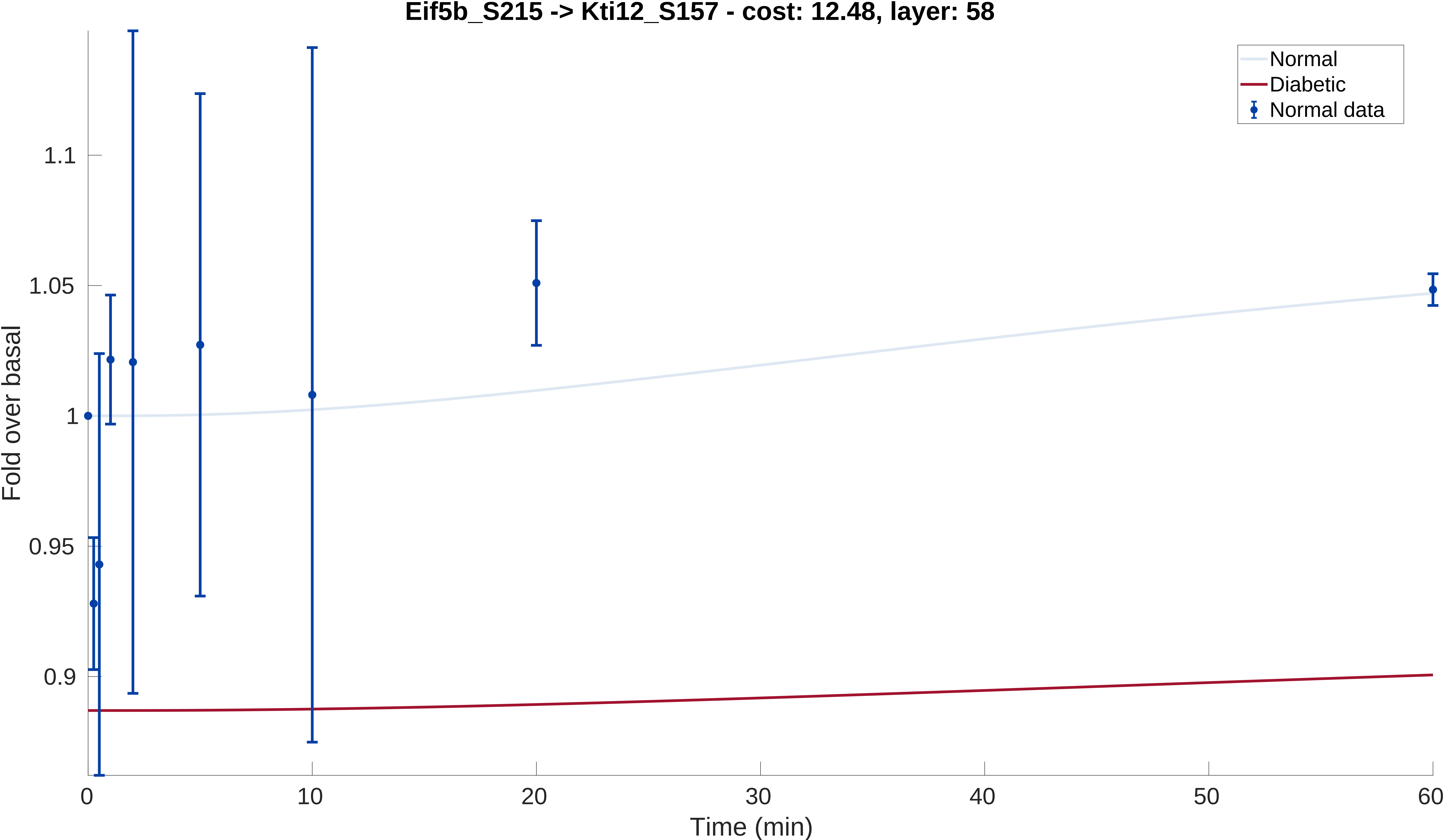
Eif5b_S215 -> Kti12_S157 - cost: 12.48, layer: 58
Normal
Diabetic
Normal data
1.1
1.05
1
0.95
0.9
Fold over basal
0
10
20
30
40
50
60
Time (min)

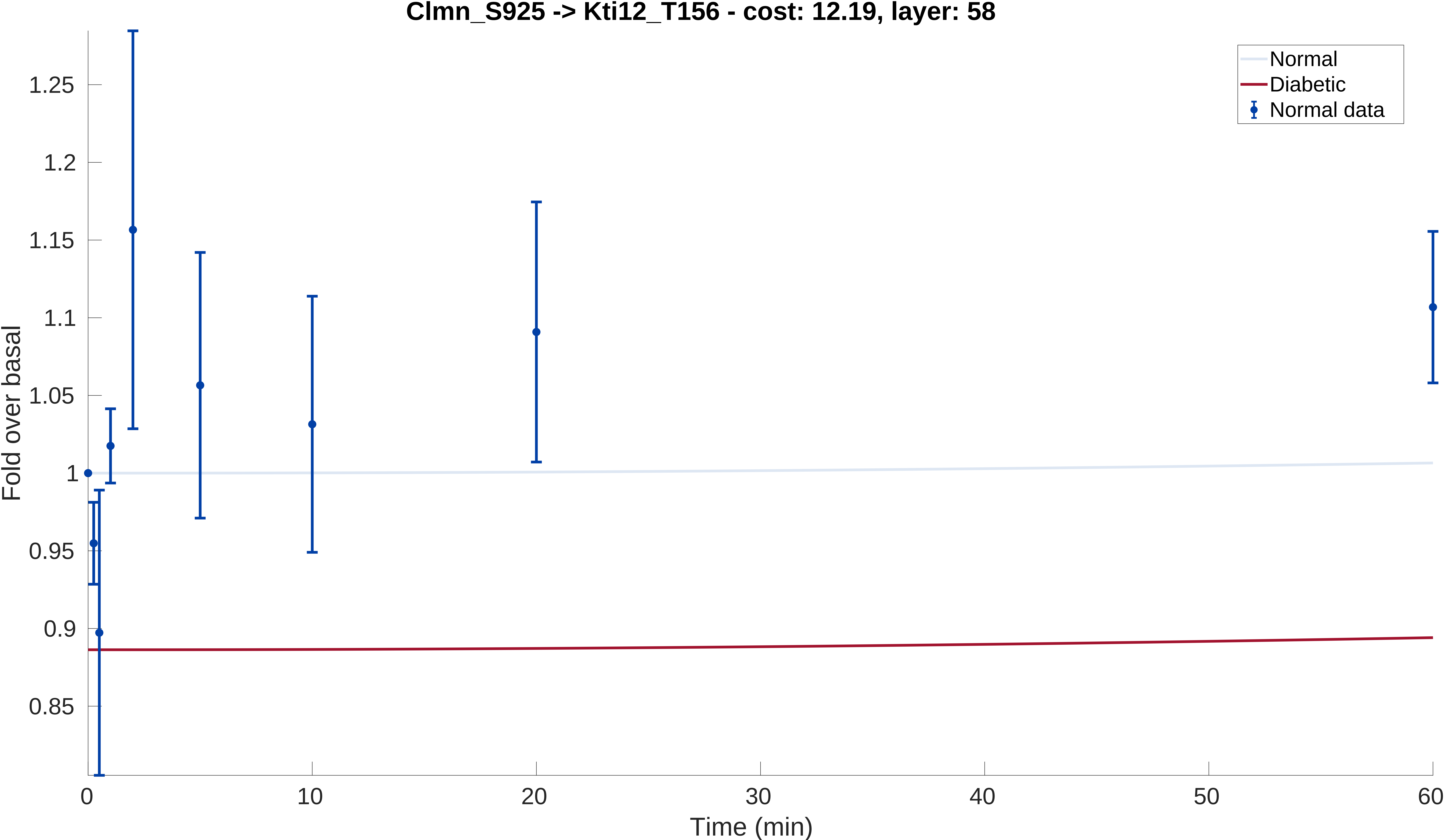

Clmn_S925 -> Kti12_T156 - cost: 12.19, layer: 58
Normal
Diabetic
Normal data
1.25
1.2
1.15
1.1
1.05
1
0.95
0.9
0.85
Fold over basal
0
10
20
30
40
50
60
Time (min)

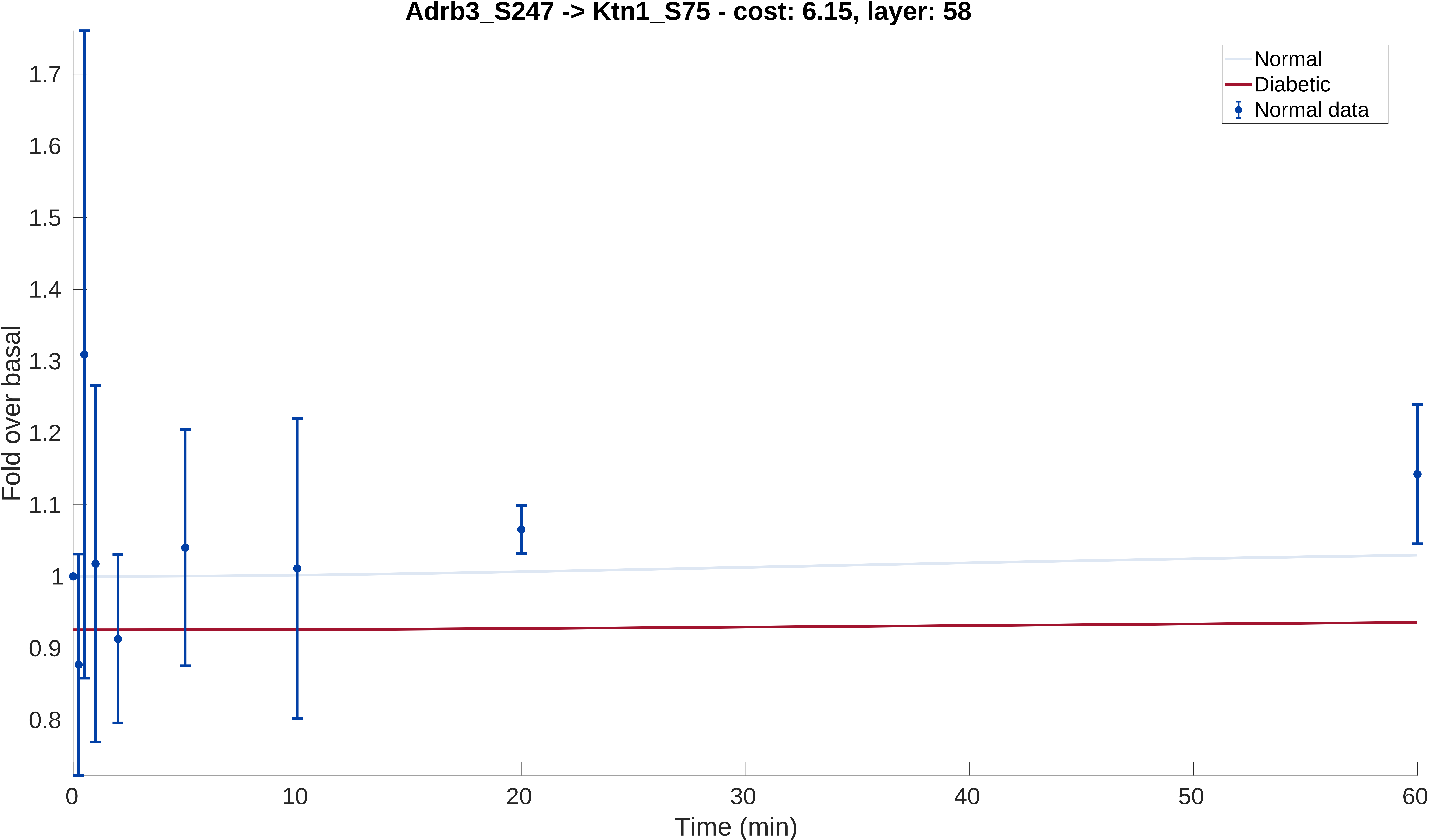

Adrb3_S247 -> Ktn1_S75 - cost: 6.15, layer: 58
Normal
Diabetic
Normal data
Fold over basal
1.7
1.6
1.5
1.4
1.3
1.2
1.1
1
0.9
0.8
0
10
20
30
40
50
60
Time (min)

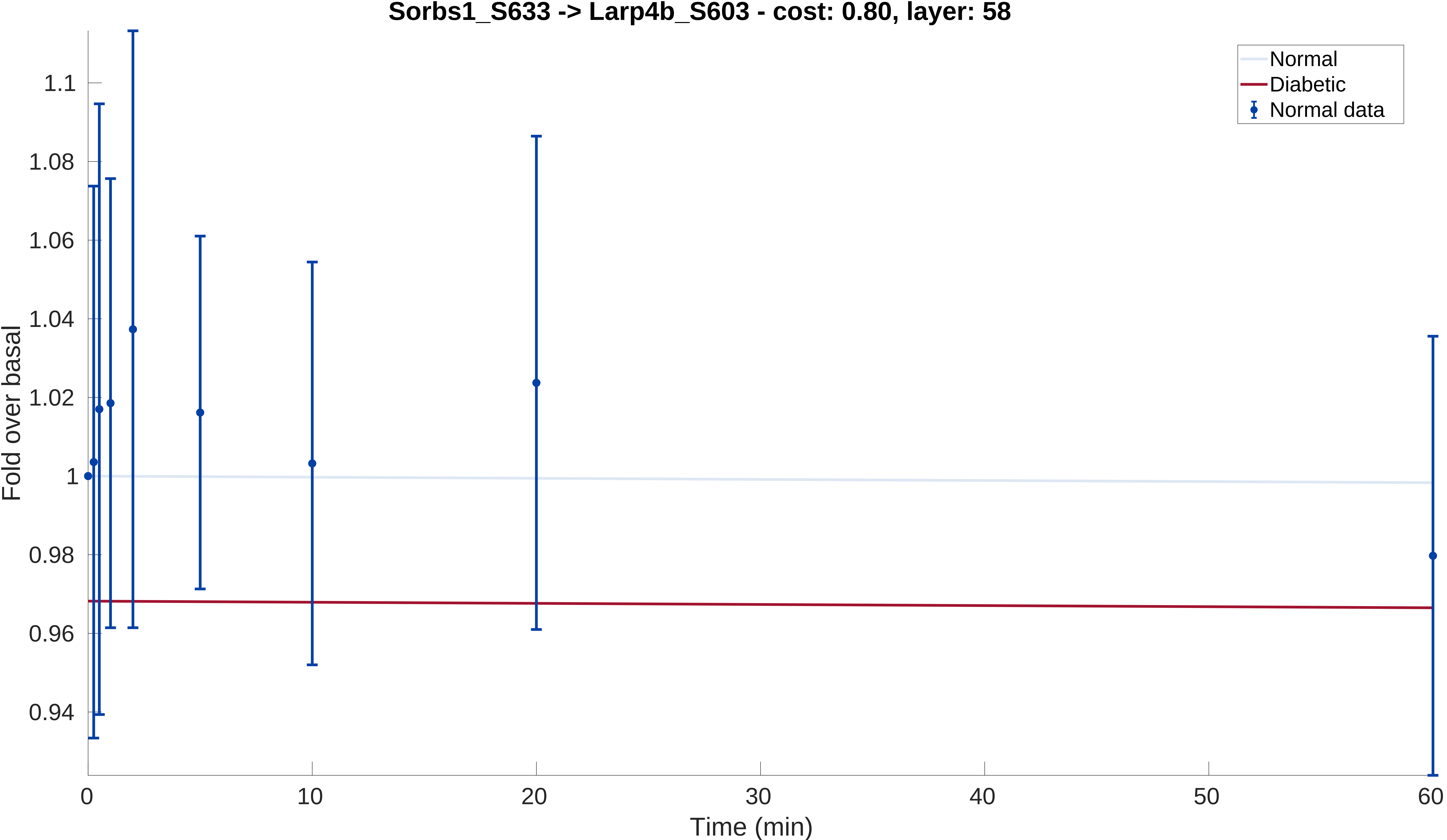
Sorbs1_S633 -> Larp4b_S603 - cost: 0.80, layer: 58
Normal
Diabetic
Normal data
Fold over basal
1.1
1.08
1.06
1.04
1.02
1
0.98
0.96
0.94
0
10
20
30
40
50
60
Time (min)

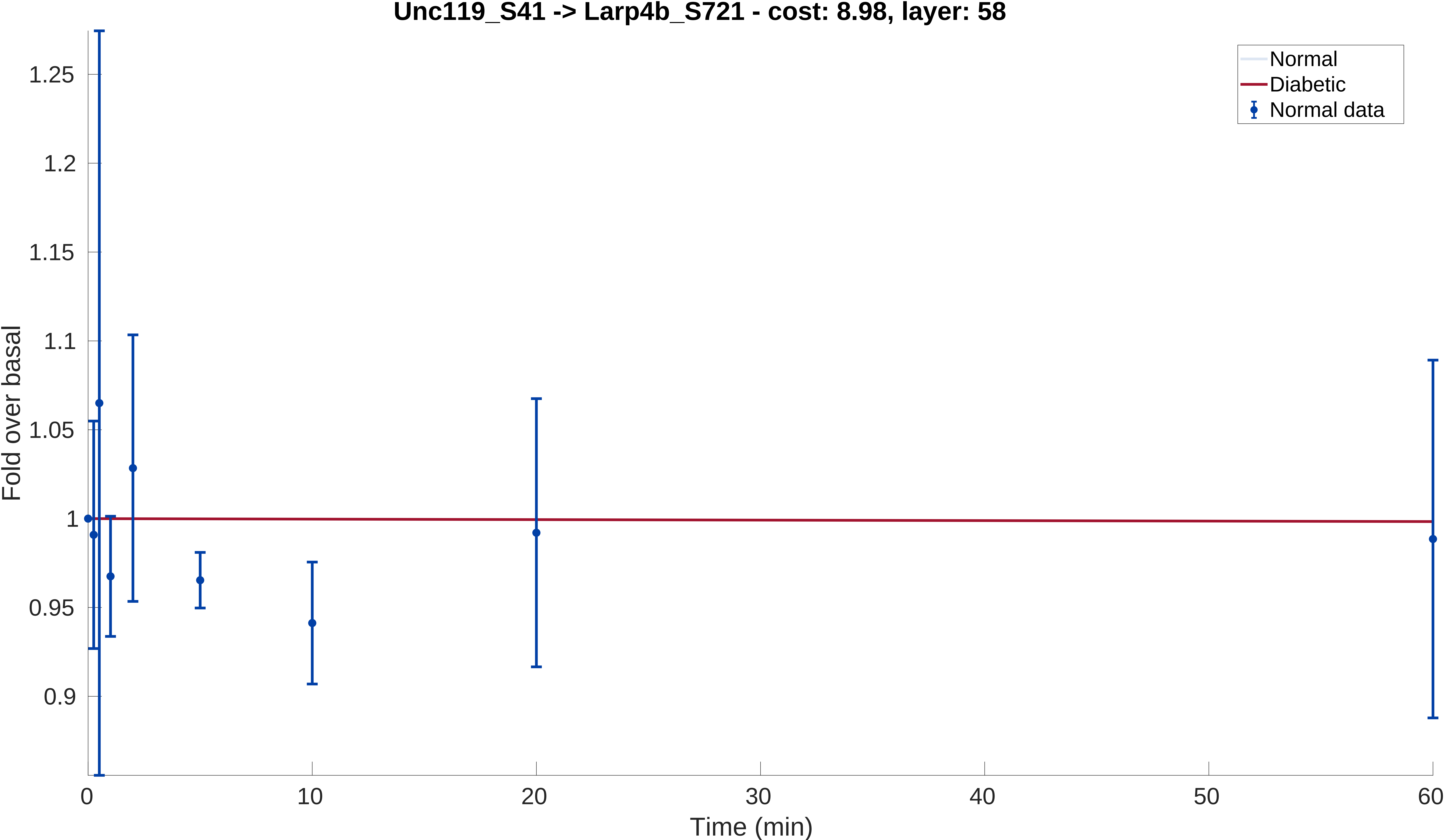
Unc119_S41 -> Larp4b_S721 - cost: 8.98, layer: 58
Normal
Diabetic
Normal data
Fold over basal
1.25
1.2
1.15
1.1
1.05
1
0.95
0.9
0
10
20
30
40
50
60
Time (min)

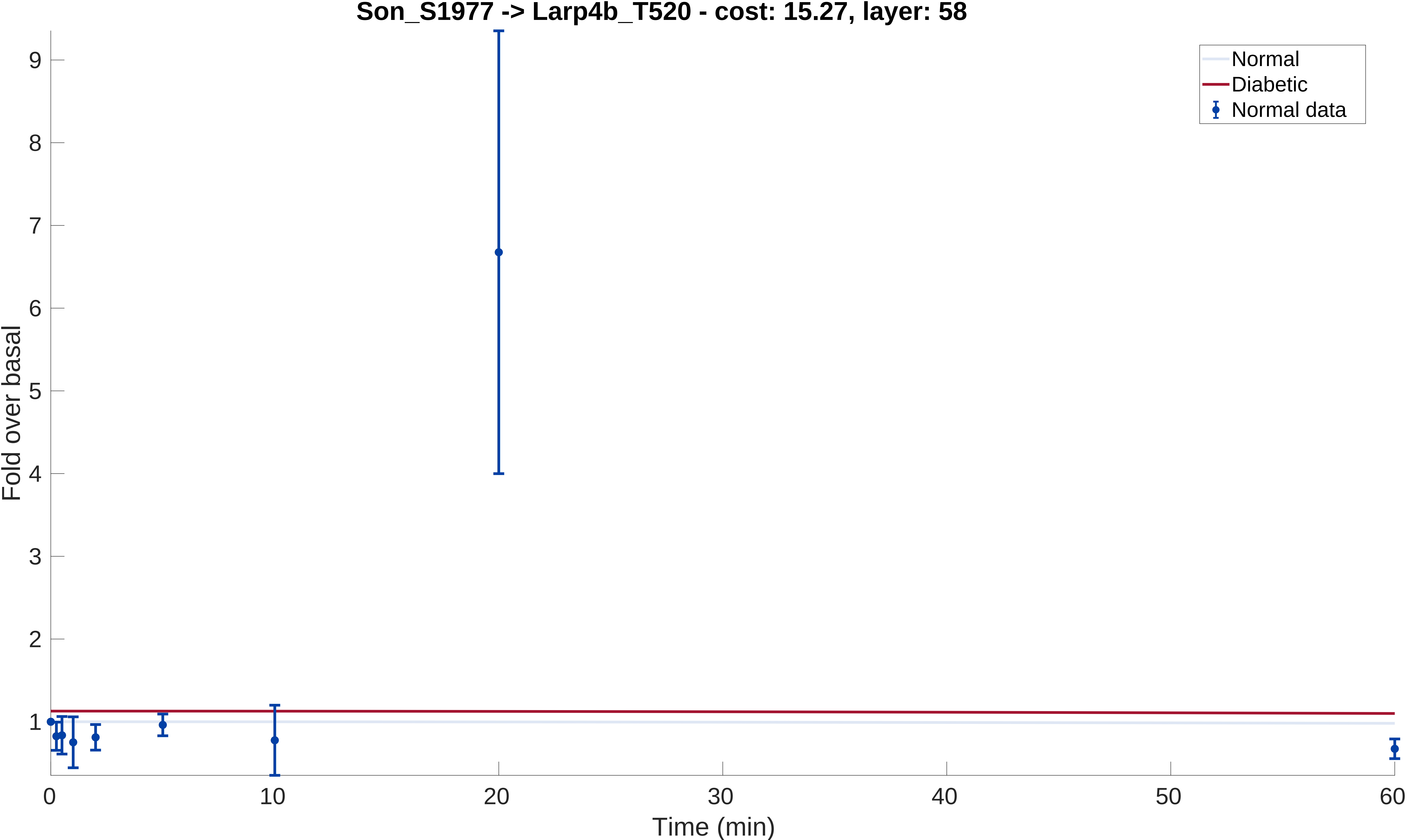

Son_S1977 -> Larp4b_T520 - cost: 15.27, layer: 58
Normal
Diabetic
Normal data
Fold over basal
Time (min)
0
10
20
30
40
50
60
1
2
3
4
5
6
7
8
9

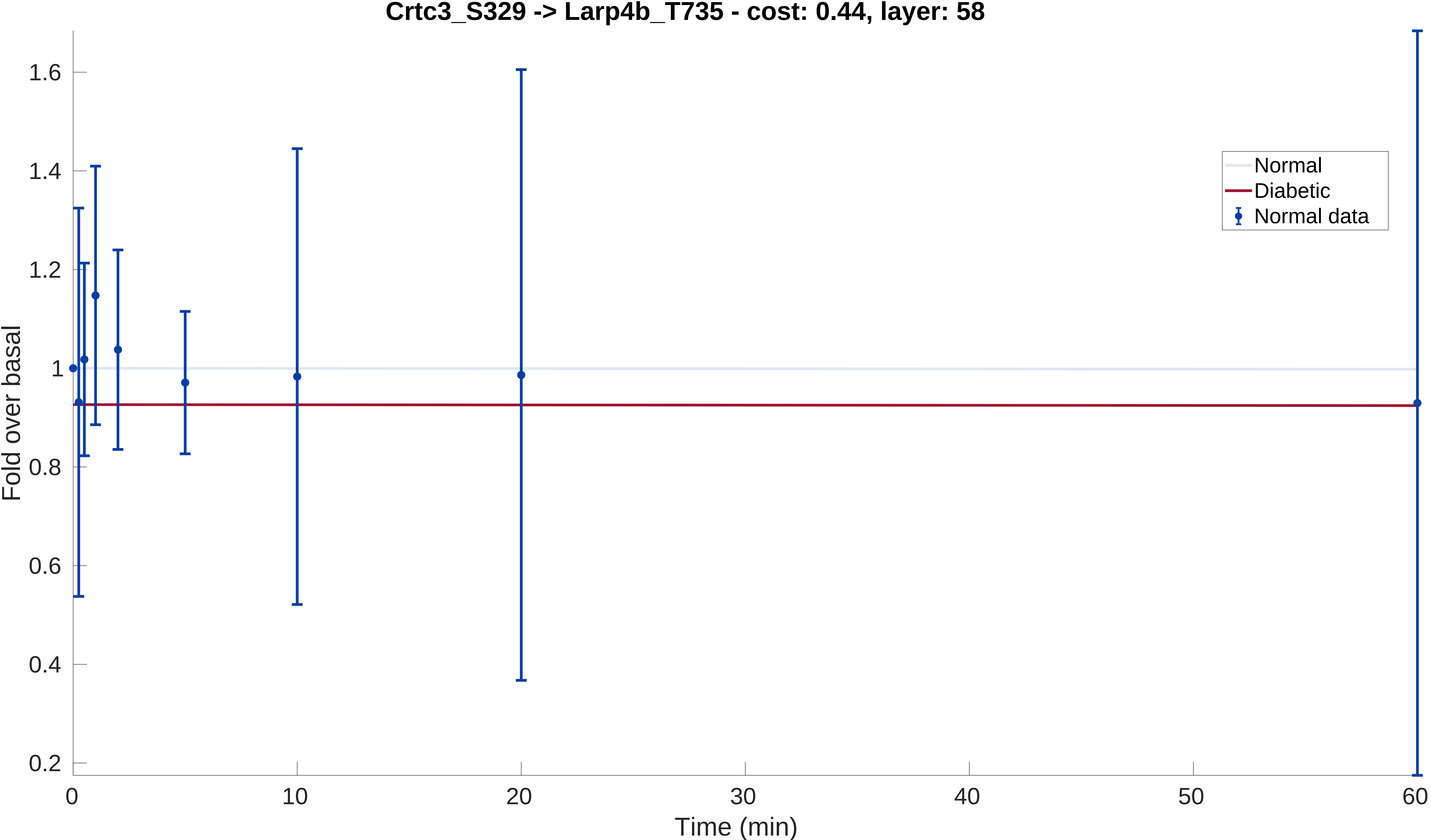

Crtc3_S329 -> Larp4b_T735 - cost: 0.44, layer: 58
Fold over basal
Time (min)
Normal
Diabetic
Normal data
1.6
1.4
1.2
1
0.8
0.6
0.4
0.2
0
10
20
30
40
50
60

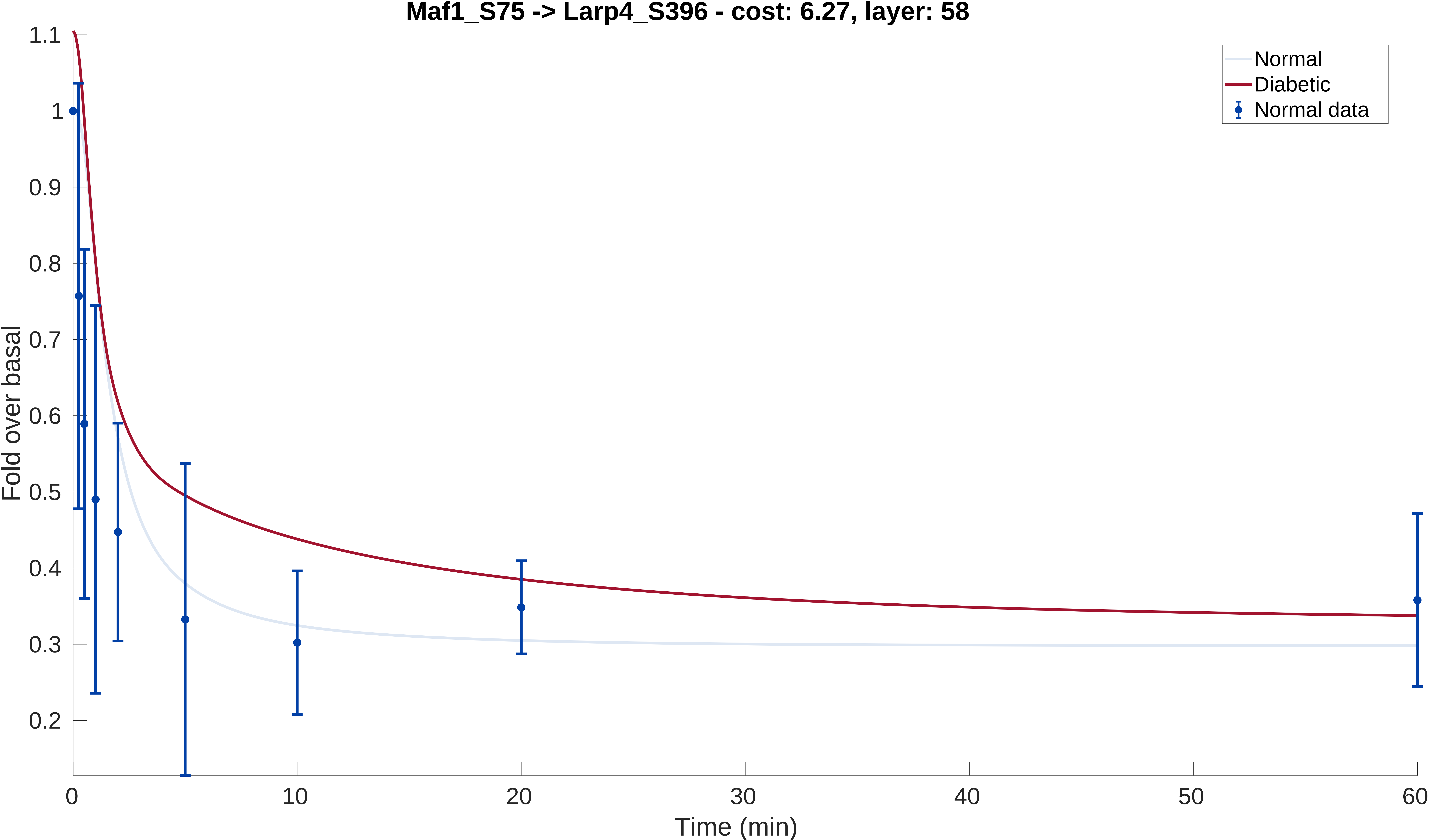

Maf1_S75 -> Larp4_S396 - cost: 6.27, layer: 58
Normal
Diabetic
Normal data
Fold over basal
Time (min)
1.1
1
0.9
0.8
0.7
0.6
0.5
0.4
0.3
0.2
0
10
20
30
40
50
60

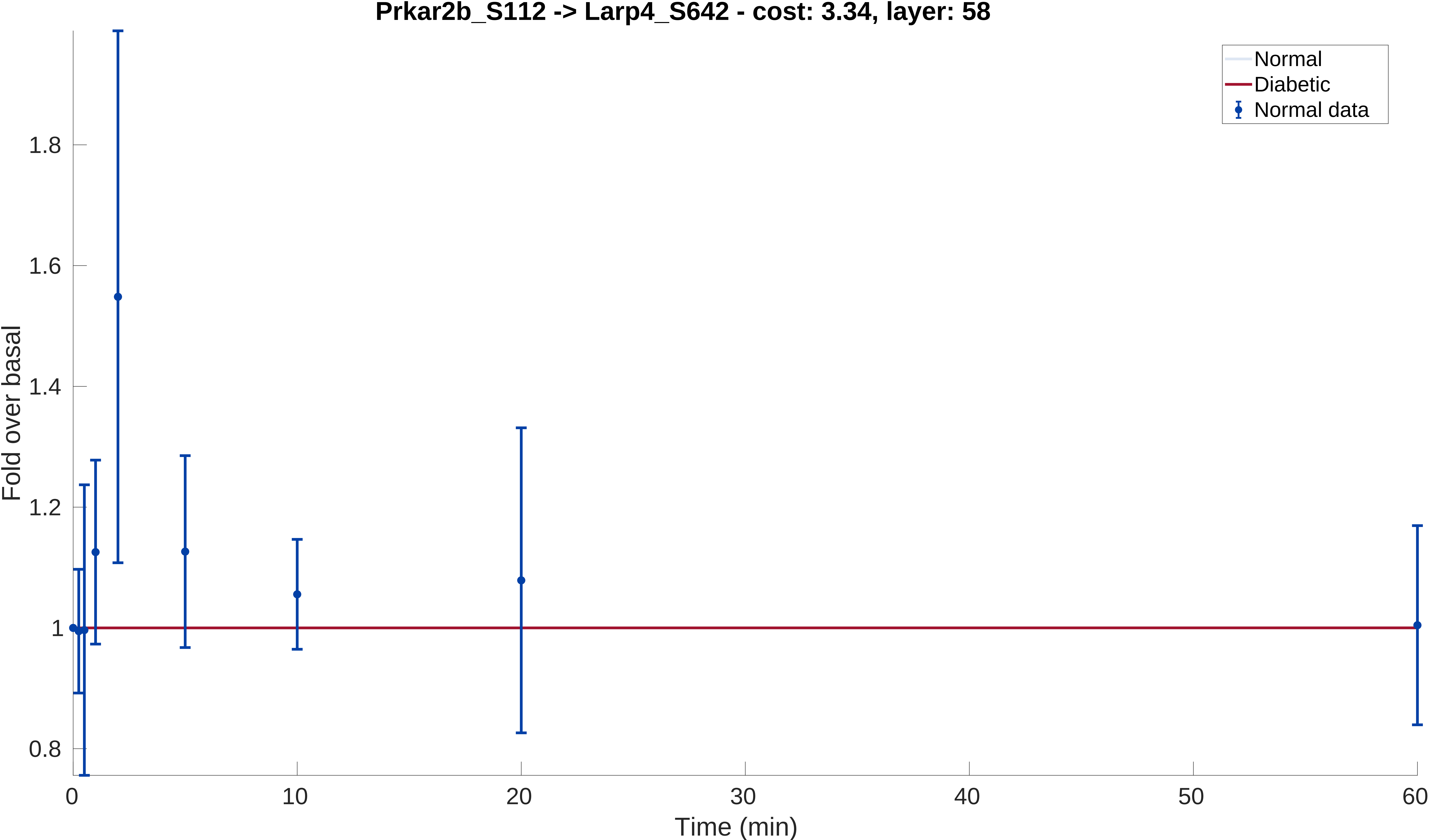

Prkar2b_S112 -> Larp4_S642 - cost: 3.34, layer: 58
Normal
Diabetic
Normal data
Fold over basal
0.8
1
1.2
1.4
1.6
1.8
0
10
20
30
40
50
60
Time (min)

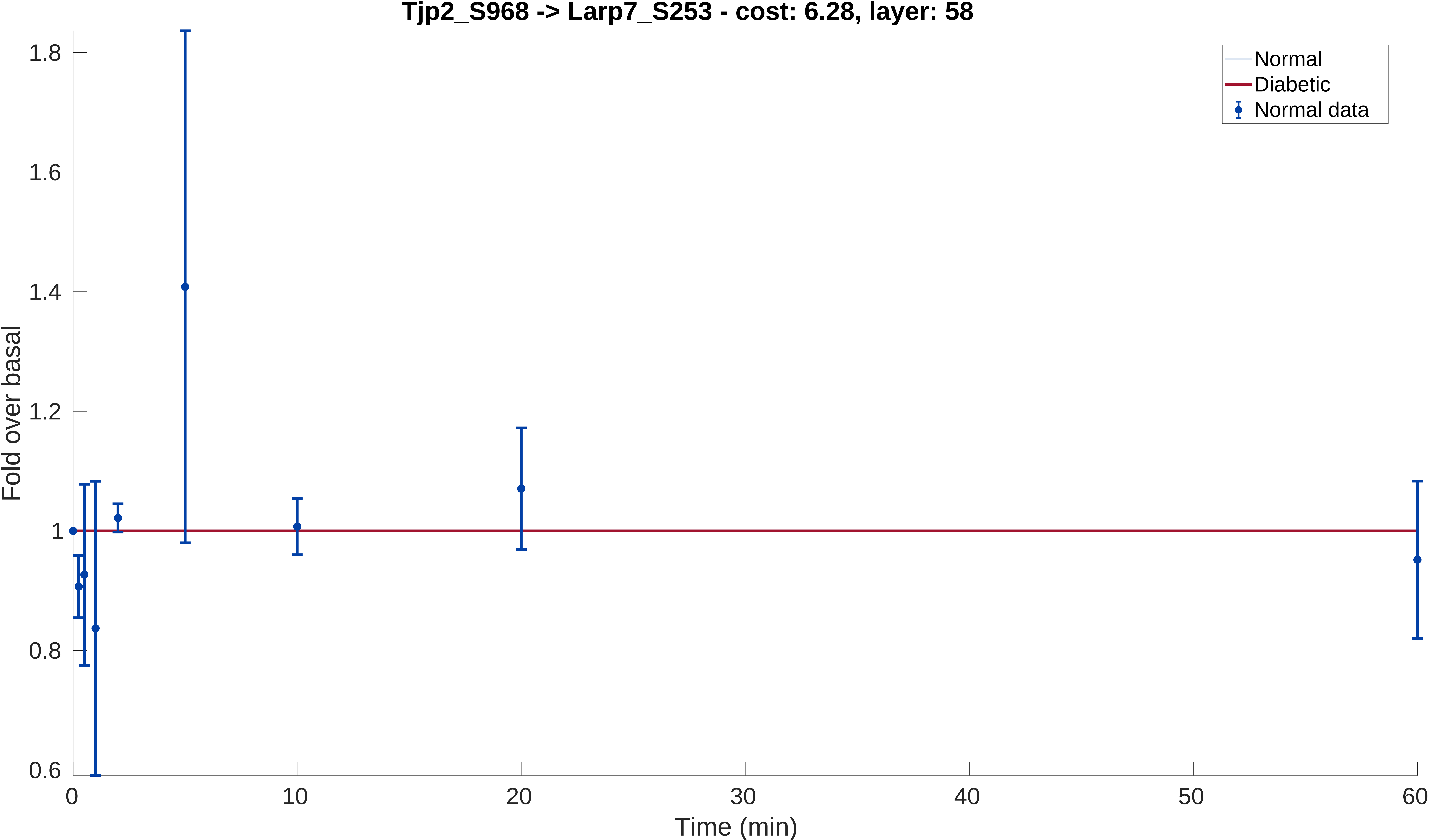

Tjp2_S968 -> Larp7_S253 - cost: 6.28, layer: 58
Normal
Diabetic
Normal data
Fold over basal
1.8
1.6
1.4
1.2
1
0.8
0.6
0
10
20
30
40
50
60
Time (min)

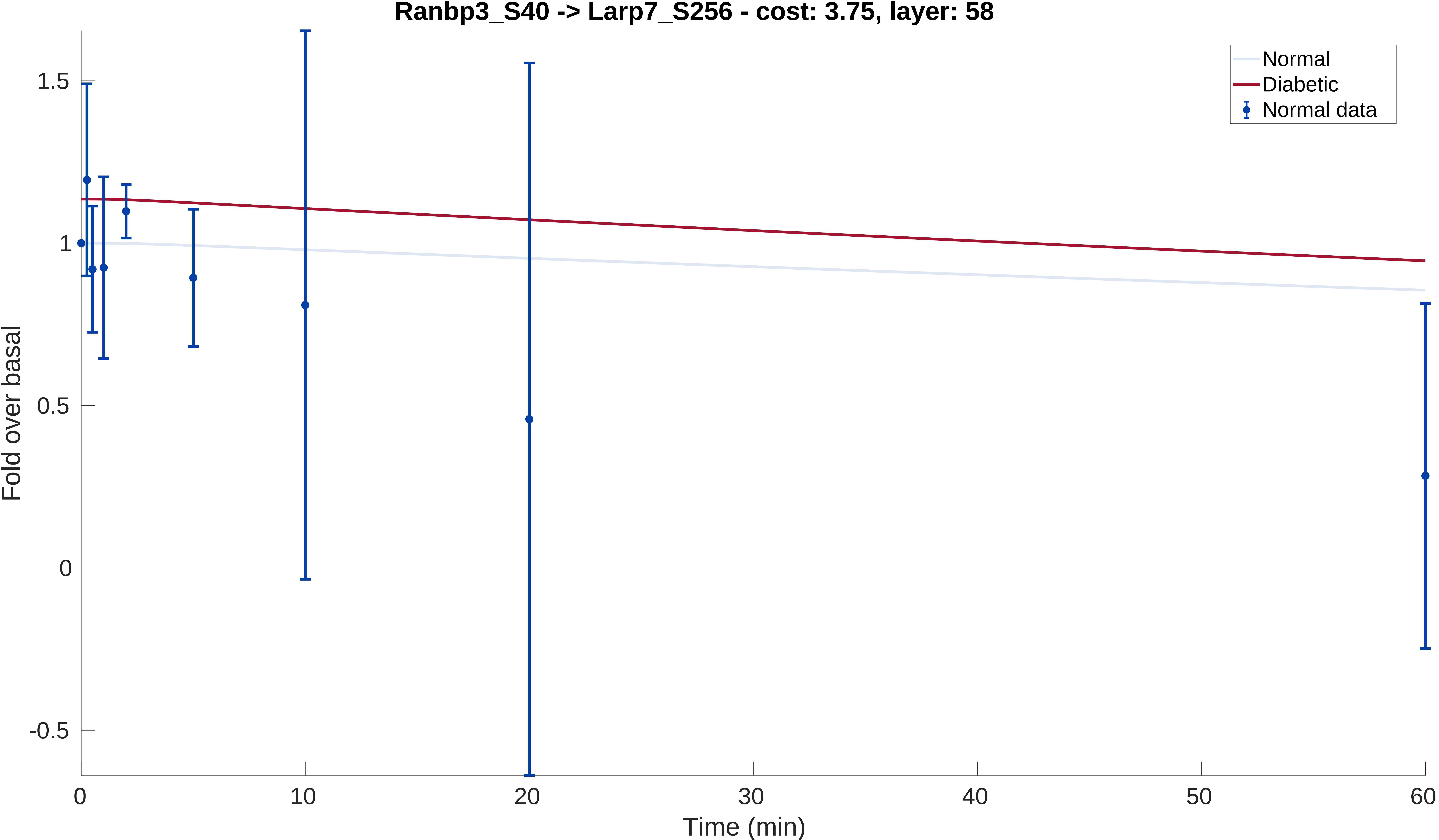

Ranbp3_S40 -> Larp7_S256 - cost: 3.75, layer: 58
Normal
Diabetic
Normal data
Fold over basal
1.5
1
0.5
0
-0.5
0
10
20
30
40
50
60
Time (min)

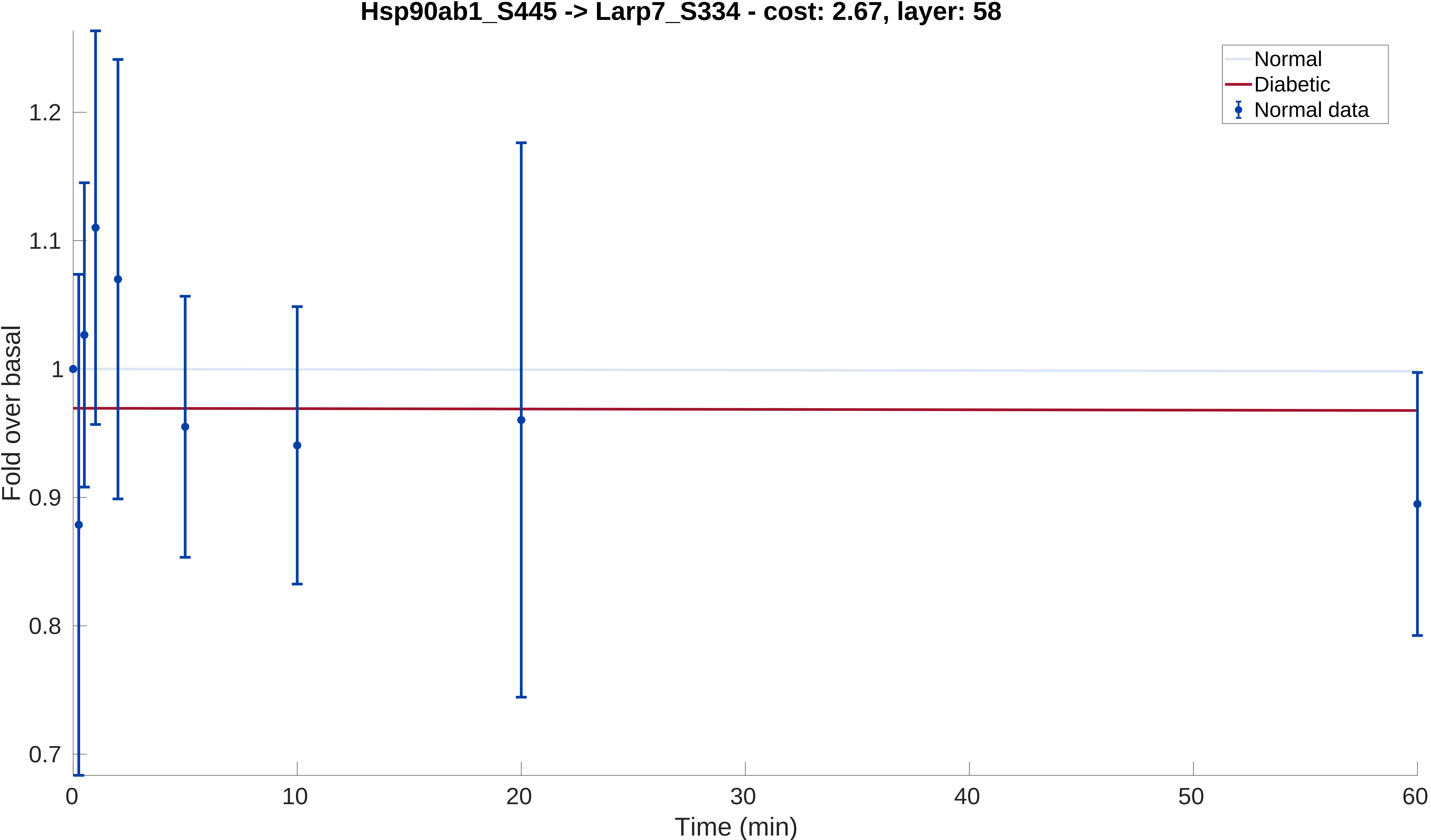

Hsp90ab1_S445 -> Larp7_S334 - cost: 2.67, layer: 58
Normal
Diabetic
Normal data
Fold over basal
1.2
1.1
1
0.9
0.8
0.7
0
10
20
30
40
50
60
Time (min)

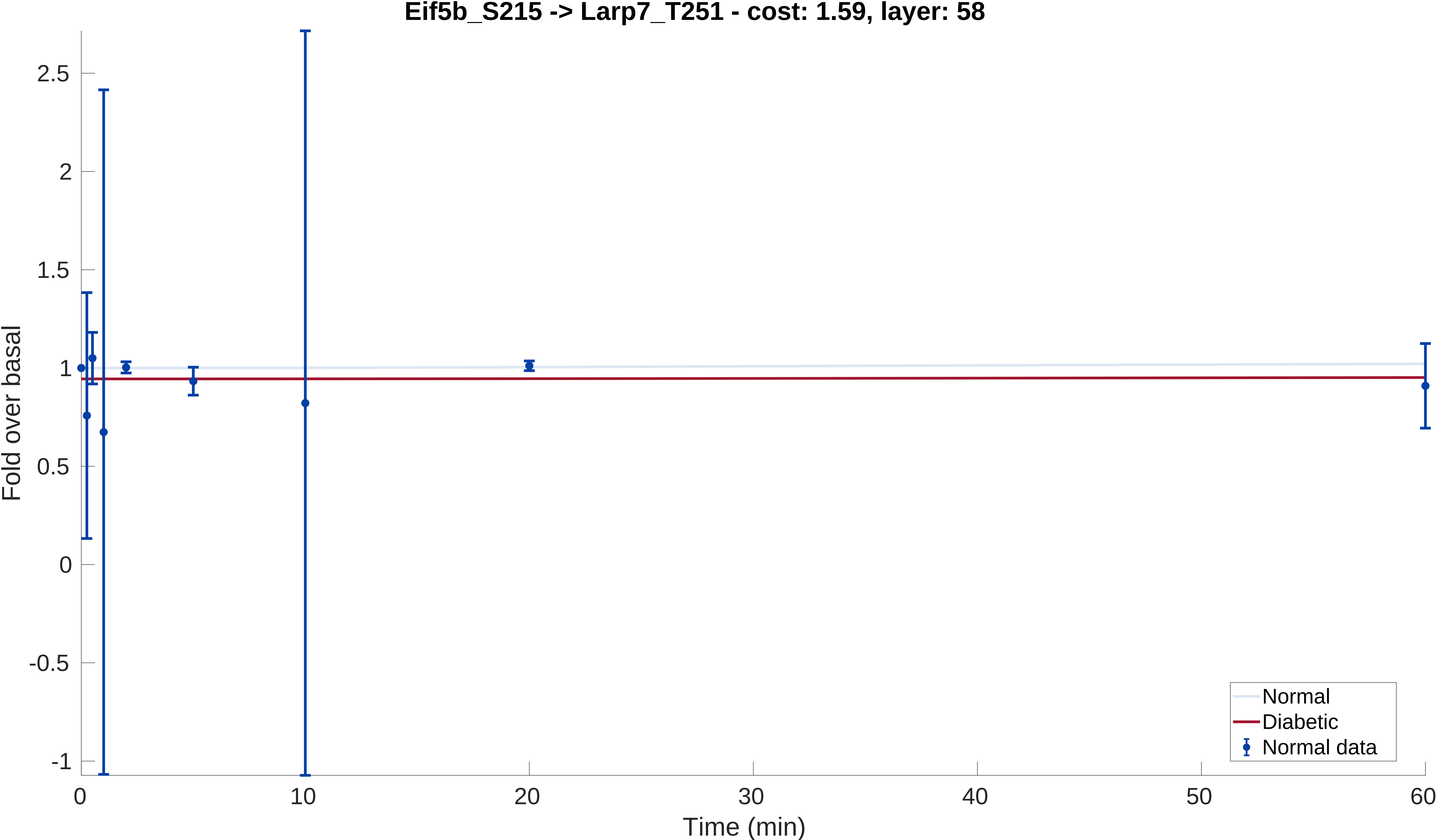

Eif5b_S215 -> Larp7_T251 - cost: 1.59, layer: 58
Fold over basal
Time (min)
2.5
2
1.5
1
0.5
0
-0.5
-1
0
10
20
30
40
50
60
Normal
Diabetic
Normal data

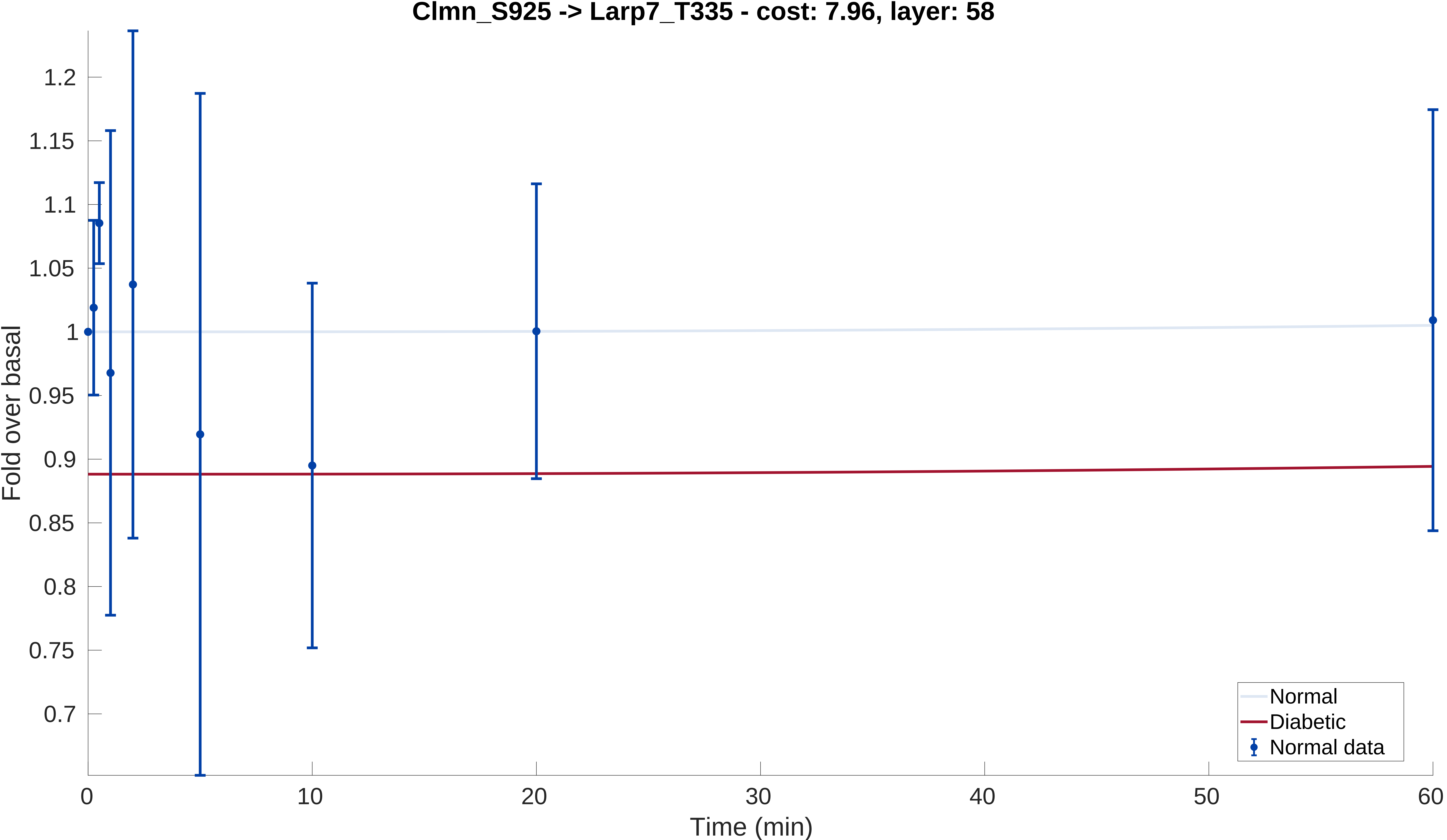
Clmn_S925 -> Larp7_T335 - cost: 7.96, layer: 58
Fold over basal
Time (min)
1.2
1.15
1.1
1.05
1
0.95
0.9
0.85
0.8
0.75
0.7
0
10
20
30
40
50
60
Normal
Diabetic
Normal data

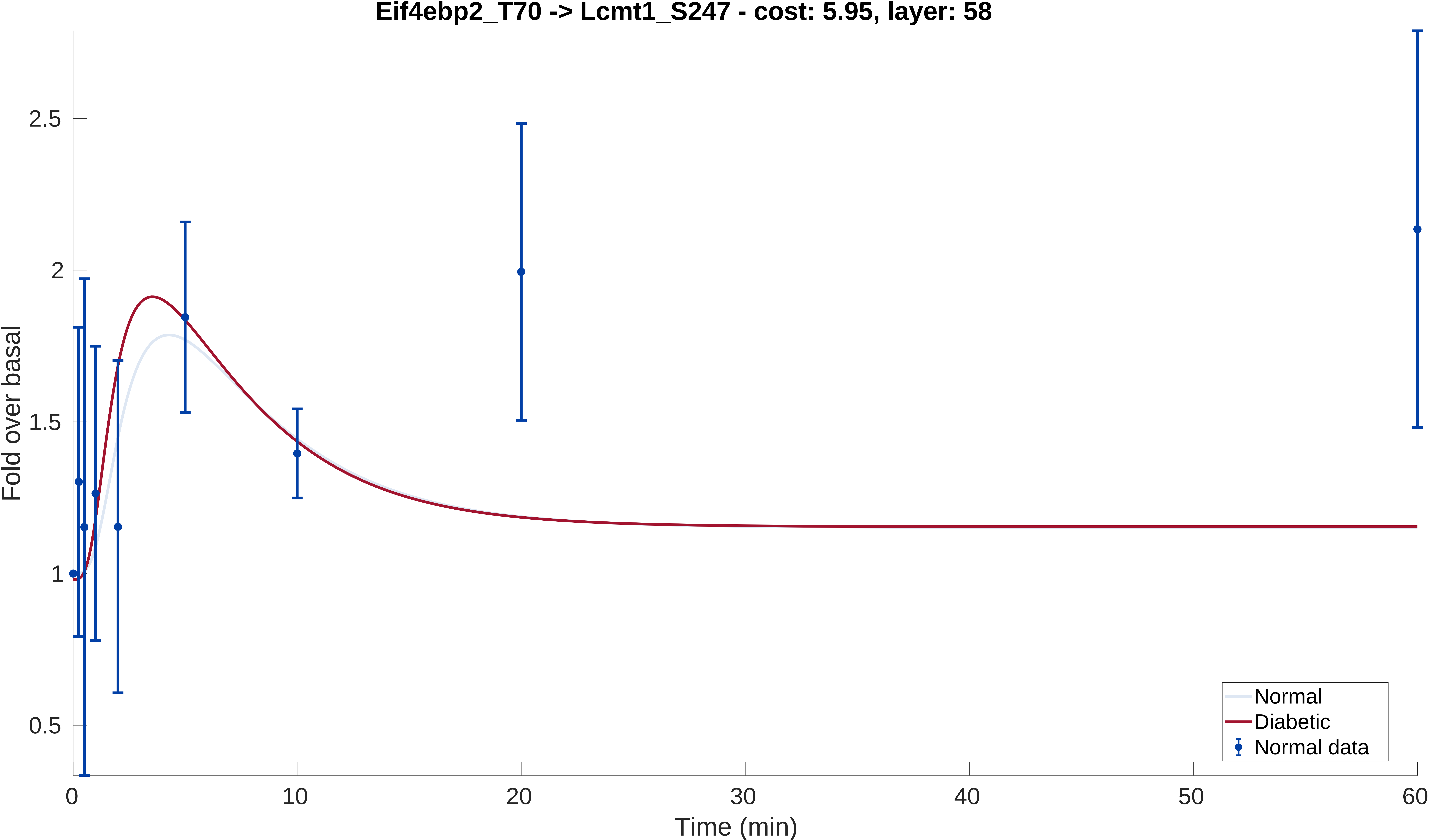

Eif4ebp2_T70 -> Lcmt1_S247 - cost: 5.95, layer: 58
Fold over basal
2.5
2
1.5
1
0.5
0
10
20
30
40
50
60
Time (min)
Normal
Diabetic
Normal data

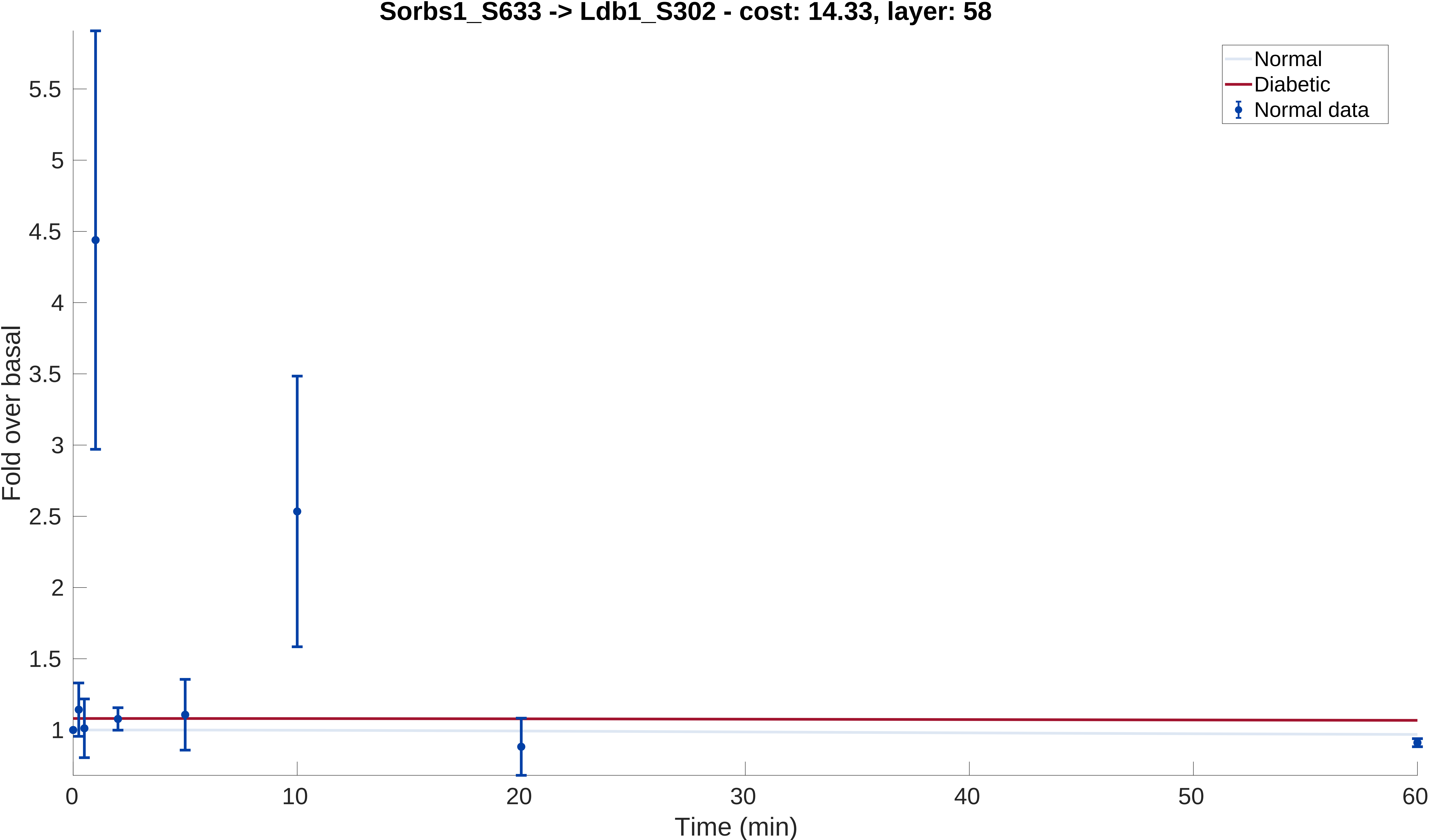

Sorbs1_S633 -> Ldb1_S302 - cost: 14.33, layer: 58
Normal
Diabetic
Normal data
Fold over basal
5.5
5
4.5
4
3.5
3
2.5
2
1.5
1
0
10
20
30
40
50
60
Time (min)

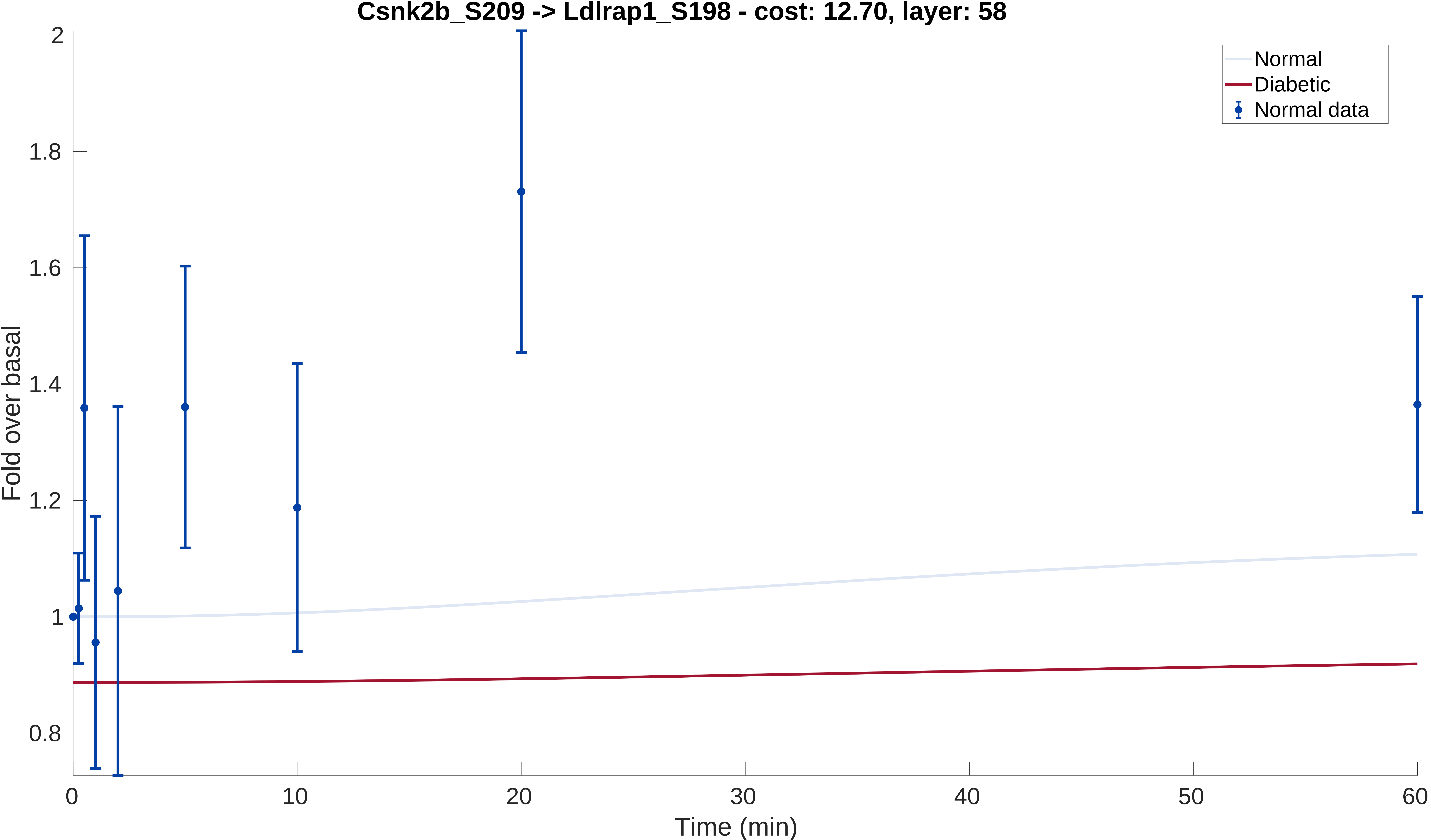

Csnk2b_S209 -> Ldlrap1_S198 - cost: 12.70, layer: 58
Normal
Diabetic
Normal data
Fold over basal
Time (min)
2
1.8
1.6
1.4
1.2
1
0.8
0
10
20
30
40
50
60

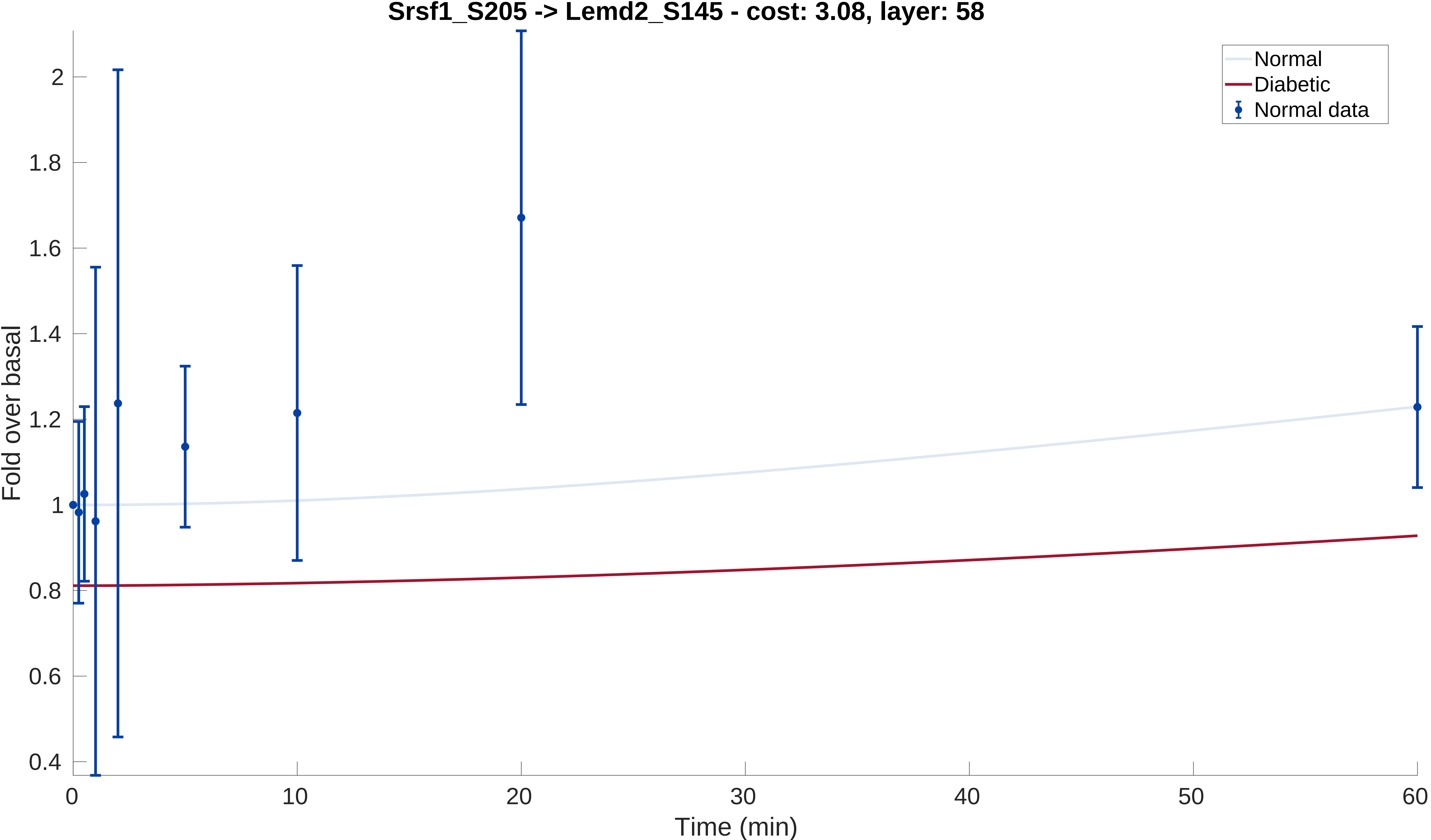
Srsf1_S205 -> Lemd2_S145 - cost: 3.08, layer: 58
Normal
Diabetic
Normal data
Fold over basal
2
1.8
1.6
1.4
1.2
1
0.8
0.6
0.4
0
10
20
30
40
50
60
Time (min)

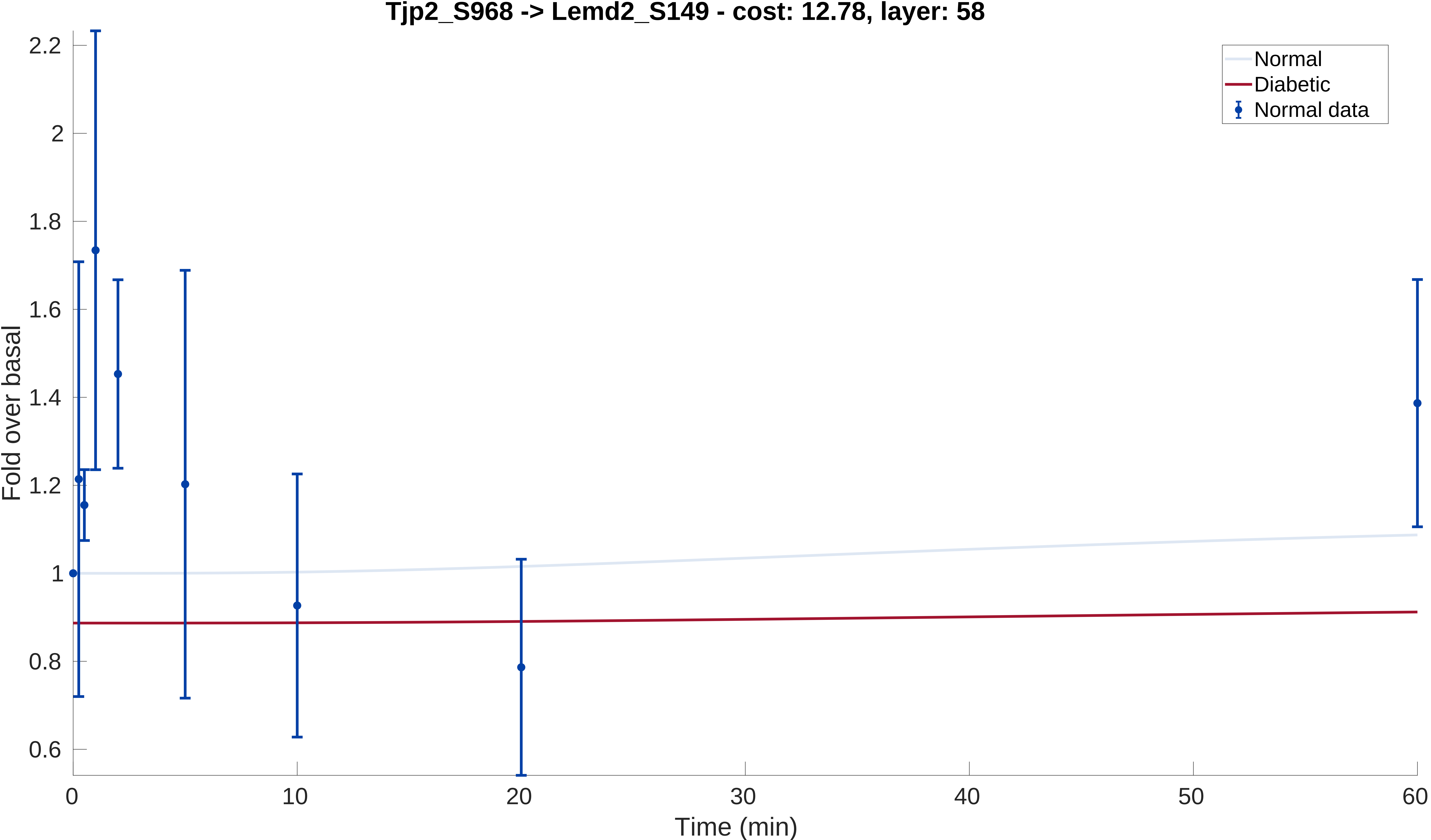
Tjp2_S968 -> Lemd2_S149 - cost: 12.78, layer: 58
Normal
Diabetic
Normal data
Fold over basal
2.2
2
1.8
1.6
1.4
1.2
1
0.8
0.6
0
10
20
30
40
50
60
Time (min)

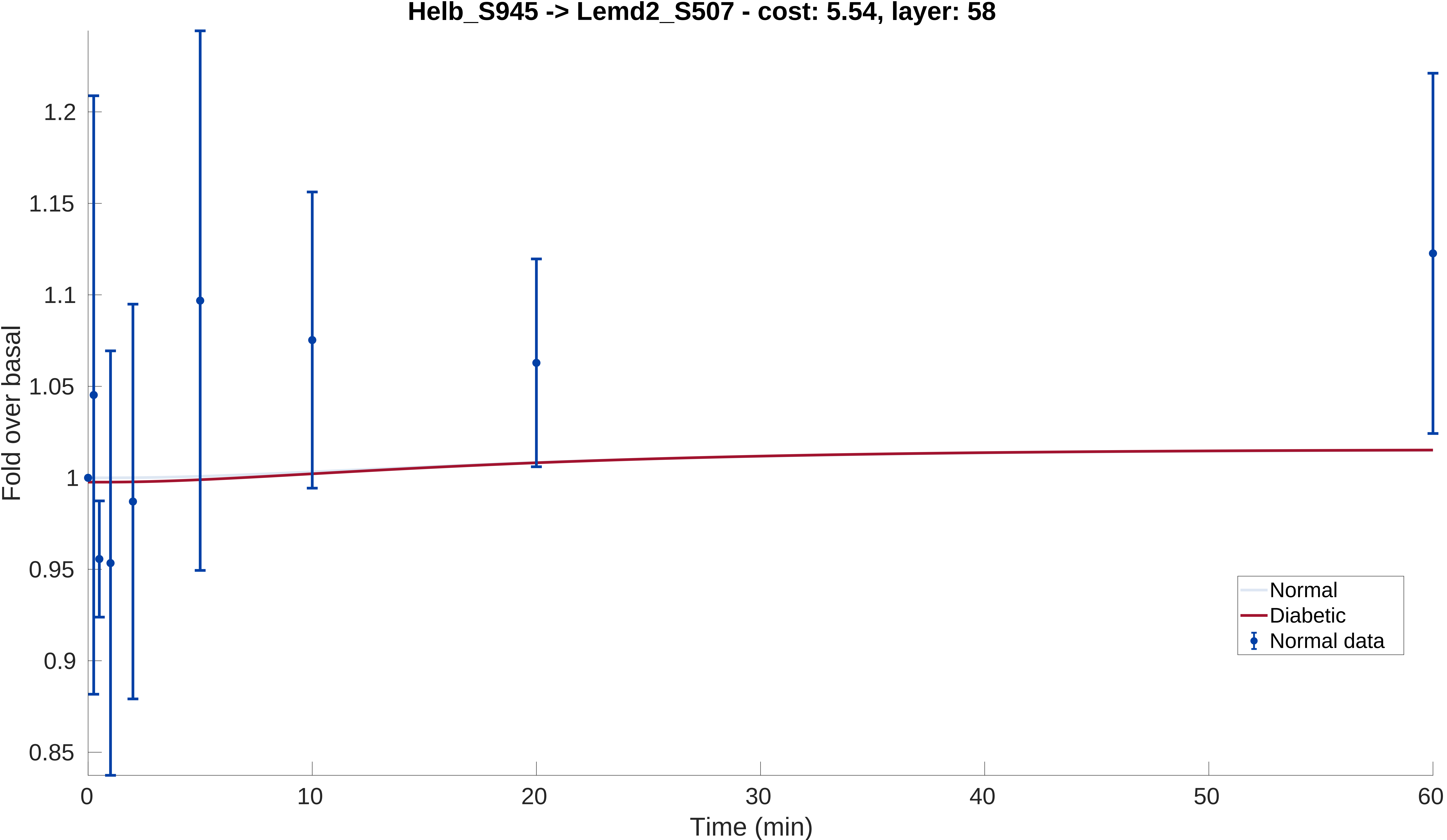
Helb_S945 -> Lemd2_S507 - cost: 5.54, layer: 58
Fold over basal
1.2
1.15
1.1
1.05
1
0.95
0.9
0.85
0
10
20
30
40
50
60
Time (min)
Normal
Diabetic
Normal data

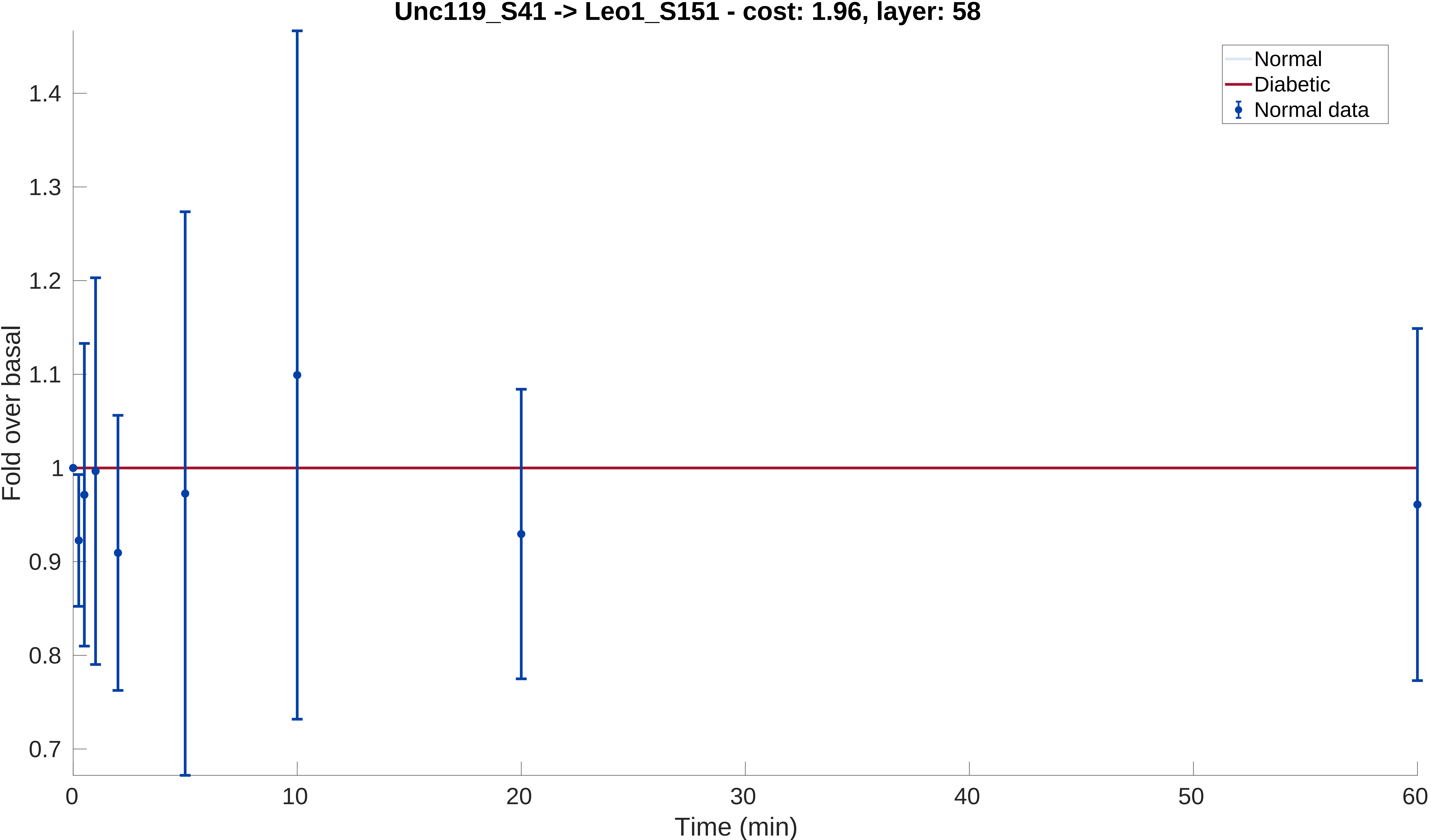

Unc119_S41 -> Leo1_S151 - cost: 1.96, layer: 58
Normal
Diabetic
Normal data
Fold over basal
1.4
1.3
1.2
1.1
1
0.9
0.8
0.7
0
10
20
30
40
50
60
Time (min)

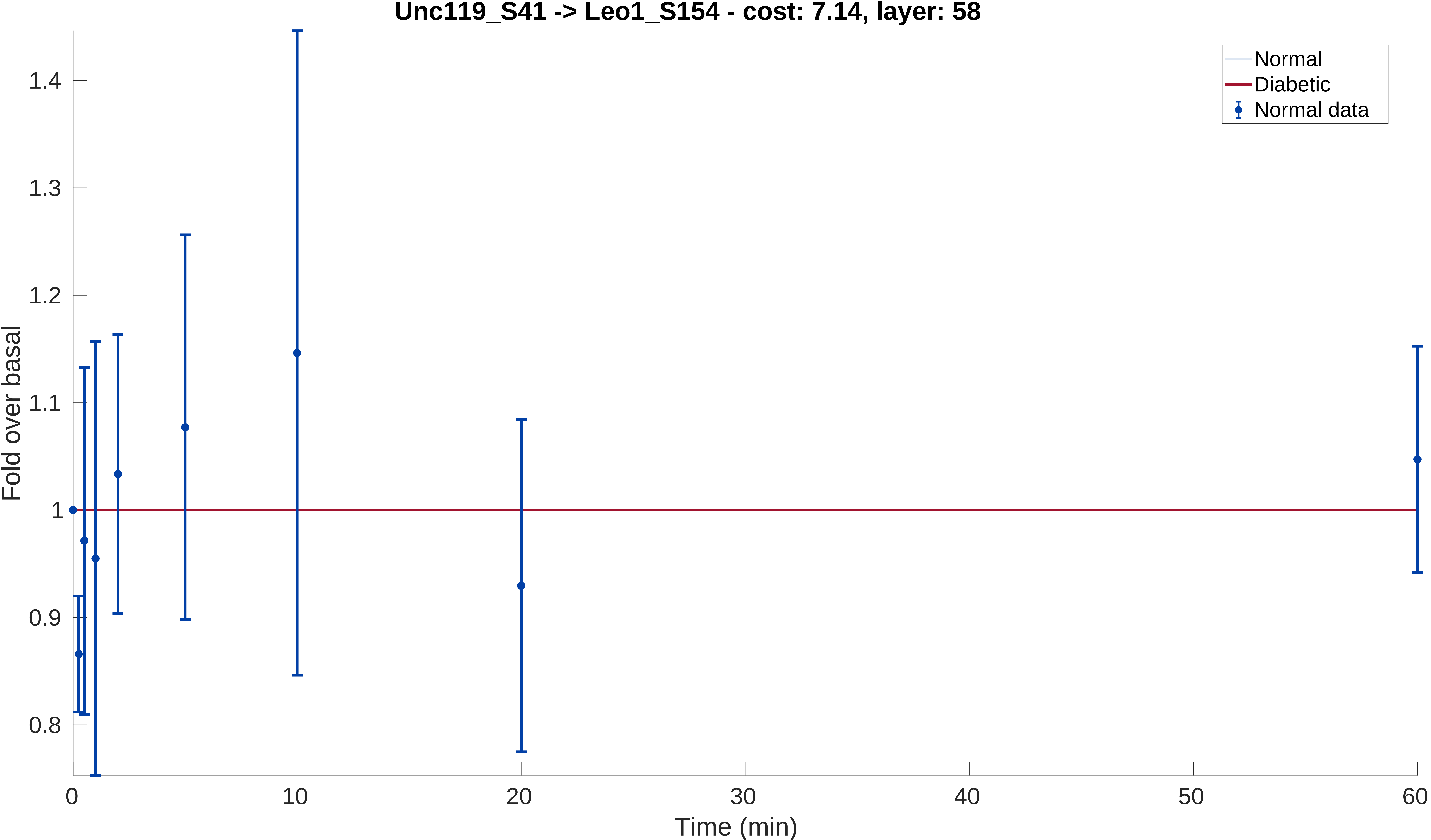

Unc119_S41 -> Leo1_S154 - cost: 7.14, layer: 58
Normal
Diabetic
Normal data
Fold over basal
Time (min)
1.4
1.3
1.2
1.1
1
0.9
0.8
0
10
20
30
40
50
60

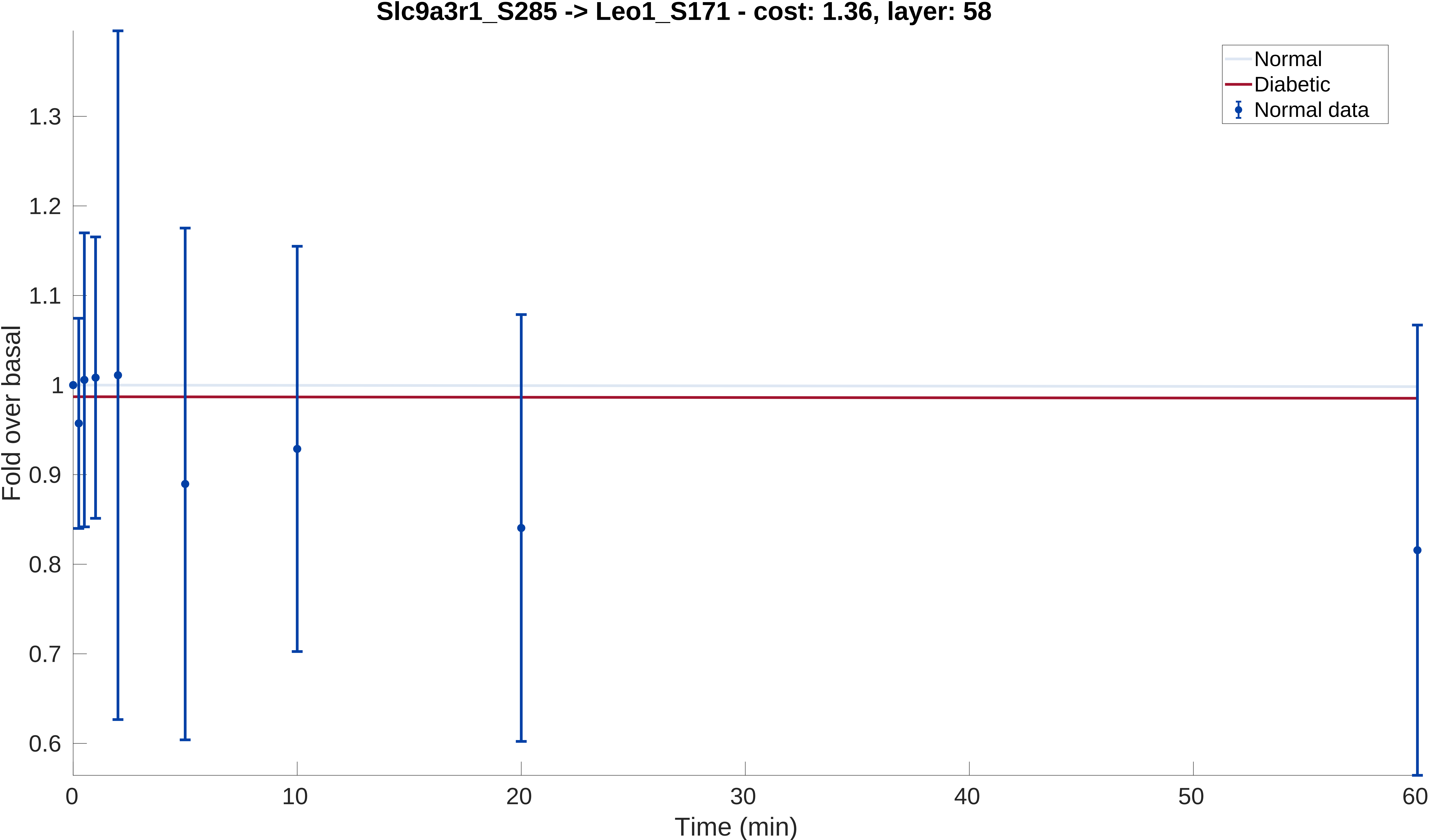

Slc9a3r1_S285 -> Leo1_S171 - cost: 1.36, layer: 58
Normal
Diabetic
Normal data
Fold over basal
Time (min)
0.6
0.7
0.8
0.9
1
1.1
1.2
1.3
0
10
20
30
40
50
60

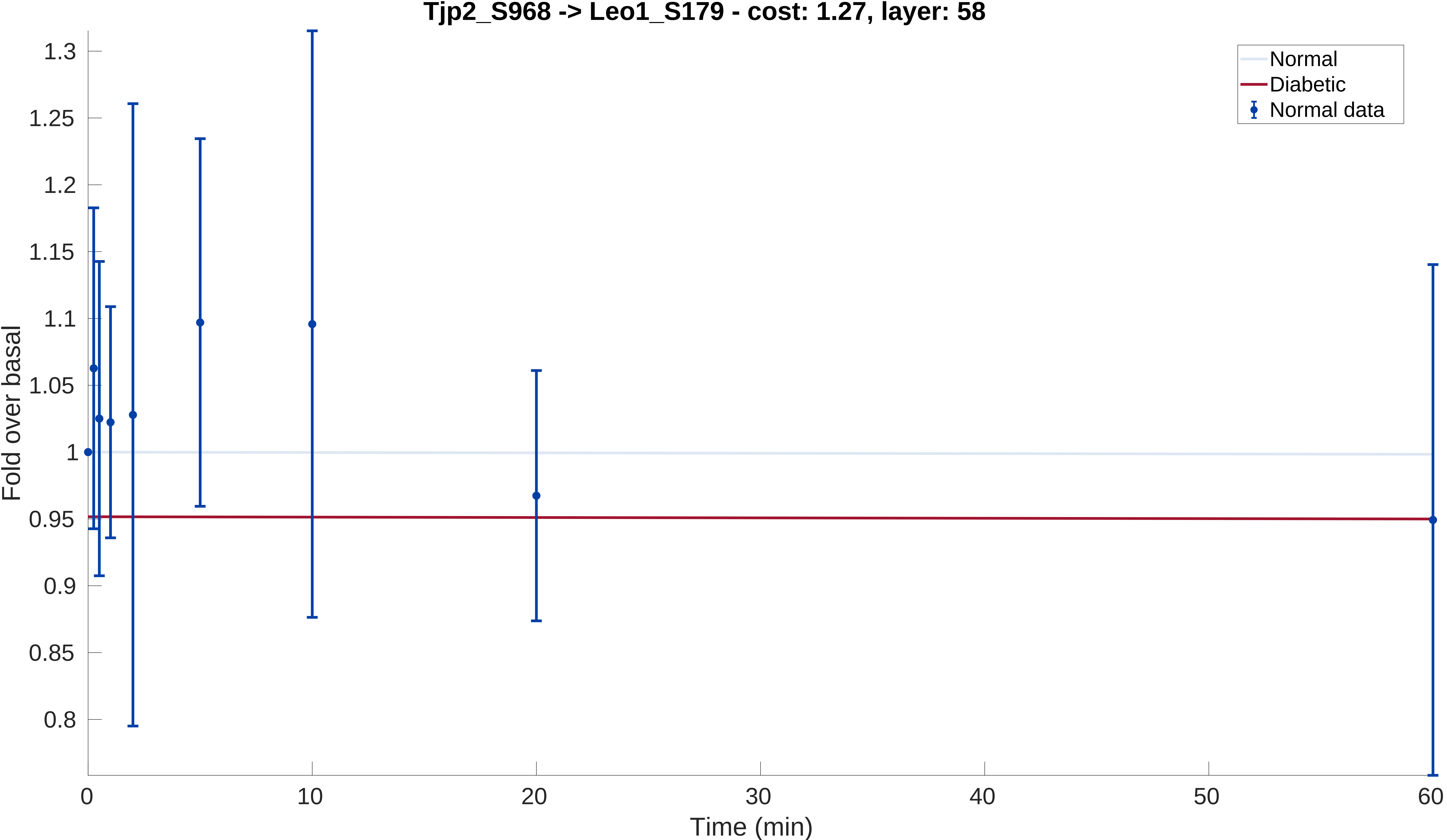

Tjp2_S968 -> Leo1_S179 - cost: 1.27, layer: 58
Normal
Diabetic
Normal data
Fold over basal
1.3
1.25
1.2
1.15
1.1
1.05
1
0.95
0.9
0.85
0.8
0
10
20
30
40
50
60
Time (min)

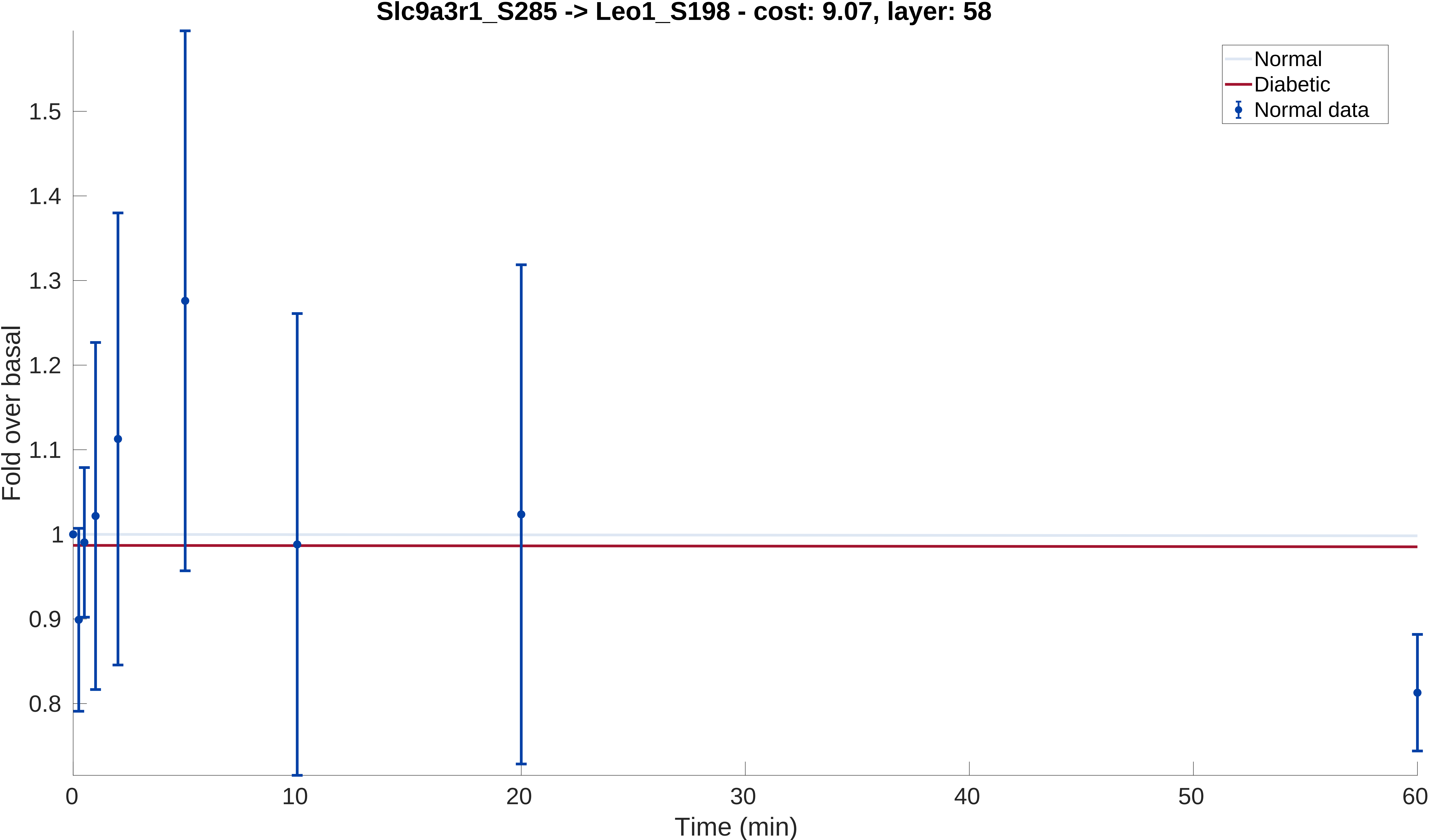

Slc9a3r1_S285 -> Leo1_S198 - cost: 9.07, layer: 58
Normal
Diabetic
Normal data
Fold over basal
1.5
1.4
1.3
1.2
1.1
1
0.9
0.8
0
10
20
30
40
50
60
Time (min)

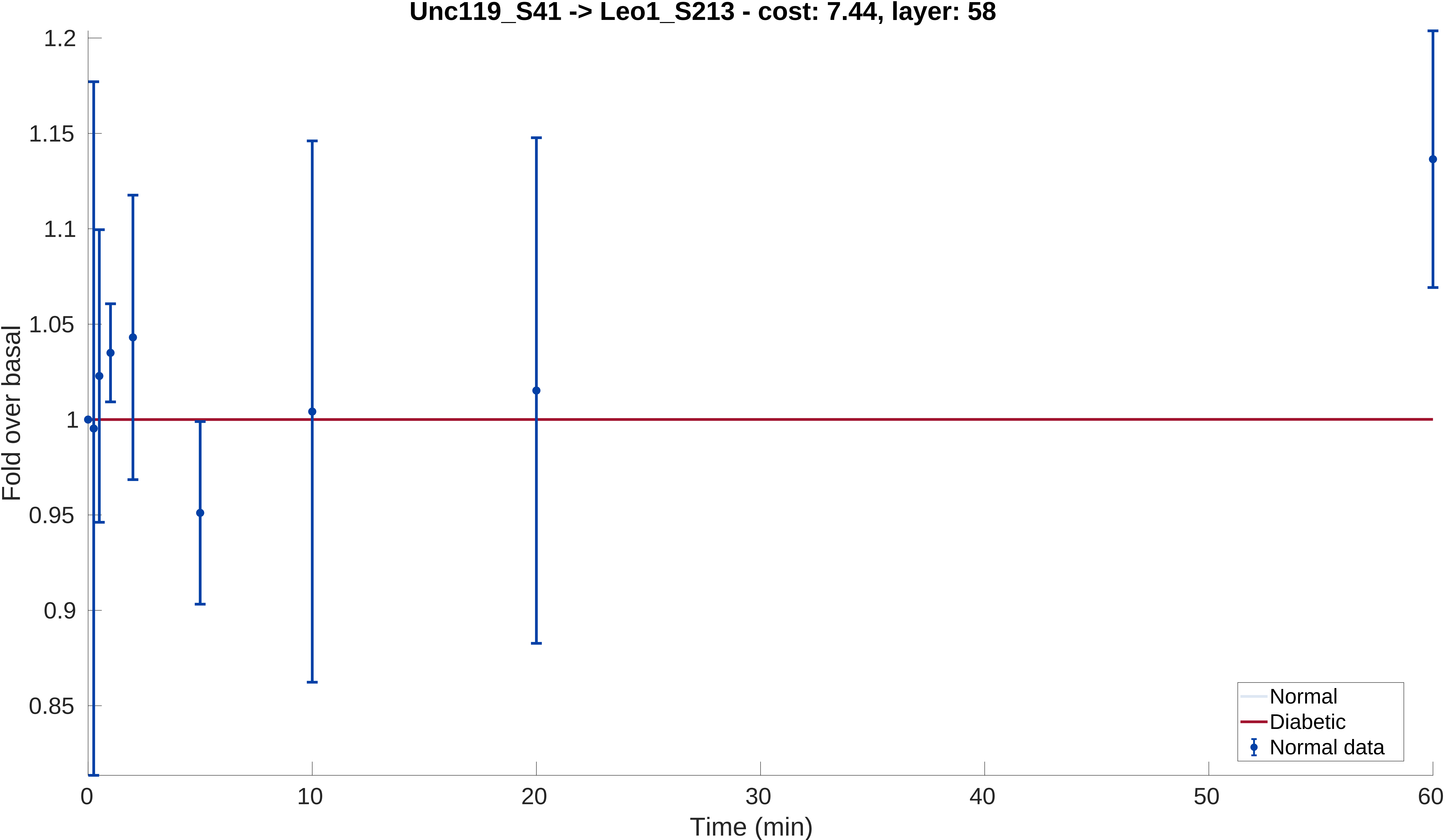

Unc119_S41 -> Leo1_S213 - cost: 7.44, layer: 58
1.2
1.15
1.1
1.05
1
0.95
0.9
0.85
Fold over basal
0
10
20
30
40
50
60
Time (min)
Normal
Diabetic
Normal data

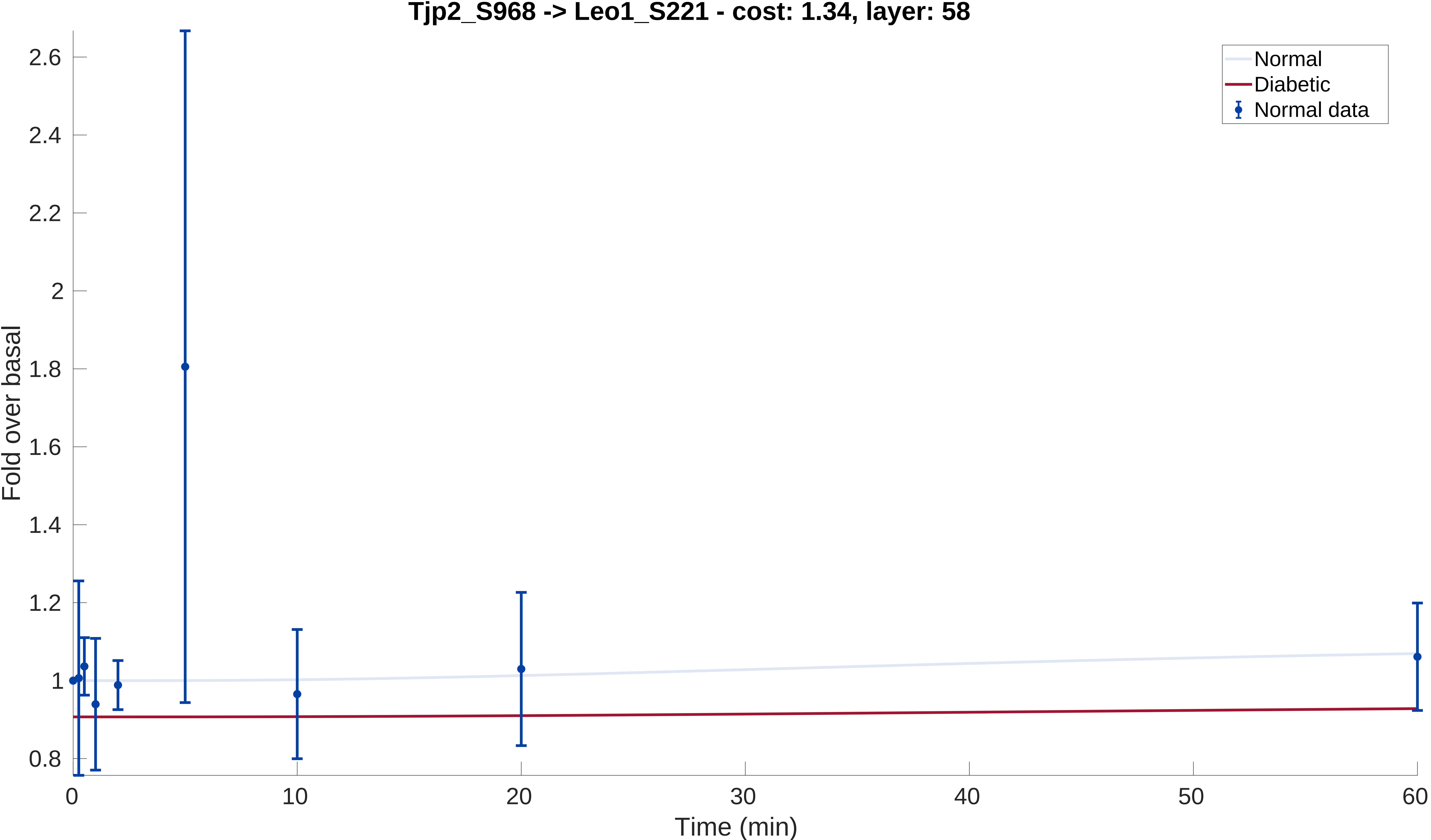
Tjp2_S968 -> Leo1_S221 - cost: 1.34, layer: 58
Normal
Diabetic
Normal data
Fold over basal
2.6
2.4
2.2
2
1.8
1.6
1.4
1.2
1
0.8
0
10
20
30
40
50
60
Time (min)

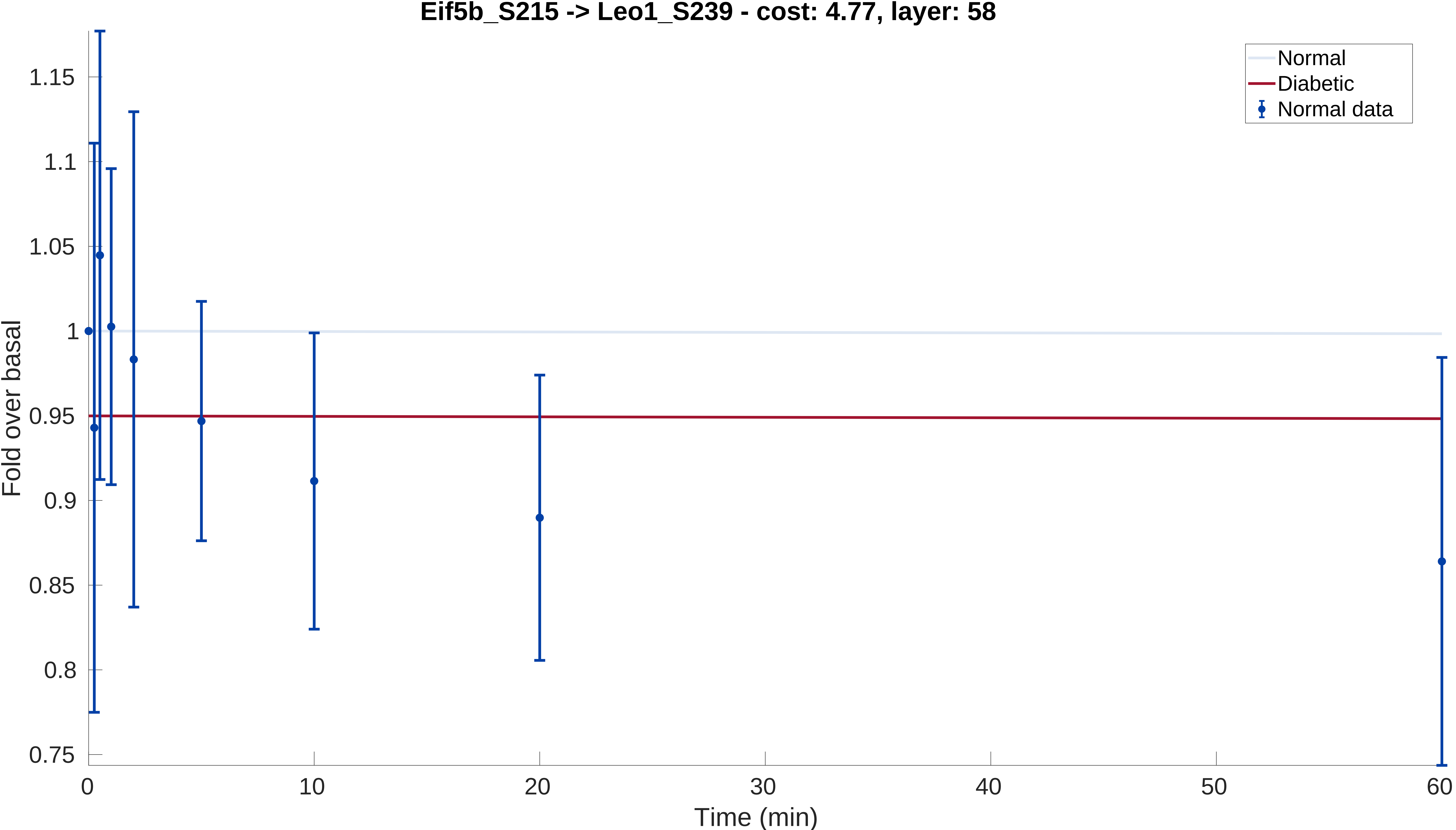
Eif5b_S215 -> Leo1_S239 - cost: 4.77, layer: 58
Normal
Diabetic
Normal data
Fold over basal
Time (min)
1.15
1.1
1.05
1
0.95
0.9
0.85
0.8
0.75
0
10
20
30
40
50
60

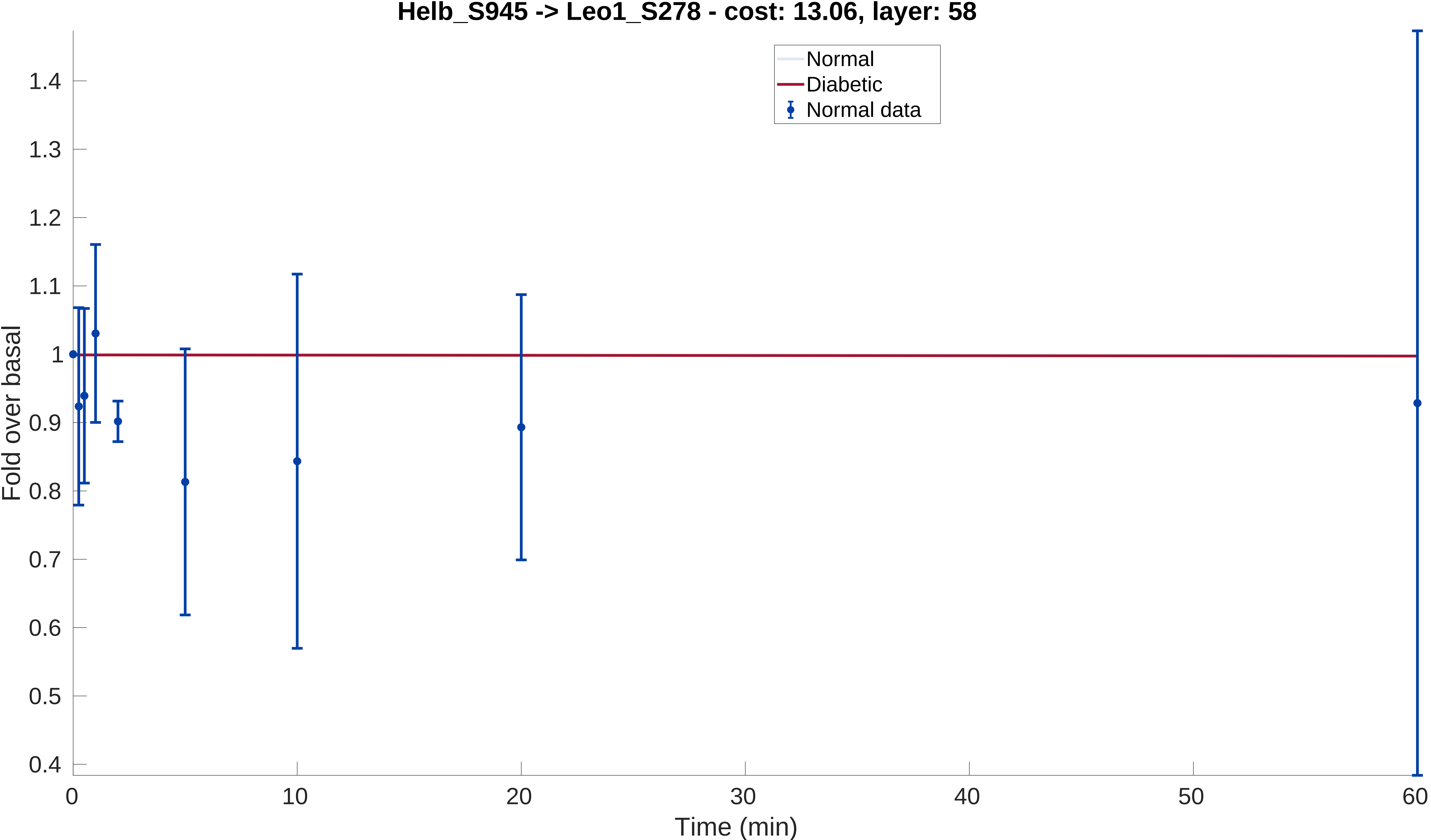

Helb_S945 -> Leo1_S278 - cost: 13.06, layer: 58
Normal
Diabetic
Normal data
Fold over basal
1.4
1.3
1.2
1.1
1
0.9
0.8
0.7
0.6
0.5
0.4
0
10
20
30
40
50
60
Time (min)

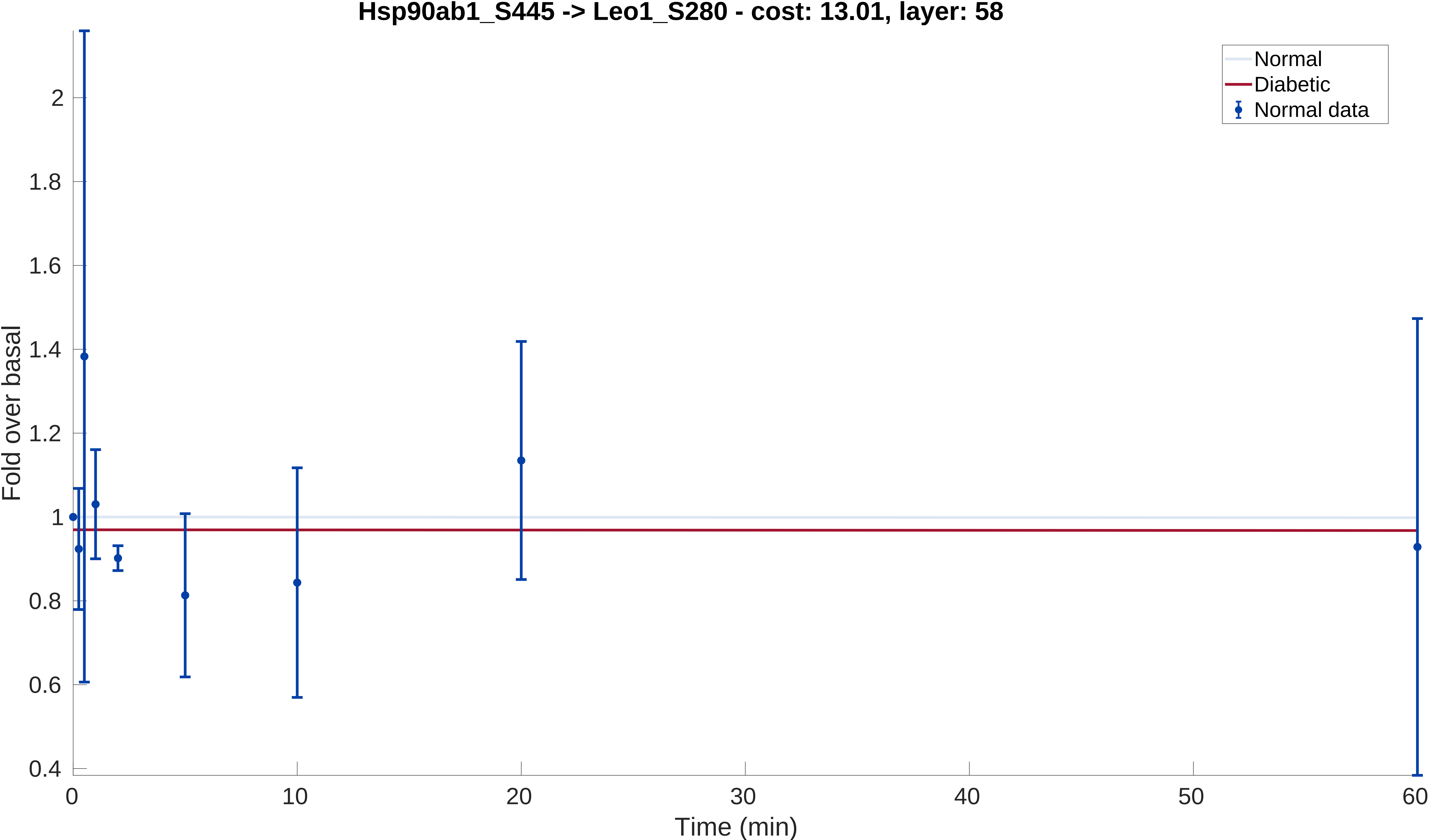

Hsp90ab1_S445 -> Leo1_S280 - cost: 13.01, layer: 58
Normal
Diabetic
Normal data
Fold over basal
2
1.8
1.6
1.4
1.2
1
0.8
0.6
0.4
0
10
20
30
40
50
60
Time (min)

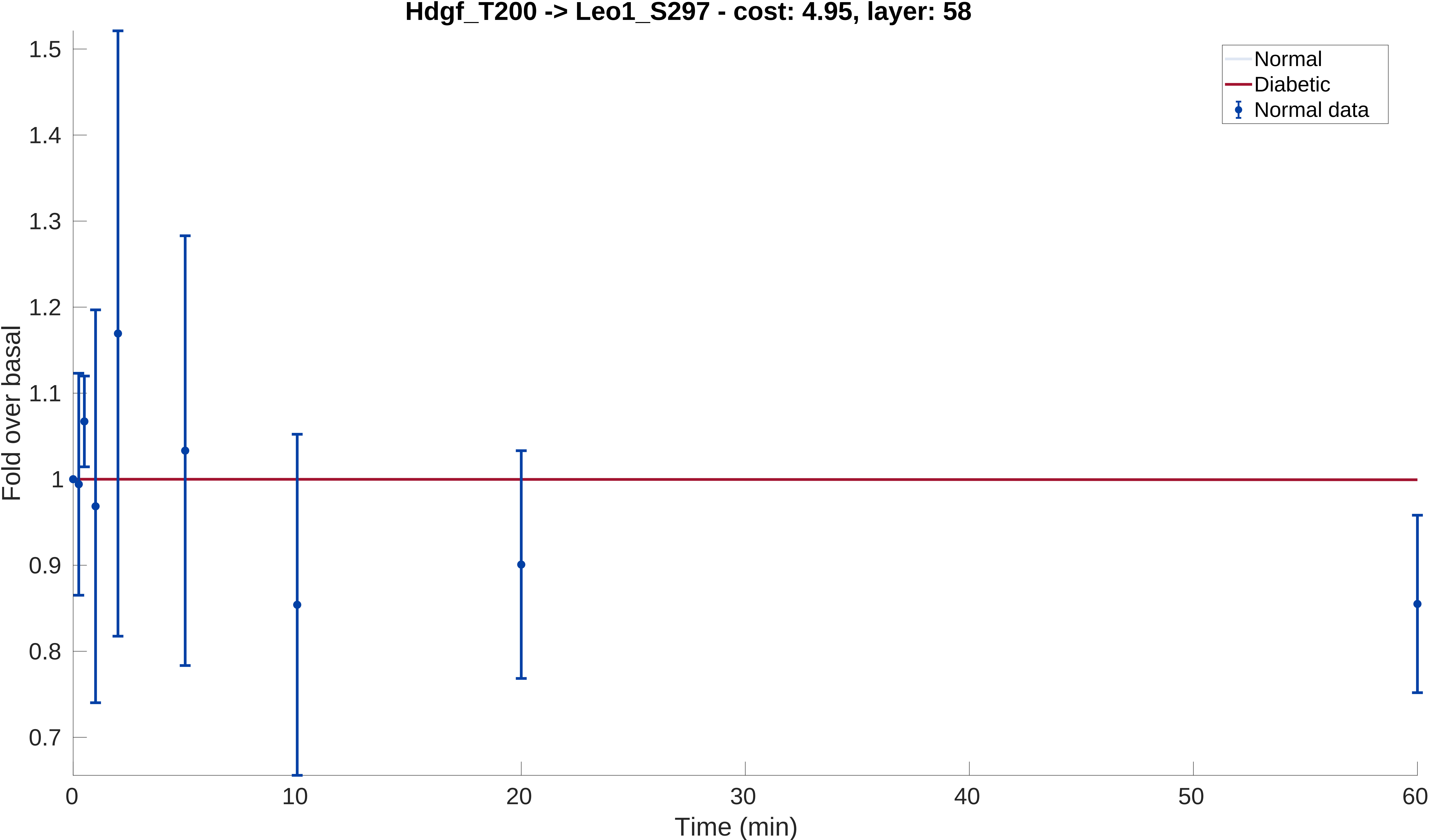

Hdgf_T200 -> Leo1_S297 - cost: 4.95, layer: 58
Normal
Diabetic
Normal data
Fold over basal
1.5
1.4
1.3
1.2
1.1
1
0.9
0.8
0.7
0
10
20
30
40
50
60
Time (min)

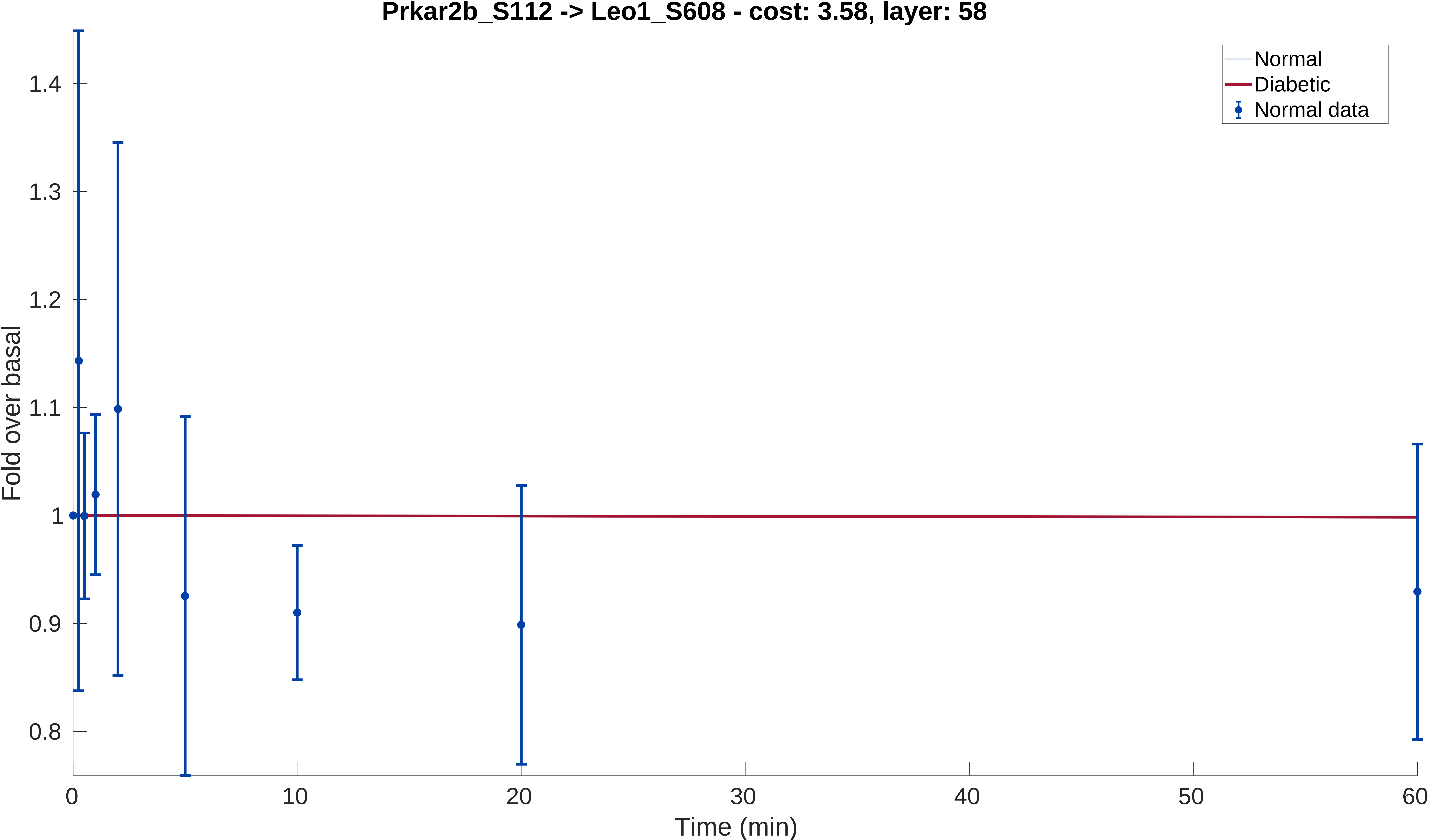

Prkar2b_S112 -> Leo1_S608 - cost: 3.58, layer: 58
Normal
Diabetic
Normal data
Fold over basal
1.4
1.3
1.2
1.1
1
0.9
0.8
0
10
20
30
40
50
60
Time (min)

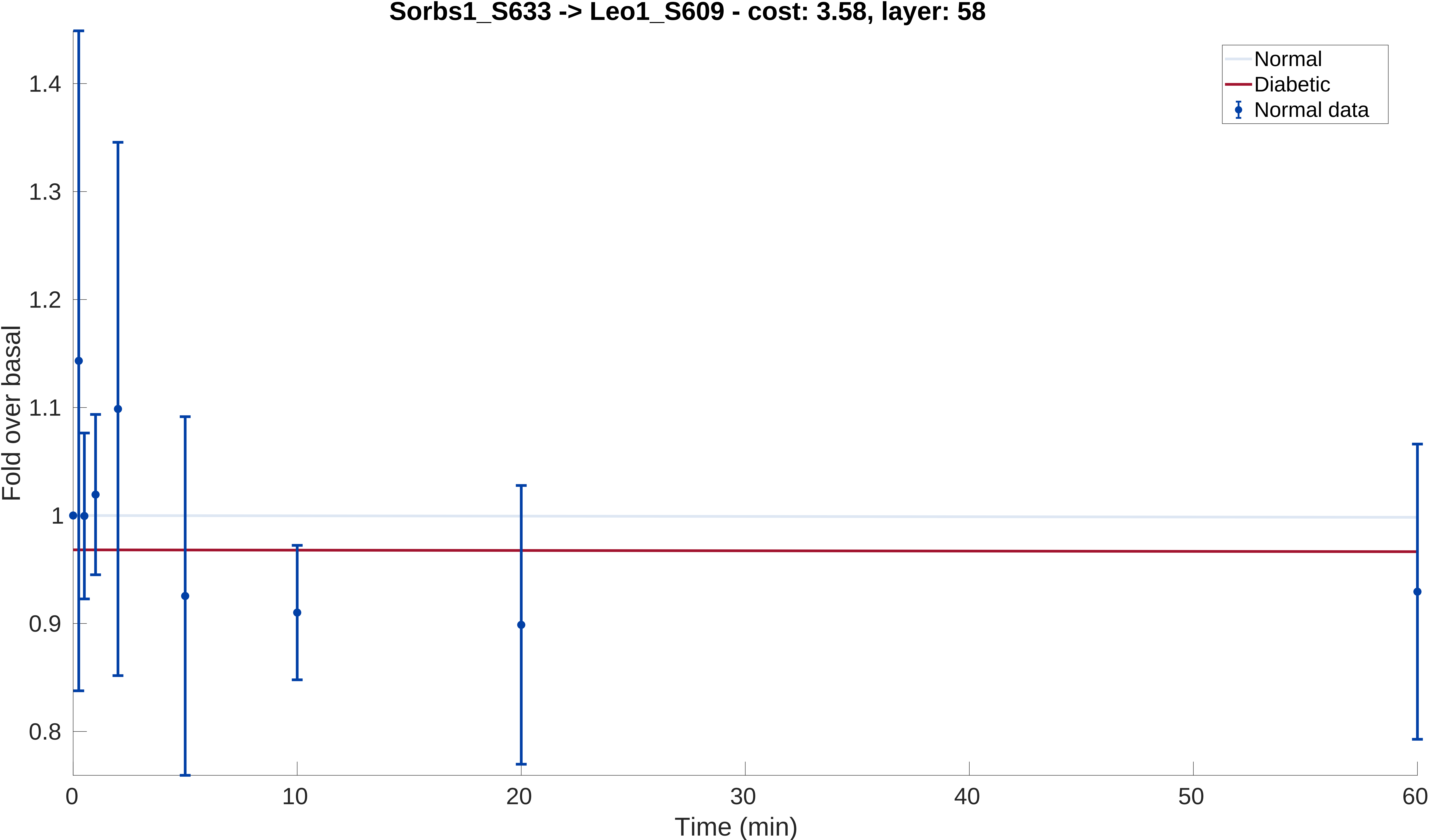
Sorbs1_S633 -> Leo1_S609 - cost: 3.58, layer: 58
Normal
Diabetic
Normal data
Fold over basal
1.4
1.3
1.2
1.1
1
0.9
0.8
0
10
20
30
40
50
60
Time (min)

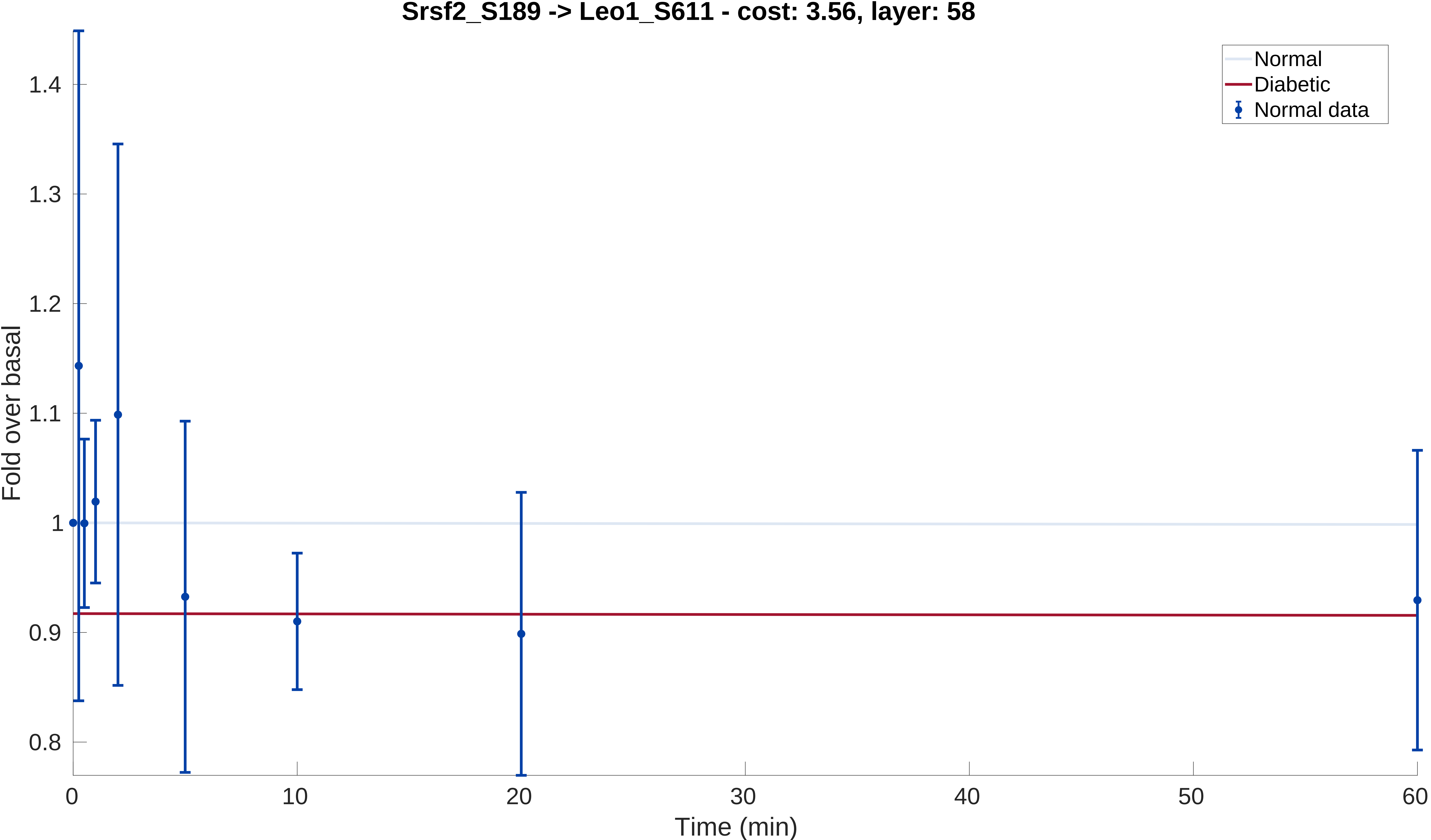

Srsf2_S189 -> Leo1_S611 - cost: 3.56, layer: 58
Normal
Diabetic
Normal data
Fold over basal
1.4
1.3
1.2
1.1
1
0.9
0.8
0
10
20
30
40
50
60
Time (min)

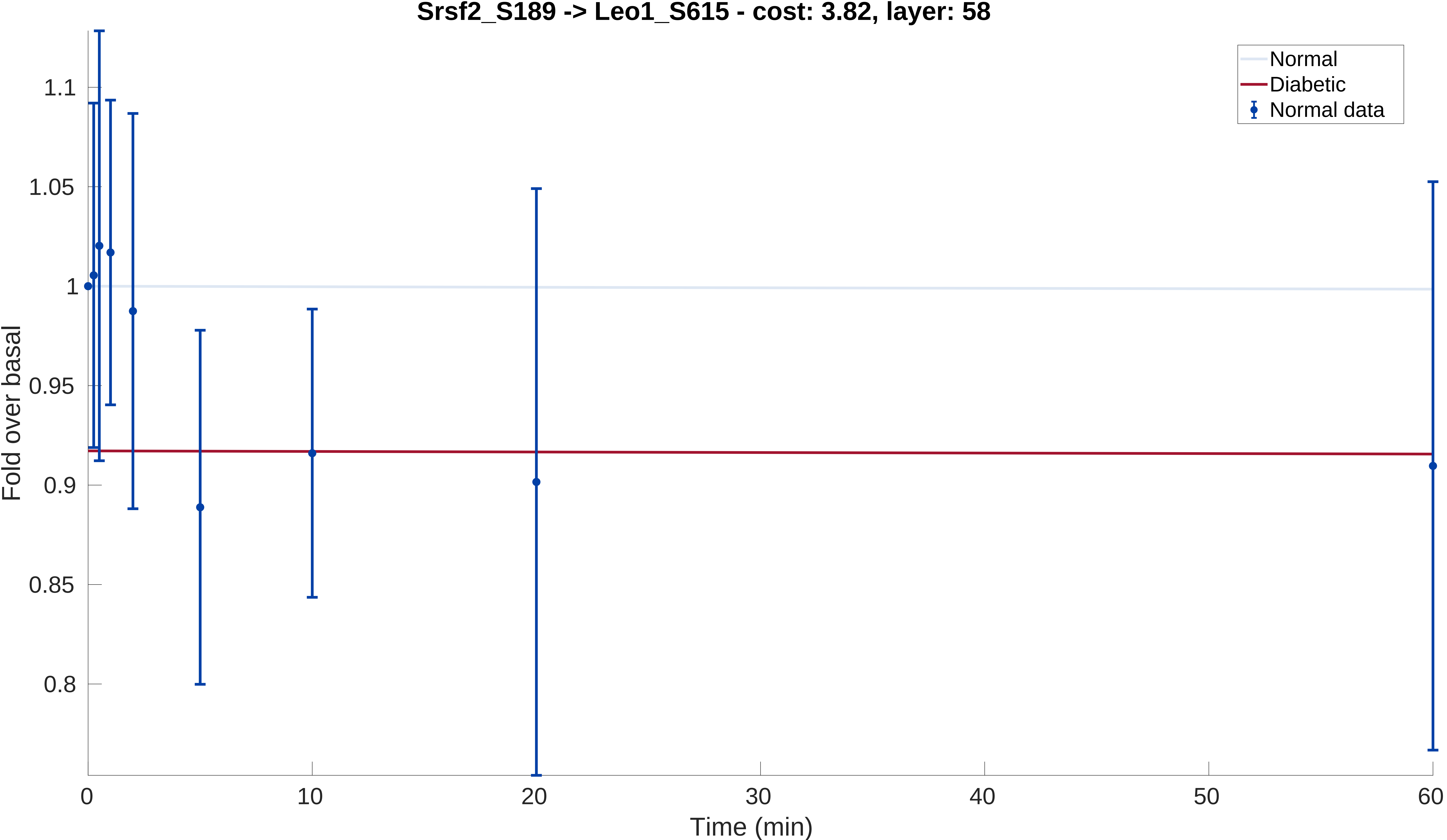
Srsf2_S189 -> Leo1_S615 - cost: 3.82, layer: 58
Normal
Diabetic
Normal data
Fold over basal
1.1
1.05
1
0.95
0.9
0.85
0.8
0
10
20
30
40
50
60
Time (min)

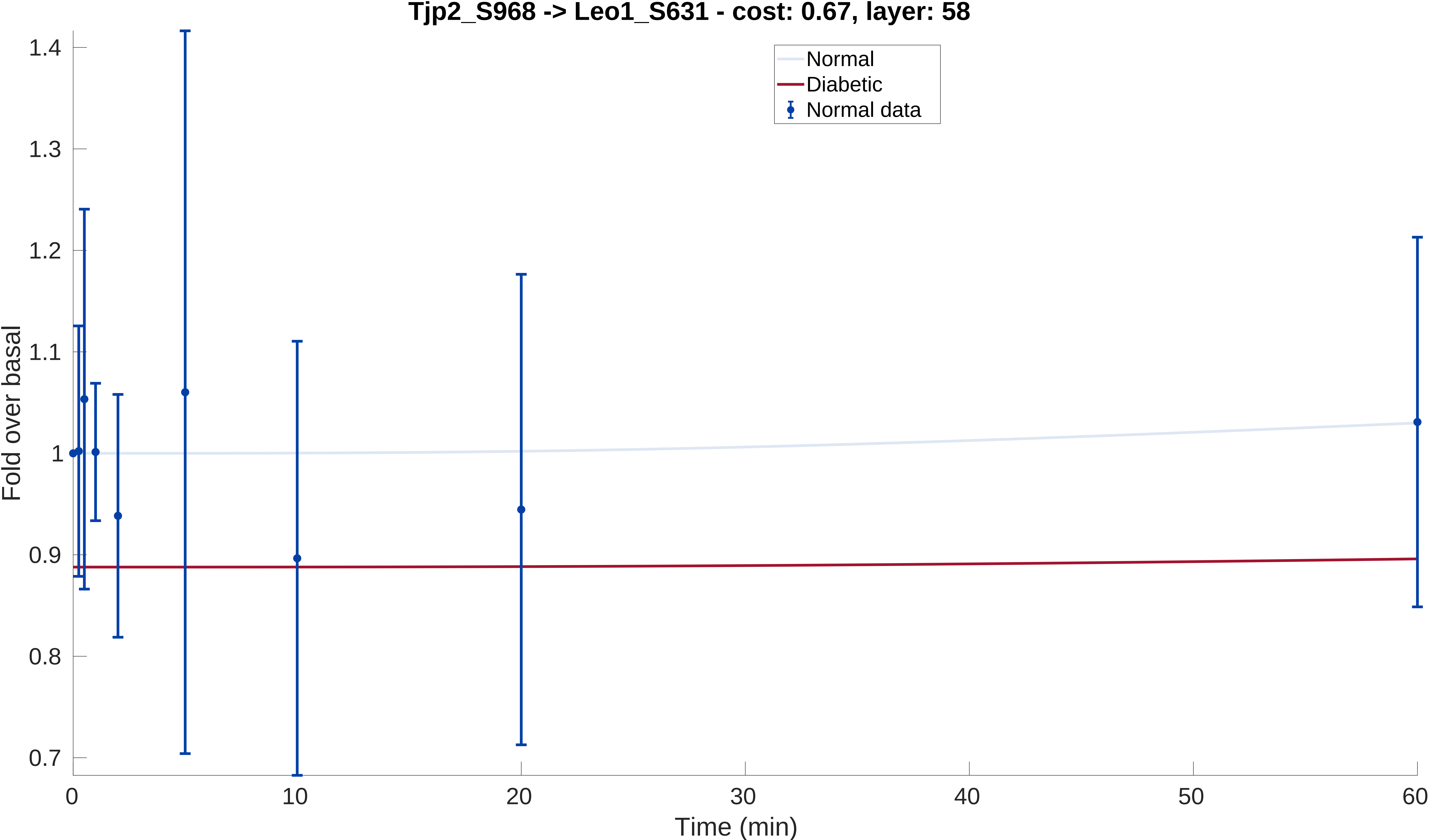

Tjp2_S968 -> Leo1_S631 - cost: 0.67, layer: 58
Normal
Diabetic
Normal data
1.4
1.3
1.2
1.1
1
0.9
0.8
0.7
Fold over basal
0
10
20
30
40
50
60
Time (min)

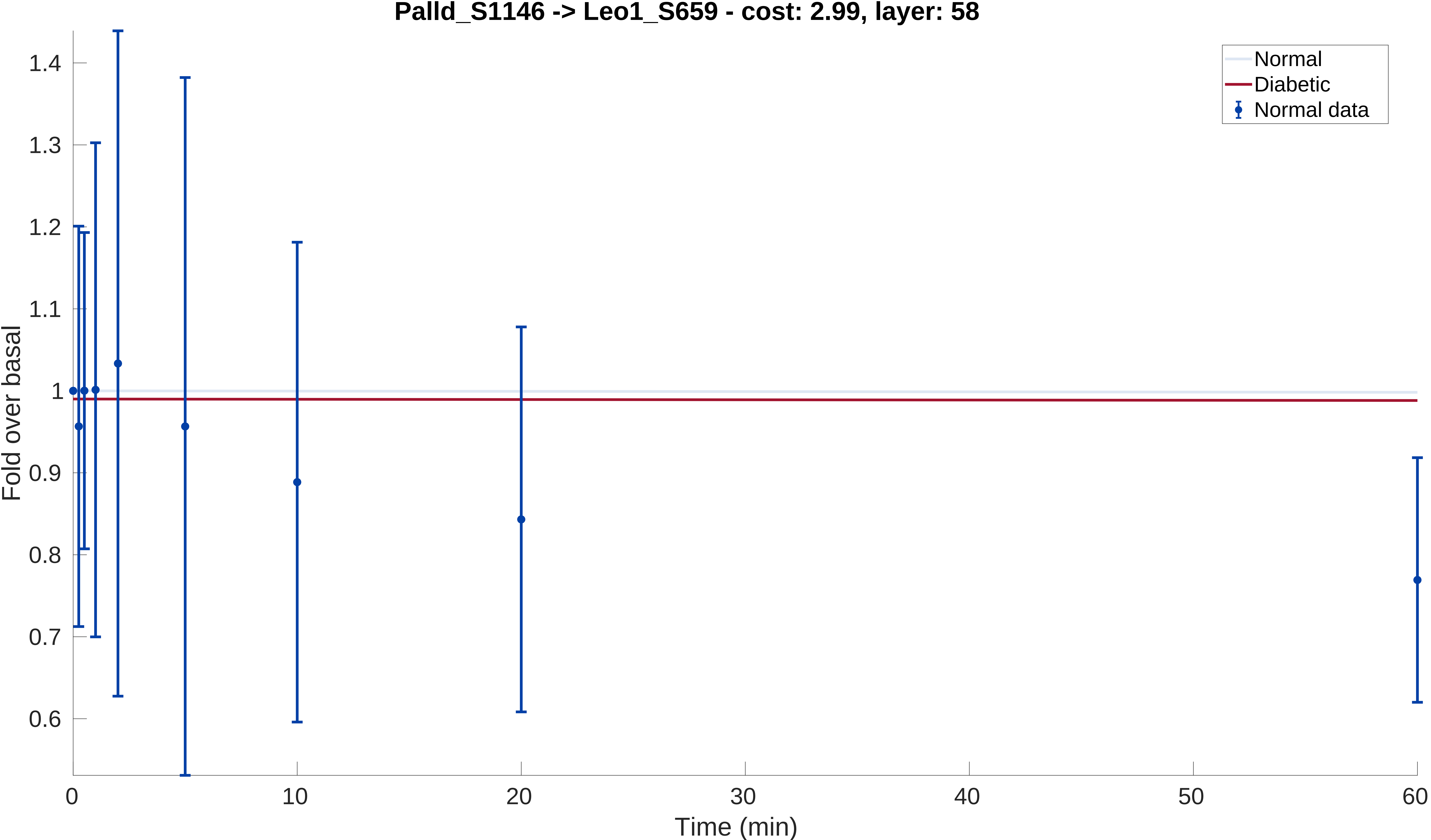
Palld_S1146 -> Leo1_S659 - cost: 2.99, layer: 58
Normal
Diabetic
Normal data
Fold over basal
1.4
1.3
1.2
1.1
1
0.9
0.8
0.7
0.6
0
10
20
30
40
50
60
Time (min)

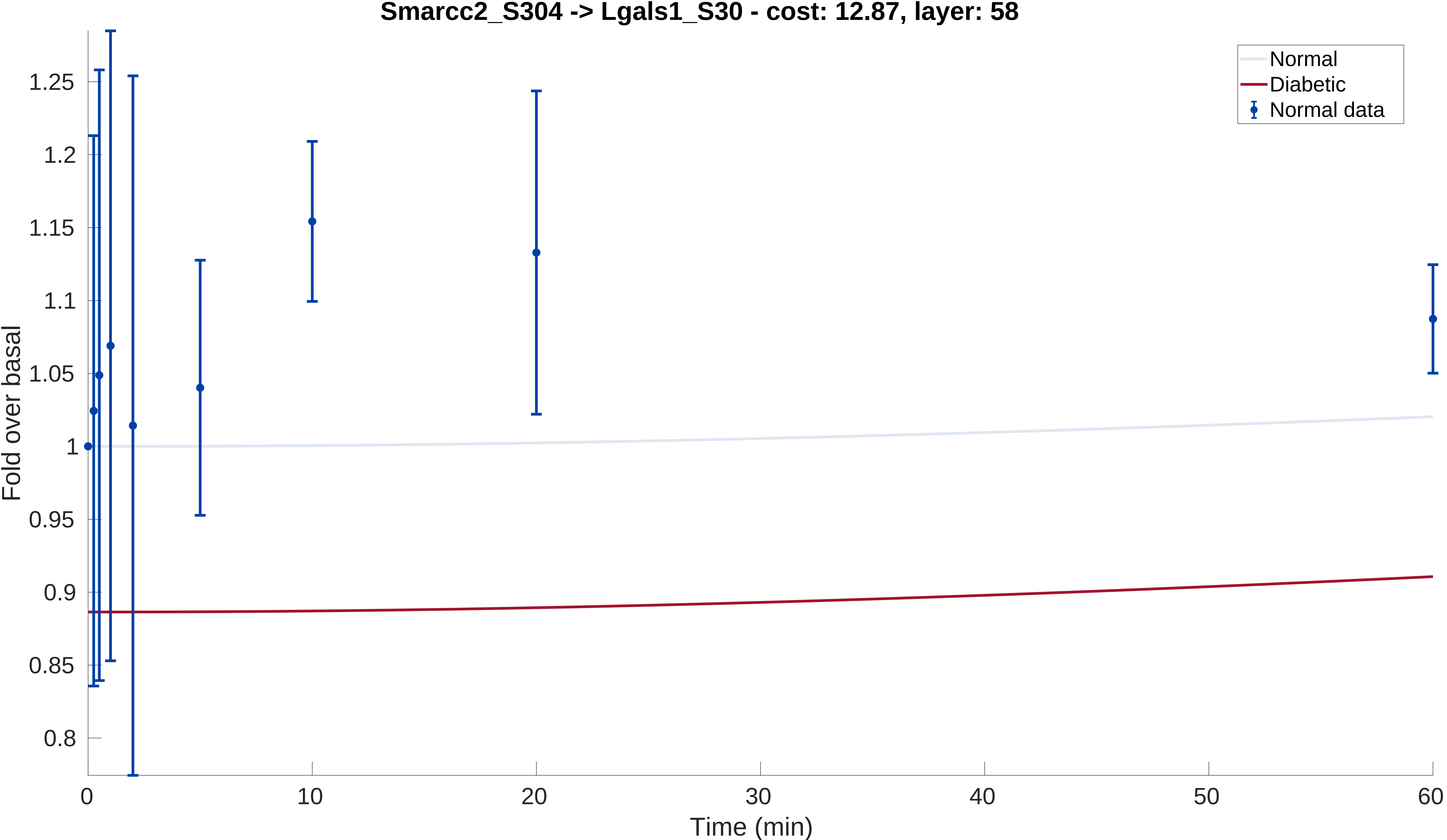

Smarcc2_S304 -> Lgals1_S30 - cost: 12.87, layer: 58
Normal
Diabetic
Normal data
1.25
1.2
1.15
1.1
1.05
1
0.95
0.9
0.85
0.8
Fold over basal
0
10
20
30
40
50
60
Time (min)

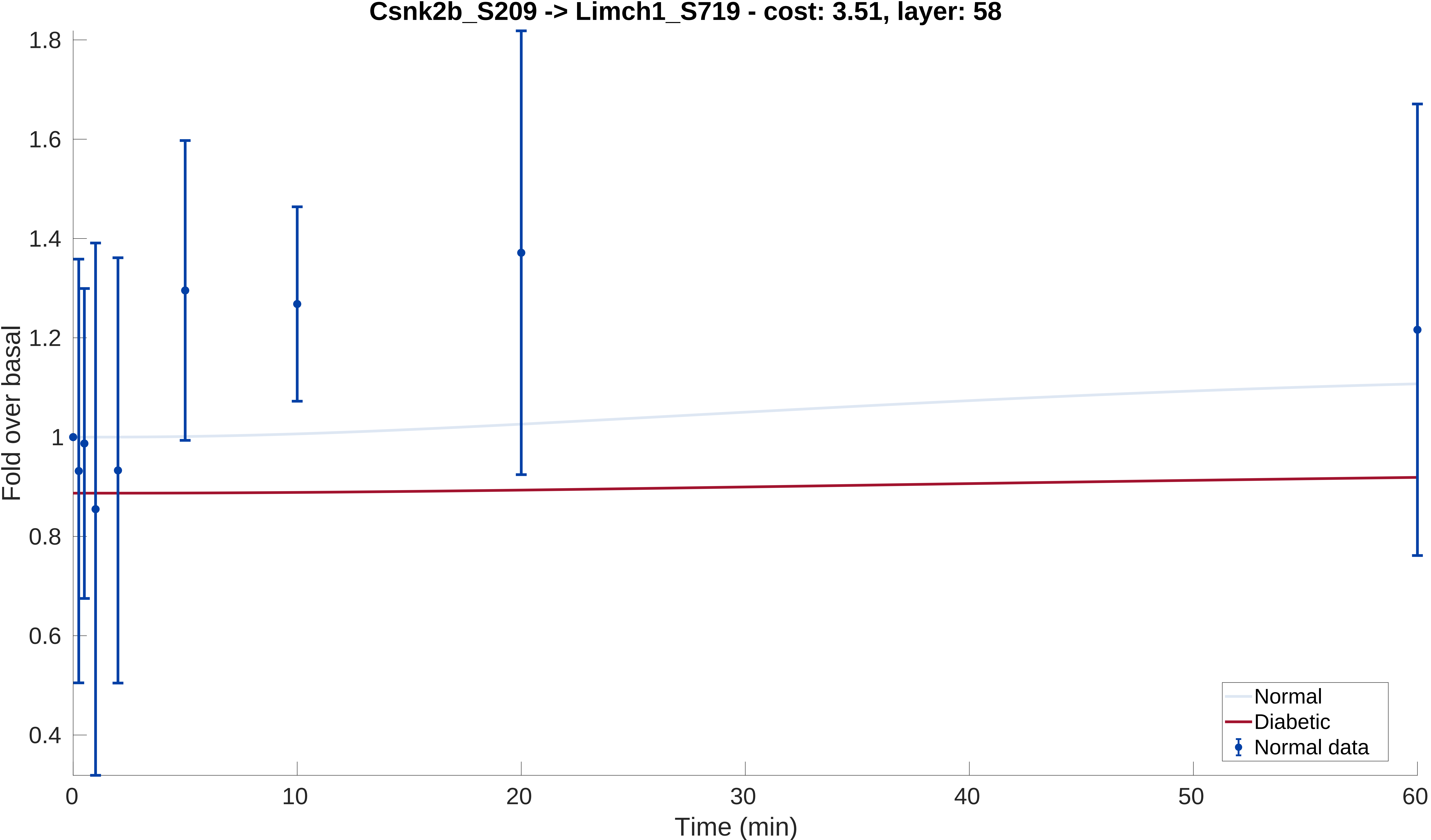

Csnk2b_S209 -> Limch1_S719 - cost: 3.51, layer: 58
Fold over basal
1.8
1.6
1.4
1.2
1
0.8
0.6
0.4
0
10
20
30
40
50
60
Time (min)
Normal
Diabetic
Normal data

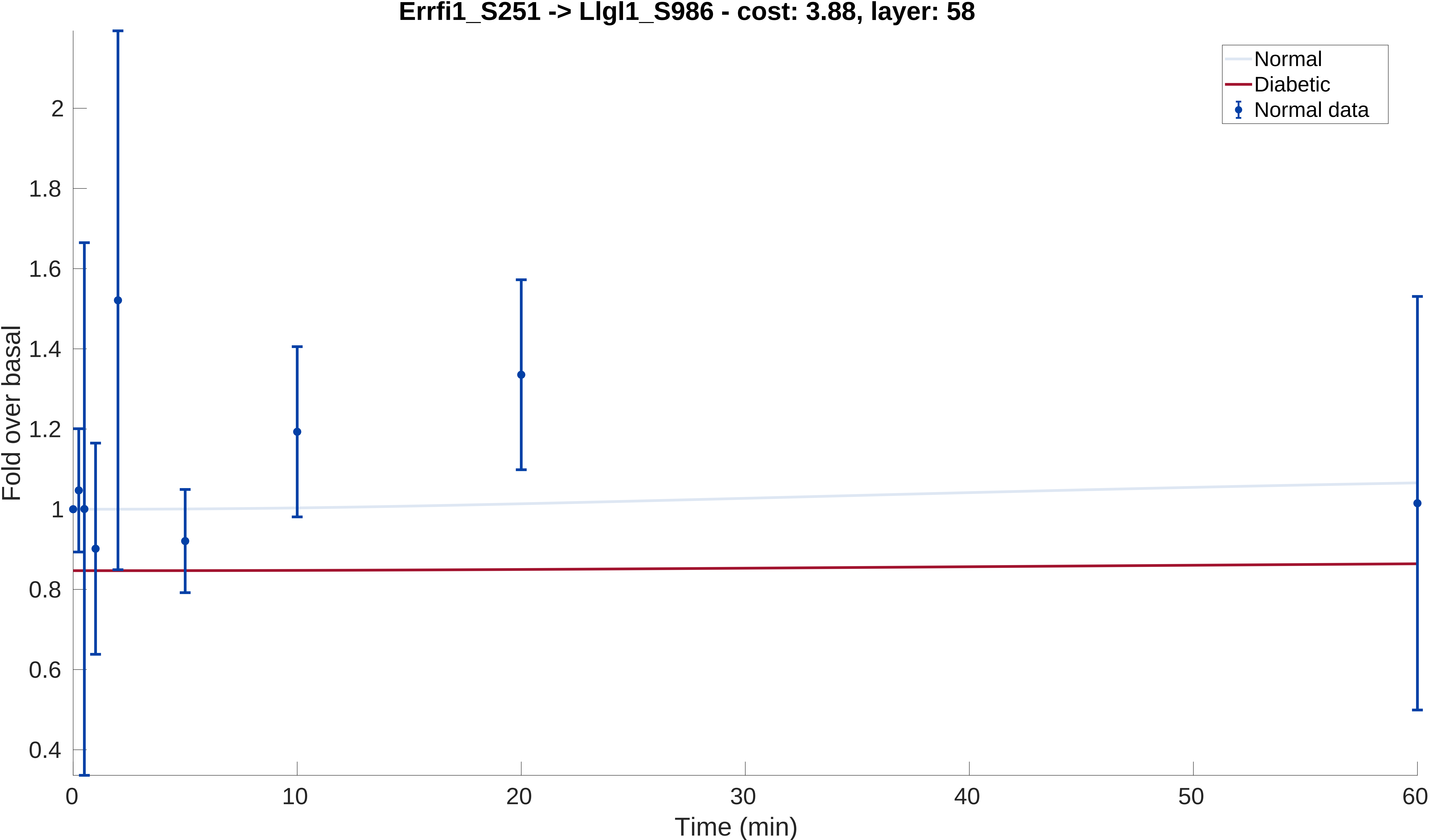

Errfi1_S251 -> Llgl1_S986 - cost: 3.88, layer: 58
Normal
Diabetic
Normal data
Fold over basal
2
1.8
1.6
1.4
1.2
1
0.8
0.6
0.4
0
10
20
30
40
50
60
Time (min)

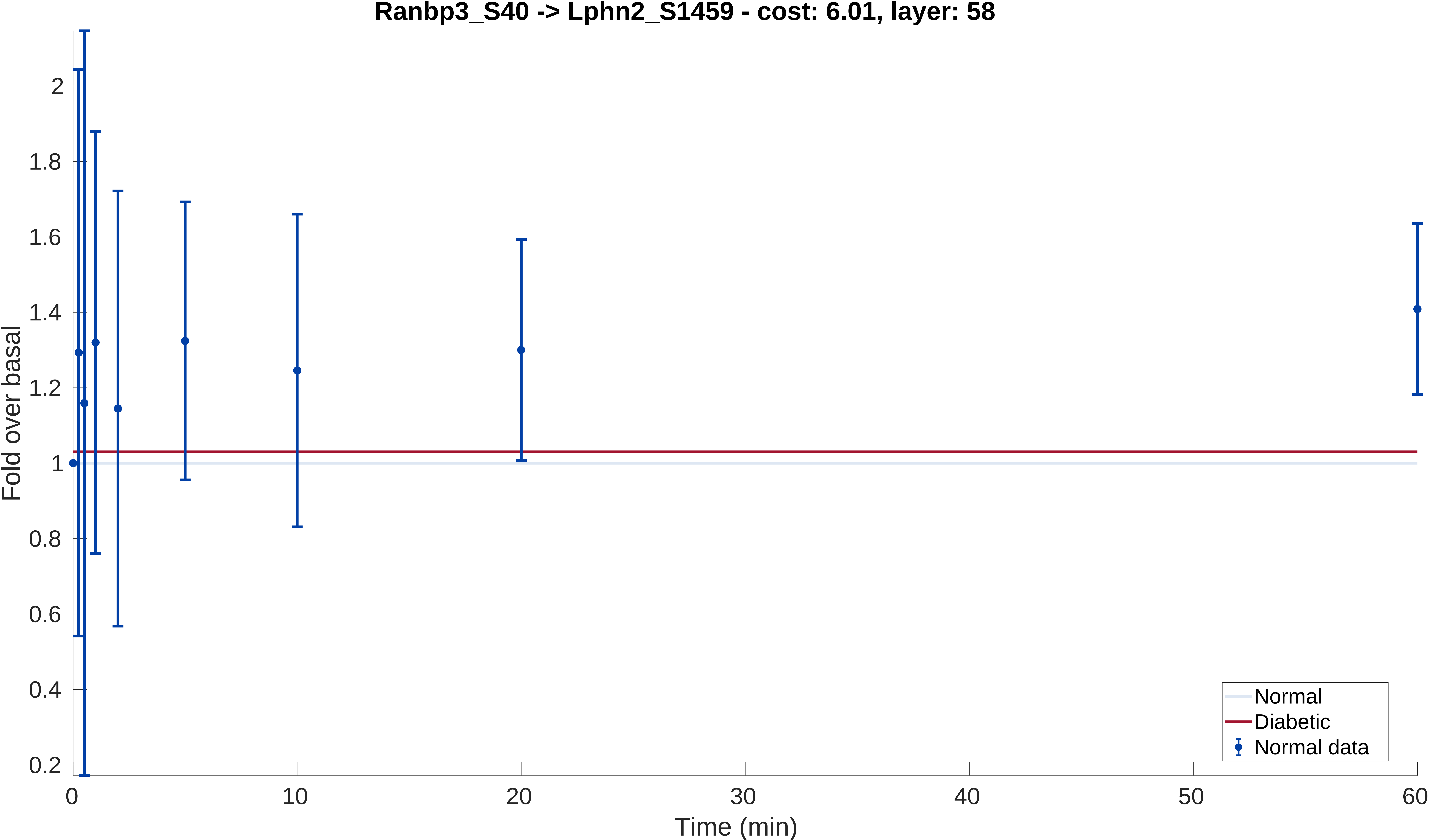
Ranbp3_S40 -> Lphn2_S1459 - cost: 6.01, layer: 58
Fold over basal
Time (min)
Normal
Diabetic
Normal data
0.2
0.4
0.6
0.8
1
1.2
1.4
1.6
1.8
2
0
10
20
30
40
50
60

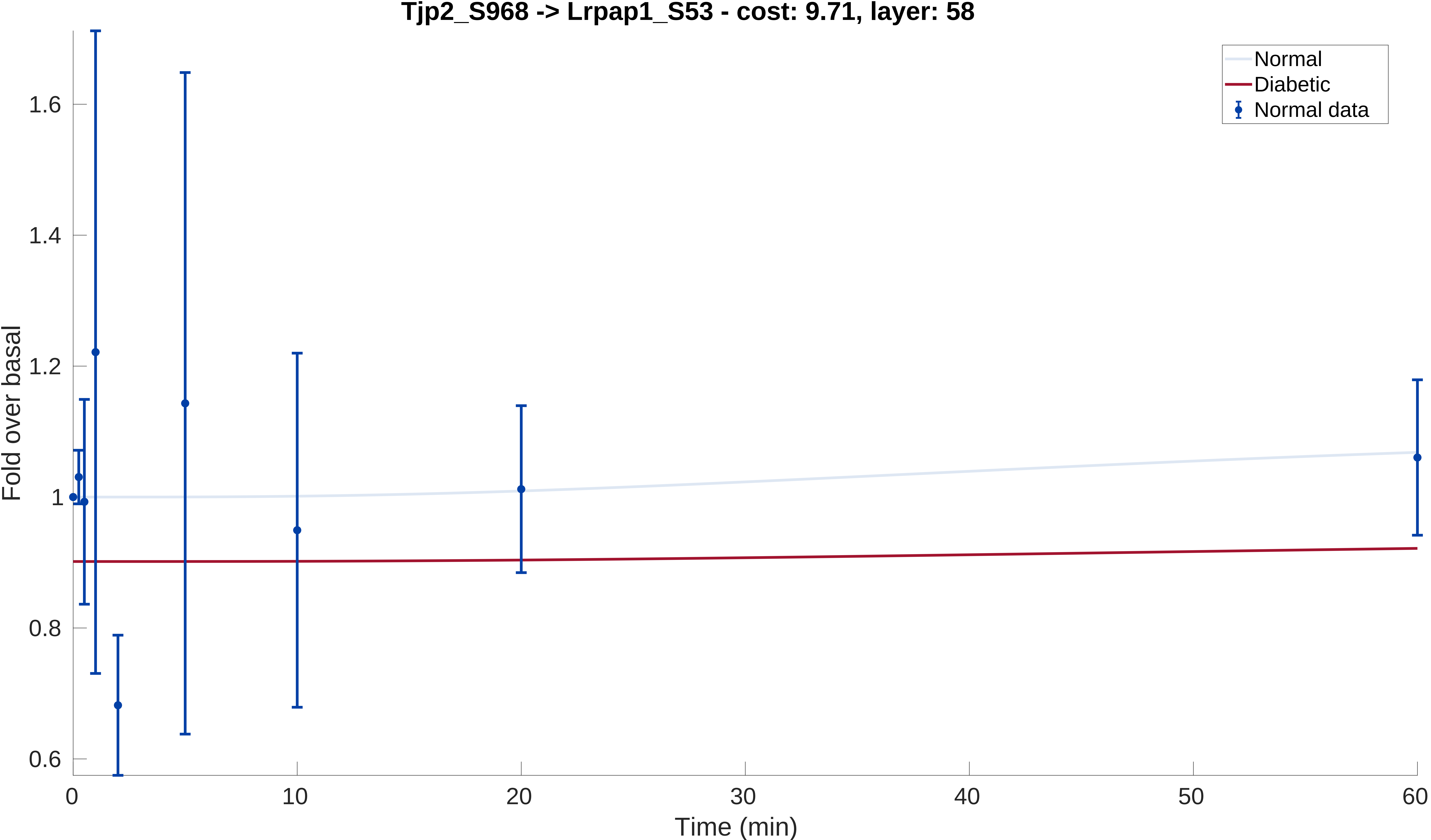

Tjp2_S968 -> Lrpap1_S53 - cost: 9.71, layer: 58
Normal
Diabetic
Normal data
Fold over basal
1.6
1.4
1.2
1
0.8
0.6
0
10
20
30
40
50
60
Time (min)

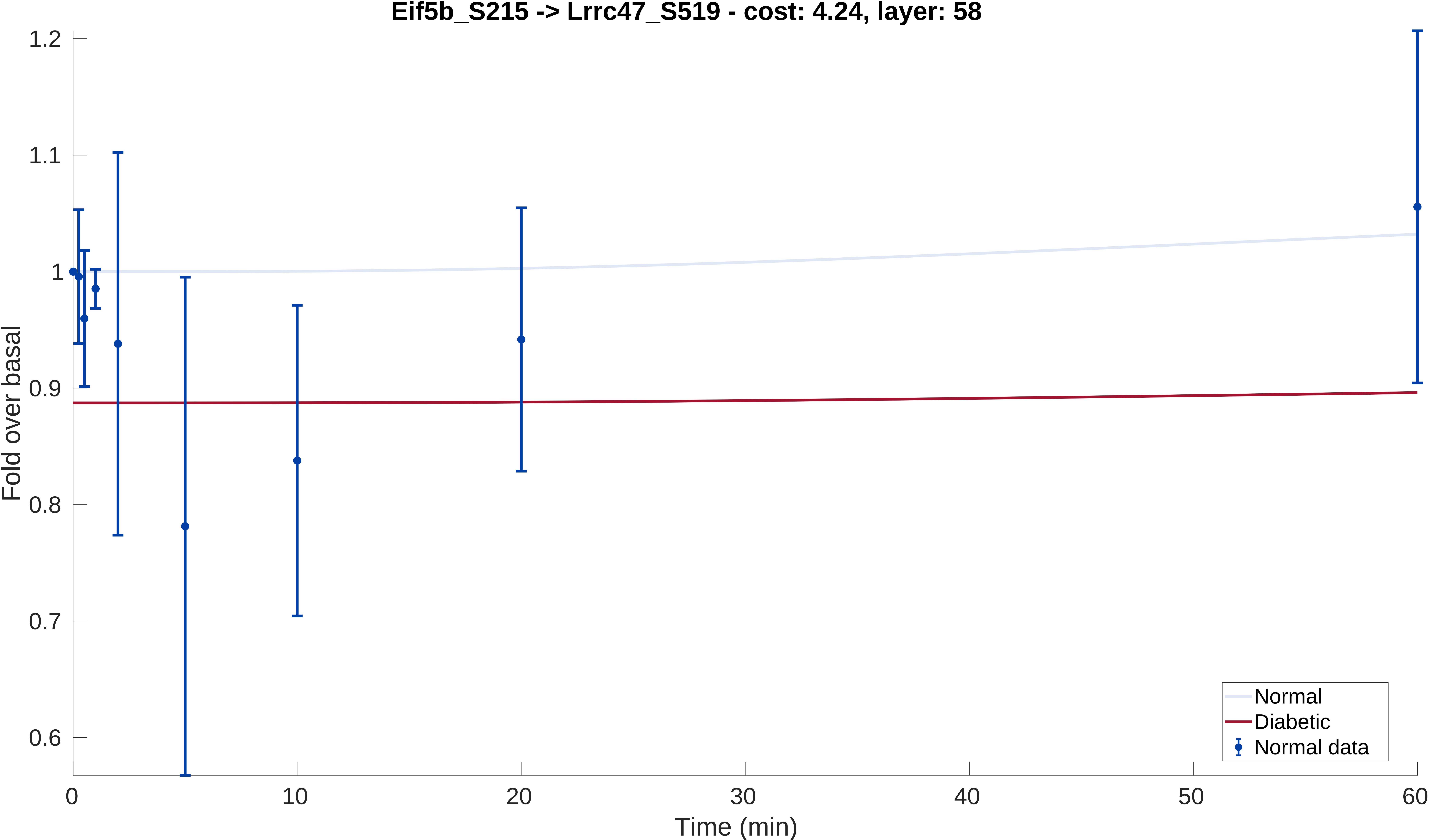

Eif5b_S215 -> Lrrc47_S519 - cost: 4.24, layer: 58
Fold over basal
1.2
1.1
1
0.9
0.8
0.7
0.6
0
10
20
30
40
50
60
Time (min)
Normal
Diabetic
Normal data

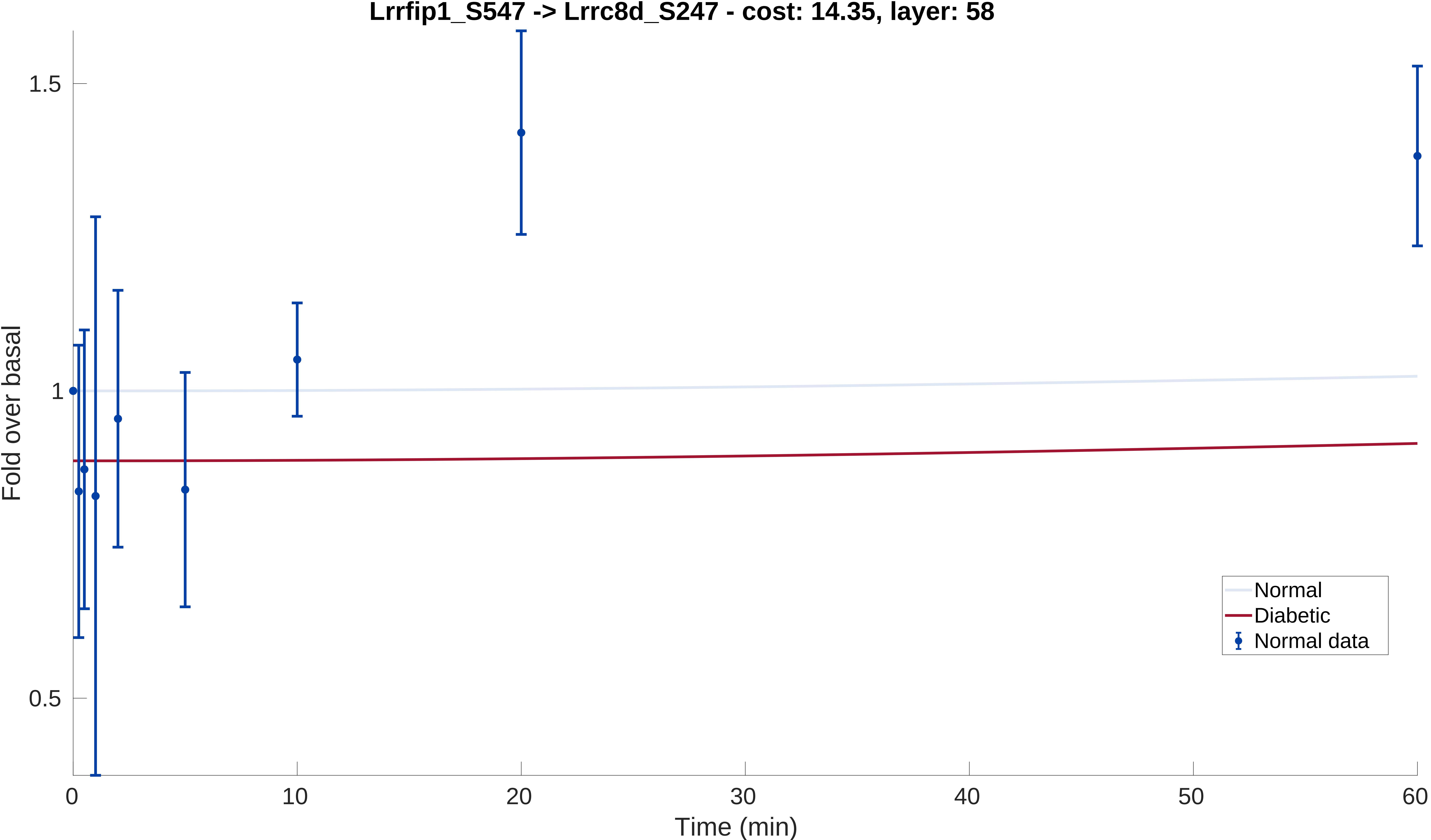

Lrrfip1_S547 -> Lrrc8d_S247 - cost: 14.35, layer: 58
Fold over basal
1.5
1
0.5
0
10
20
30
40
50
60
Time (min)
Normal
Diabetic
Normal data

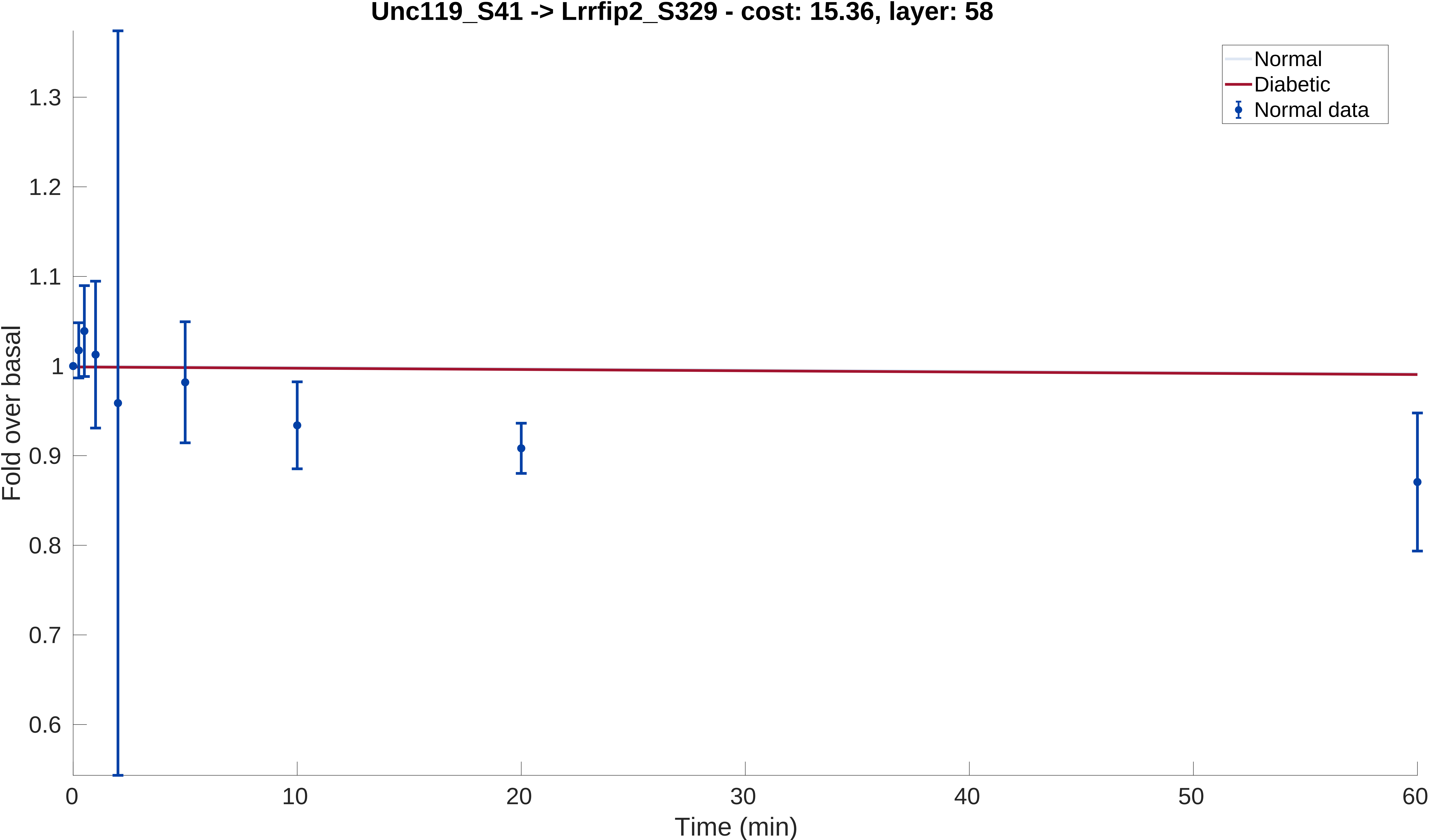

Unc119_S41 -> Lrrfip2_S329 - cost: 15.36, layer: 58
Normal
Diabetic
Normal data
Fold over basal
Time (min)
0.6
0.7
0.8
0.9
1
1.1
1.2
1.3
0
10
20
30
40
50
60

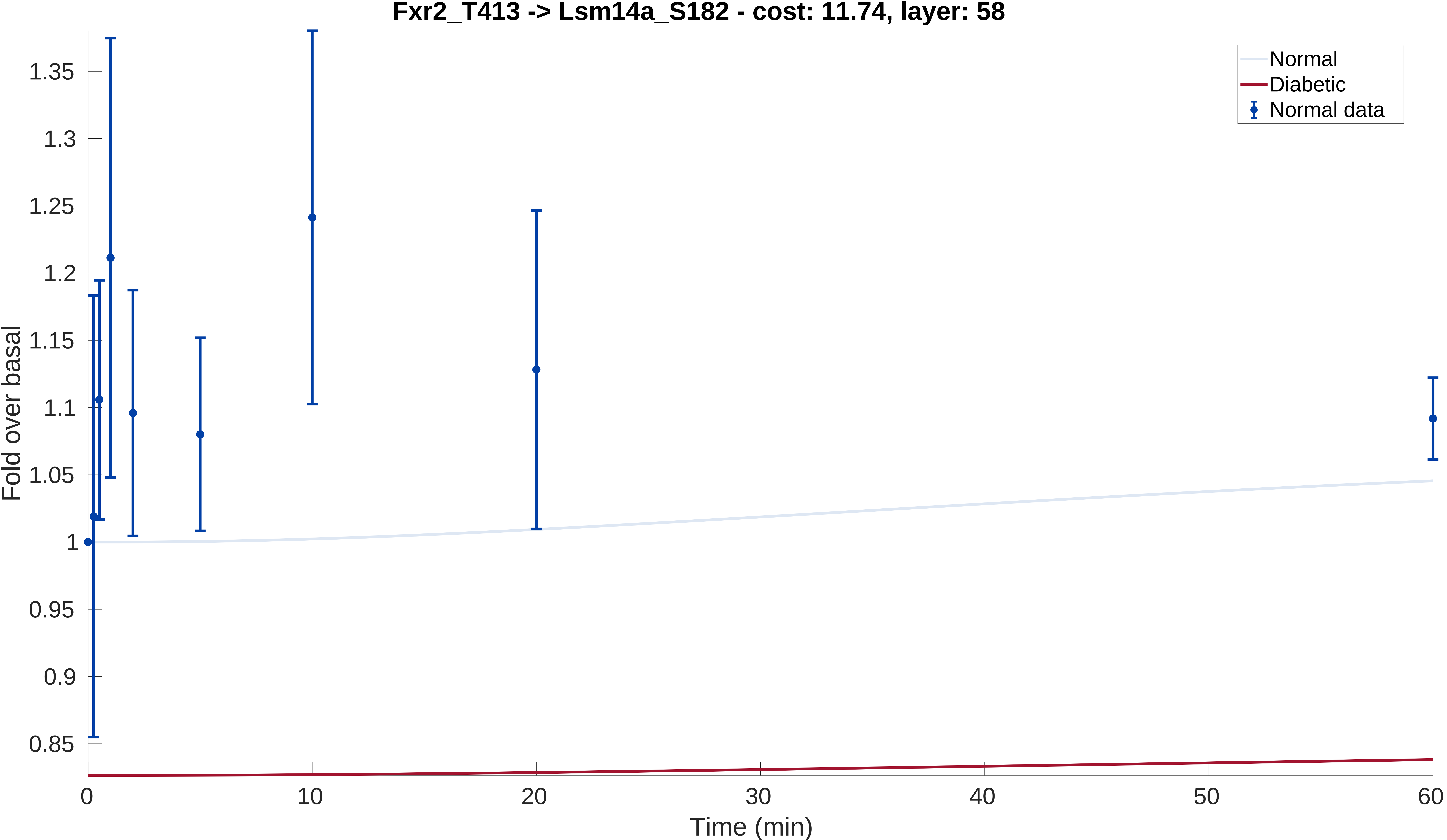
Fxr2_T413 -> Lsm14a_S182 - cost: 11.74, layer: 58
Normal
Diabetic
Normal data
Fold over basal
1.35
1.3
1.25
1.2
1.15
1.1
1.05
1
0.95
0.9
0.85
0
10
20
30
40
50
60
Time (min)

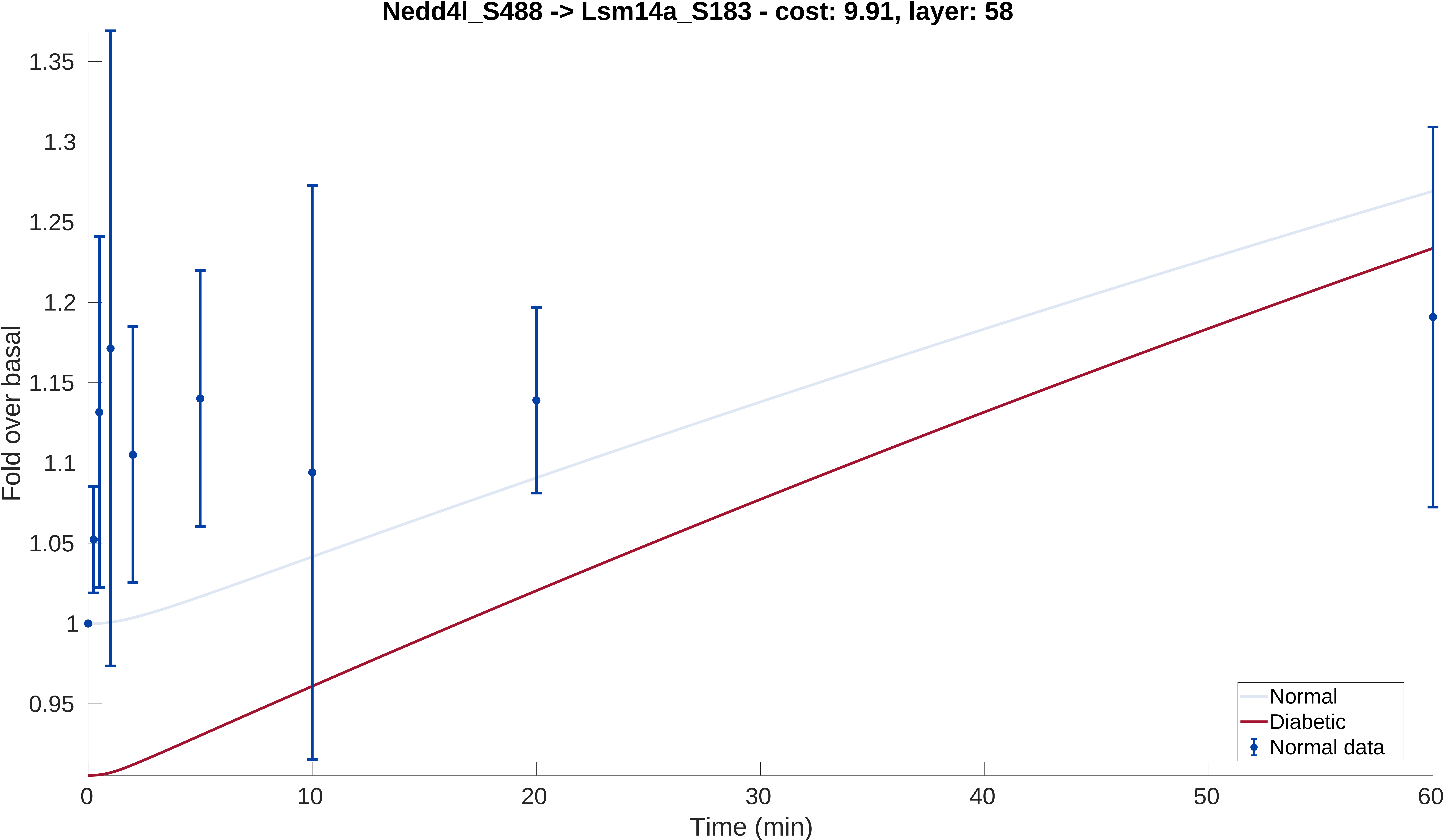

Nedd4l_S488 -> Lsm14a_S183 - cost: 9.91, layer: 58
Fold over basal
1.35
1.3
1.25
1.2
1.15
1.1
1.05
1
0.95
0
10
20
30
40
50
60
Time (min)
Normal
Diabetic
Normal data

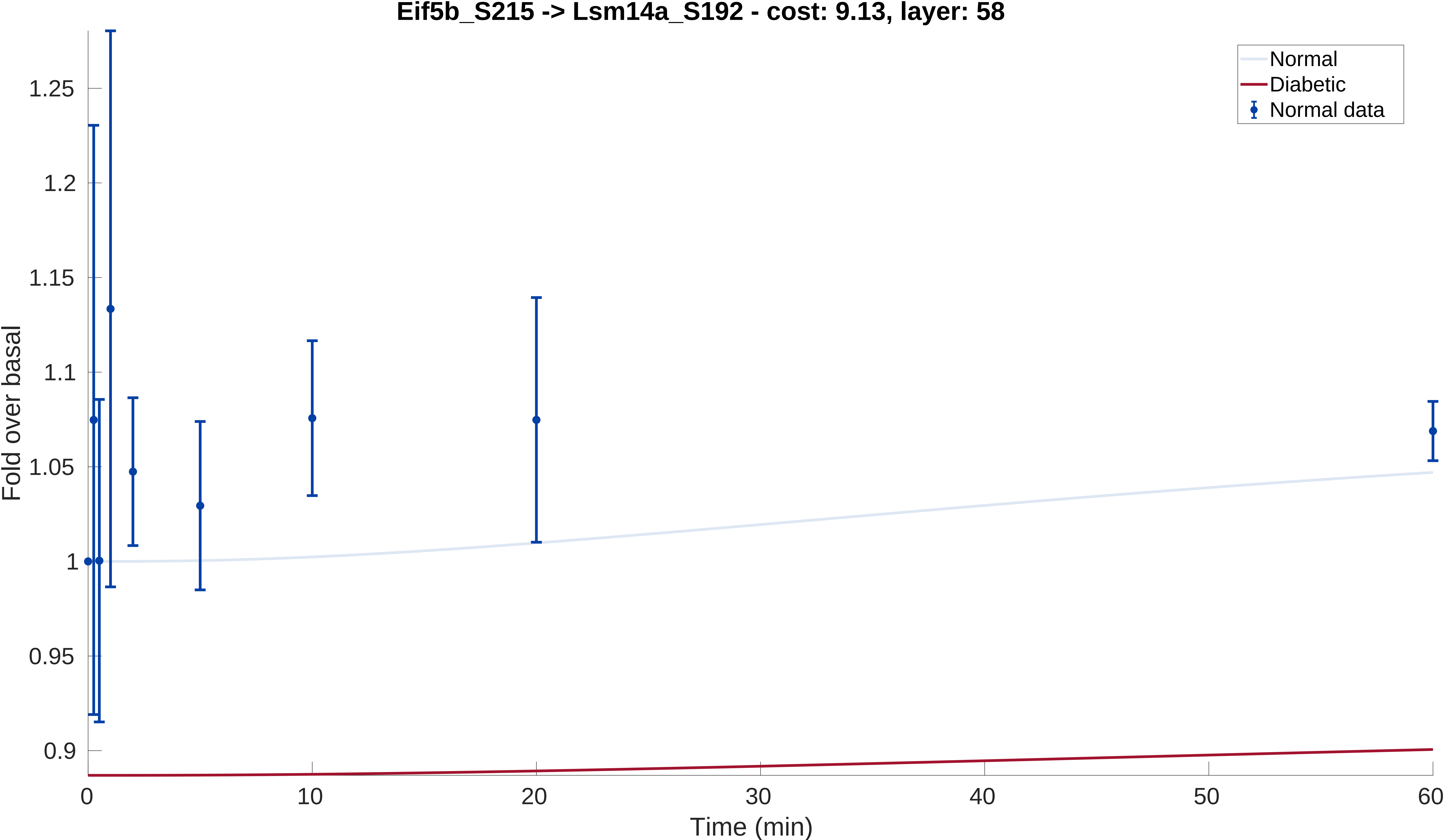

Eif5b_S215 -> Lsm14a_S192 - cost: 9.13, layer: 58
Normal
Diabetic
Normal data
Fold over basal
1.25
1.2
1.15
1.1
1.05
1
0.95
0.9
0
10
20
30
40
50
60
Time (min)

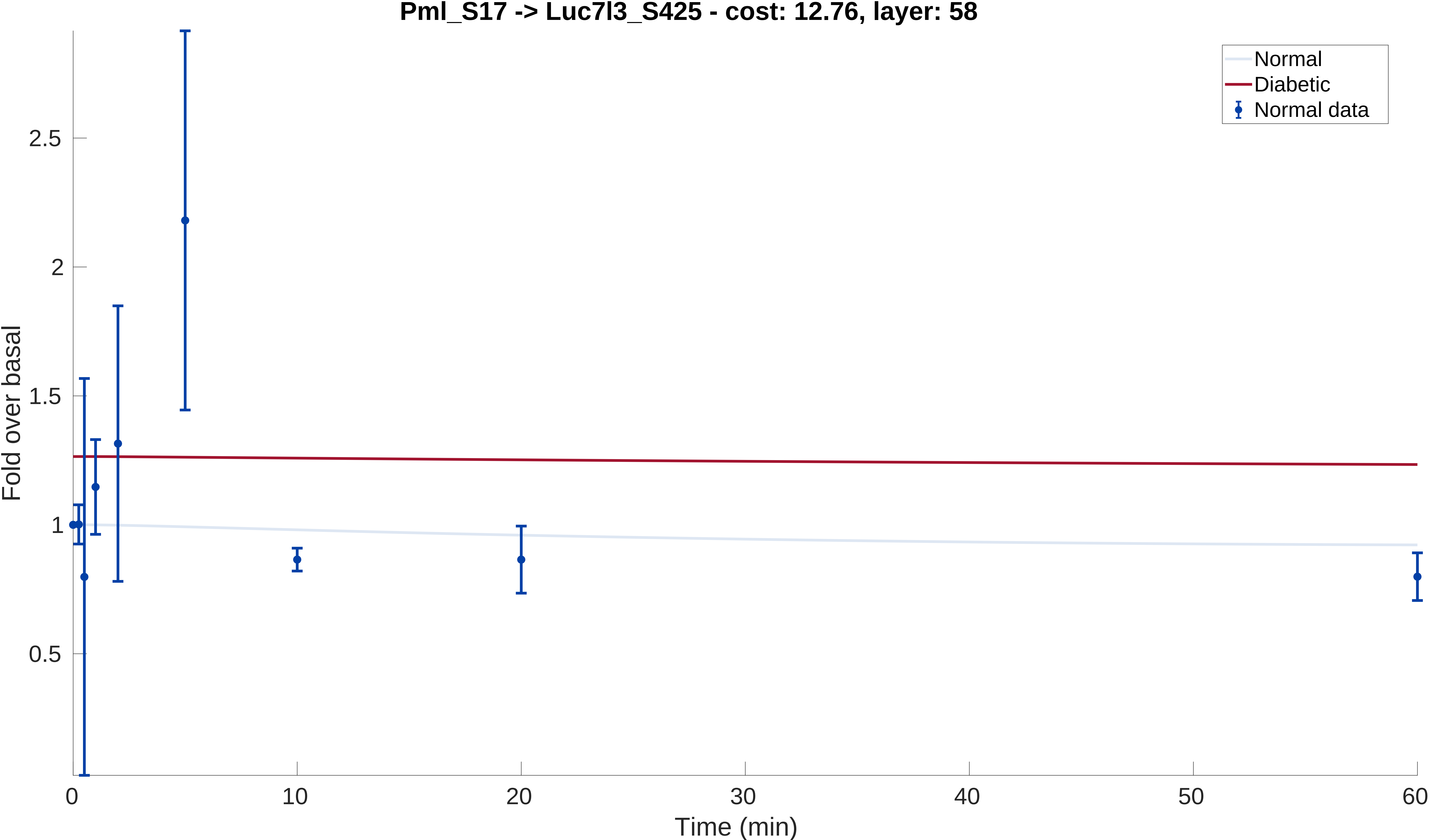

Pml_S17 -> Luc7l3_S425 - cost: 12.76, layer: 58
Normal
Diabetic
Normal data
Fold over basal
0.5
1
1.5
2
2.5
0
10
20
30
40
50
60
Time (min)

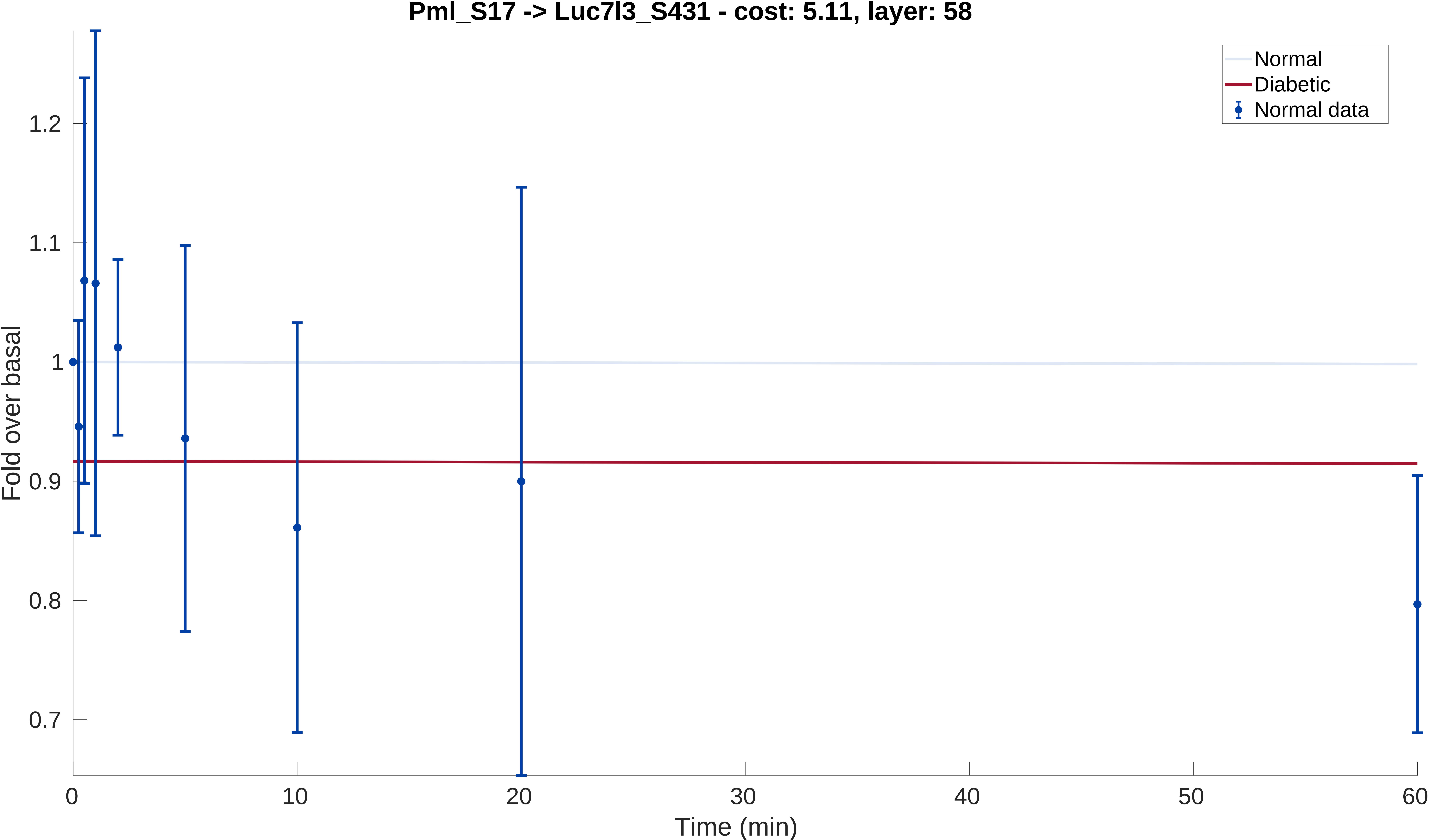

Pml_S17 -> Luc7l3_S431 - cost: 5.11, layer: 58
Normal
Diabetic
Normal data
Fold over basal
Time (min)
1.2
1.1
1
0.9
0.8
0.7
0
10
20
30
40
50
60

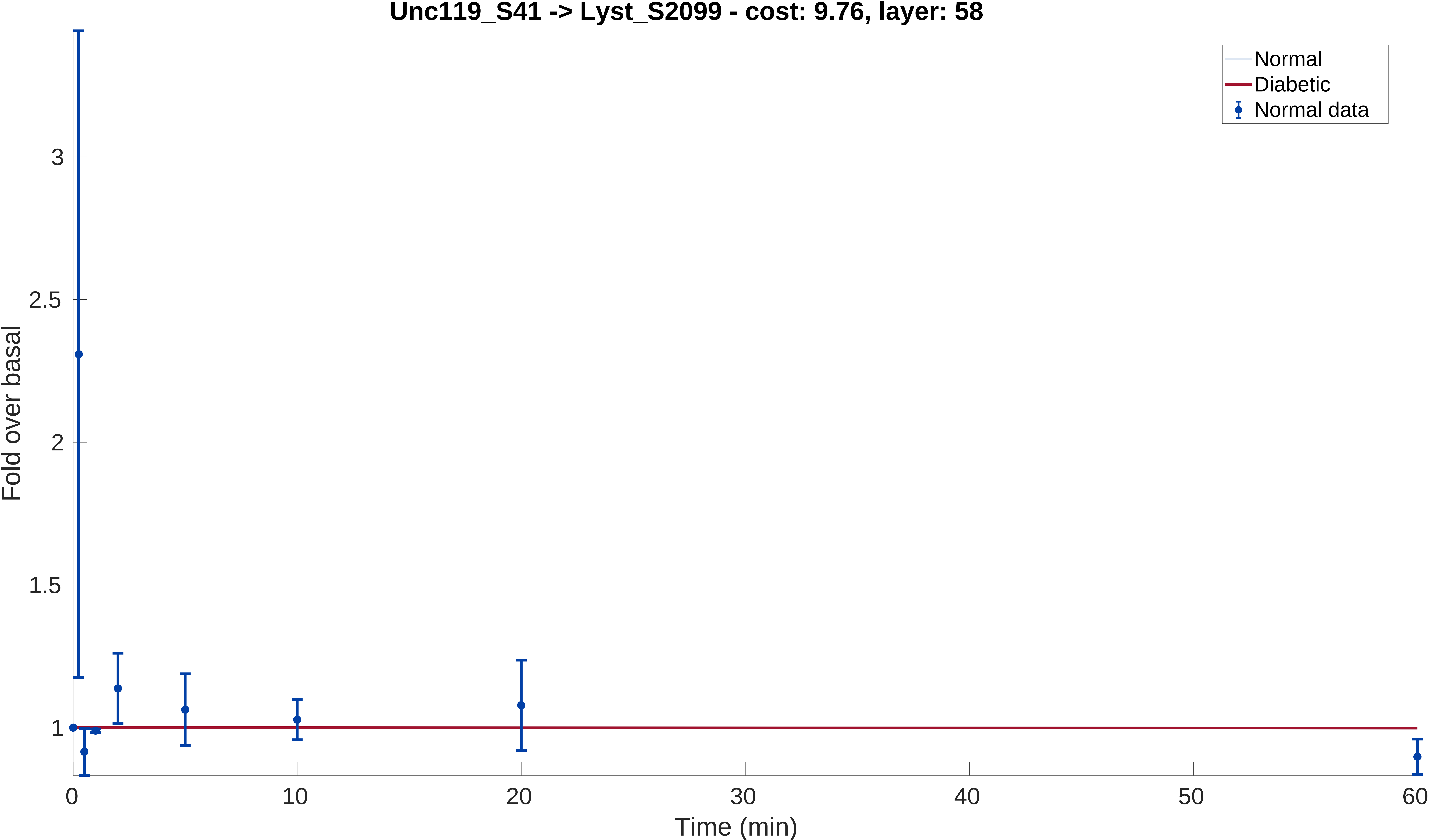

Unc119_S41 -> Lyst_S2099 - cost: 9.76, layer: 58
Normal
Diabetic
Normal data
Fold over basal
3
2.5
2
1.5
1
0
10
20
30
40
50
60
Time (min)

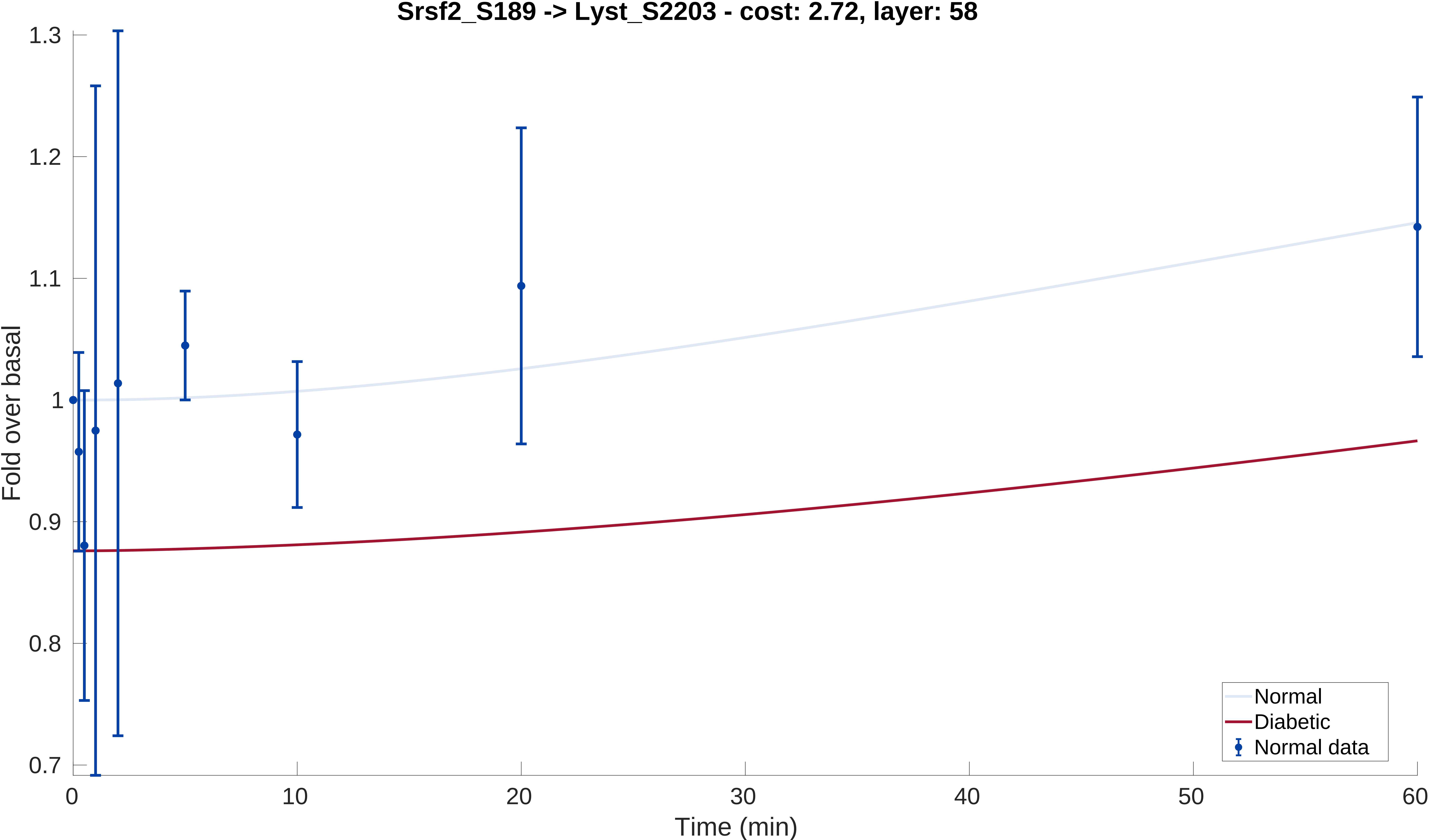

Srsf2_S189 -> Lyst_S2203 - cost: 2.72, layer: 58
Fold over basal
1.3
1.2
1.1
1
0.9
0.8
0.7
0
10
20
30
40
50
60
Time (min)
Normal
Diabetic
Normal data

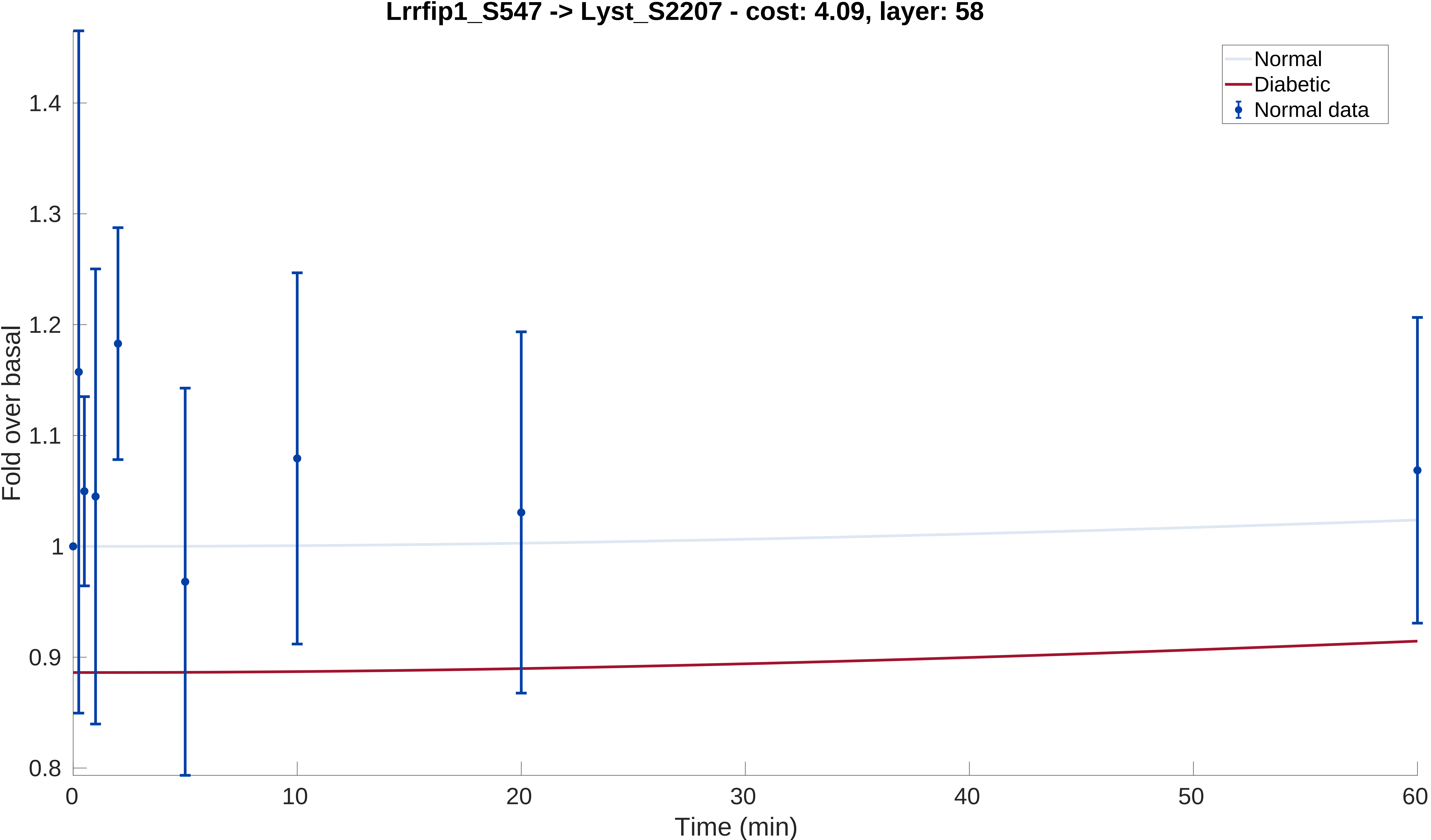

Lrrfip1_S547 -> Lyst_S2207 - cost: 4.09, layer: 58
Normal
Diabetic
Normal data
Fold over basal
1.4
1.3
1.2
1.1
1
0.9
0.8
0
10
20
30
40
50
60
Time (min)

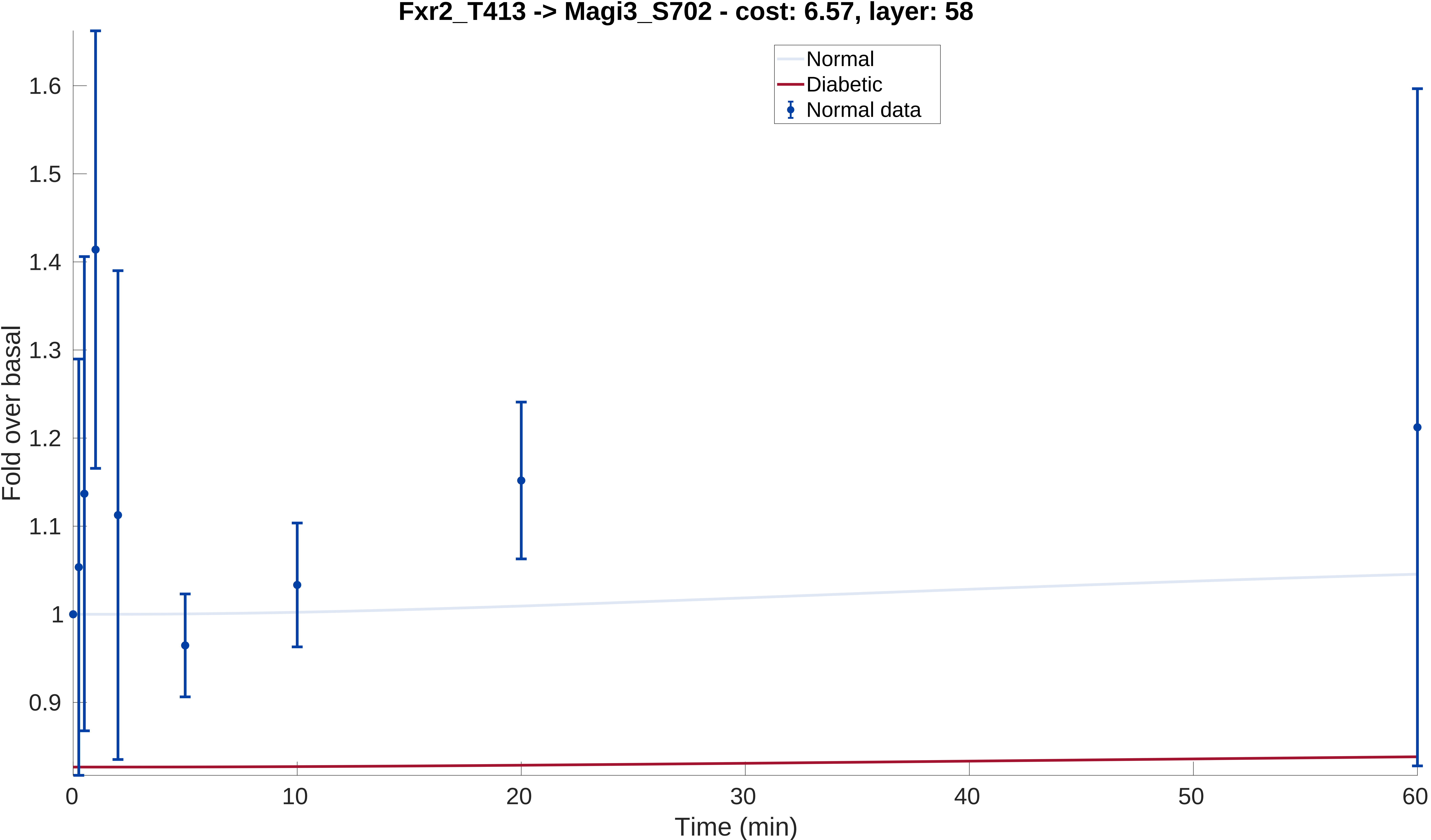

Fxr2_T413 -> Magi3_S702 - cost: 6.57, layer: 58
Normal
Diabetic
Normal data
Fold over basal
1.6
1.5
1.4
1.3
1.2
1.1
1
0.9
0
10
20
30
40
50
60
Time (min)

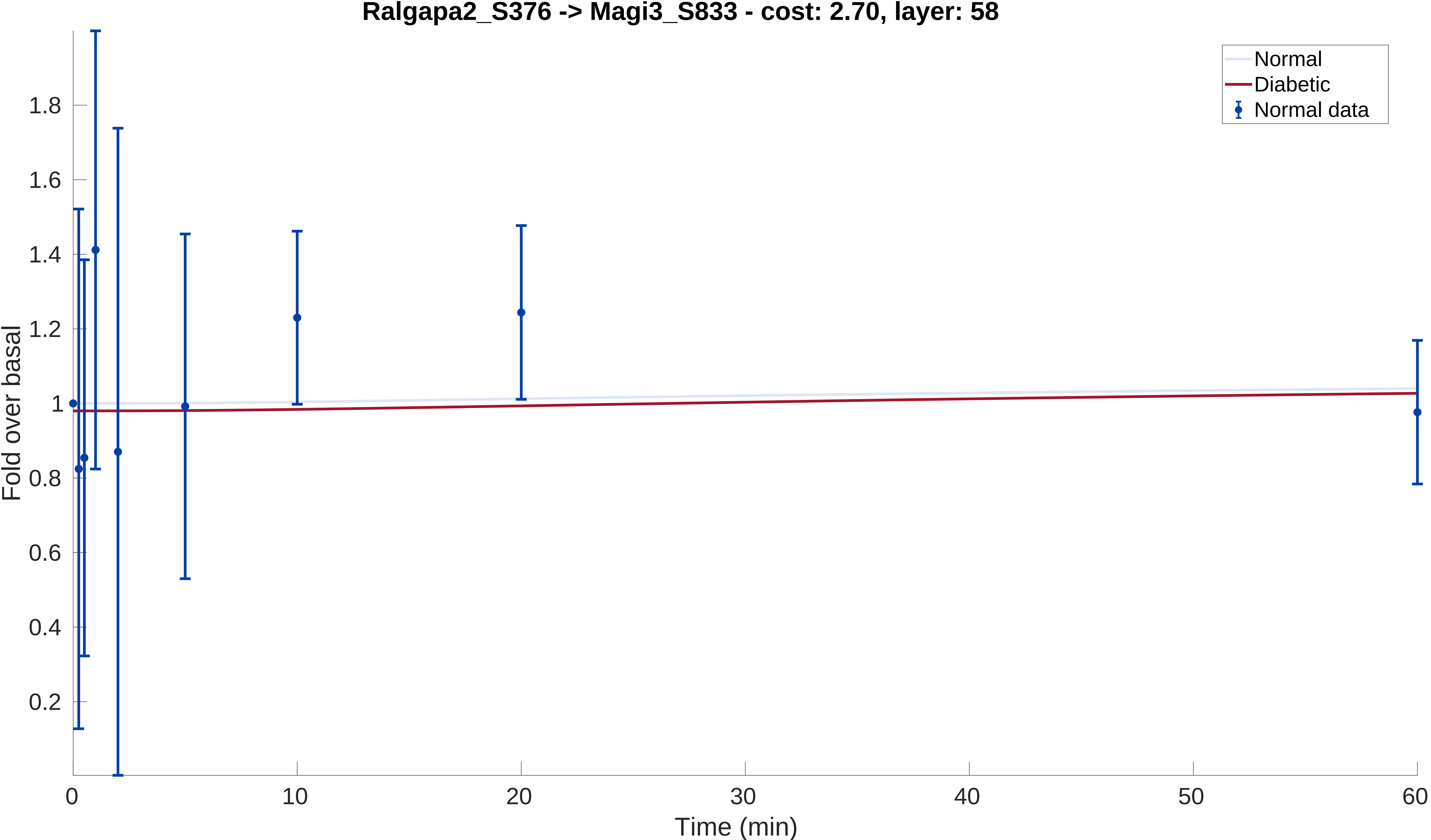

Ralgapa2_S376 -> Magi3_S833 - cost: 2.70, layer: 58
Normal
Diabetic
Normal data
Fold over basal
1.8
1.6
1.4
1.2
1
0.8
0.6
0.4
0.2
0
10
20
30
40
50
60
Time (min)

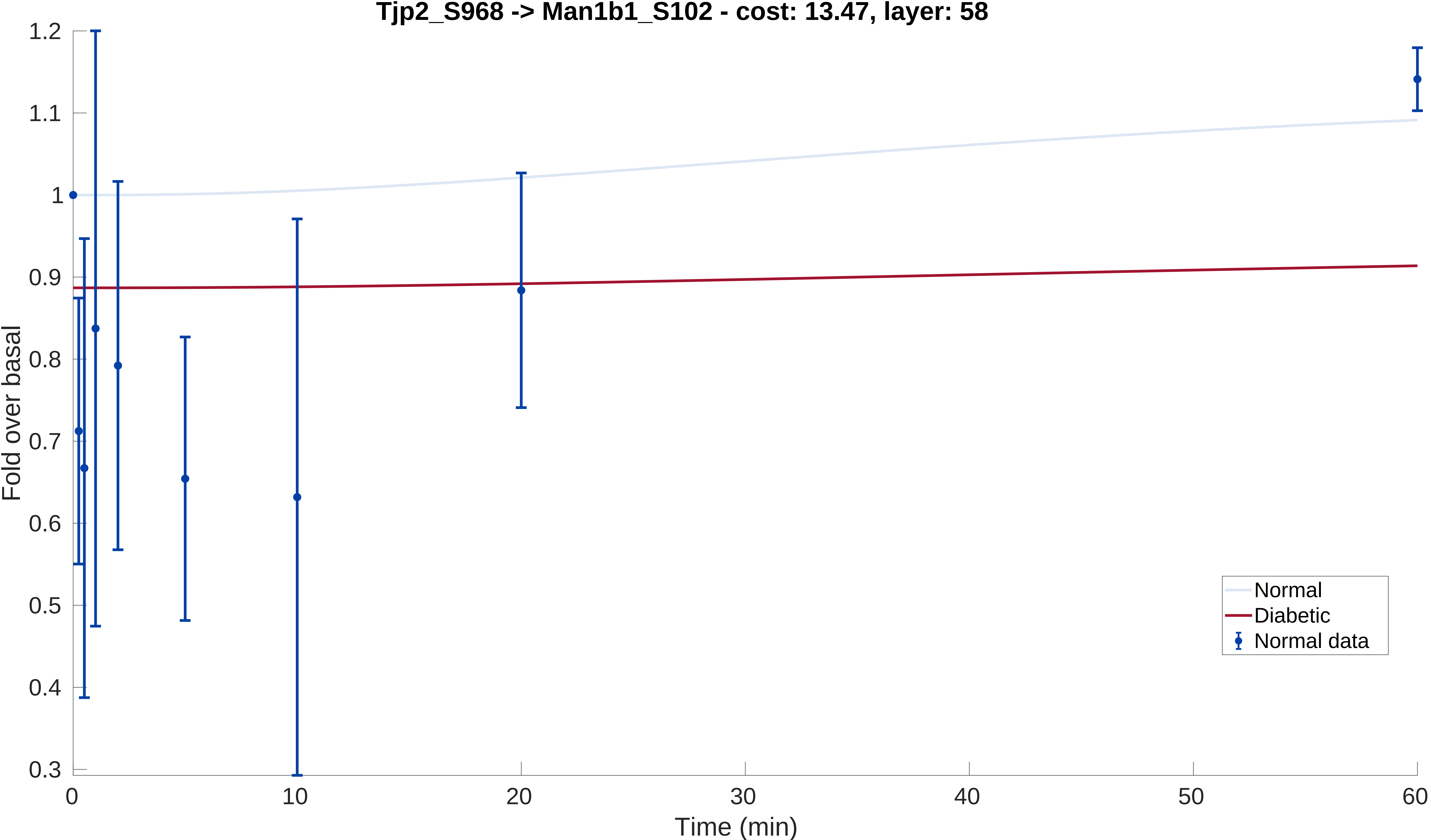

Tjp2_S968 -> Man1b1_S102 - cost: 13.47, layer: 58
Fold over basal
Time (min)
1.2
1.1
1
0.9
0.8
0.7
0.6
0.5
0.4
0.3
0
10
20
30
40
50
60
Normal
Diabetic
Normal data

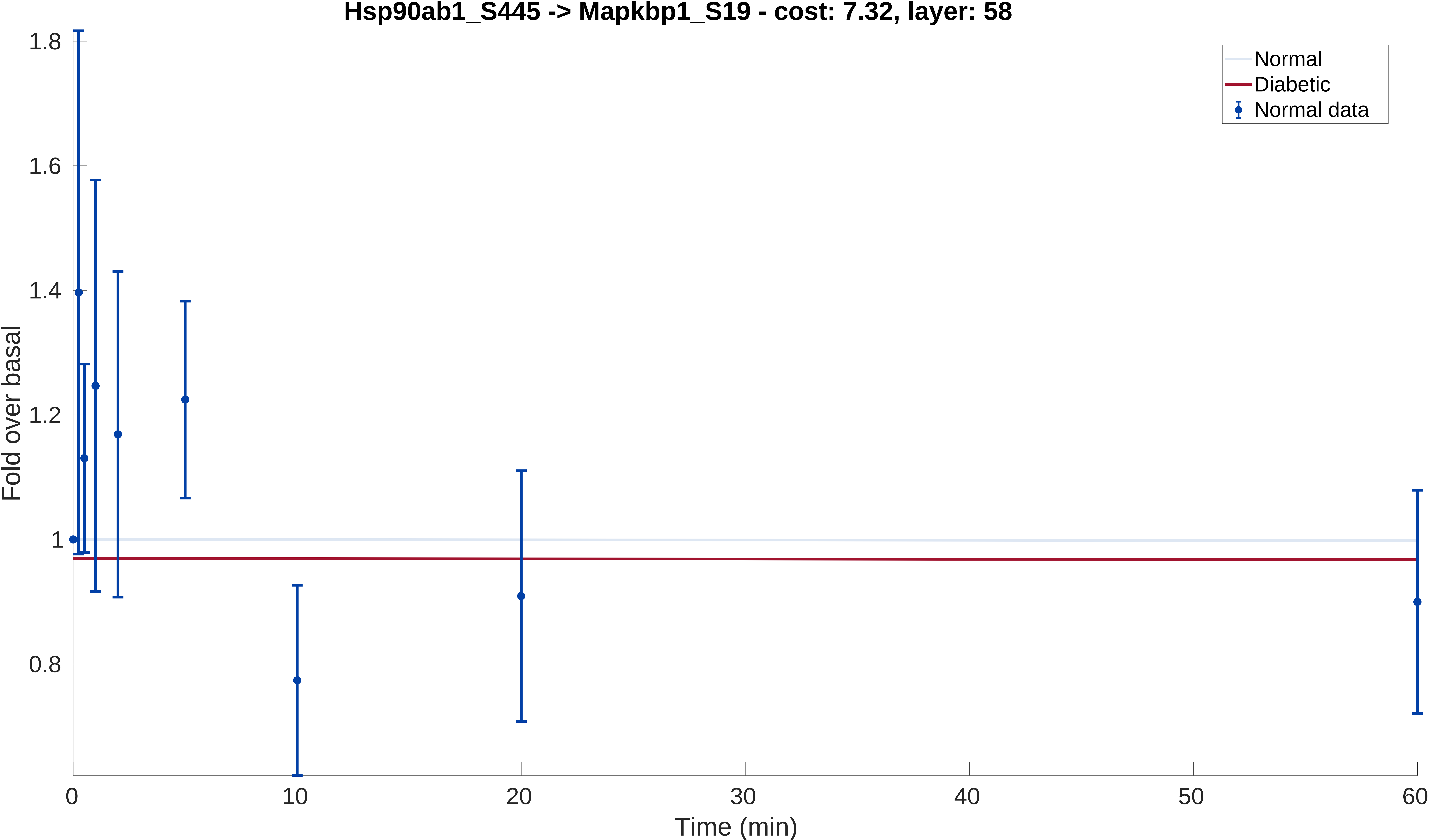

Hsp90ab1_S445 -> Mapkbp1_S19 - cost: 7.32, layer: 58
Normal
Diabetic
Normal data
Fold over basal
1.8
1.6
1.4
1.2
1
0.8
0
10
20
30
40
50
60
Time (min)

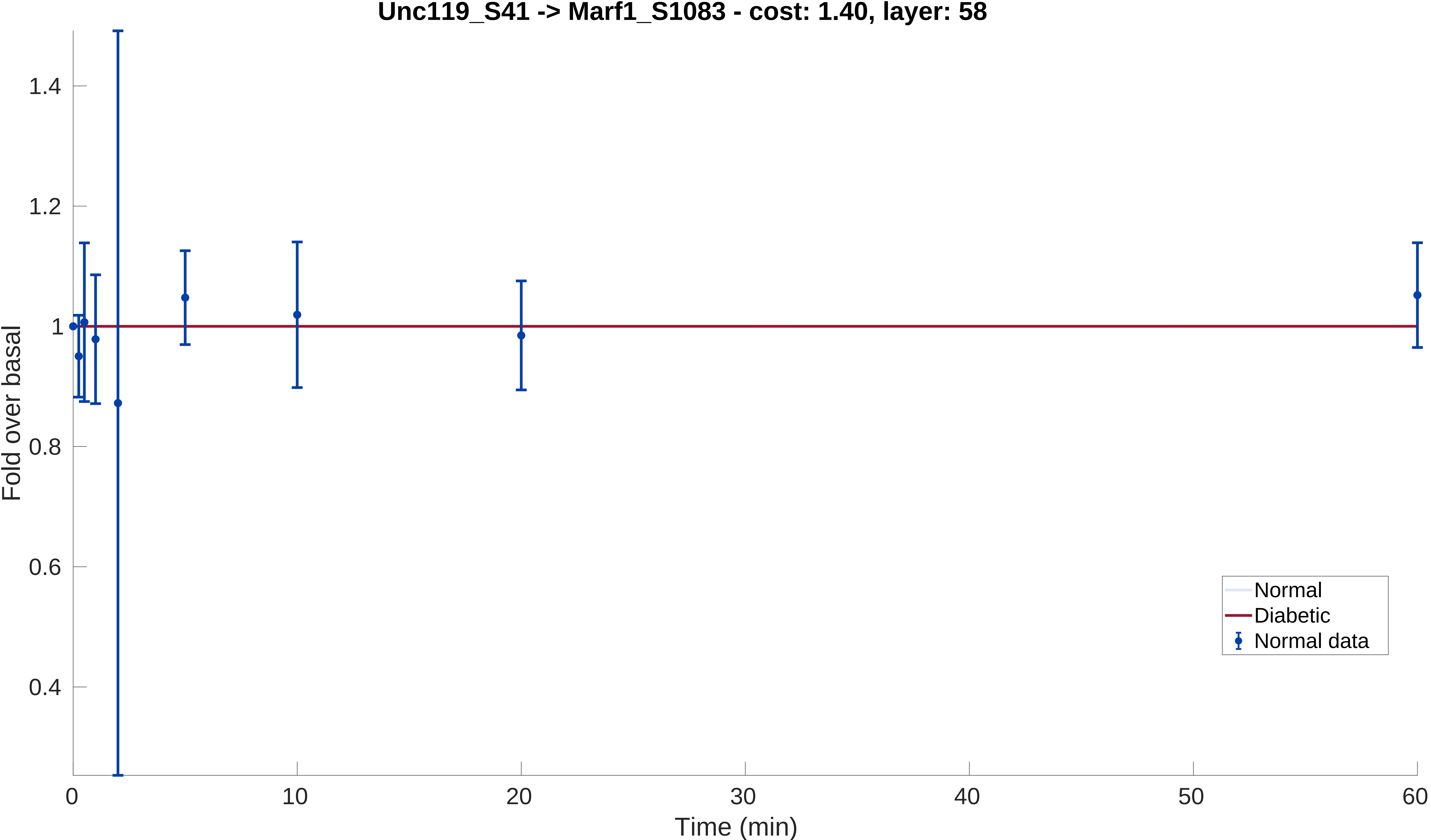

Unc119_S41 -> Marf1_S1083 - cost: 1.40, layer: 58
1.4
1.2
1
0.8
0.6
0.4
Fold over basal
0
10
20
30
40
50
60
Time (min)
Normal
Diabetic
Normal data

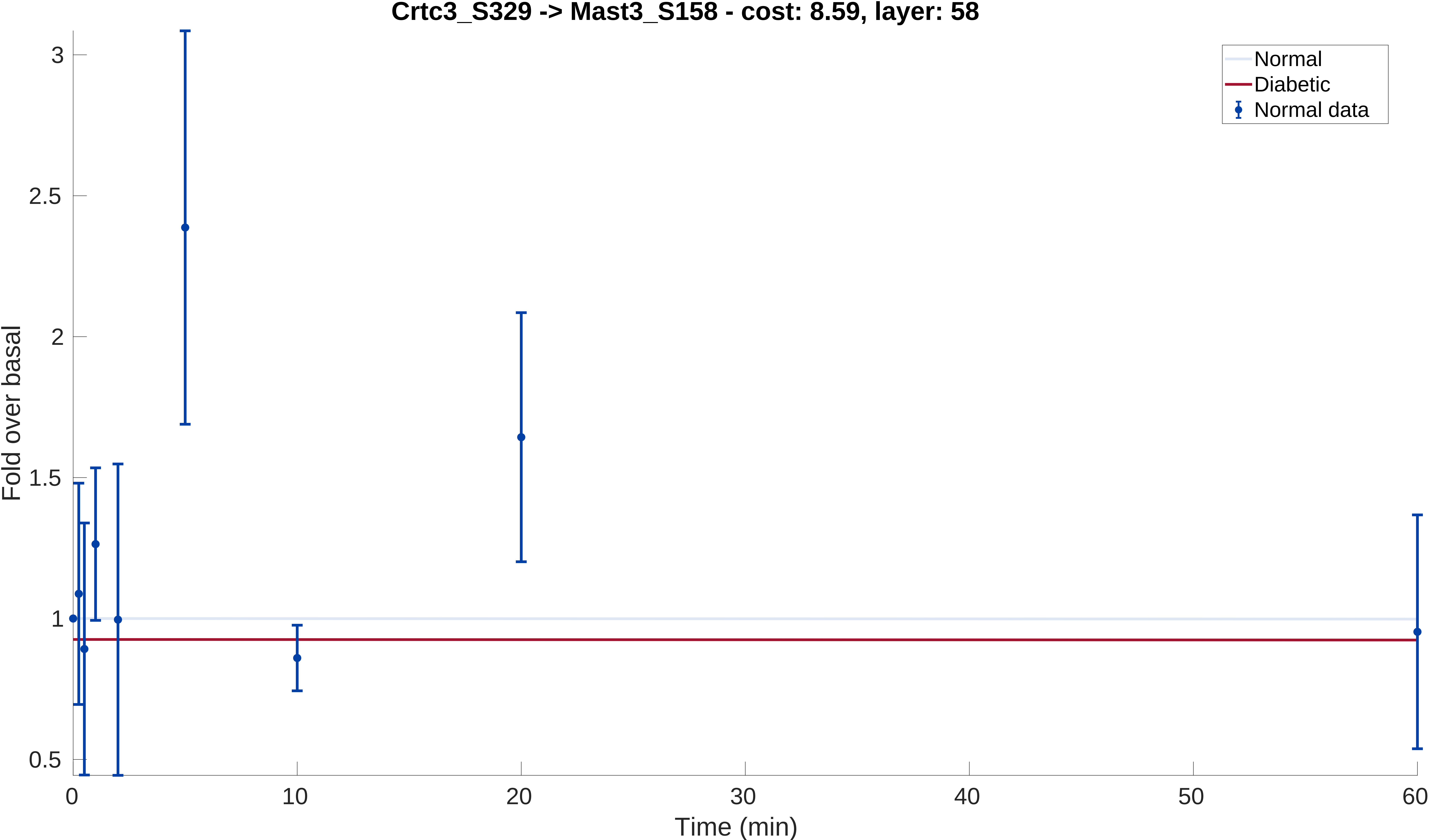

Crtc3_S329 -> Mast3_S158 - cost: 8.59, layer: 58
Normal
Diabetic
Normal data
Fold over basal
Time (min)
0
10
20
30
40
50
60
0.5
1
1.5
2
2.5
3

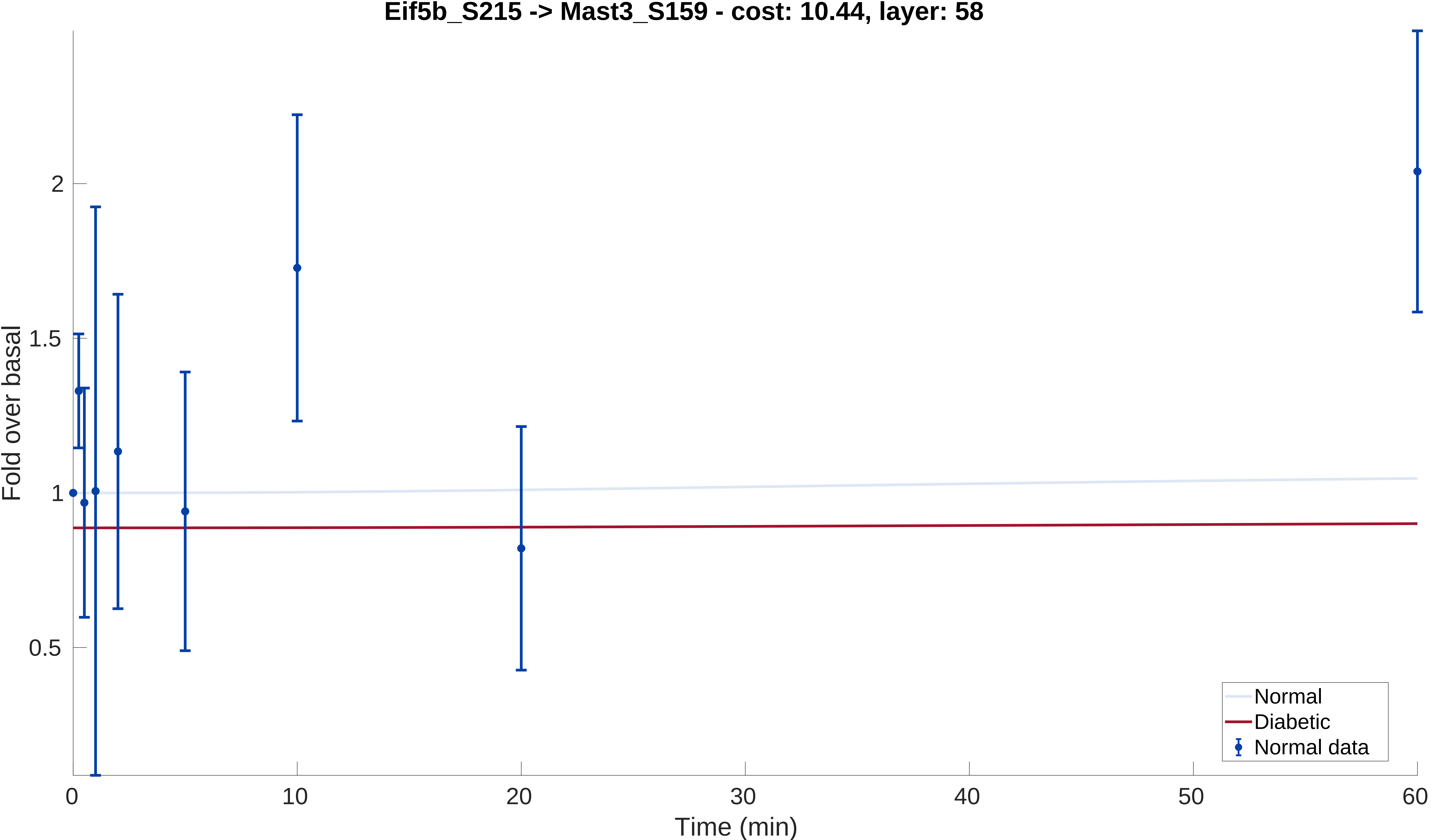
Eif5b_S215 -> Mast3_S159 - cost: 10.44, layer: 58
Fold over basal
0.5
1
1.5
2
0
10
20
30
40
50
60
Time (min)
Normal
Diabetic
Normal data

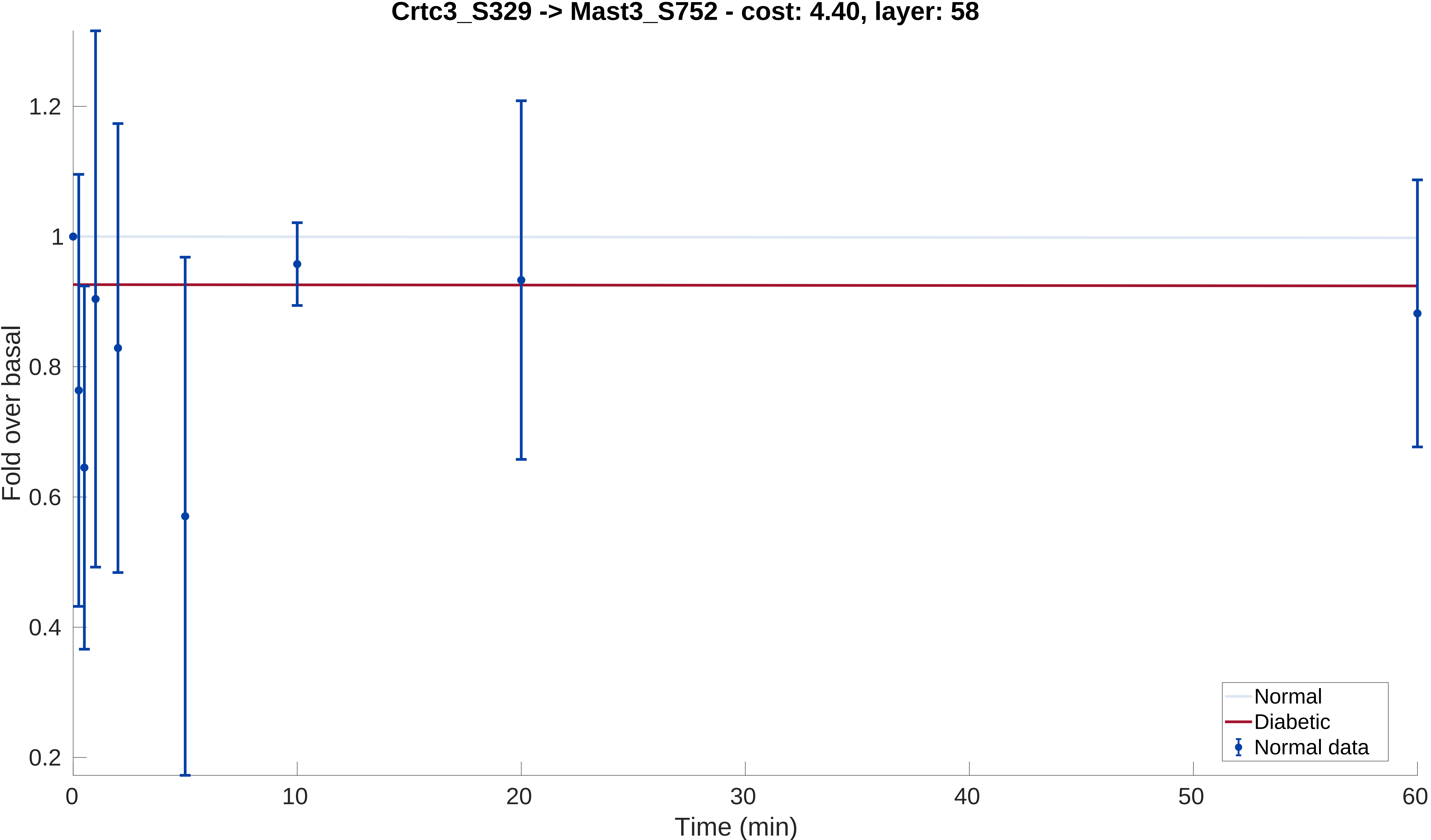

Crtc3_S329 -> Mast3_S752 - cost: 4.40, layer: 58
Fold over basal
Time (min)
0.2
0.4
0.6
0.8
1
1.2
0
10
20
30
40
50
60
Normal
Diabetic
Normal data

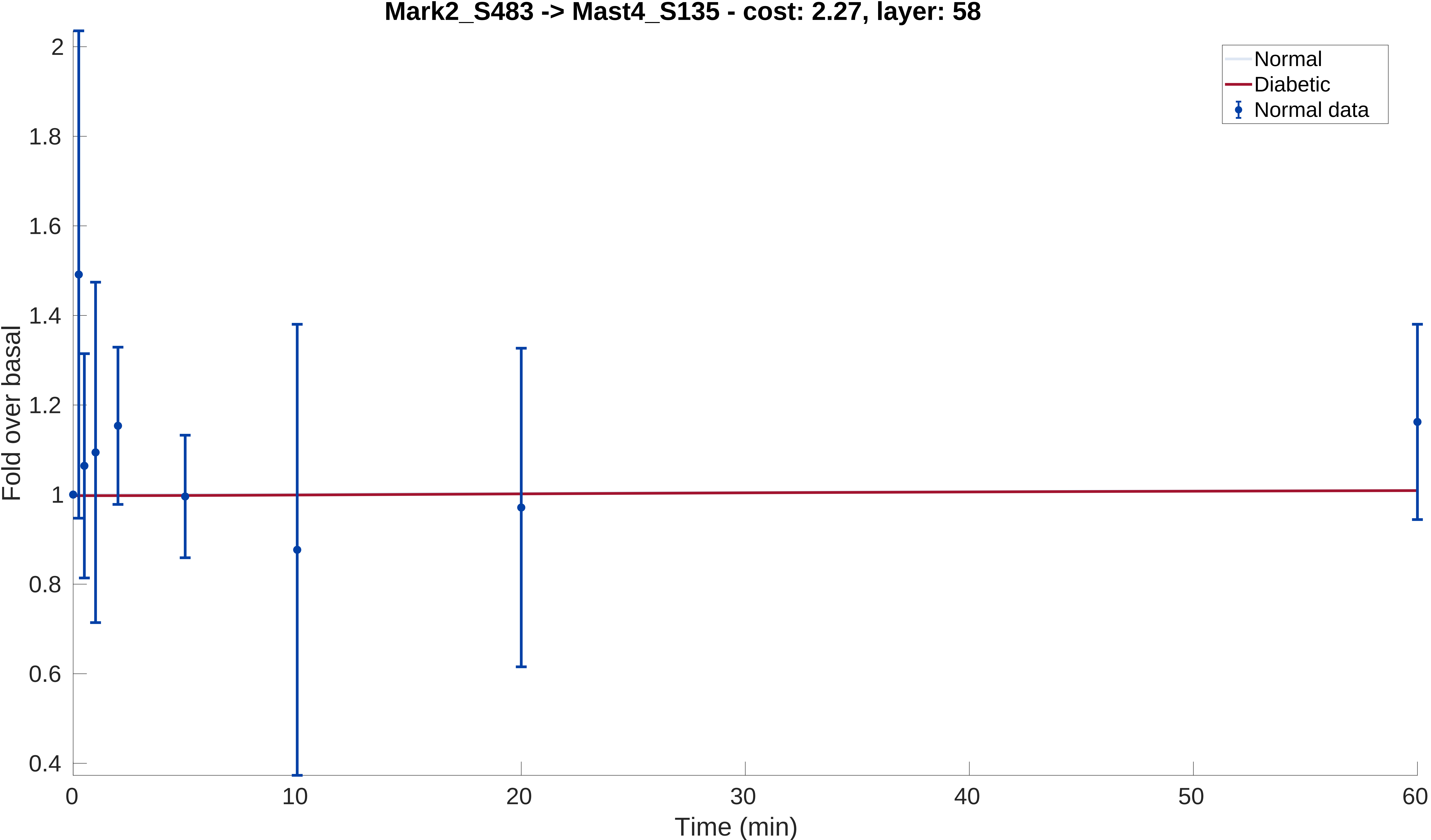

Mark2_S483 -> Mast4_S135 - cost: 2.27, layer: 58
Normal
Diabetic
Normal data
Fold over basal
Time (min)
0
10
20
30
40
50
60
0.4
0.6
0.8
1
1.2
1.4
1.6
1.8
2

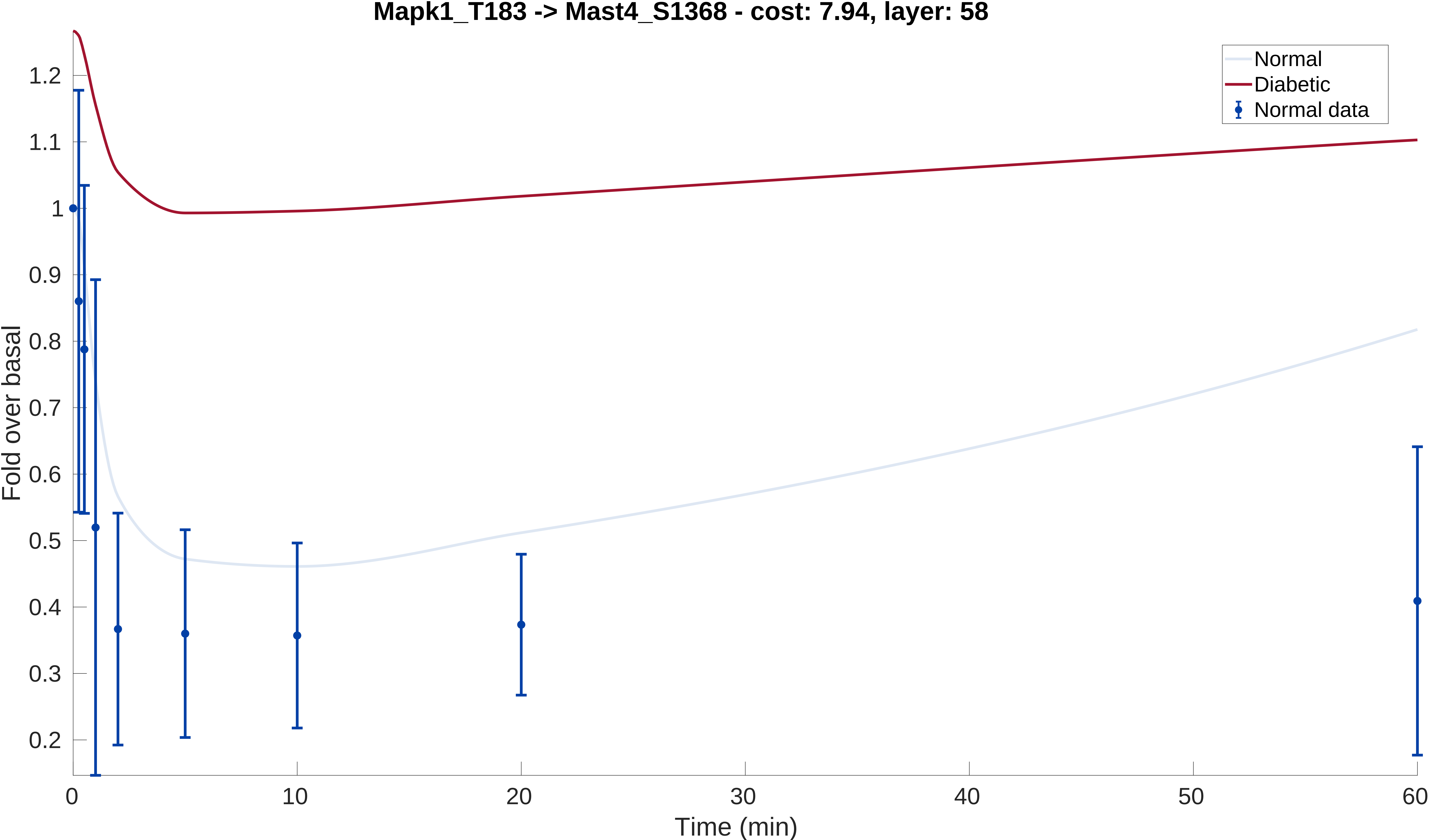
Mapk1_T183 -> Mast4_S1368 - cost: 7.94, layer: 58
Normal
Diabetic
Normal data
Fold over basal
Time (min)
0.2
0.3
0.4
0.5
0.6
0.7
0.8
0.9
1
1.1
1.2
0
10
20
30
40
50
60

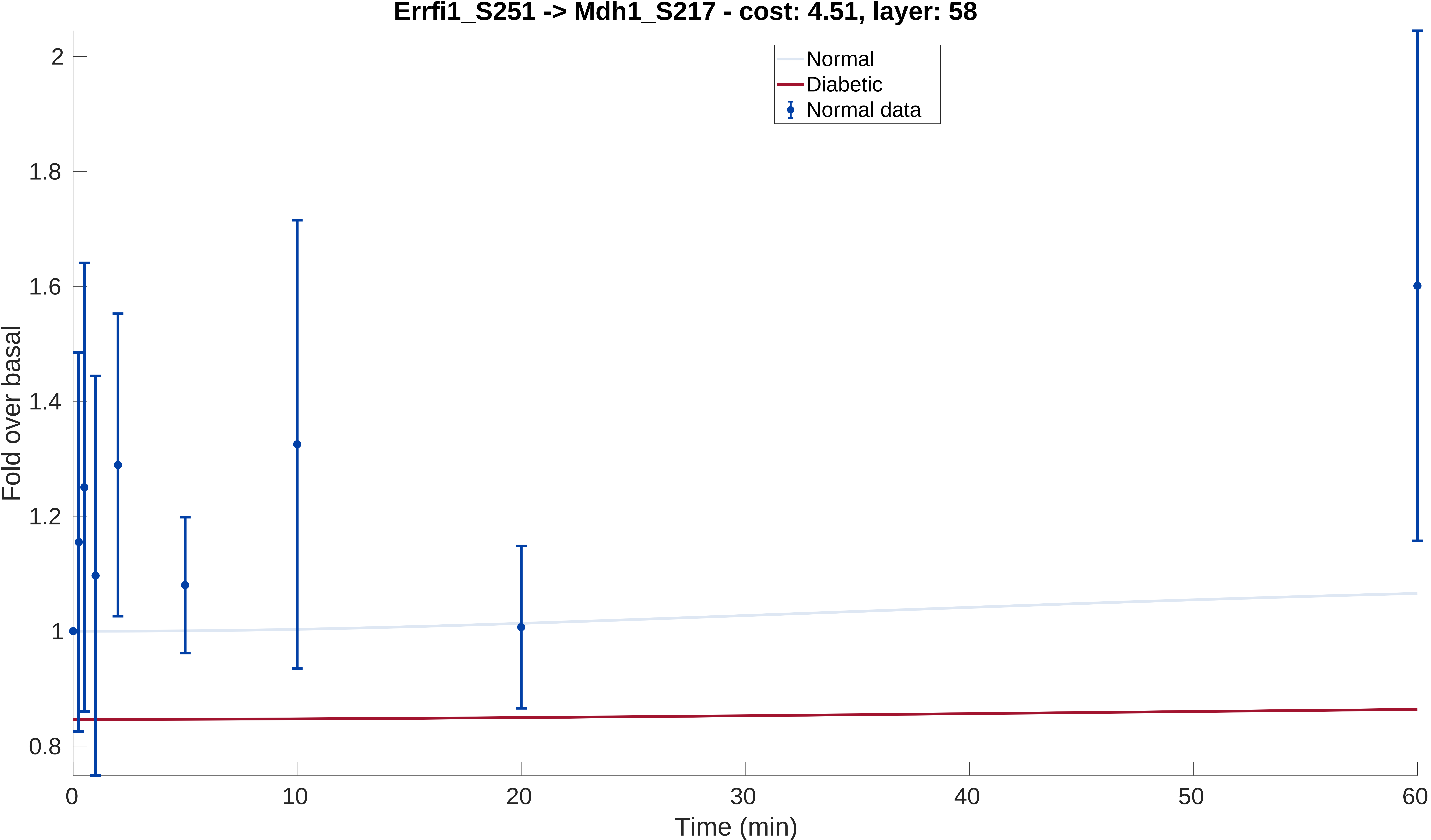

Errfi1_S251 -> Mdh1_S217 - cost: 4.51, layer: 58
Normal
Diabetic
Normal data
Fold over basal
Time (min)
2
1.8
1.6
1.4
1.2
1
0.8
0
10
20
30
40
50
60

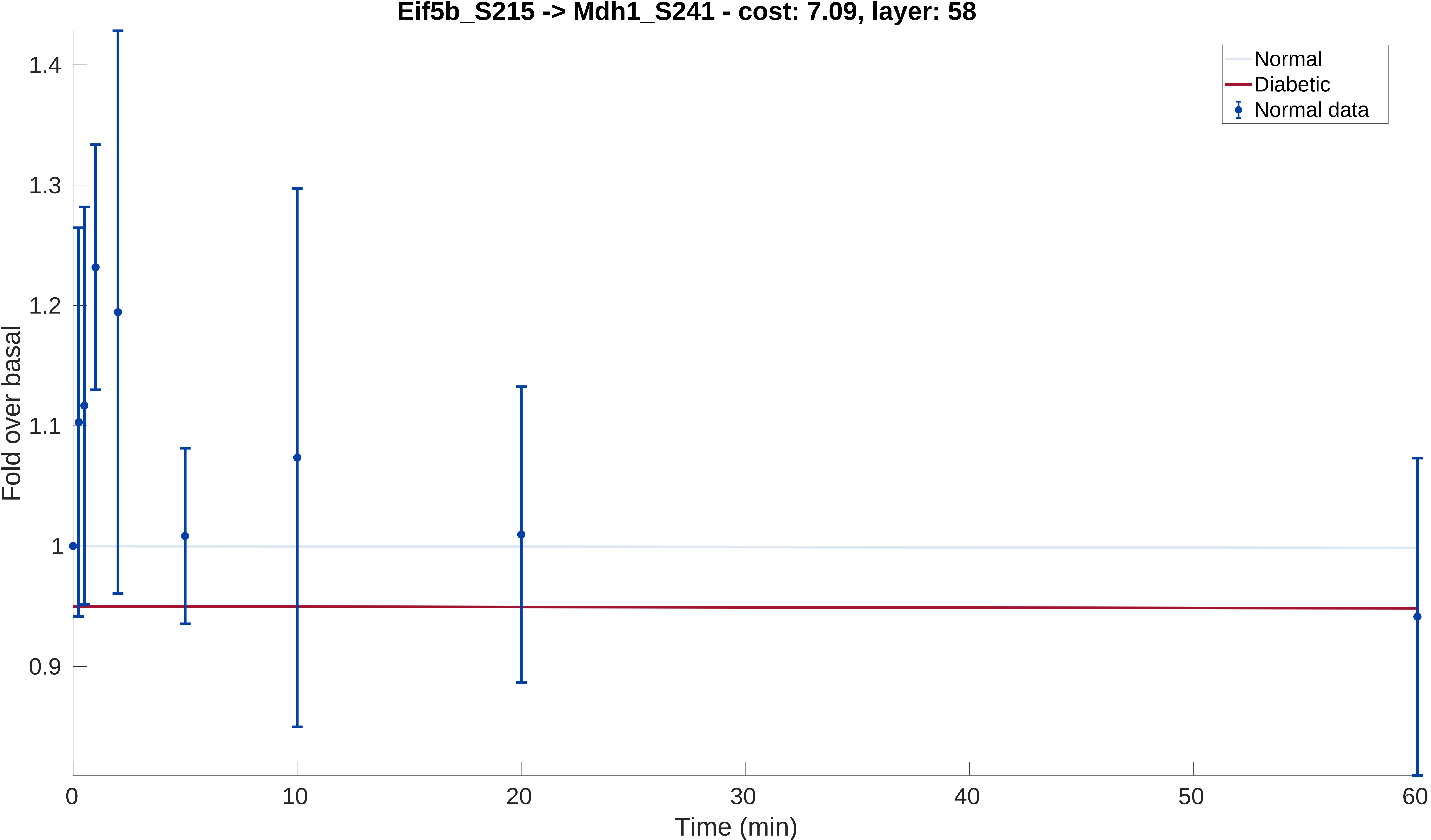

Eif5b_S215 -> Mdh1_S241 - cost: 7.09, layer: 58
Normal
Diabetic
Normal data
Fold over basal
Time (min)
1.4
1.3
1.2
1.1
1
0.9
0
10
20
30
40
50
60

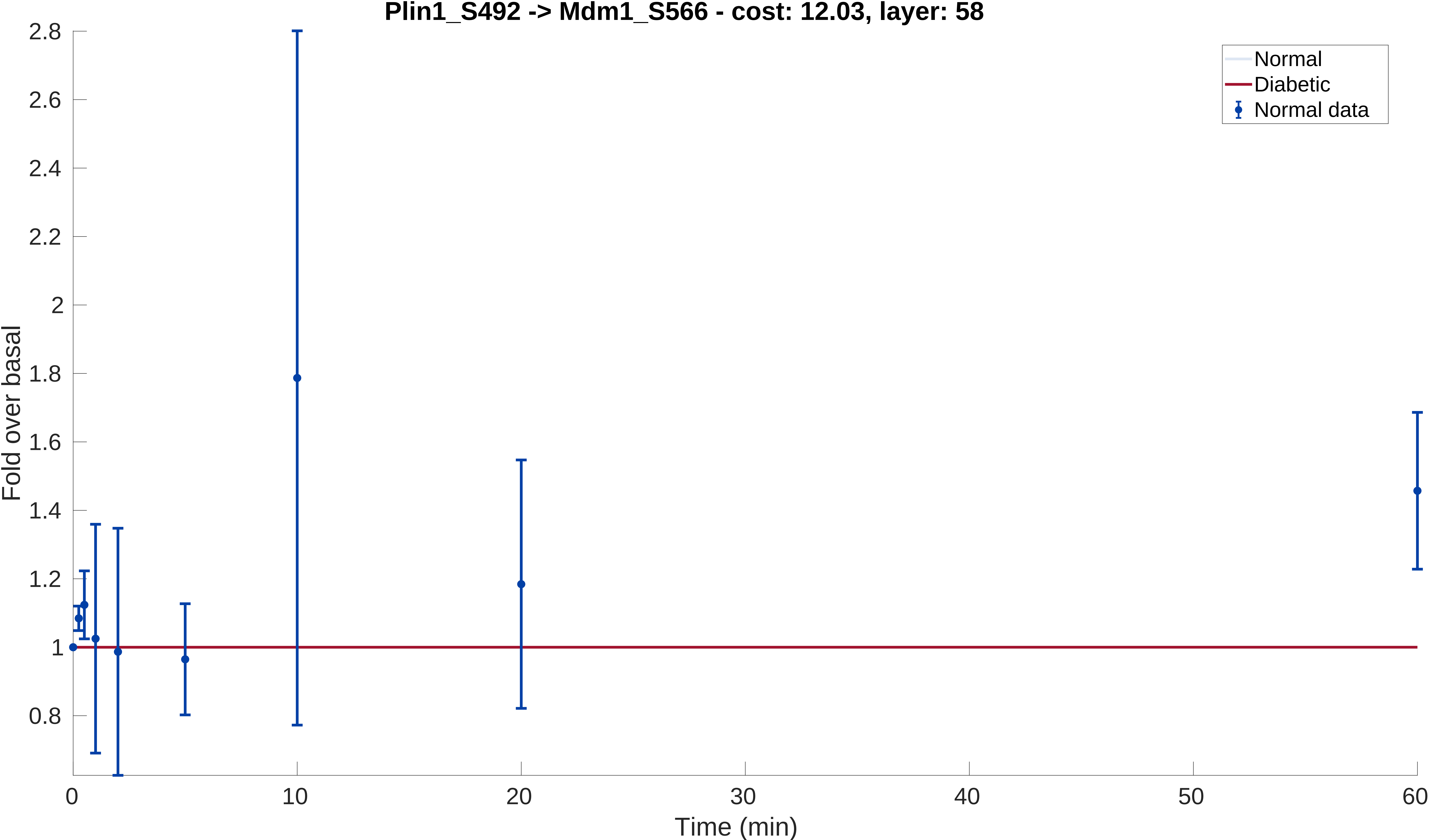

Plin1_S492 -> Mdm1_S566 - cost: 12.03, layer: 58
Normal
Diabetic
Normal data
Fold over basal
Time (min)
2.8
2.6
2.4
2.2
2
1.8
1.6
1.4
1.2
1
0.8
0
10
20
30
40
50
60

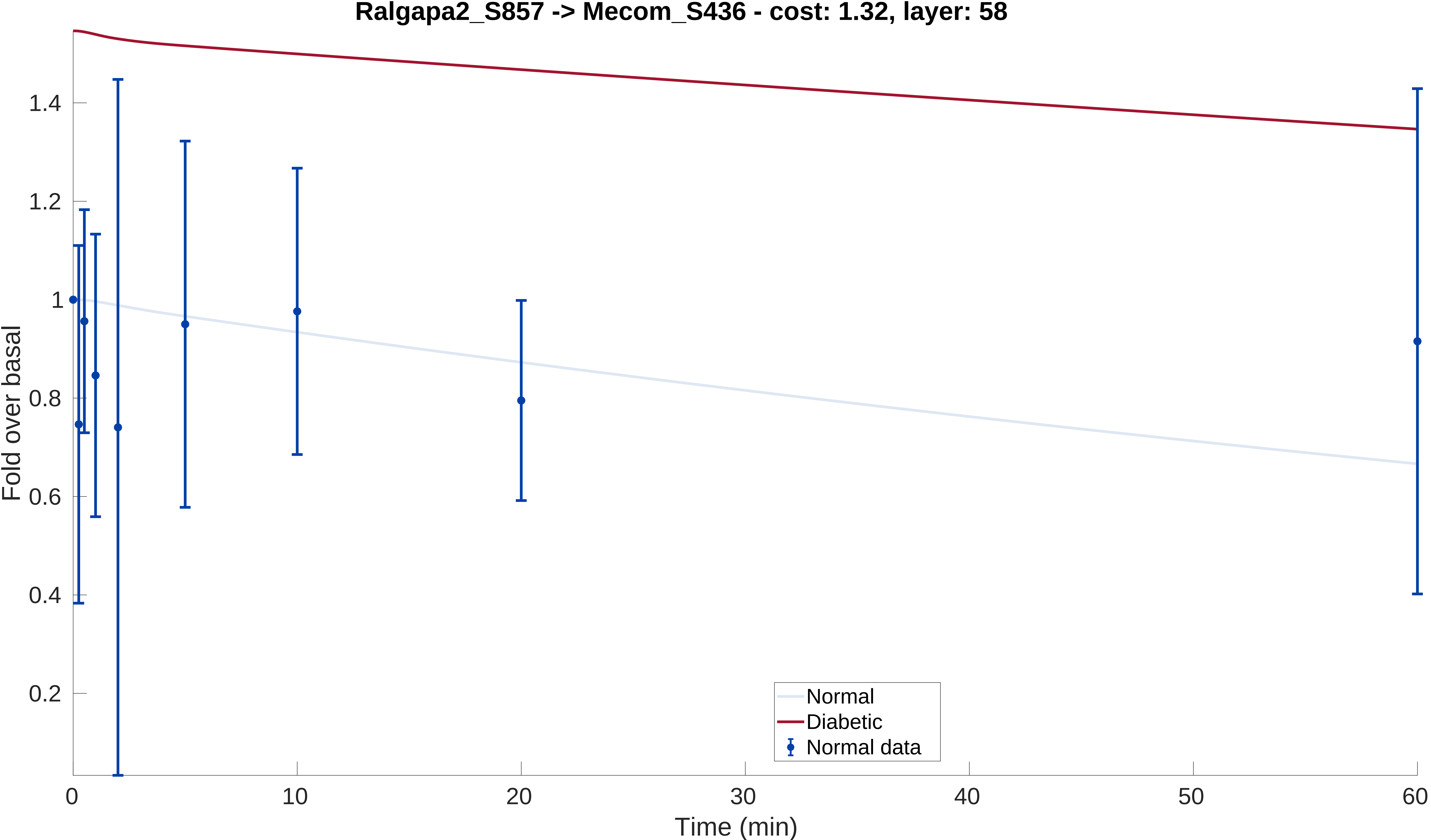

Ralgapa2_S857 -> Mecom_S436 - cost: 1.32, layer: 58
Fold over basal
Time (min)
0
10
20
30
40
50
60
0.2
0.4
0.6
0.8
1
1.2
1.4
Normal
Diabetic
Normal data

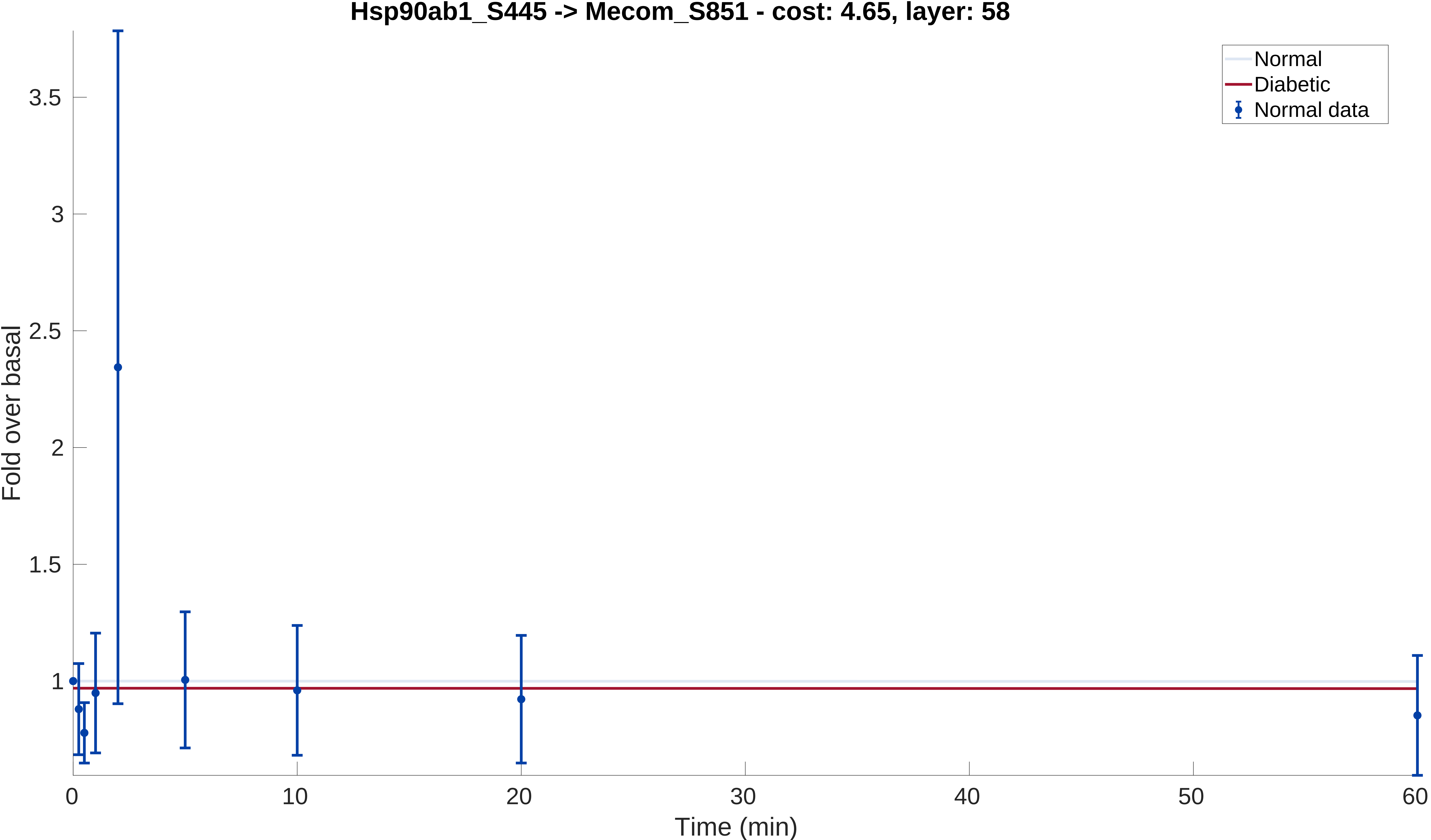
Hsp90ab1_S445 -> Mecom_S851 - cost: 4.65, layer: 58
Fold over basal
3.5
3
2.5
2
1.5
1
0
10
20
30
40
50
60
Time (min)
Normal
Diabetic
Normal data

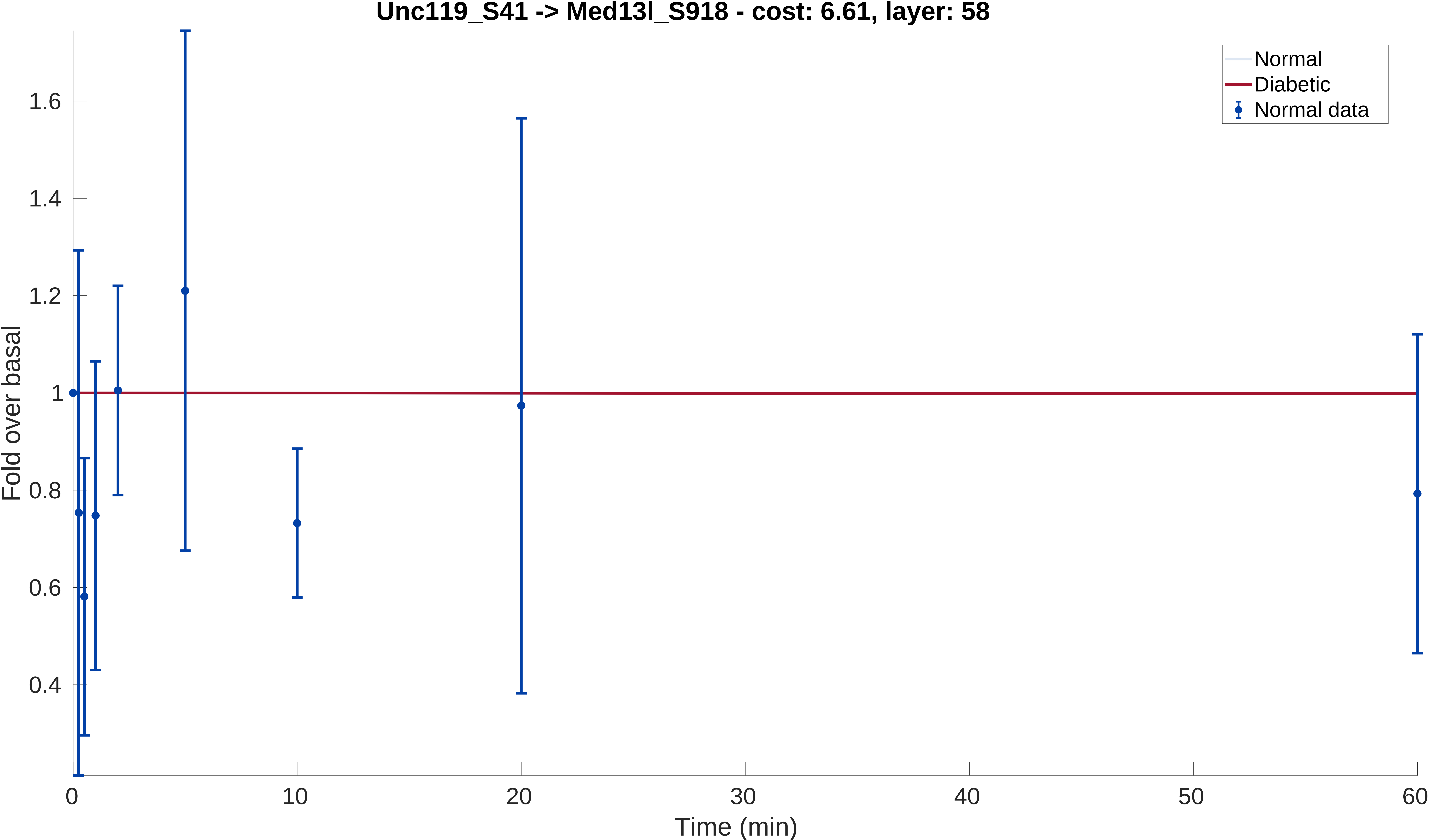

Unc119_S41 -> Med13l_S918 - cost: 6.61, layer: 58
Normal
Diabetic
Normal data
Fold over basal
Time (min)
1.6
1.4
1.2
1
0.8
0.6
0.4
0
10
20
30
40
50
60

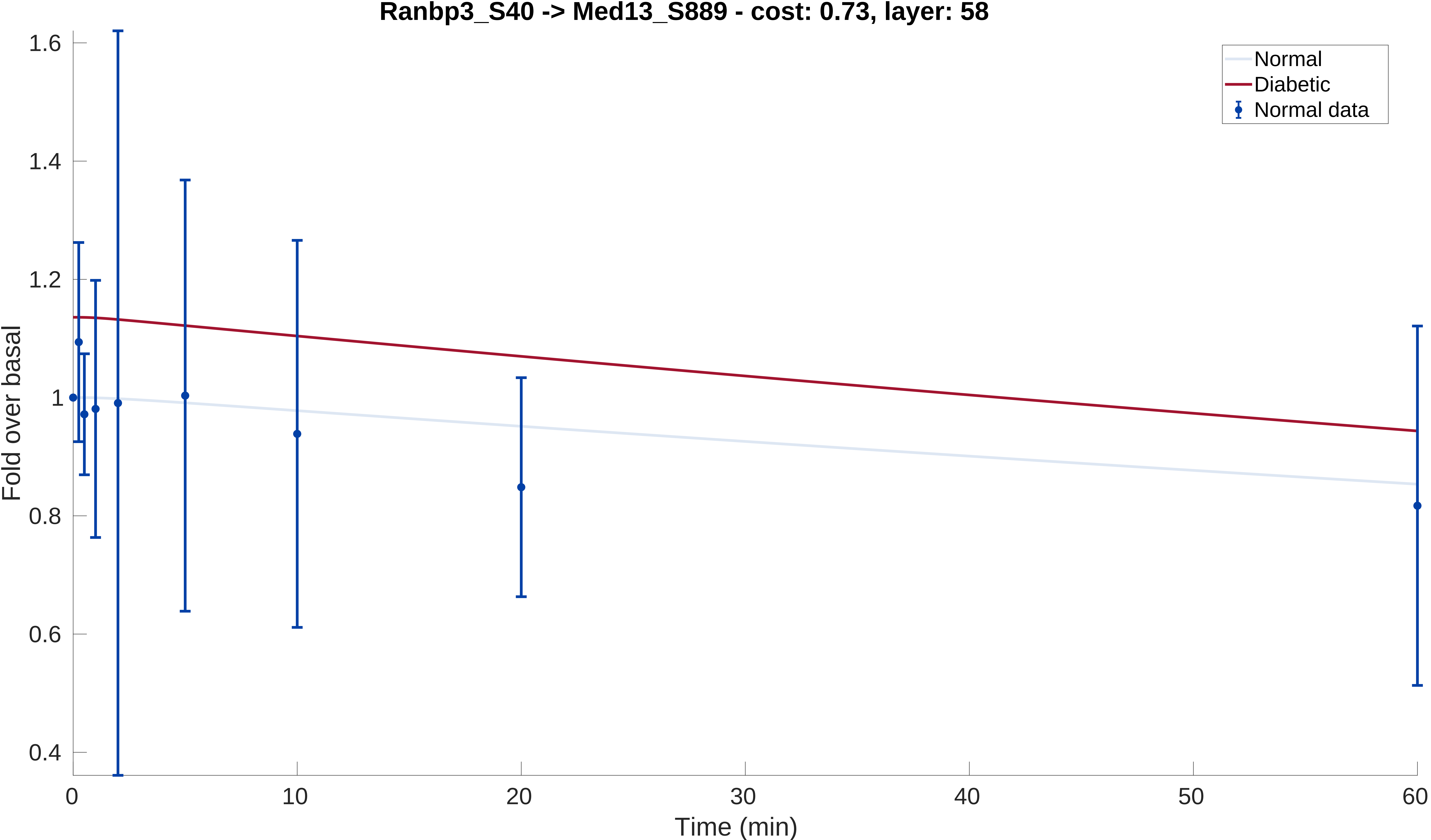

Ranbp3_S40 -> Med13_S889 - cost: 0.73, layer: 58
Normal
Diabetic
Normal data
Fold over basal
1.6
1.4
1.2
1
0.8
0.6
0.4
0
10
20
30
40
50
60
Time (min)

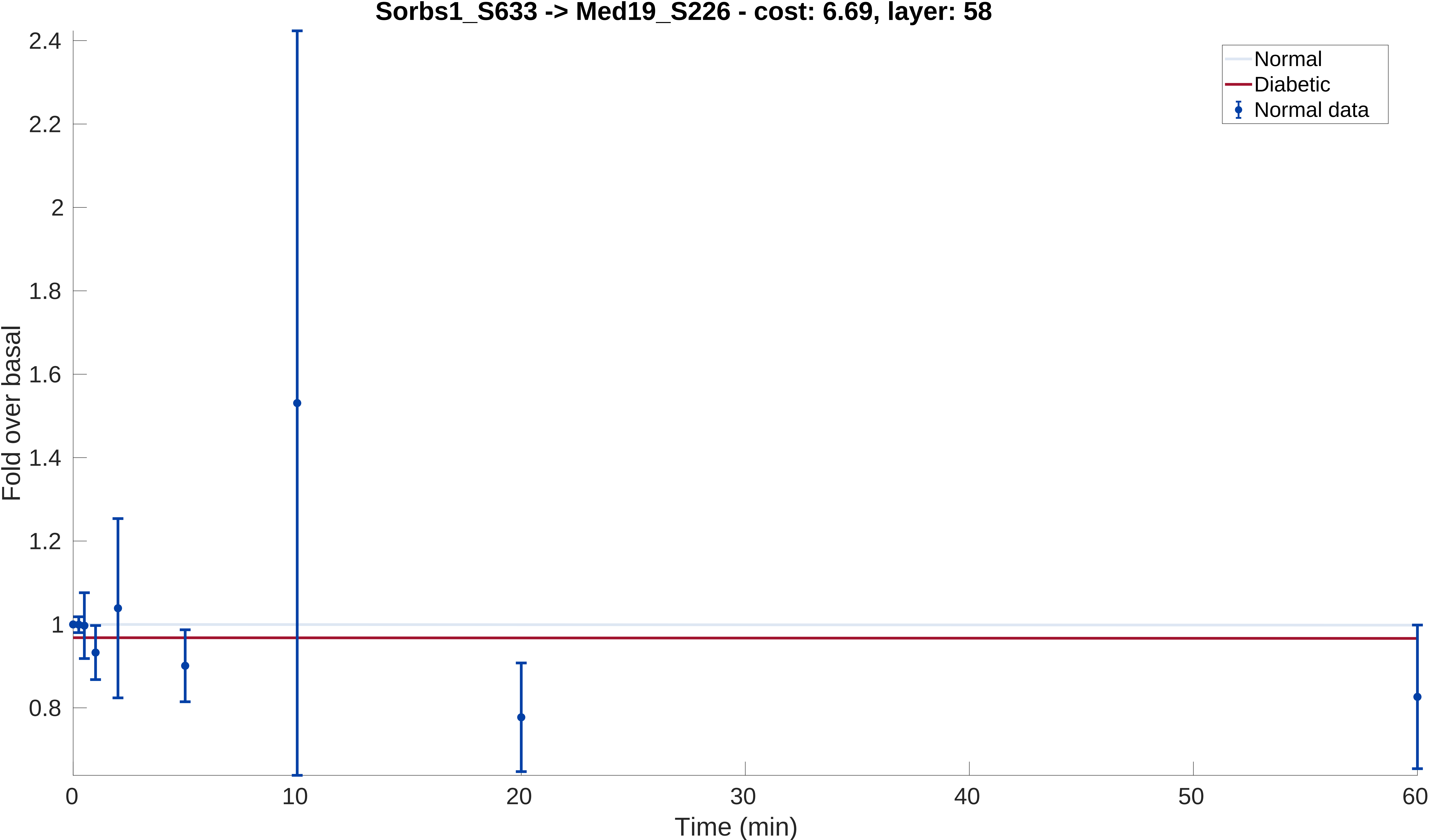

Sorbs1_S633 -> Med19_S226 - cost: 6.69, layer: 58
Normal
Diabetic
Normal data
Fold over basal
Time (min)
2.4
2.2
2
1.8
1.6
1.4
1.2
1
0.8
0
10
20
30
40
50
60

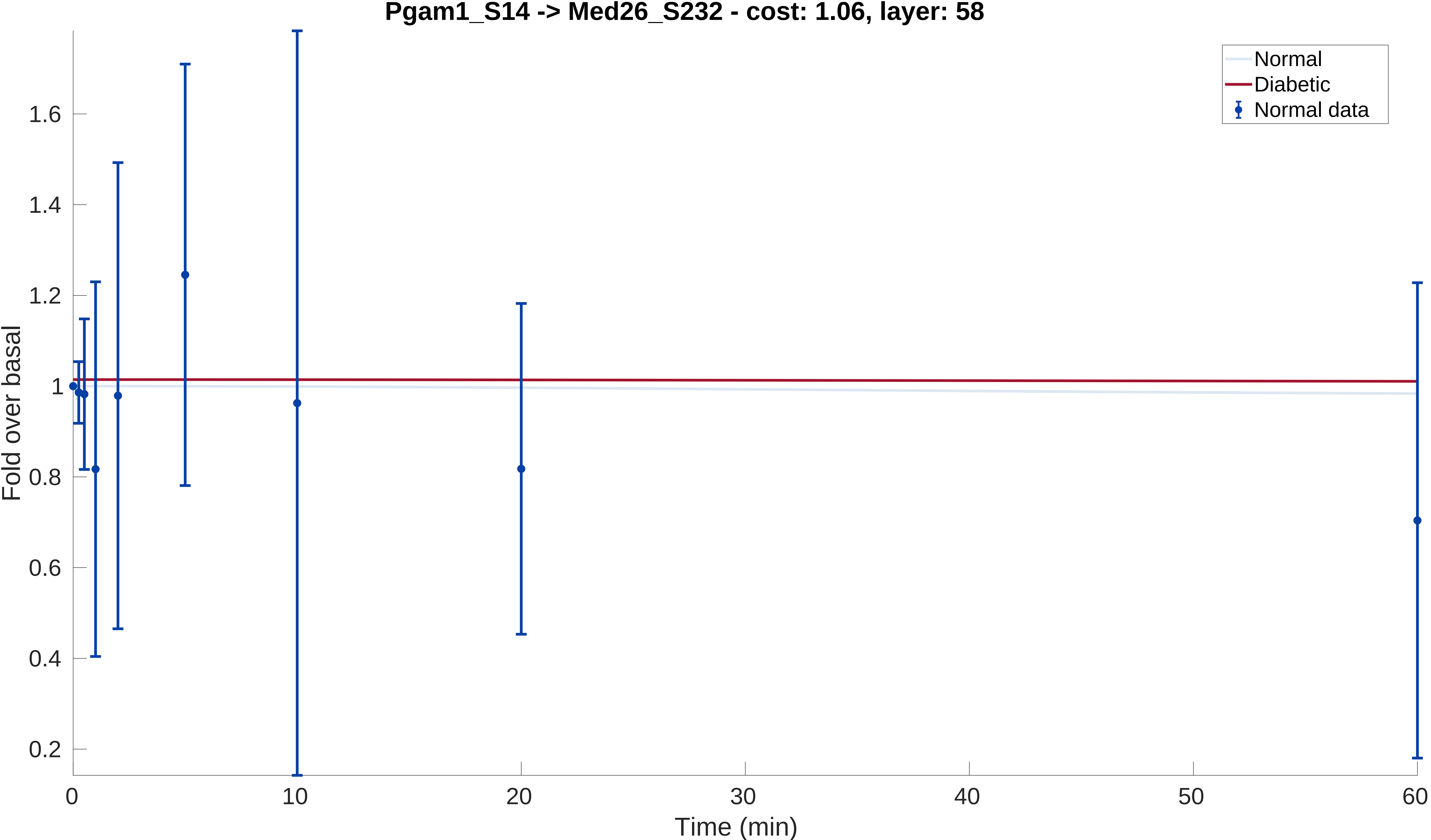

Pgam1_S14 -> Med26_S232 - cost: 1.06, layer: 58
Normal
Diabetic
Normal data
Fold over basal
1.6
1.4
1.2
1
0.8
0.6
0.4
0.2
0
10
20
30
40
50
60
Time (min)

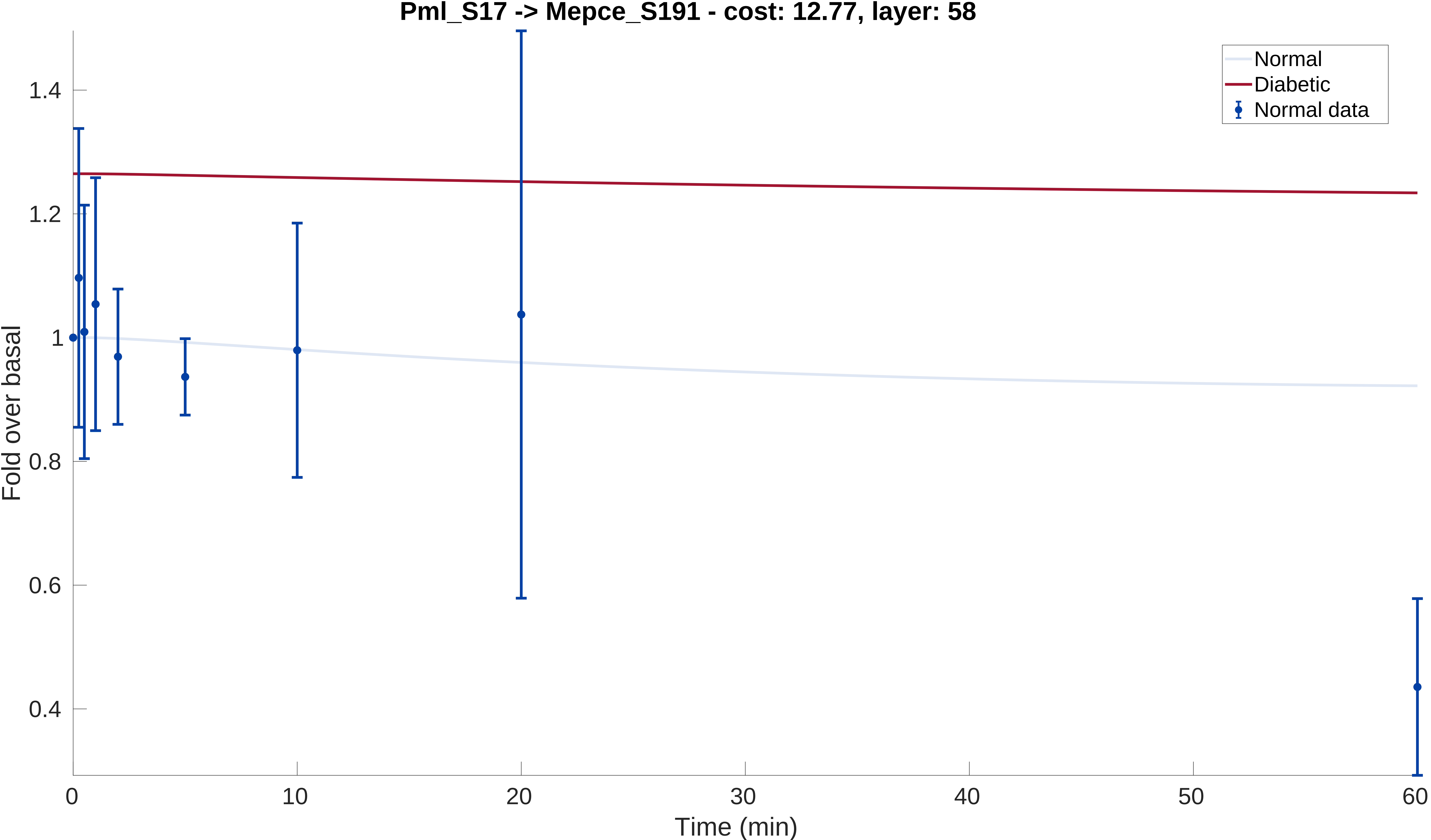

Pml_S17 -> Mepce_S191 - cost: 12.77, layer: 58
Normal
Diabetic
Normal data
Fold over basal
1.4
1.2
1
0.8
0.6
0.4
0
10
20
30
40
50
60
Time (min)

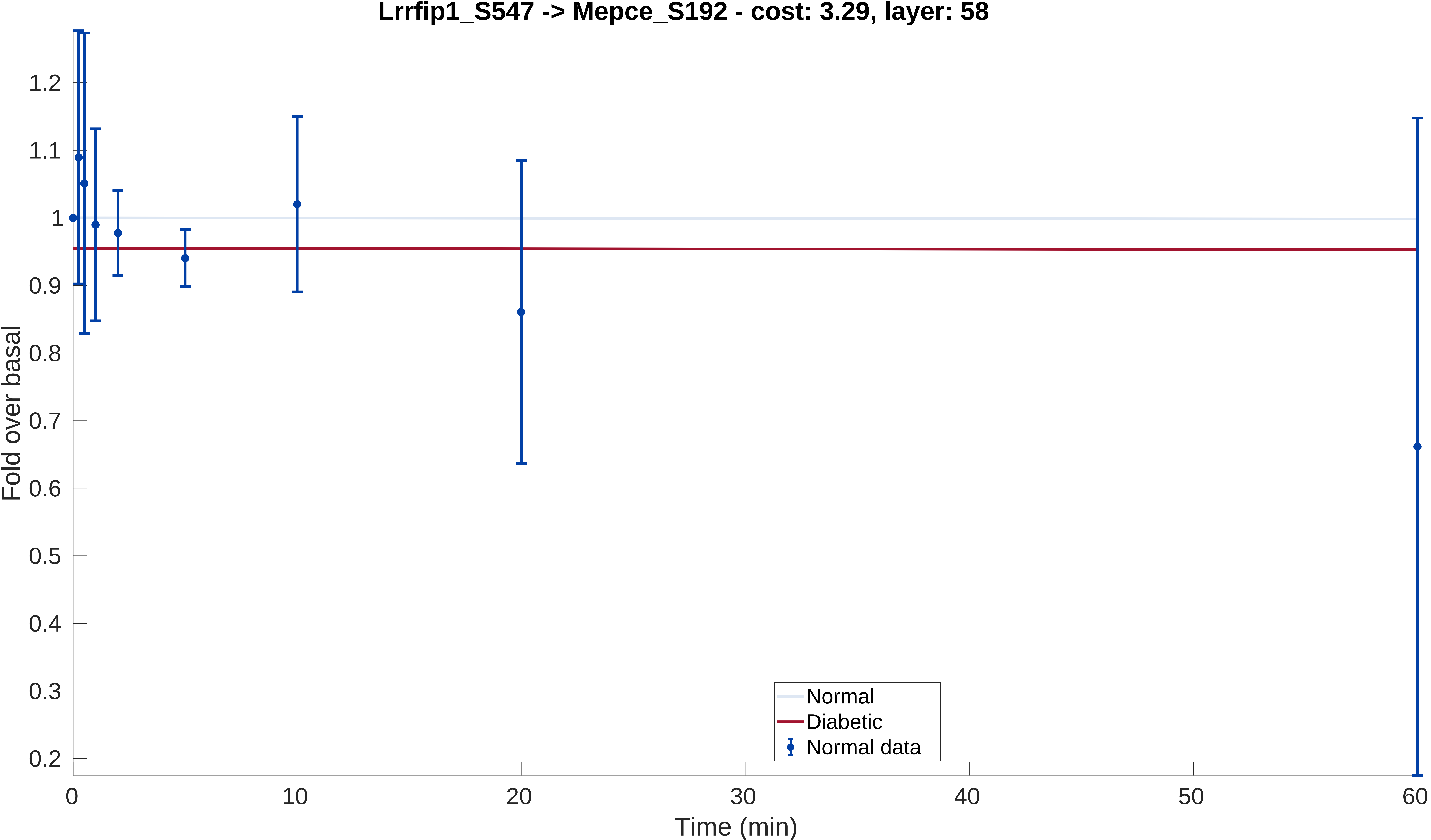

Lrrfip1_S547 -> Mepce_S192 - cost: 3.29, layer: 58
Fold over basal
1.2
1.1
1
0.9
0.8
0.7
0.6
0.5
0.4
0.3
0.2
0
10
20
30
40
50
60
Time (min)
Normal
Diabetic
Normal data

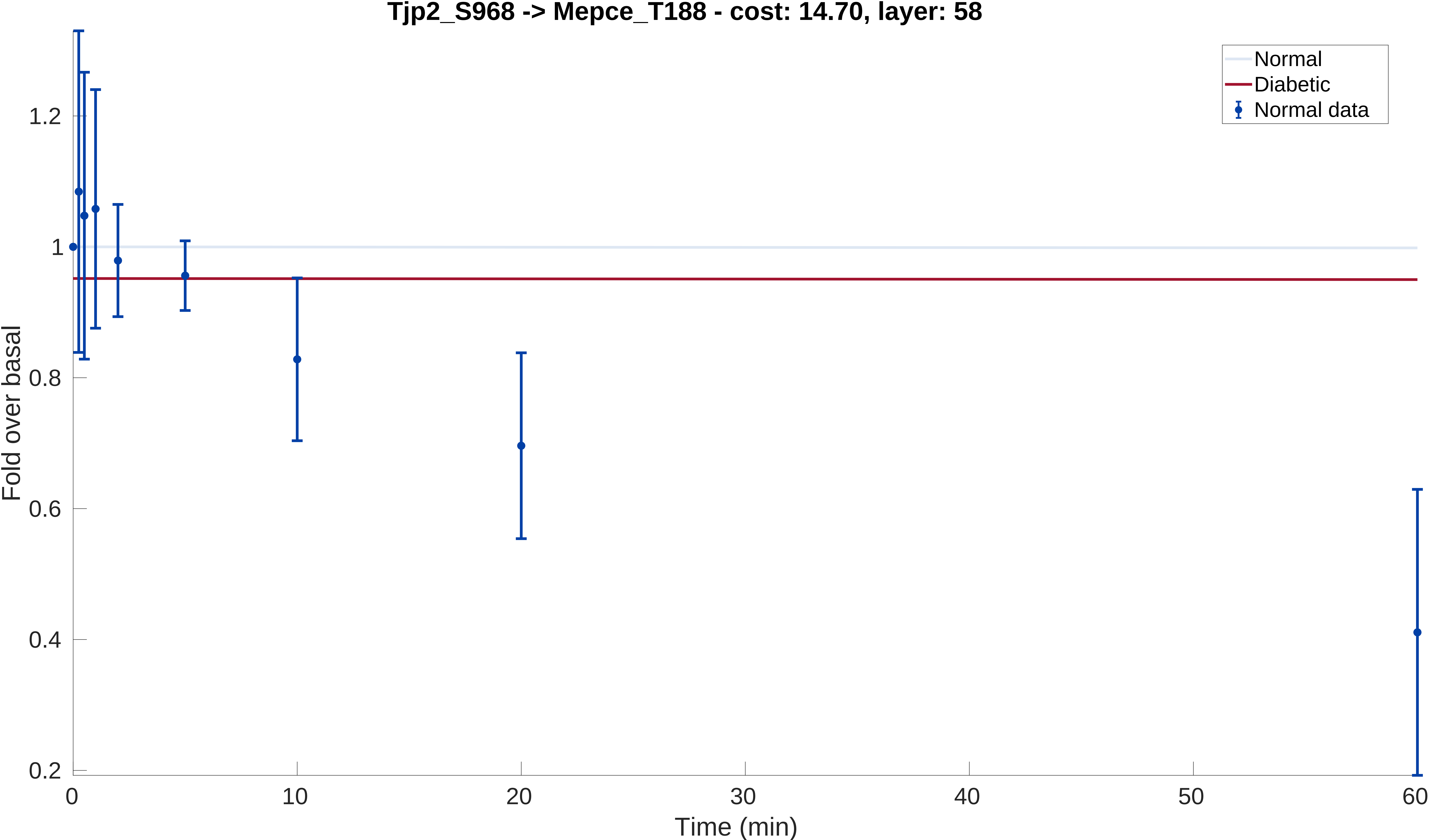

Tjp2_S968 -> Mepce_T188 - cost: 14.70, layer: 58
Normal
Diabetic
Normal data
Fold over basal
1.2
1
0.8
0.6
0.4
0.2
0
10
20
30
40
50
60
Time (min)

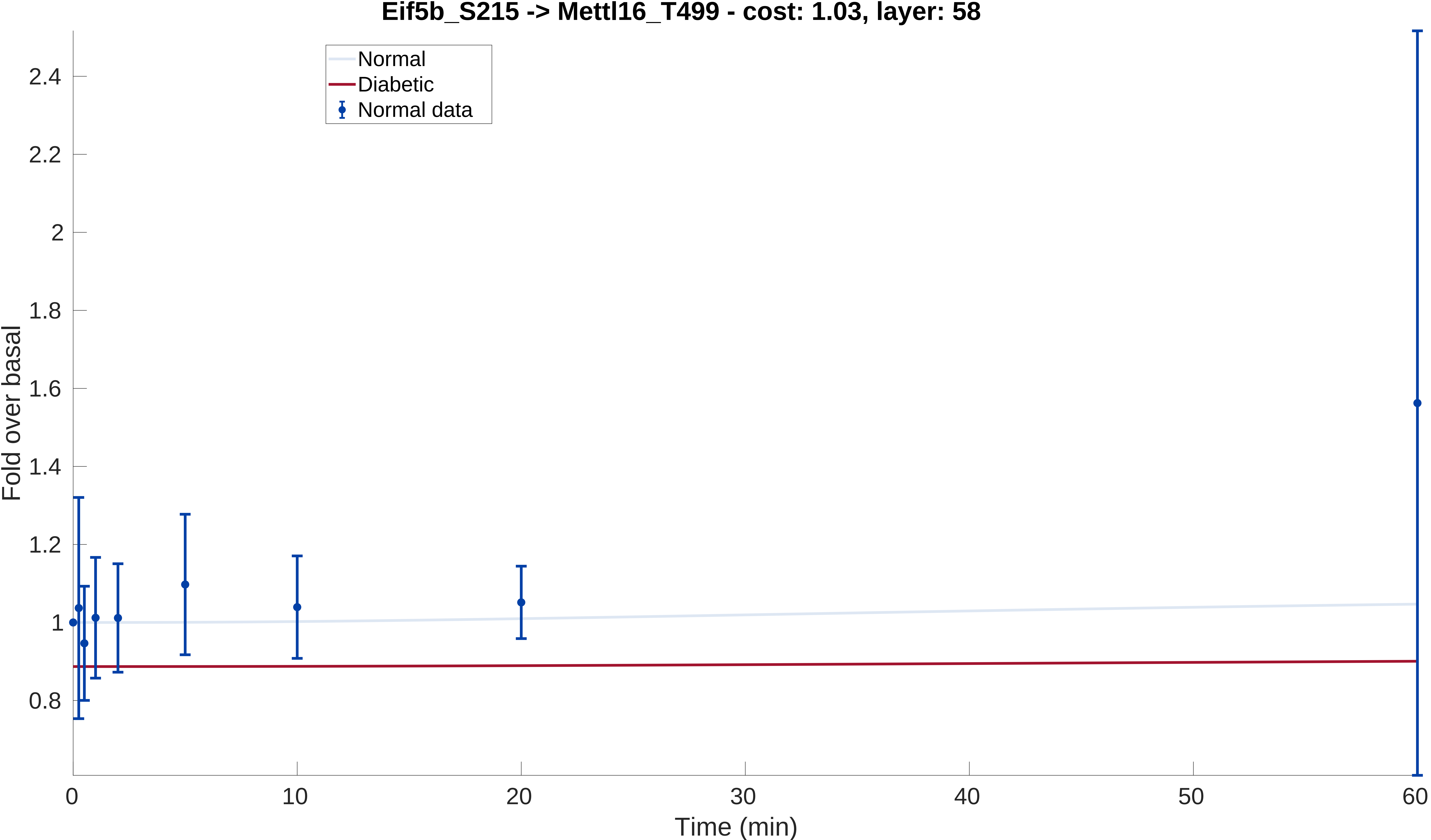
Eif5b_S215 -> Mettl16_T499 - cost: 1.03, layer: 58
Normal
Diabetic
Normal data
2.4
2.2
2
1.8
1.6
1.4
1.2
1
0.8
Fold over basal
0
10
20
30
40
50
60
Time (min)

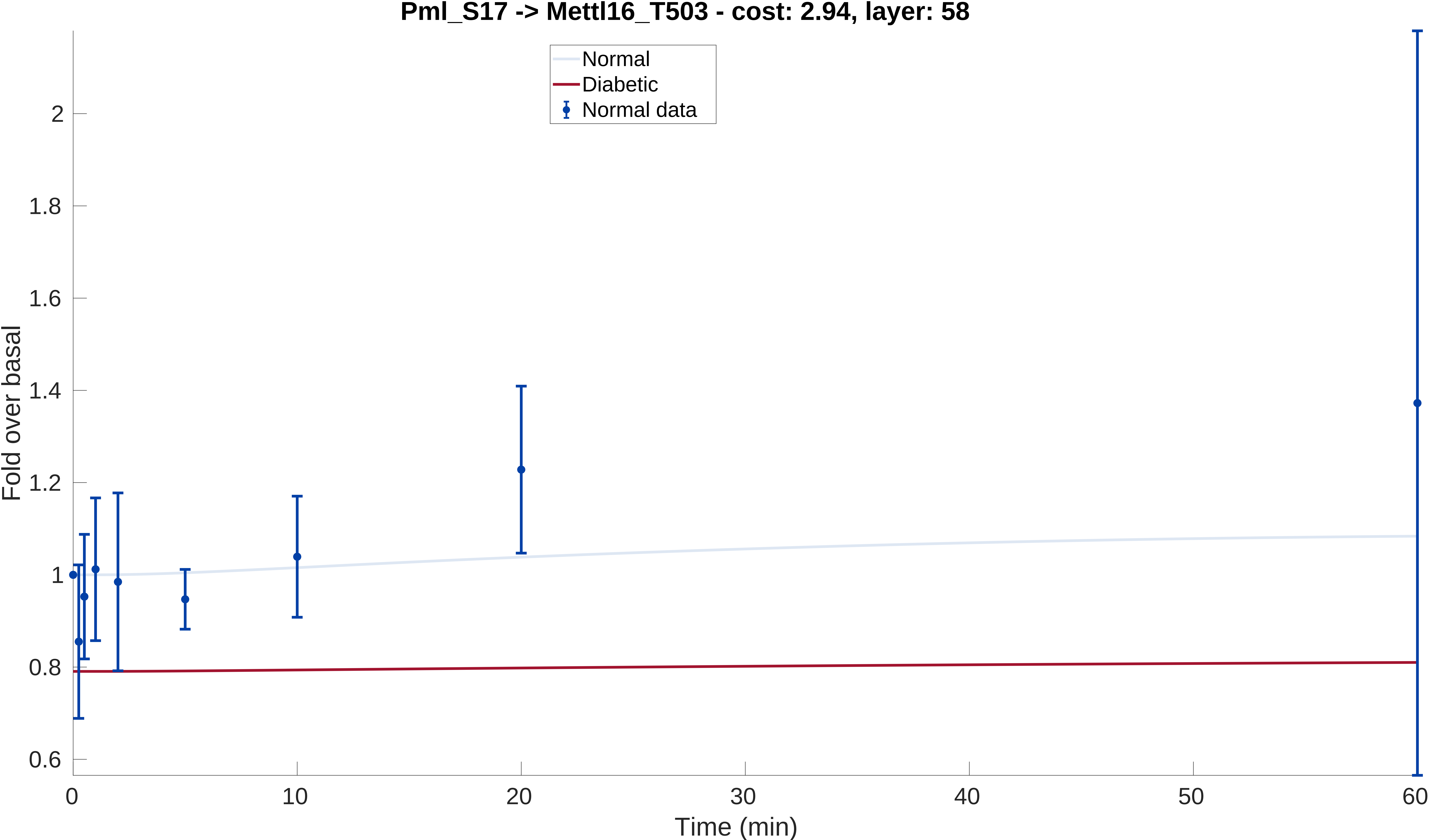

Pml_S17 -> Mettl16_T503 - cost: 2.94, layer: 58
Normal
Diabetic
Normal data
Fold over basal
2
1.8
1.6
1.4
1.2
1
0.8
0.6
0
10
20
30
40
50
60
Time (min)

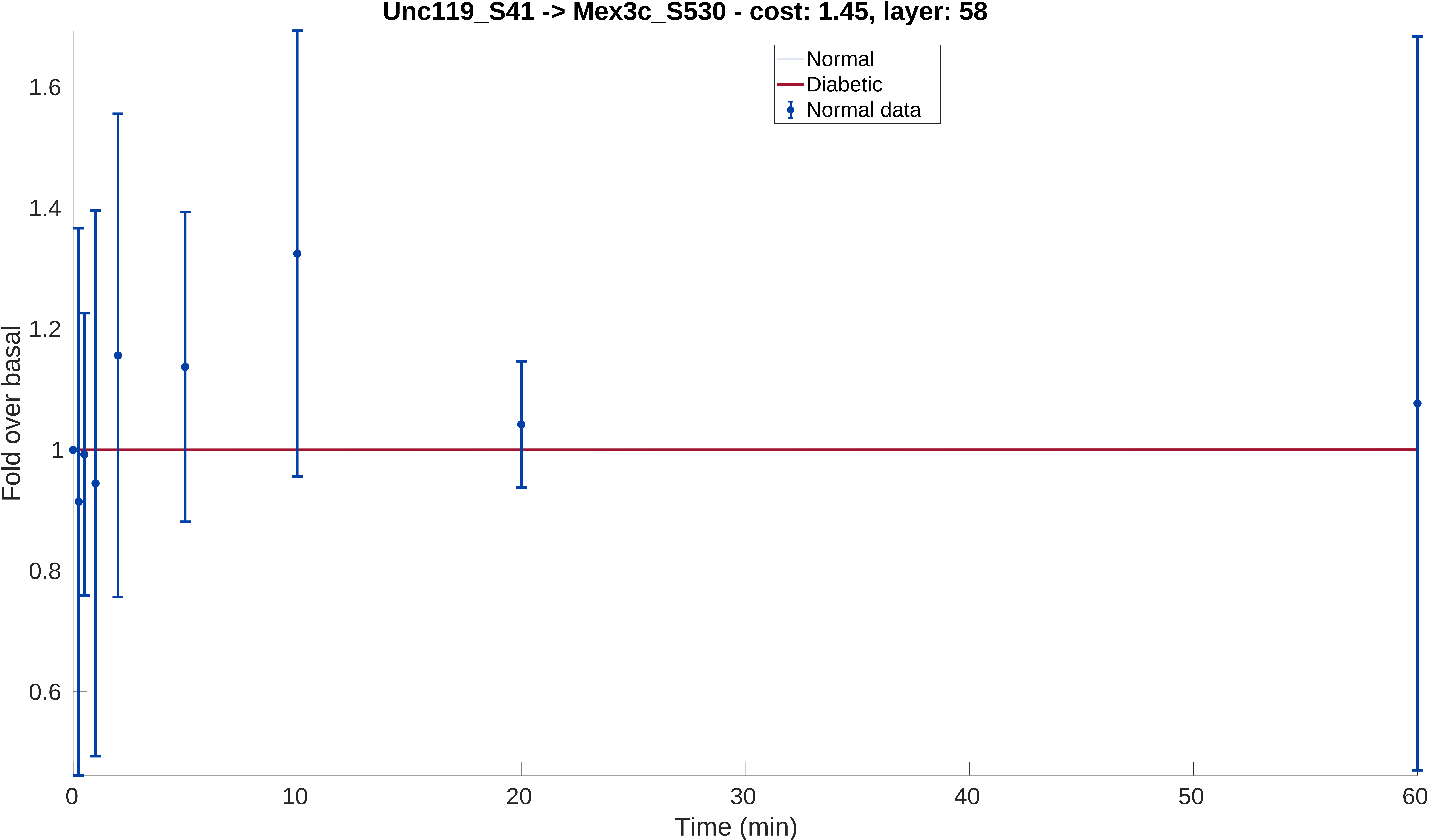

Unc119_S41 -> Mex3c_S530 - cost: 1.45, layer: 58
Normal
Diabetic
Normal data
Fold over basal
1.6
1.4
1.2
1
0.8
0.6
0
10
20
30
40
50
60
Time (min)

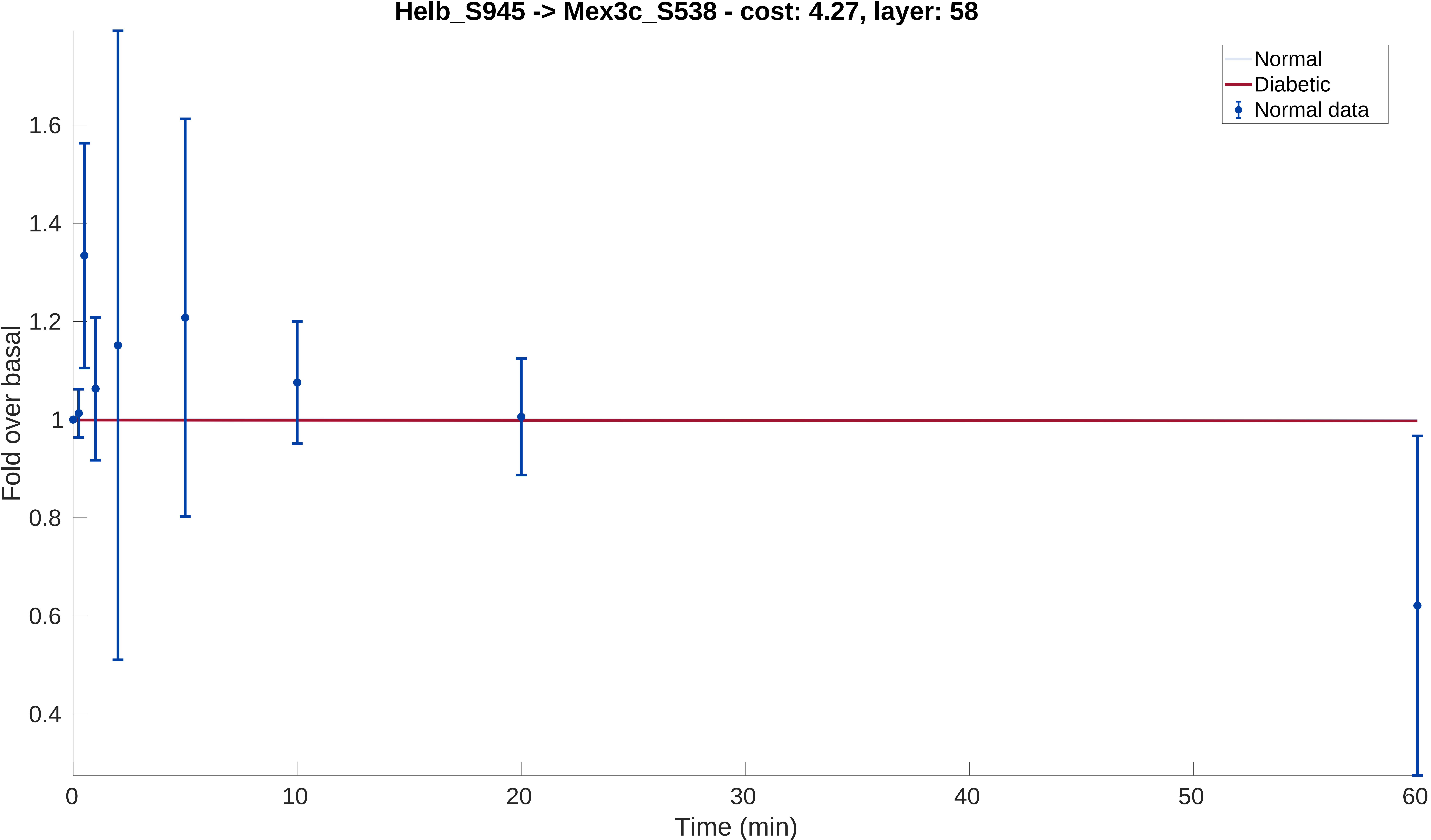

Helb_S945 -> Mex3c_S538 - cost: 4.27, layer: 58
Normal
Diabetic
Normal data
Fold over basal
1.6
1.4
1.2
1
0.8
0.6
0.4
0
10
20
30
40
50
60
Time (min)

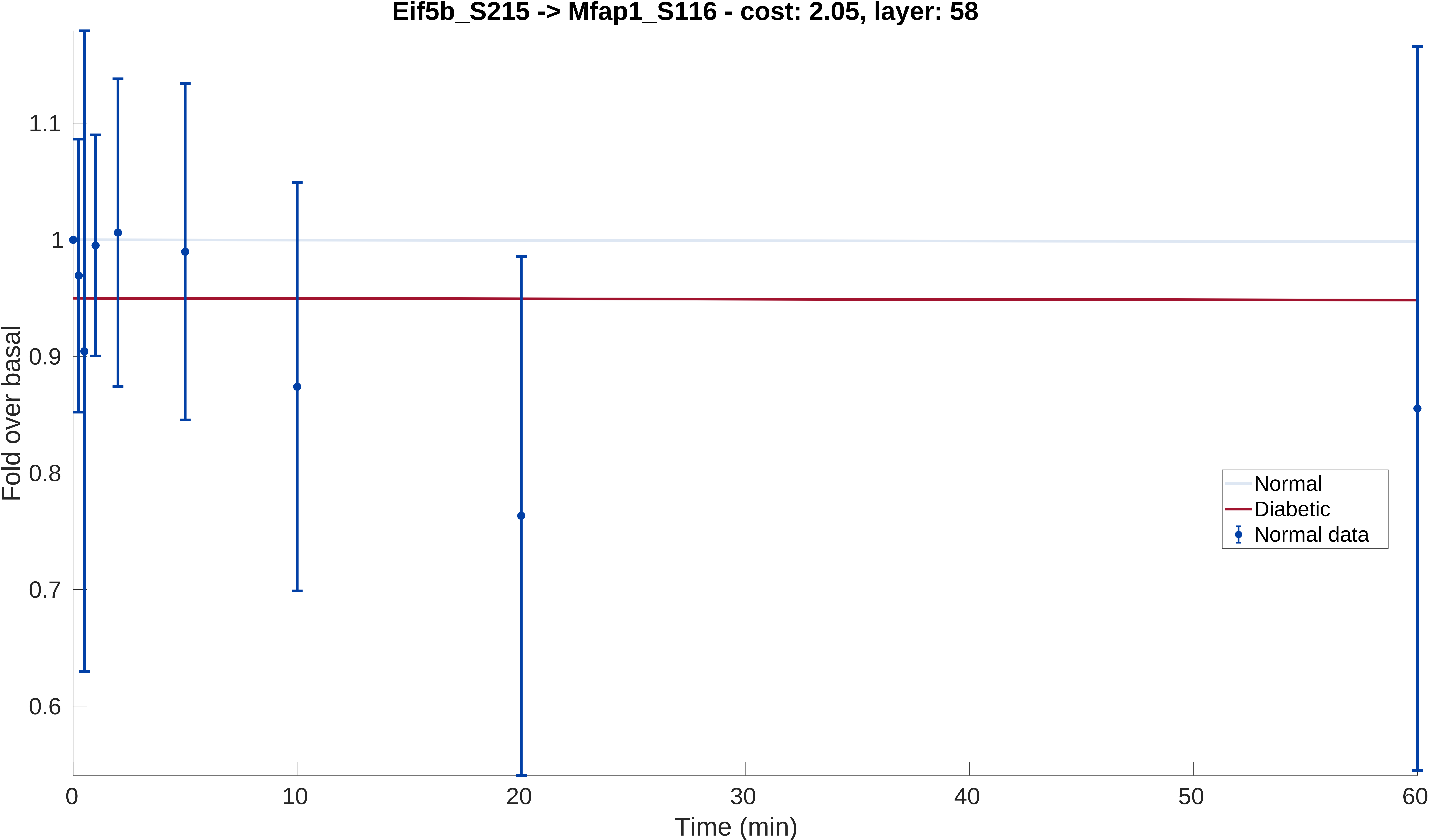

Eif5b_S215 -> Mfap1_S116 - cost: 2.05, layer: 58
Fold over basal
Time (min)
1.1
1
0.9
0.8
0.7
0.6
0
10
20
30
40
50
60
Normal
Diabetic
Normal data

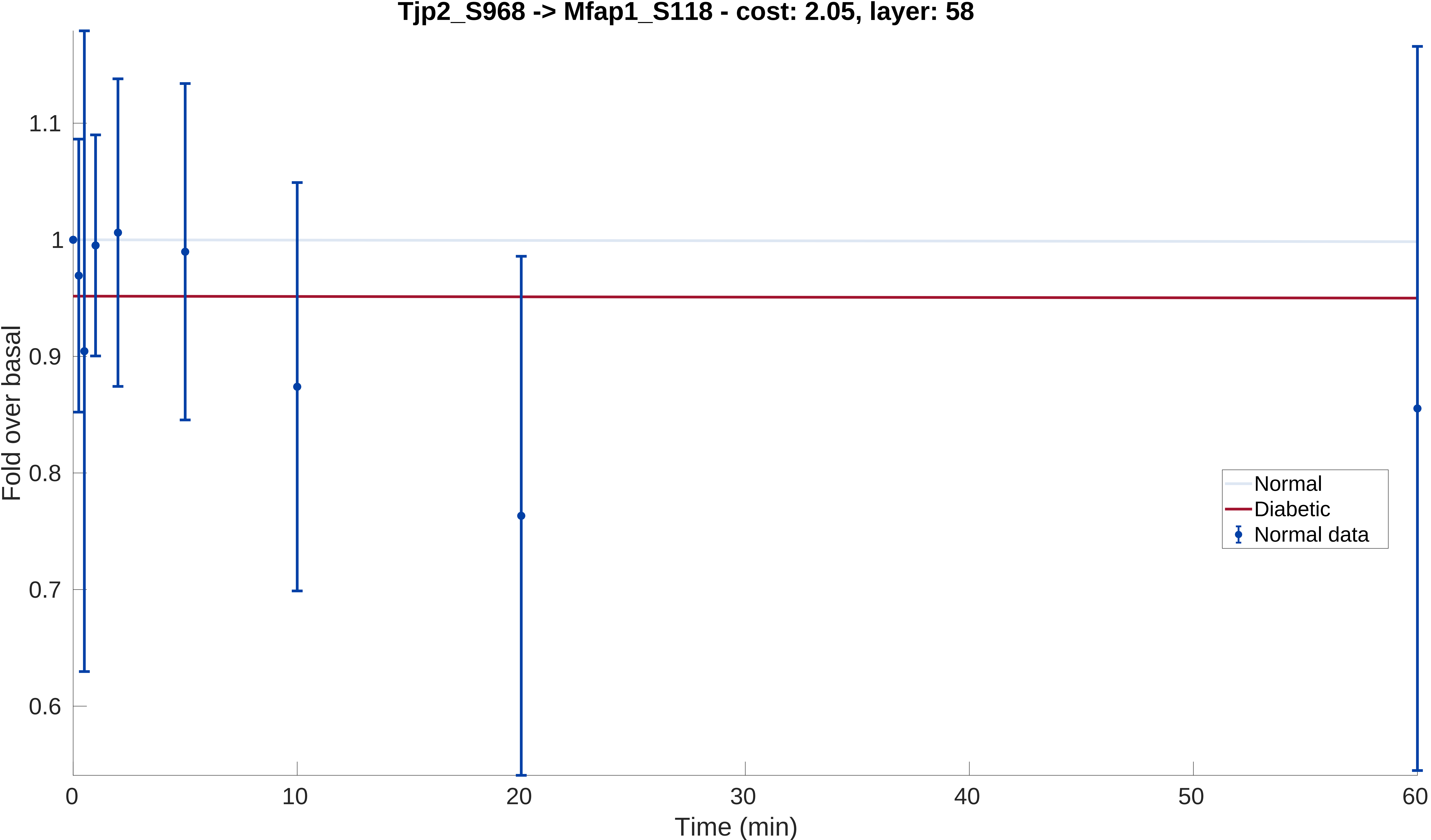
Tjp2_S968 -> Mfap1_S118 - cost: 2.05, layer: 58
Fold over basal
Time (min)
0.6
0.7
0.8
0.9
1
1.1
0
10
20
30
40
50
60
Normal
Diabetic
Normal data

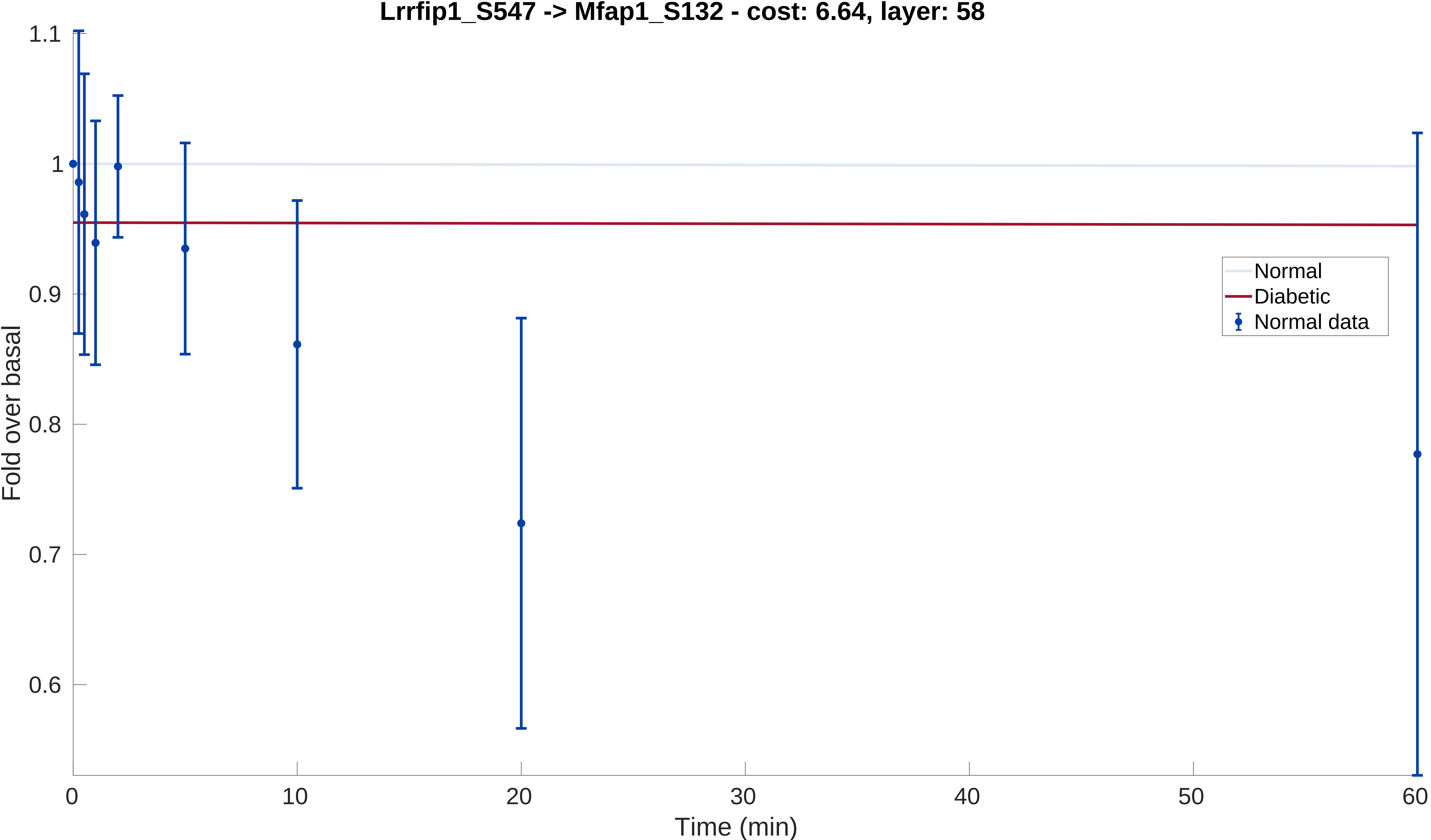
Lrrfip1_S547 -> Mfap1_S132 - cost: 6.64, layer: 58
Fold over basal
Time (min)
Normal
Diabetic
Normal data
1.1
1
0.9
0.8
0.7
0.6
0
10
20
30
40
50
60

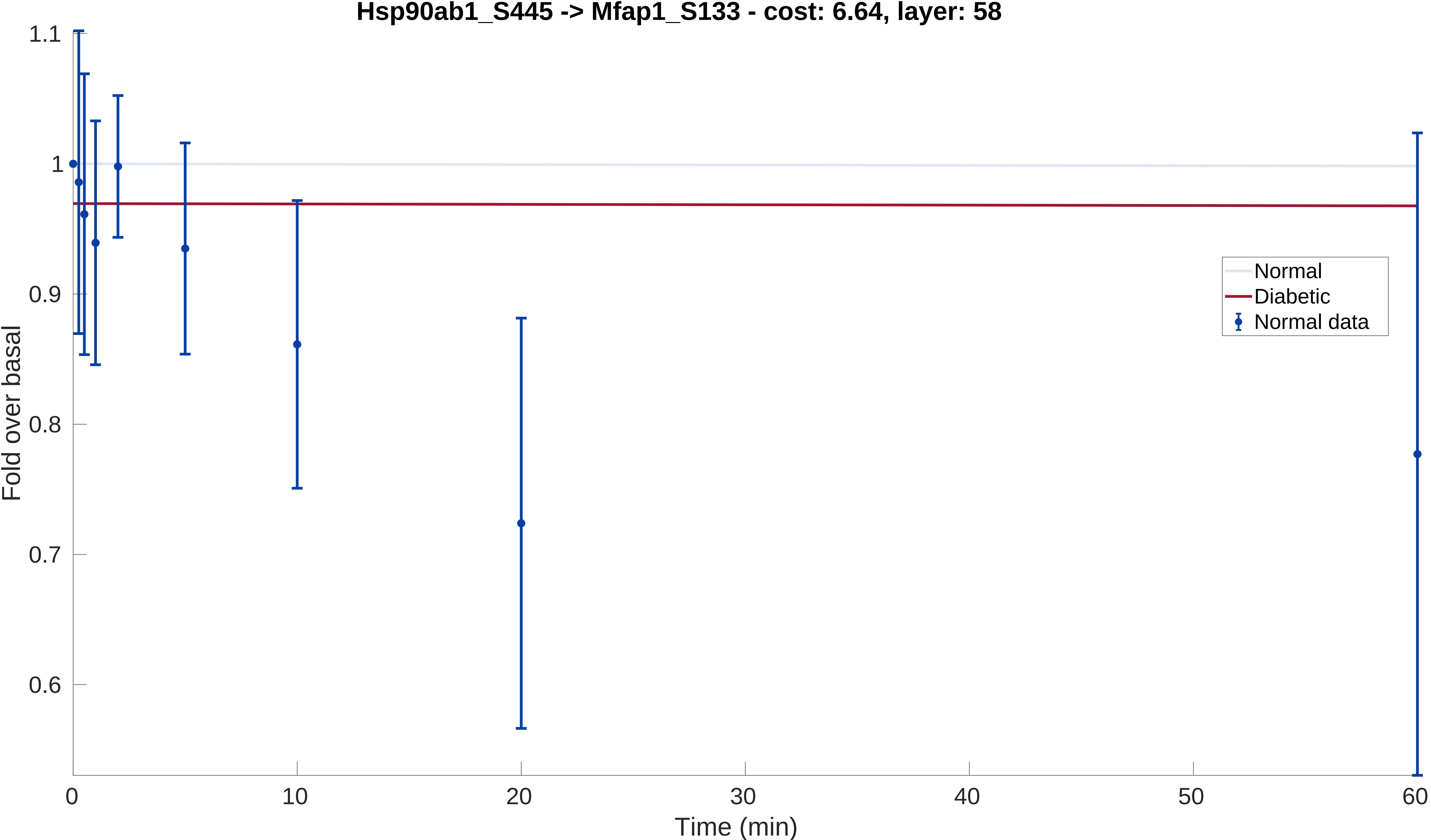

Hsp90ab1_S445 -> Mfap1_S133 - cost: 6.64, layer: 58
Fold over basal
1.1
1
0.9
0.8
0.7
0.6
0
10
20
30
40
50
60
Time (min)
Normal
Diabetic
Normal data

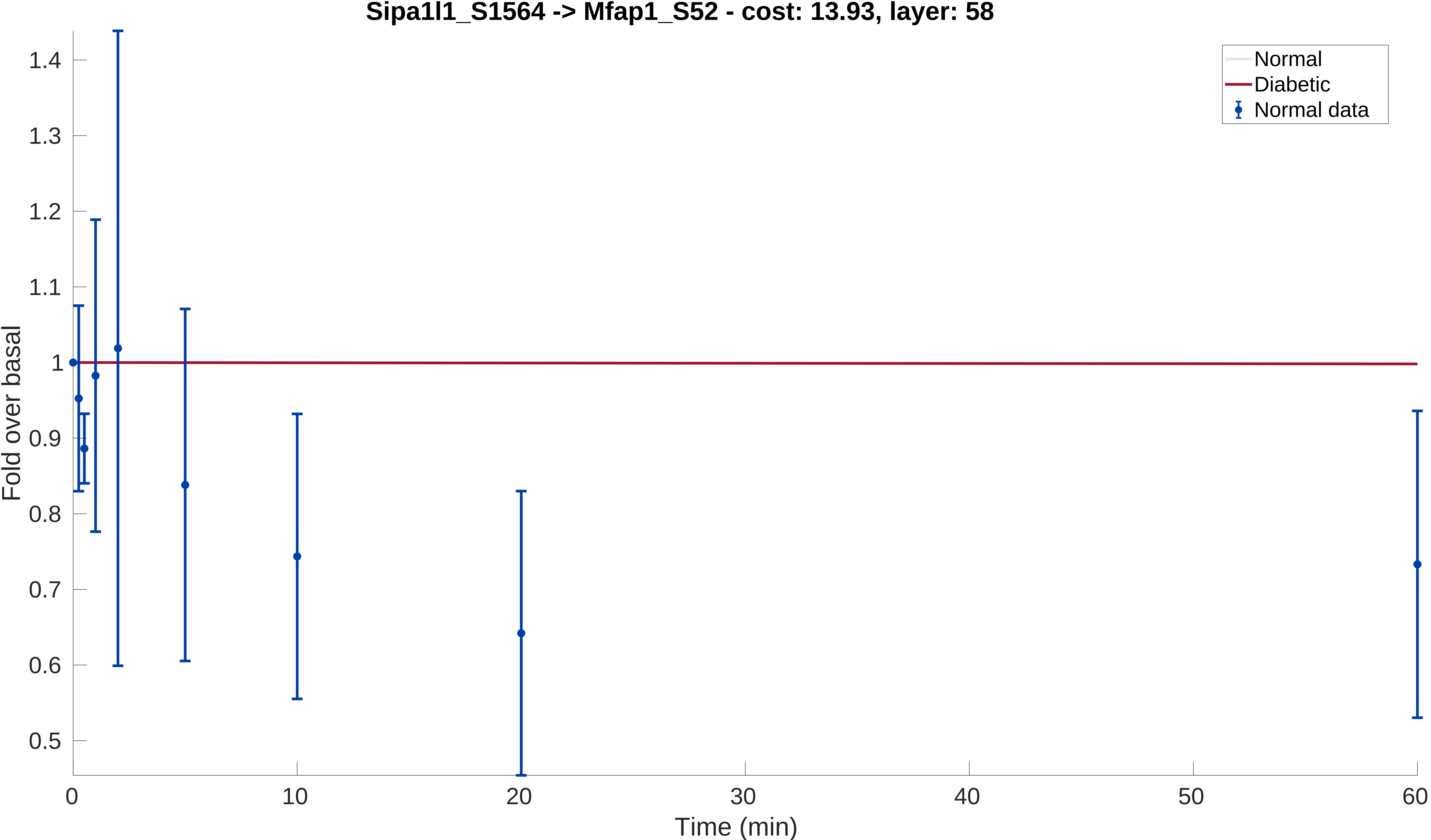
Sipa1l1_S1564 -> Mfap1_S52 - cost: 13.93, layer: 58
Normal
Diabetic
Normal data
Fold over basal
1.4
1.3
1.2
1.1
1
0.9
0.8
0.7
0.6
0.5
0
10
20
30
40
50
60
Time (min)

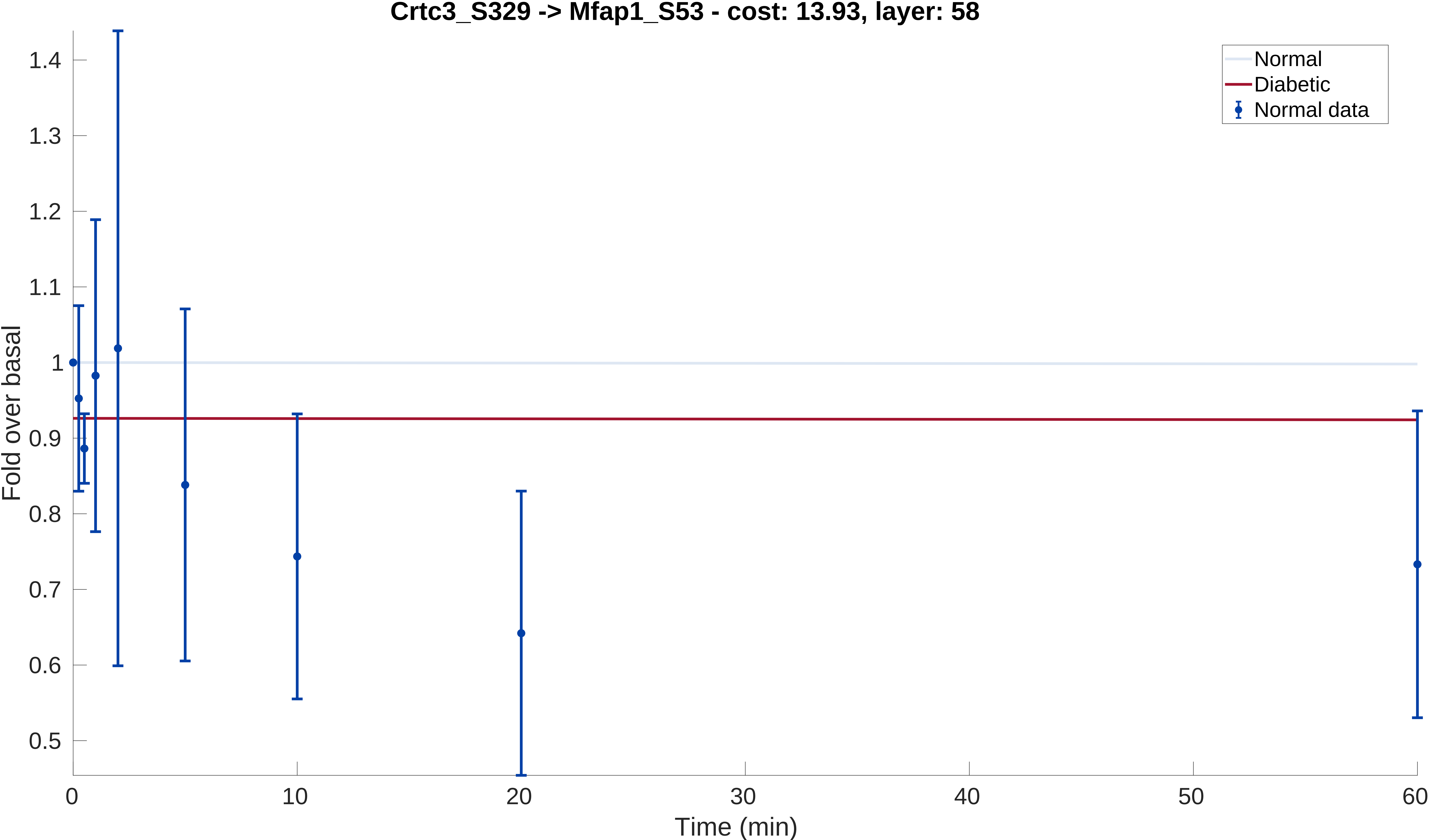

Crtc3_S329 -> Mfap1_S53 - cost: 13.93, layer: 58
Normal
Diabetic
Normal data
Fold over basal
Time (min)
1.4
1.3
1.2
1.1
1
0.9
0.8
0.7
0.6
0.5
0
10
20
30
40
50
60

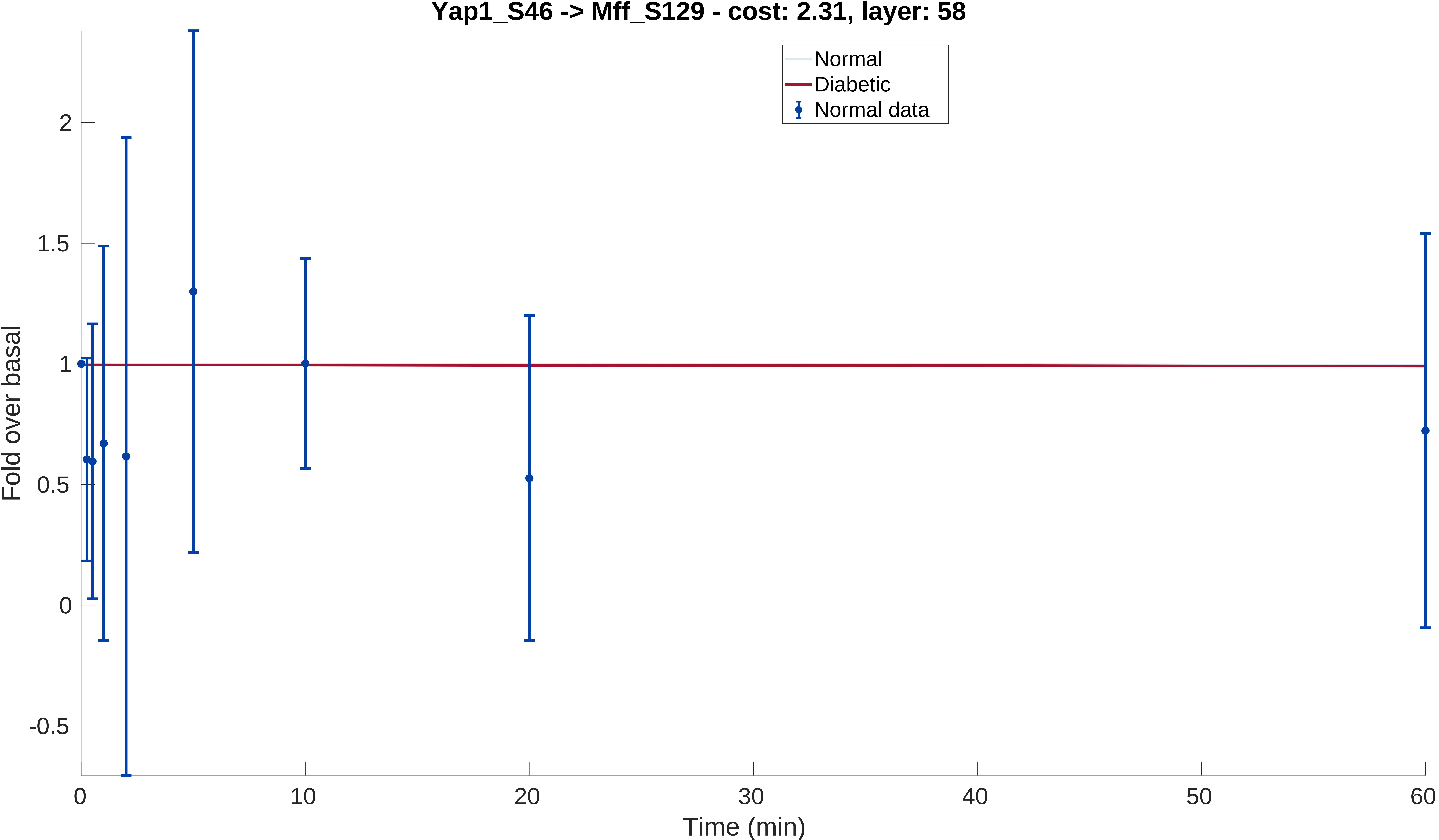

Yap1_S46 -> Mff_S129 - cost: 2.31, layer: 58
Normal
Diabetic
Normal data
Fold over basal
2
1.5
1
0.5
0
-0.5
0
10
20
30
40
50
60
Time (min)

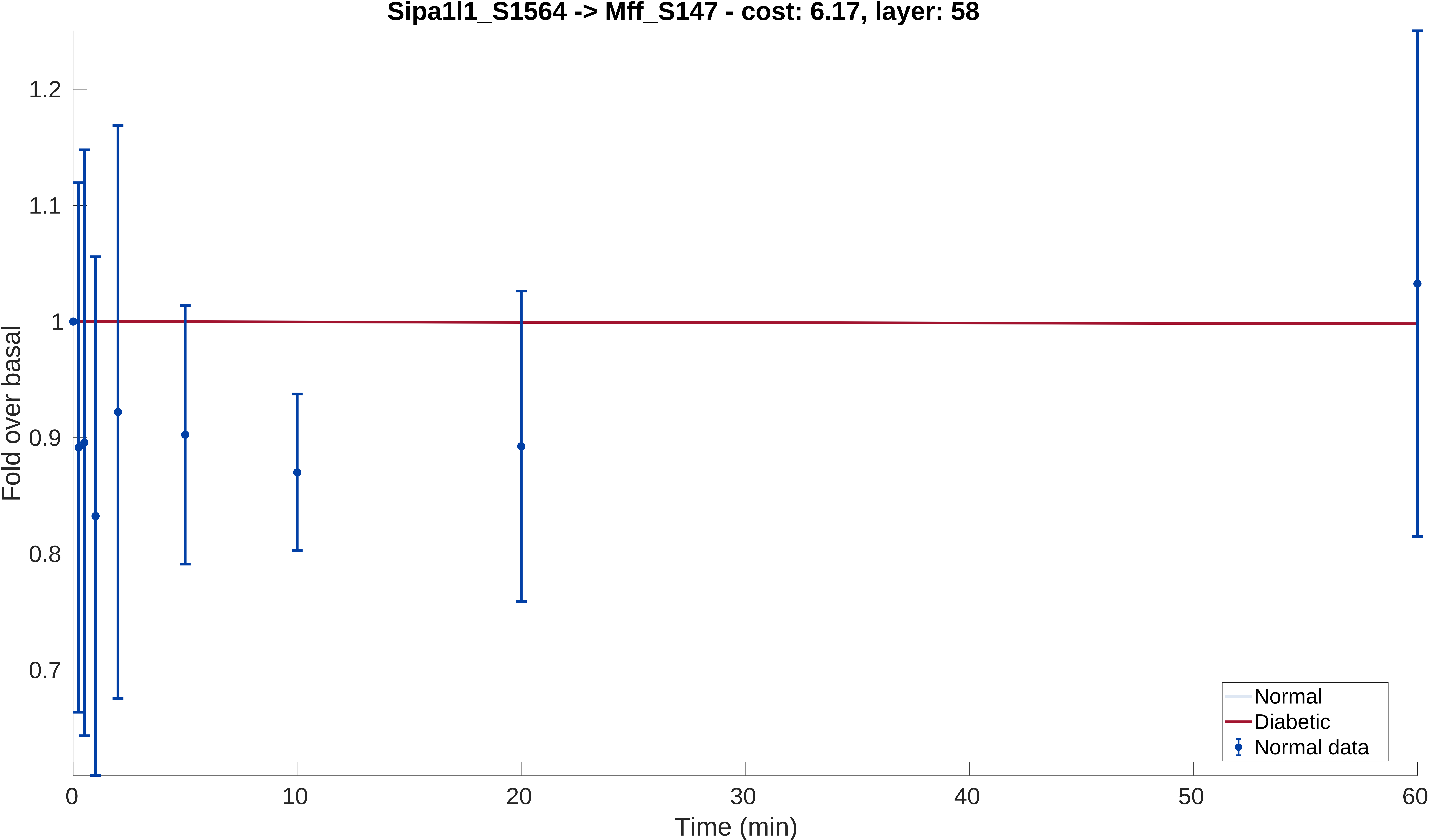

Sipa1l1_S1564 -> Mff_S147 - cost: 6.17, layer: 58
Fold over basal
1.2
1.1
1
0.9
0.8
0.7
0
10
20
30
40
50
60
Time (min)
Normal
Diabetic
Normal data

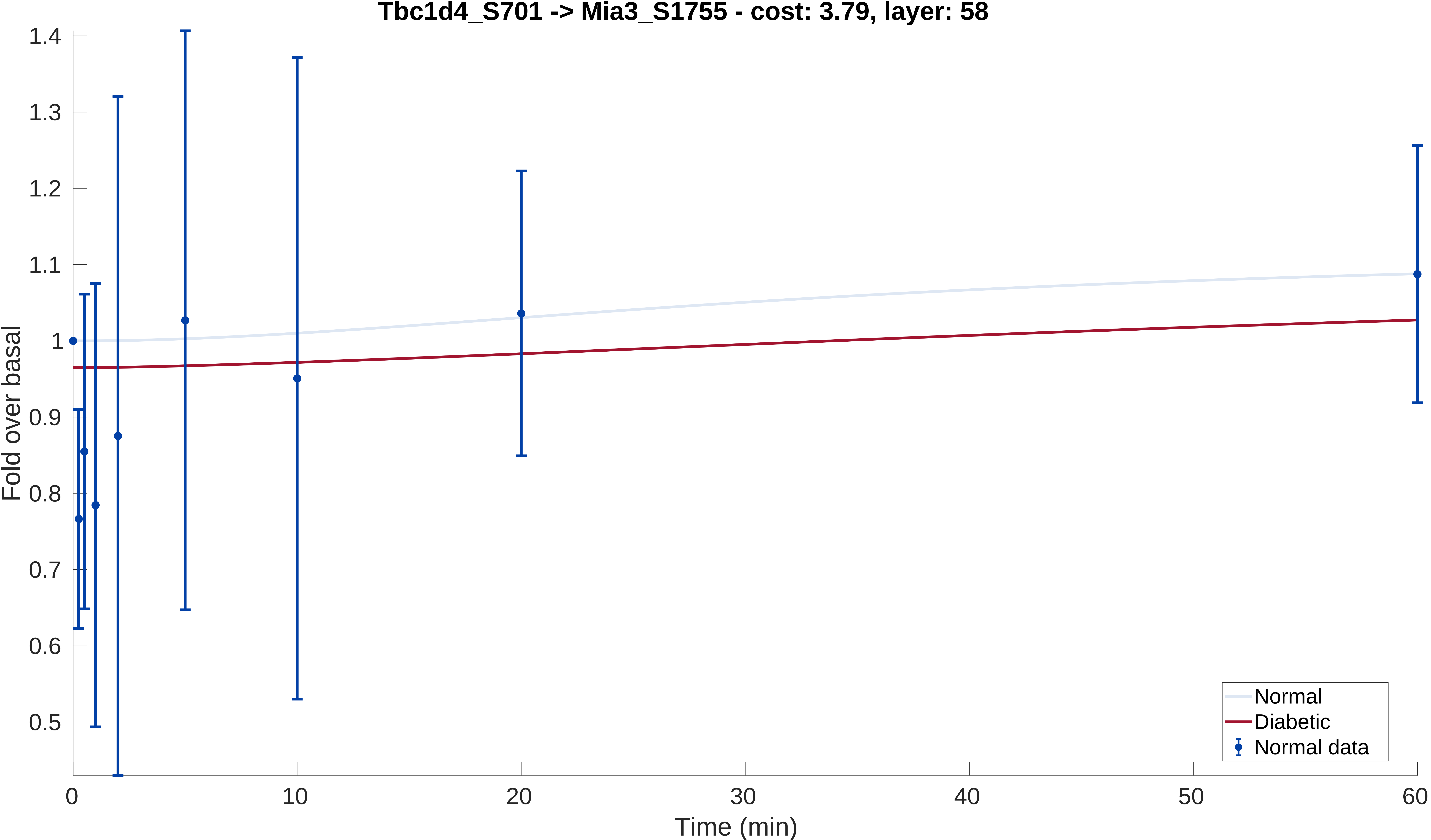

Tbc1d4_S701 -> Mia3_S1755 - cost: 3.79, layer: 58
Fold over basal
Time (min)
1.4
1.3
1.2
1.1
1
0.9
0.8
0.7
0.6
0.5
0
10
20
30
40
50
60
Normal
Diabetic
Normal data

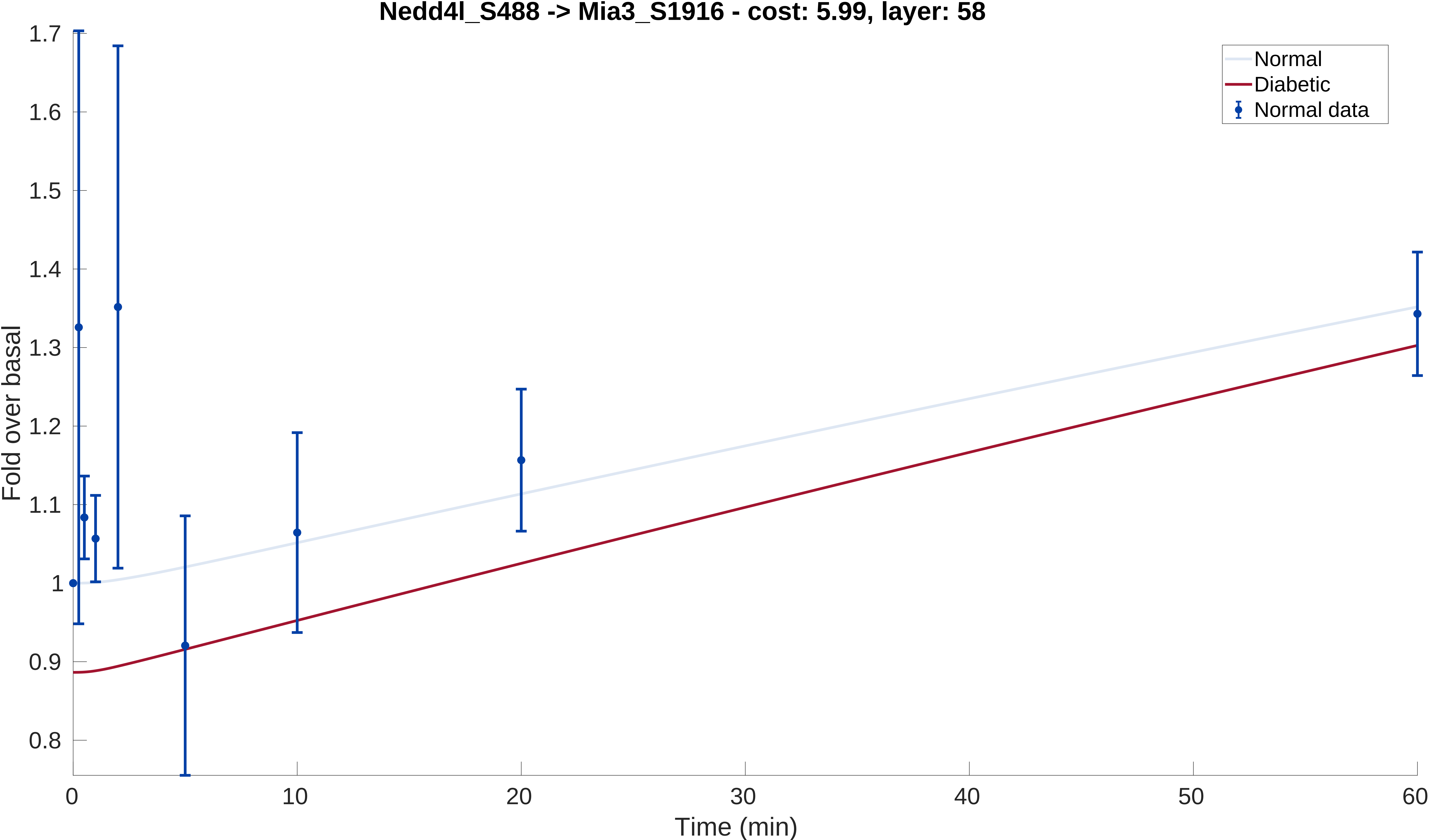
Nedd4l_S488 -> Mia3_S1916 - cost: 5.99, layer: 58
Normal
Diabetic
Normal data
Fold over basal
Time (min)
0.8
0.9
1
1.1
1.2
1.3
1.4
1.5
1.6
1.7
0
10
20
30
40
50
60

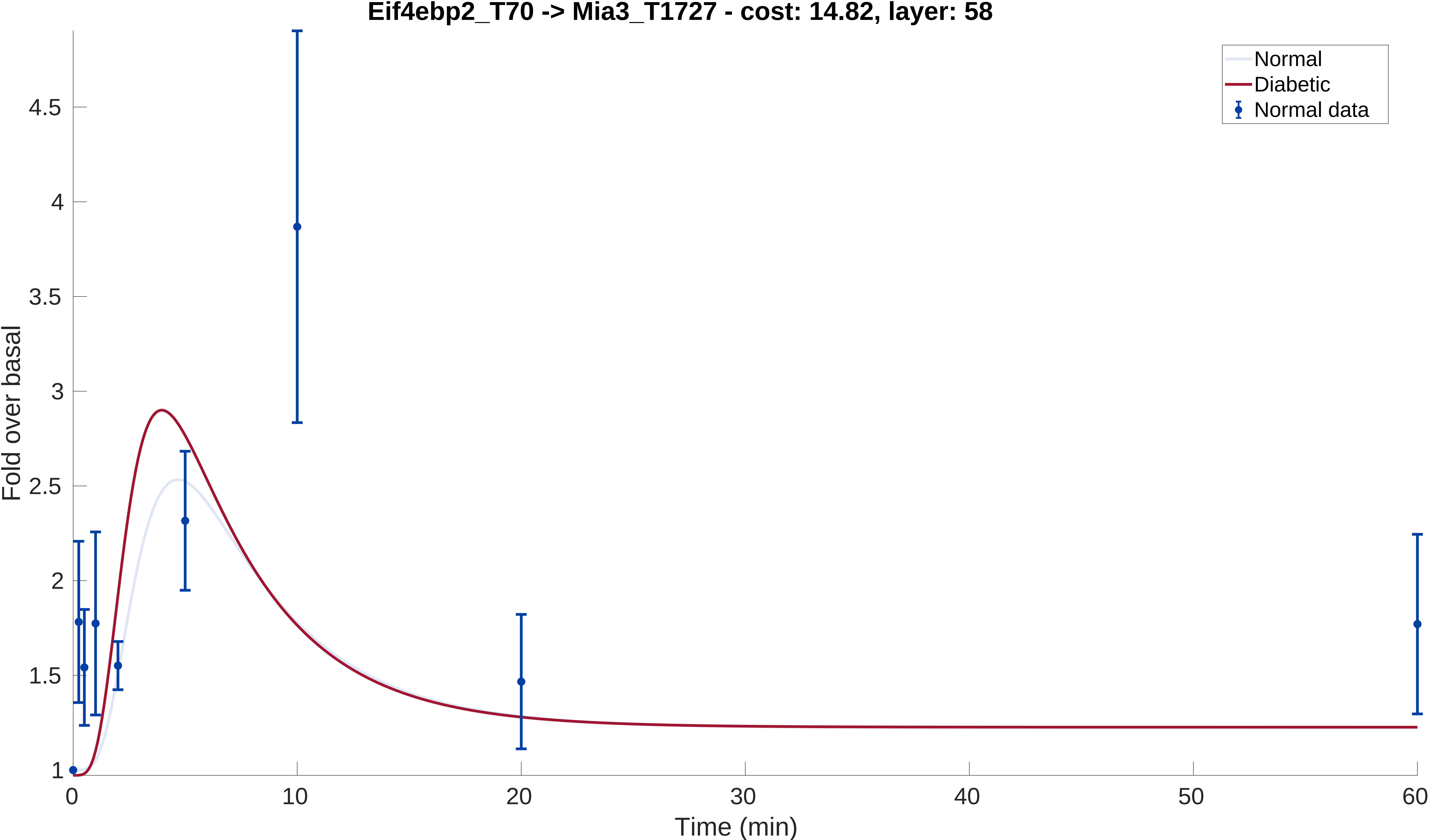
Eif4ebp2_T70 -> Mia3_T1727 - cost: 14.82, layer: 58
Normal
Diabetic
Normal data
Fold over basal
Time (min)
4.5
4
3.5
3
2.5
2
1.5
1
0
10
20
30
40
50
60

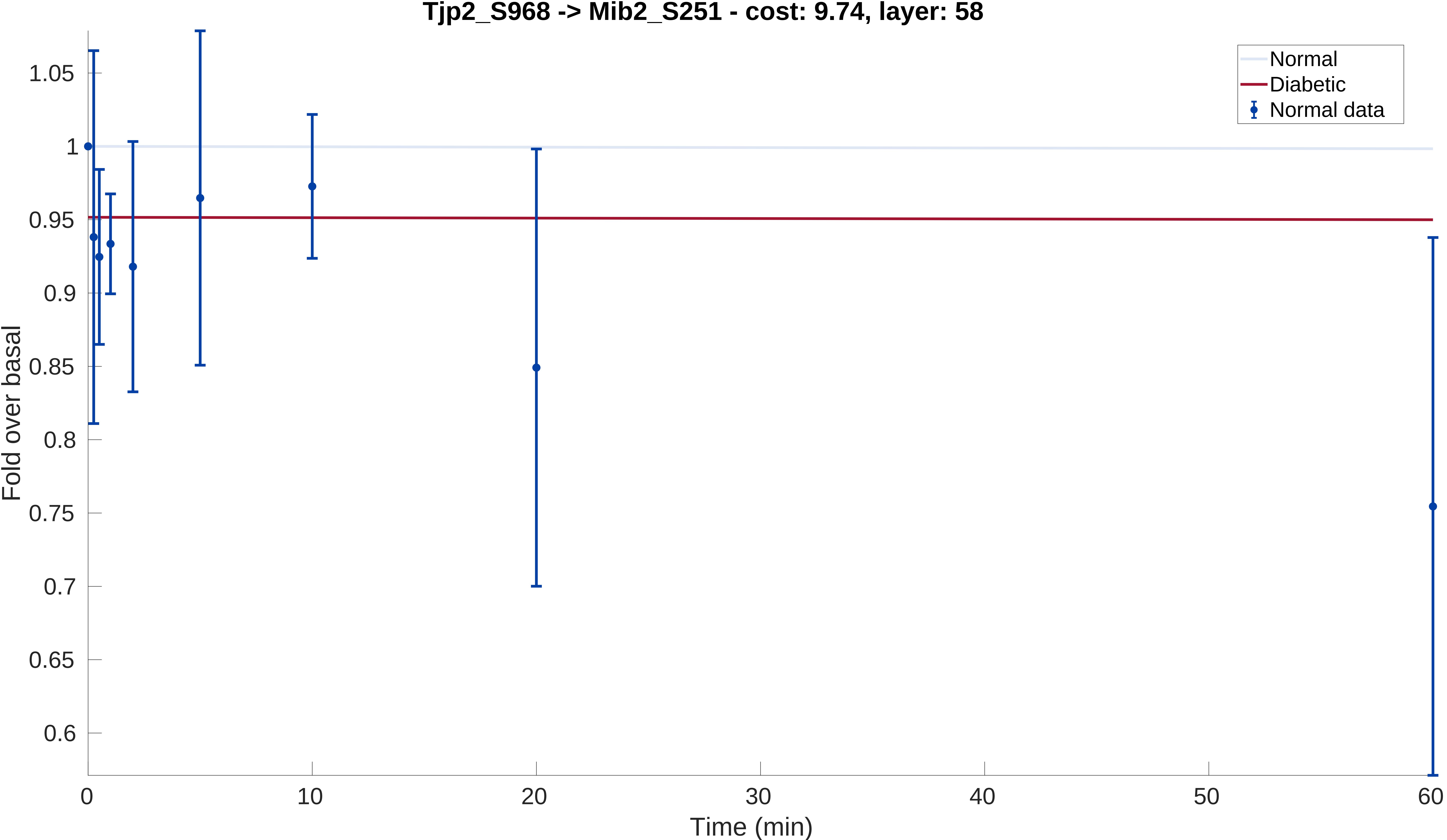

Tjp2_S968 -> Mib2_S251 - cost: 9.74, layer: 58
Normal
Diabetic
Normal data
Fold over basal
1.05
1
0.95
0.9
0.85
0.8
0.75
0.7
0.65
0.6
0
10
20
30
40
50
60
Time (min)

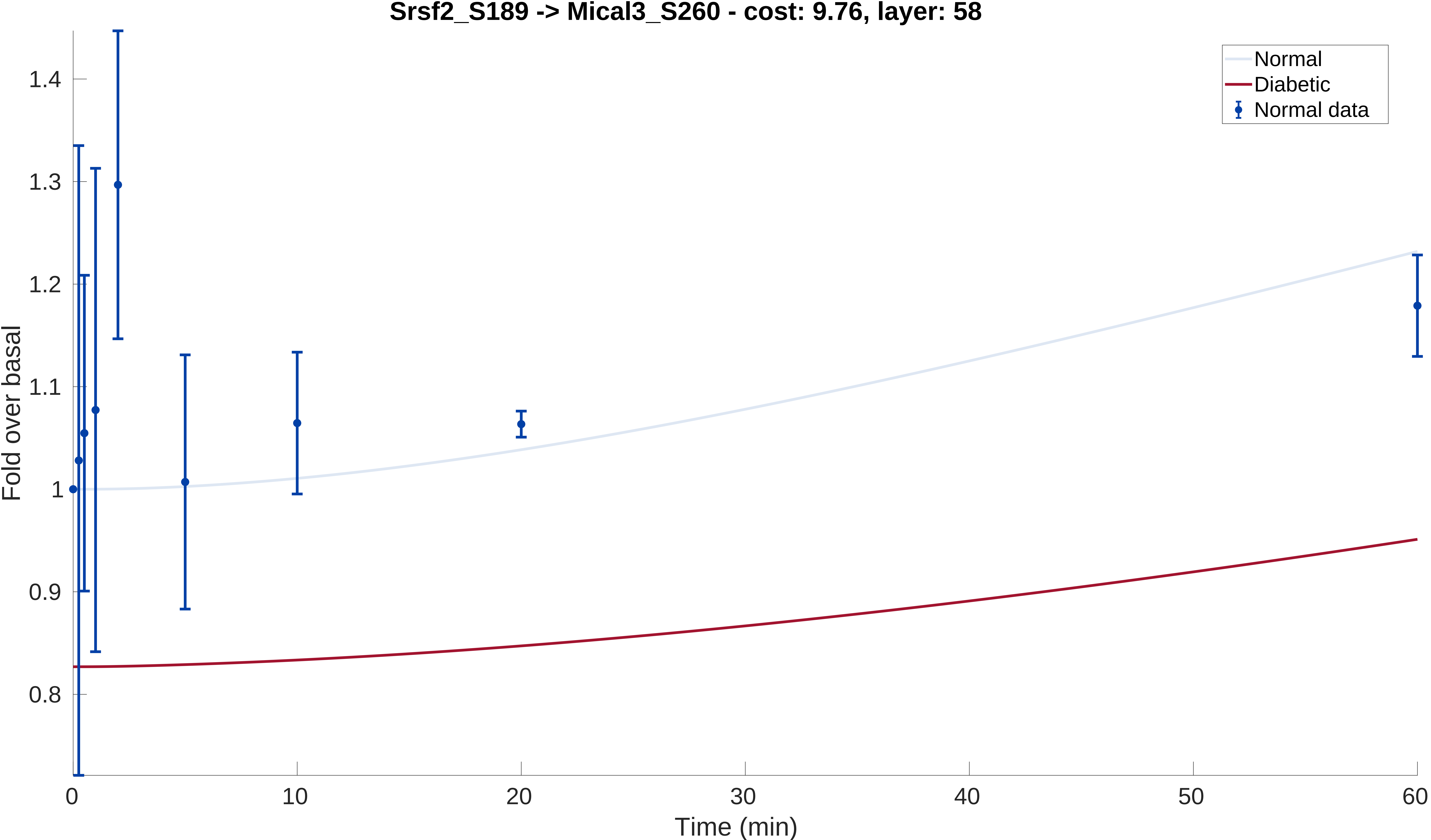

Srsf2_S189 -> Mical3_S260 - cost: 9.76, layer: 58
Normal
Diabetic
Normal data
Fold over basal
1.4
1.3
1.2
1.1
1
0.9
0.8
0
10
20
30
40
50
60
Time (min)

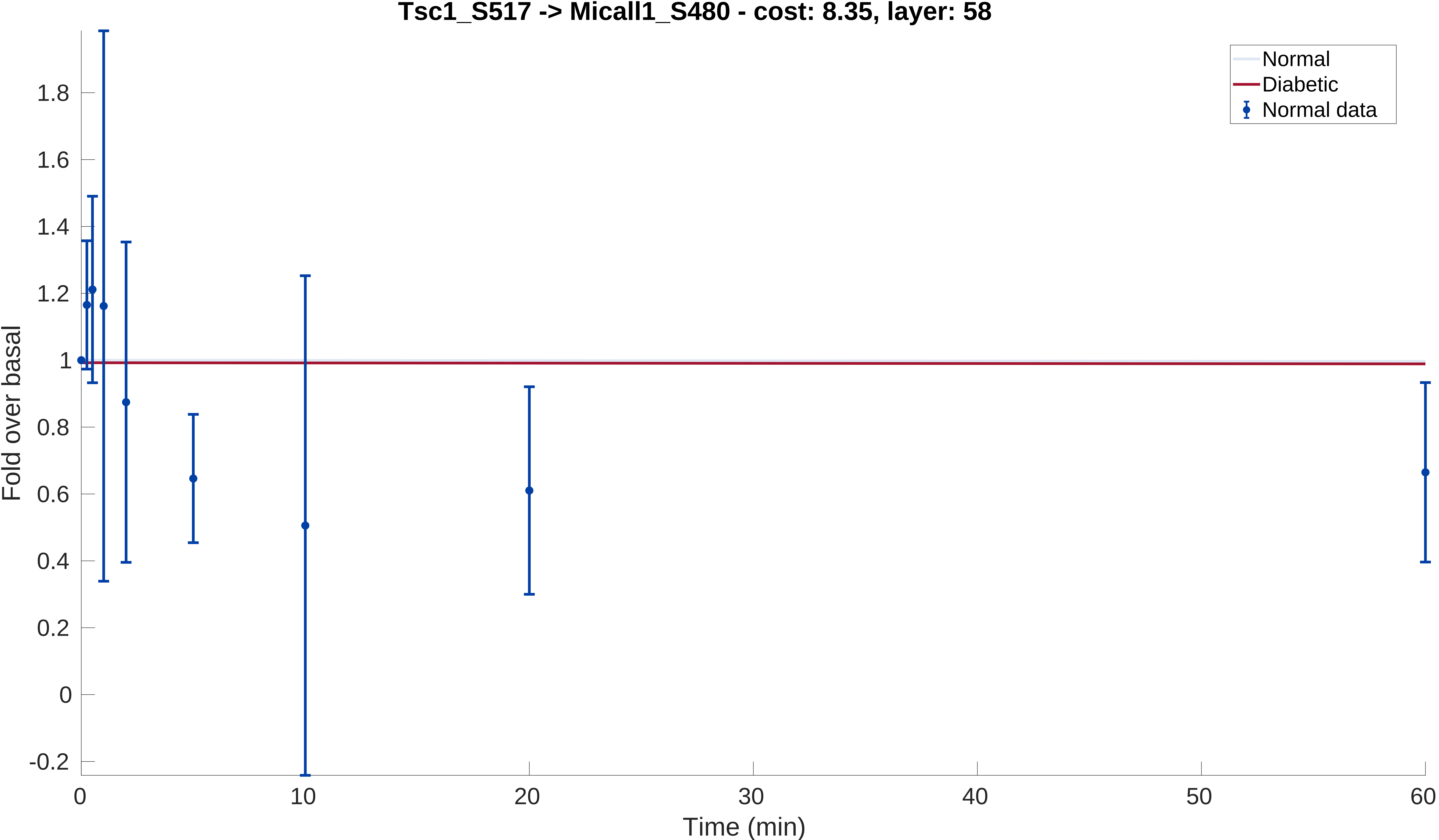
Tsc1_S517 -> Micall1_S480 - cost: 8.35, layer: 58
Normal
Diabetic
Normal data
Fold over basal
1.8
1.6
1.4
1.2
1
0.8
0.6
0.4
0.2
0
-0.2
0
10
20
30
40
50
60
Time (min)

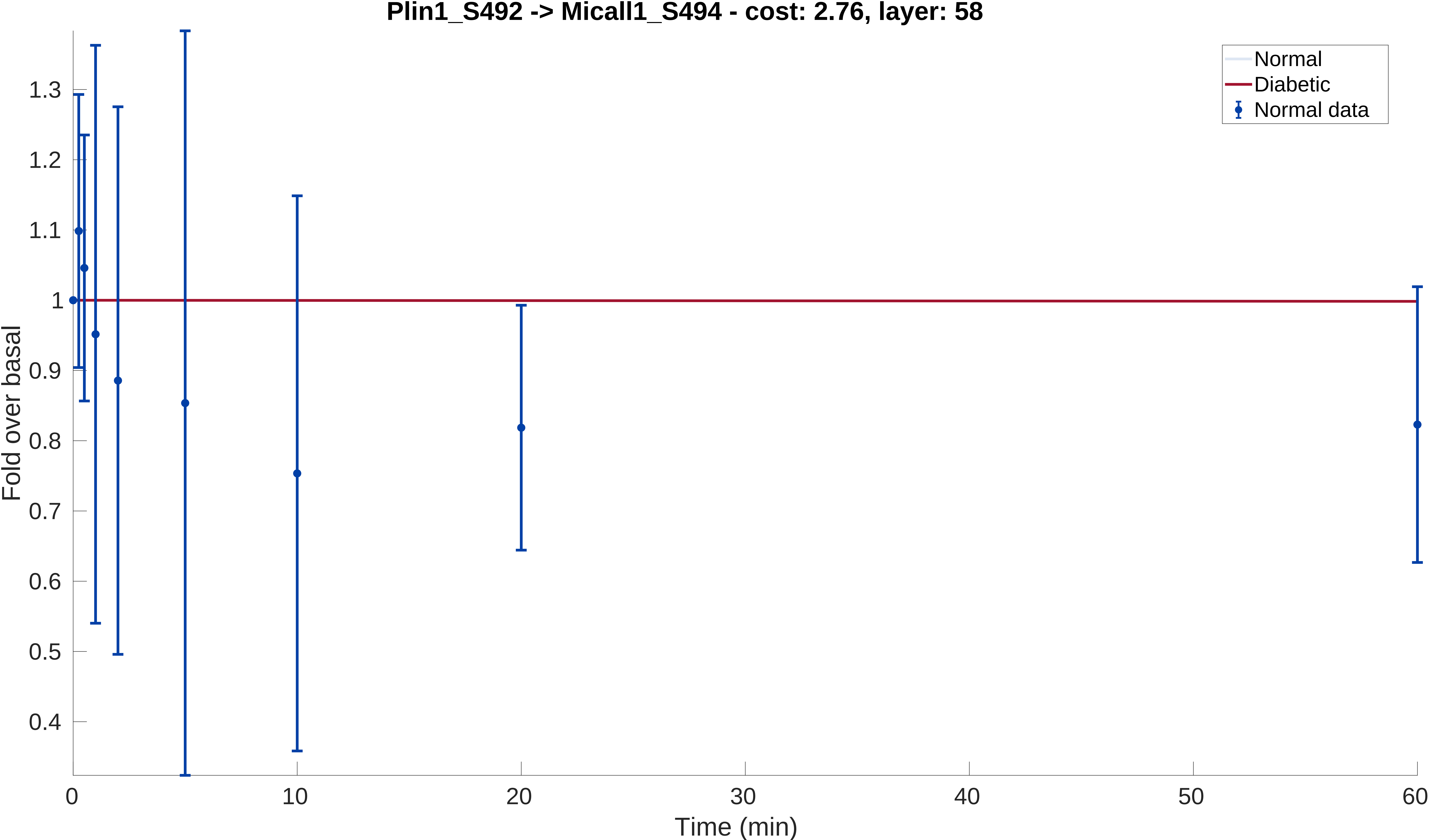

Plin1_S492 -> Micall1_S494 - cost: 2.76, layer: 58
Normal
Diabetic
Normal data
Fold over basal
1.3
1.2
1.1
1
0.9
0.8
0.7
0.6
0.5
0.4
0
10
20
30
40
50
60
Time (min)

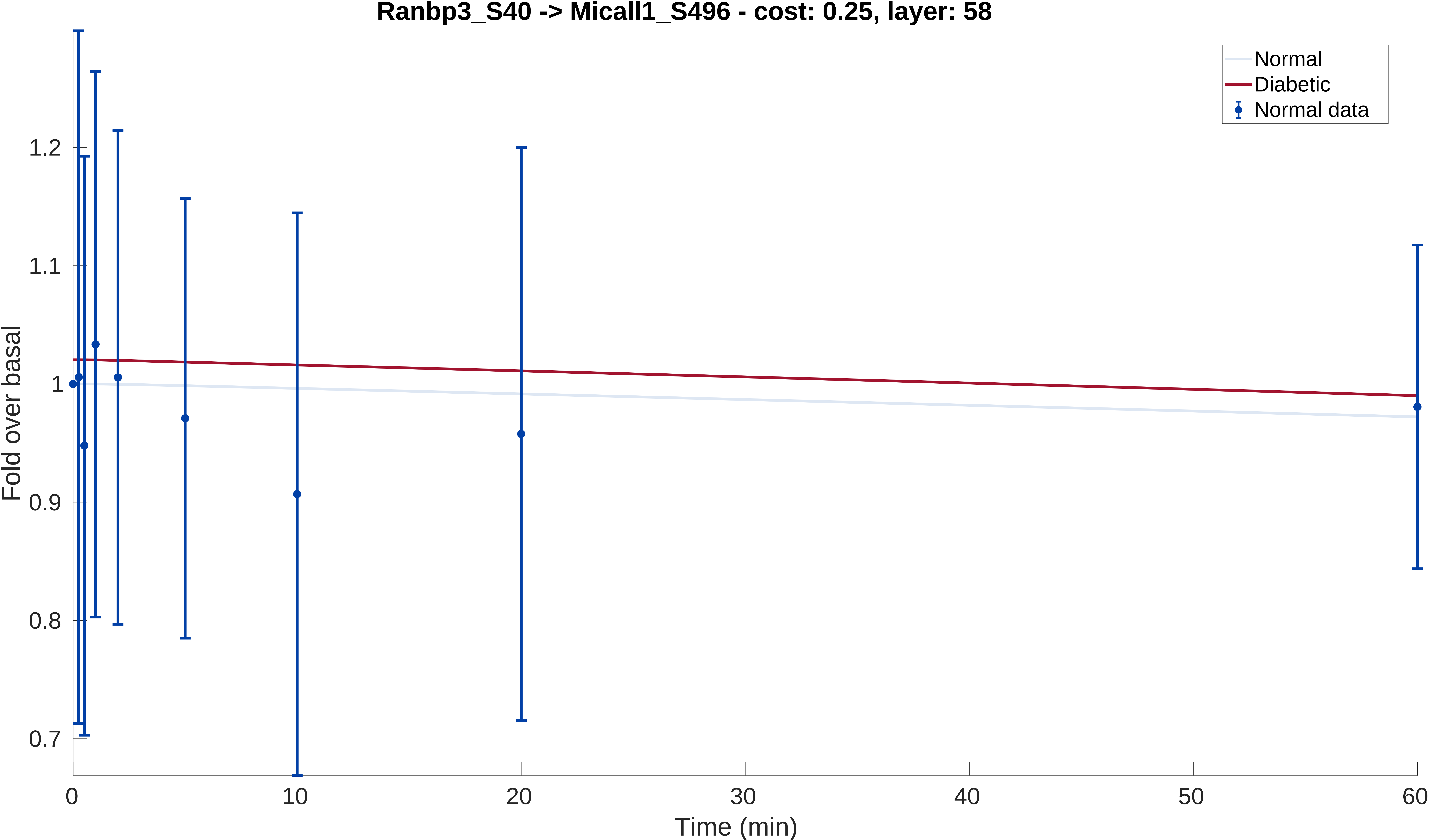

Ranbp3_S40 -> Micall1_S496 - cost: 0.25, layer: 58
Normal
Diabetic
Normal data
Fold over basal
Time (min)
0.7
0.8
0.9
1
1.1
1.2
0
10
20
30
40
50
60

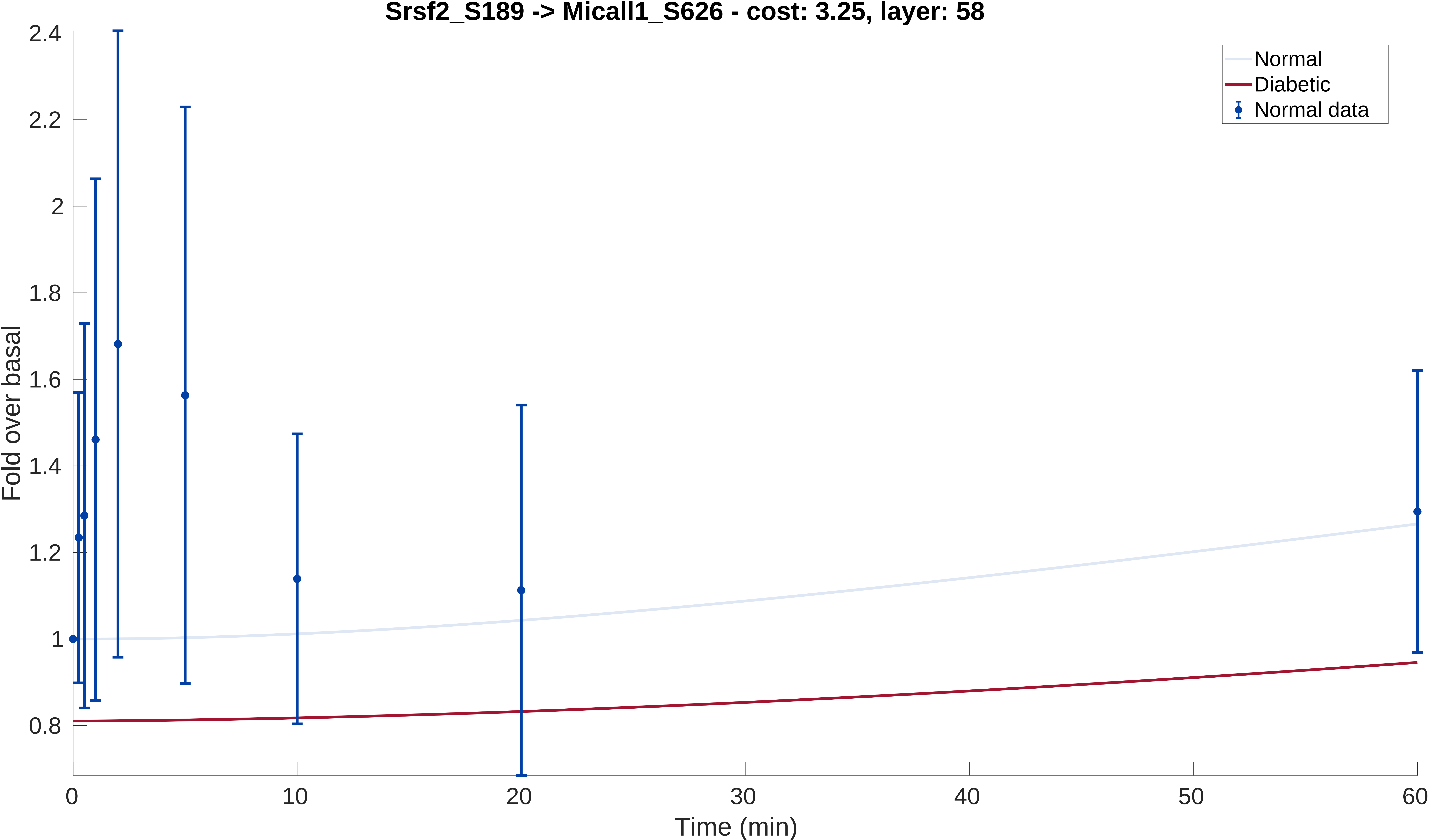

Srsf2_S189 -> Micall1_S626 - cost: 3.25, layer: 58
Normal
Diabetic
Normal data
Fold over basal
2.4
2.2
2
1.8
1.6
1.4
1.2
1
0.8
0
10
20
30
40
50
60
Time (min)

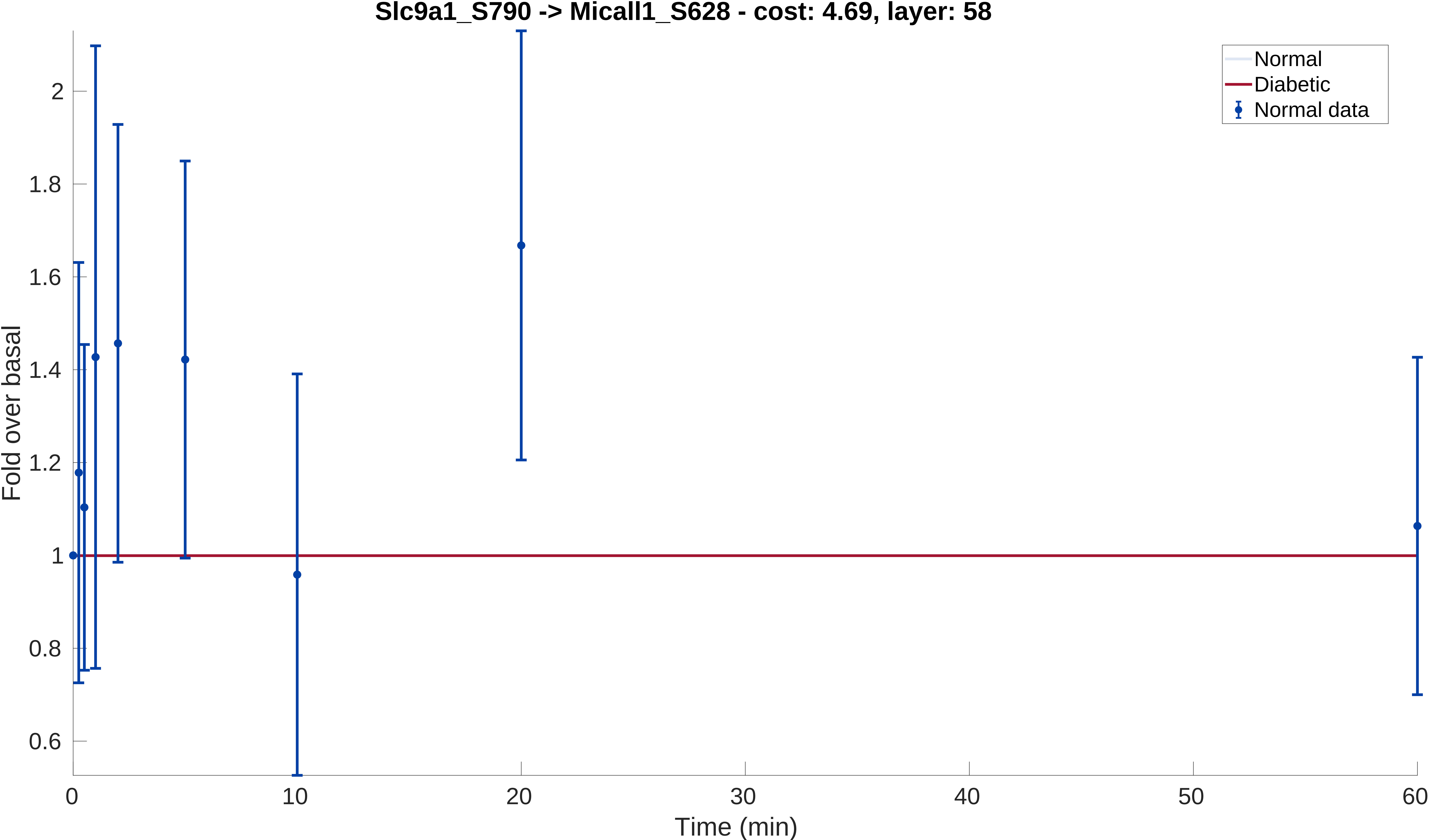
Slc9a1_S790 -> Micall1_S628 - cost: 4.69, layer: 58
Normal
Diabetic
Normal data
Fold over basal
Time (min)
0.6
0.8
1
1.2
1.4
1.6
1.8
2
0
10
20
30
40
50
60

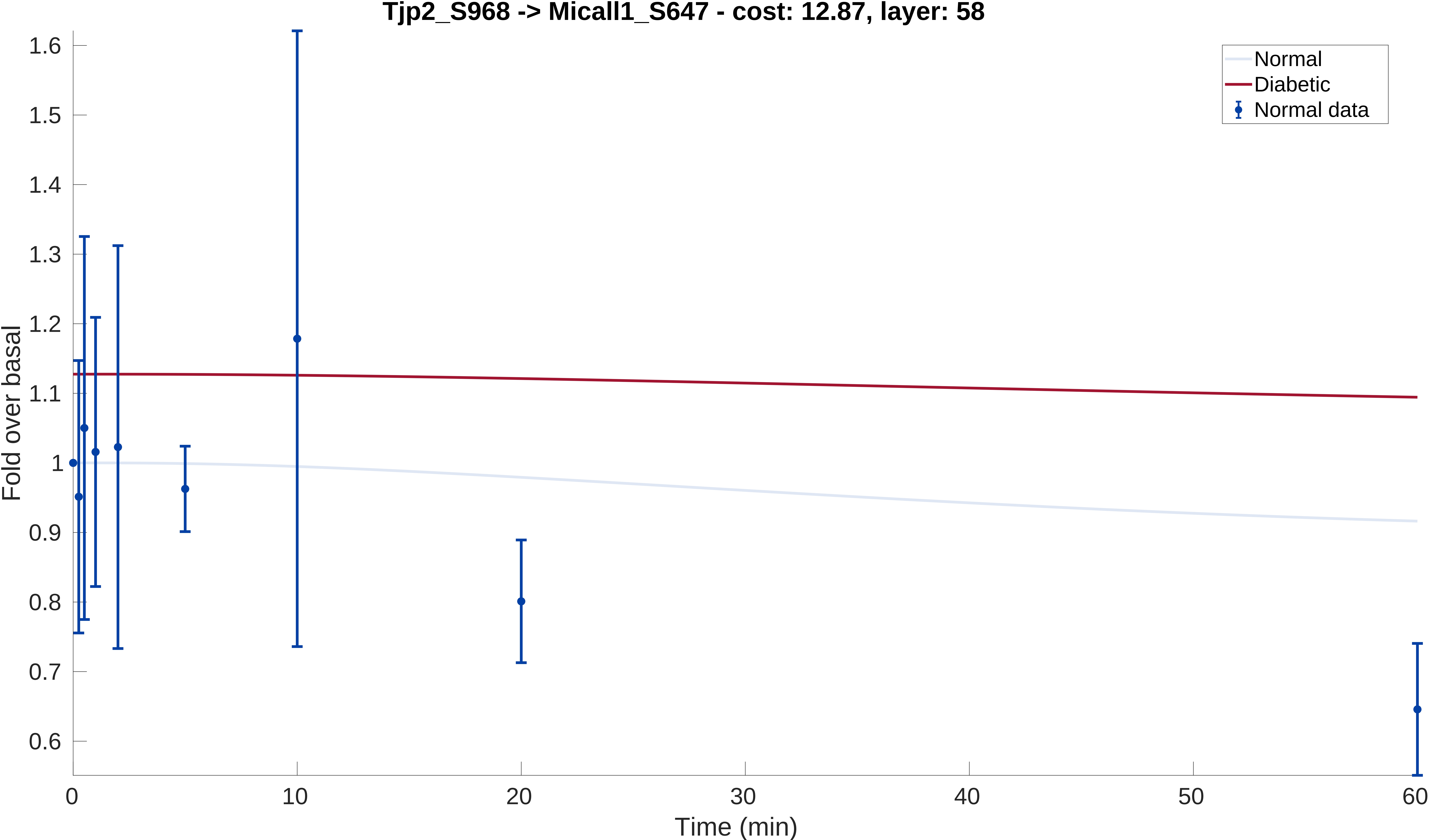
Tjp2_S968 -> Micall1_S647 - cost: 12.87, layer: 58
Fold over basal
Time (min)
Normal
Diabetic
Normal data
1.6
1.5
1.4
1.3
1.2
1.1
1
0.9
0.8
0.7
0.6
0
10
20
30
40
50
60

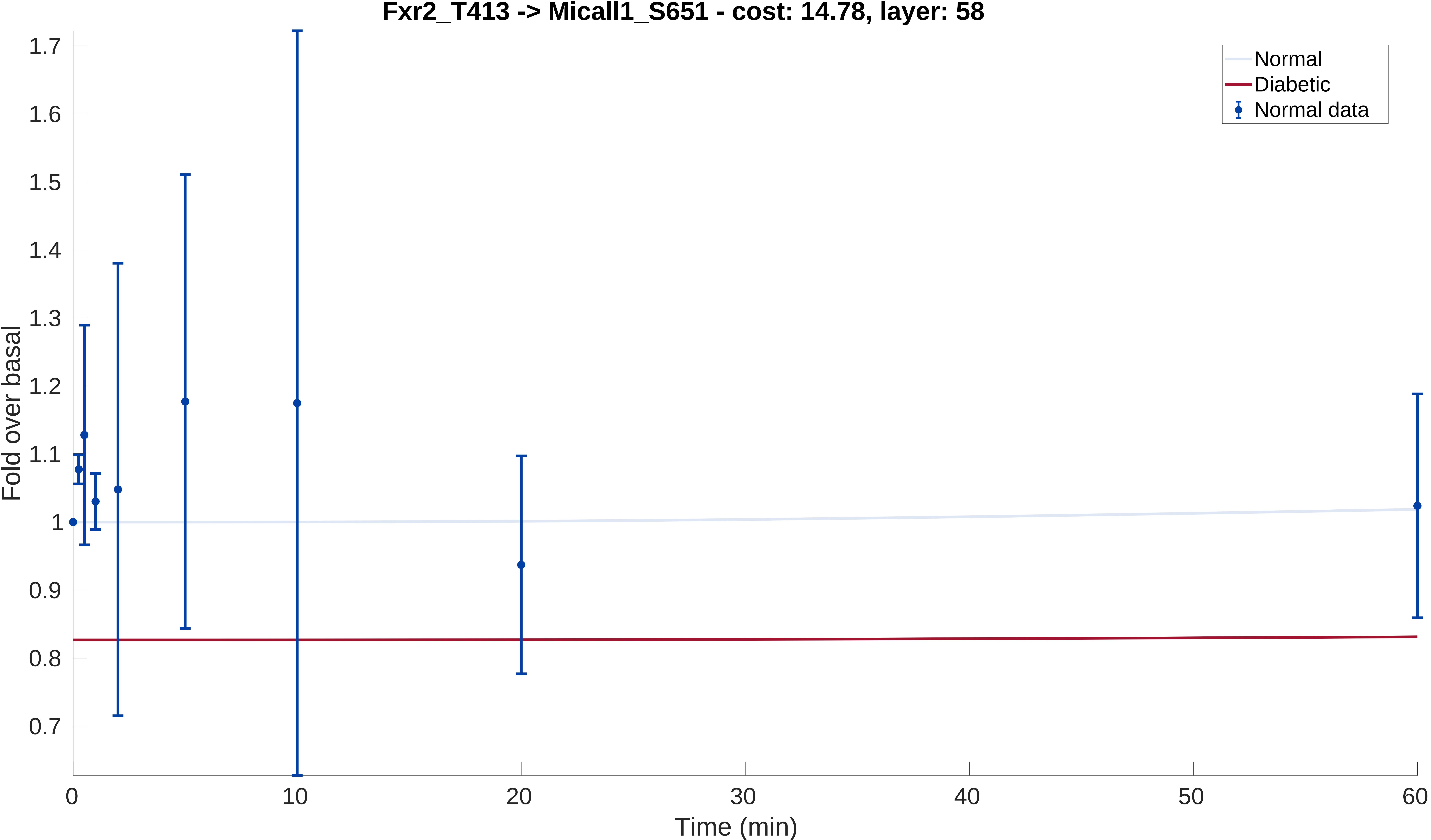
Fxr2_T413 -> Micall1_S651 - cost: 14.78, layer: 58
Normal
Diabetic
Normal data
Fold over basal
1.7
1.6
1.5
1.4
1.3
1.2
1.1
1
0.9
0.8
0.7
0
10
20
30
40
50
60
Time (min)

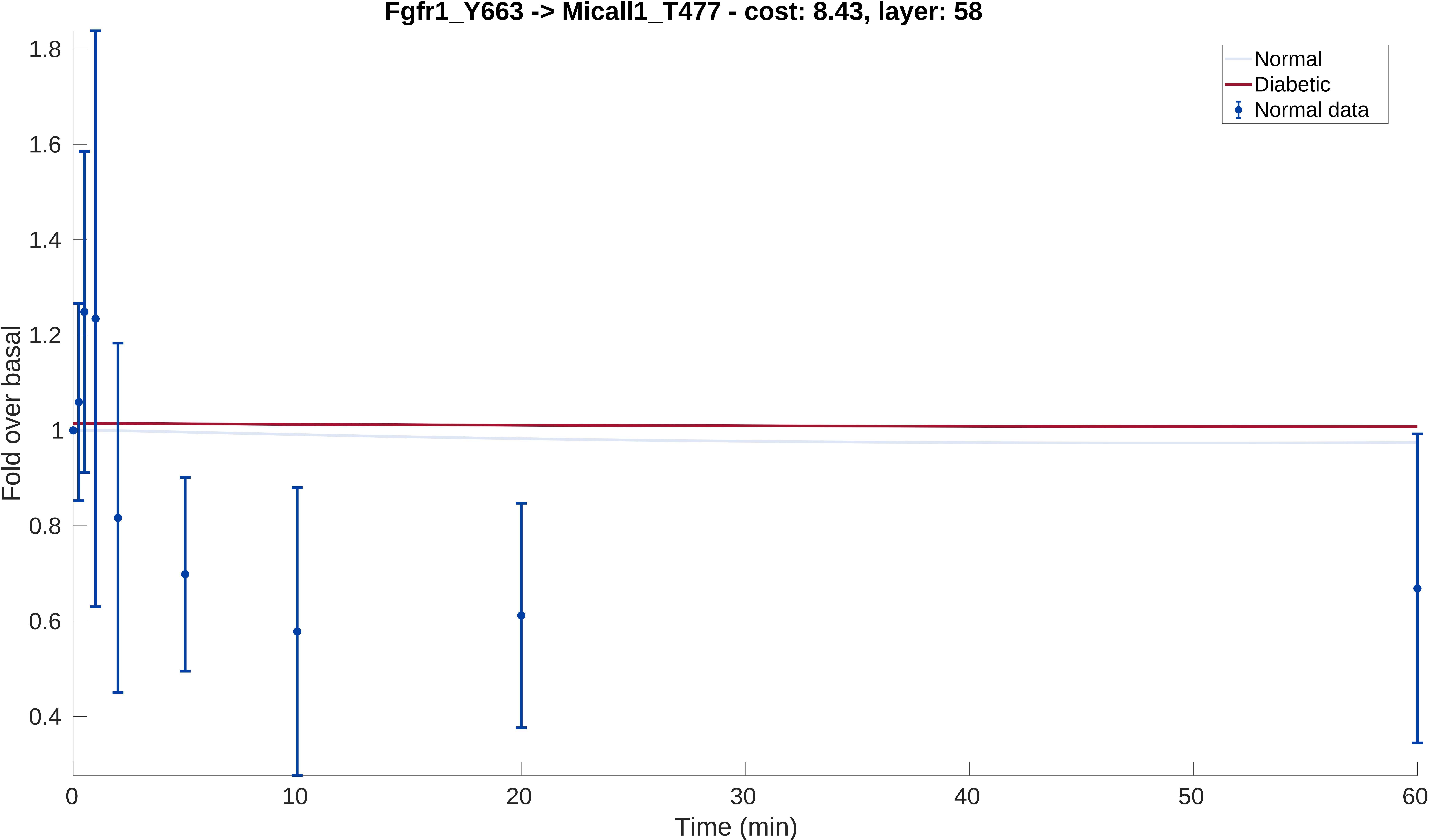

Fgfr1_Y663 -> Micall1_T477 - cost: 8.43, layer: 58
Normal
Diabetic
Normal data
Fold over basal
1.8
1.6
1.4
1.2
1
0.8
0.6
0.4
0
10
20
30
40
50
60
Time (min)

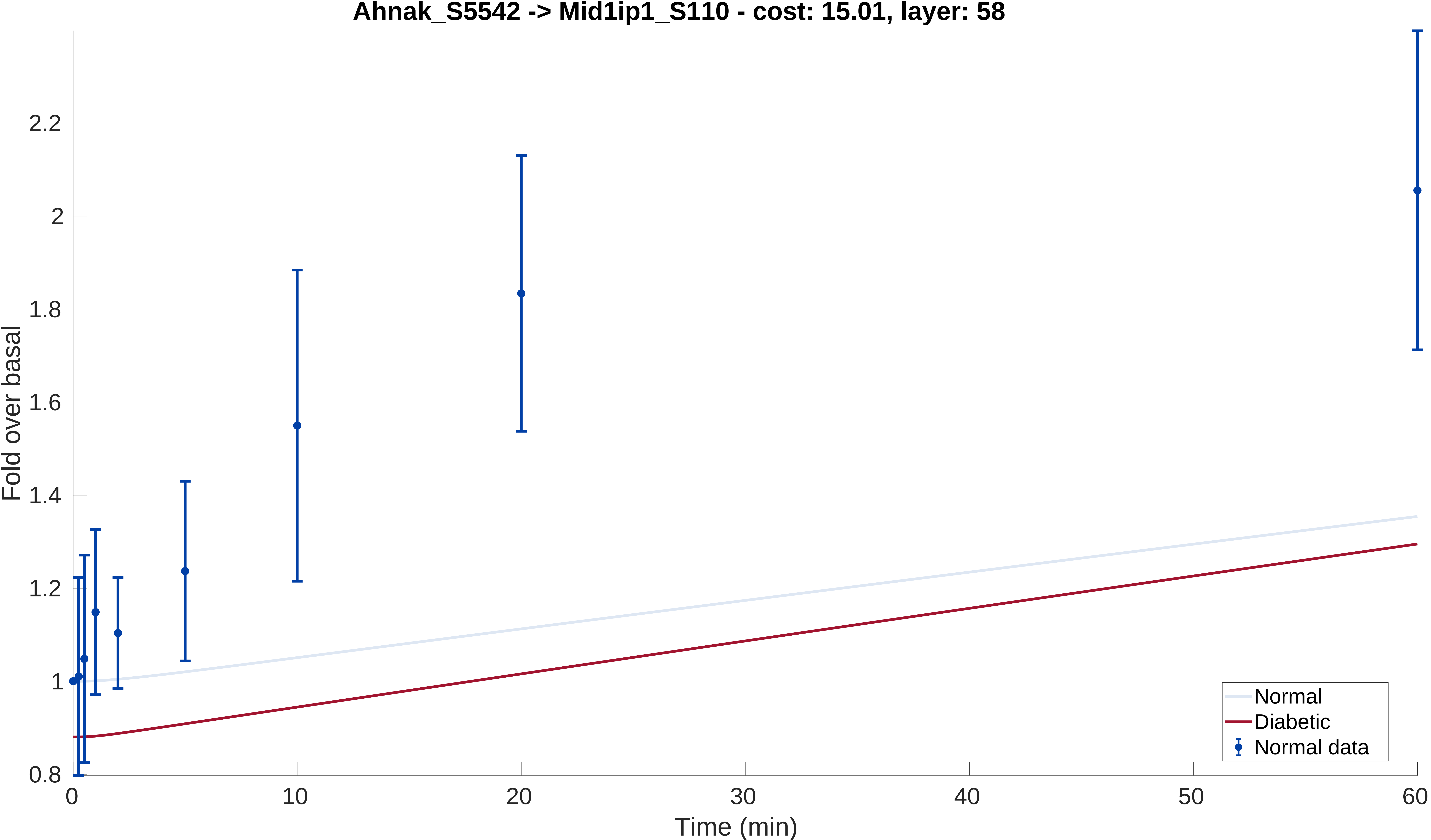
Ahnak_S5542 -> Mid1ip1_S110 - cost: 15.01, layer: 58
Fold over basal
0.8
1
1.2
1.4
1.6
1.8
2
2.2
0
10
20
30
40
50
60
Time (min)
Normal
Diabetic
Normal data

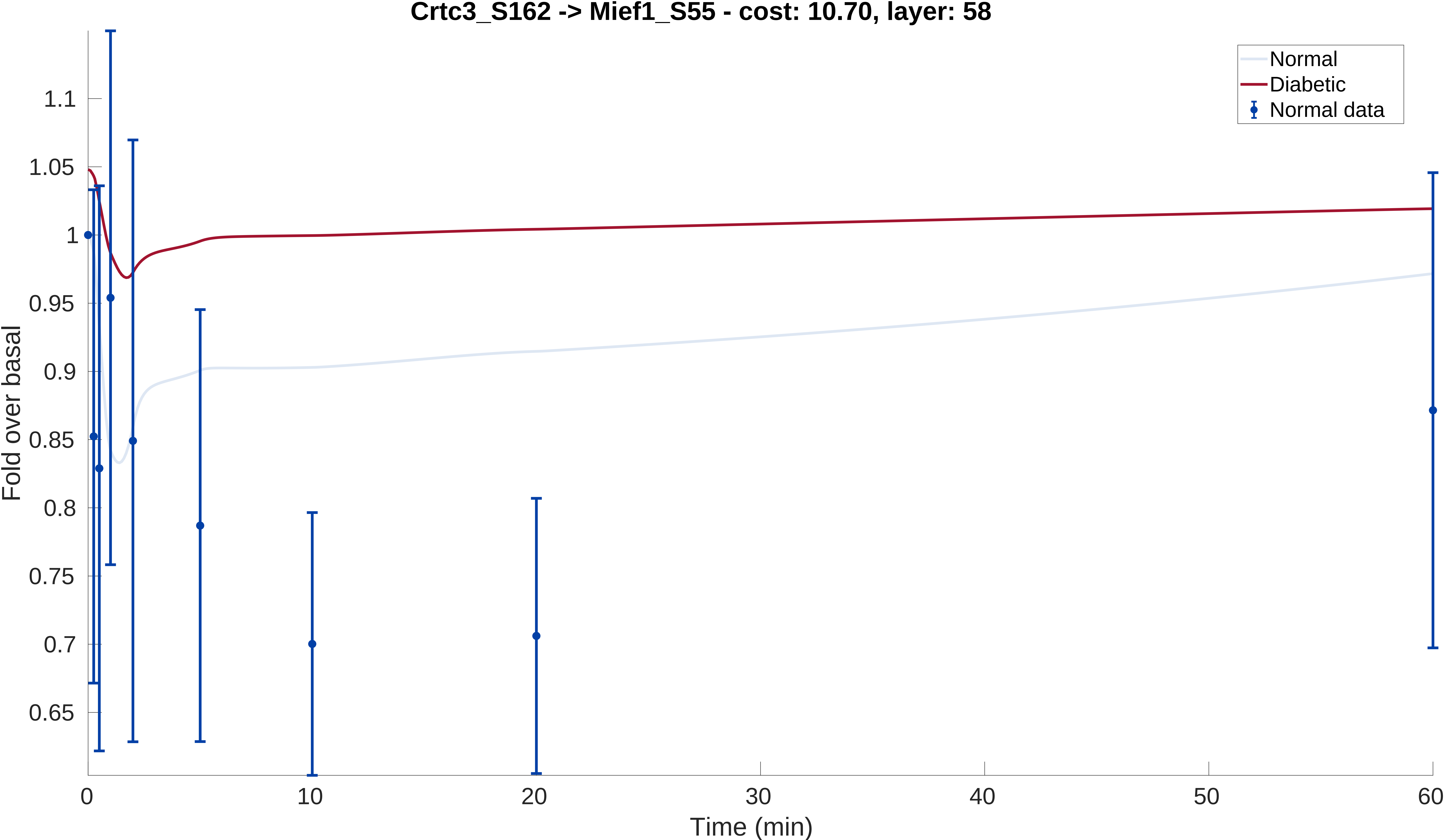

Crtc3_S162 -> Mief1_S55 - cost: 10.70, layer: 58
Normal
Diabetic
Normal data
Fold over basal
Time (min)
1.1
1.05
1
0.95
0.9
0.85
0.8
0.75
0.7
0.65
0
10
20
30
40
50
60

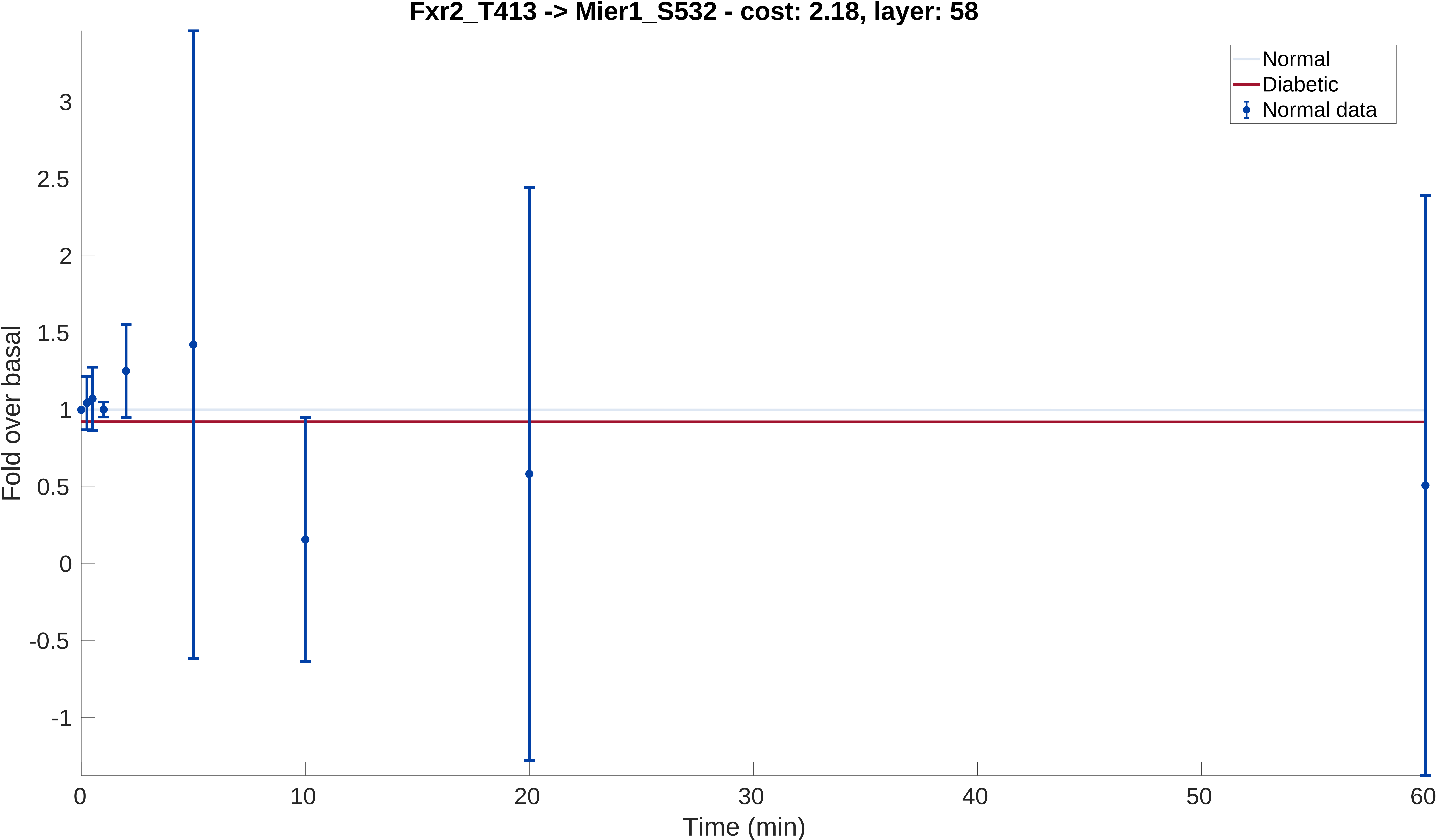

Fxr2_T413 -> Mier1_S532 - cost: 2.18, layer: 58
Normal
Diabetic
Normal data
Fold over basal
3
2.5
2
1.5
1
0.5
0
-0.5
-1
0
10
20
30
40
50
60
Time (min)

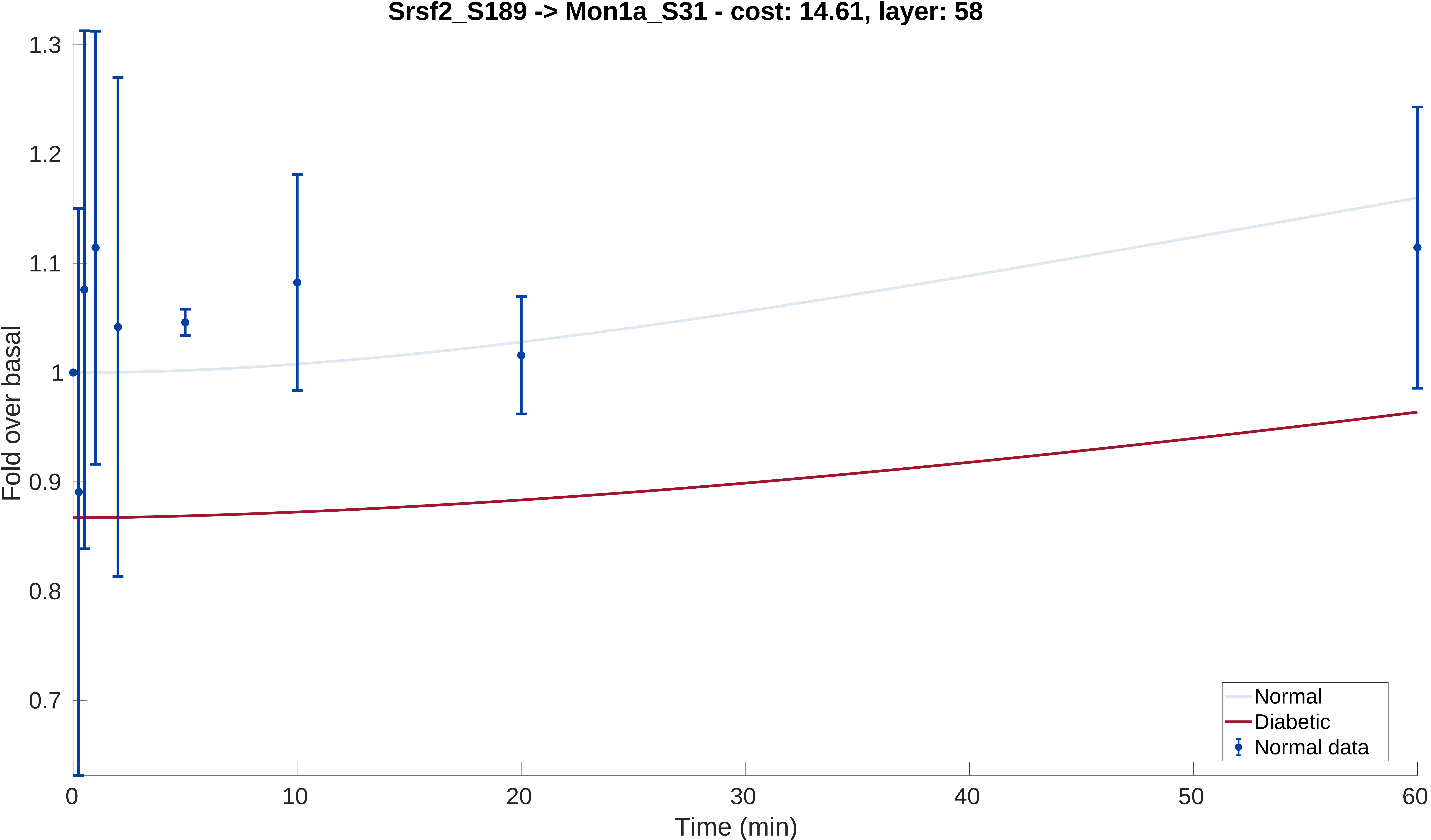
Srsf2_S189 -> Mon1a_S31 - cost: 14.61, layer: 58
Fold over basal
1.3
1.2
1.1
1
0.9
0.8
0.7
0
10
20
30
40
50
60
Time (min)
Normal
Diabetic
Normal data

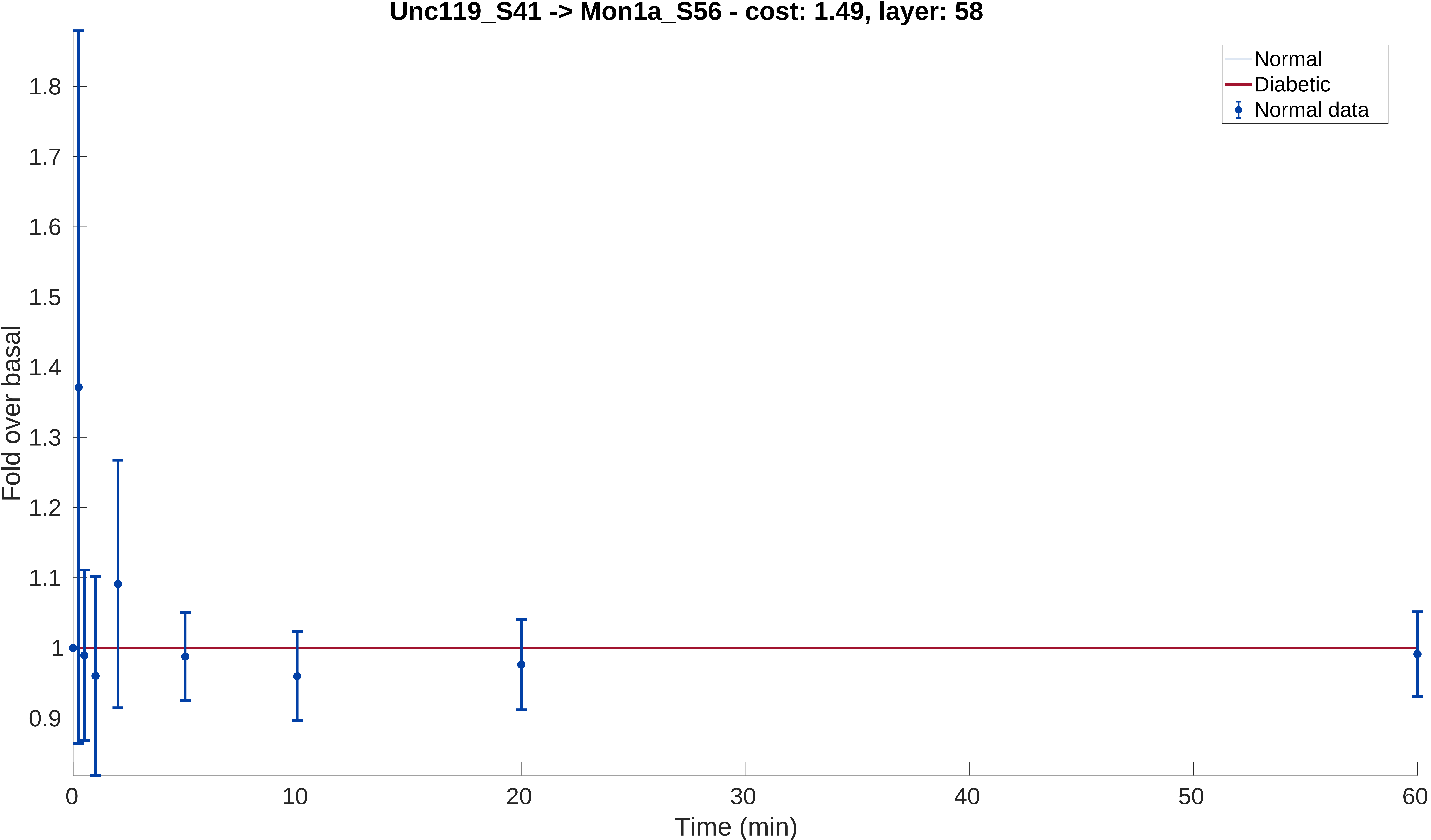

Unc119_S41 -> Mon1a_S56 - cost: 1.49, layer: 58
Normal
Diabetic
Normal data
Fold over basal
1.8
1.7
1.6
1.5
1.4
1.3
1.2
1.1
1
0.9
0
10
20
30
40
50
60
Time (min)

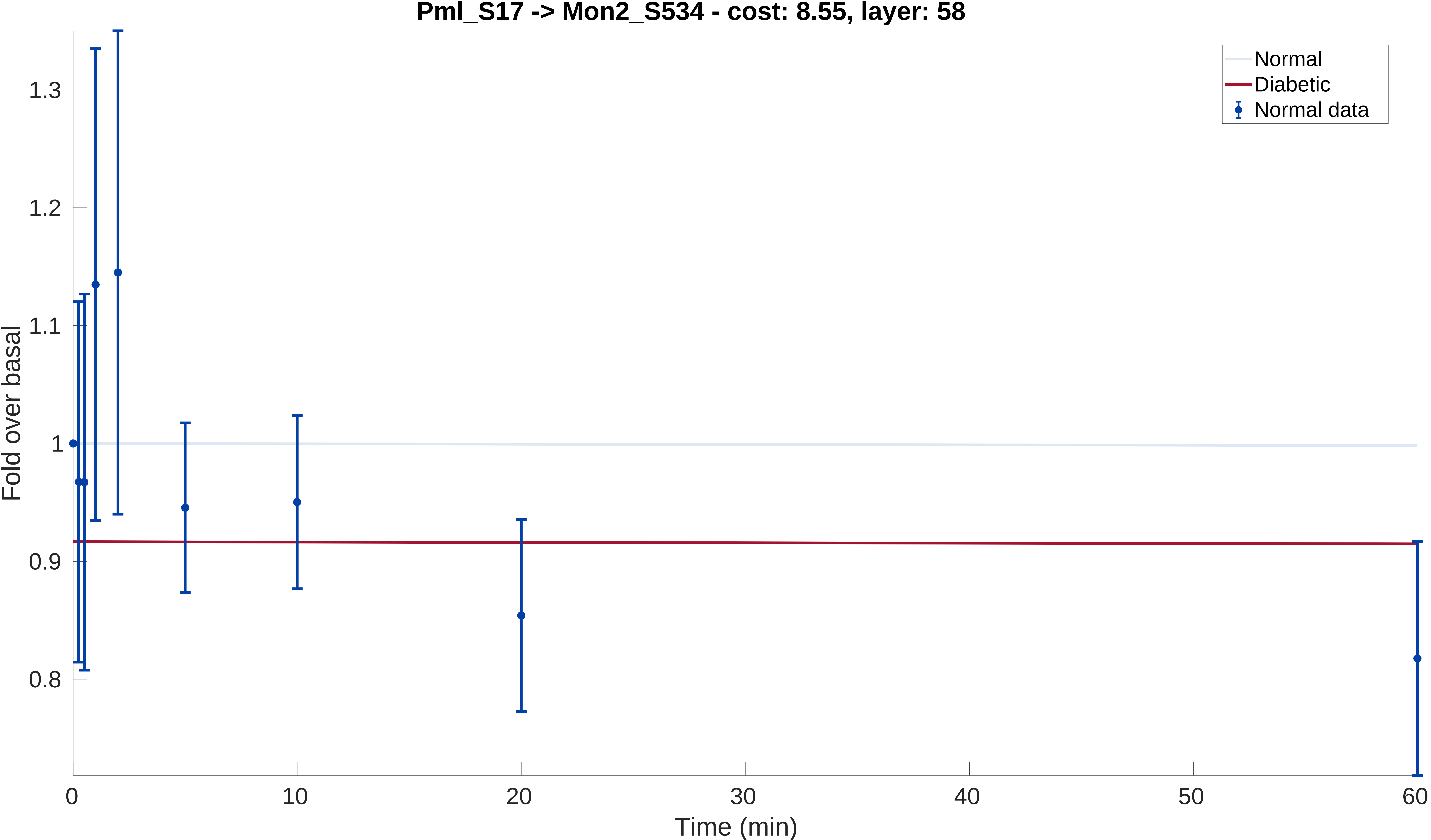
Pml_S17 -> Mon2_S534 - cost: 8.55, layer: 58
Normal
Diabetic
Normal data
Fold over basal
1.3
1.2
1.1
1
0.9
0.8
0
10
20
30
40
50
60
Time (min)

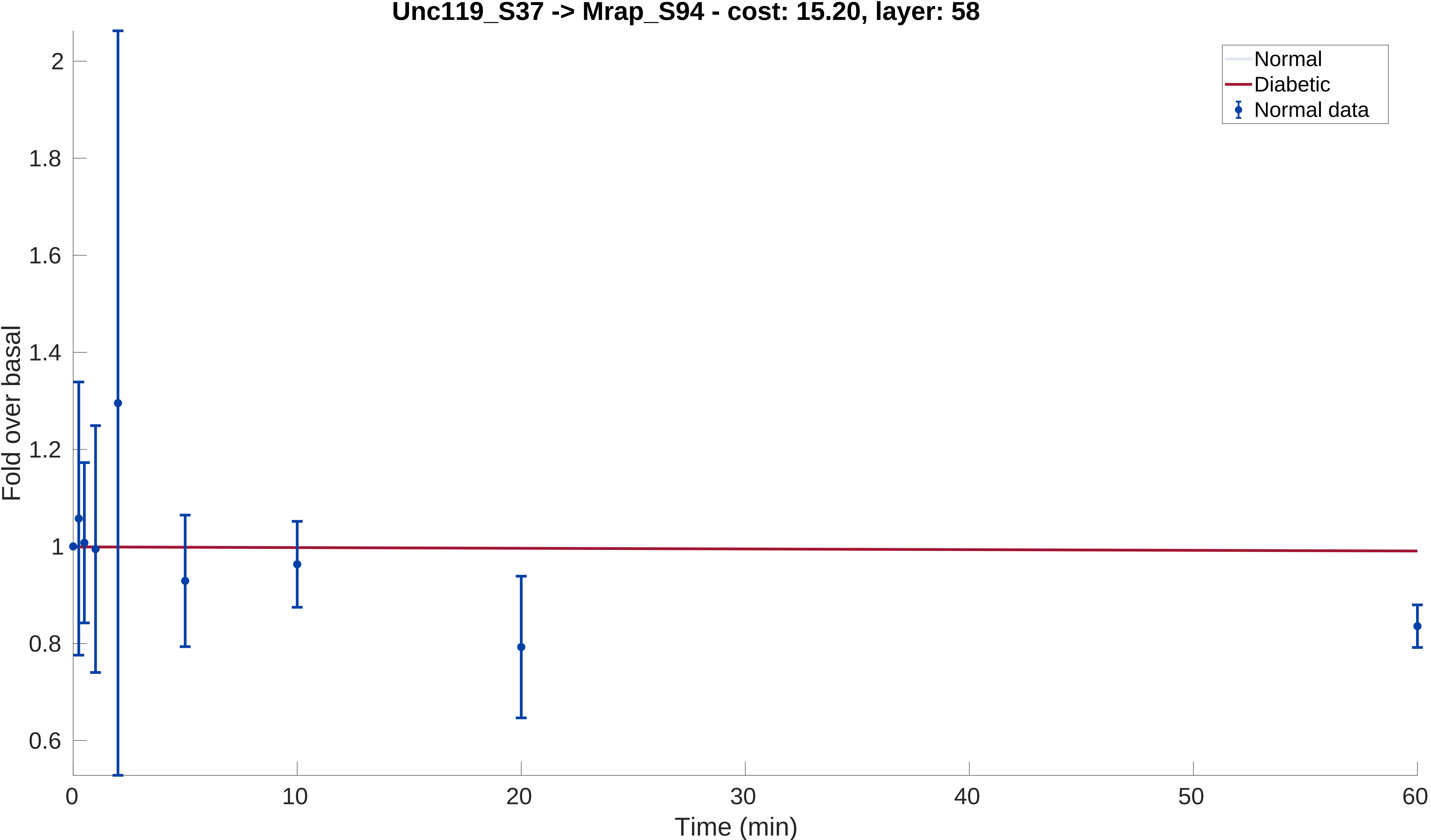

Unc119_S37 -> Mrap_S94 - cost: 15.20, layer: 58
Normal
Diabetic
Normal data
Fold over basal
2
1.8
1.6
1.4
1.2
1
0.8
0.6
0
10
20
30
40
50
60
Time (min)

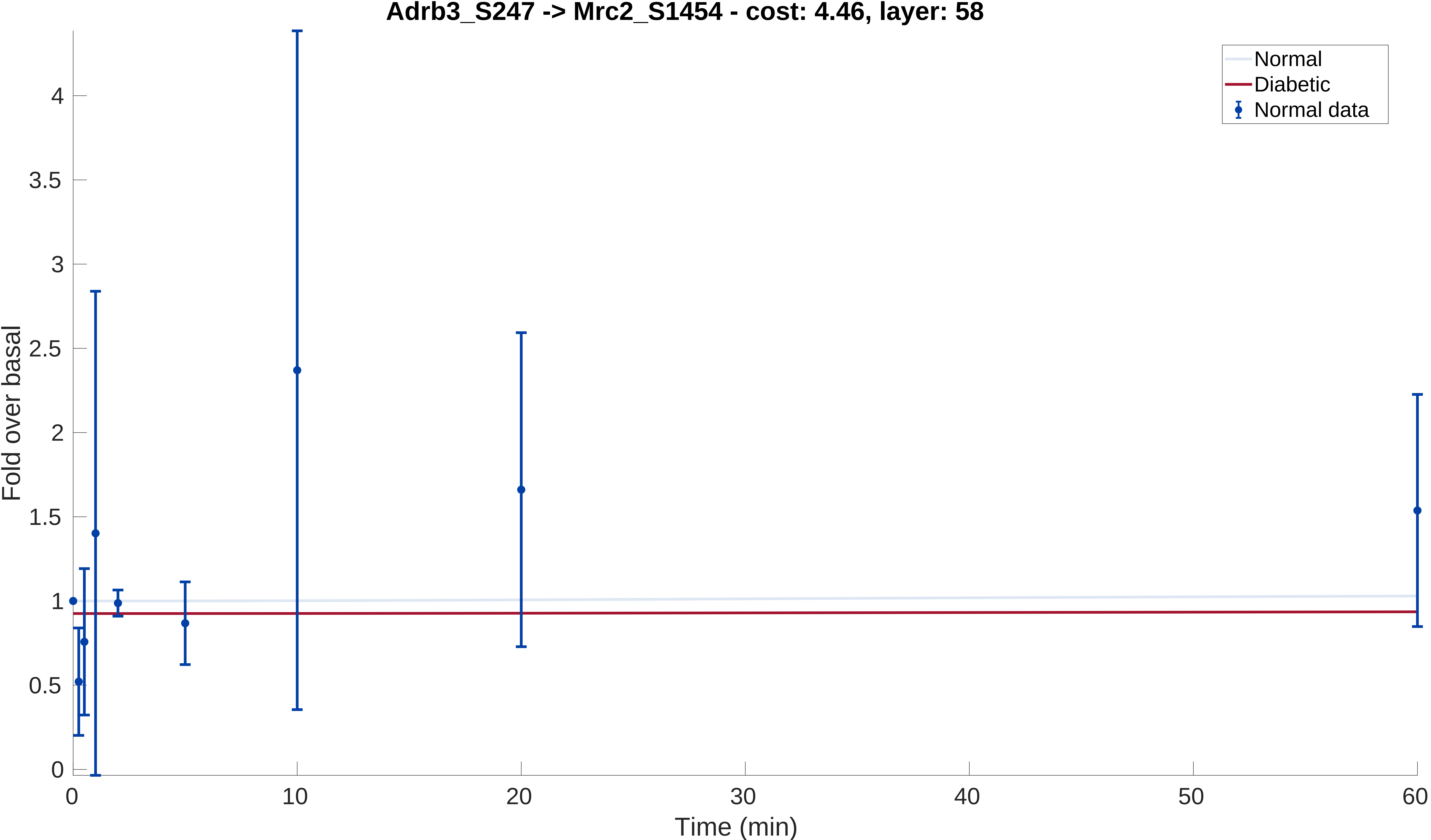

Adrb3_S247 -> Mrc2_S1454 - cost: 4.46, layer: 58
Normal
Diabetic
Normal data
Fold over basal
Time (min)
0
0.5
1
1.5
2
2.5
3
3.5
4
0
10
20
30
40
50
60

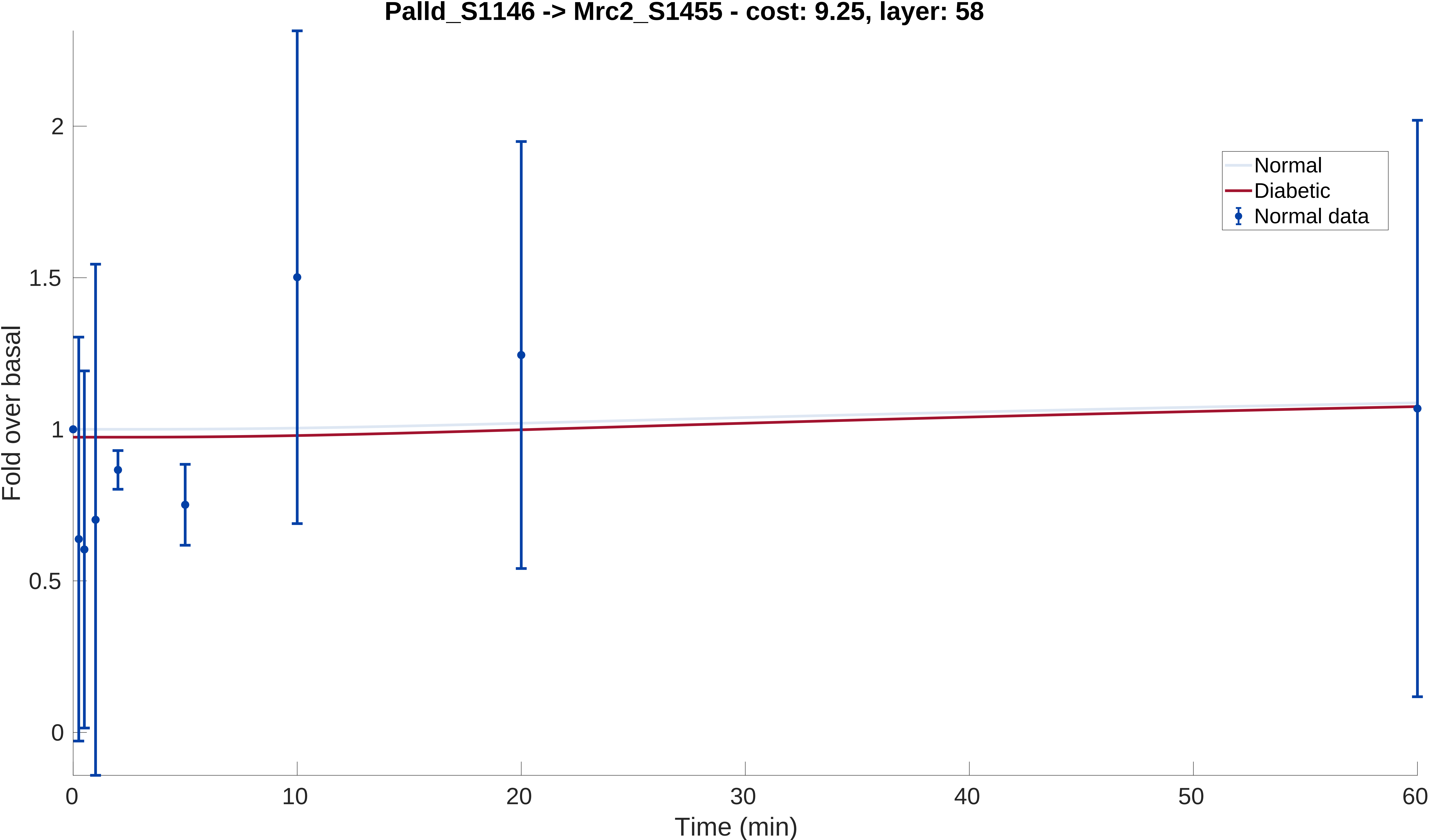

Palld_S1146 -> Mrc2_S1455 - cost: 9.25, layer: 58
Fold over basal
0
0.5
1
1.5
2
Time (min)
0
10
20
30
40
50
60
Normal
Diabetic
Normal data

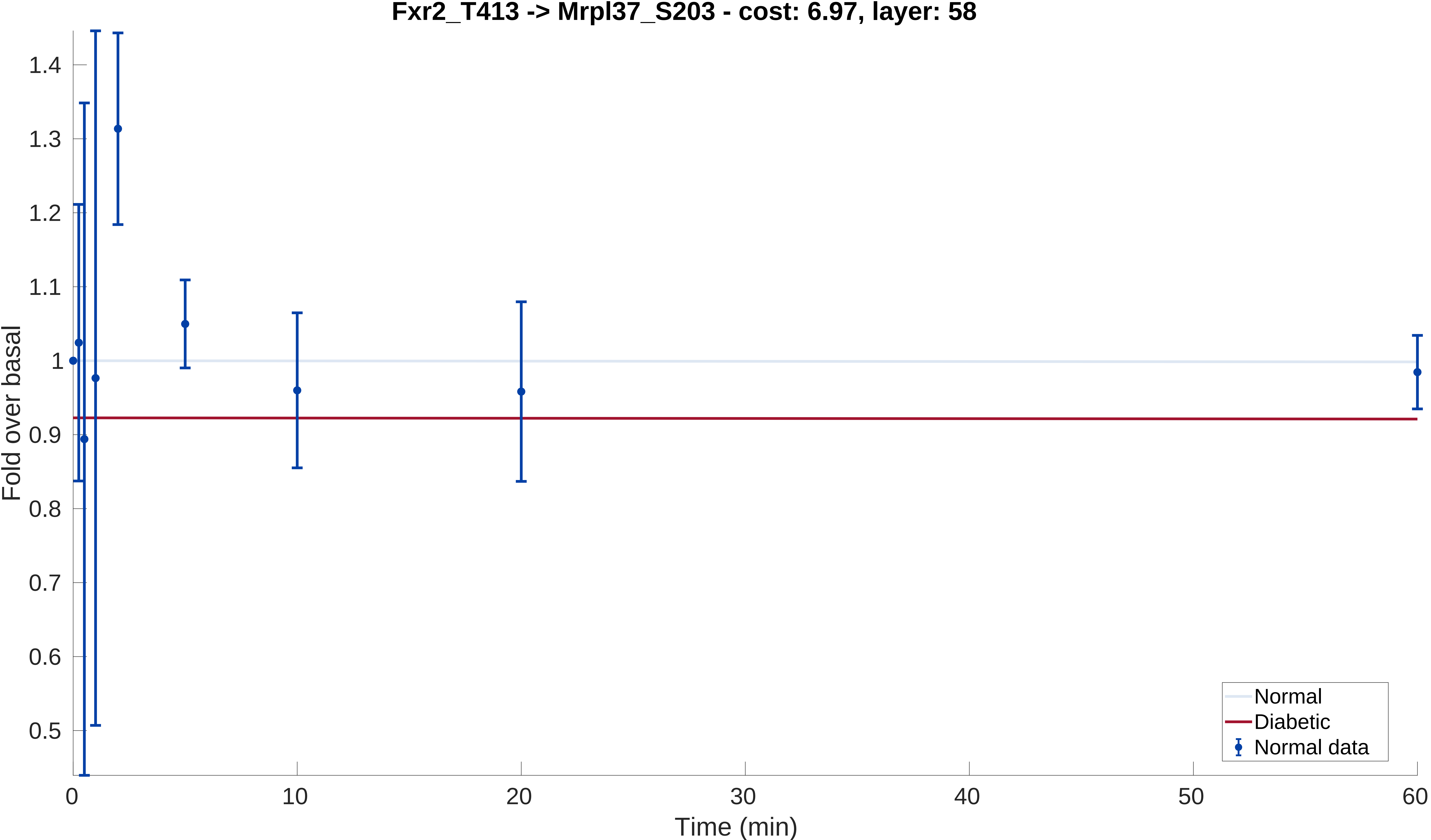
Fxr2_T413 -> Mrpl37_S203 - cost: 6.97, layer: 58
Fold over basal
Time (min)
1.4
1.3
1.2
1.1
1
0.9
0.8
0.7
0.6
0.5
0
10
20
30
40
50
60
Normal
Diabetic
Normal data

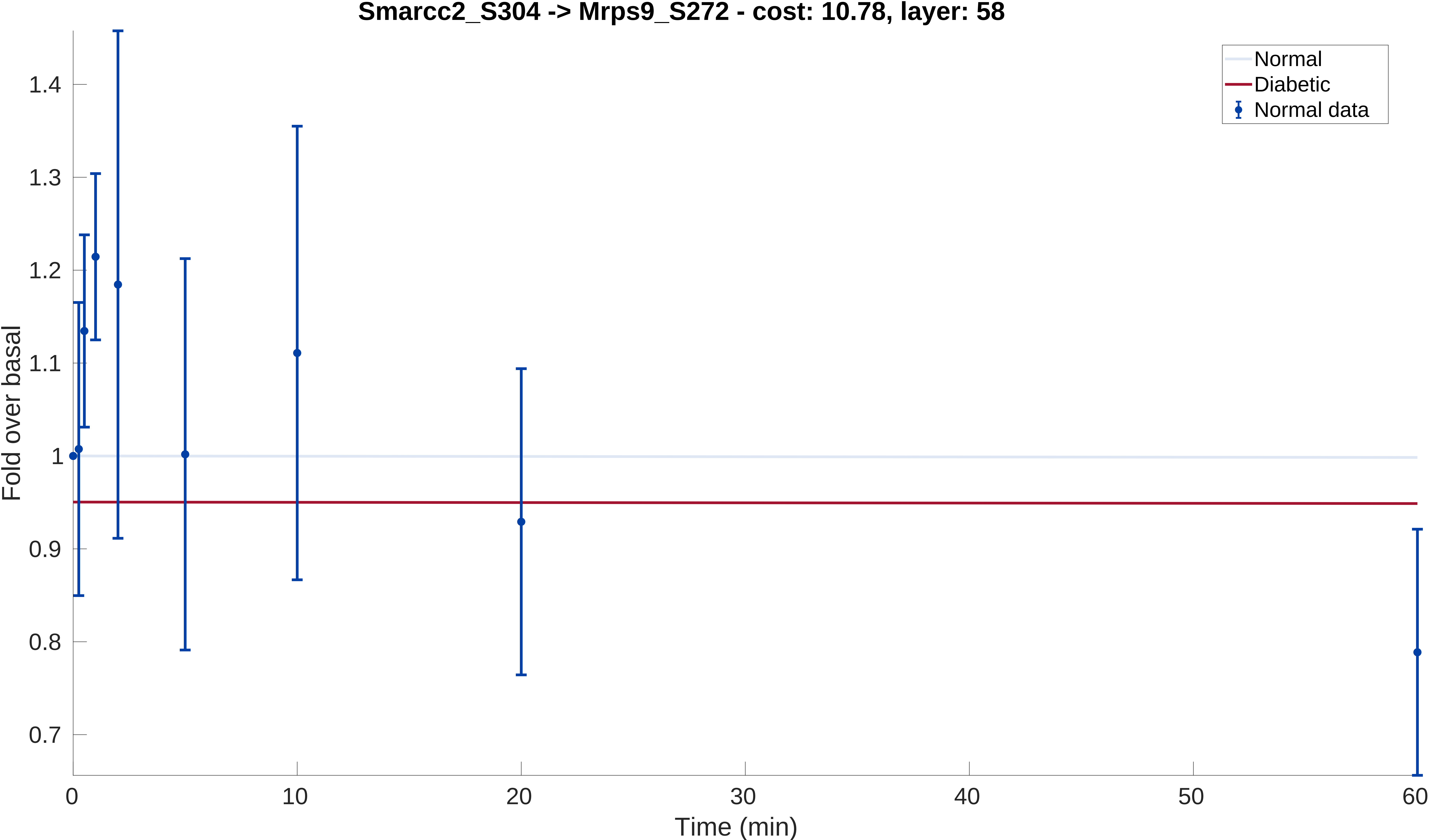

Smarcc2_S304 -> Mrps9_S272 - cost: 10.78, layer: 58
Normal
Diabetic
Normal data
1.4
1.3
1.2
1.1
1
0.9
0.8
0.7
Fold over basal
0
10
20
30
40
50
60
Time (min)

#### Sympk_S1260 -> Mta3_S370 - cost: 2.74, layer: 58

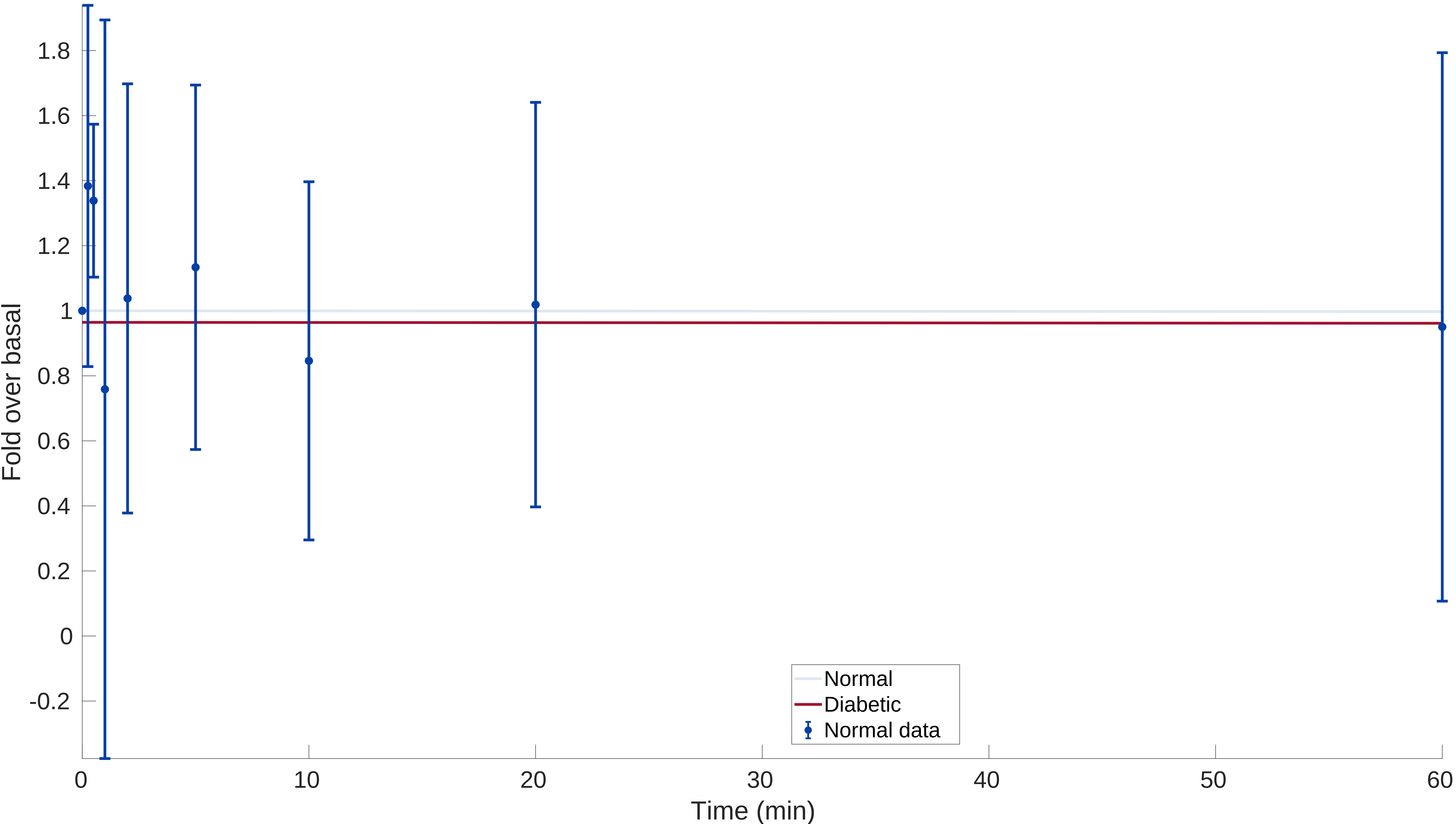

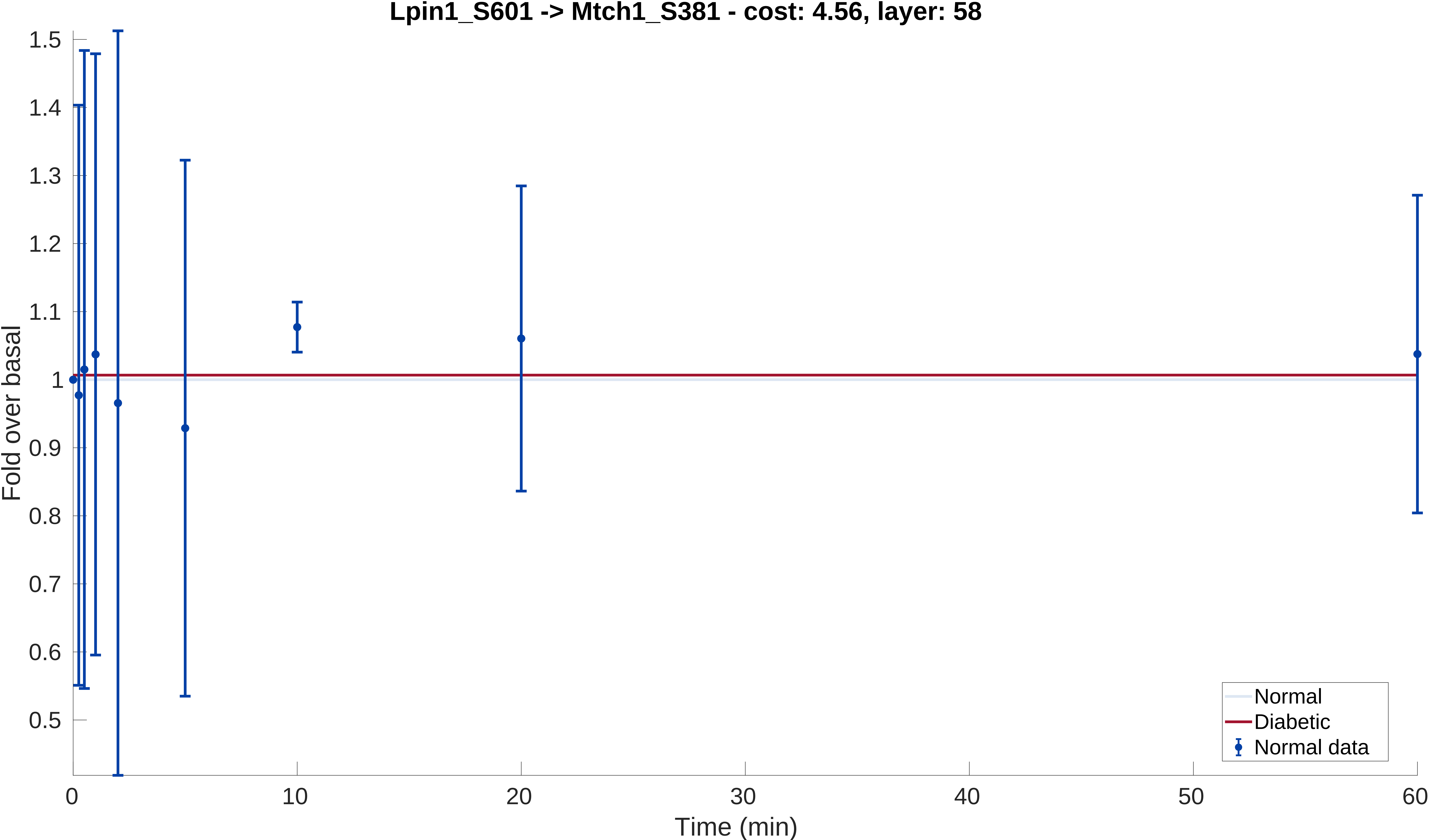

Lpin1_S601 -> Mtch1_S381 - cost: 4.56, layer: 58
Fold over basal
1.5
1.4
1.3
1.2
1.1
1
0.9
0.8
0.7
0.6
0.5
0
10
20
30
40
50
60
Time (min)
Normal
Diabetic
Normal data

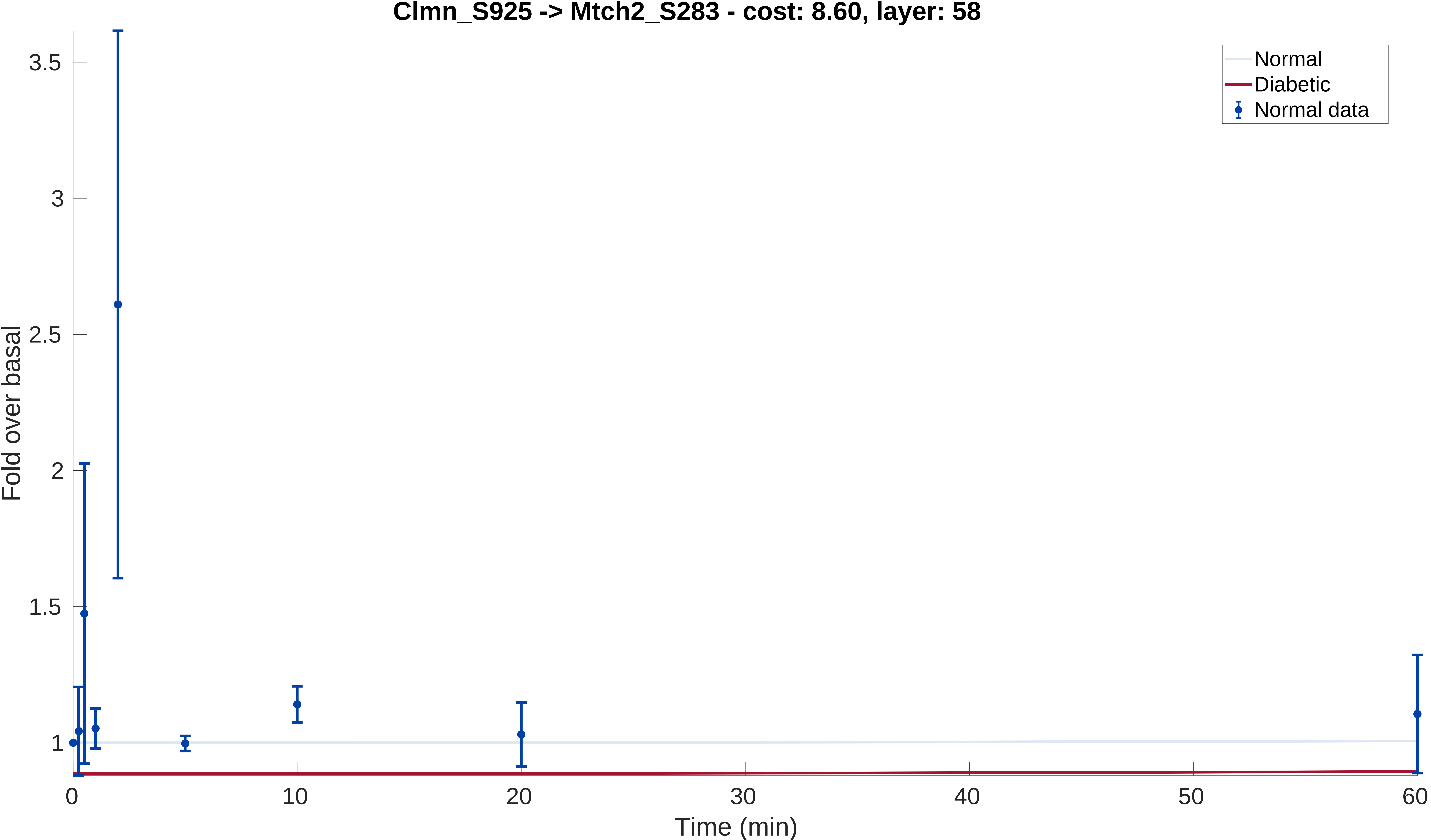

Clmn_S925 -> Mtch2_S283 - cost: 8.60, layer: 58
Fold over basal
3.5
3
2.5
2
1.5
1
0
10
20
30
40
50
60
Time (min)
Normal
Diabetic
Normal data

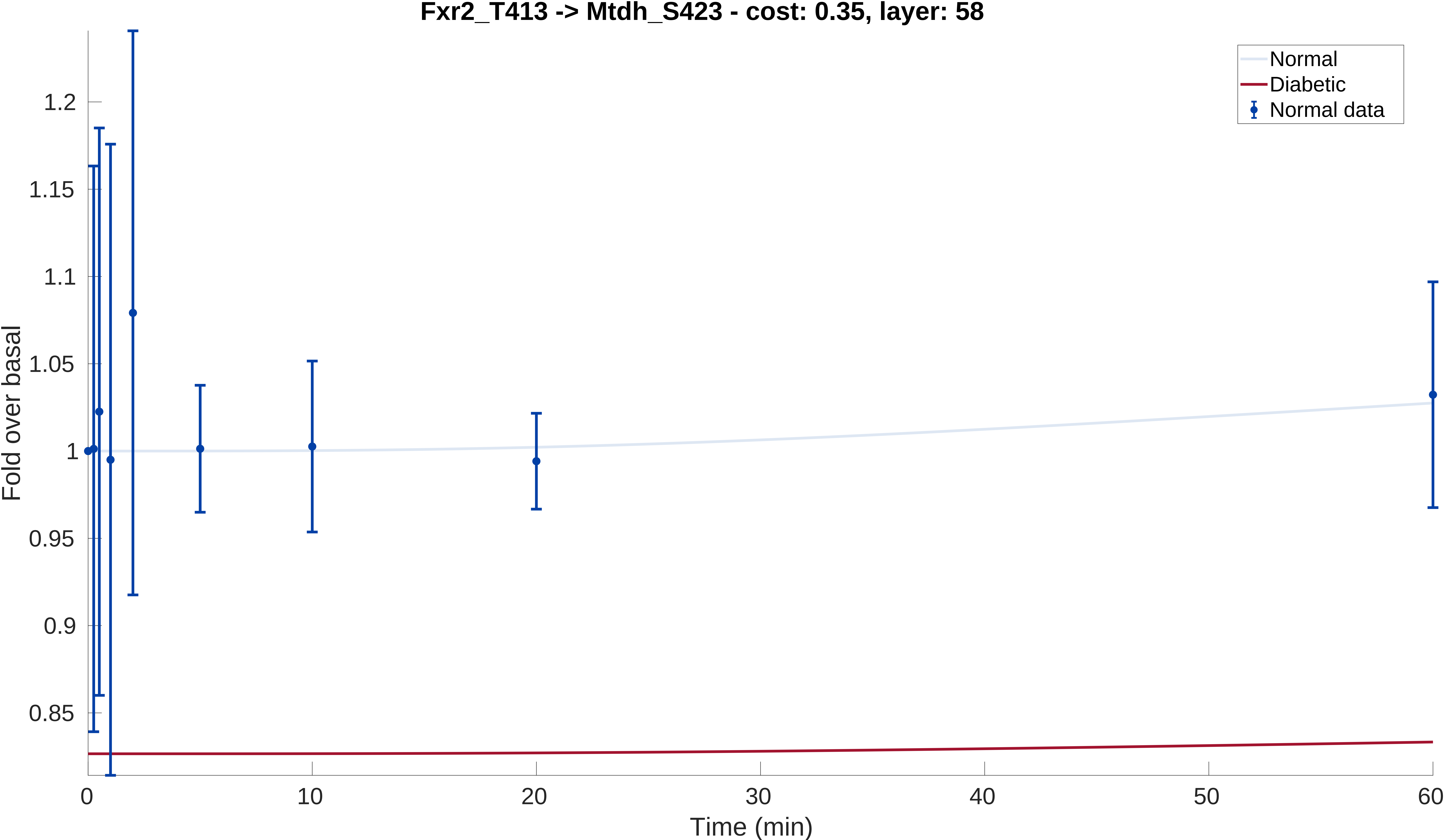

Fxr2_T413 -> Mtdh_S423 - cost: 0.35, layer: 58
Normal
Diabetic
Normal data
Fold over basal
Time (min)
1.2
1.15
1.1
1.05
1
0.95
0.9
0.85
0
10
20
30
40
50
60

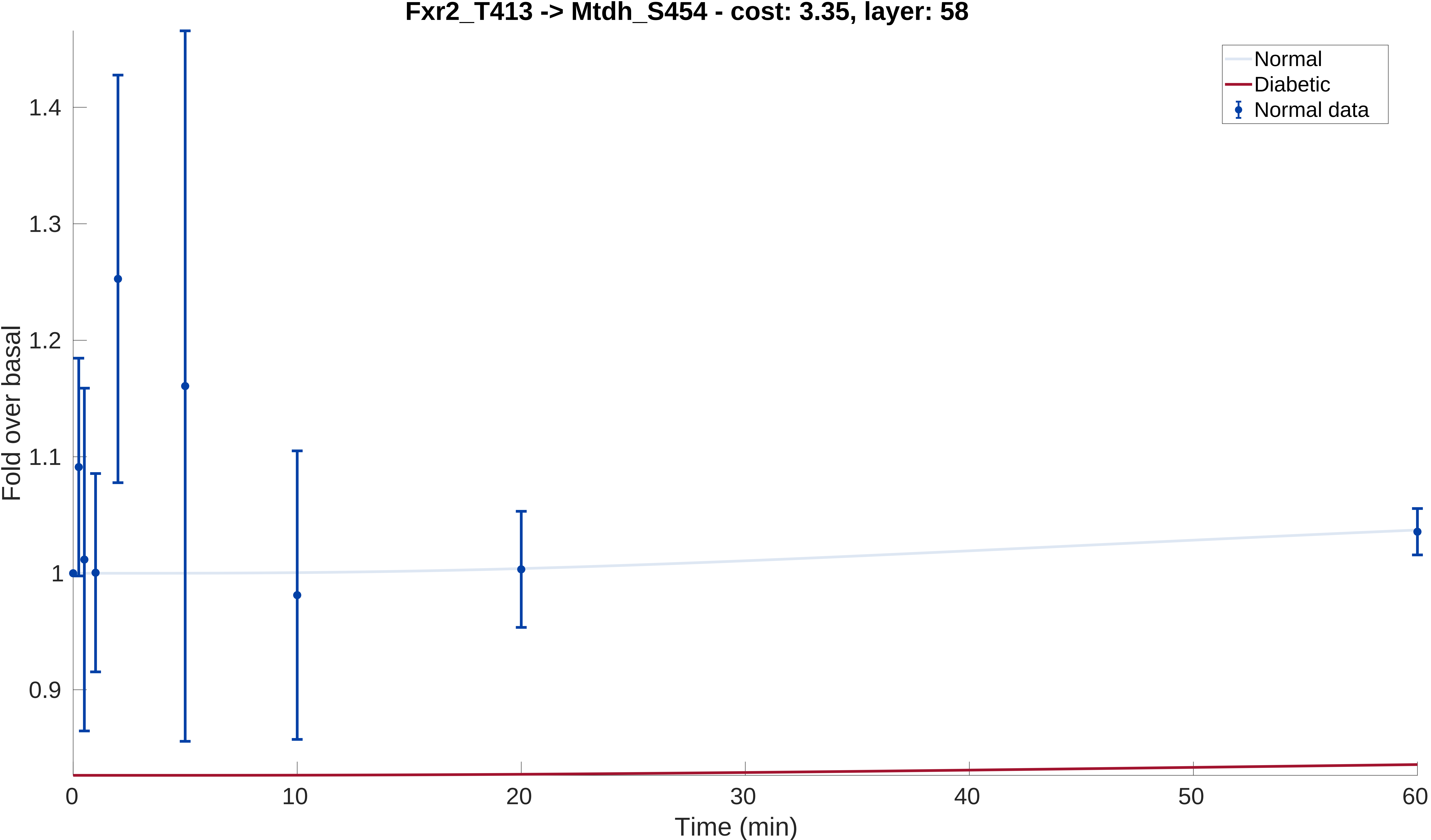
Fxr2_T413 -> Mtdh_S454 - cost: 3.35, layer: 58
Normal
Diabetic
Normal data
Fold over basal
1.4
1.3
1.2
1.1
1
0.9
0
10
20
30
40
50
60
Time (min)

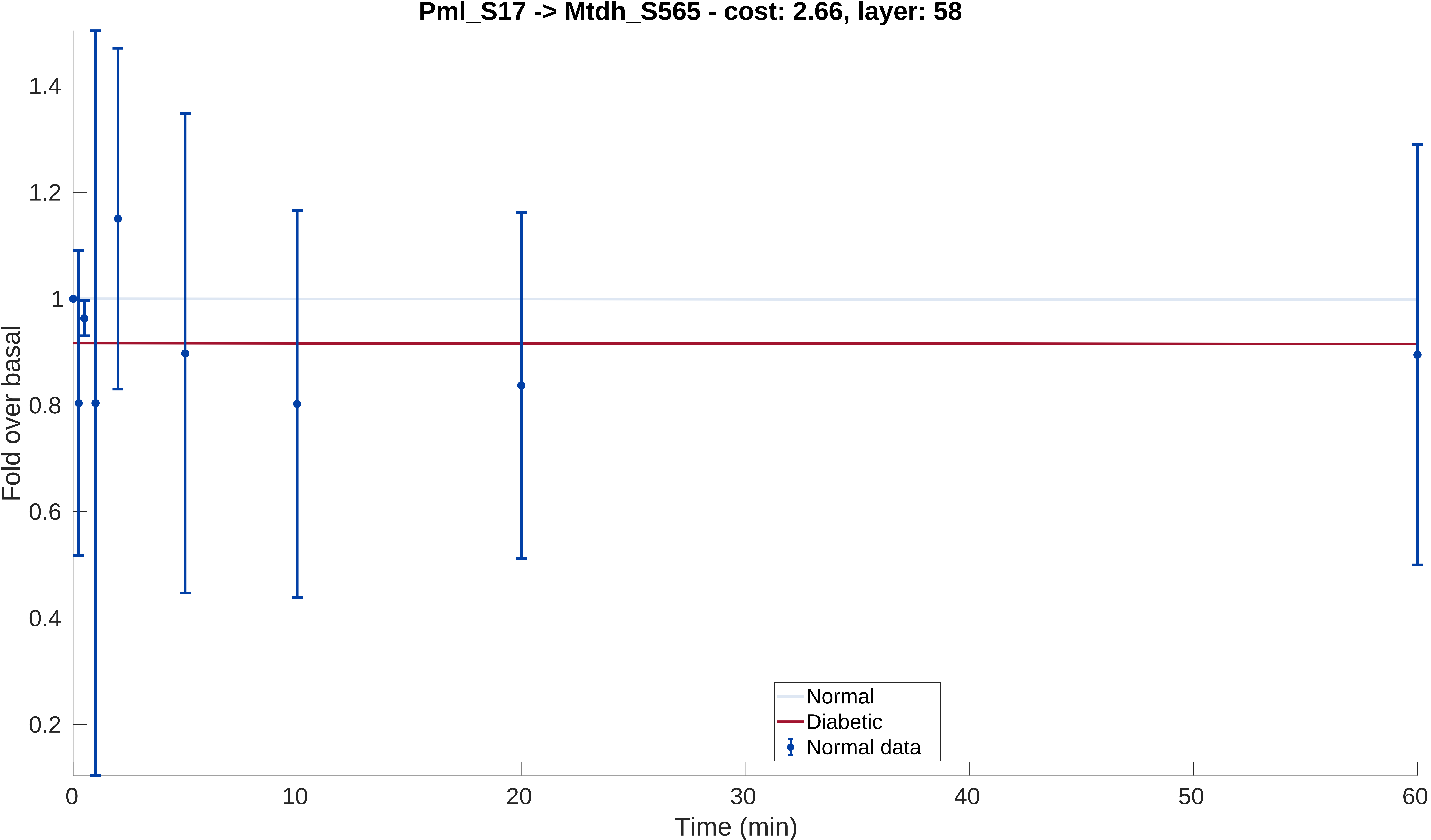

Pml_S17 -> Mtdh_S565 - cost: 2.66, layer: 58
Fold over basal
1.4
1.2
1
0.8
0.6
0.4
0.2
0
10
20
30
40
50
60
Time (min)
Normal
Diabetic
Normal data

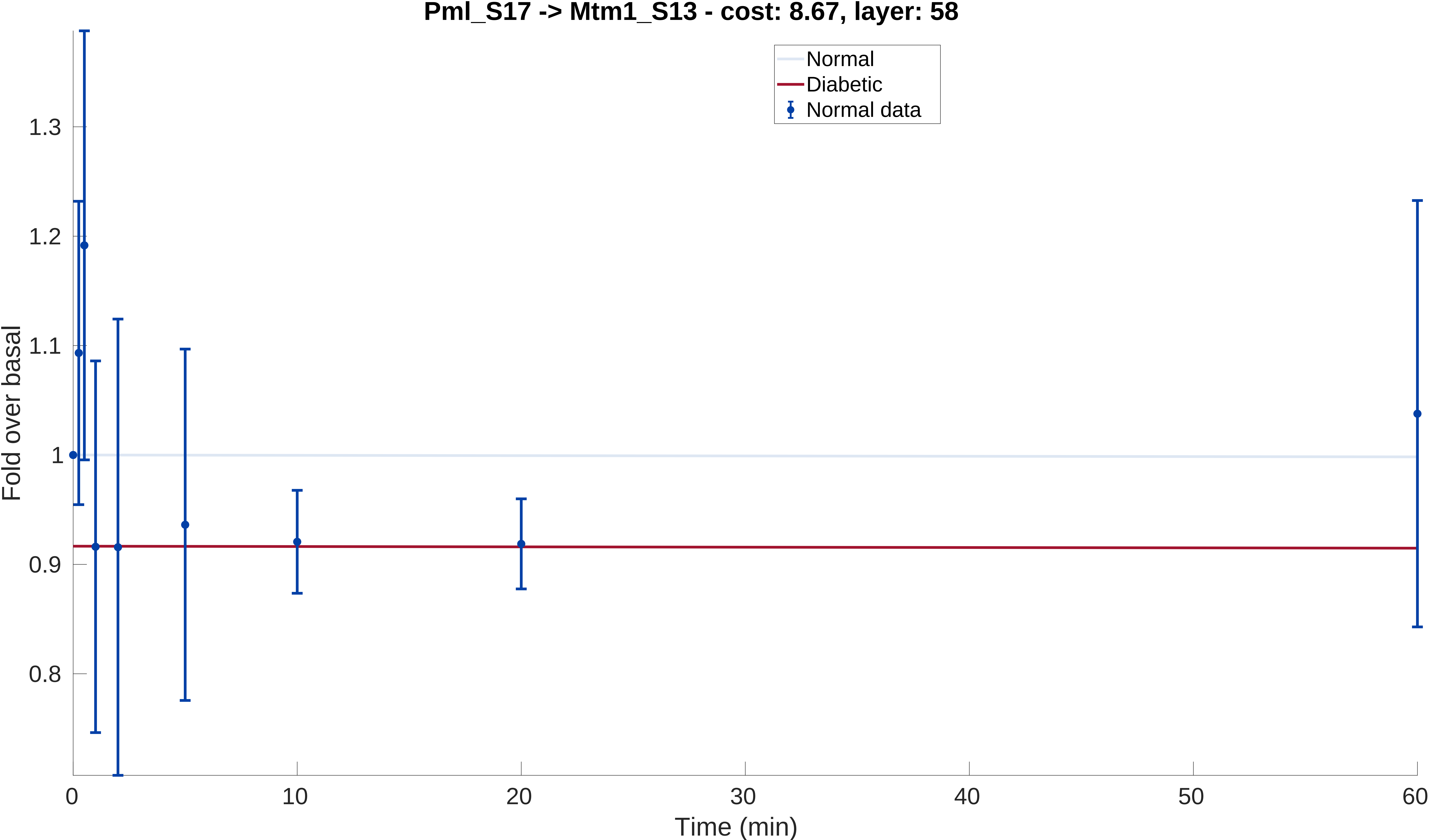

Pml_S17 -> Mtm1_S13 - cost: 8.67, layer: 58
Normal
Diabetic
Normal data
1.3
1.2
1.1
1
0.9
0.8
Fold over basal
0
10
20
30
40
50
60
Time (min)

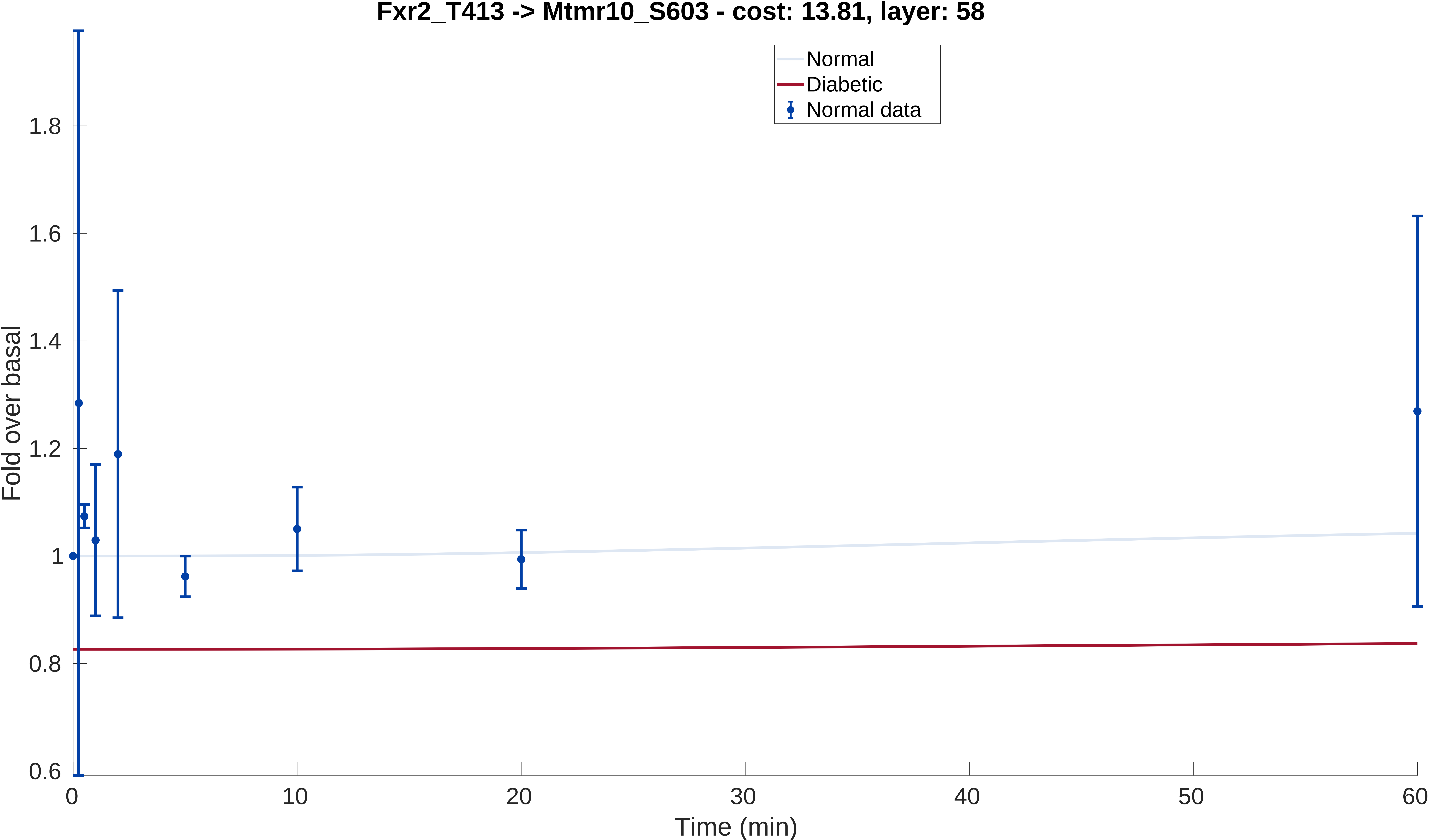

Fxr2_T413 -> Mtmr10_S603 - cost: 13.81, layer: 58
Normal
Diabetic
Normal data
Fold over basal
1.8
1.6
1.4
1.2
1
0.8
0.6
0
10
20
30
40
50
60
Time (min)

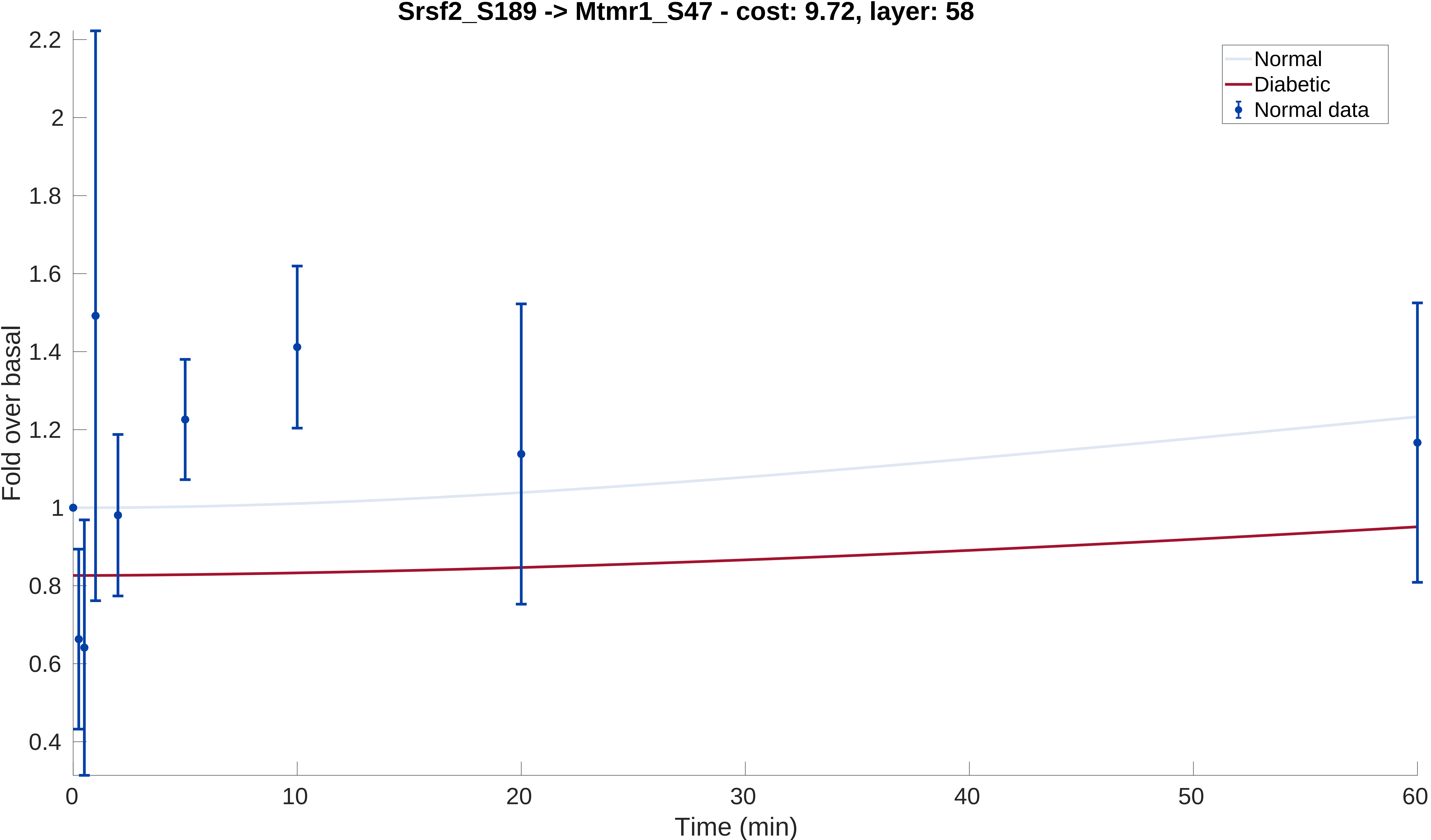

Srsf2_S189 -> Mtmr1_S47 - cost: 9.72, layer: 58
Normal
Diabetic
Normal data
Fold over basal
Time (min)
2.2
2
1.8
1.6
1.4
1.2
1
0.8
0.6
0.4
0
10
20
30
40
50
60

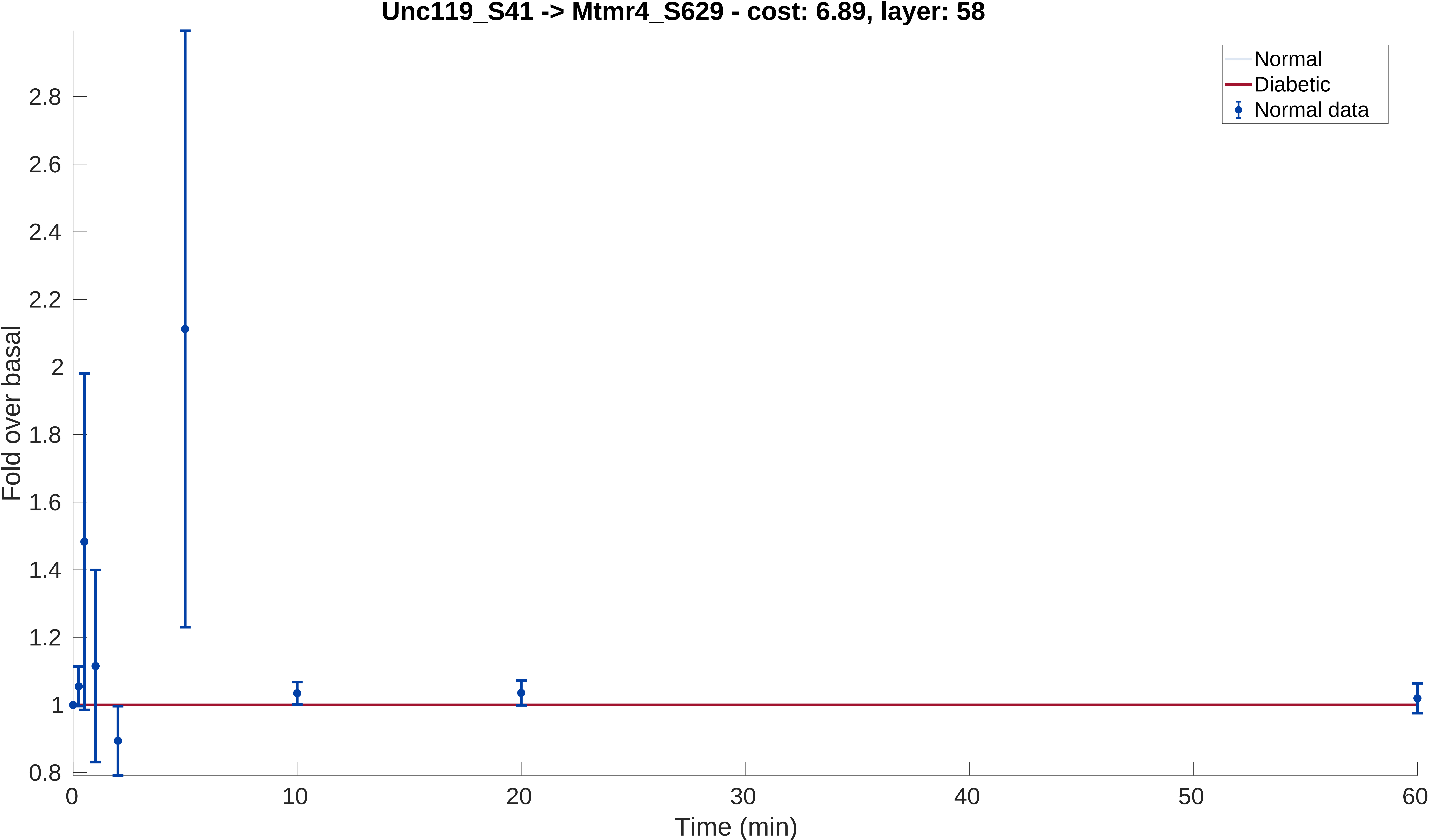

Unc119_S41 -> Mtmr4_S629 - cost: 6.89, layer: 58
Fold over basal
Time (min)
Normal
Diabetic
Normal data
0.8
1
1.2
1.4
1.6
1.8
2
2.2
2.4
2.6
2.8
0
10
20
30
40
50
60

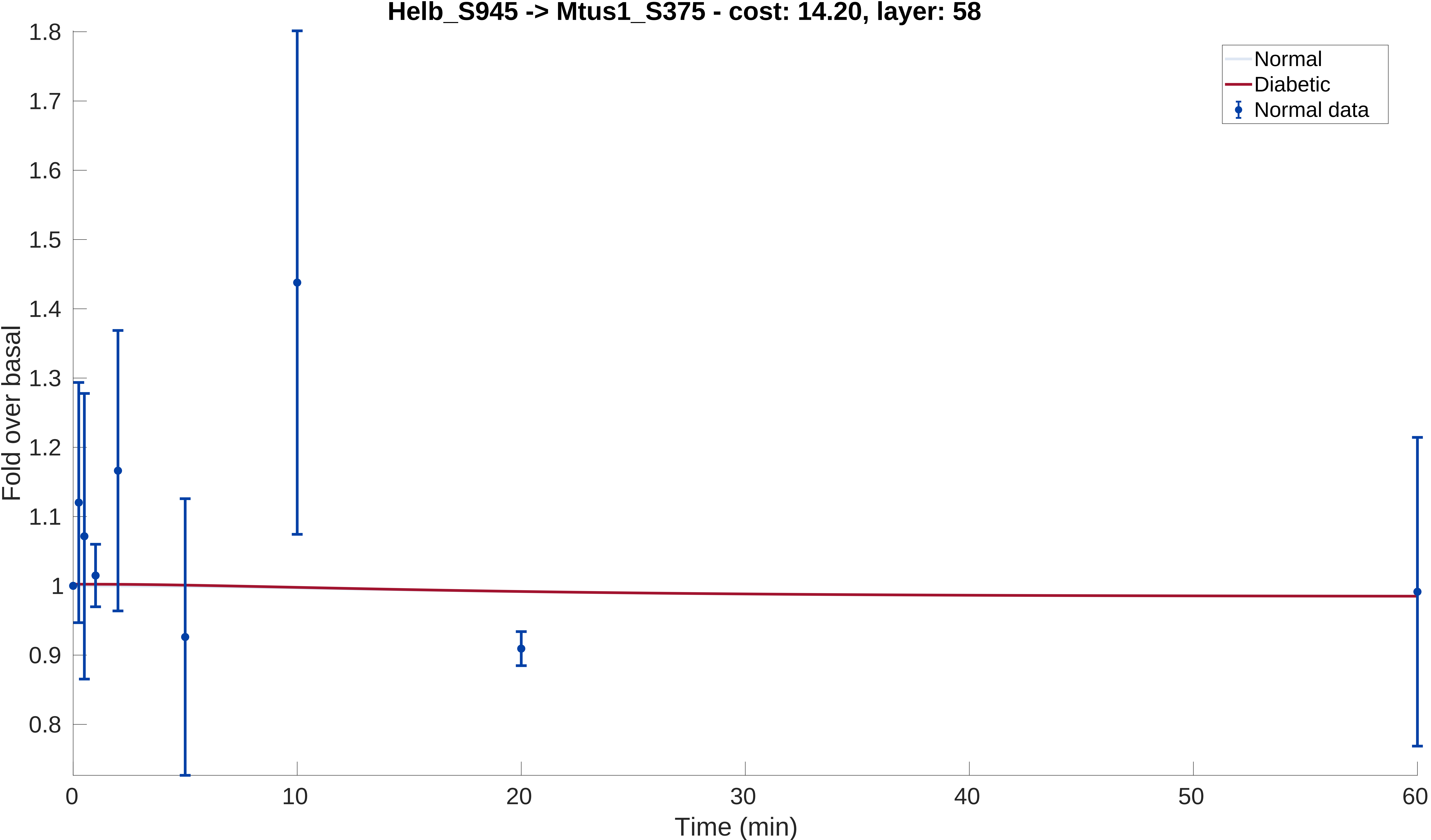

Helb_S945 -> Mtus1_S375 - cost: 14.20, layer: 58
Normal
Diabetic
Normal data
Fold over basal
Time (min)
1.8
1.7
1.6
1.5
1.4
1.3
1.2
1.1
1
0.9
0.8
0
10
20
30
40
50
60

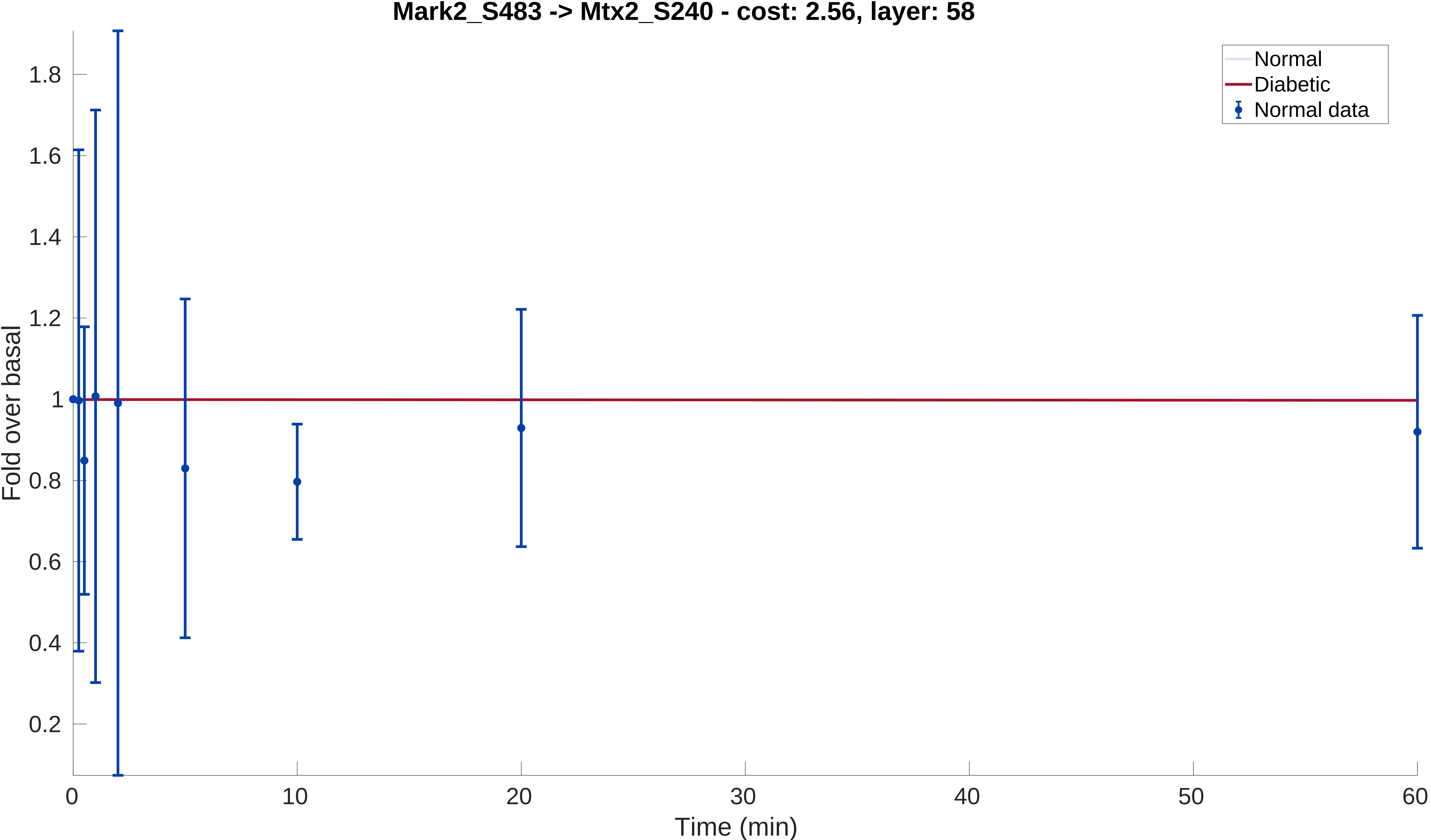

Mark2_S483 -> Mtx2_S240 - cost: 2.56, layer: 58
Normal
Diabetic
Normal data
Fold over basal
1.8
1.6
1.4
1.2
1
0.8
0.6
0.4
0.2
0
10
20
30
40
50
60
Time (min)

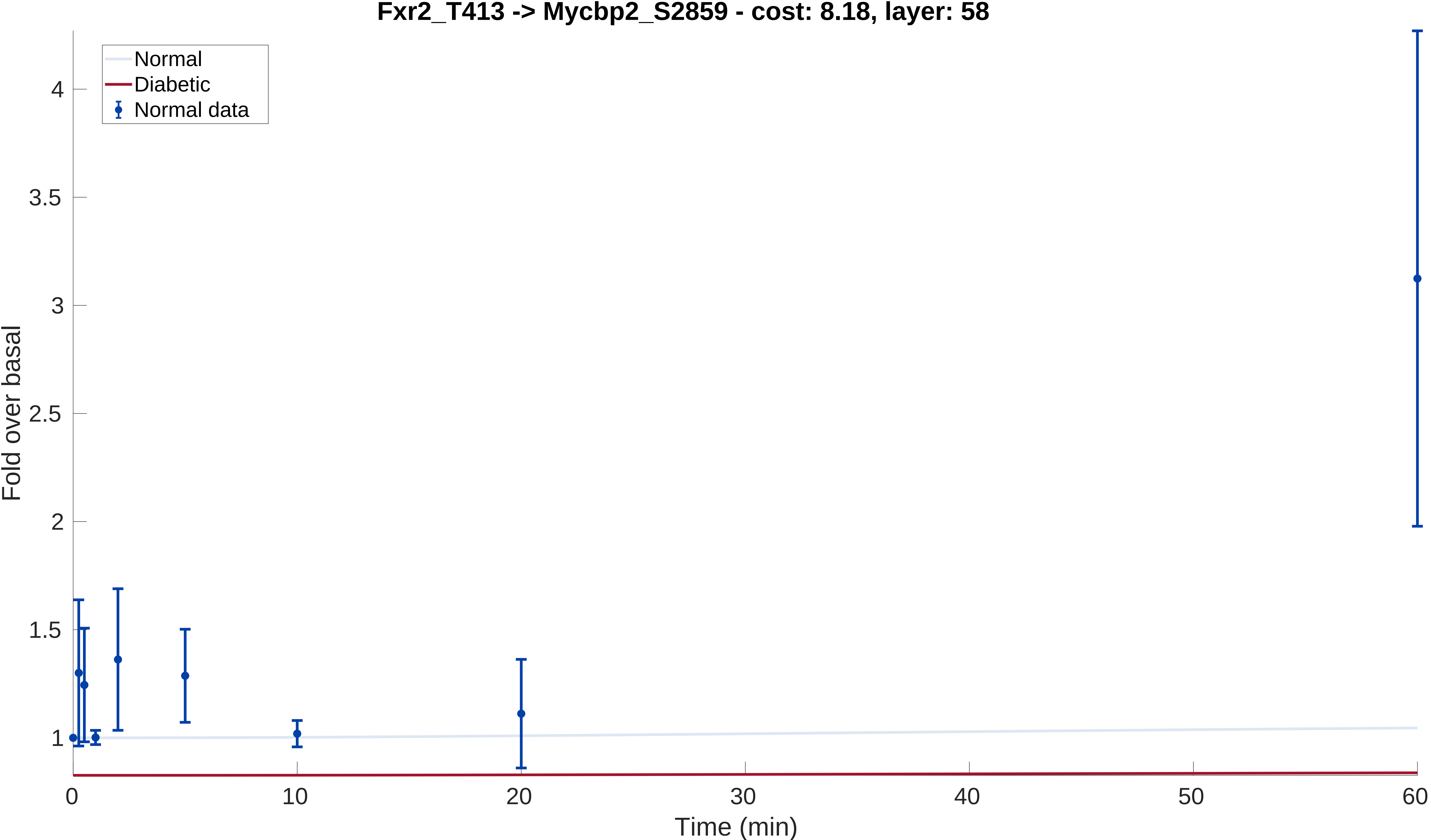
Fxr2_T413 -> Mycbp2_S2859 - cost: 8.18, layer: 58
Normal
Diabetic
Normal data
Fold over basal
1
1.5
2
2.5
3
3.5
4
0
10
20
30
40
50
60
Time (min)

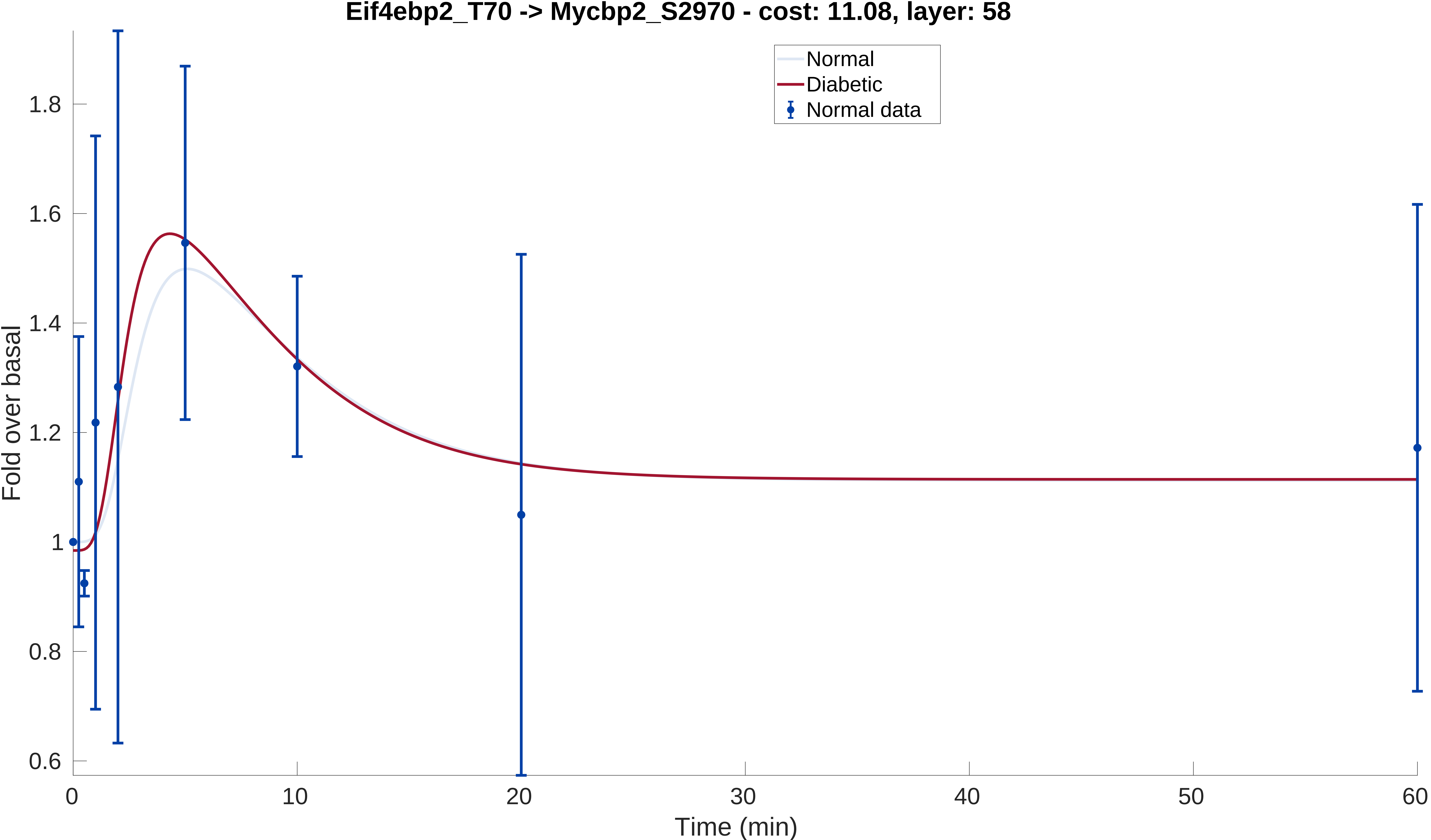

Eif4ebp2_T70 -> Mycbp2_S2970 - cost: 11.08, layer: 58
Normal
Diabetic
Normal data
Fold over basal
1.8
1.6
1.4
1.2
1
0.8
0.6
0
10
20
30
40
50
60
Time (min)

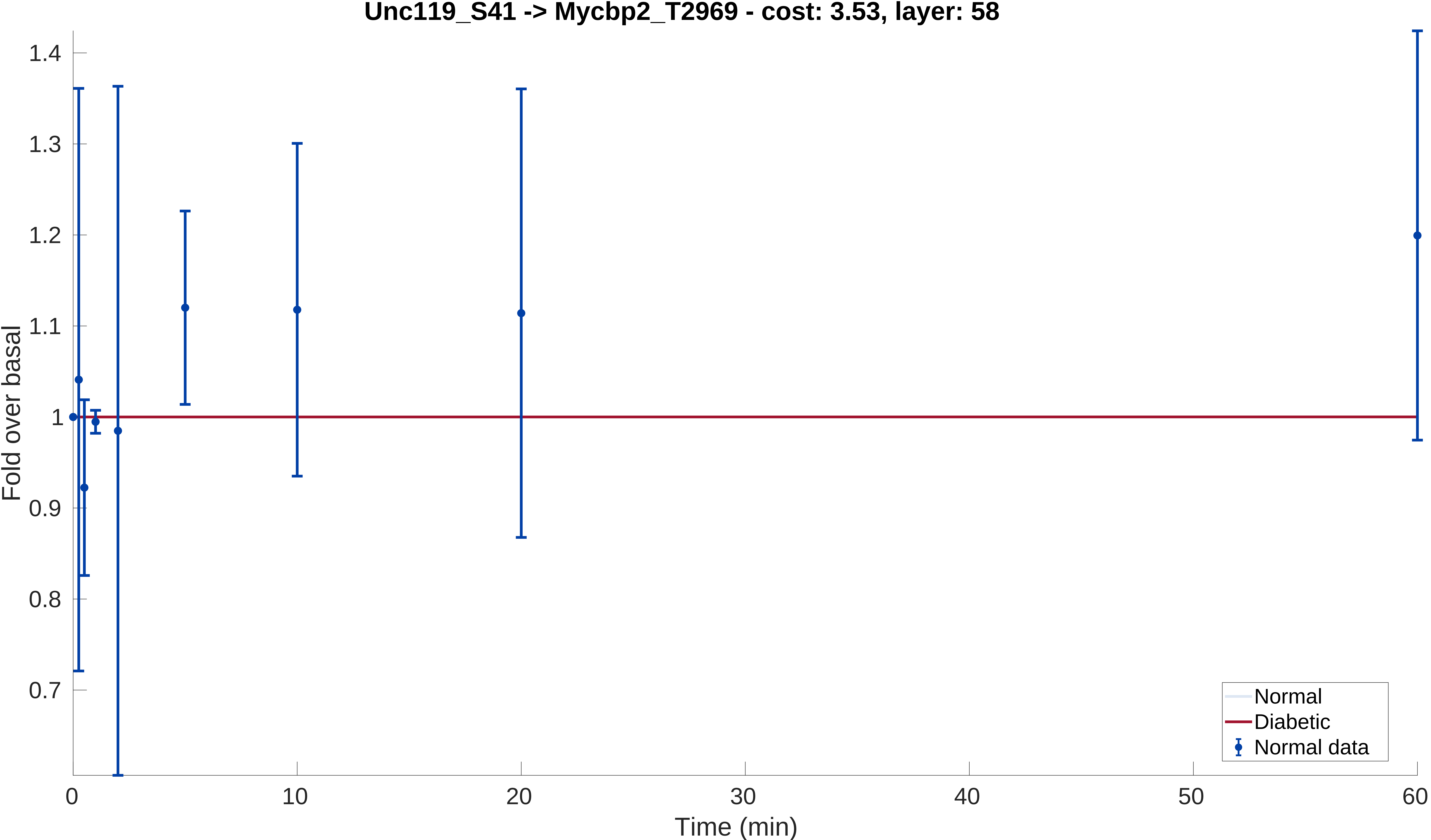

Unc119_S41 -> Mycbp2_T2969 - cost: 3.53, layer: 58
Fold over basal
1.4
1.3
1.2
1.1
1
0.9
0.8
0.7
0
10
20
30
40
50
60
Time (min)
Normal
Diabetic
Normal data

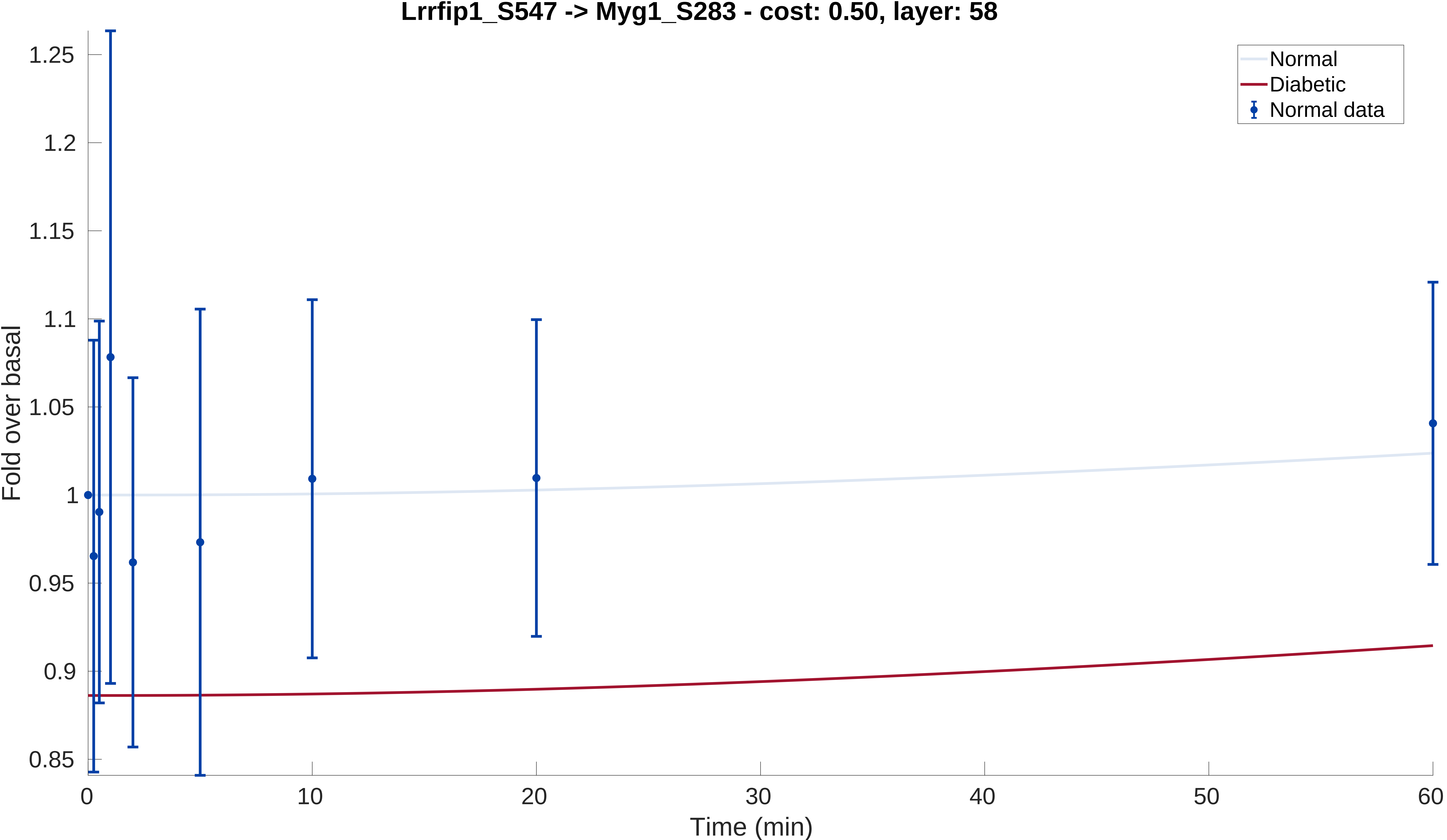

Lrrfip1_S547 -> Myg1_S283 - cost: 0.50, layer: 58
Normal
Diabetic
Normal data
Fold over basal
1.25
1.2
1.15
1.1
1.05
1
0.95
0.9
0.85
0
10
20
30
40
50
60
Time (min)

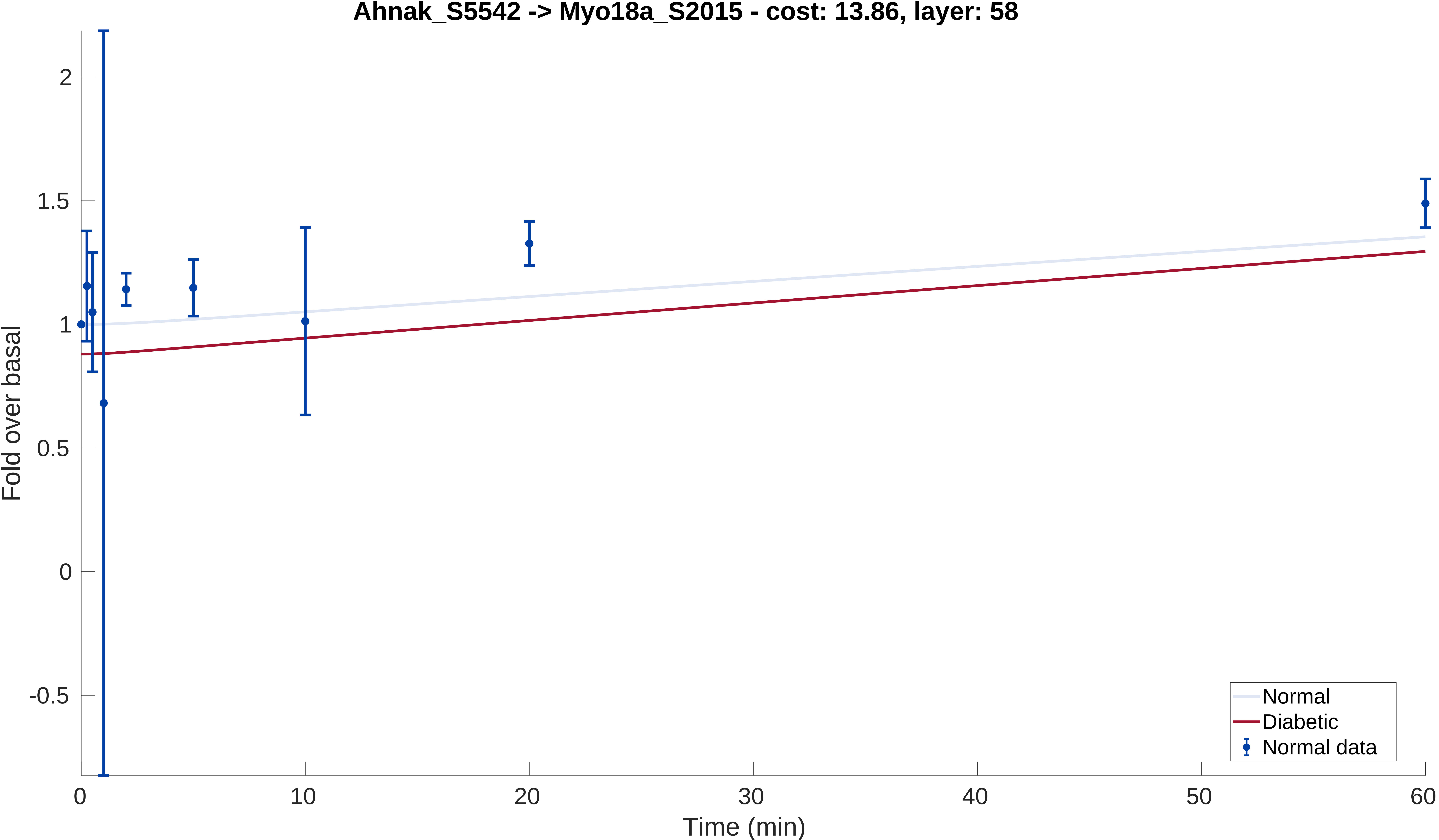

Ahnak_S5542 -> Myo18a_S2015 - cost: 13.86, layer: 58
Fold over basal
Time (min)
Normal
Diabetic
Normal data
2
1.5
1
0.5
0
-0.5
0
10
20
30
40
50
60

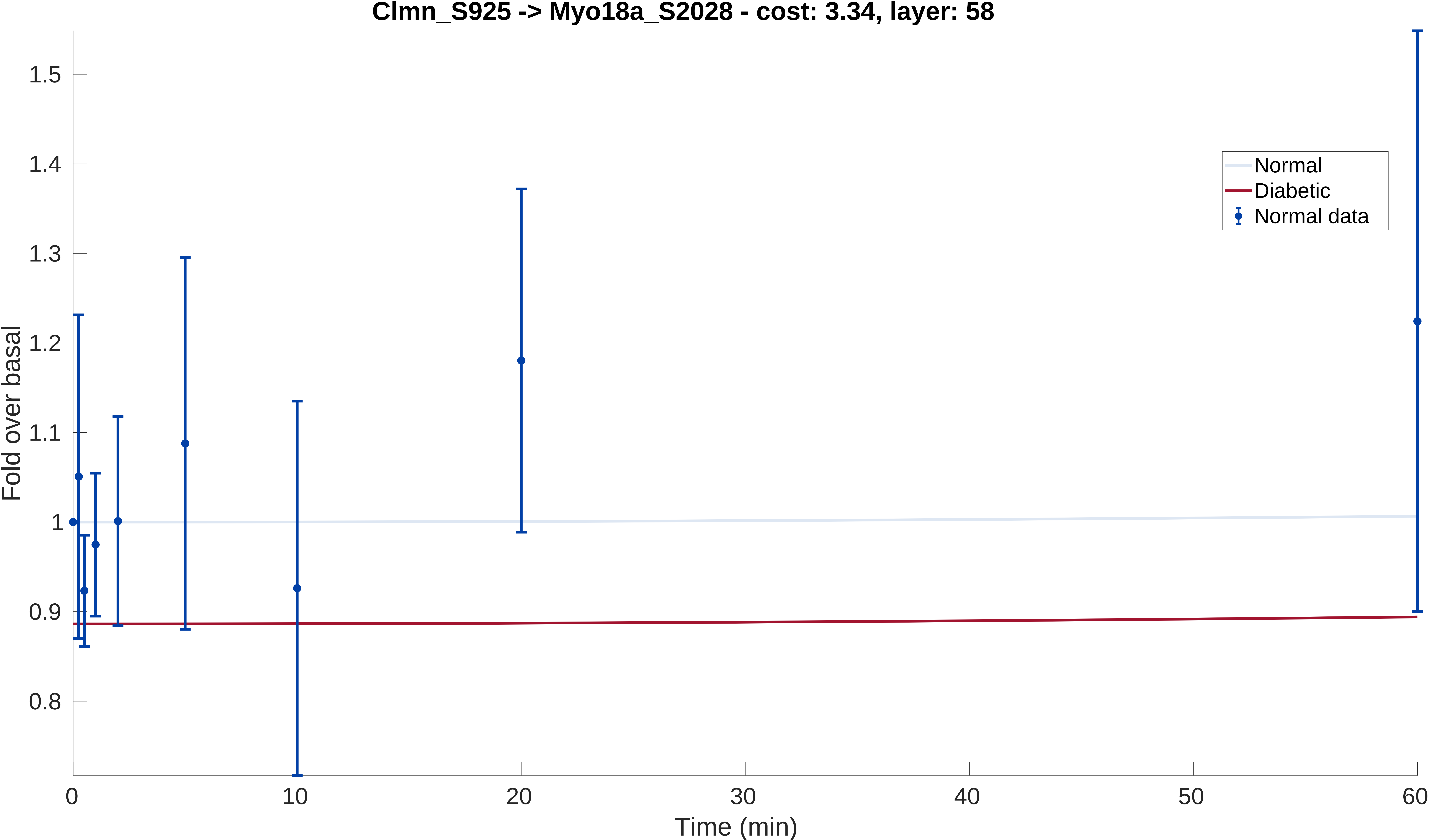
Clmn_S925 -> Myo18a_S2028 - cost: 3.34, layer: 58
Normal
Diabetic
Normal data
Fold over basal
1.5
1.4
1.3
1.2
1.1
1
0.9
0.8
0
10
20
30
40
50
60
Time (min)

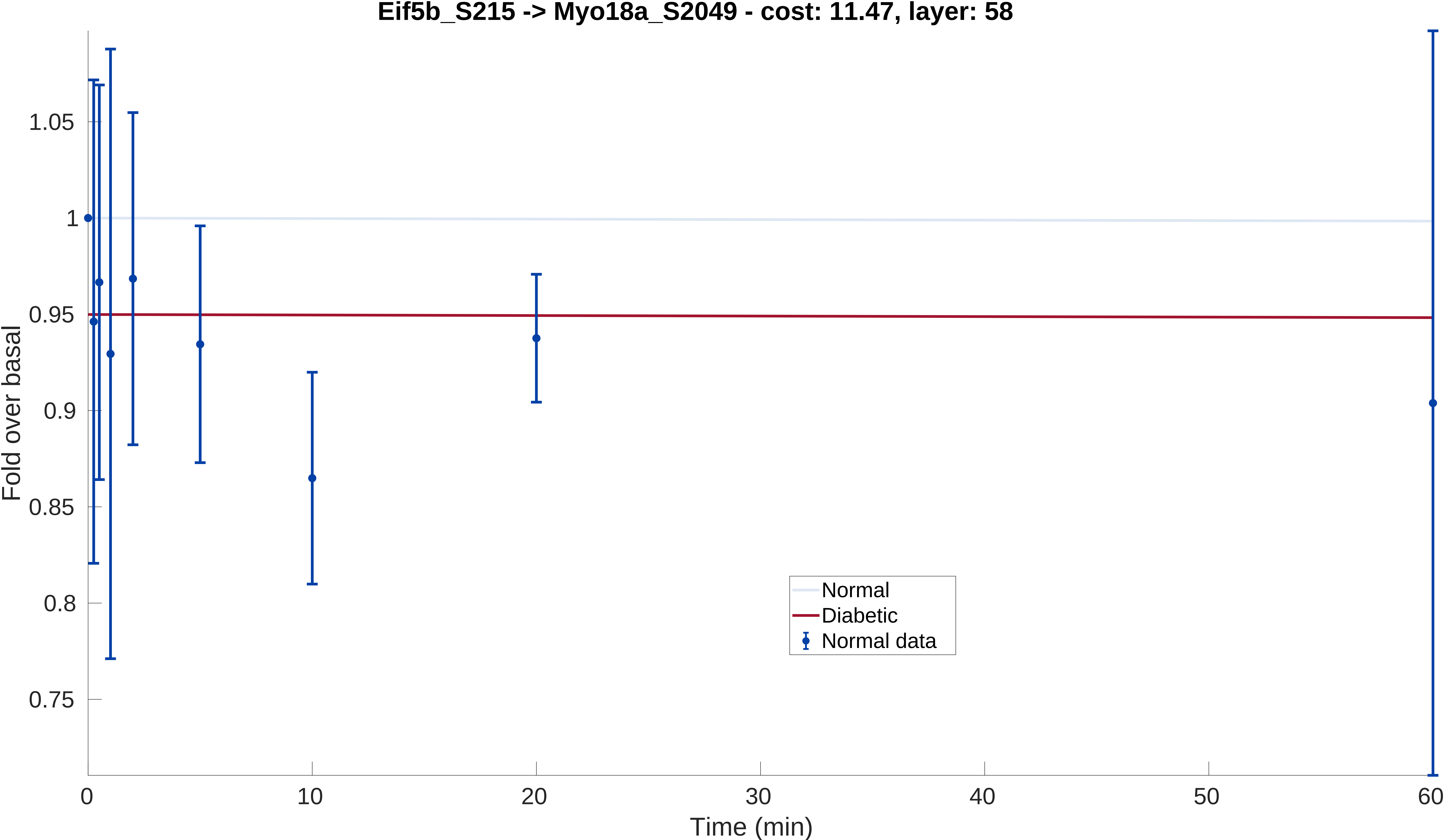

Eif5b_S215 -> Myo18a_S2049 - cost: 11.47, layer: 58
Fold over basal
1.05
1
0.95
0.9
0.85
0.8
0.75
0
10
20
30
40
50
60
Time (min)
Normal
Diabetic
Normal data

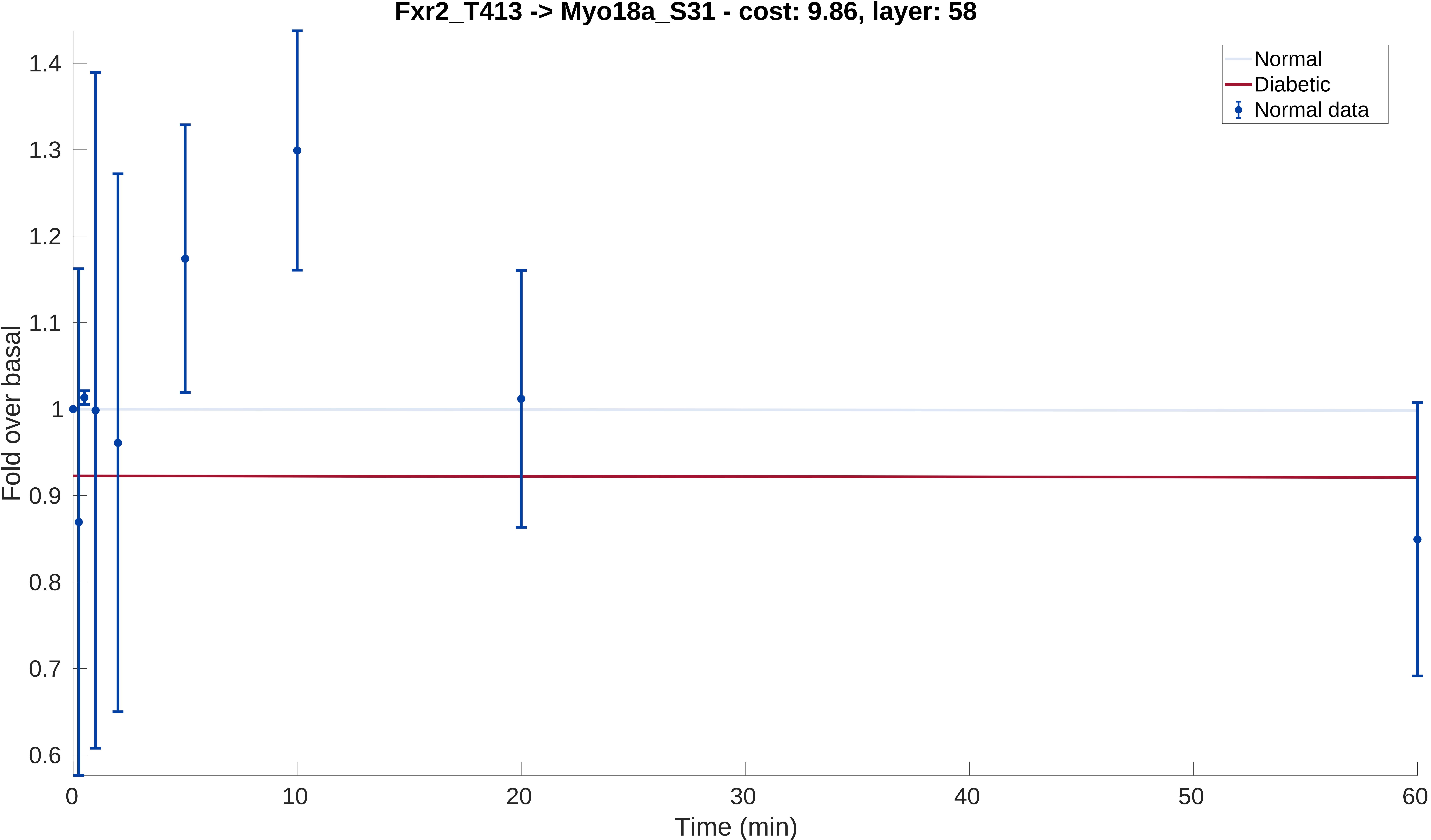

Fxr2_T413 -> Myo18a_S31 - cost: 9.86, layer: 58
Normal
Diabetic
Normal data
Fold over basal
1.4
1.3
1.2
1.1
1
0.9
0.8
0.7
0.6
0
10
20
30
40
50
60
Time (min)

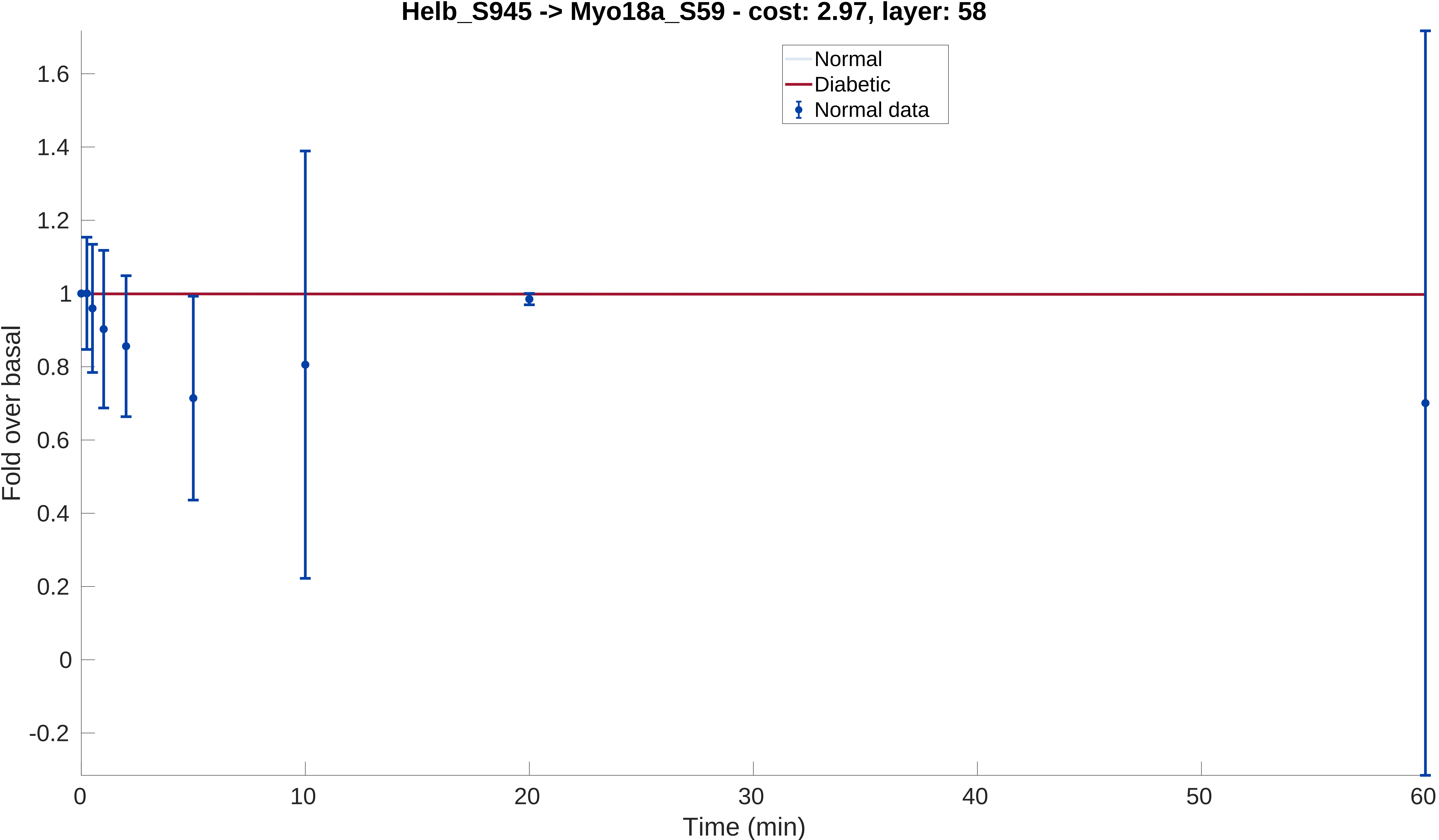

Helb_S945 -> Myo18a_S59 - cost: 2.97, layer: 58
Normal
Diabetic
Normal data
Fold over basal
1.6
1.4
1.2
1
0.8
0.6
0.4
0.2
0
-0.2
0
10
20
30
40
50
60
Time (min)

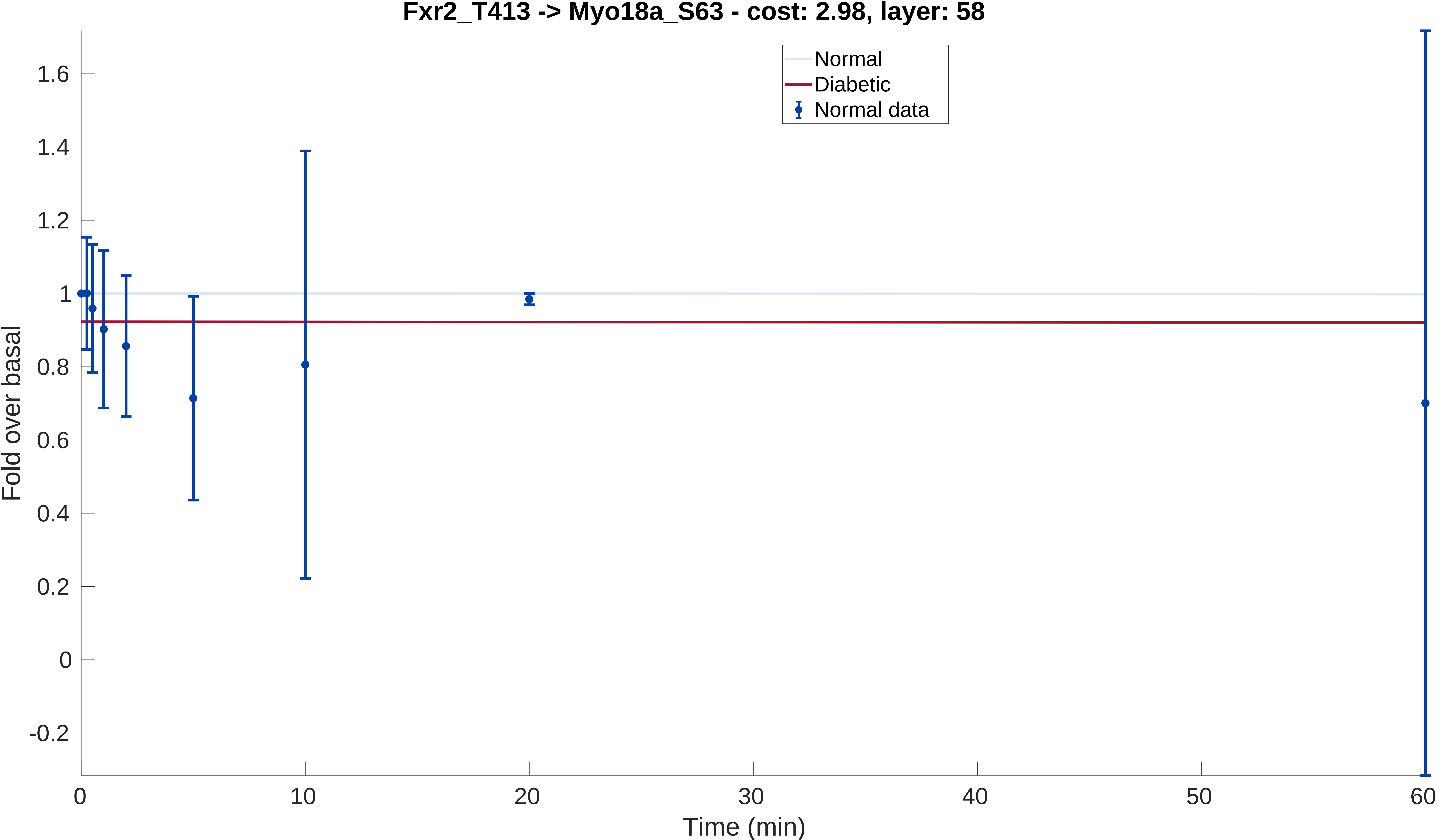
Fxr2_T413 -> Myo18a_S63 - cost: 2.98, layer: 58
Normal
Diabetic
Normal data
Fold over basal
Time (min)
1.6
1.4
1.2
1
0.8
0.6
0.4
0.2
0
-0.2
0
10
20
30
40
50
60

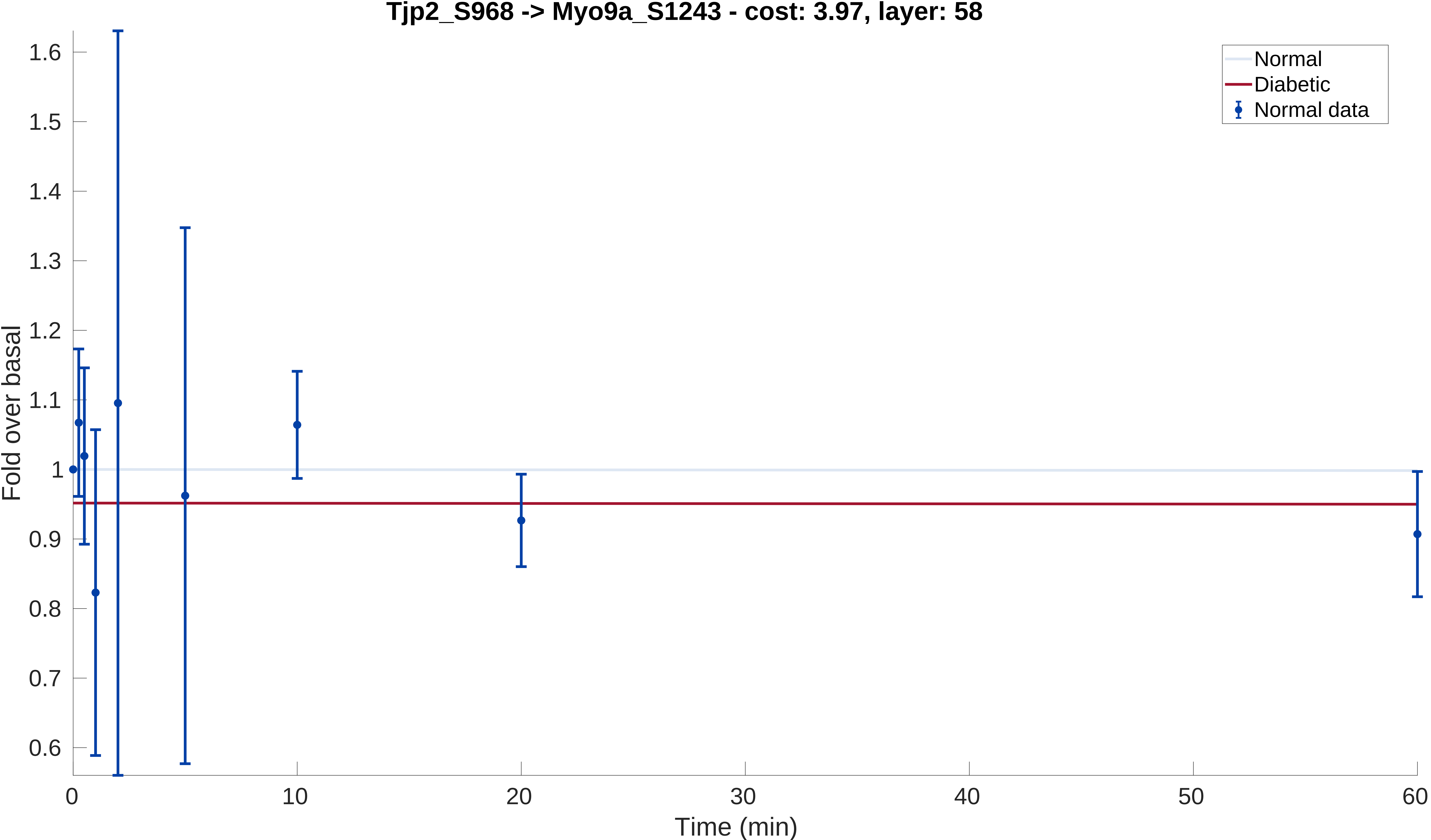
Tjp2_S968 -> Myo9a_S1243 - cost: 3.97, layer: 58
Fold over basal
Time (min)
Normal
Diabetic
Normal data
1.6
1.5
1.4
1.3
1.2
1.1
1
0.9
0.8
0.7
0.6
0
10
20
30
40
50
60

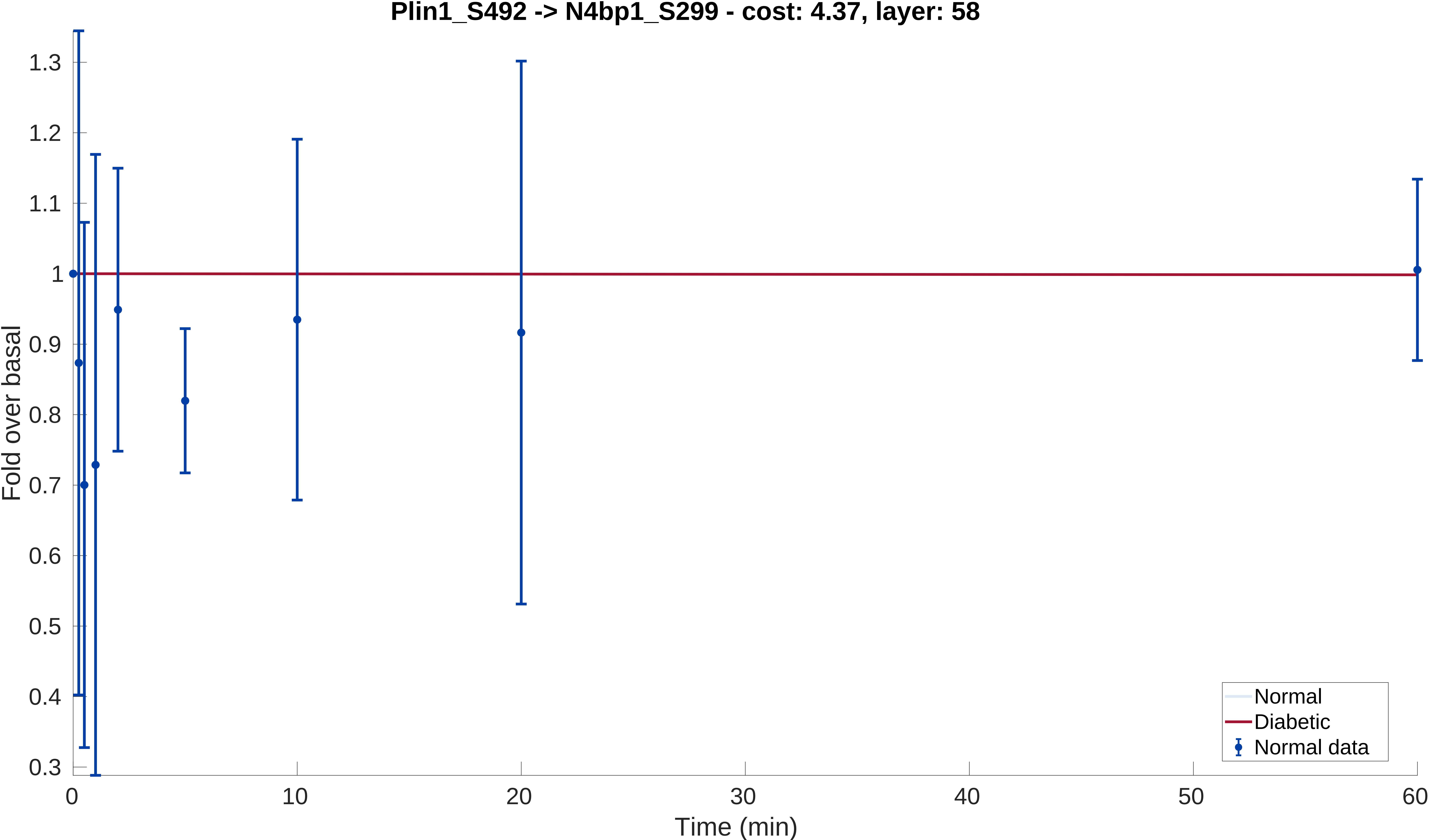
Plin1_S492 -> N4bp1_S299 - cost: 4.37, layer: 58
Fold over basal
1.3
1.2
1.1
1
0.9
0.8
0.7
0.6
0.5
0.4
0.3
0
10
20
30
40
50
60
Time (min)
Normal
Diabetic
Normal data

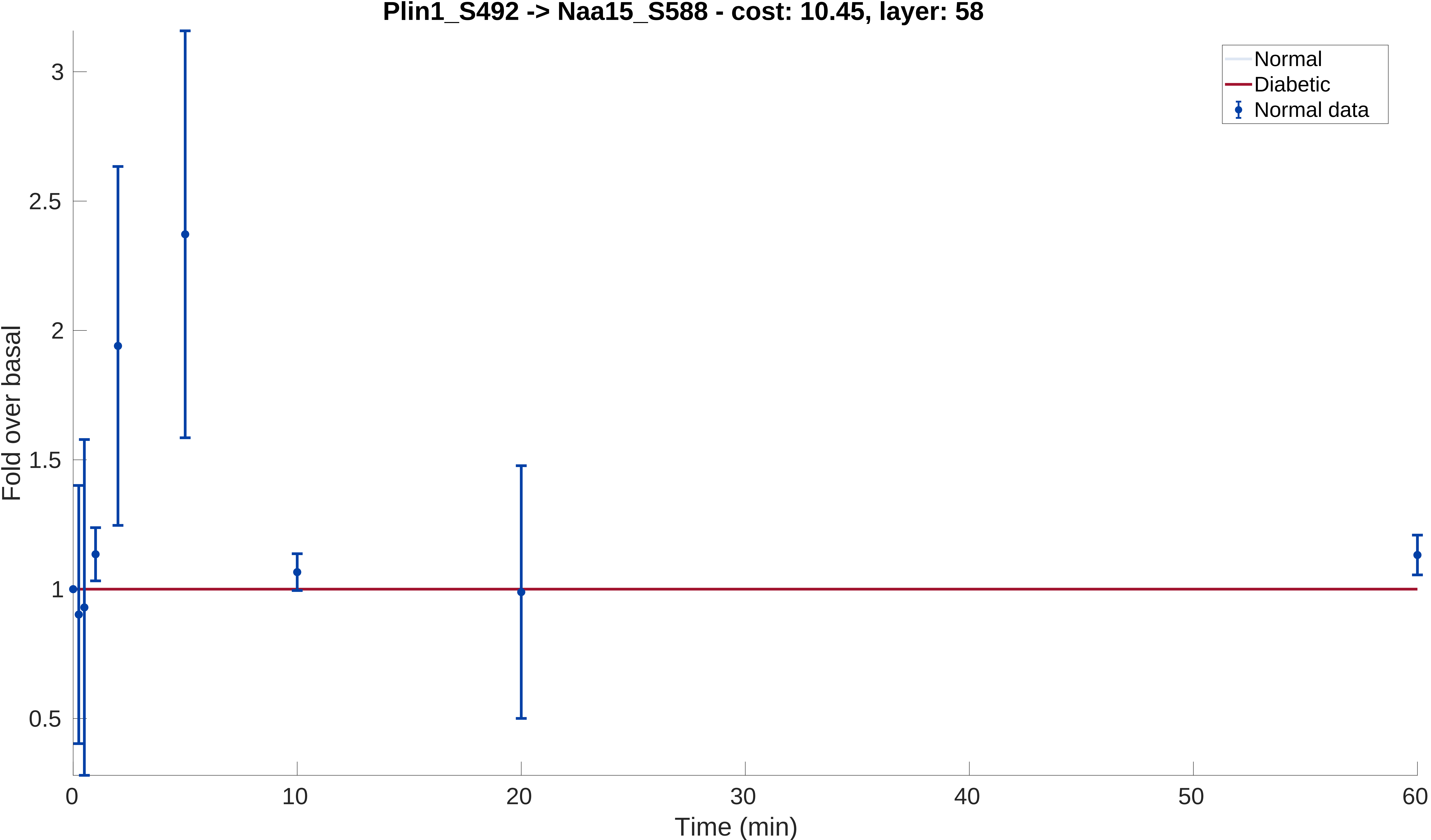
Plin1_S492 -> Naa15_S588 - cost: 10.45, layer: 58
Normal
Diabetic
Normal data
Fold over basal
Time (min)
0
10
20
30
40
50
60
0.5
1
1.5
2
2.5
3

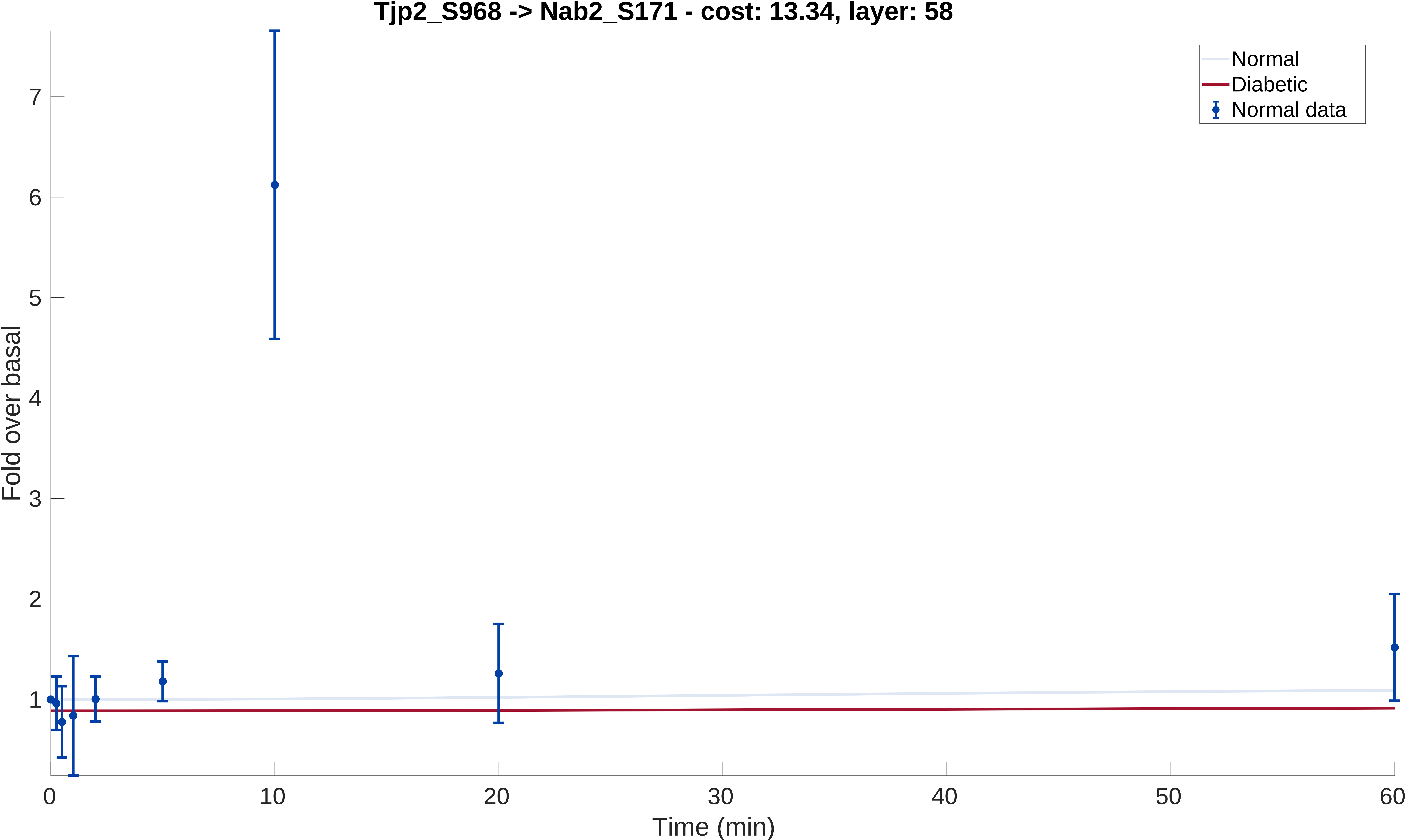

Tjp2_S968 -> Nab2_S171 - cost: 13.34, layer: 58
Normal
Diabetic
Normal data
Fold over basal
Time (min)
0
10
20
30
40
50
60
1
2
3
4
5
6
7

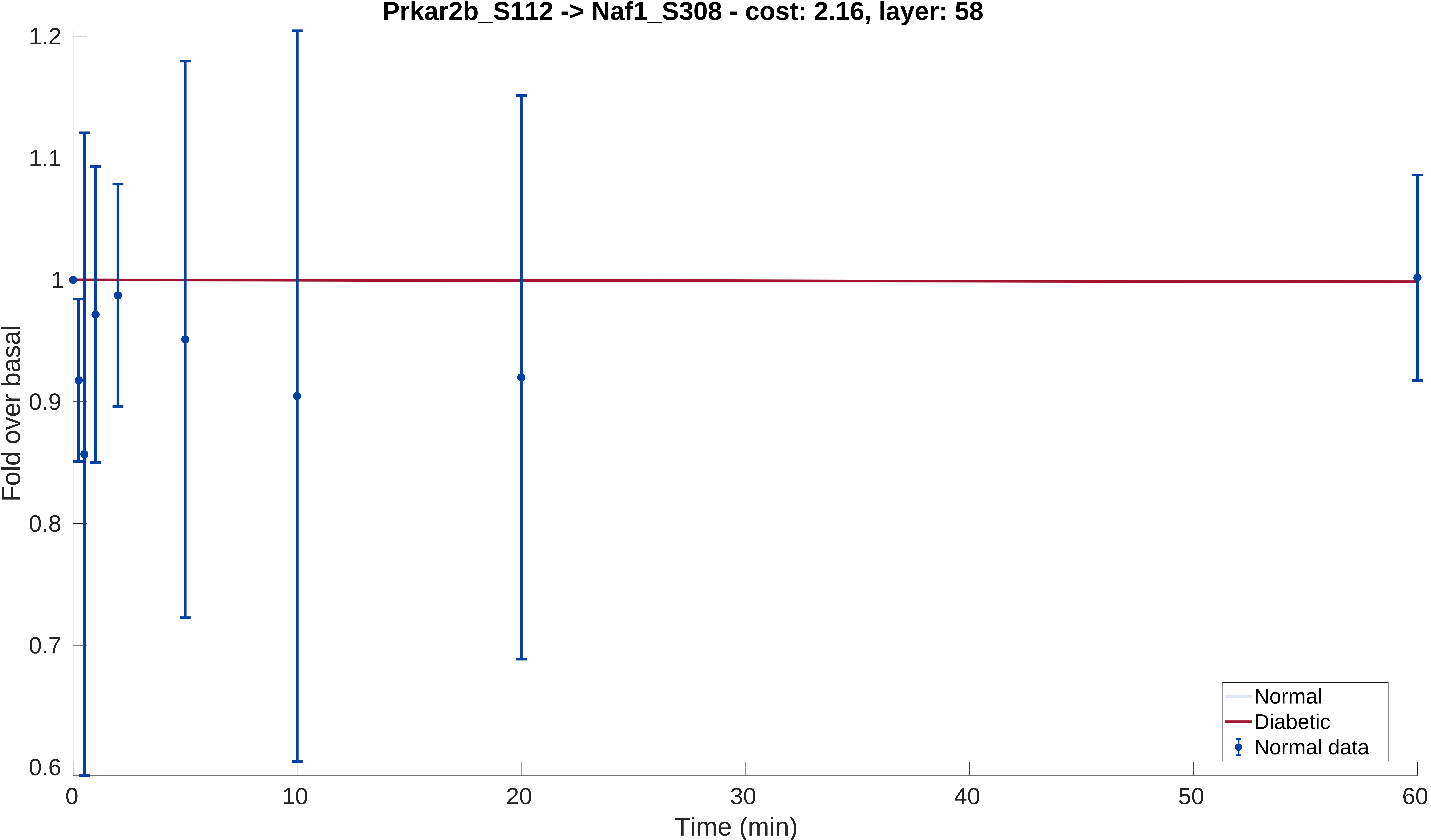

Prkar2b_S112 -> Naf1_S308 - cost: 2.16, layer: 58
Fold over basal
1.2
1.1
1
0.9
0.8
0.7
0.6
0
10
20
30
40
50
60
Time (min)
Normal
Diabetic
Normal data

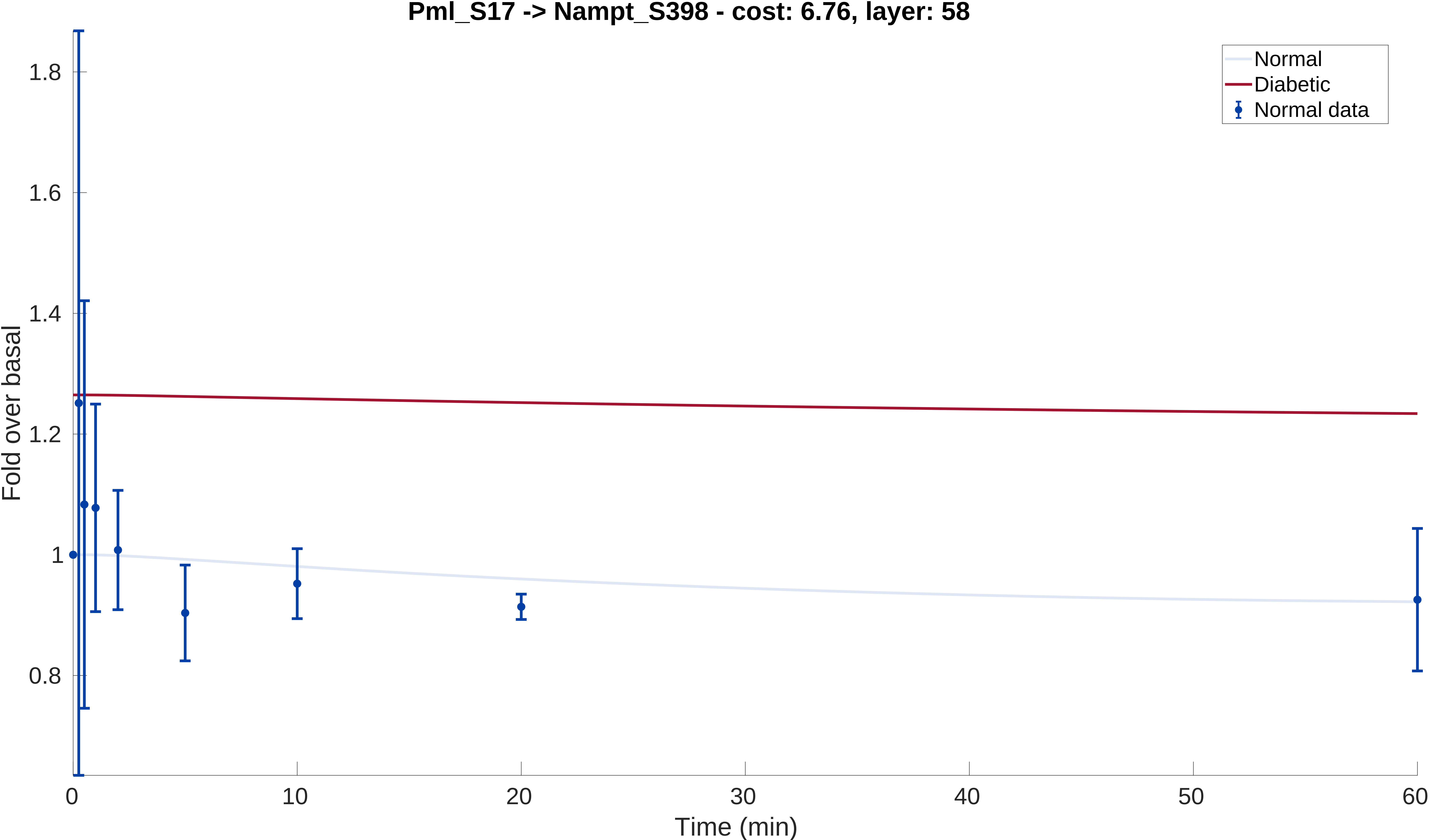
Pml_S17 -> Nampt_S398 - cost: 6.76, layer: 58
Normal
Diabetic
Normal data
Fold over basal
1.8
1.6
1.4
1.2
1
0.8
0
10
20
30
40
50
60
Time (min)

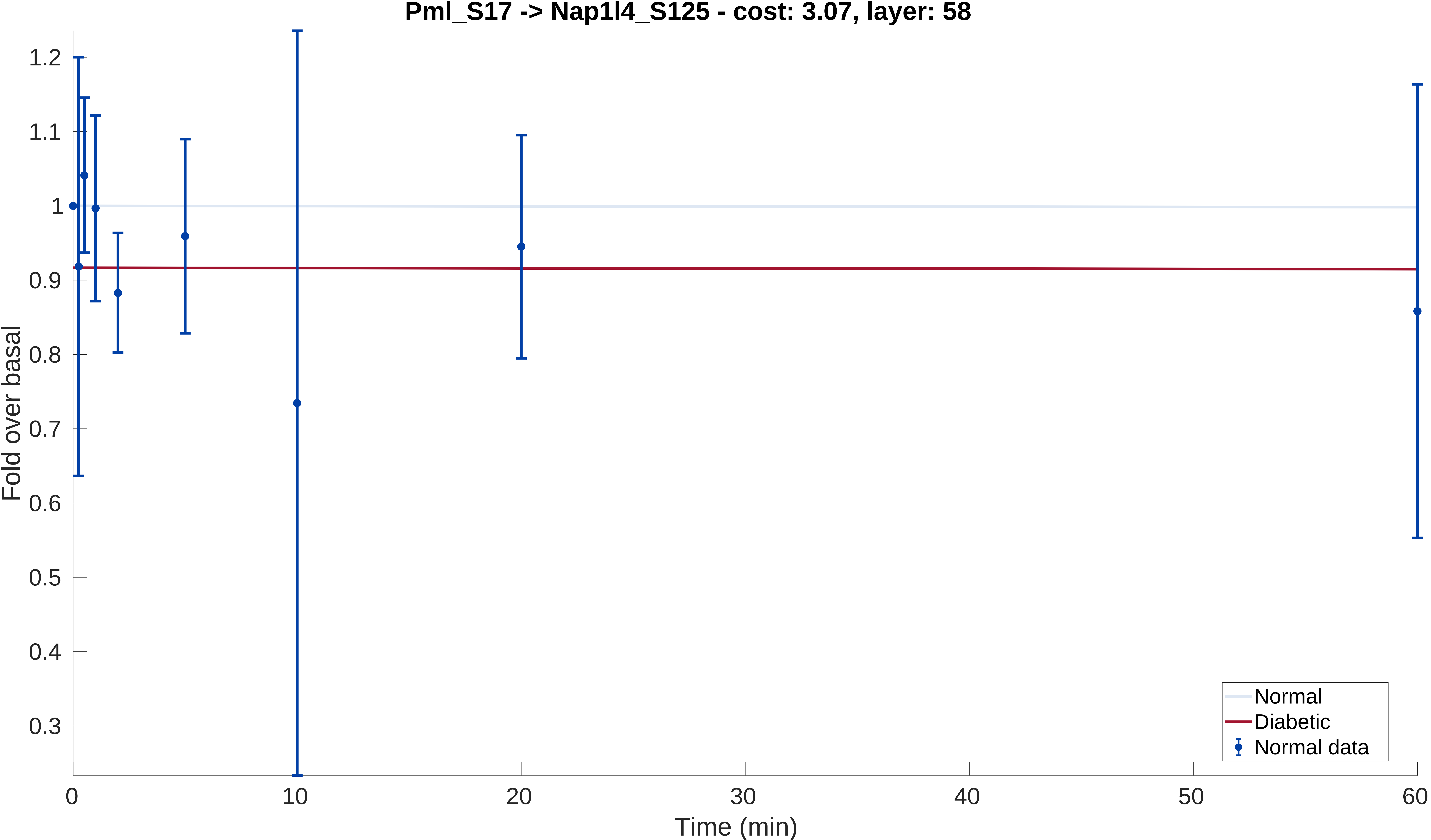

Pml_S17 -> Nap1l4_S125 - cost: 3.07, layer: 58
Fold over basal
1.2
1.1
1
0.9
0.8
0.7
0.6
0.5
0.4
0.3
0
10
20
30
40
50
60
Time (min)
Normal
Diabetic
Normal data

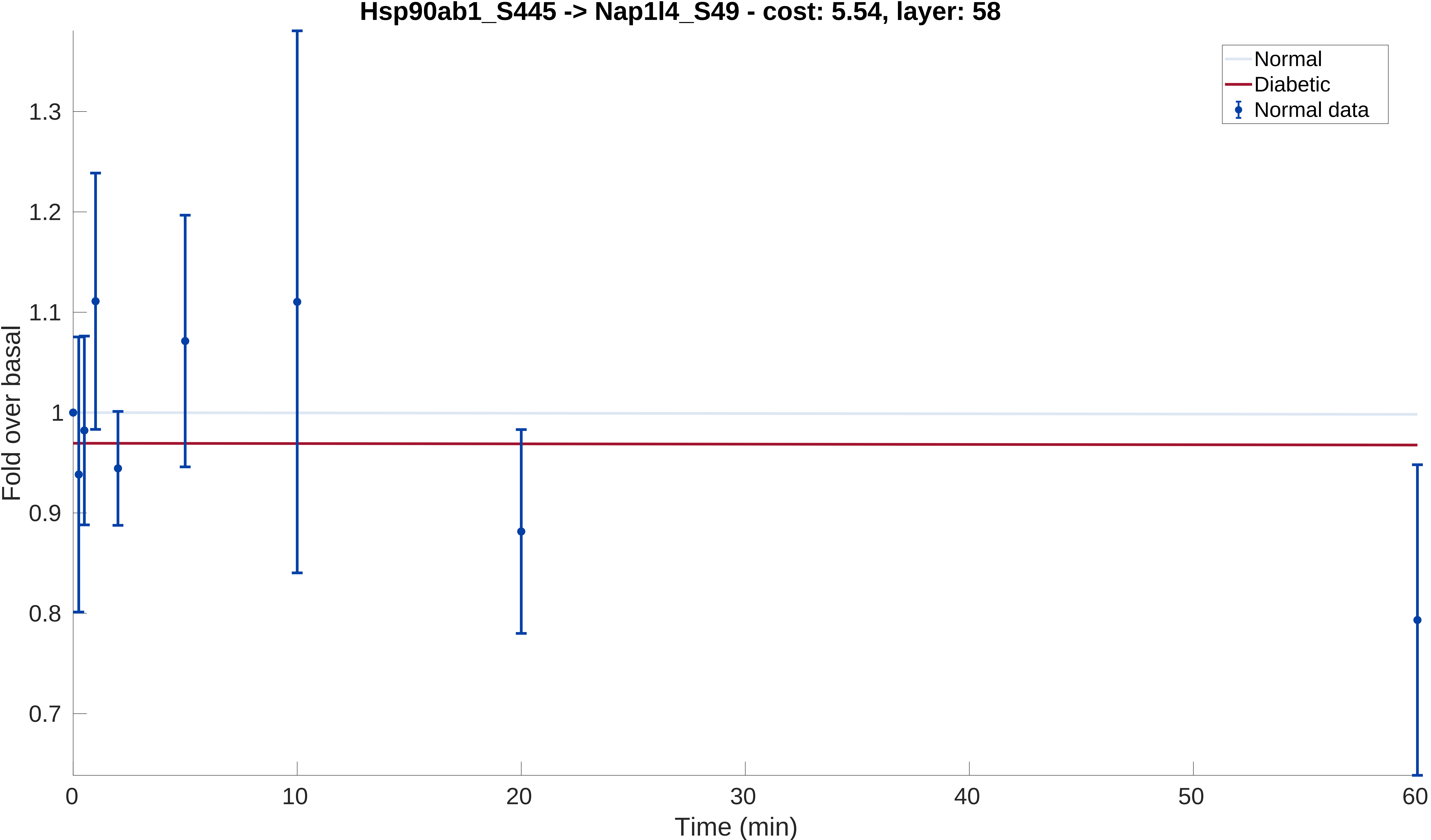

Hsp90ab1_S445 -> Nap1l4_S49 - cost: 5.54, layer: 58
Normal
Diabetic
Normal data
Fold over basal
1.3
1.2
1.1
1
0.9
0.8
0.7
0
10
20
30
40
50
60
Time (min)

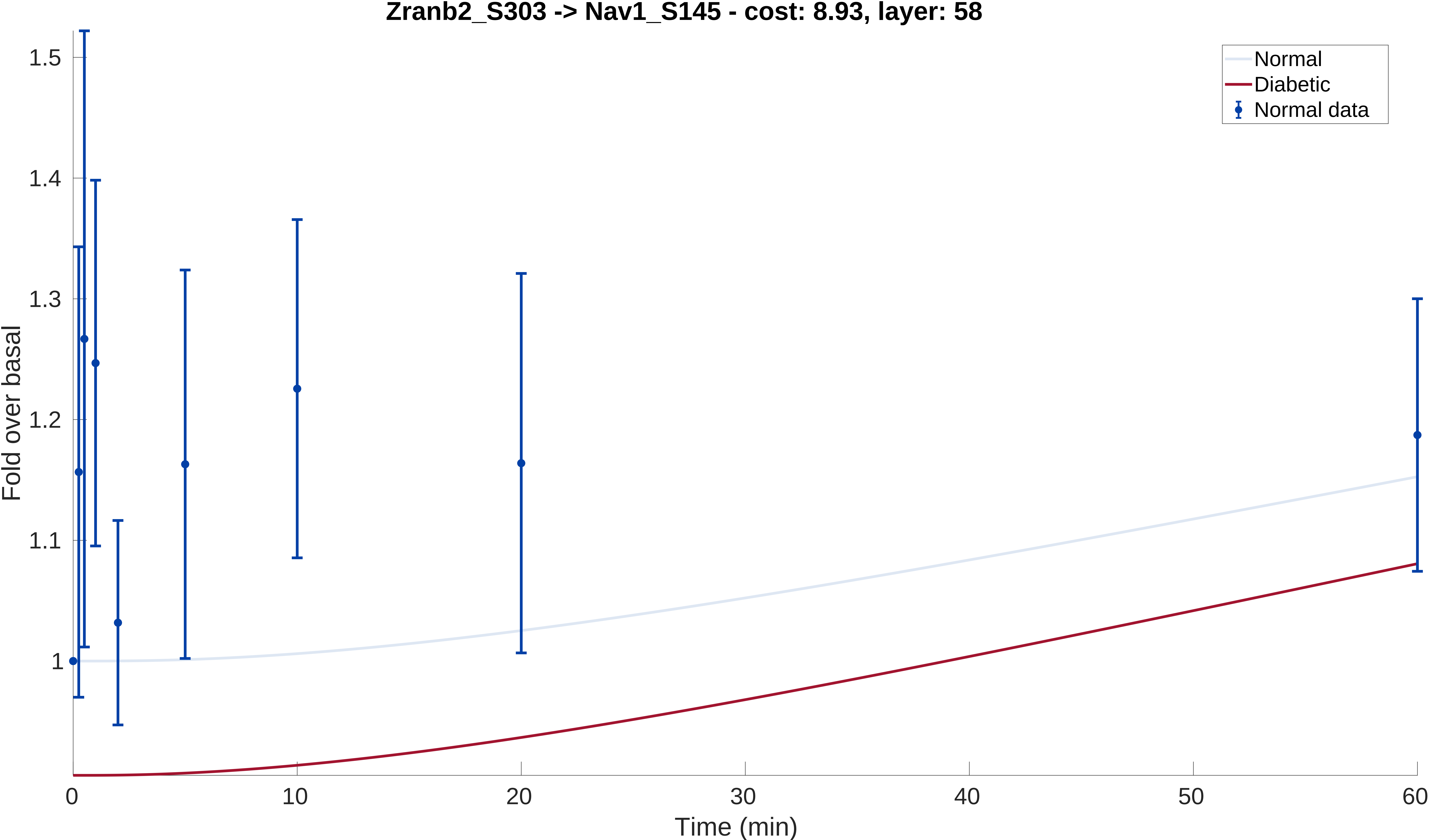

Zranb2_S303 -> Nav1_S145 - cost: 8.93, layer: 58
Normal
Diabetic
Normal data
Fold over basal
1.5
1.4
1.3
1.2
1.1
1
0
10
20
30
40
50
60
Time (min)

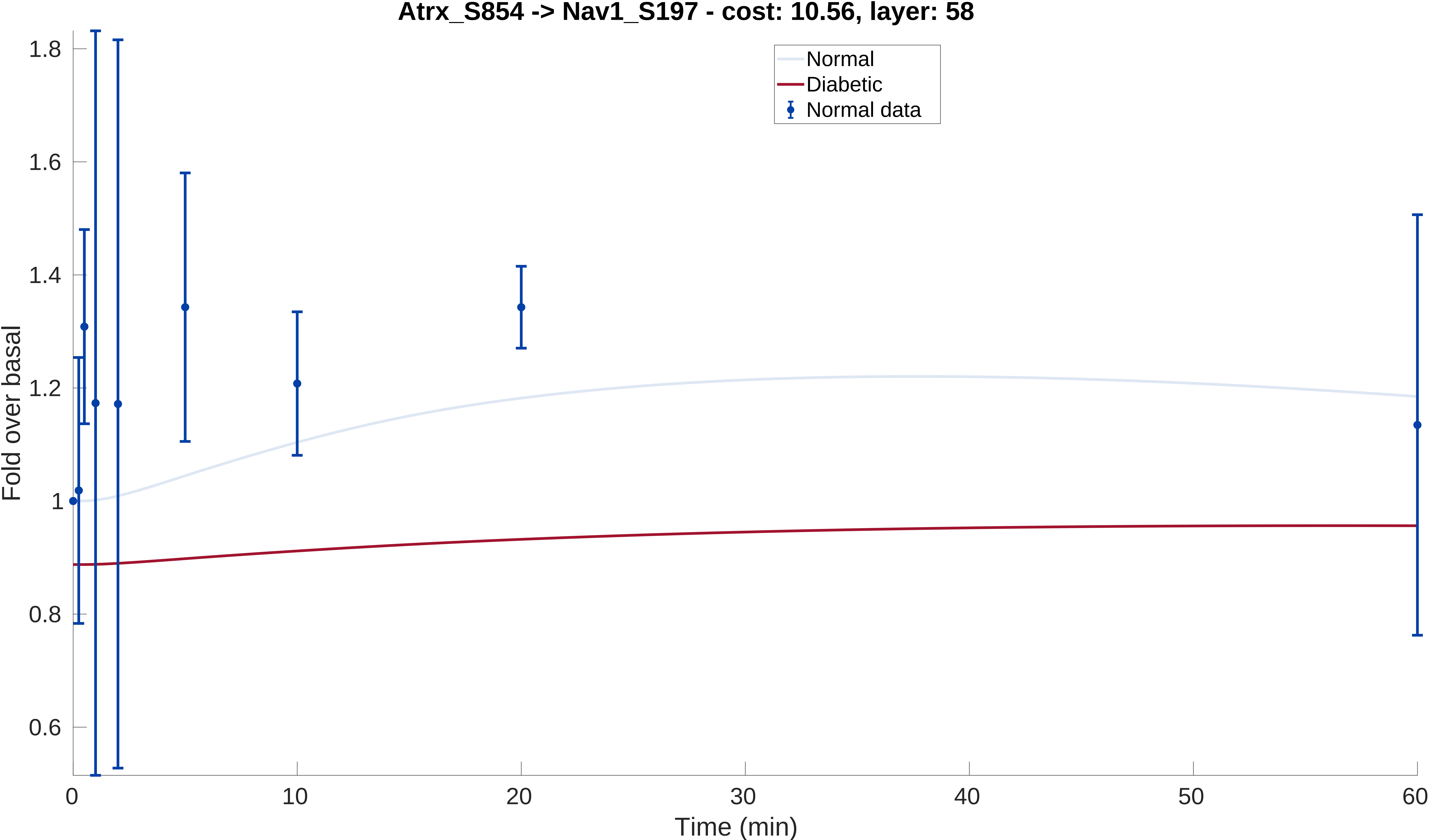

Atrx_S854 -> Nav1_S197 - cost: 10.56, layer: 58
Normal
Diabetic
Normal data
Fold over basal
Time (min)
1.8
1.6
1.4
1.2
1
0.8
0.6
0
10
20
30
40
50
60

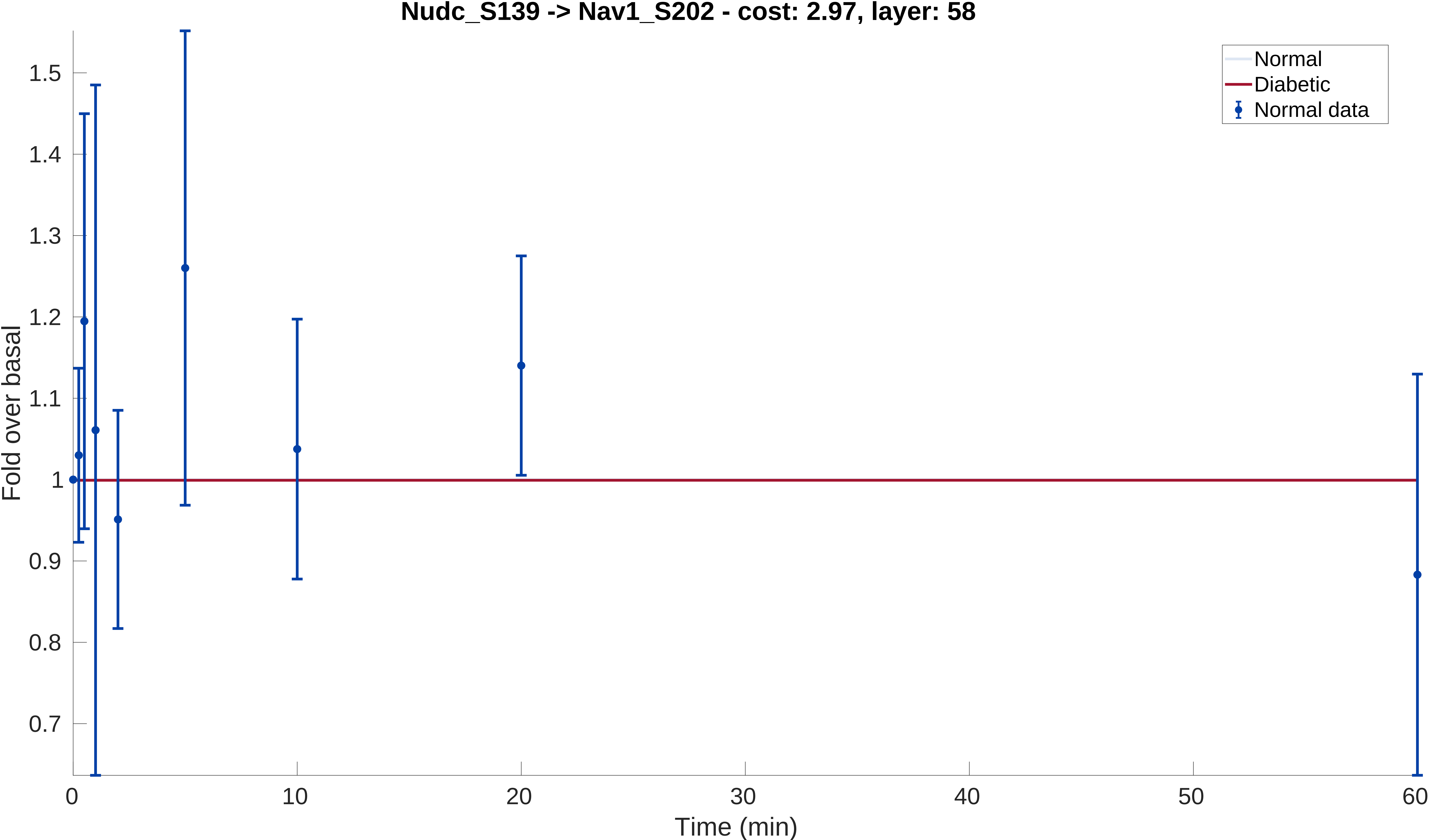

Nudc_S139 -> Nav1_S202 - cost: 2.97, layer: 58
Normal
Diabetic
Normal data
Fold over basal
1.5
1.4
1.3
1.2
1.1
1
0.9
0.8
0.7
0
10
20
30
40
50
60
Time (min)

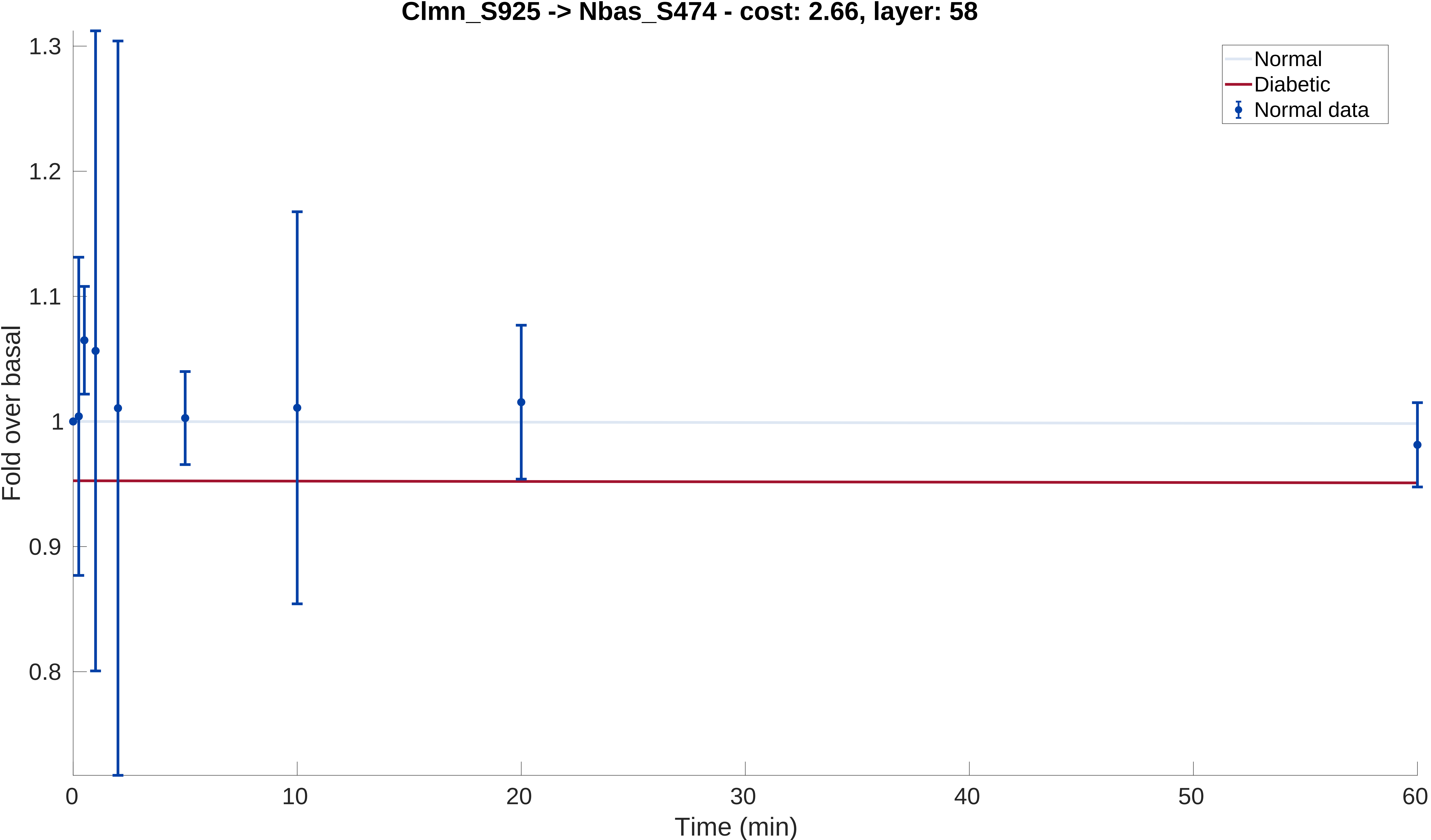

Clmn_S925 -> Nbas_S474 - cost: 2.66, layer: 58
Normal
Diabetic
Normal data
Fold over basal
Time (min)
1.3
1.2
1.1
1
0.9
0.8
0
10
20
30
40
50
60

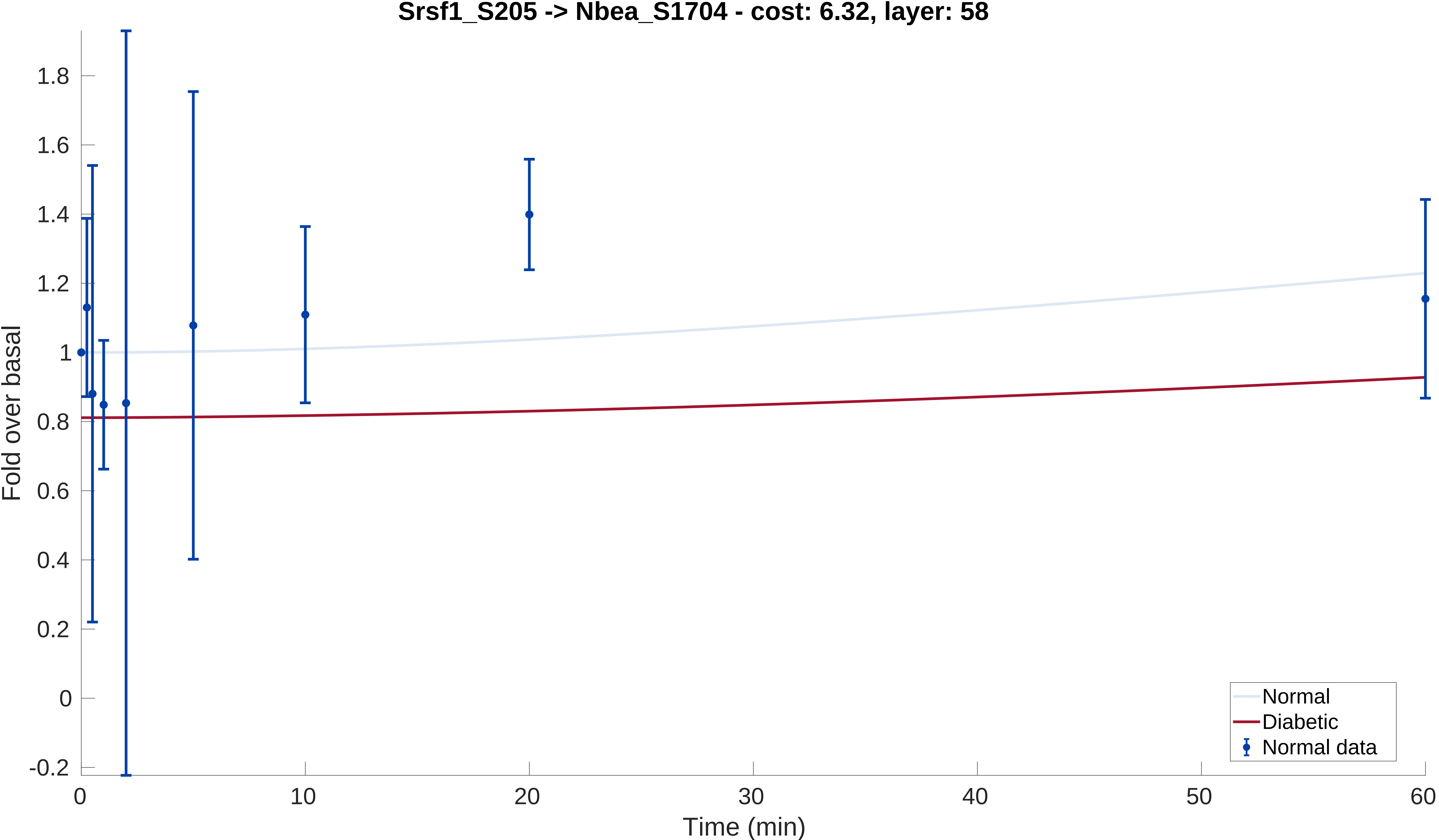

Srsf1_S205 -> Nbea_S1704 - cost: 6.32, layer: 58
Fold over basal
1.8
1.6
1.4
1.2
1
0.8
0.6
0.4
0.2
0
-0.2
0
10
20
30
40
50
60
Time (min)
Normal
Diabetic
Normal data

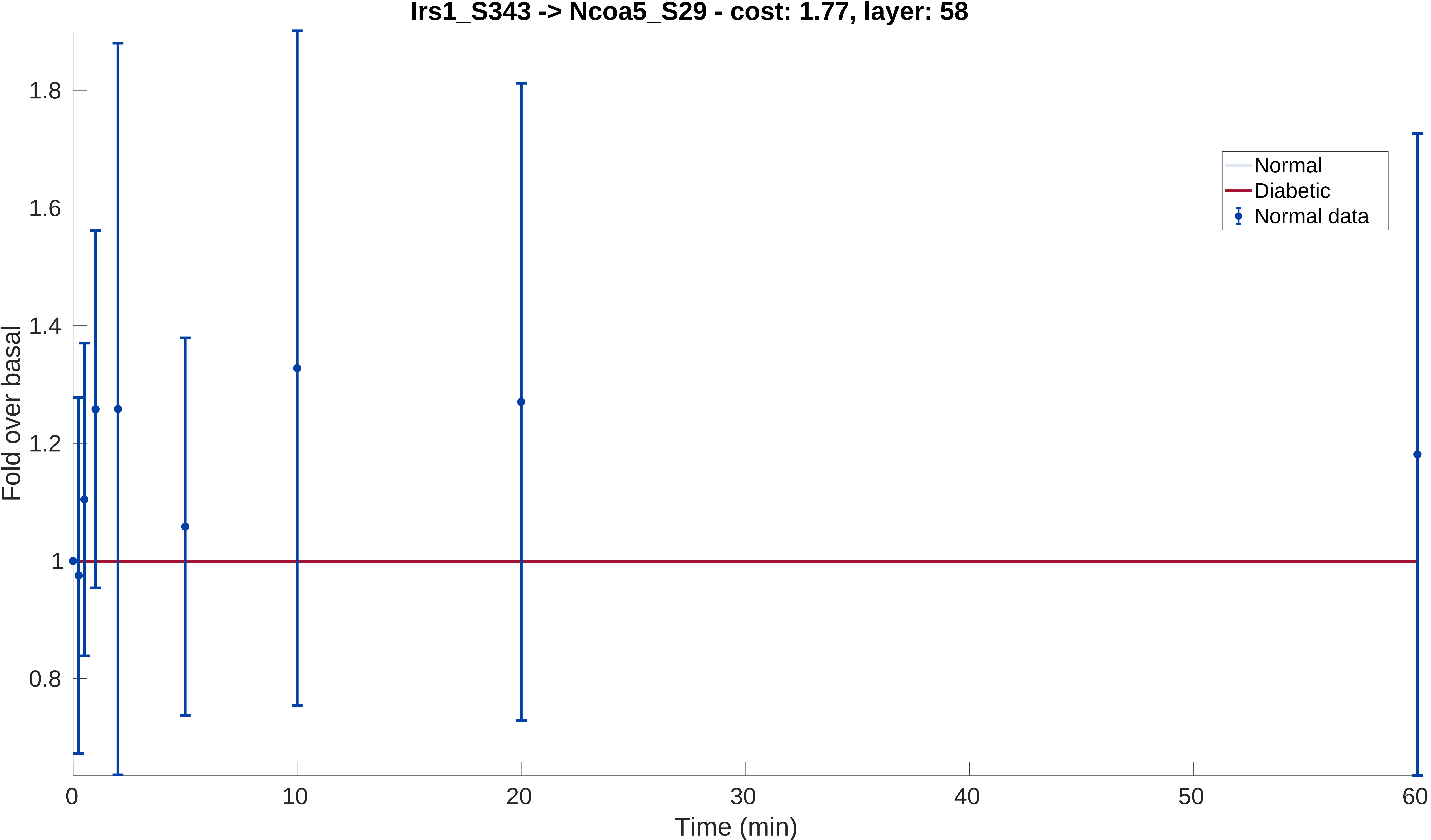
Irs1_S343 -> Ncoa5_S29 - cost: 1.77, layer: 58
Fold over basal
1.8
1.6
1.4
1.2
1
0.8
Time (min)
0
10
20
30
40
50
60
Normal
Diabetic
Normal data

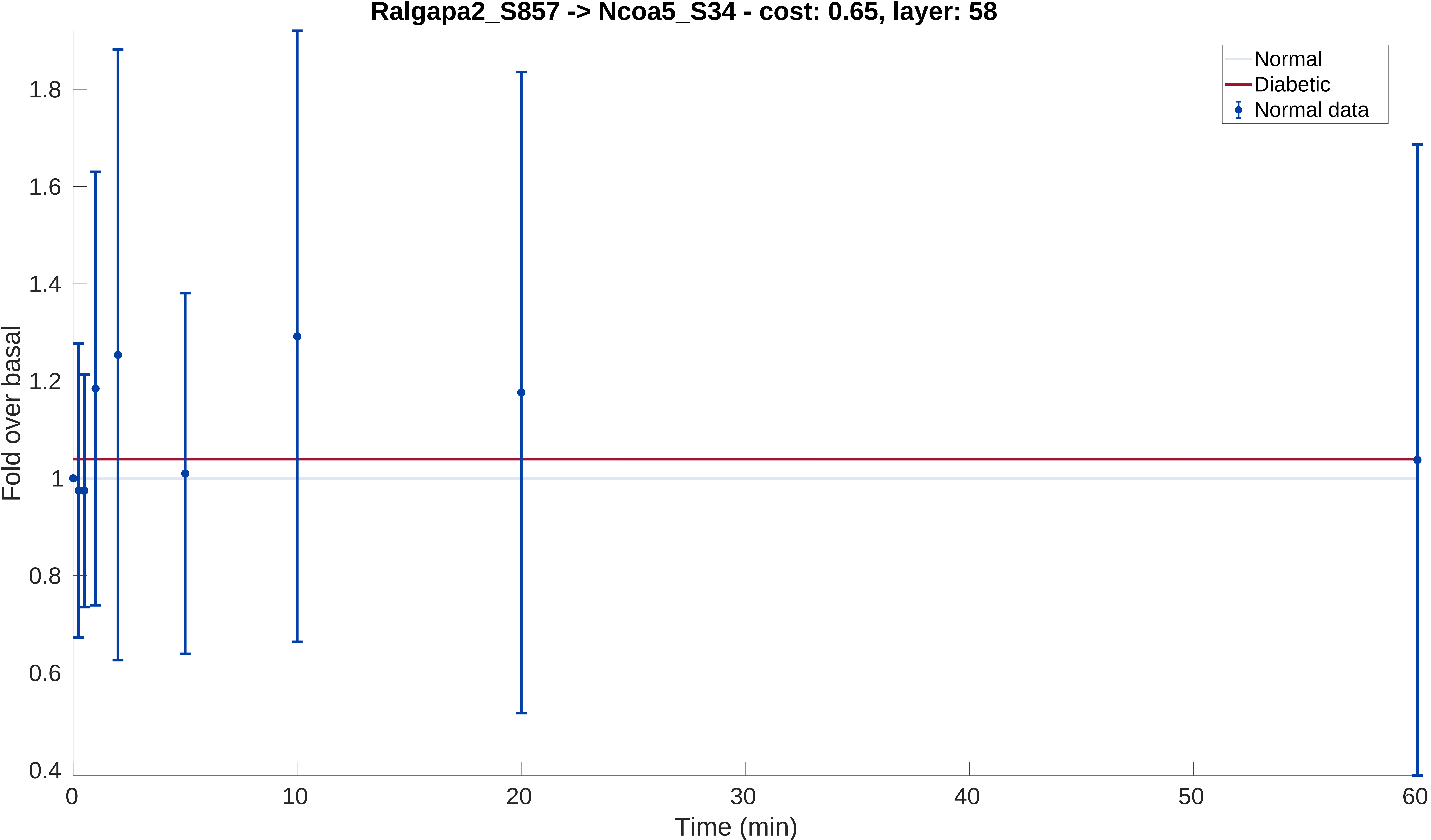

Ralgapa2_S857 -> Ncoa5_S34 - cost: 0.65, layer: 58
Normal
Diabetic
Normal data
Fold over basal
Time (min)
0.4
0.6
0.8
1
1.2
1.4
1.6
1.8
0
10
20
30
40
50
60

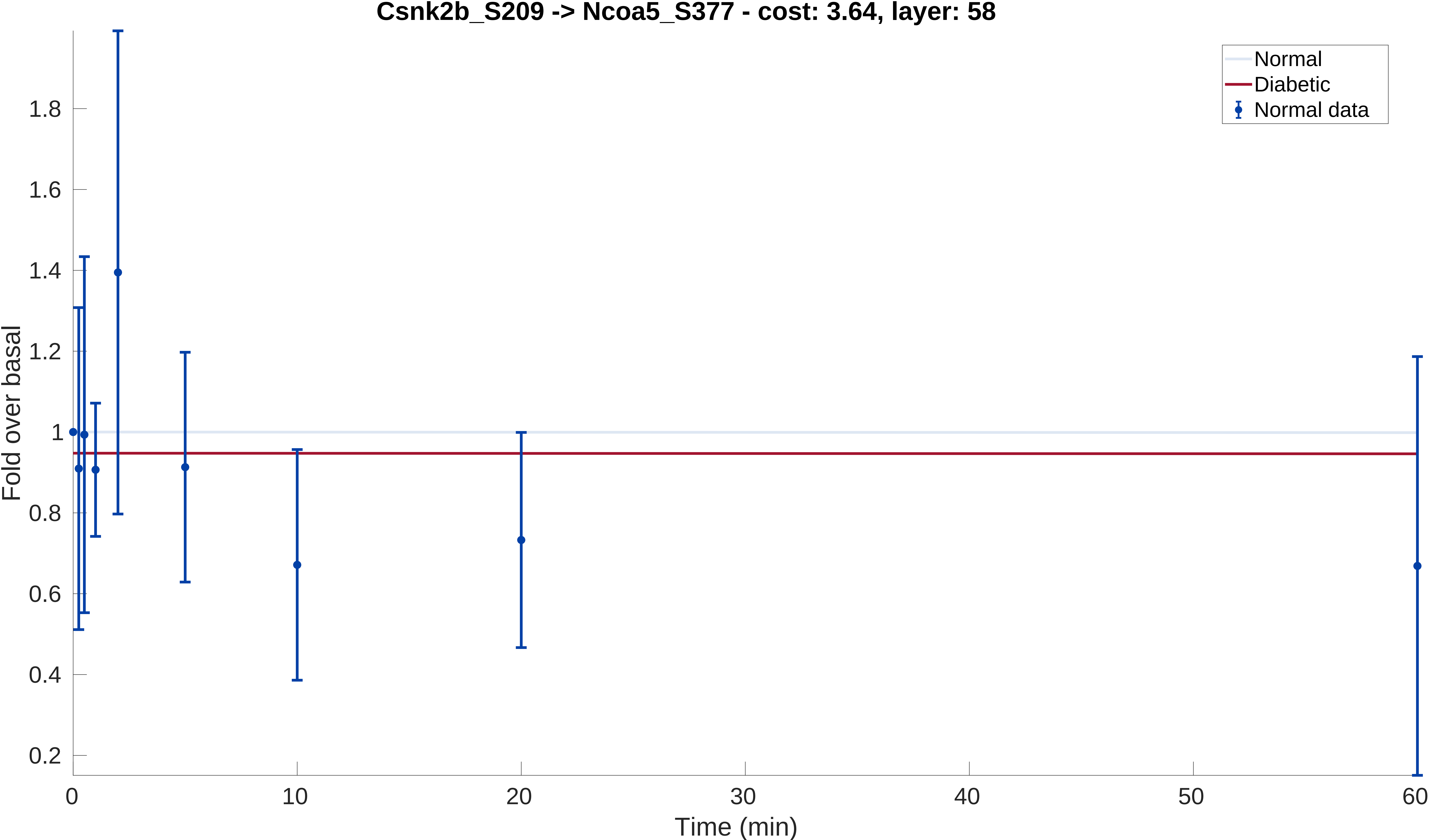

Csnk2b_S209 -> Ncoa5_S377 - cost: 3.64, layer: 58
Normal
Diabetic
Normal data
Fold over basal
1.8
1.6
1.4
1.2
1
0.8
0.6
0.4
0.2
0
10
20
30
40
50
60
Time (min)

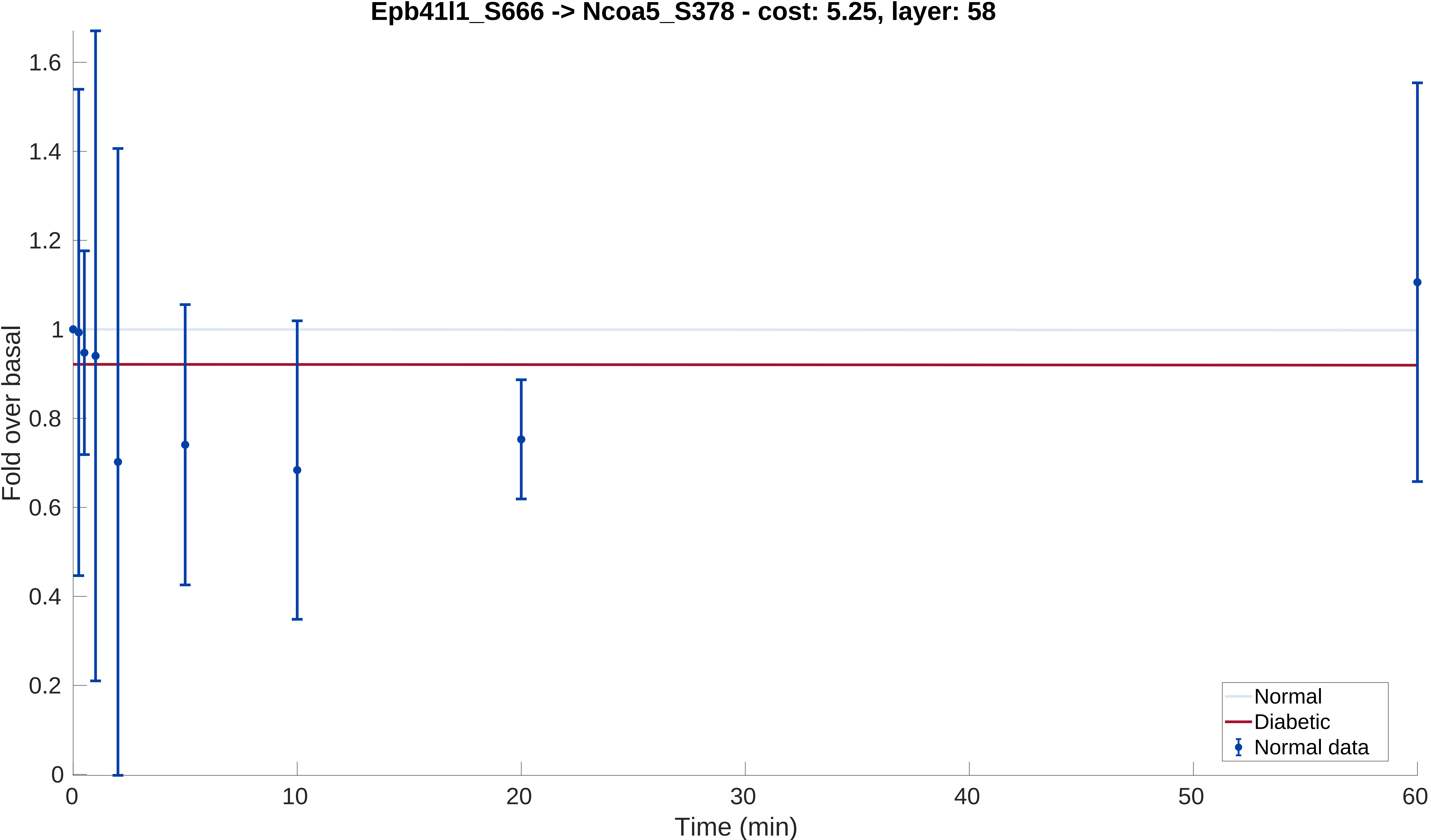

Epb41l1_S666 -> Ncoa5_S378 - cost: 5.25, layer: 58
Fold over basal
0
0.2
0.4
0.6
0.8
1
1.2
1.4
1.6
0
10
20
30
40
50
60
Time (min)
Normal
Diabetic
Normal data

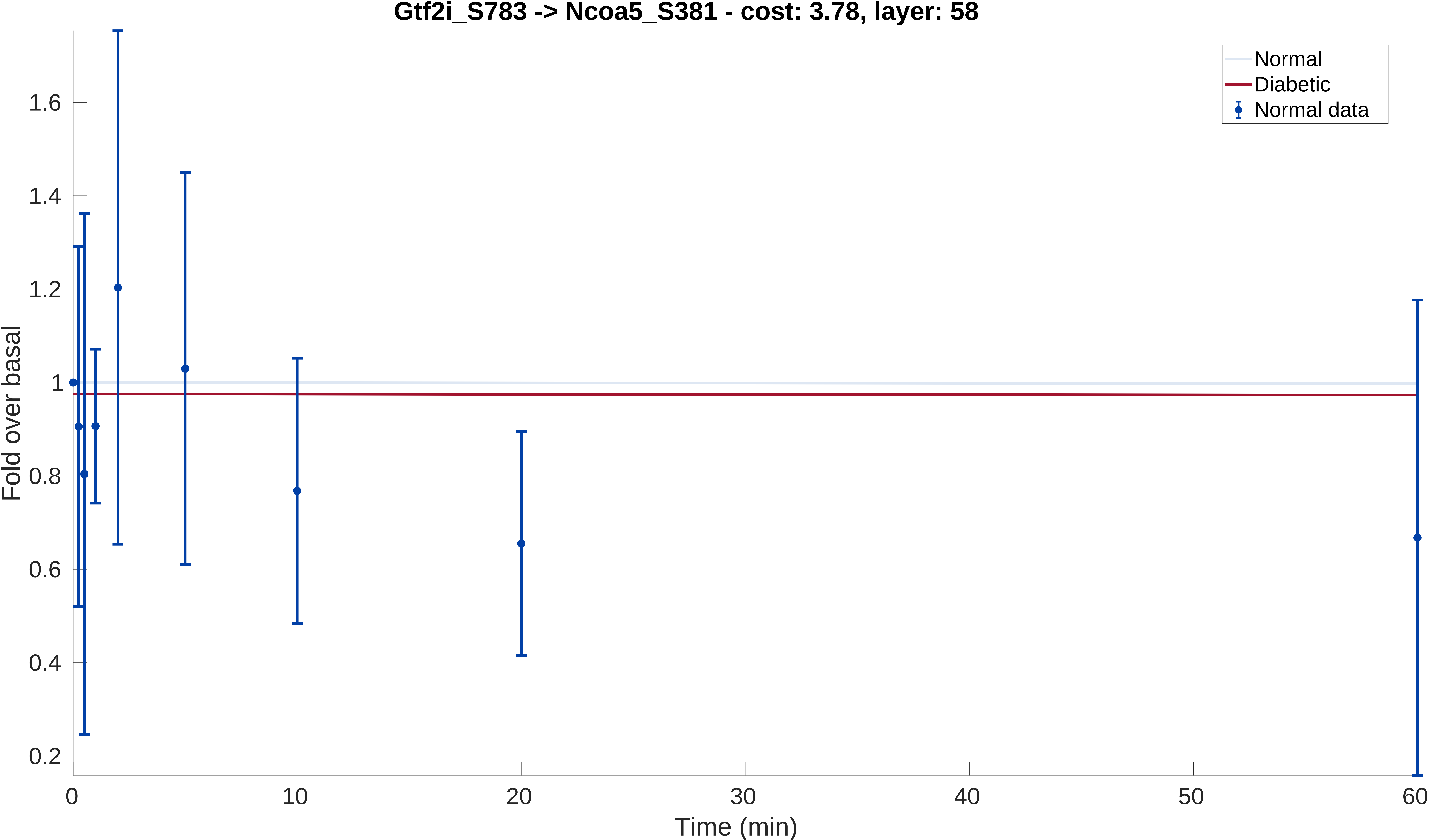

Gtf2i_S783 -> Ncoa5_S381 - cost: 3.78, layer: 58
Normal
Diabetic
Normal data
Fold over basal
1.6
1.4
1.2
1
0.8
0.6
0.4
0.2
0
10
20
30
40
50
60
Time (min)

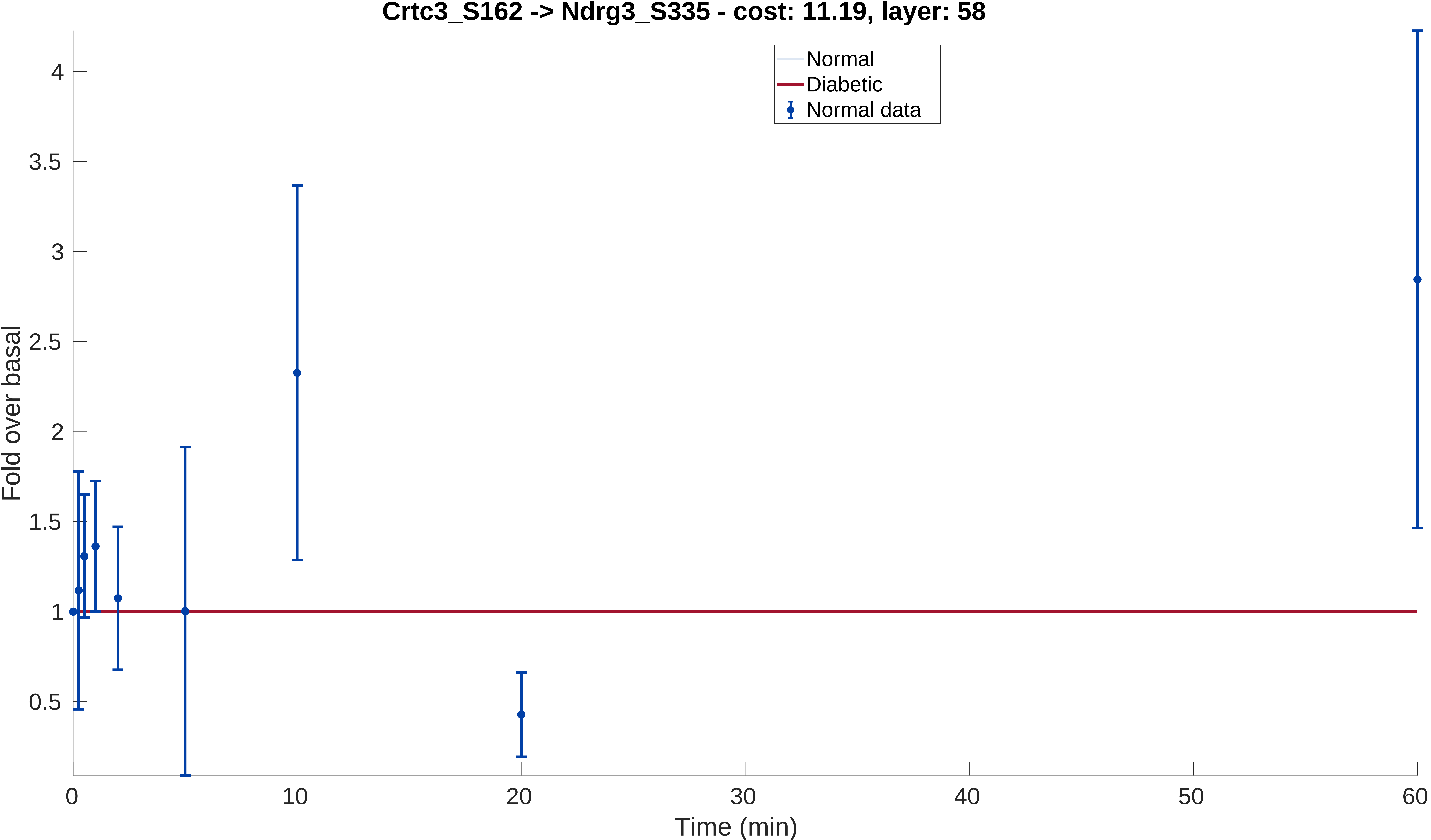
Crtc3_S162 -> Ndrg3_S335 - cost: 11.19, layer: 58
Normal
Diabetic
Normal data
Fold over basal
0.5
1
1.5
2
2.5
3
3.5
4
0
10
20
30
40
50
60
Time (min)

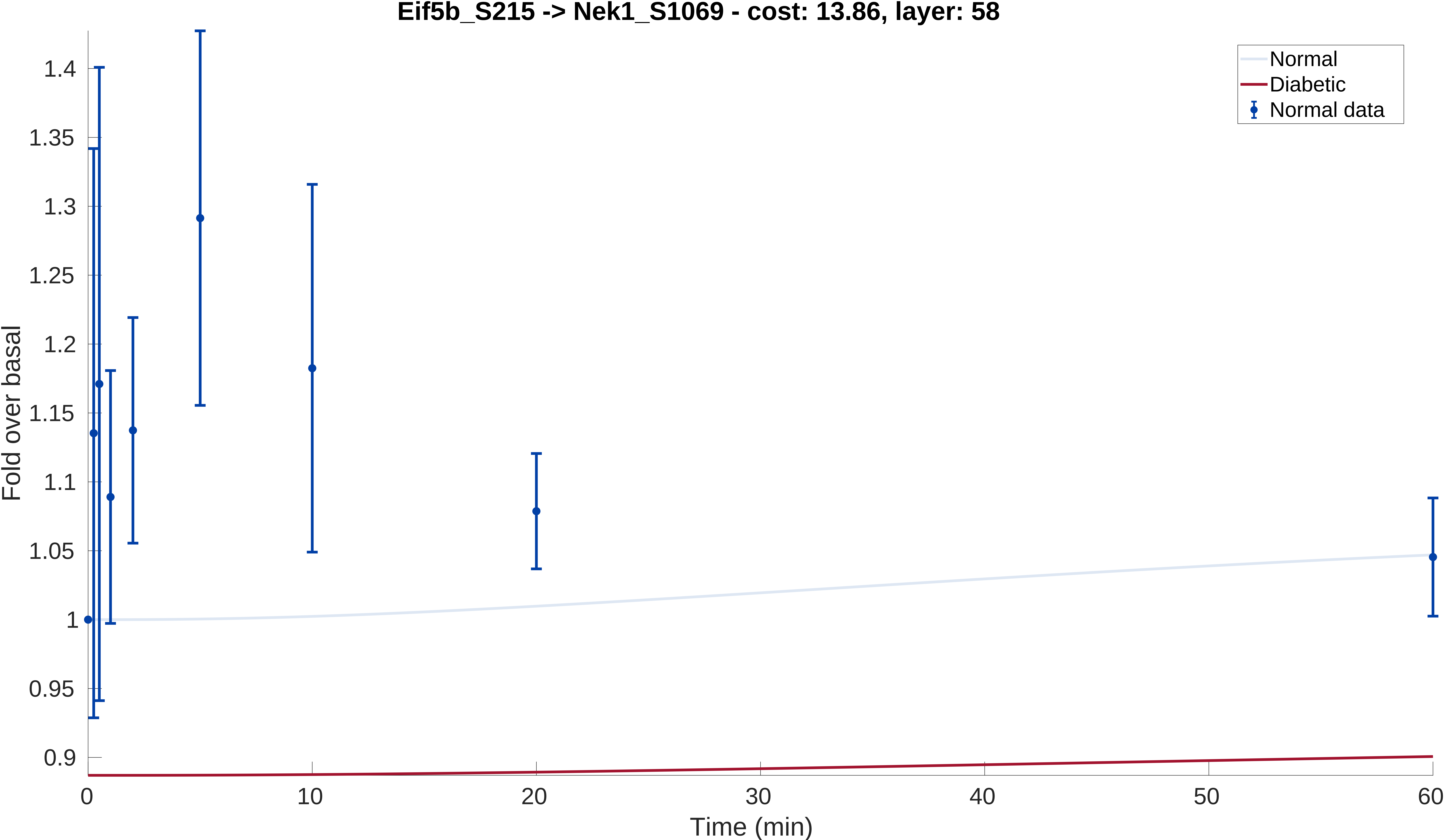

Eif5b_S215 -> Nek1_S1069 - cost: 13.86, layer: 58
Normal
Diabetic
Normal data
Fold over basal
Time (min)
1.4
1.35
1.3
1.25
1.2
1.15
1.1
1.05
1
0.95
0.9
0
10
20
30
40
50
60

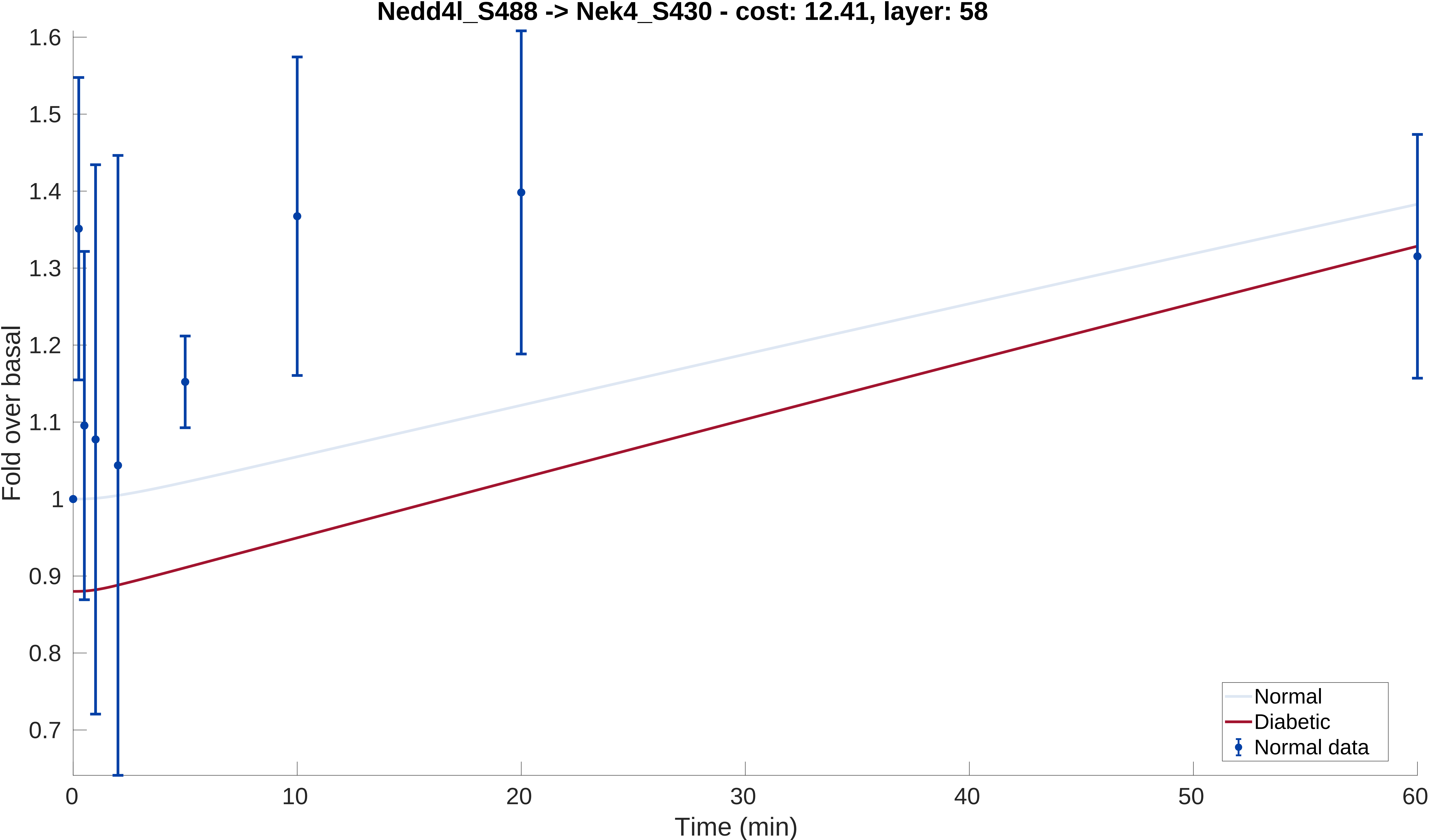

Nedd4l_S488 -> Nek4_S430 - cost: 12.41, layer: 58
1.6
1.5
1.4
1.3
1.2
1.1
1
0.9
0.8
0.7
Fold over basal
0
10
20
30
40
50
60
Time (min)
Normal
Diabetic
Normal data

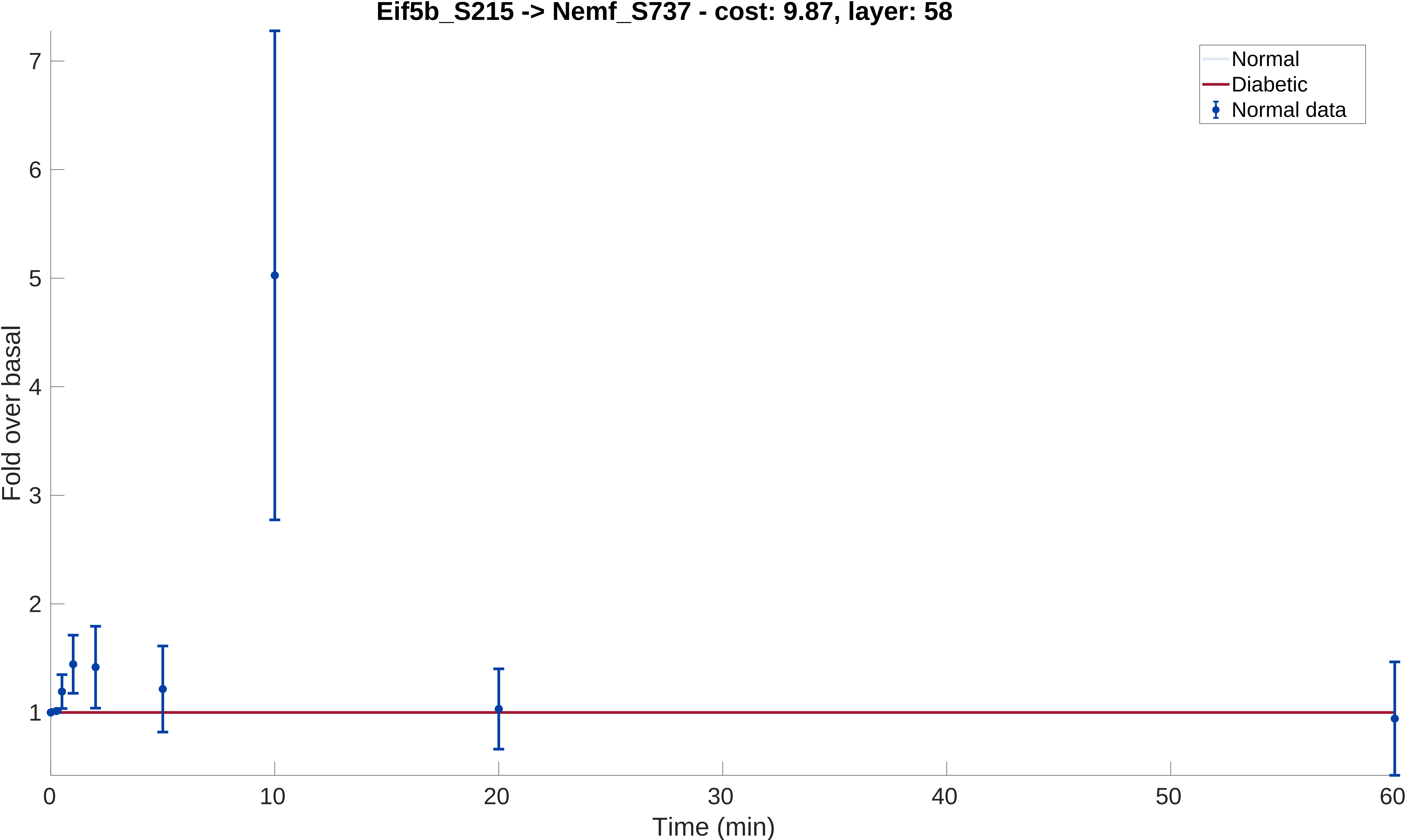
Eif5b_S215 -> Nemf_S737 - cost: 9.87, layer: 58
Fold over basal
1
2
3
4
5
6
7
0
10
20
30
40
50
60
Time (min)
Normal
Diabetic
Normal data

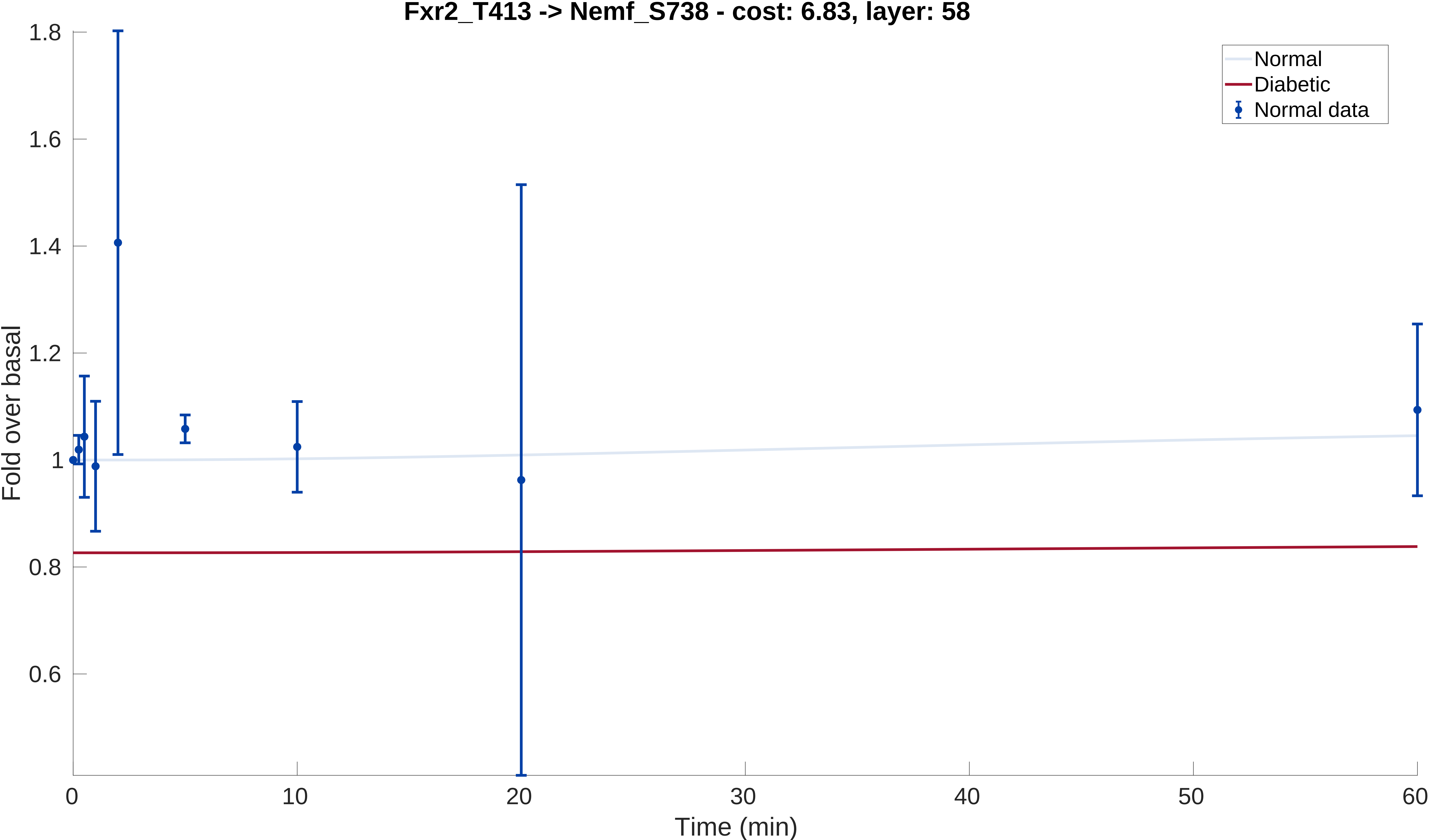

Fxr2_T413 -> Nemf_S738 - cost: 6.83, layer: 58
Normal
Diabetic
Normal data
Fold over basal
Time (min)
1.8
1.6
1.4
1.2
1
0.8
0.6
0
10
20
30
40
50
60

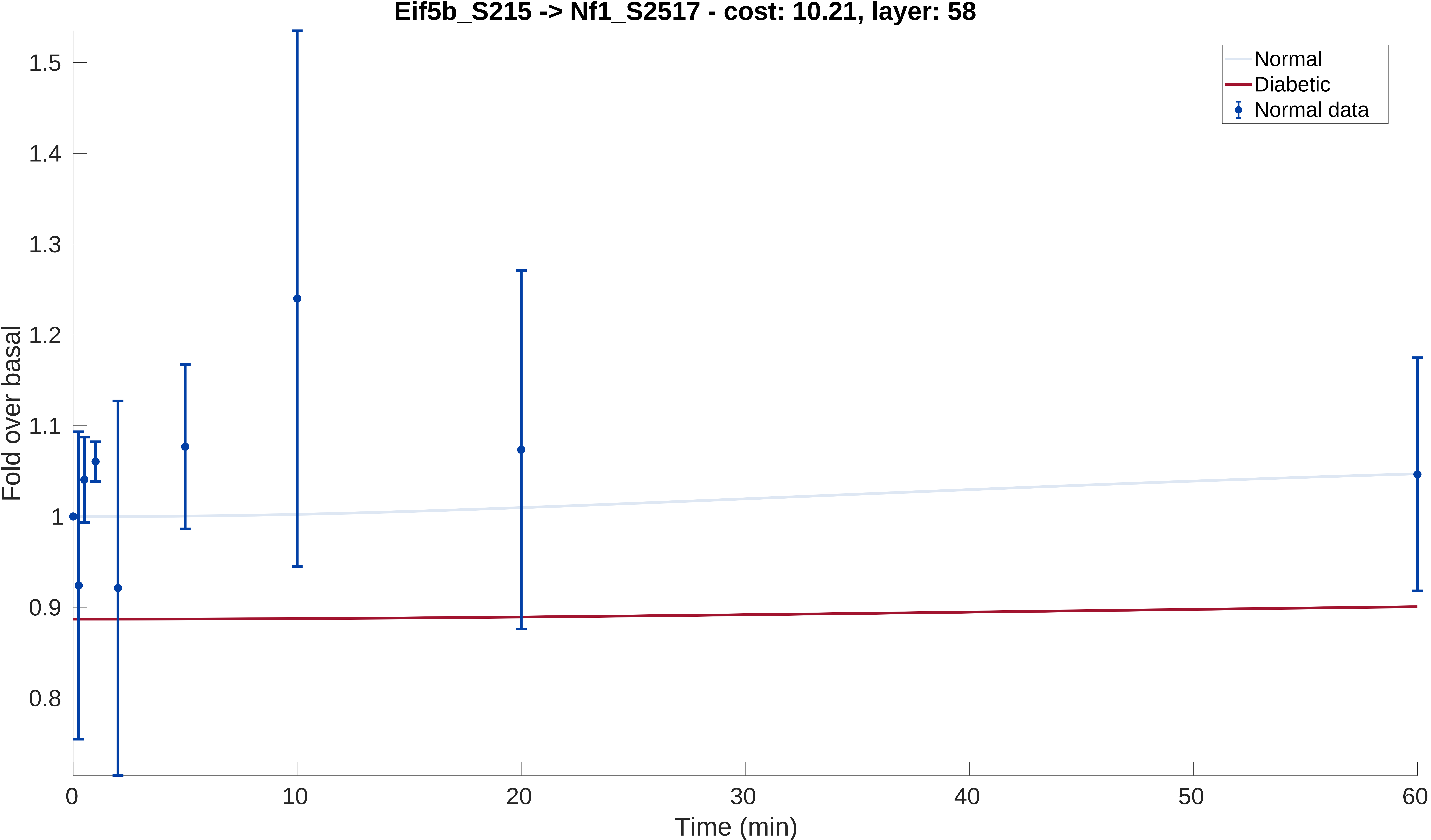
Eif5b_S215 -> Nf1_S2517 - cost: 10.21, layer: 58
Normal
Diabetic
Normal data
Fold over basal
1.5
1.4
1.3
1.2
1.1
1
0.9
0.8
0
10
20
30
40
50
60
Time (min)

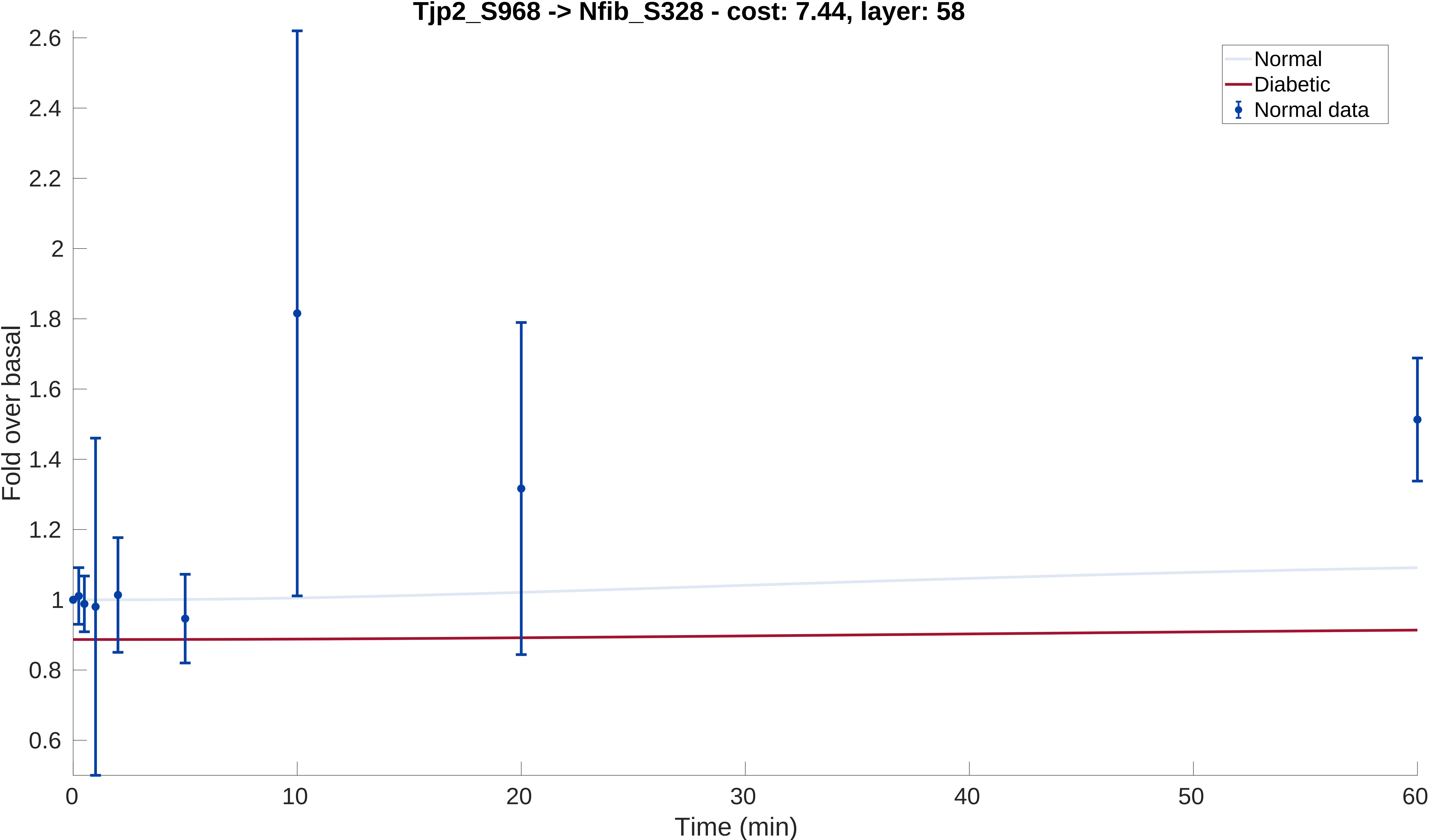
Tjp2_S968 -> Nfib_S328 - cost: 7.44, layer: 58
Fold over basal
Time (min)
Normal
Diabetic
Normal data
2.6
2.4
2.2
2
1.8
1.6
1.4
1.2
1
0.8
0.6
0
10
20
30
40
50
60

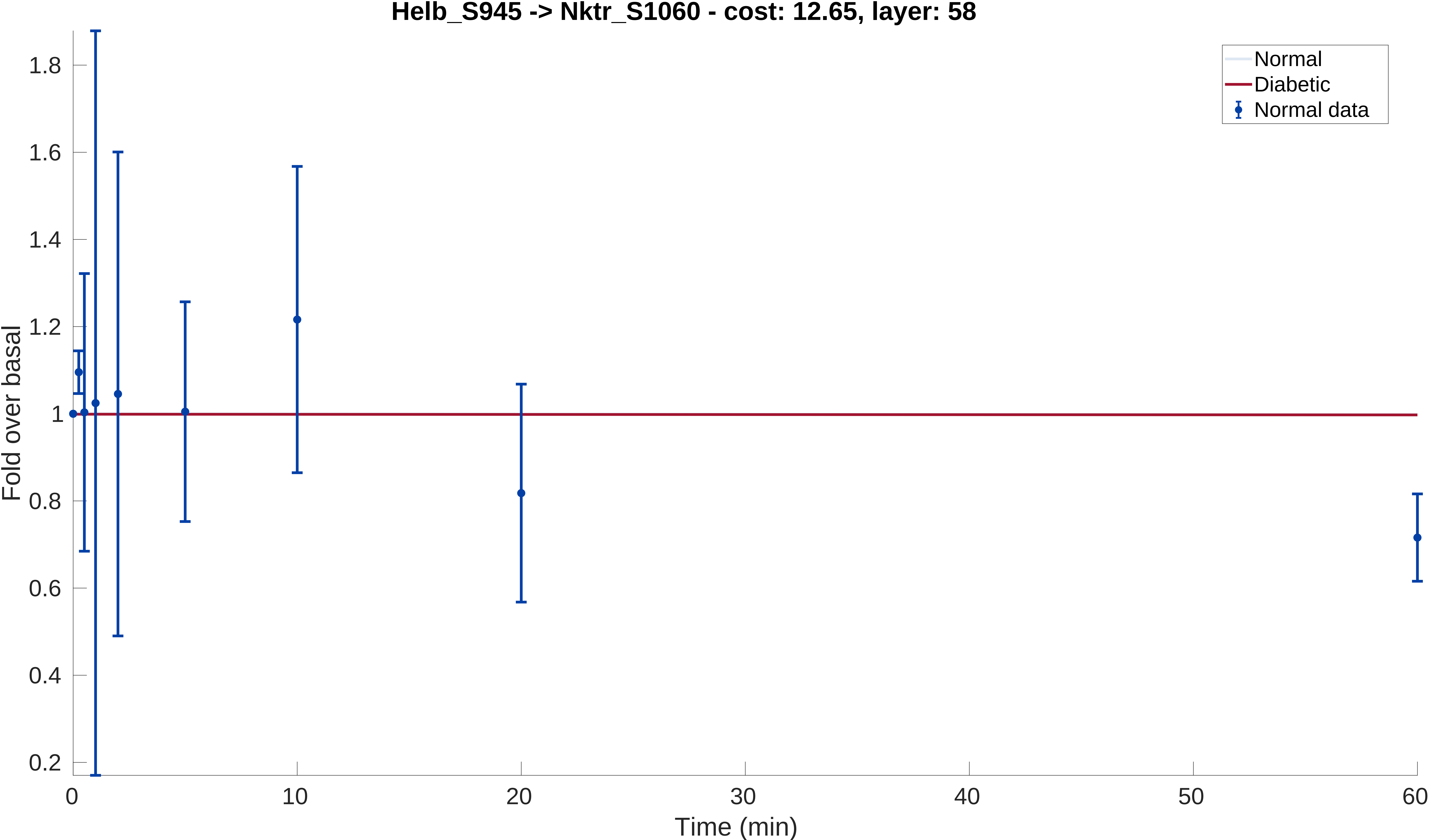

Helb_S945 -> Nktr_S1060 - cost: 12.65, layer: 58
Normal
Diabetic
Normal data
Fold over basal
0.2
0.4
0.6
0.8
1
1.2
1.4
1.6
1.8
0
10
20
30
40
50
60
Time (min)

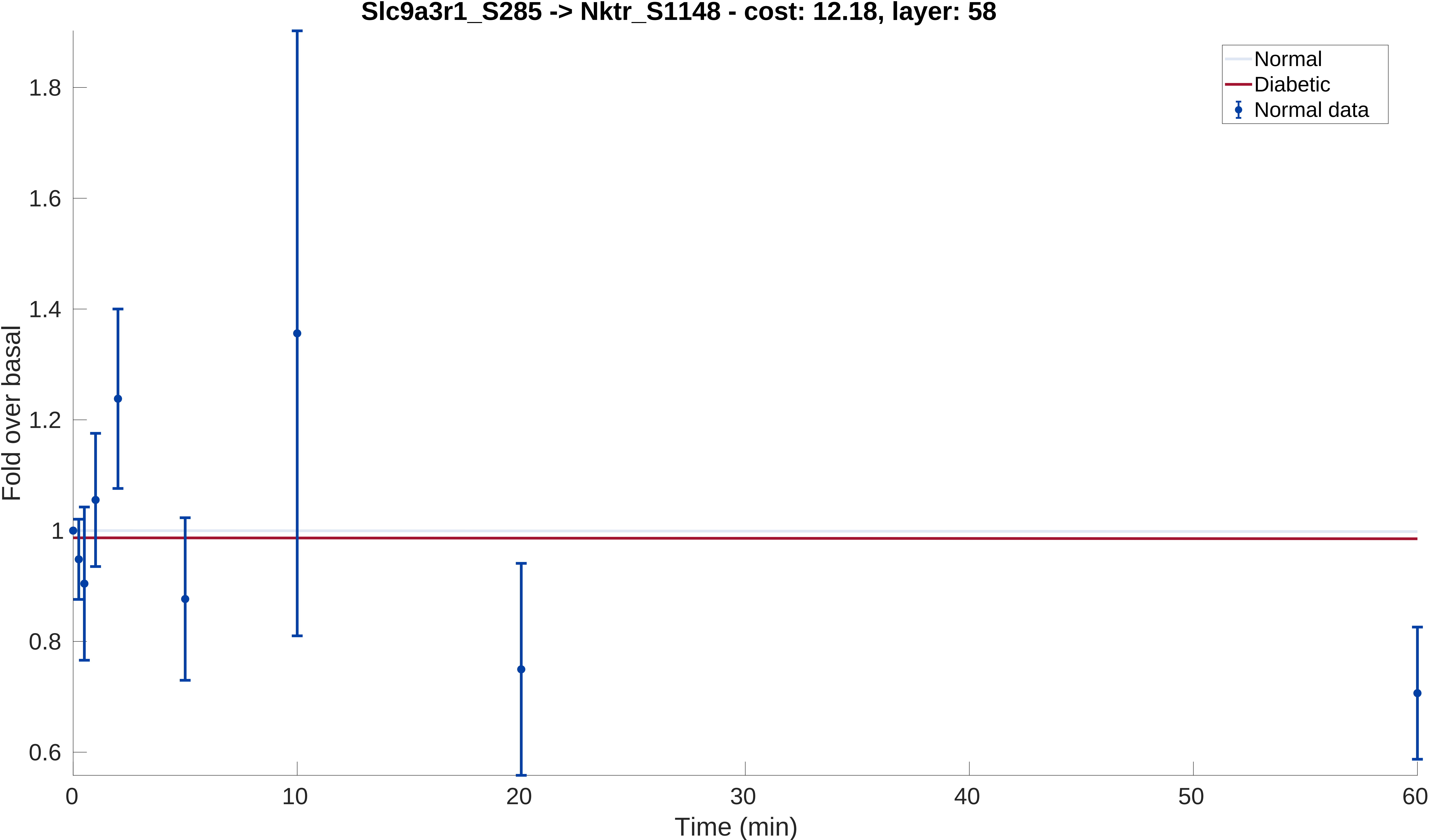

Slc9a3r1_S285 -> Nktr_S1148 - cost: 12.18, layer: 58
Normal
Diabetic
Normal data
Fold over basal
1.8
1.6
1.4
1.2
1
0.8
0.6
0
10
20
30
40
50
60
Time (min)

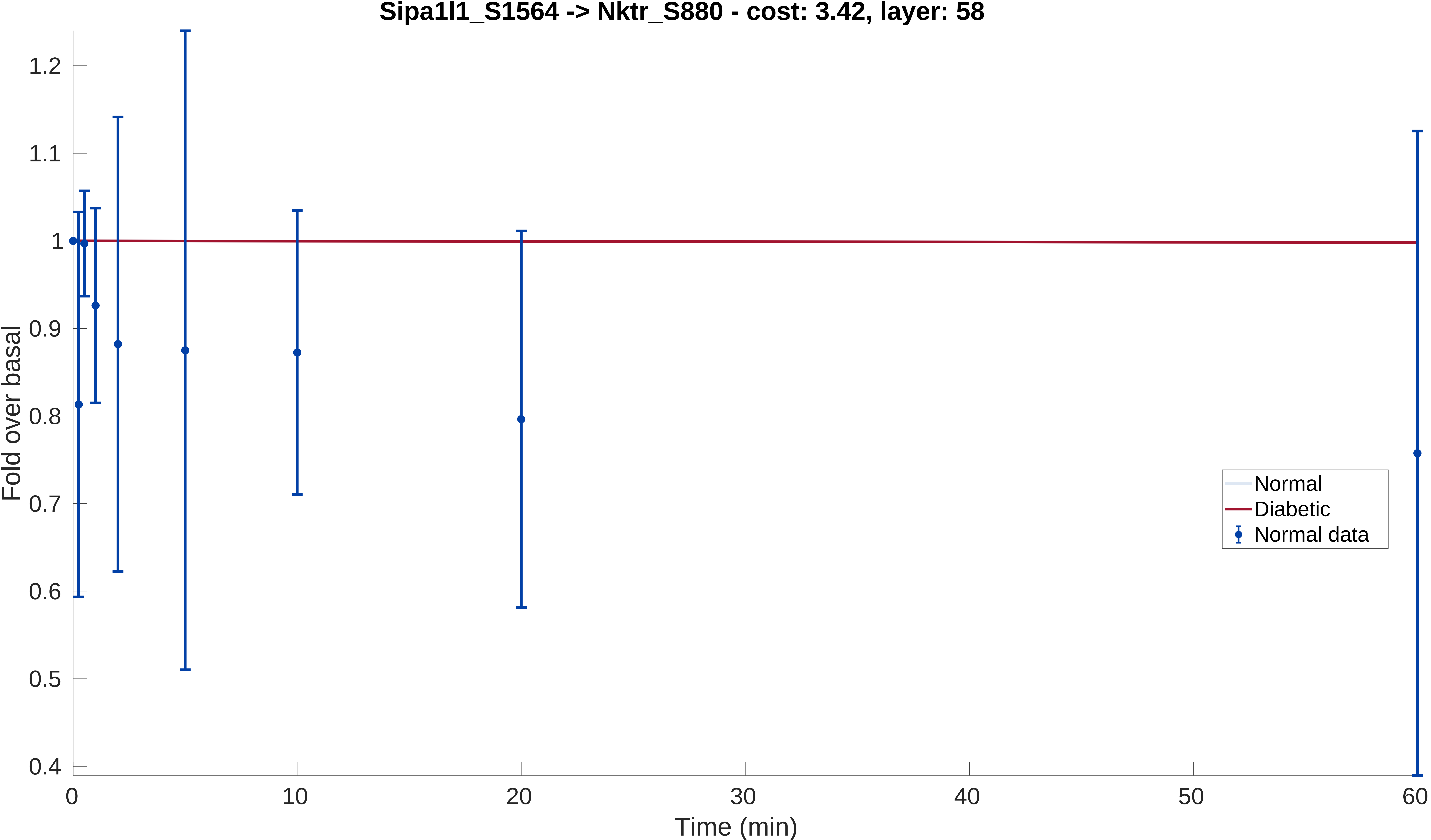

Sipa1l1_S1564 -> Nktr_S880 - cost: 3.42, layer: 58
Fold over basal
Time (min)
Normal
Diabetic
Normal data
0.4
0.5
0.6
0.7
0.8
0.9
1
1.1
1.2
0
10
20
30
40
50
60

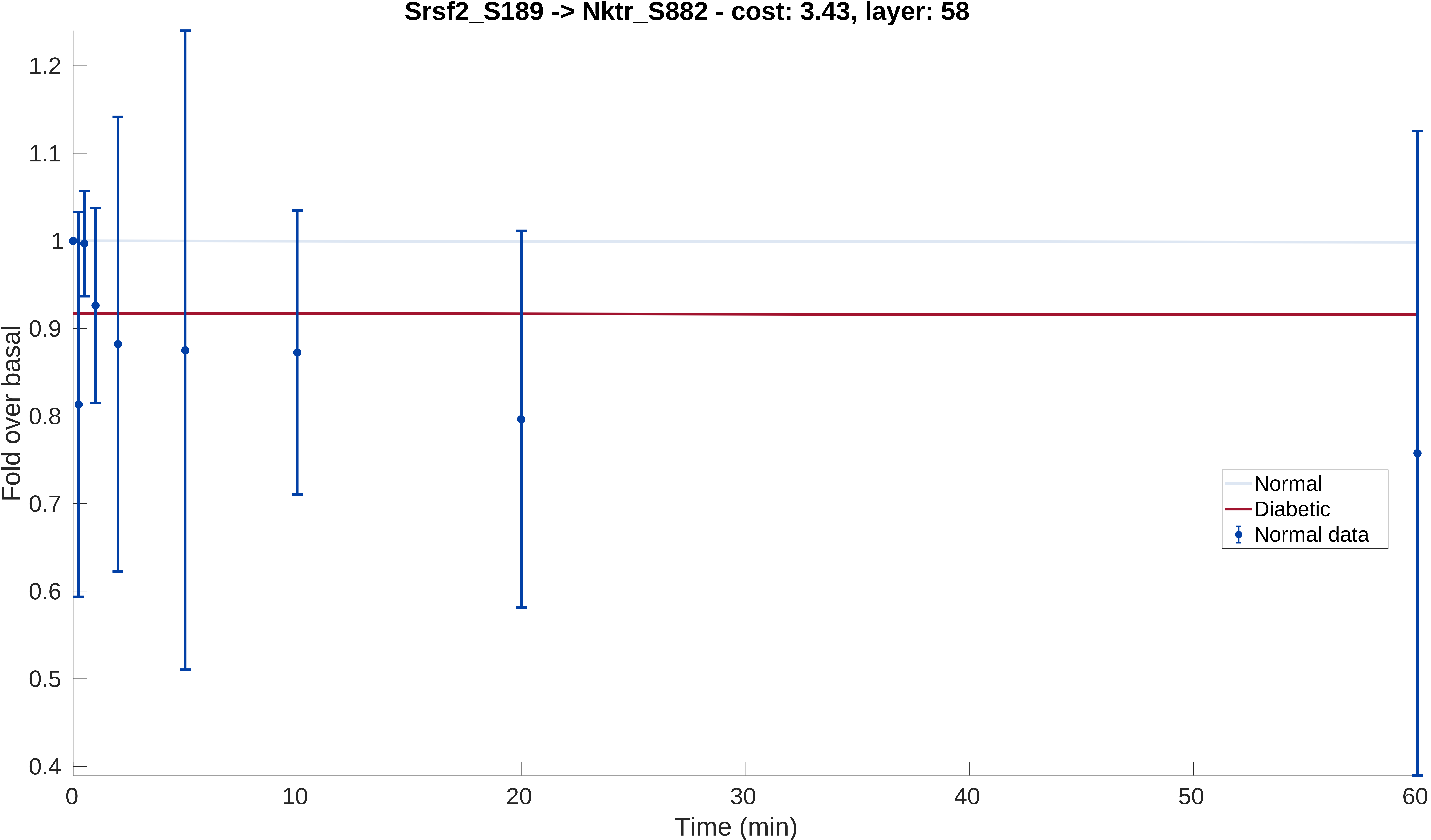
Srsf2_S189 -> Nktr_S882 - cost: 3.43, layer: 58
Fold over basal
Time (min)
1.2
1.1
1
0.9
0.8
0.7
0.6
0.5
0.4
0
10
20
30
40
50
60
Normal
Diabetic
Normal data

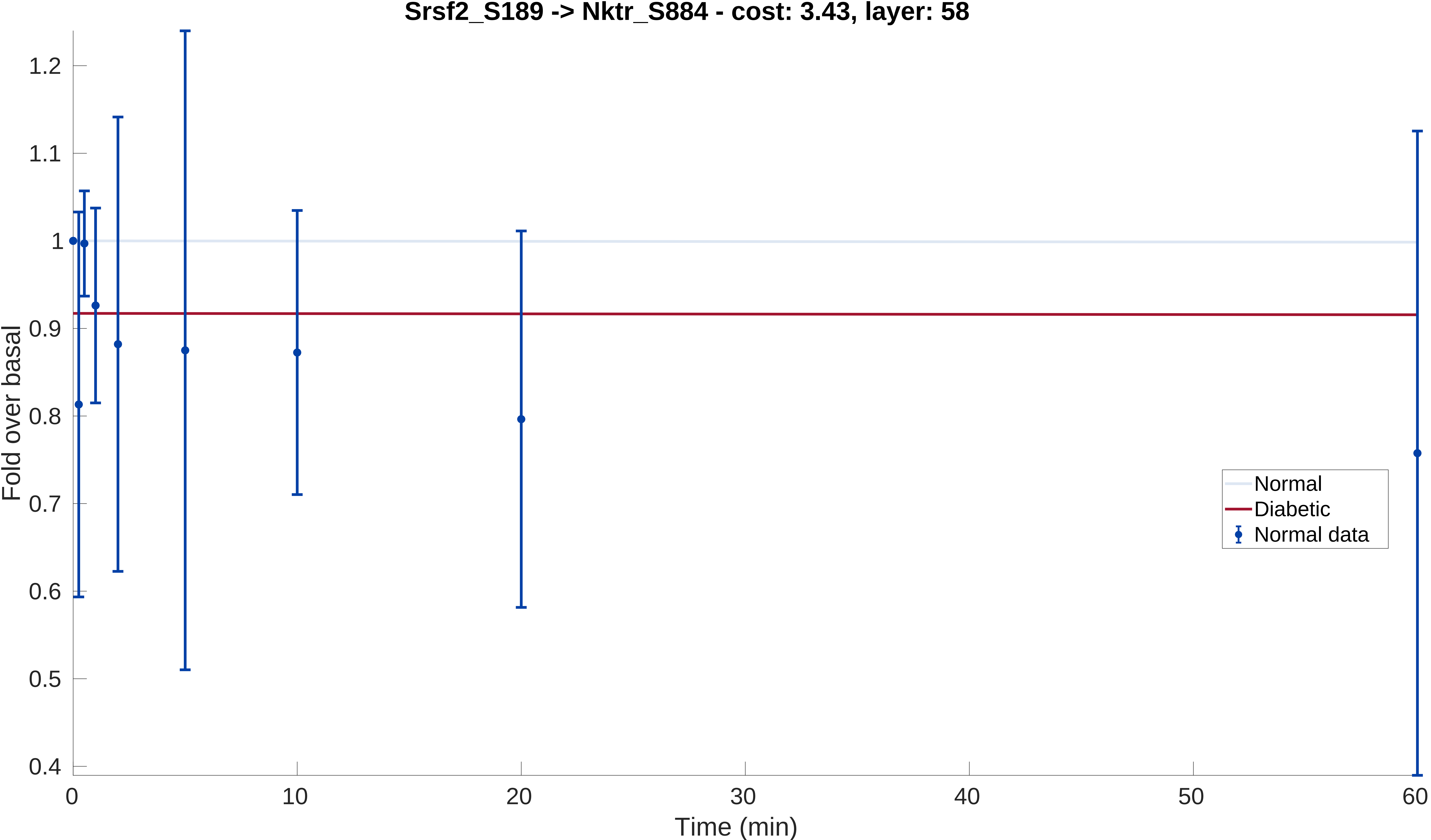
Srsf2_S189 -> Nktr_S884 - cost: 3.43, layer: 58
Fold over basal
1.2
1.1
1
0.9
0.8
0.7
0.6
0.5
0.4
0
10
20
30
40
50
60
Time (min)
Normal
Diabetic
Normal data

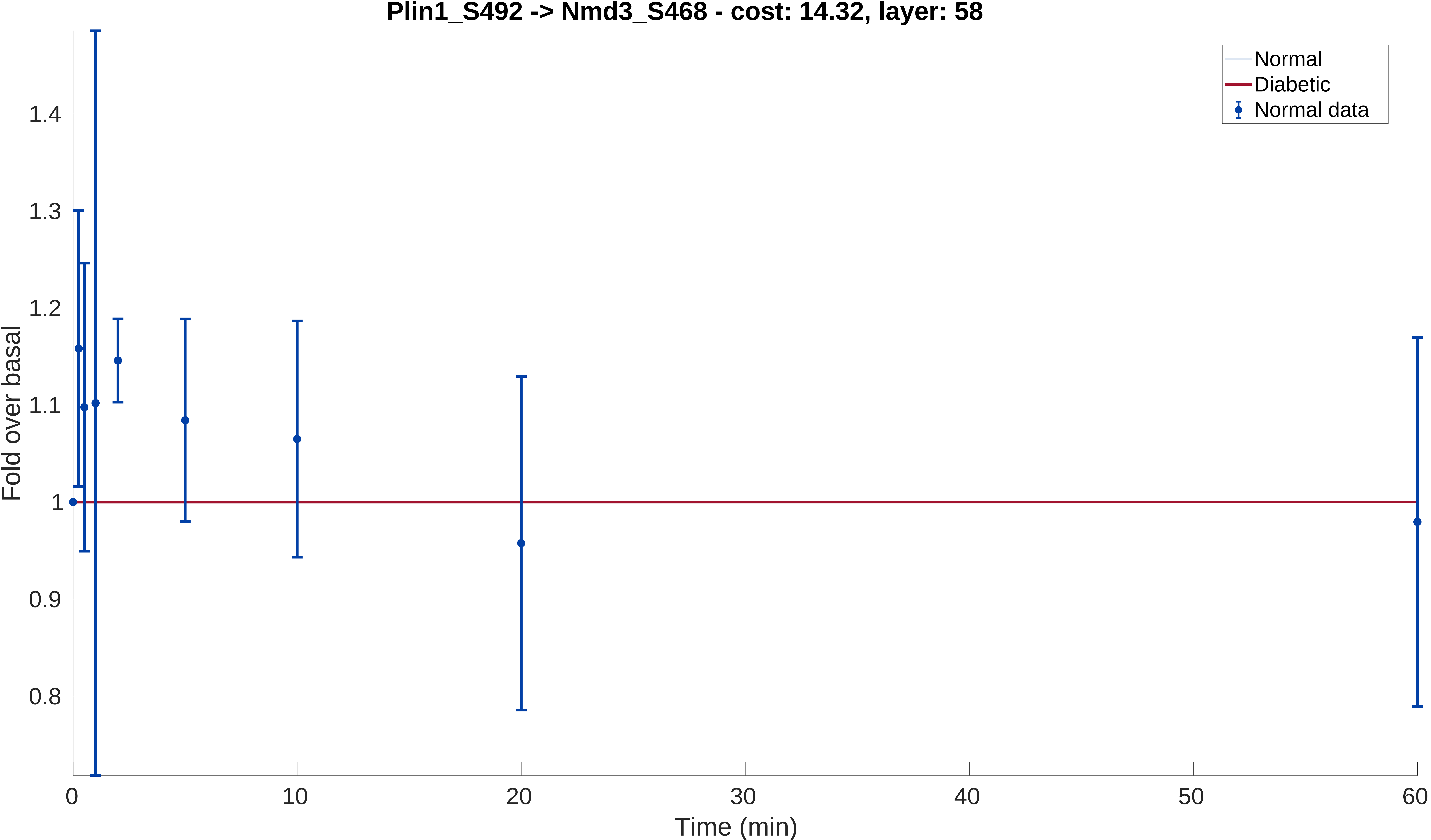

Plin1_S492 -> Nmd3_S468 - cost: 14.32, layer: 58
Normal
Diabetic
Normal data
Fold over basal
1.4
1.3
1.2
1.1
1
0.9
0.8
0
10
20
30
40
50
60
Time (min)

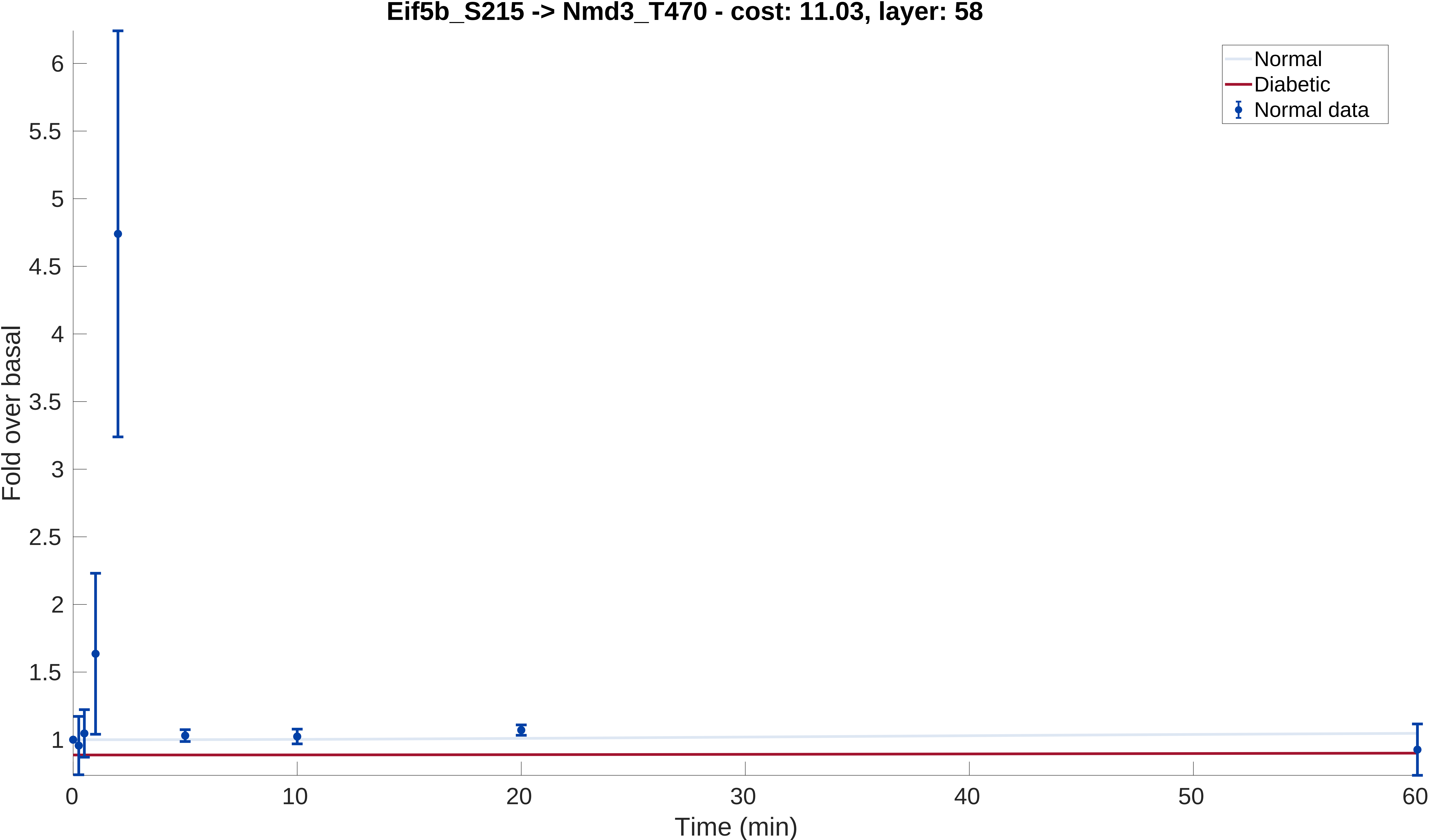
Eif5b_S215 -> Nmd3_T470 - cost: 11.03, layer: 58
Normal
Diabetic
Normal data
Fold over basal
6
5.5
5
4.5
4
3.5
3
2.5
2
1.5
1
0
10
20
30
40
50
60
Time (min)

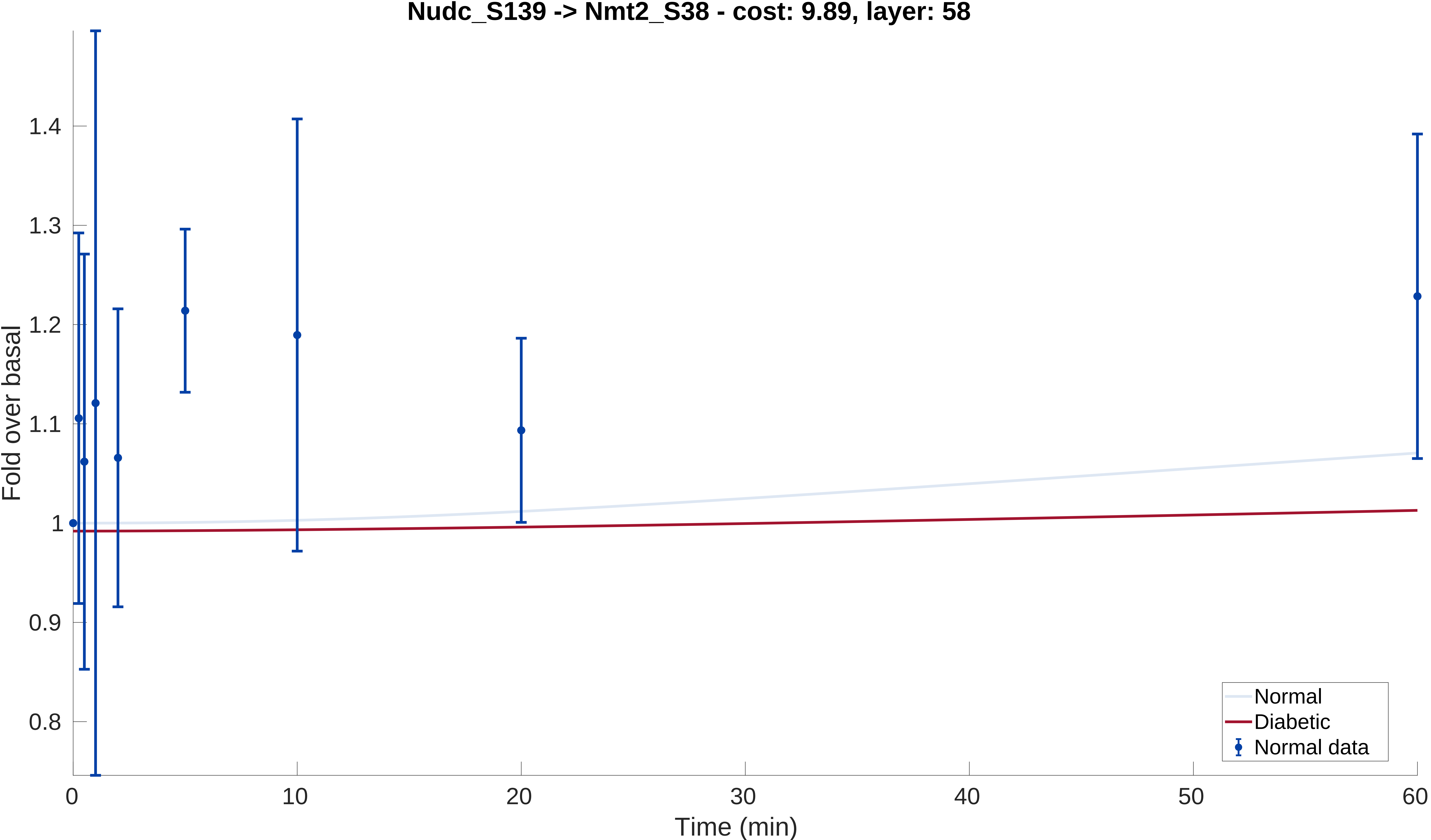

Nudc_S139 -> Nmt2_S38 - cost: 9.89, layer: 58
Fold over basal
Time (min)
0.8
0.9
1
1.1
1.2
1.3
1.4
0
10
20
30
40
50
60
Normal
Diabetic
Normal data

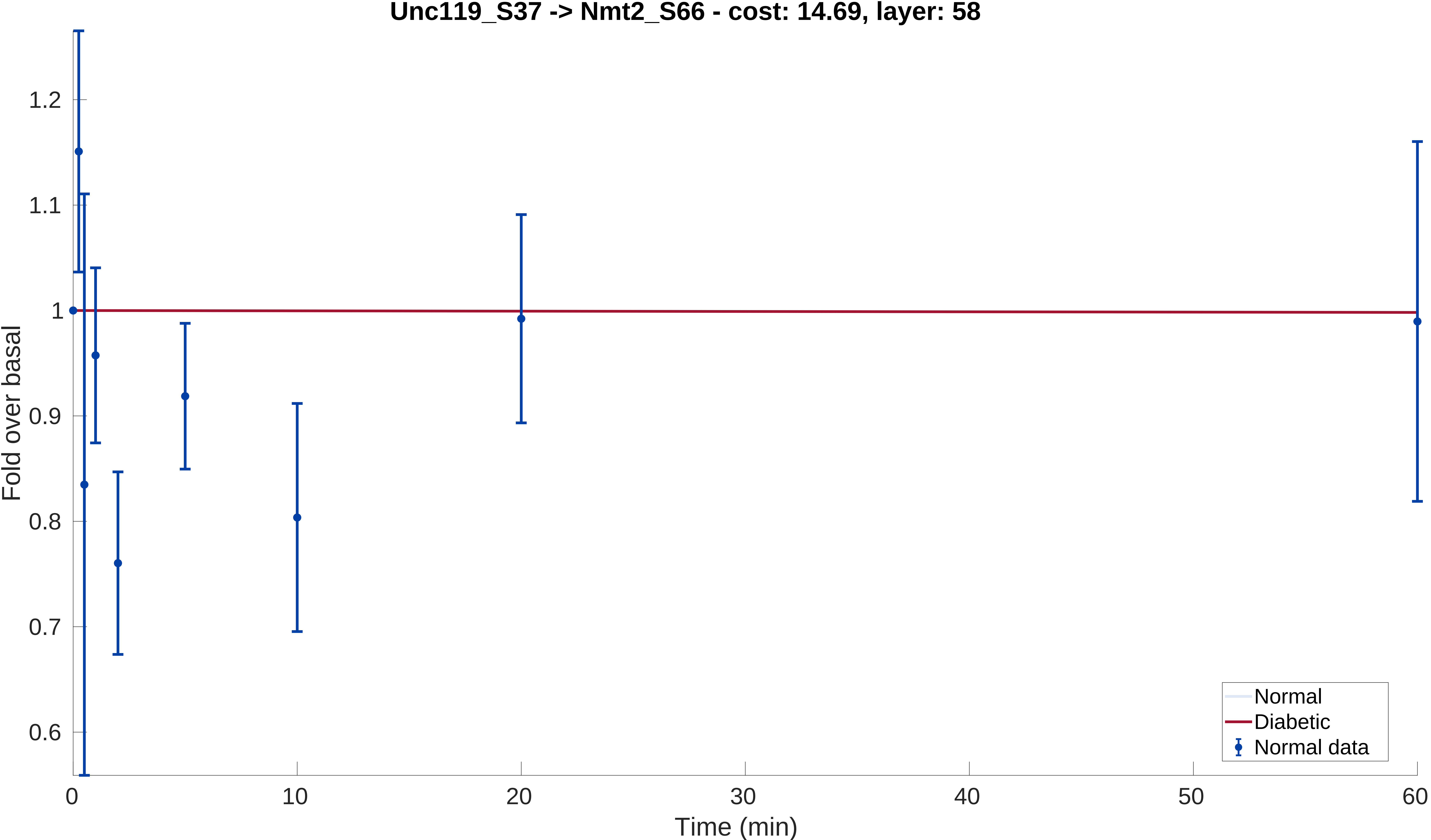

Unc119_S37 -> Nmt2_S66 - cost: 14.69, layer: 58
Fold over basal
1.2
1.1
1
0.9
0.8
0.7
0.6
0
10
20
30
40
50
60
Time (min)
Normal
Diabetic
Normal data

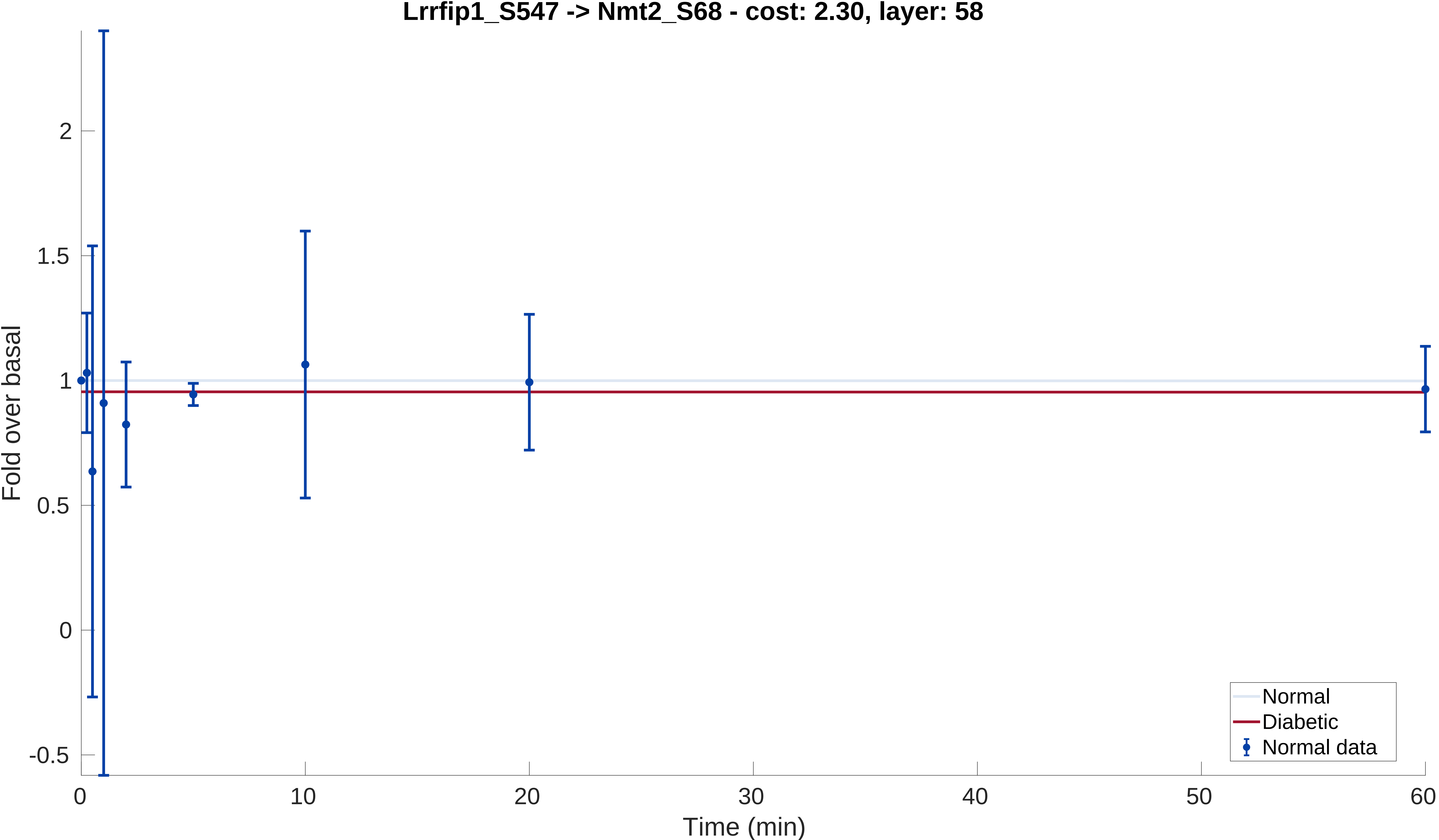

Lrrfip1_S547 -> Nmt2_S68 - cost: 2.30, layer: 58
Fold over basal
Time (min)
2
1.5
1
0.5
0
-0.5
0
10
20
30
40
50
60
Normal
Diabetic
Normal data

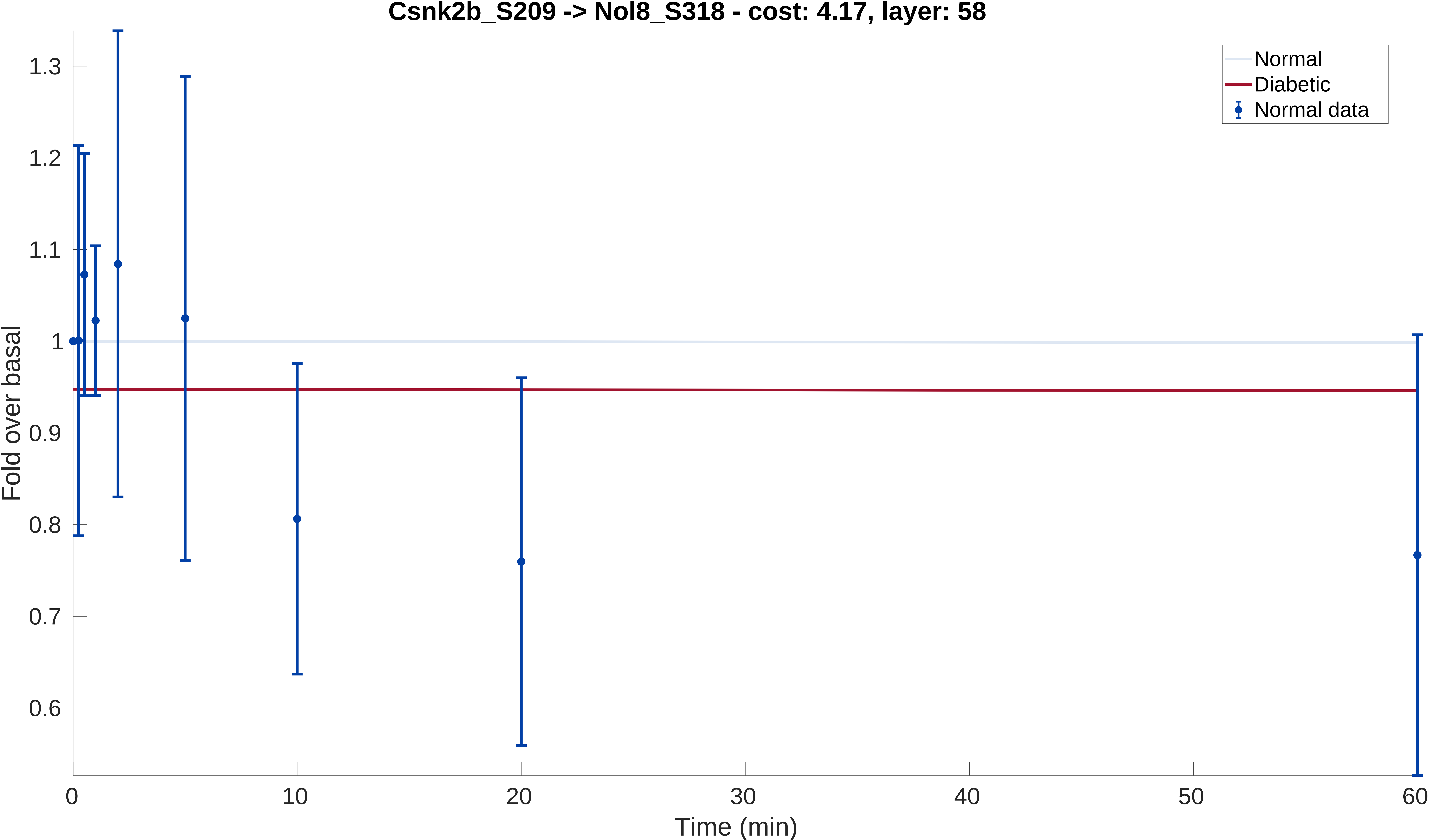

Csnk2b_S209 -> Nol8_S318 - cost: 4.17, layer: 58
Normal
Diabetic
Normal data
Fold over basal
1.3
1.2
1.1
1
0.9
0.8
0.7
0.6
0
10
20
30
40
50
60
Time (min)

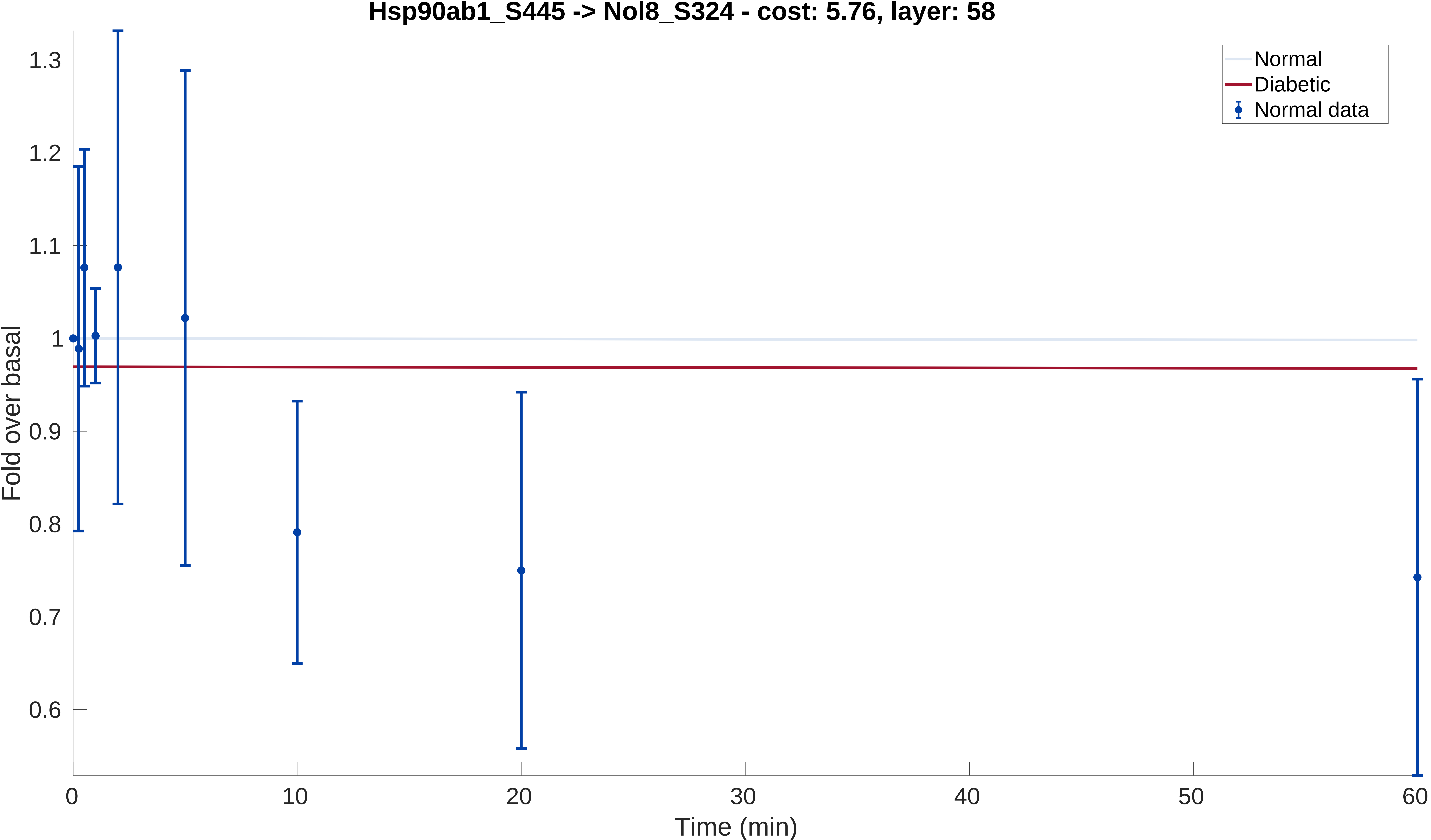

Hsp90ab1_S445 -> Nol8_S324 - cost: 5.76, layer: 58
Normal
Diabetic
Normal data
Fold over basal
Time (min)
0.6
0.7
0.8
0.9
1
1.1
1.2
1.3
0
10
20
30
40
50
60

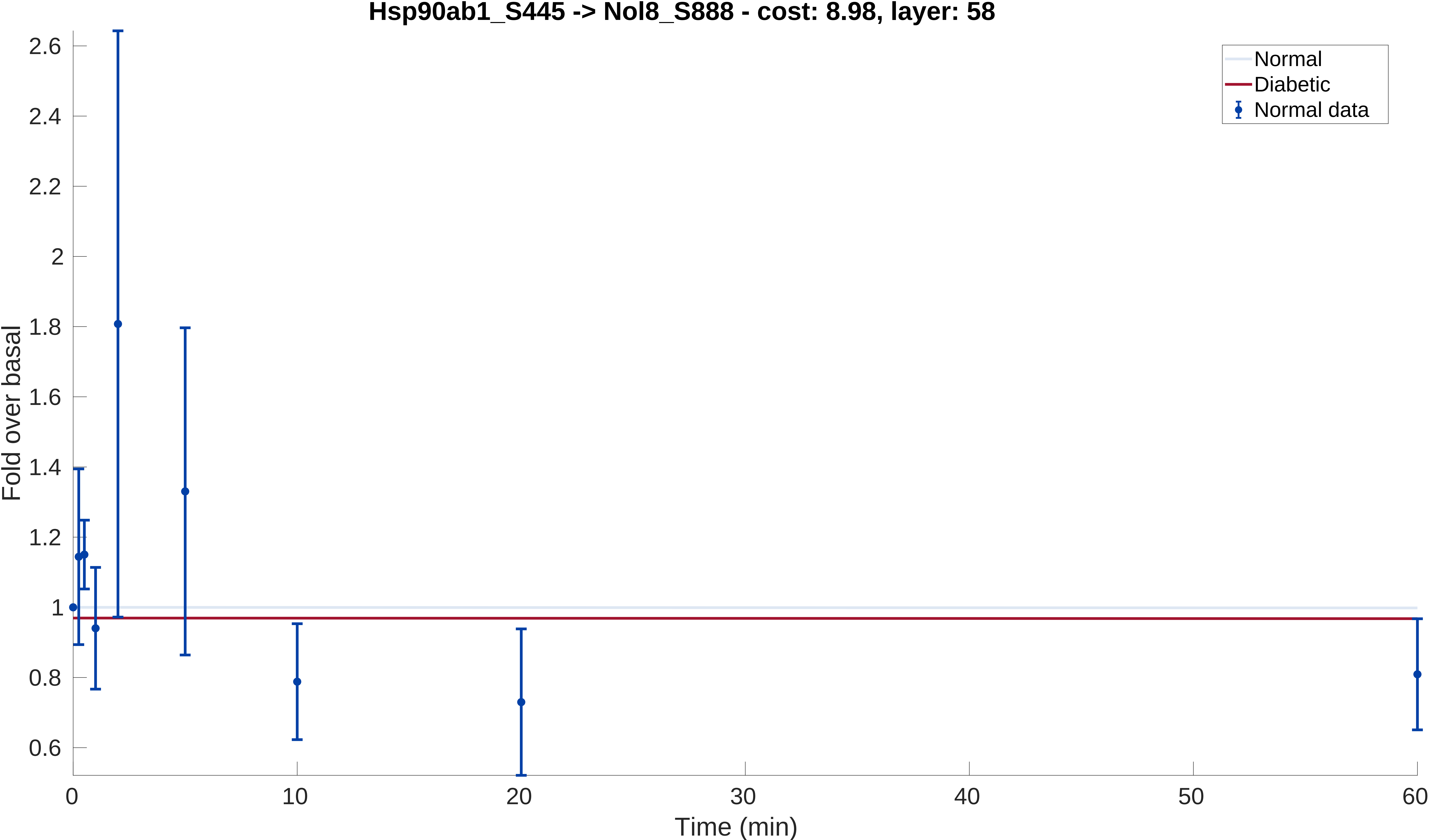
Hsp90ab1_S445 -> Nol8_S888 - cost: 8.98, layer: 58
Normal
Diabetic
Normal data
Fold over basal
2.6
2.4
2.2
2
1.8
1.6
1.4
1.2
1
0.8
0.6
0
10
20
30
40
50
60
Time (min)

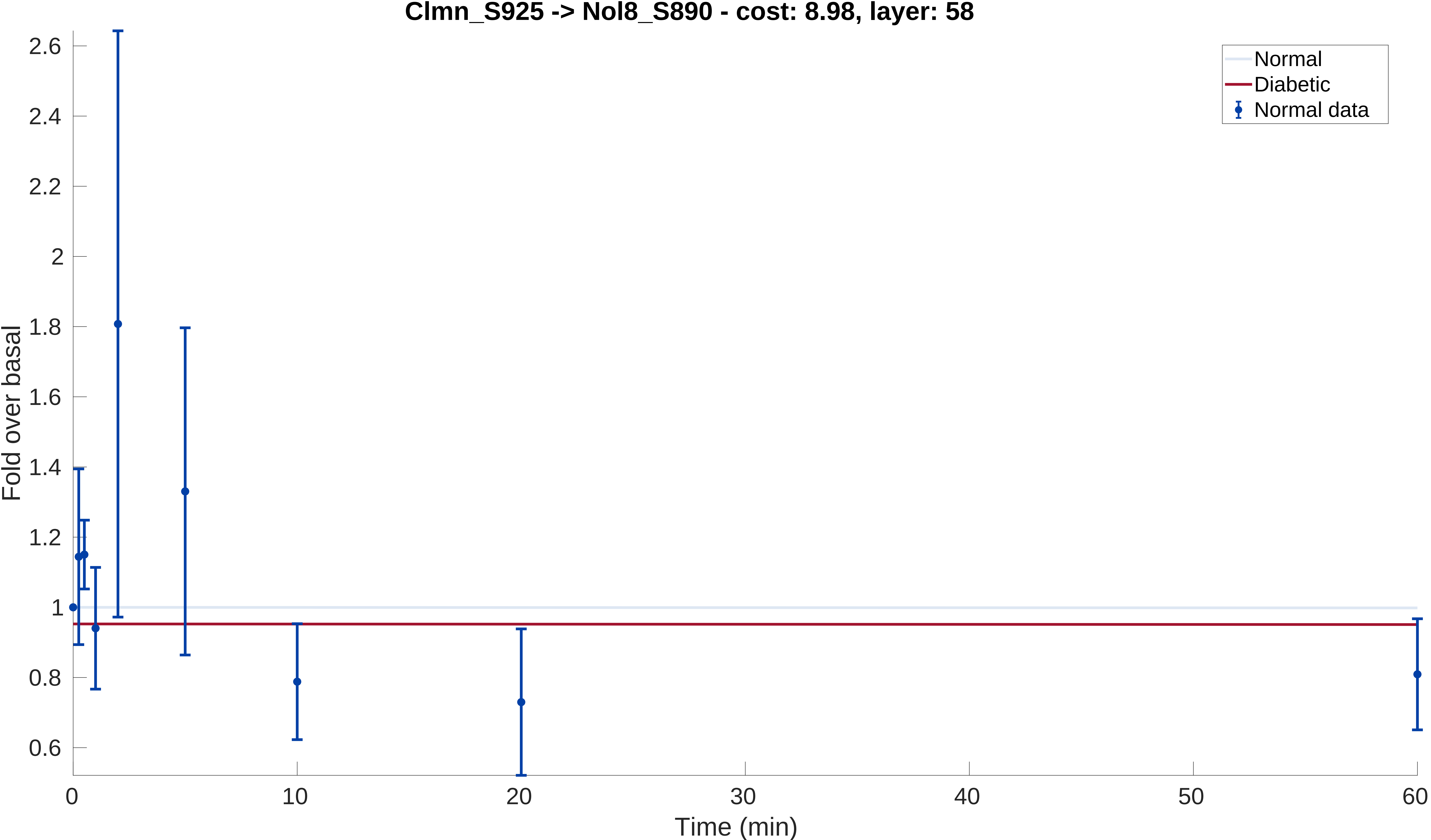

Clmn_S925 -> Nol8_S890 - cost: 8.98, layer: 58
Normal
Diabetic
Normal data
Fold over basal
2.6
2.4
2.2
2
1.8
1.6
1.4
1.2
1
0.8
0.6
0
10
20
30
40
50
60
Time (min)

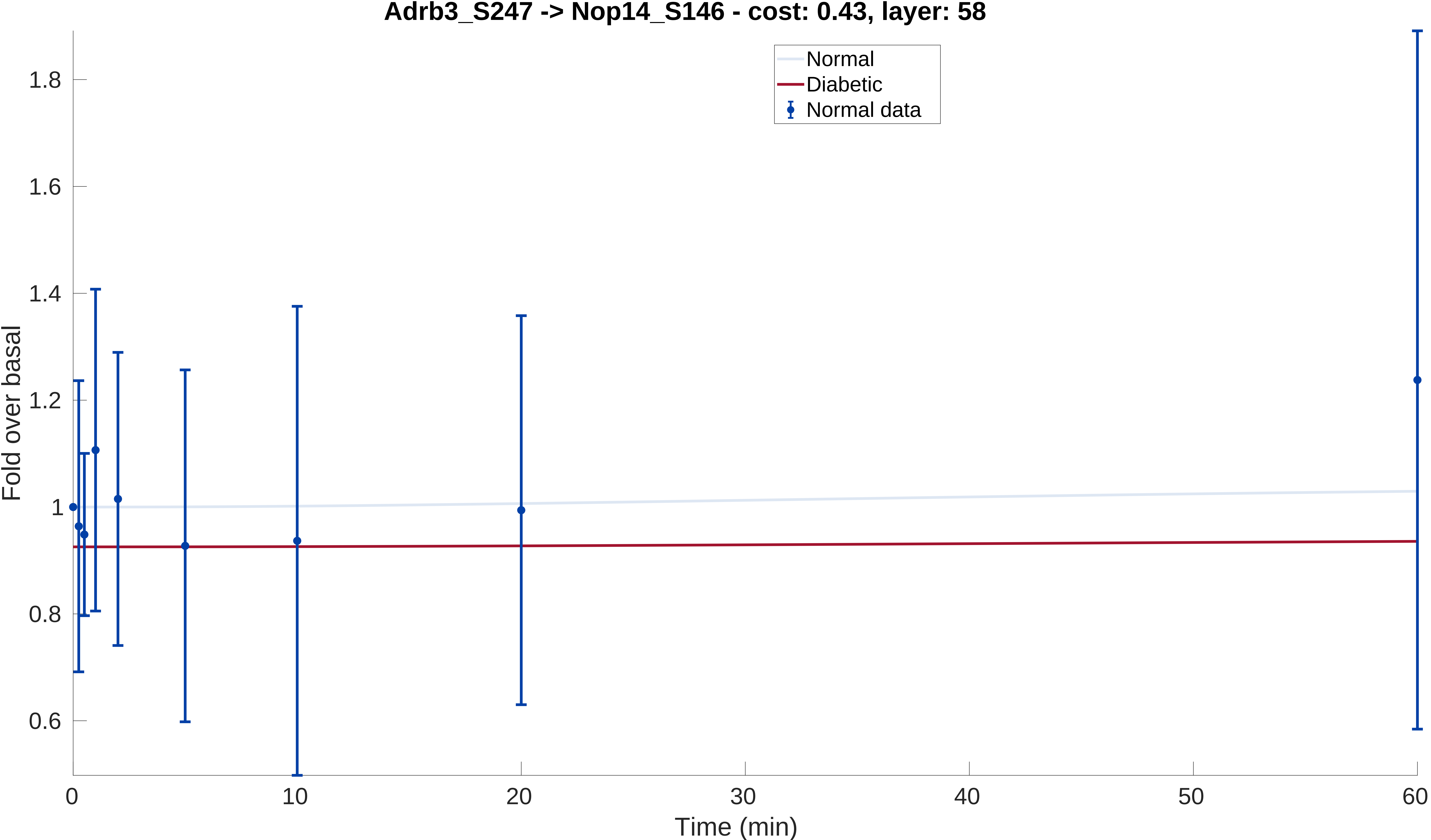

Adrb3_S247 -> Nop14_S146 - cost: 0.43, layer: 58
Normal
Diabetic
Normal data
Fold over basal
1.8
1.6
1.4
1.2
1
0.8
0.6
0
10
20
30
40
50
60
Time (min)

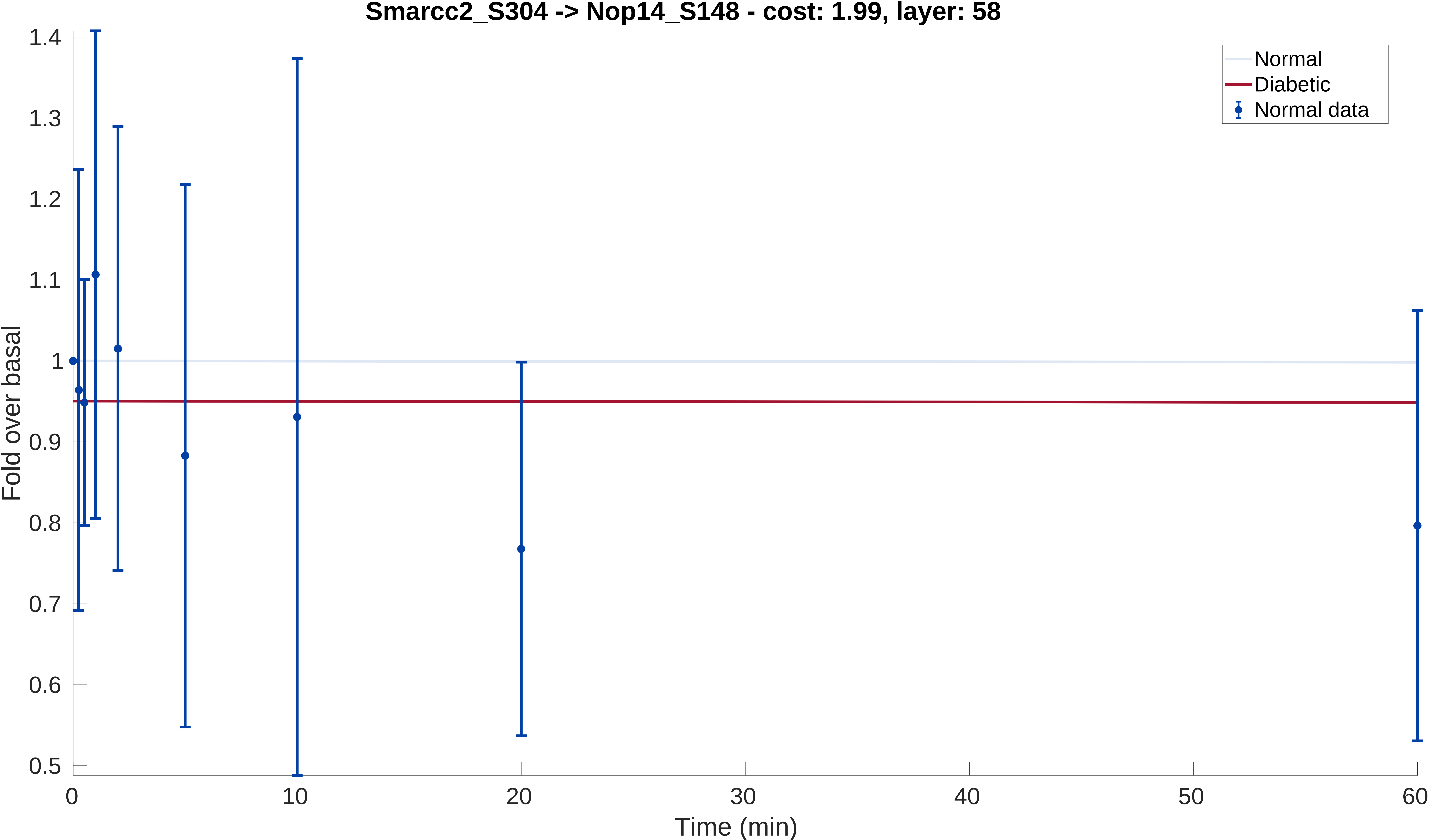
Smarcc2_S304 -> Nop14_S148 - cost: 1.99, layer: 58
Fold over basal
Time (min)
Normal
Diabetic
Normal data
1.4
1.3
1.2
1.1
1
0.9
0.8
0.7
0.6
0.5
0
10
20
30
40
50
60

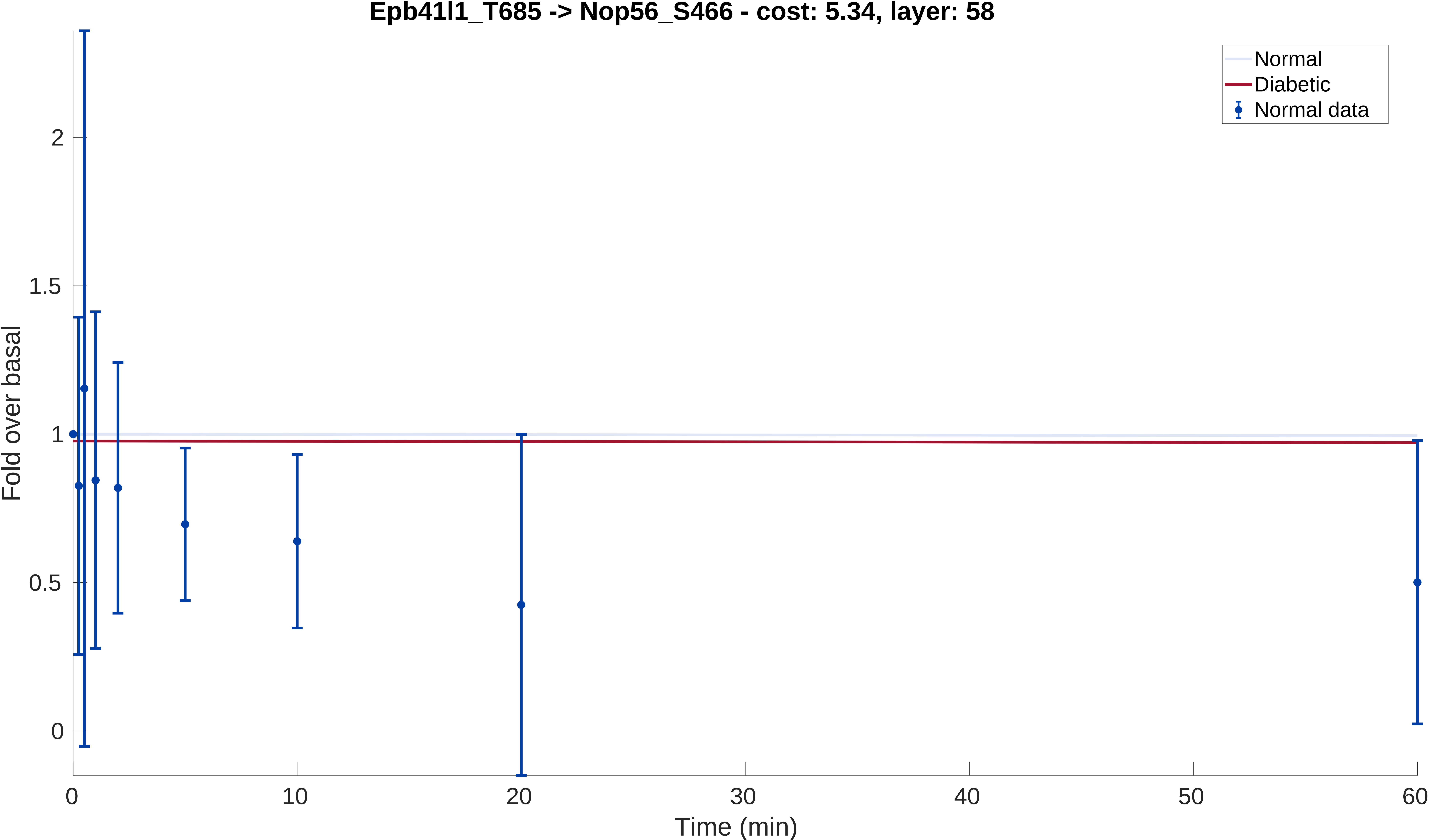

Epb41l1_T685 -> Nop56_S466 - cost: 5.34, layer: 58
Normal
Diabetic
Normal data
Fold over basal
2
1.5
1
0.5
0
0
10
20
30
40
50
60
Time (min)

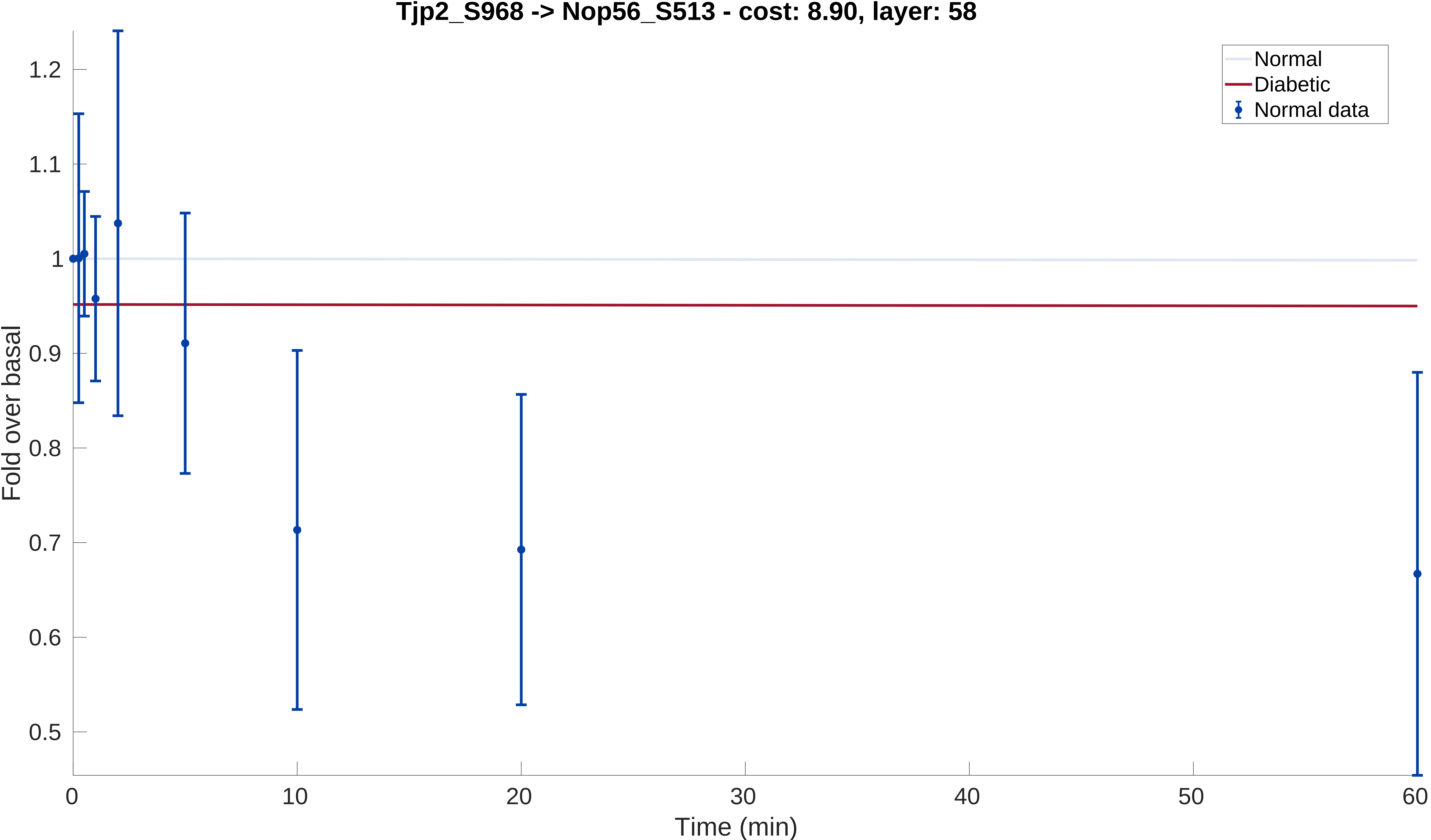

Tjp2_S968 -> Nop56_S513 - cost: 8.90, layer: 58
Normal
Diabetic
Normal data
Fold over basal
1.2
1.1
1
0.9
0.8
0.7
0.6
0.5
0
10
20
30
40
50
60
Time (min)

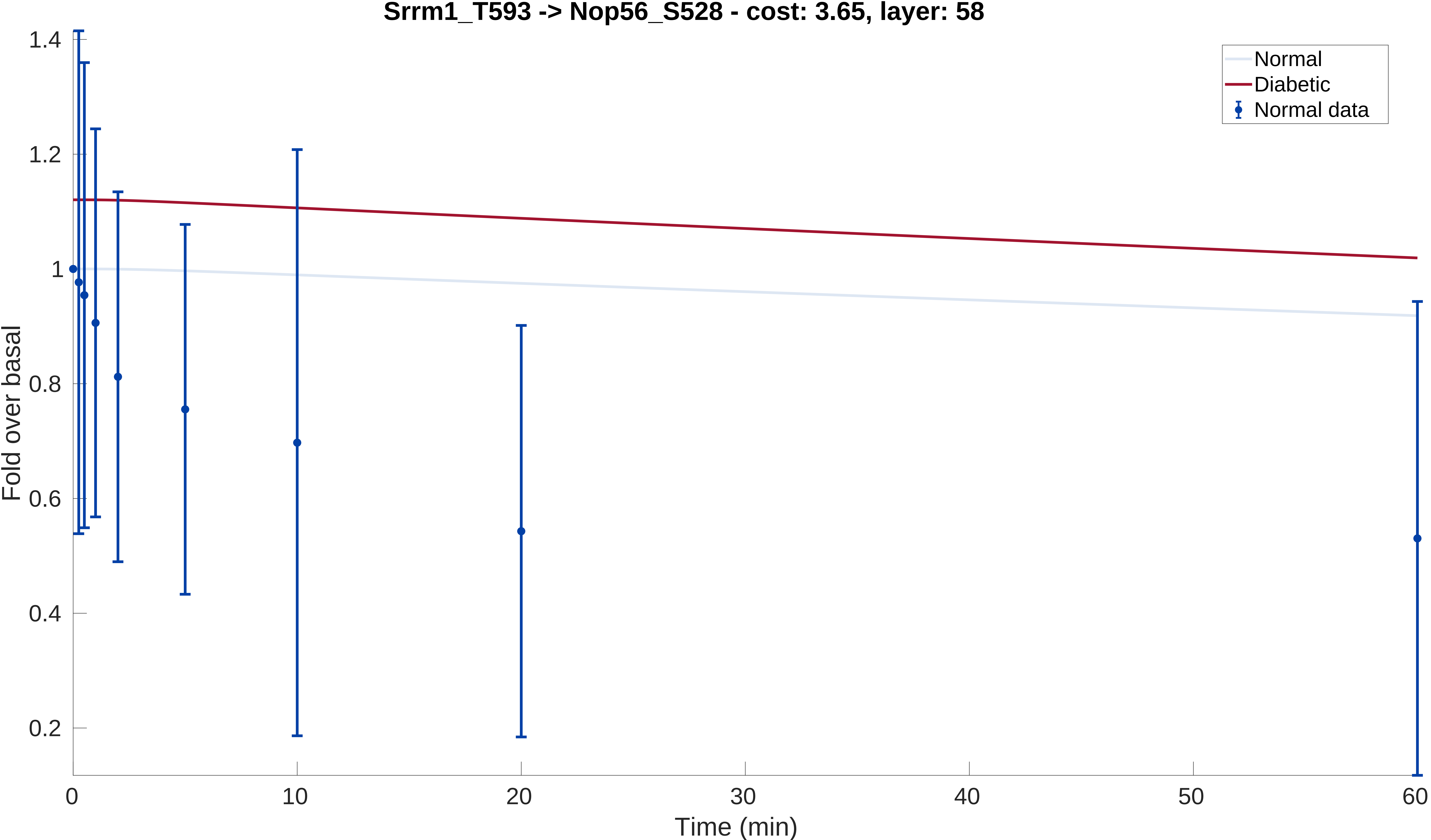
Srrm1_T593 -> Nop56_S528 - cost: 3.65, layer: 58
Normal
Diabetic
Normal data
Fold over basal
1.4
1.2
1
0.8
0.6
0.4
0.2
0
10
20
30
40
50
60
Time (min)

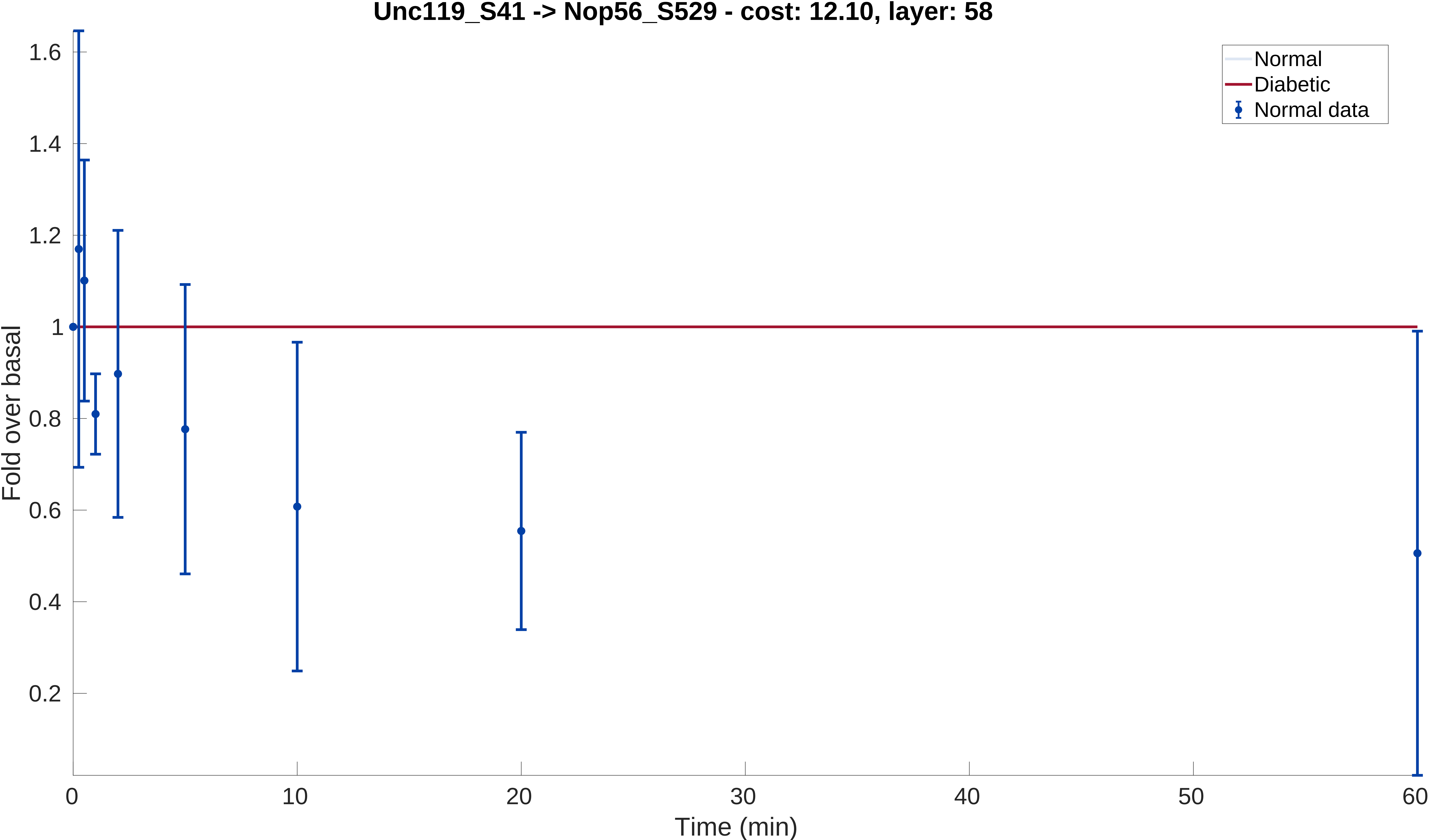

Unc119_S41 -> Nop56_S529 - cost: 12.10, layer: 58
Normal
Diabetic
Normal data
Fold over basal
1.6
1.4
1.2
1
0.8
0.6
0.4
0.2
0
10
20
30
40
50
60
Time (min)

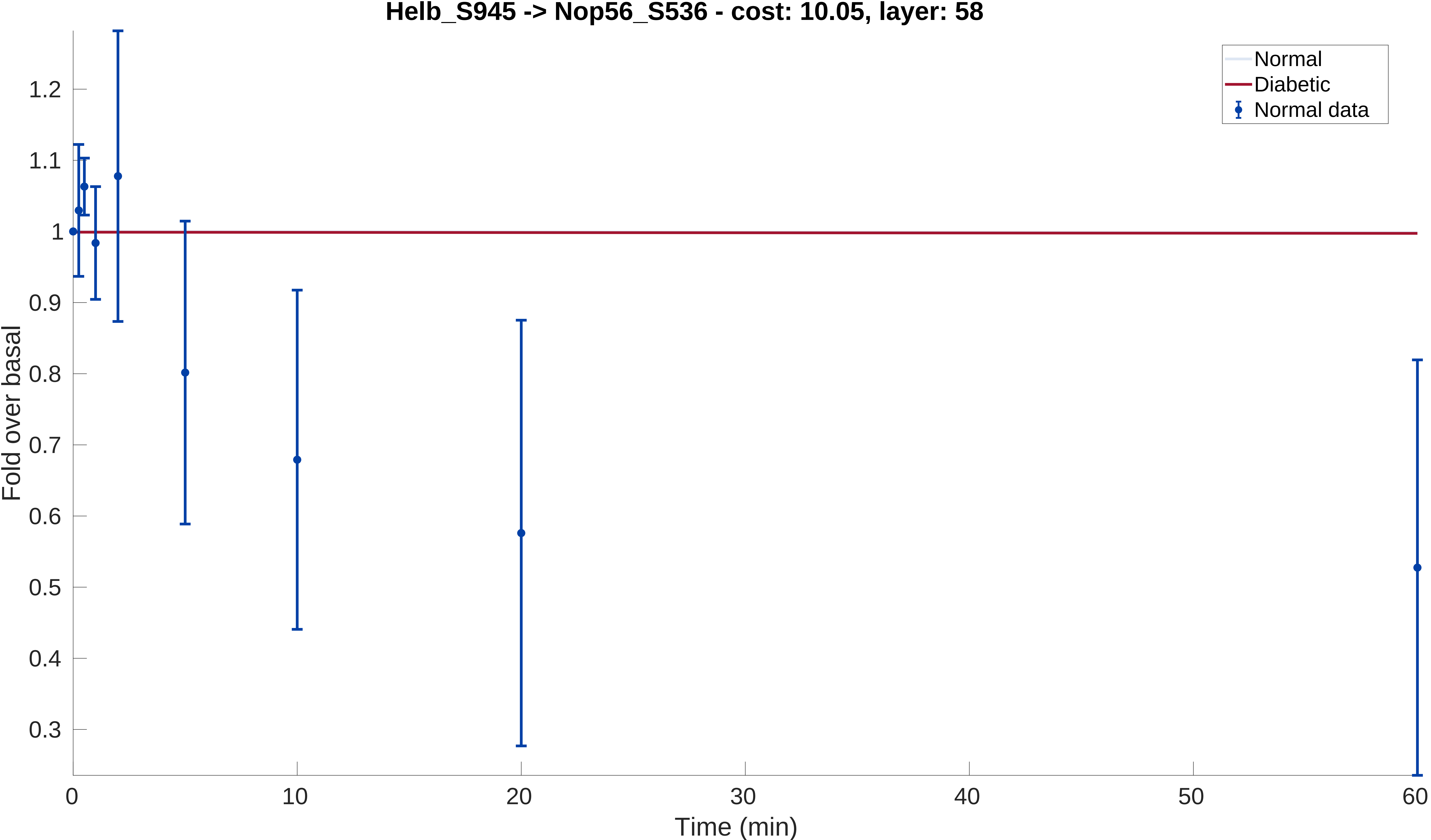

Helb_S945 -> Nop56_S536 - cost: 10.05, layer: 58
Normal
Diabetic
Normal data
Fold over basal
1.2
1.1
1
0.9
0.8
0.7
0.6
0.5
0.4
0.3
0
10
20
30
40
50
60
Time (min)

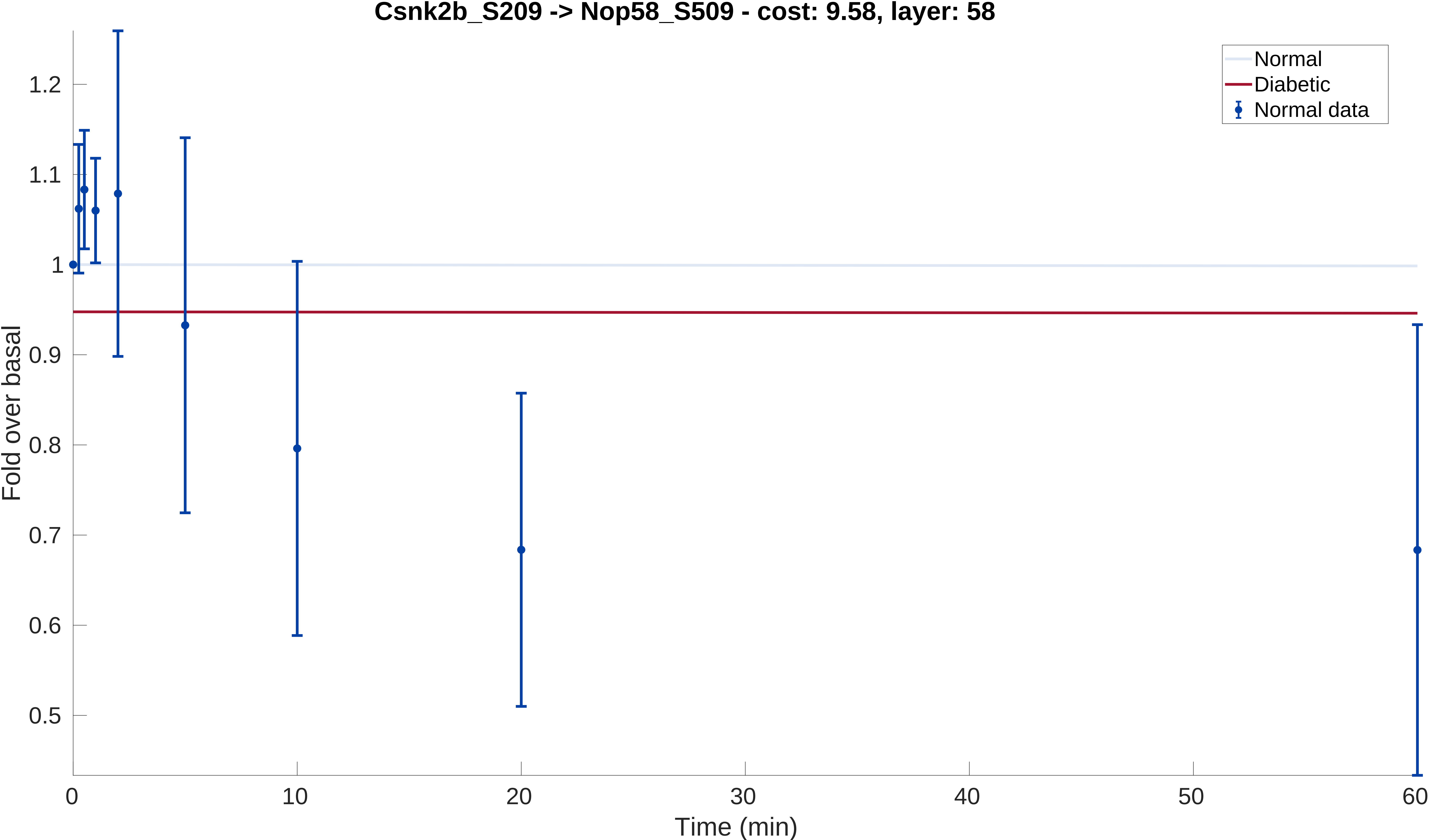
Csnk2b_S209 -> Nop58_S509 - cost: 9.58, layer: 58
Normal
Diabetic
Normal data
Fold over basal
1.2
1.1
1
0.9
0.8
0.7
0.6
0.5
0
10
20
30
40
50
60
Time (min)

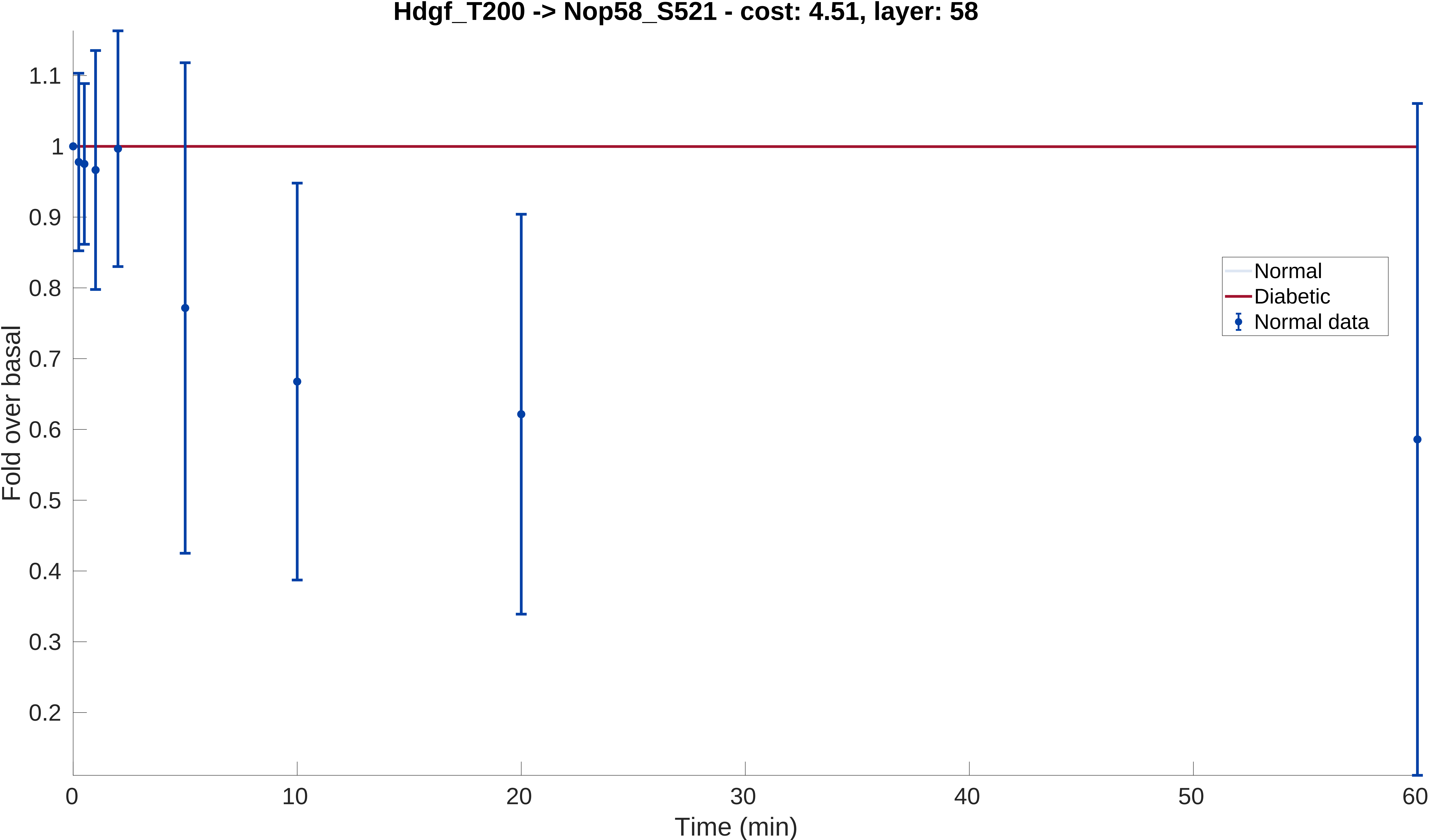

Hdgf_T200 -> Nop58_S521 - cost: 4.51, layer: 58
Fold over basal
1.1
1
0.9
0.8
0.7
0.6
0.5
0.4
0.3
0.2
0
10
20
30
40
50
60
Time (min)
Normal
Diabetic
Normal data

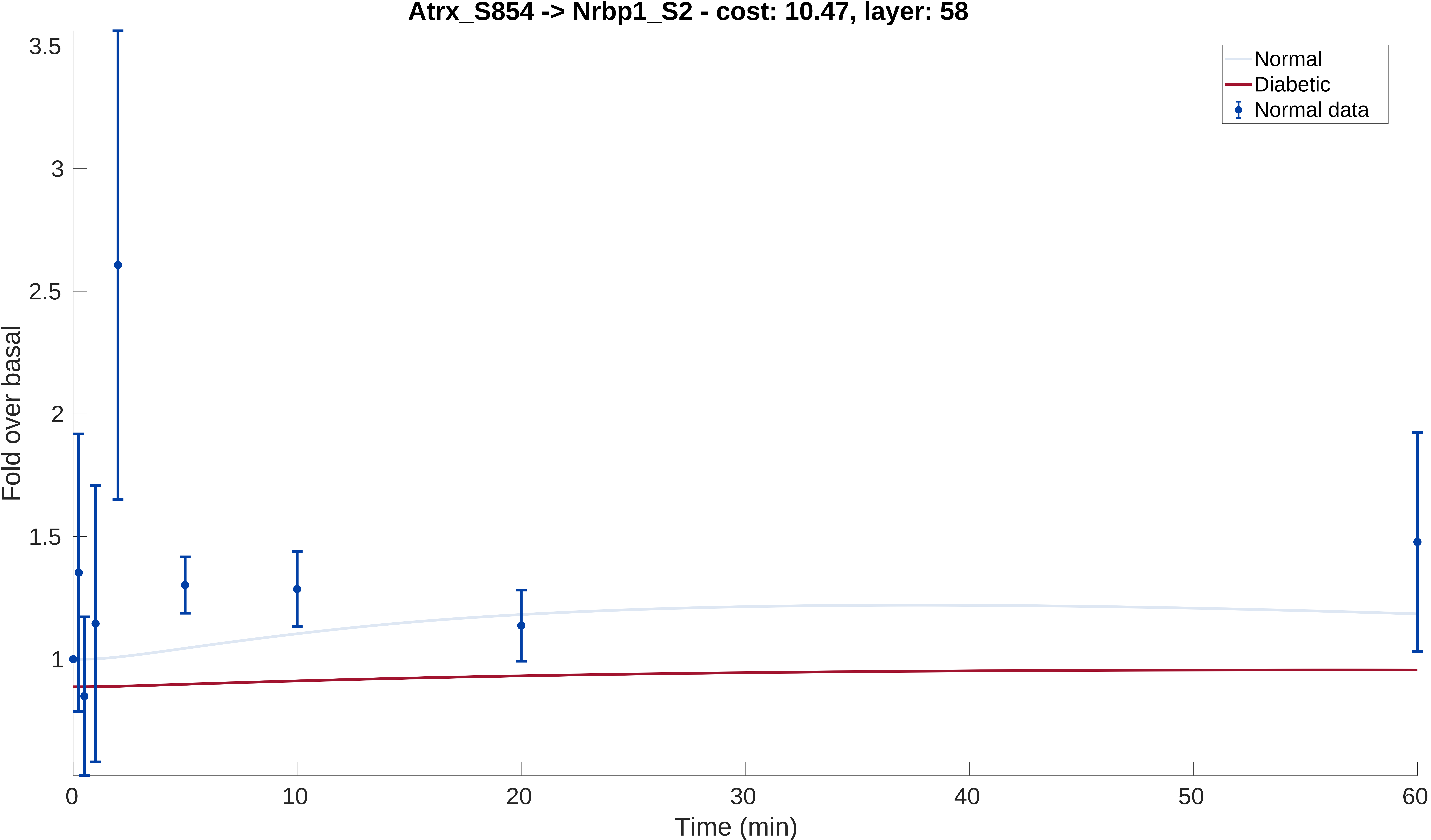
Atrx_S854 -> Nrbp1_S2 - cost: 10.47, layer: 58
Normal
Diabetic
Normal data
Fold over basal
3.5
3
2.5
2
1.5
1
0
10
20
30
40
50
60
Time (min)

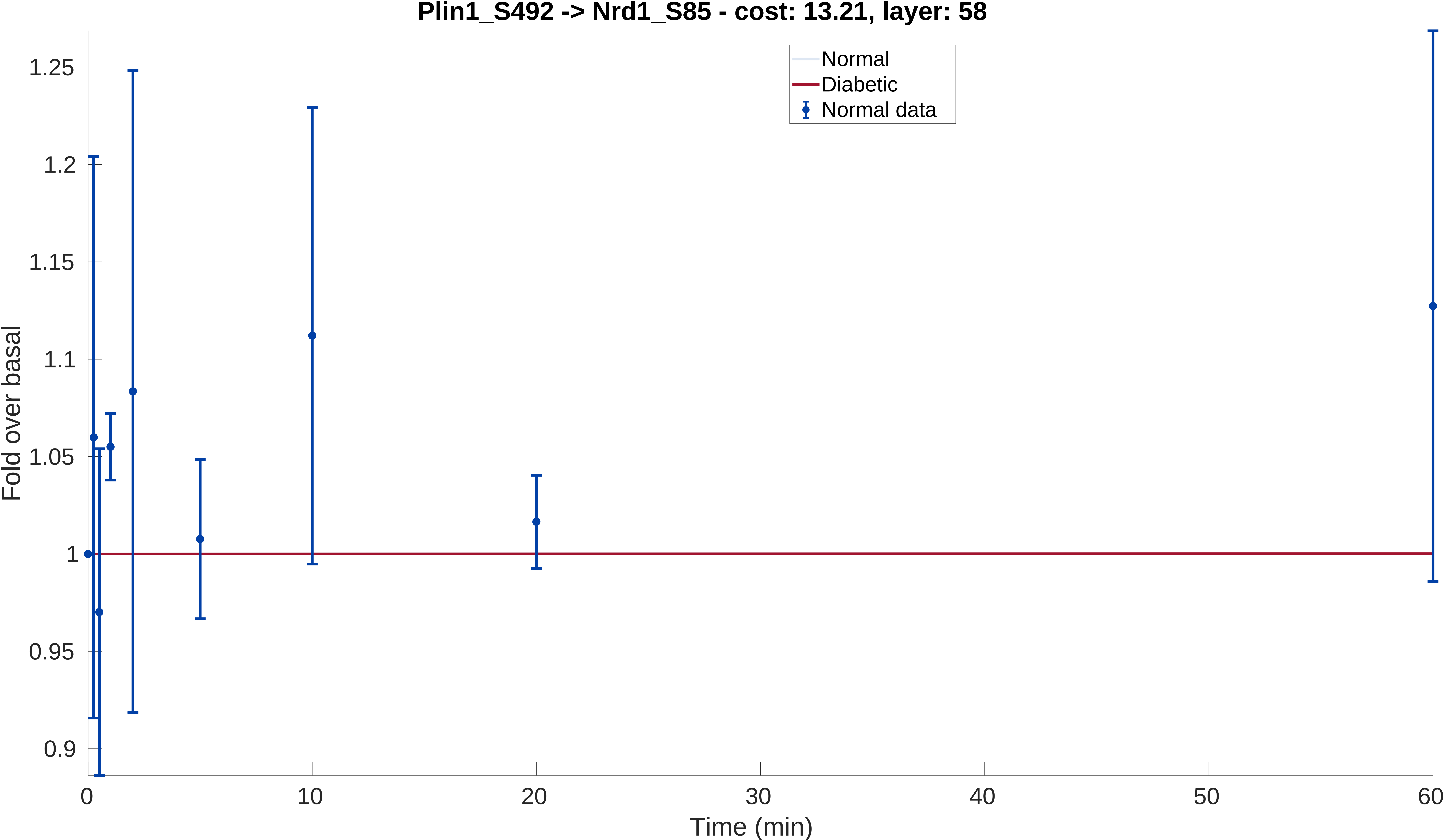
Plin1_S492 -> Nrd1_S85 - cost: 13.21, layer: 58
Normal
Diabetic
Normal data
Fold over basal
1.25
1.2
1.15
1.1
1.05
1
0.95
0.9
0
10
20
30
40
50
60
Time (min)

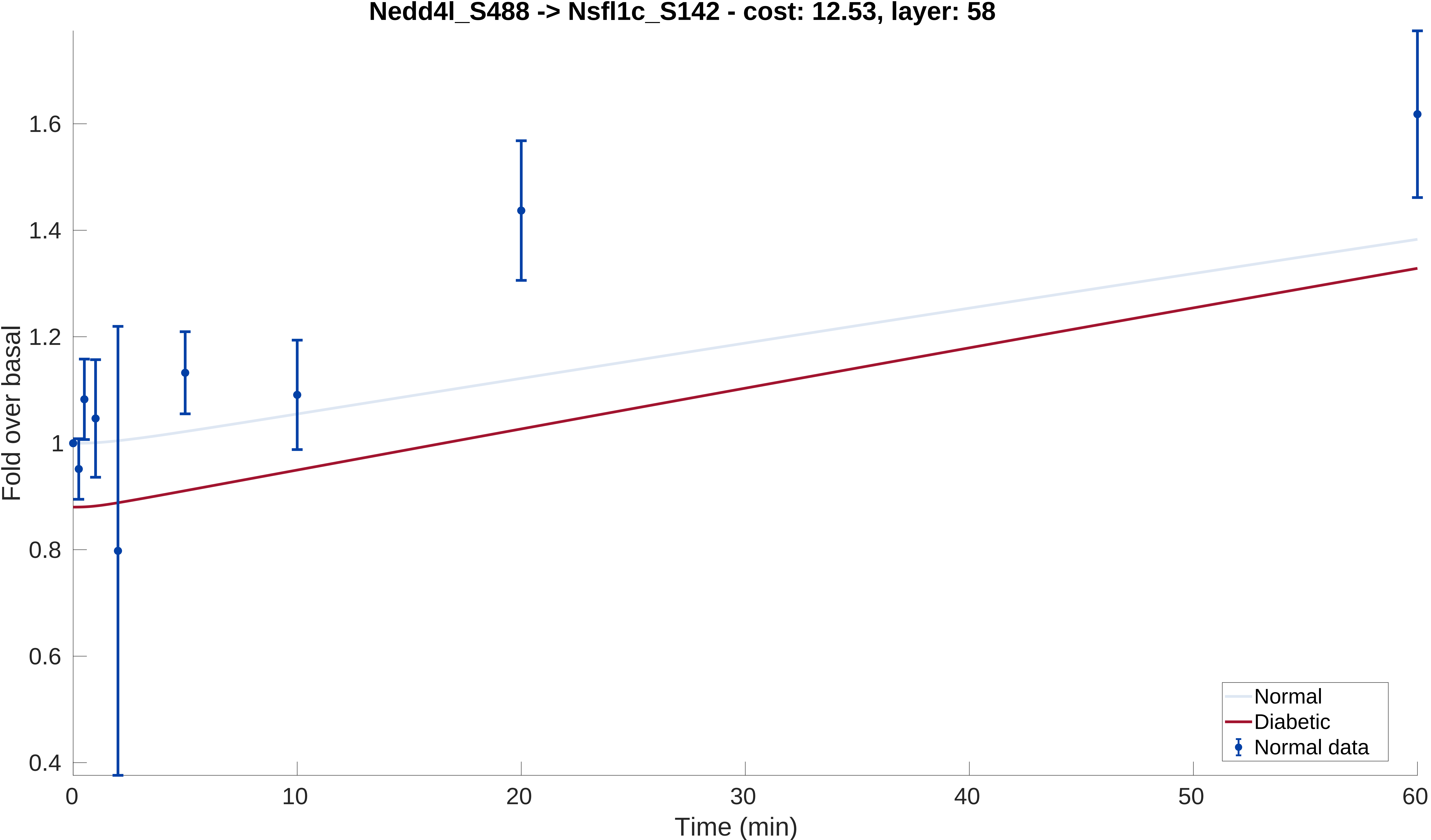

Nedd4l_S488 -> Nsfl1c_S142 - cost: 12.53, layer: 58
Fold over basal
1.6
1.4
1.2
1
0.8
0.6
0.4
0
10
20
30
40
50
60
Time (min)
Normal
Diabetic
Normal data

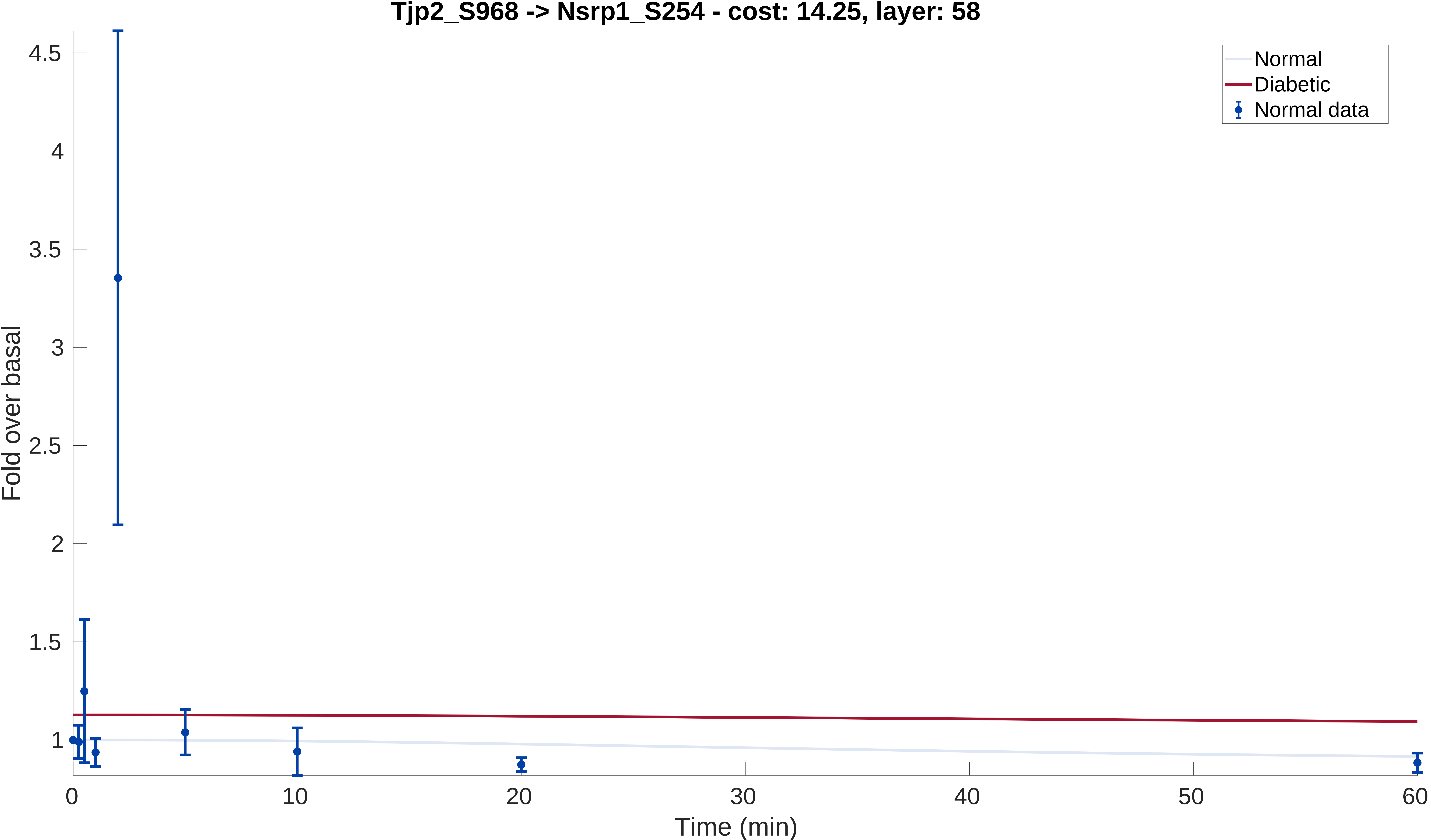

Tjp2_S968 -> Nsrp1_S254 - cost: 14.25, layer: 58
Normal
Diabetic
Normal data
Fold over basal
Time (min)
4.5
4
3.5
3
2.5
2
1.5
1
0
10
20
30
40
50
60

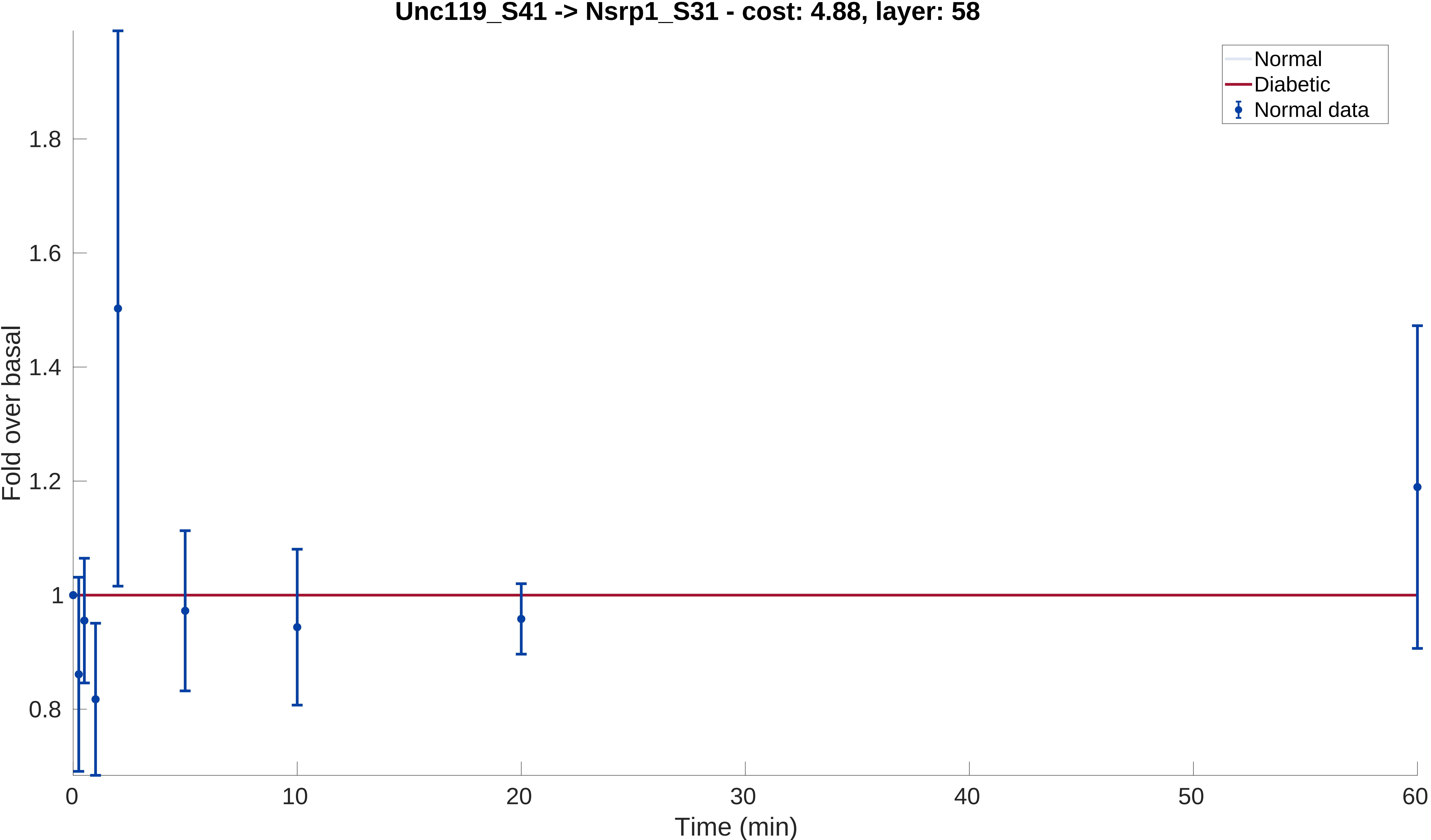
Unc119_S41 -> Nsrp1_S31 - cost: 4.88, layer: 58
Normal
Diabetic
Normal data
Fold over basal
1.8
1.6
1.4
1.2
1
0.8
0
10
20
30
40
50
60
Time (min)

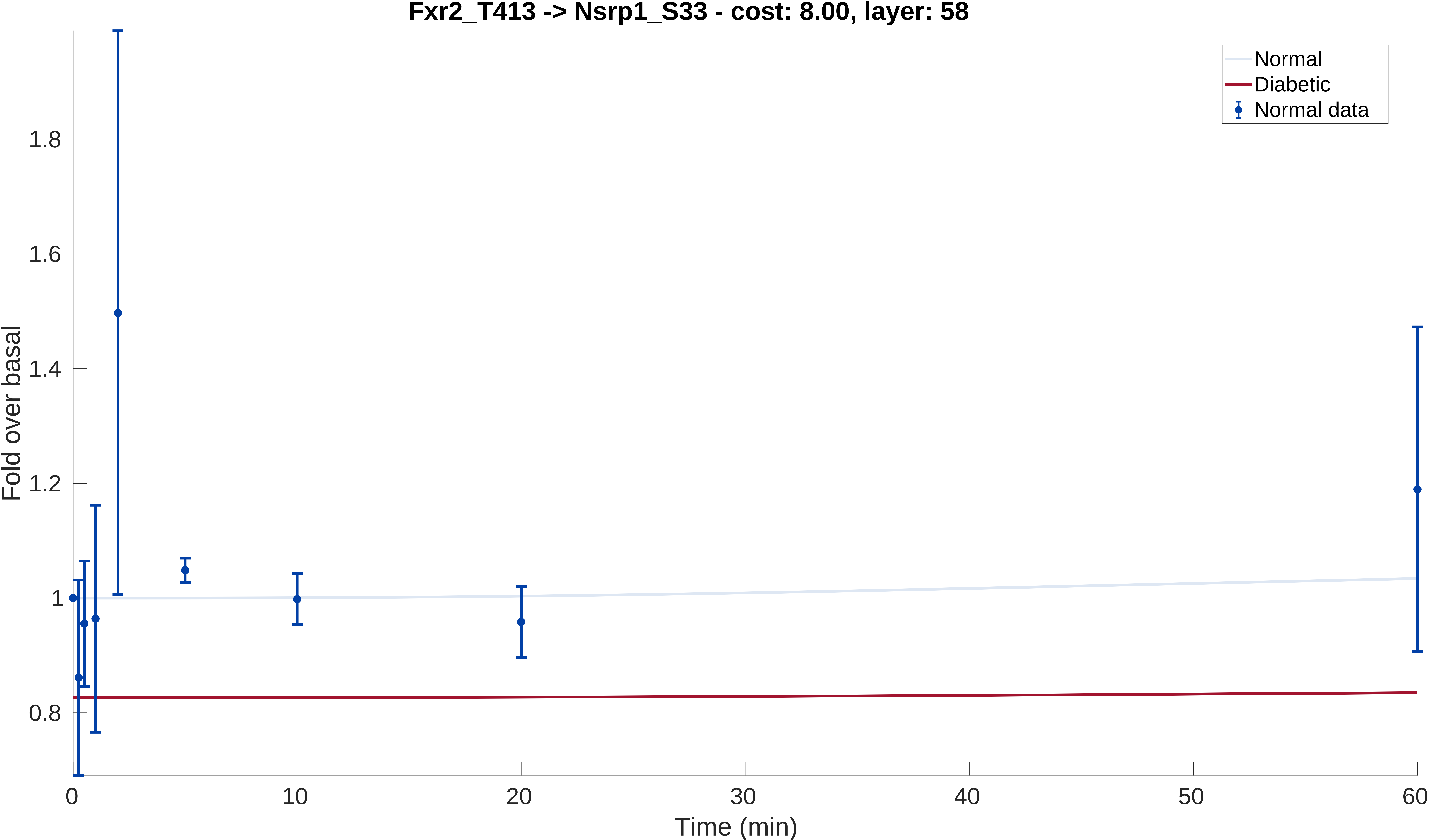

Fxr2_T413 -> Nsrp1_S33 - cost: 8.00, layer: 58
Normal
Diabetic
Normal data
Fold over basal
1.8
1.6
1.4
1.2
1
0.8
0
10
20
30
40
50
60
Time (min)

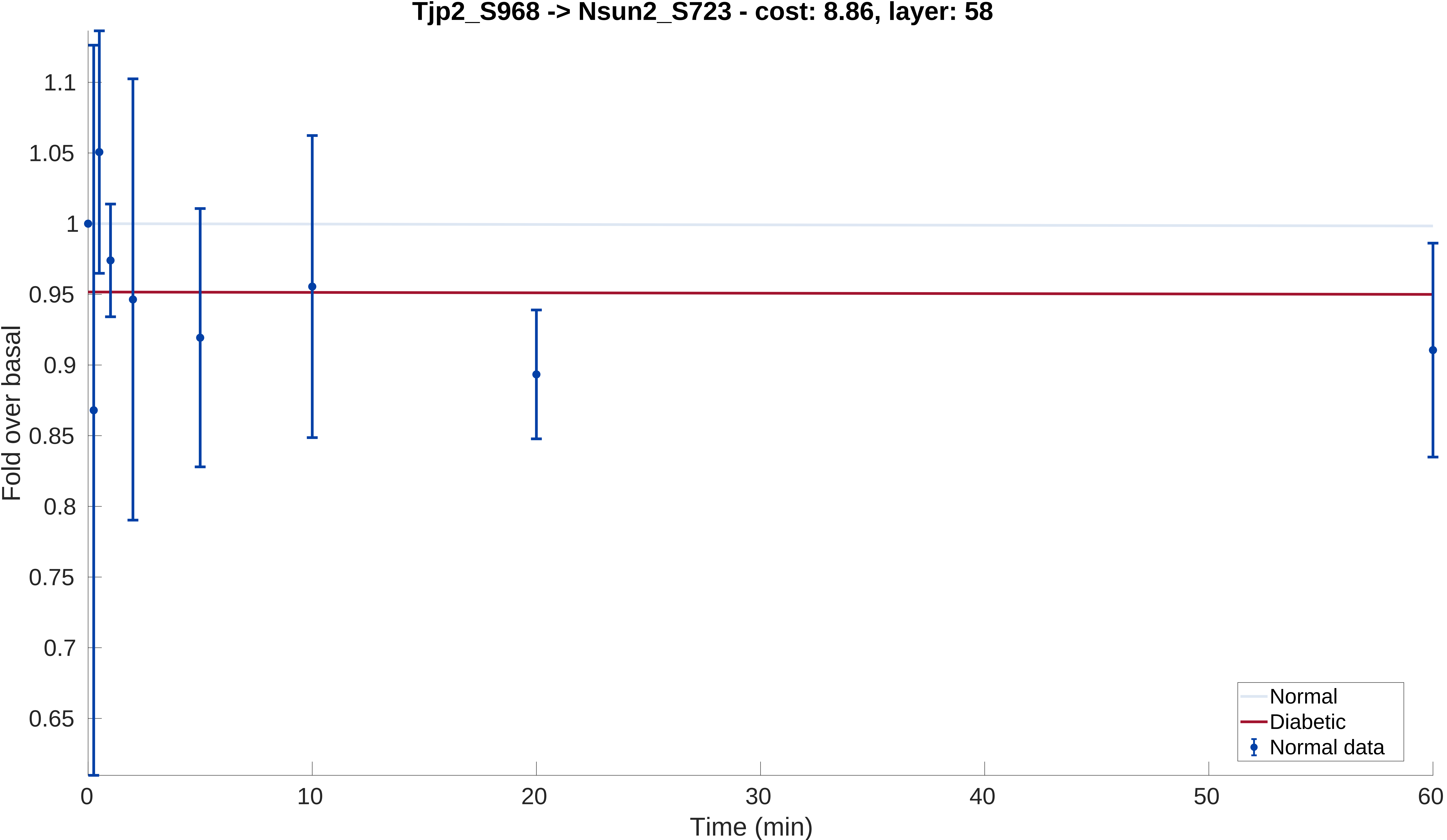

Tjp2_S968 -> Nsun2_S723 - cost: 8.86, layer: 58
Fold over basal
1.1
1.05
1
0.95
0.9
0.85
0.8
0.75
0.7
0.65
0
10
20
30
40
50
60
Time (min)
Normal
Diabetic
Normal data

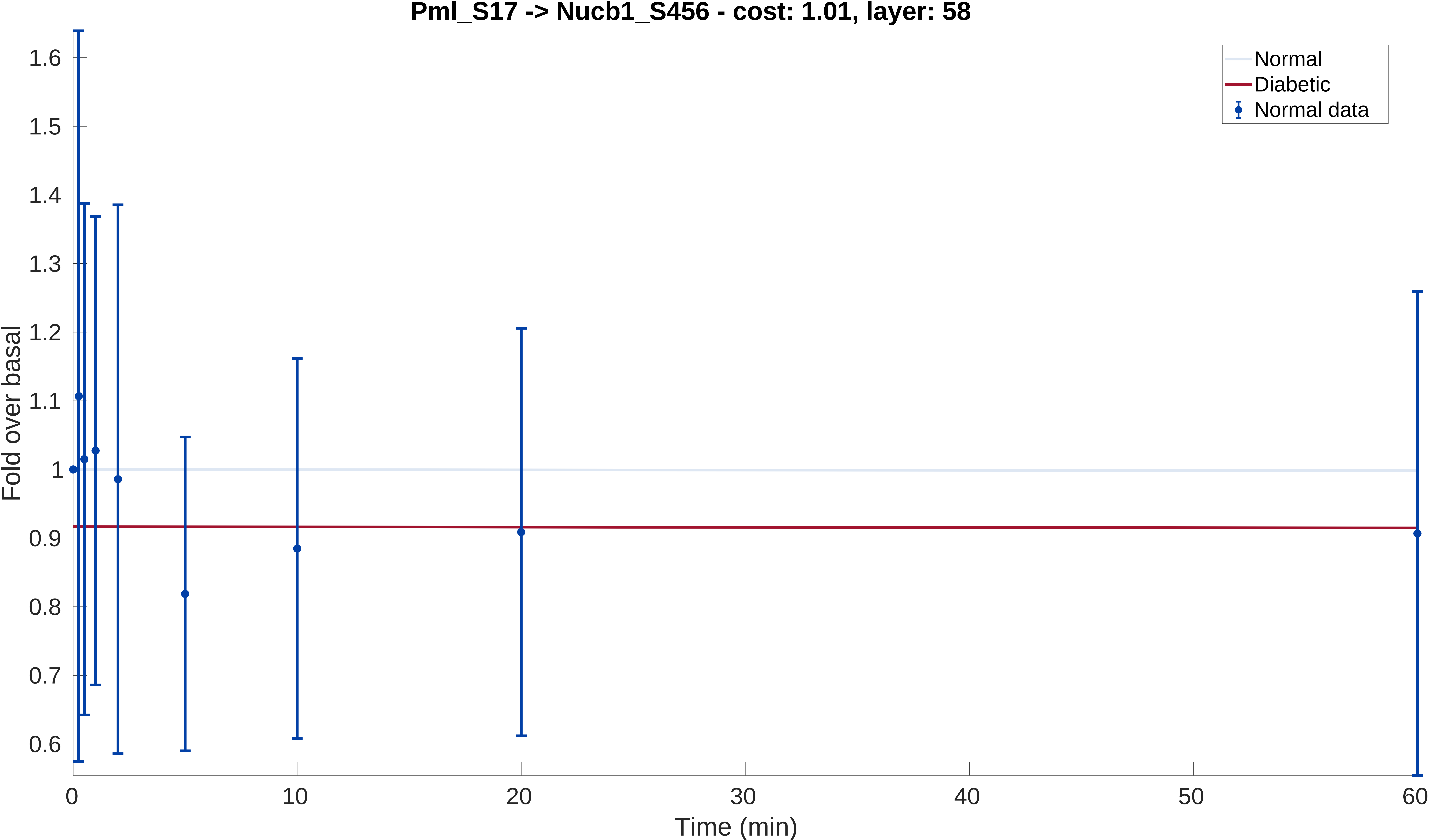

Pml_S17 -> Nucb1_S456 - cost: 1.01, layer: 58
Normal
Diabetic
Normal data
Fold over basal
Time (min)
1.6
1.5
1.4
1.3
1.2
1.1
1
0.9
0.8
0.7
0.6
0
10
20
30
40
50
60

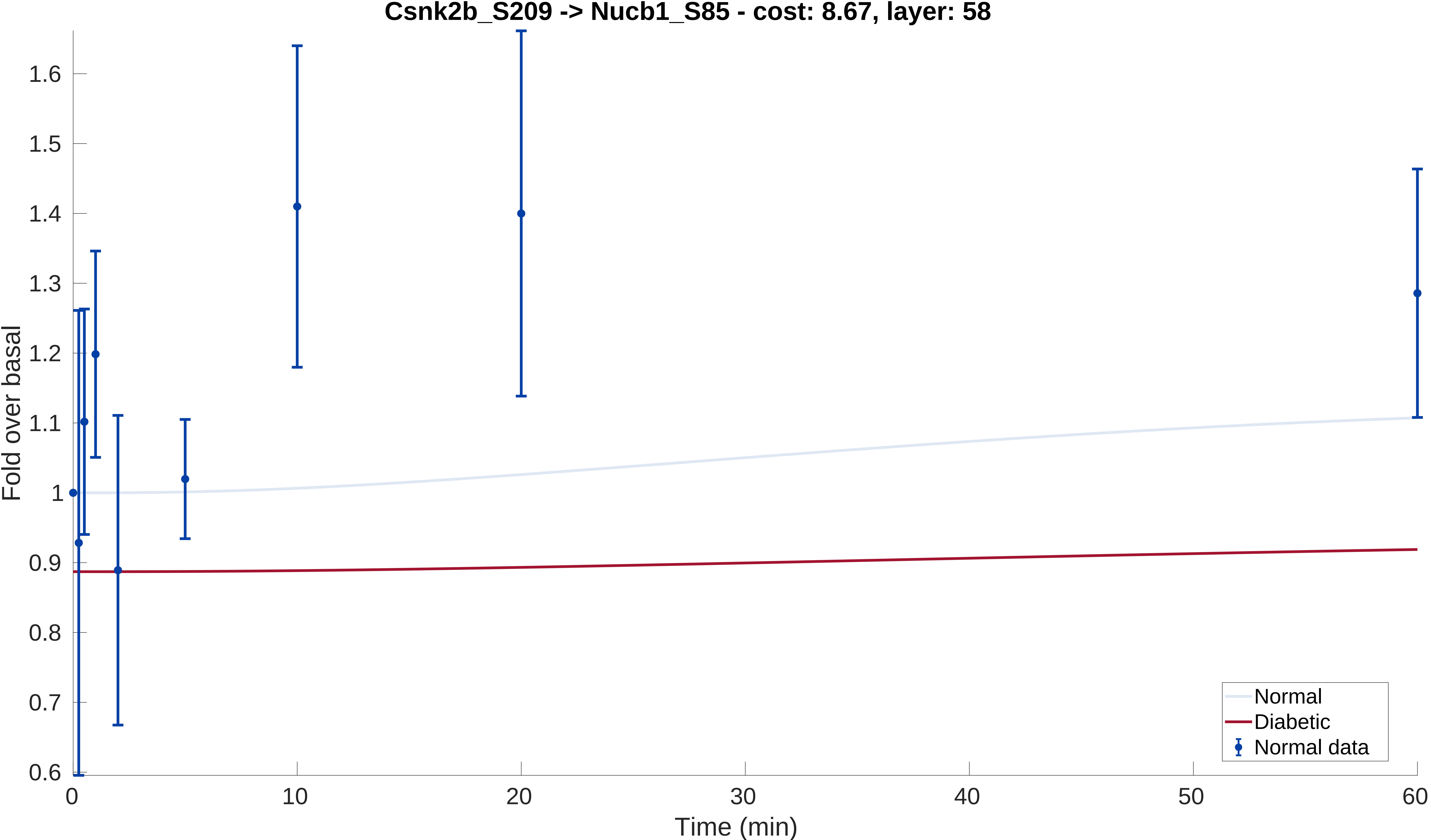

Csnk2b_S209 -> Nucb1_S85 - cost: 8.67, layer: 58
Fold over basal
1.6
1.5
1.4
1.3
1.2
1.1
1
0.9
0.8
0.7
0.6
0
10
20
30
40
50
60
Time (min)
Normal
Diabetic
Normal data

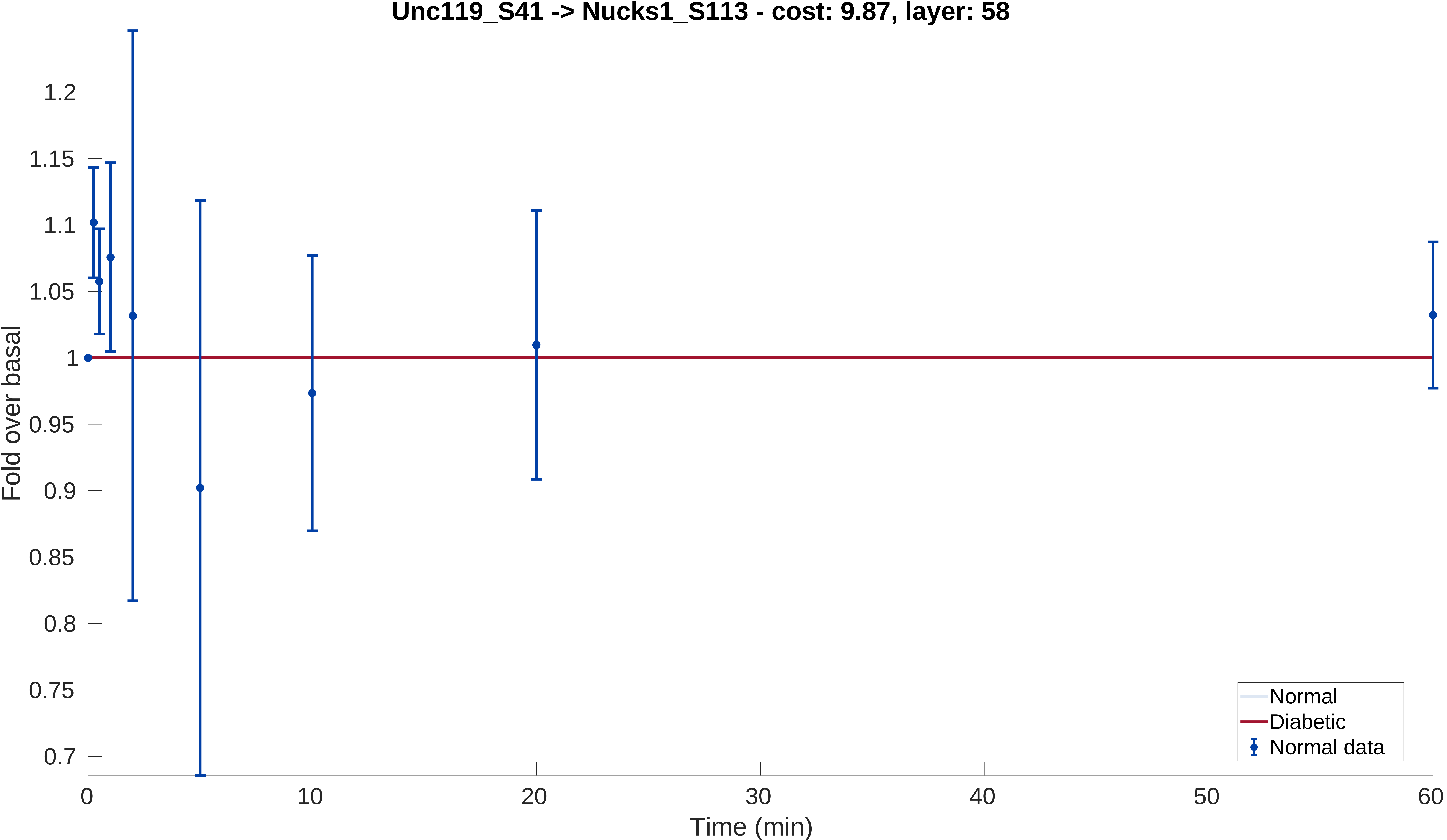

Unc119_S41 -> Nucks1_S113 - cost: 9.87, layer: 58
Fold over basal
Time (min)
1.2
1.15
1.1
1.05
1
0.95
0.9
0.85
0.8
0.75
0.7
0
10
20
30
40
50
60
Normal
Diabetic
Normal data

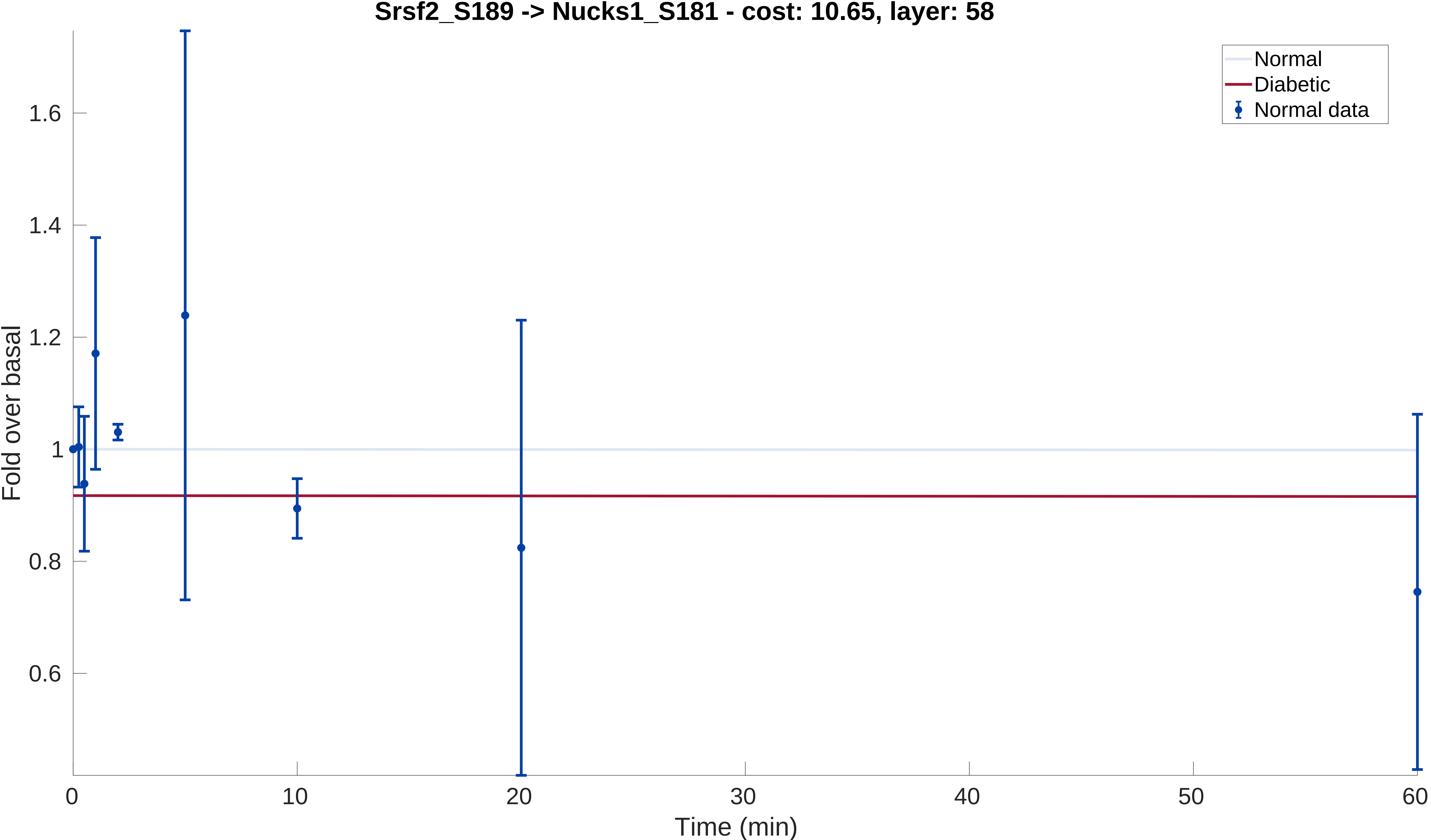

Srsf2_S189 -> Nucks1_S181 - cost: 10.65, layer: 58
Normal
Diabetic
Normal data
Fold over basal
1.6
1.4
1.2
1
0.8
0.6
0
10
20
30
40
50
60
Time (min)

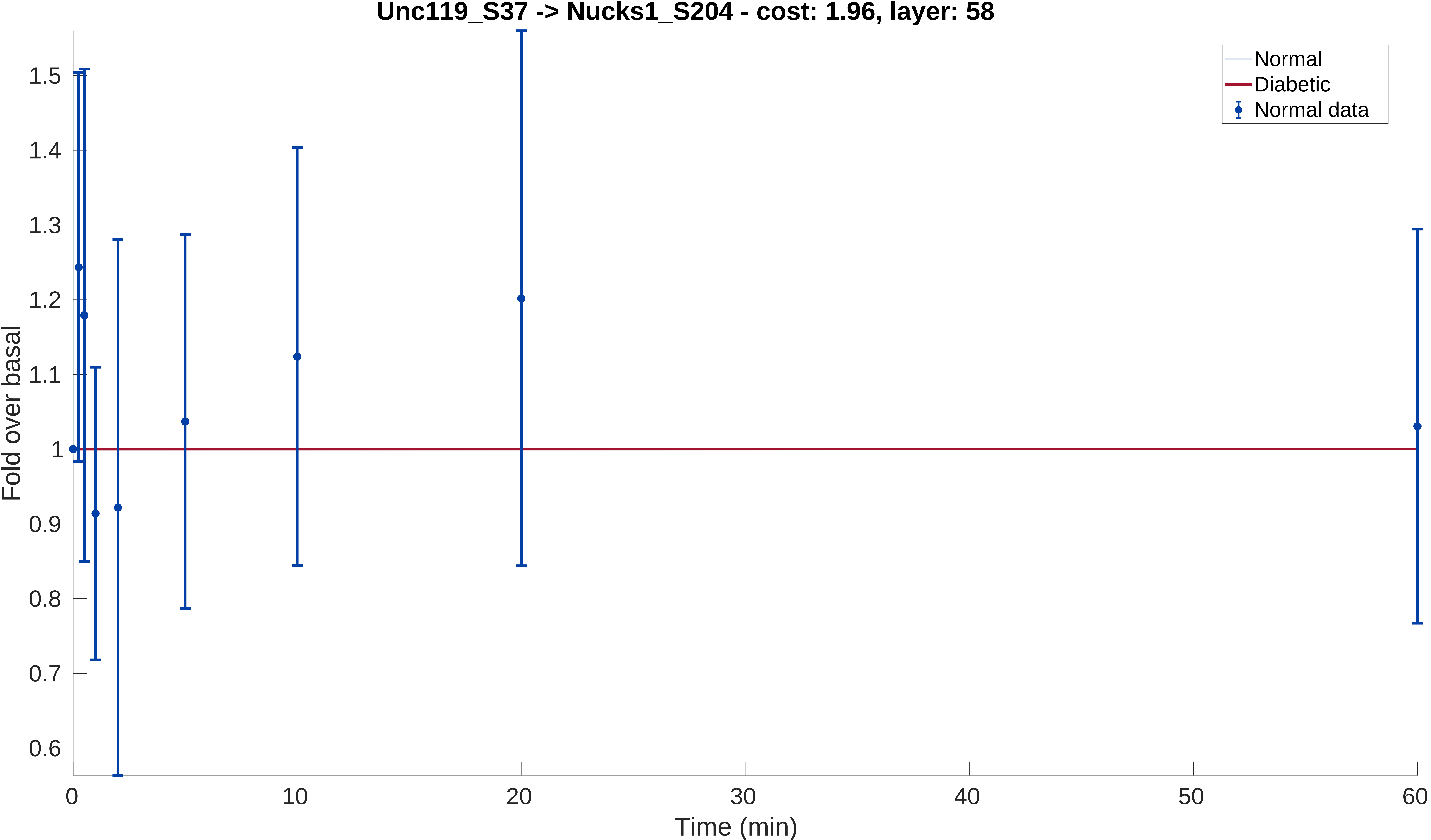
Unc119_S37 -> Nucks1_S204 - cost: 1.96, layer: 58
Normal
Diabetic
Normal data
Fold over basal
1.5
1.4
1.3
1.2
1.1
1
0.9
0.8
0.7
0.6
0
10
20
30
40
50
60
Time (min)

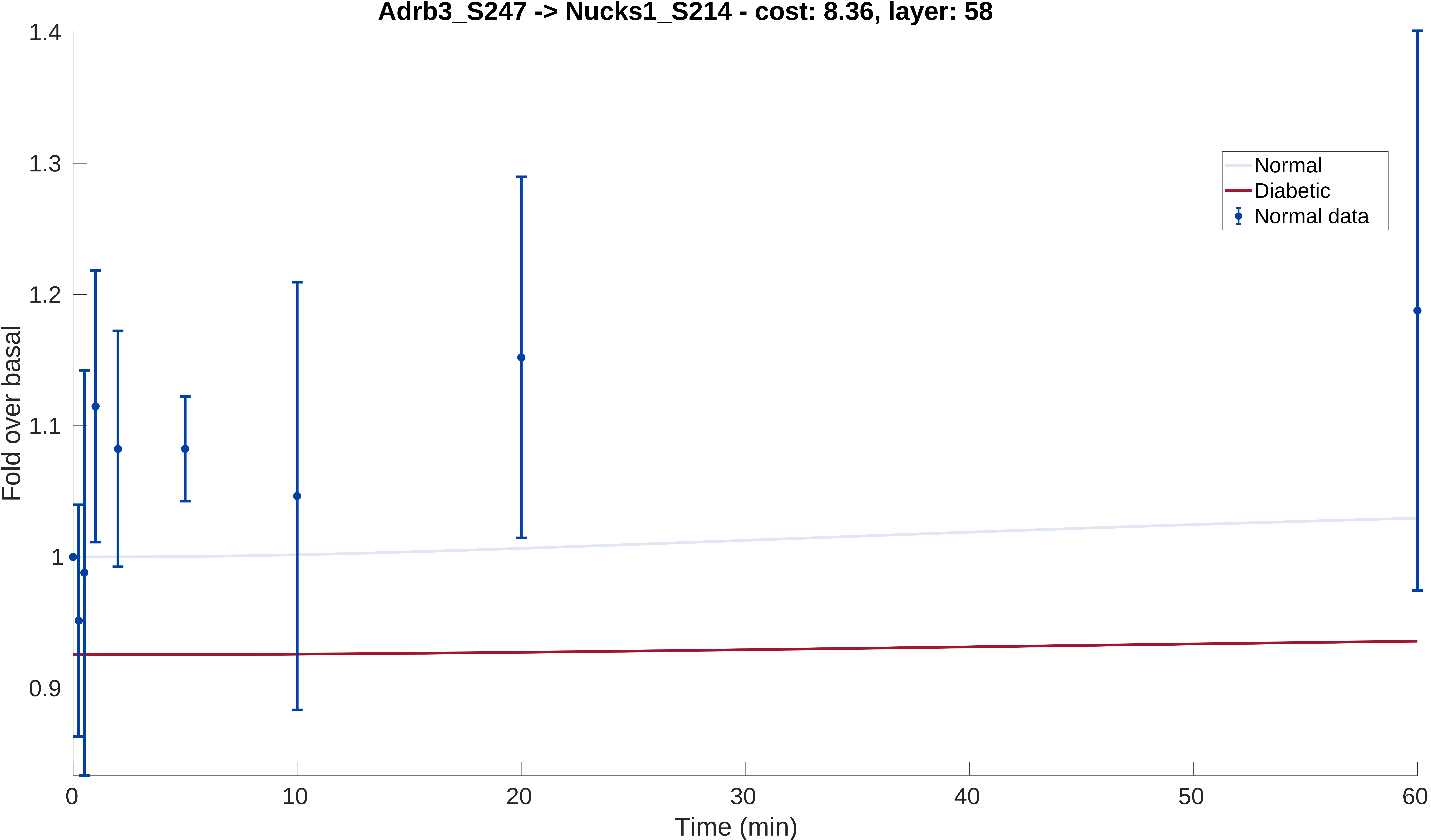
Adrb3_S247 -> Nucks1_S214 - cost: 8.36, layer: 58
1.4
1.3
1.2
1.1
1
0.9
Fold over basal
0
10
20
30
40
50
60
Time (min)
Normal
Diabetic
Normal data

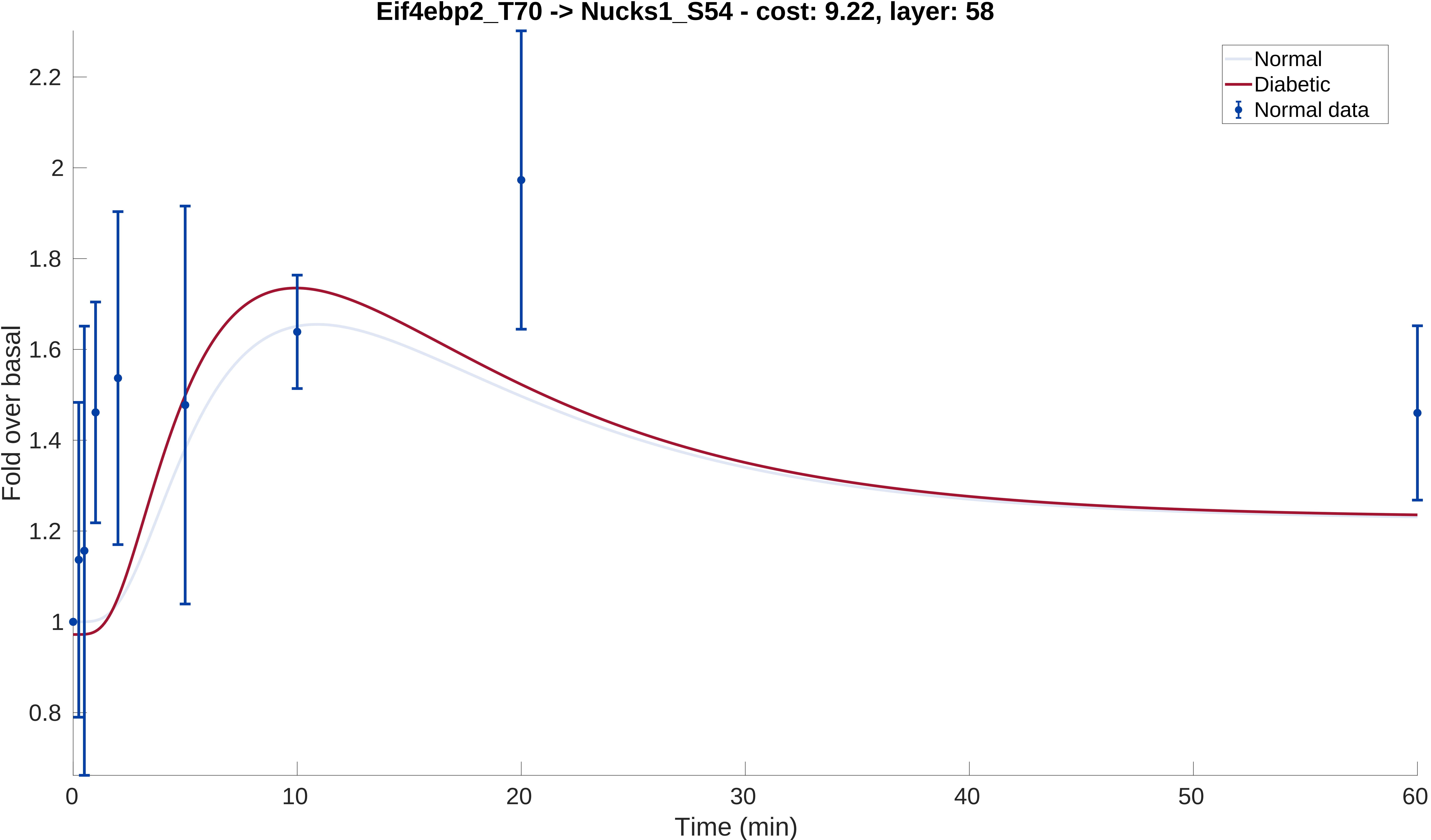

Eif4ebp2_T70 -> Nucks1_S54 - cost: 9.22, layer: 58
Normal
Diabetic
Normal data
Fold over basal
2.2
2
1.8
1.6
1.4
1.2
1
0.8
0
10
20
30
40
50
60
Time (min)

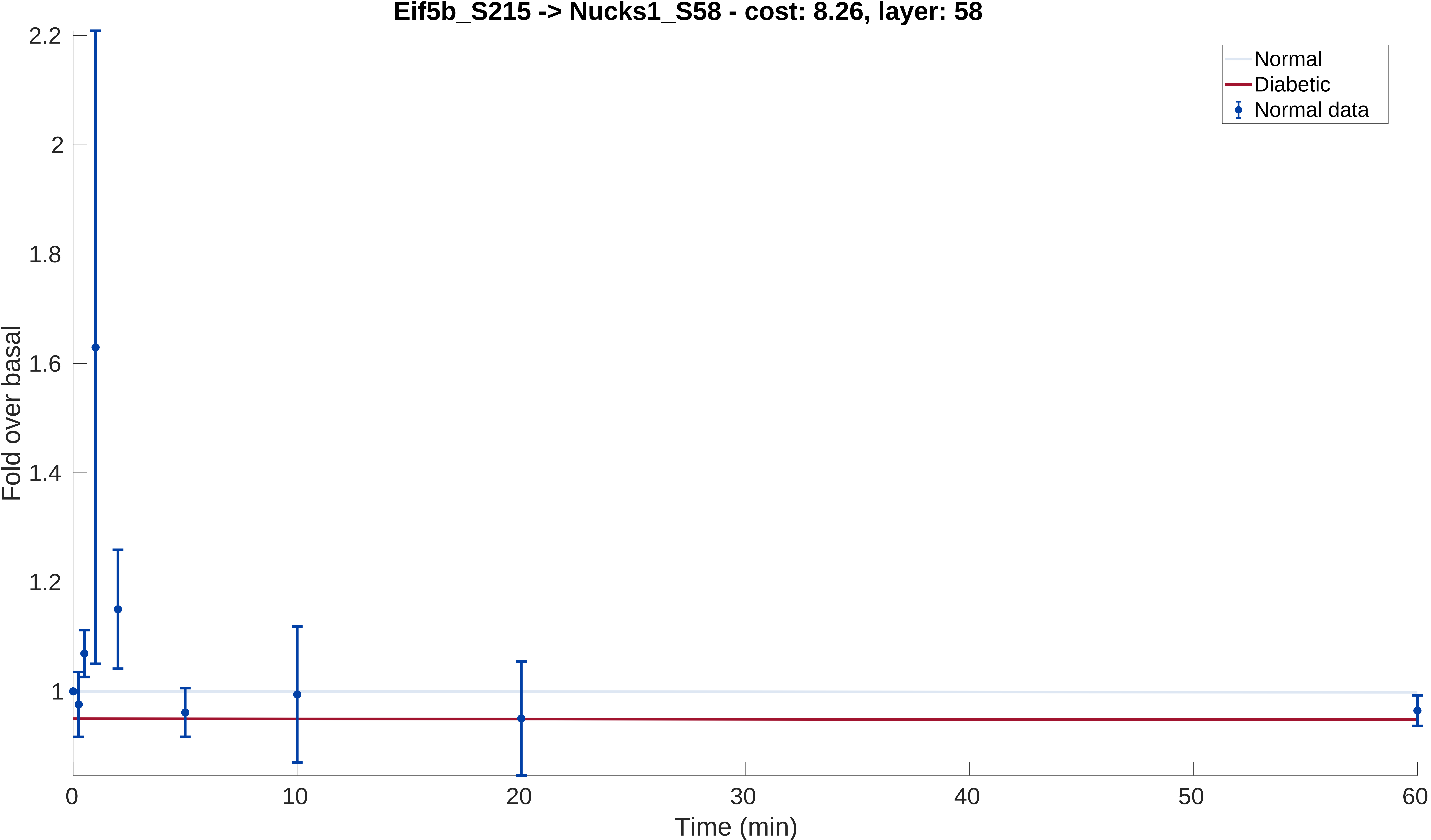

Eif5b_S215 -> Nucks1_S58 - cost: 8.26, layer: 58
Fold over basal
2.2
2
1.8
1.6
1.4
1.2
1
Time (min)
0
10
20
30
40
50
60
Normal
Diabetic
Normal data

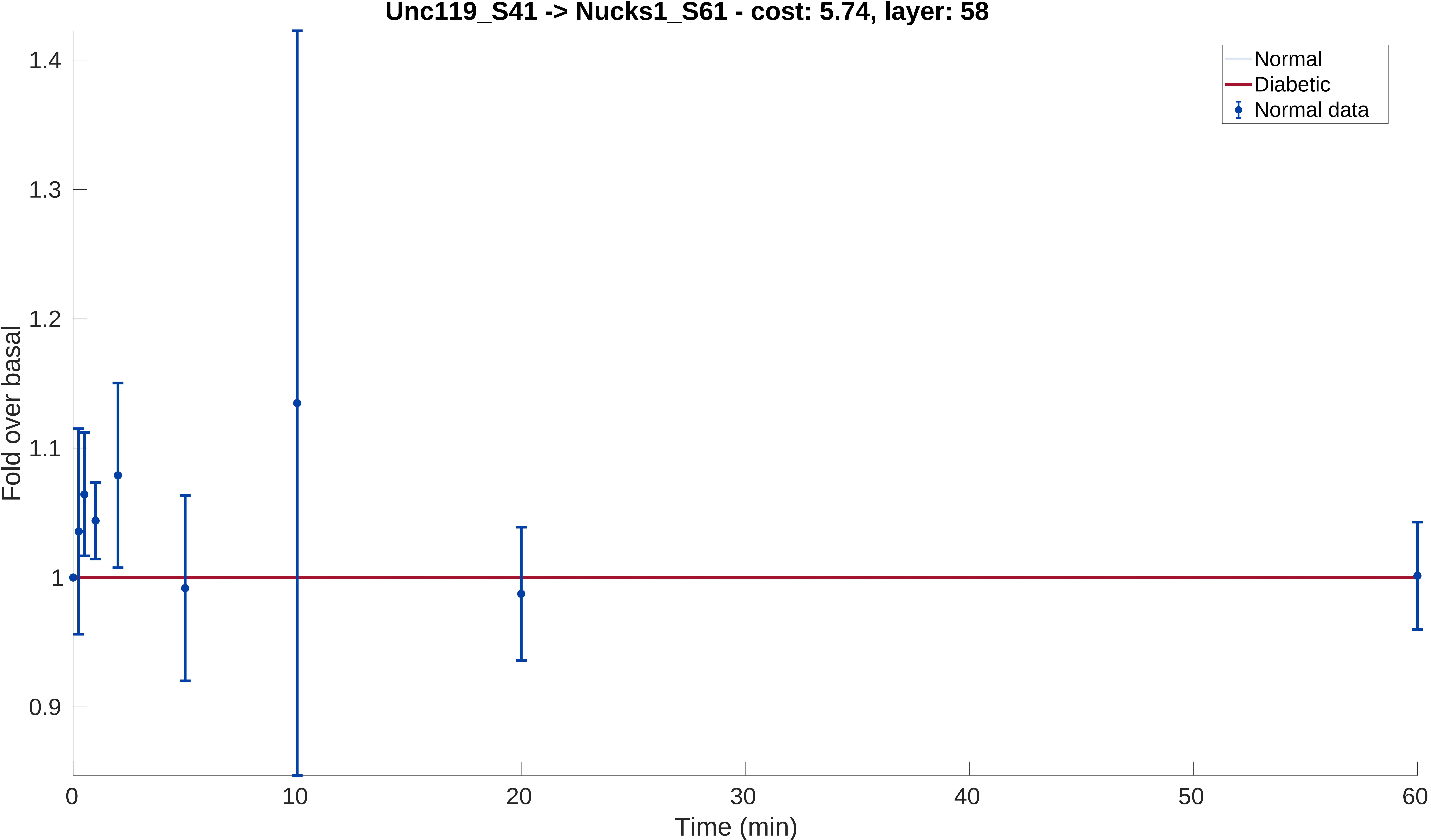

Unc119_S41 -> Nucks1_S61 - cost: 5.74, layer: 58
Normal
Diabetic
Normal data
Fold over basal
1.4
1.3
1.2
1.1
1
0.9
0
10
20
30
40
50
60
Time (min)

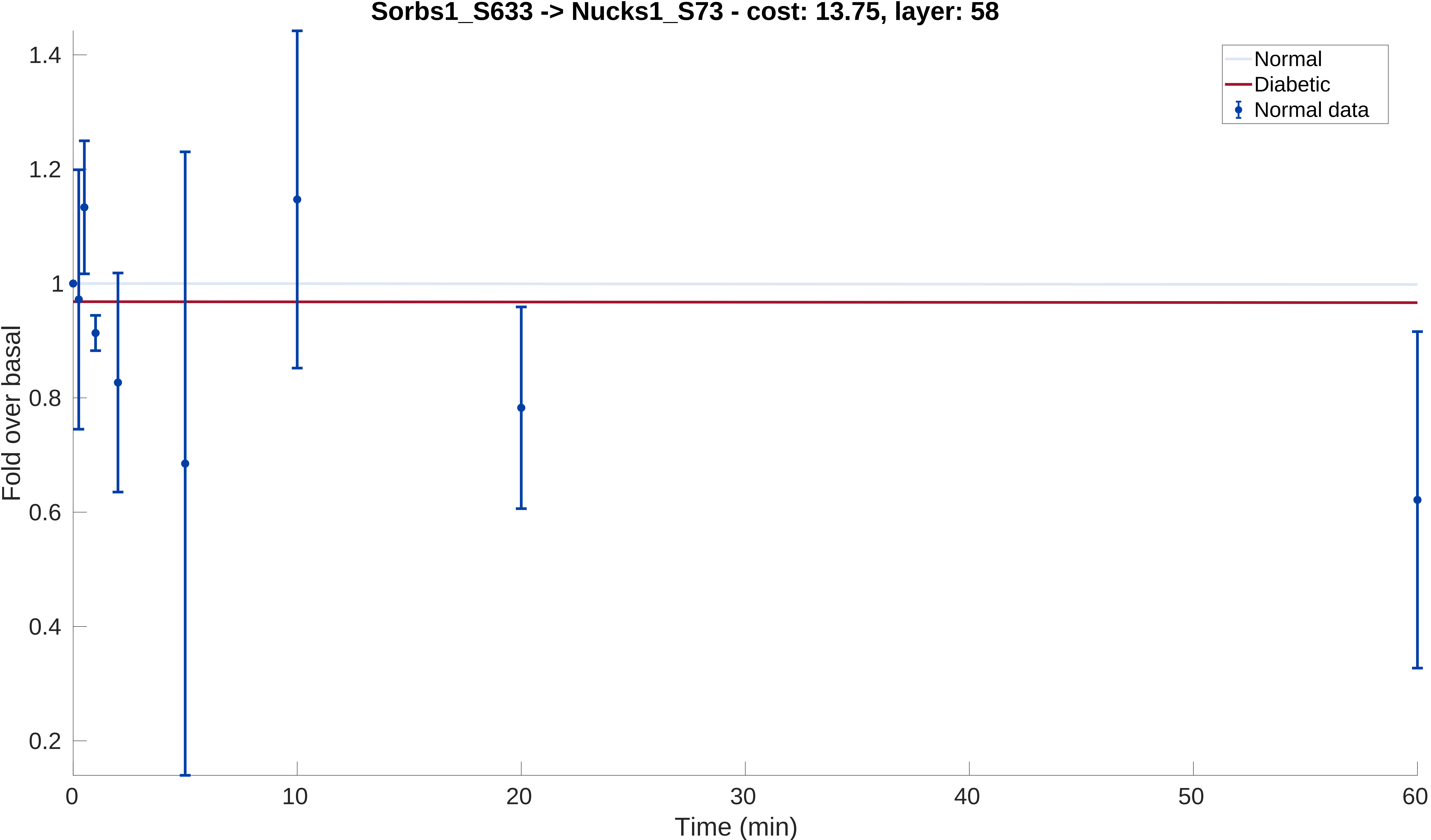

Sorbs1_S633 -> Nucks1_S73 - cost: 13.75, layer: 58
Normal
Diabetic
Normal data
Fold over basal
Time (min)
0.2
0.4
0.6
0.8
1
1.2
1.4
0
10
20
30
40
50
60

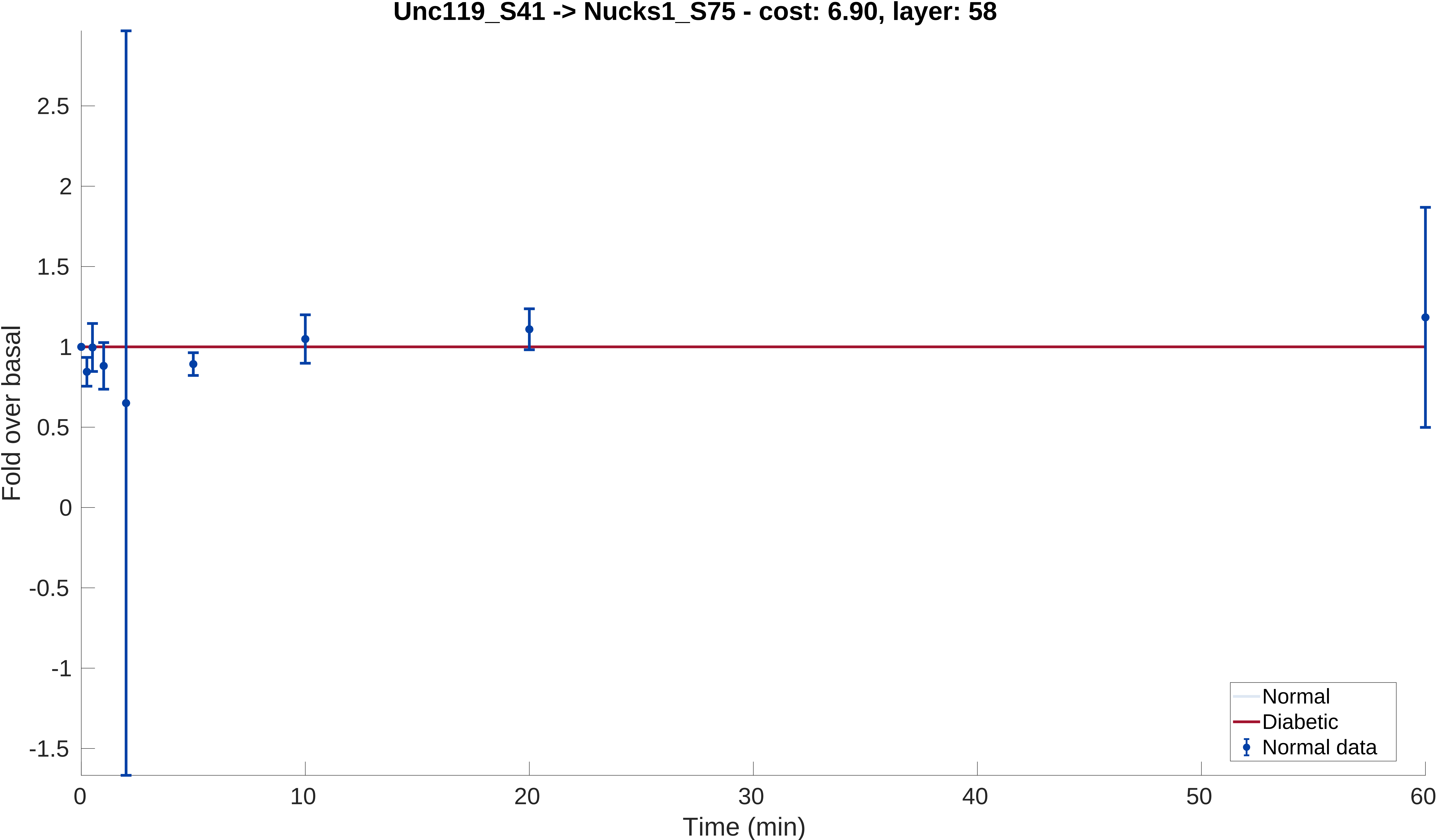

Unc119_S41 -> Nucks1_S75 - cost: 6.90, layer: 58
Fold over basal
Time (min)
2.5
2
1.5
1
0.5
0
-0.5
-1
-1.5
0
10
20
30
40
50
60
Normal
Diabetic
Normal data

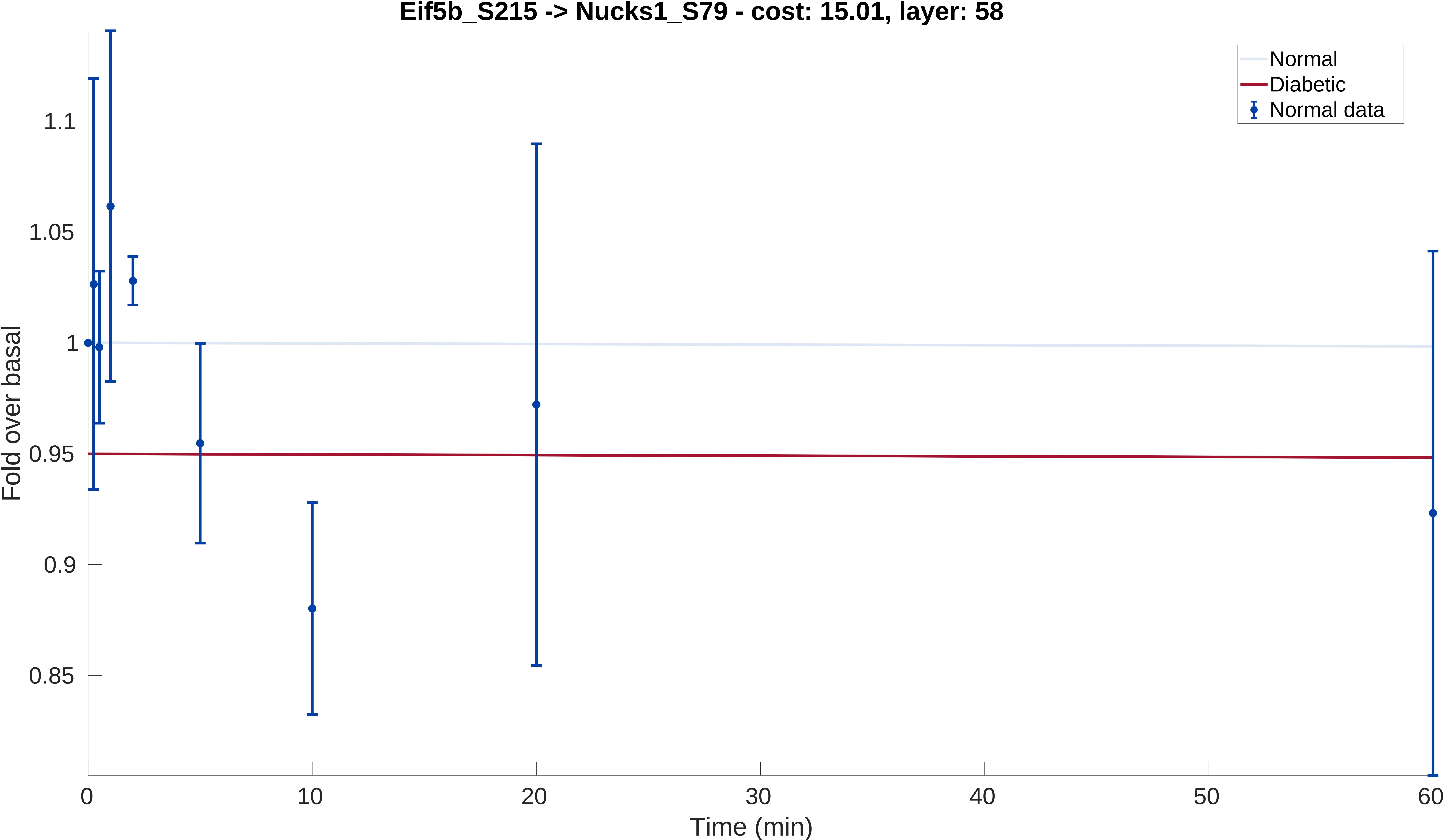
Eif5b_S215 -> Nucks1_S79 - cost: 15.01, layer: 58
Normal
Diabetic
Normal data
Fold over basal
1.1
1.05
1
0.95
0.9
0.85
0
10
20
30
40
50
60
Time (min)

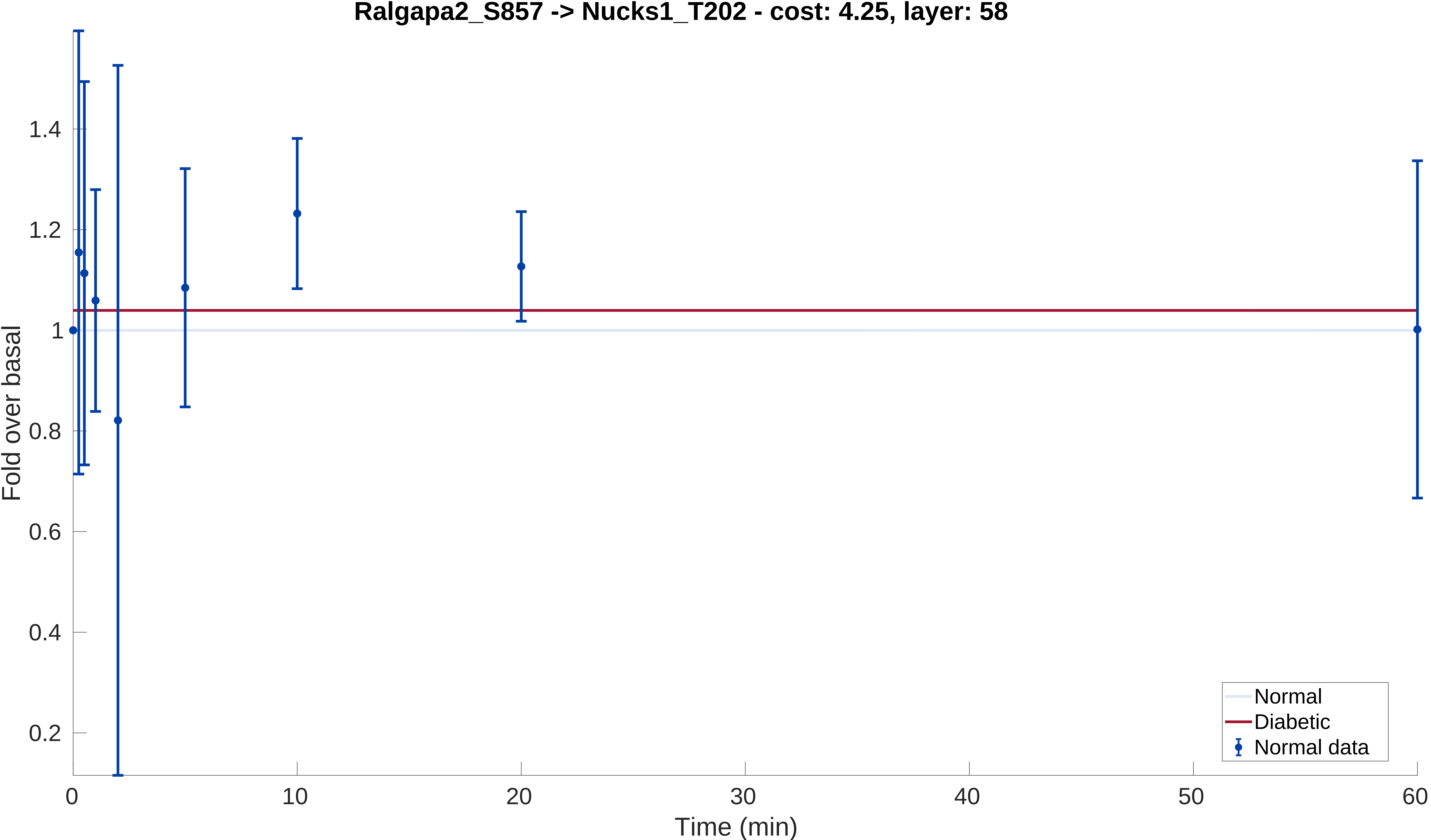

Ralgapa2_S857 -> Nucks1_T202 - cost: 4.25, layer: 58
Fold over basal
1.4
1.2
1
0.8
0.6
0.4
0.2
0
10
20
30
40
50
60
Time (min)
Normal
Diabetic
Normal data

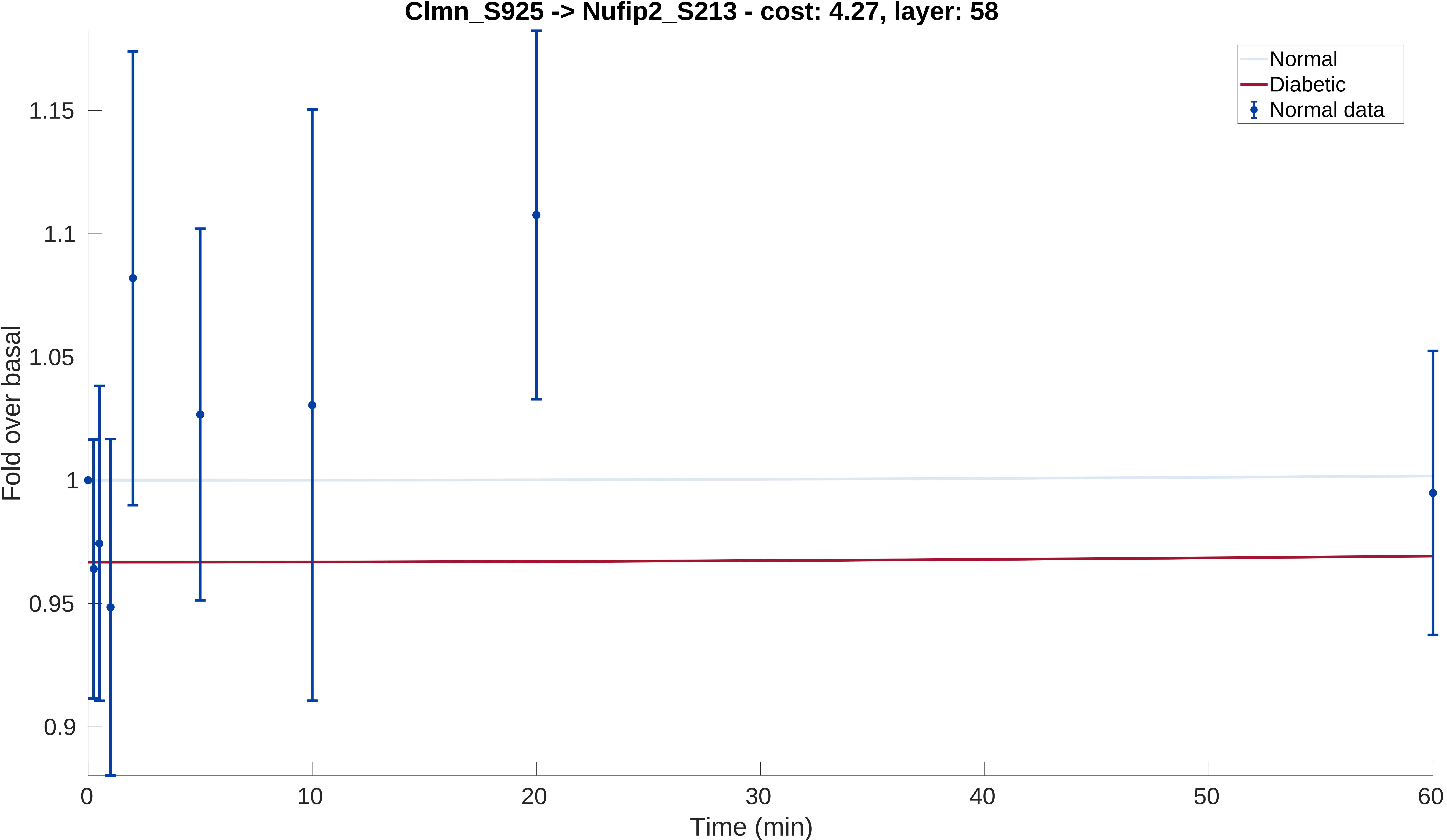
Clmn_S925 -> Nufip2_S213 - cost: 4.27, layer: 58
Normal
Diabetic
Normal data
Fold over basal
1.15
1.1
1.05
1
0.95
0.9
0
10
20
30
40
50
60
Time (min)

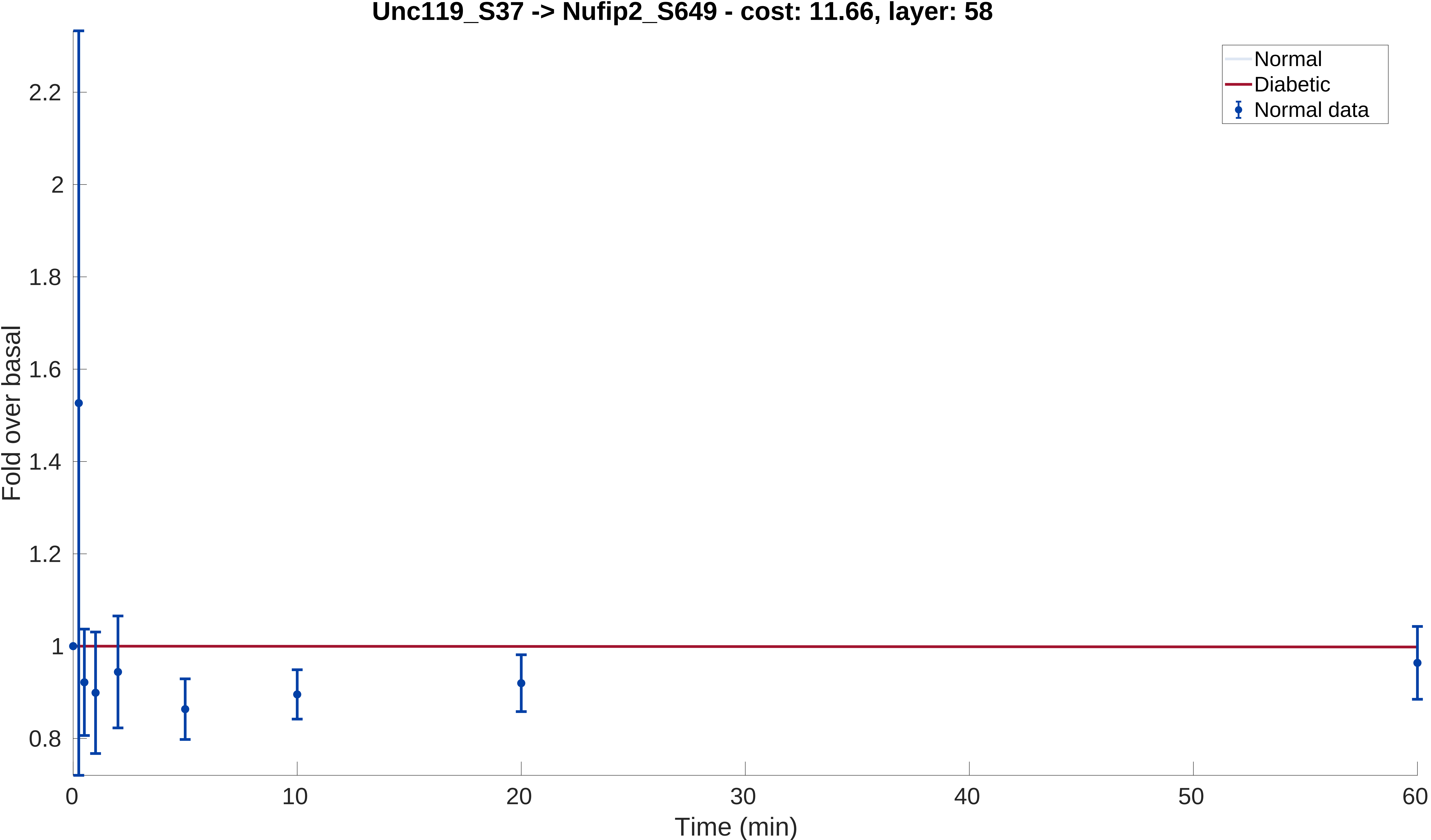

Unc119_S37 -> Nufip2_S649 - cost: 11.66, layer: 58
Normal
Diabetic
Normal data
Fold over basal
2.2
2
1.8
1.6
1.4
1.2
1
0.8
0
10
20
30
40
50
60
Time (min)

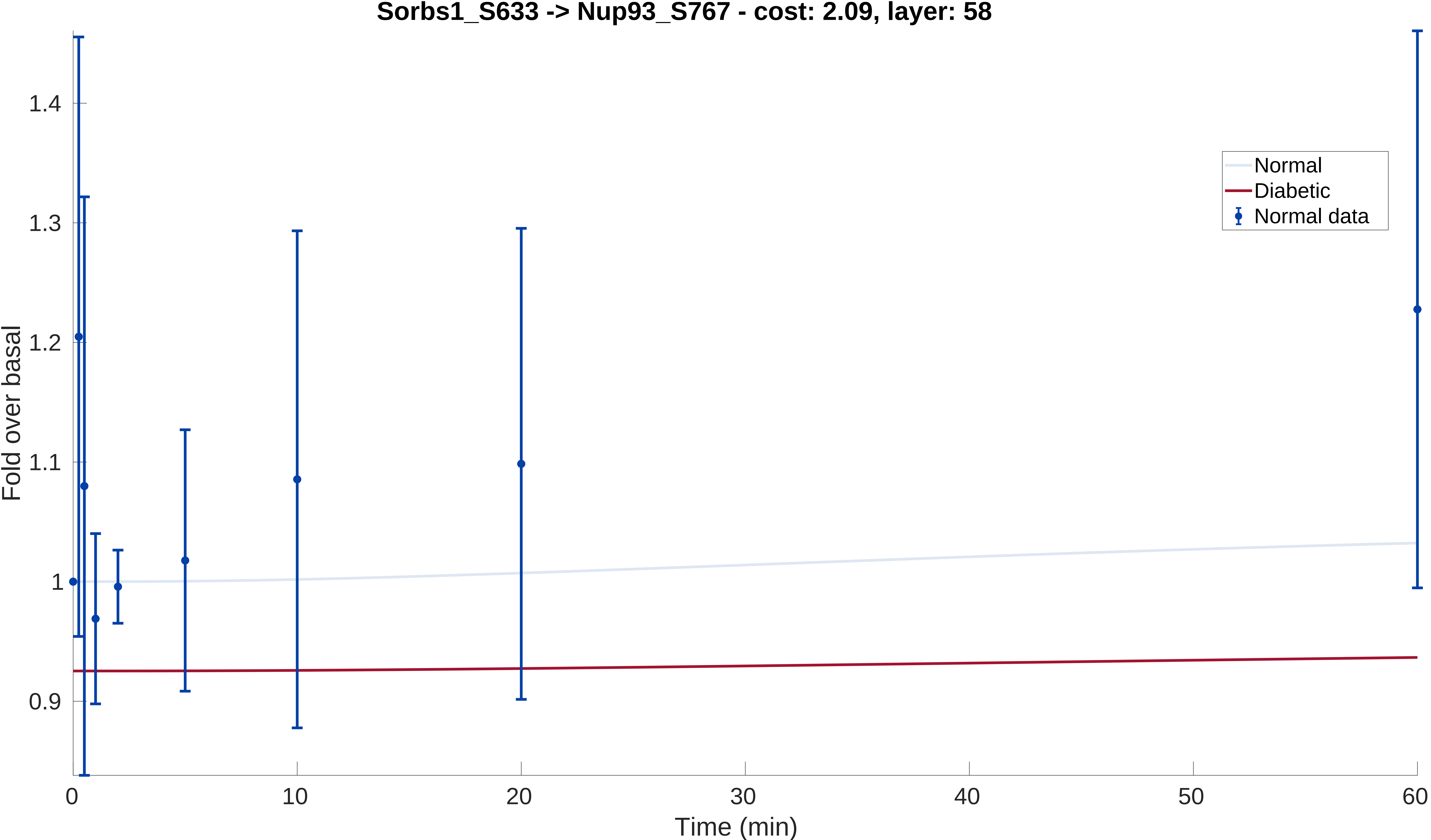

Sorbs1_S633 -> Nup93_S767 - cost: 2.09, layer: 58
Fold over basal
1.4
1.3
1.2
1.1
1
0.9
0
10
20
30
40
50
60
Time (min)
Normal
Diabetic
Normal data

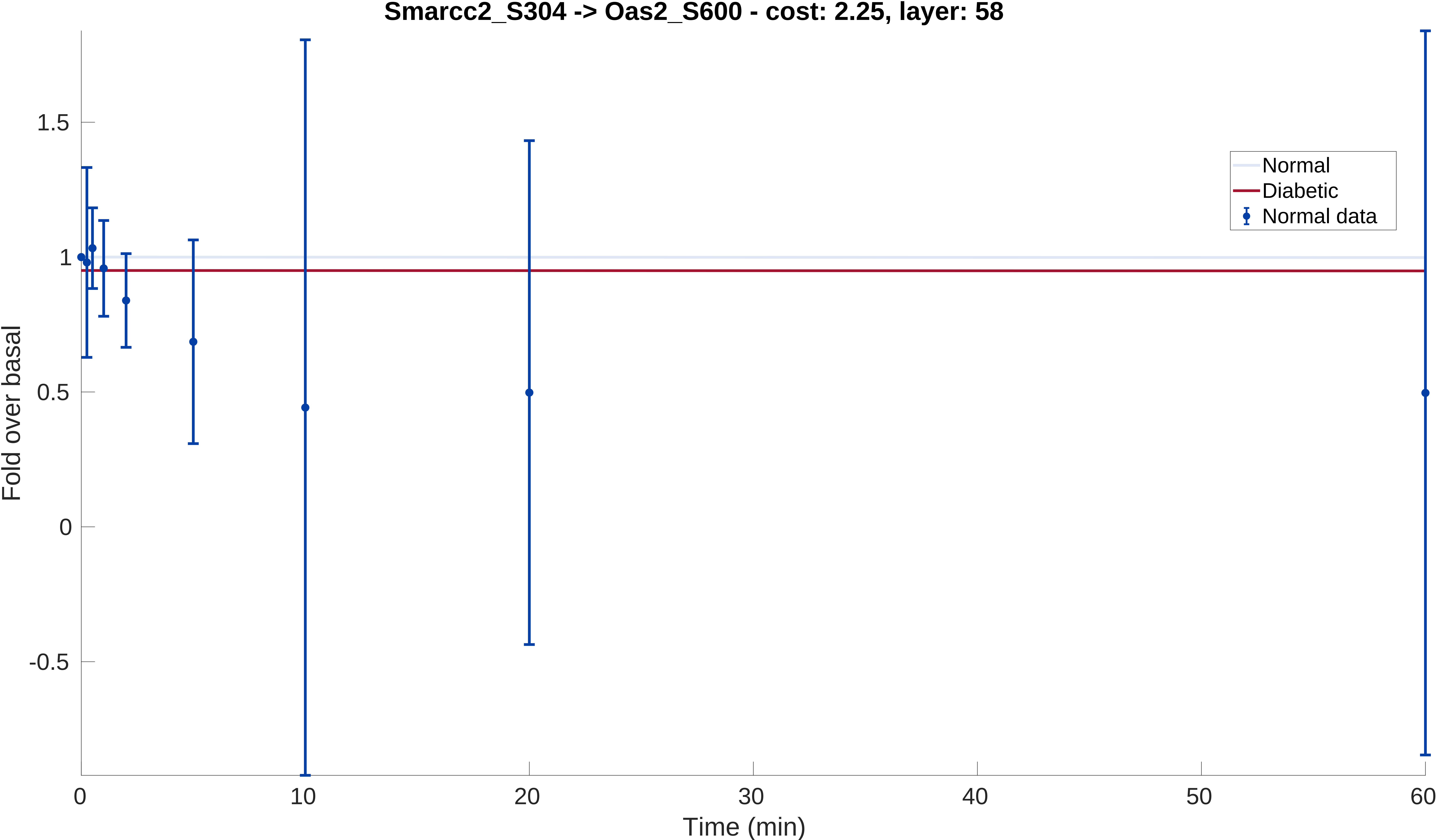
Smarcc2_S304 -> Oas2_S600 - cost: 2.25, layer: 58
Normal
Diabetic
Normal data
Fold over basal
1.5
1
0.5
0
-0.5
0
10
20
30
40
50
60
Time (min)

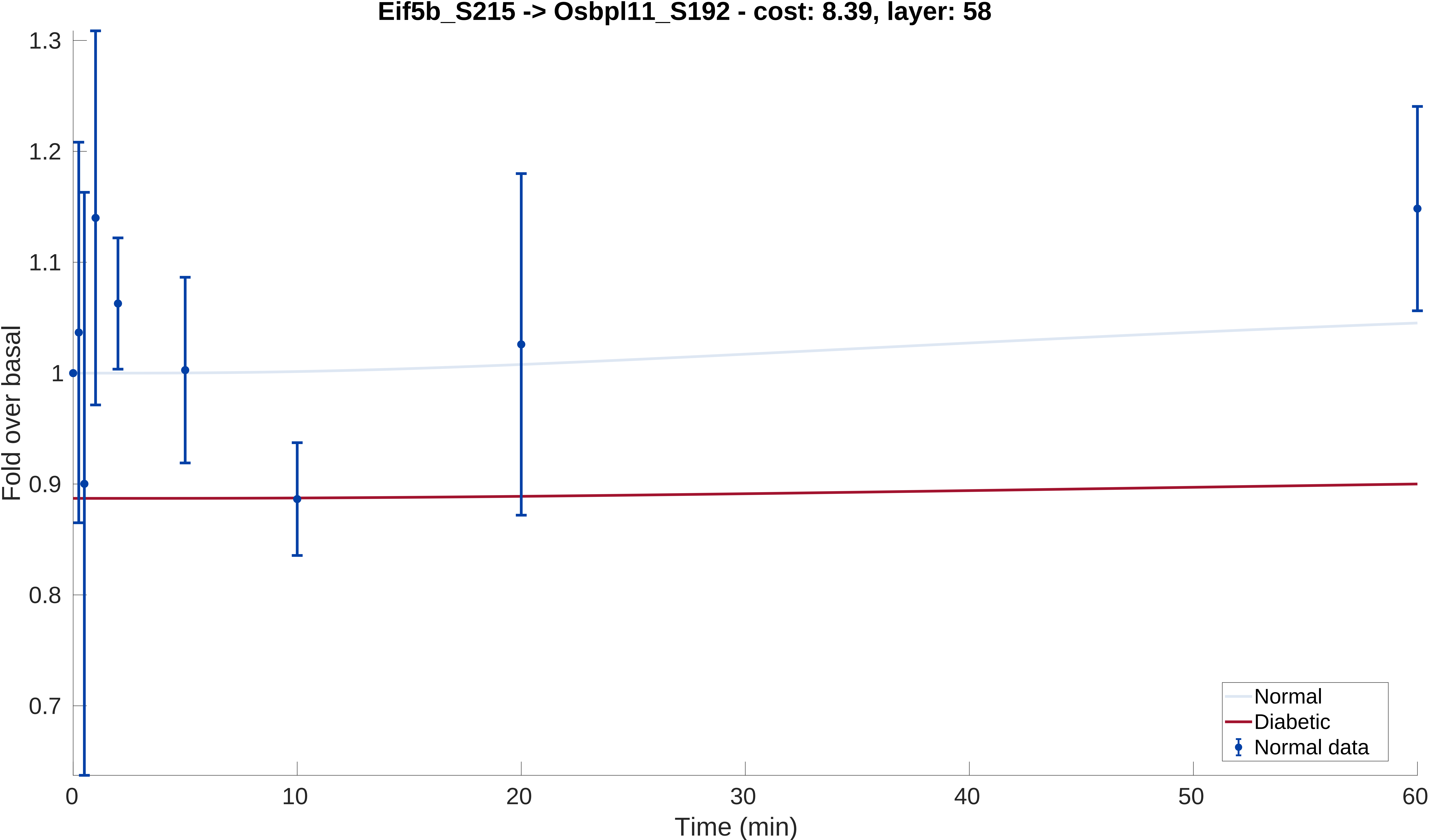

Eif5b_S215 -> Osbpl11_S192 - cost: 8.39, layer: 58
Fold over basal
1.3
1.2
1.1
1
0.9
0.8
0.7
0
10
20
30
40
50
60
Time (min)
Normal
Diabetic
Normal data

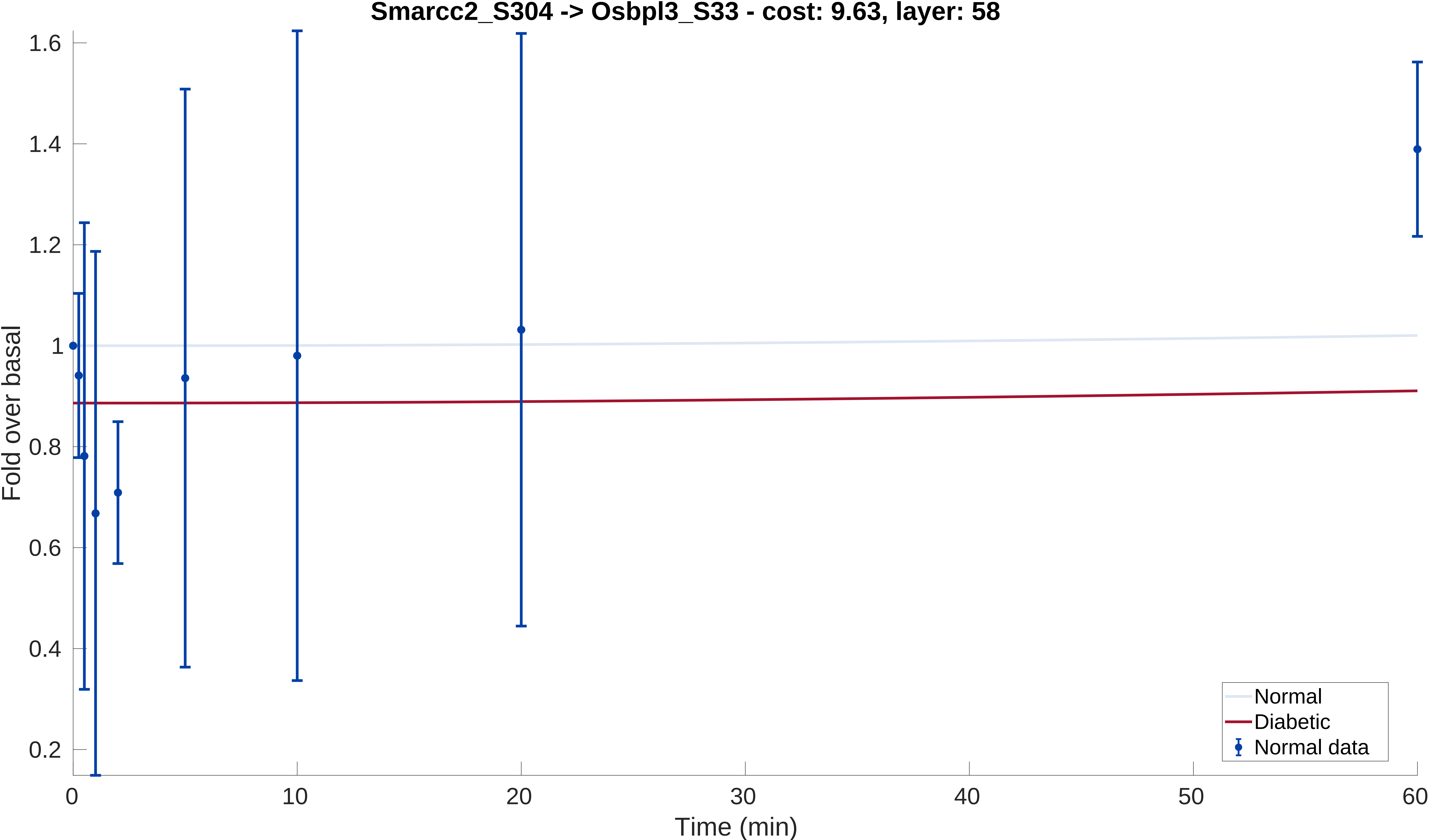

Smarcc2_S304 -> Osbpl3_S33 - cost: 9.63, layer: 58
Fold over basal
1.6
1.4
1.2
1
0.8
0.6
0.4
0.2
0
10
20
30
40
50
60
Time (min)
Normal
Diabetic
Normal data

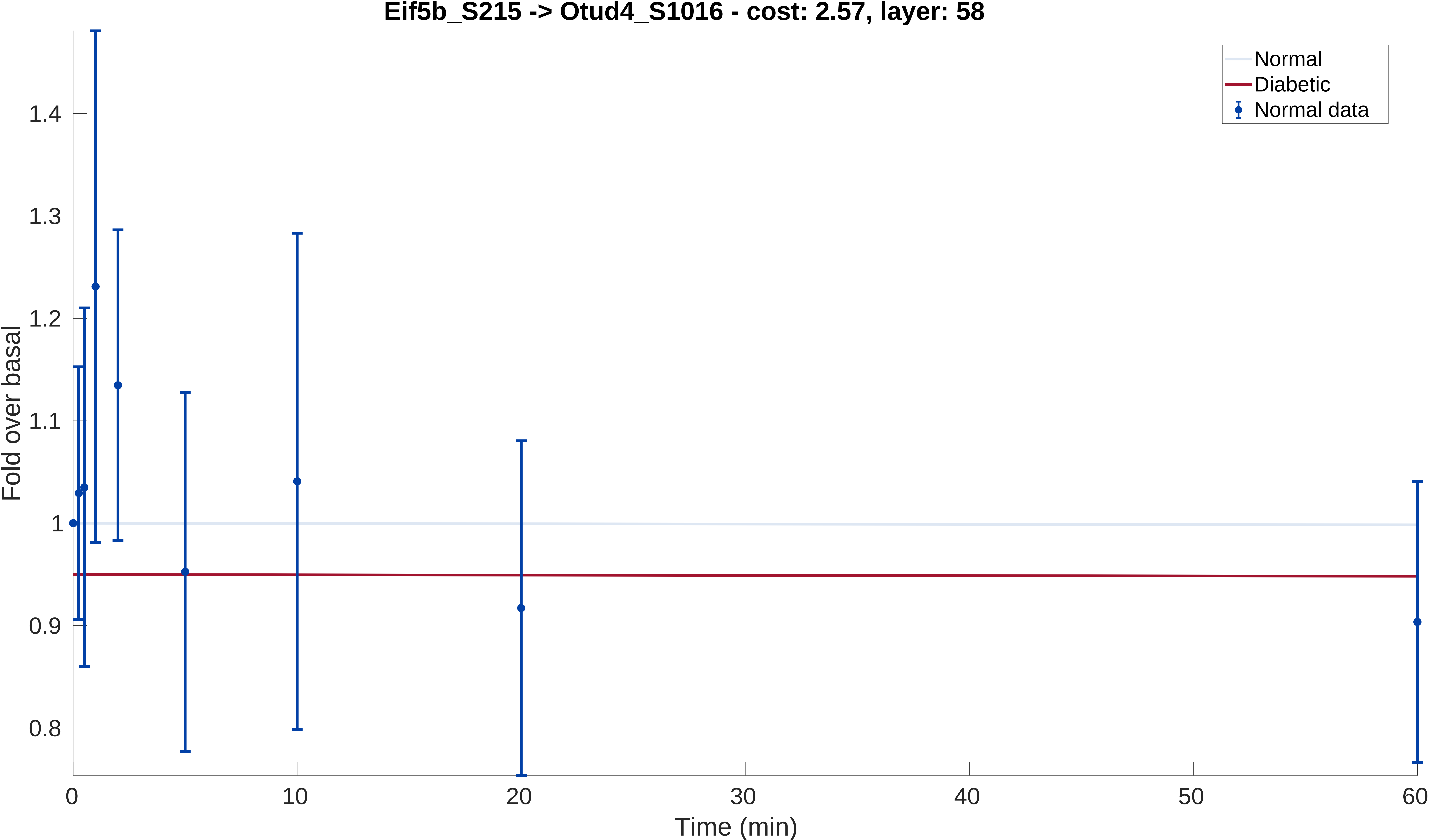
Eif5b_S215 -> Otud4_S1016 - cost: 2.57, layer: 58
Normal
Diabetic
Normal data
Fold over basal
1.4
1.3
1.2
1.1
1
0.9
0.8
0
10
20
30
40
50
60
Time (min)

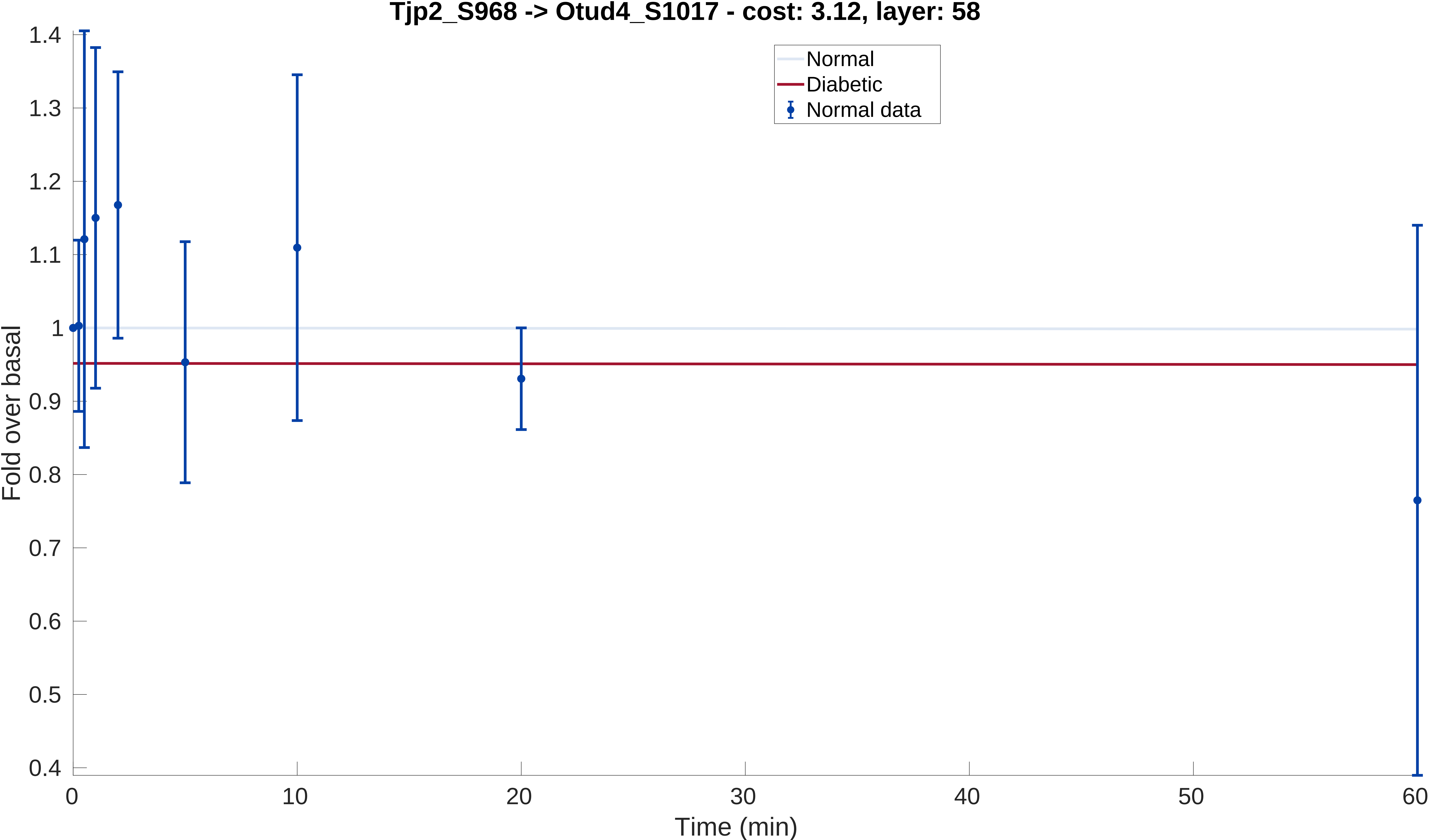

Tjp2_S968 -> Otud4_S1017 - cost: 3.12, layer: 58
Normal
Diabetic
Normal data
Fold over basal
1.4
1.3
1.2
1.1
1
0.9
0.8
0.7
0.6
0.5
0.4
0
10
20
30
40
50
60
Time (min)

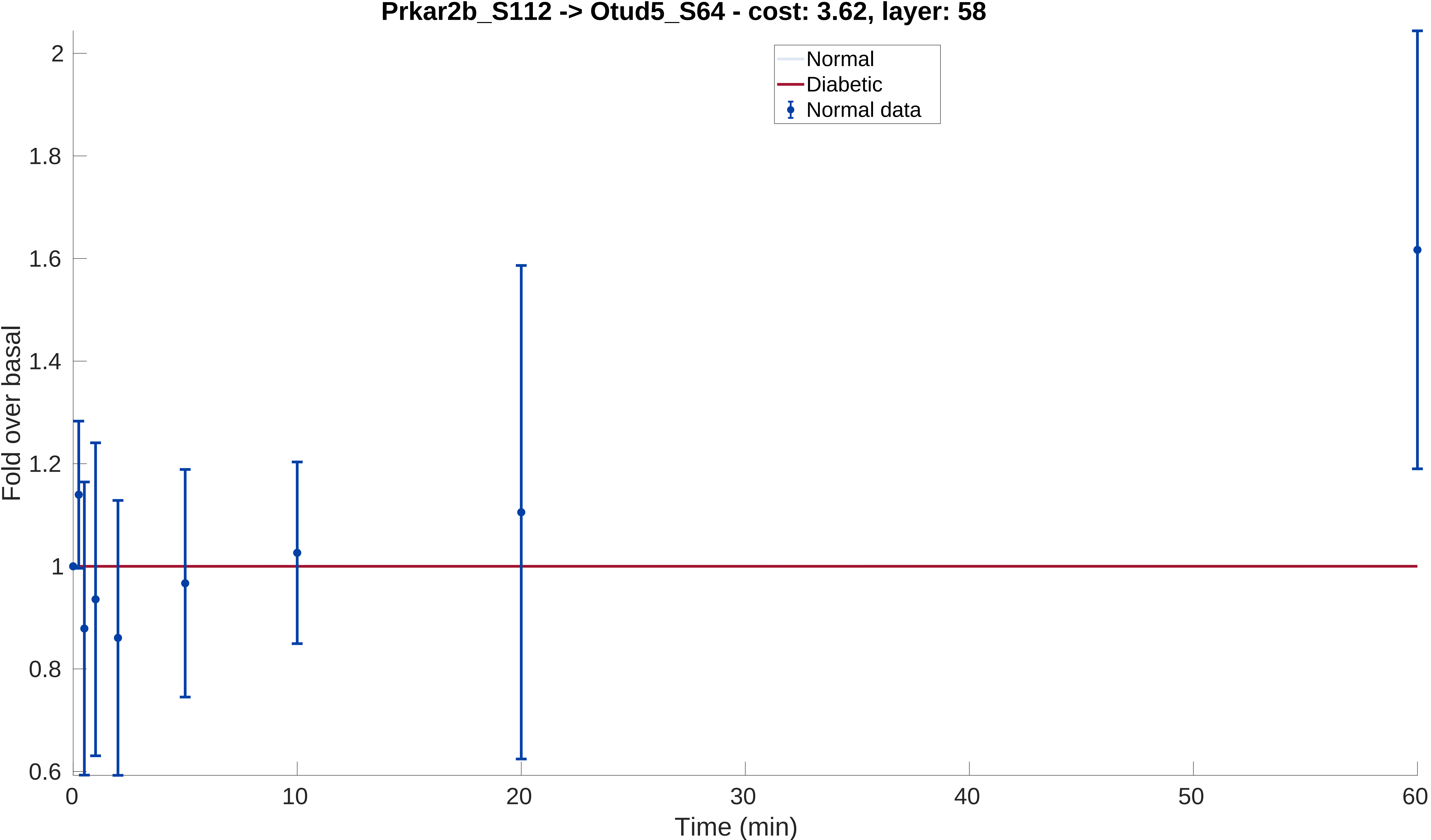

Prkar2b_S112 -> Otud5_S64 - cost: 3.62, layer: 58
Normal
Diabetic
Normal data
Fold over basal
Time (min)
2
1.8
1.6
1.4
1.2
1
0.8
0.6
0
10
20
30
40
50
60

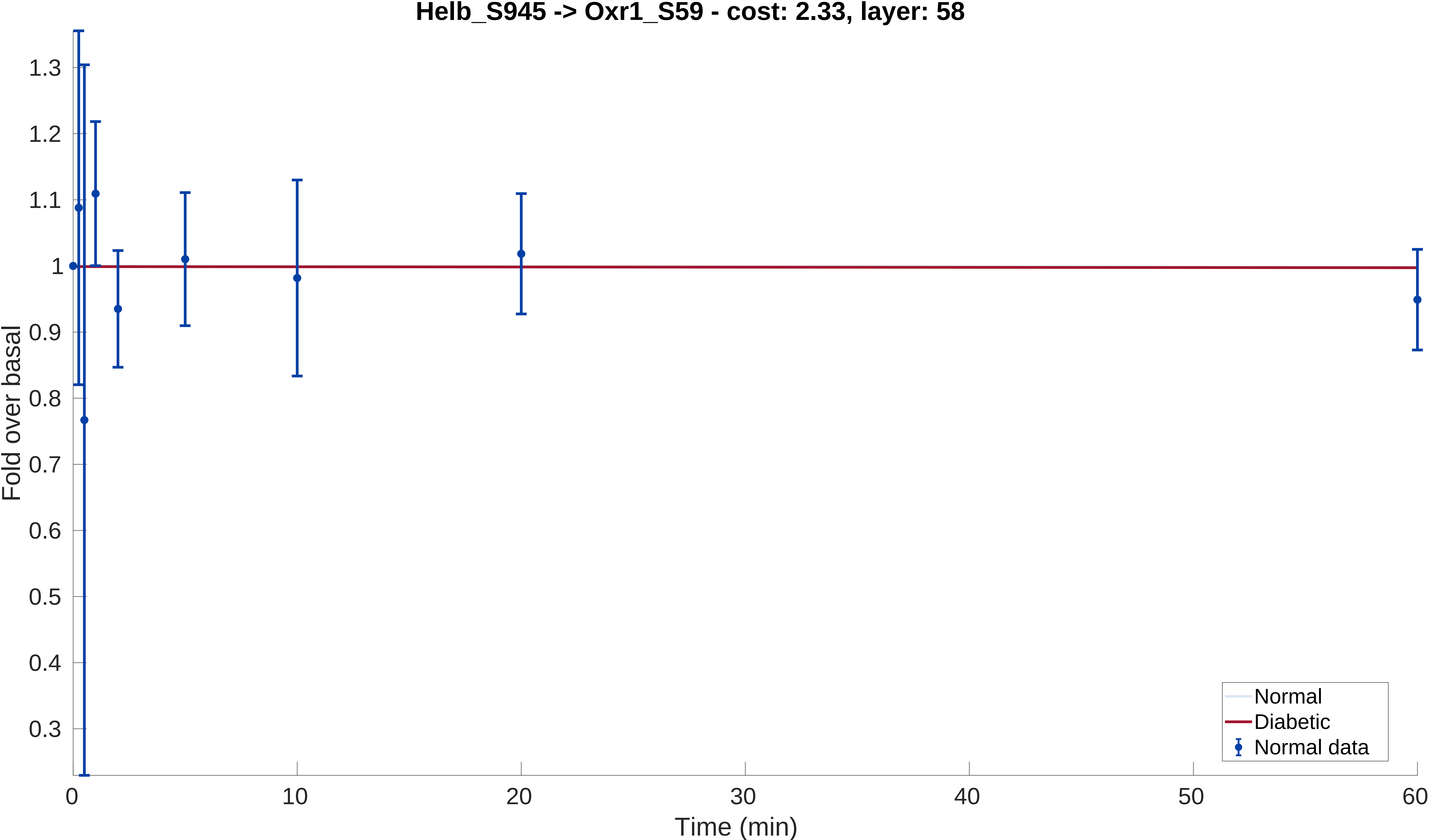

Helb_S945 -> Oxr1_S59 - cost: 2.33, layer: 58
1.3
1.2
1.1
1
0.9
0.8
0.7
0.6
0.5
0.4
0.3
Fold over basal
0
10
20
30
40
50
60
Time (min)
Normal
Diabetic
Normal data

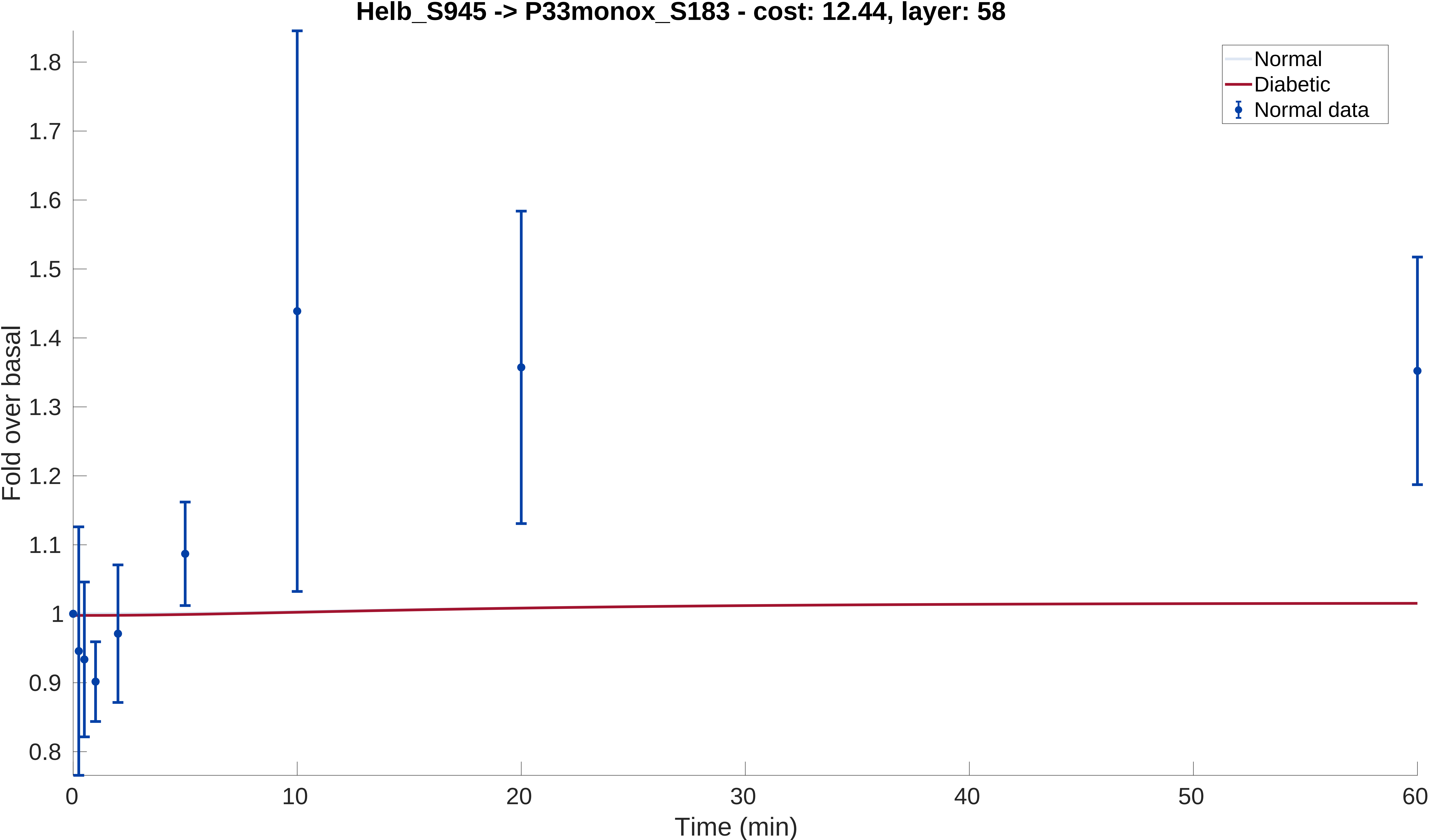

Helb_S945 -> P33monox_S183 - cost: 12.44, layer: 58
Normal
Diabetic
Normal data
Fold over basal
1.8
1.7
1.6
1.5
1.4
1.3
1.2
1.1
1
0.9
0.8
0
10
20
30
40
50
60
Time (min)

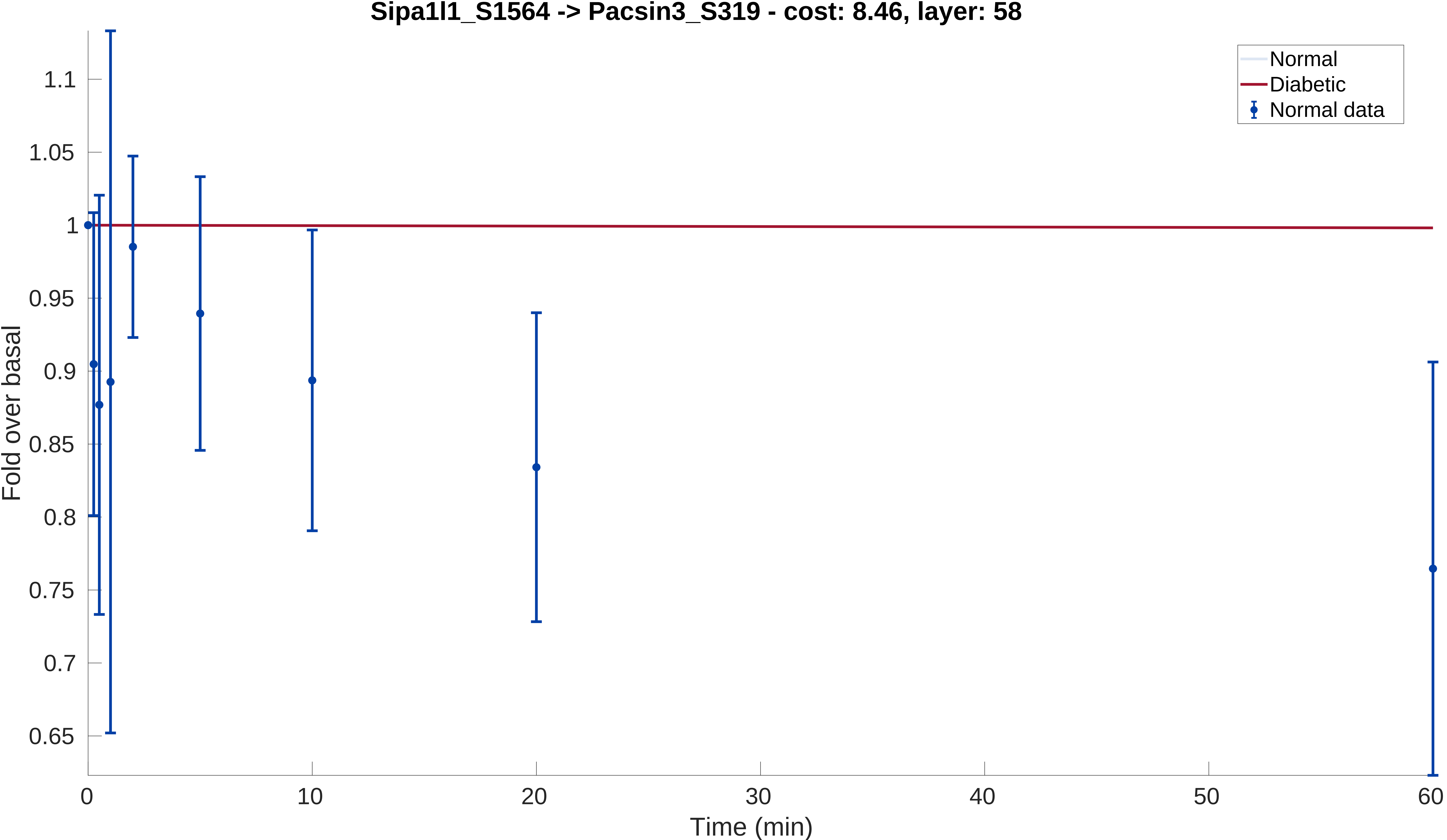

Sipa1l1_S1564 -> Pacsin3_S319 - cost: 8.46, layer: 58
Normal
Diabetic
Normal data
Fold over basal
Time (min)
1.1
1.05
1
0.95
0.9
0.85
0.8
0.75
0.7
0.65
0
10
20
30
40
50
60

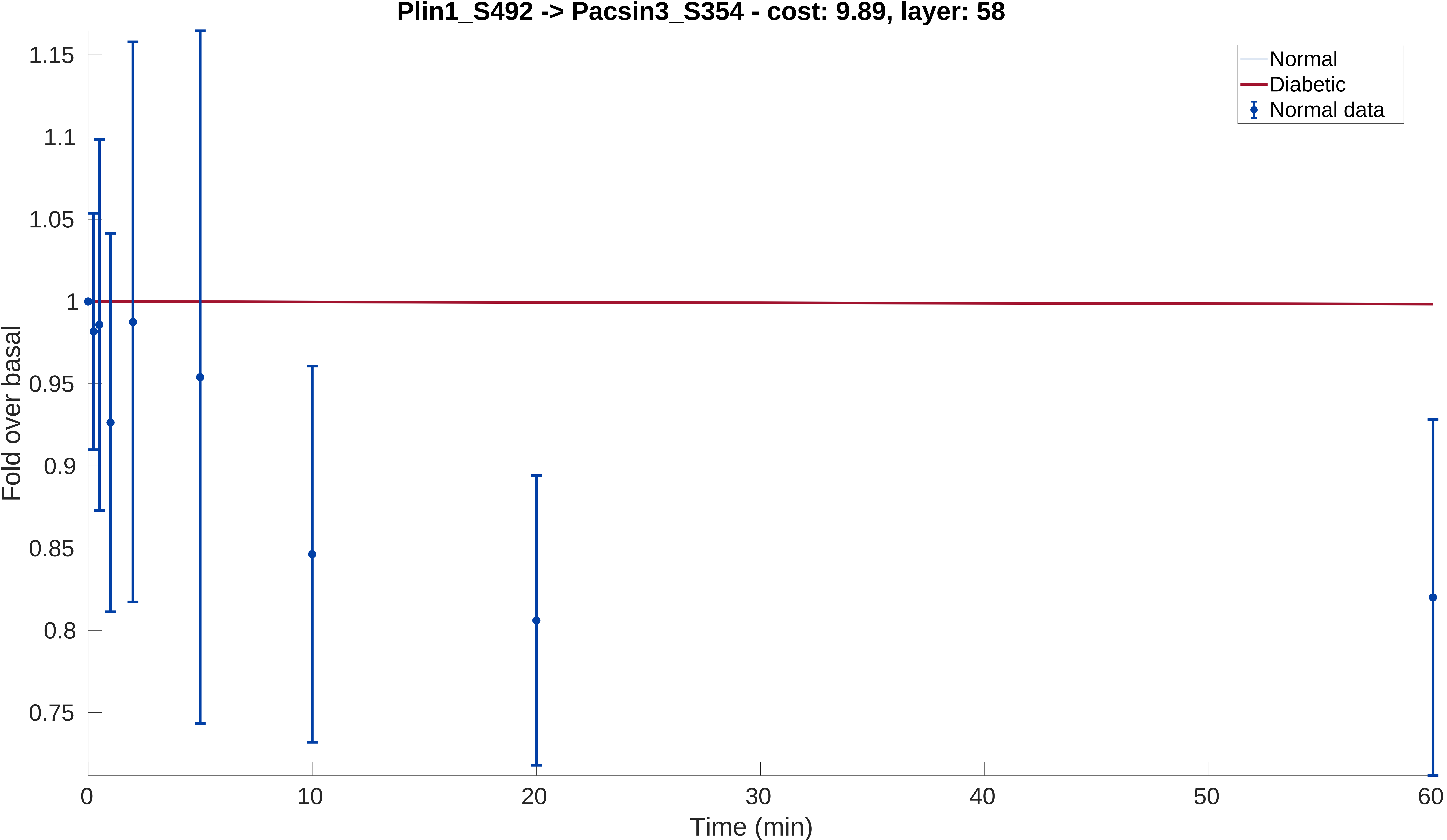

Plin1_S492 -> Pacsin3_S354 - cost: 9.89, layer: 58
Normal
Diabetic
Normal data
Fold over basal
1.15
1.1
1.05
1
0.95
0.9
0.85
0.8
0.75
0
10
20
30
40
50
60
Time (min)

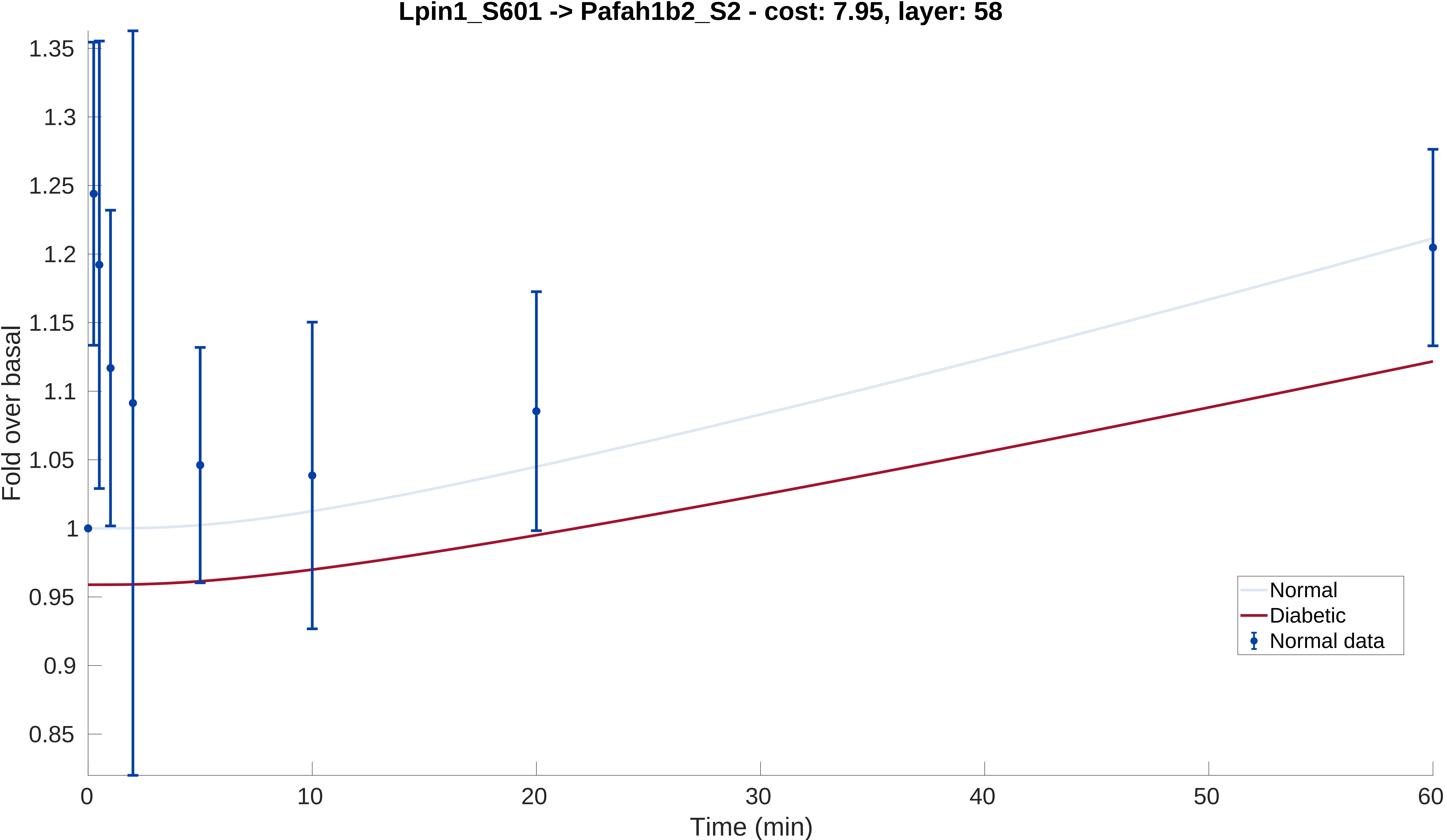
Lpin1_S601 -> Pafah1b2_S2 - cost: 7.95, layer: 58
Fold over basal
1.35
1.3
1.25
1.2
1.15
1.1
1.05
1
0.95
0.9
0.85
0
10
20
30
40
50
60
Time (min)
Normal
Diabetic
Normal data

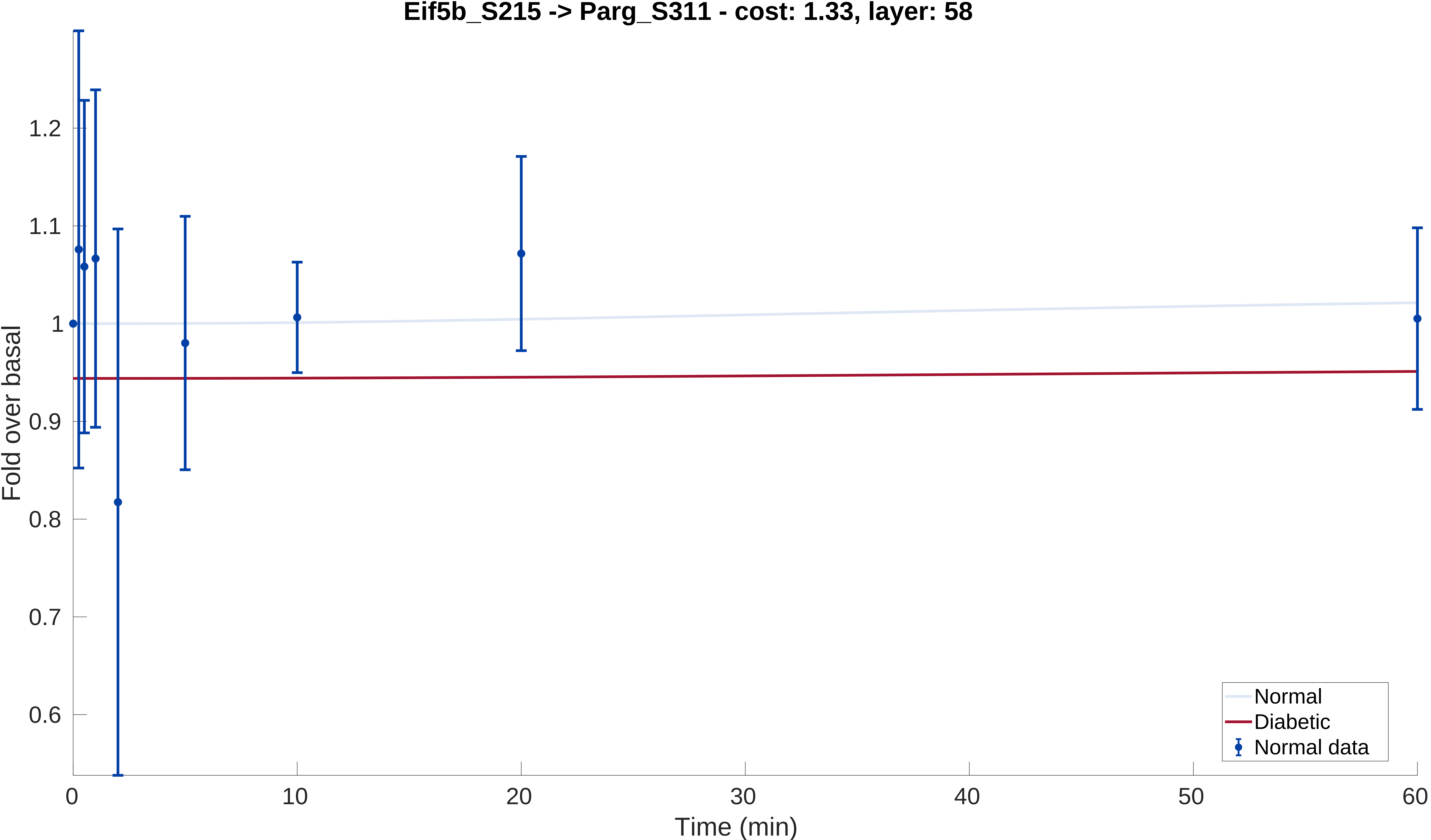

Eif5b_S215 -> Parg_S311 - cost: 1.33, layer: 58
Fold over basal
1.2
1.1
1
0.9
0.8
0.7
0.6
0
10
20
30
40
50
60
Time (min)
Normal
Diabetic
Normal data

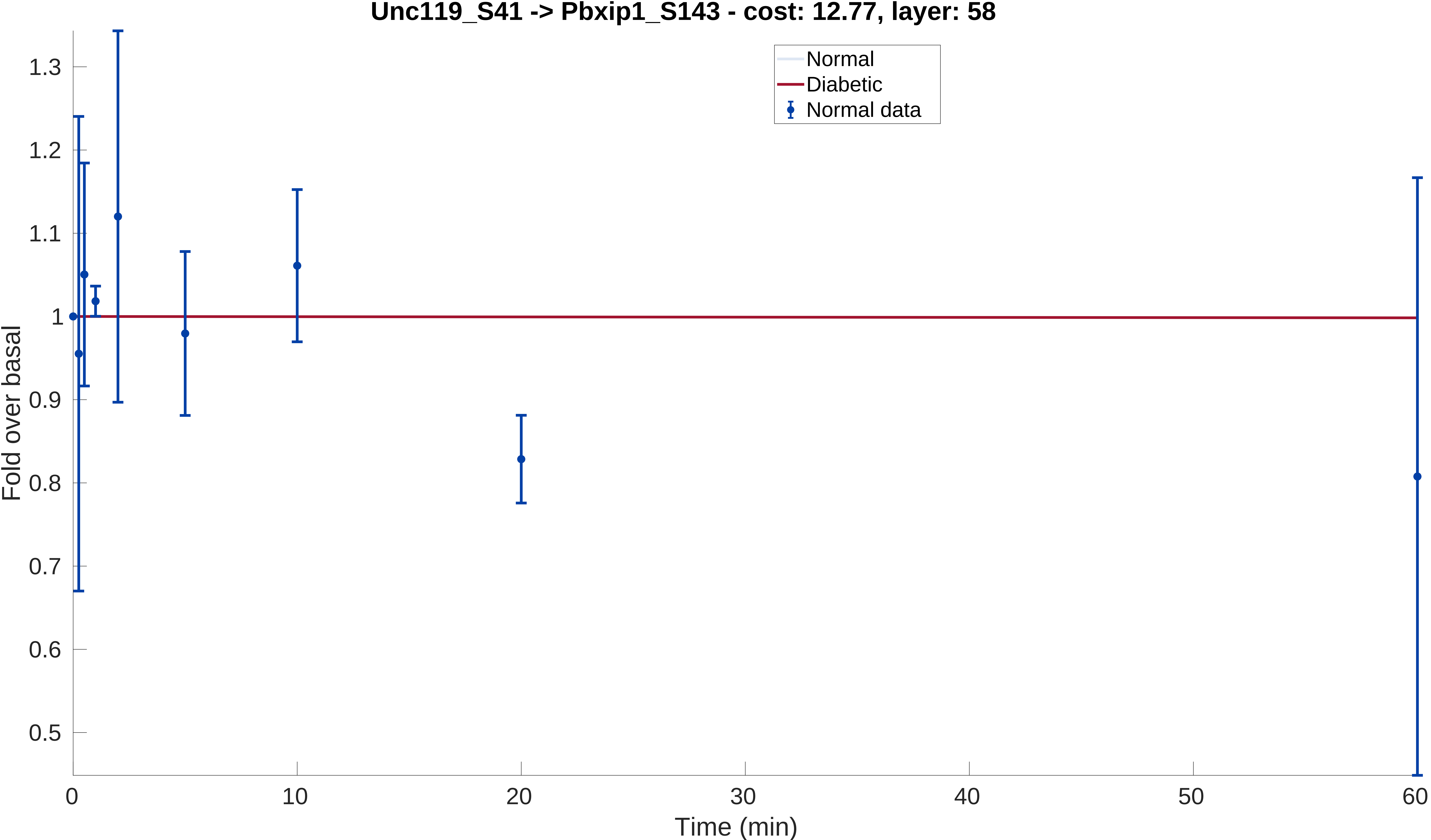

Unc119_S41 -> Pbxip1_S143 - cost: 12.77, layer: 58
Normal
Diabetic
Normal data
Fold over basal
1.3
1.2
1.1
1
0.9
0.8
0.7
0.6
0.5
0
10
20
30
40
50
60
Time (min)

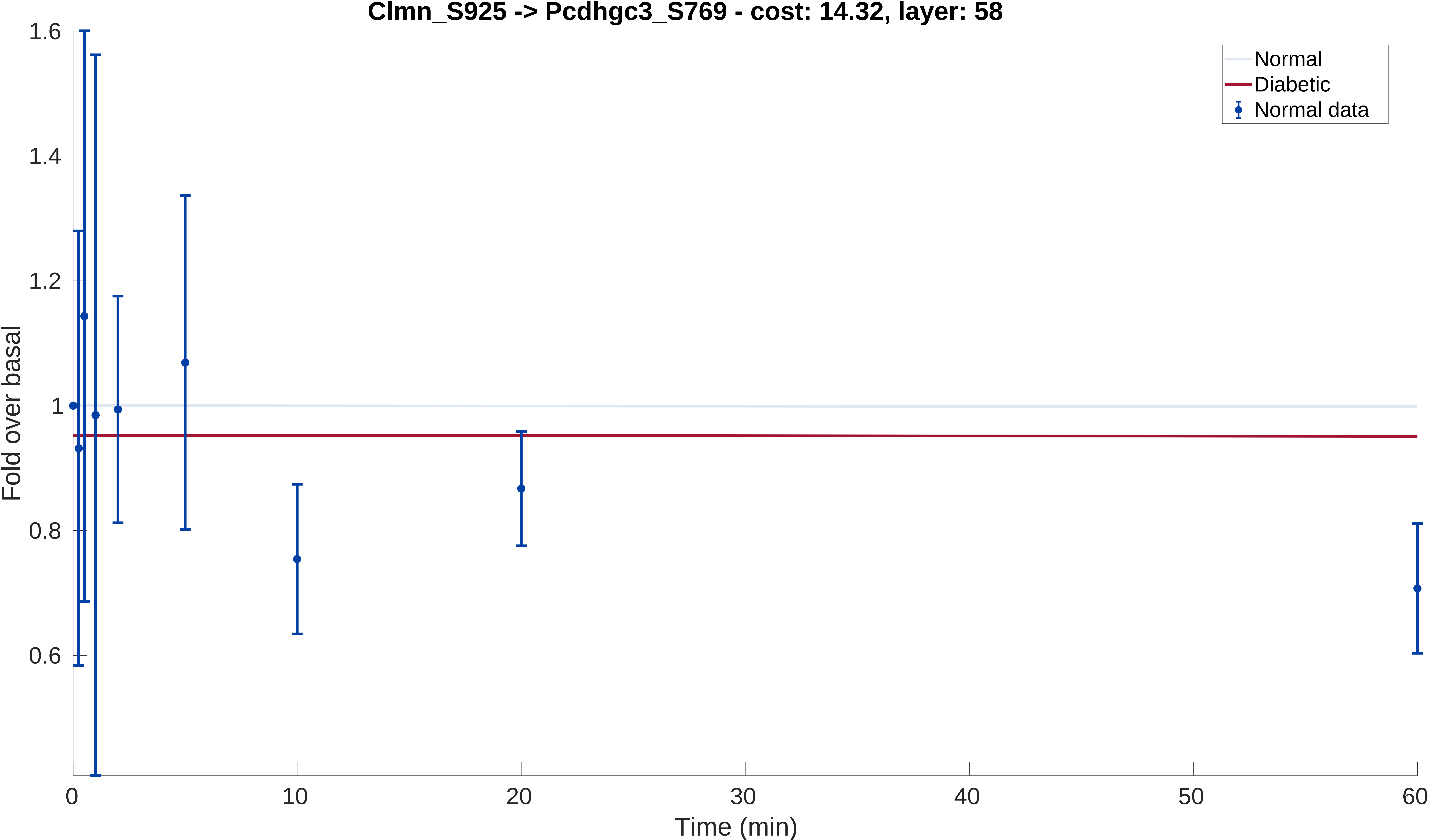

Clmn_S925 -> Pcdhgc3_S769 - cost: 14.32, layer: 58
Normal
Diabetic
Normal data
Fold over basal
1.6
1.4
1.2
1
0.8
0.6
0
10
20
30
40
50
60
Time (min)

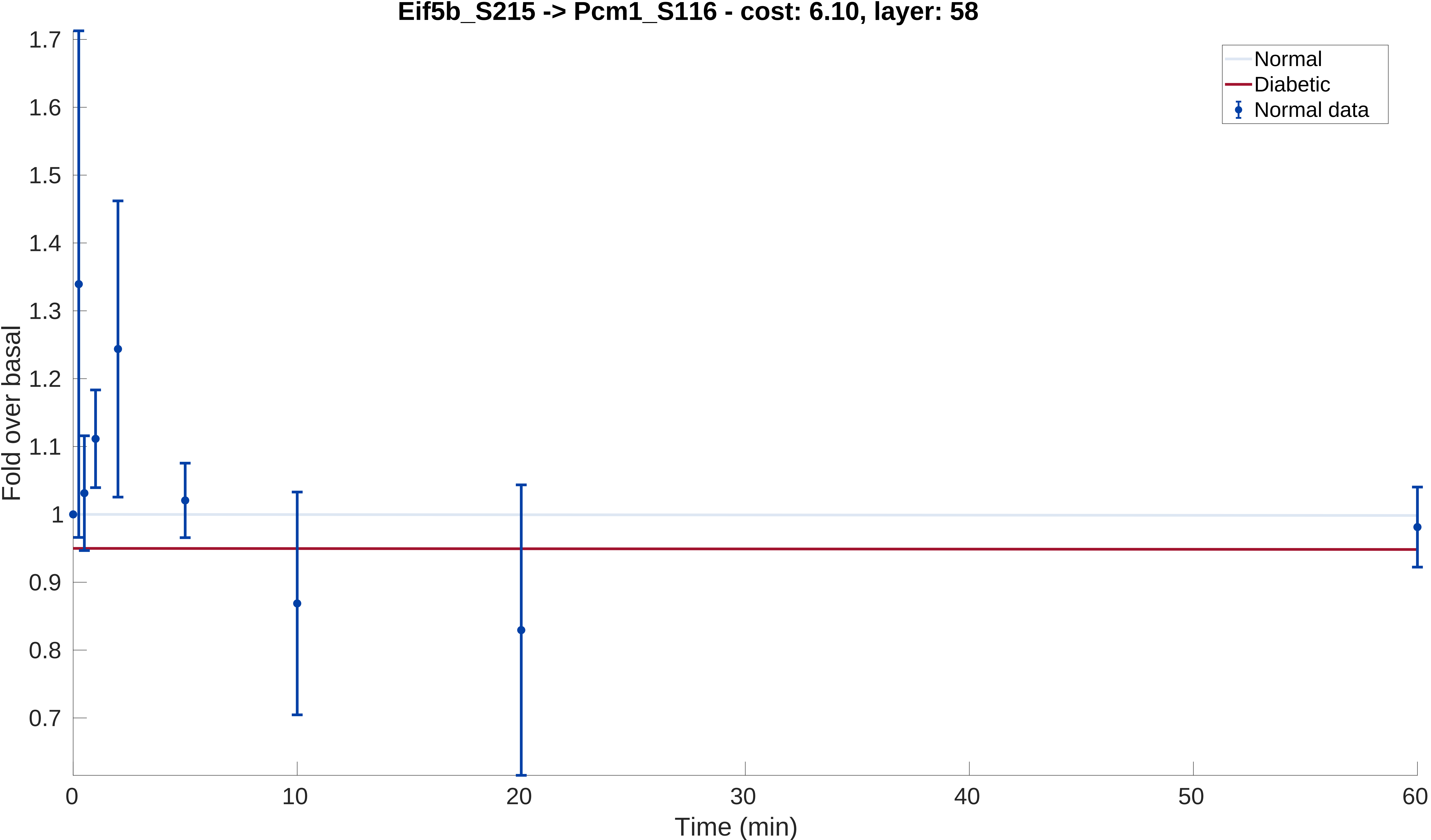

Eif5b_S215 -> Pcm1_S116 - cost: 6.10, layer: 58
Normal
Diabetic
Normal data
Fold over basal
1.7
1.6
1.5
1.4
1.3
1.2
1.1
1
0.9
0.8
0.7
0
10
20
30
40
50
60
Time (min)

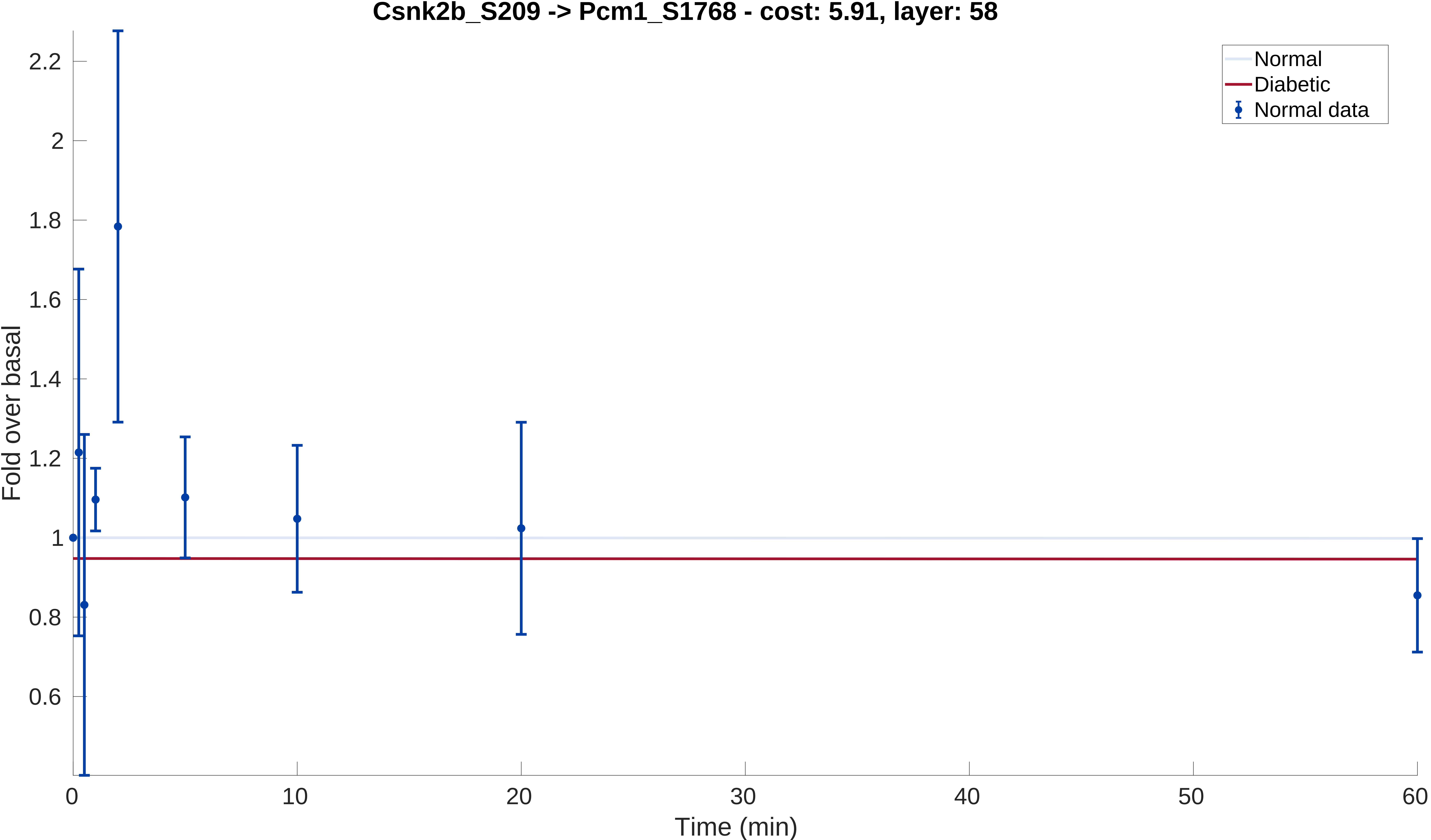

Csnk2b_S209 -> Pcm1_S1768 - cost: 5.91, layer: 58
Normal
Diabetic
Normal data
Fold over basal
2.2
2
1.8
1.6
1.4
1.2
1
0.8
0.6
0
10
20
30
40
50
60
Time (min)

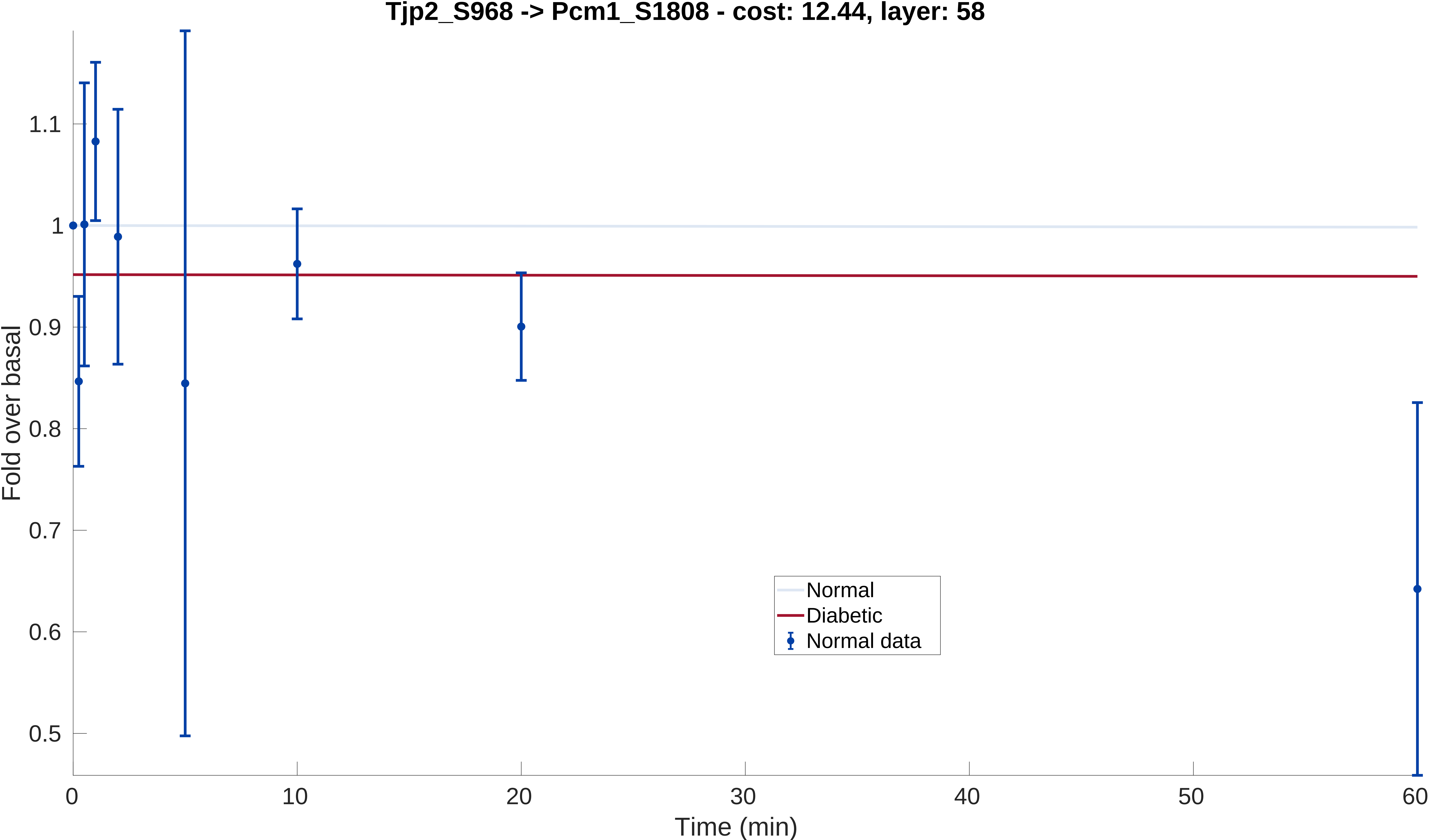

Tjp2_S968 -> Pcm1_S1808 - cost: 12.44, layer: 58
Fold over basal
1.1
1
0.9
0.8
0.7
0.6
0.5
0
10
20
30
40
50
60
Time (min)
Normal
Diabetic
Normal data

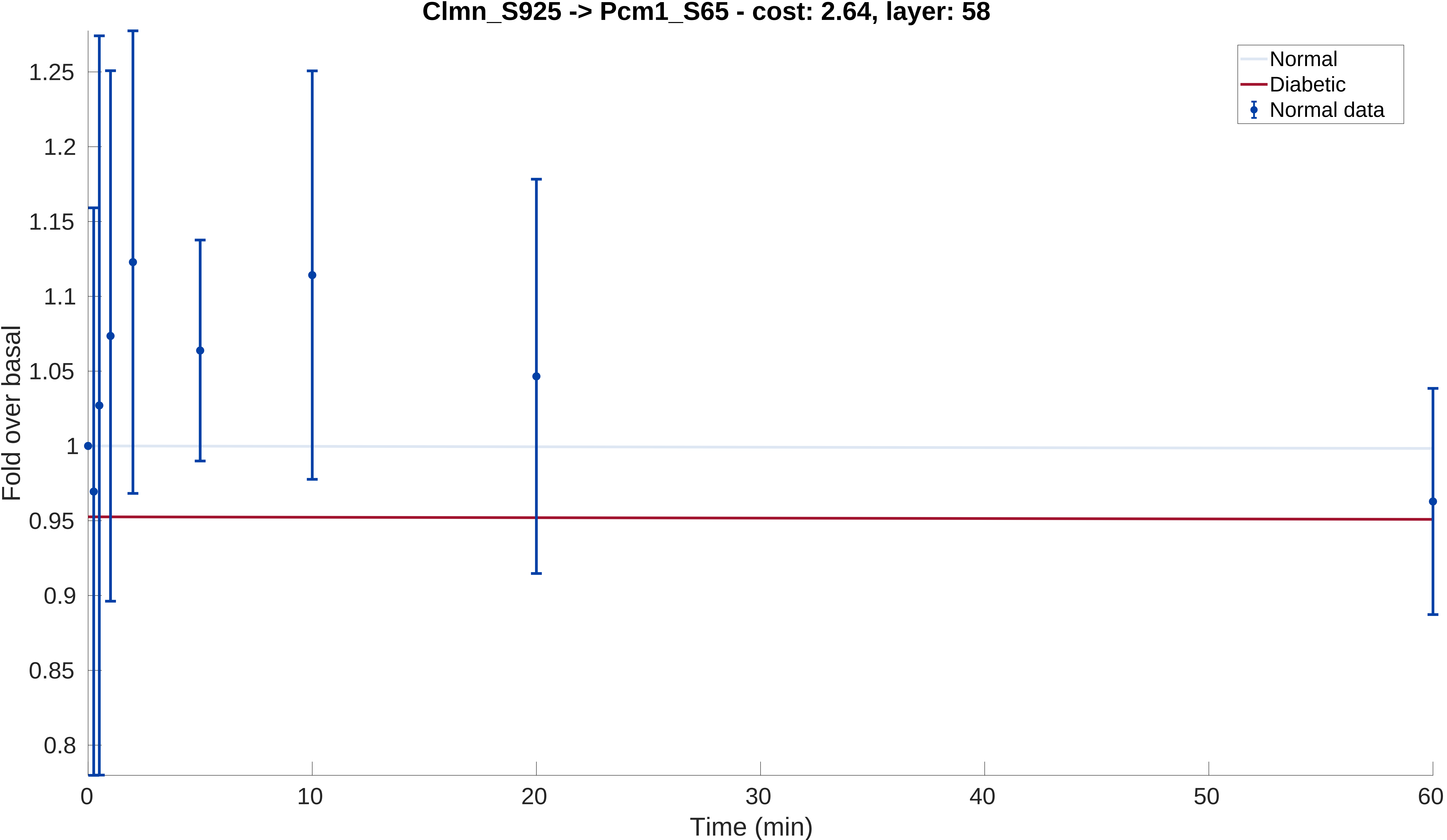

Clmn_S925 -> Pcm1_S65 - cost: 2.64, layer: 58
Normal
Diabetic
Normal data
Fold over basal
1.25
1.2
1.15
1.1
1.05
1
0.95
0.9
0.85
0.8
0
10
20
30
40
50
60
Time (min)

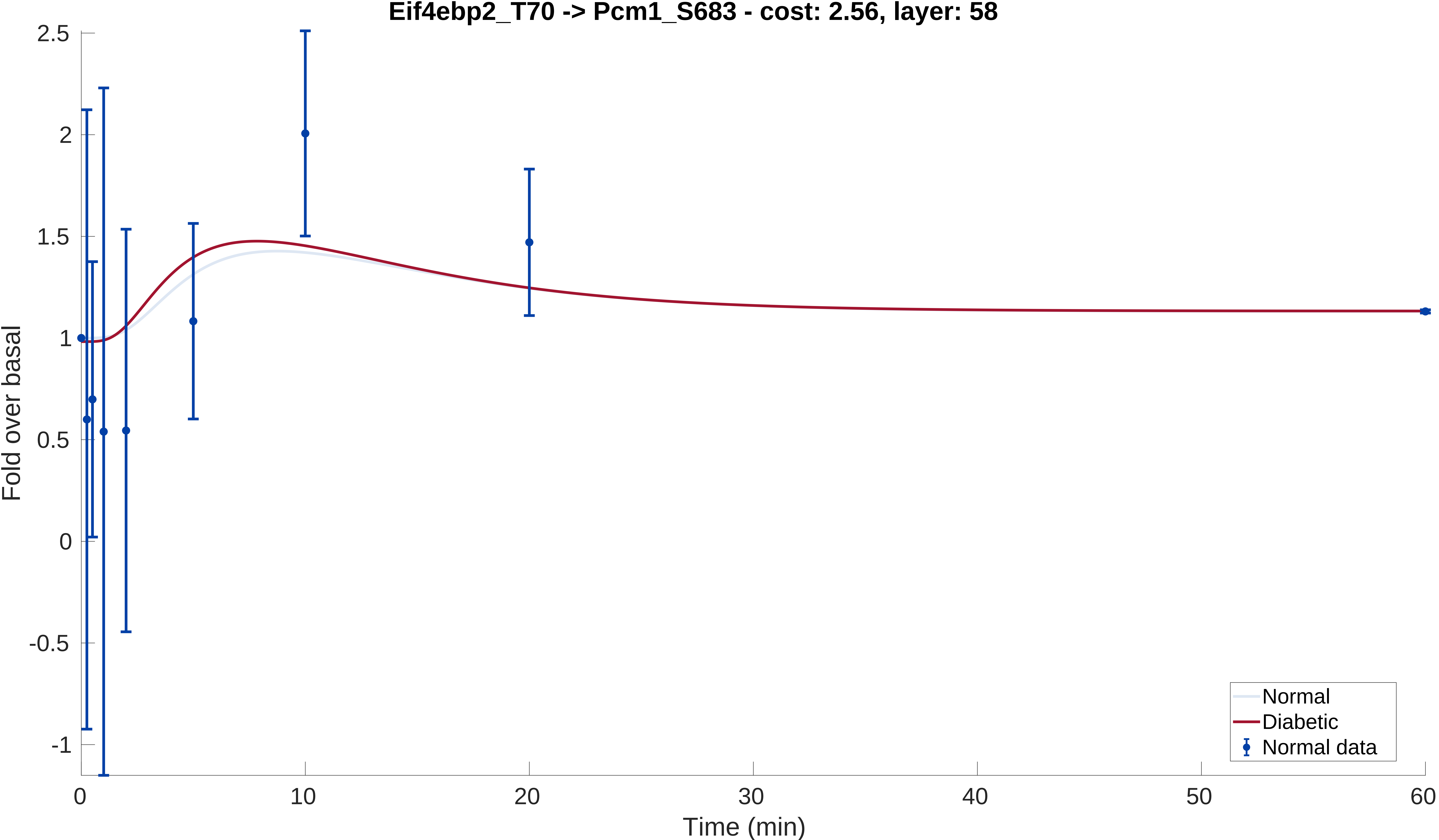

Eif4ebp2_T70 -> Pcm1_S683 - cost: 2.56, layer: 58
Fold over basal
Time (min)
2.5
2
1.5
1
0.5
0
-0.5
-1
0
10
20
30
40
50
60
Normal
Diabetic
Normal data

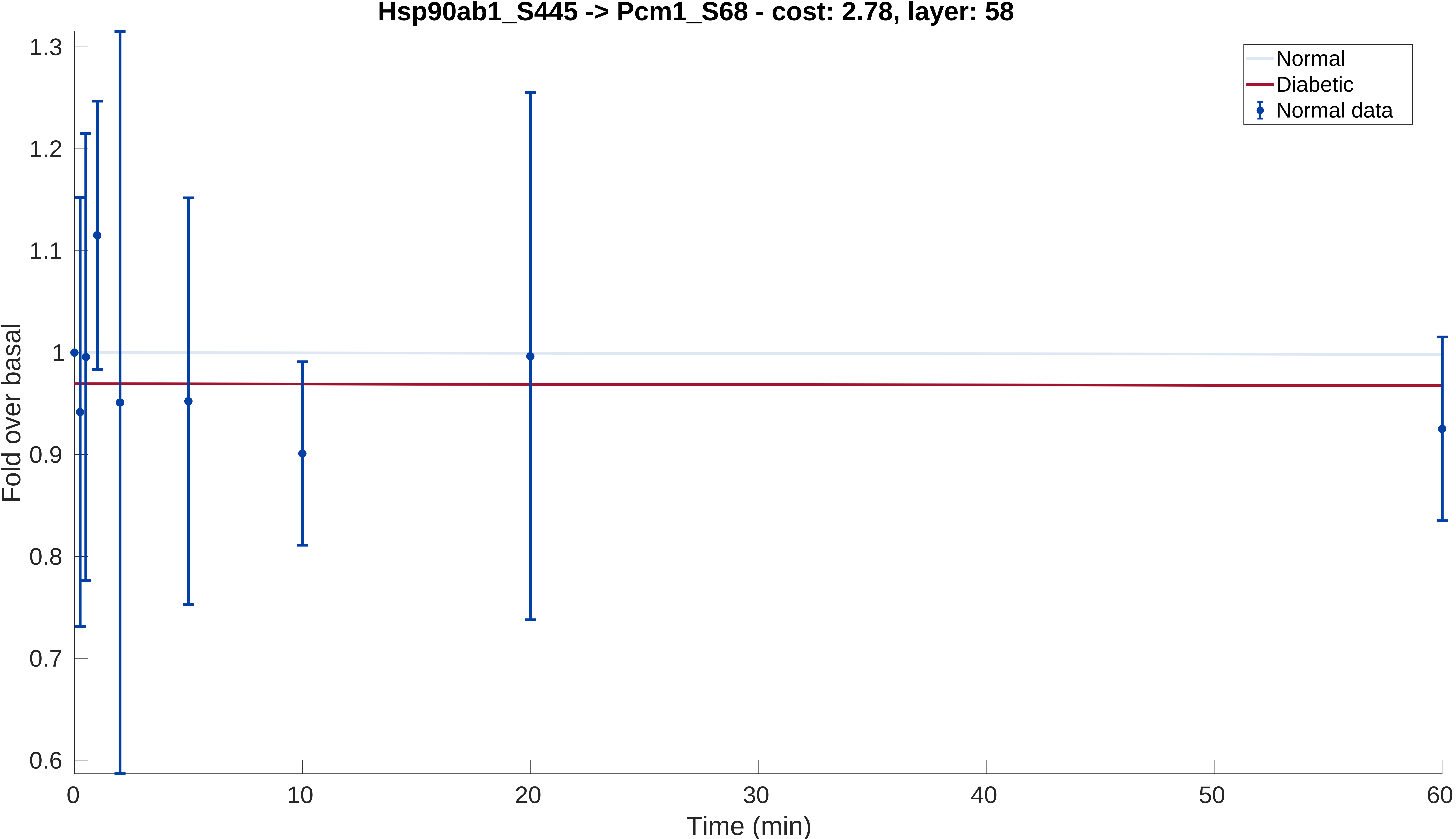
Hsp90ab1_S445 -> Pcm1_S68 - cost: 2.78, layer: 58
Normal
Diabetic
Normal data
Fold over basal
Time (min)
1.3
1.2
1.1
1
0.9
0.8
0.7
0.6
0
10
20
30
40
50
60

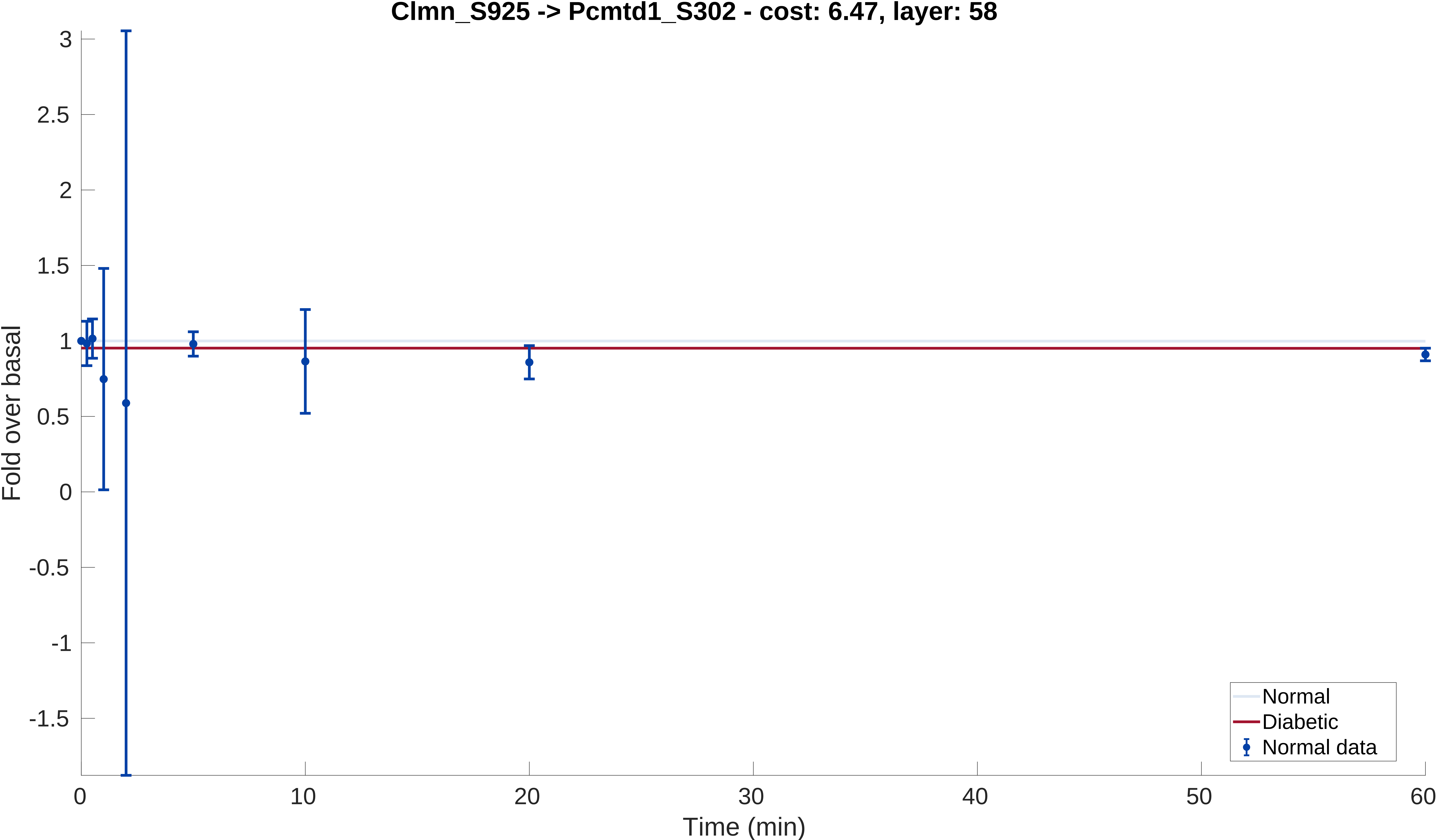

Clmn_S925 -> Pcmtd1_S302 - cost: 6.47, layer: 58
Fold over basal
Time (min)
3
2.5
2
1.5
1
0.5
0
-0.5
-1
-1.5
0
10
20
30
40
50
60
Normal
Diabetic
Normal data

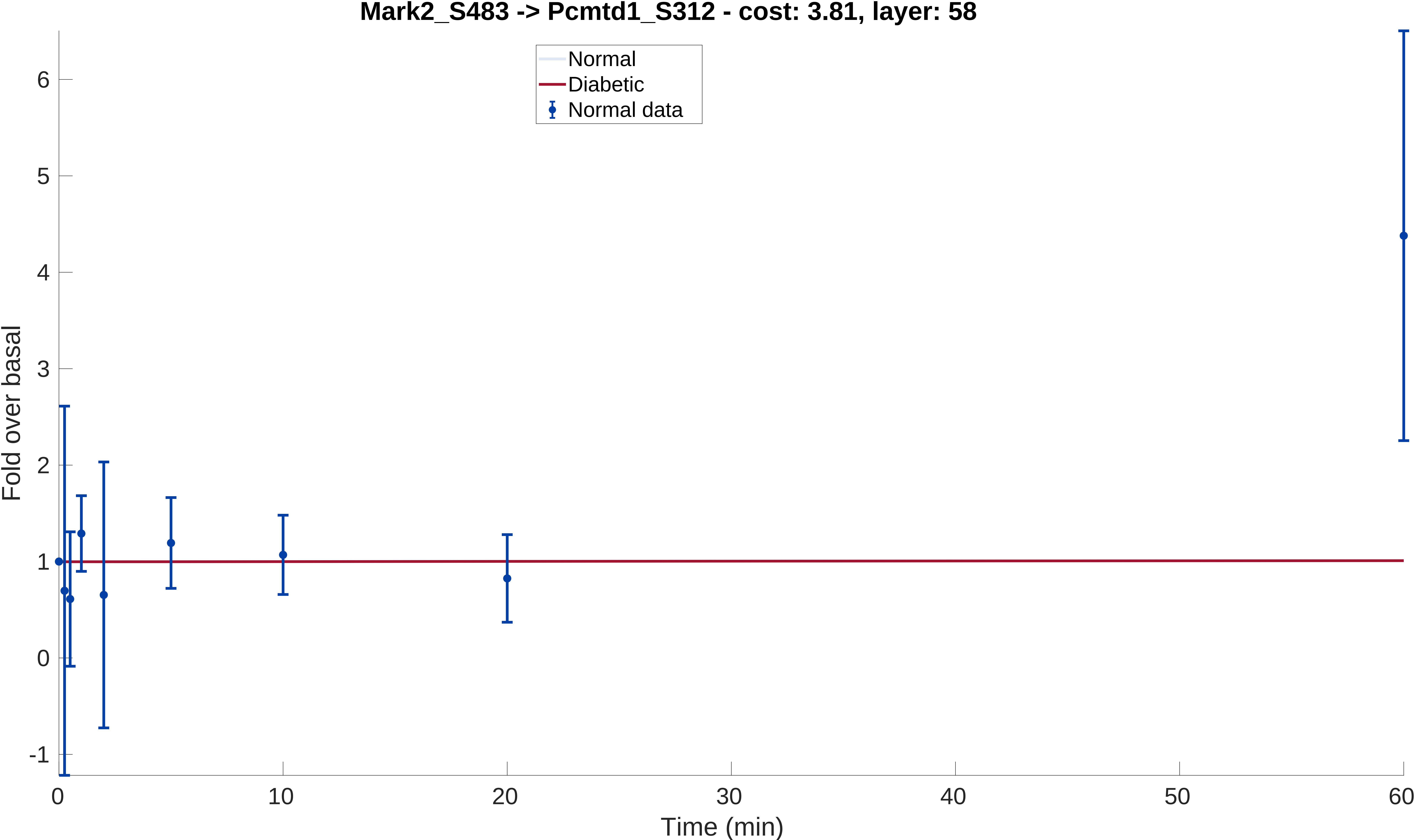

Mark2_S483 -> Pcmtd1_S312 - cost: 3.81, layer: 58
Normal
Diabetic
Normal data
Fold over basal
Time (min)
-1
0
1
2
3
4
5
6
0
10
20
30
40
50
60

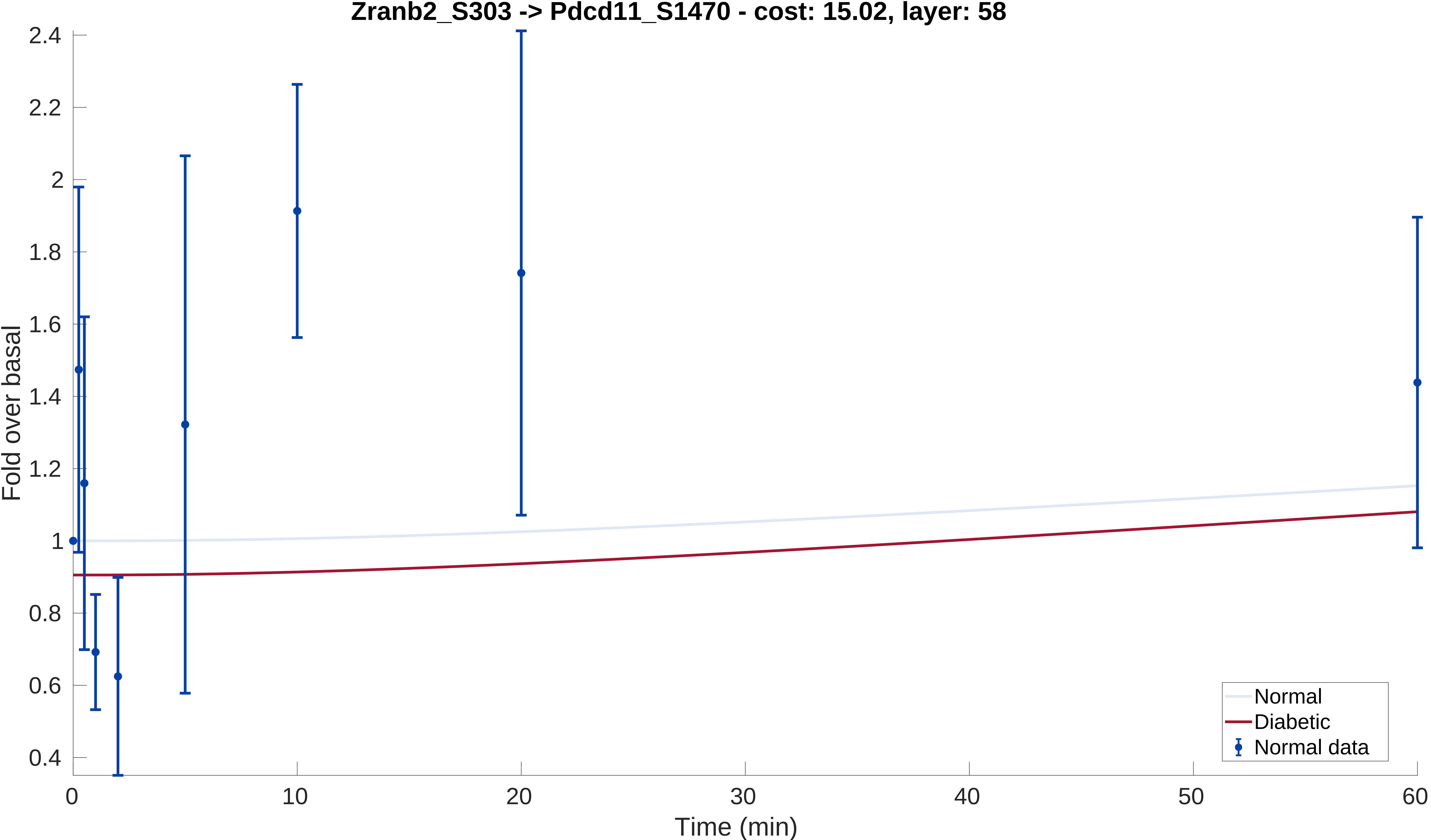

Zranb2_S303 -> Pdcd11_S1470 - cost: 15.02, layer: 58
Fold over basal
2.4
2.2
2
1.8
1.6
1.4
1.2
1
0.8
0.6
0.4
0
10
20
30
40
50
60
Time (min)
Normal
Diabetic
Normal data

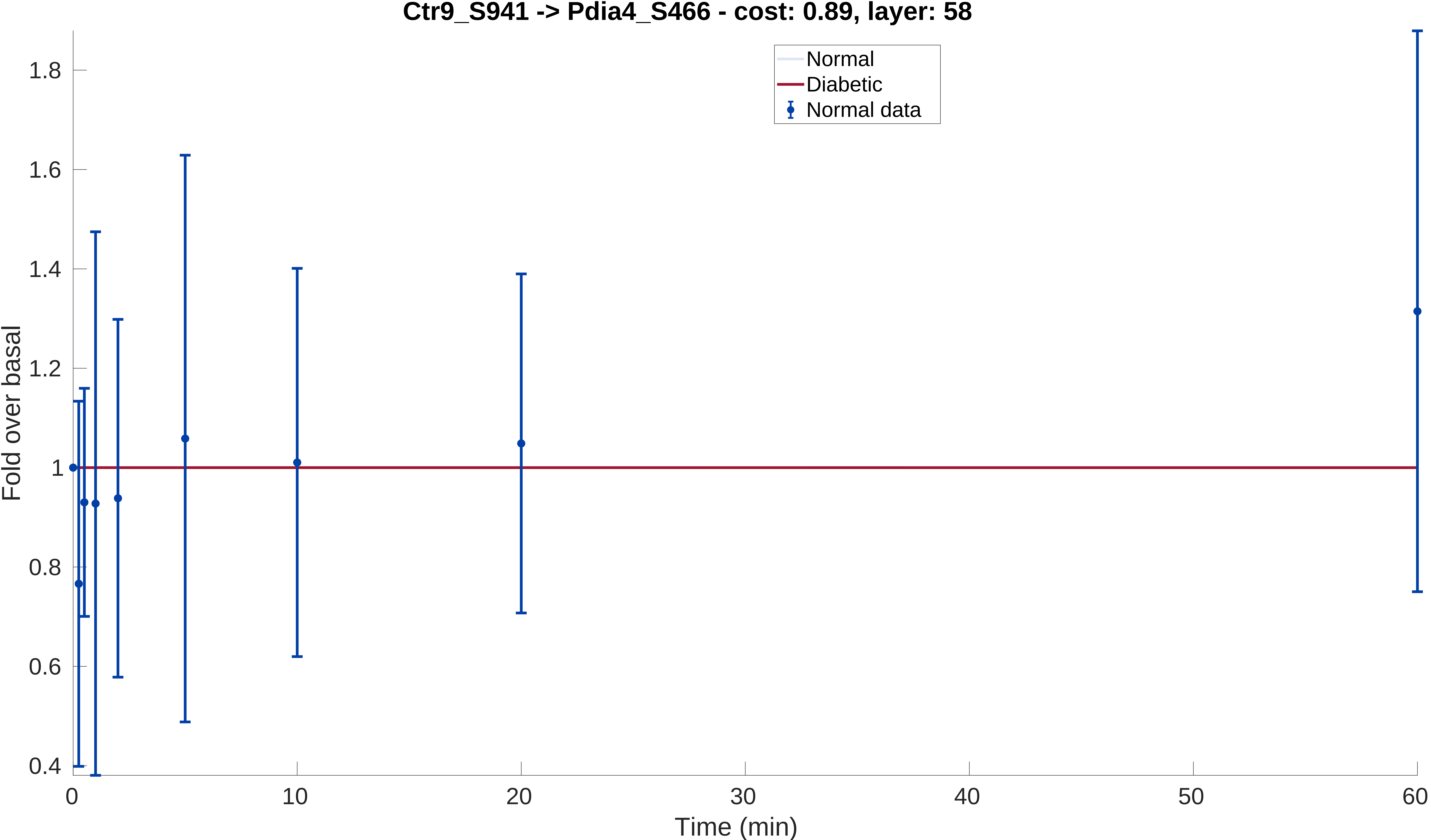
Ctr9_S941 -> Pdia4_S466 - cost: 0.89, layer: 58
Normal
Diabetic
Normal data
Fold over basal
1.8
1.6
1.4
1.2
1
0.8
0.6
0.4
0
10
20
30
40
50
60
Time (min)

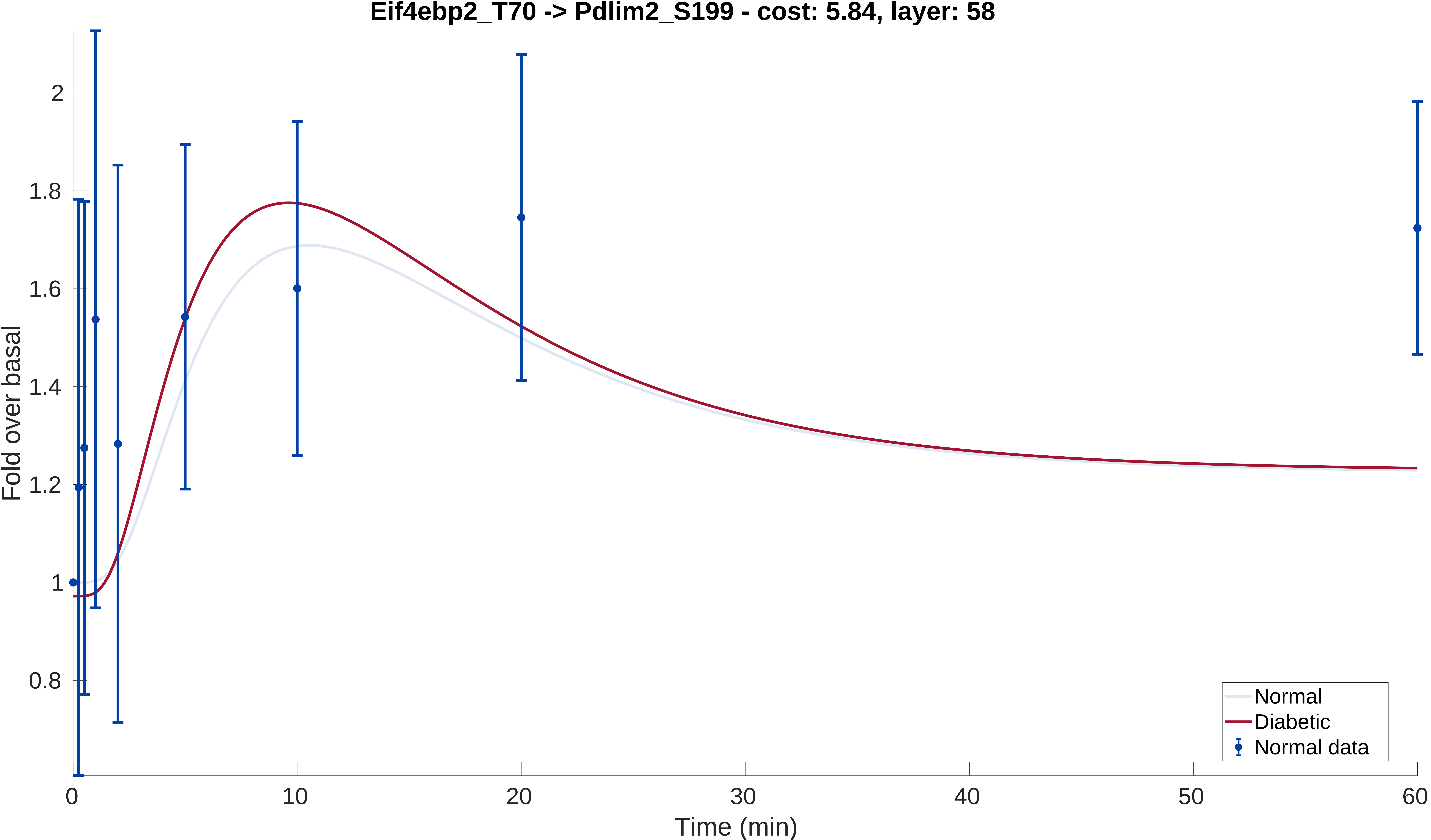
Eif4ebp2_T70 -> Pdlim2_S199 - cost: 5.84, layer: 58
Fold over basal
2
1.8
1.6
1.4
1.2
1
0.8
0
10
20
30
40
50
60
Time (min)
Normal
Diabetic
Normal data

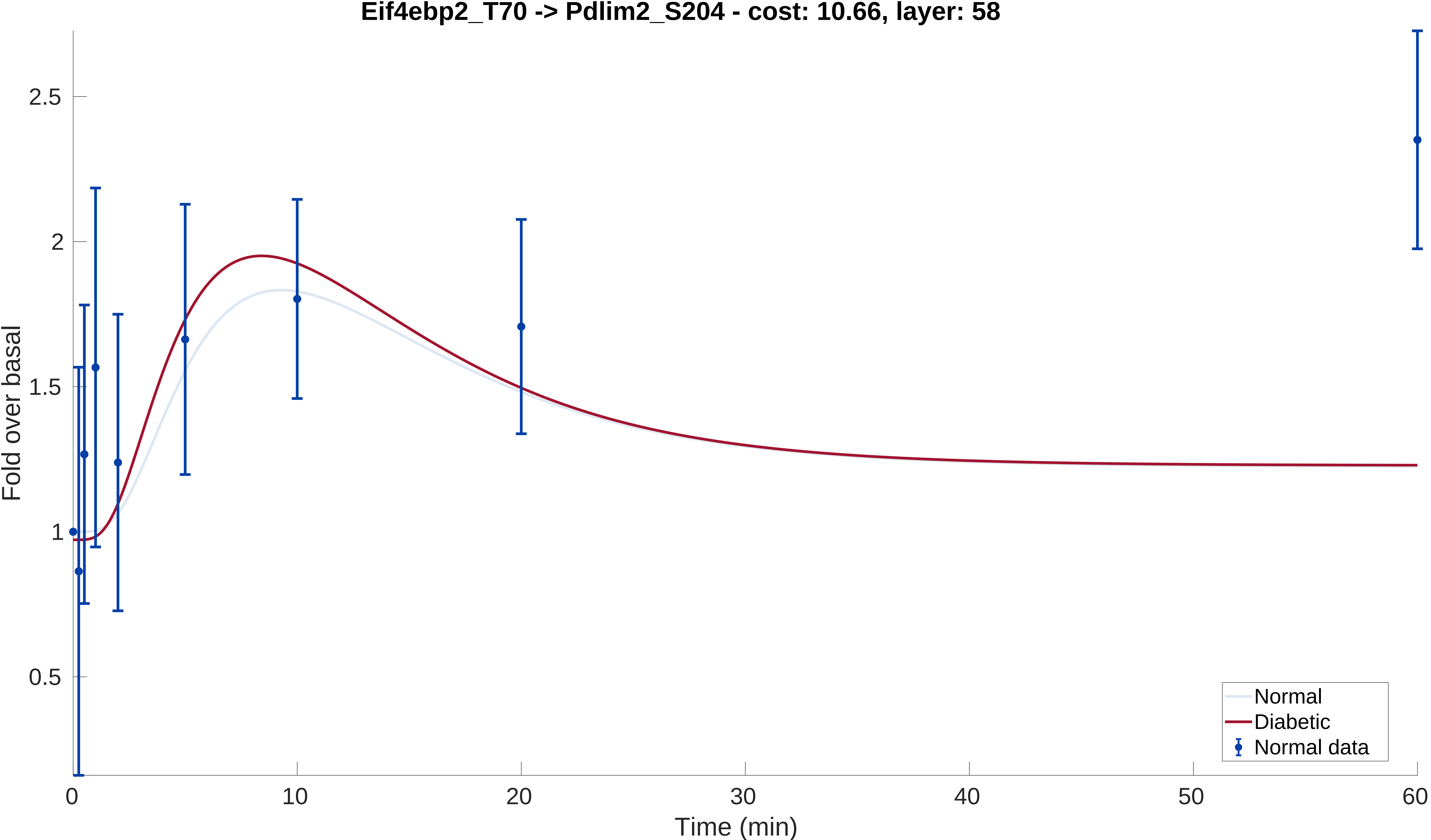
Eif4ebp2_T70 -> Pdlim2_S204 - cost: 10.66, layer: 58
Fold over basal
2.5
2
1.5
1
0.5
0
10
20
30
40
50
60
Time (min)
Normal
Diabetic
Normal data

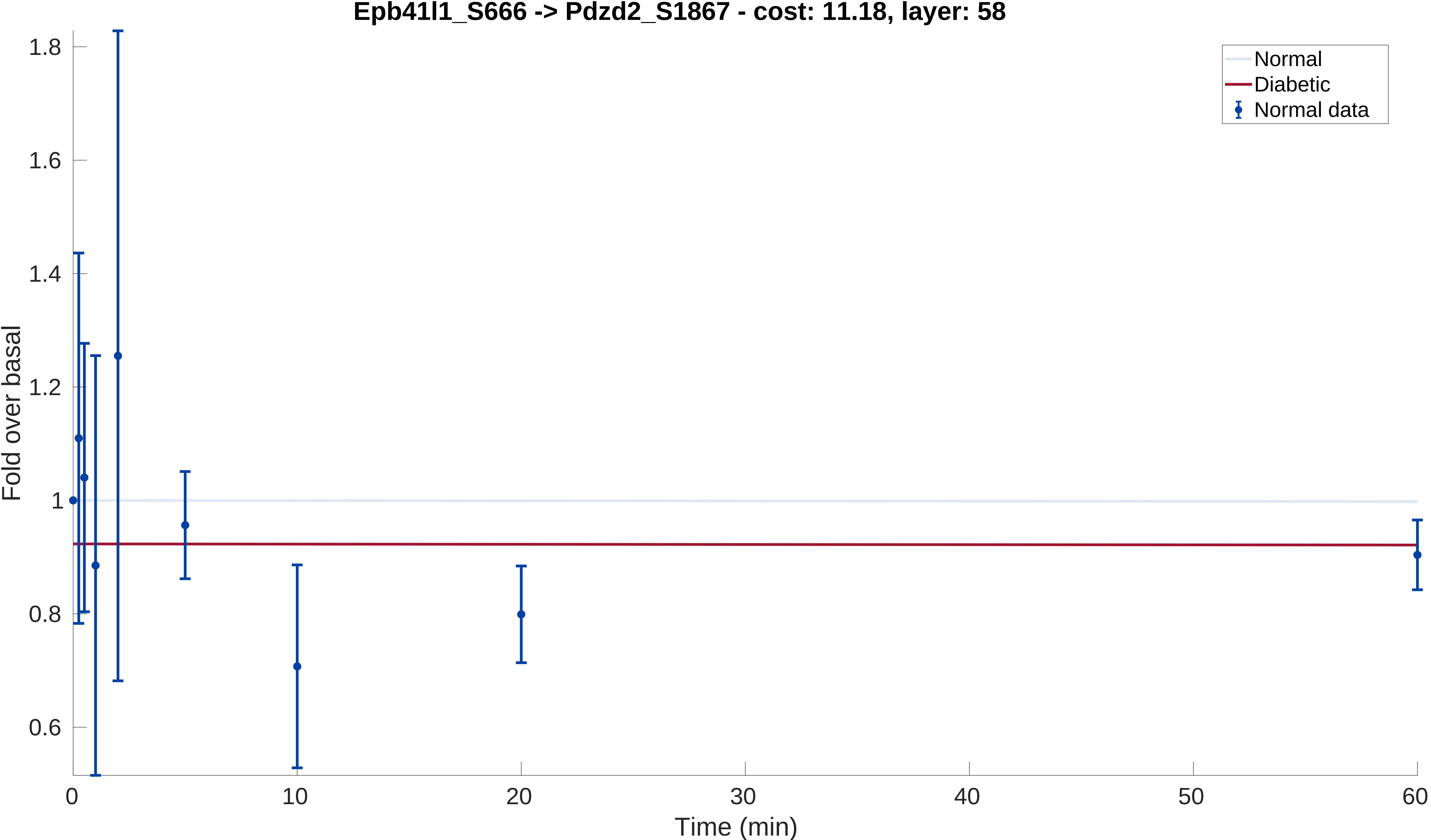
Epb41l1_S666 -> Pdzd2_S1867 - cost: 11.18, layer: 58
Normal
Diabetic
Normal data
Fold over basal
1.8
1.6
1.4
1.2
1
0.8
0.6
0
10
20
30
40
50
60
Time (min)

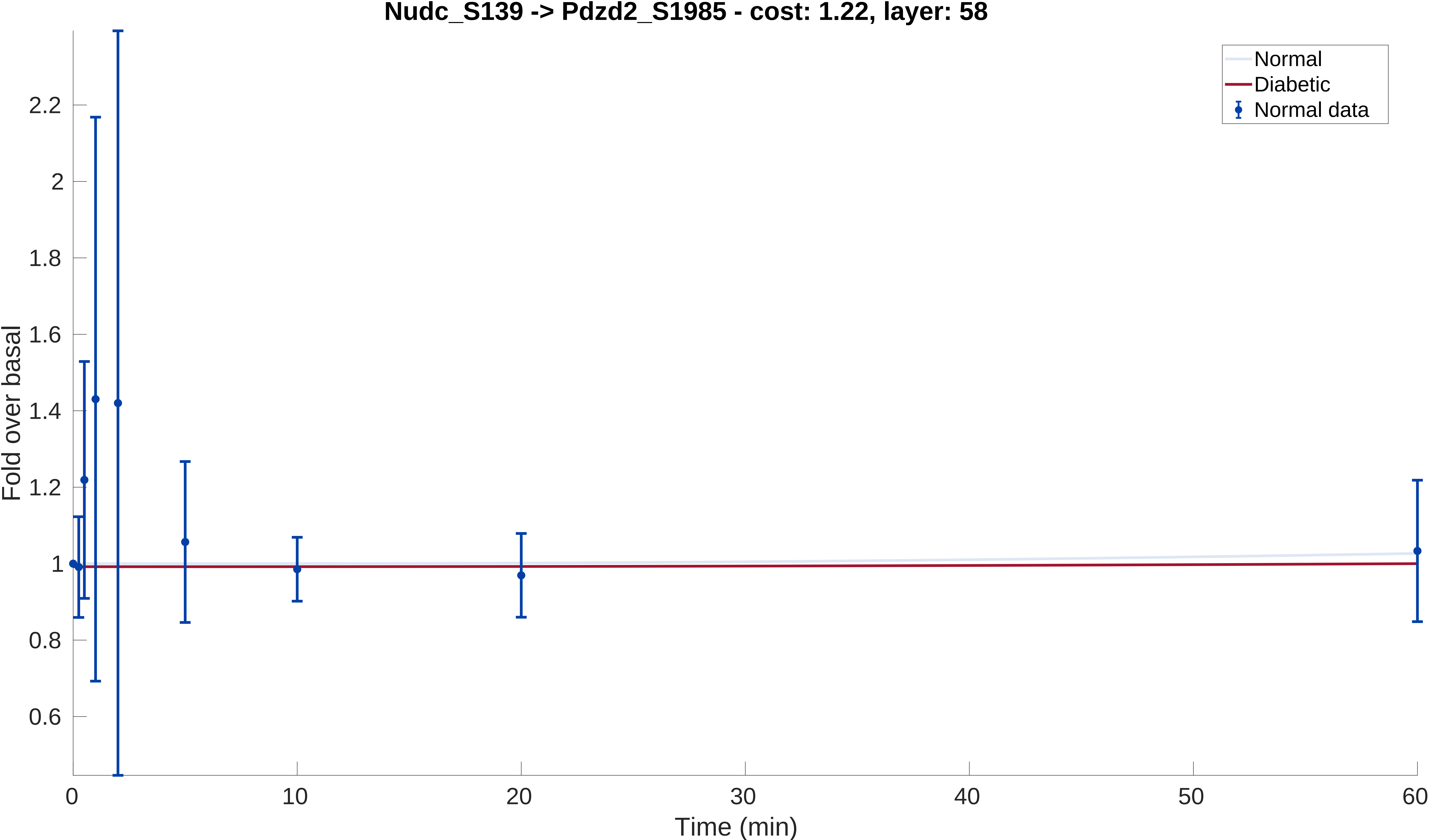

Nudc_S139 -> Pdzd2_S1985 - cost: 1.22, layer: 58
Normal
Diabetic
Normal data
Fold over basal
2.2
2
1.8
1.6
1.4
1.2
1
0.8
0.6
0
10
20
30
40
50
60
Time (min)

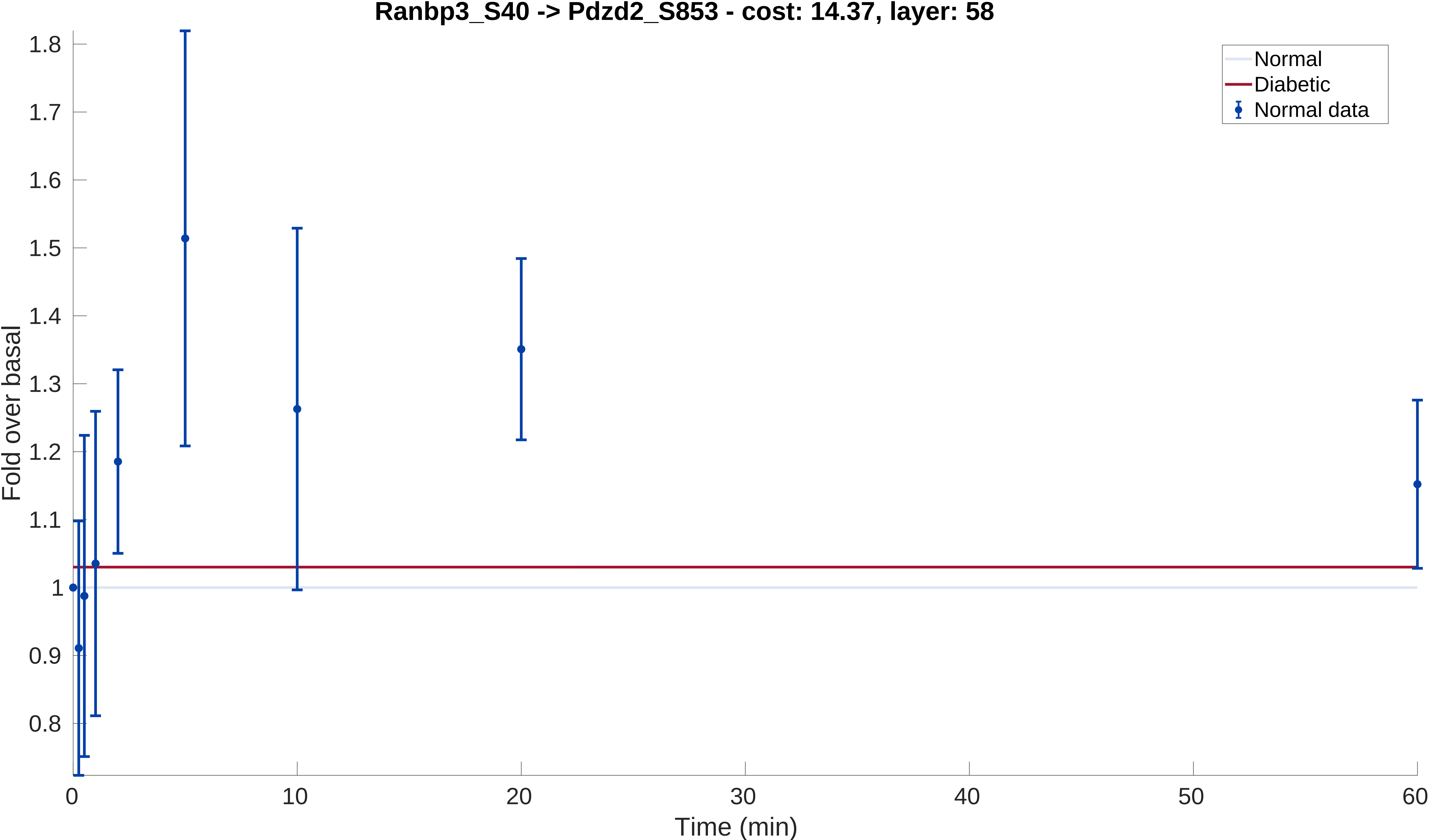

Ranbp3_S40 -> Pdzd2_S853 - cost: 14.37, layer: 58
Fold over basal
Time (min)
Normal
Diabetic
Normal data
1.8
1.7
1.6
1.5
1.4
1.3
1.2
1.1
1
0.9
0.8
0
10
20
30
40
50
60

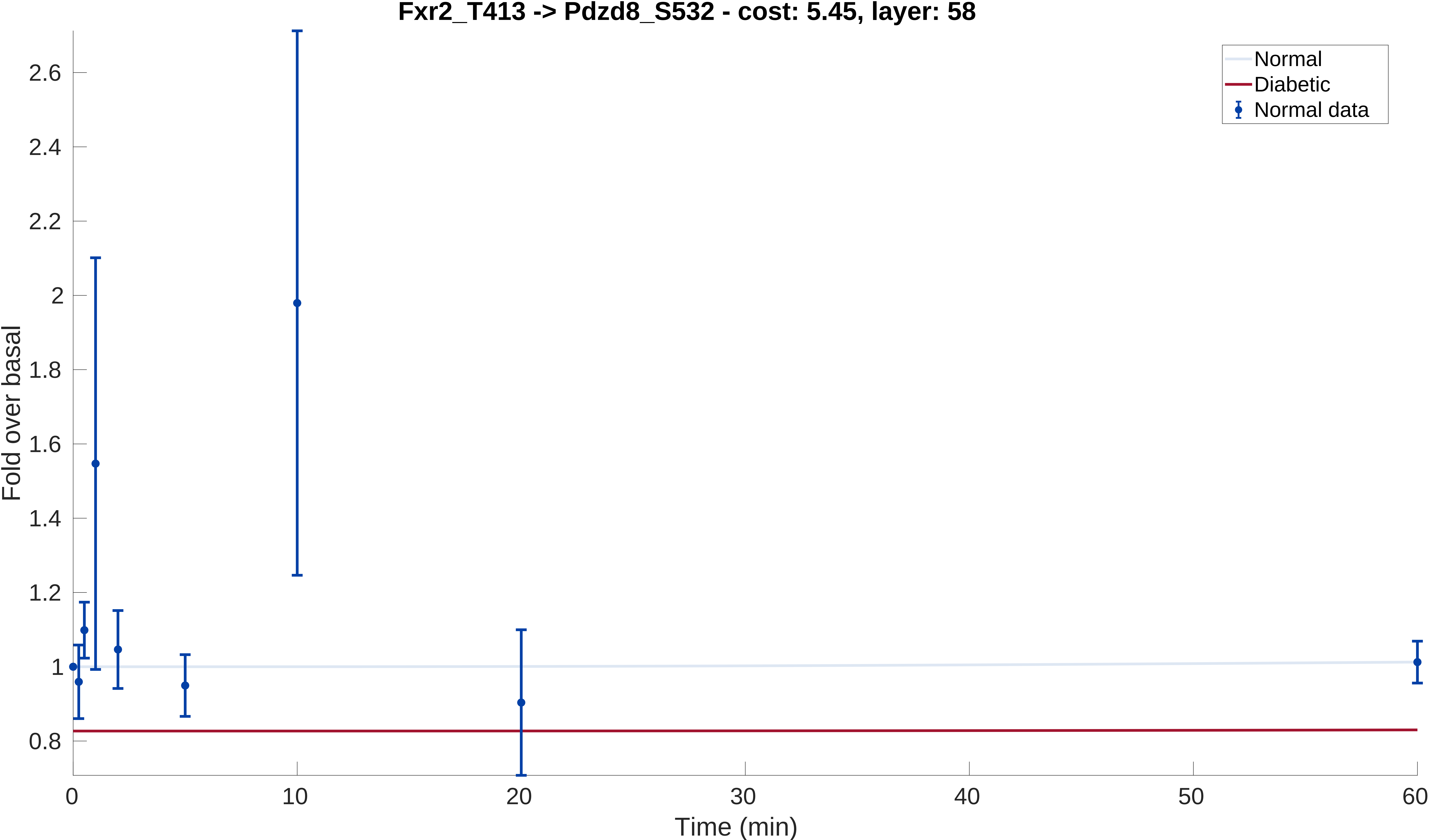

Fxr2_T413 -> Pdzd8_S532 - cost: 5.45, layer: 58
Normal
Diabetic
Normal data
Fold over basal
2.6
2.4
2.2
2
1.8
1.6
1.4
1.2
1
0.8
0
10
20
30
40
50
60
Time (min)

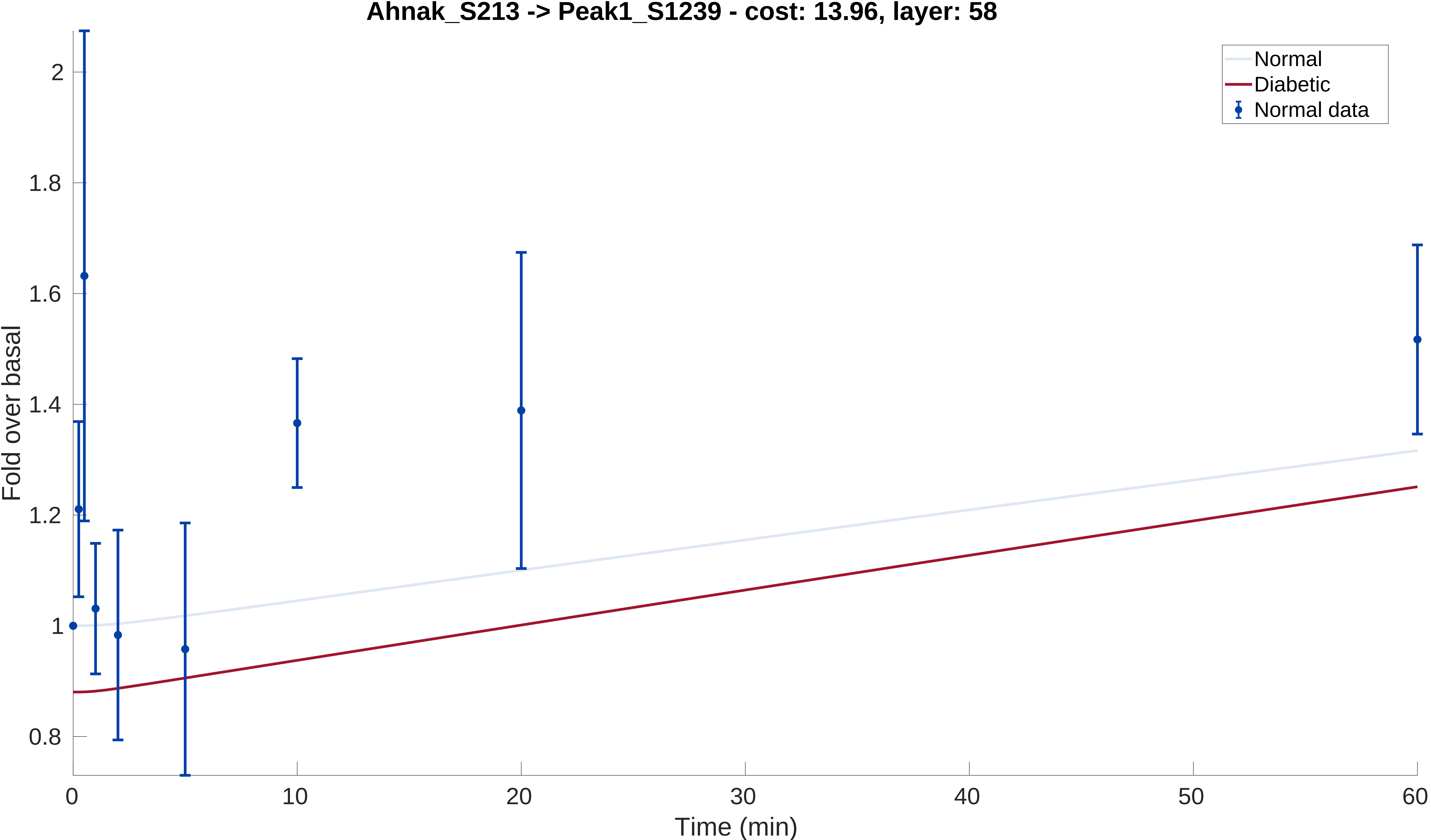

Ahnak_S213 -> Peak1_S1239 - cost: 13.96, layer: 58
Normal
Diabetic
Normal data
Fold over basal
2
1.8
1.6
1.4
1.2
1
0.8
0
10
20
30
40
50
60
Time (min)

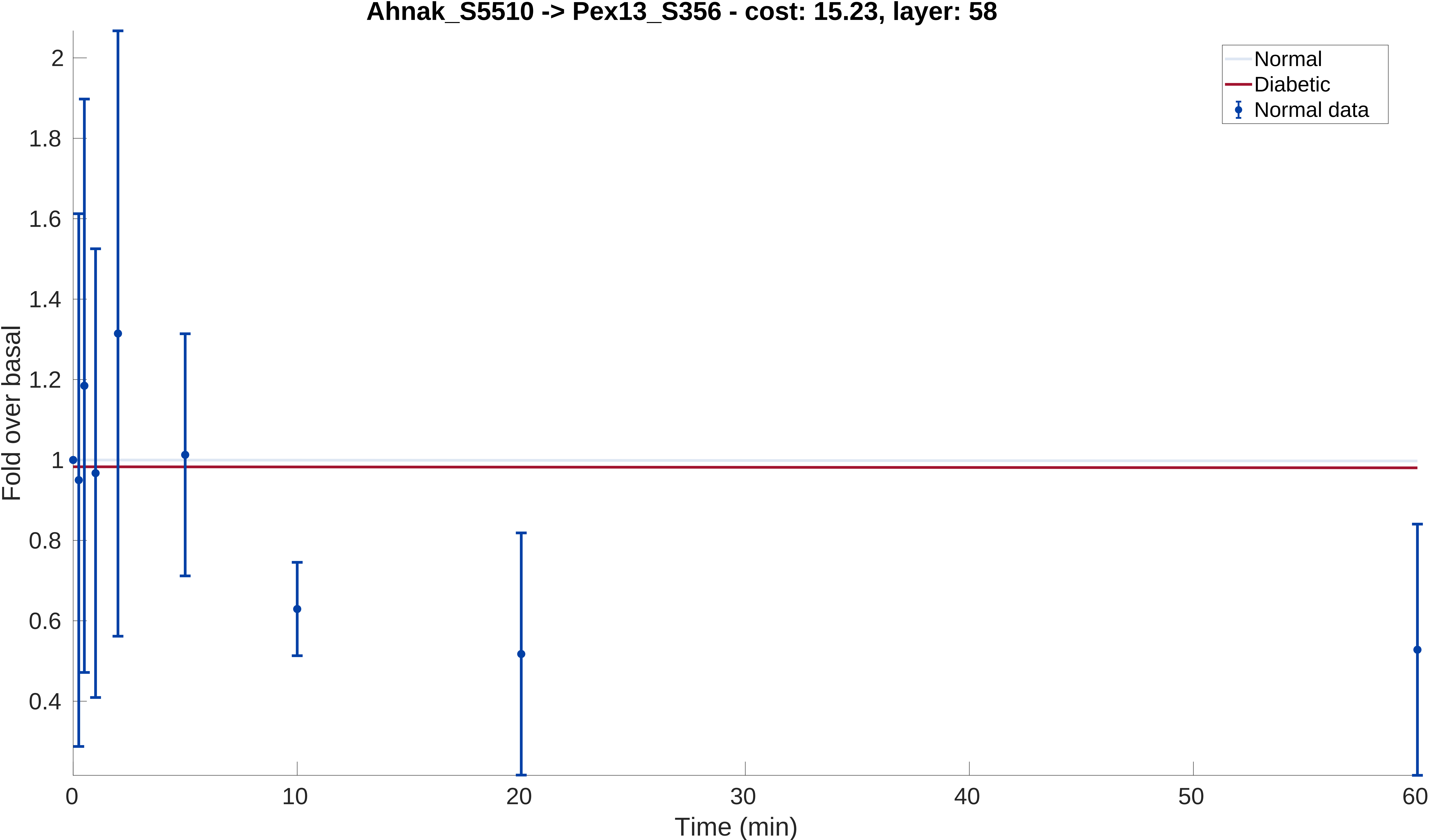

Ahnak_S5510 -> Pex13_S356 - cost: 15.23, layer: 58
Normal
Diabetic
Normal data
Fold over basal
Time (min)
2
1.8
1.6
1.4
1.2
1
0.8
0.6
0.4
0
10
20
30
40
50
60

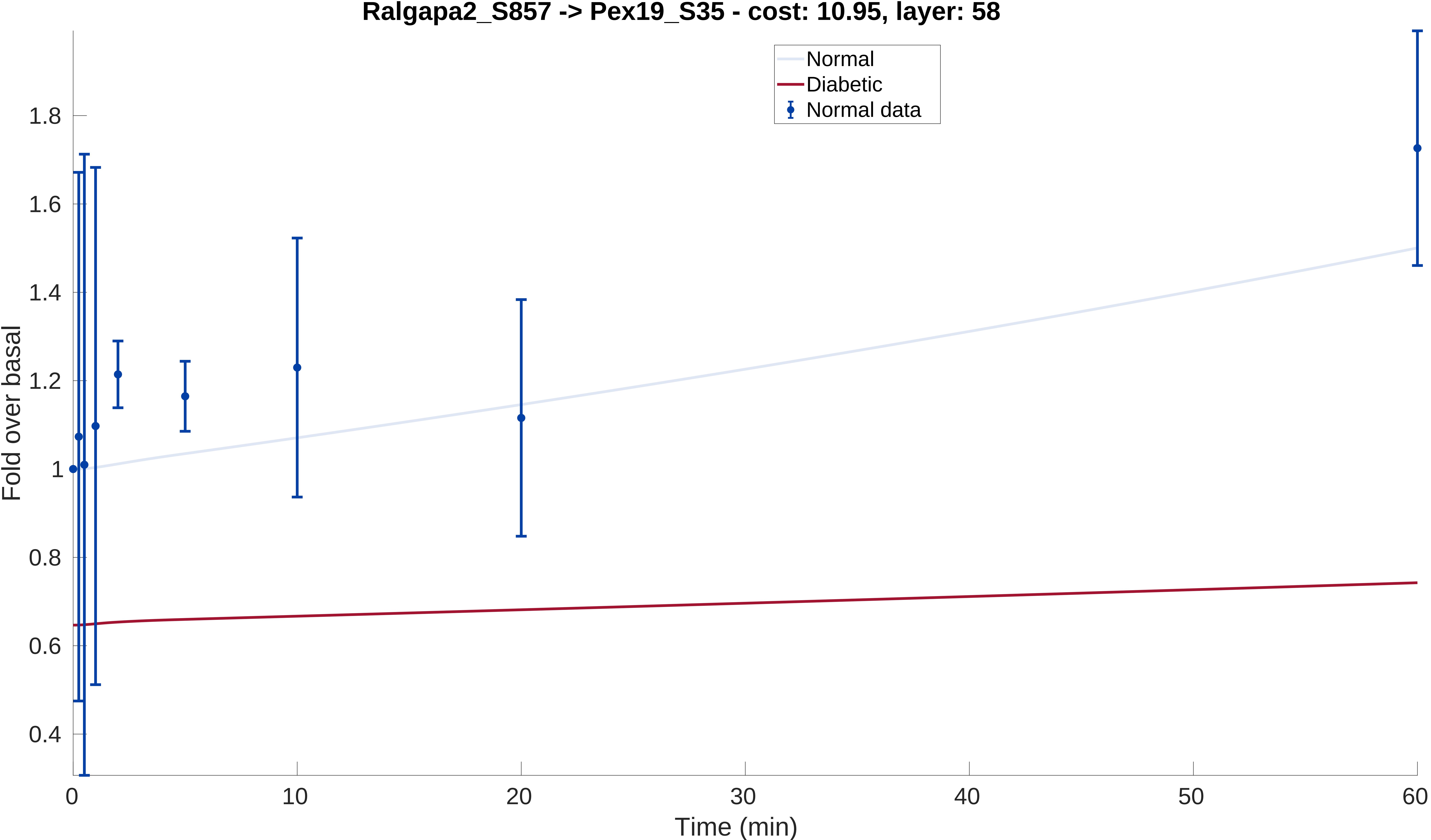

Ralgapa2_S857 -> Pex19_S35 - cost: 10.95, layer: 58
Normal
Diabetic
Normal data
Fold over basal
1.8
1.6
1.4
1.2
1
0.8
0.6
0.4
0
10
20
30
40
50
60
Time (min)

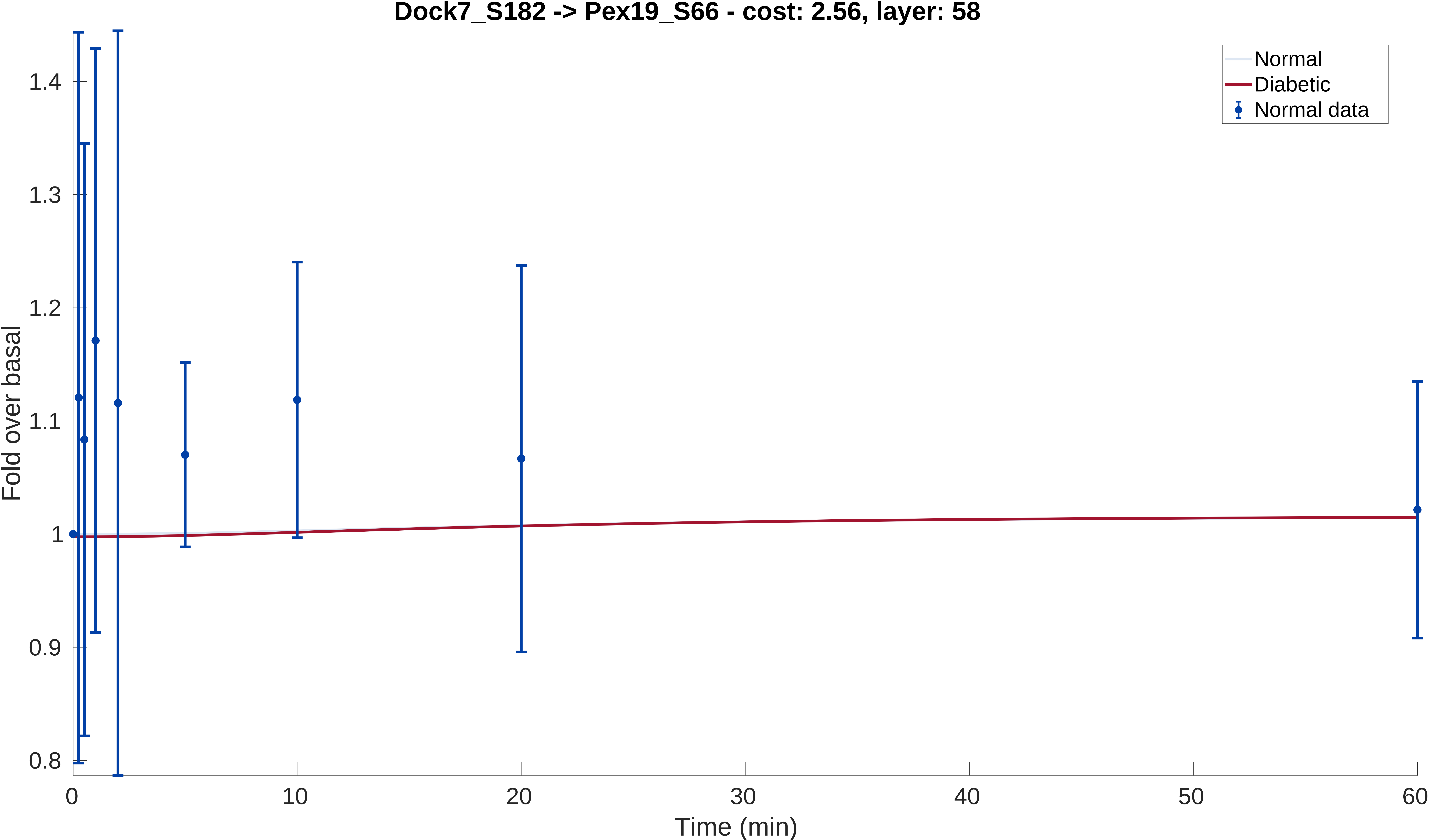

Dock7_S182 -> Pex19_S66 - cost: 2.56, layer: 58
Normal
Diabetic
Normal data
1.4
1.3
1.2
1.1
1
0.9
0.8
Fold over basal
0
10
20
30
40
50
60
Time (min)

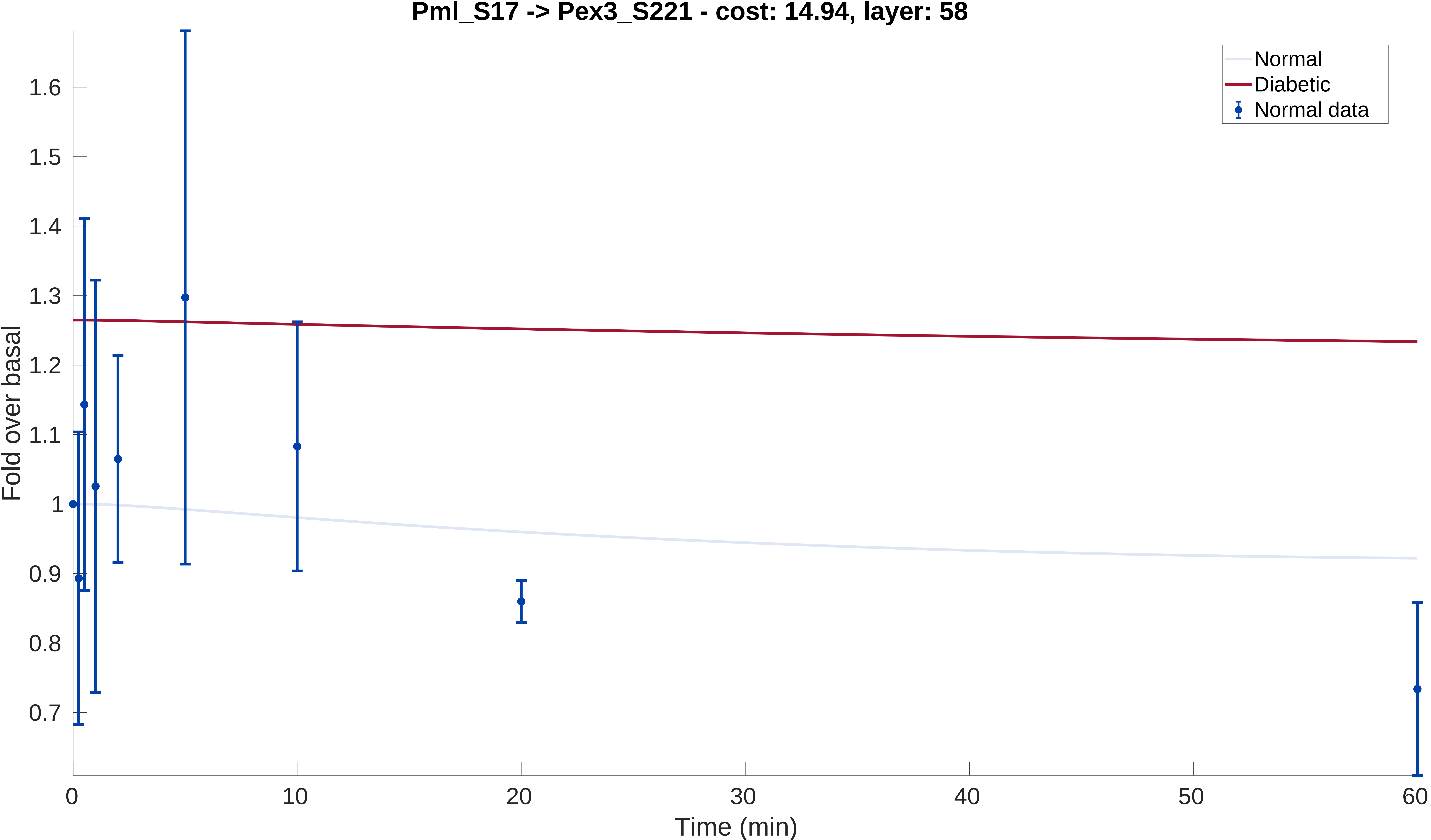

Pml_S17 -> Pex3_S221 - cost: 14.94, layer: 58
Normal
Diabetic
Normal data
Fold over basal
1.6
1.5
1.4
1.3
1.2
1.1
1
0.9
0.8
0.7
0
10
20
30
40
50
60
Time (min)

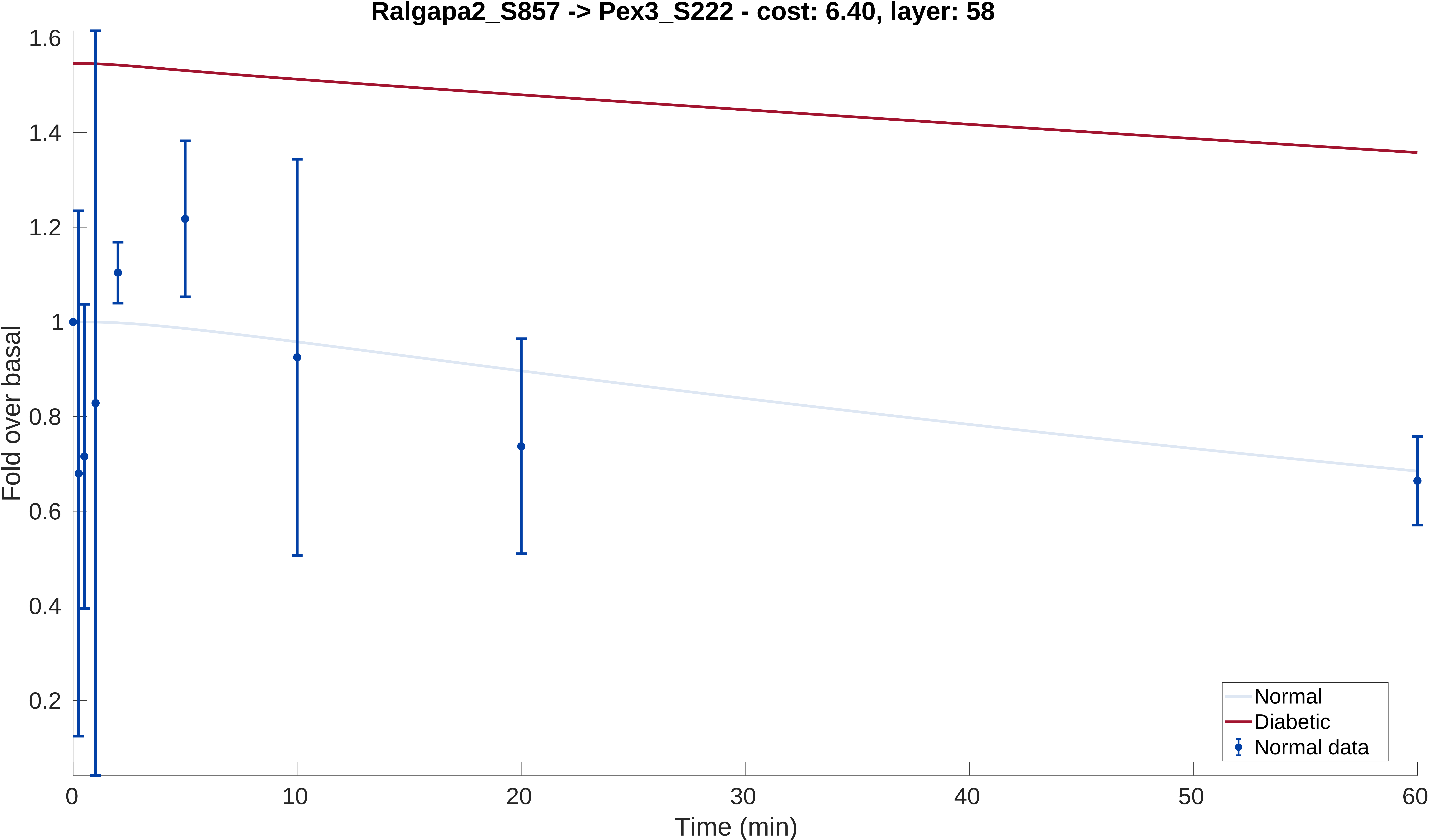

Ralgapa2_S857 -> Pex3_S222 - cost: 6.40, layer: 58
Fold over basal
1.6
1.4
1.2
1
0.8
0.6
0.4
0.2
0
10
20
30
40
50
60
Time (min)
Normal
Diabetic
Normal data

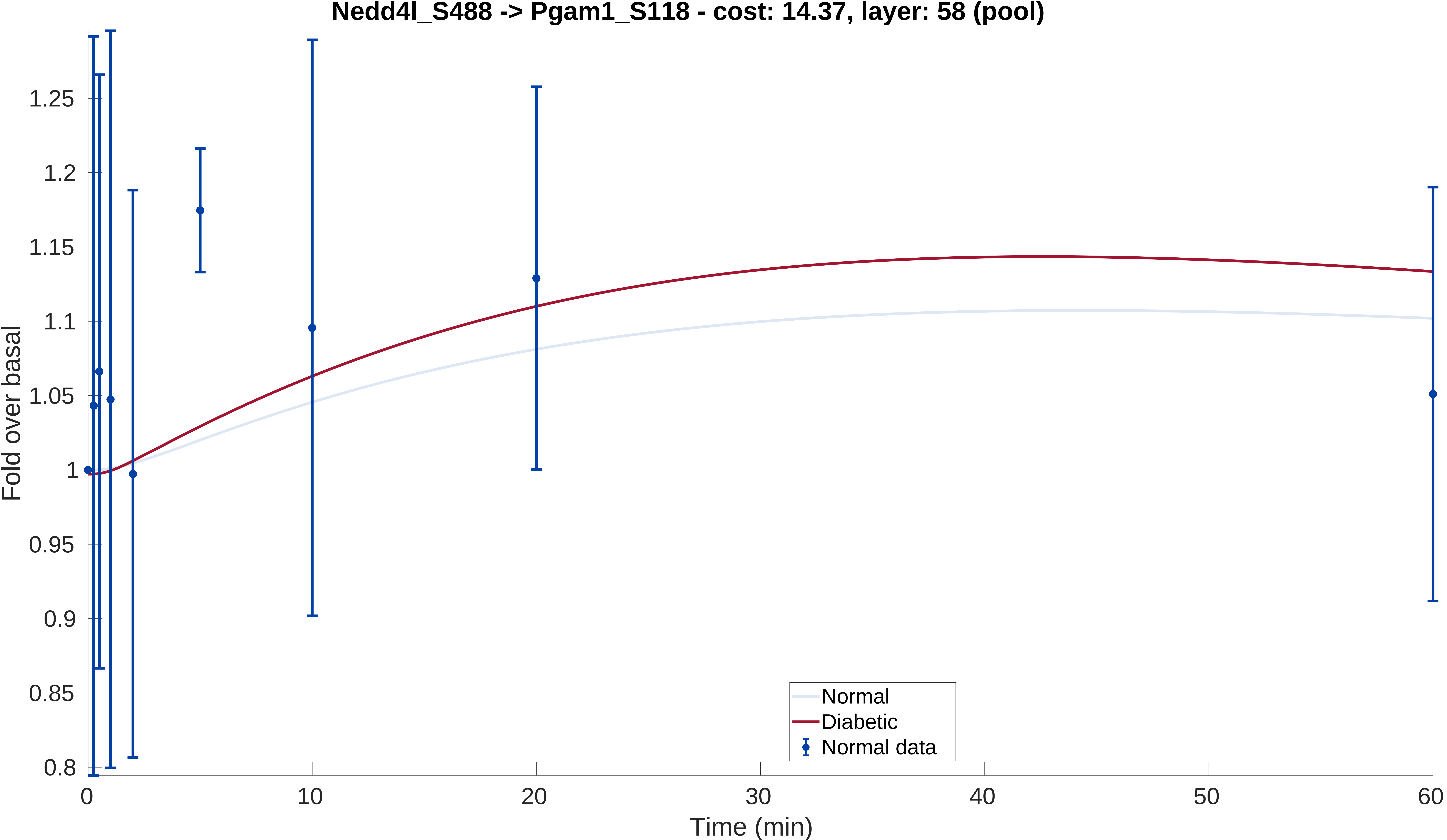

Nedd4l_S488 -> Pgam1_S118 - cost: 14.37, layer: 58 (pool)
Fold over basal
1.25
1.2
1.15
1.1
1.05
1
0.95
0.9
0.85
0.8
0
10
20
30
40
50
60
Time (min)
Normal
Diabetic
Normal data

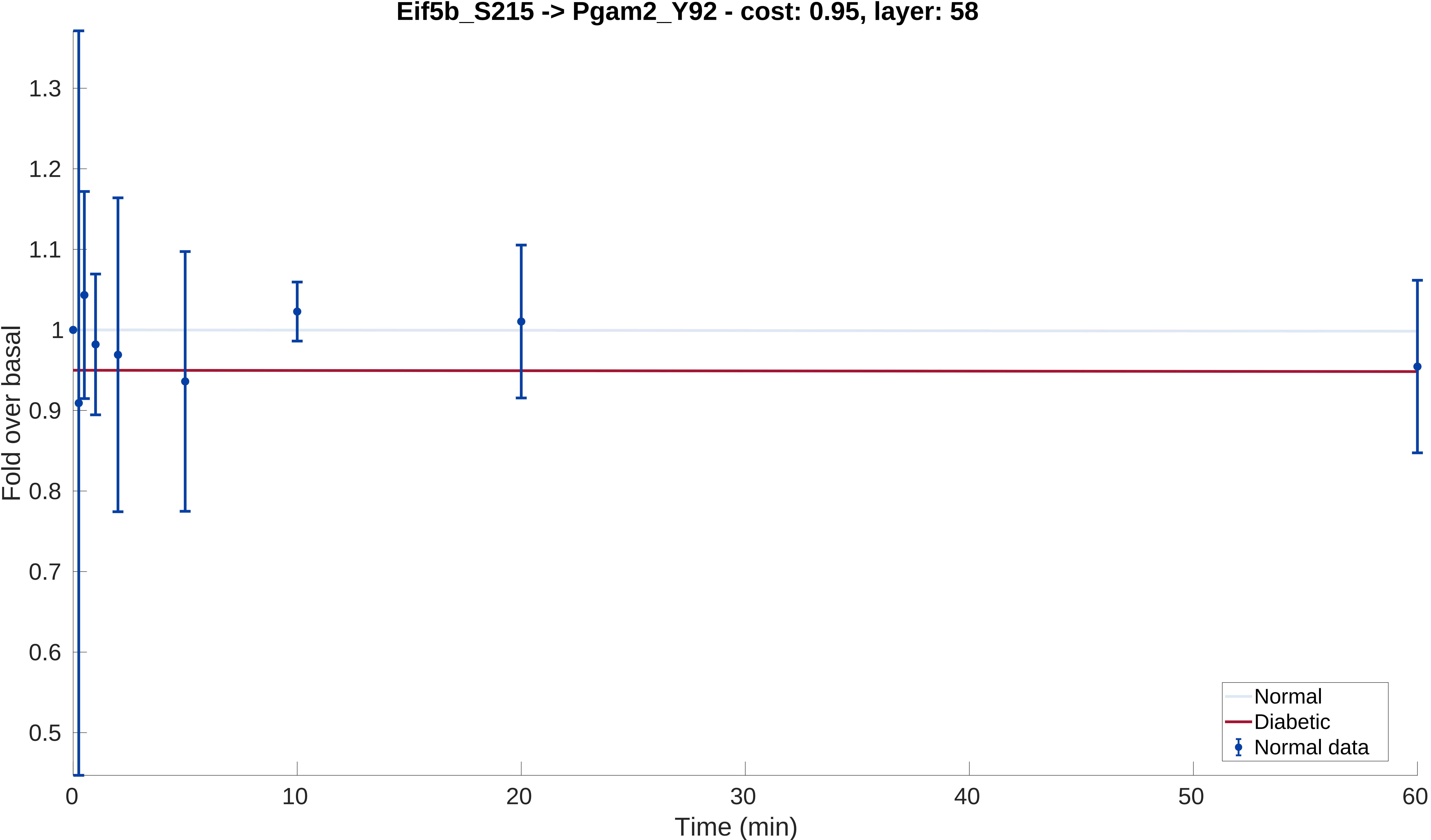
Eif5b_S215 -> Pgam2_Y92 - cost: 0.95, layer: 58
Fold over basal
Time (min)
0.5
0.6
0.7
0.8
0.9
1
1.1
1.2
1.3
0
10
20
30
40
50
60
Normal
Diabetic
Normal data

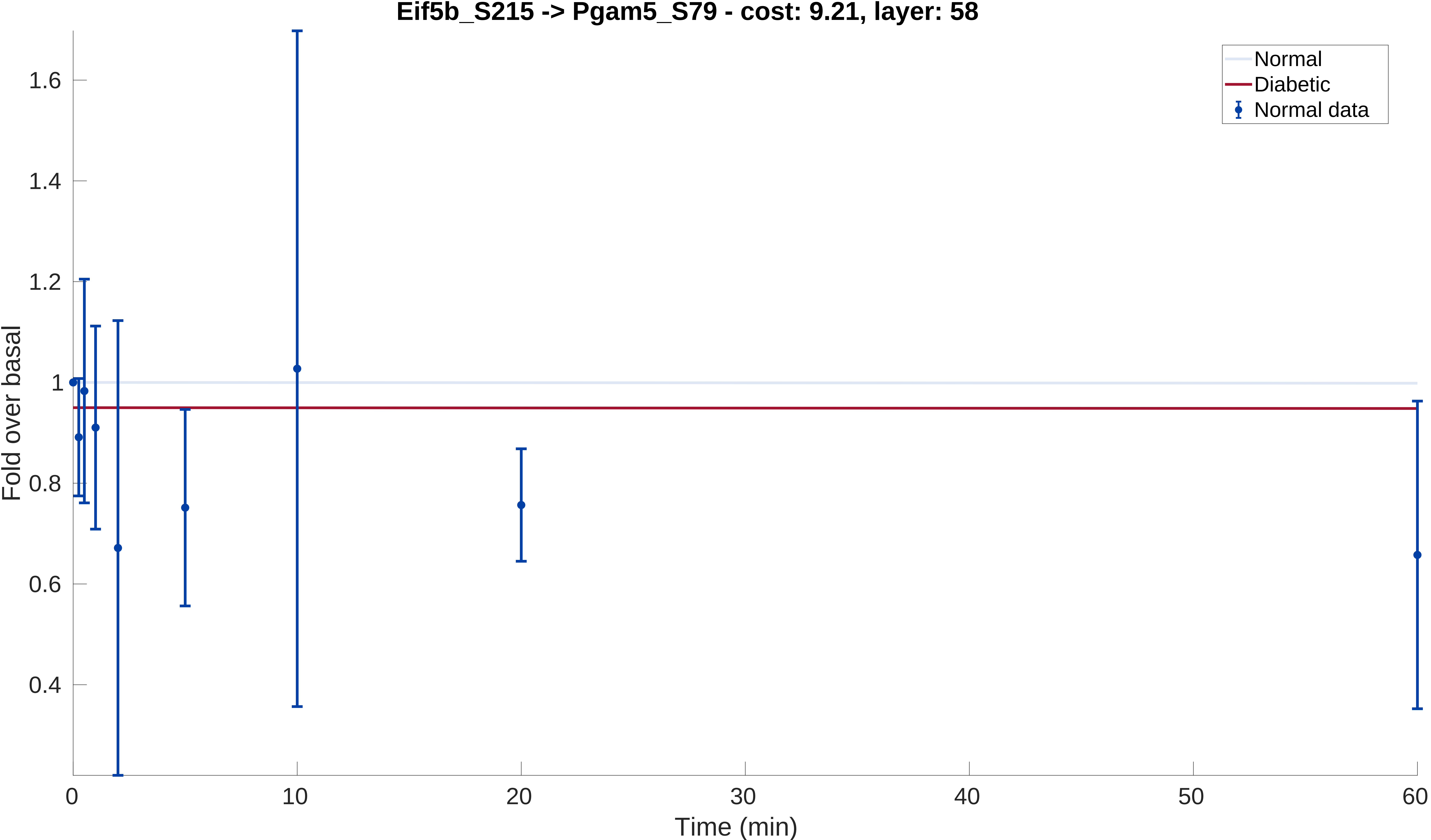

Eif5b_S215 -> Pgam5_S79 - cost: 9.21, layer: 58
Normal
Diabetic
Normal data
Fold over basal
1.6
1.4
1.2
1
0.8
0.6
0.4
0
10
20
30
40
50
60
Time (min)

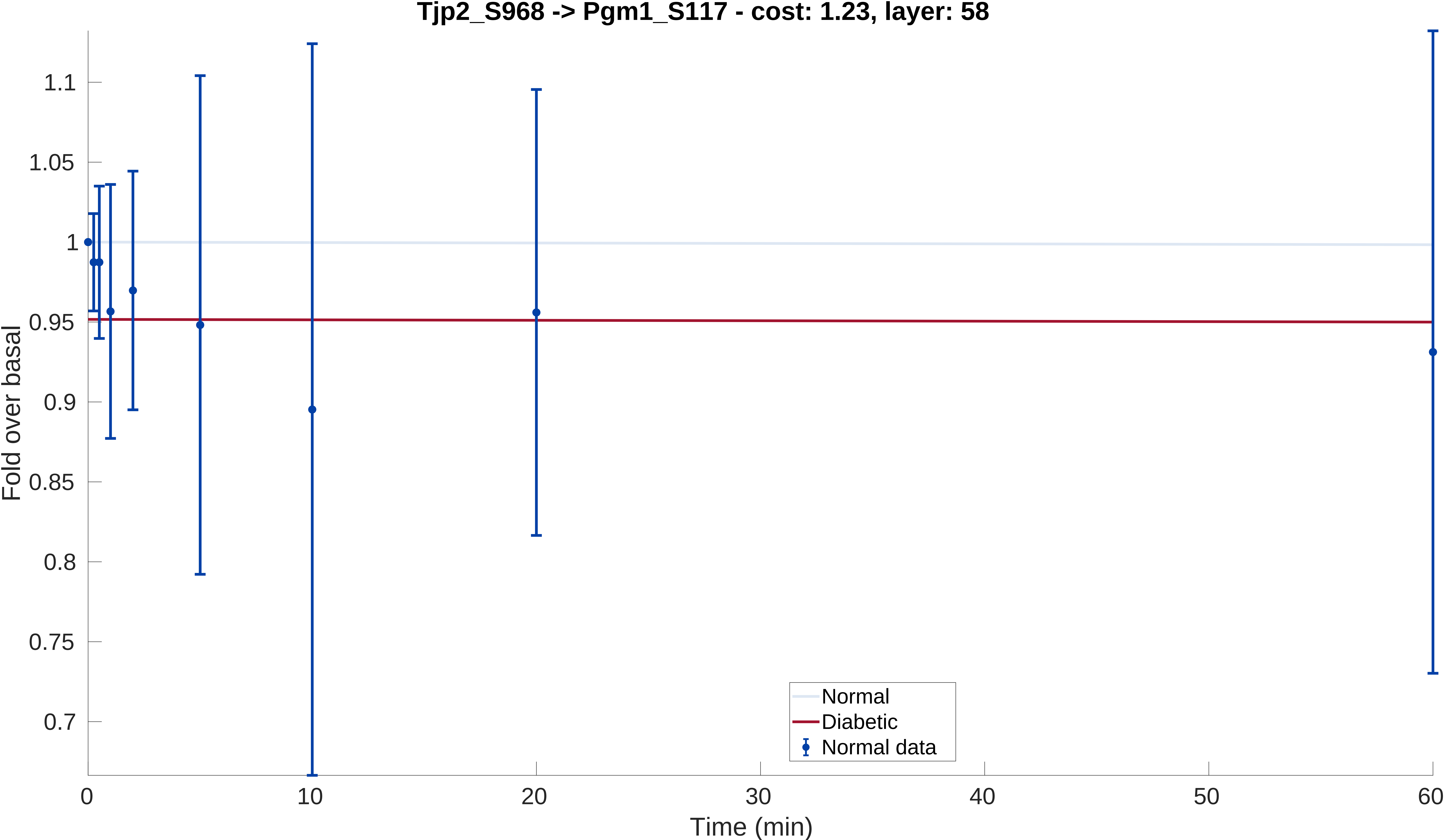

Tjp2_S968 -> Pgm1_S117 - cost: 1.23, layer: 58
Fold over basal
1.1
1.05
1
0.95
0.9
0.85
0.8
0.75
0.7
0
10
20
30
40
50
60
Time (min)
Normal
Diabetic
Normal data

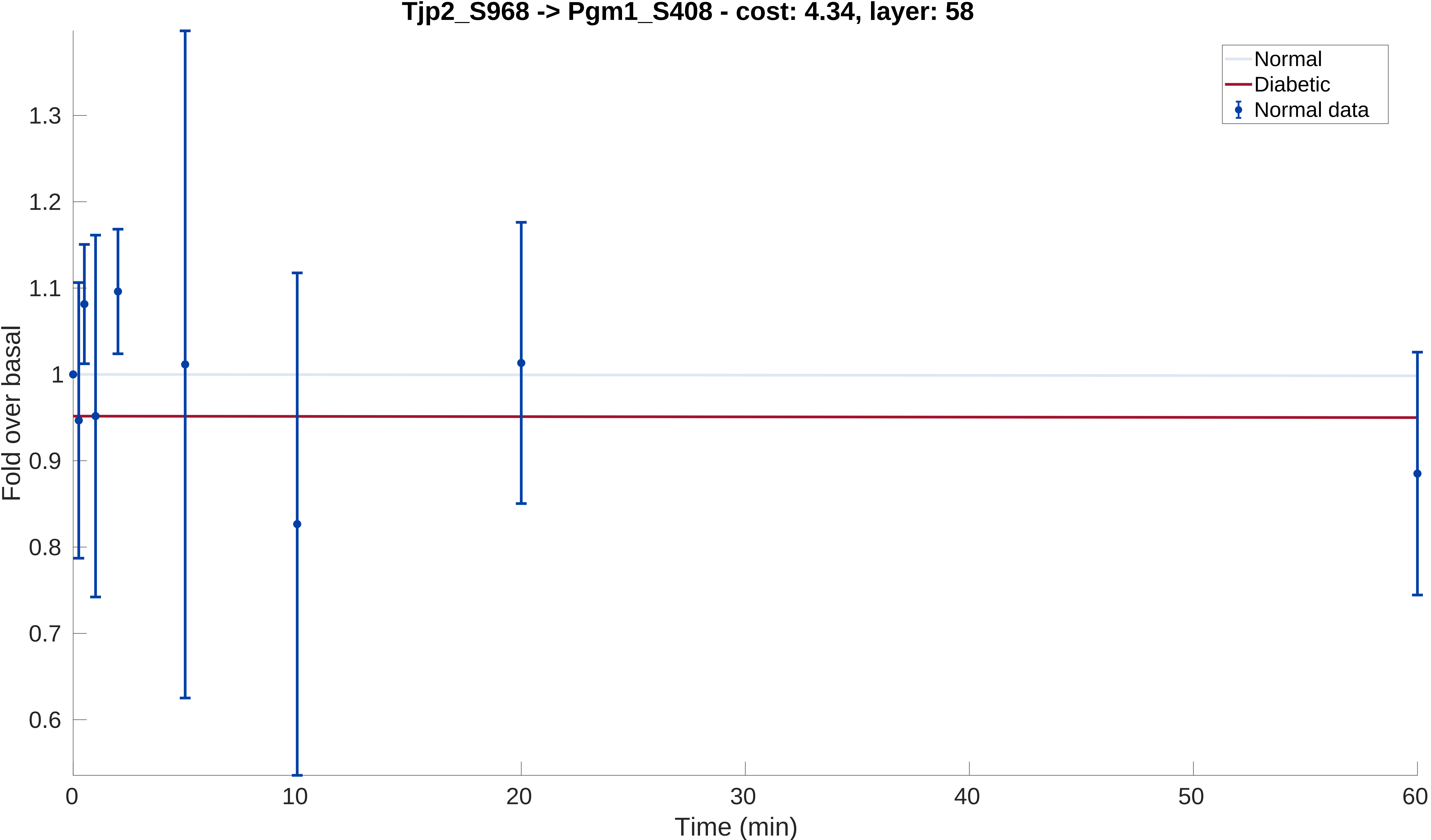

Tjp2_S968 -> Pgm1_S408 - cost: 4.34, layer: 58
Normal
Diabetic
Normal data
Fold over basal
1.3
1.2
1.1
1
0.9
0.8
0.7
0.6
0
10
20
30
40
50
60
Time (min)

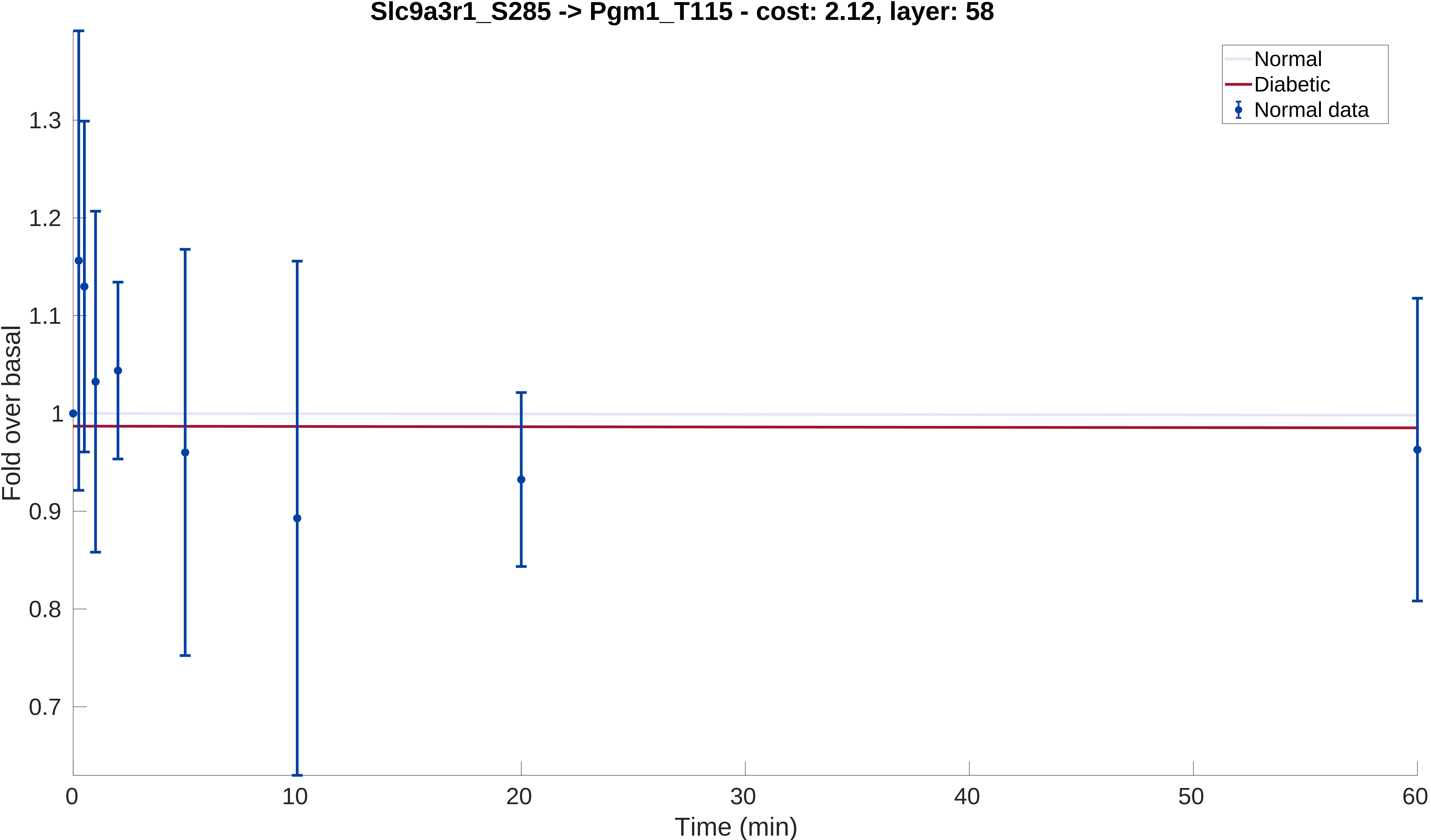

Slc9a3r1_S285 -> Pgm1_T115 - cost: 2.12, layer: 58
Normal
Diabetic
Normal data
Fold over basal
1.3
1.2
1.1
1
0.9
0.8
0.7
0
10
20
30
40
50
60
Time (min)

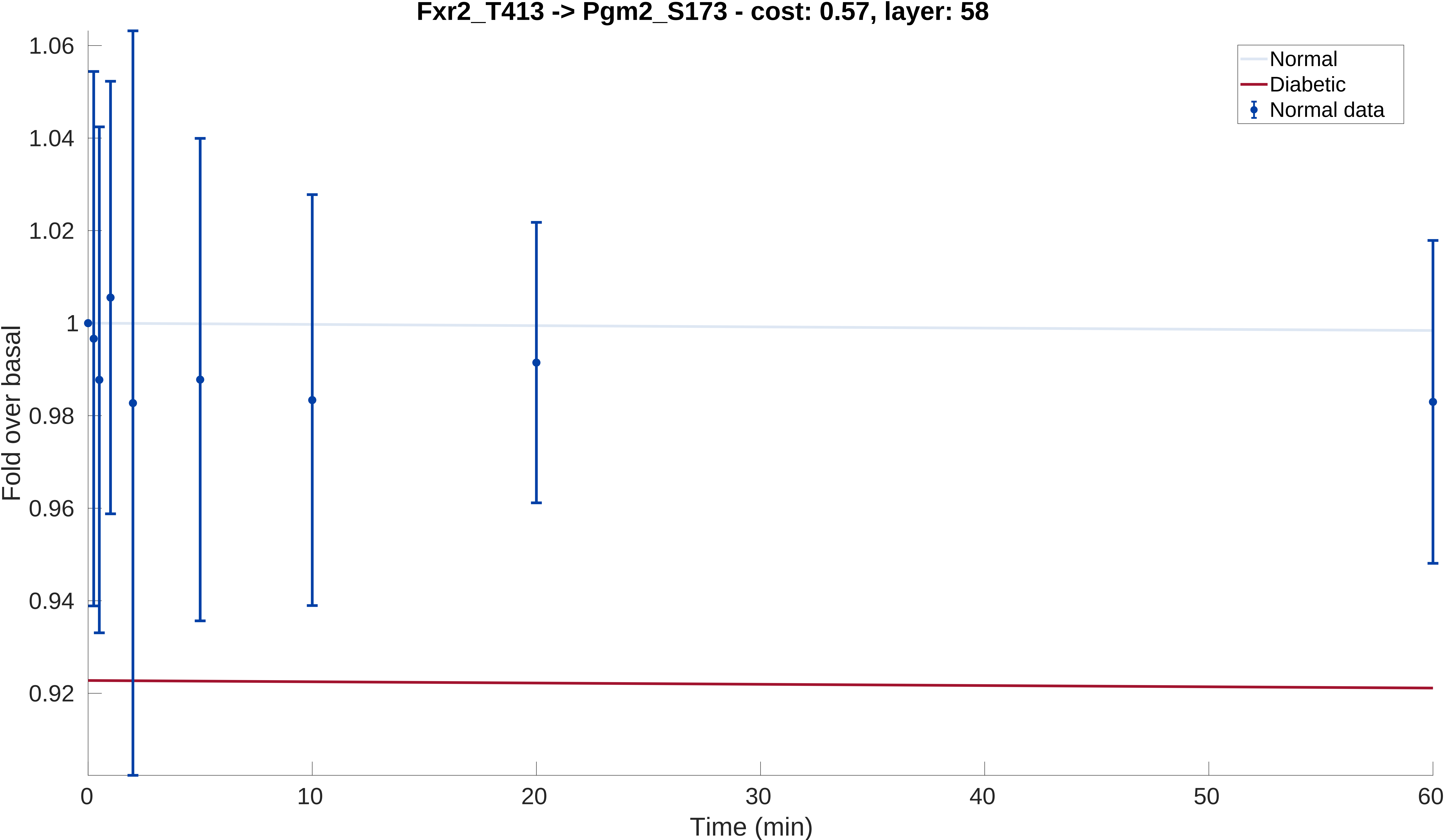

Fxr2_T413 -> Pgm2_S173 - cost: 0.57, layer: 58
Normal
Diabetic
Normal data
Fold over basal
1.06
1.04
1.02
1
0.98
0.96
0.94
0.92
0
10
20
30
40
50
60
Time (min)

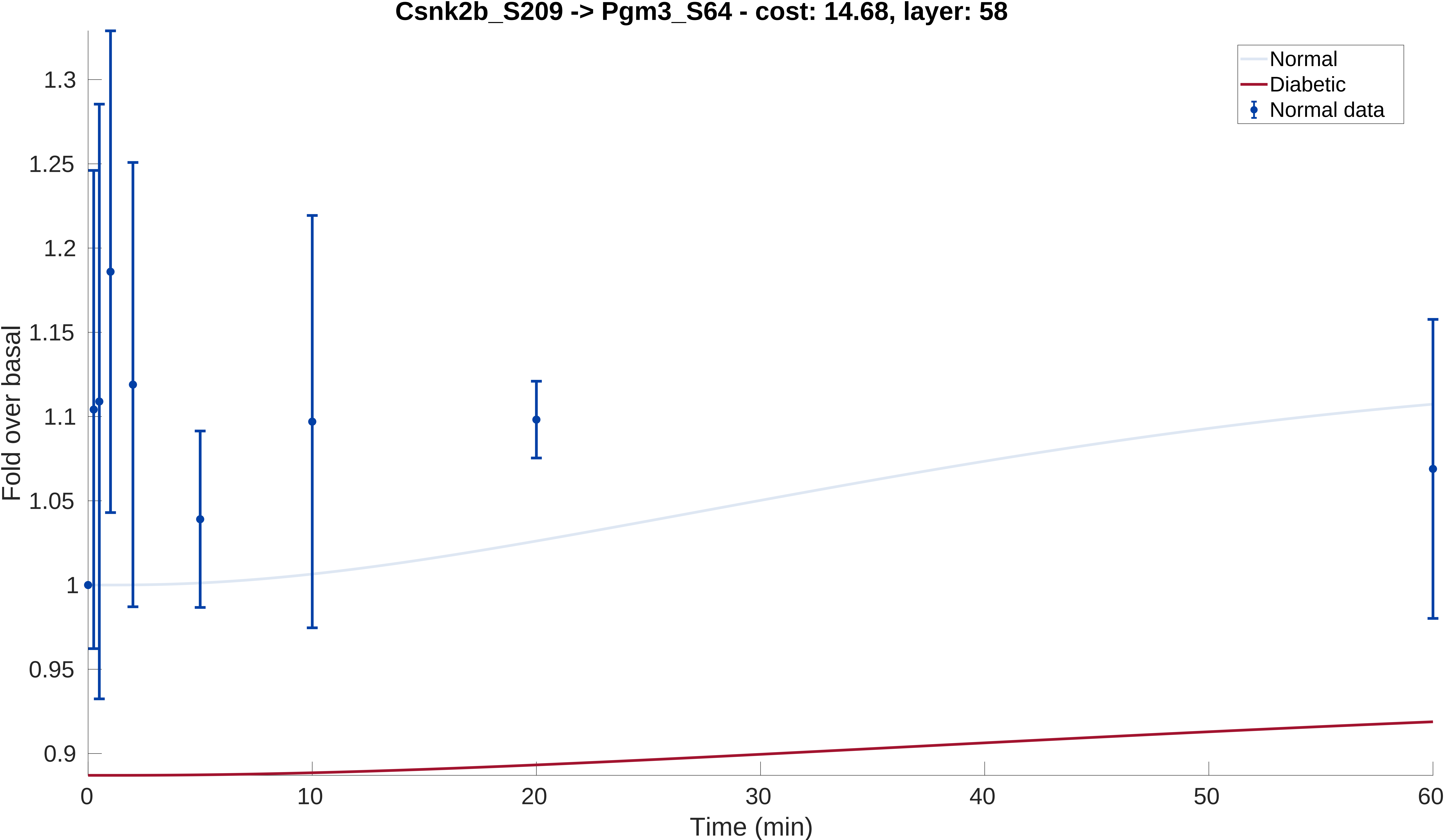

Csnk2b_S209 -> Pgm3_S64 - cost: 14.68, layer: 58
Normal
Diabetic
Normal data
Fold over basal
1.3
1.25
1.2
1.15
1.1
1.05
1
0.95
0.9
0
10
20
30
40
50
60
Time (min)

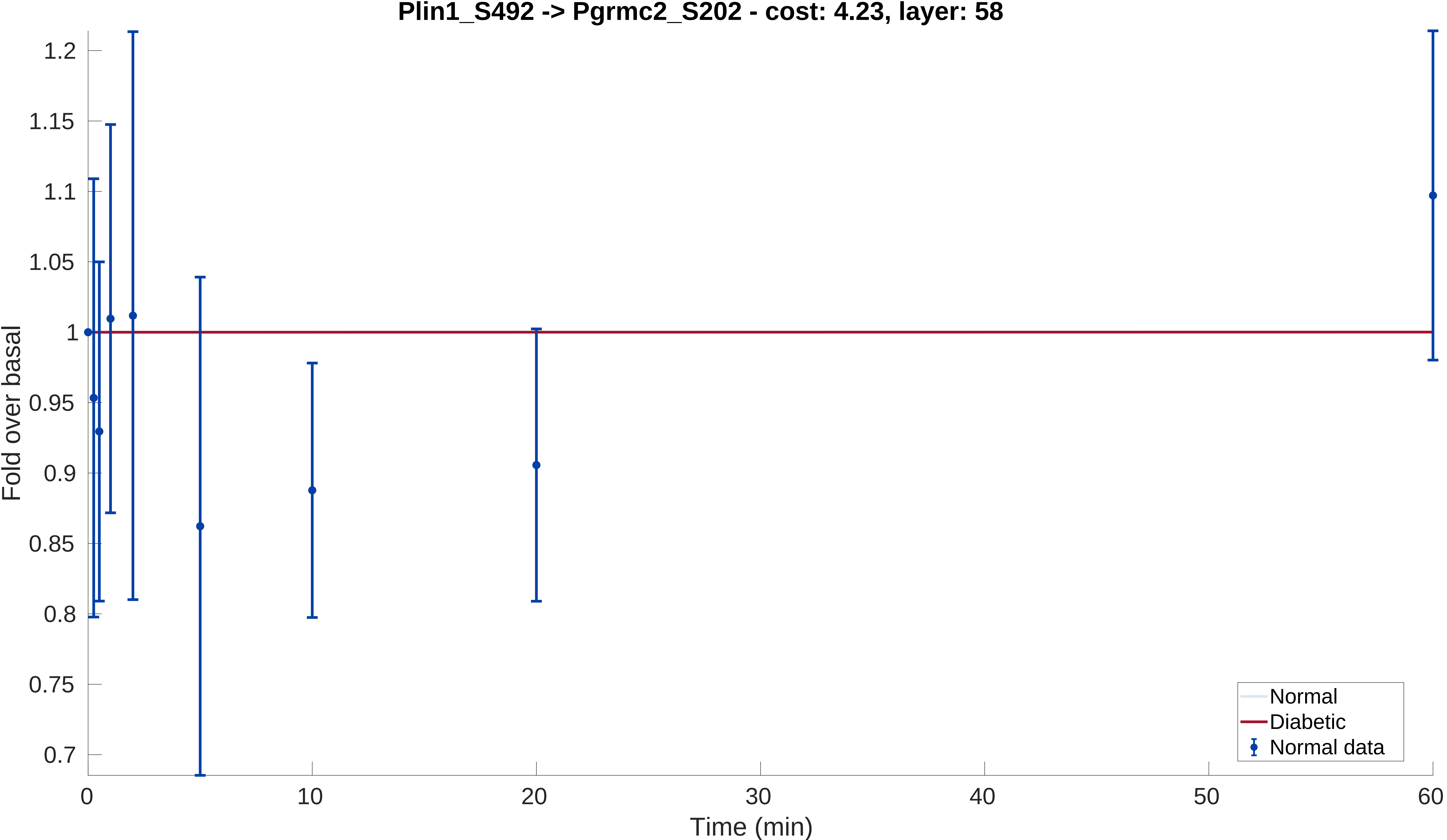

Plin1_S492 -> Pgrmc2_S202 - cost: 4.23, layer: 58
Fold over basal
1.2
1.15
1.1
1.05
1
0.95
0.9
0.85
0.8
0.75
0.7
0
10
20
30
40
50
60
Time (min)
Normal
Diabetic
Normal data

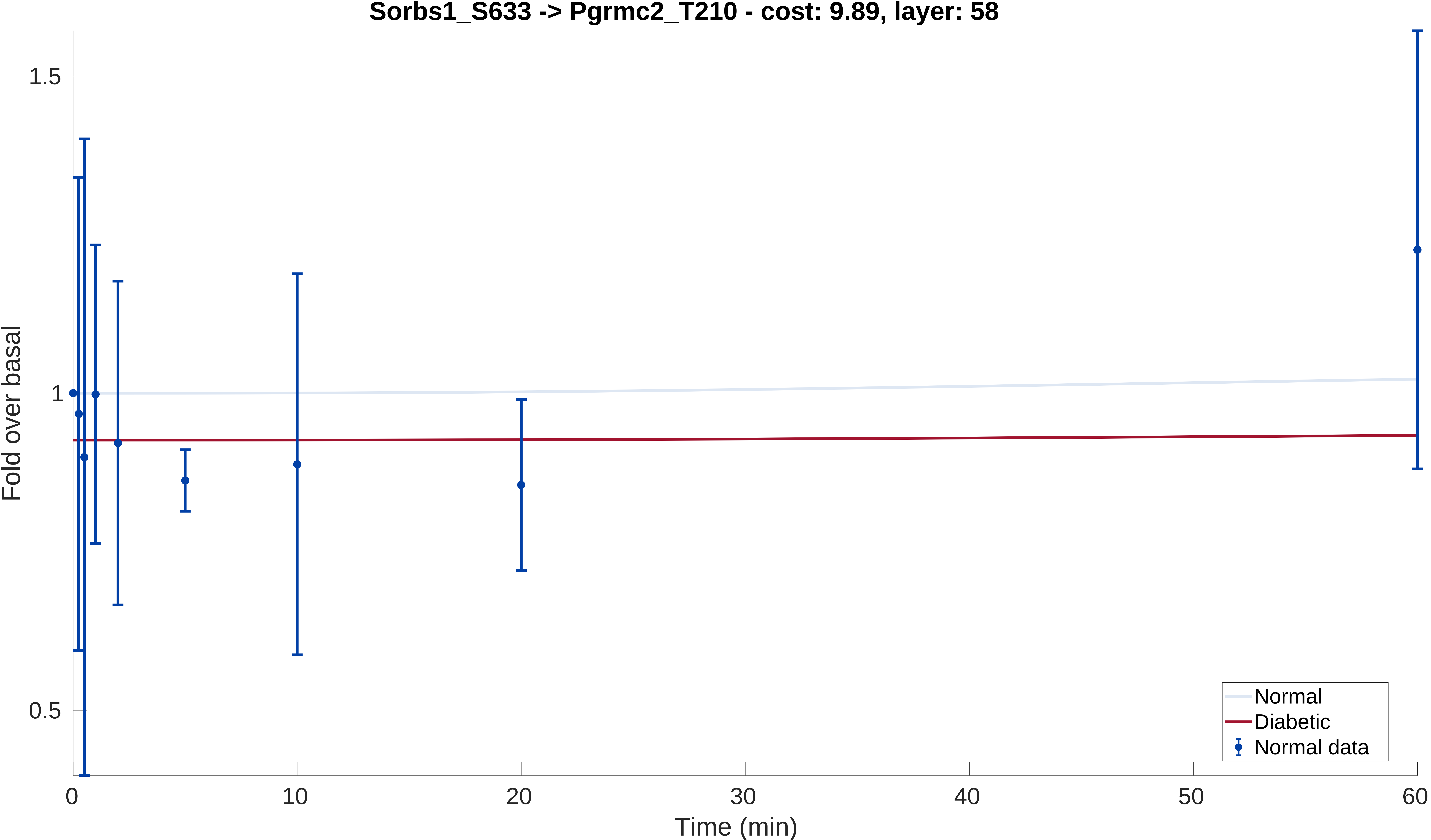

Sorbs1_S633 -> Pgrmc2_T210 - cost: 9.89, layer: 58
Fold over basal
1.5
1
0.5
0
10
20
30
40
50
60
Time (min)
Normal
Diabetic
Normal data

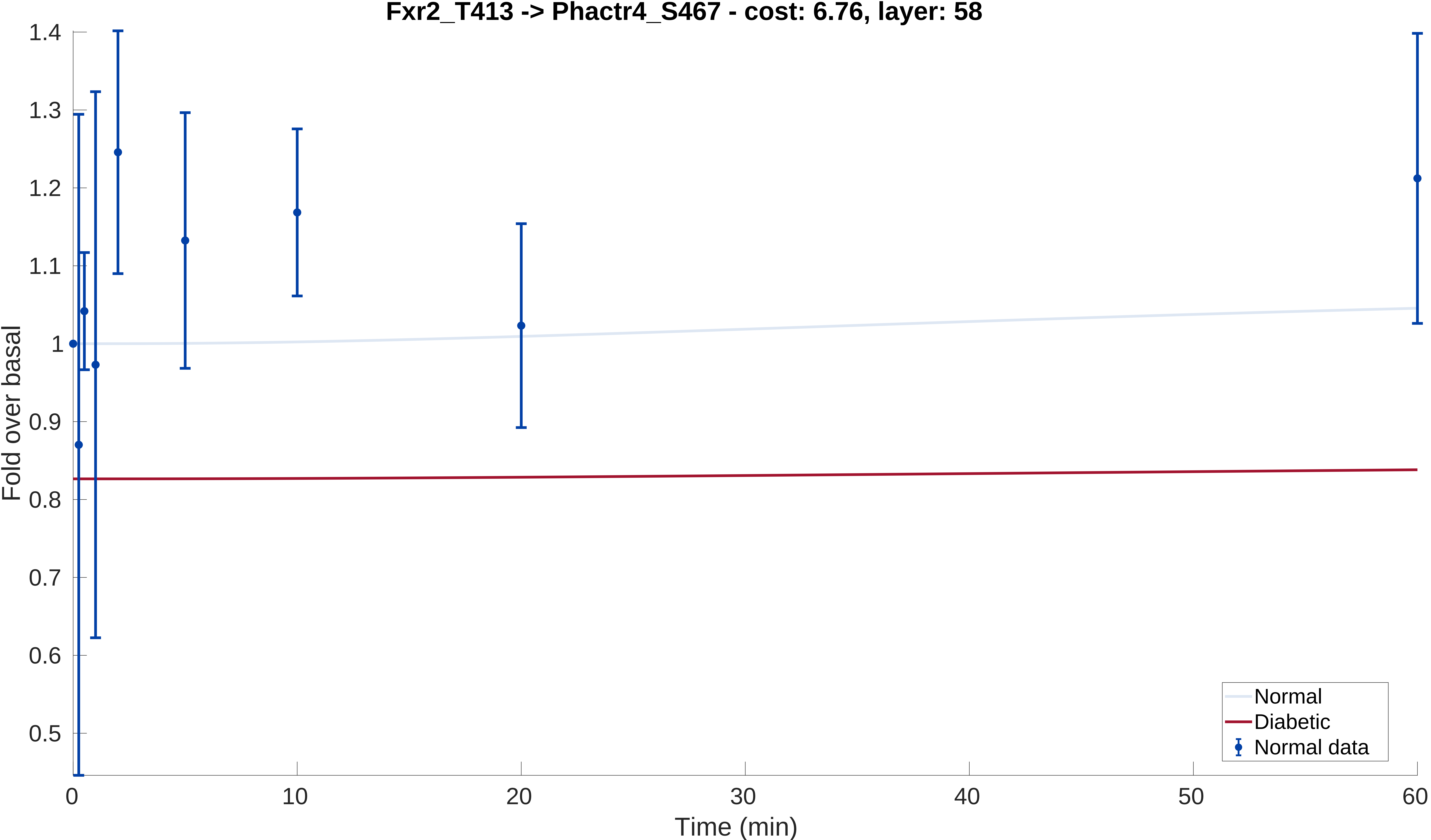

Fxr2_T413 -> Phactr4_S467 - cost: 6.76, layer: 58
Fold over basal
1.4
1.3
1.2
1.1
1
0.9
0.8
0.7
0.6
0.5
0
10
20
30
40
50
60
Time (min)
Normal
Diabetic
Normal data

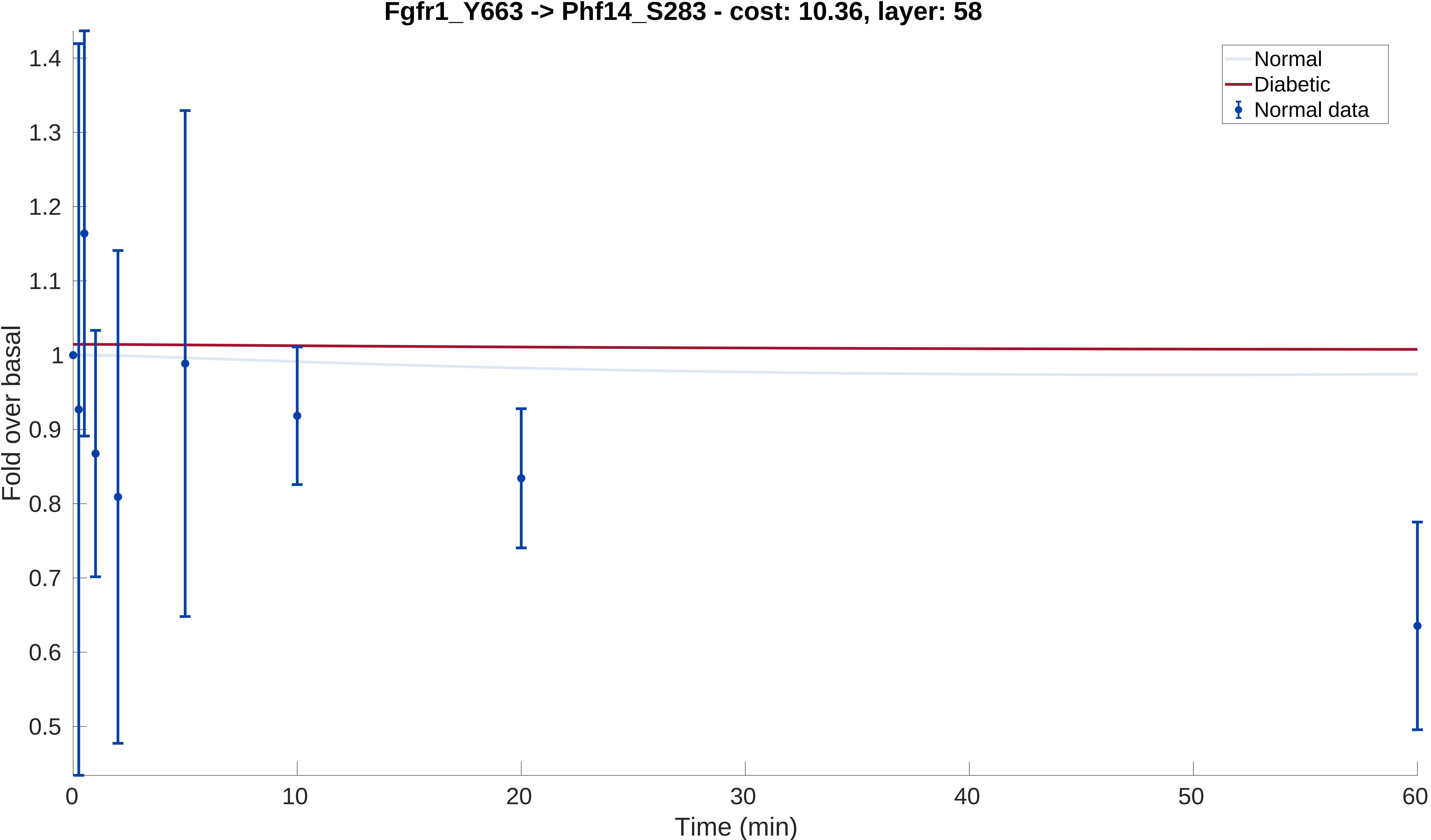
Fgfr1_Y663 -> Phf14_S283 - cost: 10.36, layer: 58
Normal
Diabetic
Normal data
Fold over basal
1.4
1.3
1.2
1.1
1
0.9
0.8
0.7
0.6
0.5
0
10
20
30
40
50
60
Time (min)

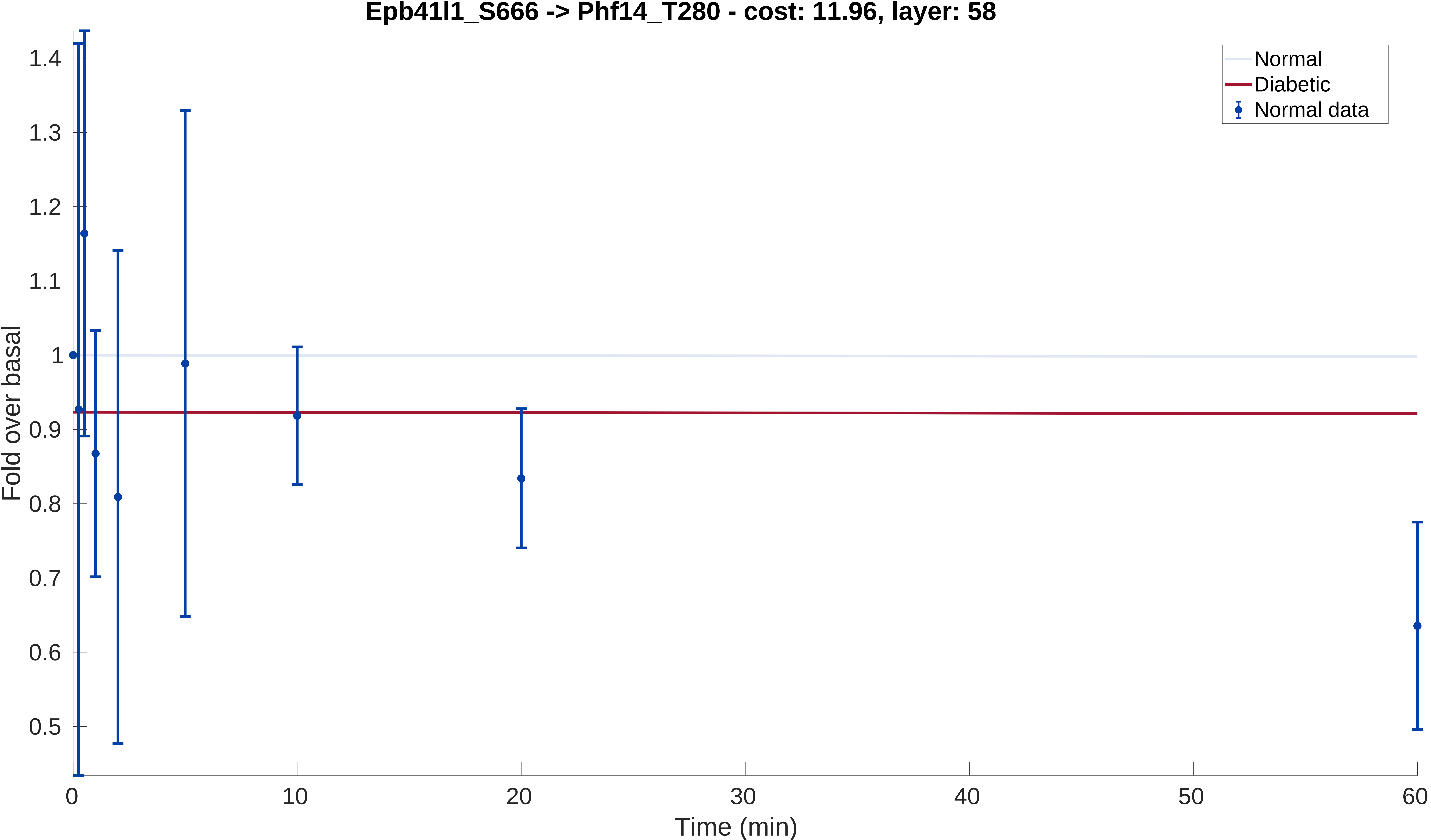

Epb41l1_S666 -> Phf14_T280 - cost: 11.96, layer: 58
Normal
Diabetic
Normal data
Fold over basal
Time (min)
1.4
1.3
1.2
1.1
1
0.9
0.8
0.7
0.6
0.5
0
10
20
30
40
50
60

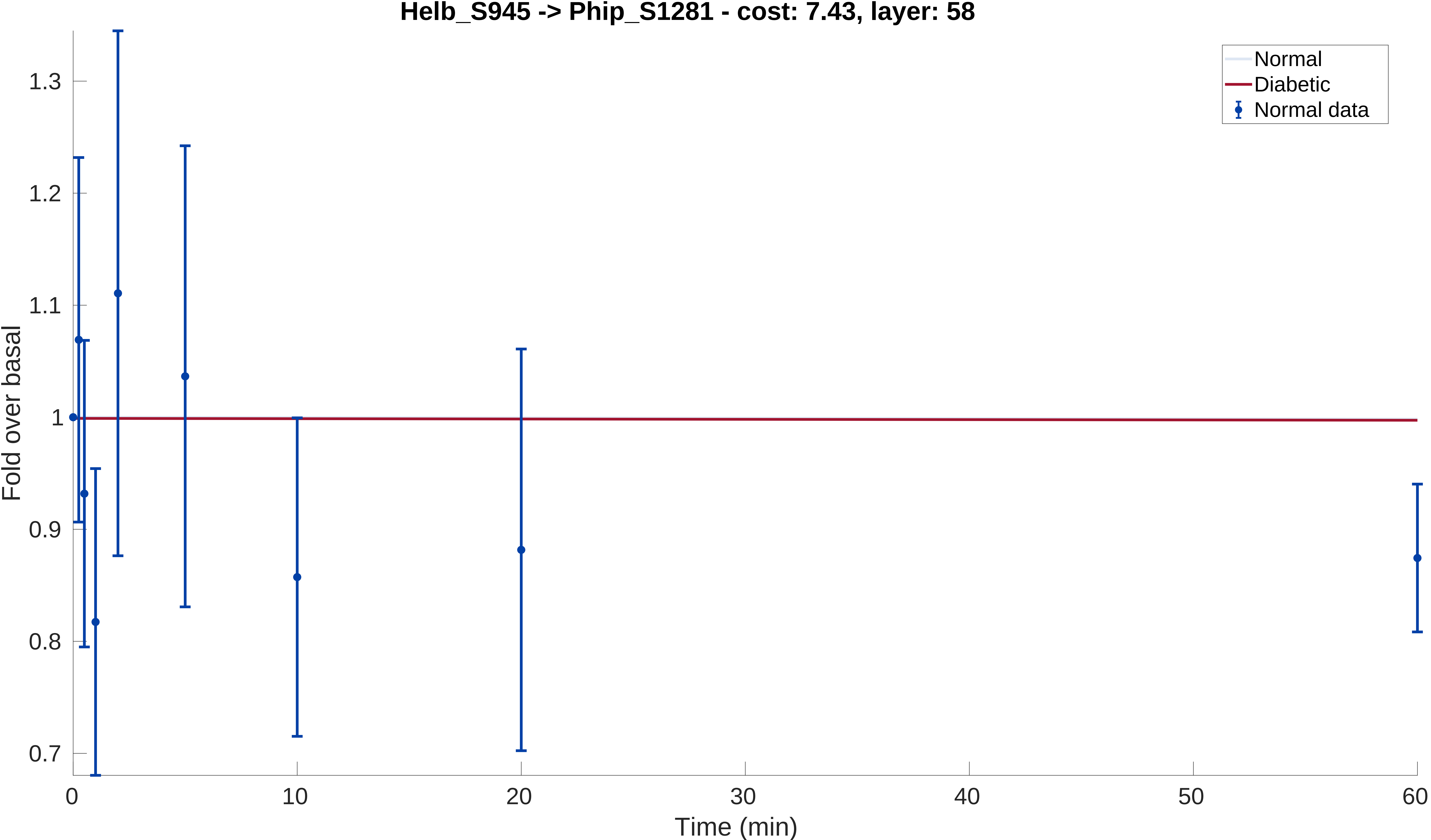

Helb_S945 -> Phip_S1281 - cost: 7.43, layer: 58
Normal
Diabetic
Normal data
Fold over basal
1.3
1.2
1.1
1
0.9
0.8
0.7
0
10
20
30
40
50
60
Time (min)

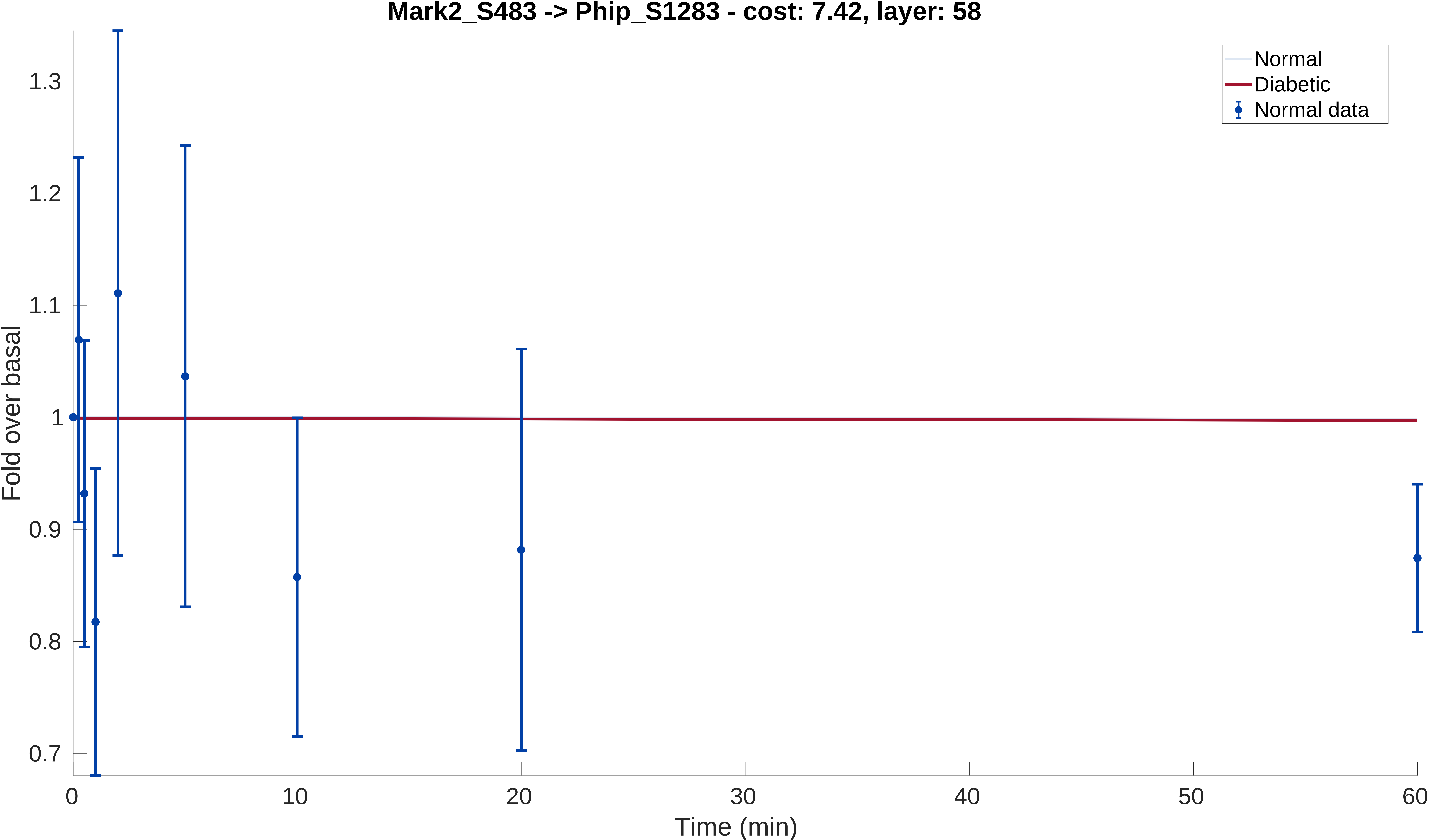

Mark2_S483 -> Phip_S1283 - cost: 7.42, layer: 58
Normal
Diabetic
Normal data
Fold over basal
1.3
1.2
1.1
1
0.9
0.8
0.7
0
10
20
30
40
50
60
Time (min)

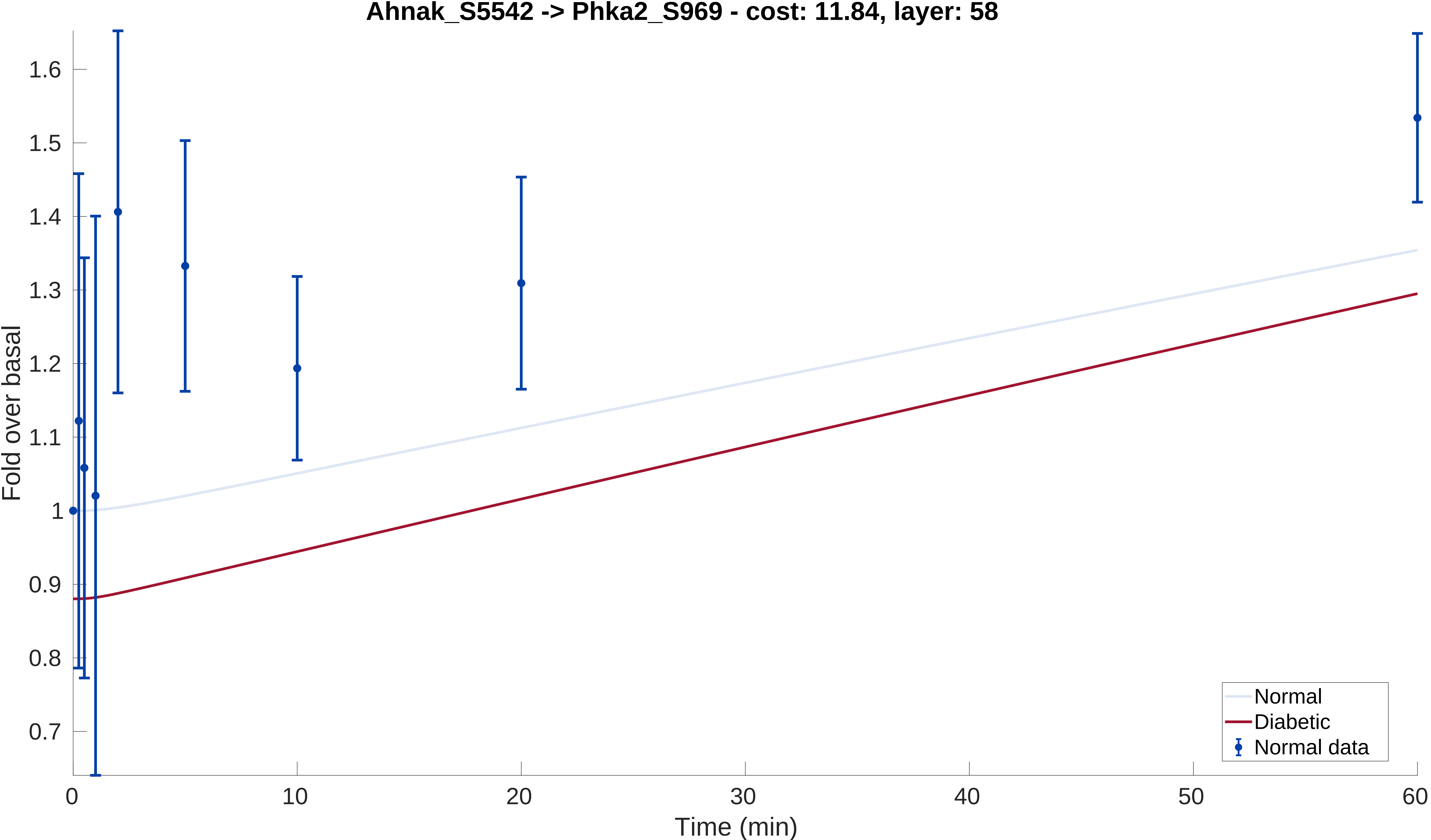

Ahnak_S5542 -> Phka2_S969 - cost: 11.84, layer: 58
Fold over basal
1.6
1.5
1.4
1.3
1.2
1.1
1
0.9
0.8
0.7
0
10
20
30
40
50
60
Time (min)
Normal
Diabetic
Normal data

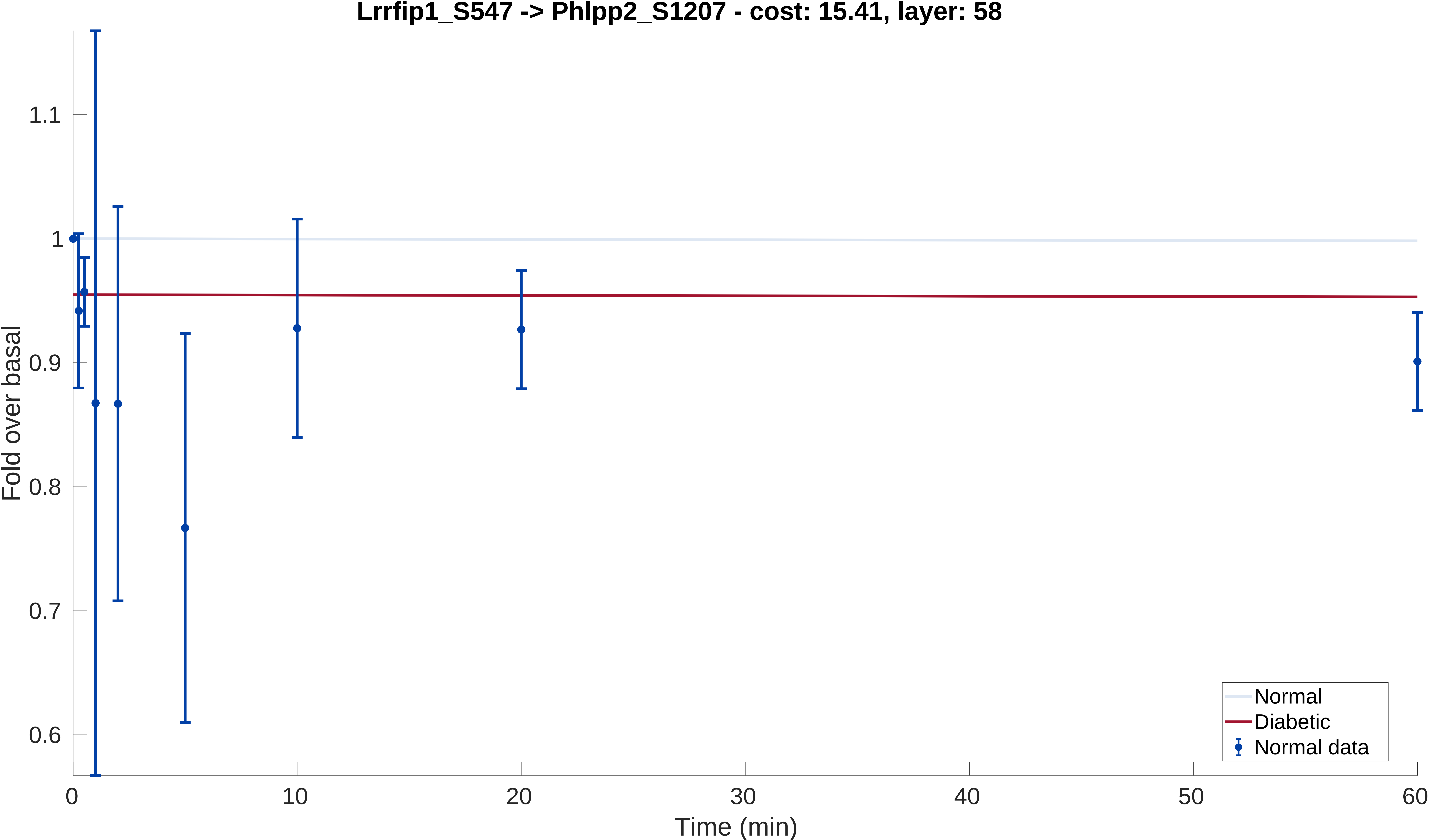
Lrrfip1_S547 -> Phlpp2_S1207 - cost: 15.41, layer: 58
Fold over basal
1.1
1
0.9
0.8
0.7
0.6
0
10
20
30
40
50
60
Time (min)
Normal
Diabetic
Normal data

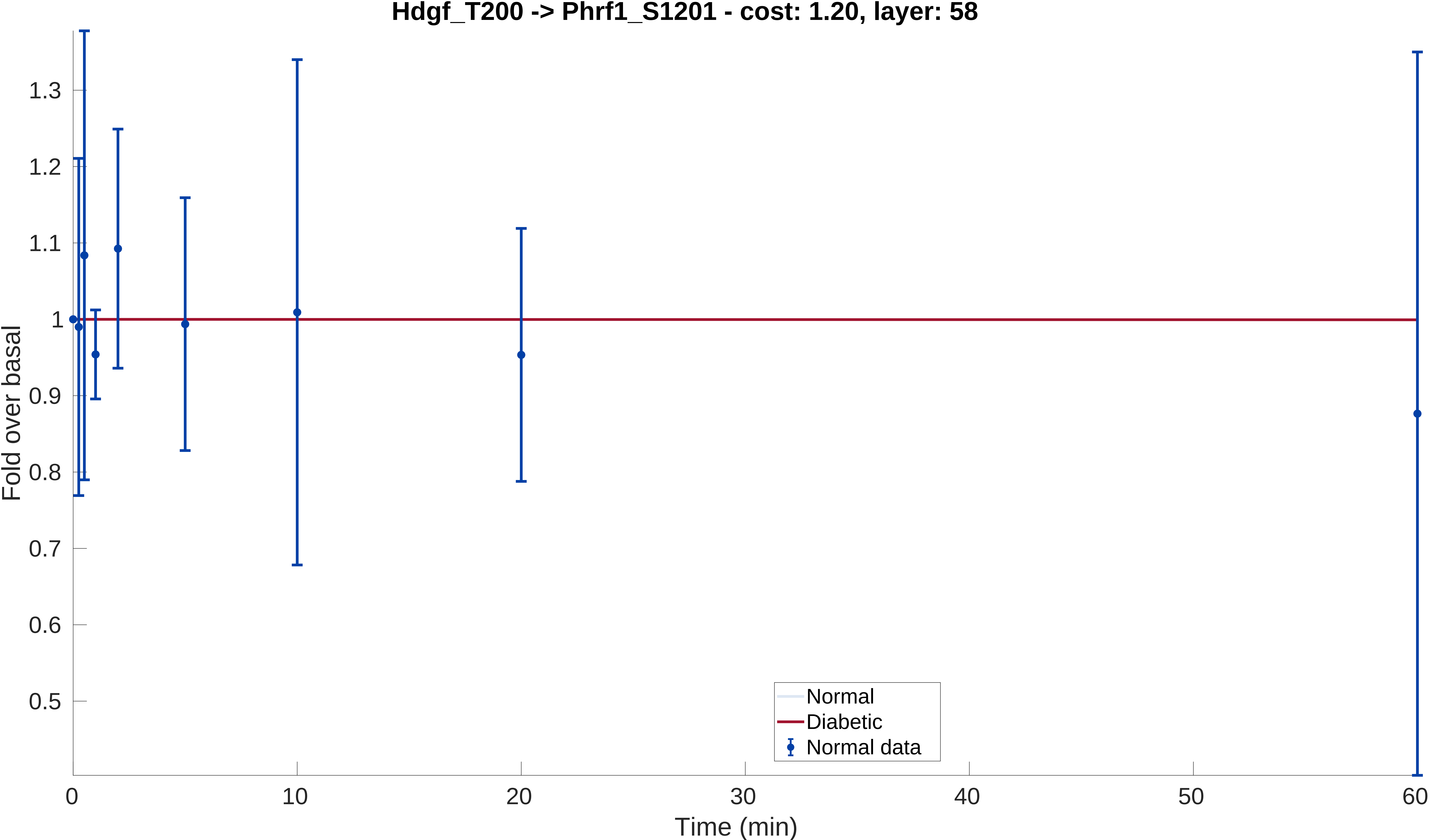
Hdgf_T200 -> Phrf1_S1201 - cost: 1.20, layer: 58
Fold over basal
1.3
1.2
1.1
1
0.9
0.8
0.7
0.6
0.5
0
10
20
30
40
50
60
Time (min)
Normal
Diabetic
Normal data

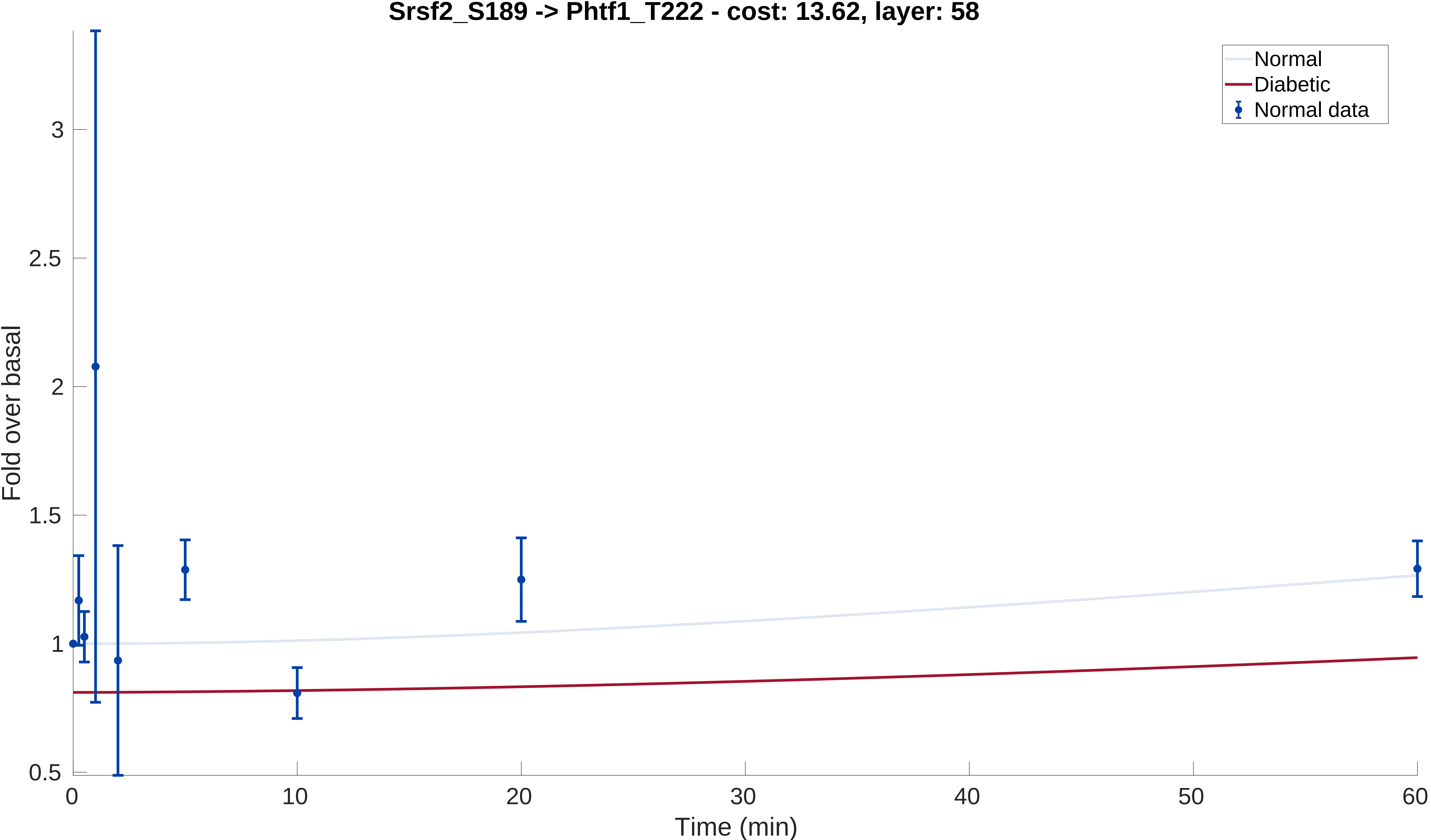

Srsf2_S189 -> Phtf1_T222 - cost: 13.62, layer: 58
Normal
Diabetic
Normal data
Fold over basal
3
2.5
2
1.5
1
0.5
0
10
20
30
40
50
60
Time (min)

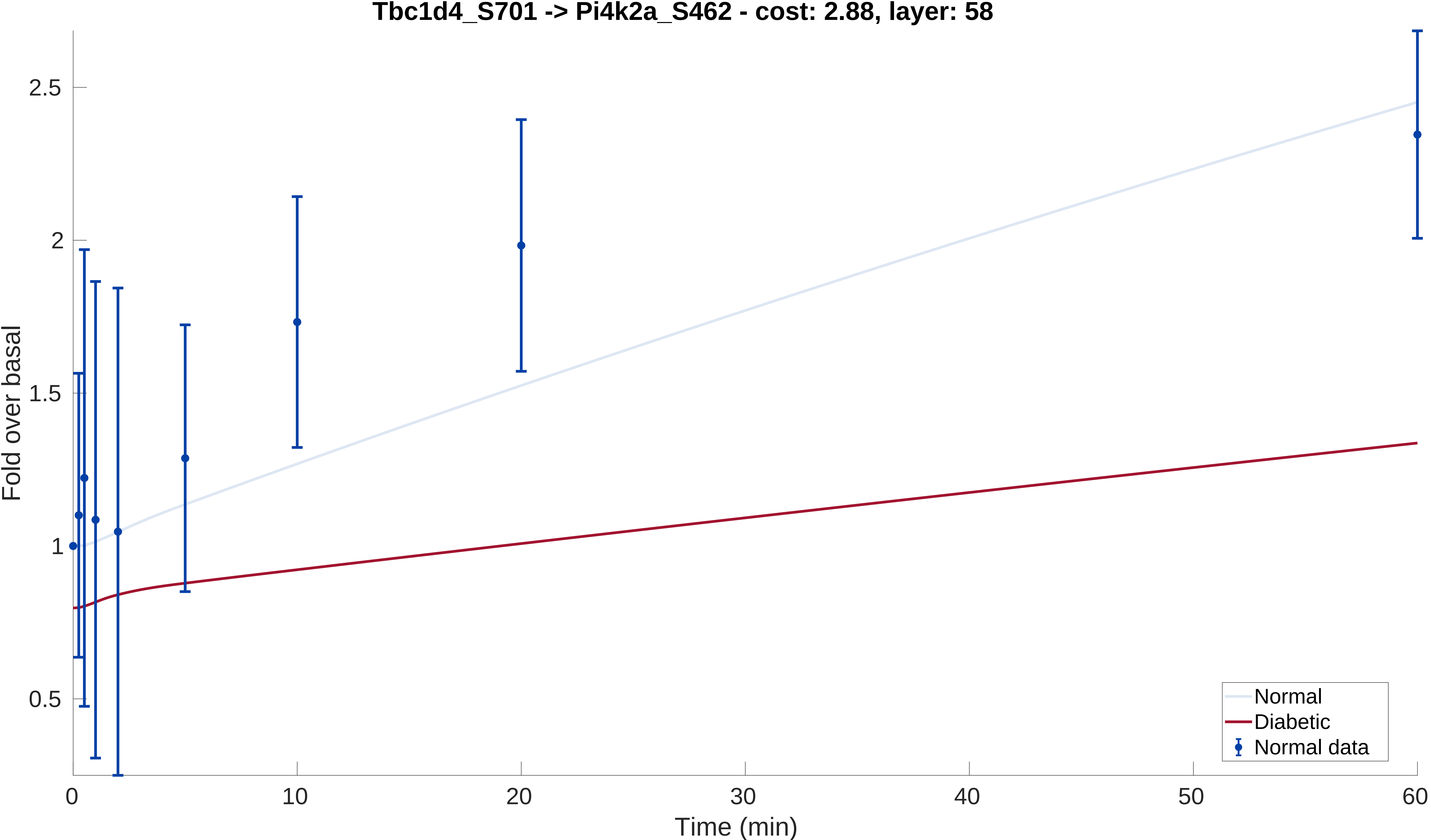

Tbc1d4_S701 -> Pi4k2a_S462 - cost: 2.88, layer: 58
Fold over basal
2.5
2
1.5
1
0.5
0
10
20
30
40
50
60
Time (min)
Normal
Diabetic
Normal data

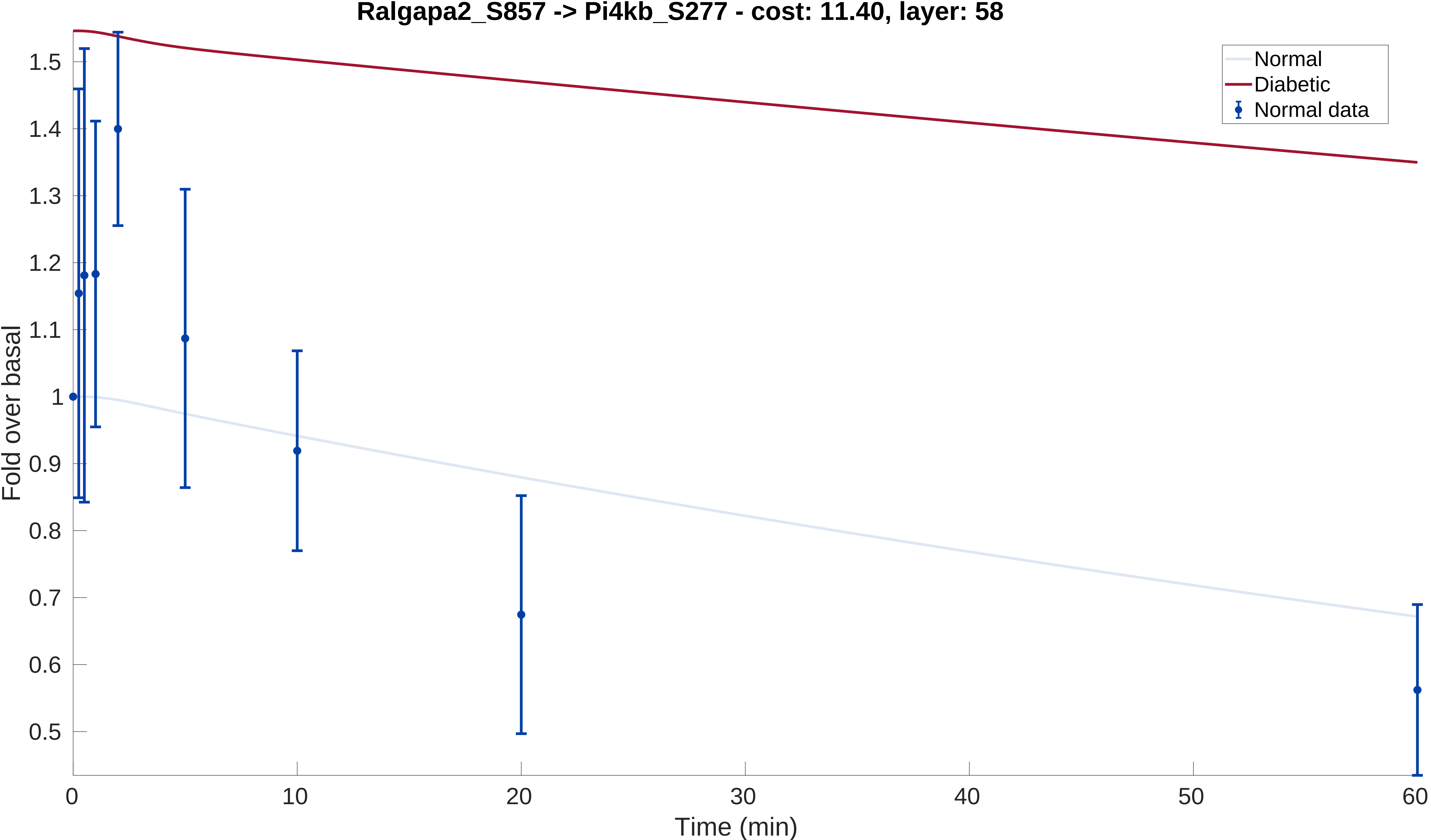
Ralgapa2_S857 -> Pi4kb_S277 - cost: 11.40, layer: 58
Normal
Diabetic
Normal data
Fold over basal
1.5
1.4
1.3
1.2
1.1
1
0.9
0.8
0.7
0.6
0.5
0
10
20
30
40
50
60
Time (min)

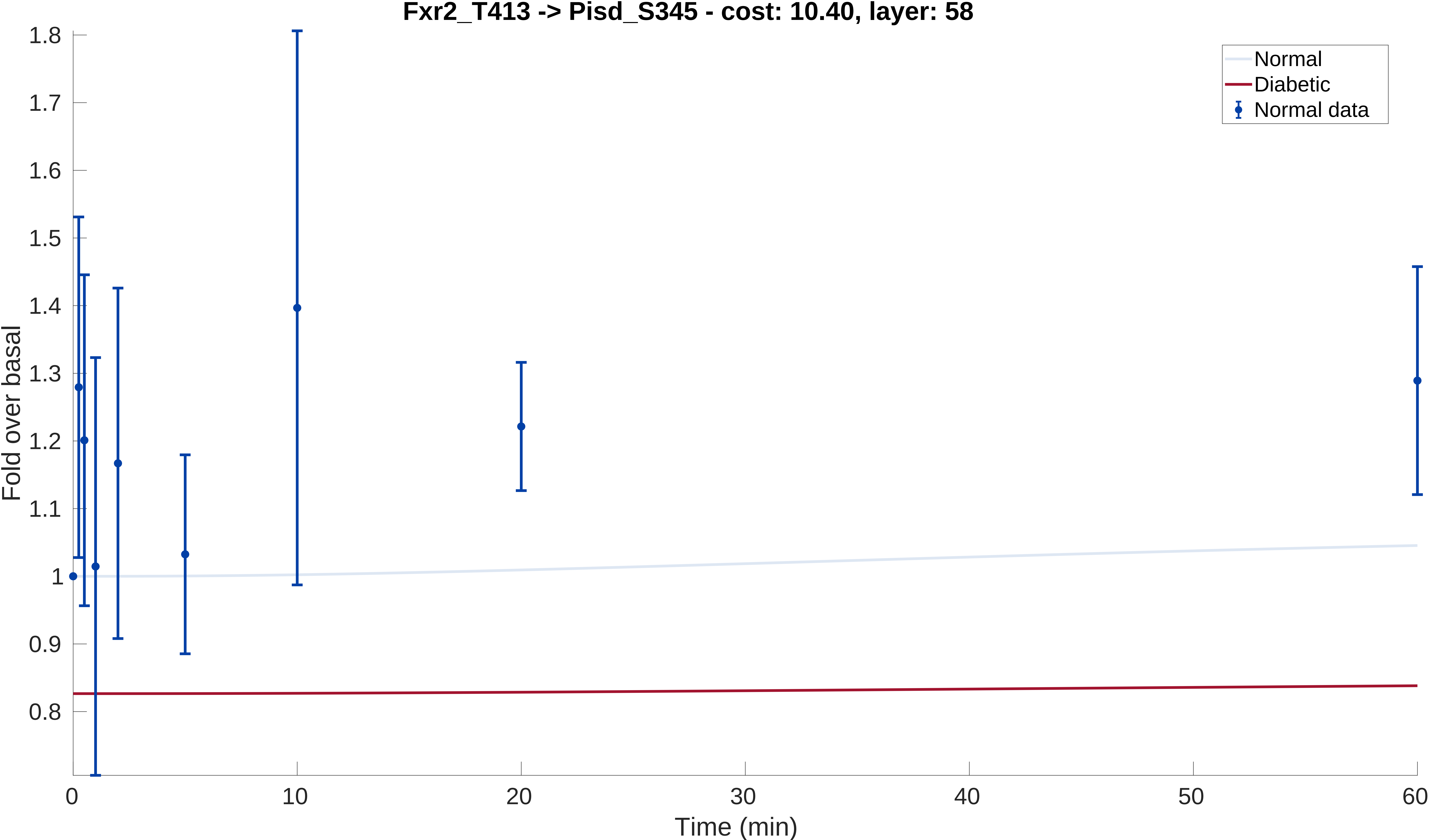

Fxr2_T413 -> Pisd_S345 - cost: 10.40, layer: 58
Normal
Diabetic
Normal data
Fold over basal
1.8
1.7
1.6
1.5
1.4
1.3
1.2
1.1
1
0.9
0.8
0
10
20
30
40
50
60
Time (min)

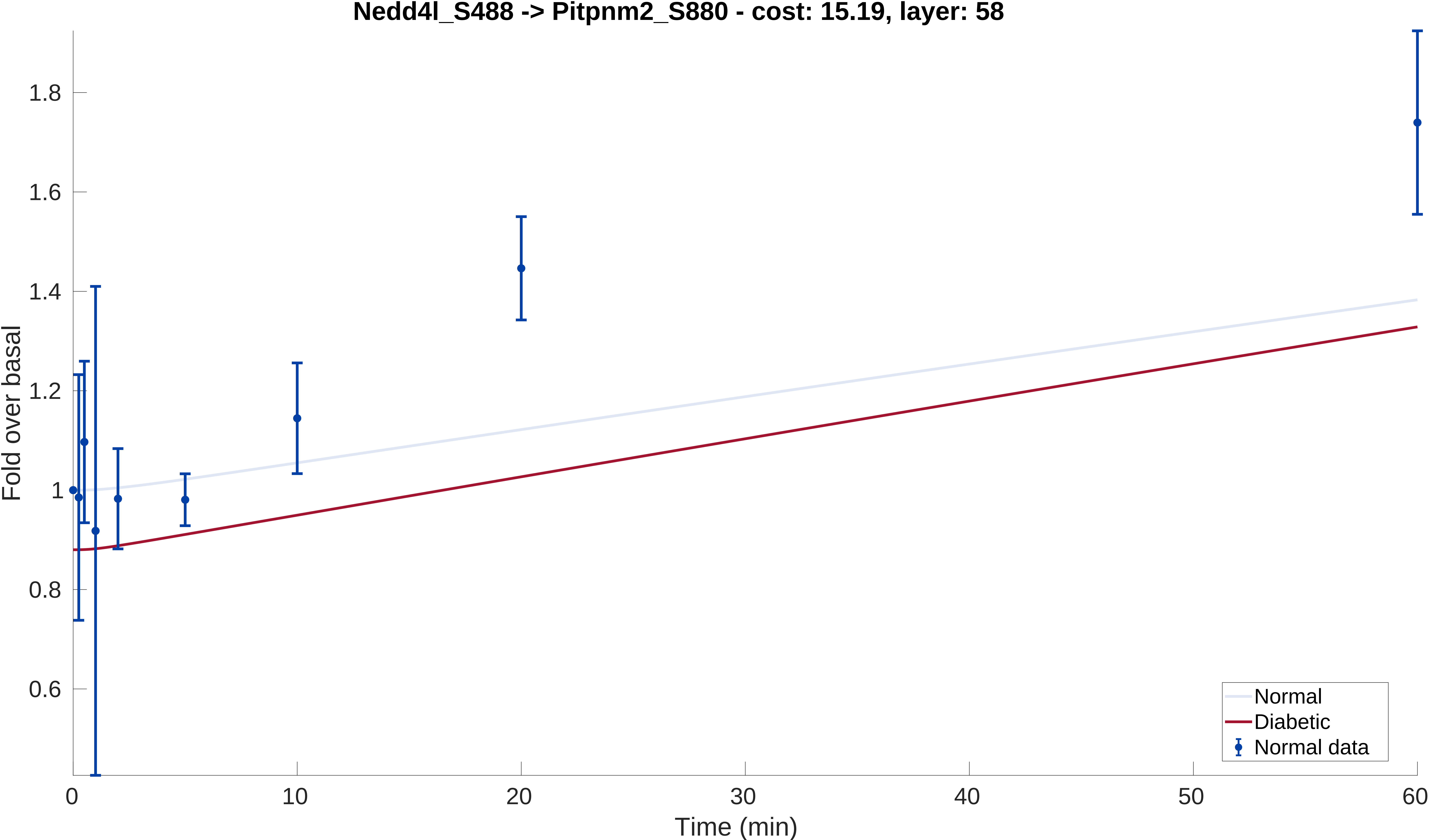

Nedd4l_S488 -> Pitpnm2_S880 - cost: 15.19, layer: 58
Fold over basal
1.8
1.6
1.4
1.2
1
0.8
0.6
0
10
20
30
40
50
60
Time (min)
Normal
Diabetic
Normal data

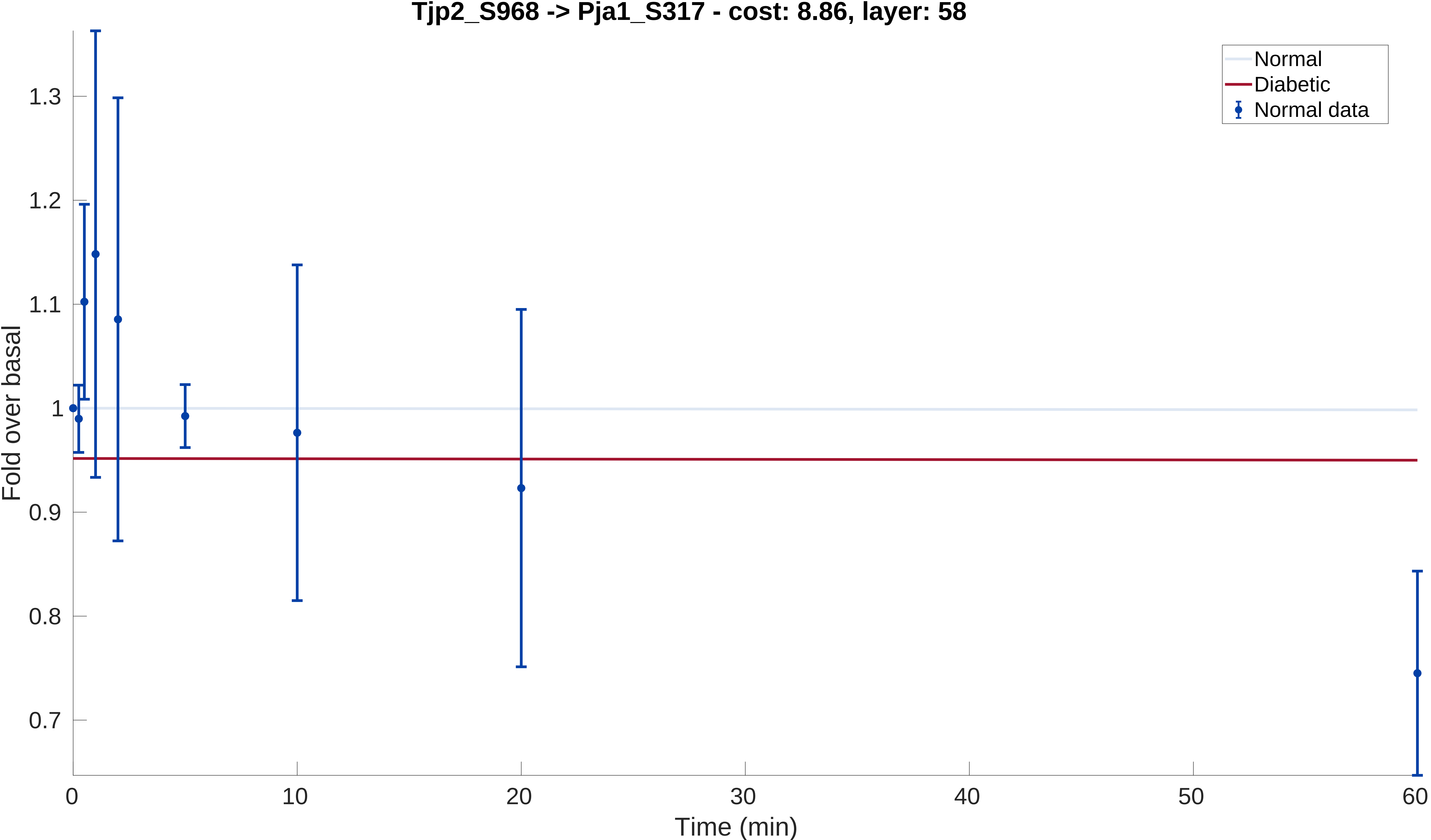

Tjp2_S968 -> Pja1_S317 - cost: 8.86, layer: 58
Normal
Diabetic
Normal data
Fold over basal
1.3
1.2
1.1
1
0.9
0.8
0.7
0
10
20
30
40
50
60
Time (min)

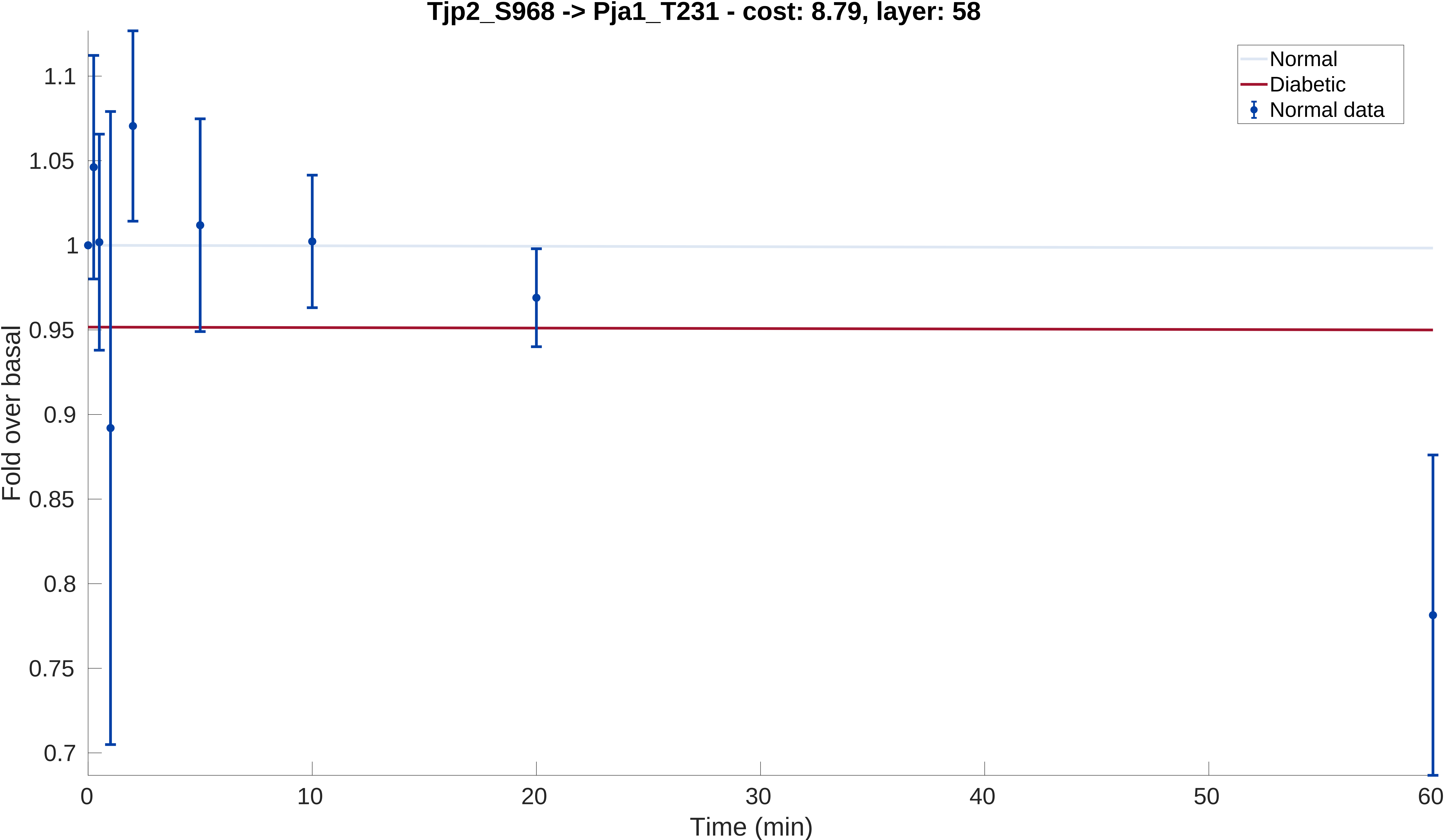

Tjp2_S968 -> Pja1_T231 - cost: 8.79, layer: 58
Normal
Diabetic
Normal data
Fold over basal
Time (min)
1.1
1.05
1
0.95
0.9
0.85
0.8
0.75
0.7
0
10
20
30
40
50
60

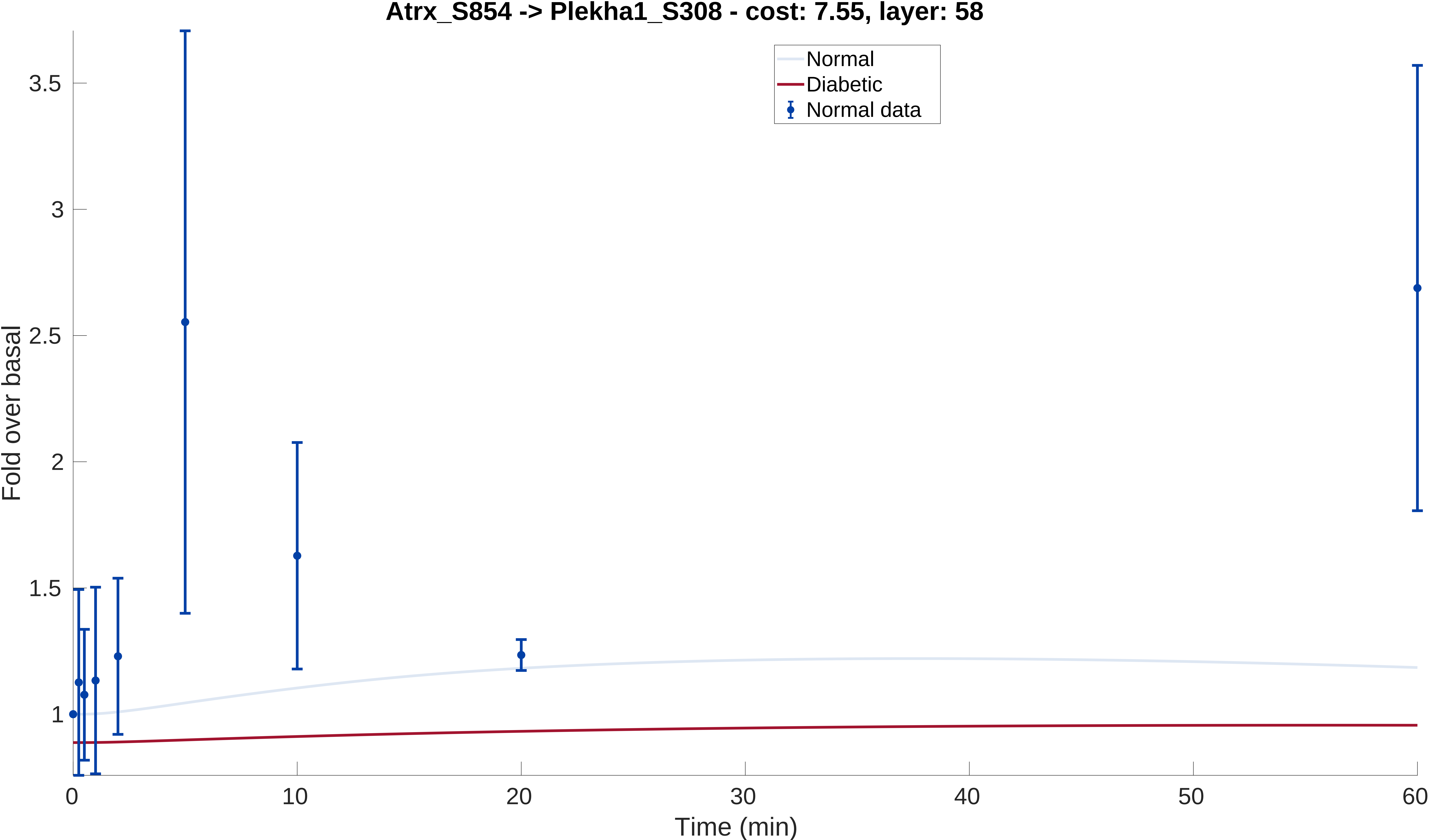
Atrx_S854 -> Plekha1_S308 - cost: 7.55, layer: 58
Normal
Diabetic
Normal data
Fold over basal
3.5
3
2.5
2
1.5
1
0
10
20
30
40
50
60
Time (min)

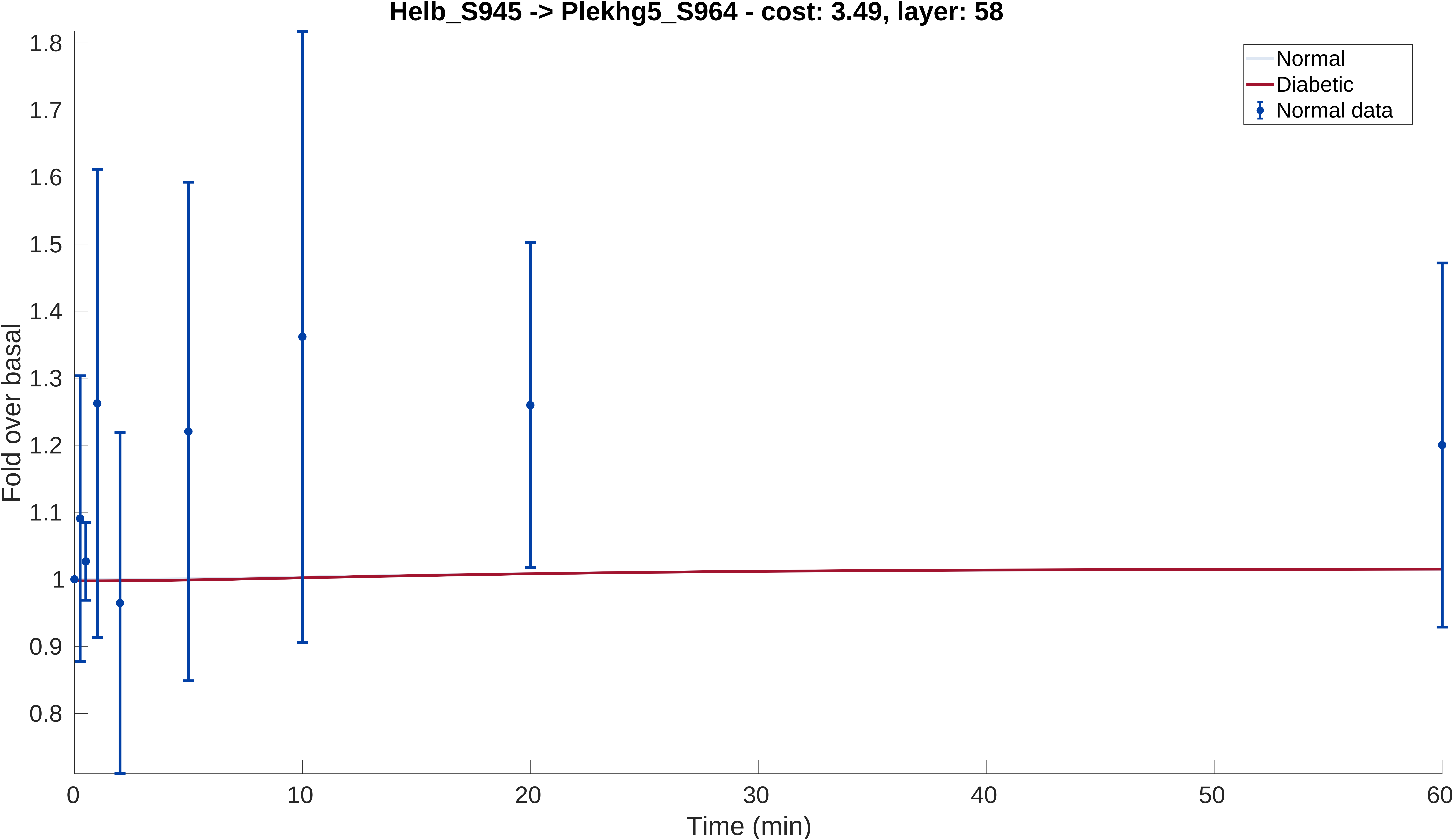

Helb_S945 -> Plekhg5_S964 - cost: 3.49, layer: 58
Normal
Diabetic
Normal data
Fold over basal
Time (min)
1.8
1.7
1.6
1.5
1.4
1.3
1.2
1.1
1
0.9
0.8
0
10
20
30
40
50
60

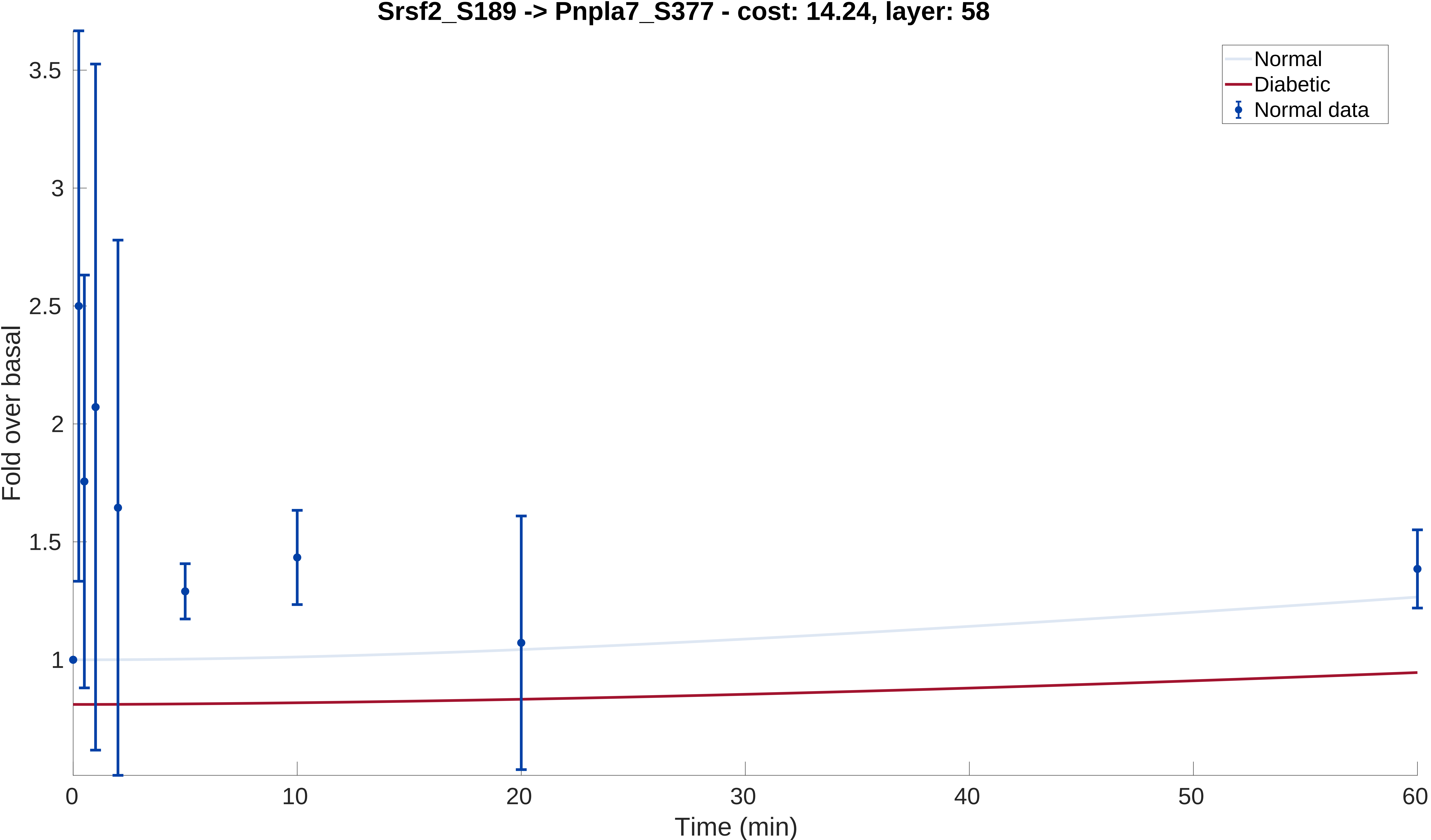

Srsf2_S189 -> Pnpla7_S377 - cost: 14.24, layer: 58
Normal
Diabetic
Normal data
Fold over basal
3.5
3
2.5
2
1.5
1
0
10
20
30
40
50
60
Time (min)

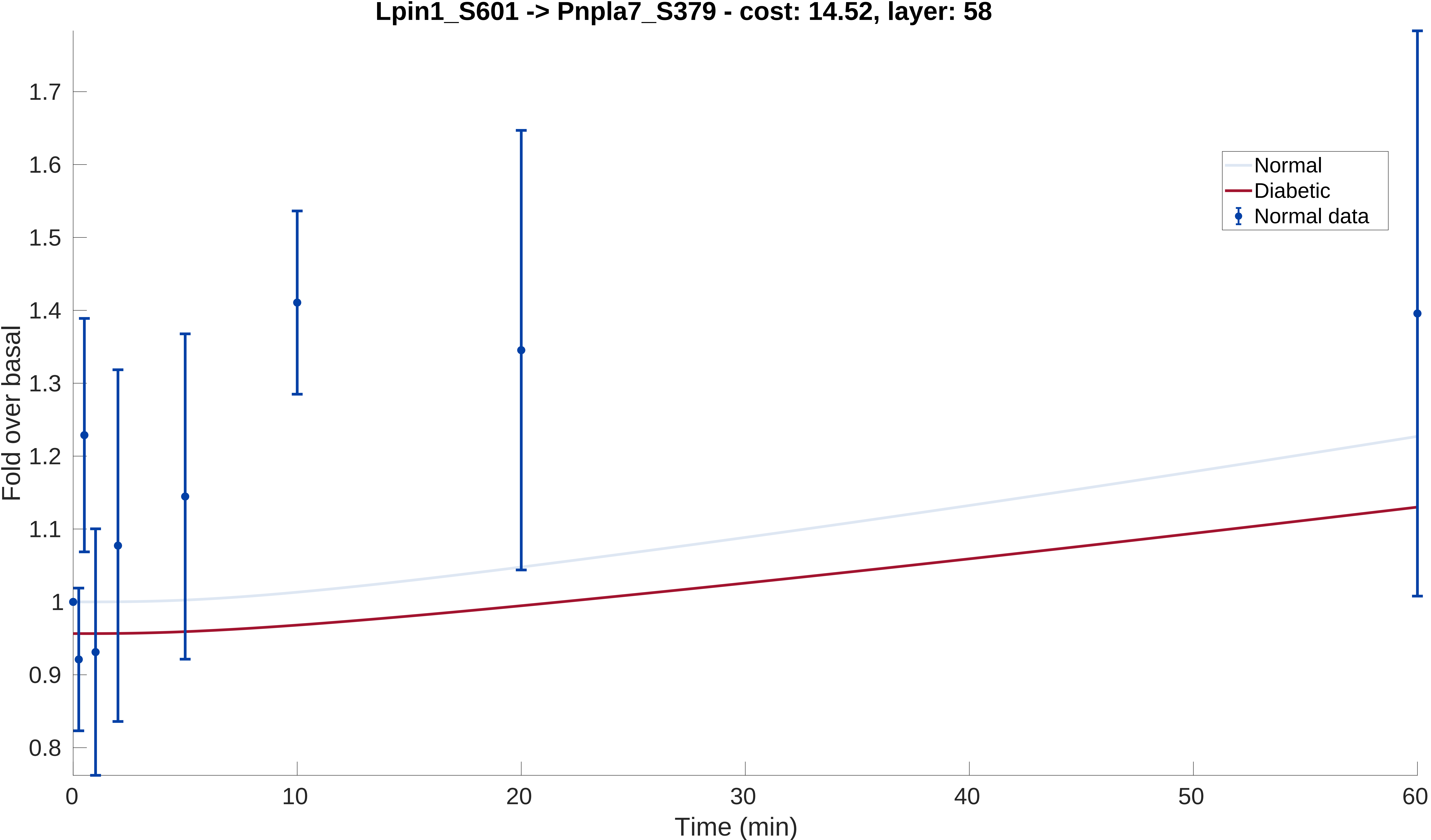

Lpin1_S601 -> Pnpla7_S379 - cost: 14.52, layer: 58
Fold over basal
1.7
1.6
1.5
1.4
1.3
1.2
1.1
1
0.9
0.8
0
10
20
30
40
50
60
Time (min)
Normal
Diabetic
Normal data

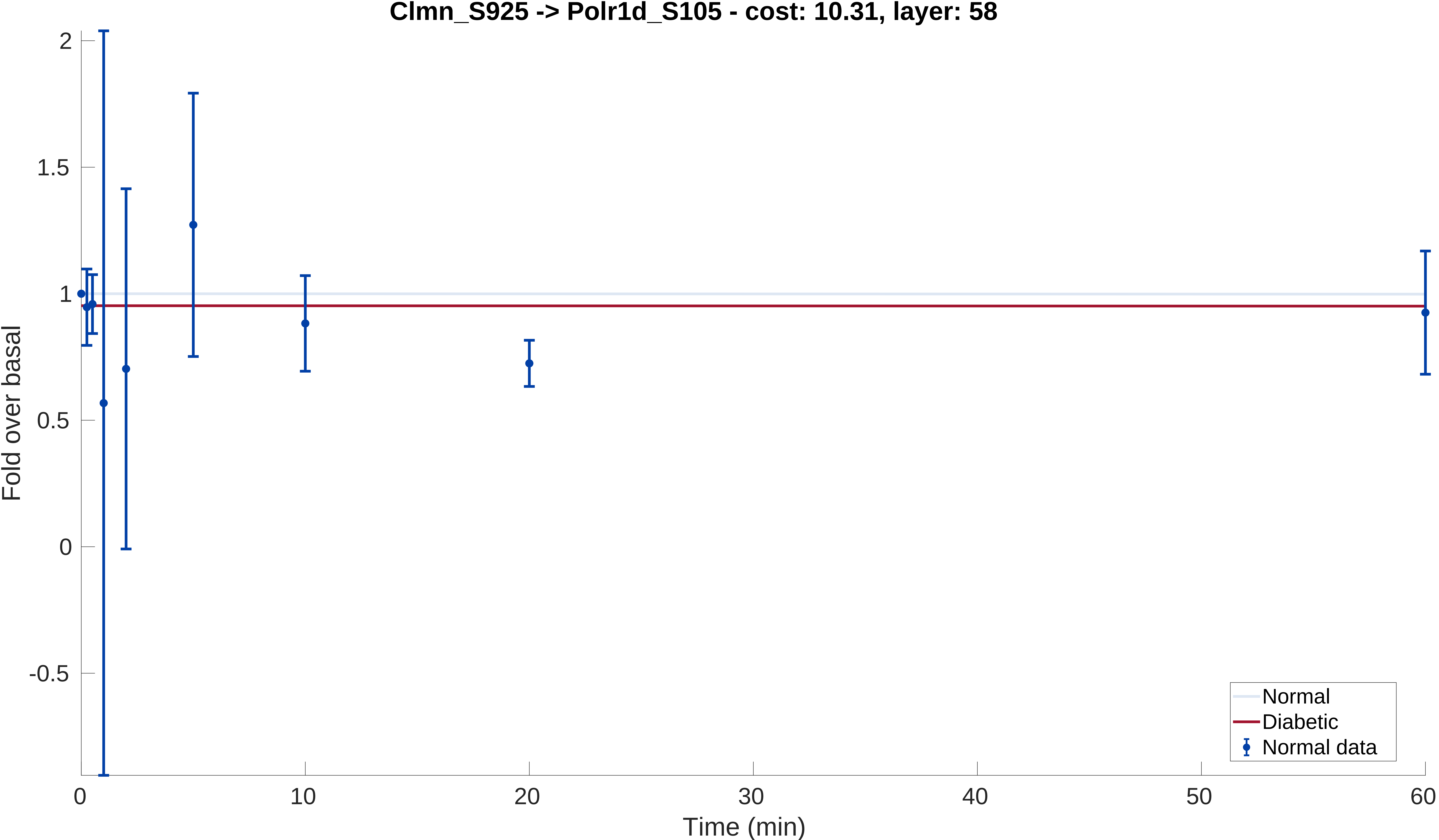

Clmn_S925 -> Polr1d_S105 - cost: 10.31, layer: 58
Fold over basal
2
1.5
1
0.5
0
-0.5
0
10
20
30
40
50
60
Time (min)
Normal
Diabetic
Normal data

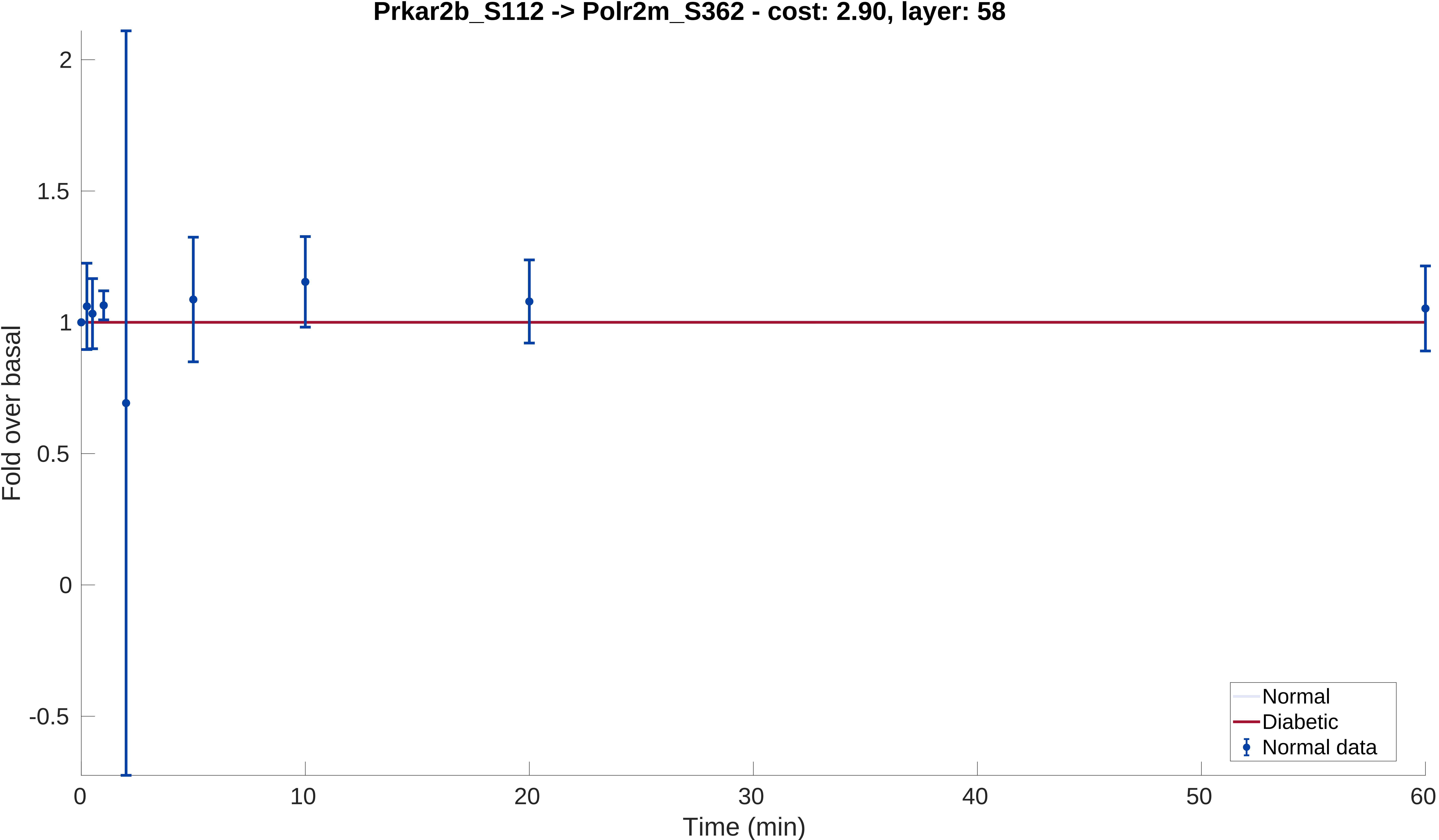
Prkar2b_S112 -> Polr2m_S362 - cost: 2.90, layer: 58
Fold over basal
2
1.5
1
0.5
0
-0.5
0
10
20
30
40
50
60
Time (min)
Normal
Diabetic
Normal data

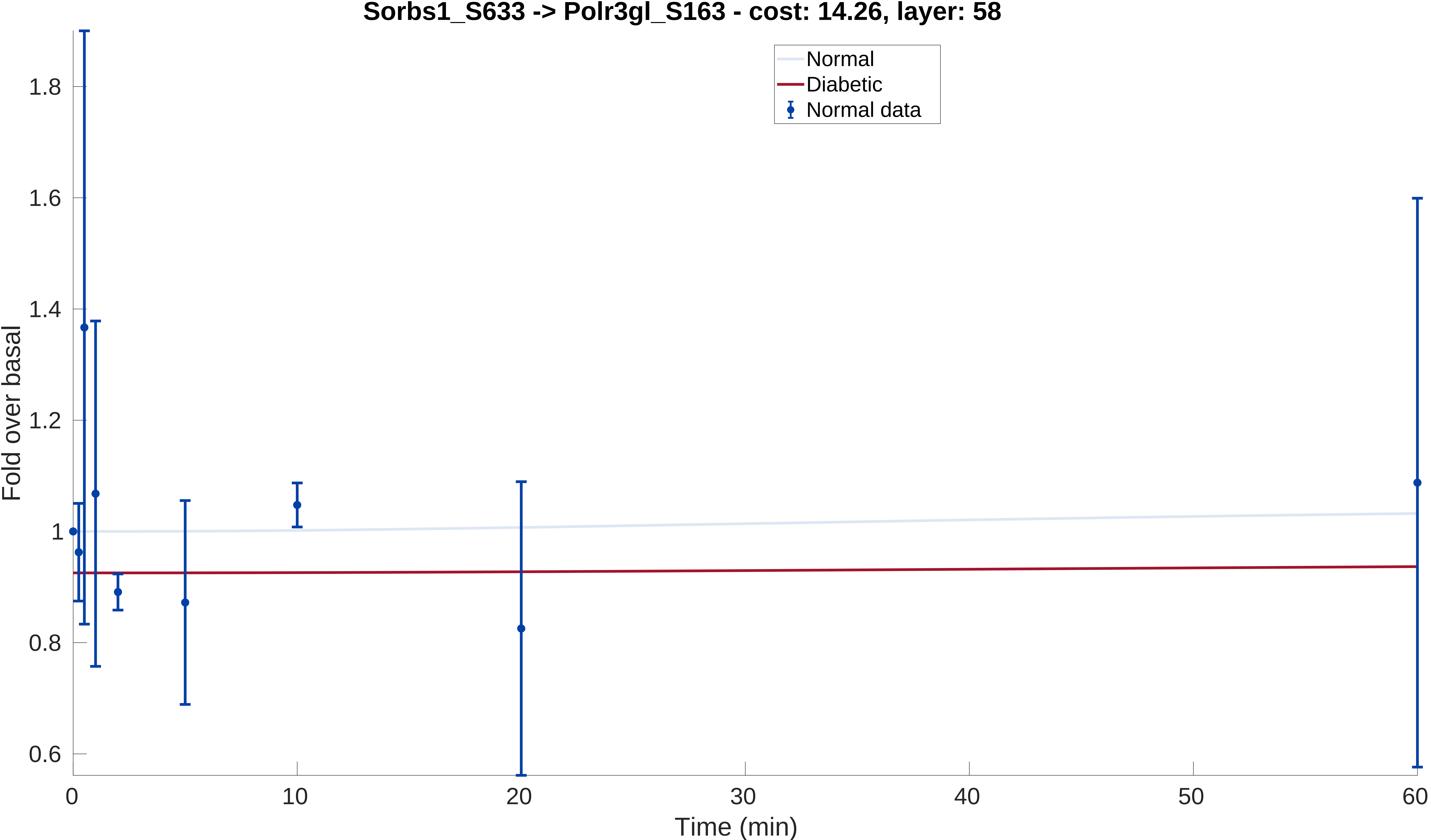

Sorbs1_S633 -> Polr3gl_S163 - cost: 14.26, layer: 58
Normal
Diabetic
Normal data
Fold over basal
1.8
1.6
1.4
1.2
1
0.8
0.6
0
10
20
30
40
50
60
Time (min)

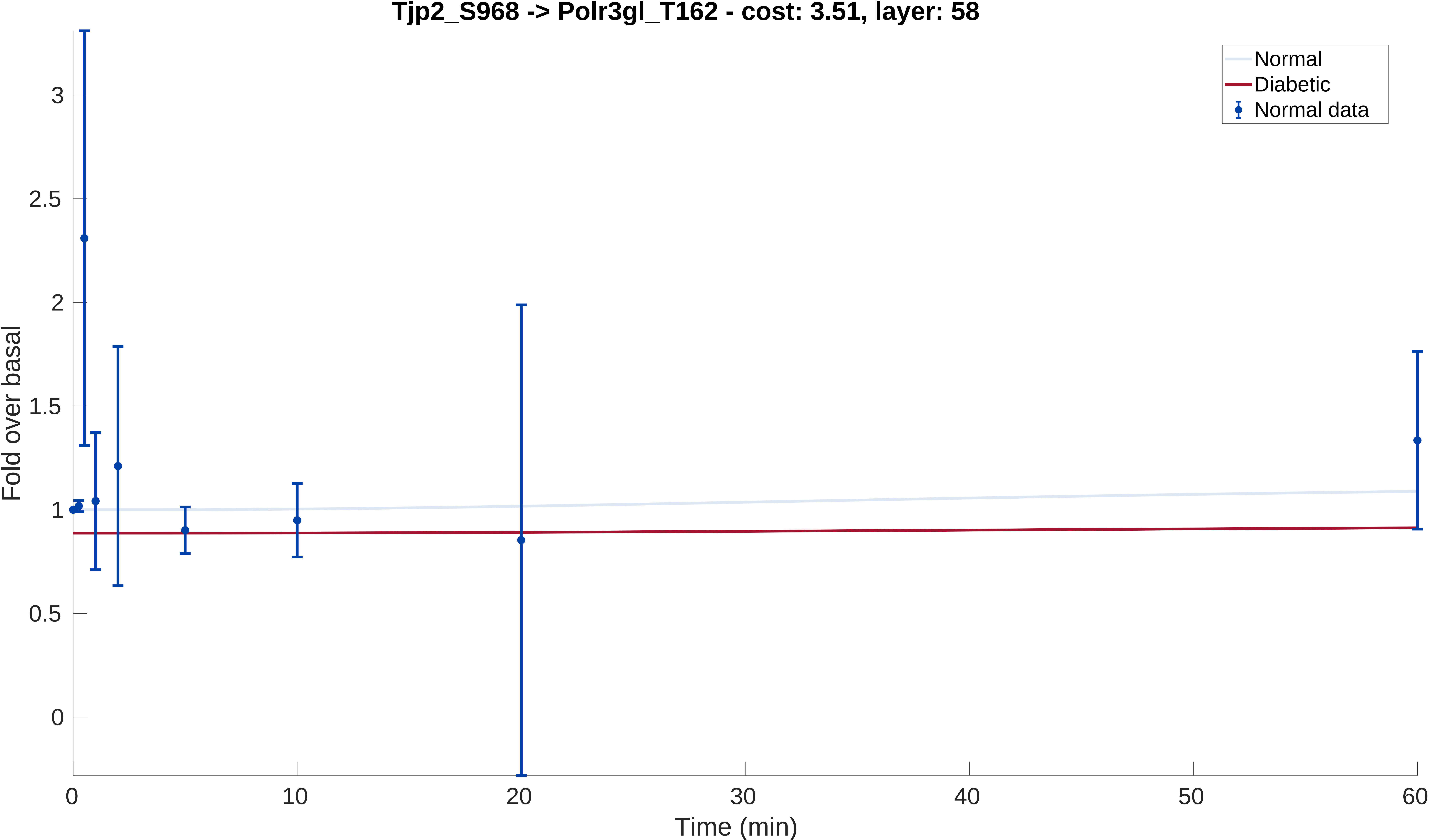

Tjp2_S968 -> Polr3gl_T162 - cost: 3.51, layer: 58
Normal
Diabetic
Normal data
Fold over basal
3
2.5
2
1.5
1
0.5
0
0
10
20
30
40
50
60
Time (min)

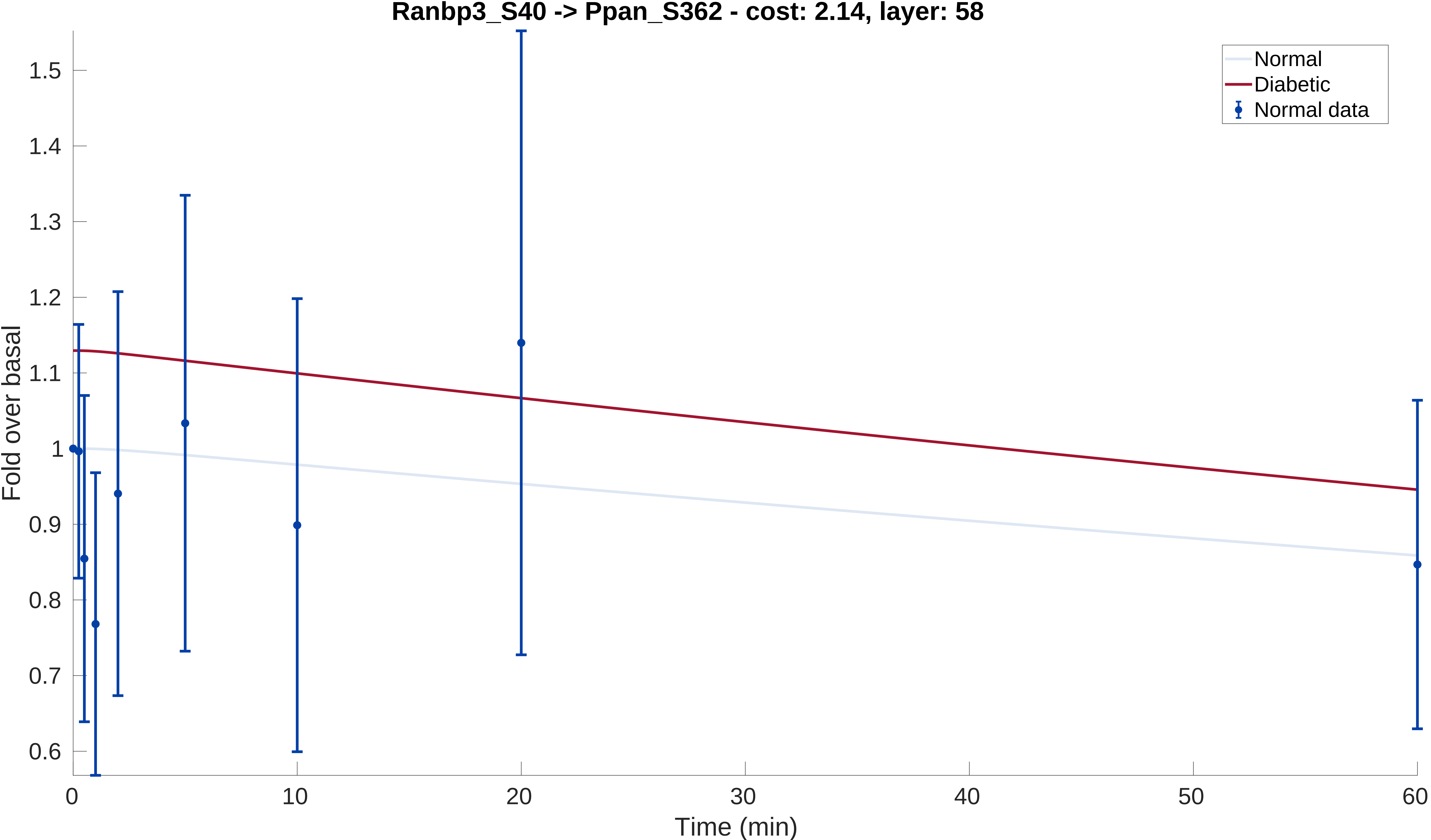

Ranbp3_S40 -> Ppan_S362 - cost: 2.14, layer: 58
Normal
Diabetic
Normal data
Fold over basal
1.5
1.4
1.3
1.2
1.1
1
0.9
0.8
0.7
0.6
0
10
20
30
40
50
60
Time (min)

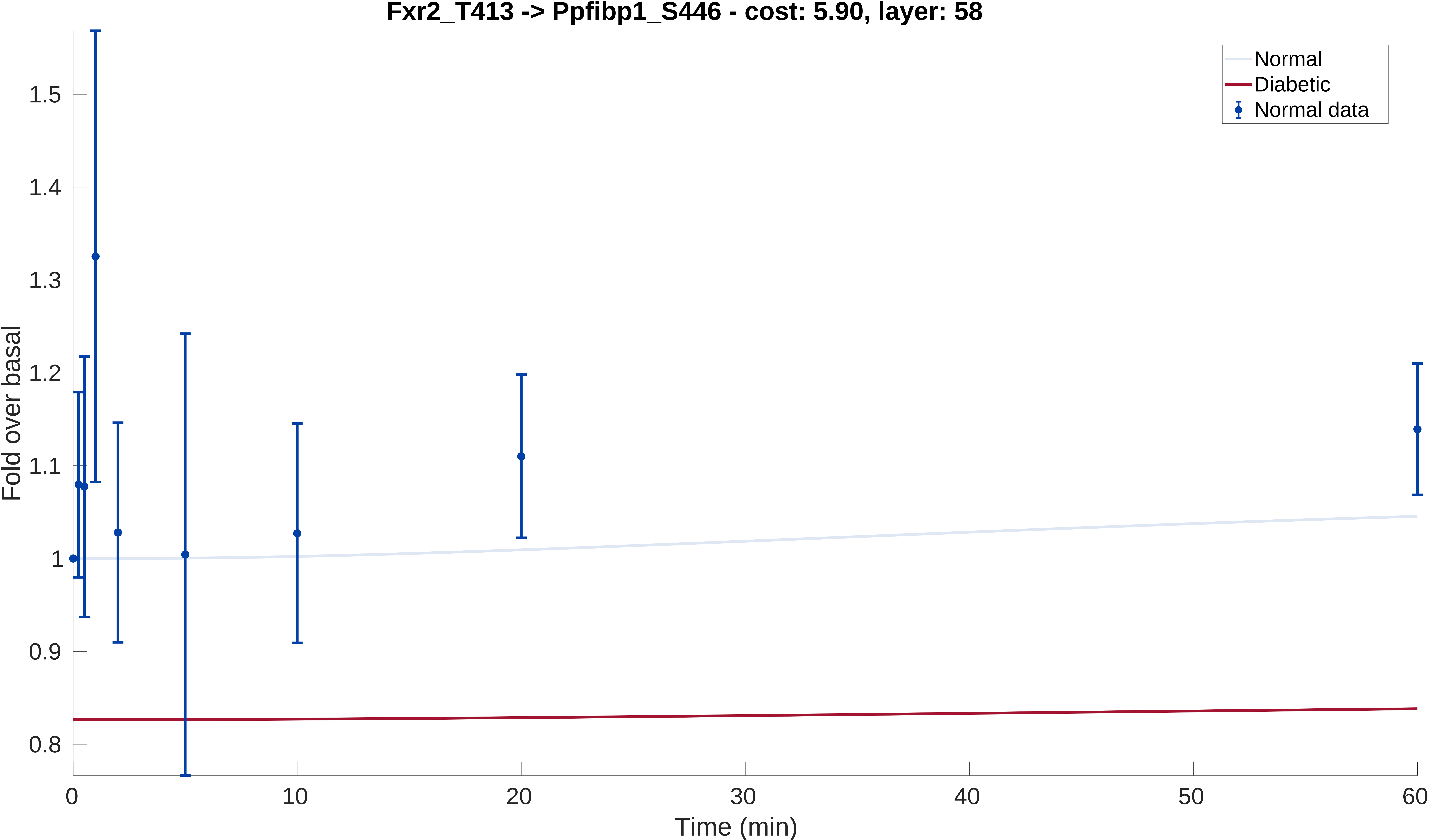

Fxr2_T413 -> Ppfibp1_S446 - cost: 5.90, layer: 58
Normal
Diabetic
Normal data
Fold over basal
1.5
1.4
1.3
1.2
1.1
1
0.9
0.8
0
10
20
30
40
50
60
Time (min)

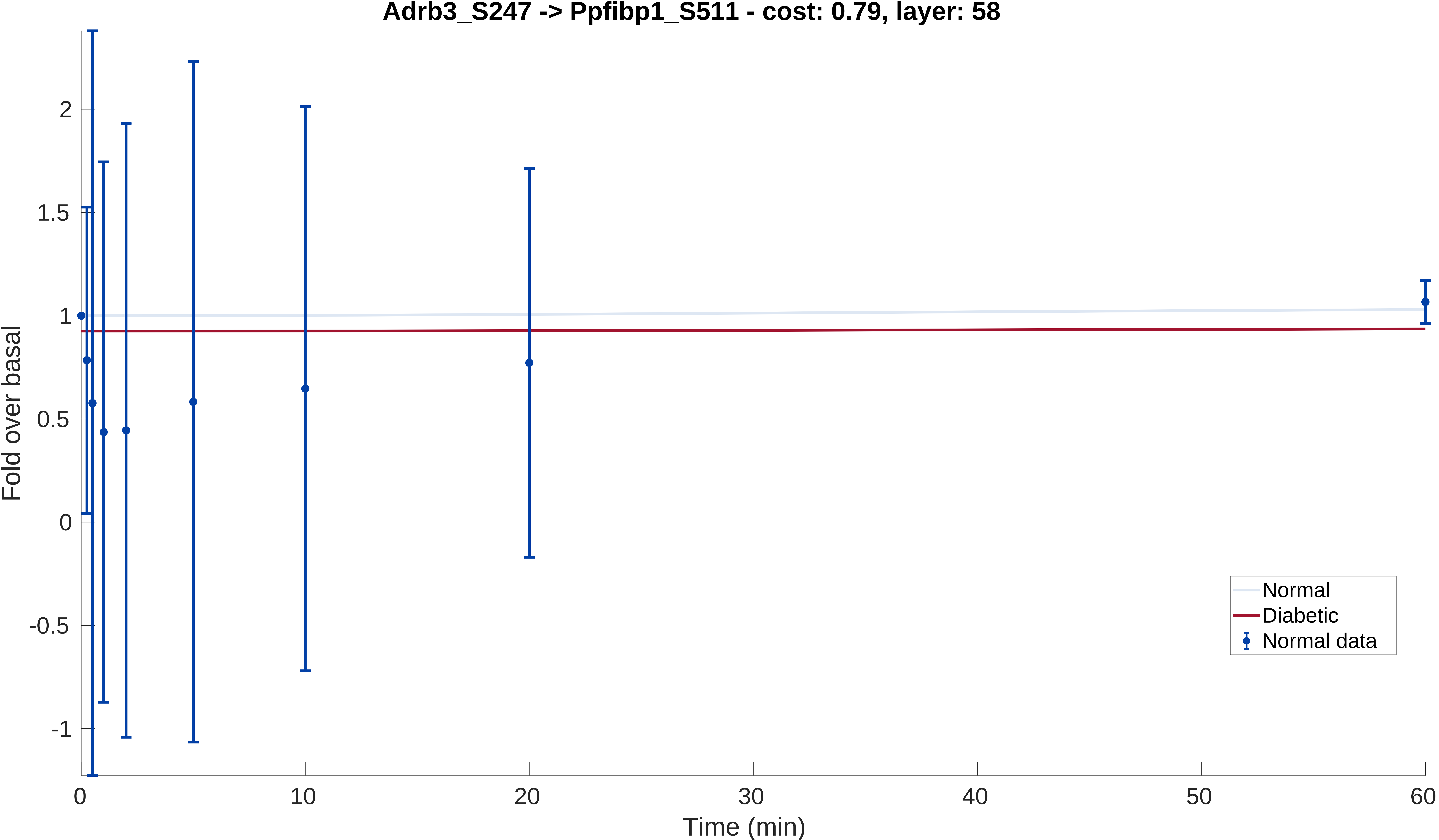
Adrb3_S247 -> Ppfibp1_S511 - cost: 0.79, layer: 58
Fold over basal
Time (min)
2
1.5
1
0.5
0
-0.5
-1
0
10
20
30
40
50
60
Normal
Diabetic
Normal data

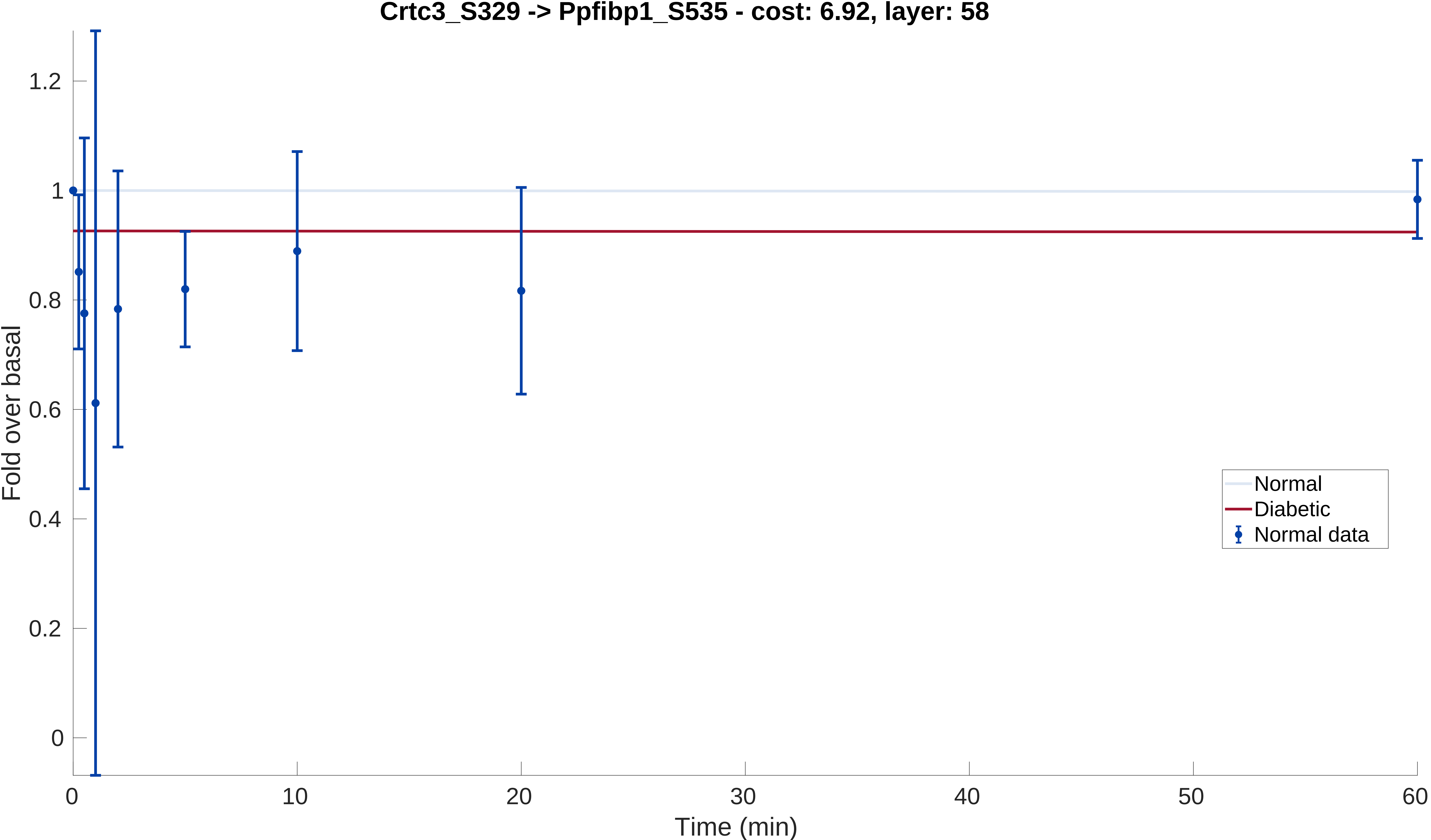

Crtc3_S329 -> Ppfibp1_S535 - cost: 6.92, layer: 58
Fold over basal
1.2
1
0.8
0.6
0.4
0.2
0
0
10
20
30
40
50
60
Time (min)
Normal
Diabetic
Normal data

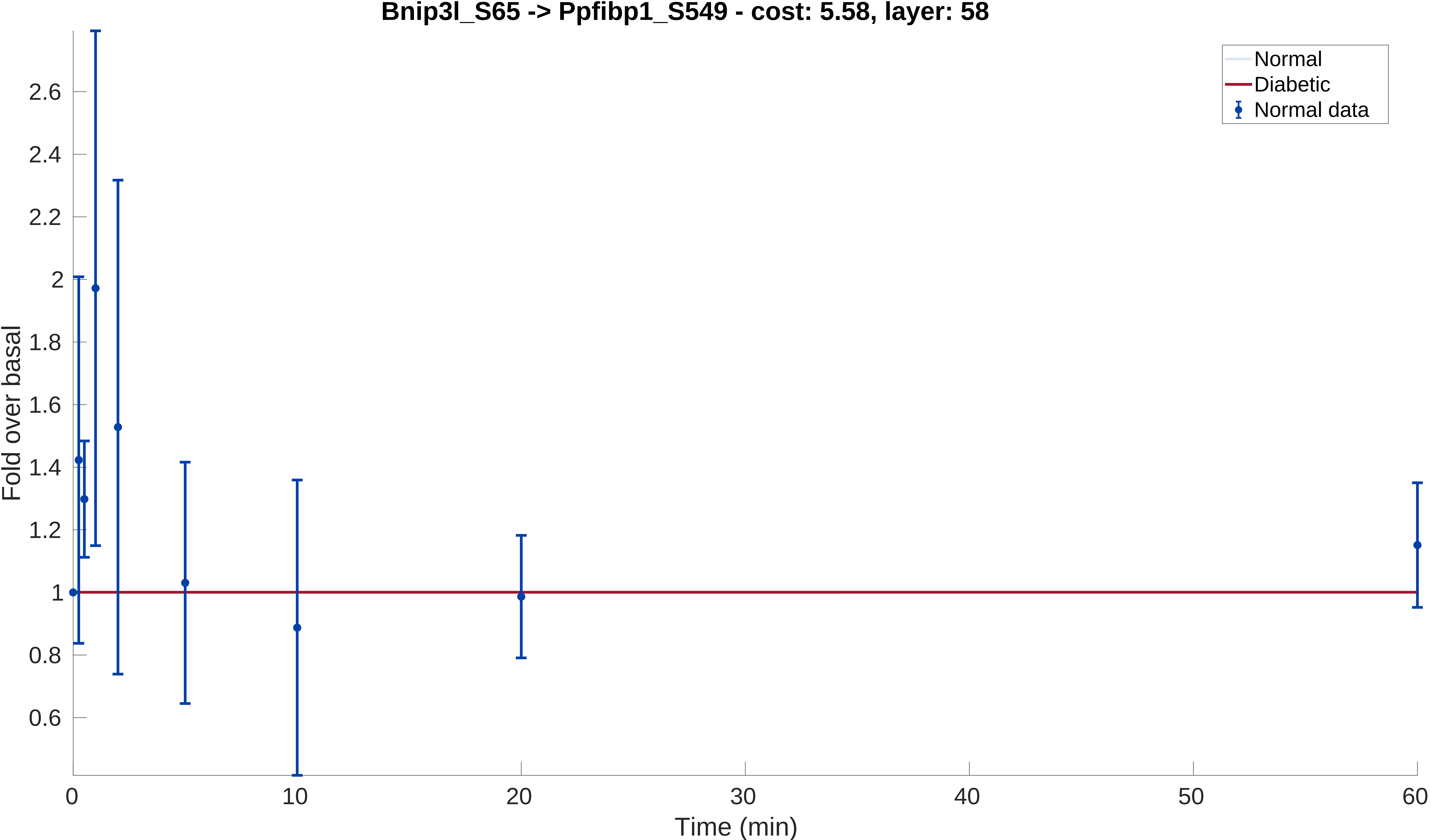
Bnip3l_S65 -> Ppfibp1_S549 - cost: 5.58, layer: 58
Normal
Diabetic
Normal data
Fold over basal
Time (min)
0.6
0.8
1
1.2
1.4
1.6
1.8
2
2.2
2.4
2.6
0
10
20
30
40
50
60

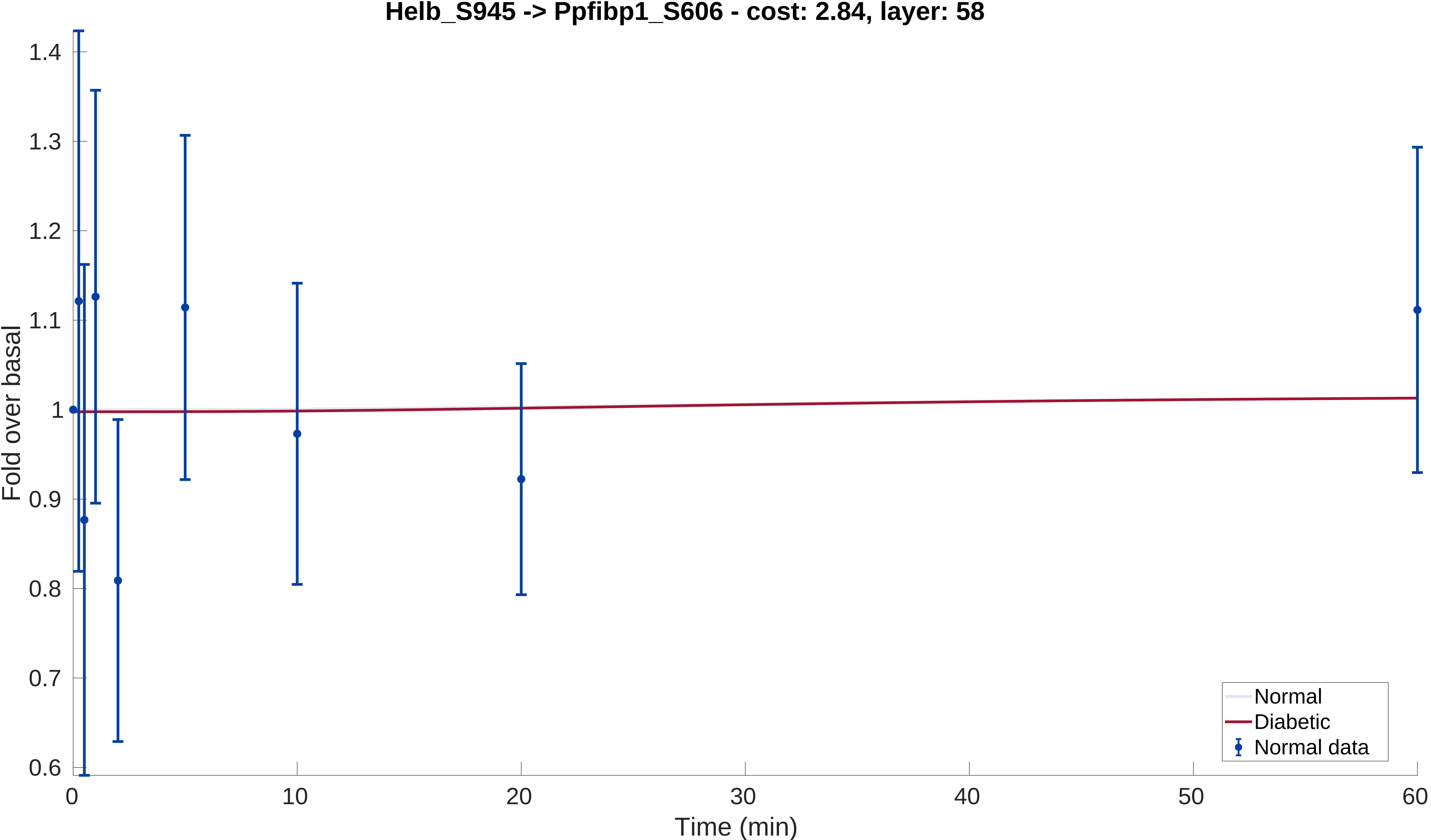

Helb_S945 -> Ppfibp1_S606 - cost: 2.84, layer: 58
Fold over basal
1.4
1.3
1.2
1.1
1
0.9
0.8
0.7
0.6
0
10
20
30
40
50
60
Time (min)
Normal
Diabetic
Normal data

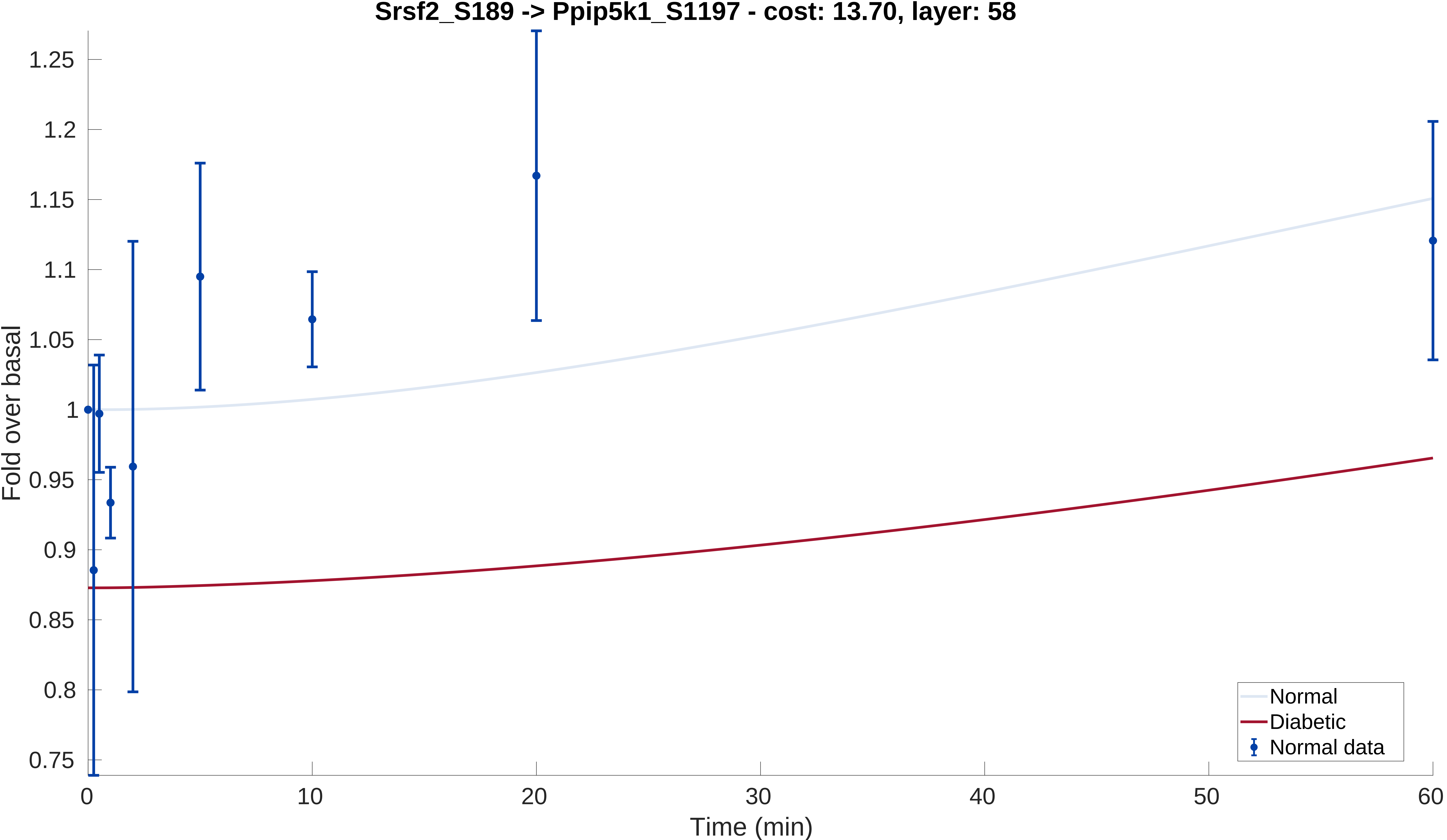
Srsf2_S189 -> Ppip5k1_S1197 - cost: 13.70, layer: 58
Fold over basal
1.25
1.2
1.15
1.1
1.05
1
0.95
0.9
0.85
0.8
0.75
0
10
20
30
40
50
60
Time (min)
Normal
Diabetic
Normal data

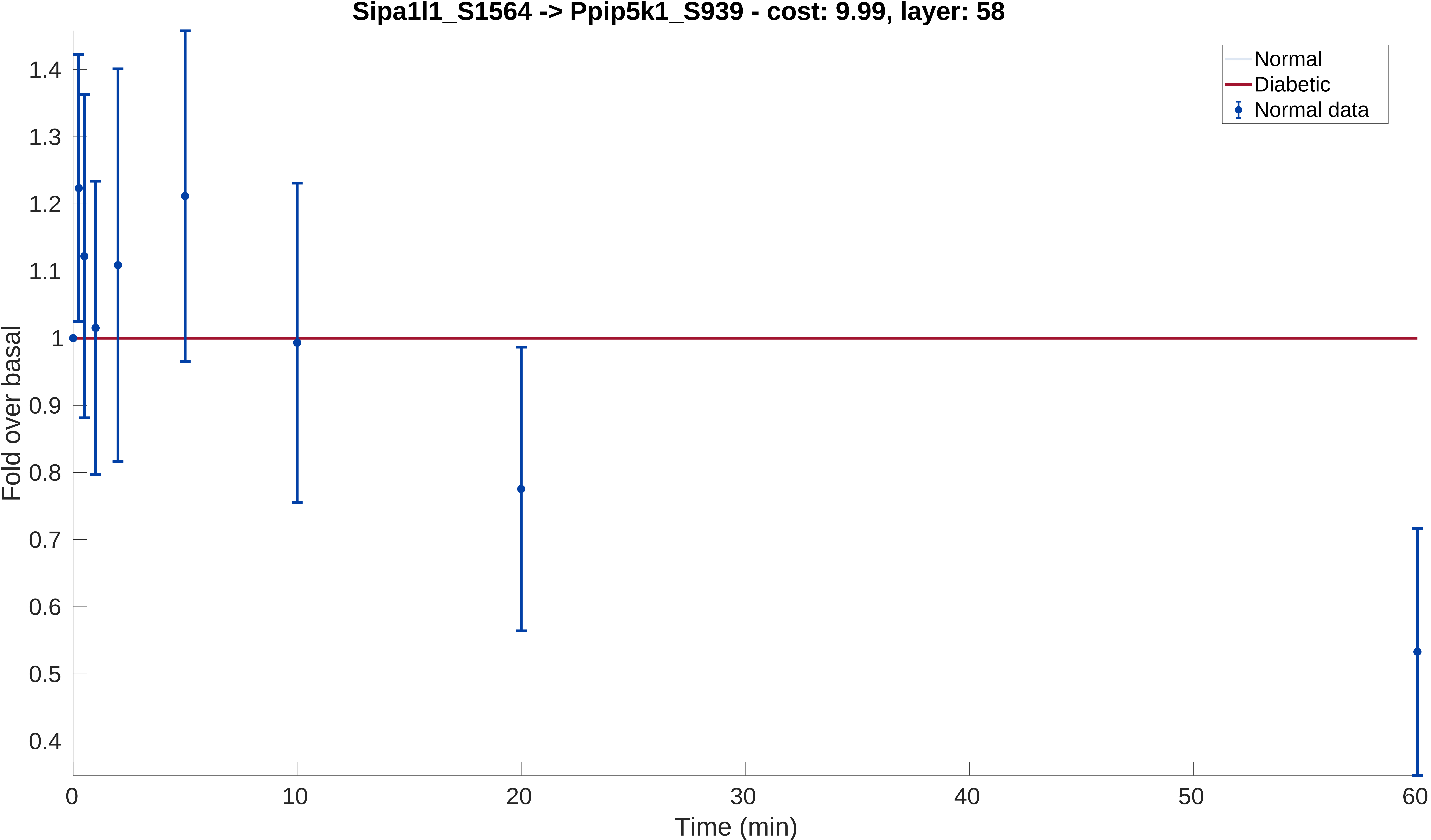

Sipa1l1_S1564 -> Ppip5k1_S939 - cost: 9.99, layer: 58
Normal
Diabetic
Normal data
Fold over basal
1.4
1.3
1.2
1.1
1
0.9
0.8
0.7
0.6
0.5
0.4
0
10
20
30
40
50
60
Time (min)

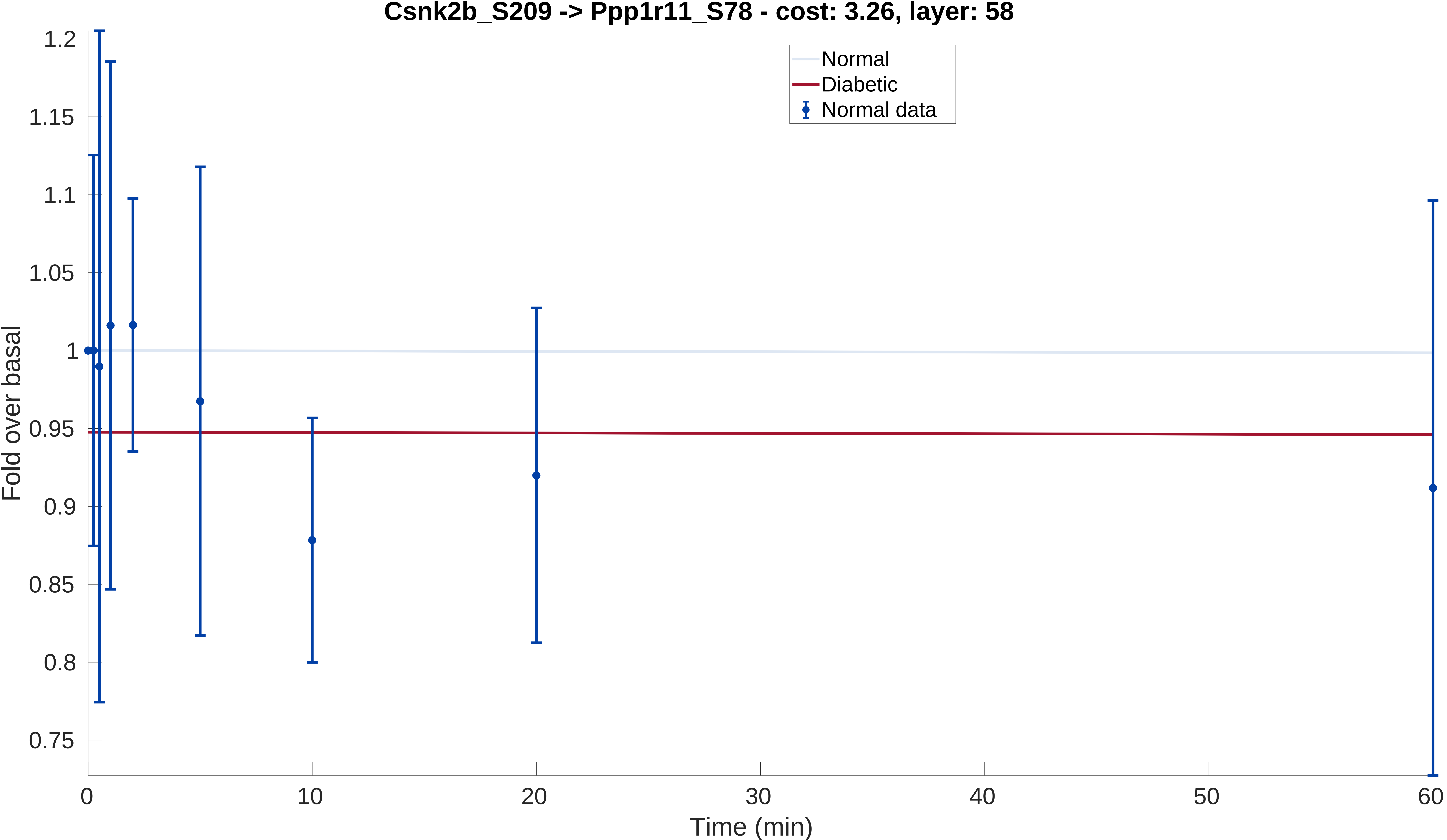

Csnk2b_S209 -> Ppp1r11_S78 - cost: 3.26, layer: 58
Normal
Diabetic
Normal data
Fold over basal
1.2
1.15
1.1
1.05
1
0.95
0.9
0.85
0.8
0.75
0
10
20
30
40
50
60
Time (min)

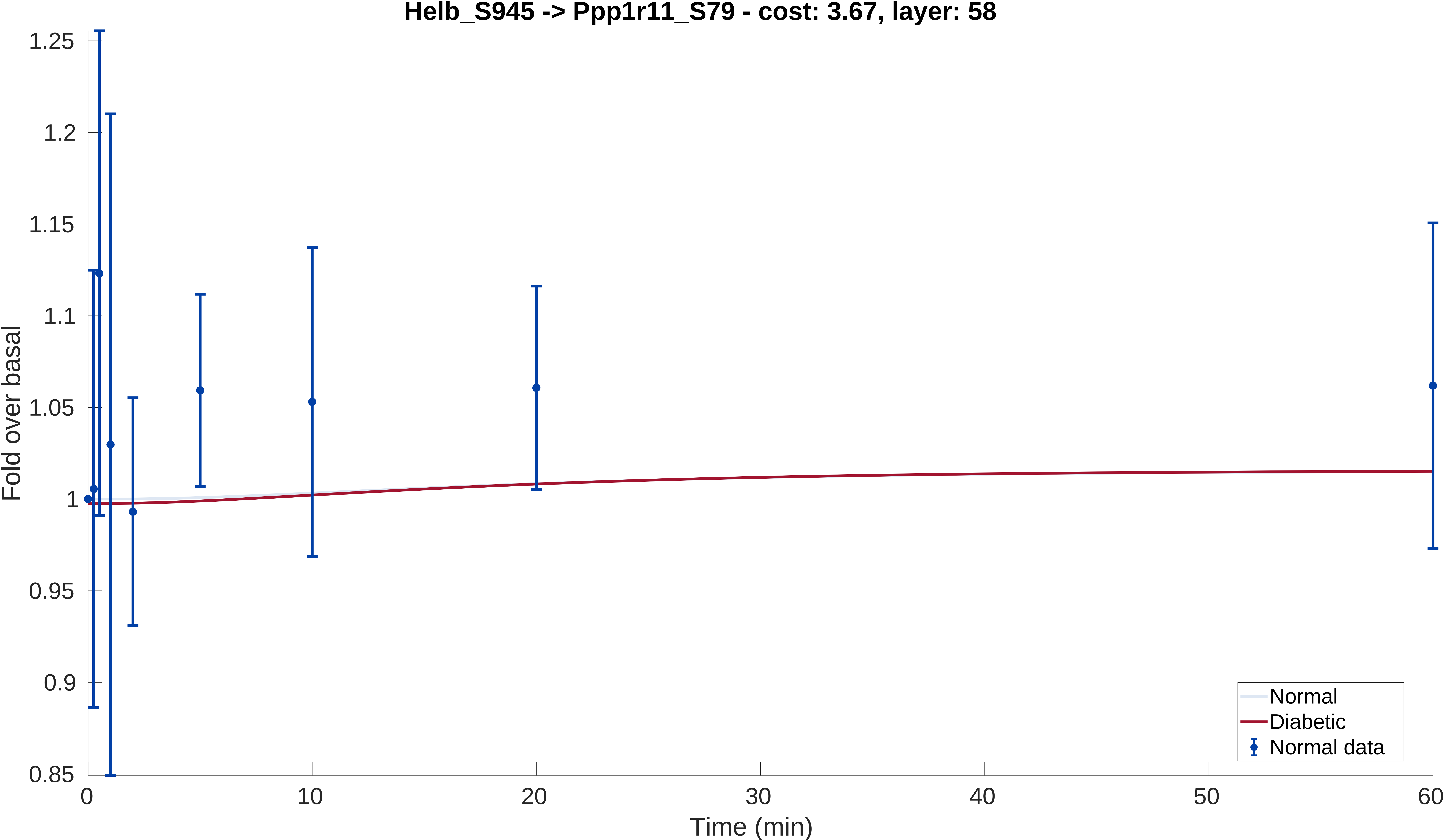

Helb_S945 -> Ppp1r11_S79 - cost: 3.67, layer: 58
Fold over basal
Time (min)
1.25
1.2
1.15
1.1
1.05
1
0.95
0.9
0.85
0
10
20
30
40
50
60
Normal
Diabetic
Normal data

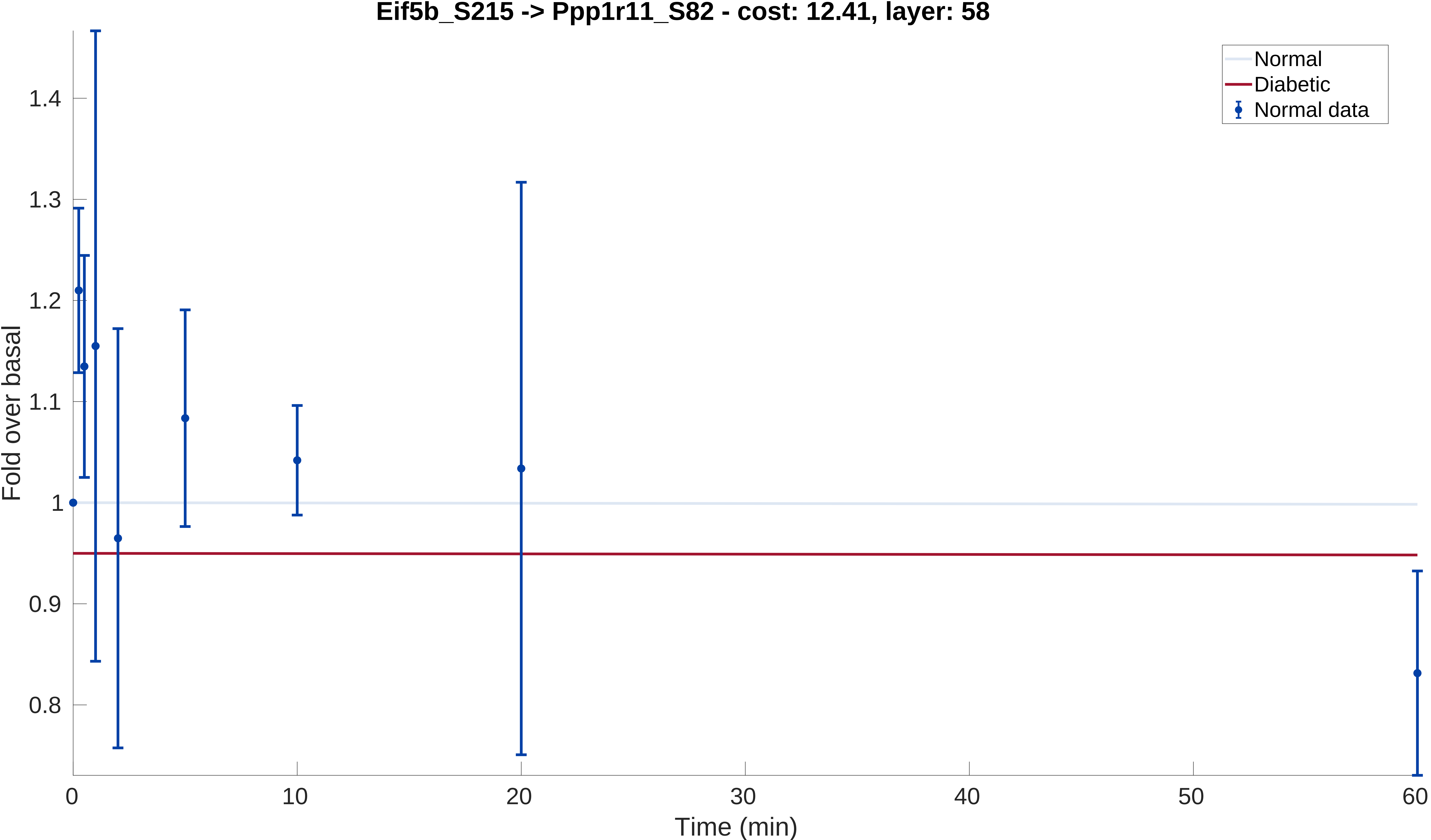

Eif5b_S215 -> Ppp1r11_S82 - cost: 12.41, layer: 58
Normal
Diabetic
Normal data
Fold over basal
1.4
1.3
1.2
1.1
1
0.9
0.8
0
10
20
30
40
50
60
Time (min)

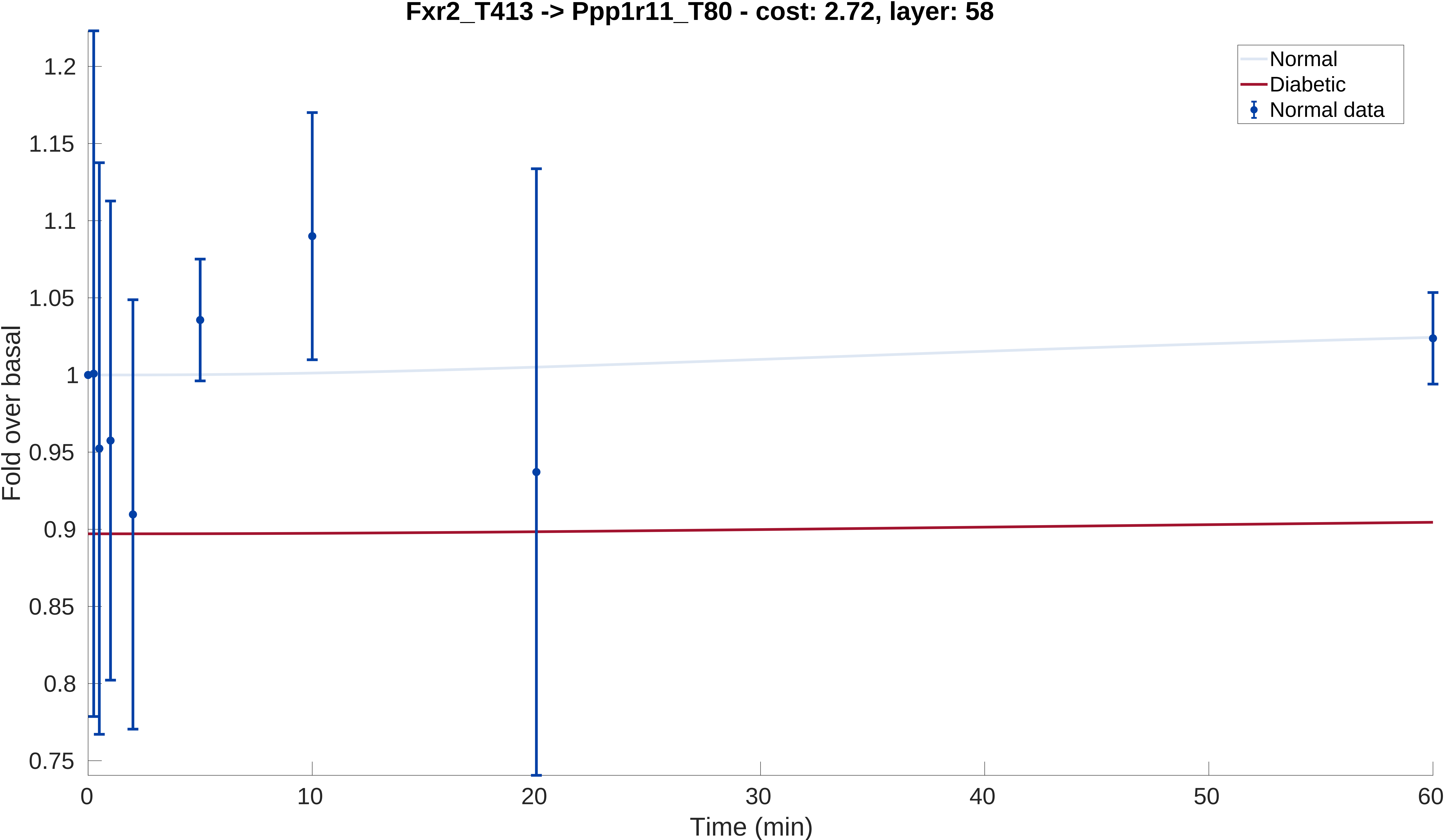
Fxr2_T413 -> Ppp1r11_T80 - cost: 2.72, layer: 58
Normal
Diabetic
Normal data
Fold over basal
1.2
1.15
1.1
1.05
1
0.95
0.9
0.85
0.8
0.75
0
10
20
30
40
50
60
Time (min)

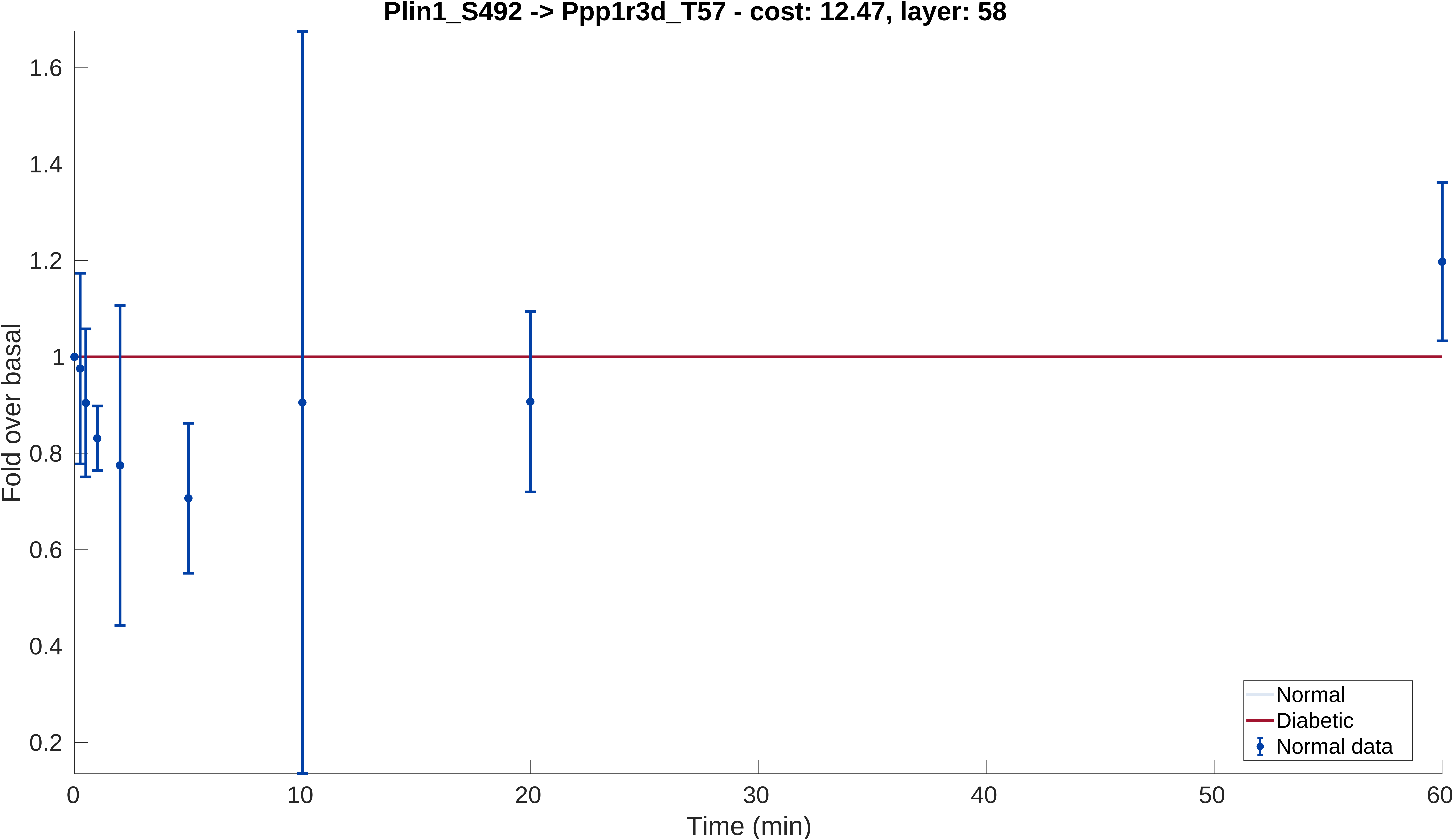
Plin1_S492 -> Ppp1r3d_T57 - cost: 12.47, layer: 58
Fold over basal
Time (min)
0
10
20
30
40
50
60
0.2
0.4
0.6
0.8
1
1.2
1.4
1.6
Normal
Diabetic
Normal data

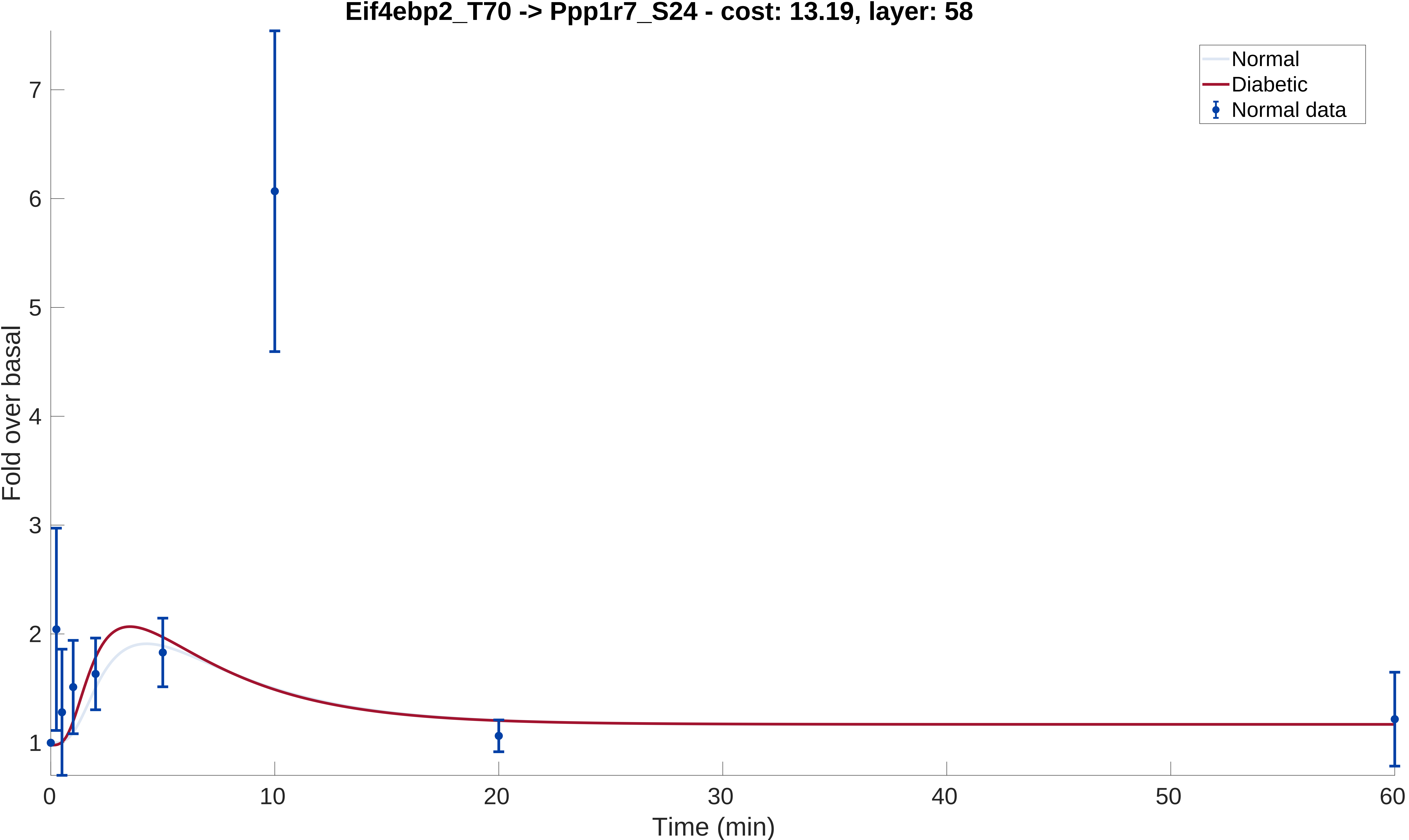
Eif4ebp2_T70 -> Ppp1r7_S24 - cost: 13.19, layer: 58
Normal
Diabetic
Normal data
Fold over basal
Time (min)
1
2
3
4
5
6
7
0
10
20
30
40
50
60

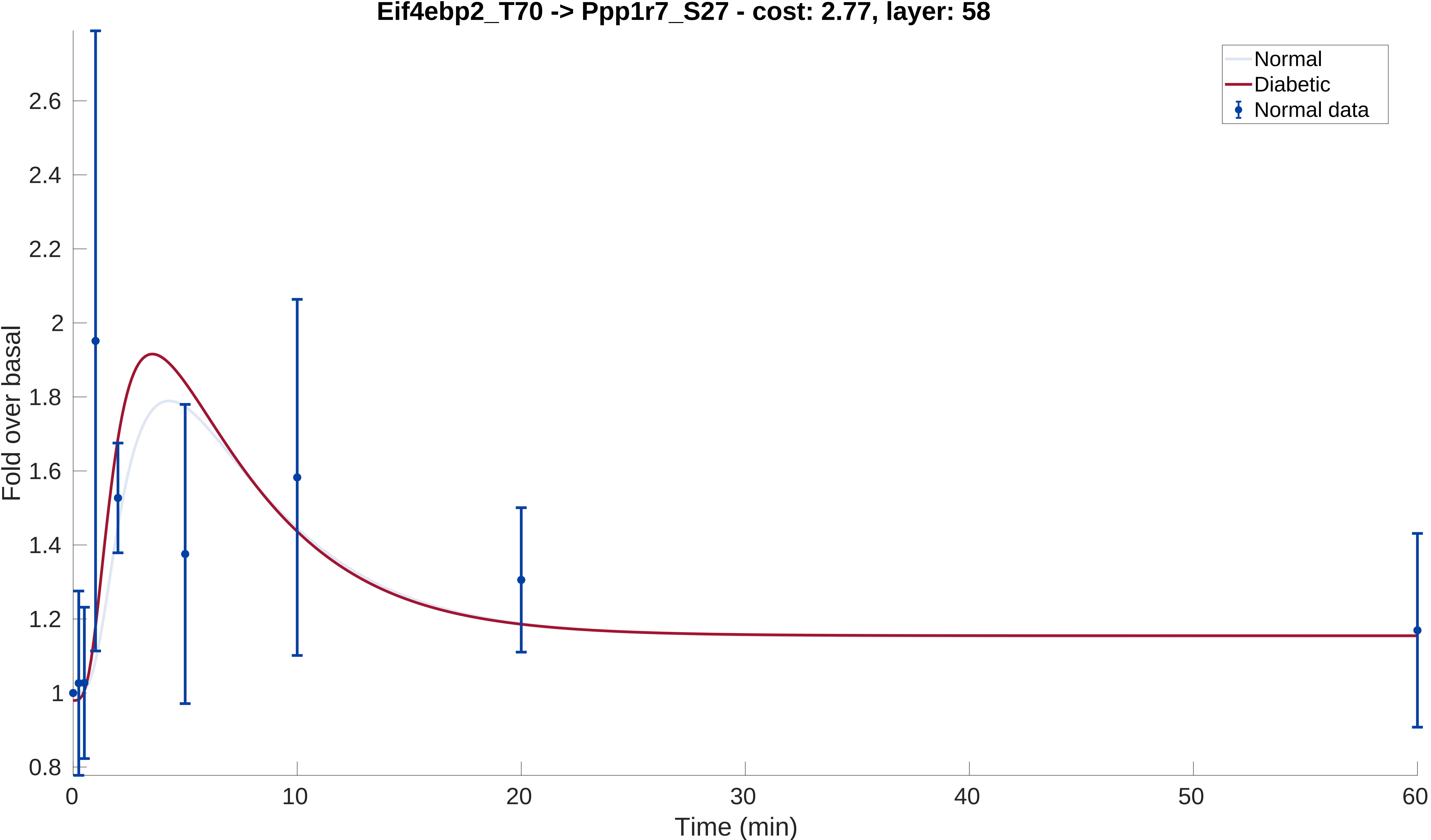

Eif4ebp2_T70 -> Ppp1r7_S27 - cost: 2.77, layer: 58
Normal
Diabetic
Normal data
Fold over basal
2.6
2.4
2.2
2
1.8
1.6
1.4
1.2
1
0.8
0
10
20
30
40
50
60
Time (min)

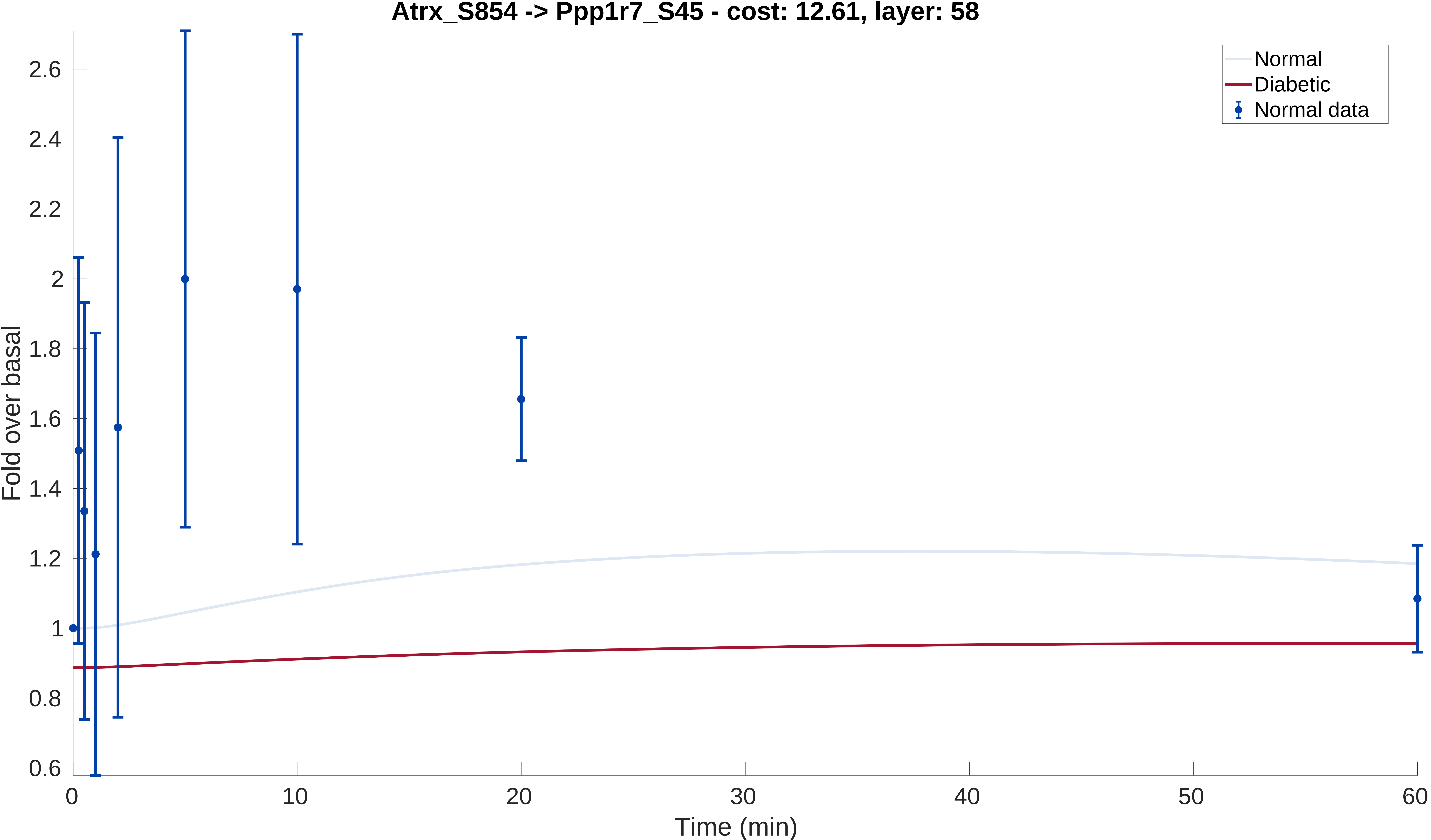
Atrx_S854 -> Ppp1r7_S45 - cost: 12.61, layer: 58
Normal
Diabetic
Normal data
Fold over basal
Time (min)
0.6
0.8
1
1.2
1.4
1.6
1.8
2
2.2
2.4
2.6
0
10
20
30
40
50
60

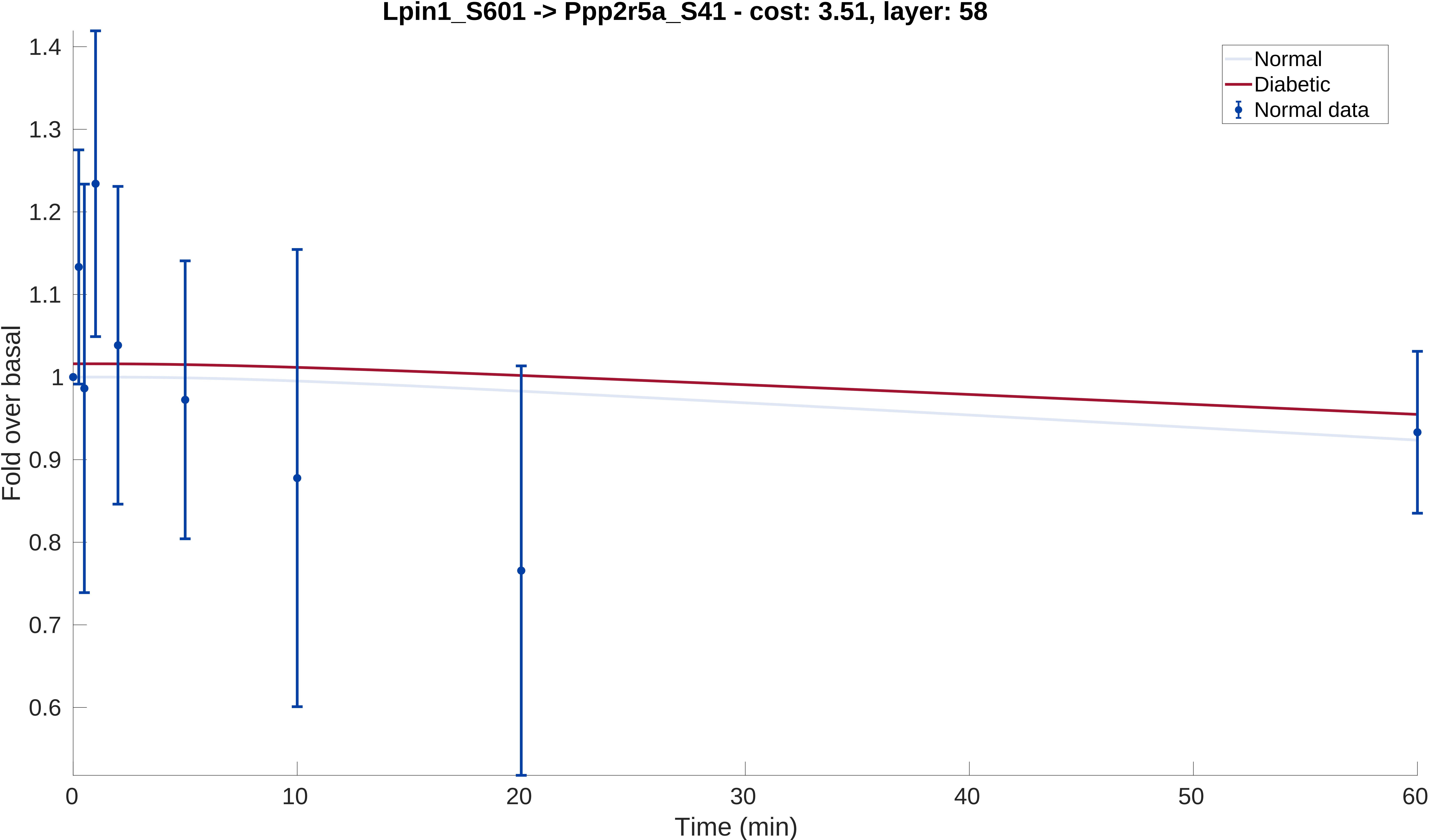
Lpin1_S601 -> Ppp2r5a_S41 - cost: 3.51, layer: 58
Normal
Diabetic
Normal data
Fold over basal
1.4
1.3
1.2
1.1
1
0.9
0.8
0.7
0.6
0
10
20
30
40
50
60
Time (min)

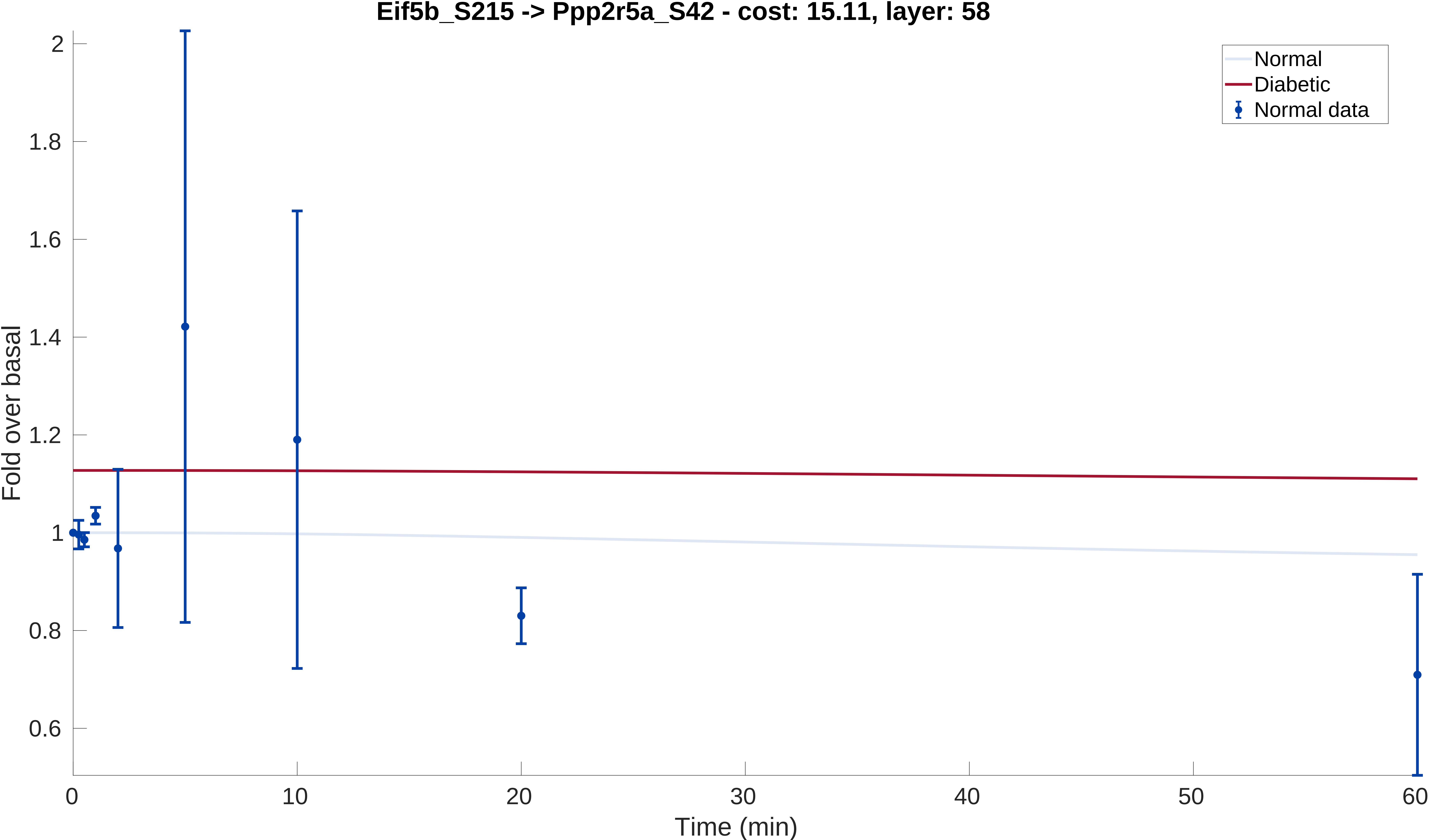

Eif5b_S215 -> Ppp2r5a_S42 - cost: 15.11, layer: 58
Normal
Diabetic
Normal data
Fold over basal
2
1.8
1.6
1.4
1.2
1
0.8
0.6
0
10
20
30
40
50
60
Time (min)

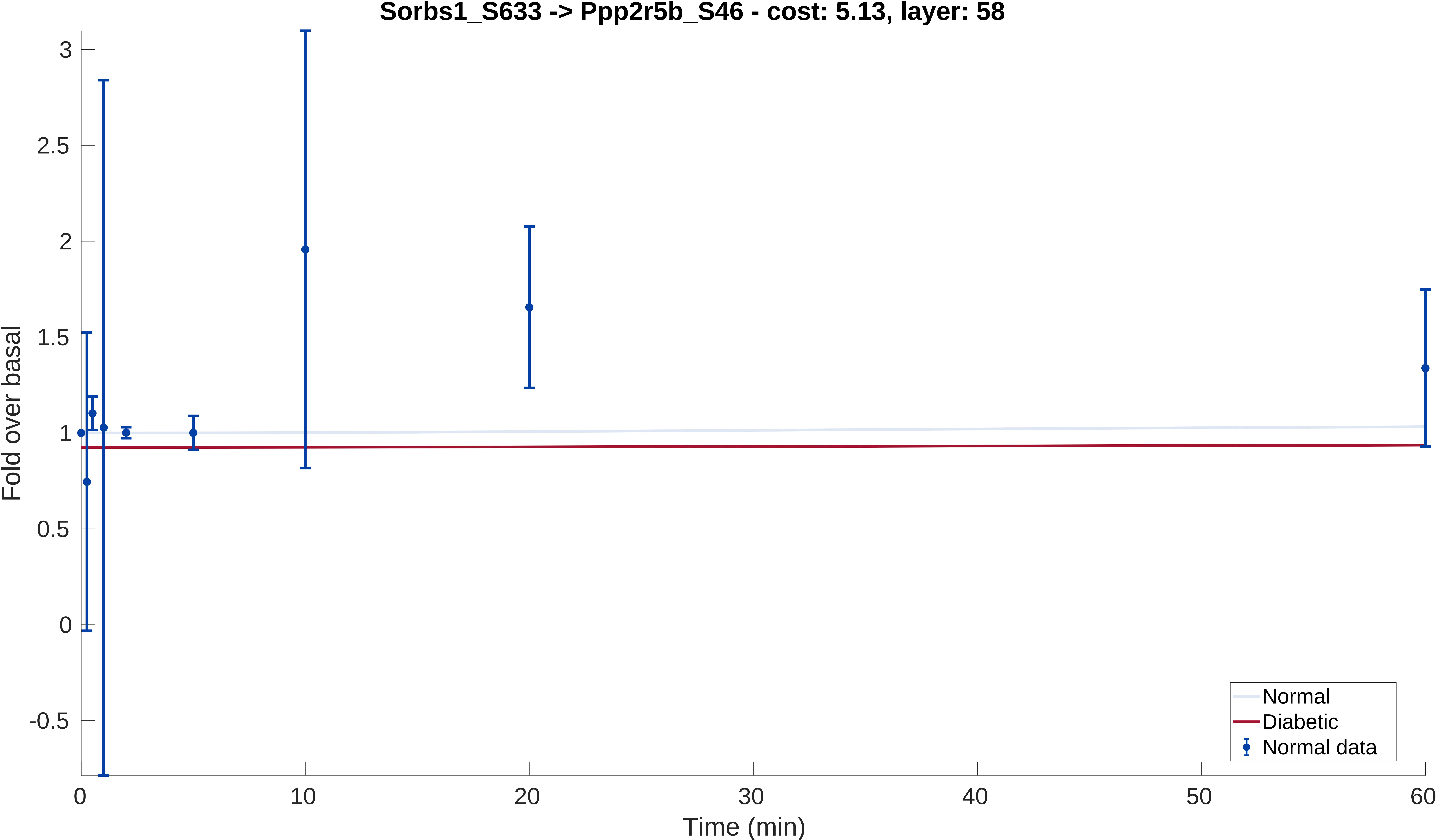
Sorbs1_S633 -> Ppp2r5b_S46 - cost: 5.13, layer: 58
Fold over basal
3
2.5
2
1.5
1
0.5
0
-0.5
0
10
20
30
40
50
60
Time (min)
Normal
Diabetic
Normal data

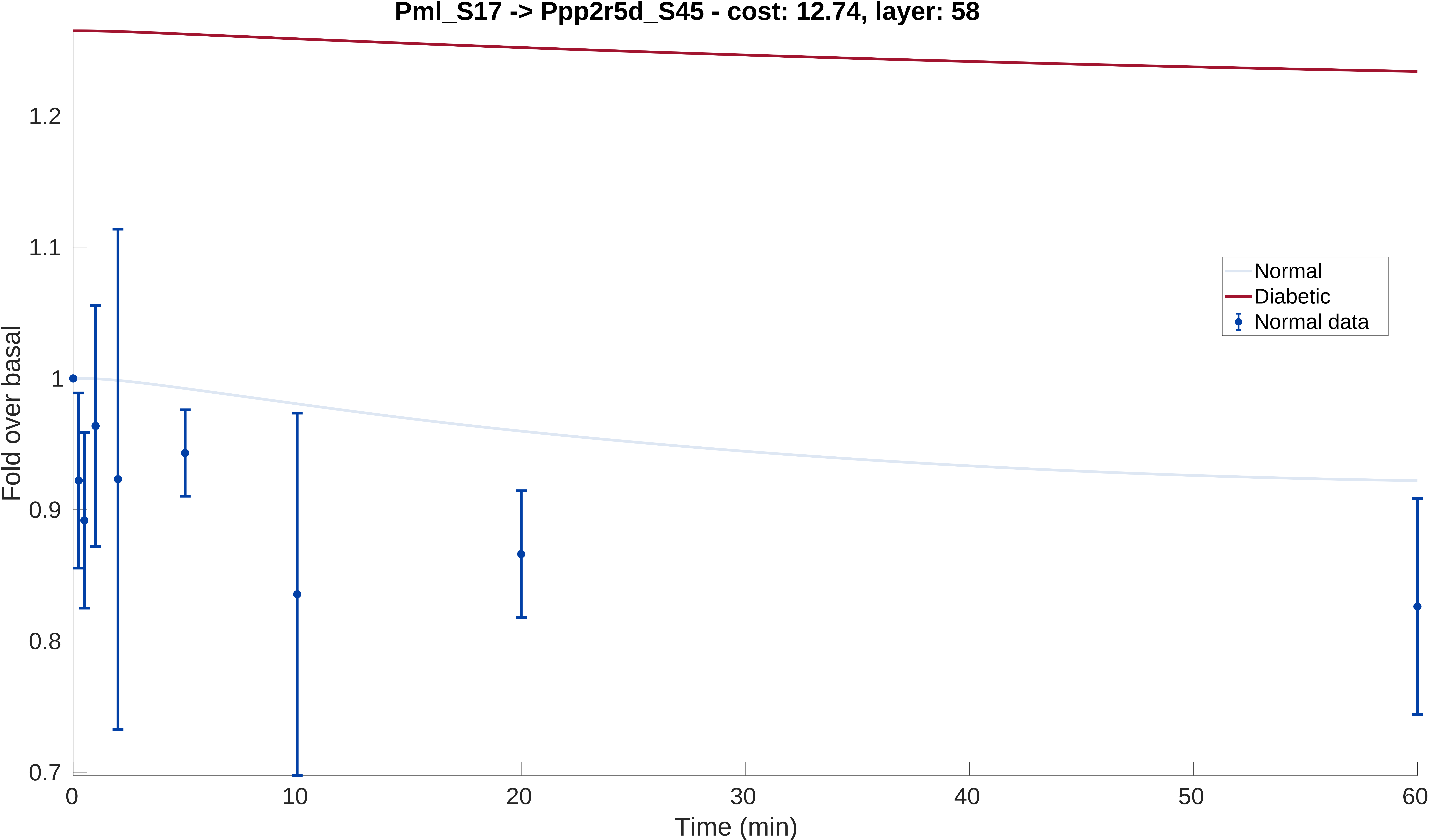

Pml_S17 -> Ppp2r5d_S45 - cost: 12.74, layer: 58
Fold over basal
Time (min)
1.2
1.1
1
0.9
0.8
0.7
0
10
20
30
40
50
60
Normal
Diabetic
Normal data

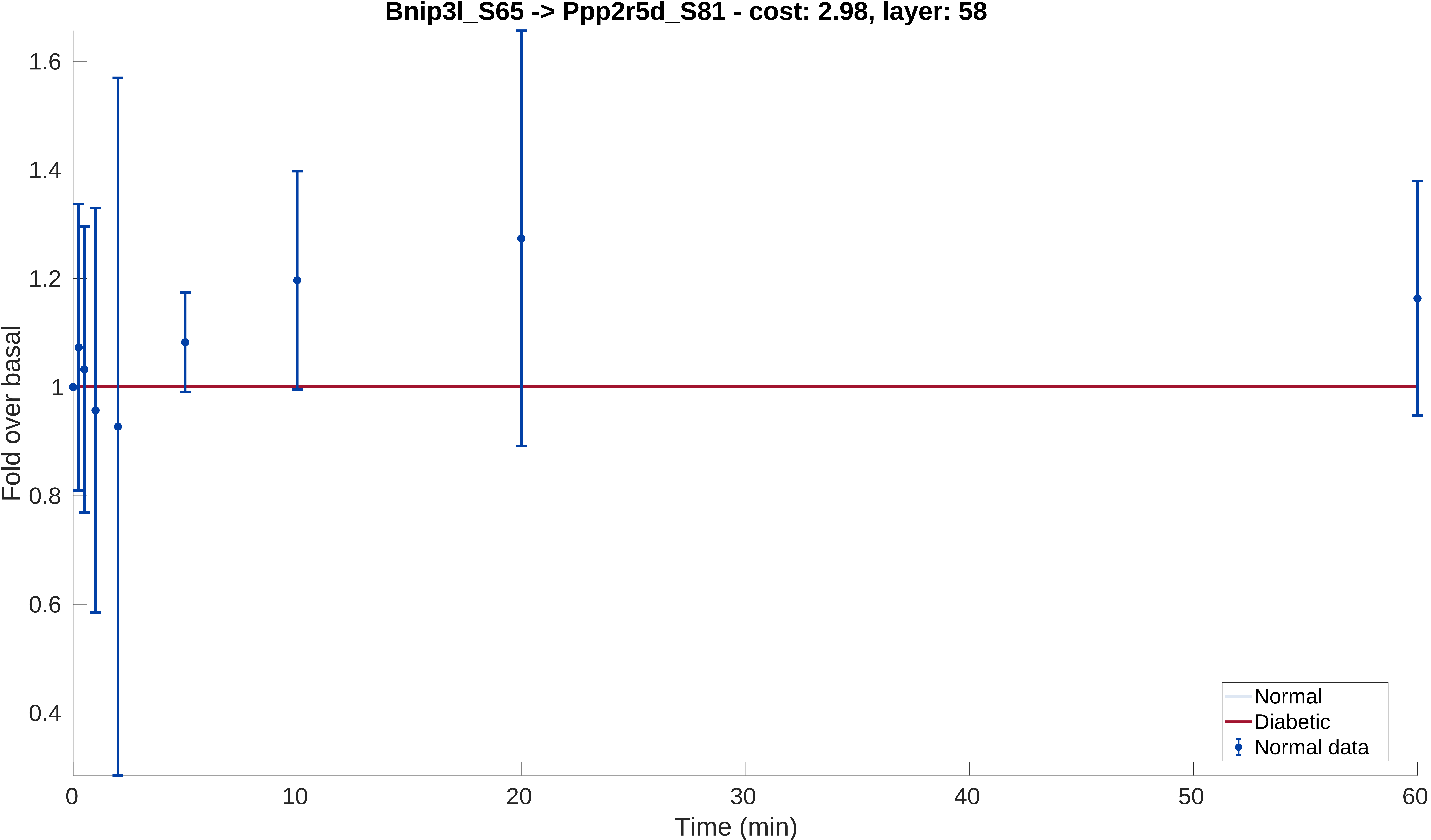

Bnip3l_S65 -> Ppp2r5d_S81 - cost: 2.98, layer: 58
Fold over basal
1.6
1.4
1.2
1
0.8
0.6
0.4
0
10
20
30
40
50
60
Time (min)
Normal
Diabetic
Normal data

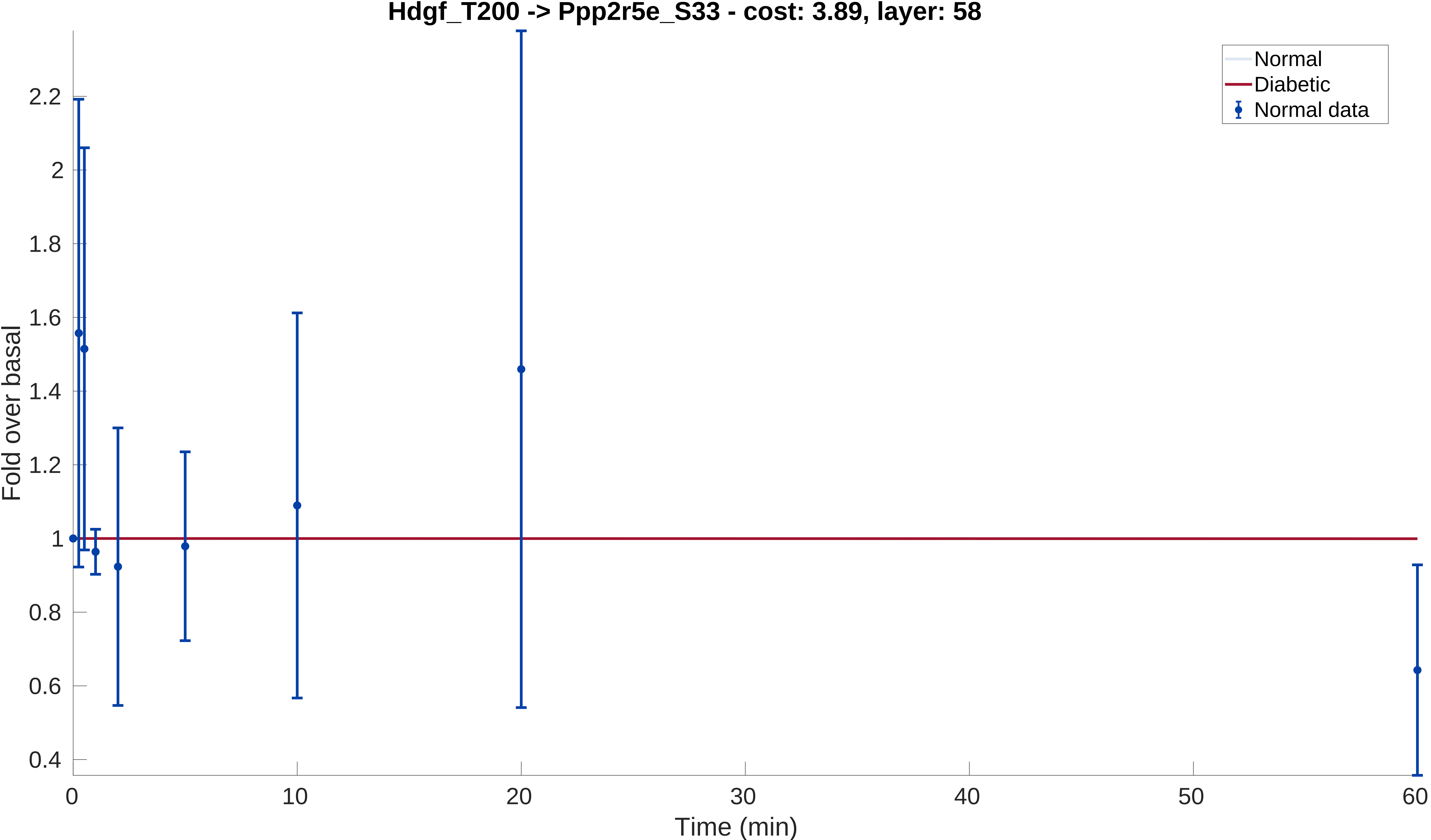
Hdgf_T200 -> Ppp2r5e_S33 - cost: 3.89, layer: 58
Normal
Diabetic
Normal data
Fold over basal
2.2
2
1.8
1.6
1.4
1.2
1
0.8
0.6
0.4
0
10
20
30
40
50
60
Time (min)

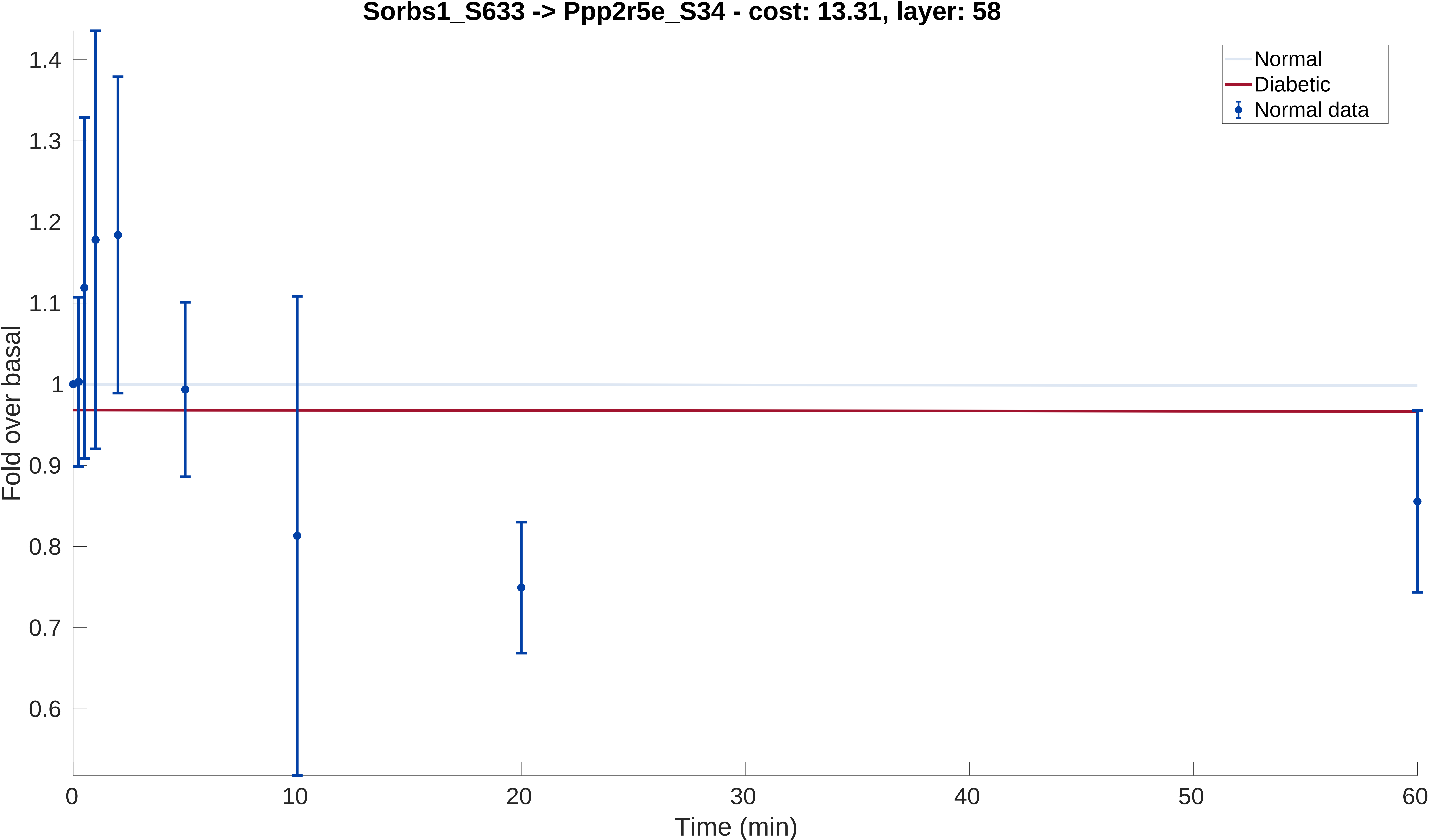
Sorbs1_S633 -> Ppp2r5e_S34 - cost: 13.31, layer: 58
Fold over basal
Time (min)
1.4
1.3
1.2
1.1
1
0.9
0.8
0.7
0.6
0
10
20
30
40
50
60
Normal
Diabetic
Normal data

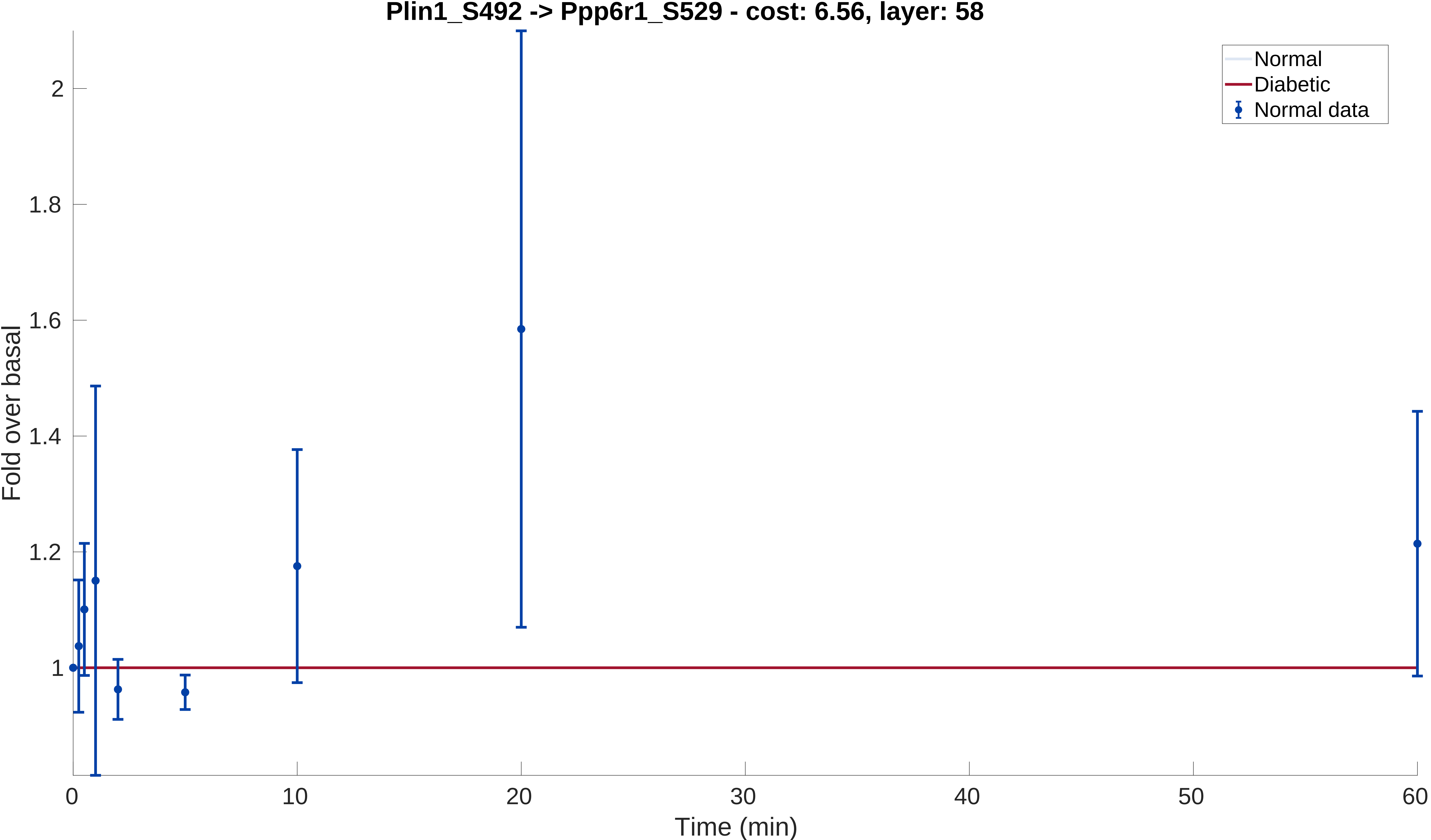

Plin1_S492 -> Ppp6r1_S529 - cost: 6.56, layer: 58
Normal
Diabetic
Normal data
Fold over basal
2
1.8
1.6
1.4
1.2
1
0
10
20
30
40
50
60
Time (min)

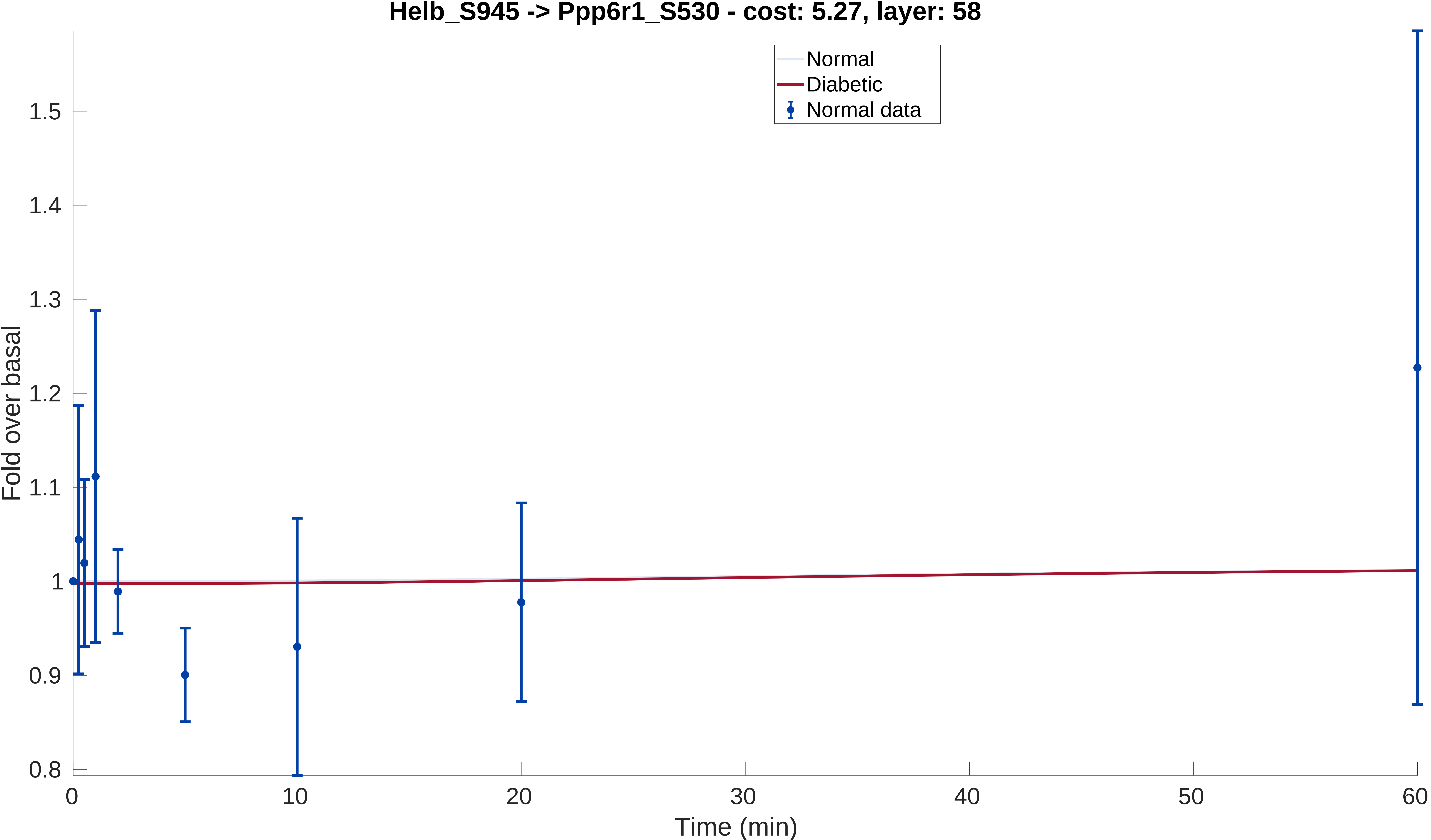

Helb_S945 -> Ppp6r1_S530 - cost: 5.27, layer: 58
Normal
Diabetic
Normal data
Fold over basal
Time (min)
0.8
0.9
1
1.1
1.2
1.3
1.4
1.5
0
10
20
30
40
50
60

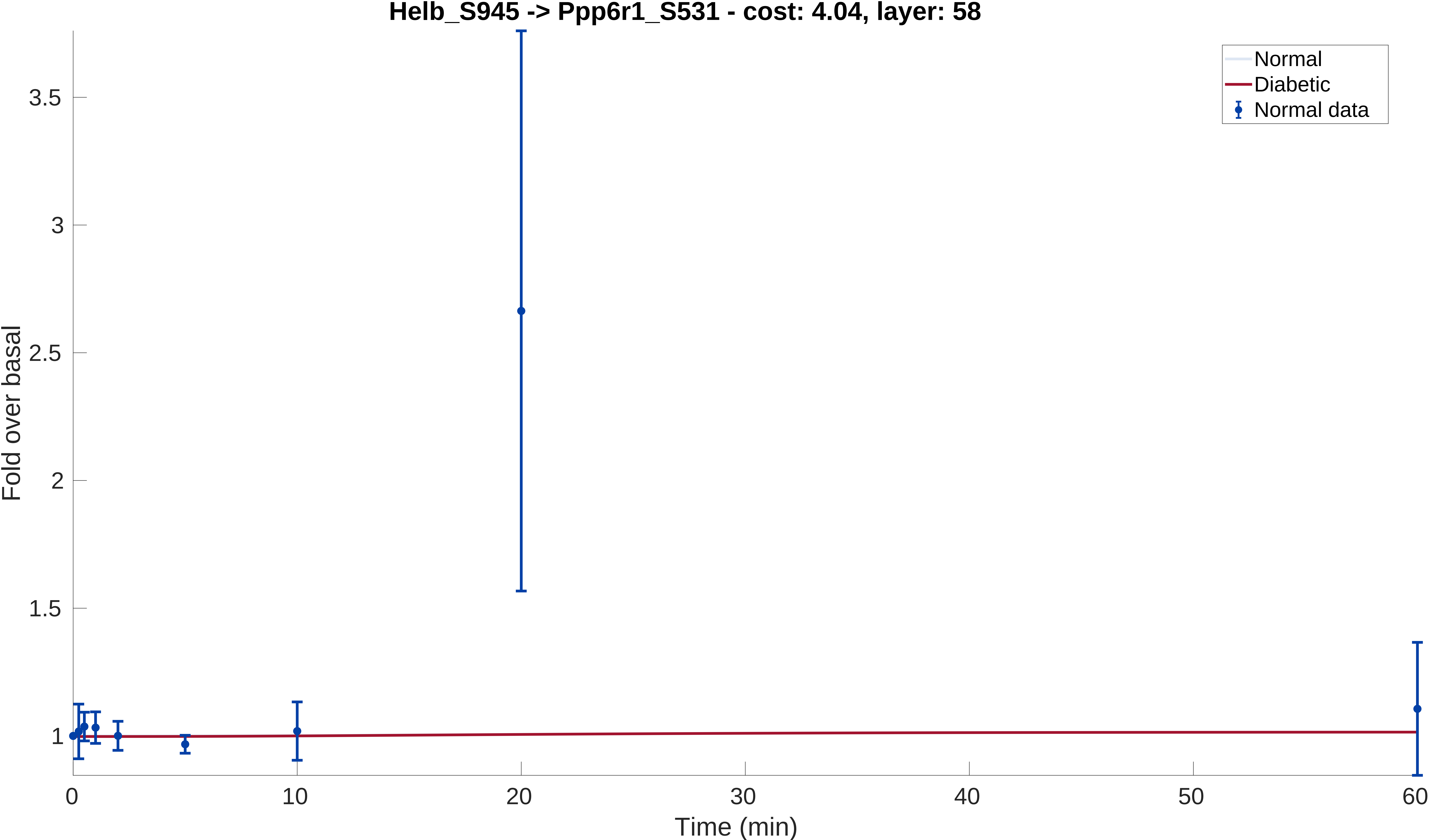
Helb_S945 -> Ppp6r1_S531 - cost: 4.04, layer: 58
Fold over basal
3.5
3
2.5
2
1.5
1
0
10
20
30
40
50
60
Time (min)
Normal
Diabetic
Normal data

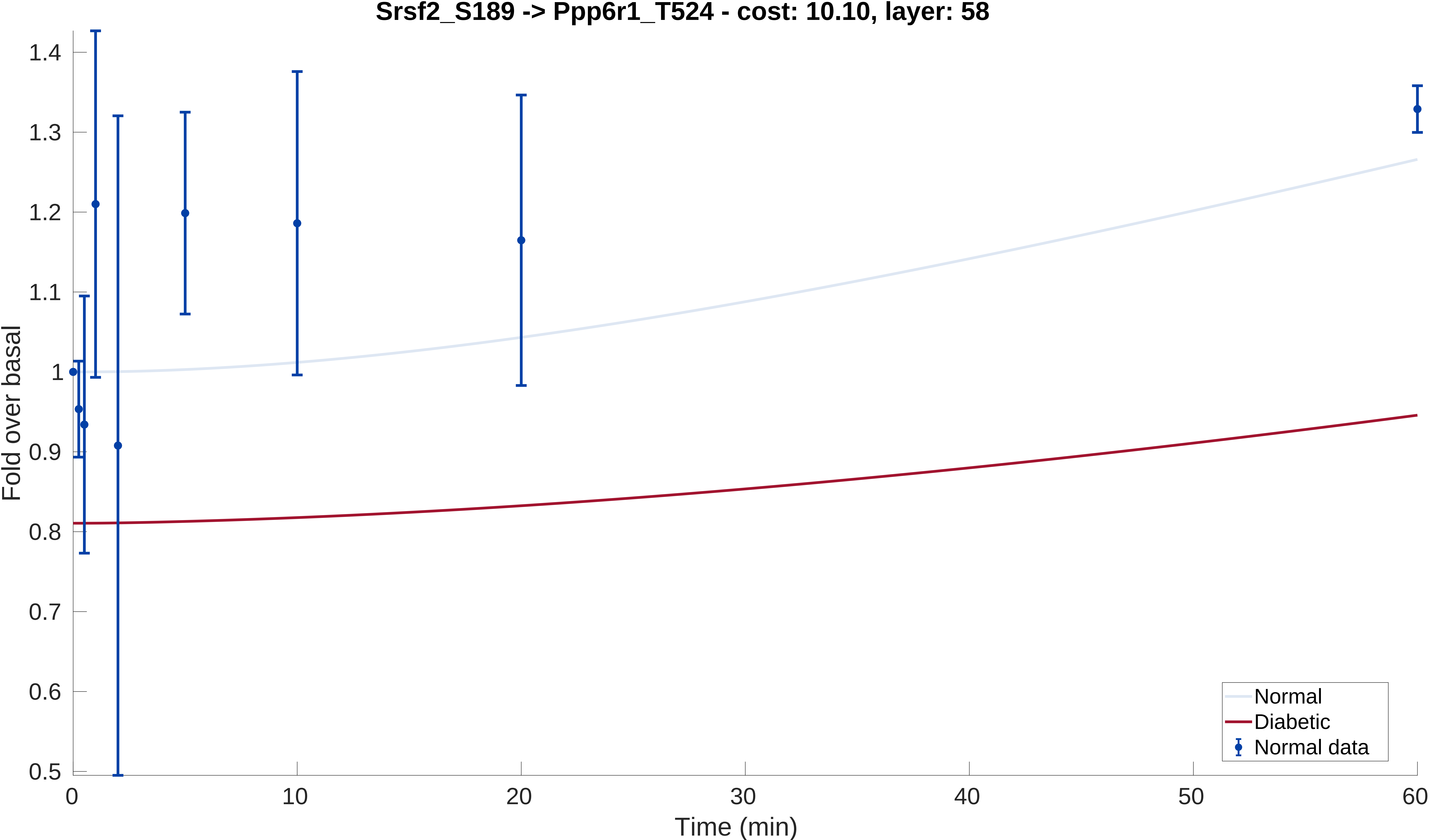

Srsf2_S189 -> Ppp6r1_T524 - cost: 10.10, layer: 58
Fold over basal
1.4
1.3
1.2
1.1
1
0.9
0.8
0.7
0.6
0.5
0
10
20
30
40
50
60
Time (min)
Normal
Diabetic
Normal data

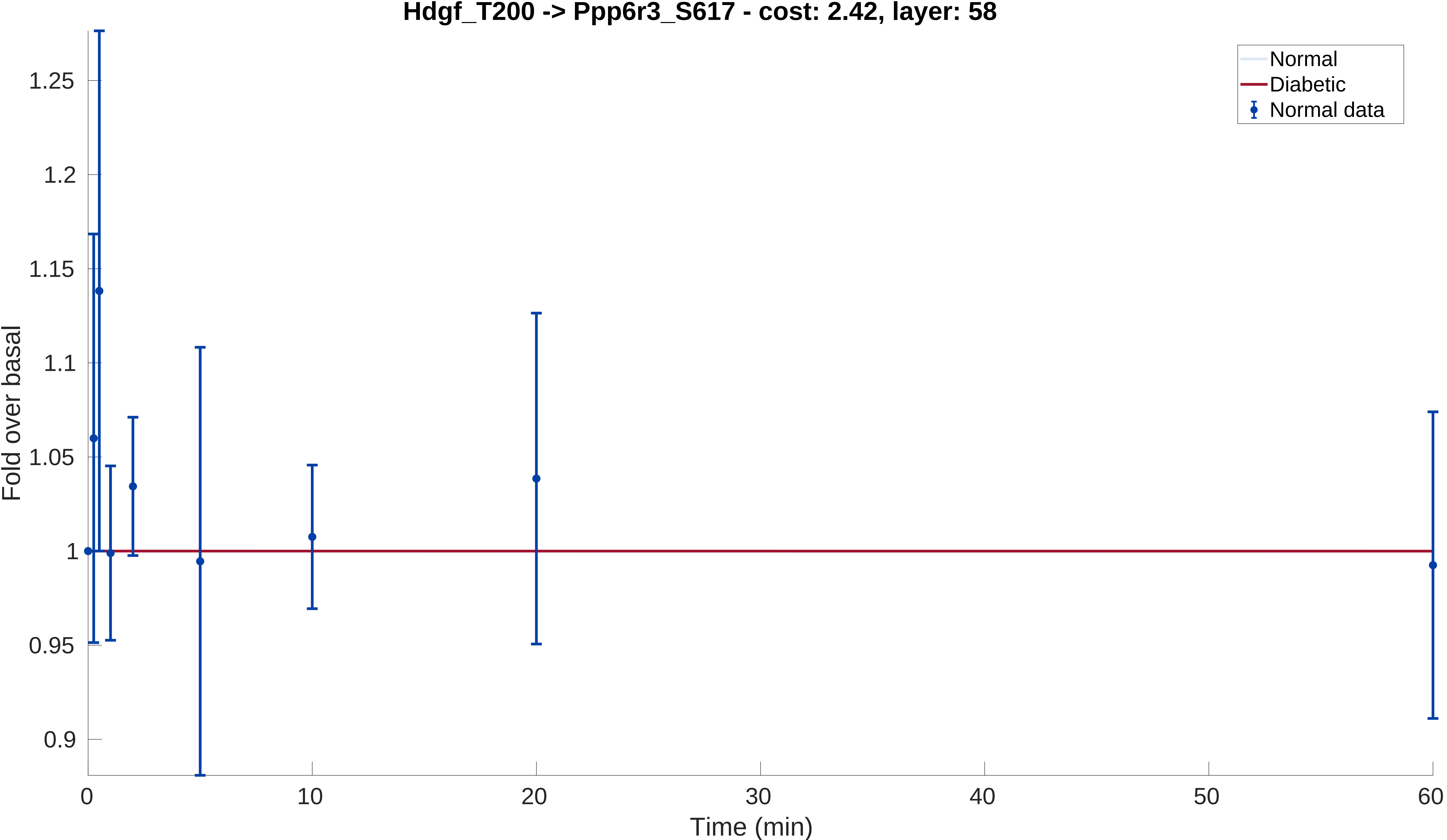

Hdgf_T200 -> Ppp6r3_S617 - cost: 2.42, layer: 58
Normal
Diabetic
Normal data
Fold over basal
Time (min)
1.25
1.2
1.15
1.1
1.05
1
0.95
0.9
0
10
20
30
40
50
60

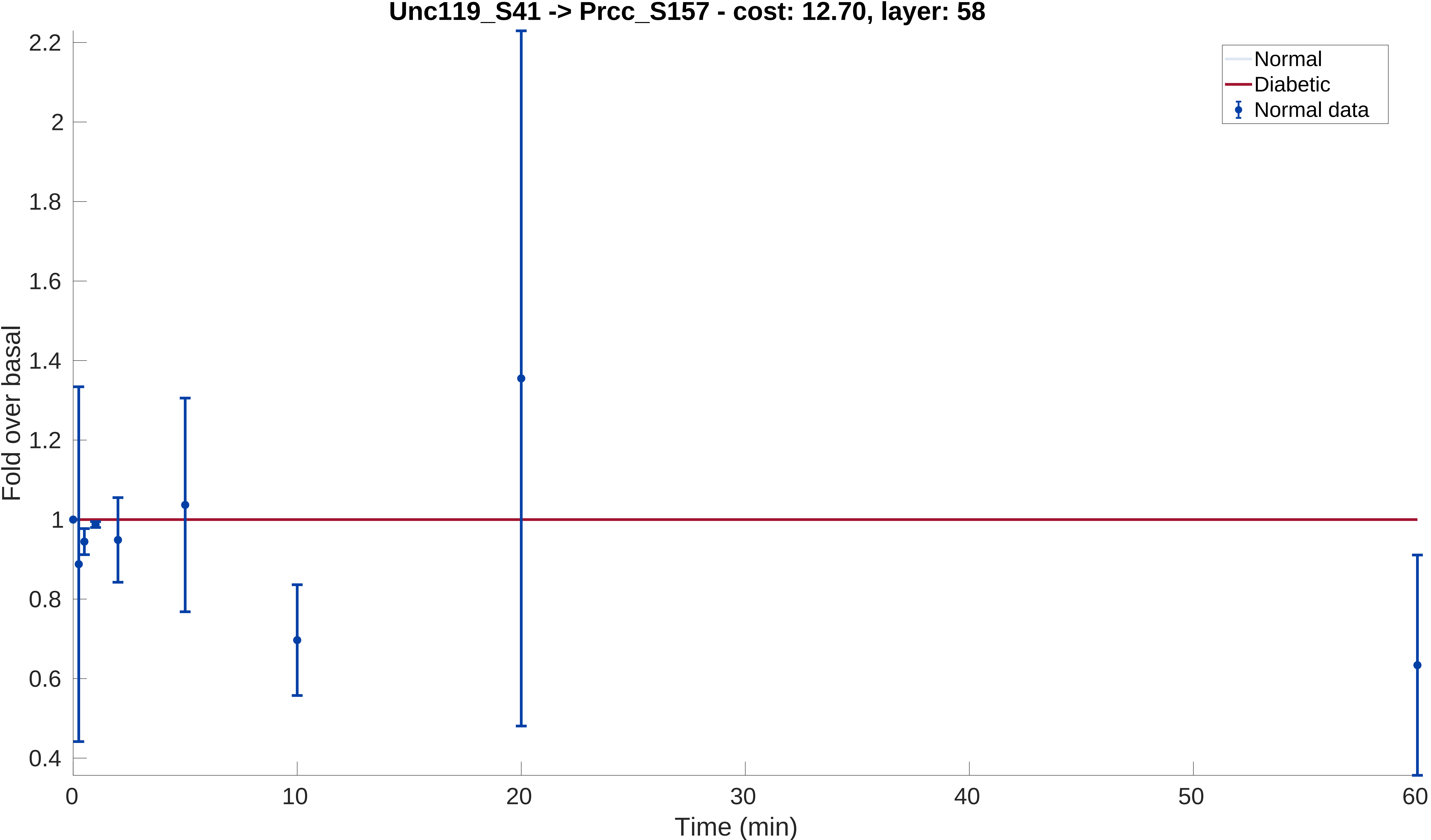

Unc119_S41 -> Prcc_S157 - cost: 12.70, layer: 58
Normal
Diabetic
Normal data
Fold over basal
Time (min)
0.4
0.6
0.8
1
1.2
1.4
1.6
1.8
2
2.2
0
10
20
30
40
50
60

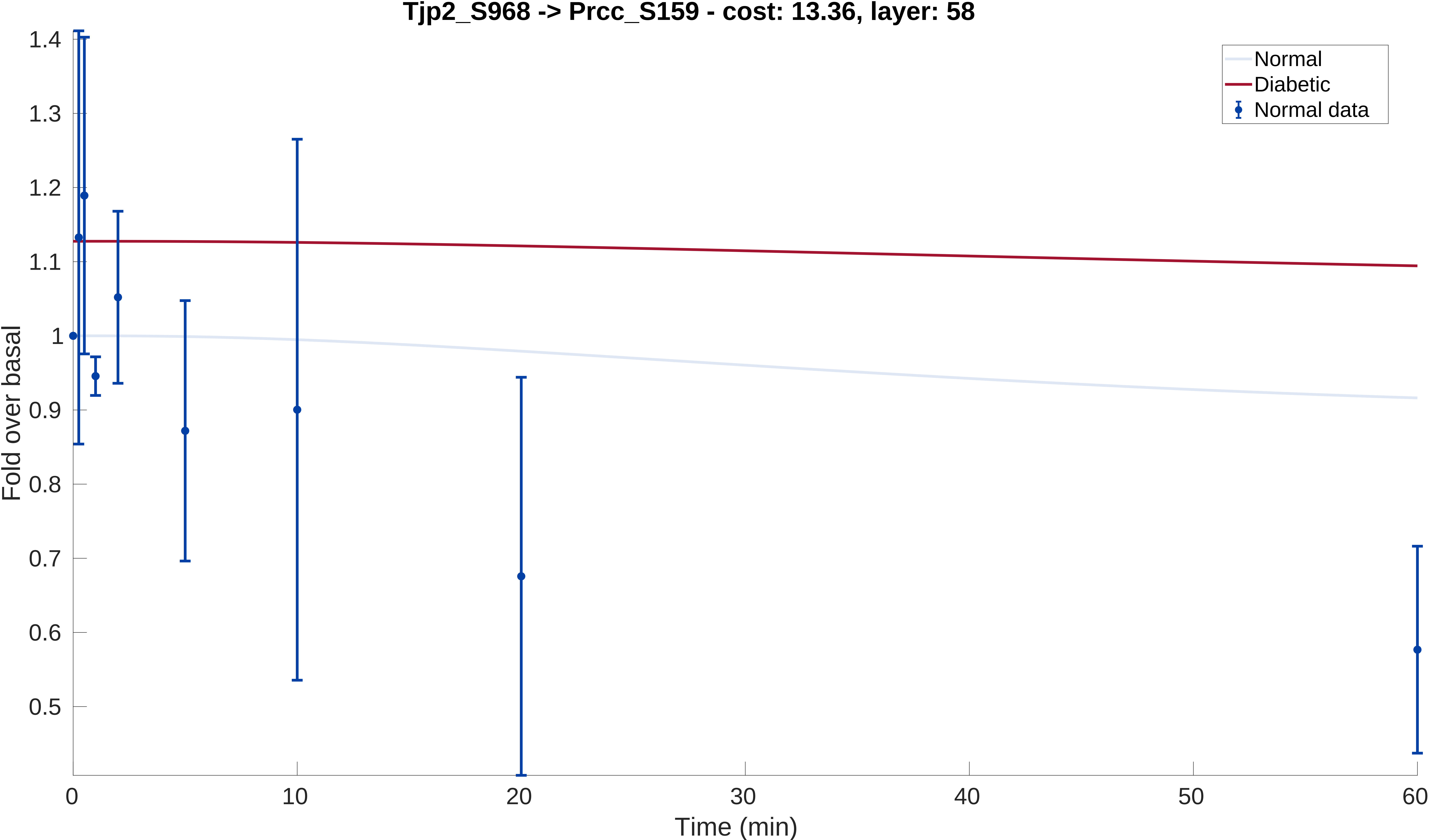

Tjp2_S968 -> Prcc_S159 - cost: 13.36, layer: 58
Normal
Diabetic
Normal data
Fold over basal
Time (min)
1.4
1.3
1.2
1.1
1
0.9
0.8
0.7
0.6
0.5
0
10
20
30
40
50
60

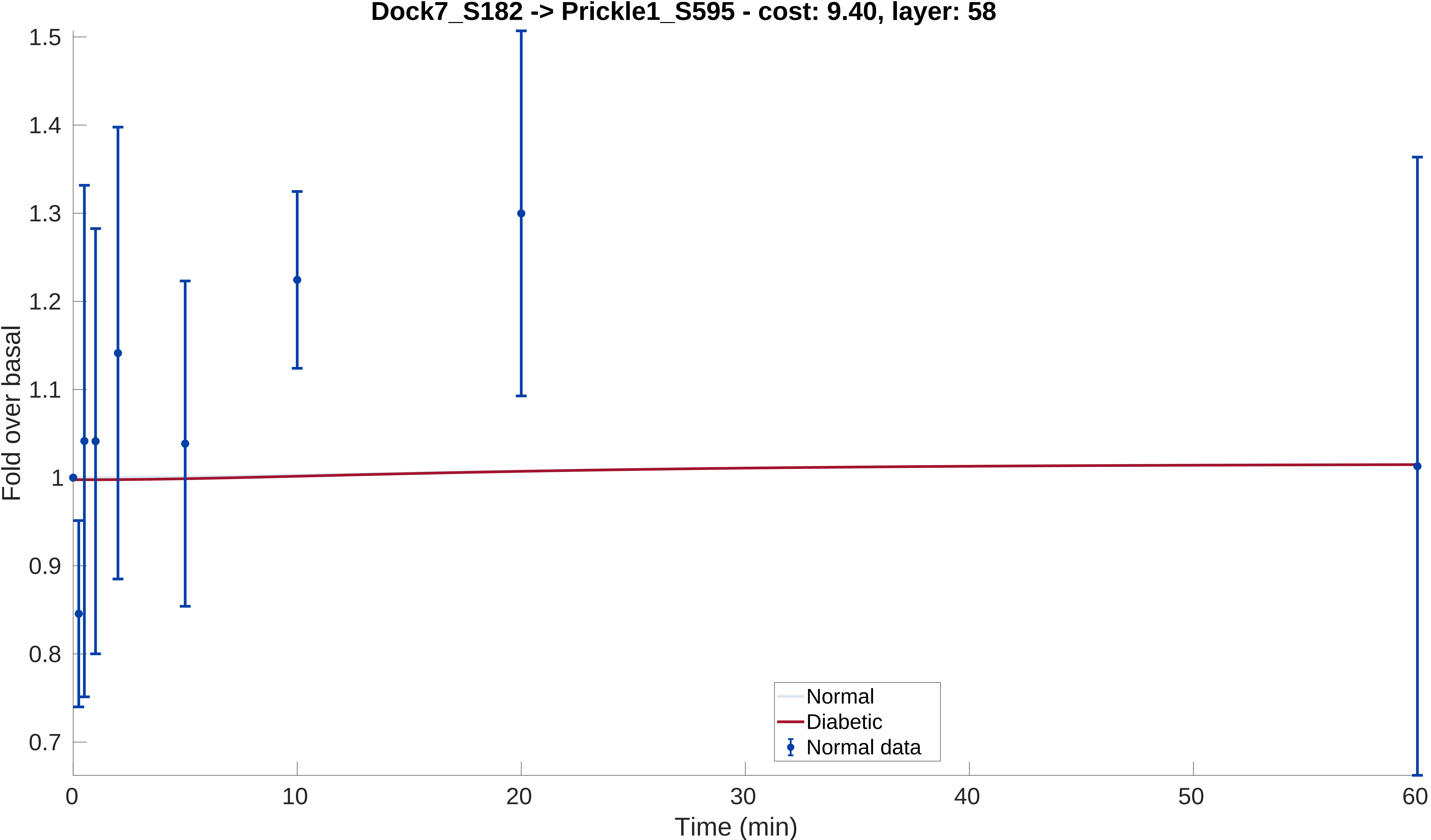
Dock7_S182 -> Prickle1_S595 - cost: 9.40, layer: 58
Fold over basal
1.5
1.4
1.3
1.2
1.1
1
0.9
0.8
0.7
0
10
20
30
40
50
60
Time (min)
Normal
Diabetic
Normal data

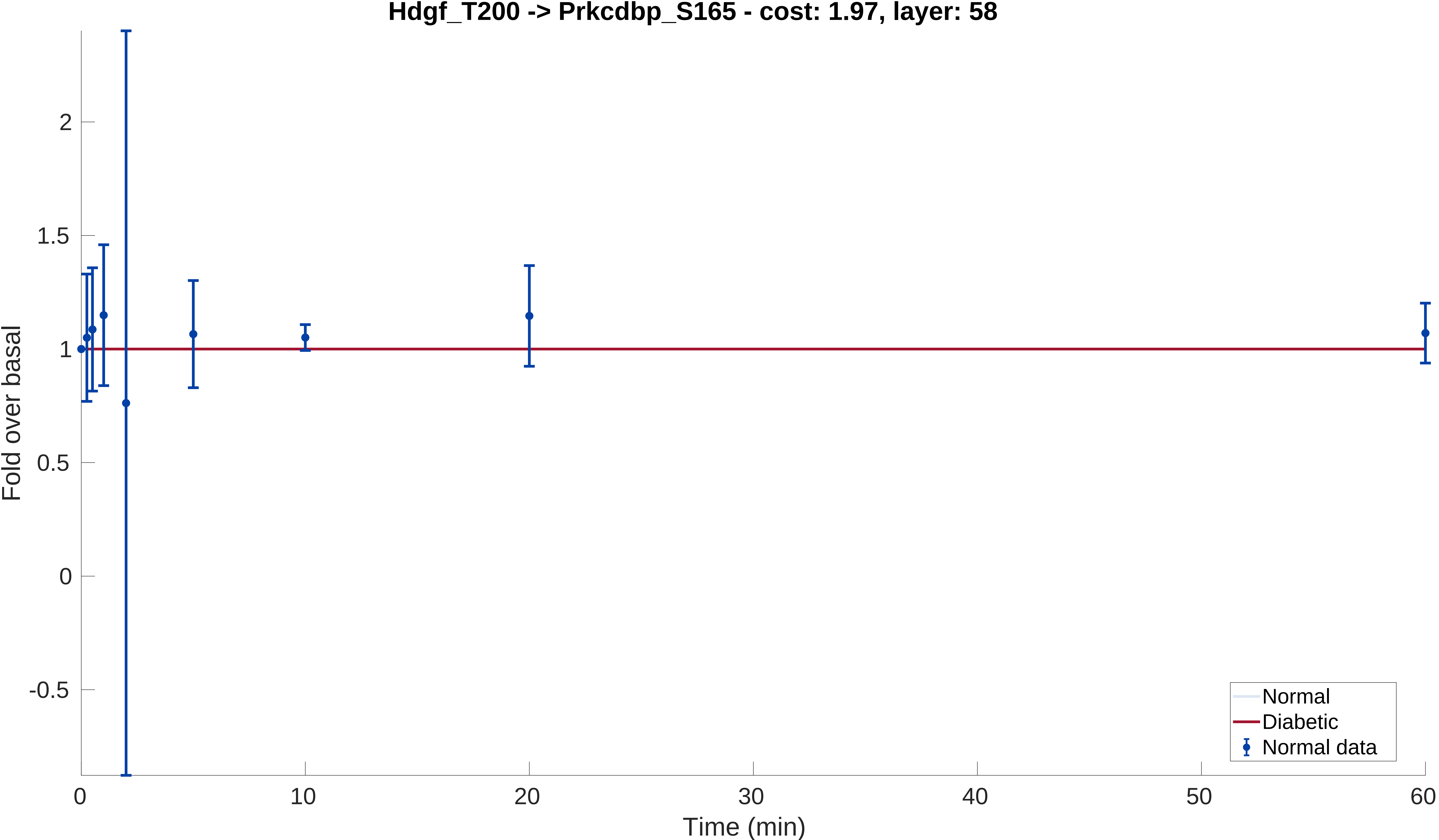
Hdgf_T200 -> Prkcdbp_S165 - cost: 1.97, layer: 58
Fold over basal
2
1.5
1
0.5
0
-0.5
0
10
20
30
40
50
60
Time (min)
Normal
Diabetic
Normal data

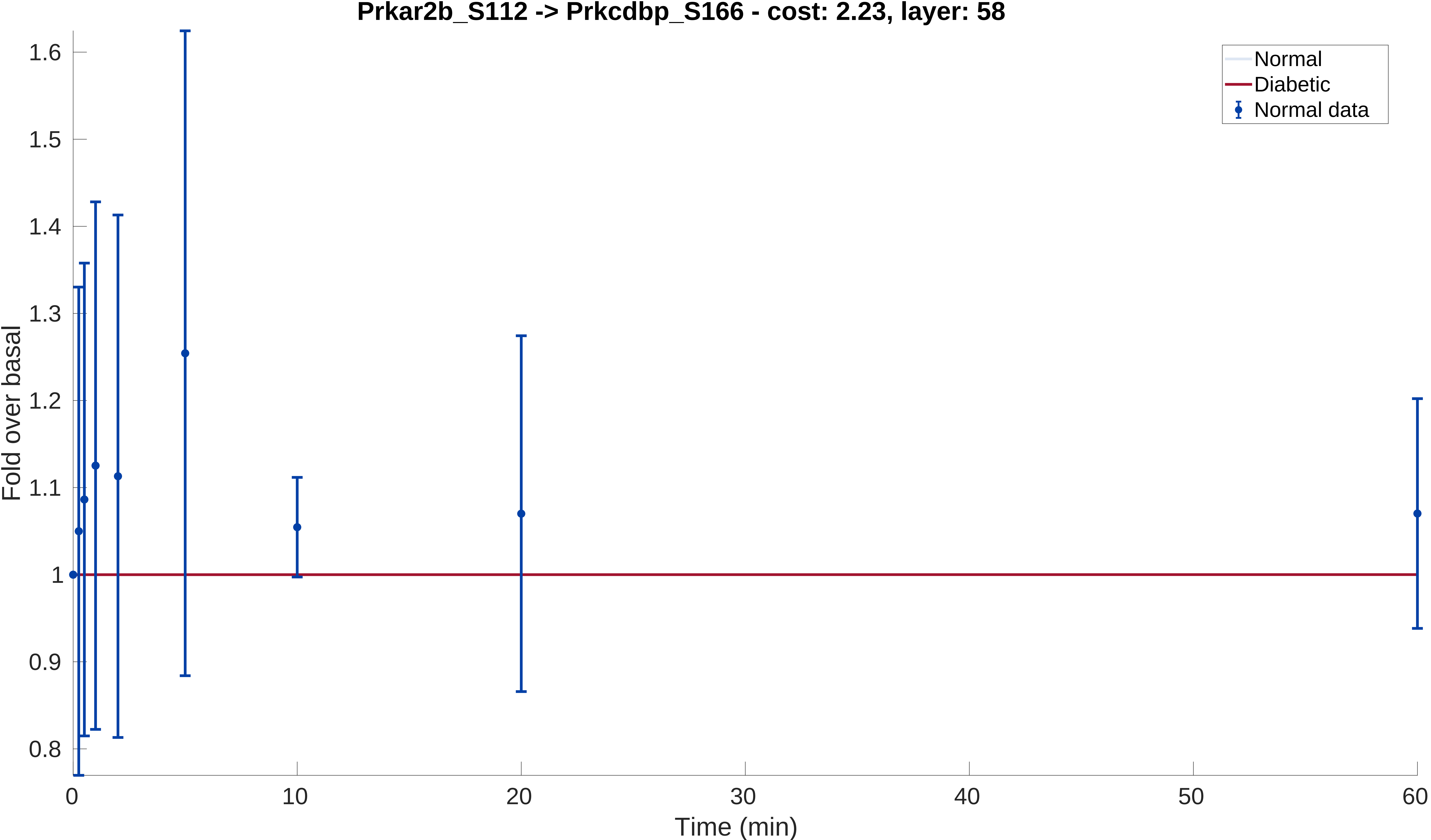

Prkar2b_S112 -> Prkcdbp_S166 - cost: 2.23, layer: 58
Normal
Diabetic
Normal data
Fold over basal
1.6
1.5
1.4
1.3
1.2
1.1
1
0.9
0.8
0
10
20
30
40
50
60
Time (min)

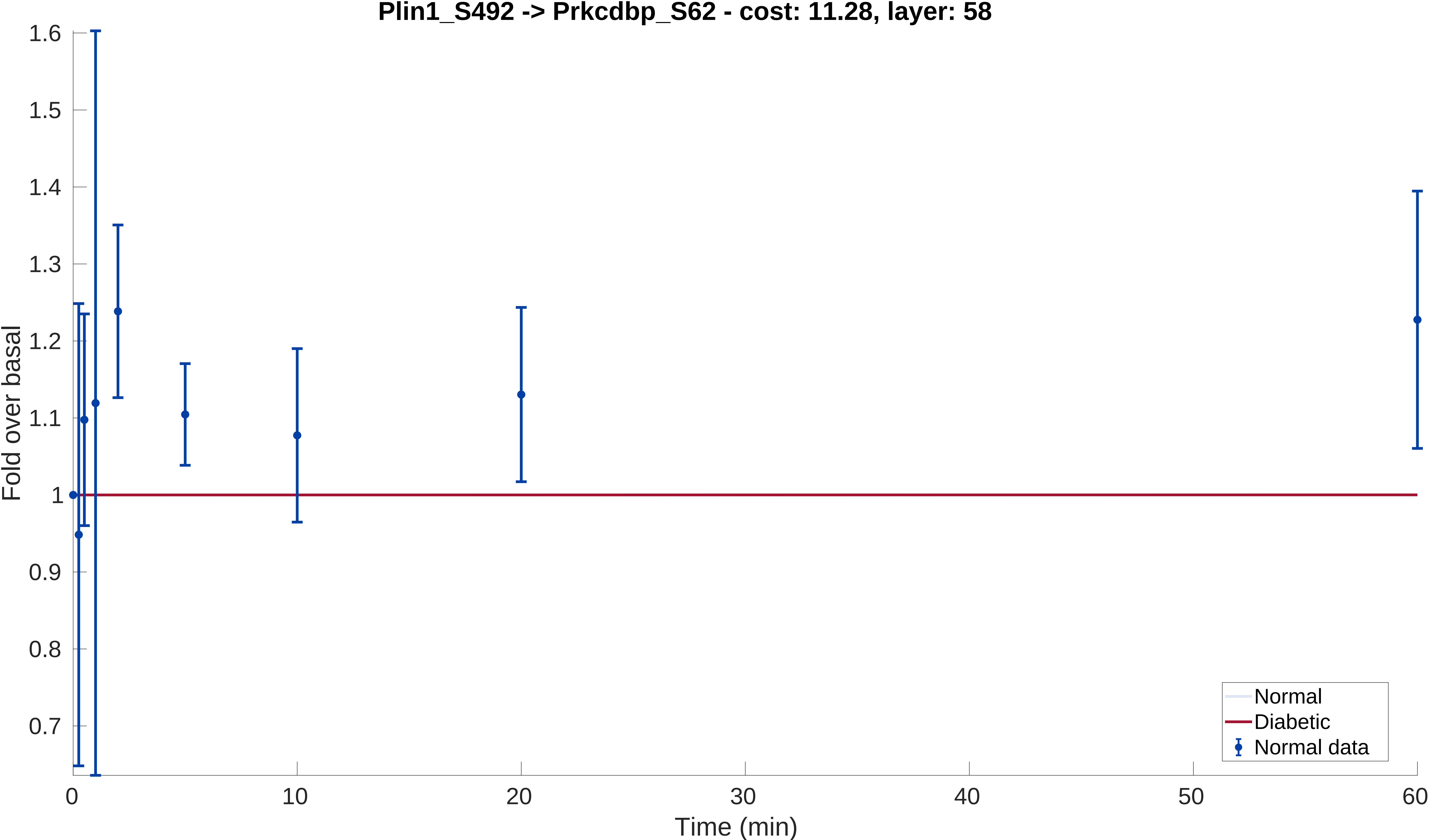

Plin1_S492 -> Prkcdbp_S62 - cost: 11.28, layer: 58
Fold over basal
1.6
1.5
1.4
1.3
1.2
1.1
1
0.9
0.8
0.7
0
10
20
30
40
50
60
Time (min)
Normal
Diabetic
Normal data

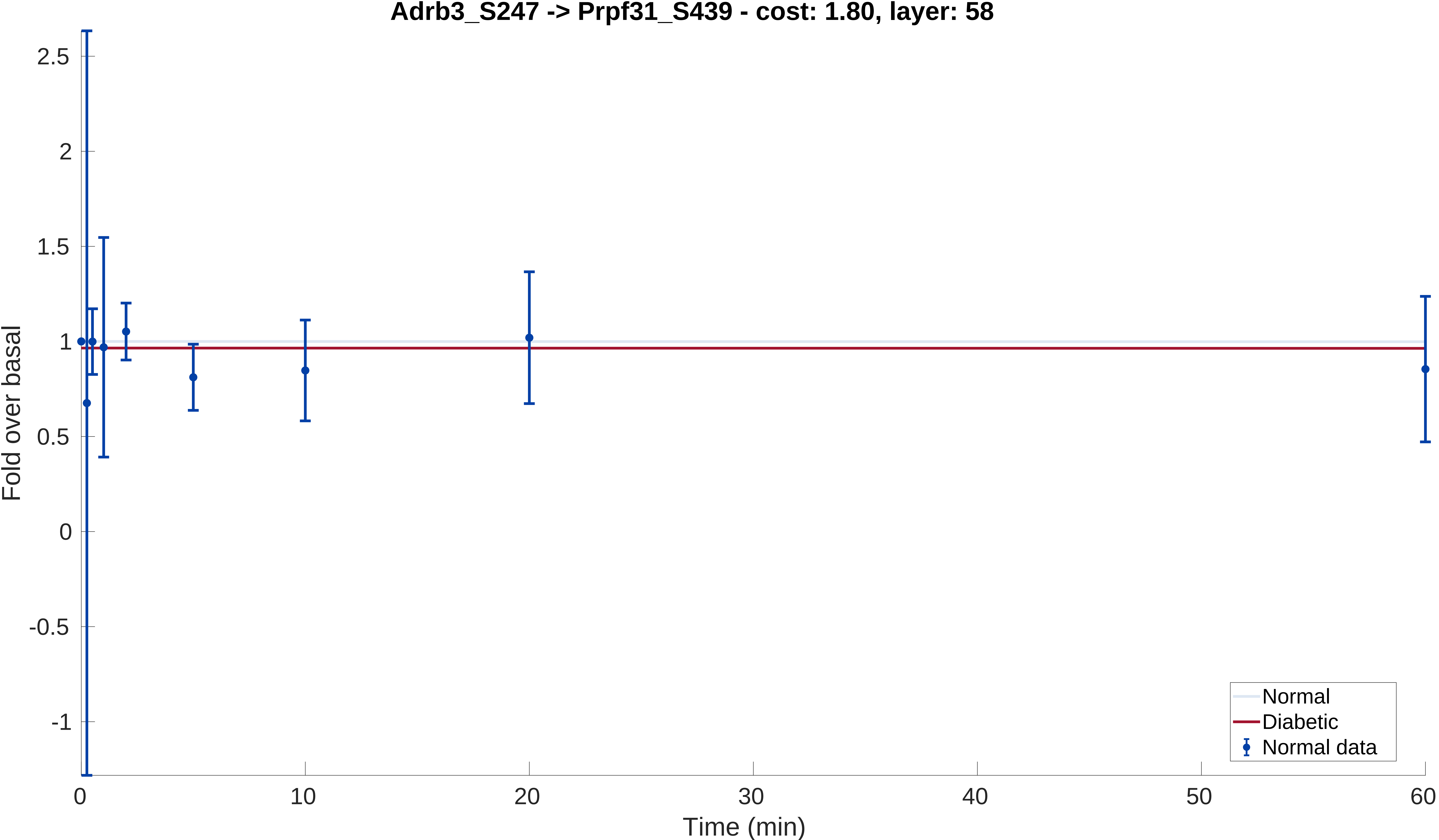

Adrb3_S247 -> Prpf31_S439 - cost: 1.80, layer: 58
Fold over basal
Time (min)
2.5
2
1.5
1
0.5
0
-0.5
-1
0
10
20
30
40
50
60
Normal
Diabetic
Normal data

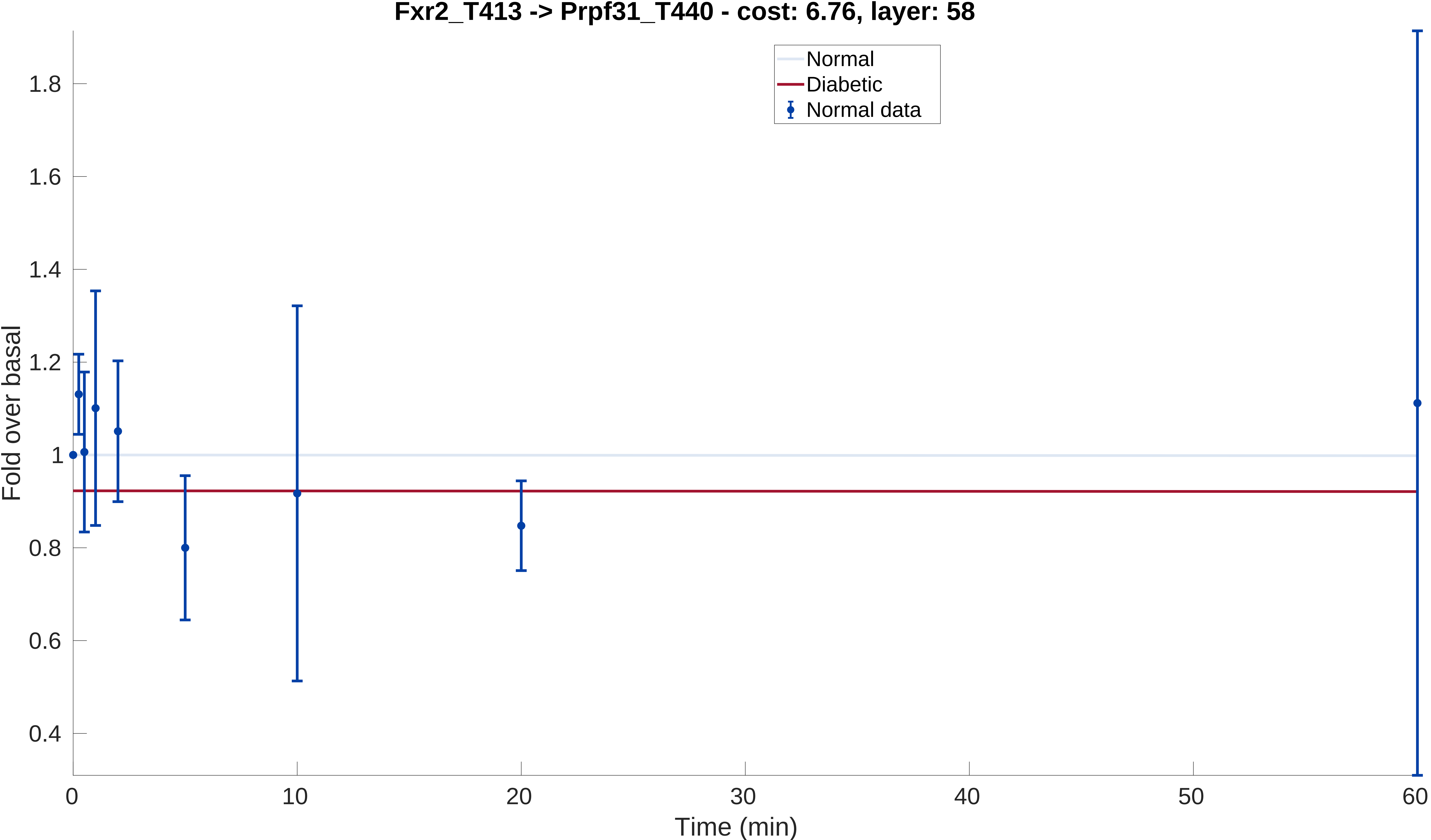
Fxr2_T413 -> Prpf31_T440 - cost: 6.76, layer: 58
Normal
Diabetic
Normal data
Fold over basal
1.8
1.6
1.4
1.2
1
0.8
0.6
0.4
0
10
20
30
40
50
60
Time (min)

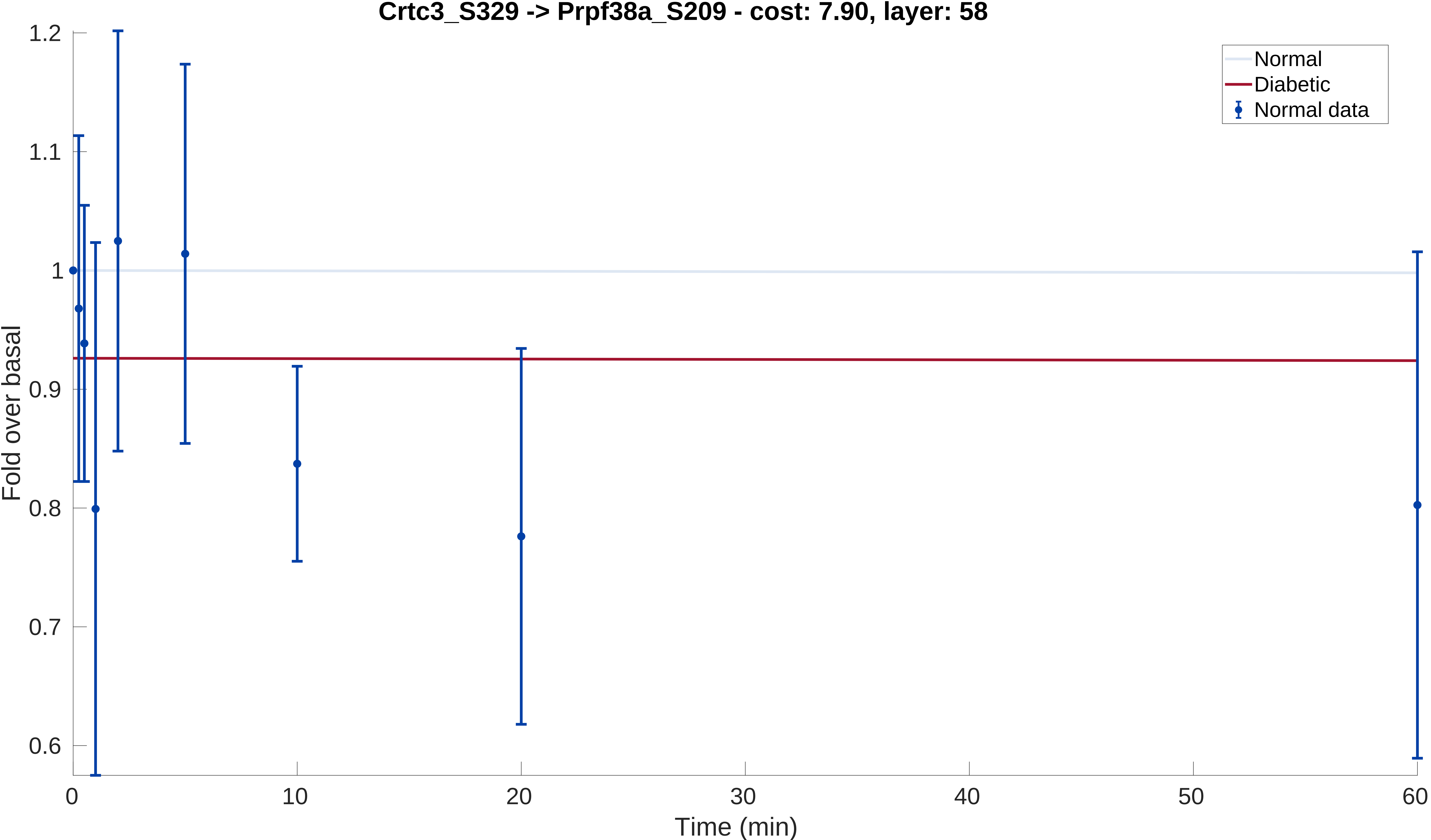

Crtc3_S329 -> Prpf38a_S209 - cost: 7.90, layer: 58
Normal
Diabetic
Normal data
Fold over basal
Time (min)
1.2
1.1
1
0.9
0.8
0.7
0.6
0
10
20
30
40
50
60

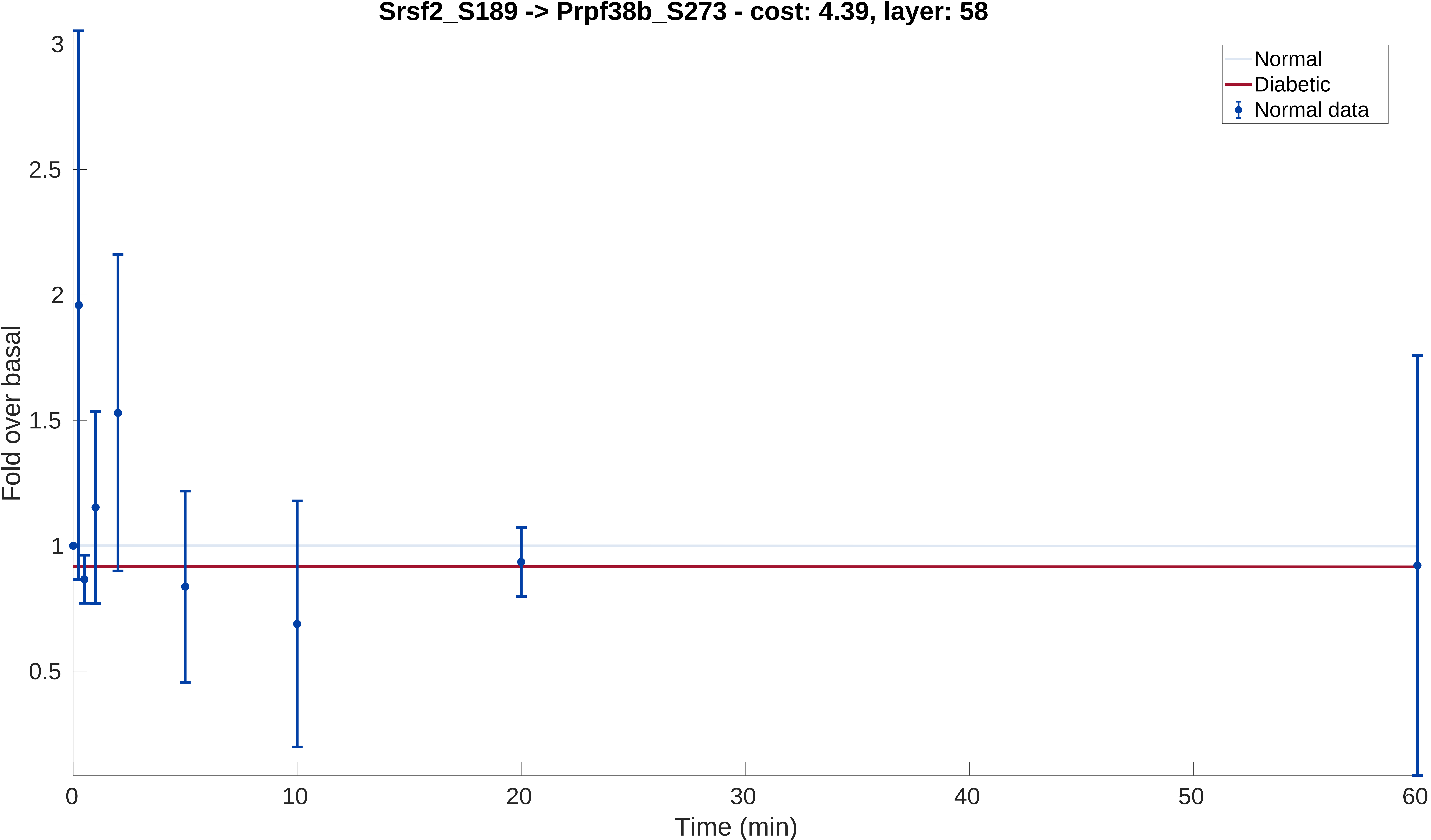

Srsf2_S189 -> Prpf38b_S273 - cost: 4.39, layer: 58
Normal
Diabetic
Normal data
Fold over basal
Time (min)
0.5
1
1.5
2
2.5
3
0
10
20
30
40
50
60

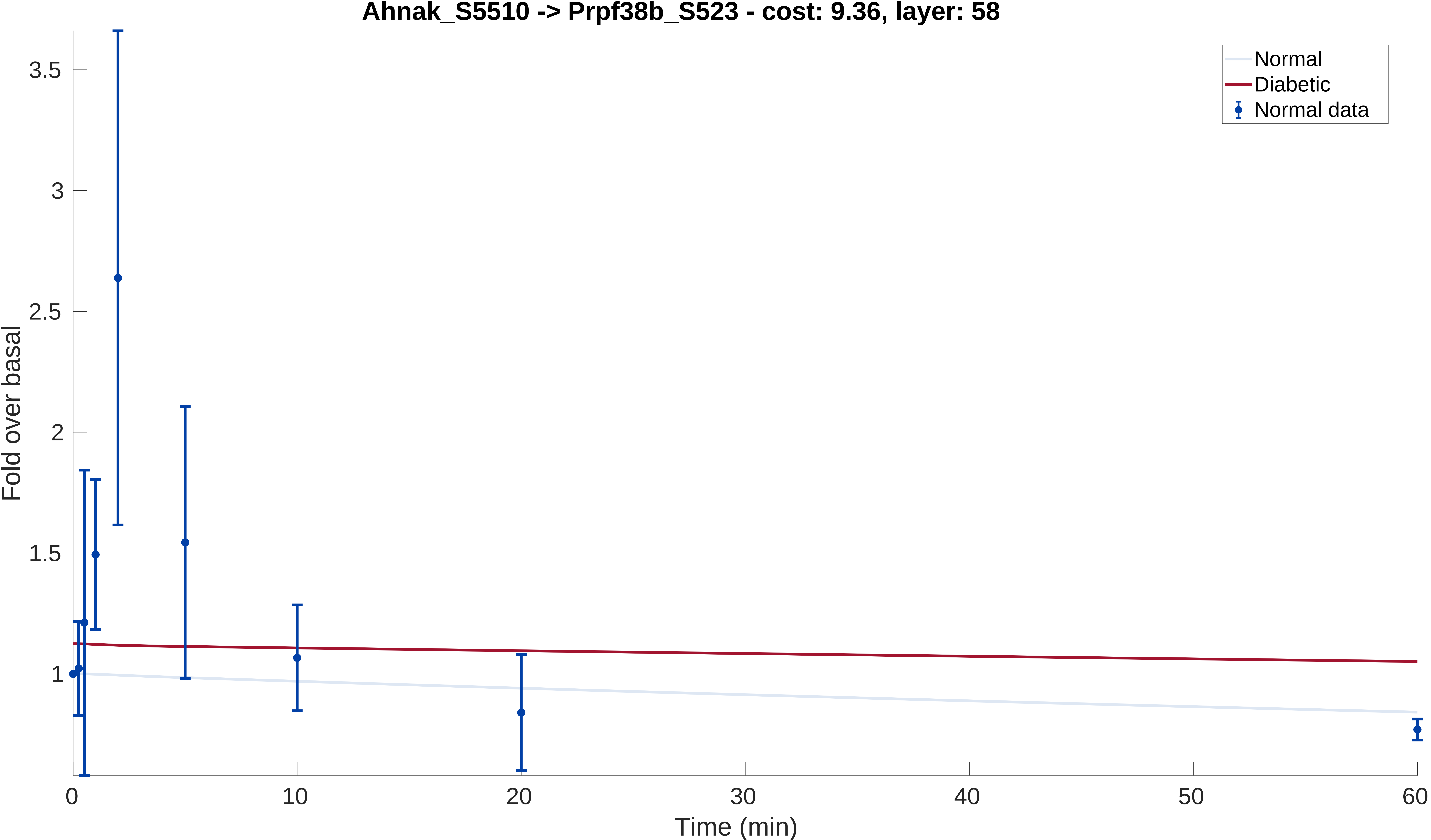

Ahnak_S5510 -> Prpf38b_S523 - cost: 9.36, layer: 58
Normal
Diabetic
Normal data
Fold over basal
3.5
3
2.5
2
1.5
1
0
10
20
30
40
50
60
Time (min)

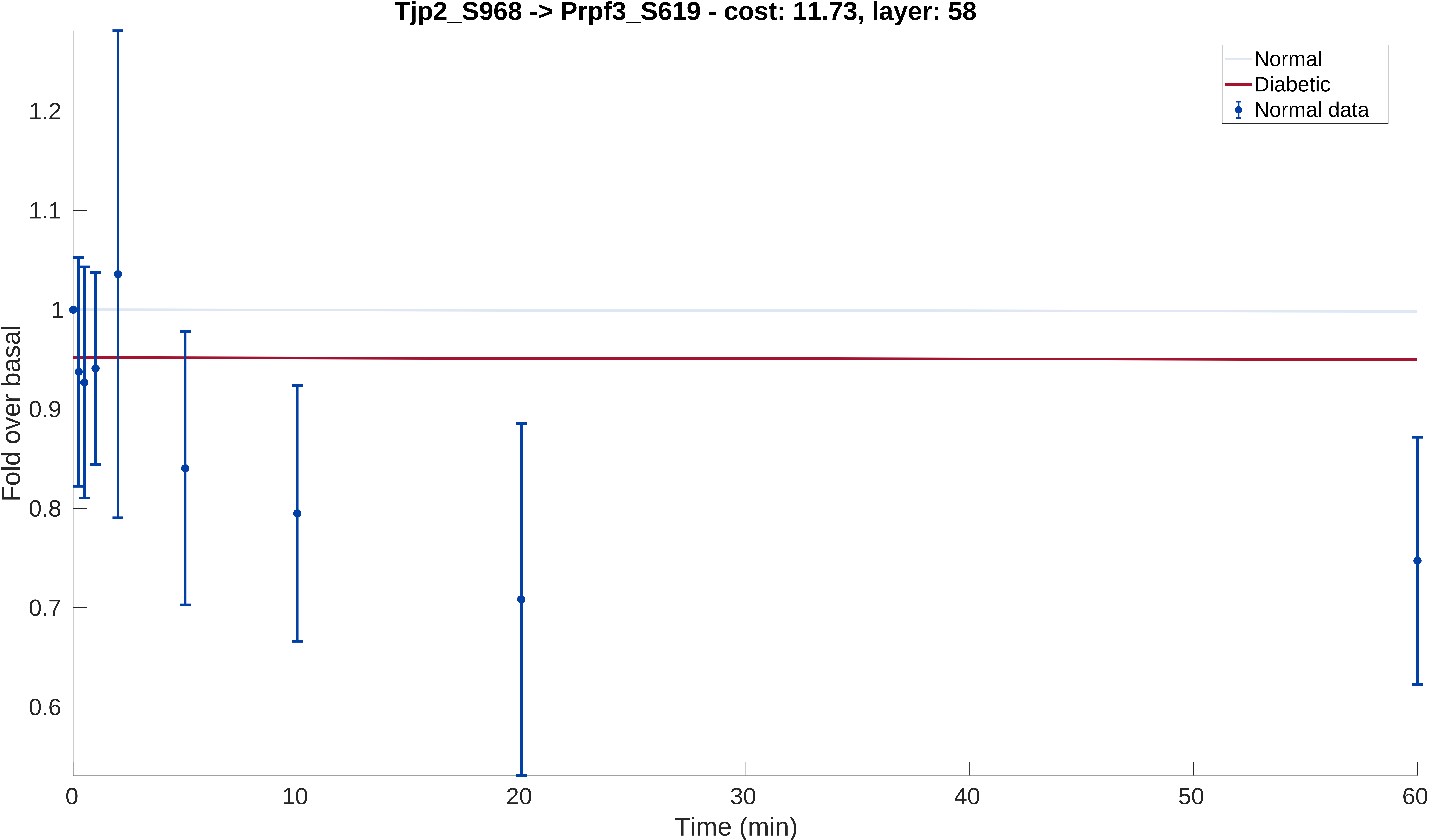

Tjp2_S968 -> Prpf3_S619 - cost: 11.73, layer: 58
Normal
Diabetic
Normal data
Fold over basal
1.2
1.1
1
0.9
0.8
0.7
0.6
0
10
20
30
40
50
60
Time (min)

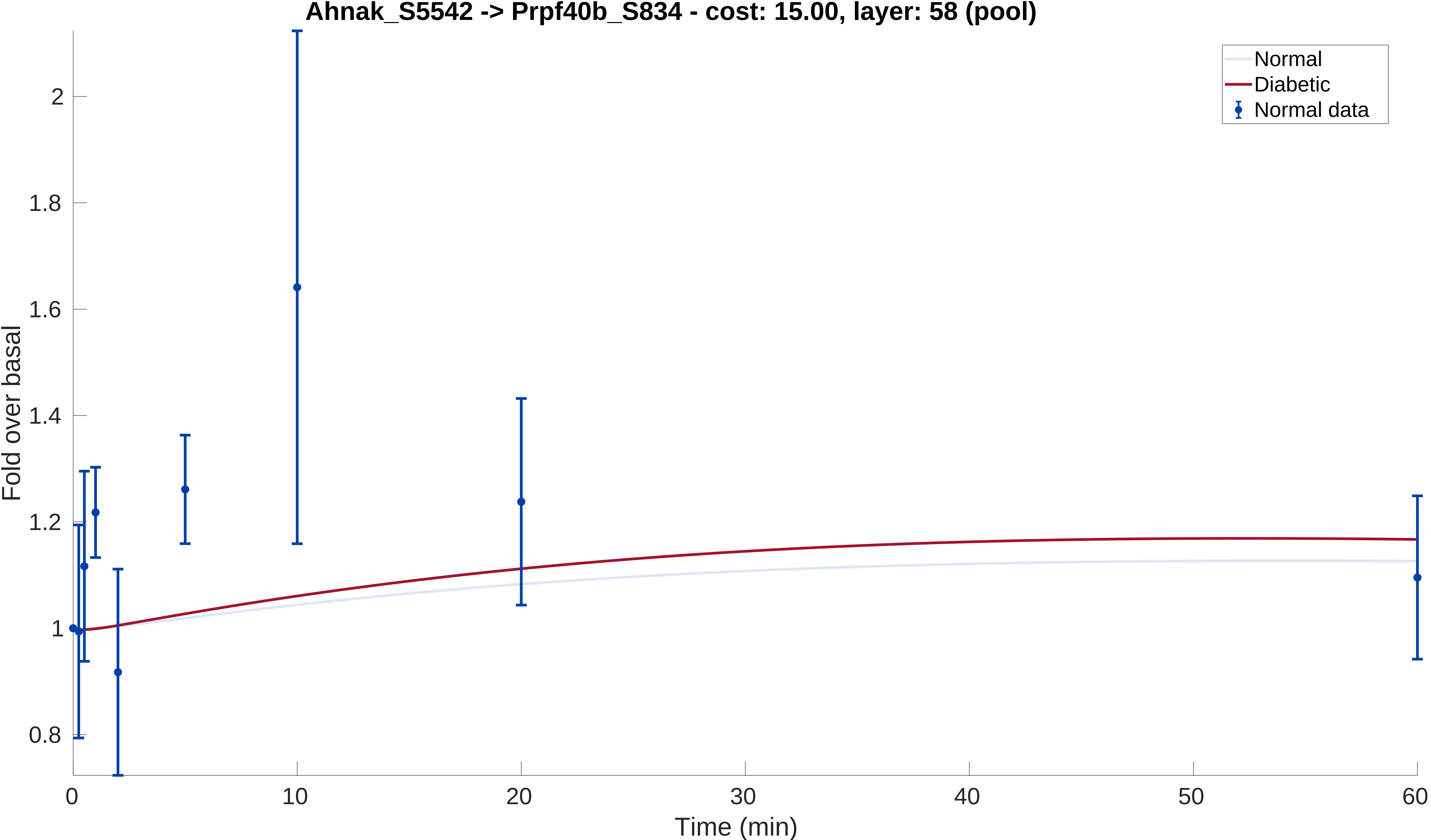
Ahnak_S5542 -> Prpf40b_S834 - cost: 15.00, layer: 58 (pool)
Normal
Diabetic
Normal data
Fold over basal
2
1.8
1.6
1.4
1.2
1
0.8
0
10
20
30
40
50
60
Time (min)

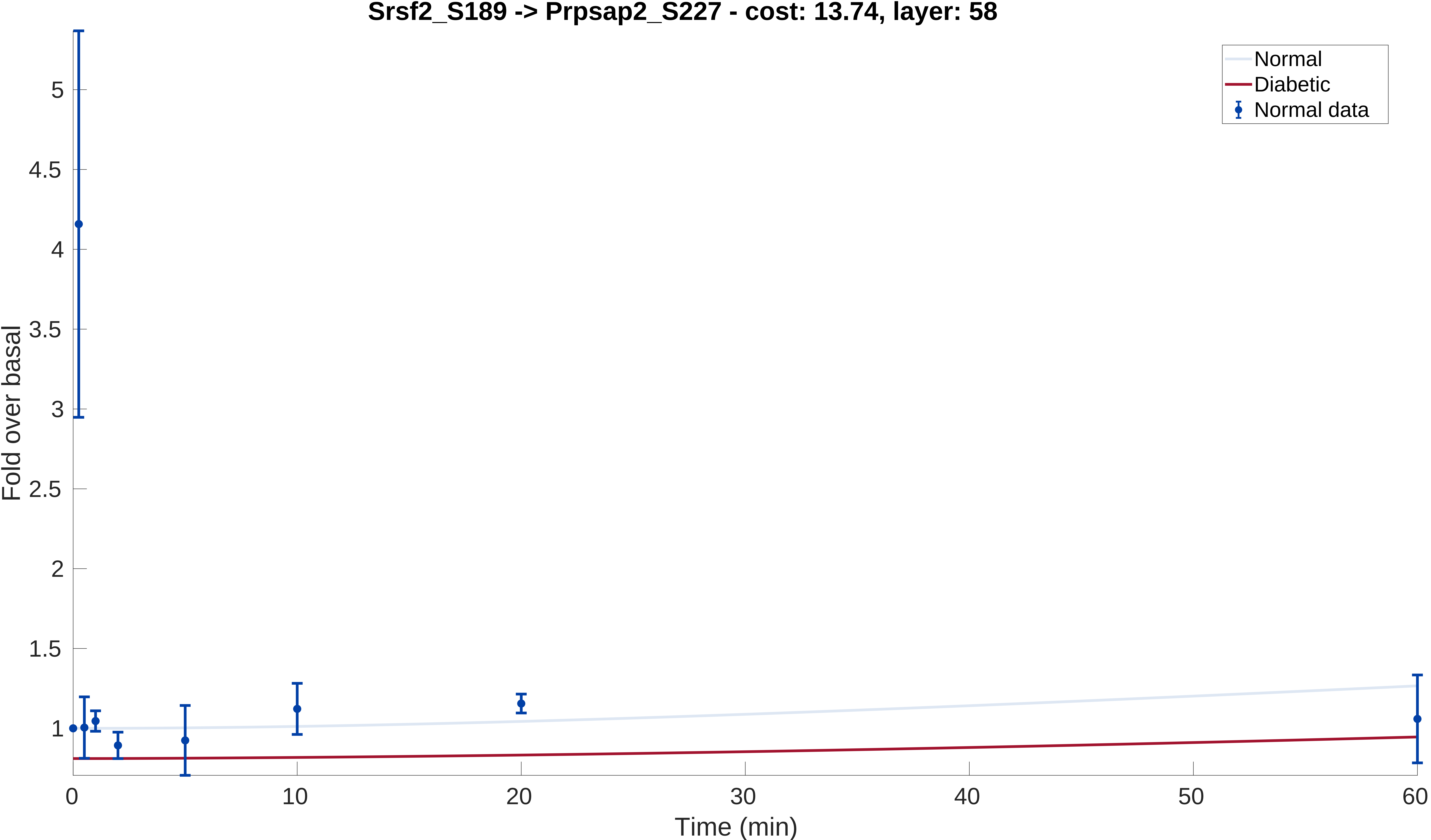
Srsf2_S189 -> Prpsap2_S227 - cost: 13.74, layer: 58
Normal
Diabetic
Normal data
Fold over basal
5
4.5
4
3.5
3
2.5
2
1.5
1
0
10
20
30
40
50
60
Time (min)

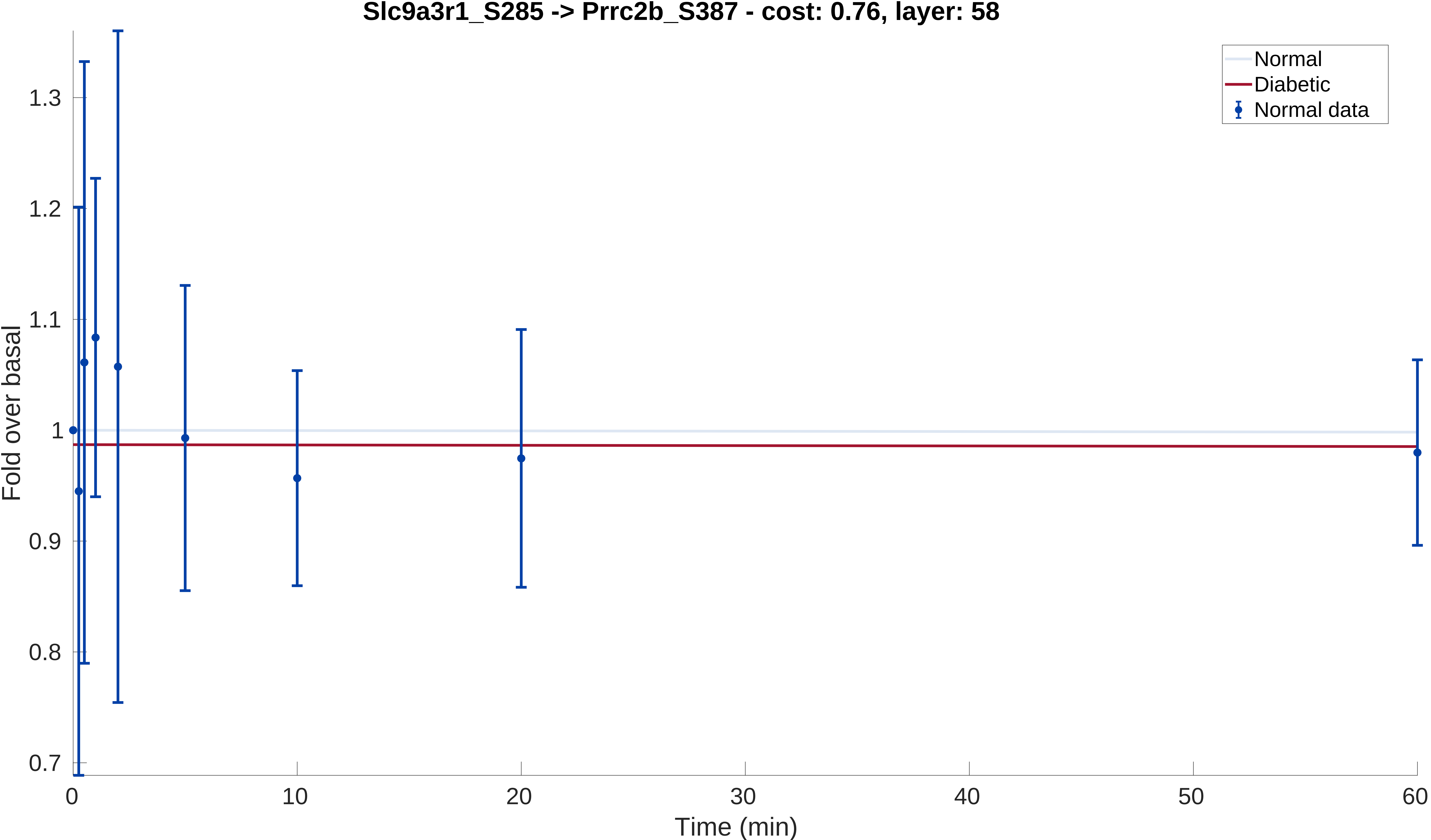

Slc9a3r1_S285 -> Prrc2b_S387 - cost: 0.76, layer: 58
Normal
Diabetic
Normal data
Fold over basal
1.3
1.2
1.1
1
0.9
0.8
0.7
0
10
20
30
40
50
60
Time (min)

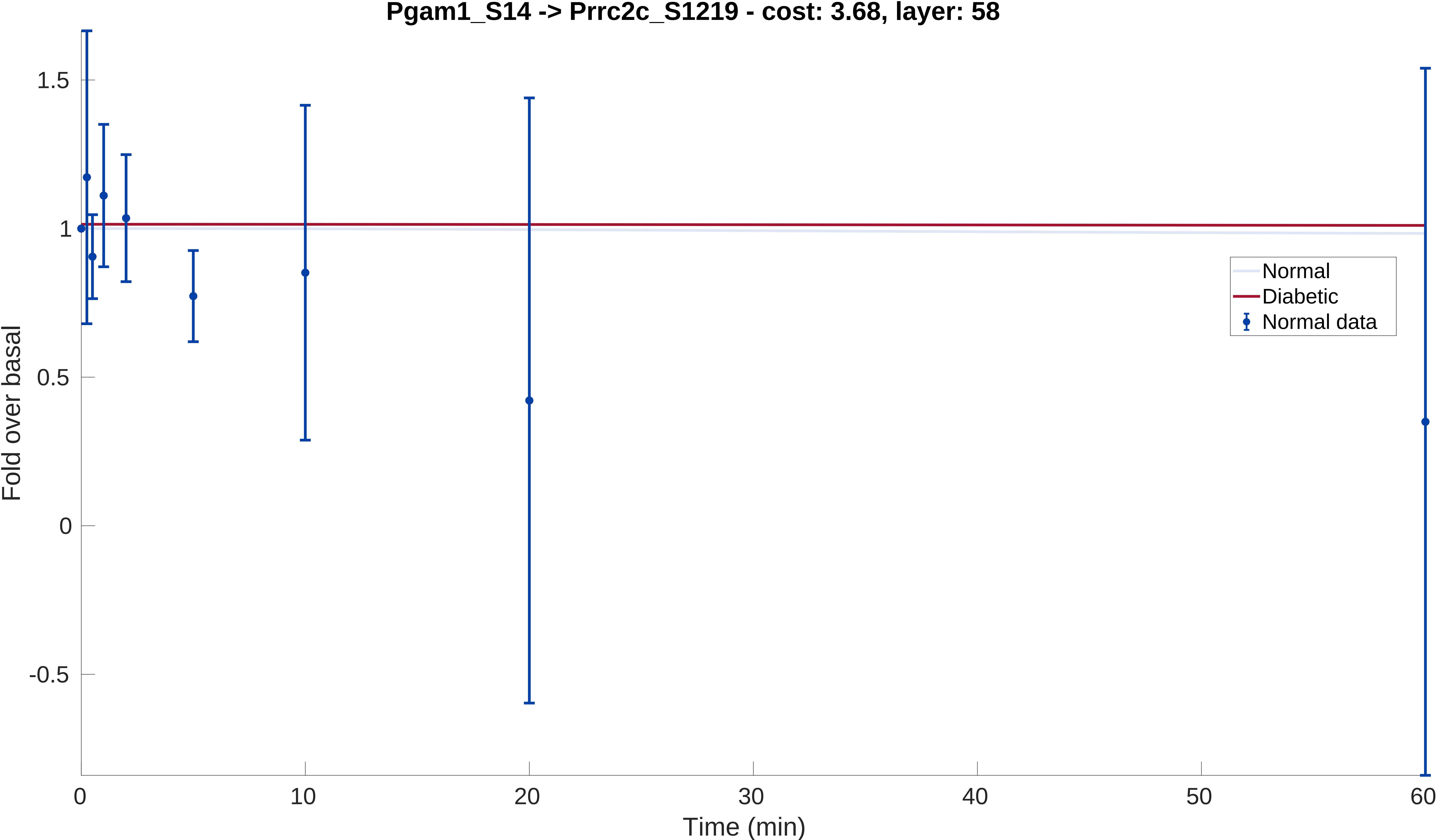
Pgam1_S14 -> Prrc2c_S1219 - cost: 3.68, layer: 58
Fold over basal
1.5
1
0.5
0
-0.5
0
10
20
30
40
50
60
Time (min)
Normal
Diabetic
Normal data

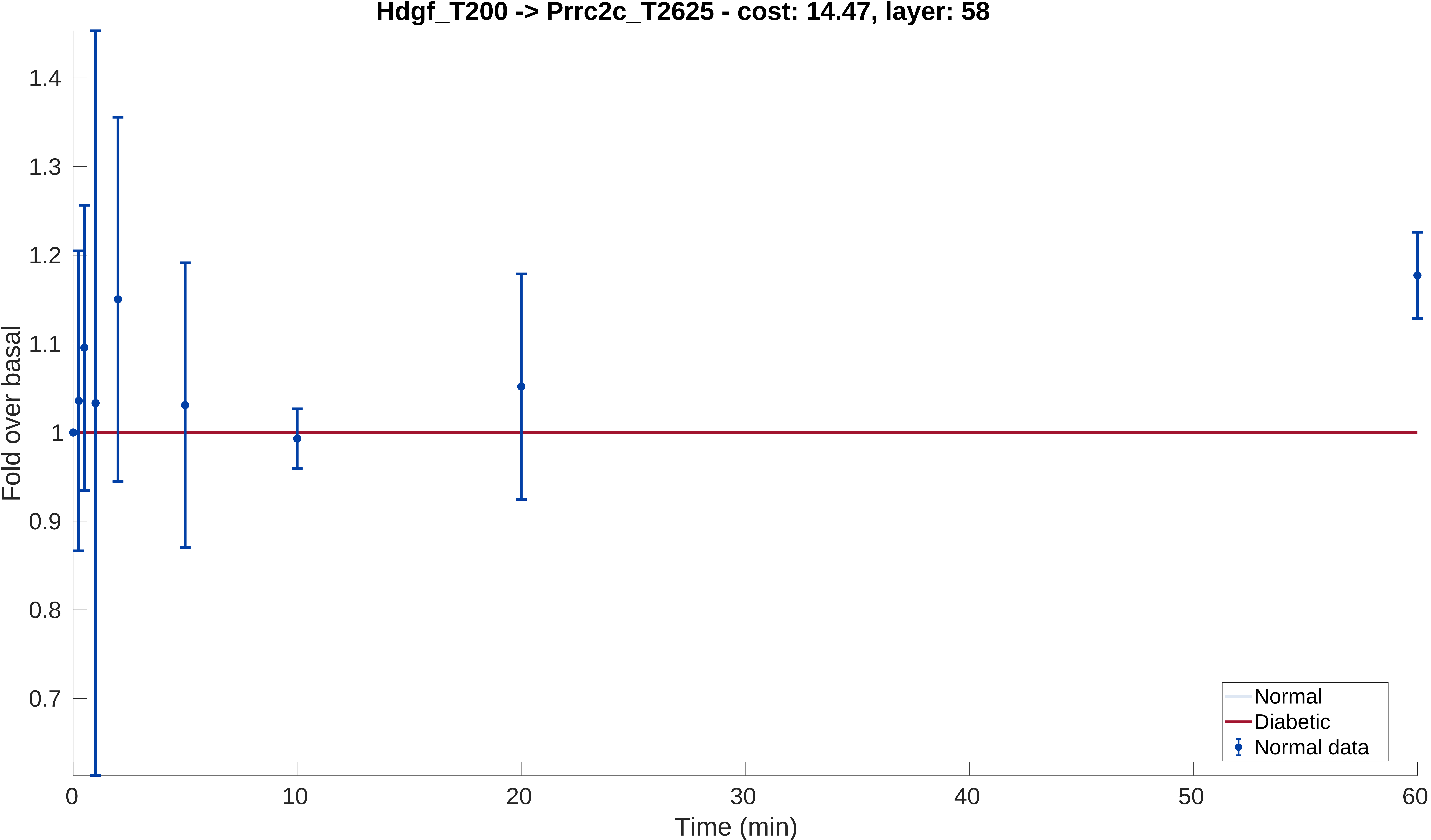

Hdgf_T200 -> Prrc2c_T2625 - cost: 14.47, layer: 58
Fold over basal
1.4
1.3
1.2
1.1
1
0.9
0.8
0.7
0
10
20
30
40
50
60
Time (min)
Normal
Diabetic
Normal data

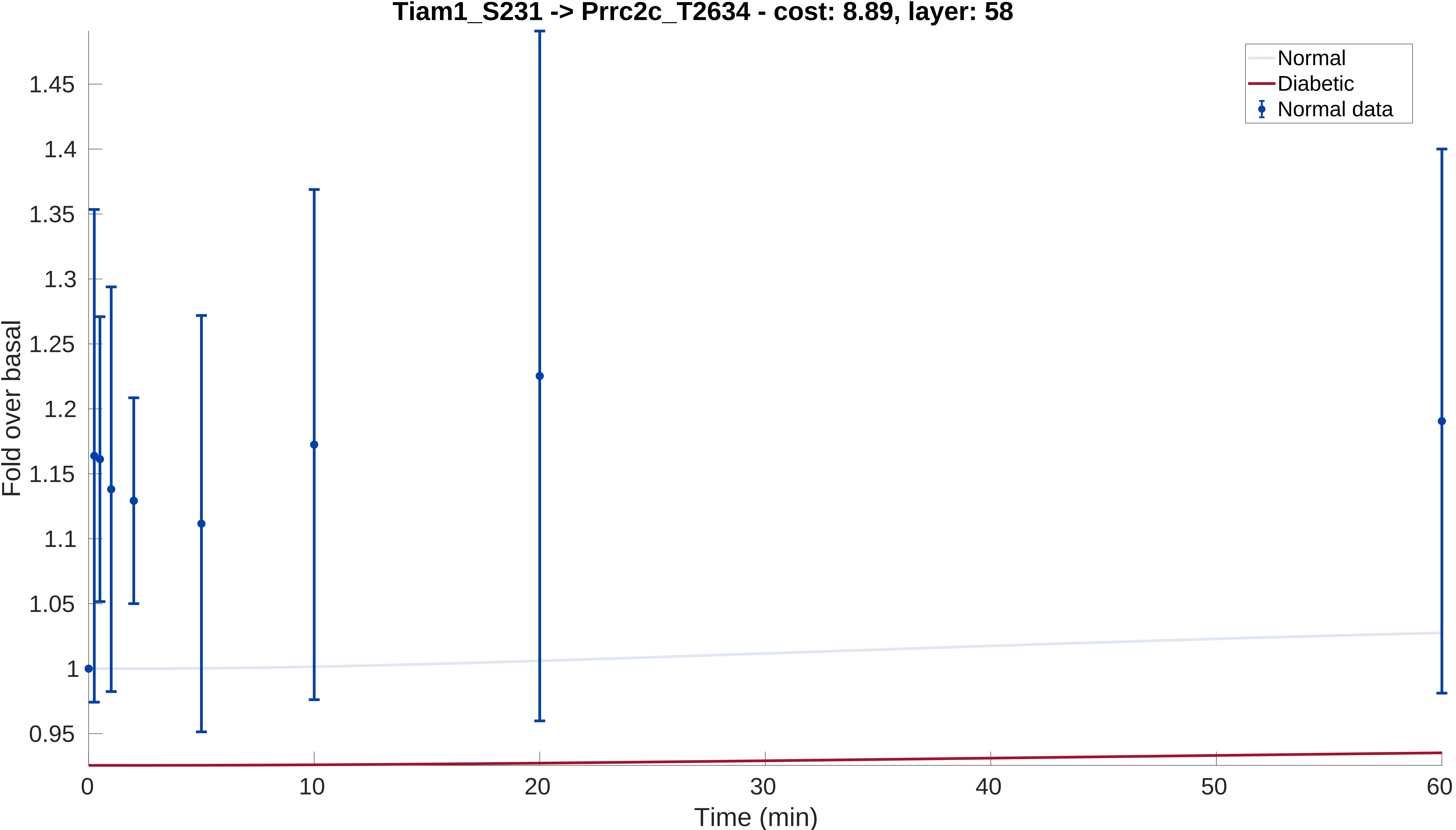

Tiam1_S231 -> Prrc2c_T2634 - cost: 8.89, layer: 58
Normal
Diabetic
Normal data
Fold over basal
1.45
1.4
1.35
1.3
1.25
1.2
1.15
1.1
1.05
1
0.95
0
10
20
30
40
50
60
Time (min)

Mark2_S483 -> Psip1_S106 - cost: 2.02, layer: 58

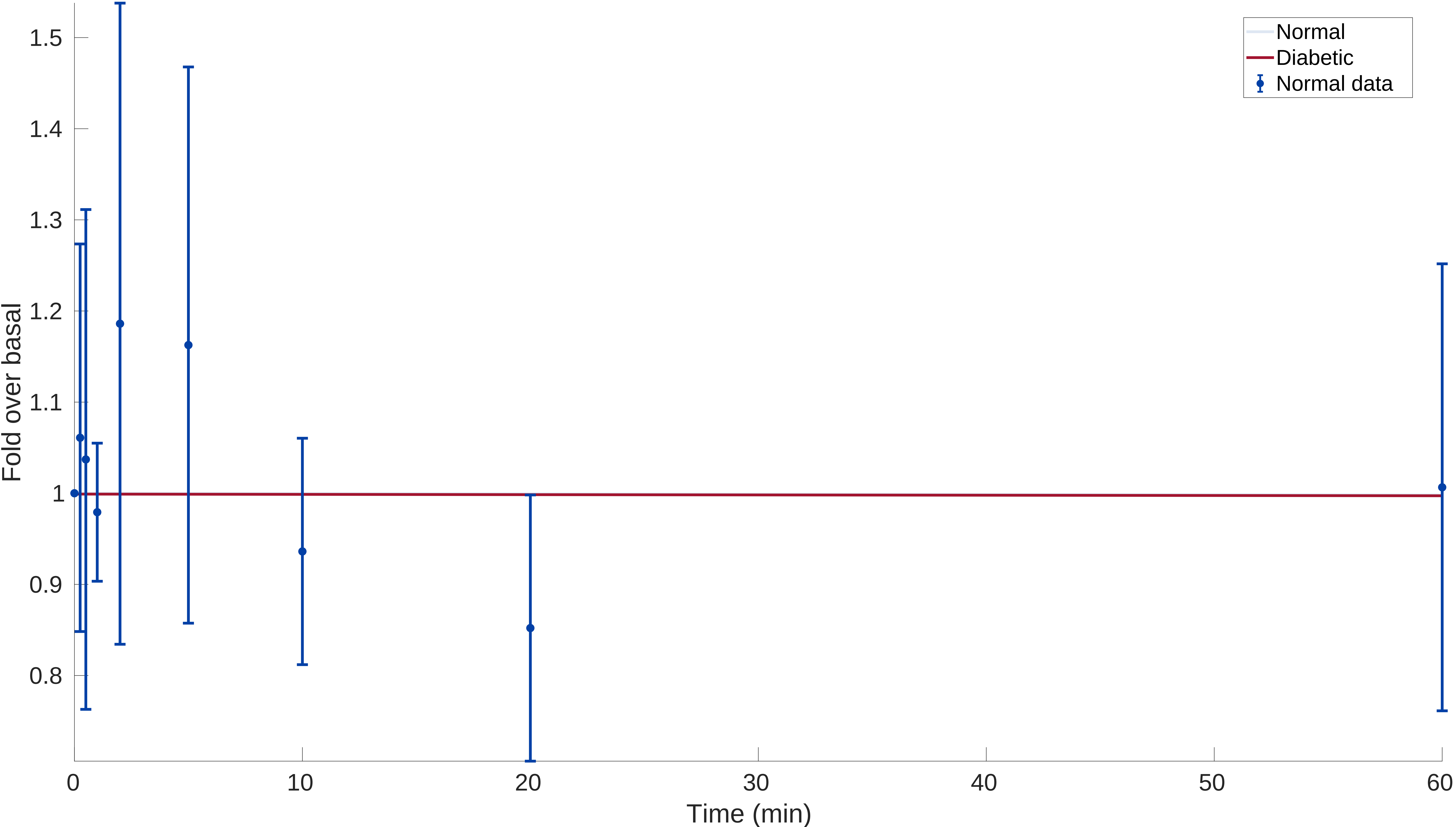

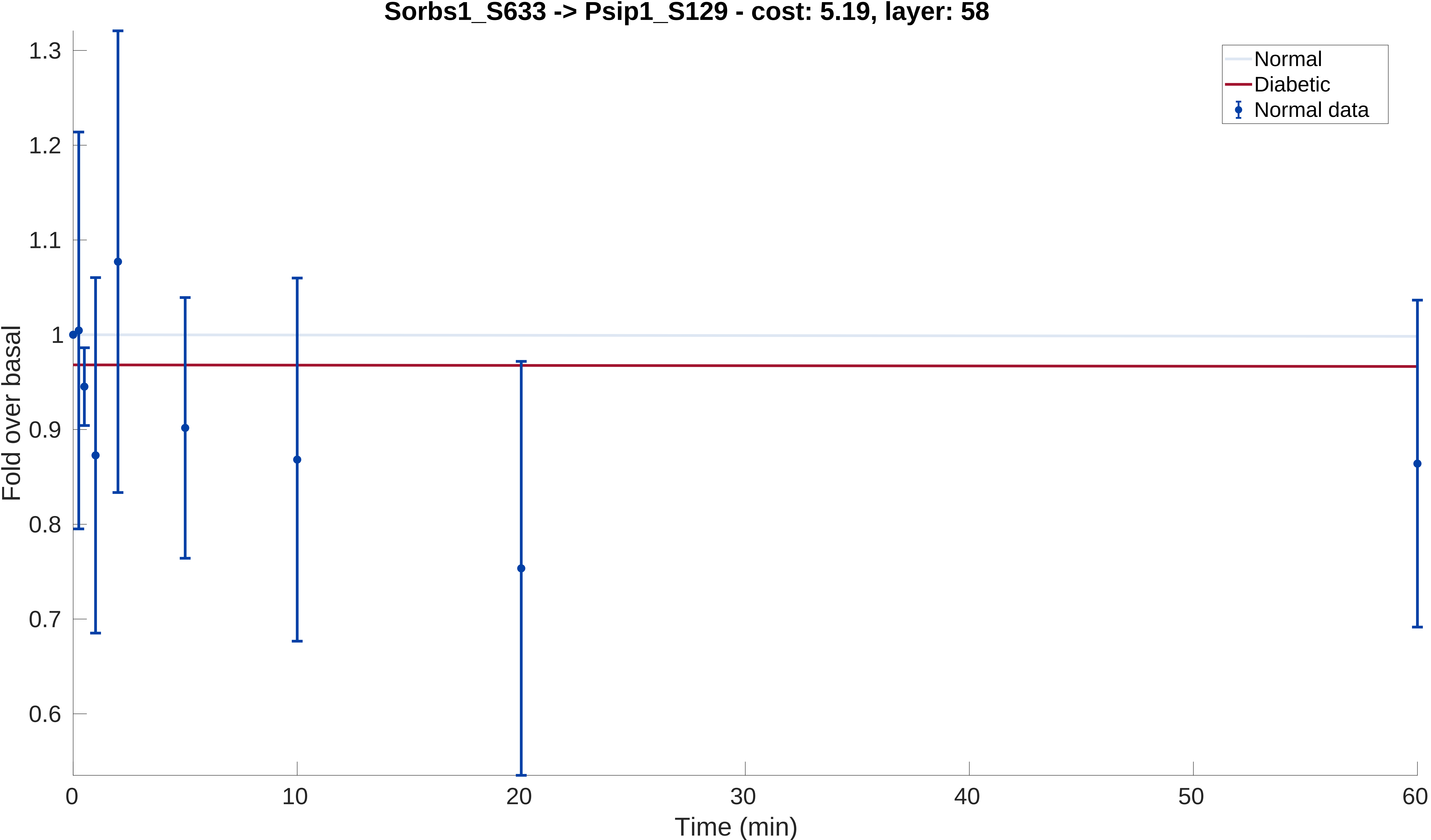

Sorbs1_S633 -> Psip1_S129 - cost: 5.19, layer: 58
Normal
Diabetic
Normal data
Fold over basal
1.3
1.2
1.1
1
0.9
0.8
0.7
0.6
0
10
20
30
40
50
60
Time (min)

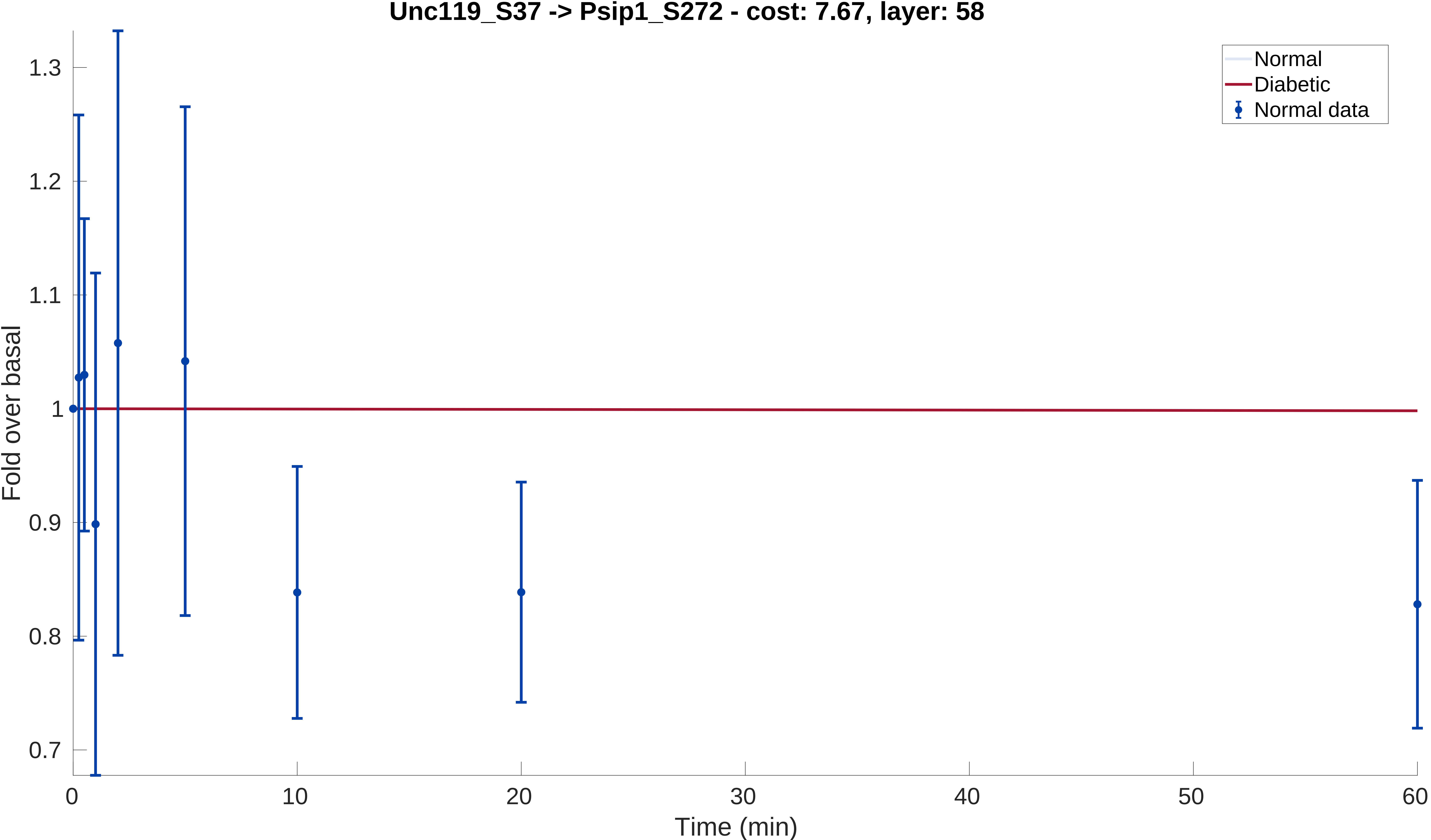
Unc119_S37 -> Psip1_S272 - cost: 7.67, layer: 58
Normal
Diabetic
Normal data
Fold over basal
Time (min)
1.3
1.2
1.1
1
0.9
0.8
0.7
0
10
20
30
40
50
60

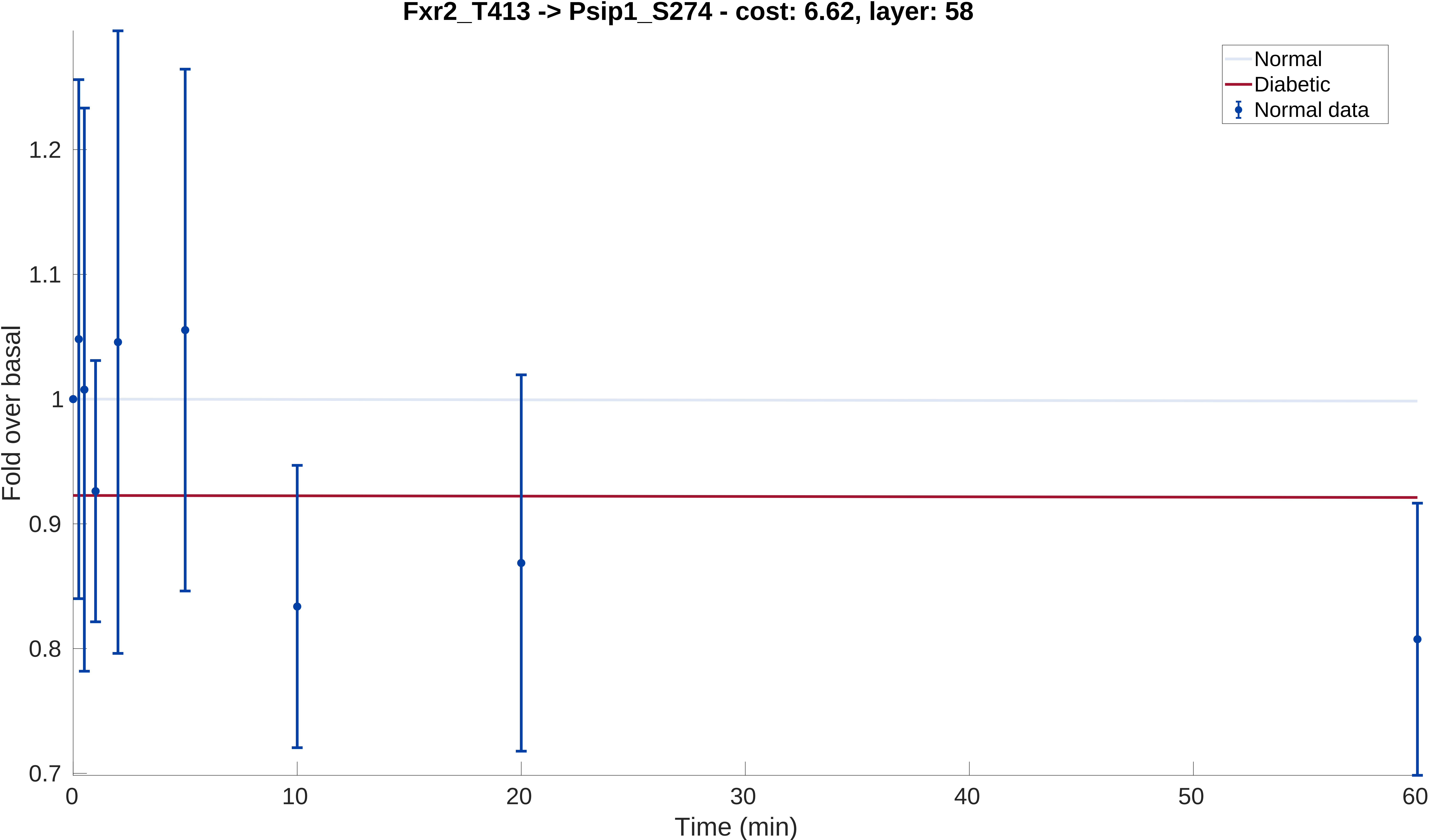

Fxr2_T413 -> Psip1_S274 - cost: 6.62, layer: 58
Normal
Diabetic
Normal data
Fold over basal
1.2
1.1
1
0.9
0.8
0.7
0
10
20
30
40
50
60
Time (min)

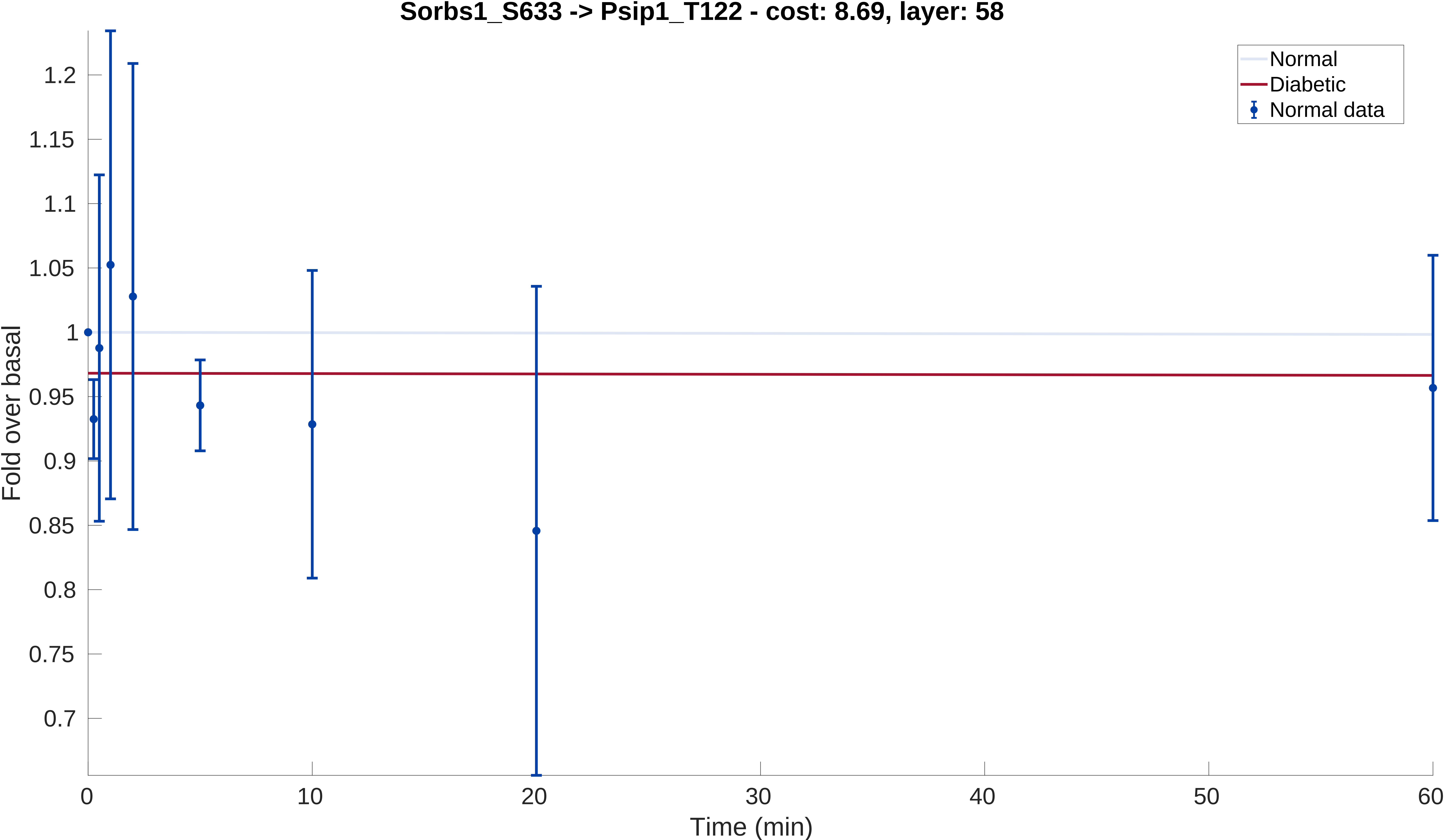

Sorbs1_S633 -> Psip1_T122 - cost: 8.69, layer: 58
Normal
Diabetic
Normal data
Fold over basal
1.2
1.15
1.1
1.05
1
0.95
0.9
0.85
0.8
0.75
0.7
0
10
20
30
40
50
60
Time (min)

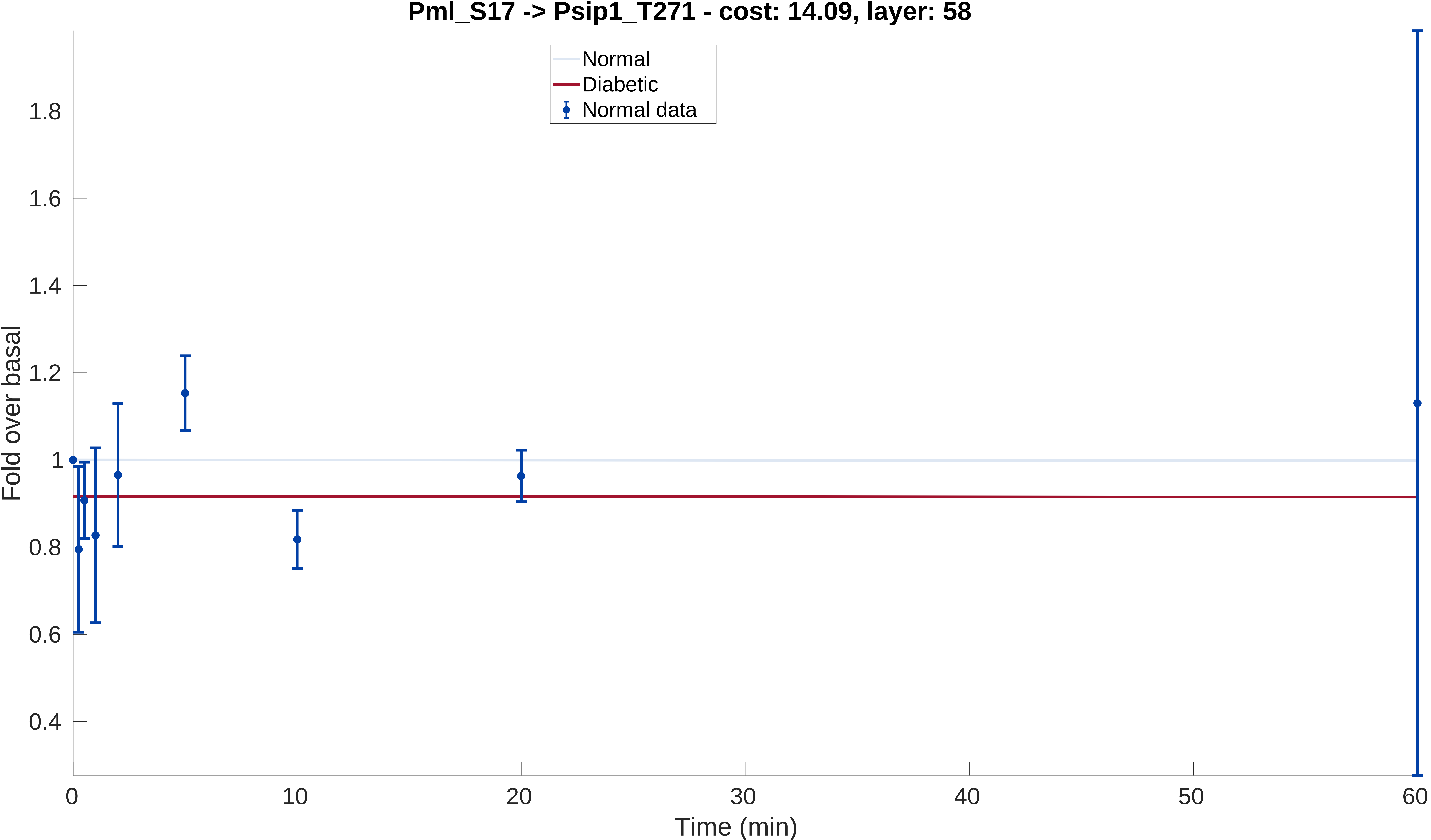

Pml_S17 -> Psip1_T271 - cost: 14.09, layer: 58
Normal
Diabetic
Normal data
1.8
1.6
1.4
1.2
1
0.8
0.6
0.4
Fold over basal
0
10
20
30
40
50
60
Time (min)

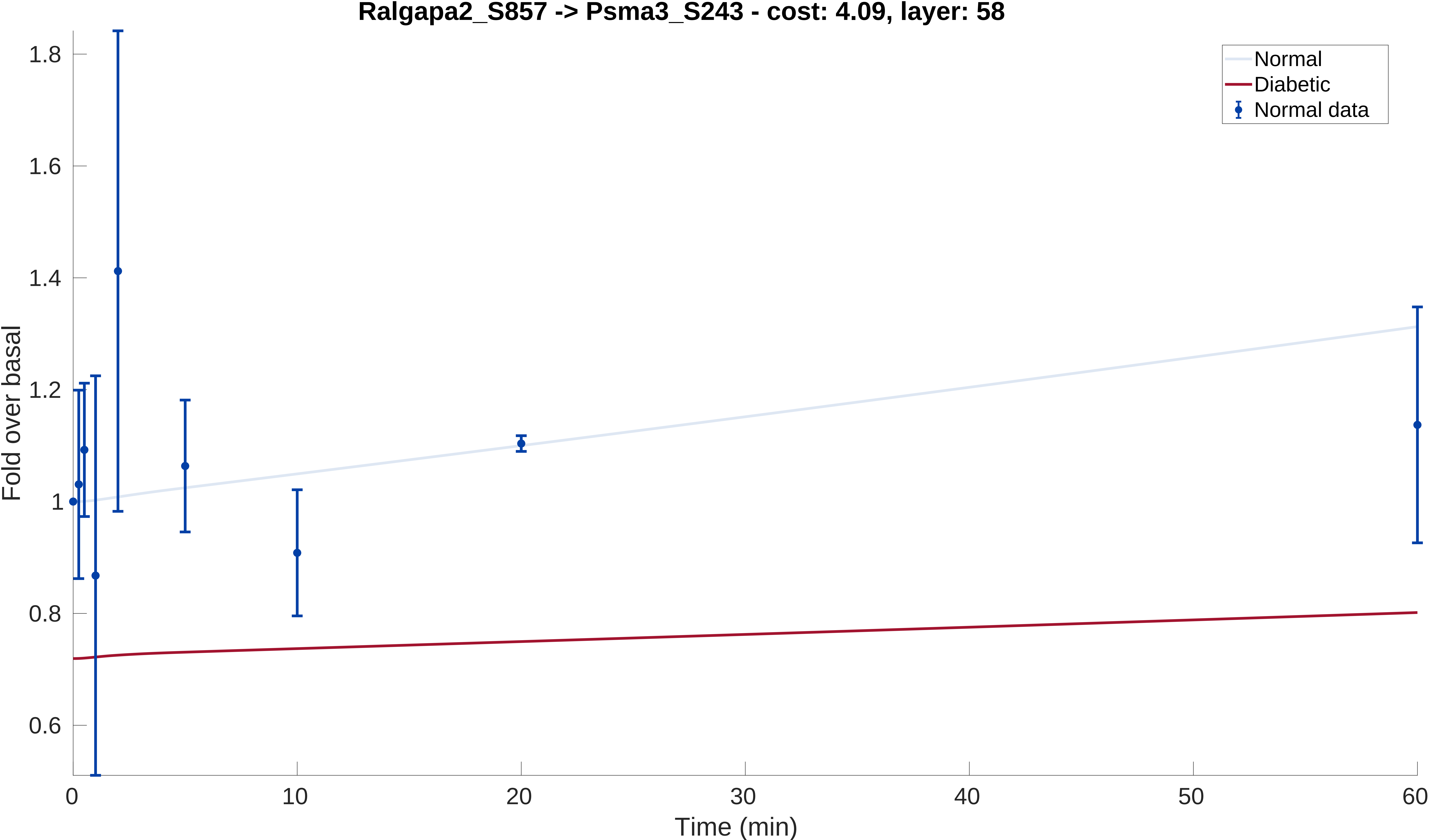

Ralgapa2_S857 -> Psma3_S243 - cost: 4.09, layer: 58
Normal
Diabetic
Normal data
Fold over basal
Time (min)
1.8
1.6
1.4
1.2
1
0.8
0.6
0
10
20
30
40
50
60

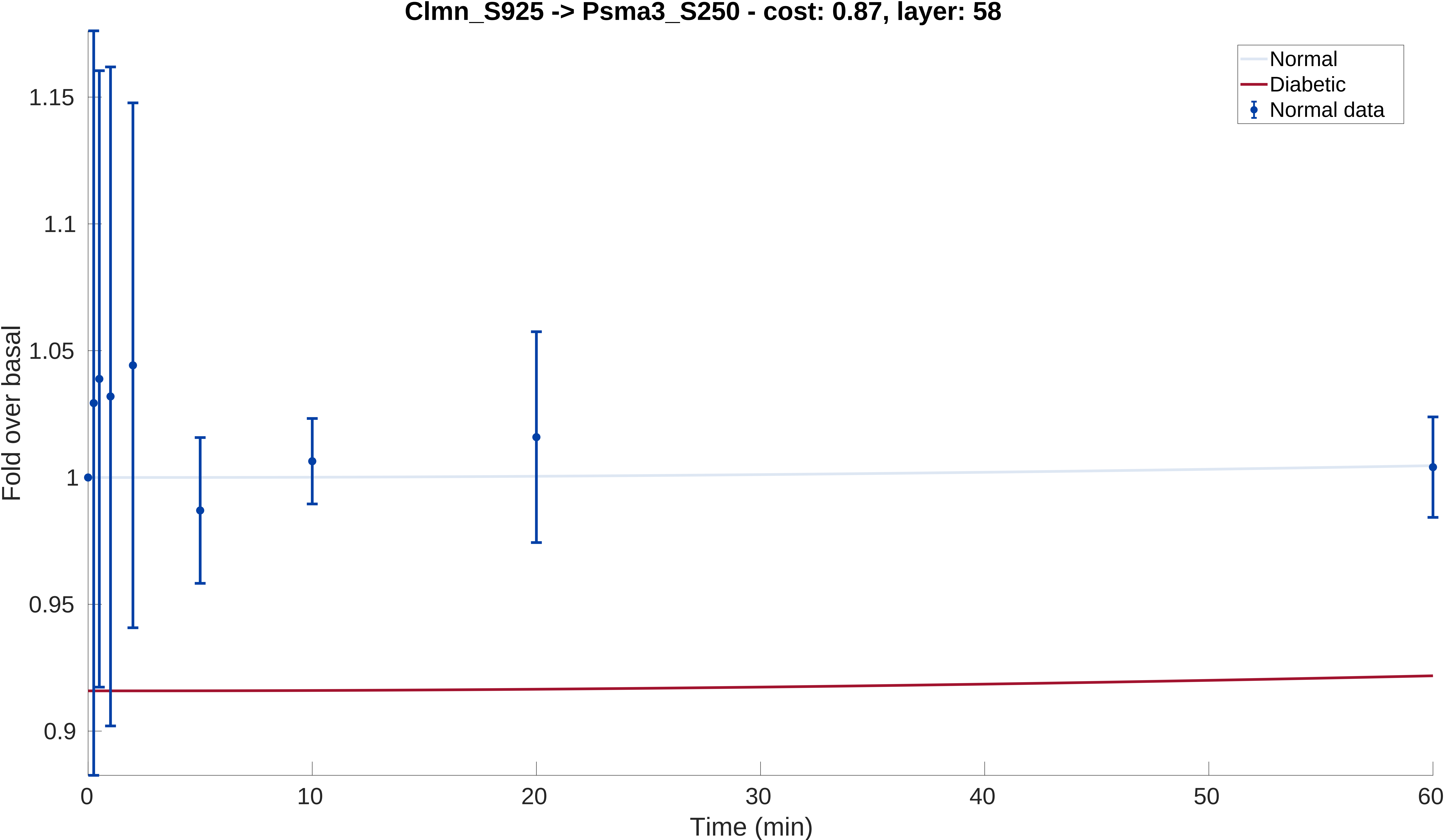
Clmn_S925 -> Psma3_S250 - cost: 0.87, layer: 58
Normal
Diabetic
Normal data
Fold over basal
1.15
1.1
1.05
1
0.95
0.9
0
10
20
30
40
50
60
Time (min)

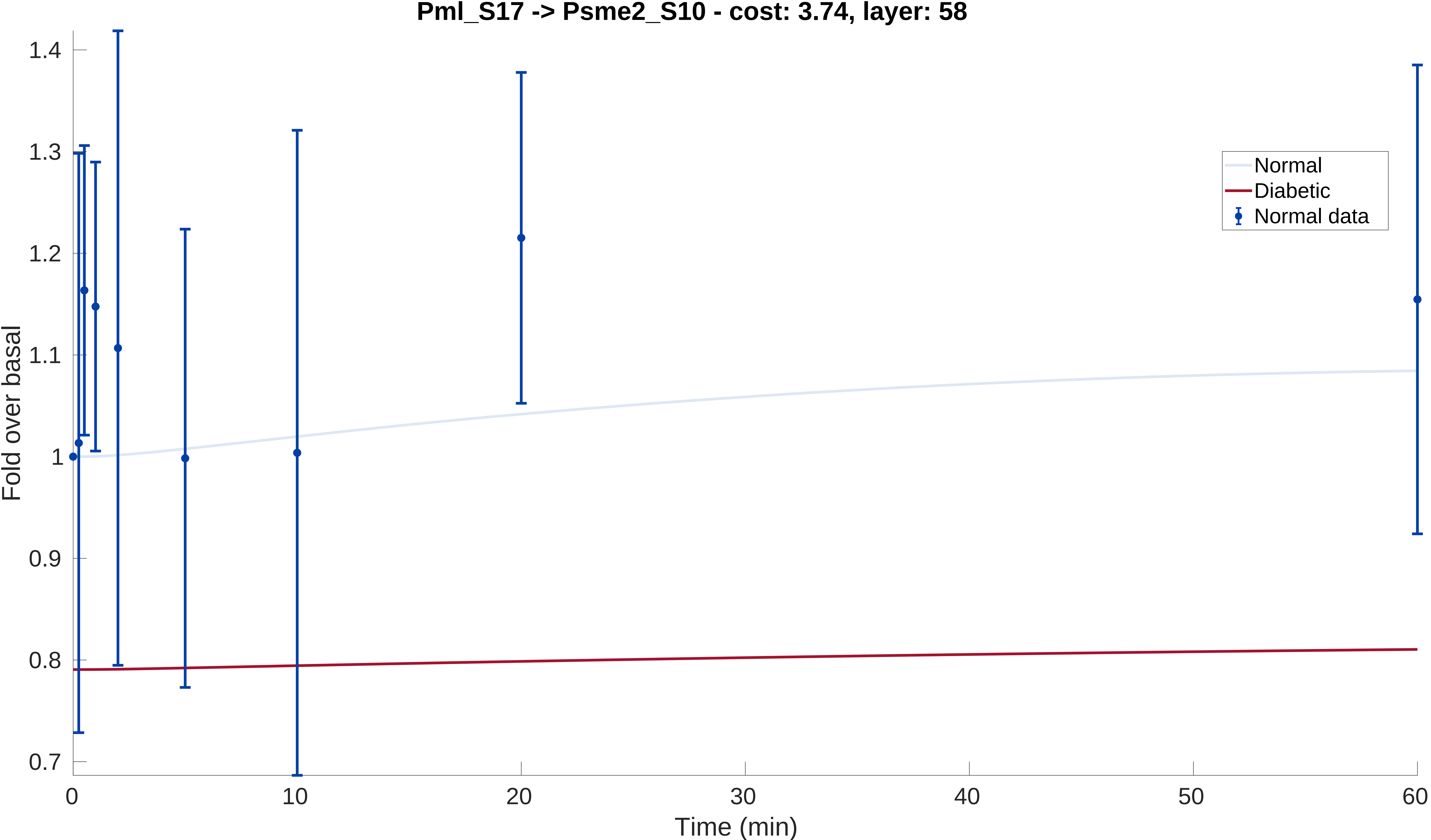

Pml_S17 -> Psme2_S10 - cost: 3.74, layer: 58
1.4
1.3
1.2
1.1
1
0.9
0.8
0.7
Fold over basal
0
10
20
30
40
50
60
Time (min)
Normal
Diabetic
Normal data

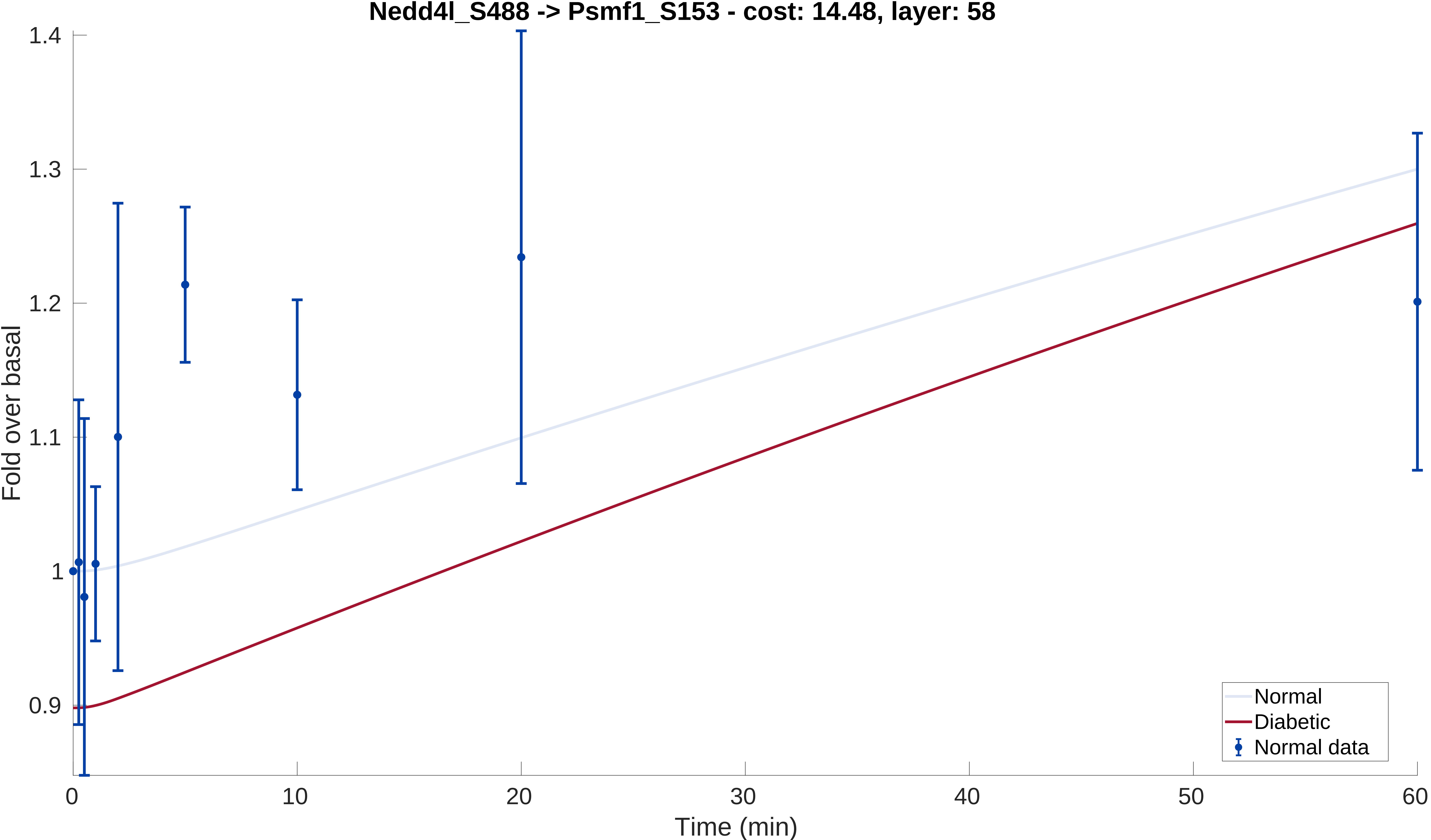
Nedd4l_S488 -> Psmf1_S153 - cost: 14.48, layer: 58
Fold over basal
1.4
1.3
1.2
1.1
1
0.9
0
10
20
30
40
50
60
Time (min)
Normal
Diabetic
Normal data

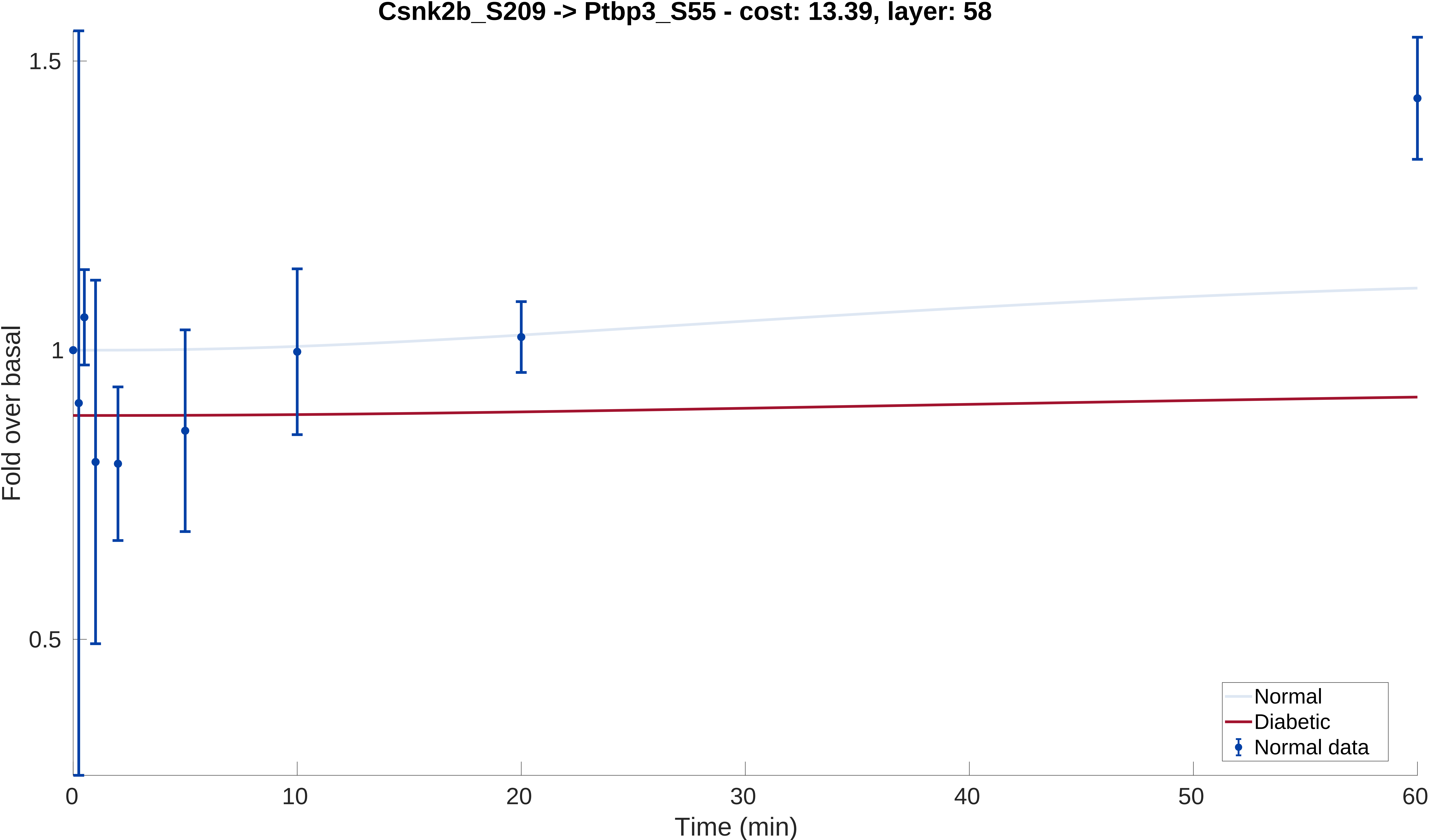

Csnk2b_S209 -> Ptbp3_S55 - cost: 13.39, layer: 58
Fold over basal
1.5
1
0.5
0
10
20
30
40
50
60
Time (min)
Normal
Diabetic
Normal data

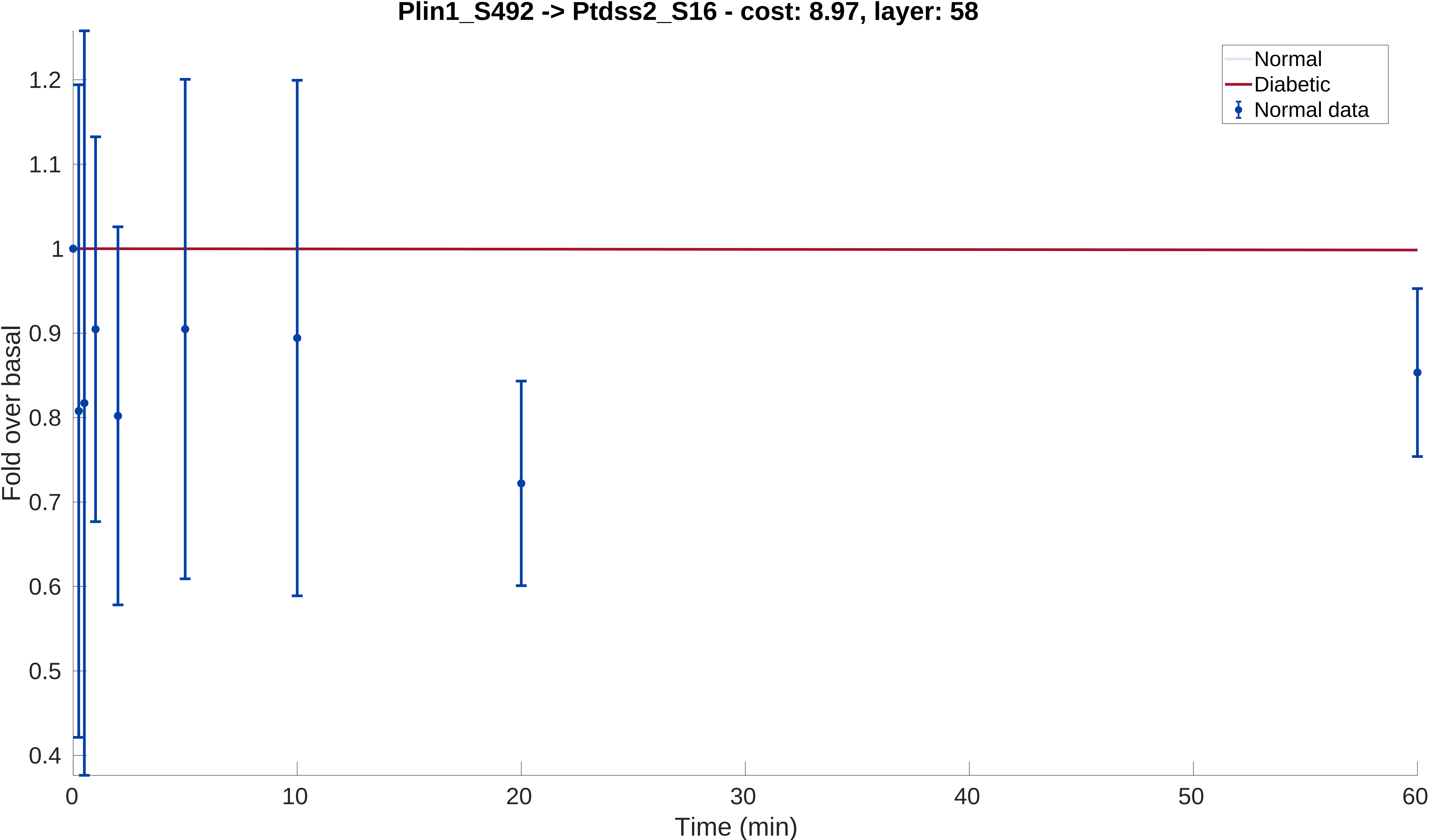

Plin1_S492 -> Ptdss2_S16 - cost: 8.97, layer: 58
Normal
Diabetic
Normal data
Fold over basal
1.2
1.1
1
0.9
0.8
0.7
0.6
0.5
0.4
0
10
20
30
40
50
60
Time (min)

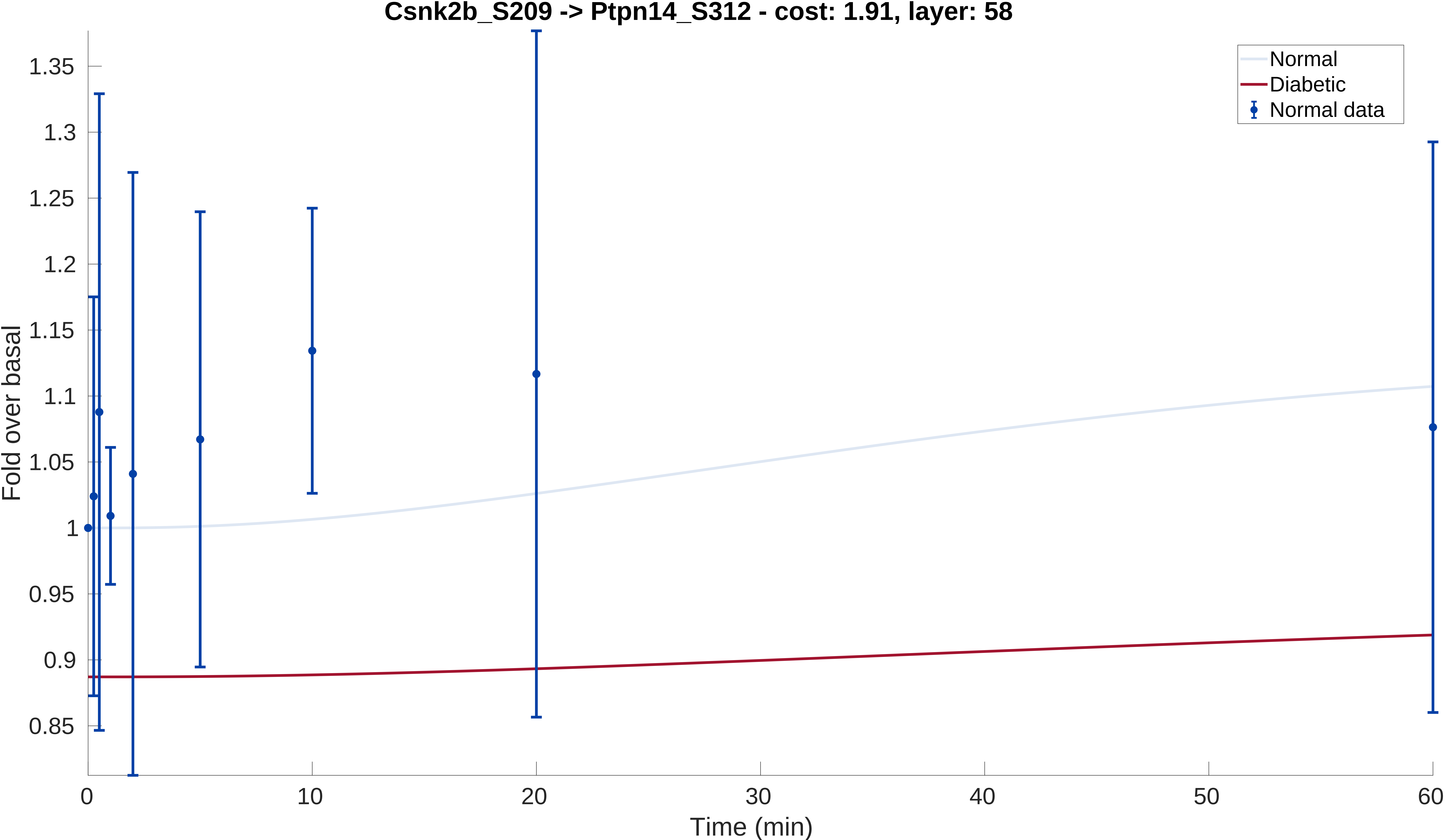

Csnk2b_S209 -> Ptpn14_S312 - cost: 1.91, layer: 58
Normal
Diabetic
Normal data
Fold over basal
1.35
1.3
1.25
1.2
1.15
1.1
1.05
1
0.95
0.9
0.85
0
10
20
30
40
50
60
Time (min)

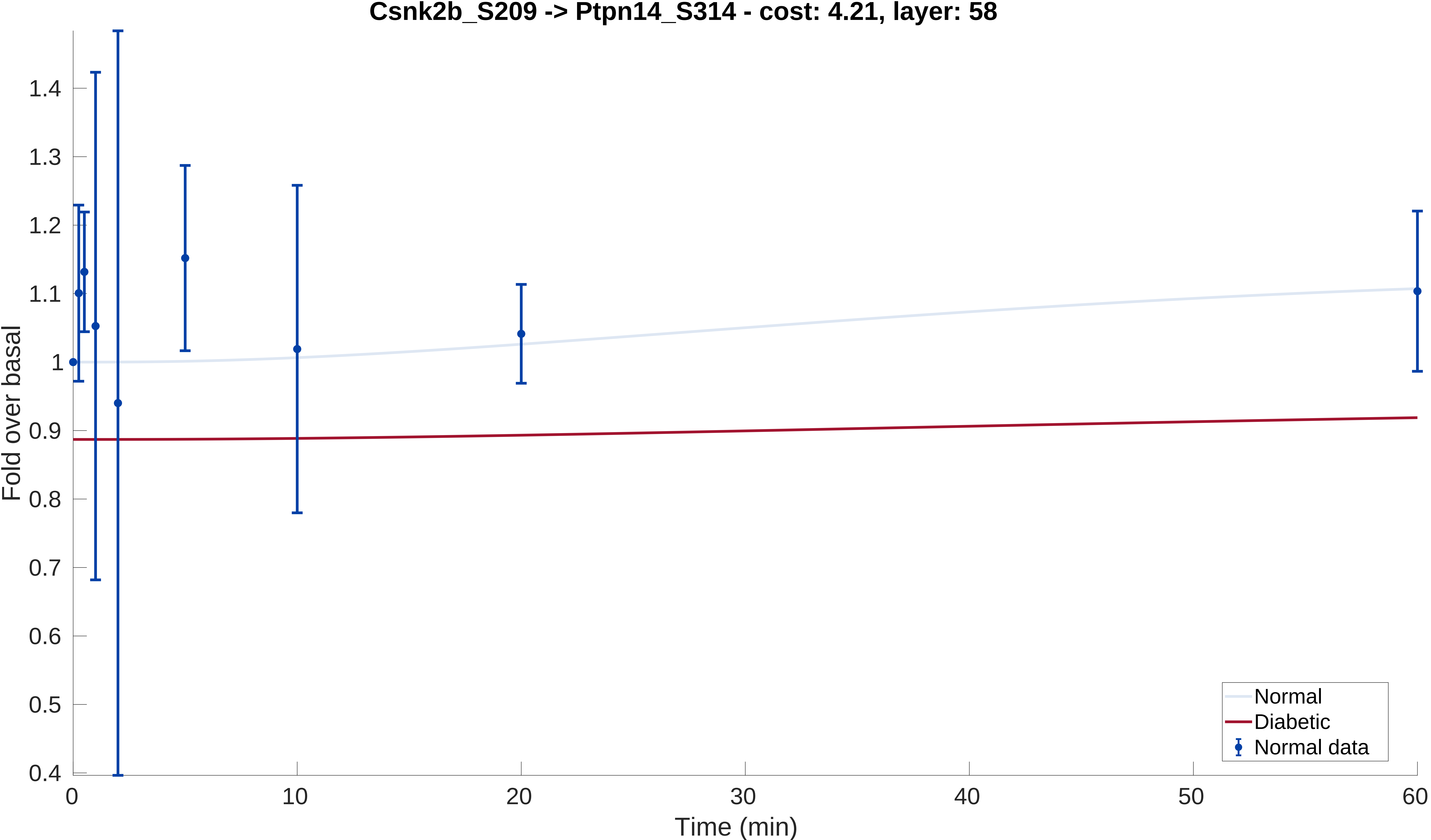

Csnk2b_S209 -> Ptpn14_S314 - cost: 4.21, layer: 58
Fold over basal
1.4
1.3
1.2
1.1
1
0.9
0.8
0.7
0.6
0.5
0.4
0
10
20
30
40
50
60
Time (min)
Normal
Diabetic
Normal data

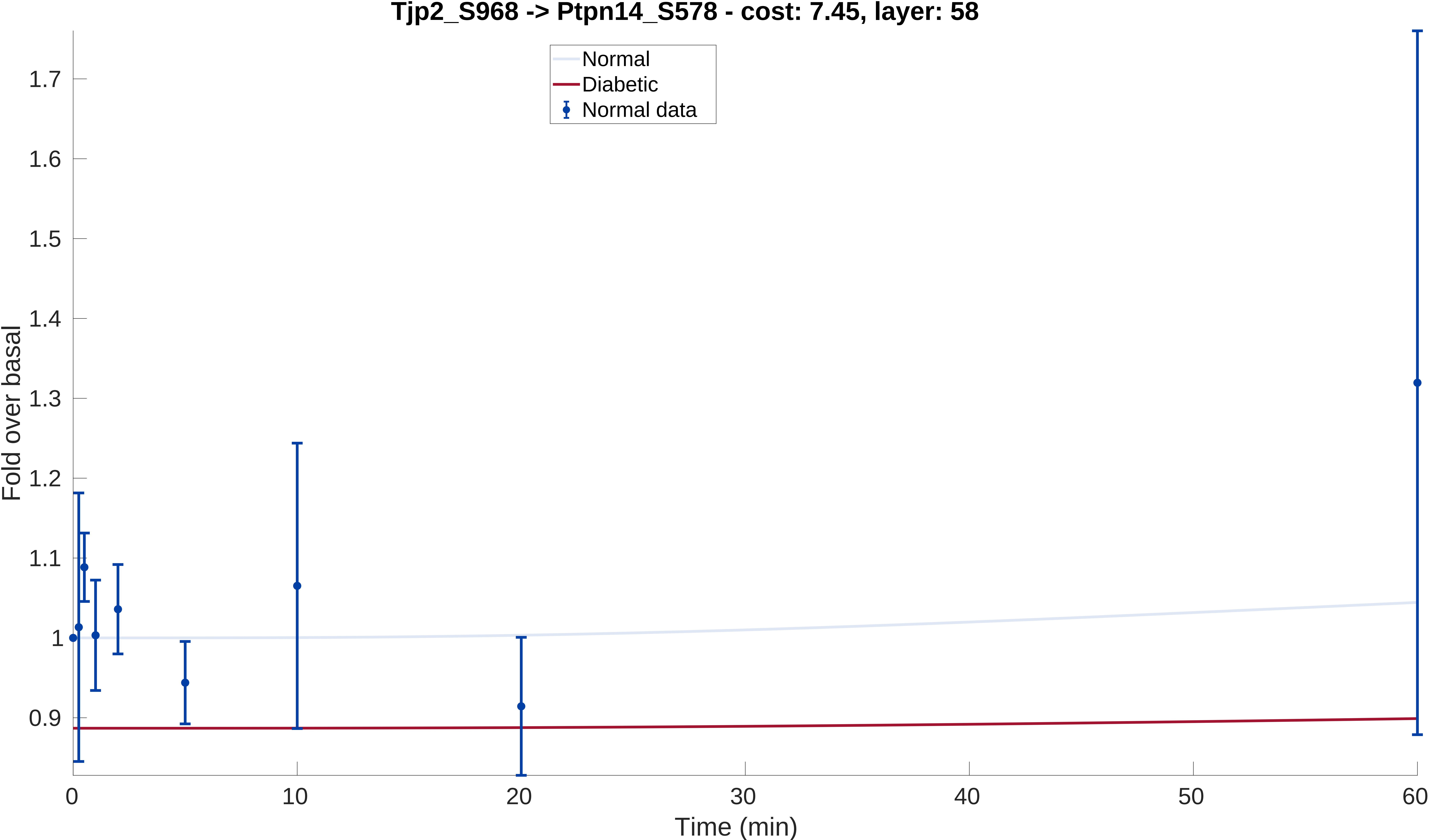

Tjp2_S968 -> Ptpn14_S578 - cost: 7.45, layer: 58
Normal
Diabetic
Normal data
Fold over basal
1.7
1.6
1.5
1.4
1.3
1.2
1.1
1
0.9
0
10
20
30
40
50
60
Time (min)

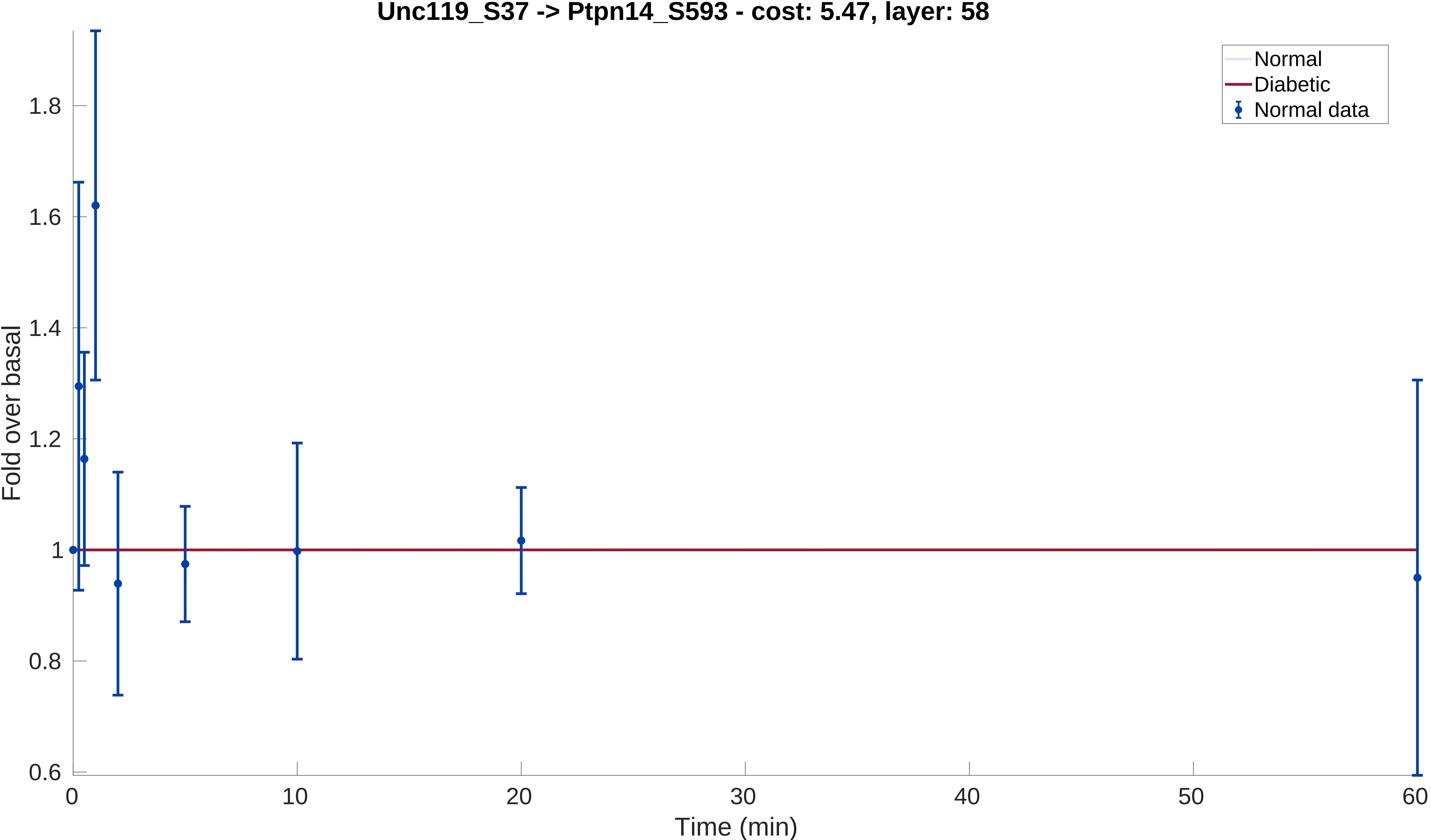

Unc119_S37 -> Ptpn14_S593 - cost: 5.47, layer: 58
Normal
Diabetic
Normal data
Fold over basal
1.8
1.6
1.4
1.2
1
0.8
0.6
0
10
20
30
40
50
60
Time (min)

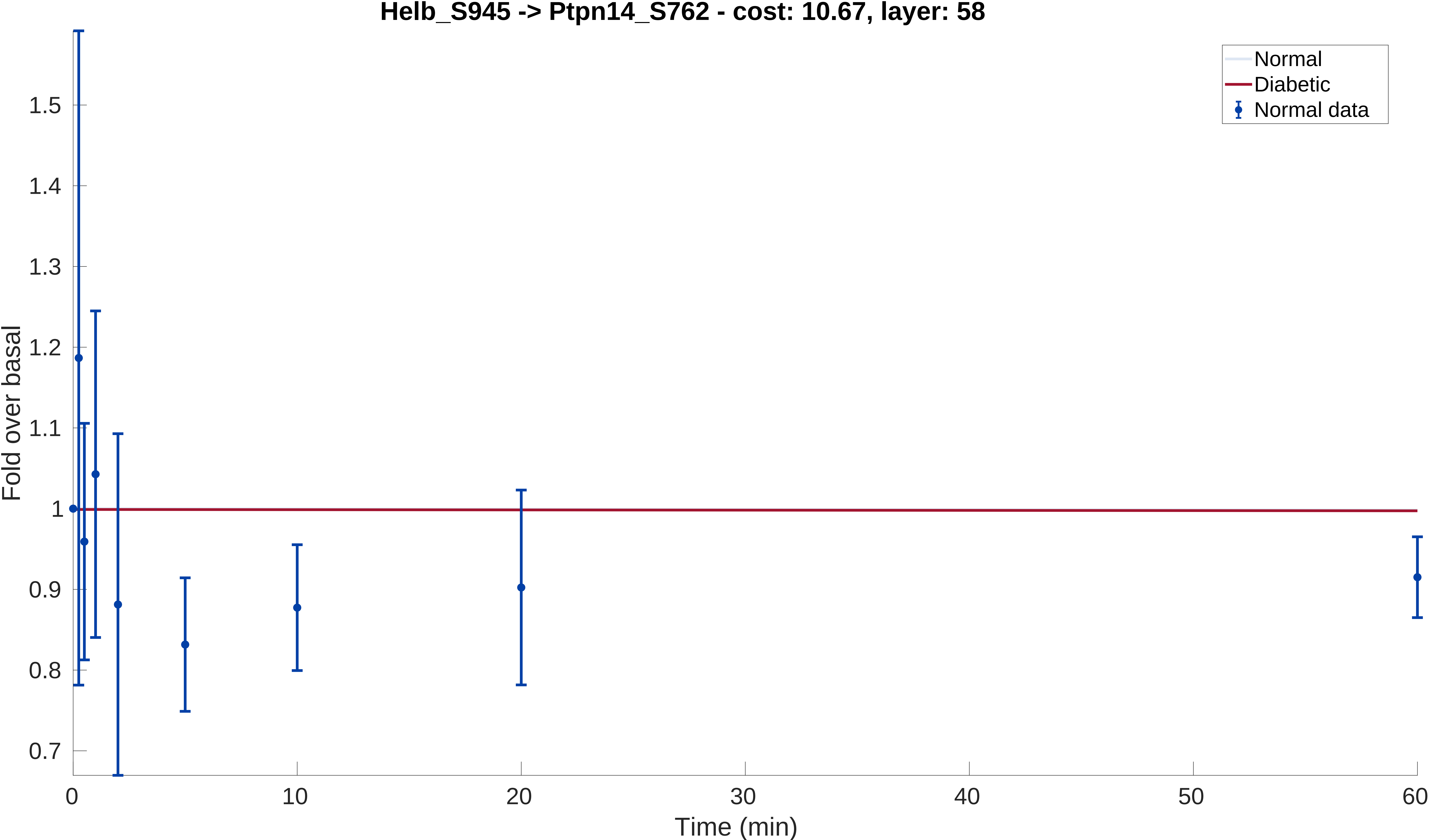

Helb_S945 -> Ptpn14_S762 - cost: 10.67, layer: 58
Normal
Diabetic
Normal data
Fold over basal
Time (min)
0.7
0.8
0.9
1
1.1
1.2
1.3
1.4
1.5
0
10
20
30
40
50
60

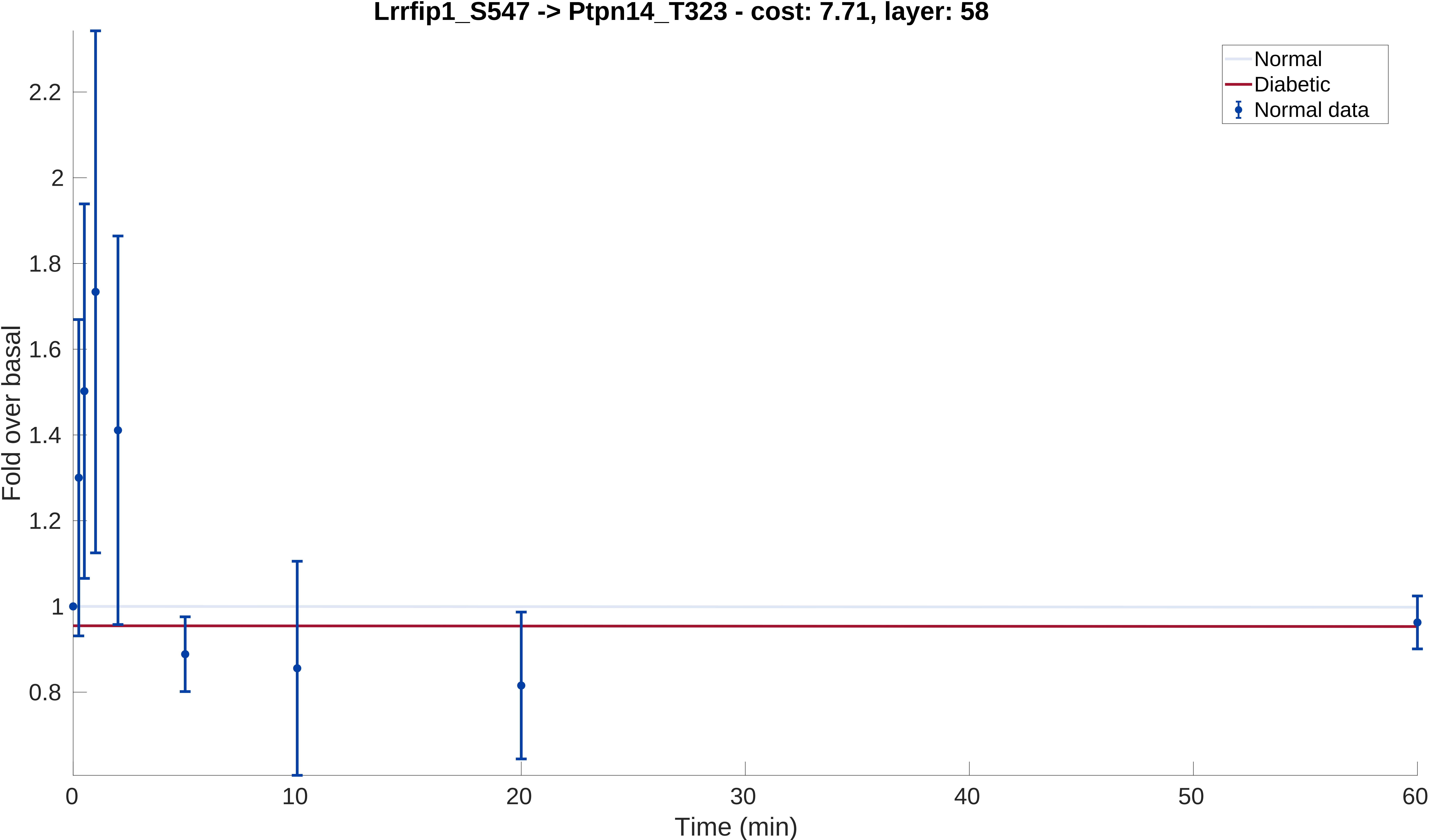

Lrrfip1_S547 -> Ptpn14_T323 - cost: 7.71, layer: 58
Normal
Diabetic
Normal data
Fold over basal
2.2
2
1.8
1.6
1.4
1.2
1
0.8
0
10
20
30
40
50
60
Time (min)

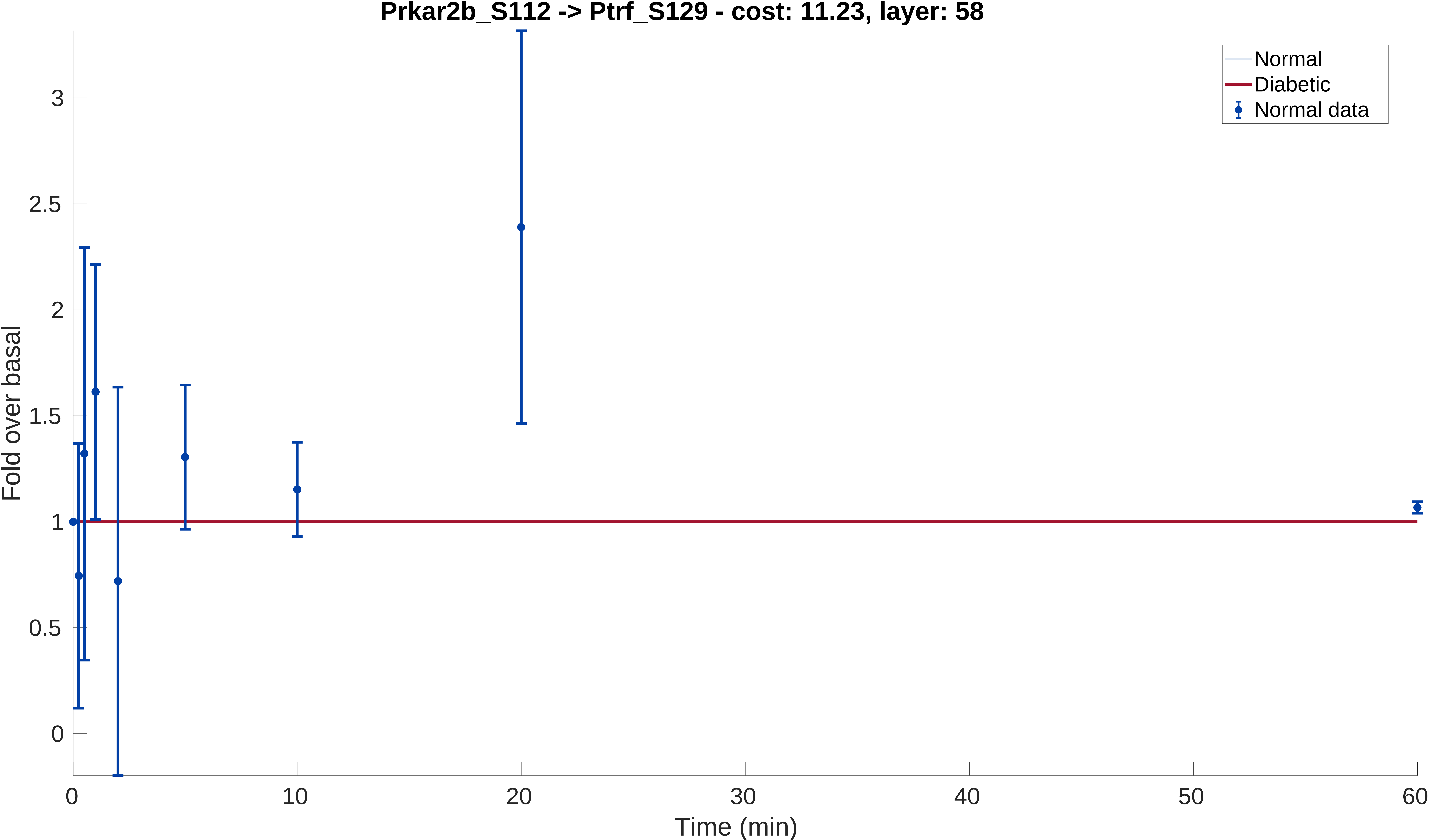

Prrkar2b_S112 -> Ptrf_S129 - cost: 11.23, layer: 58
Normal
Diabetic
Normal data
Fold over basal
Time (min)
0
0.5
1
1.5
2
2.5
3
0
10
20
30
40
50
60

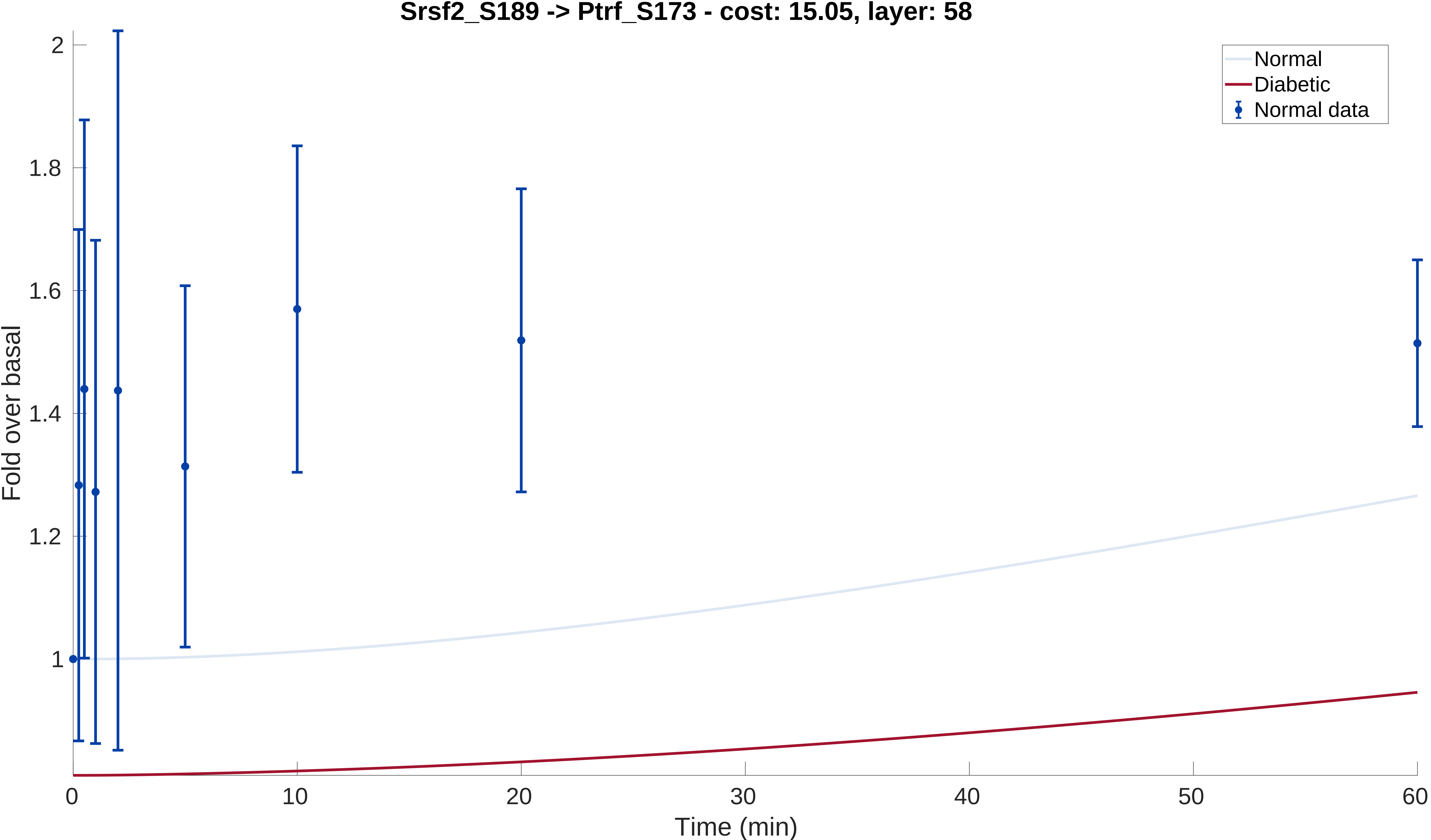

Srsf2_S189 -> Ptrf_S173 - cost: 15.05, layer: 58
Normal
Diabetic
Normal data
Fold over basal
Time (min)
2
1.8
1.6
1.4
1.2
1
0
10
20
30
40
50
60

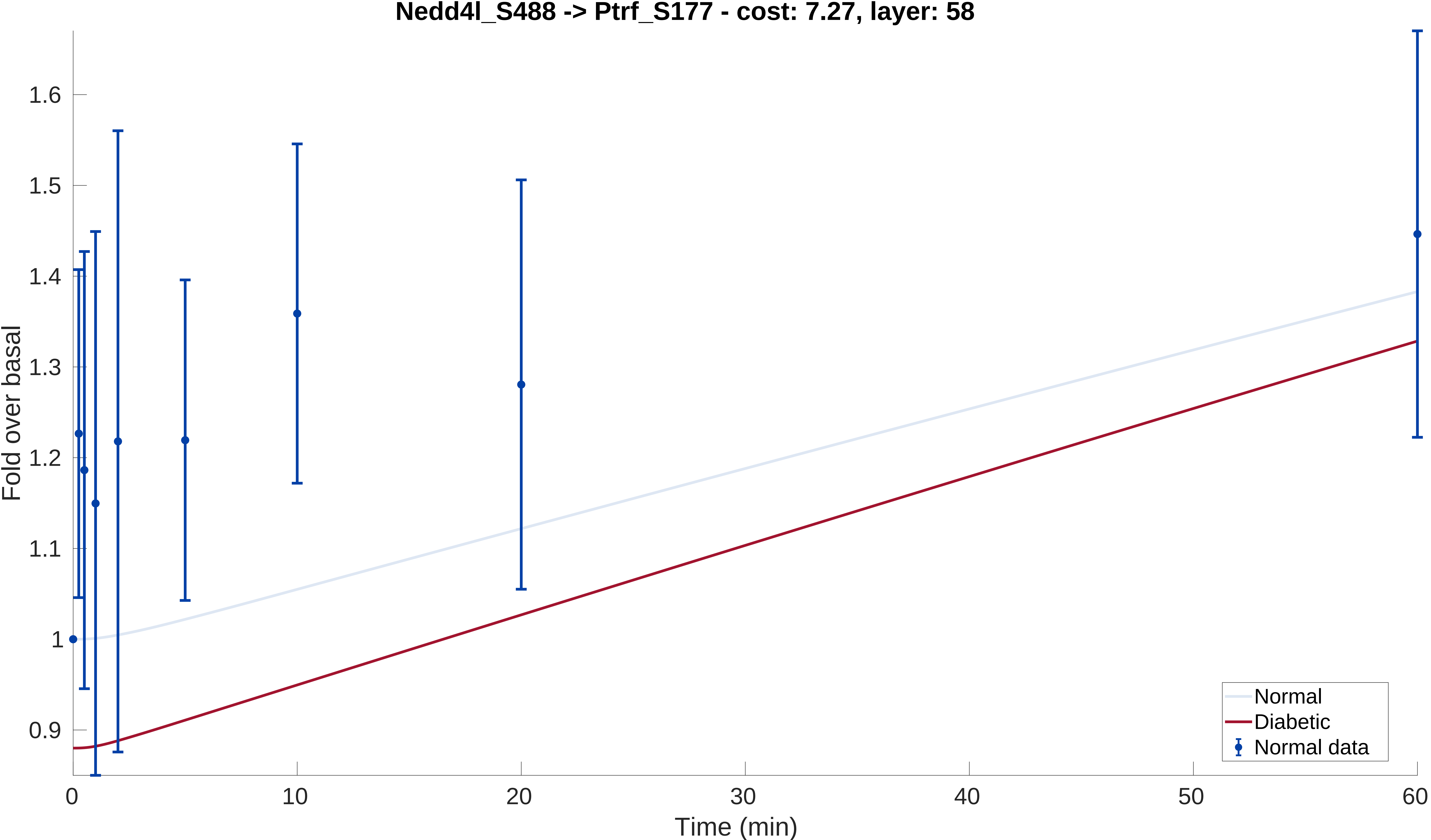
Nedd4l_S488 -> Ptrf_S177 - cost: 7.27, layer: 58
1.6
1.5
1.4
1.3
1.2
1.1
1
0.9
Fold over basal
0
10
20
30
40
50
60
Time (min)
Normal
Diabetic
Normal data

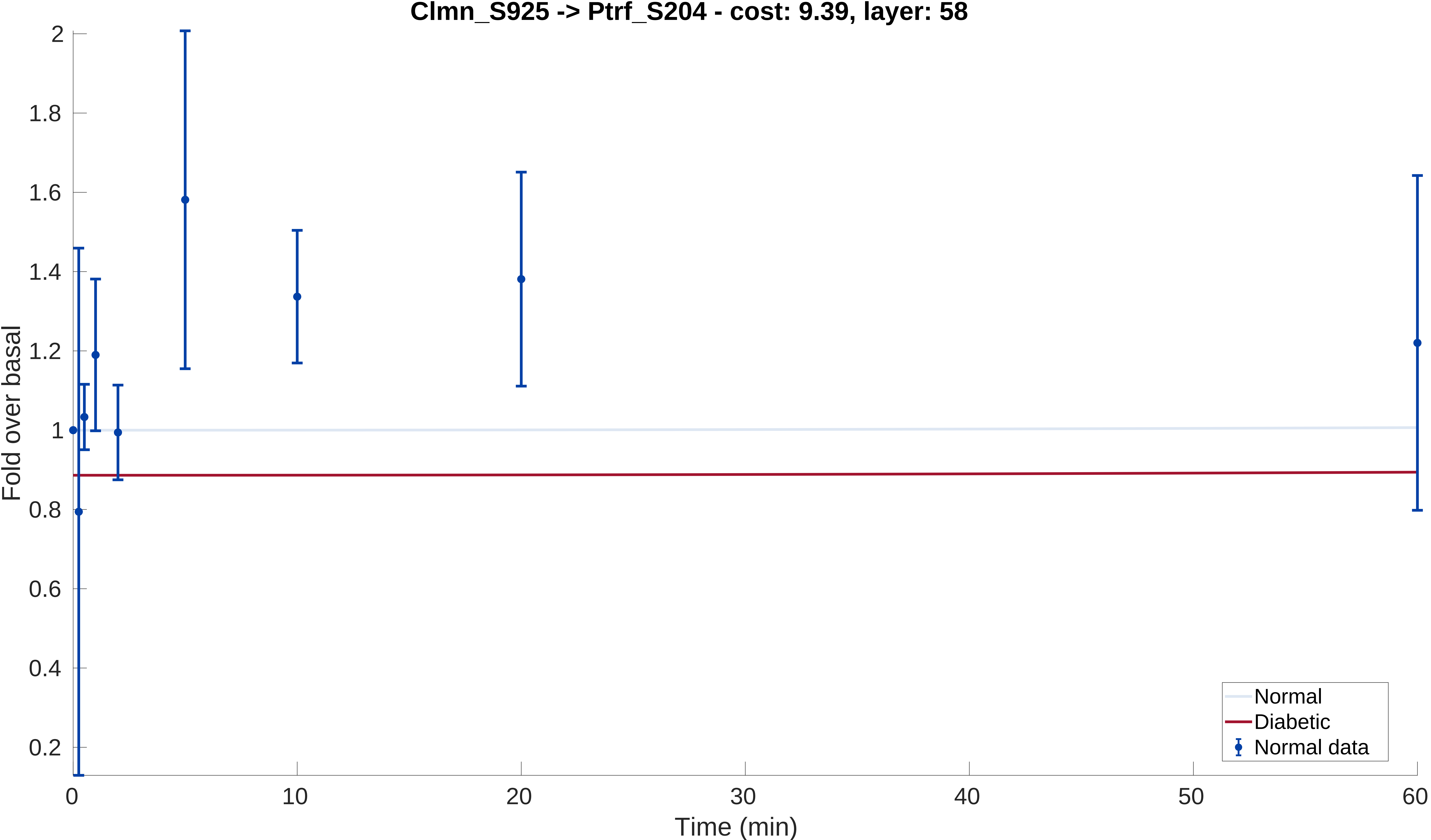
Clmn_S925 -> Ptrf_S204 - cost: 9.39, layer: 58
Fold over basal
Time (min)
2
1.8
1.6
1.4
1.2
1
0.8
0.6
0.4
0.2
0
10
20
30
40
50
60
Normal
Diabetic
Normal data

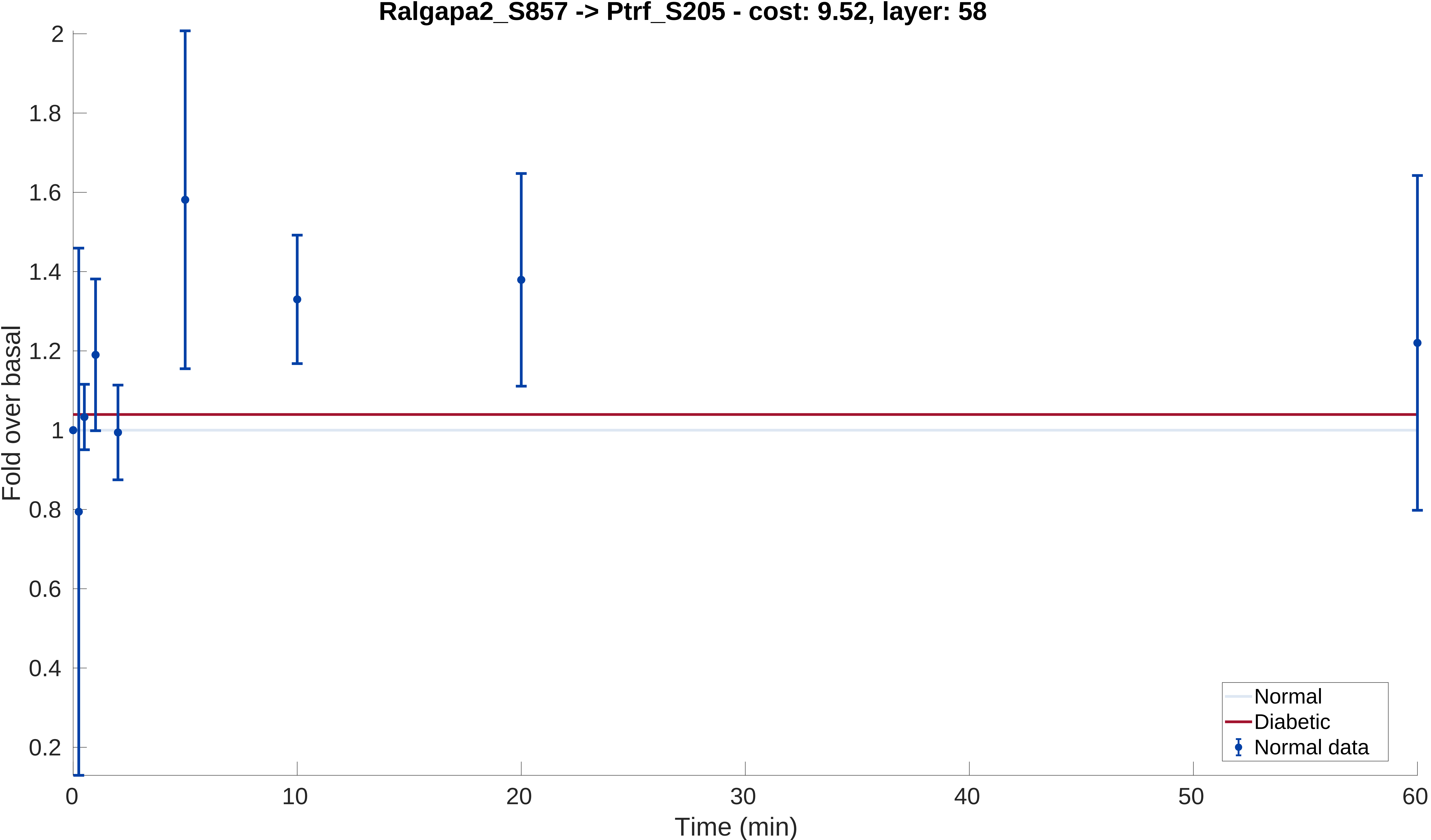

Ralgapa2_S857 -> Ptrf_S205 - cost: 9.52, layer: 58
Fold over basal
2
1.8
1.6
1.4
1.2
1
0.8
0.6
0.4
0.2
0
10
20
30
40
50
60
Time (min)
Normal
Diabetic
Normal data

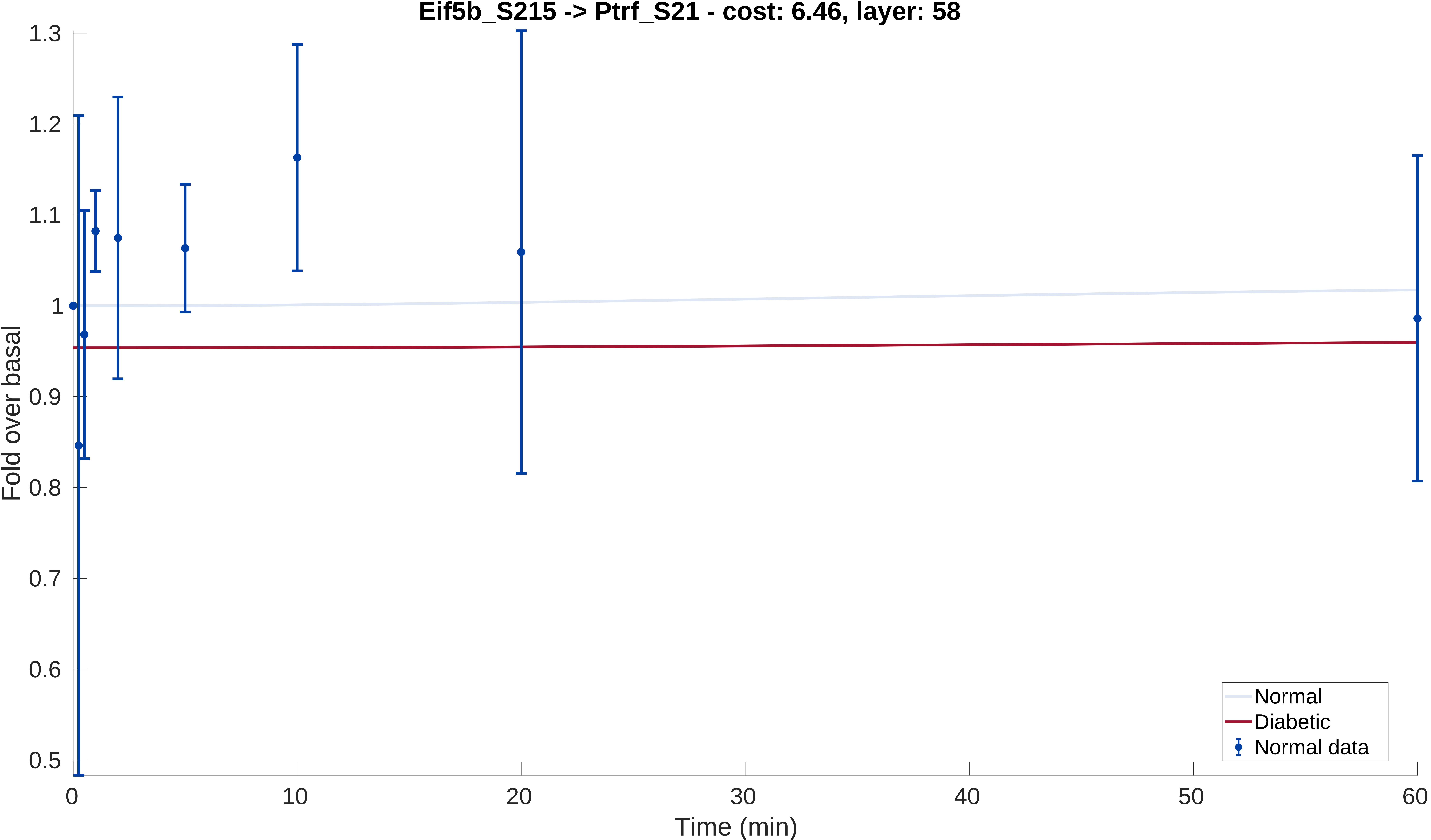
Eif5b_S215 -> Ptrf_S21 - cost: 6.46, layer: 58
Fold over basal
Time (min)
Normal
Diabetic
Normal data
1.3
1.2
1.1
1
0.9
0.8
0.7
0.6
0.5
0
10
20
30
40
50
60

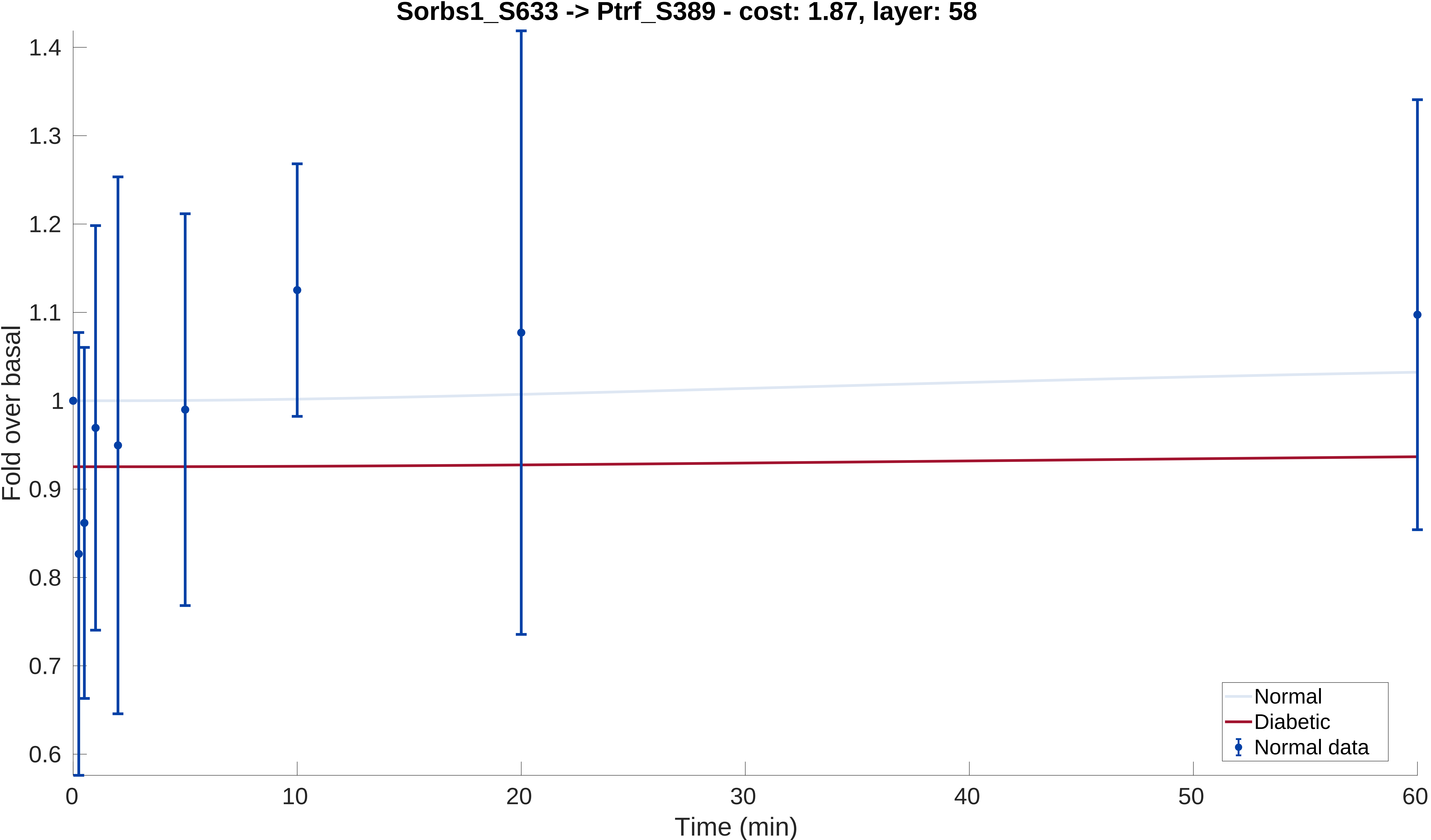

Sorbs1_S633 -> Ptrf_S389 - cost: 1.87, layer: 58
Fold over basal
1.4
1.3
1.2
1.1
1
0.9
0.8
0.7
0.6
0
10
20
30
40
50
60
Time (min)
Normal
Diabetic
Normal data

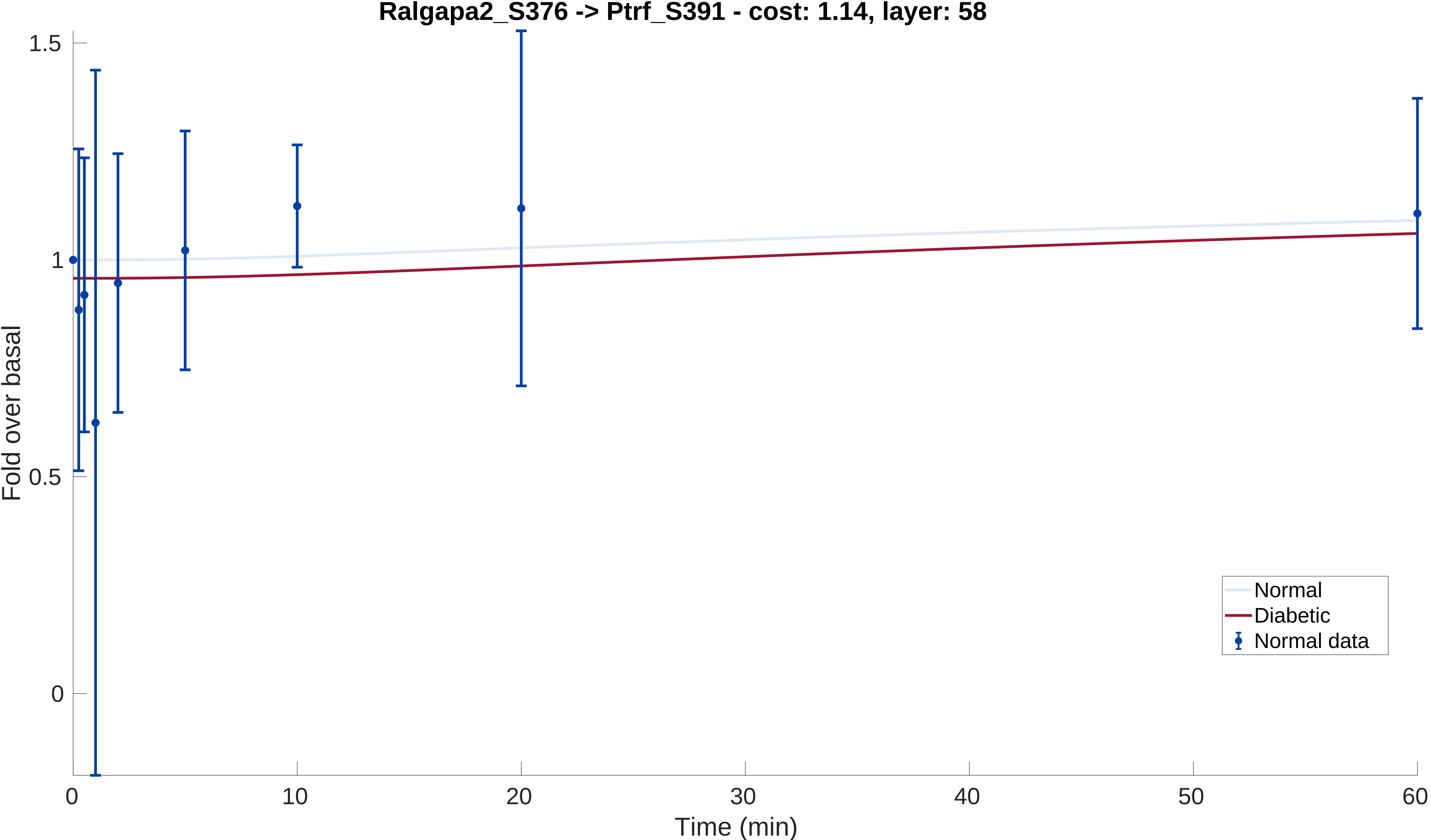

Ralgapa2_S376 -> Ptrf_S391 - cost: 1.14, layer: 58
Fold over basal
Time (min)
1.5
1
0.5
0
0
10
20
30
40
50
60
Normal
Diabetic
Normal data

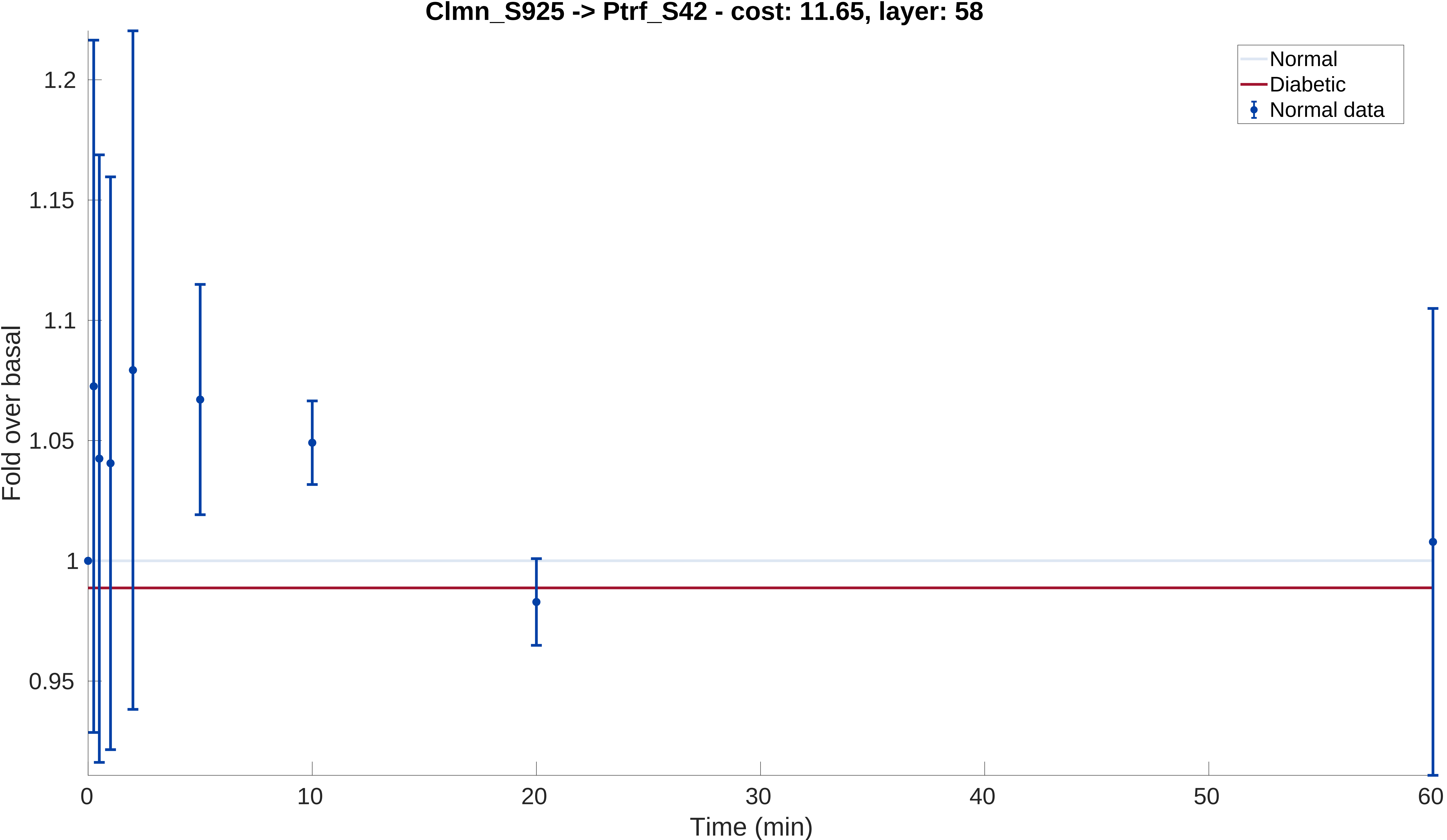

Clmn_S925 -> Ptrf_S42 - cost: 11.65, layer: 58
Normal
Diabetic
Normal data
Fold over basal
1.2
1.15
1.1
1.05
1
0.95
0
10
20
30
40
50
60
Time (min)

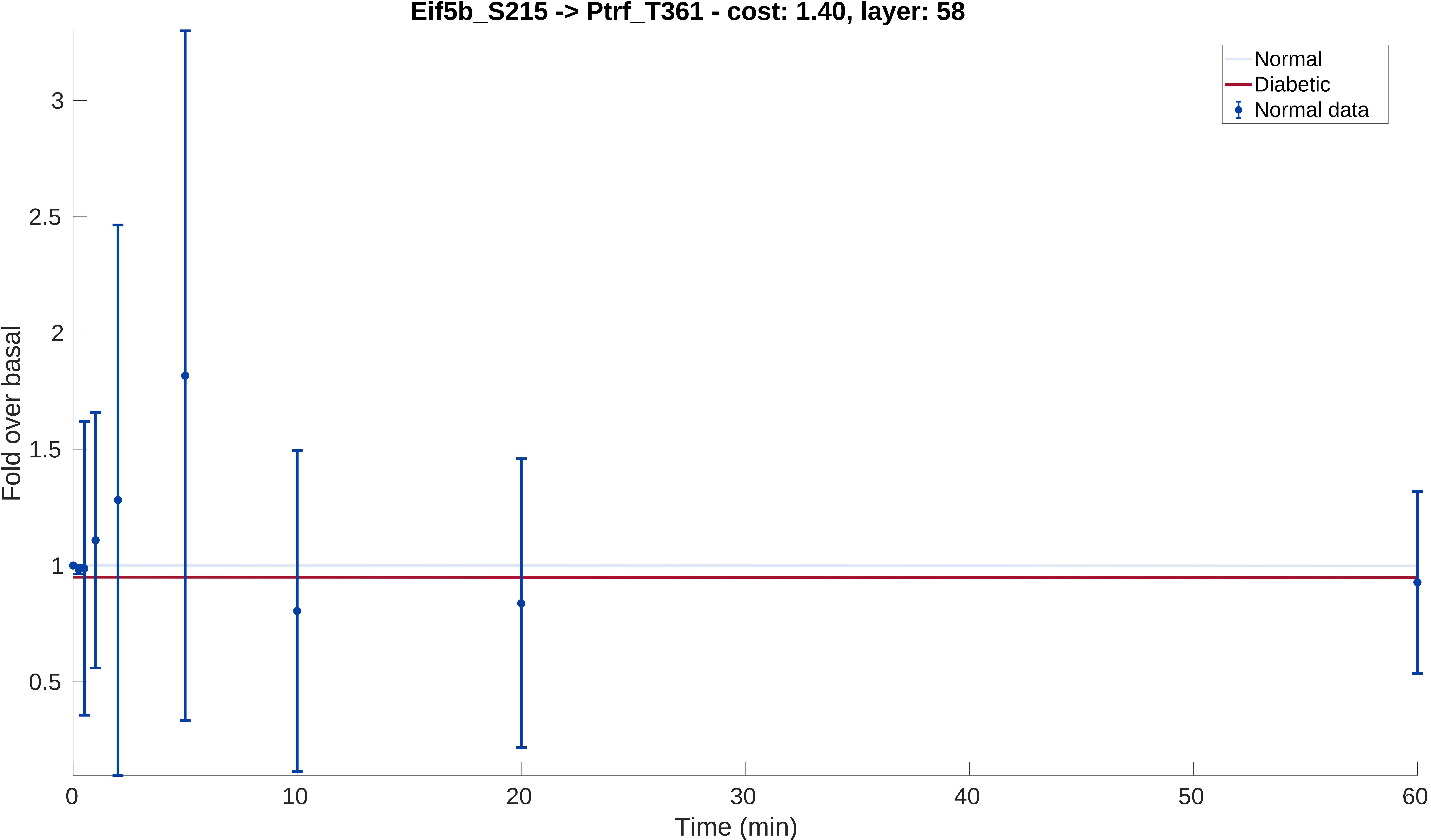

Eif5b_S215 -> Ptrf_T361 - cost: 1.40, layer: 58
Normal
Diabetic
Normal data
Fold over basal
3
2.5
2
1.5
1
0.5
0
10
20
30
40
50
60
Time (min)

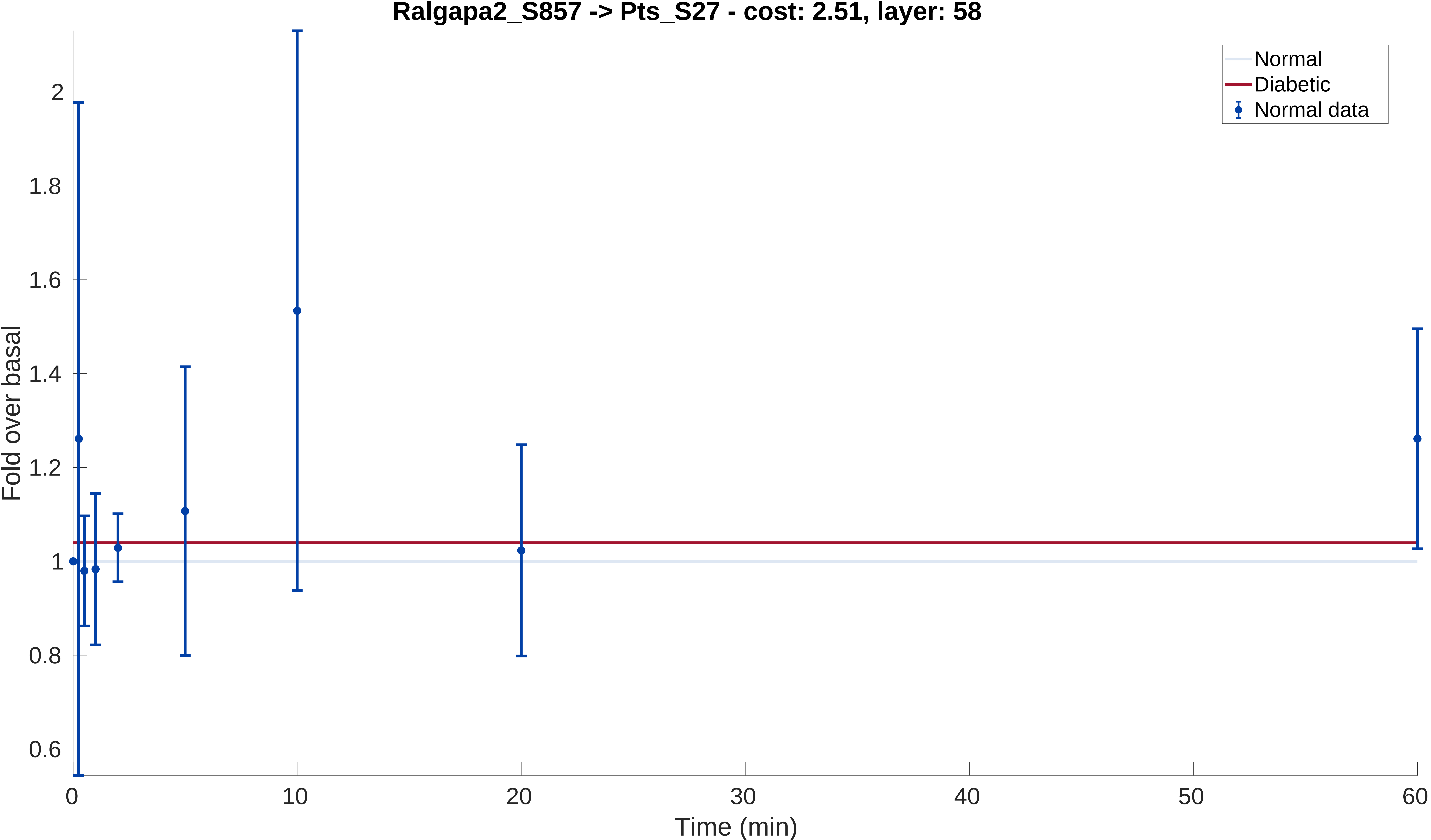

Ralgapa2_S857 -> Pts_S27 - cost: 2.51, layer: 58
Normal
Diabetic
Normal data
Fold over basal
2
1.8
1.6
1.4
1.2
1
0.8
0.6
0
10
20
30
40
50
60
Time (min)

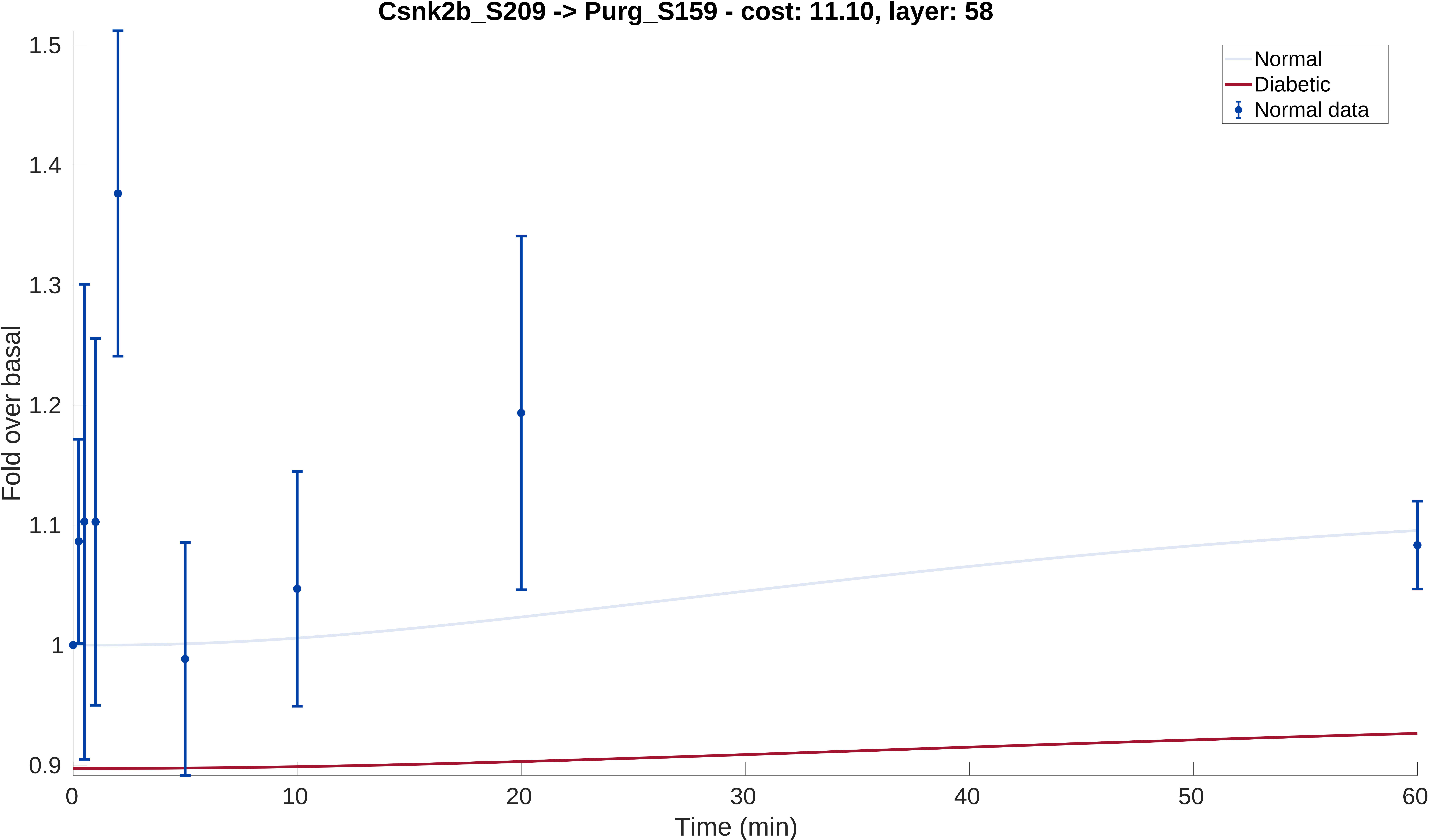

Csnk2b_S209 -> Purg_S159 - cost: 11.10, layer: 58
Fold over basal
Time (min)
Normal
Diabetic
Normal data
1.5
1.4
1.3
1.2
1.1
1
0.9
0
10
20
30
40
50
60

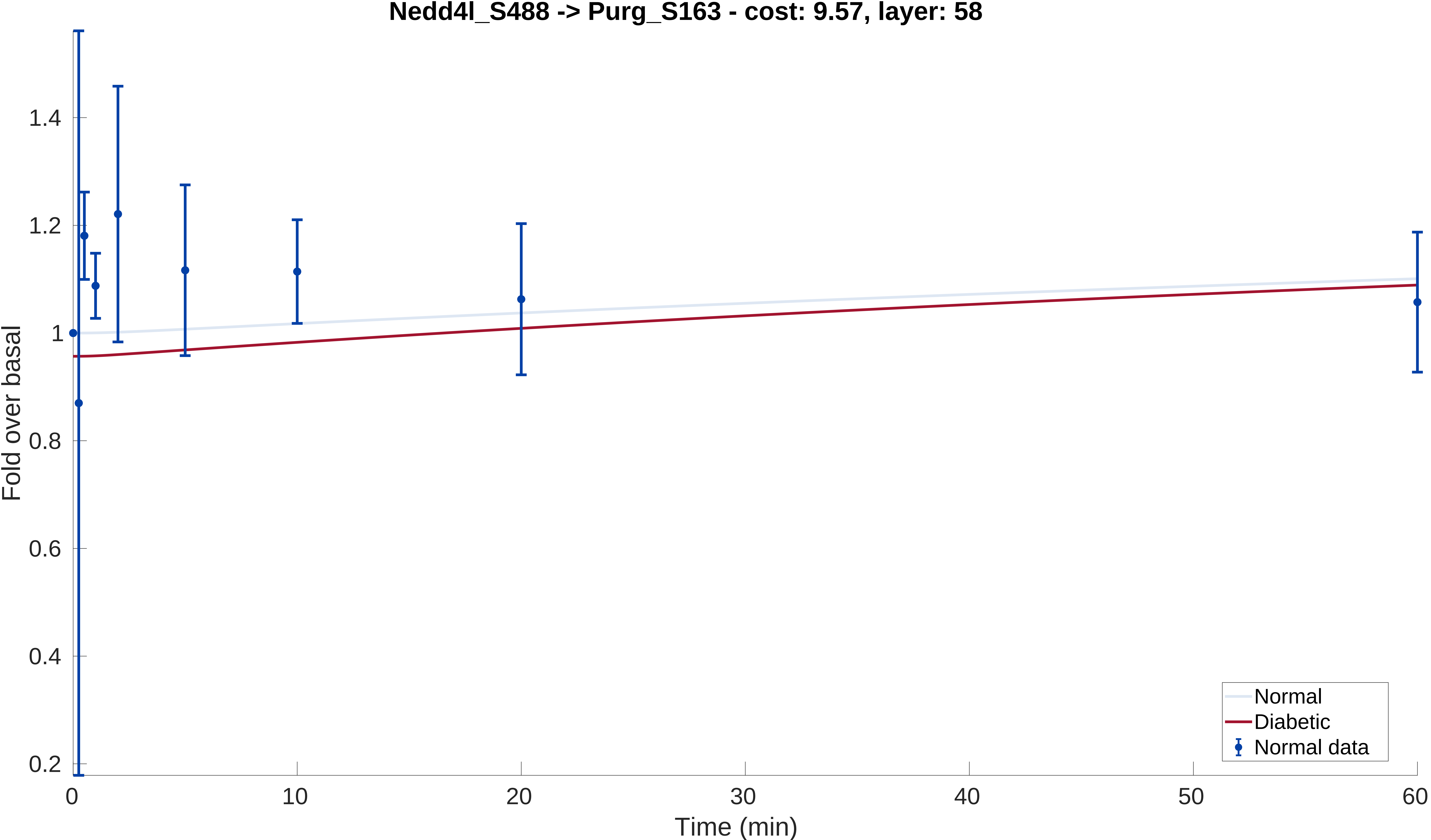

Nedd4l_S488 -> Purg_S163 - cost: 9.57, layer: 58
Fold over basal
1.4
1.2
1
0.8
0.6
0.4
0.2
0
10
20
30
40
50
60
Time (min)
Normal
Diabetic
Normal data

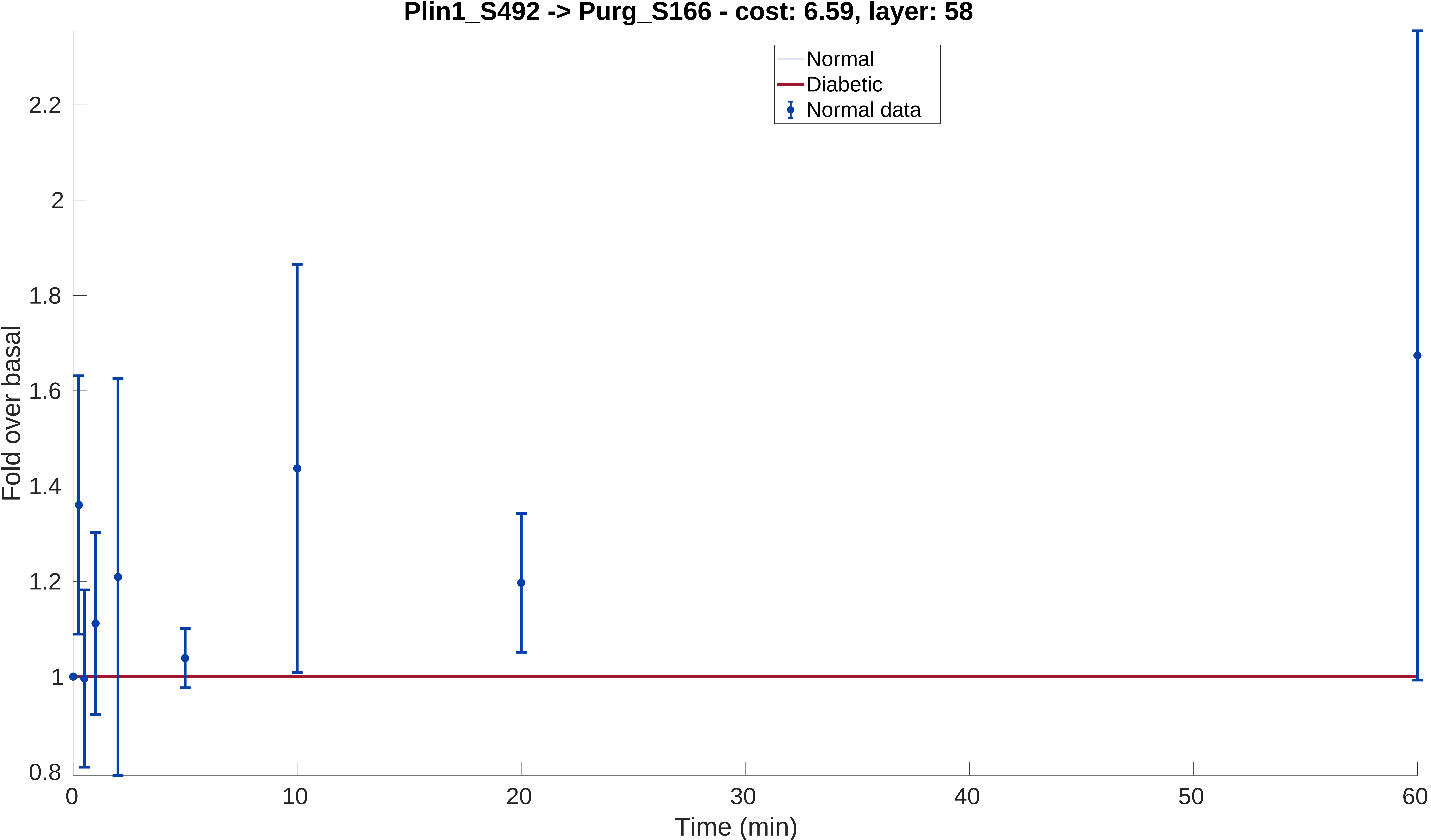
Plin1_S492 -> Purg_S166 - cost: 6.59, layer: 58
Normal
Diabetic
Normal data
Fold over basal
Time (min)
0.8
1
1.2
1.4
1.6
1.8
2
2.2
0
10
20
30
40
50
60

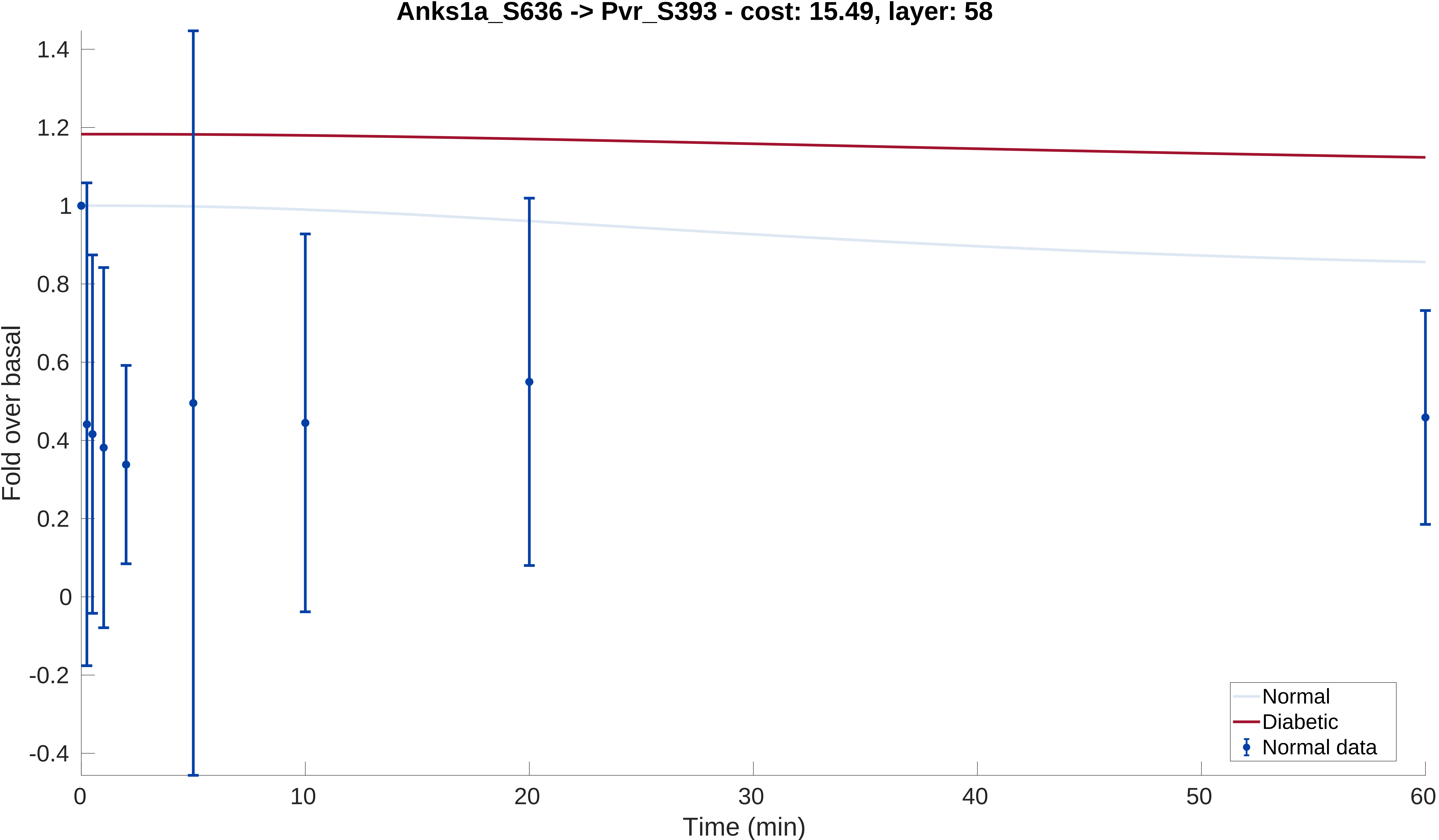

Anks1a_S636 -> Pvr_S393 - cost: 15.49, layer: 58
1.4
1.2
1
0.8
0.6
0.4
0.2
0
-0.2
-0.4
Fold over basal
0
10
20
30
40
50
60
Time (min)
Normal
Diabetic
Normal data

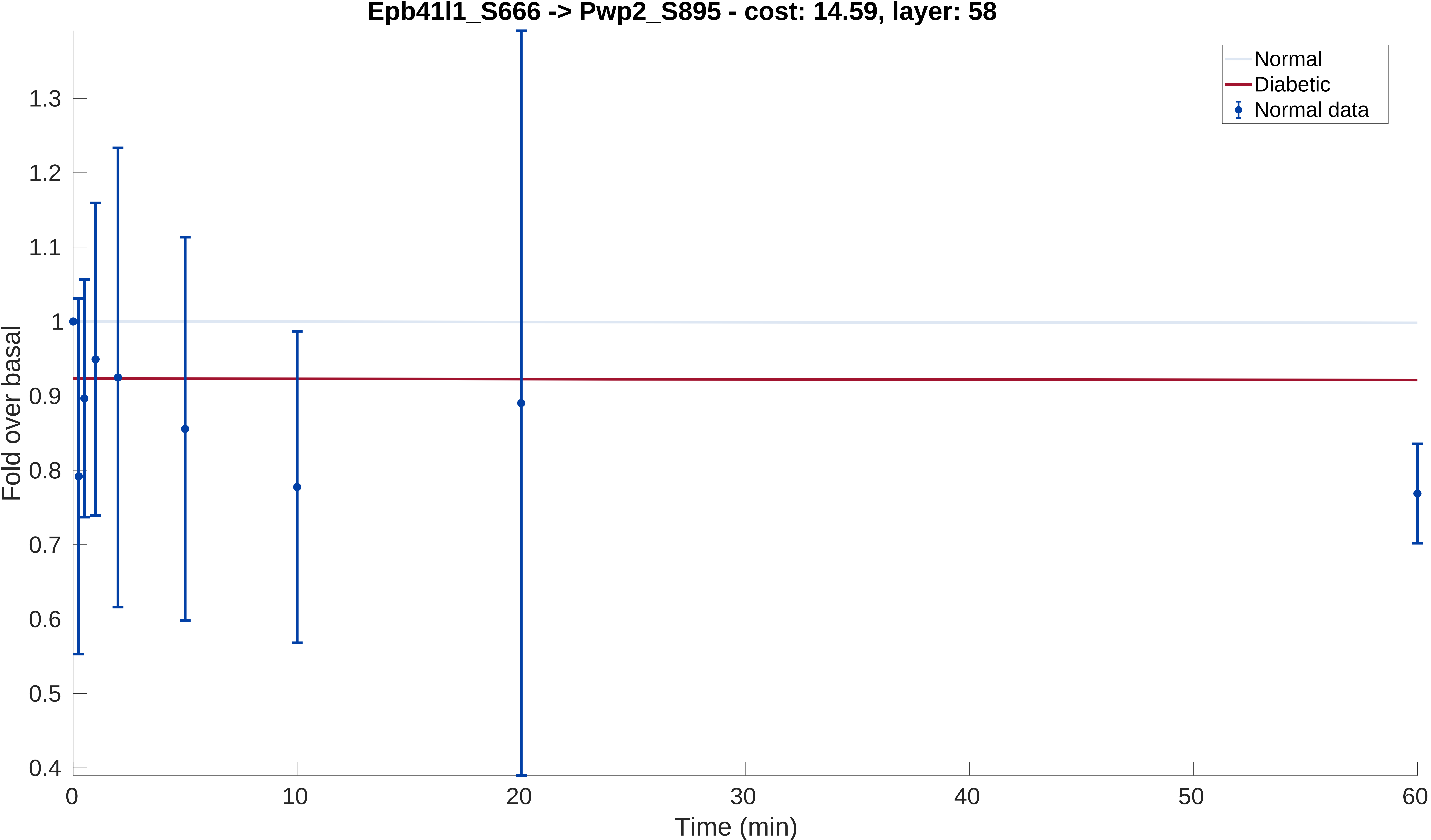

Epb41l1_S666 -> Pwp2_S895 - cost: 14.59, layer: 58
Normal
Diabetic
Normal data
1.3
1.2
1.1
1
0.9
0.8
0.7
0.6
0.5
0.4
Fold over basal
0
10
20
30
40
50
60
Time (min)

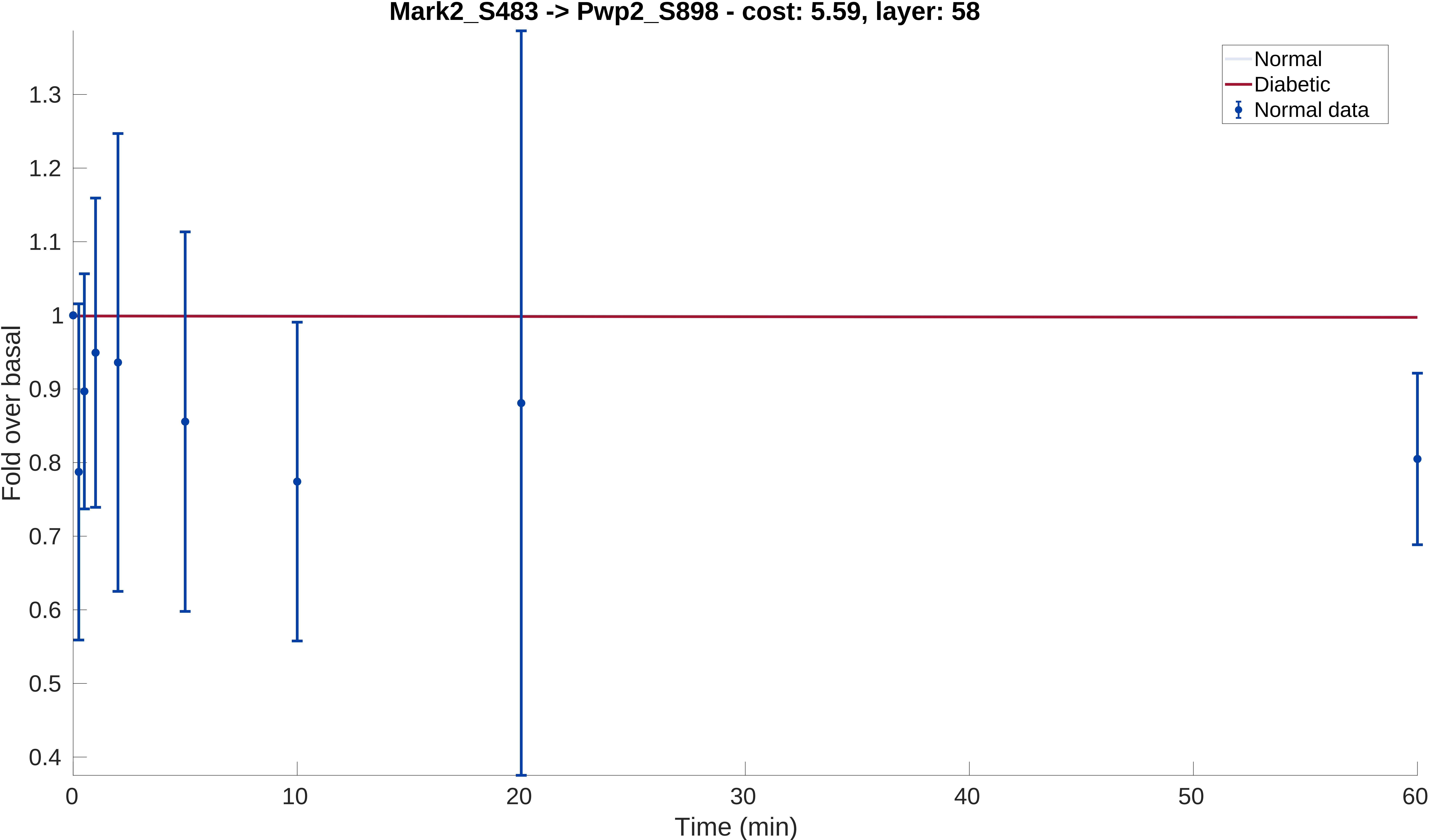

Mark2_S483 -> Pwp2_S898 - cost: 5.59, layer: 58
Normal
Diabetic
Normal data
Fold over basal
1.3
1.2
1.1
1
0.9
0.8
0.7
0.6
0.5
0.4
0
10
20
30
40
50
60
Time (min)

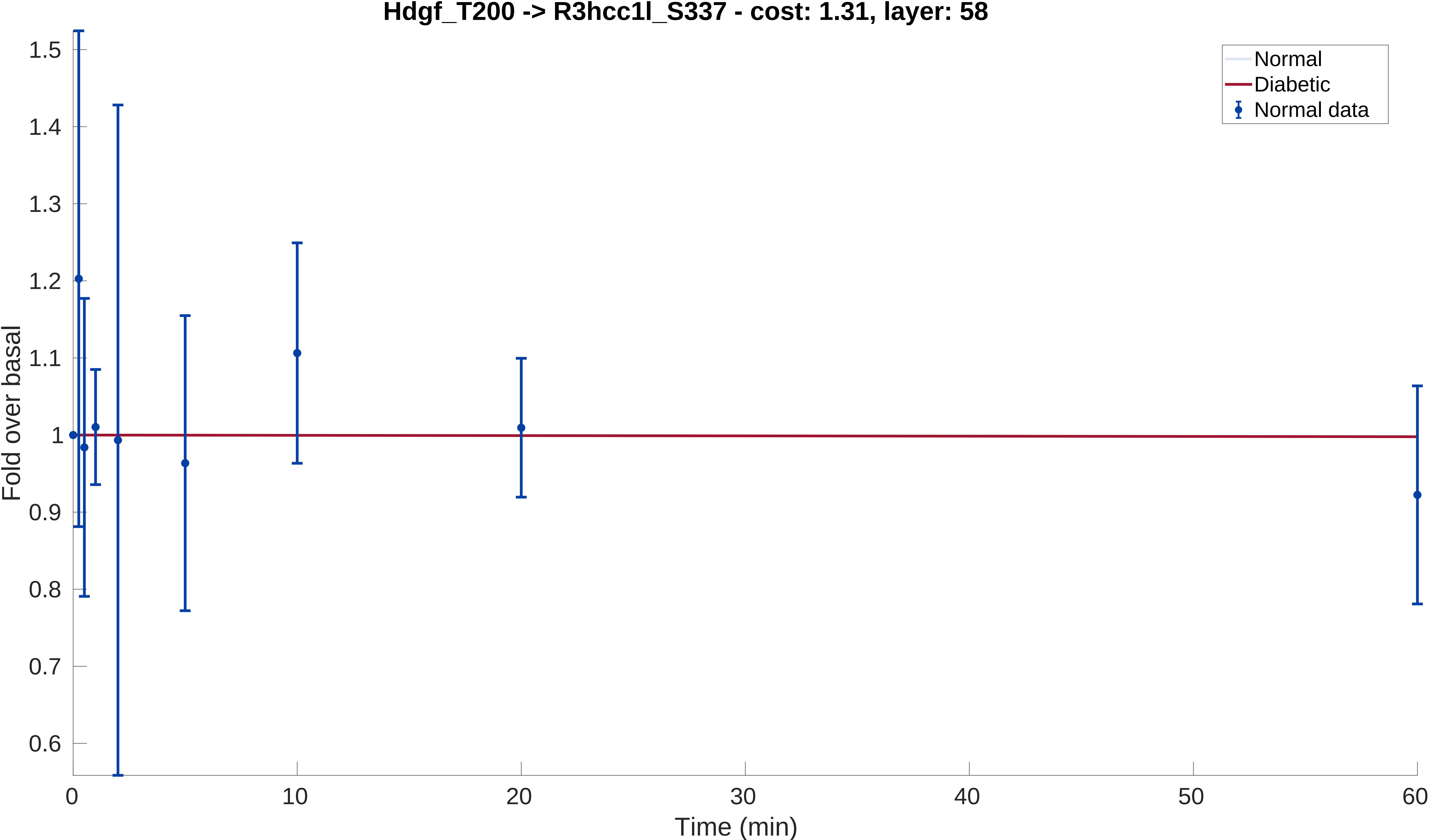
Hdgf_T200 -> R3hcc1l_S337 - cost: 1.31, layer: 58
Normal
Diabetic
Normal data
Fold over basal
1.5
1.4
1.3
1.2
1.1
1
0.9
0.8
0.7
0.6
0
10
20
30
40
50
60
Time (min)

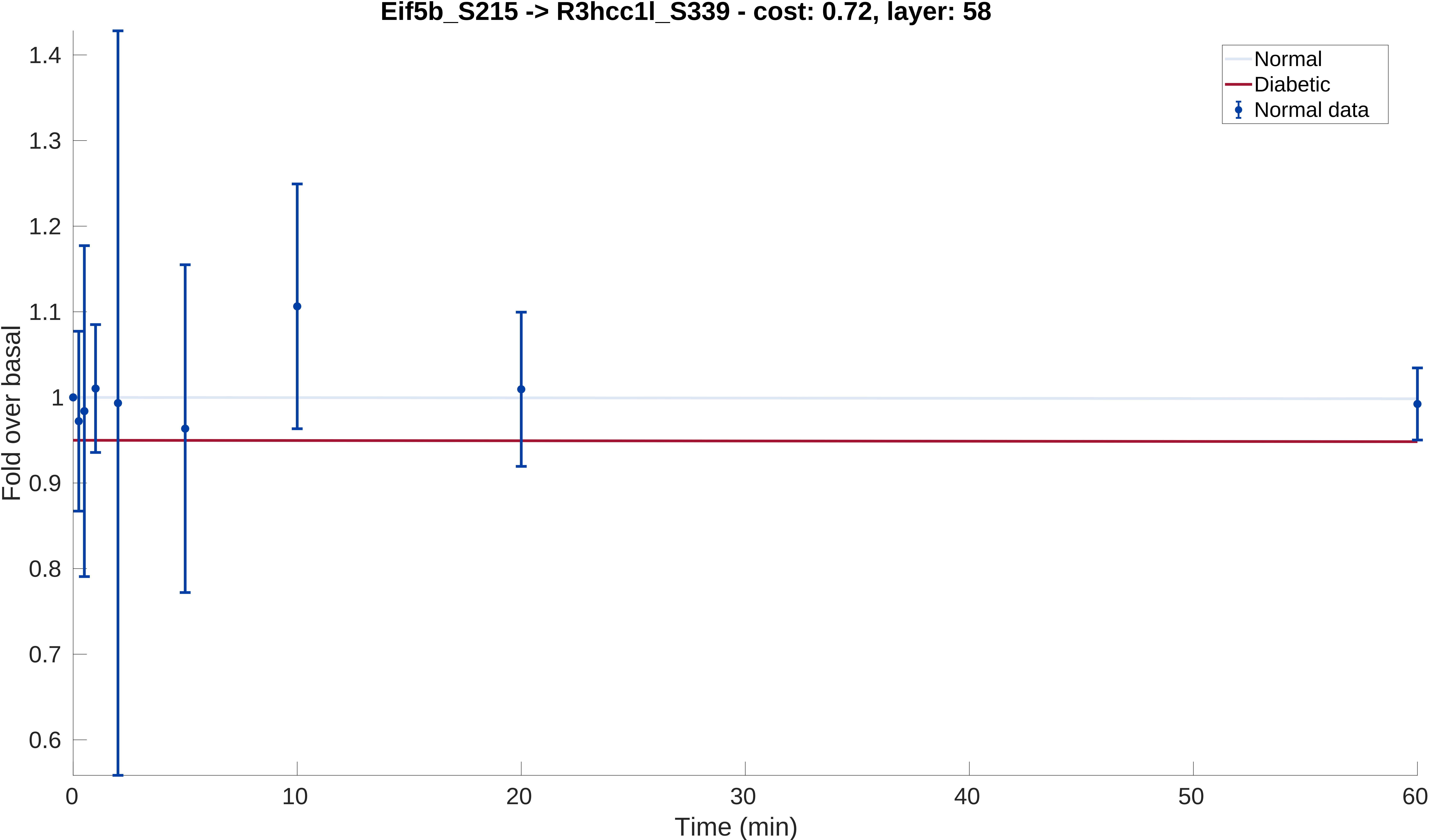

Eif5b_S215 -> R3hcc1l_S339 - cost: 0.72, layer: 58
Normal
Diabetic
Normal data
Fold over basal
1.4
1.3
1.2
1.1
1
0.9
0.8
0.7
0.6
0
10
20
30
40
50
60
Time (min)

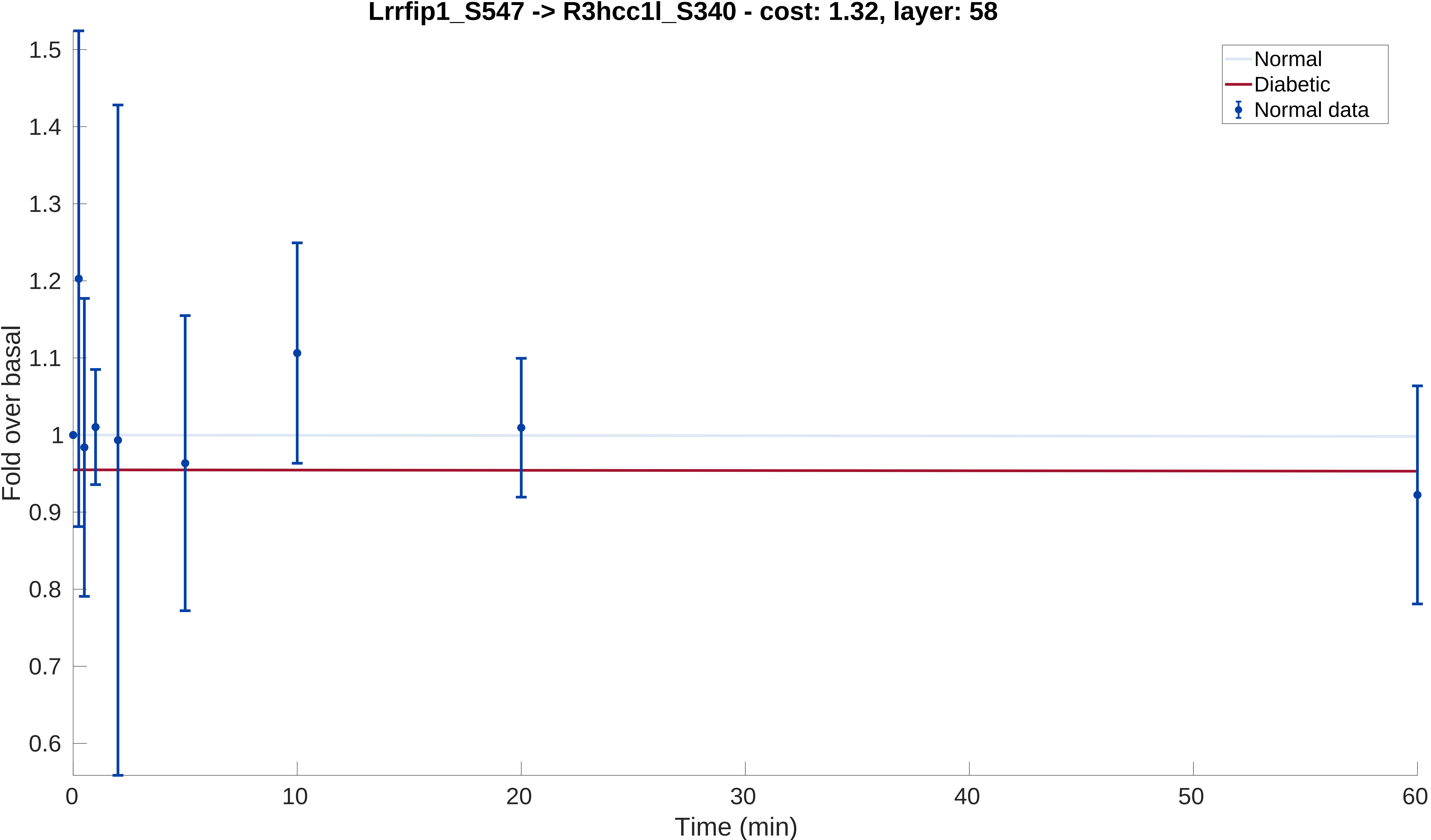

Lrrfip1_S547 -> R3hcc1l_S340 - cost: 1.32, layer: 58
Fold over basal
1.5
1.4
1.3
1.2
1.1
1
0.9
0.8
0.7
0.6
0
10
20
30
40
50
60
Time (min)
Normal
Diabetic
Normal data

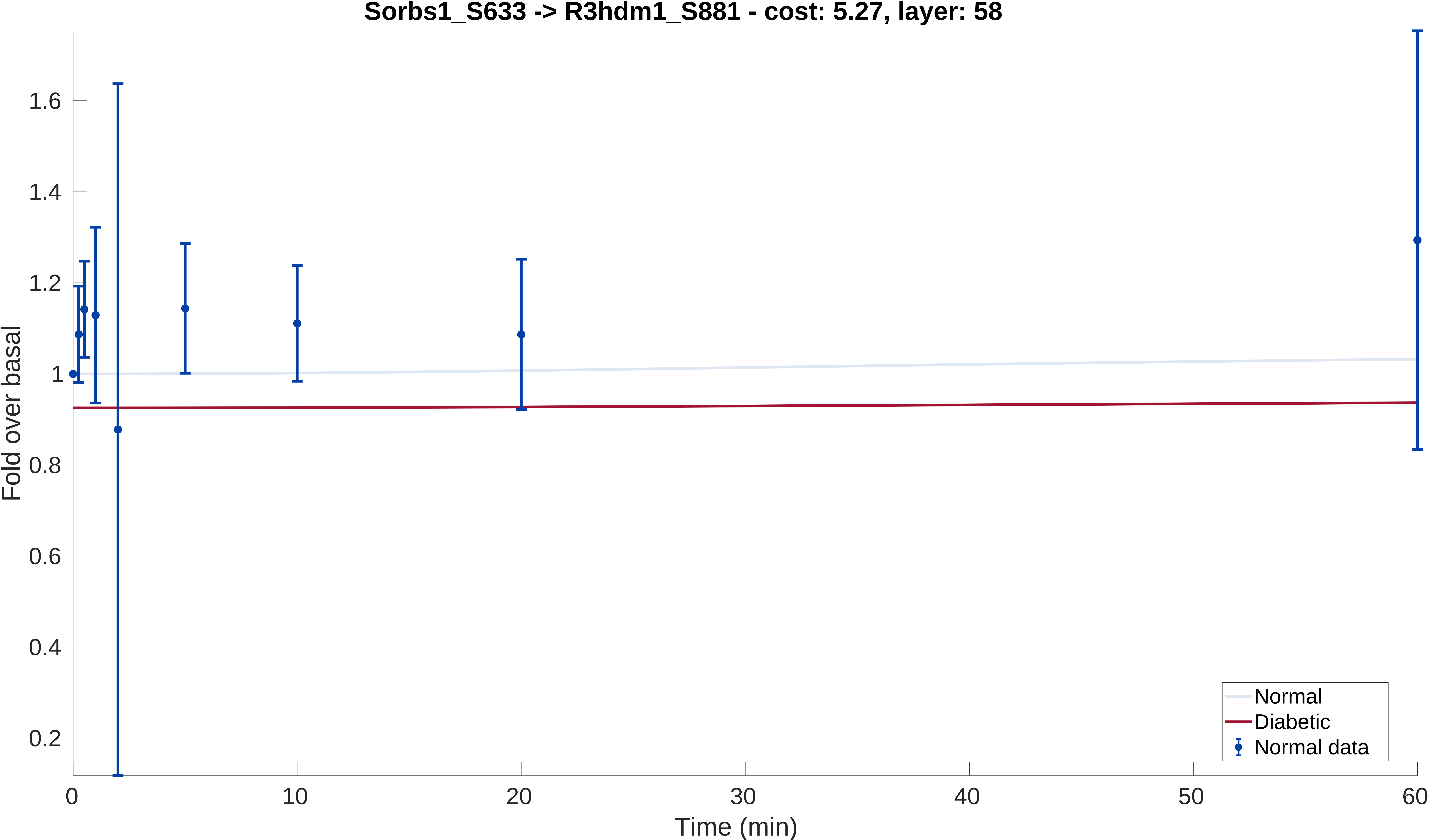

Sorbs1_S633 -> R3hdm1_S881 - cost: 5.27, layer: 58
Fold over basal
Time (min)
0.2
0.4
0.6
0.8
1
1.2
1.4
1.6
0
10
20
30
40
50
60
Normal
Diabetic
Normal data

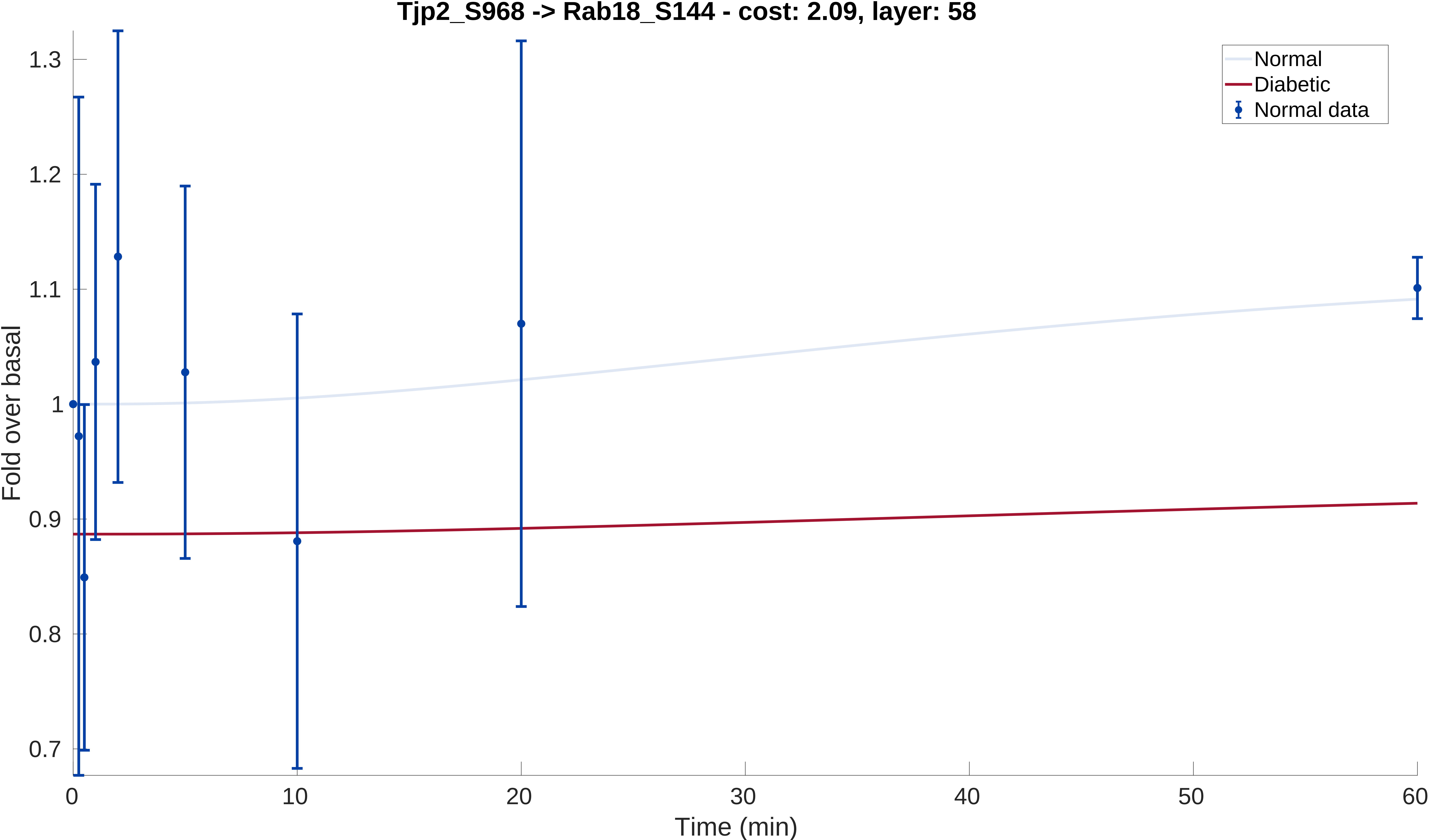
Tjp2_S968 -> Rab18_S144 - cost: 2.09, layer: 58
Normal
Diabetic
Normal data
Fold over basal
1.3
1.2
1.1
1
0.9
0.8
0.7
0
10
20
30
40
50
60
Time (min)

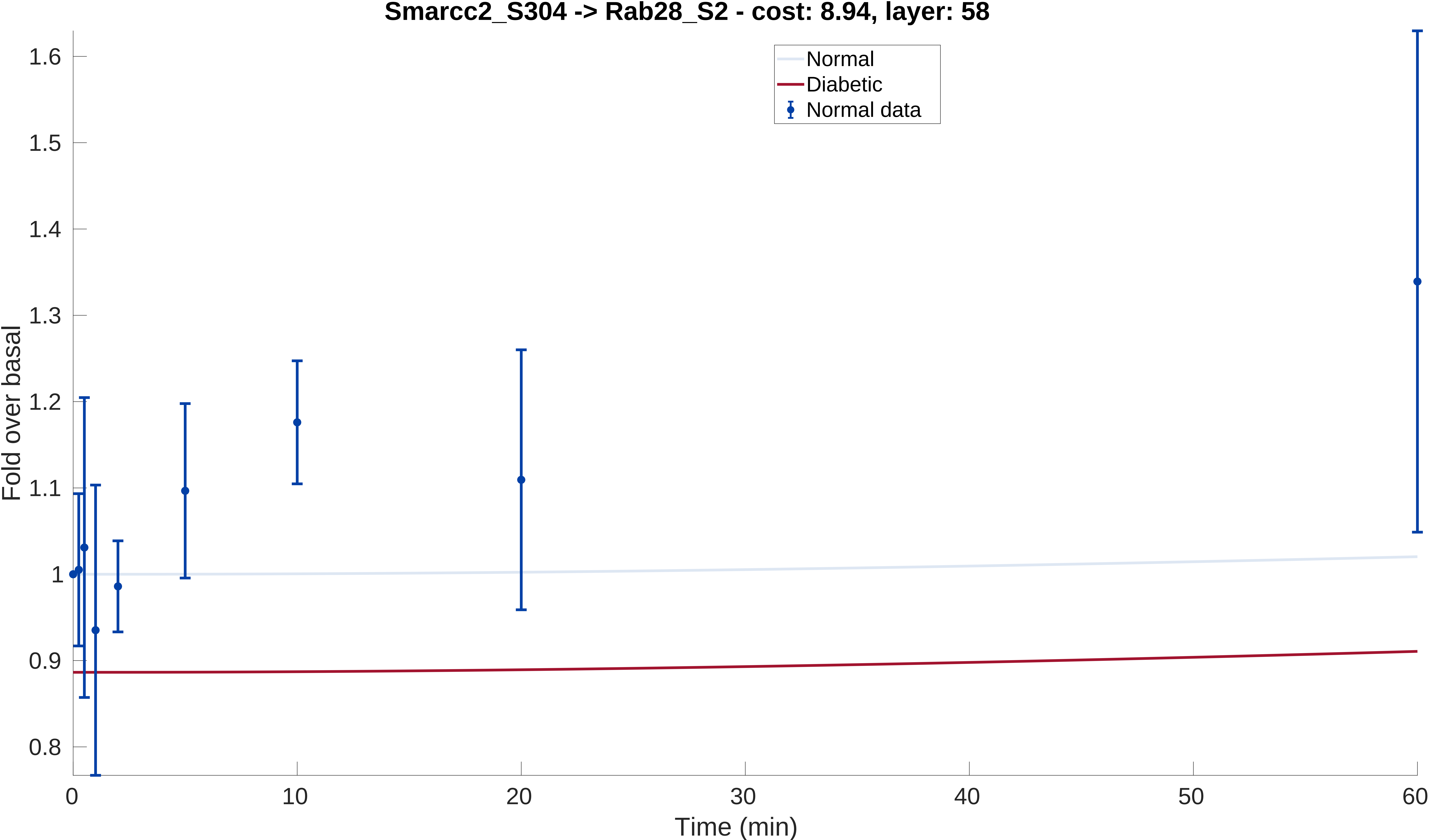

Smarcc2_S304 -> Rab28_S2 - cost: 8.94, layer: 58
Normal
Diabetic
Normal data
Fold over basal
1.6
1.5
1.4
1.3
1.2
1.1
1
0.9
0.8
0
10
20
30
40
50
60
Time (min)

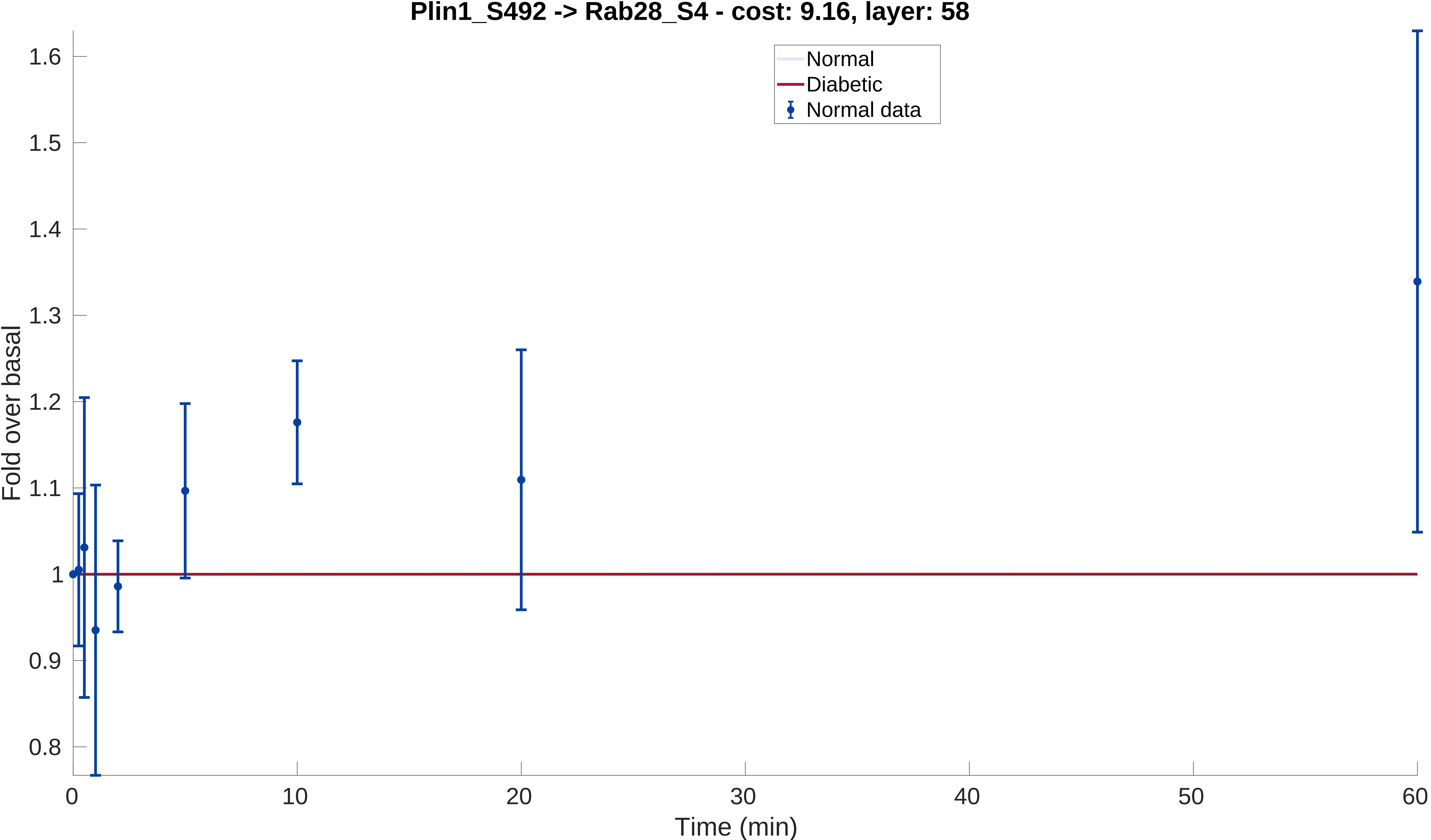
Plin1_S492 -> Rab28_S4 - cost: 9.16, layer: 58
Normal
Diabetic
Normal data
Fold over basal
1.6
1.5
1.4
1.3
1.2
1.1
1
0.9
0.8
0
10
20
30
40
50
60
Time (min)

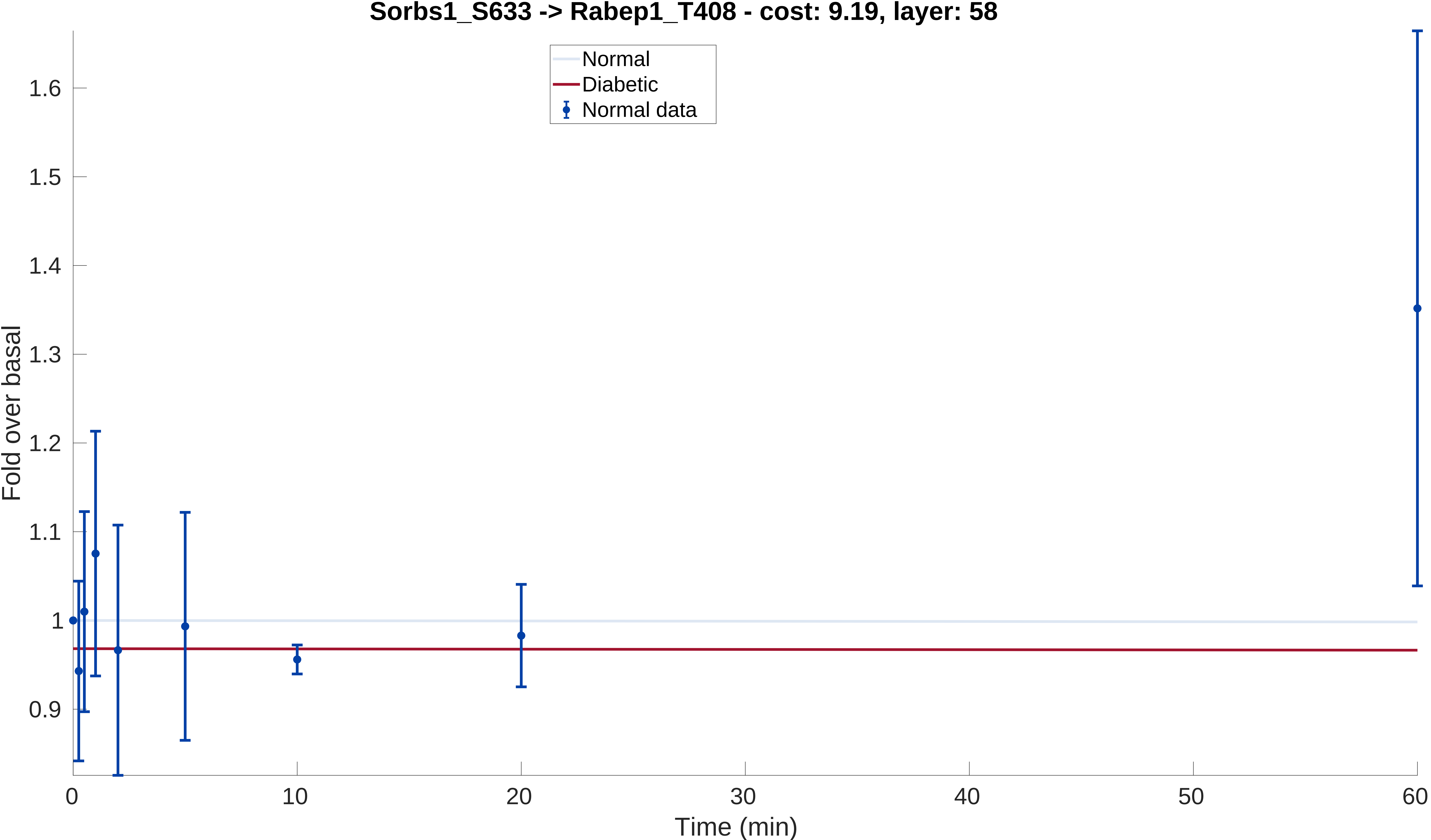
Sorbs1_S633 -> Rabep1_T408 - cost: 9.19, layer: 58
Normal
Diabetic
Normal data
Fold over basal
1.6
1.5
1.4
1.3
1.2
1.1
1
0.9
0
10
20
30
40
50
60
Time (min)

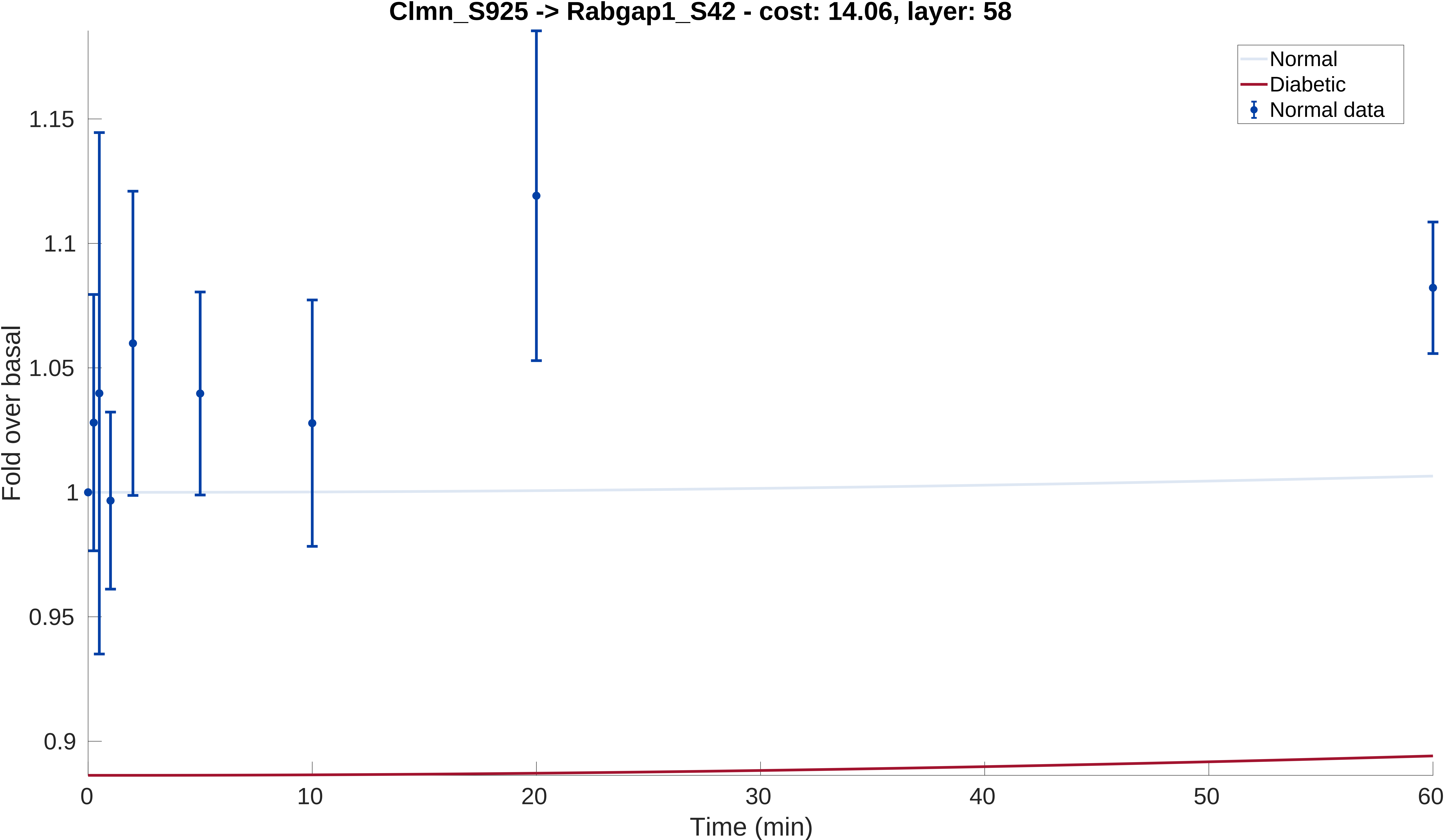

Clmn_S925 -> Rabgap1_S42 - cost: 14.06, layer: 58
Normal
Diabetic
Normal data
Fold over basal
1.15
1.1
1.05
1
0.95
0.9
0
10
20
30
40
50
60
Time (min)

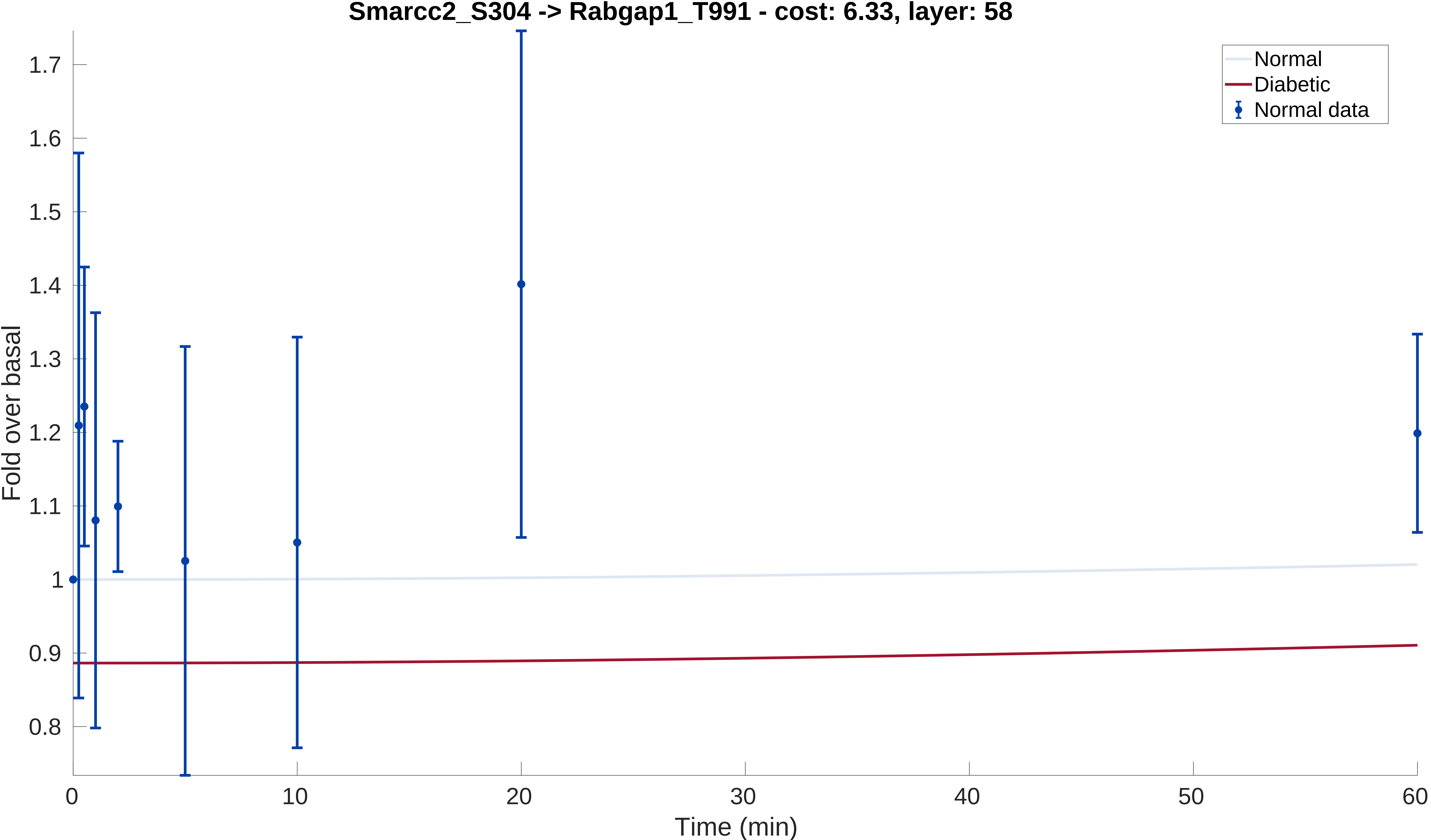

Smarcc2_S304 -> Rabgap1_T991 - cost: 6.33, layer: 58
Fold over basal
1.7
1.6
1.5
1.4
1.3
1.2
1.1
1
0.9
0.8
0
10
20
30
40
50
60
Time (min)
Normal
Diabetic
Normal data

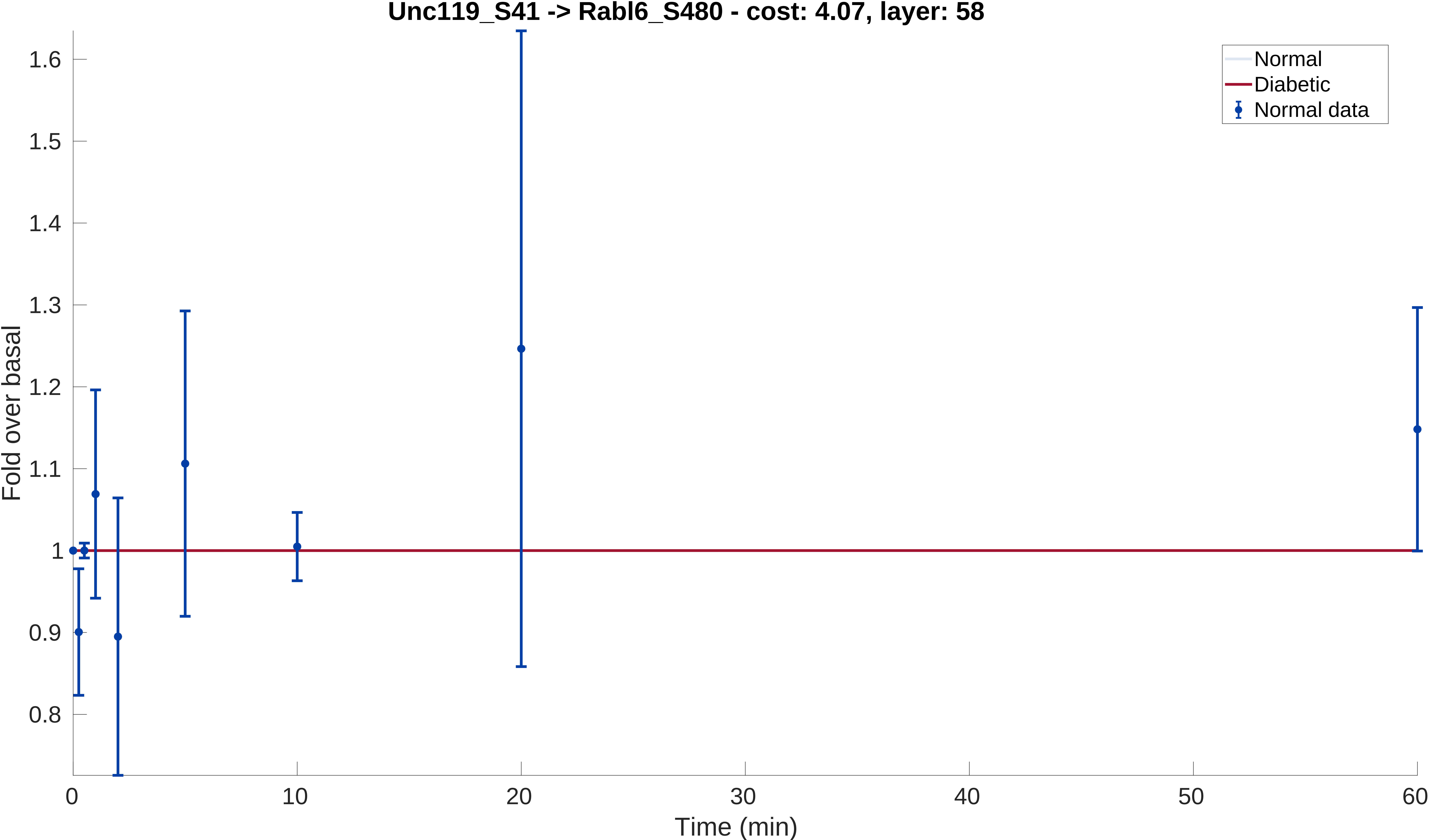

Unc119_S41 -> Rabl6_S480 - cost: 4.07, layer: 58
Normal
Diabetic
Normal data
Fold over basal
1.6
1.5
1.4
1.3
1.2
1.1
1
0.9
0.8
0
10
20
30
40
50
60
Time (min)

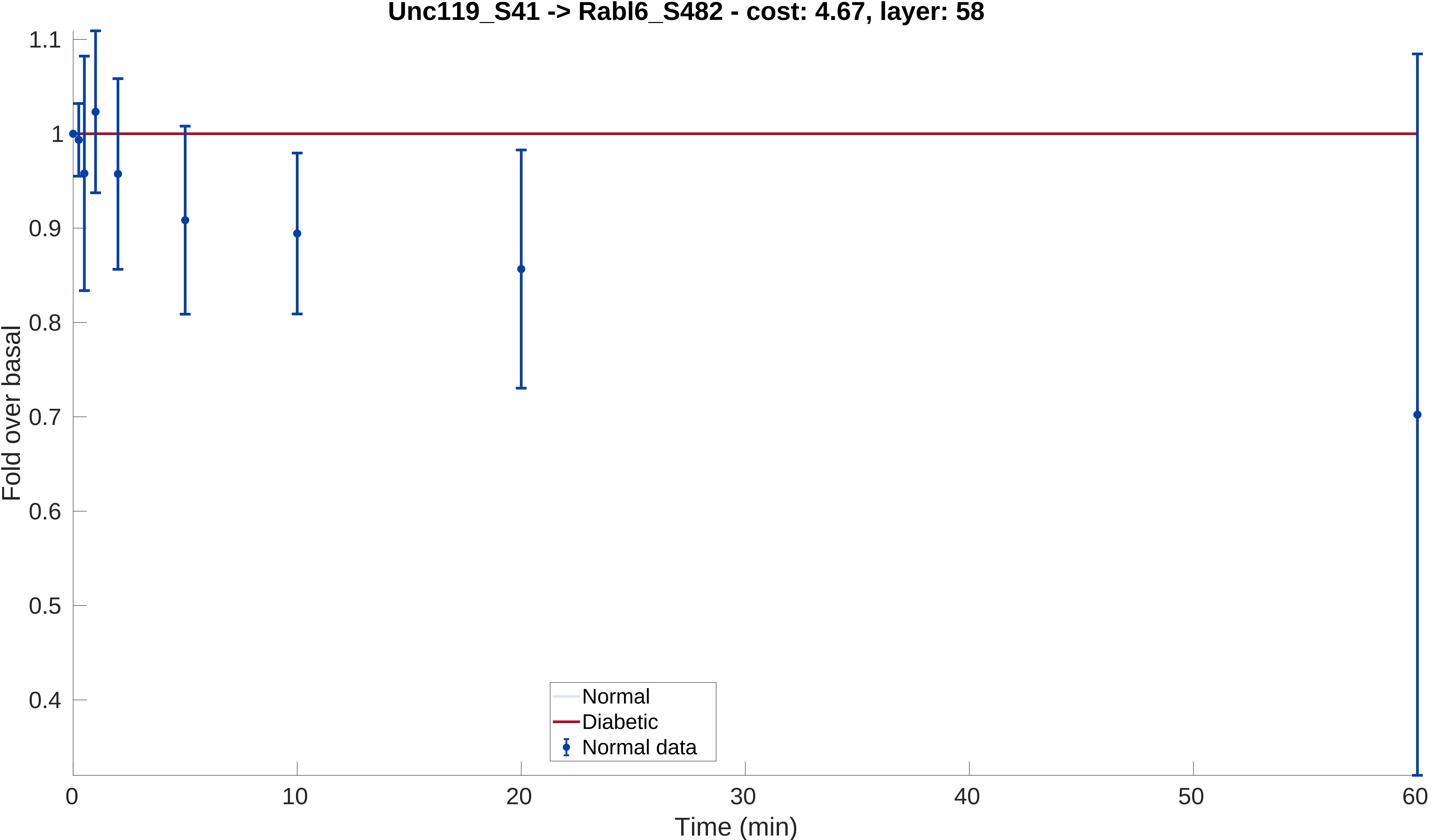

Unc119_S41 -> Rabl6_S482 - cost: 4.67, layer: 58
Fold over basal
1.1
1
0.9
0.8
0.7
0.6
0.5
0.4
0
10
20
30
40
50
60
Time (min)
Normal
Diabetic
Normal data

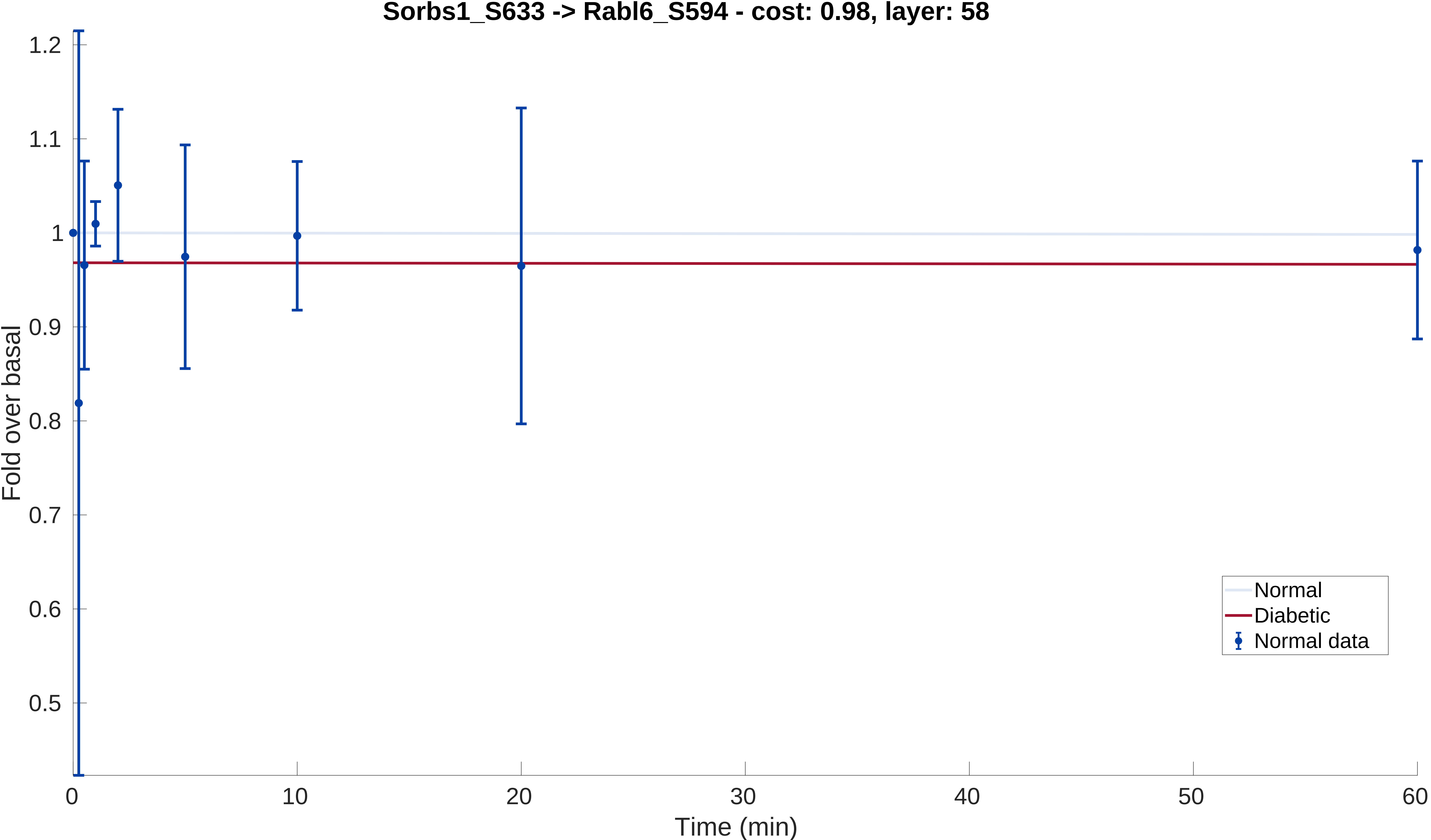

Sorbs1_S633 -> Rabl6_S594 - cost: 0.98, layer: 58
Fold over basal
1.2
1.1
1
0.9
0.8
0.7
0.6
0.5
0
10
20
30
40
50
60
Time (min)
Normal
Diabetic
Normal data

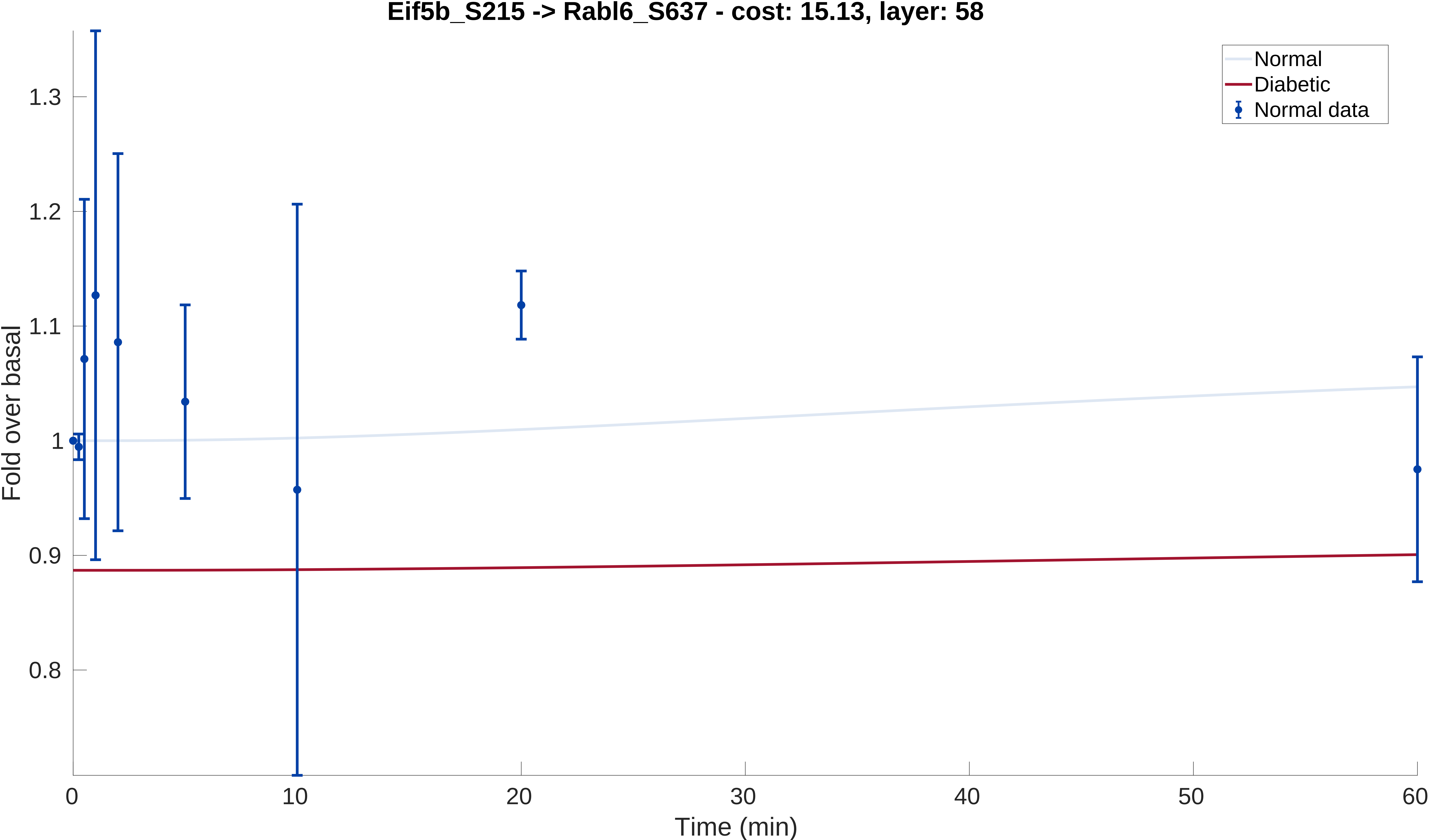
Eif5b_S215 -> Rabl6_S637 - cost: 15.13, layer: 58
Normal
Diabetic
Normal data
Fold over basal
1.3
1.2
1.1
1
0.9
0.8
0
10
20
30
40
50
60
Time (min)

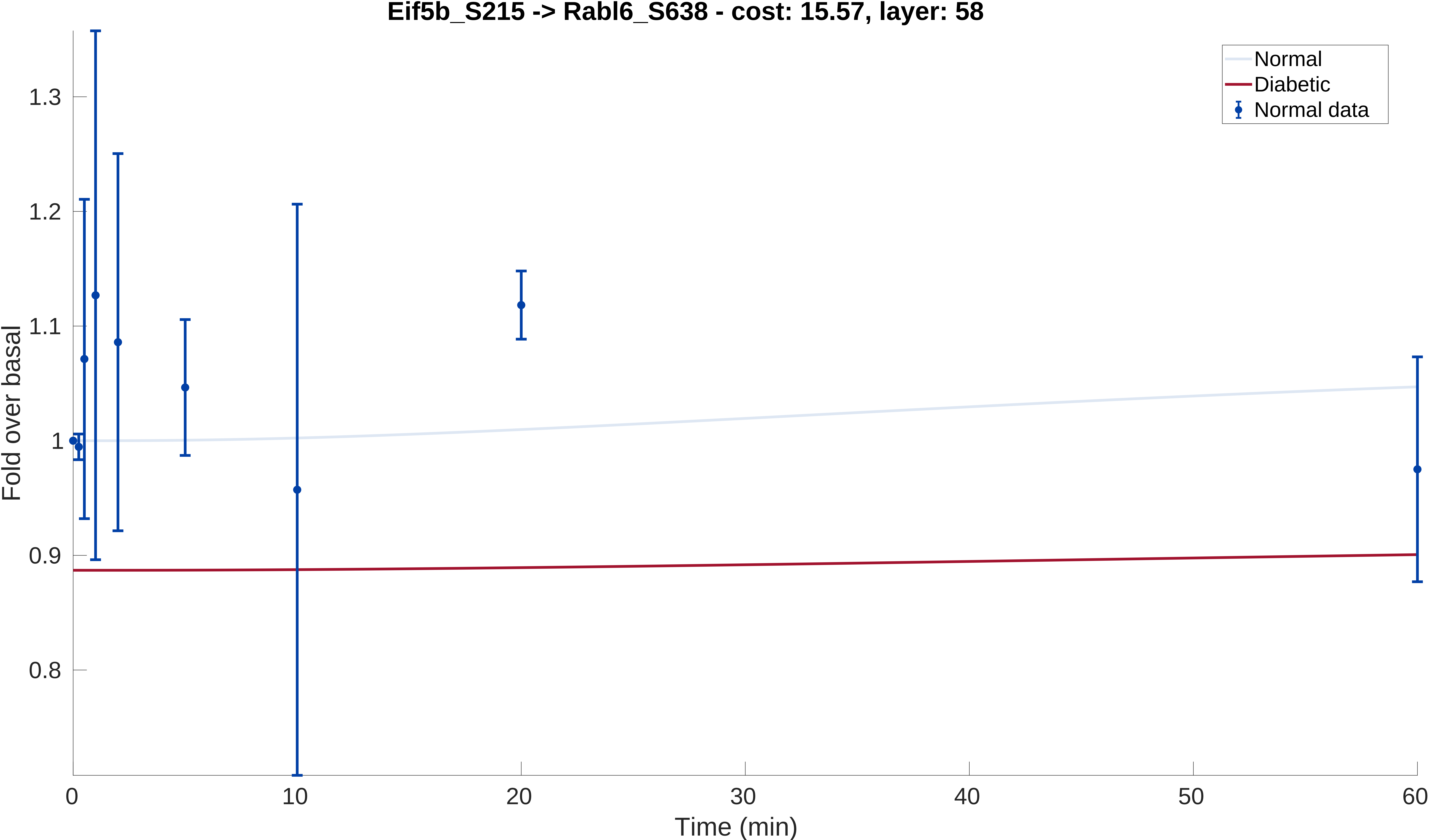

Eif5b_S215 -> Rabl6_S638 - cost: 15.57, layer: 58
Normal
Diabetic
Normal data
Fold over basal
Time (min)
1.3
1.2
1.1
1
0.9
0.8
0
10
20
30
40
50
60

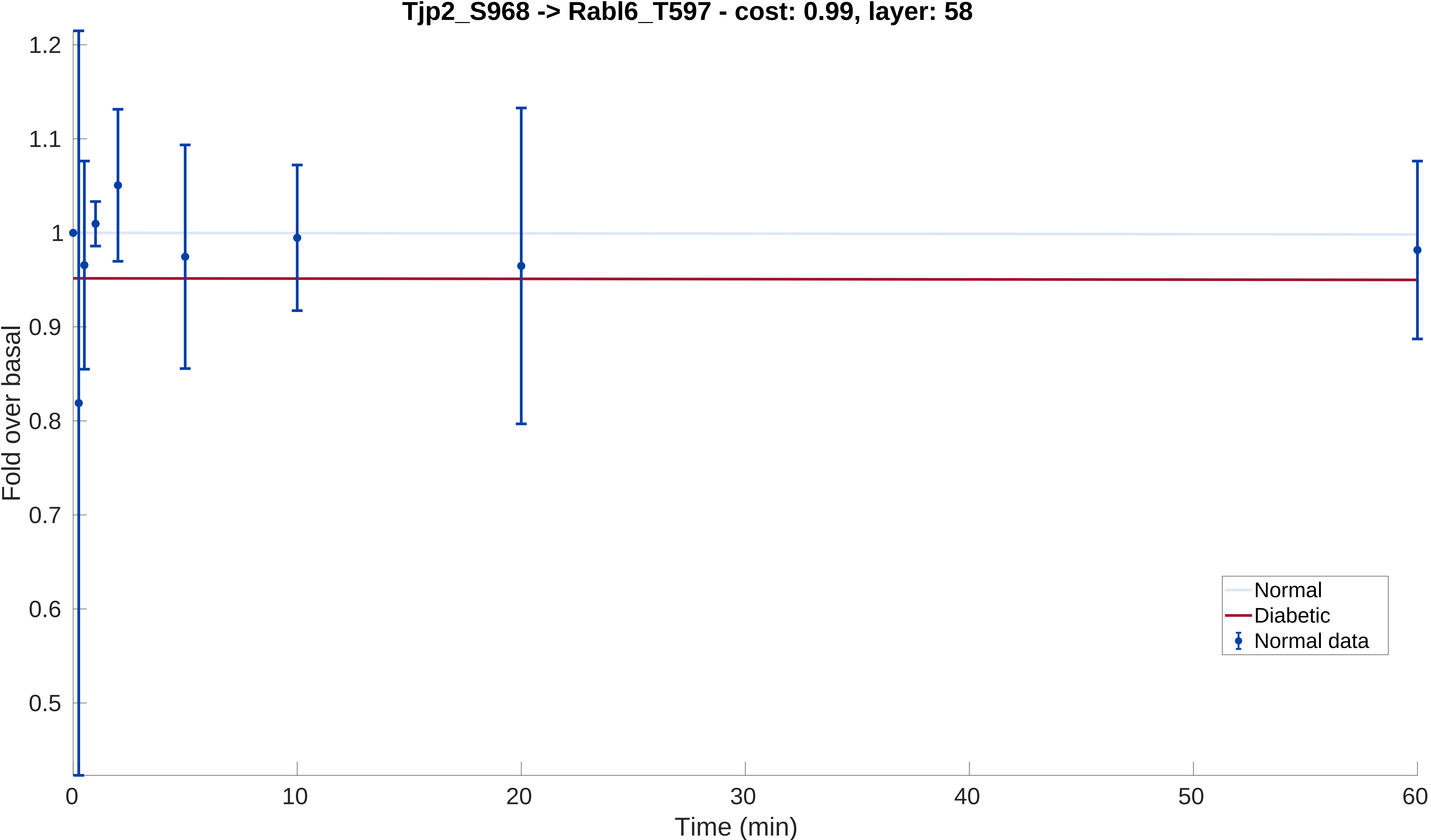

Tjp2_S968 -> Rabl6_T597 - cost: 0.99, layer: 58
Fold over basal
1.2
1.1
1
0.9
0.8
0.7
0.6
0.5
0
10
20
30
40
50
60
Time (min)
Normal
Diabetic
Normal data

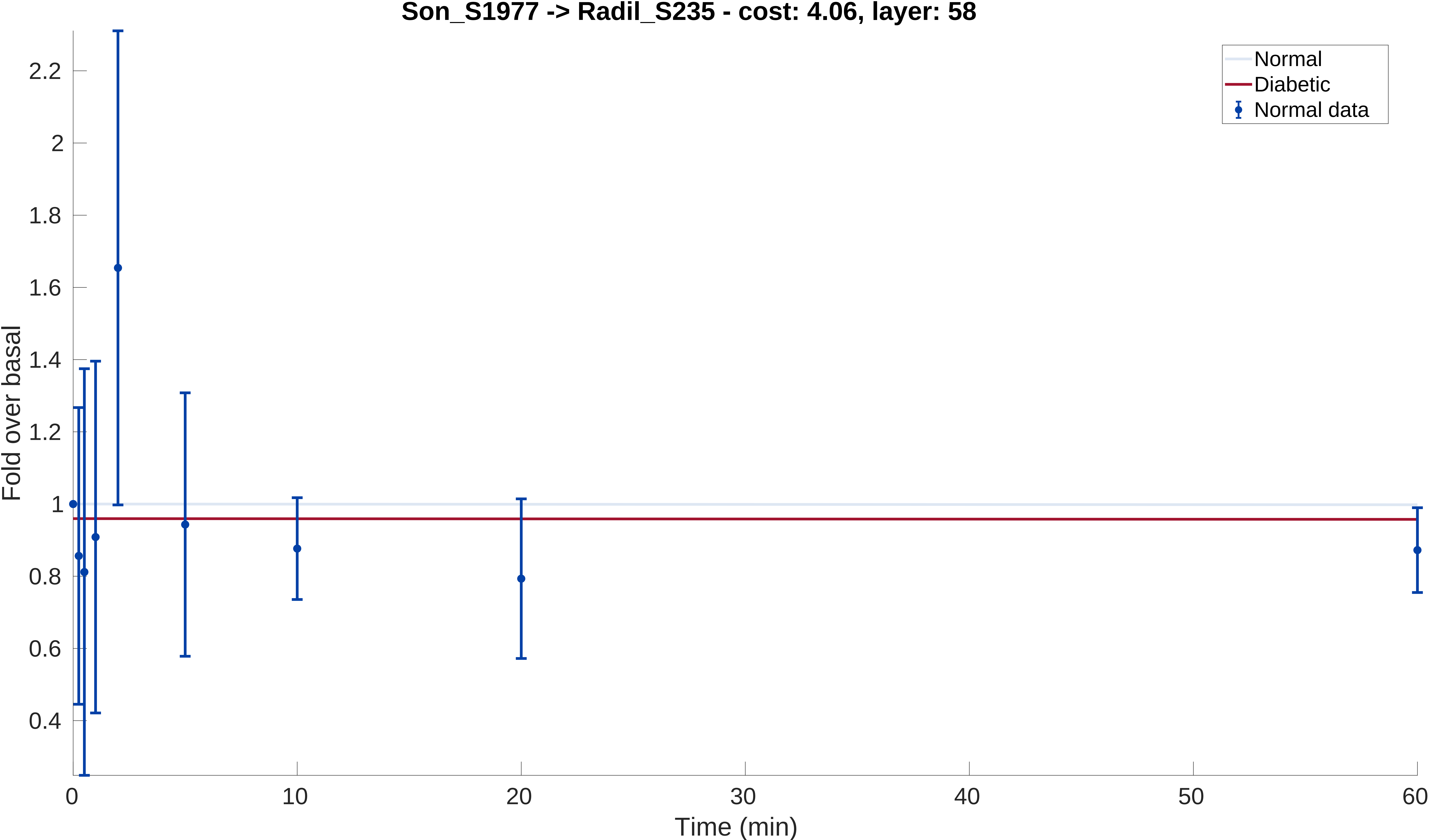

Son_S1977 -> Radil_S235 - cost: 4.06, layer: 58
Normal
Diabetic
Normal data
Fold over basal
2.2
2
1.8
1.6
1.4
1.2
1
0.8
0.6
0.4
0
10
20
30
40
50
60
Time (min)

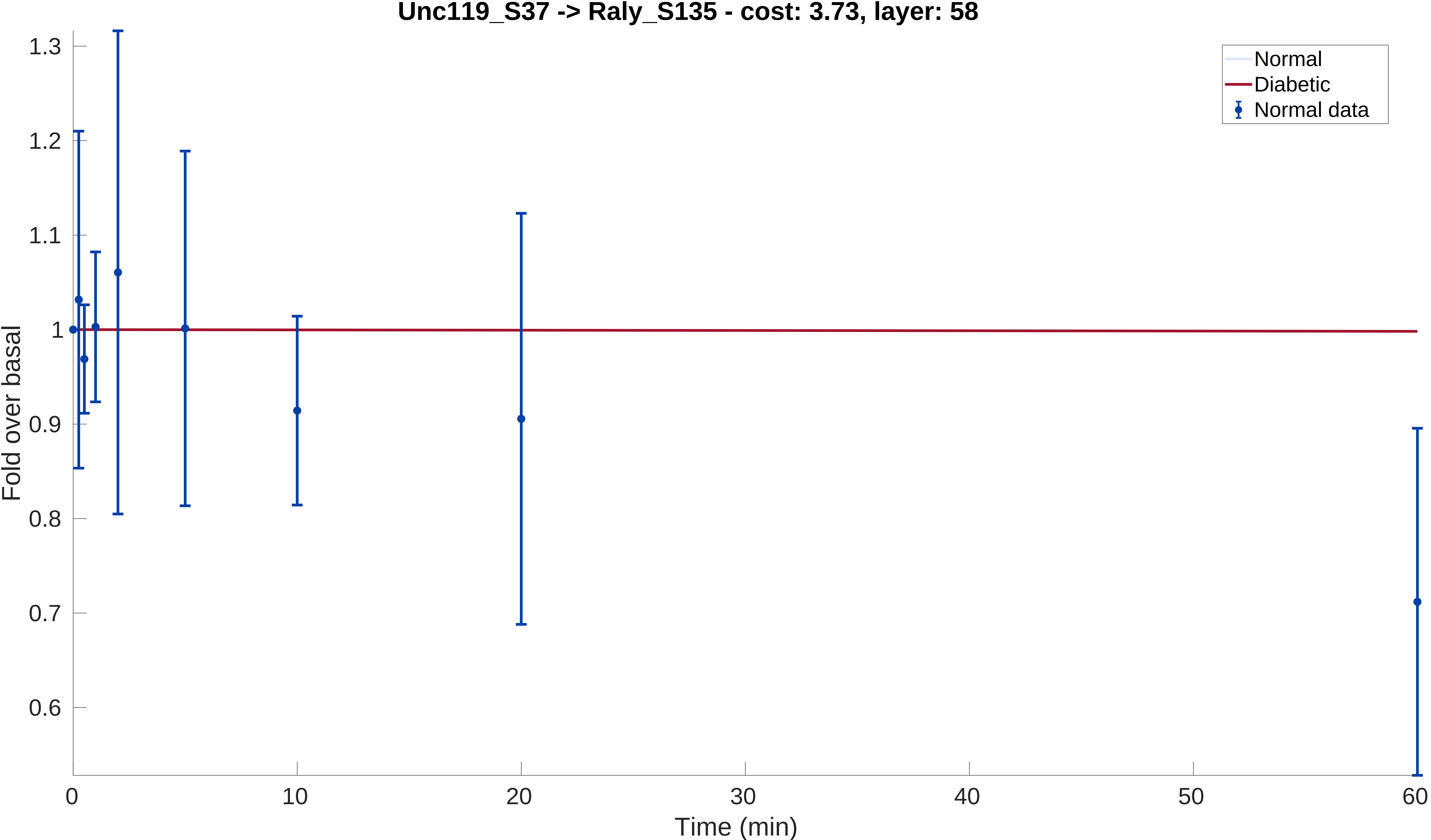

Unc119_S37 -> Raly_S135 - cost: 3.73, layer: 58
Normal
Diabetic
Normal data
Fold over basal
1.3
1.2
1.1
1
0.9
0.8
0.7
0.6
0
10
20
30
40
50
60
Time (min)

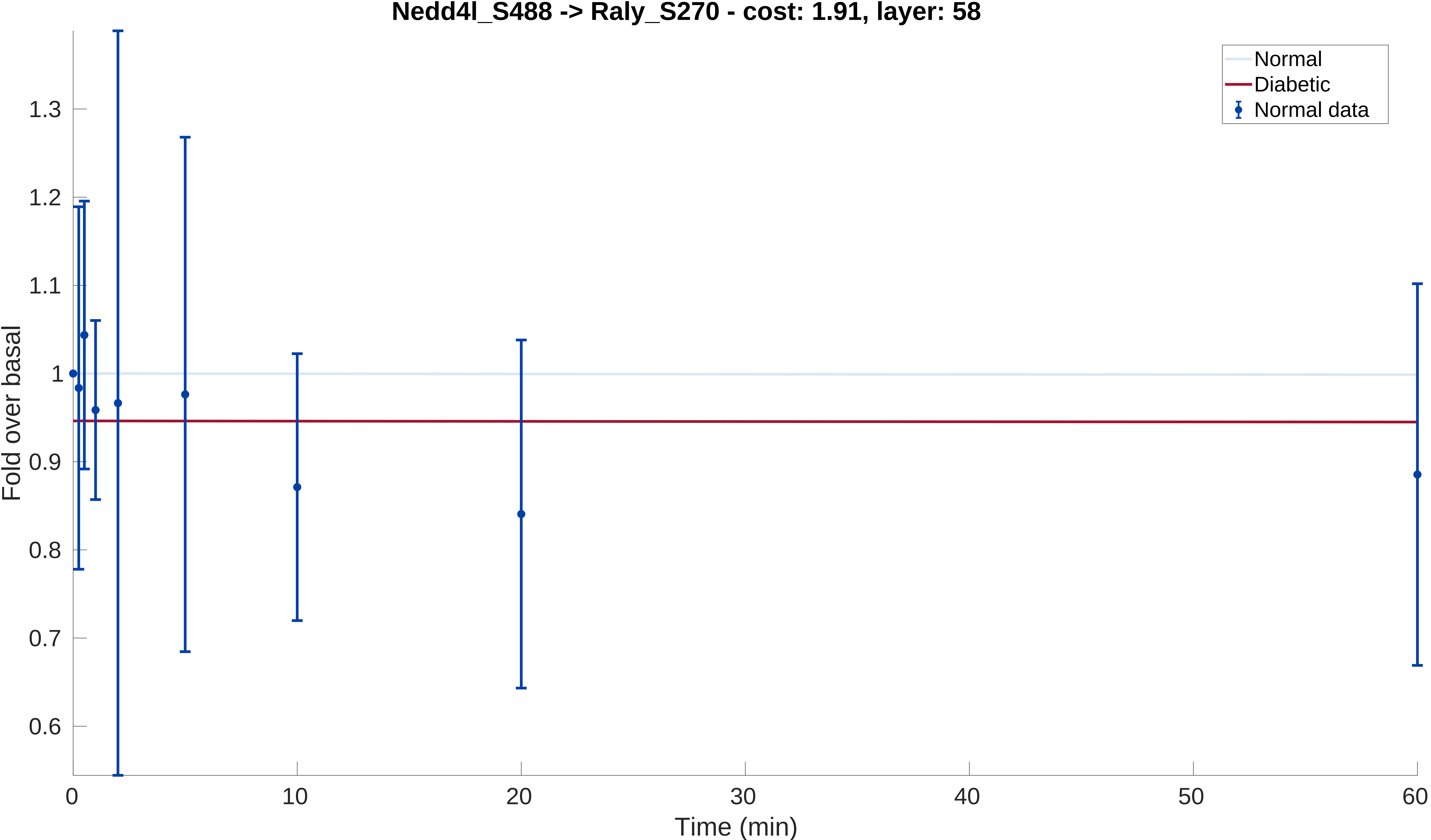

Nedd4l_S488 -> Raly_S270 - cost: 1.91, layer: 58
Normal
Diabetic
Normal data
Fold over basal
1.3
1.2
1.1
1
0.9
0.8
0.7
0.6
0
10
20
30
40
50
60
Time (min)

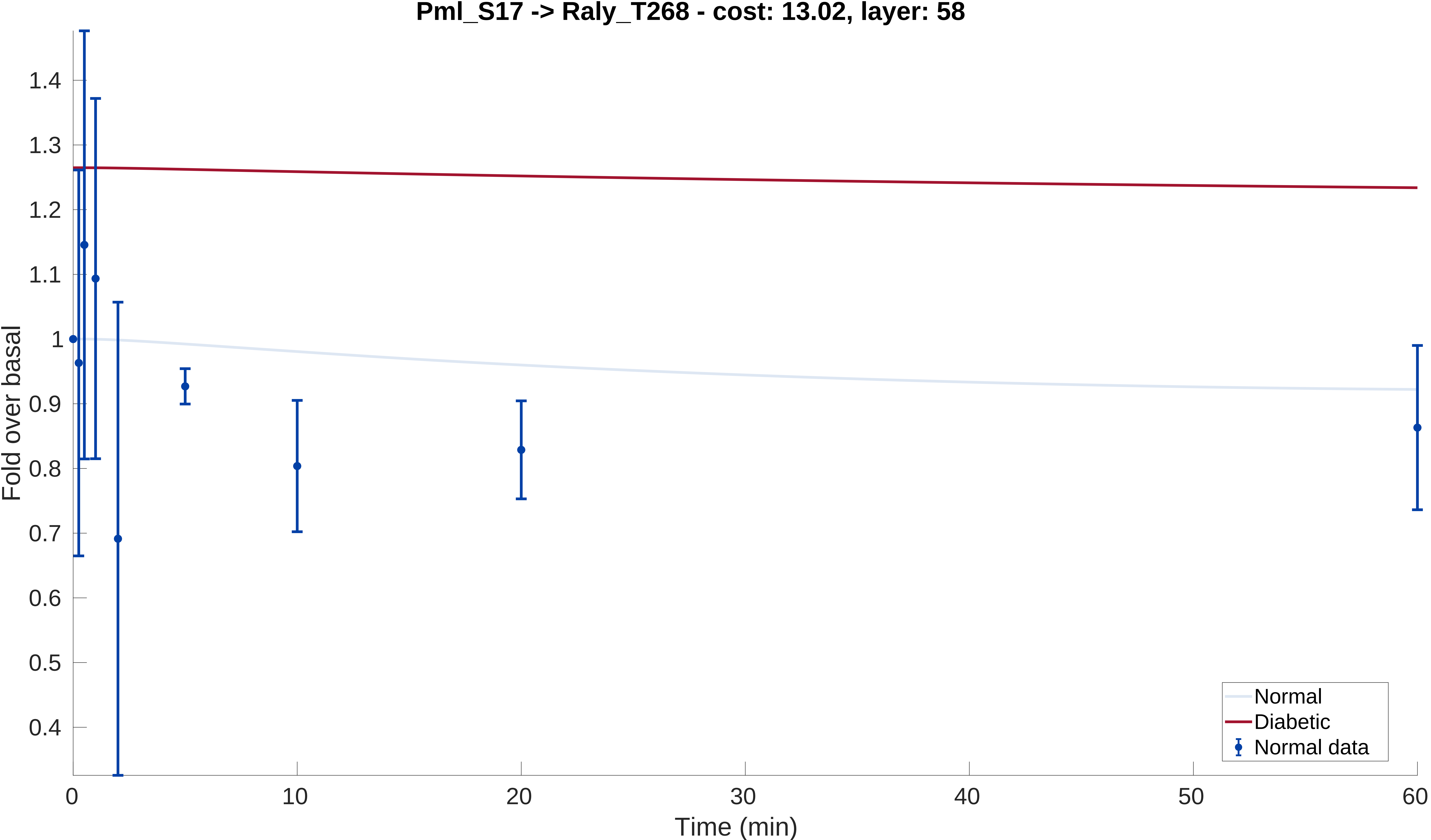

Pml_S17 -> Raly_T268 - cost: 13.02, layer: 58
Fold over basal
1.4
1.3
1.2
1.1
1
0.9
0.8
0.7
0.6
0.5
0.4
0
10
20
30
40
50
60
Time (min)
Normal
Diabetic
Normal data

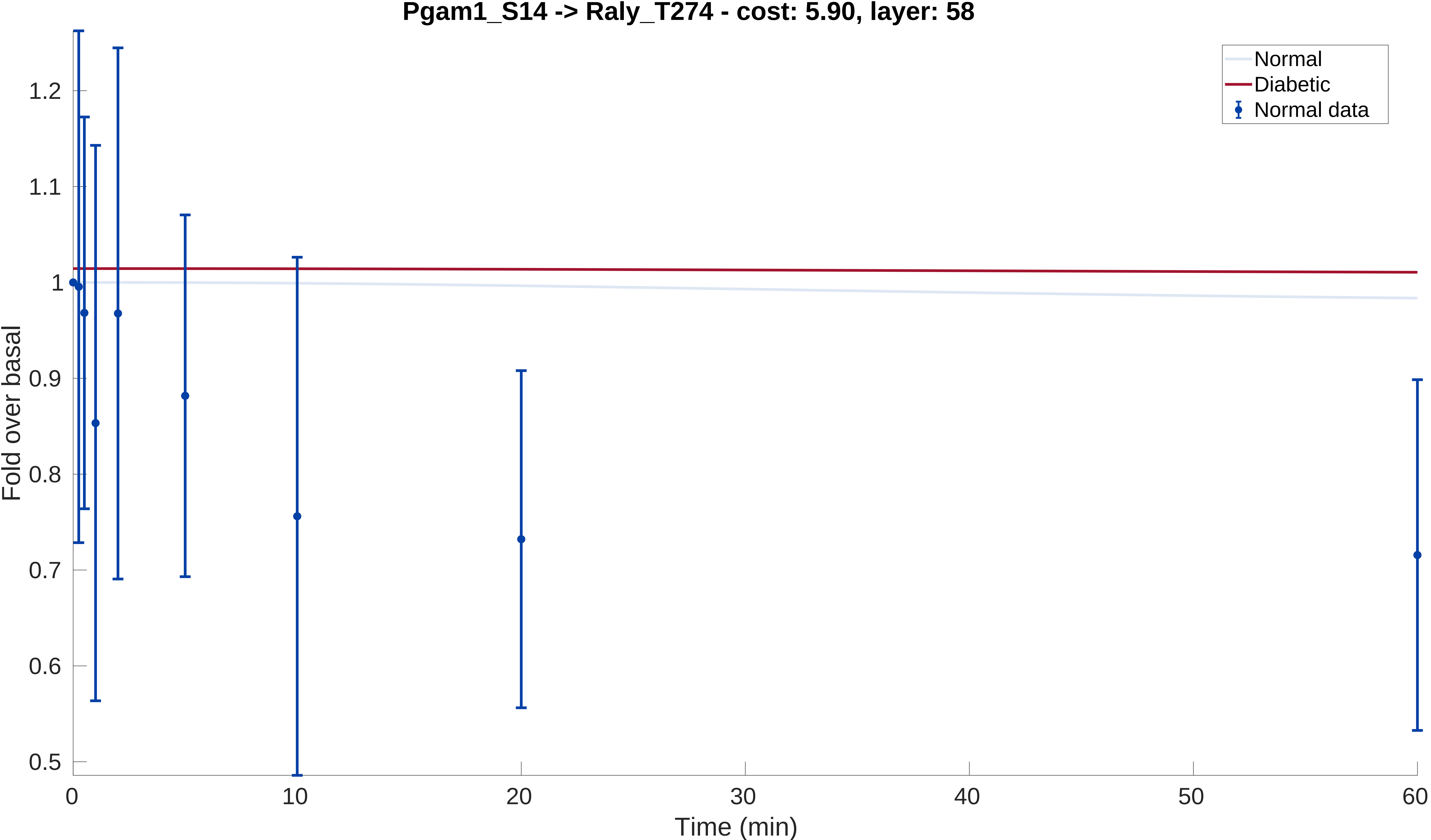
Pgam1_S14 -> Raly_T274 - cost: 5.90, layer: 58
Normal
Diabetic
Normal data
Fold over basal
1.2
1.1
1
0.9
0.8
0.7
0.6
0.5
0
10
20
30
40
50
60
Time (min)

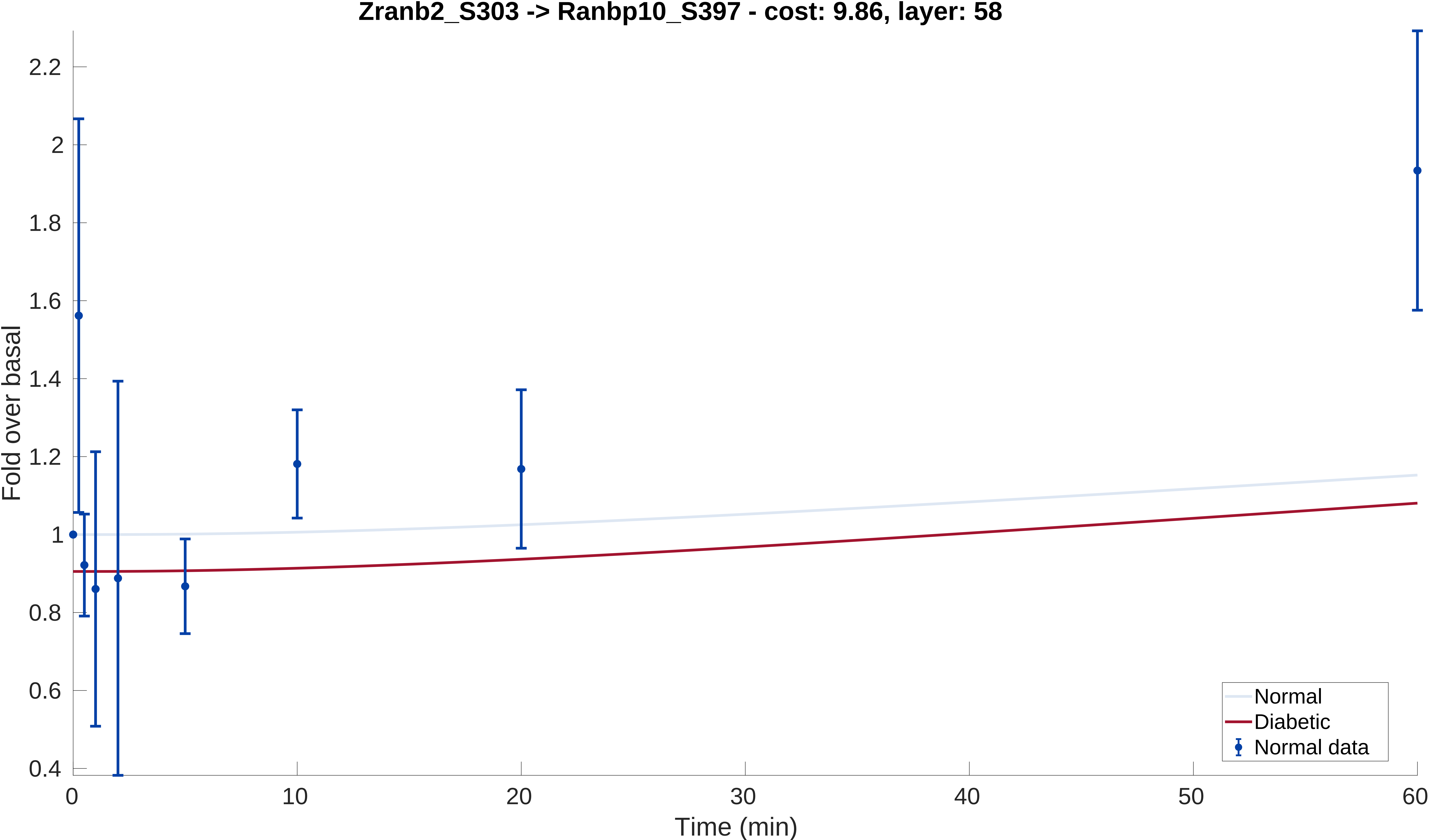

Zranb2_S303 -> Ranbp10_S397 - cost: 9.86, layer: 58
Fold over basal
Time (min)
0.4
0.6
0.8
1
1.2
1.4
1.6
1.8
2
2.2
0
10
20
30
40
50
60
Normal
Diabetic
Normal data

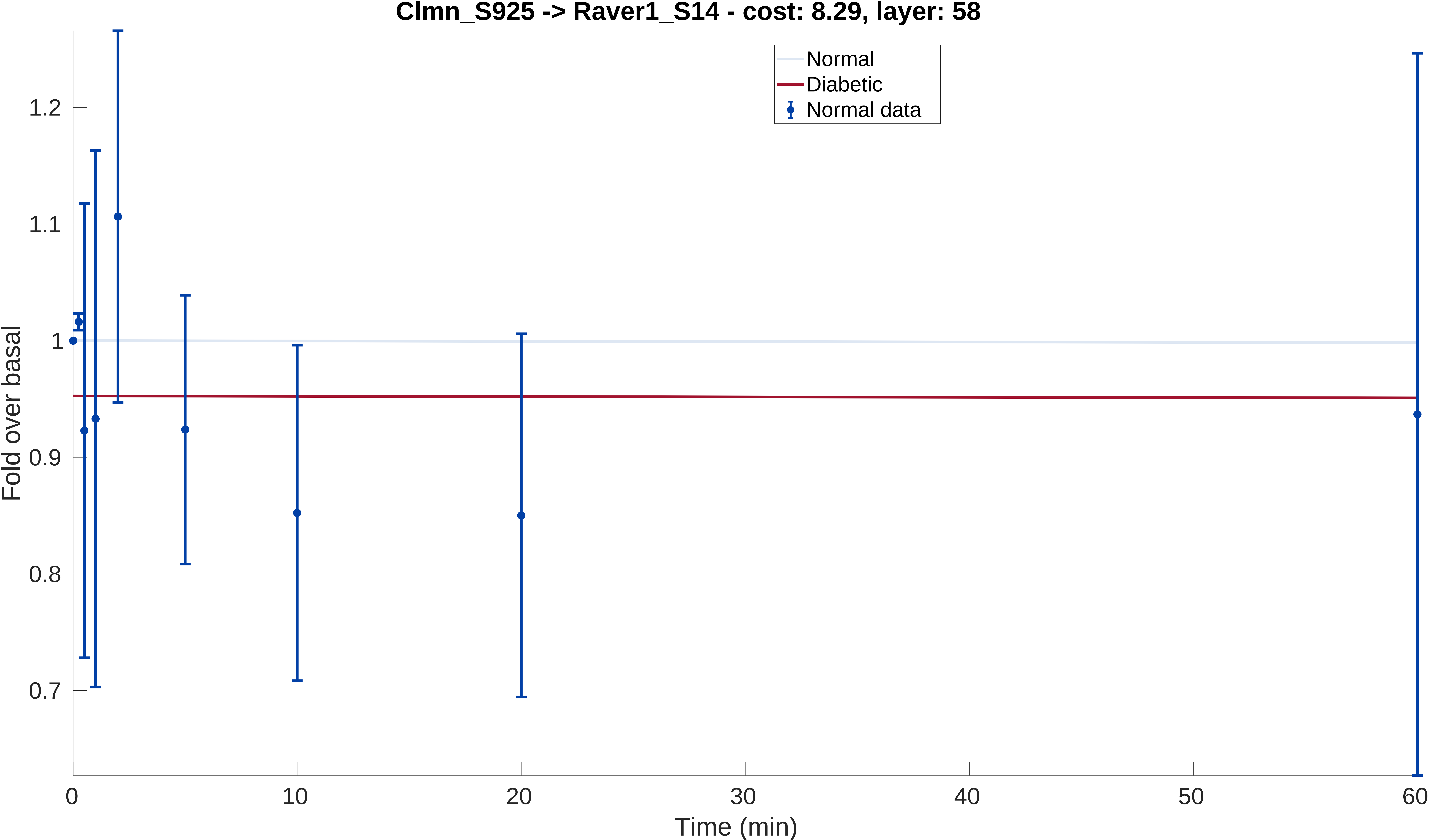

Clmn_S925 -> Raver1_S14 - cost: 8.29, layer: 58
Normal
Diabetic
Normal data
Fold over basal
1.2
1.1
1
0.9
0.8
0.7
0
10
20
30
40
50
60
Time (min)

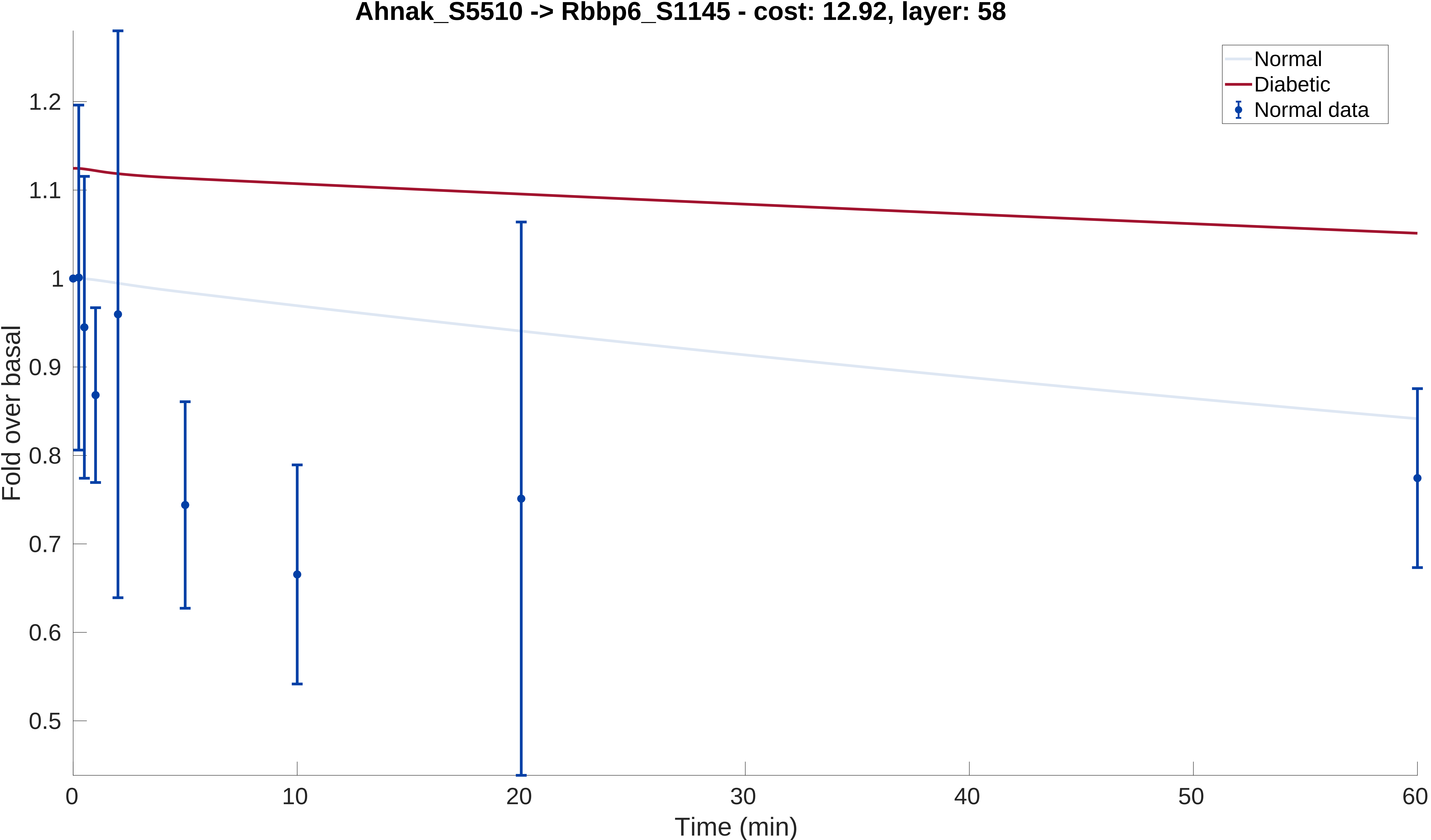
Ahnak_S5510 -> Rbbp6_S1145 - cost: 12.92, layer: 58
Normal
Diabetic
Normal data
Fold over basal
1.2
1.1
1
0.9
0.8
0.7
0.6
0.5
0
10
20
30
40
50
60
Time (min)

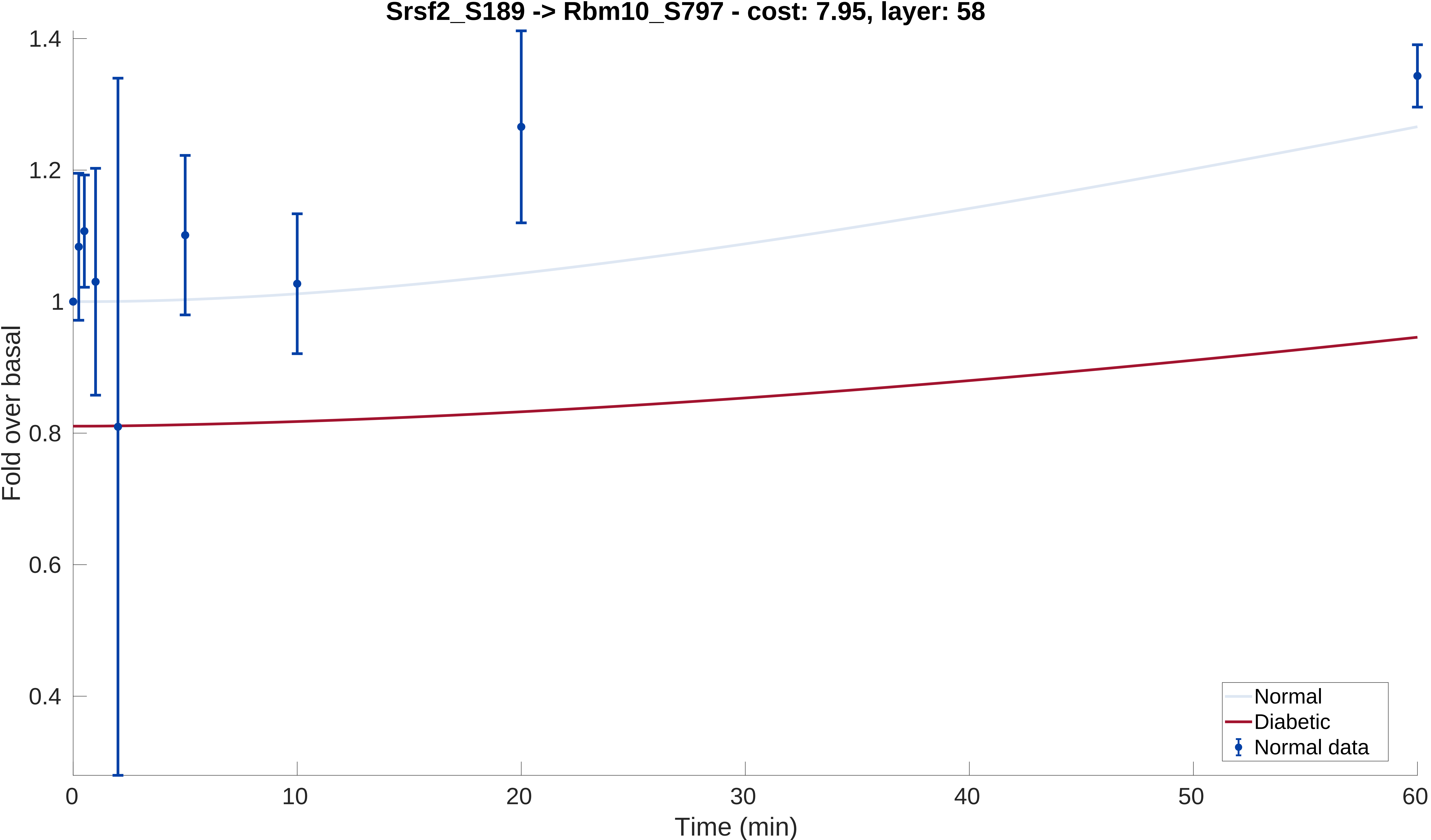
Srsf2_S189 -> Rbm10_S797 - cost: 7.95, layer: 58
Fold over basal
1.4
1.2
1
0.8
0.6
0.4
0
10
20
30
40
50
60
Time (min)
Normal
Diabetic
Normal data

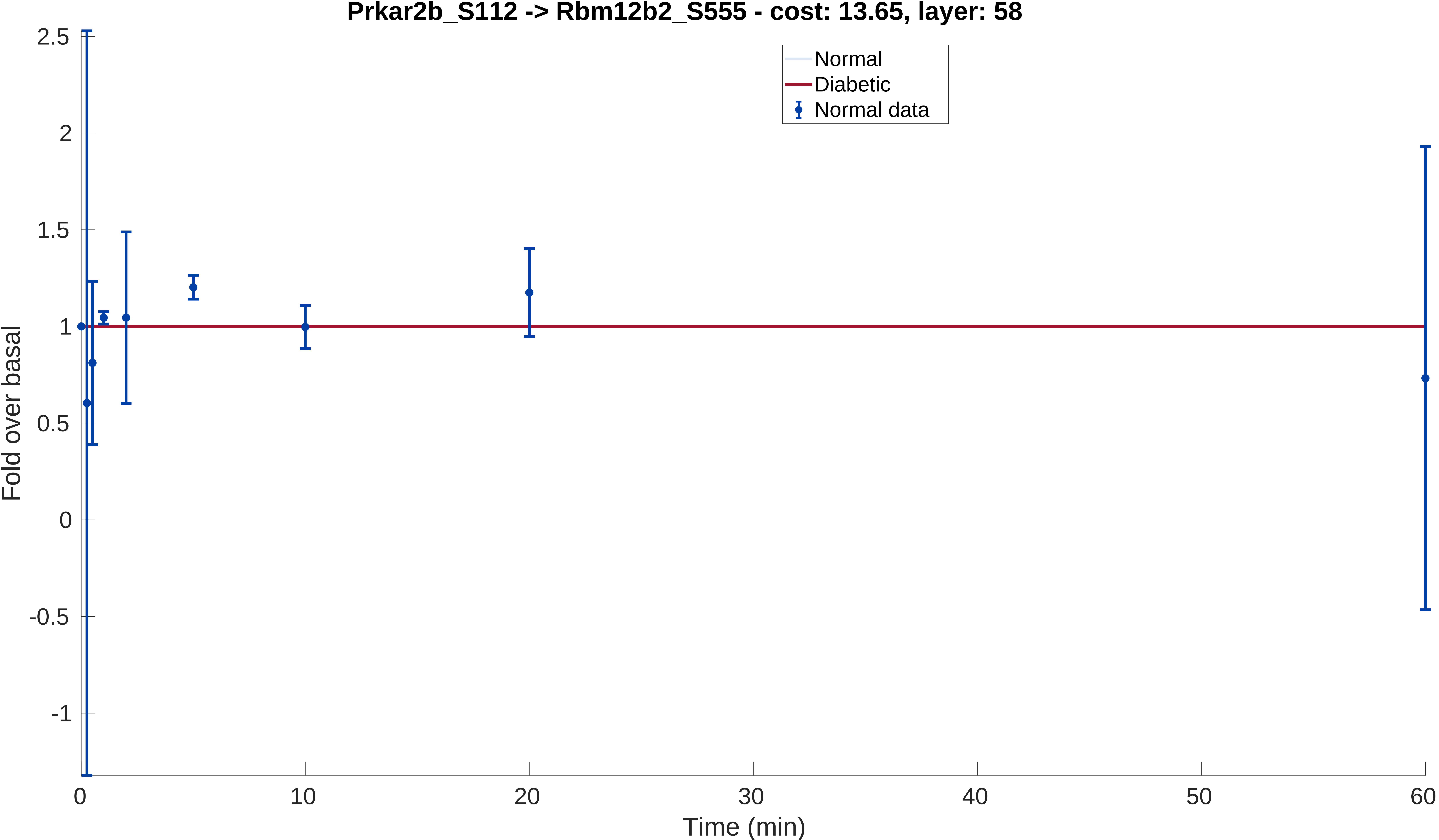

Prkar2b_S112 -> Rbm12b2_S555 - cost: 13.65, layer: 58
Normal
Diabetic
Normal data
Fold over basal
Time (min)
2.5
2
1.5
1
0.5
0
-0.5
-1
0
10
20
30
40
50
60

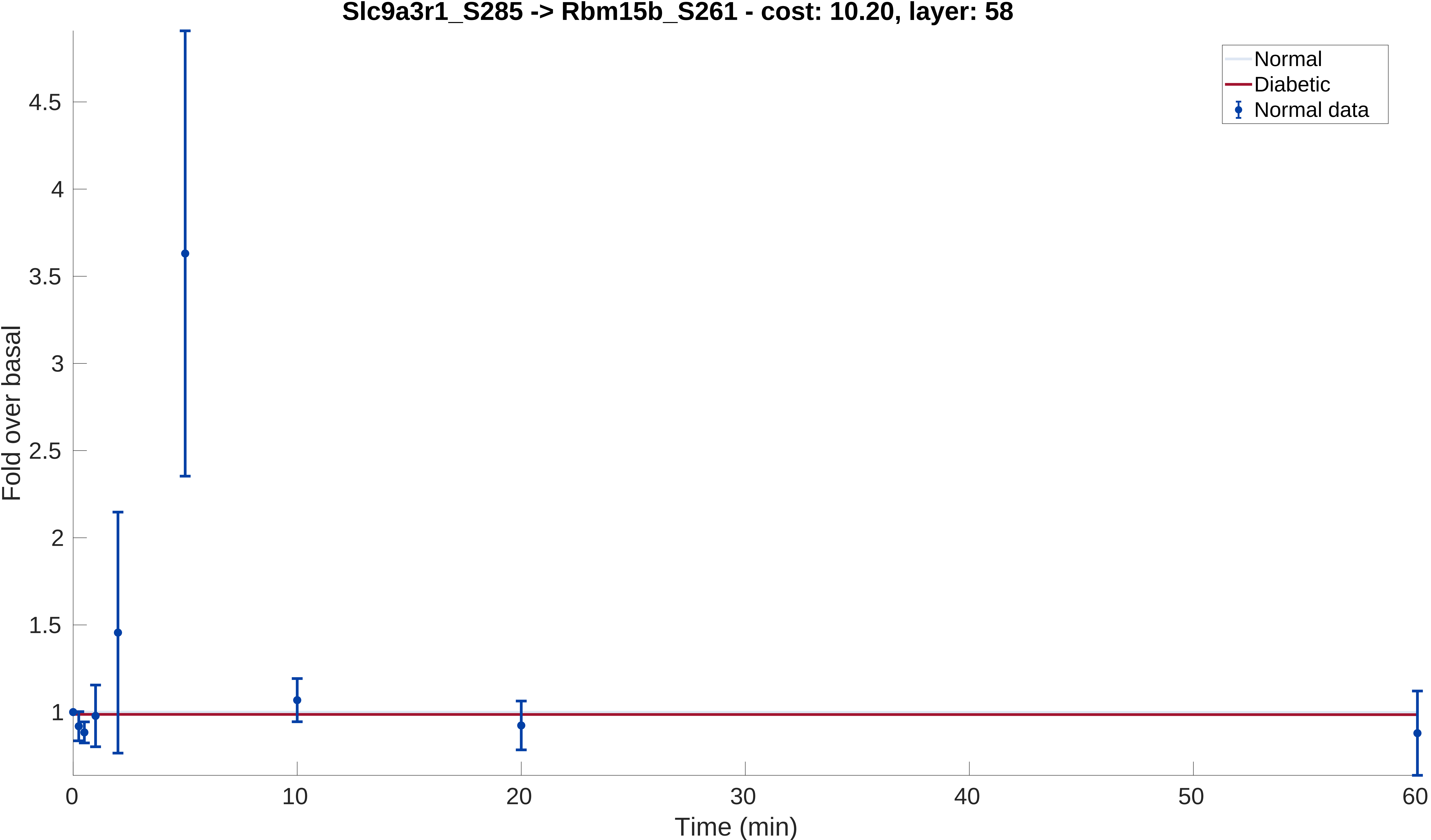

Slc9a3r1_S285 -> Rbm15b_S261 - cost: 10.20, layer: 58
Normal
Diabetic
Normal data
Fold over basal
4.5
4
3.5
3
2.5
2
1.5
1
0
10
20
30
40
50
60
Time (min)

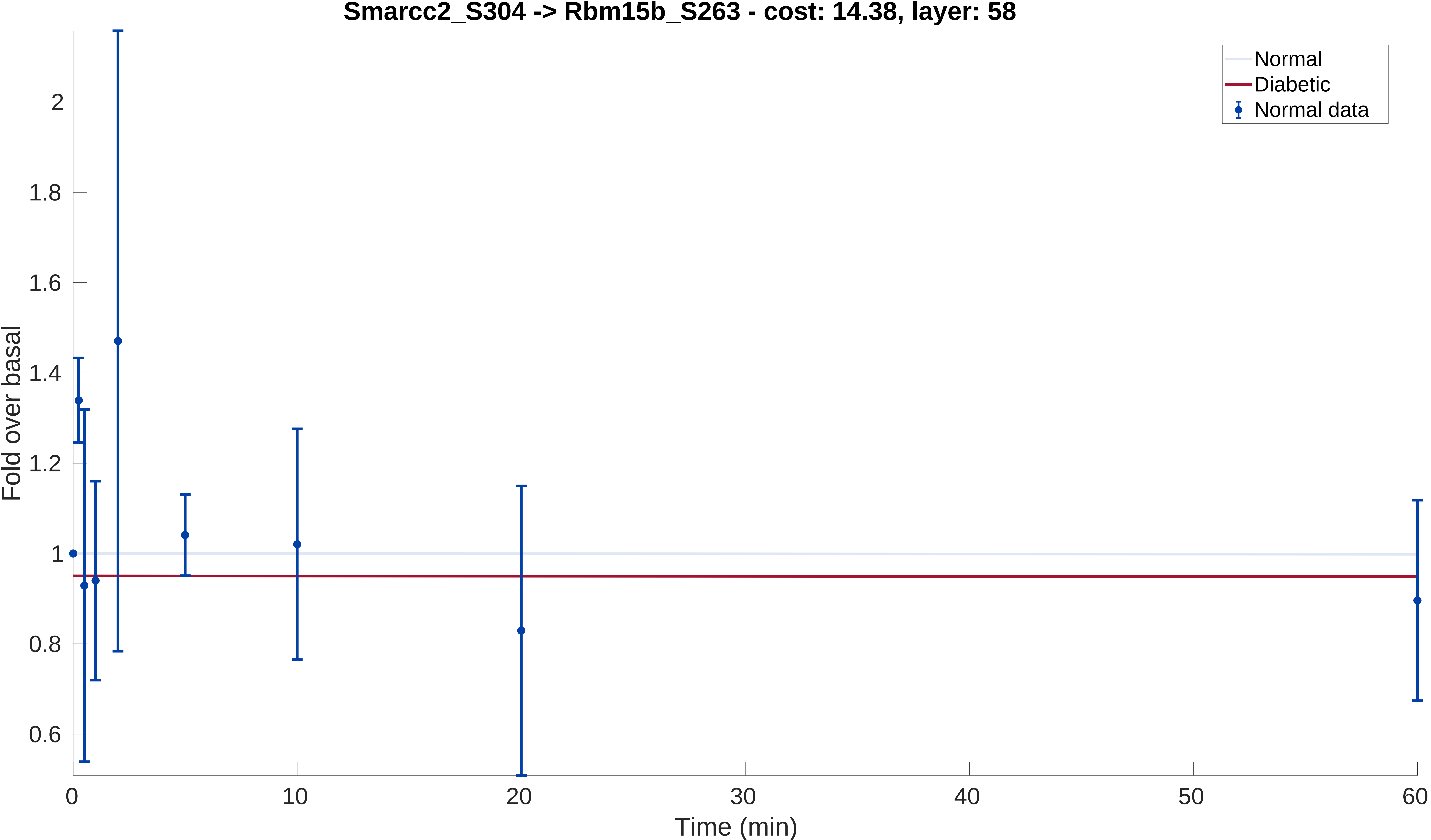

Smarcc2_S304 -> Rbm15b_S263 - cost: 14.38, layer: 58
Normal
Diabetic
Normal data
Fold over basal
Time (min)
0.6
0.8
1
1.2
1.4
1.6
1.8
2
0
10
20
30
40
50
60

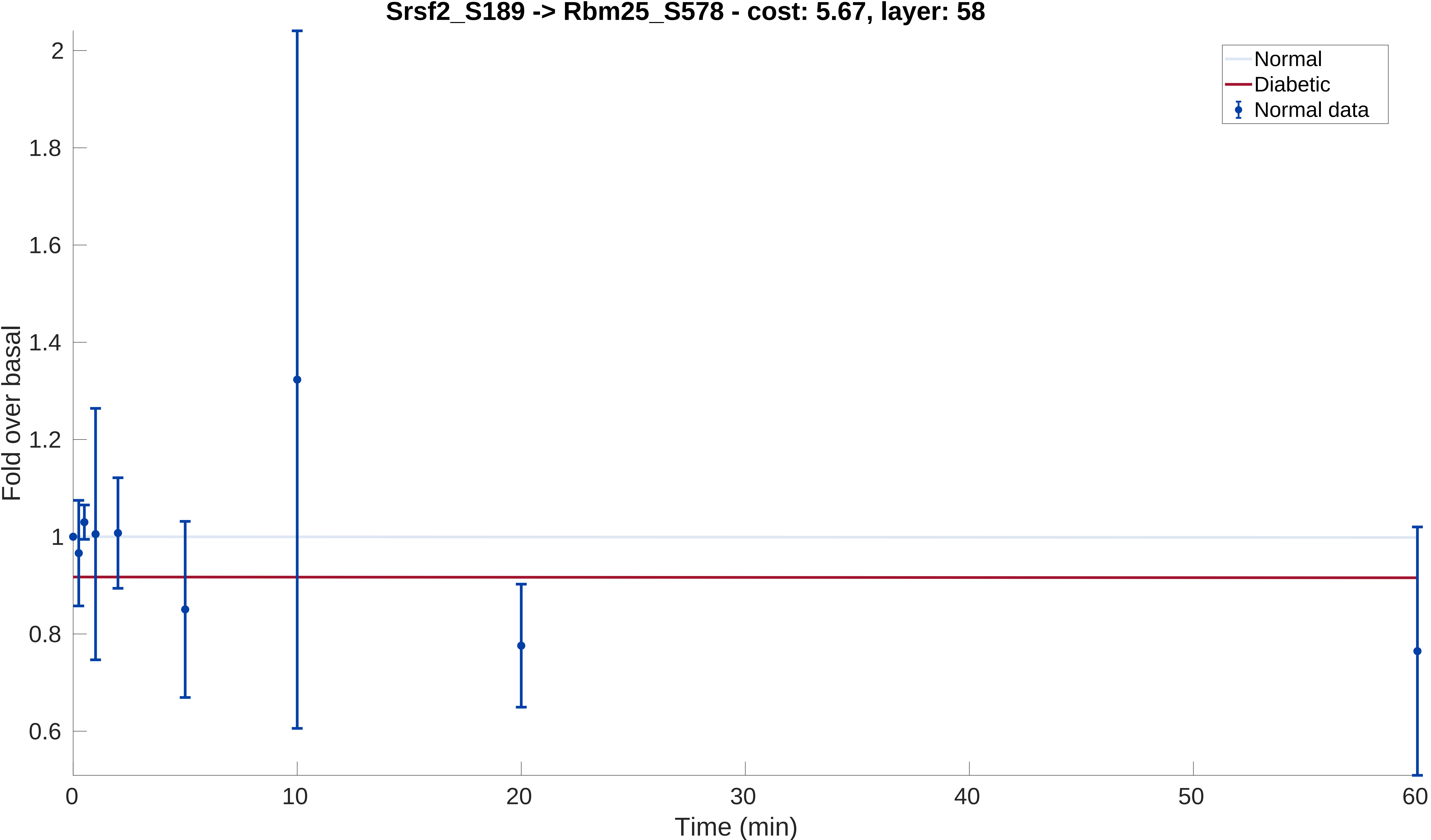

Srsf2_S189 -> Rbm25_S578 - cost: 5.67, layer: 58
Normal
Diabetic
Normal data
Fold over basal
2
1.8
1.6
1.4
1.2
1
0.8
0.6
0
10
20
30
40
50
60
Time (min)

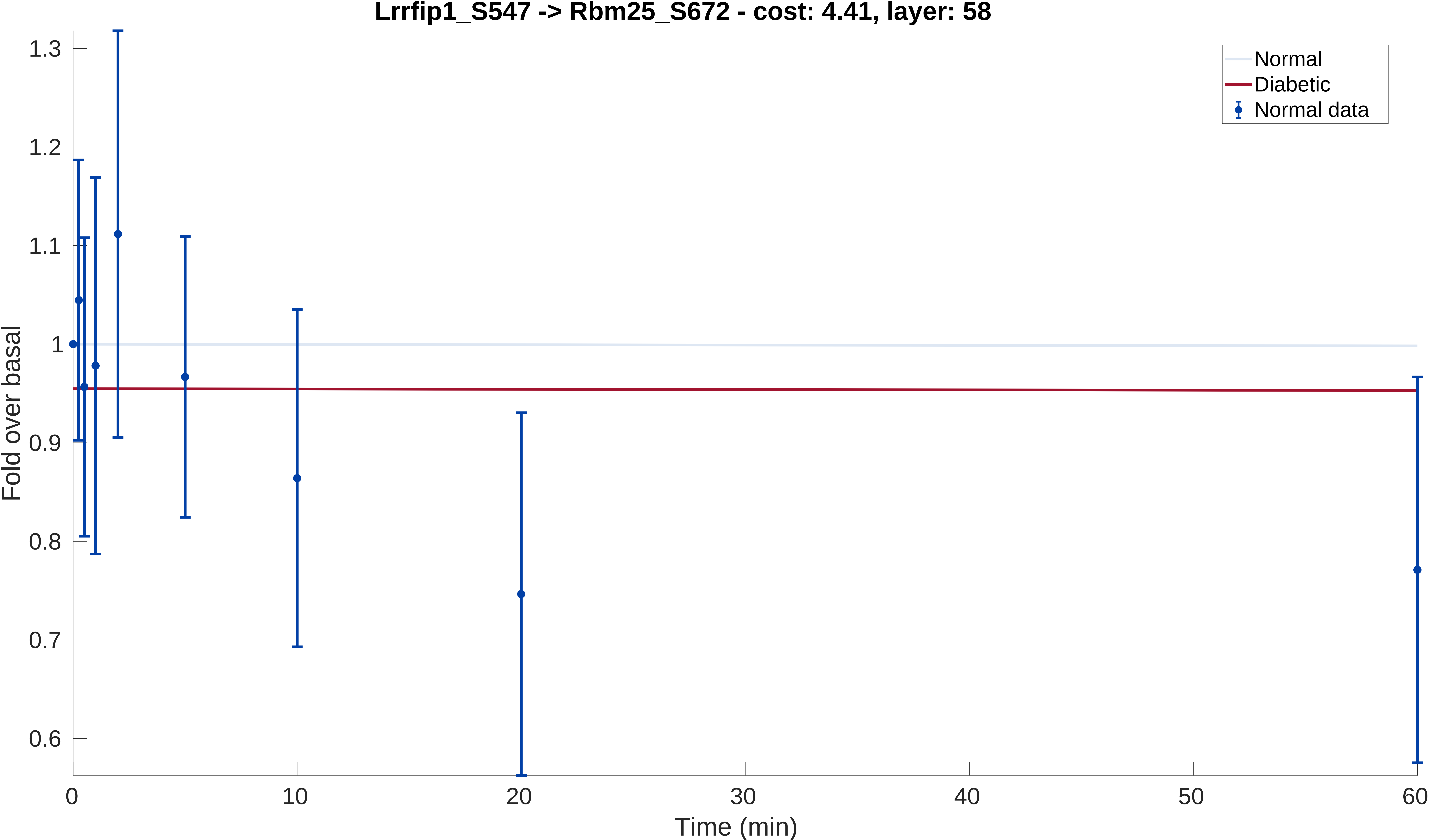

Lrrfip1_S547 -> Rbm25_S672 - cost: 4.41, layer: 58
Normal
Diabetic
Normal data
Fold over basal
Time (min)
1.3
1.2
1.1
1
0.9
0.8
0.7
0.6
0
10
20
30
40
50
60

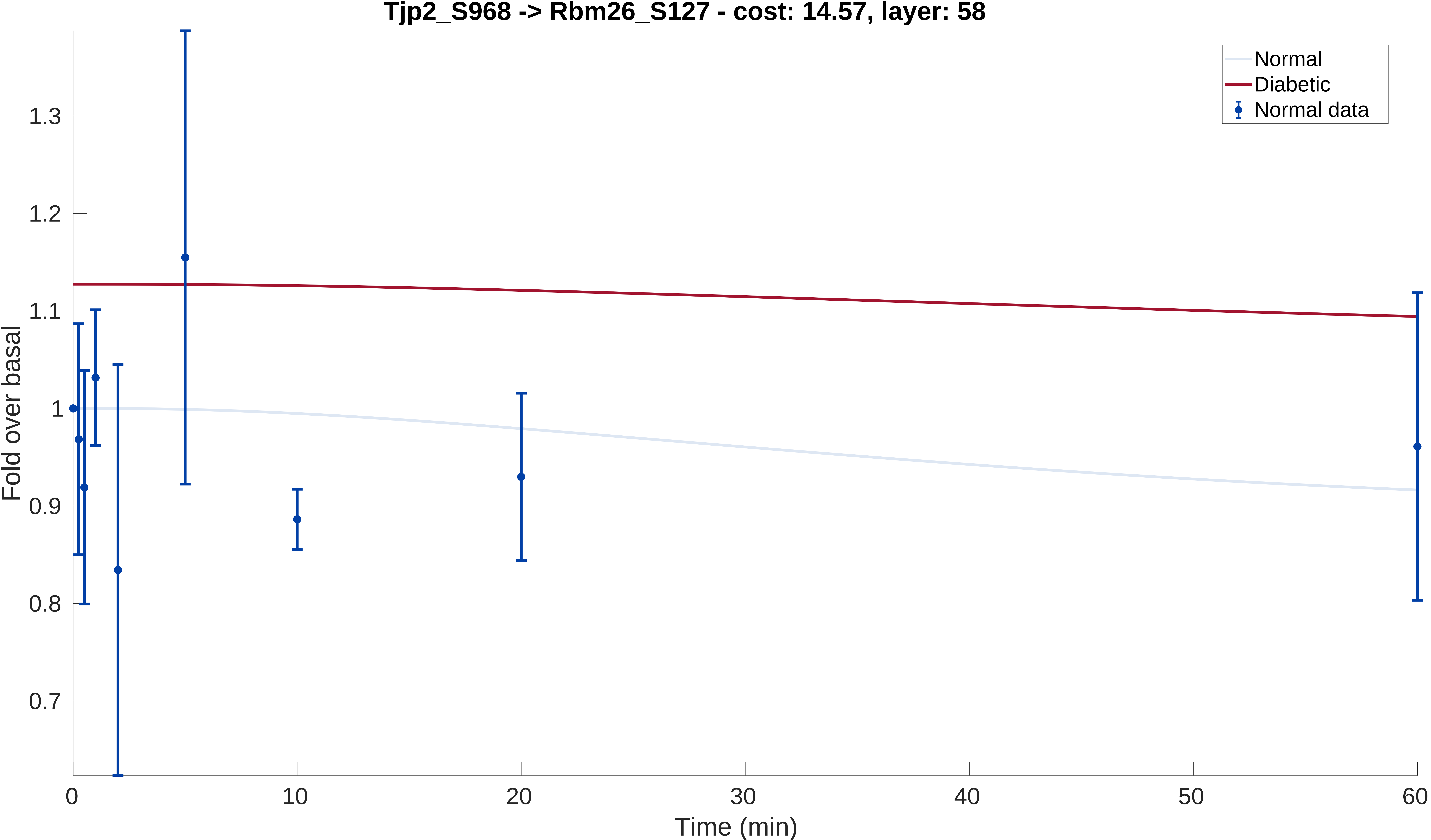
Tjp2_S968 -> Rbm26_S127 - cost: 14.57, layer: 58
Normal
Diabetic
Normal data
Fold over basal
1.3
1.2
1.1
1
0.9
0.8
0.7
0
10
20
30
40
50
60
Time (min)

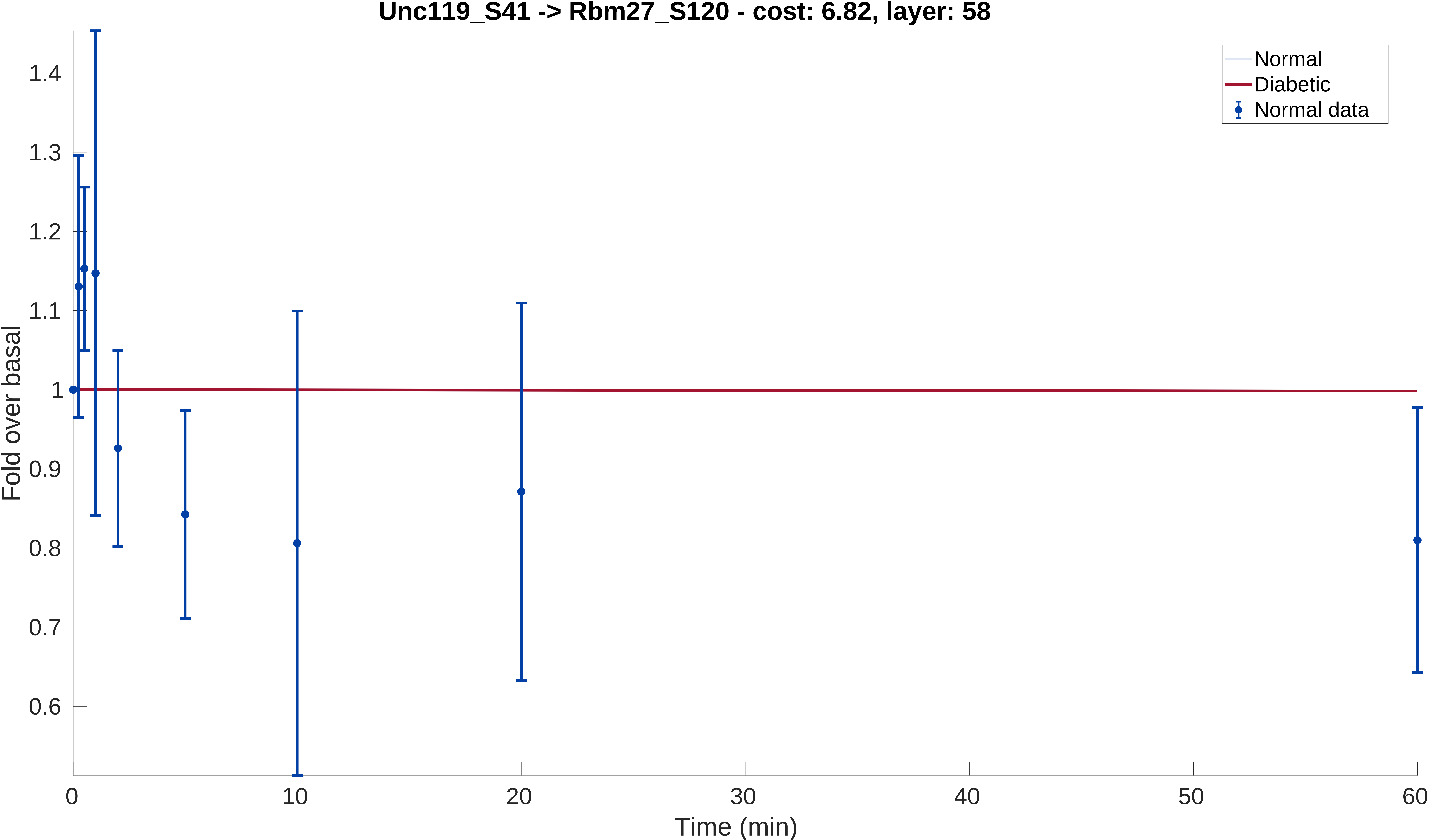

Unc119_S41 -> Rbm27_S120 - cost: 6.82, layer: 58
Normal
Diabetic
Normal data
Fold over basal
1.4
1.3
1.2
1.1
1
0.9
0.8
0.7
0.6
0
10
20
30
40
50
60
Time (min)

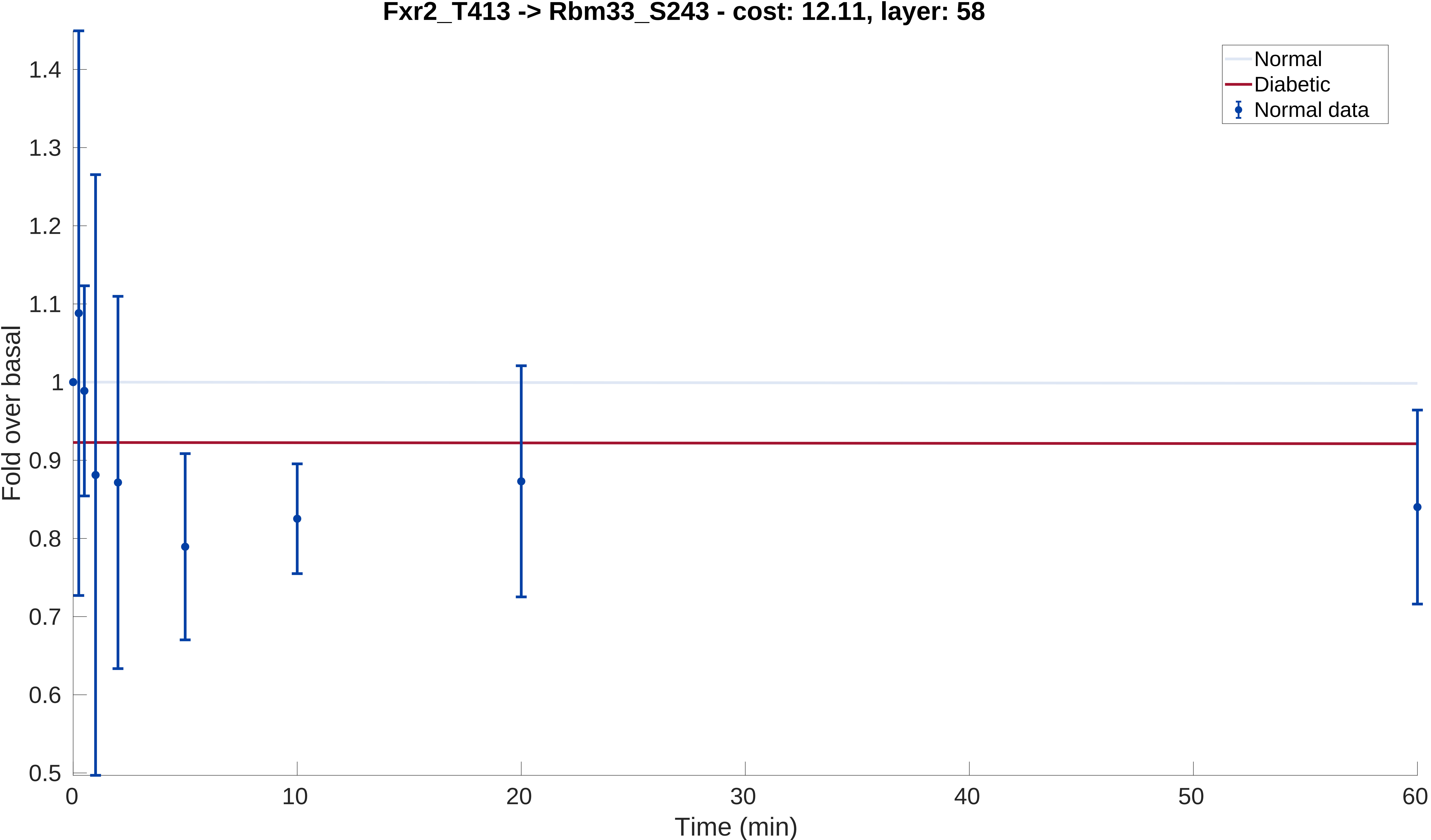
Fxr2_T413 -> Rbm33_S243 - cost: 12.11, layer: 58
Normal
Diabetic
Normal data
Fold over basal
1.4
1.3
1.2
1.1
1
0.9
0.8
0.7
0.6
0.5
0
10
20
30
40
50
60
Time (min)

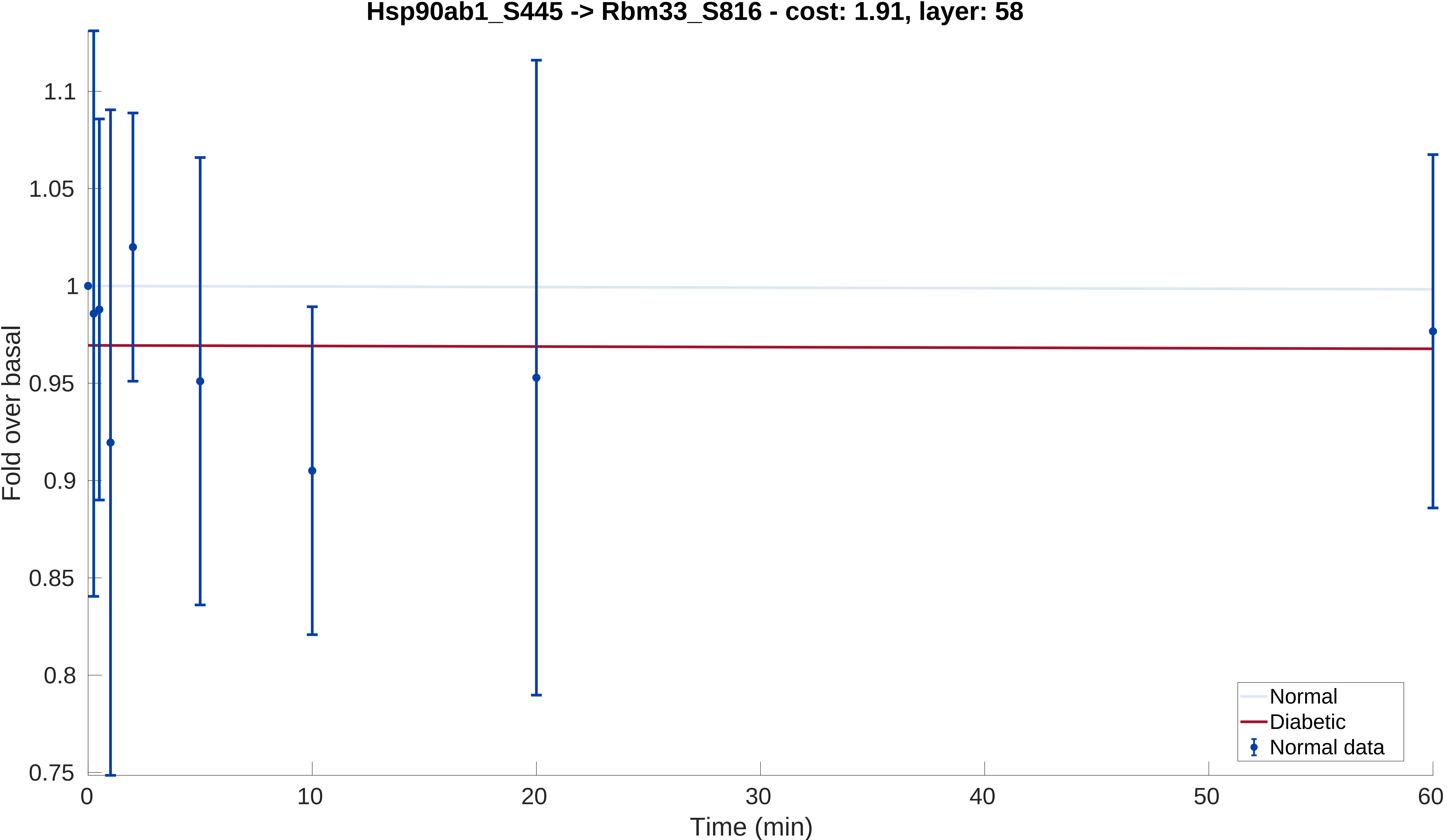

Hsp90ab1_S445 -> Rbm33_S816 - cost: 1.91, layer: 58
Fold over basal
1.1
1.05
1
0.95
0.9
0.85
0.8
0.75
0
10
20
30
40
50
60
Time (min)
Normal
Diabetic
Normal data

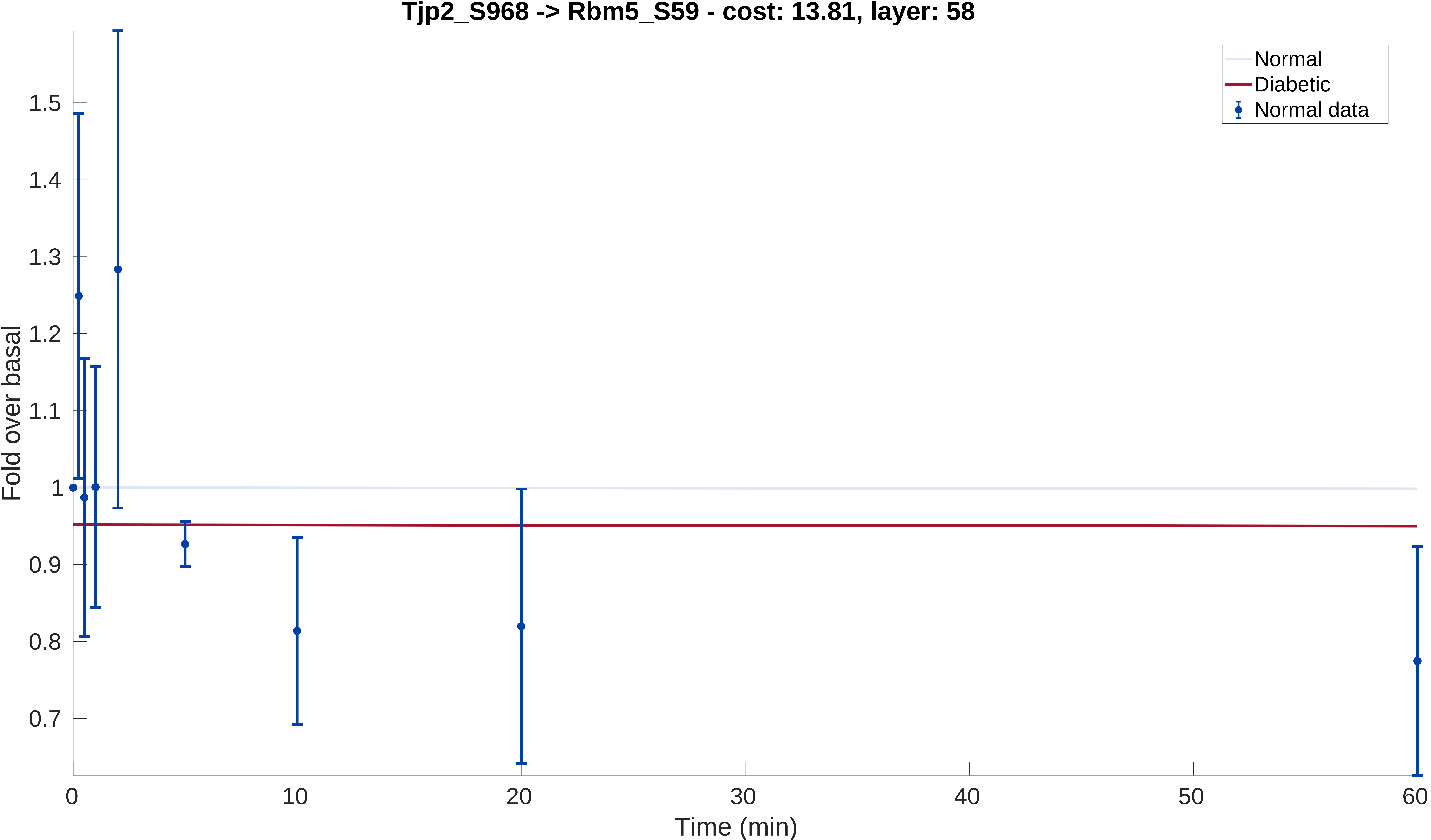

Tjp2_S968 -> Rbm5_S59 - cost: 13.81, layer: 58
Normal
Diabetic
Normal data
Fold over basal
1.5
1.4
1.3
1.2
1.1
1
0.9
0.8
0.7
0
10
20
30
40
50
60
Time (min)

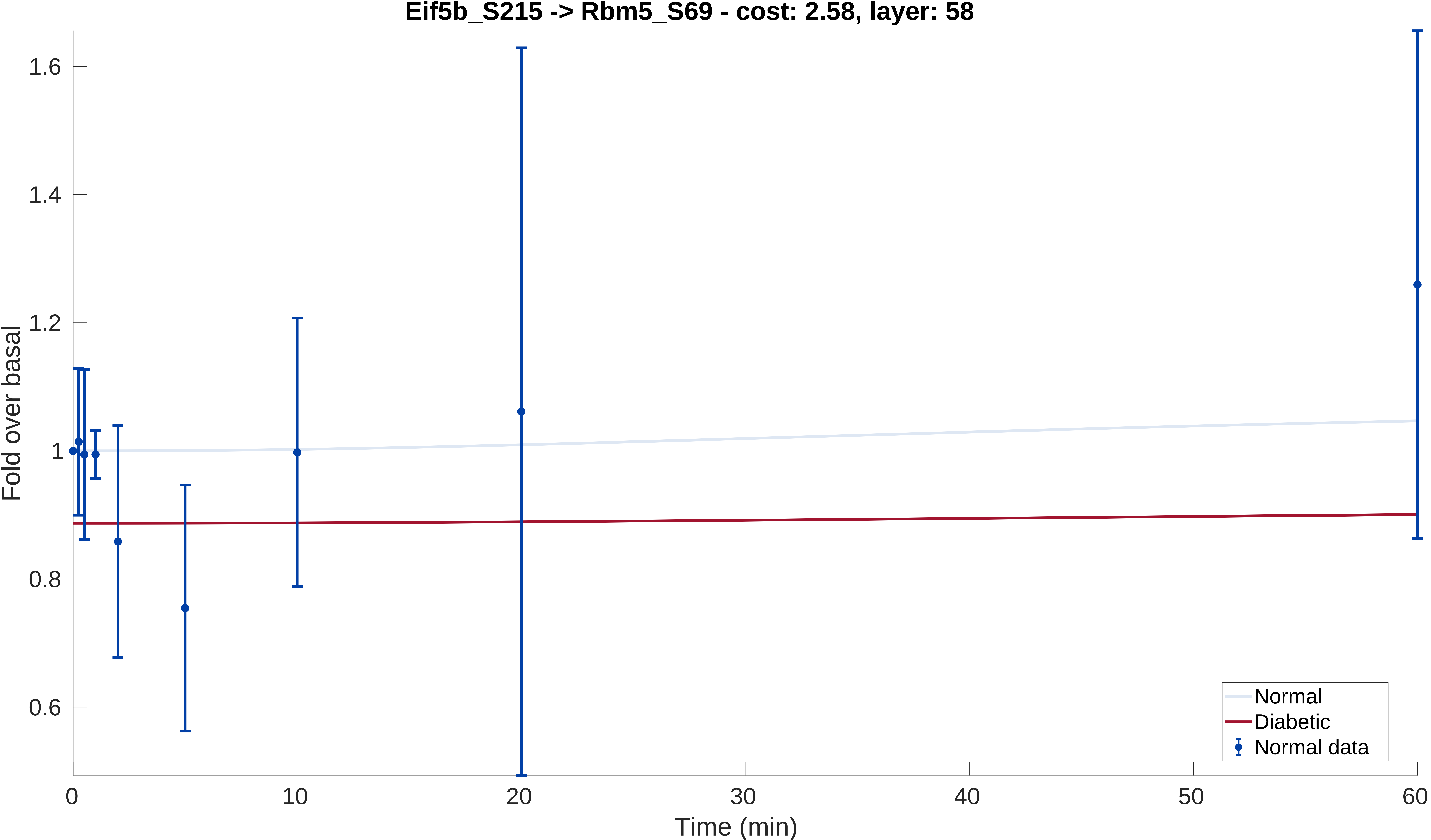

Eif5b_S215 -> Rbm5_S69 - cost: 2.58, layer: 58
Fold over basal
1.6
1.4
1.2
1
0.8
0.6
0
10
20
30
40
50
60
Time (min)
Normal
Diabetic
Normal data

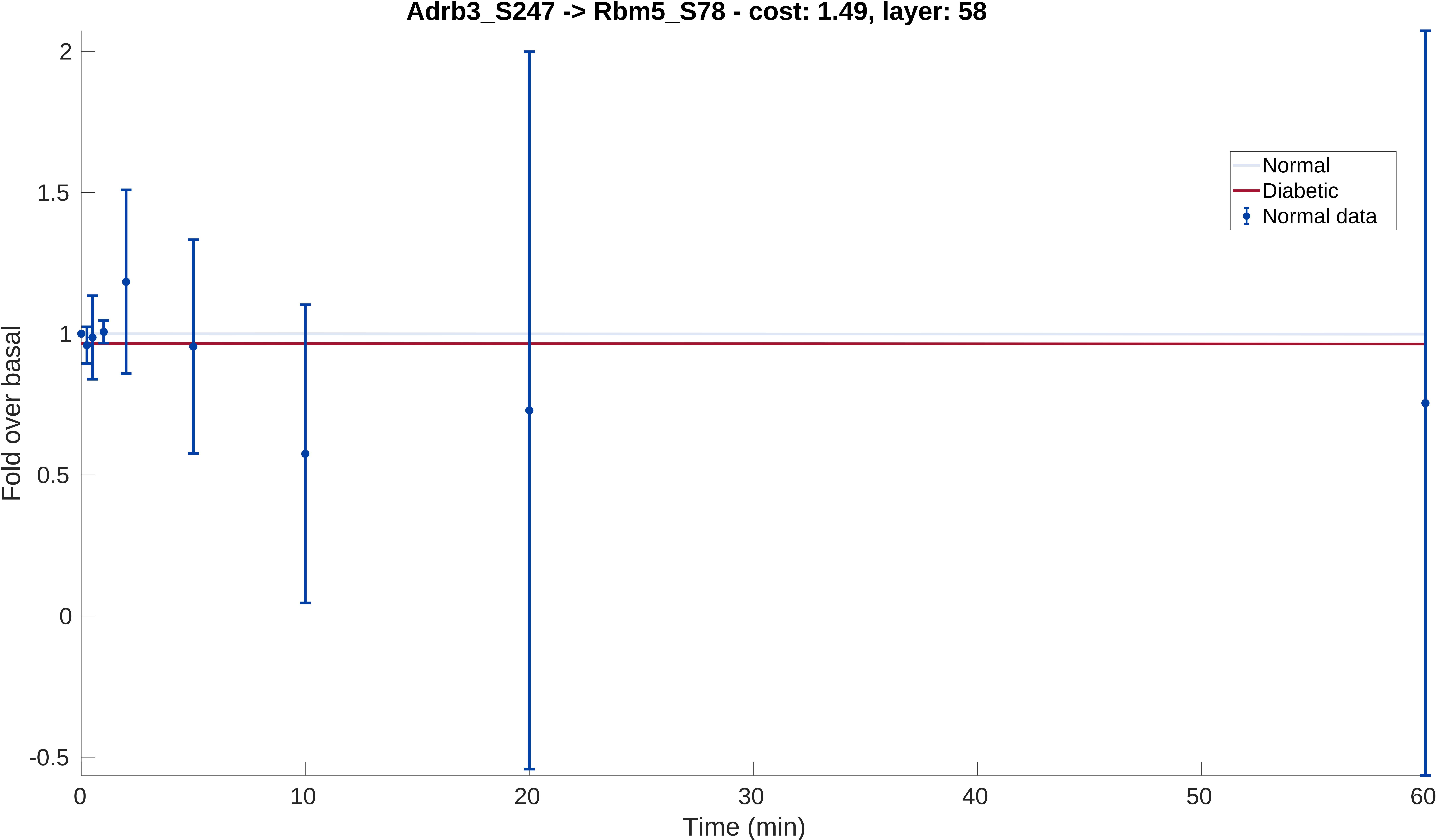
Adrb3_S247 -> Rbm5_S78 - cost: 1.49, layer: 58
Fold over basal
Time (min)
Normal
Diabetic
Normal data
2
1.5
1
0.5
0
-0.5
0
10
20
30
40
50
60

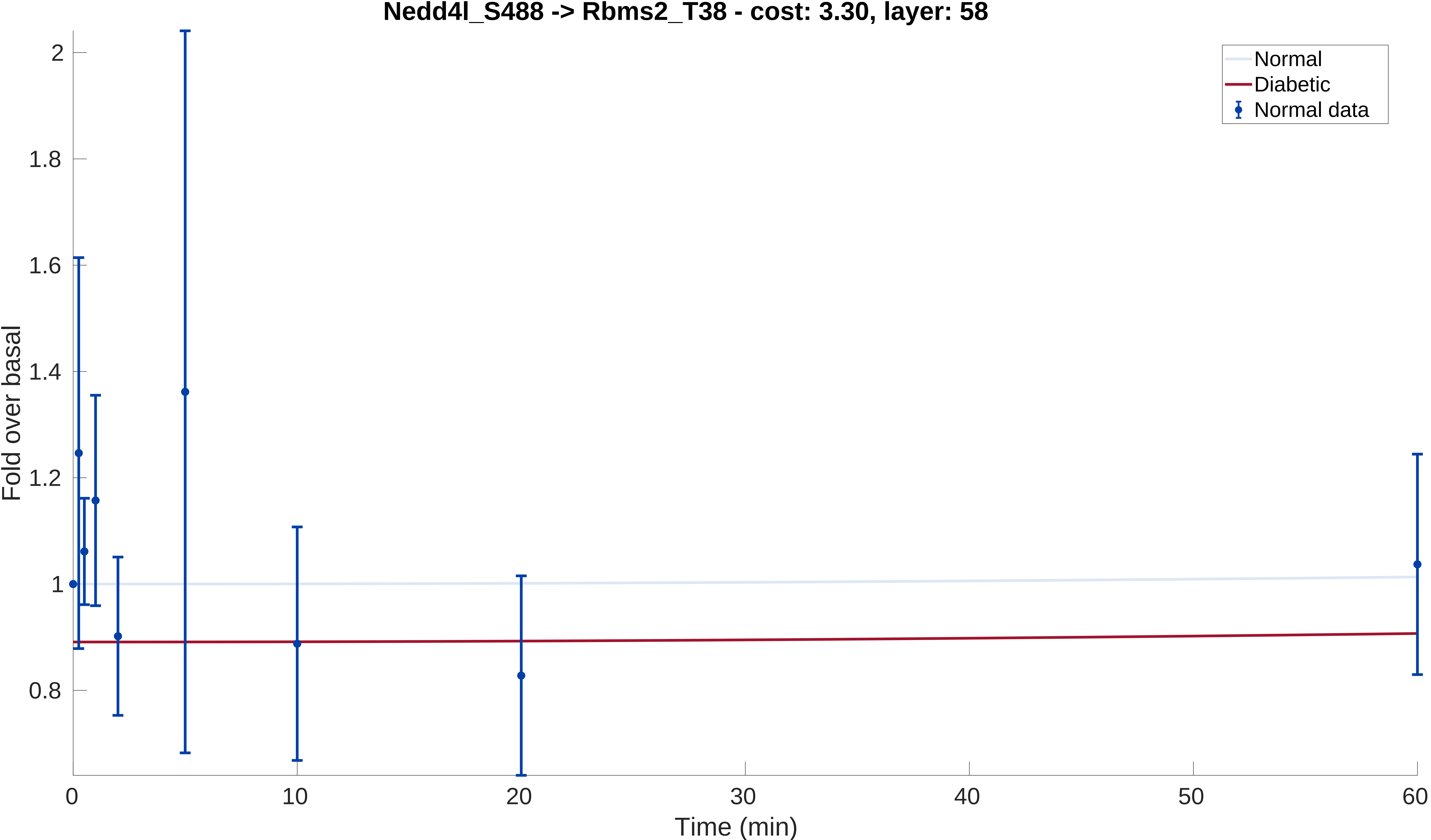
Nedd4l_S488 -> Rbms2_T38 - cost: 3.30, layer: 58
Normal
Diabetic
Normal data
Fold over basal
2
1.8
1.6
1.4
1.2
1
0.8
0
10
20
30
40
50
60
Time (min)

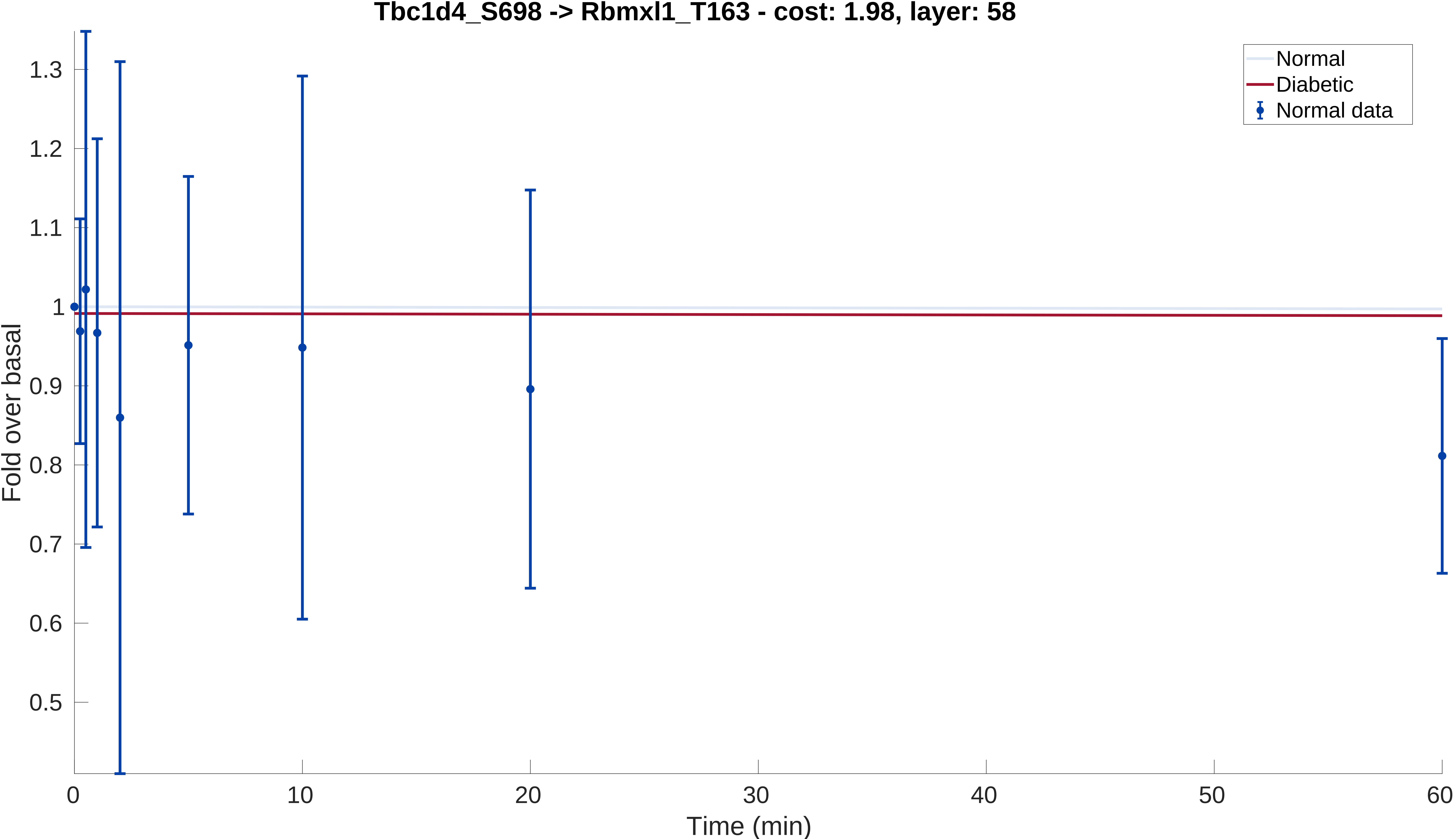
Tbc1d4_S698 -> Rbmxl1_T163 - cost: 1.98, layer: 58
Normal
Diabetic
Normal data
Fold over basal
Time (min)
1.3
1.2
1.1
1
0.9
0.8
0.7
0.6
0.5
0
10
20
30
40
50
60

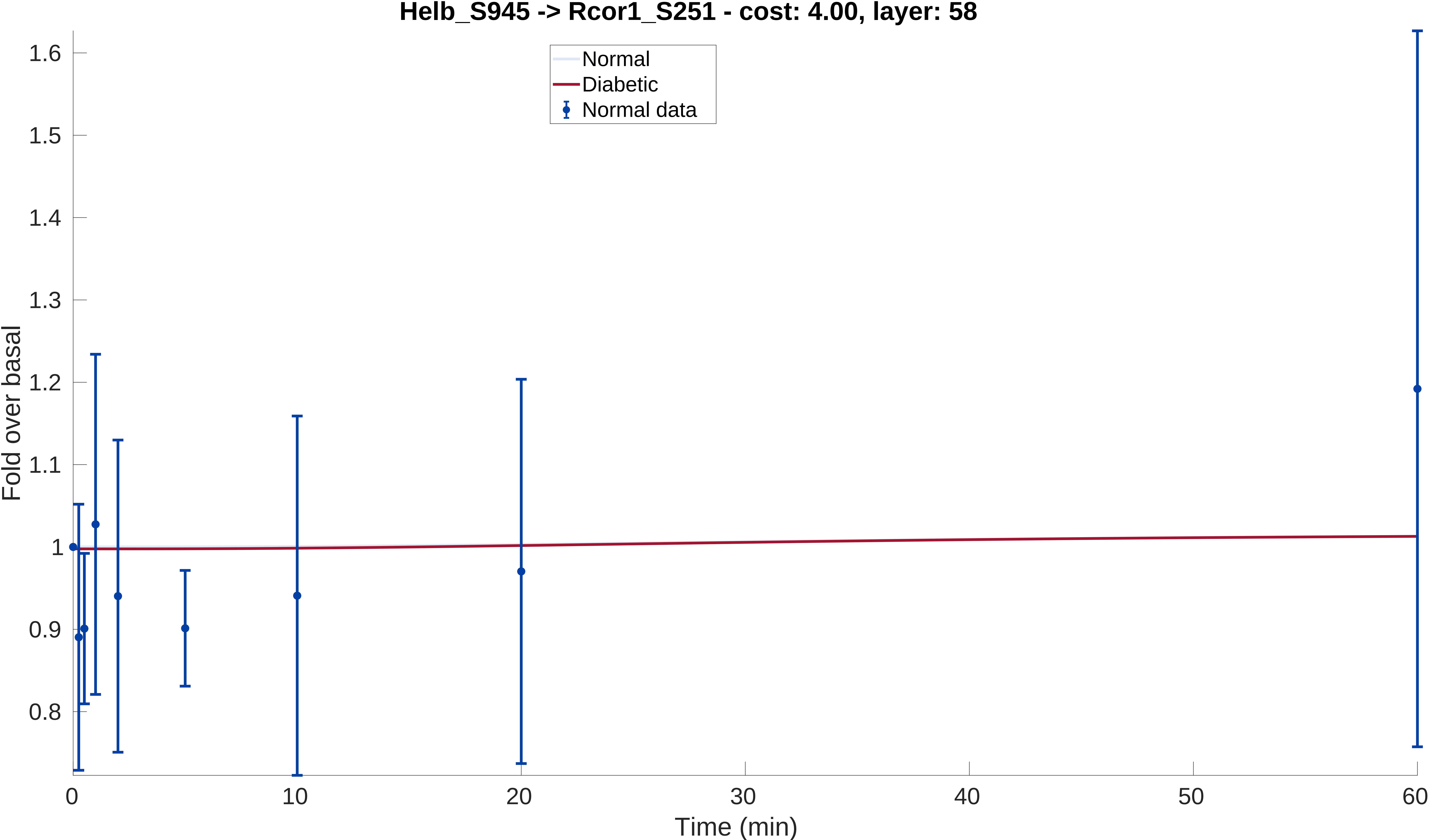

Helb_S945 -> Rcor1_S251 - cost: 4.00, layer: 58
Normal
Diabetic
Normal data
1.6
1.5
1.4
1.3
1.2
1.1
1
0.9
0.8
Fold over basal
0
10
20
30
40
50
60
Time (min)

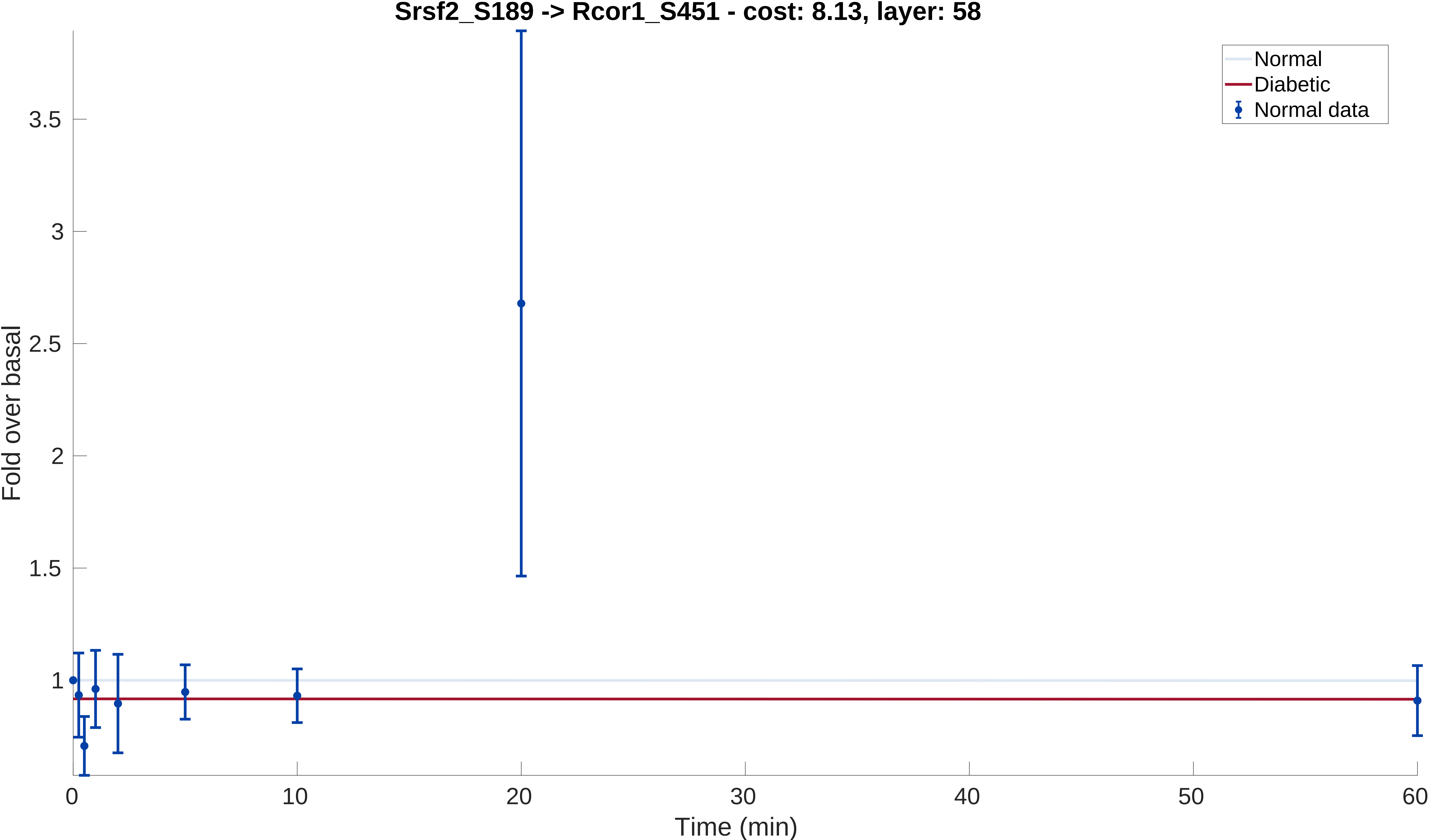
Srsf2_S189 -> Rcor1_S451 - cost: 8.13, layer: 58
Normal
Diabetic
Normal data
Fold over basal
Time (min)
3.5
3
2.5
2
1.5
1
0
10
20
30
40
50
60

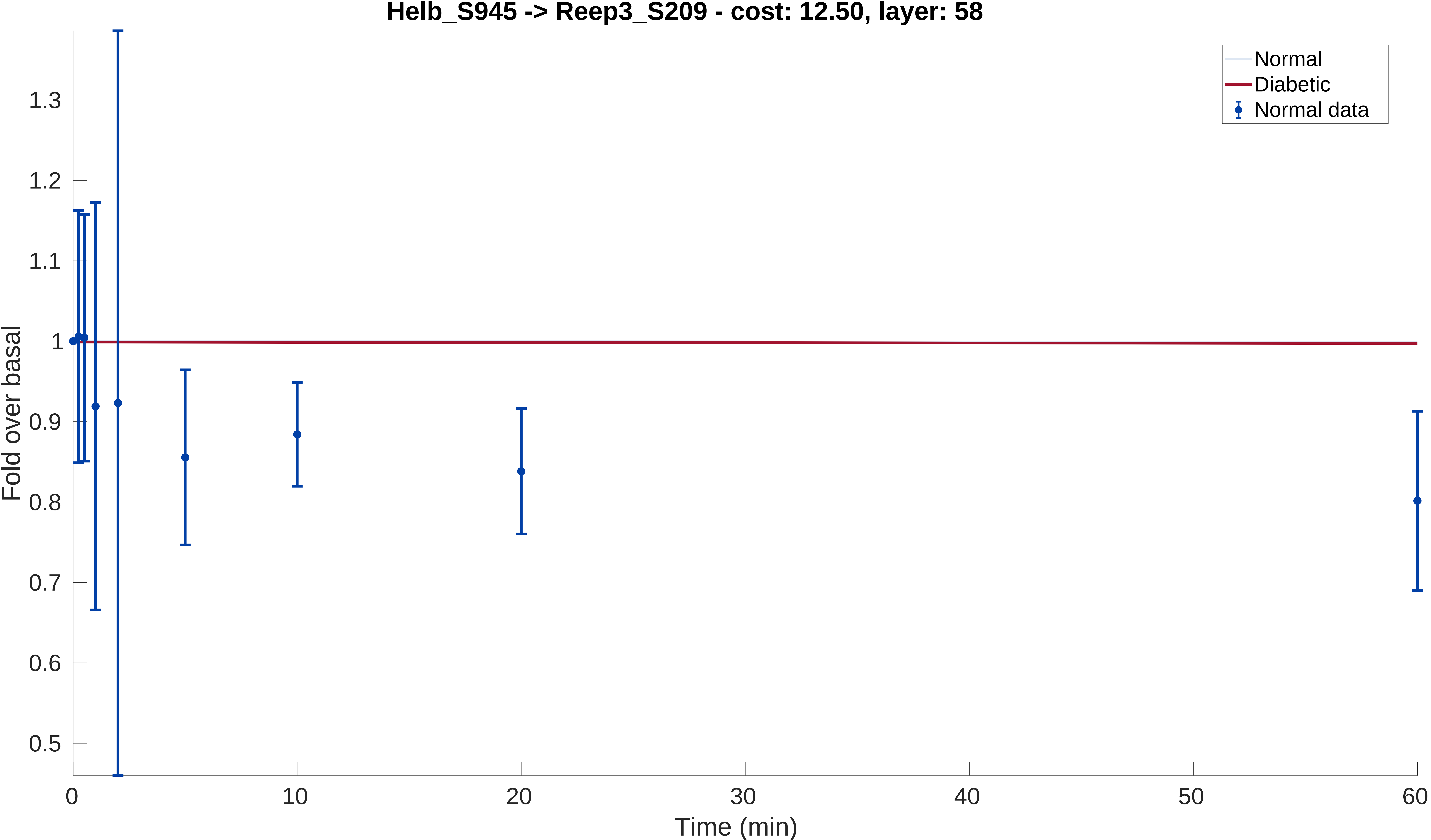

Helb_S945 -> Reep3_S209 - cost: 12.50, layer: 58
Normal
Diabetic
Normal data
Fold over basal
Time (min)
0
10
20
30
40
50
60
0.5
0.6
0.7
0.8
0.9
1
1.1
1.2
1.3

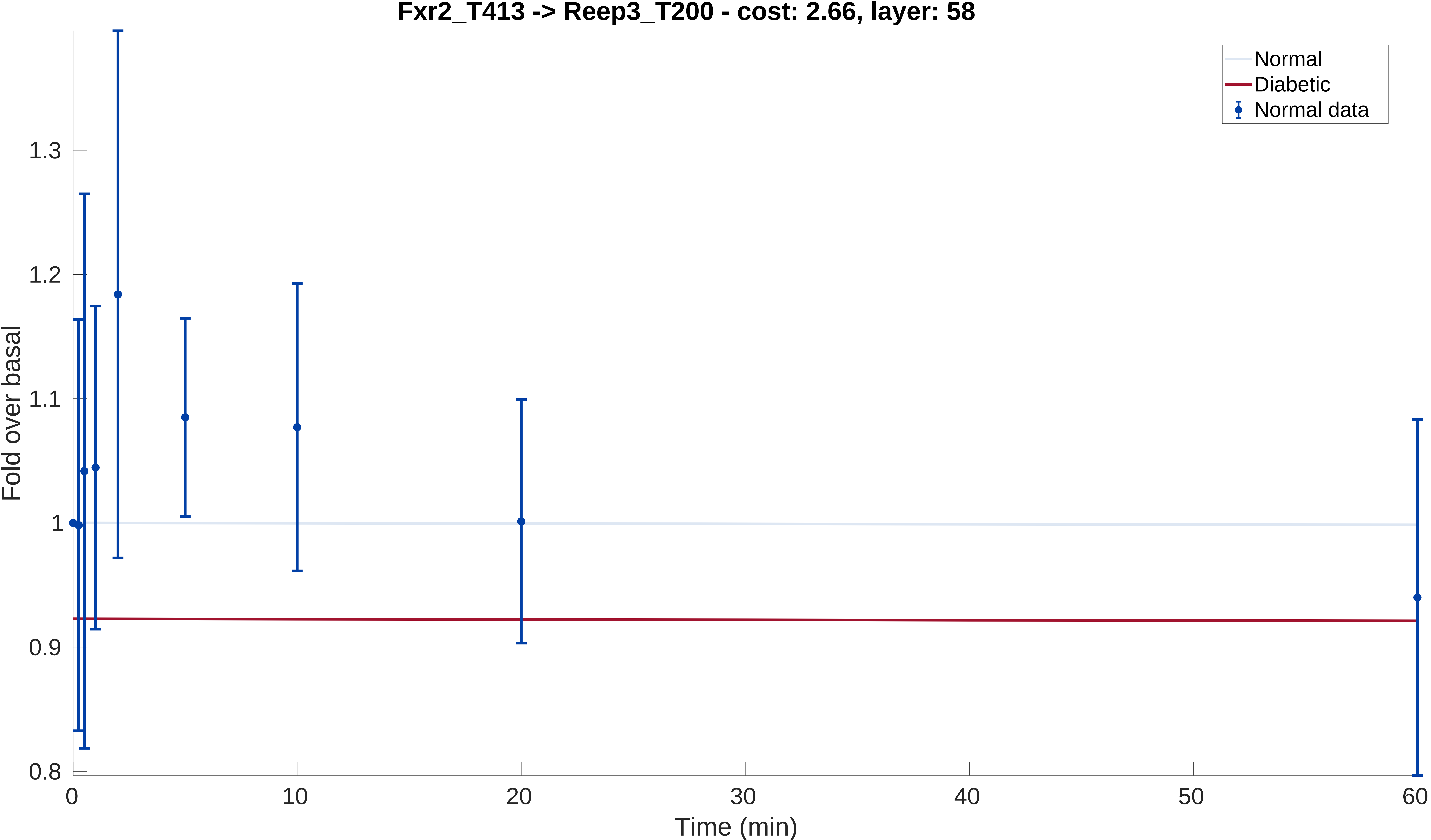

Fxr2_T413 -> Reep3_T200 - cost: 2.66, layer: 58
Normal
Diabetic
Normal data
1.3
1.2
1.1
1
0.9
0.8
Fold over basal
0
10
20
30
40
50
60
Time (min)

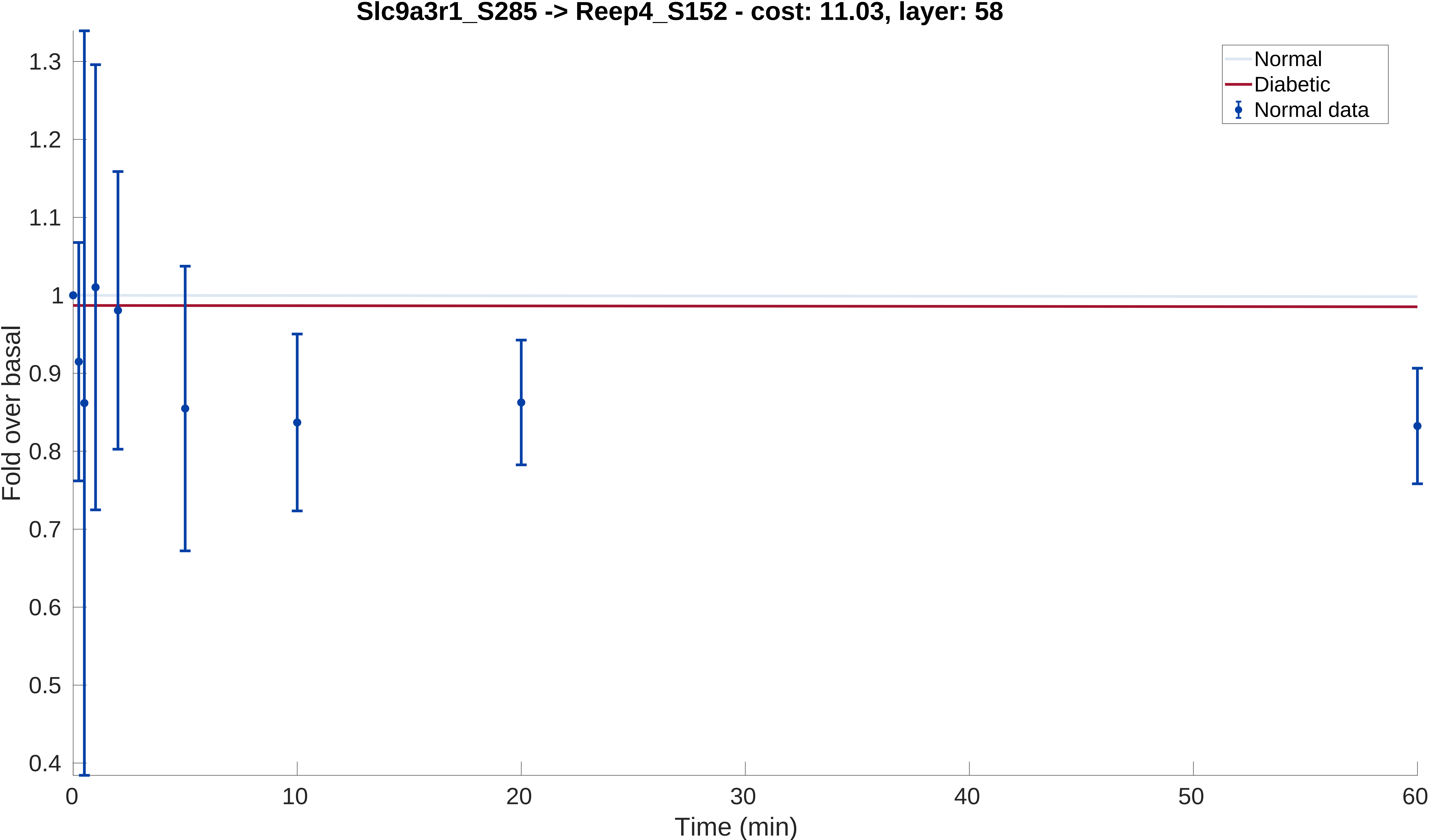

Slc9a3r1_S285 -> Reep4_S152 - cost: 11.03, layer: 58
Normal
Diabetic
Normal data
Fold over basal
Time (min)
0.4
0.5
0.6
0.7
0.8
0.9
1
1.1
1.2
1.3
0
10
20
30
40
50
60

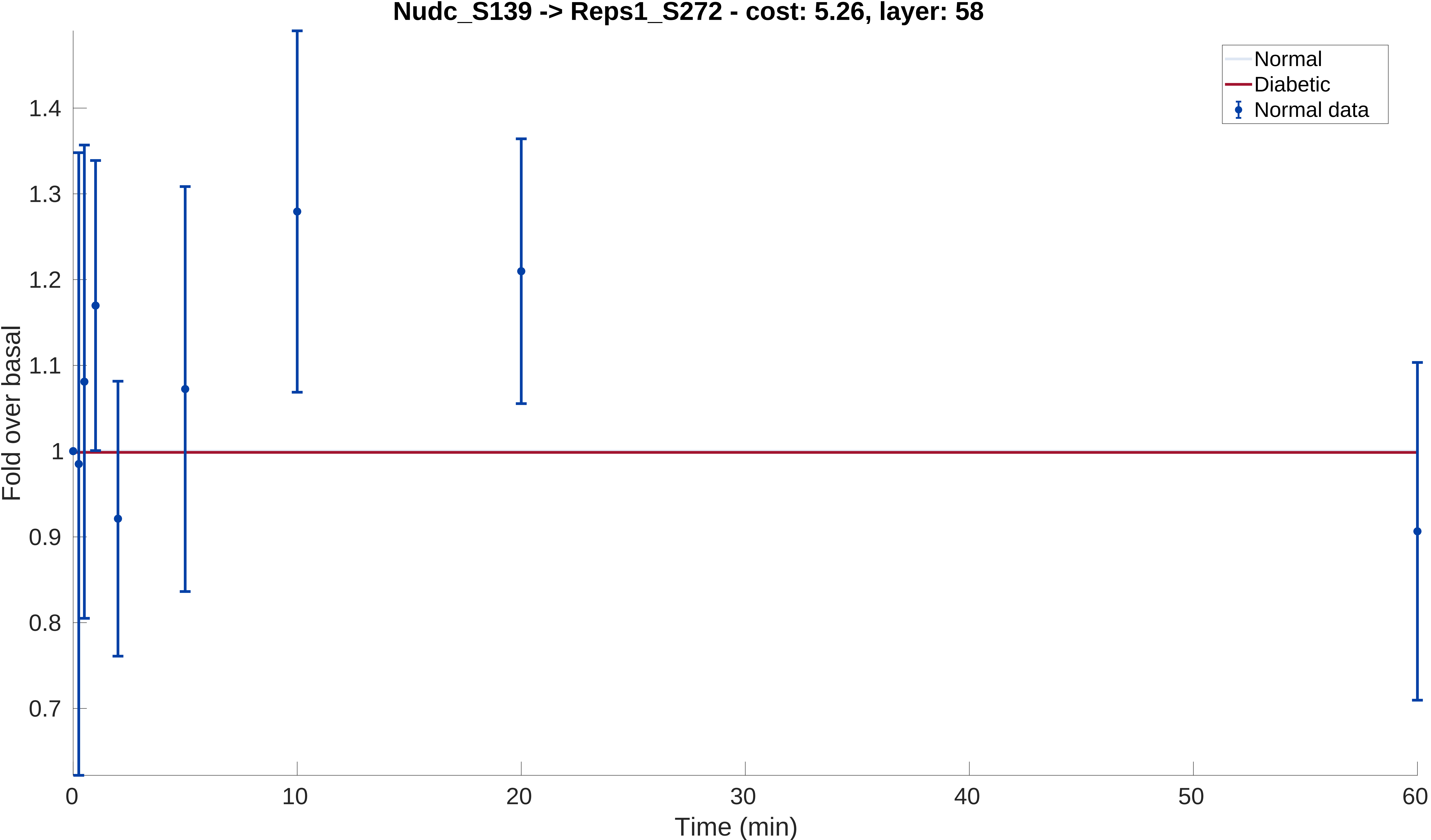
Nudc_S139 -> Reps1_S272 - cost: 5.26, layer: 58
Normal
Diabetic
Normal data
Fold over basal
1.4
1.3
1.2
1.1
1
0.9
0.8
0.7
0
10
20
30
40
50
60
Time (min)

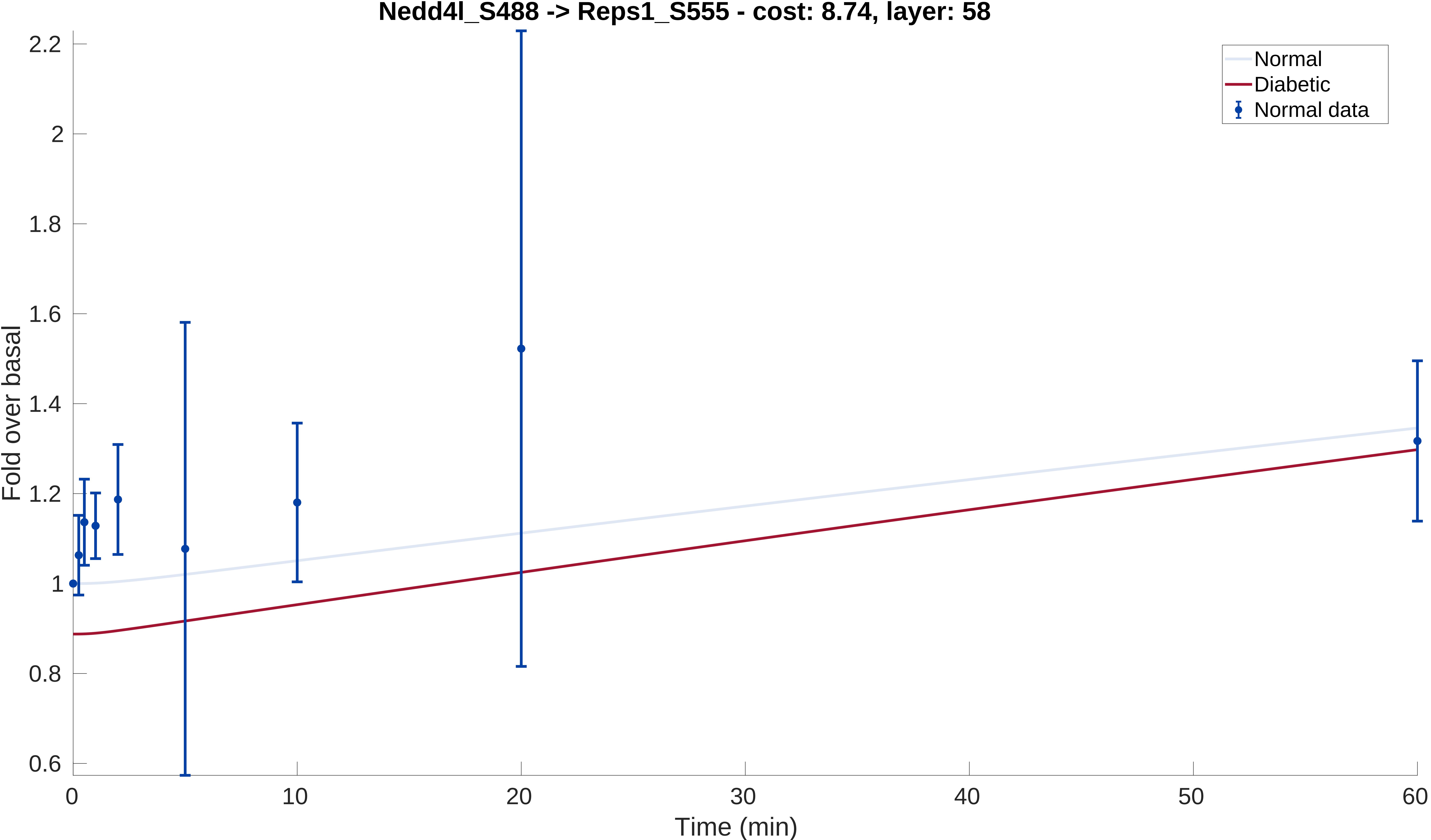

Nedd4l_S488 -> Reps1_S555 - cost: 8.74, layer: 58
Normal
Diabetic
Normal data
Fold over basal
Time (min)
0.6
0.8
1
1.2
1.4
1.6
1.8
2
2.2
0
10
20
30
40
50
60

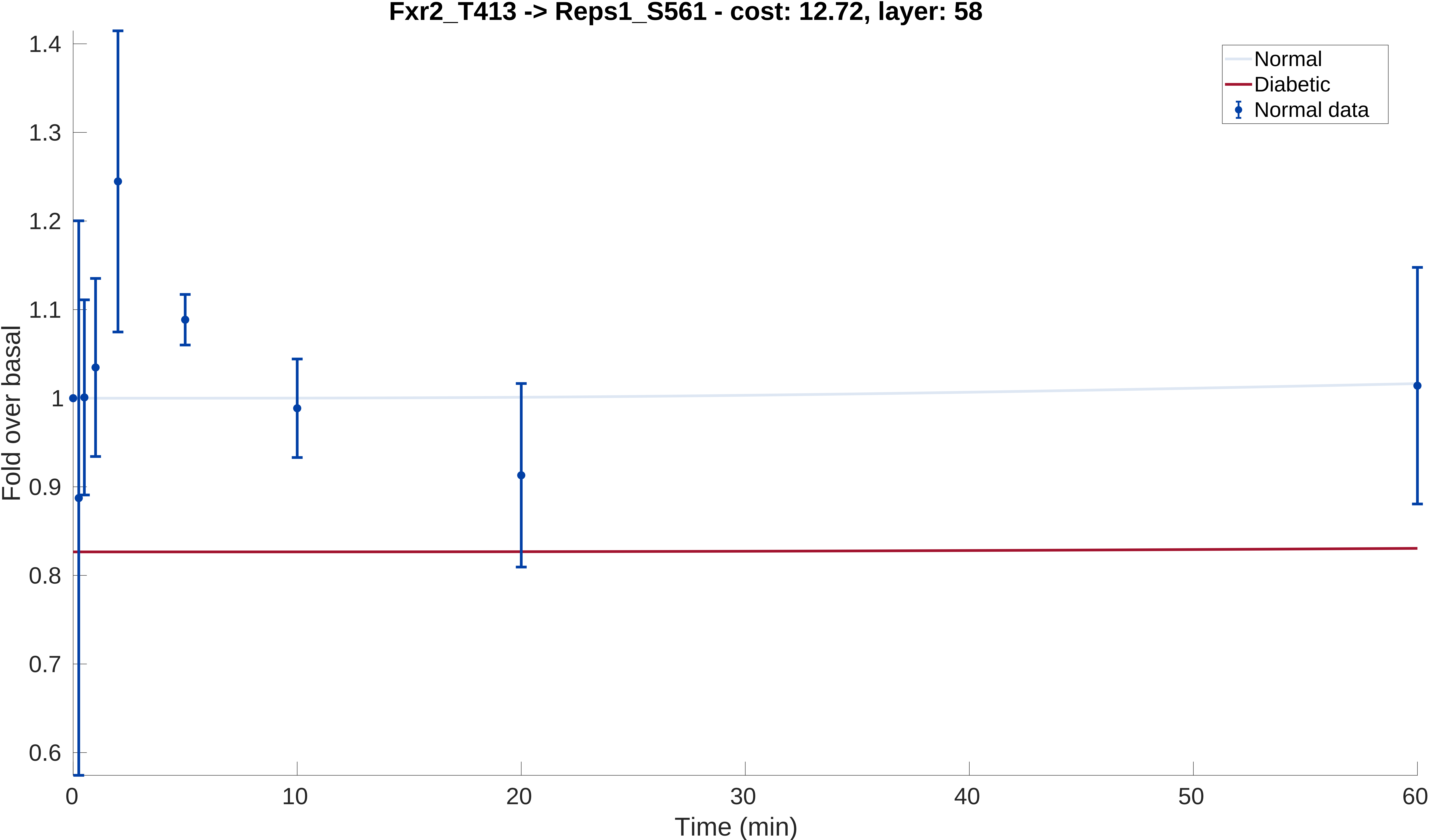

Fxr2_T413 -> Reps1_S561 - cost: 12.72, layer: 58
Normal
Diabetic
Normal data
Fold over basal
1.4
1.3
1.2
1.1
1
0.9
0.8
0.7
0.6
0
10
20
30
40
50
60
Time (min)

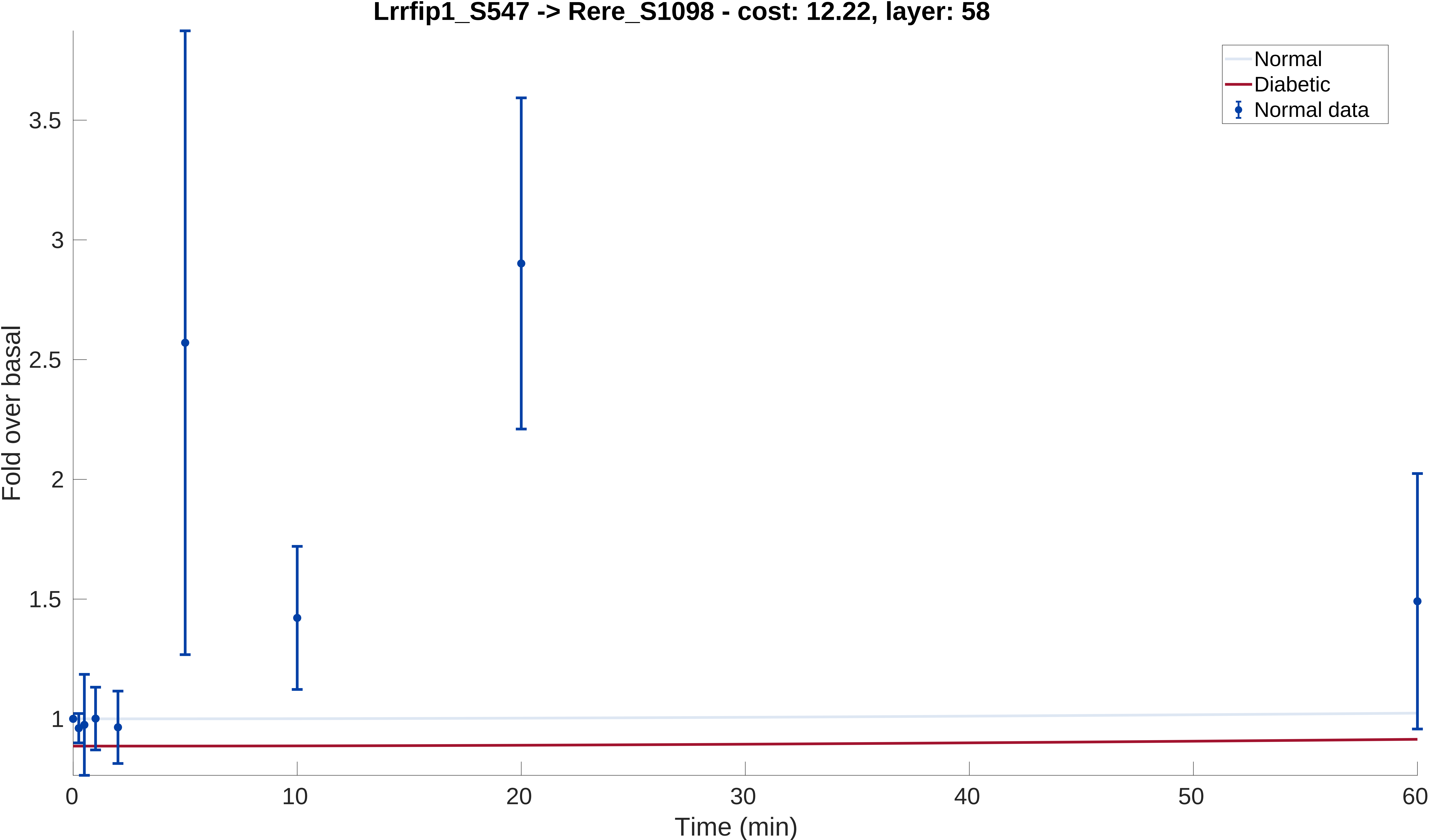
Lrrfip1_S547 -> Rere_S1098 - cost: 12.22, layer: 58
Normal
Diabetic
Normal data
Fold over basal
3.5
3
2.5
2
1.5
1
0
10
20
30
40
50
60
Time (min)

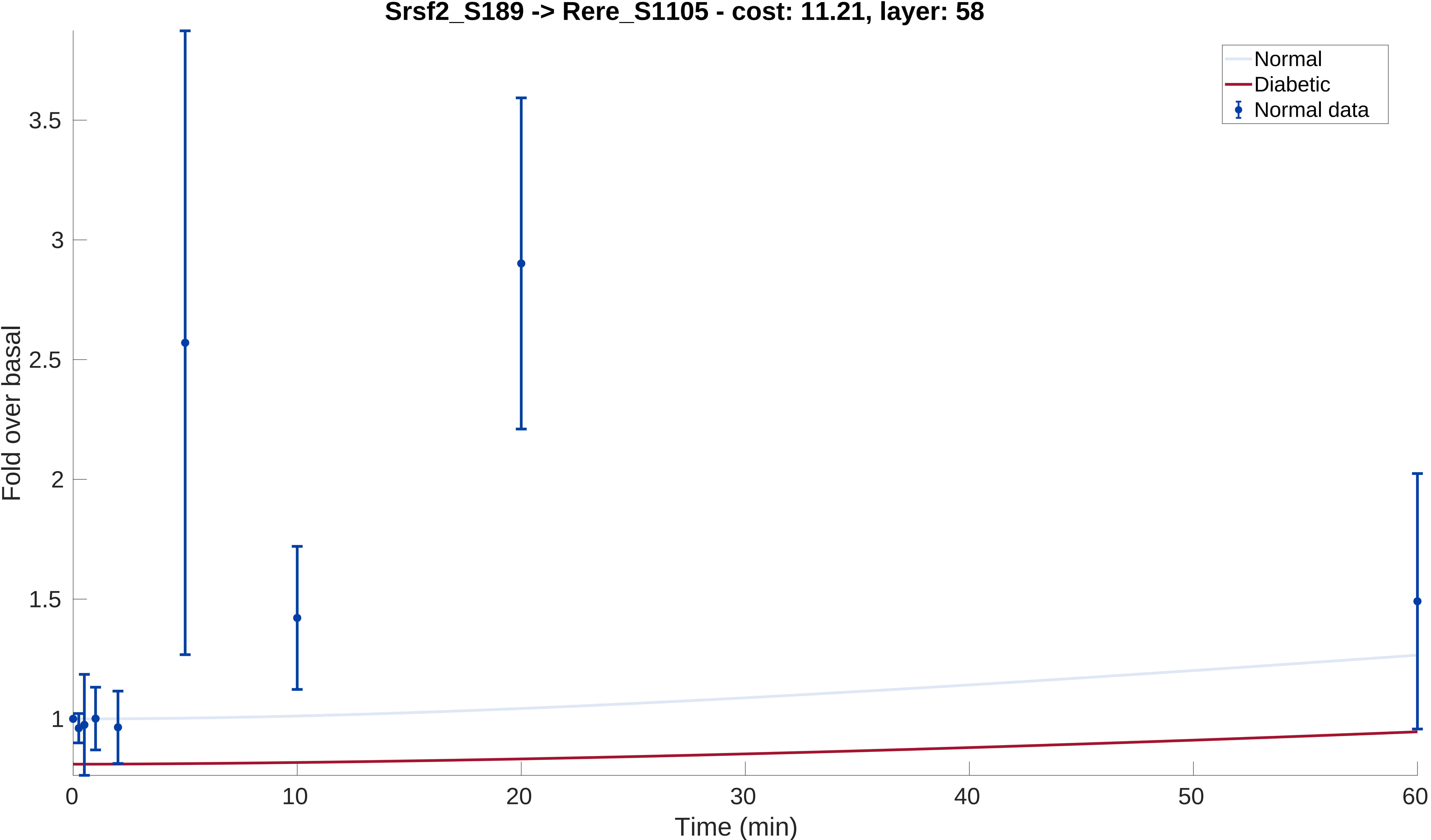

Srsf2_S189 -> Rere_S1105 - cost: 11.21, layer: 58
Normal
Diabetic
Normal data
Fold over basal
3.5
3
2.5
2
1.5
1
0
10
20
30
40
50
60
Time (min)

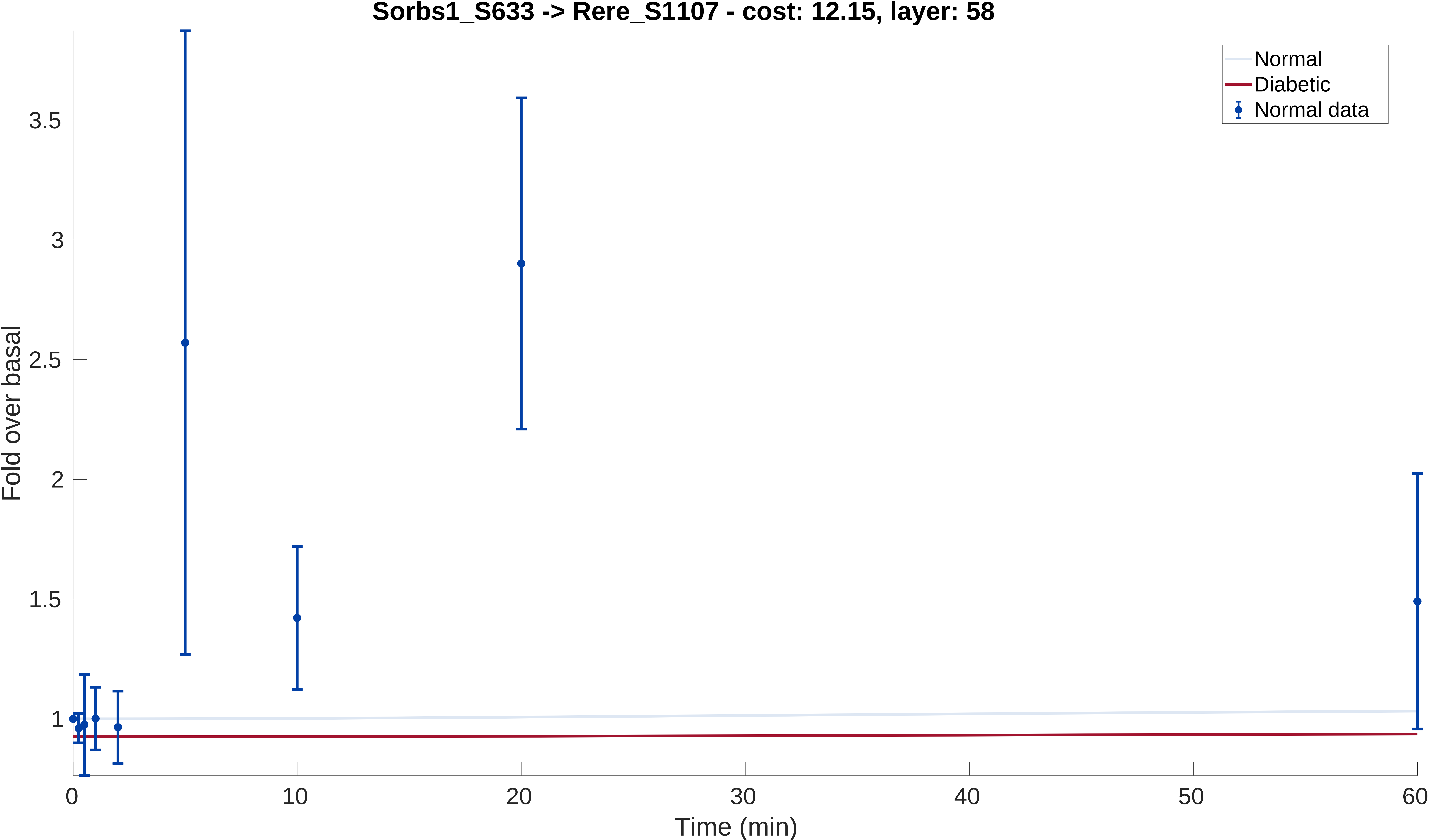

Sorbs1_S633 -> Rere_S1107 - cost: 12.15, layer: 58
Normal
Diabetic
Normal data
Fold over basal
3.5
3
2.5
2
1.5
1
0
10
20
30
40
50
60
Time (min)

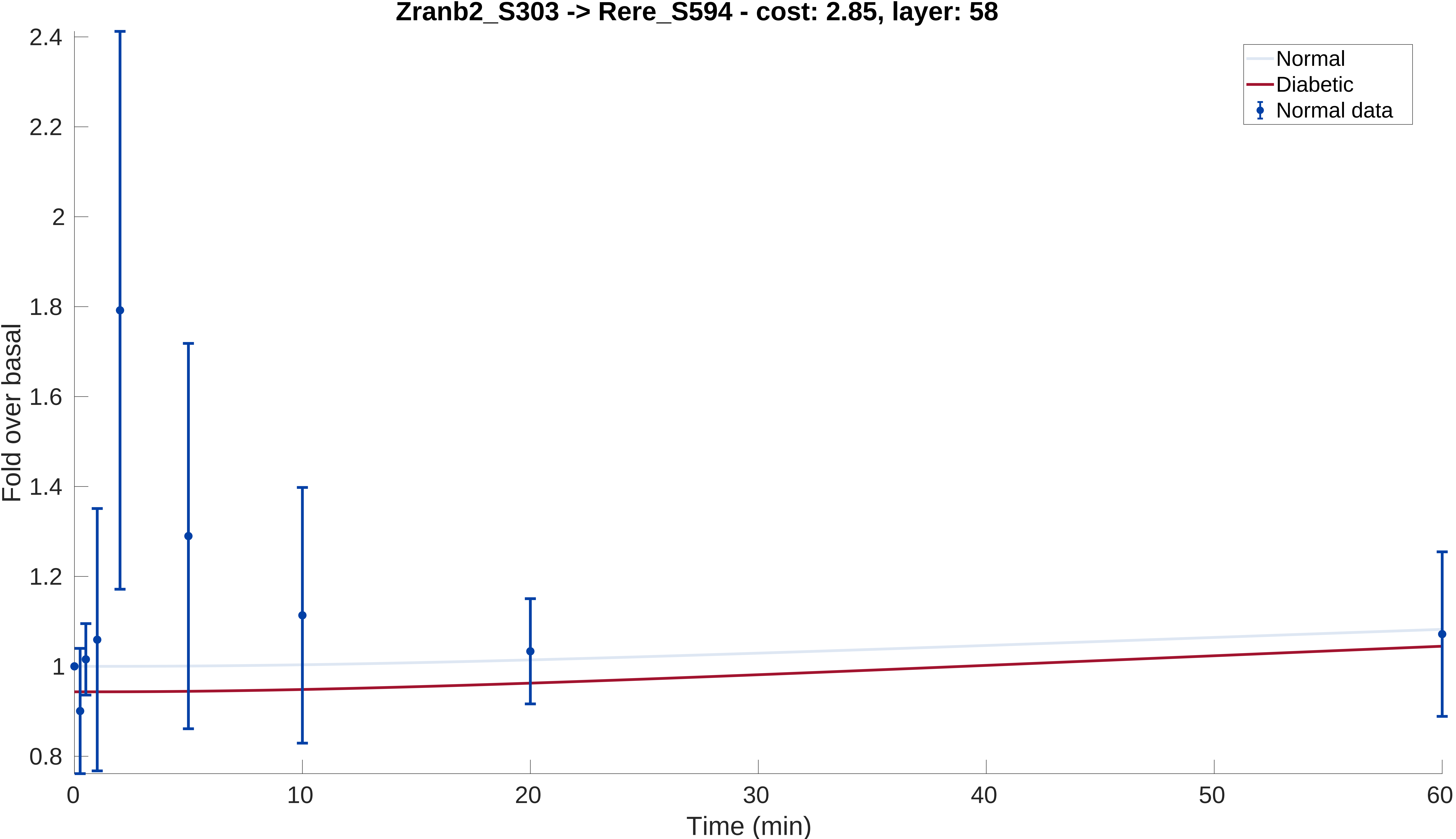
Zranb2_S303 -> Rere_S594 - cost: 2.85, layer: 58
Normal
Diabetic
Normal data
Fold over basal
Time (min)
2.4
2.2
2
1.8
1.6
1.4
1.2
1
0.8
0
10
20
30
40
50
60

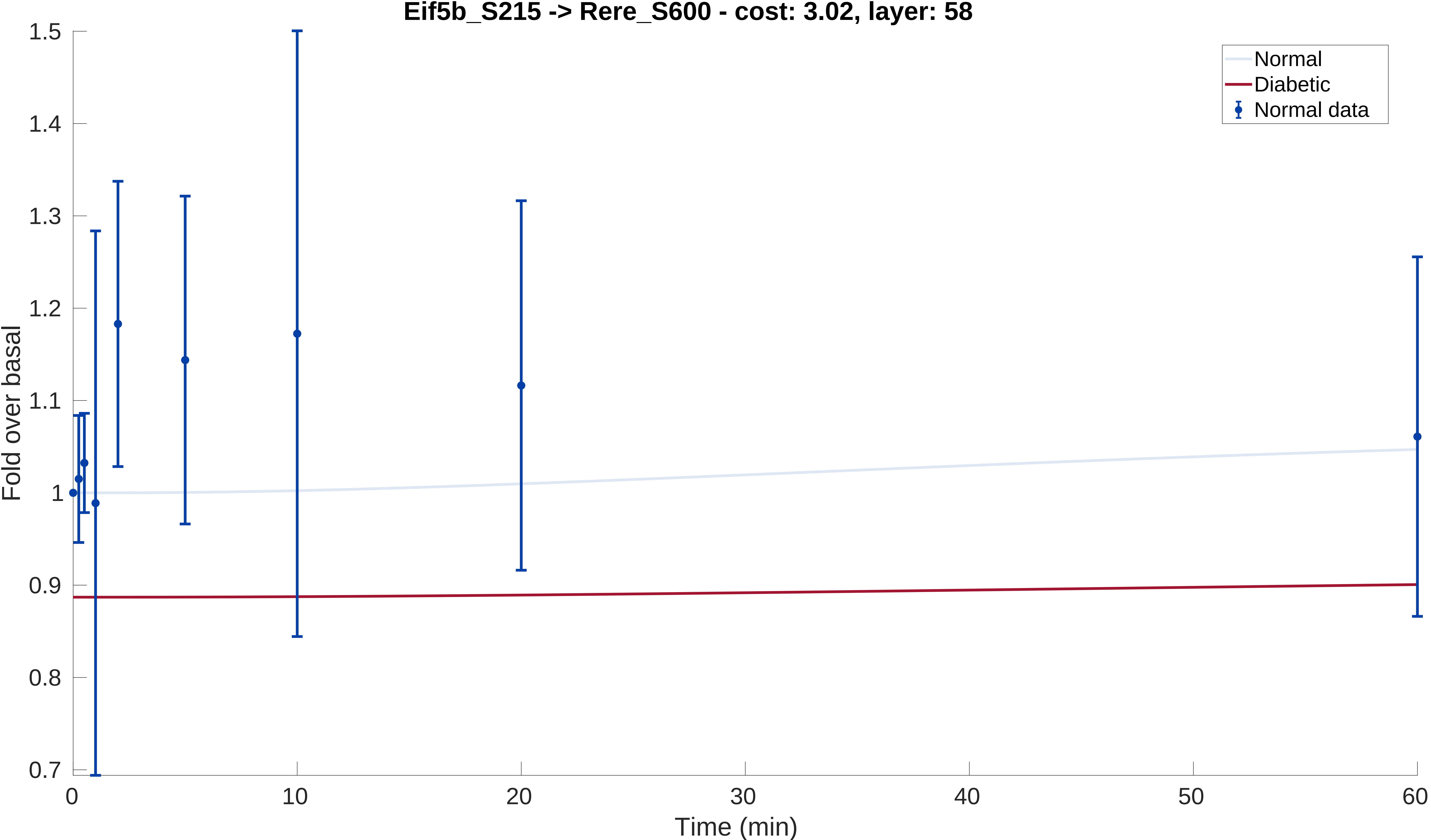

Eif5b_S215 -> Rere_S600 - cost: 3.02, layer: 58
Normal
Diabetic
Normal data
Fold over basal
Time (min)
1.5
1.4
1.3
1.2
1.1
1
0.9
0.8
0.7
0
10
20
30
40
50
60

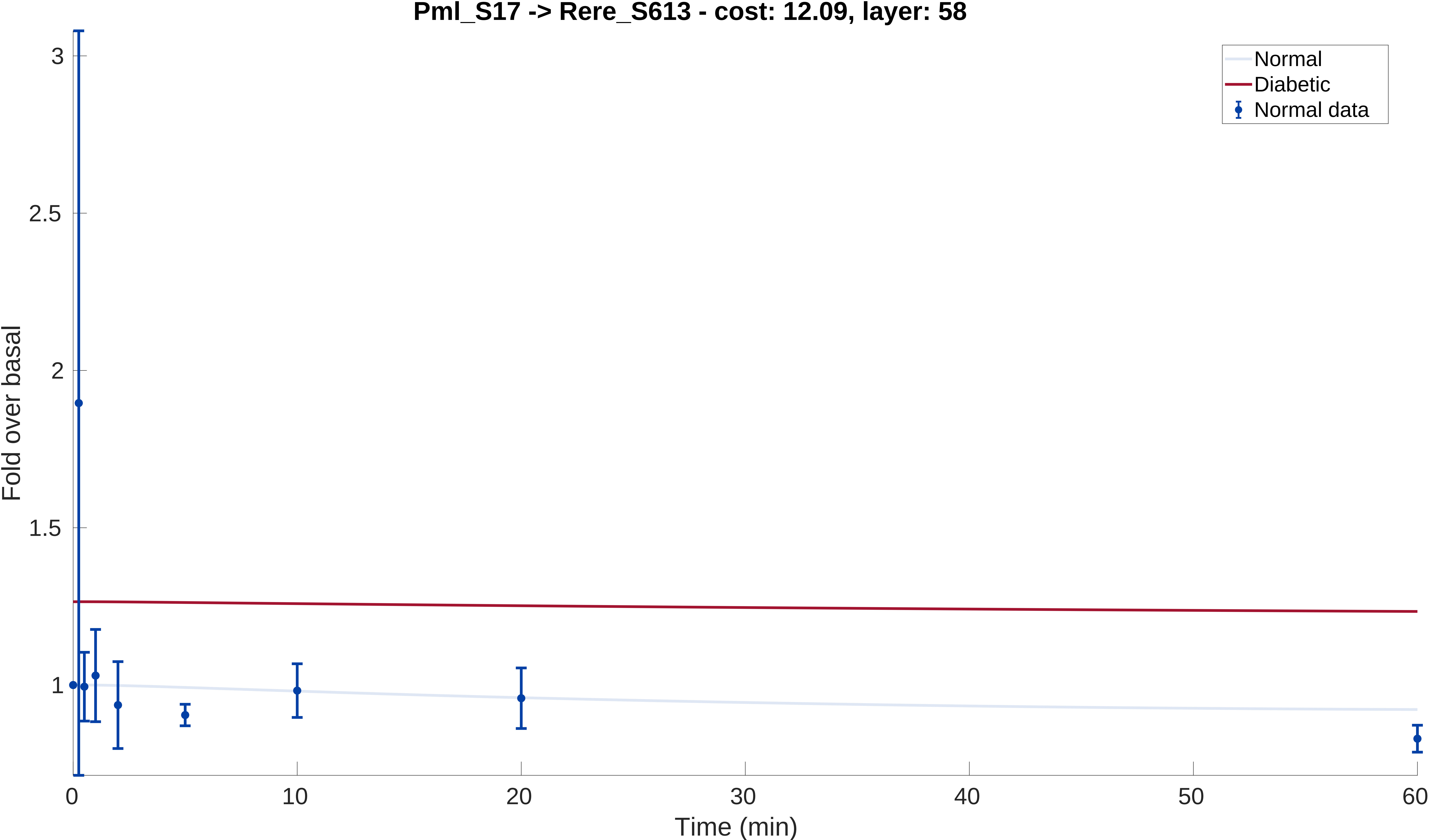

Pml_S17 -> Rere_S613 - cost: 12.09, layer: 58
Normal
Diabetic
Normal data
Fold over basal
3
2.5
2
1.5
1
0
10
20
30
40
50
60
Time (min)

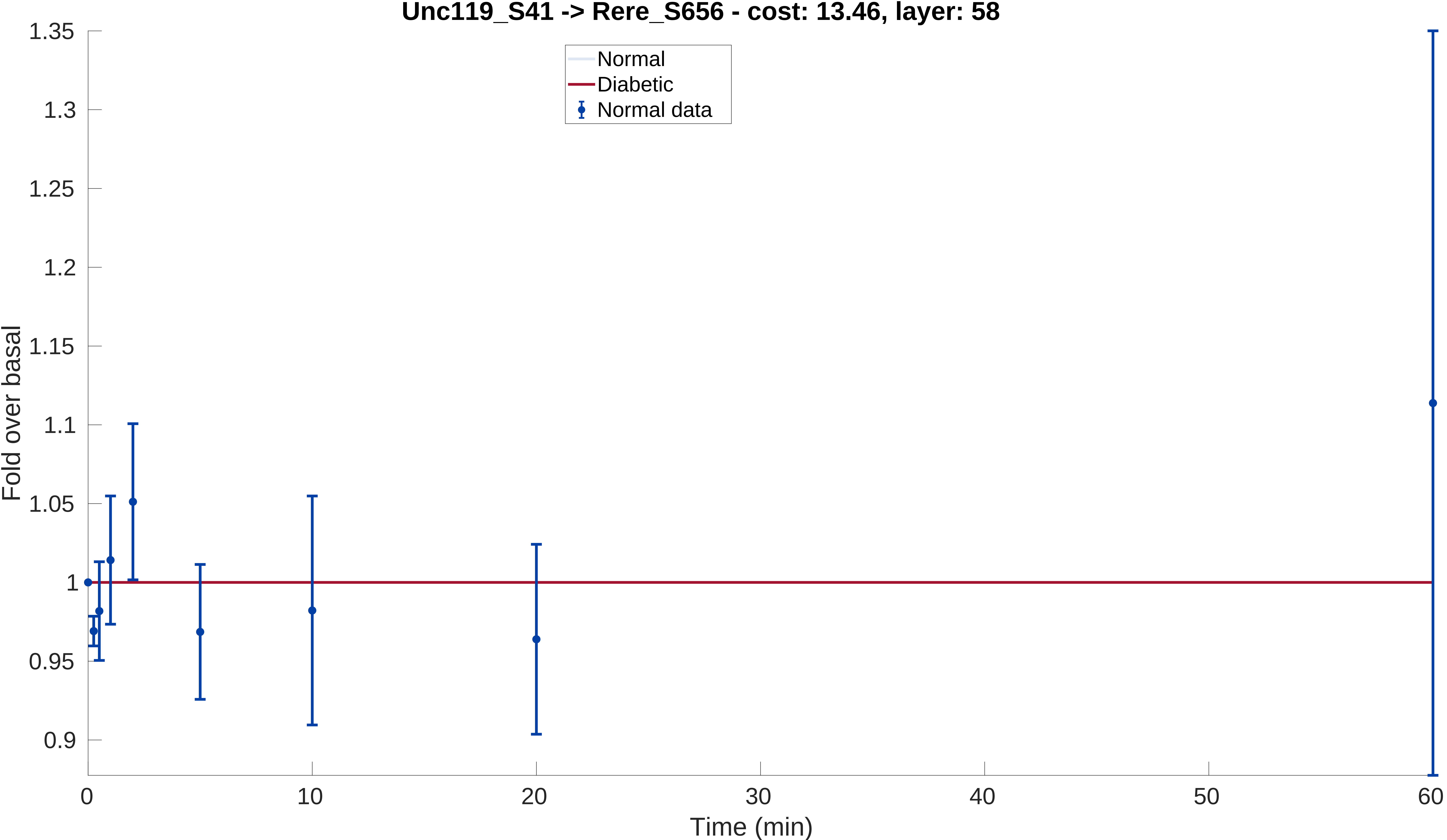
Unc119_S41 -> Rere_S656 - cost: 13.46, layer: 58
Normal
Diabetic
Normal data
Fold over basal
1.35
1.3
1.25
1.2
1.15
1.1
1.05
1
0.95
0.9
0
10
20
30
40
50
60
Time (min)

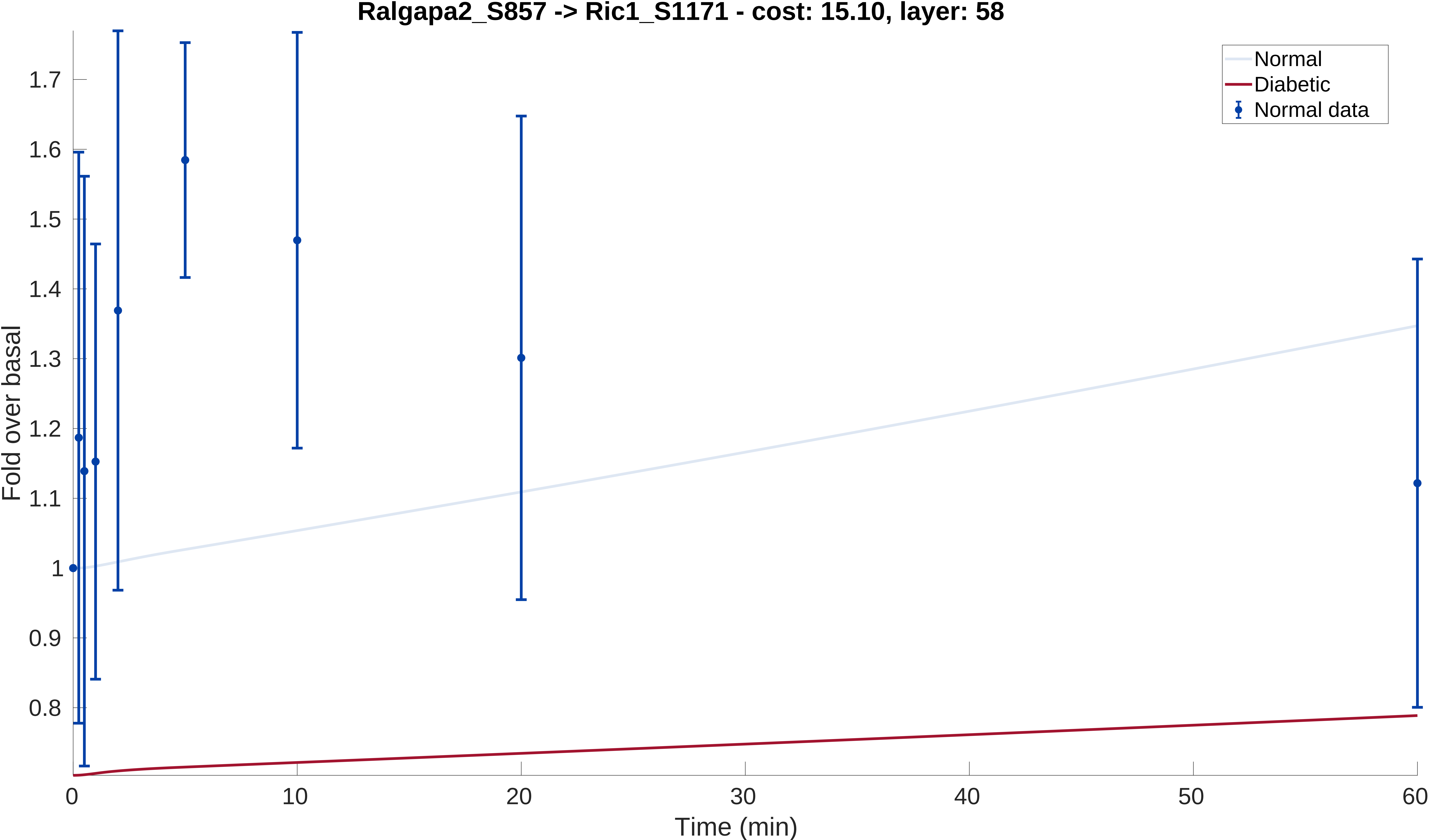

Ralgapa2_S857 -> Ric1_S1171 - cost: 15.10, layer: 58
Normal
Diabetic
Normal data
Fold over basal
Time (min)
0.8
0.9
1
1.1
1.2
1.3
1.4
1.5
1.6
1.7
0
10
20
30
40
50
60

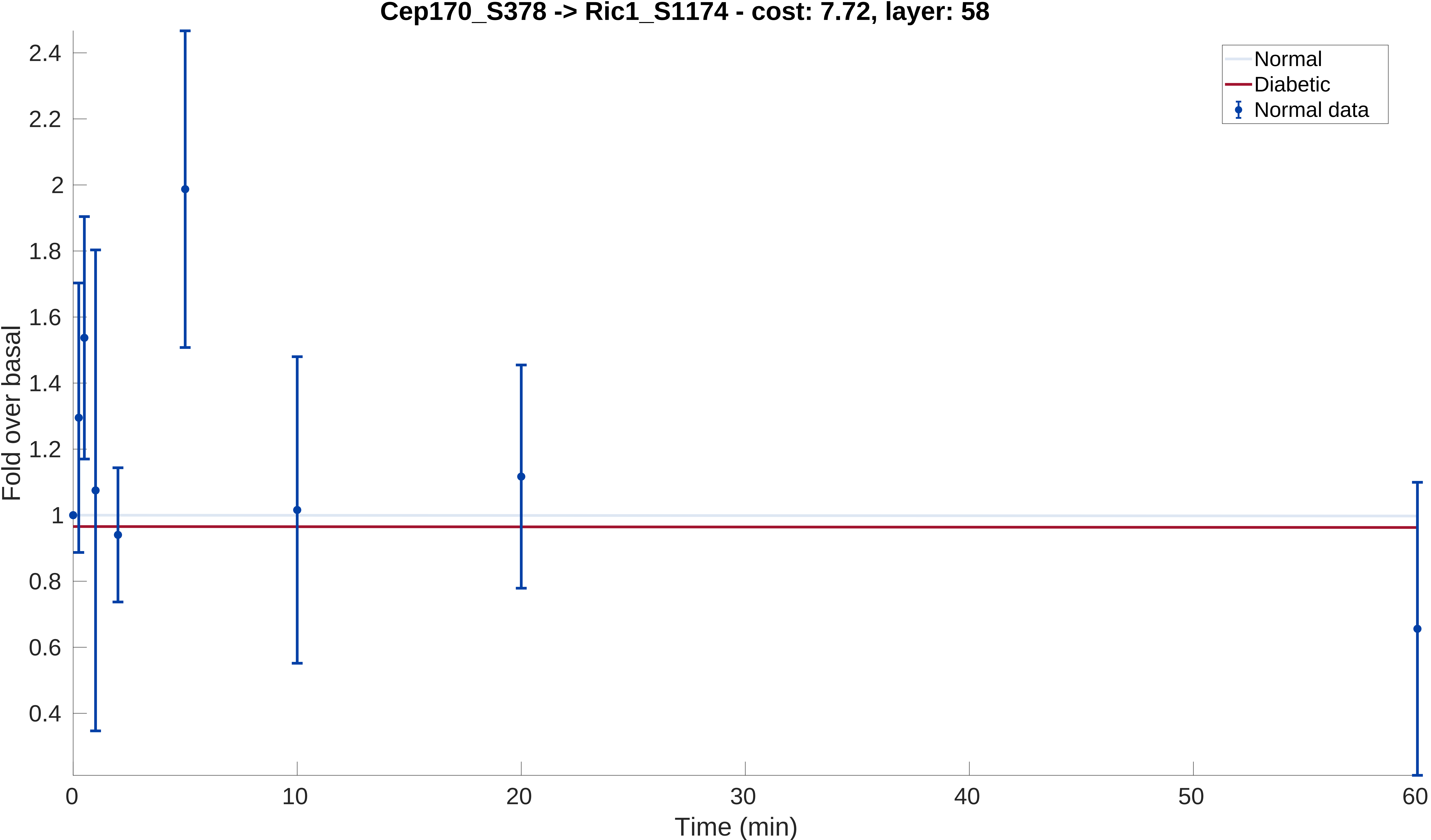

Cep170_S378 -> Ric1_S1174 - cost: 7.72, layer: 58
Normal
Diabetic
Normal data
Fold over basal
Time (min)
2.4
2.2
2
1.8
1.6
1.4
1.2
1
0.8
0.6
0.4
0
10
20
30
40
50
60

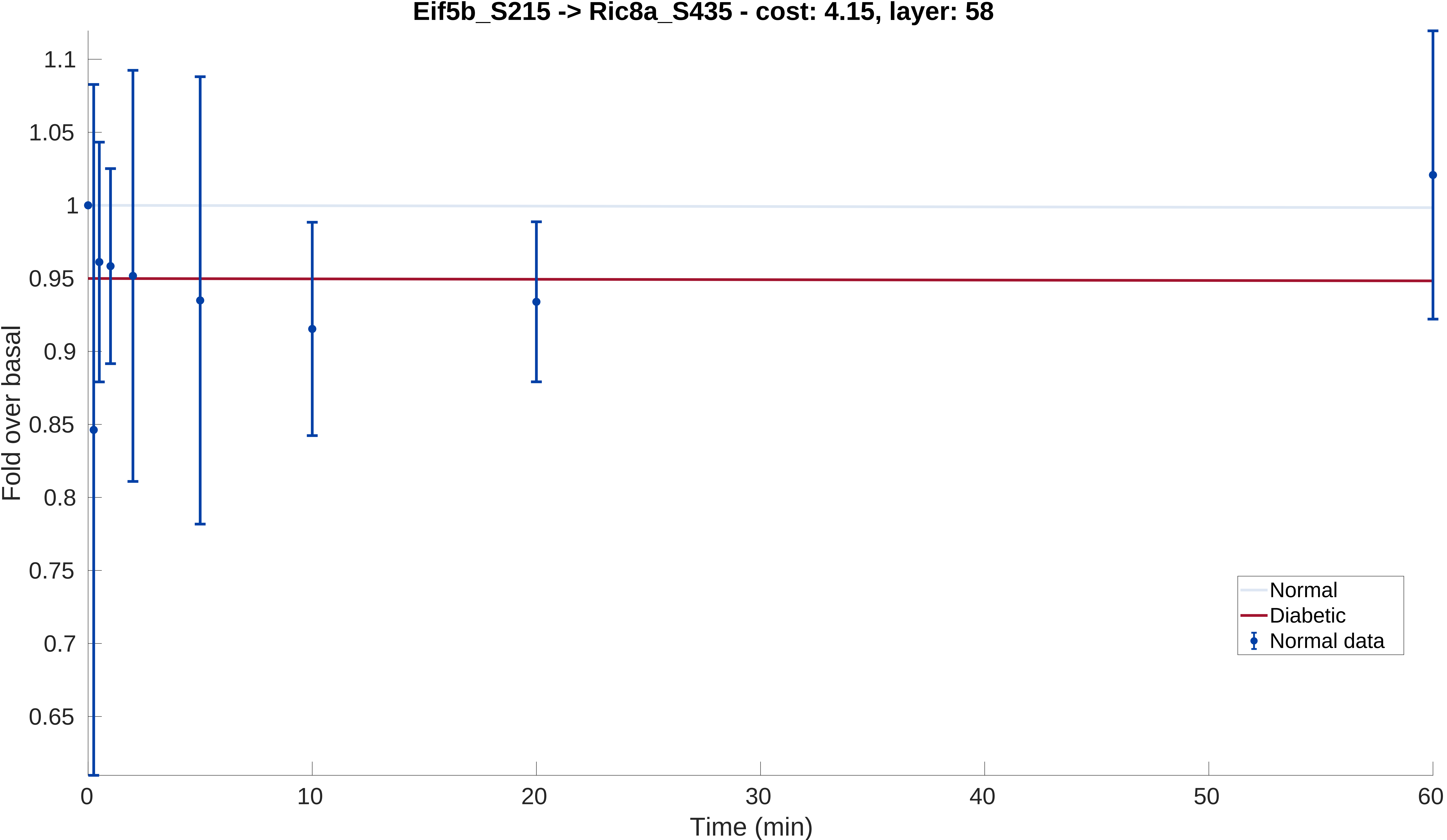

Eif5b_S215 -> Ric8a_S435 - cost: 4.15, layer: 58
Fold over basal
1.1
1.05
1
0.95
0.9
0.85
0.8
0.75
0.7
0.65
0
10
20
30
40
50
60
Time (min)
Normal
Diabetic
Normal data

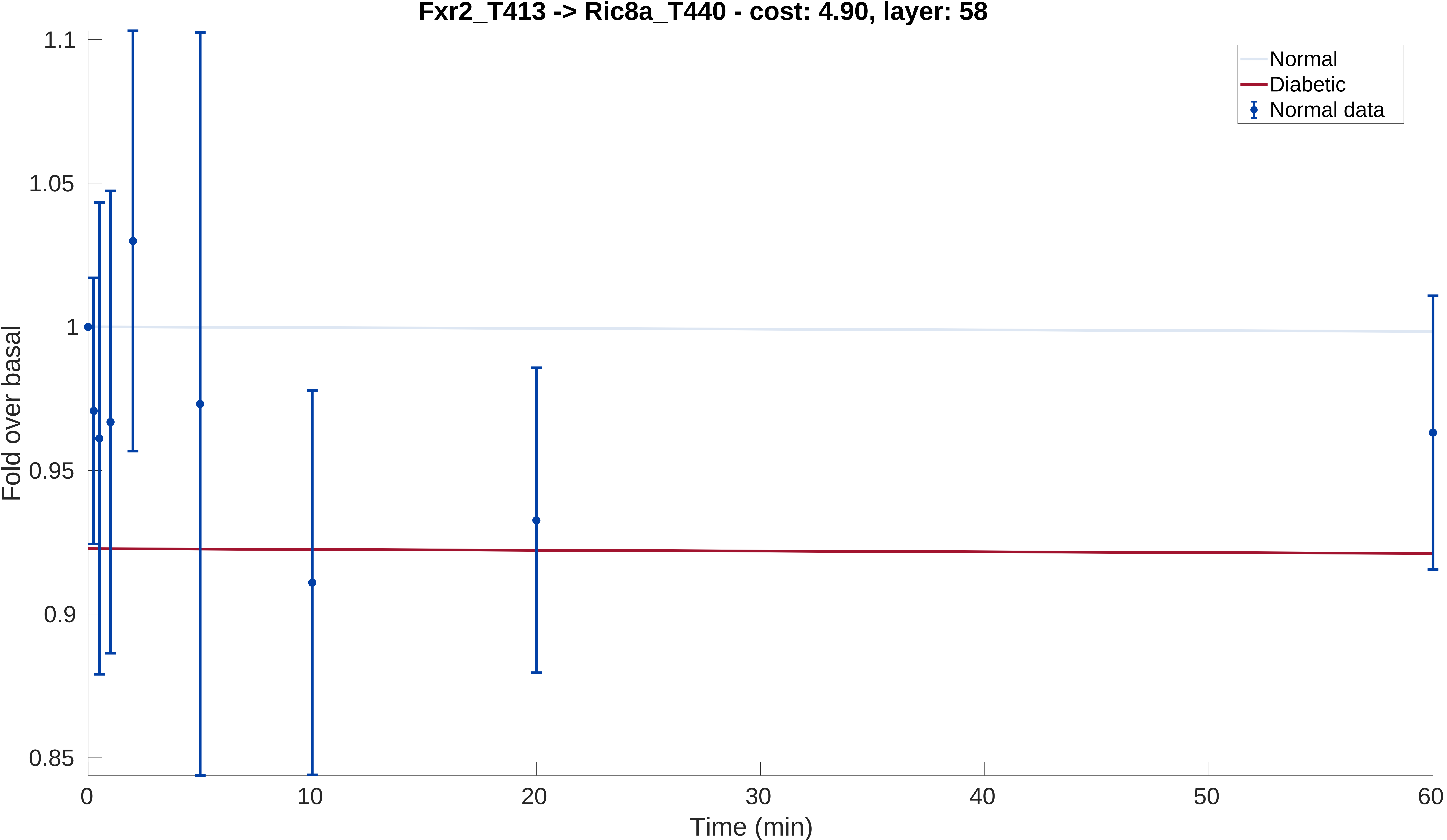

Fxr2_T413 -> Ric8a_T440 - cost: 4.90, layer: 58
Normal
Diabetic
Normal data
Fold over basal
Time (min)
1.1
1.05
1
0.95
0.9
0.85
0
10
20
30
40
50
60

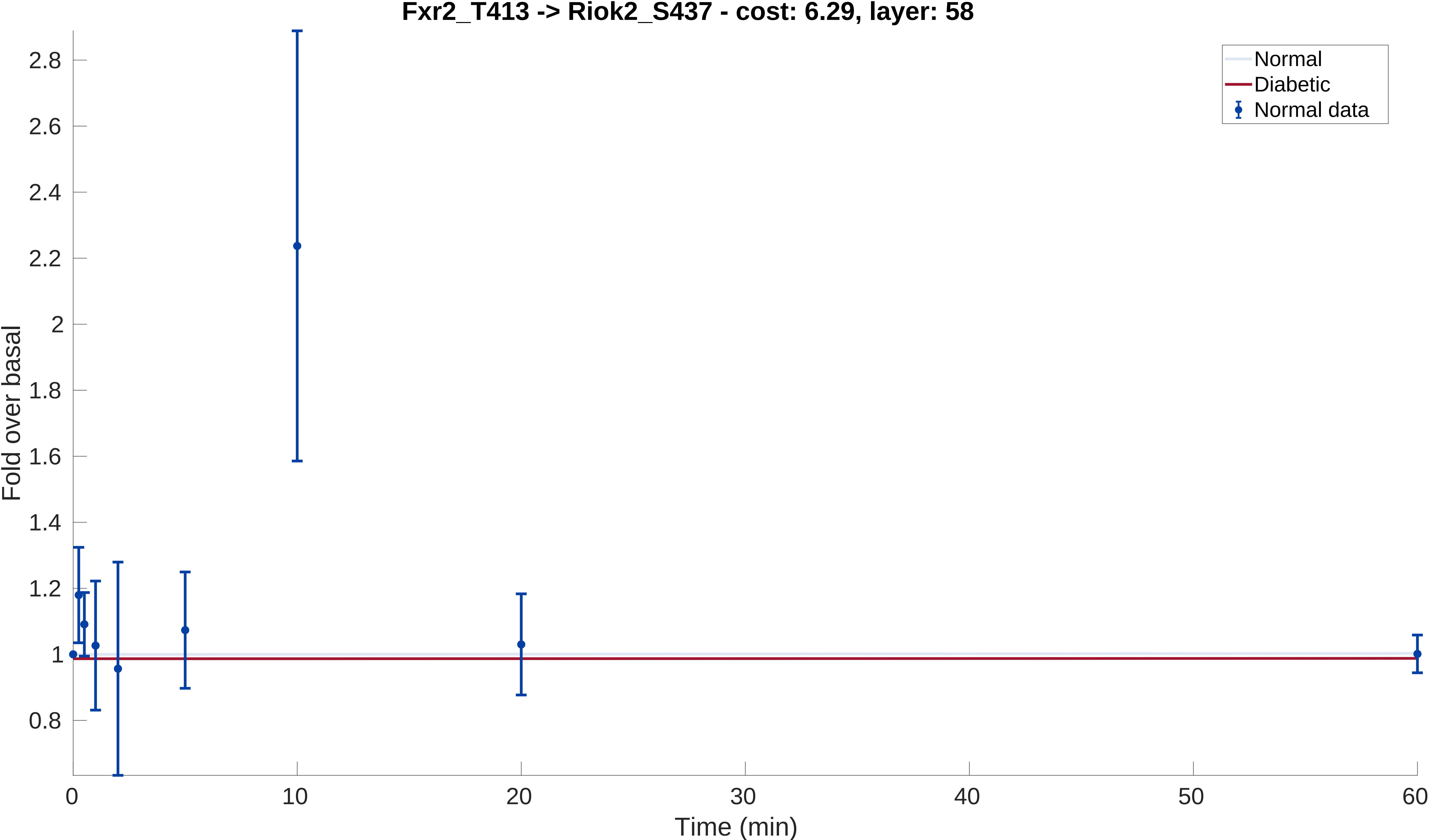

Fxr2_T413 -> Riok2_S437 - cost: 6.29, layer: 58
Normal
Diabetic
Normal data
Fold over basal
2.8
2.6
2.4
2.2
2
1.8
1.6
1.4
1.2
1
0.8
0
10
20
30
40
50
60
Time (min)

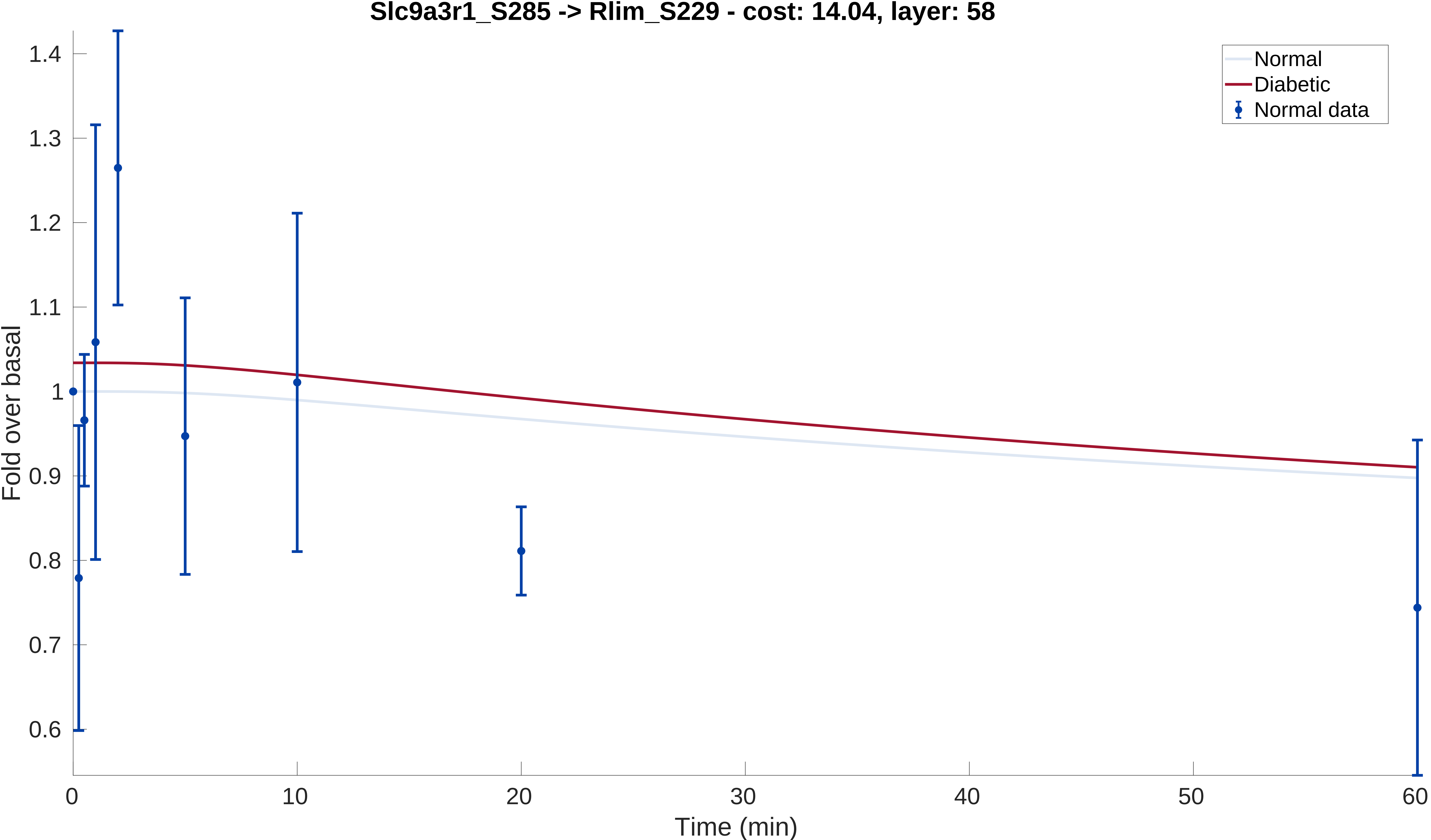

Slc9a3r1_S285 -> Rlim_S229 - cost: 14.04, layer: 58
Normal
Diabetic
Normal data
Fold over basal
Time (min)
1.4
1.3
1.2
1.1
1
0.9
0.8
0.7
0.6
0
10
20
30
40
50
60

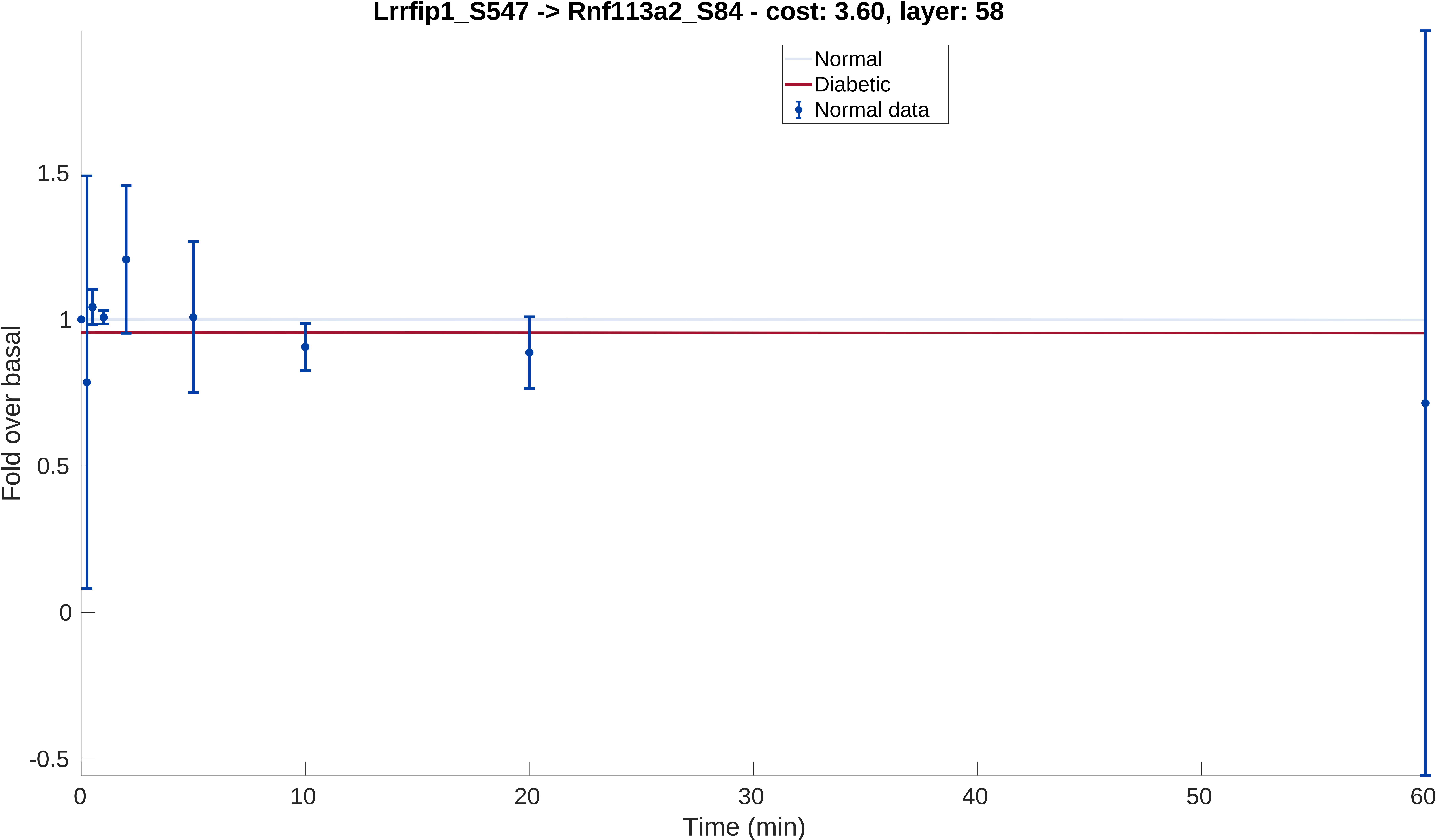

Lrrfip1_S547 -> Rnf113a2_S84 - cost: 3.60, layer: 58
Normal
Diabetic
Normal data
Fold over basal
1.5
1
0.5
0
-0.5
0
10
20
30
40
50
60
Time (min)

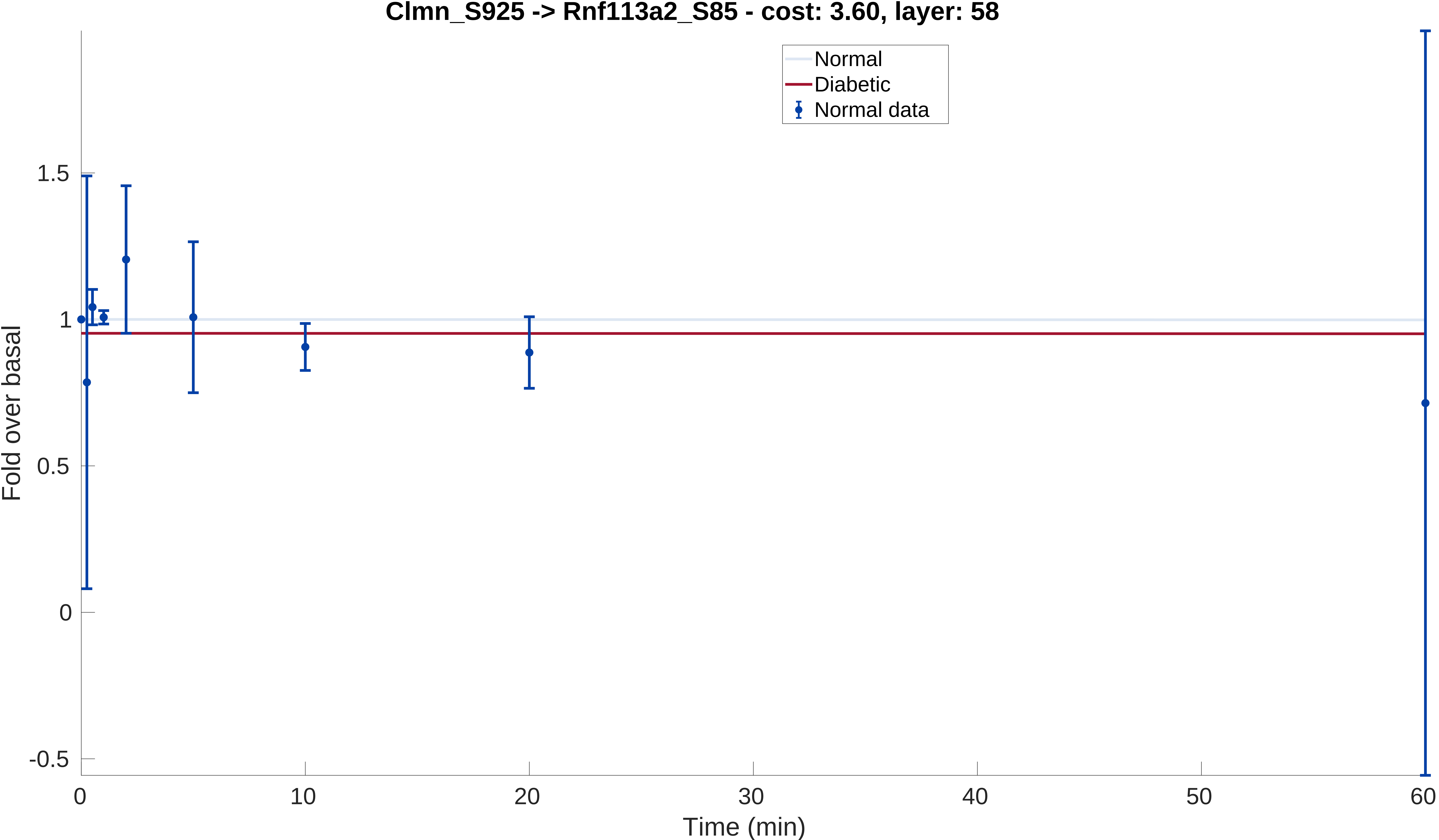

Clmn_S925 -> Rnf113a2_S85 - cost: 3.60, layer: 58
Normal
Diabetic
Normal data
Fold over basal
1.5
1
0.5
0
-0.5
0
10
20
30
40
50
60
Time (min)

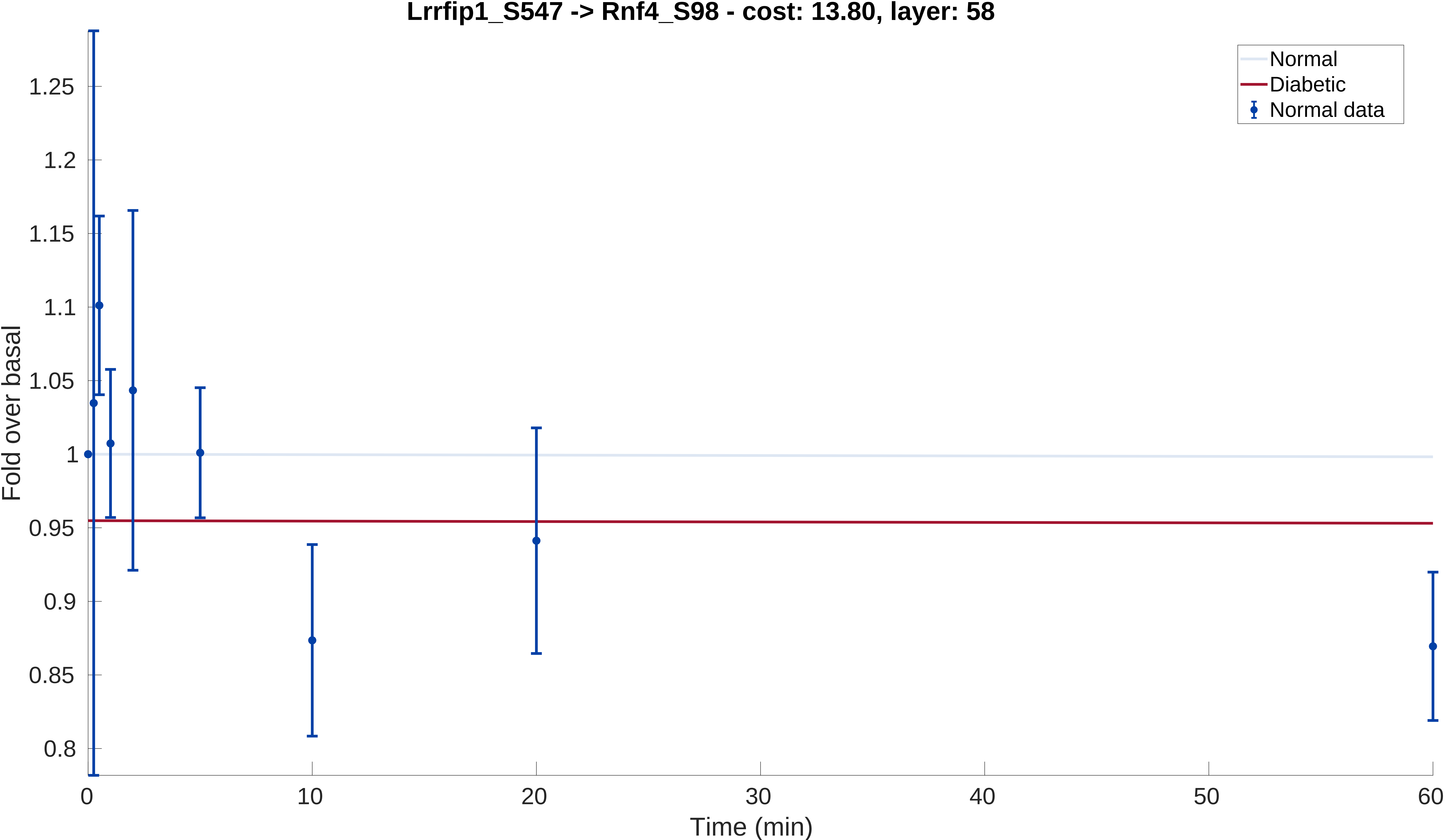

Lrrfip1_S547 -> Rnf4_S98 - cost: 13.80, layer: 58
Normal
Diabetic
Normal data
1.25
1.2
1.15
1.1
1.05
1
0.95
0.9
0.85
0.8
Fold over basal
0
10
20
30
40
50
60
Time (min)

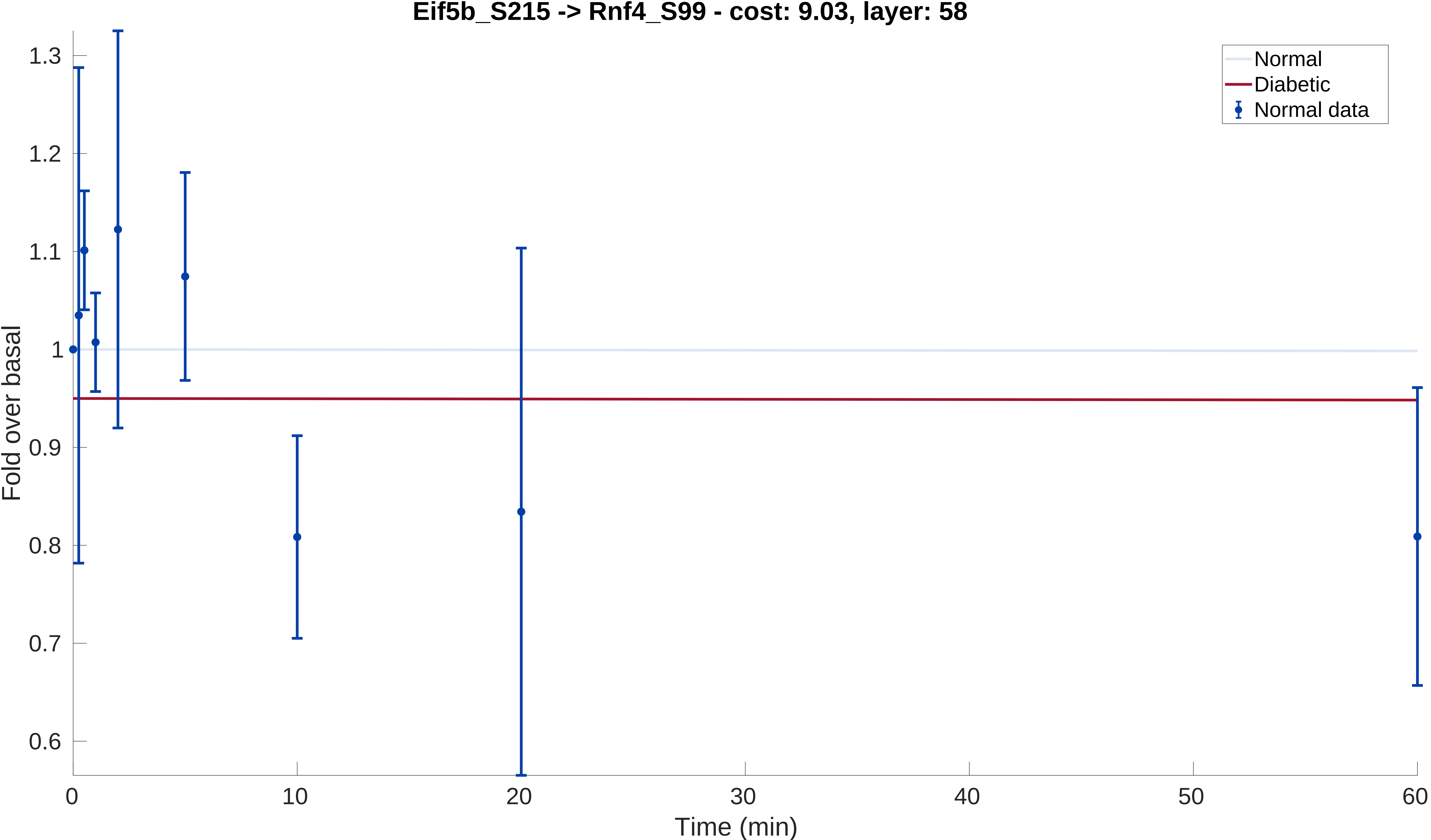
Eif5b_S215 -> Rnf4_S99 - cost: 9.03, layer: 58
Normal
Diabetic
Normal data
Fold over basal
1.3
1.2
1.1
1
0.9
0.8
0.7
0.6
0
10
20
30
40
50
60
Time (min)

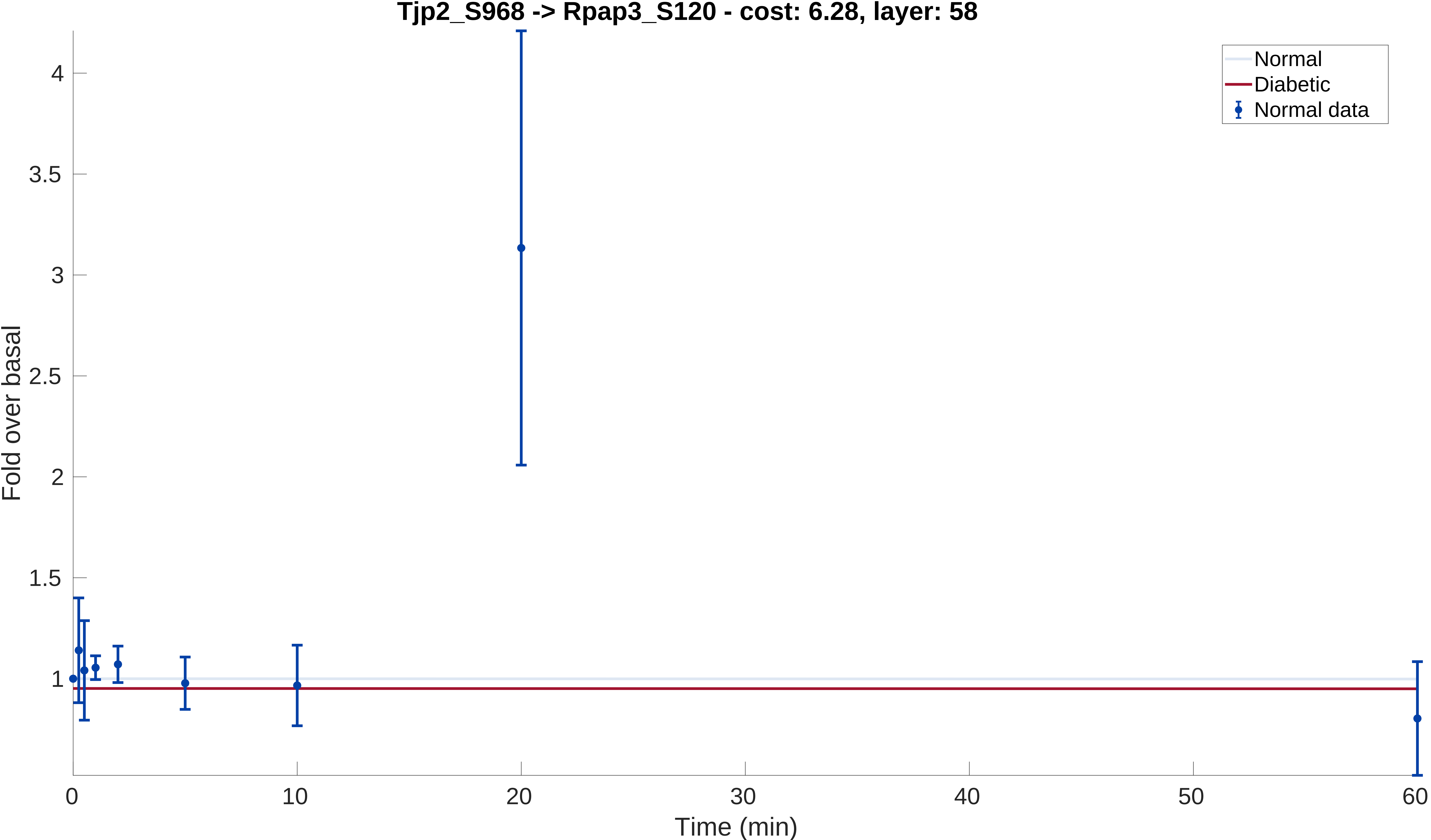

Tjp2_S968 -> Rpap3_S120 - cost: 6.28, layer: 58
Normal
Diabetic
Normal data
Fold over basal
4
3.5
3
2.5
2
1.5
1
0
10
20
30
40
50
60
Time (min)

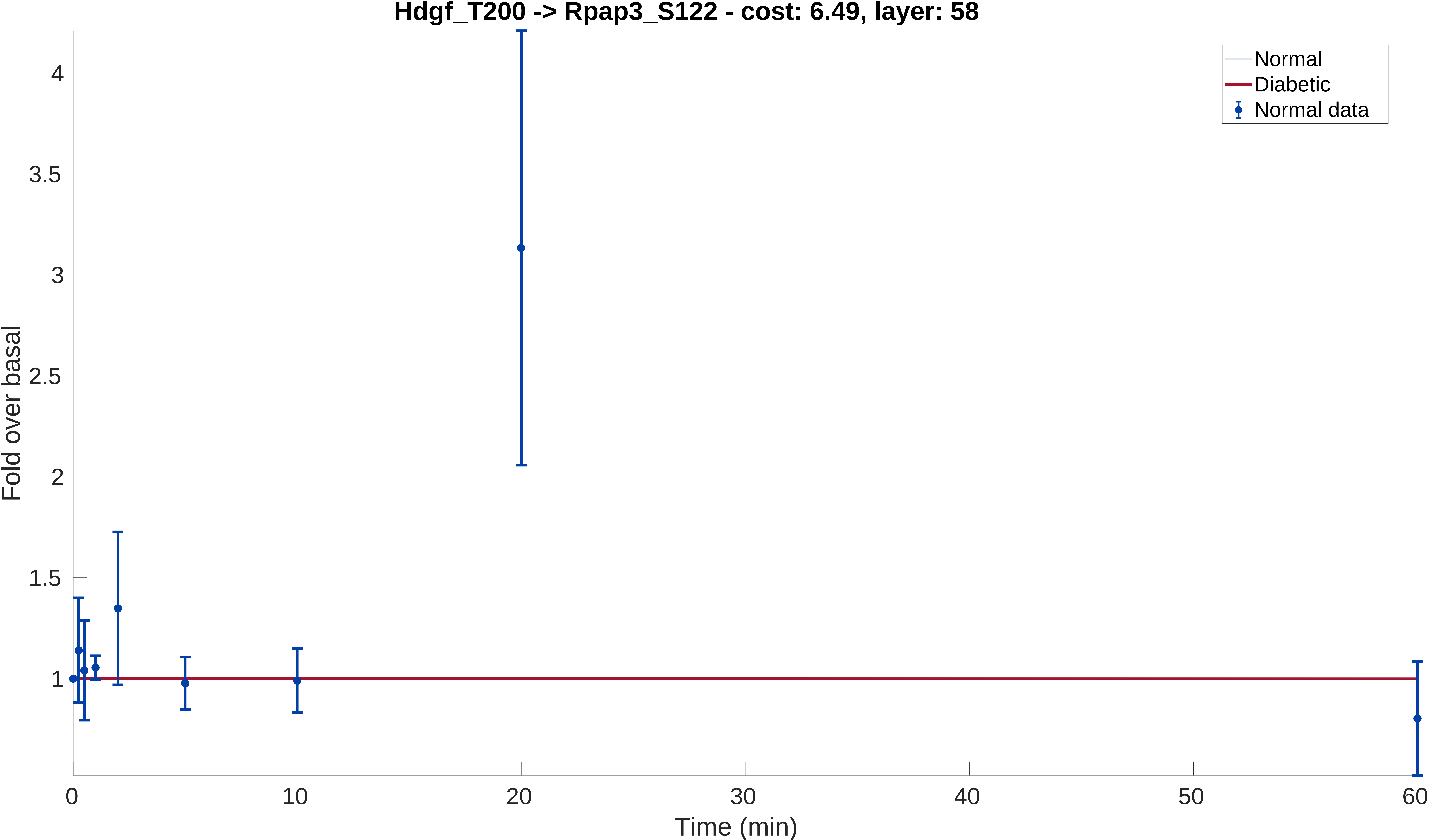

Hdgf_T200 -> Rpap3_S122 - cost: 6.49, layer: 58
Normal
Diabetic
Normal data
Fold over basal
Time (min)
4
3.5
3
2.5
2
1.5
1
0
10
20
30
40
50
60

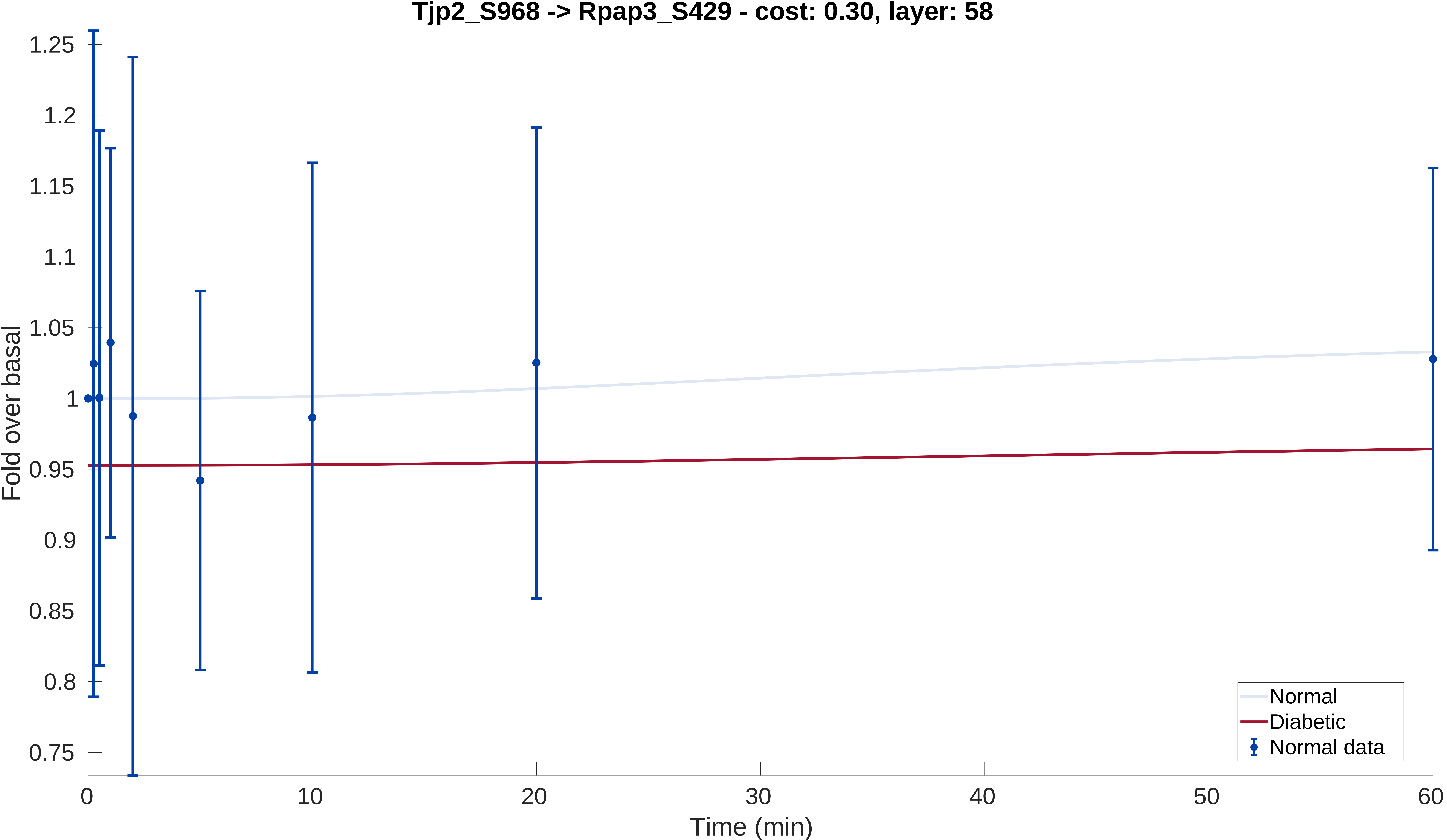
Tjp2_S968 -> Rpap3_S429 - cost: 0.30, layer: 58
Fold over basal
Time (min)
Normal
Diabetic
Normal data
1.25
1.2
1.15
1.1
1.05
1
0.95
0.9
0.85
0.8
0.75
0
10
20
30
40
50
60

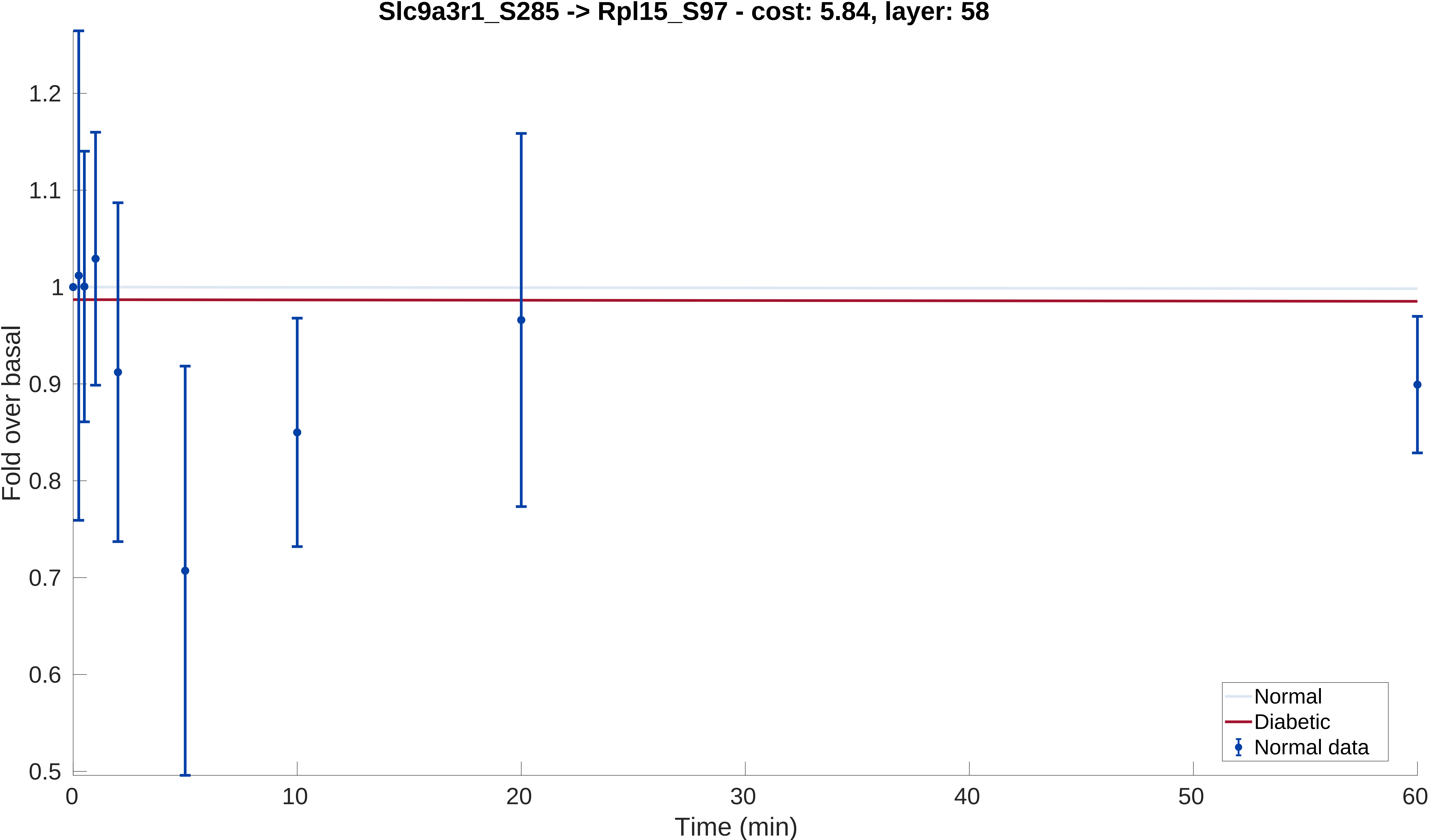
Slc9a3r1_S285 -> Rpl15_S97 - cost: 5.84, layer: 58
Fold over basal
Time (min)
1.2
1.1
1
0.9
0.8
0.7
0.6
0.5
0
10
20
30
40
50
60
Normal
Diabetic
Normal data

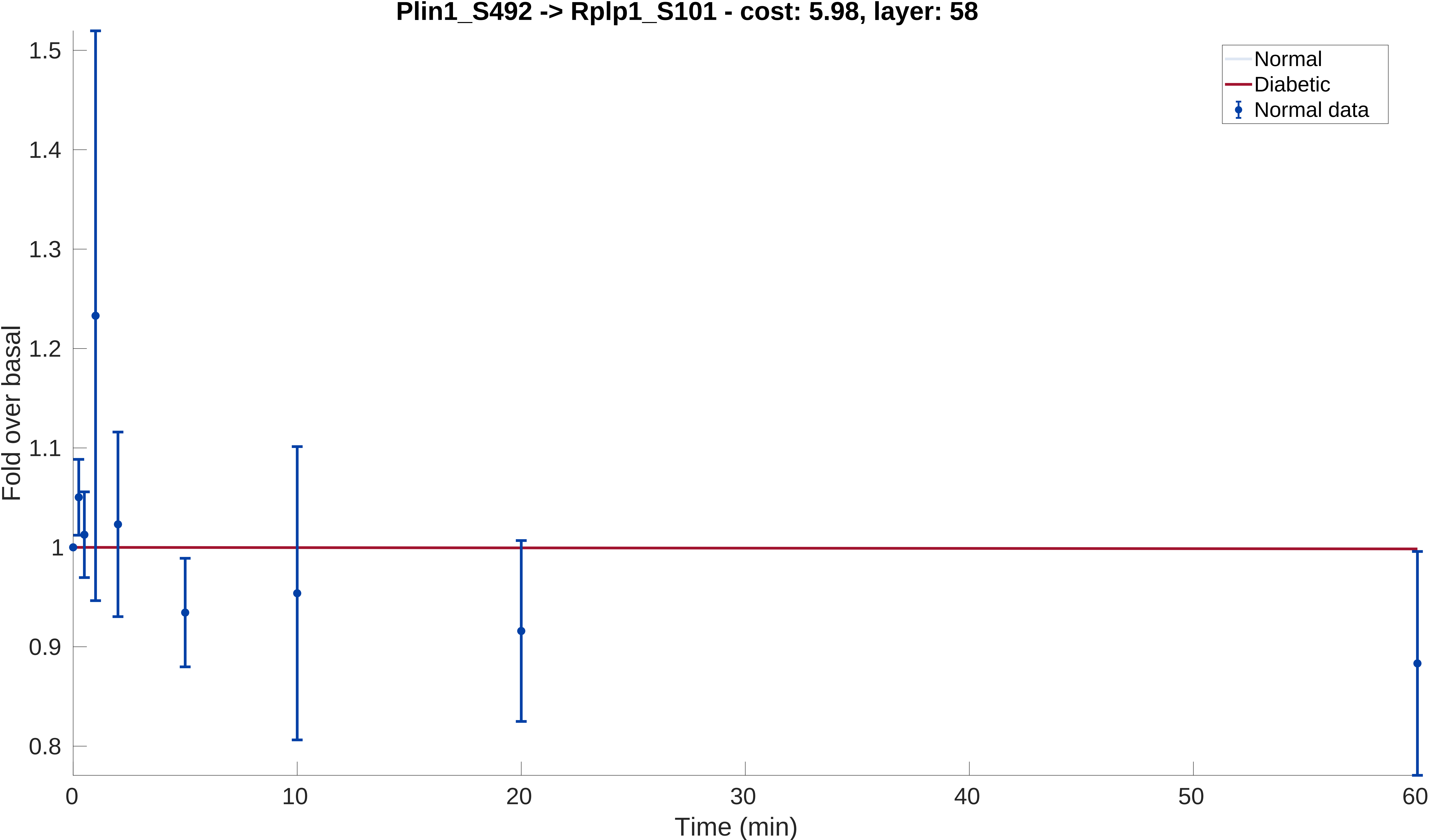

Plin1_S492 -> Rplp1_S101 - cost: 5.98, layer: 58
Normal
Diabetic
Normal data
Fold over basal
1.5
1.4
1.3
1.2
1.1
1
0.9
0.8
0
10
20
30
40
50
60
Time (min)

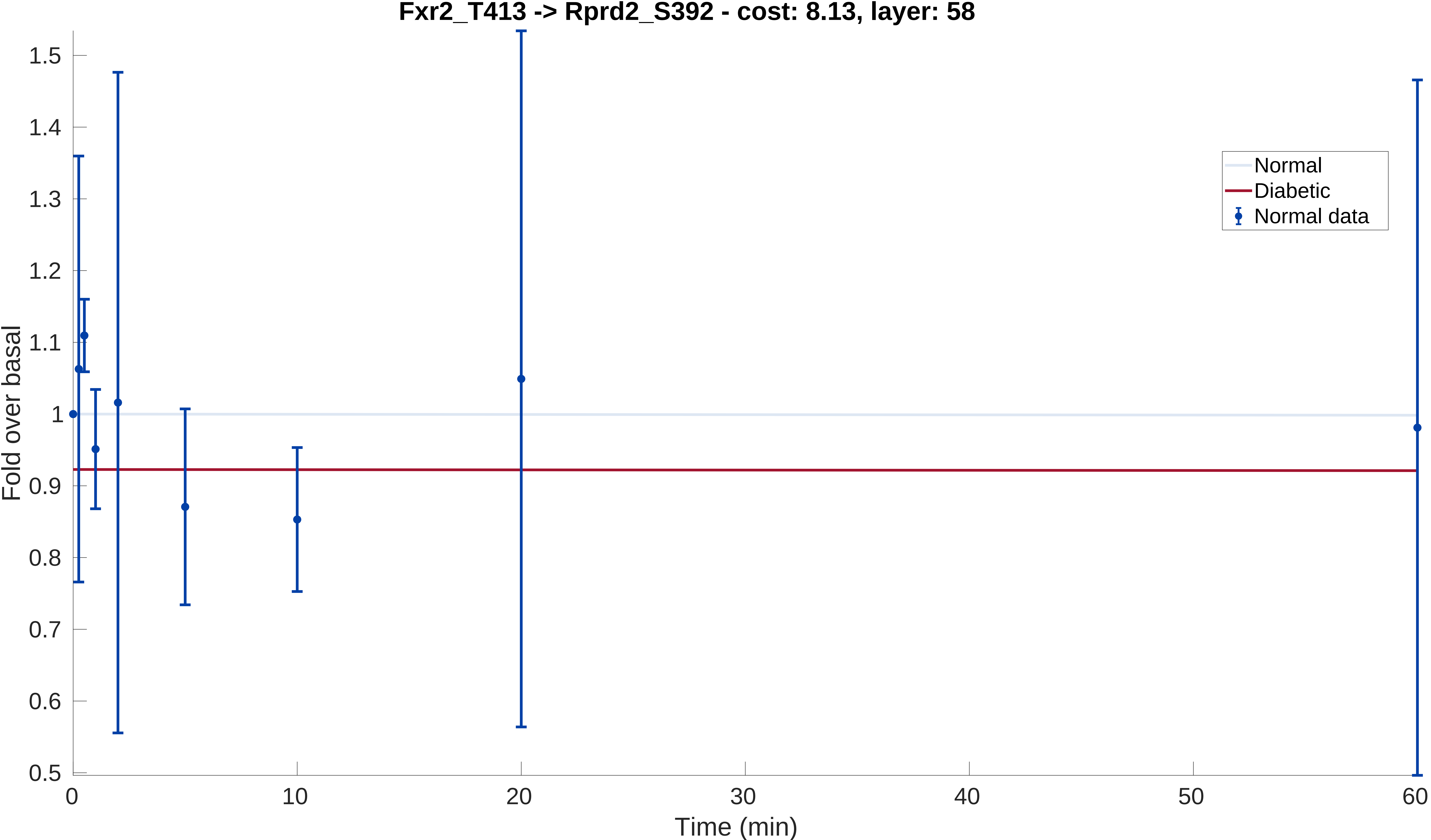
Fxr2_T413 -> Rprd2_S392 - cost: 8.13, layer: 58
Fold over basal
1.5
1.4
1.3
1.2
1.1
1
0.9
0.8
0.7
0.6
0.5
0
10
20
30
40
50
60
Time (min)
Normal
Diabetic
Normal data

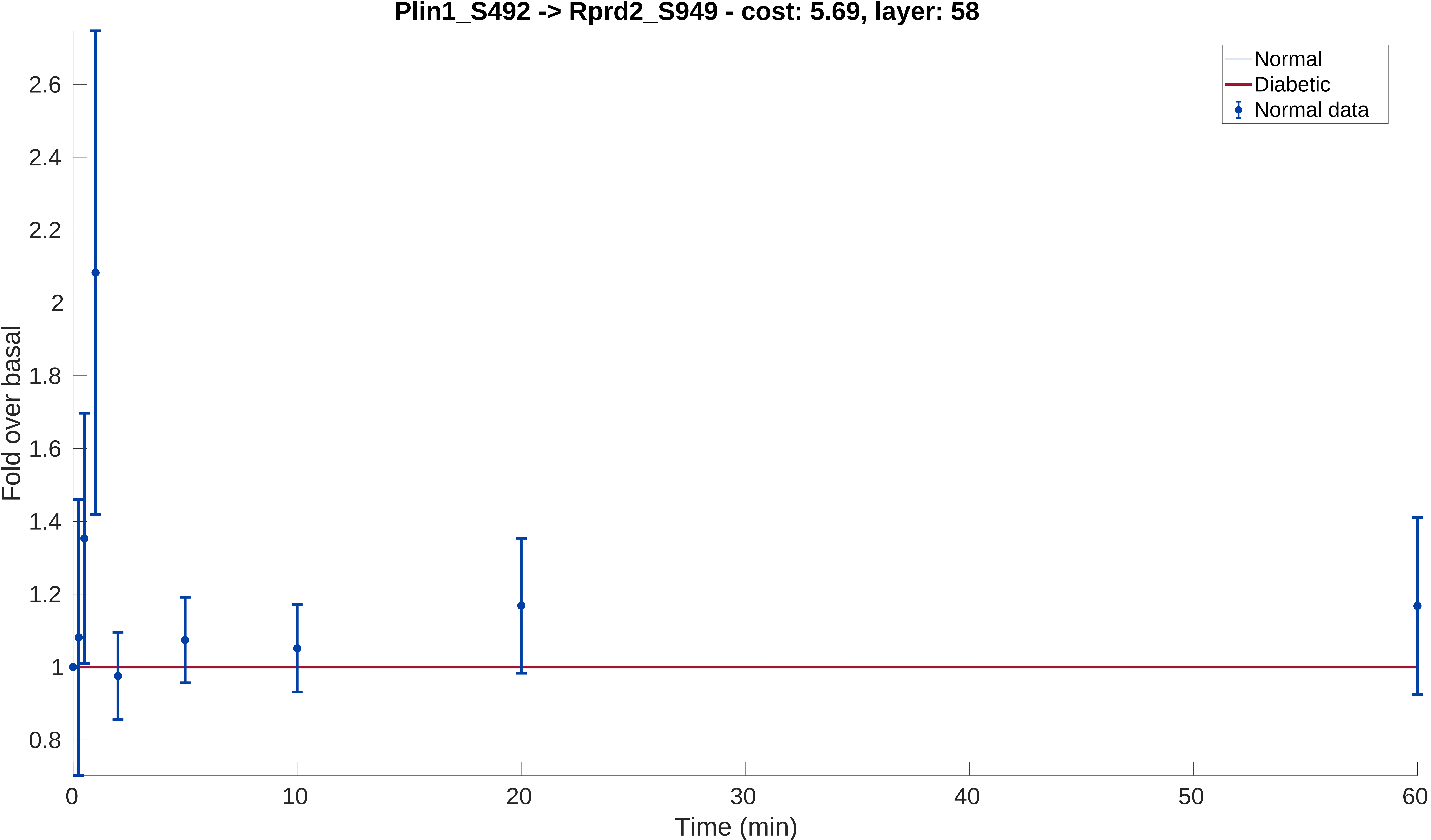

Plin1_S492 -> Rprd2_S949 - cost: 5.69, layer: 58
Normal
Diabetic
Normal data
Fold over basal
2.6
2.4
2.2
2
1.8
1.6
1.4
1.2
1
0.8
0
10
20
30
40
50
60
Time (min)

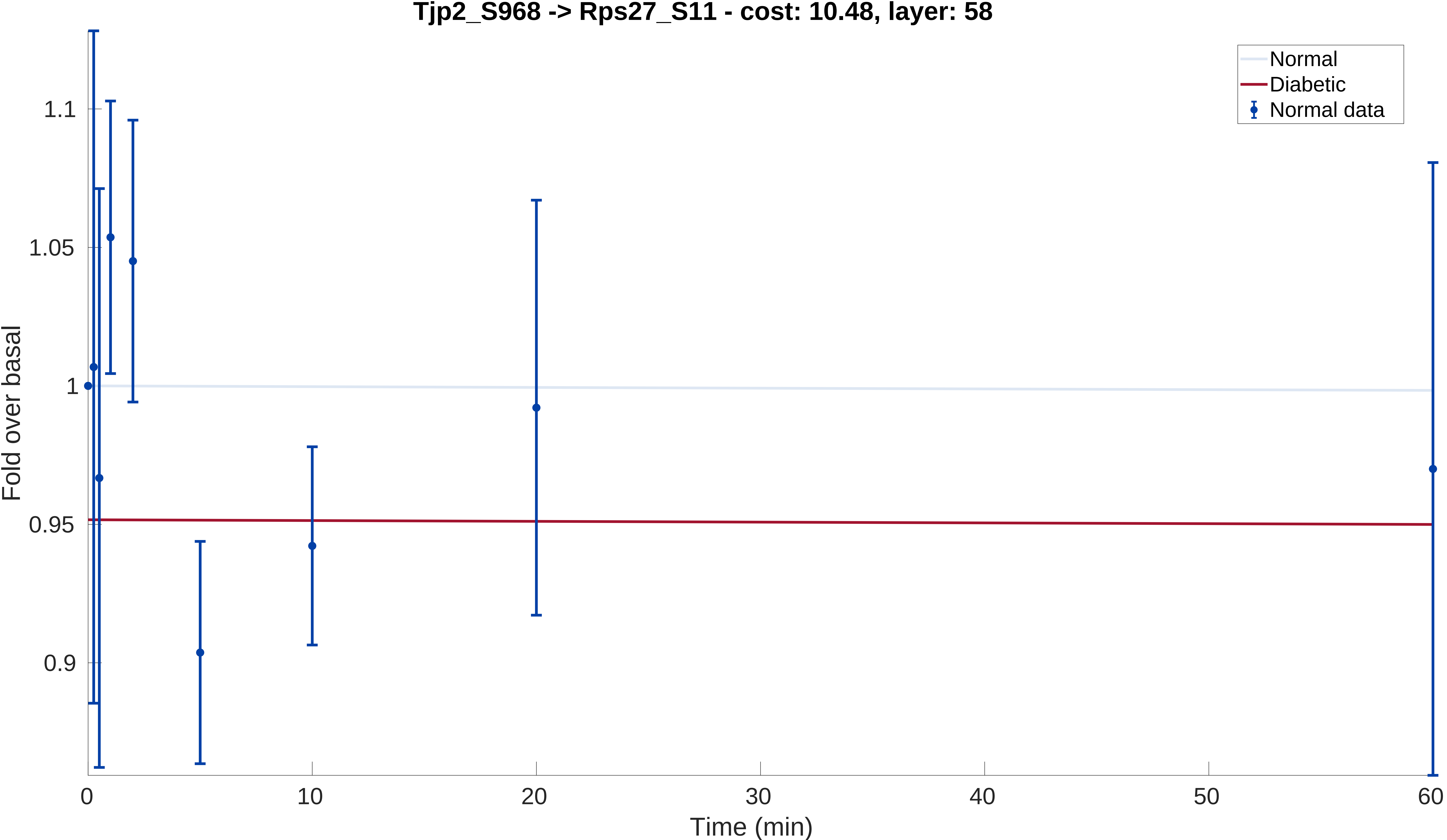
Tjp2_S968 -> Rps27_S11 - cost: 10.48, layer: 58
Normal
Diabetic
Normal data
Fold over basal
1.1
1.05
1
0.95
0.9
0
10
20
30
40
50
60
Time (min)

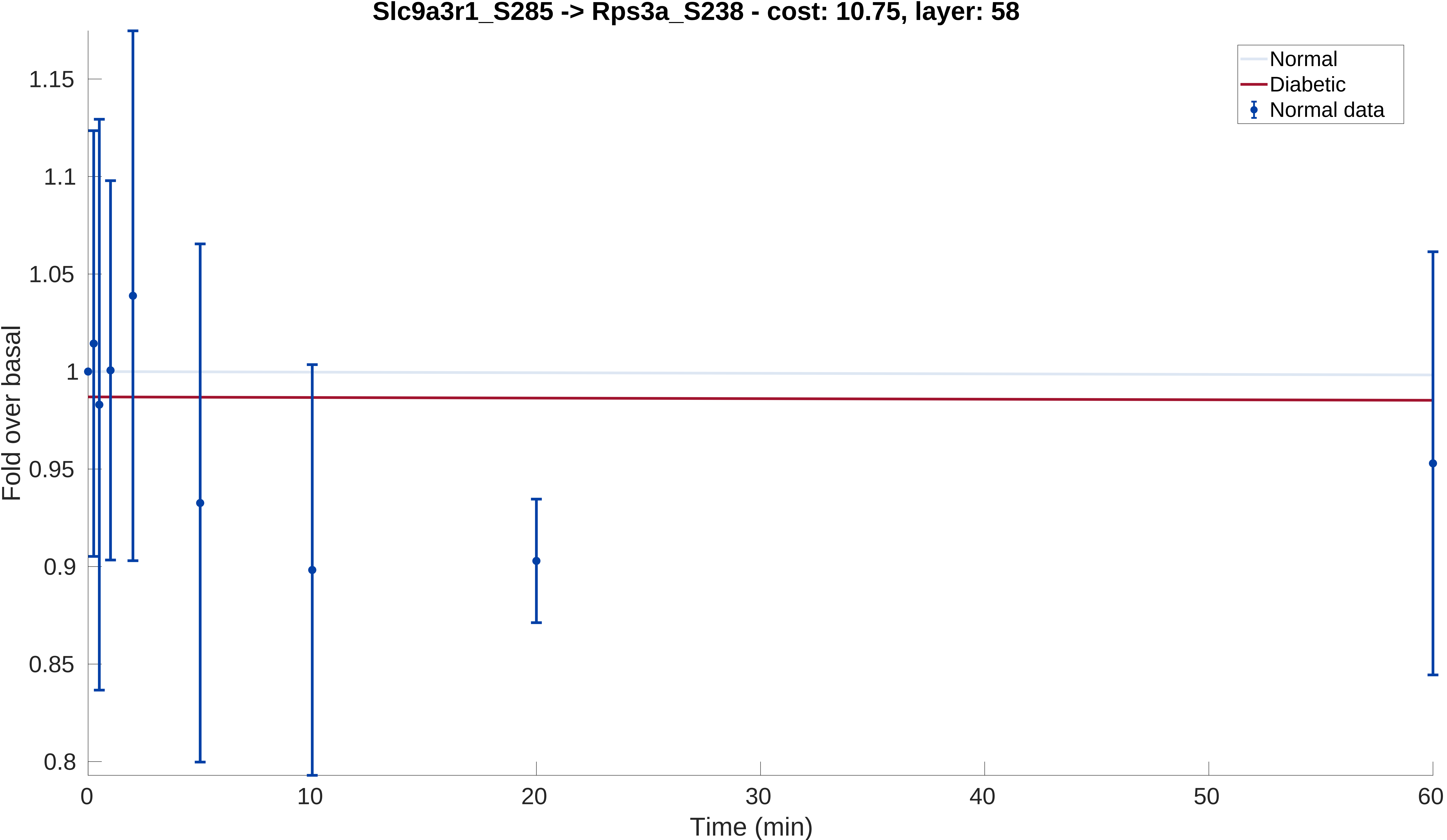
Slc9a3r1_S285 -> Rps3a_S238 - cost: 10.75, layer: 58
Normal
Diabetic
Normal data
Fold over basal
1.15
1.1
1.05
1
0.95
0.9
0.85
0.8
0
10
20
30
40
50
60
Time (min)

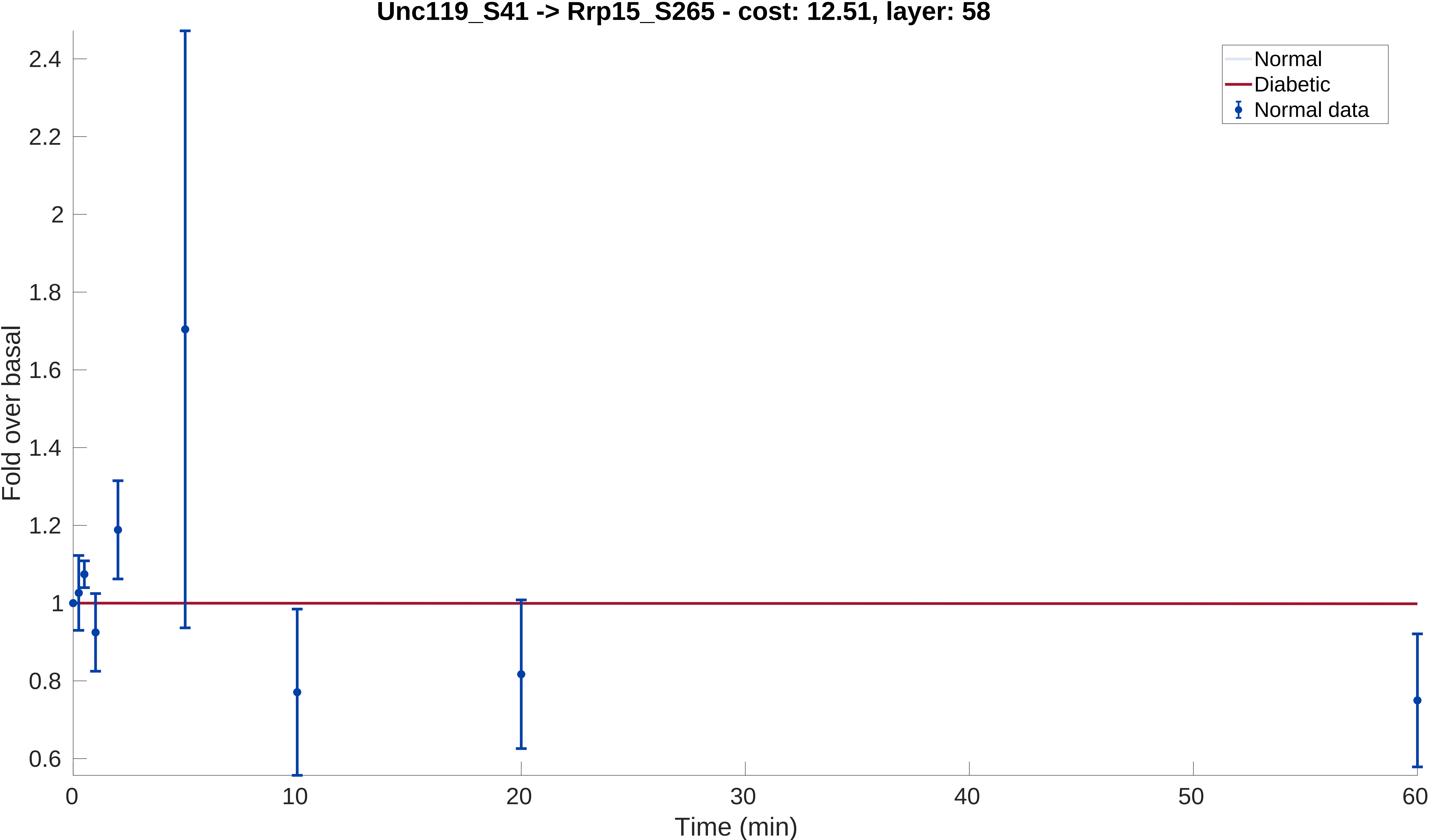

Unc119_S41 -> Rrp15_S265 - cost: 12.51, layer: 58
Normal
Diabetic
Normal data
2.4
2.2
2
1.8
1.6
1.4
1.2
1
0.8
0.6
Fold over basal
0
10
20
30
40
50
60
Time (min)

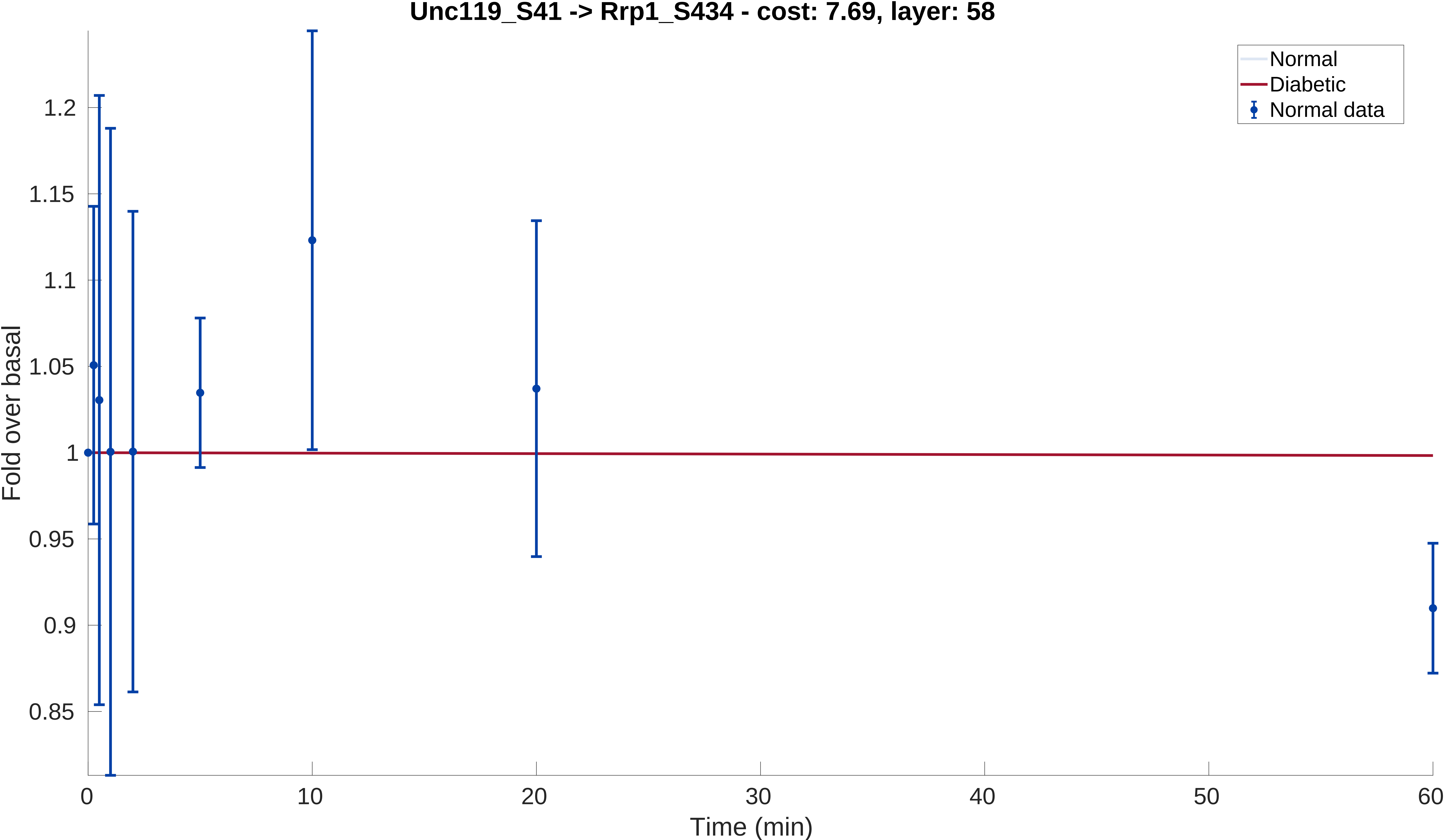

Unc119_S41 -> Rrp1_S434 - cost: 7.69, layer: 58
Normal
Diabetic
Normal data
Fold over basal
1.2
1.15
1.1
1.05
1
0.95
0.9
0.85
0
10
20
30
40
50
60
Time (min)

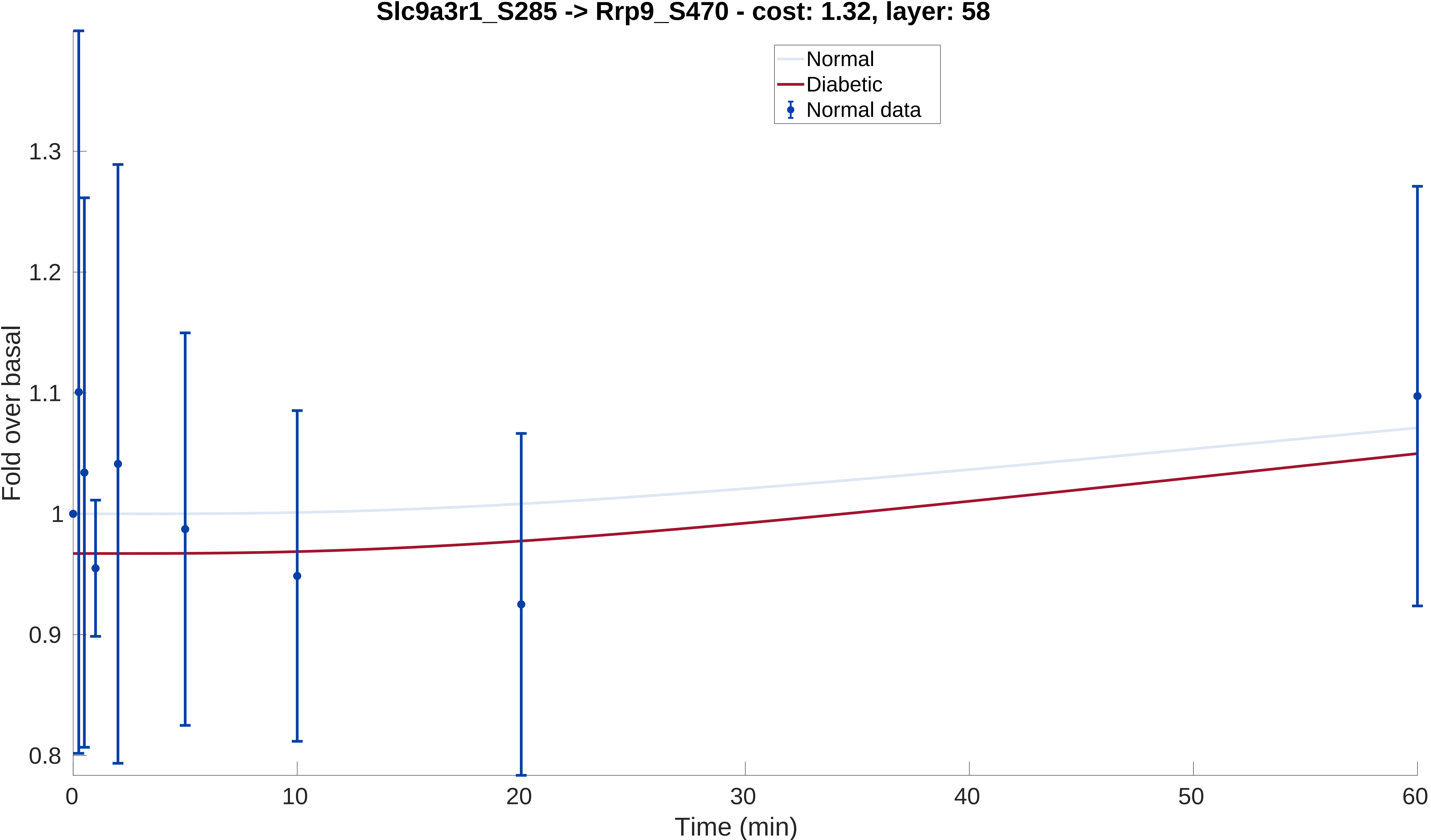

Slc9a3r1_S285 -> Rrp9_S470 - cost: 1.32, layer: 58
Normal
Diabetic
Normal data
1.3
1.2
1.1
1
0.9
0.8
Fold over basal
0
10
20
30
40
50
60
Time (min)

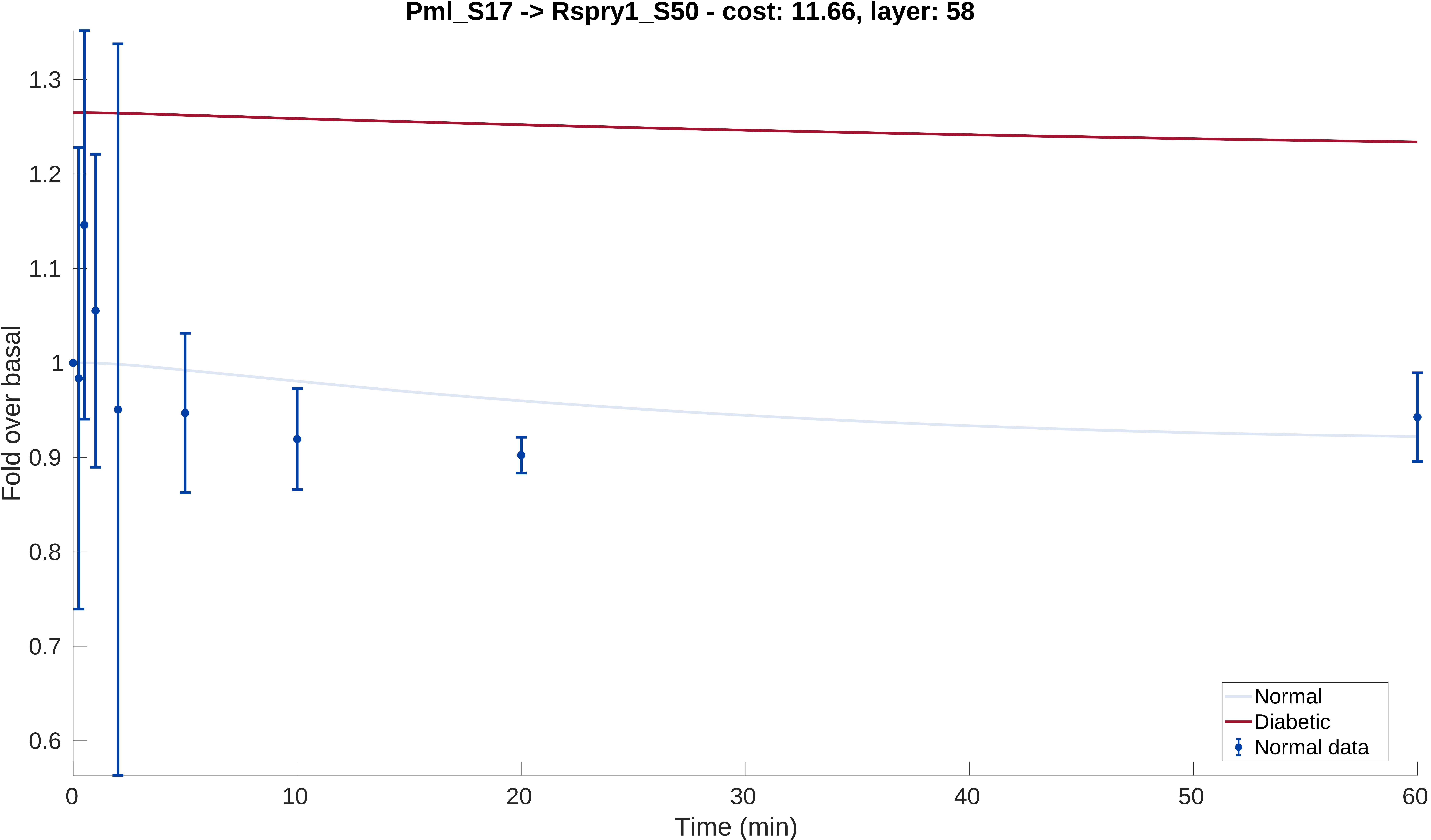

Pml_S17 -> Rspry1_S50 - cost: 11.66, layer: 58
Fold over basal
1.3
1.2
1.1
1
0.9
0.8
0.7
0.6
0
10
20
30
40
50
60
Time (min)
Normal
Diabetic
Normal data

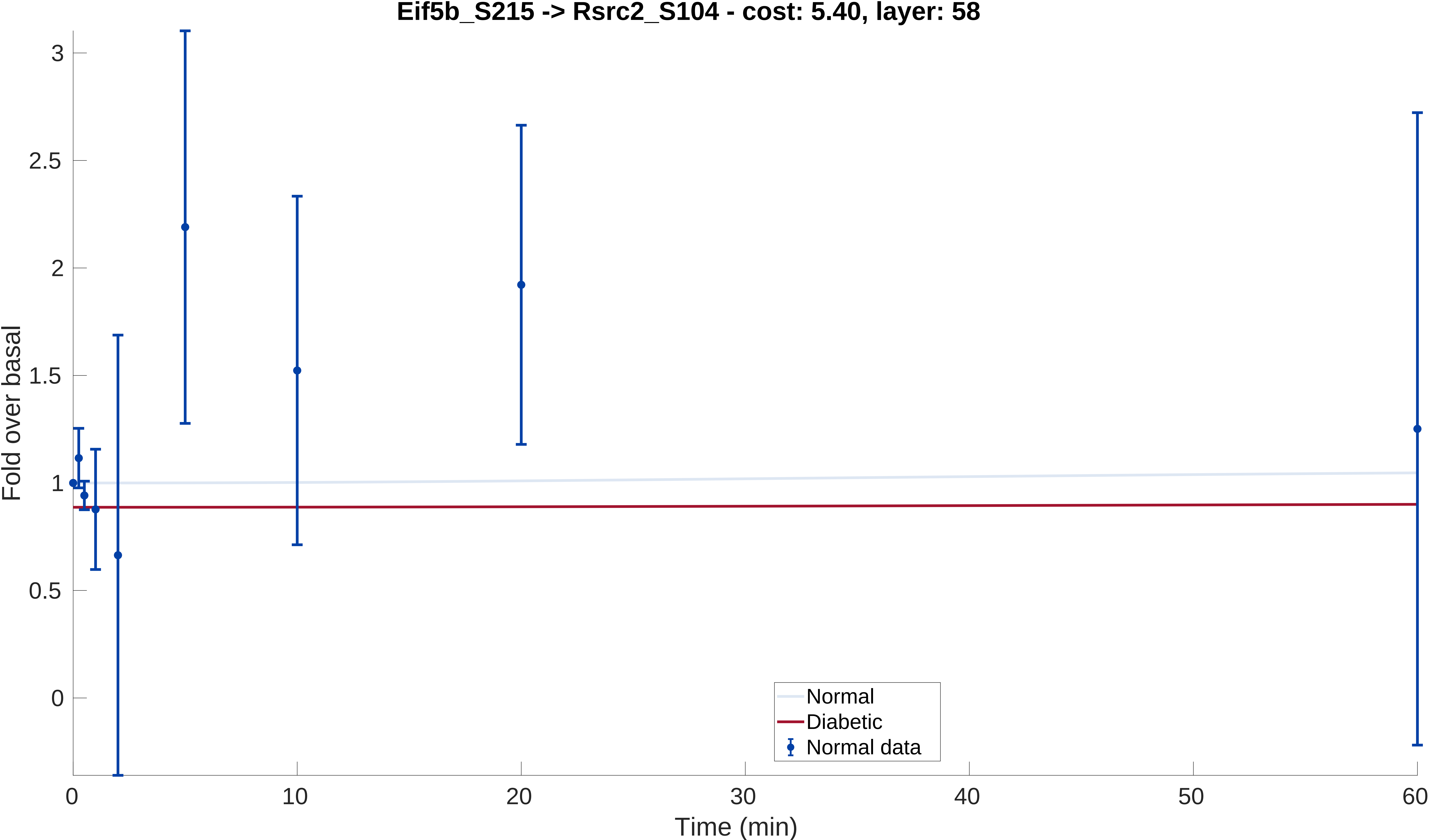

Eif5b_S215 -> Rsrc2_S104 - cost: 5.40, layer: 58
Fold over basal
3
2.5
2
1.5
1
0.5
0
0
10
20
30
40
50
60
Time (min)
Normal
Diabetic
Normal data

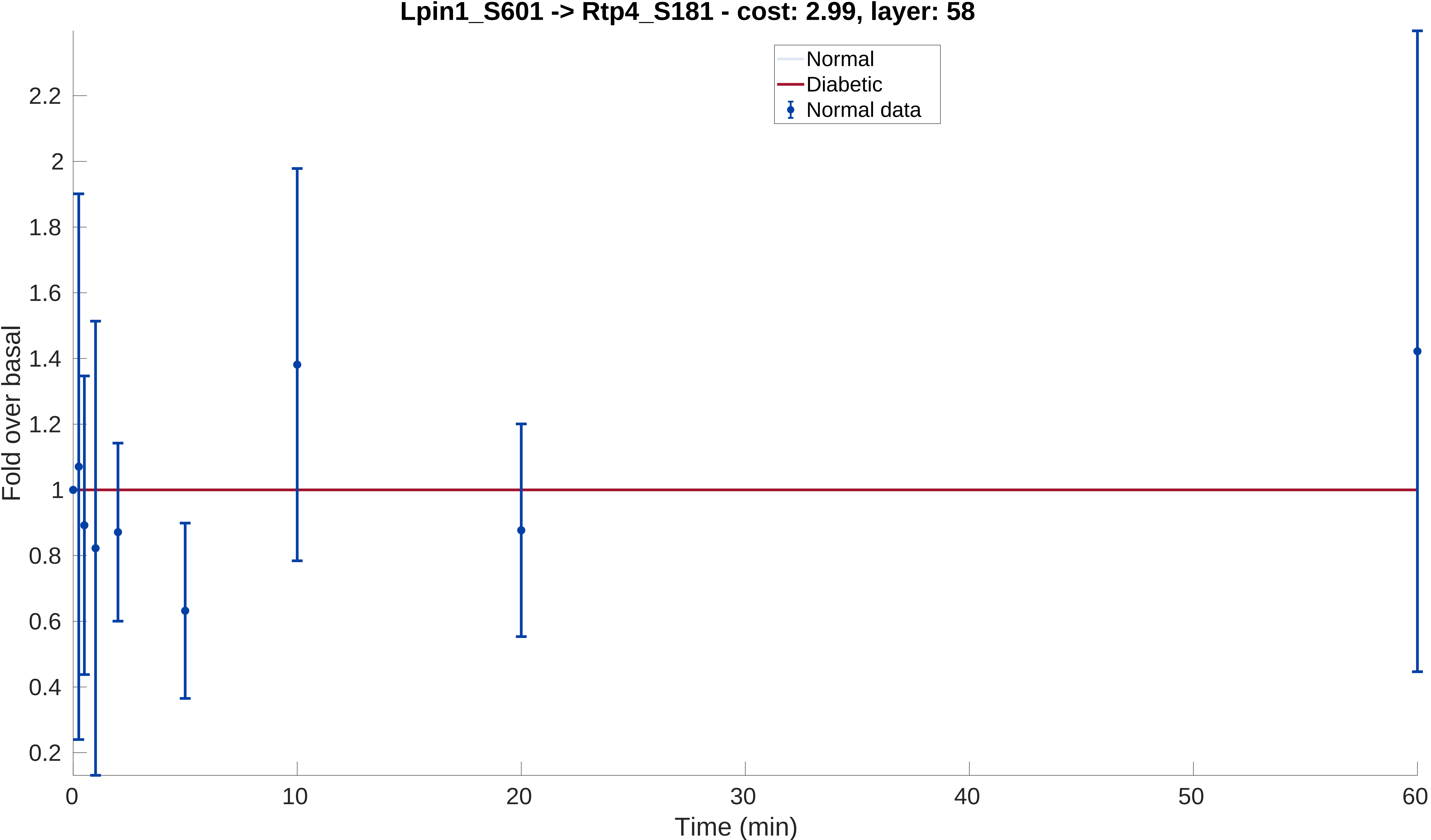

Lpin1_S601 -> Rtp4_S181 - cost: 2.99, layer: 58
Normal
Diabetic
Normal data
Fold over basal
Time (min)
0.2
0.4
0.6
0.8
1
1.2
1.4
1.6
1.8
2
2.2
0
10
20
30
40
50
60

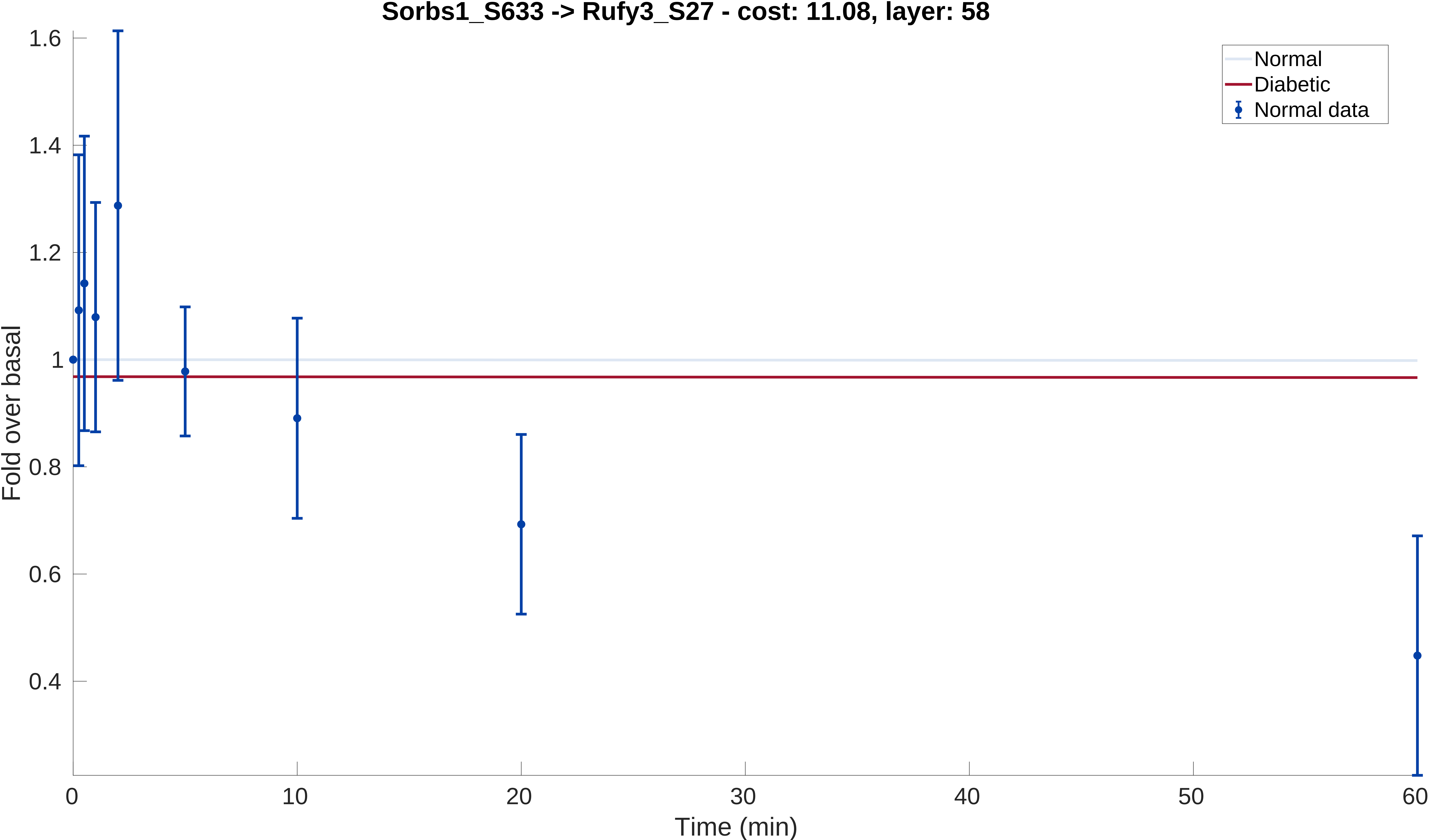

Sorbs1_S633 -> Rufy3_S27 - cost: 11.08, layer: 58
Normal
Diabetic
Normal data
Fold over basal
1.6
1.4
1.2
1
0.8
0.6
0.4
0
10
20
30
40
50
60
Time (min)

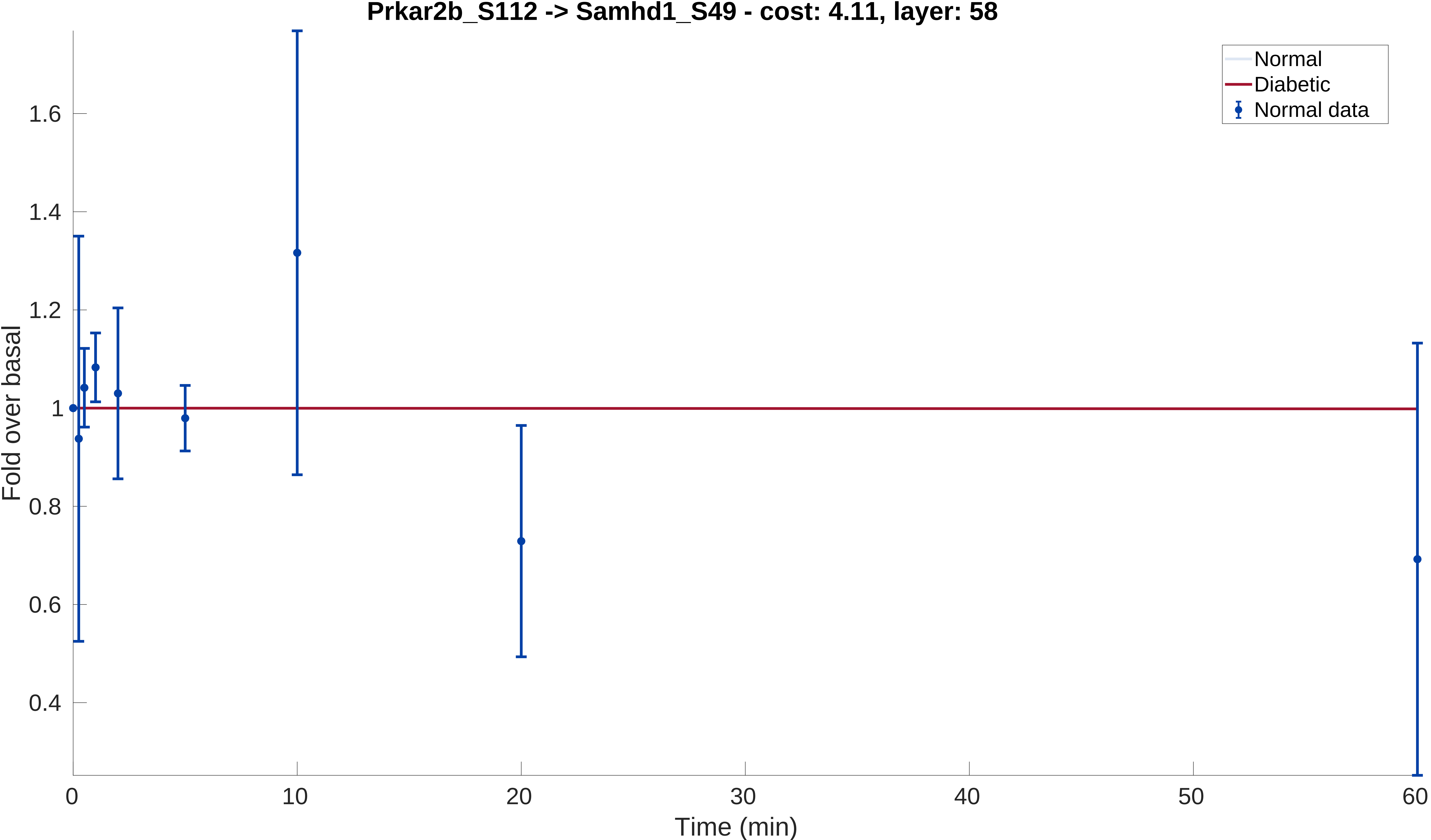

Prkar2b_S112 -> Samhd1_S49 - cost: 4.11, layer: 58
Normal
Diabetic
Normal data
Fold over basal
1.6
1.4
1.2
1
0.8
0.6
0.4
0
10
20
30
40
50
60
Time (min)

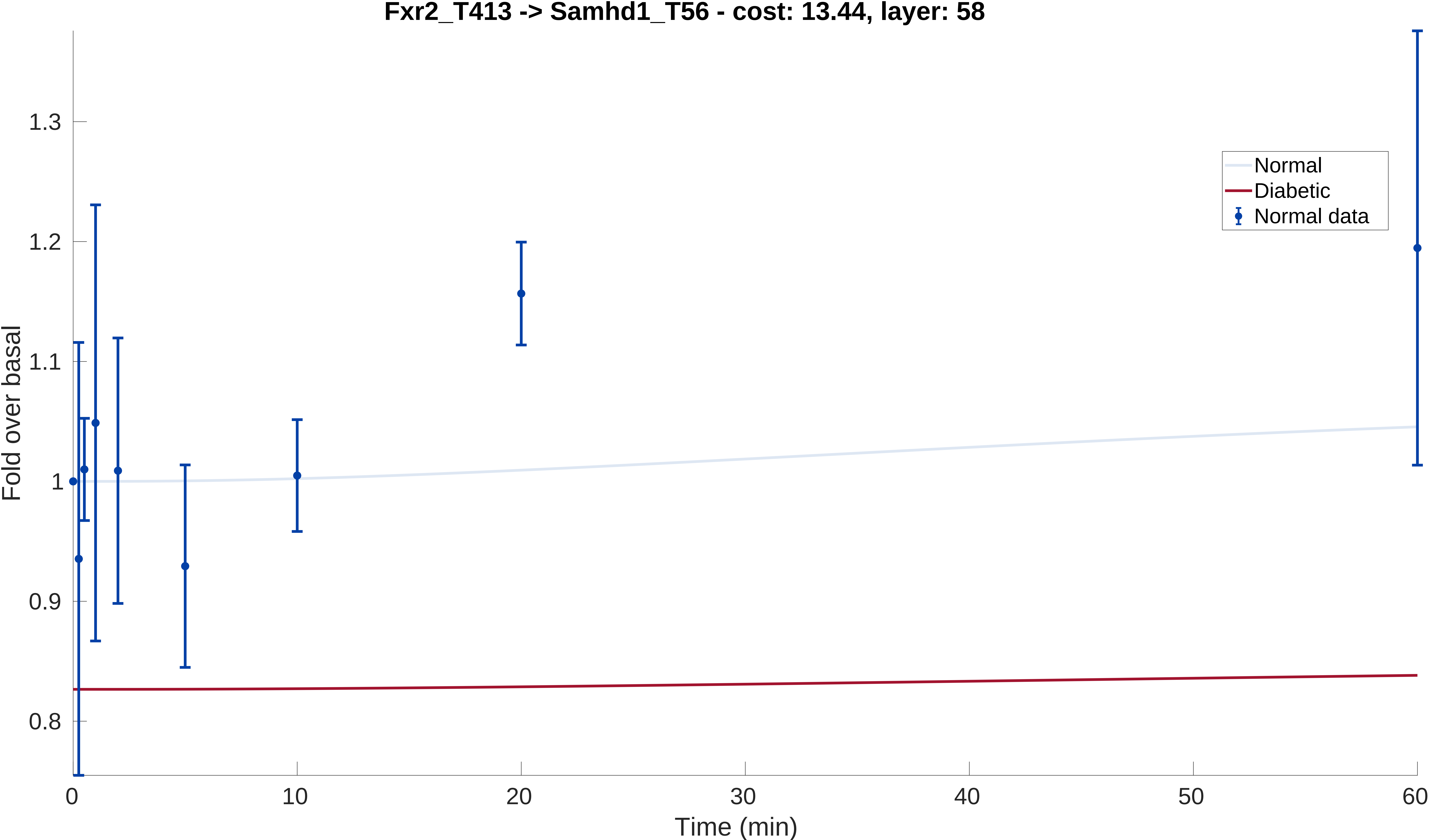
Fxr2_T413 -> Samhd1_T56 - cost: 13.44, layer: 58
Fold over basal
1.3
1.2
1.1
1
0.9
0.8
Normal
Diabetic
Normal data
0
10
20
30
40
50
60
Time (min)

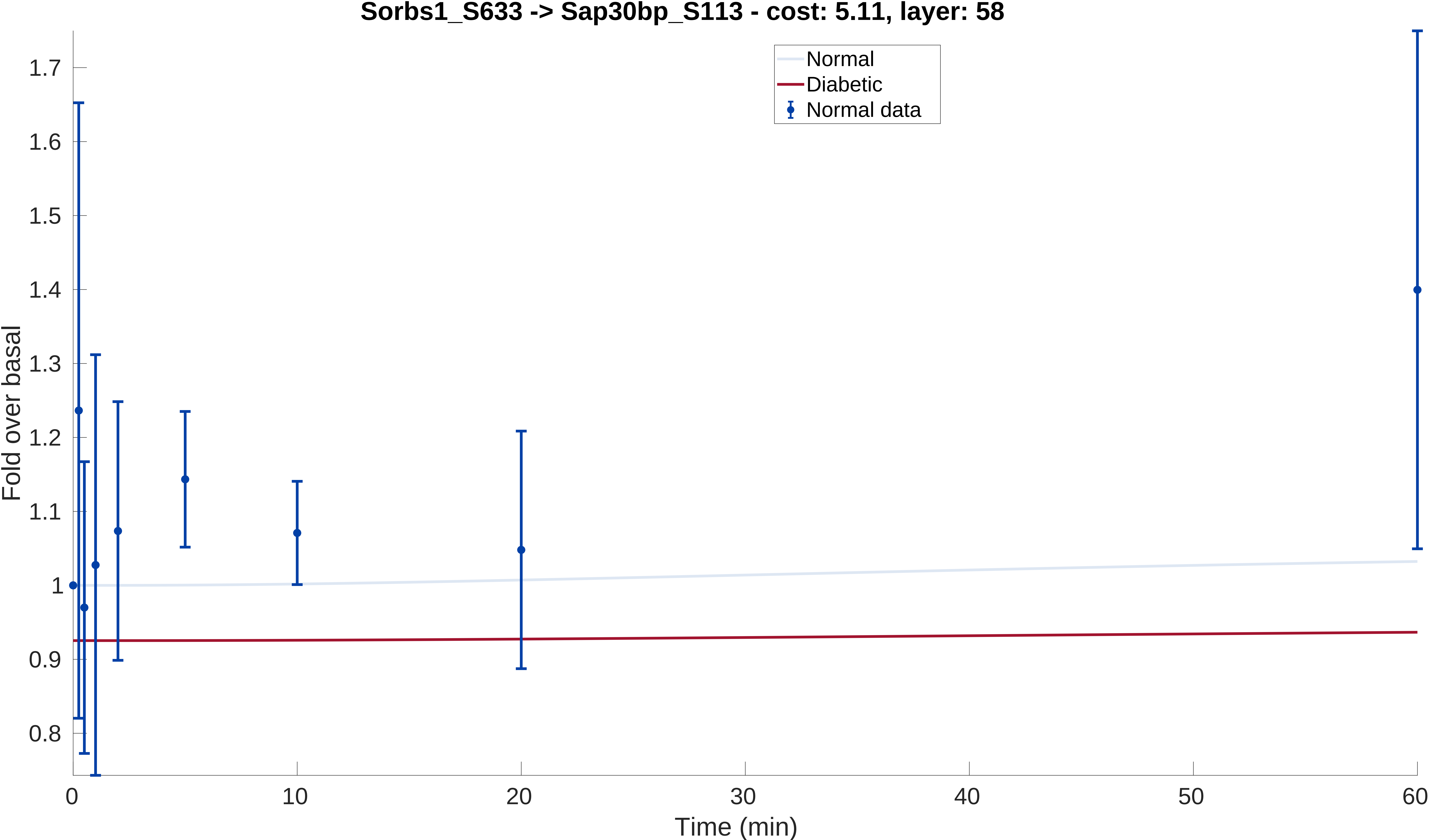

Sorbs1_S633 -> Sap30bp_S113 - cost: 5.11, layer: 58
Normal
Diabetic
Normal data
Fold over basal
1.7
1.6
1.5
1.4
1.3
1.2
1.1
1
0.9
0.8
0
10
20
30
40
50
60
Time (min)

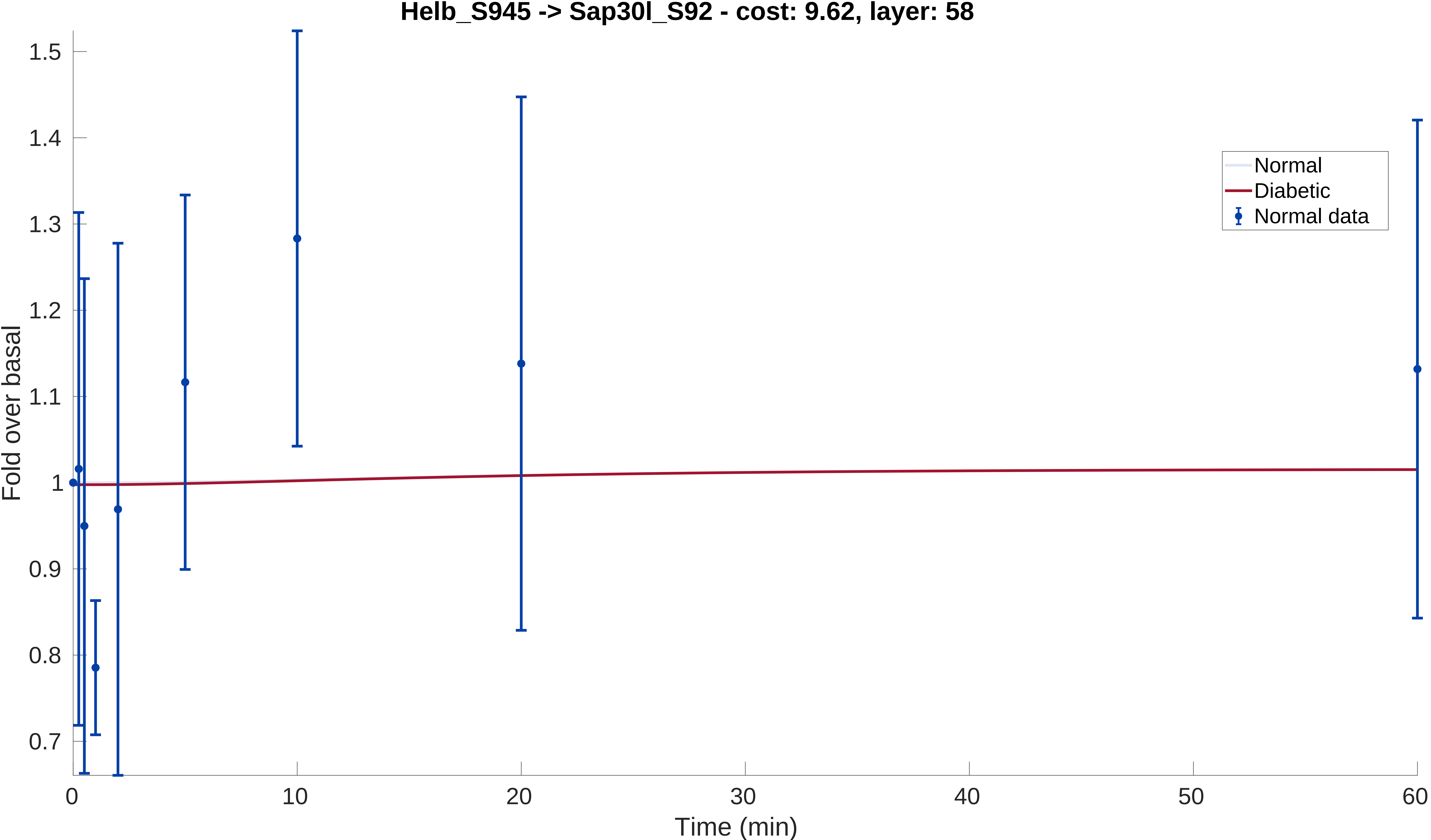

Helb_S945 -> Sap30l_S92 - cost: 9.62, layer: 58
Fold over basal
Time (min)
Normal
Diabetic
Normal data
1.5
1.4
1.3
1.2
1.1
1
0.9
0.8
0.7
0
10
20
30
40
50
60

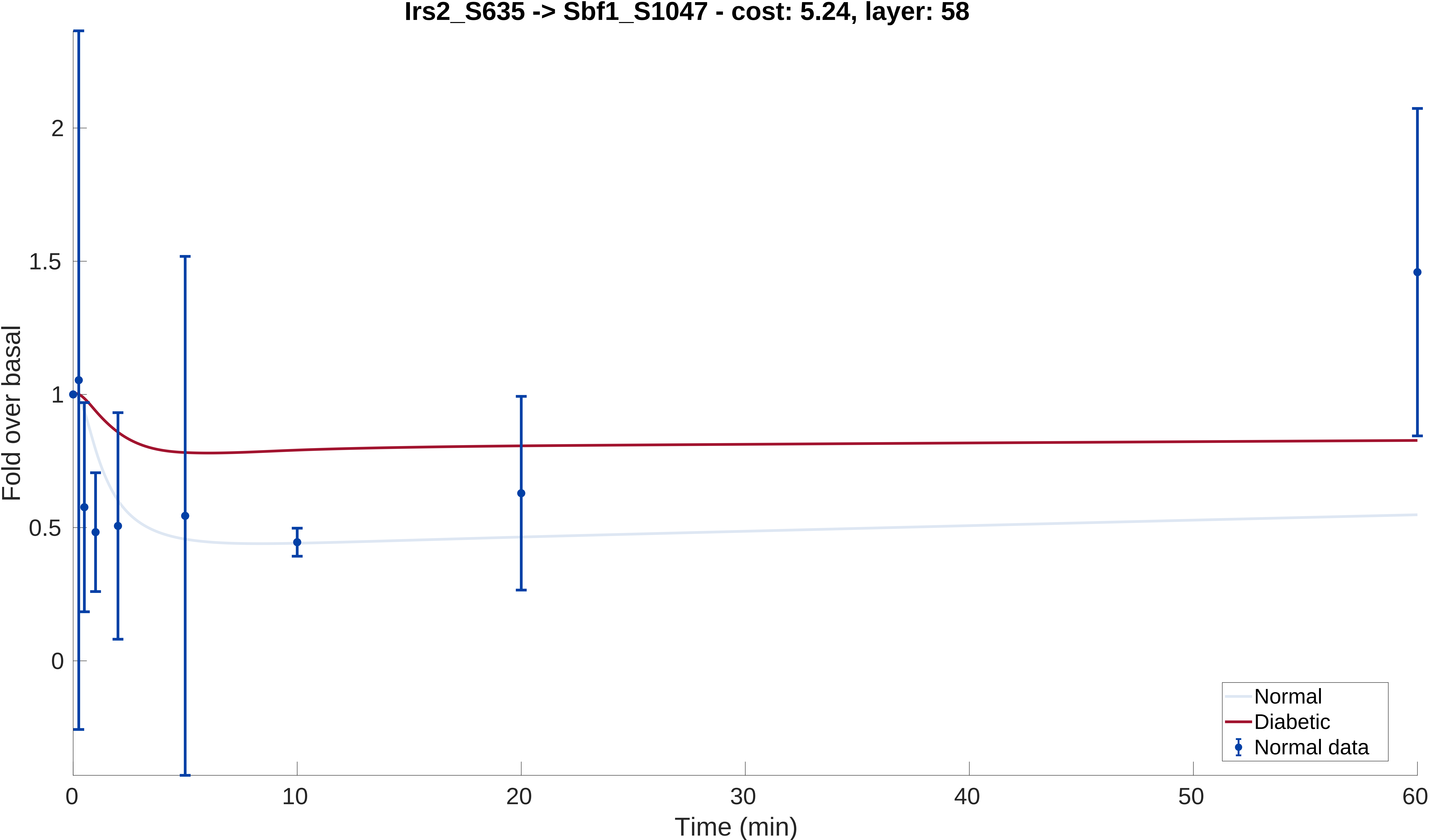

Irs2_S635 -> Sbf1_S1047 - cost: 5.24, layer: 58
Fold over basal
Time (min)
0
0.5
1
1.5
2
0
10
20
30
40
50
60
Normal
Diabetic
Normal data

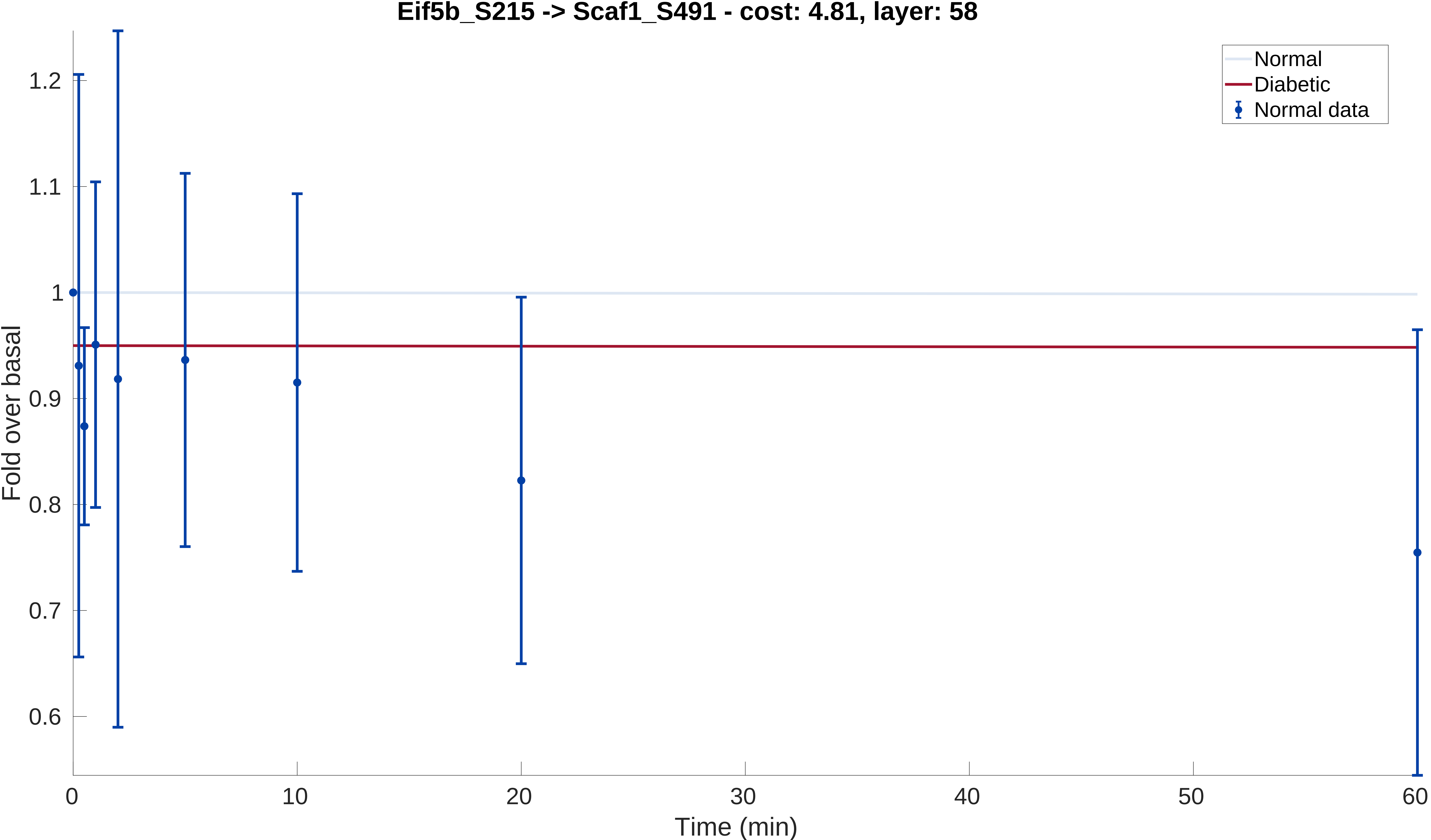

Eif5b_S215 -> Scaf1_S491 - cost: 4.81, layer: 58
Normal
Diabetic
Normal data
Fold over basal
1.2
1.1
1
0.9
0.8
0.7
0.6
0
10
20
30
40
50
60
Time (min)

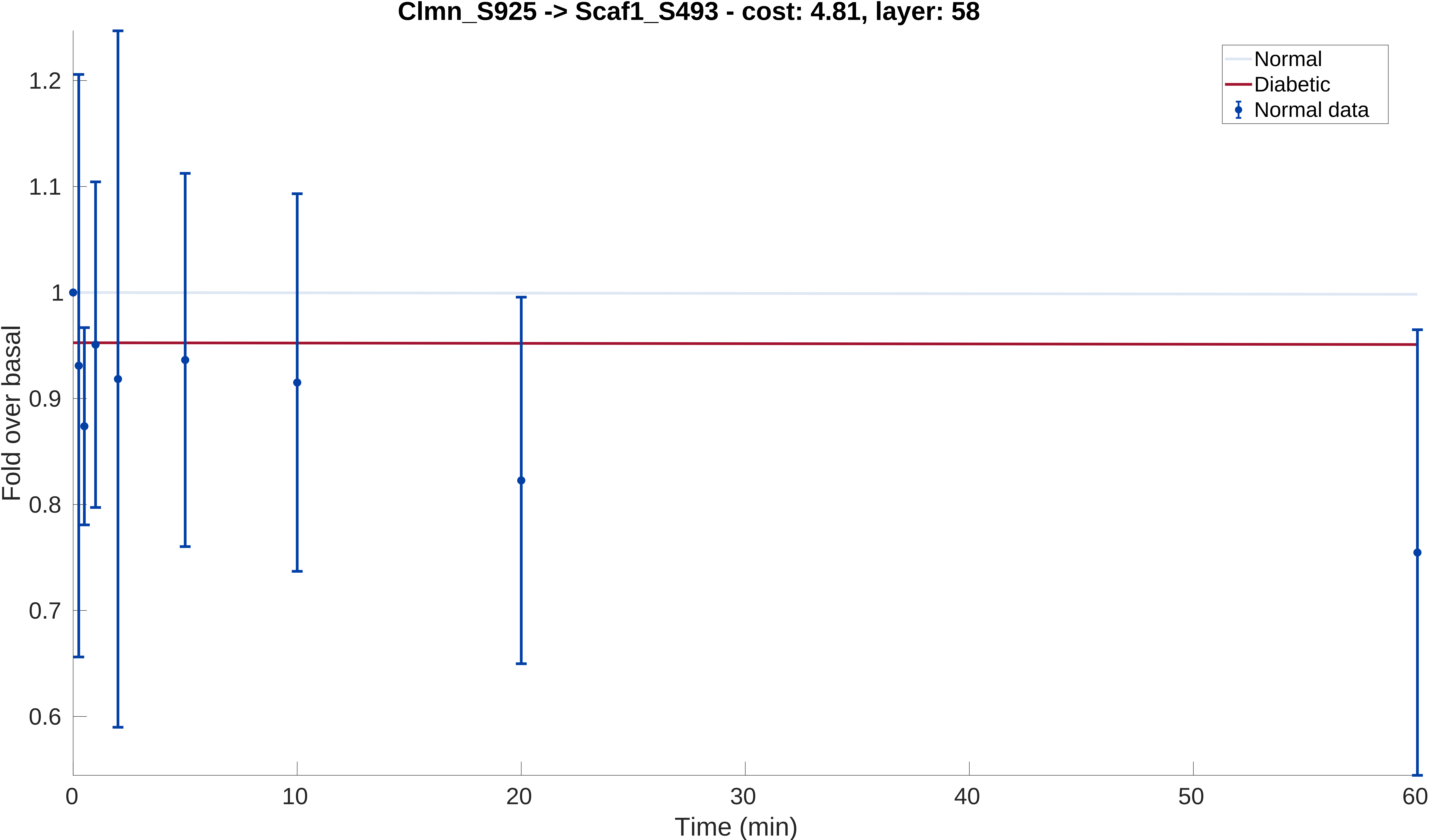

Clmn_S925 -> Scaf1_S493 - cost: 4.81, layer: 58
Normal
Diabetic
Normal data
Fold over basal
1.2
1.1
1
0.9
0.8
0.7
0.6
0
10
20
30
40
50
60
Time (min)

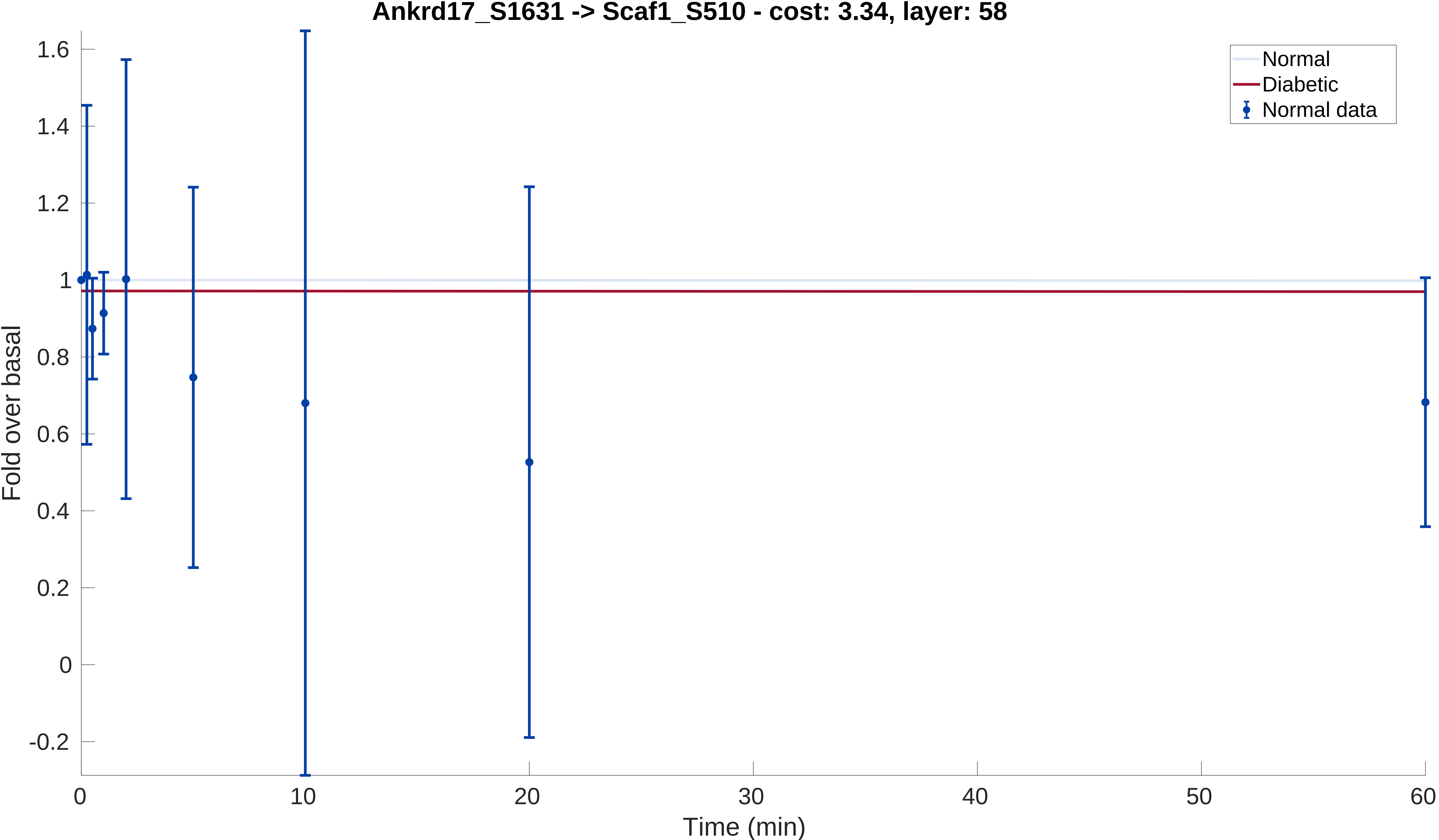

Ankrd17_S1631 -> Scaf1_S510 - cost: 3.34, layer: 58
Normal
Diabetic
Normal data
Fold over basal
Time (min)
1.6
1.4
1.2
1
0.8
0.6
0.4
0.2
0
-0.2
0
10
20
30
40
50
60

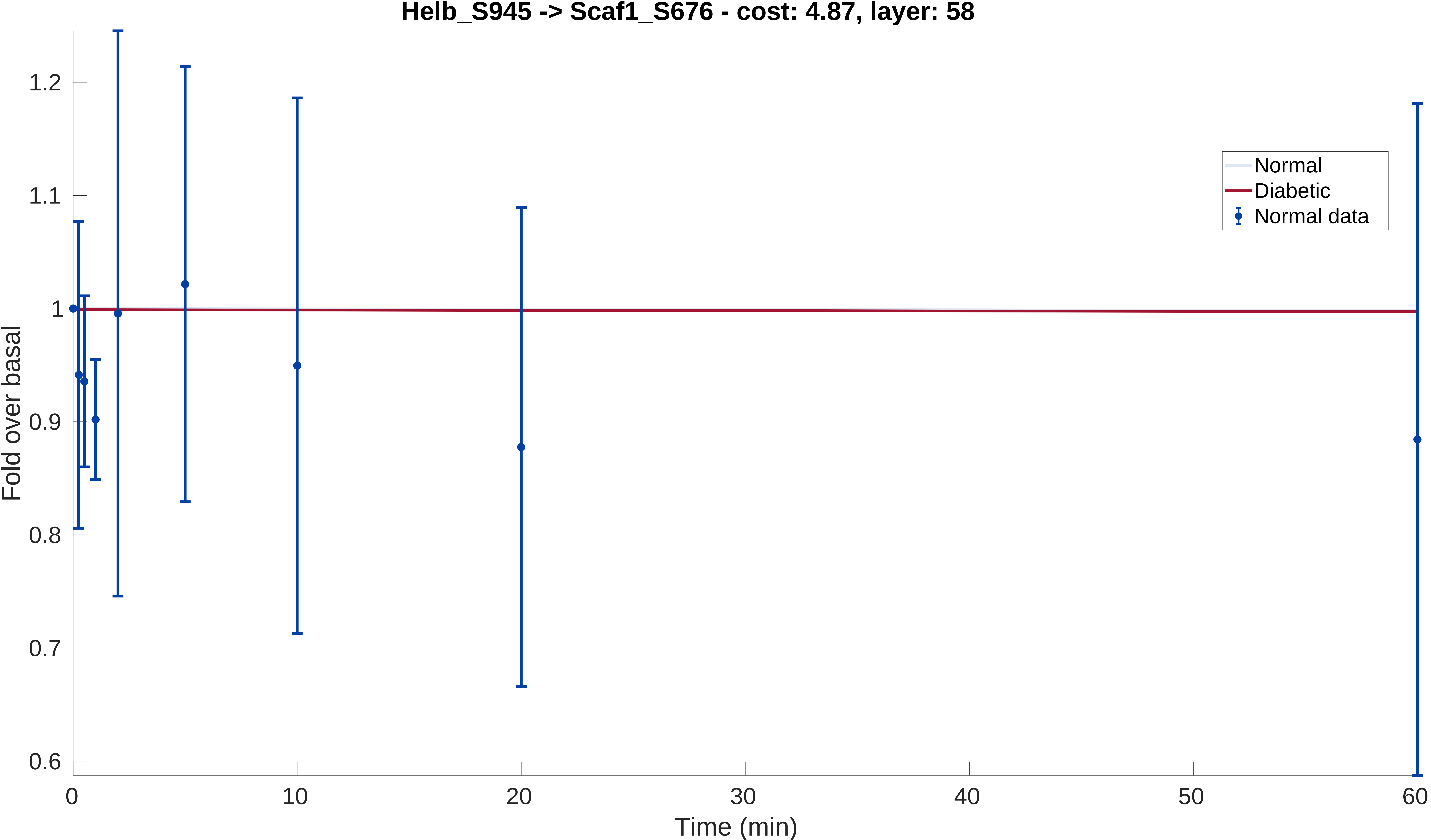

Helb_S945 -> Scaf1_S676 - cost: 4.87, layer: 58
Normal
Diabetic
Normal data
1.2
1.1
1
0.9
0.8
0.7
0.6
Fold over basal
0
10
20
30
40
50
60
Time (min)

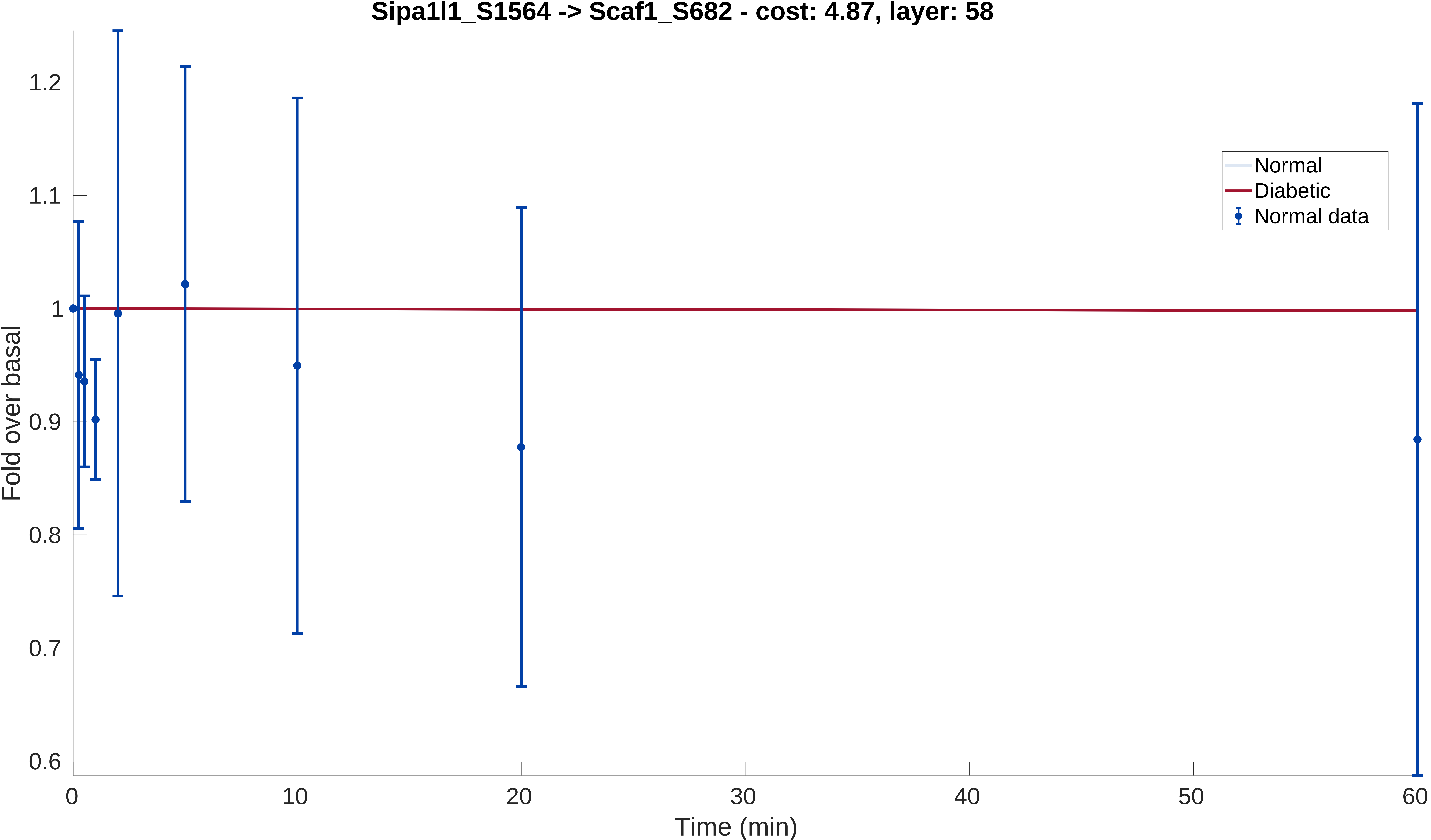

Sipa1l1_S1564 -> Scaf1_S682 - cost: 4.87, layer: 58
Fold over basal
1.2
1.1
1
0.9
0.8
0.7
0.6
0
10
20
30
40
50
60
Time (min)
Normal
Diabetic
Normal data

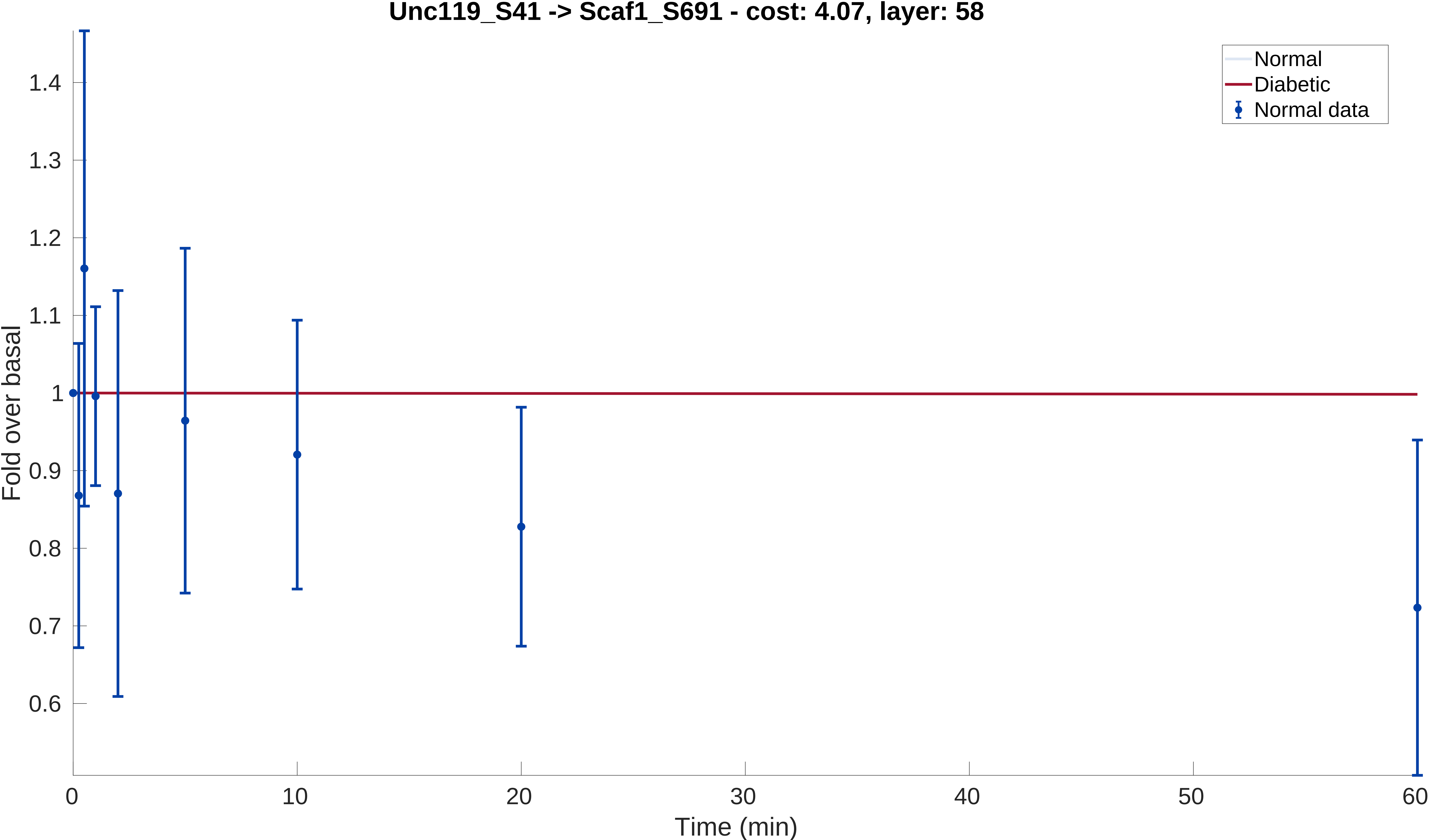

Unc119_S41 -> Scaf1_S691 - cost: 4.07, layer: 58
Normal
Diabetic
Normal data
Fold over basal
1.4
1.3
1.2
1.1
1
0.9
0.8
0.7
0.6
0
10
20
30
40
50
60
Time (min)

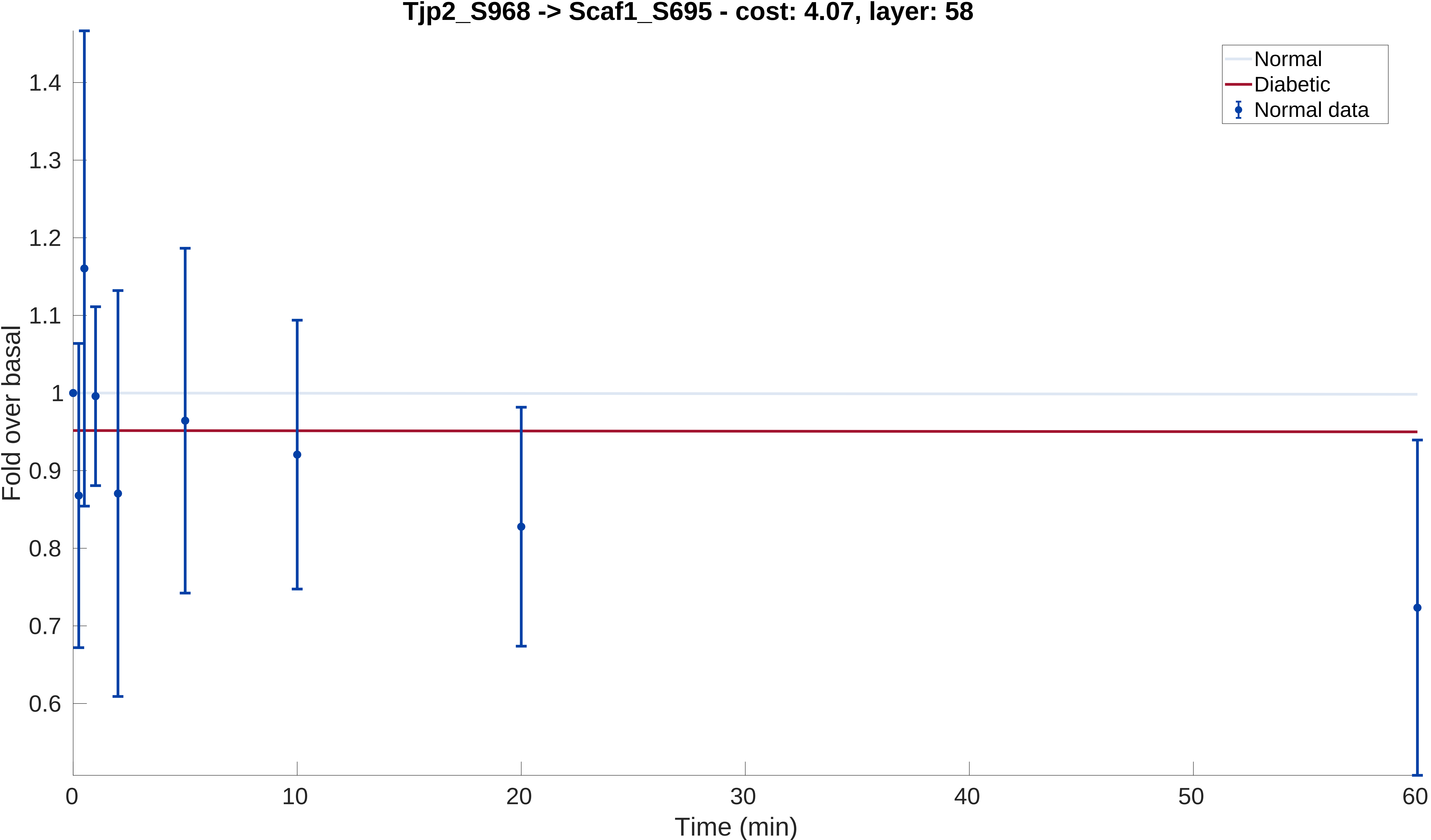
Tjp2_S968 -> Scaf1_S695 - cost: 4.07, layer: 58
Normal
Diabetic
Normal data
Fold over basal
1.4
1.3
1.2
1.1
1
0.9
0.8
0.7
0.6
0
10
20
30
40
50
60
Time (min)

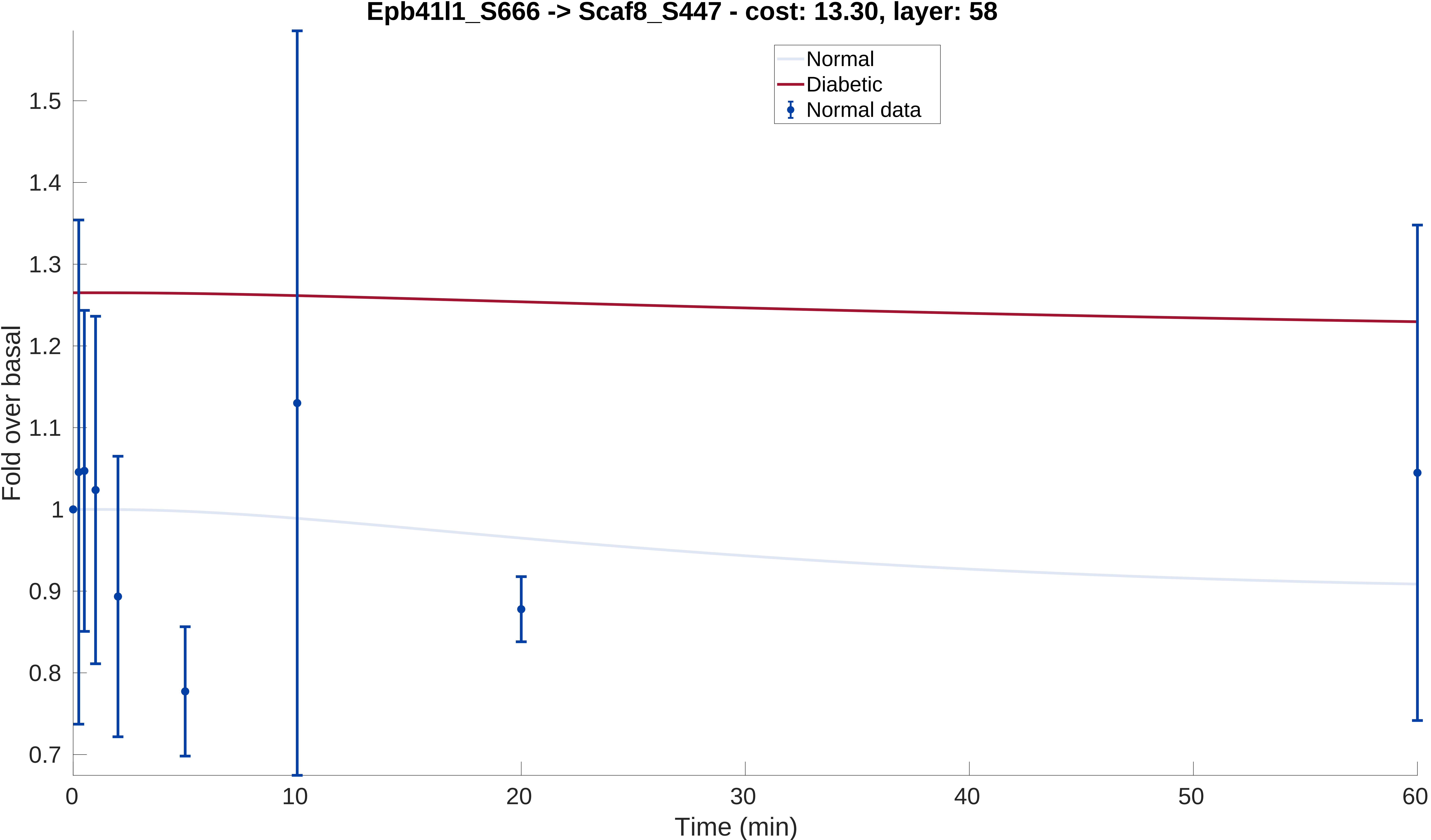
Epb41l1_S666 -> Scaf8_S447 - cost: 13.30, layer: 58
Normal
Diabetic
Normal data
Fold over basal
1.5
1.4
1.3
1.2
1.1
1
0.9
0.8
0.7
0
10
20
30
40
50
60
Time (min)

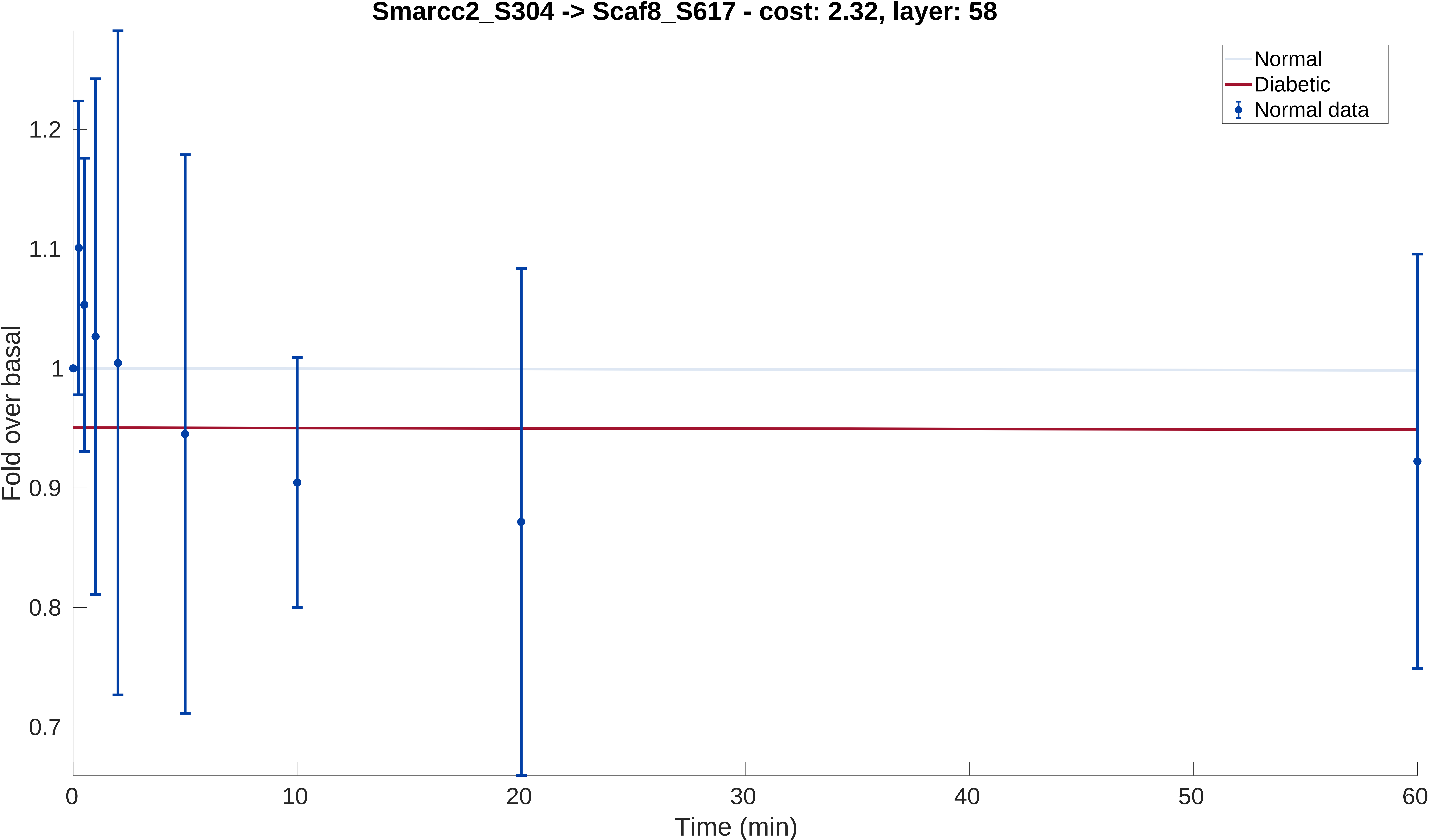

Smarcc2_S304 -> Scaf8_S617 - cost: 2.32, layer: 58
Normal
Diabetic
Normal data
Fold over basal
1.2
1.1
1
0.9
0.8
0.7
0
10
20
30
40
50
60
Time (min)

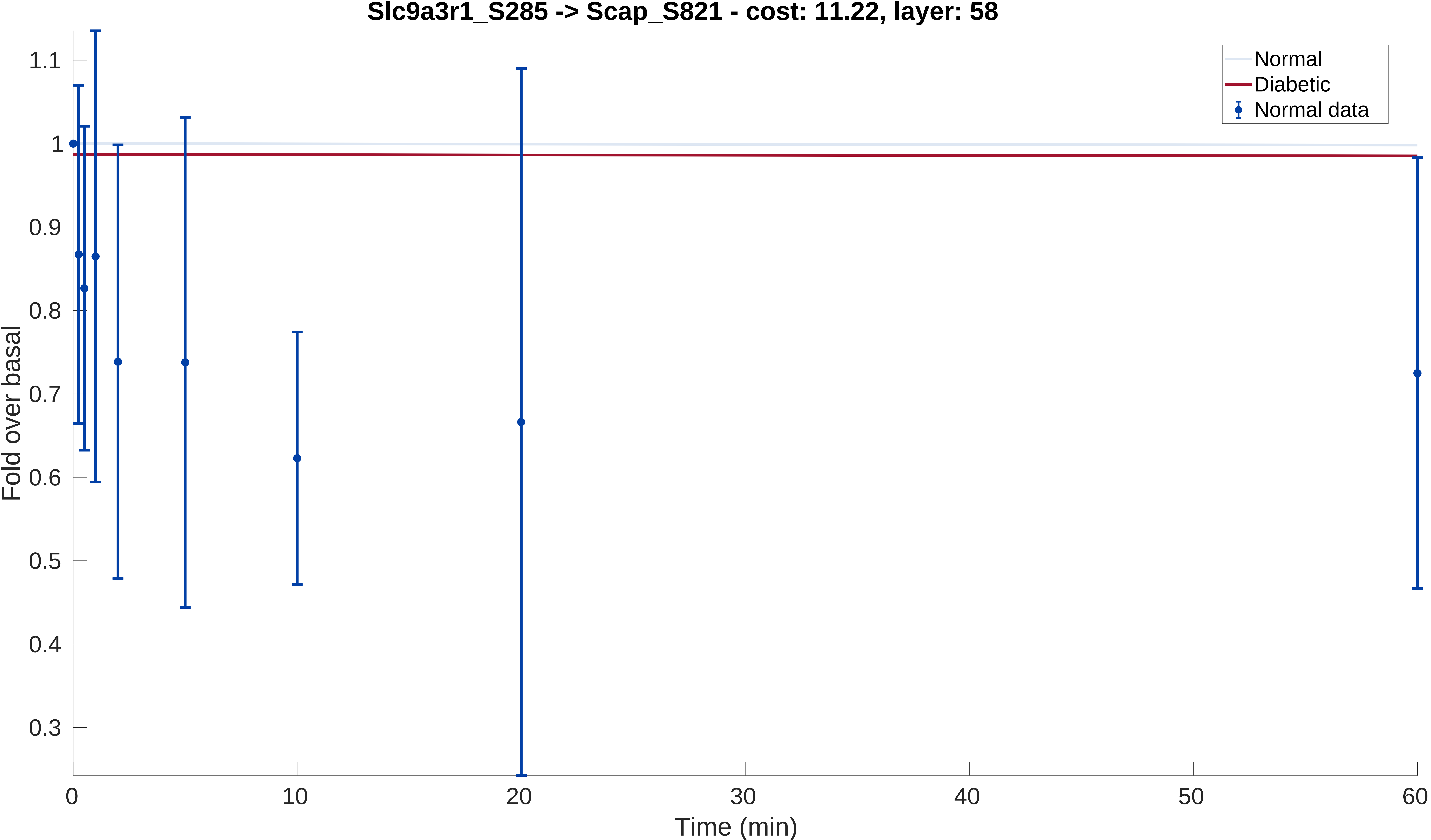

Slc9a3r1_S285 -> Scap_S821 - cost: 11.22, layer: 58
Normal
Diabetic
Normal data
Fold over basal
1.1
1
0.9
0.8
0.7
0.6
0.5
0.4
0.3
0
10
20
30
40
50
60
Time (min)

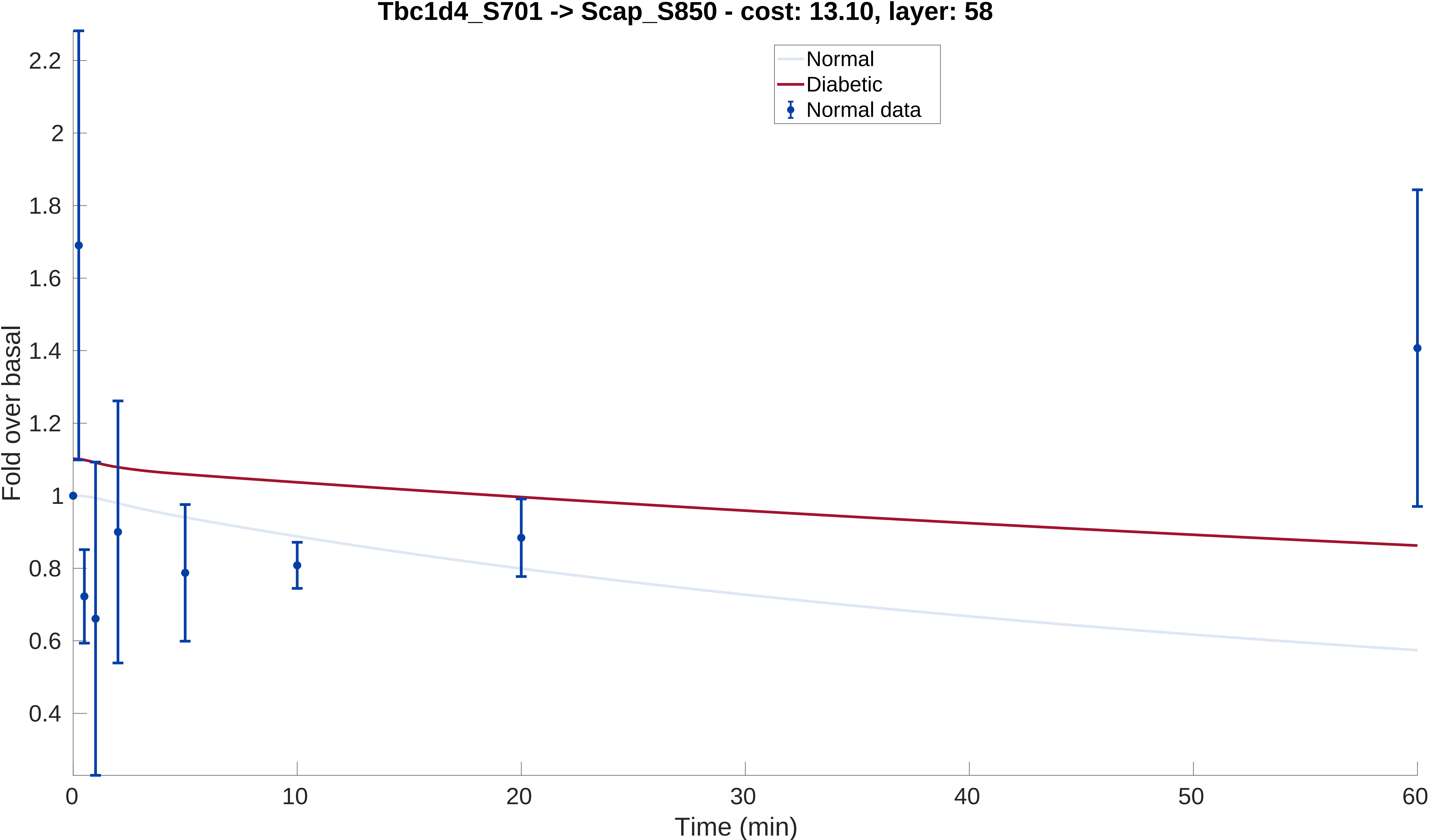

Tbc1d4_S701 -> Scap_S850 - cost: 13.10, layer: 58
Normal
Diabetic
Normal data
Fold over basal
Time (min)
0.4
0.6
0.8
1
1.2
1.4
1.6
1.8
2
2.2
0
10
20
30
40
50
60

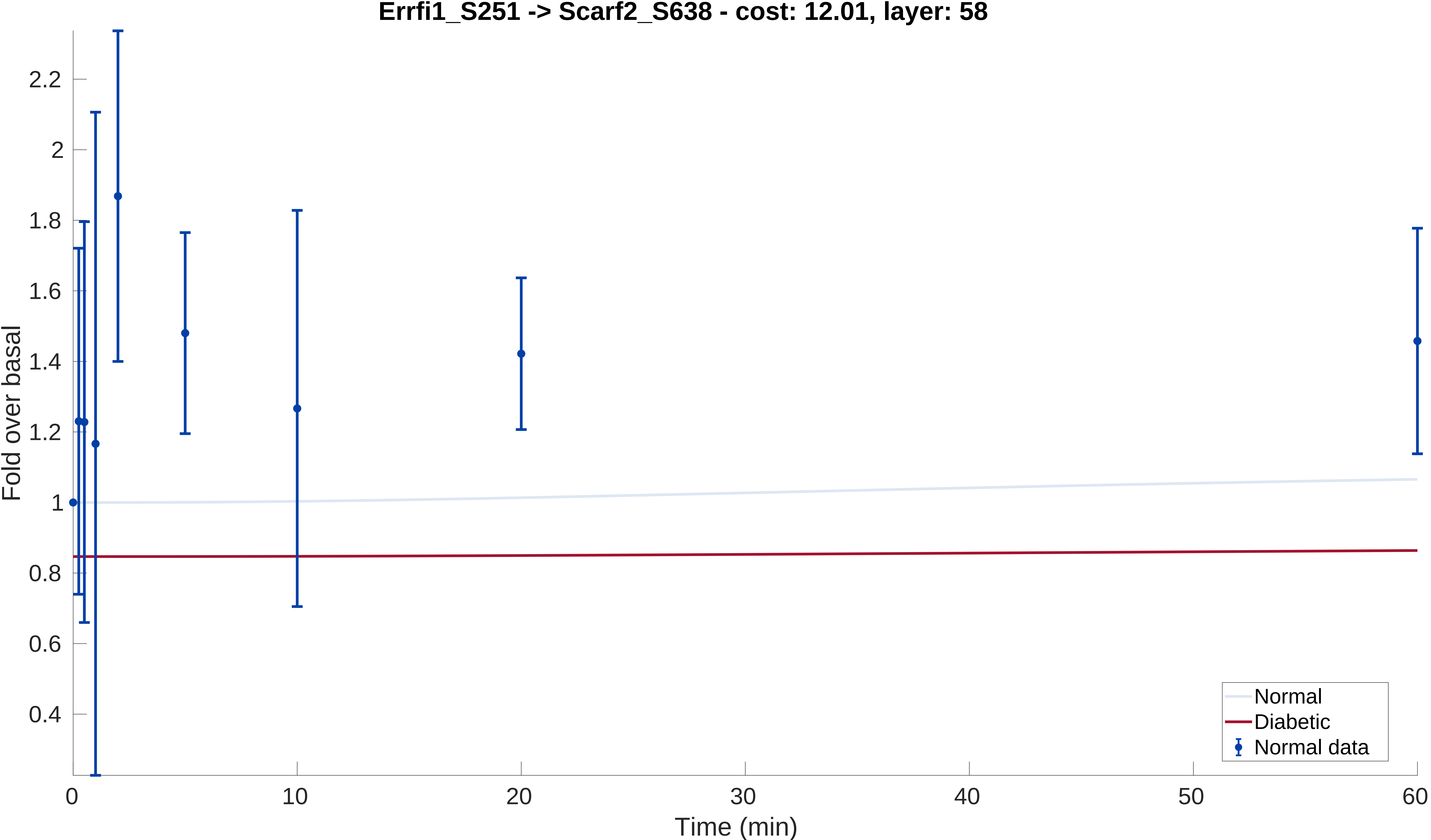

Errfi1_S251 -> Scarf2_S638 - cost: 12.01, layer: 58
Fold over basal
2.2
2
1.8
1.6
1.4
1.2
1
0.8
0.6
0.4
0
10
20
30
40
50
60
Time (min)
Normal
Diabetic
Normal data

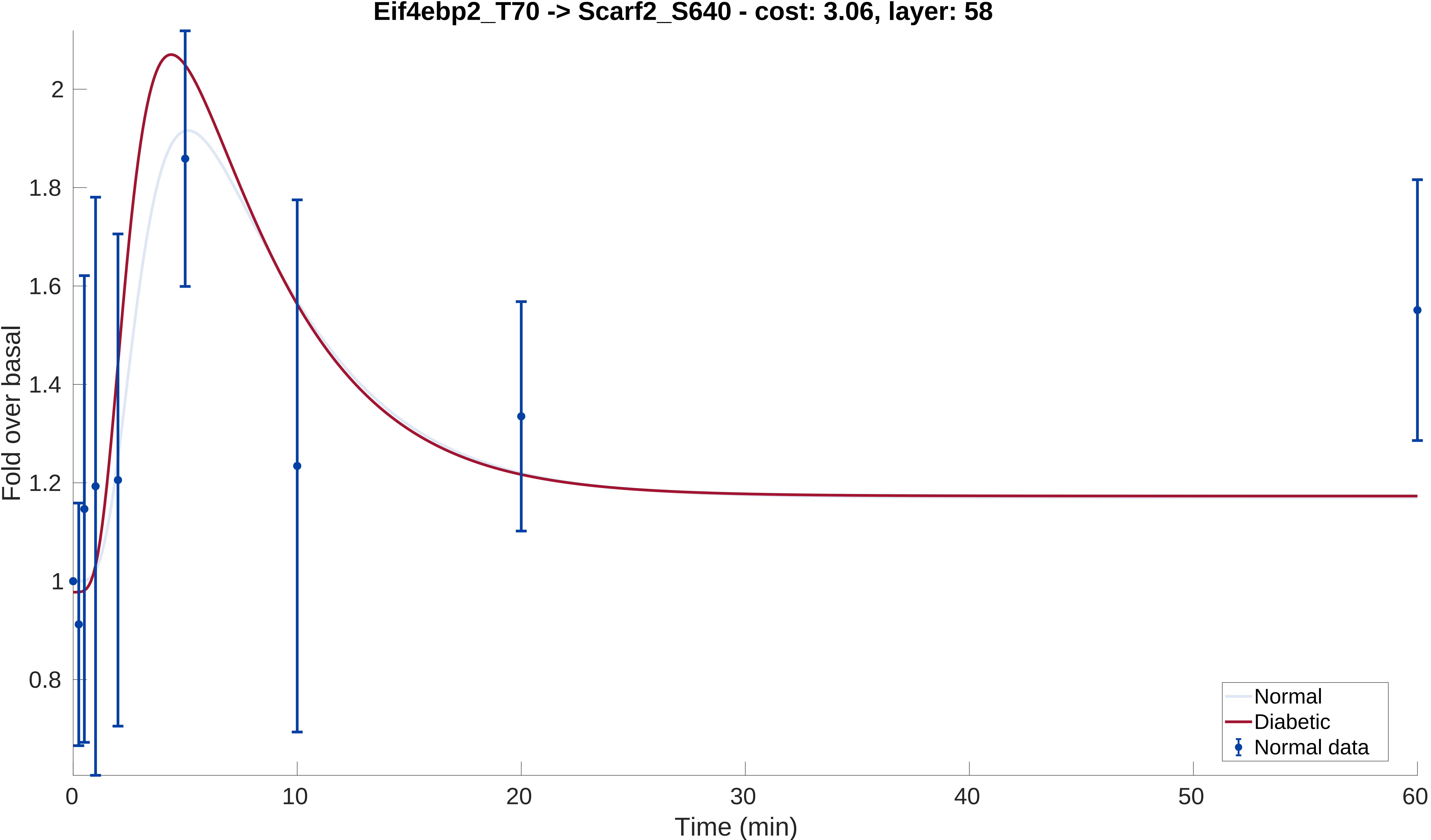
Eif4ebp2_T70 -> Scarf2_S640 - cost: 3.06, layer: 58
Fold over basal
2
1.8
1.6
1.4
1.2
1
0.8
0
10
20
30
40
50
60
Time (min)
Normal
Diabetic
Normal data

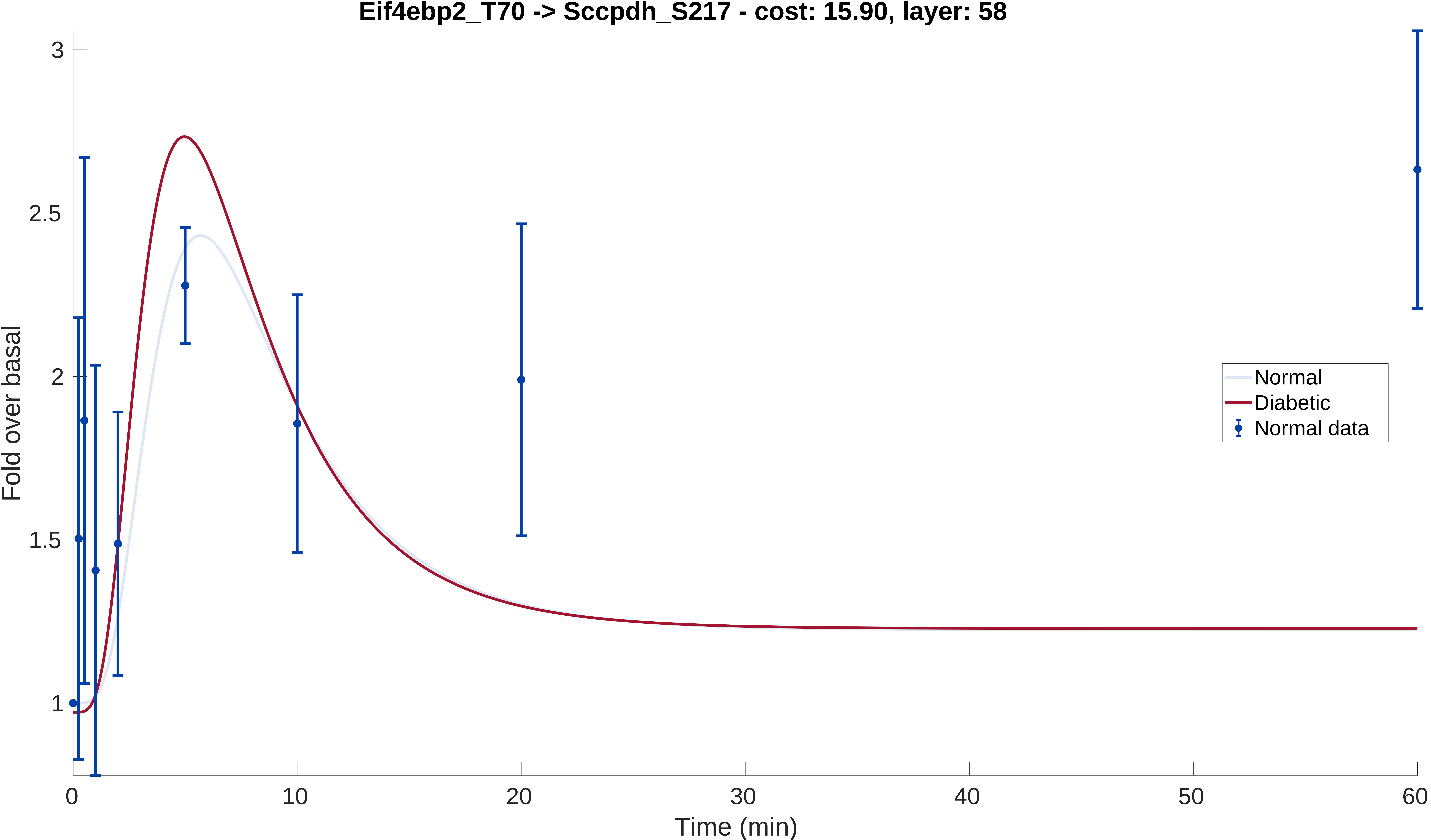

Eif4ebp2_T70 -> Sccpdh_S217 - cost: 15.90, layer: 58
Fold over basal
Time (min)
3
2.5
2
1.5
1
0
10
20
30
40
50
60
Normal
Diabetic
Normal data

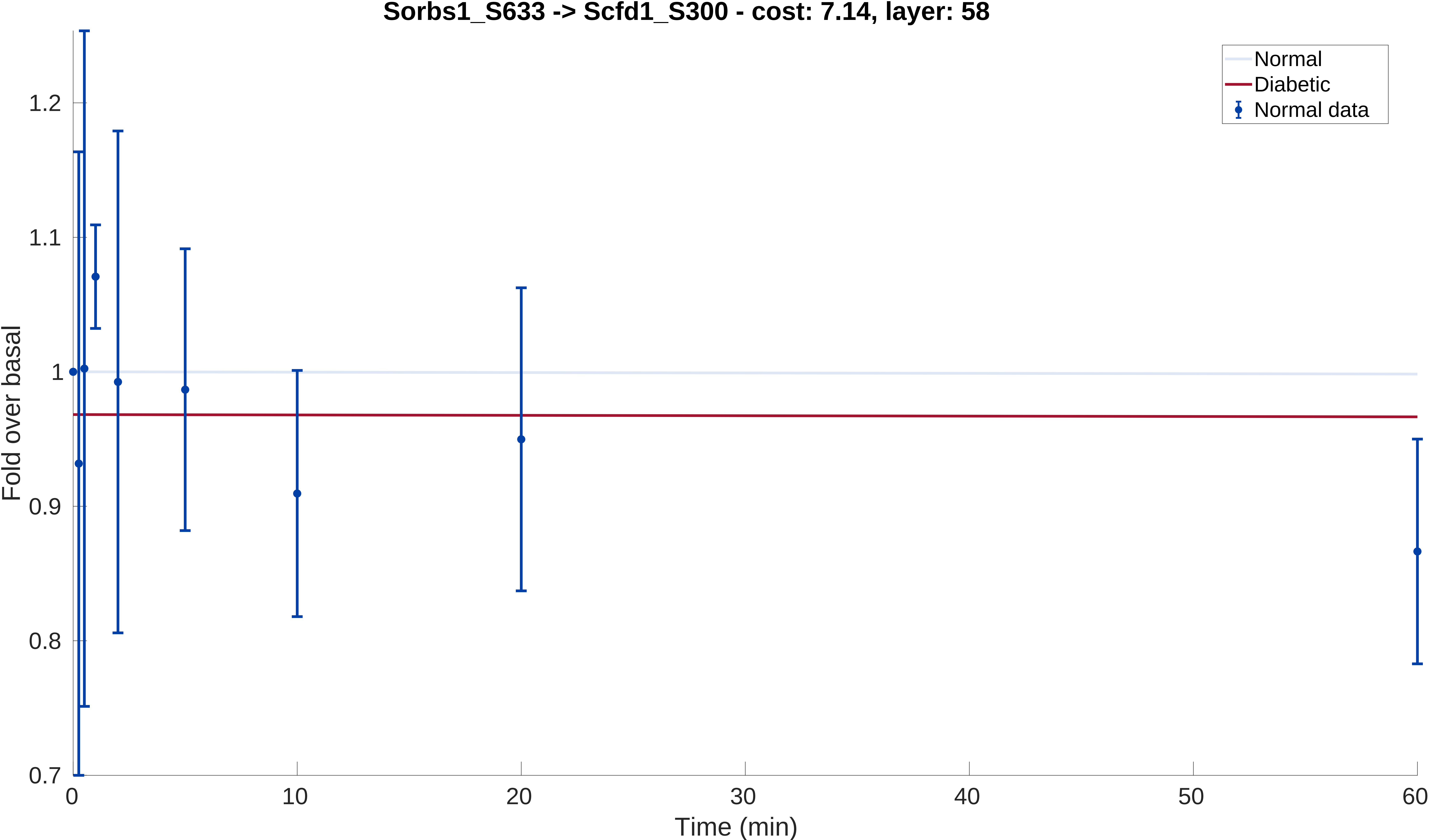
Sorbs1_S633 -> Scfd1_S300 - cost: 7.14, layer: 58
Normal
Diabetic
Normal data
Fold over basal
1.2
1.1
1
0.9
0.8
0.7
0
10
20
30
40
50
60
Time (min)

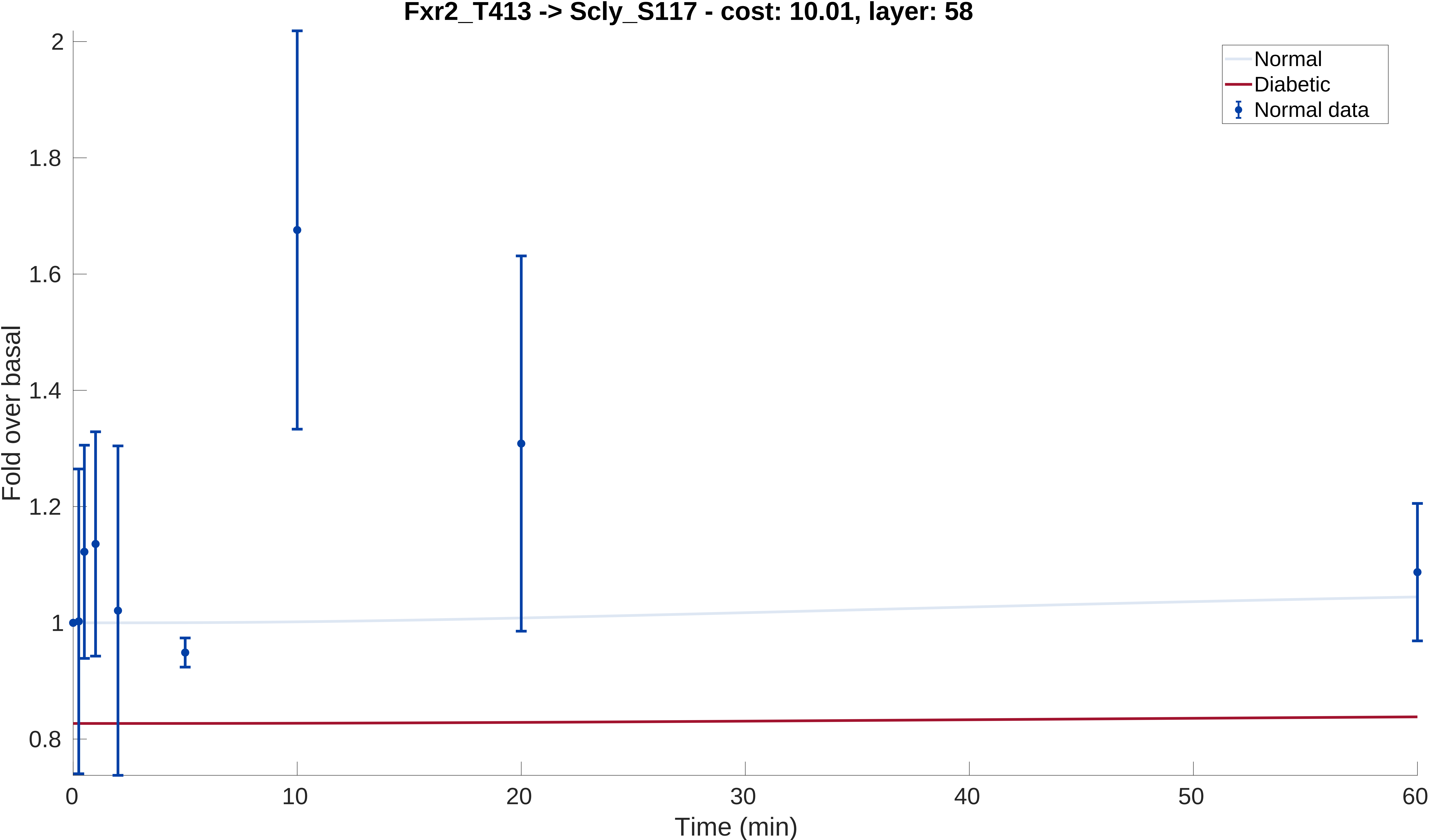

Fxr2_T413 -> Scly_S117 - cost: 10.01, layer: 58
Normal
Diabetic
Normal data
Fold over basal
Time (min)
2
1.8
1.6
1.4
1.2
1
0.8
0
10
20
30
40
50
60

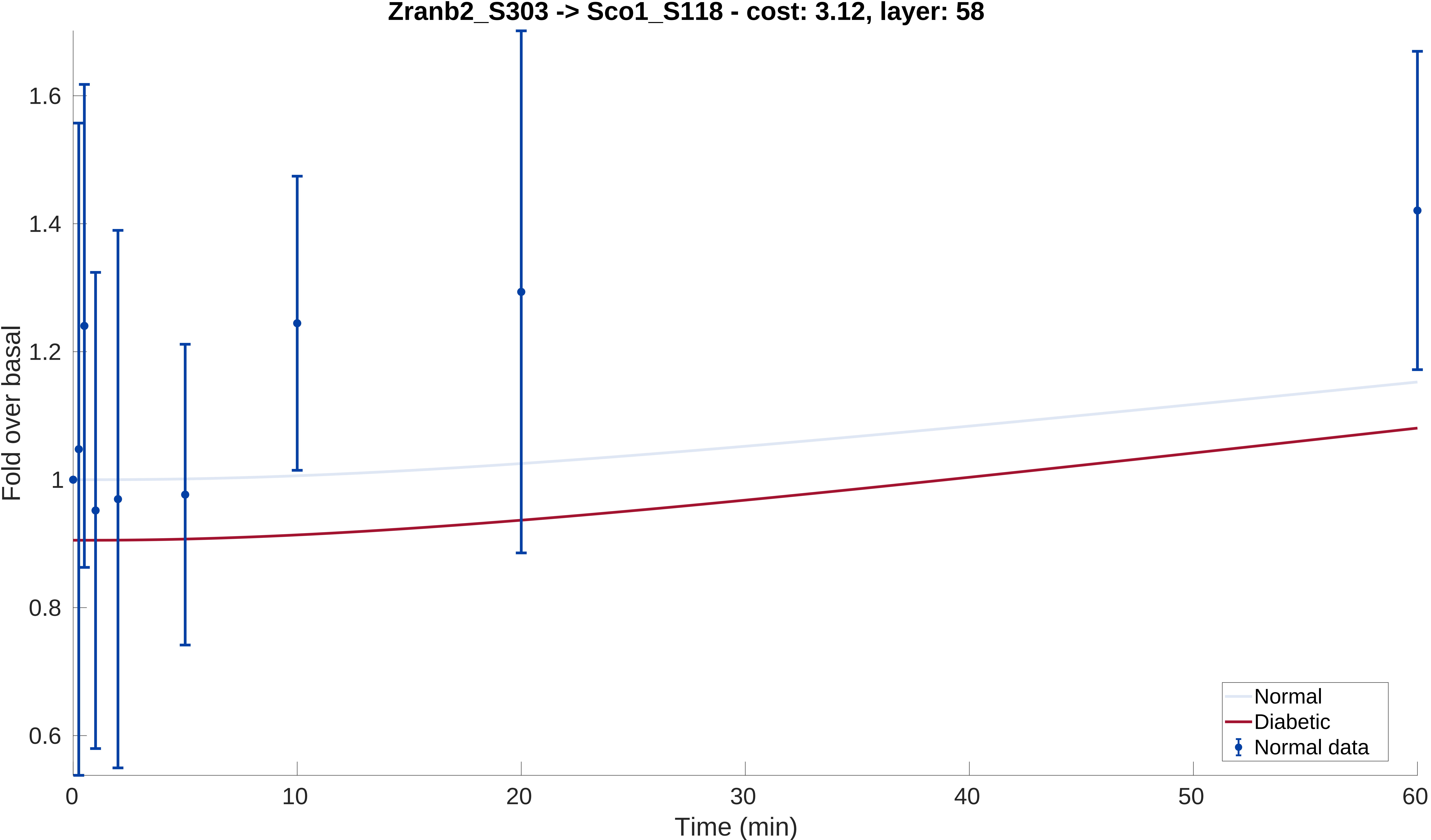

Zranb2_S303 -> Sco1_S118 - cost: 3.12, layer: 58
Fold over basal
1.6
1.4
1.2
1
0.8
0.6
0
10
20
30
40
50
60
Time (min)
Normal
Diabetic
Normal data

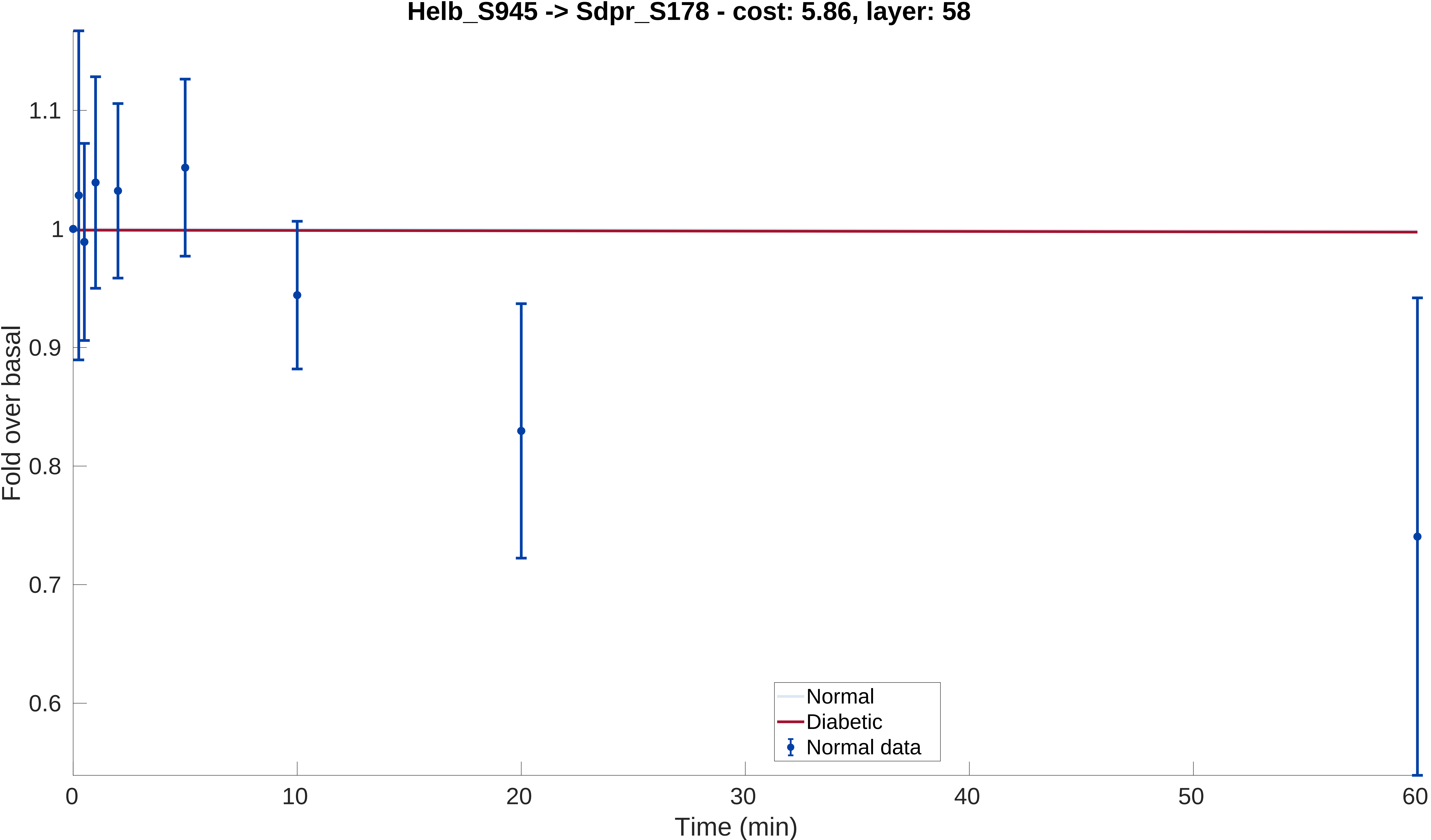

Helb_S945 -> Sdpr_S178 - cost: 5.86, layer: 58
Fold over basal
1.1
1
0.9
0.8
0.7
0.6
0
10
20
30
40
50
60
Time (min)
Normal
Diabetic
Normal data

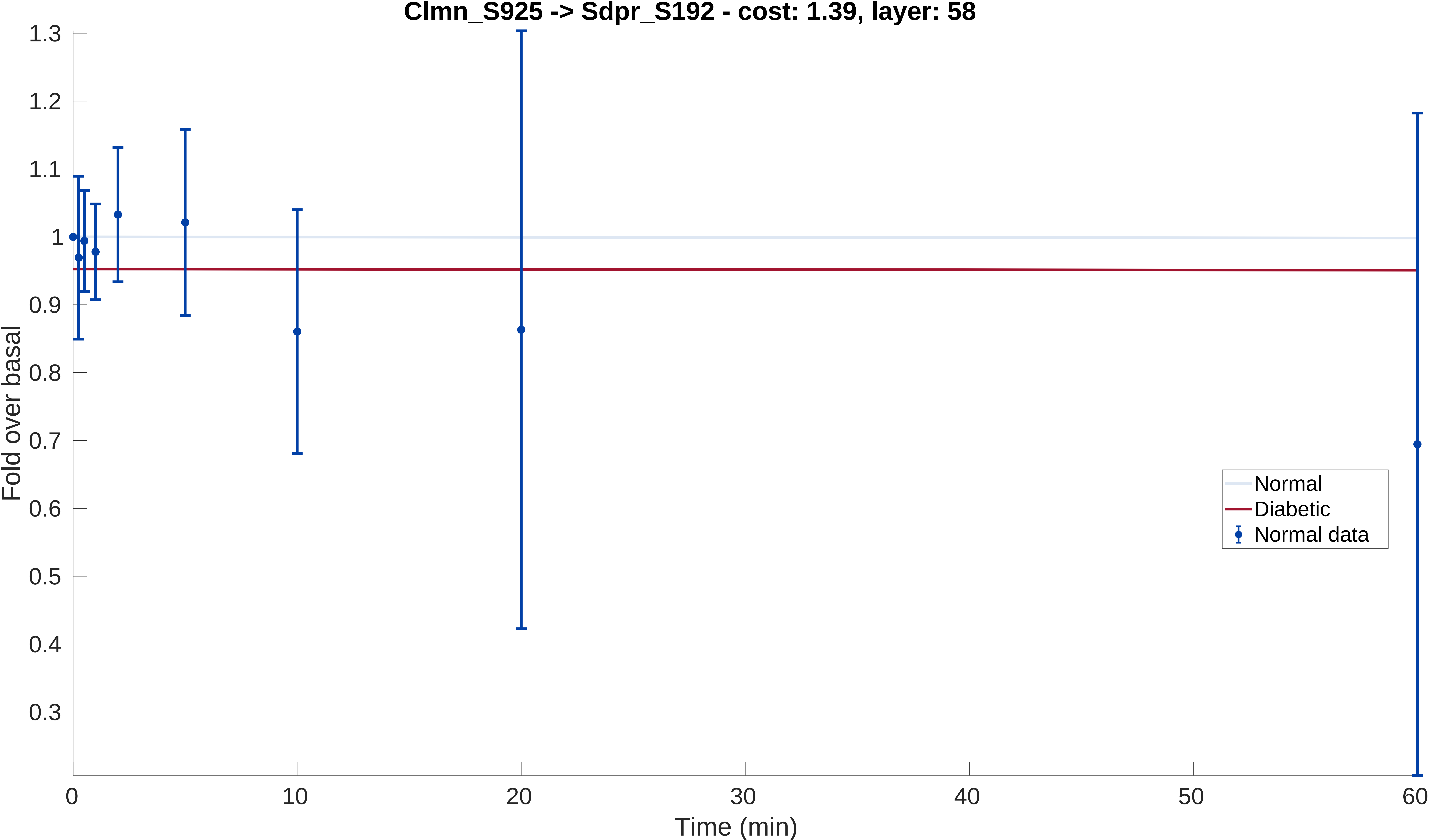

Clmn_S925 -> Sdpr_S192 - cost: 1.39, layer: 58
Fold over basal
Time (min)
Normal
Diabetic
Normal data
1.3
1.2
1.1
1
0.9
0.8
0.7
0.6
0.5
0.4
0.3
0
10
20
30
40
50
60

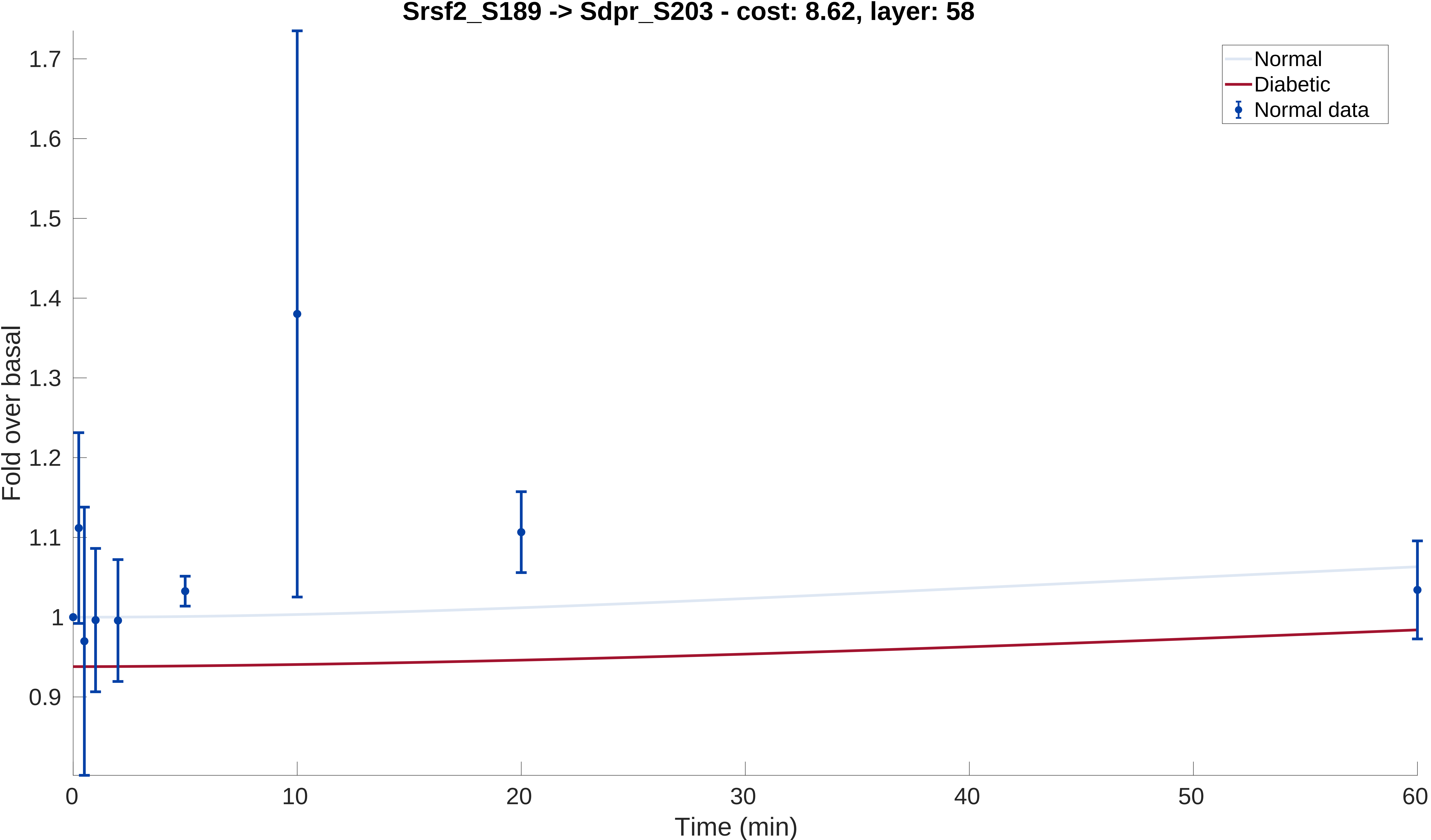

Srsf2_S189 -> Sdpr_S203 - cost: 8.62, layer: 58
Fold over basal
Time (min)
Normal
Diabetic
Normal data
1.7
1.6
1.5
1.4
1.3
1.2
1.1
1
0.9
0
10
20
30
40
50
60

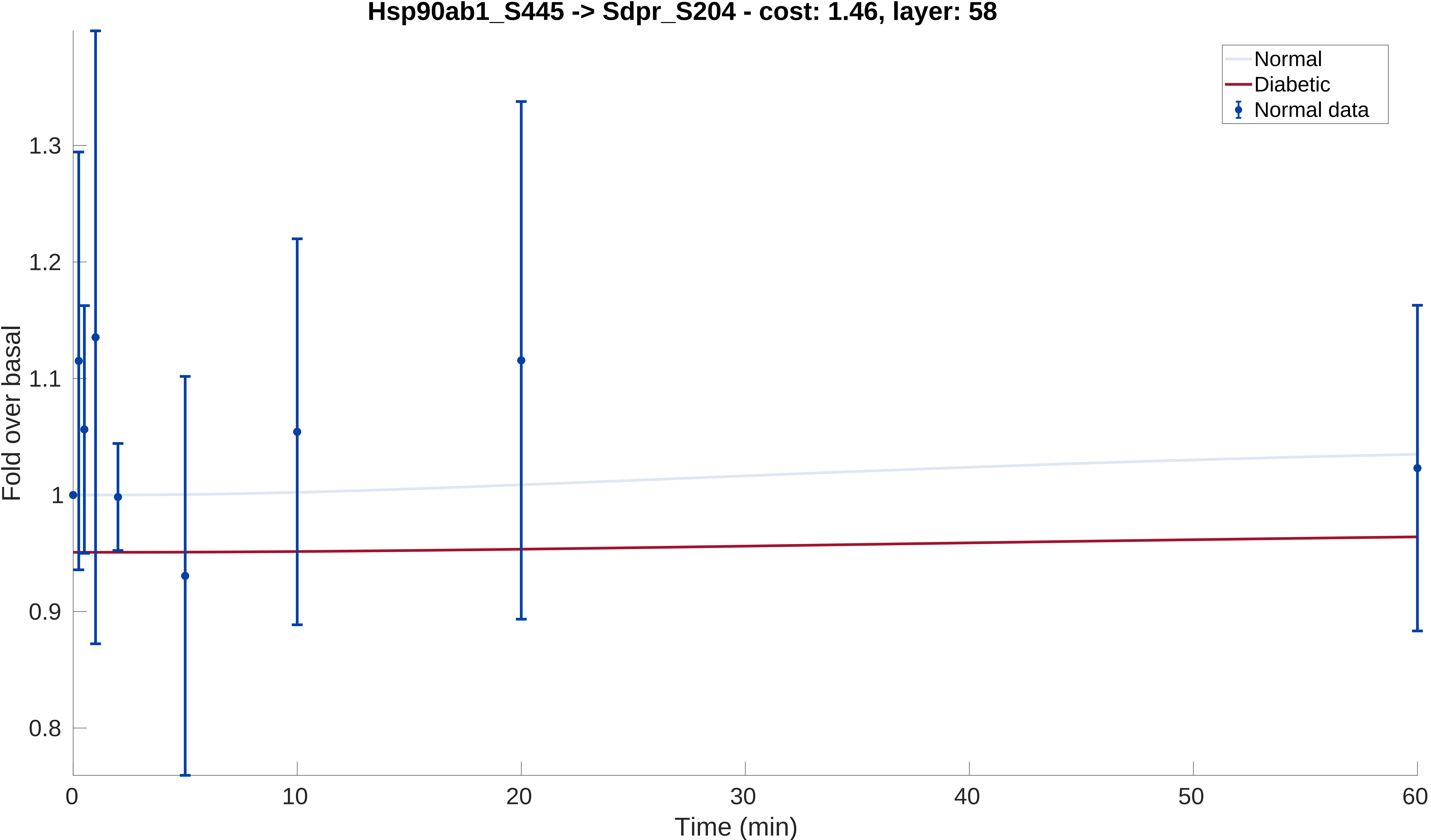

Hsp90ab1_S445 -> Sdpr_S204 - cost: 1.46, layer: 58
Normal
Diabetic
Normal data
Fold over basal
1.3
1.2
1.1
1
0.9
0.8
0
10
20
30
40
50
60
Time (min)

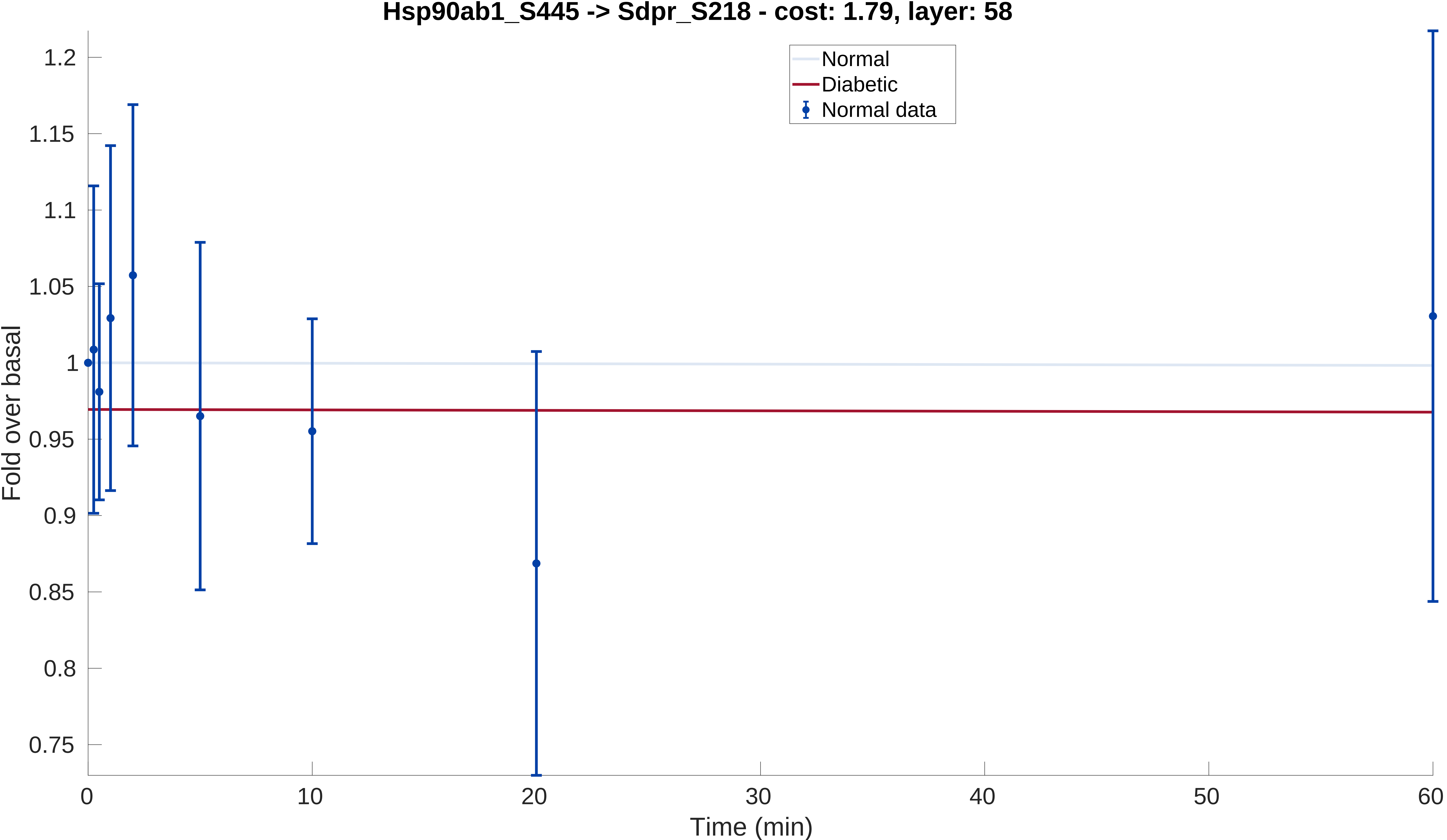
Hsp90ab1_S445 -> Sdpr_S218 - cost: 1.79, layer: 58
Normal
Diabetic
Normal data
Fold over basal
1.2
1.15
1.1
1.05
1
0.95
0.9
0.85
0.8
0.75
0
10
20
30
40
50
60
Time (min)

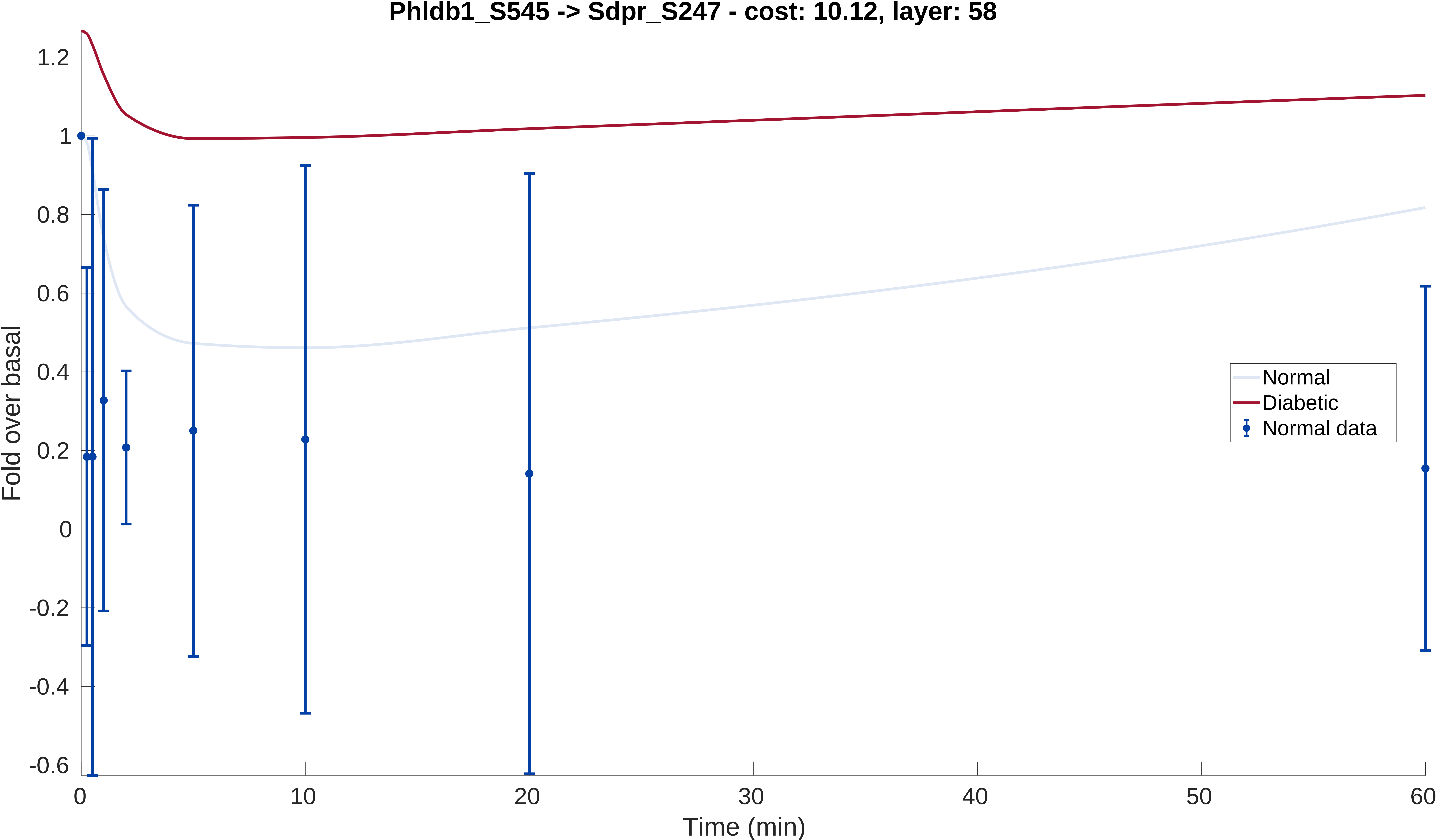
Phldb1_S545 -> Sdpr_S247 - cost: 10.12, layer: 58
Fold over basal
Time (min)
Normal
Diabetic
Normal data
1.2
1
0.8
0.6
0.4
0.2
0
-0.2
-0.4
-0.6
0
10
20
30
40
50
60

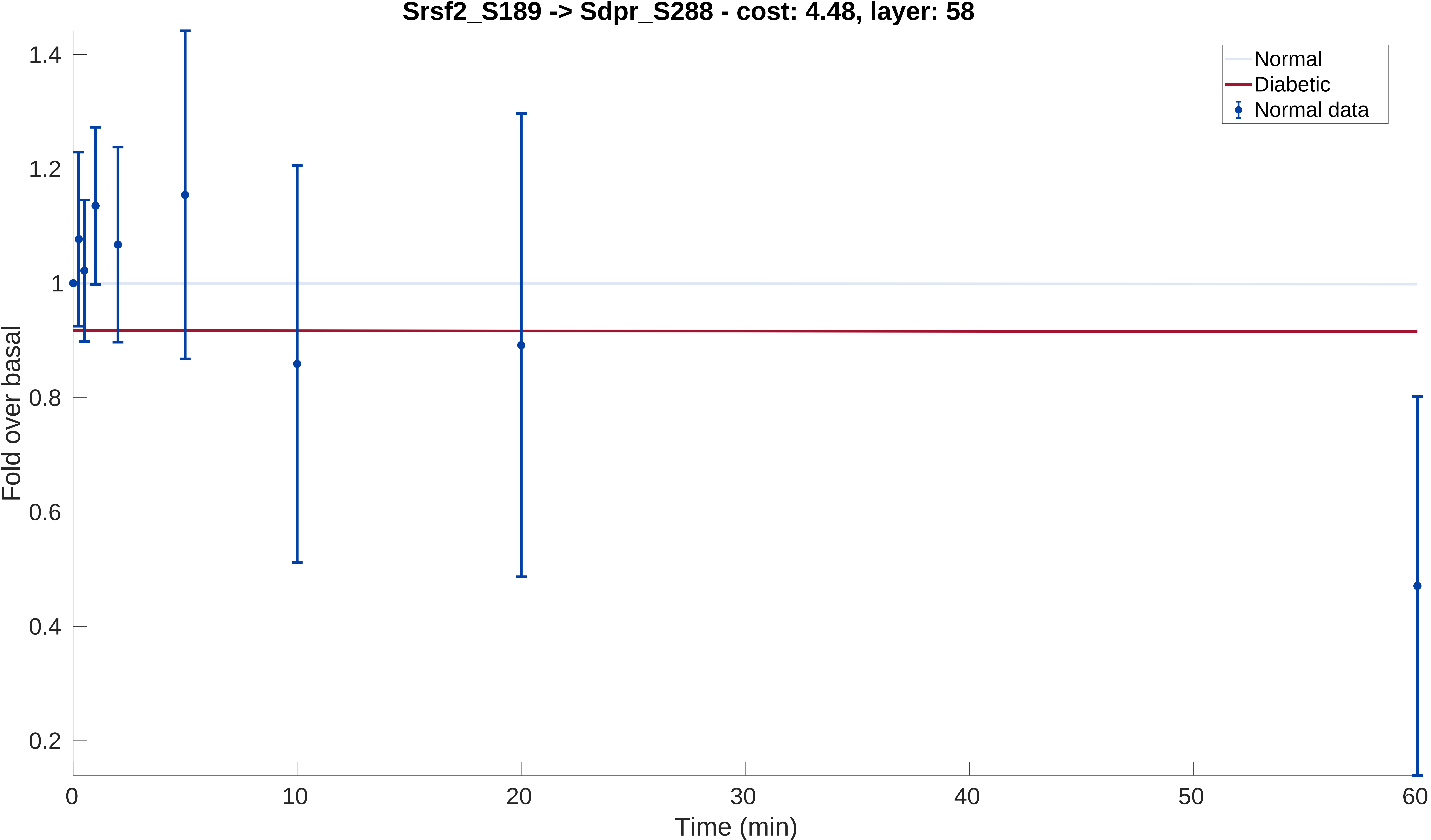

Srsf2_S189 -> Sdpr_S288 - cost: 4.48, layer: 58
Normal
Diabetic
Normal data
Fold over basal
Time (min)
1.4
1.2
1
0.8
0.6
0.4
0.2
0
10
20
30
40
50
60

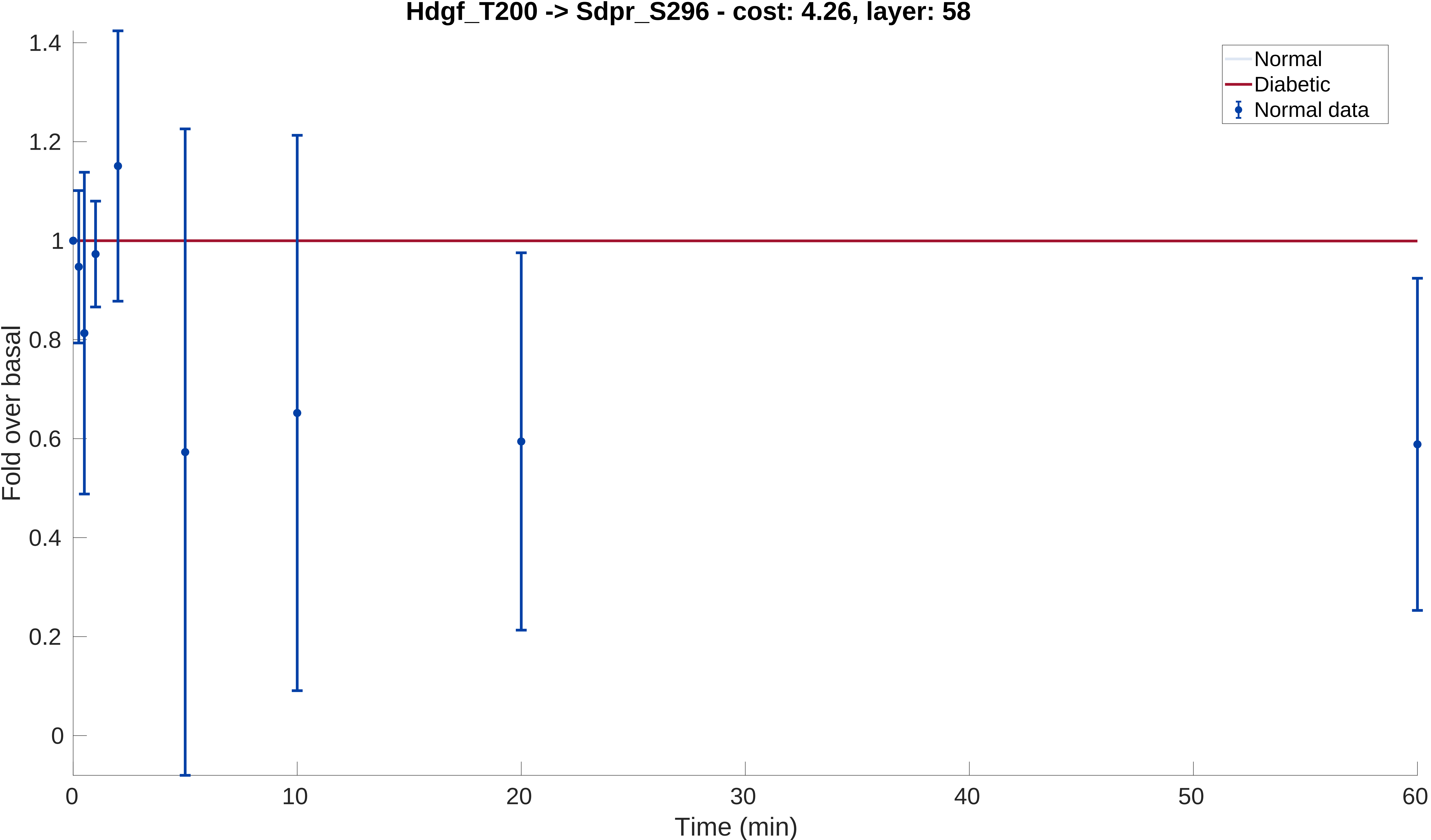

Hdgf_T200 -> Sdpr_S296 - cost: 4.26, layer: 58
Normal
Diabetic
Normal data
Fold over basal
1.4
1.2
1
0.8
0.6
0.4
0.2
0
0
10
20
30
40
50
60
Time (min)

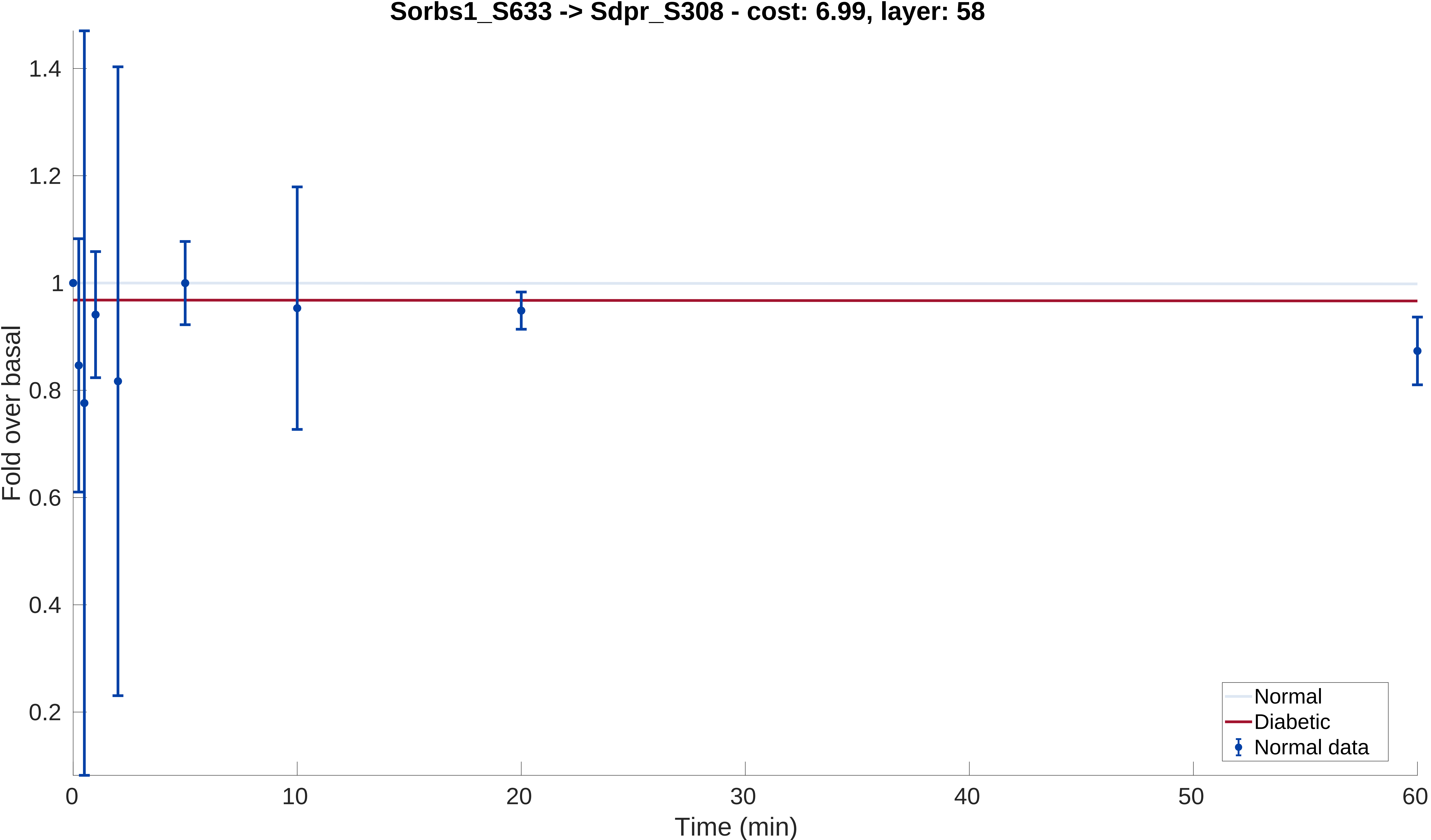
Sorbs1_S633 -> Sdpr_S308 - cost: 6.99, layer: 58
Fold over basal
1.4
1.2
1
0.8
0.6
0.4
0.2
0
10
20
30
40
50
60
Time (min)
Normal
Diabetic
Normal data

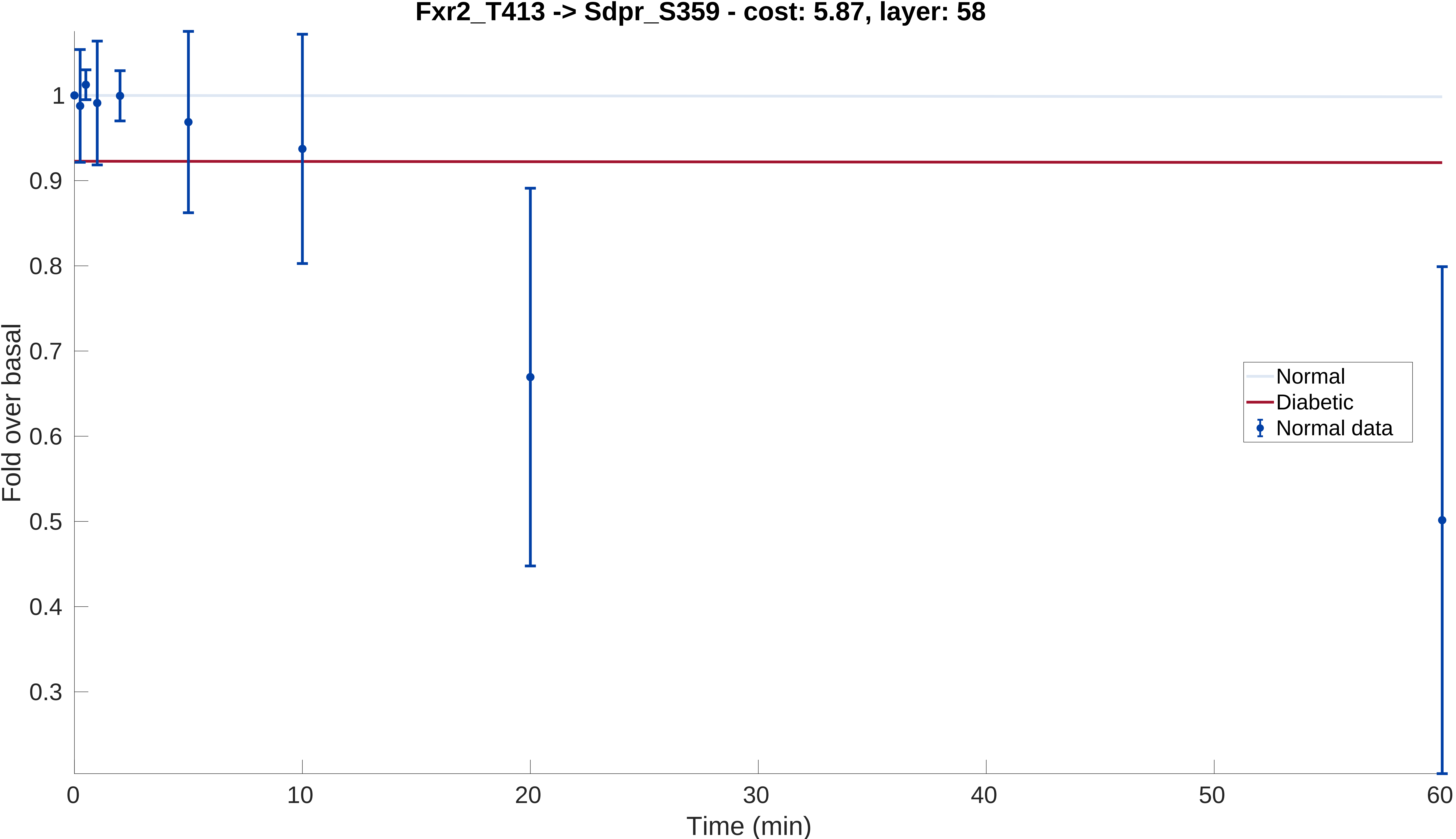

Fxr2_T413 -> Sdpr_S359 - cost: 5.87, layer: 58
Fold over basal
1
0.9
0.8
0.7
0.6
0.5
0.4
0.3
0
10
20
30
40
50
60
Time (min)
Normal
Diabetic
Normal data

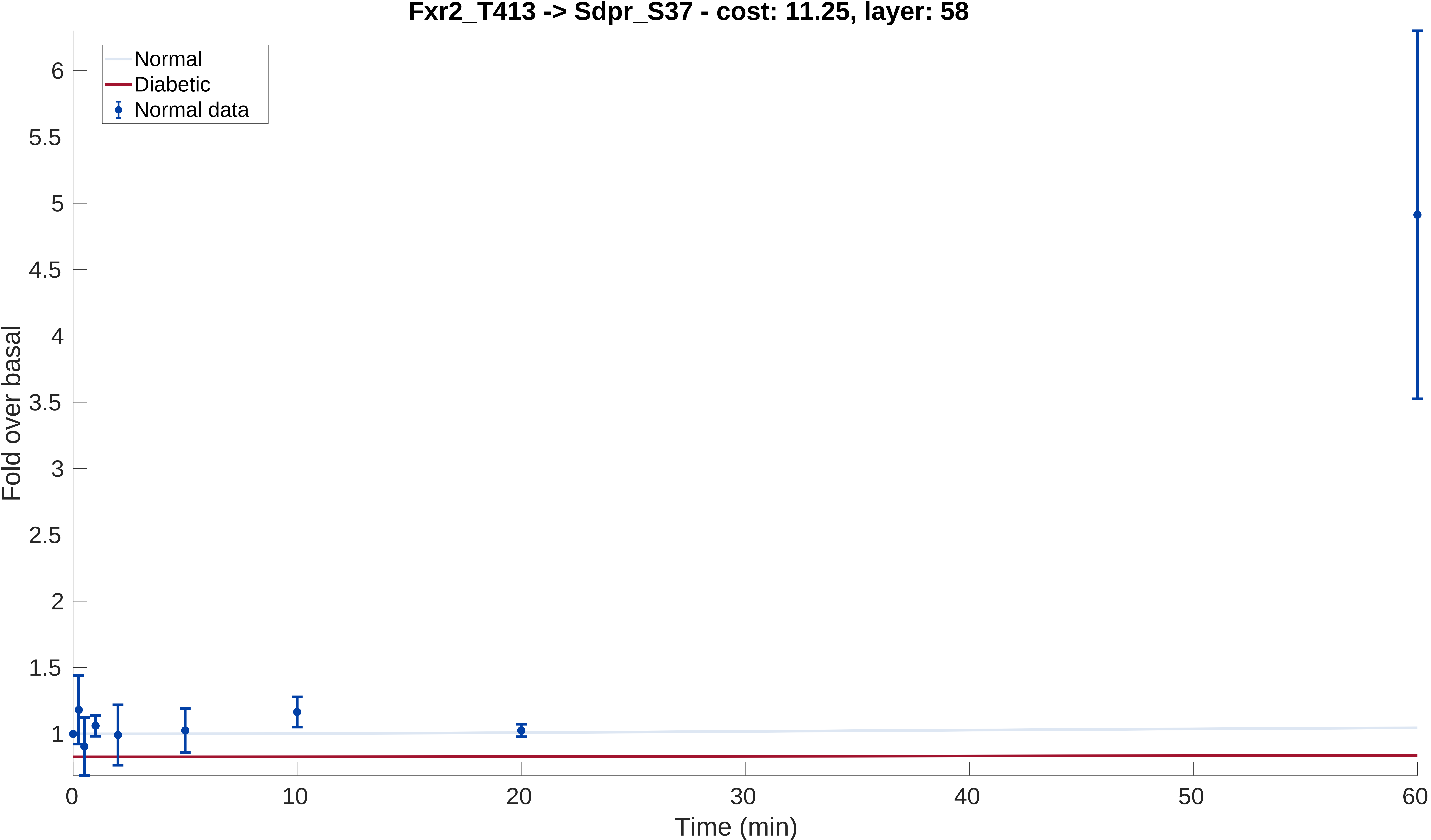

Fxr2_T413 -> Sdpr_S37 - cost: 11.25, layer: 58
Normal
Diabetic
Normal data
Fold over basal
Time (min)
0
10
20
30
40
50
60
1
1.5
2
2.5
3
3.5
4
4.5
5
5.5
6

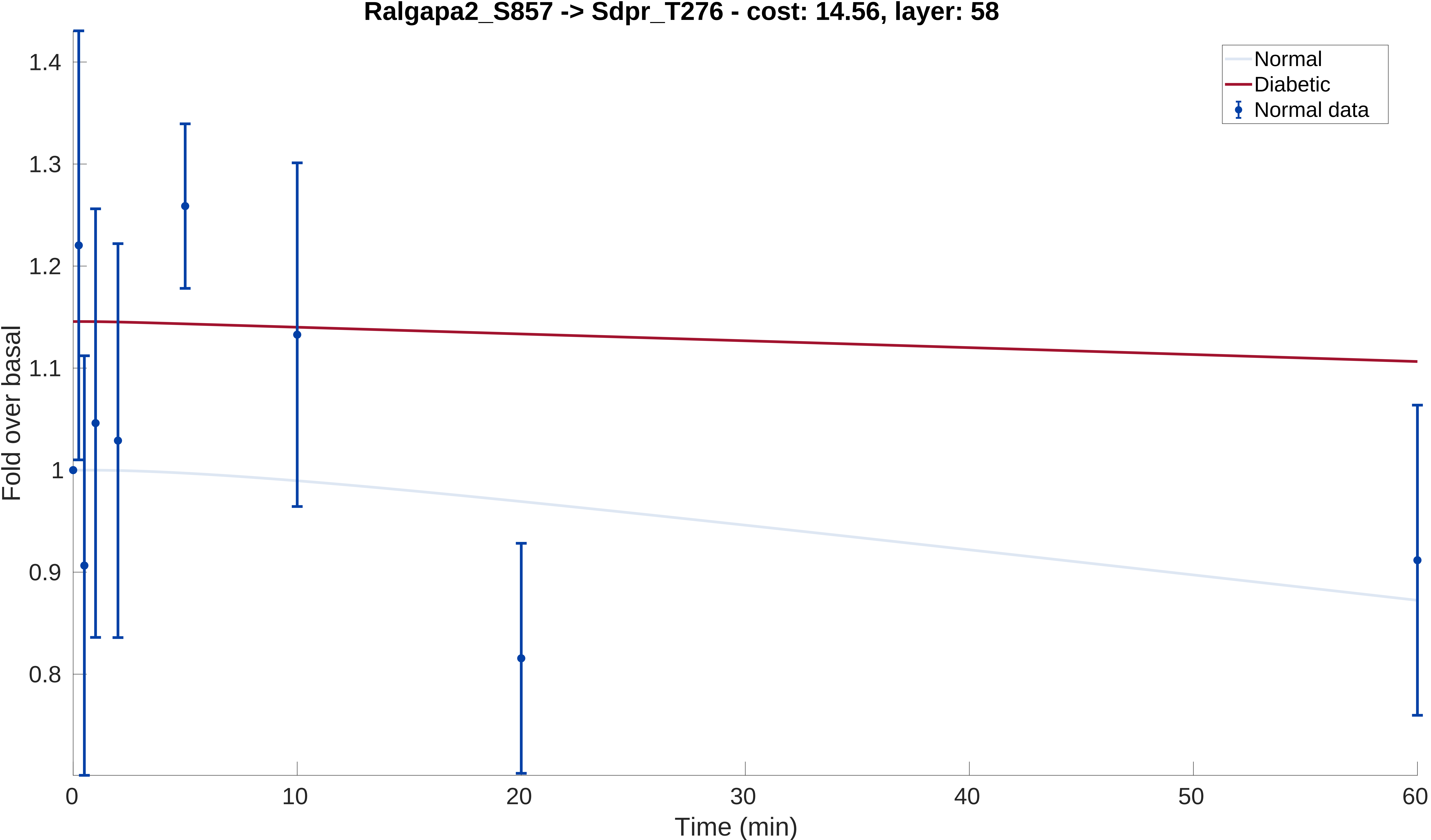

Ralgapa2_S857 -> Sdpr_T276 - cost: 14.56, layer: 58
Normal
Diabetic
Normal data
Fold over basal
1.4
1.3
1.2
1.1
1
0.9
0.8
0
10
20
30
40
50
60
Time (min)

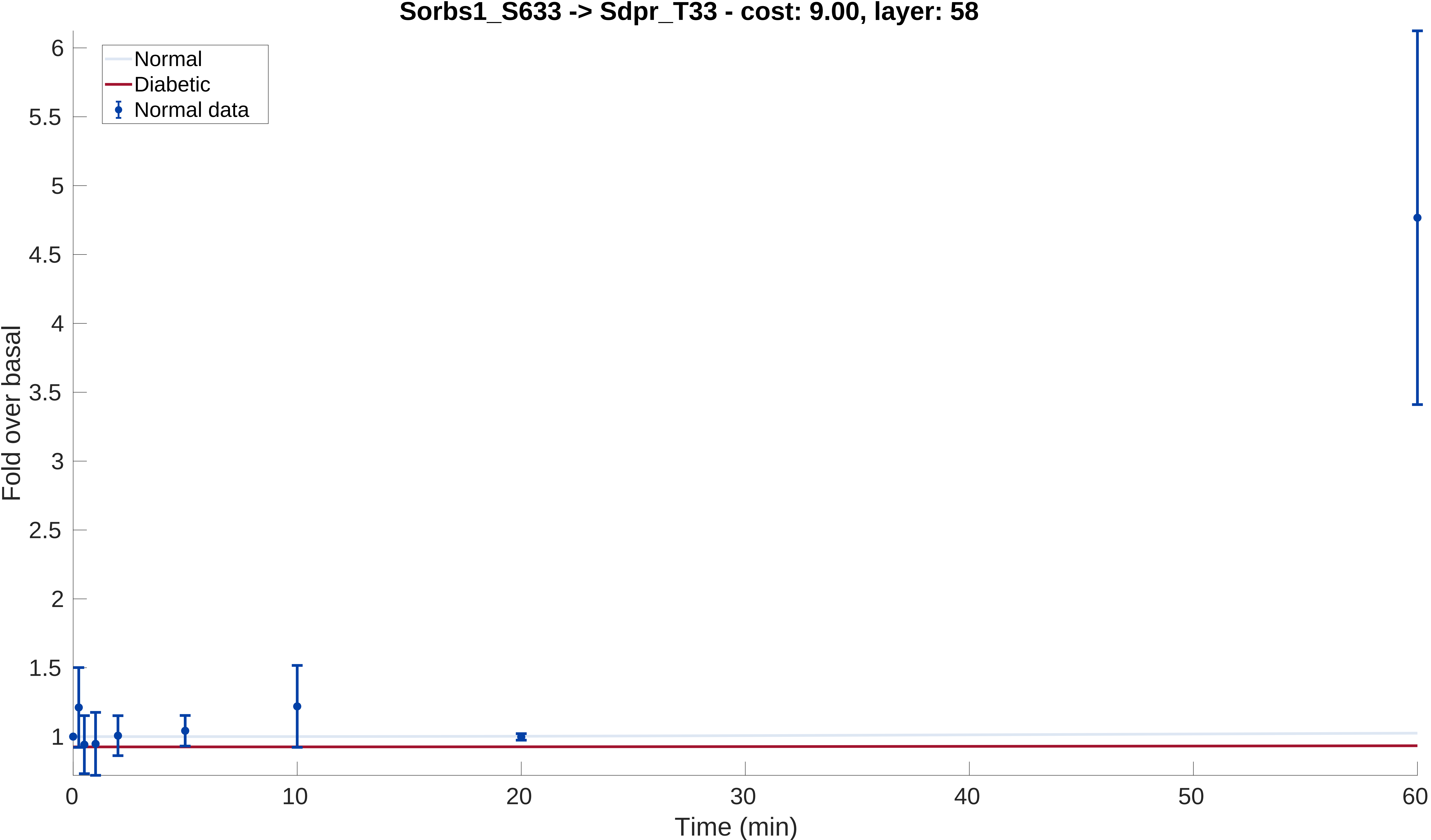
Sorbs1_S633 -> Sdpr_T33 - cost: 9.00, layer: 58
Normal
Diabetic
Normal data
Fold over basal
Time (min)
6
5.5
5
4.5
4
3.5
3
2.5
2
1.5
1
0
10
20
30
40
50
60

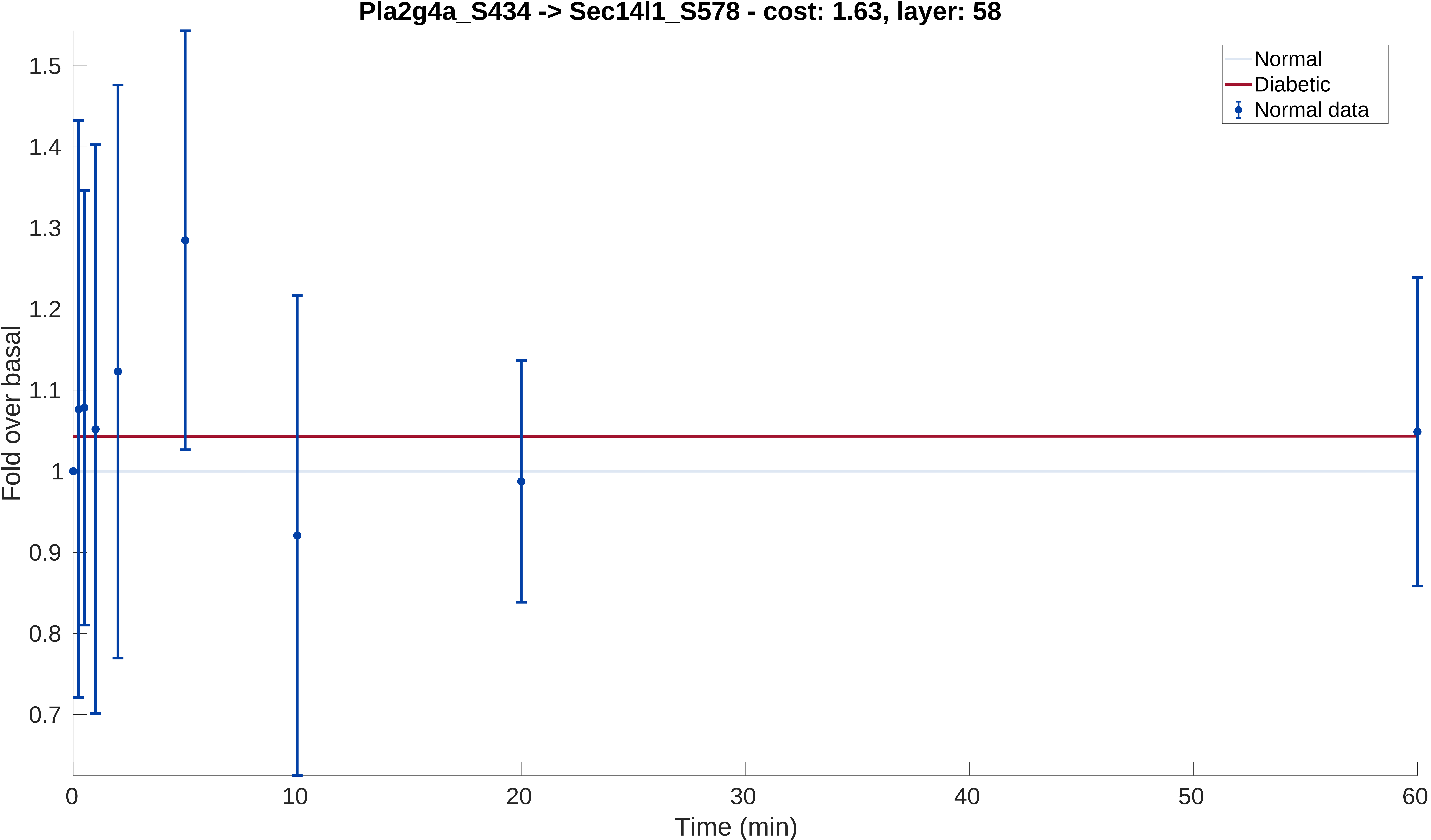

Pla2g4a_S434 -> Sec14l1_S578 - cost: 1.63, layer: 58
Normal
Diabetic
Normal data
Fold over basal
1.5
1.4
1.3
1.2
1.1
1
0.9
0.8
0.7
0
10
20
30
40
50
60
Time (min)

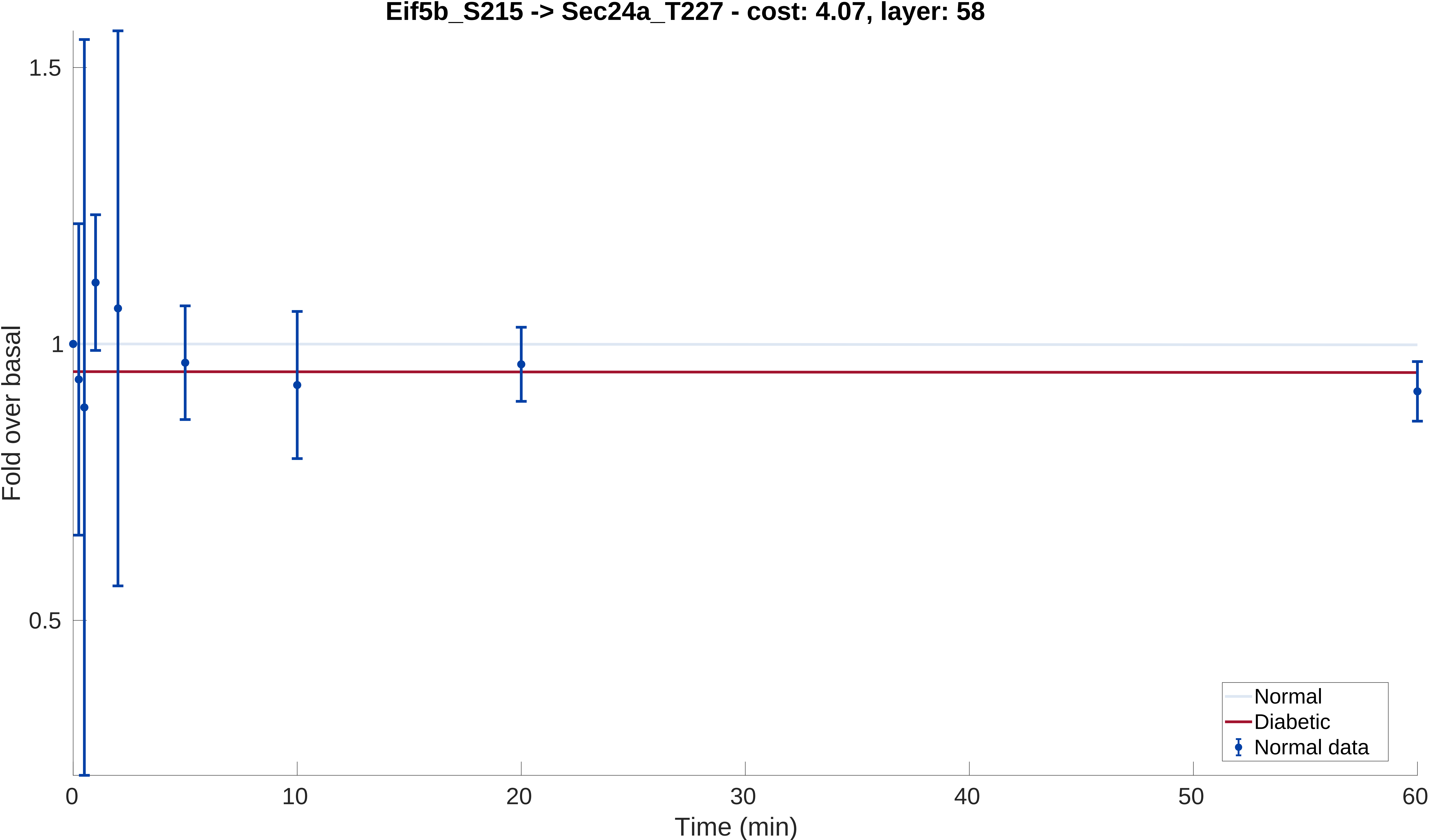

Eif5b_S215 -> Sec24a_T227 - cost: 4.07, layer: 58
Fold over basal
1.5
1
0.5
0
10
20
30
40
50
60
Time (min)
Normal
Diabetic
Normal data

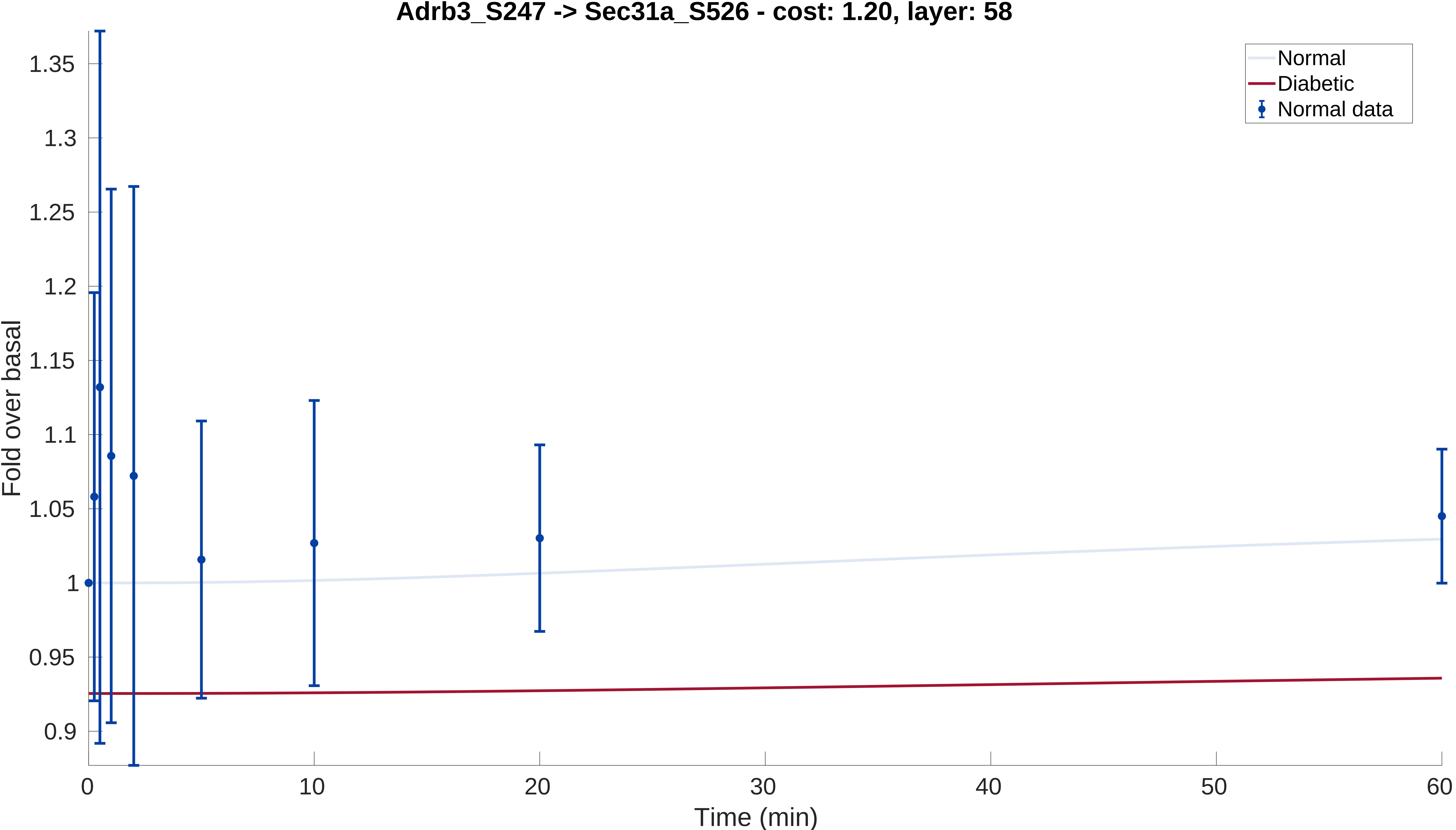

Adrb3_S247 -> Sec31a_S526 - cost: 1.20, layer: 58
Normal
Diabetic
Normal data
Fold over basal
1.35
1.3
1.25
1.2
1.15
1.1
1.05
1
0.95
0.9
0
10
20
30
40
50
60
Time (min)

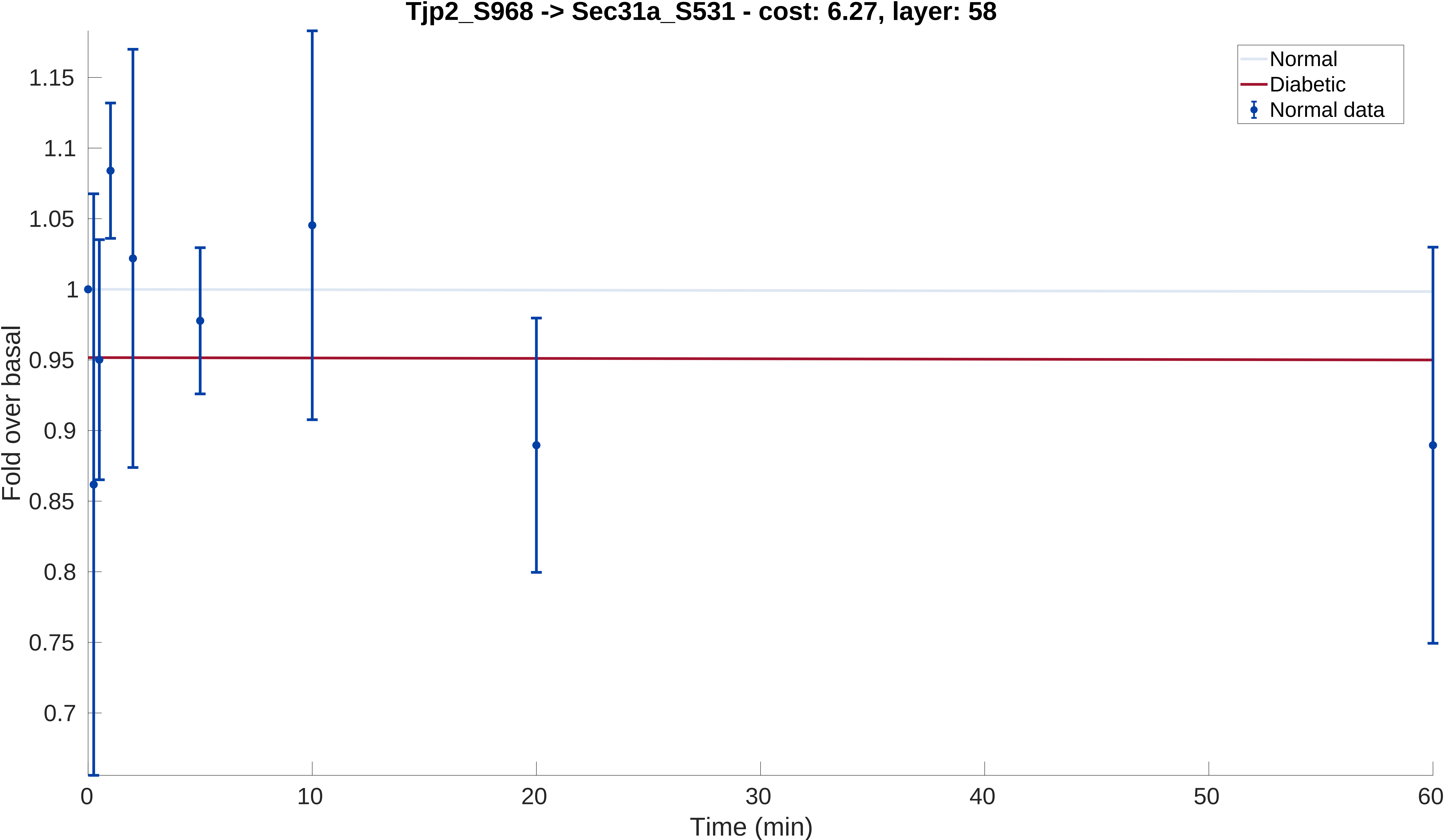

Tjp2_S968 -> Sec31a_S531 - cost: 6.27, layer: 58
Normal
Diabetic
Normal data
Fold over basal
Time (min)
1.15
1.1
1.05
1
0.95
0.9
0.85
0.8
0.75
0.7
0
10
20
30
40
50
60

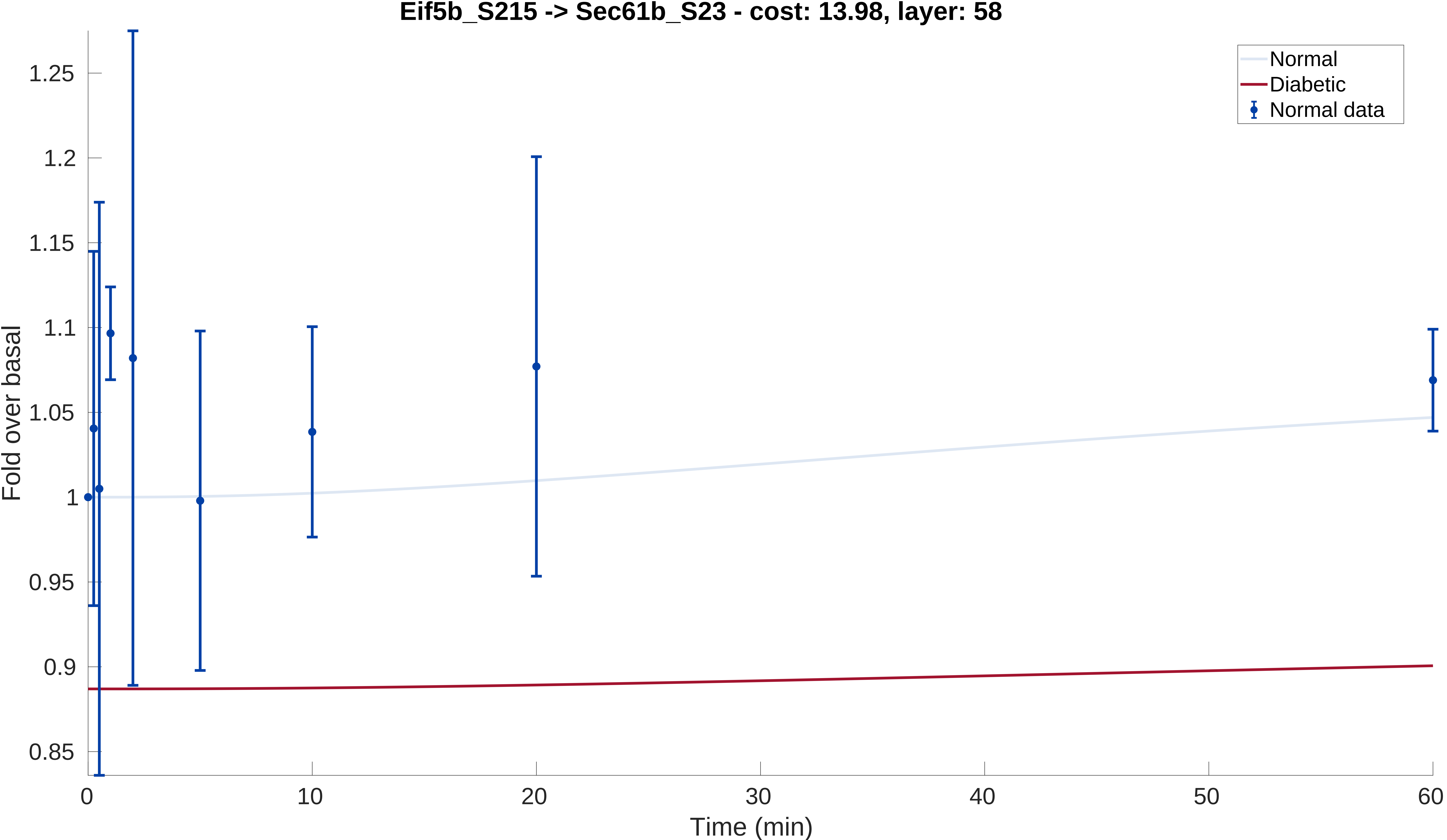

Eif5b_S215 -> Sec61b_S23 - cost: 13.98, layer: 58
Normal
Diabetic
Normal data
Fold over basal
Time (min)
1.25
1.2
1.15
1.1
1.05
1
0.95
0.9
0.85
0
10
20
30
40
50
60

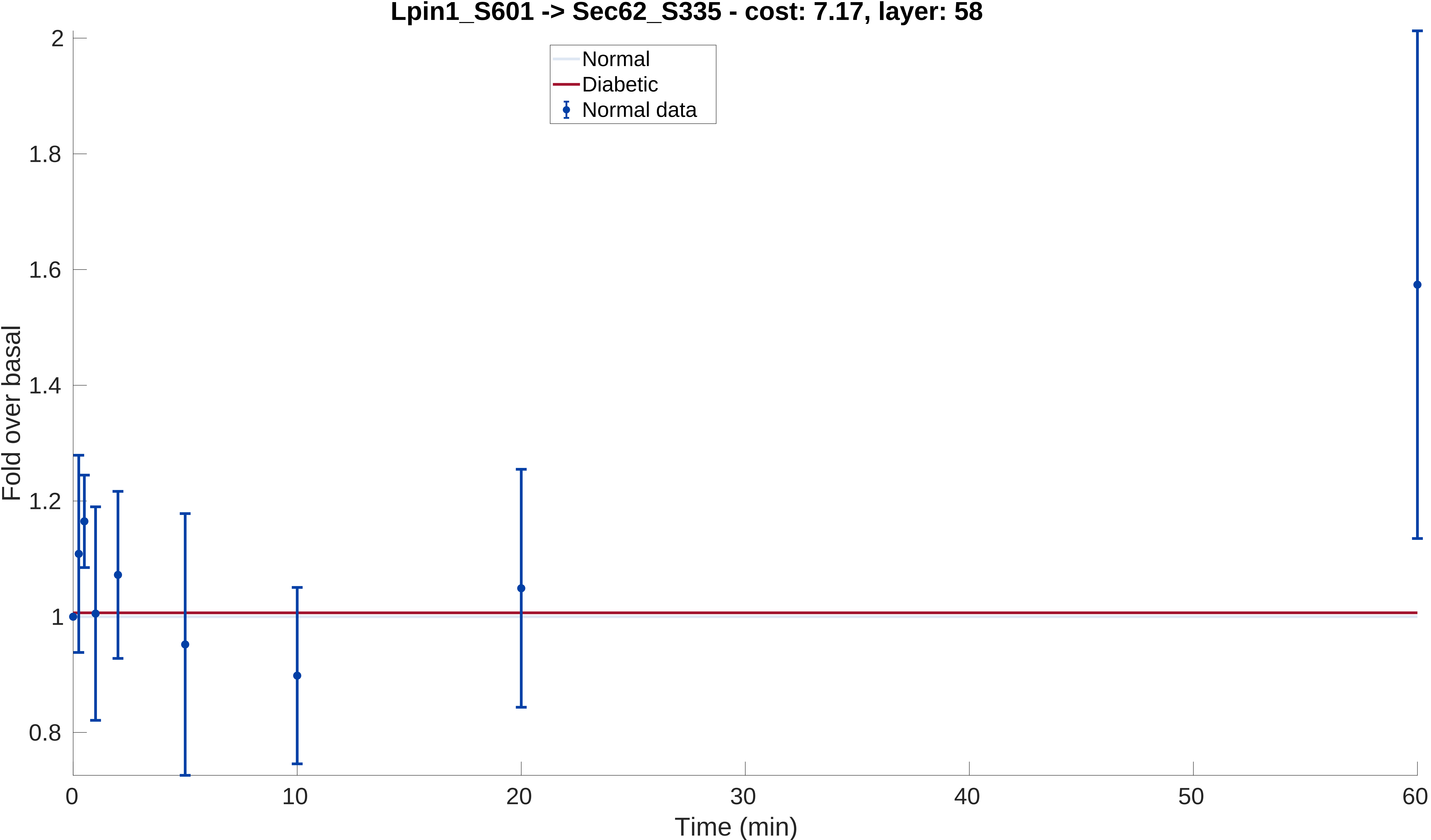
Lpin1_S601 -> Sec62_S335 - cost: 7.17, layer: 58
Normal
Diabetic
Normal data
Fold over basal
2
1.8
1.6
1.4
1.2
1
0.8
0
10
20
30
40
50
60
Time (min)

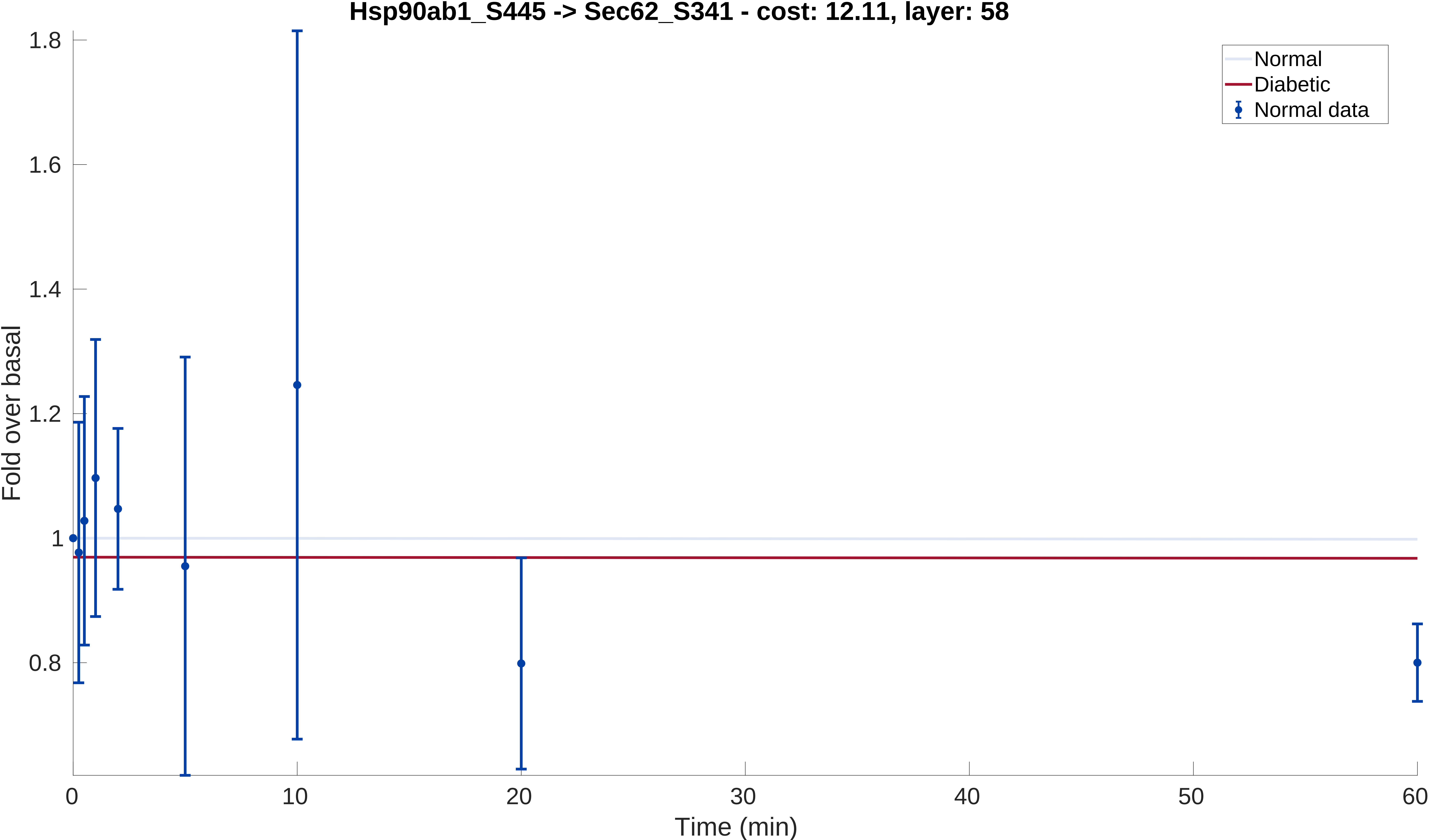

Hsp90ab1_S445 -> Sec62_S341 - cost: 12.11, layer: 58
Normal
Diabetic
Normal data
Fold over basal
1.8
1.6
1.4
1.2
1
0.8
0
10
20
30
40
50
60
Time (min)

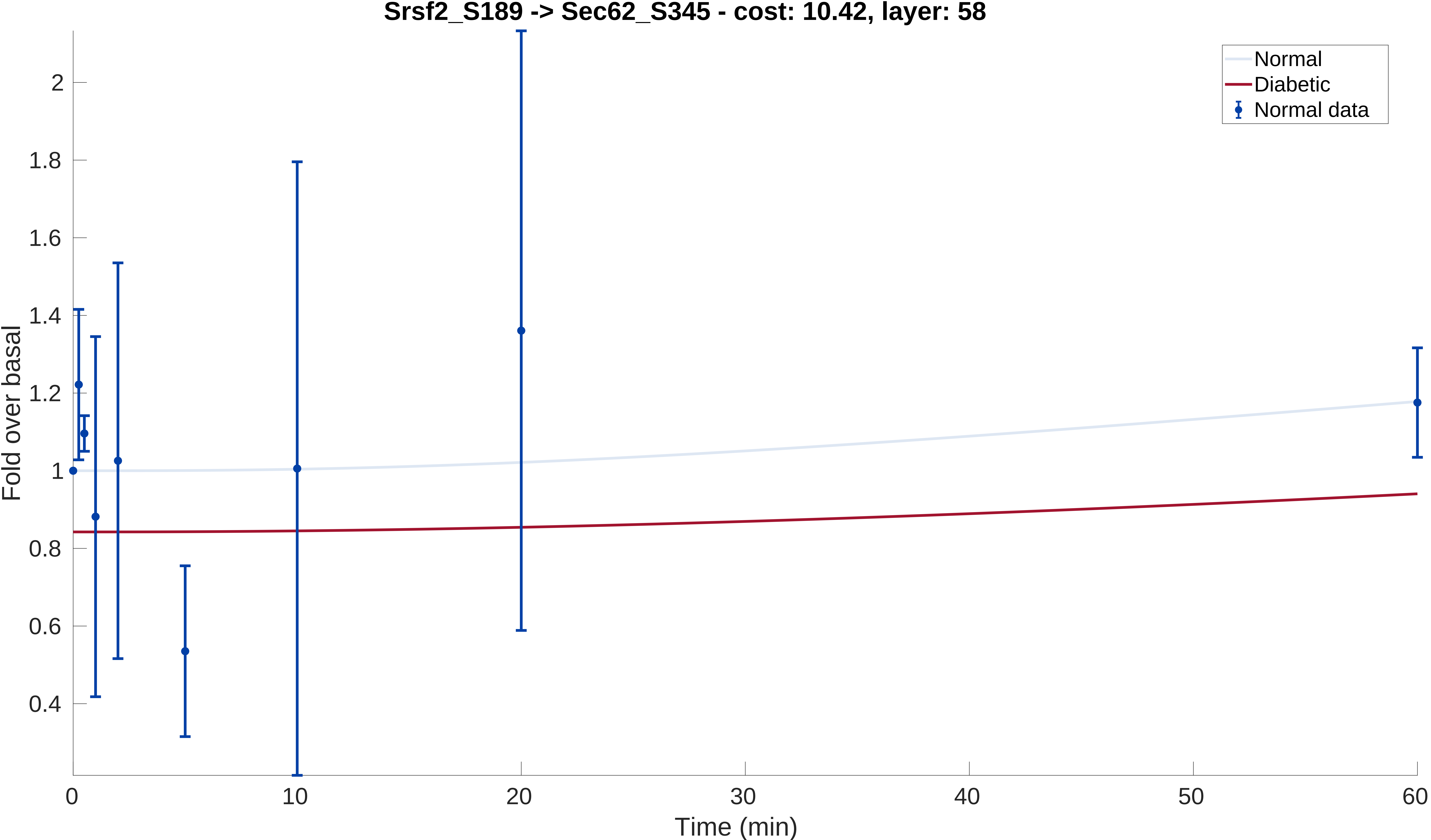
Srsf2_S189 -> Sec62_S345 - cost: 10.42, layer: 58
Normal
Diabetic
Normal data
Fold over basal
2
1.8
1.6
1.4
1.2
1
0.8
0.6
0.4
0
10
20
30
40
50
60
Time (min)

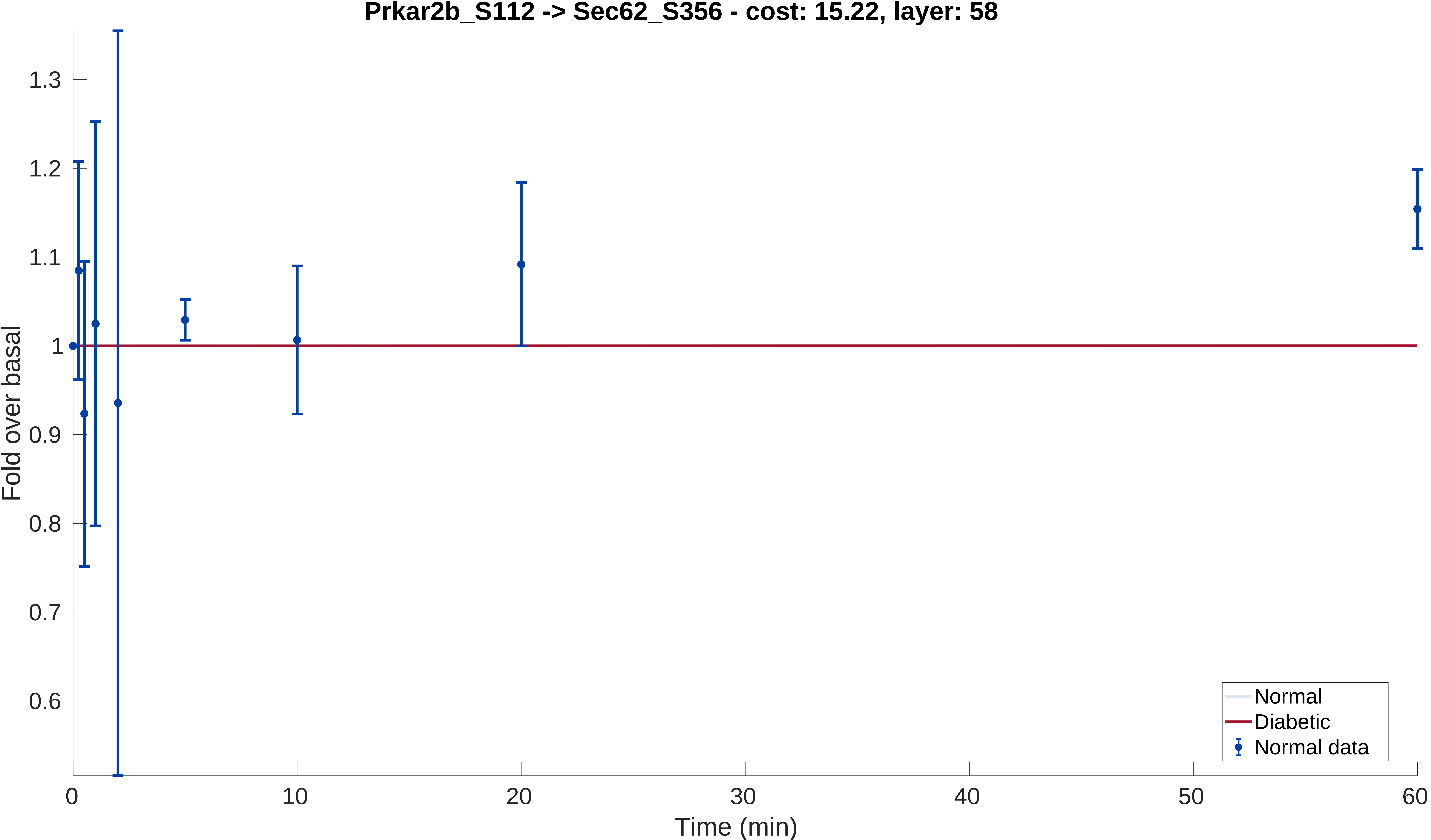
Prkar2b_S112 -> Sec62_S356 - cost: 15.22, layer: 58
Fold over basal
1.3
1.2
1.1
1
0.9
0.8
0.7
0.6
0
10
20
30
40
50
60
Time (min)
Normal
Diabetic
Normal data

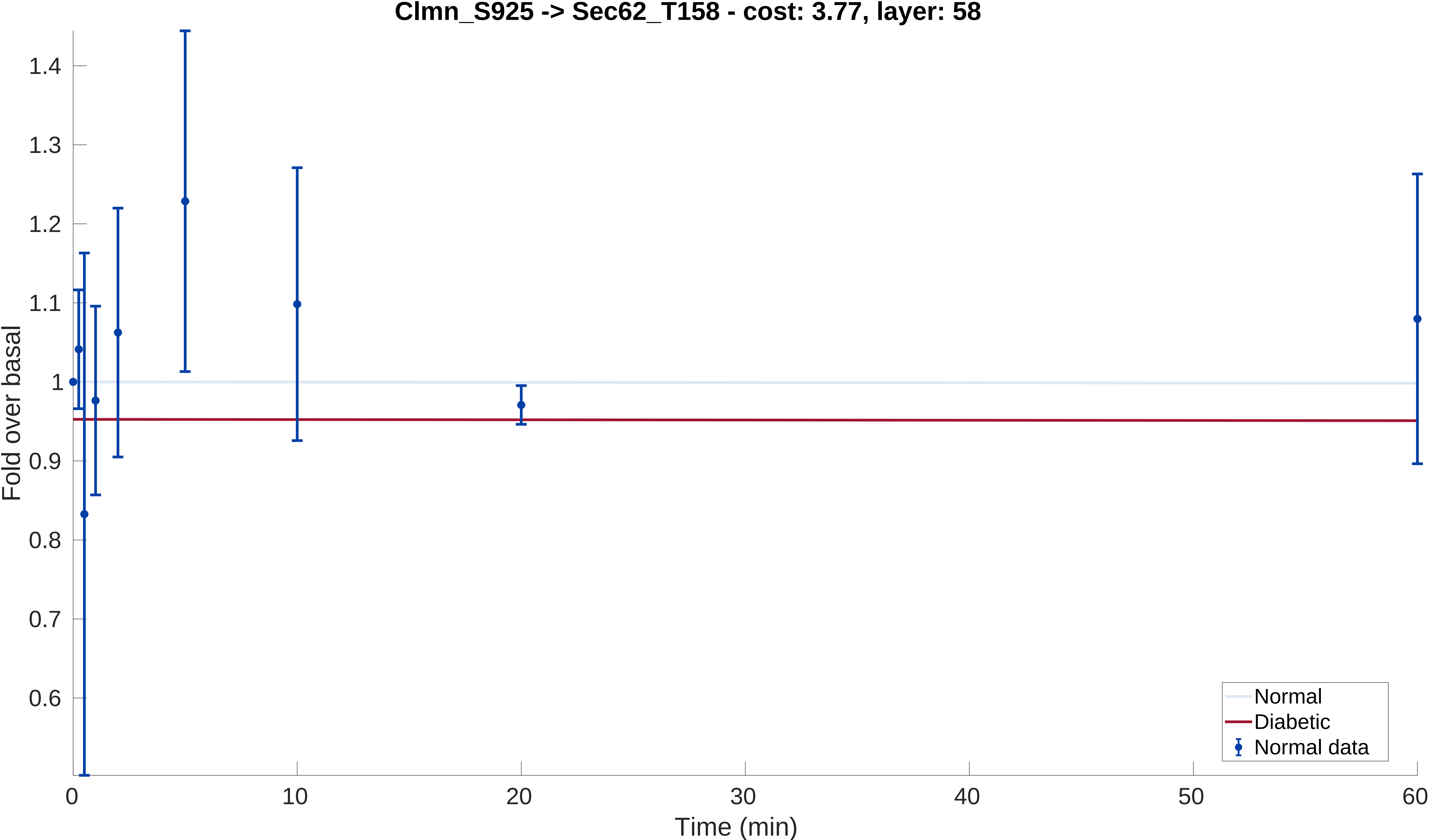
Clmn_S925 -> Sec62_T158 - cost: 3.77, layer: 58
Fold over basal
1.4
1.3
1.2
1.1
1
0.9
0.8
0.7
0.6
0
10
20
30
40
50
60
Time (min)
Normal
Diabetic
Normal data

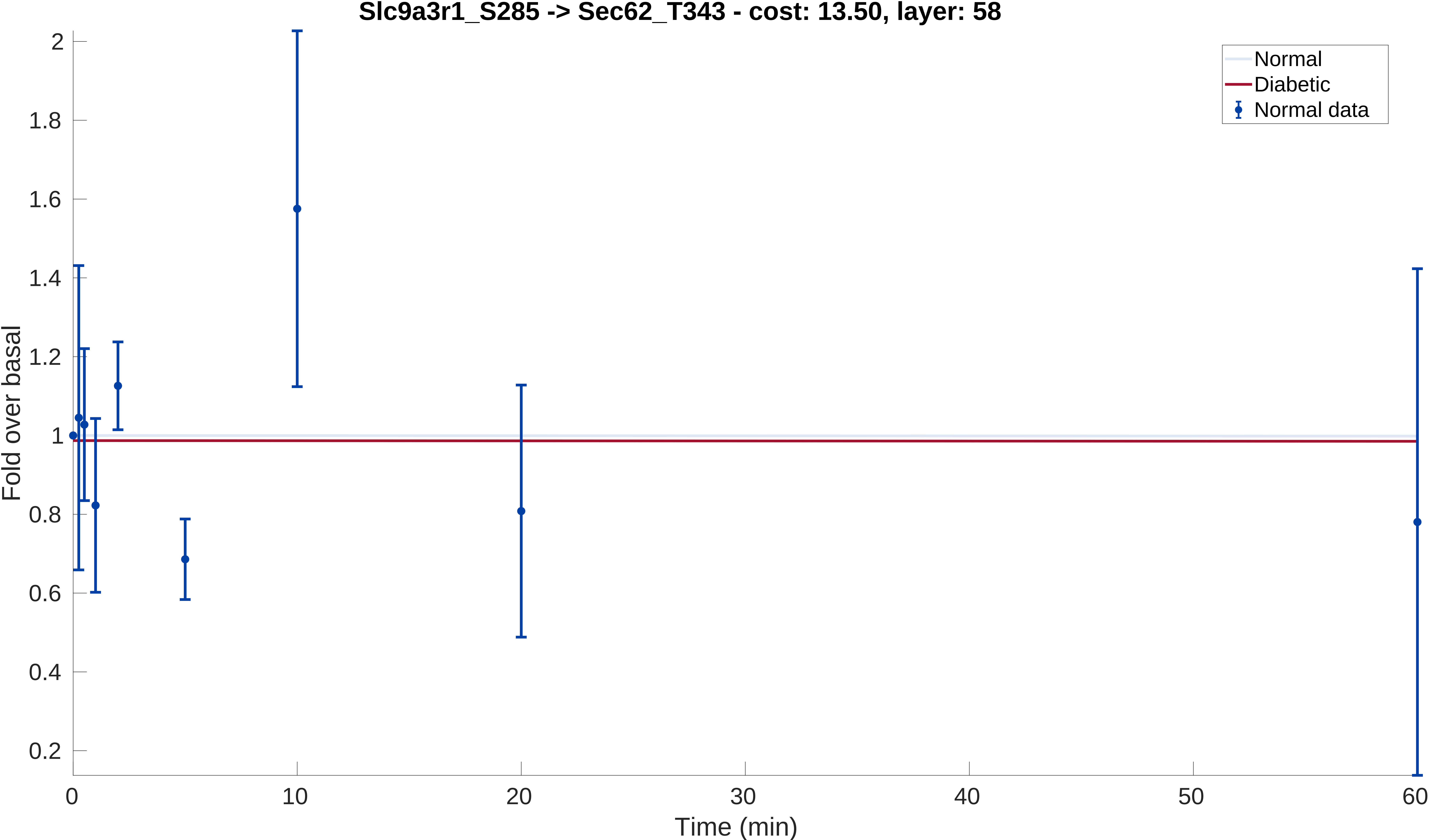

Slc9a3r1_S285 -> Sec62_T343 - cost: 13.50, layer: 58
Normal
Diabetic
Normal data
Fold over basal
Time (min)
0
10
20
30
40
50
60
0.2
0.4
0.6
0.8
1
1.2
1.4
1.6
1.8
2

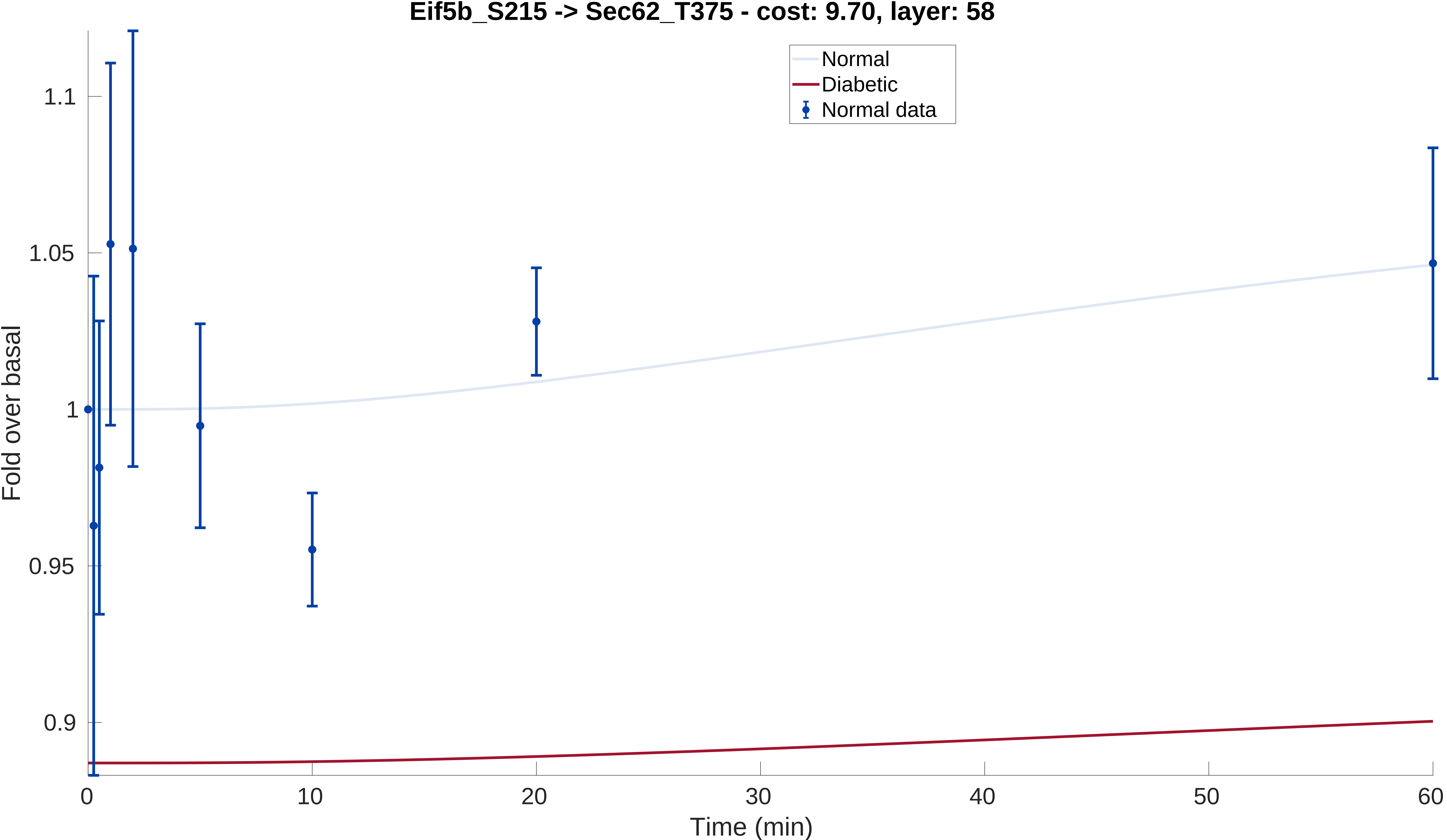

Eif5b_S215 -> Sec62_T375 - cost: 9.70, layer: 58
Normal
Diabetic
Normal data
Fold over basal
1.1
1.05
1
0.95
0.9
0
10
20
30
40
50
60
Time (min)

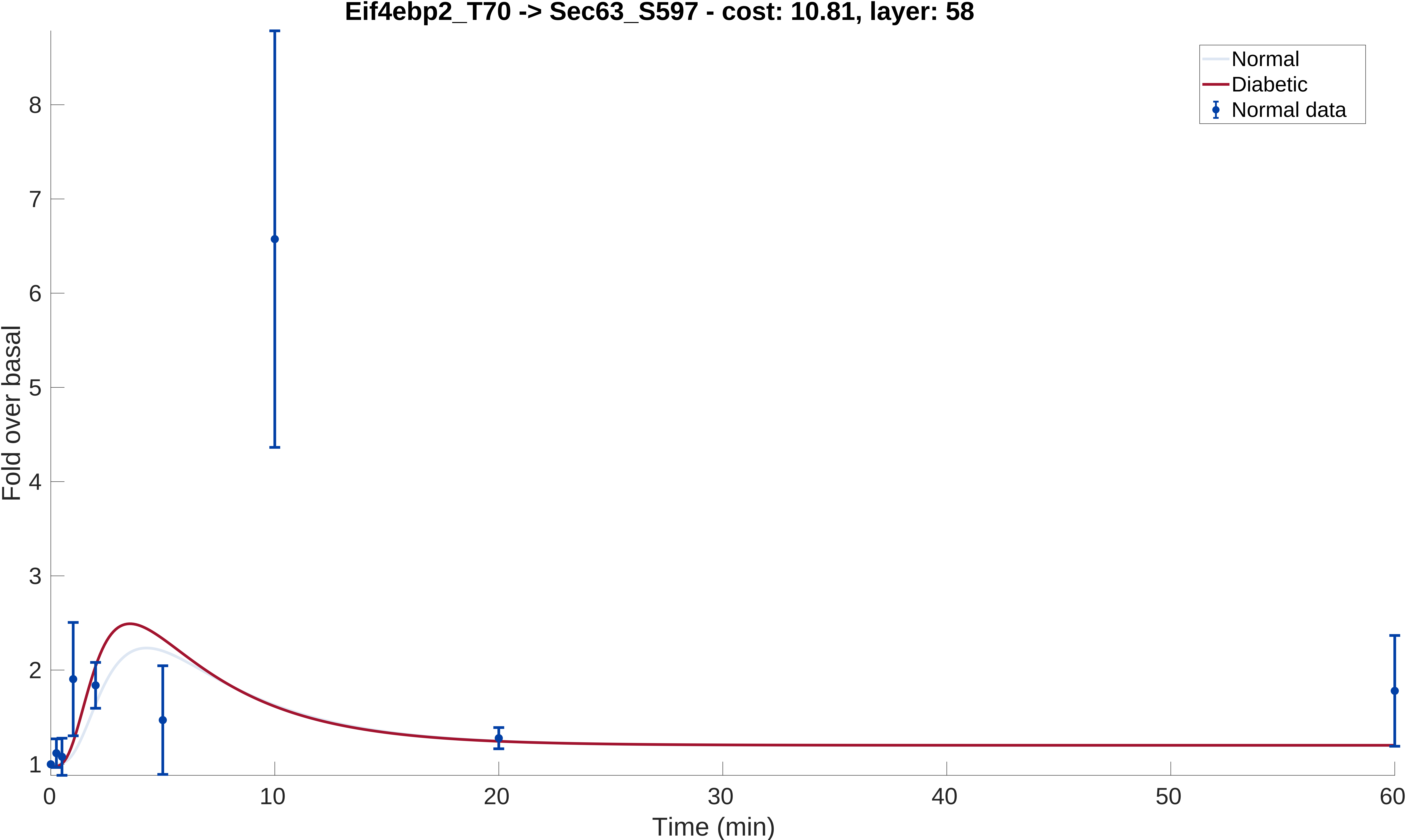

Eif4ebp2_T70 -> Sec63_S597 - cost: 10.81, layer: 58
Normal
Diabetic
Normal data
Fold over basal
Time (min)
1
2
3
4
5
6
7
8
0
10
20
30
40
50
60

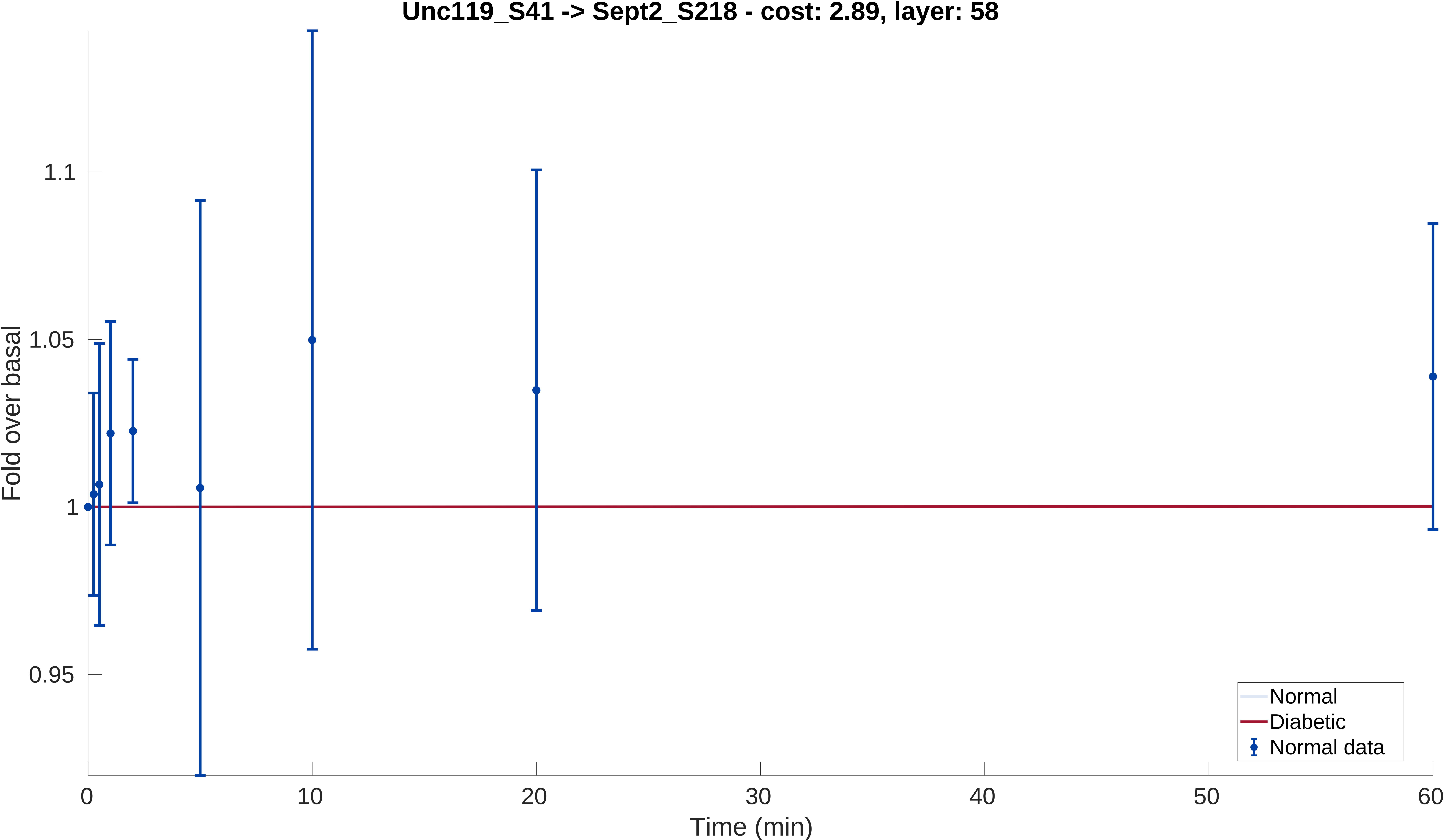

Unc119_S41 -> Sept2_S218 - cost: 2.89, layer: 58
Fold over basal
1.1
1.05
1
0.95
0
10
20
30
40
50
60
Time (min)
Normal
Diabetic
Normal data

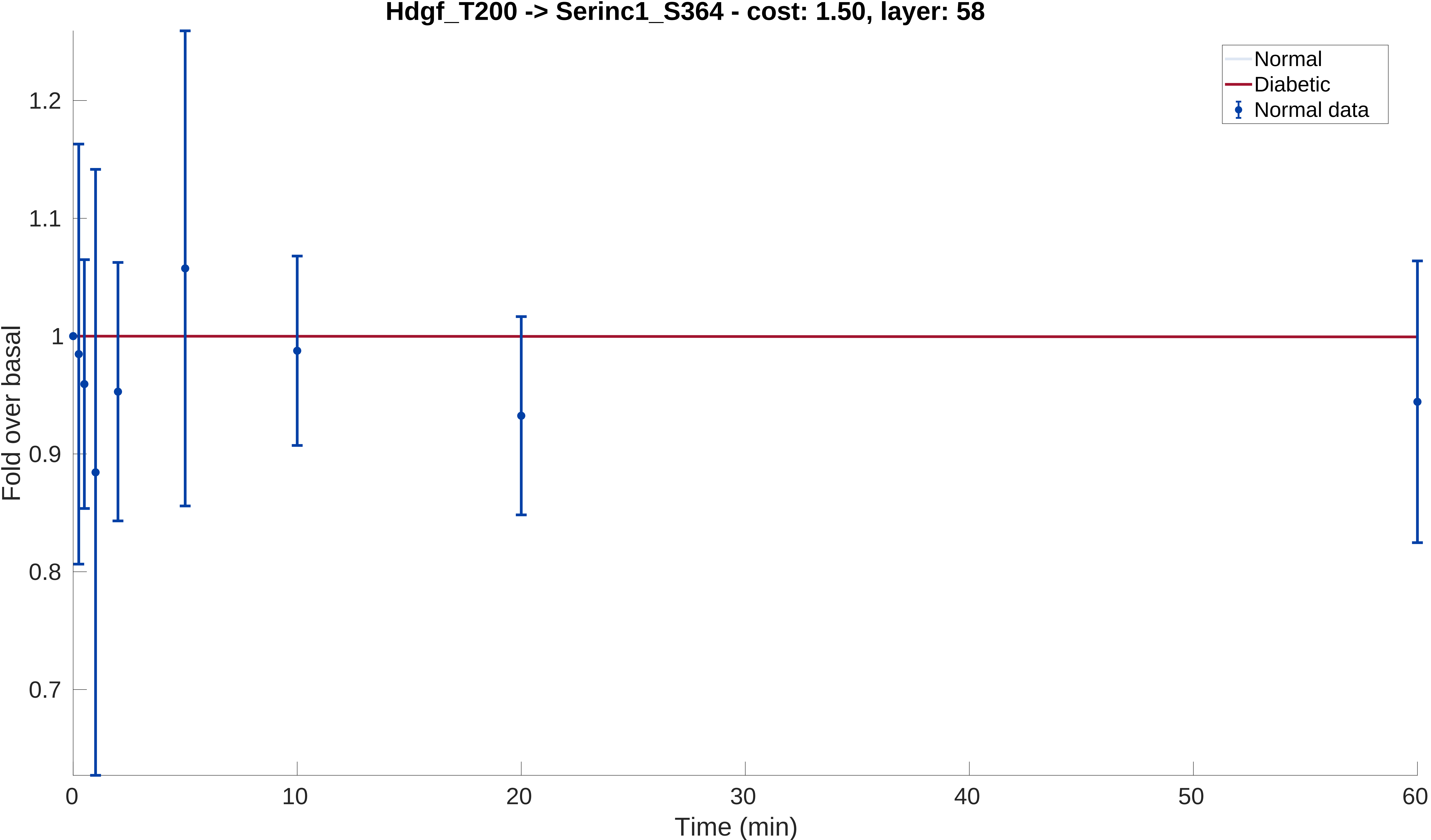

Hdgf_T200 -> Serinc1_S364 - cost: 1.50, layer: 58
Normal
Diabetic
Normal data
Fold over basal
1.2
1.1
1
0.9
0.8
0.7
0
10
20
30
40
50
60
Time (min)

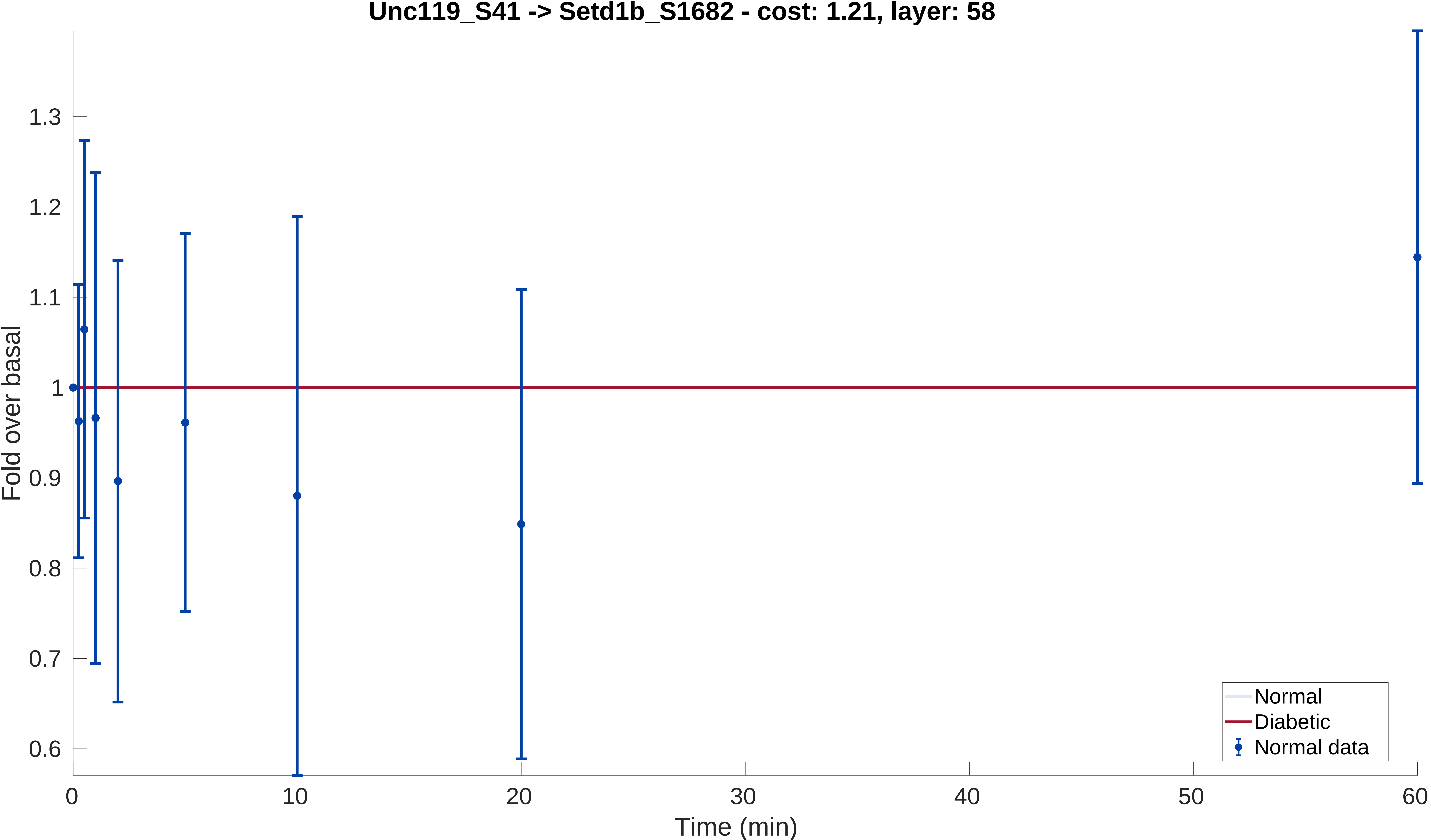

Unc119_S41 -> Setd1b_S1682 - cost: 1.21, layer: 58
Fold over basal
1.3
1.2
1.1
1
0.9
0.8
0.7
0.6
0
10
20
30
40
50
60
Time (min)
Normal
Diabetic
Normal data

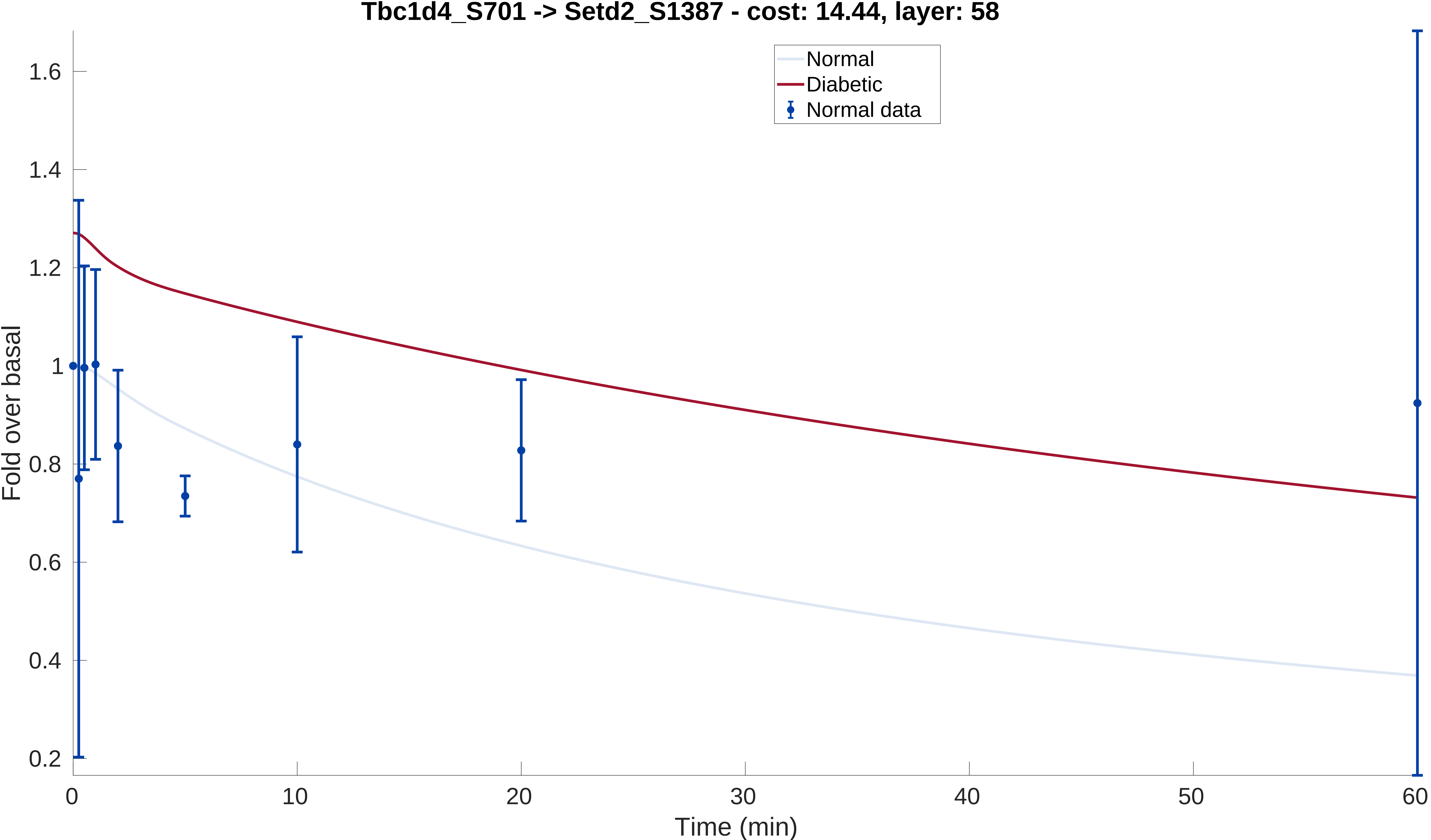

Tbc1d4_S701 -> Setd2_S1387 - cost: 14.44, layer: 58
Normal
Diabetic
Normal data
Fold over basal
Time (min)
0.2
0.4
0.6
0.8
1
1.2
1.4
1.6
0
10
20
30
40
50
60

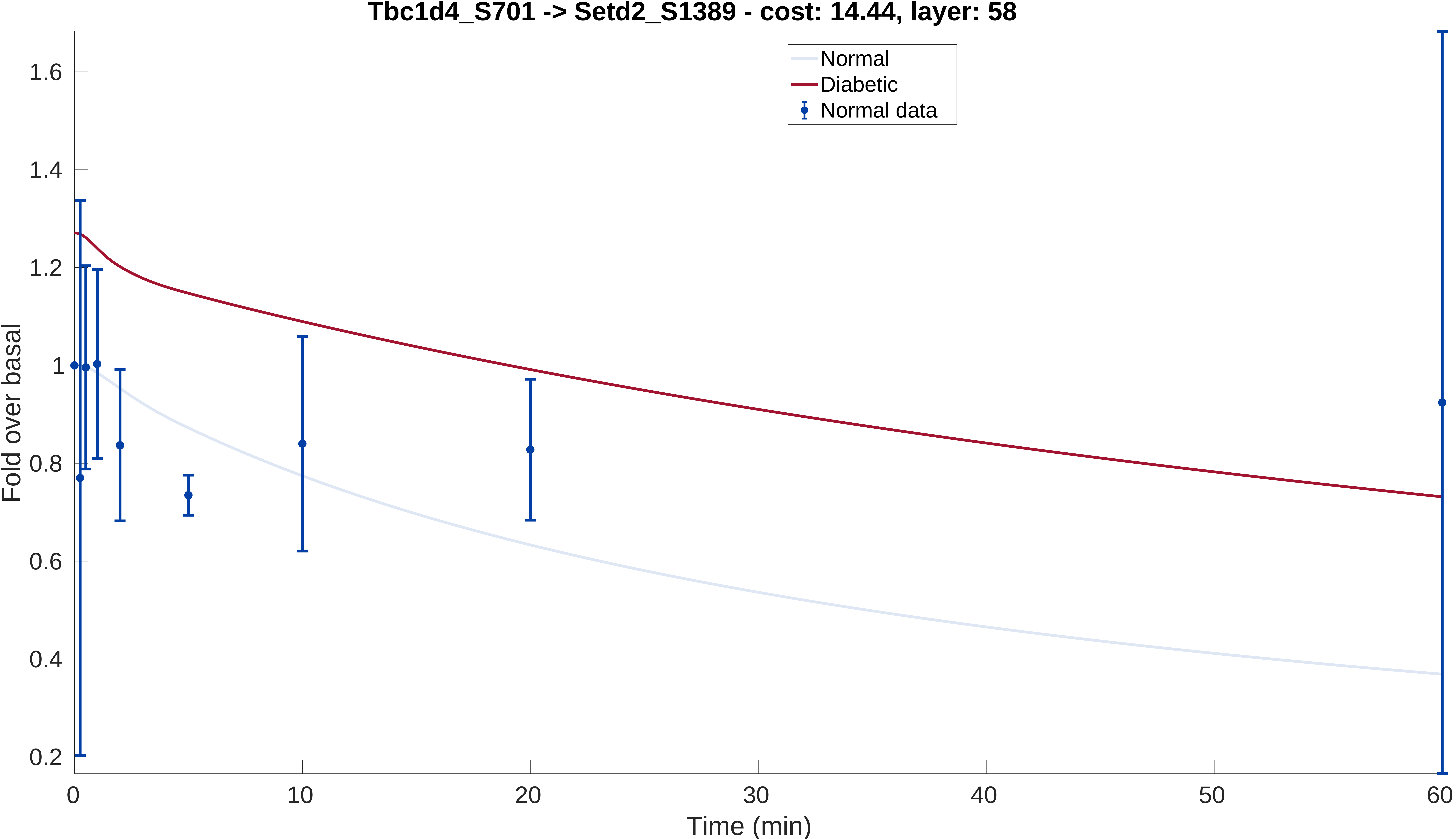
Tbc1d4_S701 -> Setd2_S1389 - cost: 14.44, layer: 58
Normal
Diabetic
Normal data
Fold over basal
1.6
1.4
1.2
1
0.8
0.6
0.4
0.2
0
10
20
30
40
50
60
Time (min)

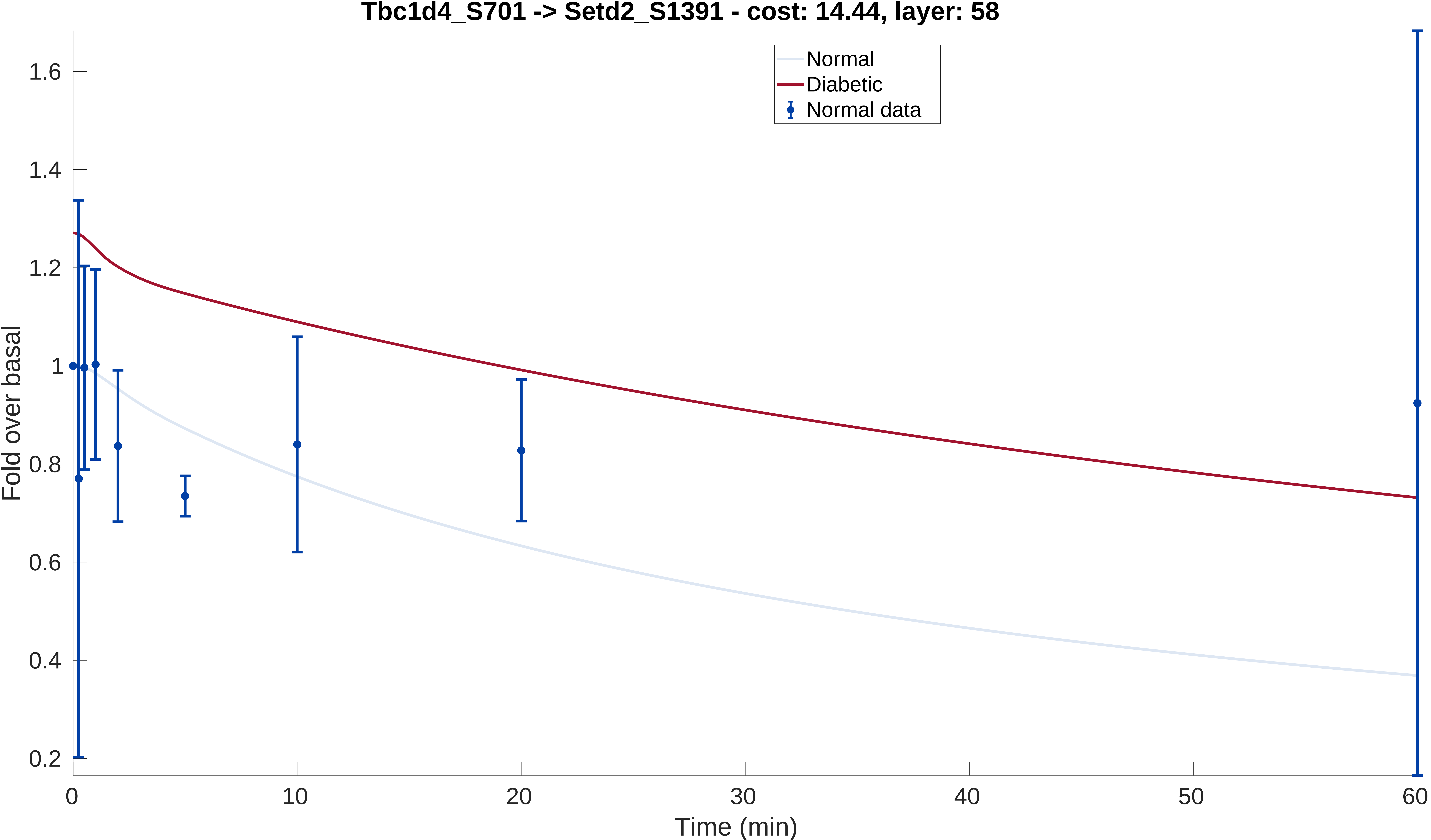

Tbc1d4_S701 -> Setd2_S1391 - cost: 14.44, layer: 58
Normal
Diabetic
Normal data
Fold over basal
1.6
1.4
1.2
1
0.8
0.6
0.4
0.2
0
10
20
30
40
50
60
Time (min)

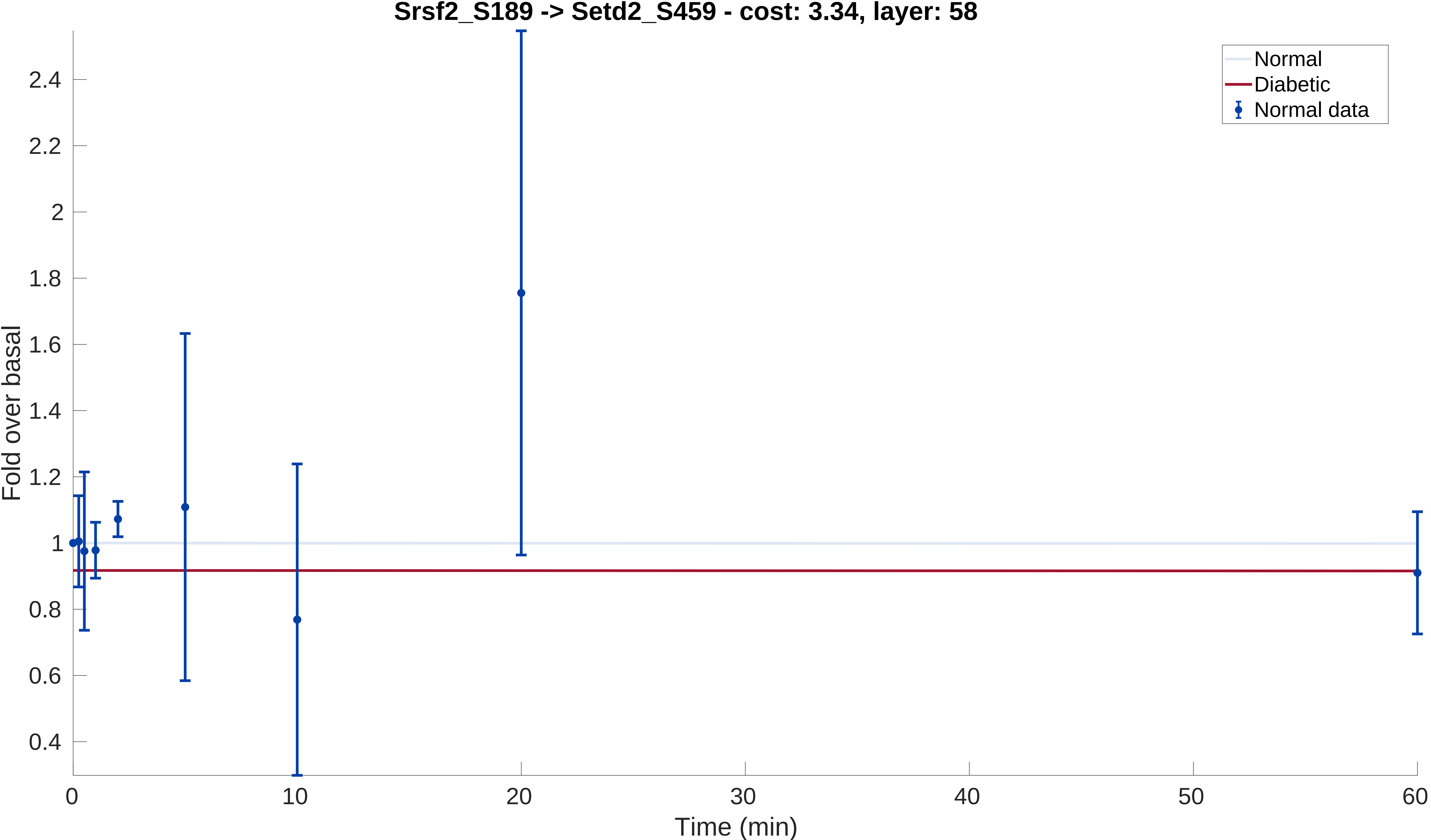

Srsf2_S189 -> Setd2_S459 - cost: 3.34, layer: 58
Normal
Diabetic
Normal data
Fold over basal
2.4
2.2
2
1.8
1.6
1.4
1.2
1
0.8
0.6
0.4
0
10
20
30
40
50
60
Time (min)

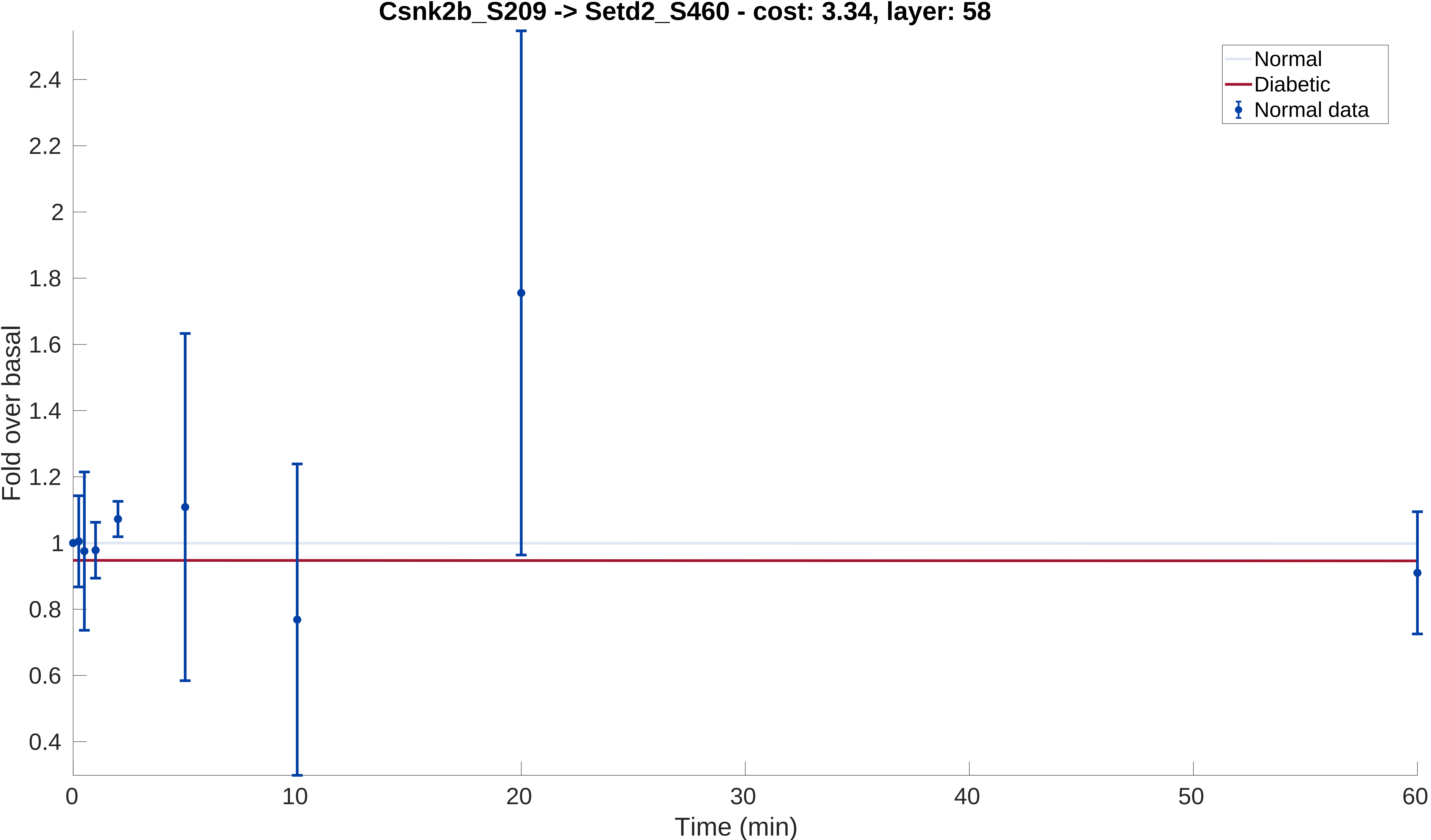

Csnk2b_S209 -> Setd2_S460 - cost: 3.34, layer: 58
Normal
Diabetic
Normal data
Fold over basal
Time (min)
2.4
2.2
2
1.8
1.6
1.4
1.2
1
0.8
0.6
0.4
0
10
20
30
40
50
60

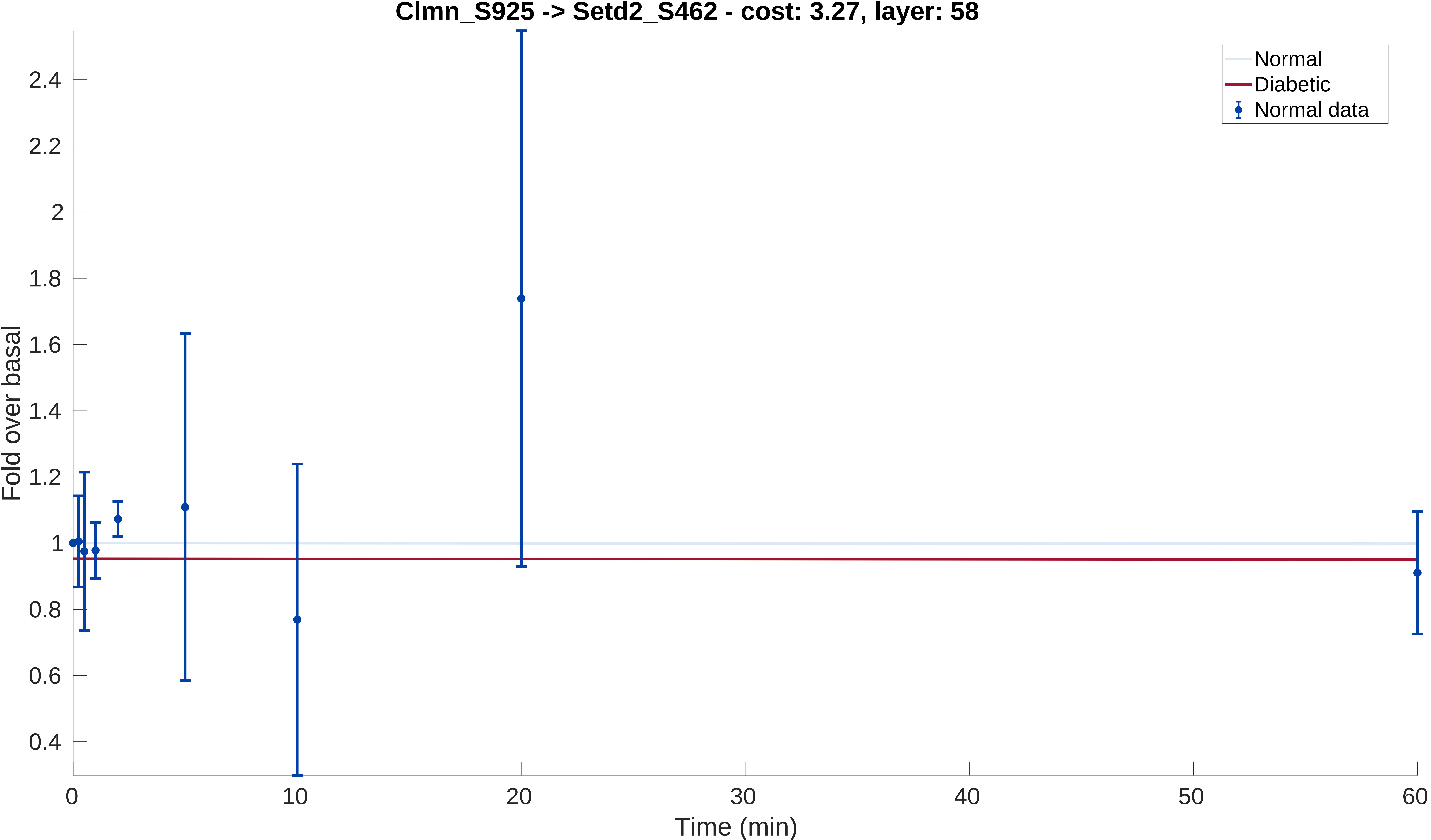

Clmn_S925 -> Setd2_S462 - cost: 3.27, layer: 58
Normal
Diabetic
Normal data
Fold over basal
2.4
2.2
2
1.8
1.6
1.4
1.2
1
0.8
0.6
0.4
0
10
20
30
40
50
60
Time (min)

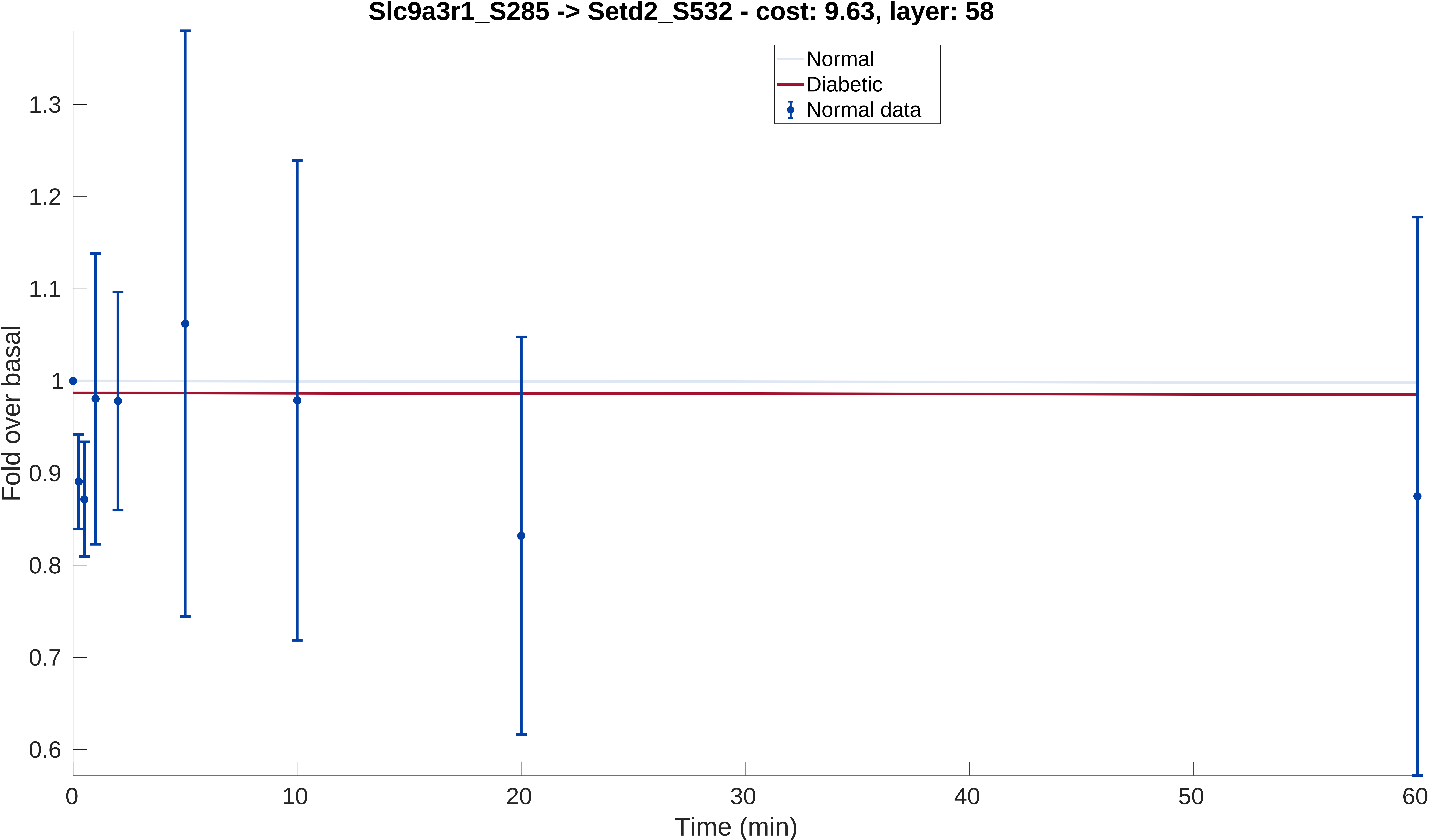

Slc9a3r1_S285 -> Setd2_S532 - cost: 9.63, layer: 58
Normal
Diabetic
Normal data
Fold over basal
1.3
1.2
1.1
1
0.9
0.8
0.7
0.6
0
10
20
30
40
50
60
Time (min)

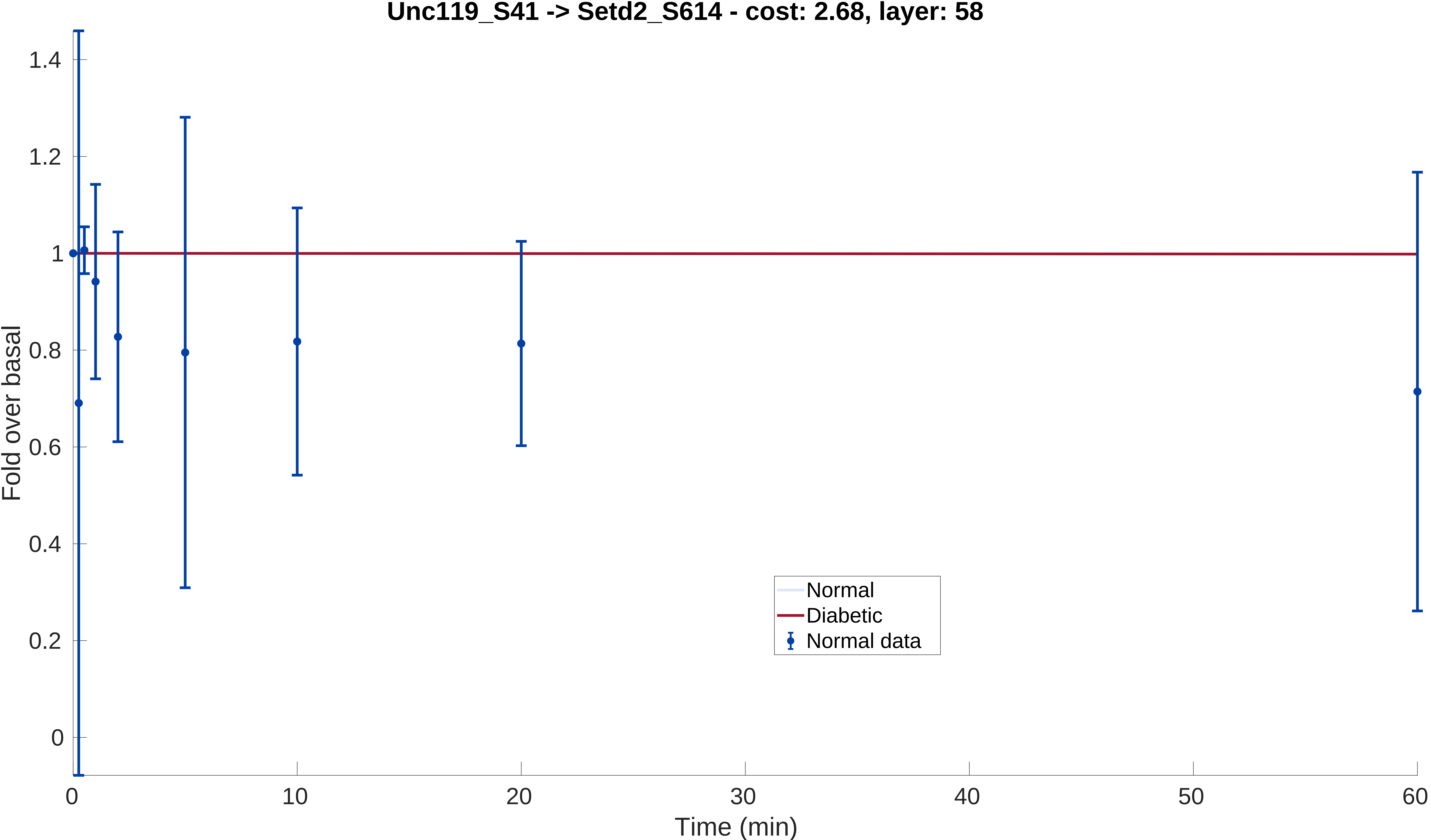
Unc119_S41 -> Setd2_S614 - cost: 2.68, layer: 58
Fold over basal
Time (min)
0
0.2
0.4
0.6
0.8
1
1.2
1.4
0
10
20
30
40
50
60
Normal
Diabetic
Normal data

#### Clmn_S925 -> Setd2_S624 - cost: 5.38, layer: 58

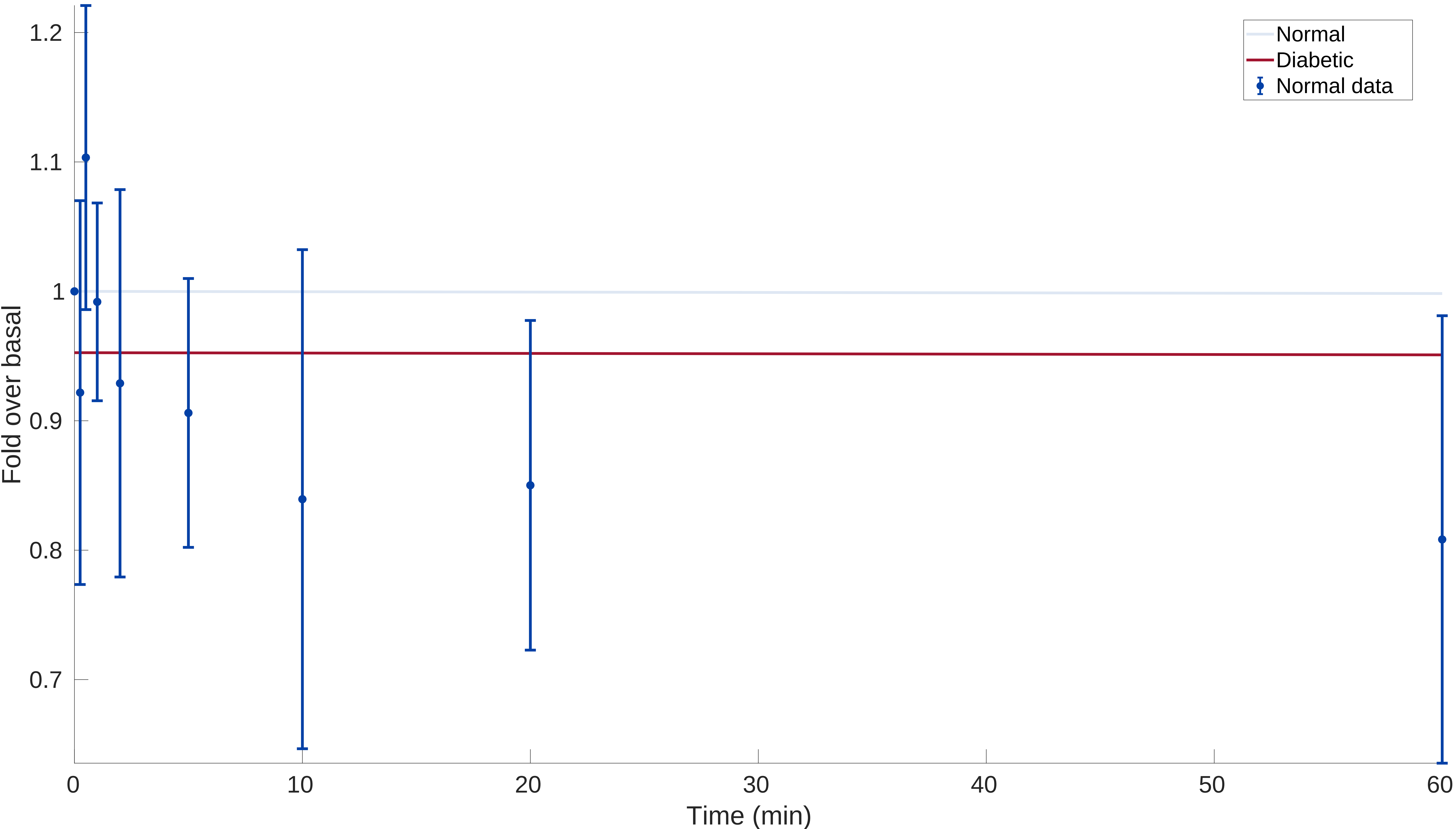

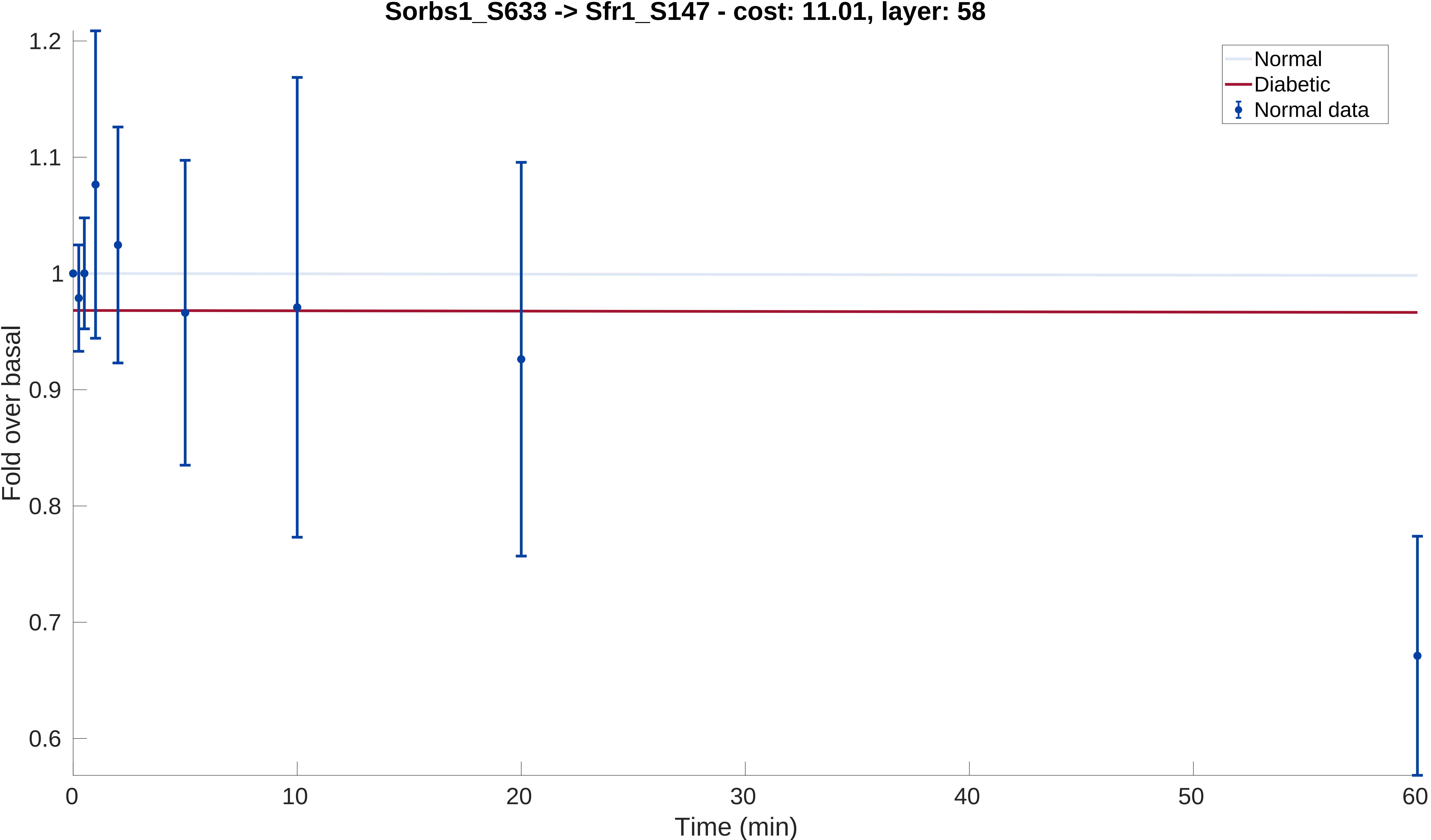
Sorbs1_S633 -> Sfr1_S147 - cost: 11.01, layer: 58
Fold over basal
Time (min)
Normal
Diabetic
Normal data
1.2
1.1
1
0.9
0.8
0.7
0.6
0
10
20
30
40
50
60

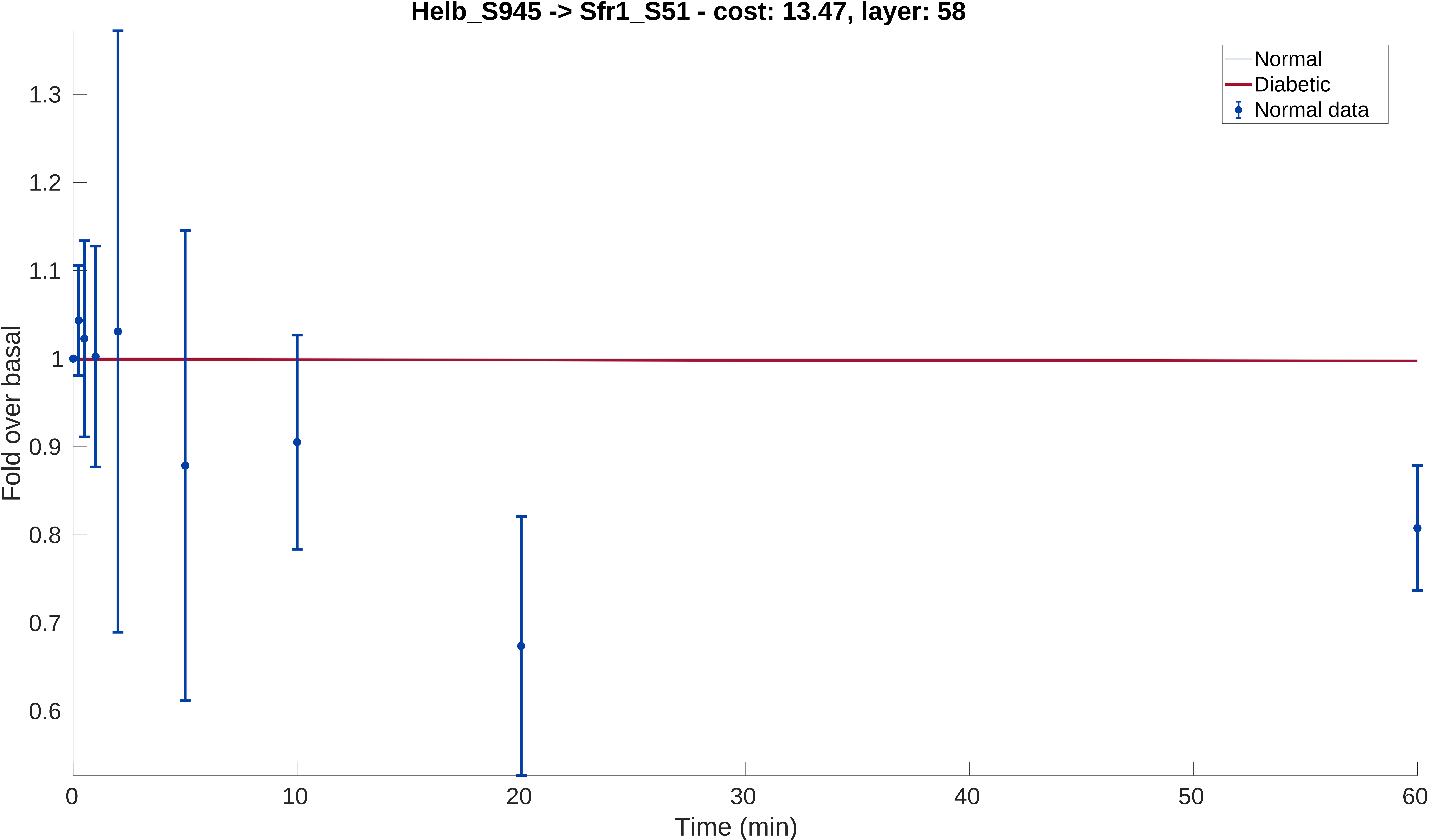
Helb_S945 -> Sfr1_S51 - cost: 13.47, layer: 58
Normal
Diabetic
Normal data
Fold over basal
Time (min)
0.6
0.7
0.8
0.9
1
1.1
1.2
1.3
0
10
20
30
40
50
60

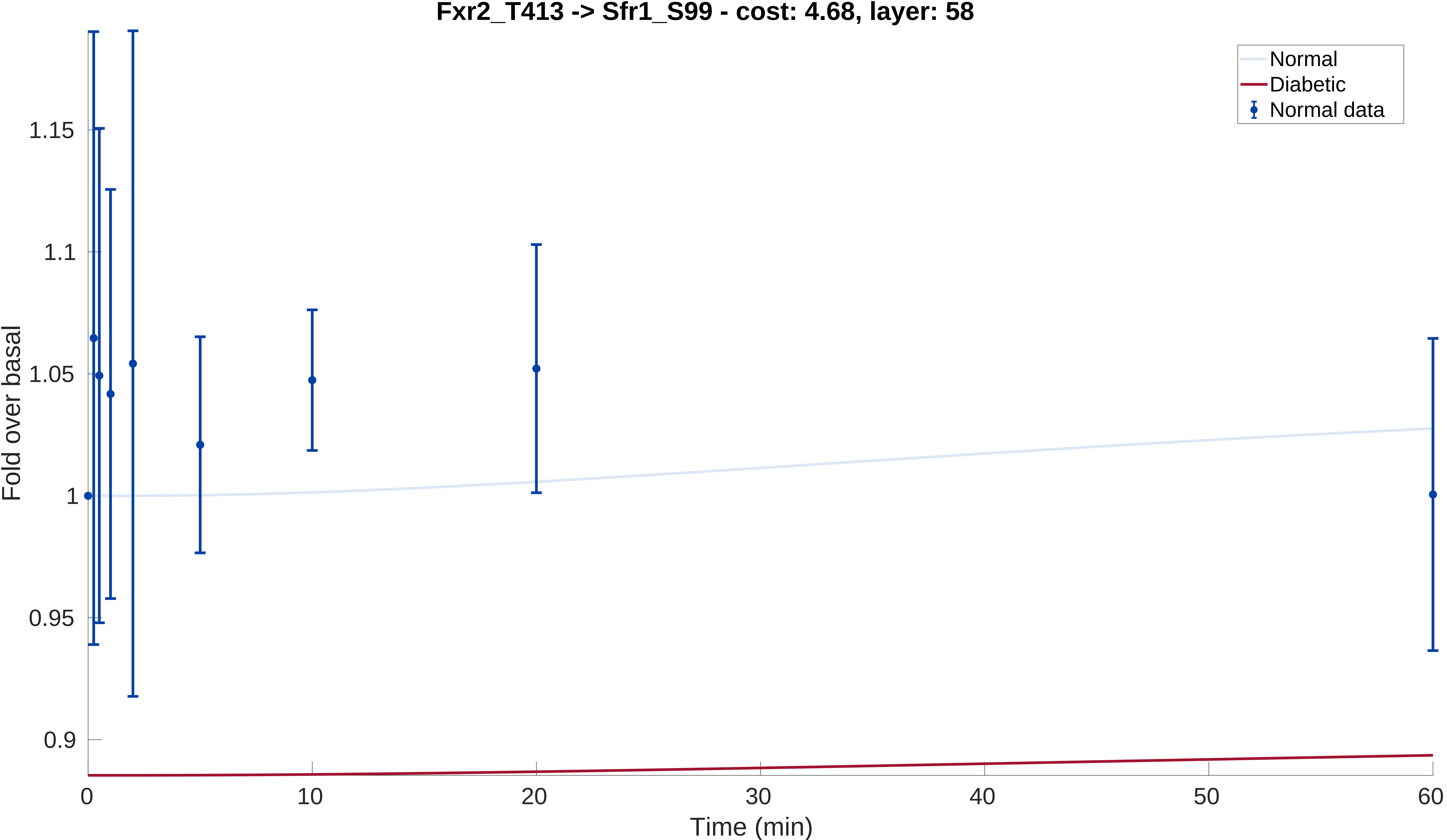

Fxr2_T413 -> Sfr1_S99 - cost: 4.68, layer: 58
Normal
Diabetic
Normal data
Fold over basal
1.15
1.1
1.05
1
0.95
0.9
0
10
20
30
40
50
60
Time (min)

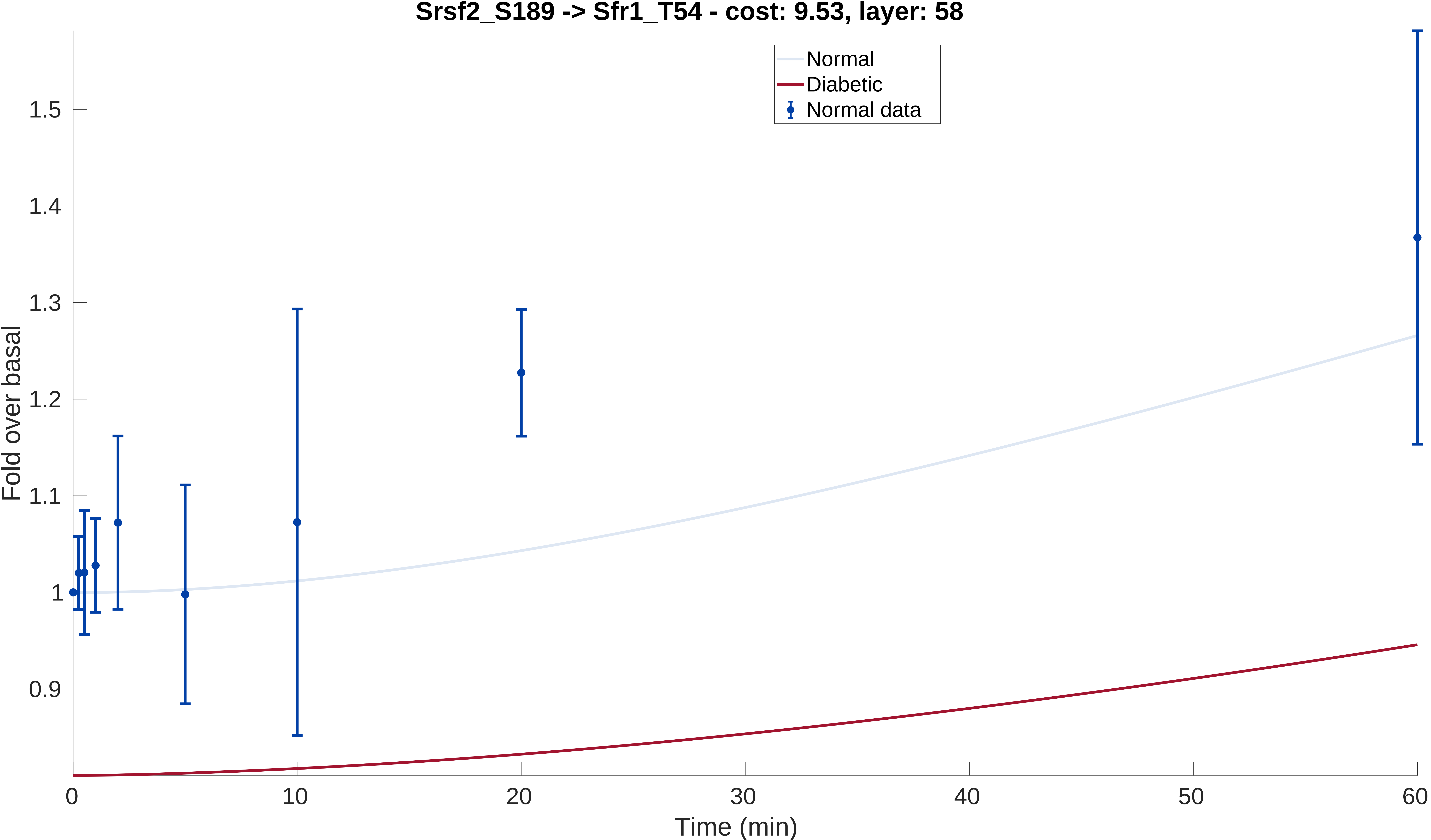
Srsf2_S189 -> Sfr1_T54 - cost: 9.53, layer: 58
Normal
Diabetic
Normal data
Fold over basal
Time (min)
0.9
1
1.1
1.2
1.3
1.4
1.5
0
10
20
30
40
50
60

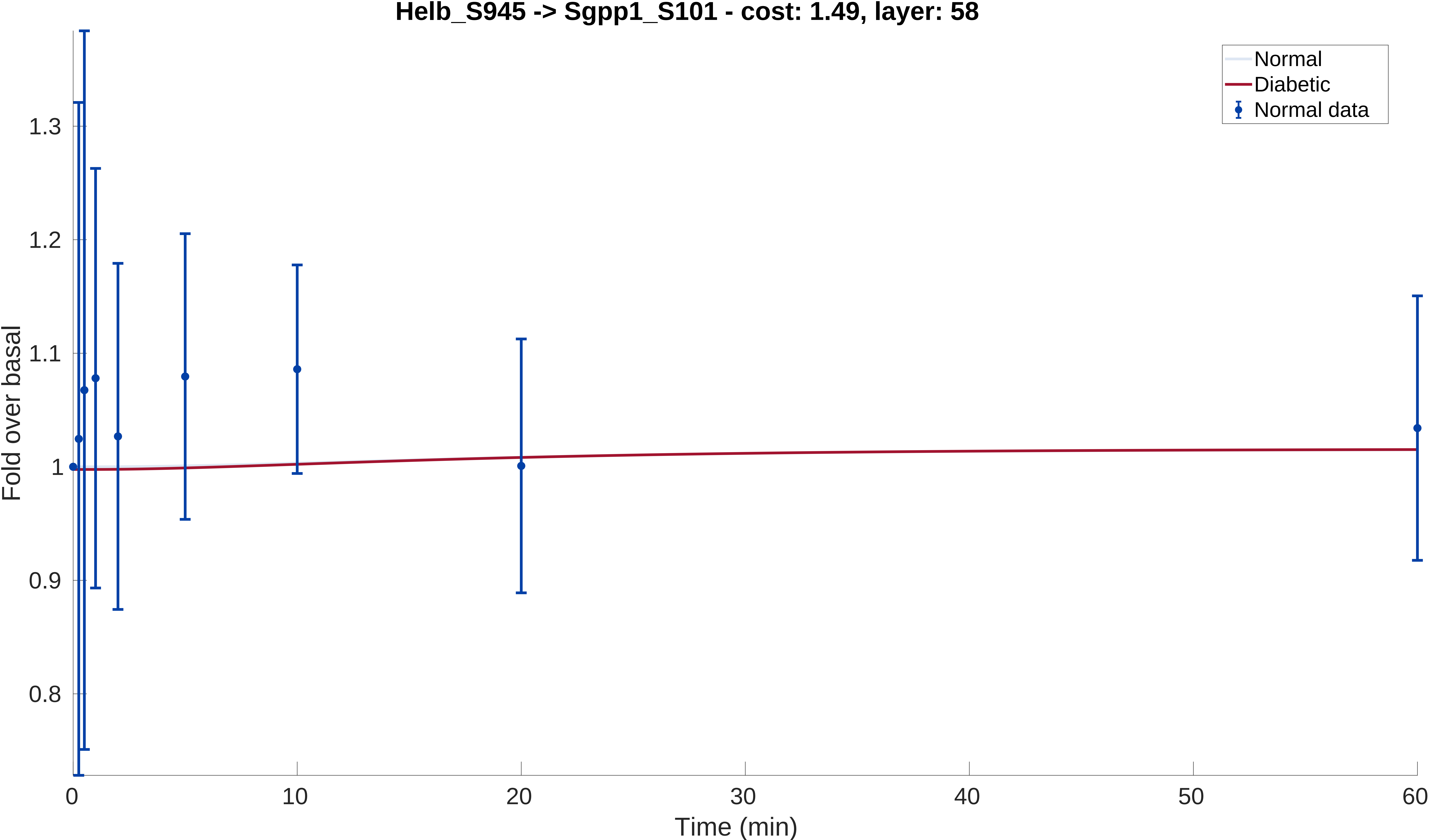

Helb_S945 -> Sgpp1_S101 - cost: 1.49, layer: 58
Normal
Diabetic
Normal data
Fold over basal
1.3
1.2
1.1
1
0.9
0.8
0
10
20
30
40
50
60
Time (min)

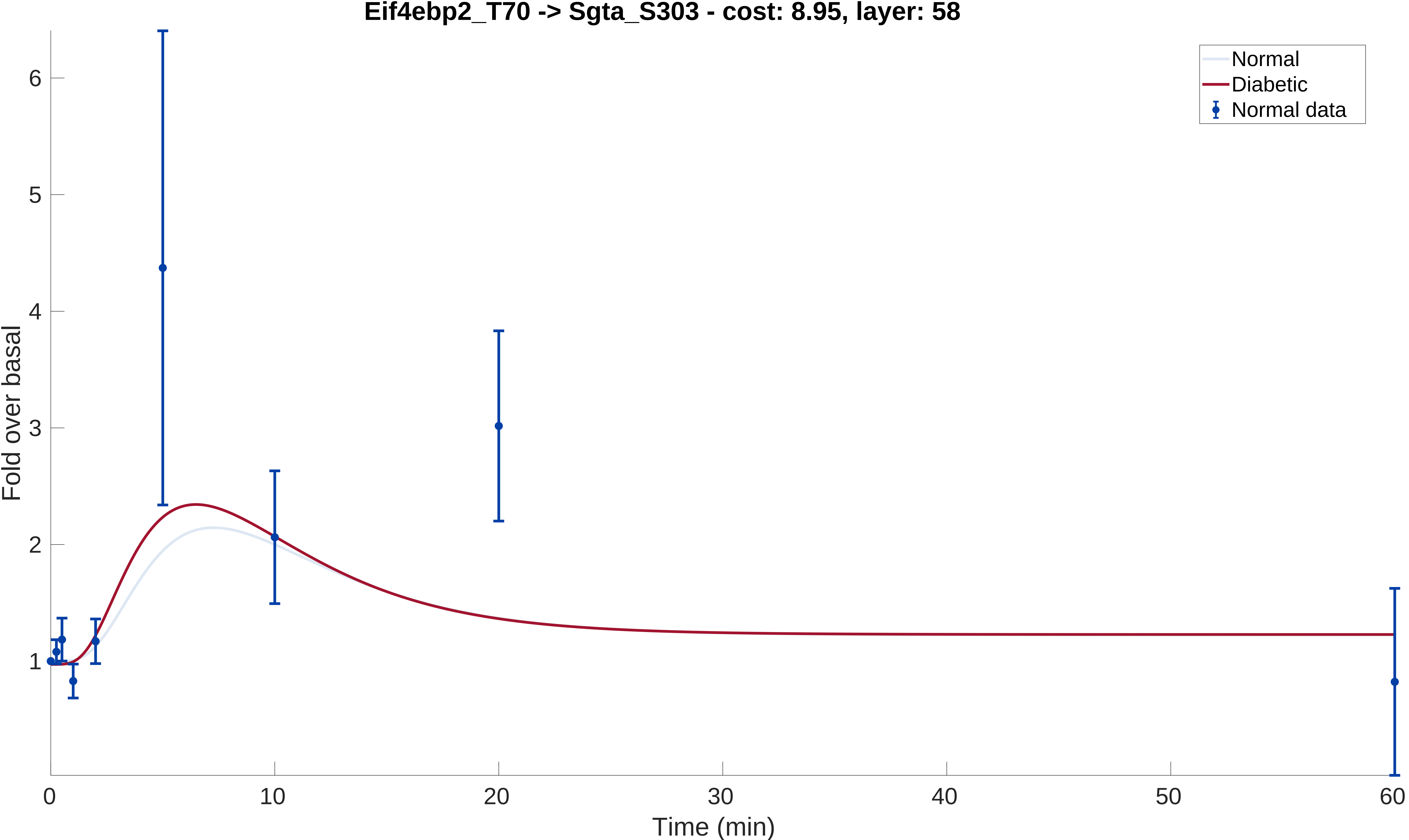
Eif4ebp2_T70 -> Sgta_S303 - cost: 8.95, layer: 58
Normal
Diabetic
Normal data
Fold over basal
Time (min)
0
10
20
30
40
50
60
1
2
3
4
5
6

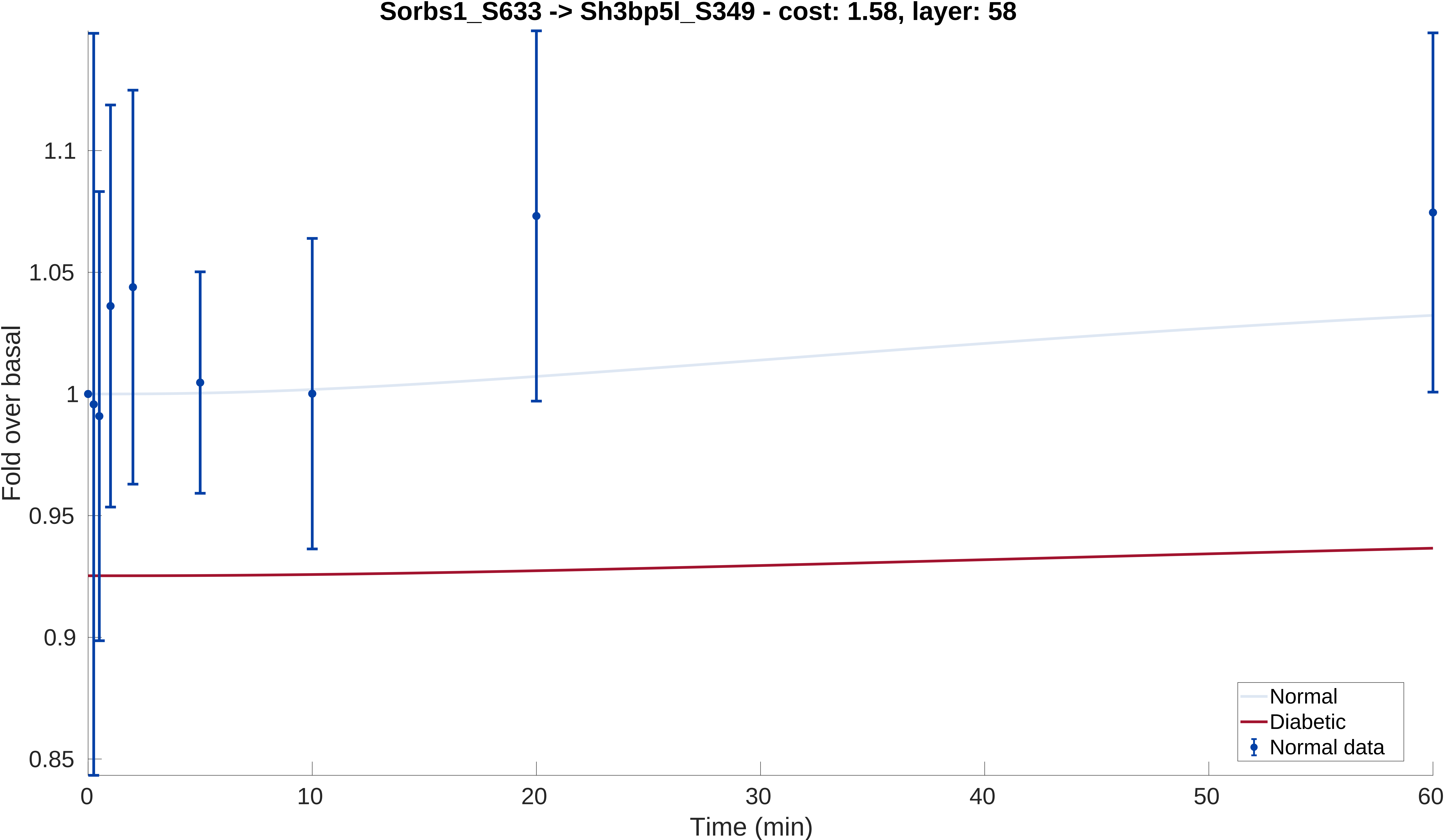

Sorbs1_S633 -> Sh3bp5l_S349 - cost: 1.58, layer: 58
Fold over basal
1.1
1.05
1
0.95
0.9
0.85
0
10
20
30
40
50
60
Time (min)
Normal
Diabetic
Normal data

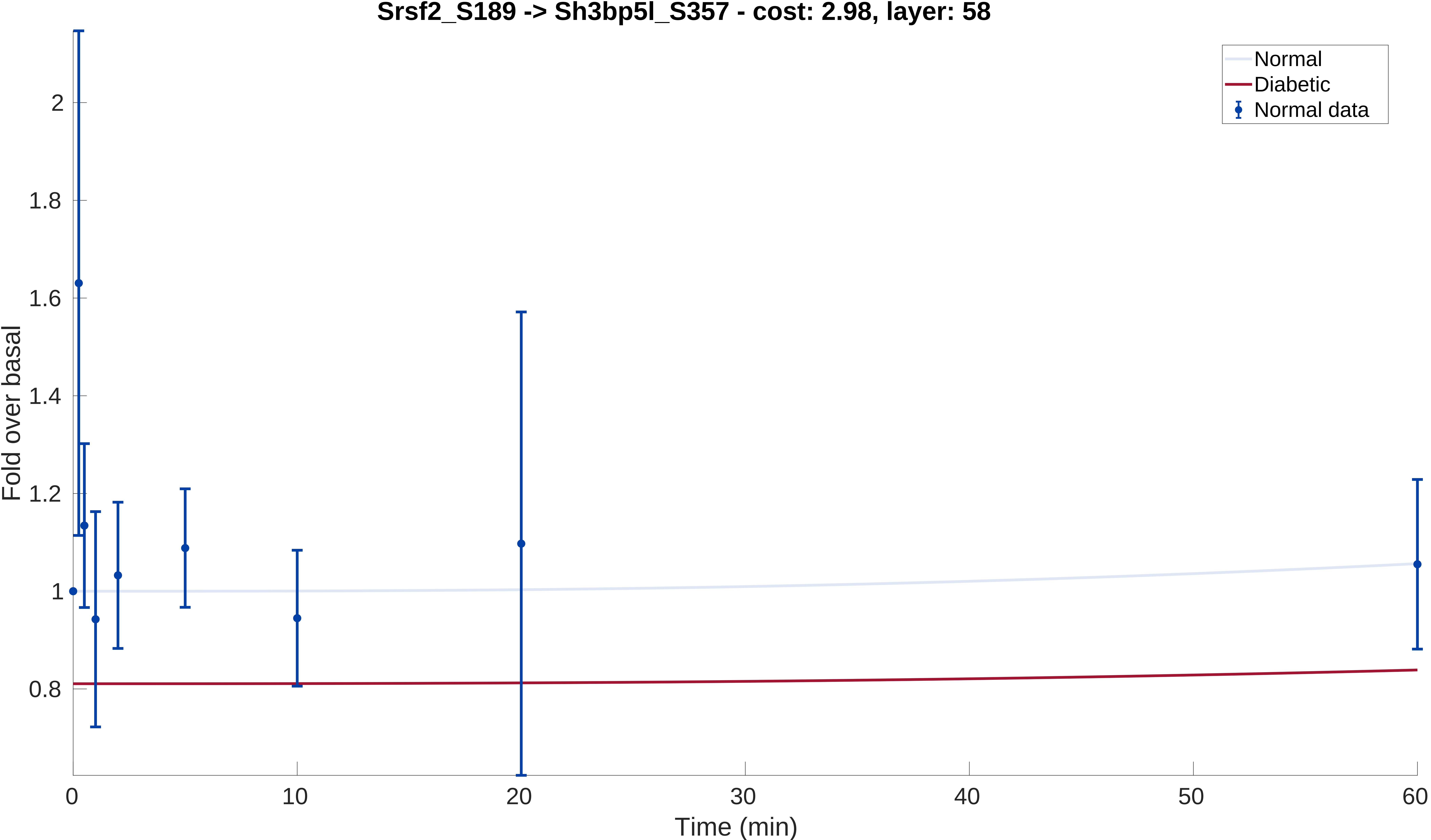

Srsf2_S189 -> Sh3bp5l_S357 - cost: 2.98, layer: 58
Normal
Diabetic
Normal data
Fold over basal
2
1.8
1.6
1.4
1.2
1
0.8
0
10
20
30
40
50
60
Time (min)

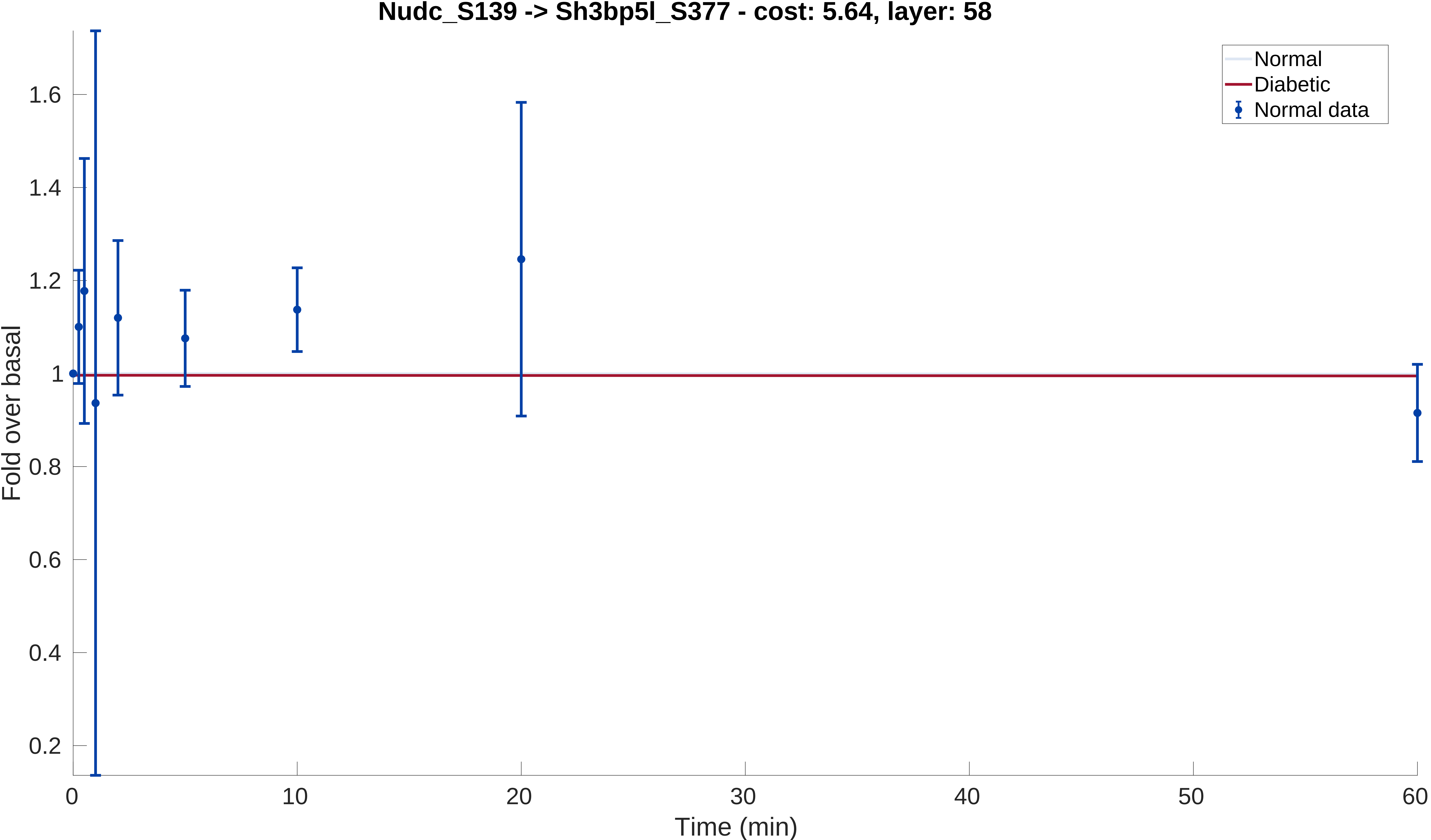

Nudc_S139 -> Sh3bp5l_S377 - cost: 5.64, layer: 58
Normal
Diabetic
Normal data
1.6
1.4
1.2
1
0.8
0.6
0.4
0.2
Fold over basal
0
10
20
30
40
50
60
Time (min)

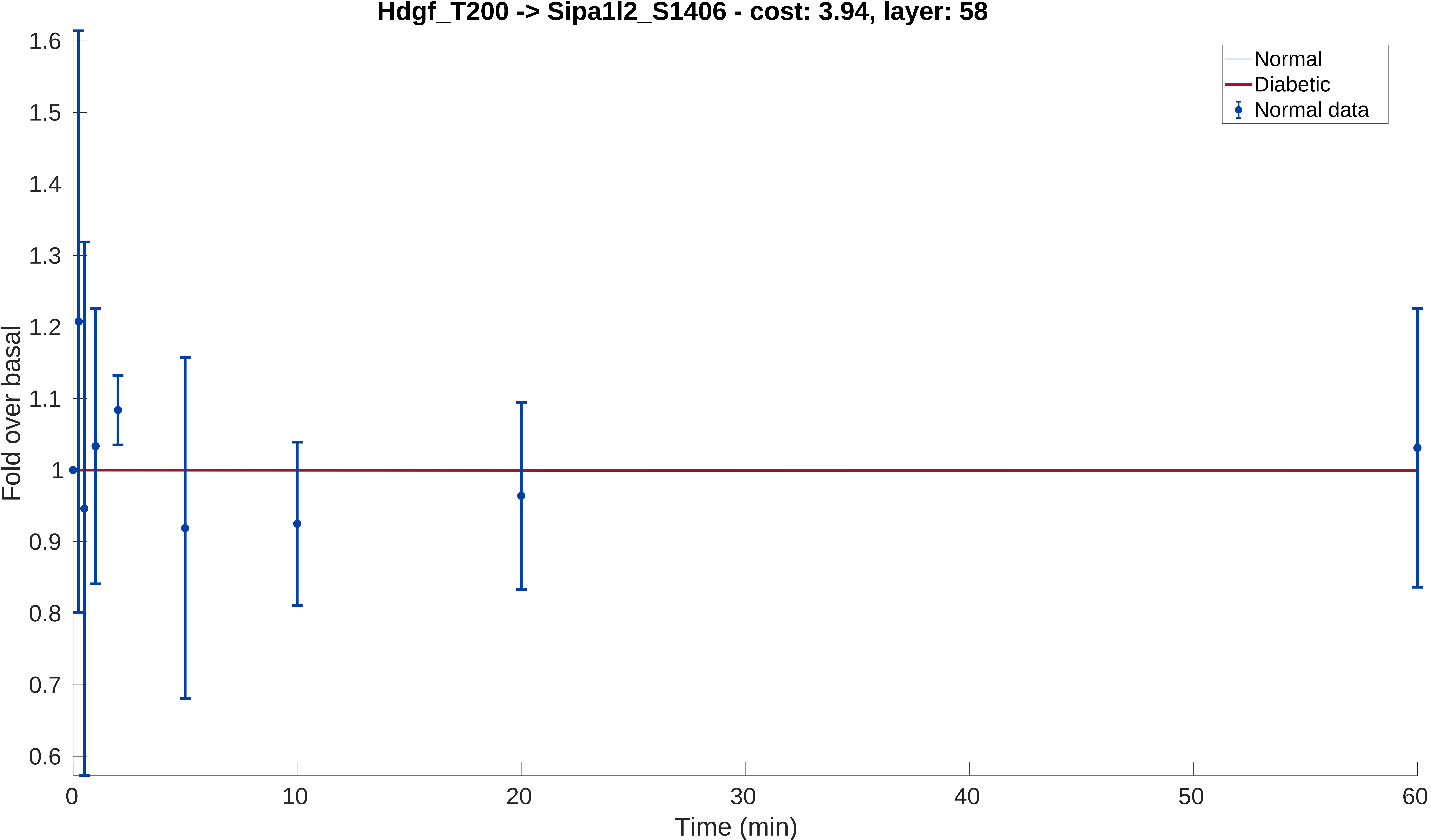

Hdgf_T200 -> Sipa1l2_S1406 - cost: 3.94, layer: 58
Fold over basal
Time (min)
0.6
0.7
0.8
0.9
1
1.1
1.2
1.3
1.4
1.5
1.6
0
10
20
30
40
50
60
Normal
Diabetic
Normal data

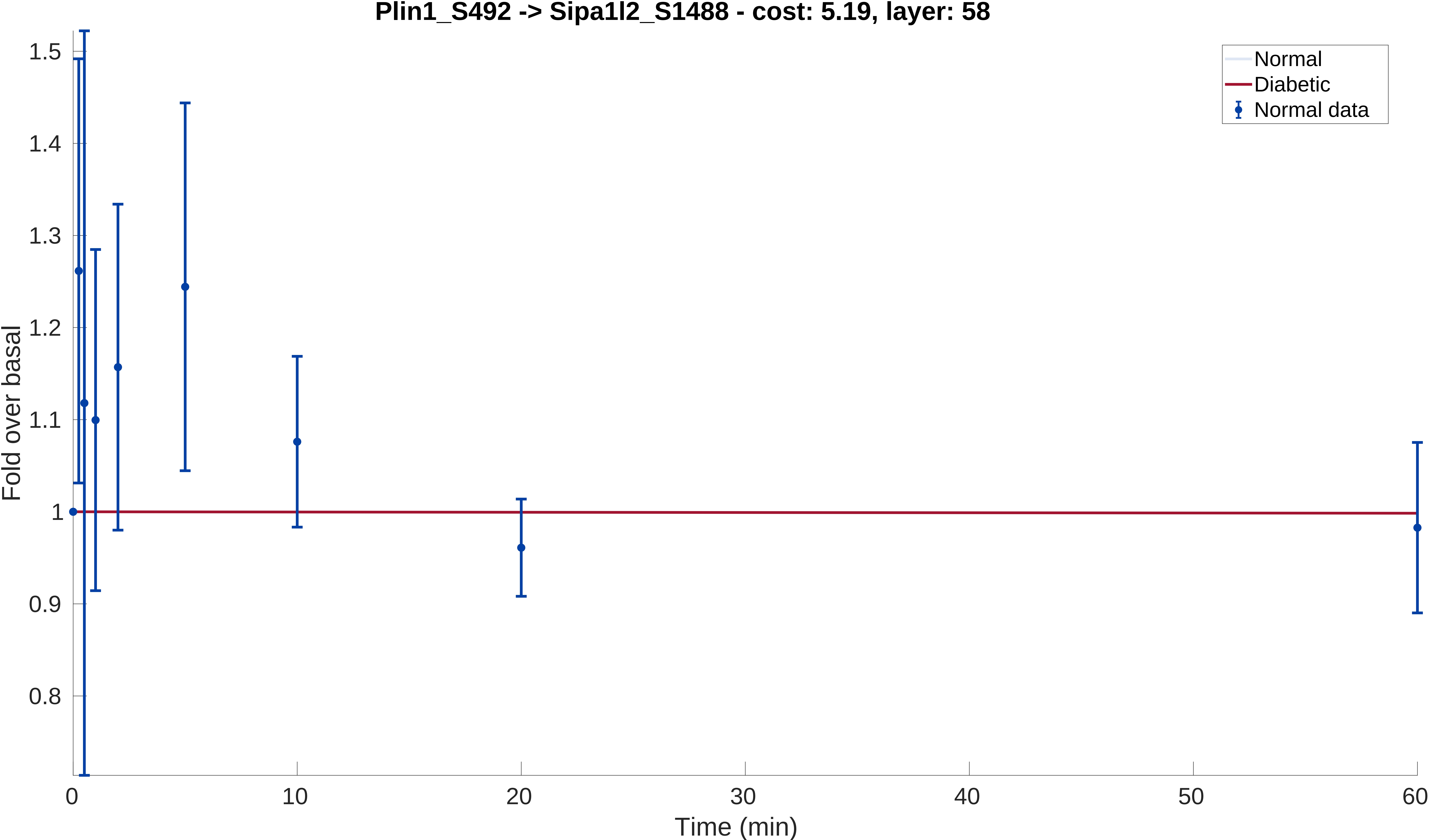
Plin1_S492 -> Sipa1l2_S1488 - cost: 5.19, layer: 58
Normal
Diabetic
Normal data
Fold over basal
1.5
1.4
1.3
1.2
1.1
1
0.9
0.8
0
10
20
30
40
50
60
Time (min)

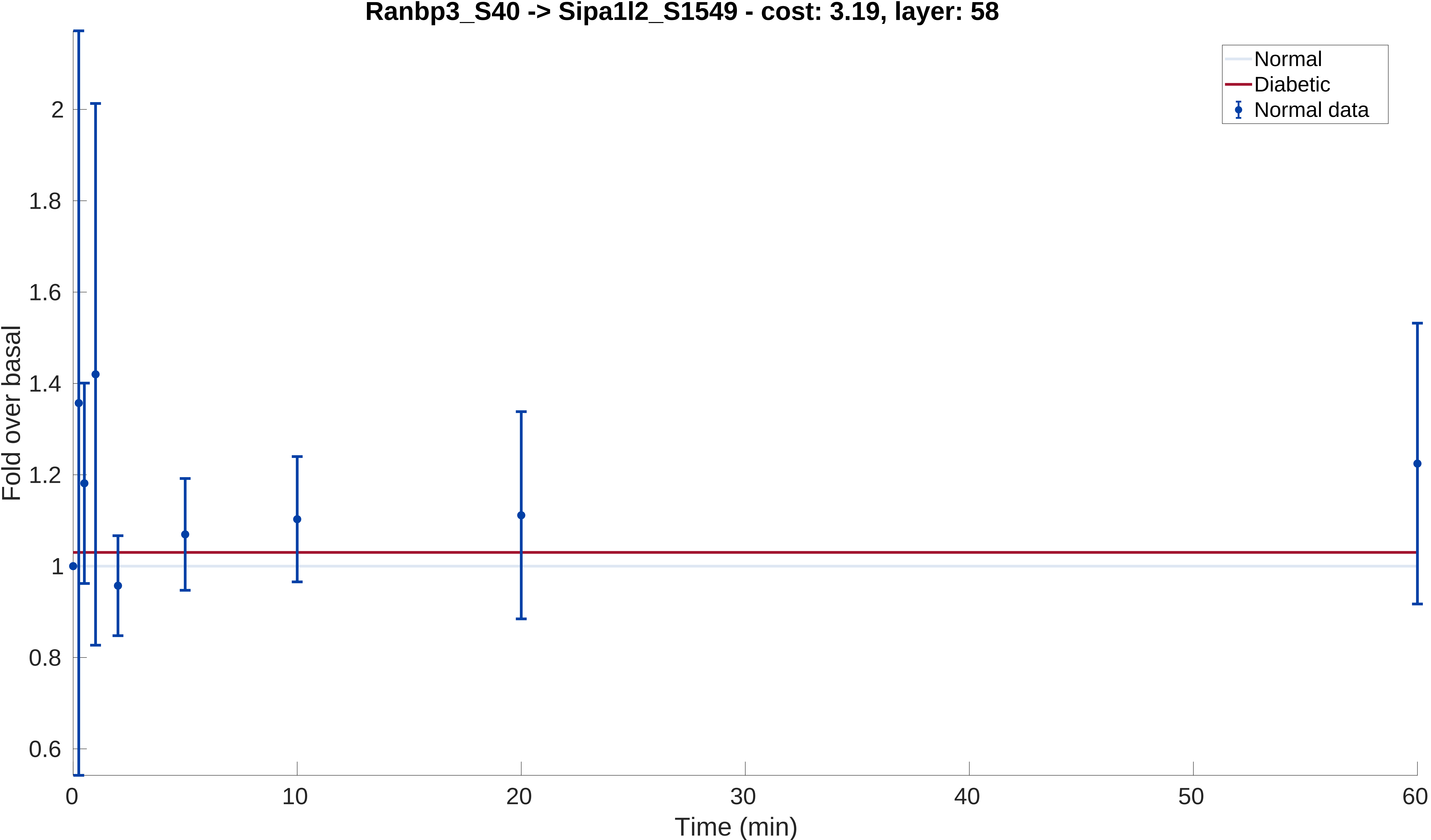
Ranbp3_S40 -> Sipa1l2_S1549 - cost: 3.19, layer: 58
Normal
Diabetic
Normal data
Fold over basal
2
1.8
1.6
1.4
1.2
1
0.8
0.6
0
10
20
30
40
50
60
Time (min)

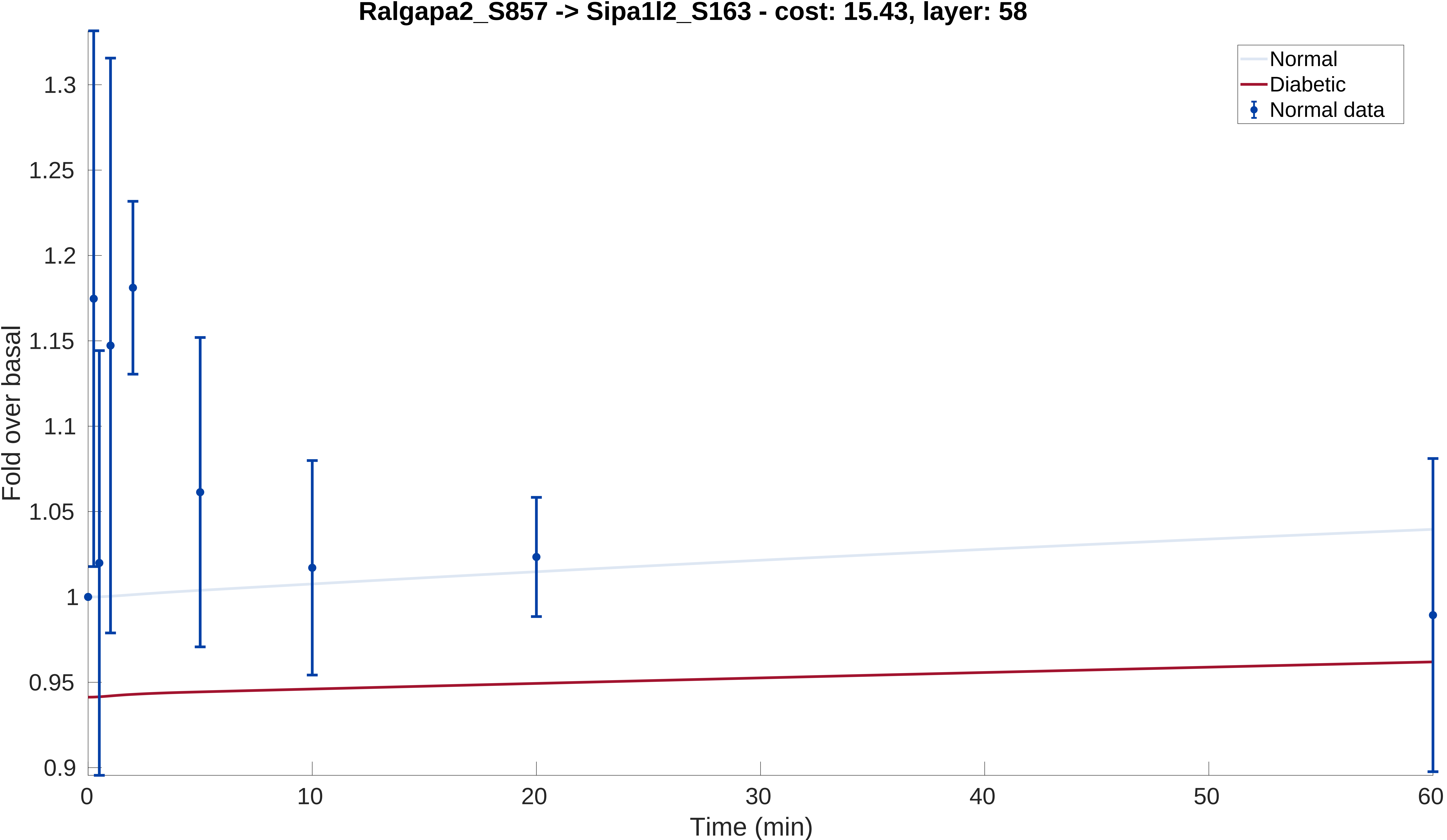

Ralgapa2_S857 -> Sipa1l2_S163 - cost: 15.43, layer: 58
Normal
Diabetic
Normal data
Fold over basal
1.3
1.25
1.2
1.15
1.1
1.05
1
0.95
0.9
0
10
20
30
40
50
60
Time (min)

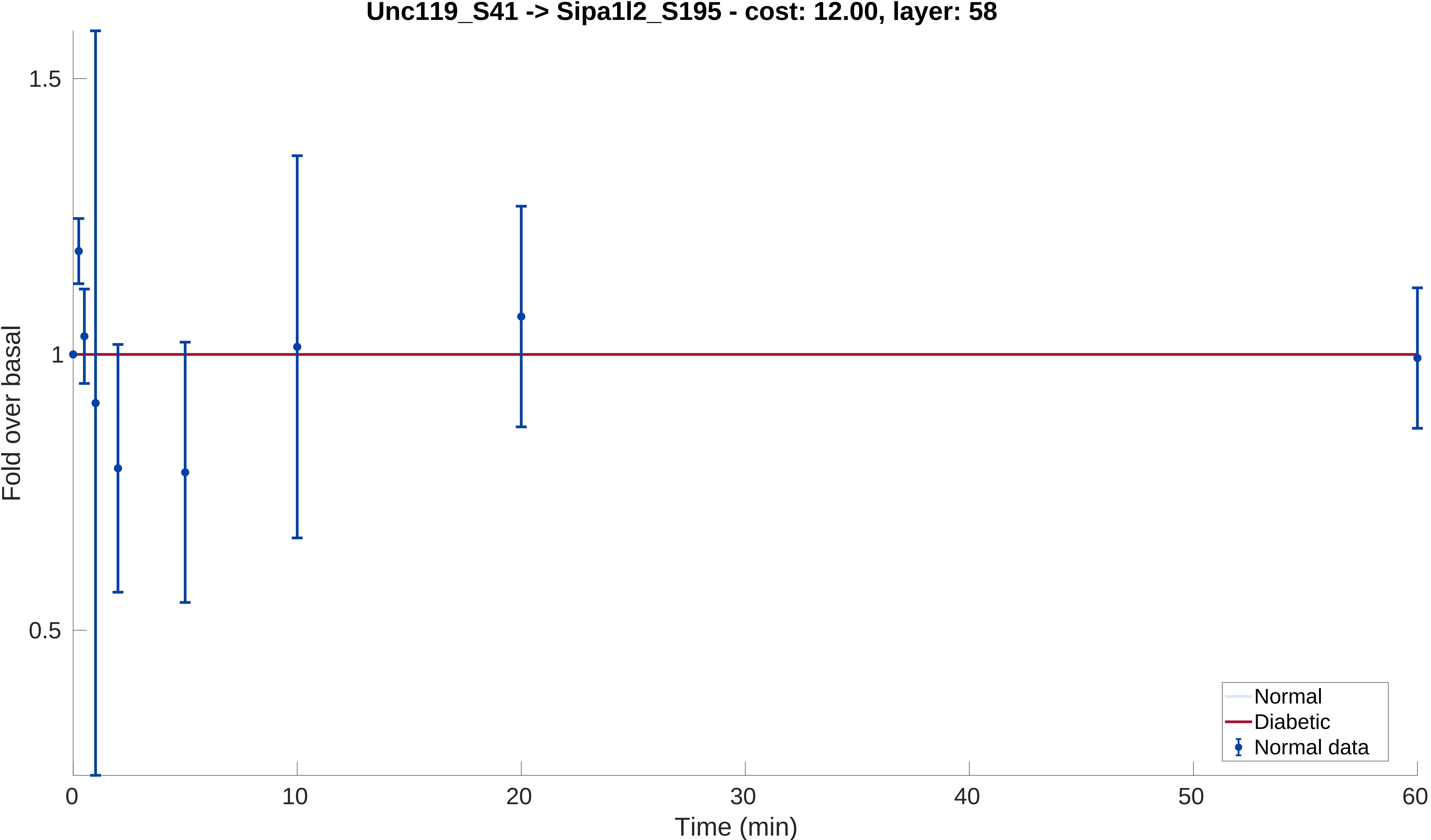
Unc119_S41 -> Sipa1l2_S195 - cost: 12.00, layer: 58
Fold over basal
1.5
1
0.5
0
10
20
30
40
50
60
Time (min)
Normal
Diabetic
Normal data

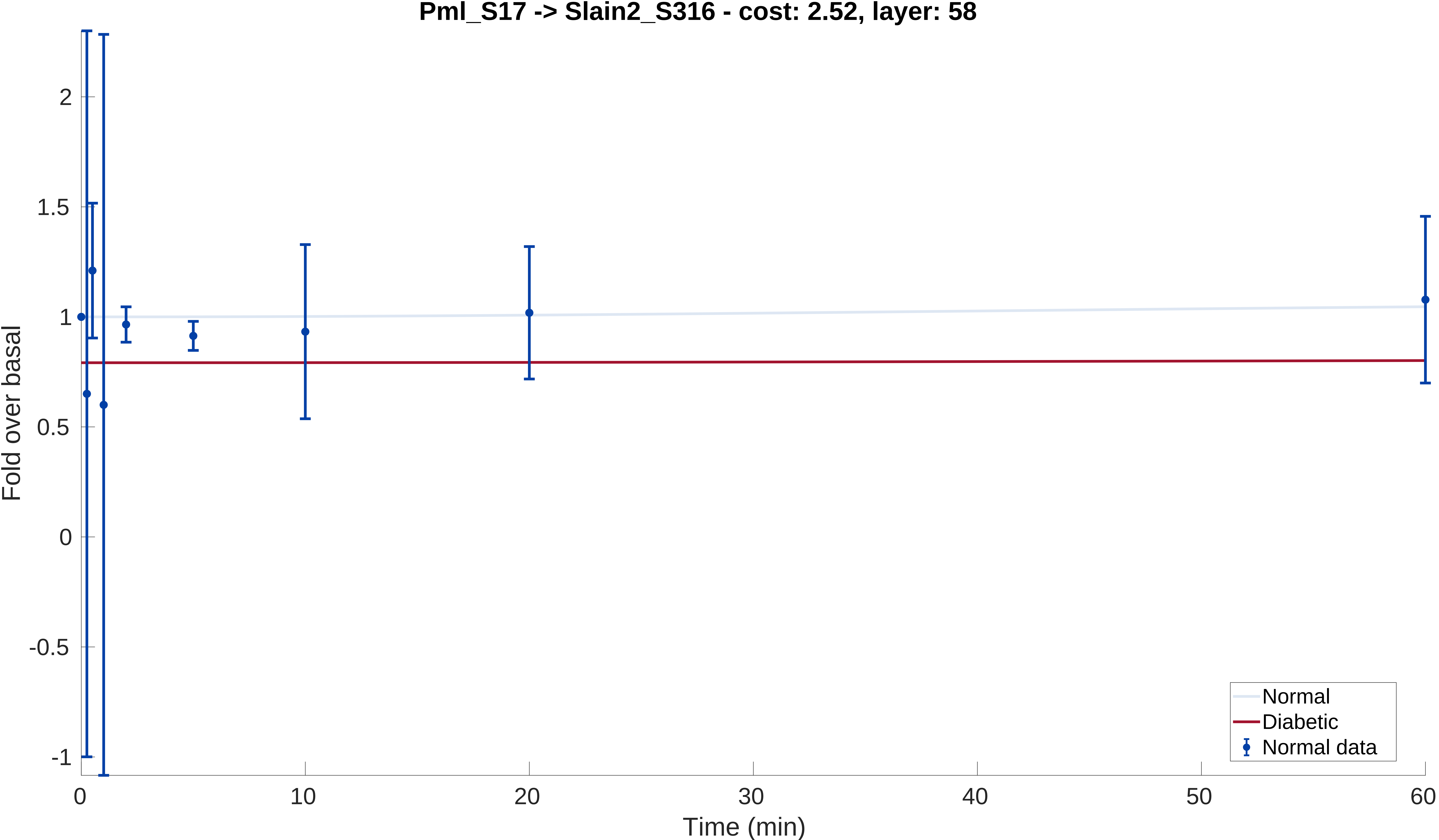

Pml_S17 -> Slain2_S316 - cost: 2.52, layer: 58
Fold over basal
Time (min)
Normal
Diabetic
Normal data
2
1.5
1
0.5
0
-0.5
-1
0
10
20
30
40
50
60

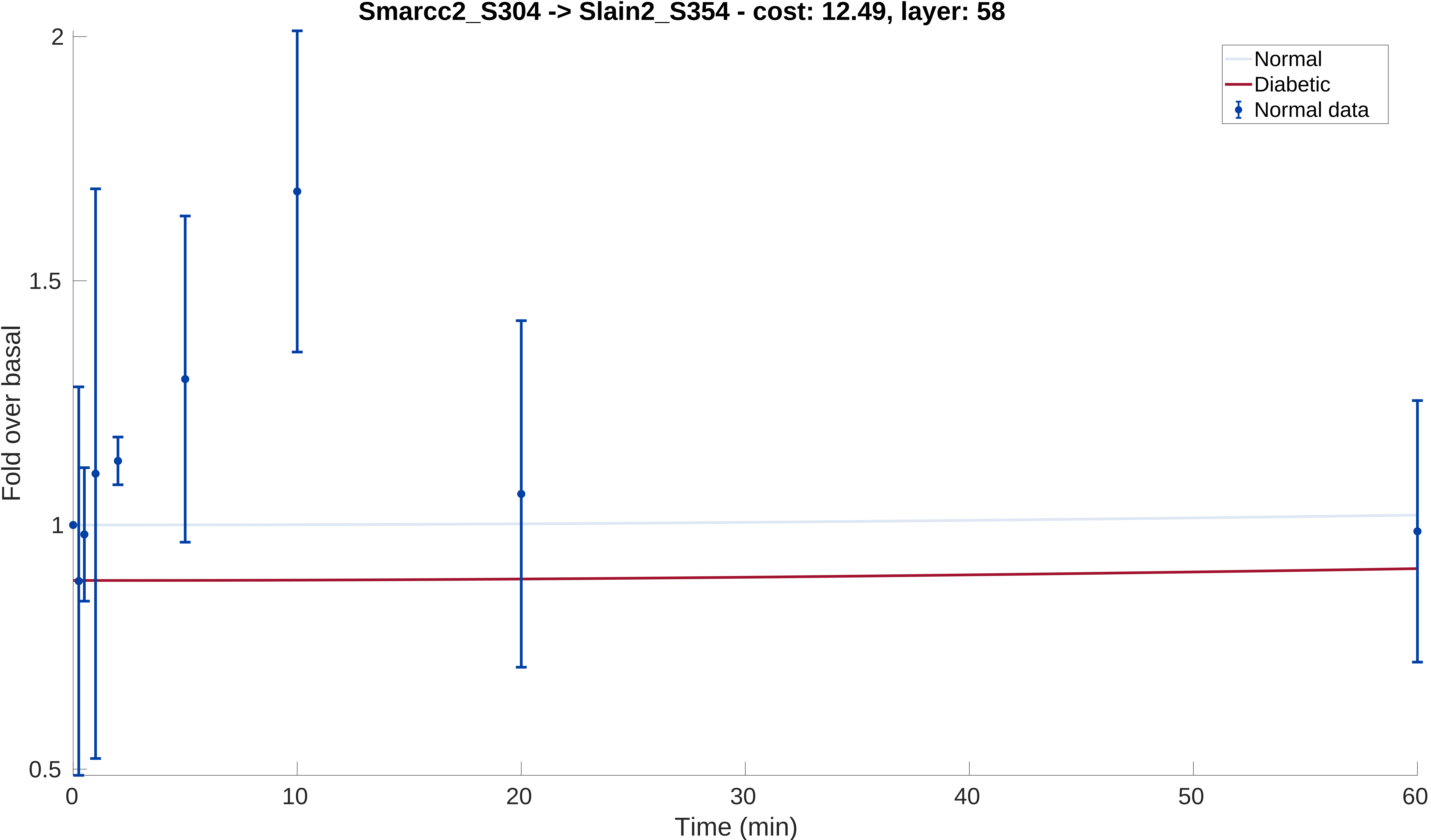

Smarcc2_S304 -> Slain2_S354 - cost: 12.49, layer: 58
Normal
Diabetic
Normal data
Fold over basal
Time (min)
2
1.5
1
0.5
0
10
20
30
40
50
60

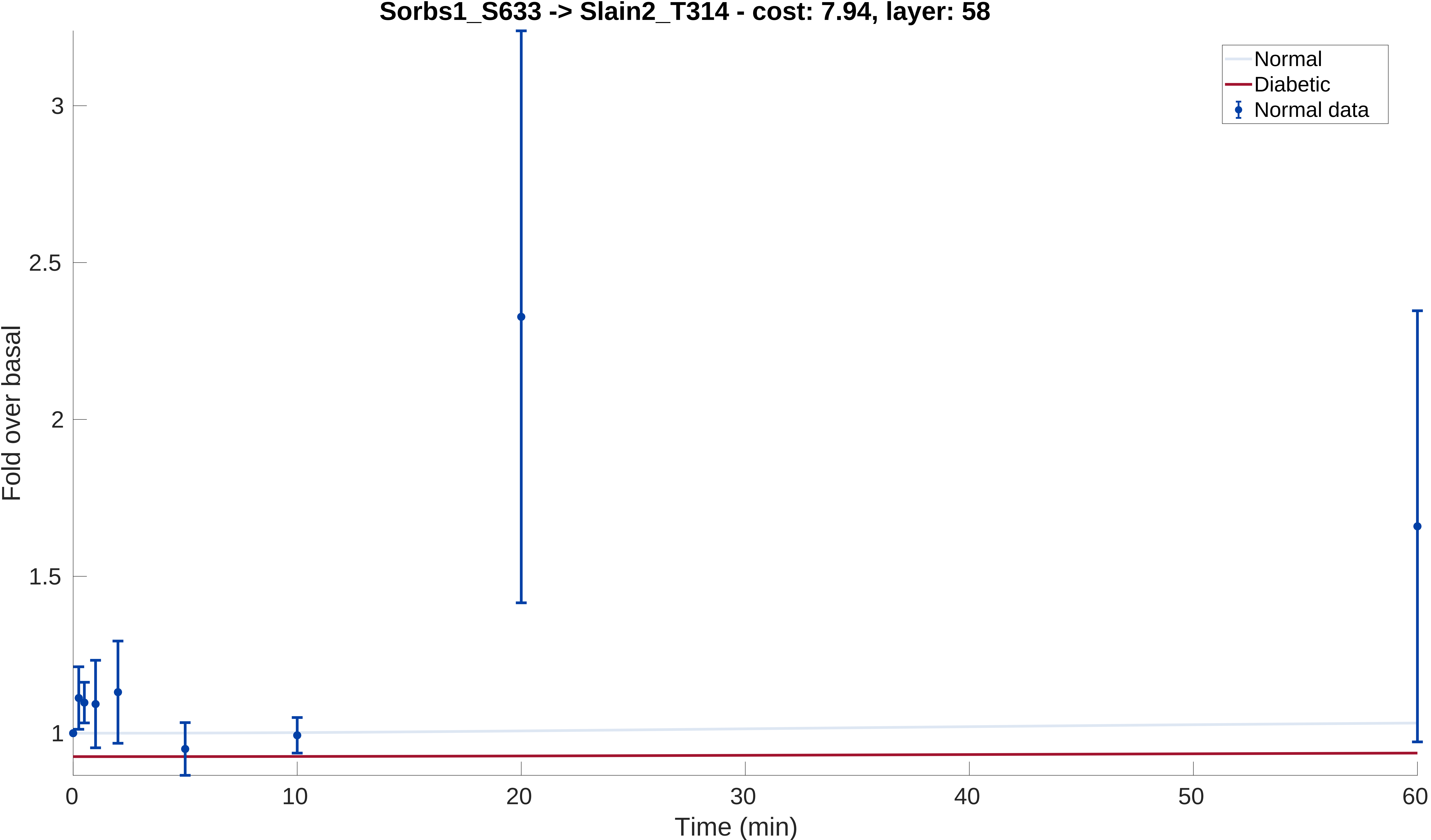
Sorbs1_S633 -> Slain2_T314 - cost: 7.94, layer: 58
Normal
Diabetic
Normal data
Fold over basal
3
2.5
2
1.5
1
0
10
20
30
40
50
60
Time (min)

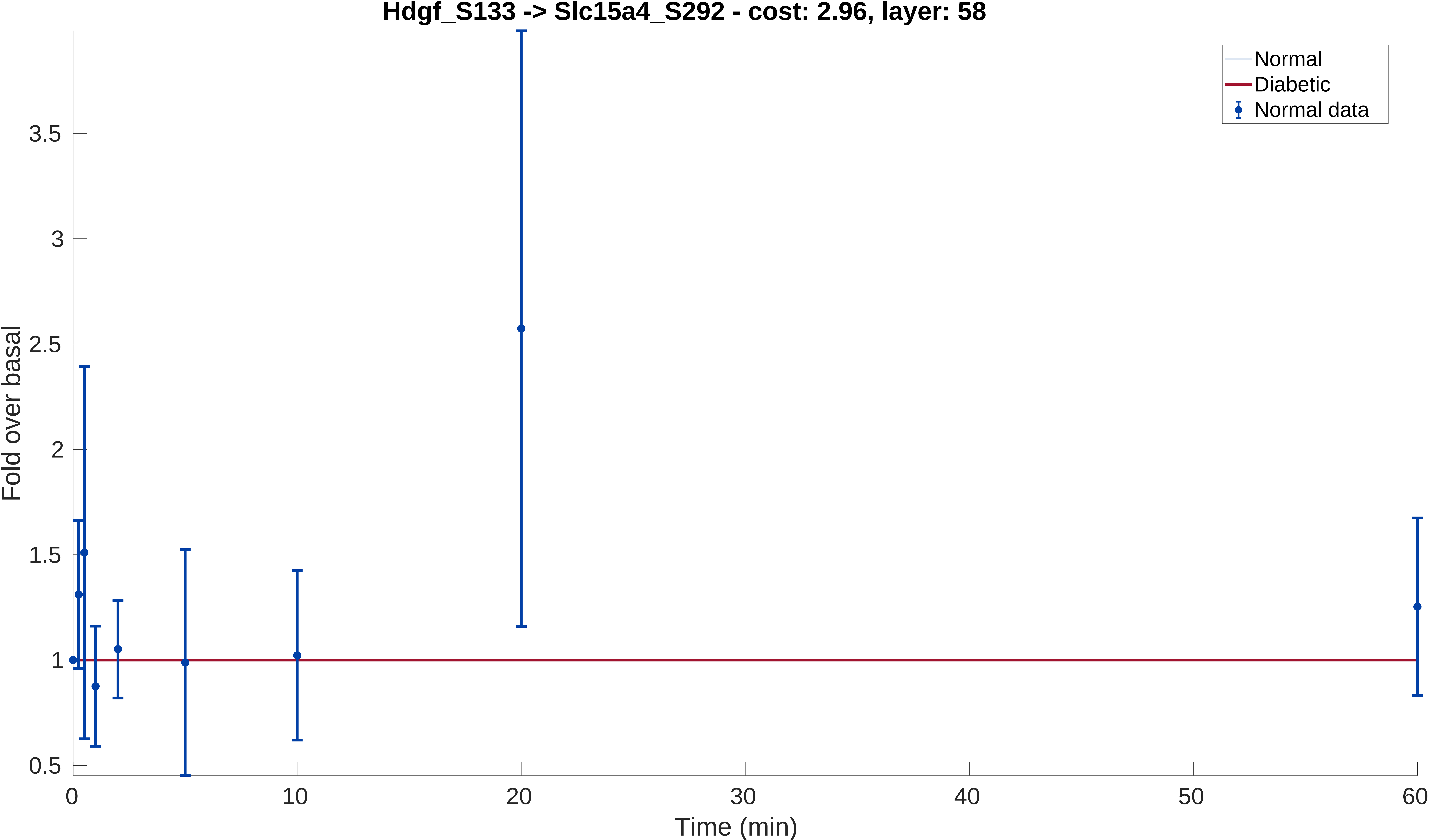

Hdgf_S133 -> Slc15a4_S292 - cost: 2.96, layer: 58
Normal
Diabetic
Normal data
Fold over basal
Time (min)
0.5
1
1.5
2
2.5
3
3.5
0
10
20
30
40
50
60

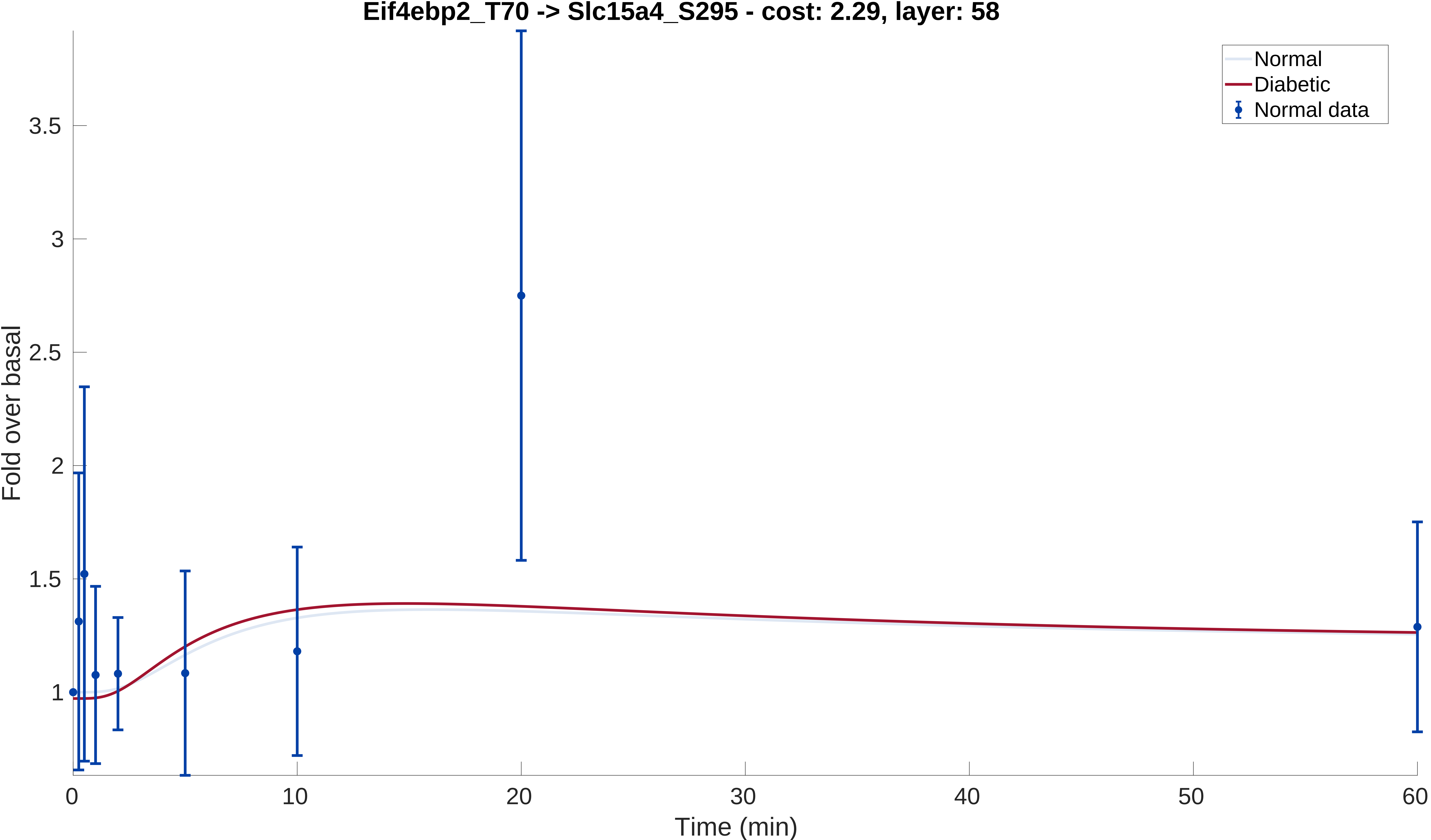

Eif4ebp2_T70 -> Slc15a4_S295 - cost: 2.29, layer: 58
Normal
Diabetic
Normal data
Fold over basal
3.5
3
2.5
2
1.5
1
0
10
20
30
40
50
60
Time (min)

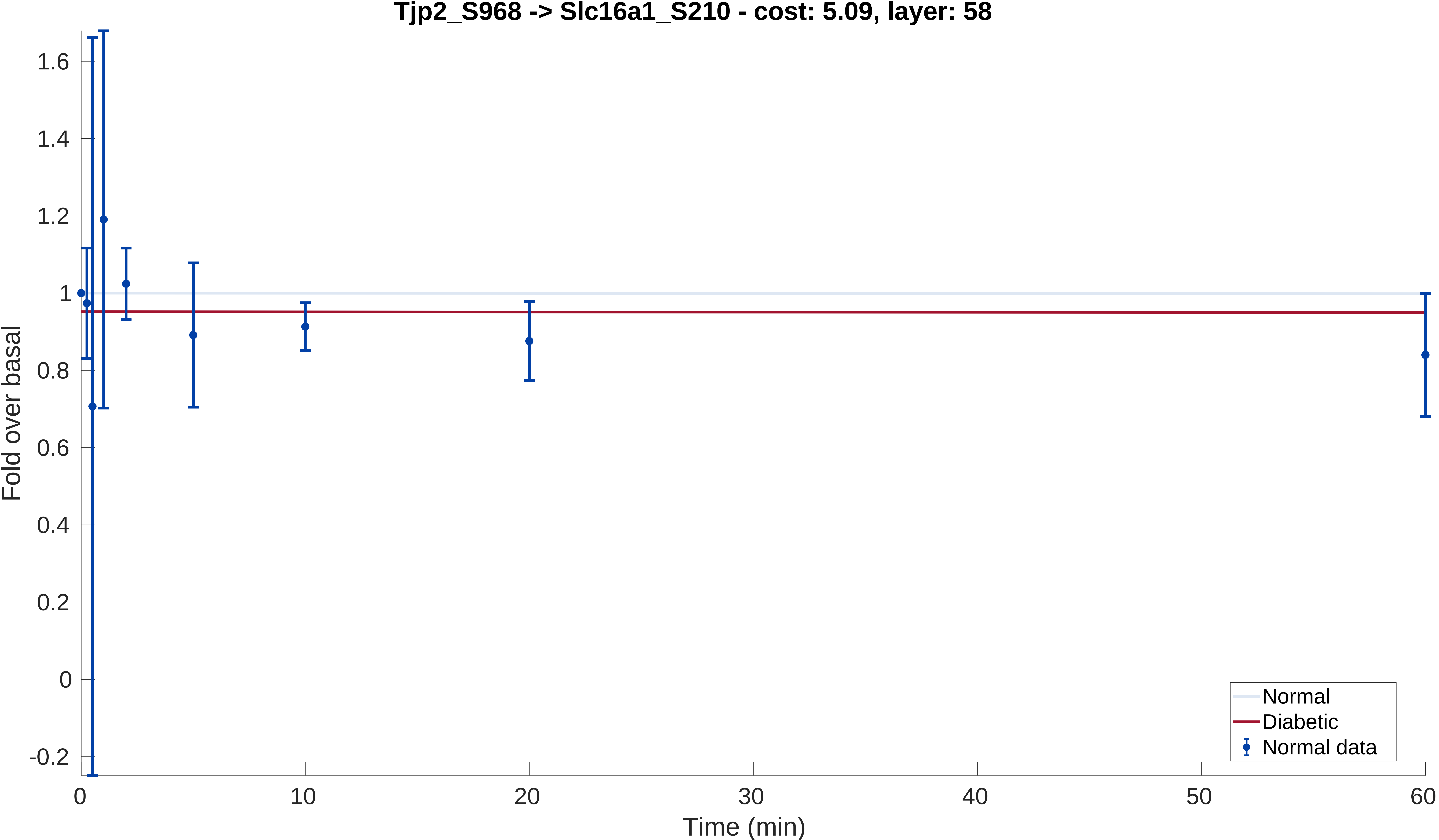

Tjp2_S968 -> Slc16a1_S210 - cost: 5.09, layer: 58
Fold over basal
1.6
1.4
1.2
1
0.8
0.6
0.4
0.2
0
-0.2
0
10
20
30
40
50
60
Time (min)
Normal
Diabetic
Normal data

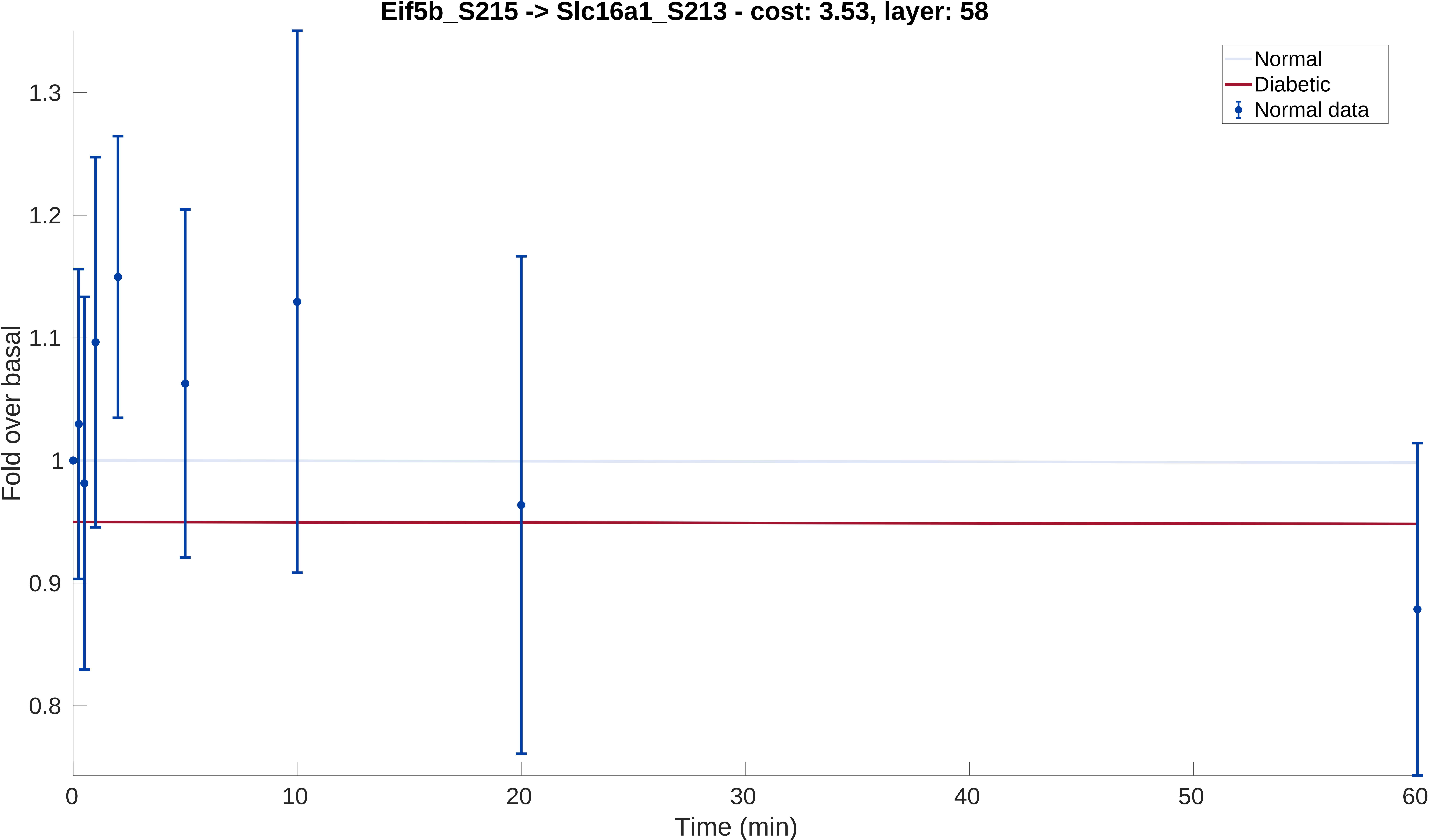
Eif5b_S215 -> Slc16a1_S213 - cost: 3.53, layer: 58
Normal
Diabetic
Normal data
Fold over basal
1.3
1.2
1.1
1
0.9
0.8
0
10
20
30
40
50
60
Time (min)

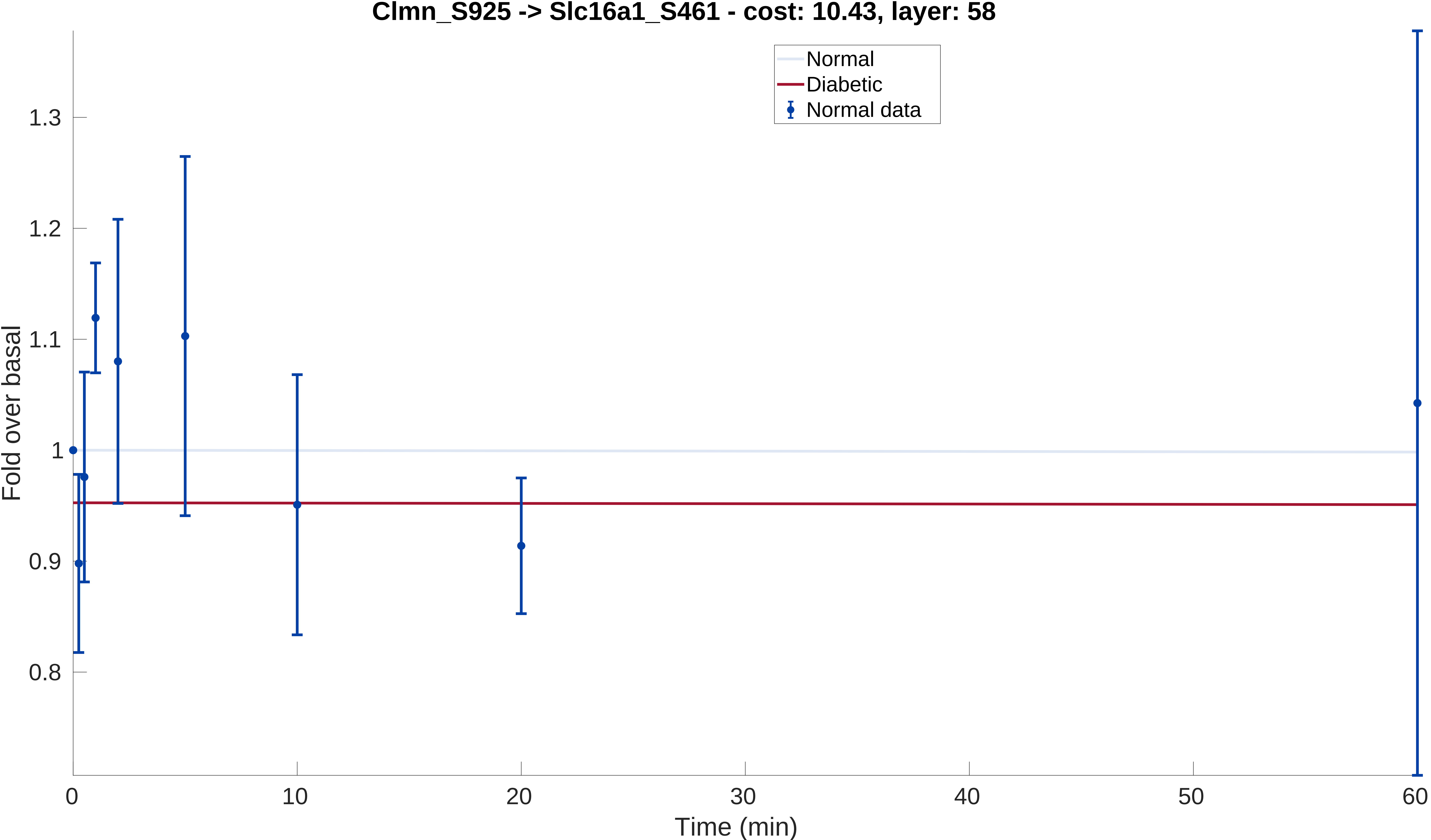

Clmn_S925 -> Slc16a1_S461 - cost: 10.43, layer: 58
Normal
Diabetic
Normal data
Fold over basal
Time (min)
1.3
1.2
1.1
1
0.9
0.8
0
10
20
30
40
50
60

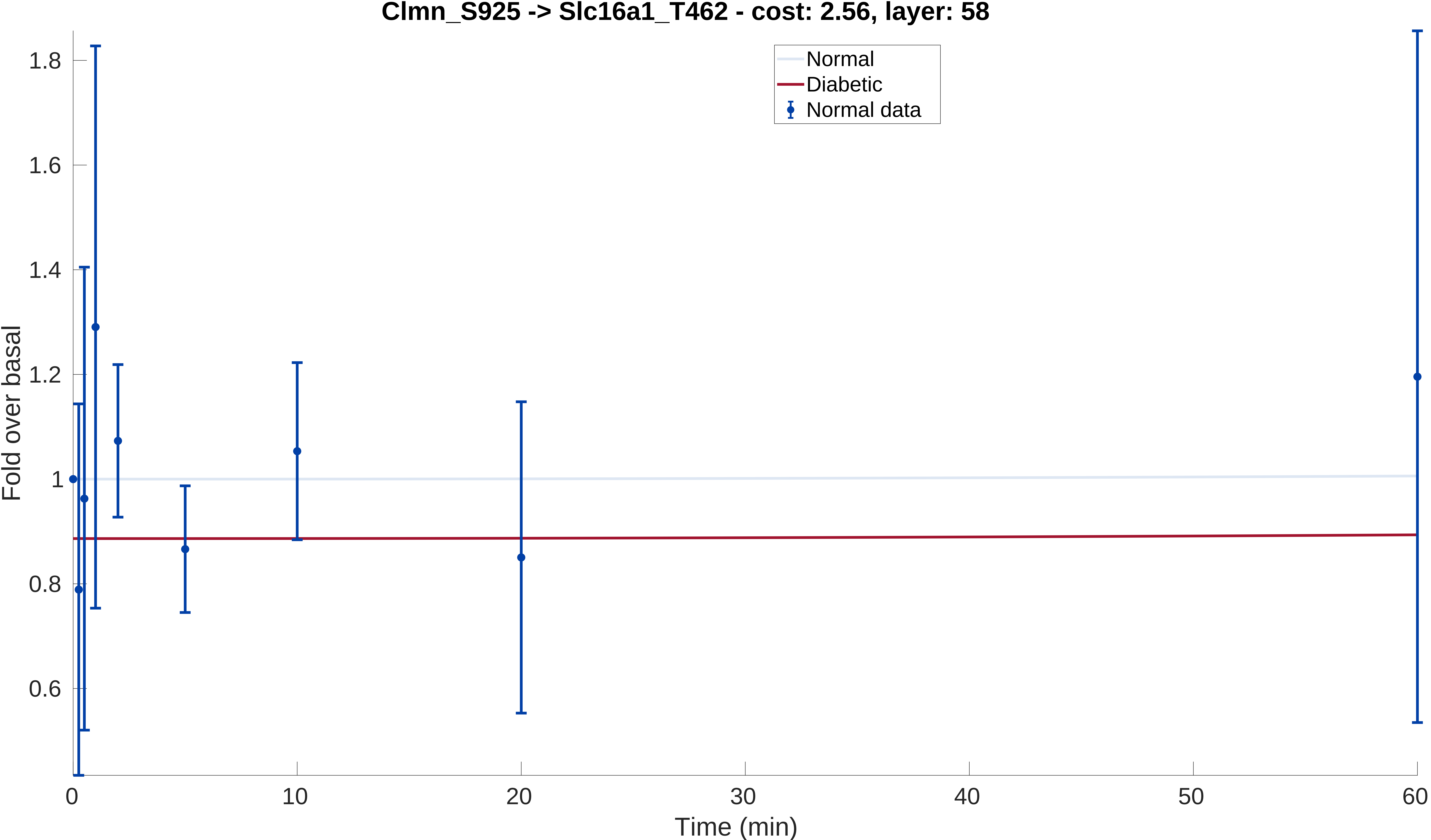

Clmn_S925 -> Slc16a1_T462 - cost: 2.56, layer: 58
Normal
Diabetic
Normal data
Fold over basal
1.8
1.6
1.4
1.2
1
0.8
0.6
0
10
20
30
40
50
60
Time (min)

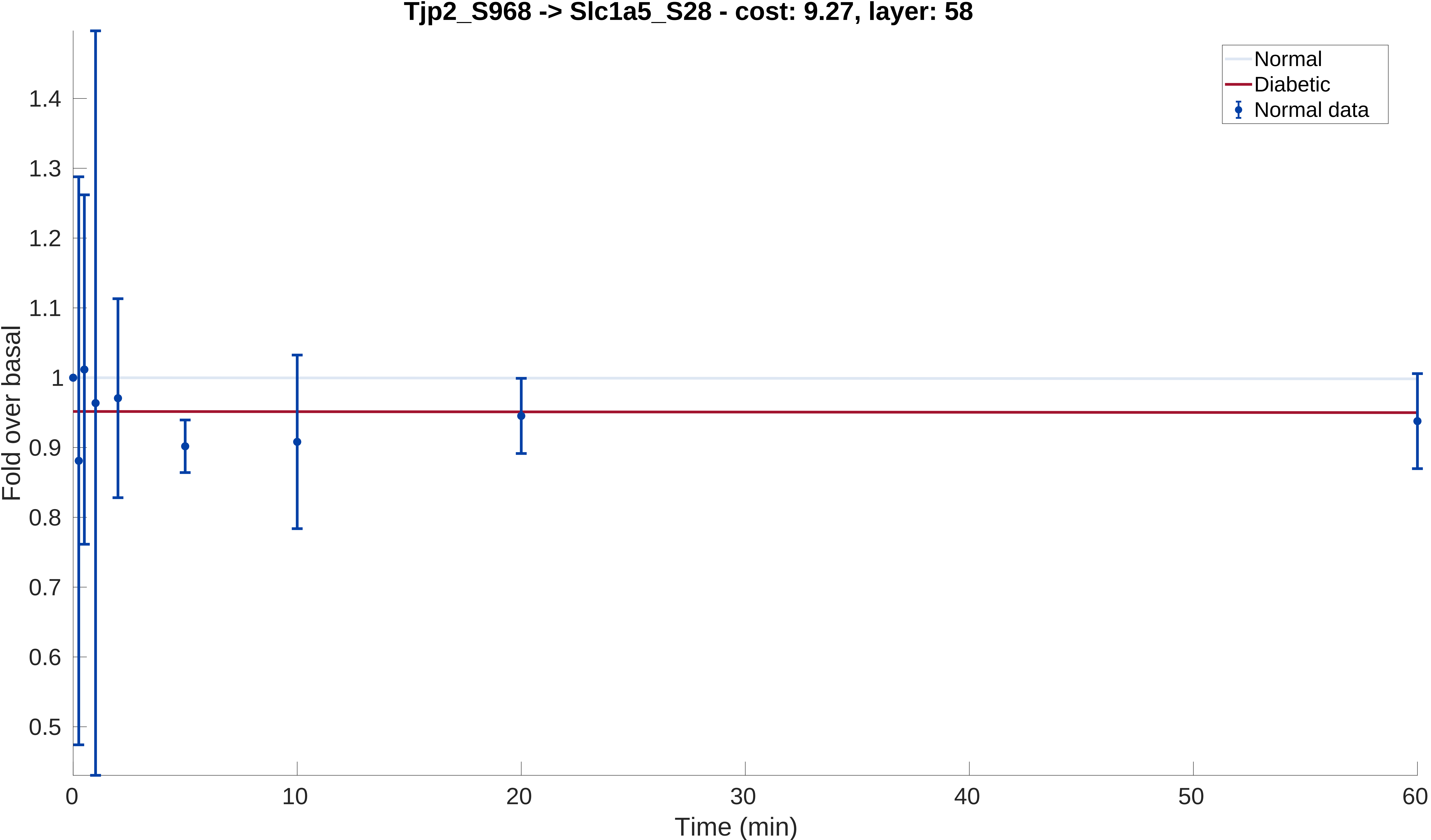
Tjp2_S968 -> Slc1a5_S28 - cost: 9.27, layer: 58
Normal
Diabetic
Normal data
Fold over basal
1.4
1.3
1.2
1.1
1
0.9
0.8
0.7
0.6
0.5
0
10
20
30
40
50
60
Time (min)

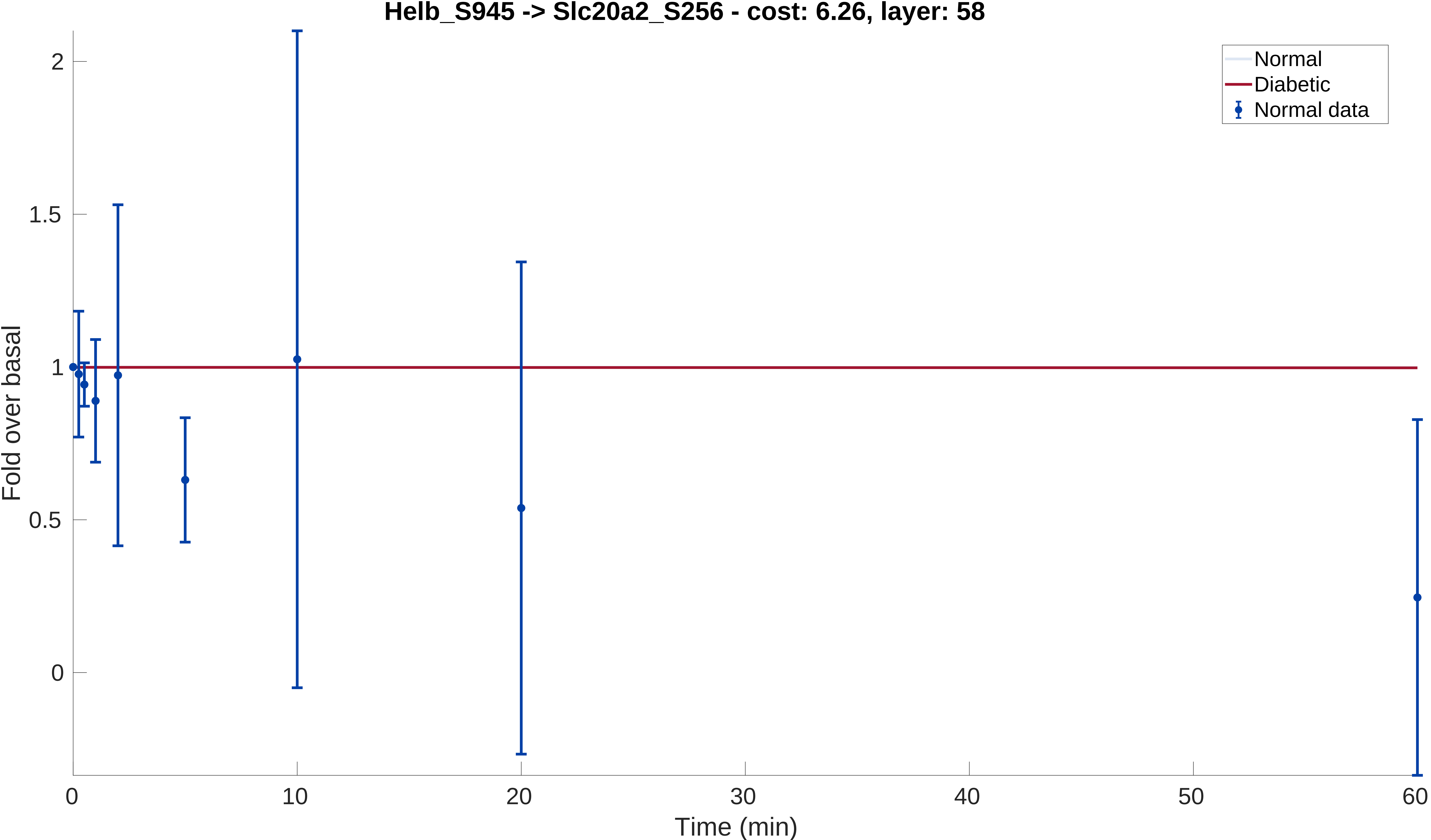

Helb_S945 -> Slc20a2_S256 - cost: 6.26, layer: 58
Normal
Diabetic
Normal data
Fold over basal
Time (min)
0
0.5
1
1.5
2
0
10
20
30
40
50
60

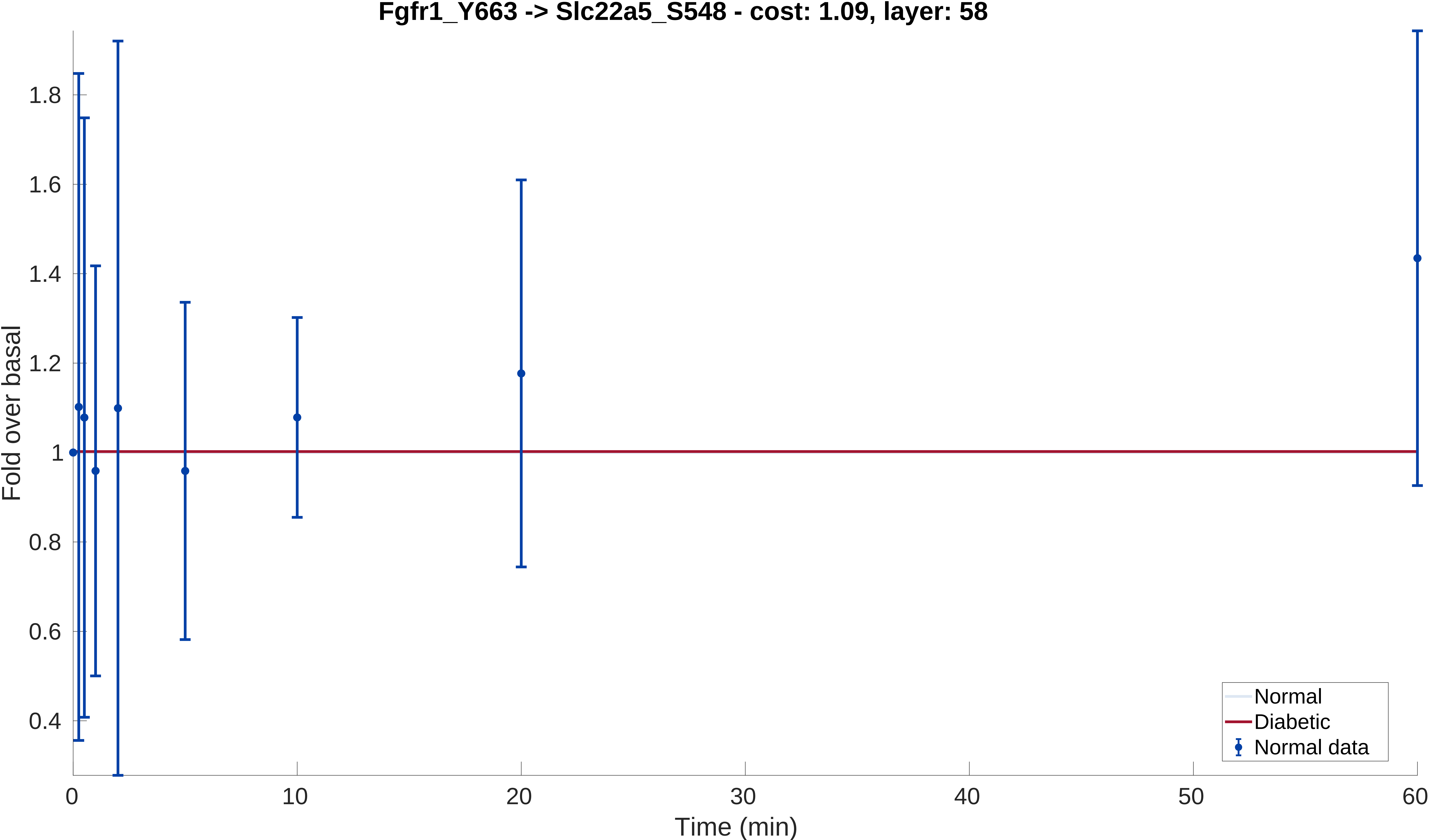

Fgfr1_Y663 -> Slc22a5_S548 - cost: 1.09, layer: 58
Fold over basal
1.8
1.6
1.4
1.2
1
0.8
0.6
0.4
0
10
20
30
40
50
60
Time (min)
Normal
Diabetic
Normal data

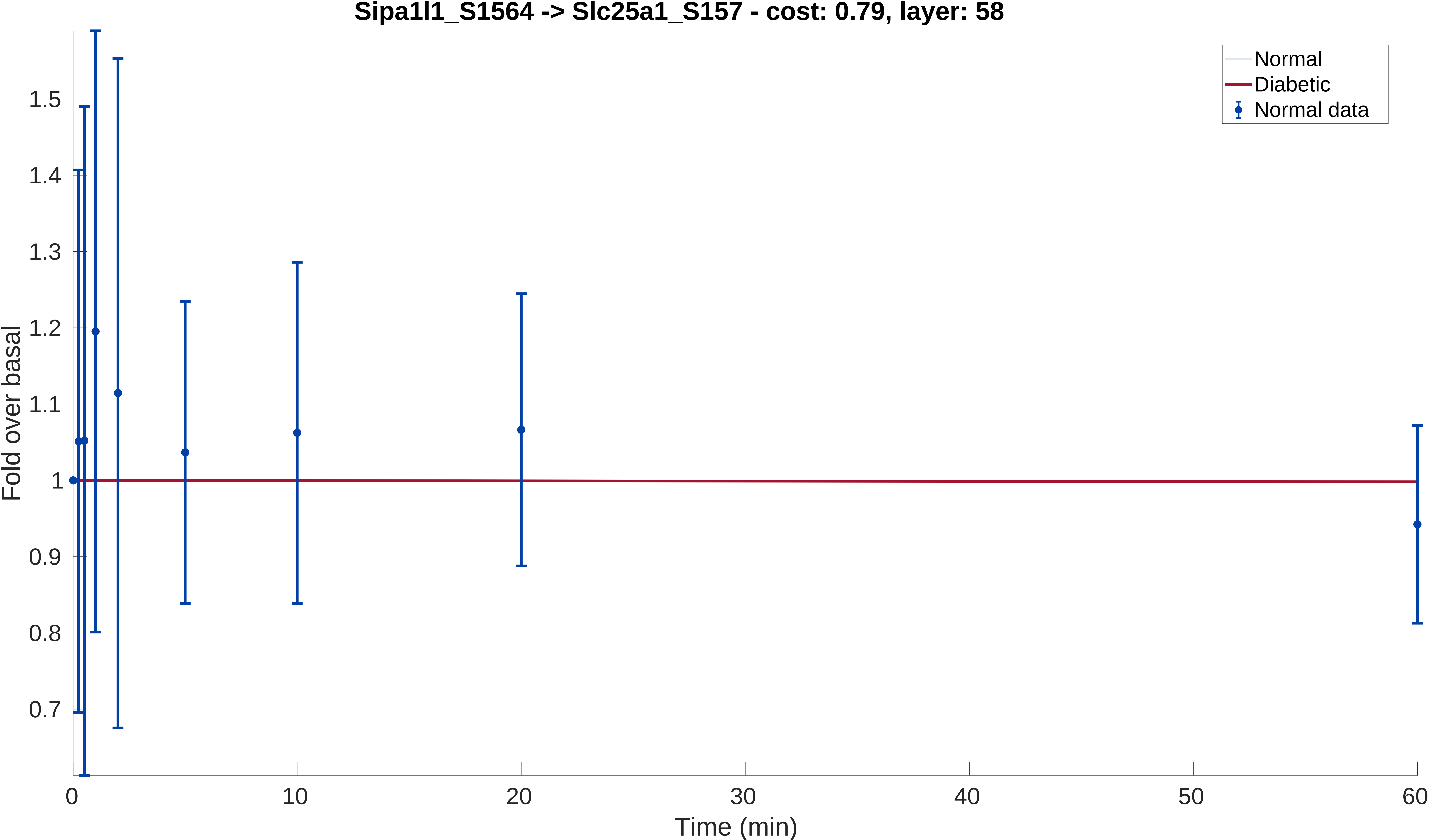

Sipa1l1_S1564 -> Slc25a1_S157 - cost: 0.79, layer: 58
Normal
Diabetic
Normal data
Fold over basal
1.5
1.4
1.3
1.2
1.1
1
0.9
0.8
0.7
0
10
20
30
40
50
60
Time (min)

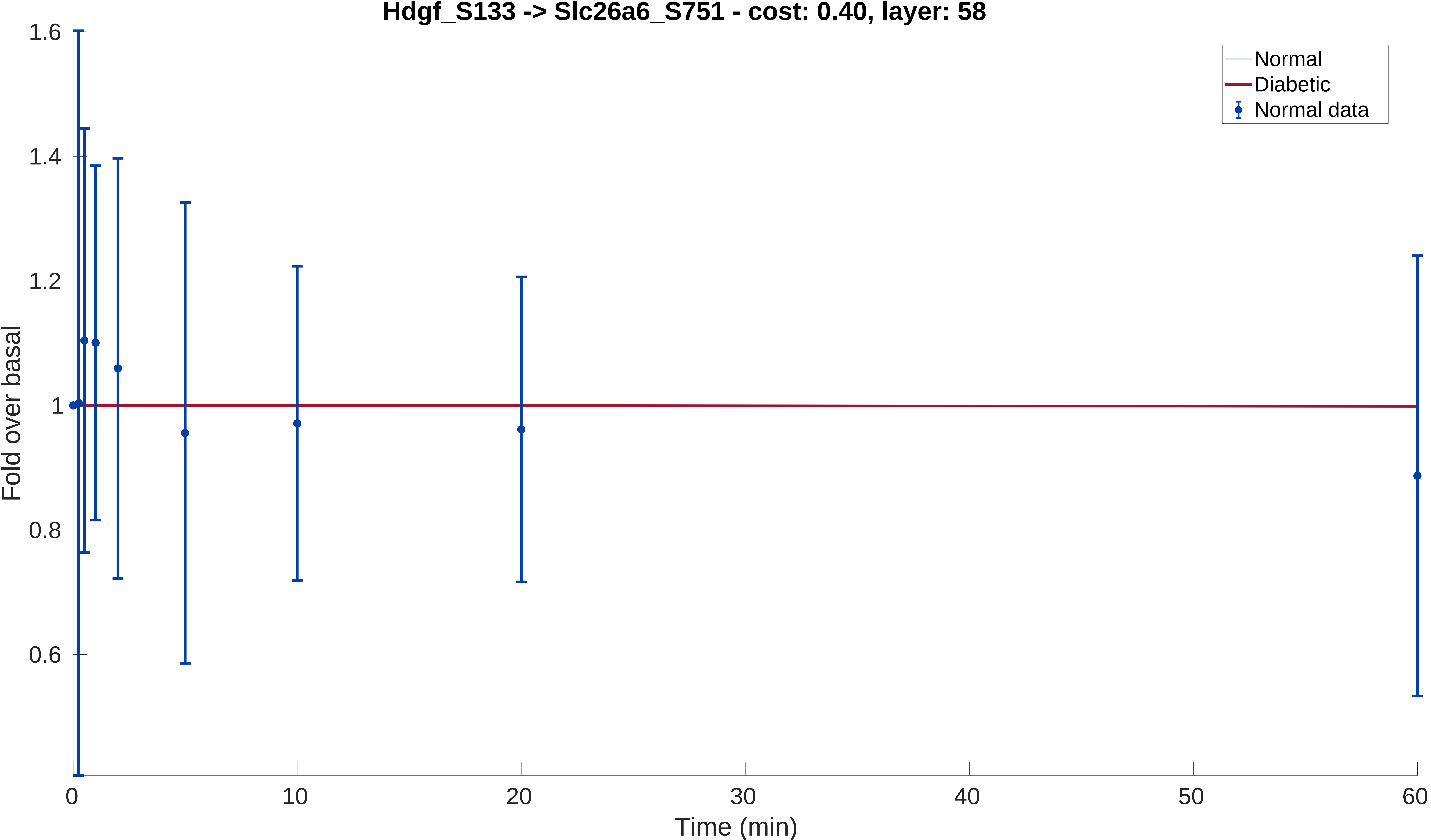

Hdgf_S133 -> Slc26a6_S751 - cost: 0.40, layer: 58
Normal
Diabetic
Normal data
Fold over basal
1.6
1.4
1.2
1
0.8
0.6
0
10
20
30
40
50
60
Time (min)

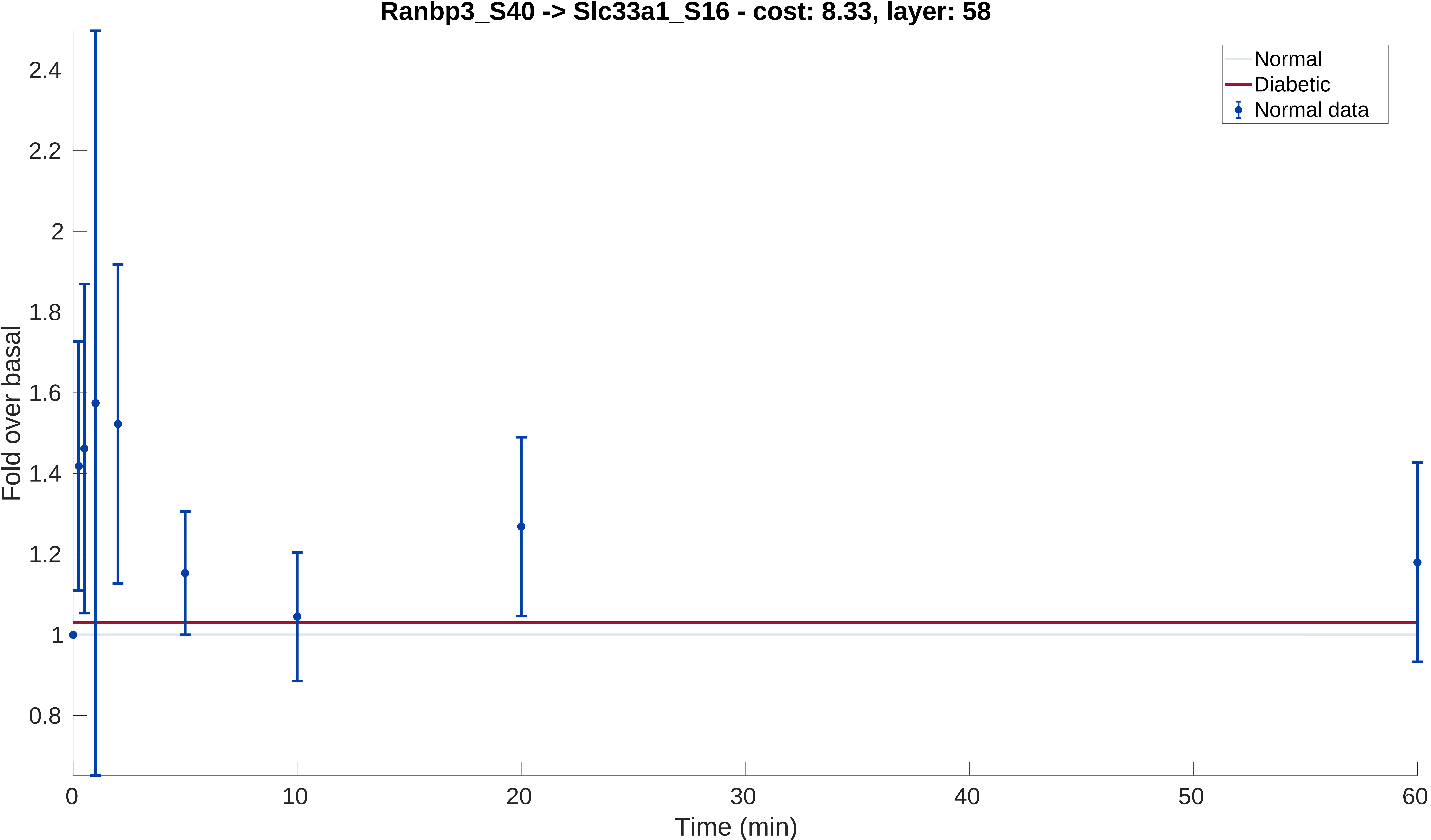

Ranbp3_S40 -> Slc33a1_S16 - cost: 8.33, layer: 58
Normal
Diabetic
Normal data
Fold over basal
Time (min)
2.4
2.2
2
1.8
1.6
1.4
1.2
1
0.8
0
10
20
30
40
50
60

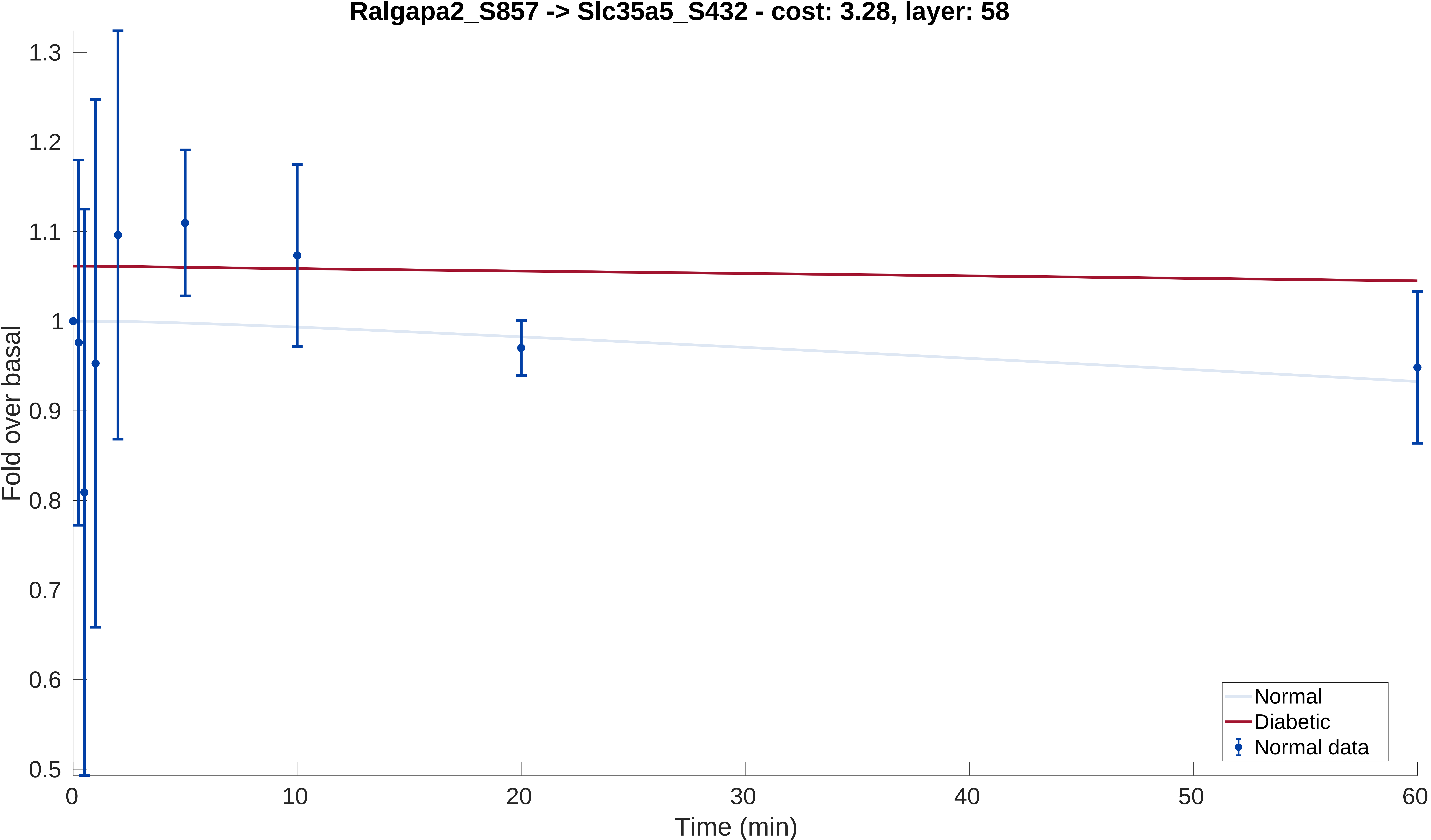
Ralgapa2_S857 -> Slc35a5_S432 - cost: 3.28, layer: 58
1.3
1.2
1.1
1
0.9
0.8
0.7
0.6
0.5
Fold over basal
0
10
20
30
40
50
60
Time (min)
Normal
Diabetic
Normal data

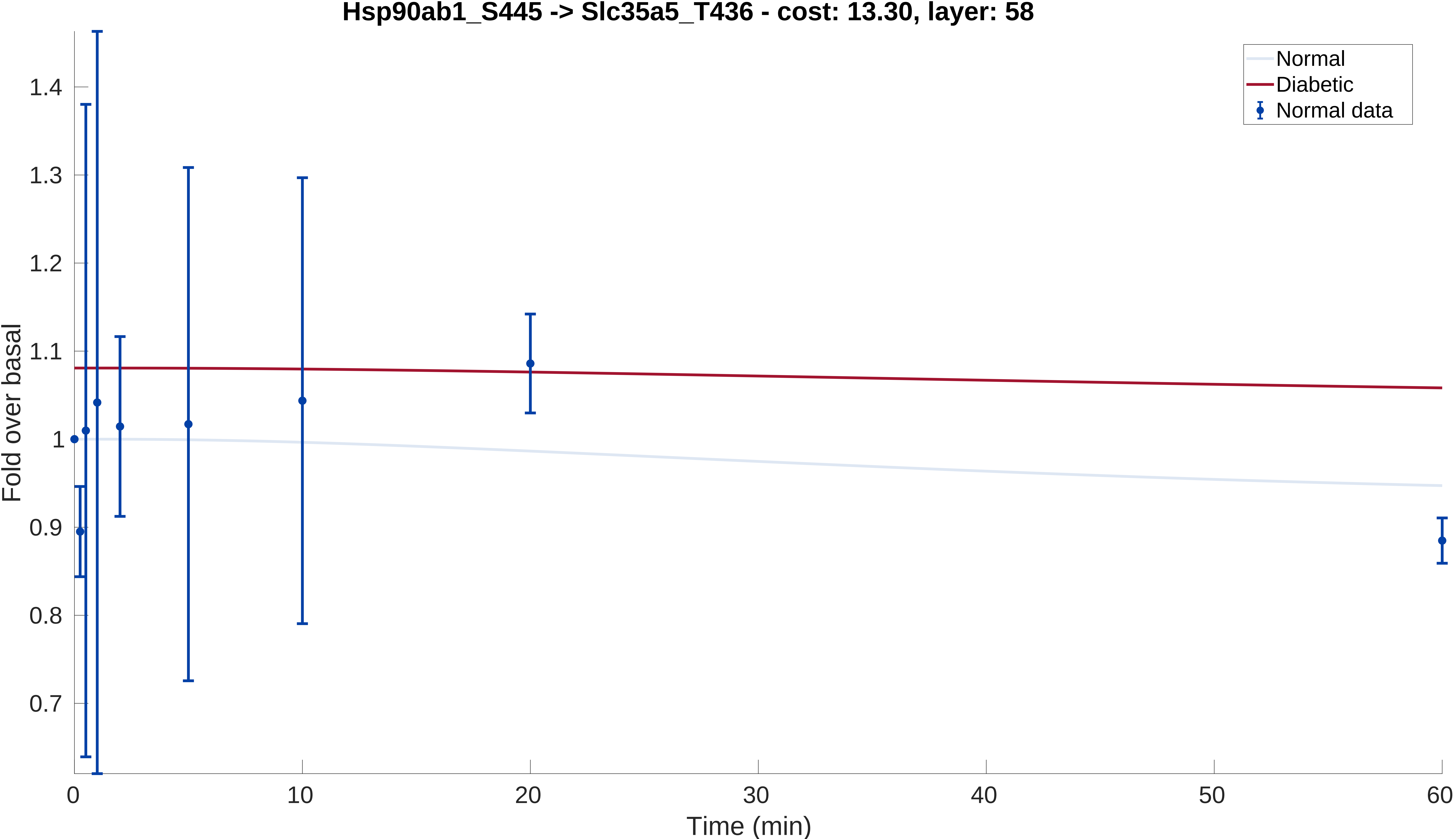

Hsp90ab1_S445 -> Slc35a5_T436 - cost: 13.30, layer: 58
Normal
Diabetic
Normal data
Fold over basal
1.4
1.3
1.2
1.1
1
0.9
0.8
0.7
0
10
20
30
40
50
60
Time (min)

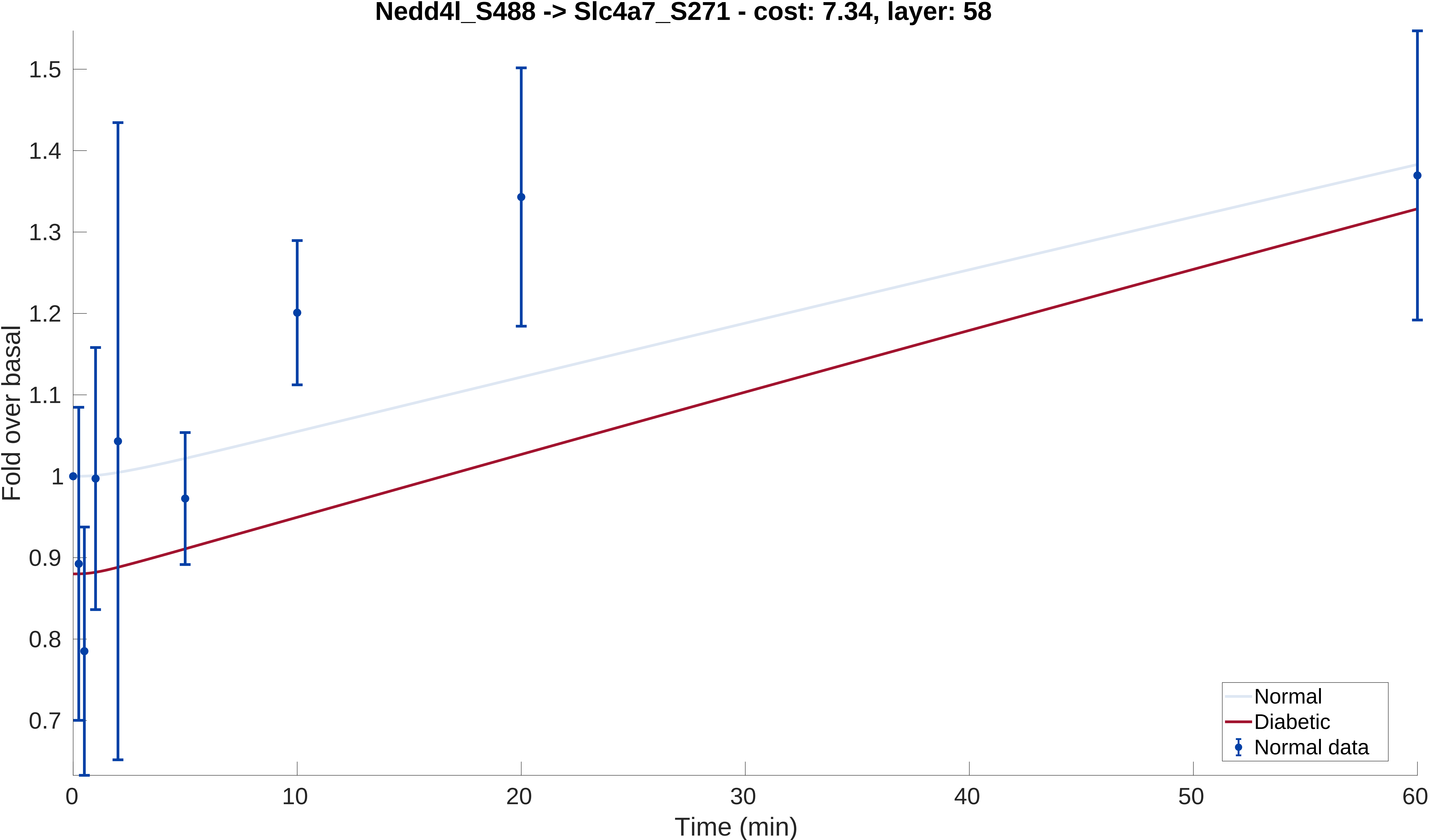
Nedd4l_S488 -> Slc4a7_S271 - cost: 7.34, layer: 58
Fold over basal
1.5
1.4
1.3
1.2
1.1
1
0.9
0.8
0.7
0
10
20
30
40
50
60
Time (min)
Normal
Diabetic
Normal data

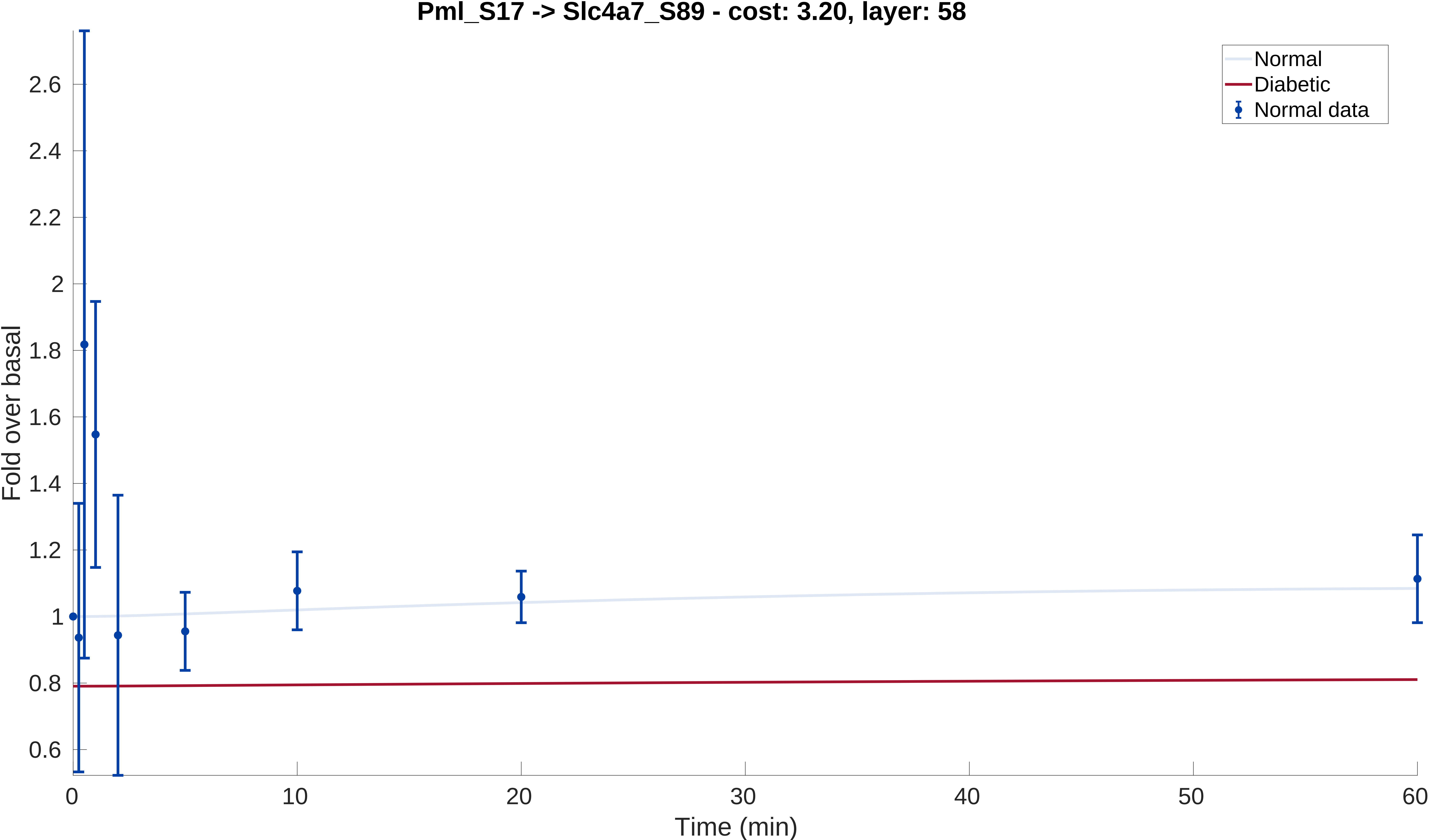
Pml_S17 -> Slc4a7_S89 - cost: 3.20, layer: 58
Normal
Diabetic
Normal data
Fold over basal
2.6
2.4
2.2
2
1.8
1.6
1.4
1.2
1
0.8
0.6
0
10
20
30
40
50
60
Time (min)

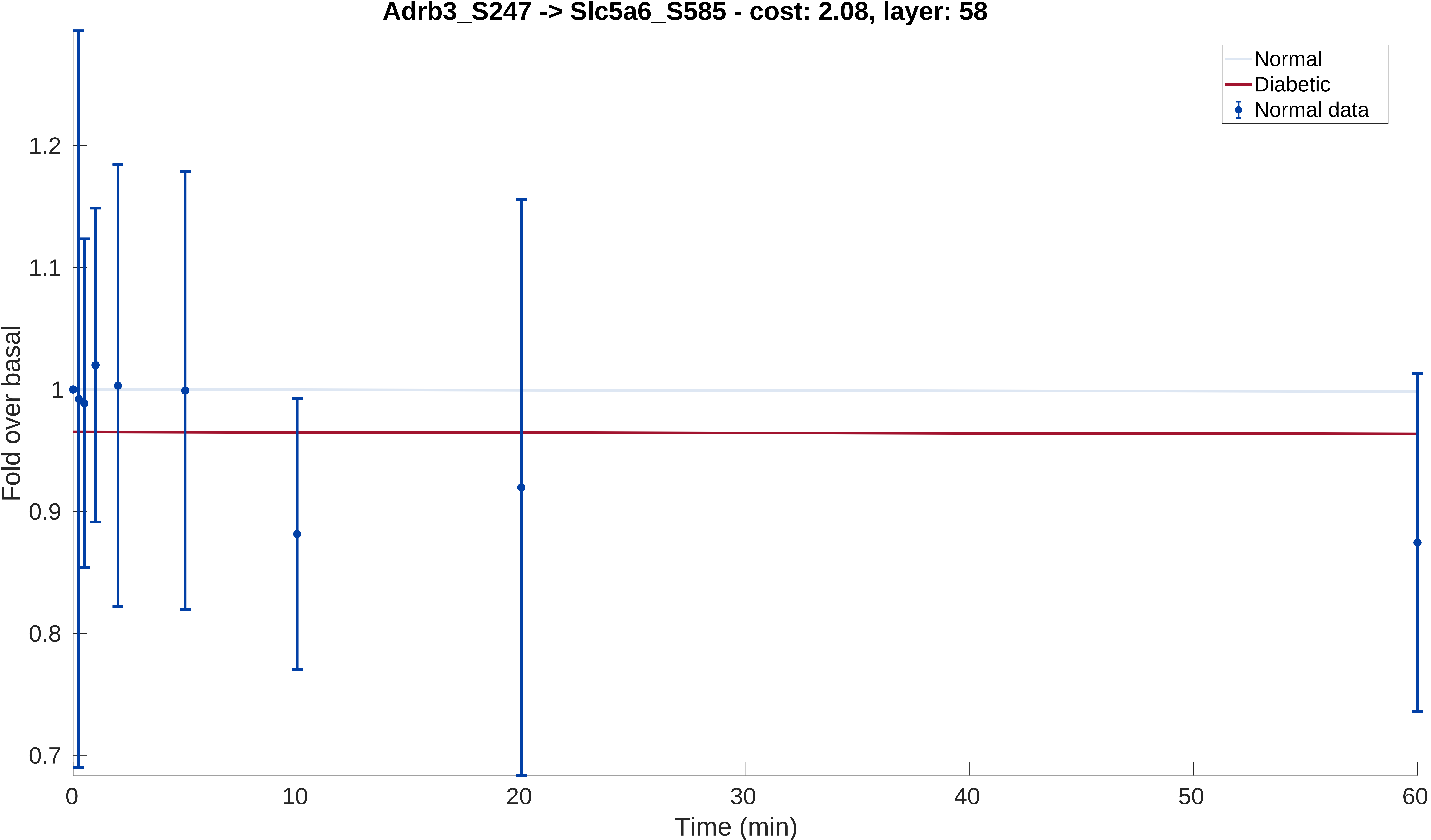
Adrb3_S247 -> Slc5a6_S585 - cost: 2.08, layer: 58
Normal
Diabetic
Normal data
1.2
1.1
1
0.9
0.8
0.7
Fold over basal
0
10
20
30
40
50
60
Time (min)

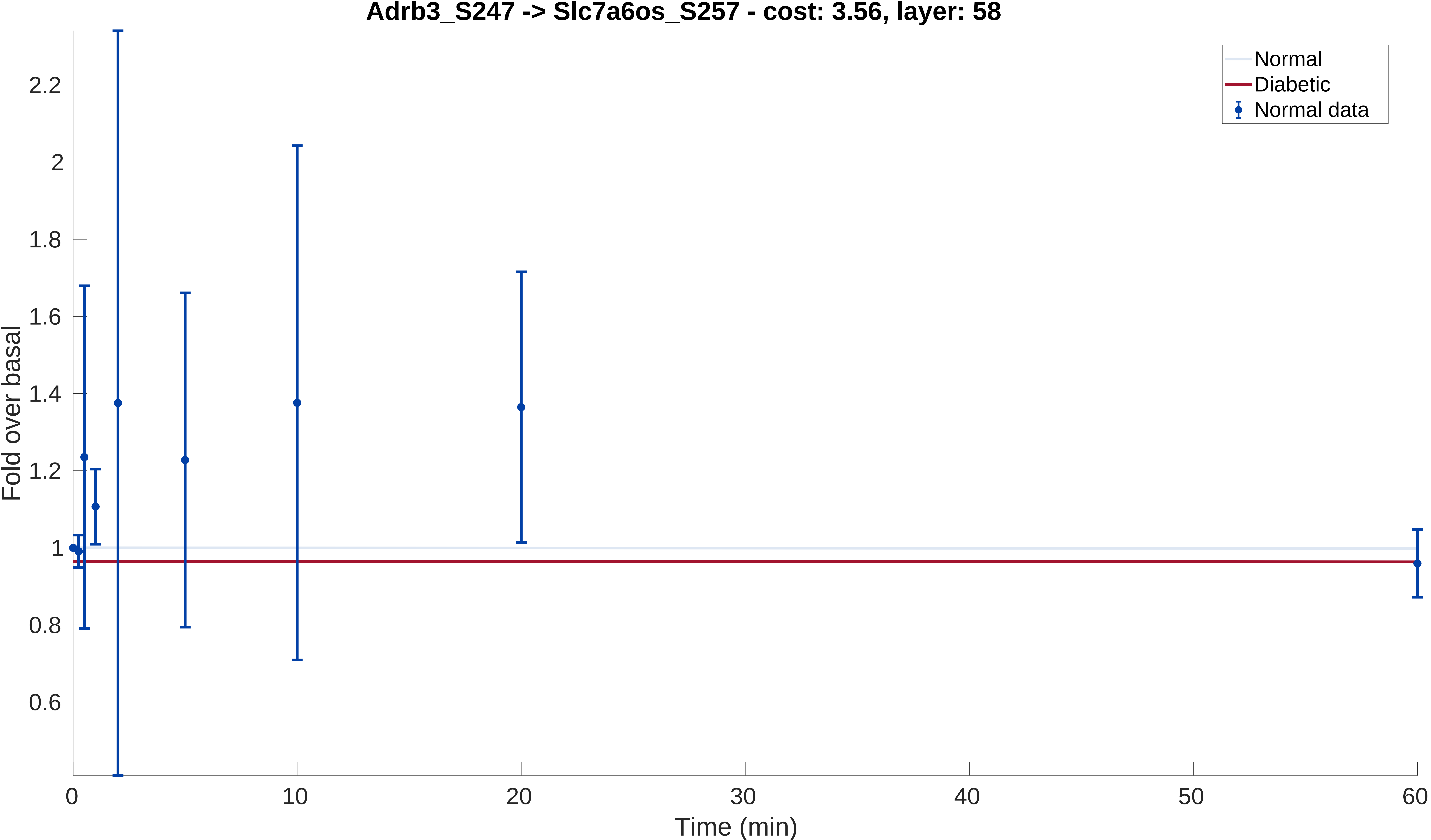
Adrb3_S247 -> Slc7a6os_S257 - cost: 3.56, layer: 58
Normal
Diabetic
Normal data
Fold over basal
2.2
2
1.8
1.6
1.4
1.2
1
0.8
0.6
0
10
20
30
40
50
60
Time (min)

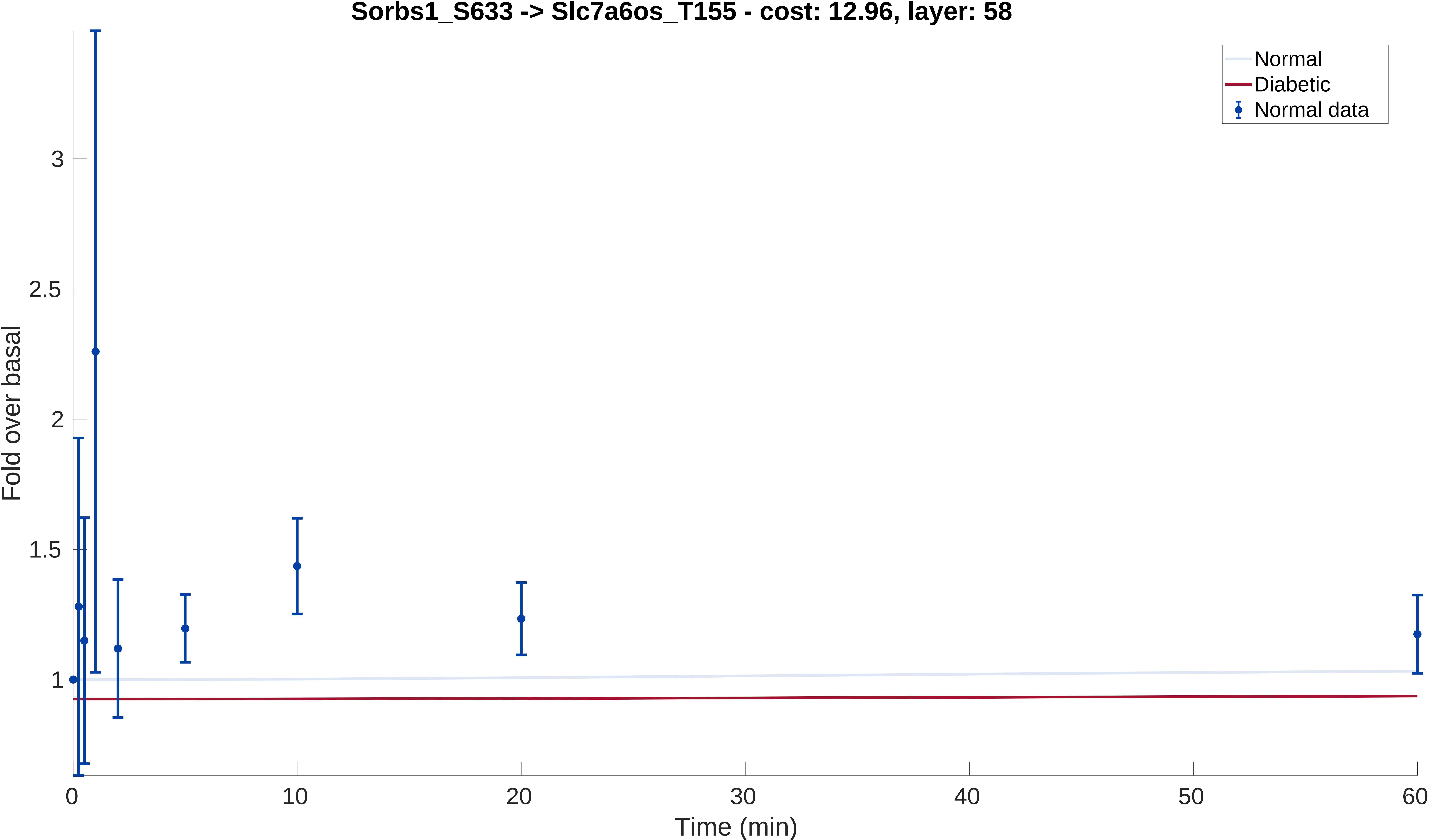

Sorbs1_S633 -> Slc7a6os_T155 - cost: 12.96, layer: 58
Normal
Diabetic
Normal data
Fold over basal
3
2.5
2
1.5
1
0
10
20
30
40
50
60
Time (min)

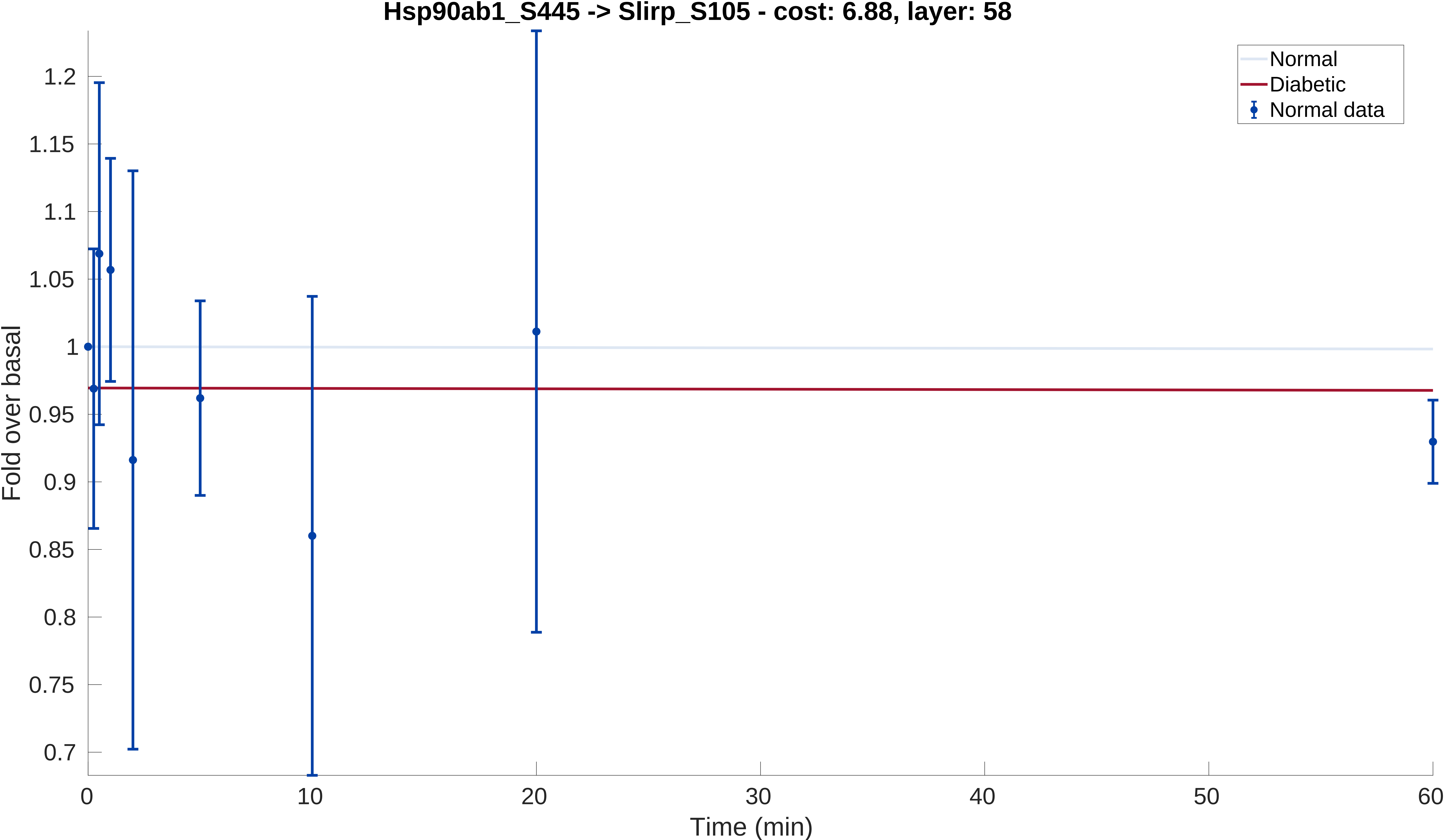

Hsp90ab1_S445 -> Slirp_S105 - cost: 6.88, layer: 58
Normal
Diabetic
Normal data
Fold over basal
1.2
1.15
1.1
1.05
1
0.95
0.9
0.85
0.8
0.75
0.7
0
10
20
30
40
50
60
Time (min)

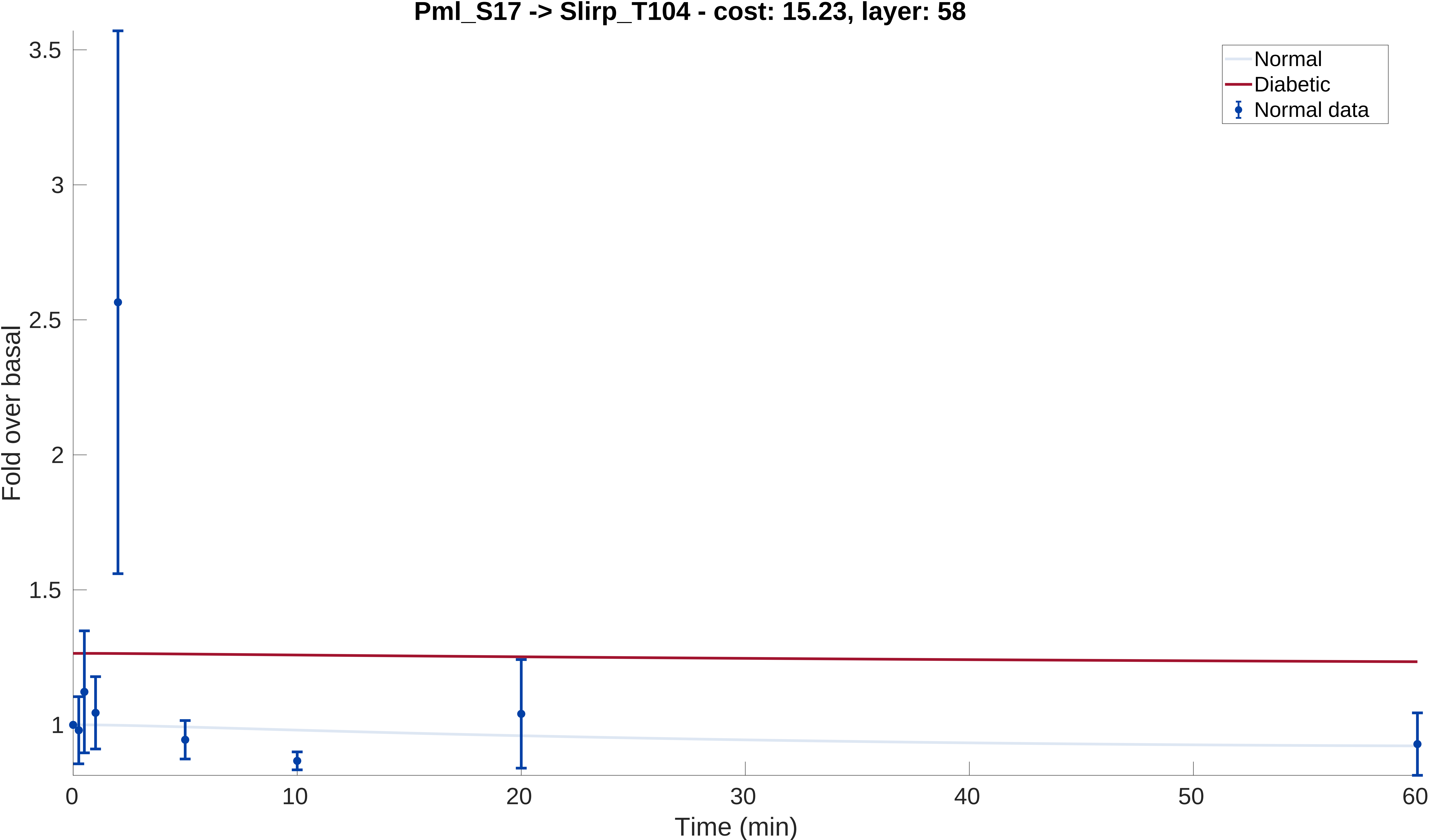
Pml_S17 -> Slirp_T104 - cost: 15.23, layer: 58
Normal
Diabetic
Normal data
Fold over basal
3.5
3
2.5
2
1.5
1
0
10
20
30
40
50
60
Time (min)

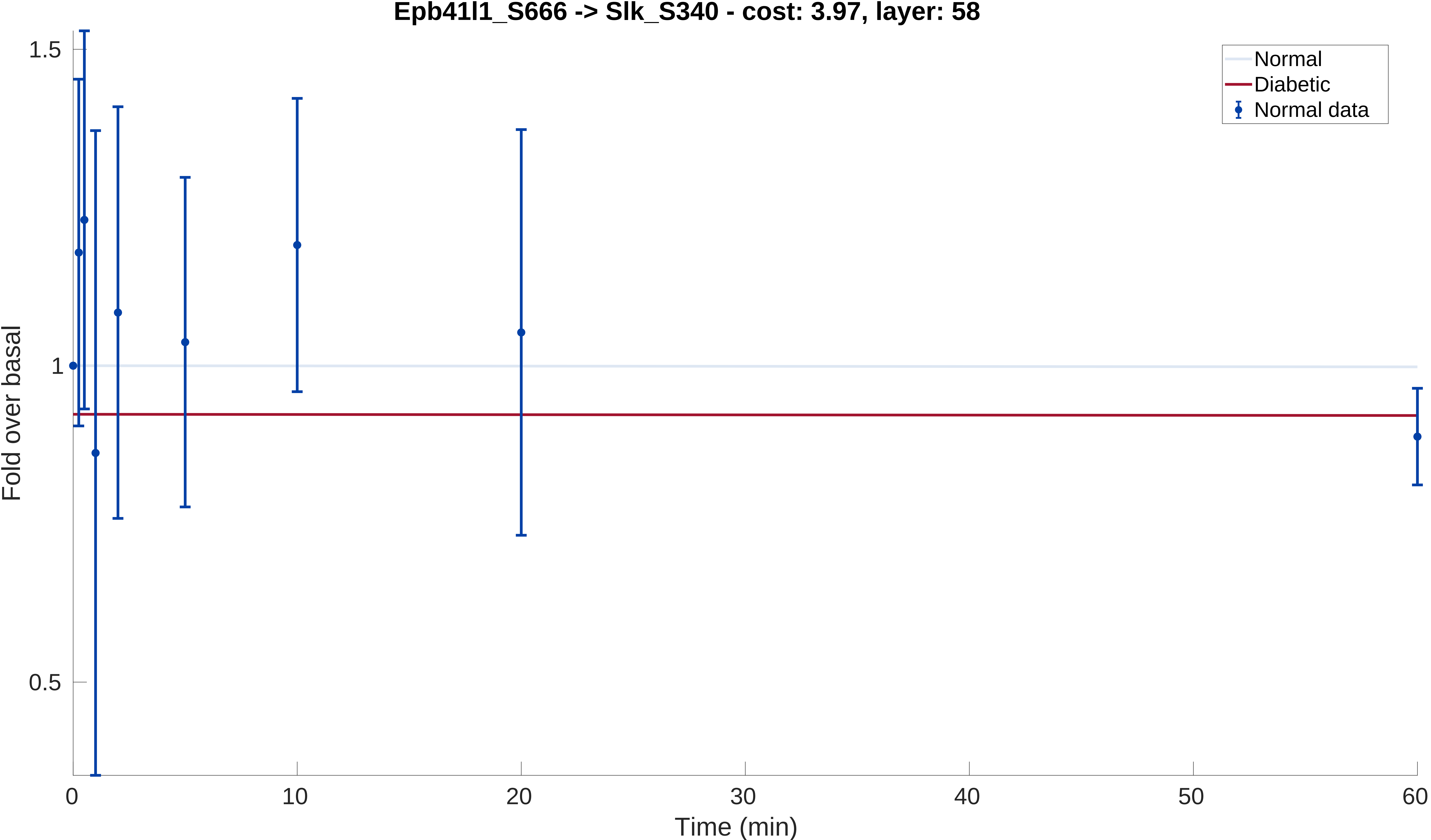

Epb41l1_S666 -> Slk_S340 - cost: 3.97, layer: 58
Normal
Diabetic
Normal data
Fold over basal
1.5
1
0.5
0
10
20
30
40
50
60
Time (min)

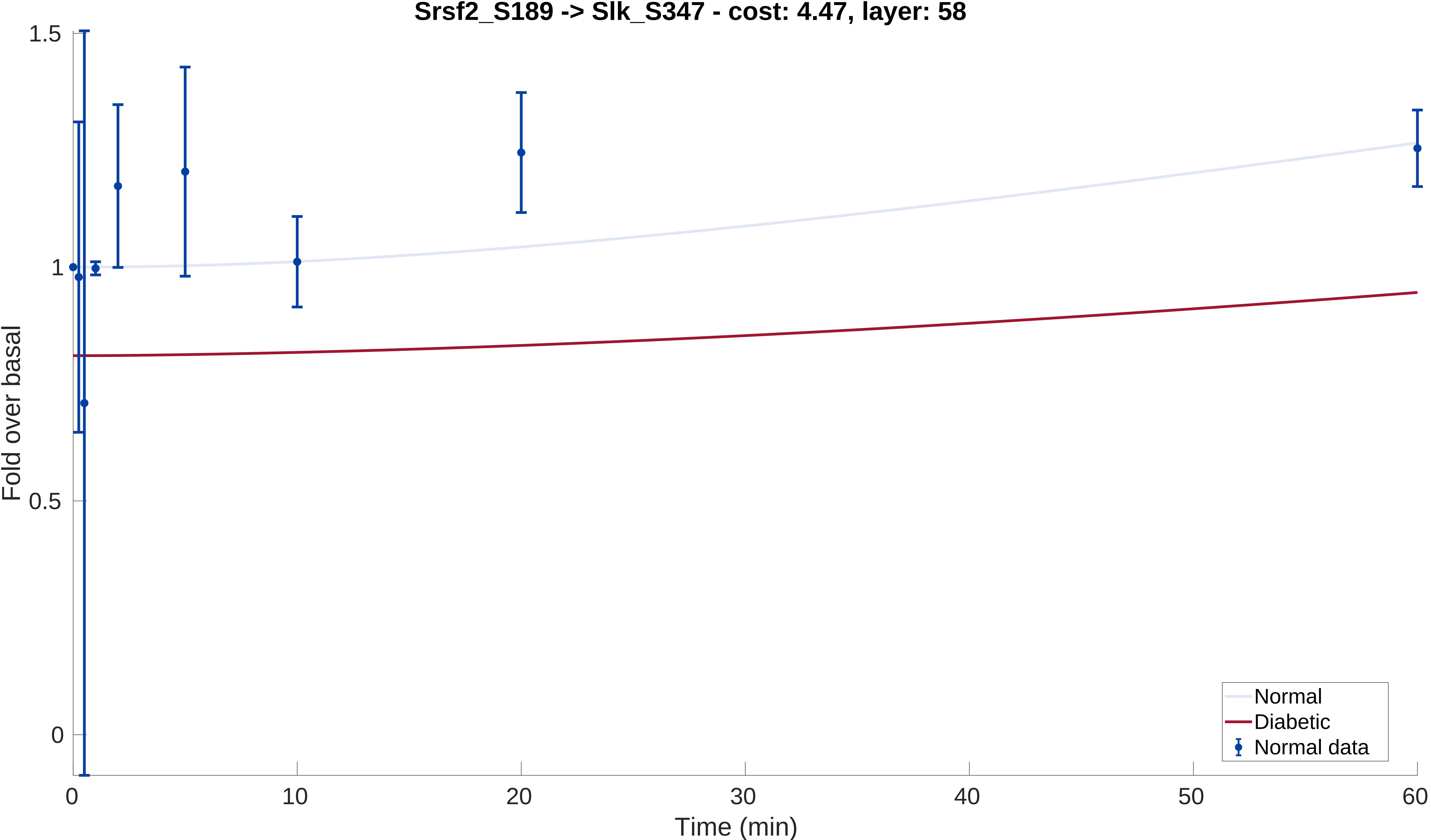

Srsf2_S189 -> Slk_S347 - cost: 4.47, layer: 58
1.5
1
0.5
0
Fold over basal
0
10
20
30
40
50
60
Time (min)
Normal
Diabetic
Normal data

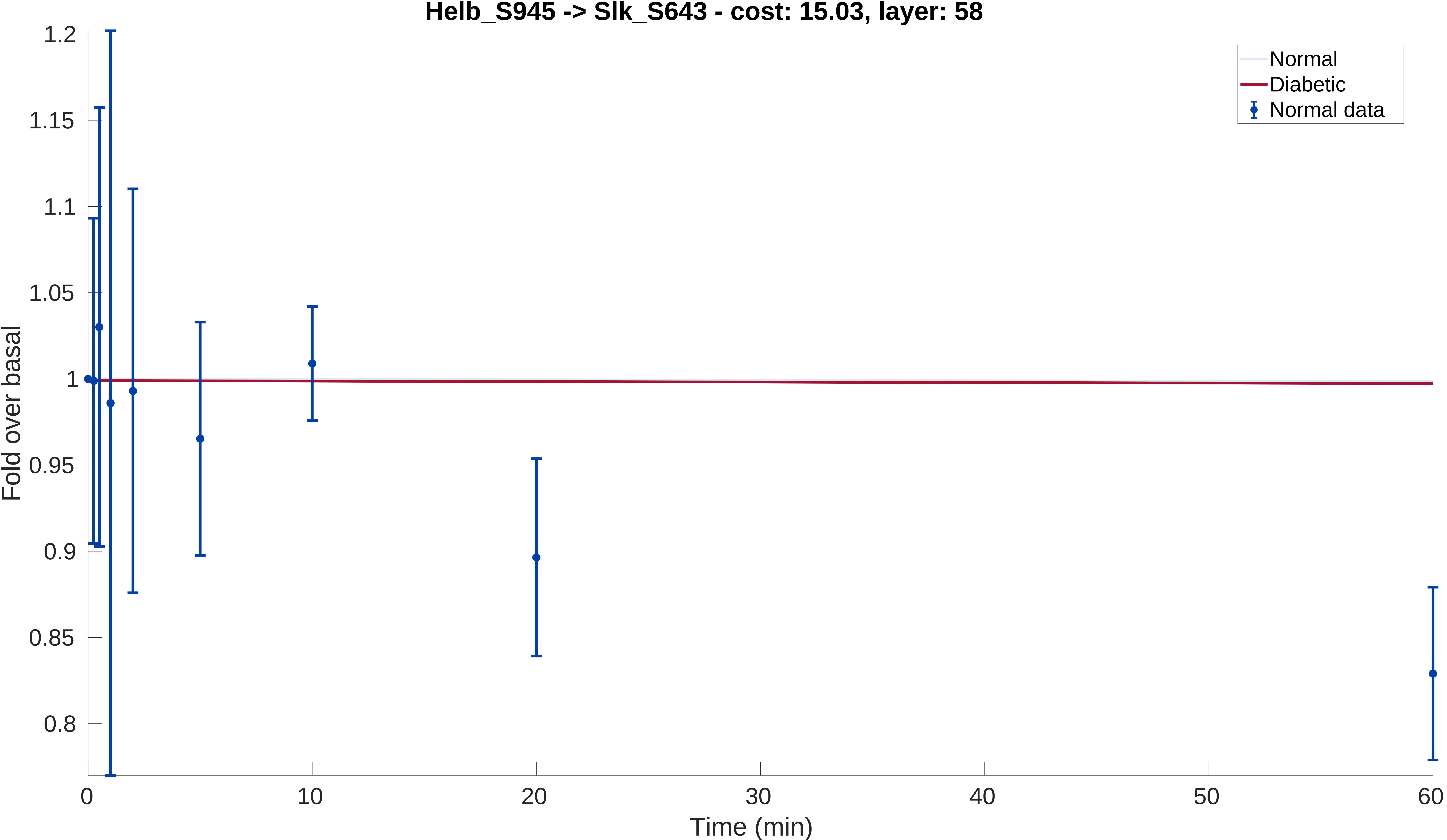

Helb_S945 -> Slk_S643 - cost: 15.03, layer: 58
Normal
Diabetic
Normal data
Fold over basal
1.2
1.15
1.1
1.05
1
0.95
0.9
0.85
0.8
0
10
20
30
40
50
60
Time (min)

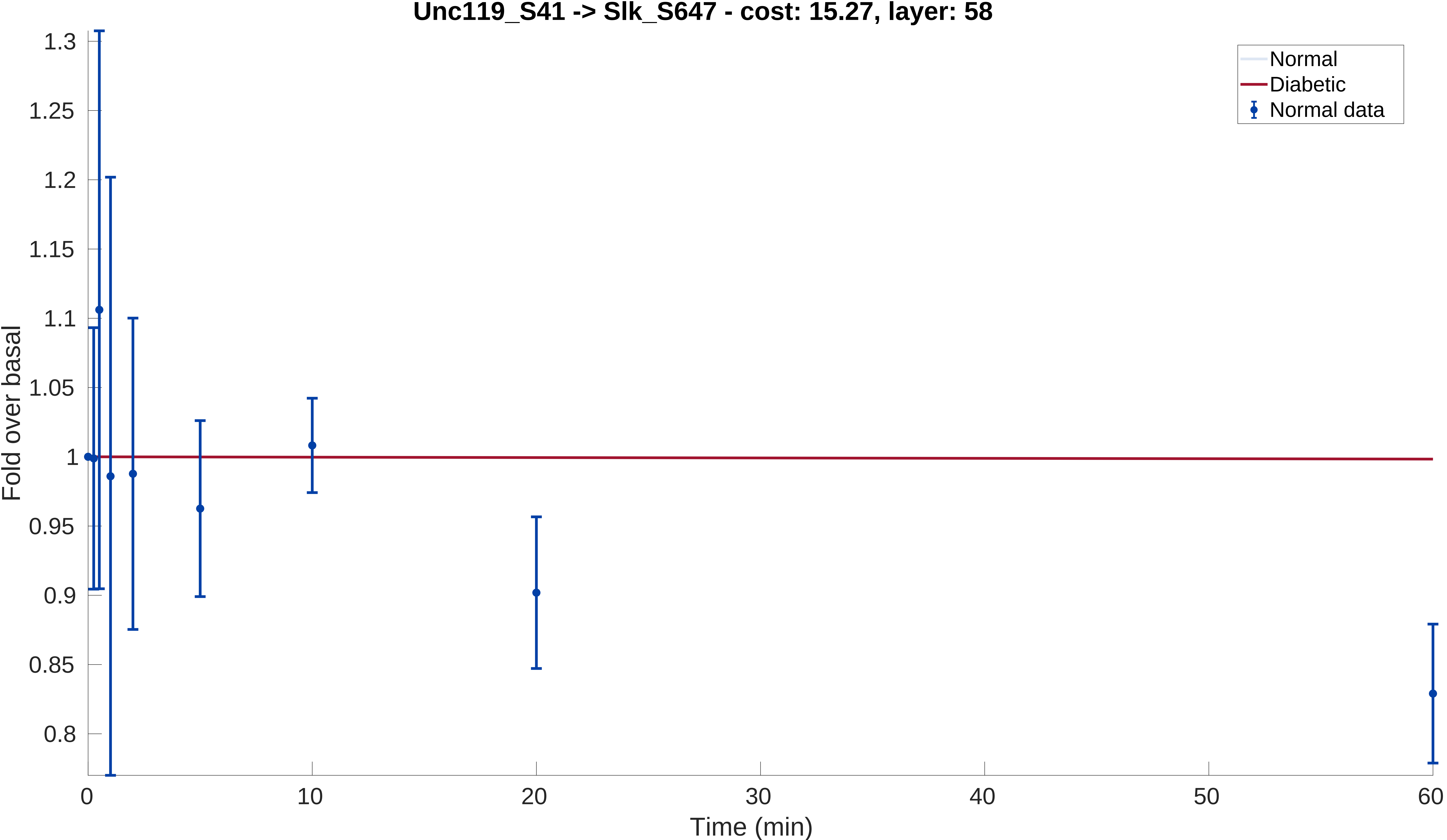
Unc119_S41 -> Slk_S647 - cost: 15.27, layer: 58
Normal
Diabetic
Normal data
Fold over basal
1.3
1.25
1.2
1.15
1.1
1.05
1
0.95
0.9
0.85
0.8
0
10
20
30
40
50
60
Time (min)

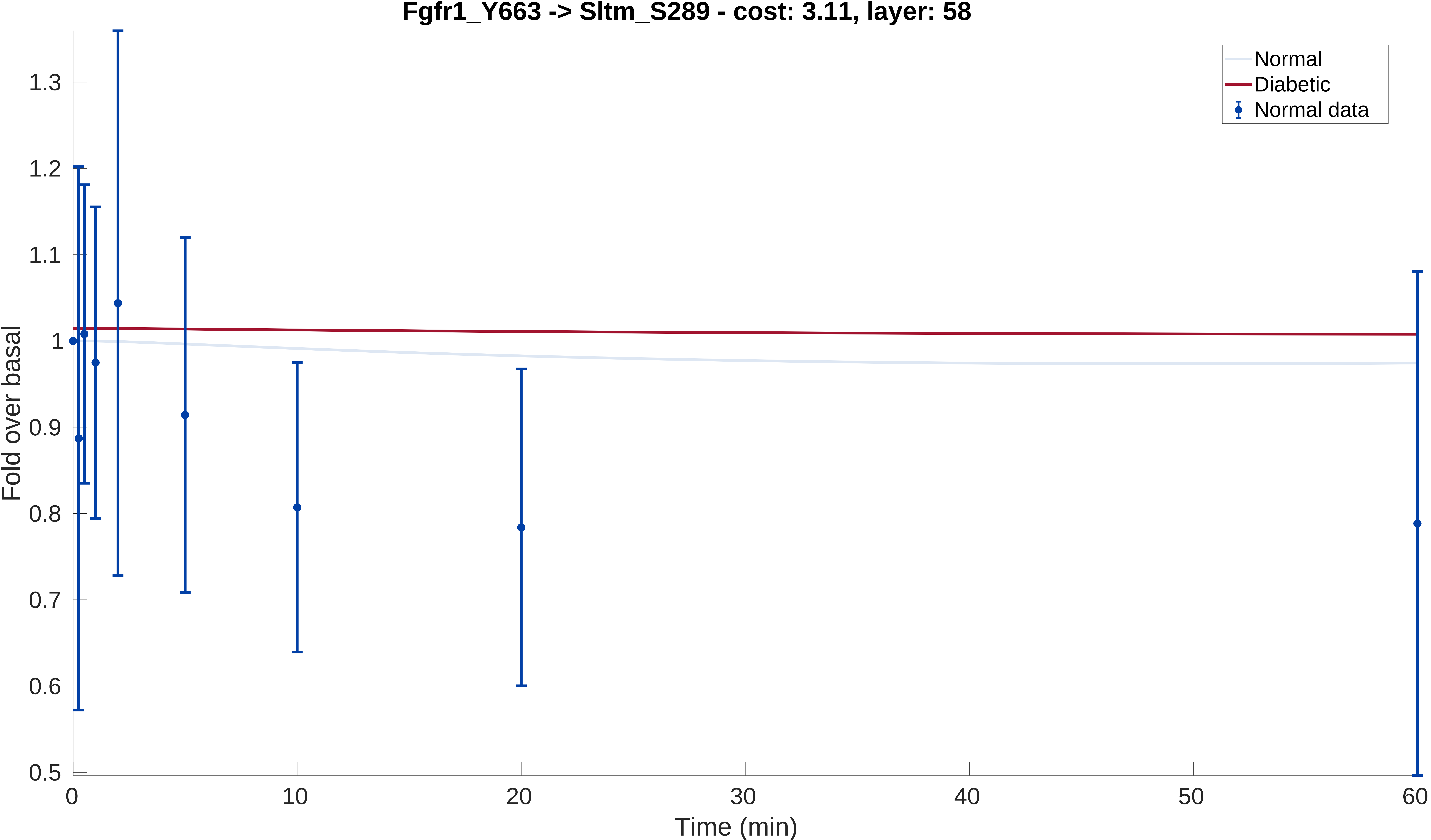

Fgfr1_Y663 -> Sltm_S289 - cost: 3.11, layer: 58
Normal
Diabetic
Normal data
Fold over basal
Time (min)
1.3
1.2
1.1
1
0.9
0.8
0.7
0.6
0.5
0
10
20
30
40
50
60

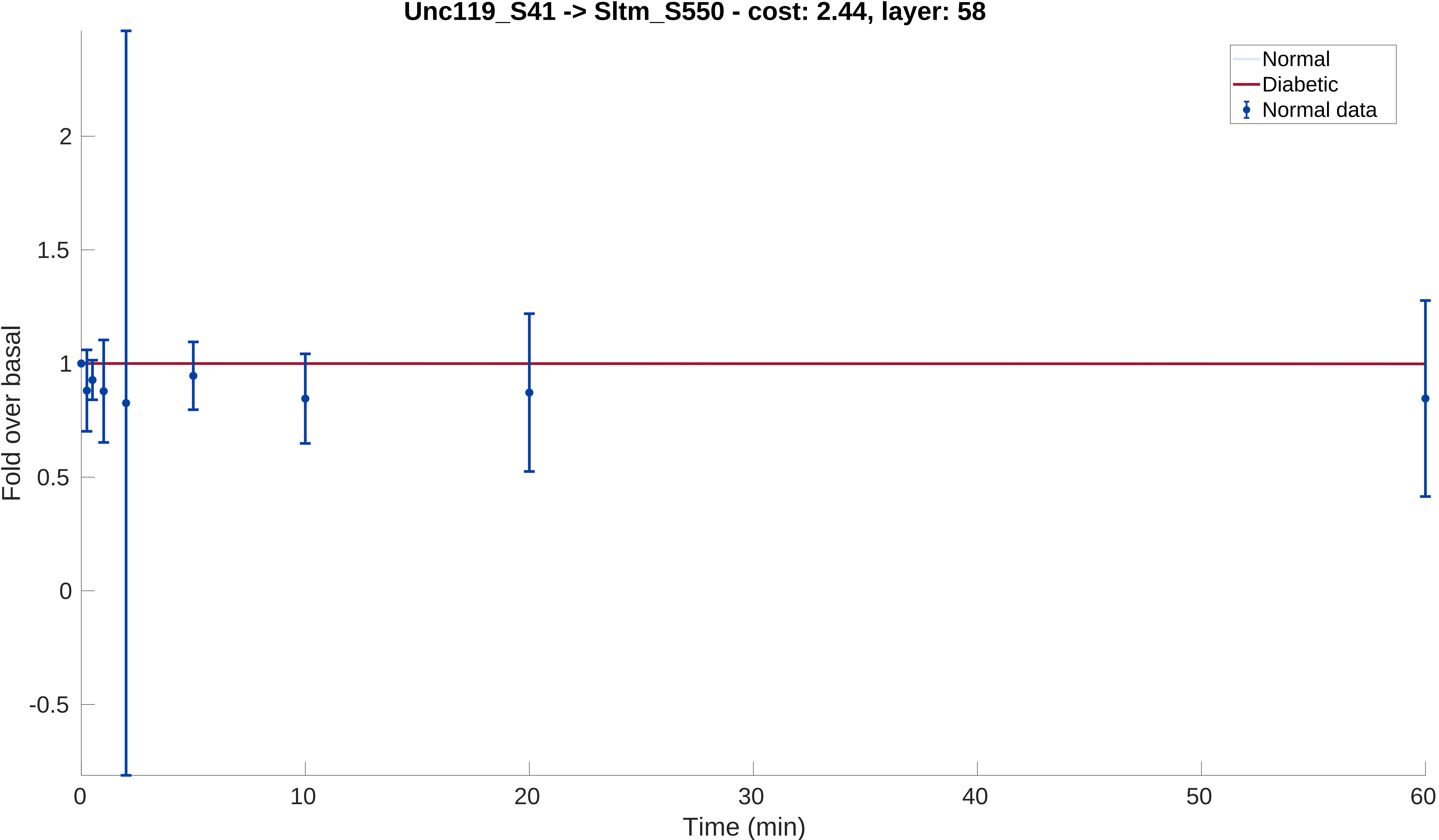

Unc119_S41 -> Sltm_S550 - cost: 2.44, layer: 58
Normal
Diabetic
Normal data
Fold over basal
2
1.5
1
0.5
0
-0.5
0
10
20
30
40
50
60
Time (min)

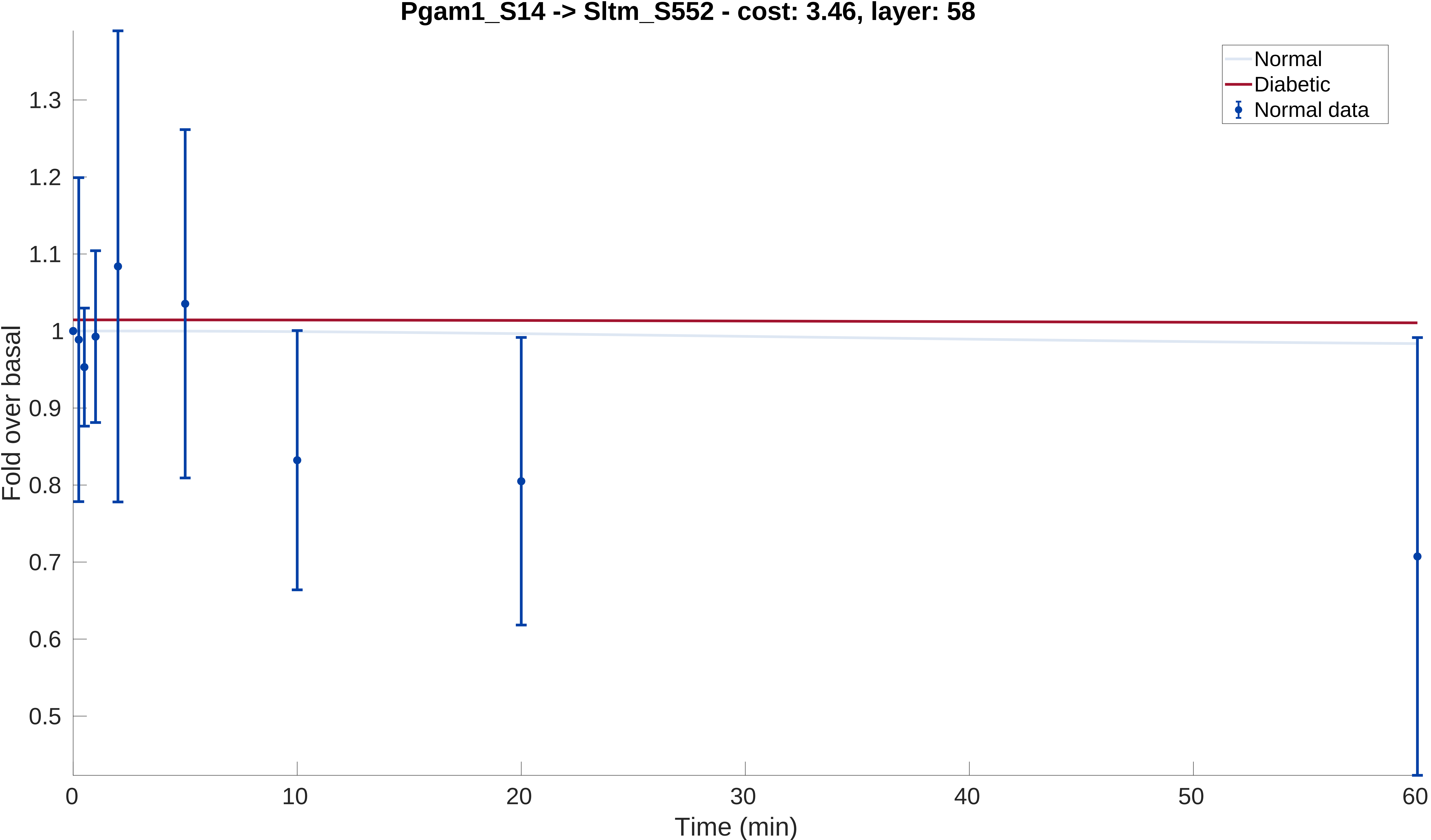

Pgam1_S14 -> Sltm_S552 - cost: 3.46, layer: 58
Normal
Diabetic
Normal data
Fold over basal
1.3
1.2
1.1
1
0.9
0.8
0.7
0.6
0.5
0
10
20
30
40
50
60
Time (min)

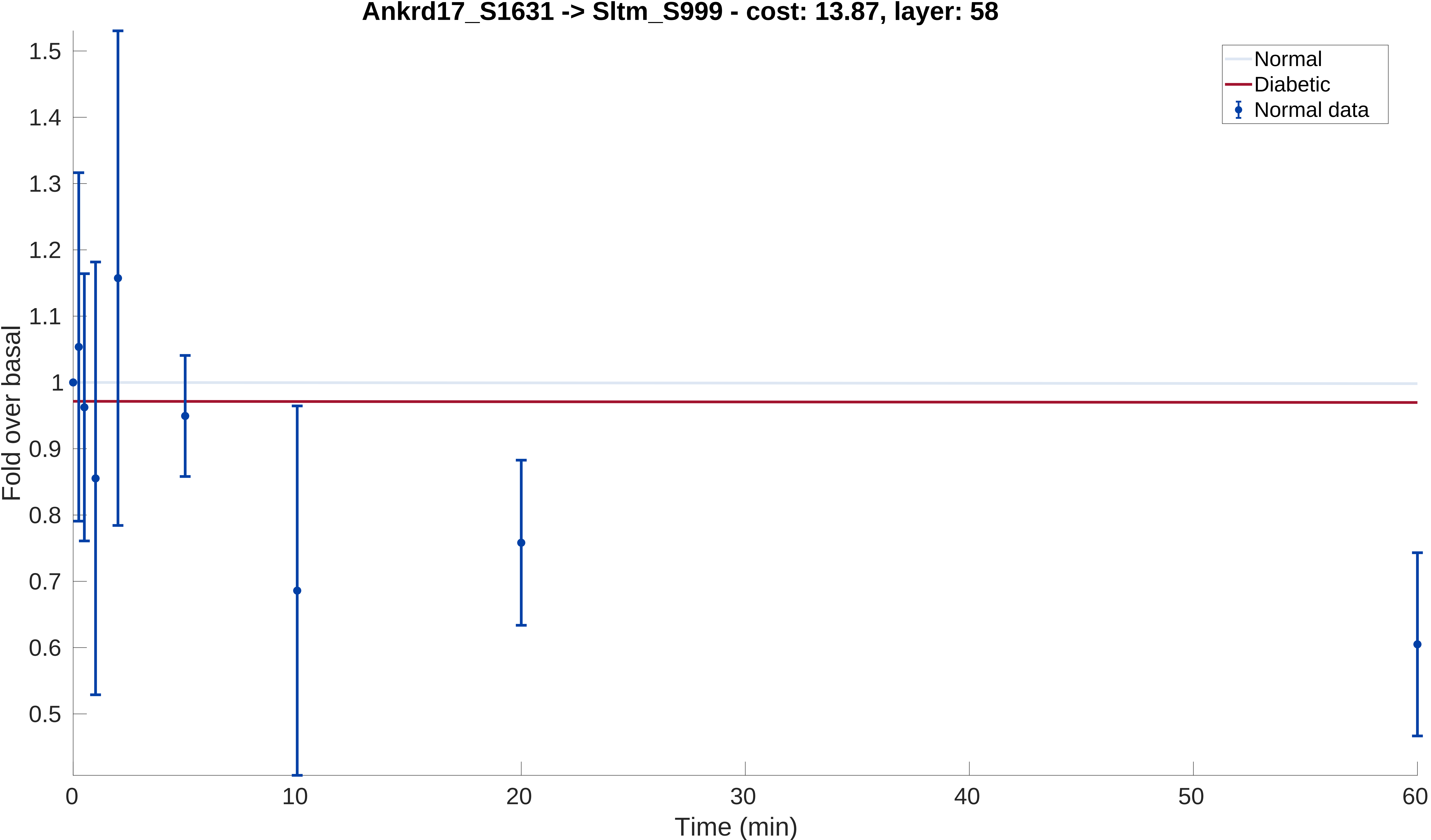

Ankrd17_S1631 -> Sltm_S999 - cost: 13.87, layer: 58
Normal
Diabetic
Normal data
Fold over basal
1.5
1.4
1.3
1.2
1.1
1
0.9
0.8
0.7
0.6
0.5
0
10
20
30
40
50
60
Time (min)

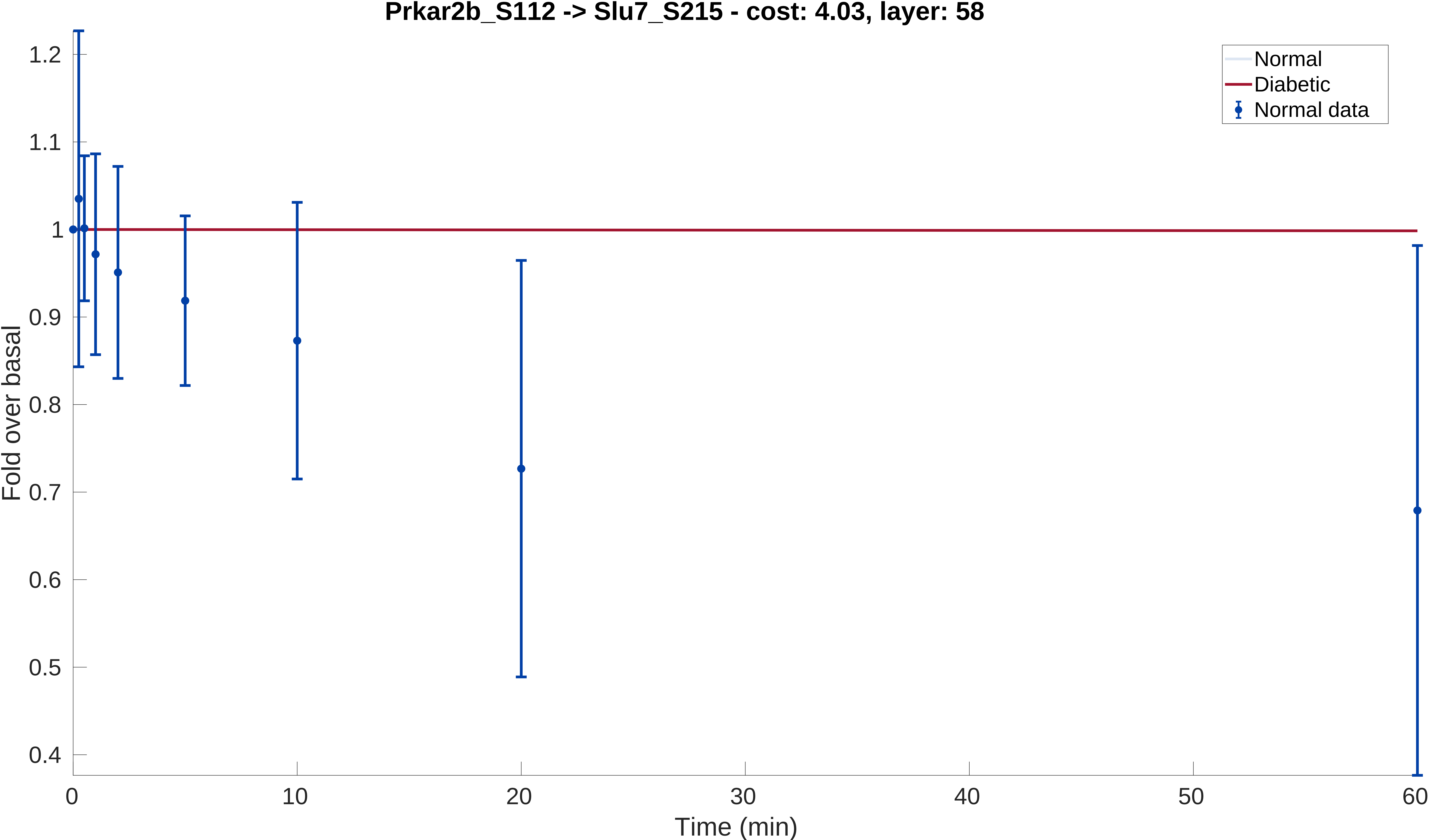

Prkar2b_S112 -> Slu7_S215 - cost: 4.03, layer: 58
Normal
Diabetic
Normal data
Fold over basal
1.2
1.1
1
0.9
0.8
0.7
0.6
0.5
0.4
0
10
20
30
40
50
60
Time (min)

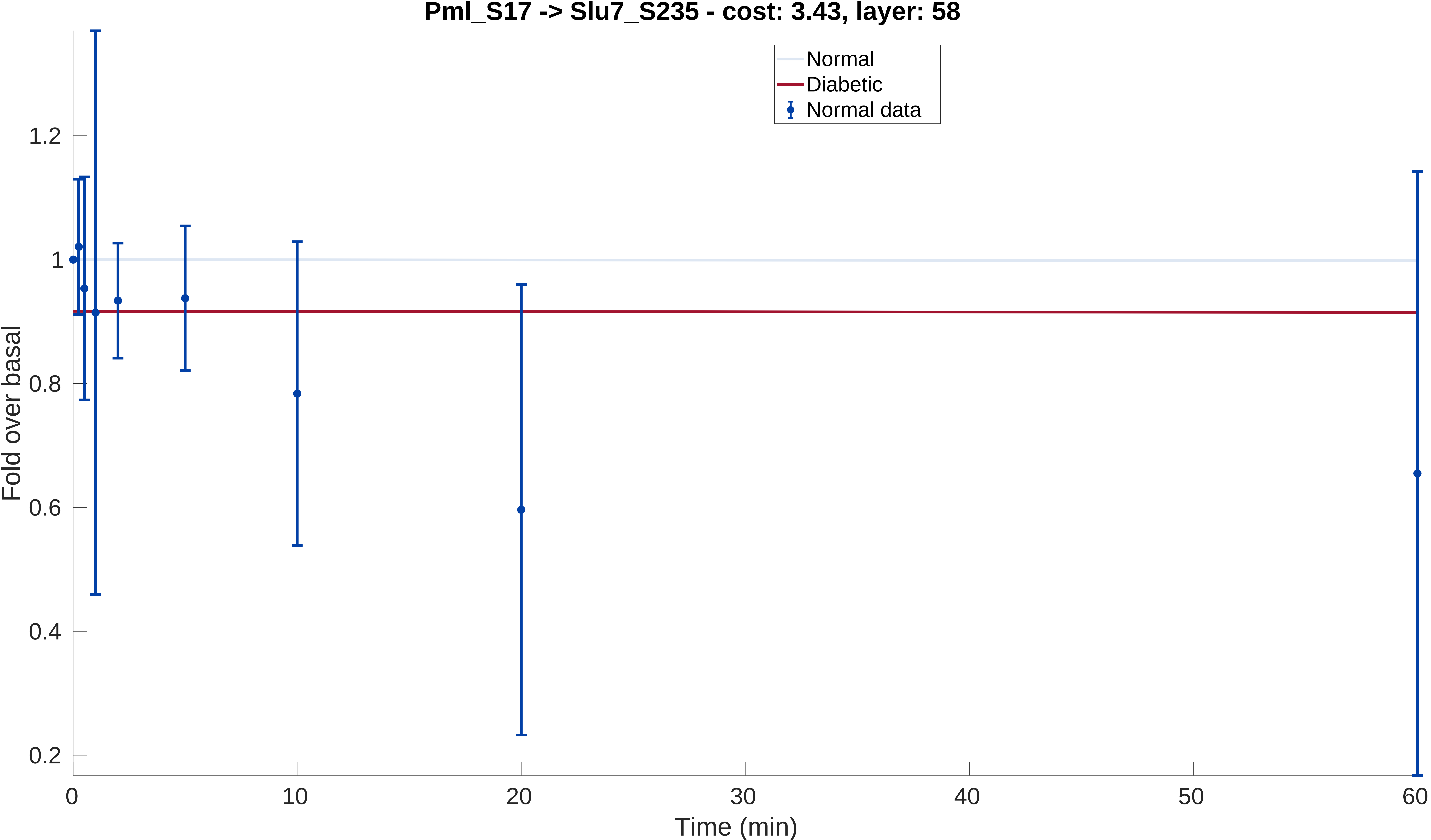

Pml_S17 -> Slu7_S235 - cost: 3.43, layer: 58
Normal
Diabetic
Normal data
Fold over basal
1.2
1
0.8
0.6
0.4
0.2
0
10
20
30
40
50
60
Time (min)

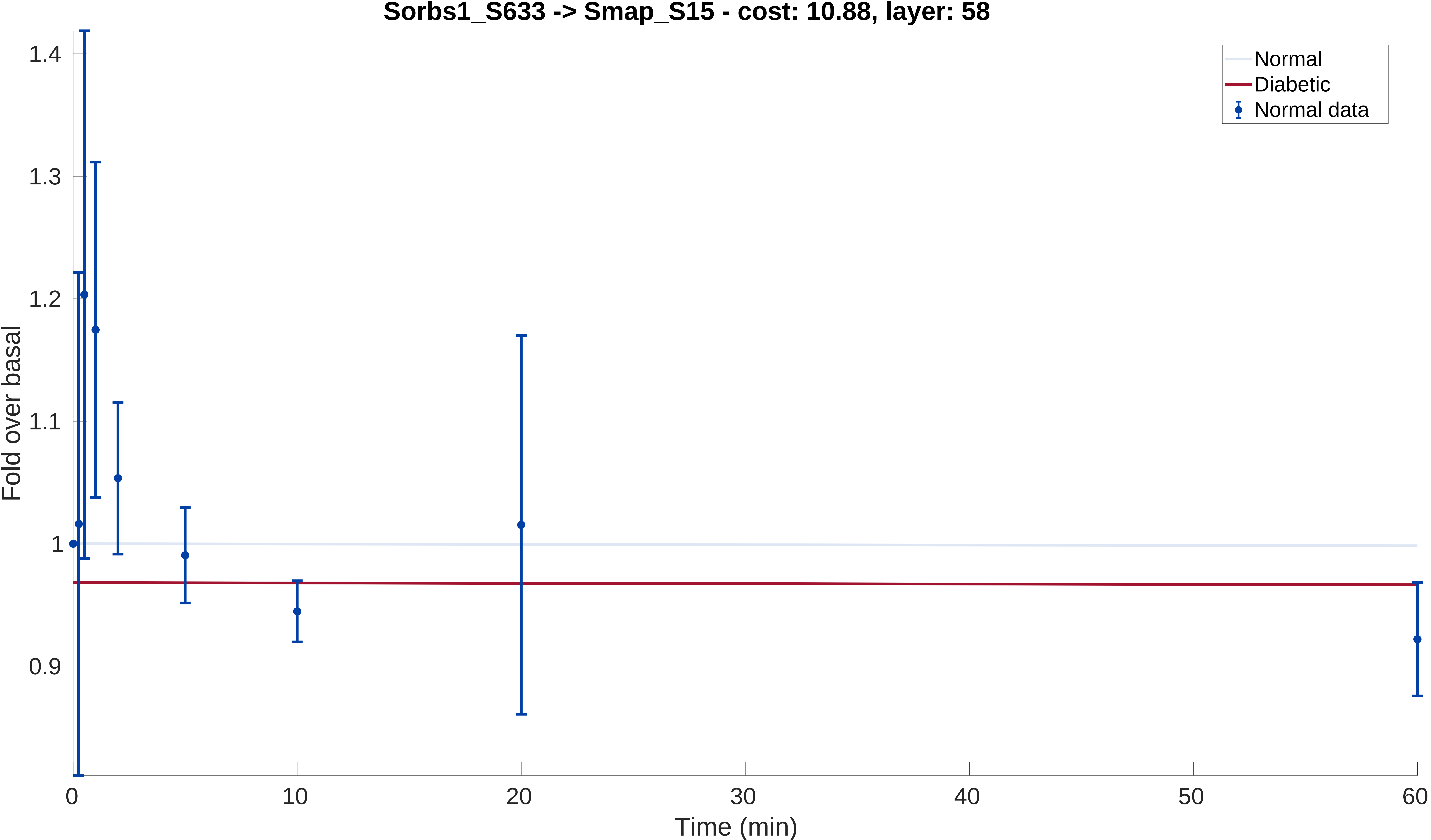

Sorbs1_S633 -> Smap_S15 - cost: 10.88, layer: 58
Normal
Diabetic
Normal data
Fold over basal
1.4
1.3
1.2
1.1
1
0.9
0
10
20
30
40
50
60
Time (min)

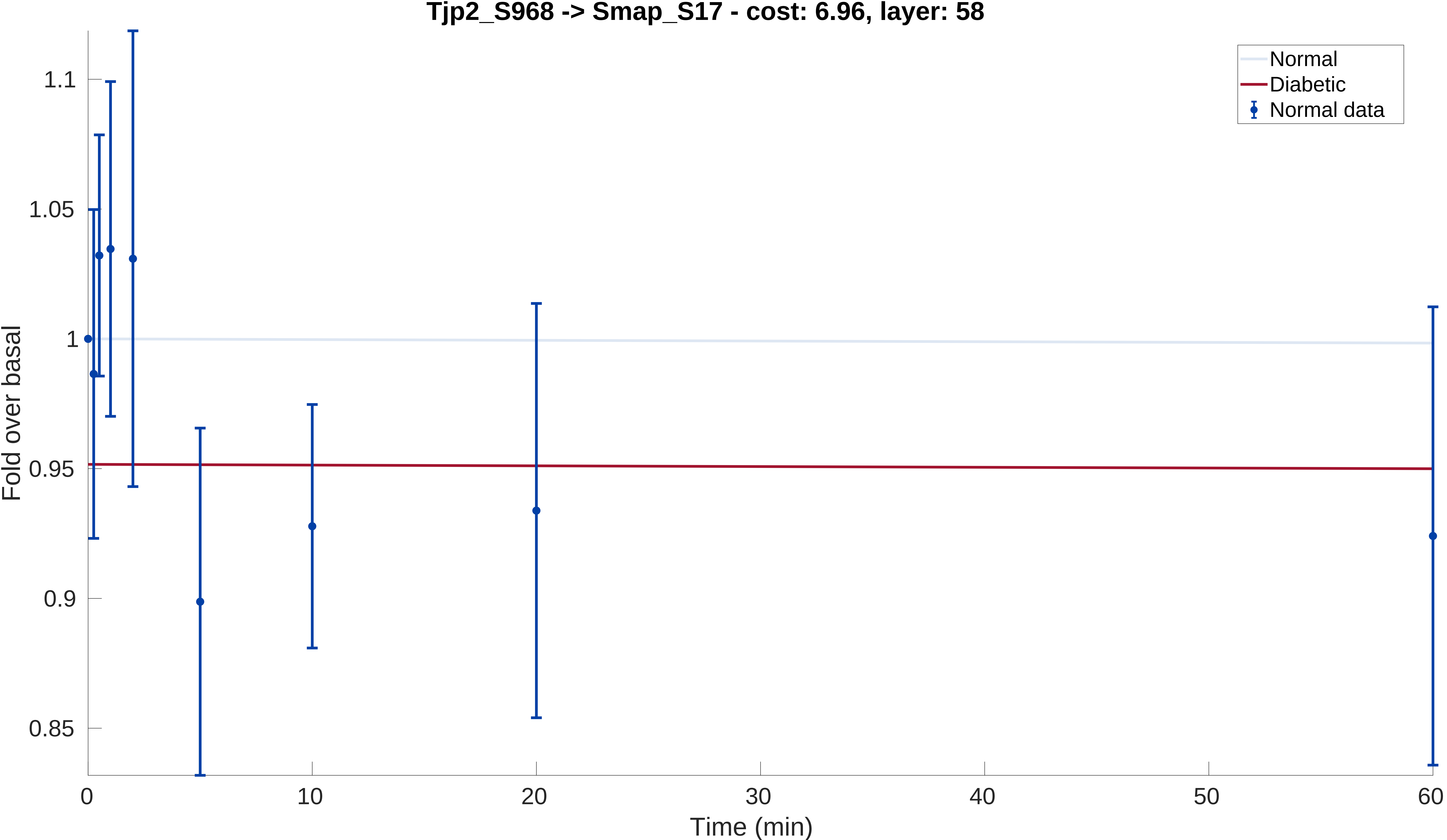
Tjp2_S968 -> Smap_S17 - cost: 6.96, layer: 58
Normal
Diabetic
Normal data
Fold over basal
1.1
1.05
1
0.95
0.9
0.85
0
10
20
30
40
50
60
Time (min)

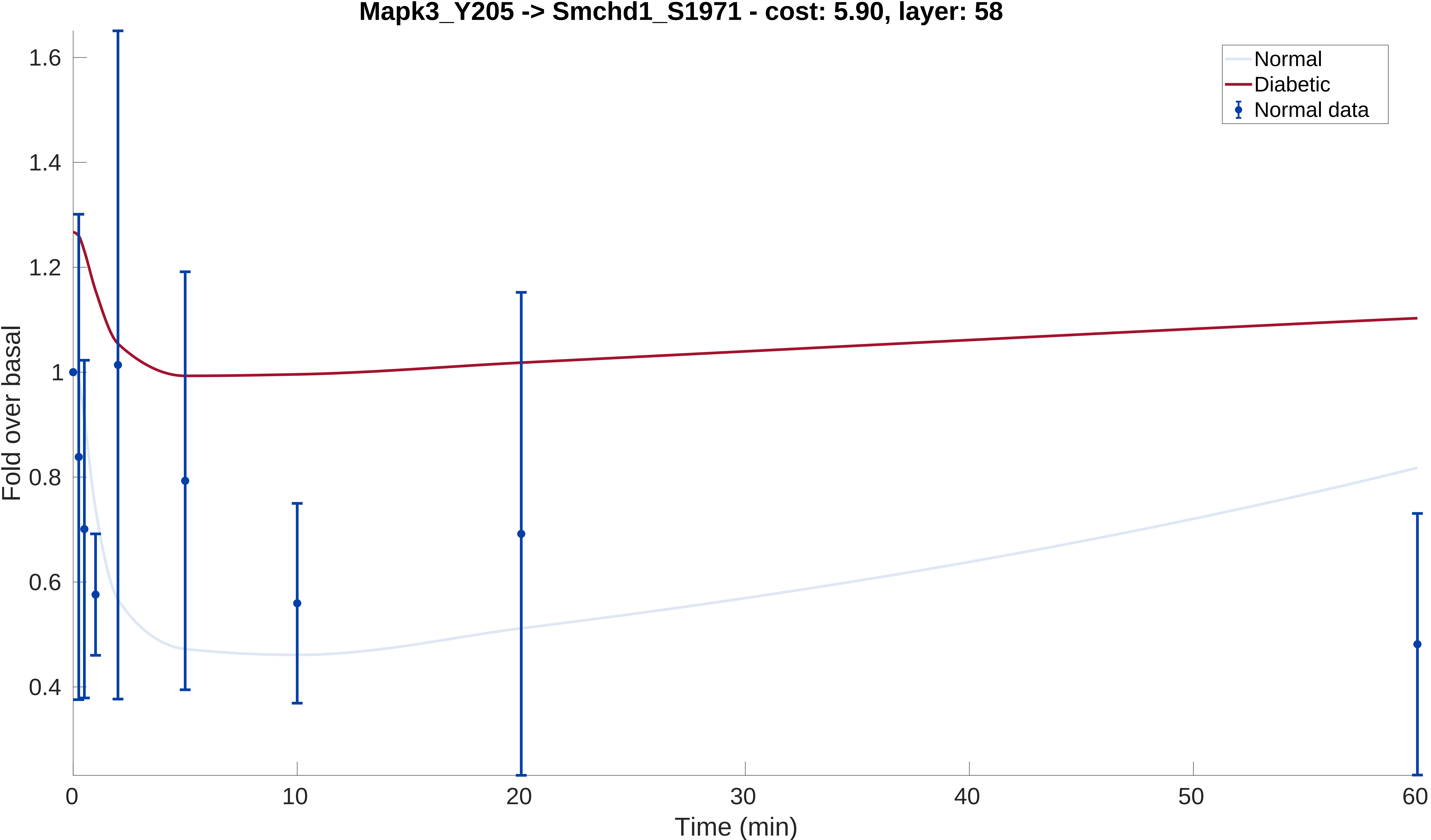

Mapk3_Y205 -> Smchd1_S1971 - cost: 5.90, layer: 58
Normal
Diabetic
Normal data
Fold over basal
1.6
1.4
1.2
1
0.8
0.6
0.4
0
10
20
30
40
50
60
Time (min)

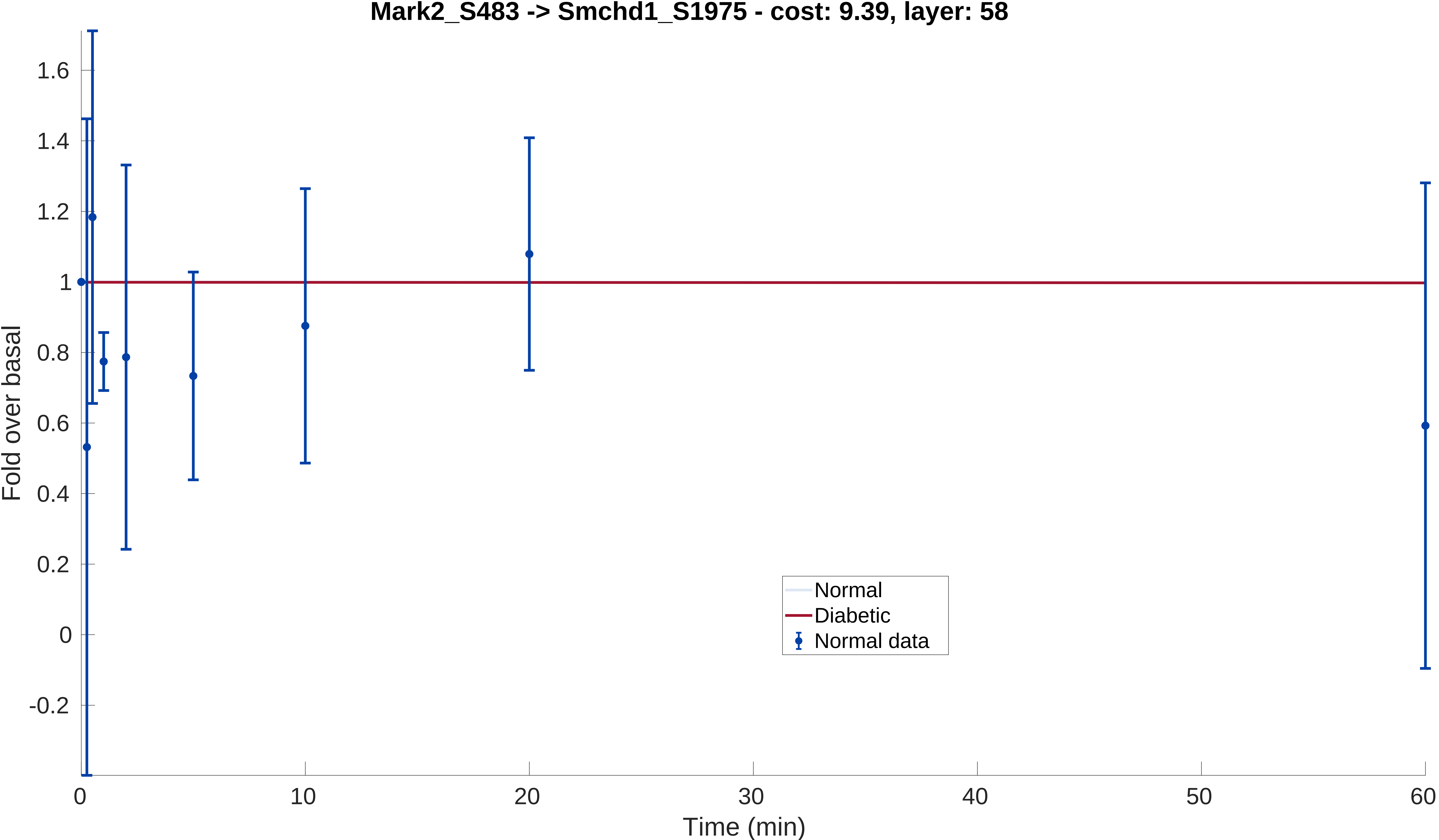

Mark2_S483 -> Smchd1_S1975 - cost: 9.39, layer: 58
Fold over basal
Time (min)
1.6
1.4
1.2
1
0.8
0.6
0.4
0.2
0
-0.2
0
10
20
30
40
50
60
Normal
Diabetic
Normal data

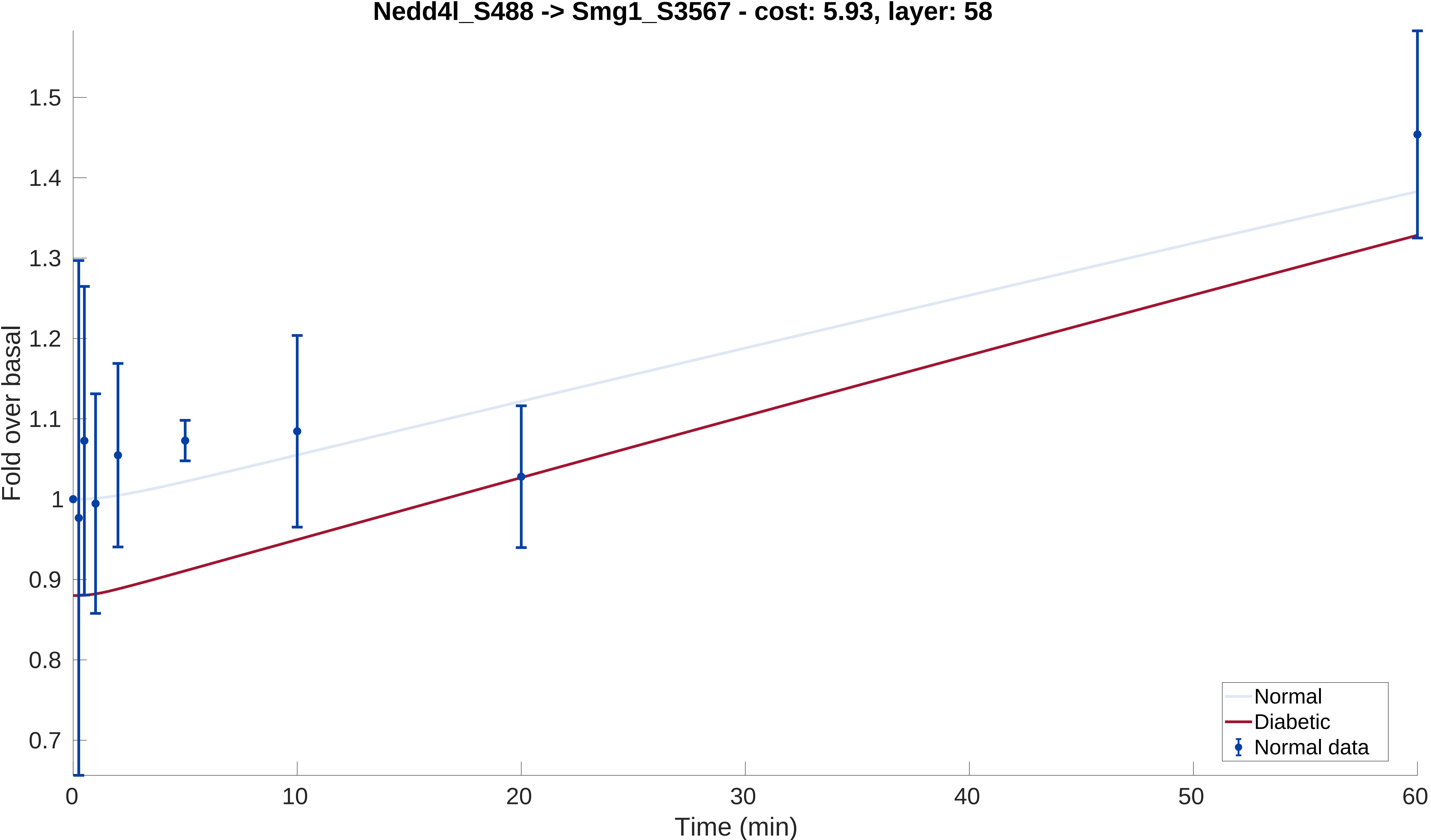
Nedd4l_S488 -> Smg1_S3567 - cost: 5.93, layer: 58
Fold over basal
Time (min)
1.5
1.4
1.3
1.2
1.1
1
0.9
0.8
0.7
0
10
20
30
40
50
60
Normal
Diabetic
Normal data

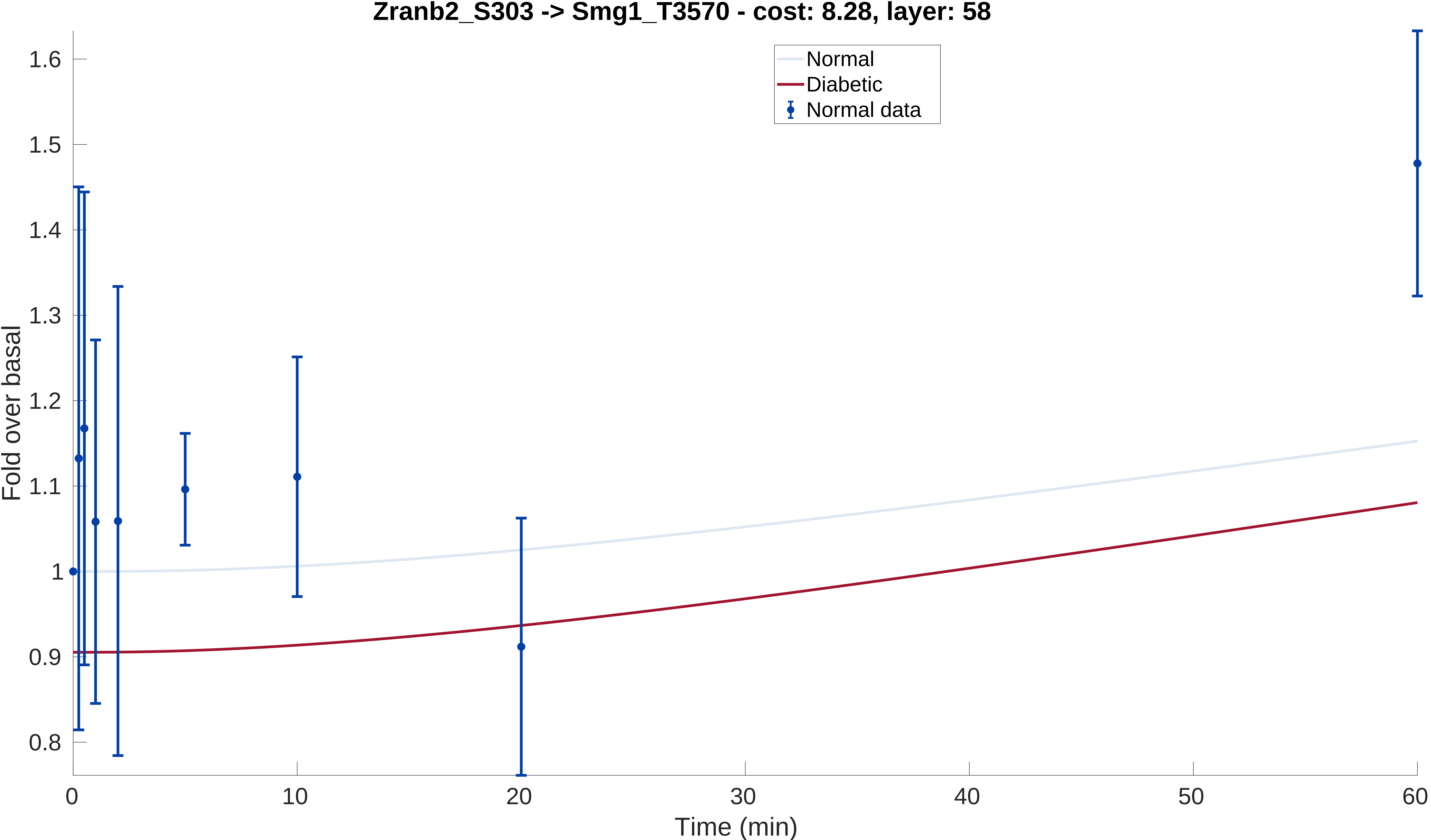

Zranb2_S303 -> Smg1_T3570 - cost: 8.28, layer: 58
Normal
Diabetic
Normal data
Fold over basal
1.6
1.5
1.4
1.3
1.2
1.1
1
0.9
0.8
0
10
20
30
40
50
60
Time (min)

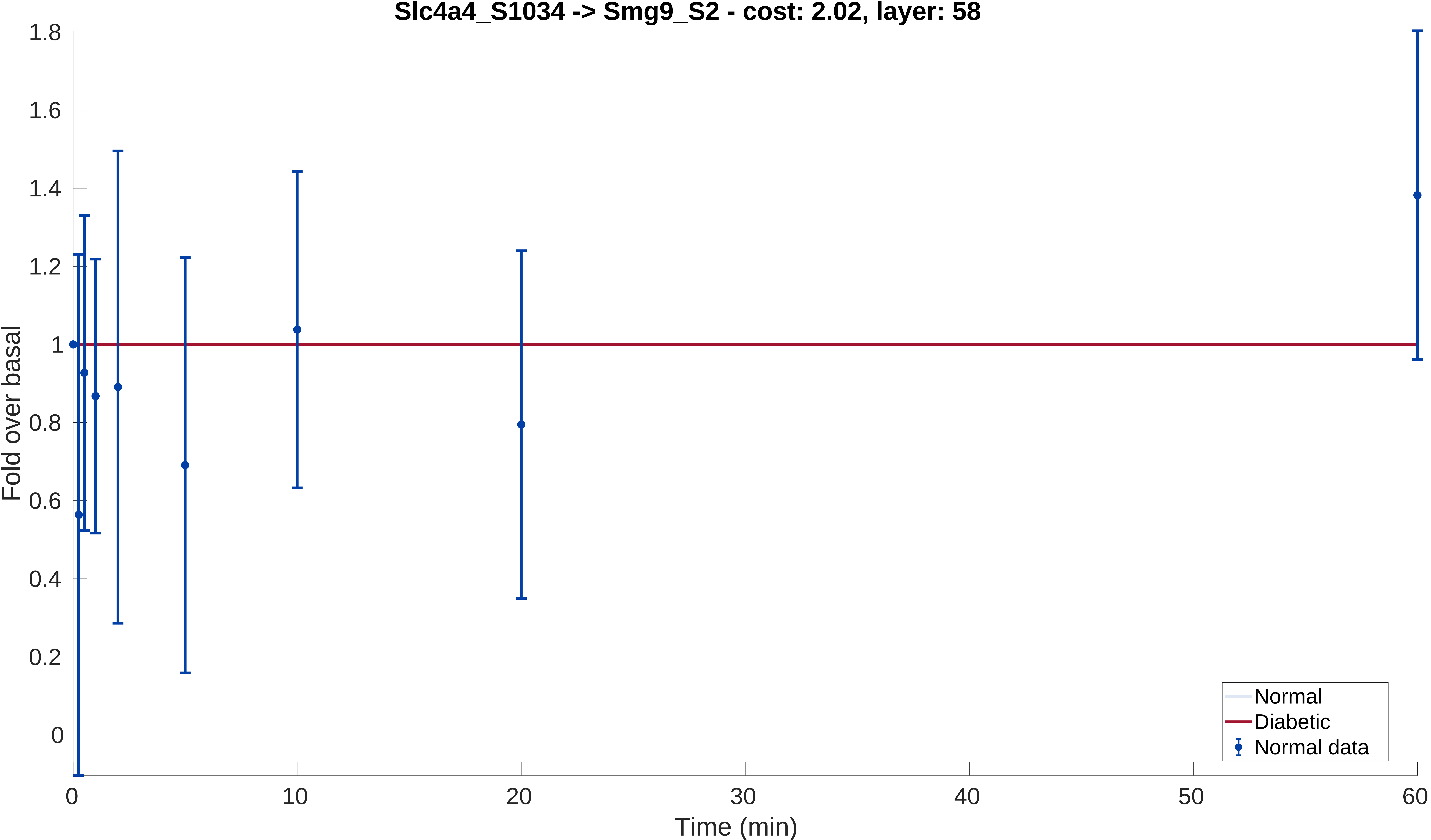
Slc4a4_S1034 -> Smg9_S2 - cost: 2.02, layer: 58
Fold over basal
Time (min)
0
0.2
0.4
0.6
0.8
1
1.2
1.4
1.6
1.8
0
10
20
30
40
50
60
Normal
Diabetic
Normal data

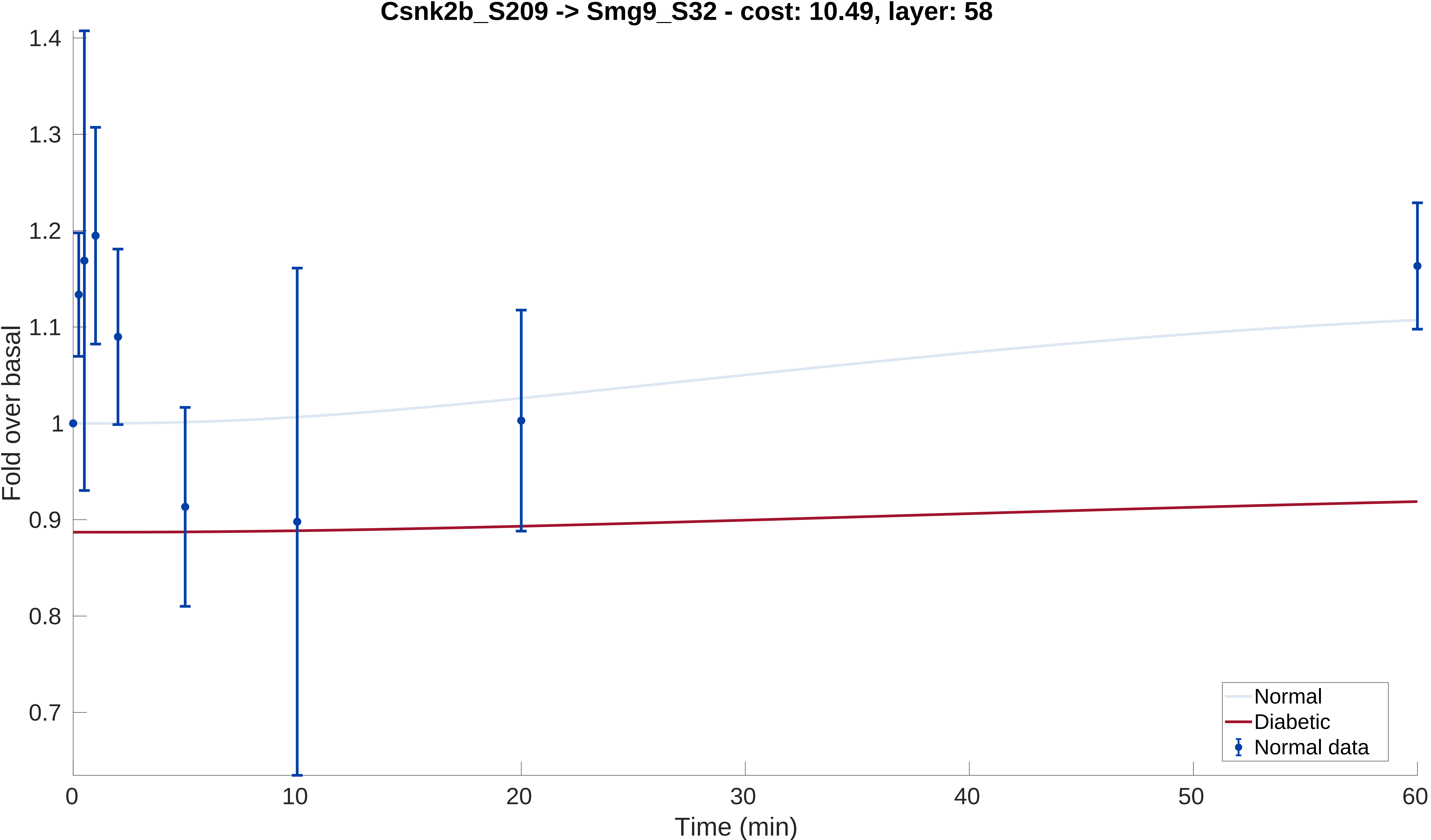

Csnk2b_S209 -> Smg9_S32 - cost: 10.49, layer: 58
Fold over basal
1.4
1.3
1.2
1.1
1
0.9
0.8
0.7
0
10
20
30
40
50
60
Time (min)
Normal
Diabetic
Normal data

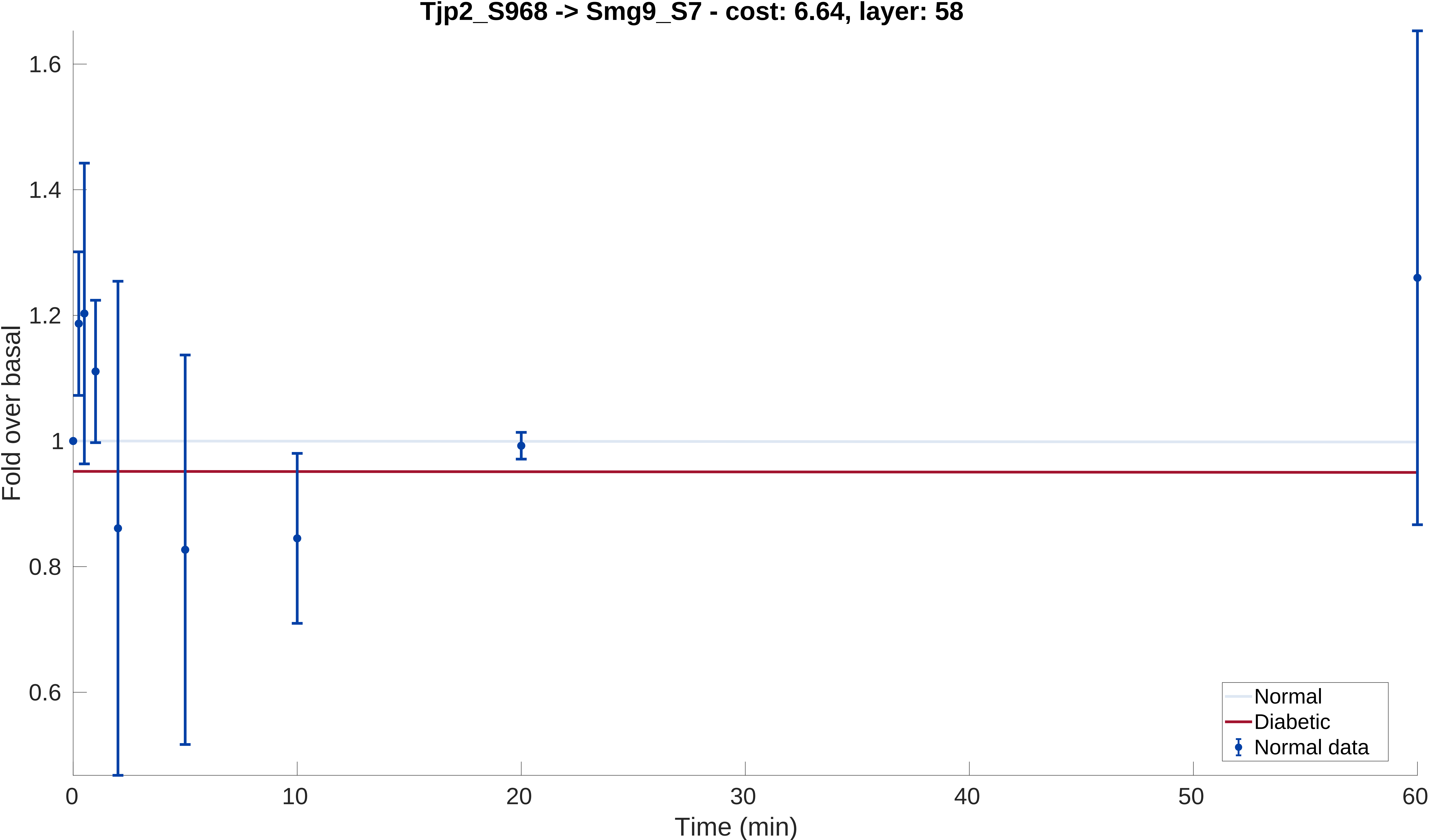
Tjp2_S968 -> Smg9_S7 - cost: 6.64, layer: 58
Fold over basal
Time (min)
Normal
Diabetic
Normal data
0.6
0.8
1
1.2
1.4
1.6
0
10
20
30
40
50
60

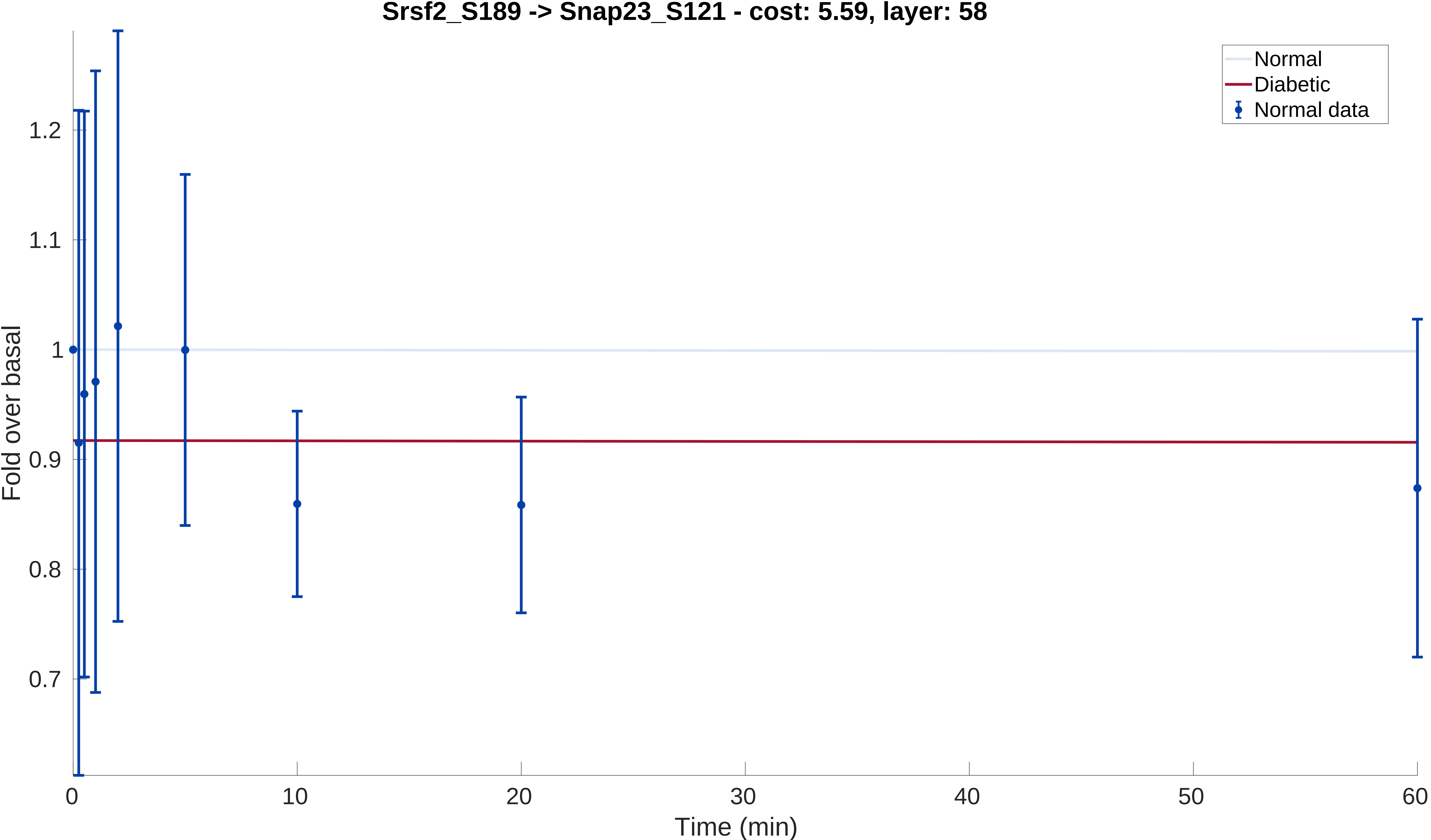

Srsf2_S189 -> Snap23_S121 - cost: 5.59, layer: 58
Normal
Diabetic
Normal data
Fold over basal
1.2
1.1
1
0.9
0.8
0.7
0
10
20
30
40
50
60
Time (min)

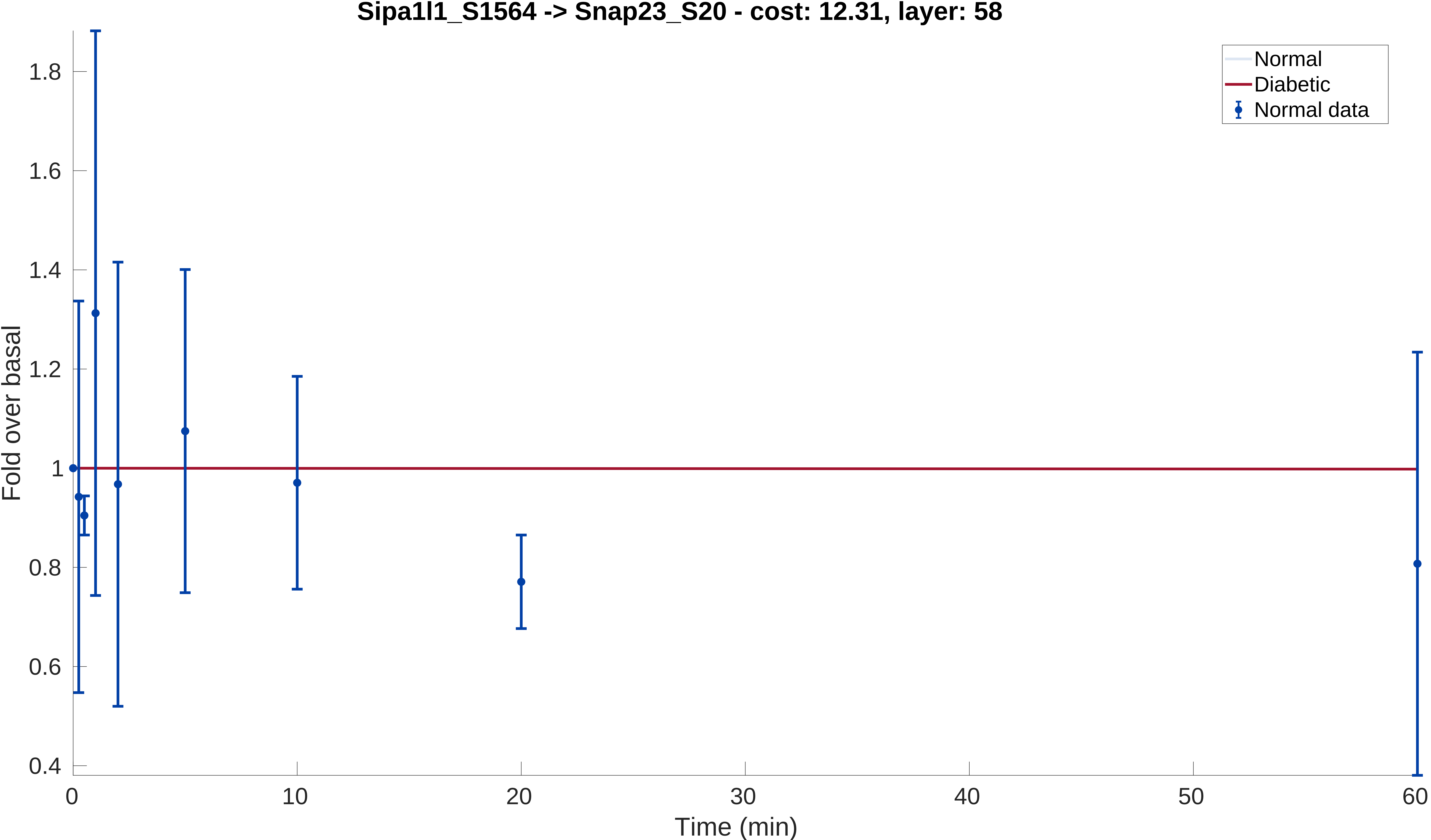
Sipa1l1_S1564 -> Snap23_S20 - cost: 12.31, layer: 58
Normal
Diabetic
Normal data
Fold over basal
1.8
1.6
1.4
1.2
1
0.8
0.6
0.4
0
10
20
30
40
50
60
Time (min)

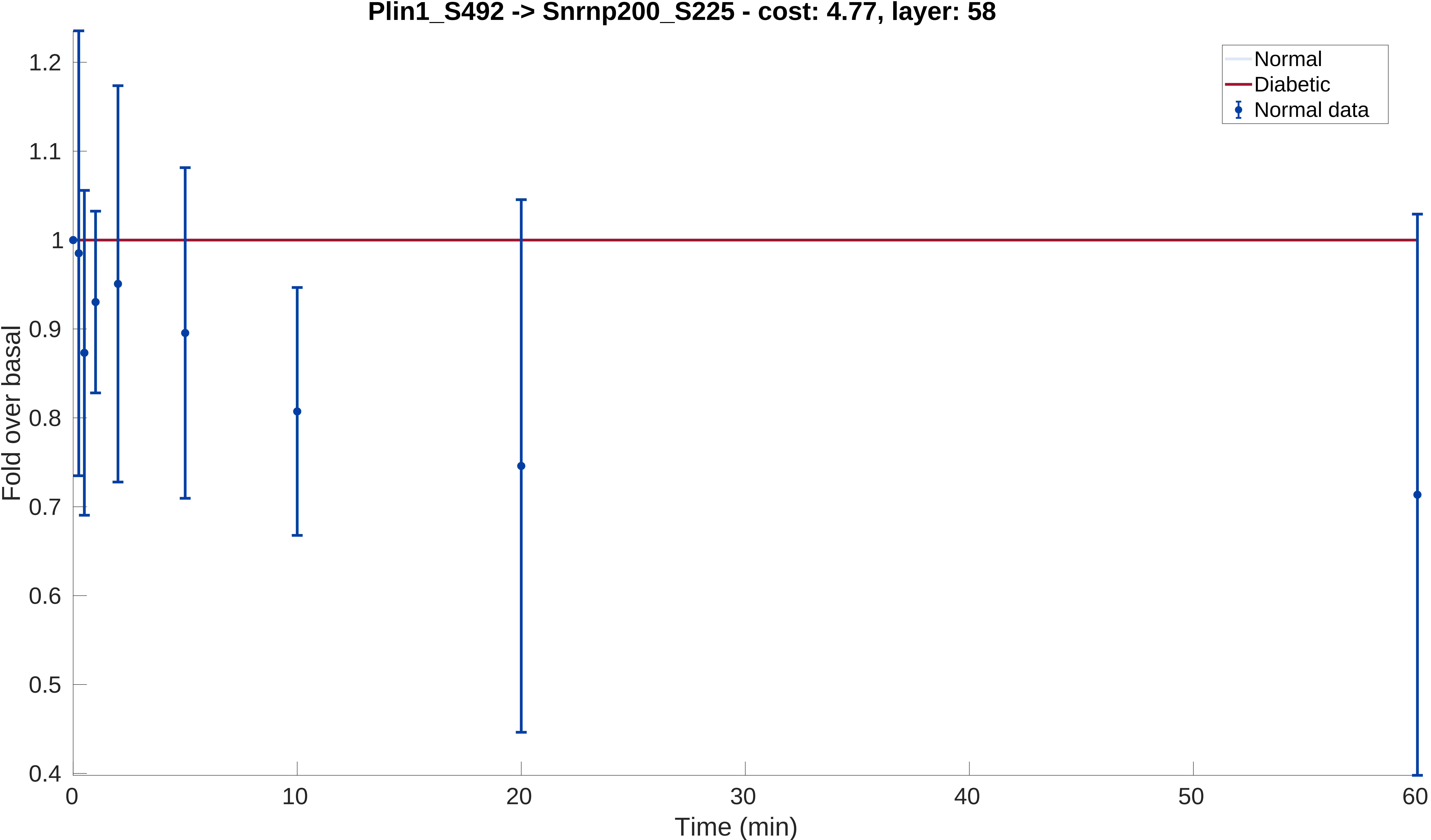

Plin1_S492 -> Snrnp200_S225 - cost: 4.77, layer: 58
Normal
Diabetic
Normal data
Fold over basal
1.2
1.1
1
0.9
0.8
0.7
0.6
0.5
0.4
0
10
20
30
40
50
60
Time (min)

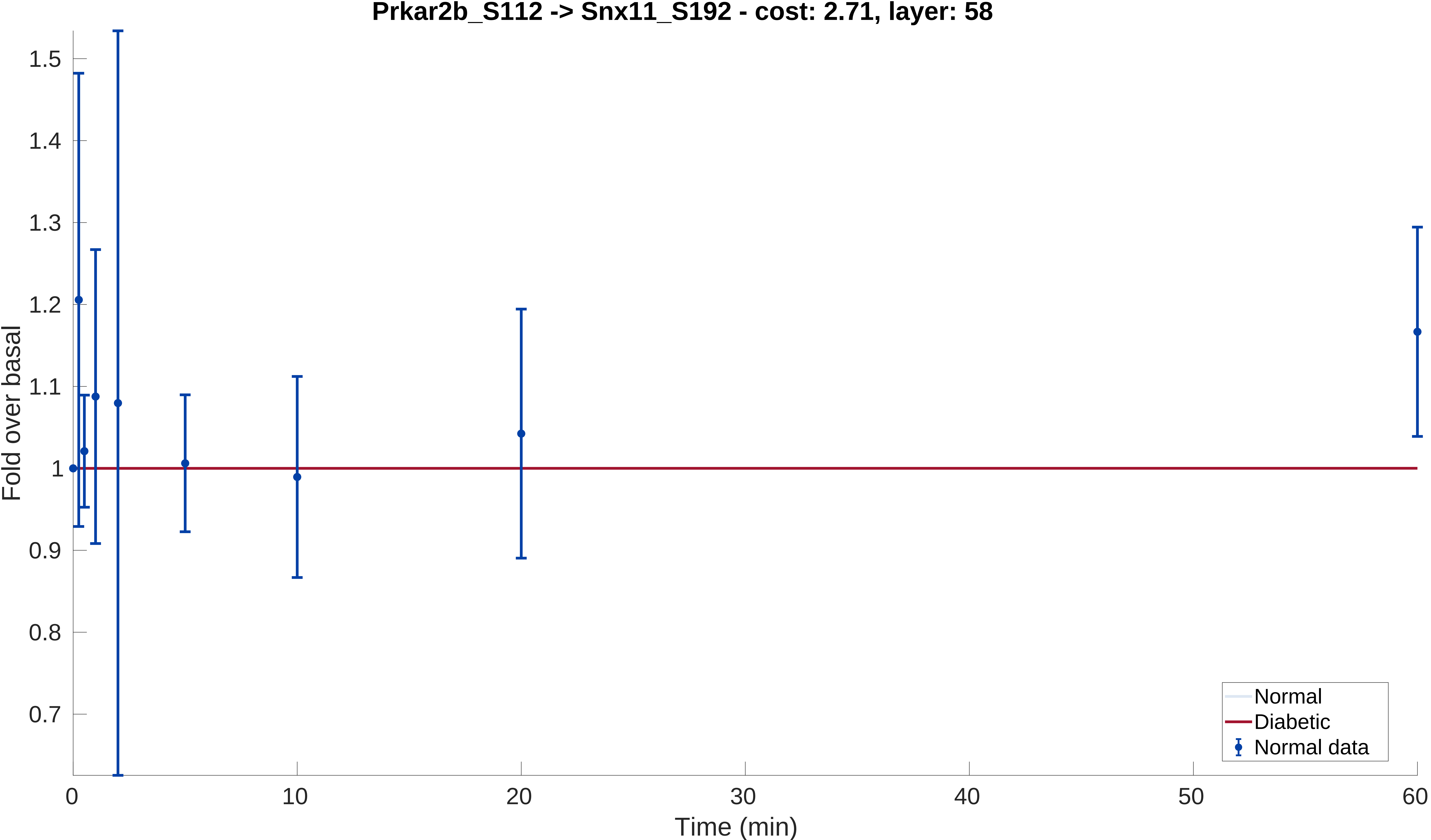
Prkar2b_S112 -> Snx11_S192 - cost: 2.71, layer: 58
Fold over basal
1.5
1.4
1.3
1.2
1.1
1
0.9
0.8
0.7
0
10
20
30
40
50
60
Time (min)
Normal
Diabetic
Normal data

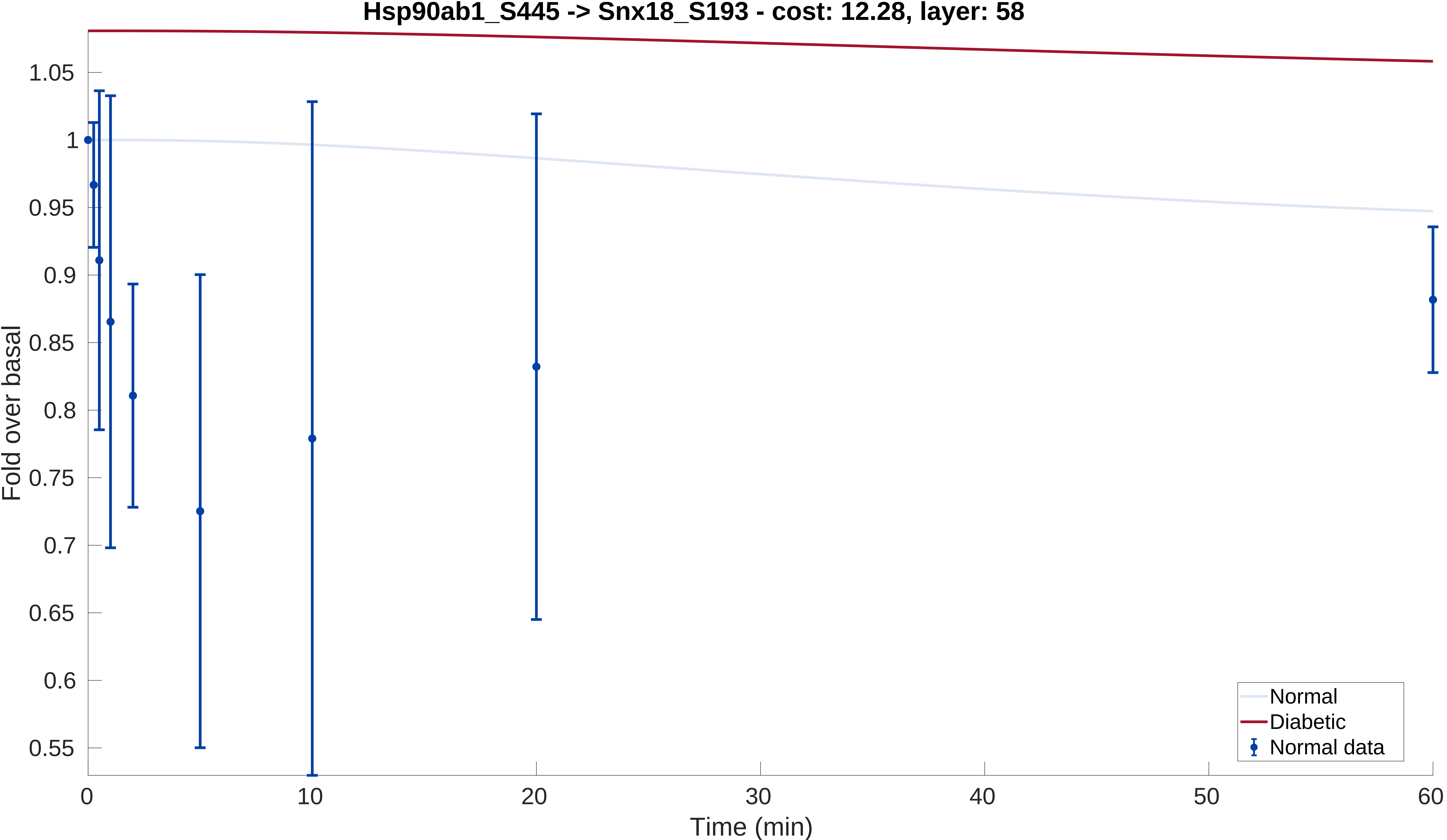

Hsp90ab1_S445 -> Snx18_S193 - cost: 12.28, layer: 58
Fold over basal
1.05
1
0.95
0.9
0.85
0.8
0.75
0.7
0.65
0.6
0.55
0
10
20
30
40
50
60
Time (min)
Normal
Diabetic
Normal data

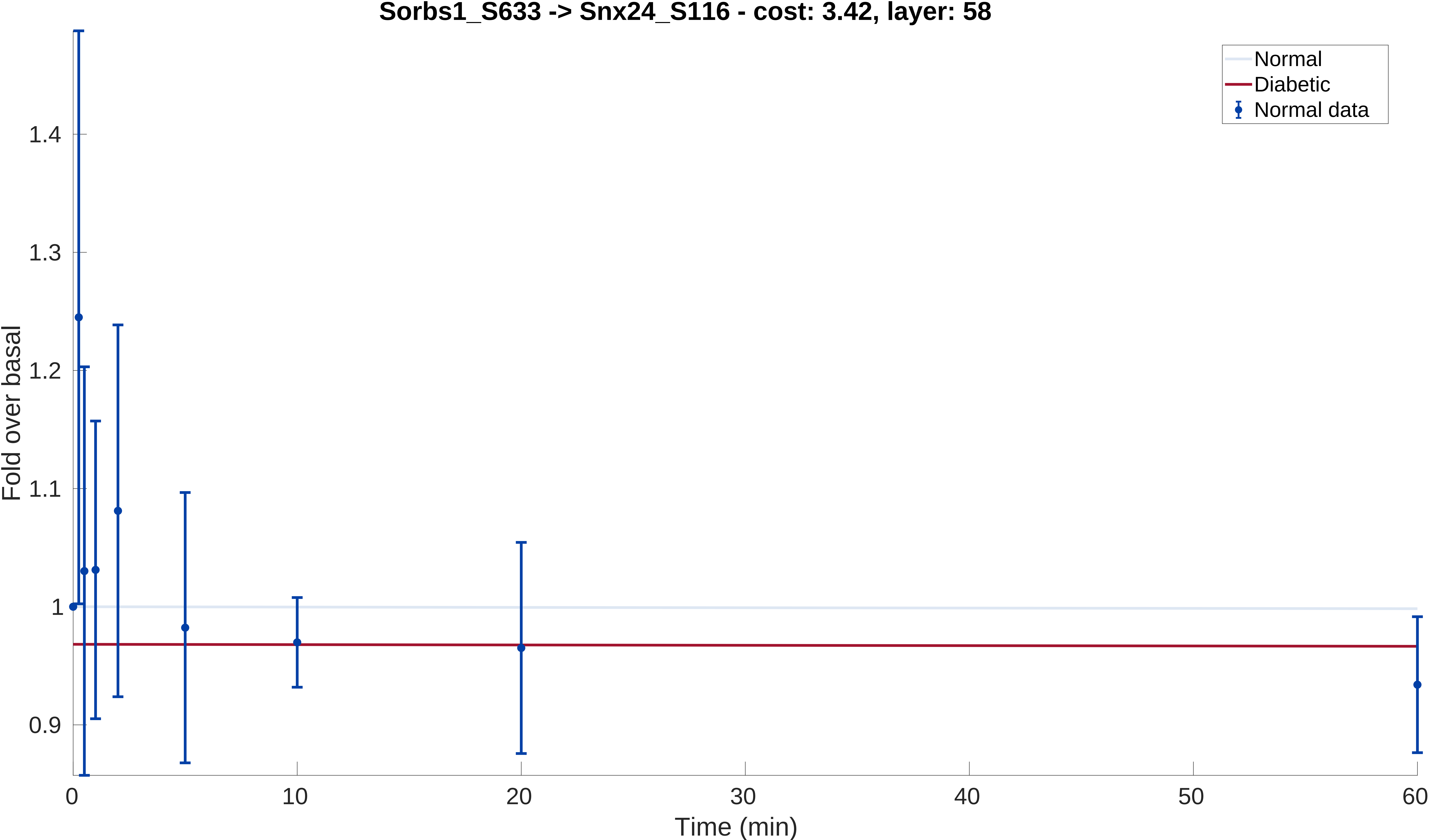

Sorbs1_S633 -> Snx24_S116 - cost: 3.42, layer: 58
Normal
Diabetic
Normal data
Fold over basal
1.4
1.3
1.2
1.1
1
0.9
0
10
20
30
40
50
60
Time (min)

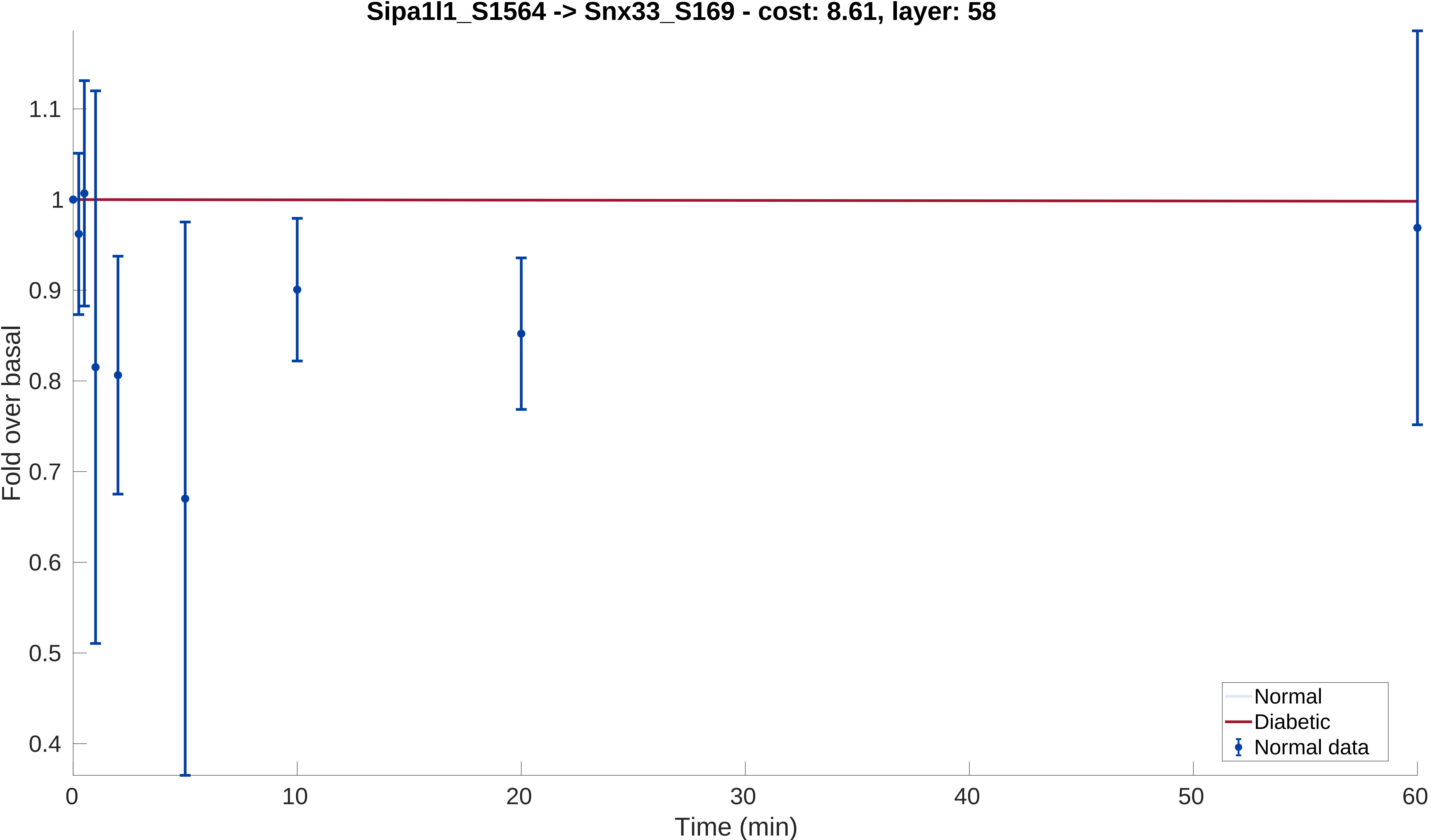

Sipa1l1_S1564 -> Snx33_S169 - cost: 8.61, layer: 58
Fold over basal
1.1
1
0.9
0.8
0.7
0.6
0.5
0.4
0
10
20
30
40
50
60
Time (min)
Normal
Diabetic
Normal data

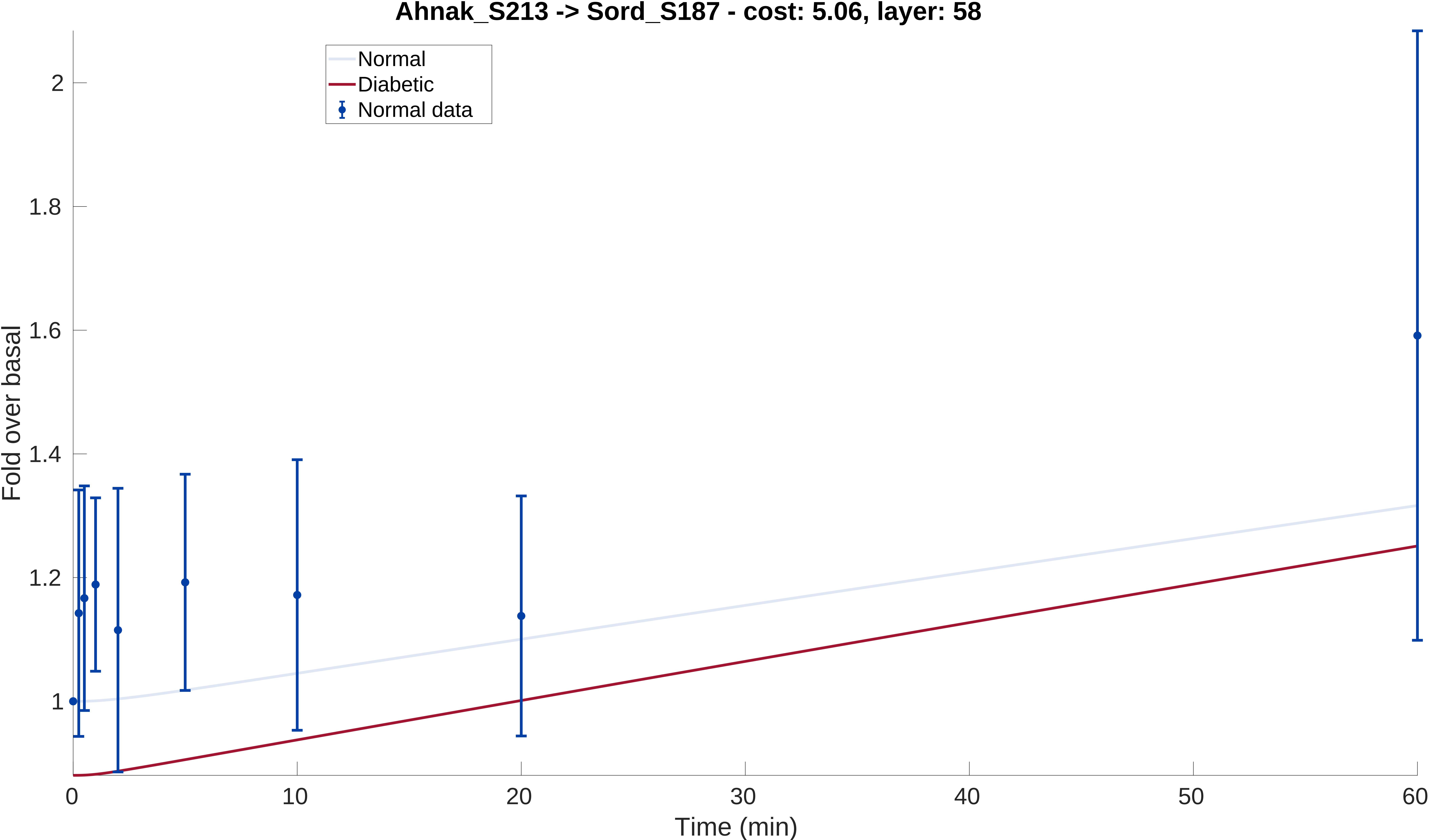

Ahnak_S213 -> Sord_S187 - cost: 5.06, layer: 58
Normal
Diabetic
Normal data
Fold over basal
2
1.8
1.6
1.4
1.2
1
0
10
20
30
40
50
60
Time (min)

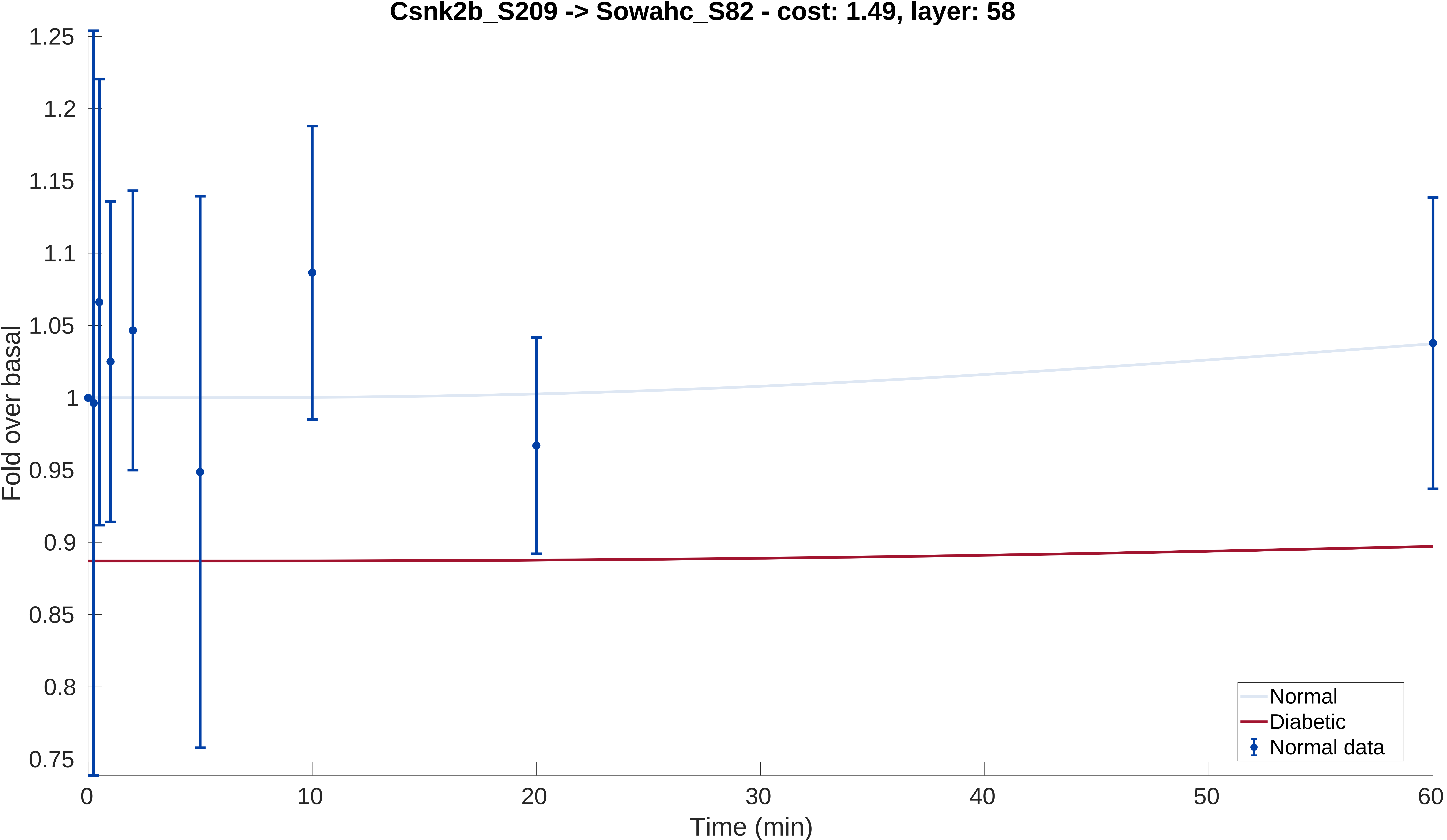

Csnk2b_S209 -> Sowahc_S82 - cost: 1.49, layer: 58
Fold over basal
1.25
1.2
1.15
1.1
1.05
1
0.95
0.9
0.85
0.8
0.75
0
10
20
30
40
50
60
Time (min)
Normal
Diabetic
Normal data

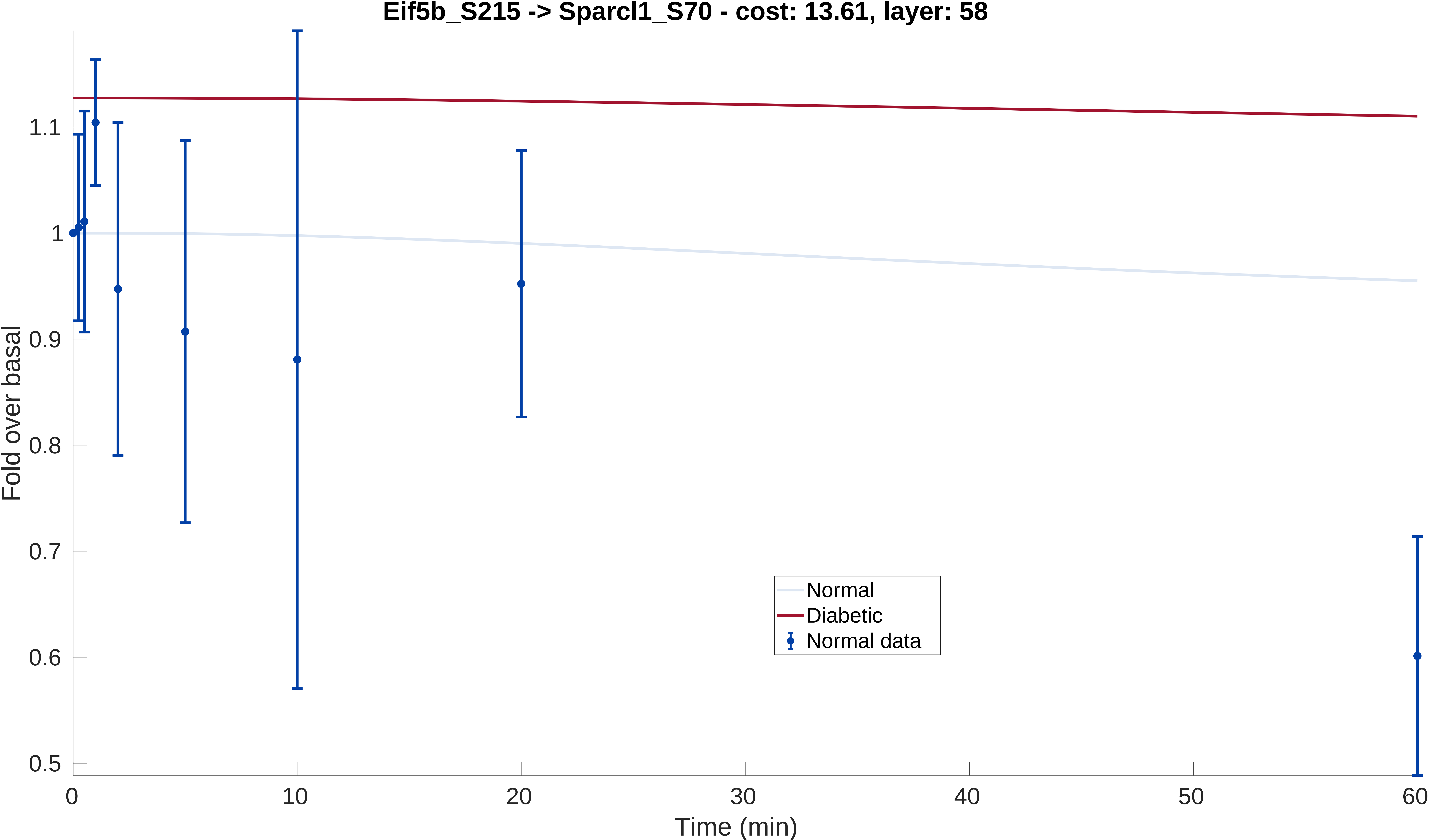

Eif5b_S215 -> Sparcl1_S70 - cost: 13.61, layer: 58
Fold over basal
1.1
1
0.9
0.8
0.7
0.6
0.5
0
10
20
30
40
50
60
Time (min)
Normal
Diabetic
Normal data

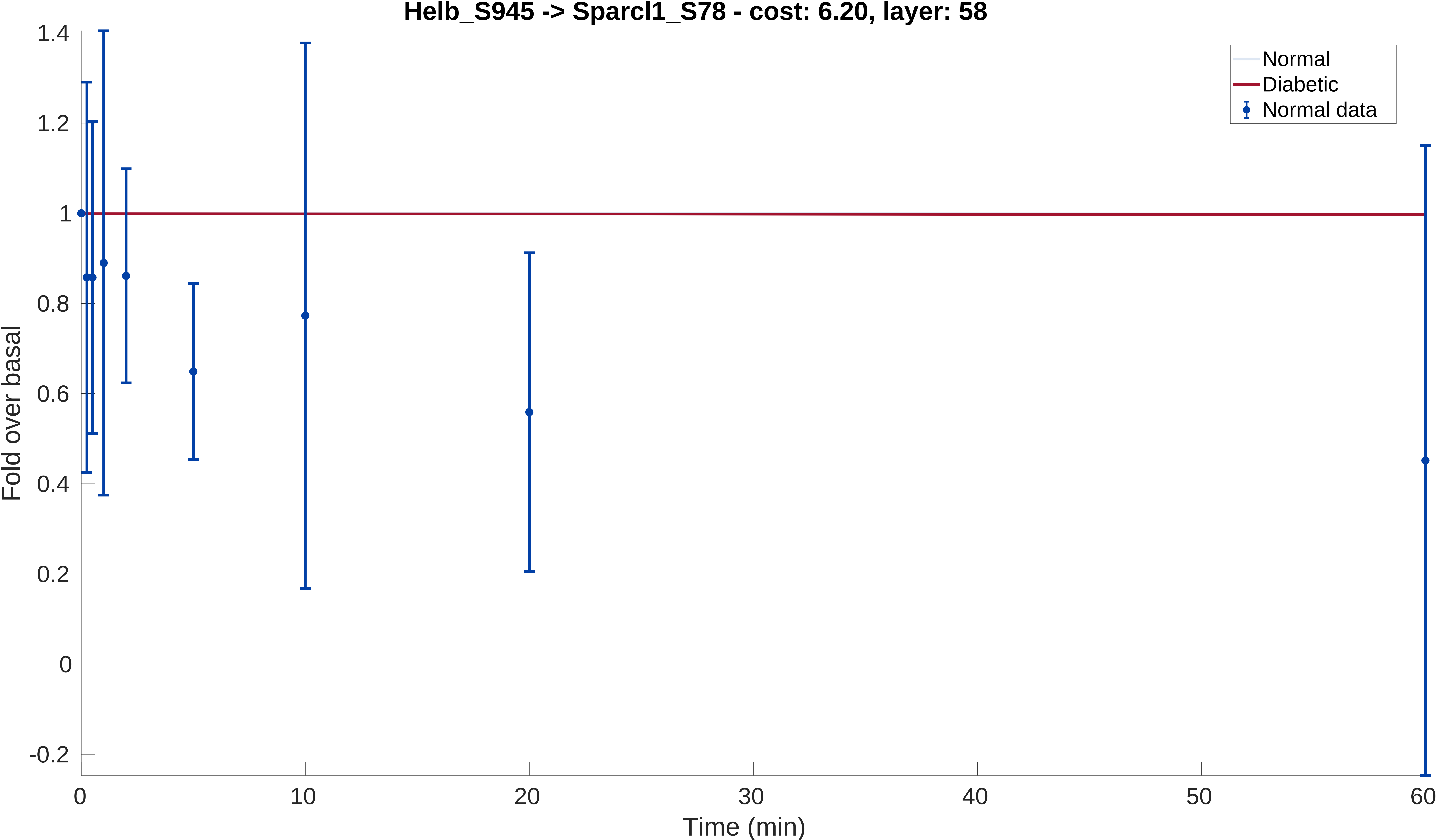

Helb_S945 -> Sparcl1_S78 - cost: 6.20, layer: 58
Normal
Diabetic
Normal data
Fold over basal
Time (min)
1.4
1.2
1
0.8
0.6
0.4
0.2
0
-0.2
0
10
20
30
40
50
60

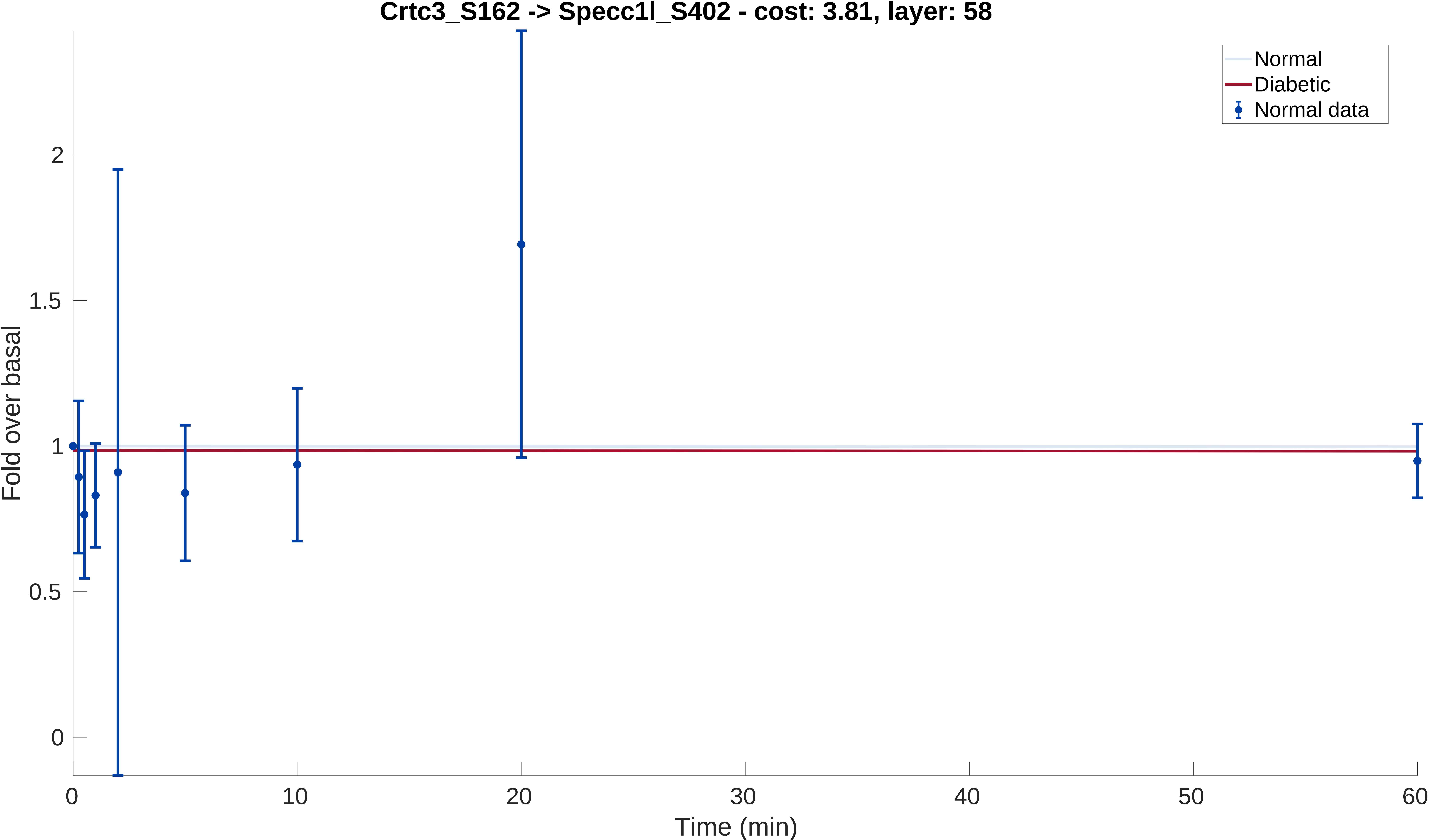

Crtc3_S162 -> Specc1l_S402 - cost: 3.81, layer: 58
Normal
Diabetic
Normal data
Fold over basal
Time (min)
0
0.5
1
1.5
2
0
10
20
30
40
50
60

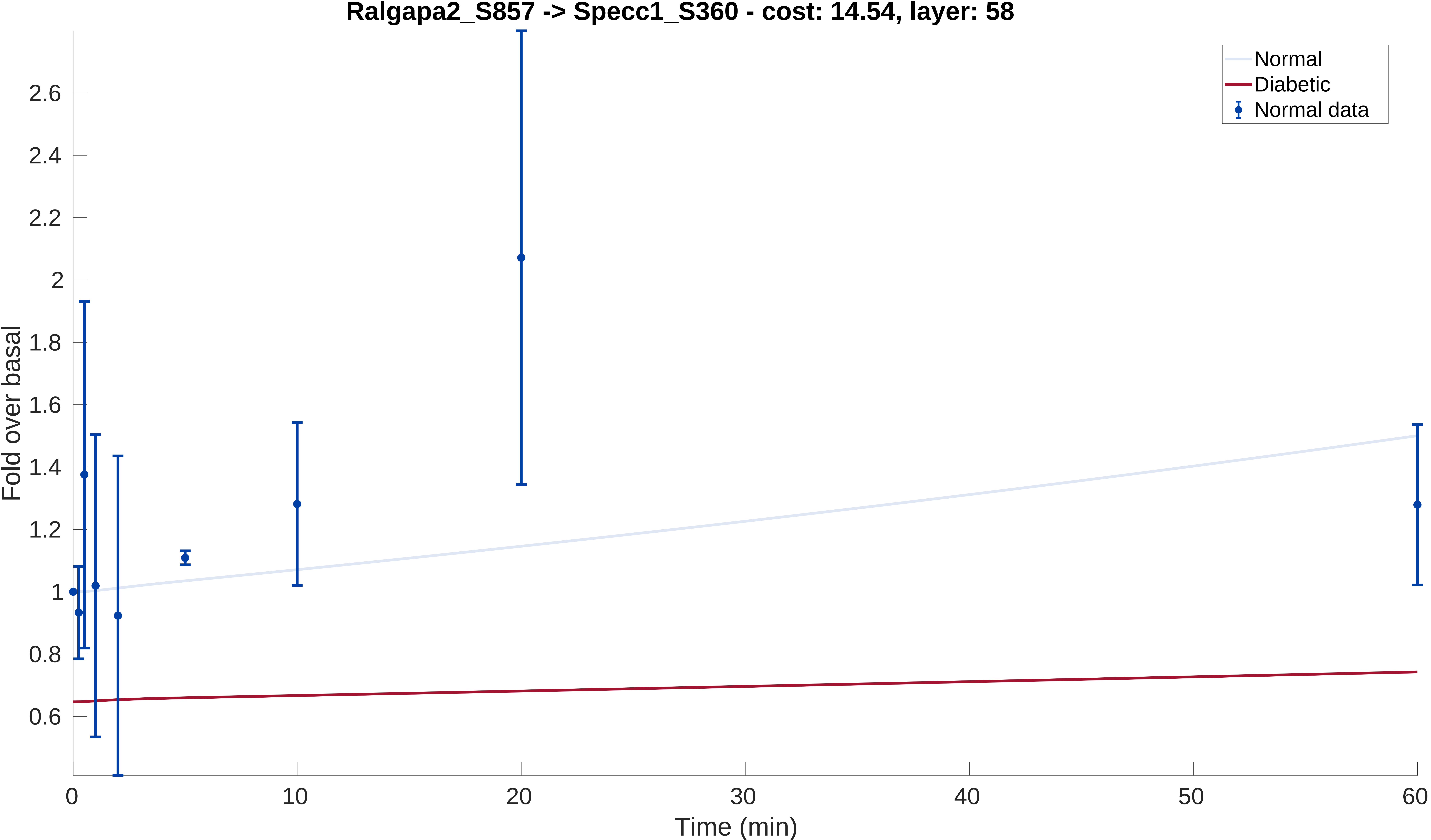

Ralgapa2_S857 -> Specc1_S360 - cost: 14.54, layer: 58
Normal
Diabetic
Normal data
Fold over basal
Time (min)
2.6
2.4
2.2
2
1.8
1.6
1.4
1.2
1
0.8
0.6
0
10
20
30
40
50
60

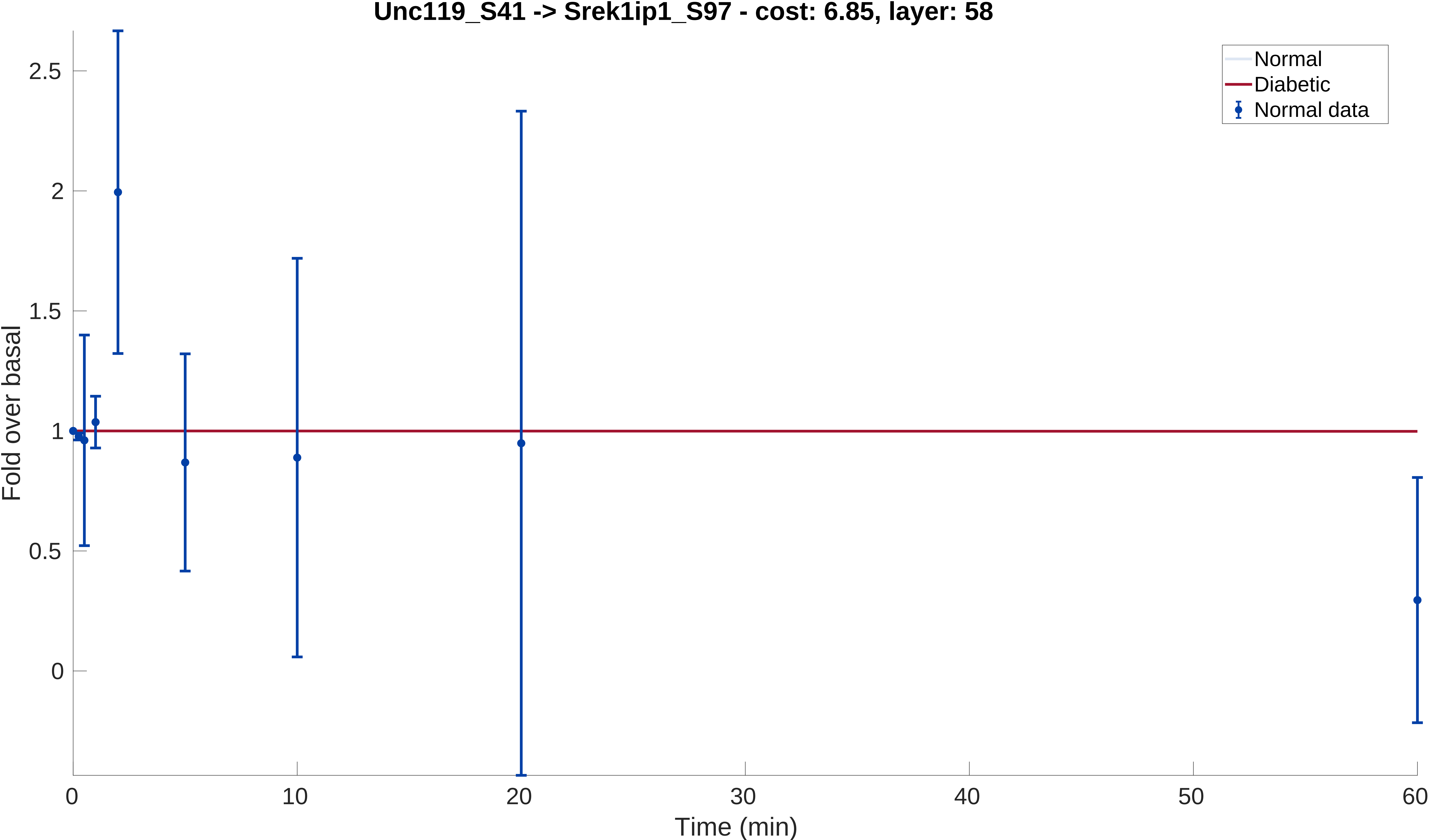
Unc119_S41 -> Srek1ip1_S97 - cost: 6.85, layer: 58
Normal
Diabetic
Normal data
Fold over basal
Time (min)
0
0.5
1
1.5
2
2.5
0
10
20
30
40
50
60

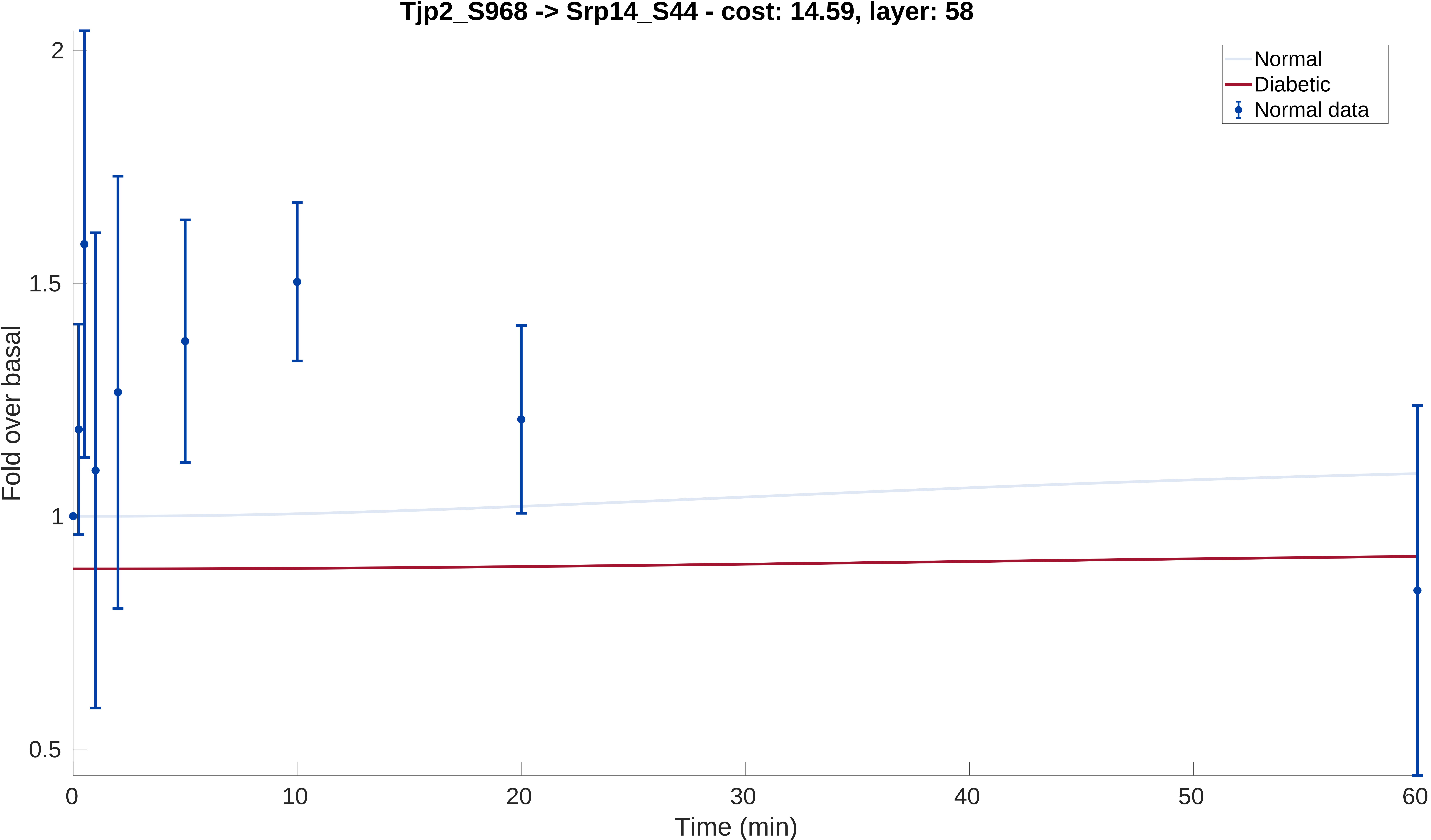

Tjp2_S968 -> Srp14_S44 - cost: 14.59, layer: 58
Normal
Diabetic
Normal data
2
1.5
1
0.5
Fold over basal
0
10
20
30
40
50
60
Time (min)

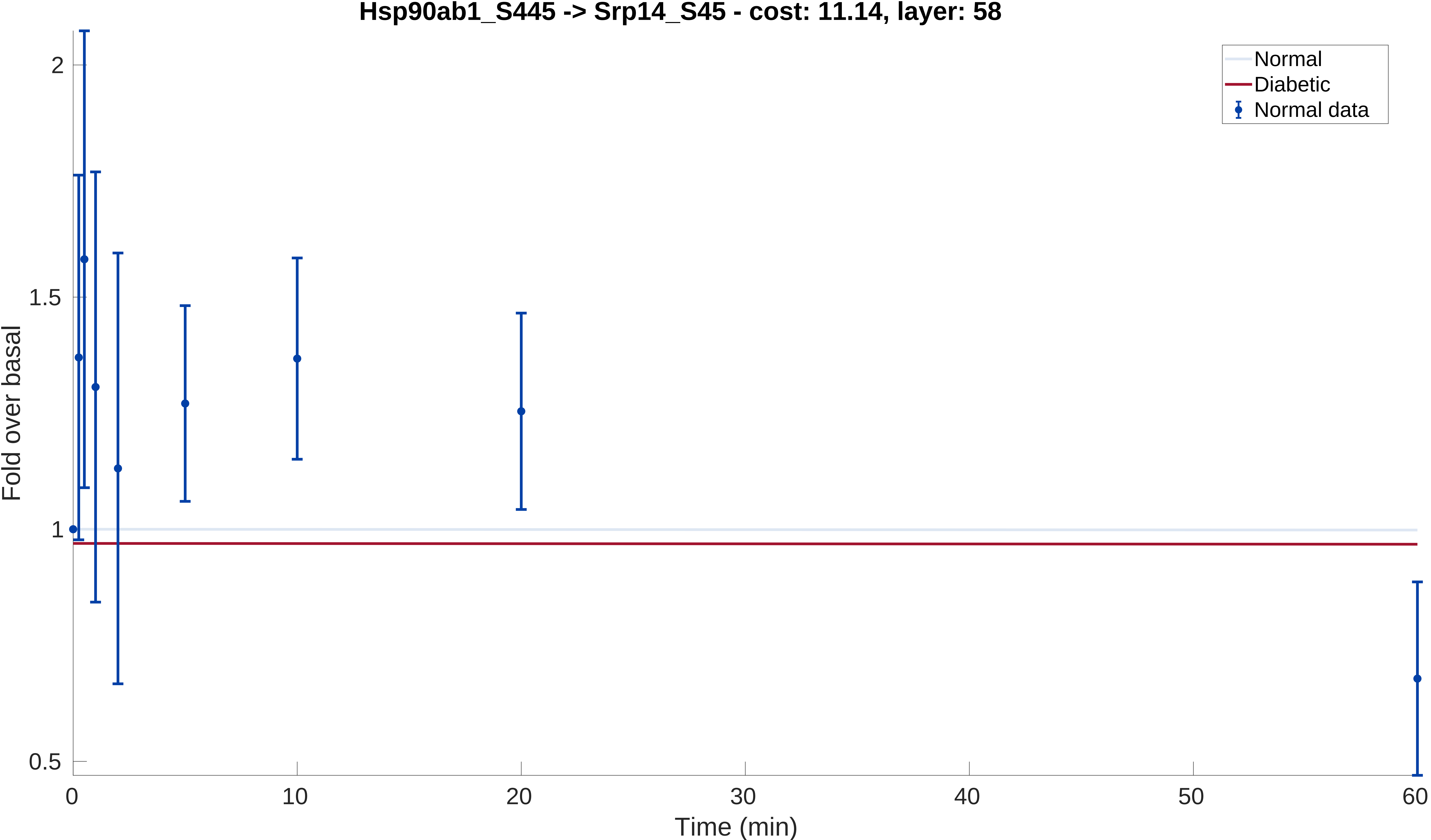
Hsp90ab1_S445 -> Srp14_S45 - cost: 11.14, layer: 58
Normal
Diabetic
Normal data
Fold over basal
2
1.5
1
0.5
0
10
20
30
40
50
60
Time (min)

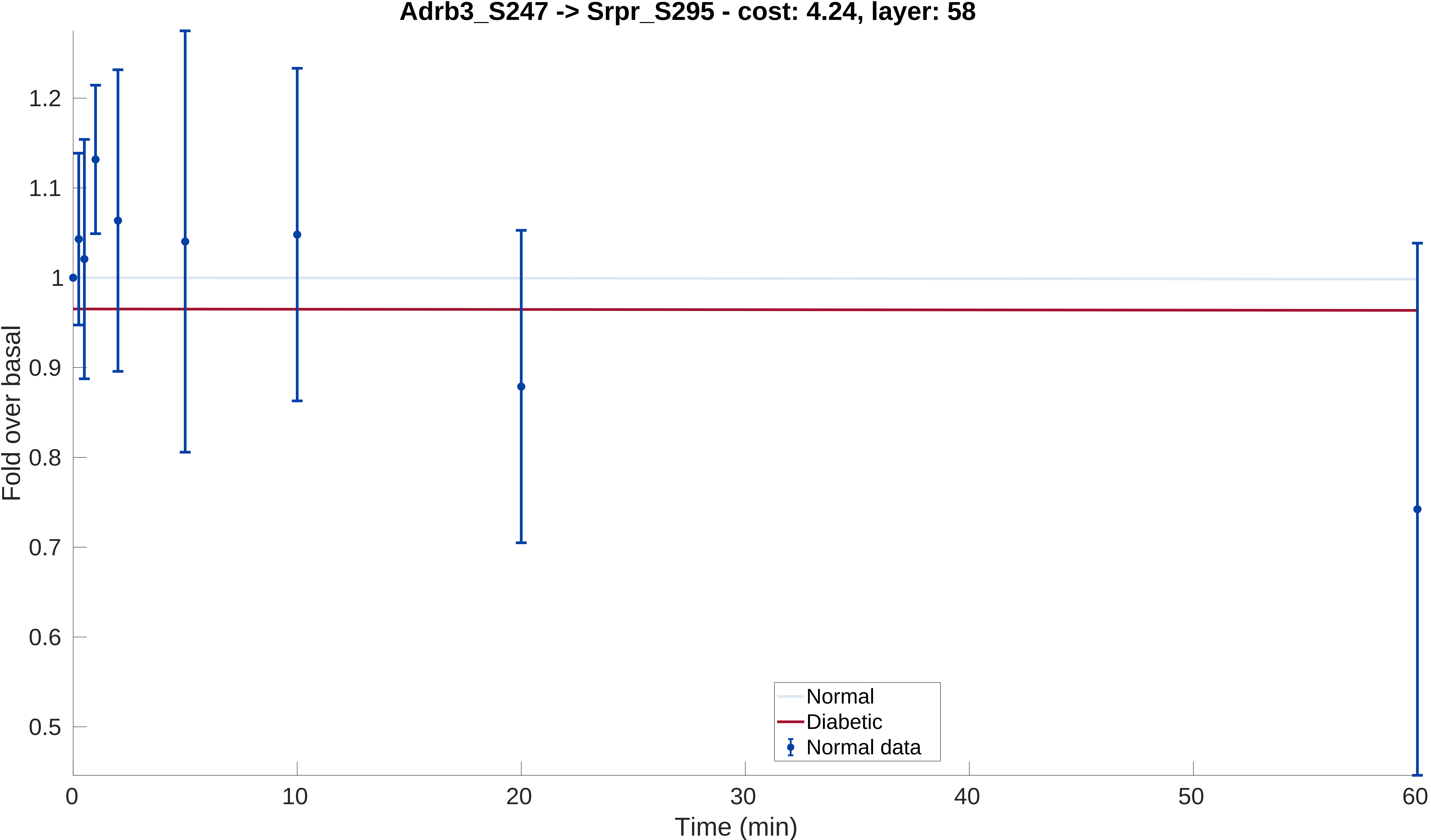
Adrb3_S247 -> Srpr_S295 - cost: 4.24, layer: 58
Fold over basal
1.2
1.1
1
0.9
0.8
0.7
0.6
0.5
0
10
20
30
40
50
60
Time (min)
Normal
Diabetic
Normal data

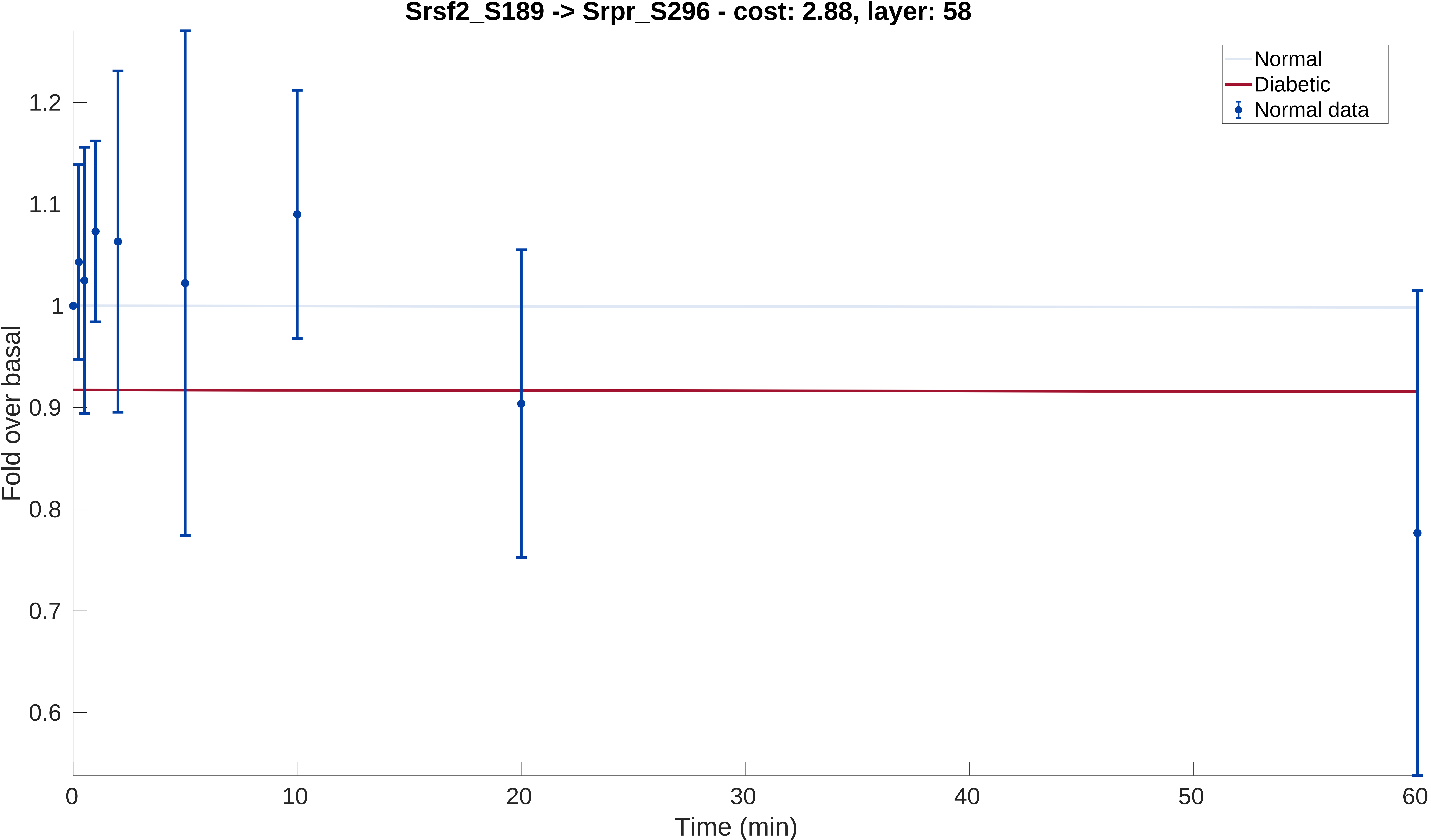
Srsf2_S189 -> Srpr_S296 - cost: 2.88, layer: 58
Normal
Diabetic
Normal data
Fold over basal
1.2
1.1
1
0.9
0.8
0.7
0.6
0
10
20
30
40
50
60
Time (min)

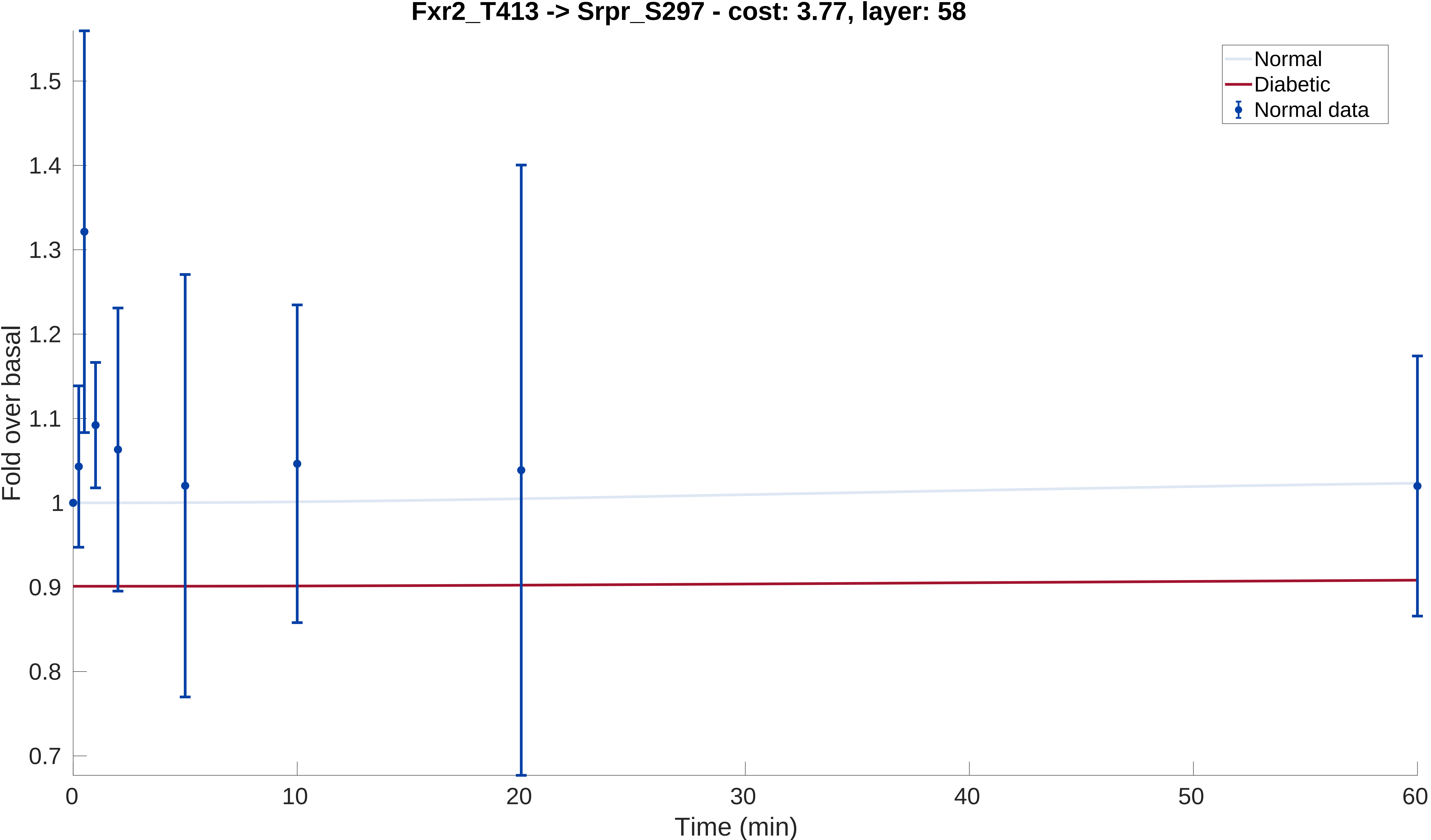

Fxr2_T413 -> Srpr_S297 - cost: 3.77, layer: 58
Normal
Diabetic
Normal data
Fold over basal
1.5
1.4
1.3
1.2
1.1
1
0.9
0.8
0.7
0
10
20
30
40
50
60
Time (min)

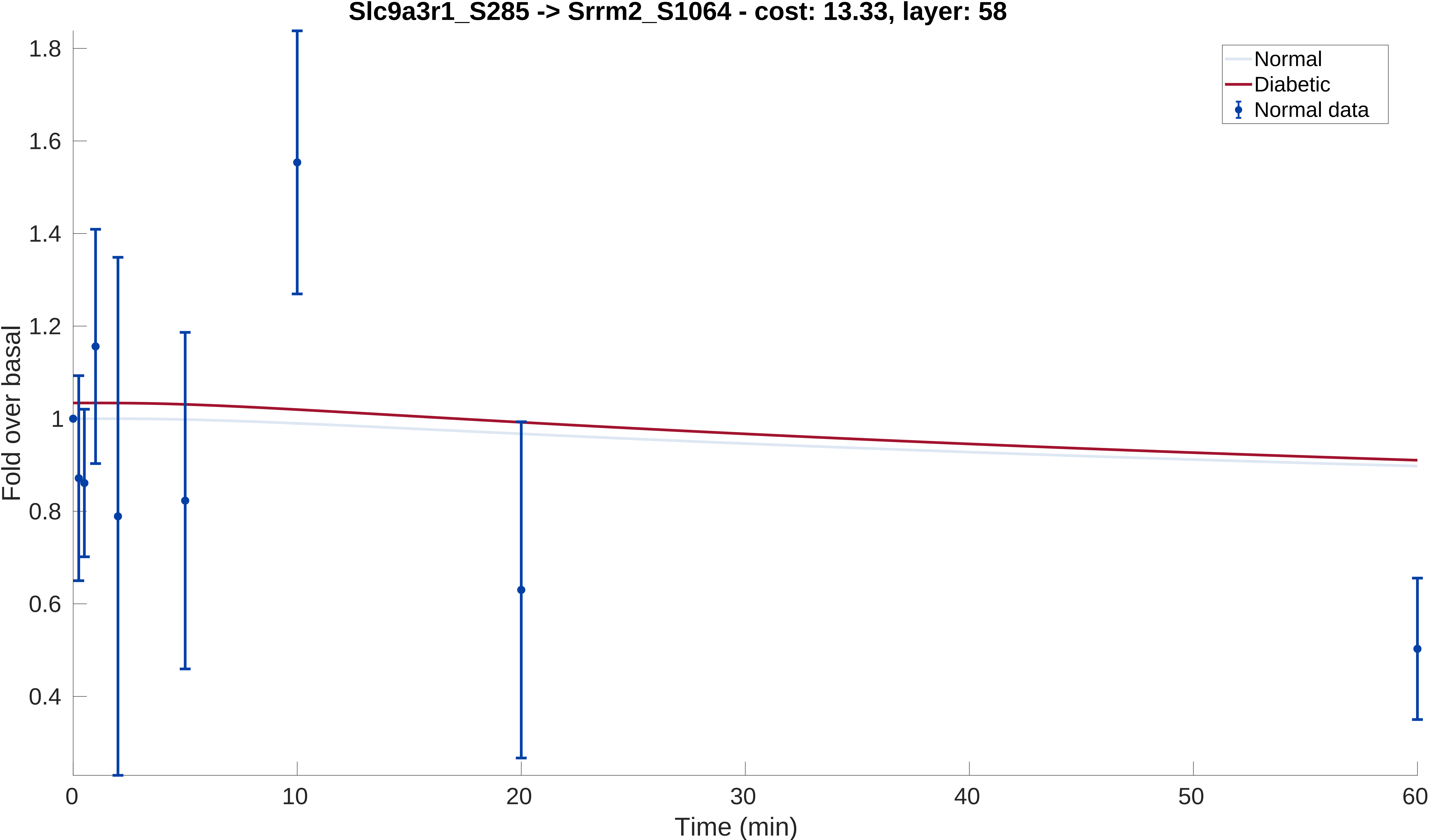
Slc9a3r1_S285 -> Srrm2_S1064 - cost: 13.33, layer: 58
Normal
Diabetic
Normal data
Fold over basal
1.8
1.6
1.4
1.2
1
0.8
0.6
0.4
0
10
20
30
40
50
60
Time (min)

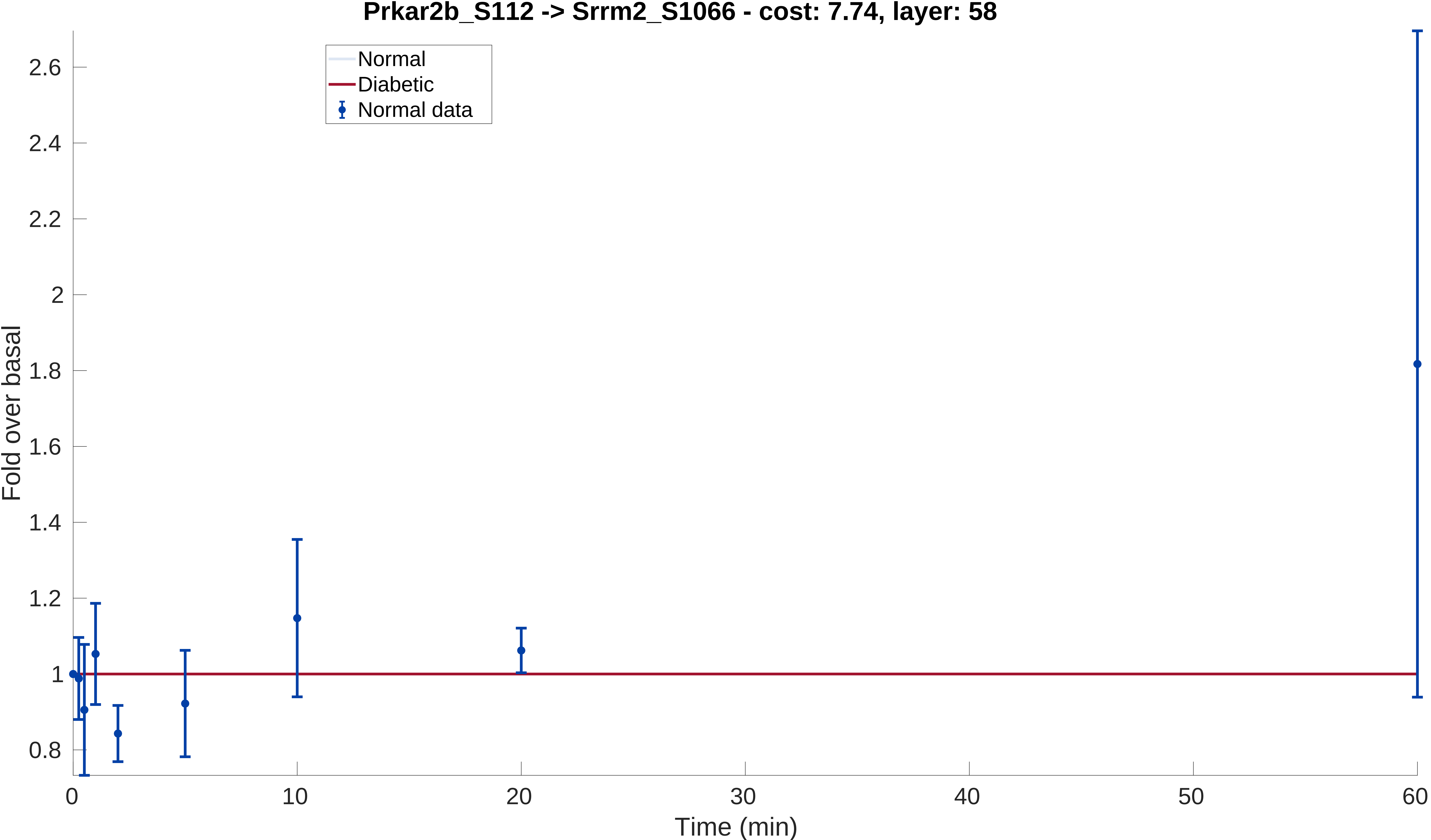
Prkar2b_S112 -> Srrm2_S1066 - cost: 7.74, layer: 58
Normal
Diabetic
Normal data
Fold over basal
2.6
2.4
2.2
2
1.8
1.6
1.4
1.2
1
0.8
0
10
20
30
40
50
60
Time (min)

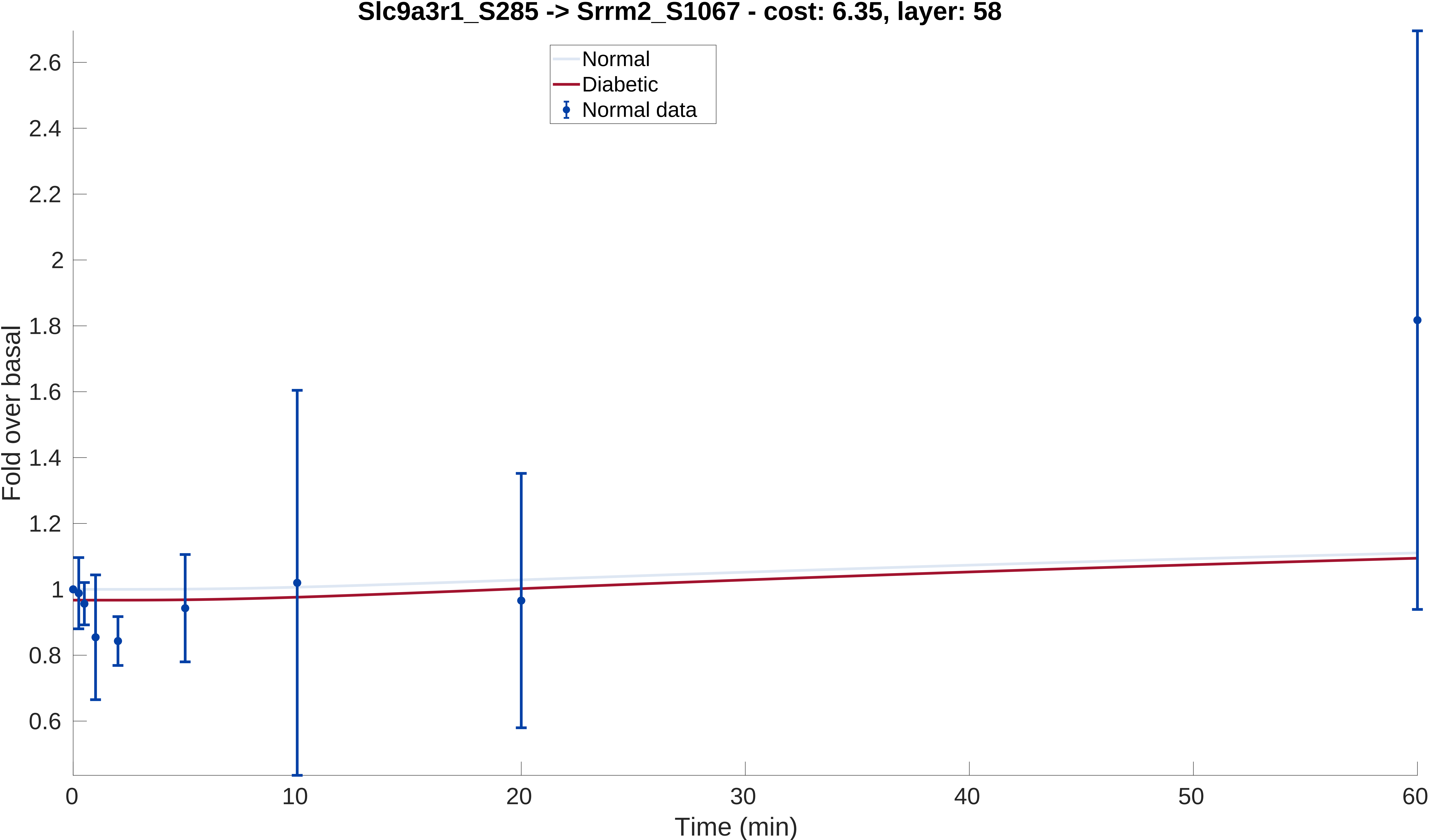

Slc9a3r1_S285 -> Srrm2_S1067 - cost: 6.35, layer: 58
Normal
Diabetic
Normal data
Fold over basal
2.6
2.4
2.2
2
1.8
1.6
1.4
1.2
1
0.8
0.6
0
10
20
30
40
50
60
Time (min)

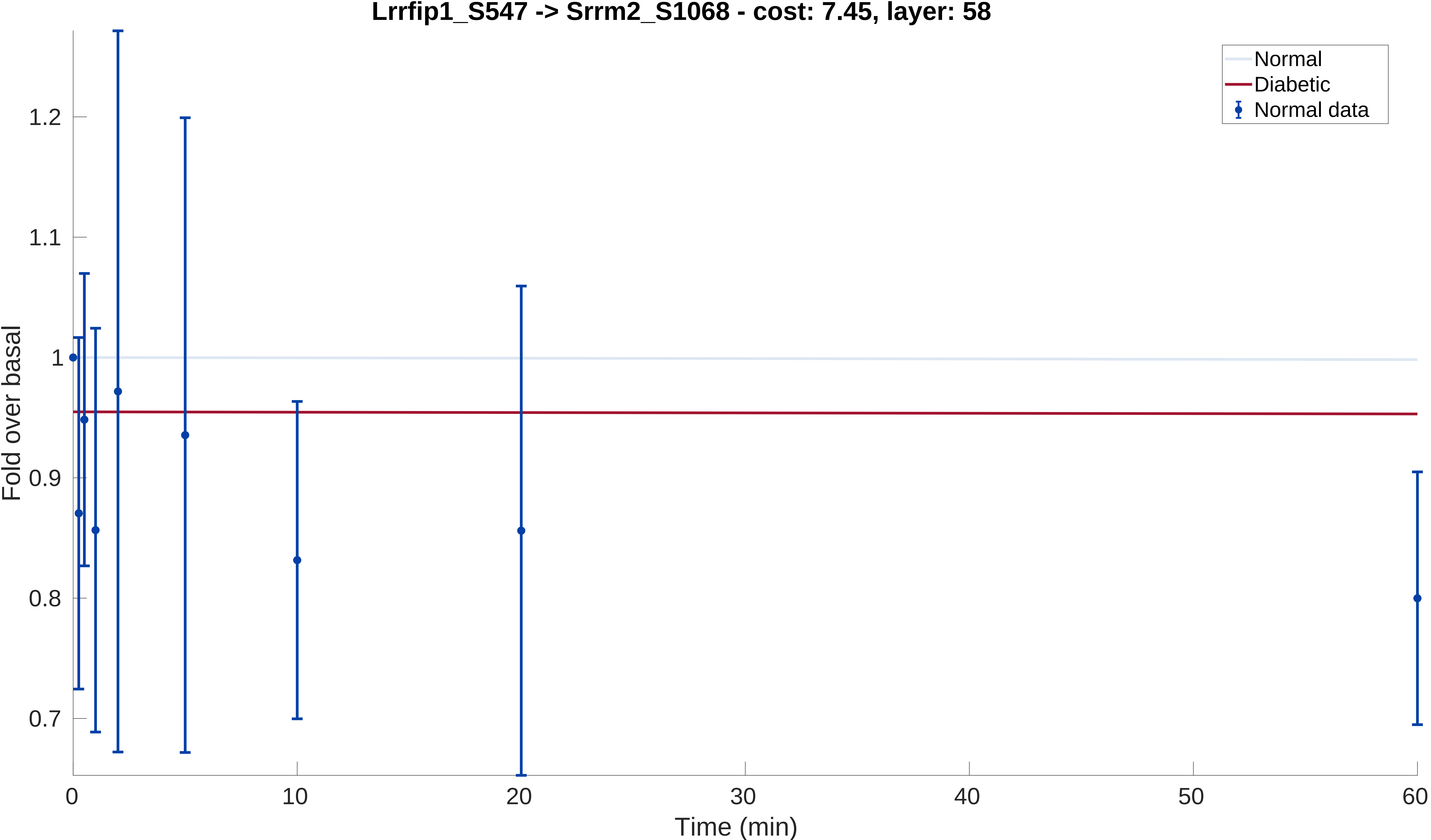

Lrrfip1_S547 -> Srrm2_S1068 - cost: 7.45, layer: 58
Normal
Diabetic
Normal data
Fold over basal
1.2
1.1
1
0.9
0.8
0.7
0
10
20
30
40
50
60
Time (min)

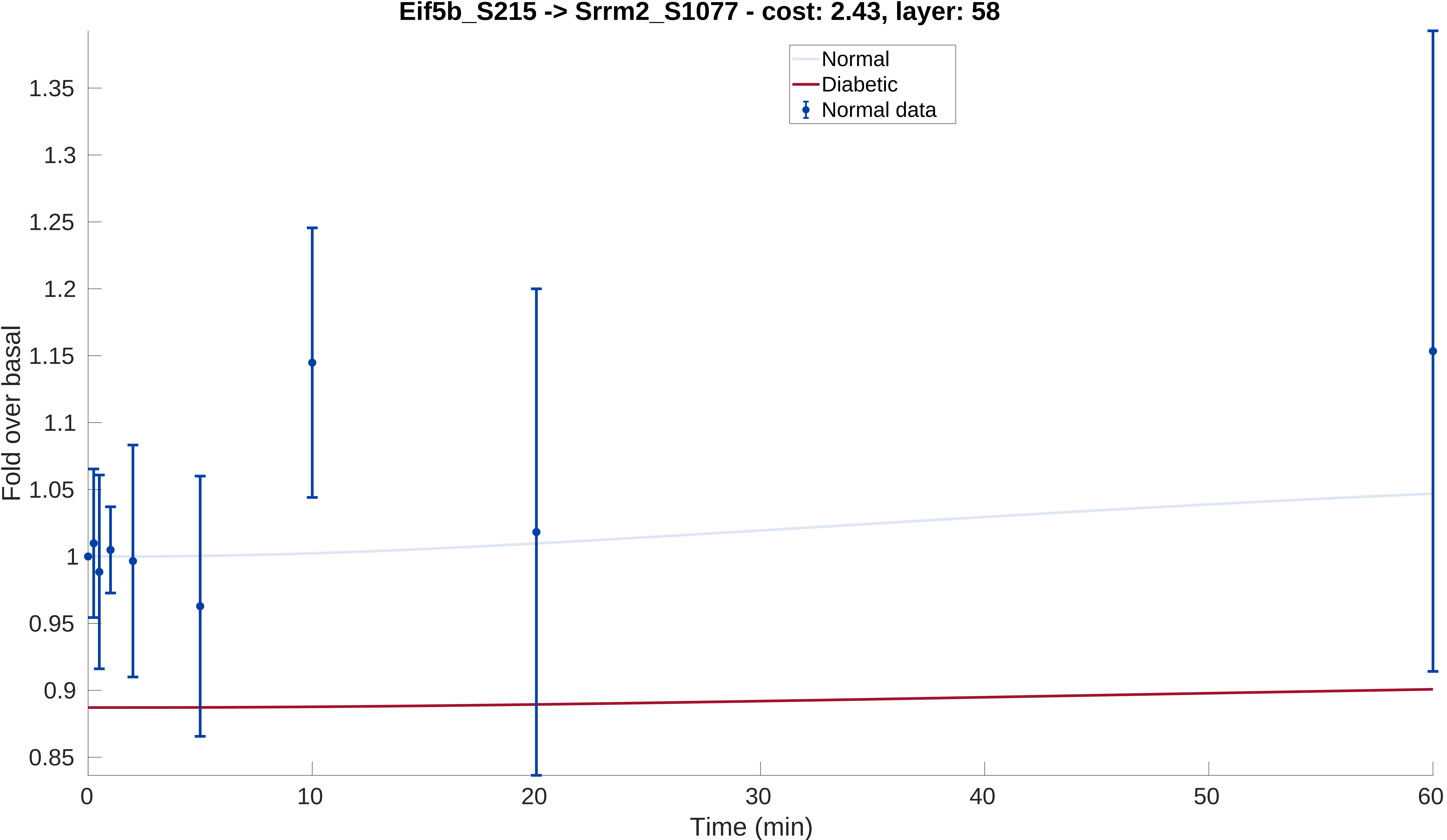
Eif5b_S215 -> Srrm2_S1077 - cost: 2.43, layer: 58
Normal
Diabetic
Normal data
Fold over basal
1.35
1.3
1.25
1.2
1.15
1.1
1.05
1
0.95
0.9
0.85
0
10
20
30
40
50
60
Time (min)

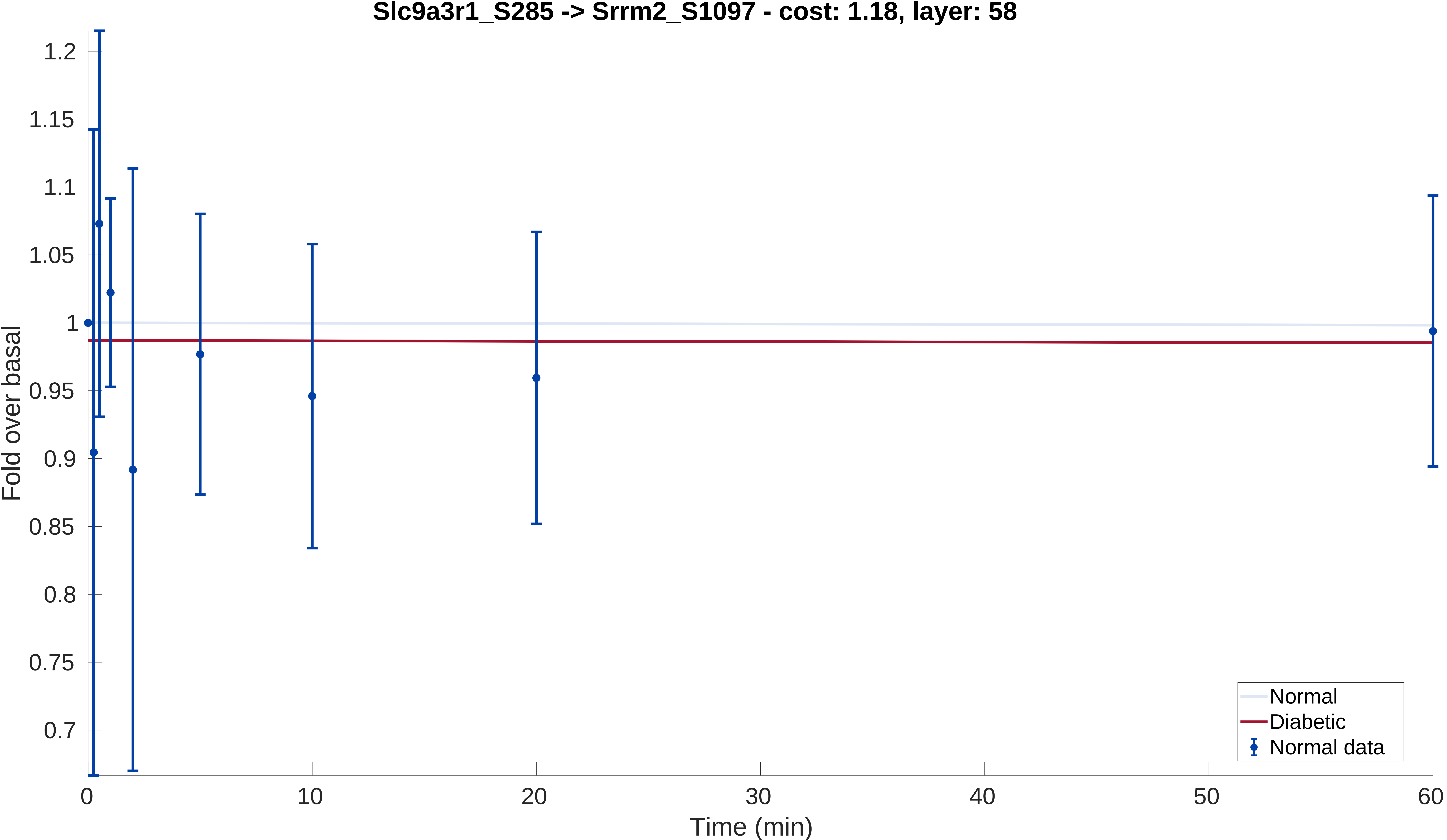

Slc9a3r1_S285 -> Srrm2_S1097 - cost: 1.18, layer: 58
Fold over basal
1.2
1.15
1.1
1.05
1
0.95
0.9
0.85
0.8
0.75
0.7
0
10
20
30
40
50
60
Time (min)
Normal
Diabetic
Normal data

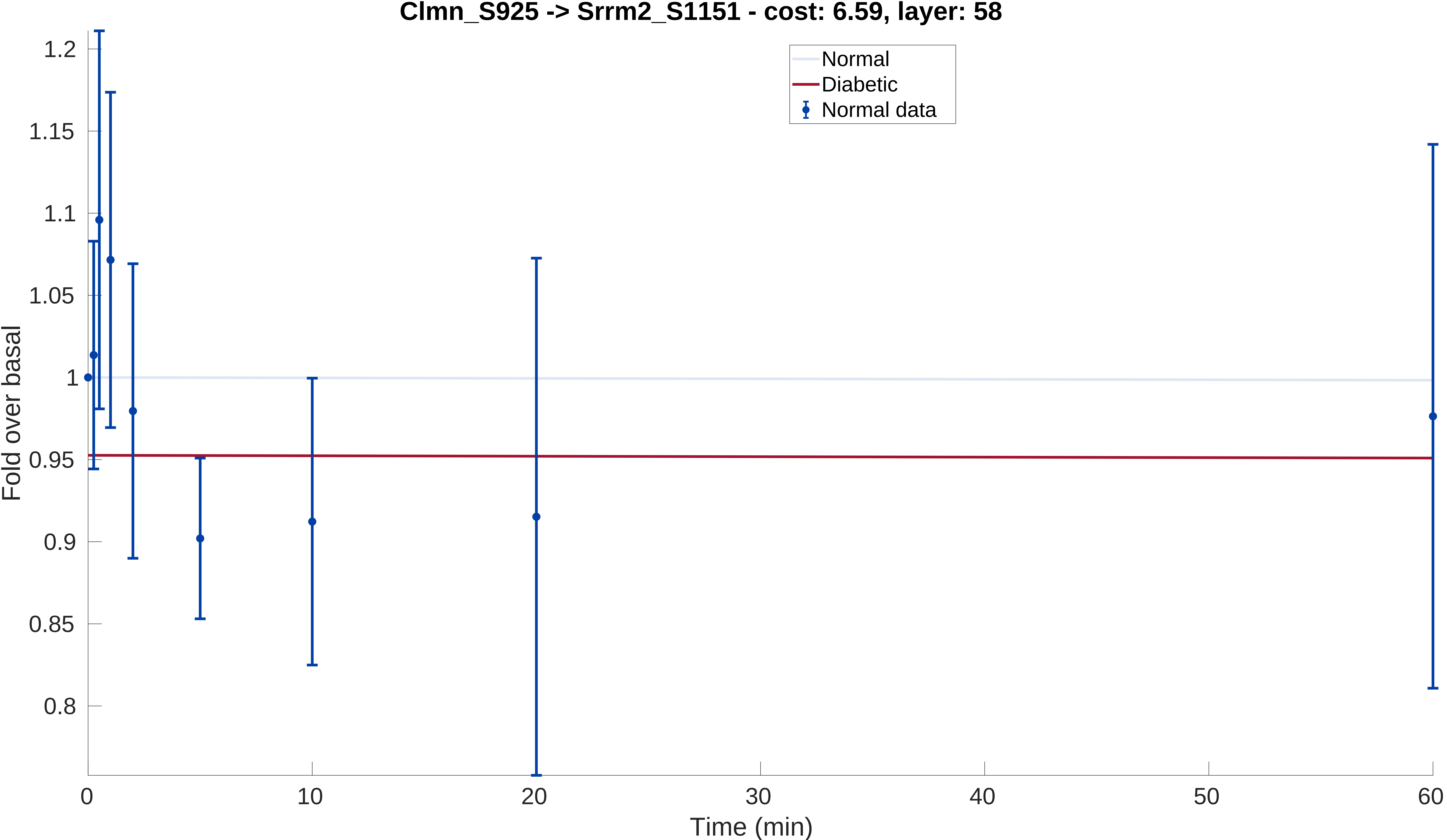

Clmn_S925 -> Srrm2_S1151 - cost: 6.59, layer: 58
Normal
Diabetic
Normal data
Fold over basal
1.2
1.15
1.1
1.05
1
0.95
0.9
0.85
0.8
0
10
20
30
40
50
60
Time (min)

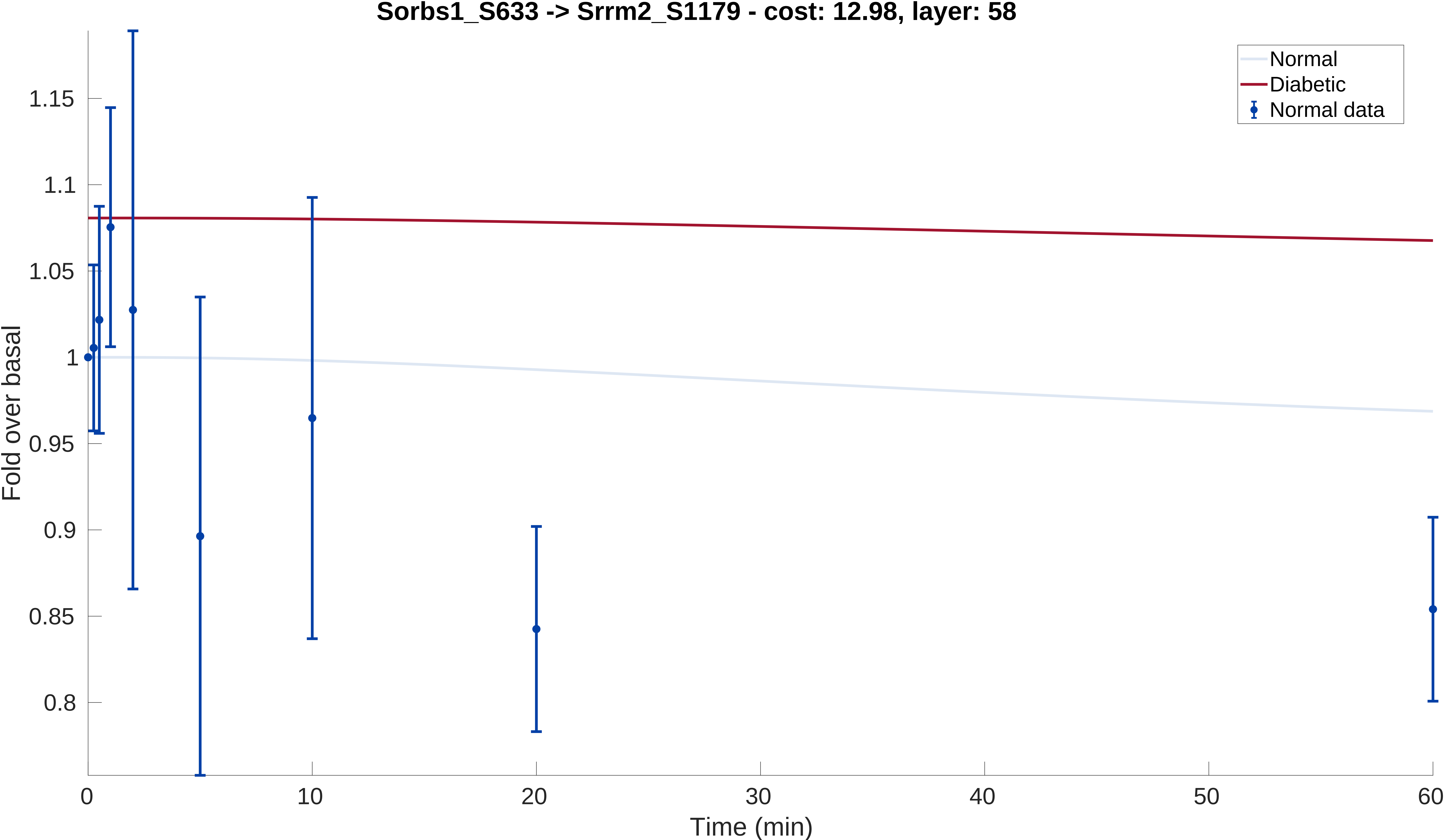
Sorbs1_S633 -> Srrm2_S1179 - cost: 12.98, layer: 58
Normal
Diabetic
Normal data
Fold over basal
1.15
1.1
1.05
1
0.95
0.9
0.85
0.8
0
10
20
30
40
50
60
Time (min)

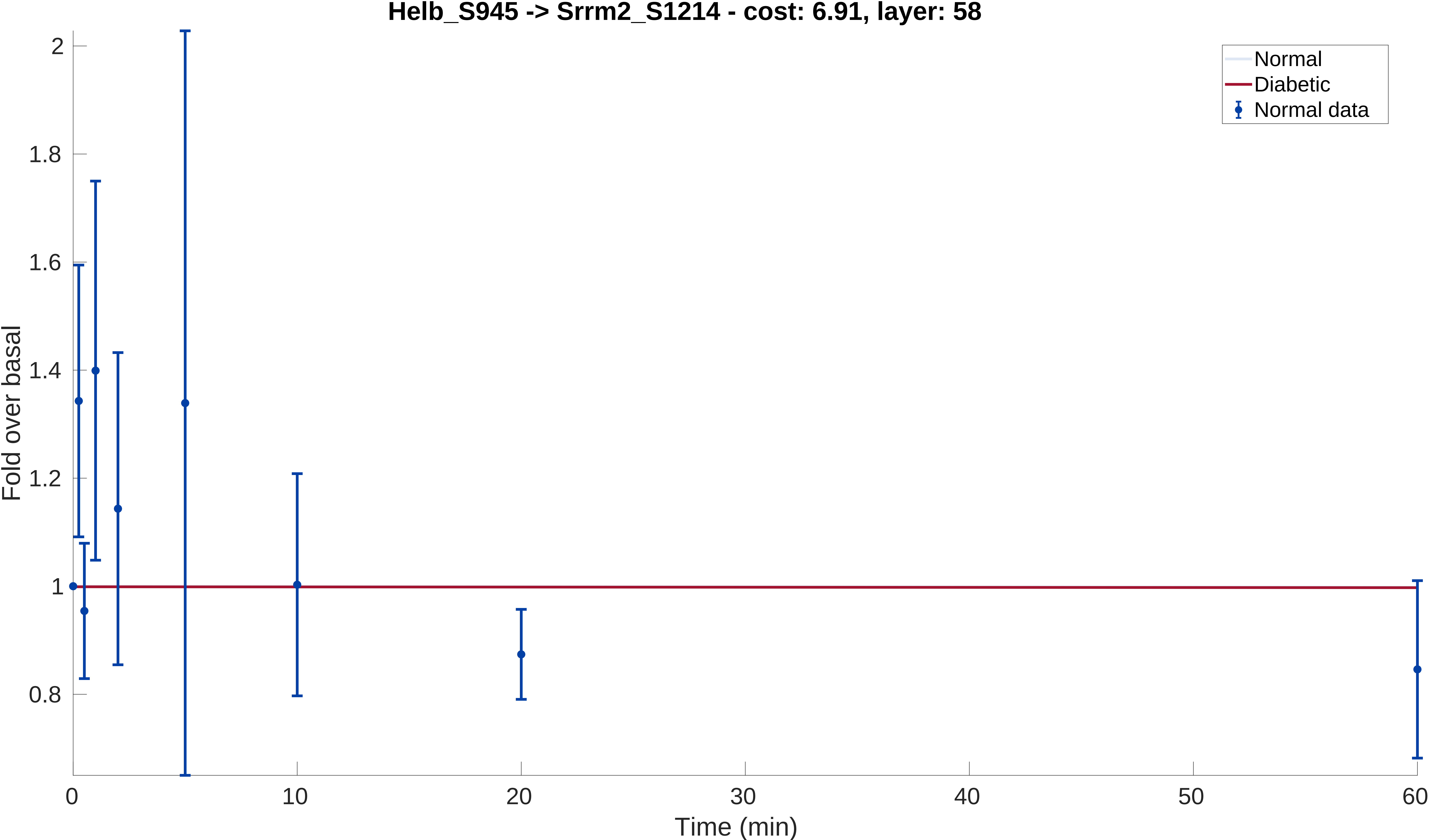

Helb_S945 -> Srrm2_S1214 - cost: 6.91, layer: 58
Normal
Diabetic
Normal data
Fold over basal
Time (min)
0.8
1
1.2
1.4
1.6
1.8
2
0
10
20
30
40
50
60

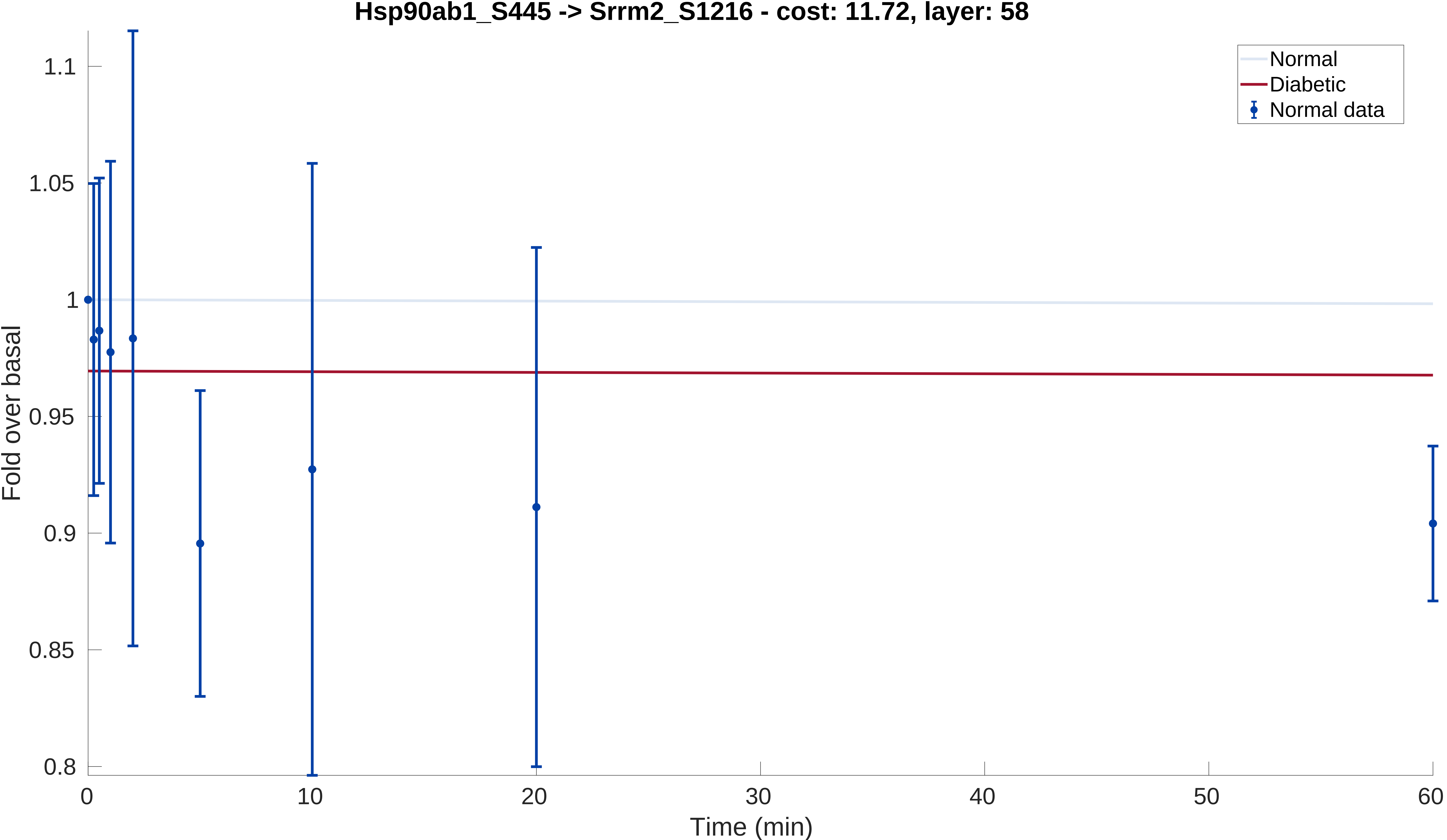

Hsp90ab1_S445 -> Srrm2_S1216 - cost: 11.72, layer: 58
Normal
Diabetic
Normal data
Fold over basal
Time (min)
1.1
1.05
1
0.95
0.9
0.85
0.8
0
10
20
30
40
50
60

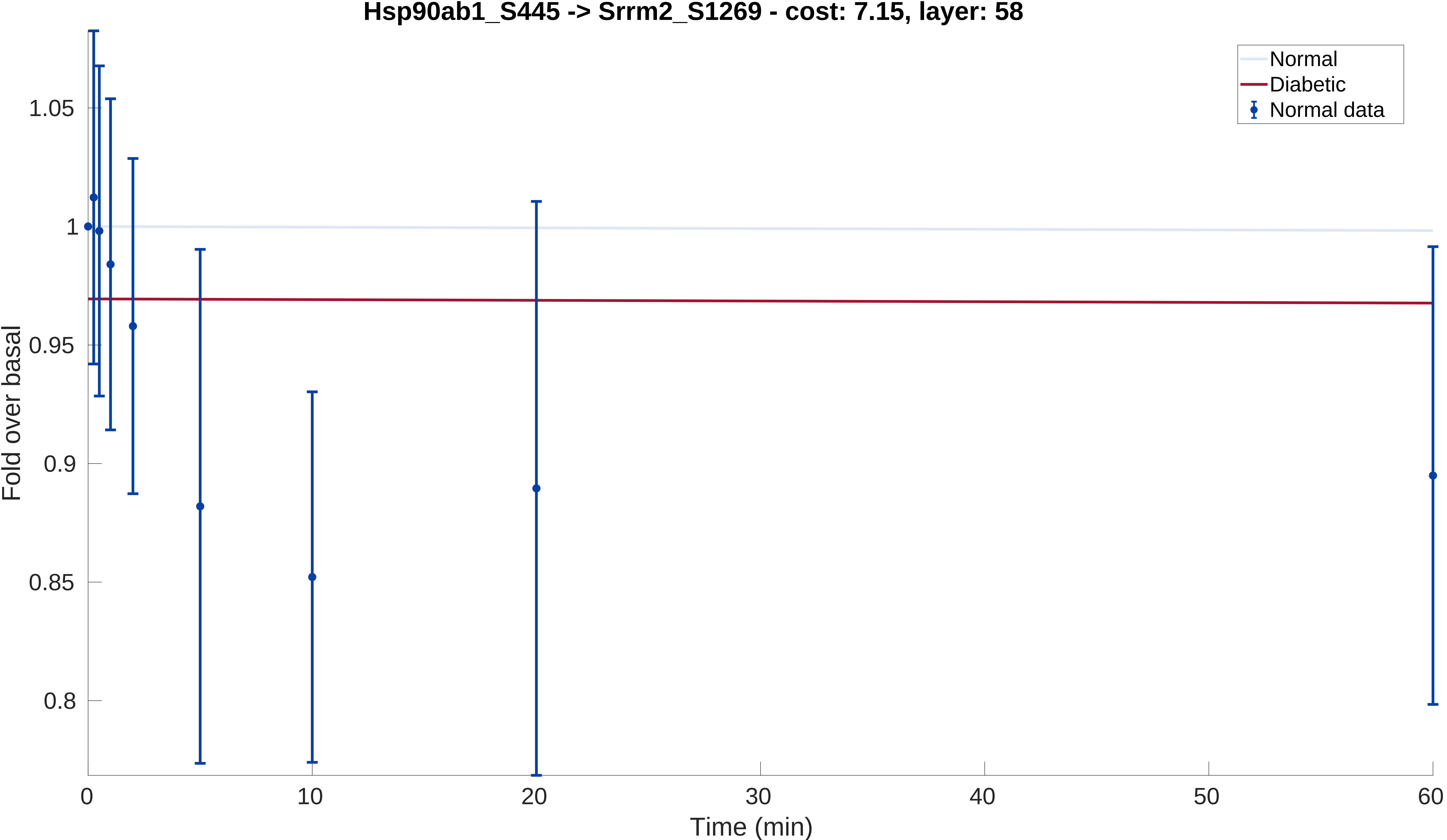

Hsp90ab1_S445 -> Srrm2_S1269 - cost: 7.15, layer: 58
Normal
Diabetic
Normal data
Fold over basal
1.05
1
0.95
0.9
0.85
0.8
0
10
20
30
40
50
60
Time (min)

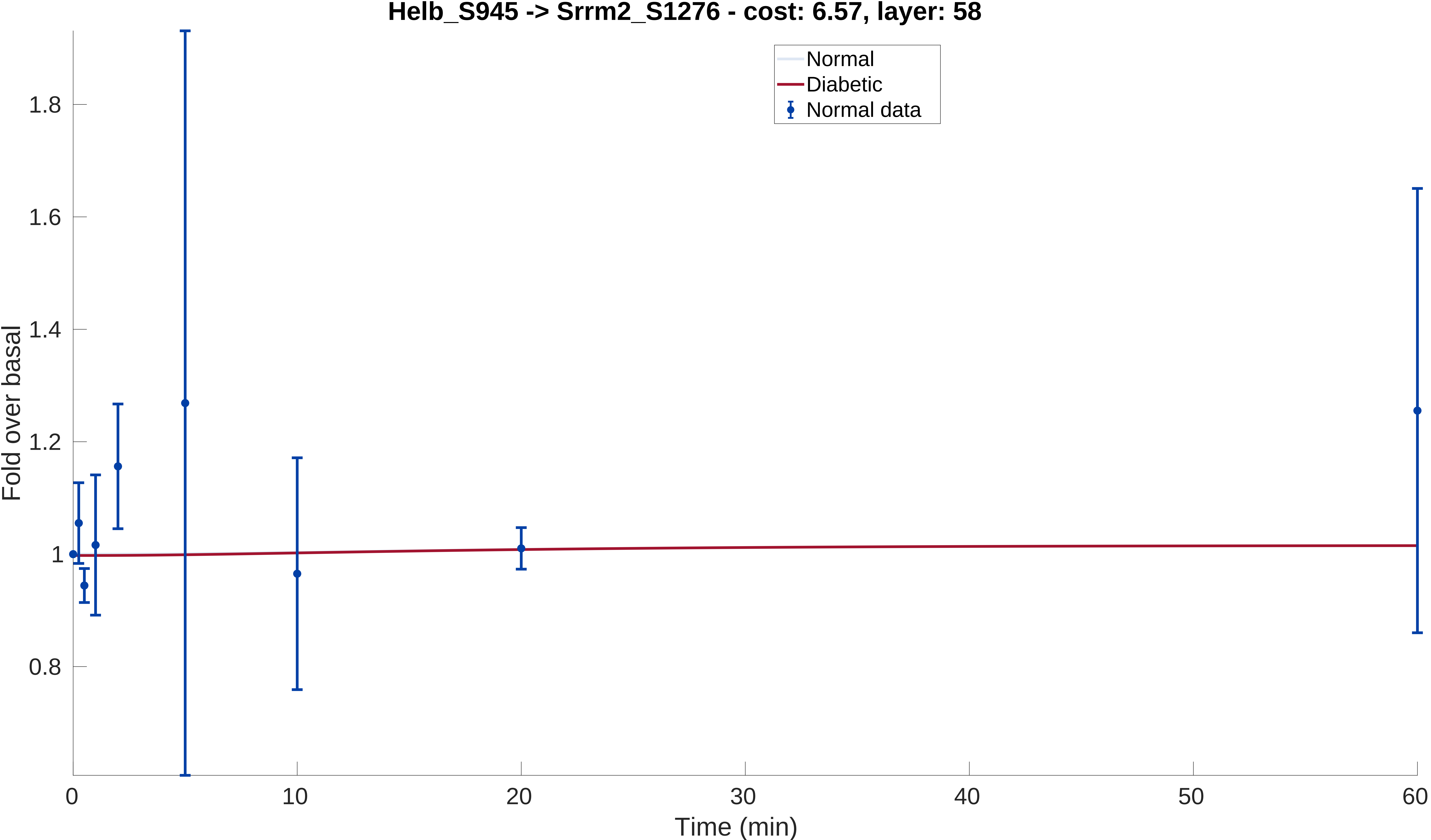

Helb_S945 -> Srrm2_S1276 - cost: 6.57, layer: 58
Normal
Diabetic
Normal data
Fold over basal
Time (min)
0.8
1
1.2
1.4
1.6
1.8
0
10
20
30
40
50
60

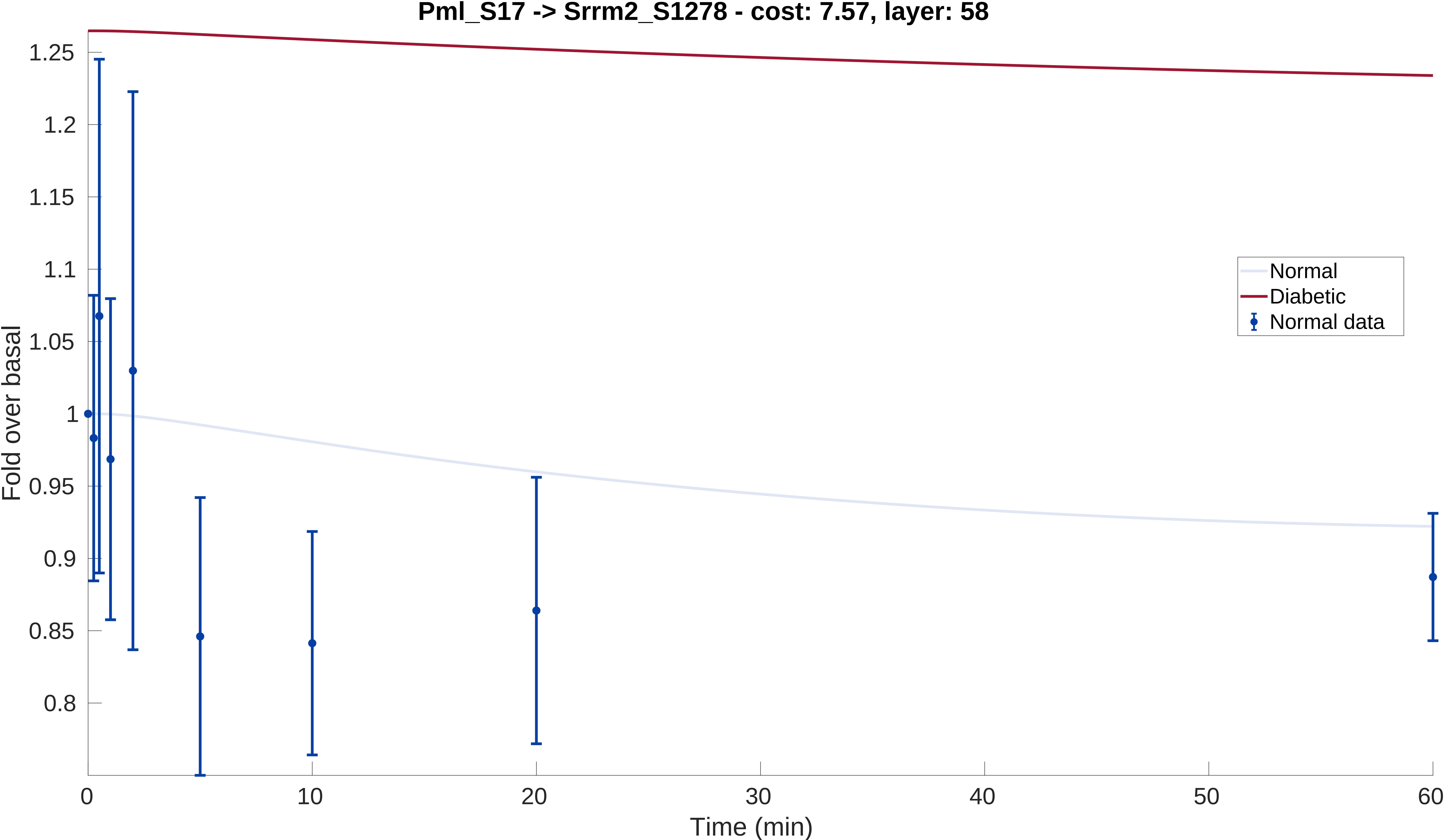

Pml_S17 -> Srrm2_S1278 - cost: 7.57, layer: 58
Fold over basal
Time (min)
Normal
Diabetic
Normal data
1.25
1.2
1.15
1.1
1.05
1
0.95
0.9
0.85
0.8
0
10
20
30
40
50
60

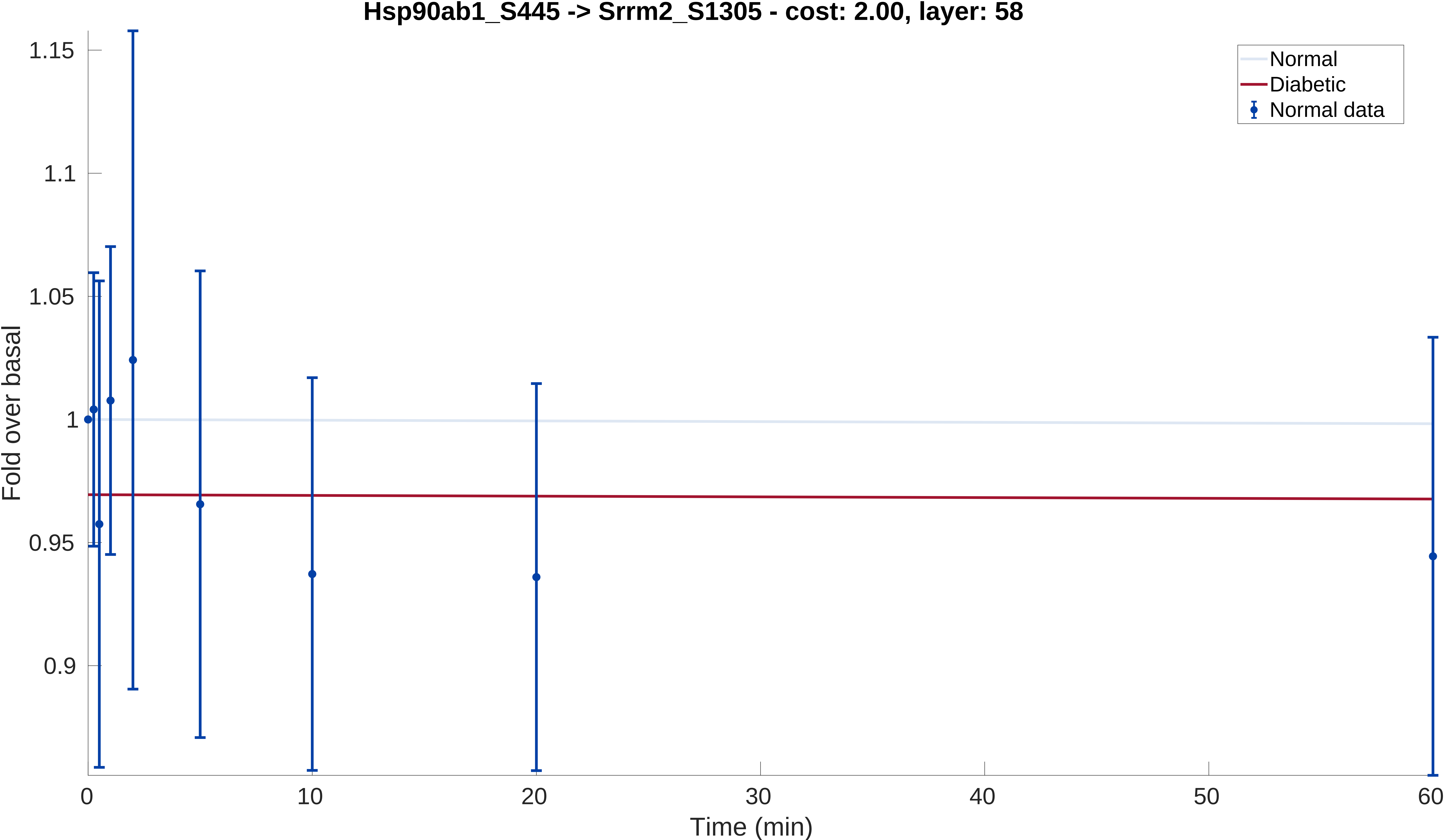
Hsp90ab1_S445 -> Srrm2_S1305 - cost: 2.00, layer: 58
Fold over basal
1.15
1.1
1.05
1
0.95
0.9
0
10
20
30
40
50
60
Time (min)
Normal
Diabetic
Normal data

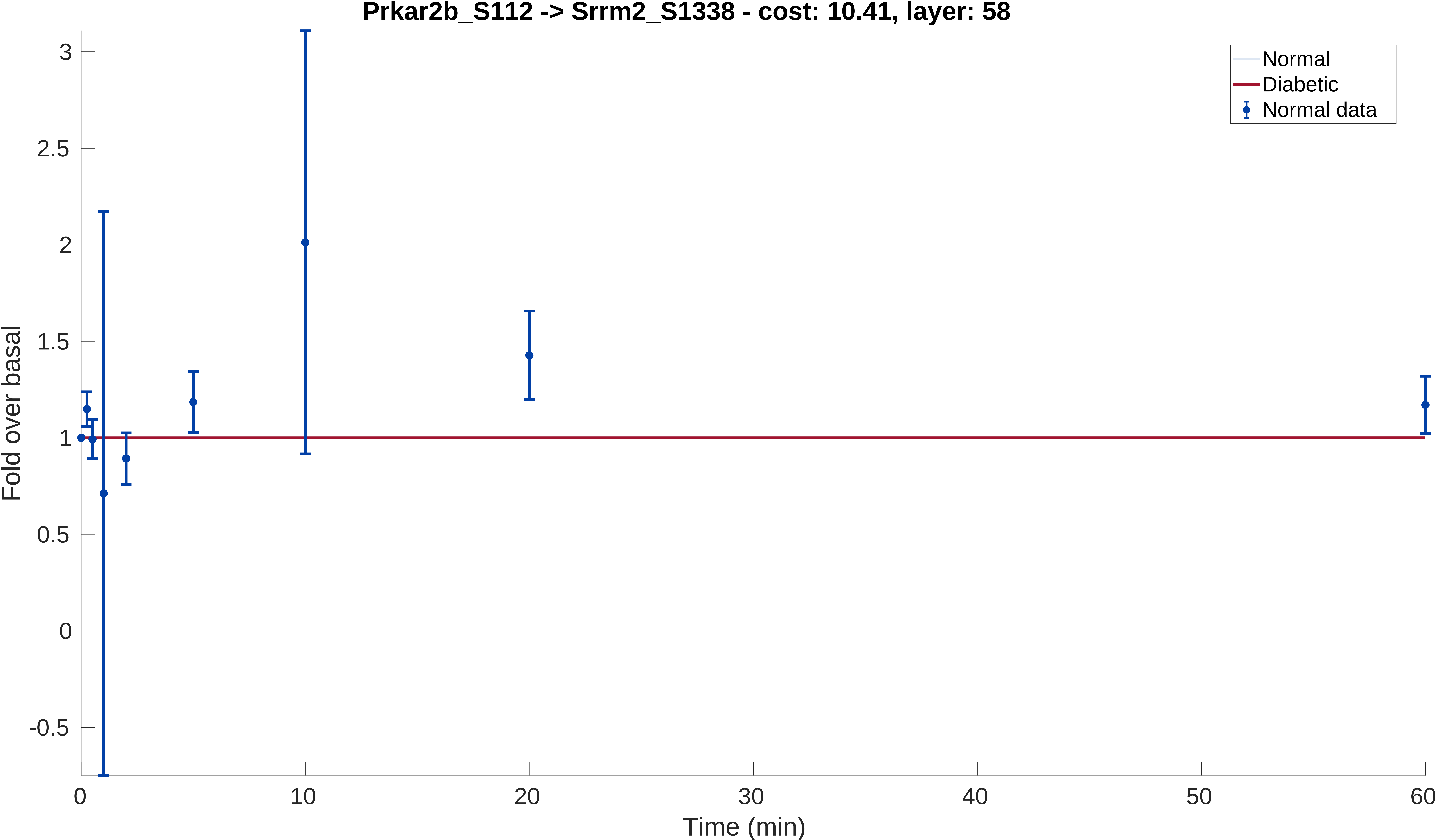

Prkar2b_S112 -> Srrm2_S1338 - cost: 10.41, layer: 58
Normal
Diabetic
Normal data
Fold over basal
3
2.5
2
1.5
1
0.5
0
-0.5
0
10
20
30
40
50
60
Time (min)

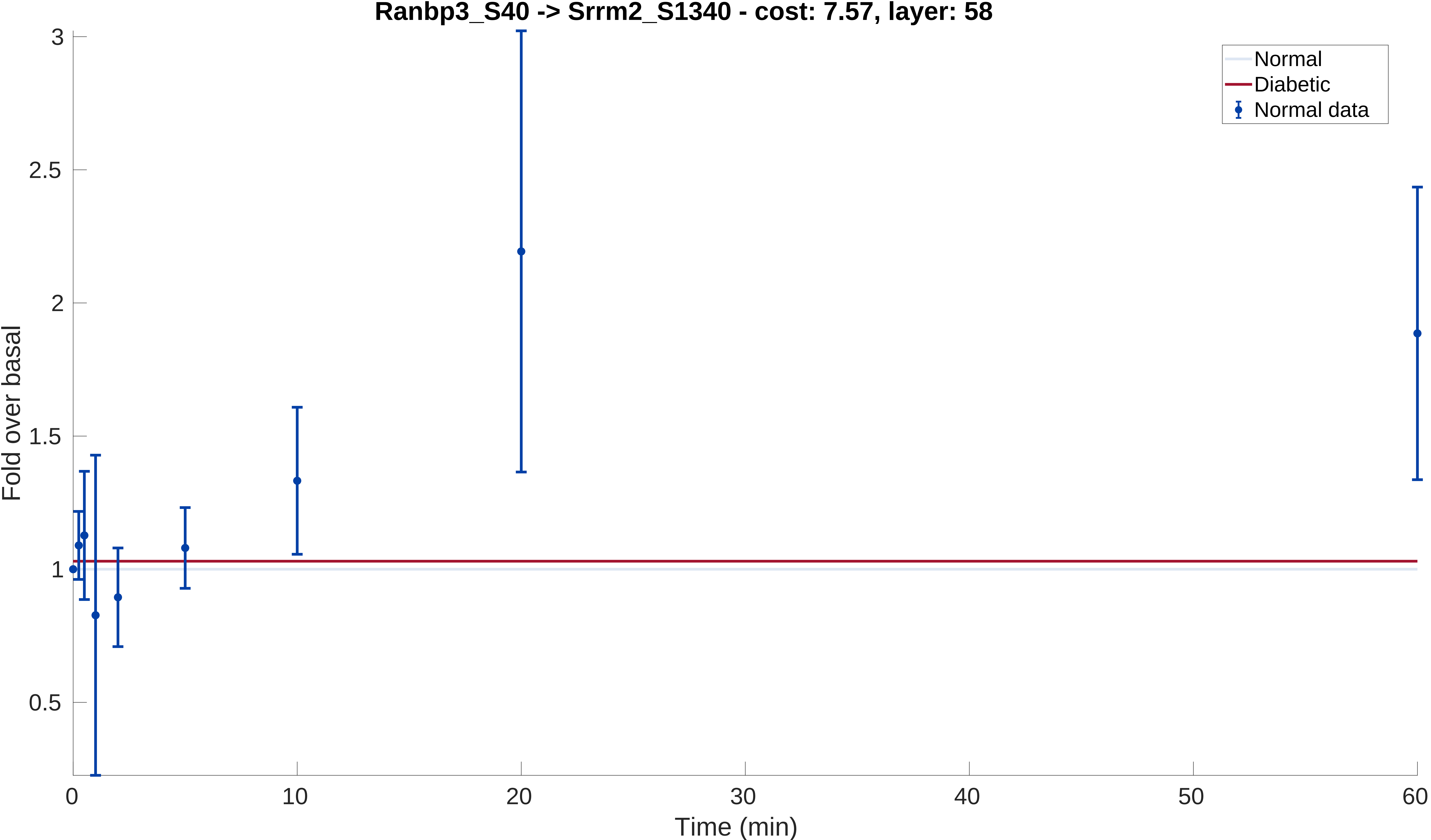

Ranbp3_S40 -> Srrm2_S1340 - cost: 7.57, layer: 58
Normal
Diabetic
Normal data
Fold over basal
Time (min)
0.5
1
1.5
2
2.5
3
0
10
20
30
40
50
60

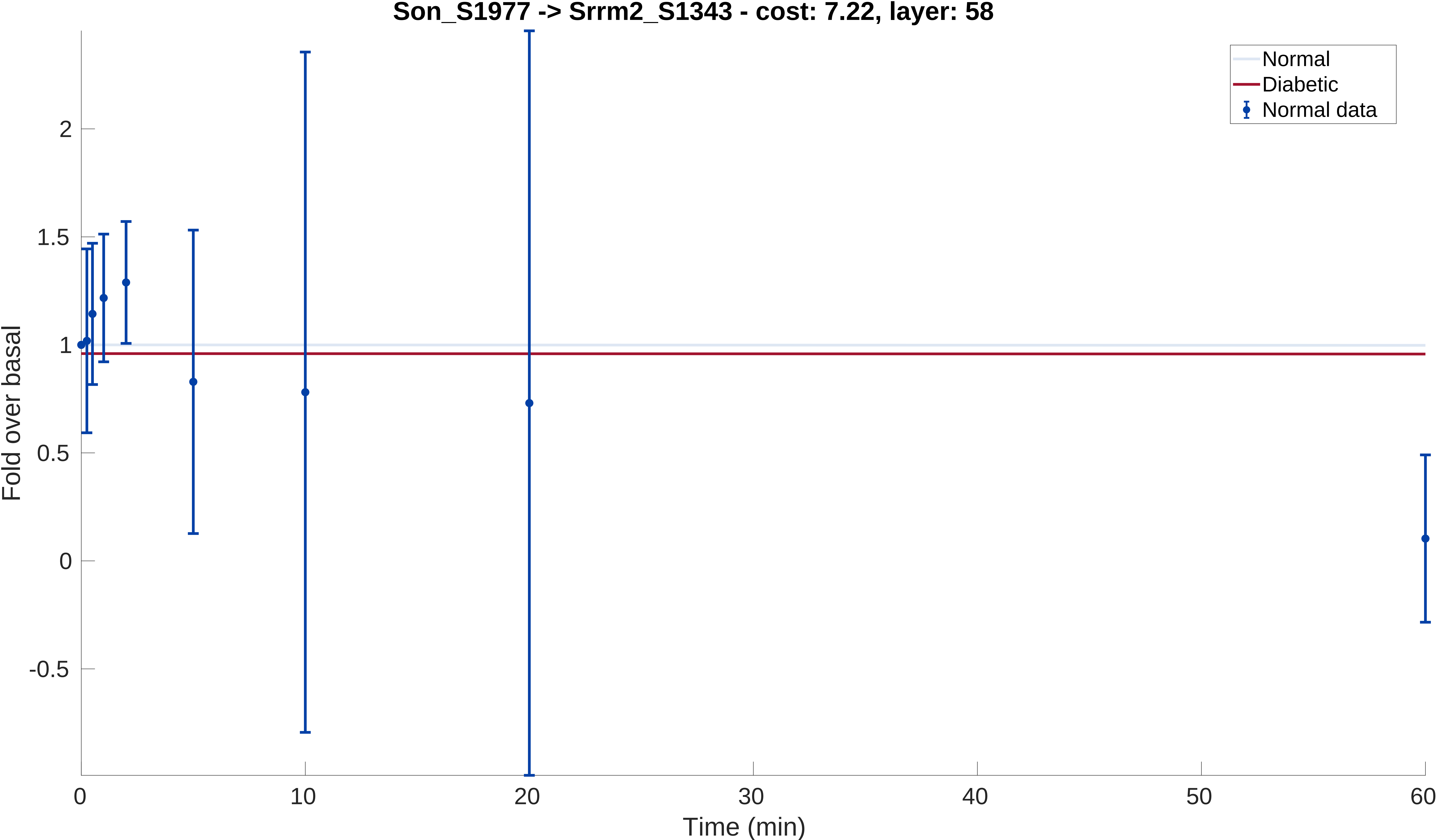
Son_S1977 -> Srrm2_S1343 - cost: 7.22, layer: 58
Normal
Diabetic
Normal data
Fold over basal
Time (min)
2
1.5
1
0.5
0
-0.5
0
10
20
30
40
50
60

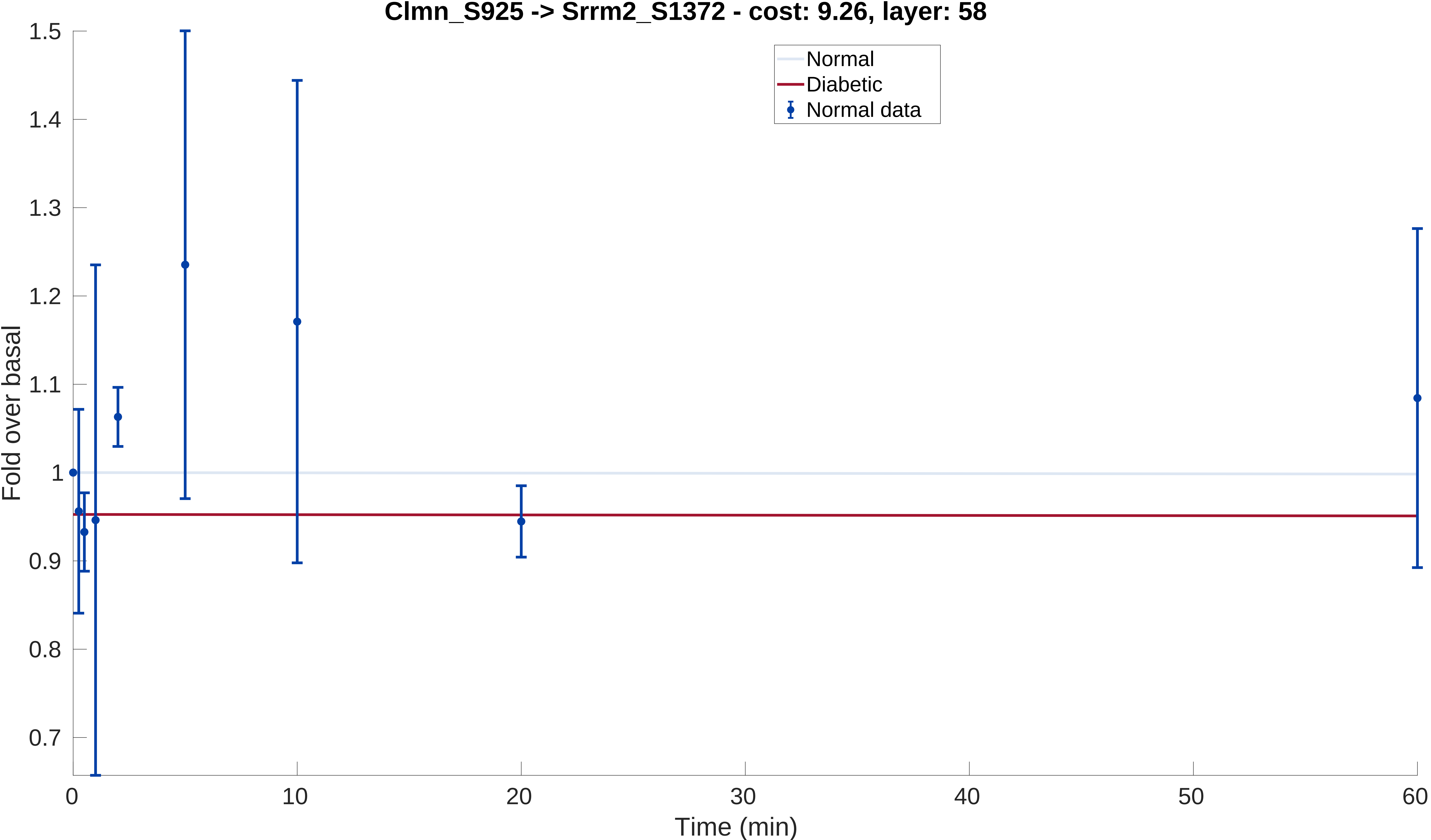

Clmn_S925 -> Srrm2_S1372 - cost: 9.26, layer: 58
Normal
Diabetic
Normal data
Fold over basal
Time (min)
1.5
1.4
1.3
1.2
1.1
1
0.9
0.8
0.7
0
10
20
30
40
50
60

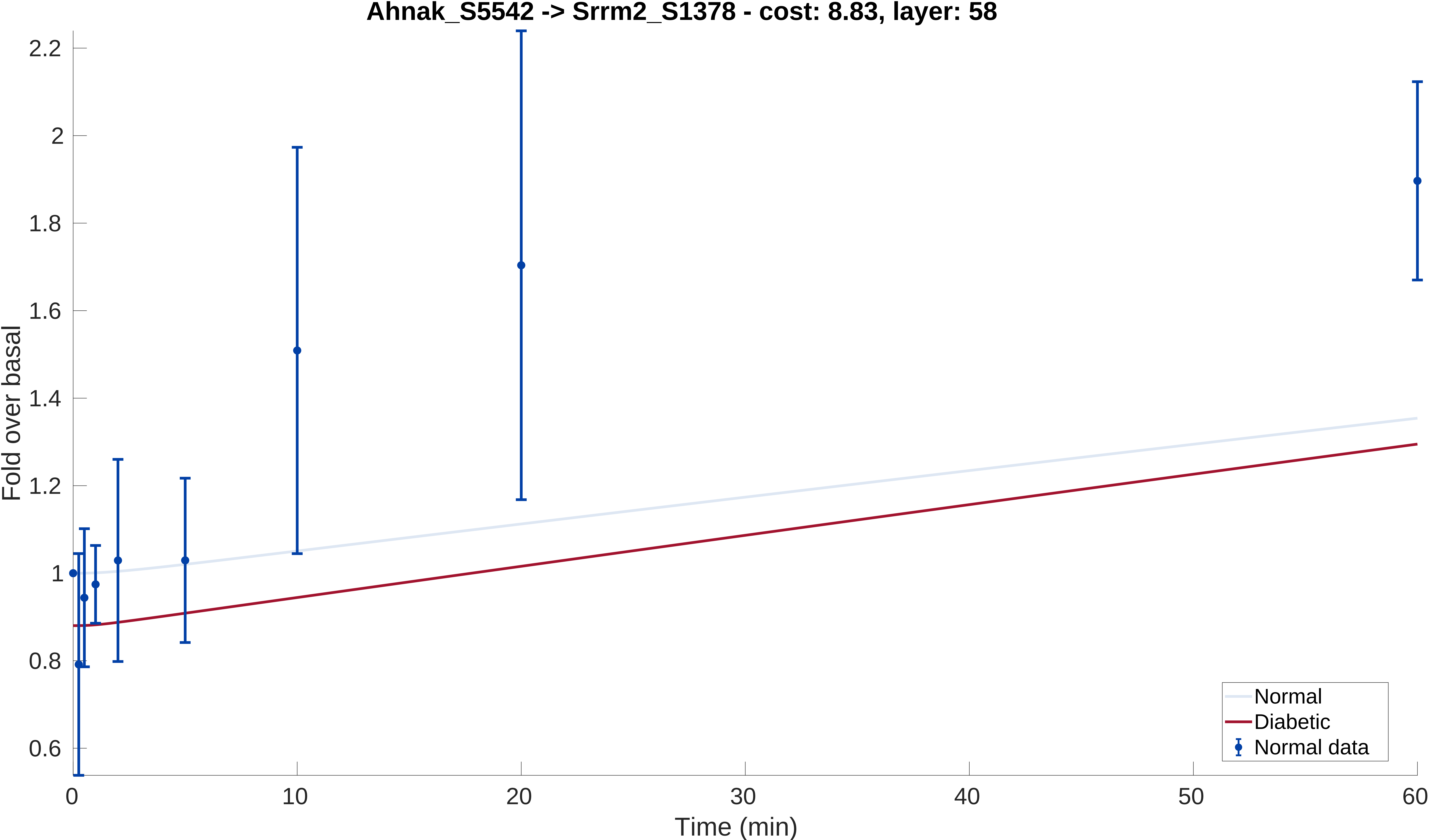

Ahnak_S5542 -> Srrm2_S1378 - cost: 8.83, layer: 58
Fold over basal
2.2
2
1.8
1.6
1.4
1.2
1
0.8
0.6
0
10
20
30
40
50
60
Time (min)
Normal
Diabetic
Normal data

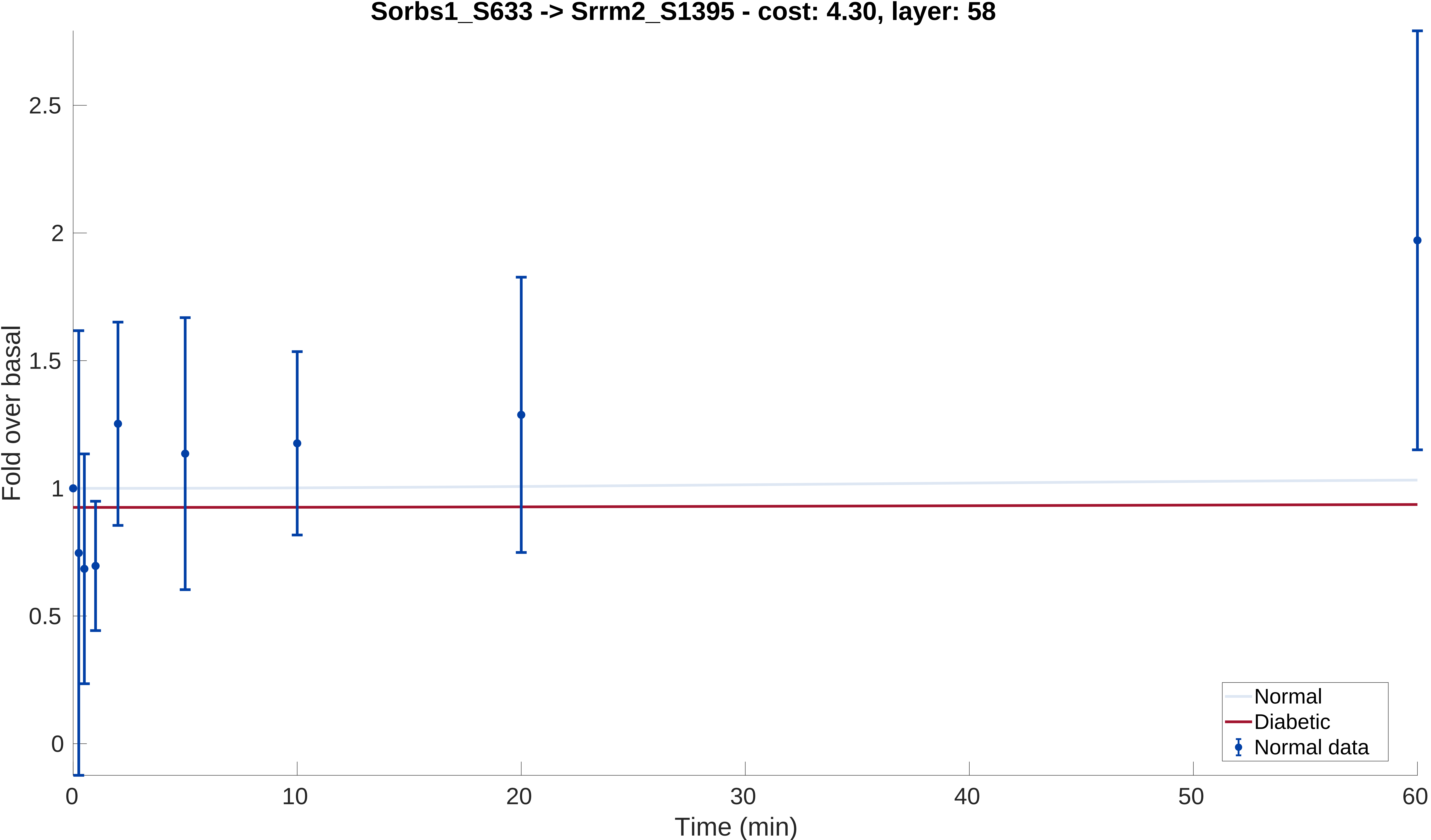

Sorbs1_S633 -> Srrm2_S1395 - cost: 4.30, layer: 58
Fold over basal
2.5
2
1.5
1
0.5
0
0
10
20
30
40
50
60
Time (min)
Normal
Diabetic
Normal data

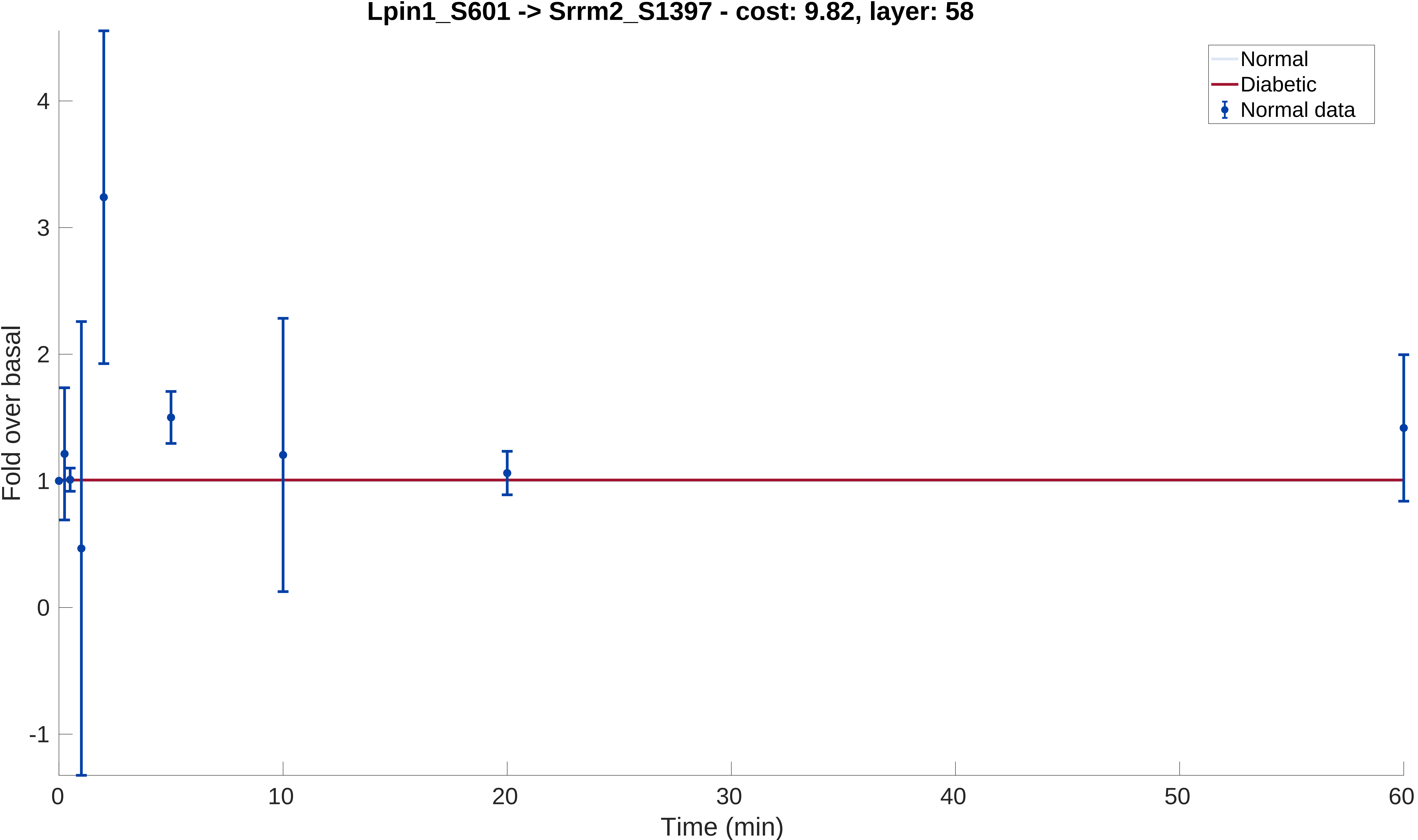

Lpin1_S601 -> Srrm2_S1397 - cost: 9.82, layer: 58
Normal
Diabetic
Normal data
Fold over basal
Time (min)
4
3
2
1
0
-1
0
10
20
30
40
50
60

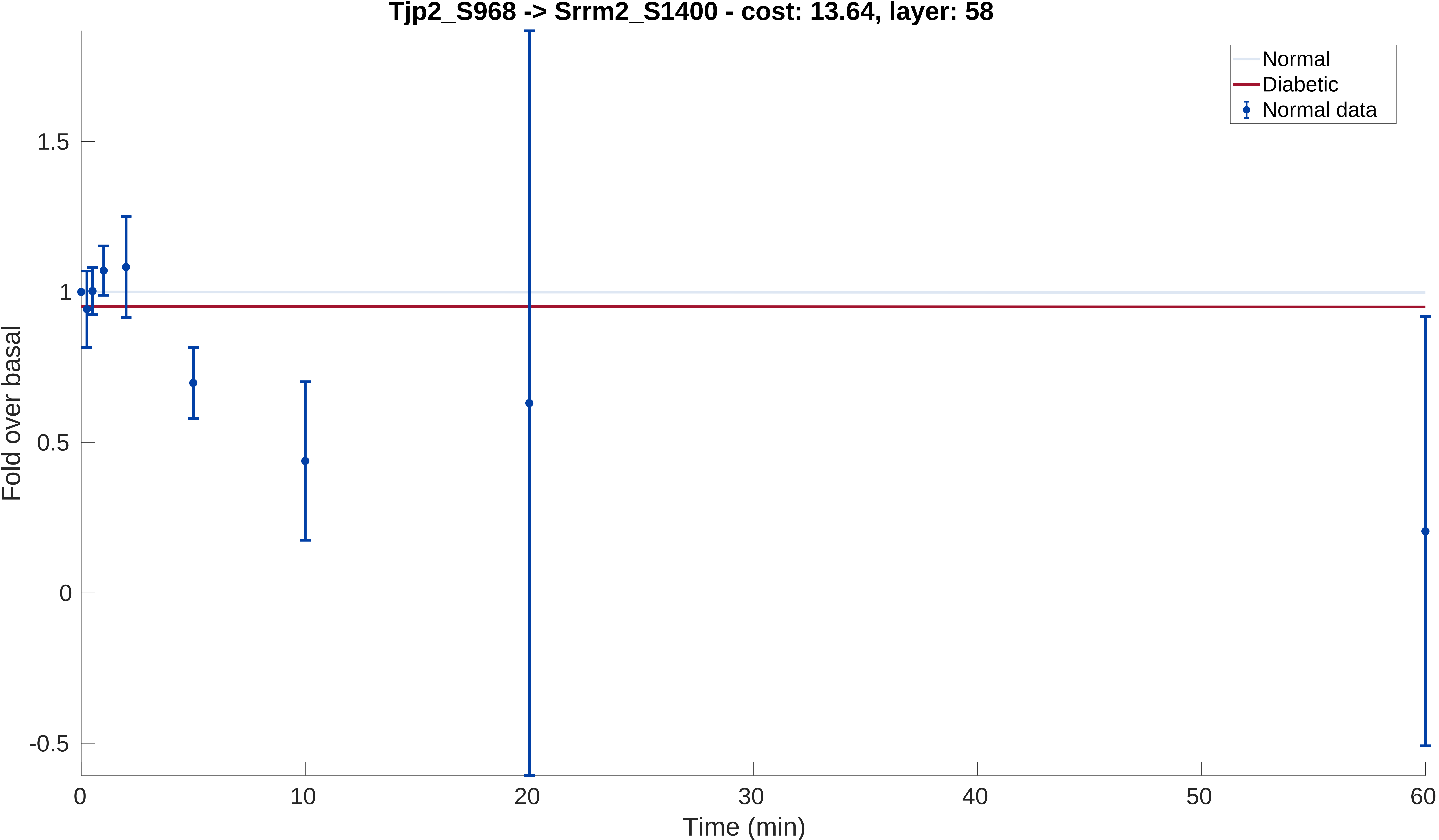
Tjp2_S968 -> Srrm2_S1400 - cost: 13.64, layer: 58
Normal
Diabetic
Normal data
Fold over basal
1.5
1
0.5
0
-0.5
0
10
20
30
40
50
60
Time (min)

#### Unc119_S41 -> Srrm2_S1418 - cost: 11.47, layer: 58

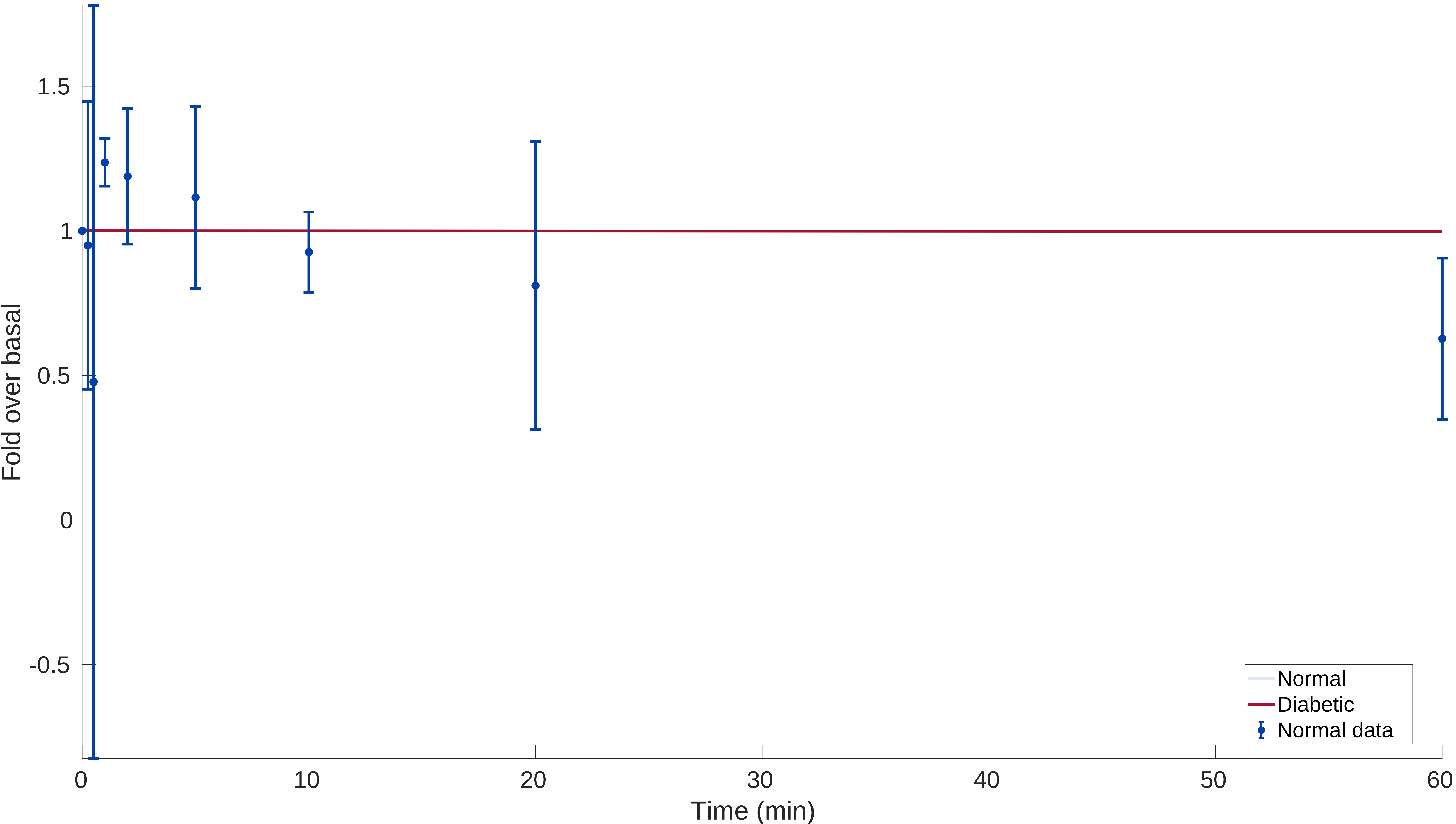

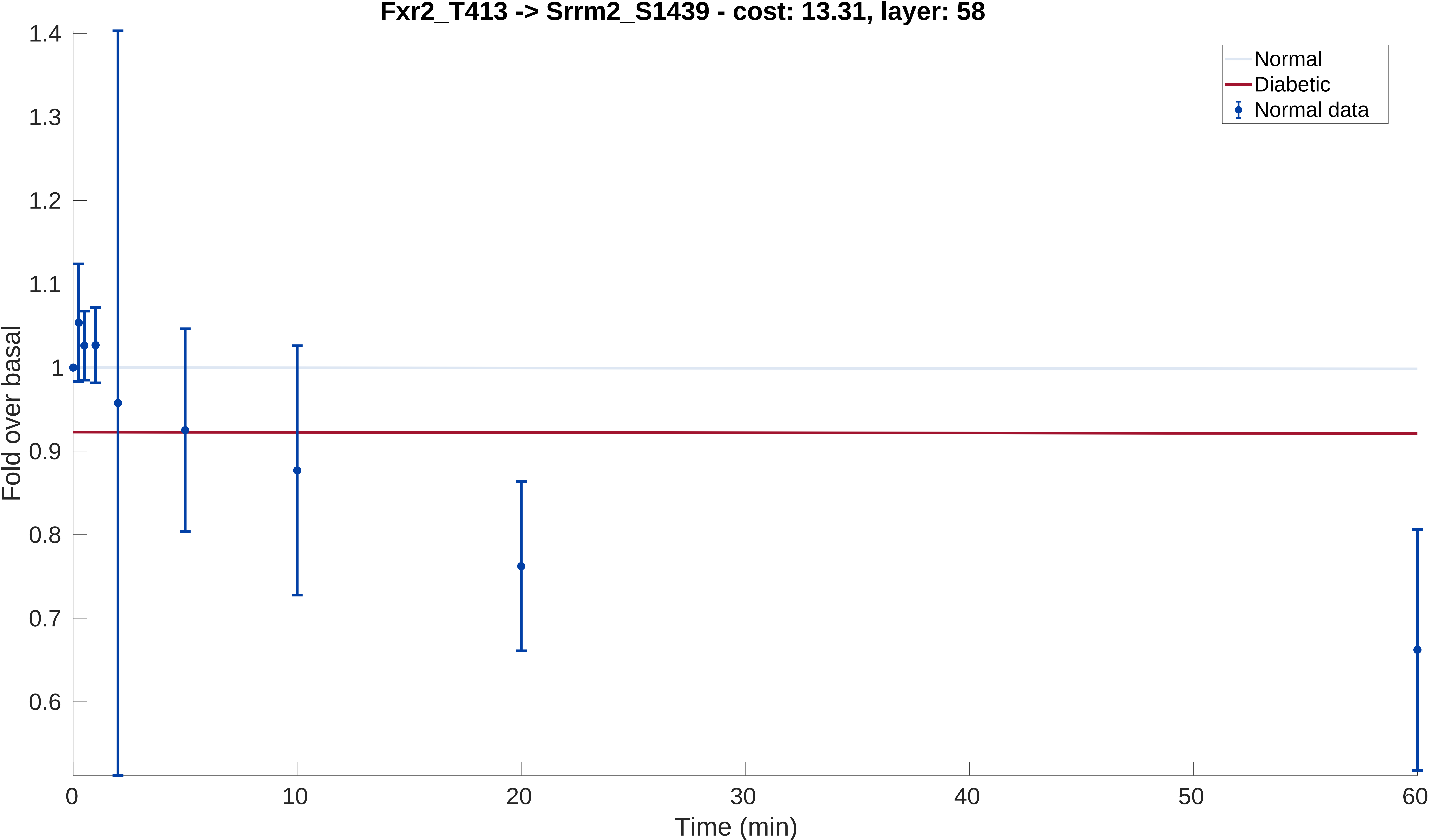

Fxr2_T413 -> Srrm2_S1439 - cost: 13.31, layer: 58
Normal
Diabetic
Normal data
Fold over basal
Time (min)
1.4
1.3
1.2
1.1
1
0.9
0.8
0.7
0.6
0
10
20
30
40
50
60

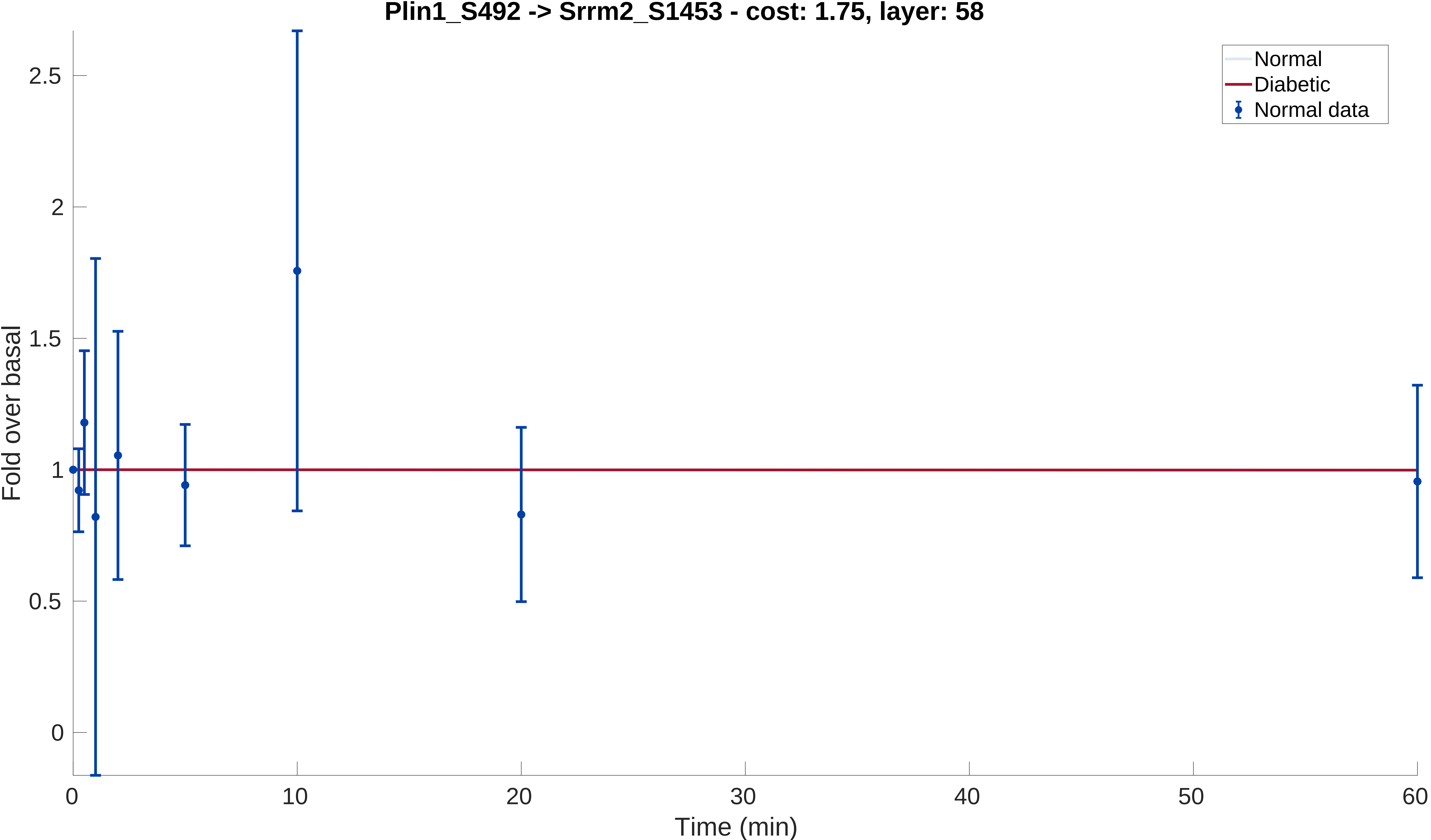

Plin1_S492 -> Srrm2_S1453 - cost: 1.75, layer: 58
Normal
Diabetic
Normal data
Fold over basal
Time (min)
0
0.5
1
1.5
2
2.5
0
10
20
30
40
50
60

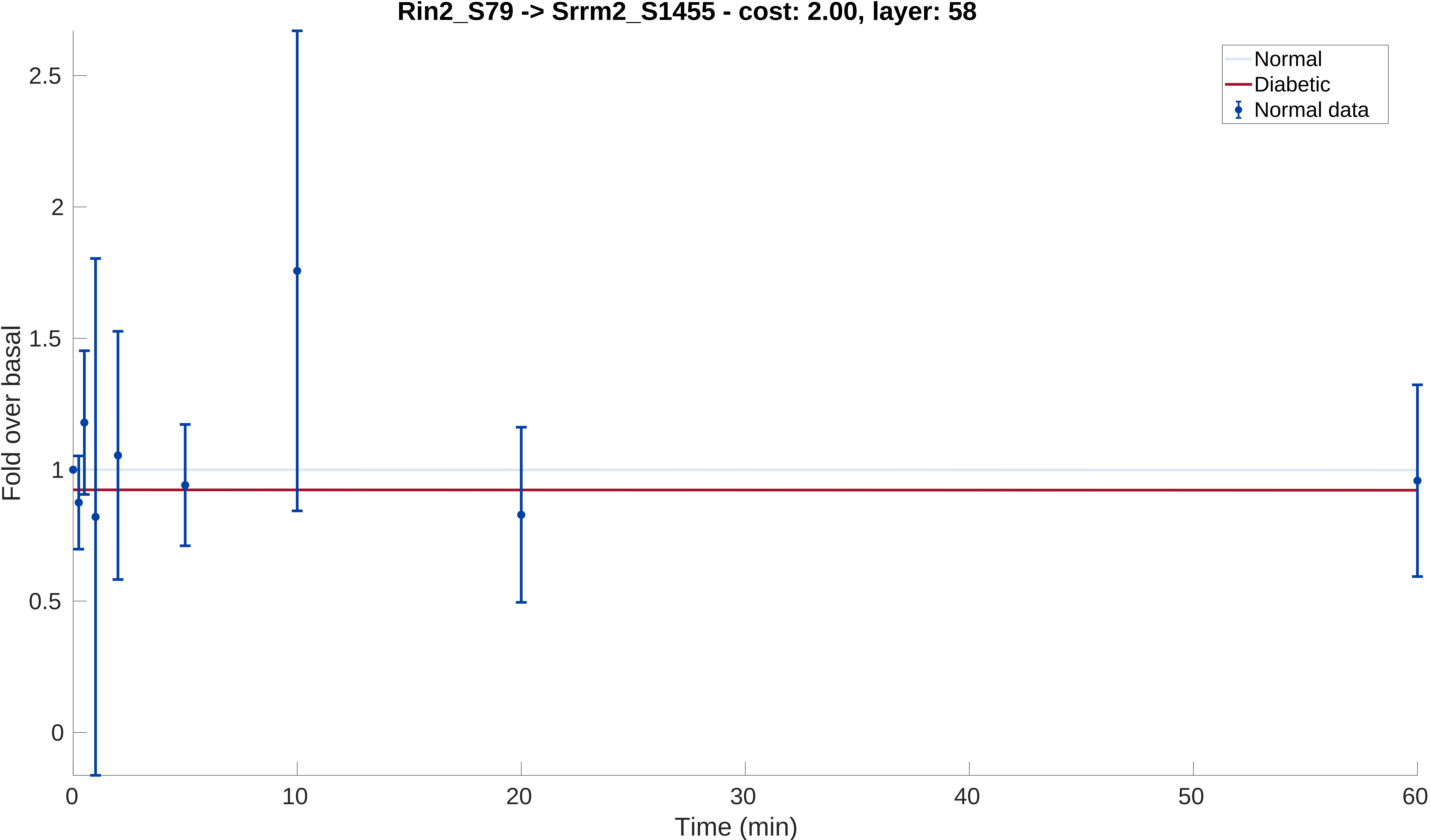

Rin2_S79 -> Srrm2_S1455 - cost: 2.00, layer: 58
Normal
Diabetic
Normal data
Fold over basal
Time (min)
0
0.5
1
1.5
2
2.5
0
10
20
30
40
50
60

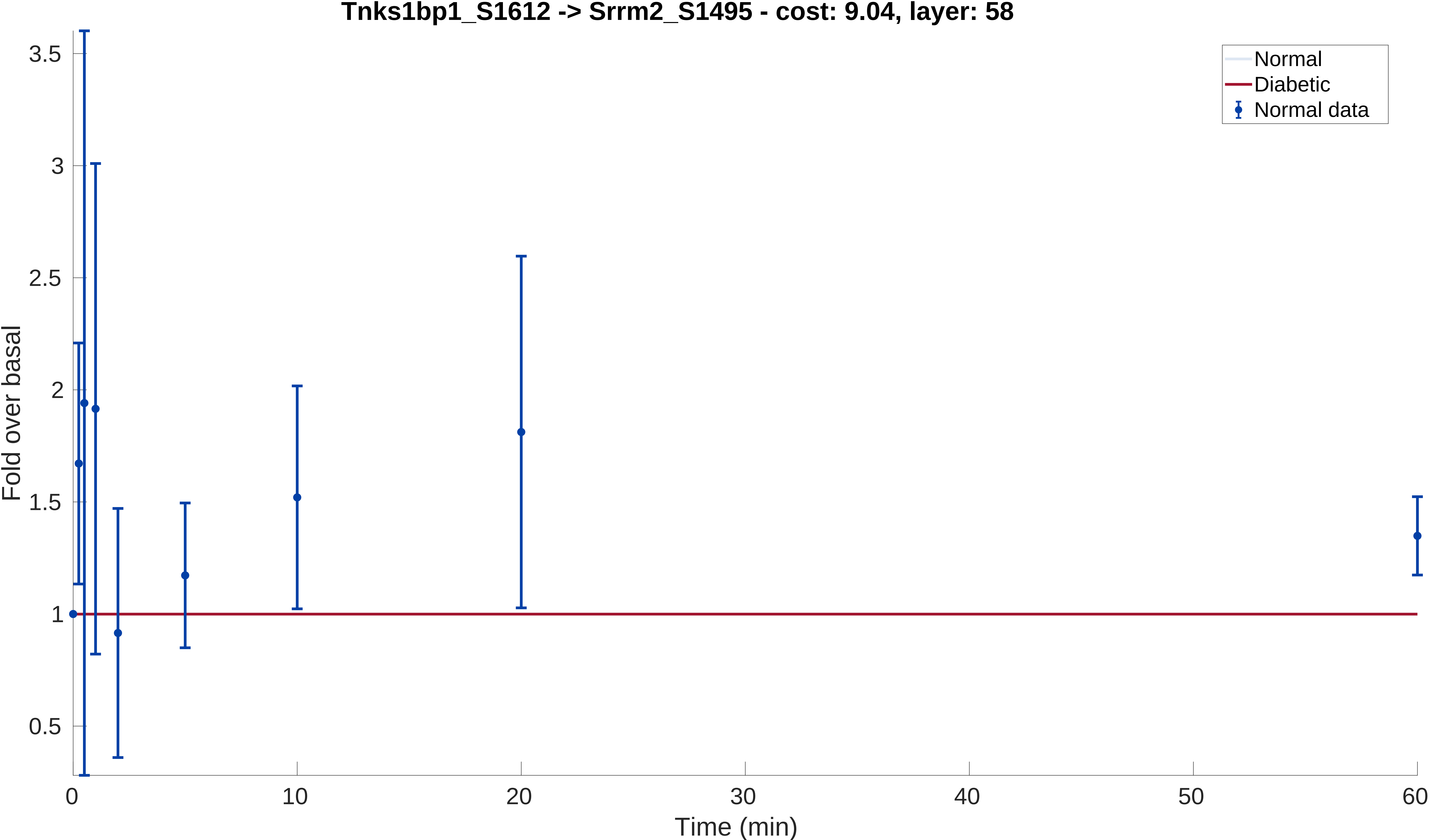

Tnks1bp1_S1612 -> Srrm2_S1495 - cost: 9.04, layer: 58
Normal
Diabetic
Normal data
Fold over basal
3.5
3
2.5
2
1.5
1
0.5
0
10
20
30
40
50
60
Time (min)

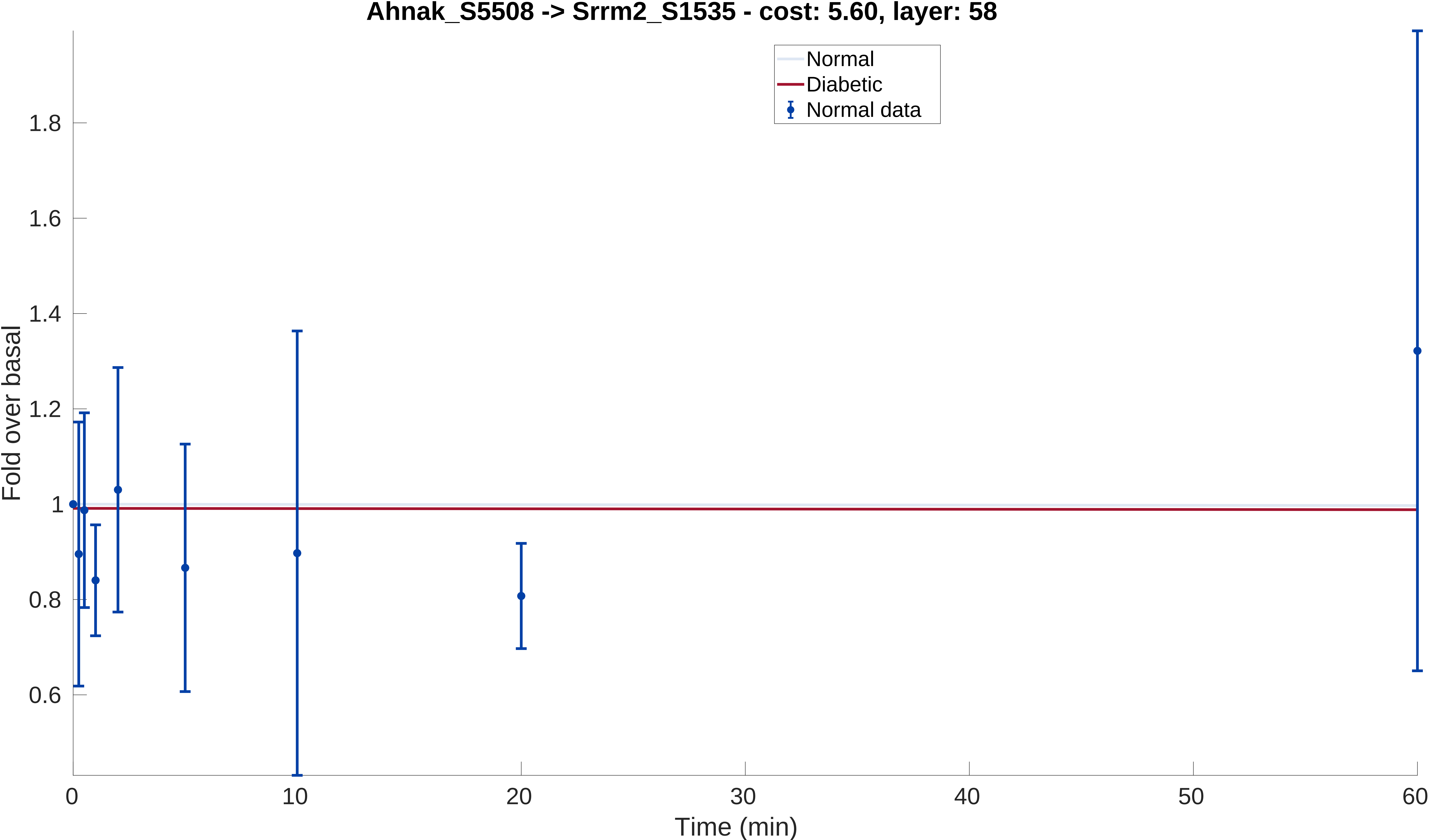
Ahnak_S5508 -> Srrm2_S1535 - cost: 5.60, layer: 58
Normal
Diabetic
Normal data
Fold over basal
1.8
1.6
1.4
1.2
1
0.8
0.6
0
10
20
30
40
50
60
Time (min)

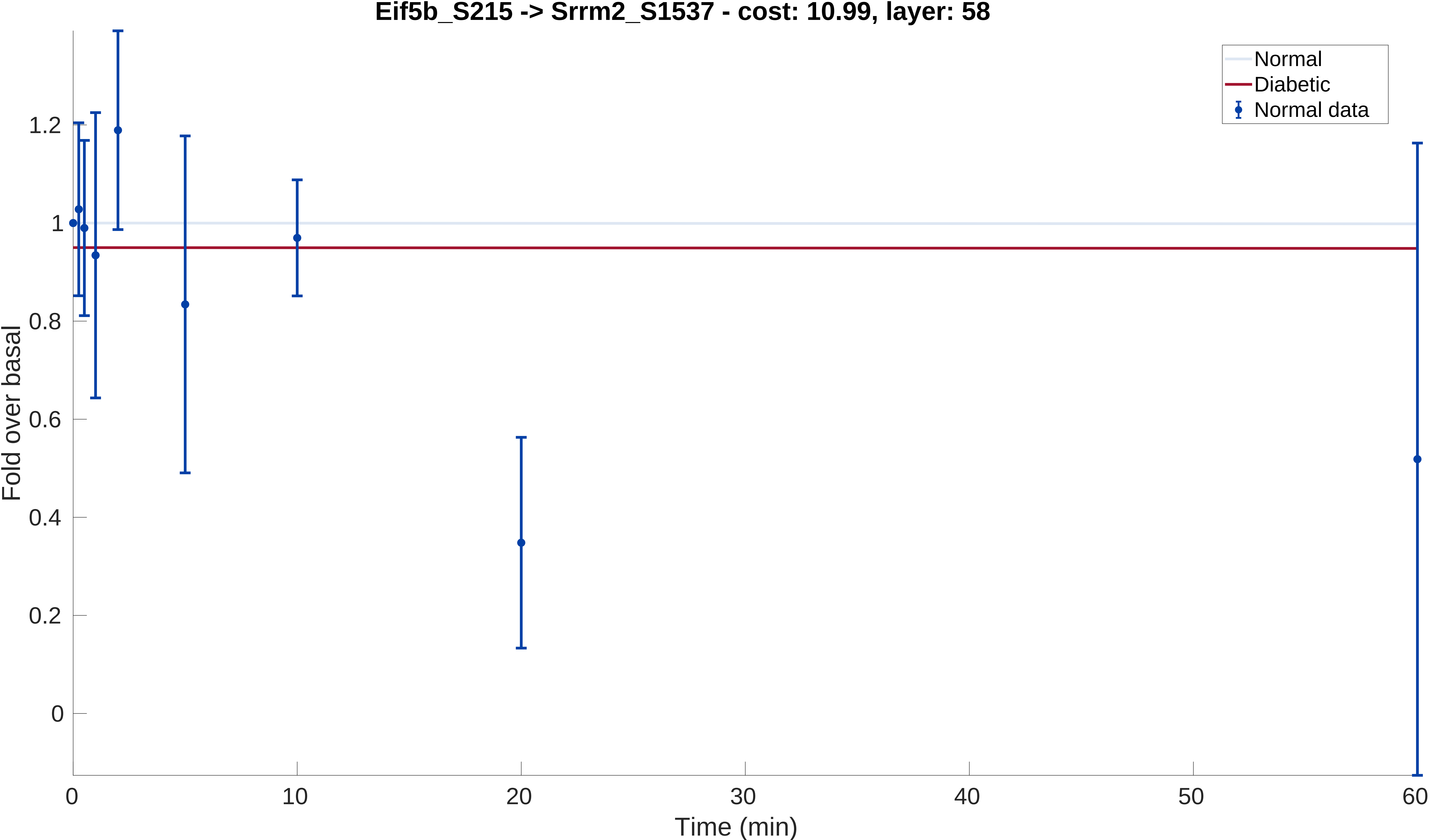

Eif5b_S215 -> Srrm2_S1537 - cost: 10.99, layer: 58
Normal
Diabetic
Normal data
Fold over basal
Time (min)
1.2
1
0.8
0.6
0.4
0.2
0
0
10
20
30
40
50
60

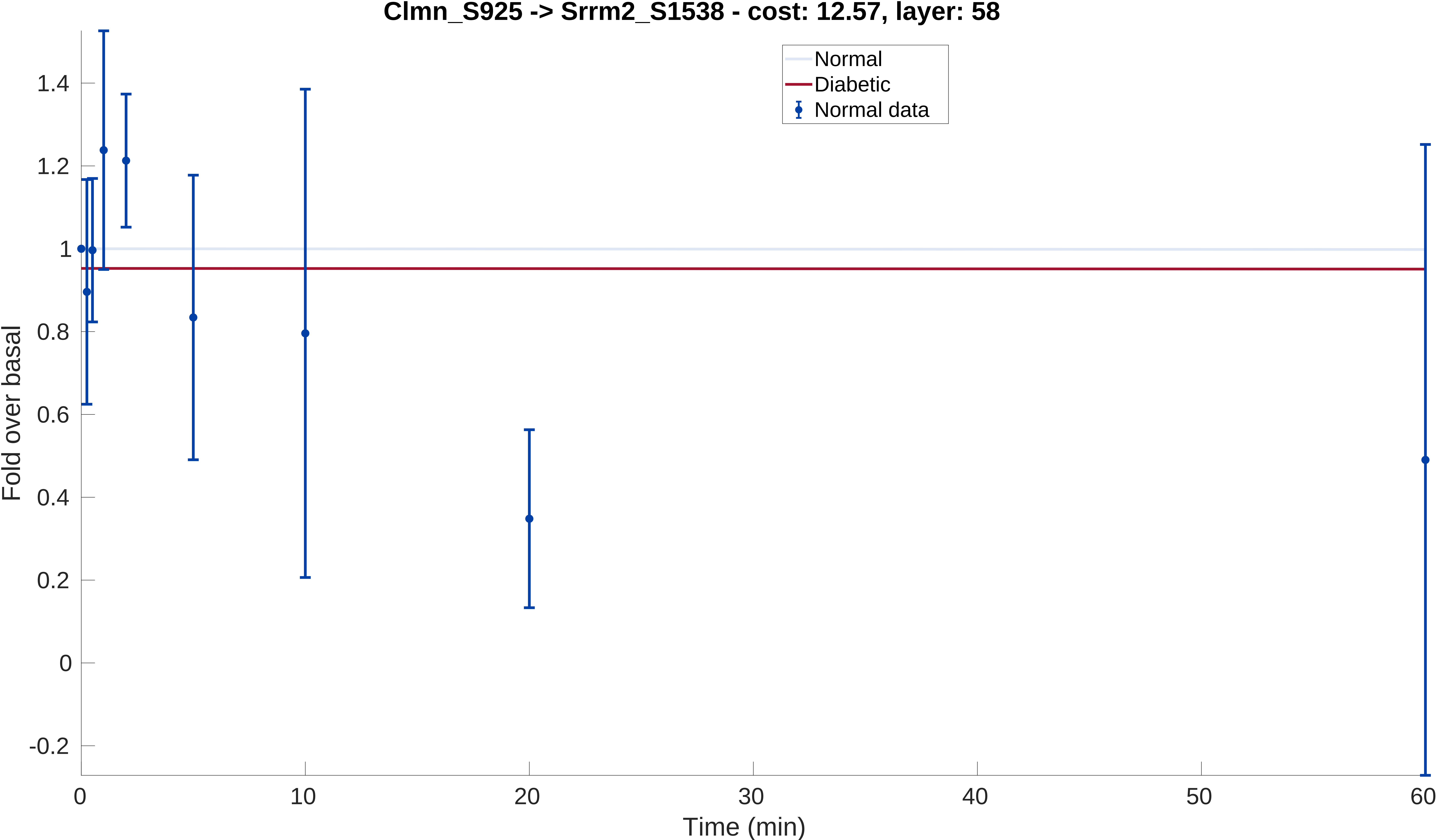
Clmn_S925 -> Srrm2_S1538 - cost: 12.57, layer: 58
Normal
Diabetic
Normal data
1.4
1.2
1
0.8
0.6
0.4
0.2
0
-0.2
Fold over basal
0
10
20
30
40
50
60
Time (min)

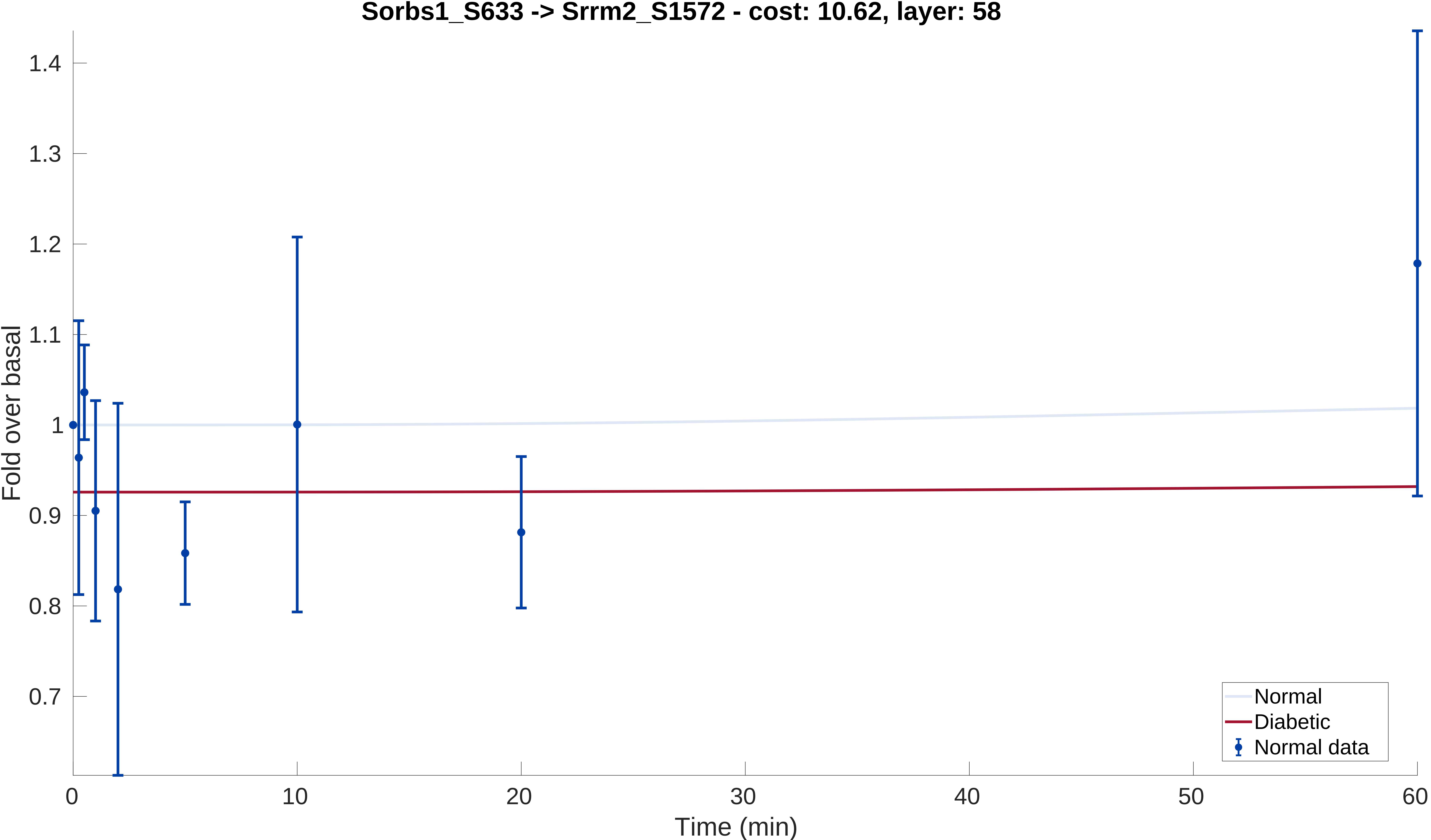

Sorbs1_S633 -> Srrm2_S1572 - cost: 10.62, layer: 58
Fold over basal
1.4
1.3
1.2
1.1
1
0.9
0.8
0.7
0
10
20
30
40
50
60
Time (min)
Normal
Diabetic
Normal data

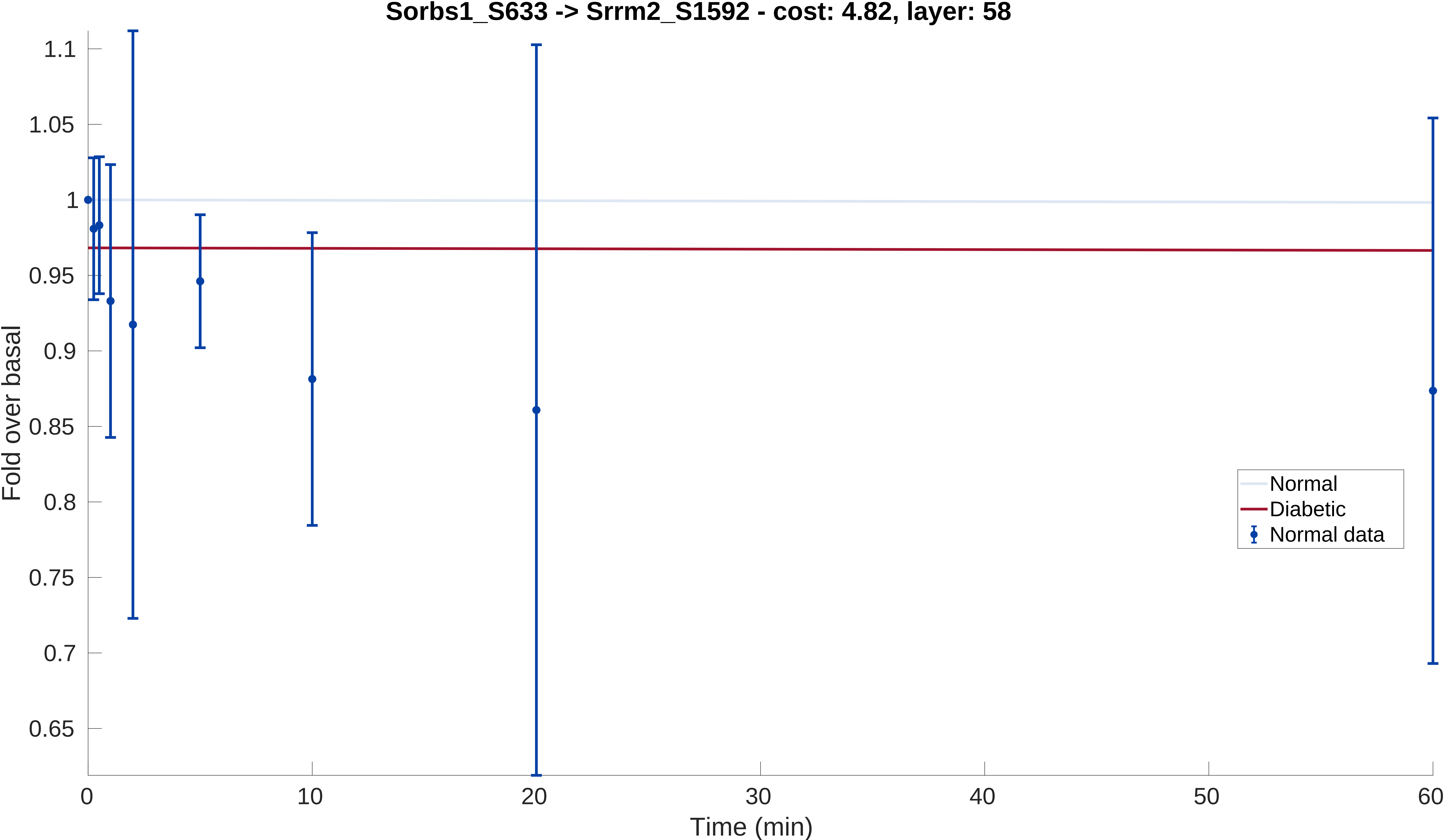

Sorbs1_S633 -> Srrm2_S1592 - cost: 4.82, layer: 58
Fold over basal
1.1
1.05
1
0.95
0.9
0.85
0.8
0.75
0.7
0.65
0
10
20
30
40
50
60
Time (min)
Normal
Diabetic
Normal data

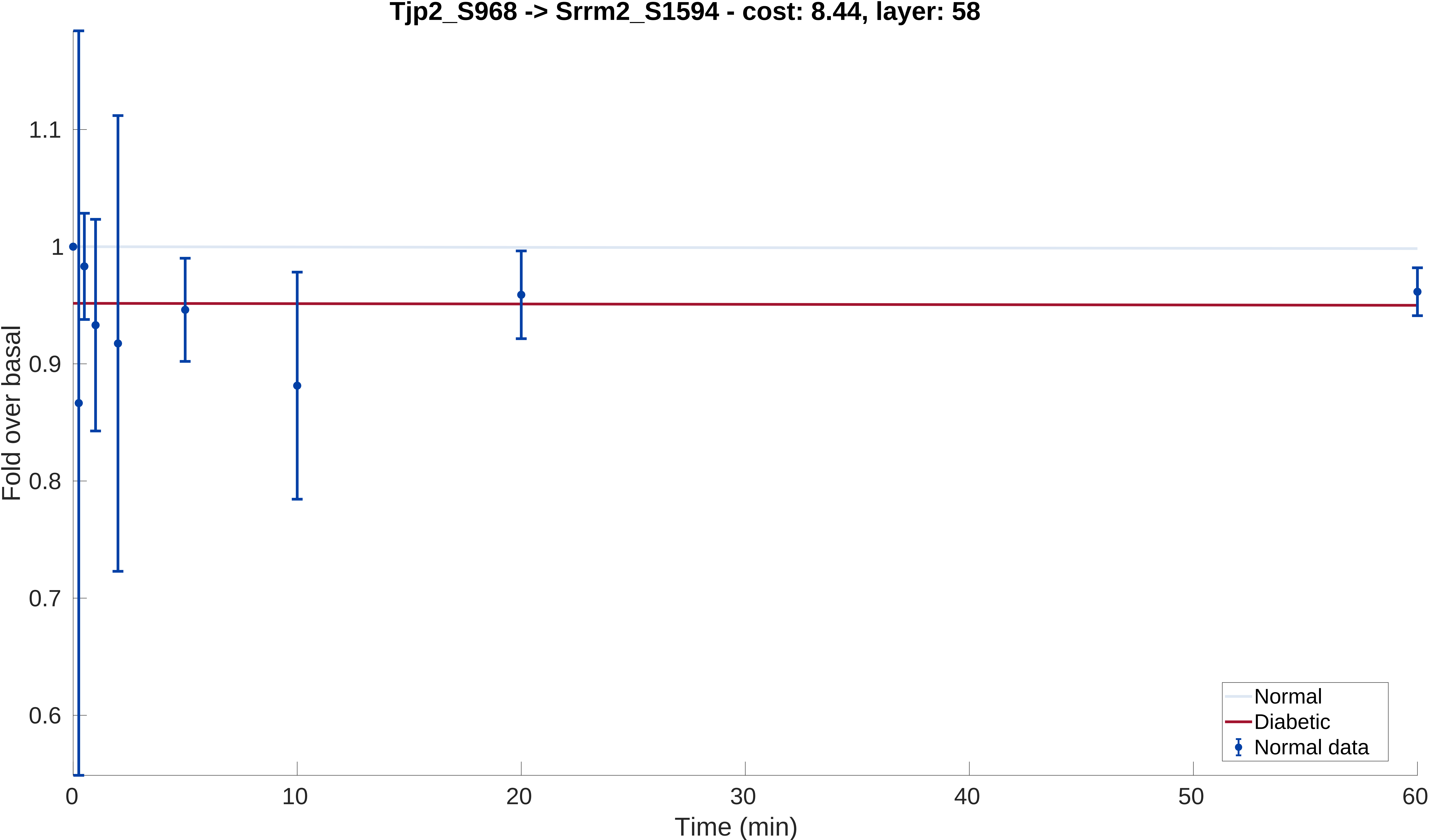

Tjp2_S968 -> Srrm2_S1594 - cost: 8.44, layer: 58
Fold over basal
1.1
1
0.9
0.8
0.7
0.6
0
10
20
30
40
50
60
Time (min)
Normal
Diabetic
Normal data

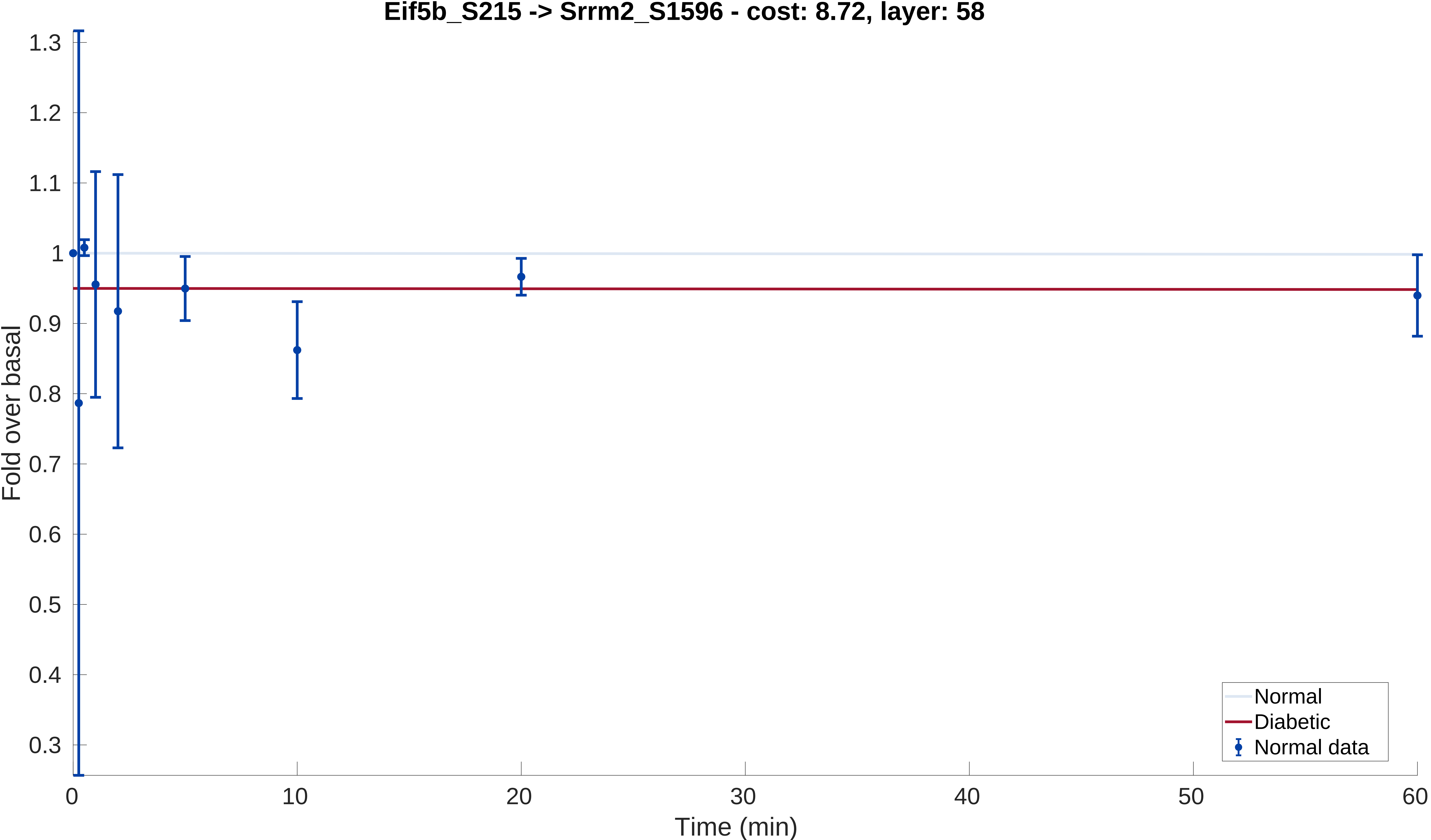

Eif5b_S215 -> Srrm2_S1596 - cost: 8.72, layer: 58
Fold over basal
1.3
1.2
1.1
1
0.9
0.8
0.7
0.6
0.5
0.4
0.3
0
10
20
30
40
50
60
Time (min)
Normal
Diabetic
Normal data

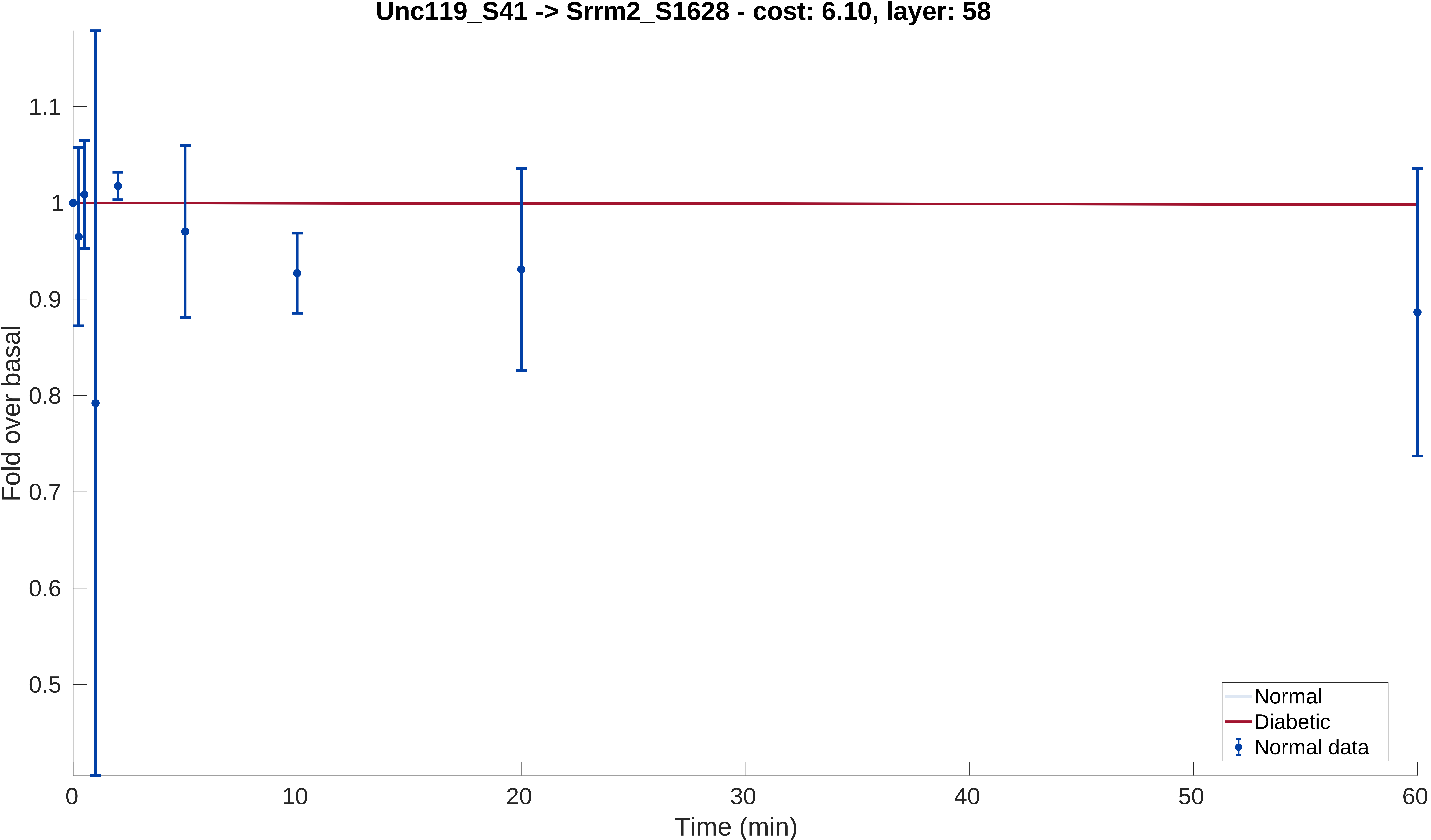
Unc119_S41 -> Srrm2_S1628 - cost: 6.10, layer: 58
Fold over basal
1.1
1
0.9
0.8
0.7
0.6
0.5
0
10
20
30
40
50
60
Time (min)
Normal
Diabetic
Normal data

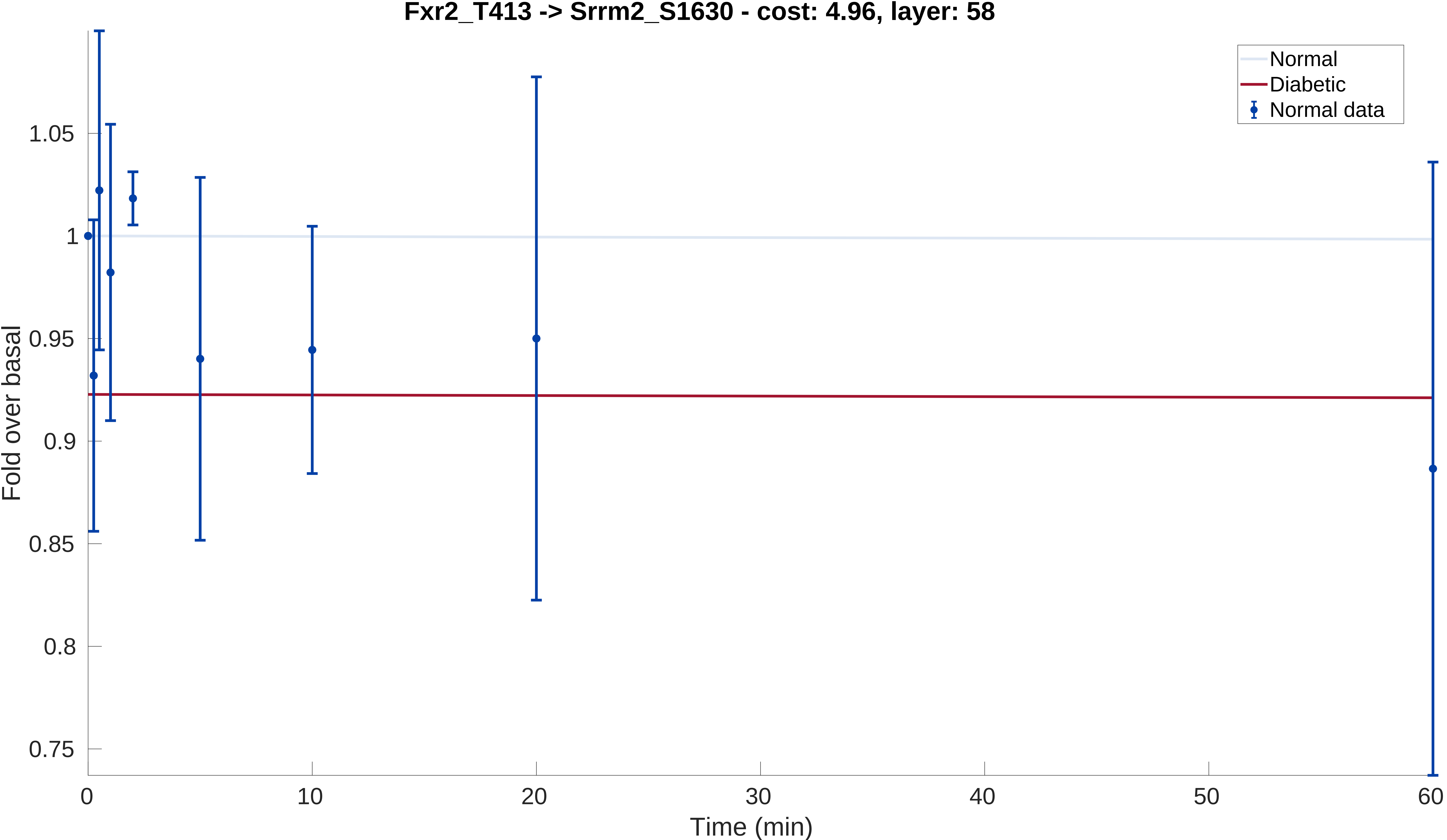
Fxr2_T413 -> Srrm2_S1630 - cost: 4.96, layer: 58
Normal
Diabetic
Normal data
Fold over basal
1.05
1
0.95
0.9
0.85
0.8
0.75
0
10
20
30
40
50
60
Time (min)

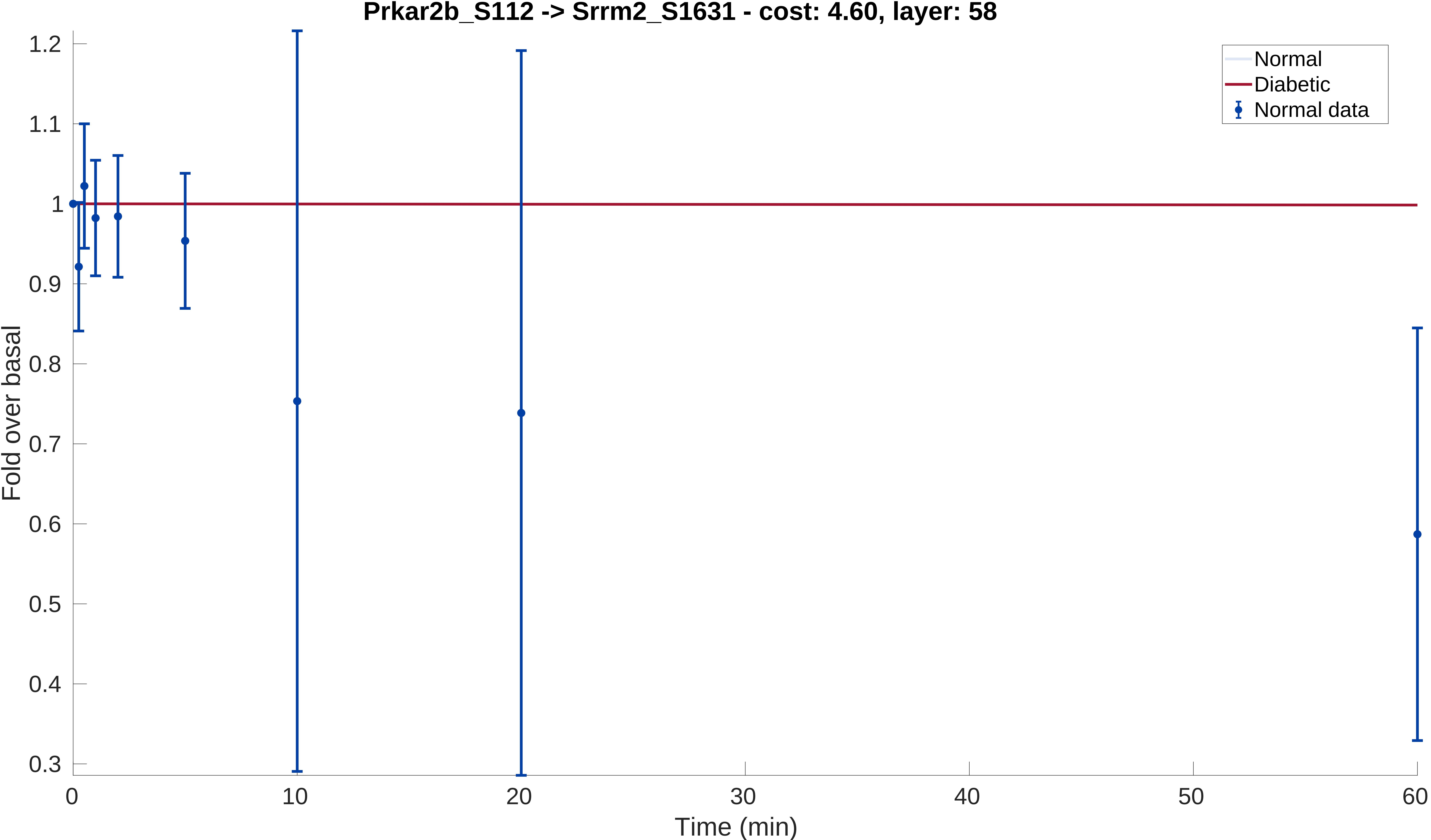

Prkar2b_S112 -> Srrm2_S1631 - cost: 4.60, layer: 58
Normal
Diabetic
Normal data
Fold over basal
Time (min)
1.2
1.1
1
0.9
0.8
0.7
0.6
0.5
0.4
0.3
0
10
20
30
40
50
60

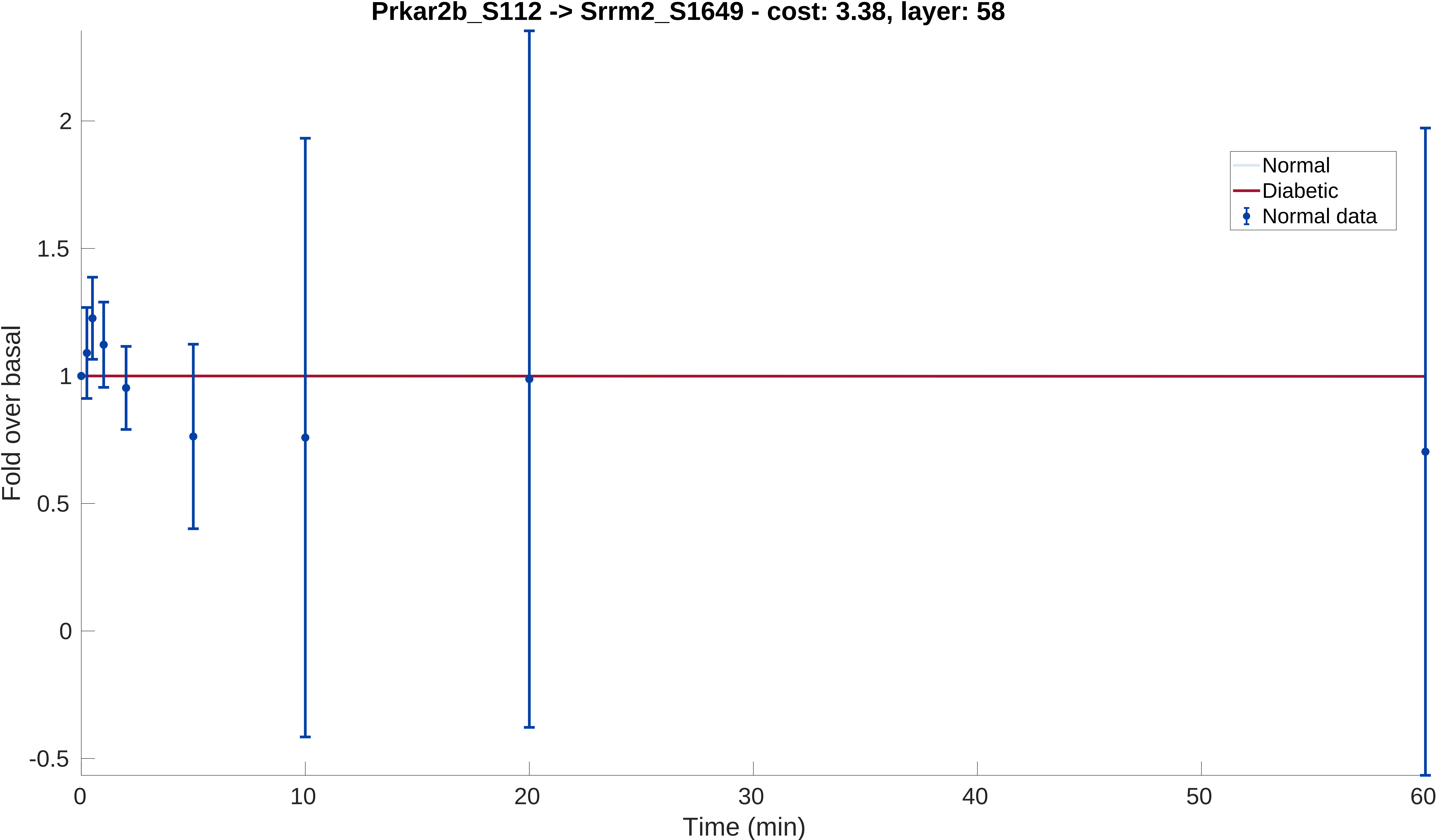

Prkar2b_S112 -> Srrm2_S1649 - cost: 3.38, layer: 58
Normal
Diabetic
Normal data
Fold over basal
Time (min)
-0.5
0
0.5
1
1.5
2
0
10
20
30
40
50
60

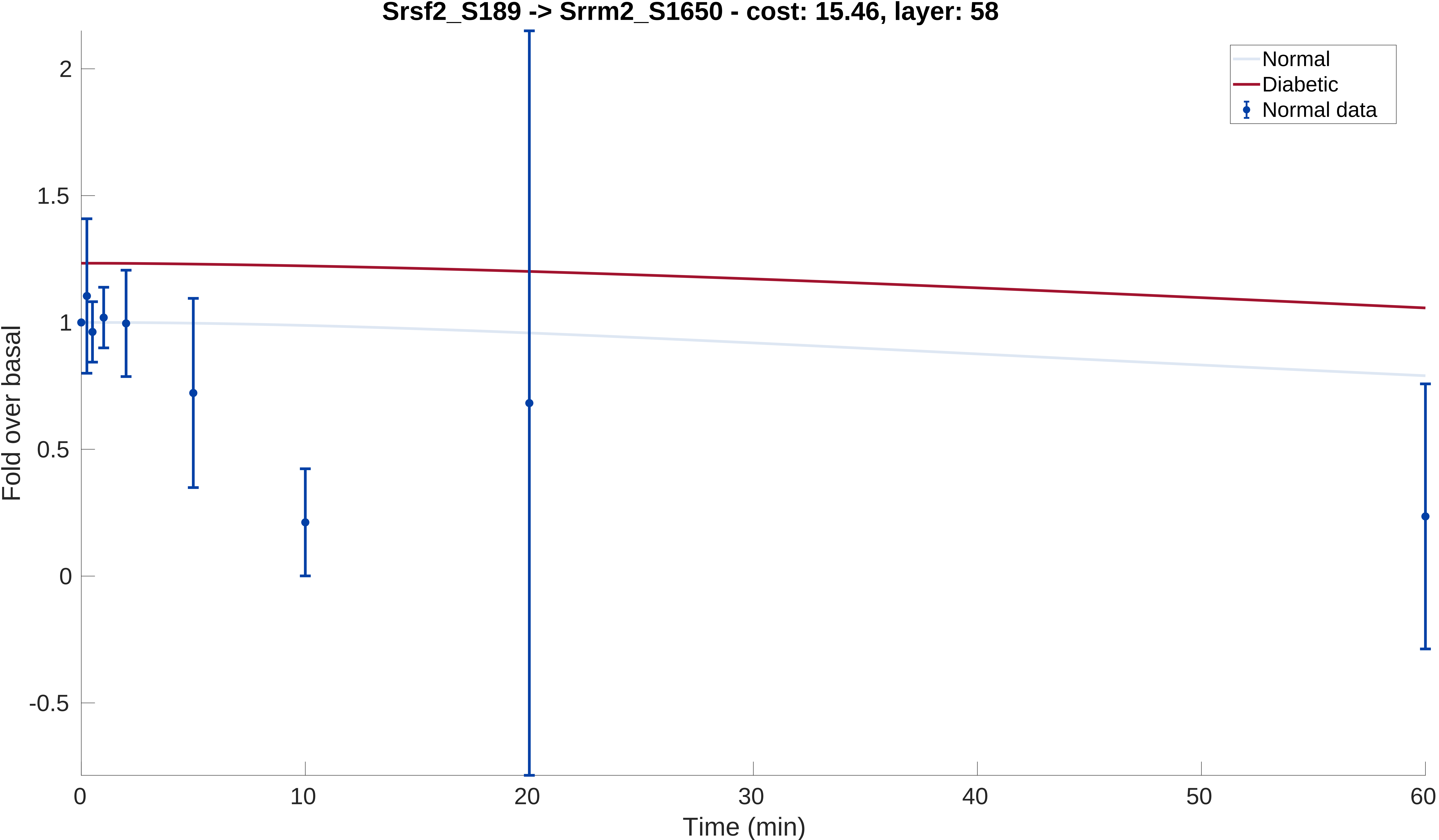

Srsf2_S189 -> Srrm2_S1650 - cost: 15.46, layer: 58
Normal
Diabetic
Normal data
Fold over basal
2
1.5
1
0.5
0
-0.5
0
10
20
30
40
50
60
Time (min)

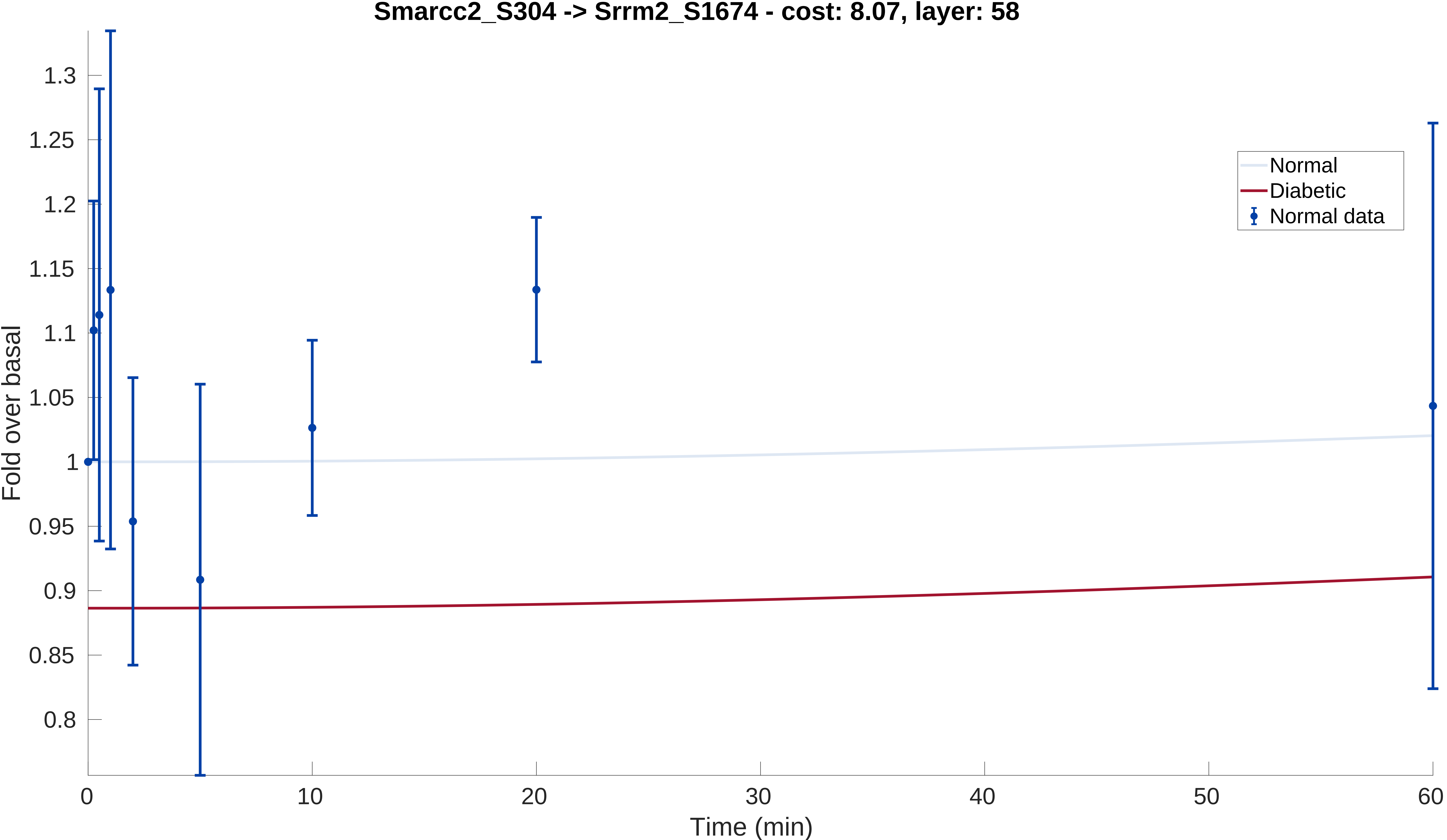

Smarcc2_S304 -> Srrm2_S1674 - cost: 8.07, layer: 58
Fold over basal
Time (min)
Normal
Diabetic
Normal data
1.3
1.25
1.2
1.15
1.1
1.05
1
0.95
0.9
0.85
0.8
0
10
20
30
40
50
60

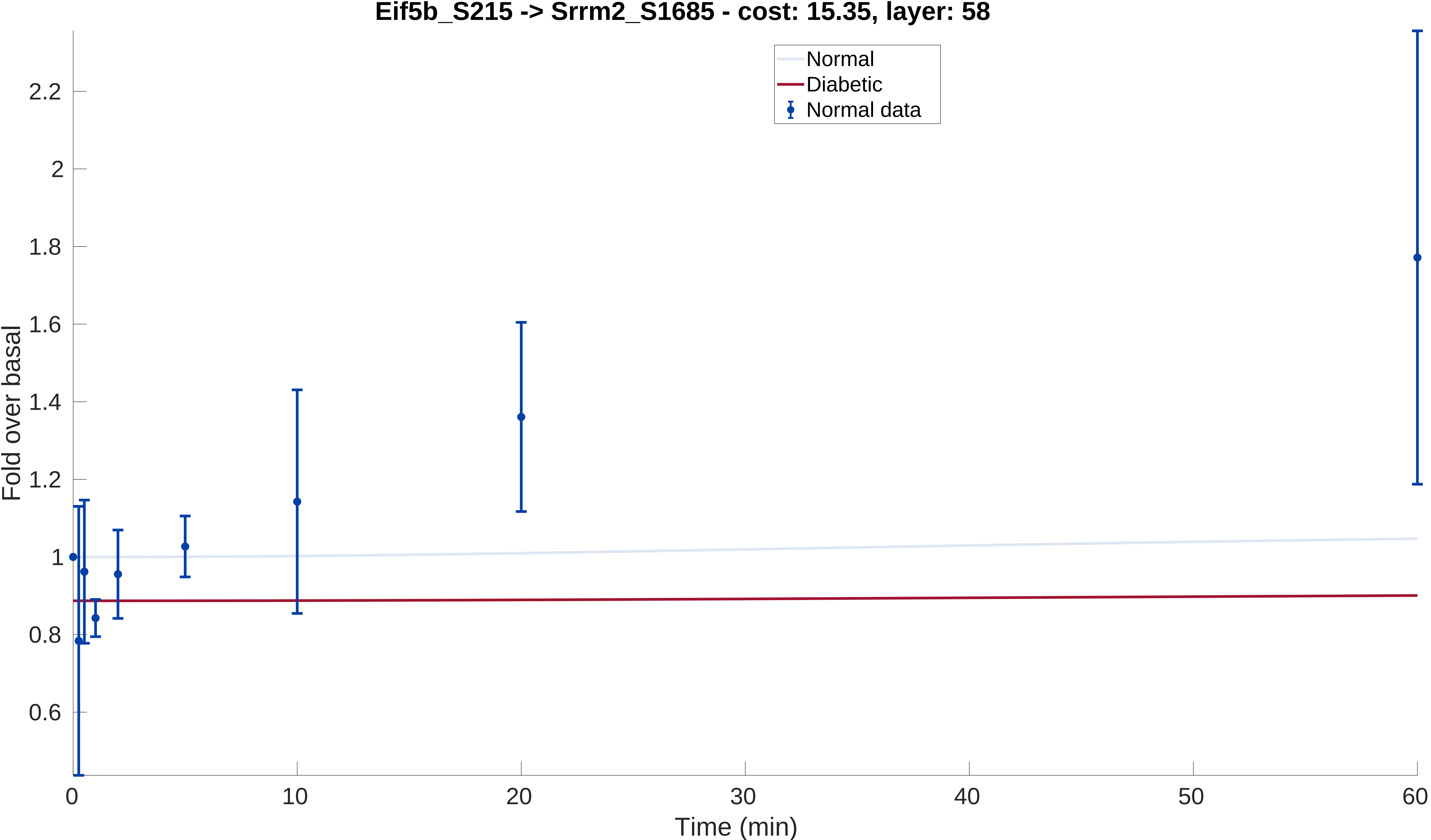

Eif5b_S215 -> Srrm2_S1685 - cost: 15.35, layer: 58
Normal
Diabetic
Normal data
Fold over basal
2.2
2
1.8
1.6
1.4
1.2
1
0.8
0.6
0
10
20
30
40
50
60
Time (min)

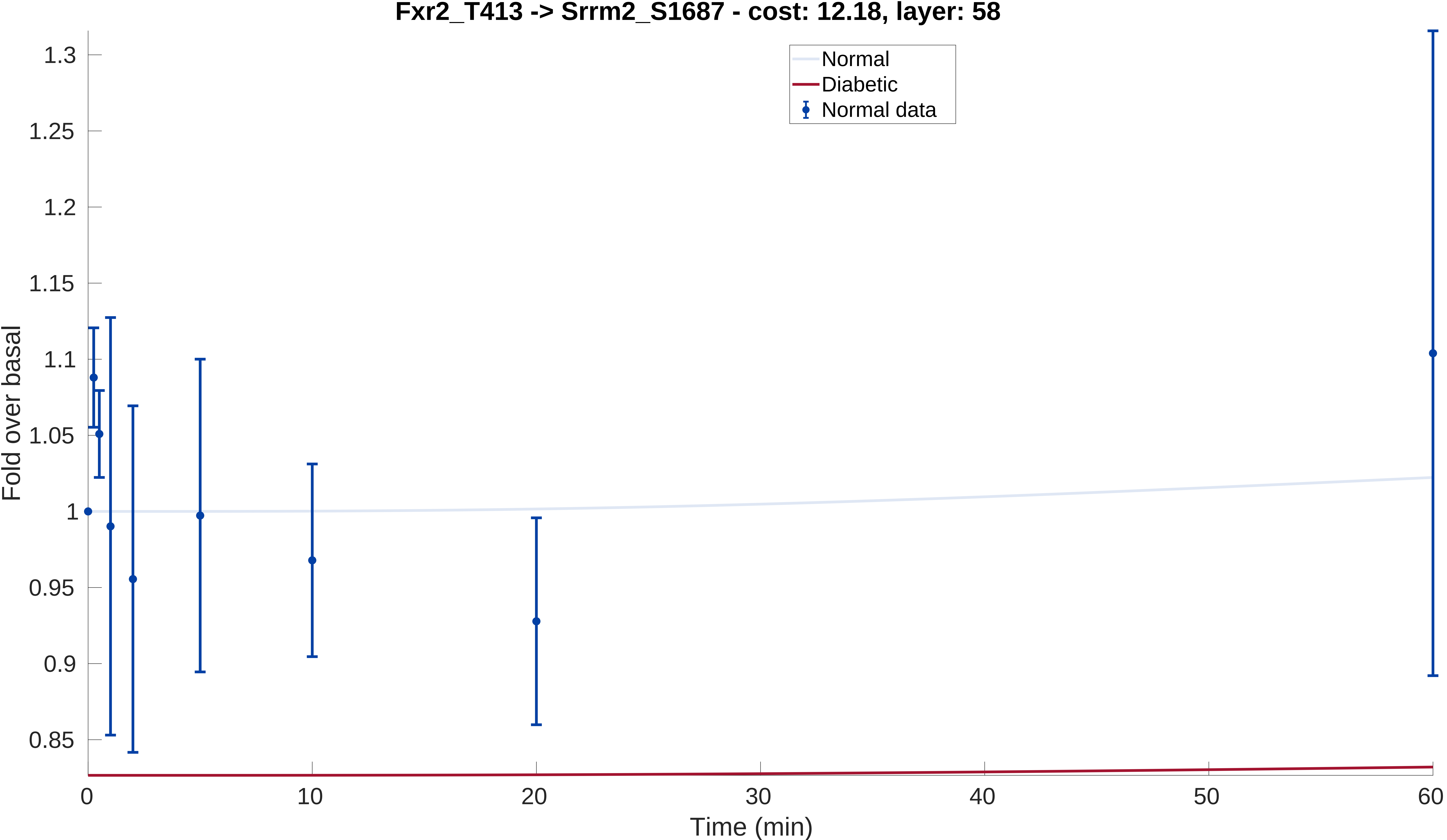

Fxr2_T413 -> Srrm2_S1687 - cost: 12.18, layer: 58
Normal
Diabetic
Normal data
Fold over basal
1.3
1.25
1.2
1.15
1.1
1.05
1
0.95
0.9
0.85
0
10
20
30
40
50
60
Time (min)

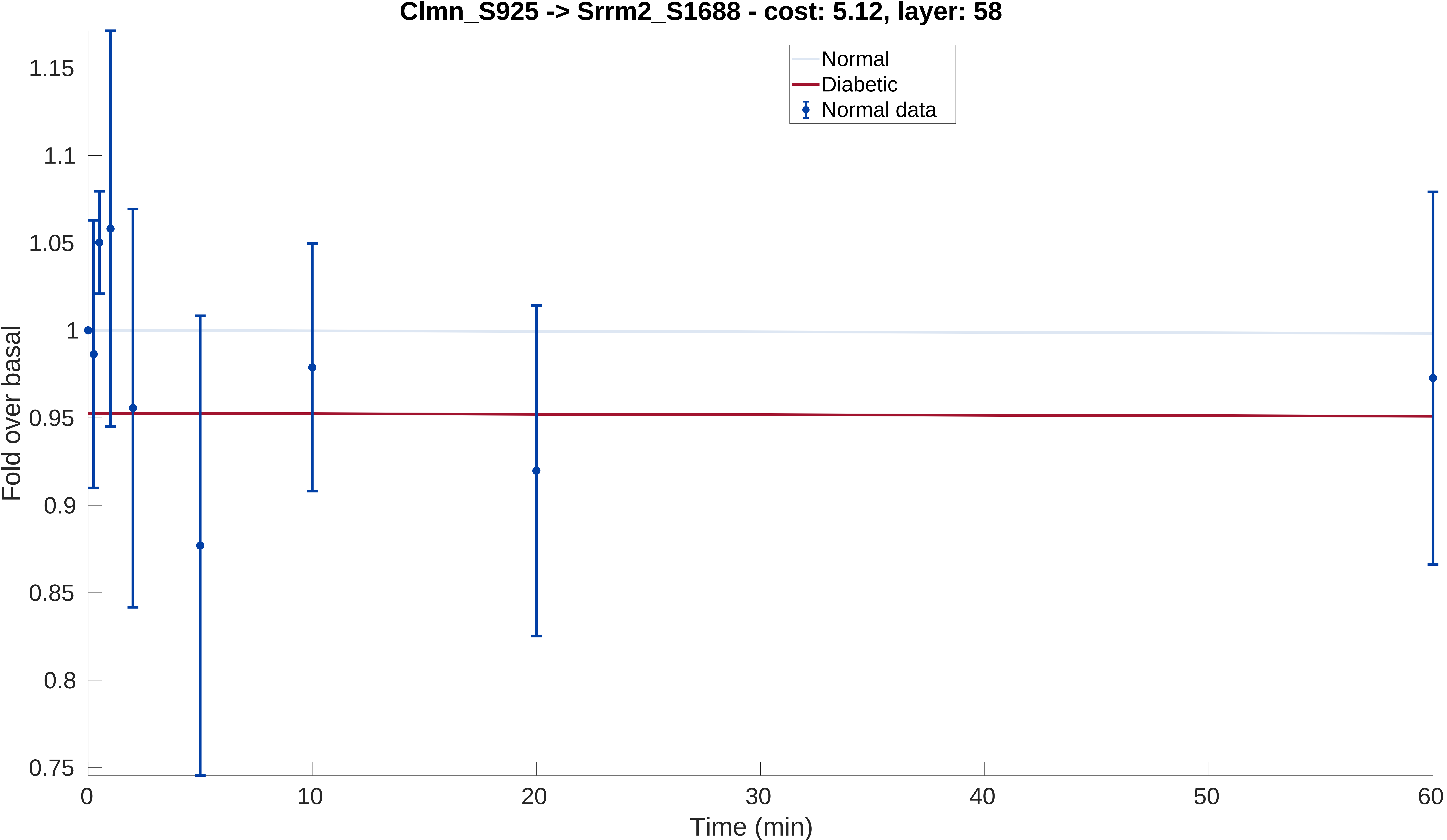

Clmn_S925 -> Srrm2_S1688 - cost: 5.12, layer: 58
Normal
Diabetic
Normal data
Fold over basal
1.15
1.1
1.05
1
0.95
0.9
0.85
0.8
0.75
0
10
20
30
40
50
60
Time (min)

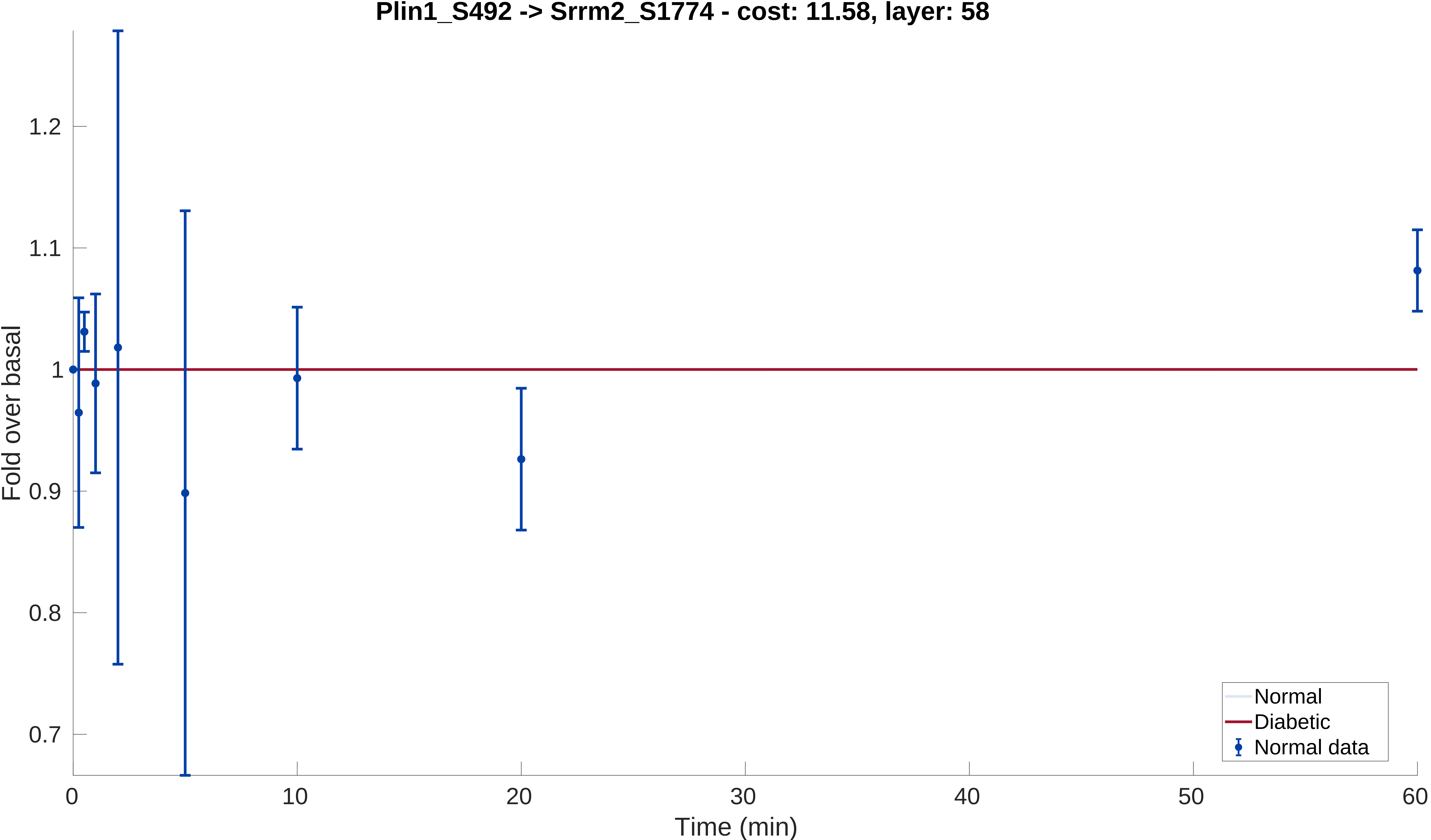
Plin1_S492 -> Srrm2_S1774 - cost: 11.58, layer: 58
Fold over basal
1.2
1.1
1
0.9
0.8
0.7
0
10
20
30
40
50
60
Time (min)
Normal
Diabetic
Normal data

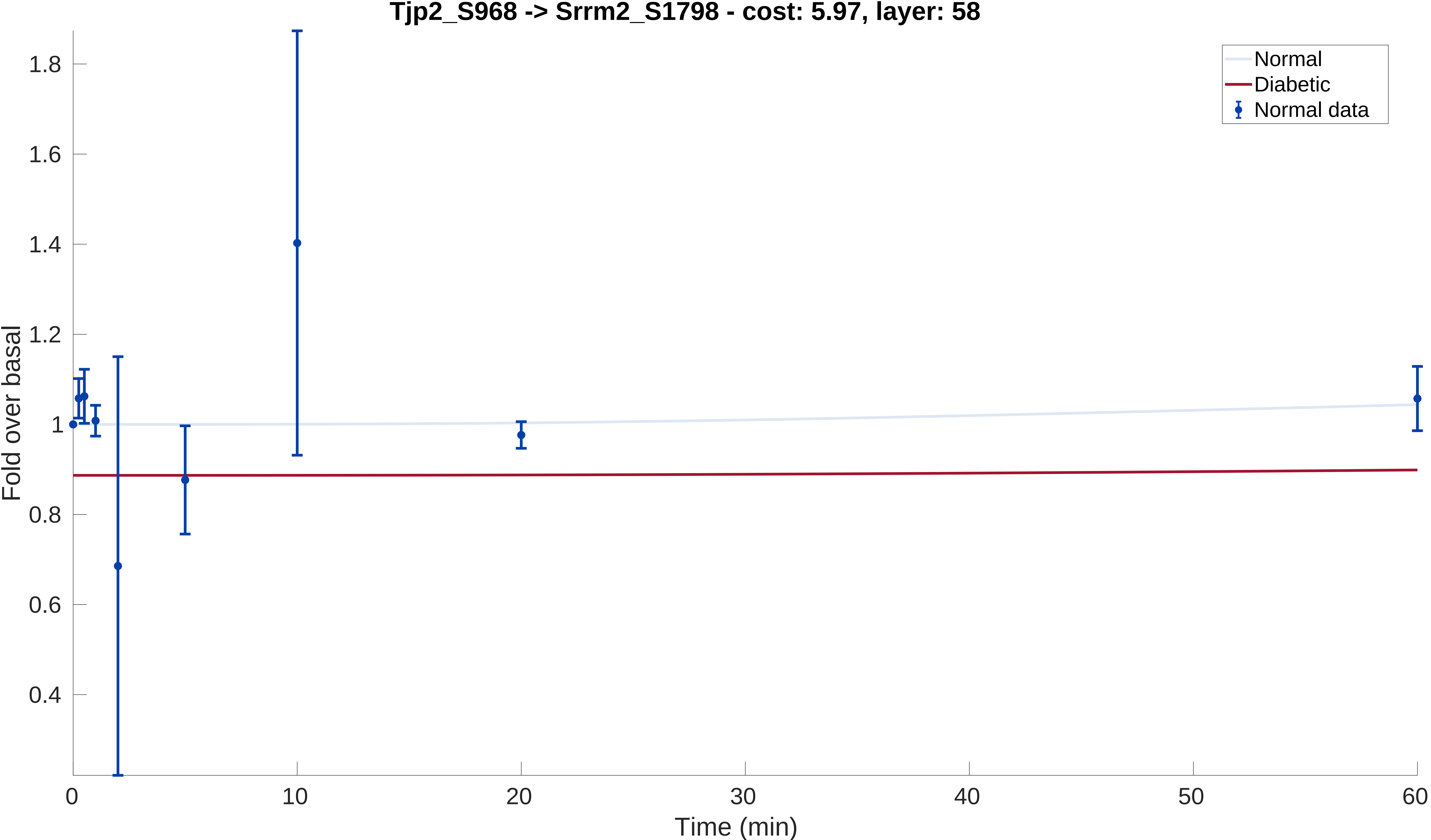

Tjp2_S968 -> Srrm2_S1798 - cost: 5.97, layer: 58
Normal
Diabetic
Normal data
Fold over basal
1.8
1.6
1.4
1.2
1
0.8
0.6
0.4
0
10
20
30
40
50
60
Time (min)

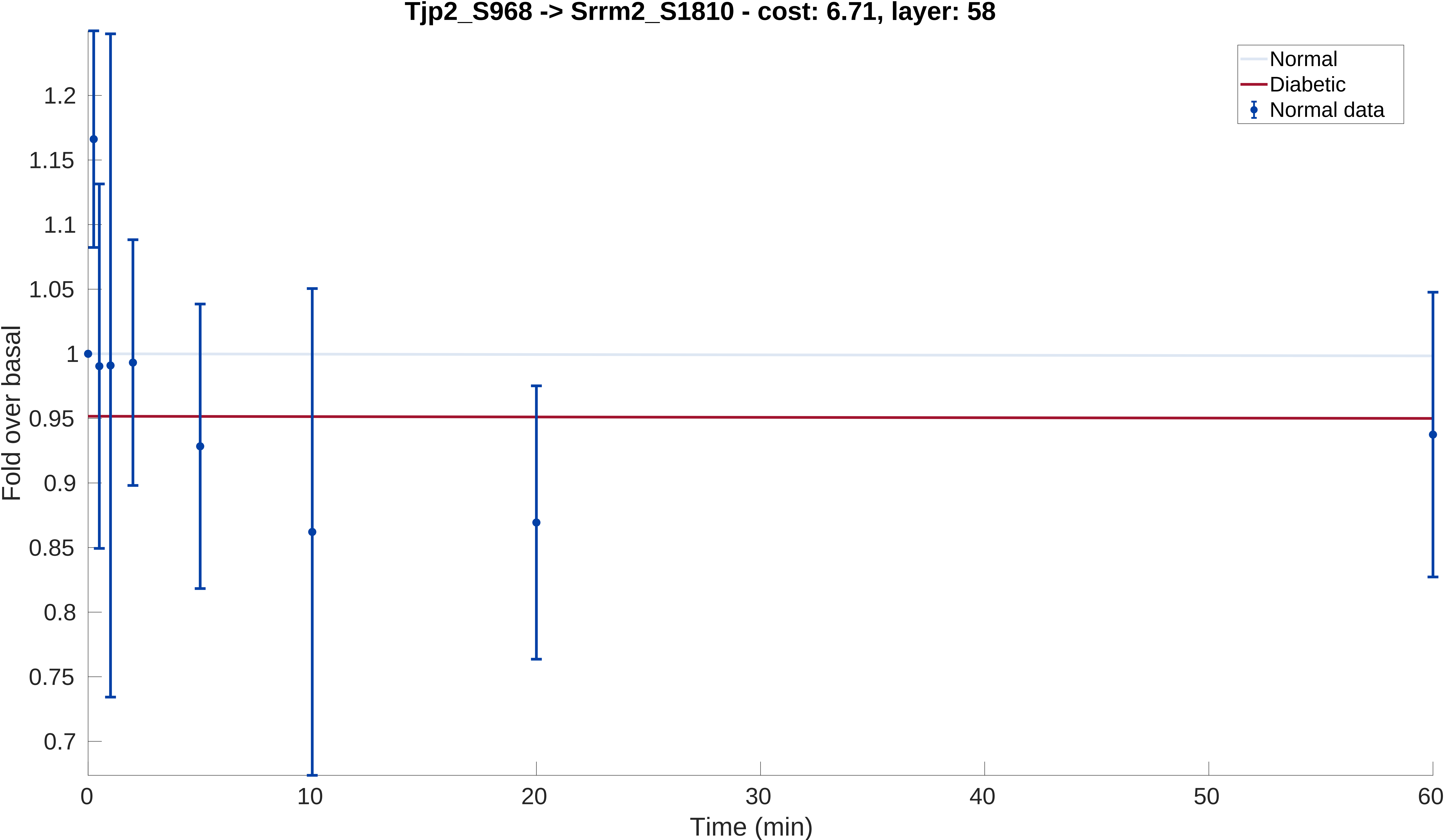

Tjp2_S968 -> Srrm2_S1810 - cost: 6.71, layer: 58
Normal
Diabetic
Normal data
Fold over basal
1.2
1.15
1.1
1.05
1
0.95
0.9
0.85
0.8
0.75
0.7
0
10
20
30
40
50
60
Time (min)

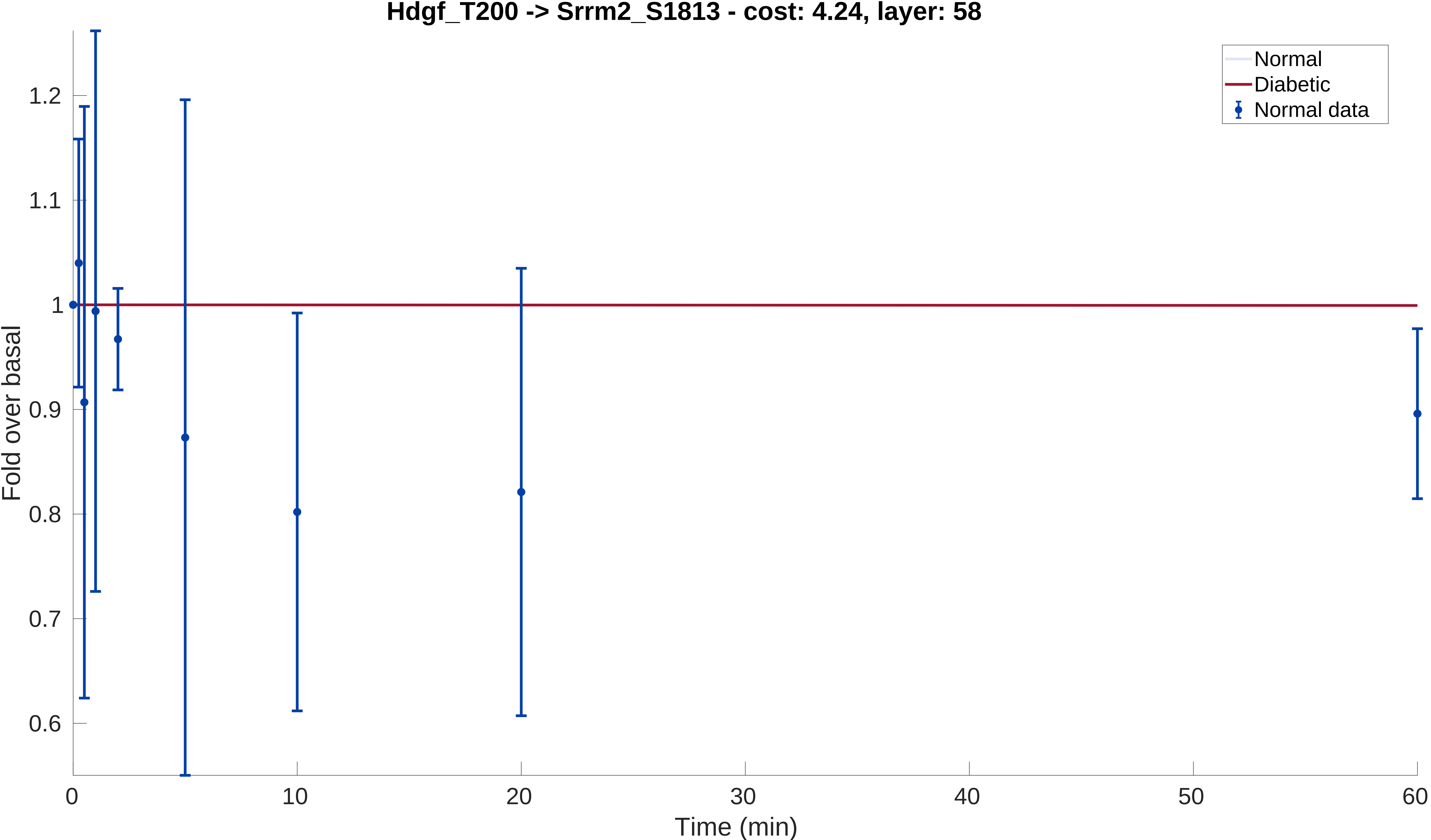

Hdgf_T200 -> Srrm2_S1813 - cost: 4.24, layer: 58
Normal
Diabetic
Normal data
Fold over basal
1.2
1.1
1
0.9
0.8
0.7
0.6
0
10
20
30
40
50
60
Time (min)

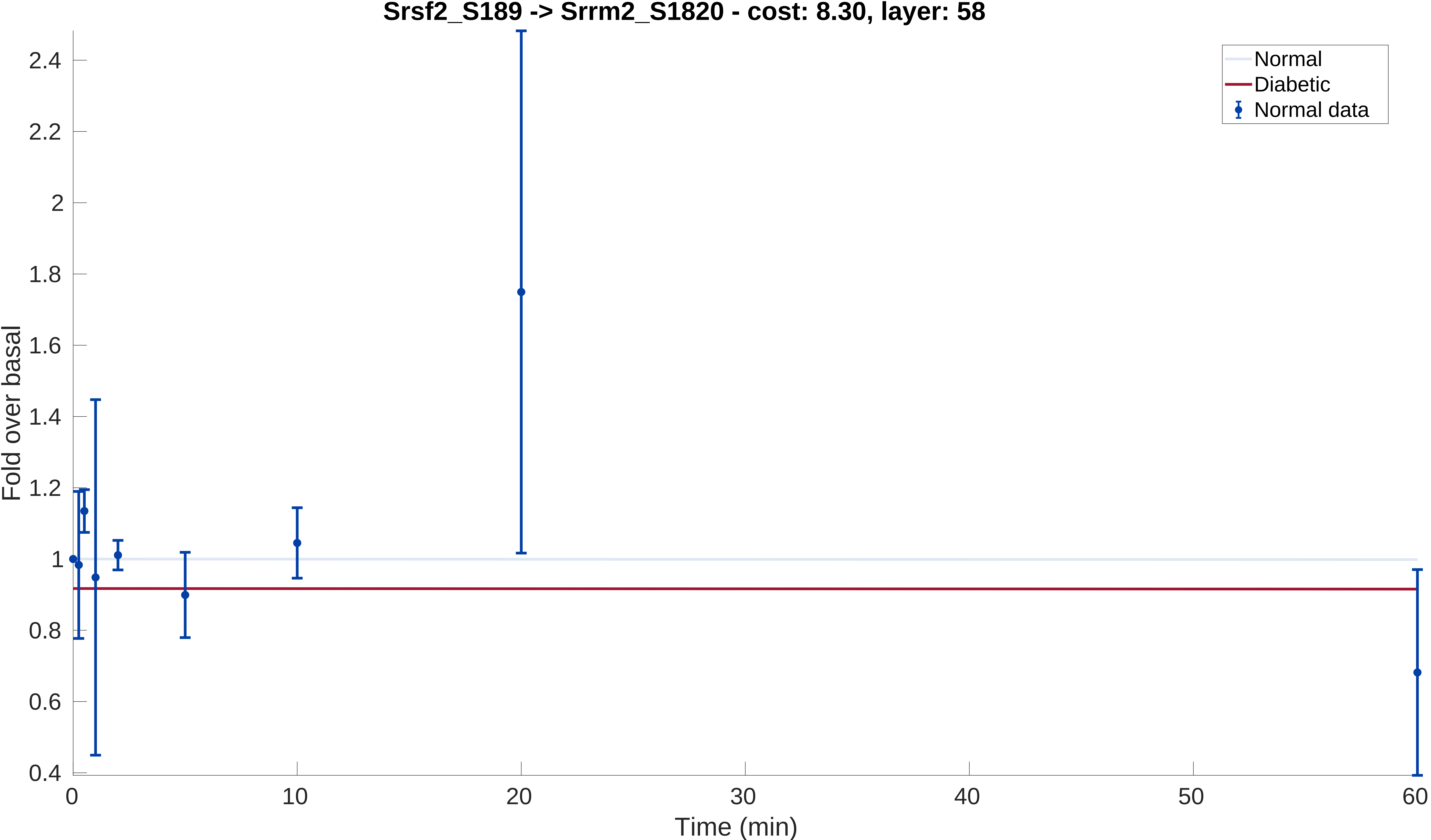
Srsf2_S189 -> Srrm2_S1820 - cost: 8.30, layer: 58
Normal
Diabetic
Normal data
Fold over basal
2.4
2.2
2
1.8
1.6
1.4
1.2
1
0.8
0.6
0.4
0
10
20
30
40
50
60
Time (min)

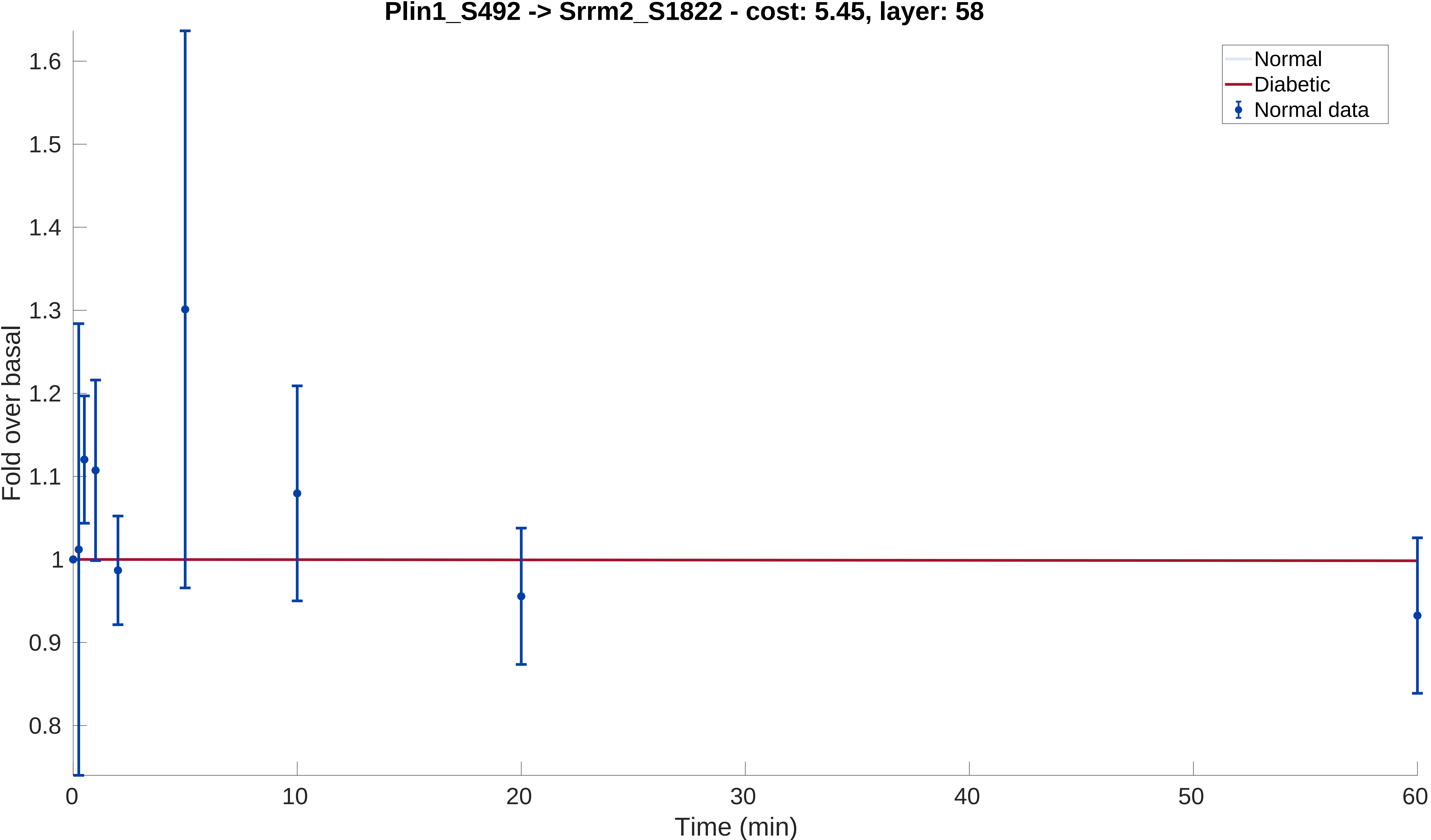

Plin1_S492 -> Srrm2_S1822 - cost: 5.45, layer: 58
Normal
Diabetic
Normal data
Fold over basal
1.6
1.5
1.4
1.3
1.2
1.1
1
0.9
0.8
0
10
20
30
40
50
60
Time (min)

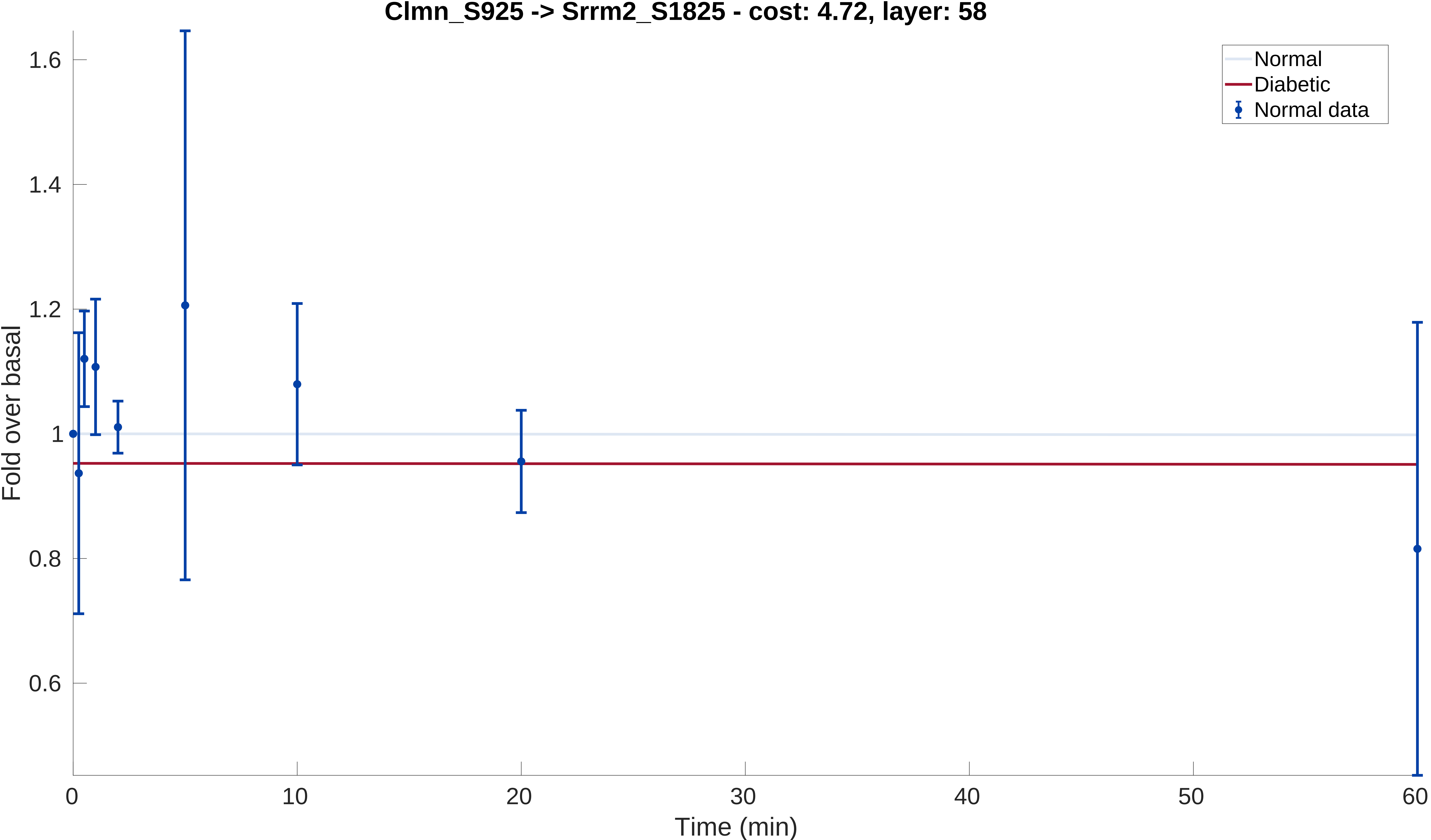
Clmn_S925 -> Srrm2_S1825 - cost: 4.72, layer: 58
Normal
Diabetic
Normal data
Fold over basal
1.6
1.4
1.2
1
0.8
0.6
0
10
20
30
40
50
60
Time (min)

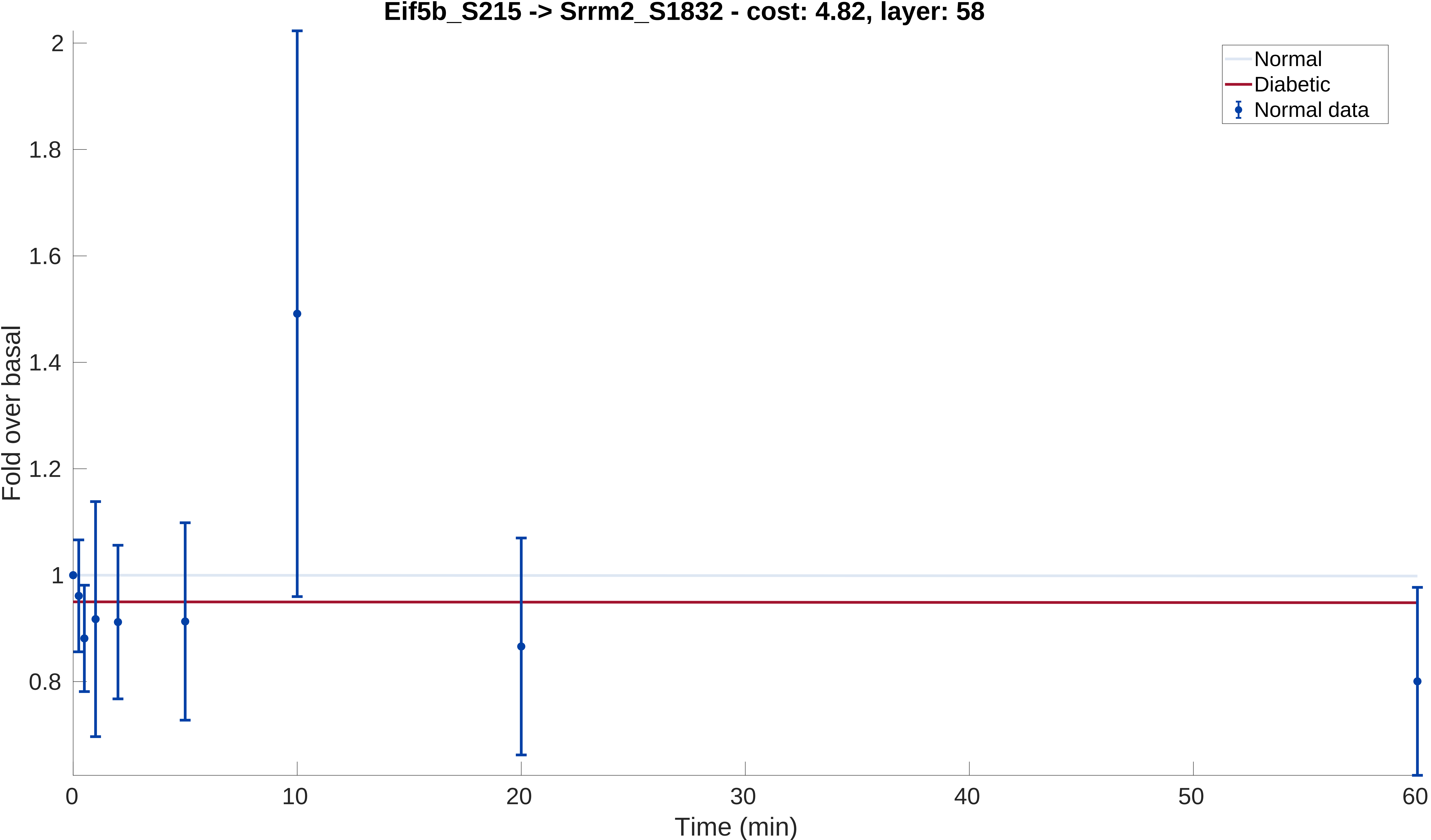

Eif5b_S215 -> Srrm2_S1832 - cost: 4.82, layer: 58
Normal
Diabetic
Normal data
Fold over basal
2
1.8
1.6
1.4
1.2
1
0.8
0
10
20
30
40
50
60
Time (min)

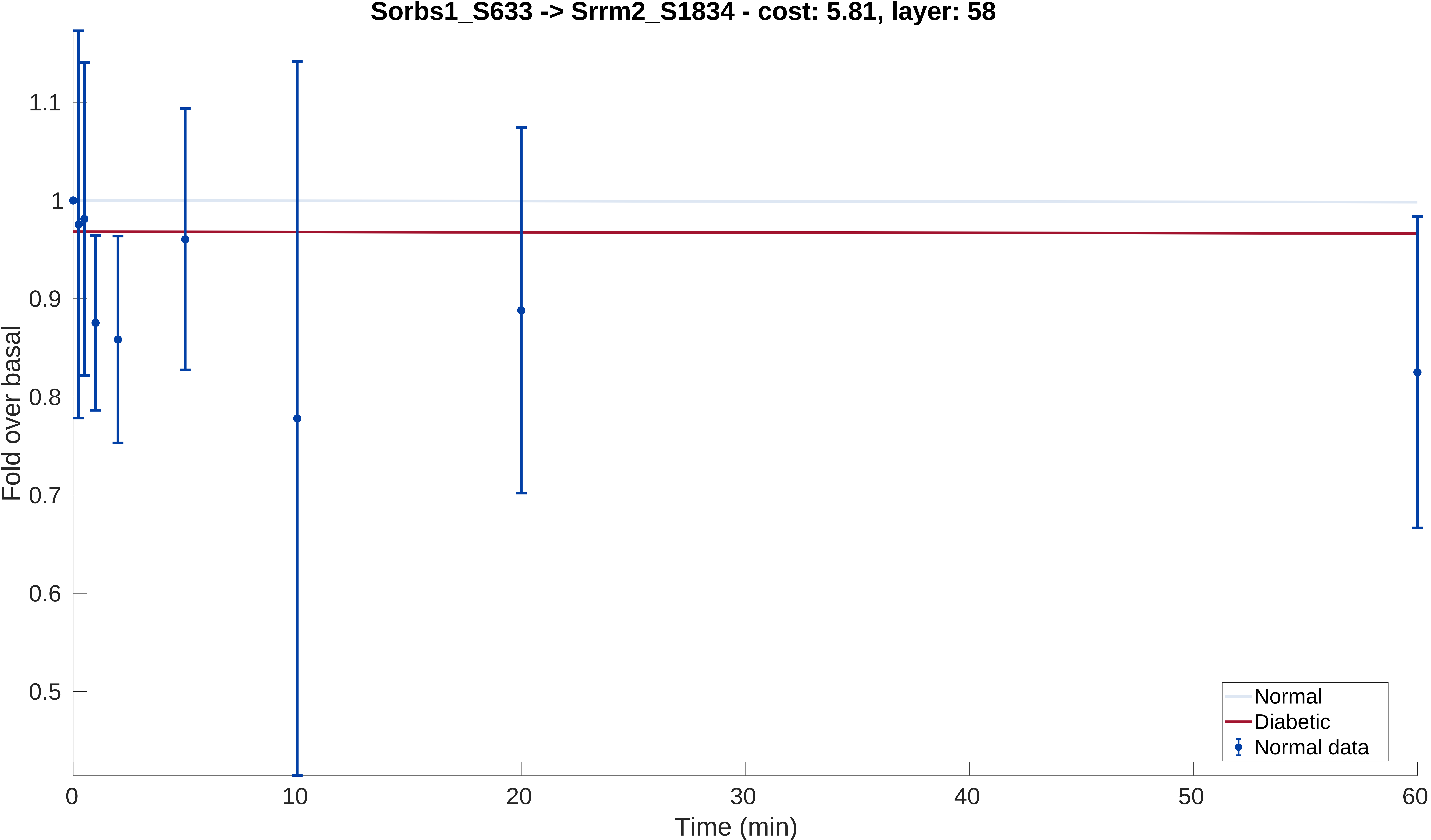

Sorbs1_S633 -> Srrm2_S1834 - cost: 5.81, layer: 58
Fold over basal
Time (min)
0.5
0.6
0.7
0.8
0.9
1
1.1
0
10
20
30
40
50
60
Normal
Diabetic
Normal data

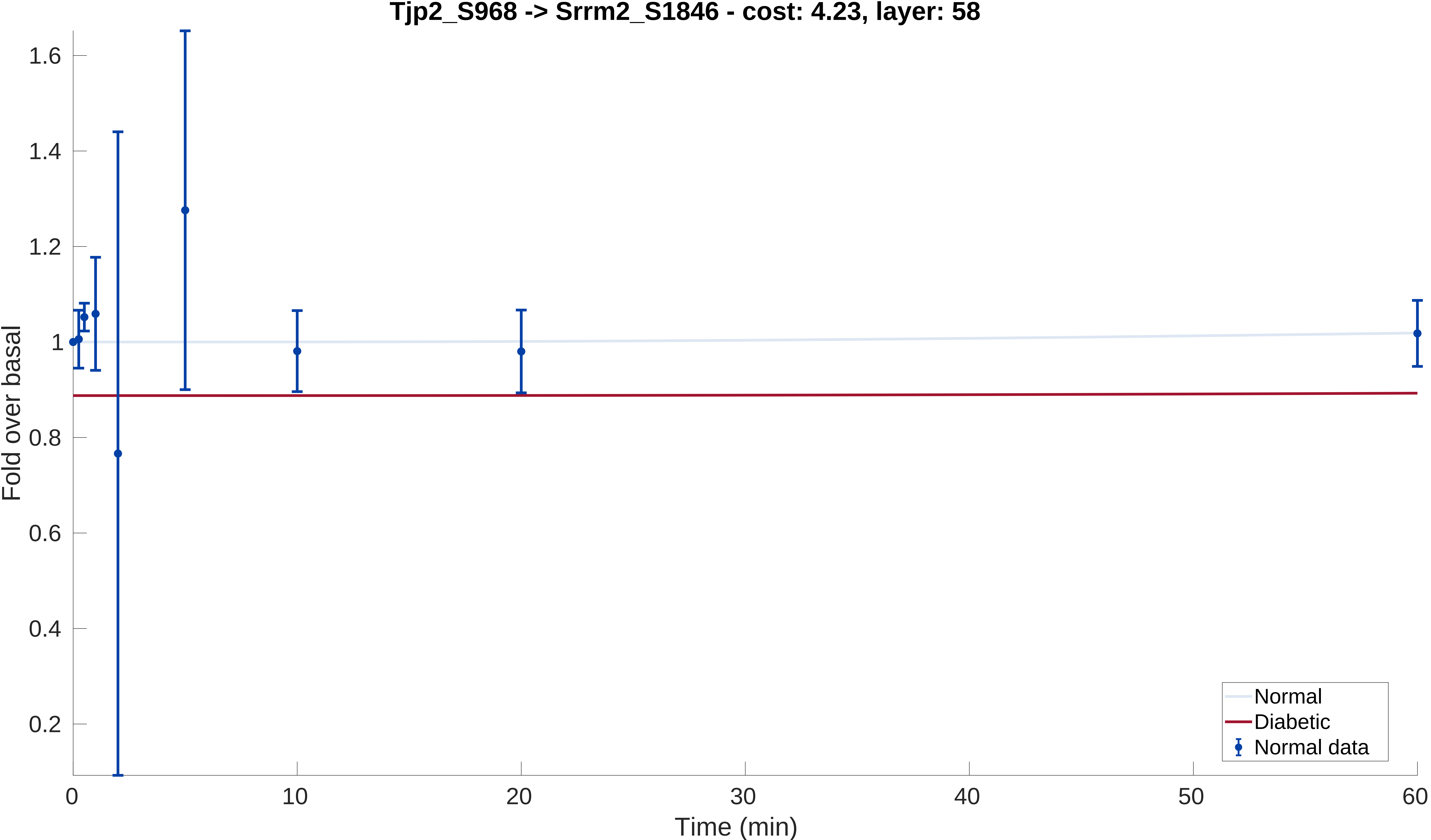
Tjp2_S968 -> Srrm2_S1846 - cost: 4.23, layer: 58
Fold over basal
1.6
1.4
1.2
1
0.8
0.6
0.4
0.2
0
10
20
30
40
50
60
Time (min)
Normal
Diabetic
Normal data

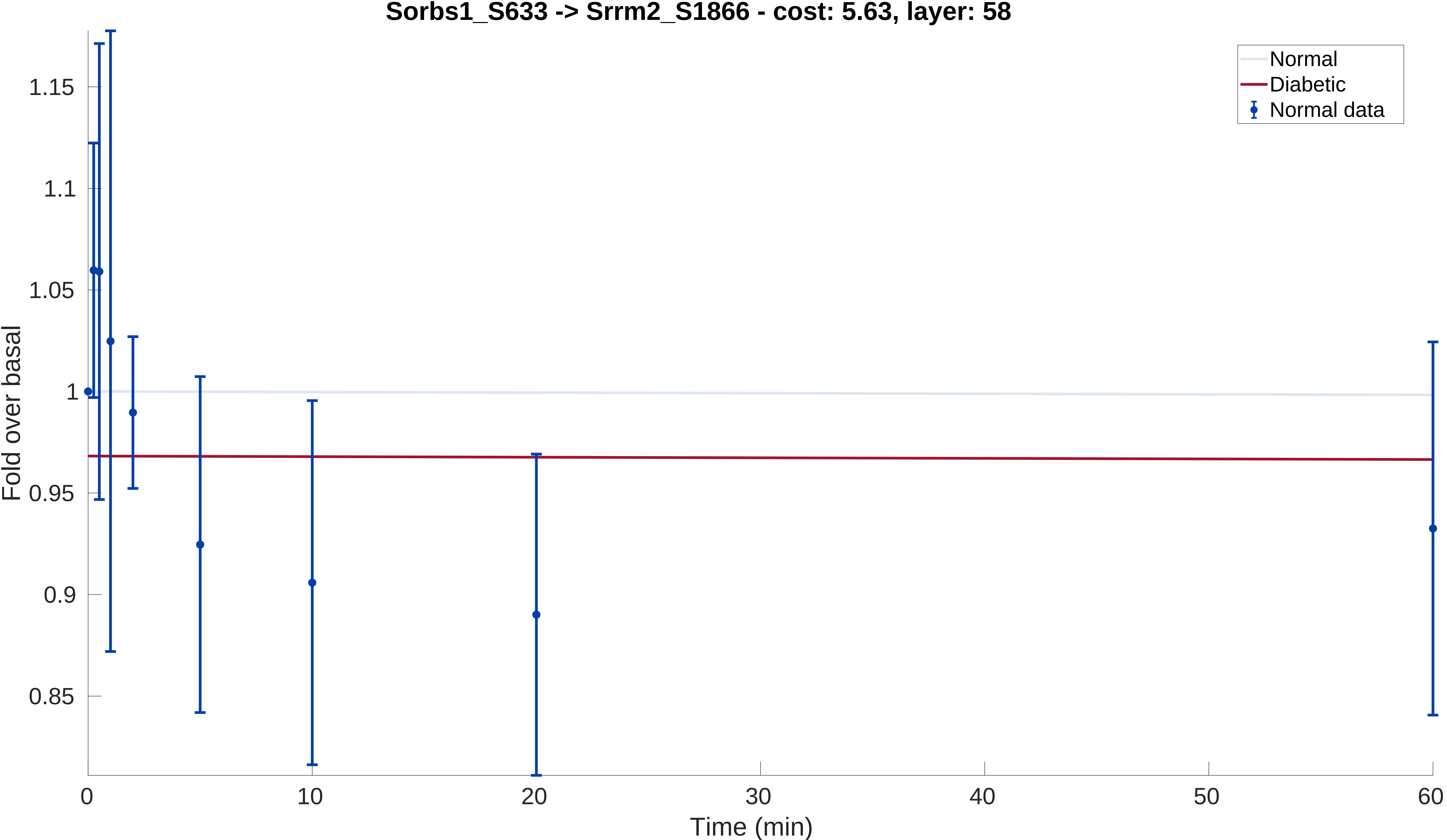

Sorbs1_S633 -> Srrm2_S1866 - cost: 5.63, layer: 58
Normal
Diabetic
Normal data
Fold over basal
1.15
1.1
1.05
1
0.95
0.9
0.85
0
10
20
30
40
50
60
Time (min)

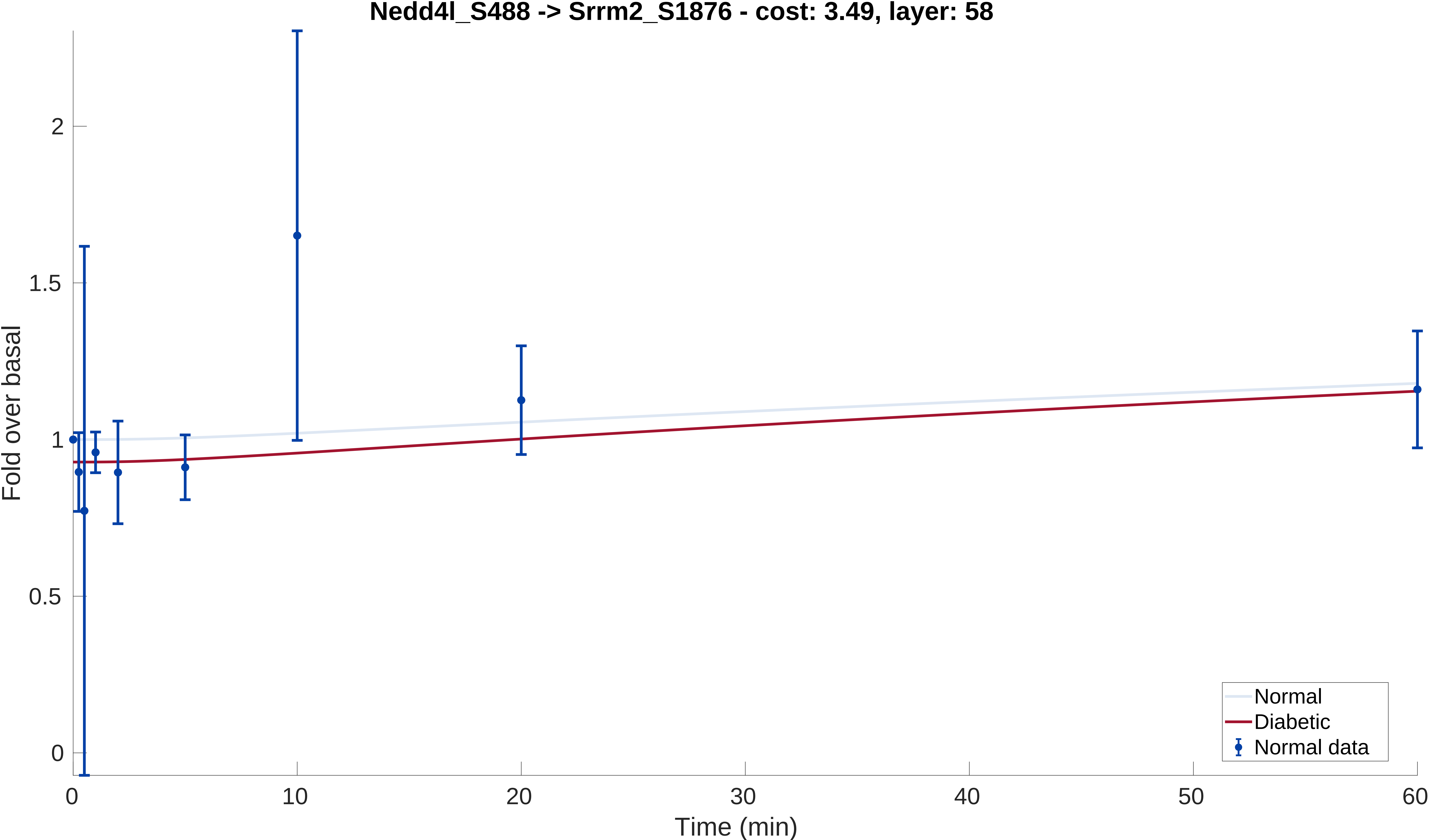

Nedd4l_S488 -> Srrm2_S1876 - cost: 3.49, layer: 58
Fold over basal
Time (min)
0
0.5
1
1.5
2
0
10
20
30
40
50
60
Normal
Diabetic
Normal data

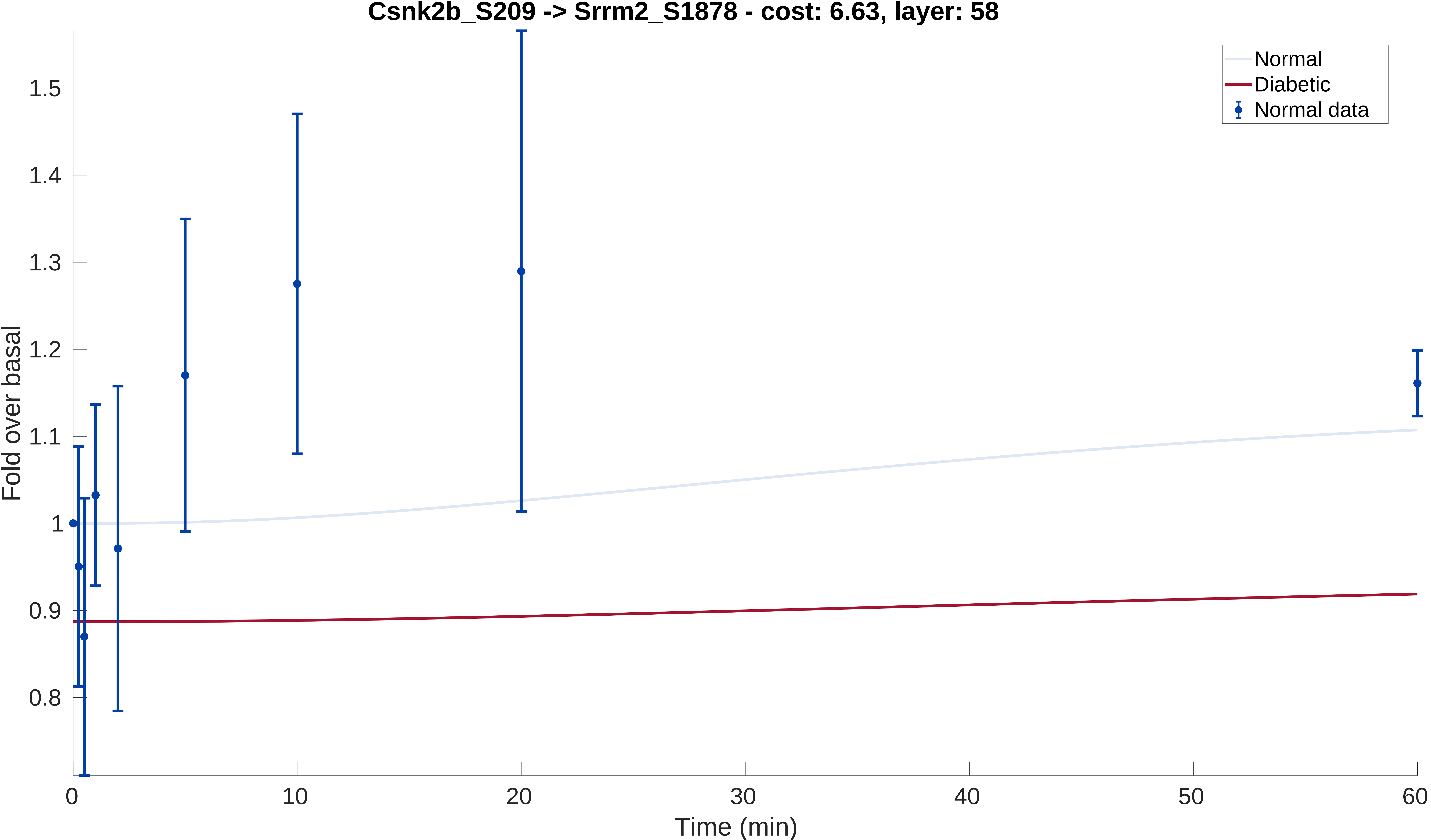

Csnk2b_S209 -> Srrm2_S1878 - cost: 6.63, layer: 58
Normal
Diabetic
Normal data
Fold over basal
1.5
1.4
1.3
1.2
1.1
1
0.9
0.8
0
10
20
30
40
50
60
Time (min)

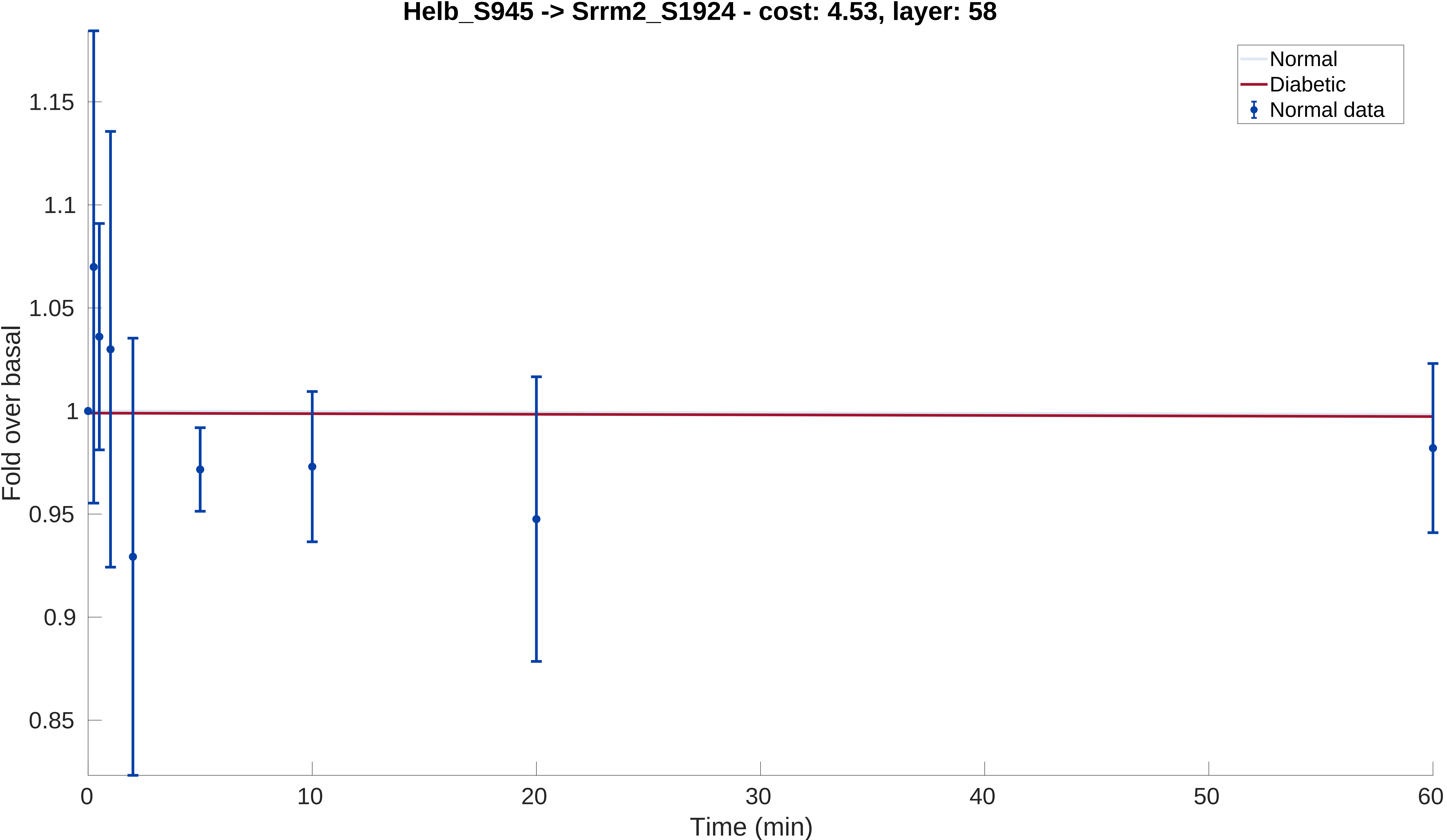
Helb_S945 -> Srrm2_S1924 - cost: 4.53, layer: 58
Normal
Diabetic
Normal data
Fold over basal
1.15
1.1
1.05
1
0.95
0.9
0.85
0
10
20
30
40
50
60
Time (min)

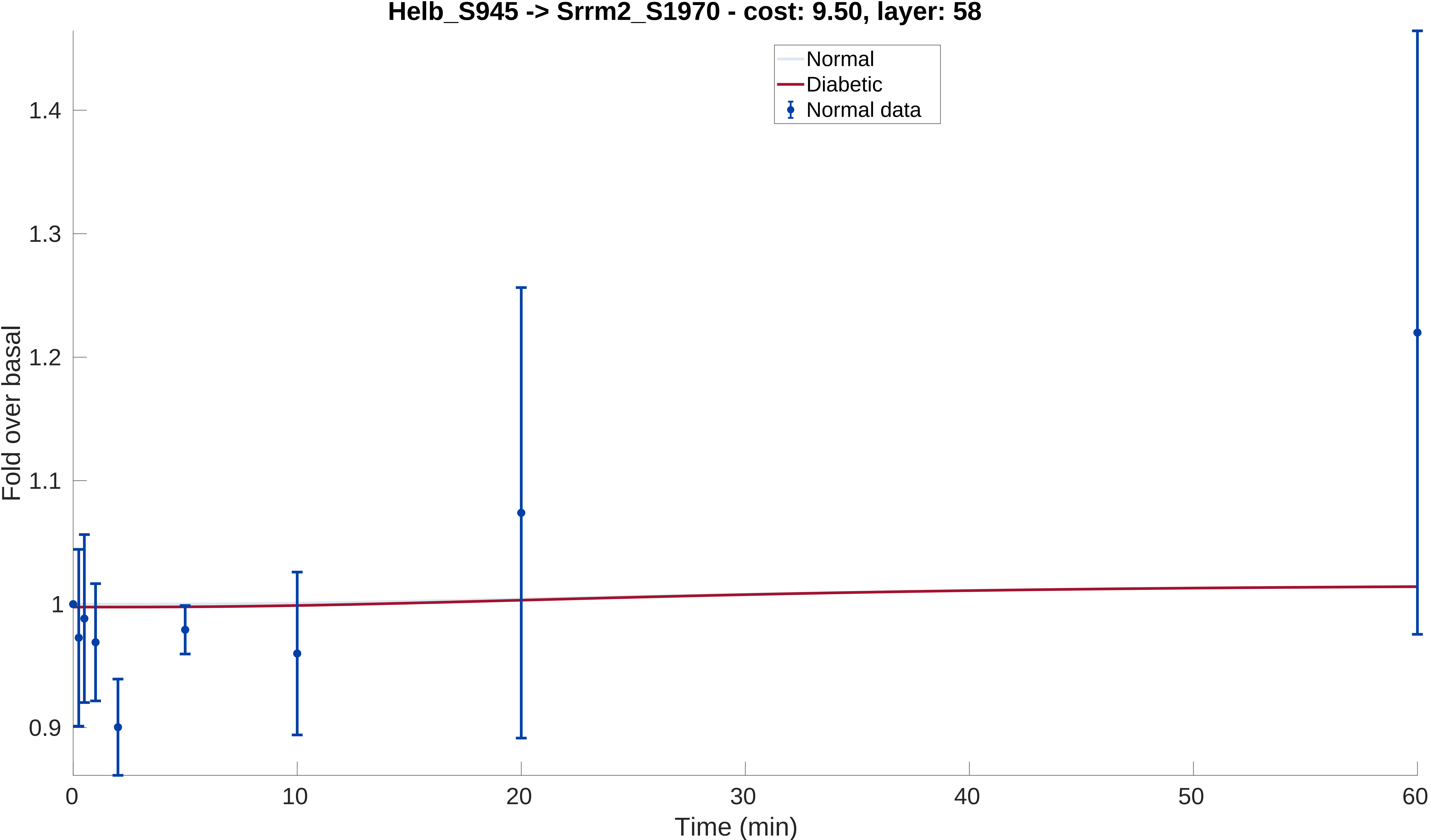

Helb_S945 -> Srrm2_S1970 - cost: 9.50, layer: 58
Normal
Diabetic
Normal data
Fold over basal
1.4
1.3
1.2
1.1
1
0.9
0
10
20
30
40
50
60
Time (min)

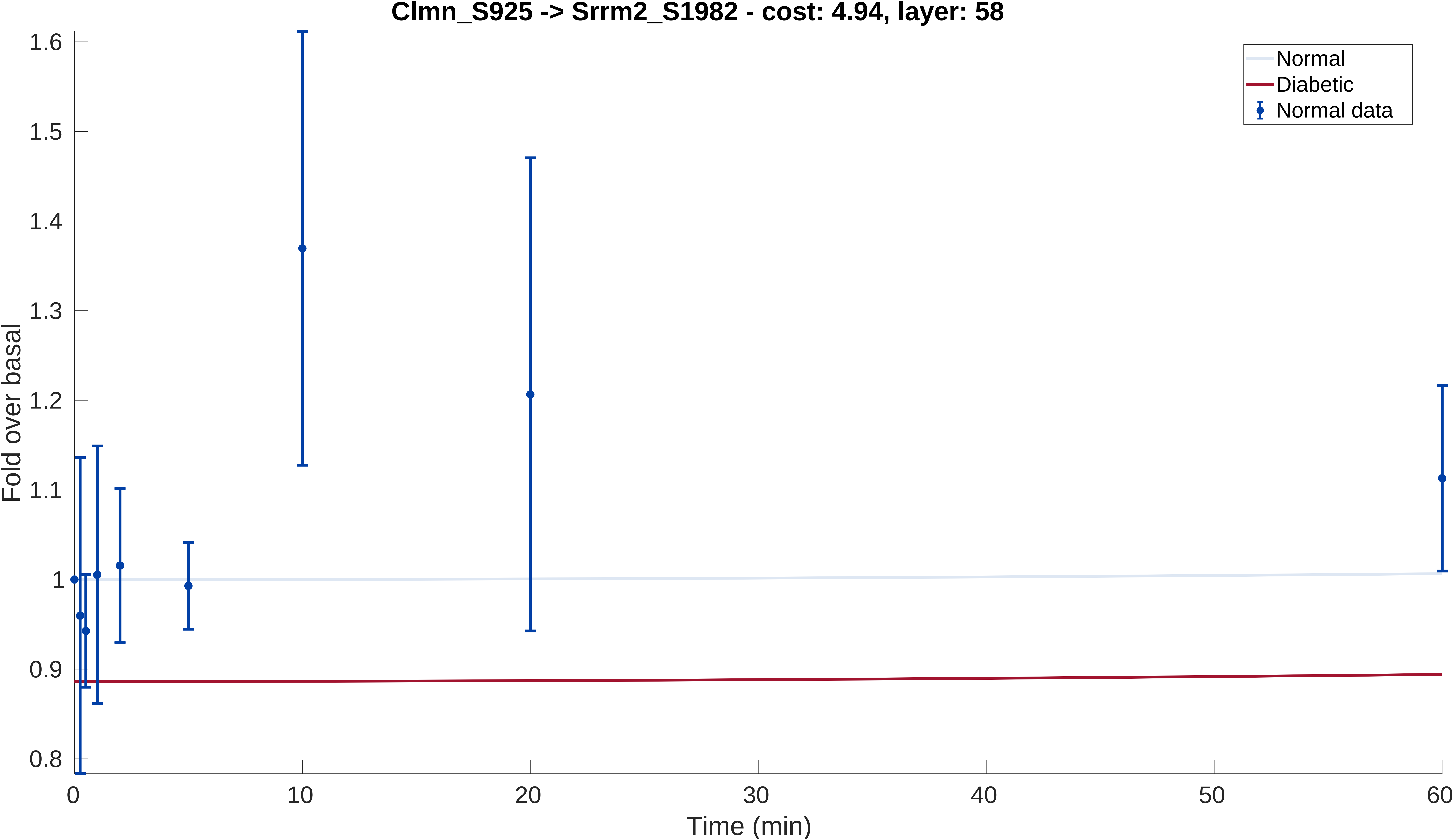

Clmn_S925 -> Srrm2_S1982 - cost: 4.94, layer: 58
Normal
Diabetic
Normal data
Fold over basal
Time (min)
1.6
1.5
1.4
1.3
1.2
1.1
1
0.9
0.8
0
10
20
30
40
50
60

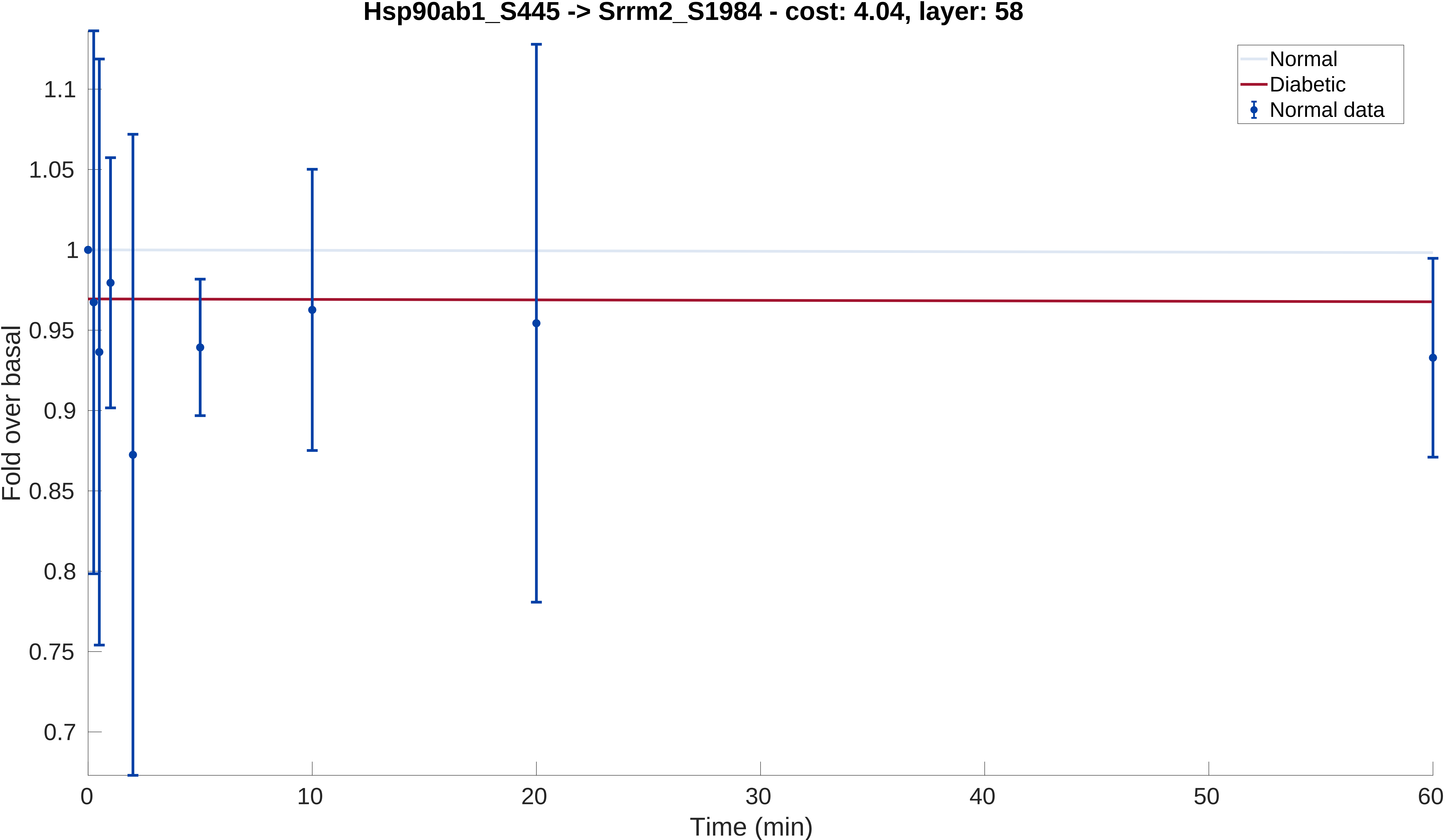

Hsp90ab1_S445 -> Srrm2_S1984 - cost: 4.04, layer: 58
Normal
Diabetic
Normal data
Fold over basal
1.1
1.05
1
0.95
0.9
0.85
0.8
0.75
0.7
0
10
20
30
40
50
60
Time (min)

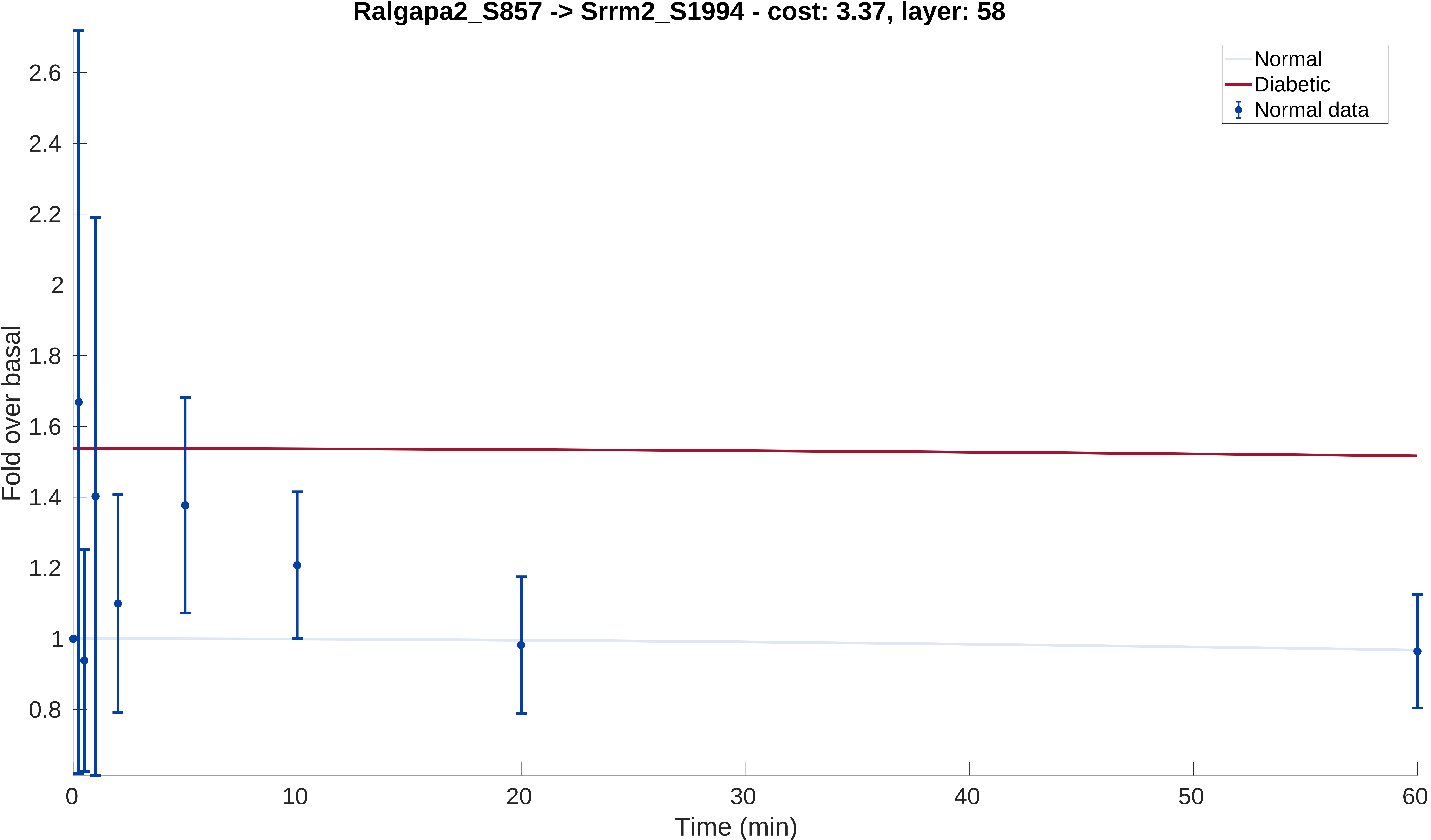

Ralgapa2_S857 -> Srrm2_S1994 - cost: 3.37, layer: 58
Normal
Diabetic
Normal data
Fold over basal
Time (min)
0.8
1
1.2
1.4
1.6
1.8
2
2.2
2.4
2.6
0
10
20
30
40
50
60

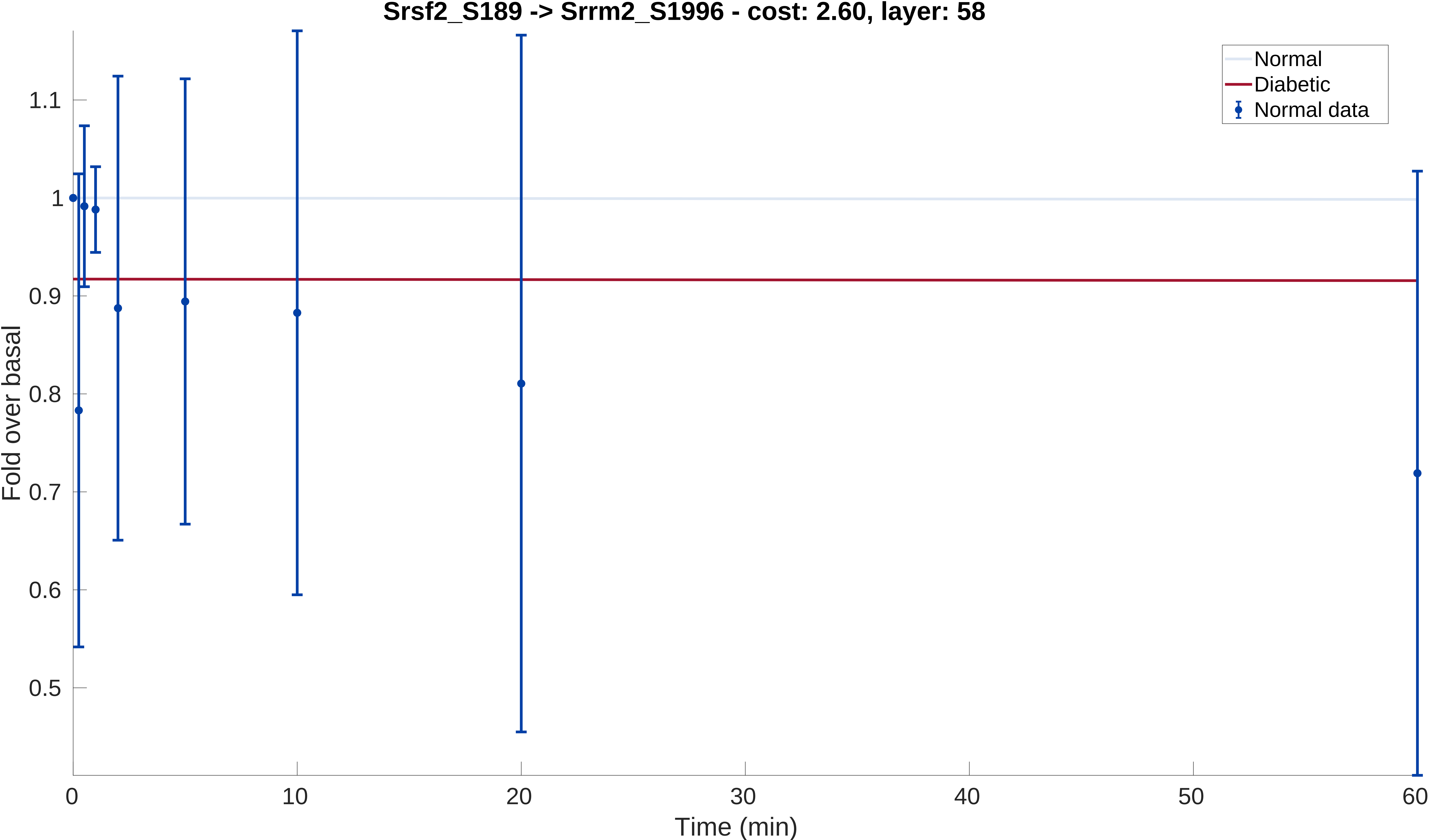
Srsf2_S189 -> Srrm2_S1996 - cost: 2.60, layer: 58
Normal
Diabetic
Normal data
Fold over basal
1.1
1
0.9
0.8
0.7
0.6
0.5
0
10
20
30
40
50
60
Time (min)

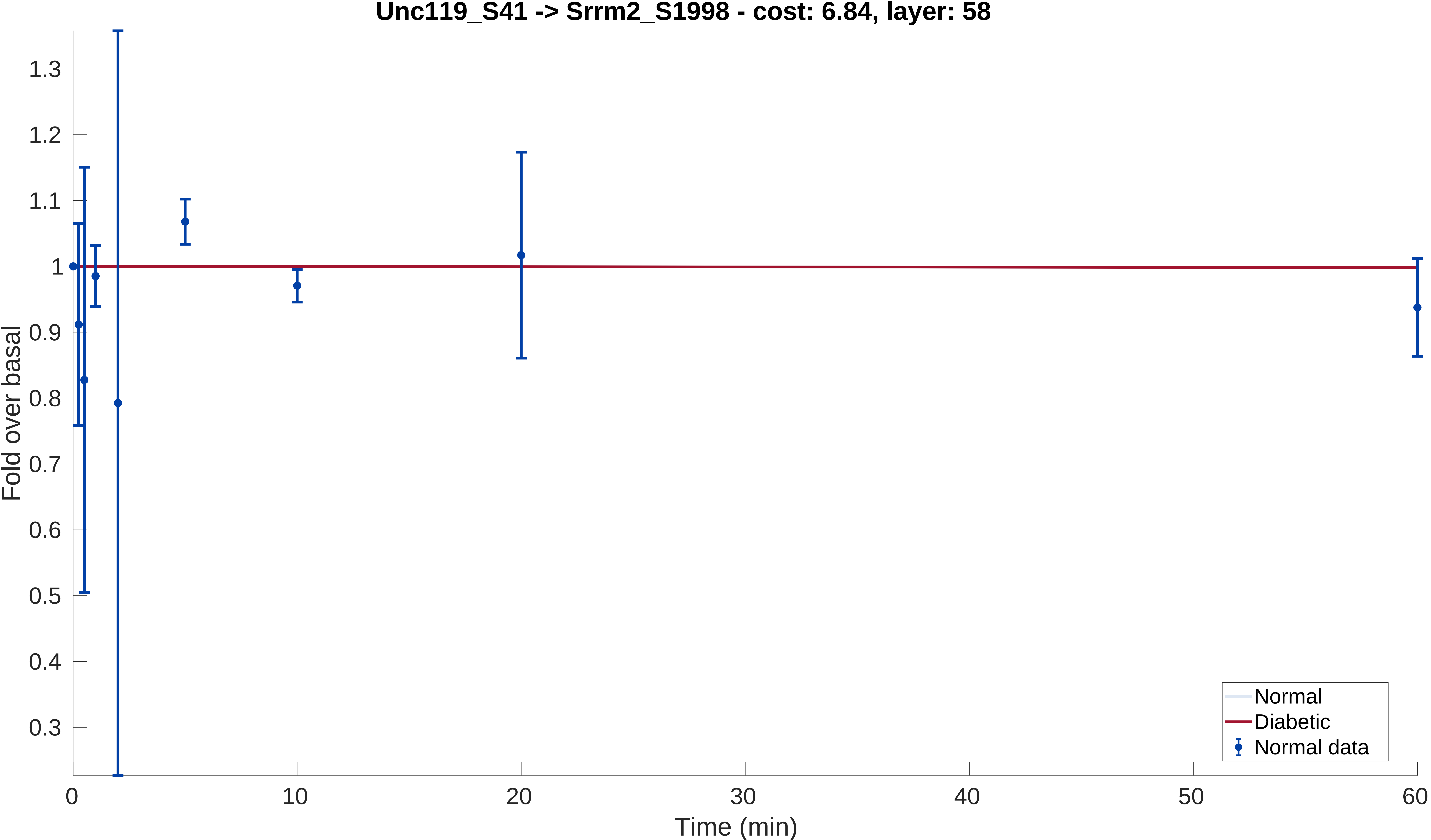

Unc119_S41 -> Srrm2_S1998 - cost: 6.84, layer: 58
Fold over basal
1.3
1.2
1.1
1
0.9
0.8
0.7
0.6
0.5
0.4
0.3
0
10
20
30
40
50
60
Time (min)
Normal
Diabetic
Normal data

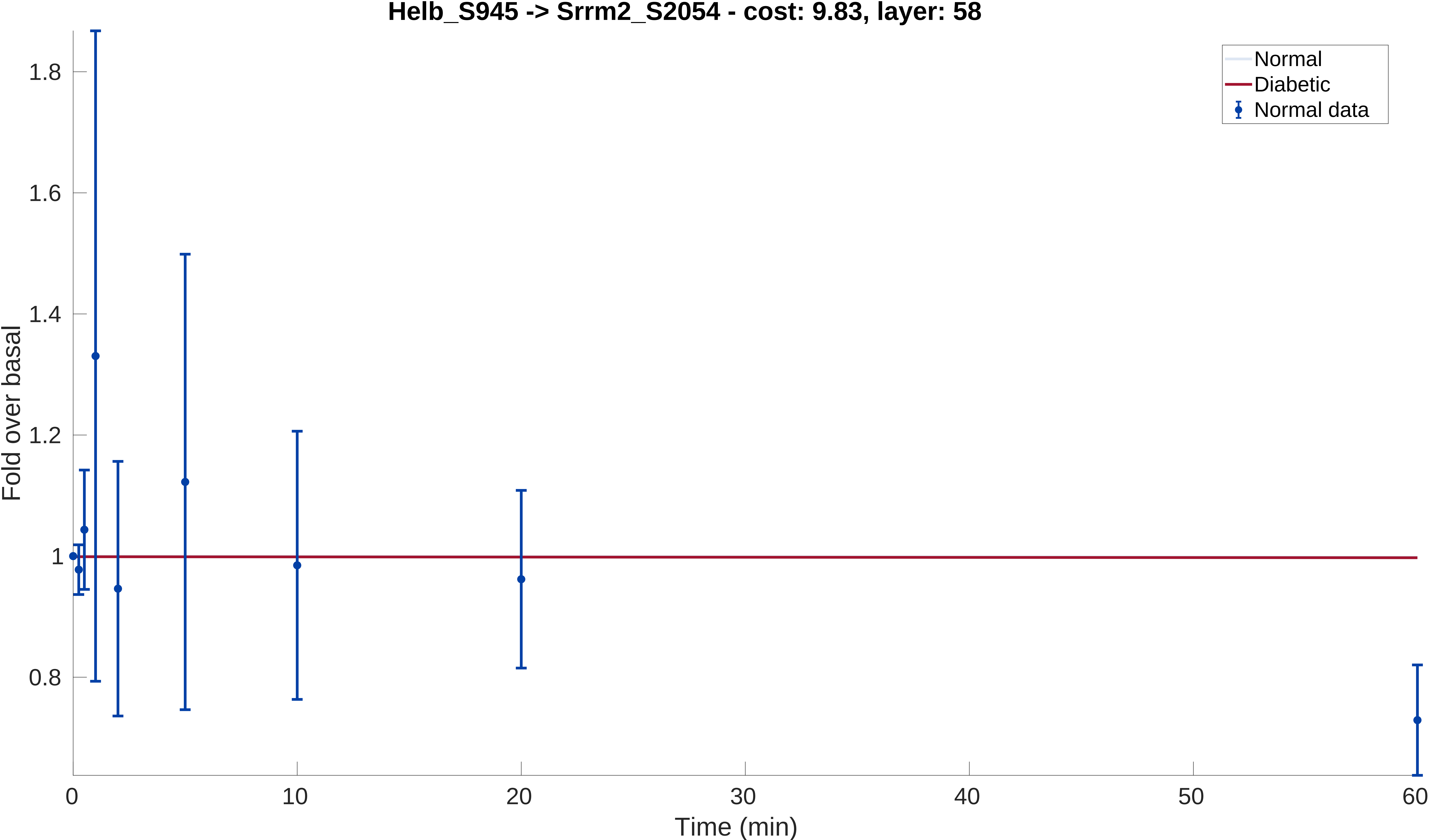
Helb_S945 -> Srrm2_S2054 - cost: 9.83, layer: 58
Normal
Diabetic
Normal data
Fold over basal
1.8
1.6
1.4
1.2
1
0.8
0
10
20
30
40
50
60
Time (min)

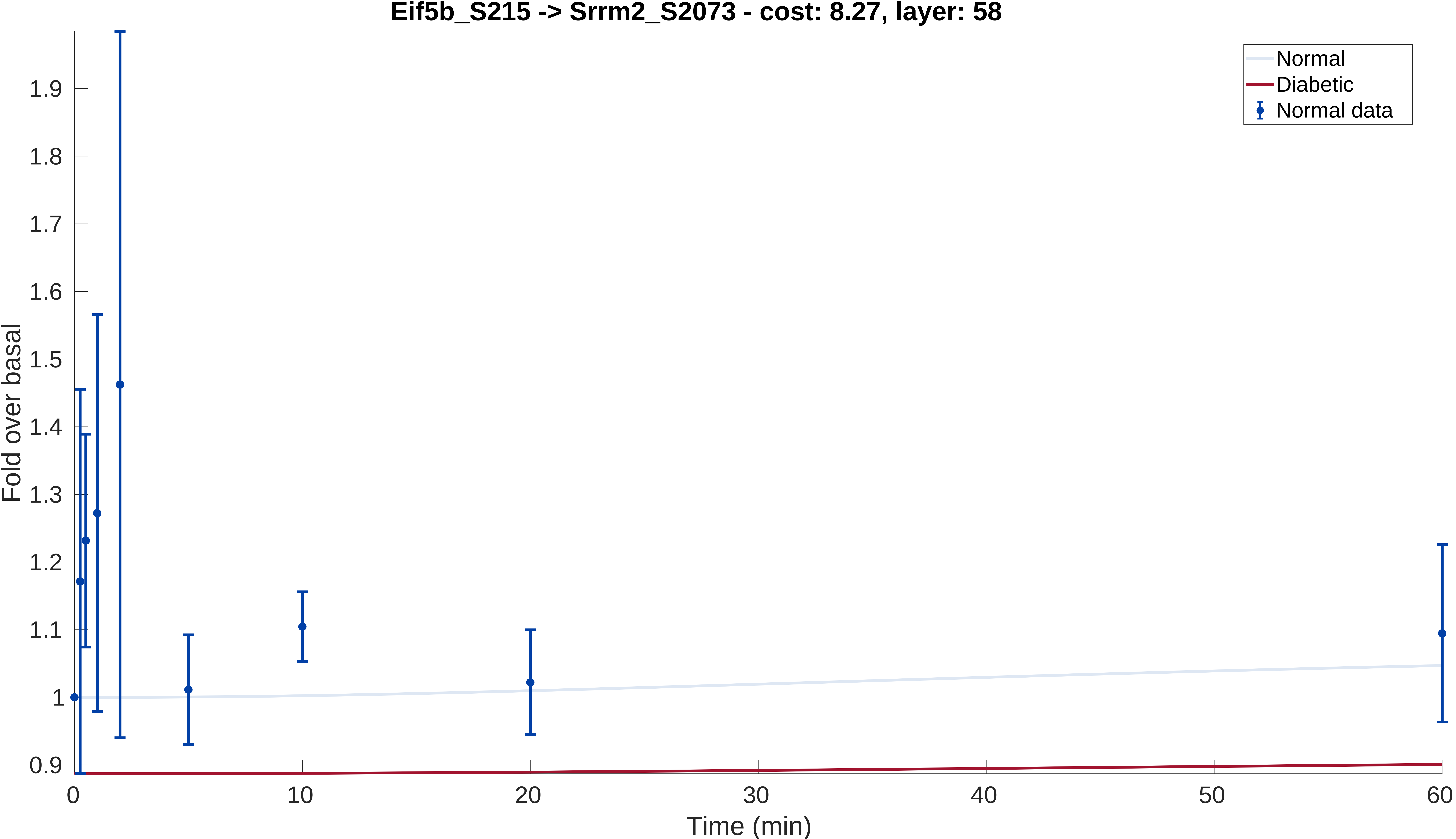
Eif5b_S215 -> Srrm2_S2073 - cost: 8.27, layer: 58
Normal
Diabetic
Normal data
Fold over basal
1.9
1.8
1.7
1.6
1.5
1.4
1.3
1.2
1.1
1
0.9
0
10
20
30
40
50
60
Time (min)

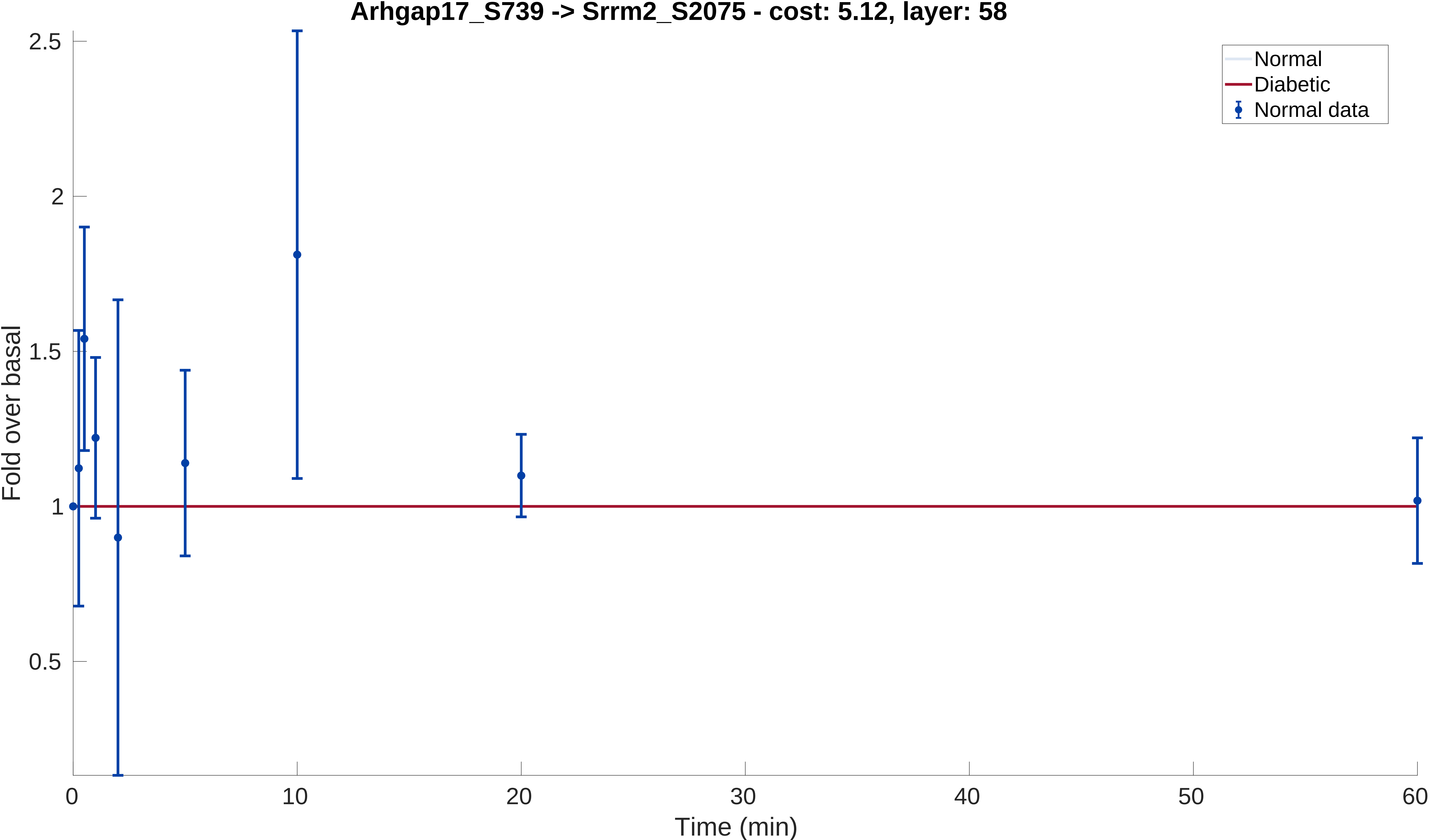
Arhgap17_S739 -> Srrm2_S2075 - cost: 5.12, layer: 58
Normal
Diabetic
Normal data
Fold over basal
Time (min)
0
10
20
30
40
50
60
0.5
1
1.5
2
2.5

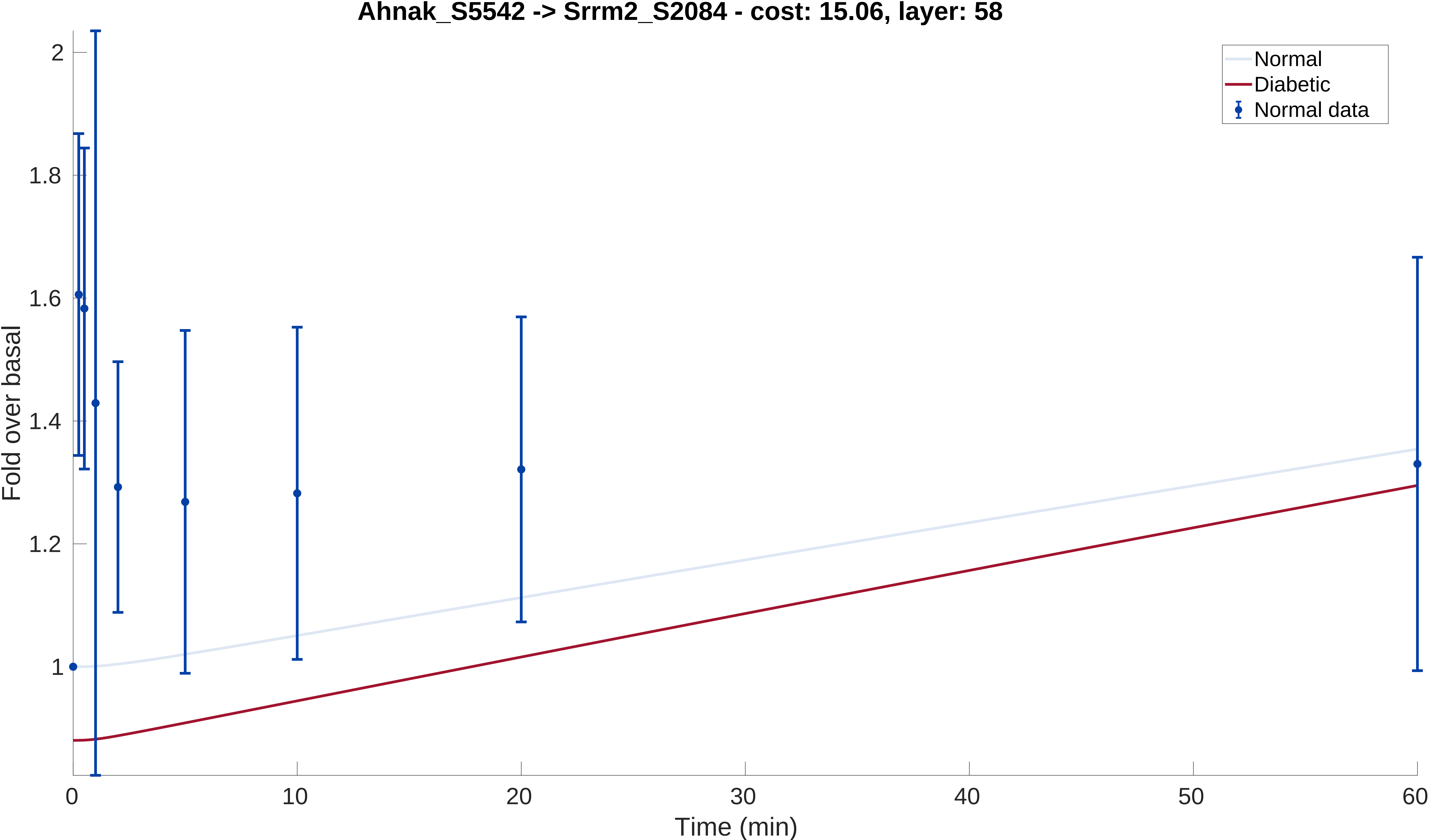

Ahnak_S5542 -> Srrm2_S2084 - cost: 15.06, layer: 58
Normal
Diabetic
Normal data
Fold over basal
2
1.8
1.6
1.4
1.2
1
0
10
20
30
40
50
60
Time (min)

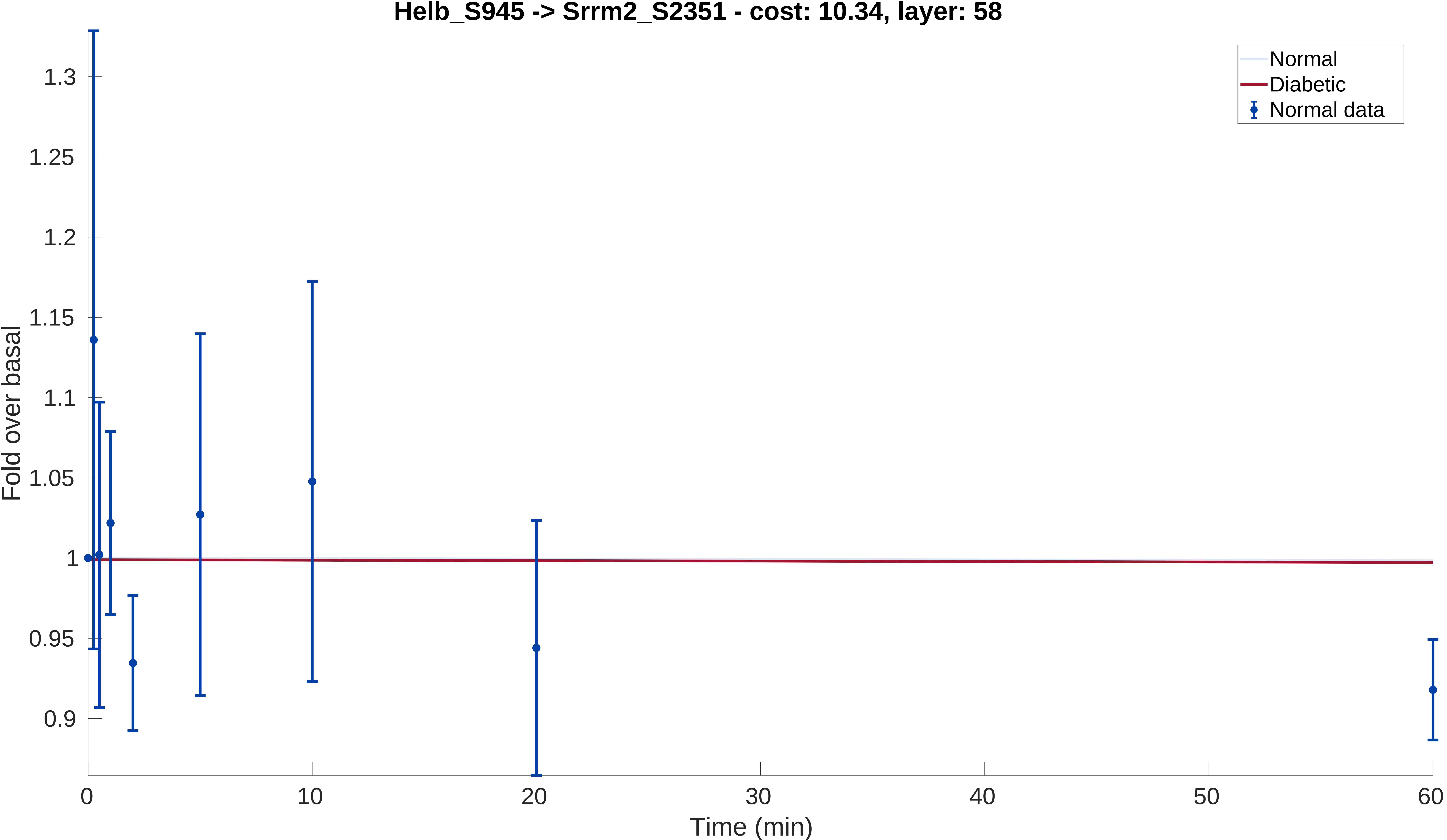

Helb_S945 -> Srrm2_S2351 - cost: 10.34, layer: 58
Normal
Diabetic
Normal data
Fold over basal
1.3
1.25
1.2
1.15
1.1
1.05
1
0.95
0.9
0
10
20
30
40
50
60
Time (min)

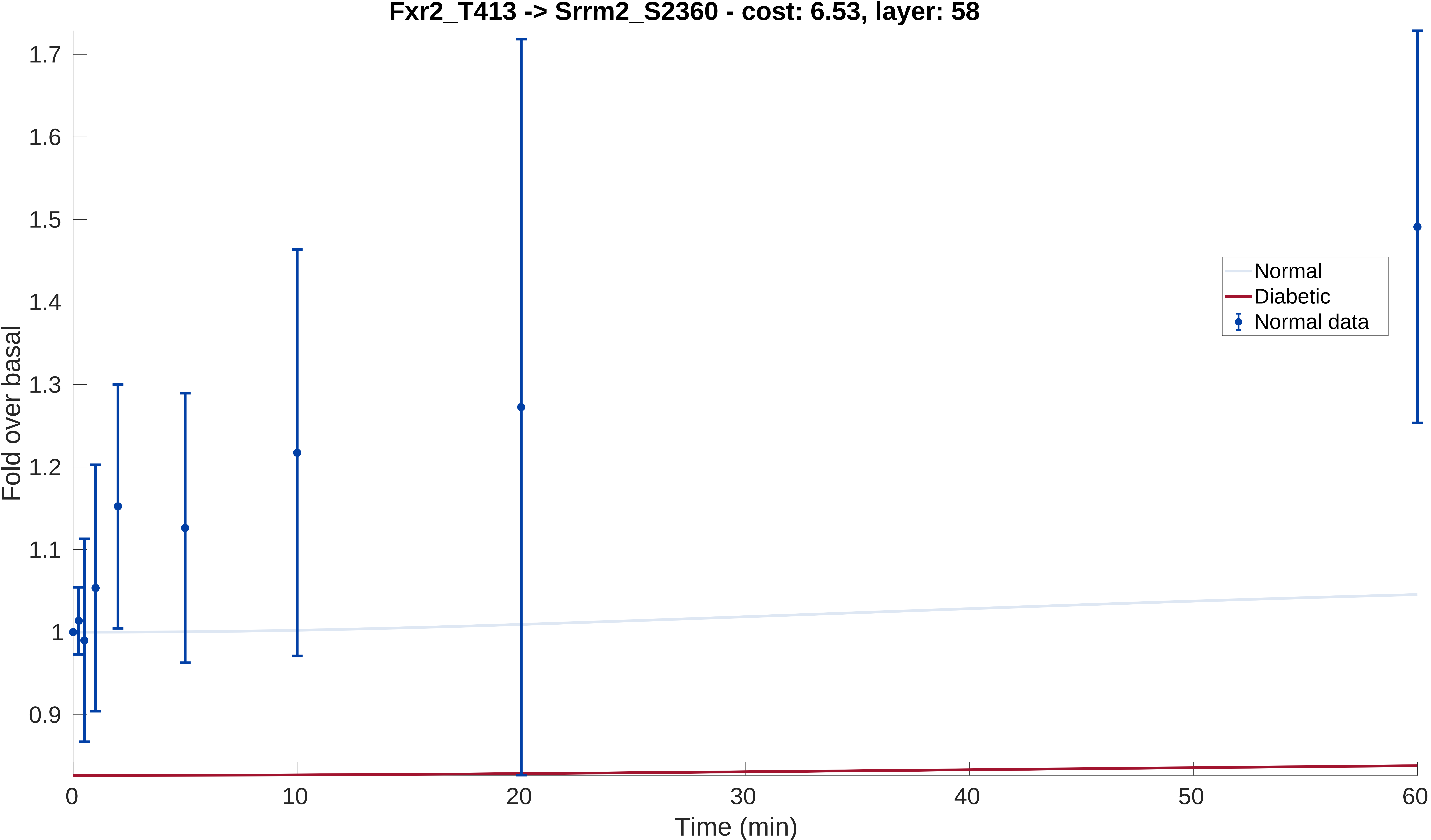

Fxr2_T413 -> Srrm2_S2360 - cost: 6.53, layer: 58
Fold over basal
1.7
1.6
1.5
1.4
1.3
1.2
1.1
1
0.9
0
10
20
30
40
50
60
Time (min)
Normal
Diabetic
Normal data

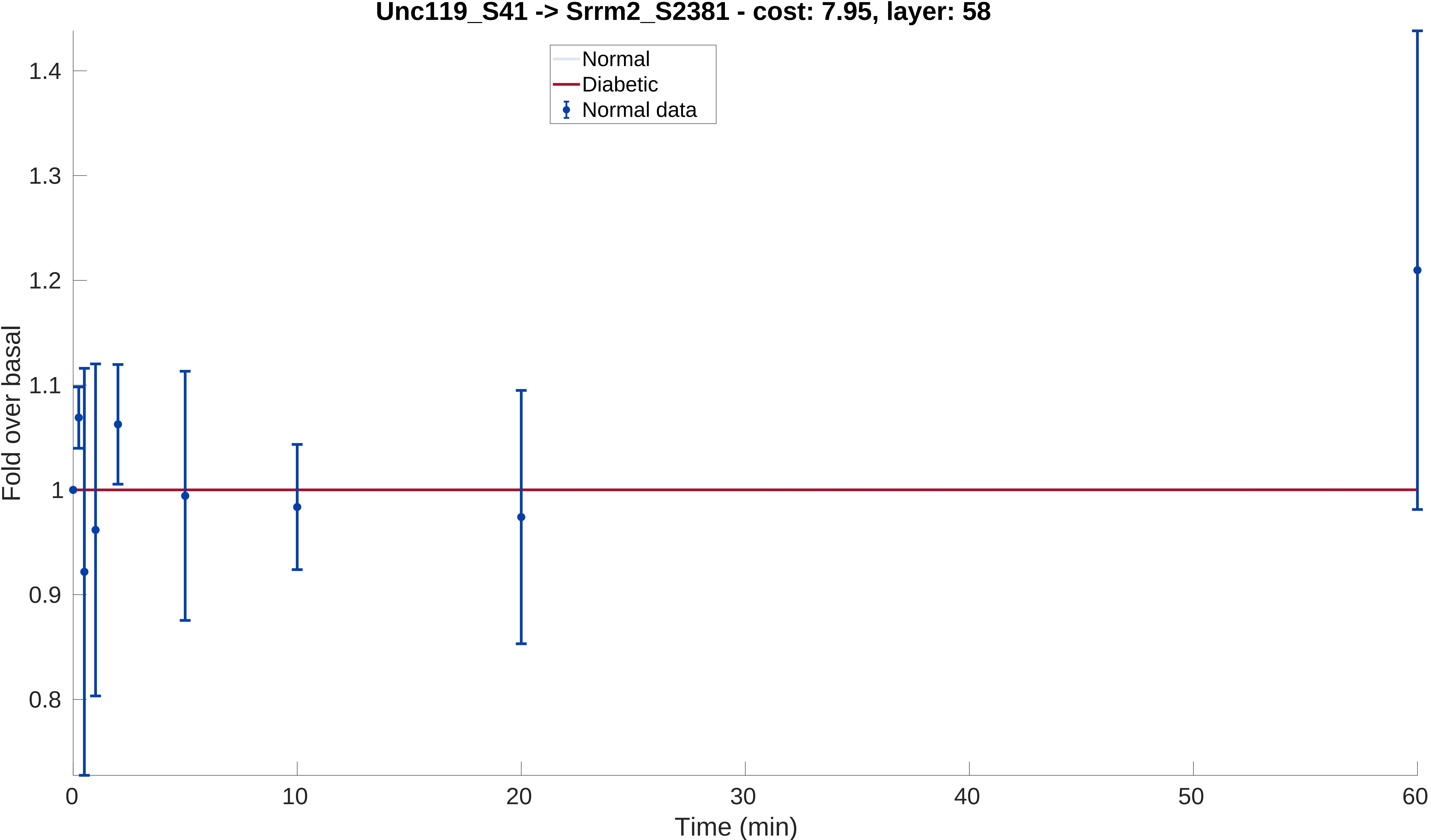

Unc119_S41 -> Srrm2_S2381 - cost: 7.95, layer: 58
Normal
Diabetic
Normal data
Fold over basal
1.4
1.3
1.2
1.1
1
0.9
0.8
0
10
20
30
40
50
60
Time (min)

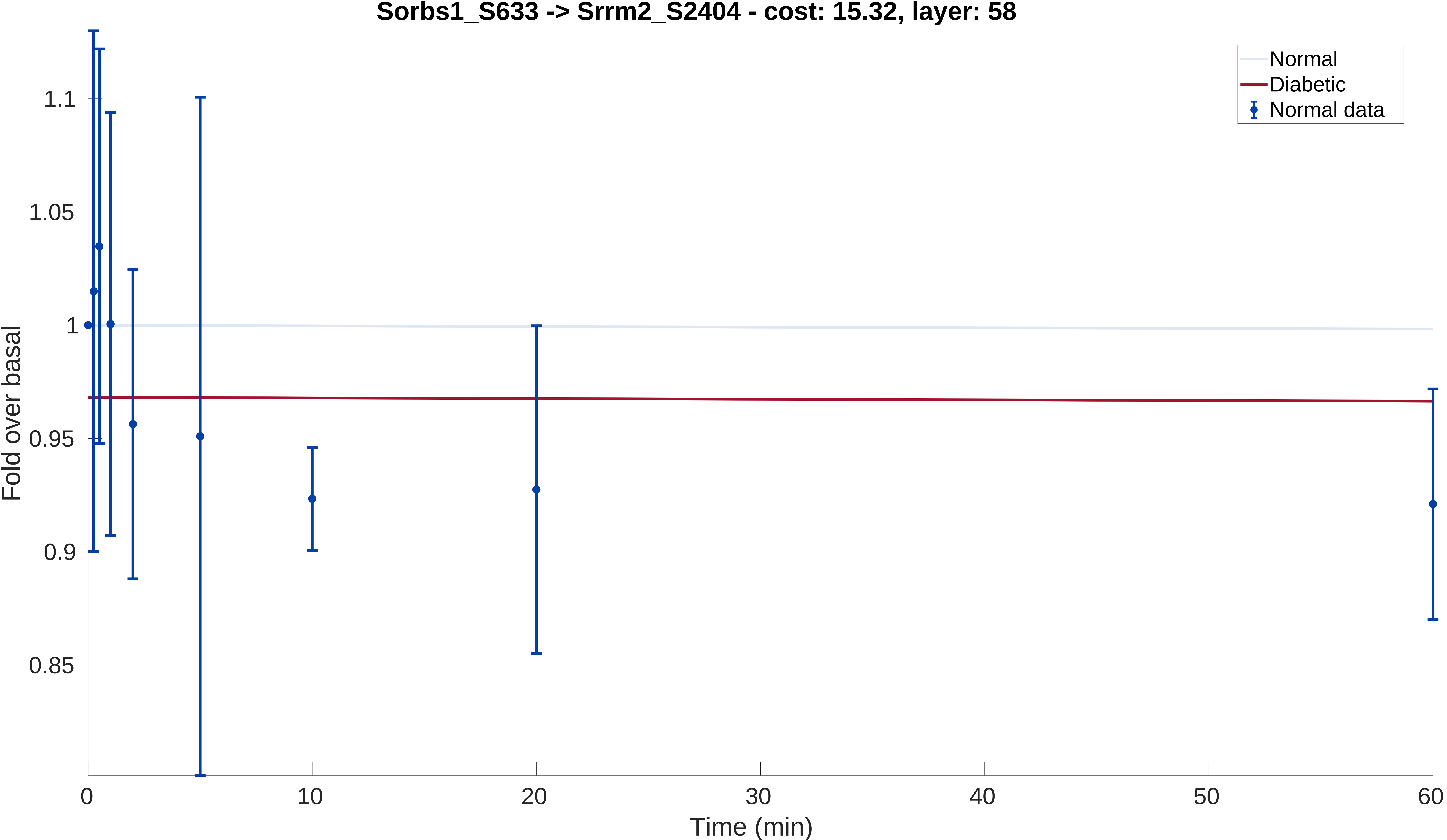
Sorbs1_S633 -> Srrm2_S2404 - cost: 15.32, layer: 58
Normal
Diabetic
Normal data
Fold over basal
1.1
1.05
1
0.95
0.9
0.85
0
10
20
30
40
50
60
Time (min)

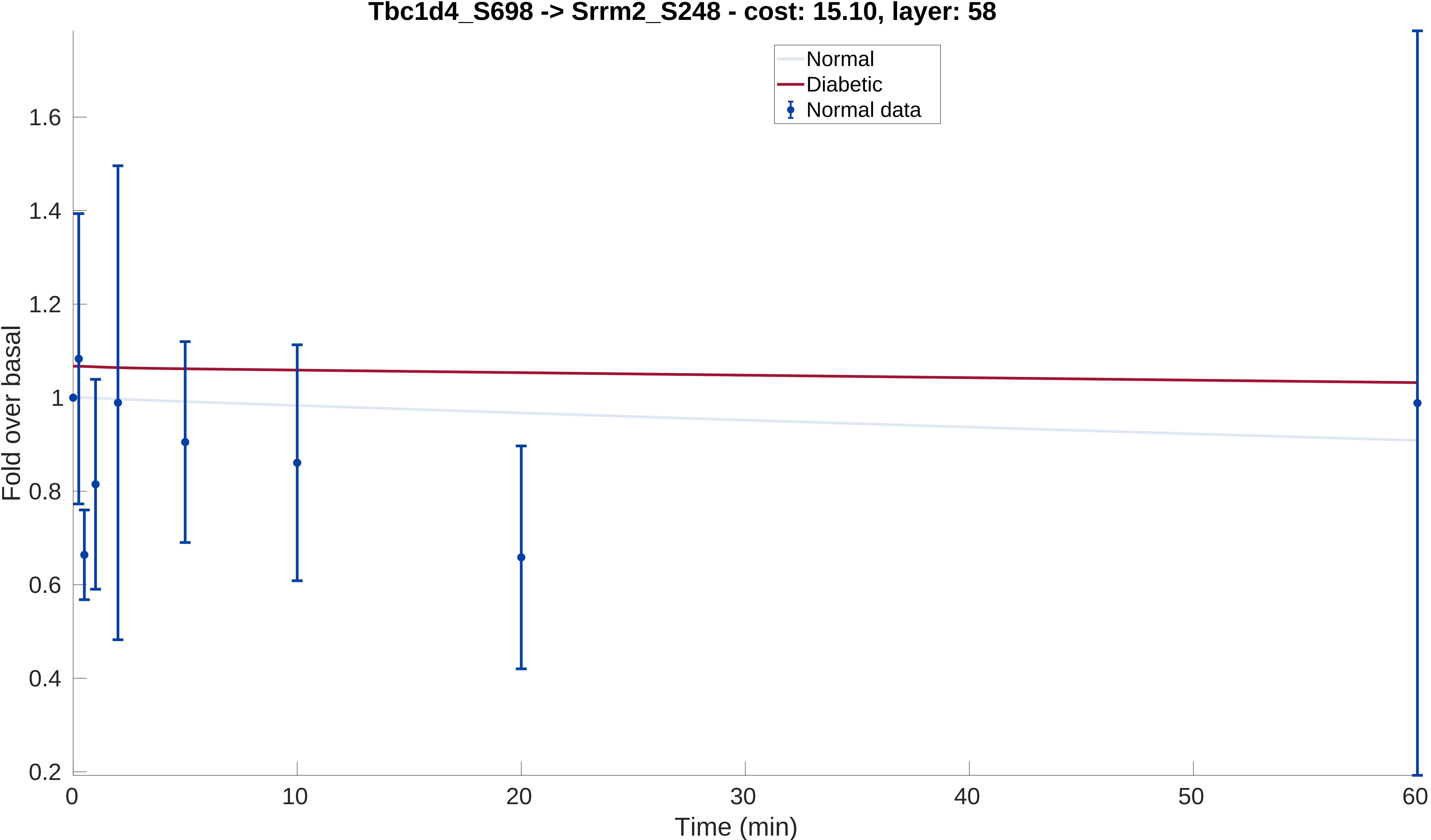

Tbc1d4_S698 -> Srrm2_S248 - cost: 15.10, layer: 58
Normal
Diabetic
Normal data
Fold over basal
0.2
0.4
0.6
0.8
1
1.2
1.4
1.6
0
10
20
30
40
50
60
Time (min)

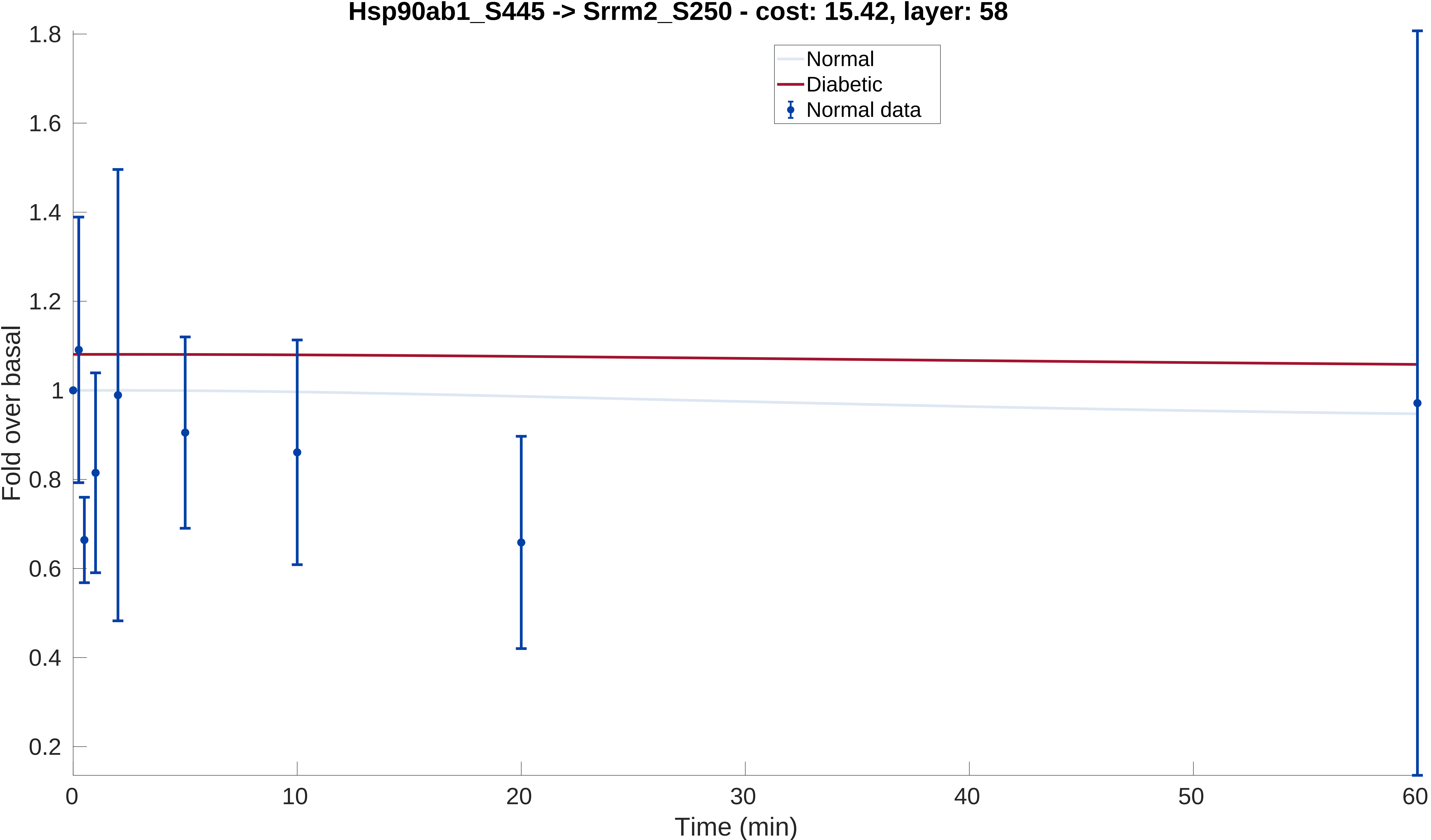

Hsp90ab1_S445 -> Srrm2_S250 - cost: 15.42, layer: 58
Normal
Diabetic
Normal data
Fold over basal
Time (min)
1.8
1.6
1.4
1.2
1
0.8
0.6
0.4
0.2
0
10
20
30
40
50
60

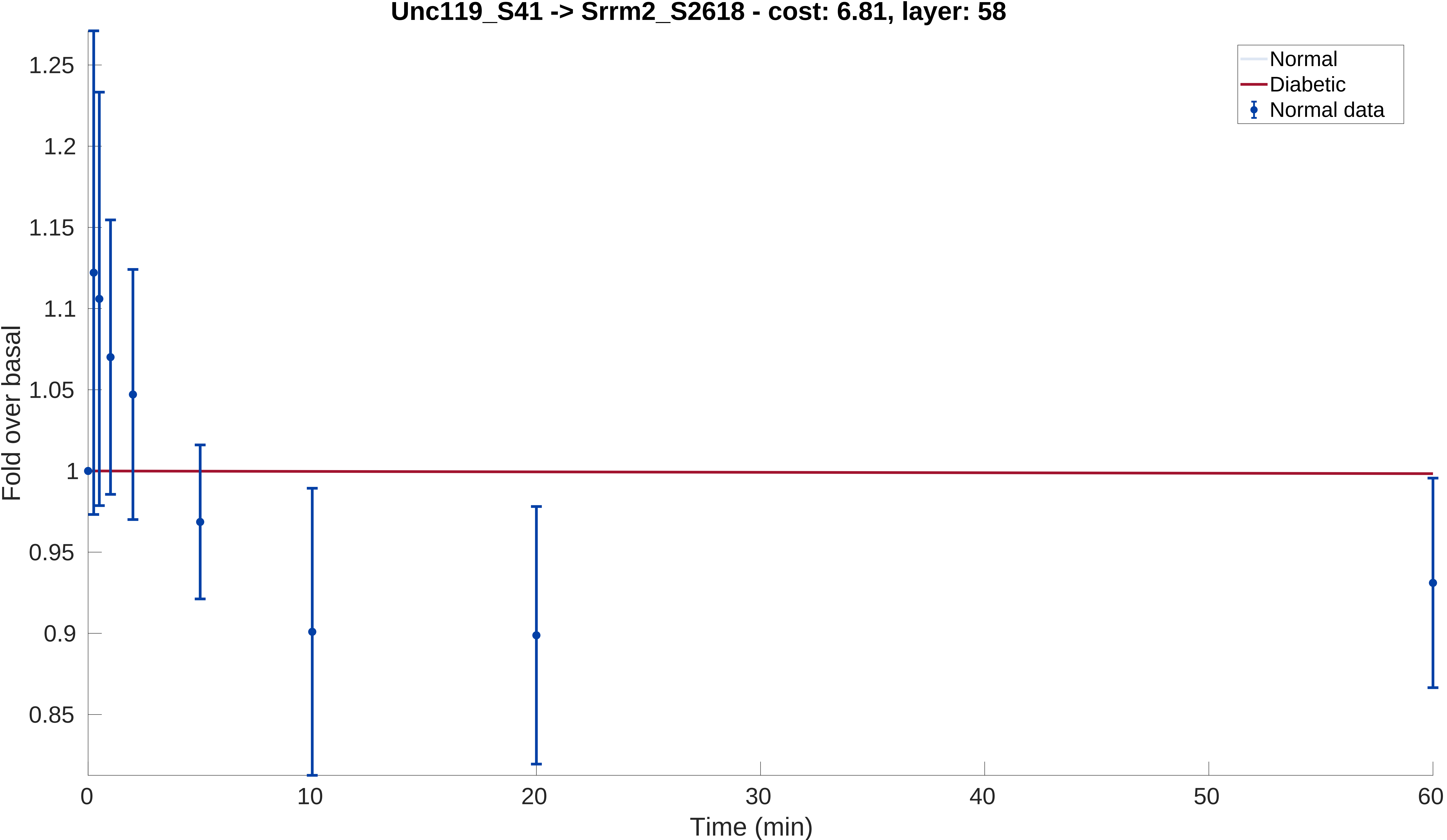

Unc119_S41 -> Srrm2_S2618 - cost: 6.81, layer: 58
Normal
Diabetic
Normal data
Fold over basal
1.25
1.2
1.15
1.1
1.05
1
0.95
0.9
0.85
Time (min)
0
10
20
30
40
50
60

Prkar2b_S112 -> Srrm2_S2642 - cost: 13.73, layer: 58

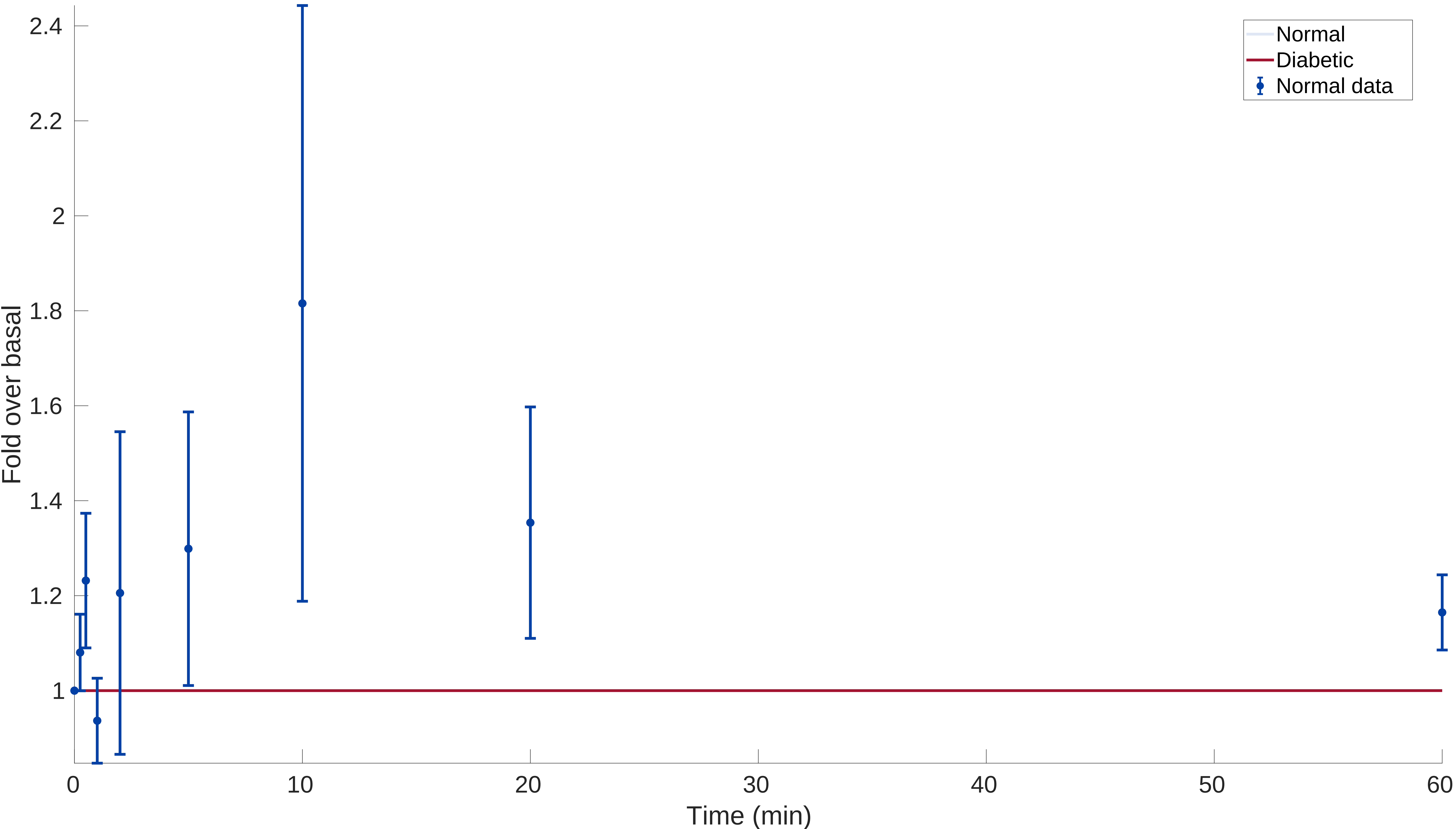

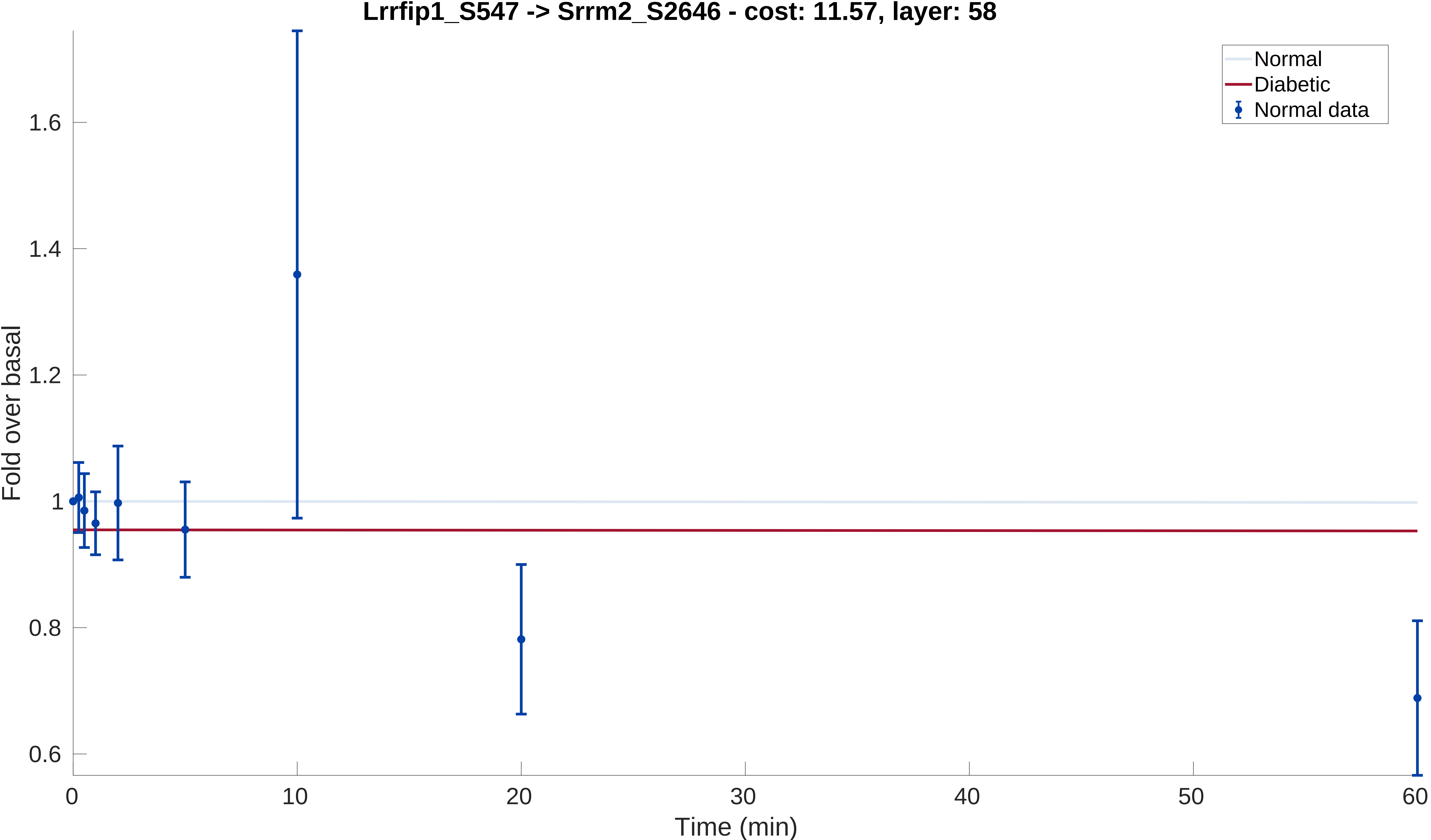

Lrrfip1_S547 -> Srrm2_S2646 - cost: 11.57, layer: 58
Normal
Diabetic
Normal data
Fold over basal
1.6
1.4
1.2
1
0.8
0.6
0
10
20
30
40
50
60
Time (min)

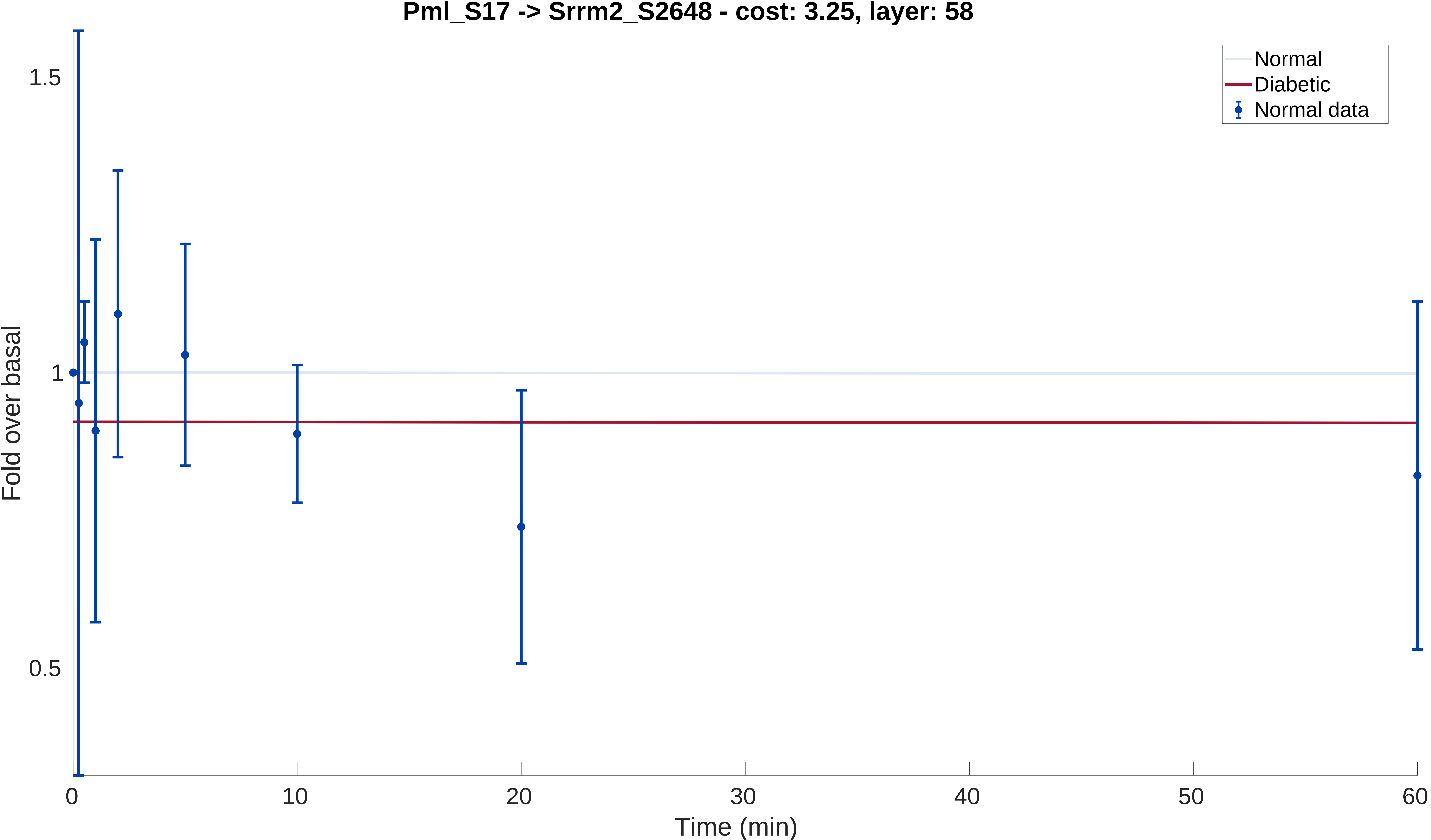
Pml_S17 -> Srrm2_S2648 - cost: 3.25, layer: 58
Normal
Diabetic
Normal data
1.5
1
0.5
Fold over basal
0
10
20
30
40
50
60
Time (min)

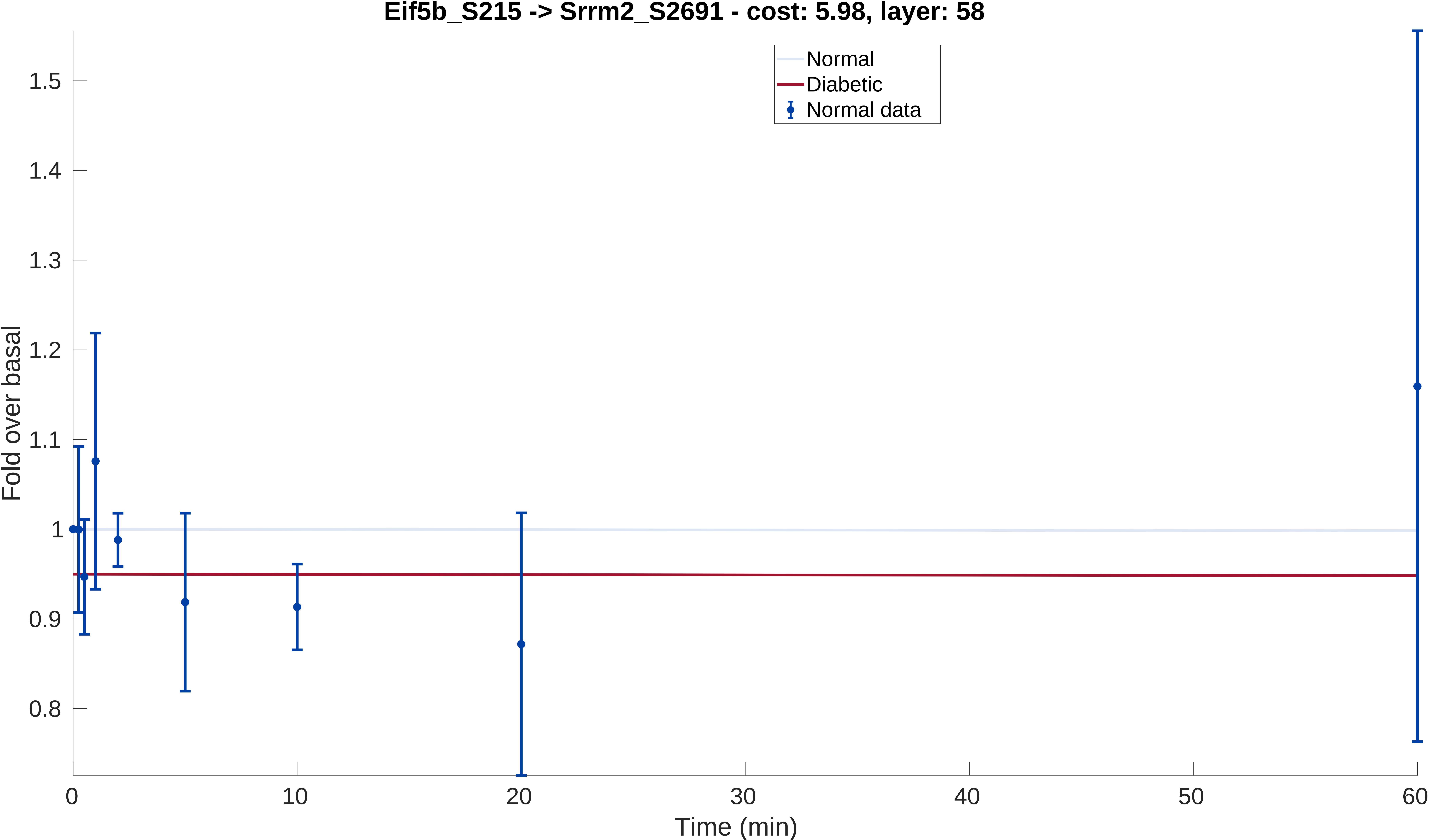

Eif5b_S215 -> Srrm2_S2691 - cost: 5.98, layer: 58
Normal
Diabetic
Normal data
Fold over basal
1.5
1.4
1.3
1.2
1.1
1
0.9
0.8
0
10
20
30
40
50
60
Time (min)

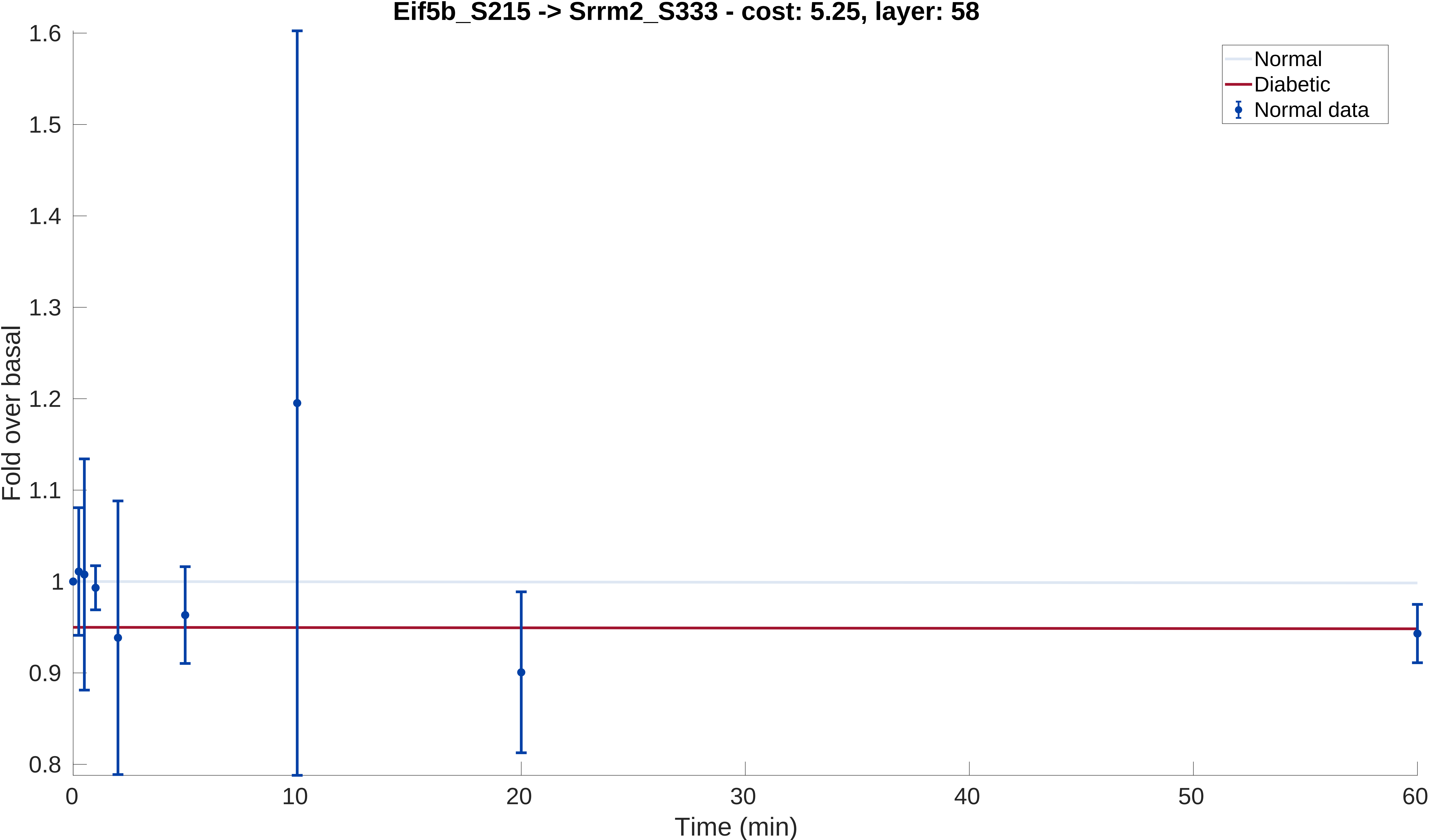

Eif5b_S215 -> Srrm2_S333 - cost: 5.25, layer: 58
Normal
Diabetic
Normal data
Fold over basal
Time (min)
1.6
1.5
1.4
1.3
1.2
1.1
1
0.9
0.8
0
10
20
30
40
50
60

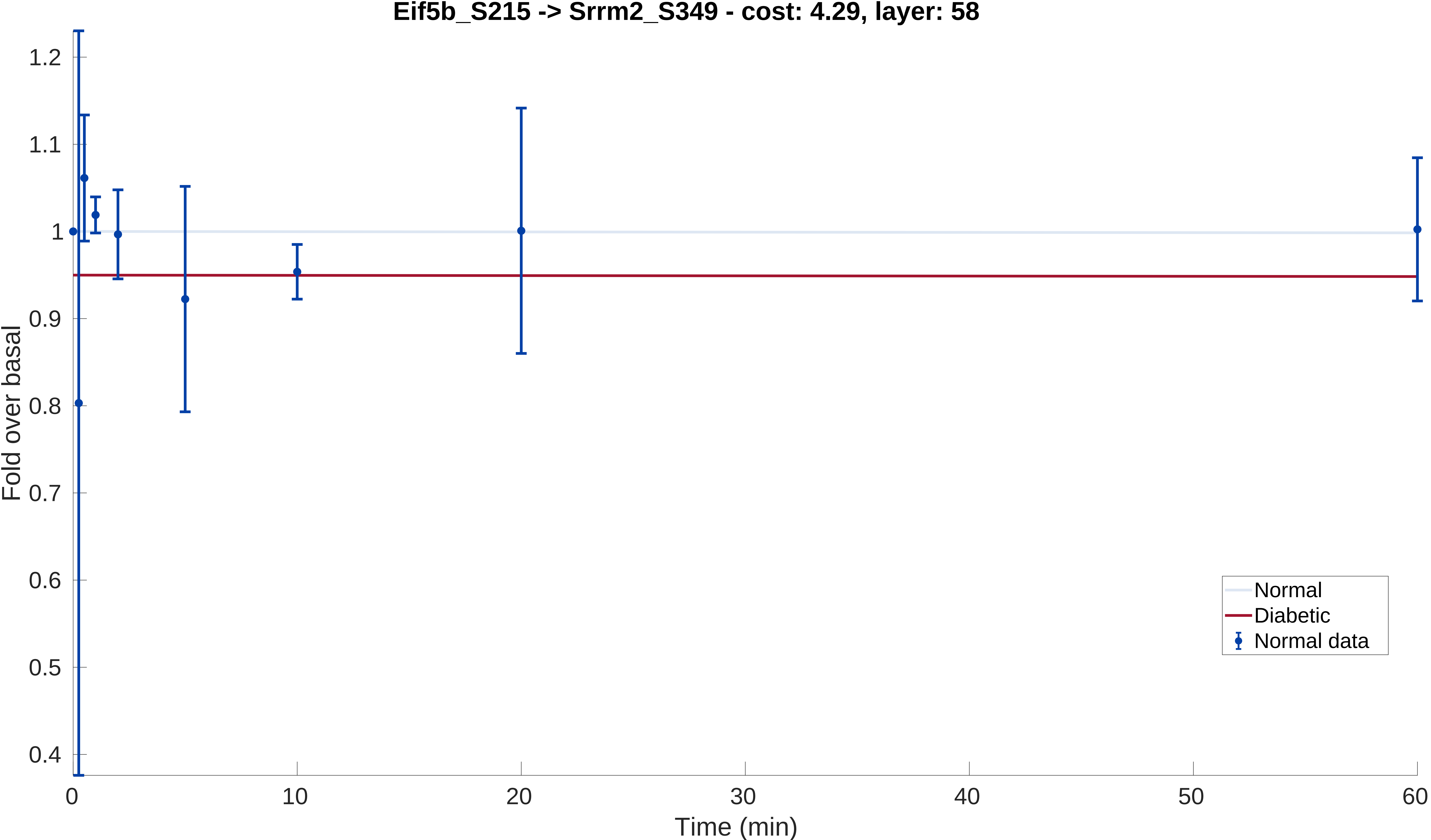

Eif5b_S215 -> Srrm2_S349 - cost: 4.29, layer: 58
Fold over basal
1.2
1.1
1
0.9
0.8
0.7
0.6
0.5
0.4
0
10
20
30
40
50
60
Time (min)
Normal
Diabetic
Normal data

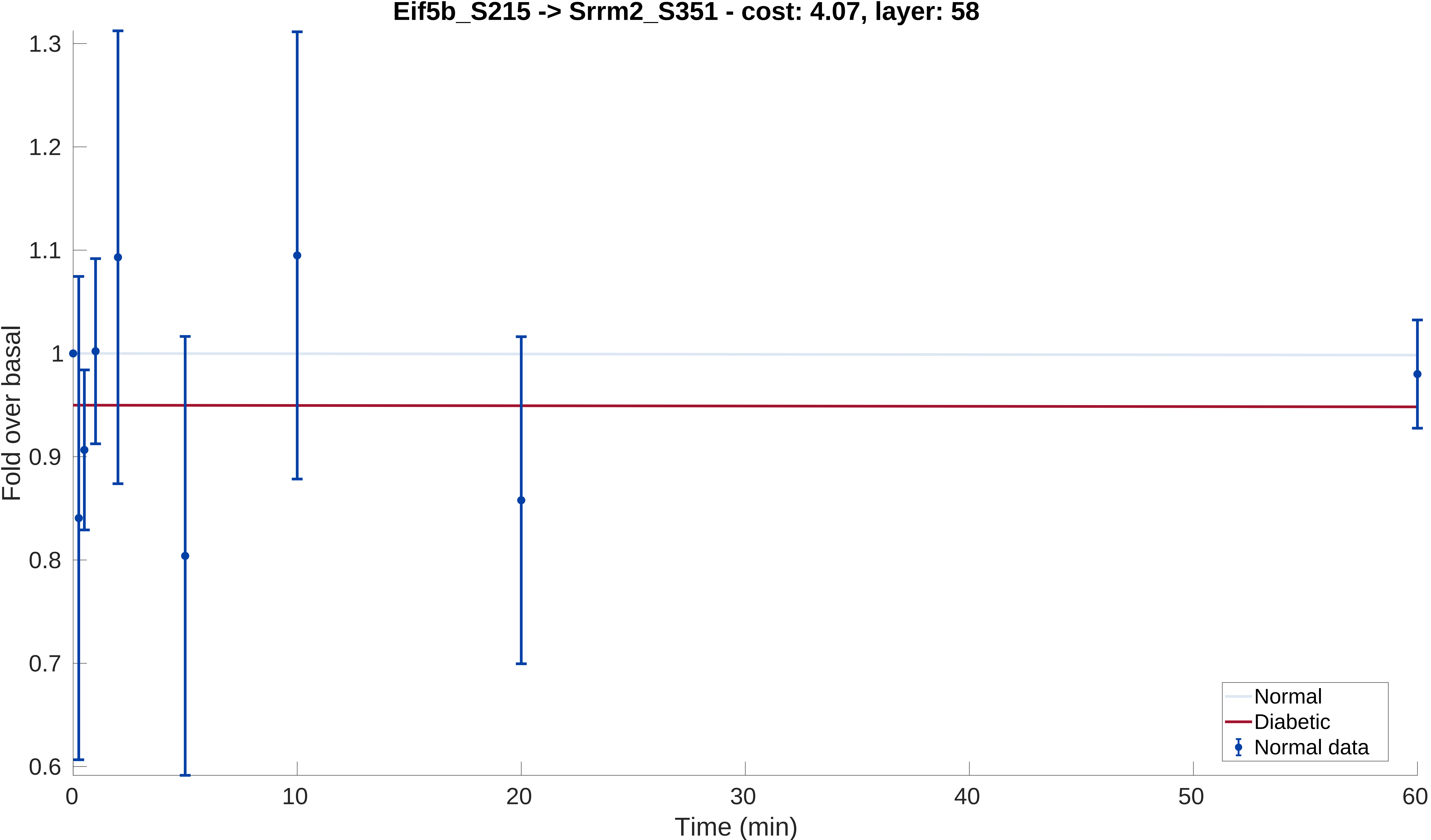

Eif5b_S215 -> Srrm2_S351 - cost: 4.07, layer: 58
1.3
1.2
1.1
1
0.9
0.8
0.7
0.6
Fold over basal
0
10
20
30
40
50
60
Time (min)
Normal
Diabetic
Normal data

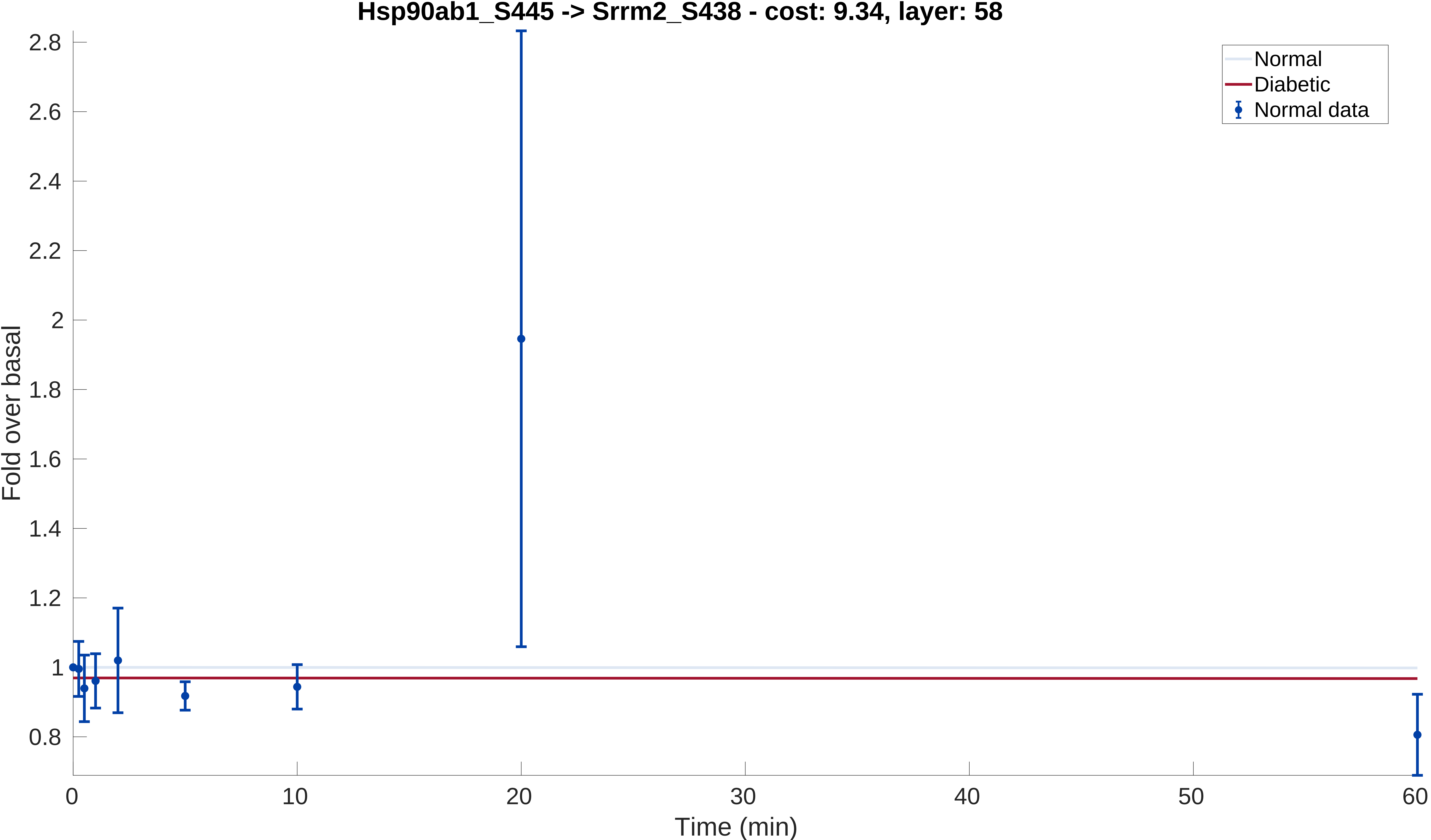
Hsp90ab1_S445 -> Srrm2_S438 - cost: 9.34, layer: 58
Normal
Diabetic
Normal data
Fold over basal
2.8
2.6
2.4
2.2
2
1.8
1.6
1.4
1.2
1
0.8
0
10
20
30
40
50
60
Time (min)

#### Sorbs1_S633 -> Srrm2_S453 - cost: 2.45, layer: 58

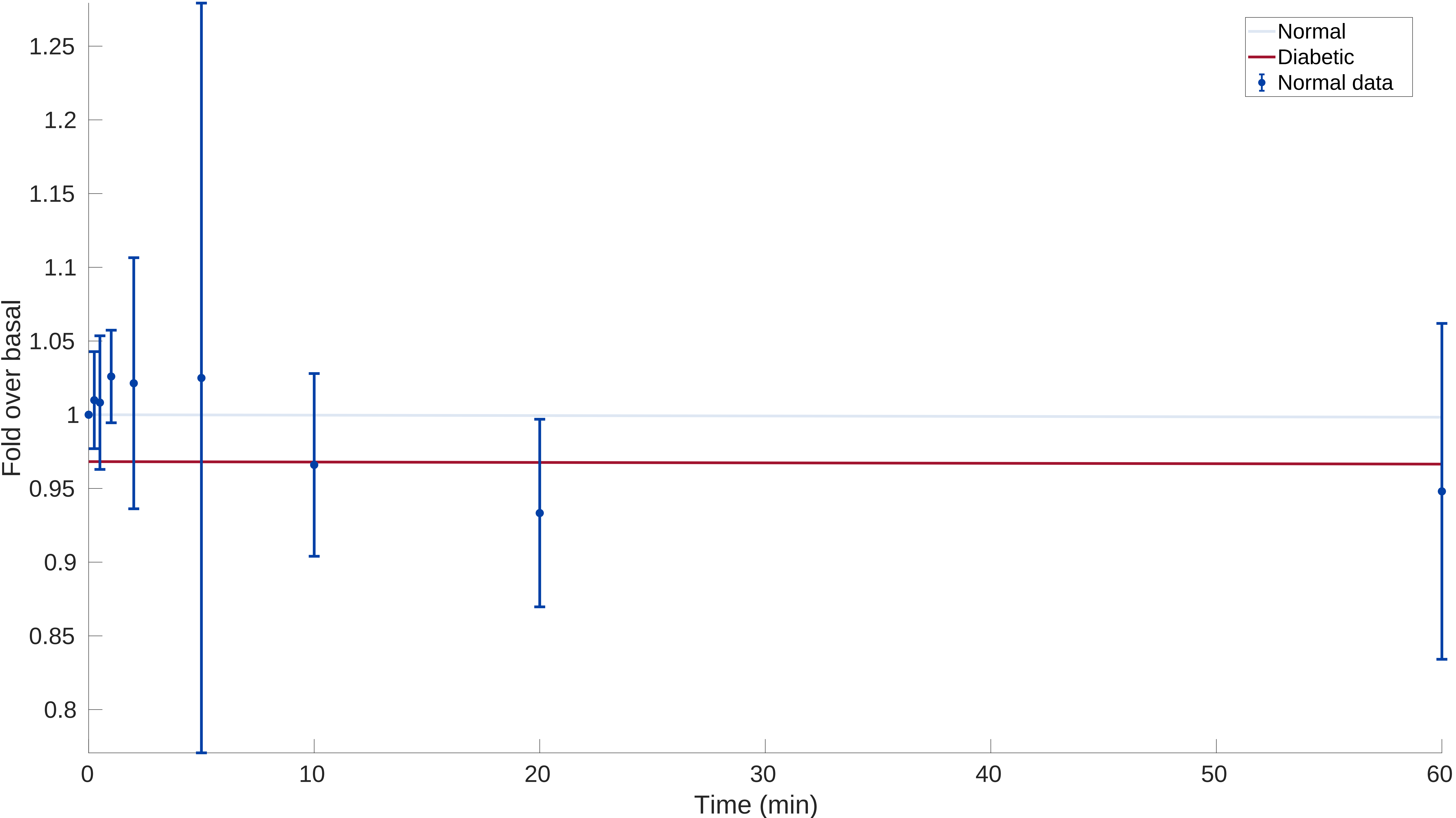

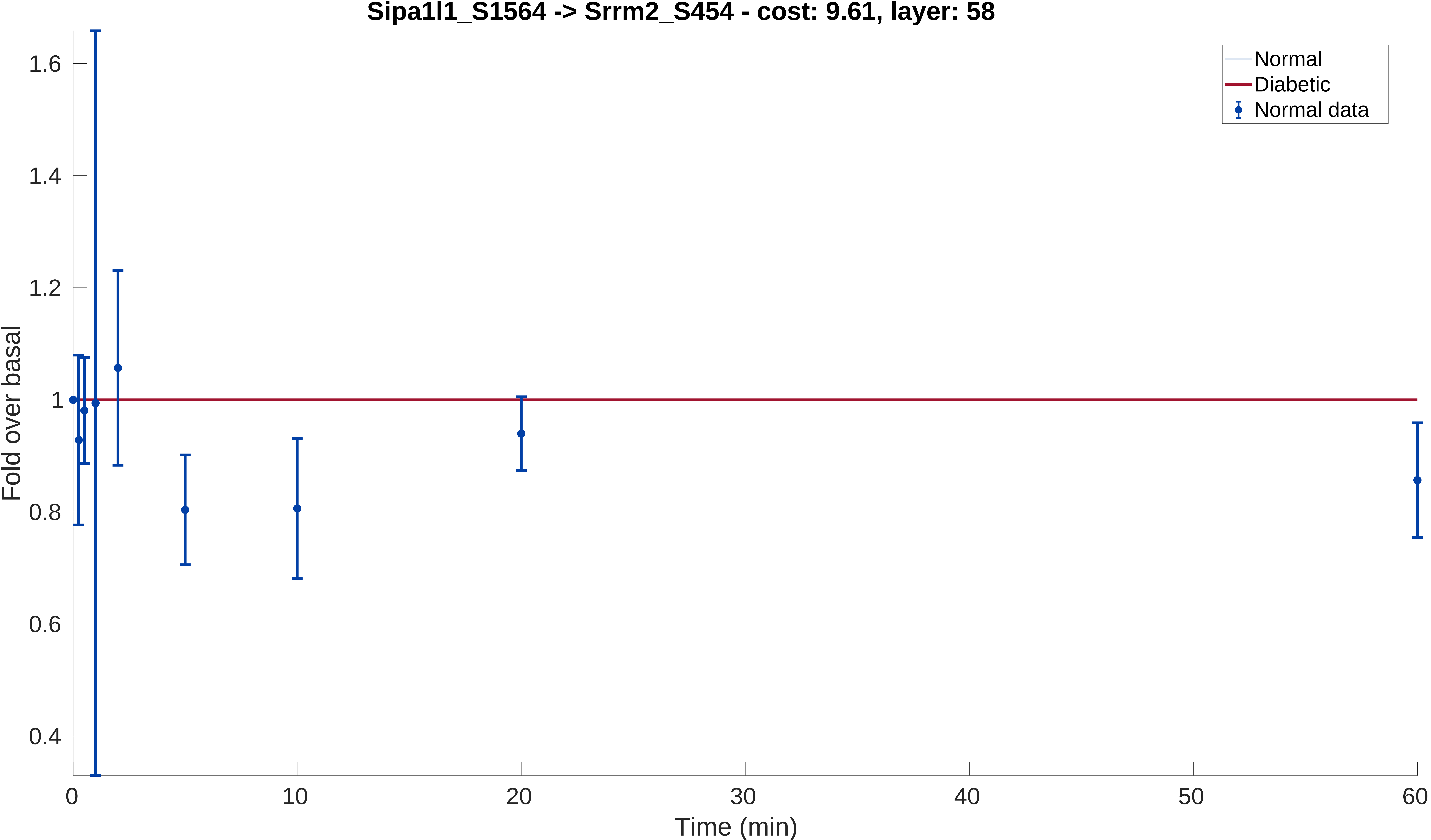
Sipa1l1_S1564 -> Srrm2_S454 - cost: 9.61, layer: 58
Normal
Diabetic
Normal data
Fold over basal
1.6
1.4
1.2
1
0.8
0.6
0.4
0
10
20
30
40
50
60
Time (min)

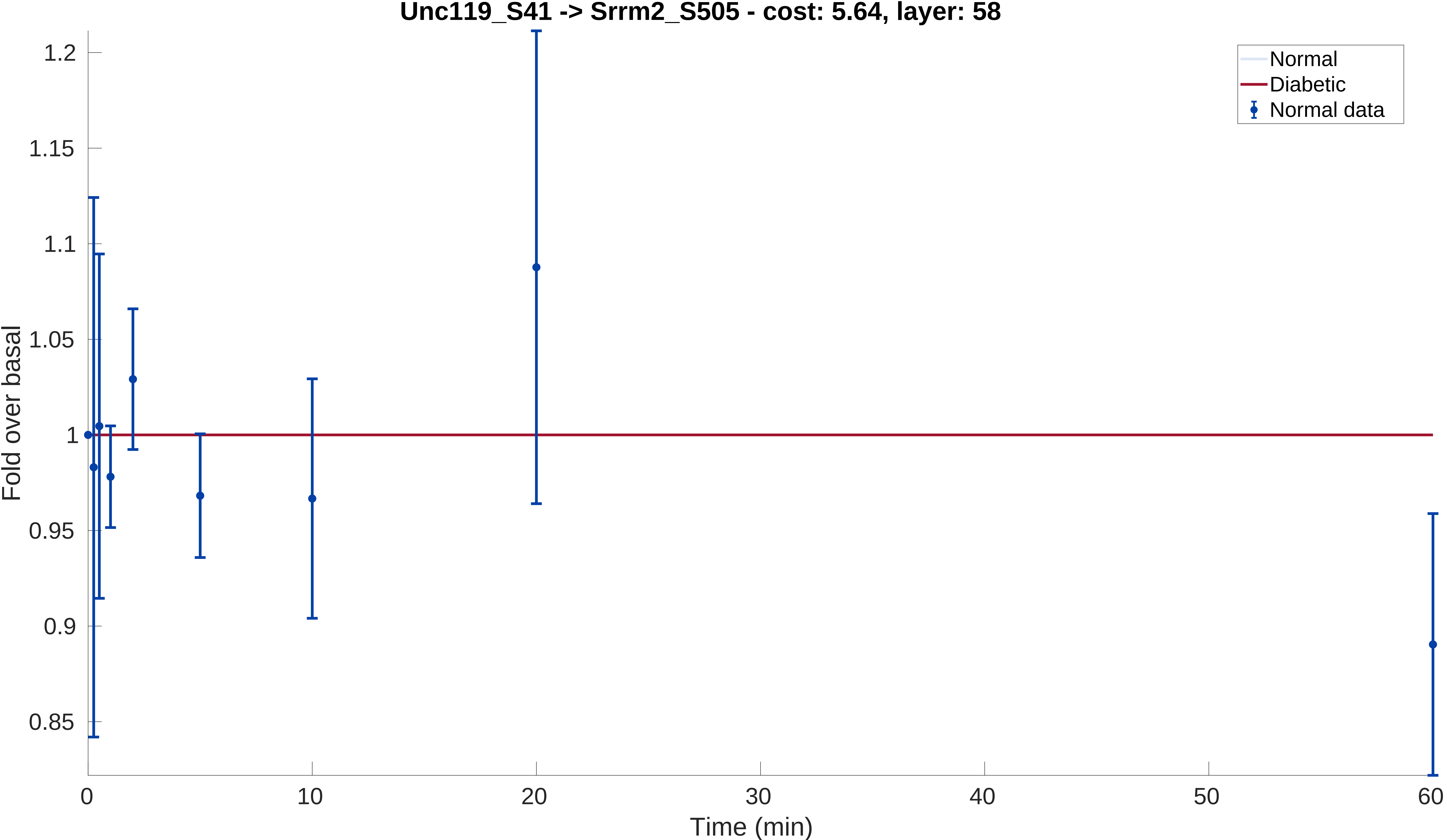

Unc119_S41 -> Srrm2_S505 - cost: 5.64, layer: 58
Normal
Diabetic
Normal data
Fold over basal
Time (min)
1.2
1.15
1.1
1.05
1
0.95
0.9
0.85
0
10
20
30
40
50
60

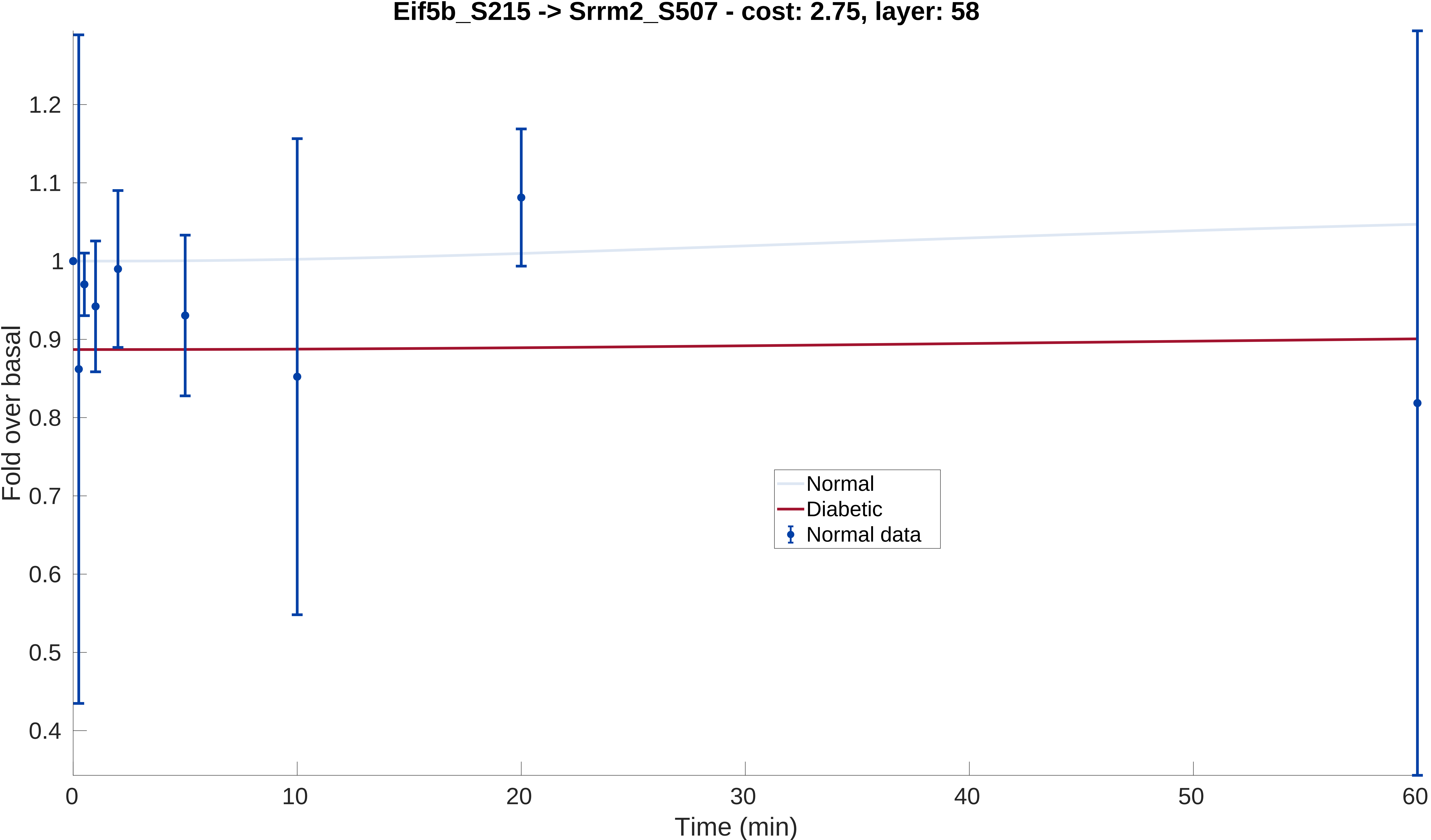

Eif5b_S215 -> Srrm2_S507 - cost: 2.75, layer: 58
1.2
1.1
1
0.9
0.8
0.7
0.6
0.5
0.4
Fold over basal
0
10
20
30
40
50
60
Time (min)
Normal
Diabetic
Normal data

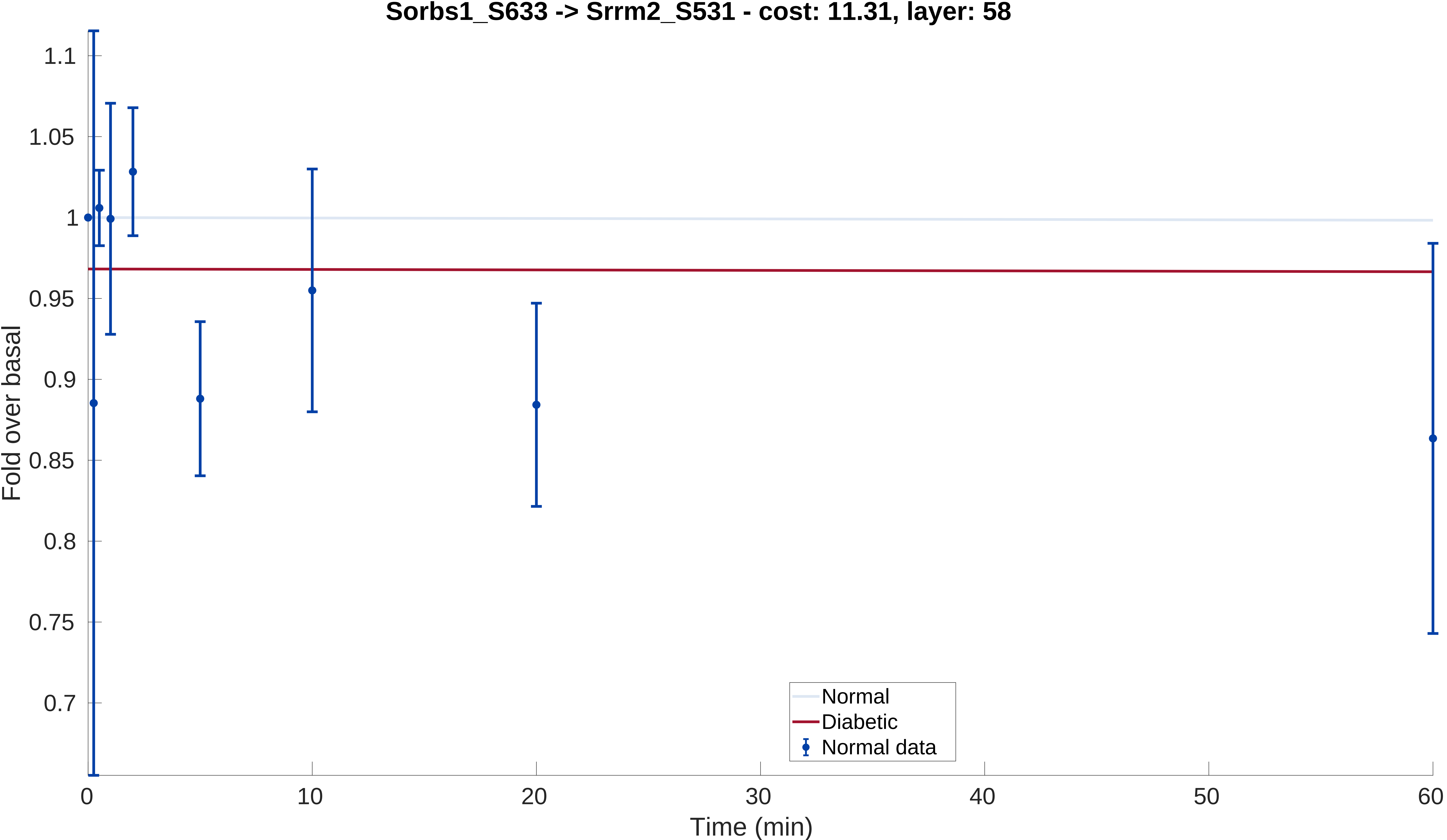

Sorbs1_S633 -> Srrm2_S531 - cost: 11.31, layer: 58
1.1
1.05
1
0.95
0.9
0.85
0.8
0.75
0.7
Fold over basal
0
10
20
30
40
50
60
Time (min)
Normal
Diabetic
Normal data

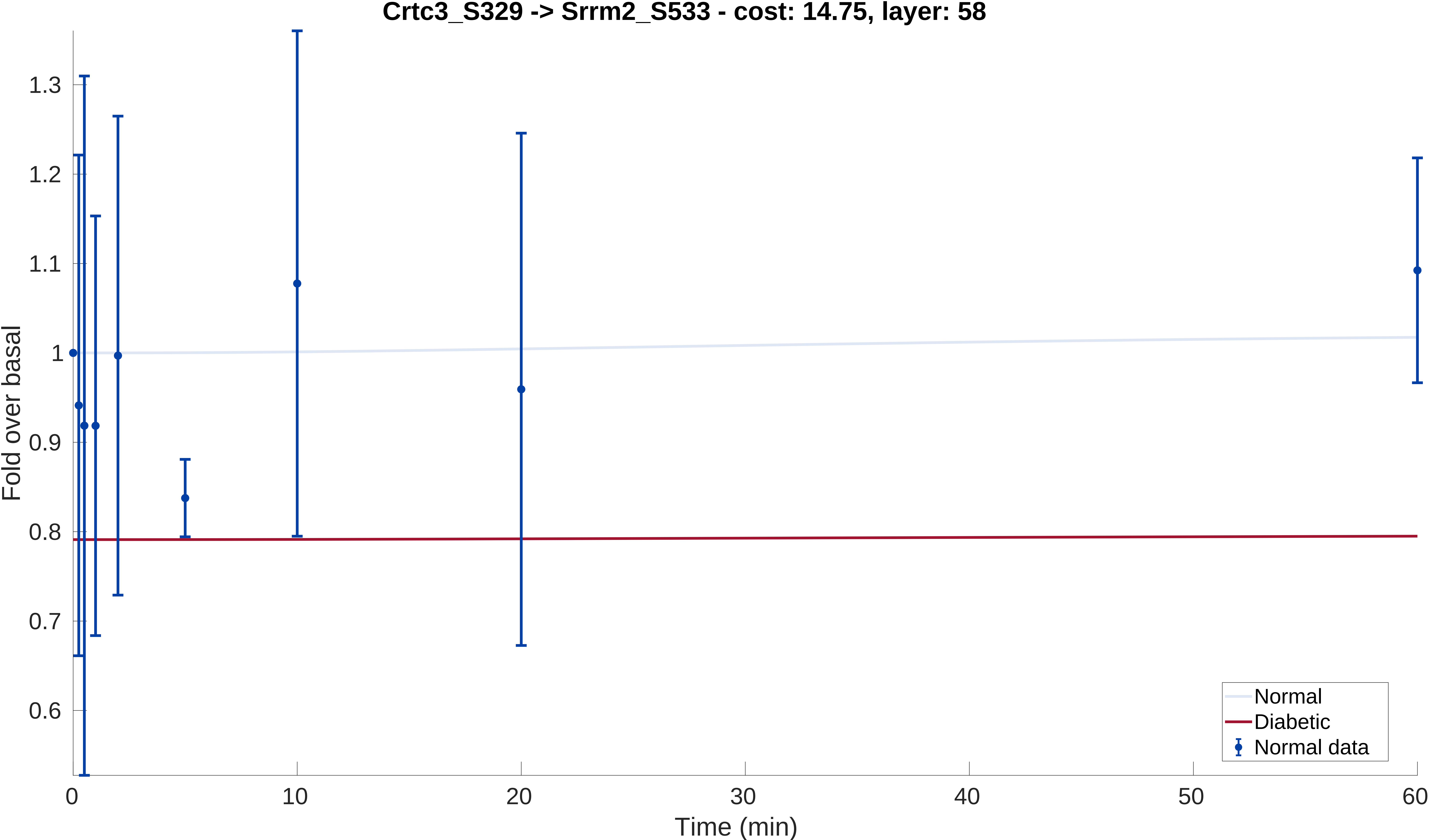
Crtc3_S329 -> Srrm2_S533 - cost: 14.75, layer: 58
Fold over basal
Time (min)
1.3
1.2
1.1
1
0.9
0.8
0.7
0.6
0
10
20
30
40
50
60
Normal
Diabetic
Normal data

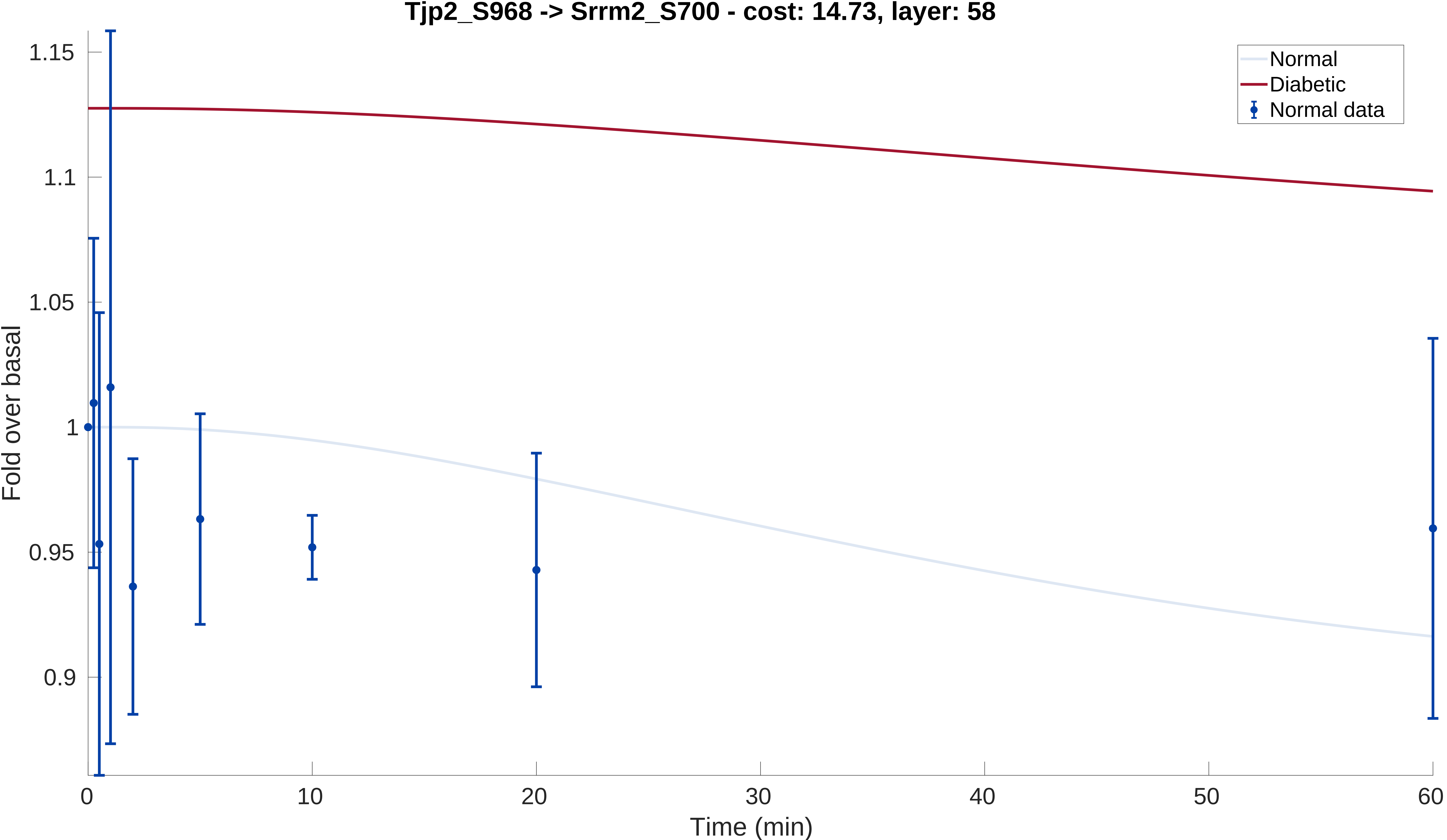

Tjp2_S968 -> Srrm2_S700 - cost: 14.73, layer: 58
Normal
Diabetic
Normal data
1.15
1.1
1.05
1
0.95
0.9
Fold over basal
0
10
20
30
40
50
60
Time (min)

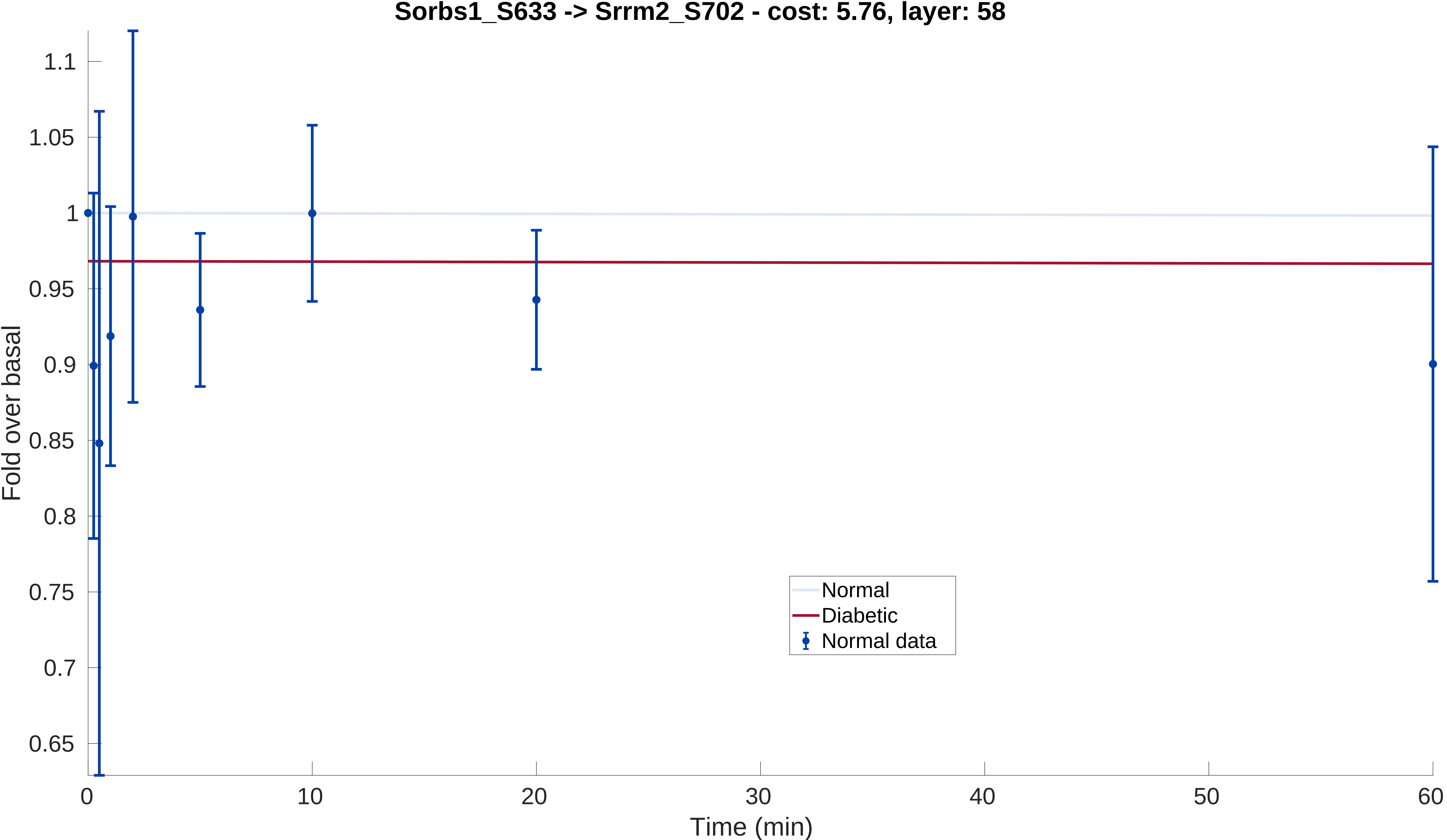

Sorbs1_S633 -> Srrm2_S702 - cost: 5.76, layer: 58
Fold over basal
1.1
1.05
1
0.95
0.9
0.85
0.8
0.75
0.7
0.65
0
10
20
30
40
50
60
Time (min)
Normal
Diabetic
Normal data

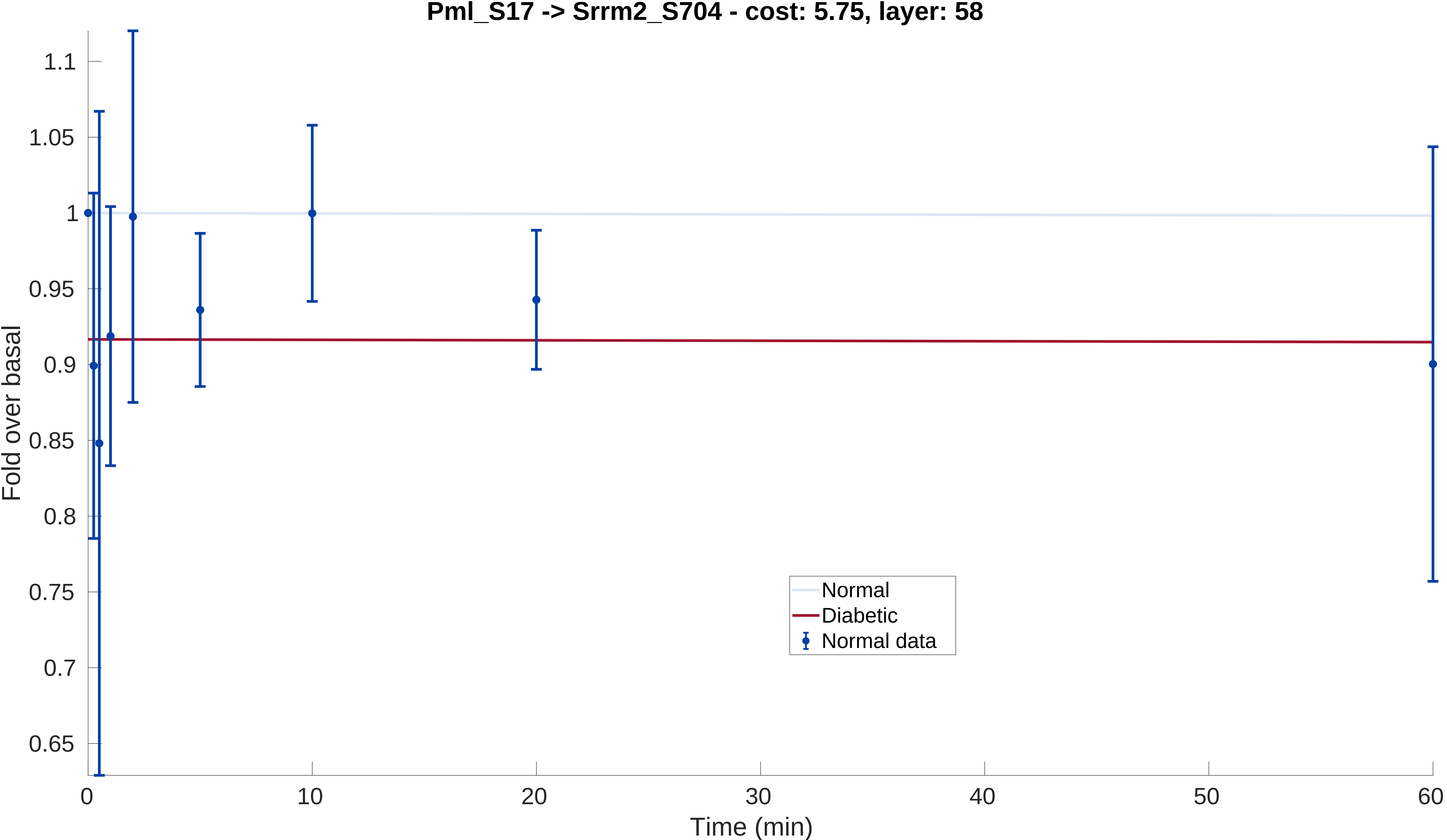

Pml_S17 -> Srrm2_S704 - cost: 5.75, layer: 58
Fold over basal
1.1
1.05
1
0.95
0.9
0.85
0.8
0.75
0.7
0.65
0
10
20
30
40
50
60
Time (min)
Normal
Diabetic
Normal data

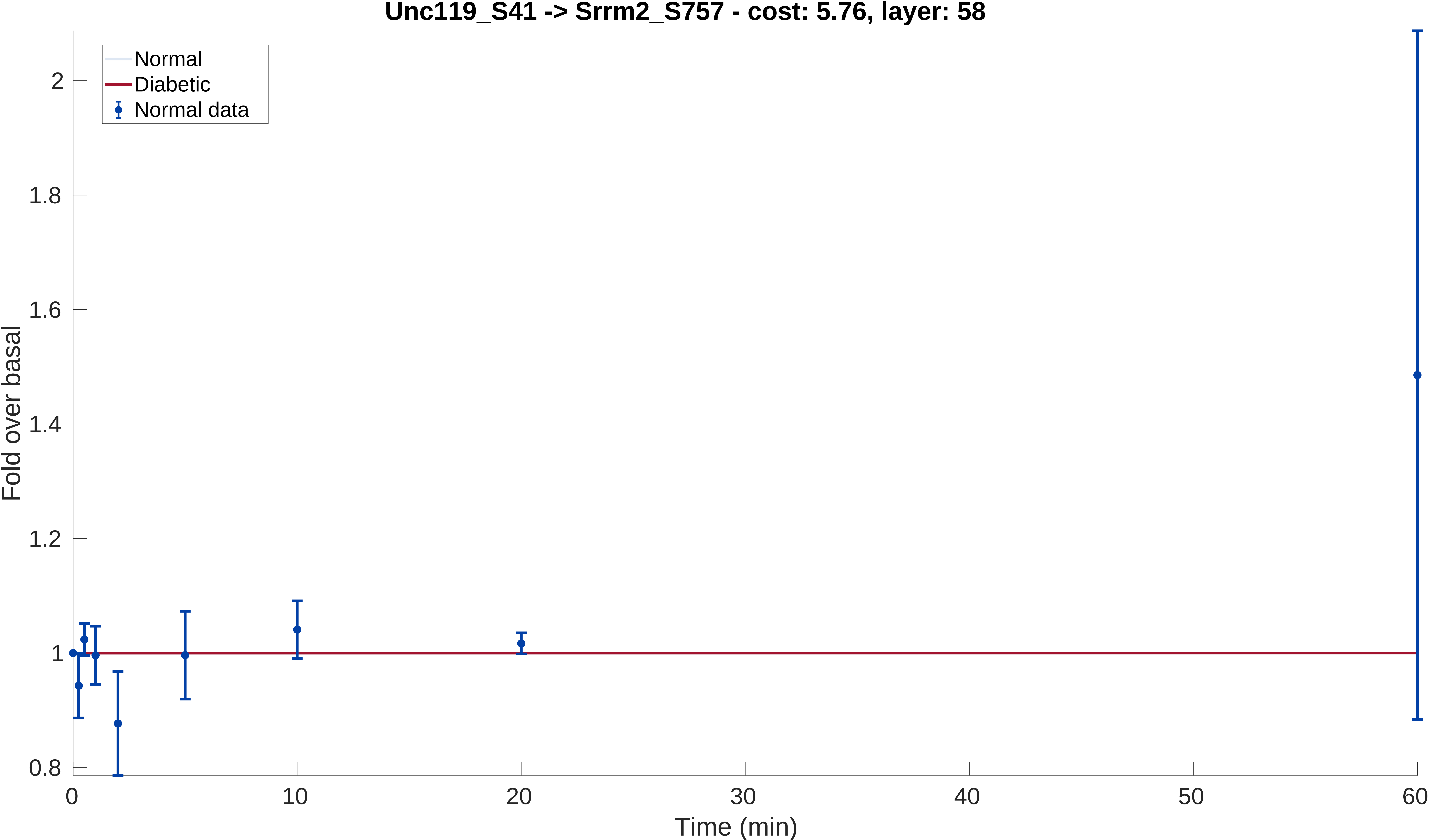

Unc119_S41 -> Srrm2_S757 - cost: 5.76, layer: 58
Normal
Diabetic
Normal data
Fold over basal
Time (min)
0.8
1
1.2
1.4
1.6
1.8
2
0
10
20
30
40
50
60

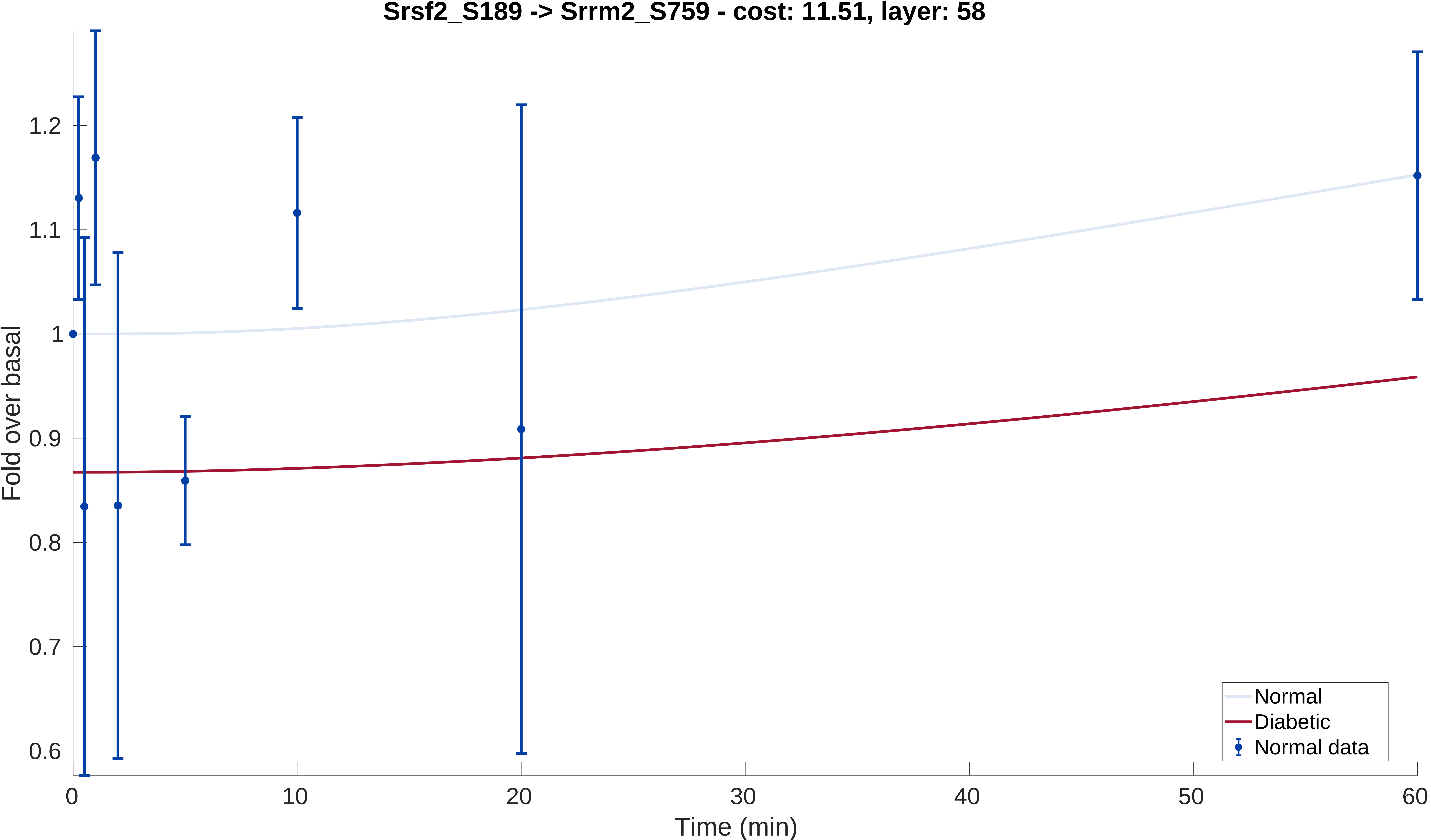

Srsf2_S189 -> Srrm2_S759 - cost: 11.51, layer: 58
Fold over basal
1.2
1.1
1
0.9
0.8
0.7
0.6
0
10
20
30
40
50
60
Time (min)
Normal
Diabetic
Normal data

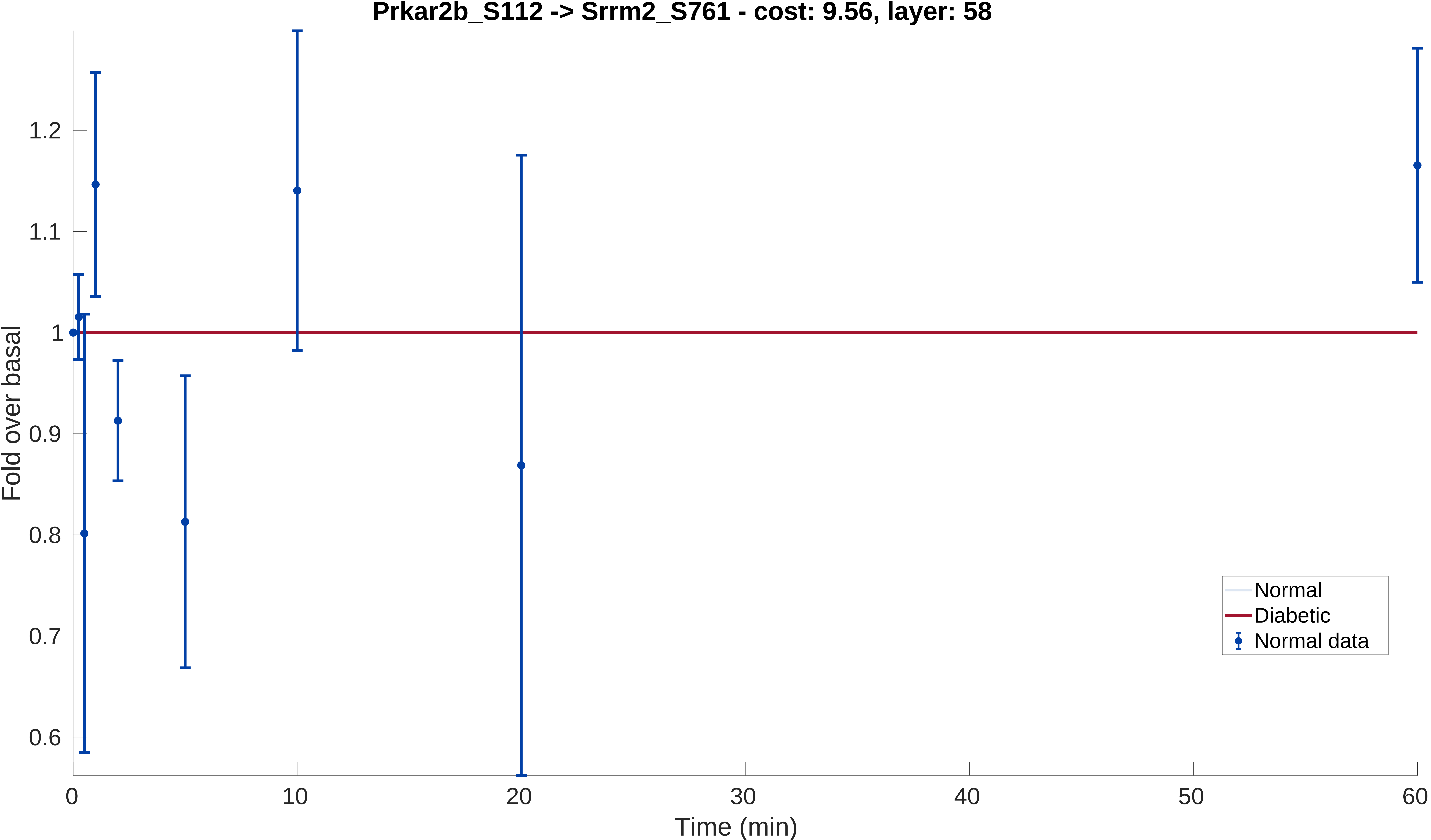

Prkar2b_S112 -> Srrm2_S761 - cost: 9.56, layer: 58
Fold over basal
1.2
1.1
1
0.9
0.8
0.7
0.6
0
10
20
30
40
50
60
Time (min)
Normal
Diabetic
Normal data

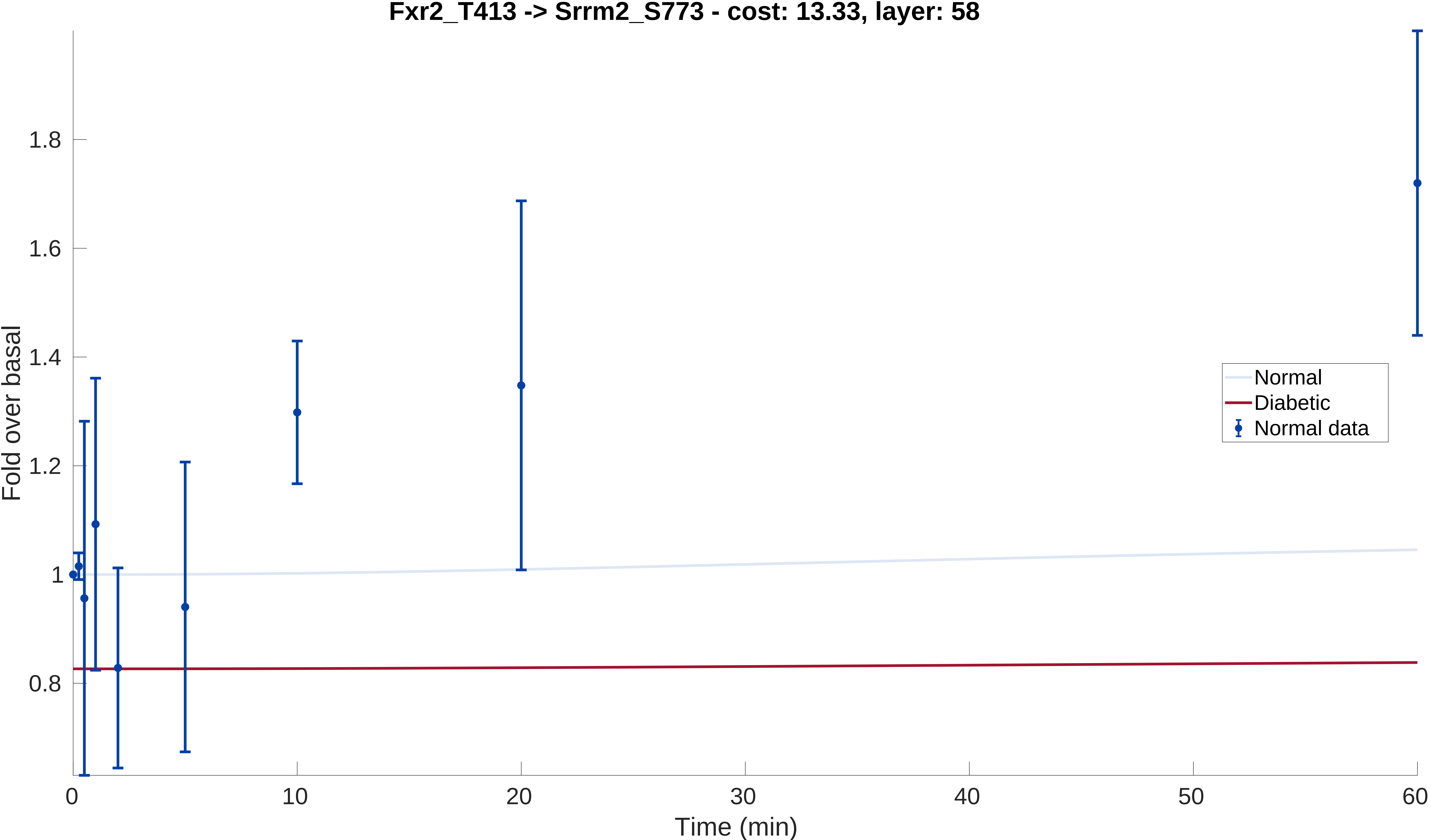

Fxr2_T413 -> Srrm2_S773 - cost: 13.33, layer: 58
Fold over basal
1.8
1.6
1.4
1.2
1
0.8
0
10
20
30
40
50
60
Time (min)
Normal
Diabetic
Normal data

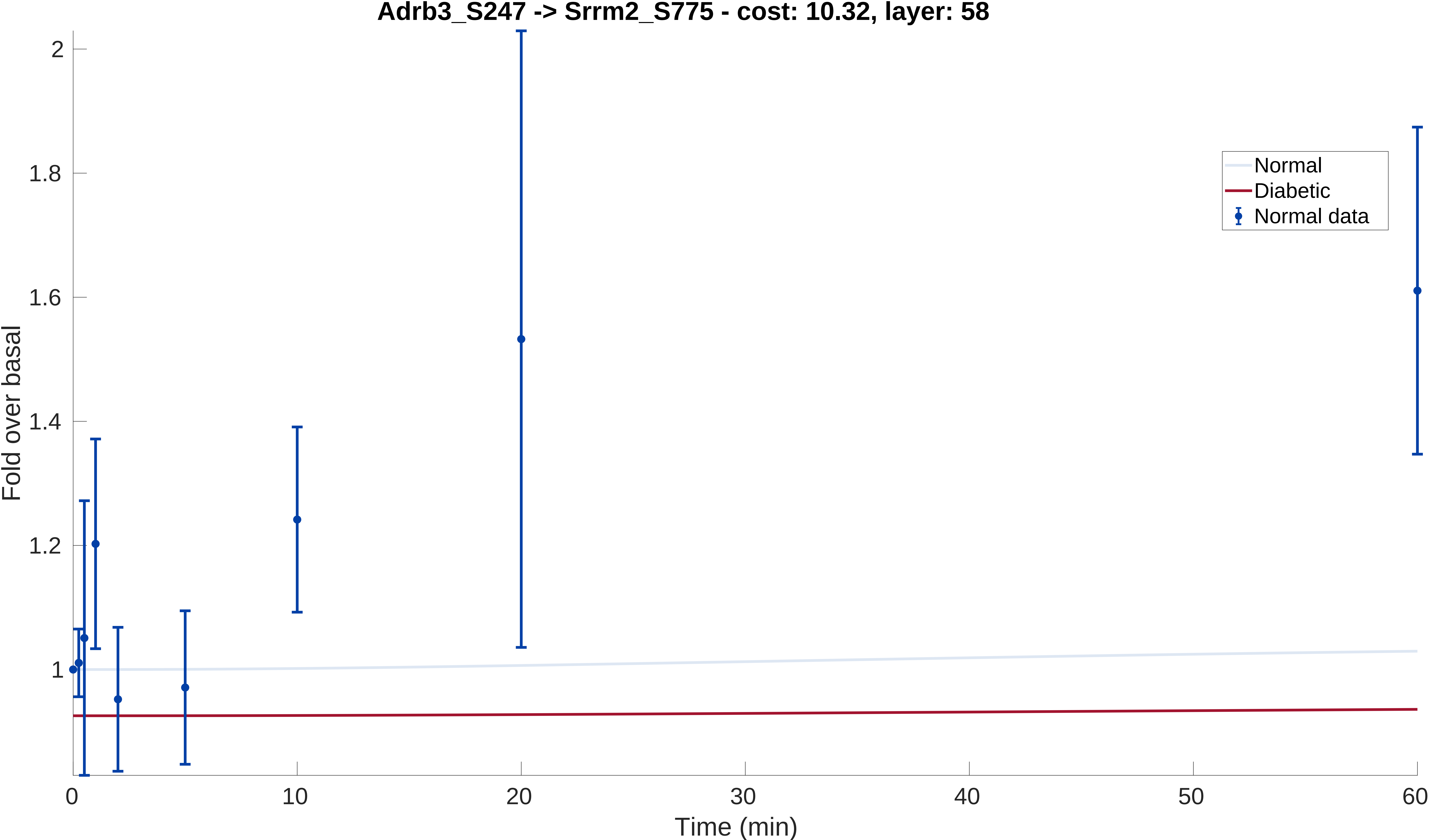

Adrb3_S247 -> Srrm2_S775 - cost: 10.32, layer: 58
Fold over basal
2
1.8
1.6
1.4
1.2
1
Time (min)
0
10
20
30
40
50
60
Normal
Diabetic
Normal data

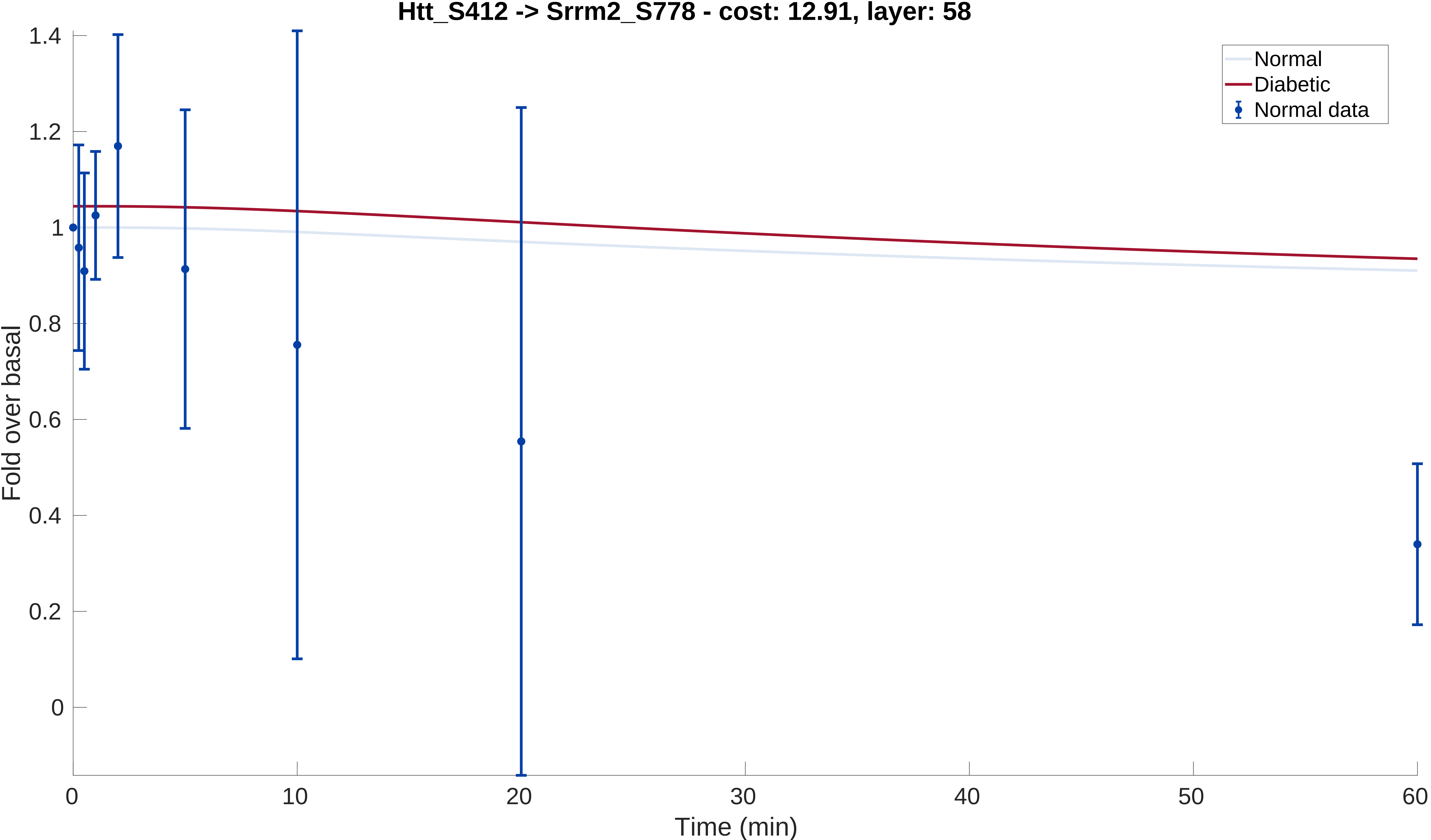
Htt_S412 -> Srrm2_S778 - cost: 12.91, layer: 58
Normal
Diabetic
Normal data
Fold over basal
Time (min)
1.4
1.2
1
0.8
0.6
0.4
0.2
0
0
10
20
30
40
50
60

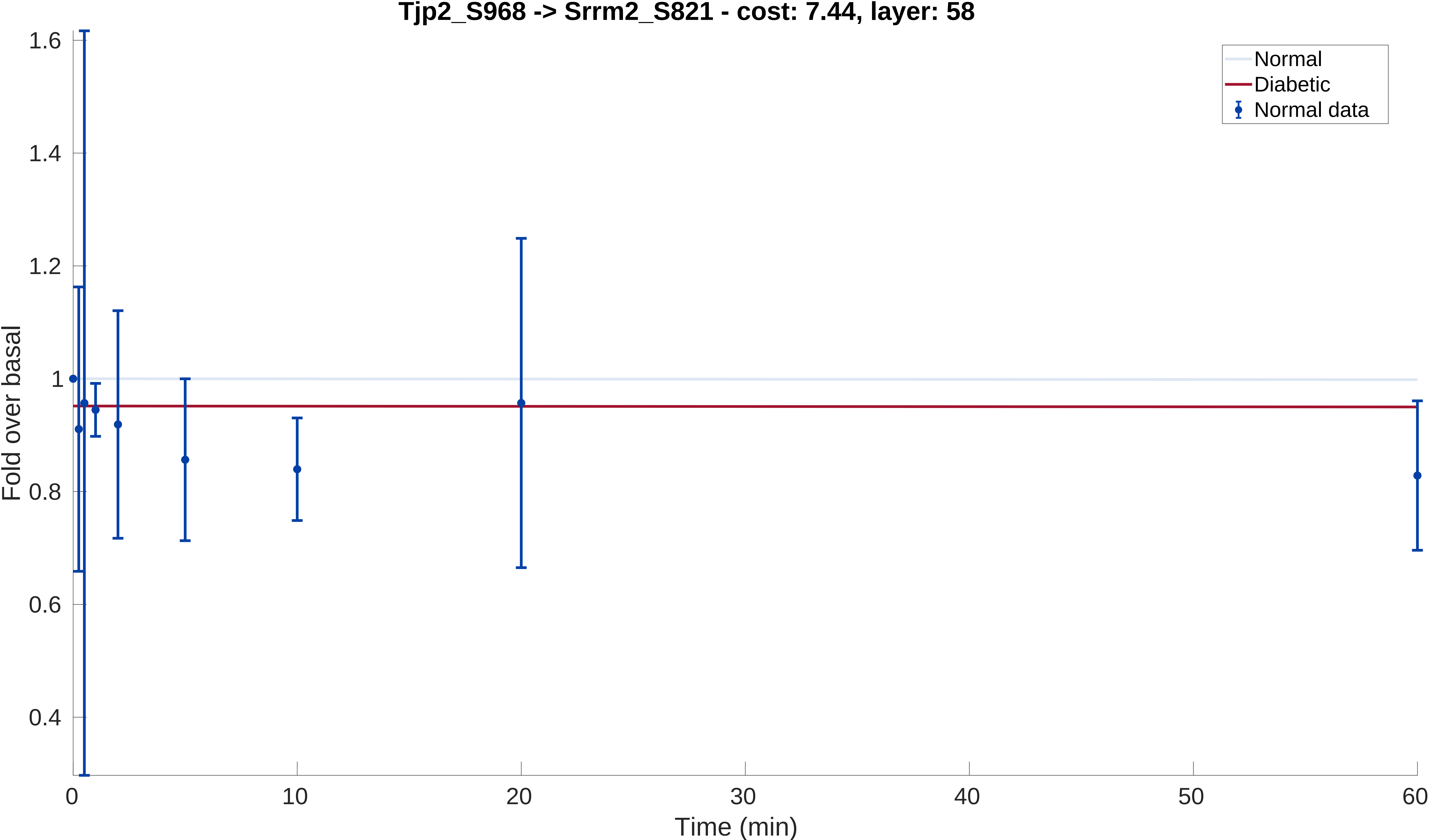

Tjp2_S968 -> Srrm2_S821 - cost: 7.44, layer: 58
Normal
Diabetic
Normal data
Fold over basal
1.6
1.4
1.2
1
0.8
0.6
0.4
0
10
20
30
40
50
60
Time (min)

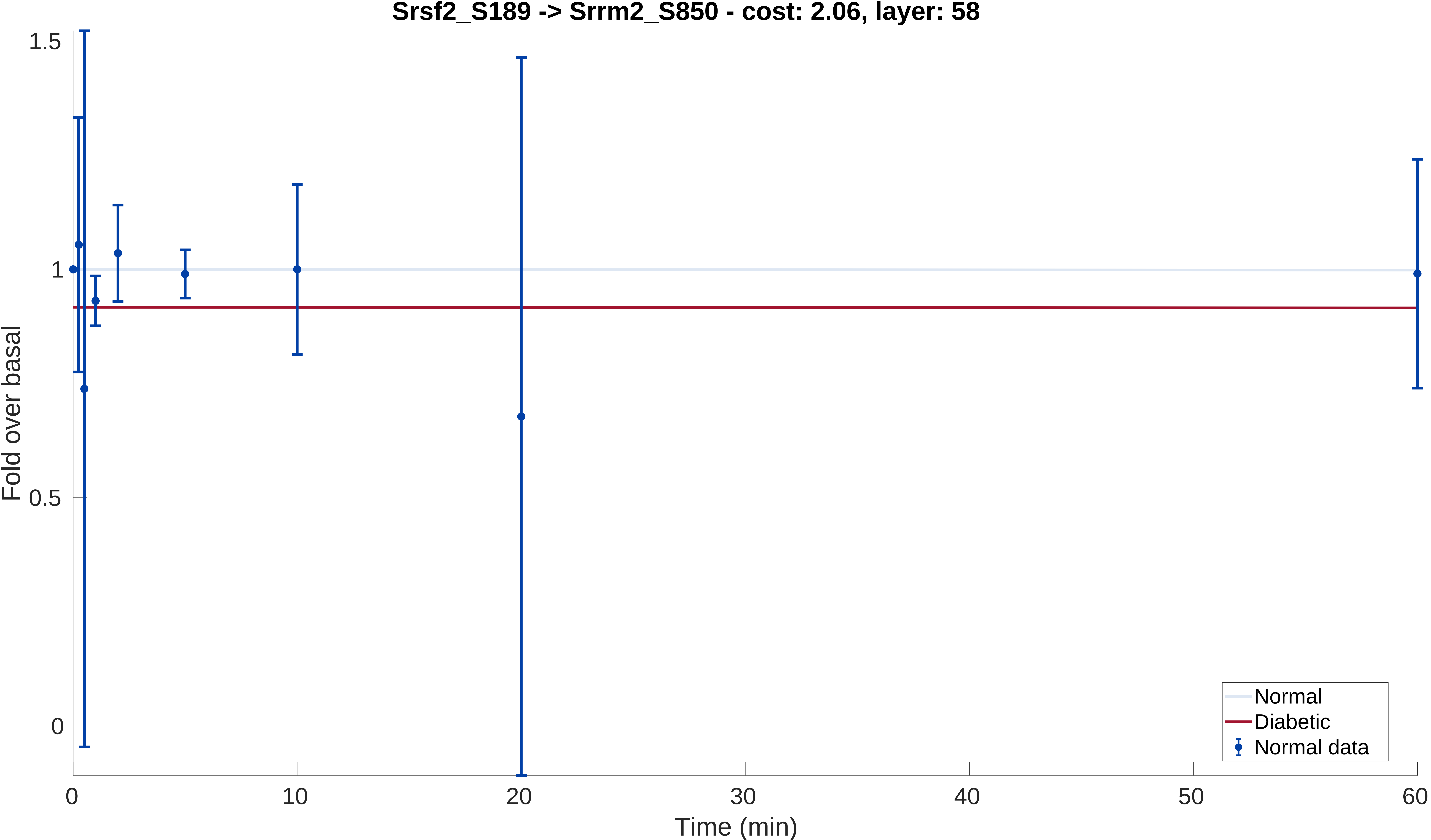
Srsf2_S189 -> Srrm2_S850 - cost: 2.06, layer: 58
Fold over basal
1.5
1
0.5
0
0
10
20
30
40
50
60
Time (min)
Normal
Diabetic
Normal data

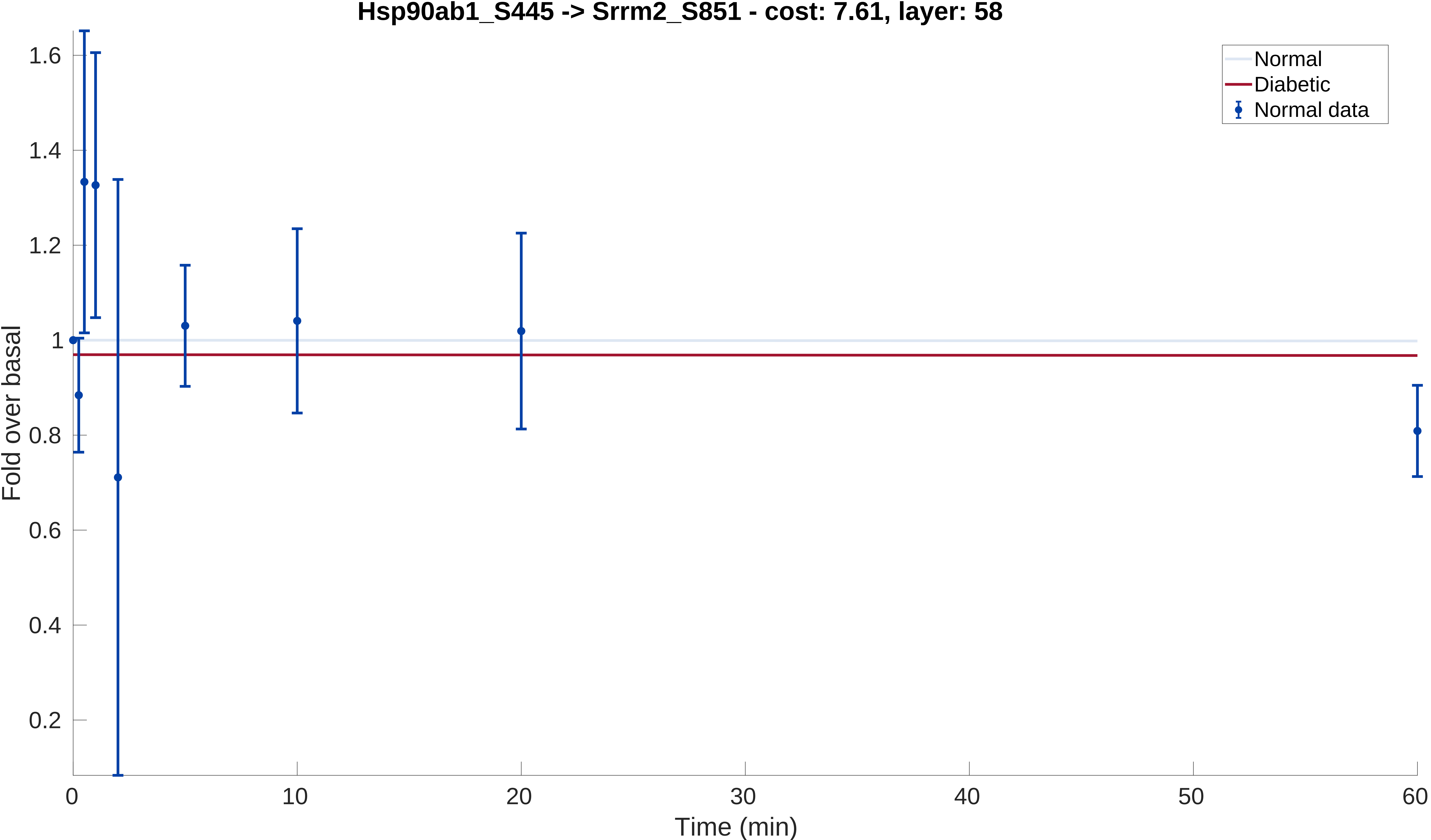

Hsp90ab1_S445 -> Srrm2_S851 - cost: 7.61, layer: 58
Normal
Diabetic
Normal data
Fold over basal
1.6
1.4
1.2
1
0.8
0.6
0.4
0.2
0
10
20
30
40
50
60
Time (min)

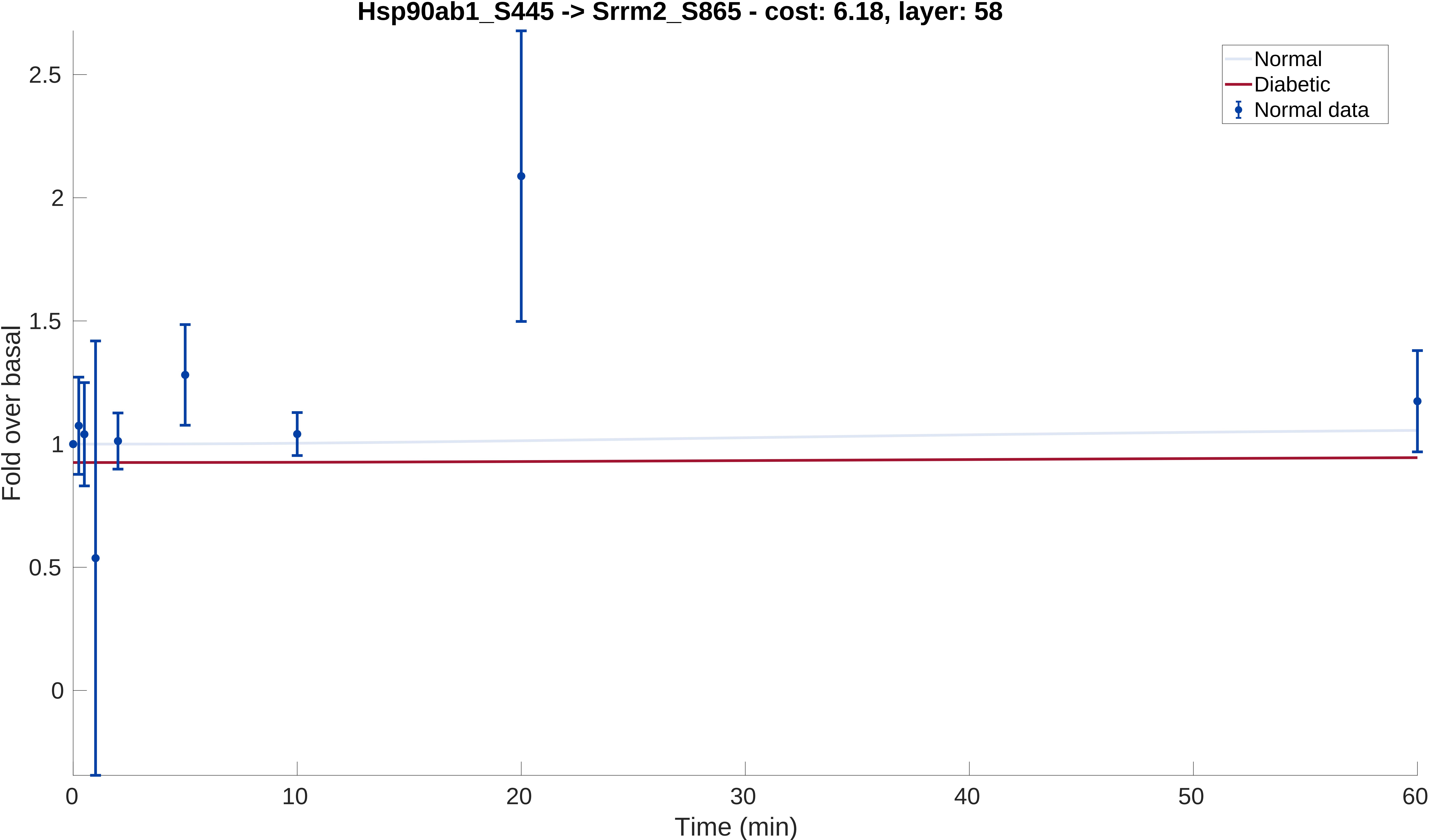
Hsp90ab1_S445 -> Srrm2_S865 - cost: 6.18, layer: 58
Normal
Diabetic
Normal data
Fold over basal
2.5
2
1.5
1
0.5
0
0
10
20
30
40
50
60
Time (min)

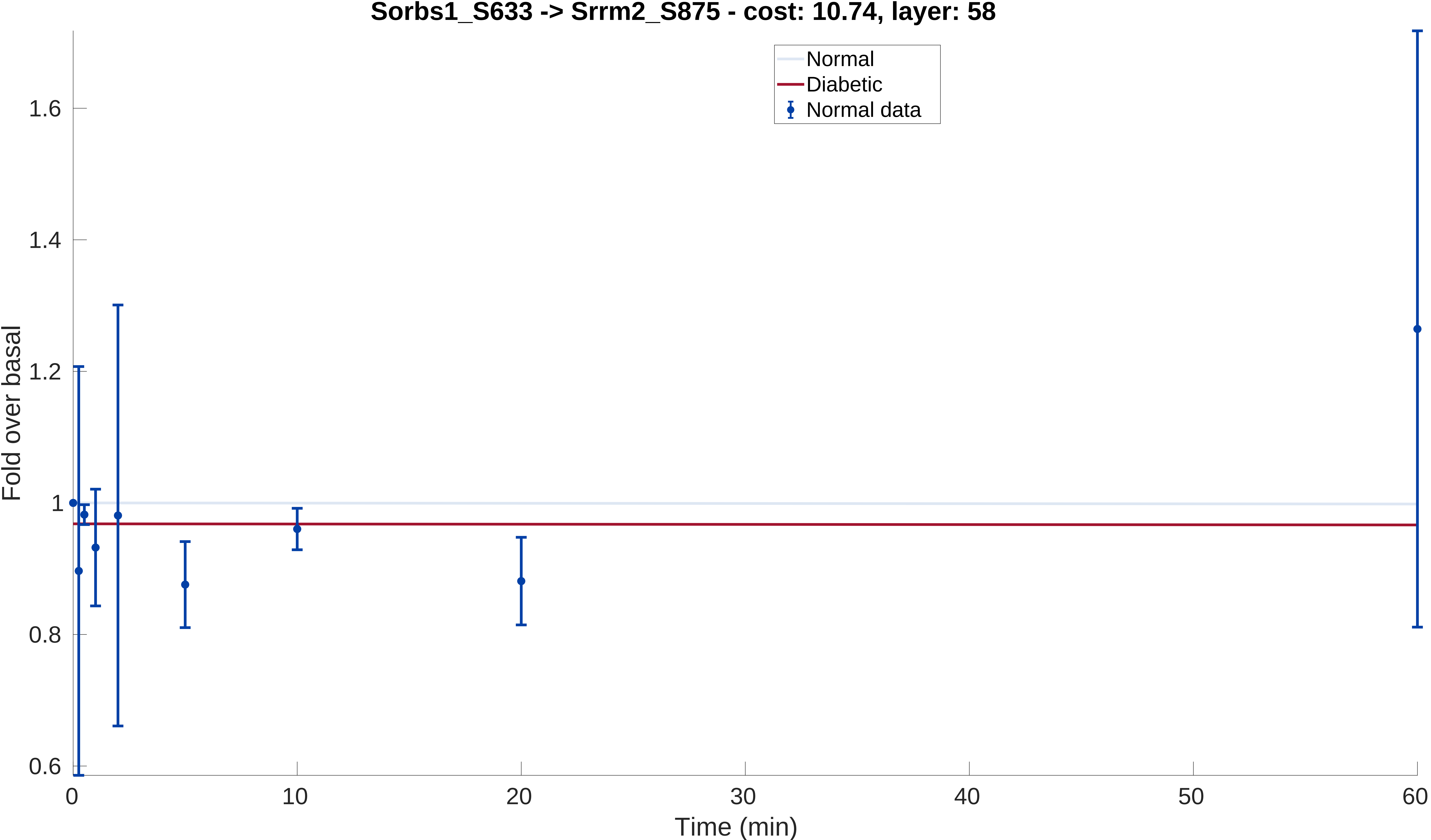
Sorbs1_S633 -> Srrm2_S875 - cost: 10.74, layer: 58
Normal
Diabetic
Normal data
Fold over basal
1.6
1.4
1.2
1
0.8
0.6
0
10
20
30
40
50
60
Time (min)

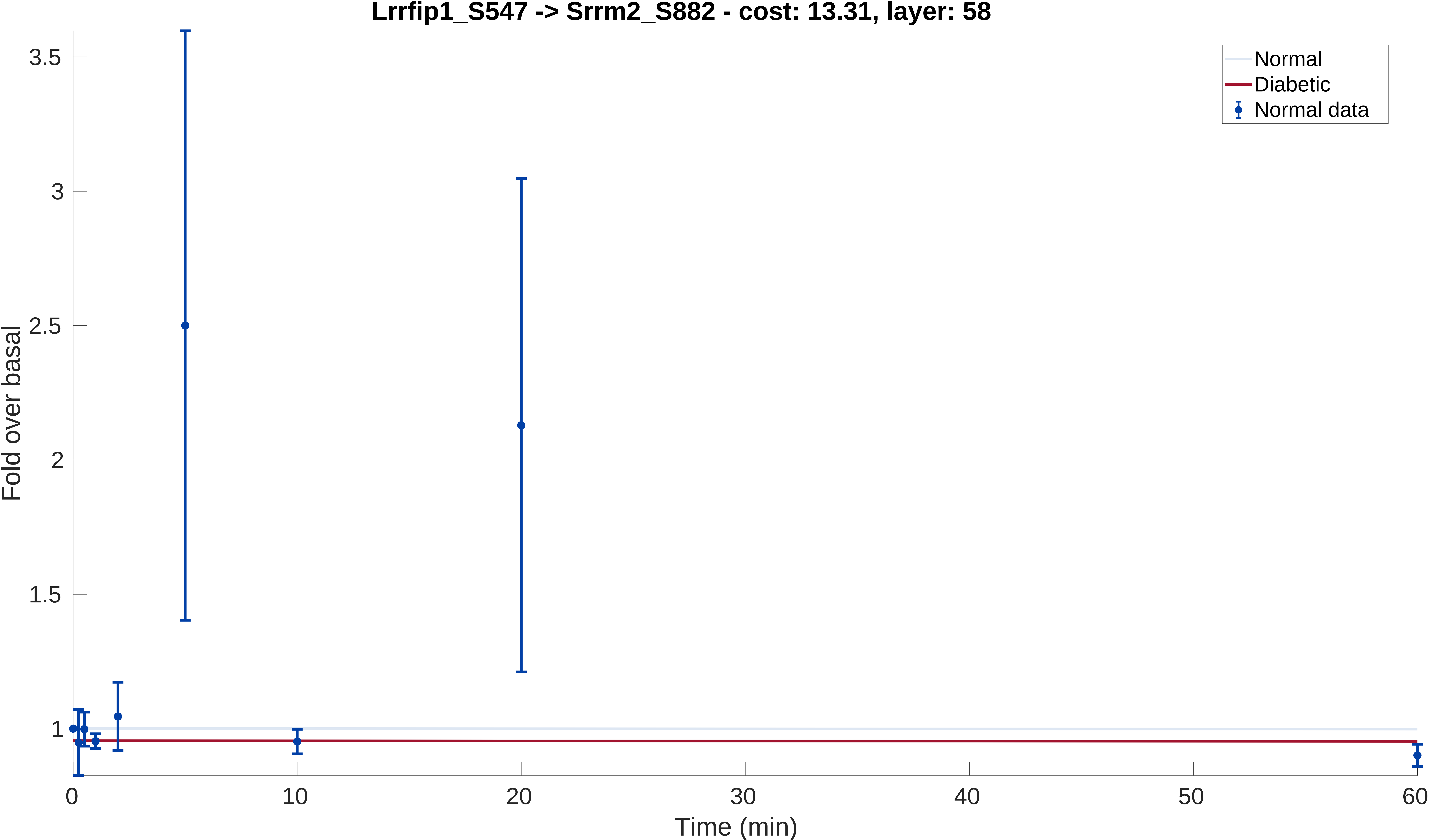
Lrrfip1_S547 -> Srrm2_S882 - cost: 13.31, layer: 58
Fold over basal
Time (min)
Normal
Diabetic
Normal data
3.5
3
2.5
2
1.5
1
0
10
20
30
40
50
60

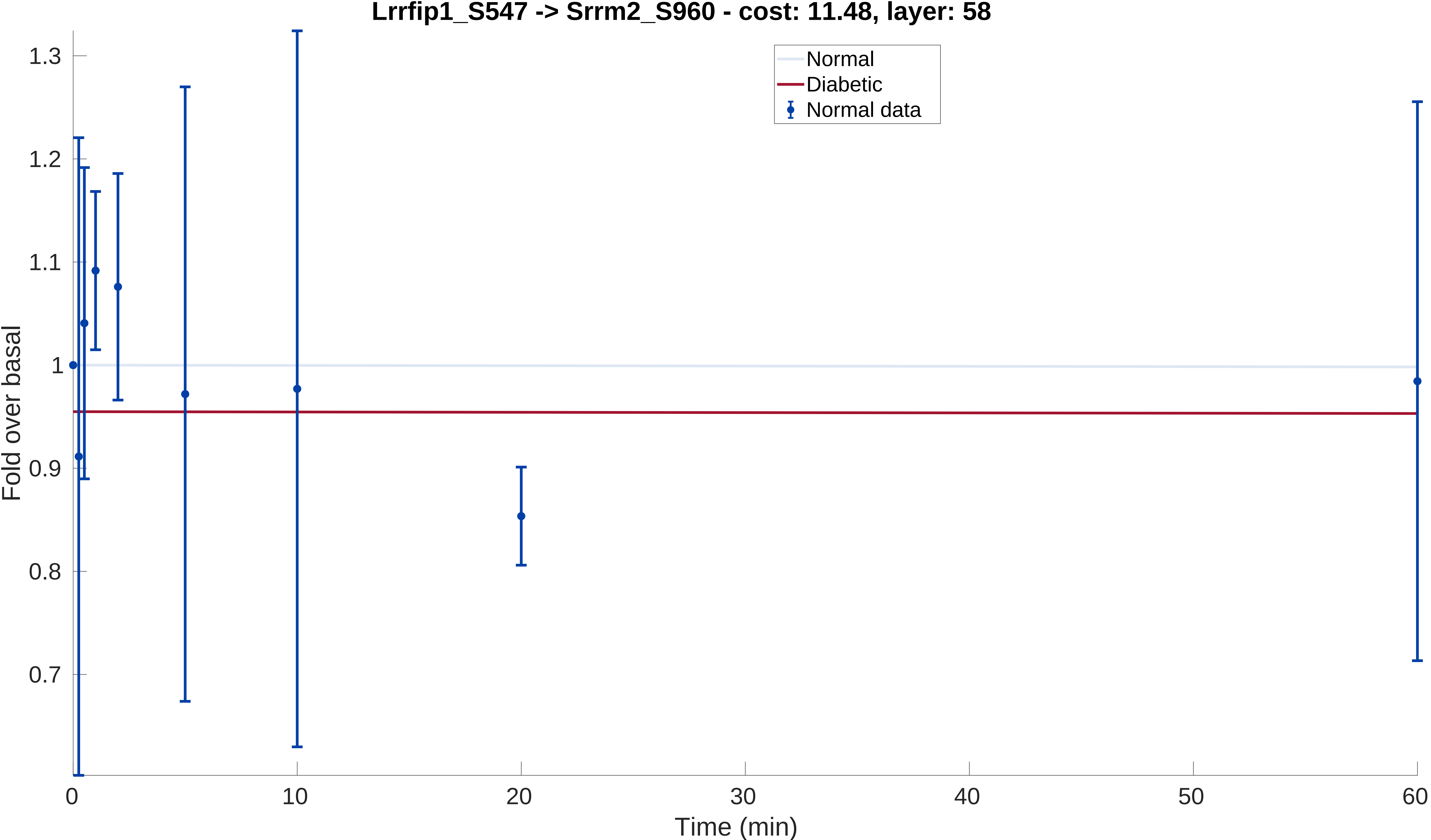

Lrrfip1_S547 -> Srrm2_S960 - cost: 11.48, layer: 58
Normal
Diabetic
Normal data
Fold over basal
1.3
1.2
1.1
1
0.9
0.8
0.7
0
10
20
30
40
50
60
Time (min)

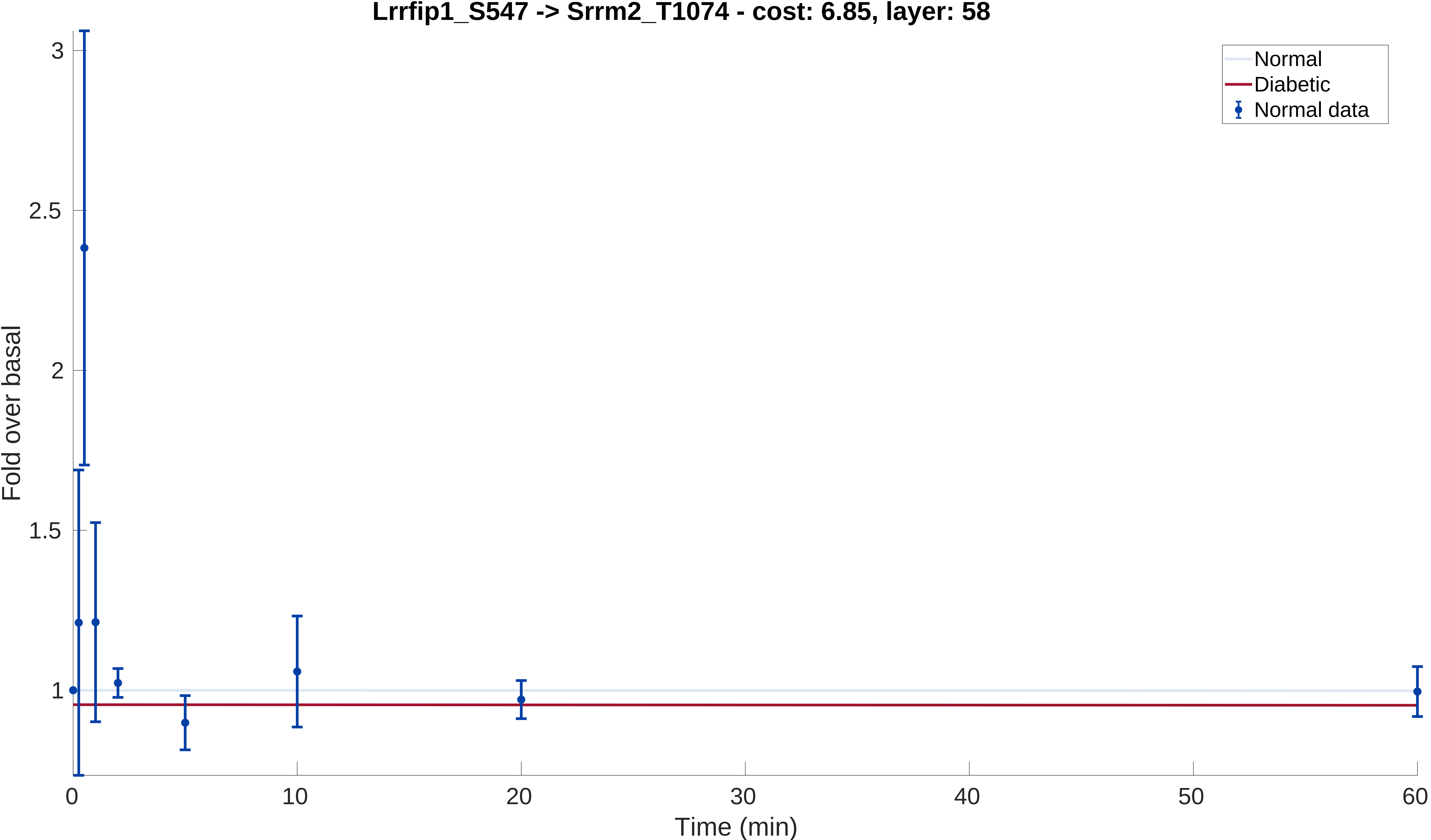
Lrrfip1_S547 -> Srrm2_T1074 - cost: 6.85, layer: 58
Normal
Diabetic
Normal data
Fold over basal
3
2.5
2
1.5
1
0
10
20
30
40
50
60
Time (min)

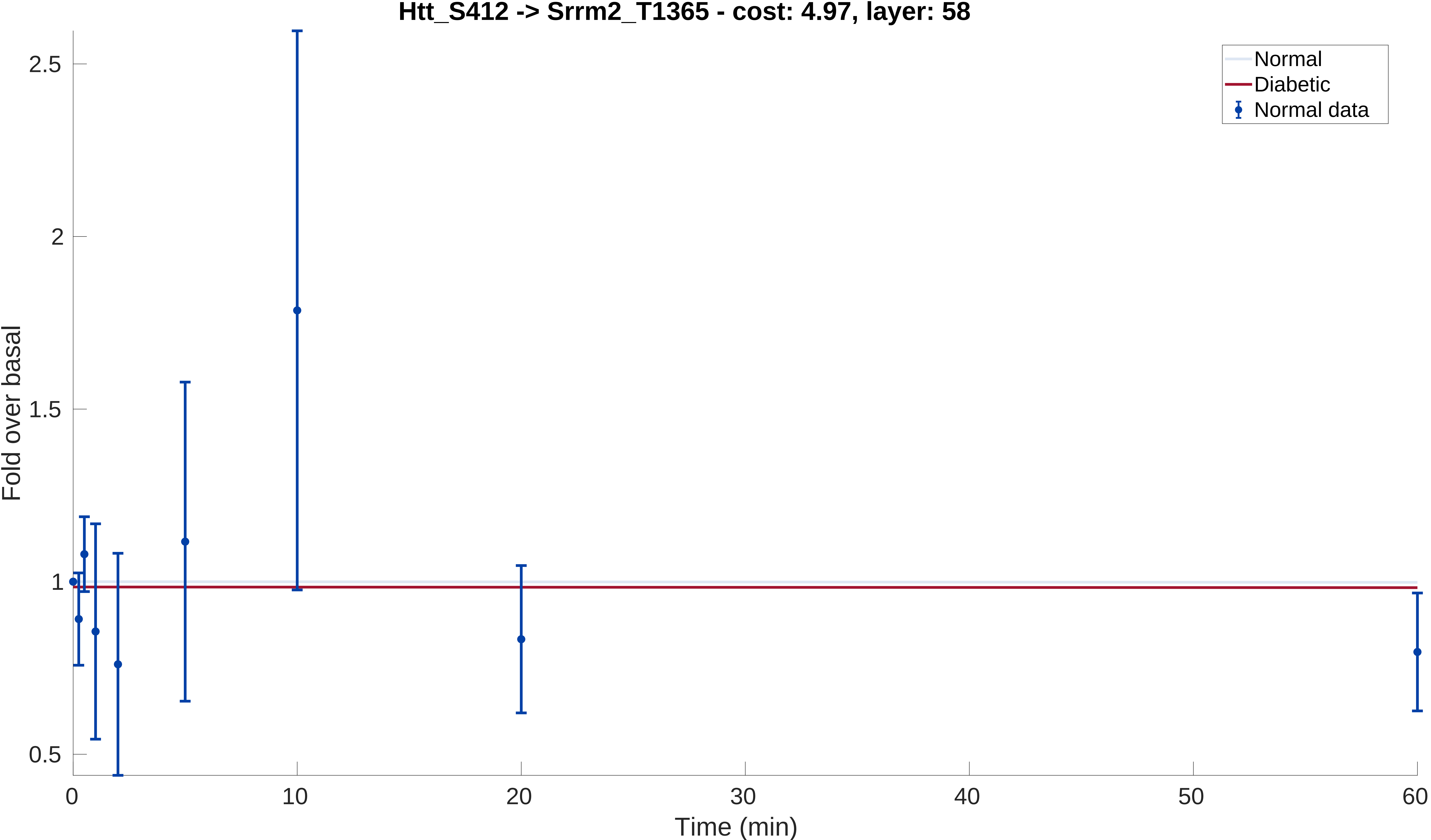

Htt_S412 -> Srrm2_T1365 - cost: 4.97, layer: 58
Normal
Diabetic
Normal data
Fold over basal
2.5
2
1.5
1
0.5
0
10
20
30
40
50
60
Time (min)

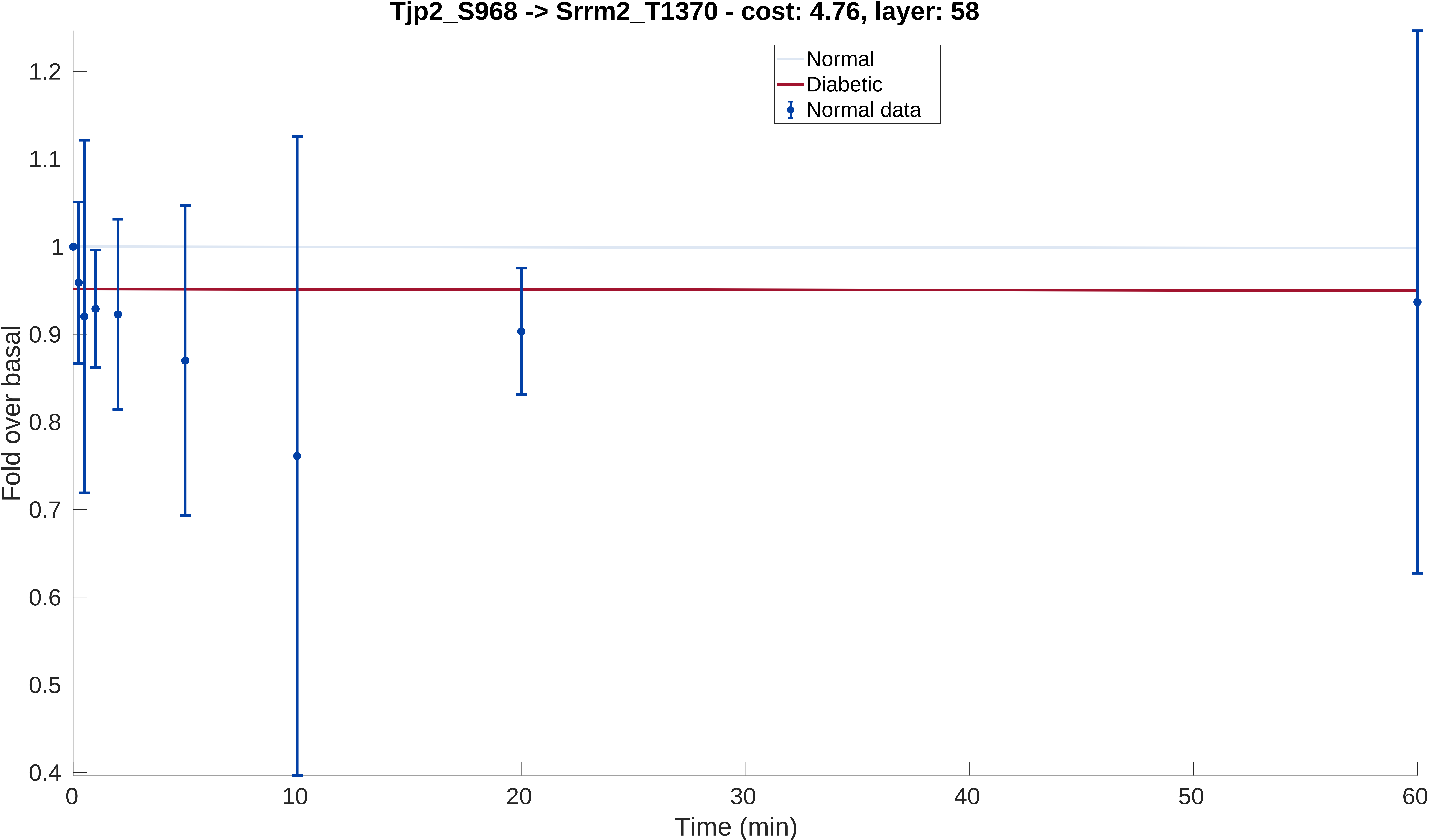

Tjp2_S968 -> Srrm2_T1370 - cost: 4.76, layer: 58
Normal
Diabetic
Normal data
Fold over basal
Time (min)
0.4
0.5
0.6
0.7
0.8
0.9
1
1.1
1.2
0
10
20
30
40
50
60

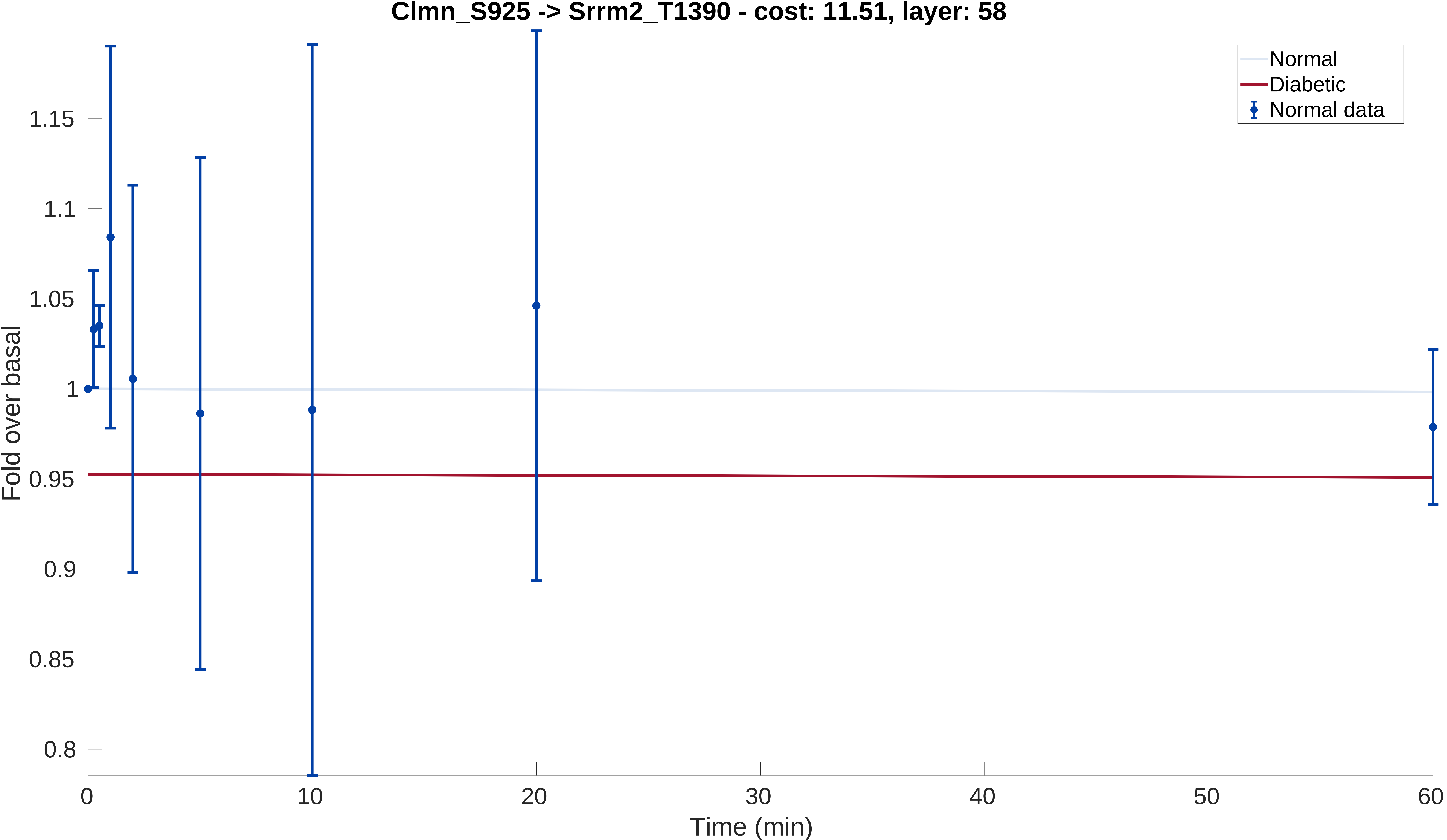
Clmn_S925 -> Srrm2_T1390 - cost: 11.51, layer: 58
Normal
Diabetic
Normal data
1.15
1.1
1.05
1
0.95
0.9
0.85
0.8
Fold over basal
0
10
20
30
40
50
60
Time (min)

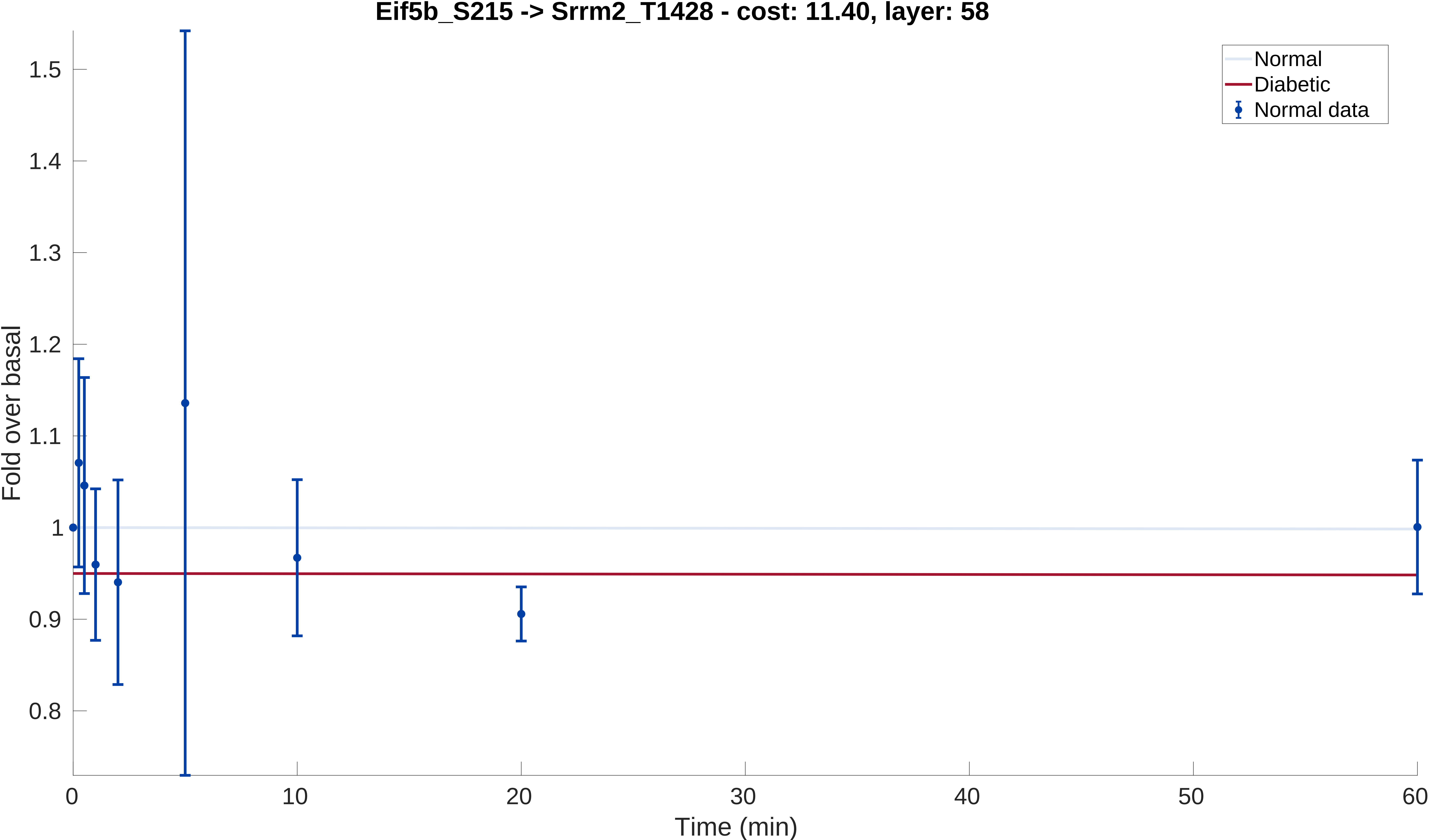
Eif5b_S215 -> Srrm2_T1428 - cost: 11.40, layer: 58
Normal
Diabetic
Normal data
Fold over basal
1.5
1.4
1.3
1.2
1.1
1
0.9
0.8
0
10
20
30
40
50
60
Time (min)

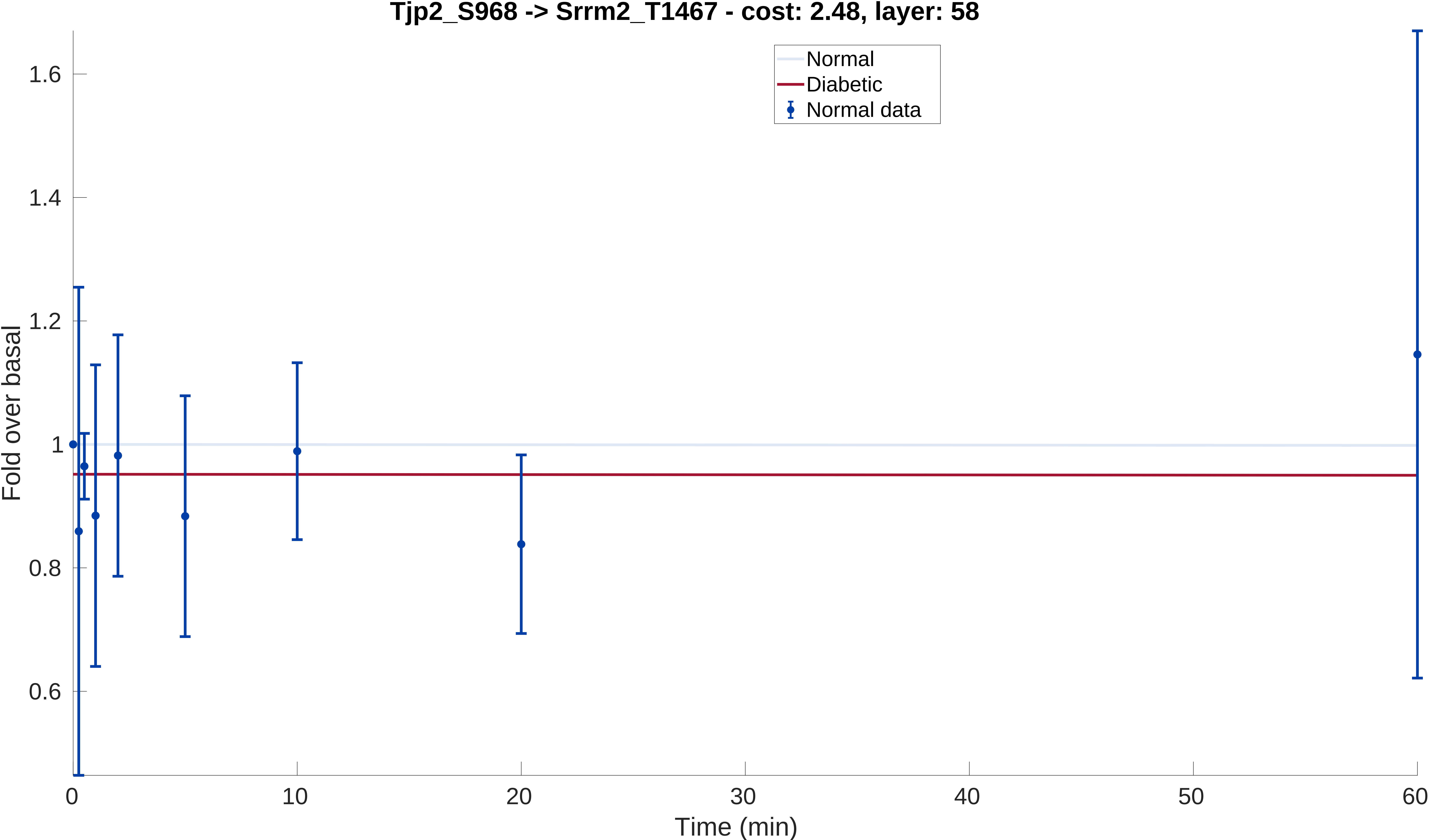

Tjp2_S968 -> Srrm2_T1467 - cost: 2.48, layer: 58
Normal
Diabetic
Normal data
Fold over basal
1.6
1.4
1.2
1
0.8
0.6
0
10
20
30
40
50
60
Time (min)

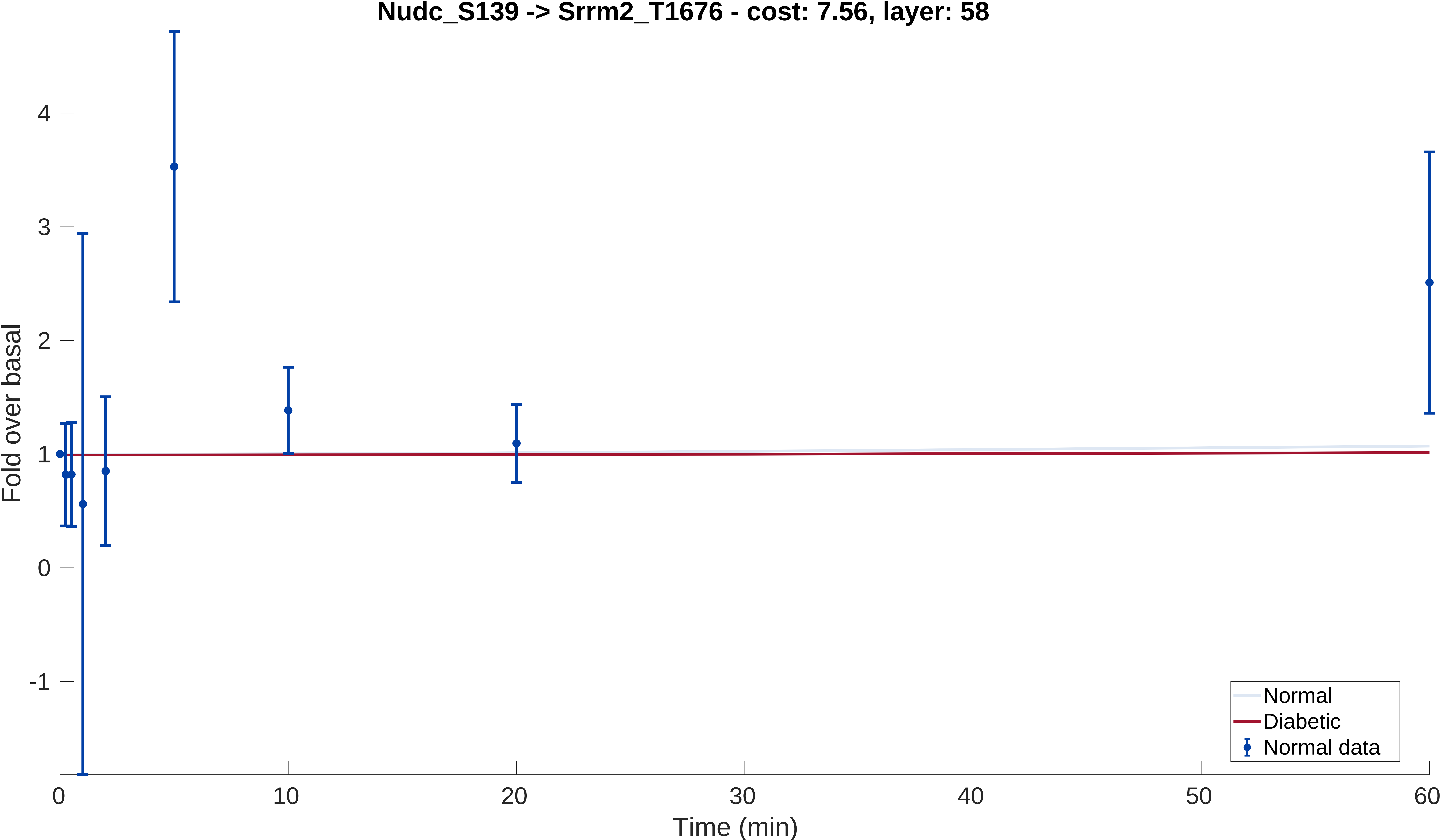
Nudc_S139 -> Srrm2_T1676 - cost: 7.56, layer: 58
Fold over basal
Time (min)
4
3
2
1
0
-1
0
10
20
30
40
50
60
Normal
Diabetic
Normal data

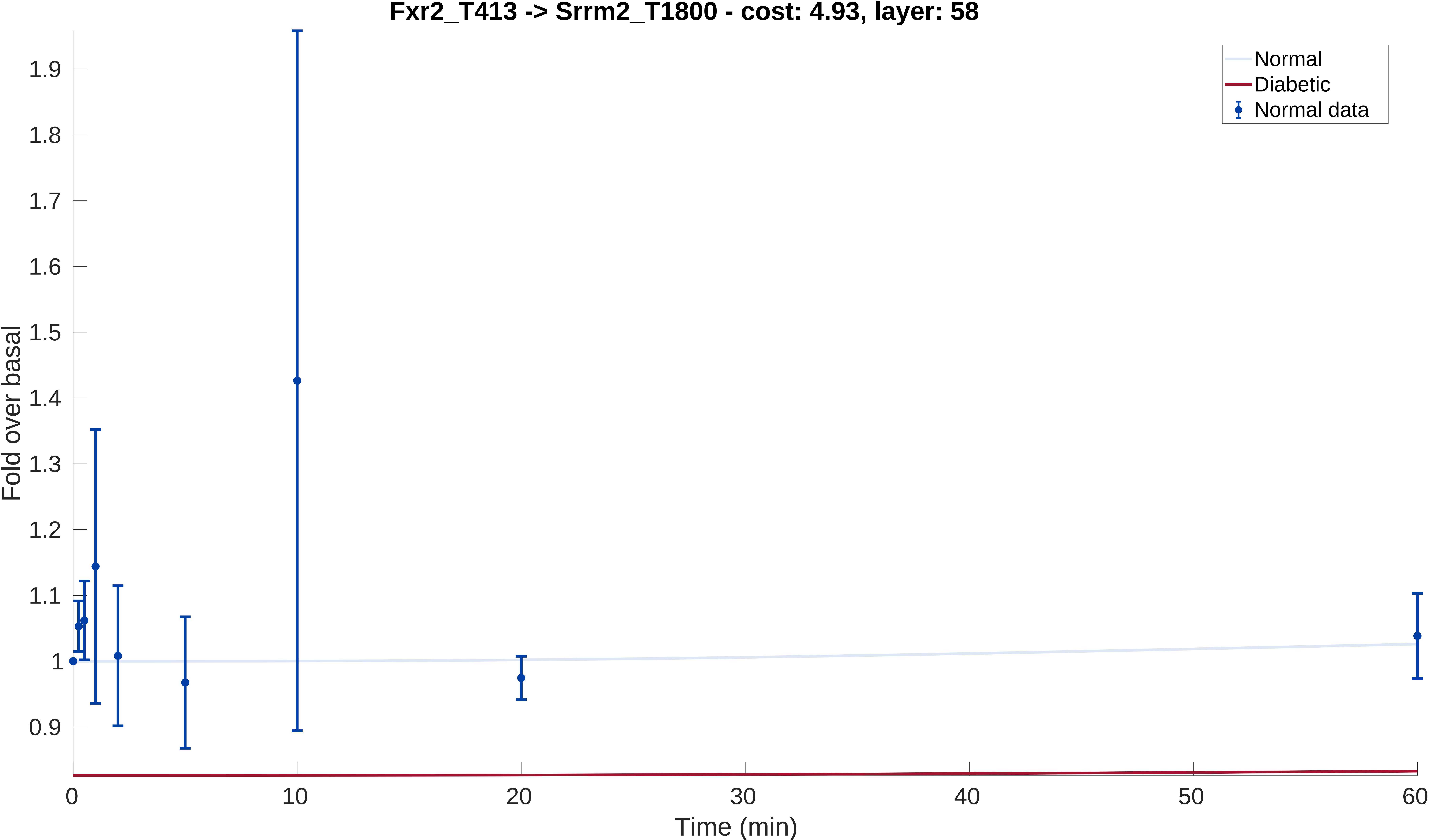

Fxr2_T413 -> Srrm2_T1800 - cost: 4.93, layer: 58
Fold over basal
1.9
1.8
1.7
1.6
1.5
1.4
1.3
1.2
1.1
1
0.9
0
10
20
30
40
50
60
Time (min)
Normal
Diabetic
Normal data

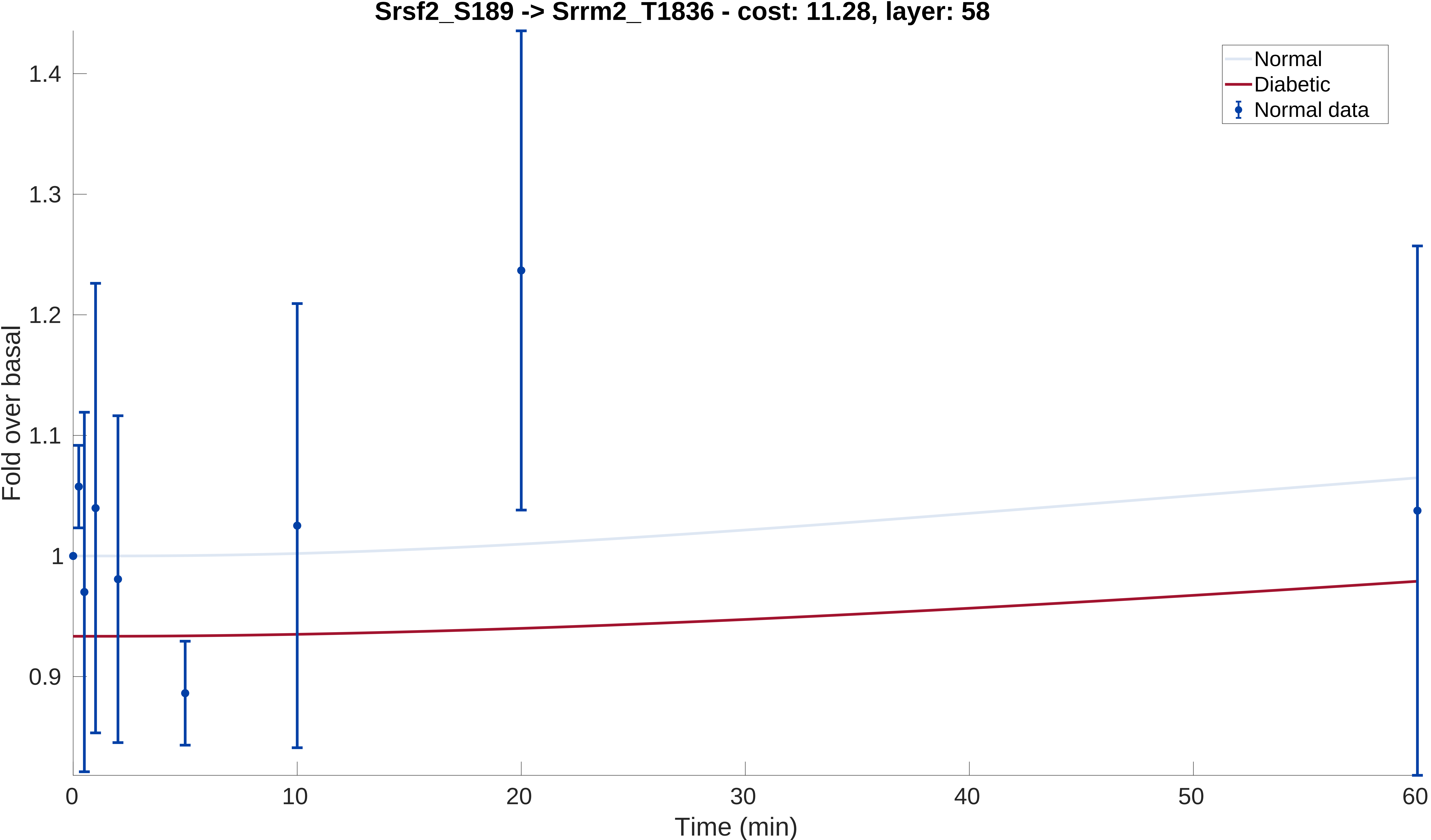

Srsf2_S189 -> Srrm2_T1836 - cost: 11.28, layer: 58
Normal
Diabetic
Normal data
1.4
1.3
1.2
1.1
1
0.9
Fold over basal
0
10
20
30
40
50
60
Time (min)

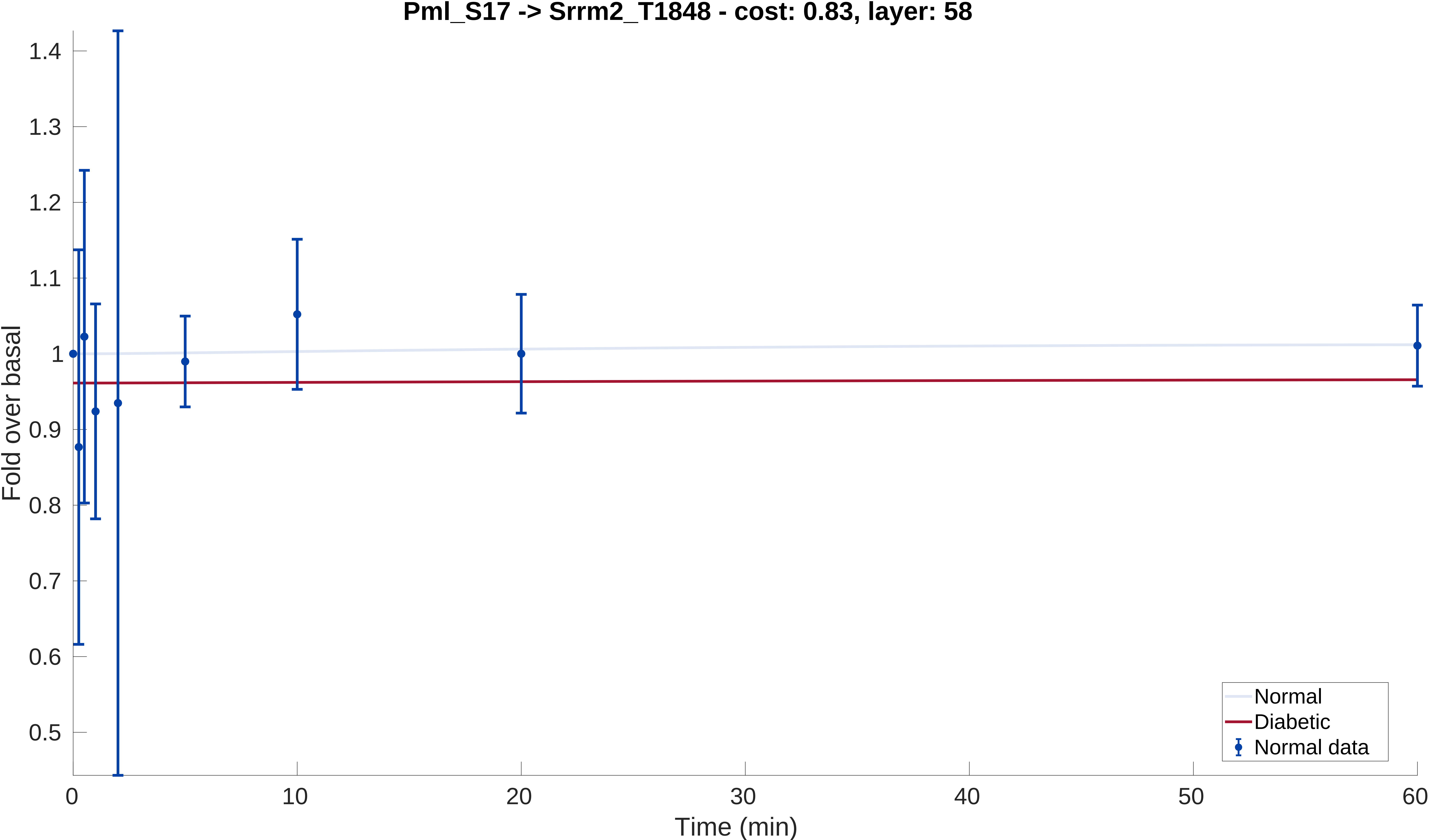
Pml_S17 -> Srrm2_T1848 - cost: 0.83, layer: 58
Fold over basal
1.4
1.3
1.2
1.1
1
0.9
0.8
0.7
0.6
0.5
0
10
20
30
40
50
60
Time (min)
Normal
Diabetic
Normal data

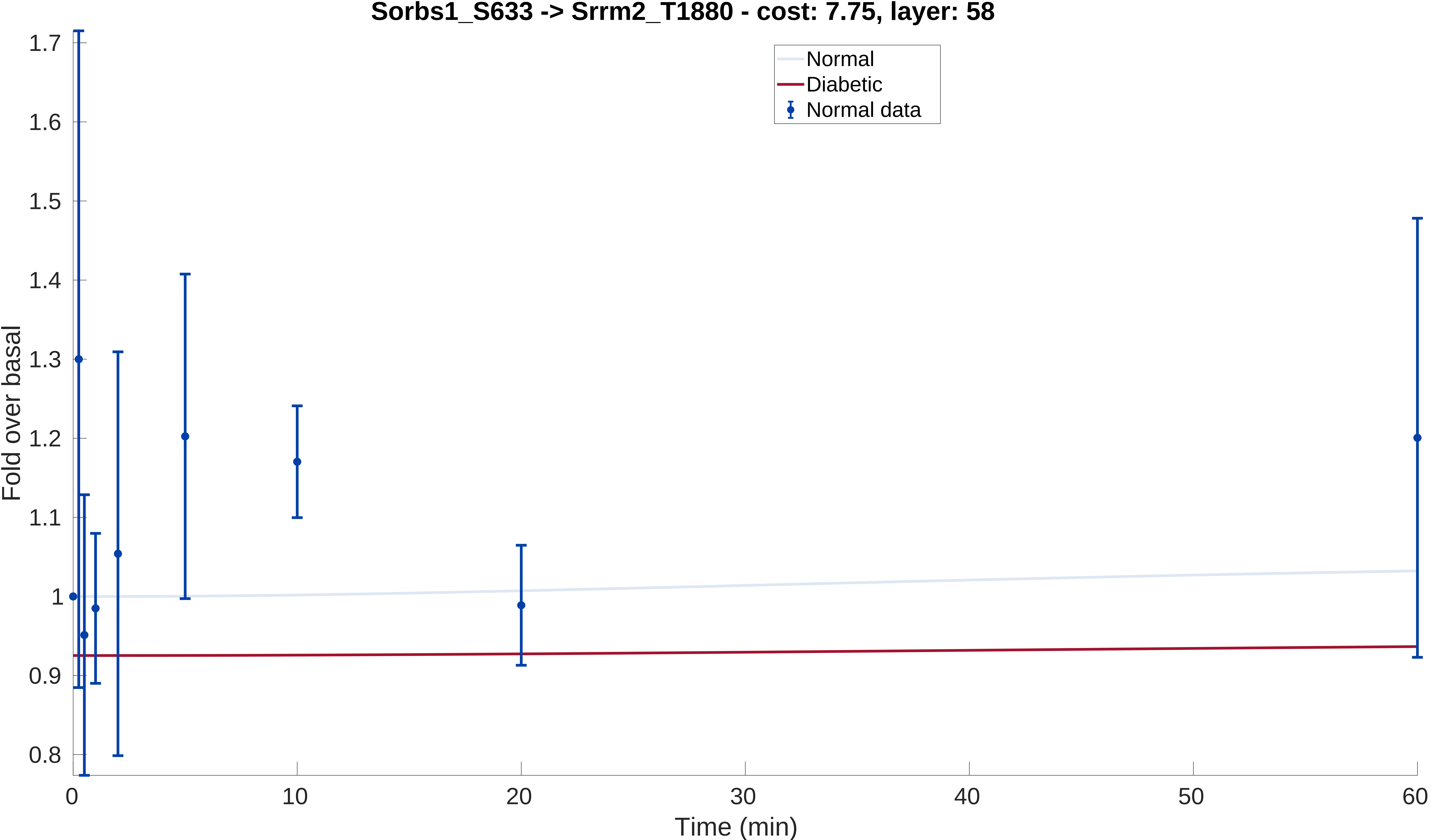

Sorbs1_S633 -> Srrm2_T1880 - cost: 7.75, layer: 58
Normal
Diabetic
Normal data
Fold over basal
Time (min)
0.8
0.9
1
1.1
1.2
1.3
1.4
1.5
1.6
1.7
0
10
20
30
40
50
60

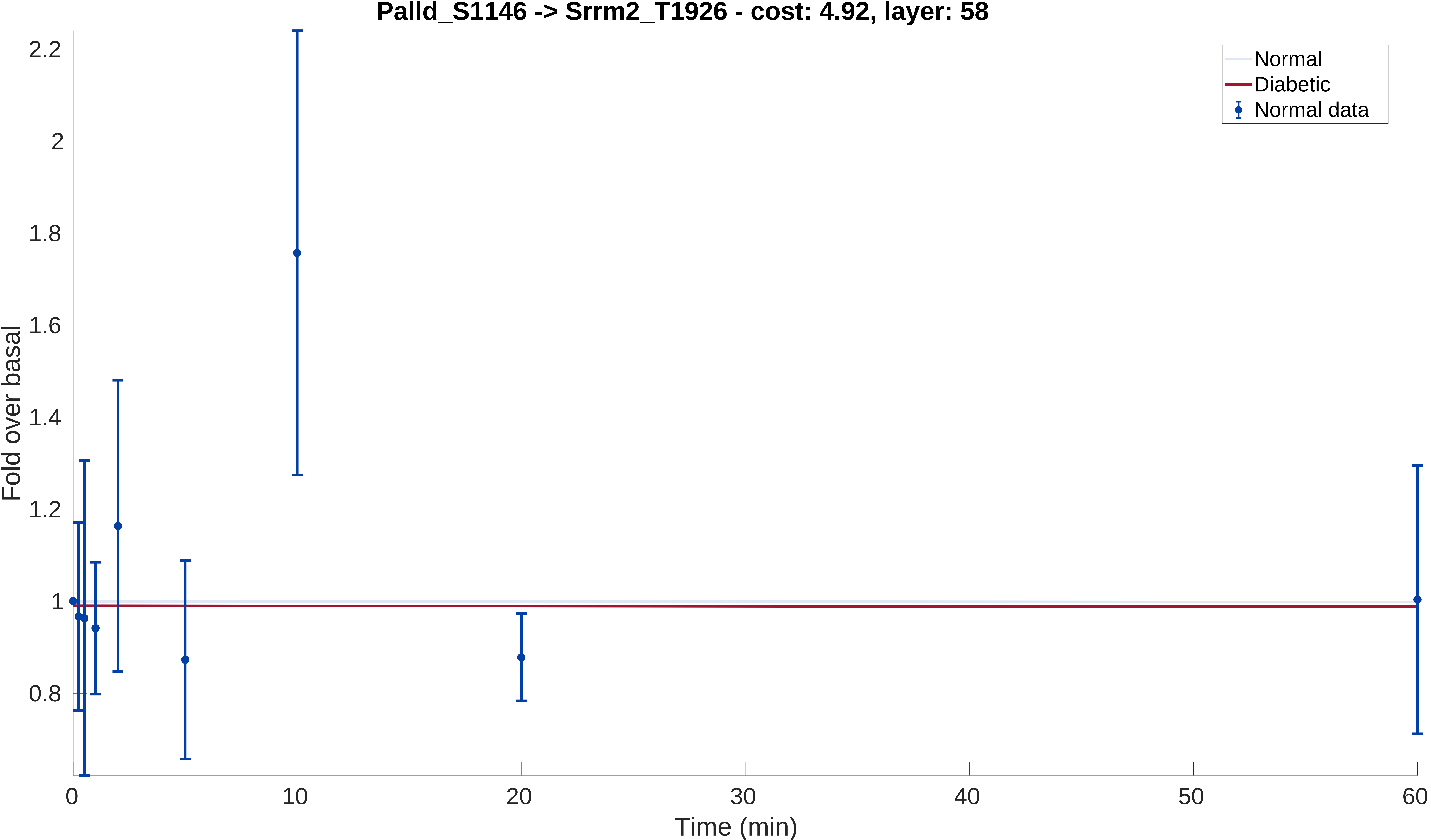

Palld_S1146 -> Srrm2_T1926 - cost: 4.92, layer: 58
Normal
Diabetic
Normal data
Fold over basal
2.2
2
1.8
1.6
1.4
1.2
1
0.8
0
10
20
30
40
50
60
Time (min)

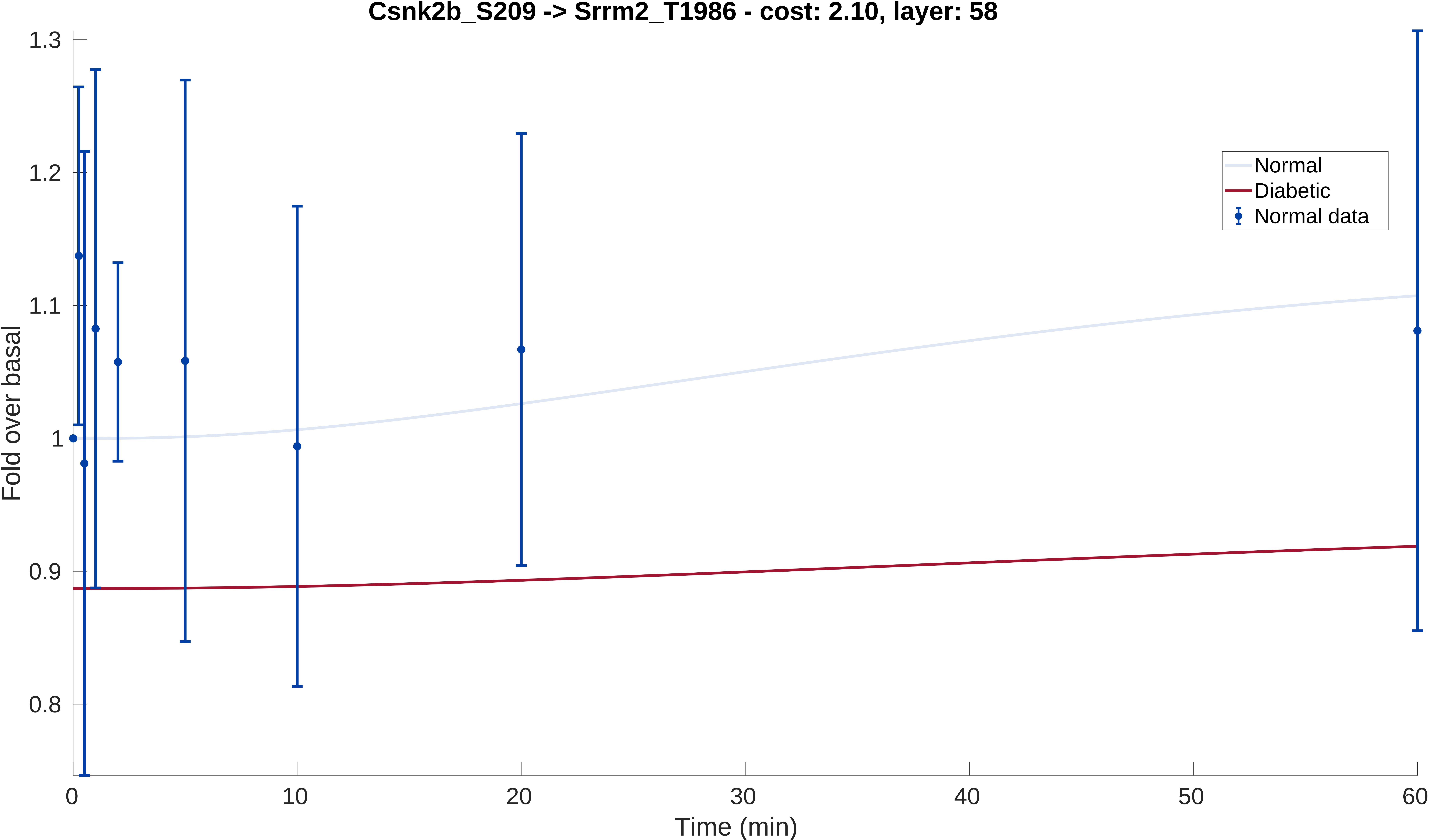
Csnk2b_S209 -> Srrm2_T1986 - cost: 2.10, layer: 58
Fold over basal
1.3
1.2
1.1
1
0.9
0.8
0
10
20
30
40
50
60
Time (min)
Normal
Diabetic
Normal data

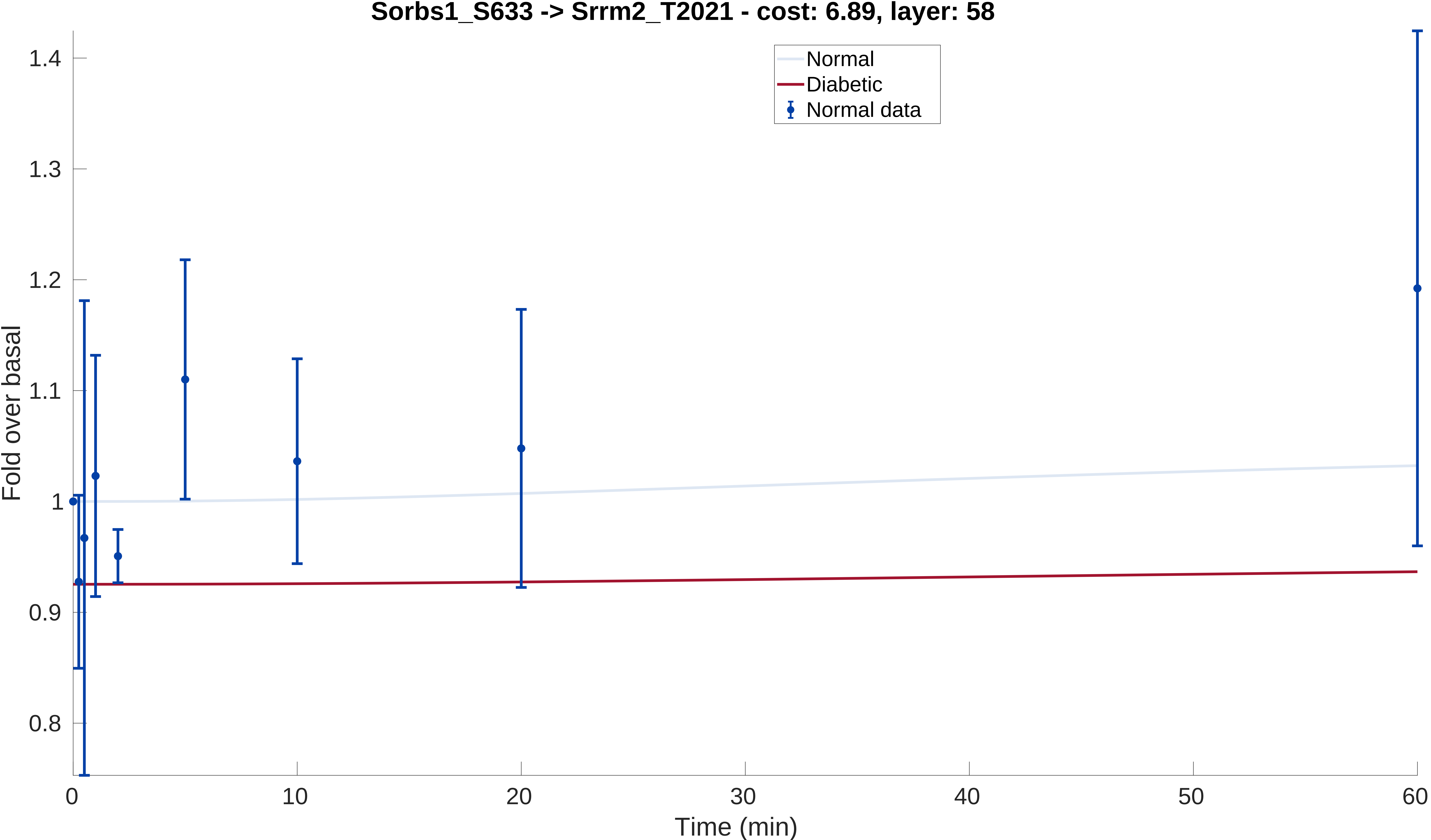
Sorbs1_S633 -> Srrm2_T2021 - cost: 6.89, layer: 58
Normal
Diabetic
Normal data
Fold over basal
1.4
1.3
1.2
1.1
1
0.9
0.8
0
10
20
30
40
50
60
Time (min)

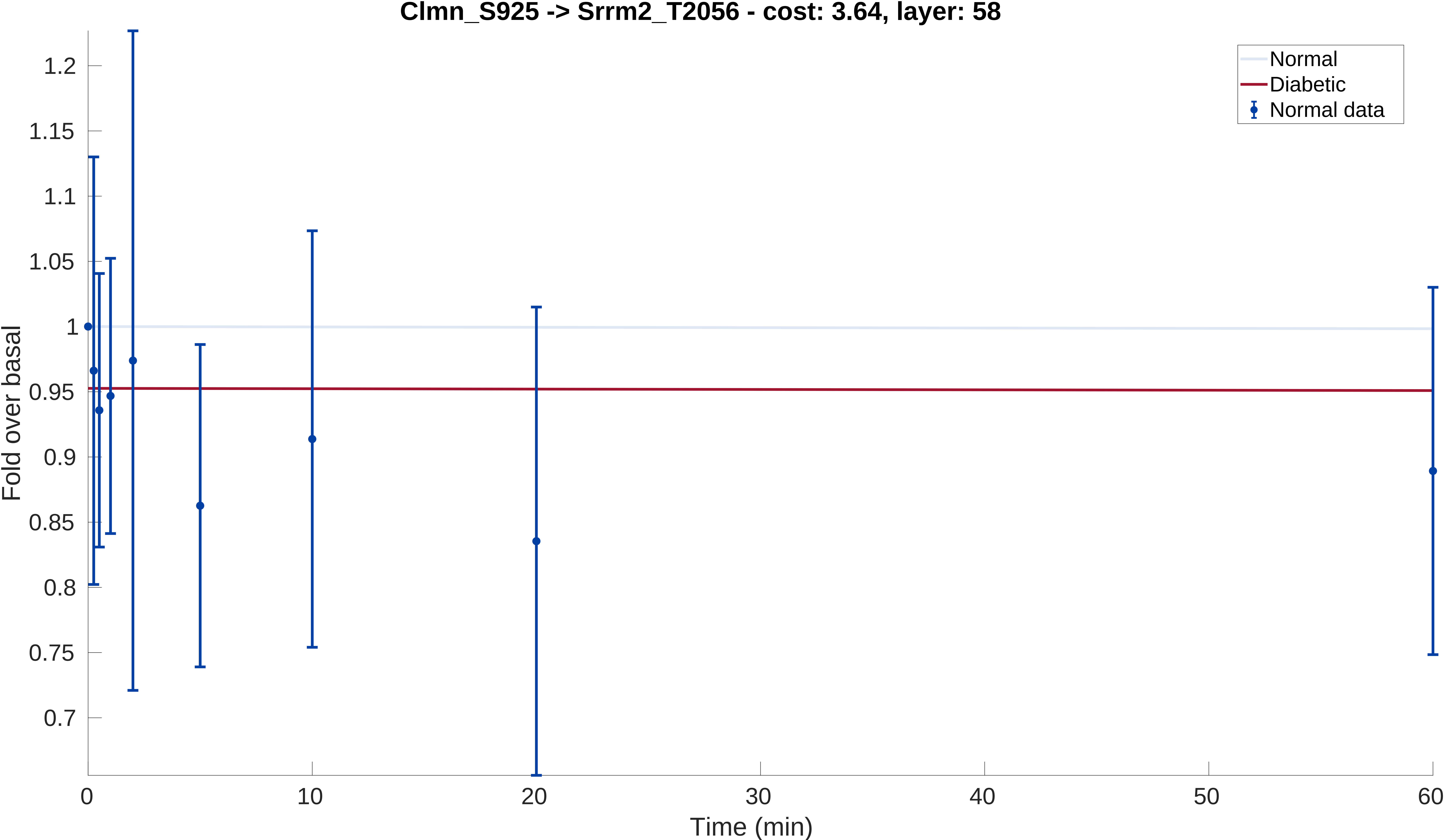
Clmn_S925 -> Srrm2_T2056 - cost: 3.64, layer: 58
Fold over basal
Time (min)
Normal
Diabetic
Normal data
1.2
1.15
1.1
1.05
1
0.95
0.9
0.85
0.8
0.75
0.7
0
10
20
30
40
50
60

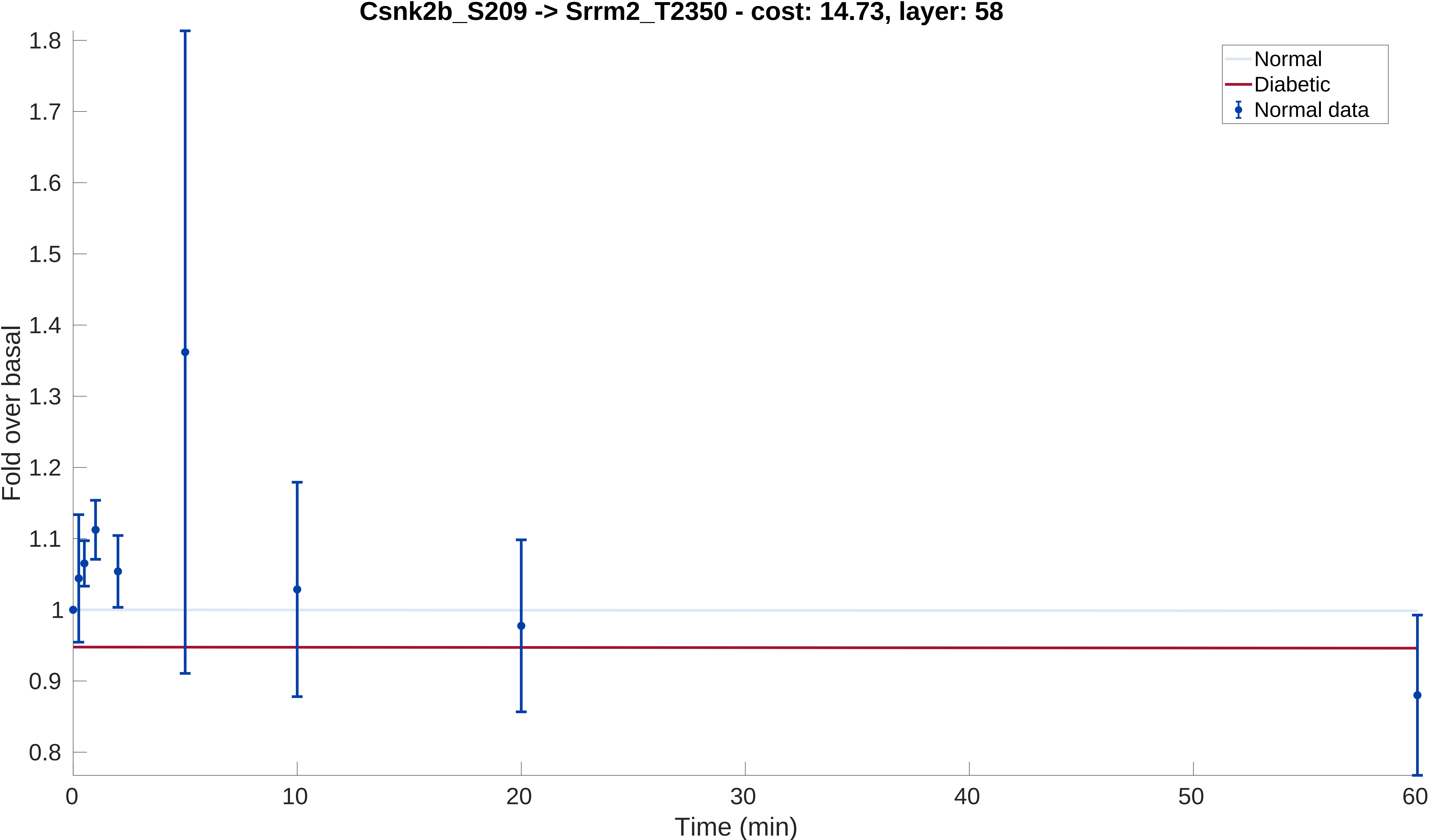

Csnk2b_S209 -> Srrm2_T2350 - cost: 14.73, layer: 58
Normal
Diabetic
Normal data
Fold over basal
Time (min)
1.8
1.7
1.6
1.5
1.4
1.3
1.2
1.1
1
0.9
0.8
0
10
20
30
40
50
60

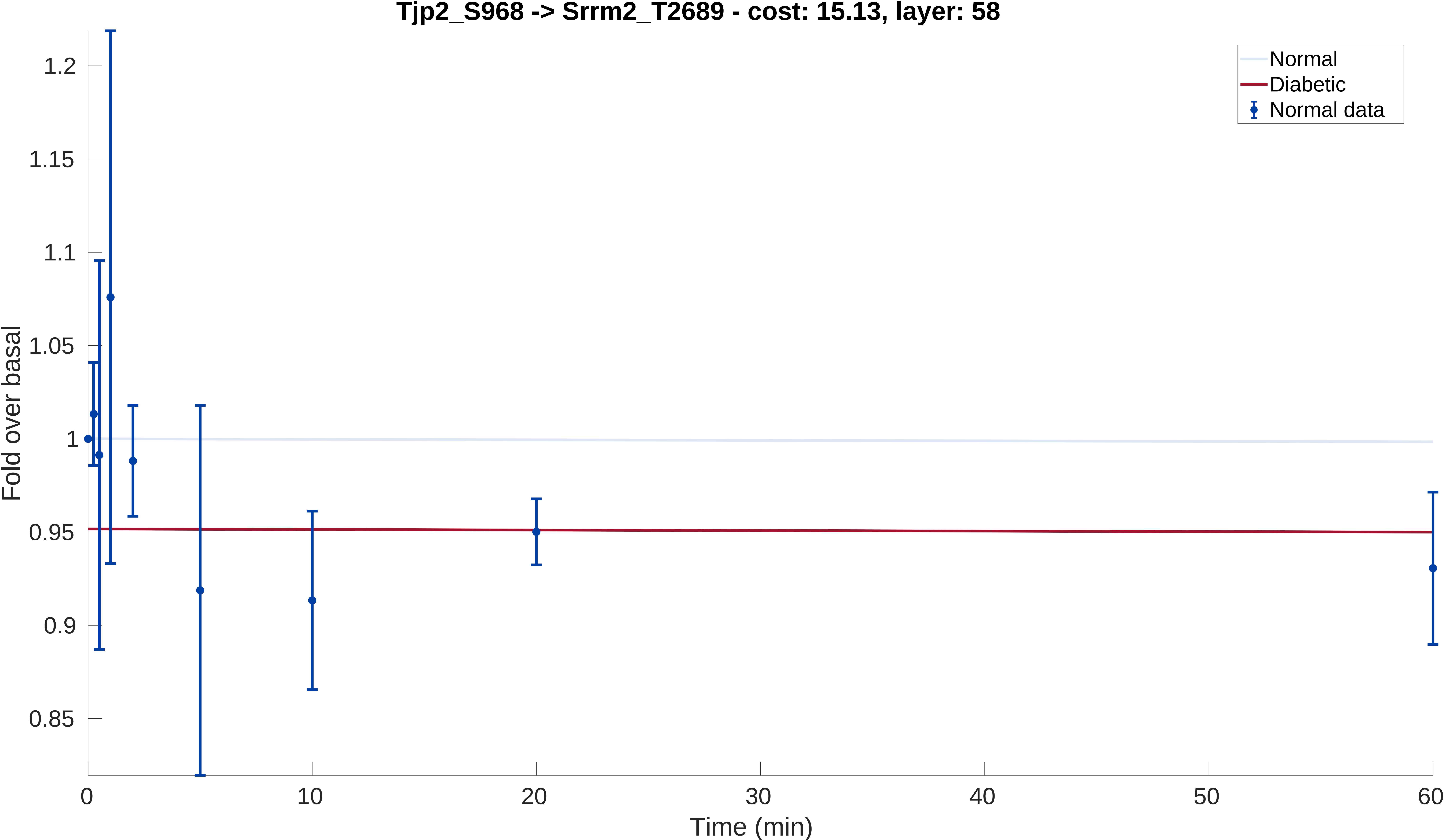
Tjp2_S968 -> Srrm2_T2689 - cost: 15.13, layer: 58
Normal
Diabetic
Normal data
Fold over basal
1.2
1.15
1.1
1.05
1
0.95
0.9
0.85
0
10
20
30
40
50
60
Time (min)

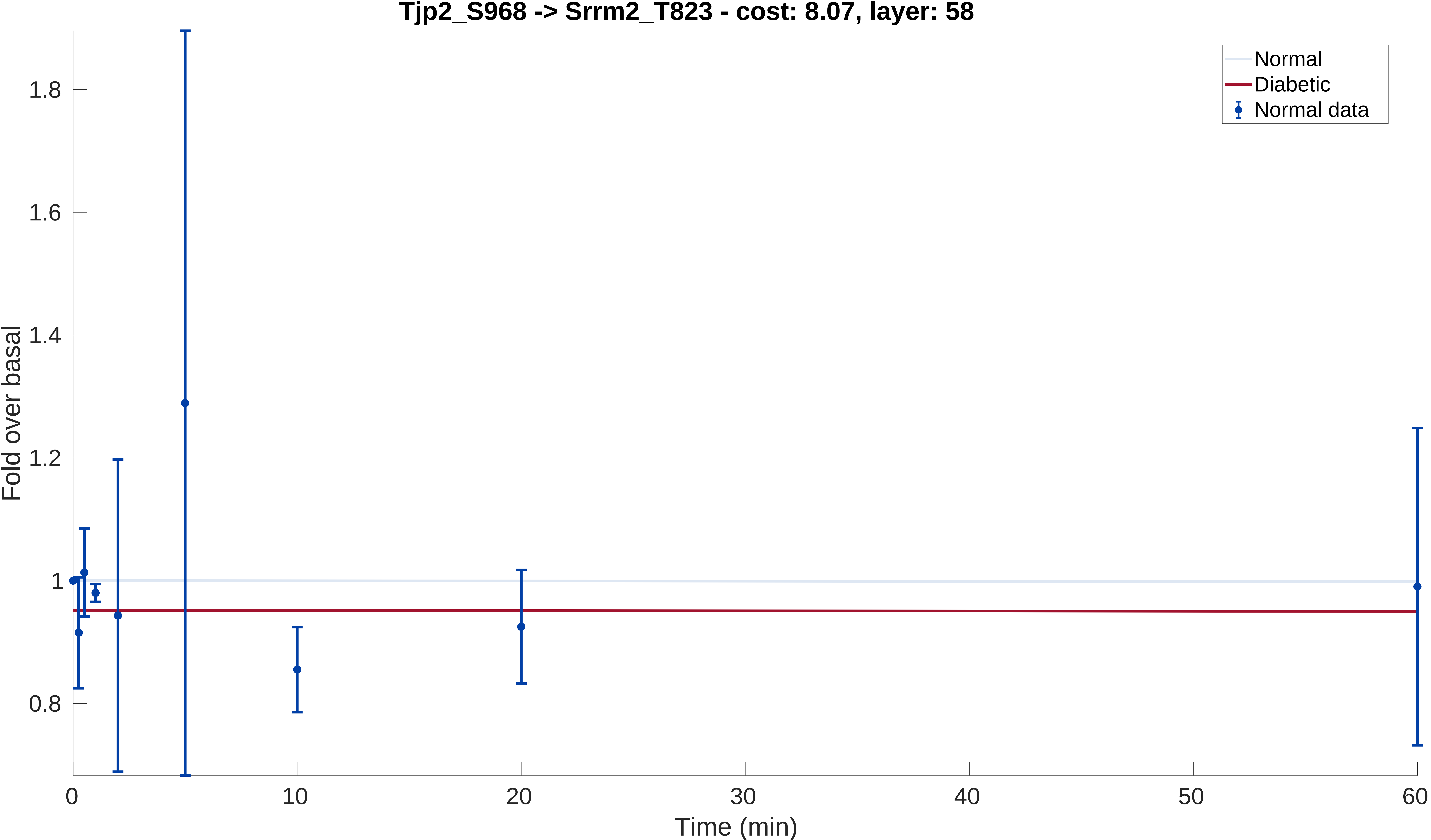
Tjp2_S968 -> Srrm2_T823 - cost: 8.07, layer: 58
Normal
Diabetic
Normal data
Fold over basal
1.8
1.6
1.4
1.2
1
0.8
0
10
20
30
40
50
60
Time (min)

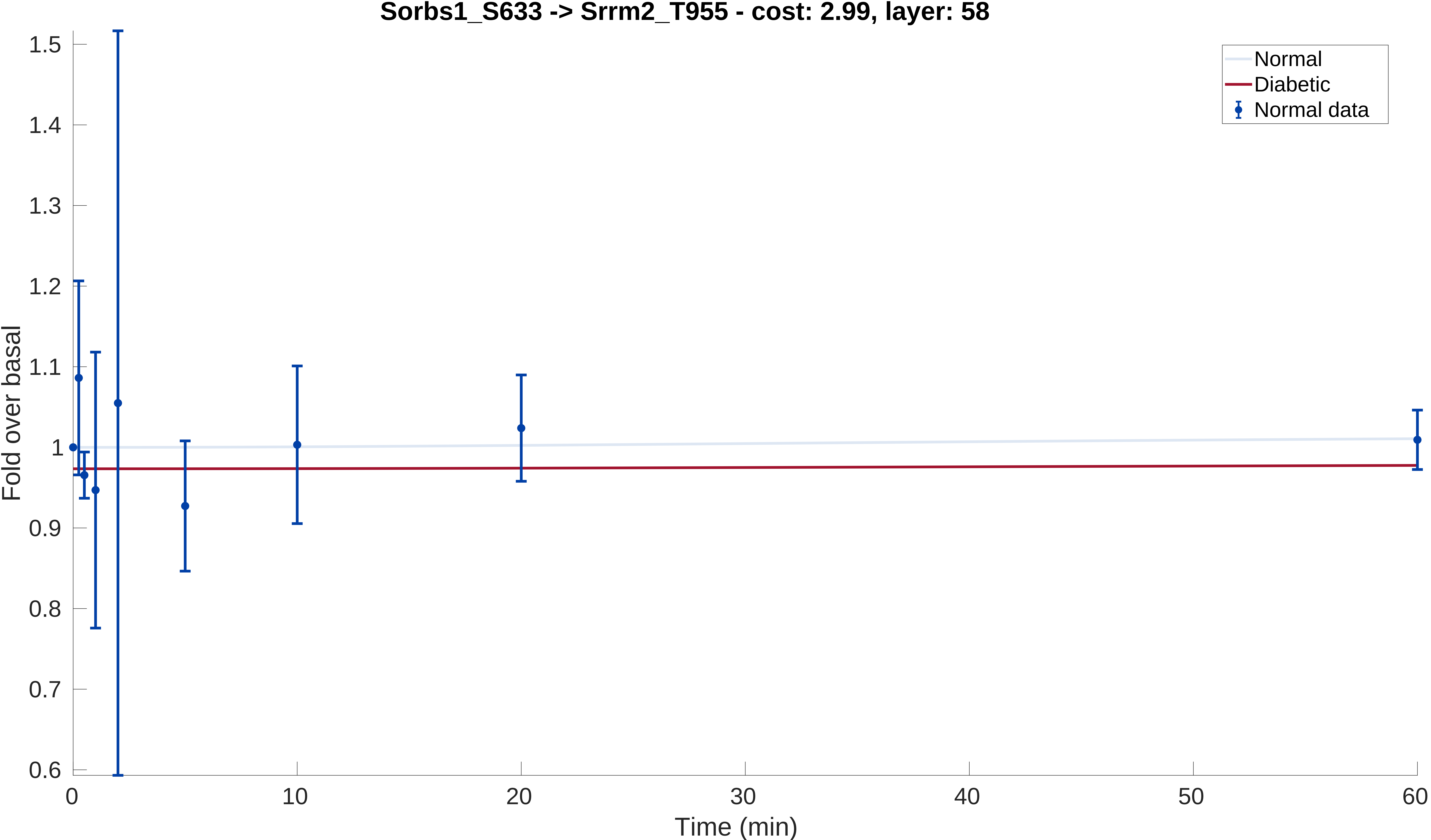

Sorbs1_S633 -> Srrm2_T955 - cost: 2.99, layer: 58
Normal
Diabetic
Normal data
Fold over basal
1.5
1.4
1.3
1.2
1.1
1
0.9
0.8
0.7
0.6
0
10
20
30
40
50
60
Time (min)

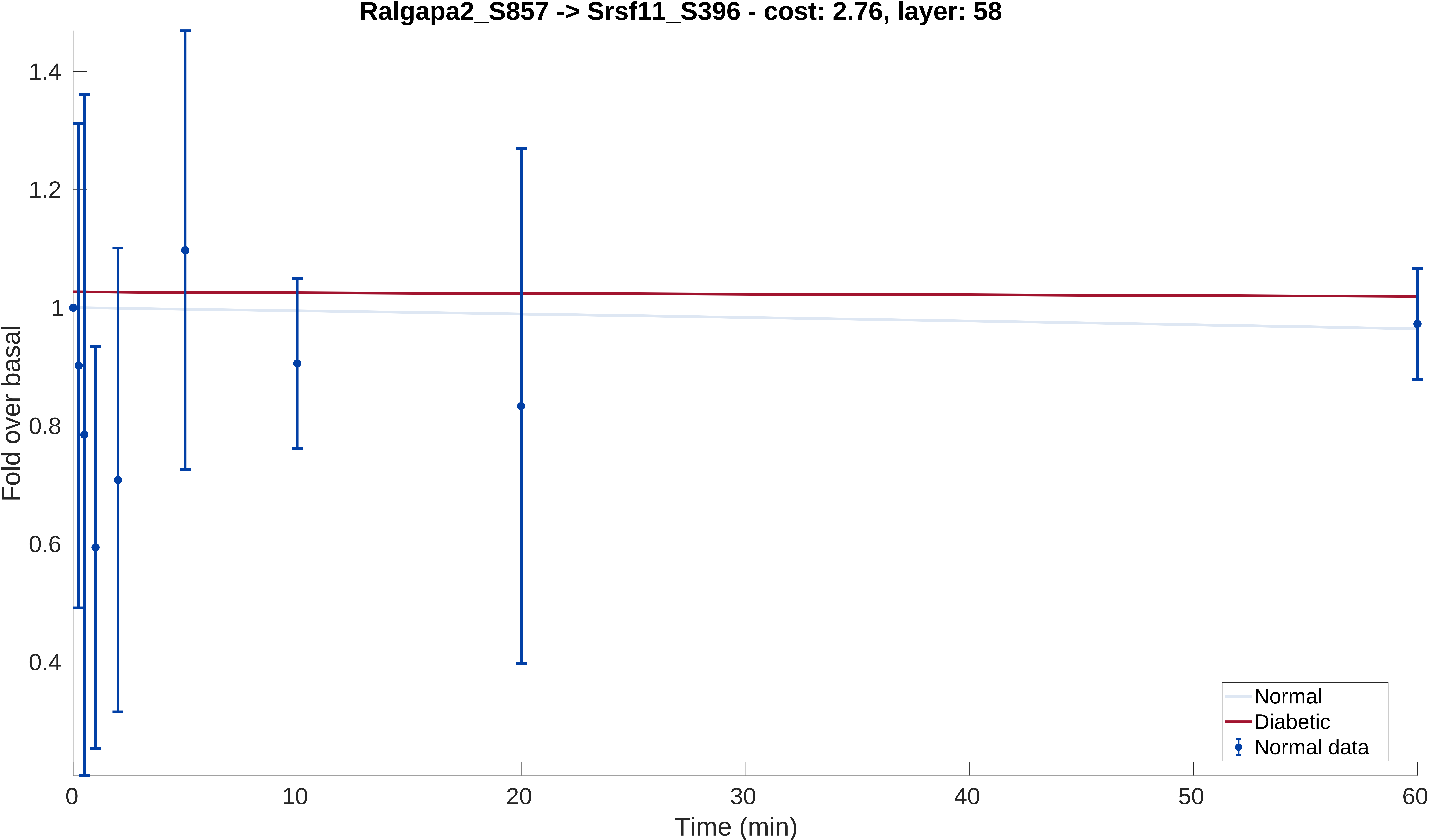
Ralgapa2_S857 -> Srsf11_S396 - cost: 2.76, layer: 58
1.4
1.2
1
0.8
0.6
0.4
Fold over basal
0
10
20
30
40
50
60
Time (min)
Normal
Diabetic
Normal data

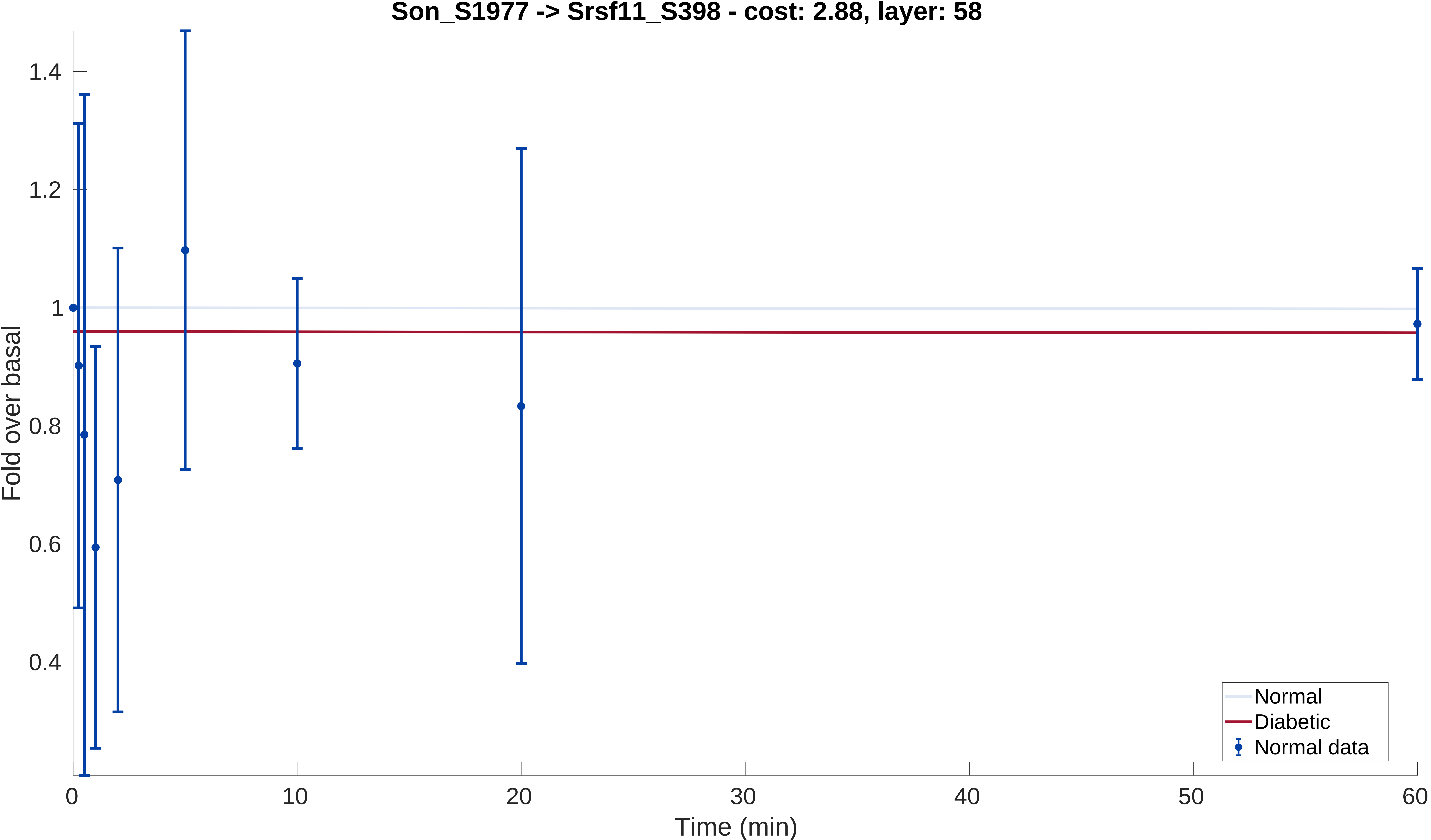

Son_S1977 -> Srsf11_S398 - cost: 2.88, layer: 58
Fold over basal
1.4
1.2
1
0.8
0.6
0.4
0
10
20
30
40
50
60
Time (min)
Normal
Diabetic
Normal data

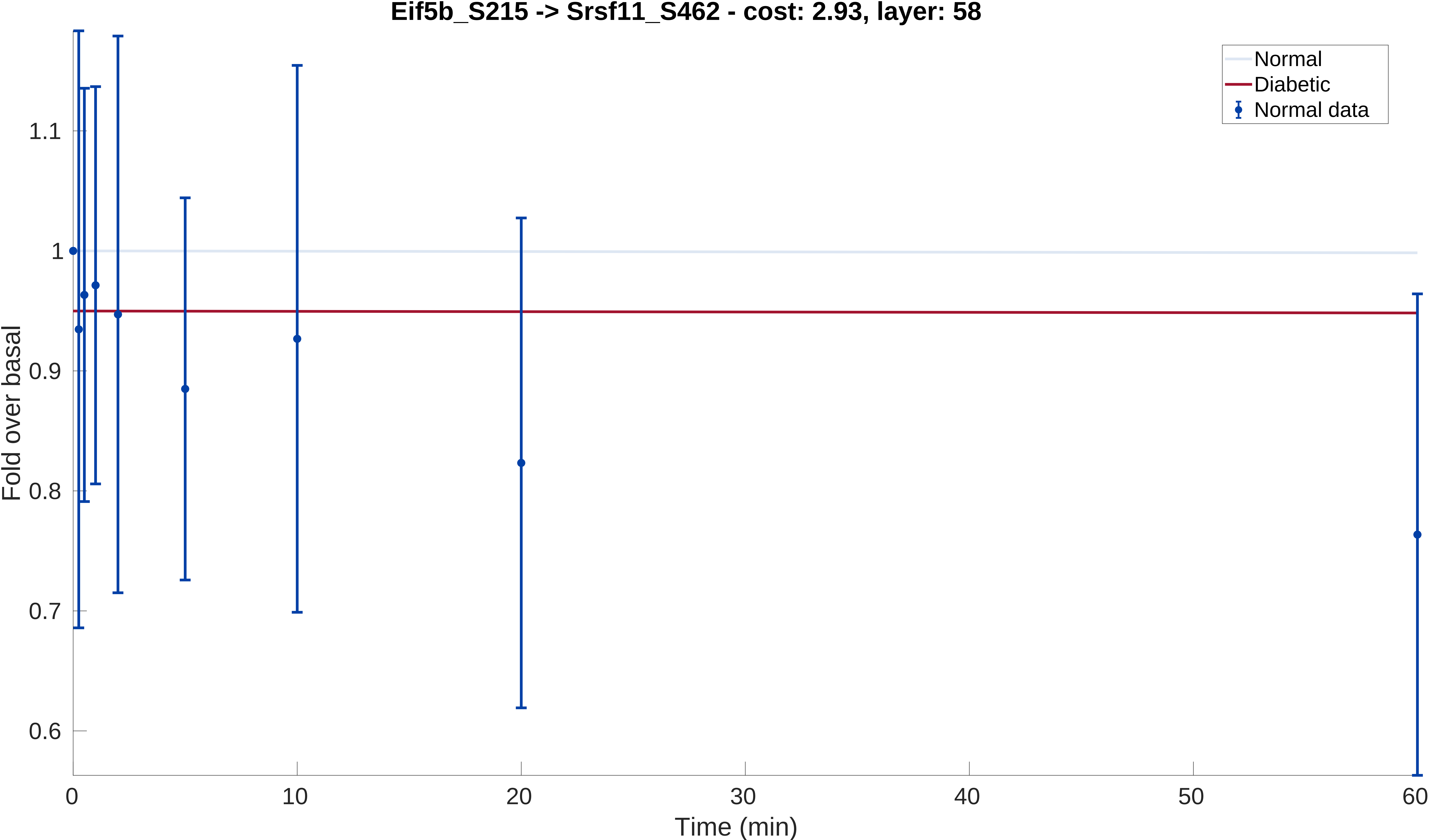
Eif5b_S215 -> Srsf11_S462 - cost: 2.93, layer: 58
Normal
Diabetic
Normal data
Fold over basal
1.1
1
0.9
0.8
0.7
0.6
0
10
20
30
40
50
60
Time (min)

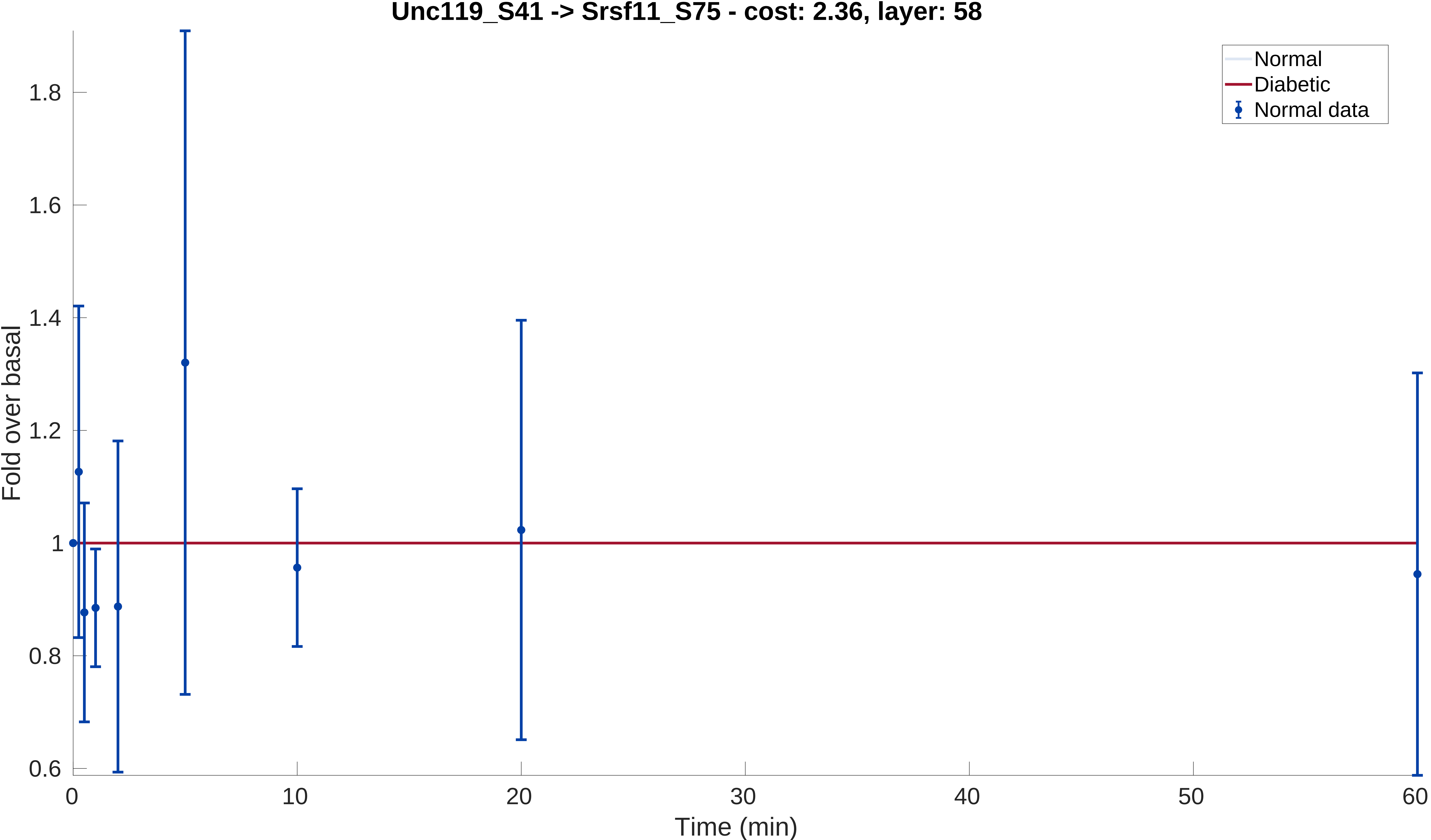
Unc119_S41 -> Srsf11_S75 - cost: 2.36, layer: 58
Normal
Diabetic
Normal data
Fold over basal
Time (min)
0.6
0.8
1
1.2
1.4
1.6
1.8
0
10
20
30
40
50
60

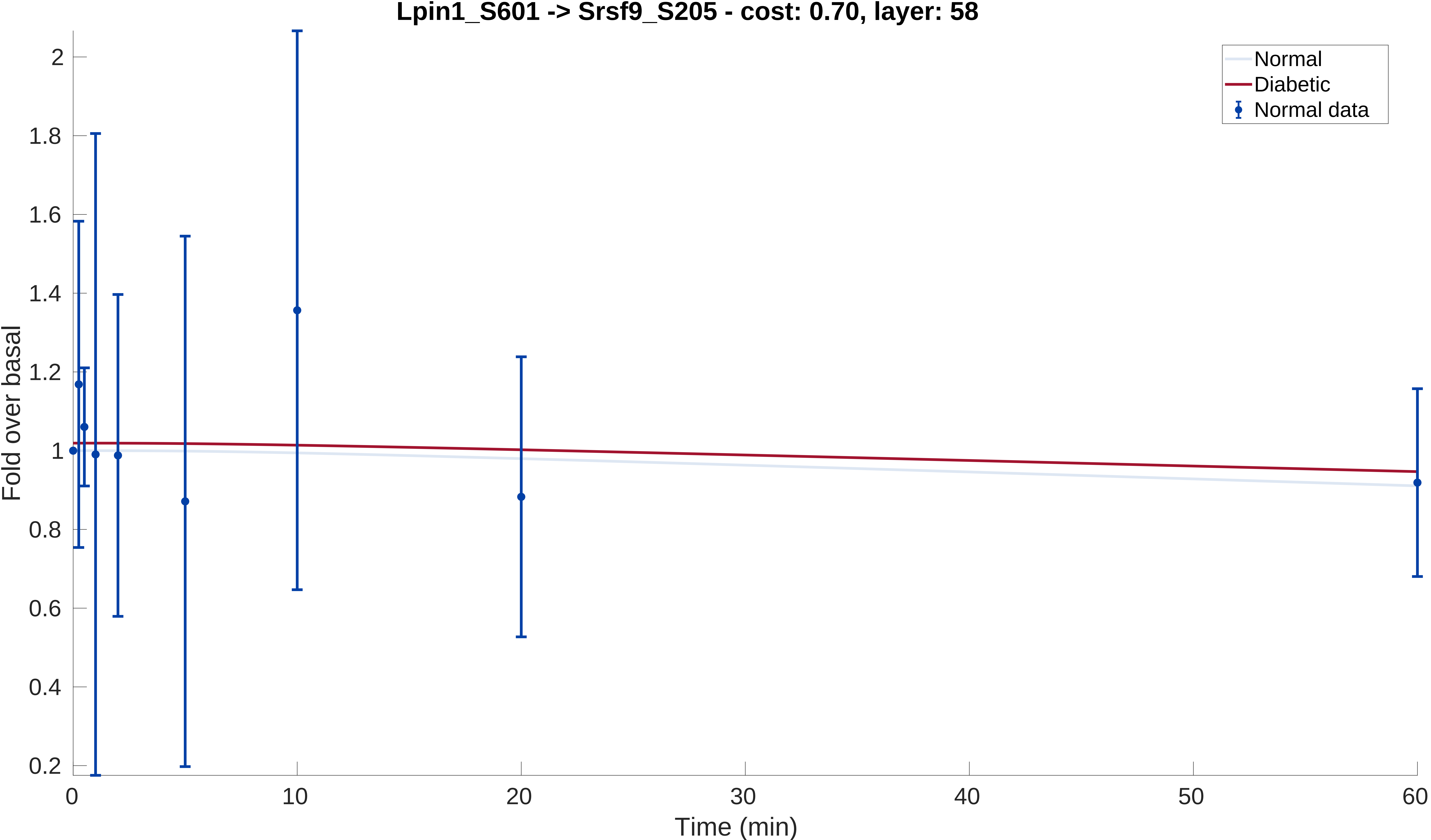

Lpin1_S601 -> Srsf9_S205 - cost: 0.70, layer: 58
Normal
Diabetic
Normal data
Fold over basal
Time (min)
0.2
0.4
0.6
0.8
1
1.2
1.4
1.6
1.8
2
0
10
20
30
40
50
60

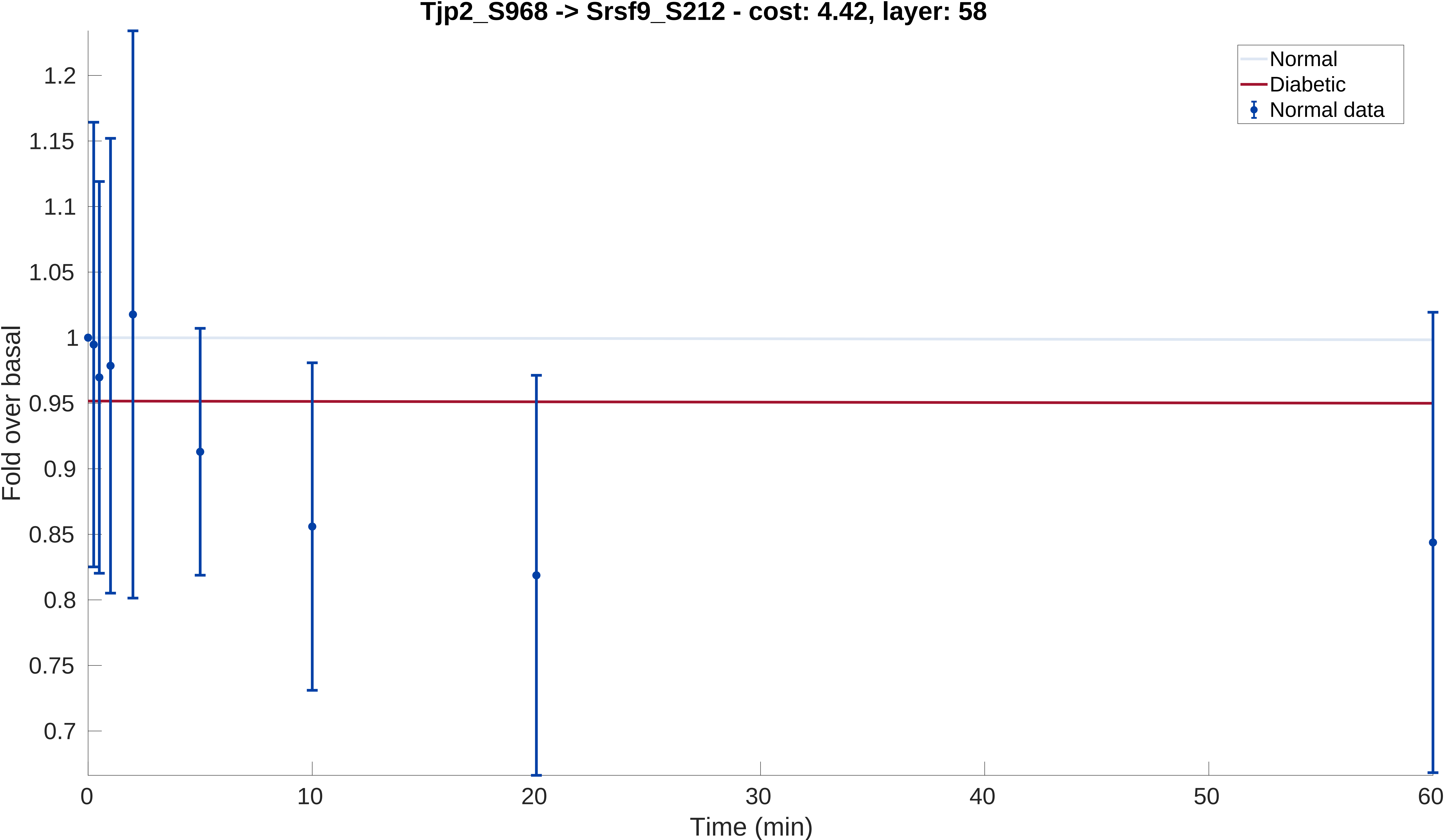

Tjp2_S968 -> Srsf9_S212 - cost: 4.42, layer: 58
Normal
Diabetic
Normal data
Fold over basal
1.2
1.15
1.1
1.05
1
0.95
0.9
0.85
0.8
0.75
0.7
0
10
20
30
40
50
60
Time (min)

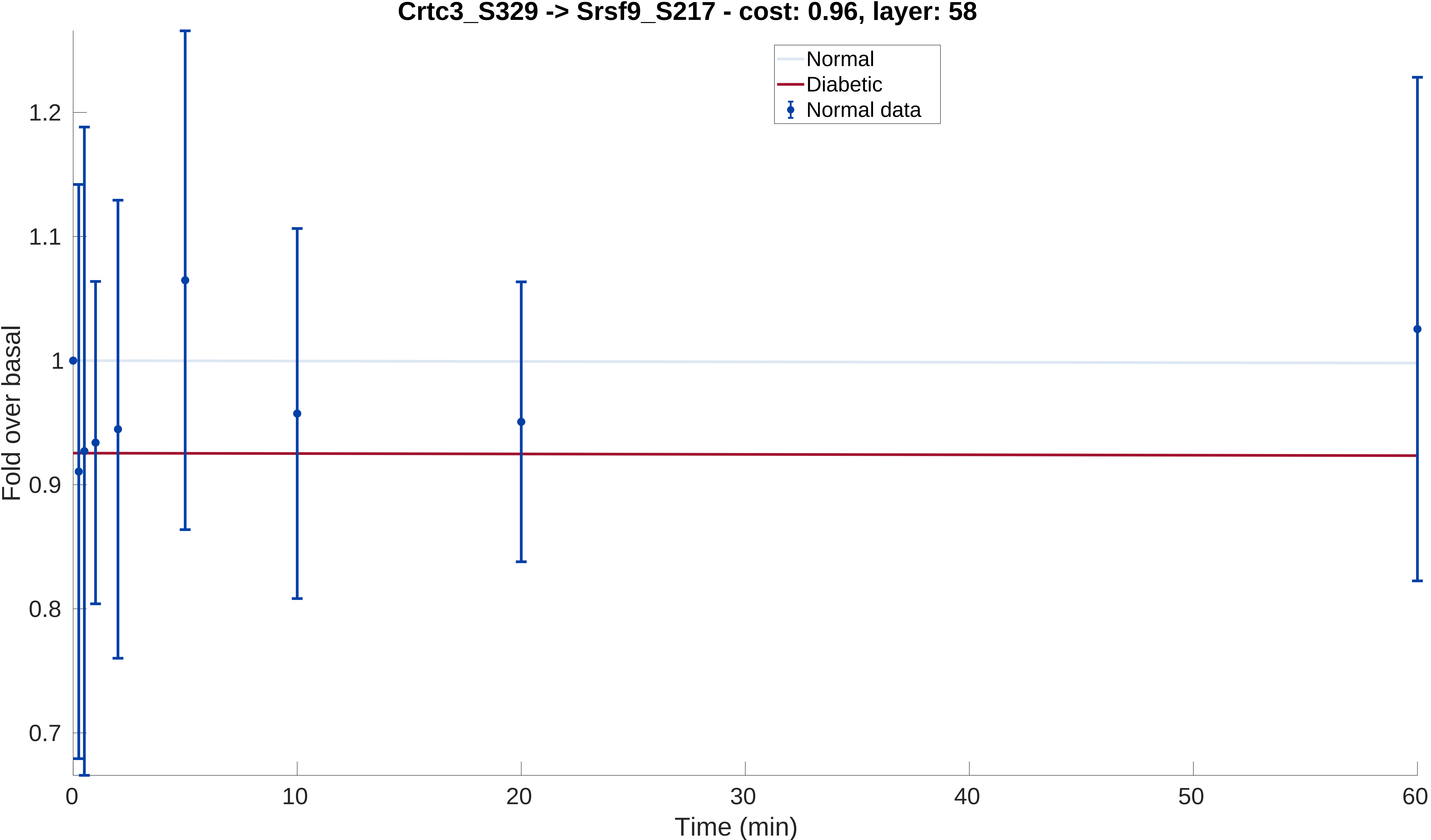
Crtc3_S329 -> Srsf9_S217 - cost: 0.96, layer: 58
Normal
Diabetic
Normal data
Fold over basal
Time (min)
1.2
1.1
1
0.9
0.8
0.7
0
10
20
30
40
50
60

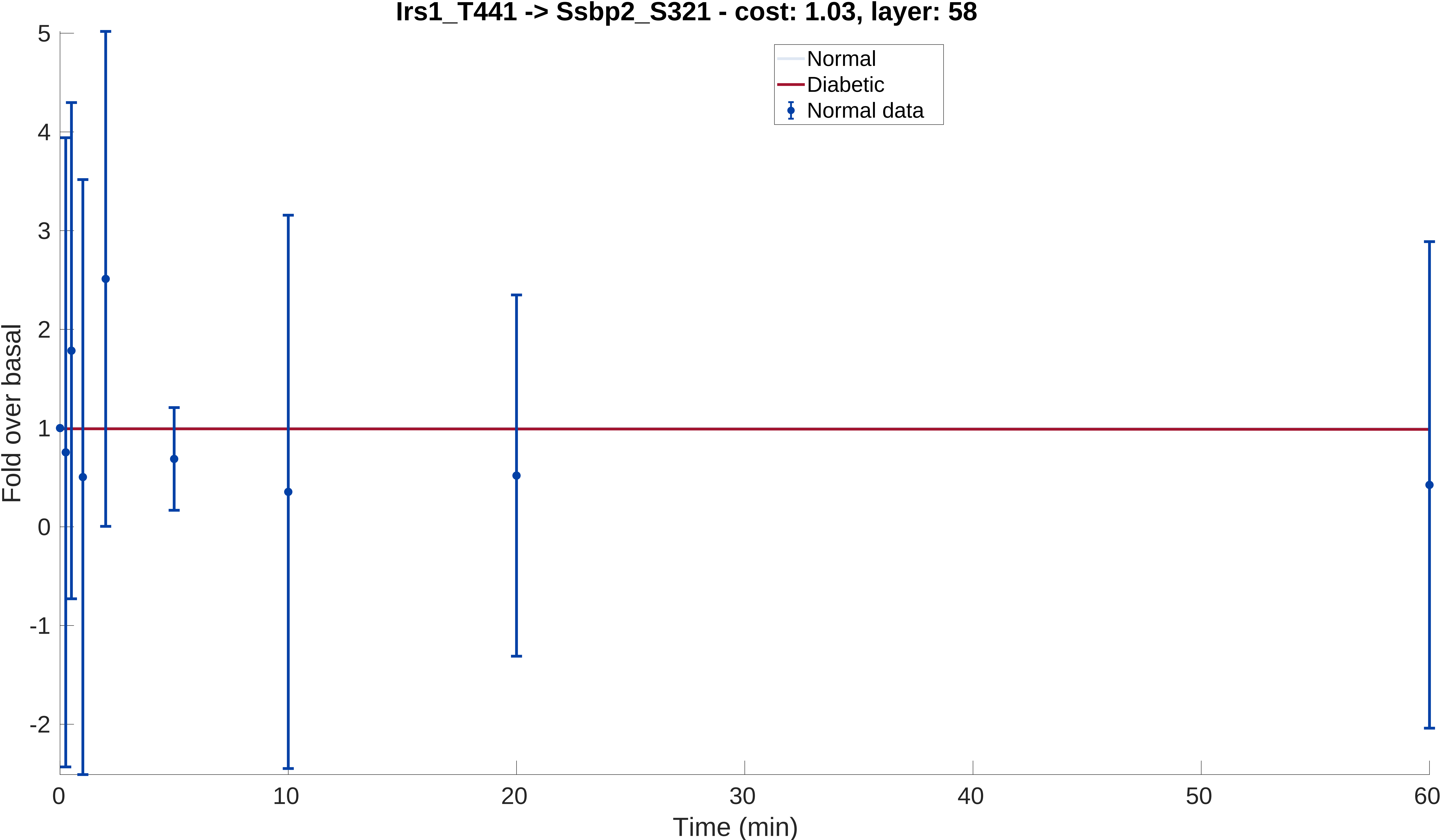
Irs1_T441 -> Ssbp2_S321 - cost: 1.03, layer: 58
Normal
Diabetic
Normal data
Fold over basal
Time (min)
5
4
3
2
1
0
-1
-2
0
10
20
30
40
50
60

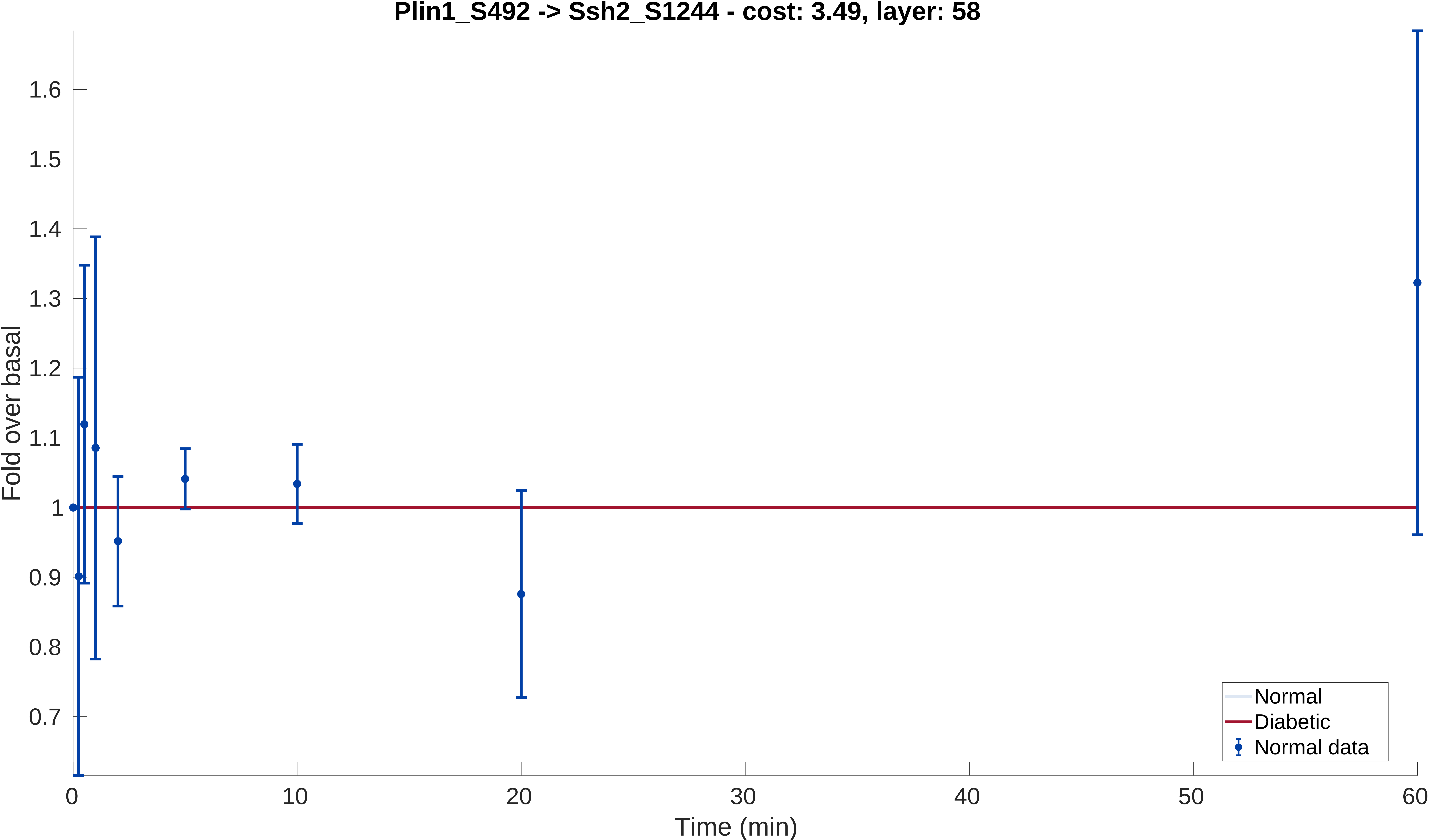

Plin1_S492 -> Ssh2_S1244 - cost: 3.49, layer: 58
Fold over basal
1.6
1.5
1.4
1.3
1.2
1.1
1
0.9
0.8
0.7
0
10
20
30
40
50
60
Time (min)
Normal
Diabetic
Normal data

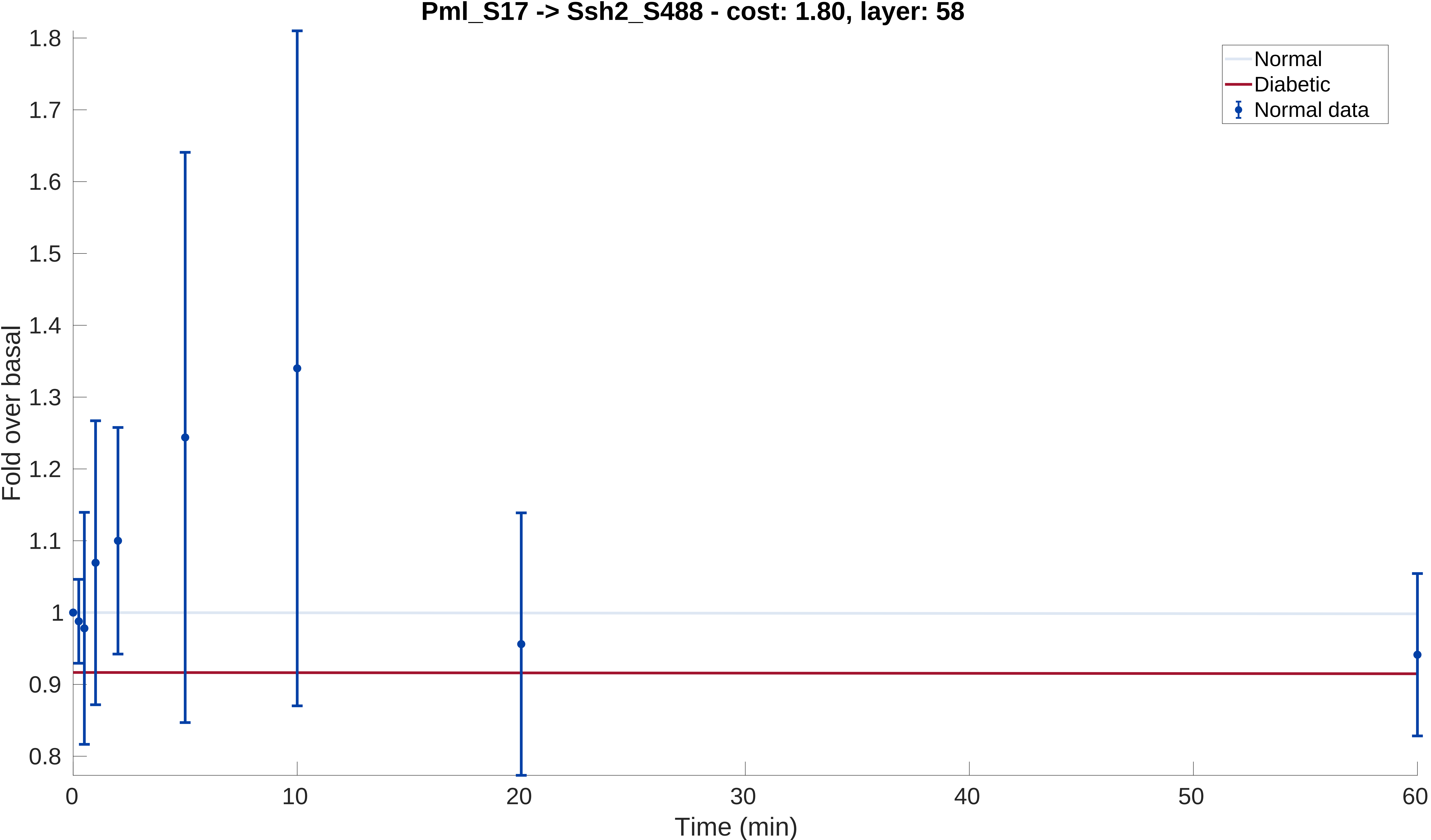
Pml_S17 -> Ssh2_S488 - cost: 1.80, layer: 58
Normal
Diabetic
Normal data
Fold over basal
Time (min)
1.8
1.7
1.6
1.5
1.4
1.3
1.2
1.1
1
0.9
0.8
0
10
20
30
40
50
60

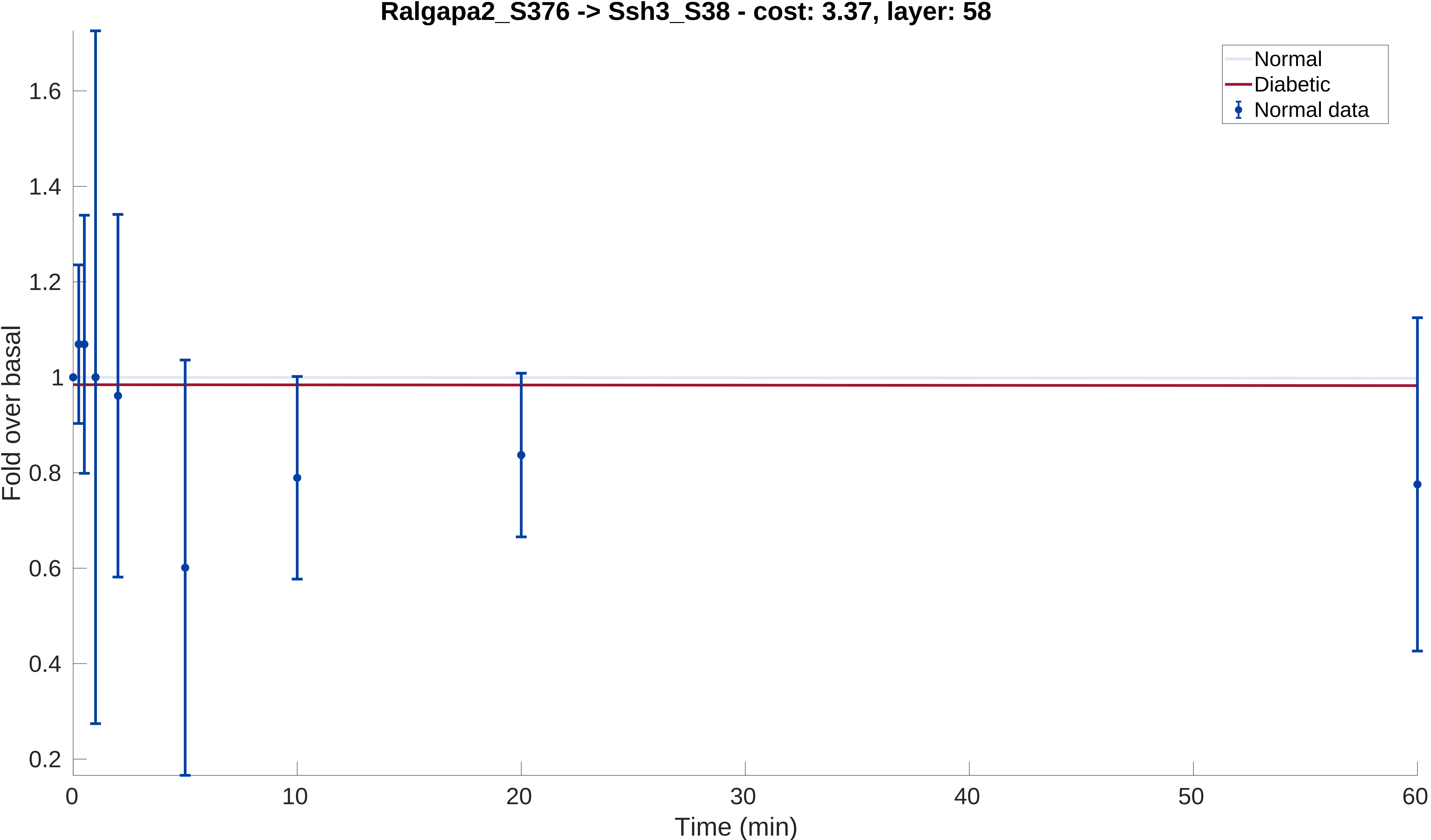

Ralgapa2_S376 -> Ssh3_S38 - cost: 3.37, layer: 58
Normal
Diabetic
Normal data
Fold over basal
1.6
1.4
1.2
1
0.8
0.6
0.4
0.2
0
10
20
30
40
50
60
Time (min)

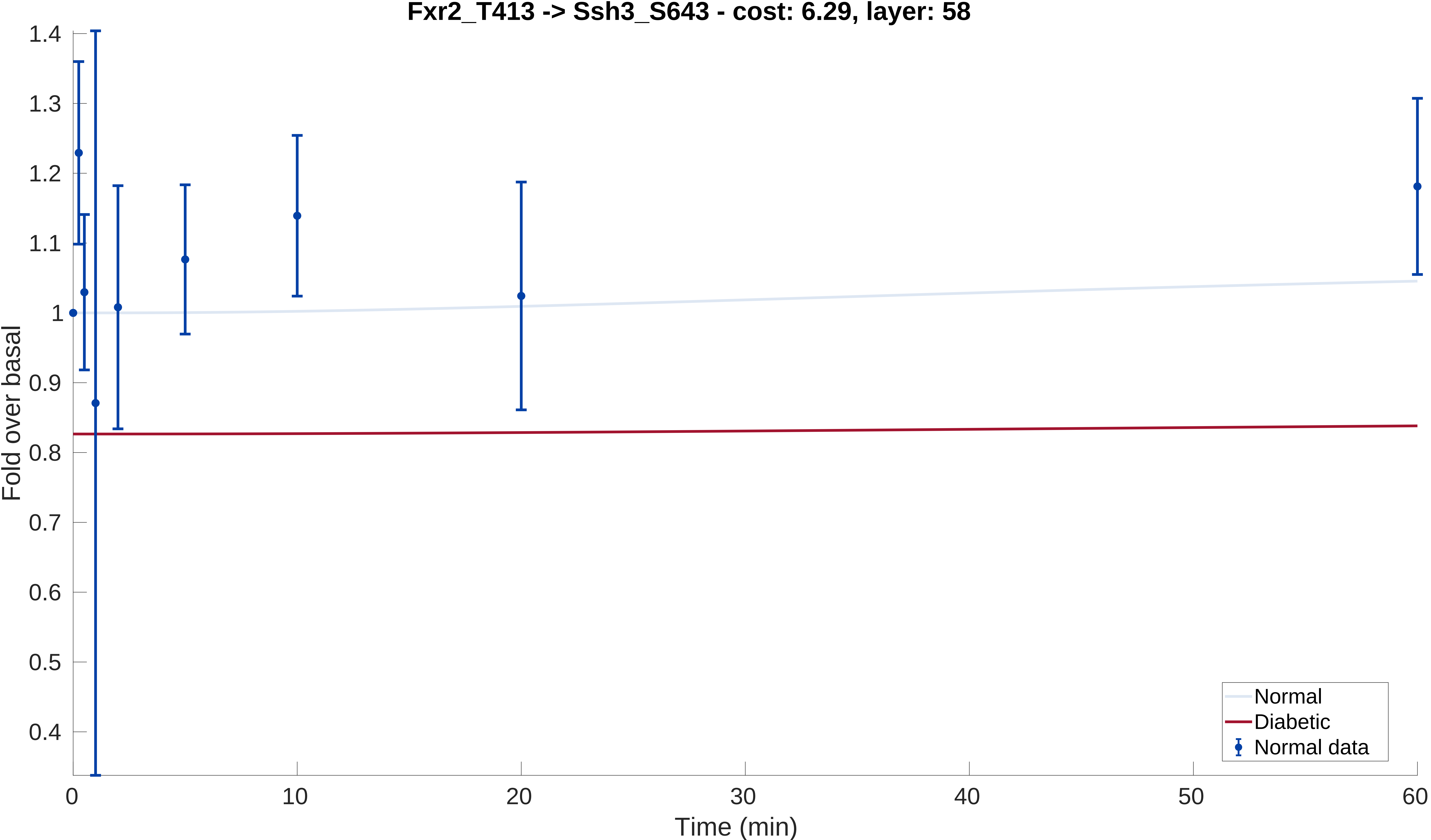

Fxr2_T413 -> Ssh3_S643 - cost: 6.29, layer: 58
Fold over basal
Time (min)
1.4
1.3
1.2
1.1
1
0.9
0.8
0.7
0.6
0.5
0.4
0
10
20
30
40
50
60
Normal
Diabetic
Normal data

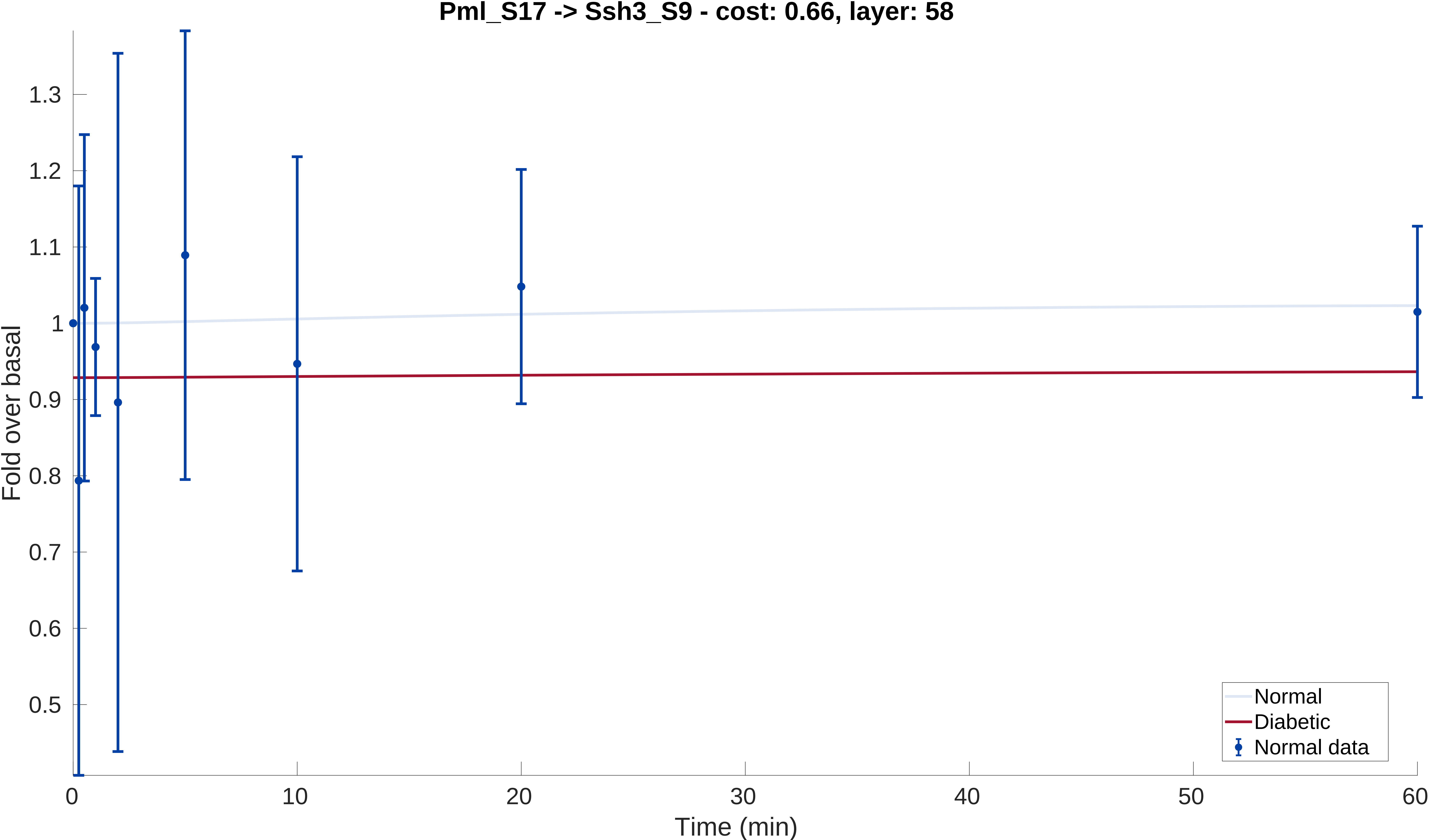

Pml_S17 -> Ssh3_S9 - cost: 0.66, layer: 58
Fold over basal
1.3
1.2
1.1
1
0.9
0.8
0.7
0.6
0.5
0
10
20
30
40
50
60
Time (min)
Normal
Diabetic
Normal data

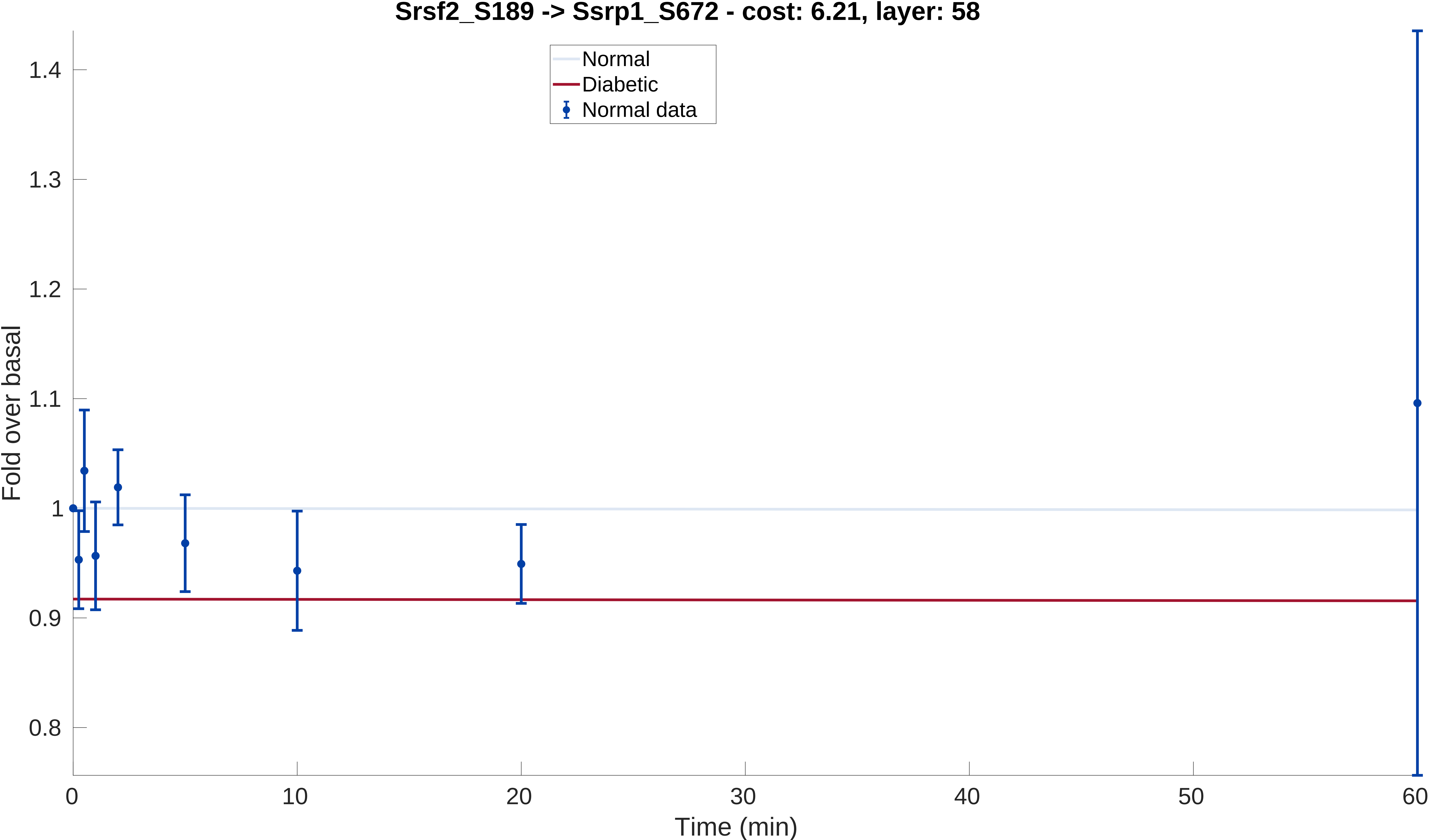

Srsf2_S189 -> Ssrp1_S672 - cost: 6.21, layer: 58
Normal
Diabetic
Normal data
Fold over basal
1.4
1.3
1.2
1.1
1
0.9
0.8
0
10
20
30
40
50
60
Time (min)

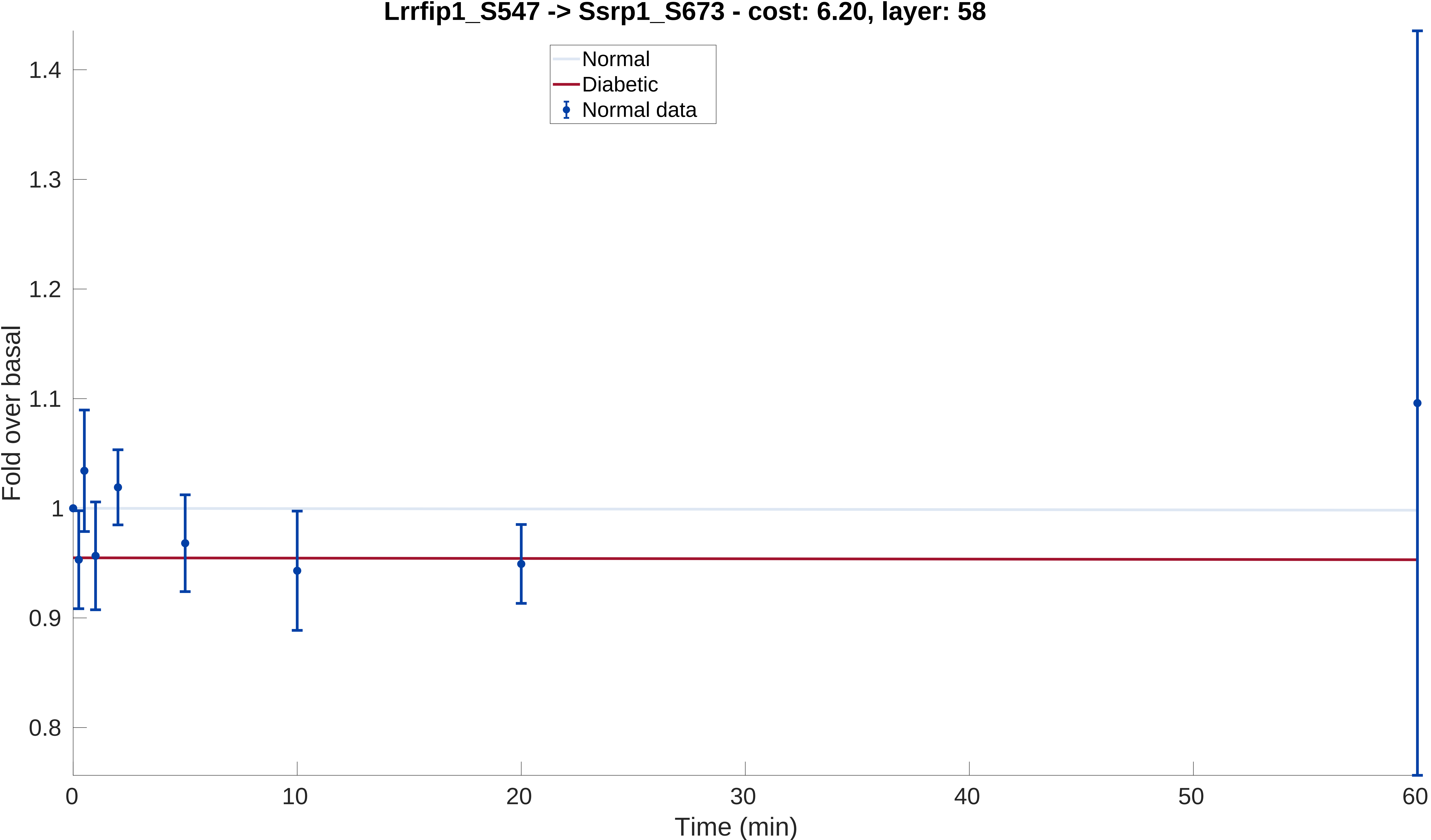

Lrrfip1_S547 -> Ssrp1_S673 - cost: 6.20, layer: 58
Normal
Diabetic
Normal data
Fold over basal
Time (min)
1.4
1.3
1.2
1.1
1
0.9
0.8
0
10
20
30
40
50
60

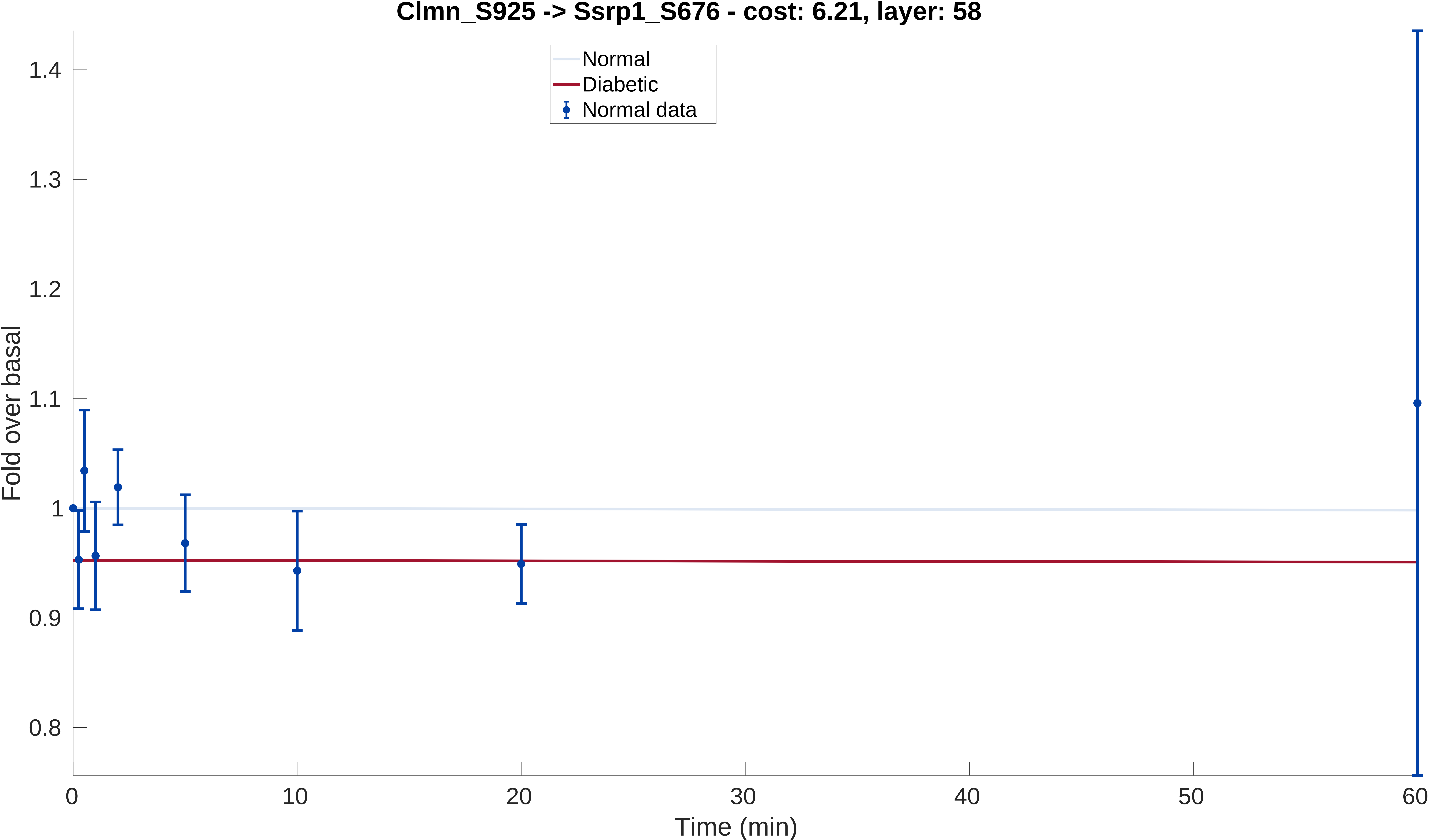
Clmn_S925 -> Ssrp1_S676 - cost: 6.21, layer: 58
Normal
Diabetic
Normal data
Fold over basal
1.4
1.3
1.2
1.1
1
0.9
0.8
0
10
20
30
40
50
60
Time (min)

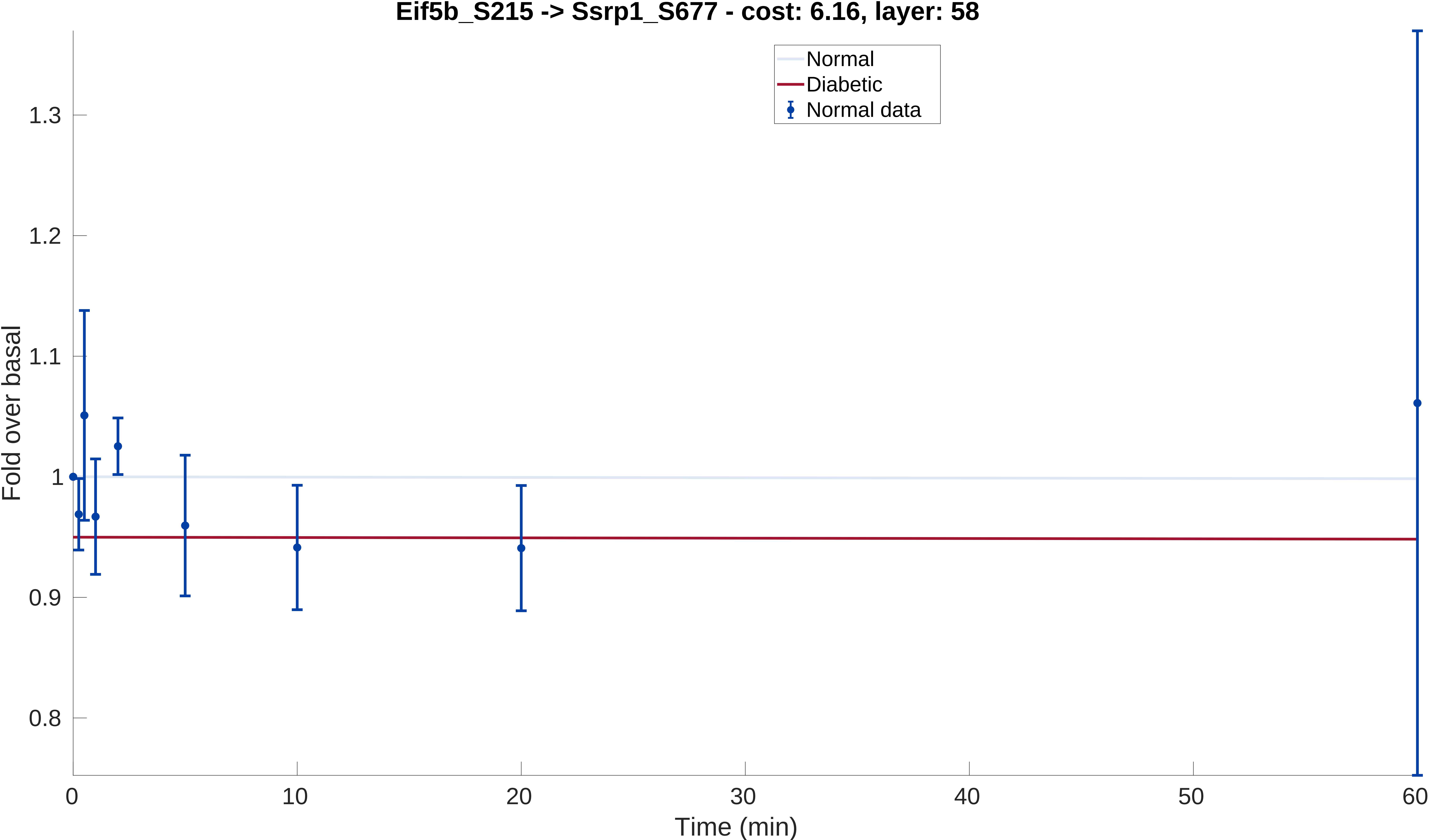

Eif5b_S215 -> Ssrp1_S677 - cost: 6.16, layer: 58
Normal
Diabetic
Normal data
Fold over basal
1.3
1.2
1.1
1
0.9
0.8
0
10
20
30
40
50
60
Time (min)

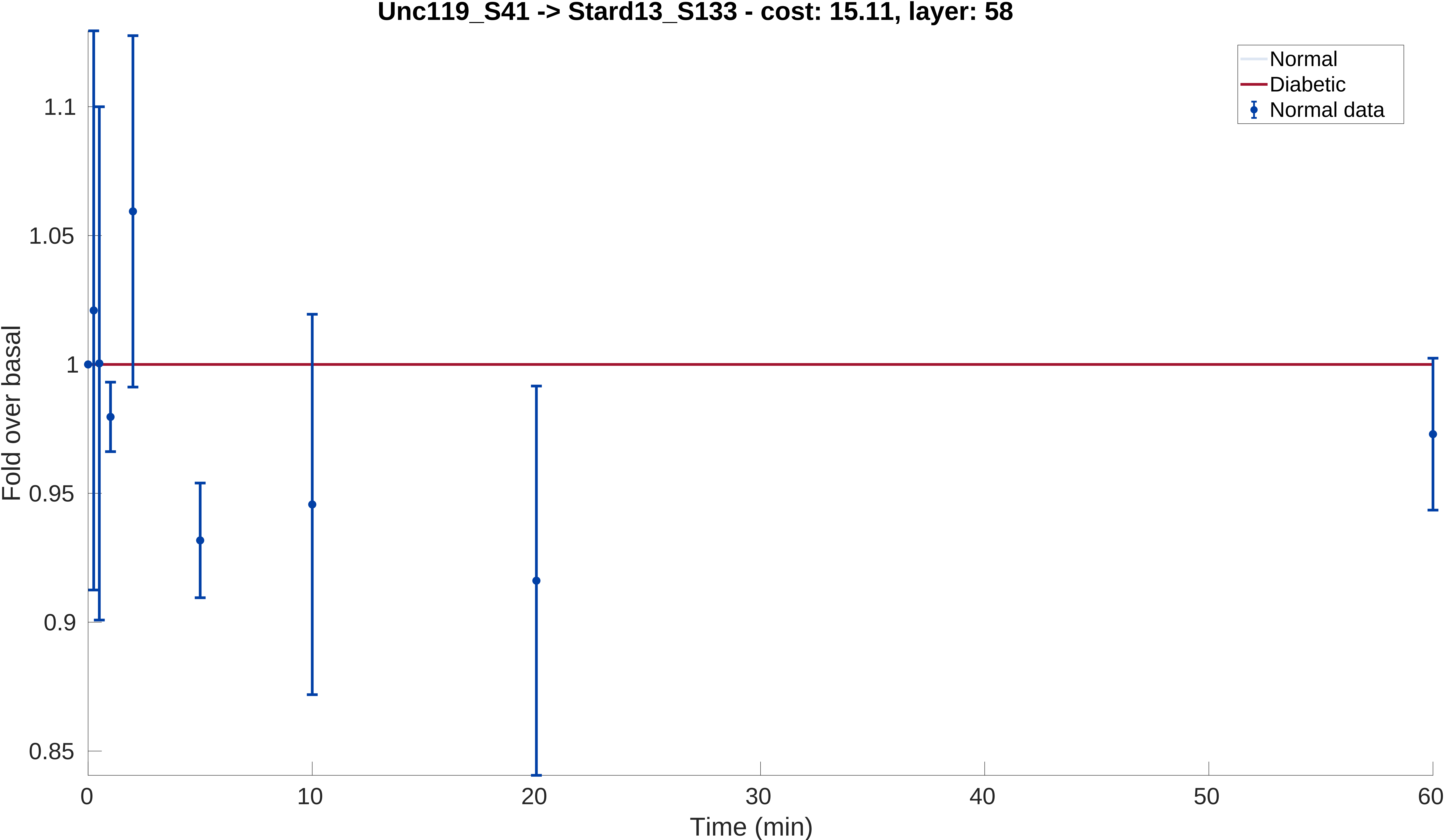
Unc119_S41 -> Stard13_S133 - cost: 15.11, layer: 58
Normal
Diabetic
Normal data
Fold over basal
1.1
1.05
1
0.95
0.9
0.85
0
10
20
30
40
50
60
Time (min)

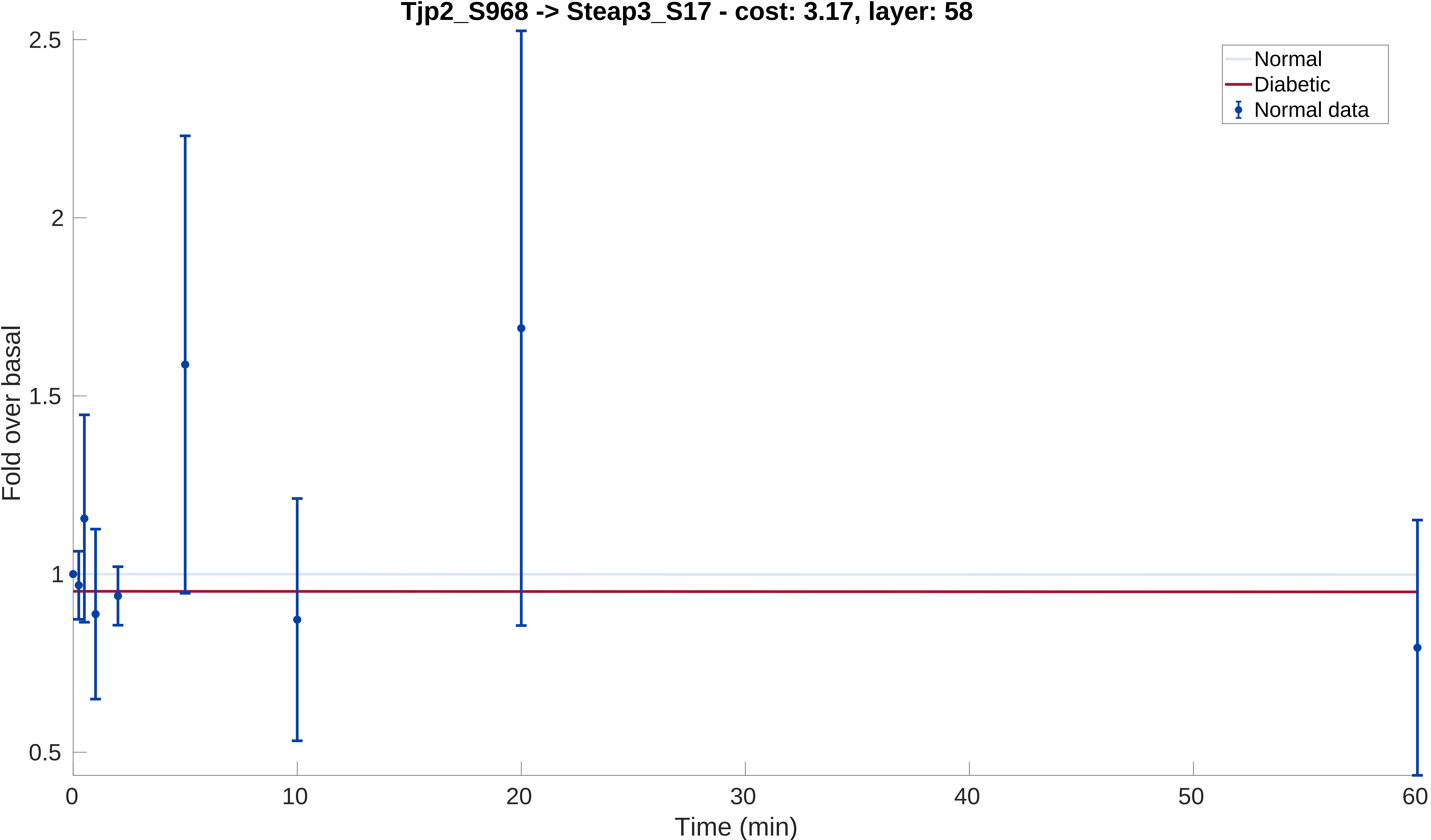

Tjp2_S968 -> Steap3_S17 - cost: 3.17, layer: 58
Normal
Diabetic
Normal data
Fold over basal
2.5
2
1.5
1
0.5
0
10
20
30
40
50
60
Time (min)

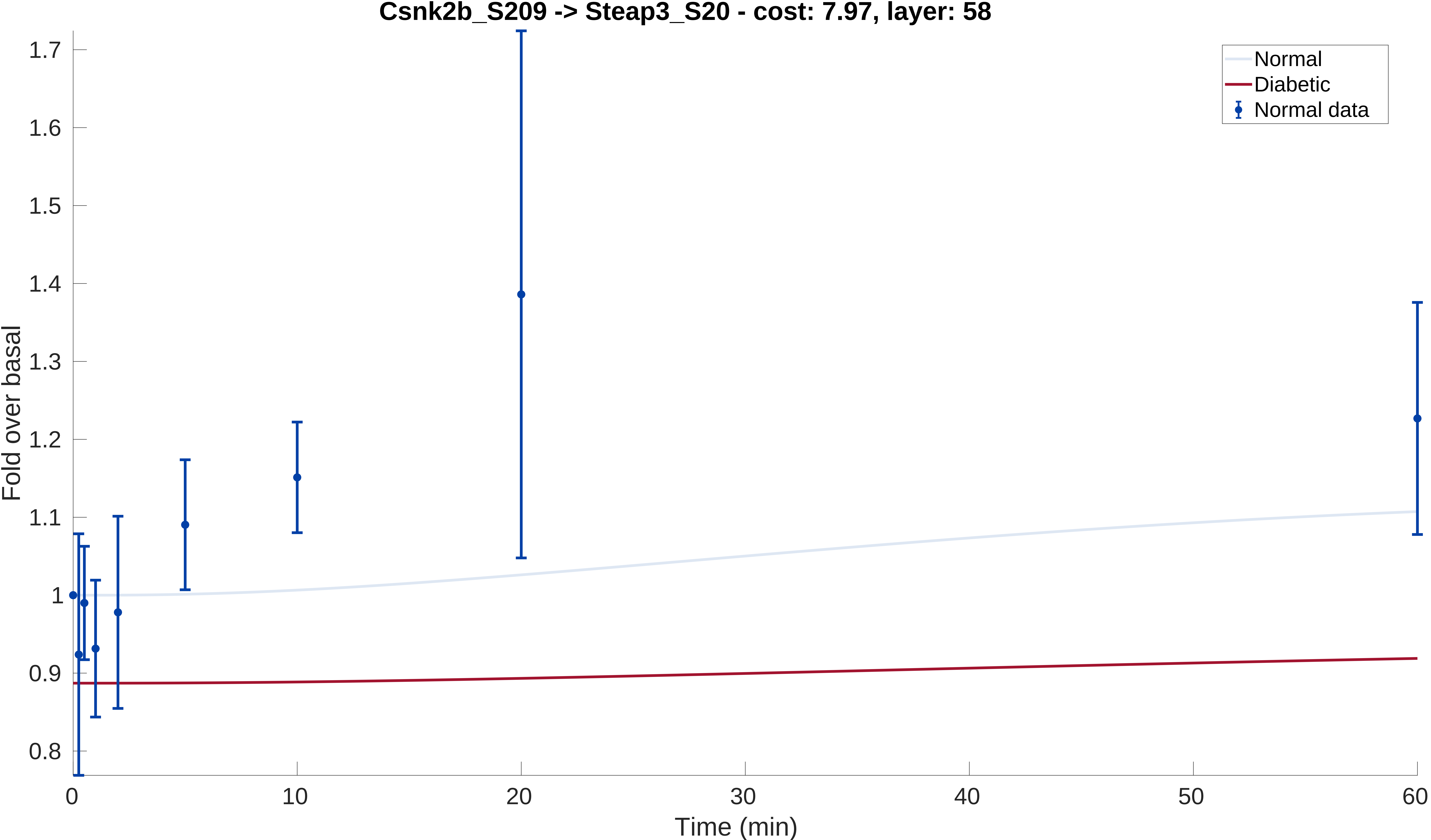

Csnk2b_S209 -> Steap3_S20 - cost: 7.97, layer: 58
Normal
Diabetic
Normal data
Fold over basal
Time (min)
1.7
1.6
1.5
1.4
1.3
1.2
1.1
1
0.9
0.8
0
10
20
30
40
50
60

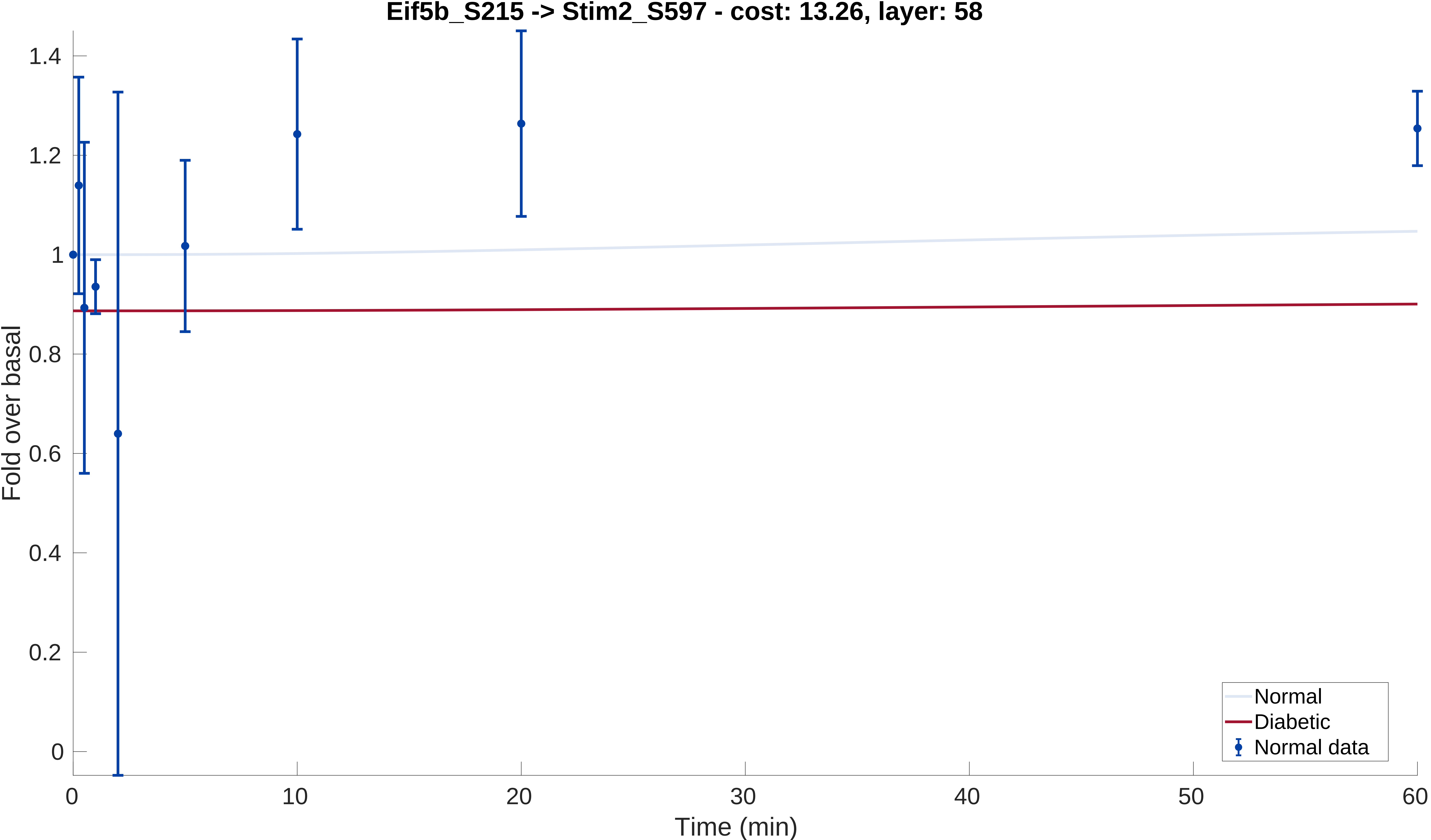
Eif5b_S215 -> Stim2_S597 - cost: 13.26, layer: 58
Fold over basal
1.4
1.2
1
0.8
0.6
0.4
0.2
0
Time (min)
0
10
20
30
40
50
60
Normal
Diabetic
Normal data

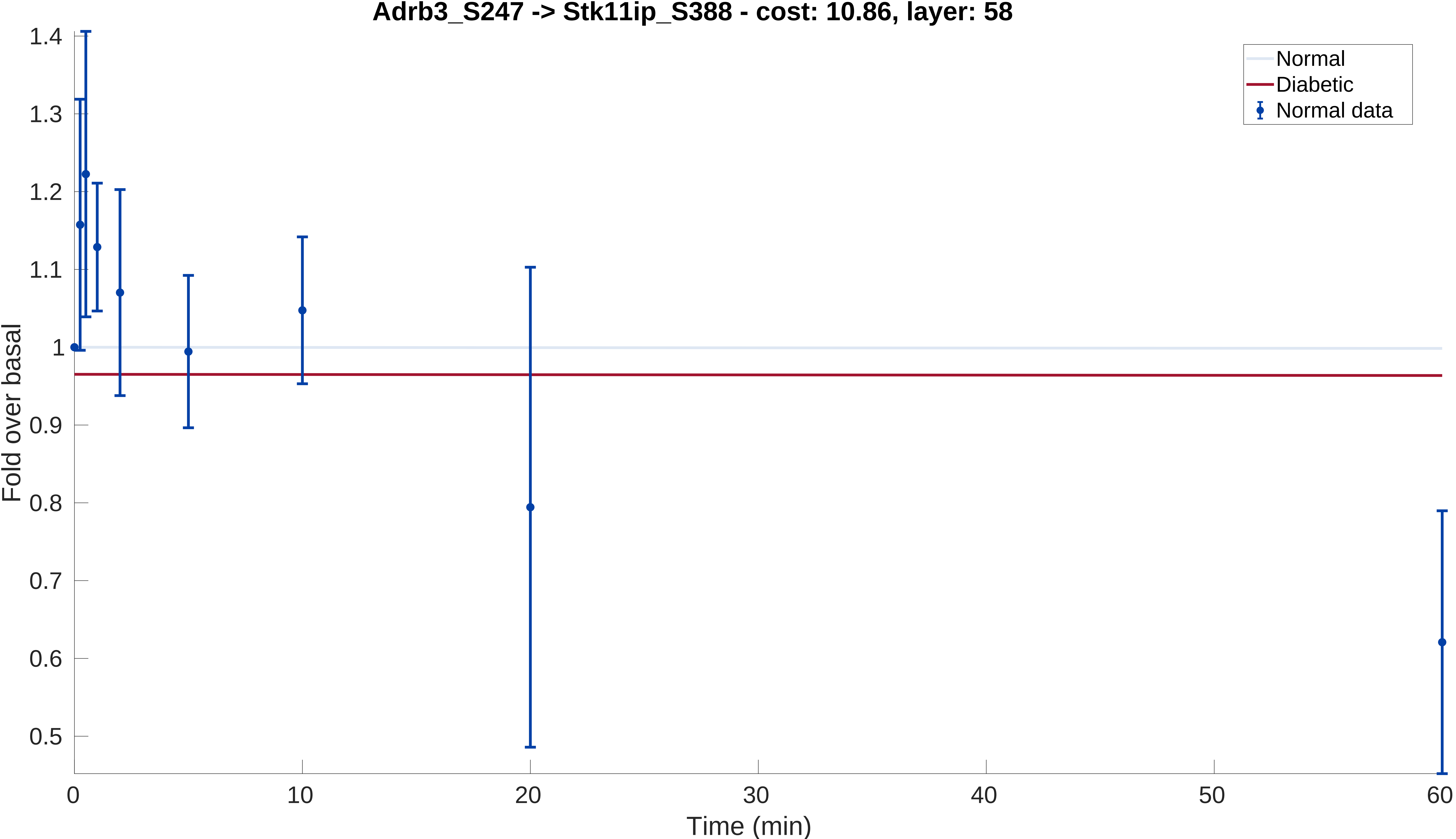

Adrb3_S247 -> Stk11ip_S388 - cost: 10.86, layer: 58
Fold over basal
Time (min)
Normal
Diabetic
Normal data
1.4
1.3
1.2
1.1
1
0.9
0.8
0.7
0.6
0.5
0
10
20
30
40
50
60

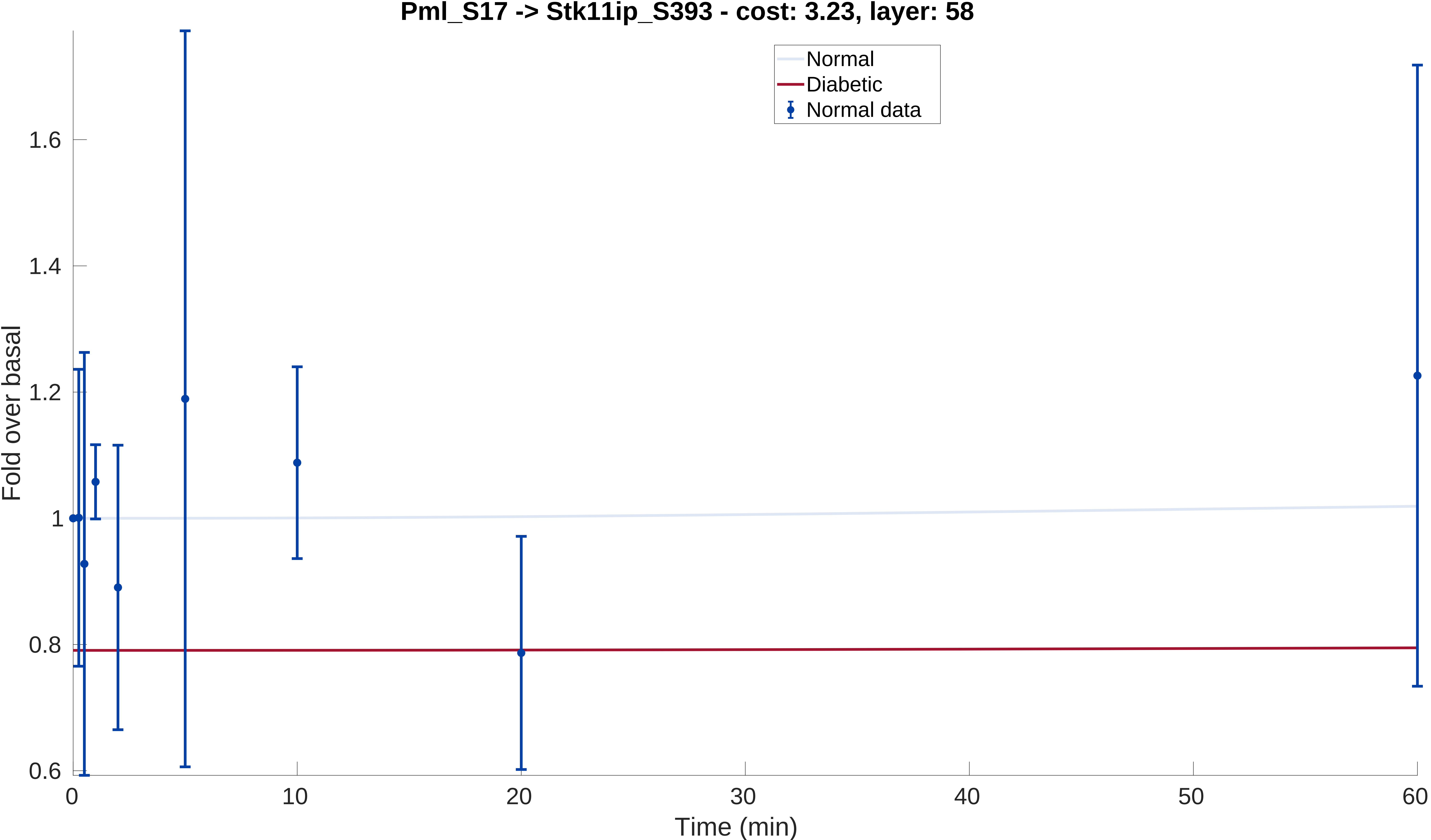

Pml_S17 -> Stk11ip_S393 - cost: 3.23, layer: 58
Normal
Diabetic
Normal data
Fold over basal
1.6
1.4
1.2
1
0.8
0.6
0
10
20
30
40
50
60
Time (min)

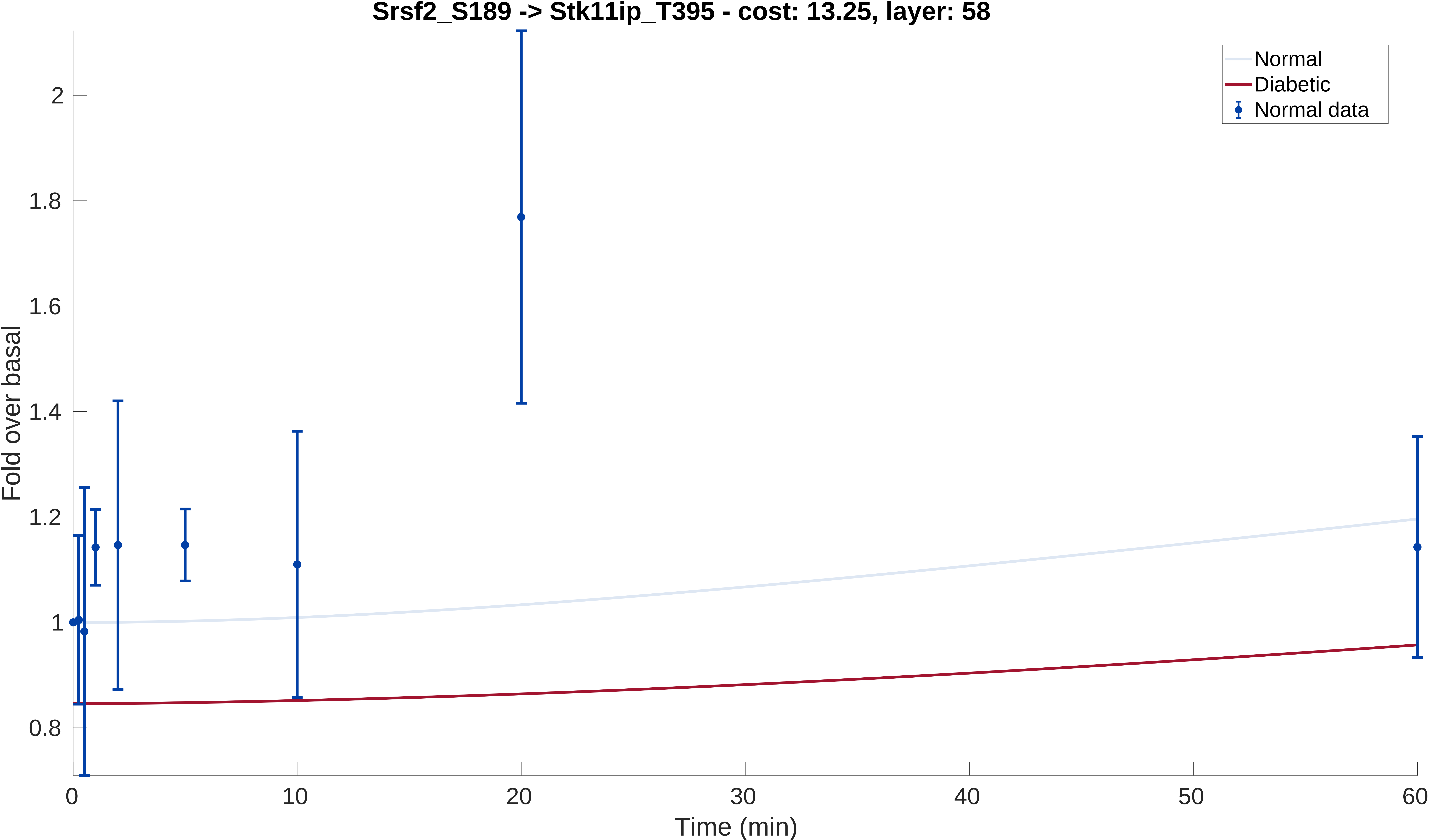
Srsf2_S189 -> Stk11ip_T395 - cost: 13.25, layer: 58
Normal
Diabetic
Normal data
Fold over basal
2
1.8
1.6
1.4
1.2
1
0.8
0
10
20
30
40
50
60
Time (min)

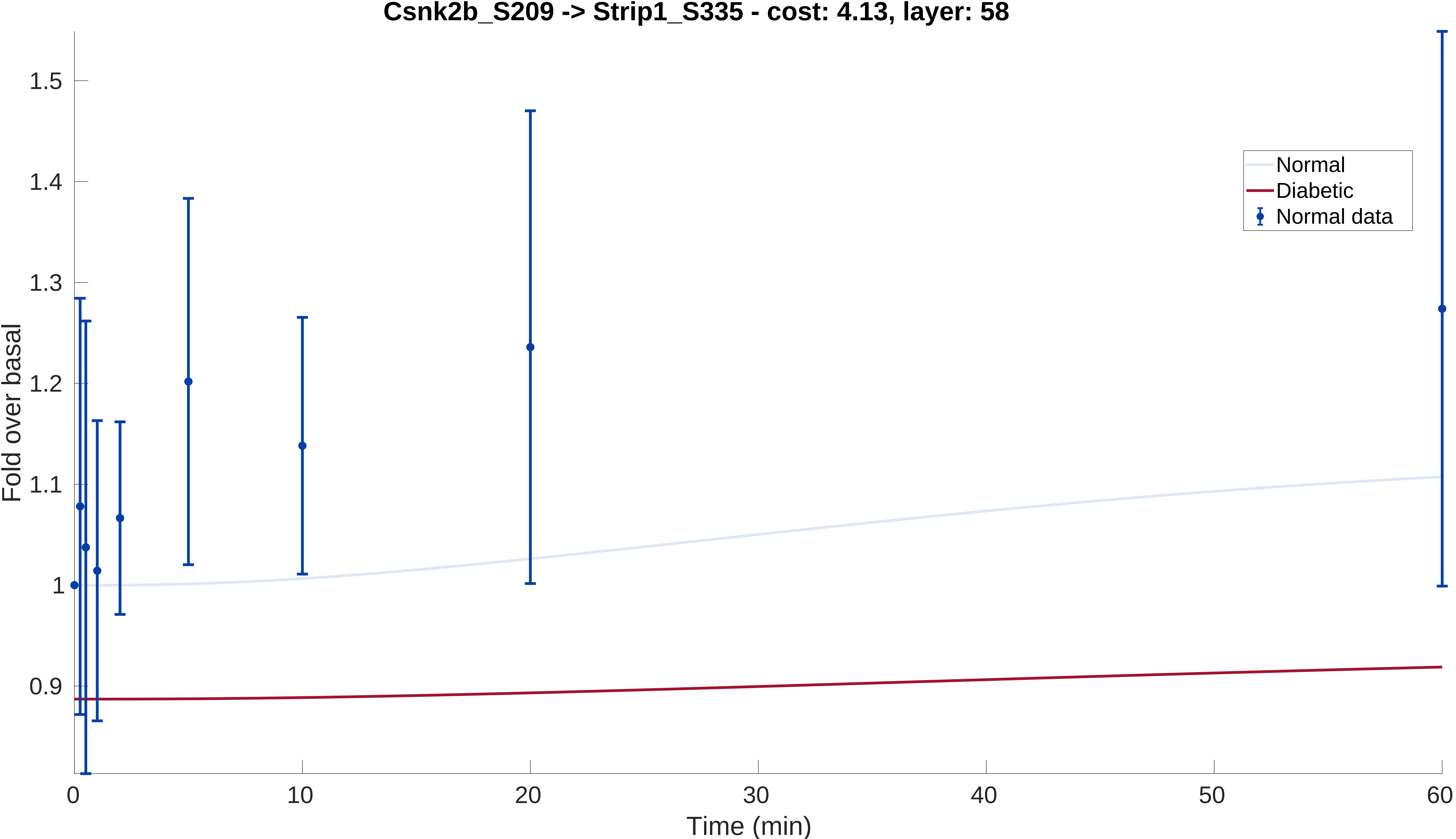
Csnk2b_S209 -> Strip1_S335 - cost: 4.13, layer: 58
Fold over basal
1.5
1.4
1.3
1.2
1.1
1
0.9
0
10
20
30
40
50
60
Time (min)
Normal
Diabetic
Normal data

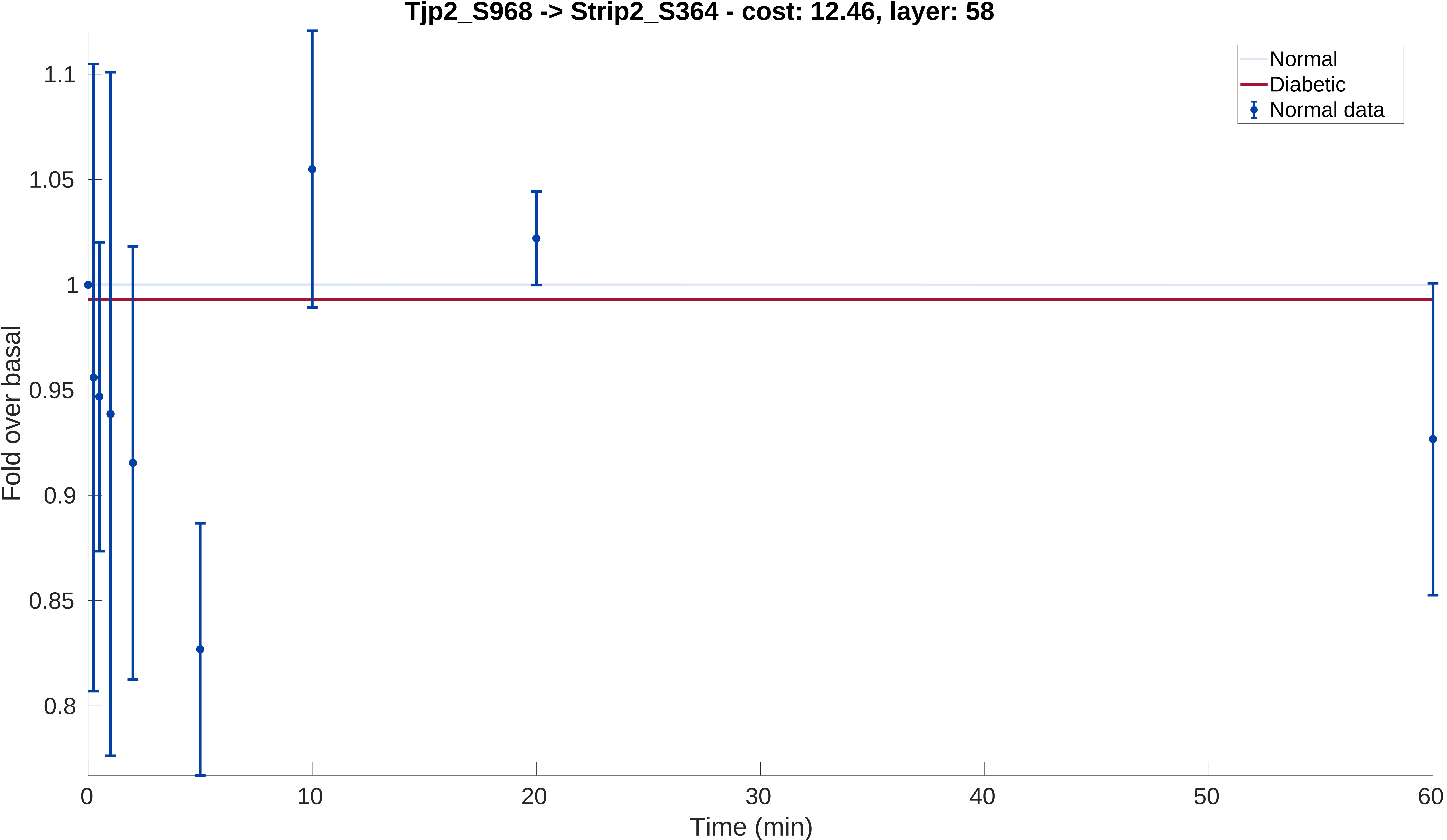

Tjp2_S968 -> Strip2_S364 - cost: 12.46, layer: 58
Normal
Diabetic
Normal data
Fold over basal
1.1
1.05
1
0.95
0.9
0.85
0.8
0
10
20
30
40
50
60
Time (min)

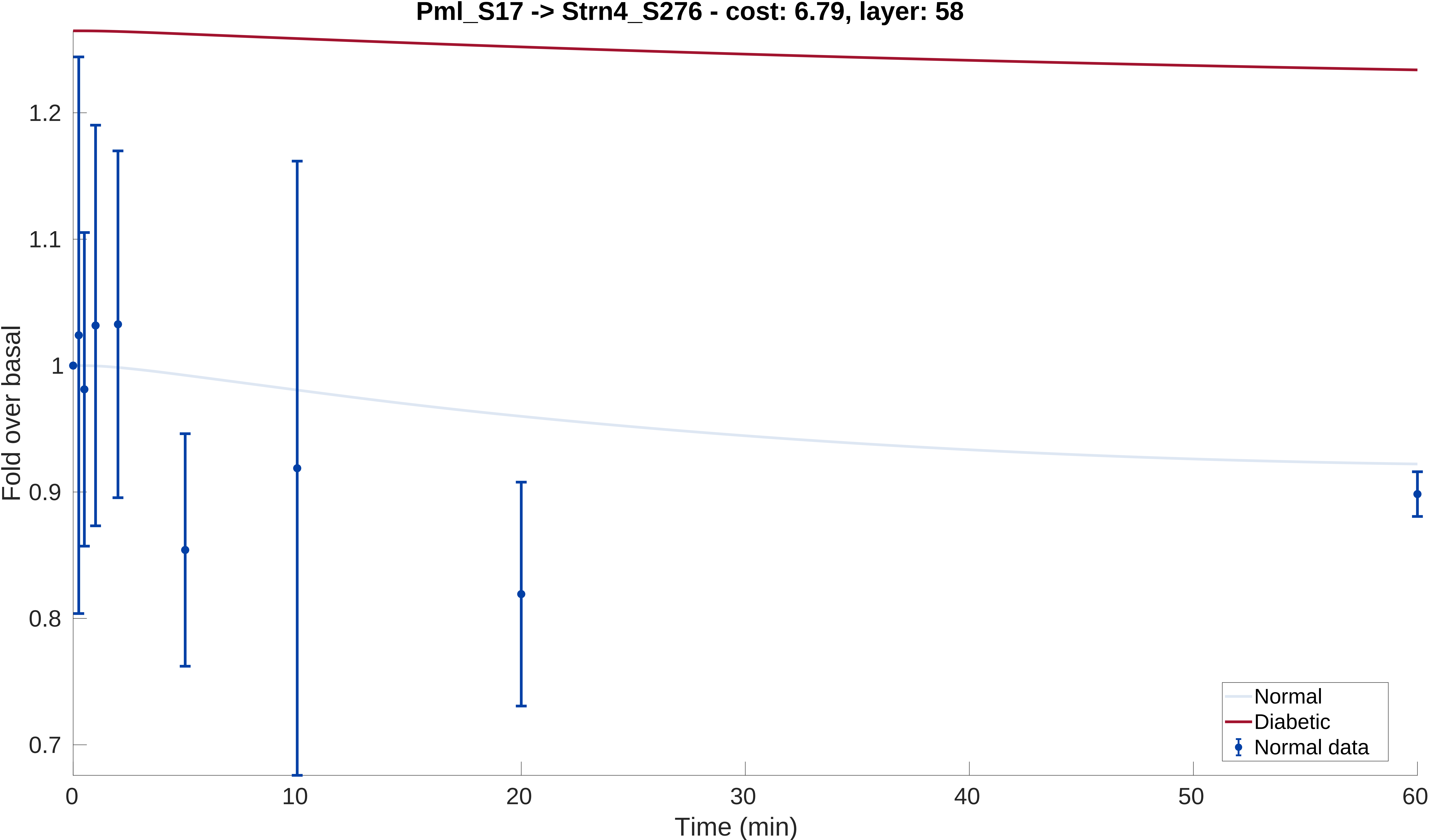
Pml_S17 -> Strn4_S276 - cost: 6.79, layer: 58
Fold over basal
1.2
1.1
1
0.9
0.8
0.7
0
10
20
30
40
50
60
Time (min)
Normal
Diabetic
Normal data

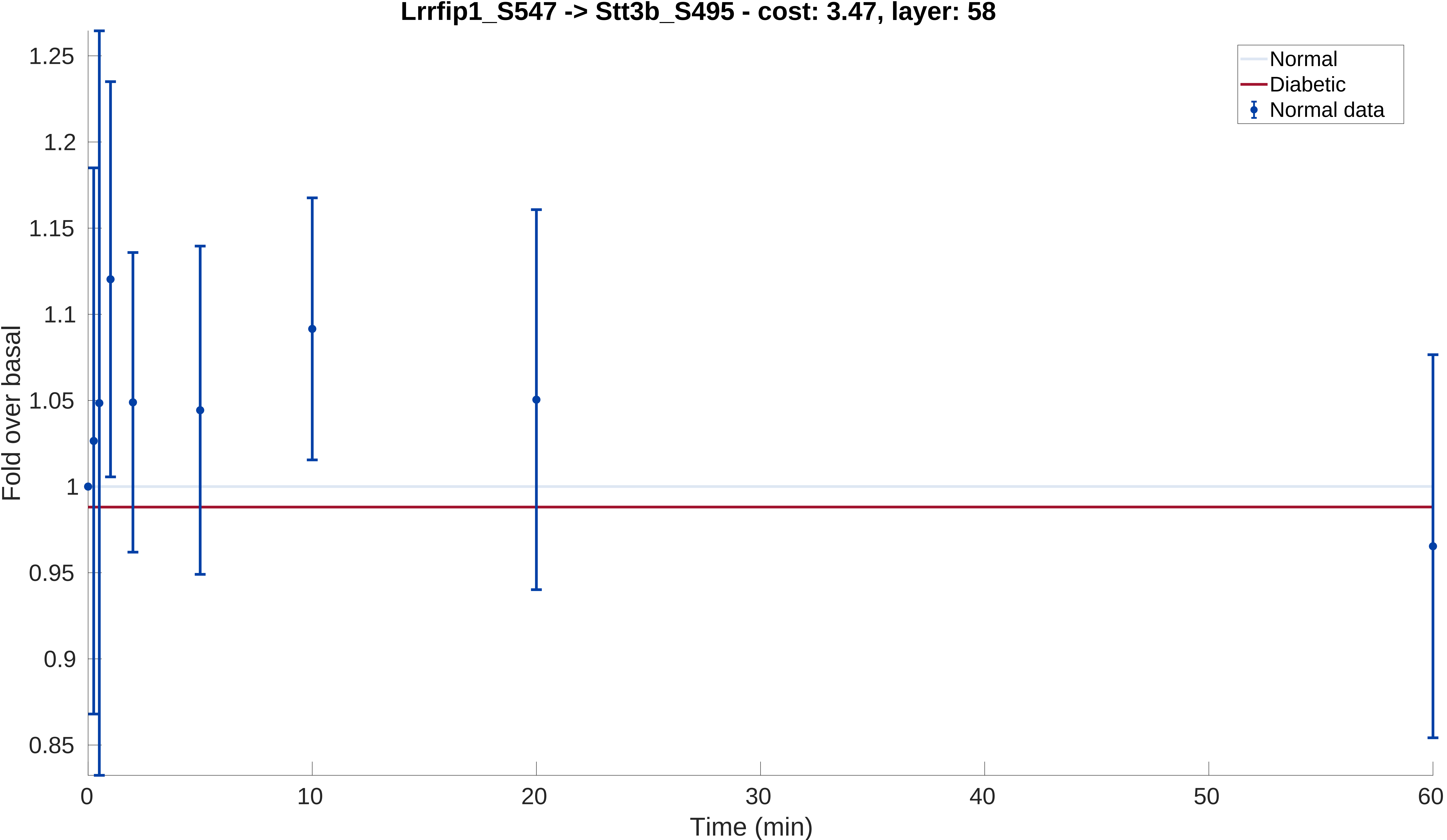

Lrrfip1_S547 -> Stt3b_S495 - cost: 3.47, layer: 58
Normal
Diabetic
Normal data
Fold over basal
Time (min)
1.25
1.2
1.15
1.1
1.05
1
0.95
0.9
0.85
0
10
20
30
40
50
60

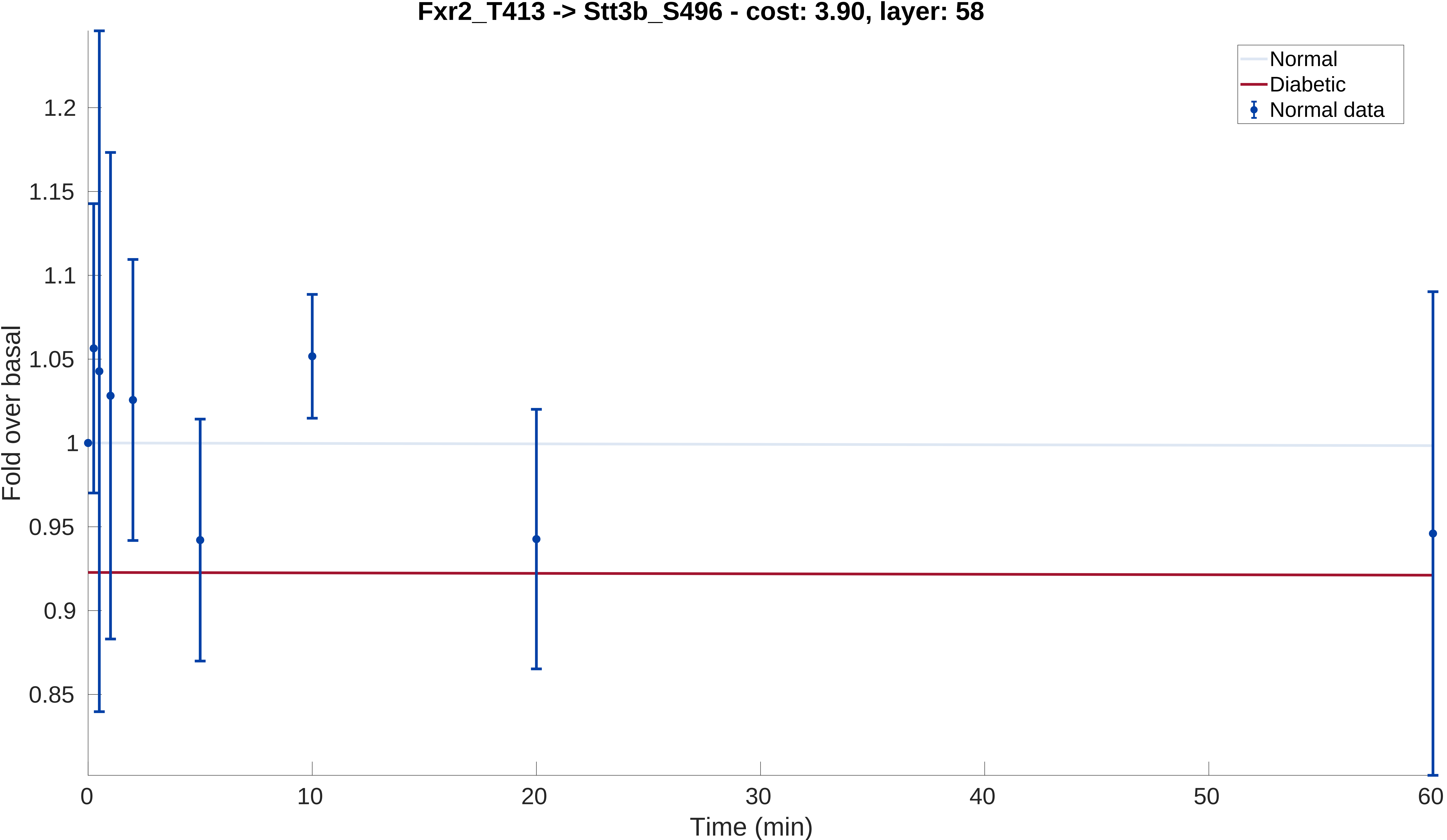
Fxr2_T413 -> Stt3b_S496 - cost: 3.90, layer: 58
Normal
Diabetic
Normal data
1.2
1.15
1.1
1.05
1
0.95
0.9
0.85
Fold over basal
0
10
20
30
40
50
60
Time (min)

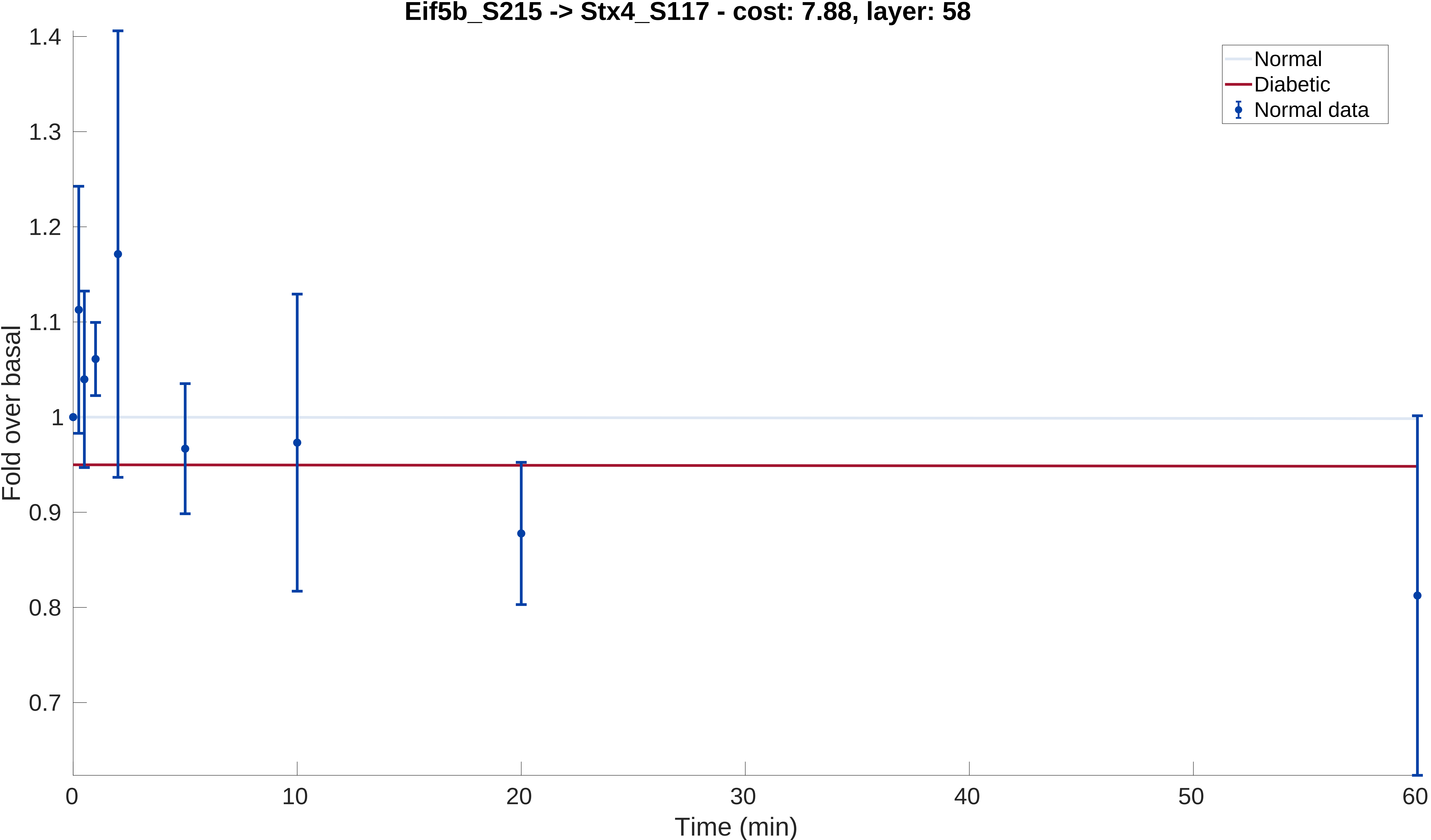

Eif5b_S215 -> Stx4_S117 - cost: 7.88, layer: 58
Normal
Diabetic
Normal data
Fold over basal
1.4
1.3
1.2
1.1
1
0.9
0.8
0.7
0
10
20
30
40
50
60
Time (min)

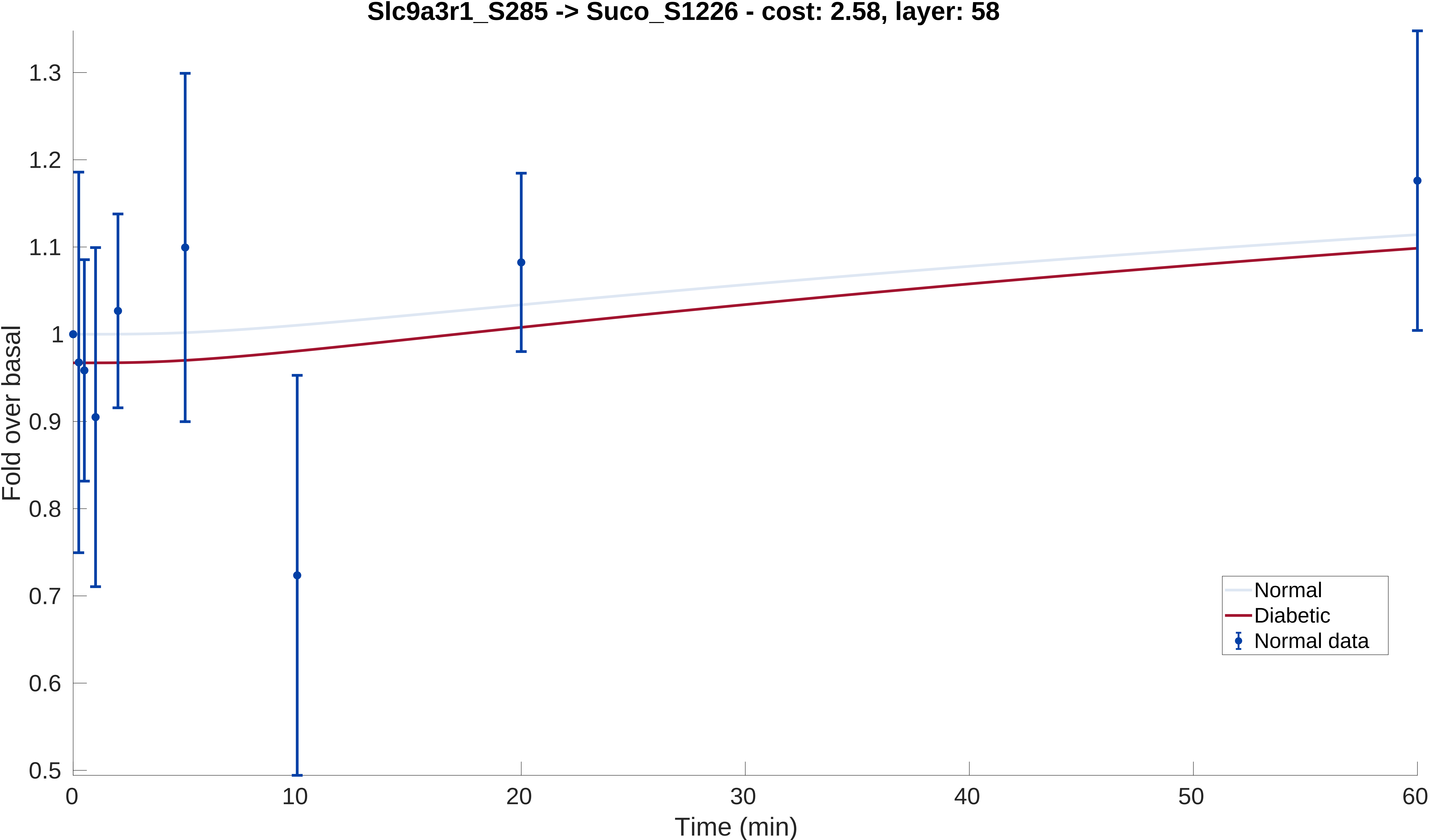
Slc9a3r1_S285 -> Suco_S1226 - cost: 2.58, layer: 58
Fold over basal
Time (min)
1.3
1.2
1.1
1
0.9
0.8
0.7
0.6
0.5
0
10
20
30
40
50
60
Normal
Diabetic
Normal data

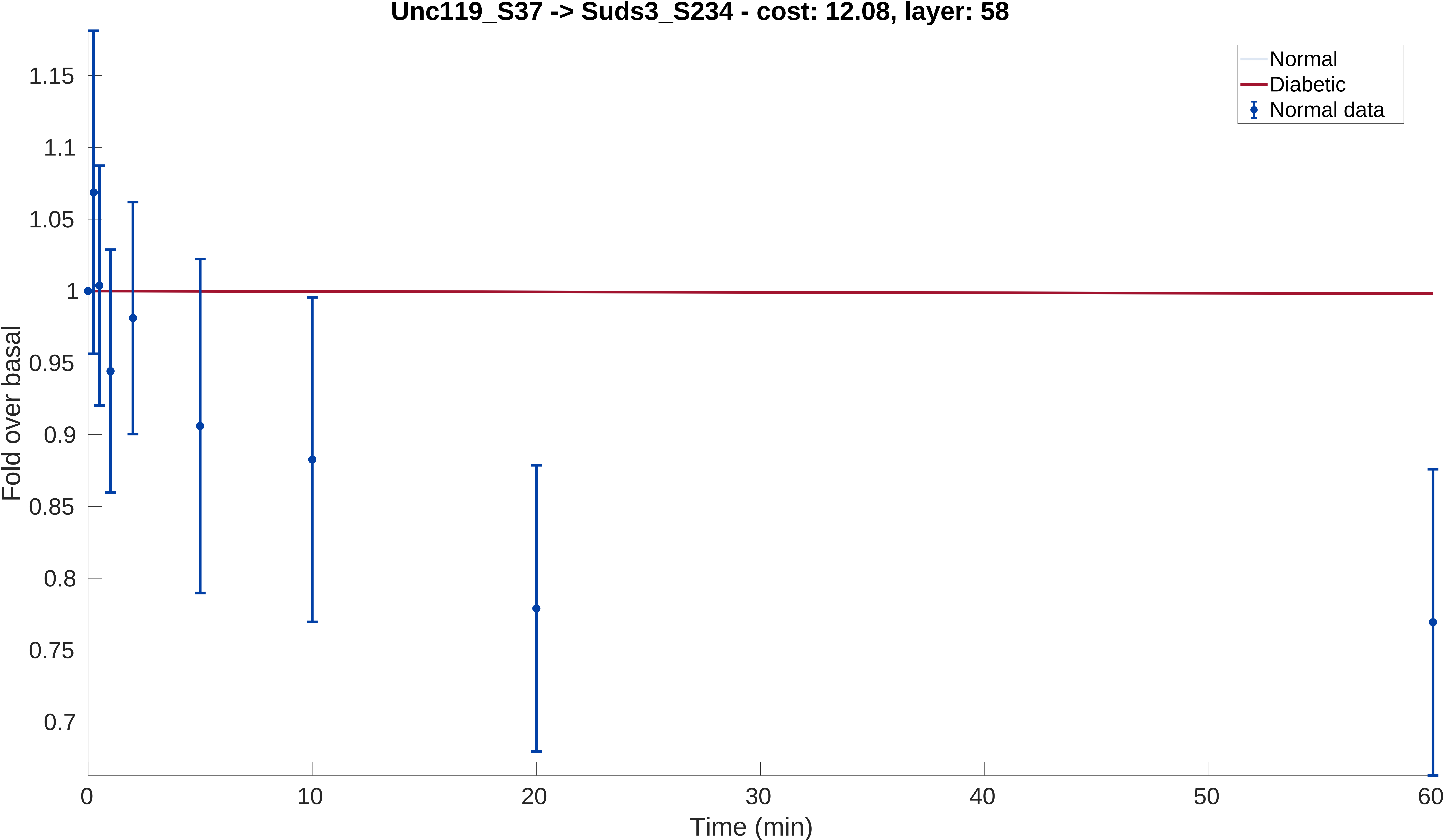
Unc119_S37 -> Suds3_S234 - cost: 12.08, layer: 58
Normal
Diabetic
Normal data
Fold over basal
1.15
1.1
1.05
1
0.95
0.9
0.85
0.8
0.75
0.7
0
10
20
30
40
50
60
Time (min)

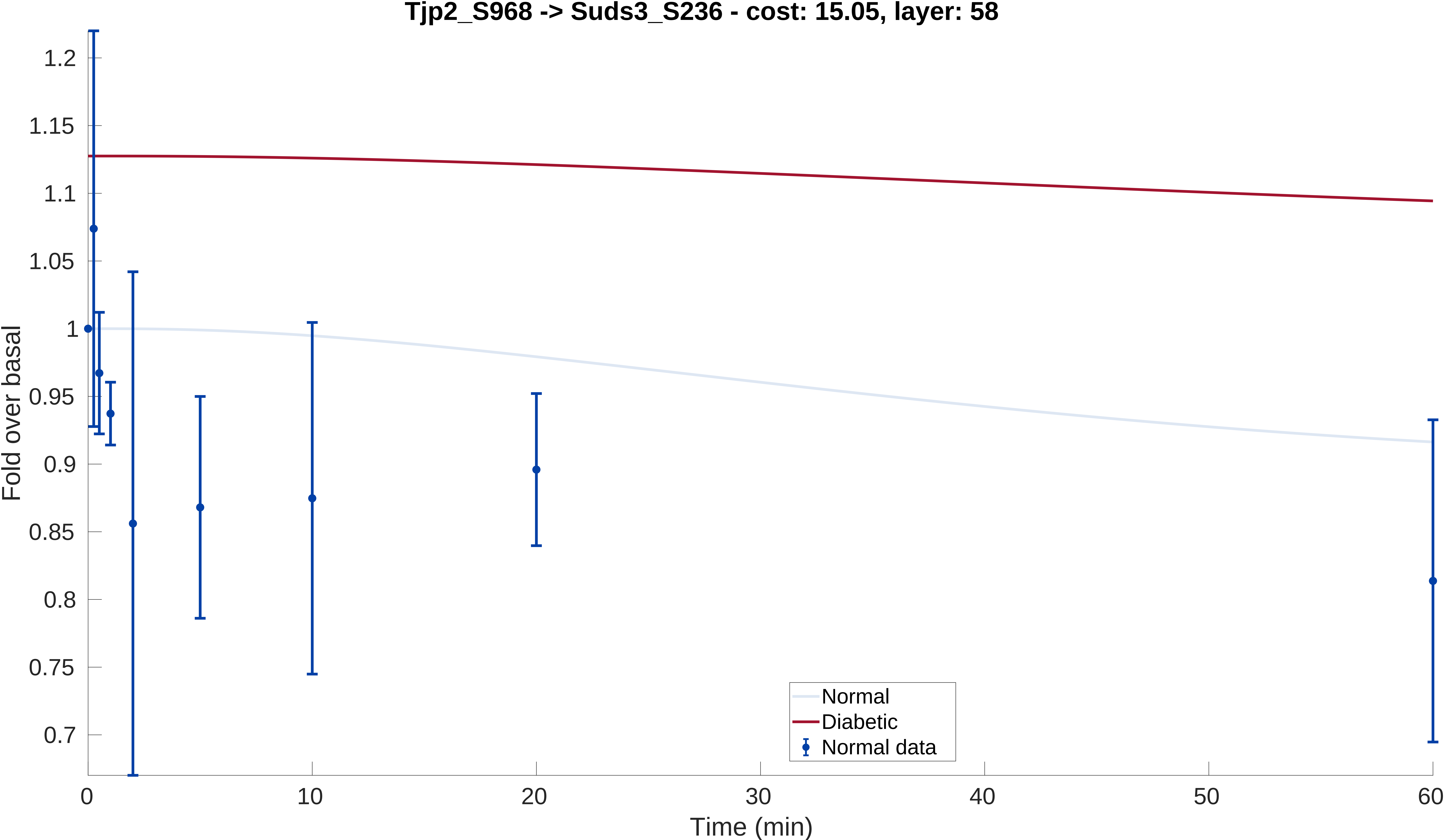

Tjp2_S968 -> Suds3_S236 - cost: 15.05, layer: 58
Fold over basal
1.2
1.15
1.1
1.05
1
0.95
0.9
0.85
0.8
0.75
0.7
0
10
20
30
40
50
60
Time (min)
Normal
Diabetic
Normal data

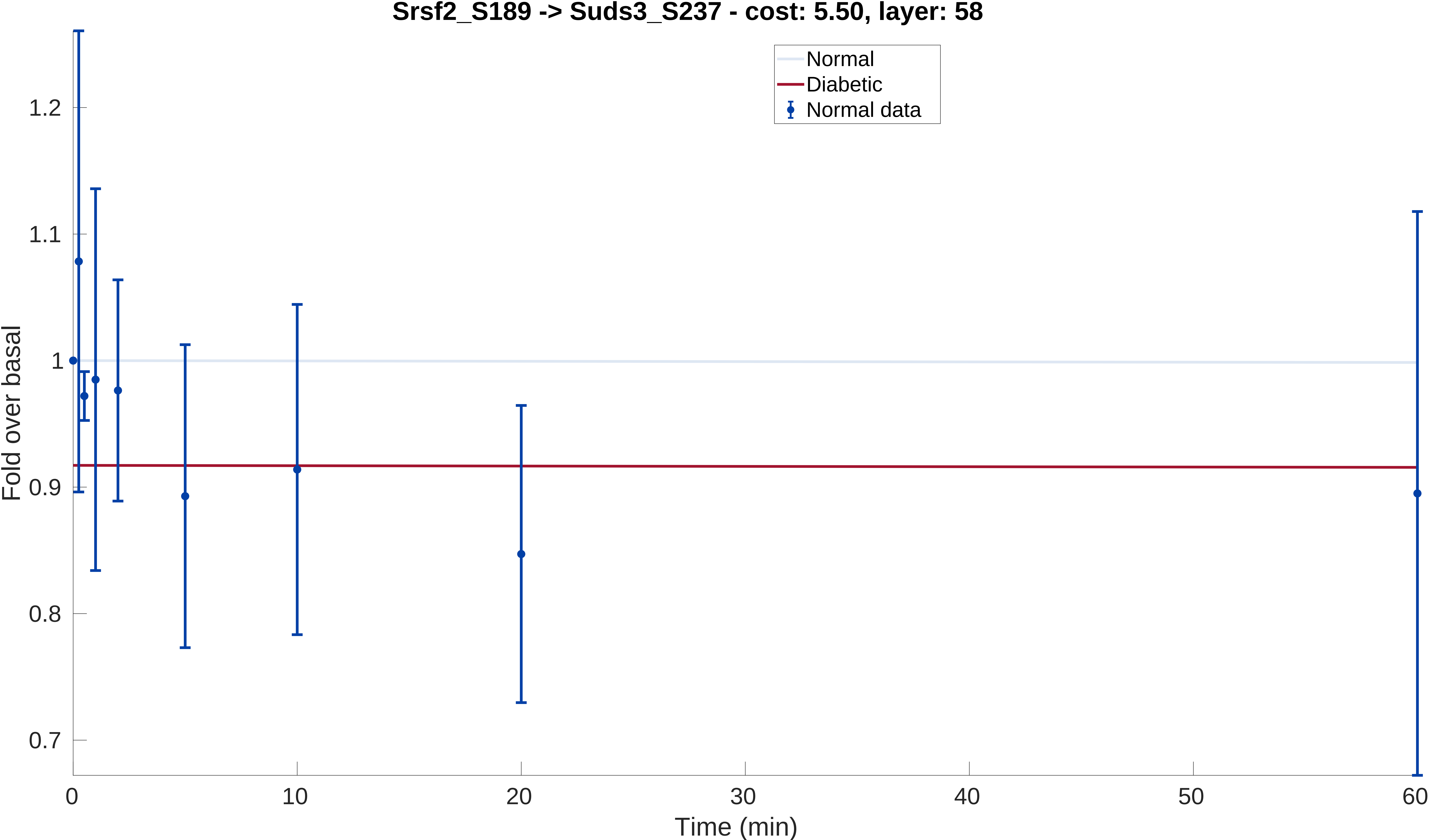
Srsf2_S189 -> Suds3_S237 - cost: 5.50, layer: 58
Normal
Diabetic
Normal data
Fold over basal
Time (min)
0.7
0.8
0.9
1
1.1
1.2
0
10
20
30
40
50
60

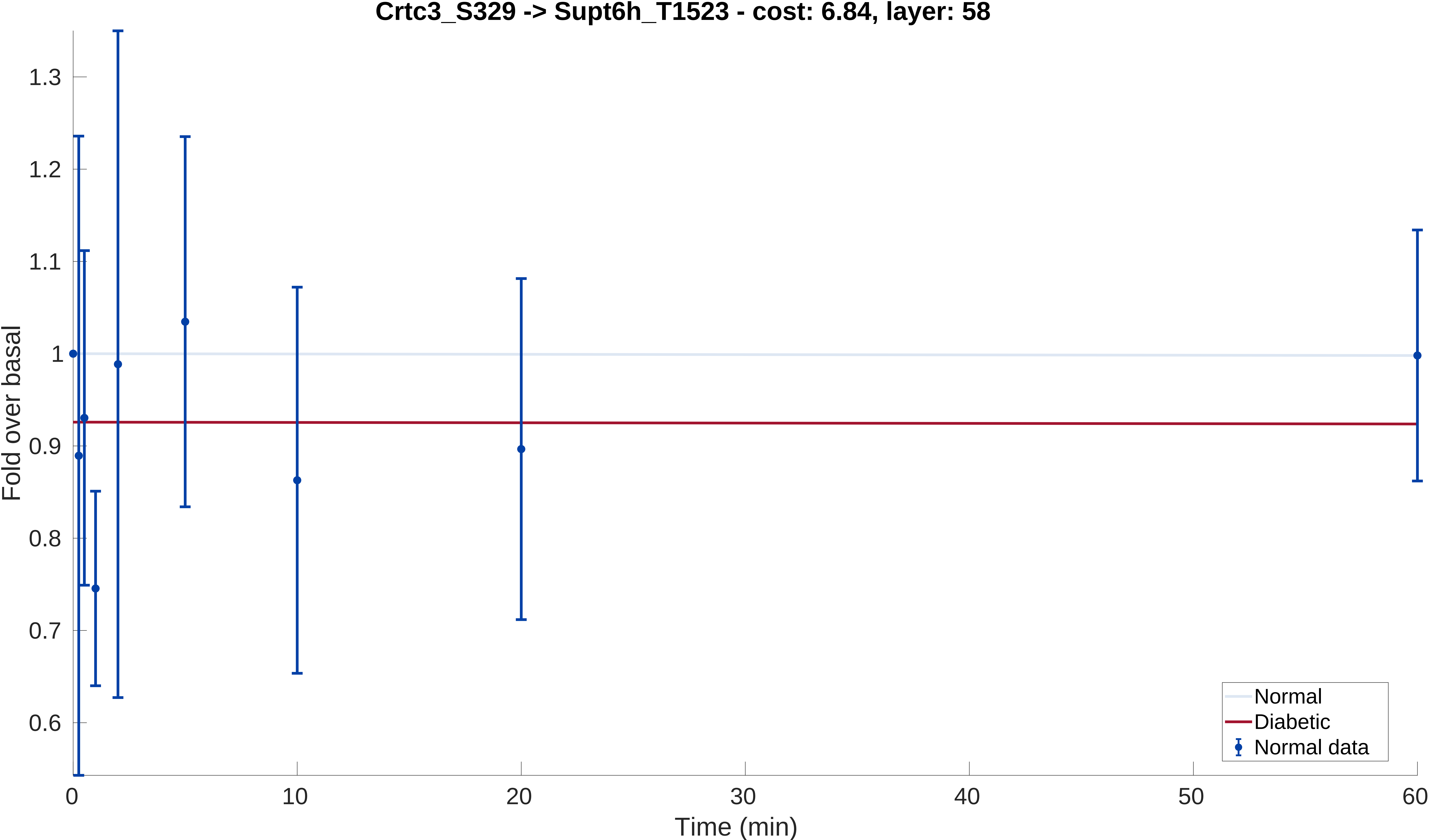

Crtc3_S329 -> Supt6h_T1523 - cost: 6.84, layer: 58
Fold over basal
1.3
1.2
1.1
1
0.9
0.8
0.7
0.6
0
10
20
30
40
50
60
Time (min)
Normal
Diabetic
Normal data

Unc119_S41 -> Svil_S241 - cost: 4.89, layer: 58

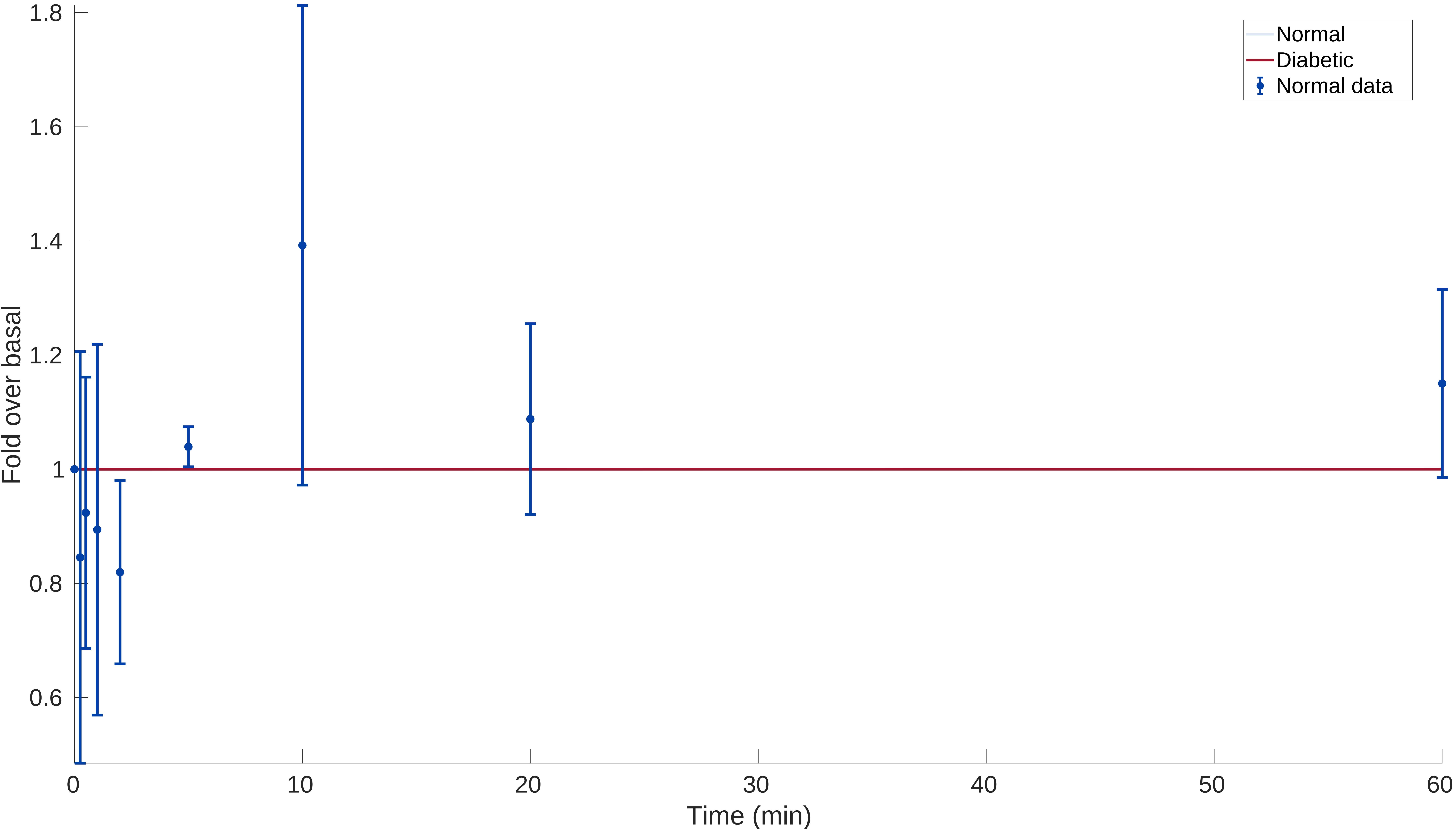

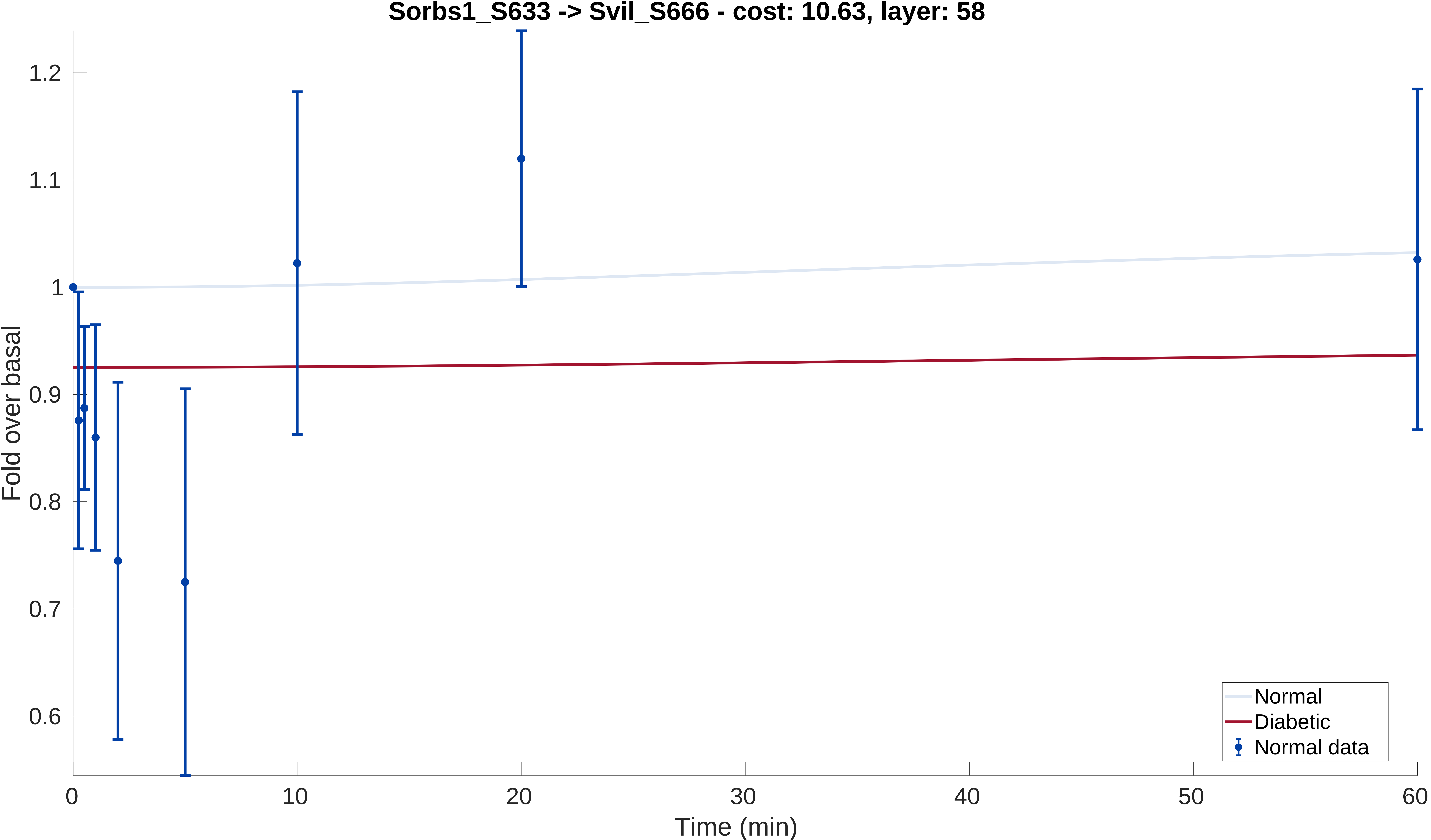

Sorbs1_S633 -> Svil_S666 - cost: 10.63, layer: 58
1.2
1.1
1
0.9
0.8
0.7
0.6
Fold over basal
0
10
20
30
40
50
60
Time (min)
Normal
Diabetic
Normal data

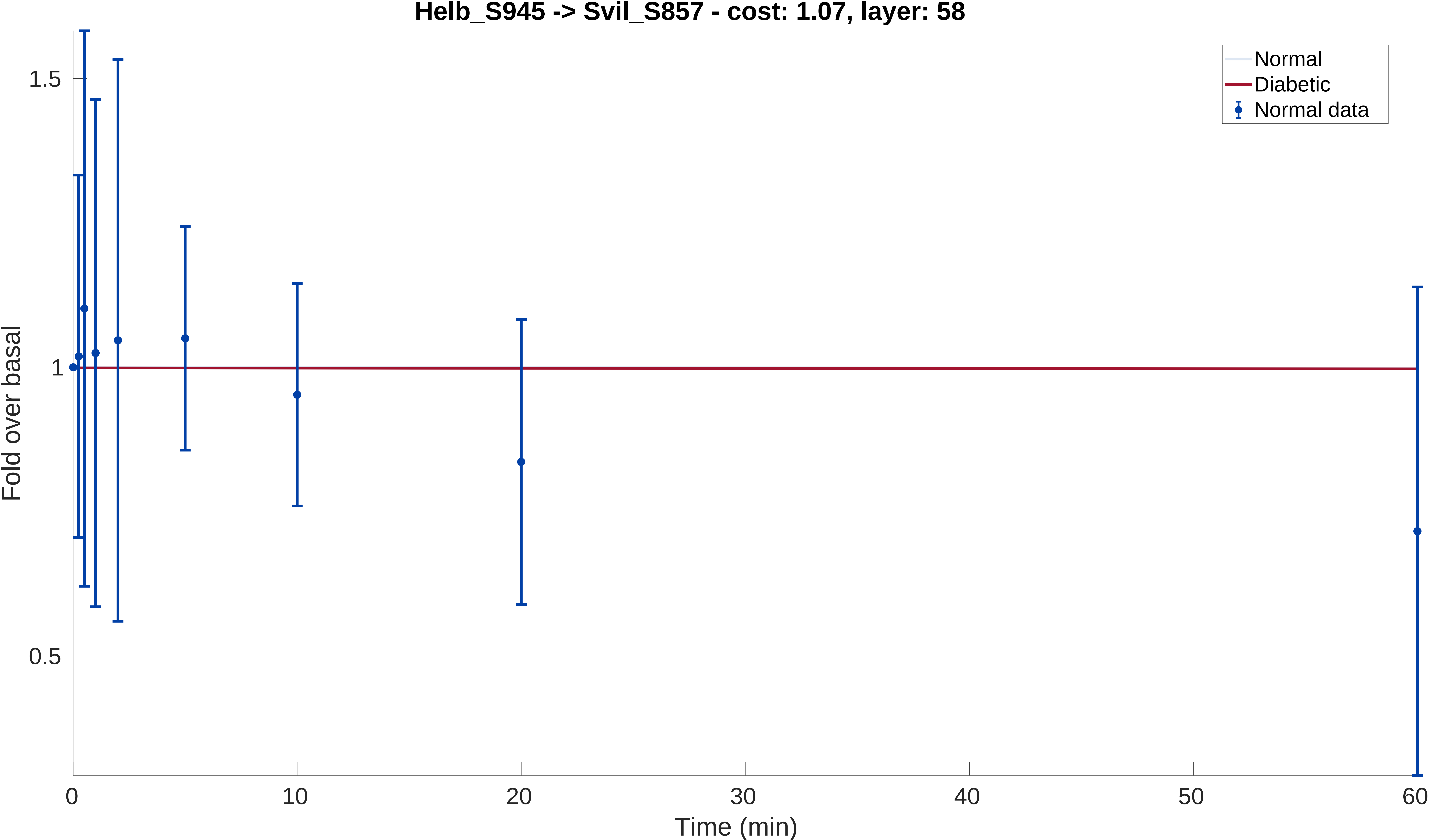

Helb_S945 -> Svil_S857 - cost: 1.07, layer: 58
Normal
Diabetic
Normal data
Fold over basal
1.5
1
0.5
0
10
20
30
40
50
60
Time (min)

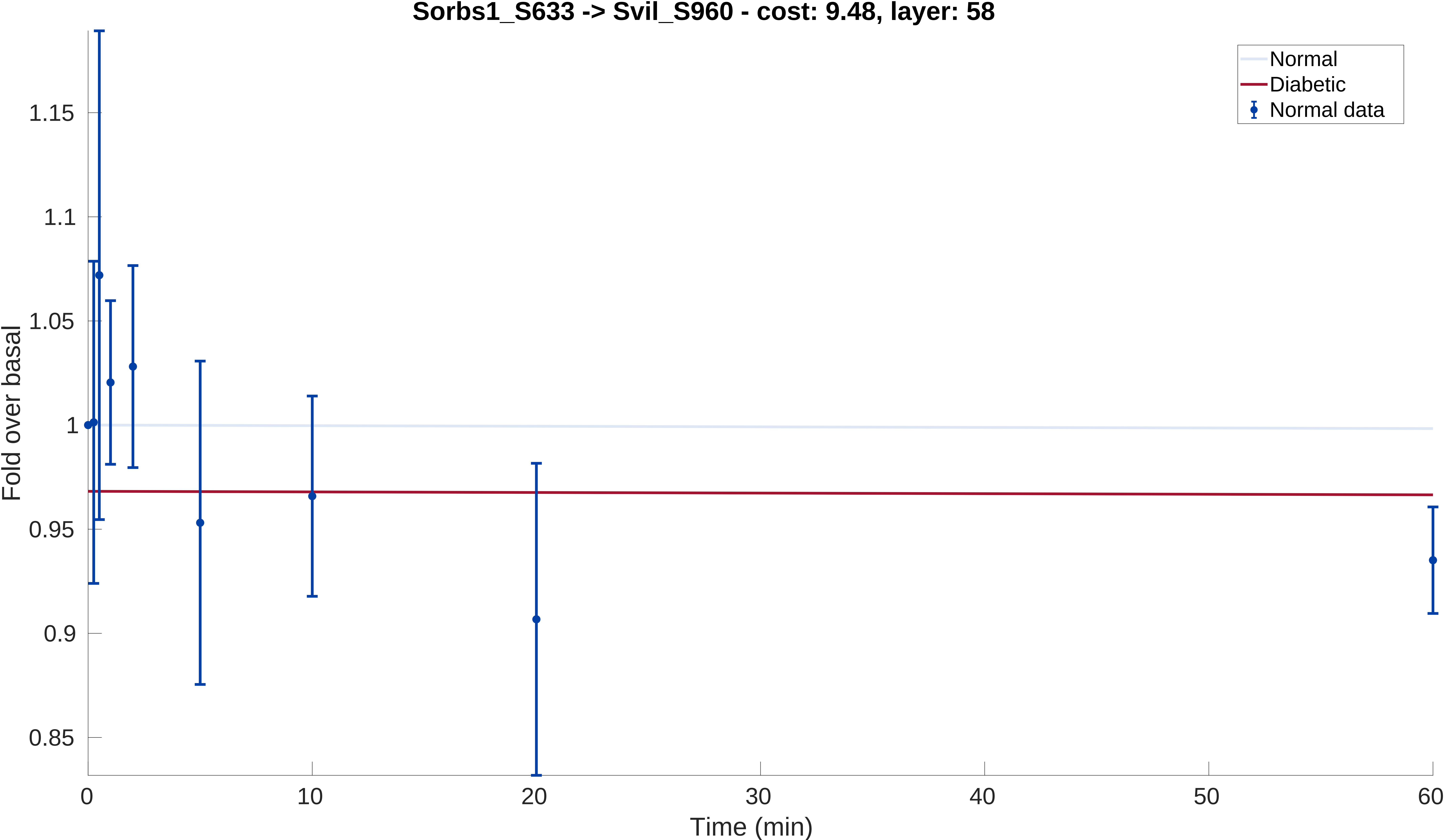

Sorbs1_S633 -> Svil_S960 - cost: 9.48, layer: 58
Normal
Diabetic
Normal data
1.15
1.1
1.05
1
0.95
0.9
0.85
Fold over basal
0
10
20
30
40
50
60
Time (min)

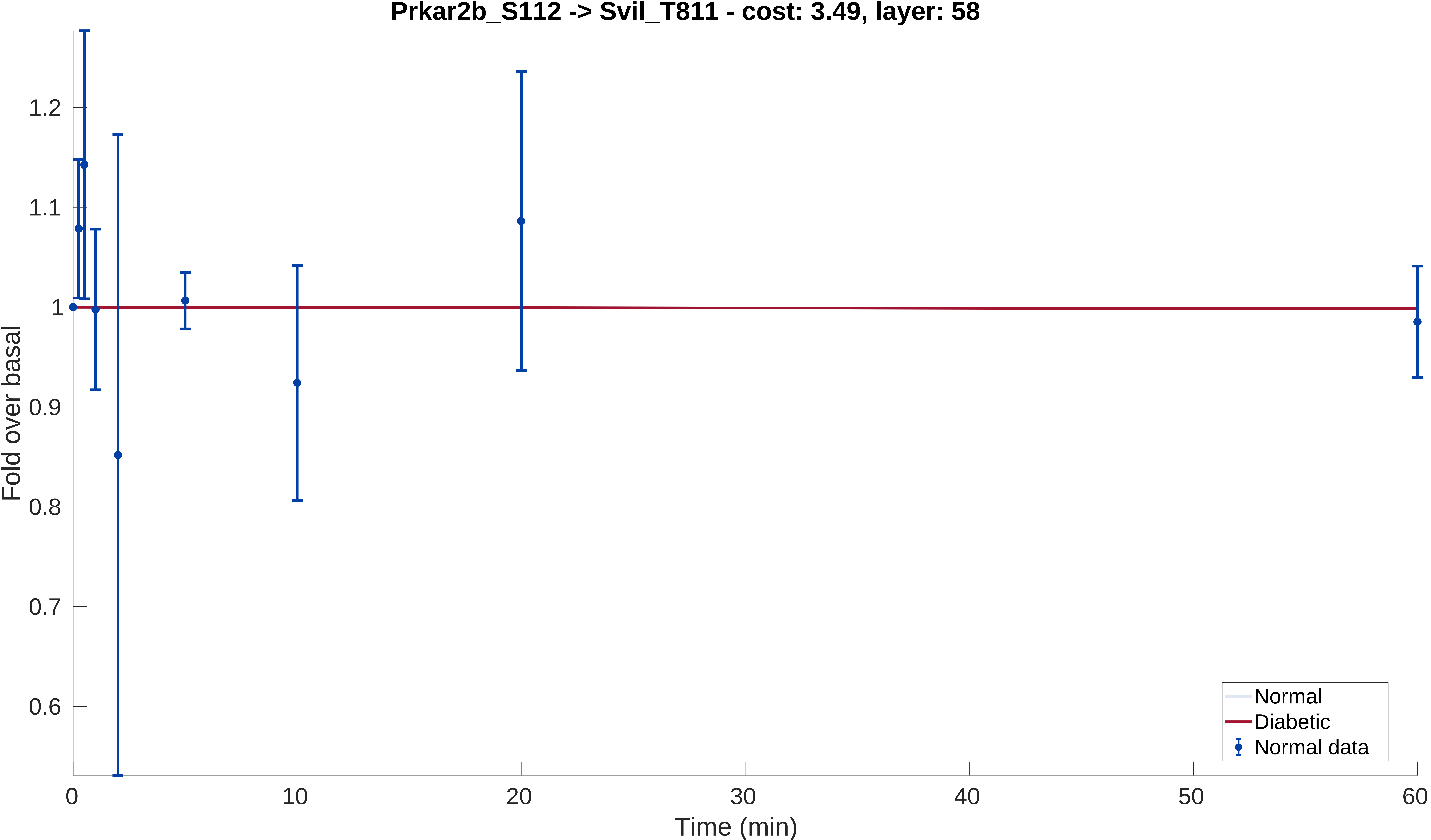

Prkar2b_S112 -> Svil_T811 - cost: 3.49, layer: 58
Fold over basal
1.2
1.1
1
0.9
0.8
0.7
0.6
0
10
20
30
40
50
60
Time (min)
Normal
Diabetic
Normal data

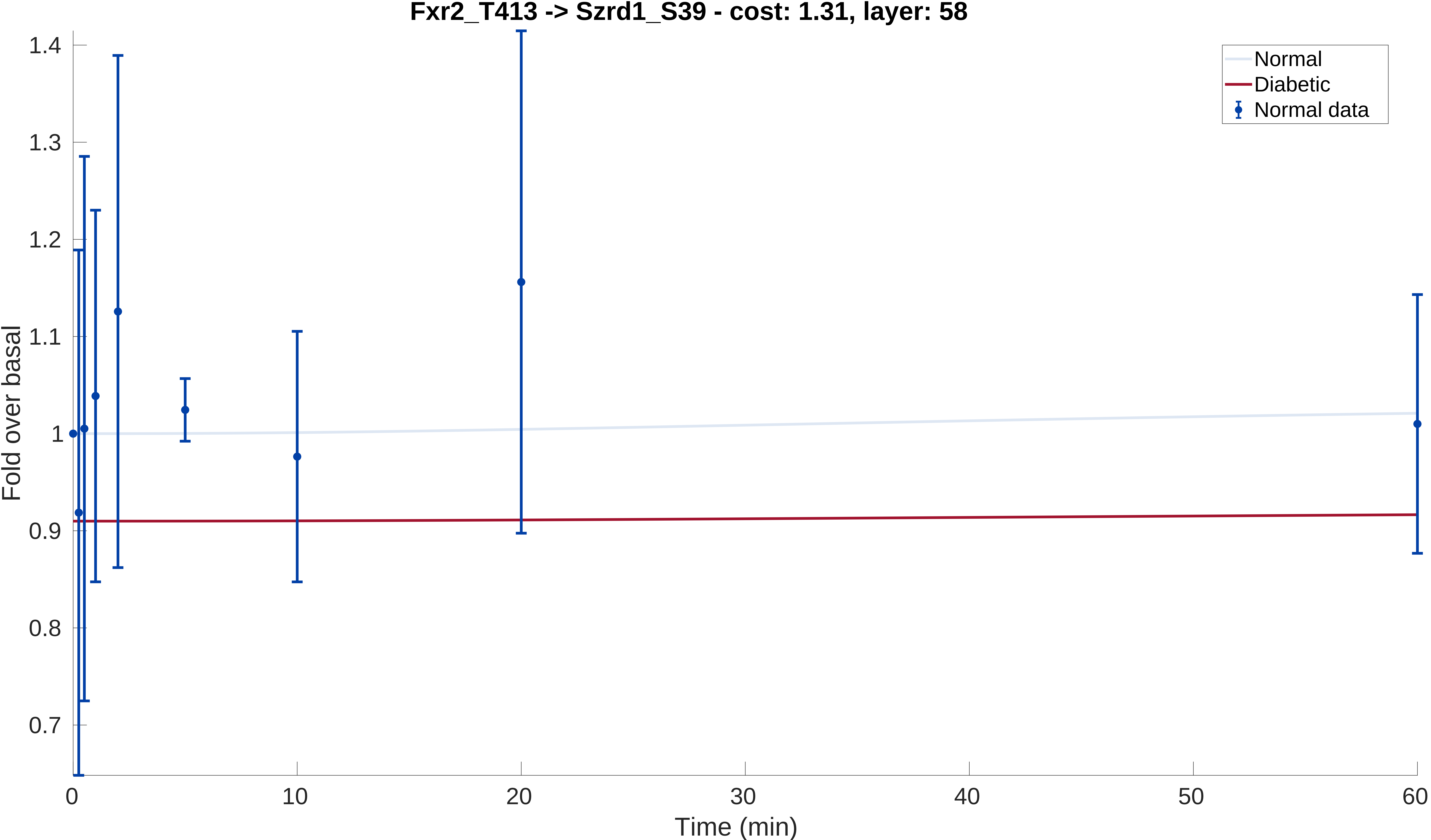

Fxr2_T413 -> Szrd1_S39 - cost: 1.31, layer: 58
Normal
Diabetic
Normal data
Fold over basal
Time (min)
1.4
1.3
1.2
1.1
1
0.9
0.8
0.7
0
10
20
30
40
50
60

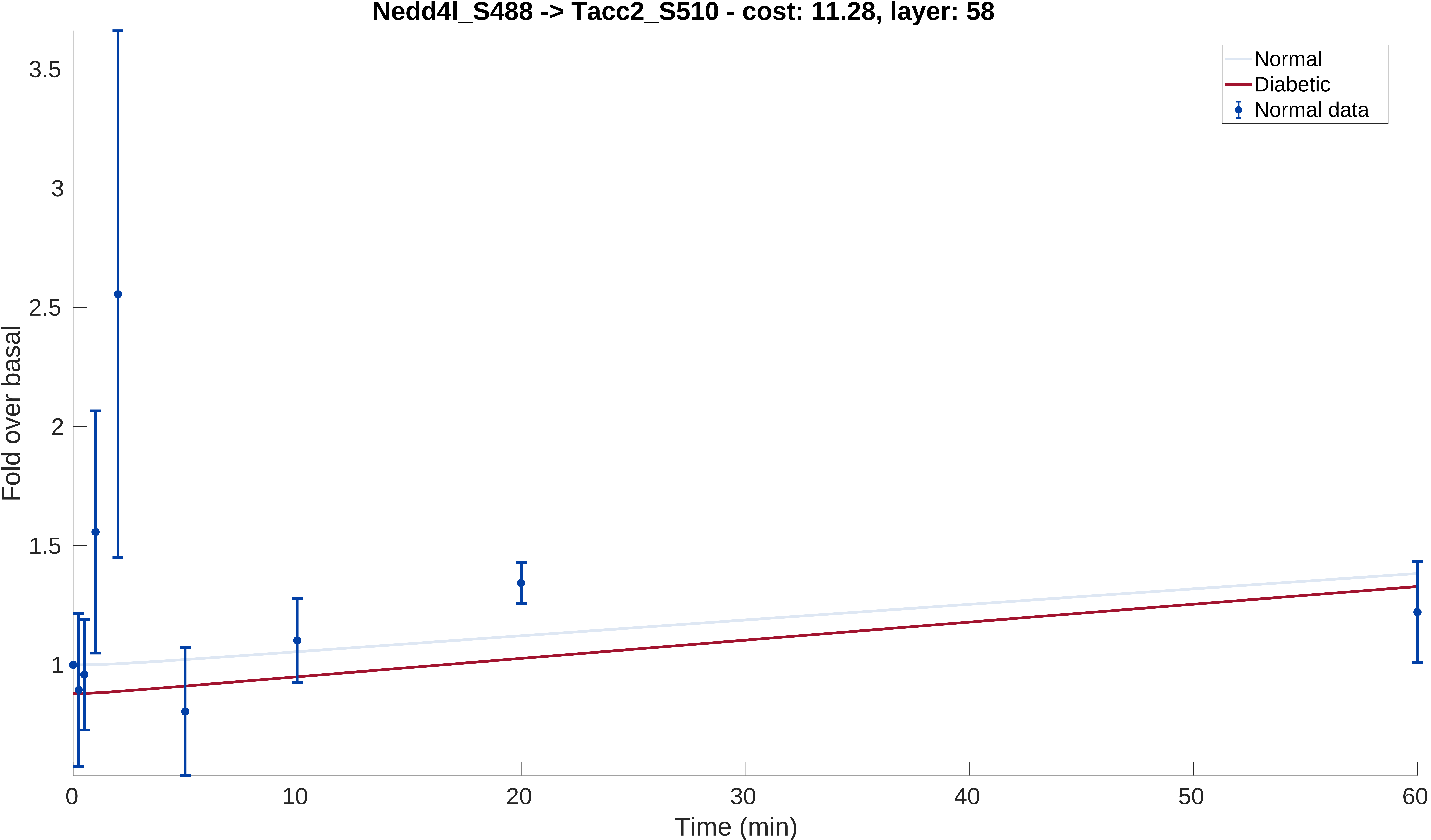
Nedd4l_S488 -> Tacc2_S510 - cost: 11.28, layer: 58
Normal
Diabetic
Normal data
Fold over basal
3.5
3
2.5
2
1.5
1
0
10
20
30
40
50
60
Time (min)

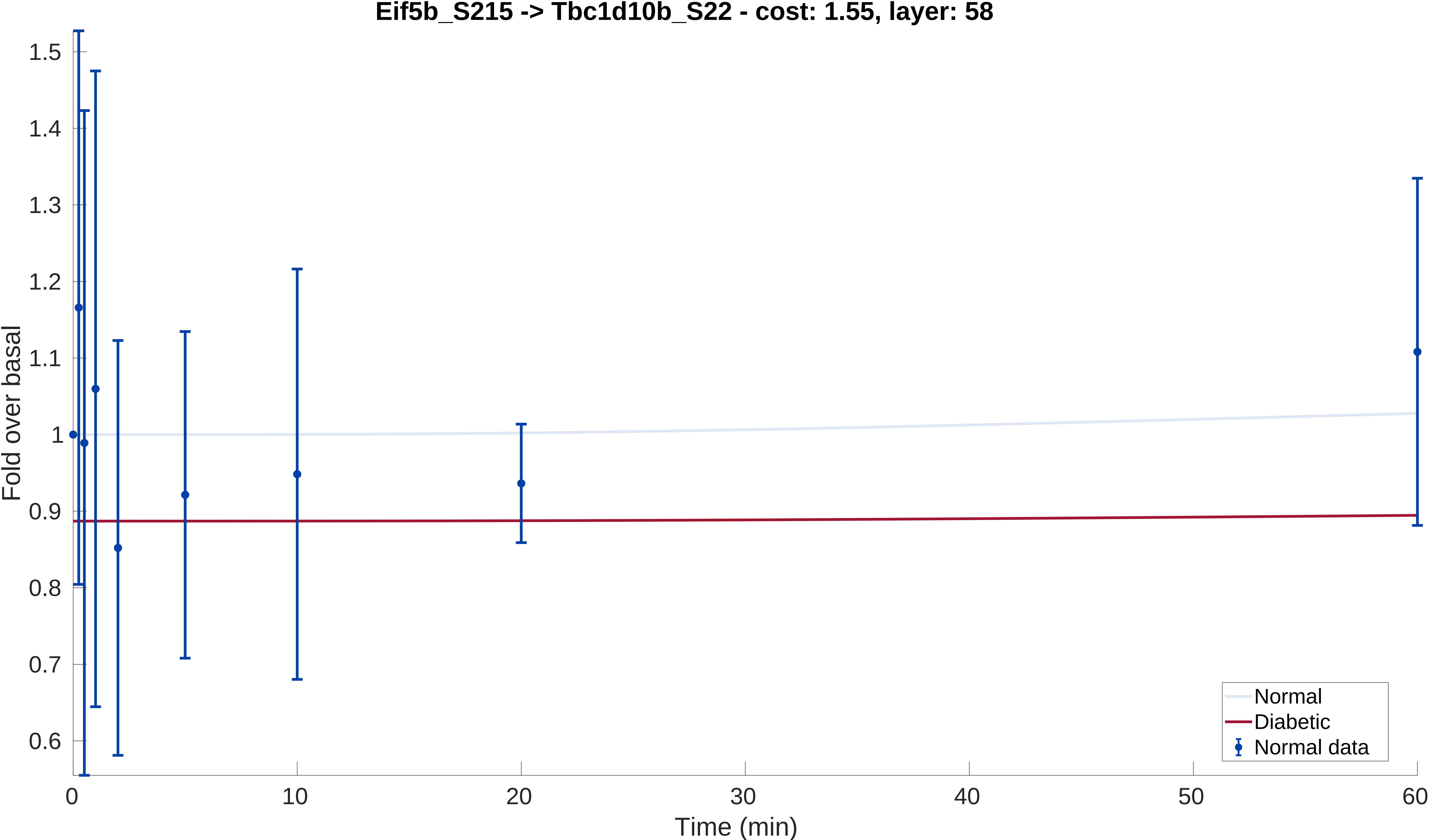
Eif5b_S215 -> Tbc1d10b_S22 - cost: 1.55, layer: 58
Fold over basal
Time (min)
Normal
Diabetic
Normal data
0
10
20
30
40
50
60
0.6
0.7
0.8
0.9
1
1.1
1.2
1.3
1.4
1.5

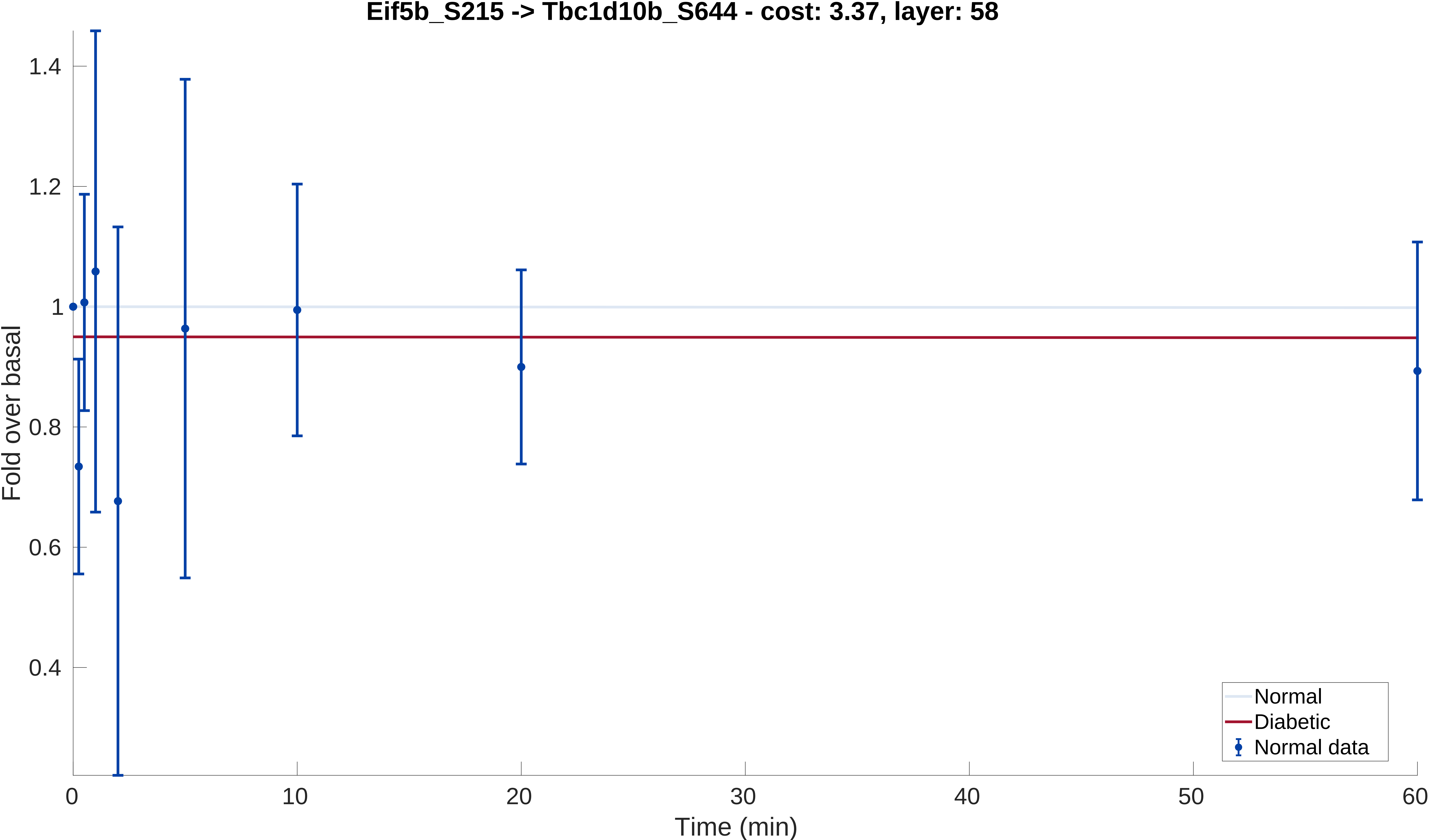

Eif5b_S215 -> Tbc1d10b_S644 - cost: 3.37, layer: 58
1.4
1.2
1
0.8
0.6
0.4
Fold over basal
0
10
20
30
40
50
60
Time (min)
Normal
Diabetic
Normal data

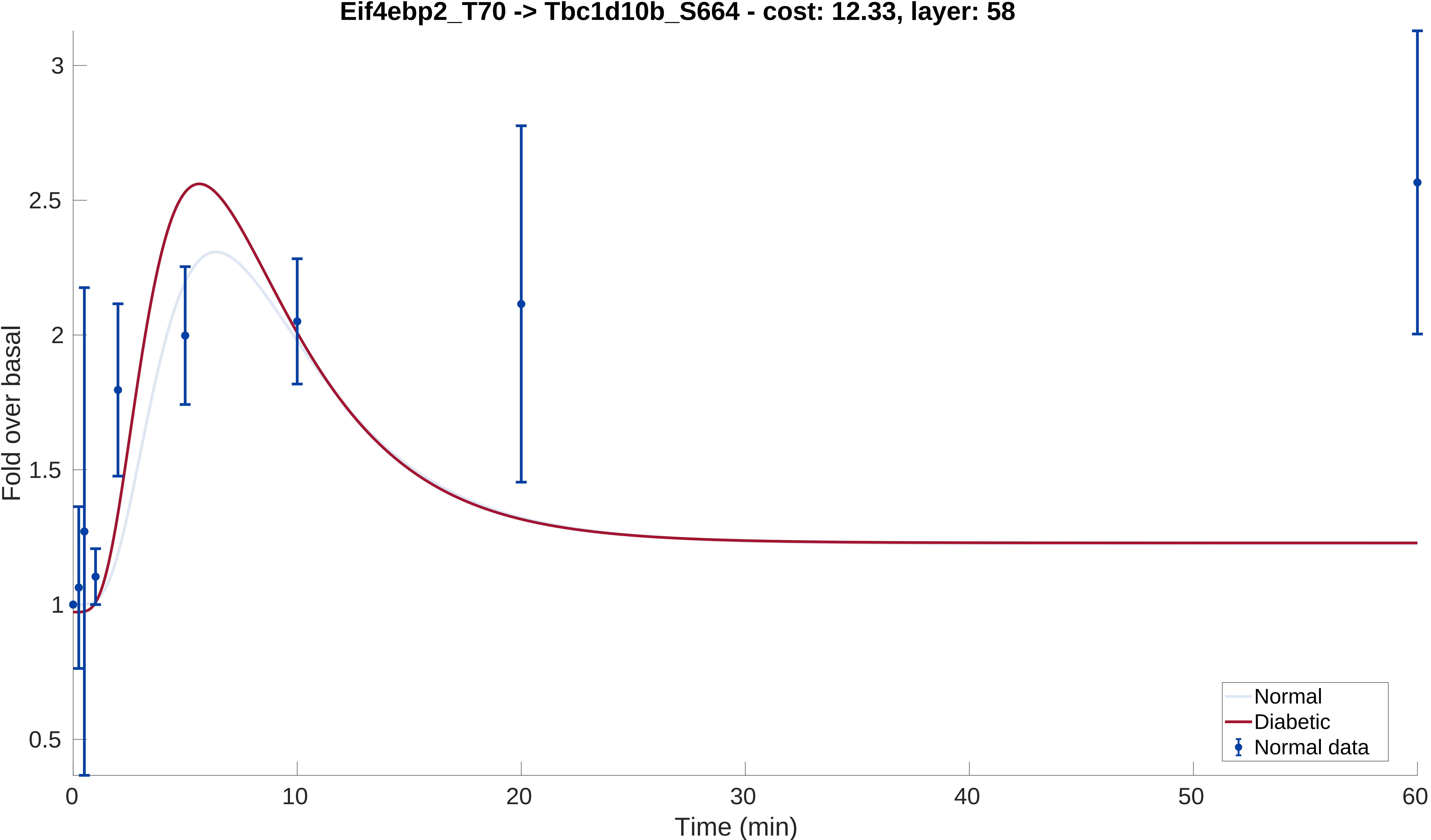

Eif4ebp2_T70 -> Tbc1d10b_S664 - cost: 12.33, layer: 58
Fold over basal
3
2.5
2
1.5
1
0.5
0
10
20
30
40
50
60
Time (min)
Normal
Diabetic
Normal data

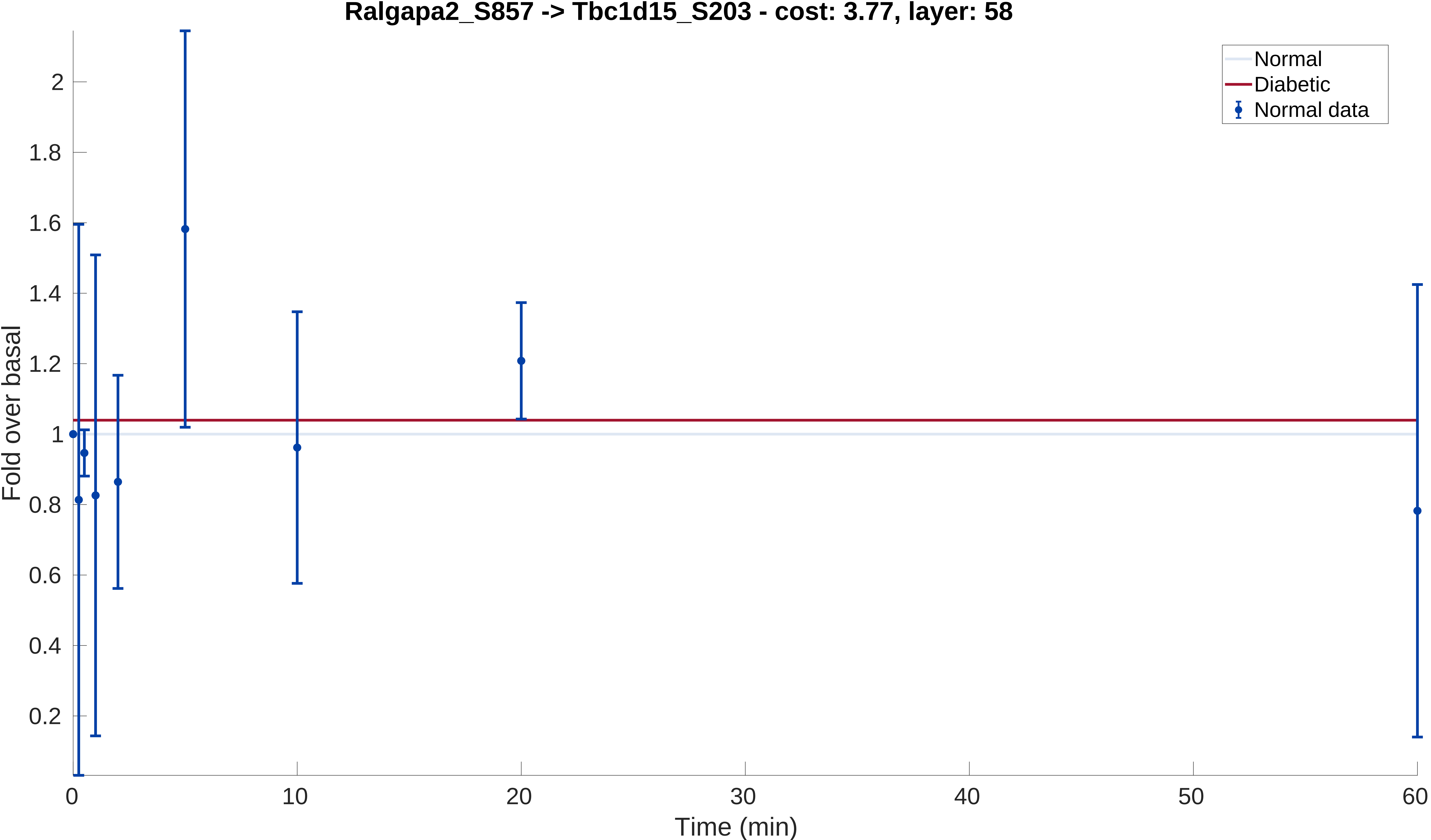

Ralgapa2_S857 -> Tbc1d15_S203 - cost: 3.77, layer: 58
Normal
Diabetic
Normal data
Fold over basal
2
1.8
1.6
1.4
1.2
1
0.8
0.6
0.4
0.2
0
10
20
30
40
50
60
Time (min)

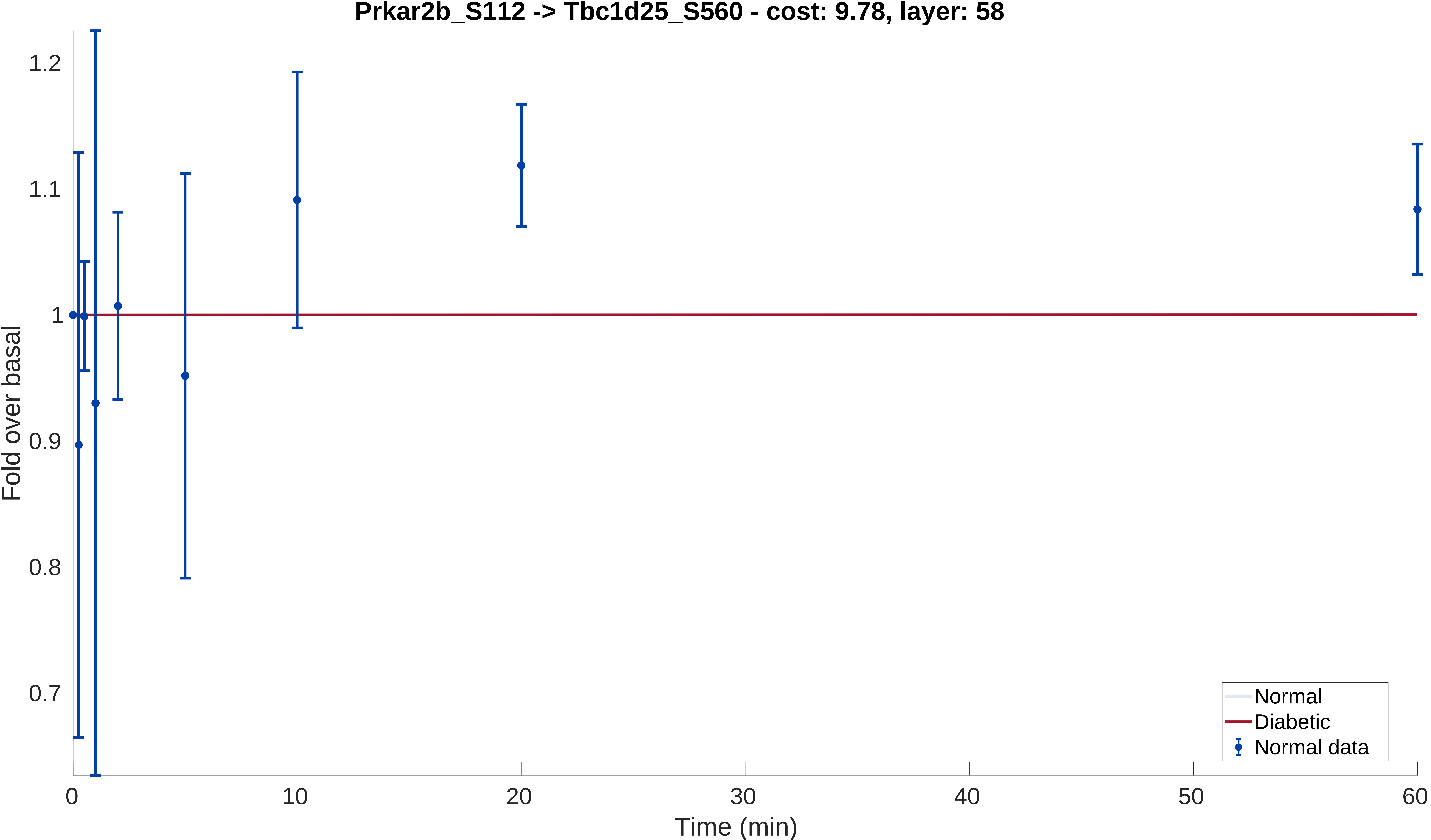
Prkar2b_S112 -> Tbc1d25_S560 - cost: 9.78, layer: 58
Fold over basal
1.2
1.1
1
0.9
0.8
0.7
Time (min)
0
10
20
30
40
50
60
Normal
Diabetic
Normal data

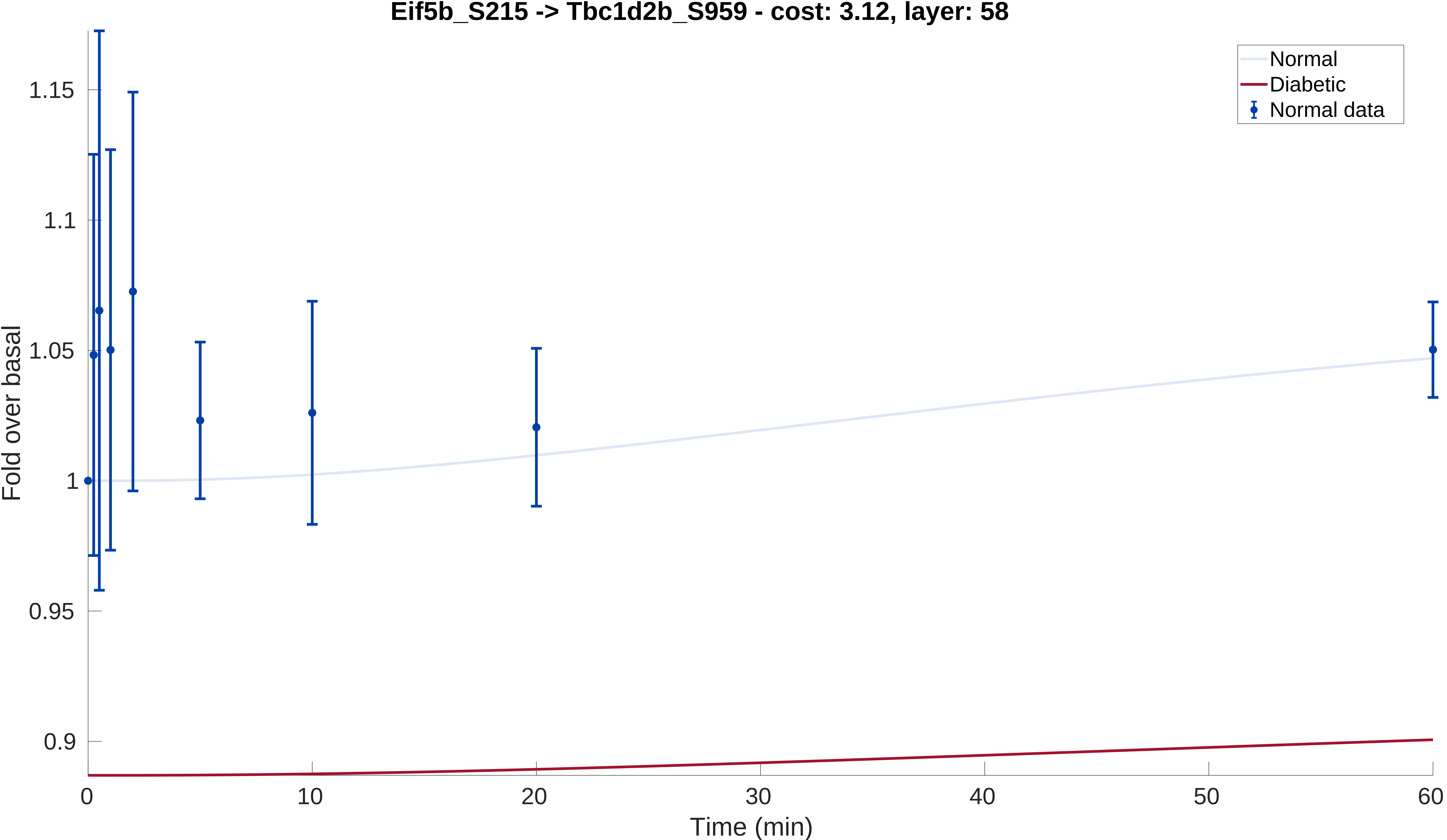
Eif5b_S215 -> Tbc1d2b_S959 - cost: 3.12, layer: 58
Normal
Diabetic
Normal data
Fold over basal
1.15
1.1
1.05
1
0.95
0.9
0
10
20
30
40
50
60
Time (min)

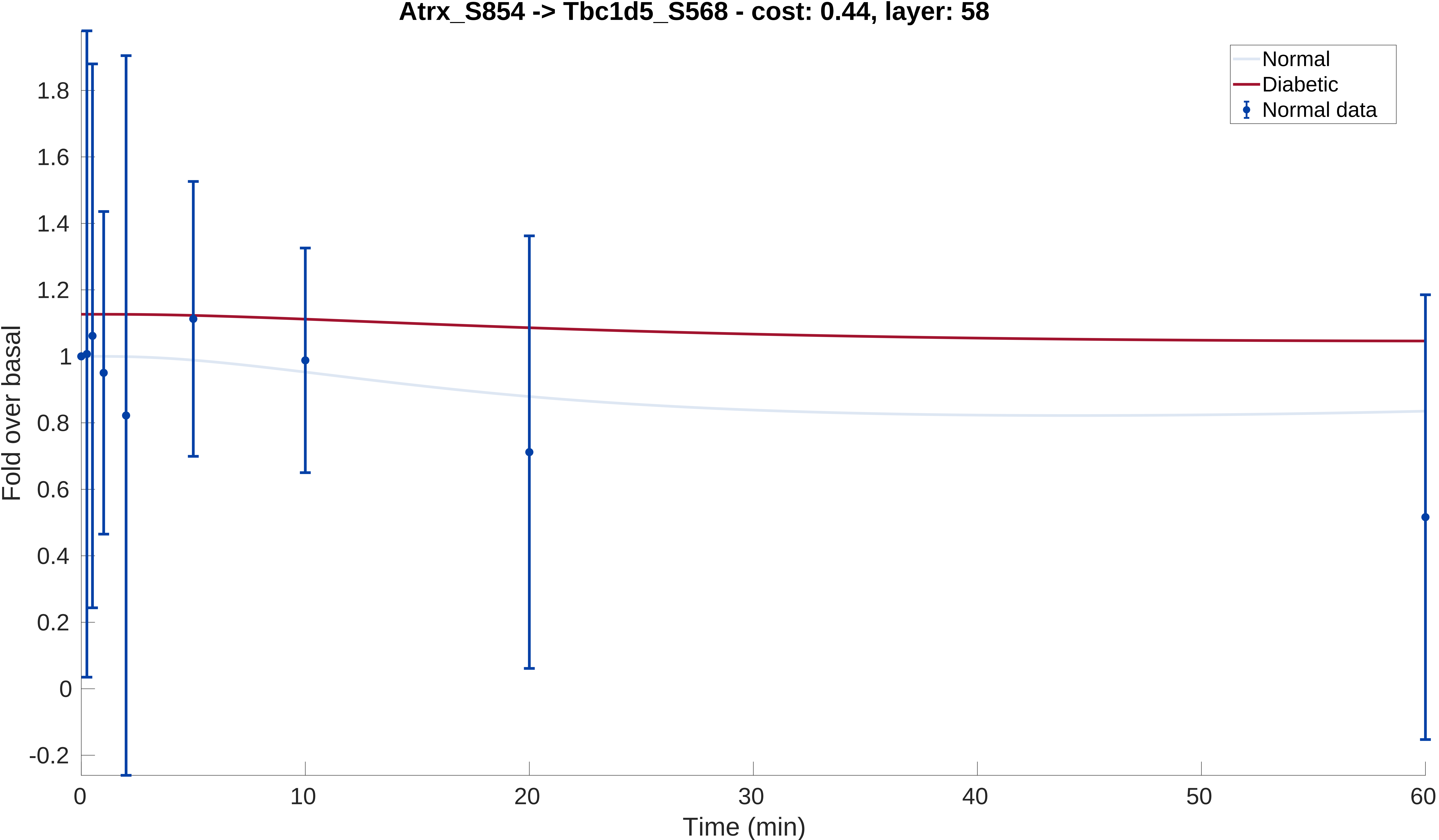
Atrx_S854 -> Tbc1d5_S568 - cost: 0.44, layer: 58
Normal
Diabetic
Normal data
Fold over basal
1.8
1.6
1.4
1.2
1
0.8
0.6
0.4
0.2
0
-0.2
0
10
20
30
40
50
60
Time (min)

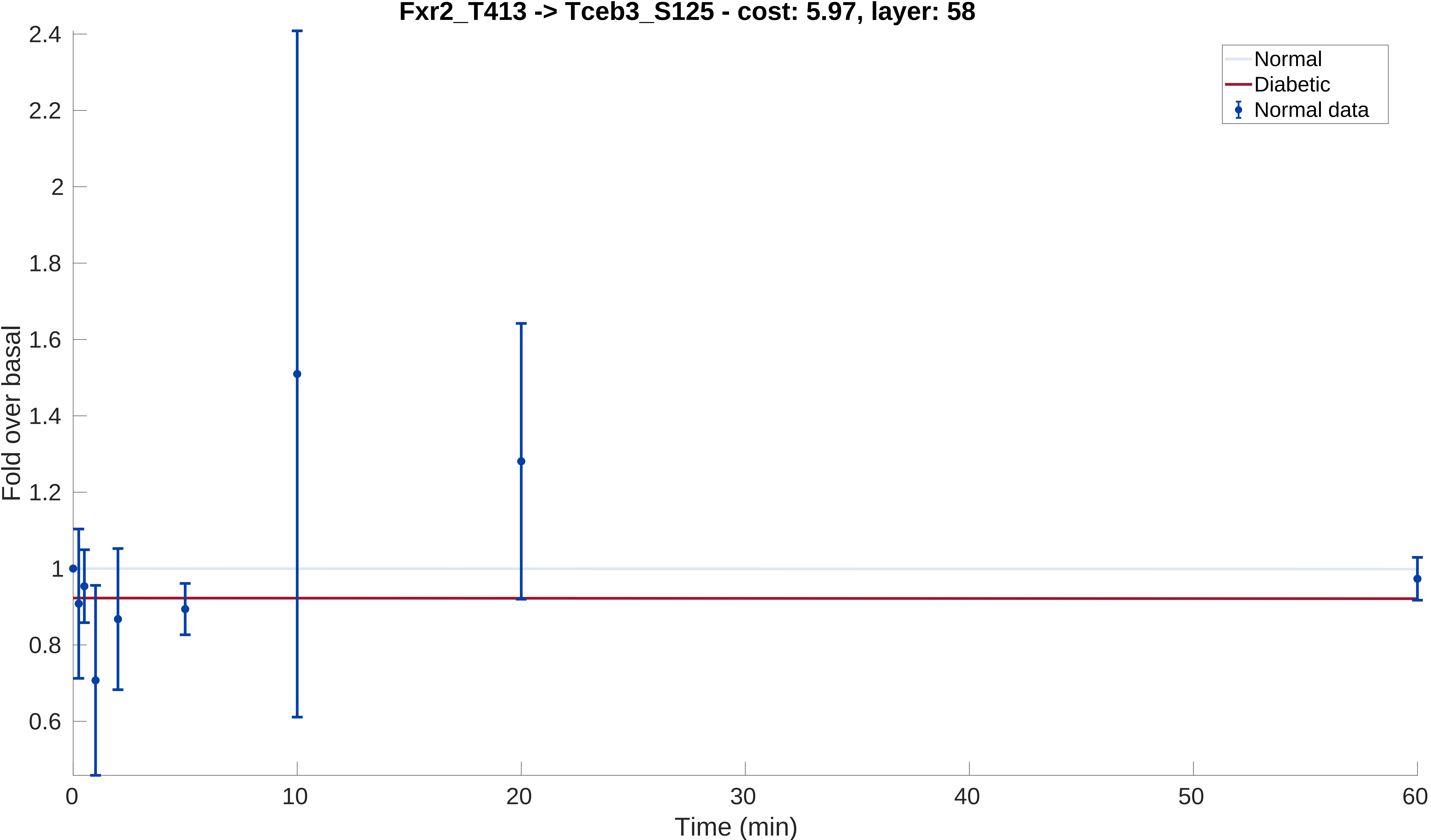
Fxr2_T413 -> Tceb3_S125 - cost: 5.97, layer: 58
Normal
Diabetic
Normal data
Fold over basal
2.4
2.2
2
1.8
1.6
1.4
1.2
1
0.8
0.6
0
10
20
30
40
50
60
Time (min)

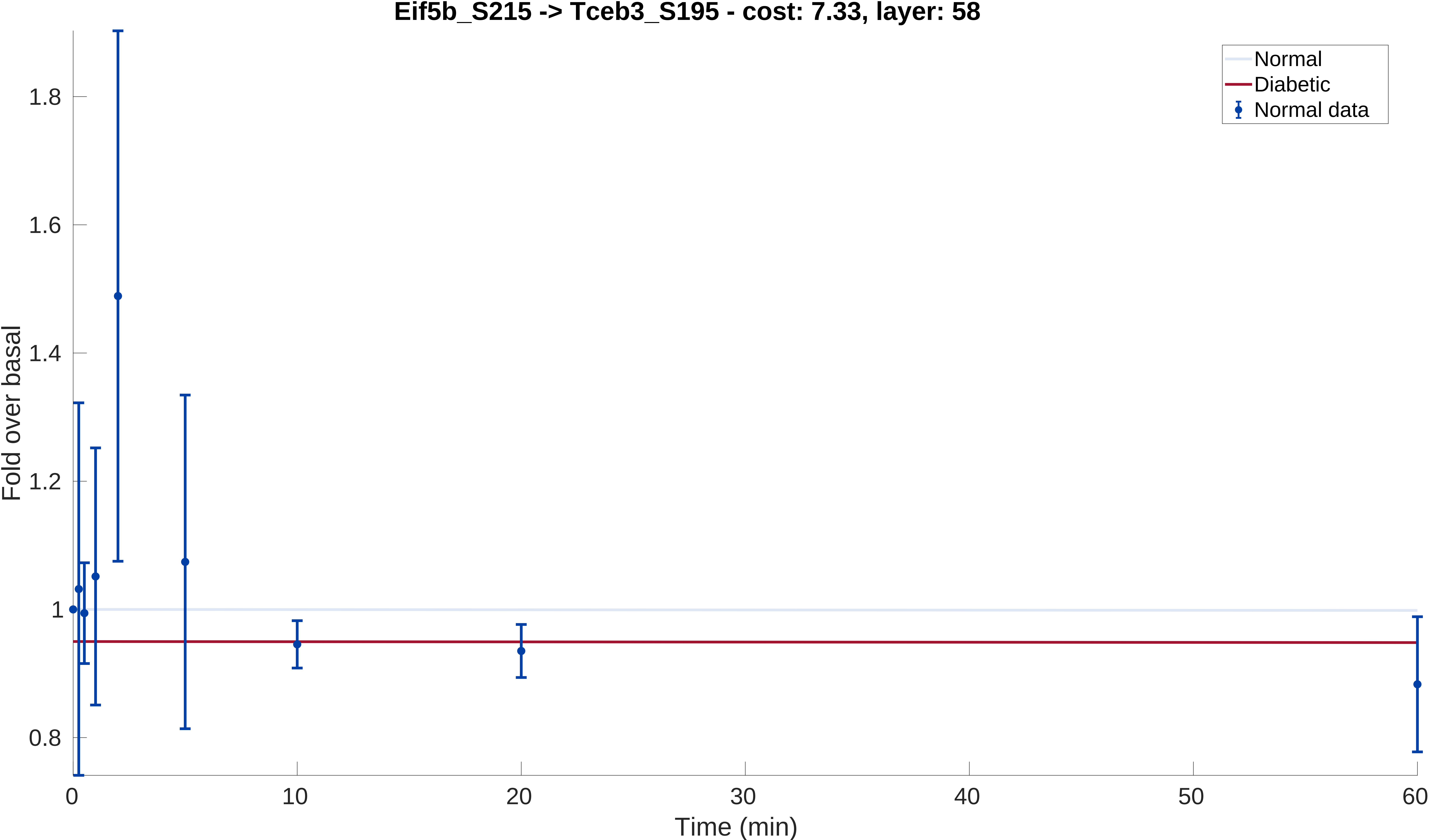

Eif5b_S215 -> Tceb3_S195 - cost: 7.33, layer: 58
Normal
Diabetic
Normal data
1.8
1.6
1.4
1.2
1
0.8
Fold over basal
0
10
20
30
40
50
60
Time (min)

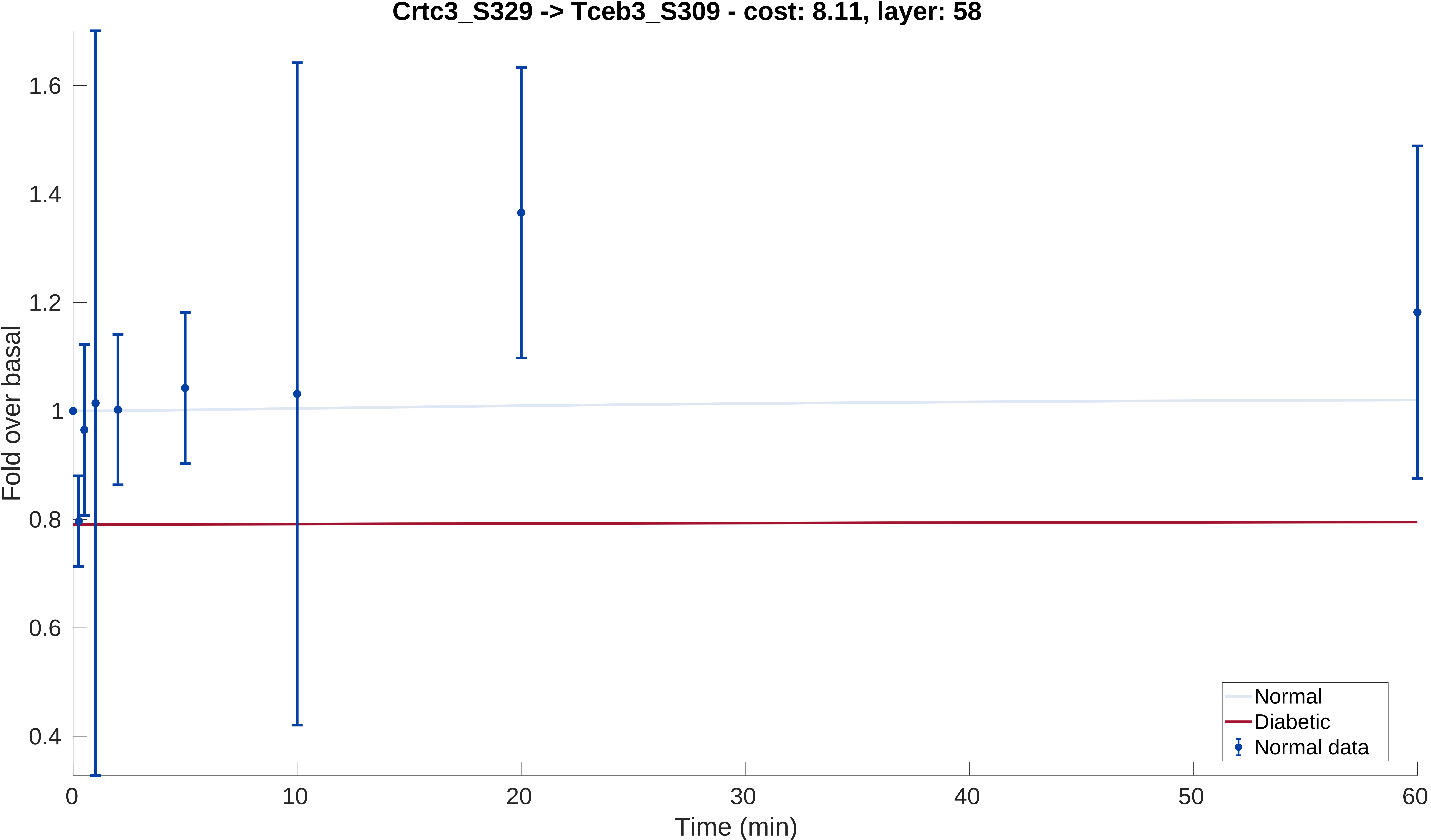
Crtc3_S329 -> Tceb3_S309 - cost: 8.11, layer: 58
Fold over basal
1.6
1.4
1.2
1
0.8
0.6
0.4
0
10
20
30
40
50
60
Time (min)
Normal
Diabetic
Normal data

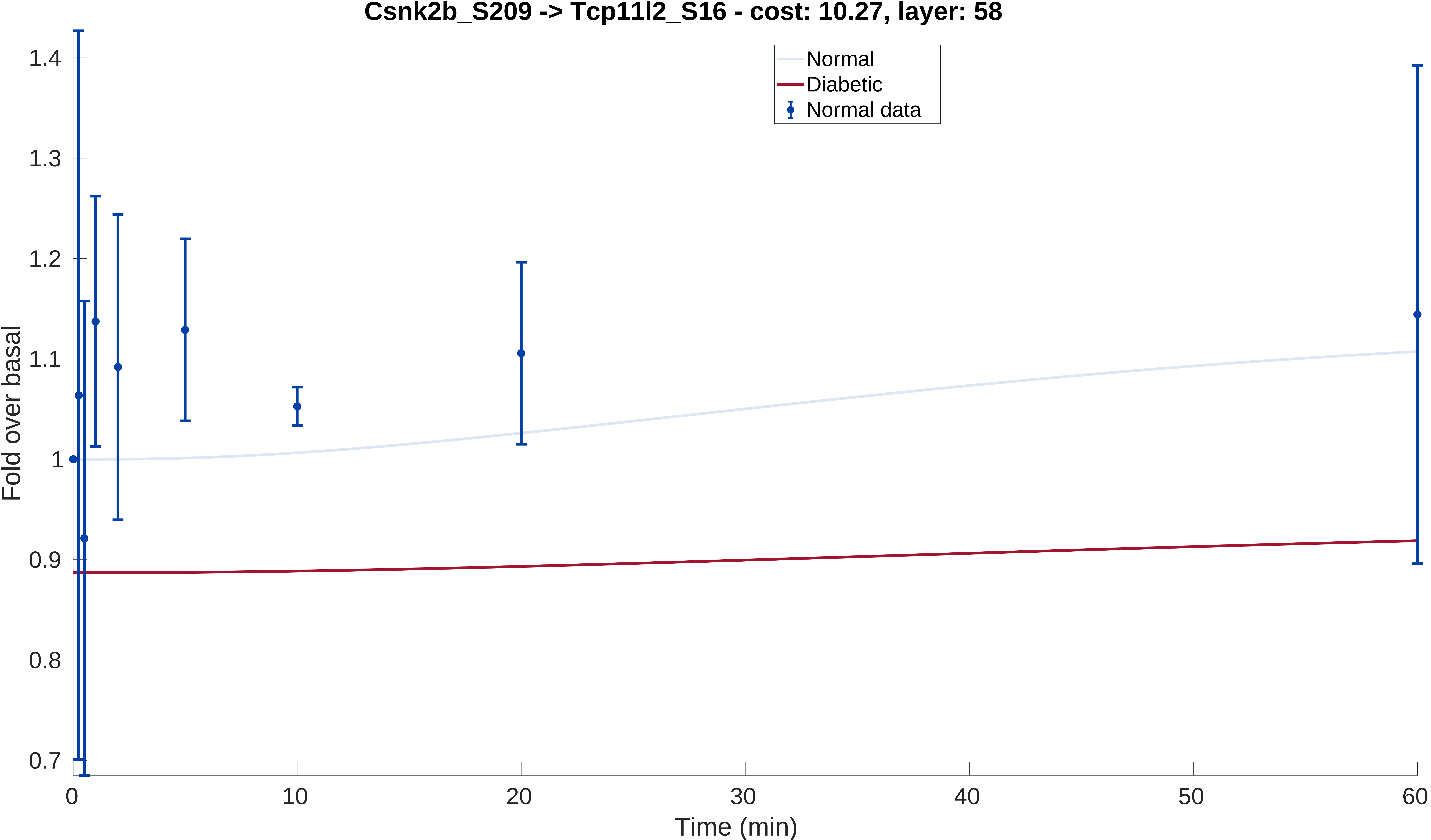

Csnk2b_S209 -> Tcp11l2_S16 - cost: 10.27, layer: 58
Normal
Diabetic
Normal data
Fold over basal
1.4
1.3
1.2
1.1
1
0.9
0.8
0.7
0
10
20
30
40
50
60
Time (min)

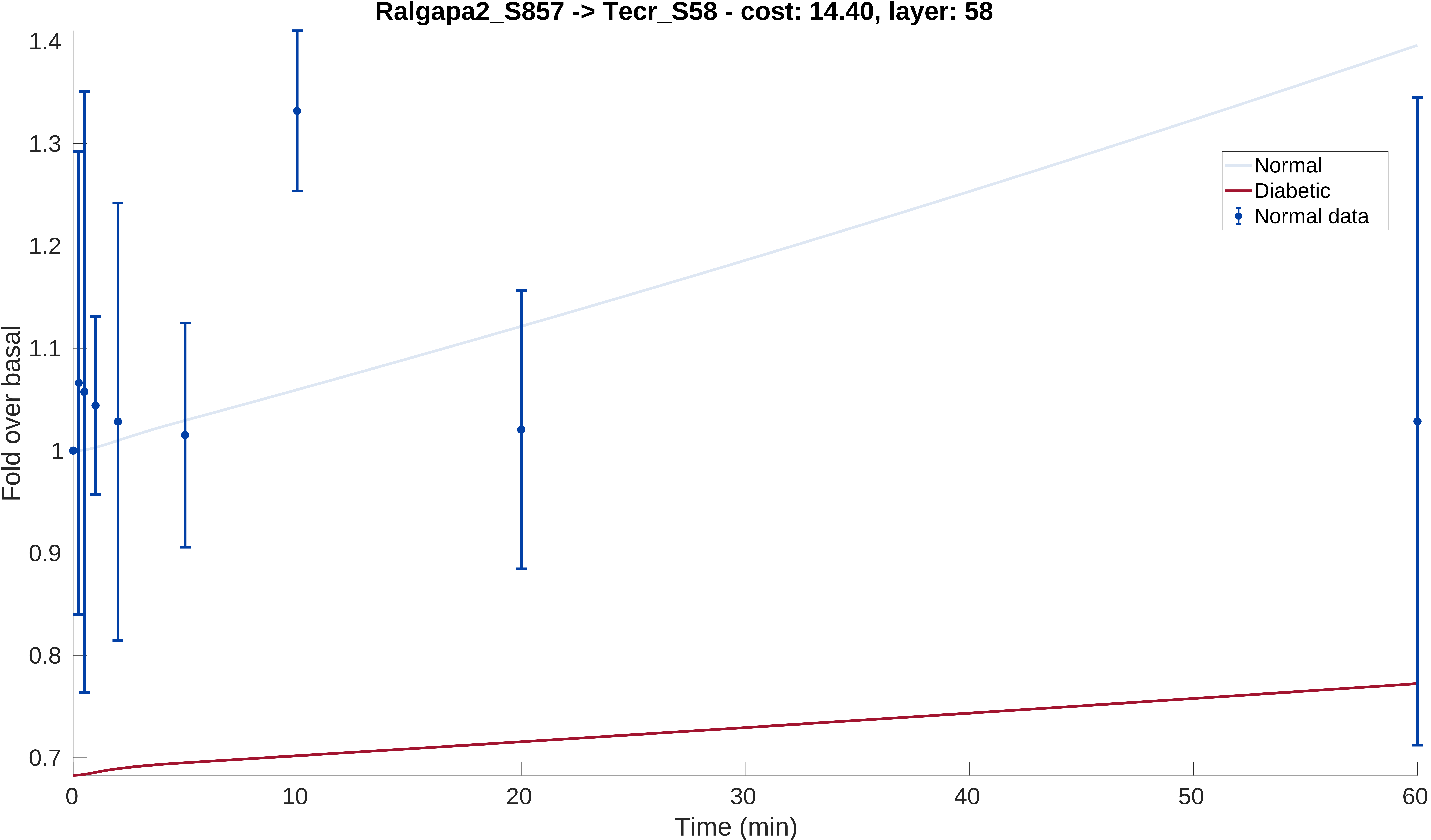

Ralgapa2_S857 -> Tecr_S58 - cost: 14.40, layer: 58
Fold over basal
Time (min)
1.4
1.3
1.2
1.1
1
0.9
0.8
0.7
0
10
20
30
40
50
60
Normal
Diabetic
Normal data

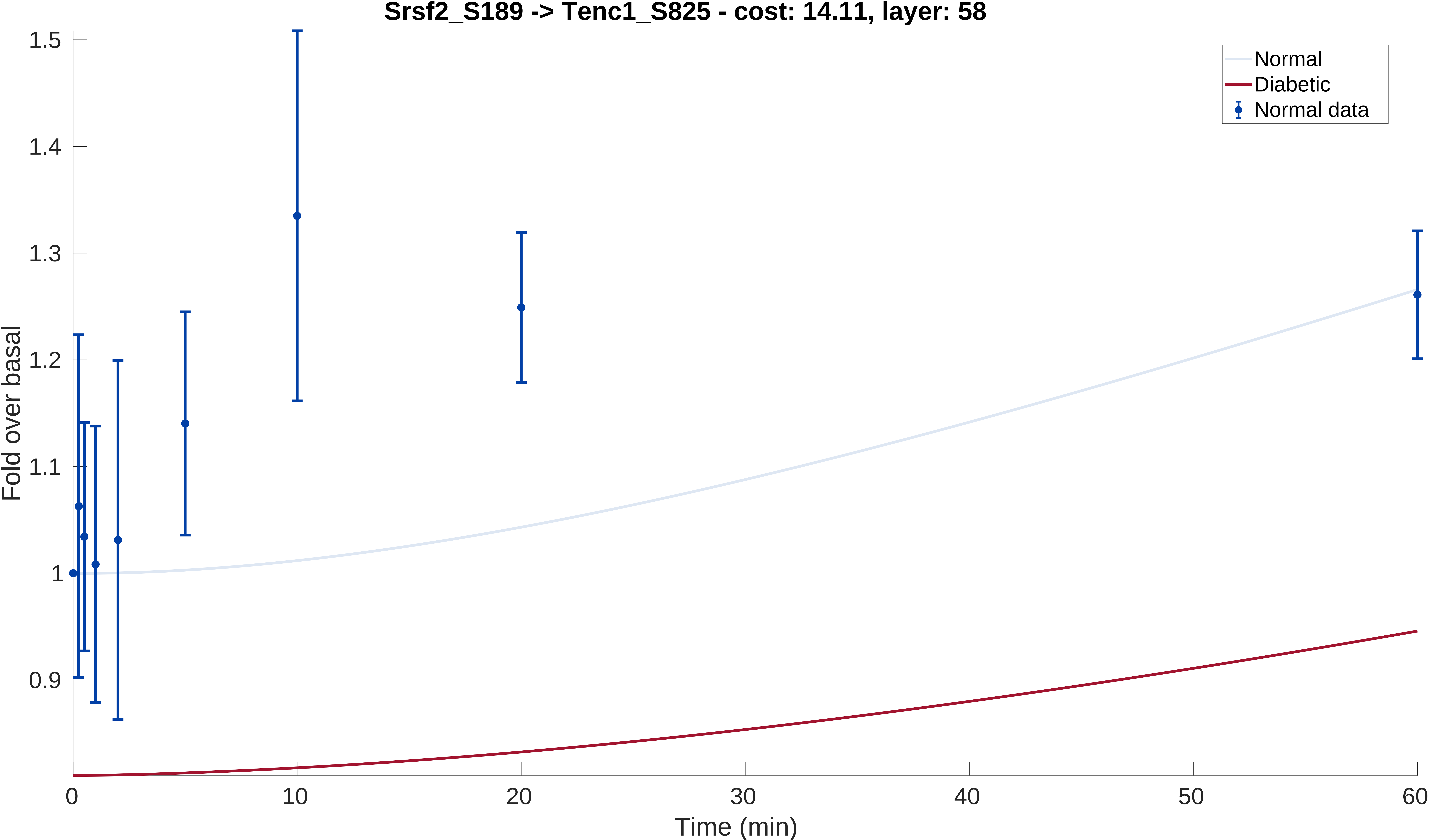
Srsf2_S189 -> Tenc1_S825 - cost: 14.11, layer: 58
Normal
Diabetic
Normal data
Fold over basal
1.5
1.4
1.3
1.2
1.1
1
0.9
0
10
20
30
40
50
60
Time (min)

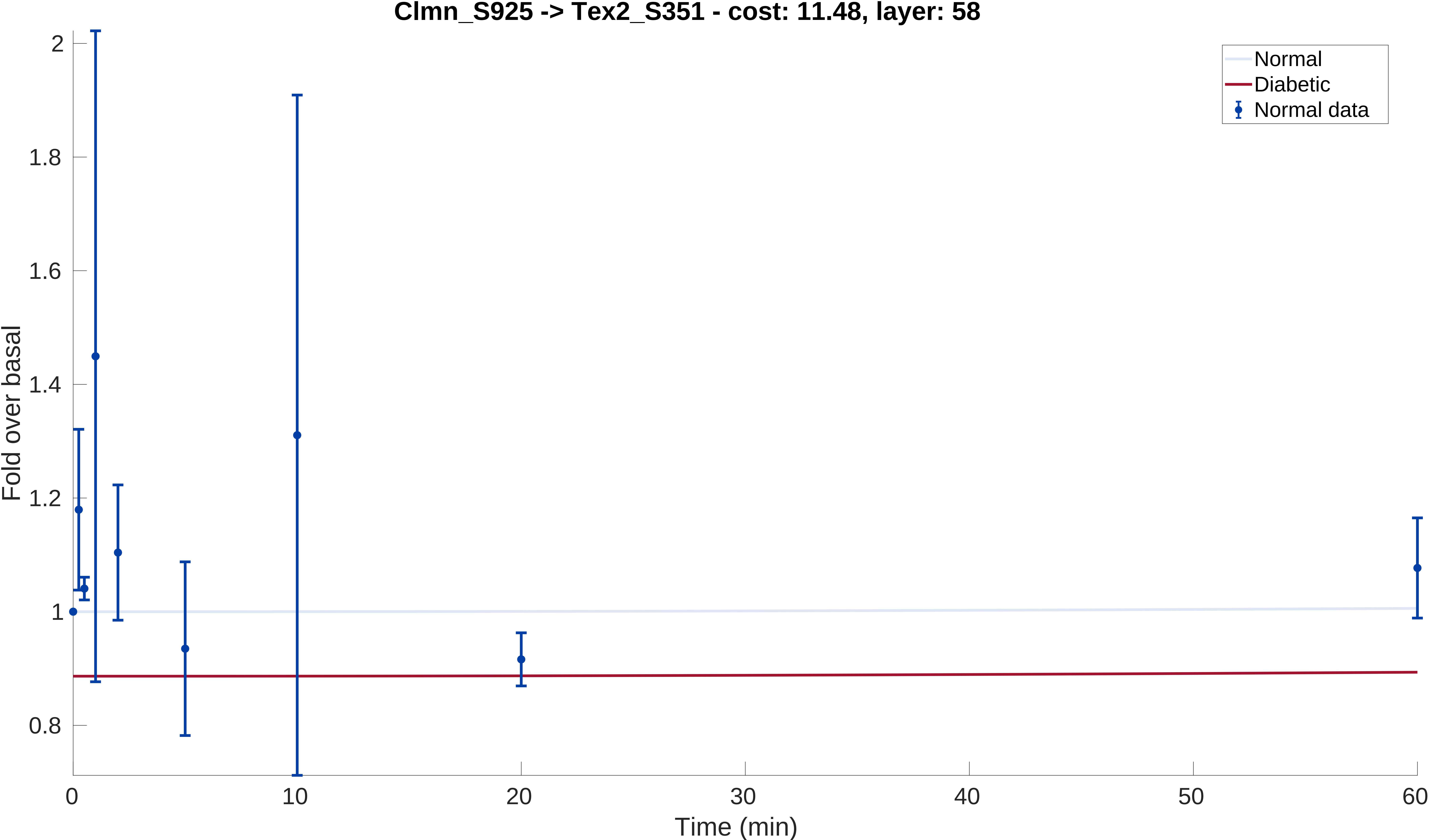

Clmn_S925 -> Tex2_S351 - cost: 11.48, layer: 58
Normal
Diabetic
Normal data
Fold over basal
2
1.8
1.6
1.4
1.2
1
0.8
0
10
20
30
40
50
60
Time (min)

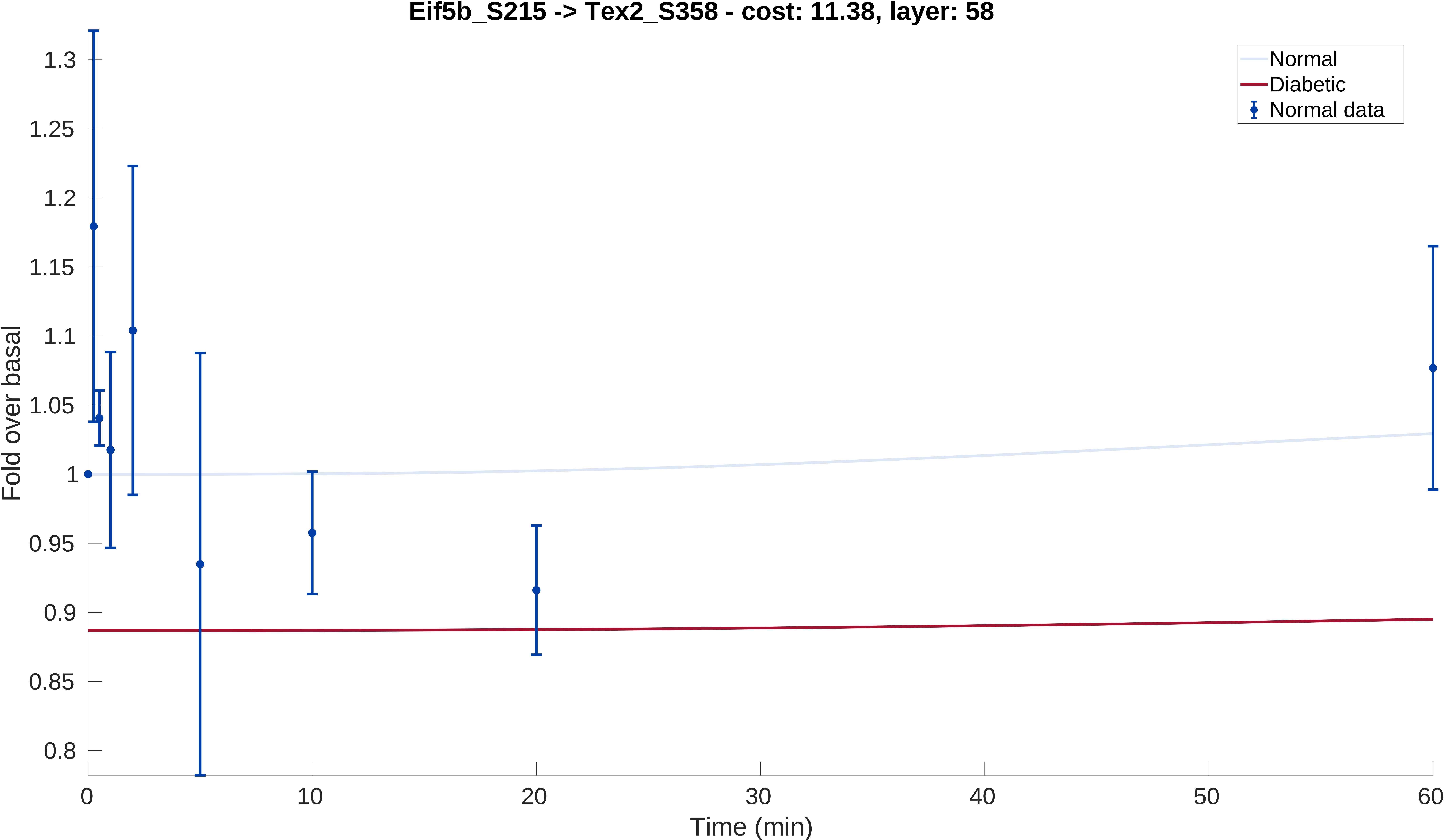

Eif5b_S215 -> Tex2_S358 - cost: 11.38, layer: 58
Normal
Diabetic
Normal data
Fold over basal
Time (min)
1.3
1.25
1.2
1.15
1.1
1.05
1
0.95
0.9
0.85
0.8
0
10
20
30
40
50
60

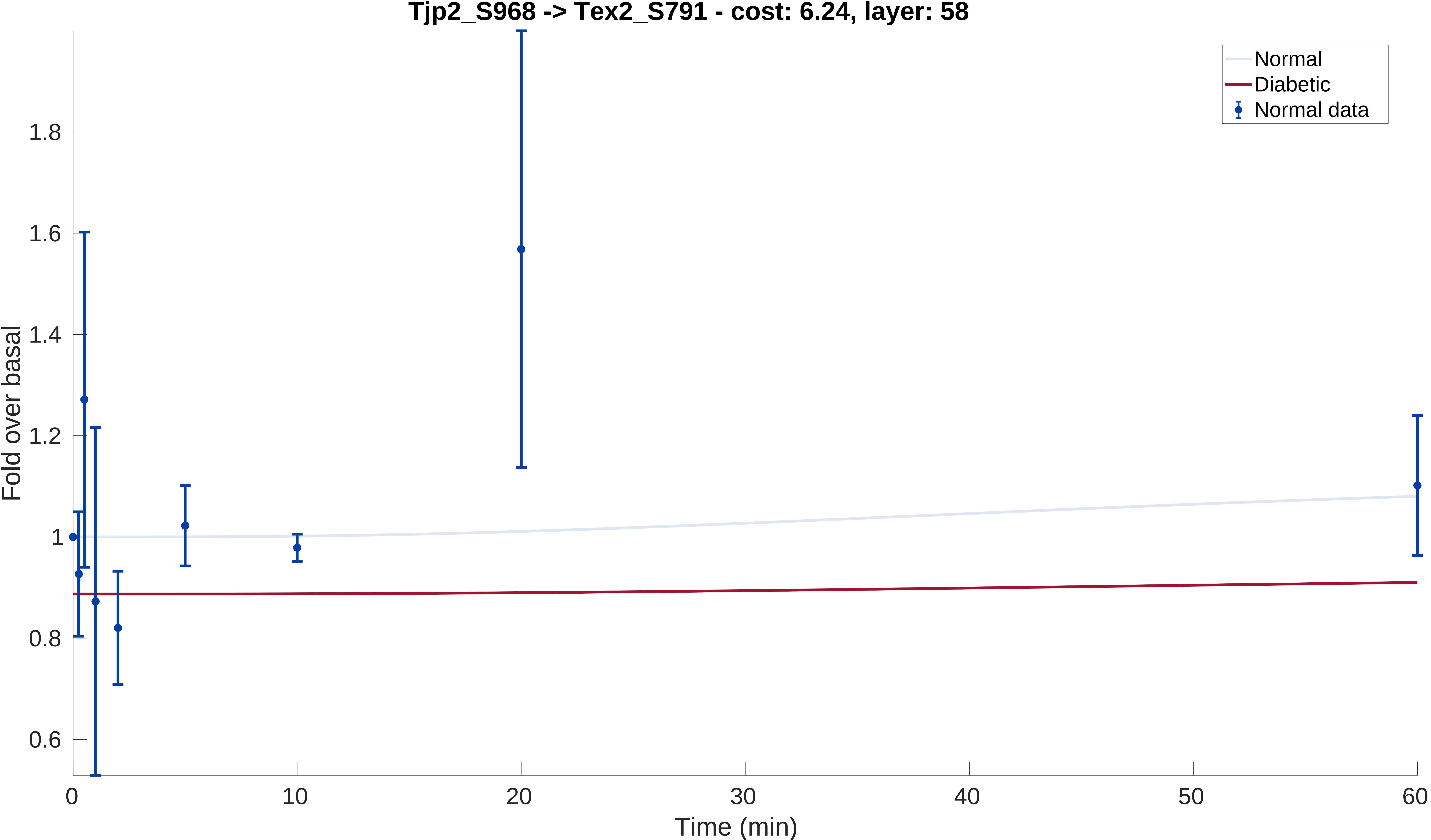

Tjp2_S968 -> Tex2_S791 - cost: 6.24, layer: 58
Normal
Diabetic
Normal data
Fold over basal
1.8
1.6
1.4
1.2
1
0.8
0.6
0
10
20
30
40
50
60
Time (min)

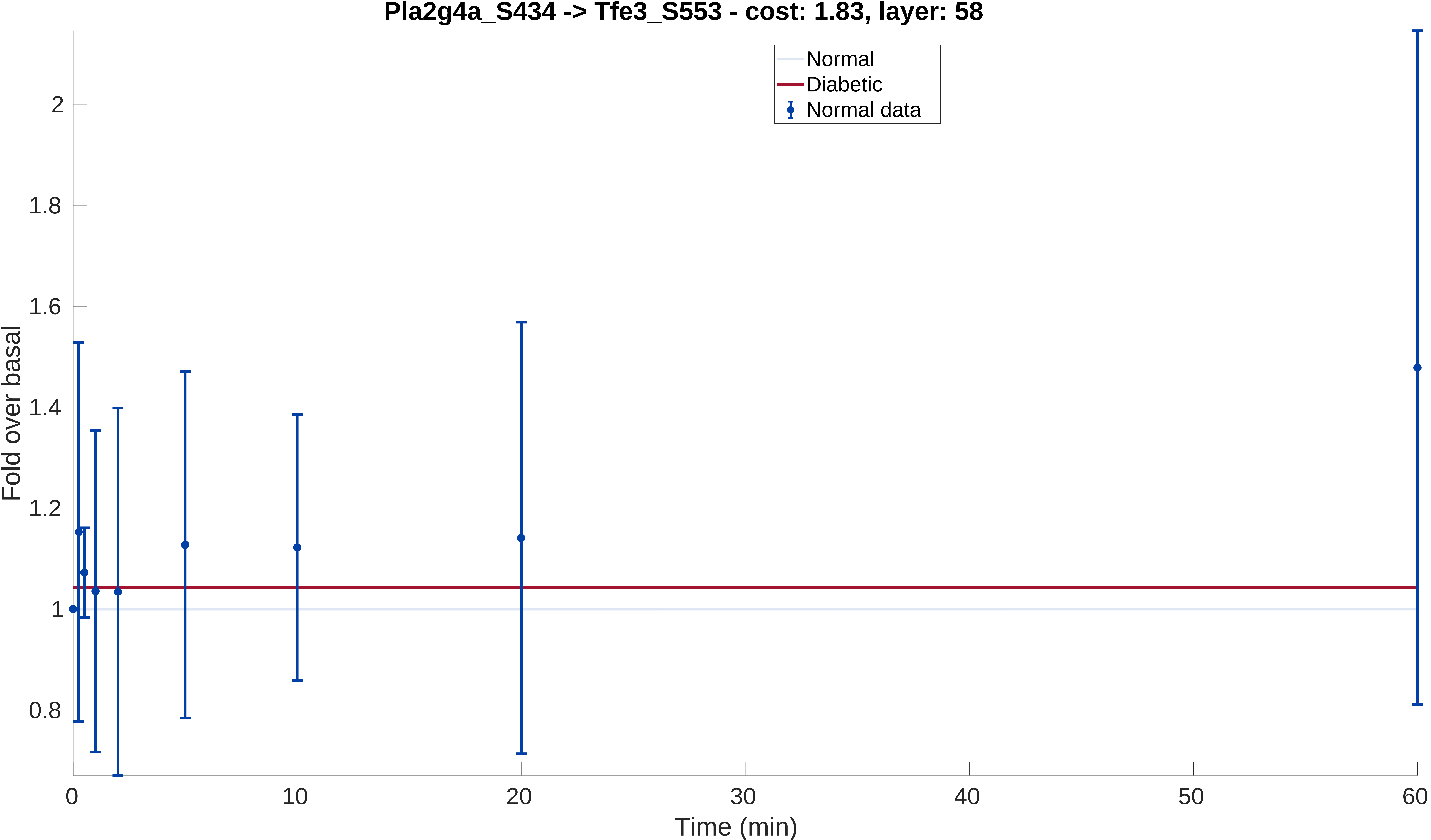

Pla2g4a_S434 -> Tfe3_S553 - cost: 1.83, layer: 58
Normal
Diabetic
Normal data
Fold over basal
Time (min)
0.8
1
1.2
1.4
1.6
1.8
2
0
10
20
30
40
50
60

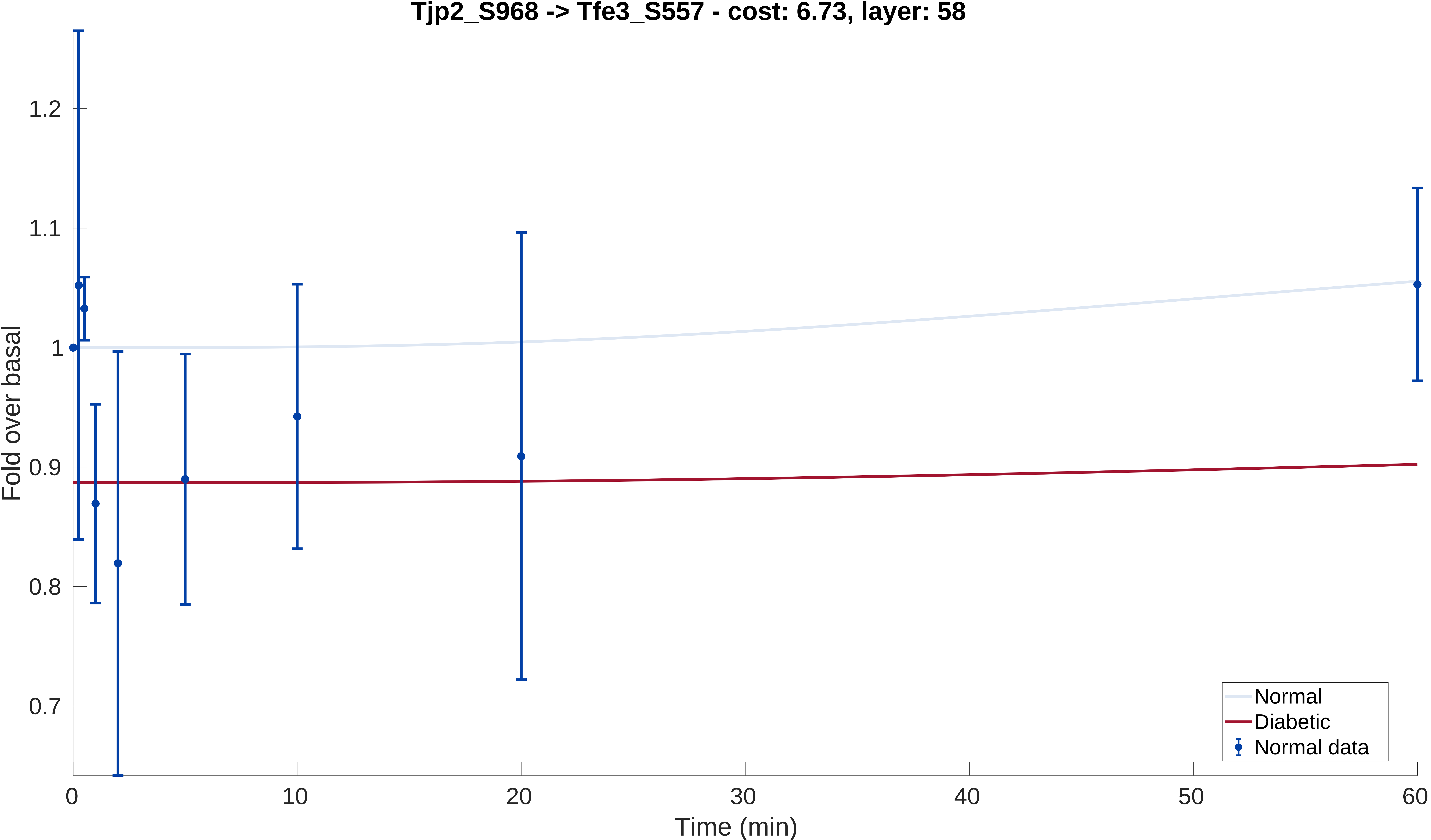
Tjp2_S968 -> Tfe3_S557 - cost: 6.73, layer: 58
Fold over basal
1.2
1.1
1
0.9
0.8
0.7
0
10
20
30
40
50
60
Time (min)
Normal
Diabetic
Normal data

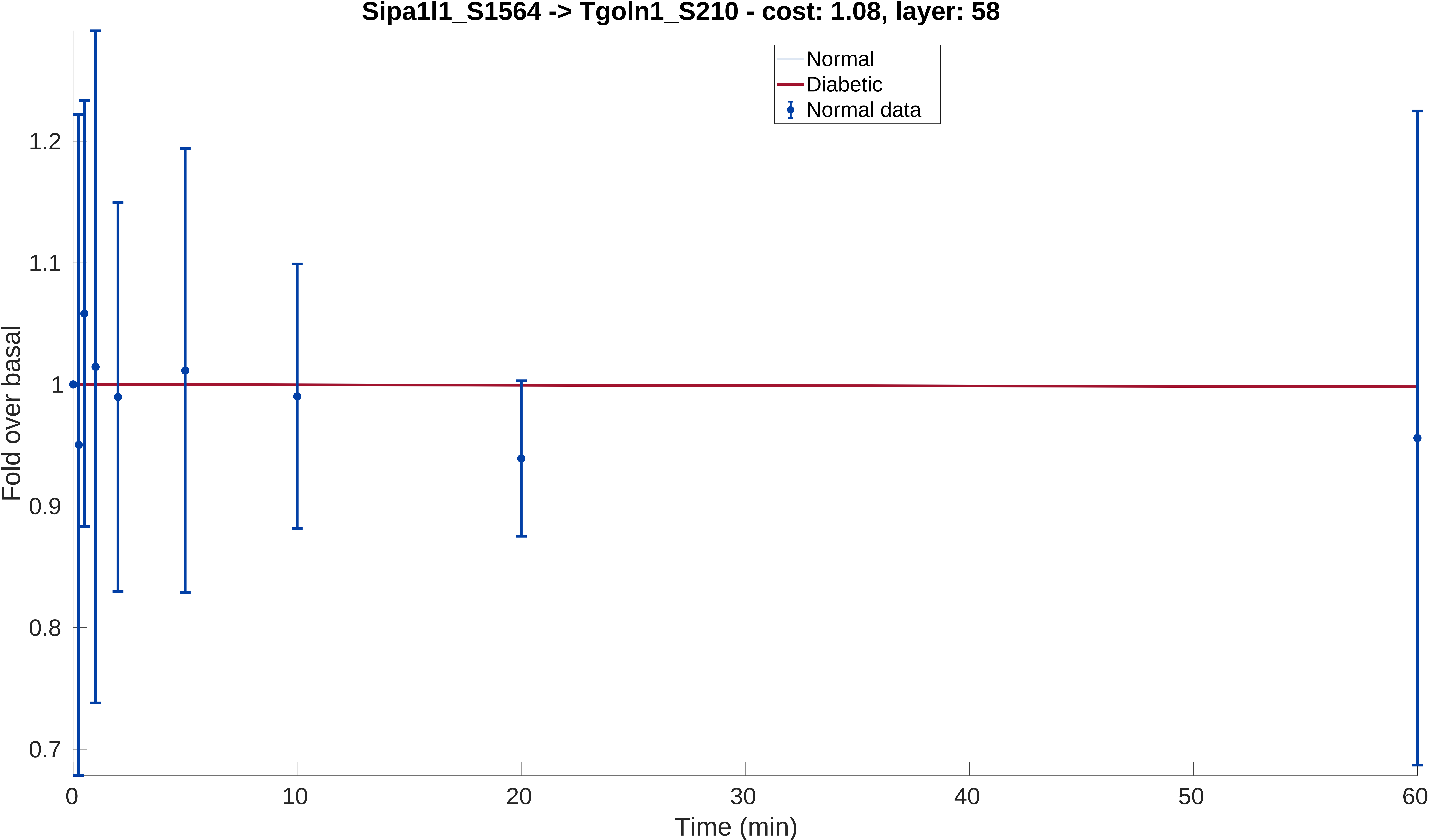

Sipa1l1_S1564 -> Tgoln1_S210 - cost: 1.08, layer: 58
Normal
Diabetic
Normal data
Fold over basal
1.2
1.1
1
0.9
0.8
0.7
0
10
20
30
40
50
60
Time (min)

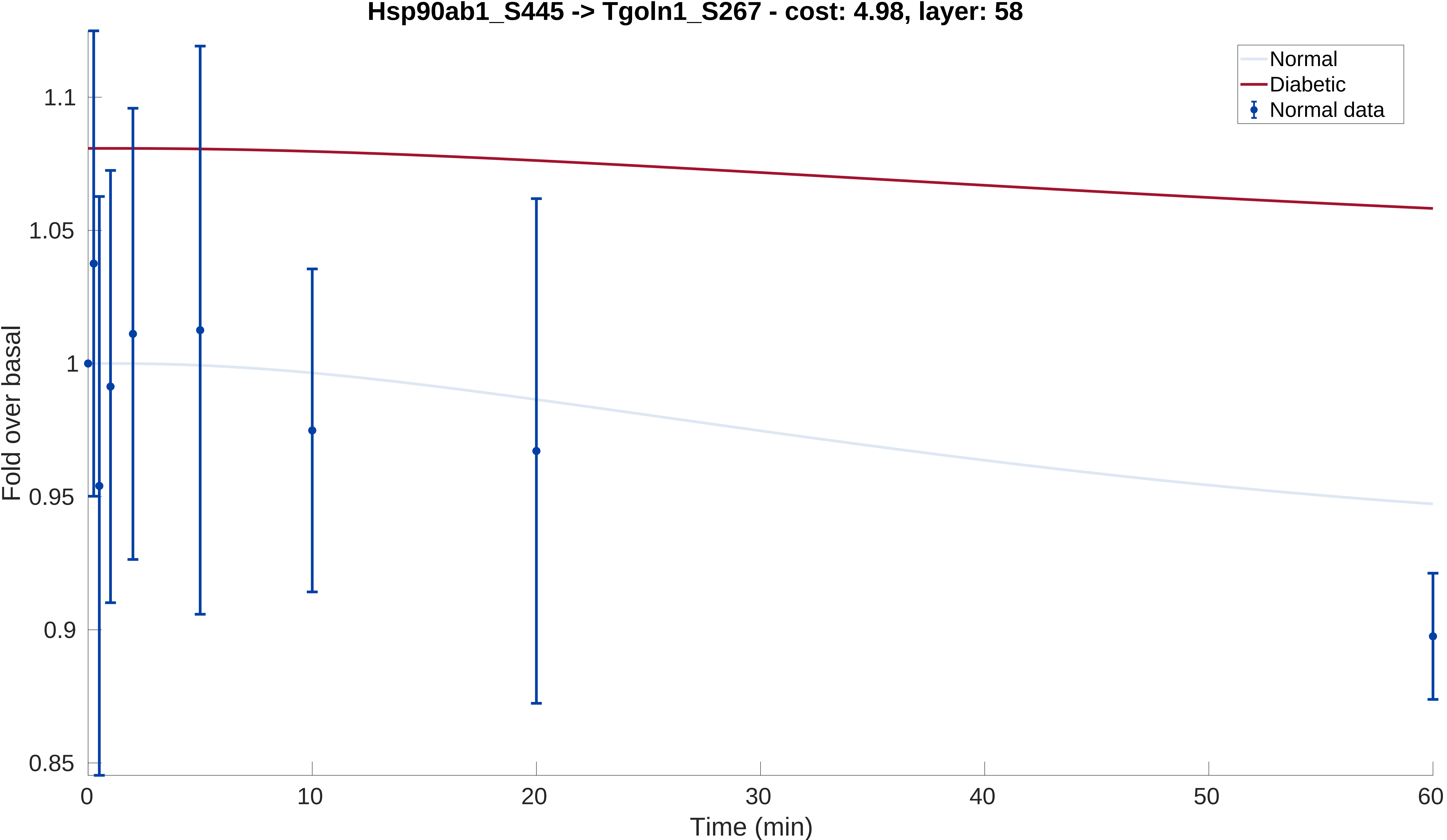
Hsp90ab1_S445 -> Tgoln1_S267 - cost: 4.98, layer: 58
Normal
Diabetic
Normal data
1.1
1.05
1
0.95
0.9
0.85
Fold over basal
0
10
20
30
40
50
60
Time (min)

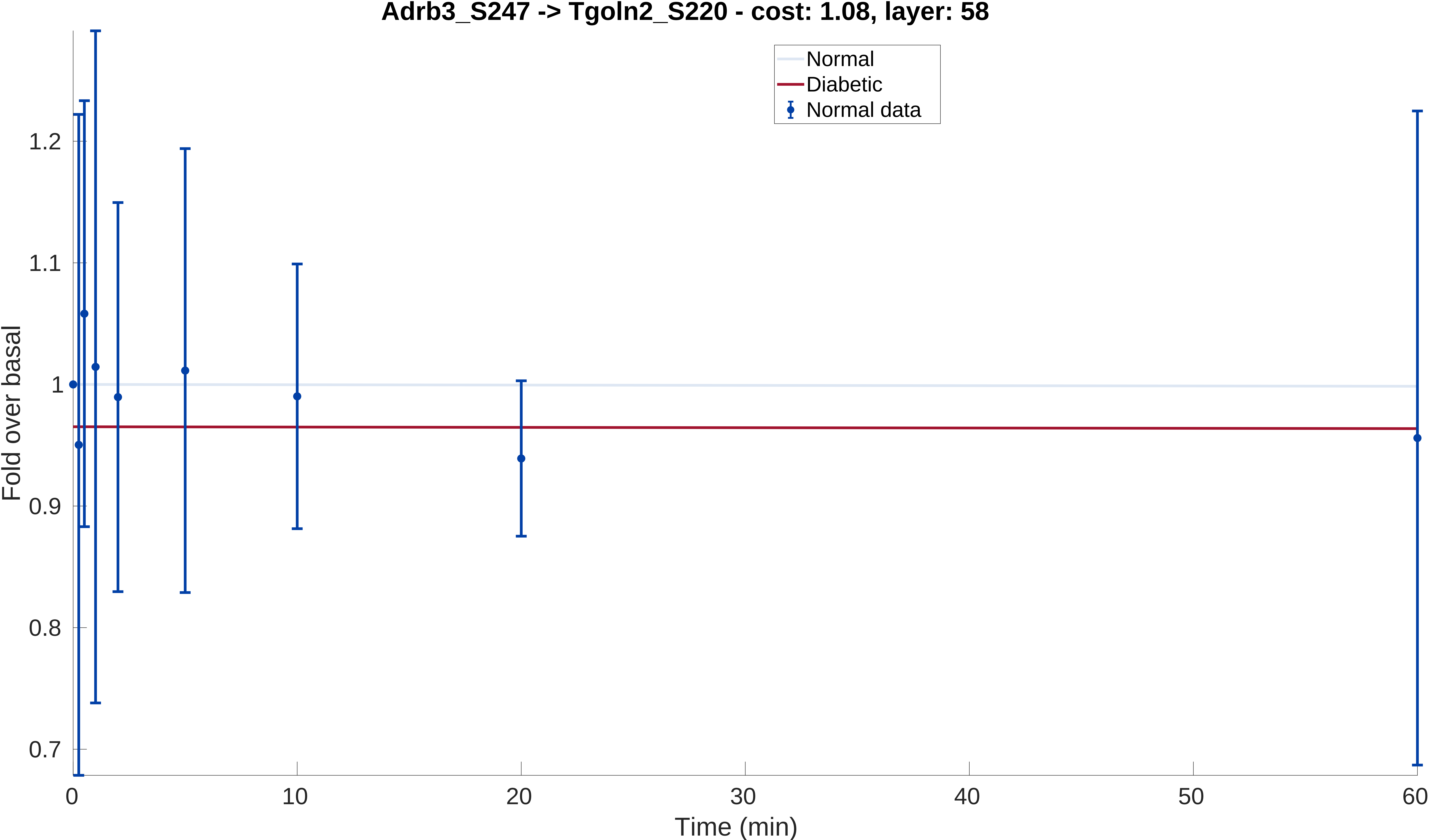

Adrb3_S247 -> Tgoln2_S220 - cost: 1.08, layer: 58
Normal
Diabetic
Normal data
Fold over basal
1.2
1.1
1
0.9
0.8
0.7
0
10
20
30
40
50
60
Time (min)

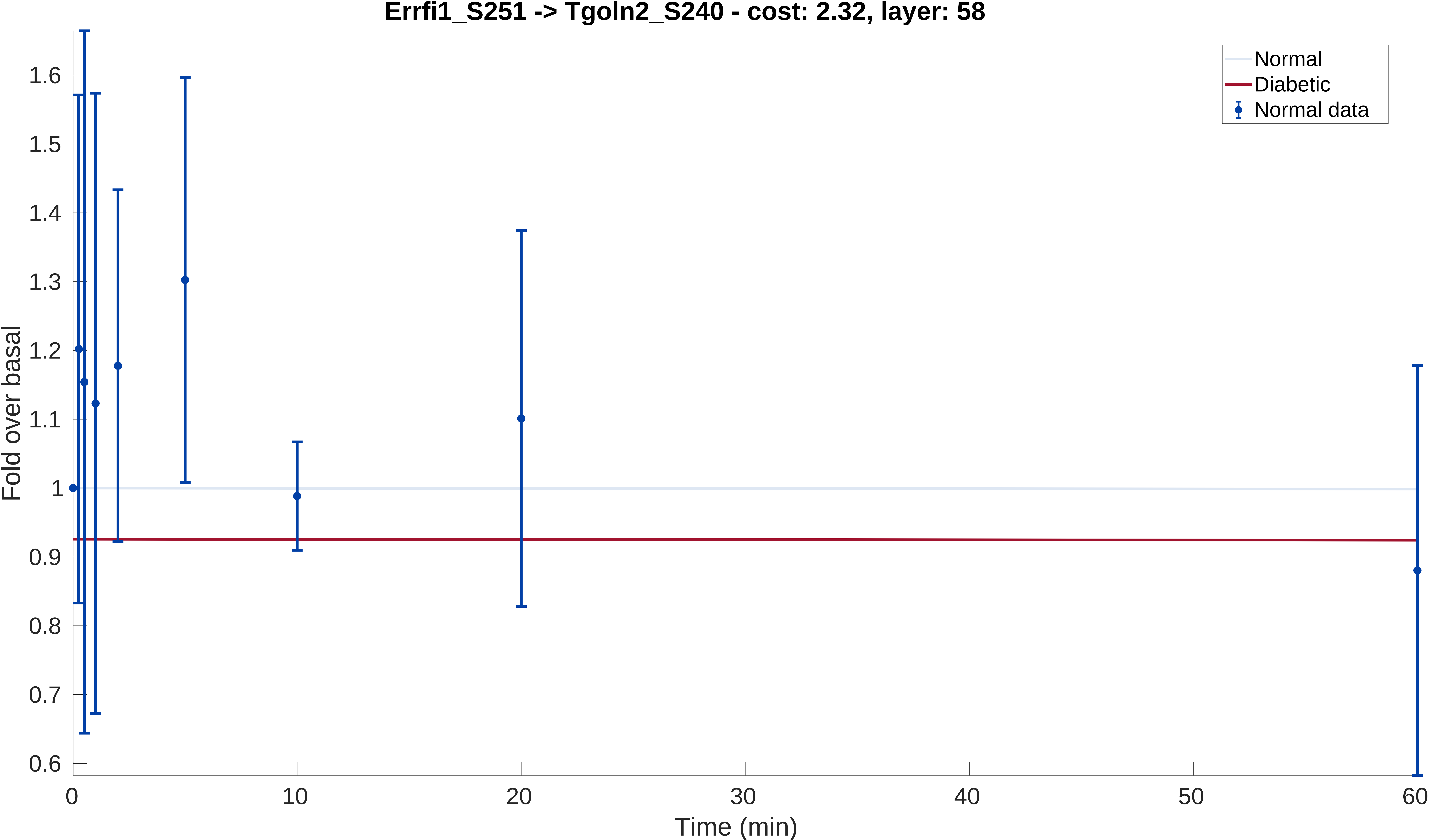

Errfi1_S251 -> Tgoln2_S240 - cost: 2.32, layer: 58
Normal
Diabetic
Normal data
Fold over basal
1.6
1.5
1.4
1.3
1.2
1.1
1
0.9
0.8
0.7
0.6
0
10
20
30
40
50
60
Time (min)

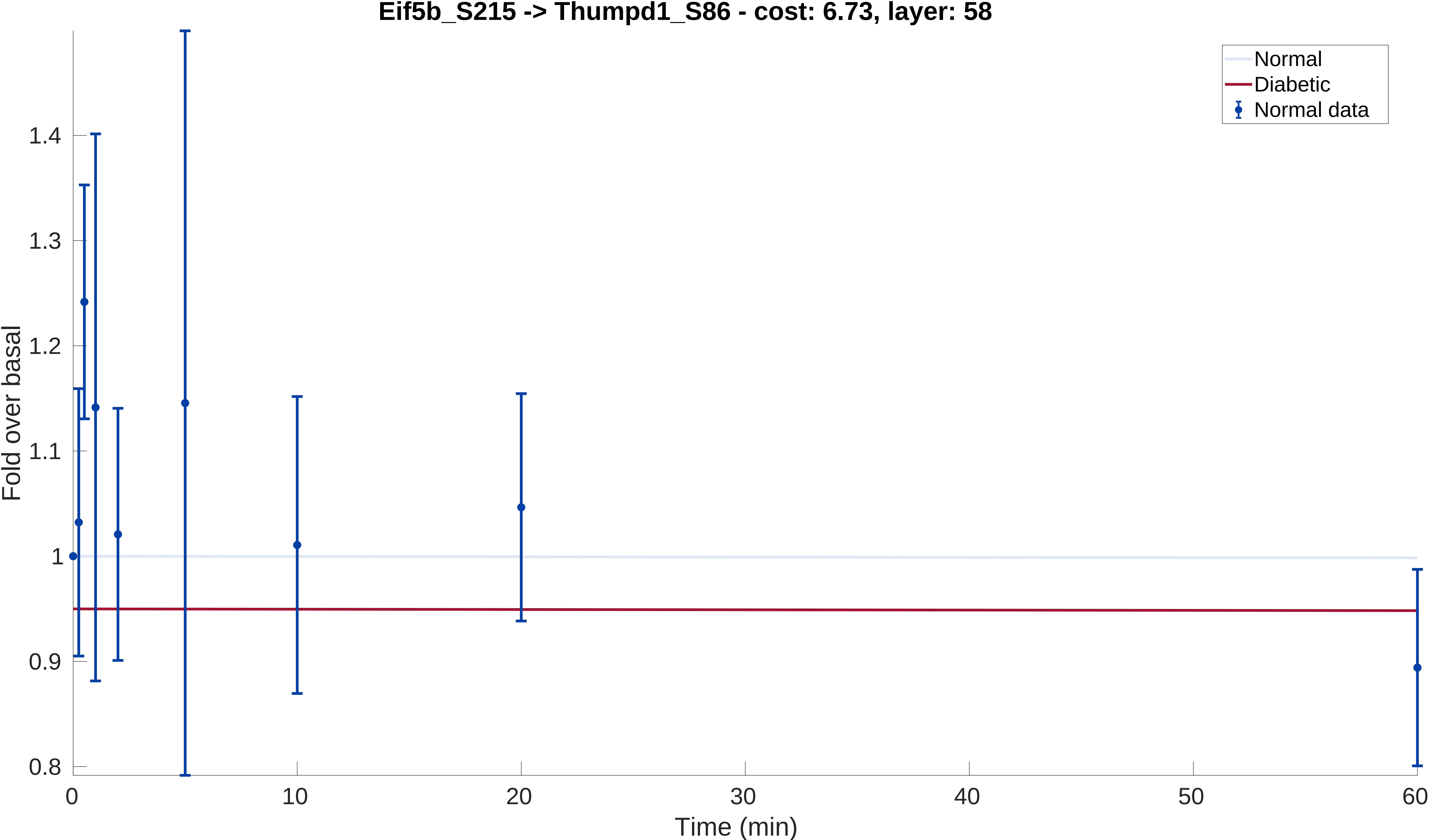

Eif5b_S215 -> Thumpd1_S86 - cost: 6.73, layer: 58
Normal
Diabetic
Normal data
Fold over basal
1.4
1.3
1.2
1.1
1
0.9
0.8
0
10
20
30
40
50
60
Time (min)

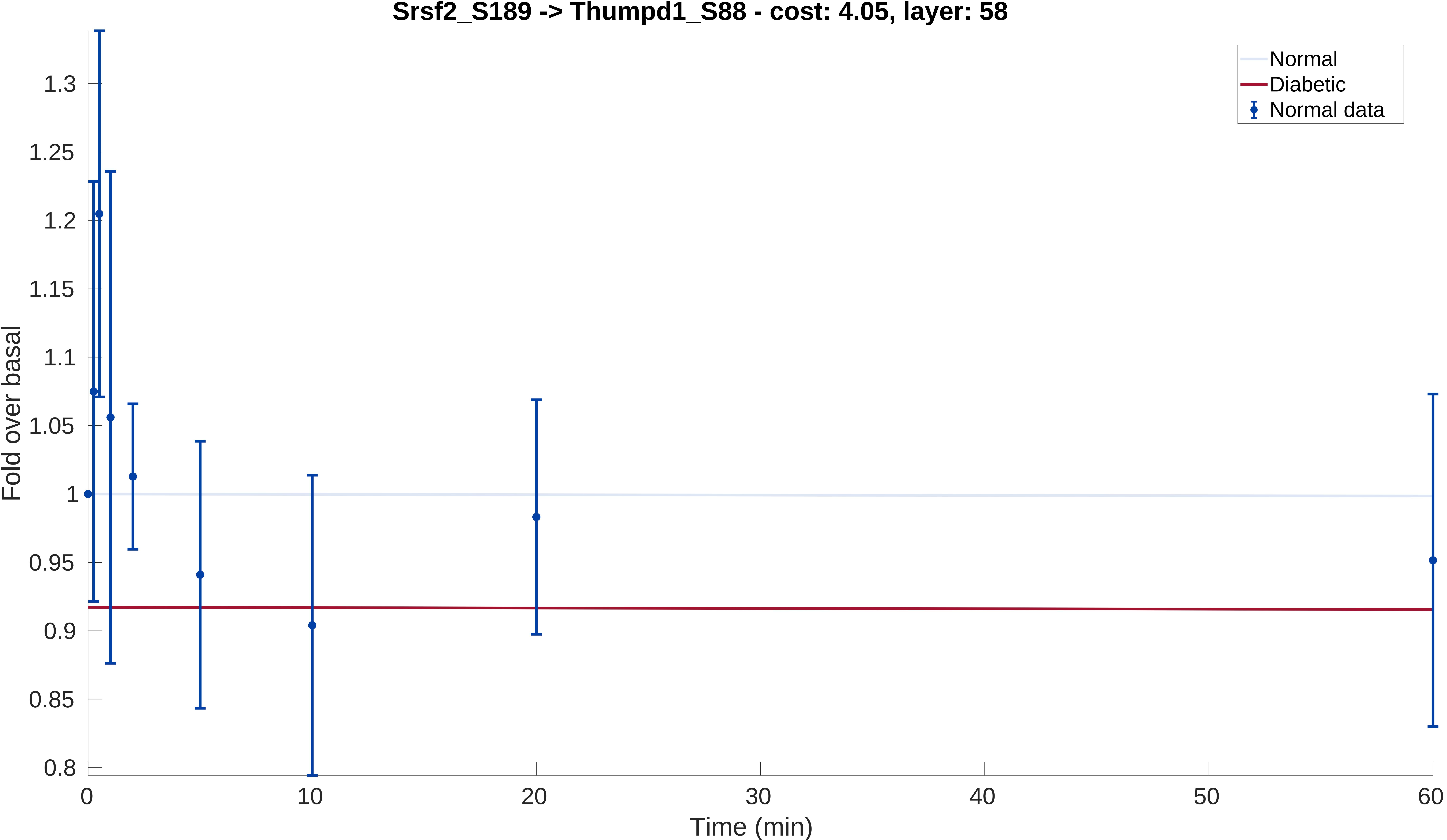

Srsf2_S189 -> Thumpd1_S88 - cost: 4.05, layer: 58
Normal
Diabetic
Normal data
Fold over basal
1.3
1.25
1.2
1.15
1.1
1.05
1
0.95
0.9
0.85
0.8
0
10
20
30
40
50
60
Time (min)

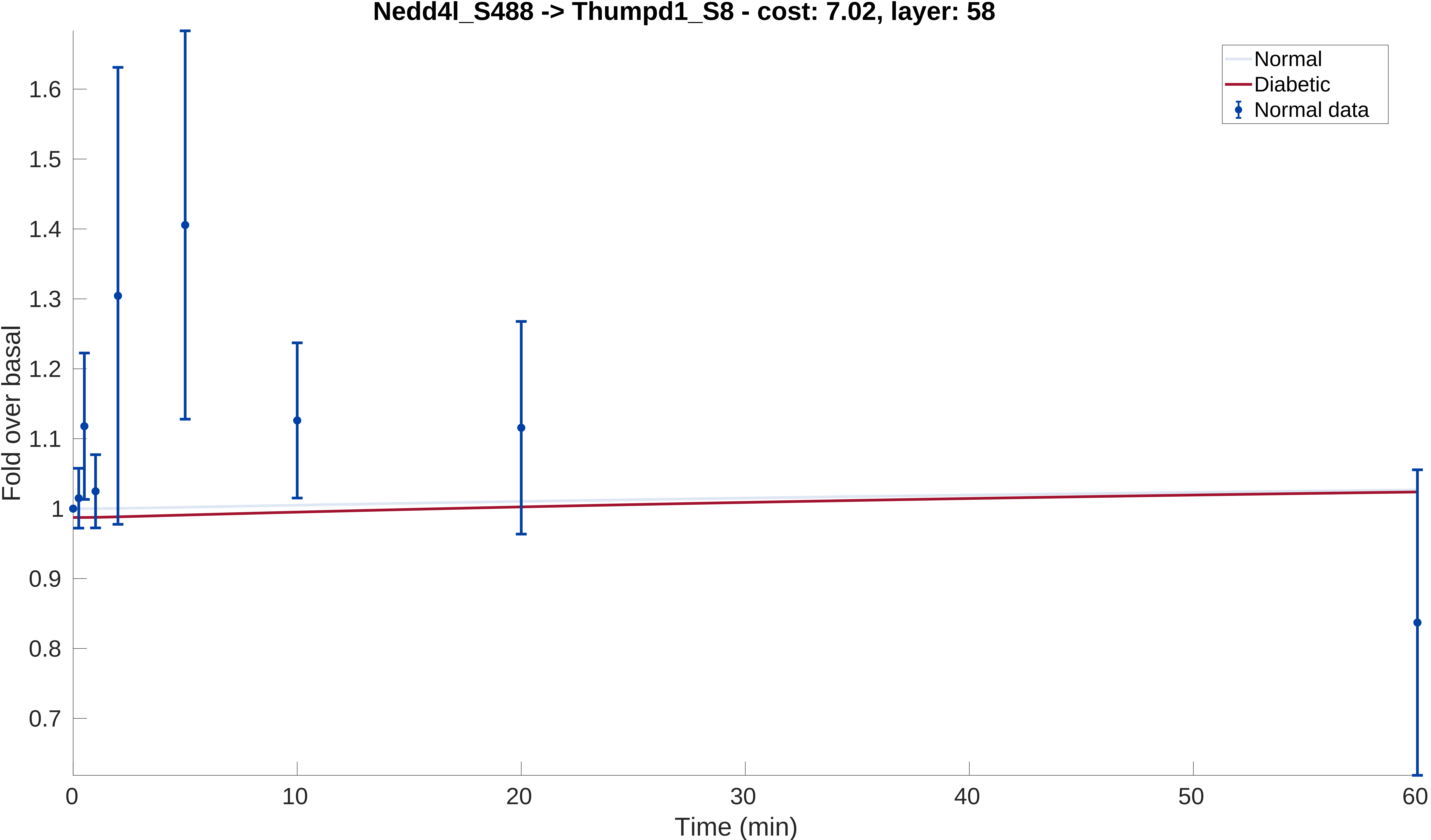

Nedd4l_S488 -> Thumpd1_S8 - cost: 7.02, layer: 58
Normal
Diabetic
Normal data
Fold over basal
Time (min)
1.6
1.5
1.4
1.3
1.2
1.1
1
0.9
0.8
0.7
0
10
20
30
40
50
60

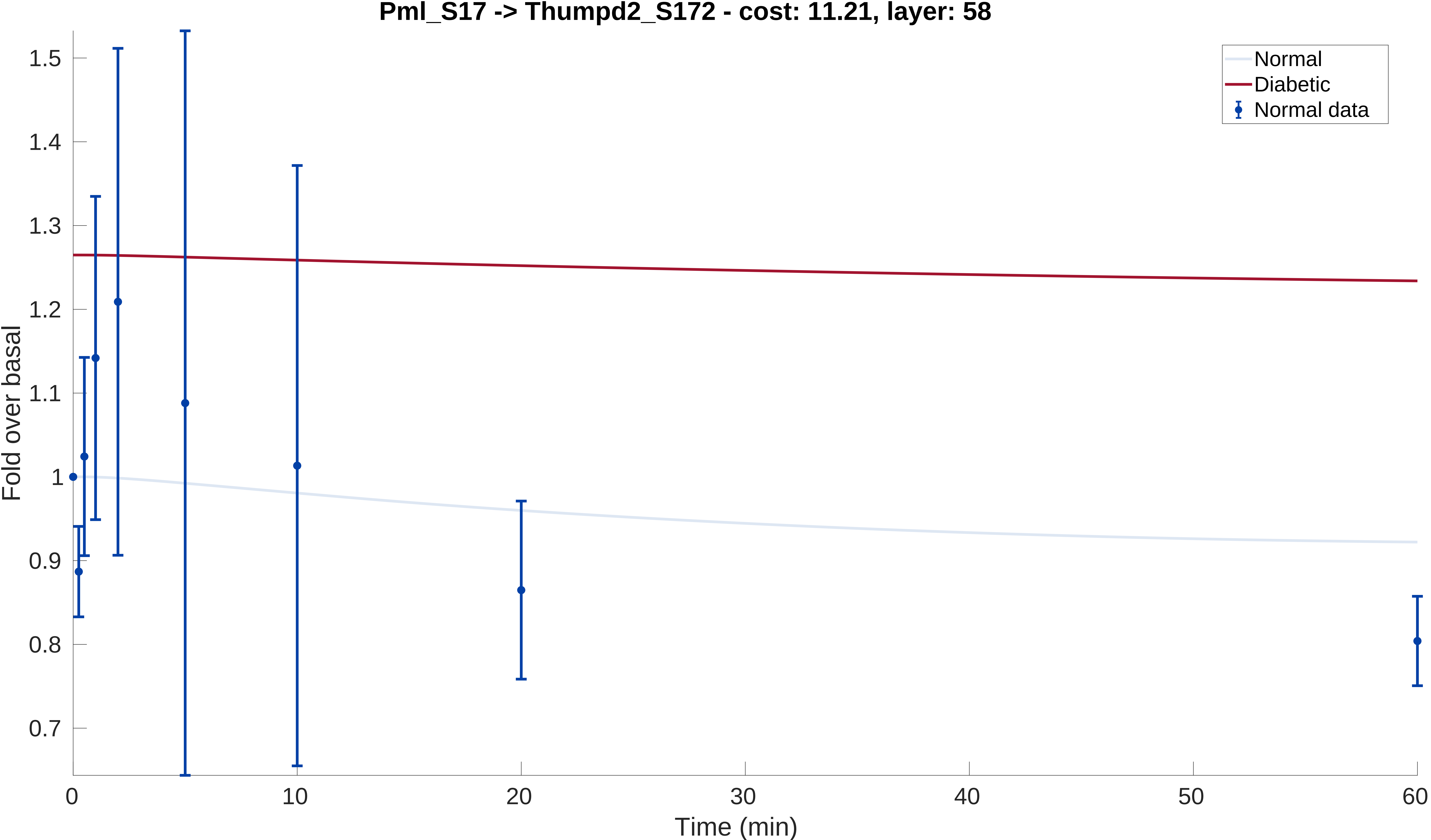
Pml_S17 -> Thumpd2_S172 - cost: 11.21, layer: 58
Normal
Diabetic
Normal data
Fold over basal
Time (min)
0.7
0.8
0.9
1
1.1
1.2
1.3
1.4
1.5
0
10
20
30
40
50
60

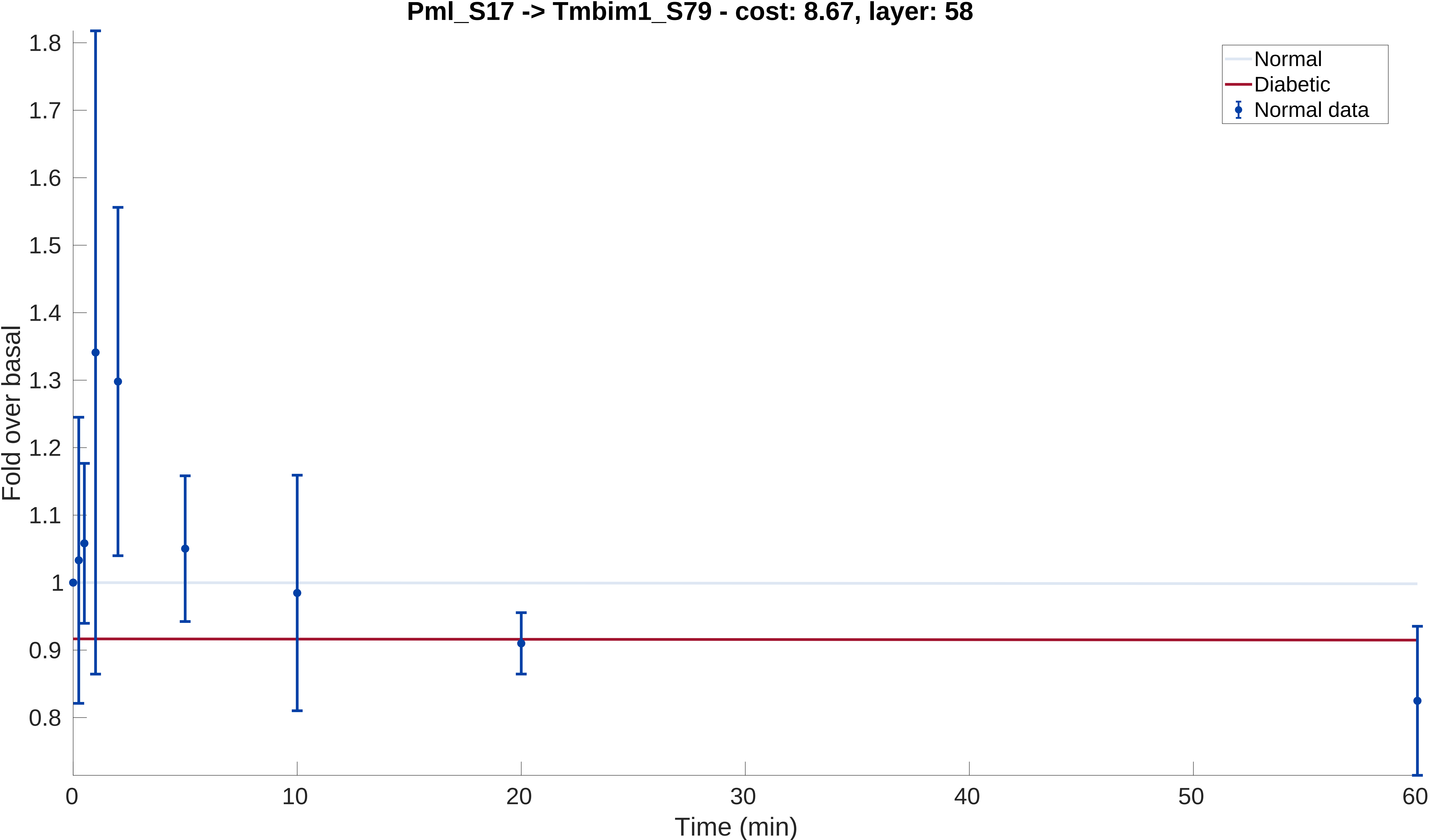

Pml_S17 -> Tmbim1_S79 - cost: 8.67, layer: 58
Normal
Diabetic
Normal data
Fold over basal
1.8
1.7
1.6
1.5
1.4
1.3
1.2
1.1
1
0.9
0.8
0
10
20
30
40
50
60
Time (min)

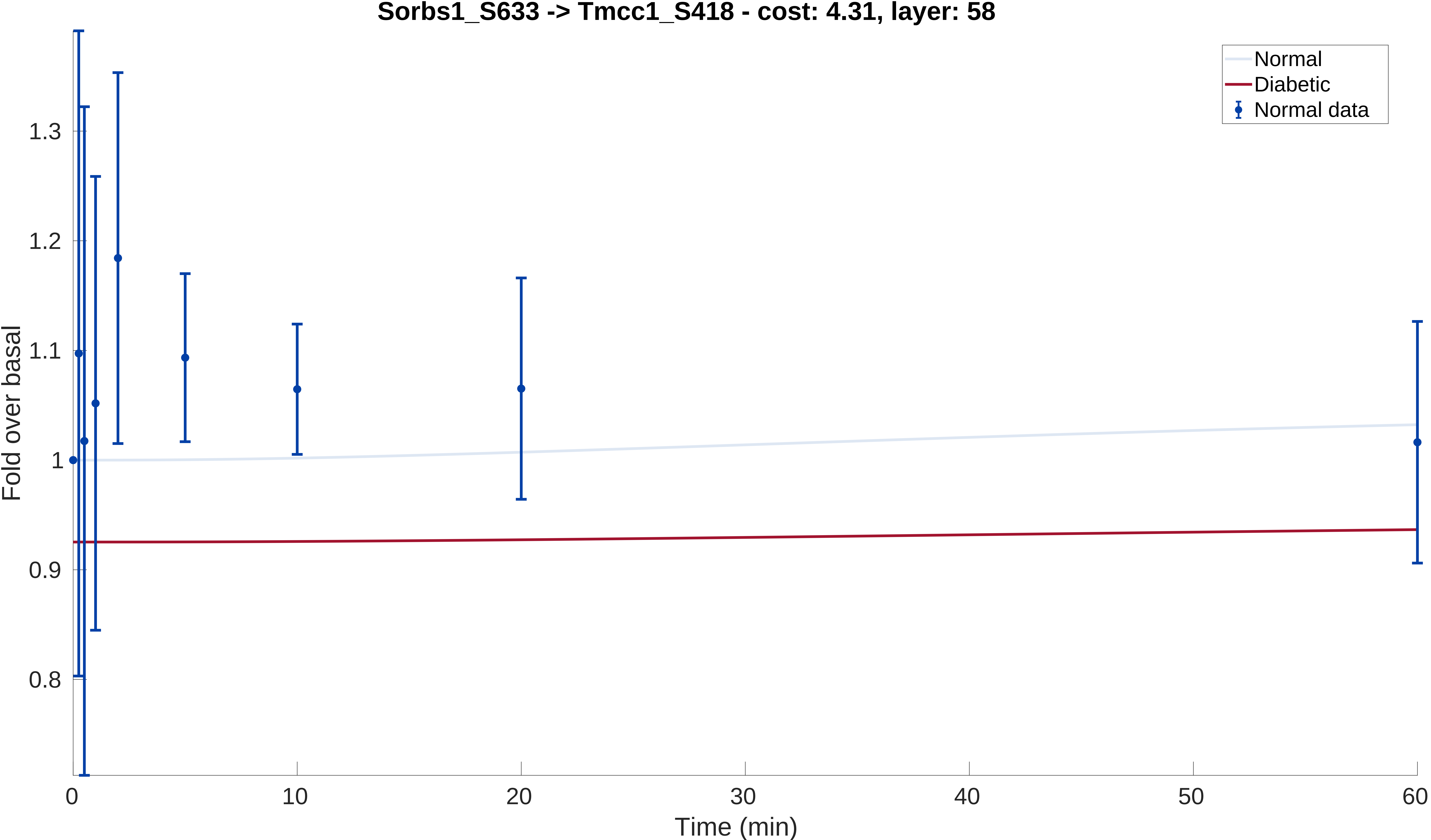

Sorbs1_S633 -> Tmcc1_S418 - cost: 4.31, layer: 58
Normal
Diabetic
Normal data
Fold over basal
Time (min)
1.3
1.2
1.1
1
0.9
0.8
0
10
20
30
40
50
60

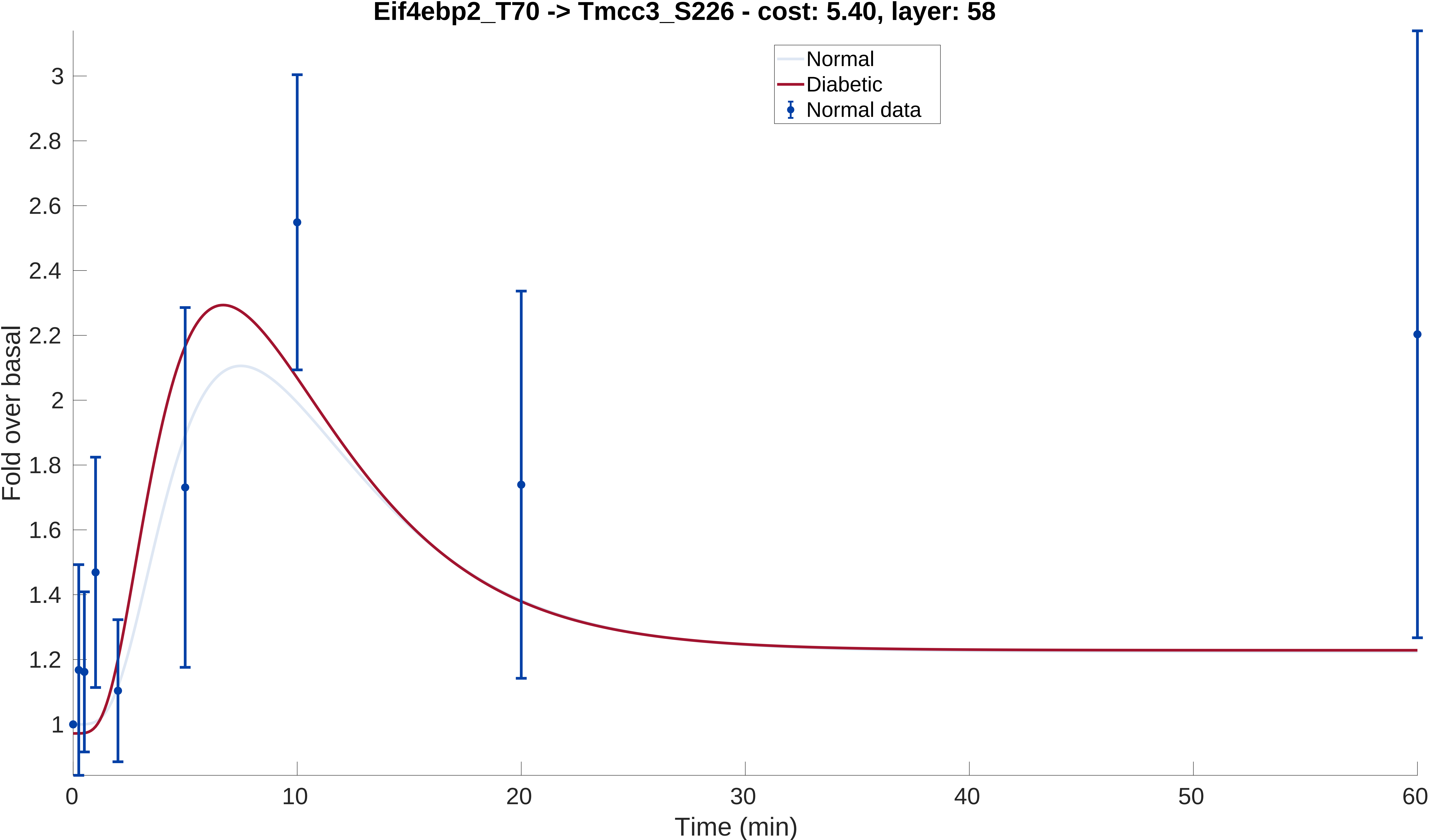

Eif4ebp2_T70 -> Tmcc3_S226 - cost: 5.40, layer: 58
Normal
Diabetic
Normal data
Fold over basal
3
2.8
2.6
2.4
2.2
2
1.8
1.6
1.4
1.2
1
0
10
20
30
40
50
60
Time (min)

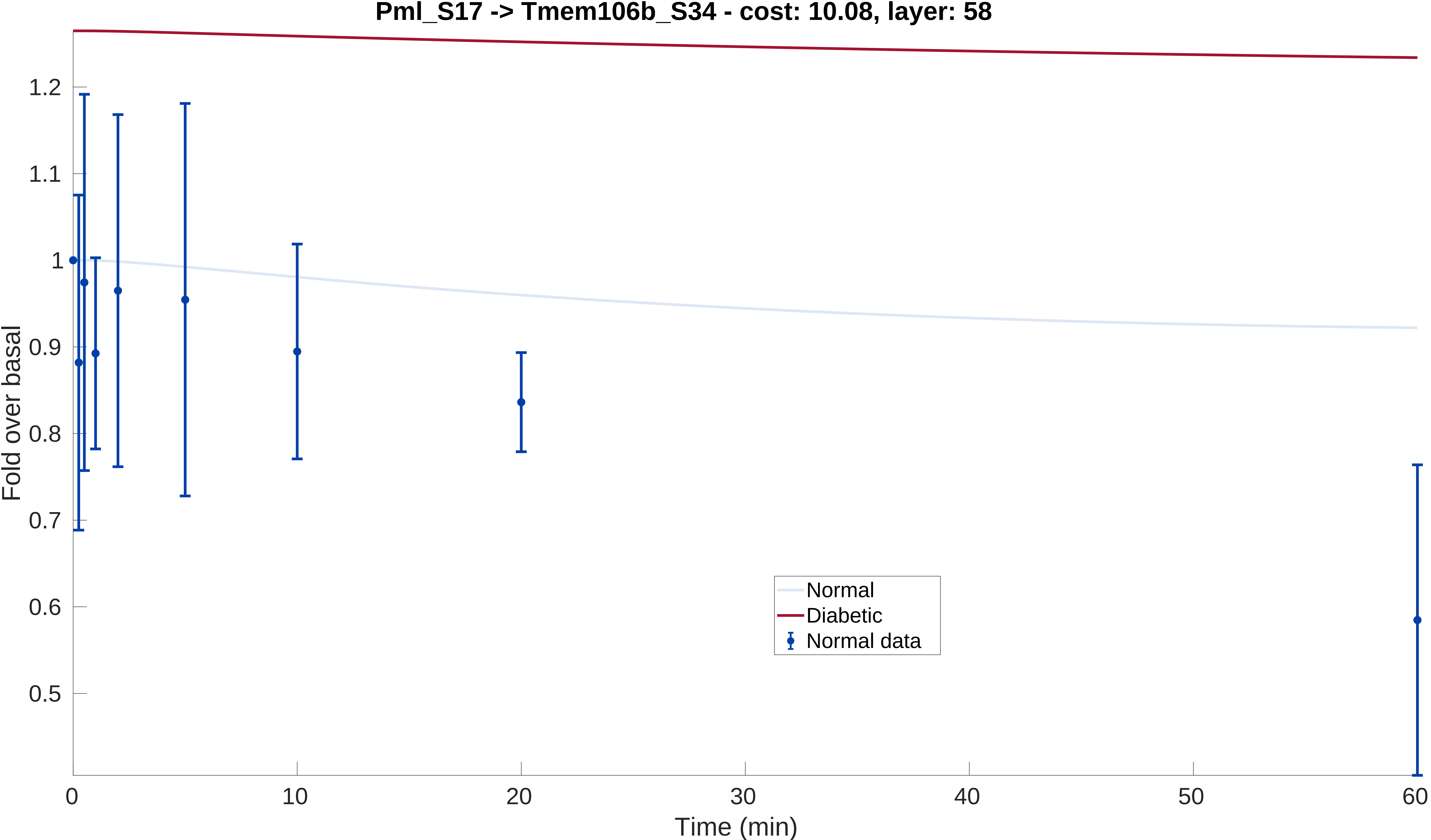

Pml_S17 -> Tmem106b_S34 - cost: 10.08, layer: 58
Fold over basal
1.2
1.1
1
0.9
0.8
0.7
0.6
0.5
0
10
20
30
40
50
60
Time (min)
Normal
Diabetic
Normal data

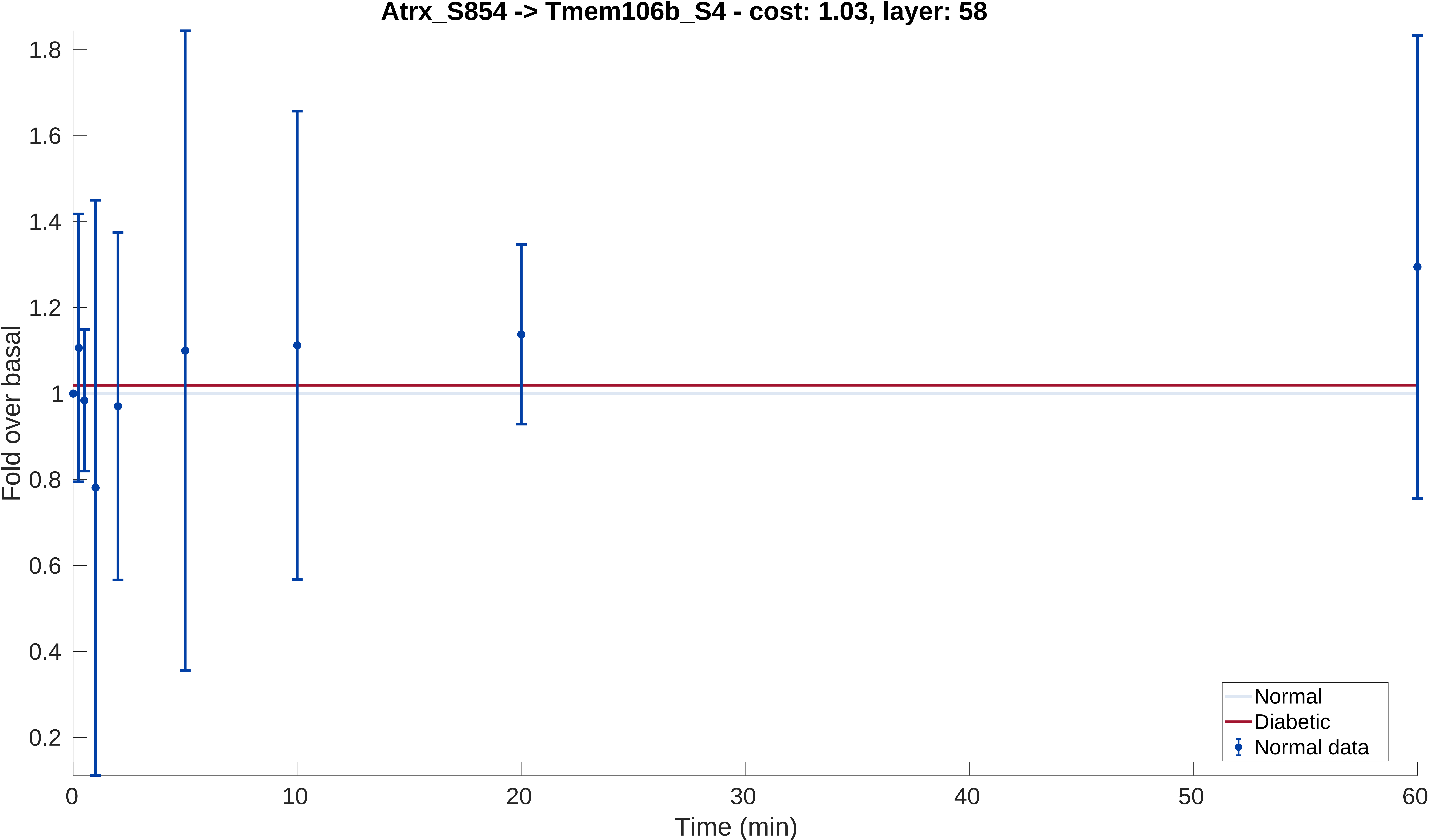

Atrx_S854 -> Tmem106b_S4 - cost: 1.03, layer: 58
Fold over basal
1.8
1.6
1.4
1.2
1
0.8
0.6
0.4
0.2
0
10
20
30
40
50
60
Time (min)
Normal
Diabetic
Normal data

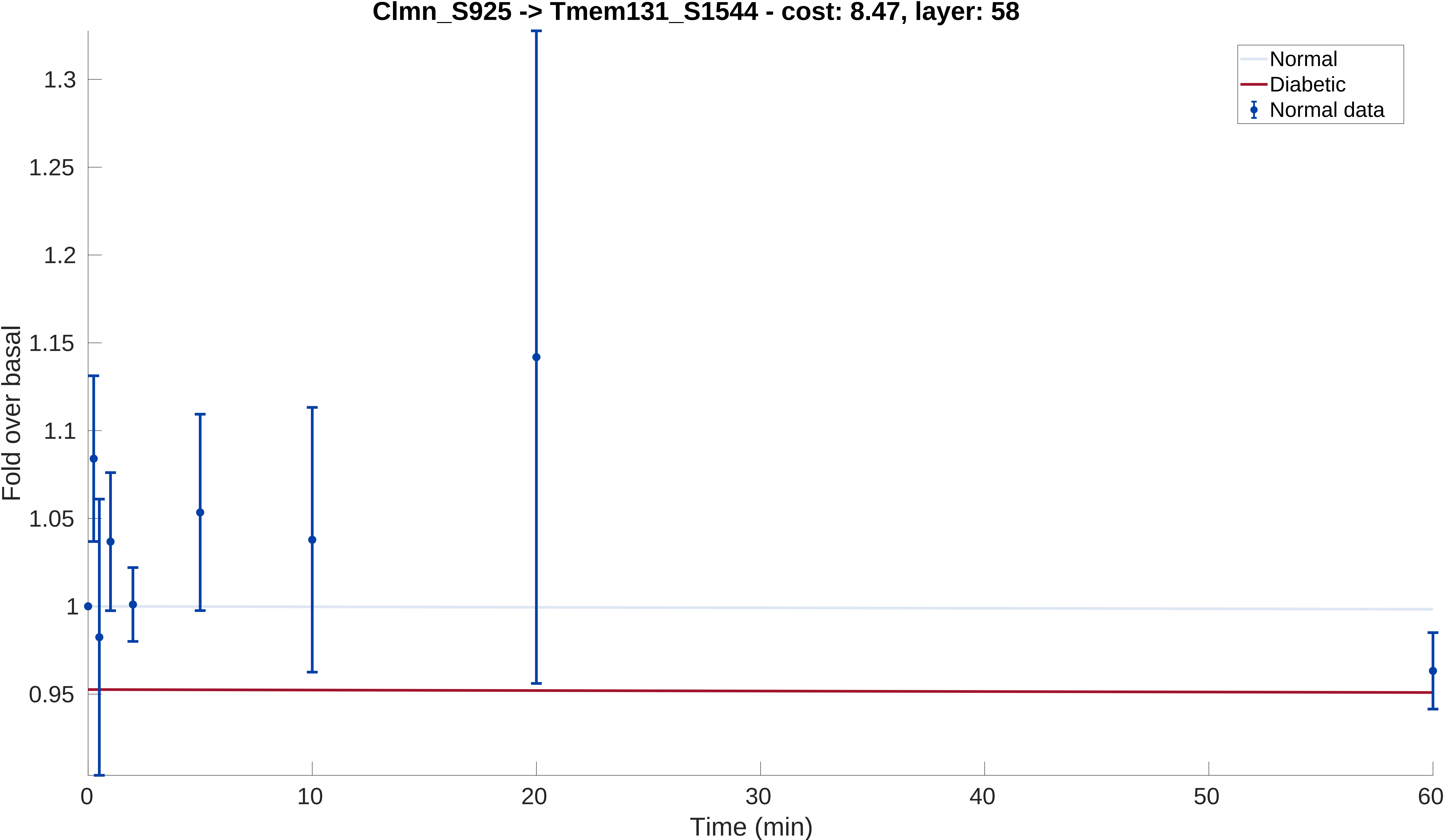

Clmn_S925 -> Tmem131_S1544 - cost: 8.47, layer: 58
Normal
Diabetic
Normal data
Fold over basal
1.3
1.25
1.2
1.15
1.1
1.05
1
0.95
0
10
20
30
40
50
60
Time (min)

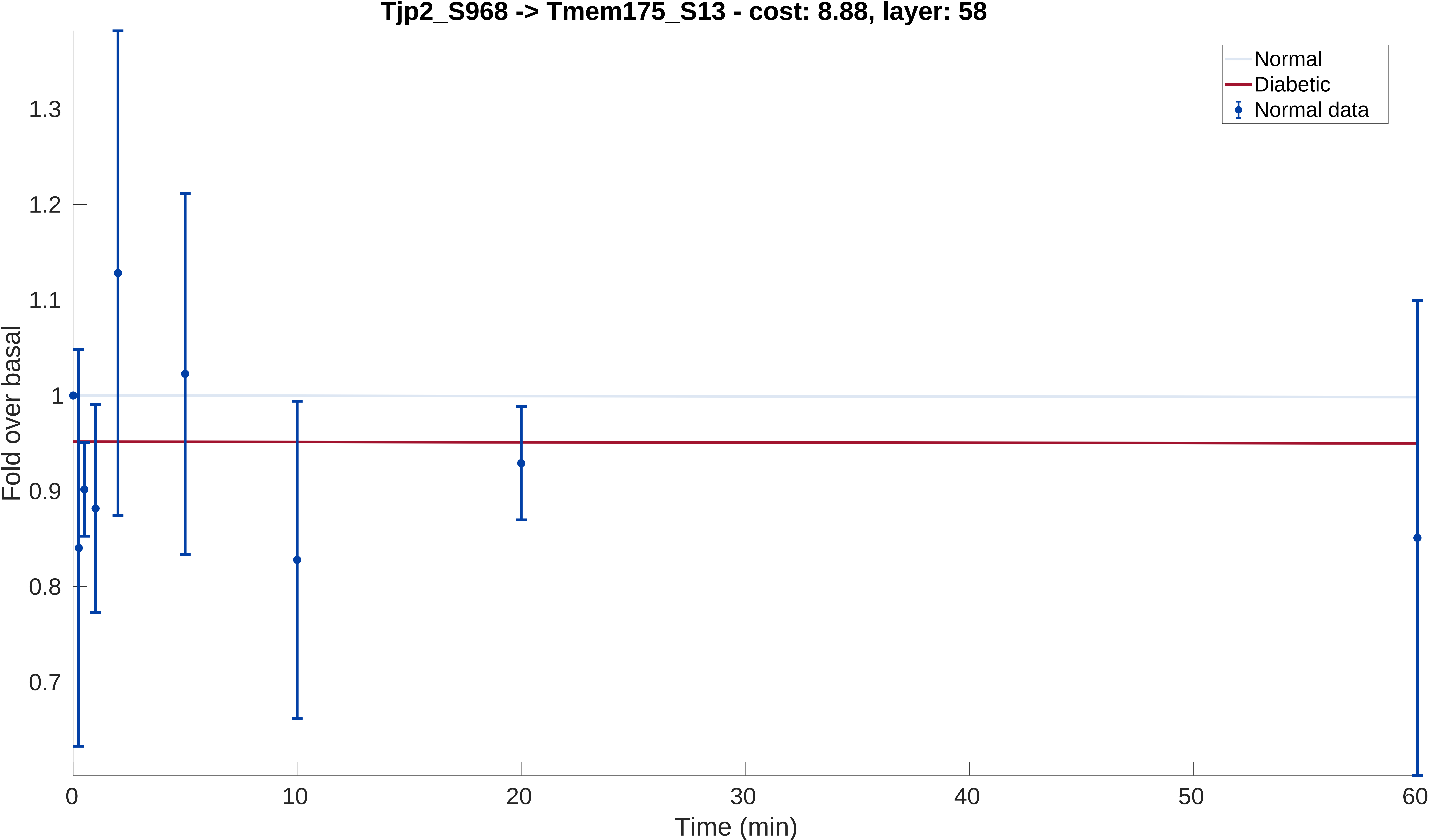

Tjp2_S968 -> Tmem175_S13 - cost: 8.88, layer: 58
Normal
Diabetic
Normal data
Fold over basal
1.3
1.2
1.1
1
0.9
0.8
0.7
0
10
20
30
40
50
60
Time (min)

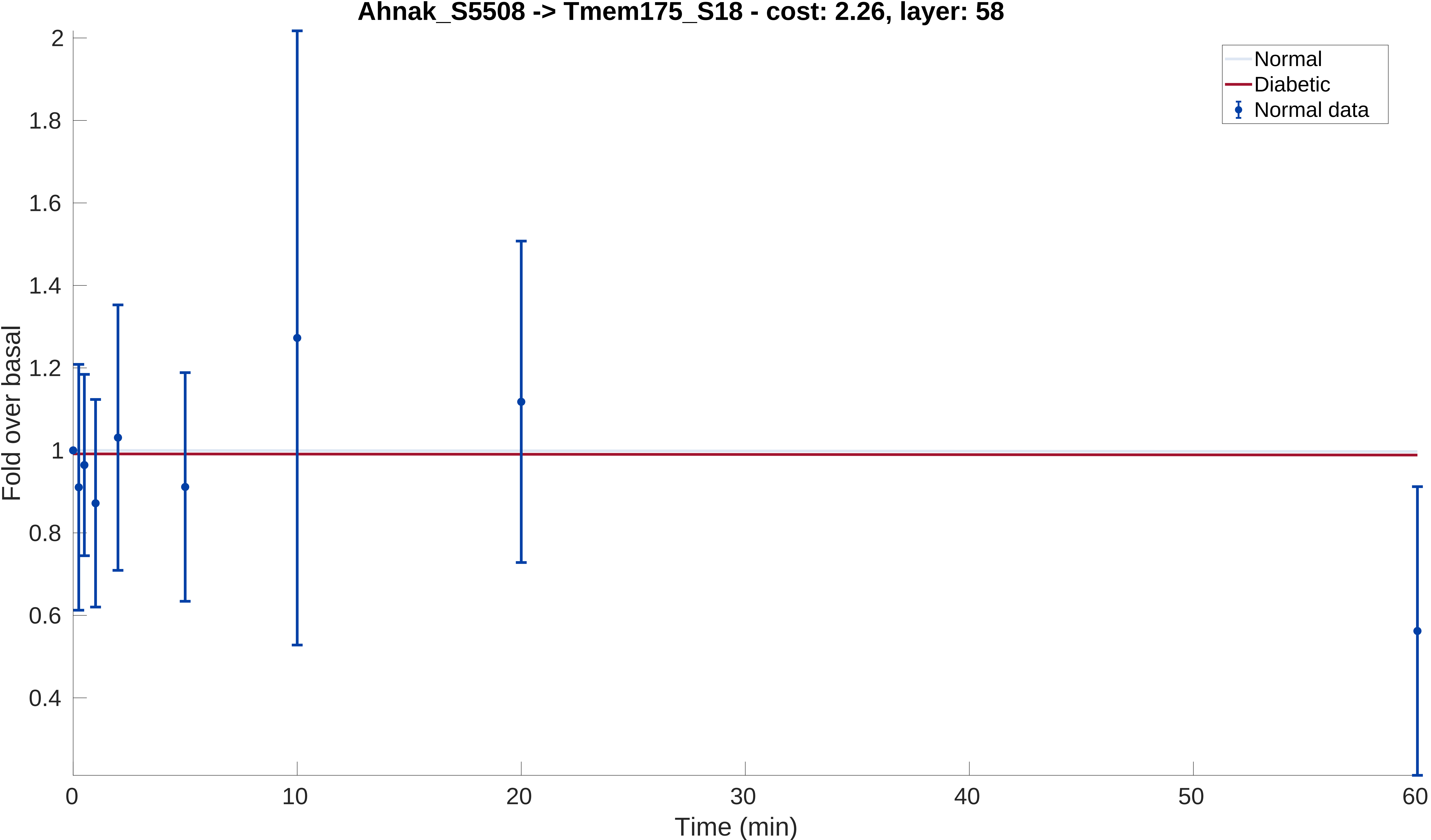

Ahnak_S5508 -> Tmem175_S18 - cost: 2.26, layer: 58
Normal
Diabetic
Normal data
Fold over basal
Time (min)
2
1.8
1.6
1.4
1.2
1
0.8
0.6
0.4
0
10
20
30
40
50
60

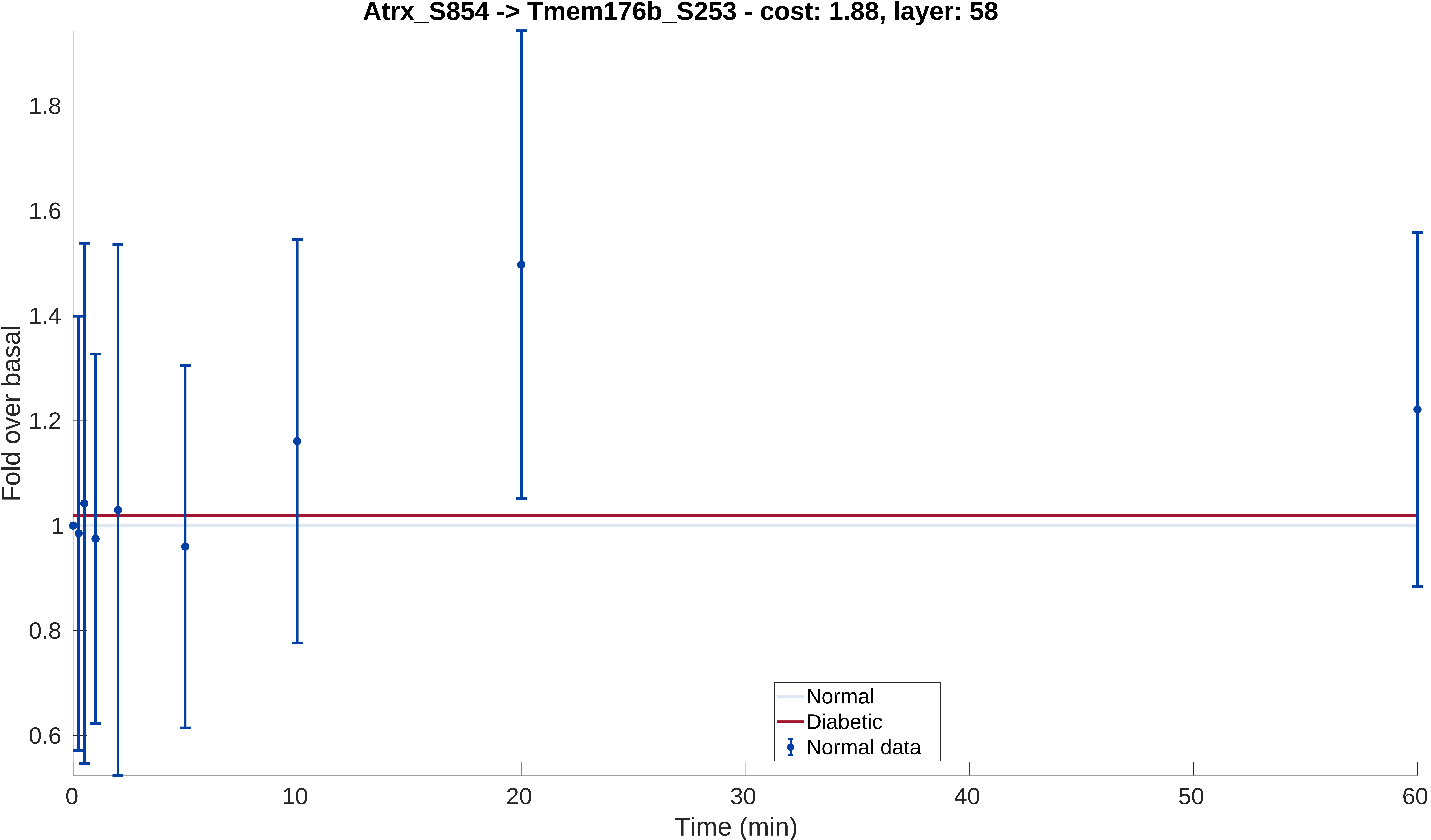

Atrx_S854 -> Tmem176b_S253 - cost: 1.88, layer: 58
Fold over basal
Time (min)
0.6
0.8
1
1.2
1.4
1.6
1.8
0
10
20
30
40
50
60
Normal
Diabetic
Normal data

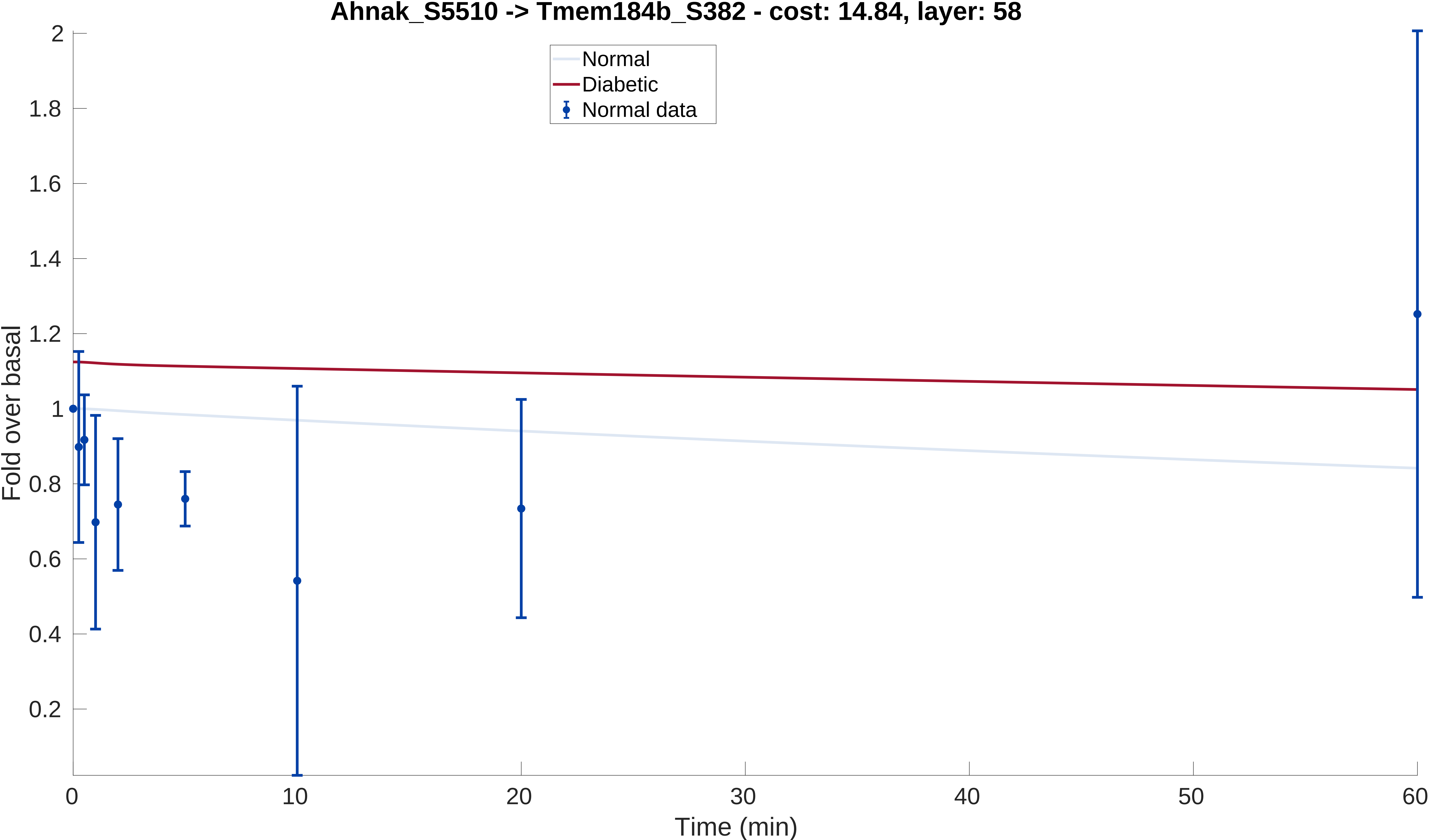

Ahnak_S5510 -> Tmem184b_S382 - cost: 14.84, layer: 58
Normal
Diabetic
Normal data
Fold over basal
2
1.8
1.6
1.4
1.2
1
0.8
0.6
0.4
0.2
0
10
20
30
40
50
60
Time (min)

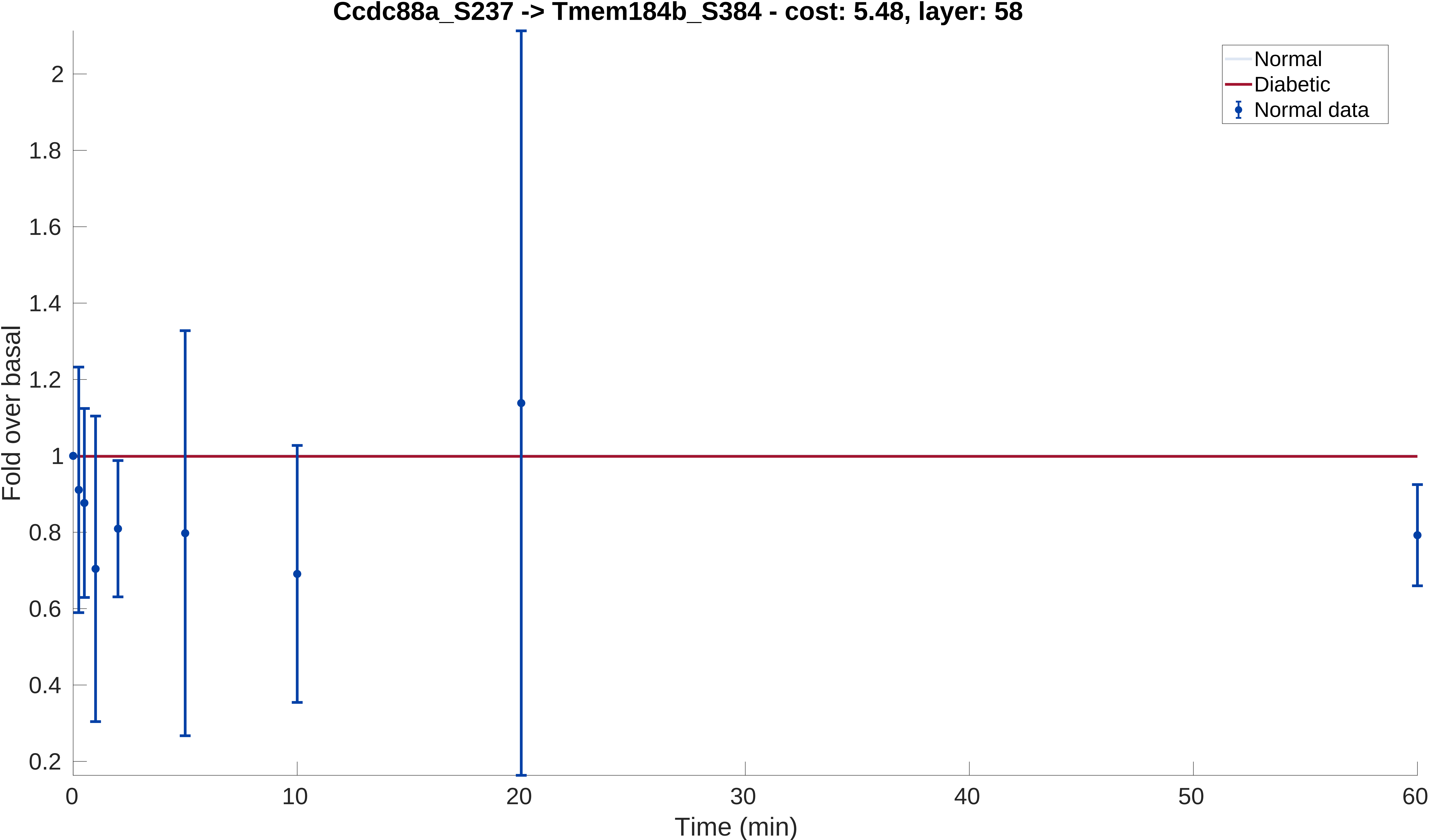

Ccdc88a_S237 -> Tmem184b_S384 - cost: 5.48, layer: 58
Normal
Diabetic
Normal data
Fold over basal
Time (min)
0
10
20
30
40
50
60
0.2
0.4
0.6
0.8
1
1.2
1.4
1.6
1.8
2

#### Helb_S945 -> Tmem186_S48 - cost: 3.89, layer: 58

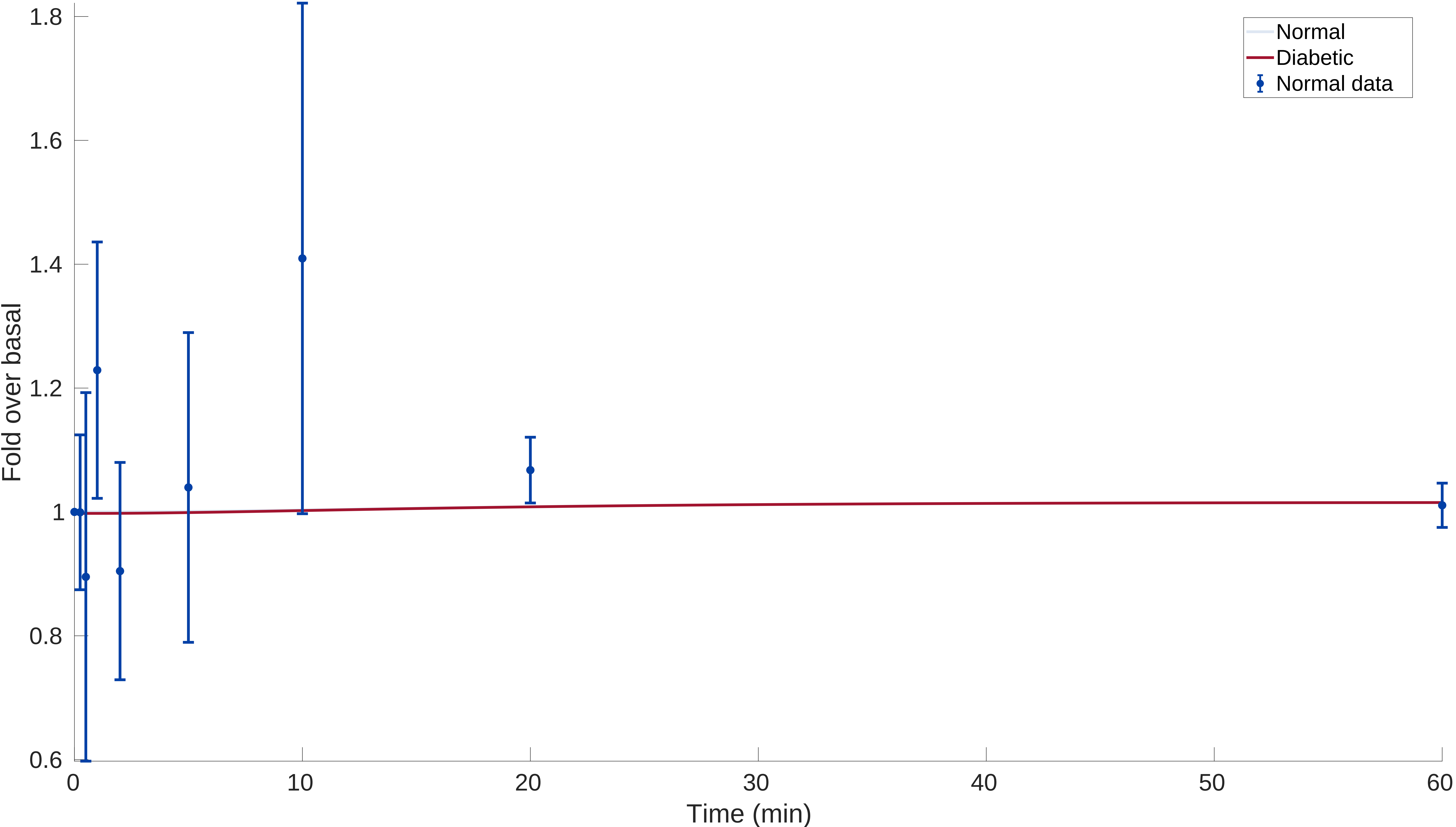

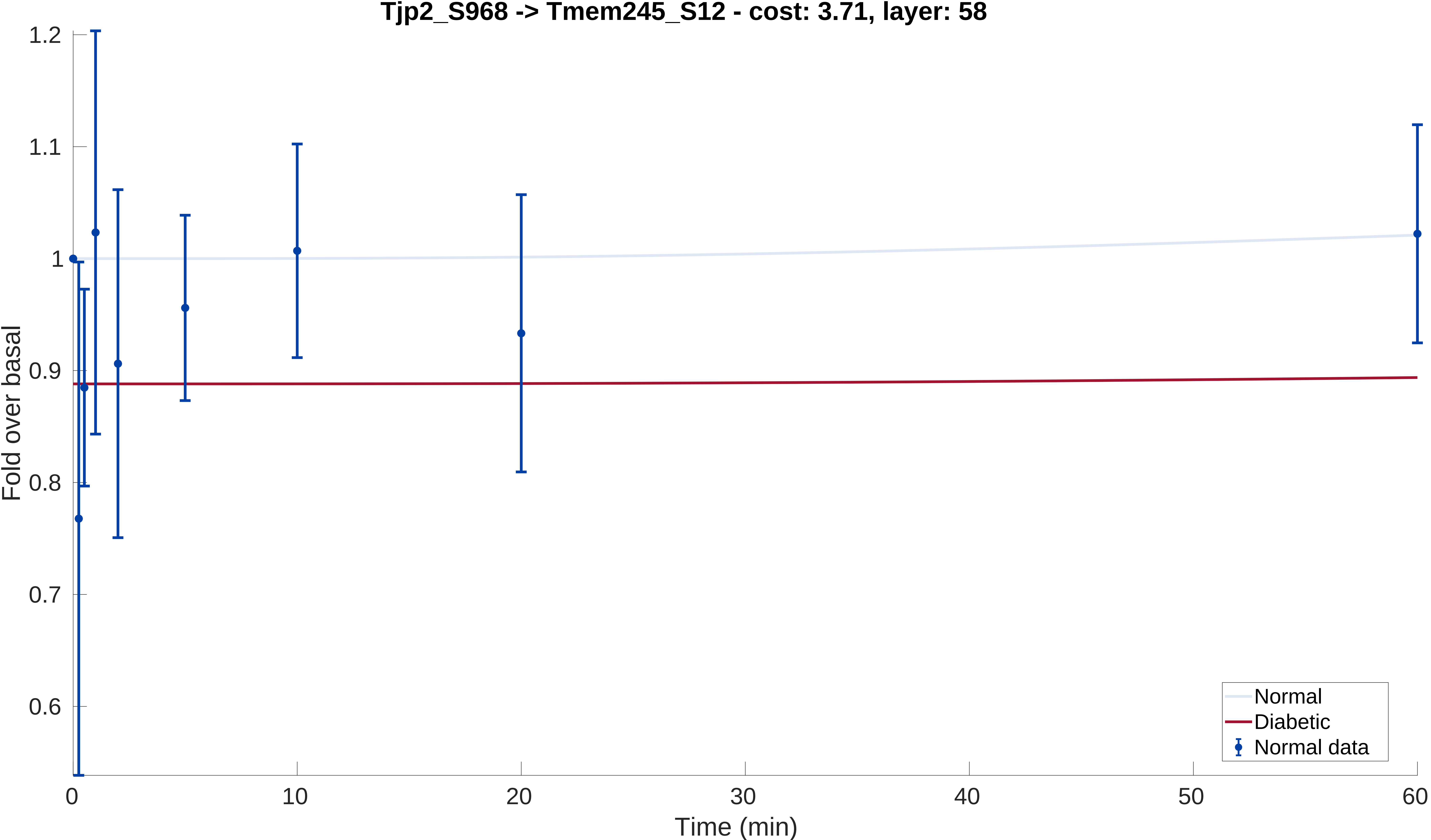

Tjp2_S968 -> Tmem245_S12 - cost: 3.71, layer: 58
Fold over basal
1.2
1.1
1
0.9
0.8
0.7
0.6
0
10
20
30
40
50
60
Time (min)
Normal
Diabetic
Normal data

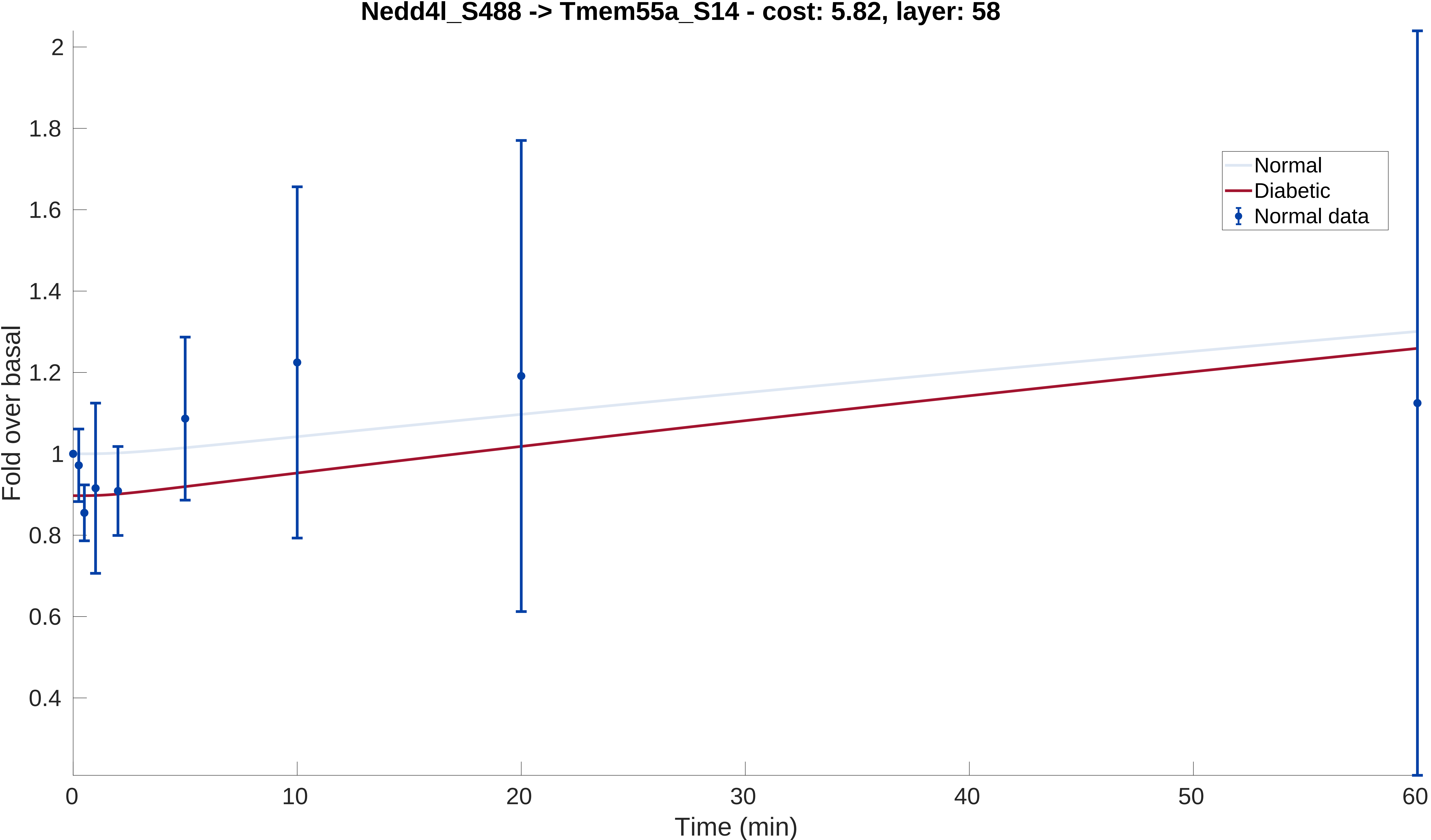

Nedd4l_S488 -> Tmem55a_S14 - cost: 5.82, layer: 58
Fold over basal
2
1.8
1.6
1.4
1.2
1
0.8
0.6
0.4
0
10
20
30
40
50
60
Time (min)
Normal
Diabetic
Normal data

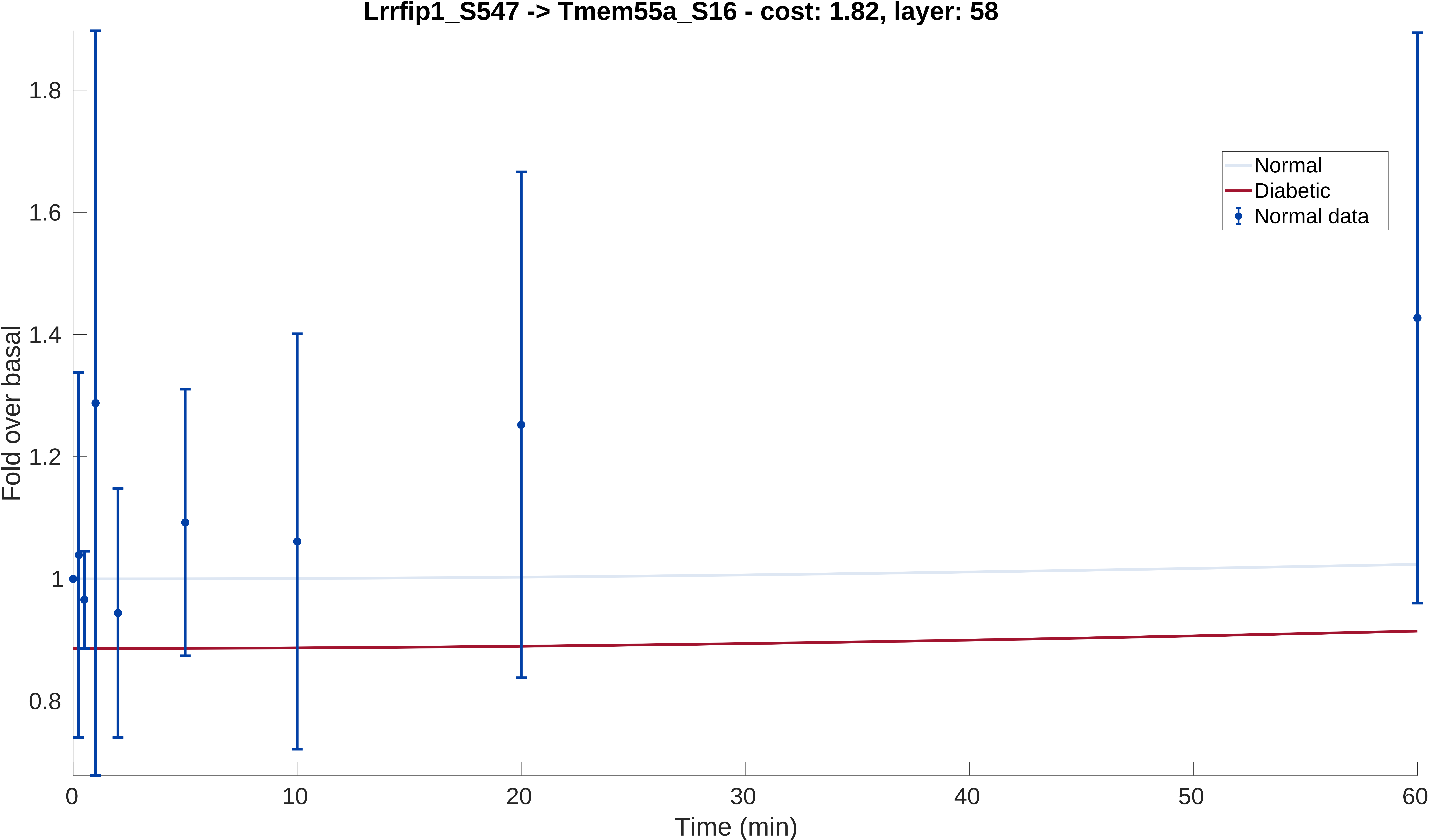
Lrrfip1_S547 -> Tmem55a_S16 - cost: 1.82, layer: 58
Fold over basal
Time (min)
Normal
Diabetic
Normal data
1.8
1.6
1.4
1.2
1
0.8
0
10
20
30
40
50
60

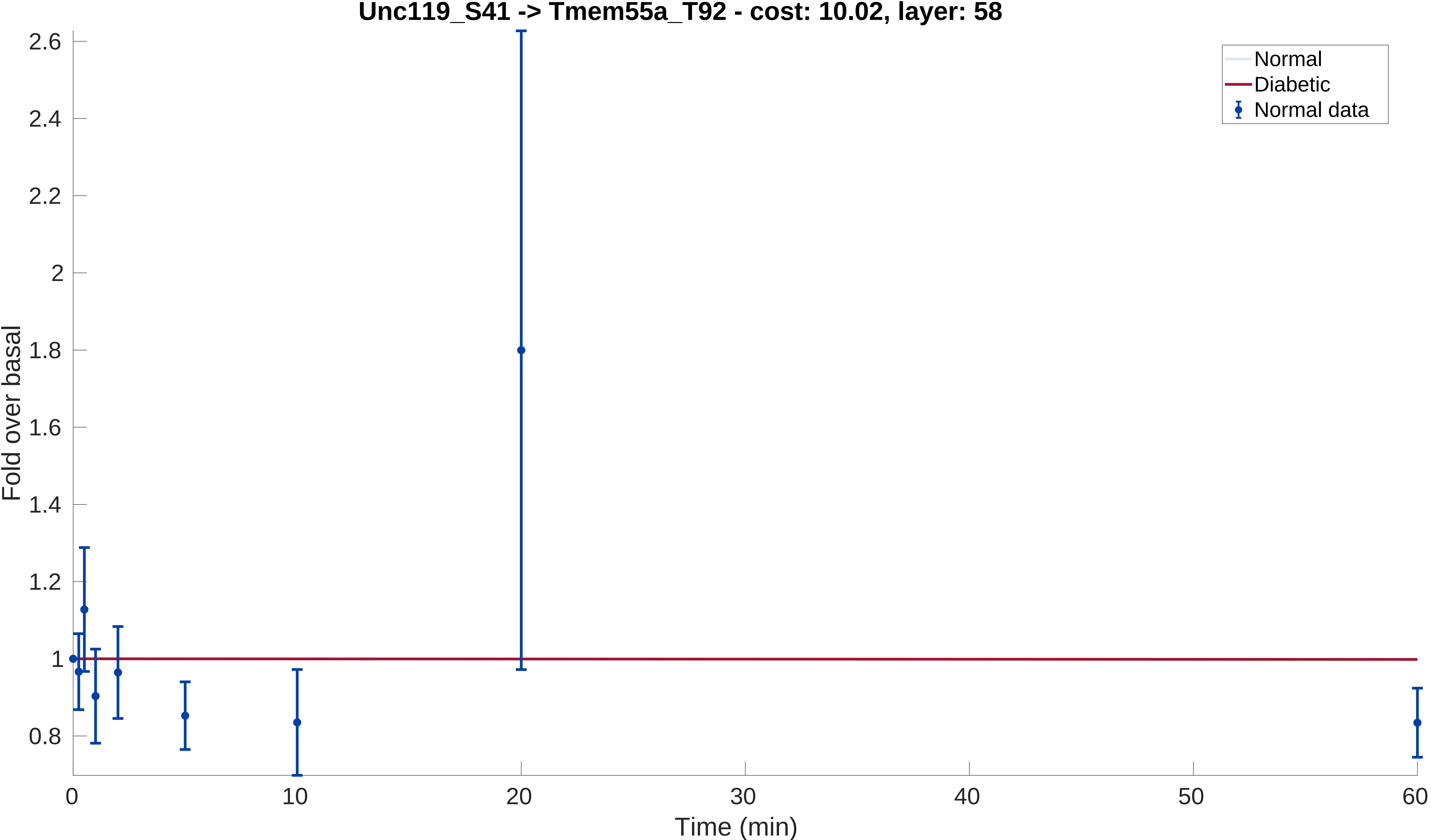

Unc119_S41 -> Tmem55a_T92 - cost: 10.02, layer: 58
Normal
Diabetic
Normal data
Fold over basal
Time (min)
2.6
2.4
2.2
2
1.8
1.6
1.4
1.2
1
0.8
0
10
20
30
40
50
60

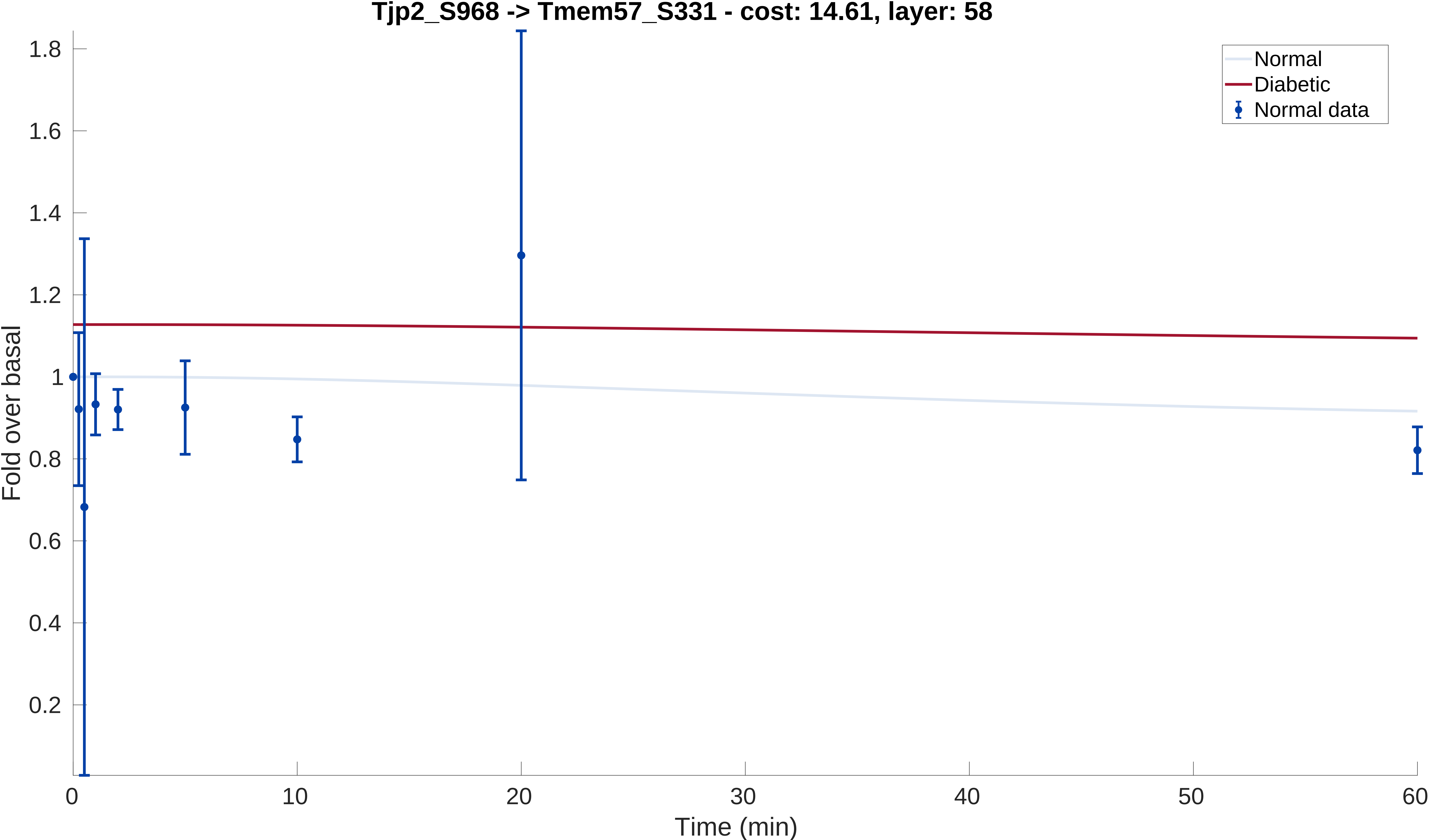

Tjp2_S968 -> Tmem57_S331 - cost: 14.61, layer: 58
Fold over basal
1.8
1.6
1.4
1.2
1
0.8
0.6
0.4
0.2
0
10
20
30
40
50
60
Time (min)
Normal
Diabetic
Normal data

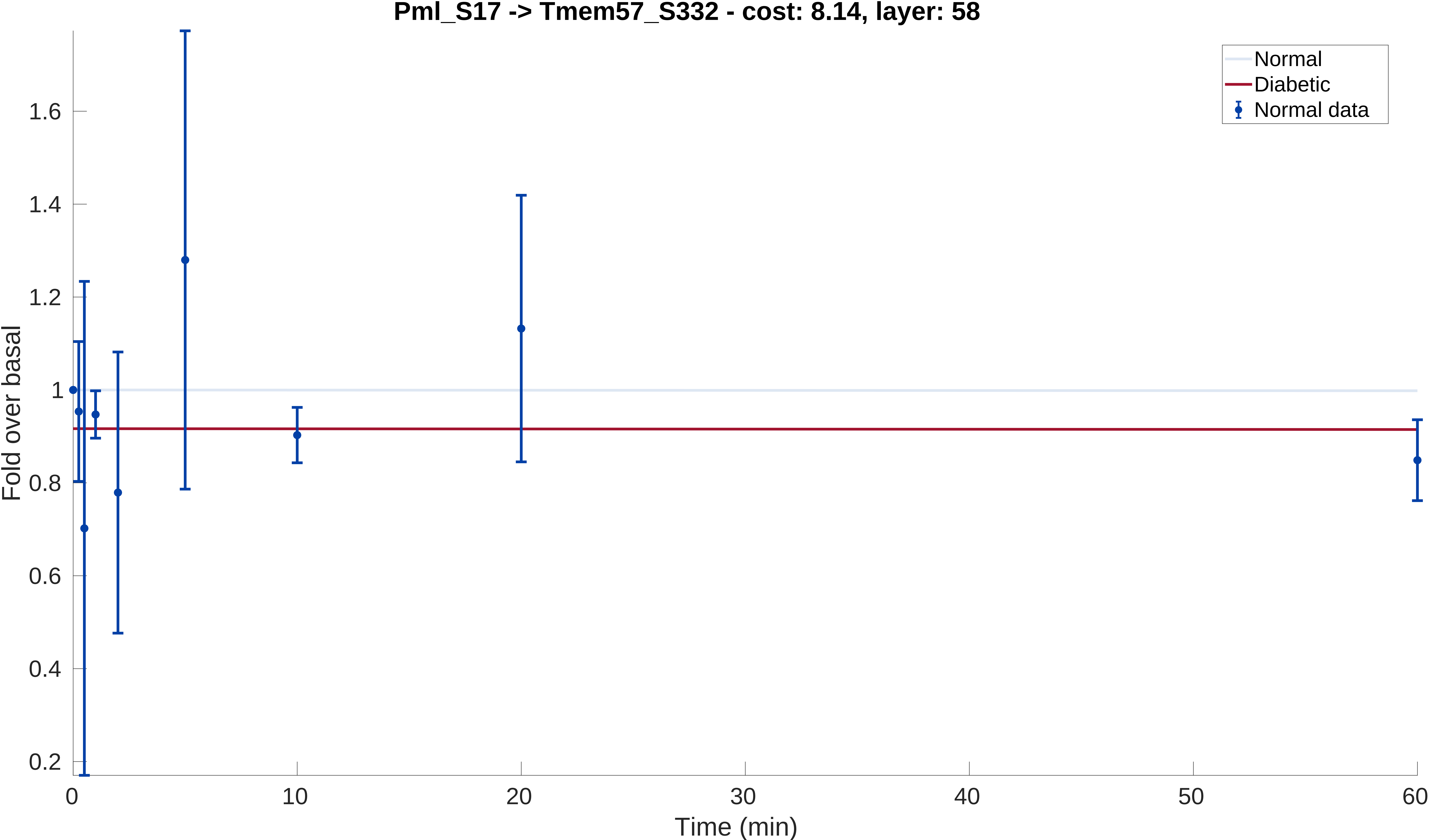

Pml_S17 -> Tmem57_S332 - cost: 8.14, layer: 58
Normal
Diabetic
Normal data
Fold over basal
Time (min)
1.6
1.4
1.2
1
0.8
0.6
0.4
0.2
0
10
20
30
40
50
60

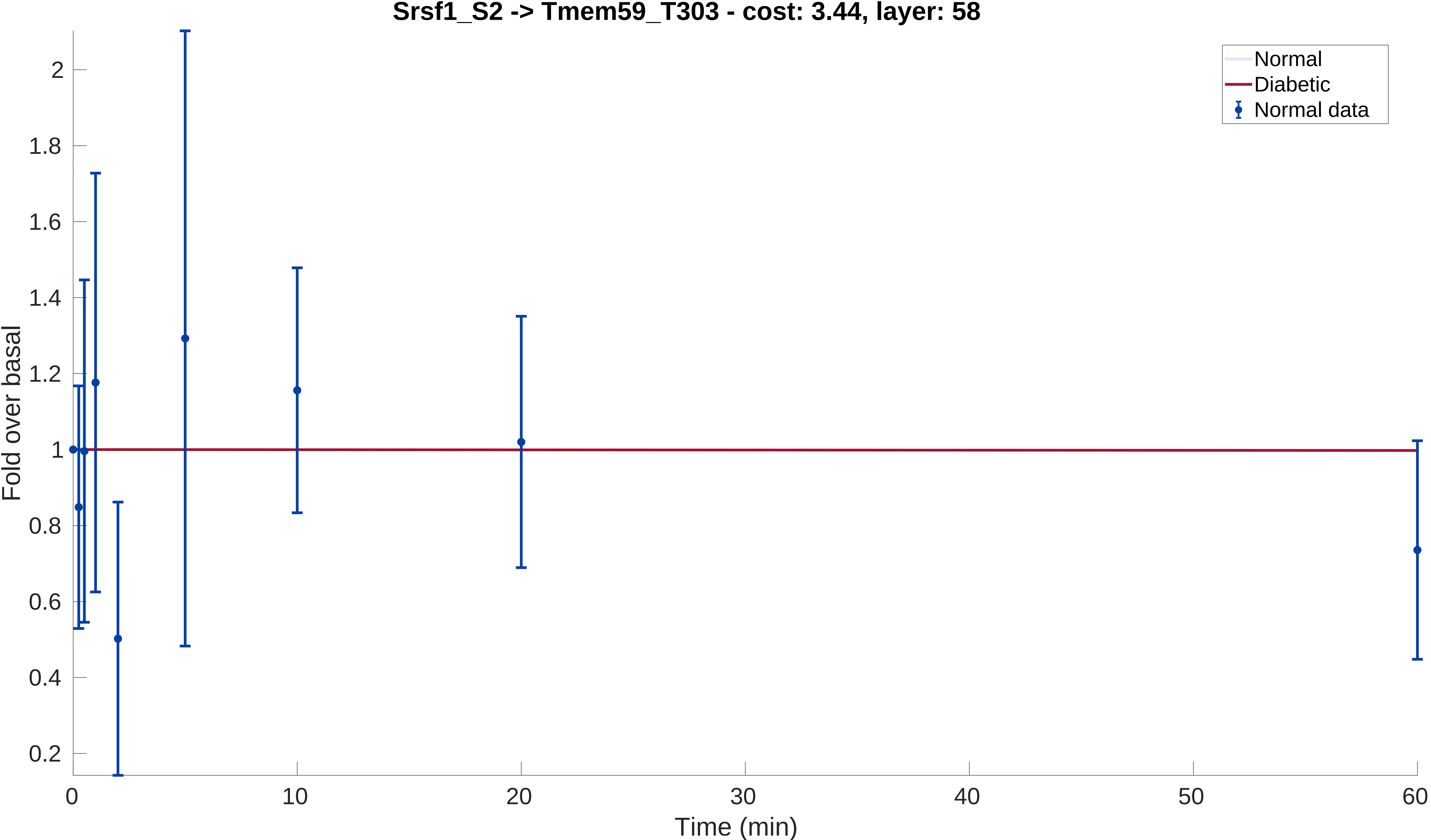
Srsf1_S2 -> Tmem59_T303 - cost: 3.44, layer: 58
Normal
Diabetic
Normal data
Fold over basal
Time (min)
2
1.8
1.6
1.4
1.2
1
0.8
0.6
0.4
0.2
0
10
20
30
40
50
60

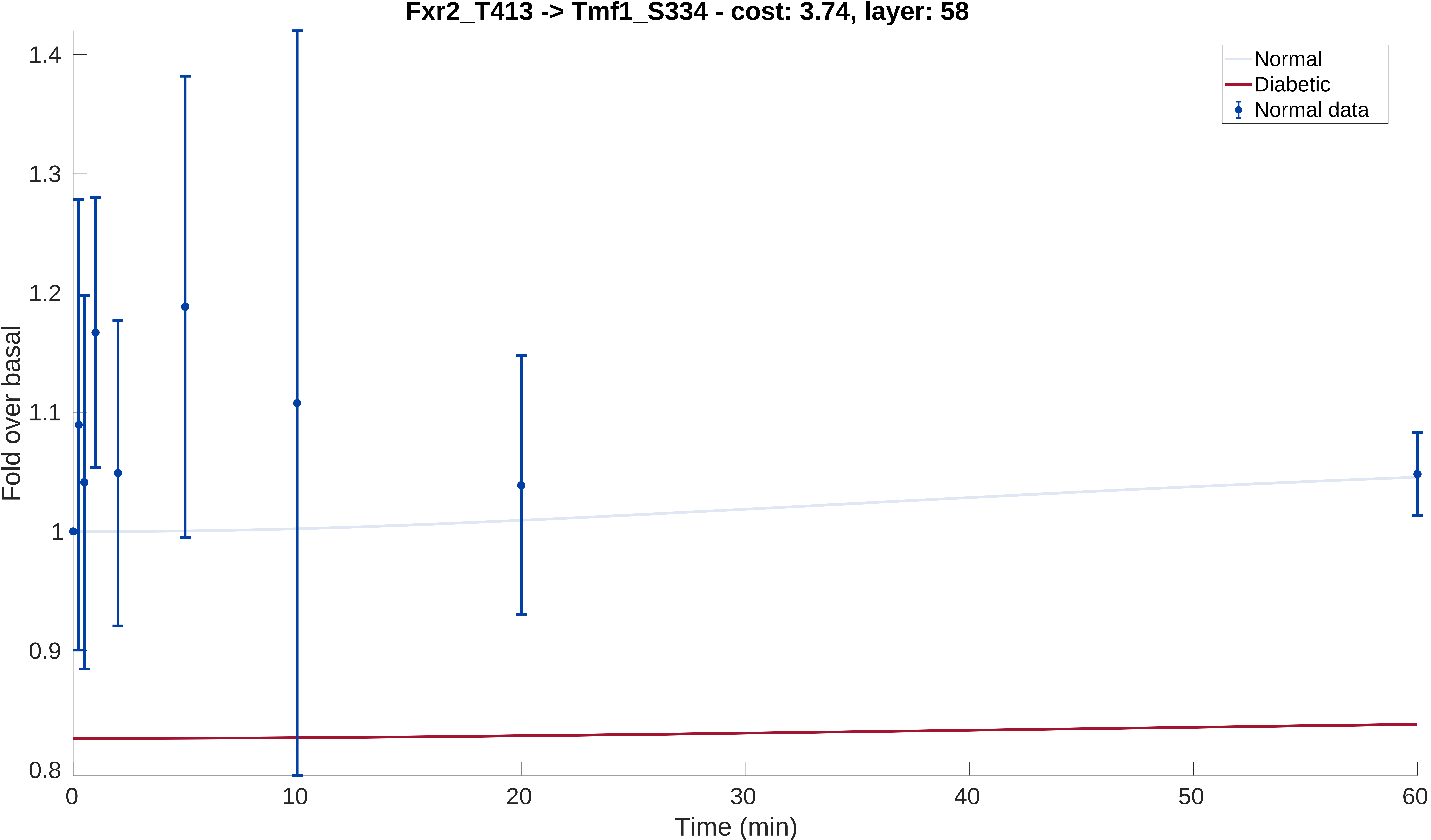
Fxr2_T413 -> Tmf1_S334 - cost: 3.74, layer: 58
Normal
Diabetic
Normal data
Fold over basal
Time (min)
0.8
0.9
1
1.1
1.2
1.3
1.4
0
10
20
30
40
50
60

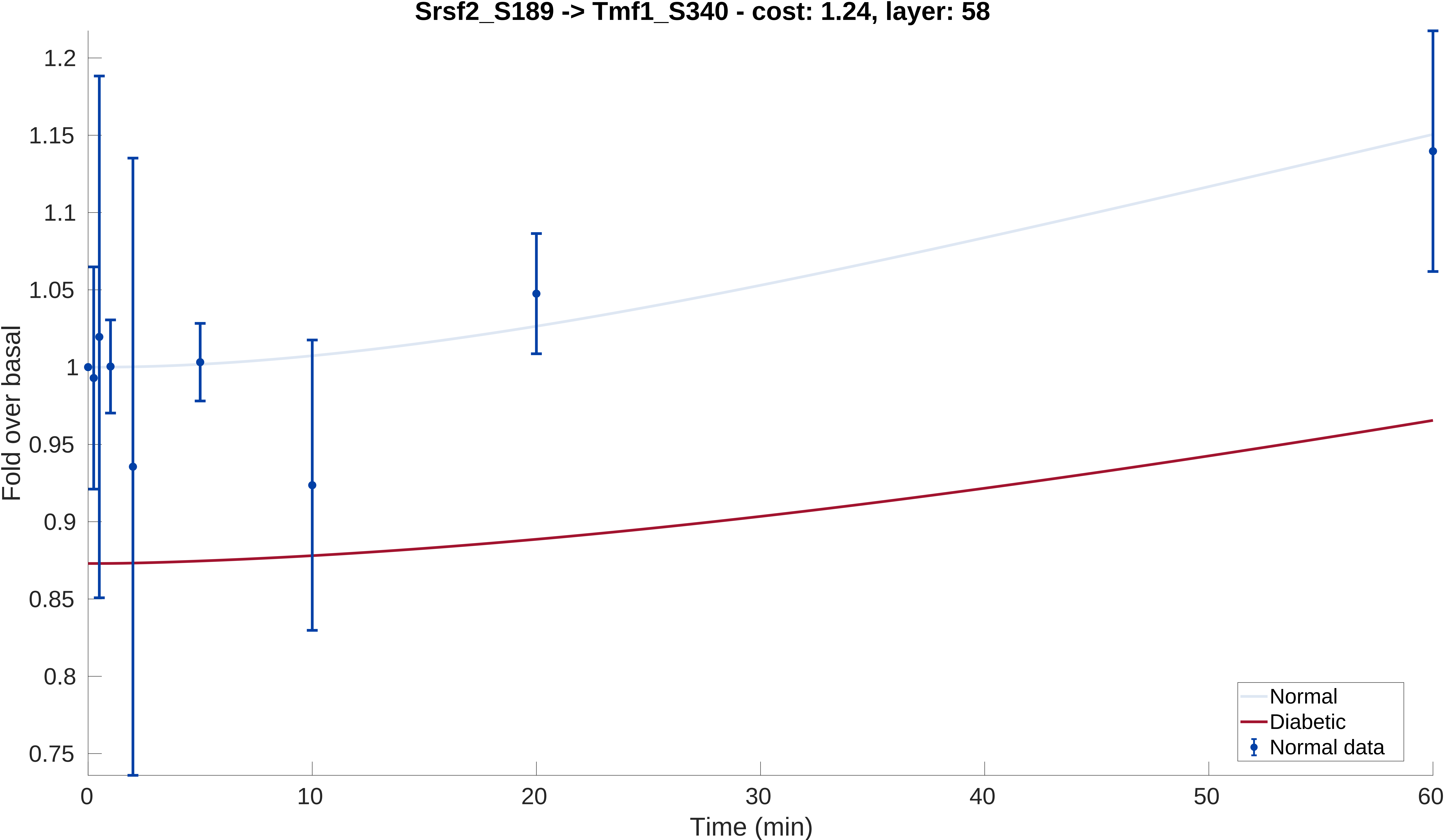

Srsf2_S189 -> Tmf1_S340 - cost: 1.24, layer: 58
Fold over basal
1.2
1.15
1.1
1.05
1
0.95
0.9
0.85
0.8
0.75
0
10
20
30
40
50
60
Time (min)
Normal
Diabetic
Normal data

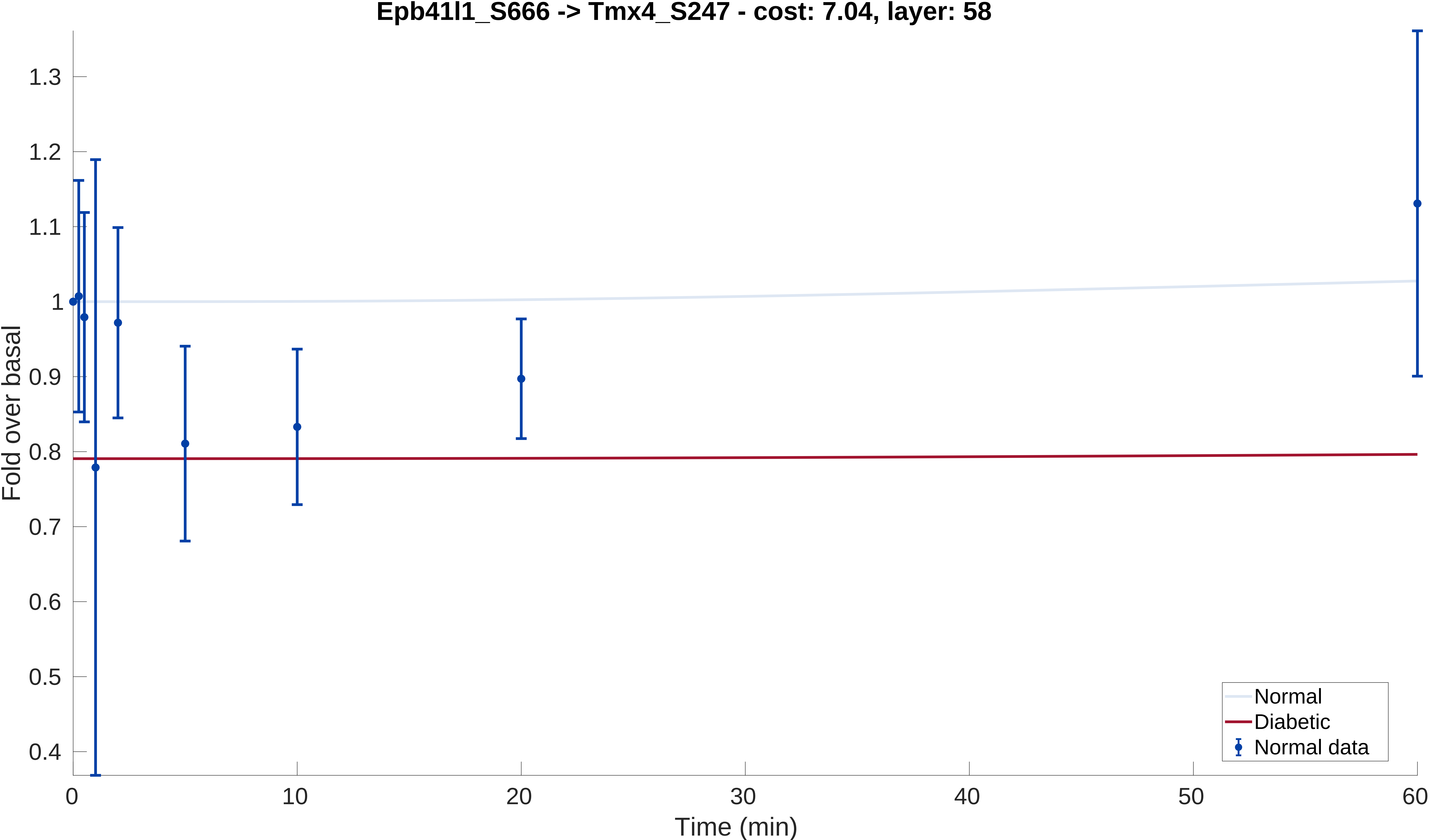
Epb41l1_S666 -> Tmx4_S247 - cost: 7.04, layer: 58
Fold over basal
1.3
1.2
1.1
1
0.9
0.8
0.7
0.6
0.5
0.4
0
10
20
30
40
50
60
Time (min)
Normal
Diabetic
Normal data

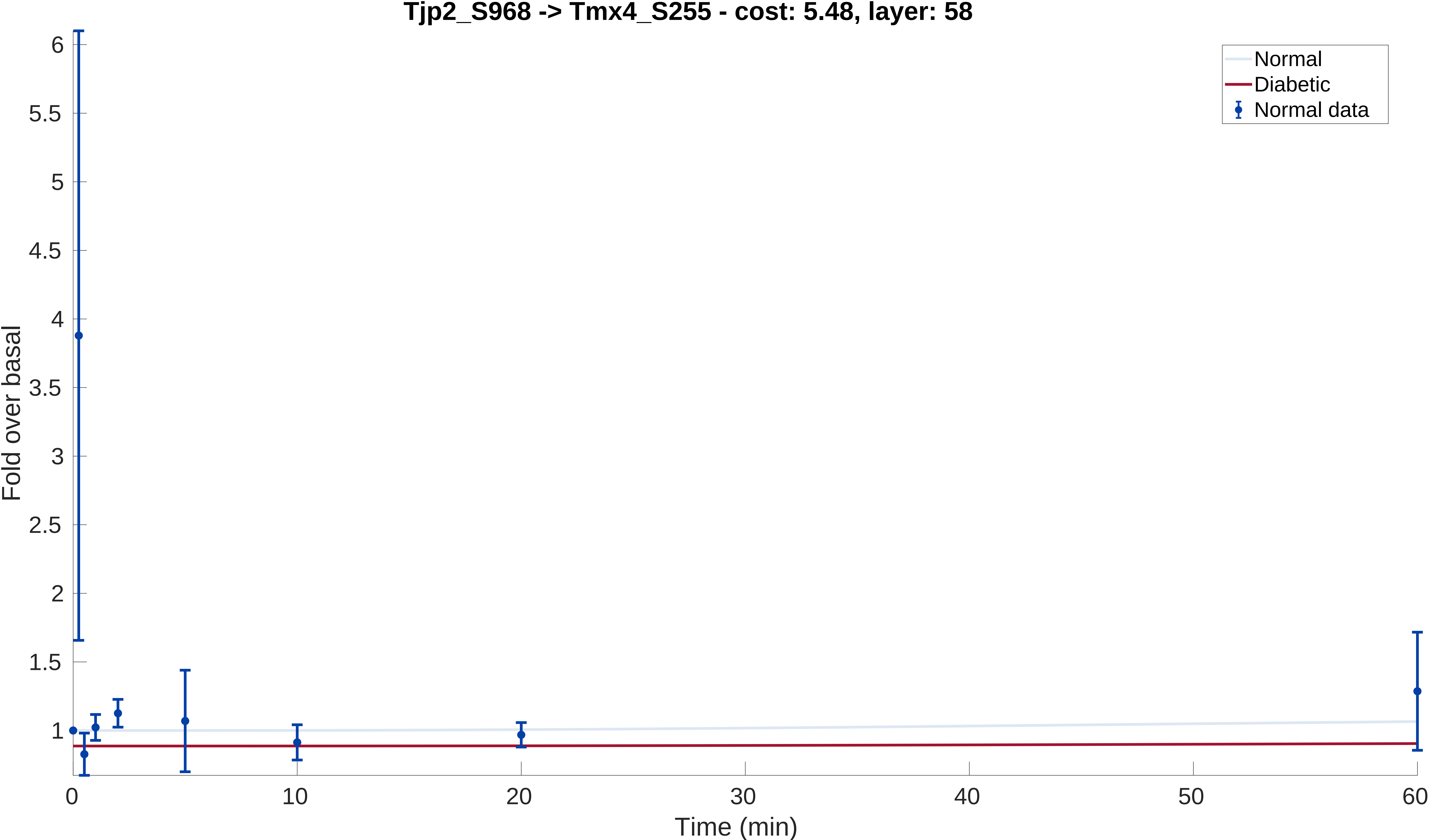
Tjp2_S968 -> Tmx4_S255 - cost: 5.48, layer: 58
Fold over basal
6
5.5
5
4.5
4
3.5
3
2.5
2
1.5
1
0
10
20
30
40
50
60
Time (min)
Normal
Diabetic
Normal data

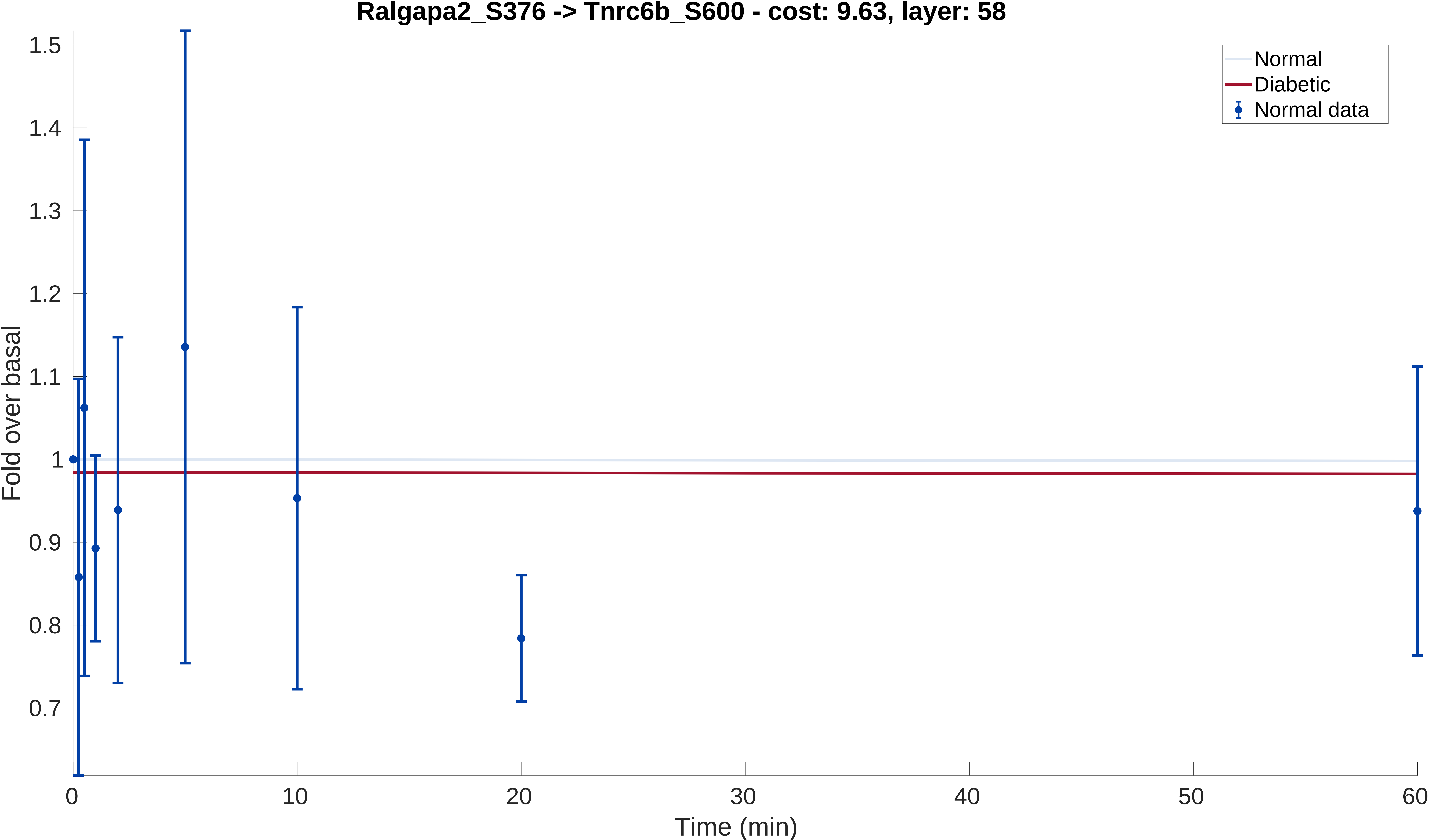

Ralgapa2_S376 -> Tnrc6b_S600 - cost: 9.63, layer: 58
Normal
Diabetic
Normal data
Fold over basal
Time (min)
1.5
1.4
1.3
1.2
1.1
1
0.9
0.8
0.7
0
10
20
30
40
50
60

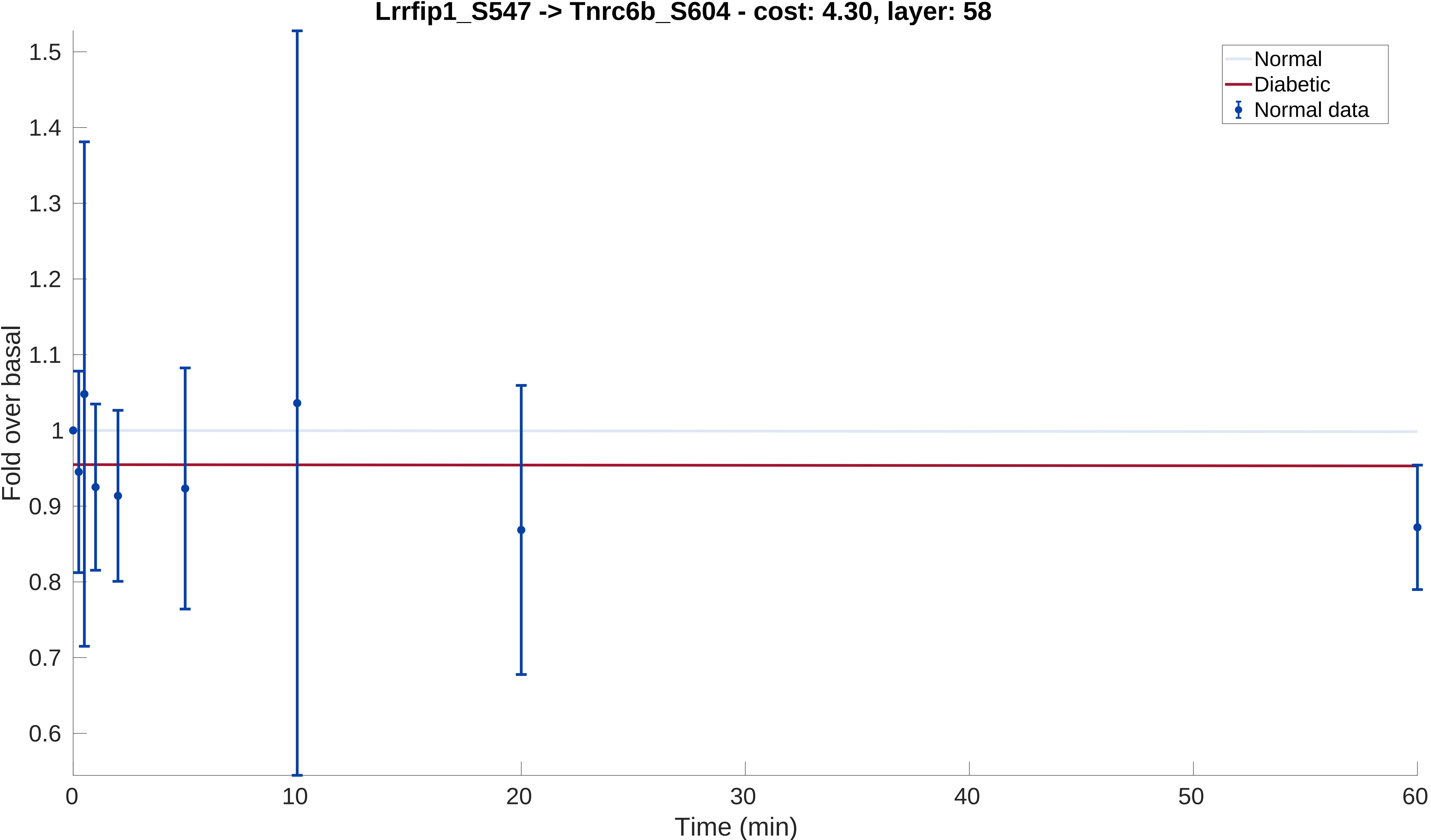

Lrrfip1_S547 -> Tnrc6b_S604 - cost: 4.30, layer: 58
Normal
Diabetic
Normal data
Fold over basal
Time (min)
1.5
1.4
1.3
1.2
1.1
1
0.9
0.8
0.7
0.6
0
10
20
30
40
50
60

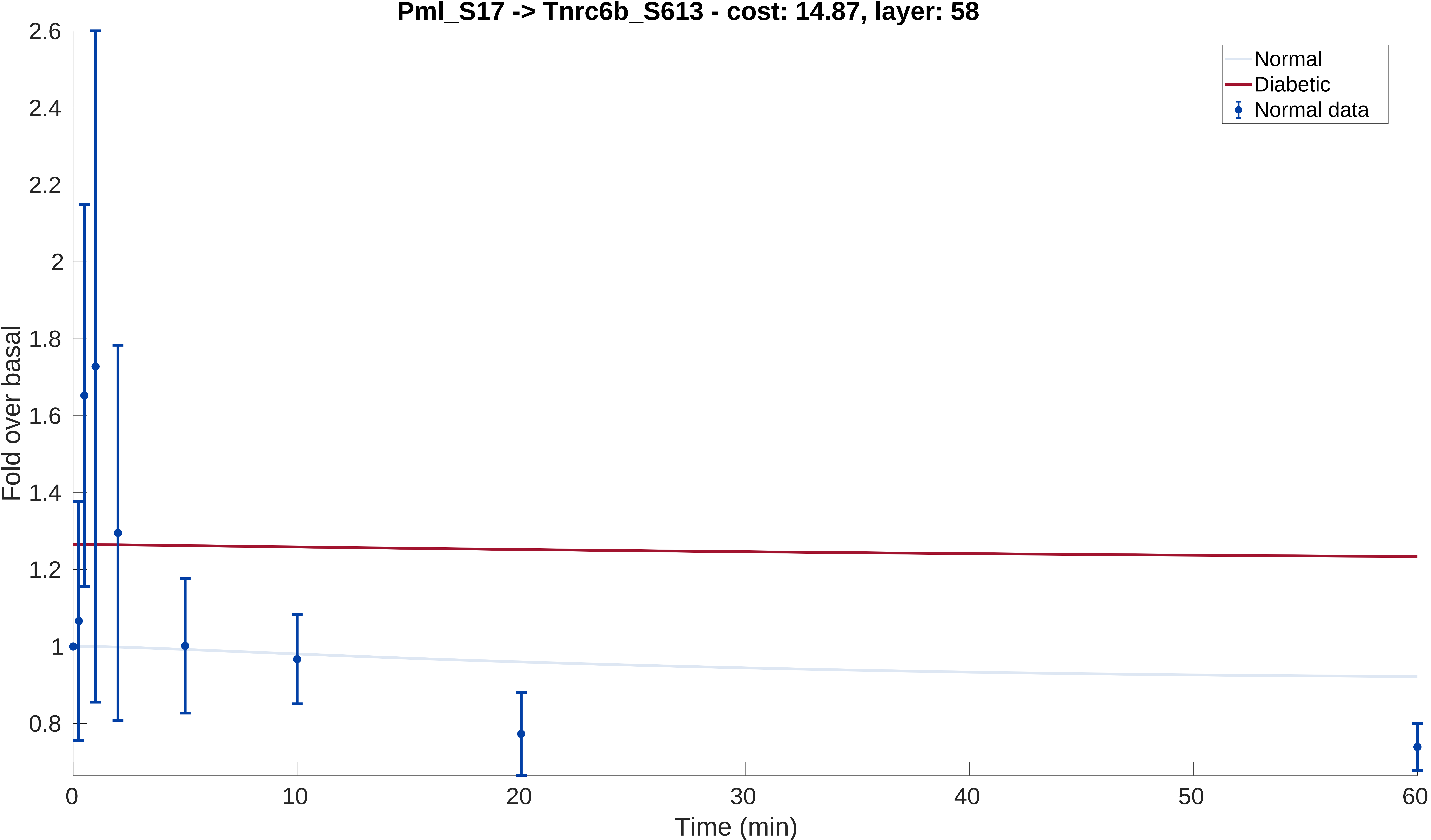

Pml_S17 -> Tnrc6b_S613 - cost: 14.87, layer: 58
Normal
Diabetic
Normal data
Fold over basal
Time (min)
2.6
2.4
2.2
2
1.8
1.6
1.4
1.2
1
0.8
0
10
20
30
40
50
60

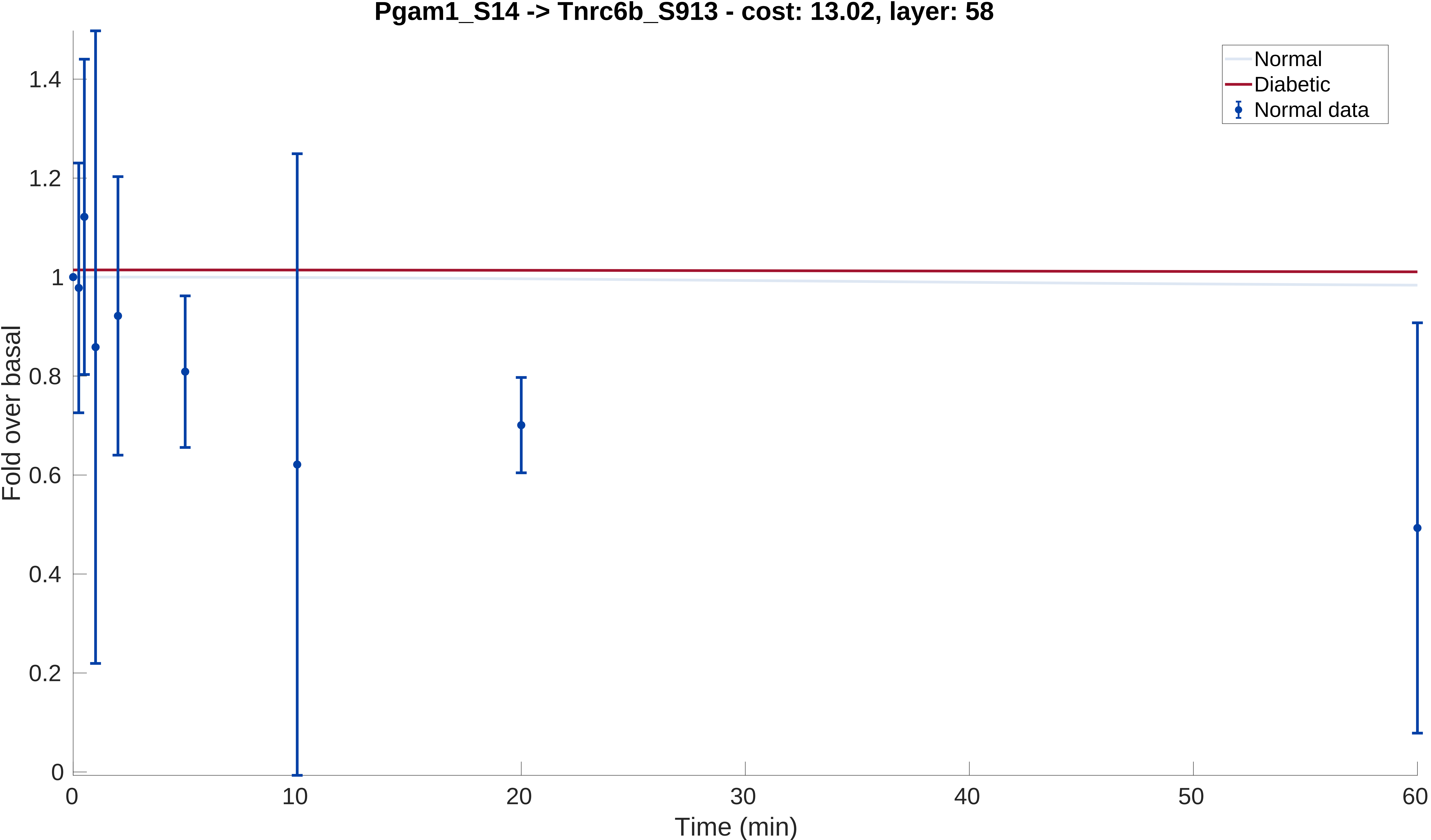

Pgam1_S14 -> Tnrc6b_S913 - cost: 13.02, layer: 58
Normal
Diabetic
Normal data
Fold over basal
1.4
1.2
1
0.8
0.6
0.4
0.2
0
0
10
20
30
40
50
60
Time (min)

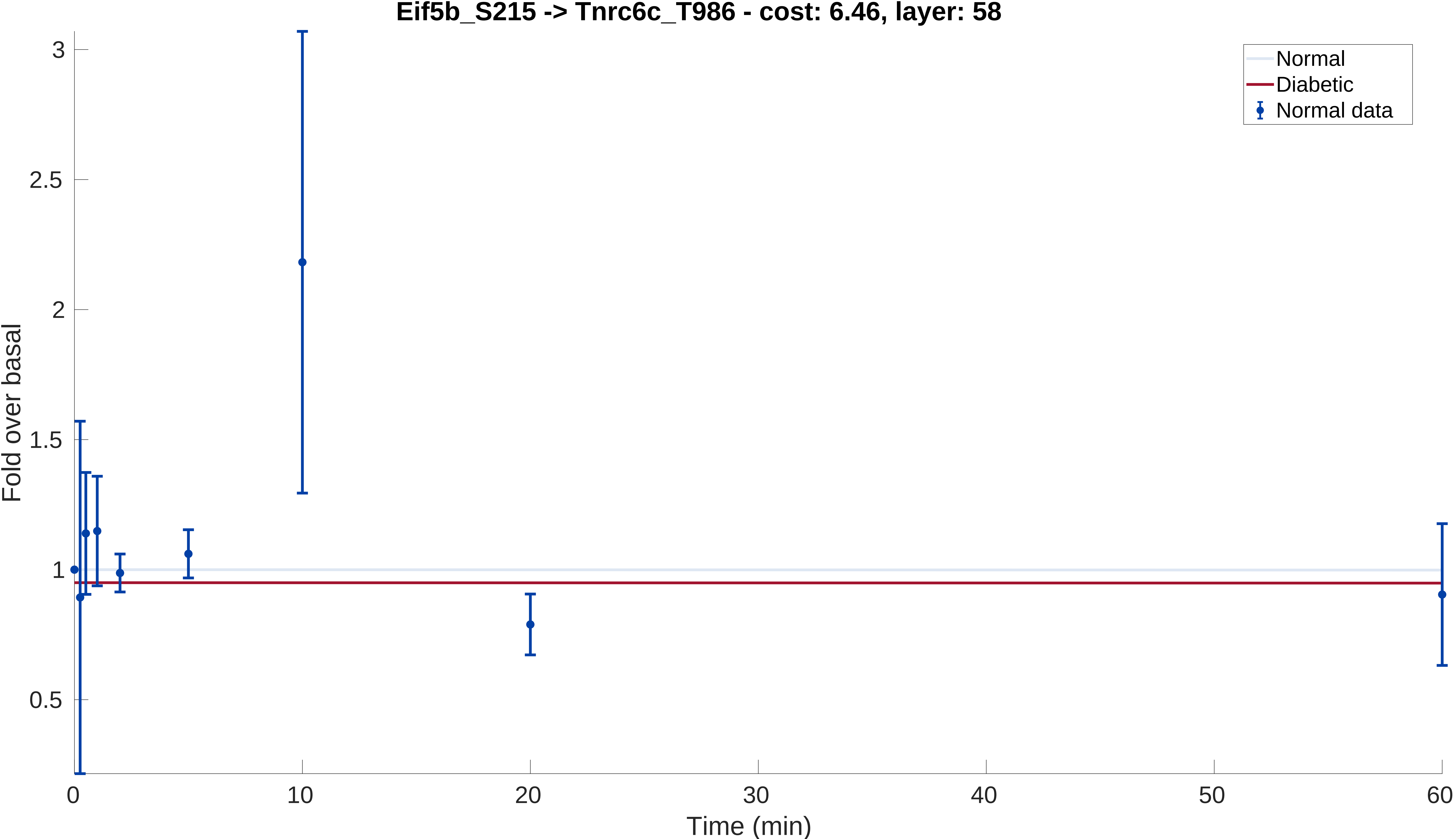
Eif5b_S215 -> Tnrc6c_T986 - cost: 6.46, layer: 58
Normal
Diabetic
Normal data
Fold over basal
3
2.5
2
1.5
1
0.5
0
10
20
30
40
50
60
Time (min)

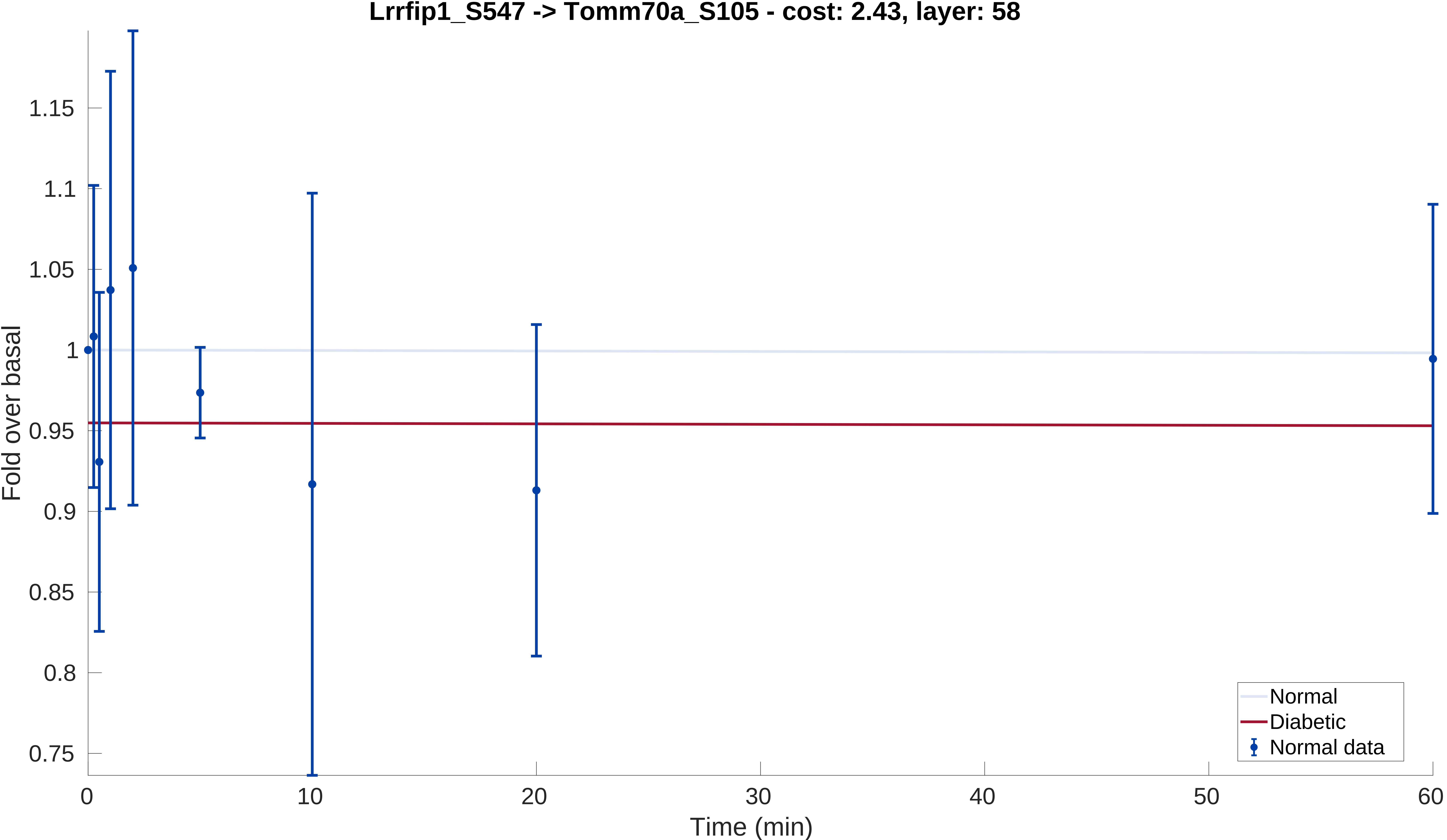

Lrrfip1_S547 -> Tomm70a_S105 - cost: 2.43, layer: 58
Fold over basal
1.15
1.1
1.05
1
0.95
0.9
0.85
0.8
0.75
0
10
20
30
40
50
60
Time (min)
Normal
Diabetic
Normal data

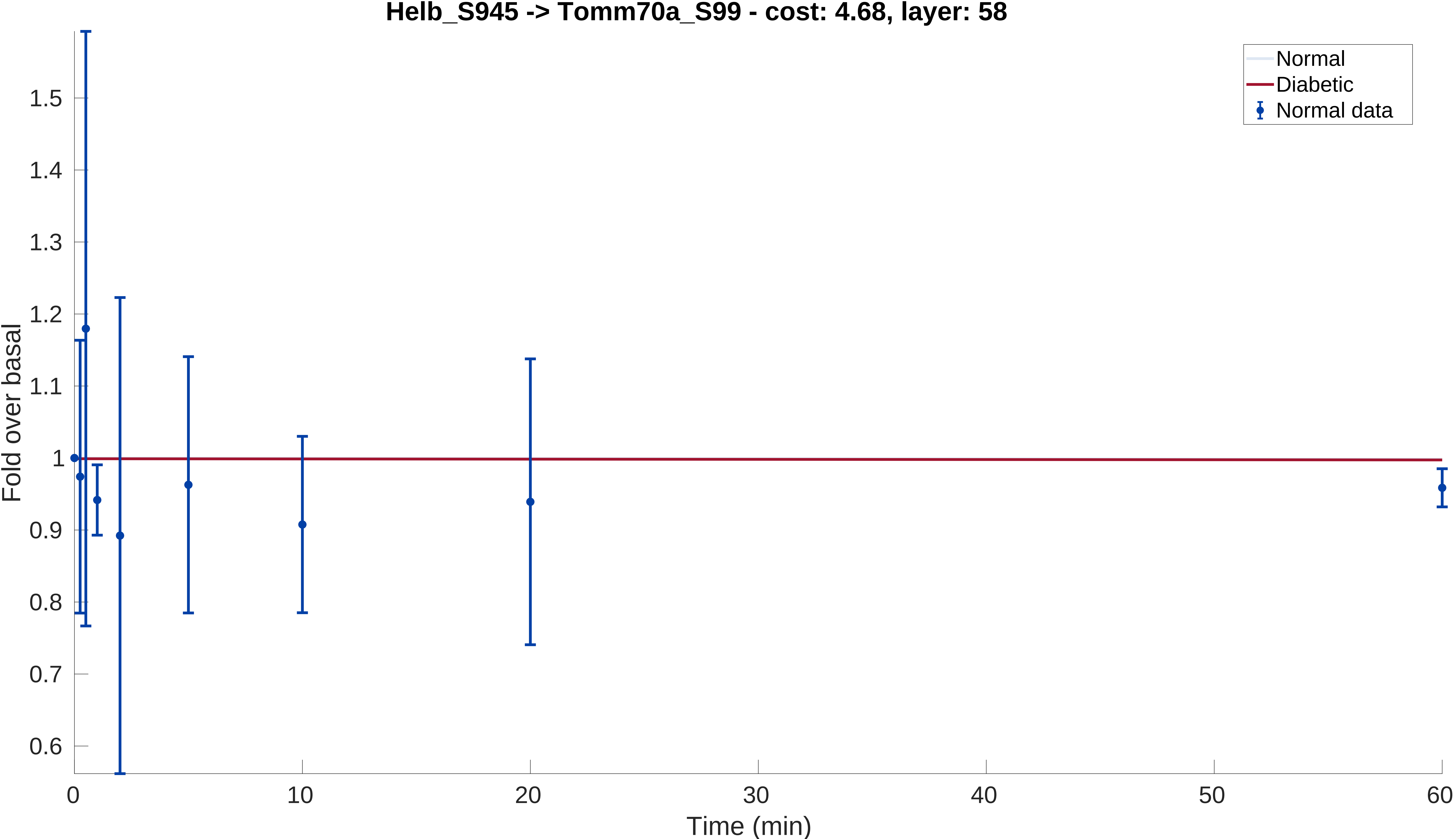
Helb_S945 -> Tomm70a_S99 - cost: 4.68, layer: 58
Normal
Diabetic
Normal data
1.5
1.4
1.3
1.2
1.1
1
0.9
0.8
0.7
0.6
Fold over basal
0
10
20
30
40
50
60
Time (min)

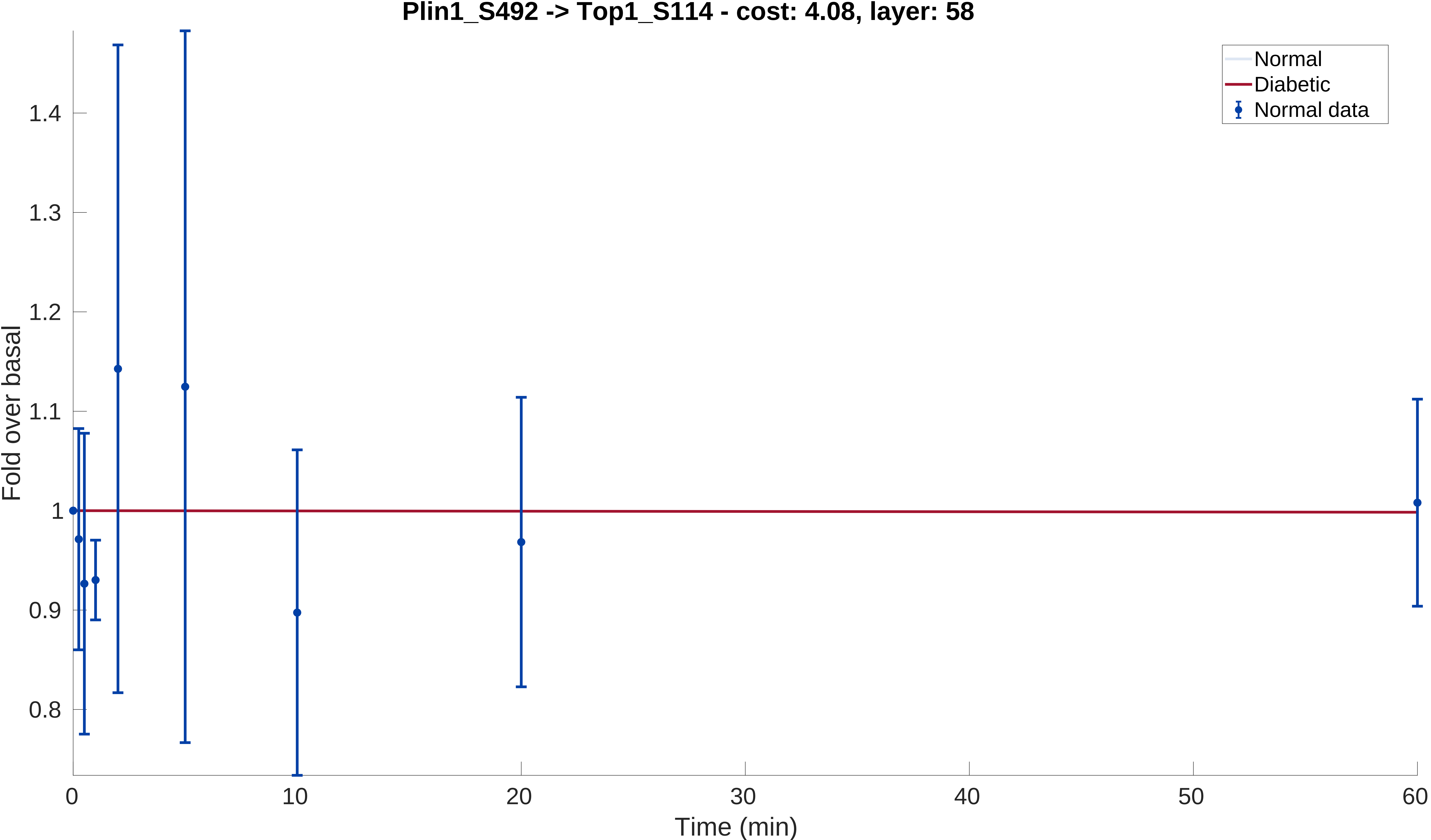

Plin1_S492 -> Top1_S114 - cost: 4.08, layer: 58
Normal
Diabetic
Normal data
Fold over basal
1.4
1.3
1.2
1.1
1
0.9
0.8
0
10
20
30
40
50
60
Time (min)

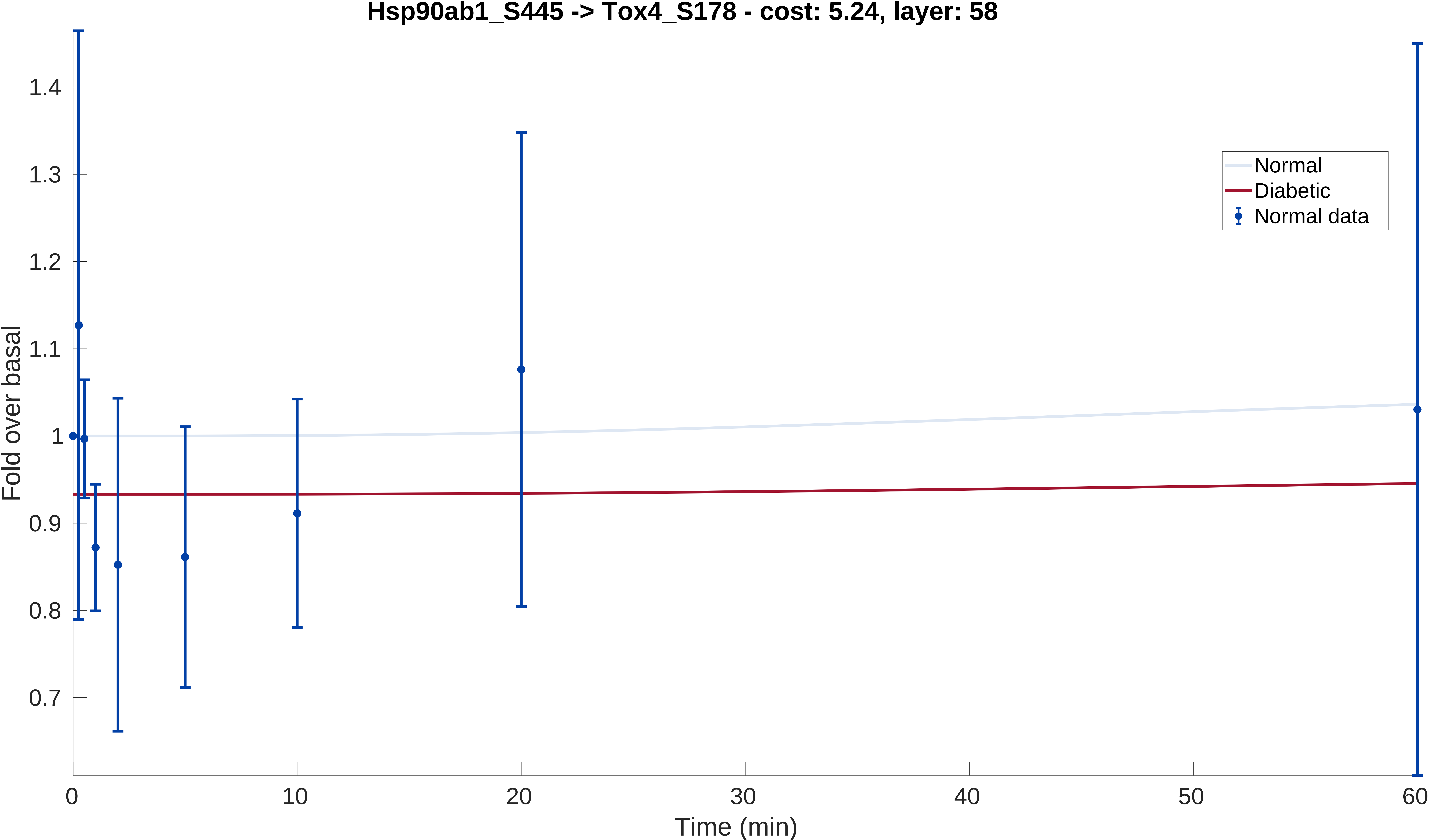
Hsp90ab1_S445 -> Tox4_S178 - cost: 5.24, layer: 58
Normal
Diabetic
Normal data
Fold over basal
Time (min)
0.7
0.8
0.9
1
1.1
1.2
1.3
1.4
0
10
20
30
40
50
60

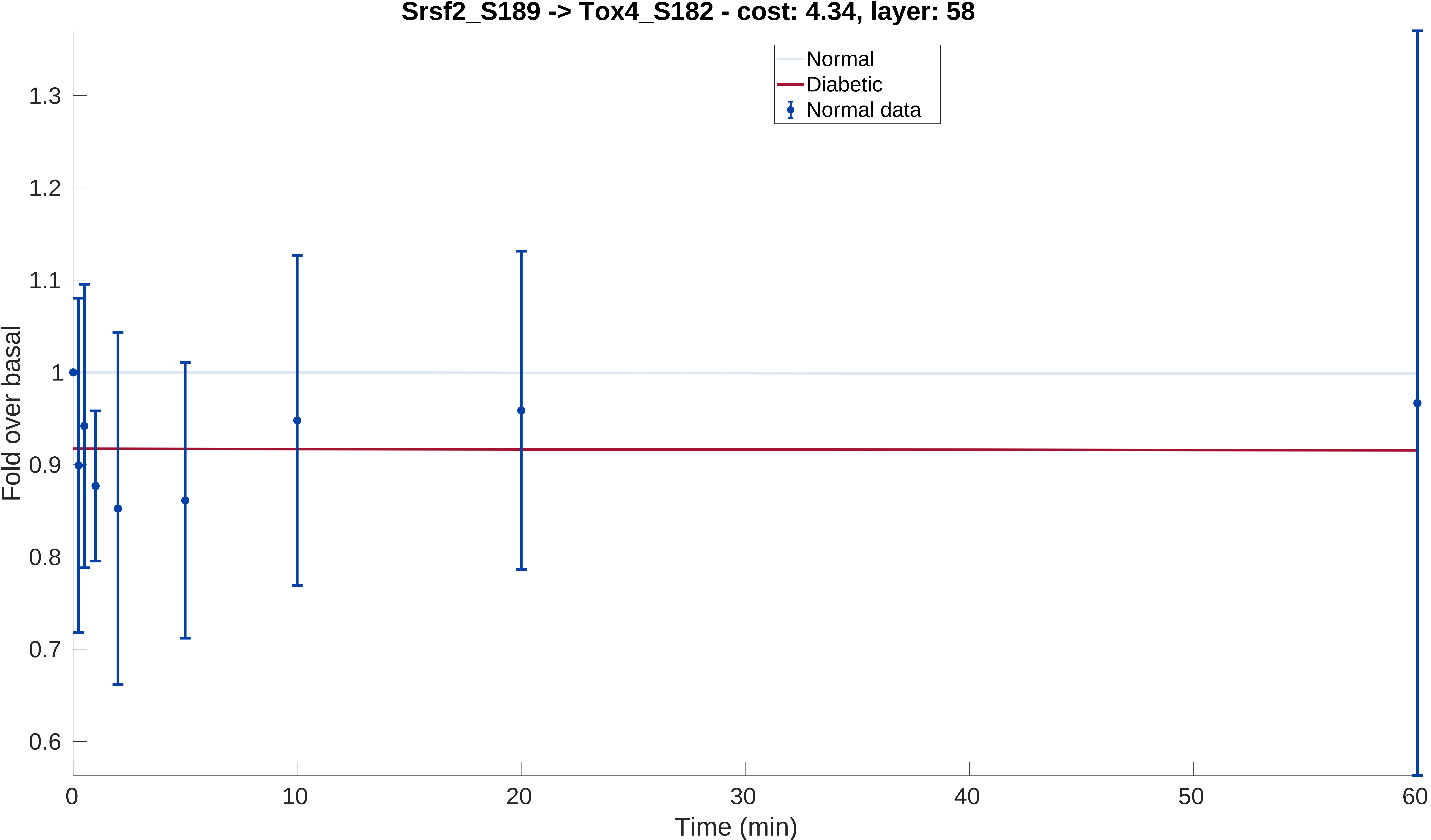

Srsf2_S189 -> Tox4_S182 - cost: 4.34, layer: 58
Normal
Diabetic
Normal data
Fold over basal
Time (min)
1.3
1.2
1.1
1
0.9
0.8
0.7
0.6
0
10
20
30
40
50
60

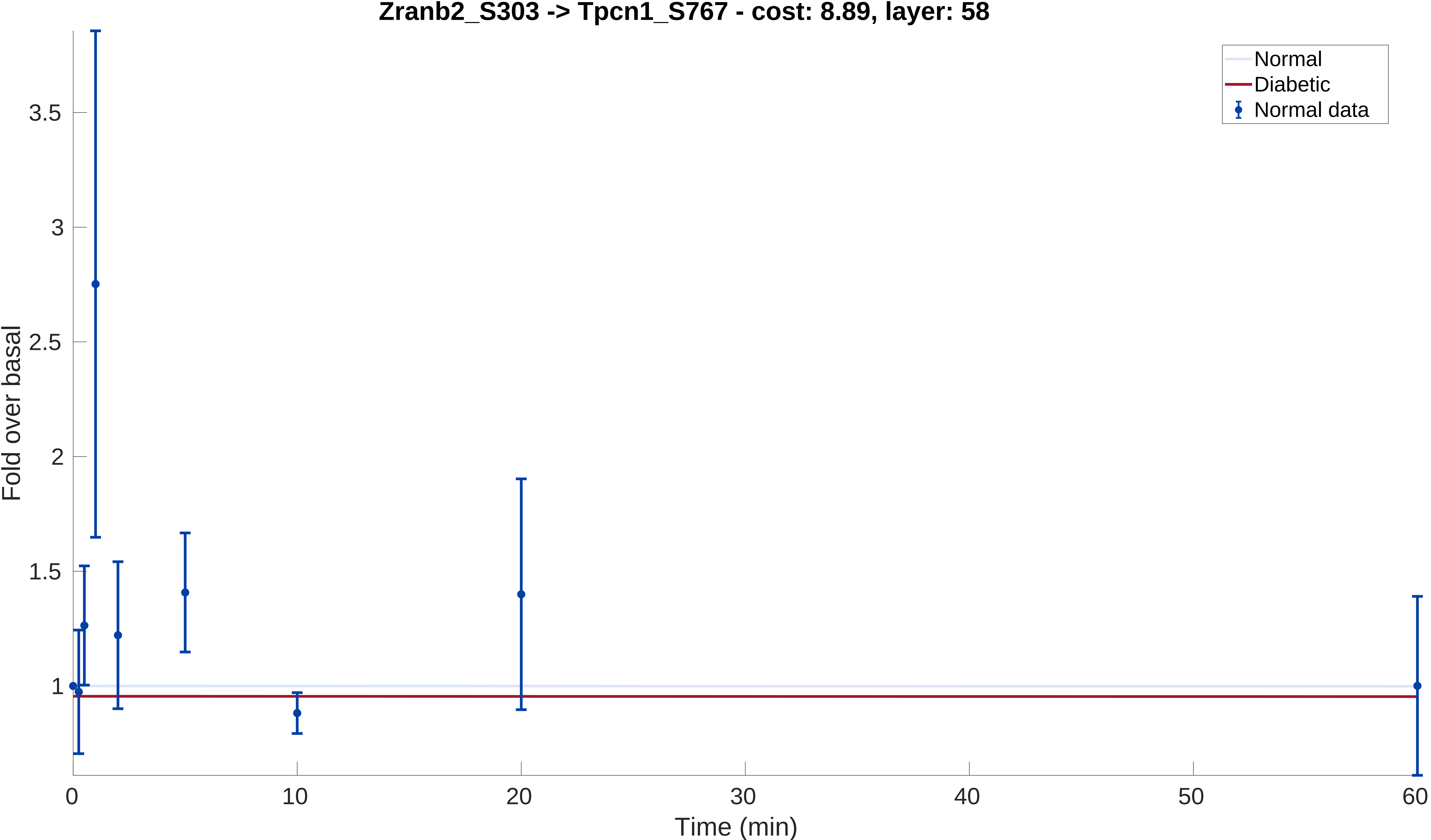

Zranb2_S303 -> Tpcn1_S767 - cost: 8.89, layer: 58
Normal
Diabetic
Normal data
Fold over basal
3.5
3
2.5
2
1.5
1
0
10
20
30
40
50
60
Time (min)

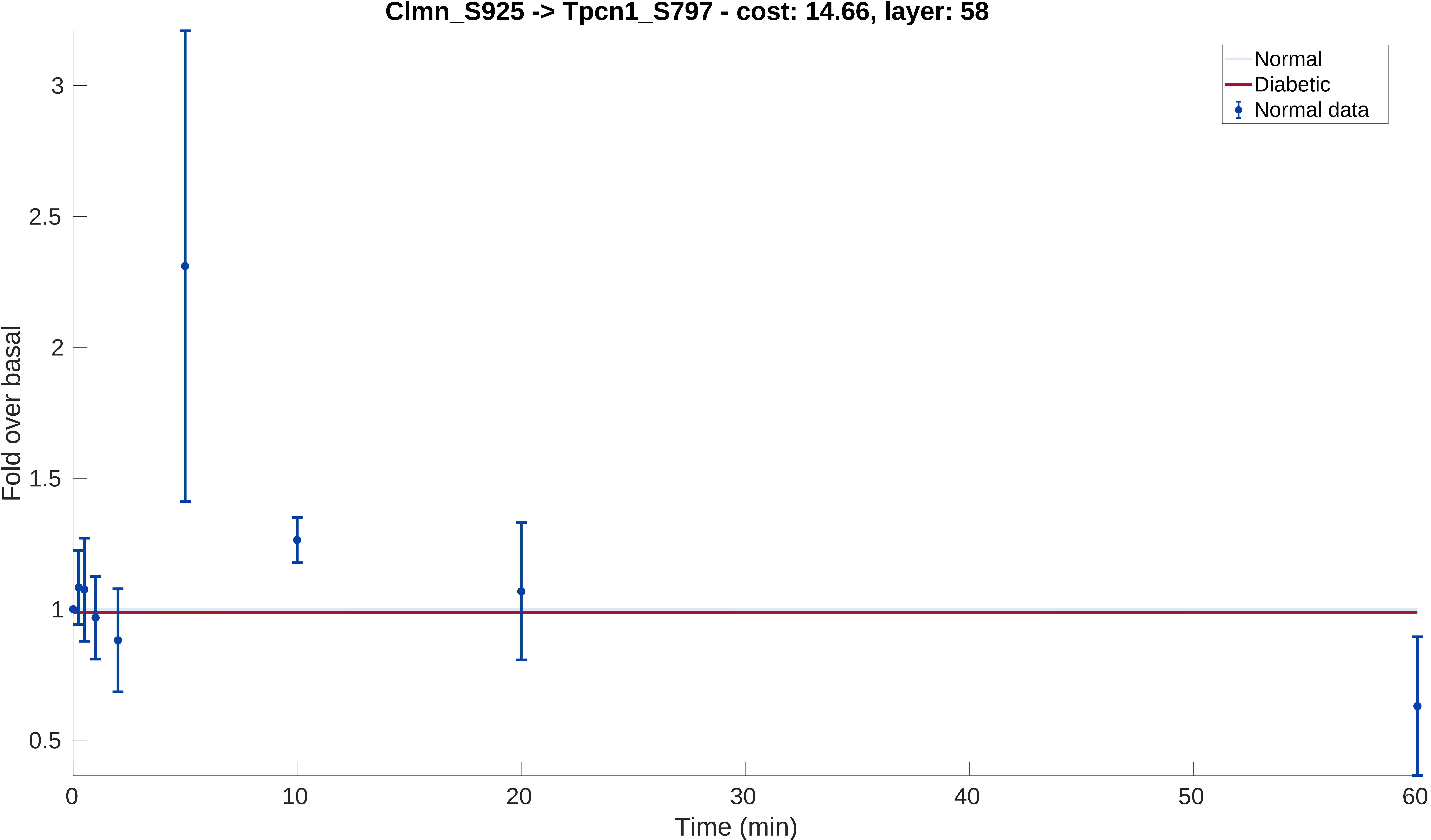

Clmn_S925 -> Tpcn1_S797 - cost: 14.66, layer: 58
Normal
Diabetic
Normal data
Fold over basal
Time (min)
3
2.5
2
1.5
1
0.5
0
10
20
30
40
50
60

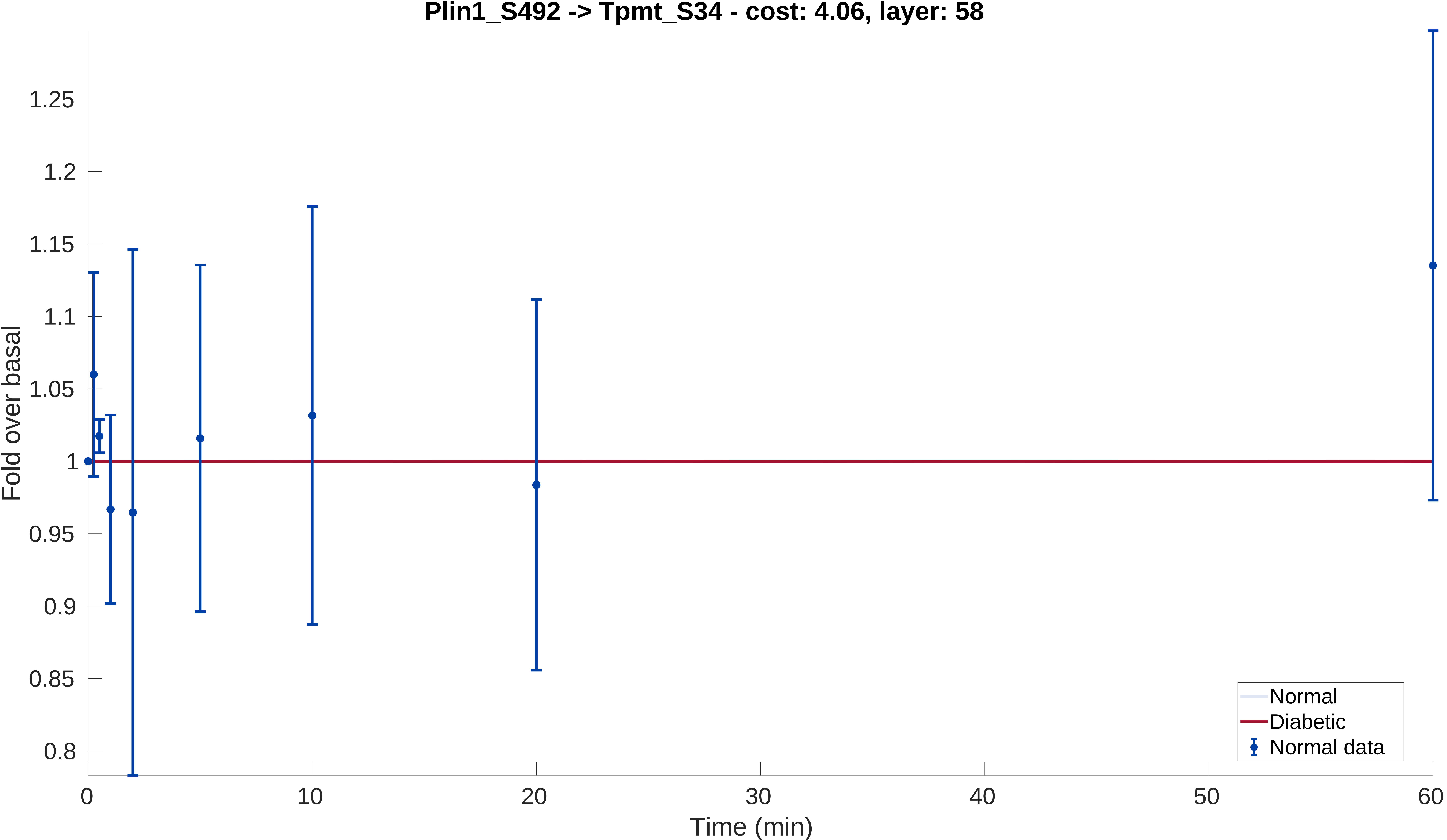

Plin1_S492 -> Tpmt_S34 - cost: 4.06, layer: 58
Fold over basal
1.25
1.2
1.15
1.1
1.05
1
0.95
0.9
0.85
0.8
0
10
20
30
40
50
60
Time (min)
Normal
Diabetic
Normal data

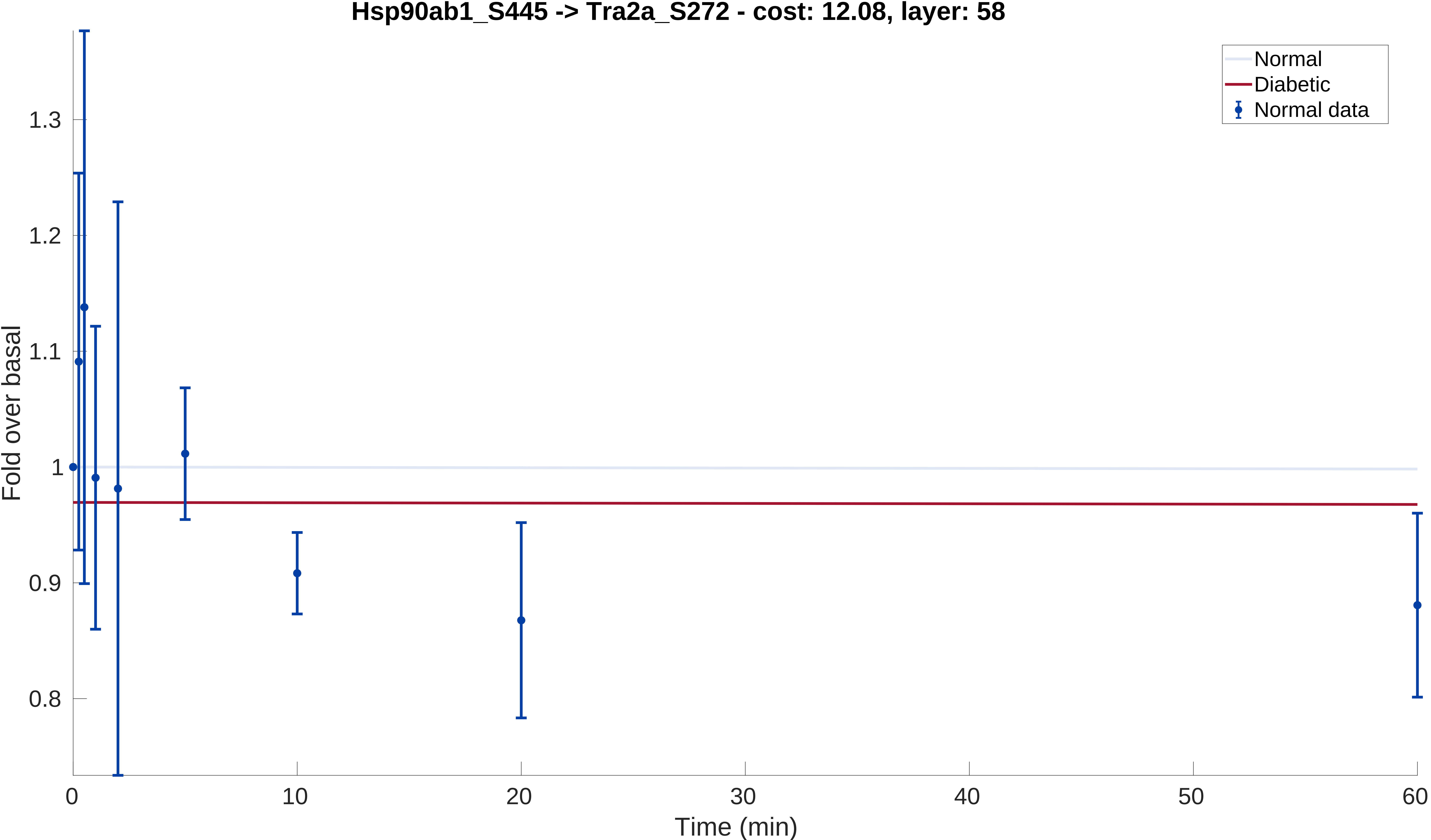
Hsp90ab1_S445 -> Tra2a_S272 - cost: 12.08, layer: 58
Normal
Diabetic
Normal data
Fold over basal
1.3
1.2
1.1
1
0.9
0.8
0
10
20
30
40
50
60
Time (min)

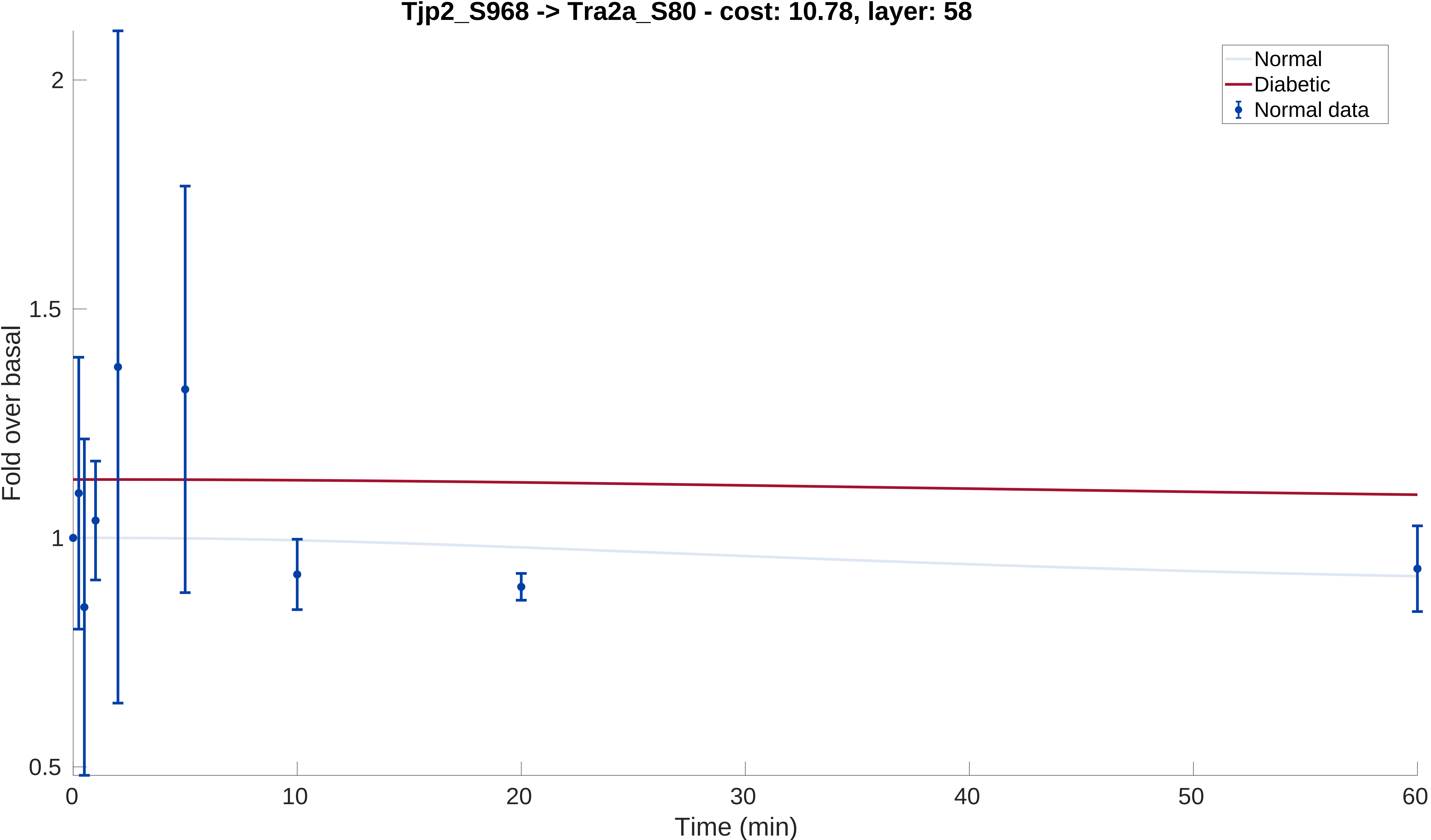
Tjp2_S968 -> Tra2a_S80 - cost: 10.78, layer: 58
Normal
Diabetic
Normal data
Fold over basal
Time (min)
0.5
1
1.5
2
0
10
20
30
40
50
60

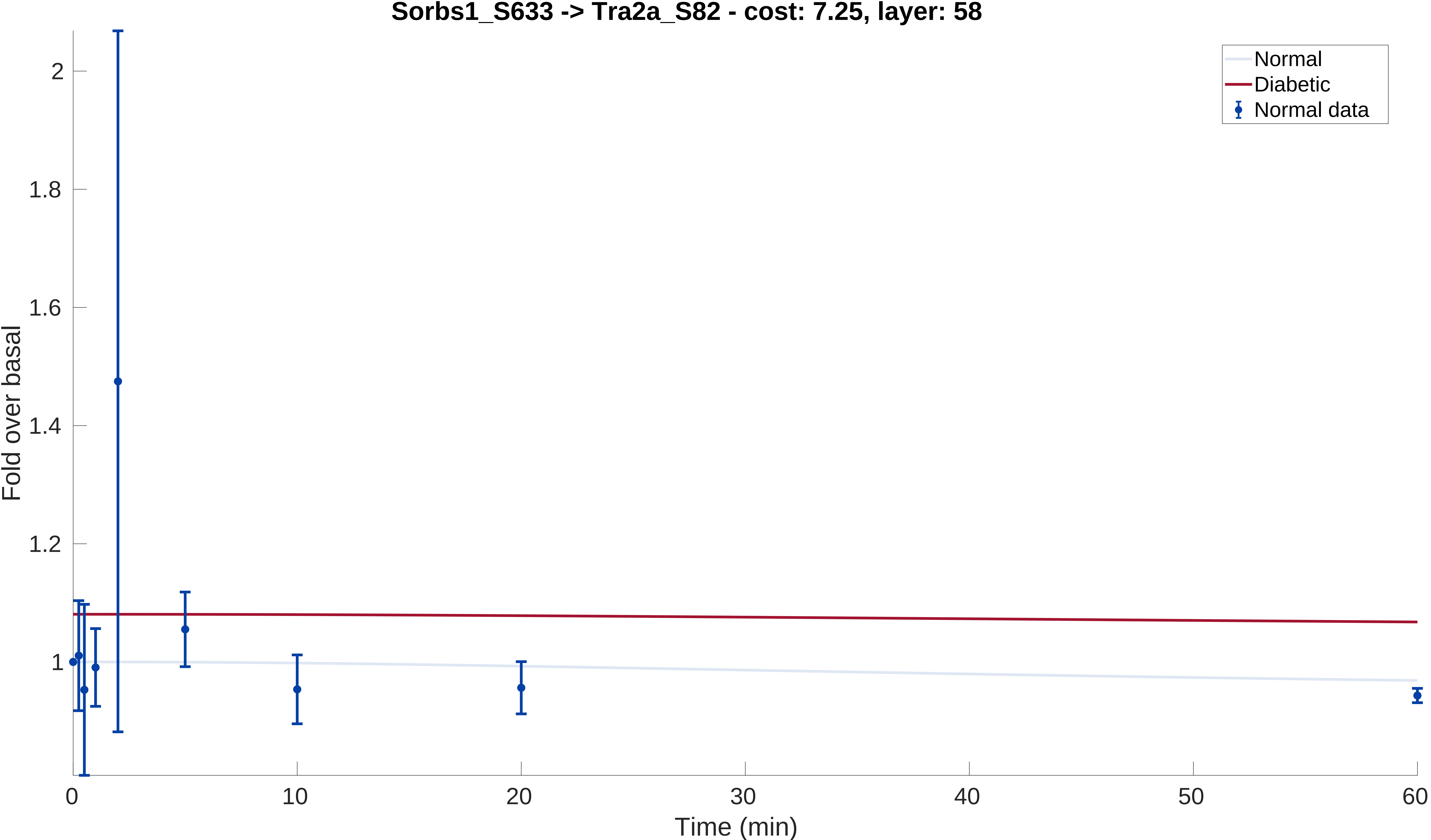

Sorbs1_S633 -> Tra2a_S82 - cost: 7.25, layer: 58
Normal
Diabetic
Normal data
Fold over basal
2
1.8
1.6
1.4
1.2
1
0
10
20
30
40
50
60
Time (min)

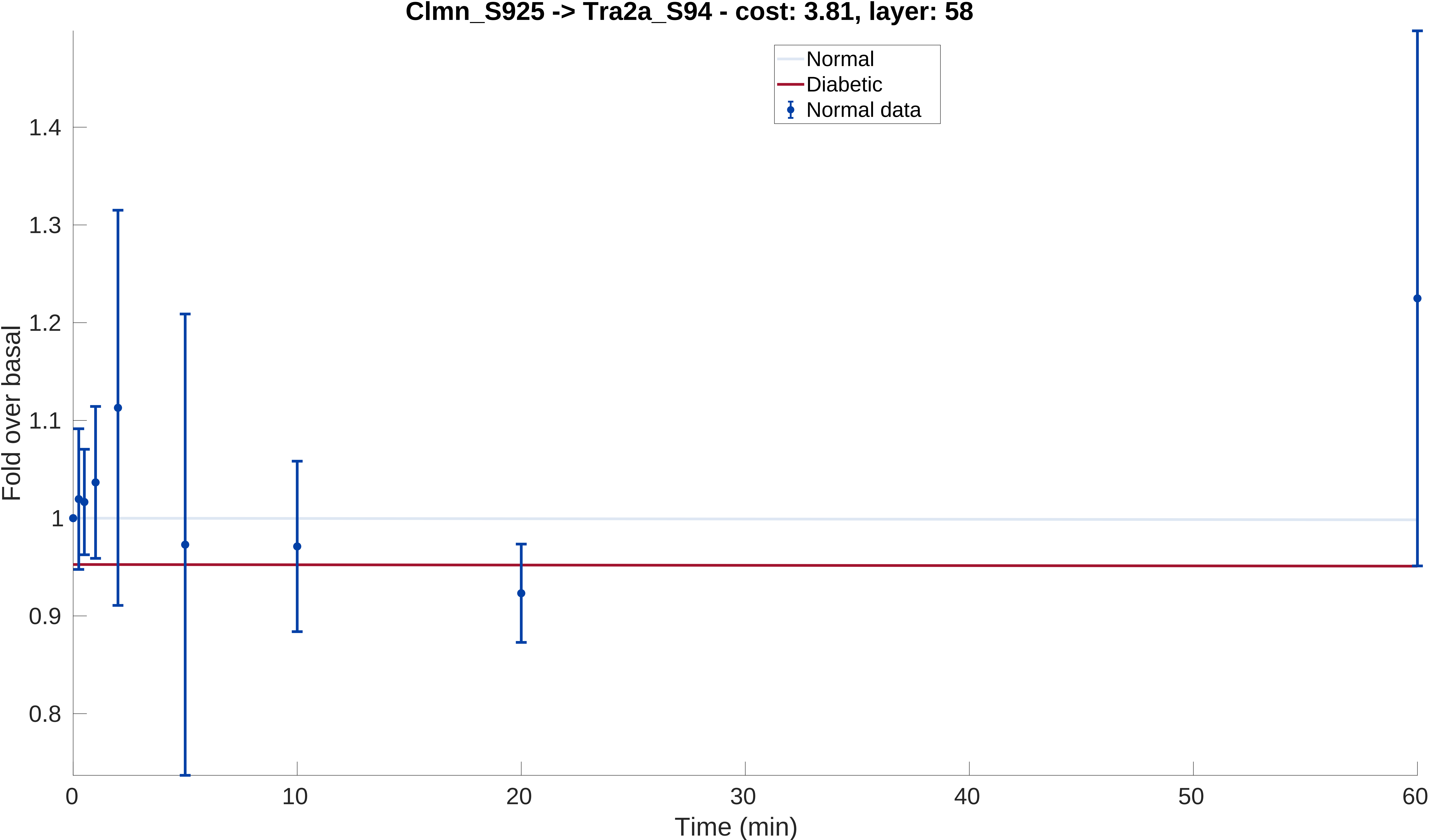

Clmn_S925 -> Tra2a_S94 - cost: 3.81, layer: 58
Normal
Diabetic
Normal data
1.4
1.3
1.2
1.1
1
0.9
0.8
Fold over basal
0
10
20
30
40
50
60
Time (min)

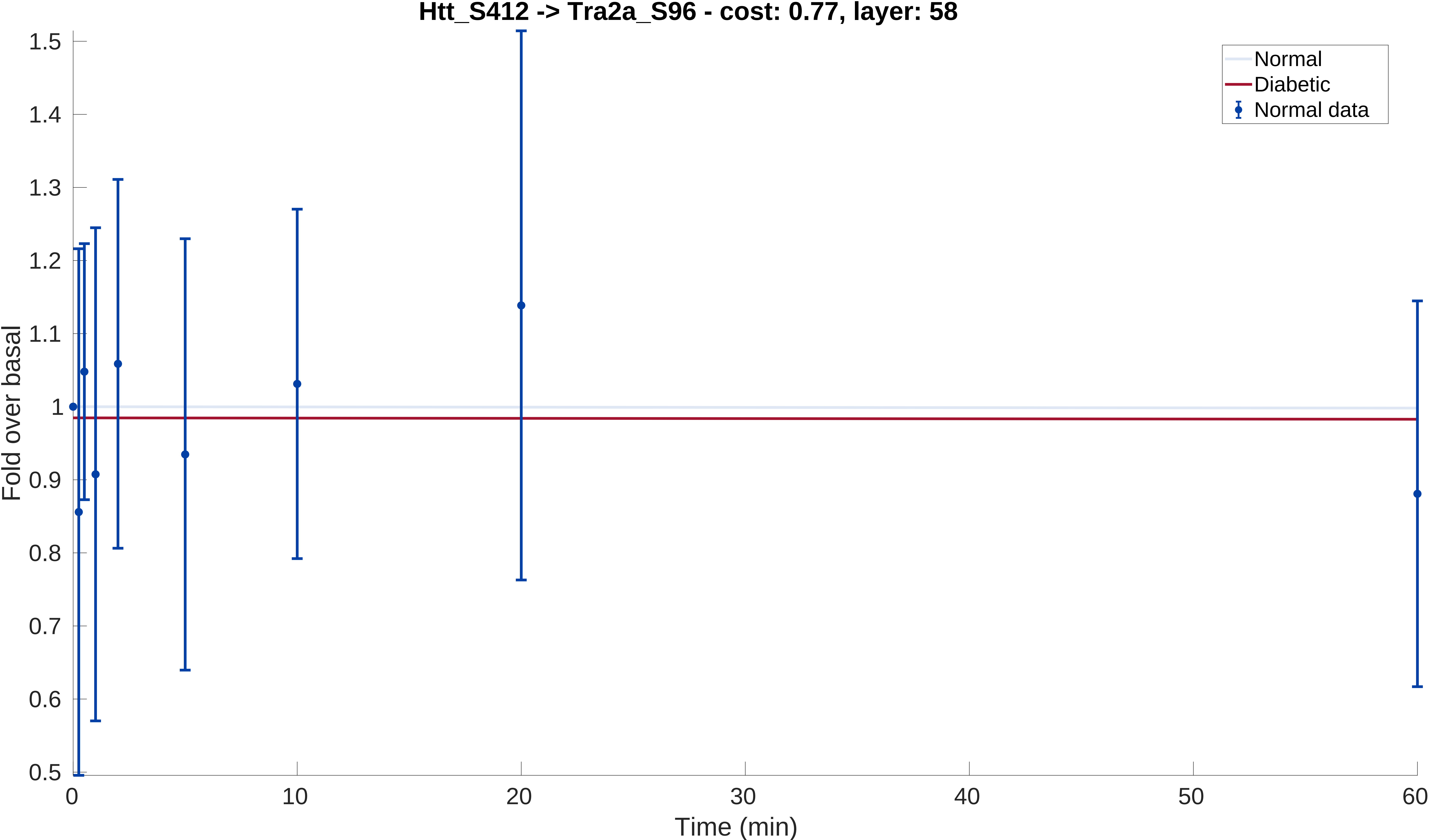

Htt_S412 -> Tra2a_S96 - cost: 0.77, layer: 58
Normal
Diabetic
Normal data
Fold over basal
Time (min)
0
10
20
30
40
50
60
0.5
0.6
0.7
0.8
0.9
1
1.1
1.2
1.3
1.4
1.5

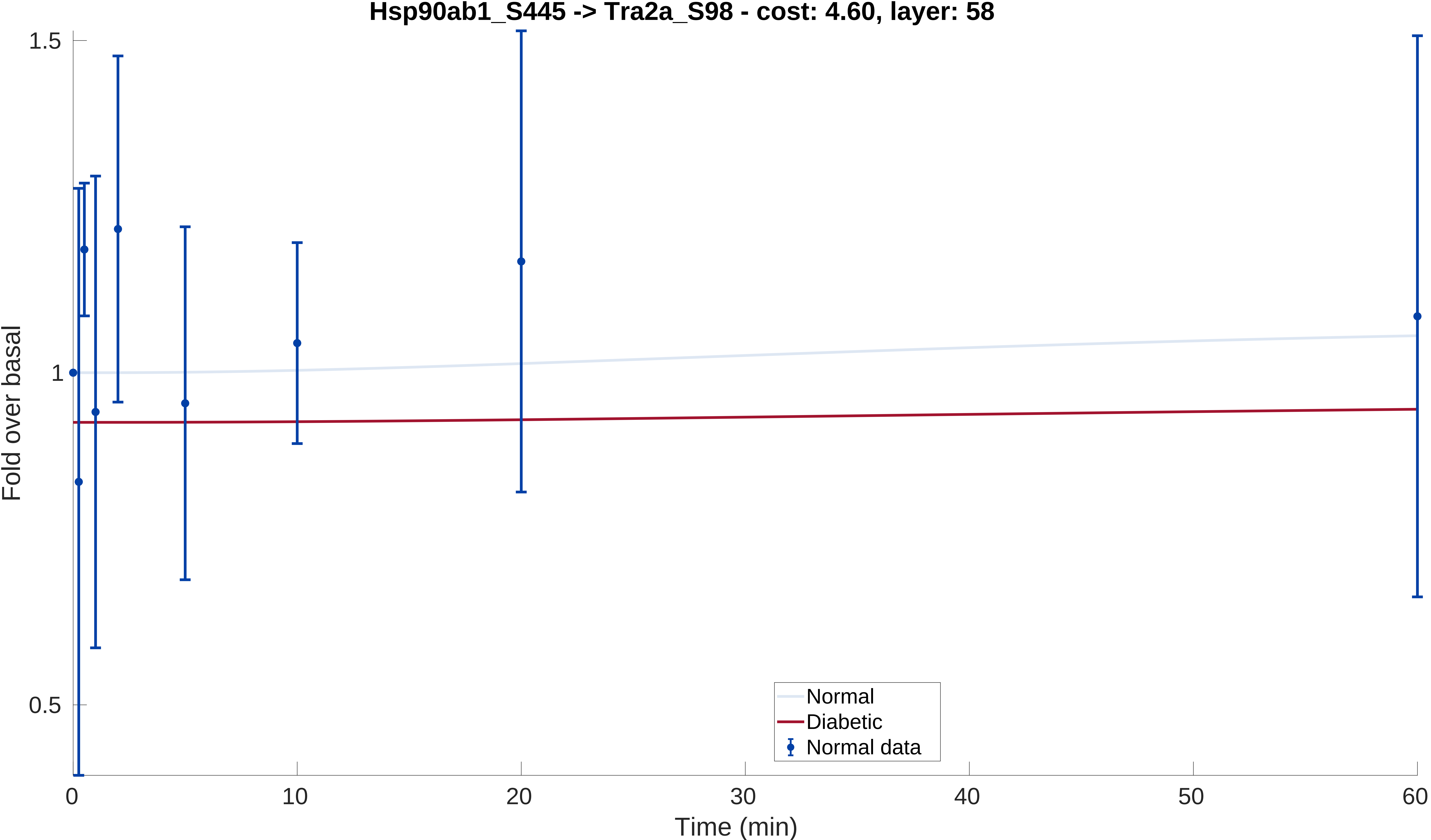
Hsp90ab1_S445 -> Tra2a_S98 - cost: 4.60, layer: 58
1.5
1
0.5
Fold over basal
0
10
20
30
40
50
60
Time (min)
Normal
Diabetic
Normal data

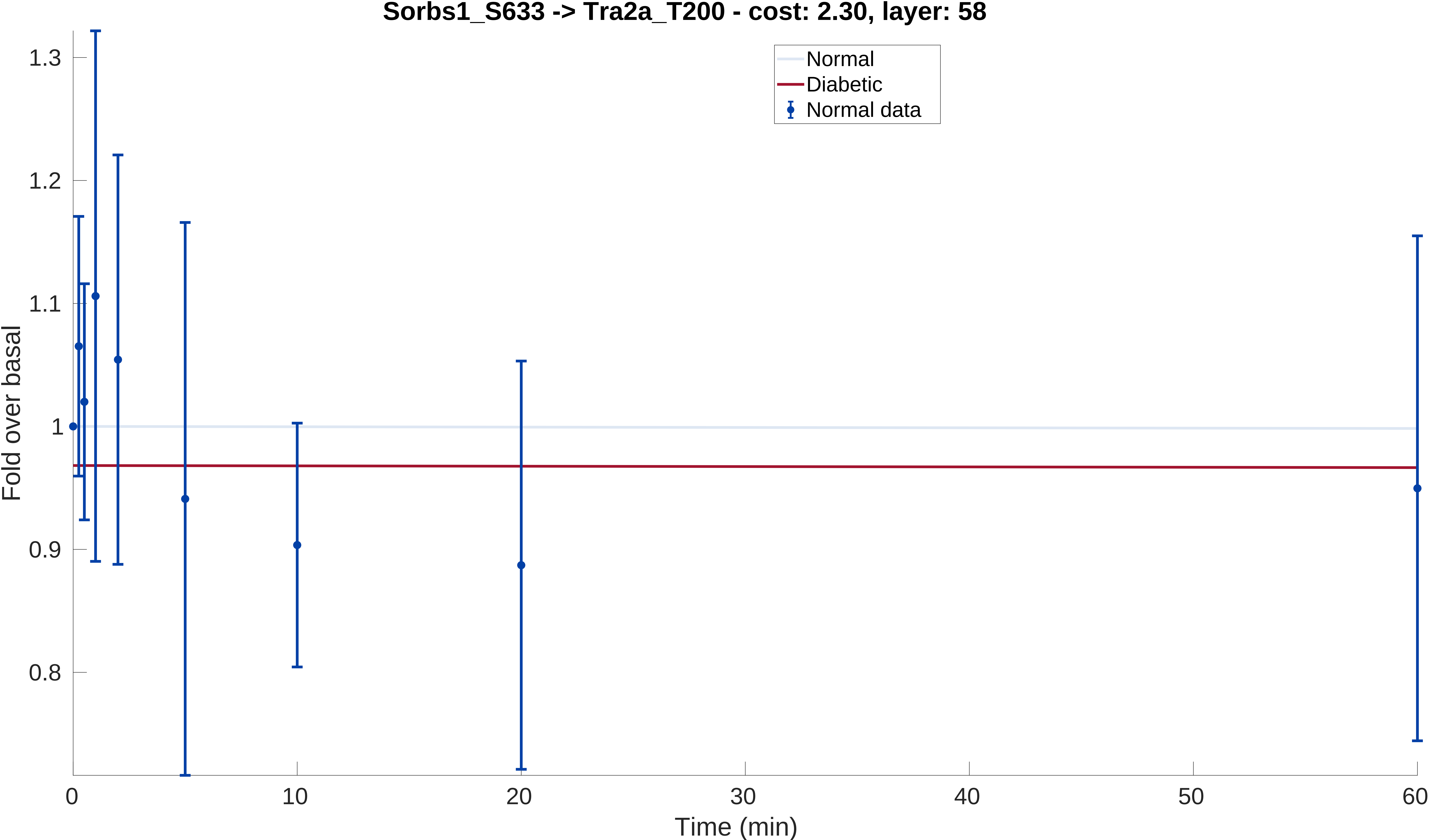

Sorbs1_S633 -> Tra2a_T200 - cost: 2.30, layer: 58
Normal
Diabetic
Normal data
Fold over basal
1.3
1.2
1.1
1
0.9
0.8
0
10
20
30
40
50
60
Time (min)

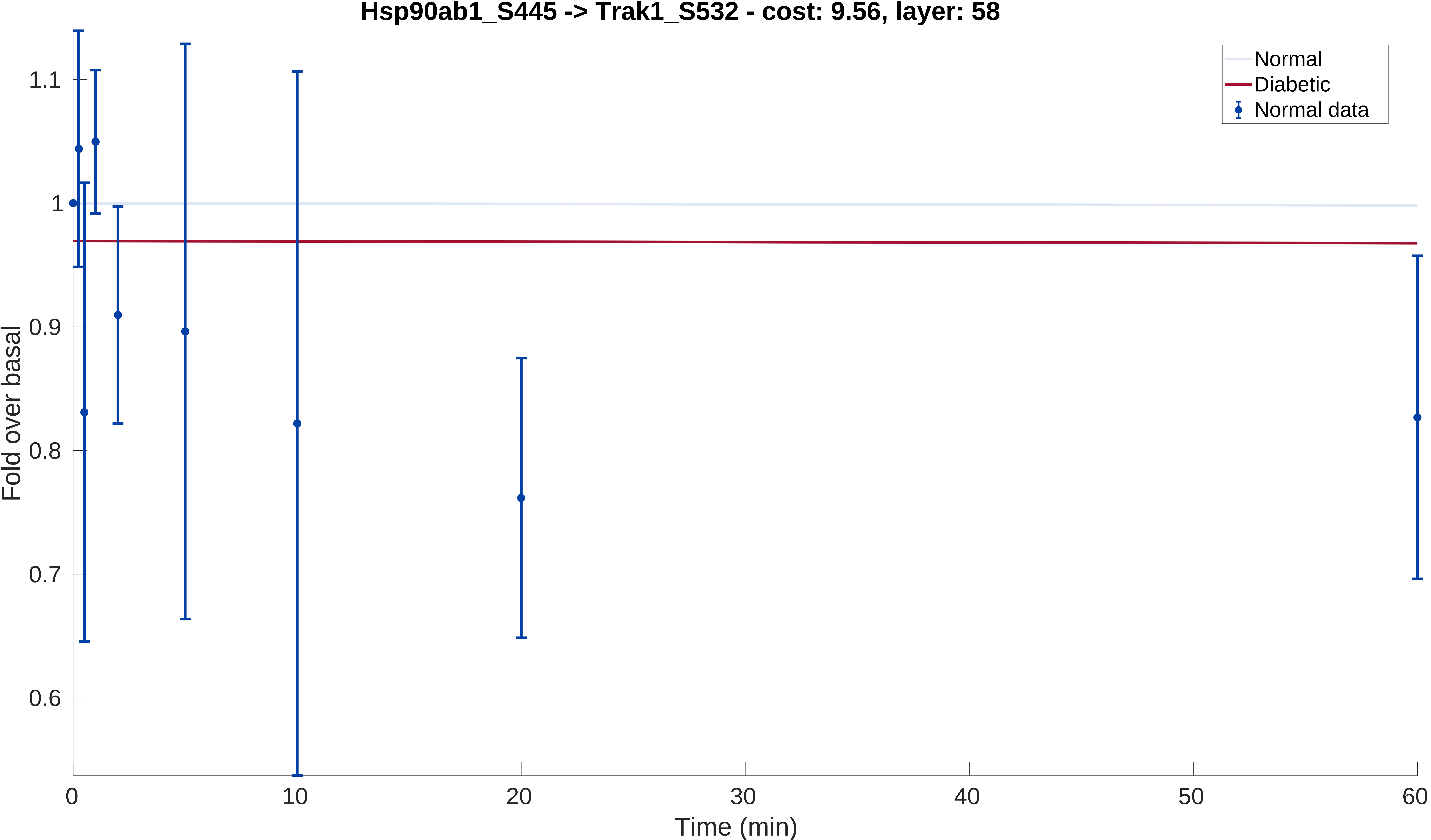

Hsp90ab1_S445 -> Trak1_S532 - cost: 9.56, layer: 58
Normal
Diabetic
Normal data
1.1
1
0.9
0.8
0.7
0.6
Fold over basal
0
10
20
30
40
50
60
Time (min)

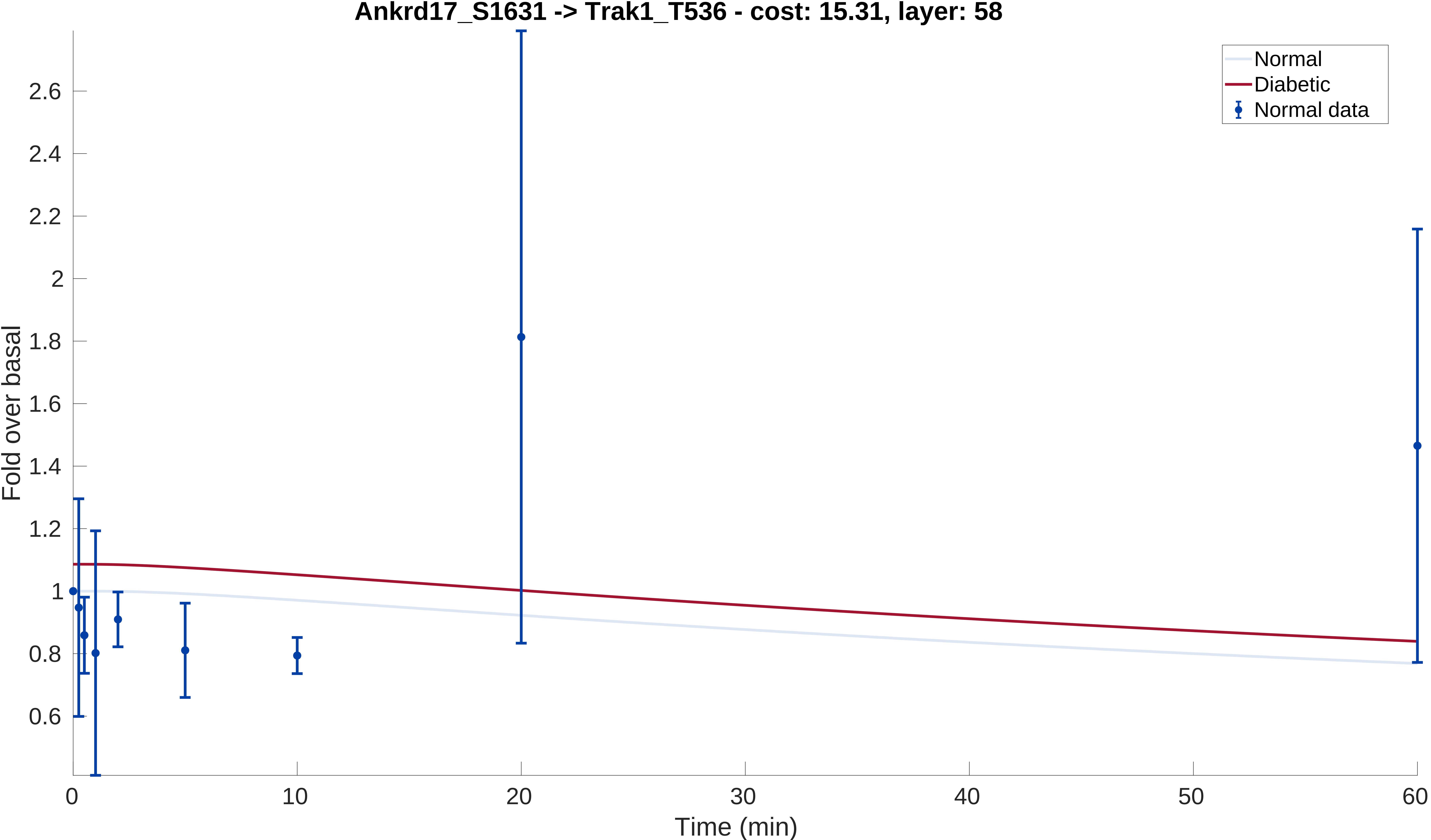

Ankrd17_S1631 -> Trak1_T536 - cost: 15.31, layer: 58
Normal
Diabetic
Normal data
Fold over basal
2.6
2.4
2.2
2
1.8
1.6
1.4
1.2
1
0.8
0.6
0
10
20
30
40
50
60
Time (min)

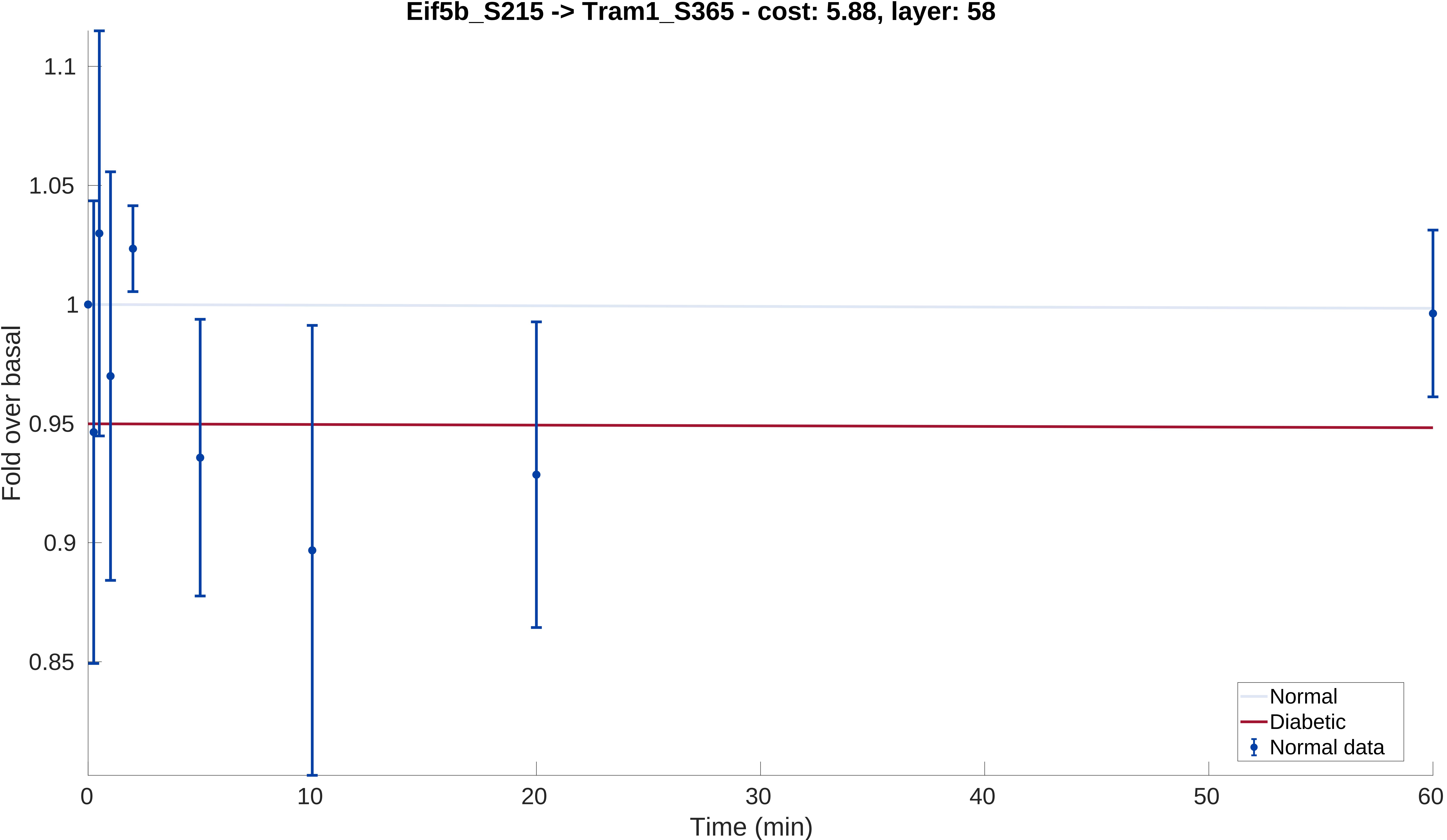
Eif5b_S215 -> Tram1_S365 - cost: 5.88, layer: 58
Fold over basal
1.1
1.05
1
0.95
0.9
0.85
0
10
20
30
40
50
60
Time (min)
Normal
Diabetic
Normal data

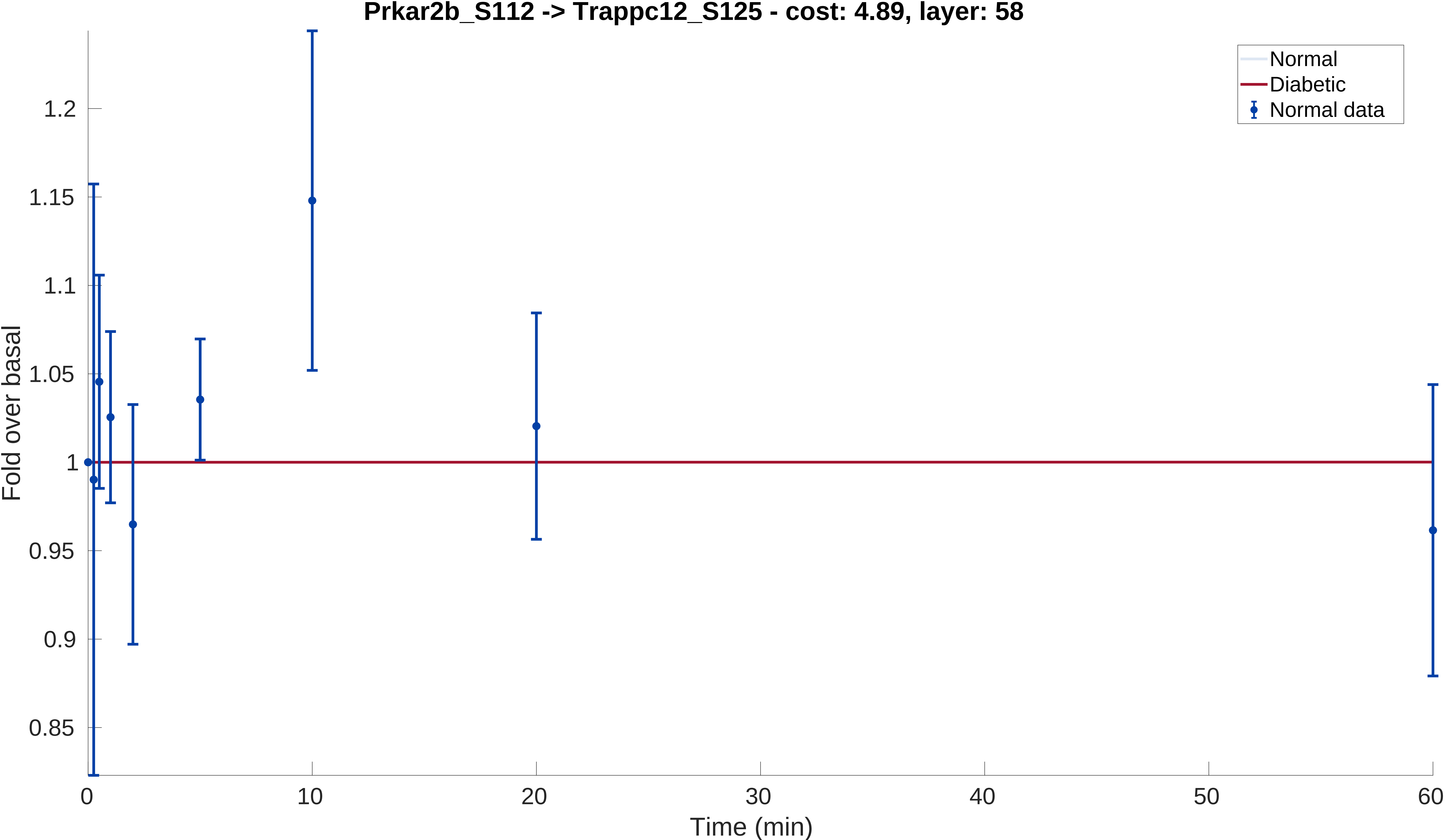

Prkar2b_S112 -> Trappc12_S125 - cost: 4.89, layer: 58
Normal
Diabetic
Normal data
Fold over basal
1.2
1.15
1.1
1.05
1
0.95
0.9
0.85
0
10
20
30
40
50
60
Time (min)

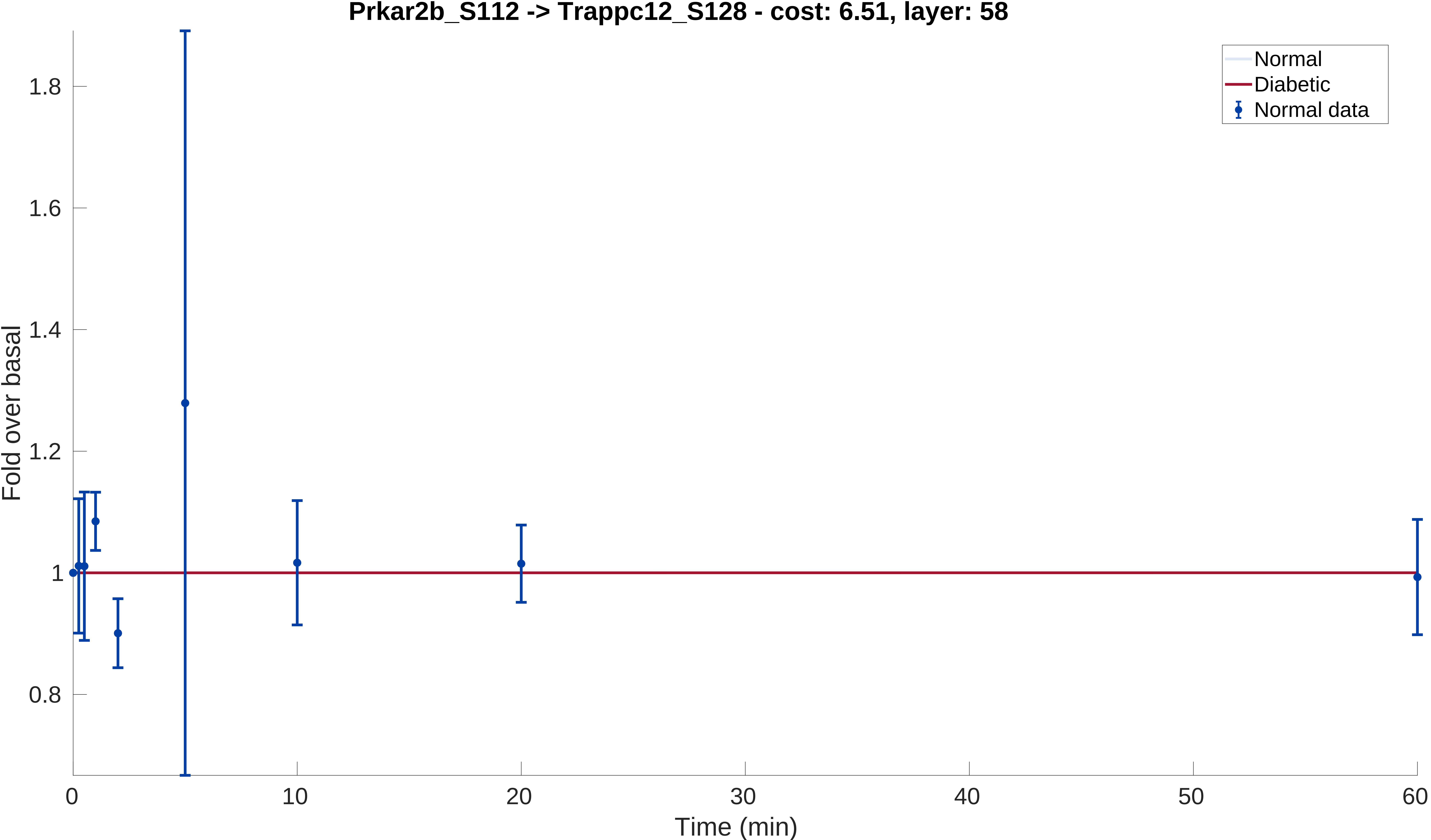

Prkar2b_S112 -> Trappc12_S128 - cost: 6.51, layer: 58
Normal
Diabetic
Normal data
Fold over basal
1.8
1.6
1.4
1.2
1
0.8
0
10
20
30
40
50
60
Time (min)

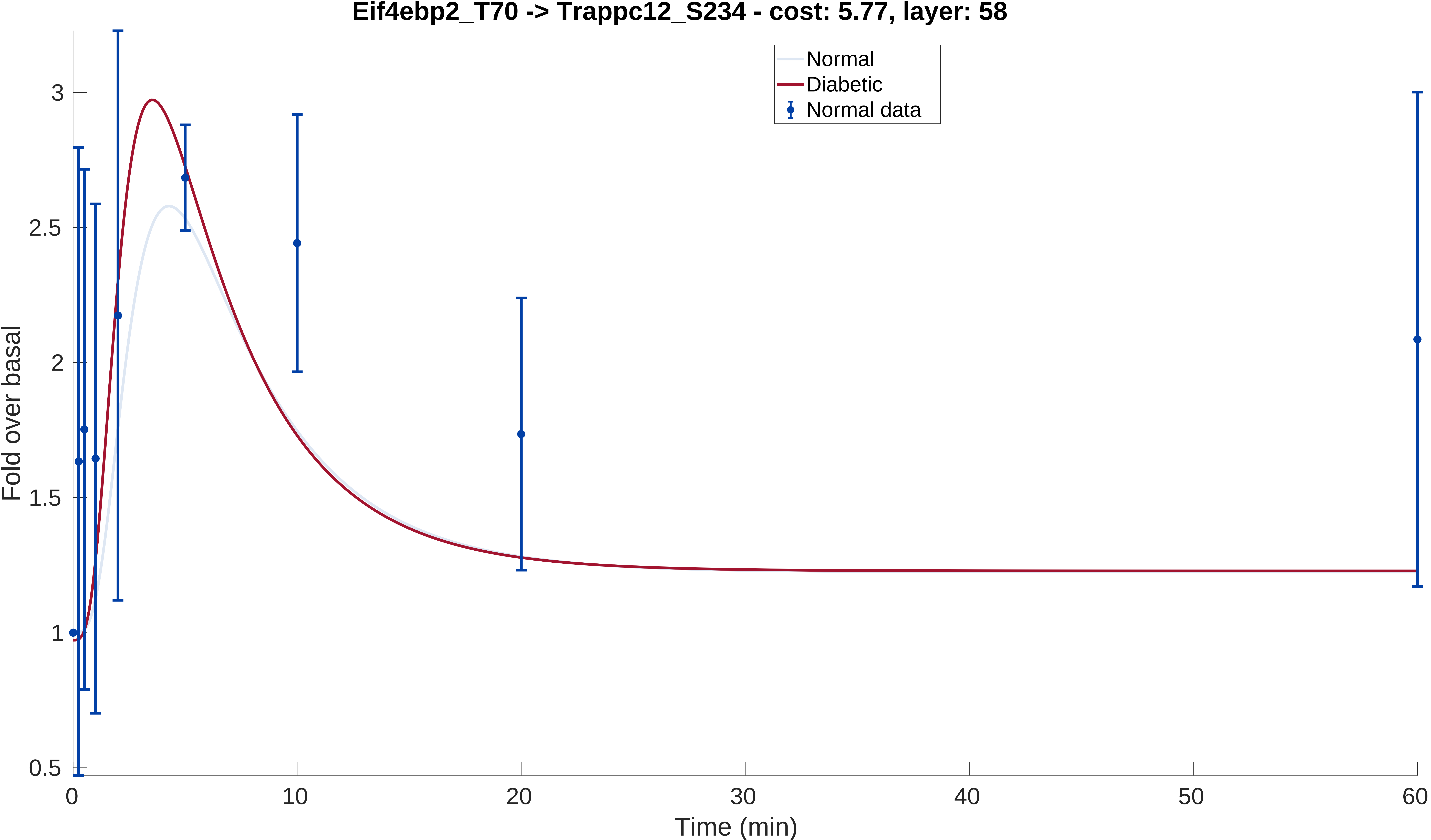

Eif4ebp2_T70 -> Trappc12_S234 - cost: 5.77, layer: 58
Normal
Diabetic
Normal data
Fold over basal
3
2.5
2
1.5
1
0.5
0
10
20
30
40
50
60
Time (min)

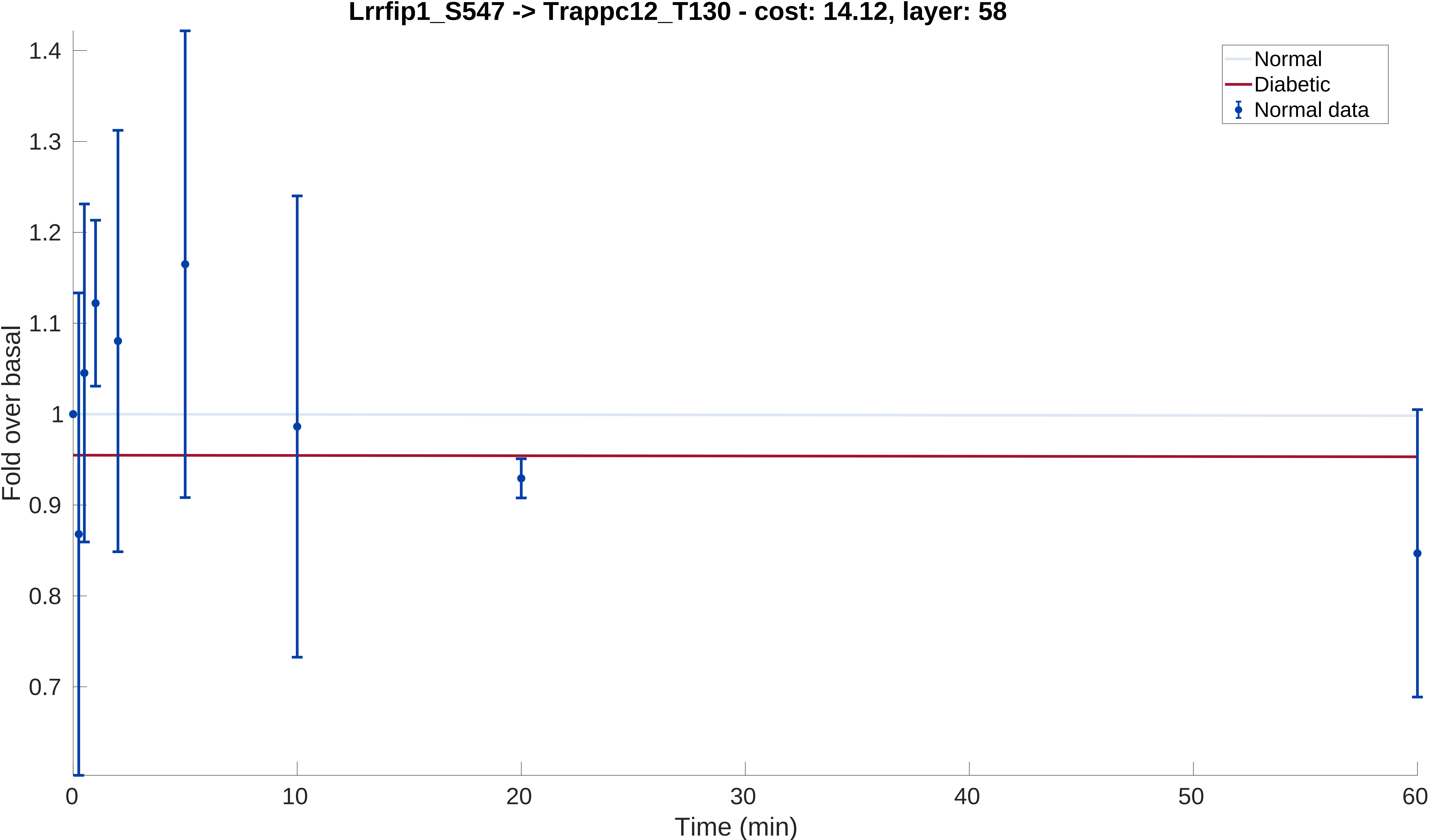
Lrrfip1_S547 -> Trappc12_T130 - cost: 14.12, layer: 58
Normal
Diabetic
Normal data
Fold over basal
Time (min)
1.4
1.3
1.2
1.1
1
0.9
0.8
0.7
0
10
20
30
40
50
60

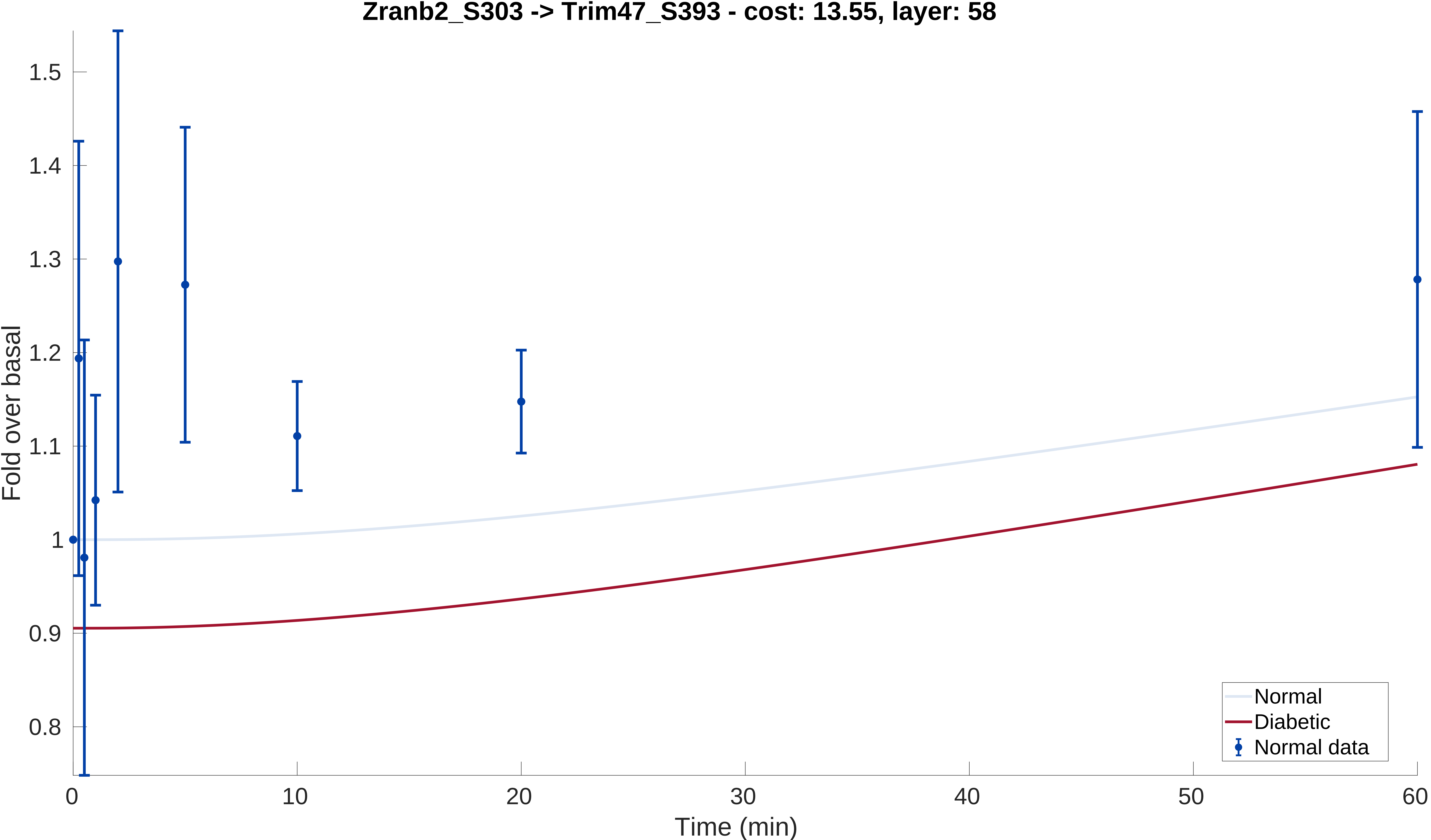

Zranb2_S303 -> Trim47_S393 - cost: 13.55, layer: 58
Fold over basal
1.5
1.4
1.3
1.2
1.1
1
0.9
0.8
0
10
20
30
40
50
60
Time (min)
Normal
Diabetic
Normal data

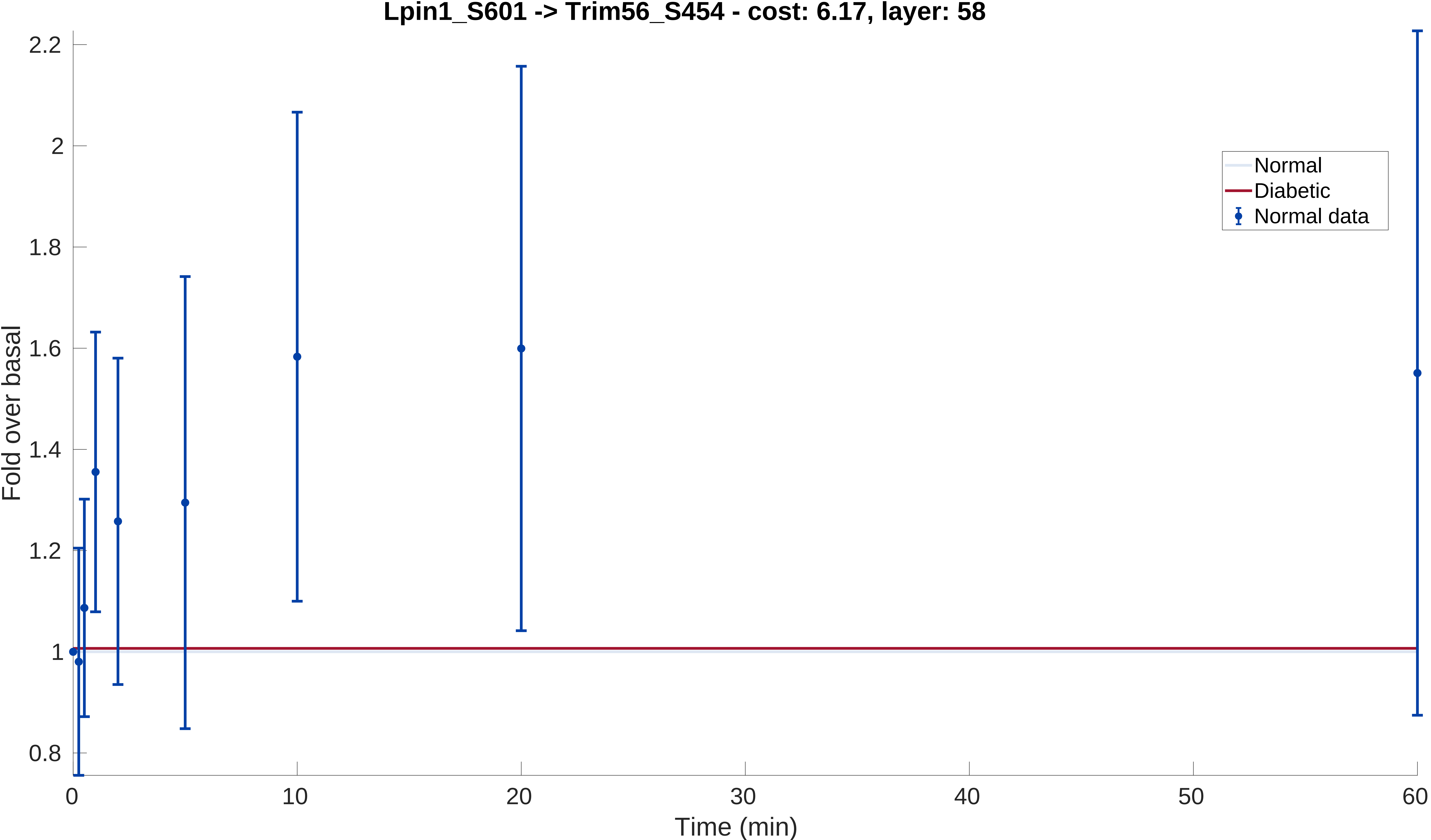

Lpin1_S601 -> Trim56_S454 - cost: 6.17, layer: 58
Fold over basal
2.2
2
1.8
1.6
1.4
1.2
1
0.8
0
10
20
30
40
50
60
Time (min)
Normal
Diabetic
Normal data

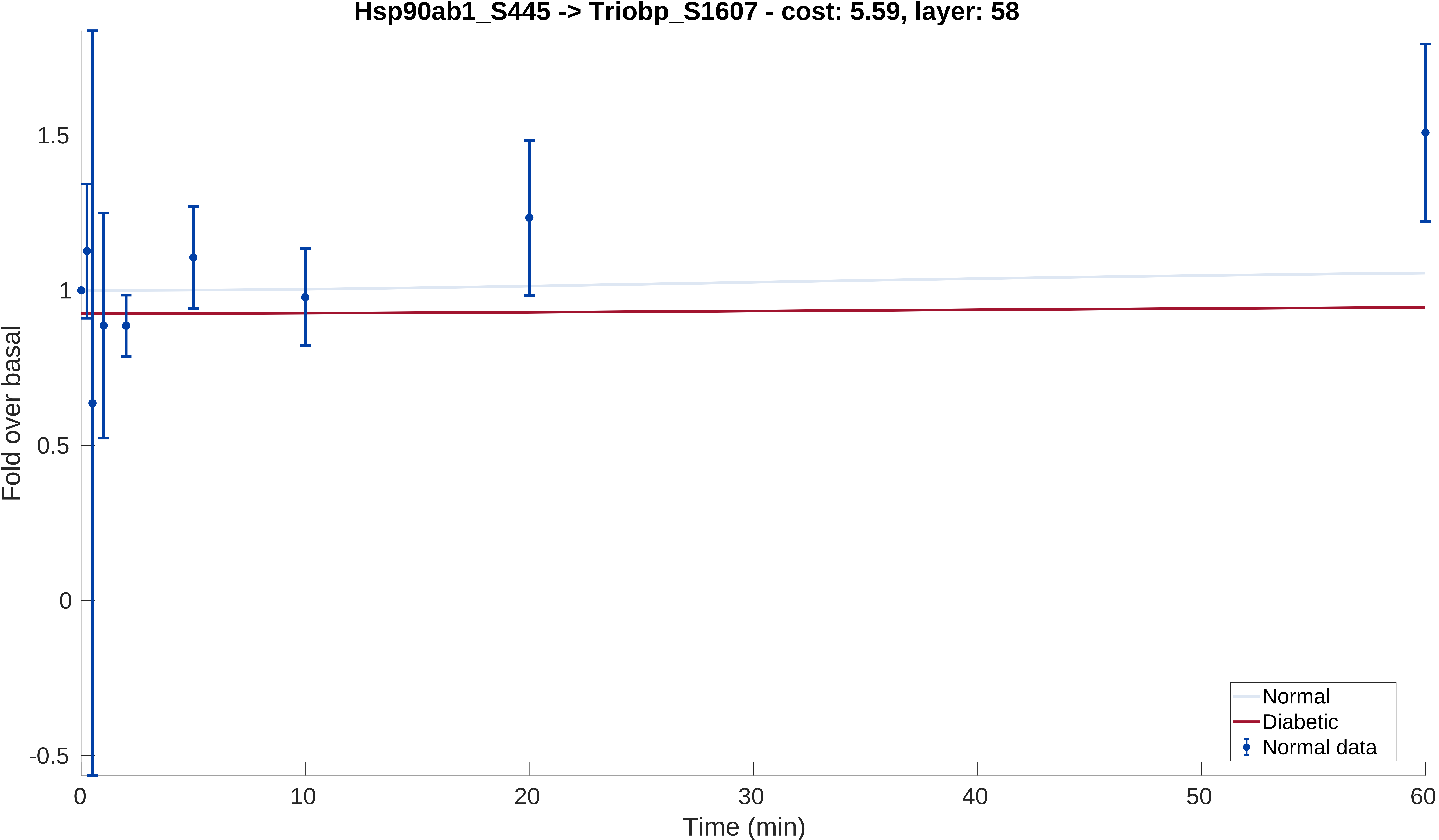

Hsp90ab1_S445 -> Triobp_S1607 - cost: 5.59, layer: 58
Fold over basal
1.5
1
0.5
0
-0.5
0
10
20
30
40
50
60
Time (min)
Normal
Diabetic
Normal data

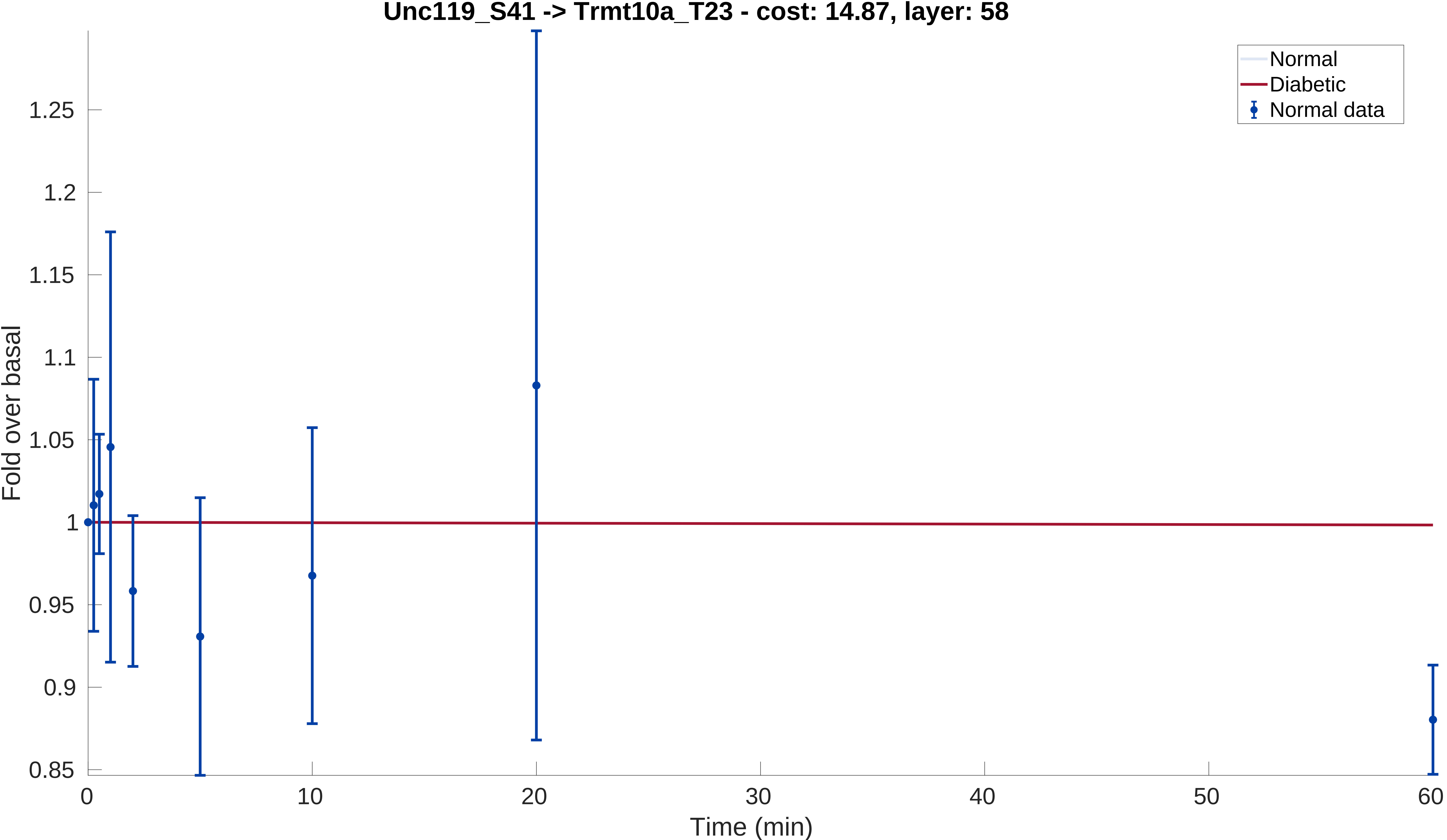
Unc119_S41 -> Trmt10a_T23 - cost: 14.87, layer: 58
Normal
Diabetic
Normal data
Fold over basal
Time (min)
1.25
1.2
1.15
1.1
1.05
1
0.95
0.9
0.85
0
10
20
30
40
50
60

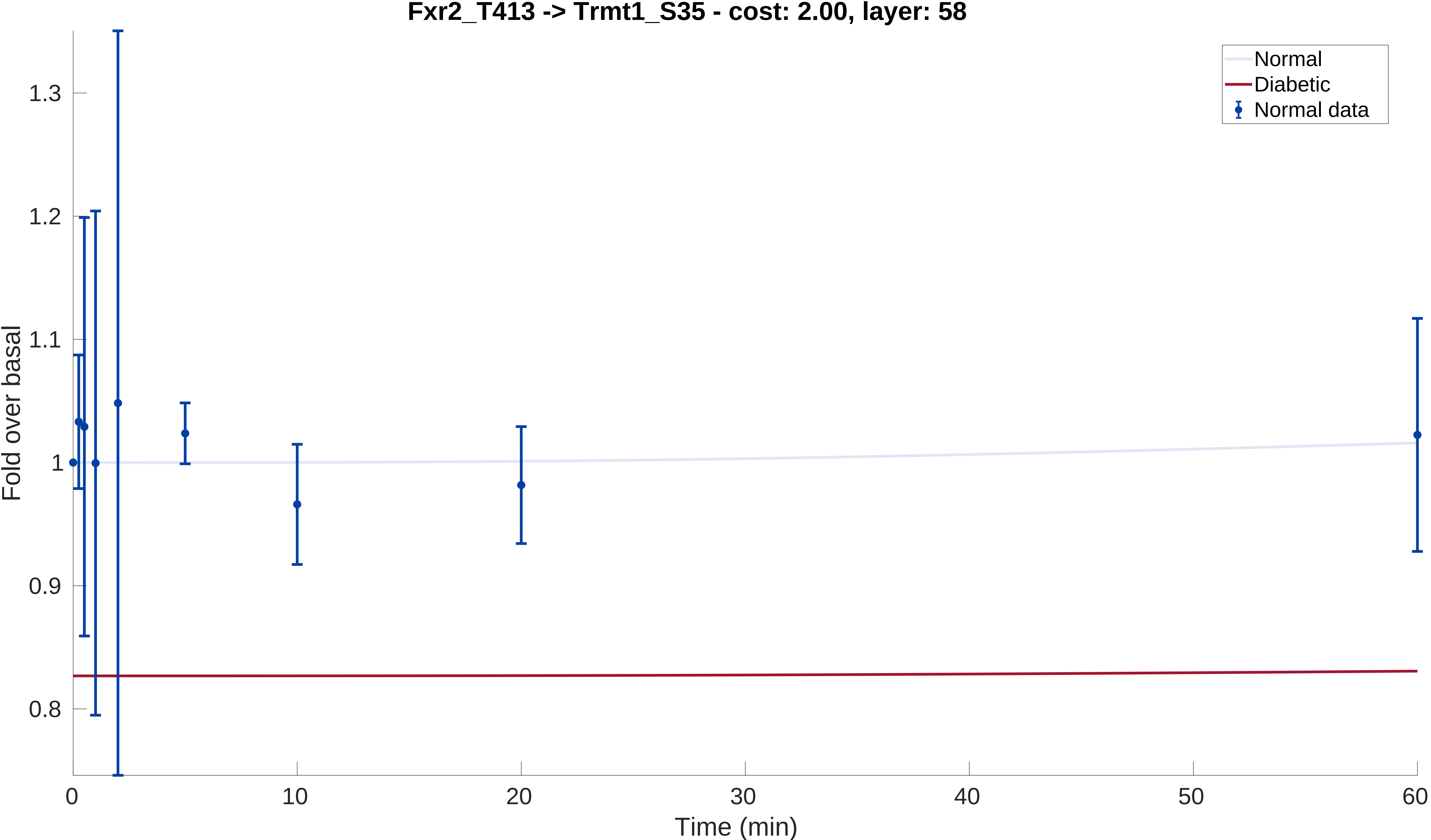

Fxr2_T413 -> Trmt1_S35 - cost: 2.00, layer: 58
Normal
Diabetic
Normal data
1.3
1.2
1.1
1
0.9
0.8
Fold over basal
0
10
20
30
40
50
60
Time (min)

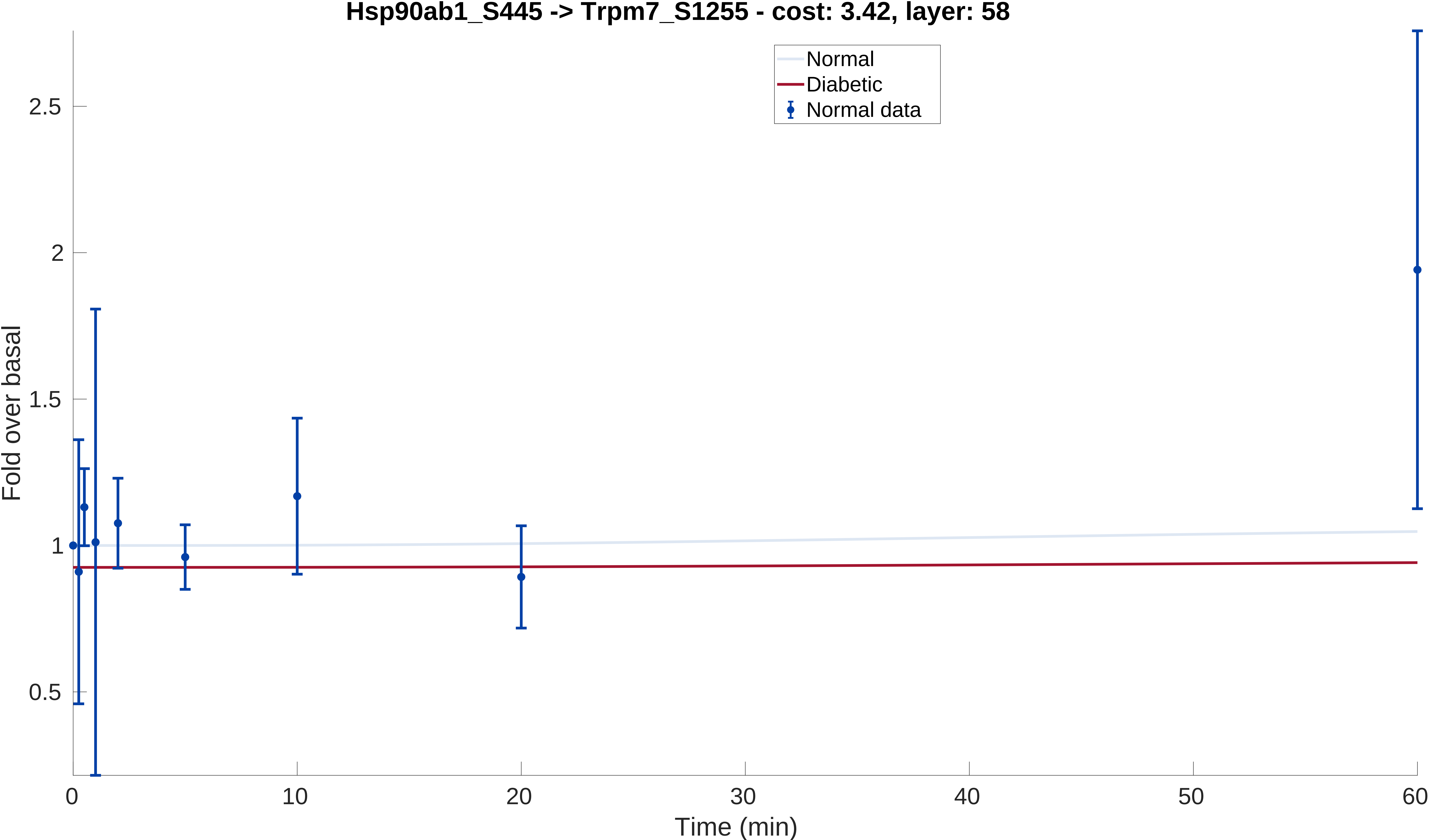

Hsp90ab1_S445 -> Trpm7_S1255 - cost: 3.42, layer: 58
Normal
Diabetic
Normal data
Fold over basal
2.5
2
1.5
1
0.5
0
10
20
30
40
50
60
Time (min)

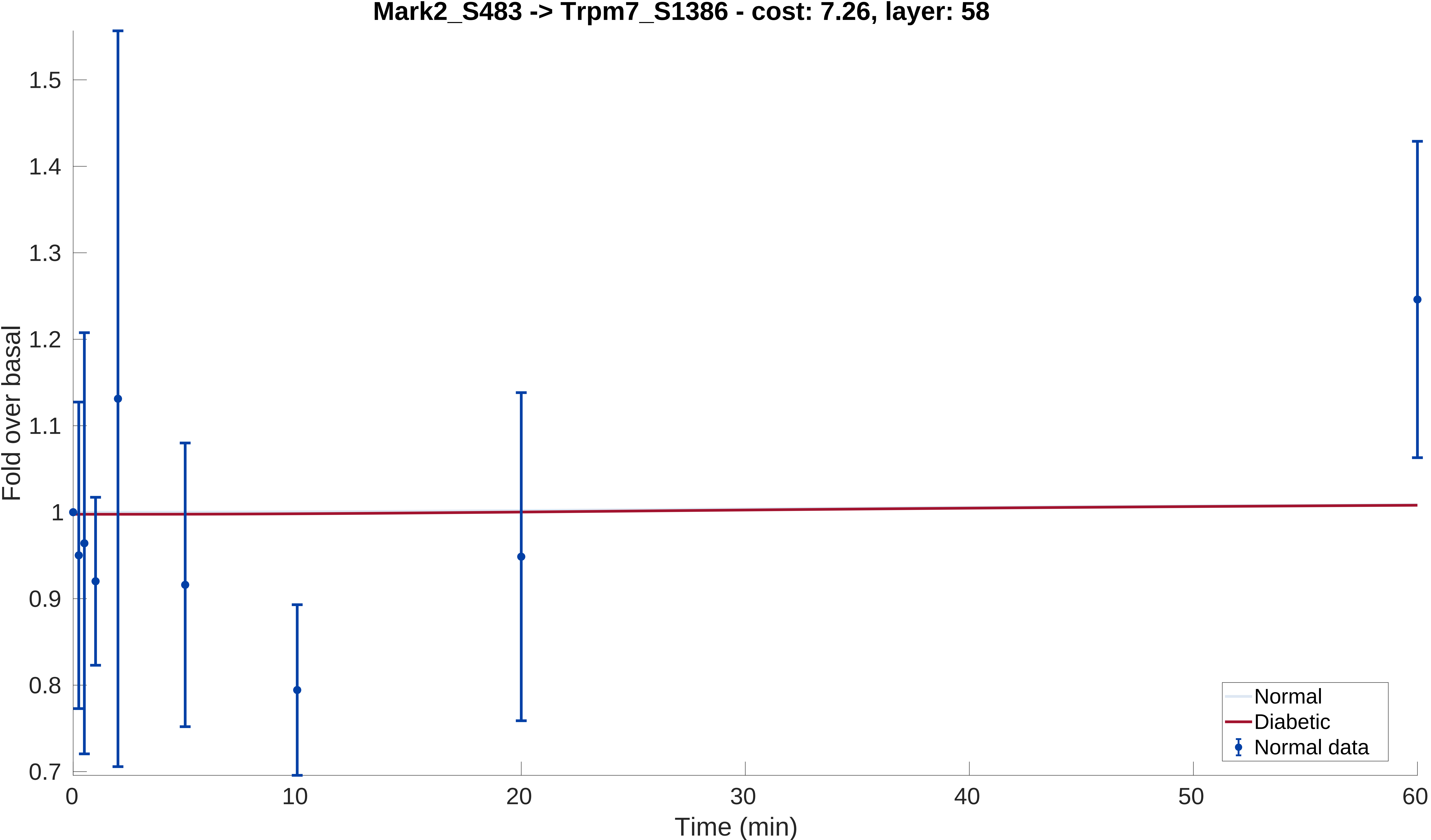
Mark2_S483 -> Trpm7_S1386 - cost: 7.26, layer: 58
Fold over basal
1.5
1.4
1.3
1.2
1.1
1
0.9
0.8
0.7
0
10
20
30
40
50
60
Time (min)
Normal
Diabetic
Normal data

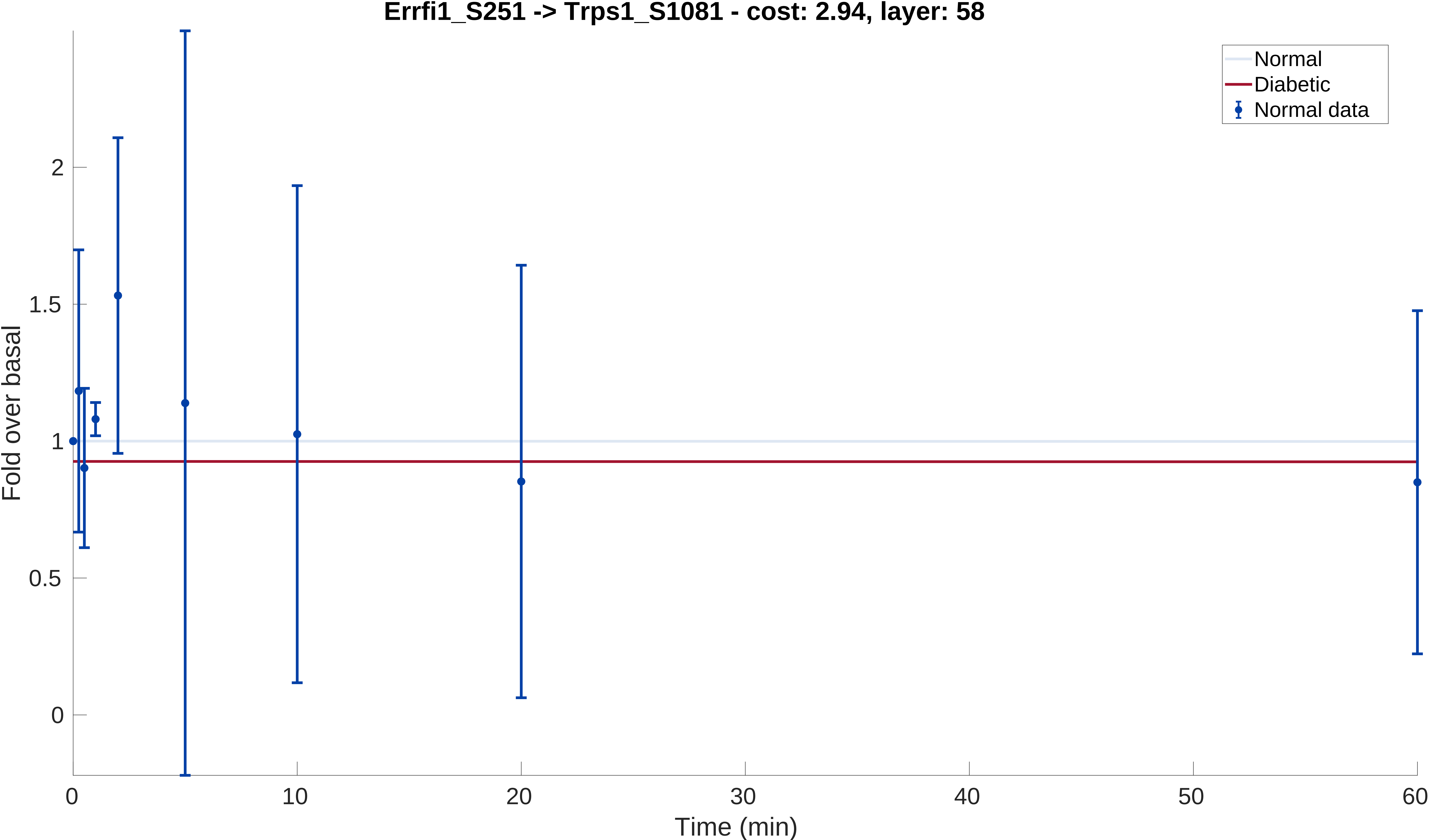

Errfi1_S251 -> Trps1_S1081 - cost: 2.94, layer: 58
Normal
Diabetic
Normal data
Fold over basal
Time (min)
0
0.5
1
1.5
2
0
10
20
30
40
50
60

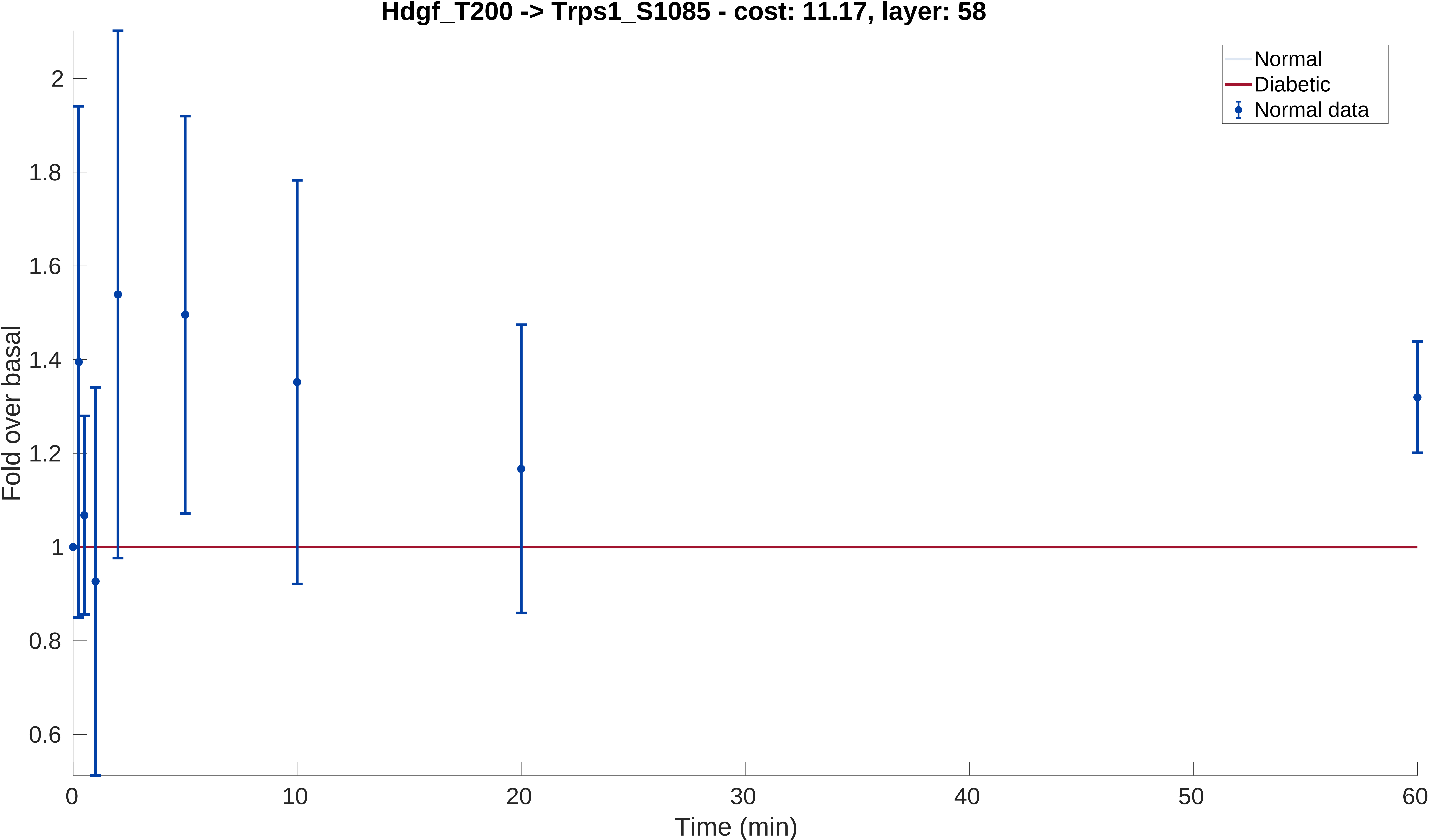

Hdgf_T200 -> Trps1_S1085 - cost: 11.17, layer: 58
Normal
Diabetic
Normal data
Fold over basal
2
1.8
1.6
1.4
1.2
1
0.8
0.6
0
10
20
30
40
50
60
Time (min)

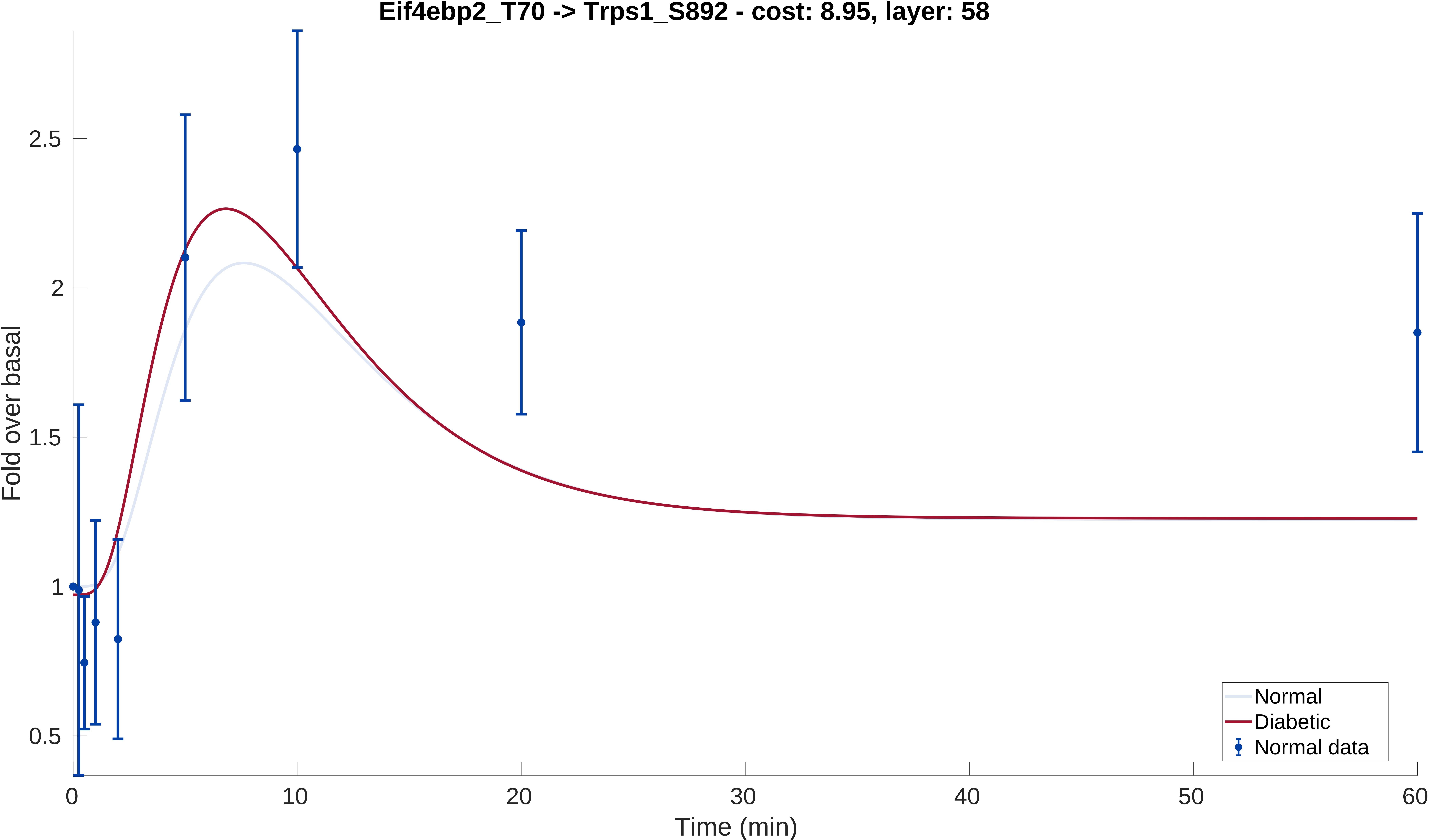
Eif4ebp2_T70 -> Trps1_S892 - cost: 8.95, layer: 58
Fold over basal
Time (min)
0.5
1
1.5
2
2.5
0
10
20
30
40
50
60
Normal
Diabetic
Normal data

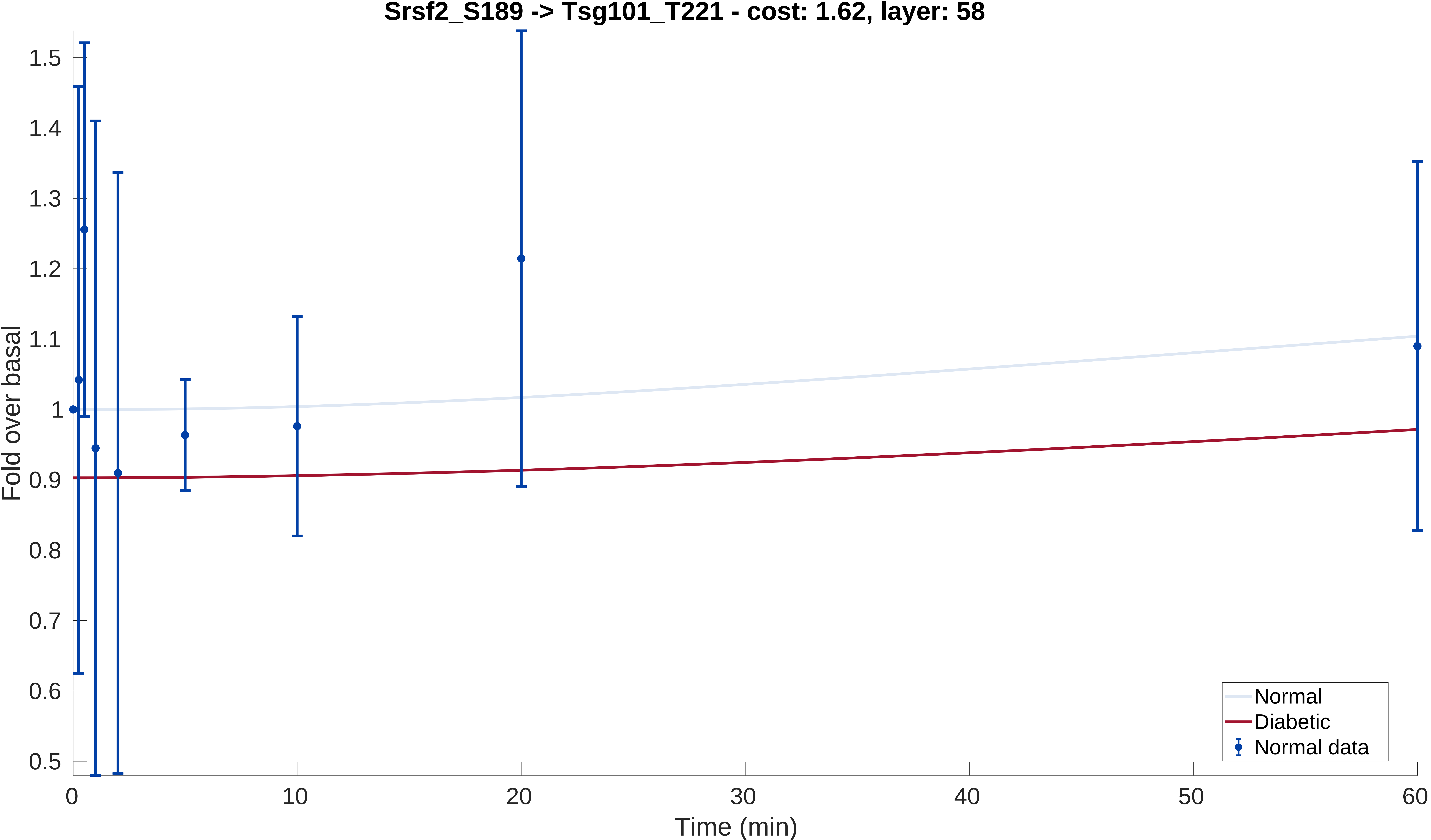

Srsf2_S189 -> Tsg101_T221 - cost: 1.62, layer: 58
Fold over basal
1.5
1.4
1.3
1.2
1.1
1
0.9
0.8
0.7
0.6
0.5
0
10
20
30
40
50
60
Time (min)
Normal
Diabetic
Normal data

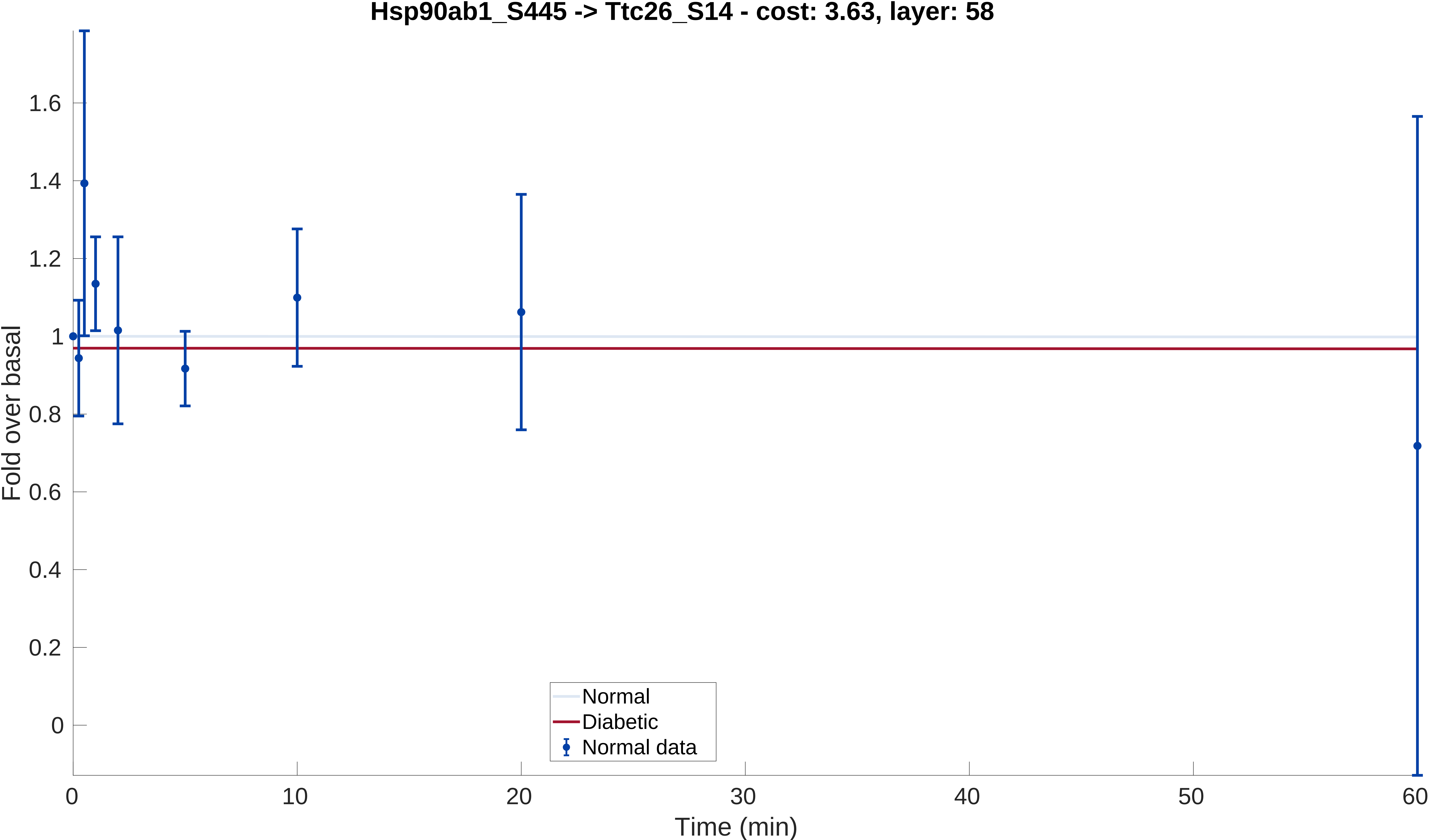

Hsp90ab1_S445 -> Ttc26_S14 - cost: 3.63, layer: 58
Fold over basal
1.6
1.4
1.2
1
0.8
0.6
0.4
0.2
0
Normal
Diabetic
Normal data
0
10
20
30
40
50
60
Time (min)

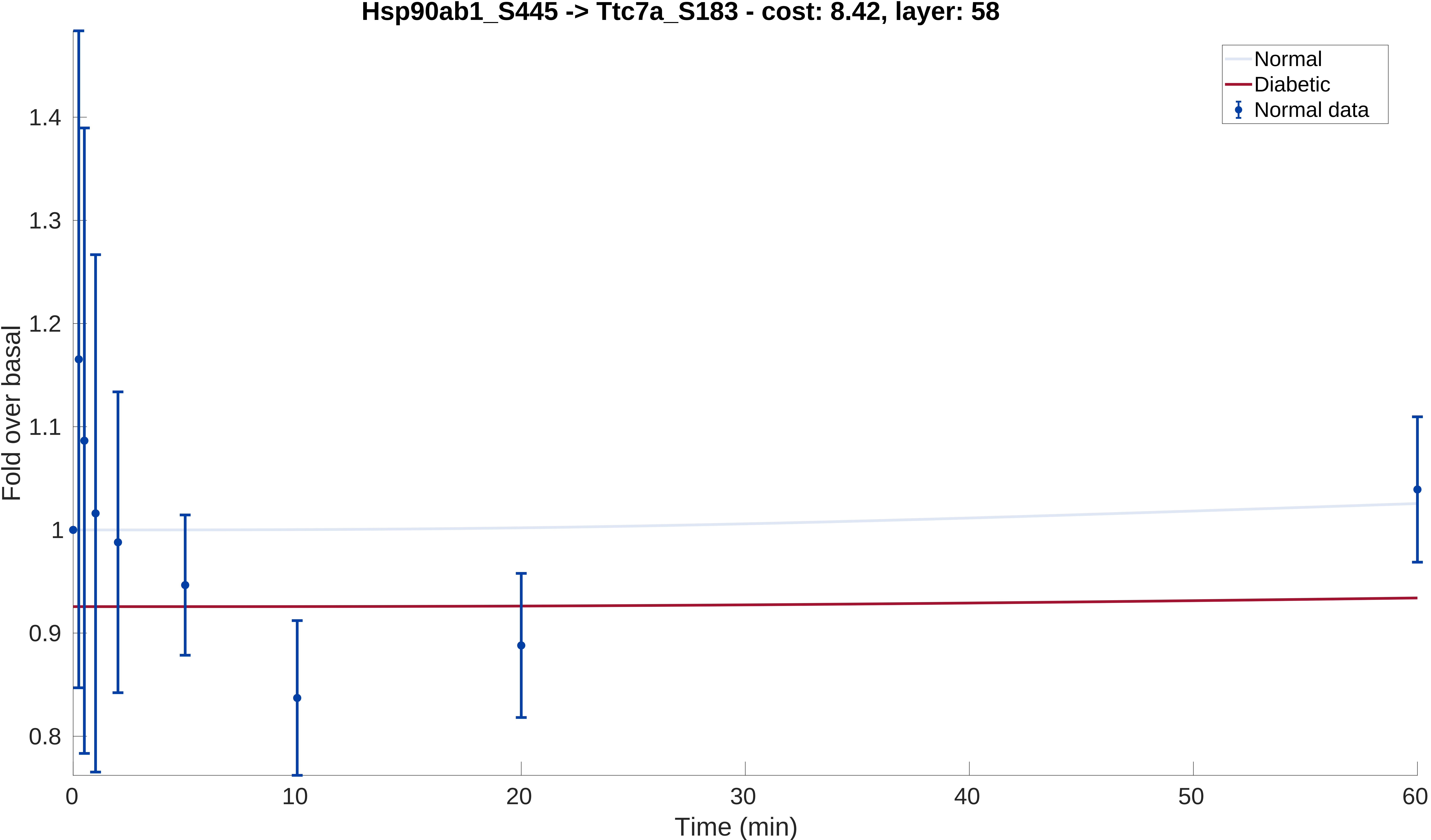
Hsp90ab1_S445 -> Ttc7a_S183 - cost: 8.42, layer: 58
Normal
Diabetic
Normal data
Fold over basal
1.4
1.3
1.2
1.1
1
0.9
0.8
0
10
20
30
40
50
60
Time (min)

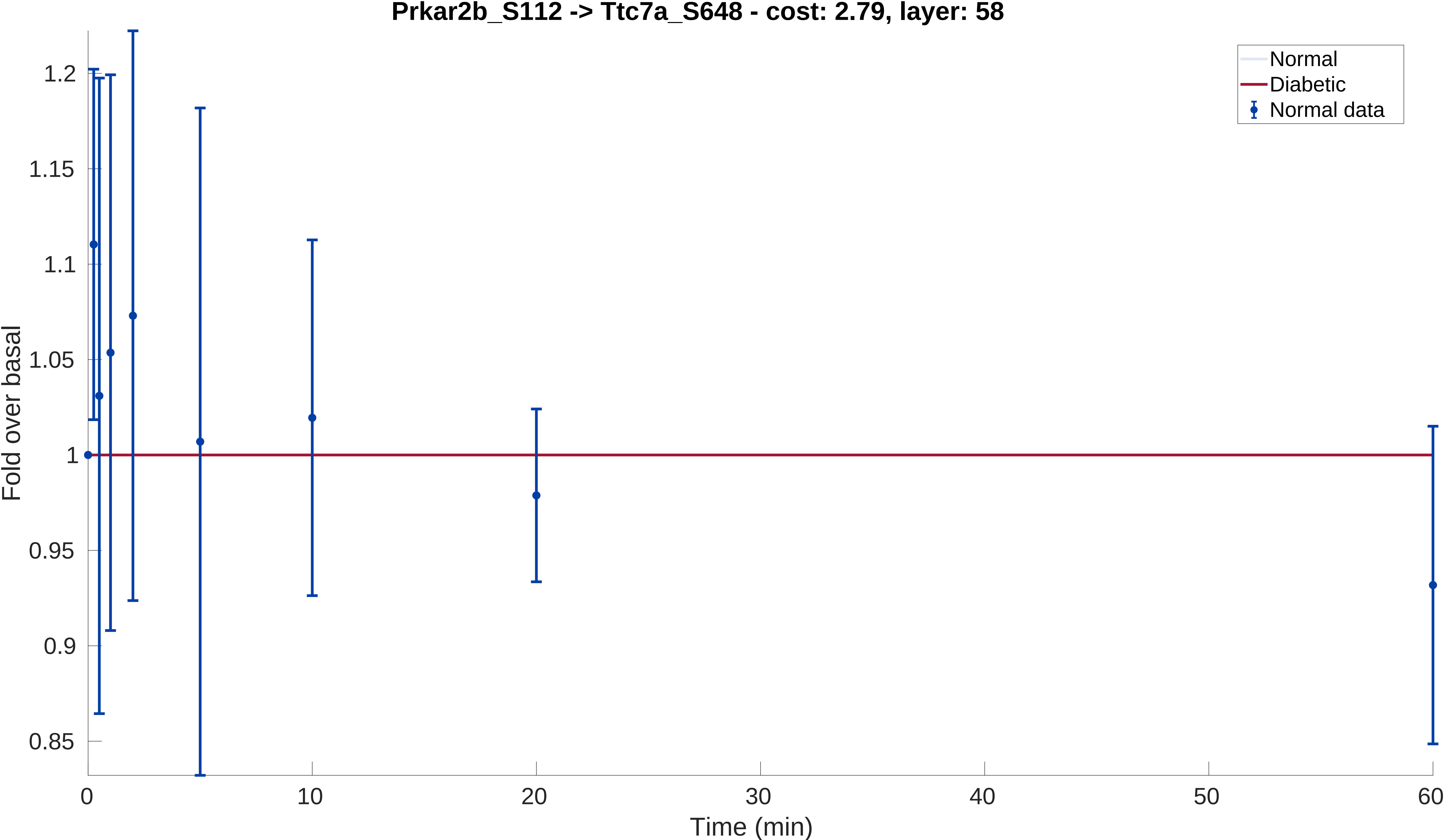

Prkar2b_S112 -> Ttc7a_S648 - cost: 2.79, layer: 58
Normal
Diabetic
Normal data
Fold over basal
Time (min)
1.2
1.15
1.1
1.05
1
0.95
0.9
0.85
0
10
20
30
40
50
60

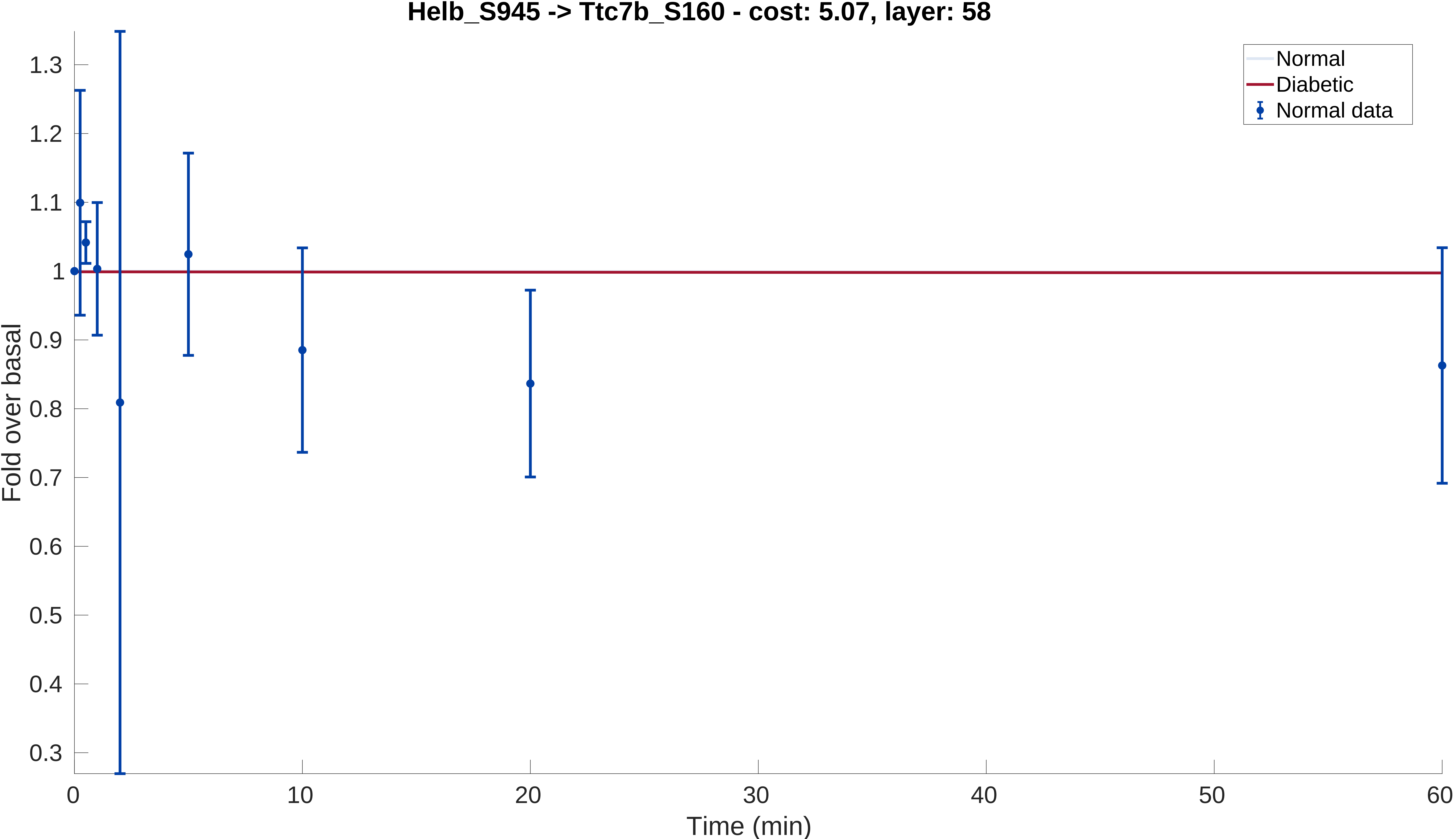

Helb_S945 -> Ttc7b_S160 - cost: 5.07, layer: 58
Normal
Diabetic
Normal data
Fold over basal
Time (min)
1.3
1.2
1.1
1
0.9
0.8
0.7
0.6
0.5
0.4
0.3
0
10
20
30
40
50
60

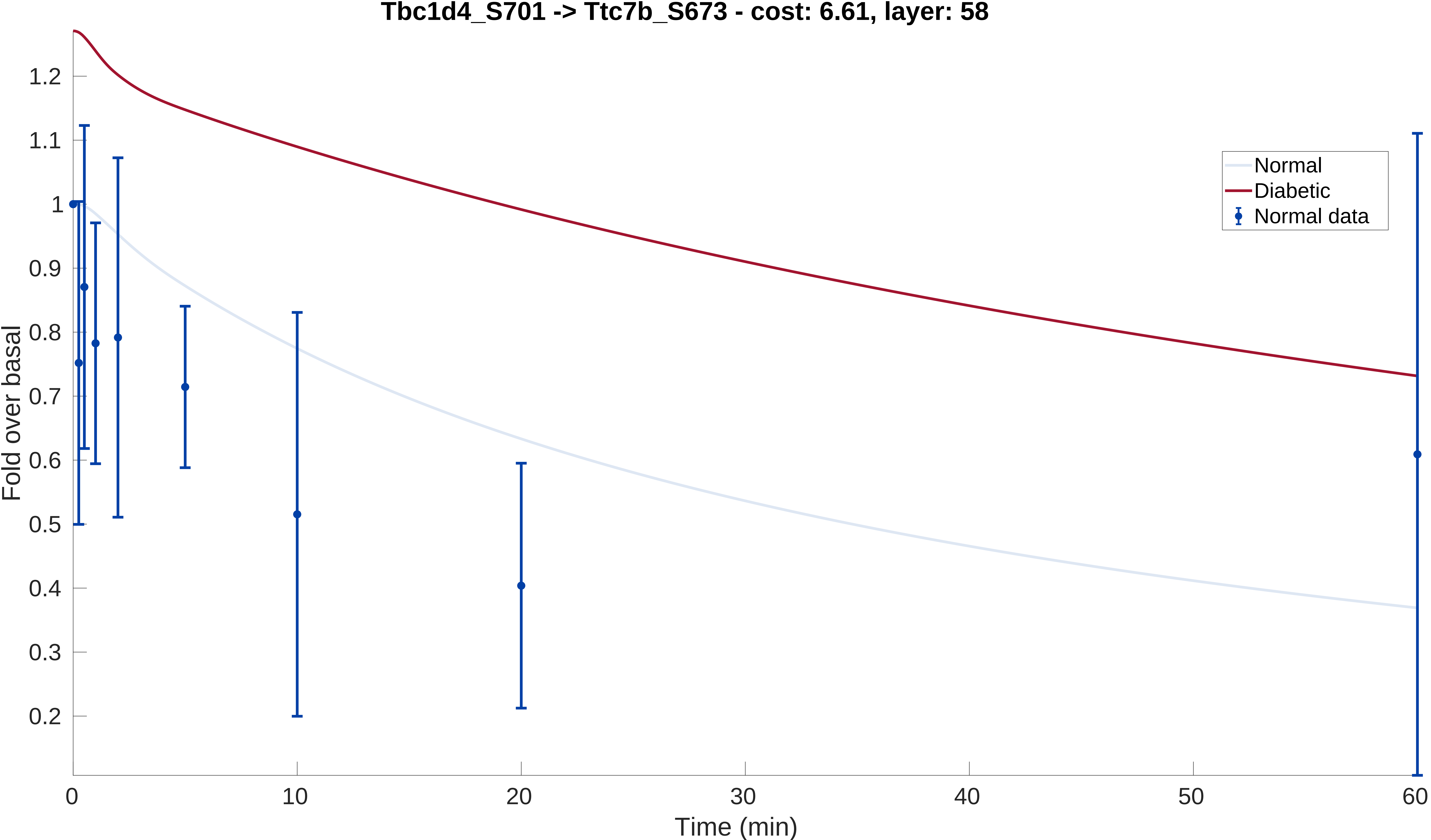

Tbc1d4_S701 -> Ttc7b_S673 - cost: 6.61, layer: 58
Fold over basal
1.2
1.1
1
0.9
0.8
0.7
0.6
0.5
0.4
0.3
0.2
0
10
20
30
40
50
60
Time (min)
Normal
Diabetic
Normal data

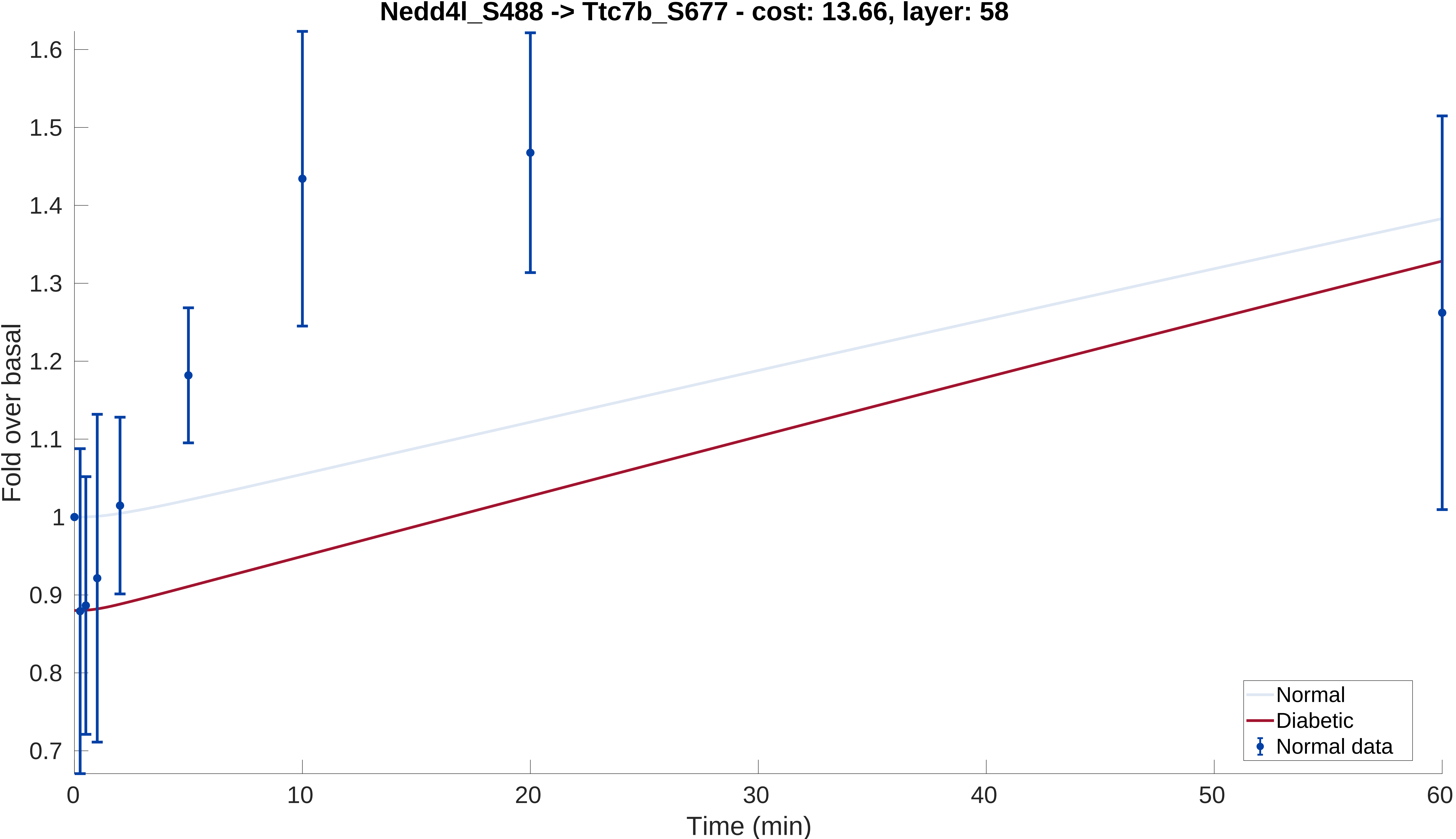
Nedd4l_S488 -> Ttc7b_S677 - cost: 13.66, layer: 58
Fold over basal
Time (min)
Normal
Diabetic
Normal data
0.7
0.8
0.9
1
1.1
1.2
1.3
1.4
1.5
1.6
0
10
20
30
40
50
60

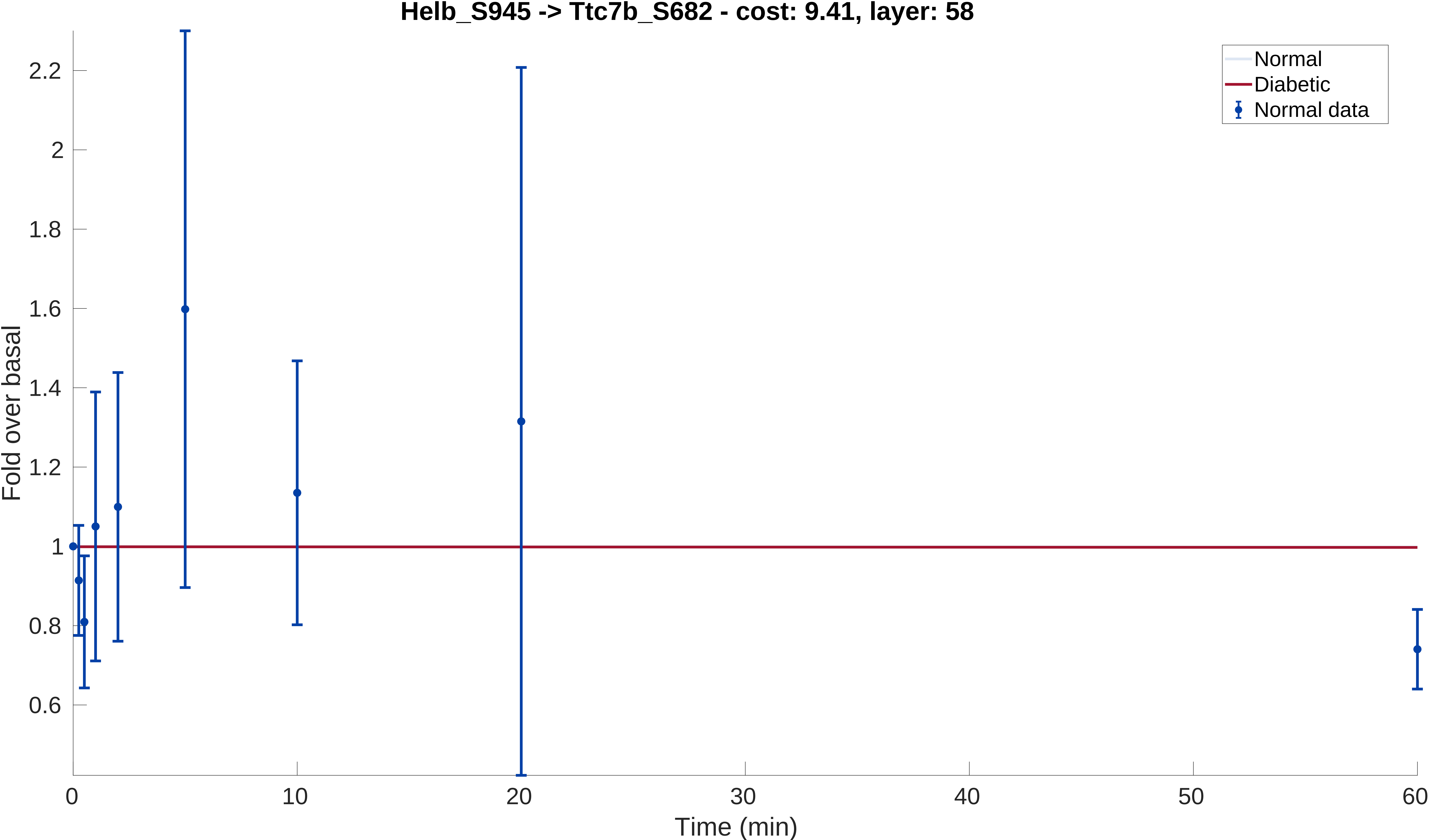

Helb_S945 -> Ttc7b_S682 - cost: 9.41, layer: 58
Normal
Diabetic
Normal data
Fold over basal
2.2
2
1.8
1.6
1.4
1.2
1
0.8
0.6
0
10
20
30
40
50
60
Time (min)

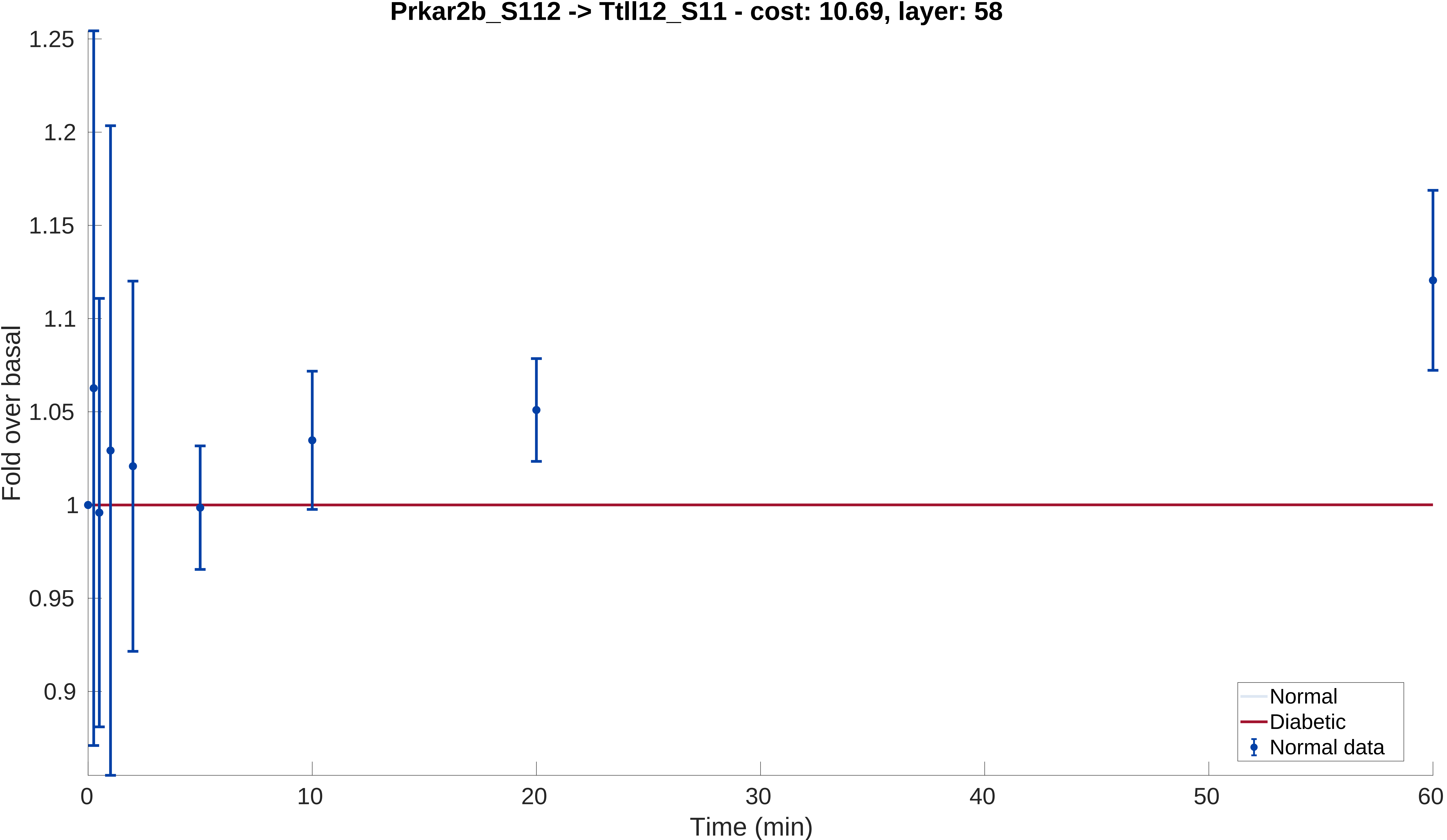

Prkar2b_S112 -> Ttll12_S11 - cost: 10.69, layer: 58
Fold over basal
1.25
1.2
1.15
1.1
1.05
1
0.95
0.9
0
10
20
30
40
50
60
Time (min)
Normal
Diabetic
Normal data

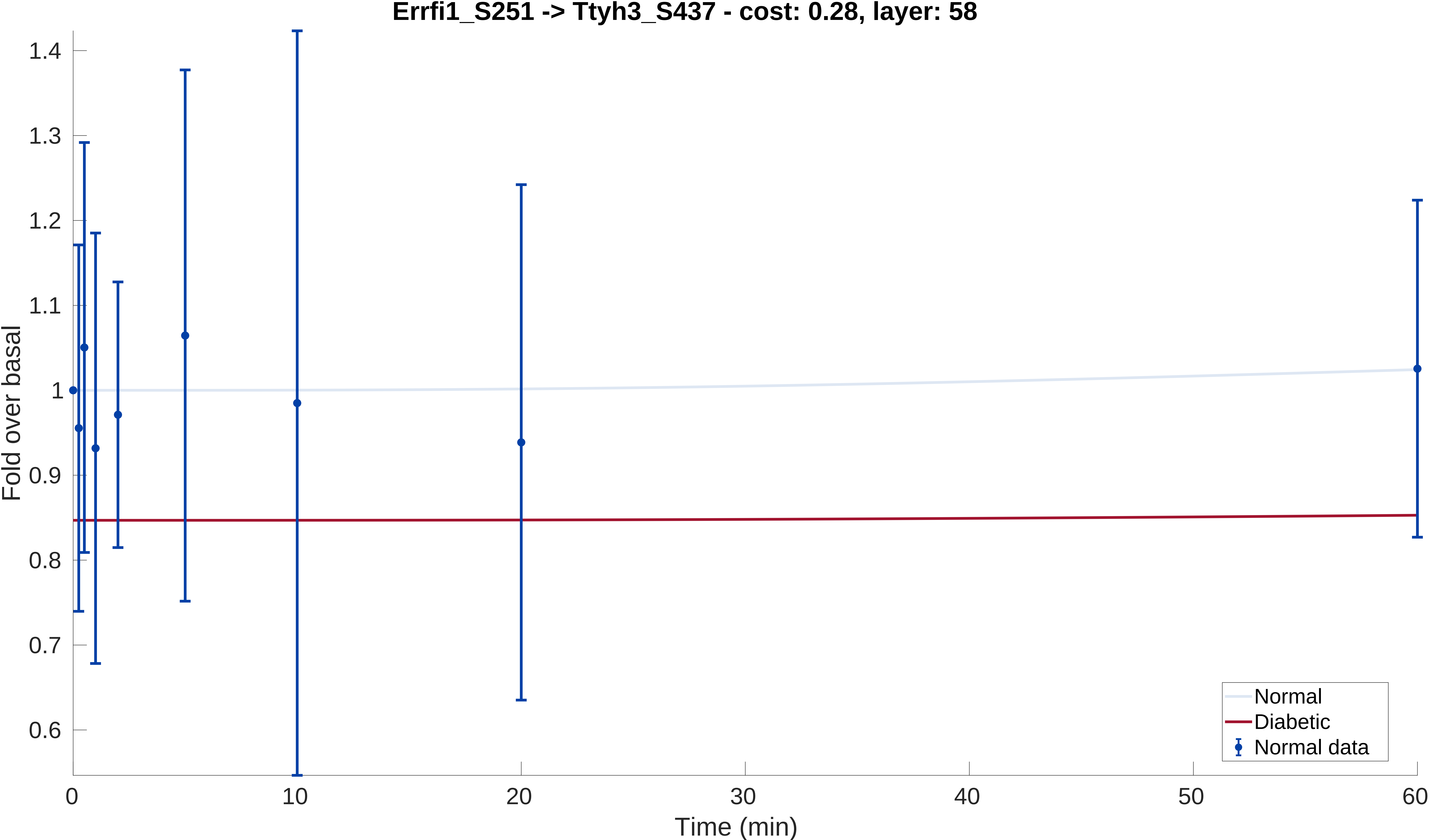
Errfi1_S251 -> Ttyh3_S437 - cost: 0.28, layer: 58
Fold over basal
Time (min)
1.4
1.3
1.2
1.1
1
0.9
0.8
0.7
0.6
0
10
20
30
40
50
60
Normal
Diabetic
Normal data

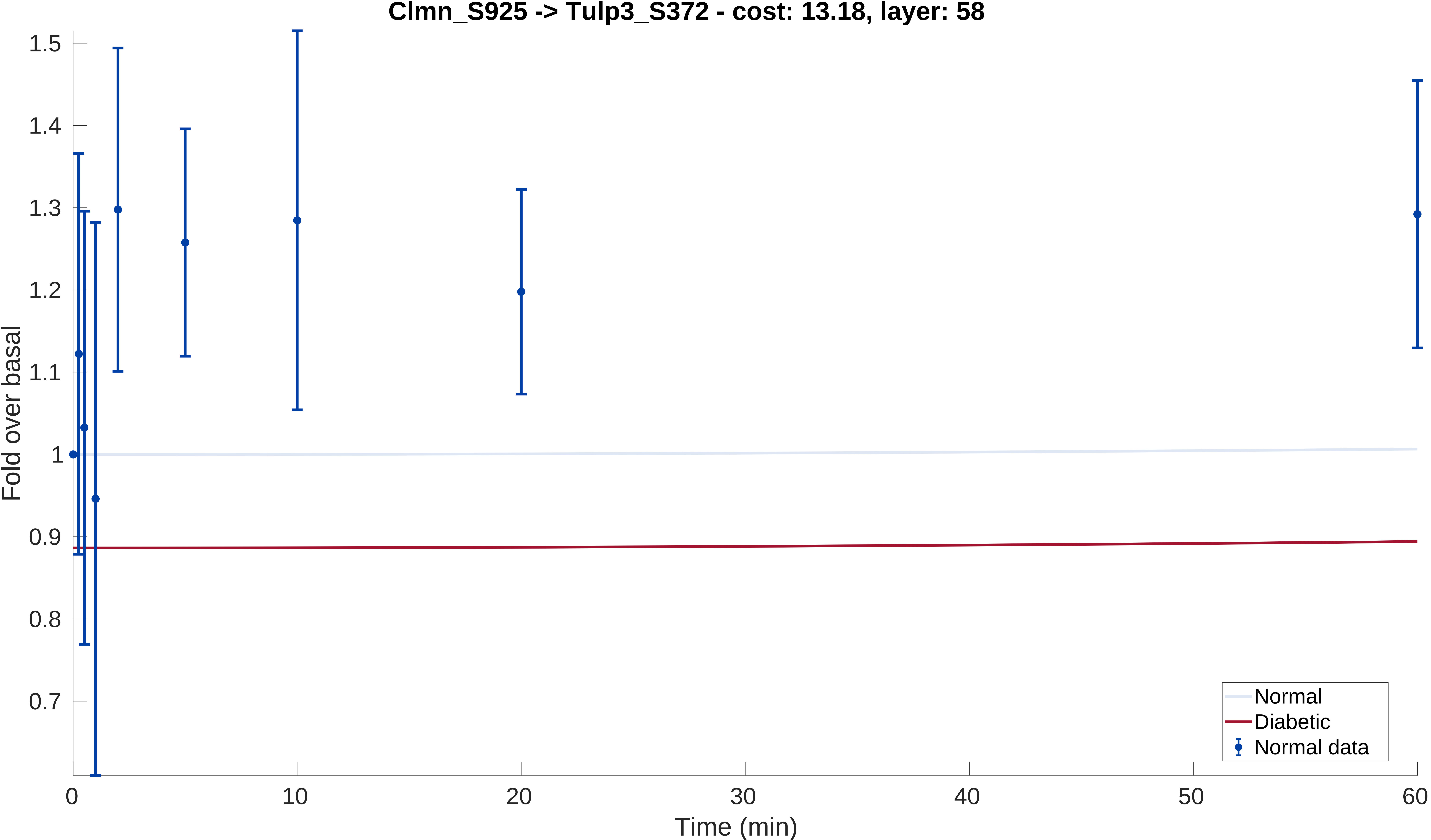

Clmn_S925 -> Tulp3_S372 - cost: 13.18, layer: 58
Fold over basal
1.5
1.4
1.3
1.2
1.1
1
0.9
0.8
0.7
0
10
20
30
40
50
60
Time (min)
Normal
Diabetic
Normal data

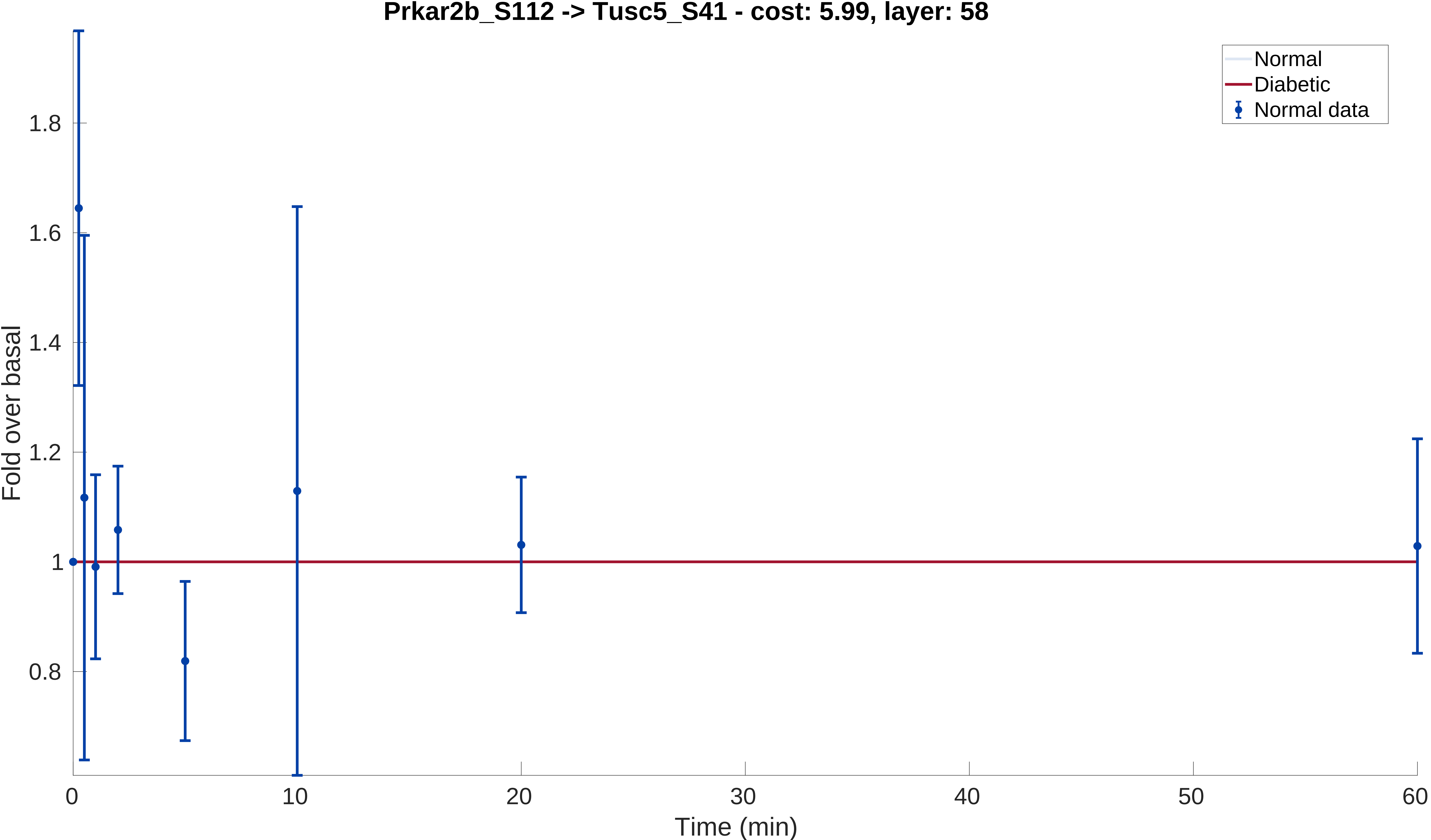

Prkar2b_S112 -> Tusc5_S41 - cost: 5.99, layer: 58
Normal
Diabetic
Normal data
Fold over basal
1.8
1.6
1.4
1.2
1
0.8
0
10
20
30
40
50
60
Time (min)

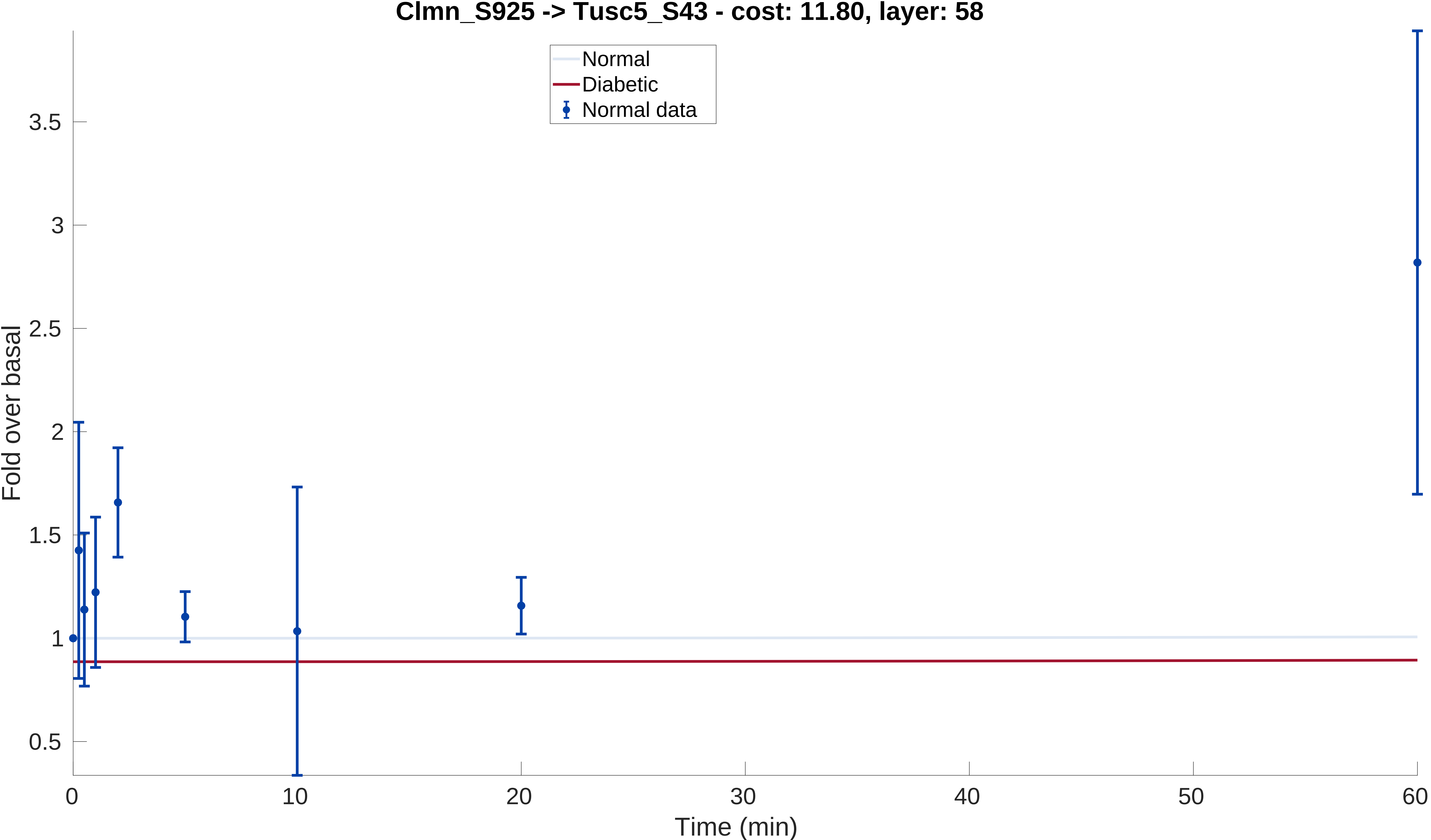

Clmn_S925 -> Tusc5_S43 - cost: 11.80, layer: 58
Normal
Diabetic
Normal data
Fold over basal
0.5
1
1.5
2
2.5
3
3.5
0
10
20
30
40
50
60
Time (min)

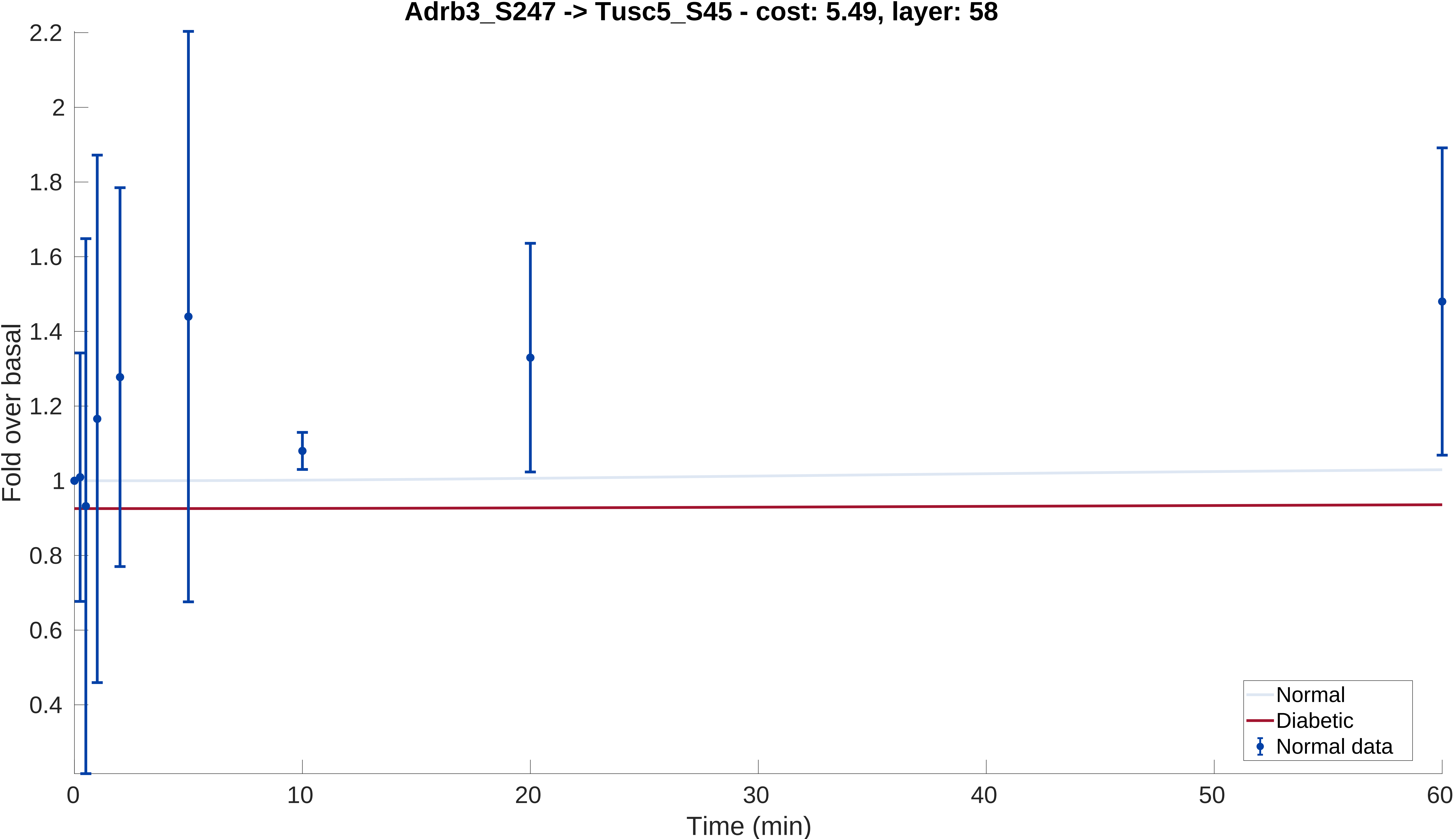

Adrb3_S247 -> Tusc5_S45 - cost: 5.49, layer: 58
Fold over basal
Time (min)
2.2
2
1.8
1.6
1.4
1.2
1
0.8
0.6
0.4
0
10
20
30
40
50
60
Normal
Diabetic
Normal data

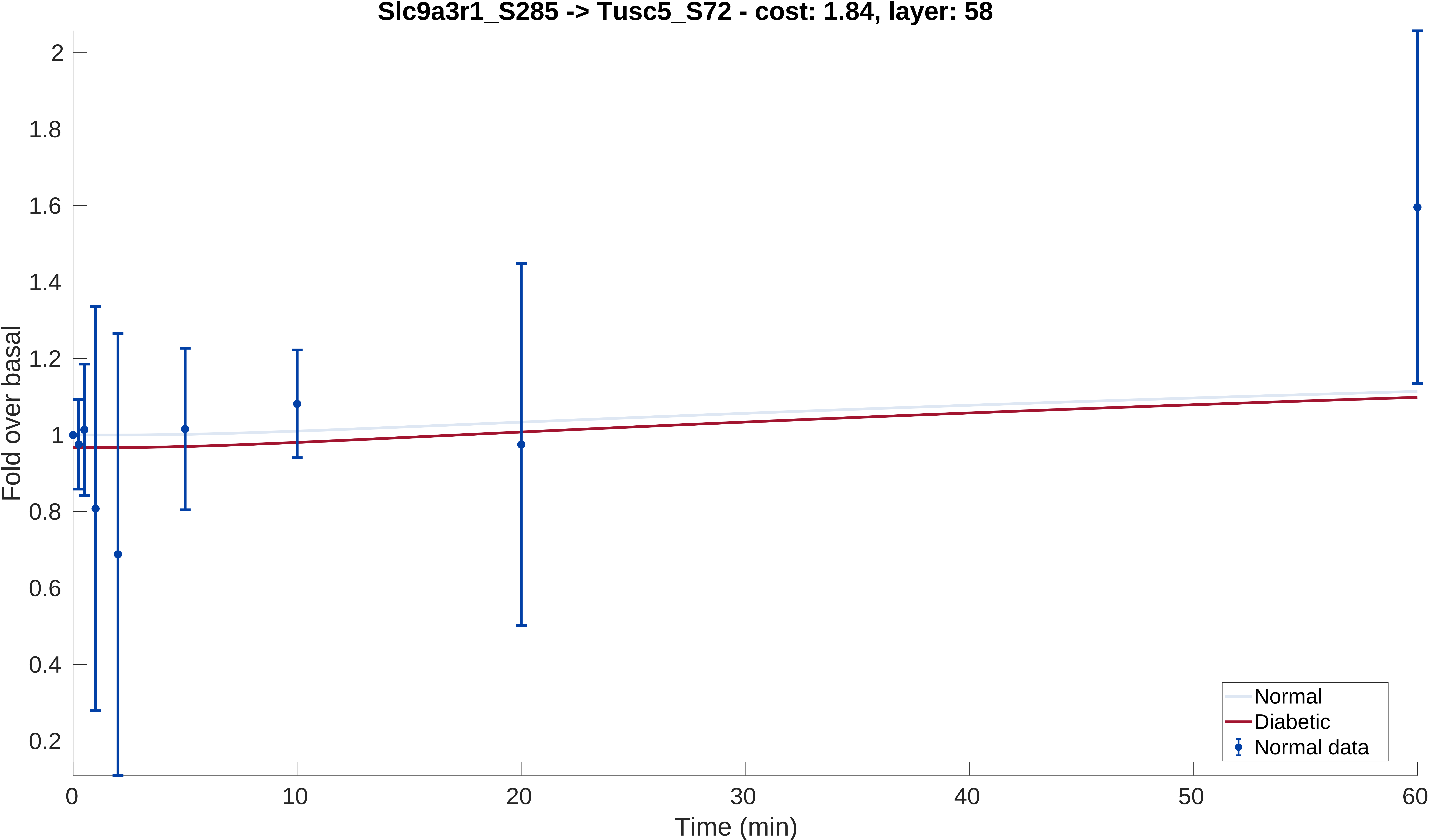
Slc9a3r1_S285 -> Tusc5_S72 - cost: 1.84, layer: 58
Fold over basal
Time (min)
2
1.8
1.6
1.4
1.2
1
0.8
0.6
0.4
0.2
0
10
20
30
40
50
60
Normal
Diabetic
Normal data

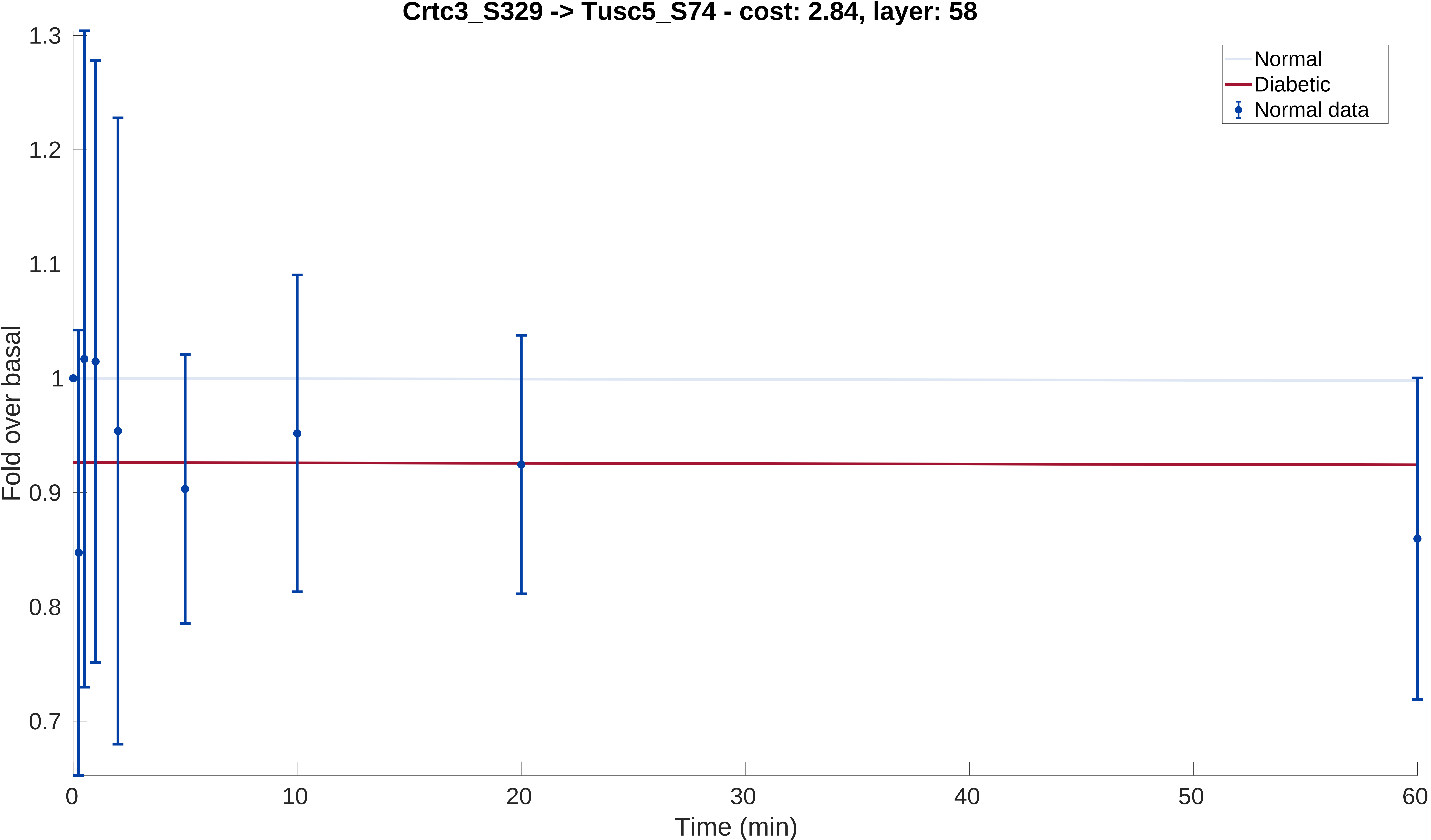

Crtc3_S329 -> Tusc5_S74 - cost: 2.84, layer: 58
Normal
Diabetic
Normal data
Fold over basal
1.3
1.2
1.1
1
0.9
0.8
0.7
0
10
20
30
40
50
60
Time (min)

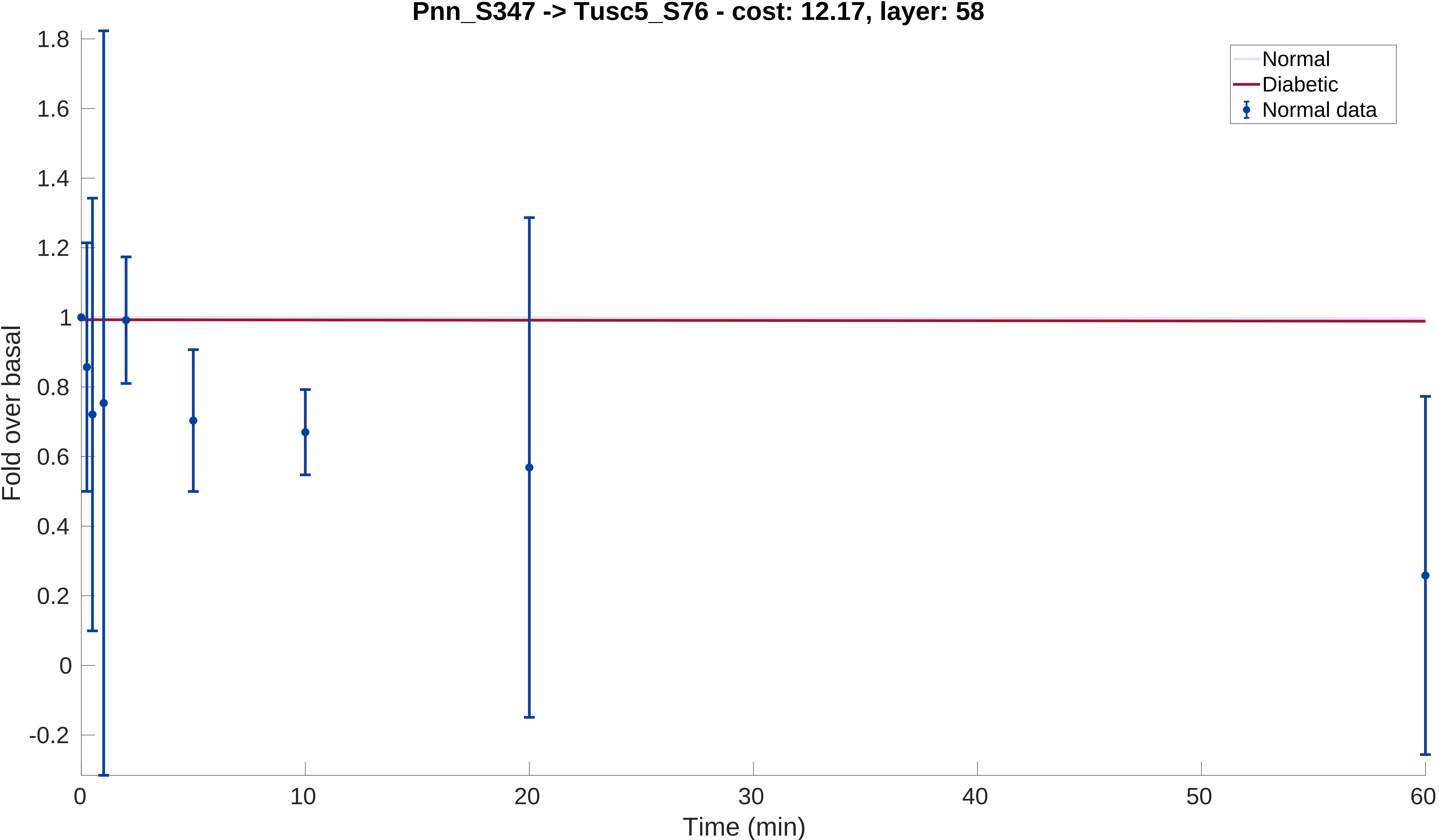

Pnn_S347 -> Tusc5_S76 - cost: 12.17, layer: 58
Normal
Diabetic
Normal data
Fold over basal
Time (min)
1.8
1.6
1.4
1.2
1
0.8
0.6
0.4
0.2
0
-0.2
0
10
20
30
40
50
60

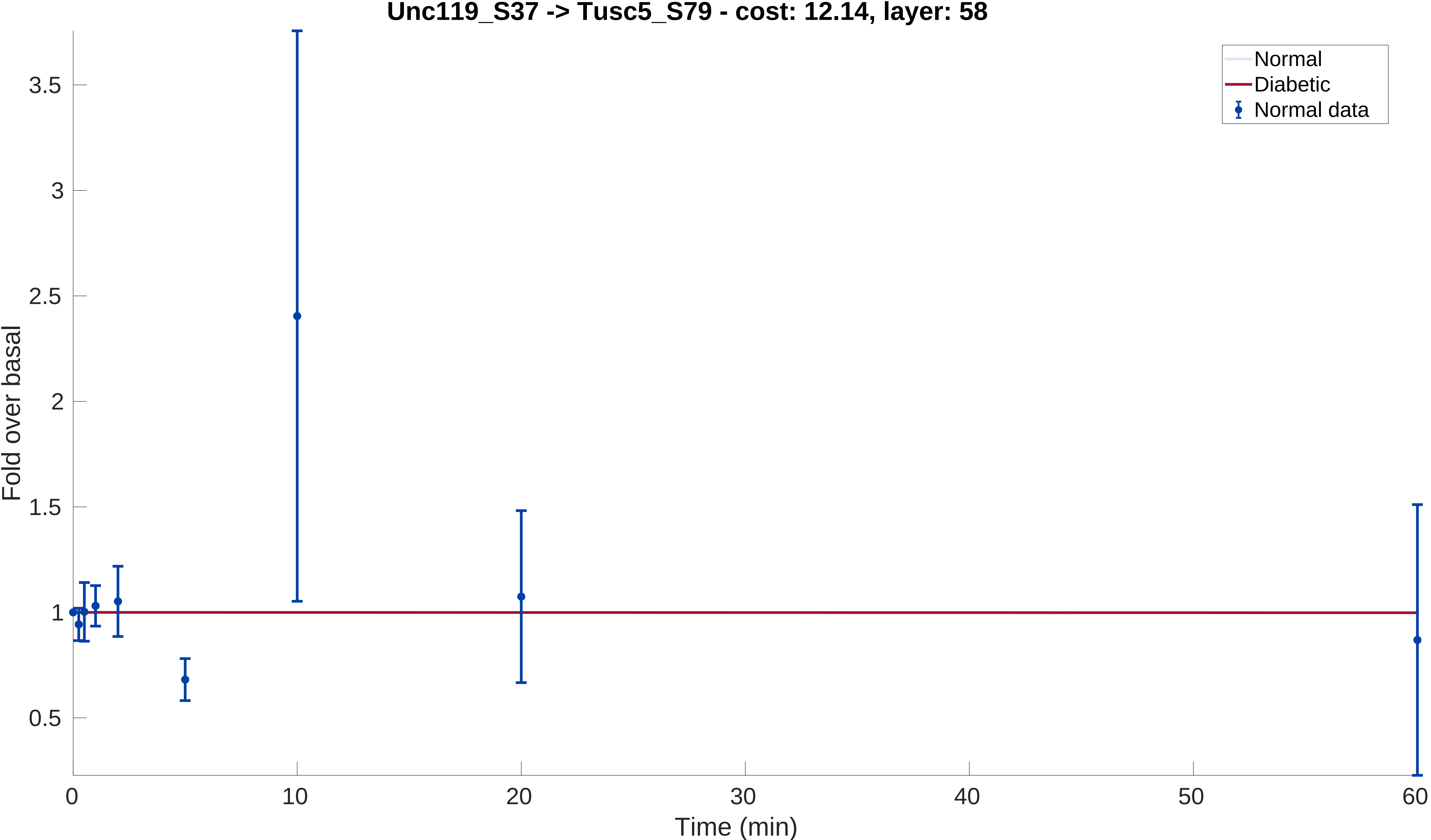
Unc119_S37 -> Tusc5_S79 - cost: 12.14, layer: 58
Normal
Diabetic
Normal data
Fold over basal
3.5
3
2.5
2
1.5
1
0.5
0
10
20
30
40
50
60
Time (min)

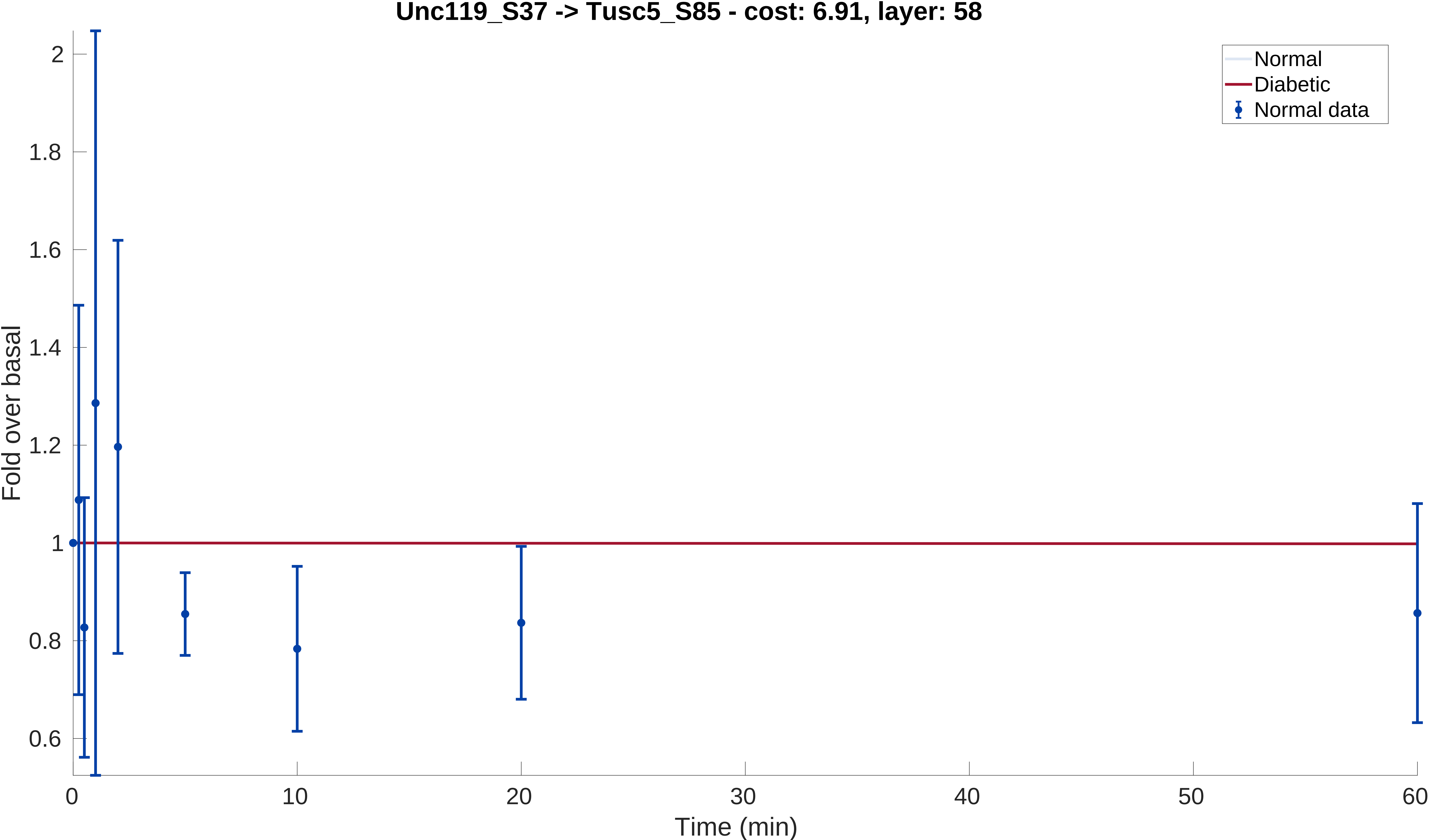

Unc119_S37 -> Tusc5_S85 - cost: 6.91, layer: 58
Normal
Diabetic
Normal data
Fold over basal
2
1.8
1.6
1.4
1.2
1
0.8
0.6
0
10
20
30
40
50
60
Time (min)

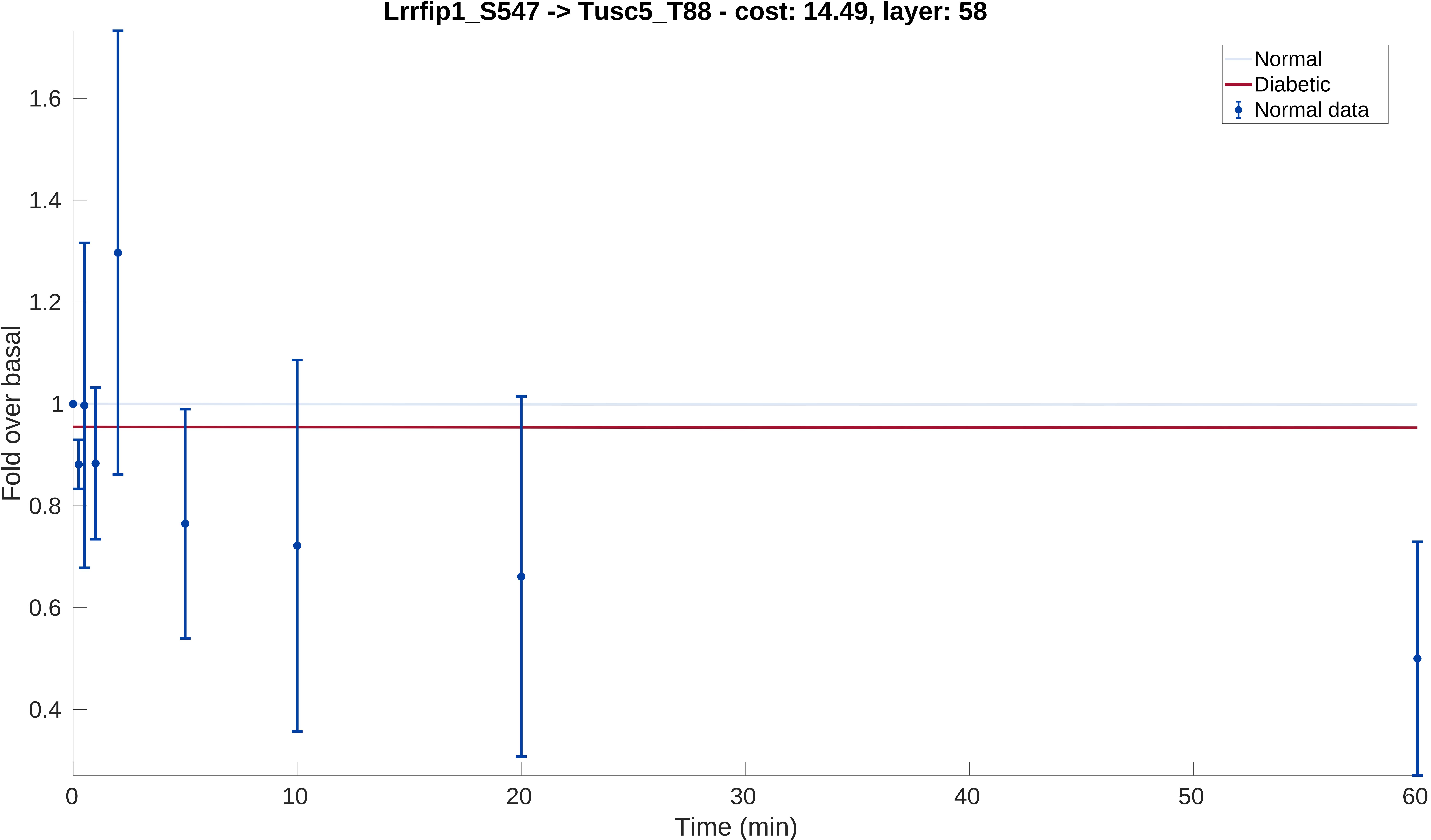

Lrrfip1_S547 -> Tusc5_T88 - cost: 14.49, layer: 58
Normal
Diabetic
Normal data
1.6
1.4
1.2
1
0.8
0.6
0.4
Fold over basal
0
10
20
30
40
50
60
Time (min)

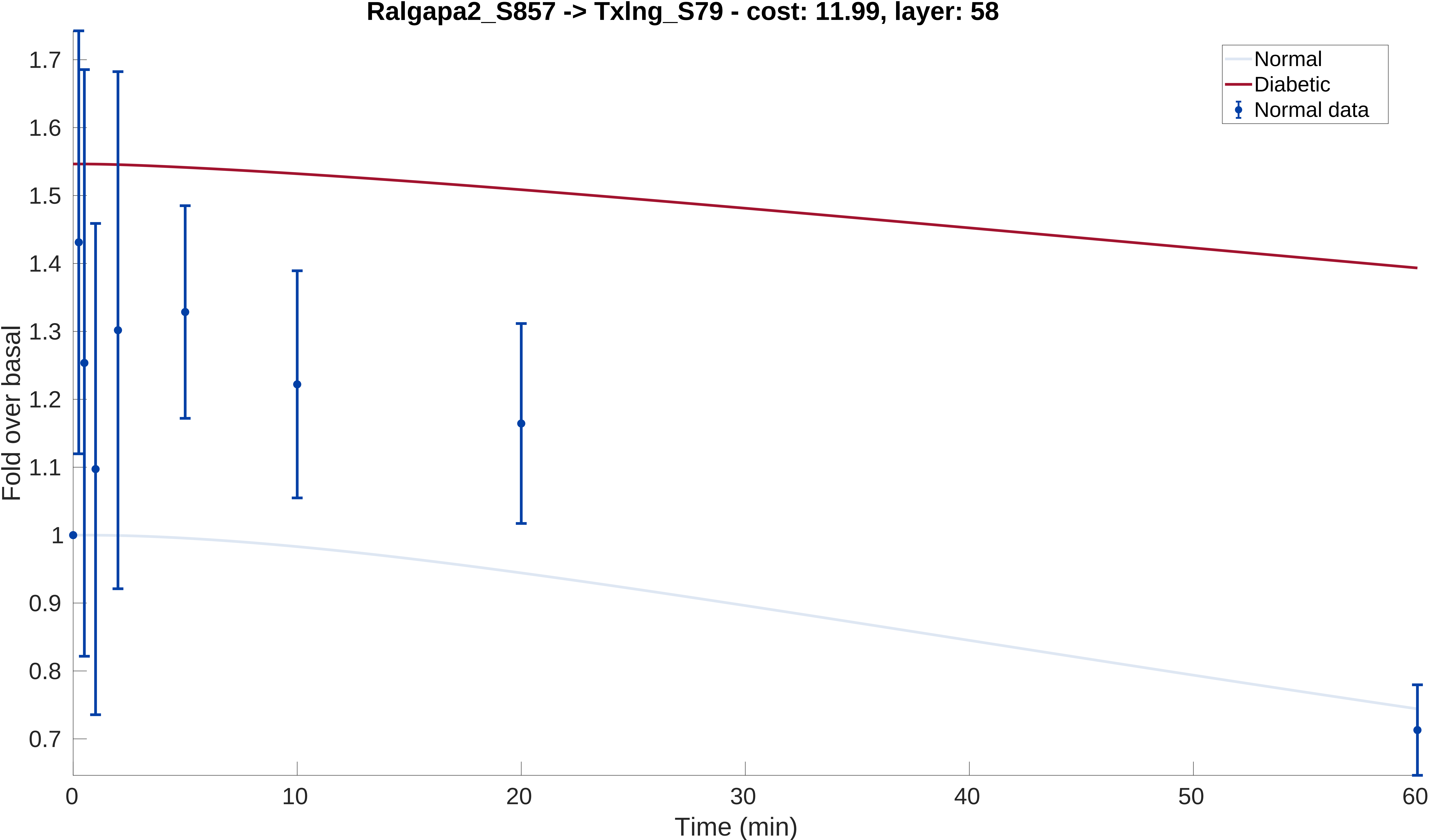

Ralgapa2_S857 -> Txlng_S79 - cost: 11.99, layer: 58
Normal
Diabetic
Normal data
Fold over basal
1.7
1.6
1.5
1.4
1.3
1.2
1.1
1
0.9
0.8
0.7
0
10
20
30
40
50
60
Time (min)

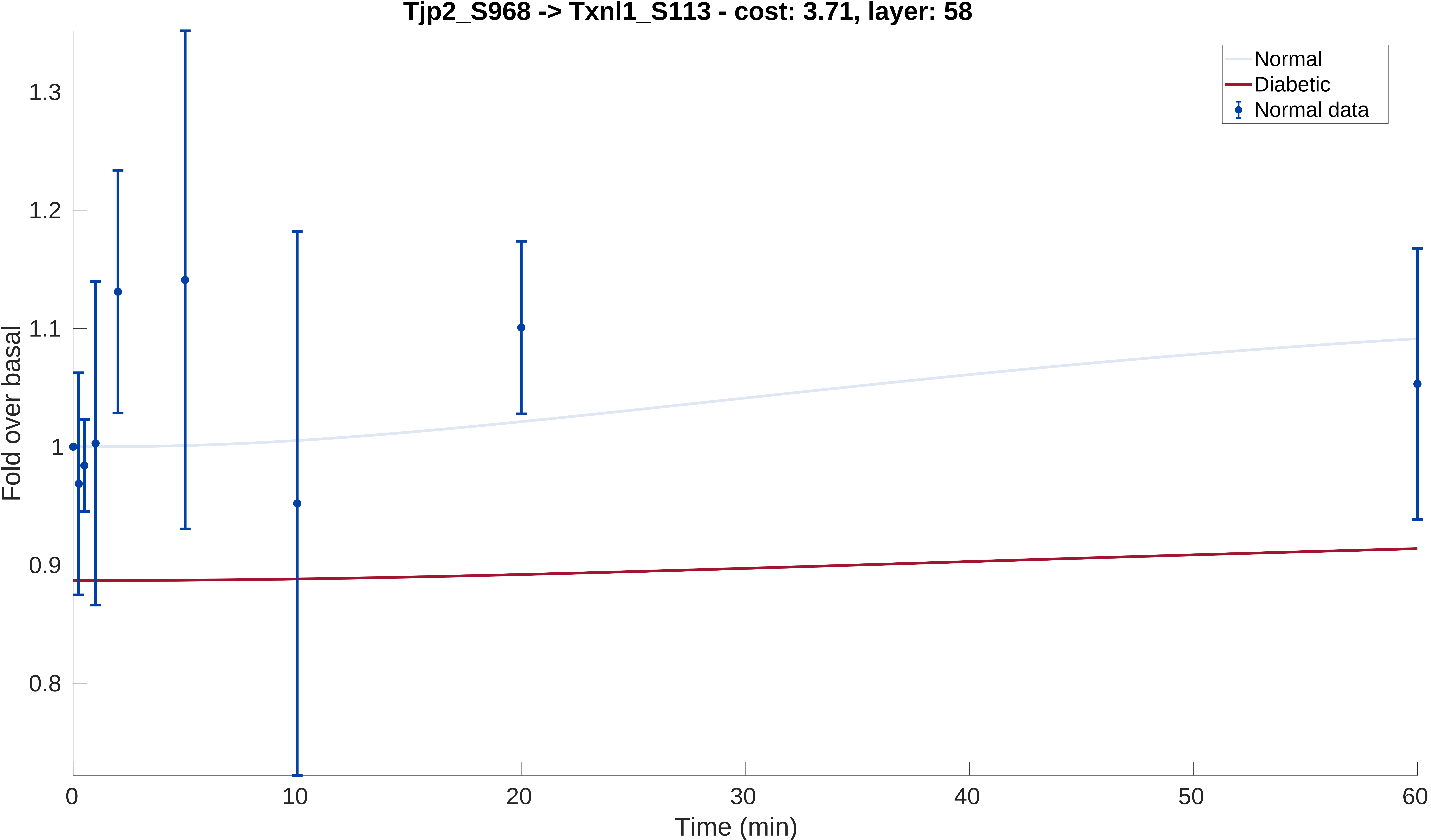

Tjp2_S968 -> Txnl1_S113 - cost: 3.71, layer: 58
Normal
Diabetic
Normal data
Fold over basal
Time (min)
1.3
1.2
1.1
1
0.9
0.8
0
10
20
30
40
50
60

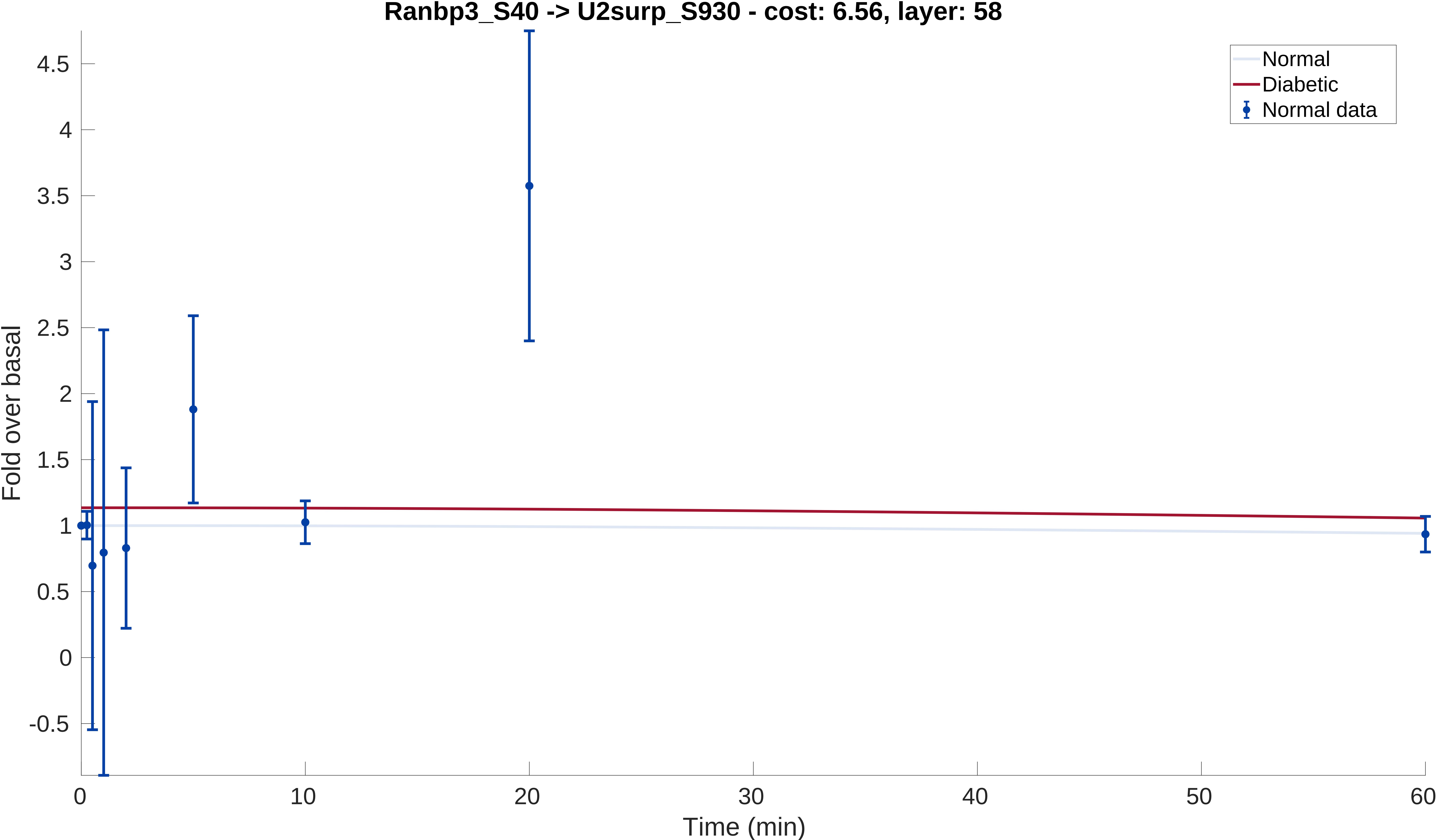

Ranbp3_S40 -> U2surp_S930 - cost: 6.56, layer: 58
Normal
Diabetic
Normal data
Fold over basal
Time (min)
4.5
4
3.5
3
2.5
2
1.5
1
0.5
0
-0.5
0
10
20
30
40
50
60

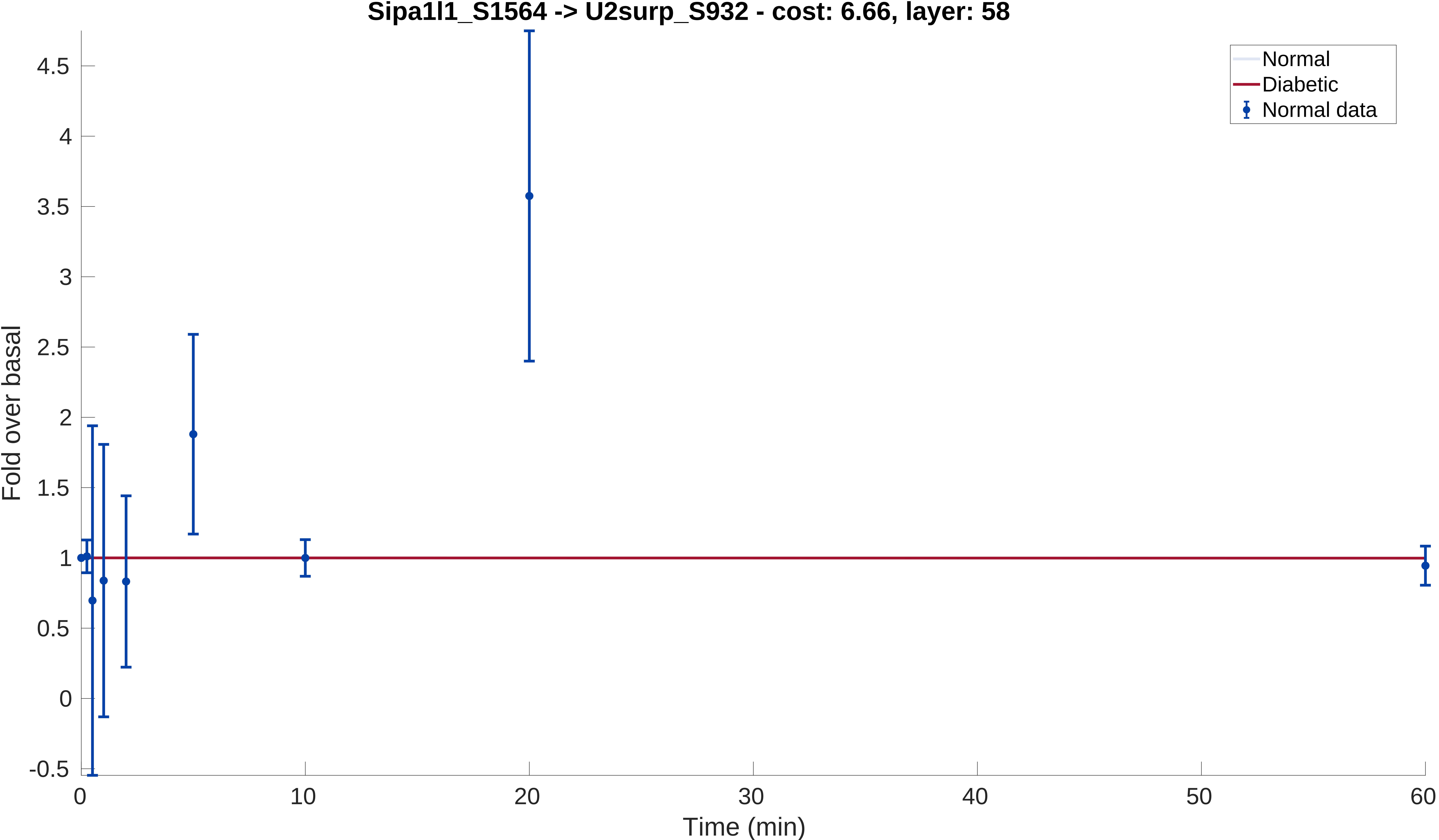
Sipa1l1_S1564 -> U2surp_S932 - cost: 6.66, layer: 58
Normal
Diabetic
Normal data
Fold over basal
Time (min)
4.5
4
3.5
3
2.5
2
1.5
1
0.5
0
-0.5
0
10
20
30
40
50
60

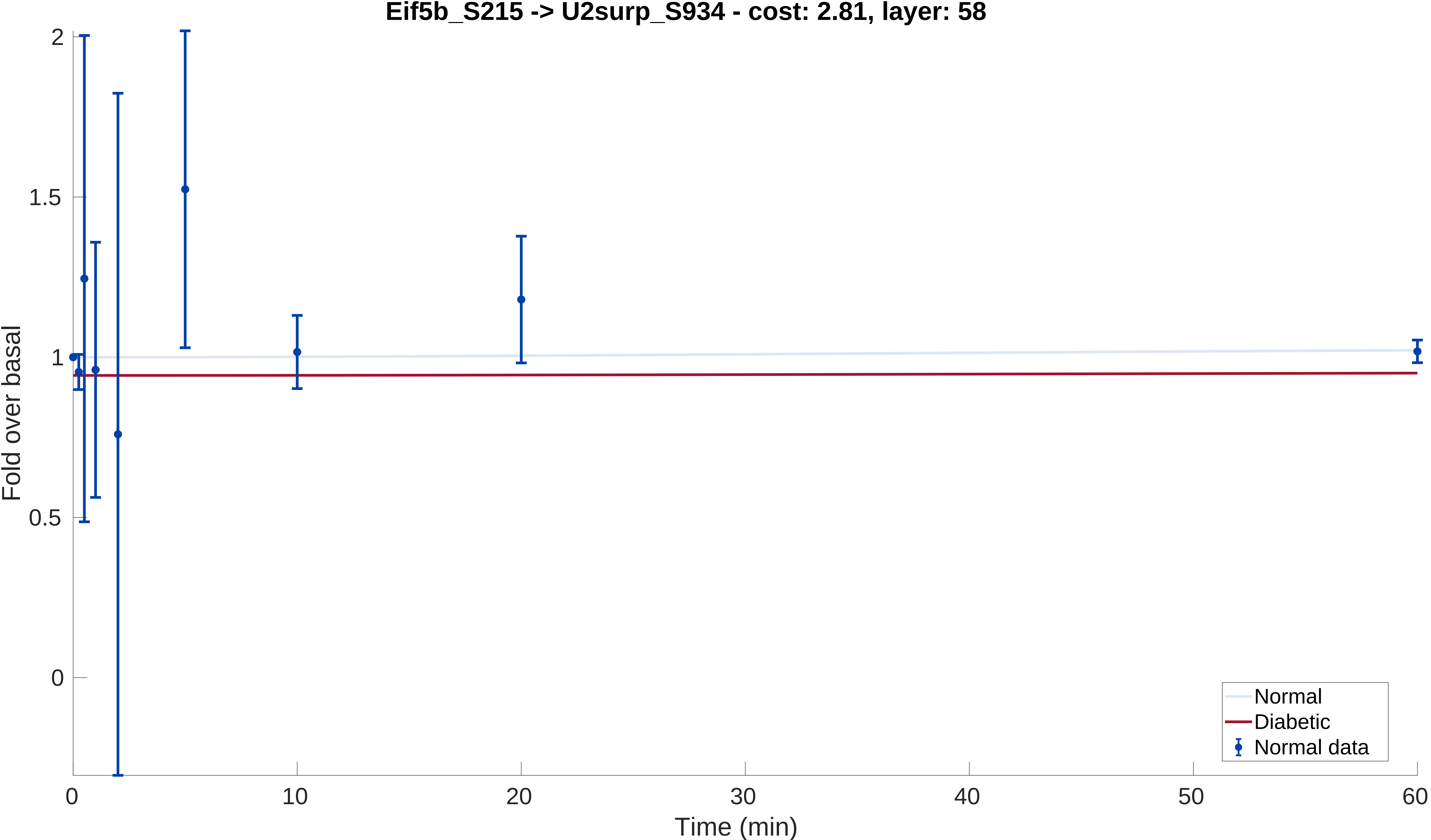

Eif5b_S215 -> U2surp_S934 - cost: 2.81, layer: 58
Fold over basal
Time (min)
0
0.5
1
1.5
2
0
10
20
30
40
50
60
Normal
Diabetic
Normal data

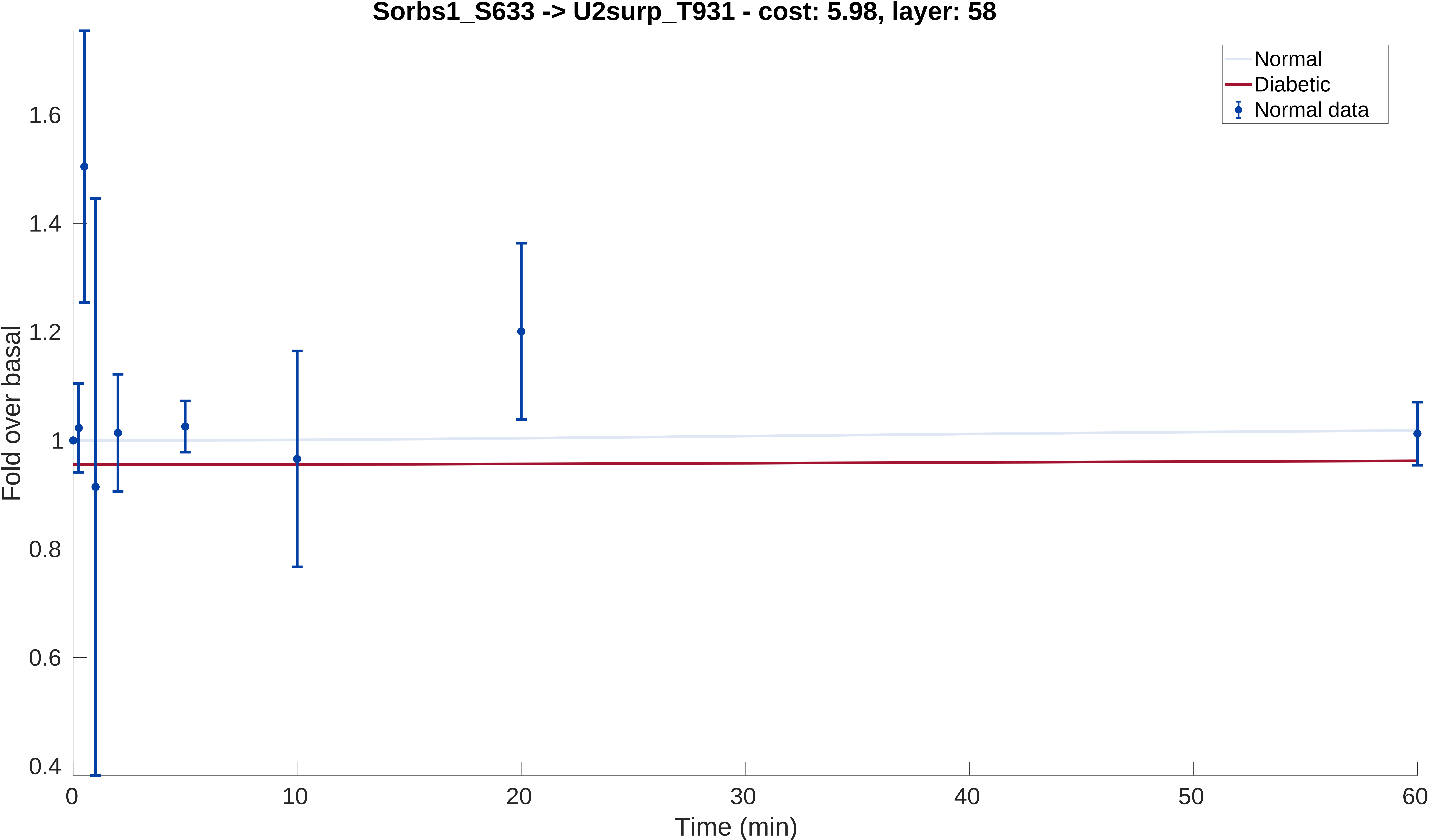

Sorbs1_S633 -> U2surp_T931 - cost: 5.98, layer: 58
Normal
Diabetic
Normal data
Fold over basal
1.6
1.4
1.2
1
0.8
0.6
0.4
0
10
20
30
40
50
60
Time (min)

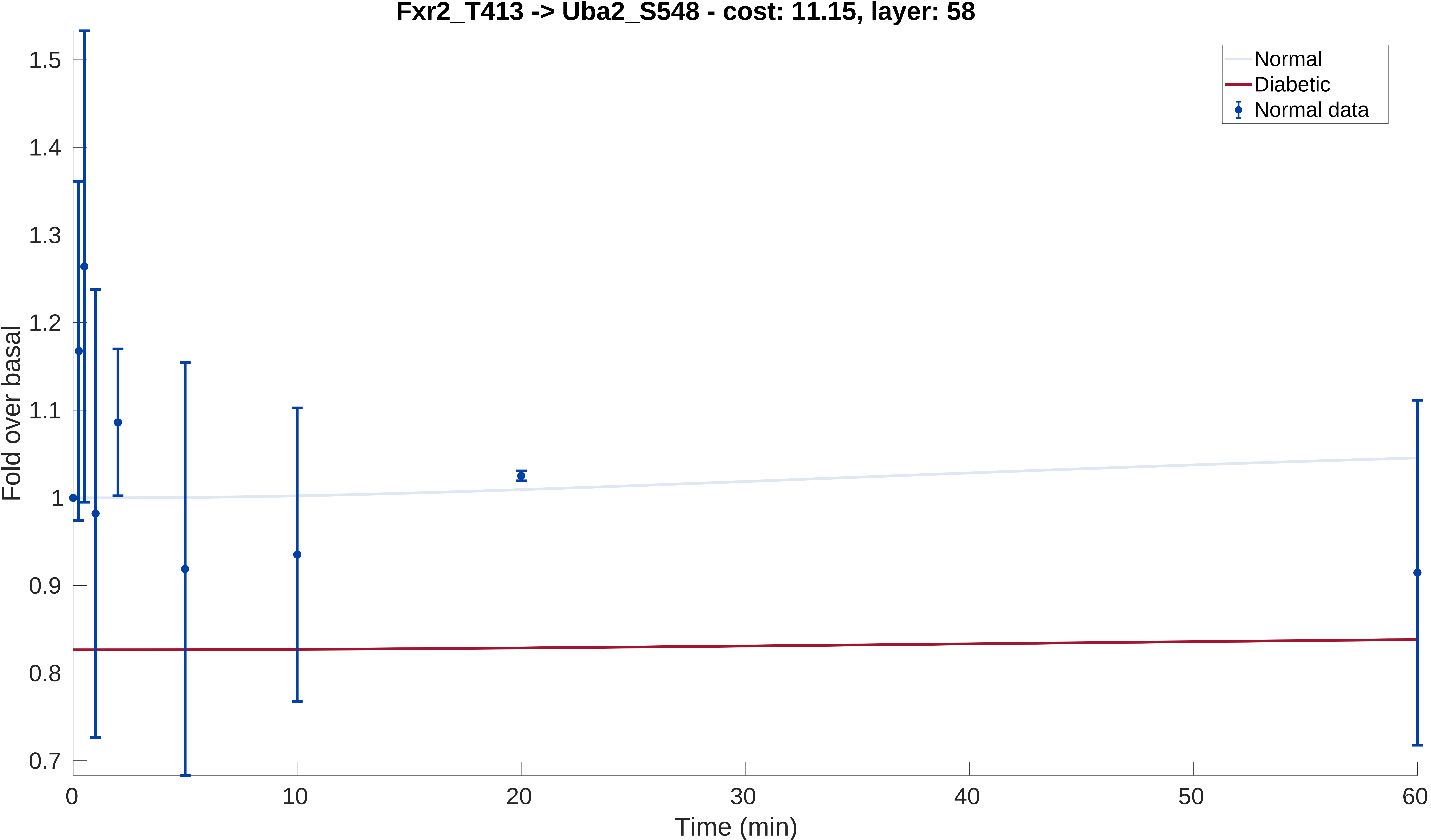
Fxr2_T413 -> Uba2_S548 - cost: 11.15, layer: 58
Normal
Diabetic
Normal data
Fold over basal
1.5
1.4
1.3
1.2
1.1
1
0.9
0.8
0.7
0
10
20
30
40
50
60
Time (min)

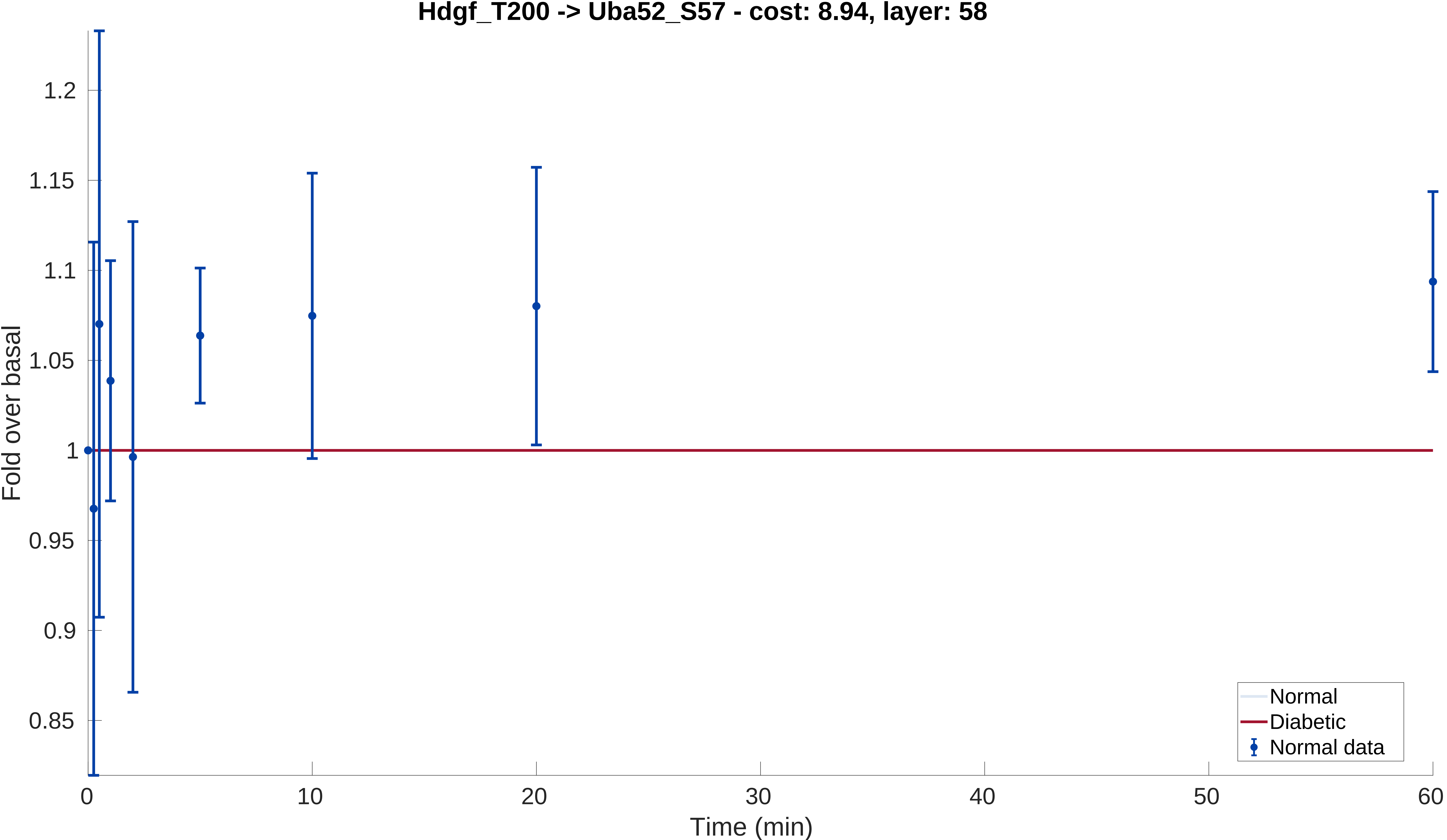

Hdgf_T200 -> Uba52_S57 - cost: 8.94, layer: 58
1.2
1.15
1.1
1.05
1
0.95
0.9
0.85
Fold over basal
0
10
20
30
40
50
60
Time (min)
Normal
Diabetic
Normal data

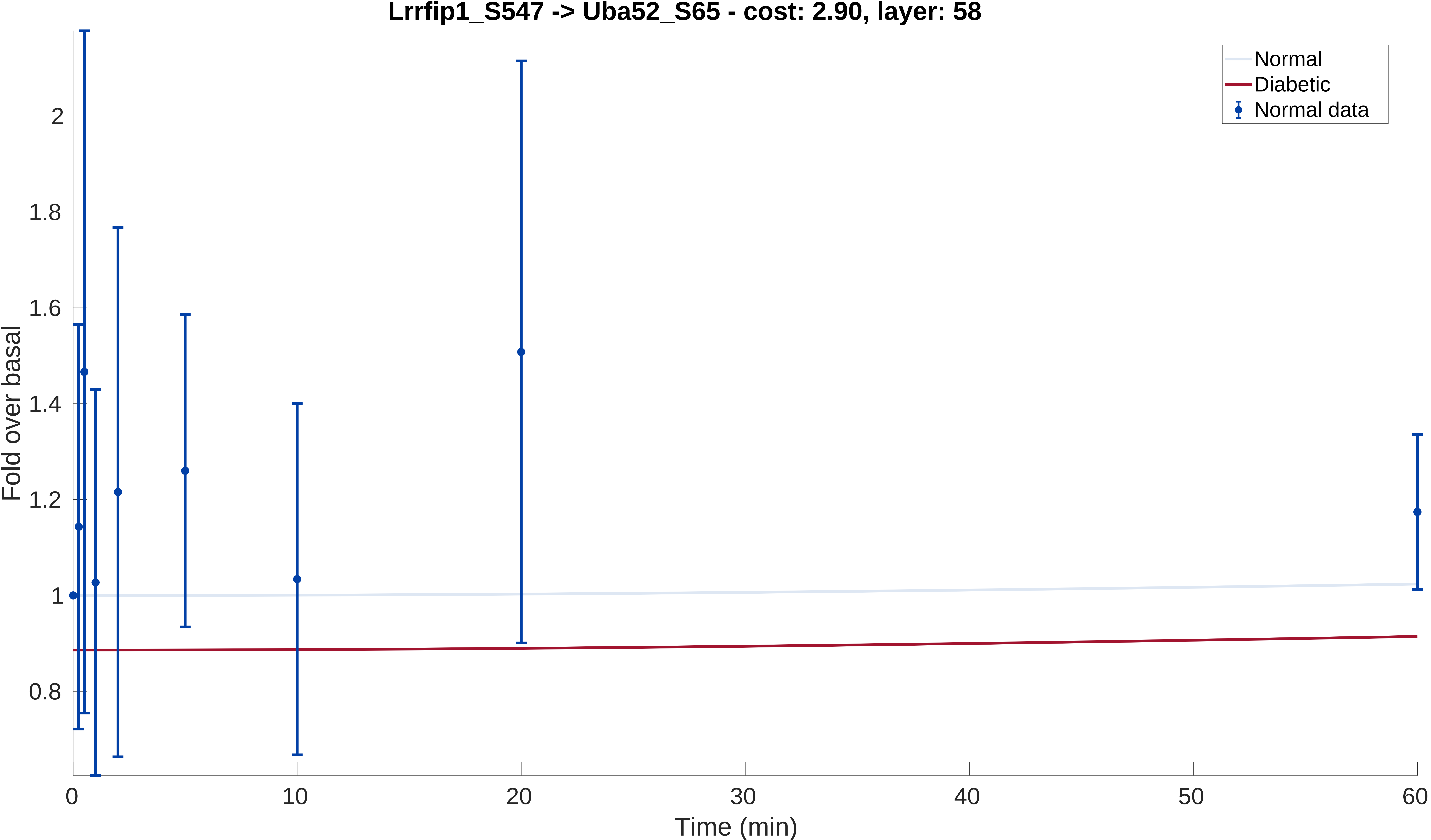
Lrrfip1_S547 -> Uba52_S65 - cost: 2.90, layer: 58
Normal
Diabetic
Normal data
Fold over basal
2
1.8
1.6
1.4
1.2
1
0.8
0
10
20
30
40
50
60
Time (min)

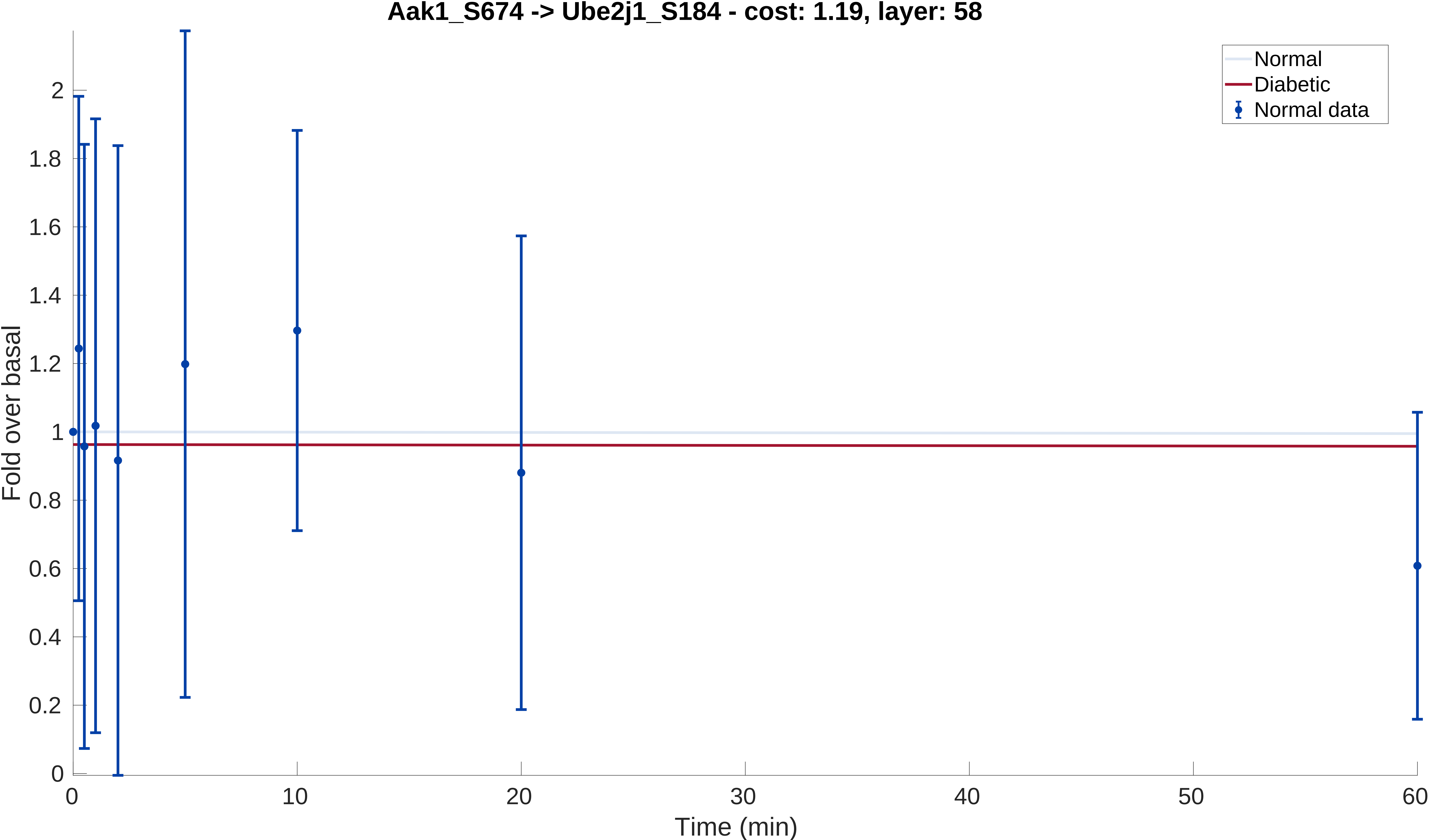
Aak1_S674 -> Ube2j1_S184 - cost: 1.19, layer: 58
Normal
Diabetic
Normal data
Fold over basal
Time (min)
0
0.2
0.4
0.6
0.8
1
1.2
1.4
1.6
1.8
2
0
10
20
30
40
50
60

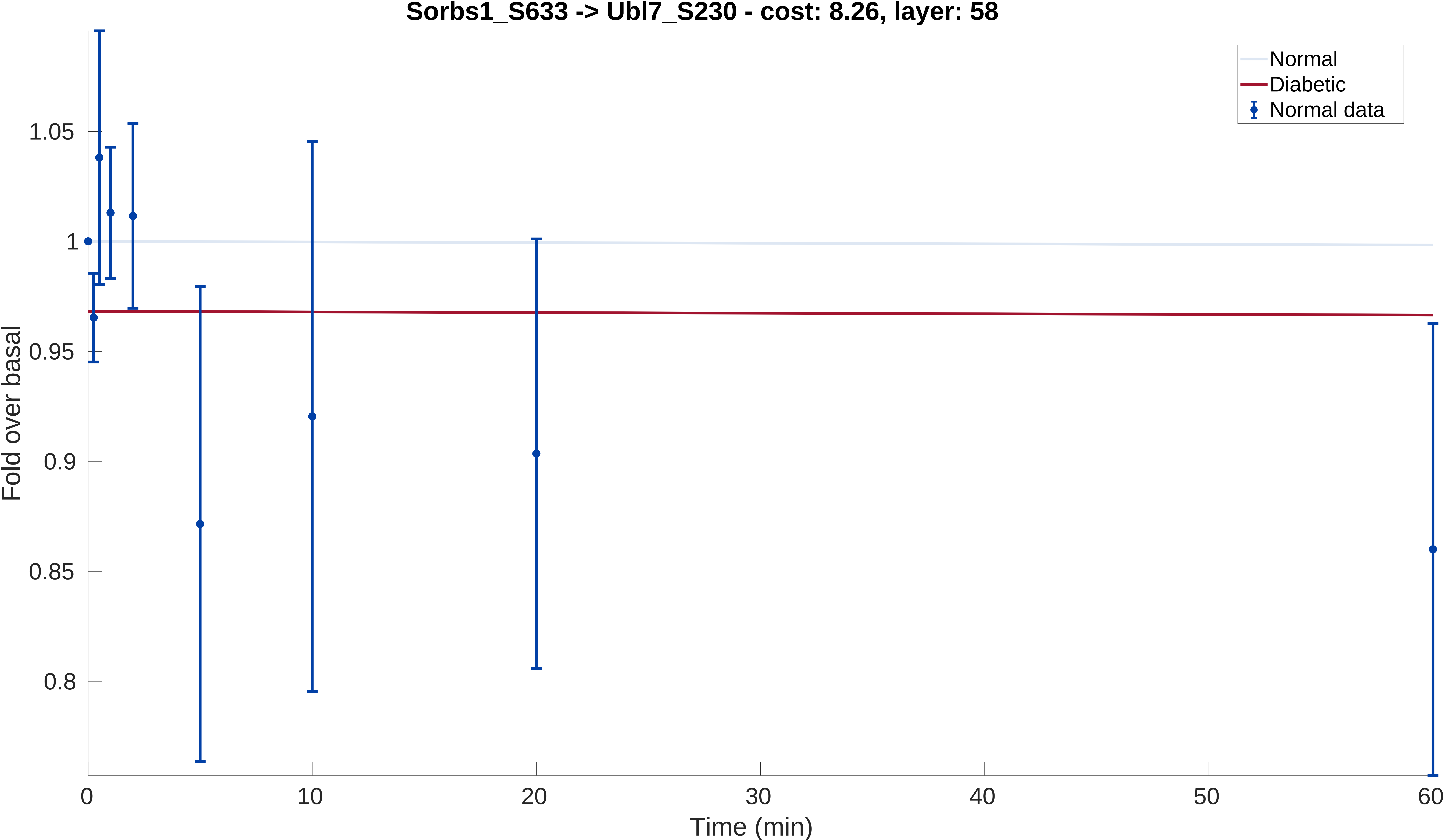
Sorbs1_S633 -> Ubl7_S230 - cost: 8.26, layer: 58
Normal
Diabetic
Normal data
Fold over basal
1.05
1
0.95
0.9
0.85
0.8
0
10
20
30
40
50
60
Time (min)

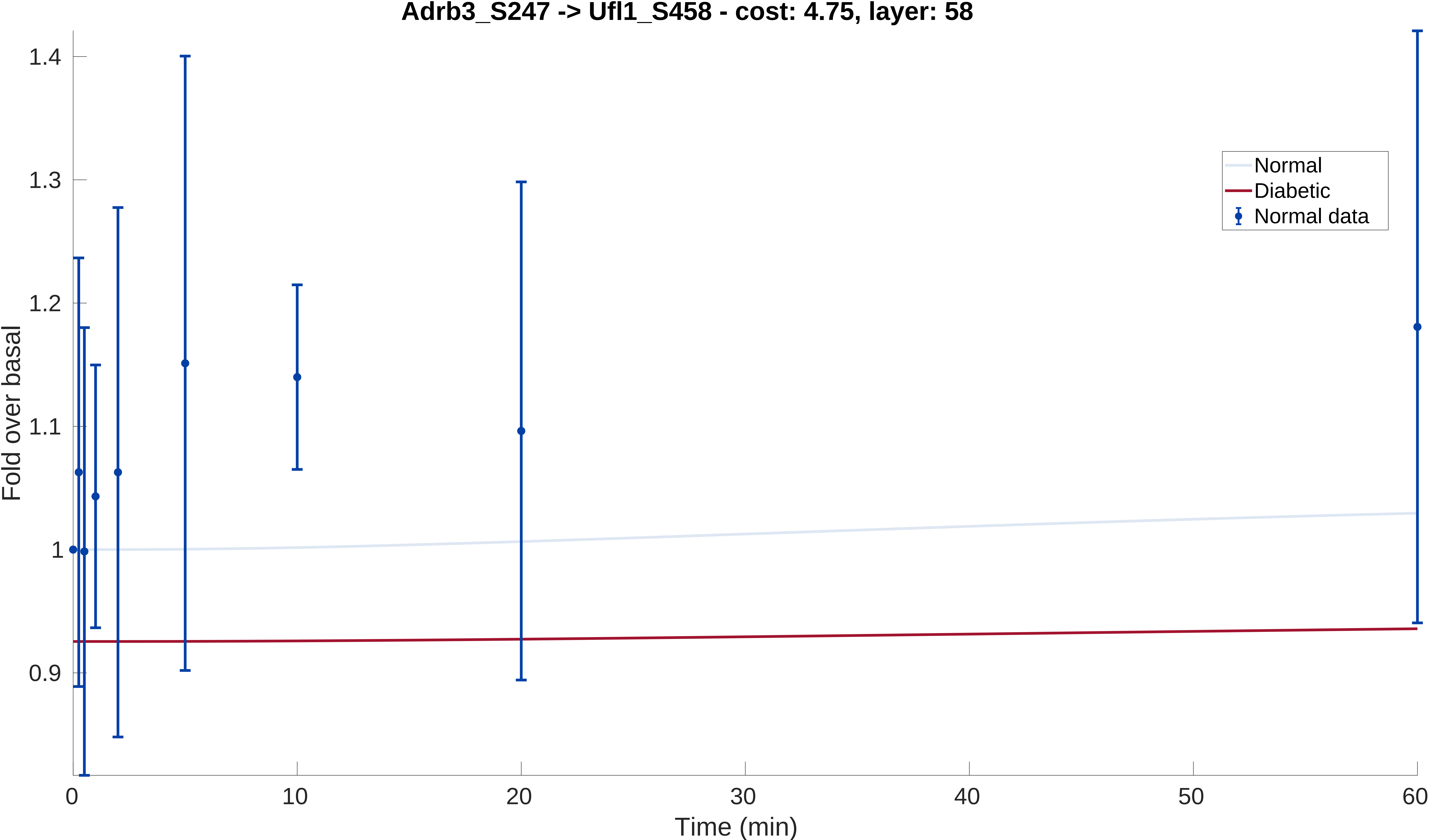

Adrb3_S247 -> Ufl1_S458 - cost: 4.75, layer: 58
Fold over basal
1.4
1.3
1.2
1.1
1
0.9
0
10
20
30
40
50
60
Time (min)
Normal
Diabetic
Normal data

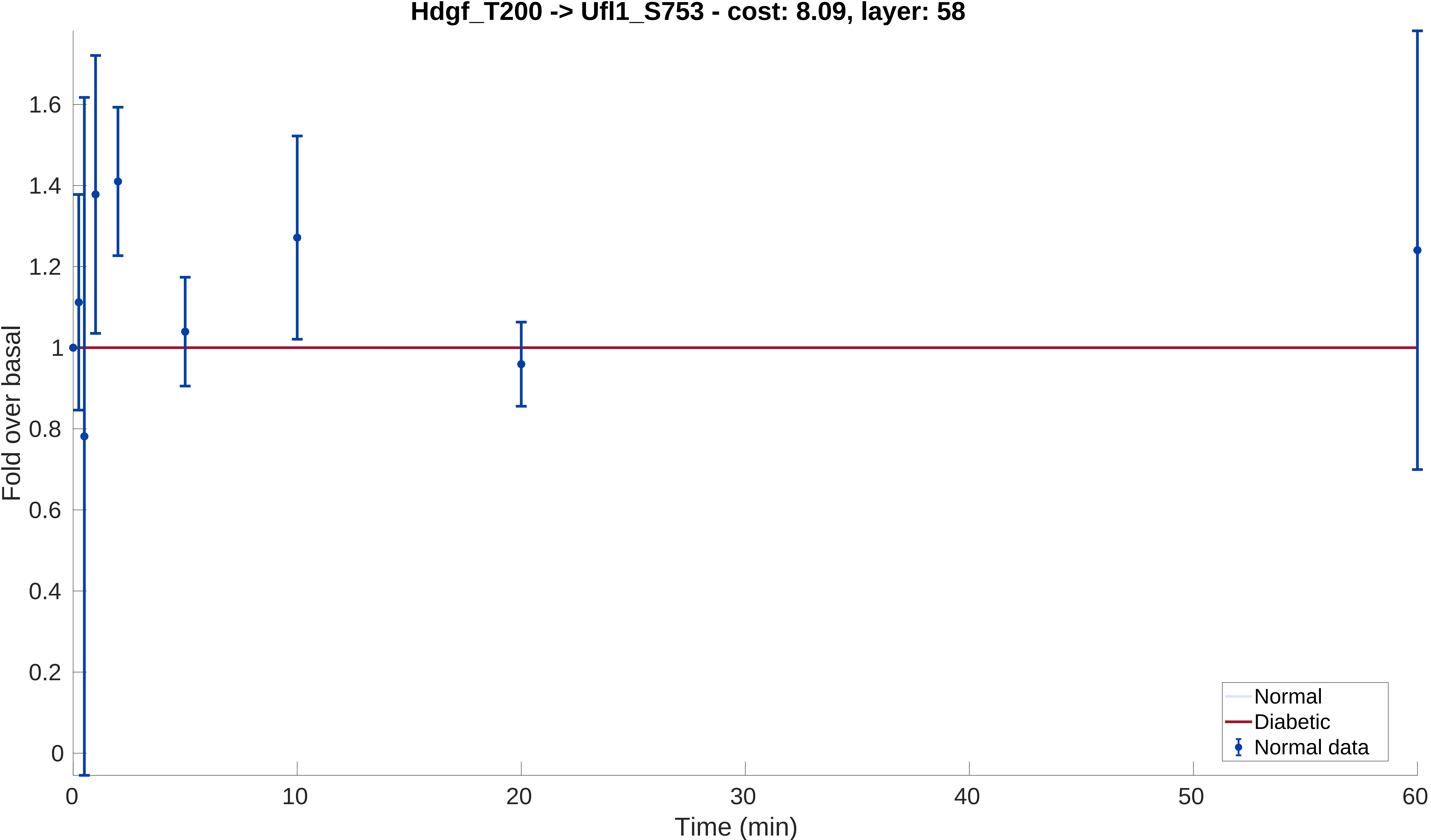

Hdgf_T200 -> Ufl1_S753 - cost: 8.09, layer: 58
Fold over basal
0
0.2
0.4
0.6
0.8
1
1.2
1.4
1.6
0
10
20
30
40
50
60
Time (min)
Normal
Diabetic
Normal data

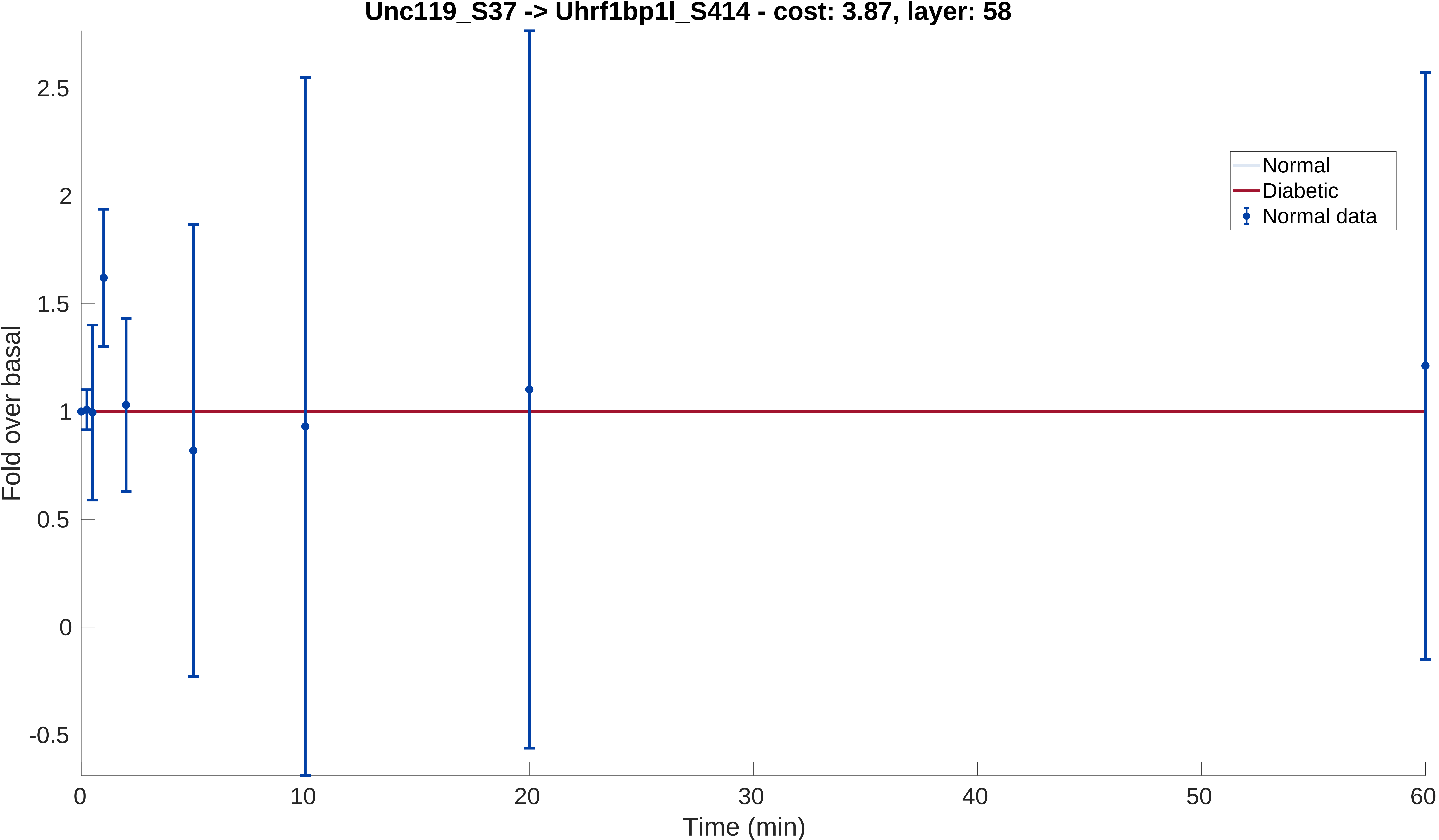

Unc119_S37 -> Uhrf1bp1l_S414 - cost: 3.87, layer: 58
Normal
Diabetic
Normal data
Fold over basal
Time (min)
2.5
2
1.5
1
0.5
0
-0.5
0
10
20
30
40
50
60

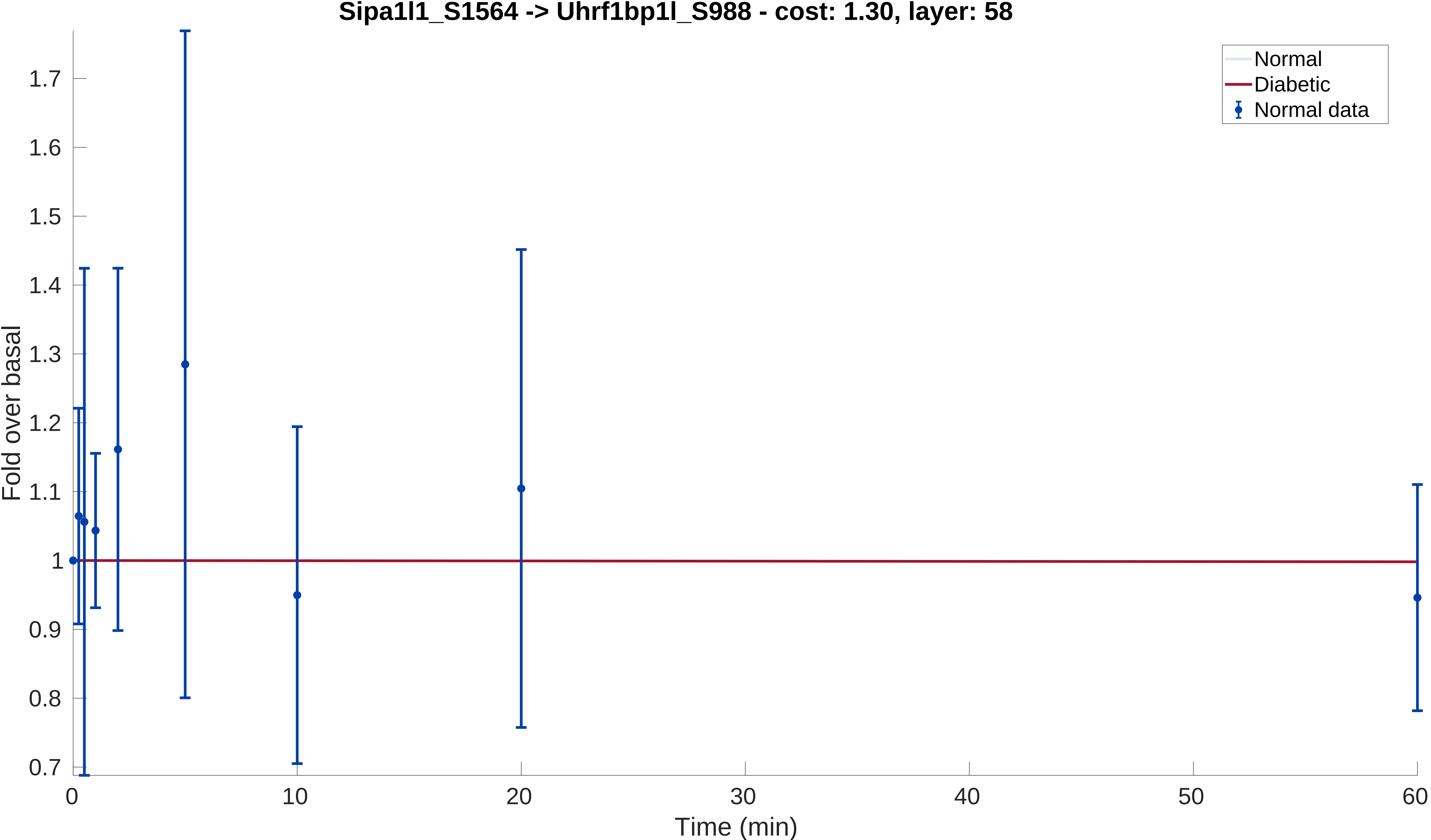

Sipa1l1_S1564 -> Uhrf1bp1l_S988 - cost: 1.30, layer: 58
Normal
Diabetic
Normal data
Fold over basal
Time (min)
1.7
1.6
1.5
1.4
1.3
1.2
1.1
1
0.9
0.8
0.7
0
10
20
30
40
50
60

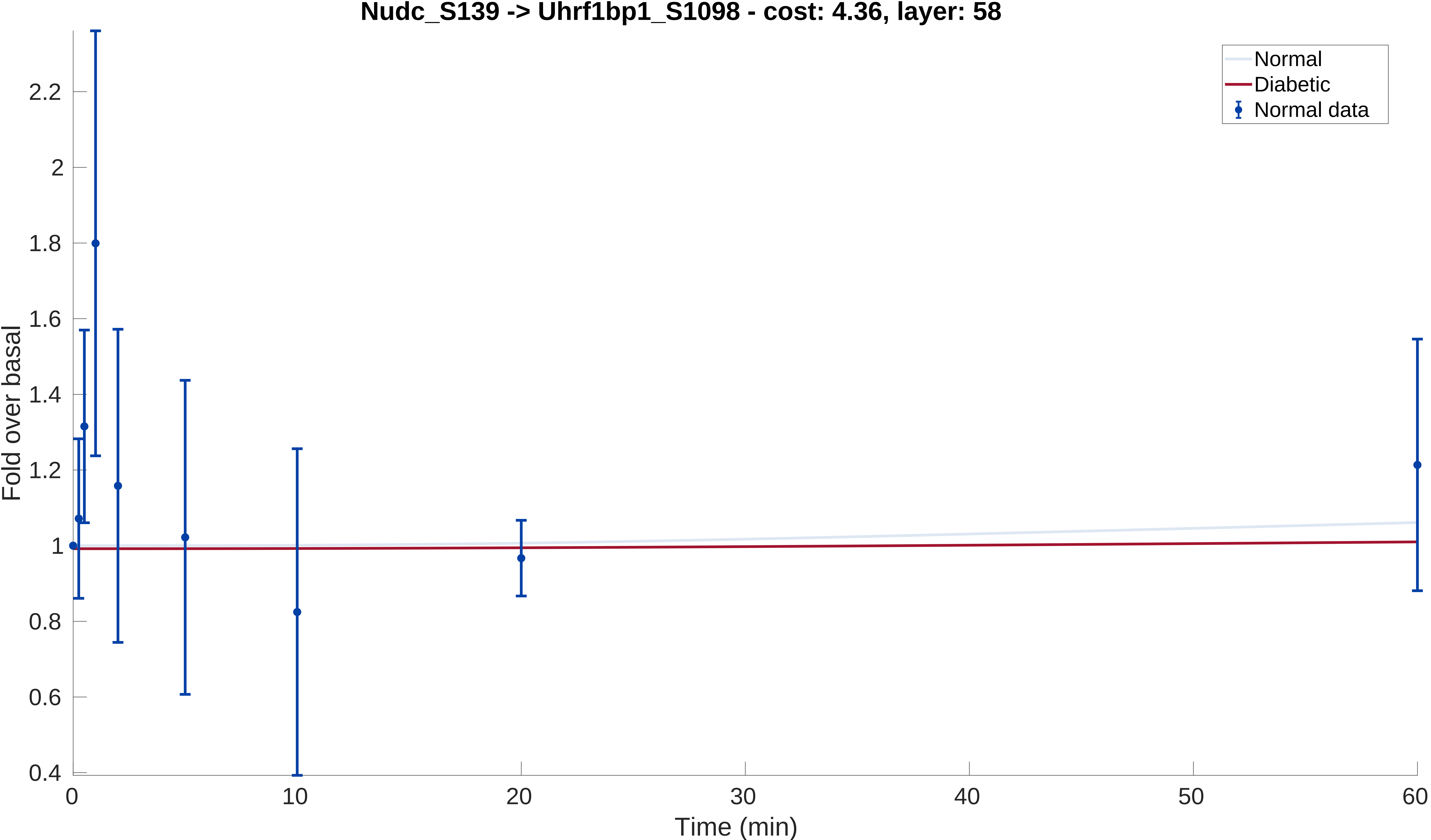

Nudc_S139 -> Uhrf1bp1_S1098 - cost: 4.36, layer: 58
Normal
Diabetic
Normal data
Fold over basal
Time (min)
0.4
0.6
0.8
1
1.2
1.4
1.6
1.8
2
2.2
0
10
20
30
40
50
60

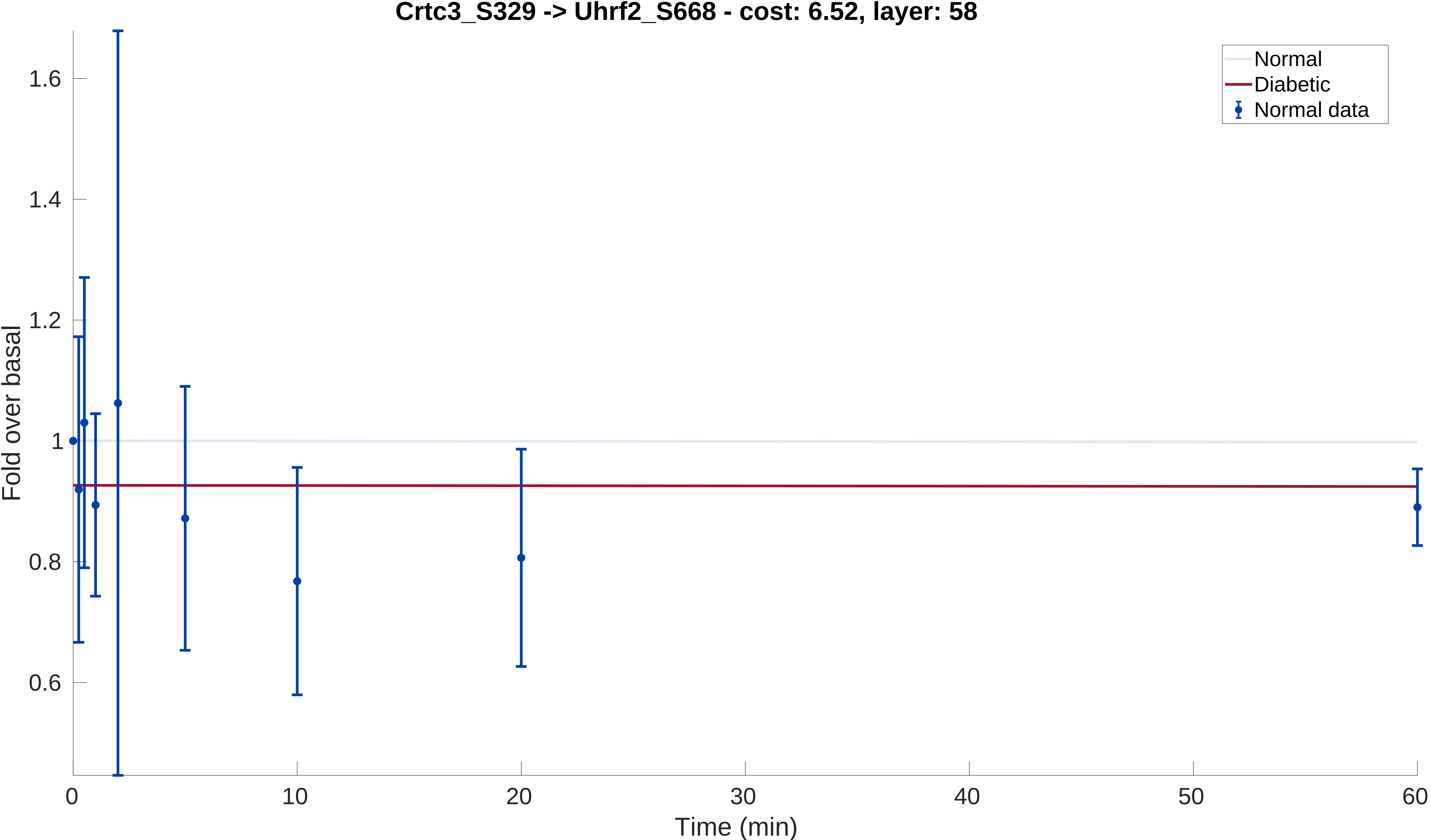

Crtc3_S329 -> Uhrf2_S668 - cost: 6.52, layer: 58
Normal
Diabetic
Normal data
Fold over basal
1.6
1.4
1.2
1
0.8
0.6
0
10
20
30
40
50
60
Time (min)

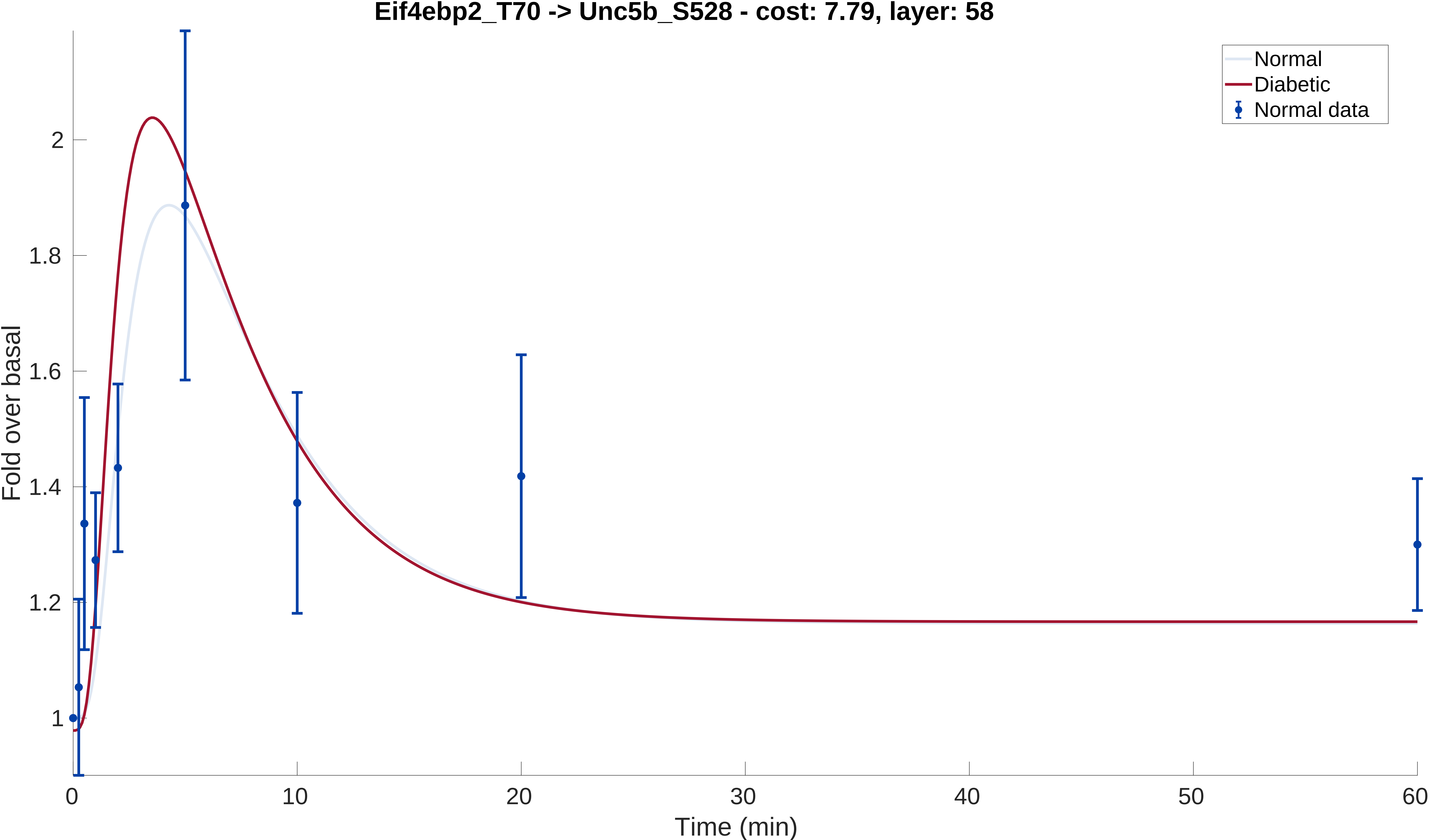

Eif4ebp2_T70 -> Unc5b_S528 - cost: 7.79, layer: 58
Normal
Diabetic
Normal data
Fold over basal
1
1.2
1.4
1.6
1.8
2
0
10
20
30
40
50
60
Time (min)

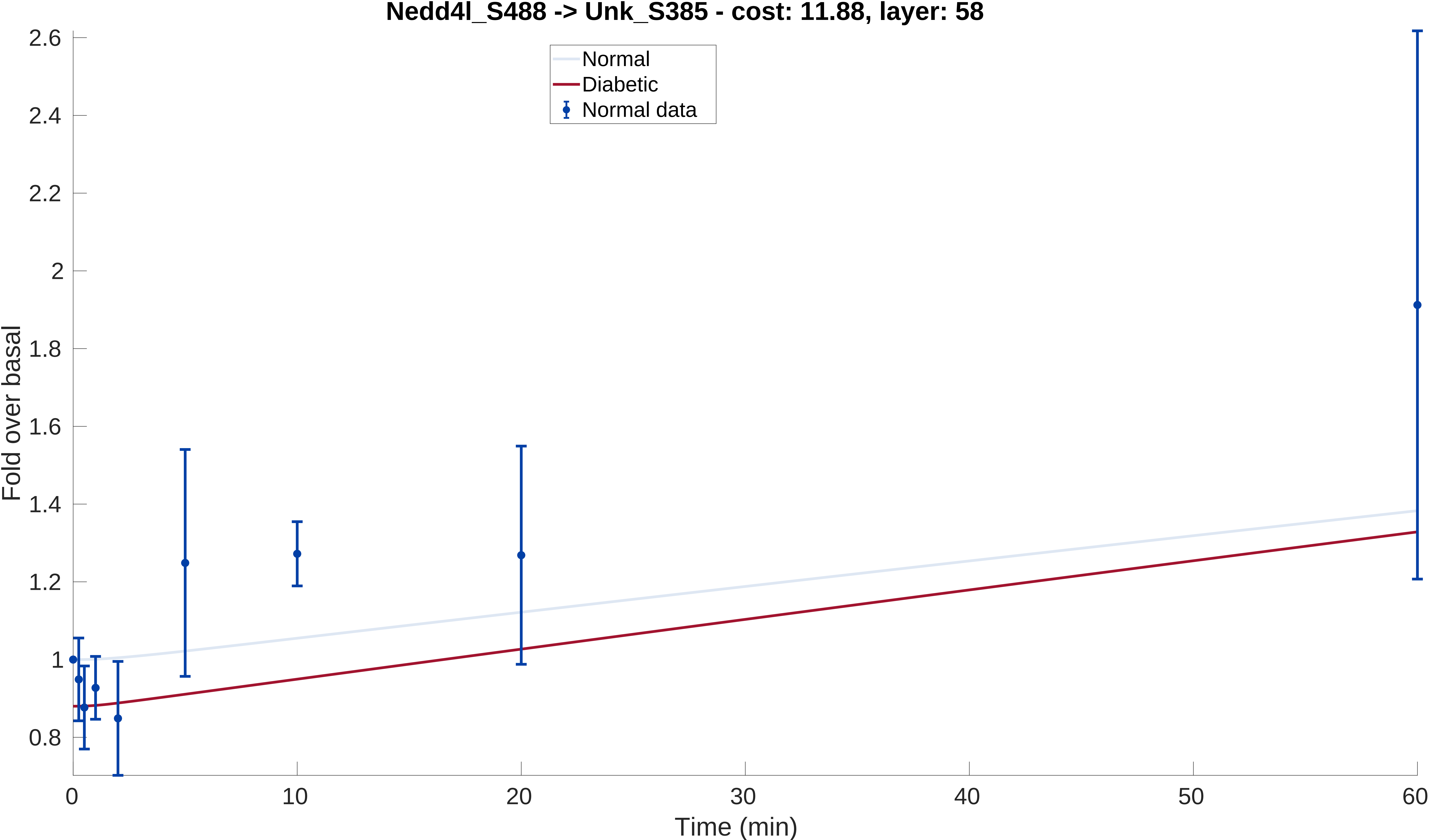
Nedd4l_S488 -> Unk_S385 - cost: 11.88, layer: 58
Normal
Diabetic
Normal data
Fold over basal
2.6
2.4
2.2
2
1.8
1.6
1.4
1.2
1
0.8
0
10
20
30
40
50
60
Time (min)

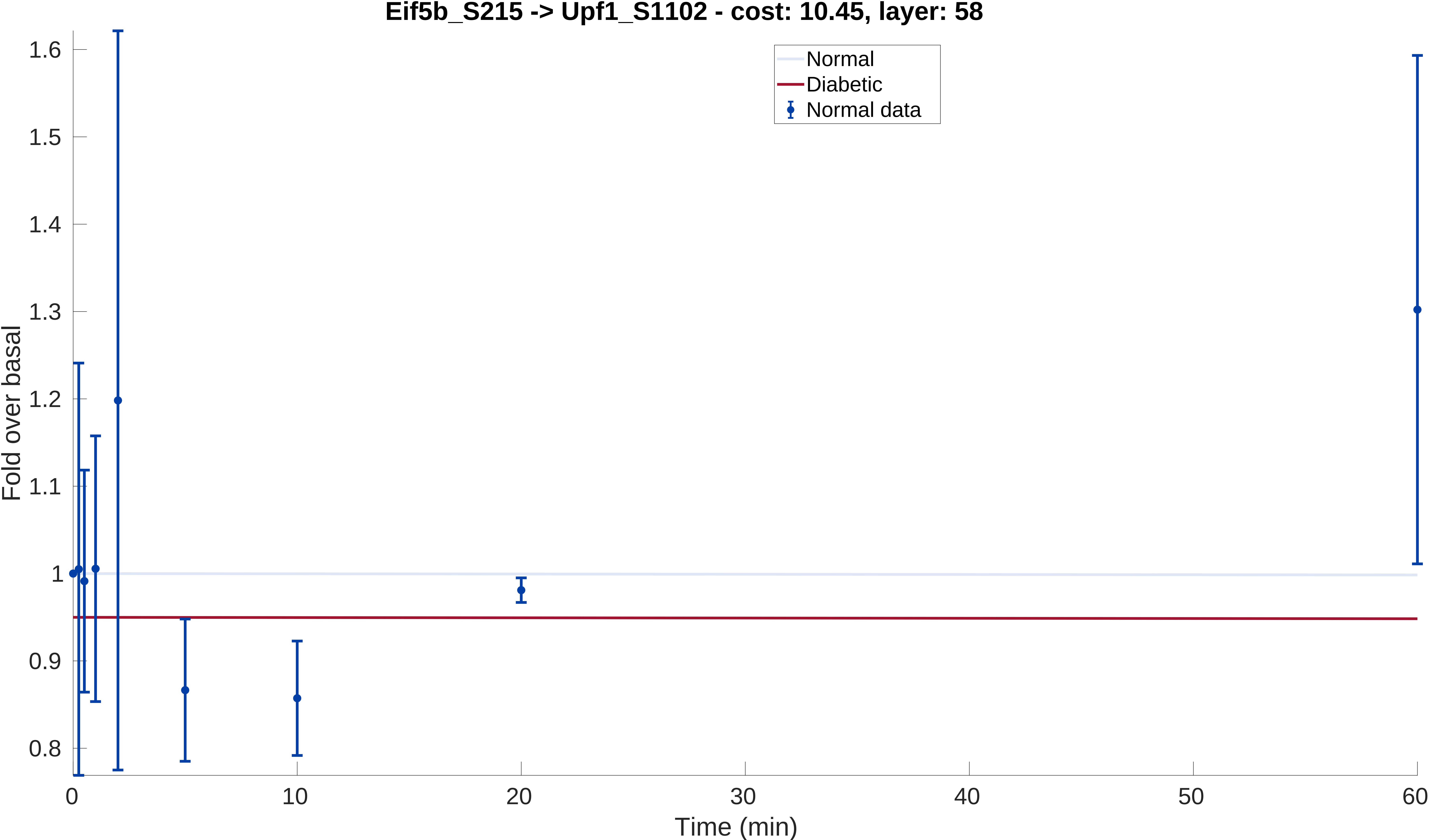
Eif5b_S215 -> Upf1_S1102 - cost: 10.45, layer: 58
Normal
Diabetic
Normal data
Fold over basal
1.6
1.5
1.4
1.3
1.2
1.1
1
0.9
0.8
0
10
20
30
40
50
60
Time (min)

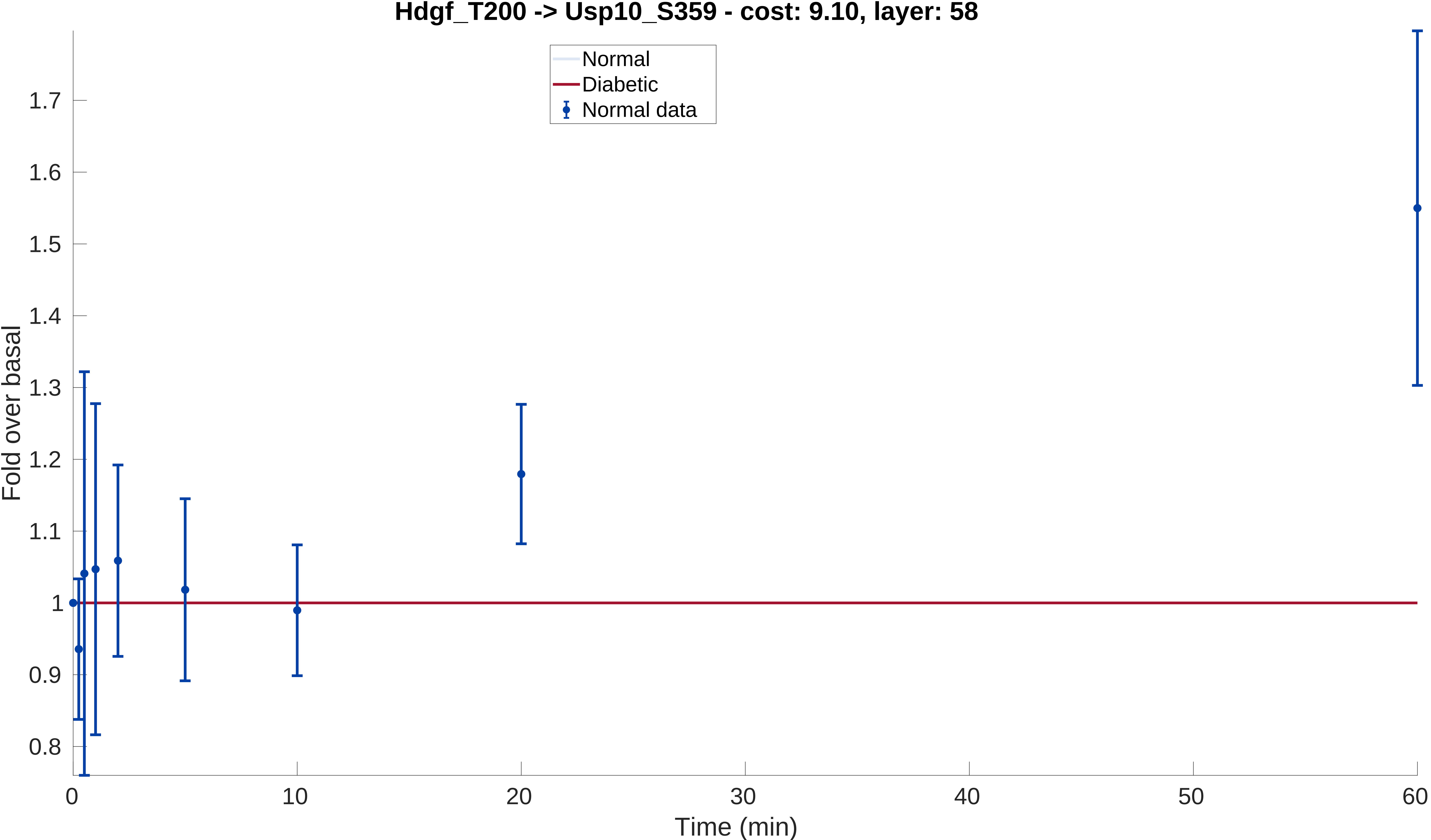

Hdgf_T200 -> Usp10_S359 - cost: 9.10, layer: 58
Normal
Diabetic
Normal data
Fold over basal
1.7
1.6
1.5
1.4
1.3
1.2
1.1
1
0.9
0.8
0
10
20
30
40
50
60
Time (min)

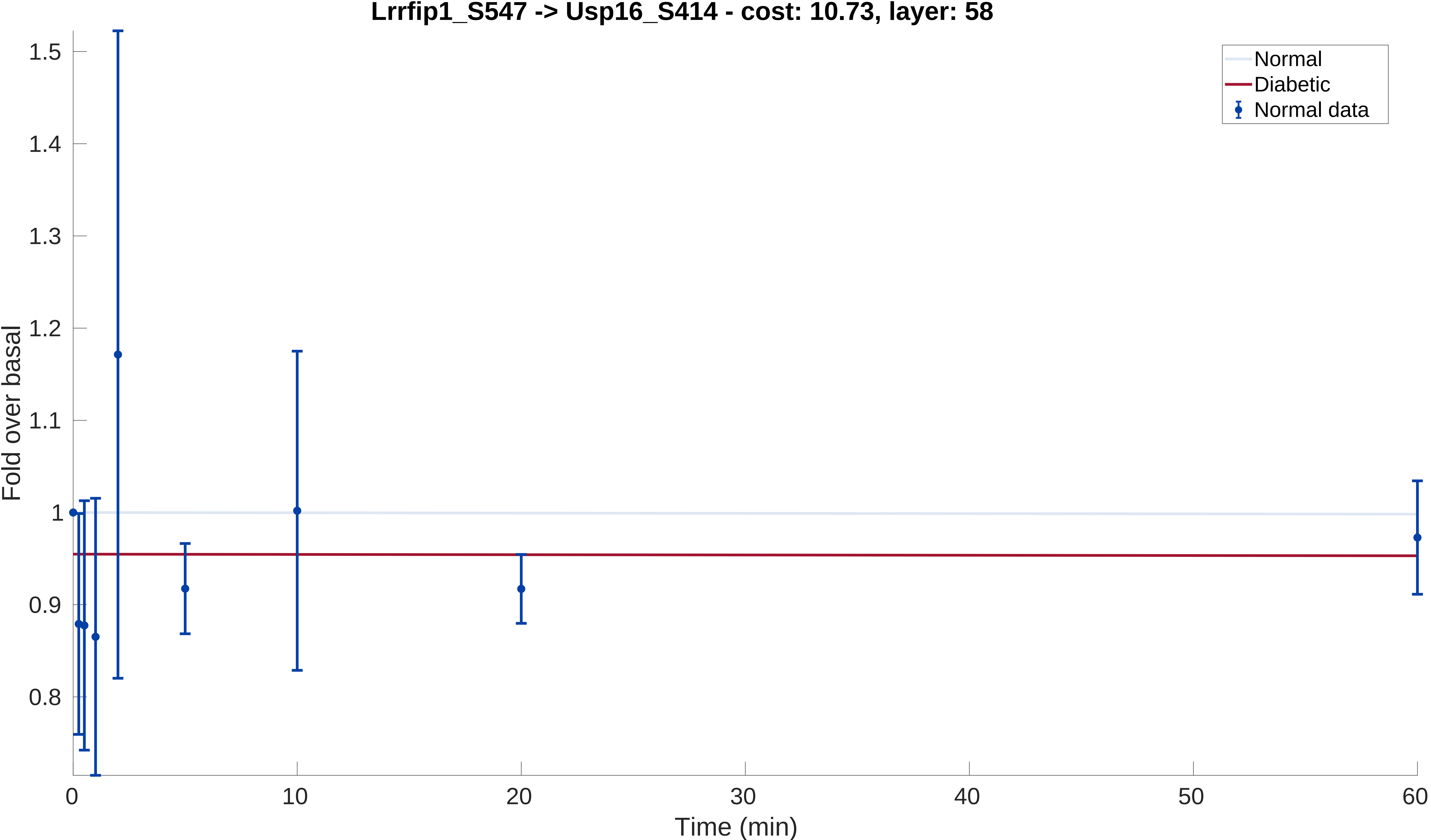

Lrrfip1_S547 -> Usp16_S414 - cost: 10.73, layer: 58
Normal
Diabetic
Normal data
Fold over basal
Time (min)
1.5
1.4
1.3
1.2
1.1
1
0.9
0.8
0
10
20
30
40
50
60

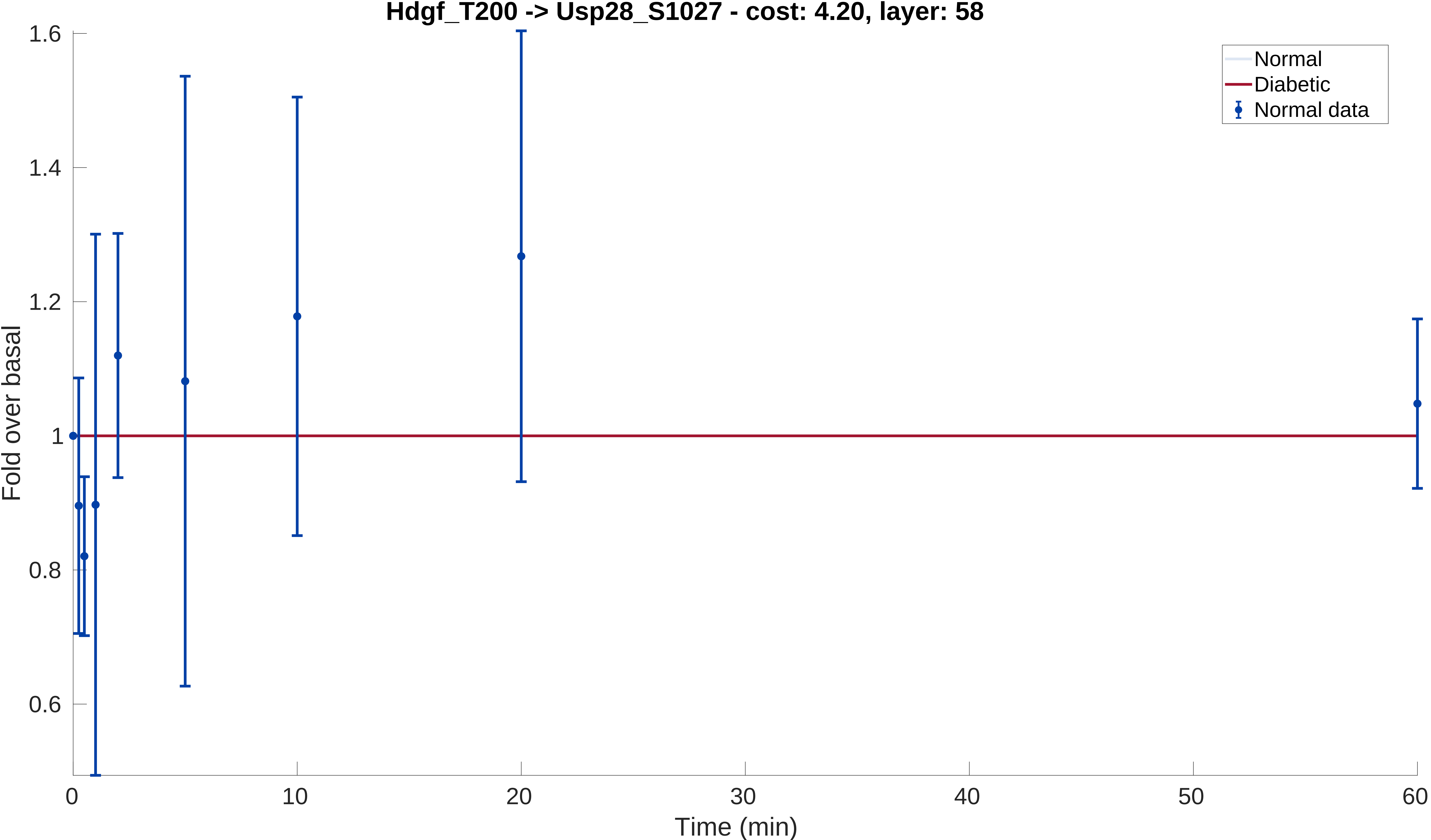

Hdgf_T200 -> Usp28_S1027 - cost: 4.20, layer: 58
Normal
Diabetic
Normal data
Fold over basal
Time (min)
1.6
1.4
1.2
1
0.8
0.6
0
10
20
30
40
50
60

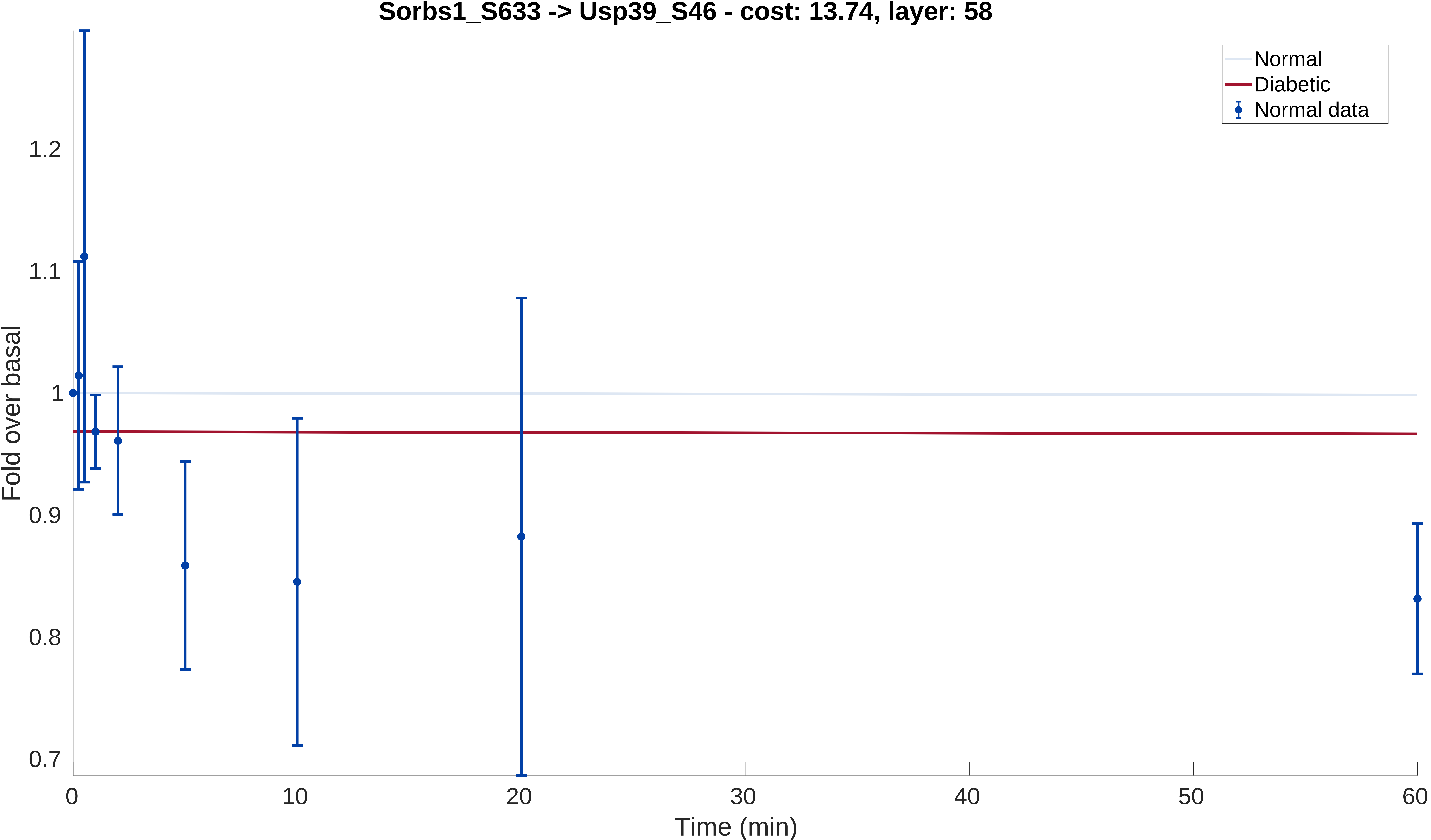
Sorbs1_S633 -> Usp39_S46 - cost: 13.74, layer: 58
Normal
Diabetic
Normal data
Fold over basal
1.2
1.1
1
0.9
0.8
0.7
0
10
20
30
40
50
60
Time (min)

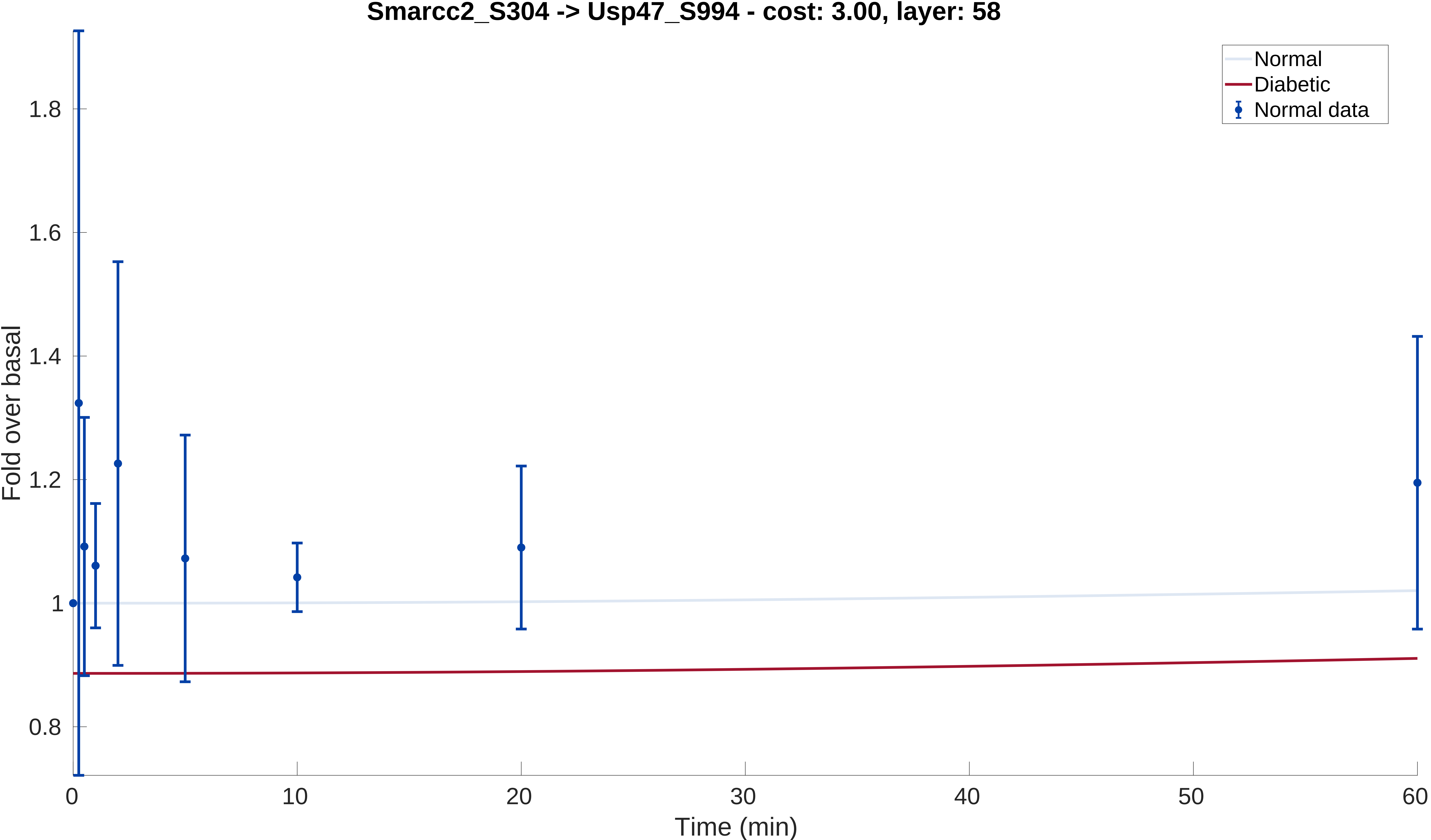
Smarcc2_S304 -> Usp47_S994 - cost: 3.00, layer: 58
Normal
Diabetic
Normal data
Fold over basal
1.8
1.6
1.4
1.2
1
0.8
0
10
20
30
40
50
60
Time (min)

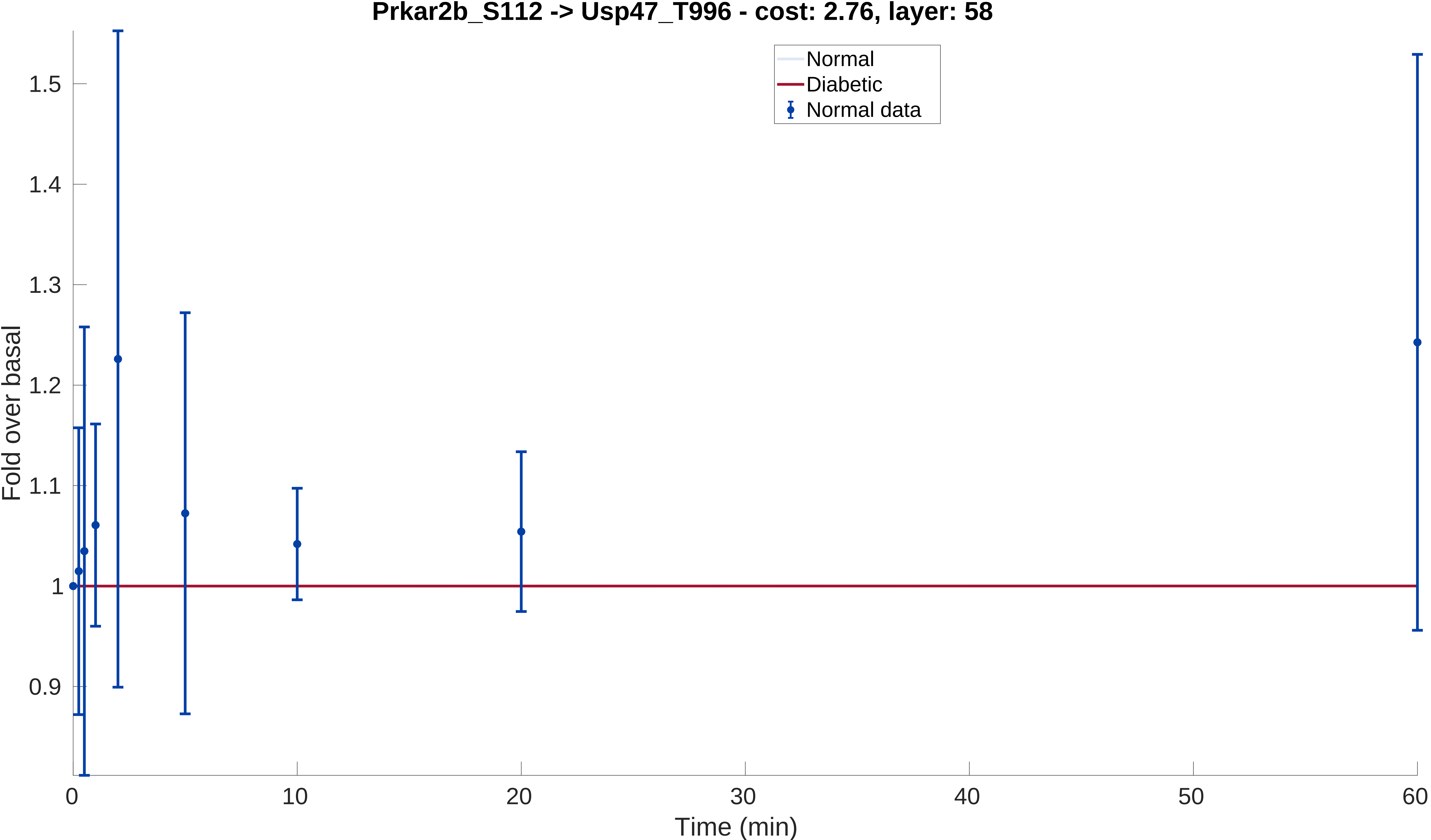

Prkar2b_S112 -> Usp47_T996 - cost: 2.76, layer: 58
Normal
Diabetic
Normal data
Fold over basal
1.5
1.4
1.3
1.2
1.1
1
0.9
0
10
20
30
40
50
60
Time (min)

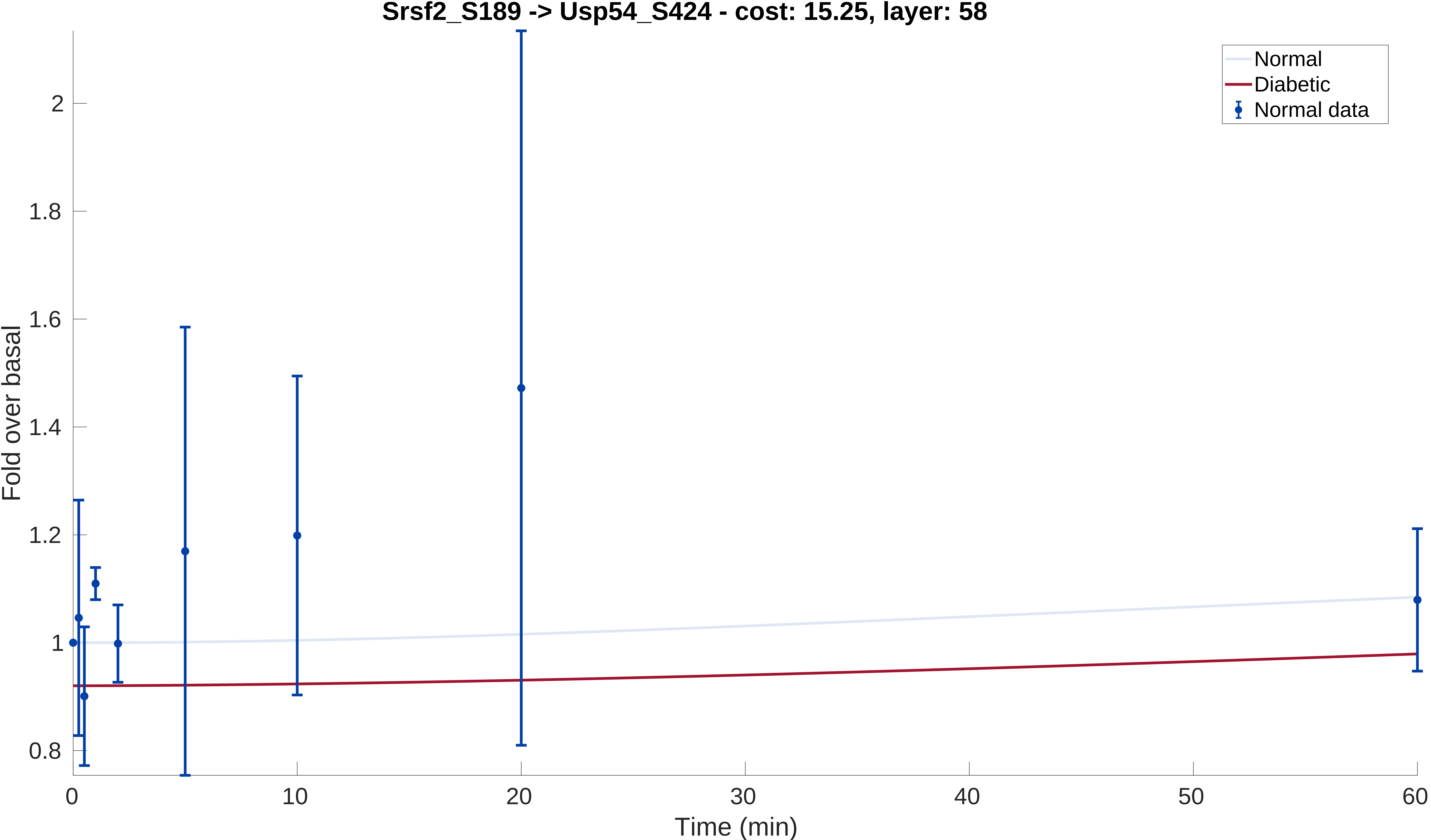
Srsf2_S189 -> Usp54_S424 - cost: 15.25, layer: 58
Normal
Diabetic
Normal data
Fold over basal
2
1.8
1.6
1.4
1.2
1
0.8
0
10
20
30
40
50
60
Time (min)

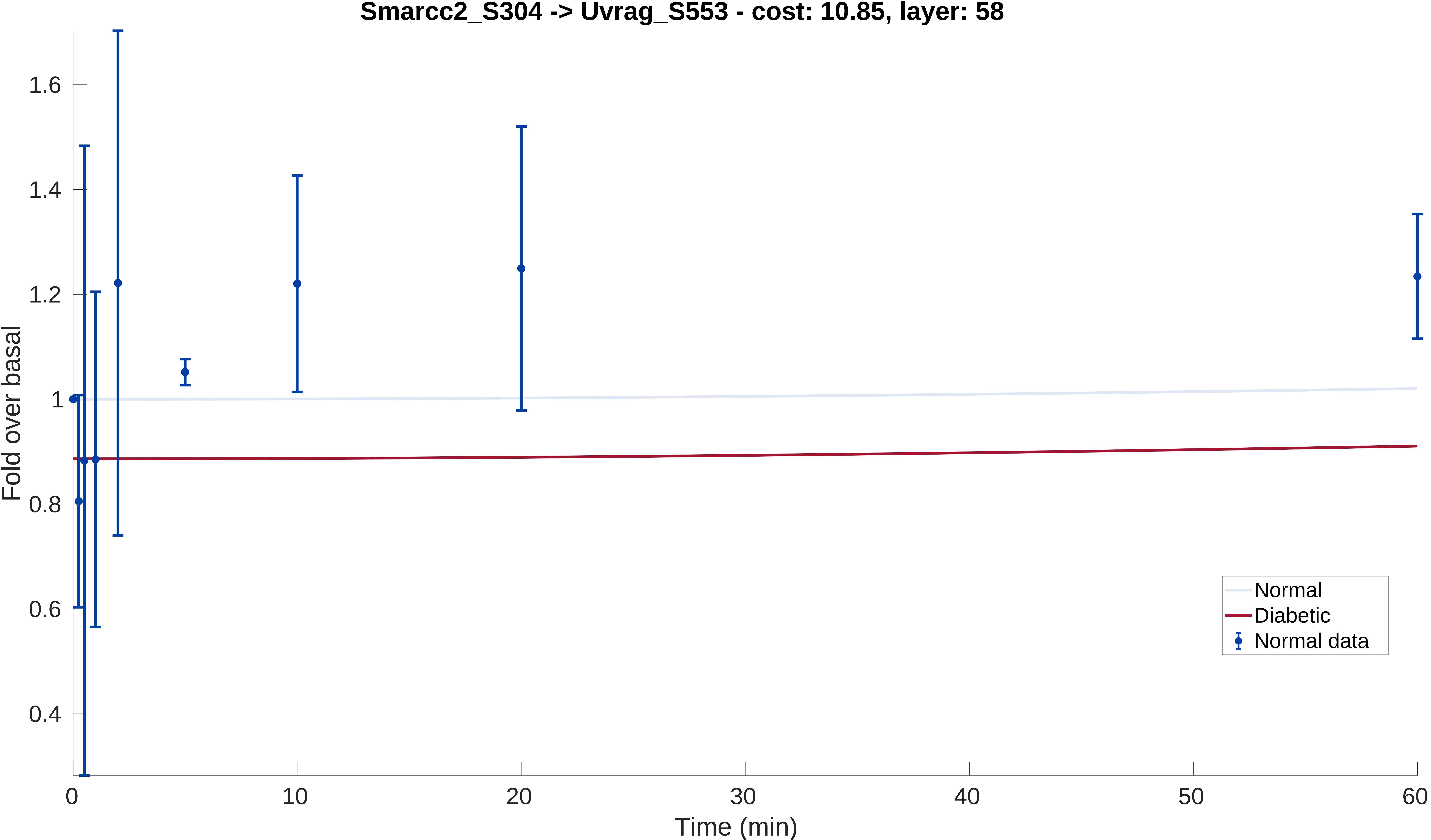

Smarcc2_S304 -> Uvrag_S553 - cost: 10.85, layer: 58
Fold over basal
1.6
1.4
1.2
1
0.8
0.6
0.4
0
10
20
30
40
50
60
Time (min)
Normal
Diabetic
Normal data

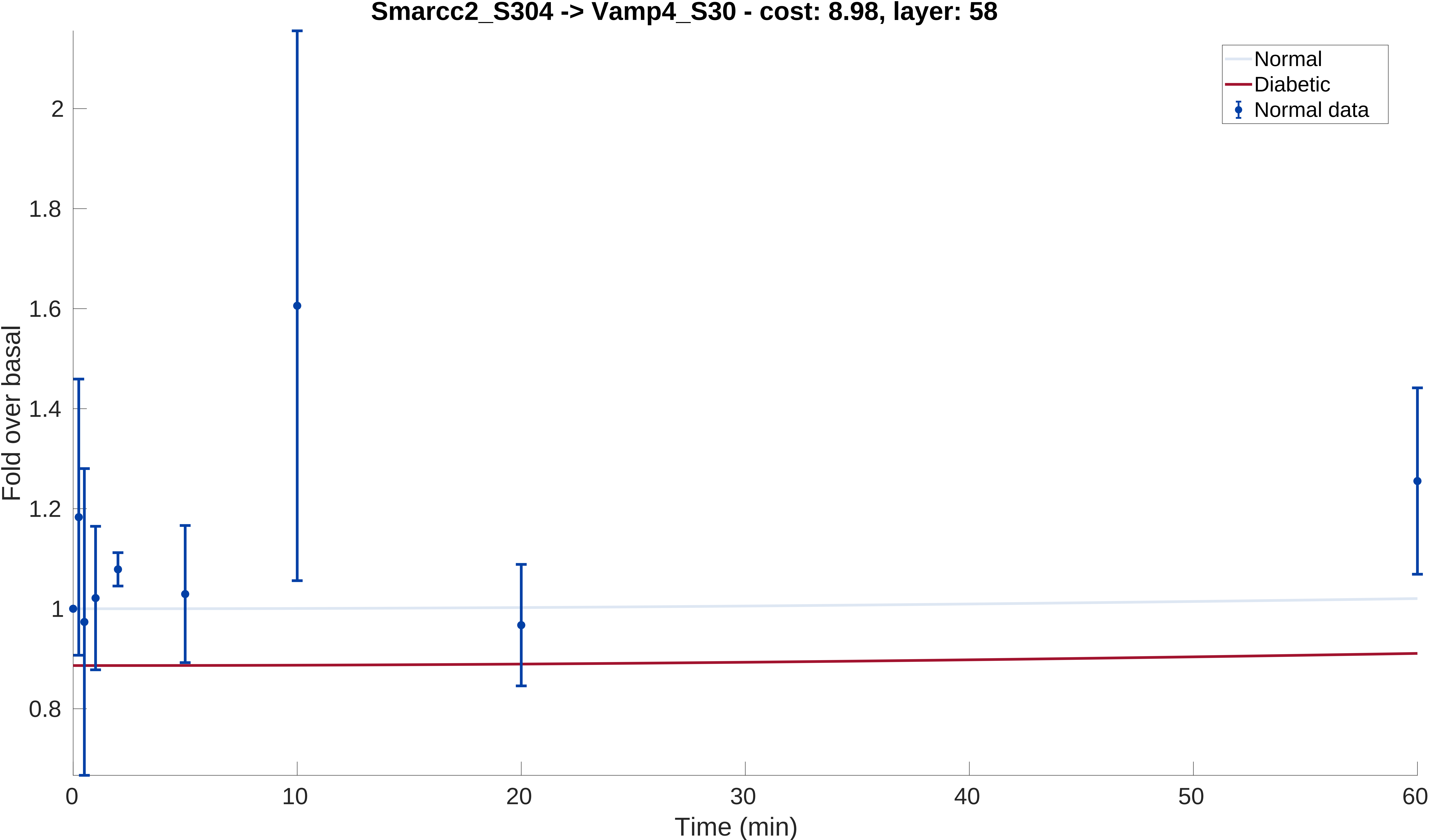

Smarcc2_S304 -> Vamp4_S30 - cost: 8.98, layer: 58
Normal
Diabetic
Normal data
Fold over basal
2
1.8
1.6
1.4
1.2
1
0.8
0
10
20
30
40
50
60
Time (min)

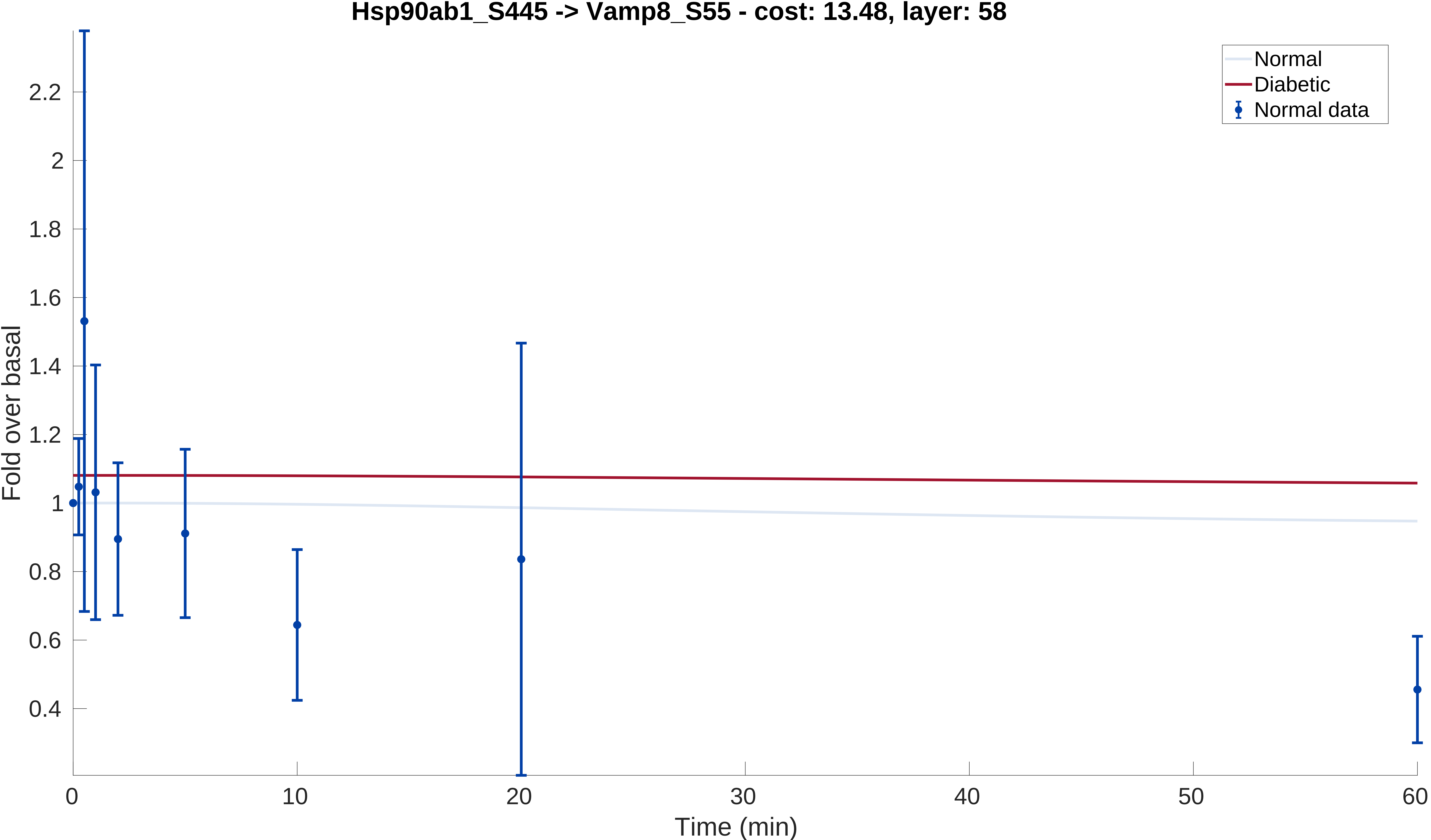

Hsp90ab1_S445 -> Vamp8_S55 - cost: 13.48, layer: 58
Normal
Diabetic
Normal data
Fold over basal
2.2
2
1.8
1.6
1.4
1.2
1
0.8
0.6
0.4
0
10
20
30
40
50
60
Time (min)

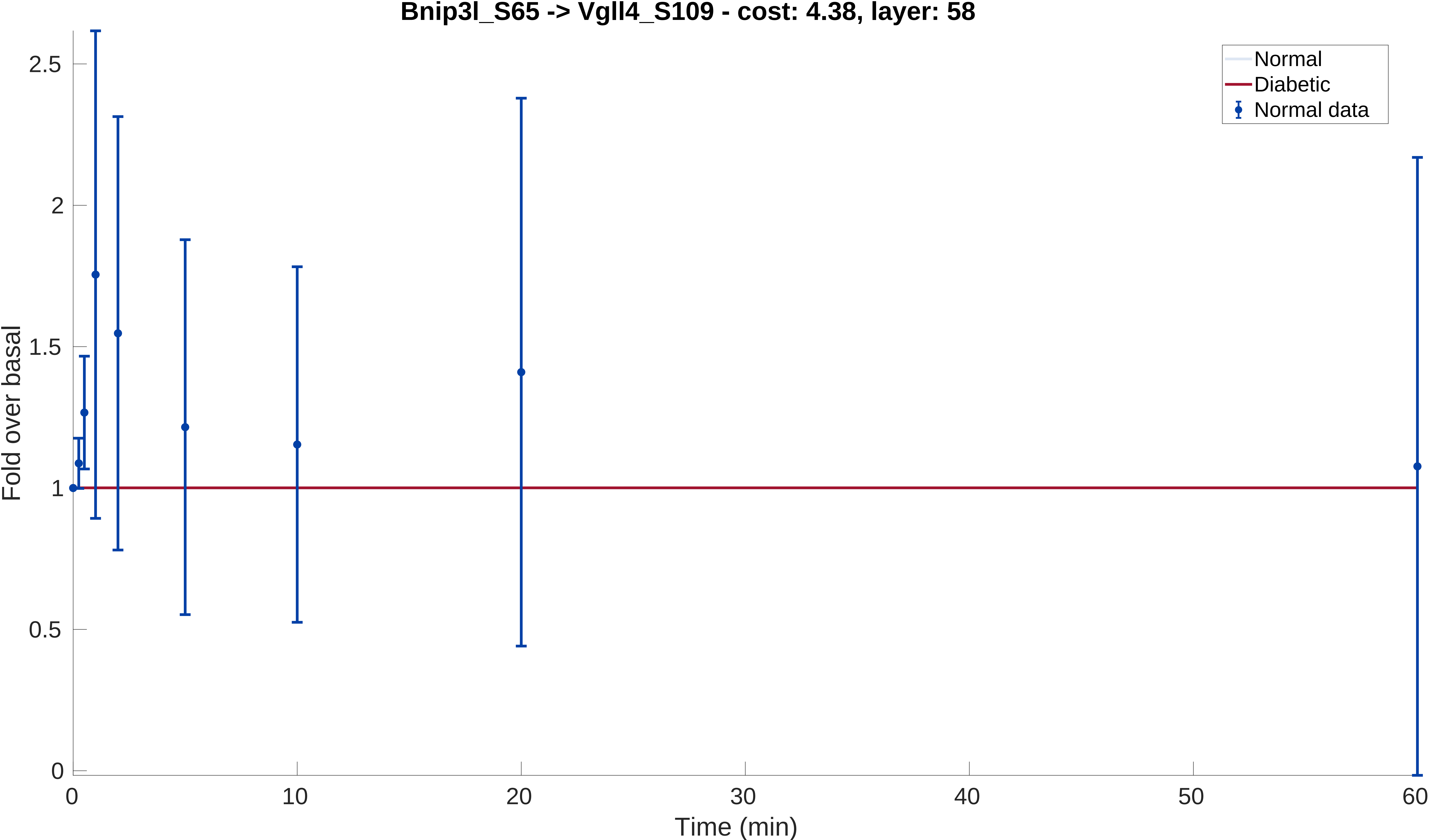

Bnip3l_S65 -> Vgll4_S109 - cost: 4.38, layer: 58
Normal
Diabetic
Normal data
Fold over basal
Time (min)
0
0.5
1
1.5
2
2.5
0
10
20
30
40
50
60

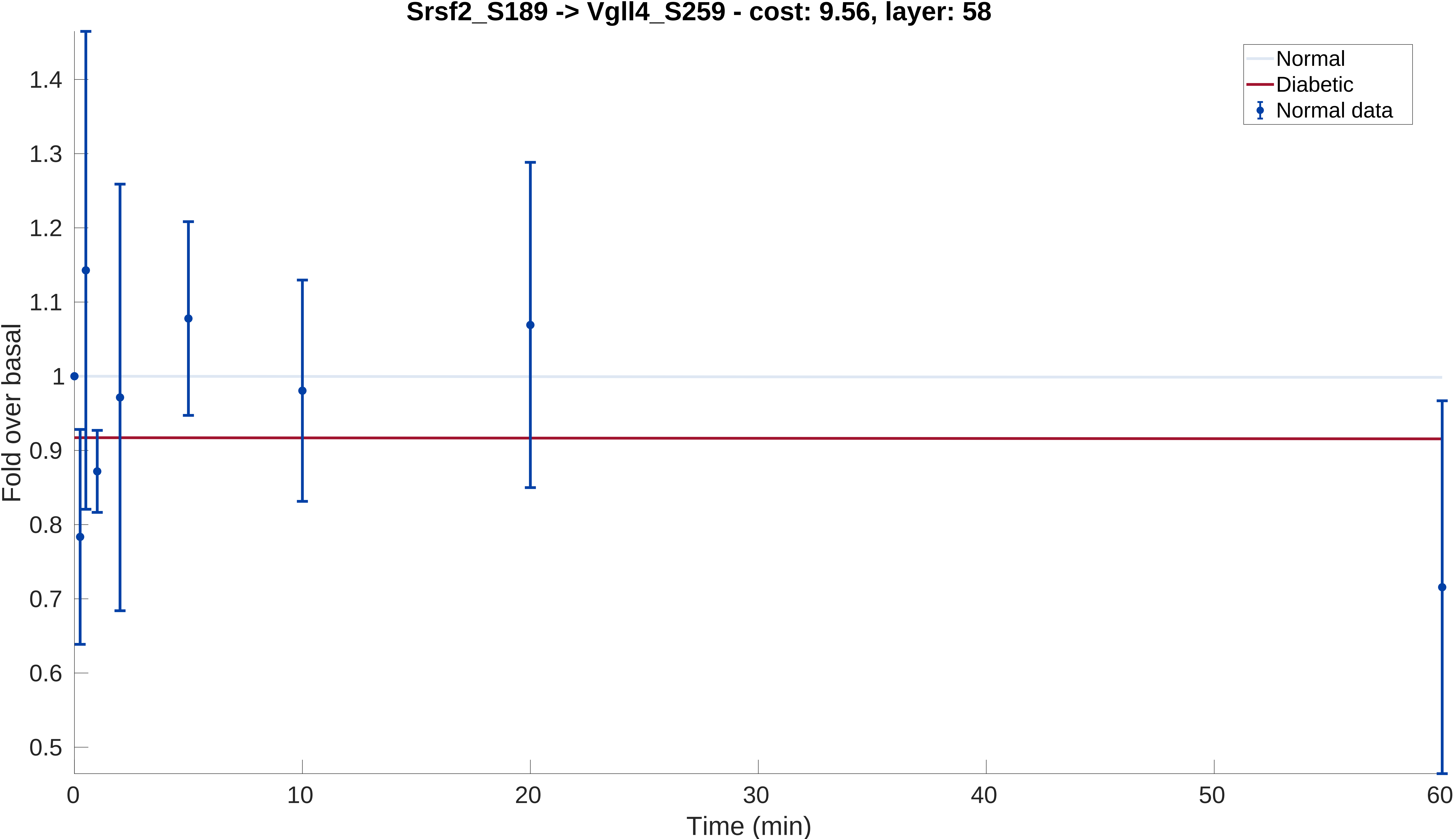
Srsf2_S189 -> Vgll4_S259 - cost: 9.56, layer: 58
Normal
Diabetic
Normal data
Fold over basal
Time (min)
1.4
1.3
1.2
1.1
1
0.9
0.8
0.7
0.6
0.5
0
10
20
30
40
50
60

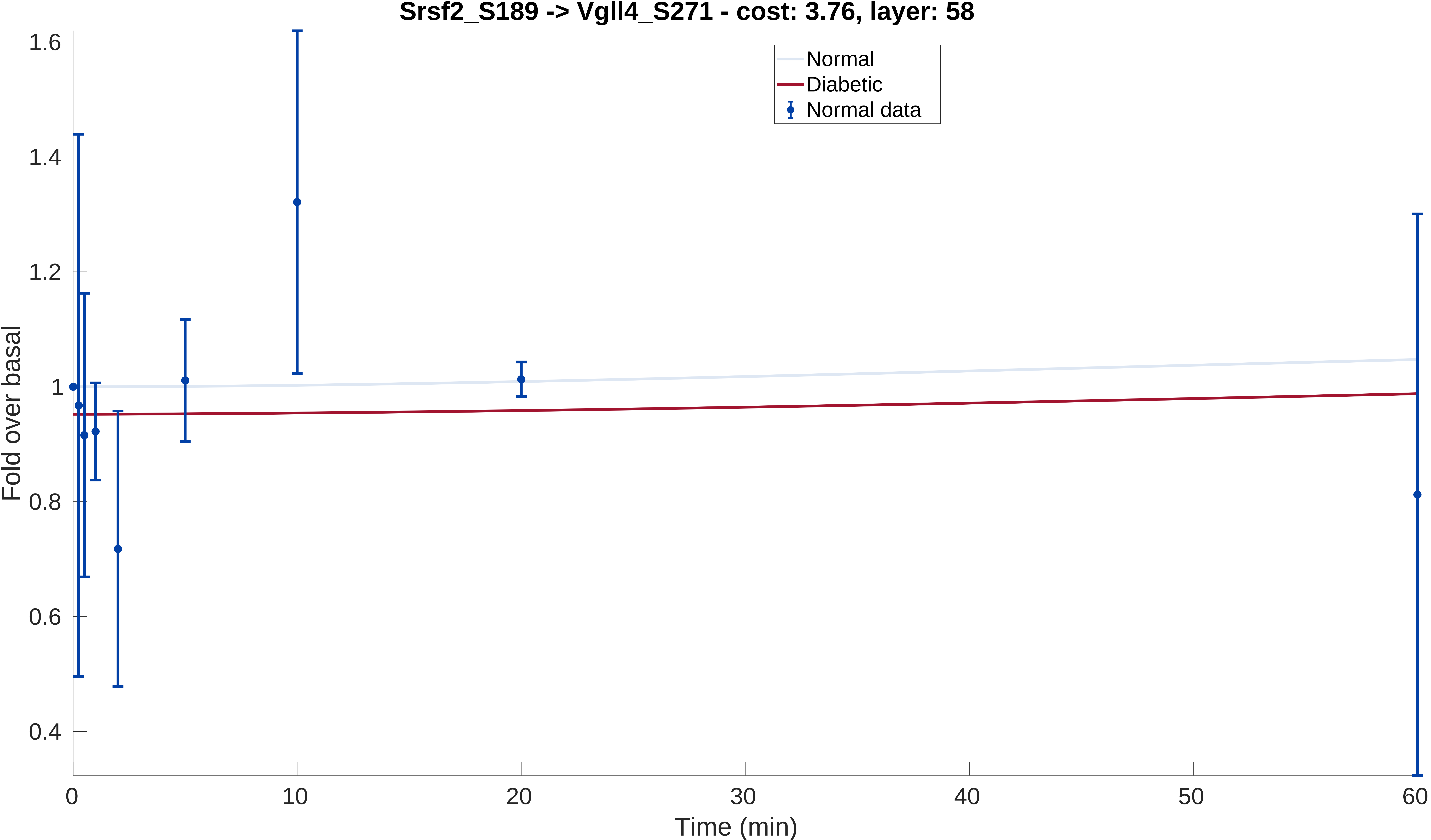

Srsf2_S189 -> Vgll4_S271 - cost: 3.76, layer: 58
Normal
Diabetic
Normal data
Fold over basal
1.6
1.4
1.2
1
0.8
0.6
0.4
0
10
20
30
40
50
60
Time (min)

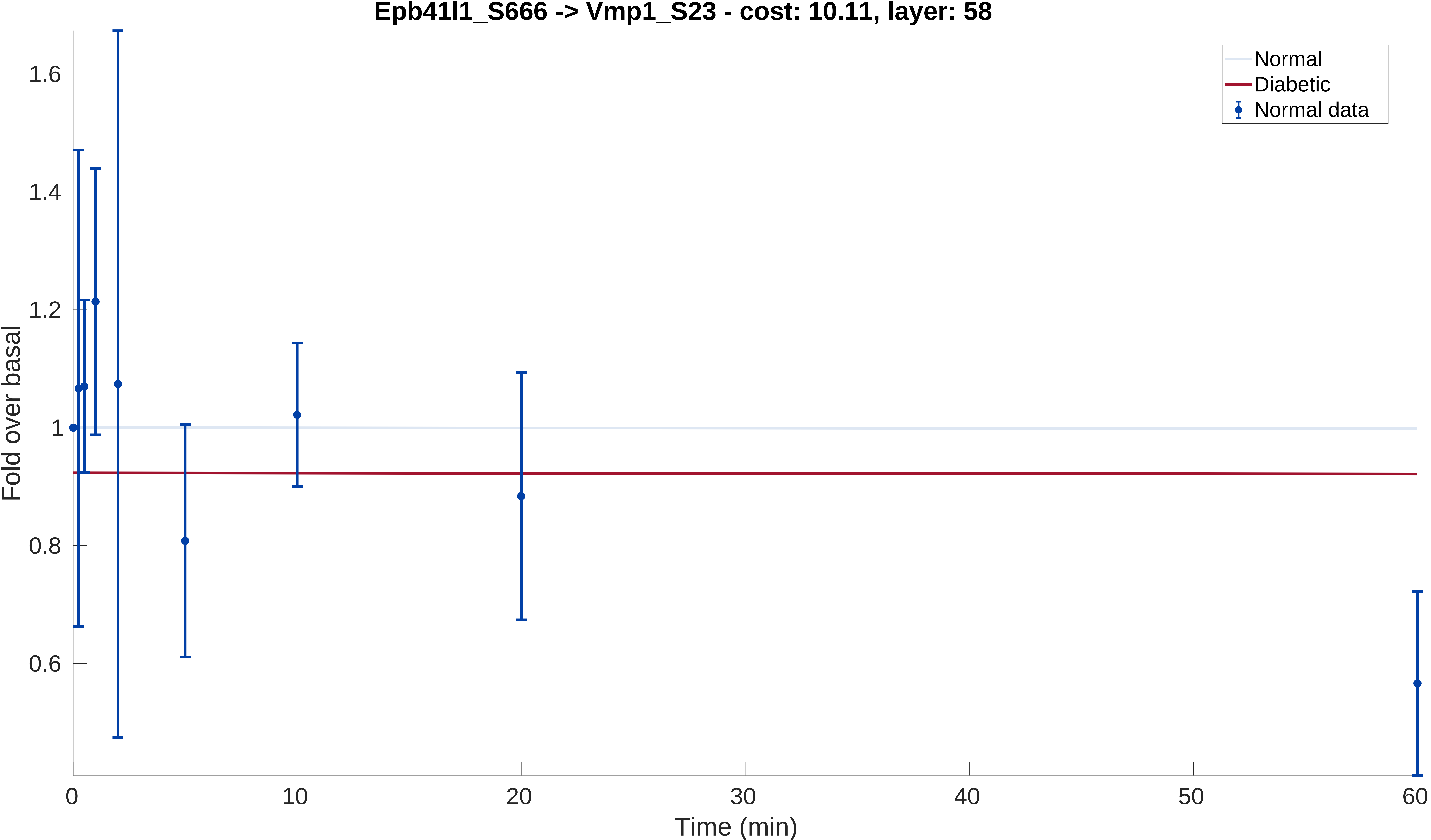

Epb41l1_S666 -> Vmp1_S23 - cost: 10.11, layer: 58
Normal
Diabetic
Normal data
Fold over basal
1.6
1.4
1.2
1
0.8
0.6
0
10
20
30
40
50
60
Time (min)

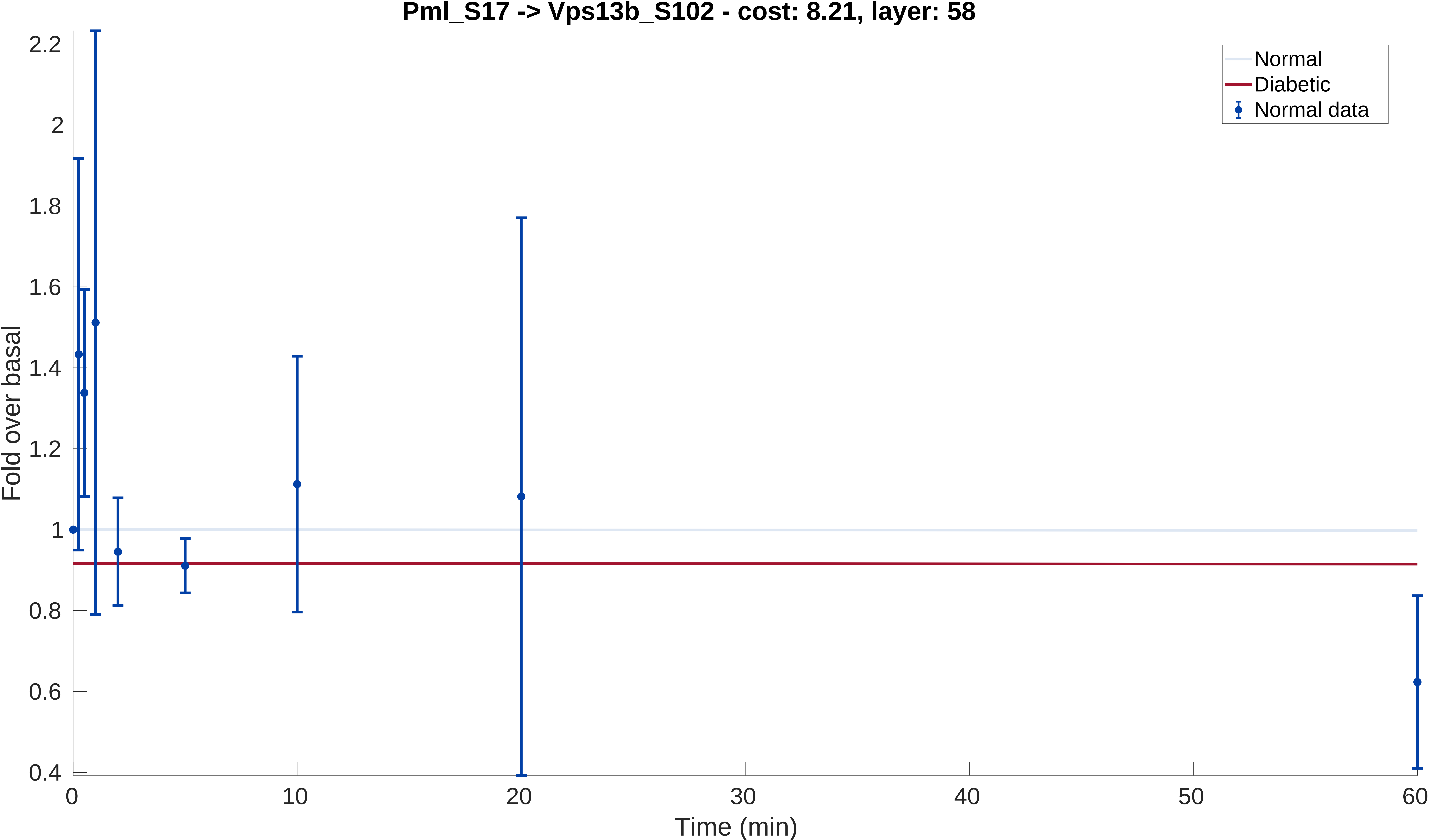
Pml_S17 -> Vps13b_S102 - cost: 8.21, layer: 58
Normal
Diabetic
Normal data
Fold over basal
Time (min)
0.4
0.6
0.8
1
1.2
1.4
1.6
1.8
2
2.2
0
10
20
30
40
50
60

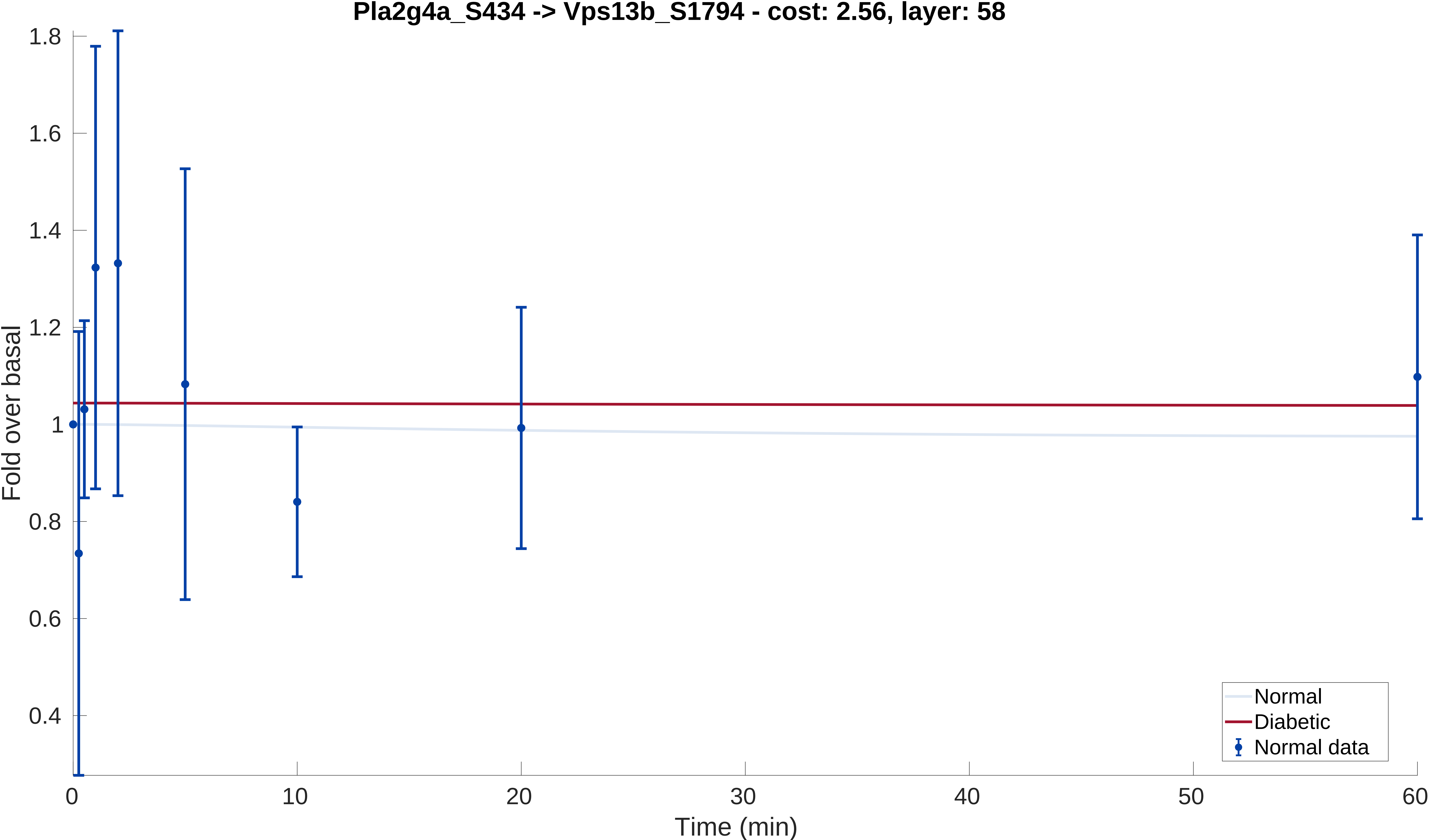

Pla2g4a_S434 -> Vps13b_S1794 - cost: 2.56, layer: 58
Fold over basal
Time (min)
1.8
1.6
1.4
1.2
1
0.8
0.6
0.4
0
10
20
30
40
50
60
Normal
Diabetic
Normal data

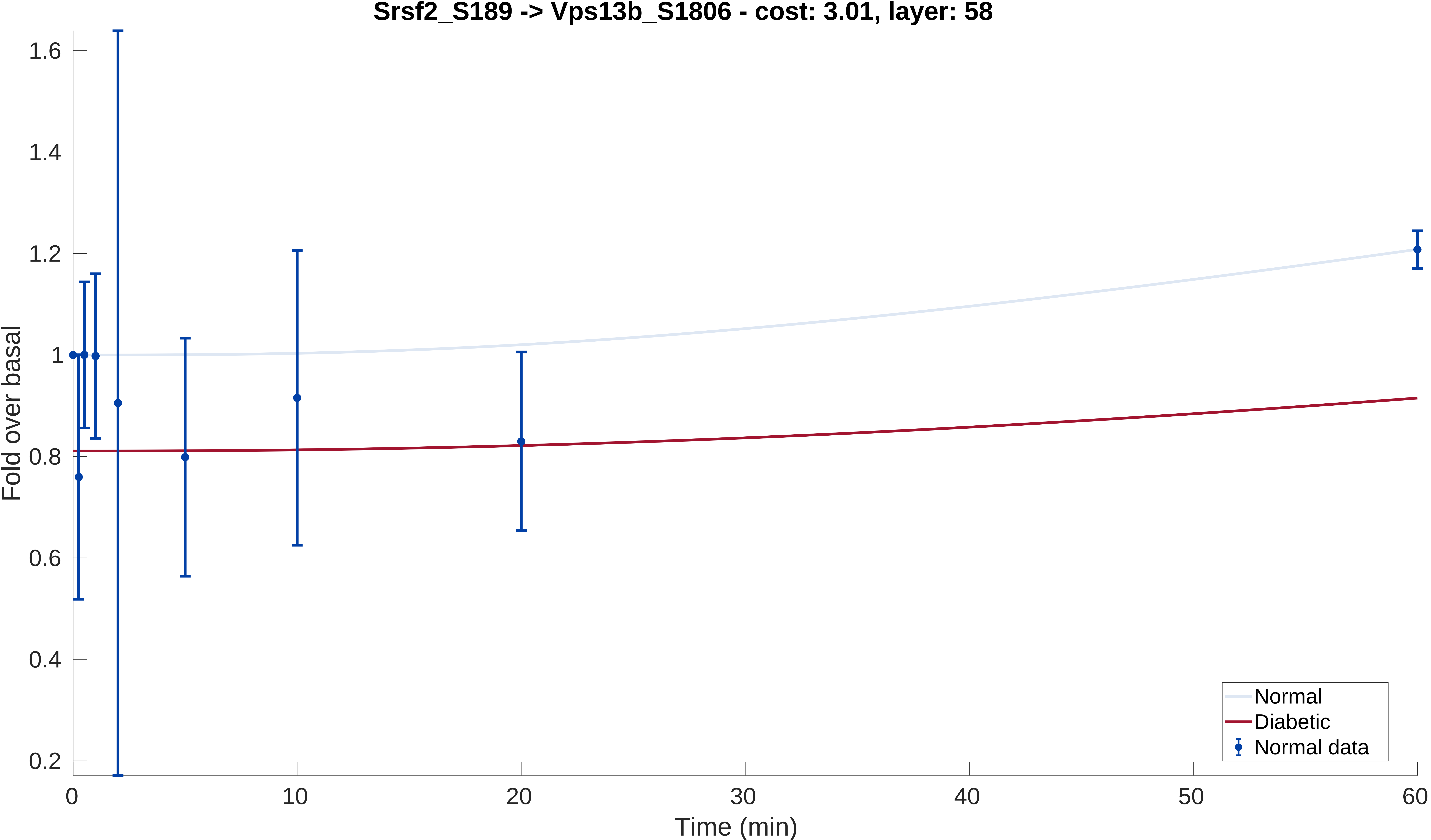

Srsf2_S189 -> Vps13b_S1806 - cost: 3.01, layer: 58
Fold over basal
1.6
1.4
1.2
1
0.8
0.6
0.4
0.2
0
10
20
30
40
50
60
Time (min)
Normal
Diabetic
Normal data

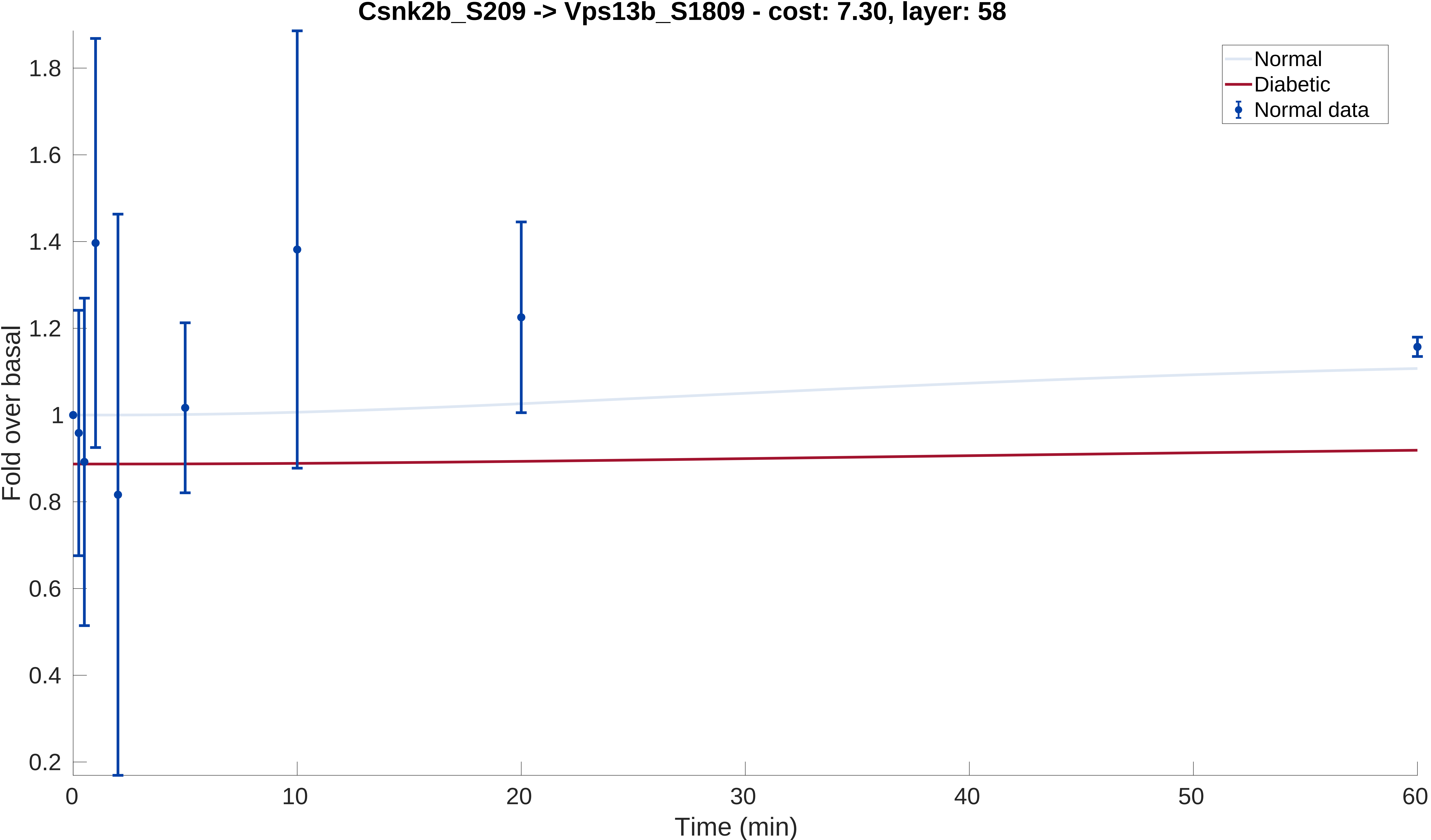
Csnk2b_S209 -> Vps13b_S1809 - cost: 7.30, layer: 58
Normal
Diabetic
Normal data
Fold over basal
0.2
0.4
0.6
0.8
1
1.2
1.4
1.6
1.8
0
10
20
30
40
50
60
Time (min)

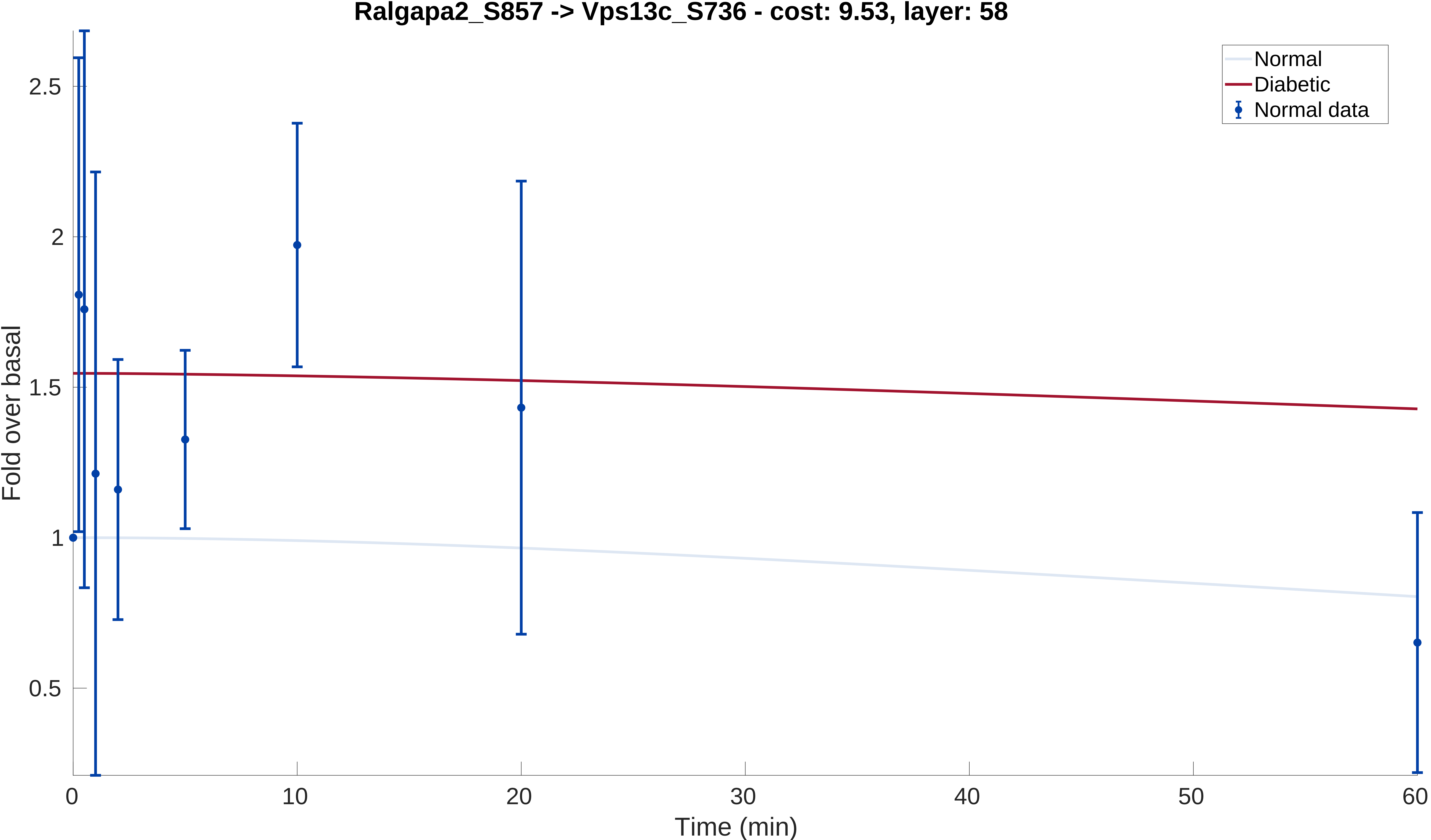

Ralgapa2_S857 -> Vps13c_S736 - cost: 9.53, layer: 58
Normal
Diabetic
Normal data
Fold over basal
2.5
2
1.5
1
0.5
0
10
20
30
40
50
60
Time (min)

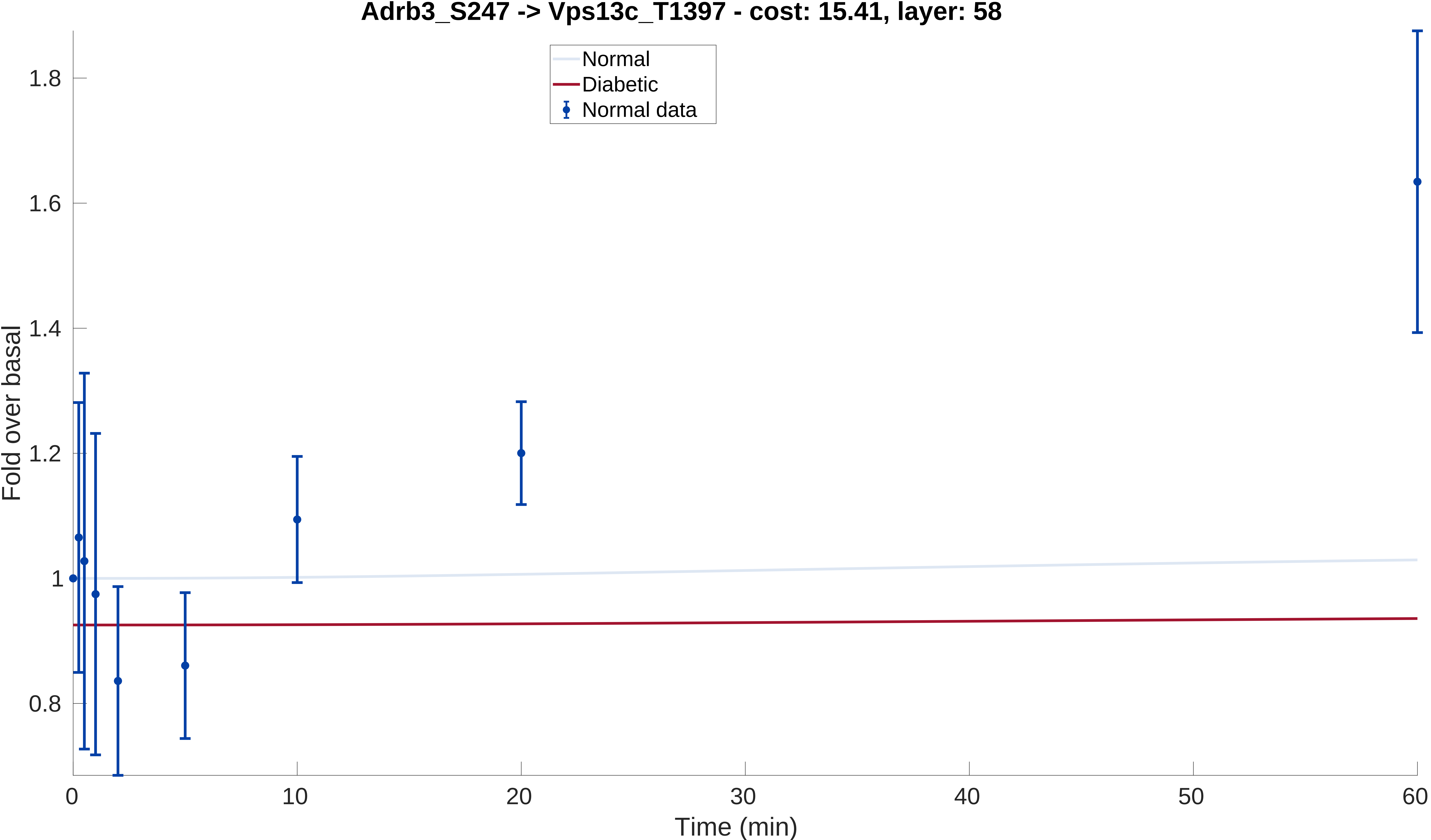
Adrb3_S247 -> Vps13c_T1397 - cost: 15.41, layer: 58
Normal
Diabetic
Normal data
Fold over basal
1.8
1.6
1.4
1.2
1
0.8
0
10
20
30
40
50
60
Time (min)

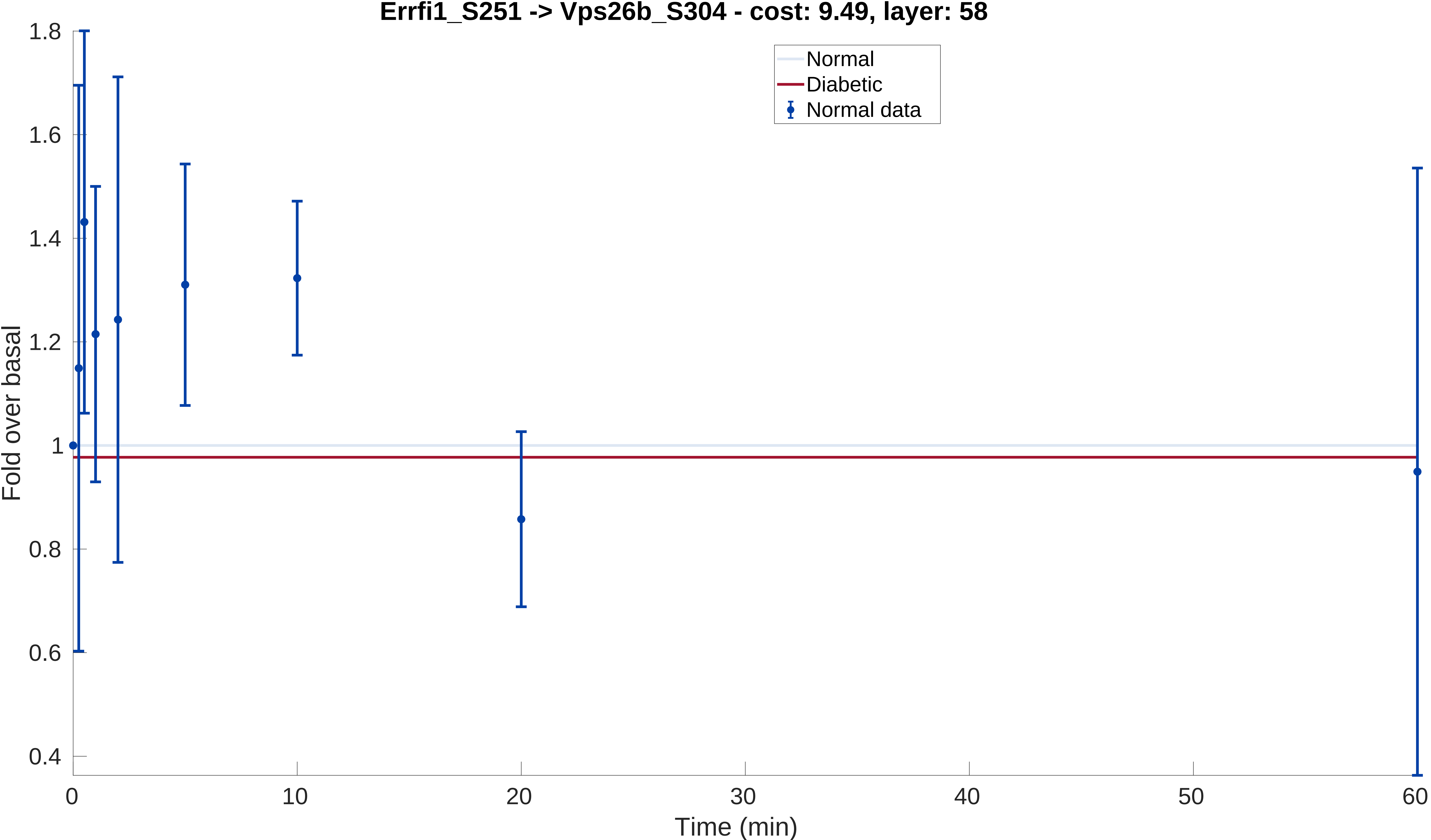
Errfi1_S251 -> Vps26b_S304 - cost: 9.49, layer: 58
Normal
Diabetic
Normal data
Fold over basal
Time (min)
1.8
1.6
1.4
1.2
1
0.8
0.6
0.4
0
10
20
30
40
50
60

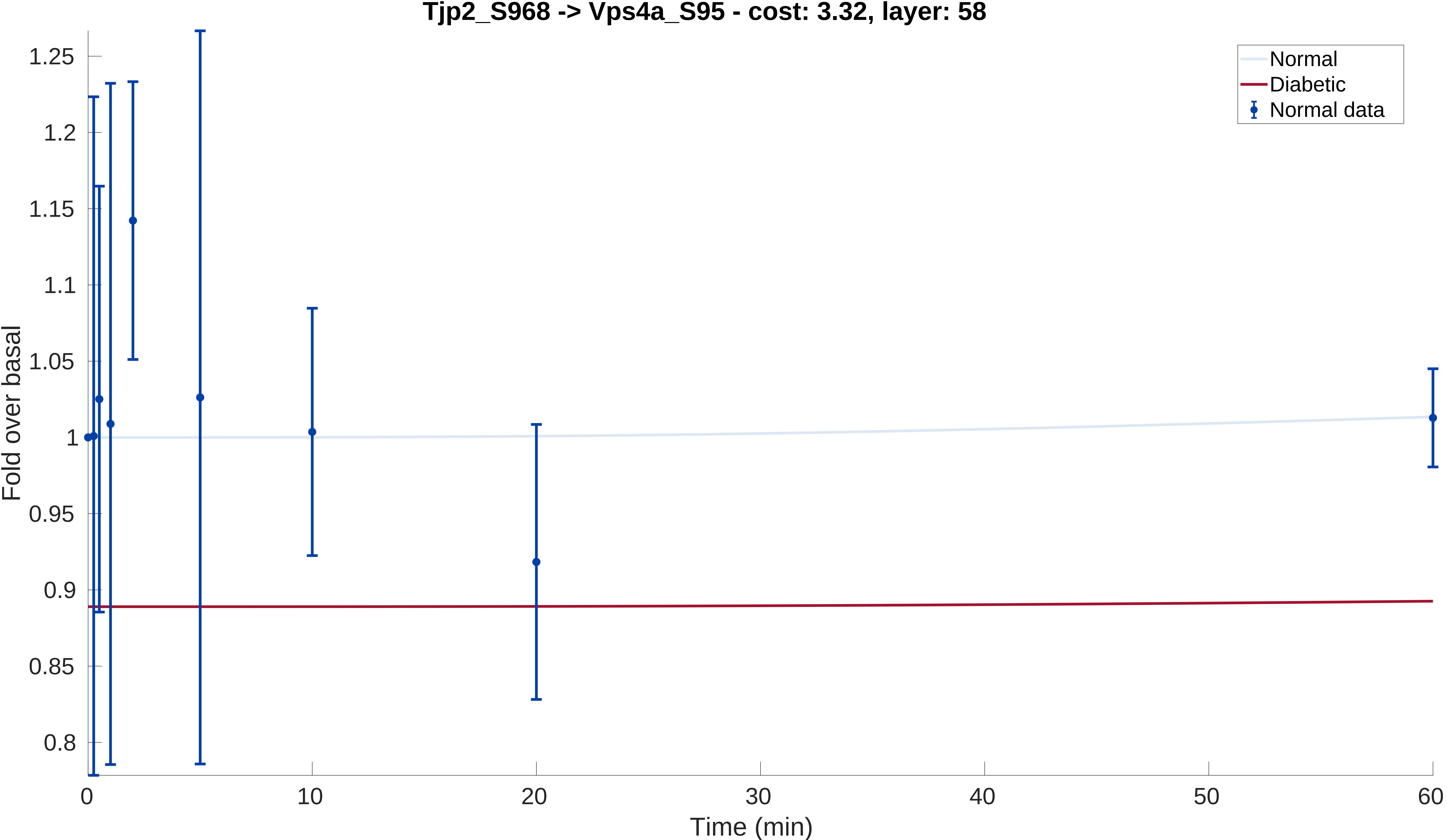

Tjp2_S968 -> Vps4a_S95 - cost: 3.32, layer: 58
Normal
Diabetic
Normal data
Fold over basal
1.25
1.2
1.15
1.1
1.05
1
0.95
0.9
0.85
0.8
0
10
20
30
40
50
60
Time (min)

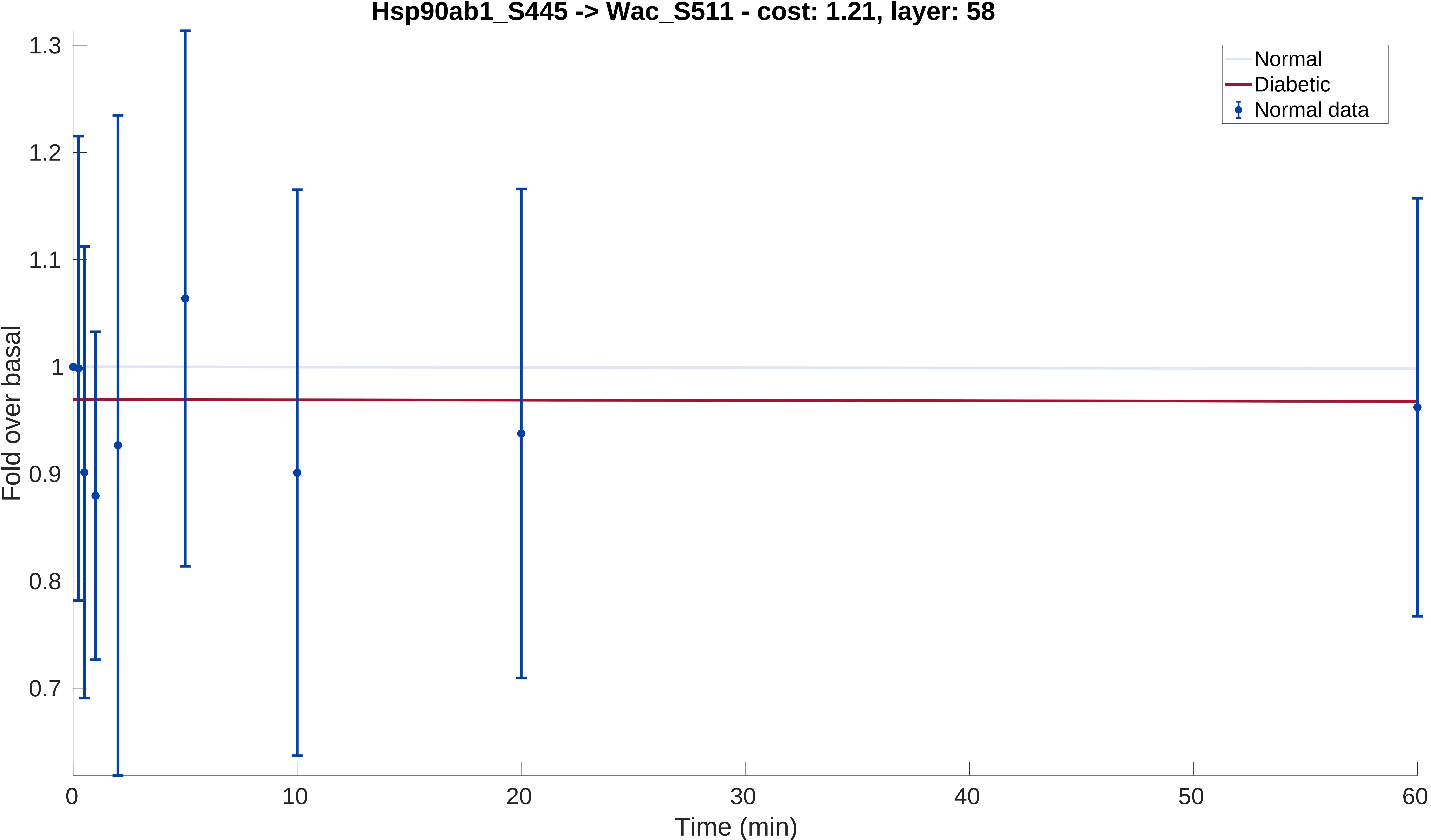

Hsp90ab1_S445 -> Wac_S511 - cost: 1.21, layer: 58
Normal
Diabetic
Normal data
Fold over basal
Time (min)
1.3
1.2
1.1
1
0.9
0.8
0.7
0
10
20
30
40
50
60

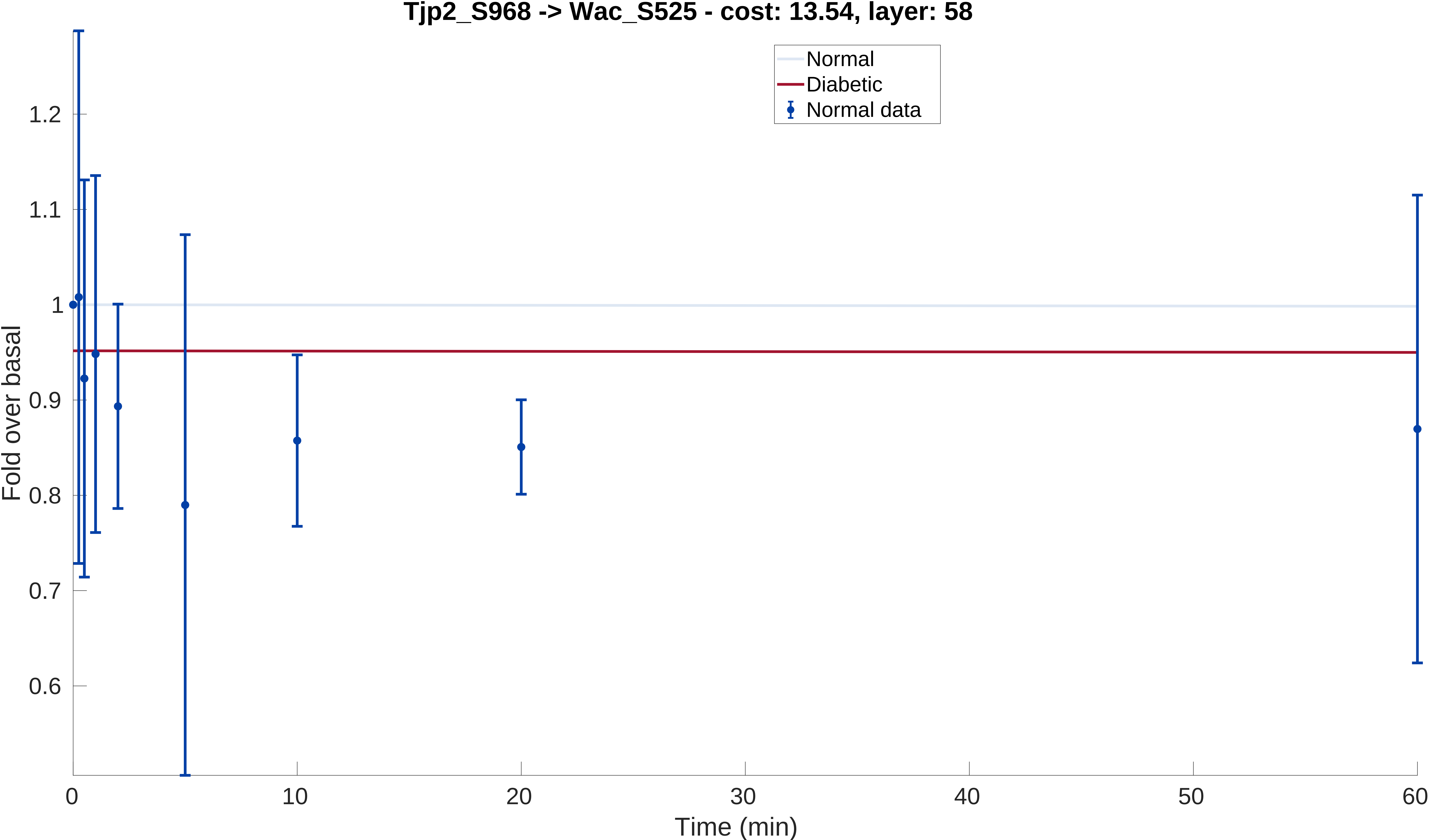

Tjp2_S968 -> Wac_S525 - cost: 13.54, layer: 58
Normal
Diabetic
Normal data
Fold over basal
1.2
1.1
1
0.9
0.8
0.7
0.6
0
10
20
30
40
50
60
Time (min)

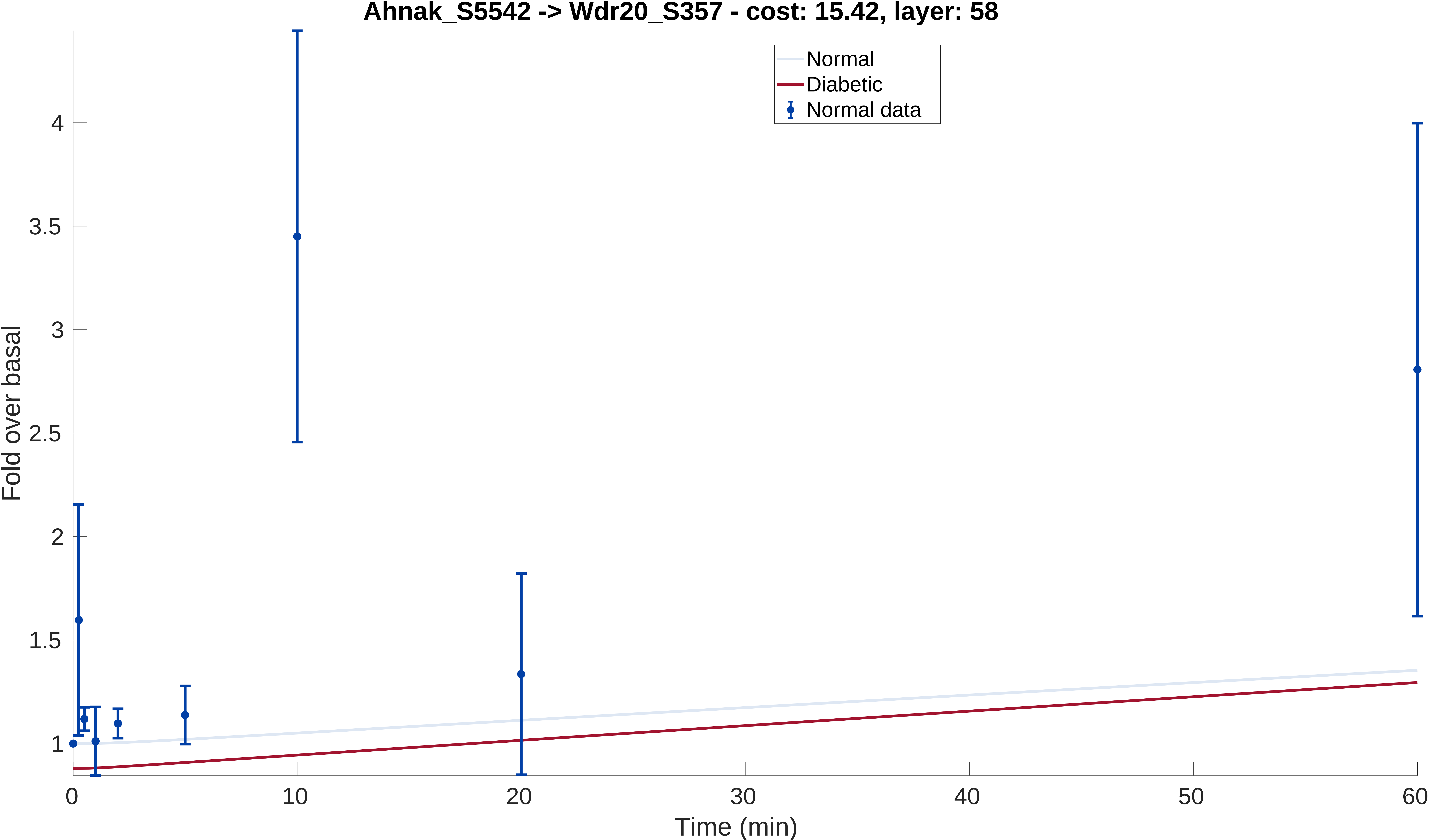

Ahnak_S5542 -> Wdr20_S357 - cost: 15.42, layer: 58
Normal
Diabetic
Normal data
Fold over basal
Time (min)
4
3.5
3
2.5
2
1.5
1
0
10
20
30
40
50
60

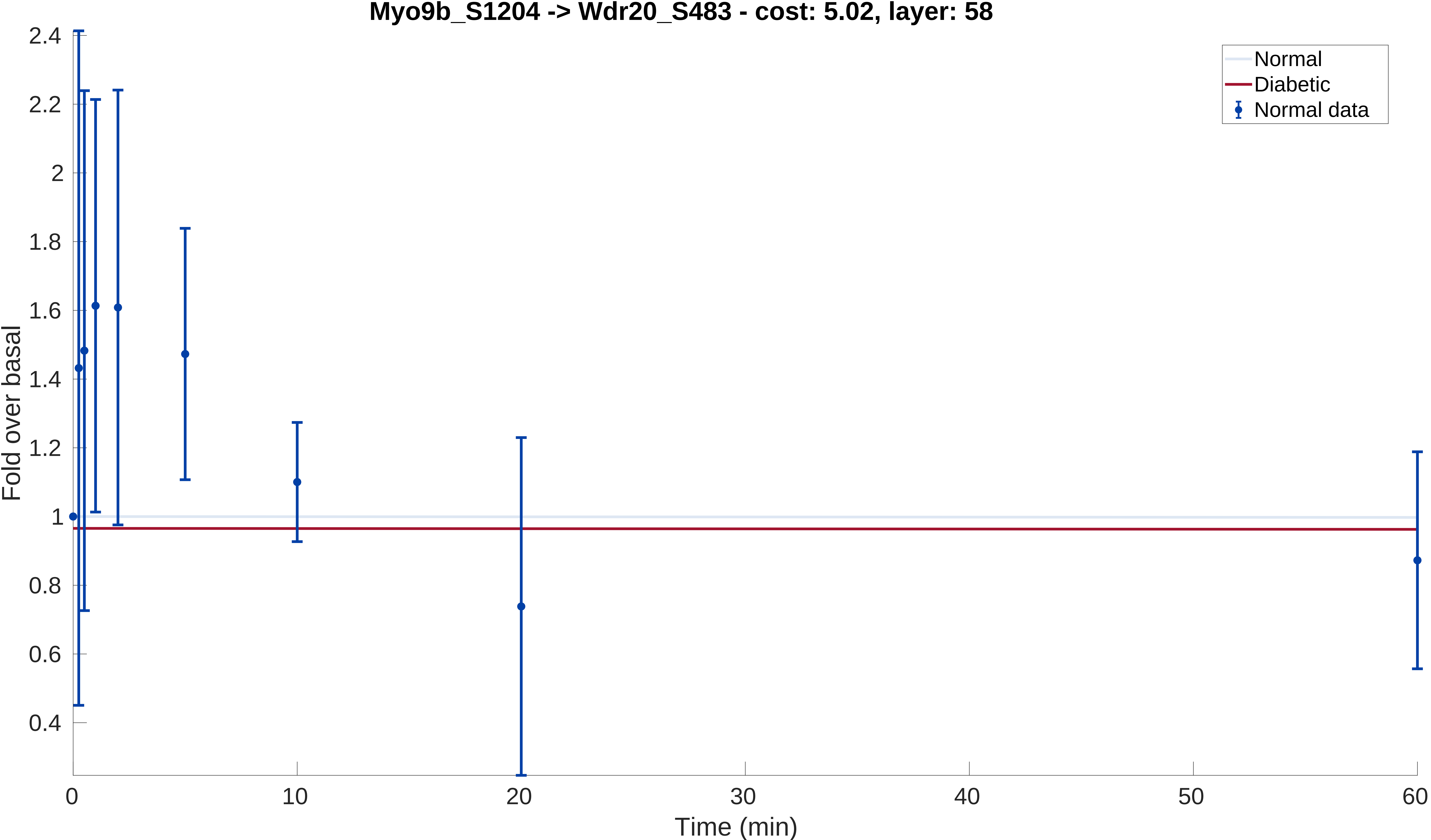

Myo9b_S1204 -> Wdr20_S483 - cost: 5.02, layer: 58
Normal
Diabetic
Normal data
Fold over basal
Time (min)
2.4
2.2
2
1.8
1.6
1.4
1.2
1
0.8
0.6
0.4
0
10
20
30
40
50
60

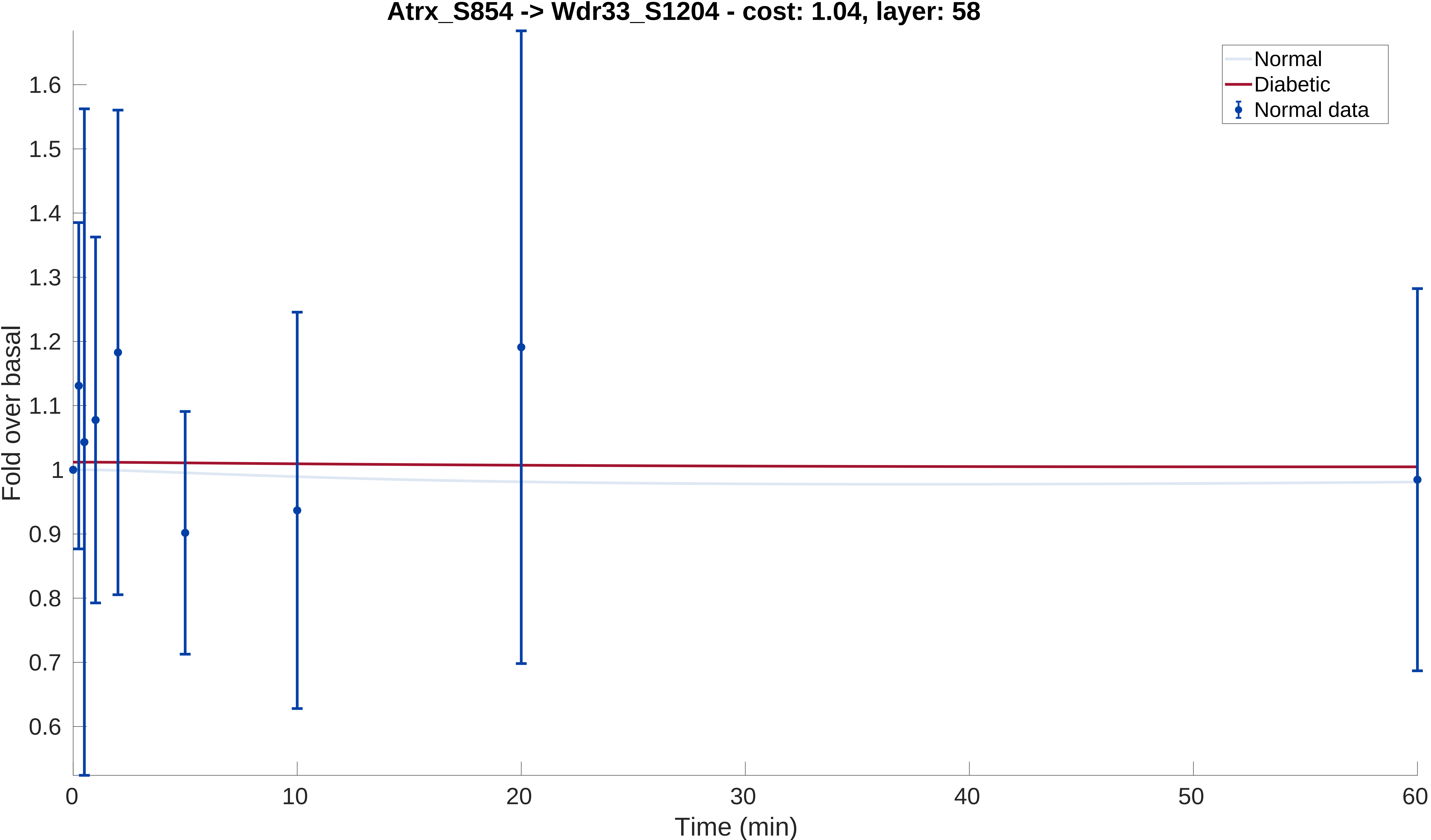

Atrx_S854 -> Wdr33_S1204 - cost: 1.04, layer: 58
Normal
Diabetic
Normal data
Fold over basal
1.6
1.5
1.4
1.3
1.2
1.1
1
0.9
0.8
0.7
0.6
0
10
20
30
40
50
60
Time (min)

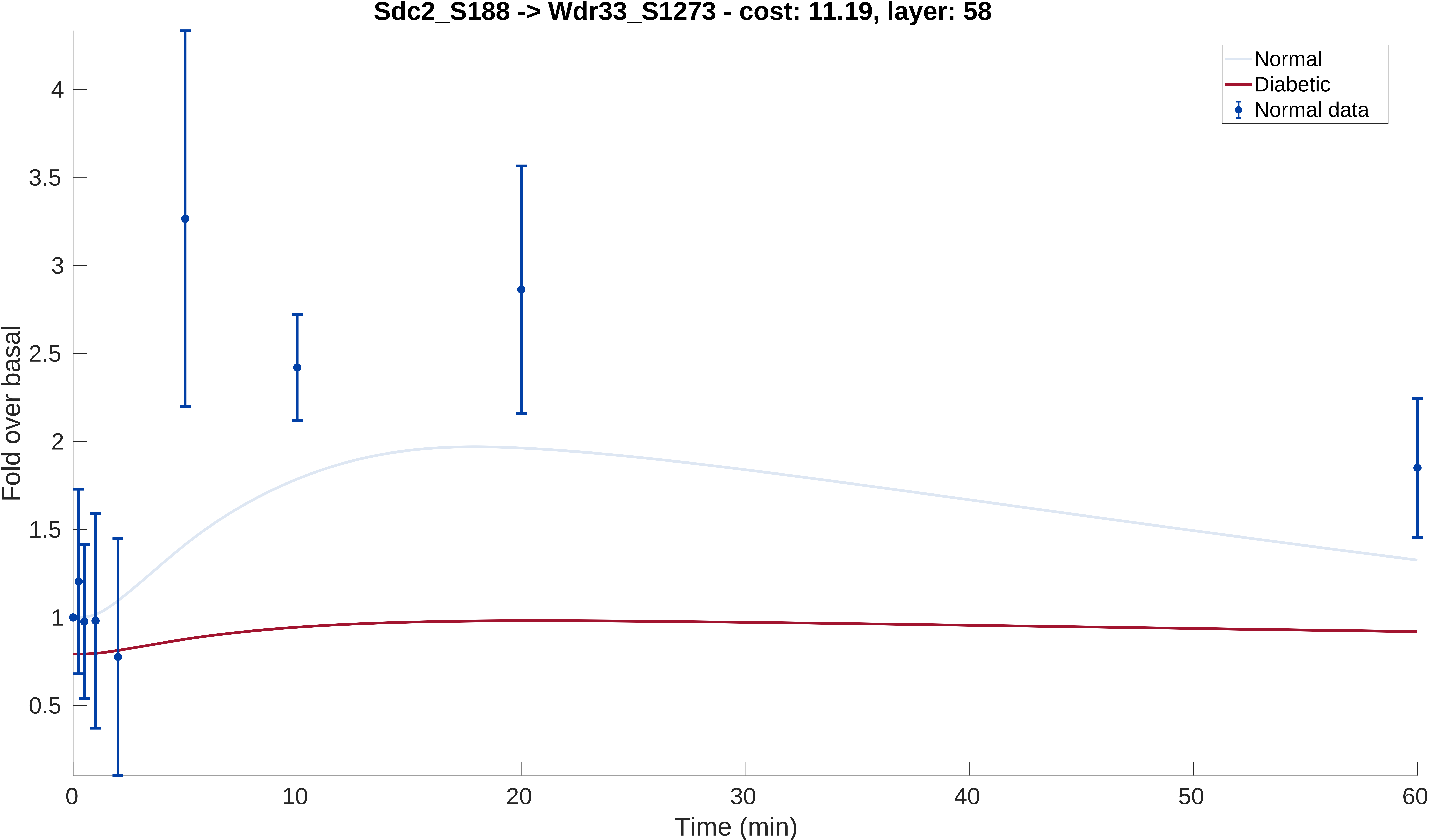

Sdc2_S188 -> Wdr33_S1273 - cost: 11.19, layer: 58
Normal
Diabetic
Normal data
Fold over basal
Time (min)
0.5
1
1.5
2
2.5
3
3.5
4
0
10
20
30
40
50
60

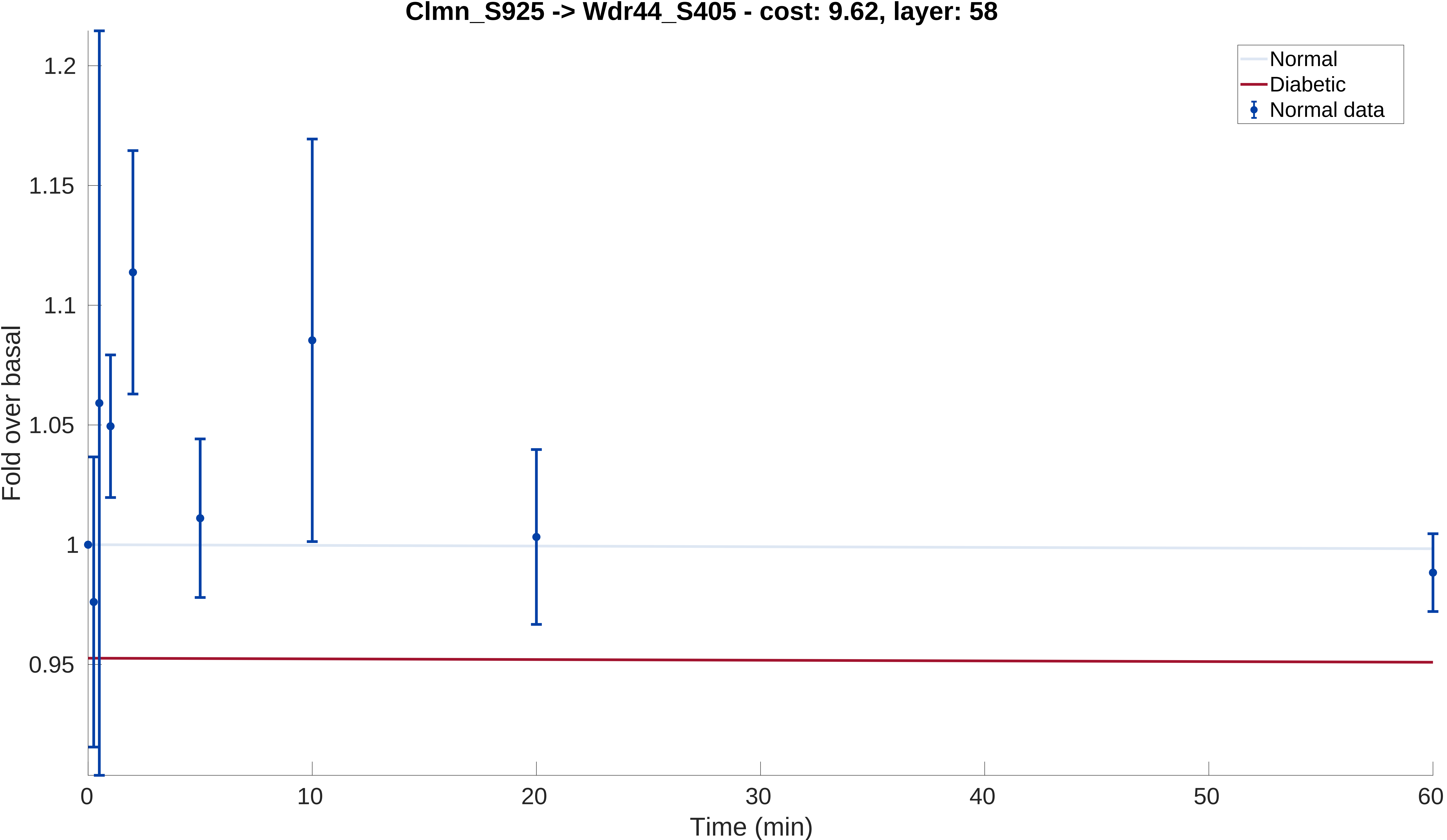
Clmn_S925 -> Wdr44_S405 - cost: 9.62, layer: 58
Normal
Diabetic
Normal data
Fold over basal
1.2
1.15
1.1
1.05
1
0.95
0
10
20
30
40
50
60
Time (min)

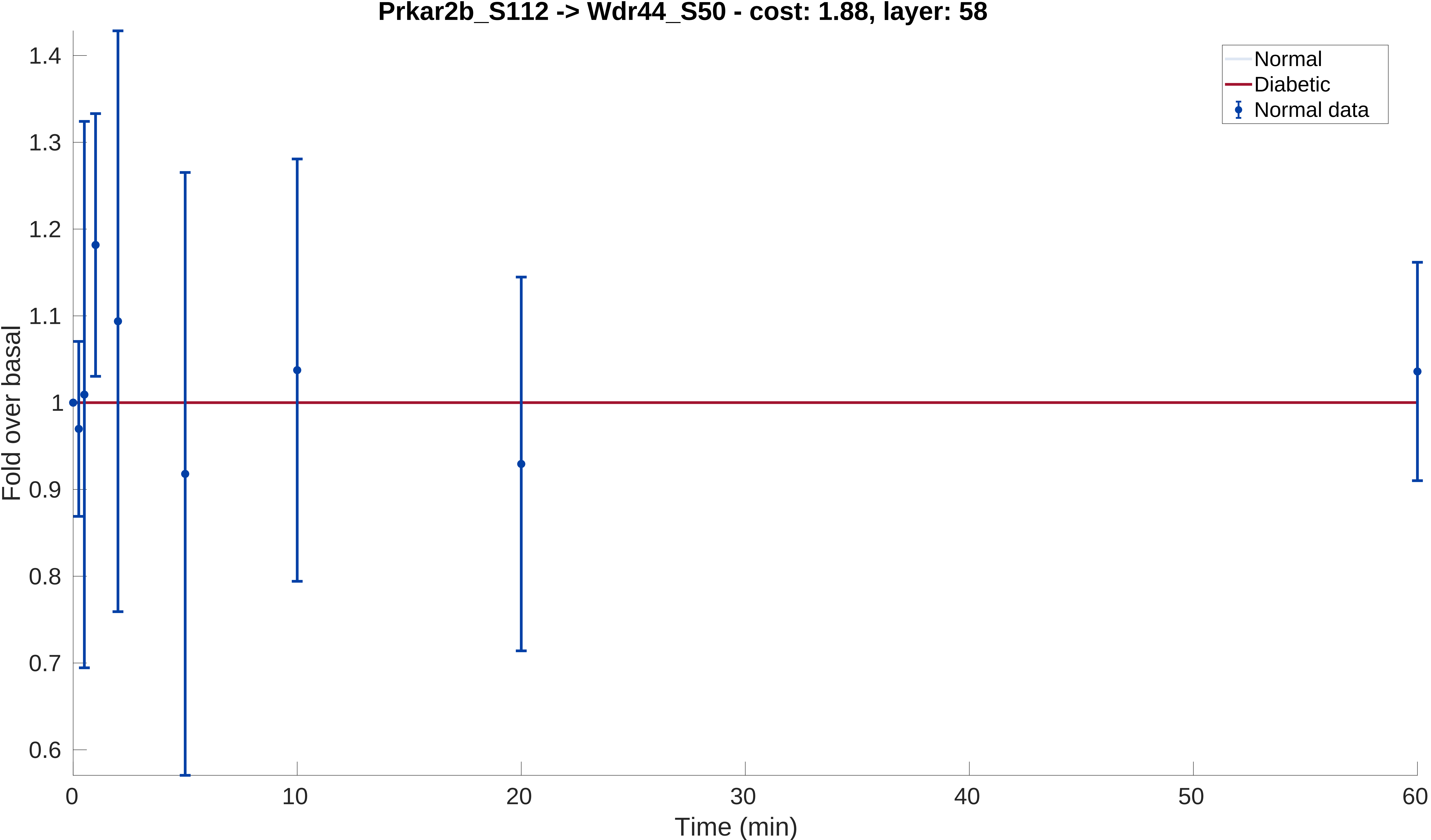

Prkar2b_S112 -> Wdr44_S50 - cost: 1.88, layer: 58
Normal
Diabetic
Normal data
Fold over basal
1.4
1.3
1.2
1.1
1
0.9
0.8
0.7
0.6
0
10
20
30
40
50
60
Time (min)

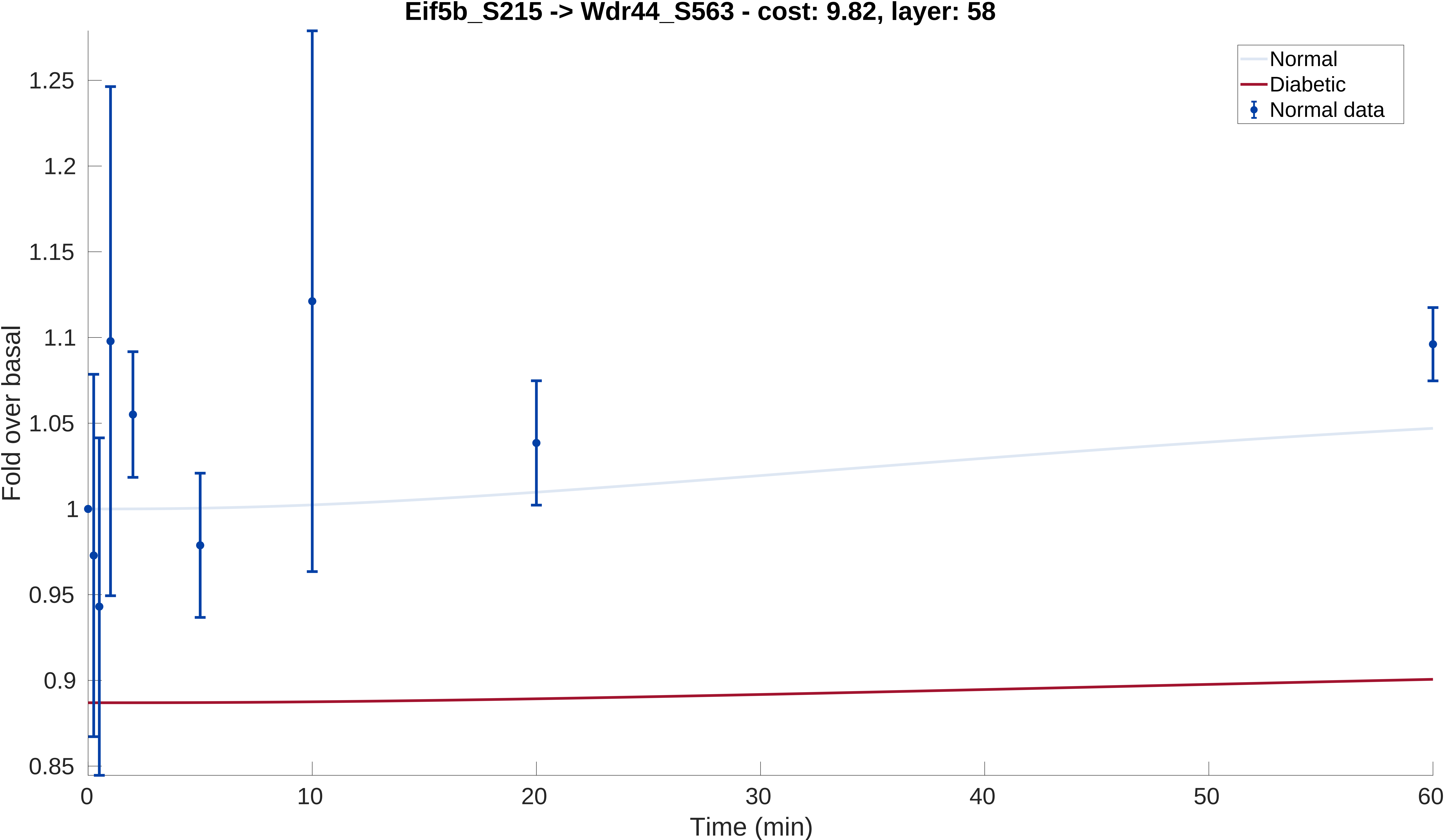

Eif5b_S215 -> Wdr44_S563 - cost: 9.82, layer: 58
Normal
Diabetic
Normal data
Fold over basal
1.25
1.2
1.15
1.1
1.05
1
0.95
0.9
0.85
0
10
20
30
40
50
60
Time (min)

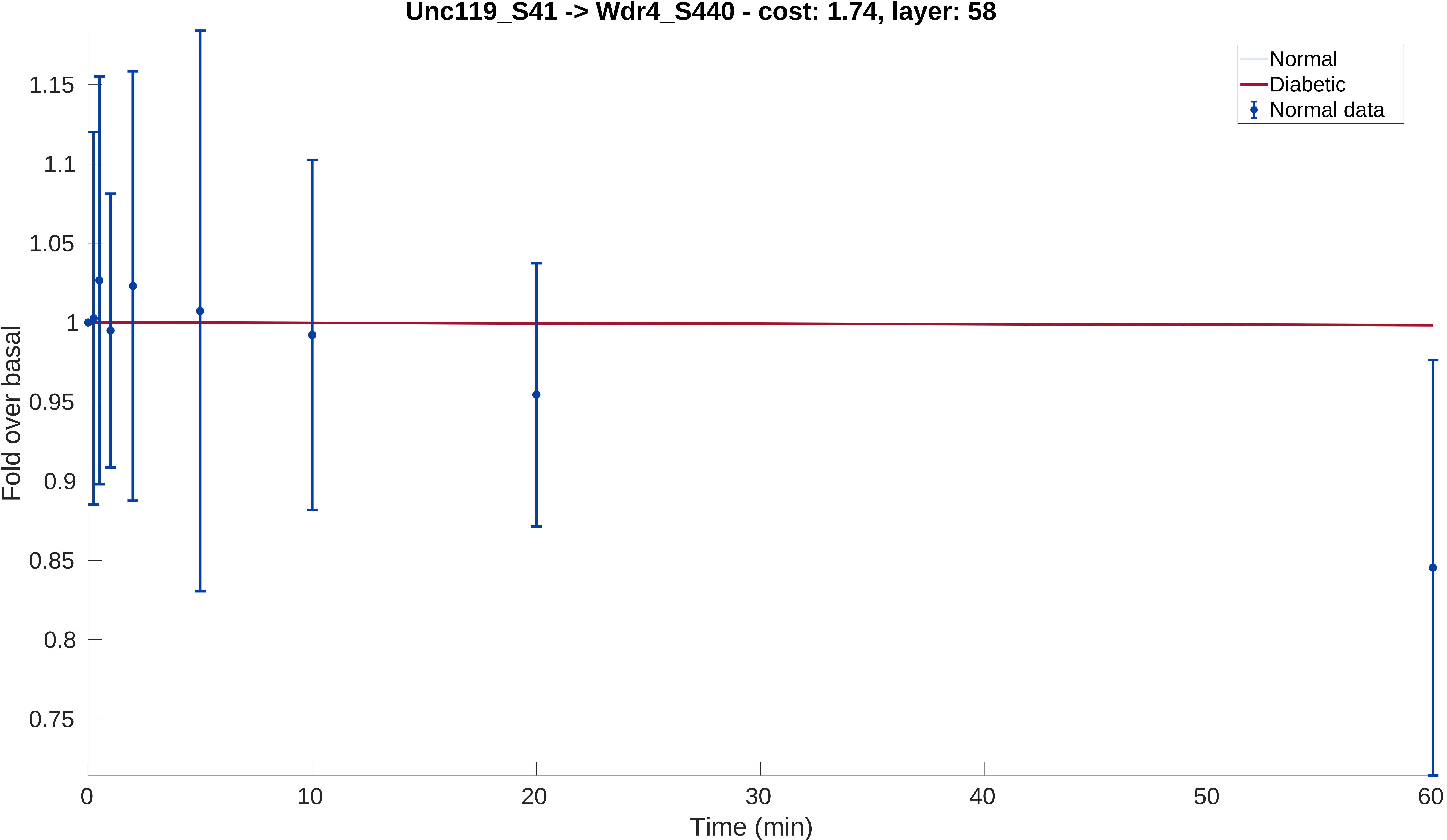
Unc119_S41 -> Wdr4_S440 - cost: 1.74, layer: 58
Normal
Diabetic
Normal data
Fold over basal
1.15
1.1
1.05
1
0.95
0.9
0.85
0.8
0.75
0
10
20
30
40
50
60
Time (min)

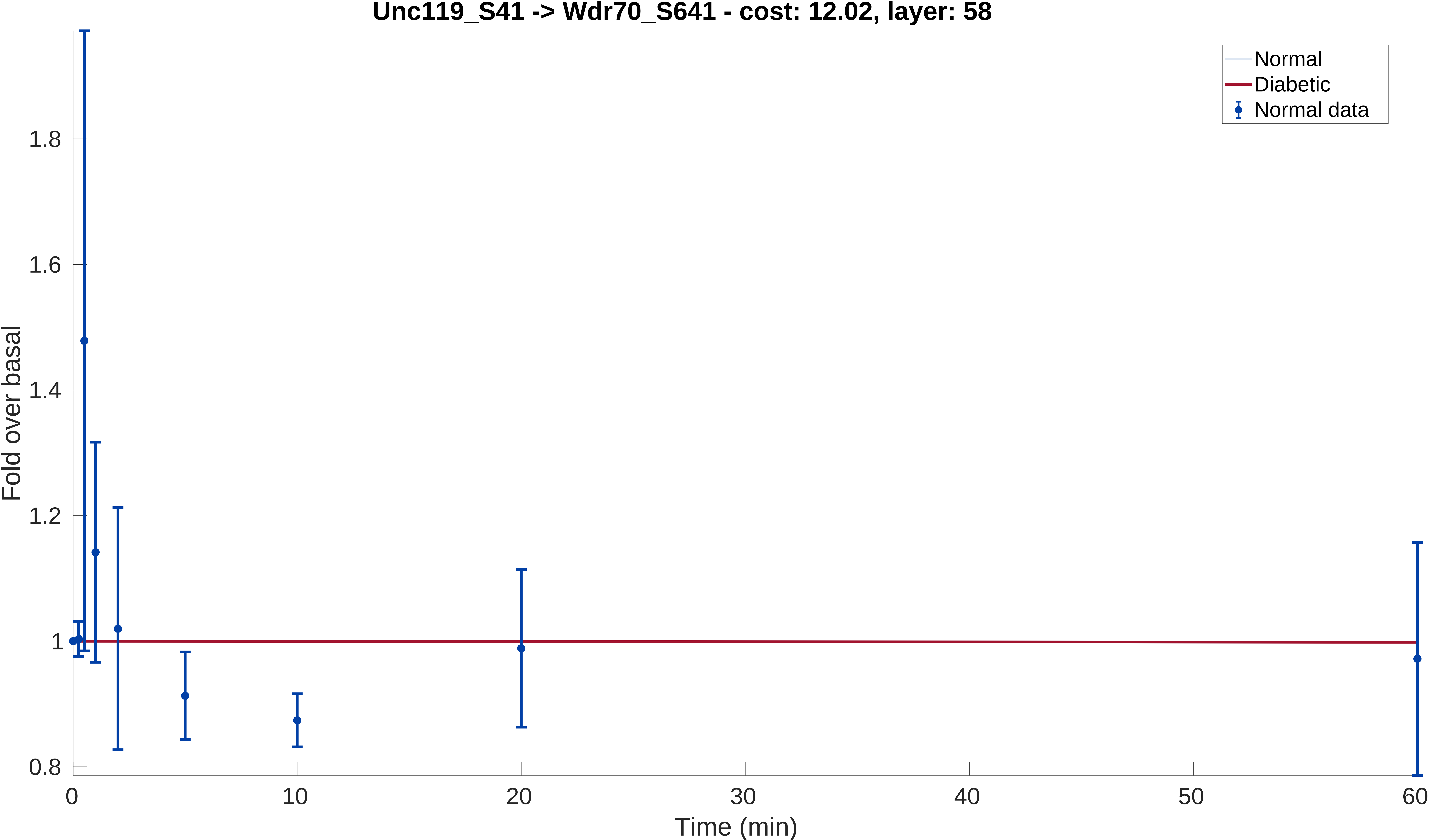
Unc119_S41 -> Wdr70_S641 - cost: 12.02, layer: 58
Normal
Diabetic
Normal data
Fold over basal
1.8
1.6
1.4
1.2
1
0.8
0
10
20
30
40
50
60
Time (min)

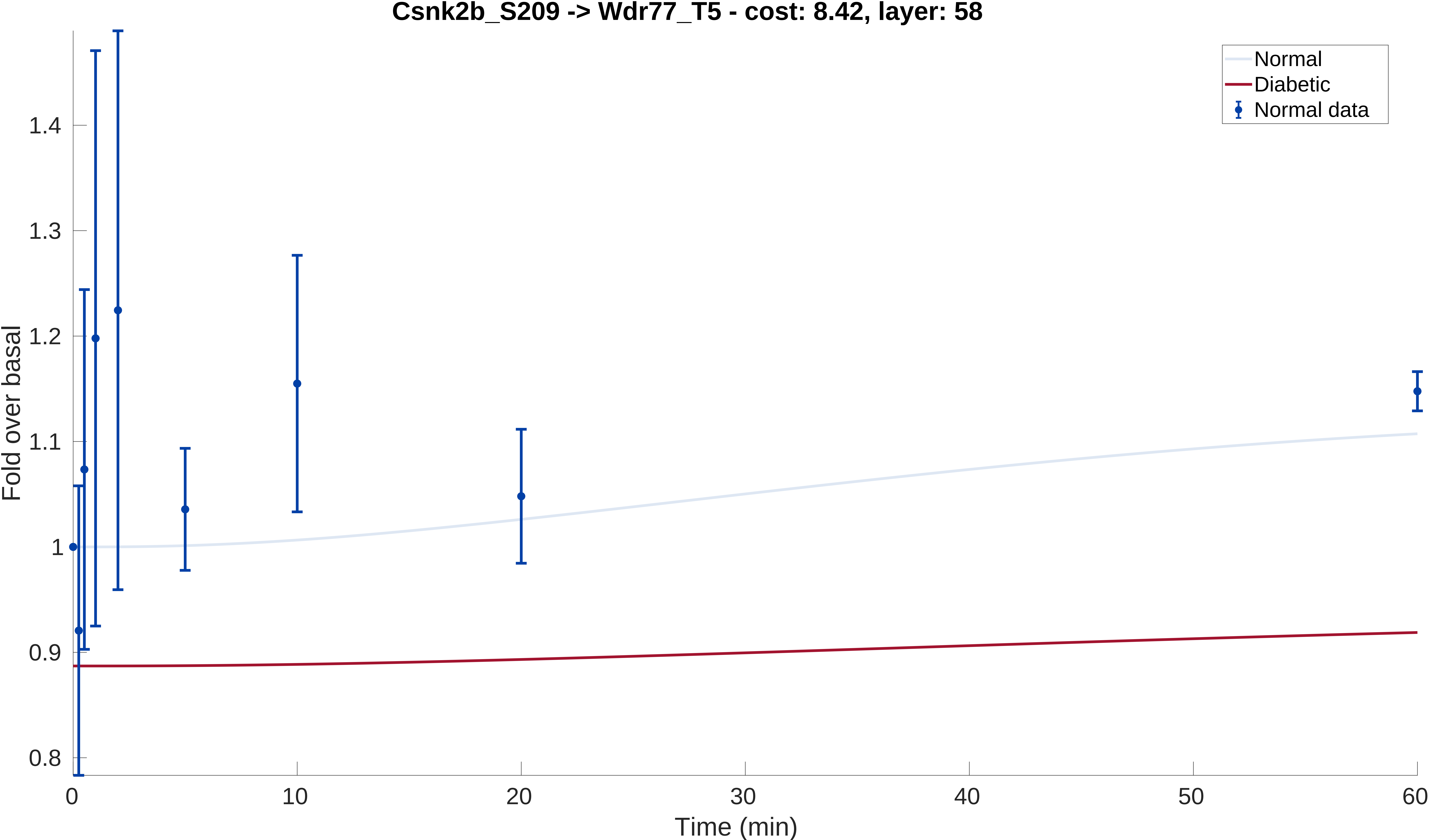
Csnk2b_S209 -> Wdr77_T5 - cost: 8.42, layer: 58
Normal
Diabetic
Normal data
Fold over basal
1.4
1.3
1.2
1.1
1
0.9
0.8
0
10
20
30
40
50
60
Time (min)

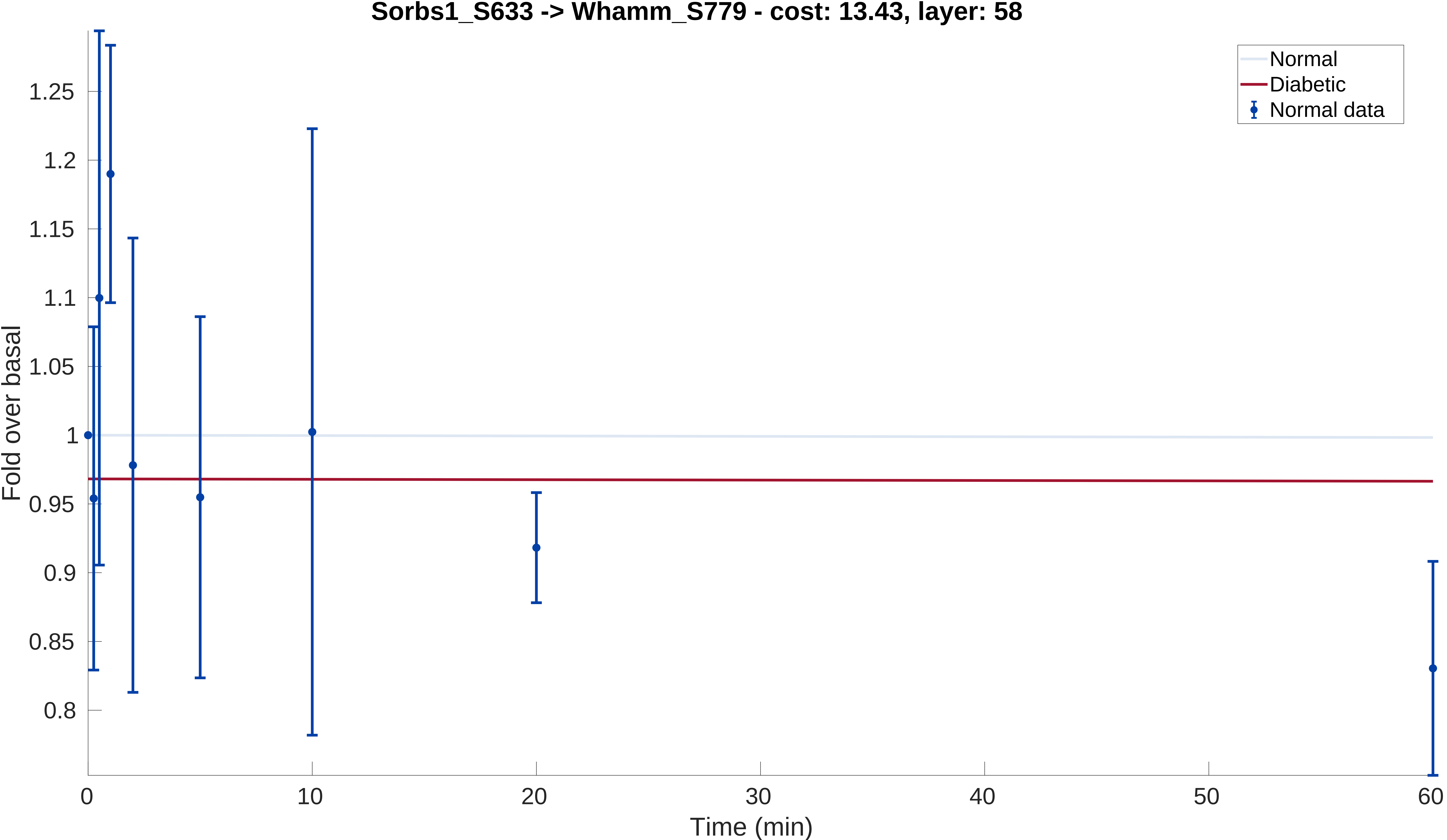

Sorbs1_S633 -> Whamm_S779 - cost: 13.43, layer: 58
Normal
Diabetic
Normal data
Fold over basal
1.25
1.2
1.15
1.1
1.05
1
0.95
0.9
0.85
0.8
0
10
20
30
40
50
60
Time (min)

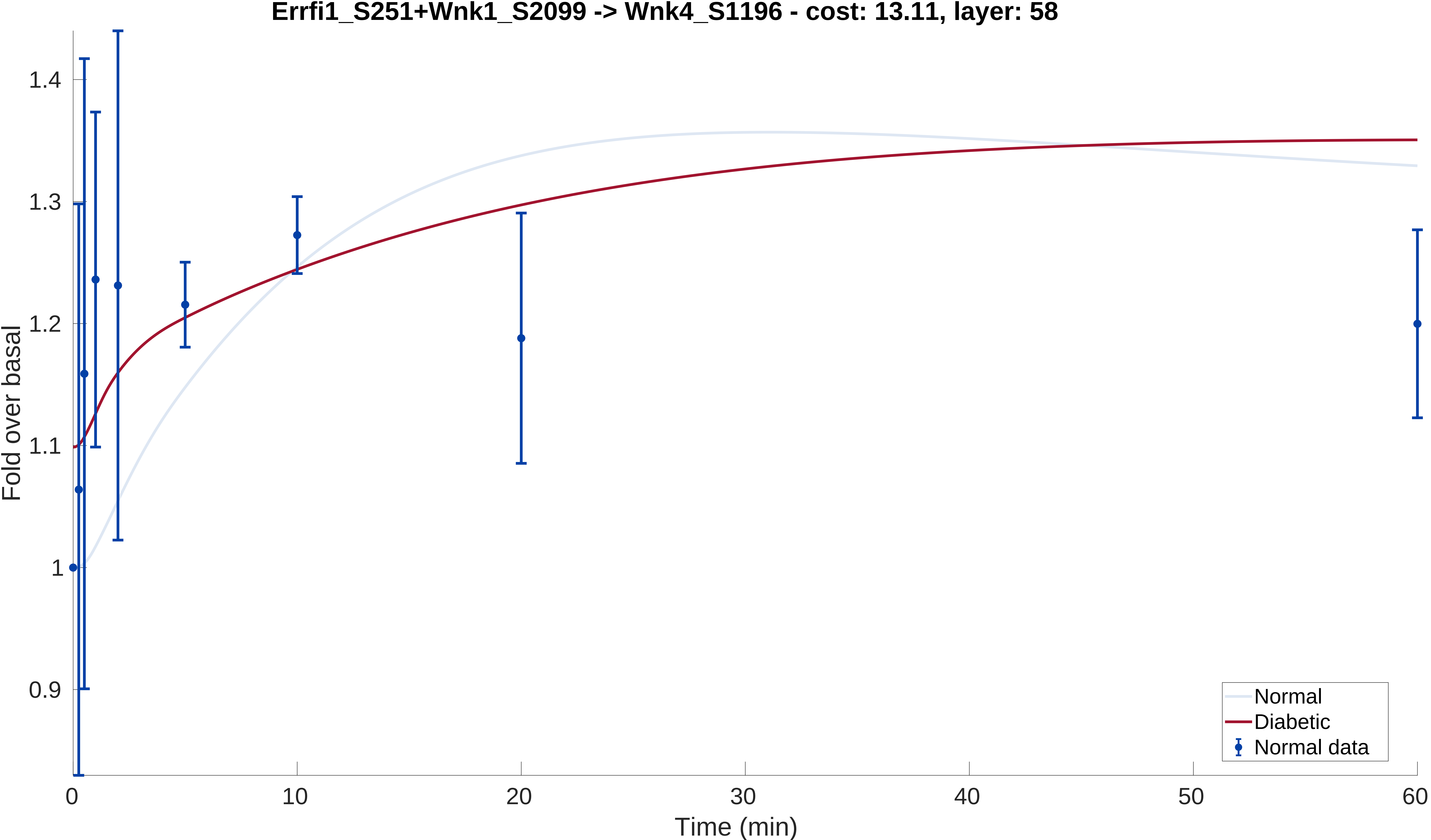

Errfi1_S251+Wnk1_S2099 -> Wnk4_S1196 - cost: 13.11, layer: 58
Fold over basal
1.4
1.3
1.2
1.1
1
0.9
0
10
20
30
40
50
60
Time (min)
Normal
Diabetic
Normal data

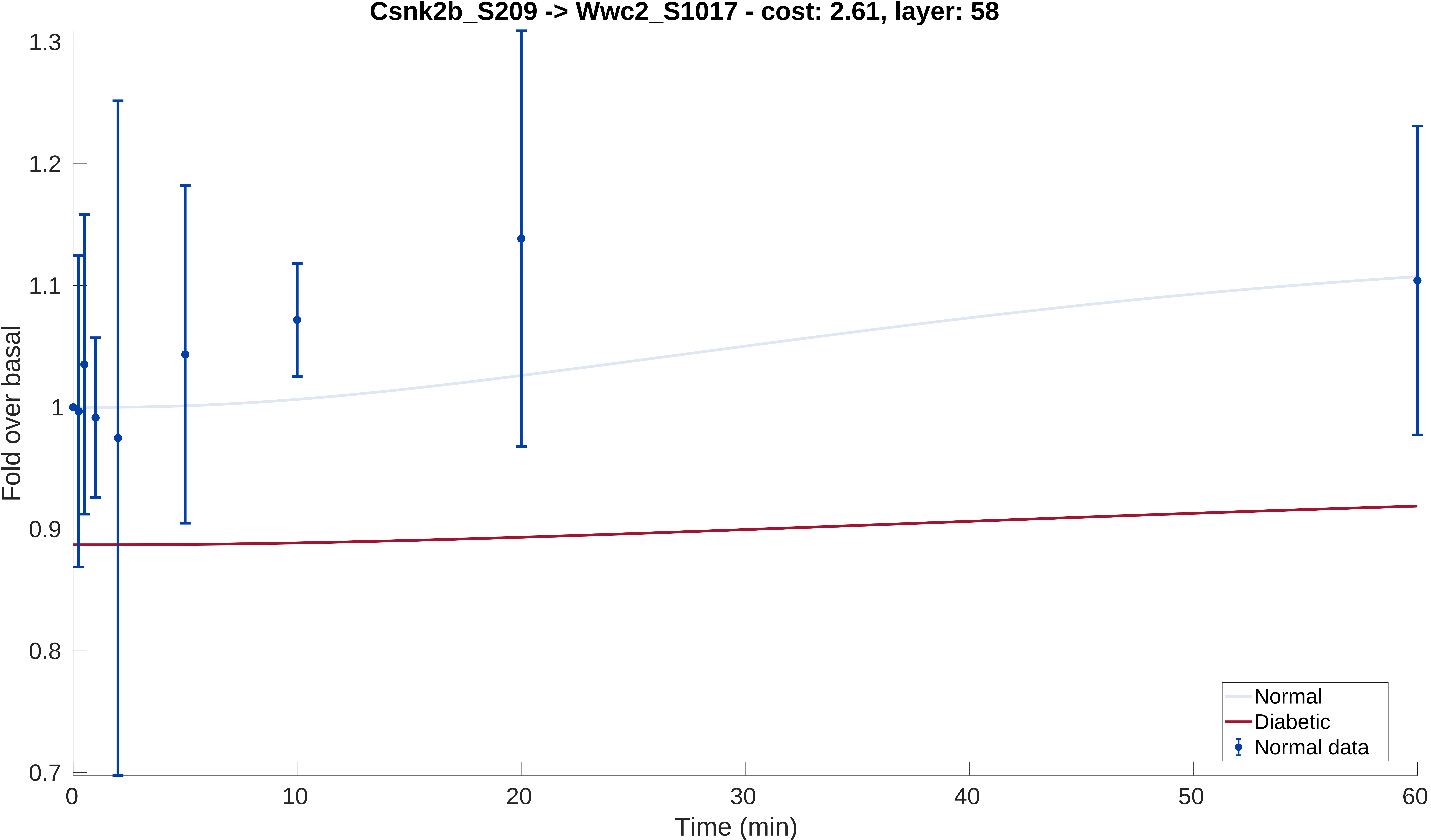
Csnk2b_S209 -> Wwc2_S1017 - cost: 2.61, layer: 58
Fold over basal
1.3
1.2
1.1
1
0.9
0.8
0.7
0
10
20
30
40
50
60
Time (min)
Normal
Diabetic
Normal data

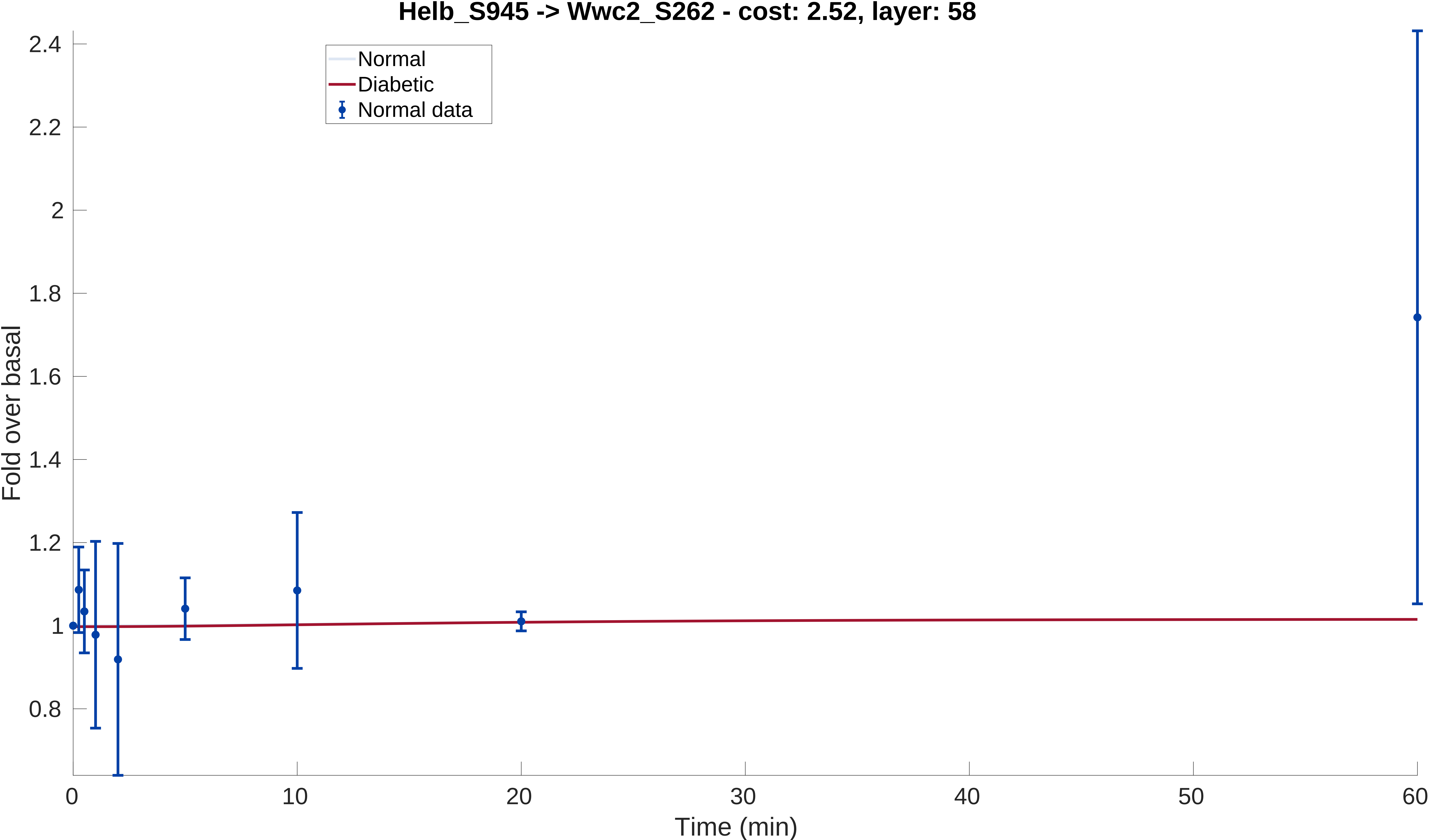

Helb_S945 -> Wwc2_S262 - cost: 2.52, layer: 58
Normal
Diabetic
Normal data
Fold over basal
Time (min)
0
10
20
30
40
50
60
0.8
1
1.2
1.4
1.6
1.8
2
2.2
2.4

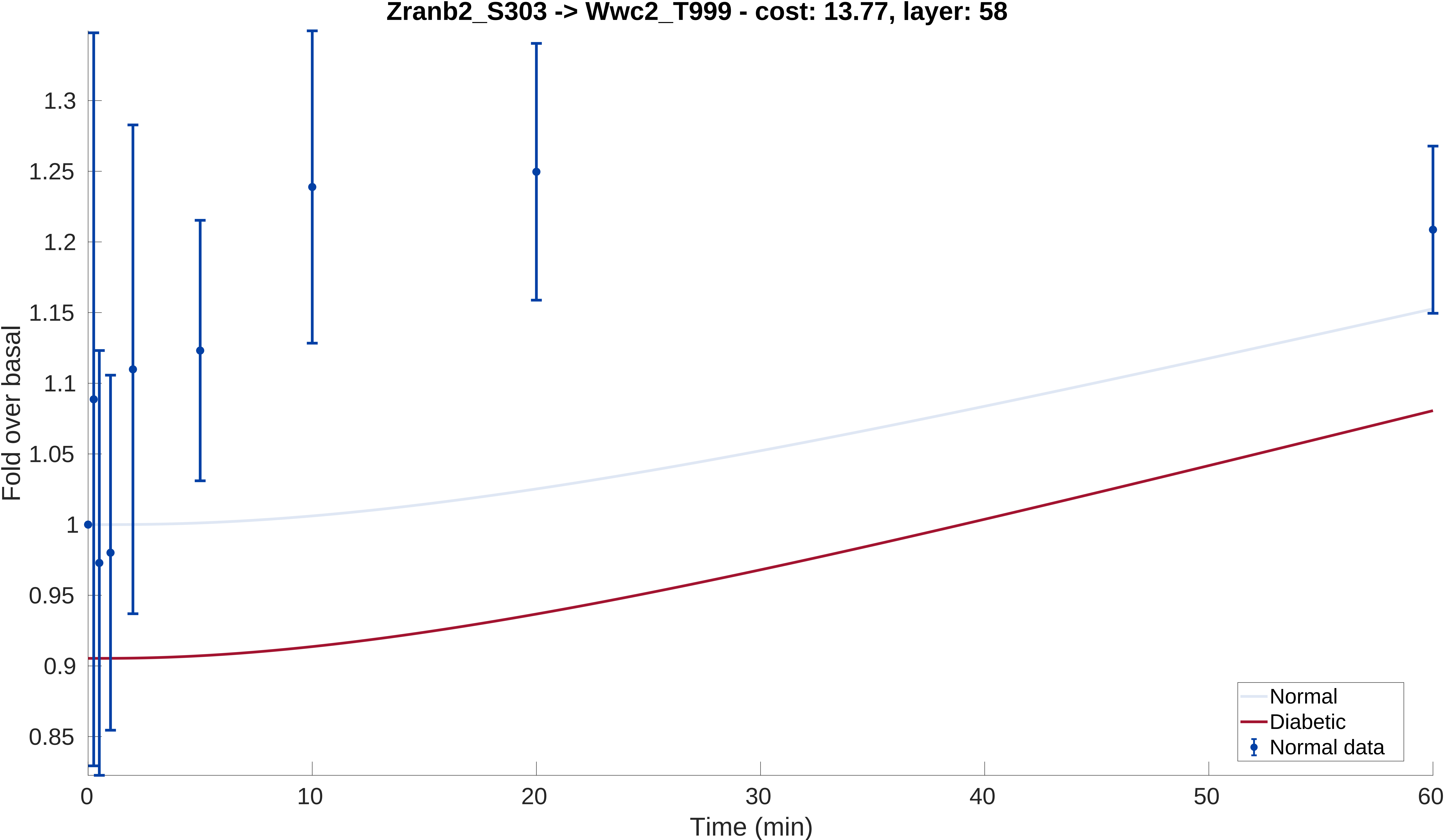

Zranb2_S303 -> Wwc2_T999 - cost: 13.77, layer: 58
Fold over basal
Time (min)
1.3
1.25
1.2
1.15
1.1
1.05
1
0.95
0.9
0.85
0
10
20
30
40
50
60
Normal
Diabetic
Normal data

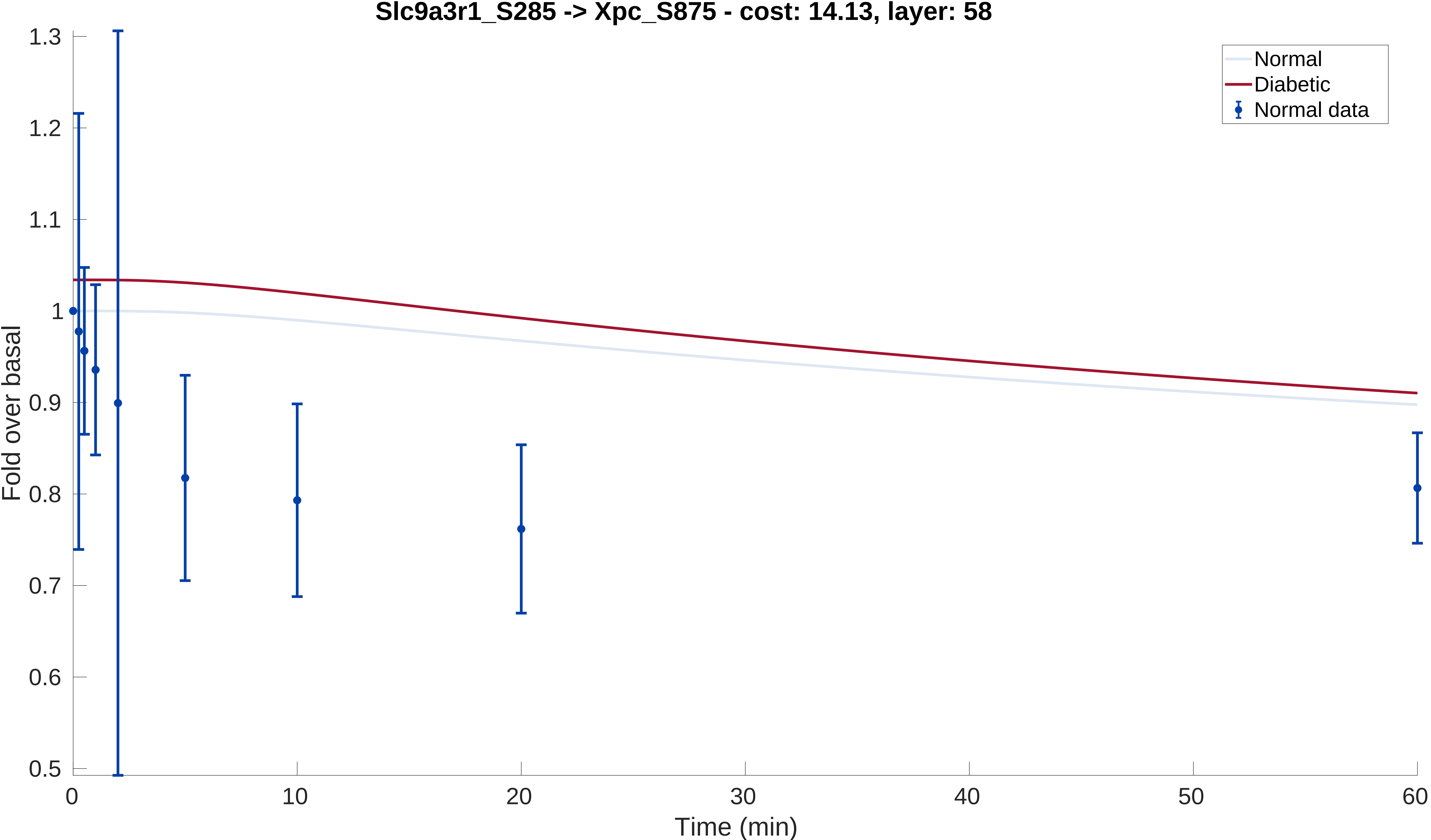
Slc9a3r1_S285 -> Xpc_S875 - cost: 14.13, layer: 58
Normal
Diabetic
Normal data
Fold over basal
Time (min)
1.3
1.2
1.1
1
0.9
0.8
0.7
0.6
0.5
0
10
20
30
40
50
60

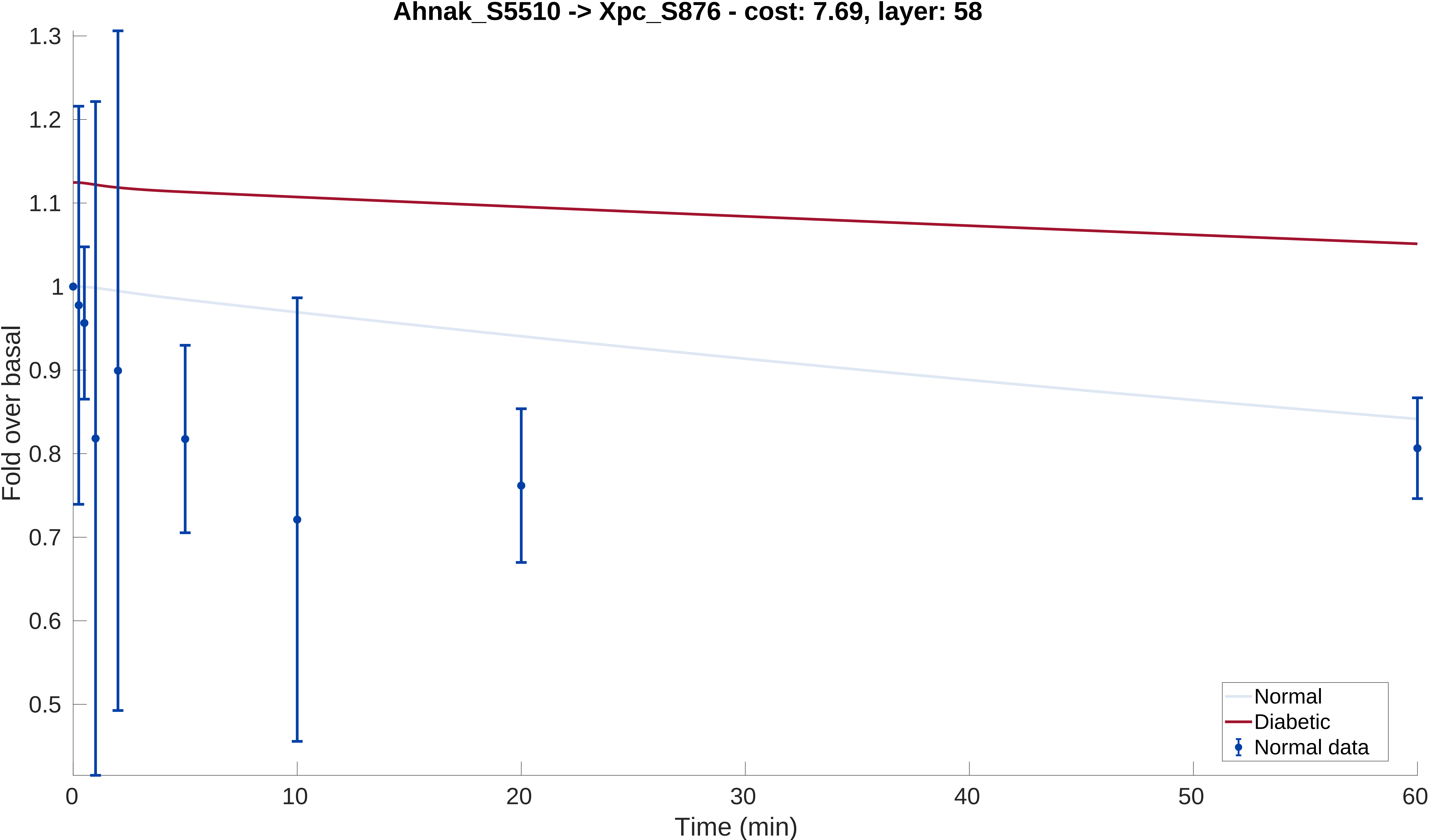
Ahnak_S5510 -> Xpc_S876 - cost: 7.69, layer: 58
Fold over basal
Time (min)
0
10
20
30
40
50
60
0.5
0.6
0.7
0.8
0.9
1
1.1
1.2
1.3
Normal
Diabetic
Normal data

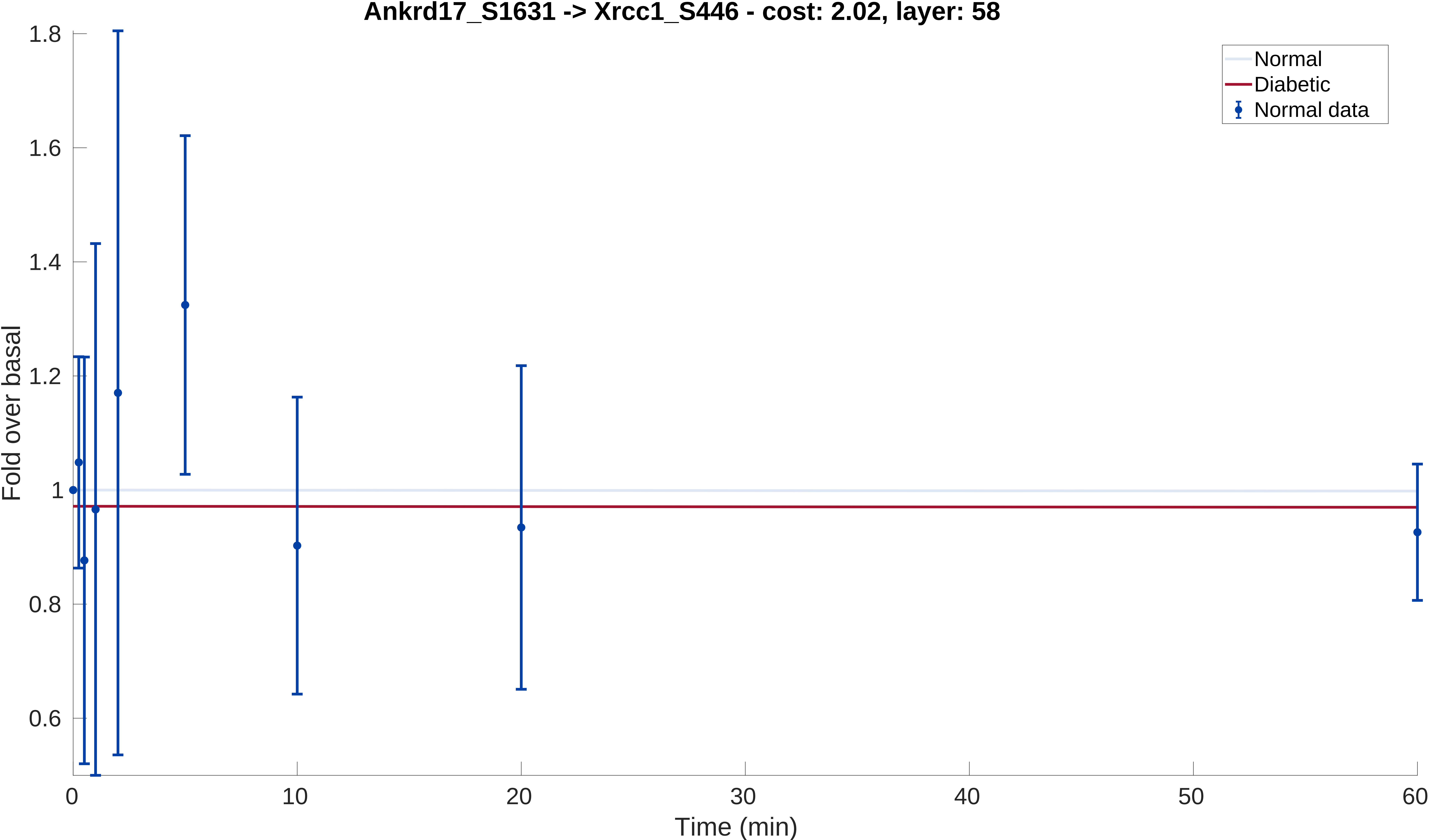

Ankrd17_S1631 -> Xrcc1_S446 - cost: 2.02, layer: 58
Normal
Diabetic
Normal data
Fold over basal
1.8
1.6
1.4
1.2
1
0.8
0.6
0
10
20
30
40
50
60
Time (min)

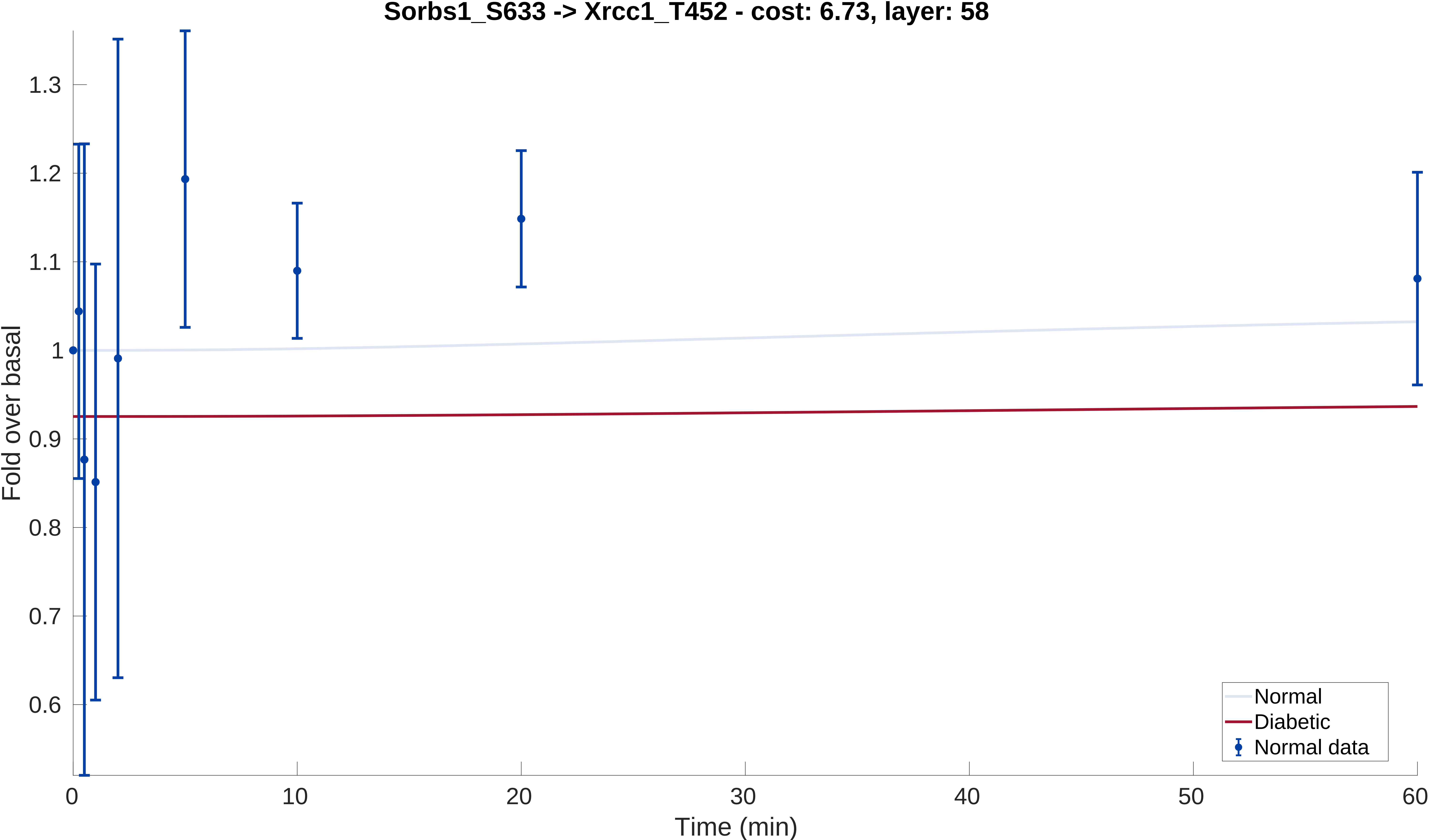

Sorbs1_S633 -> Xrcc1_T452 - cost: 6.73, layer: 58
Fold over basal
1.3
1.2
1.1
1
0.9
0.8
0.7
0.6
0
10
20
30
40
50
60
Time (min)
Normal
Diabetic
Normal data

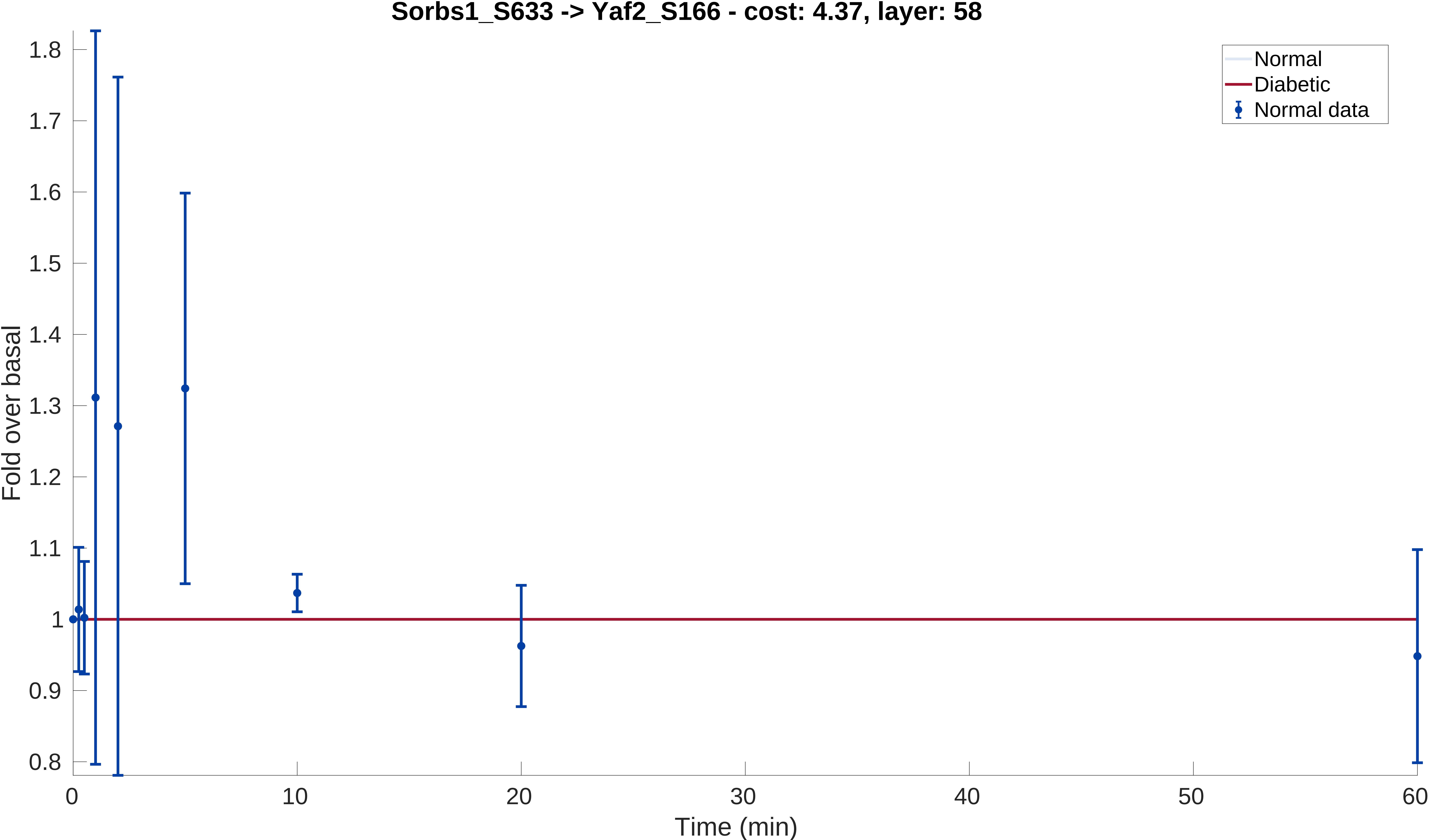

Sorbs1_S633 -> Yaf2_S166 - cost: 4.37, layer: 58
Normal
Diabetic
Normal data
Fold over basal
1.8
1.7
1.6
1.5
1.4
1.3
1.2
1.1
1
0.9
0.8
0
10
20
30
40
50
60
Time (min)

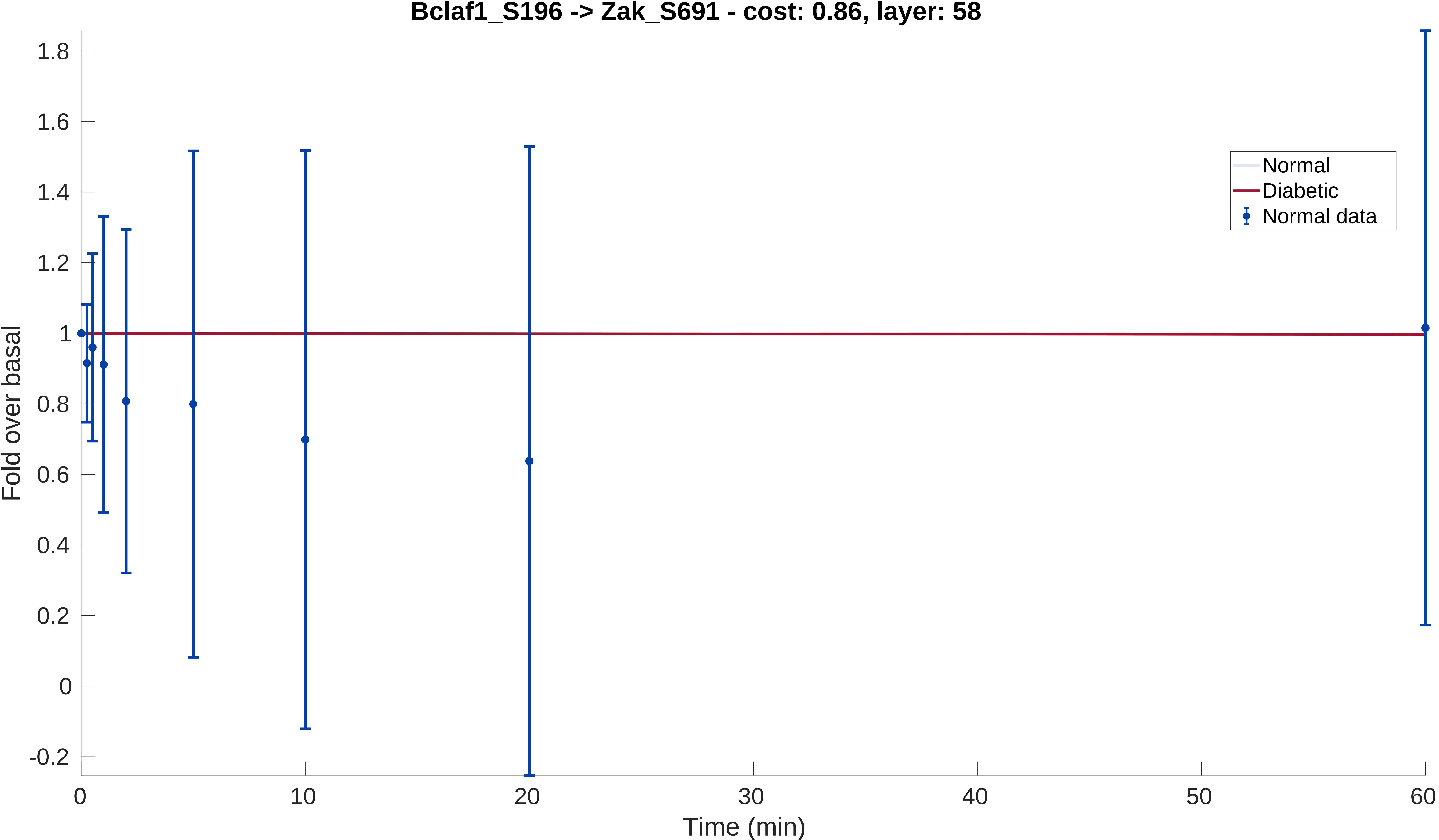

Bclaf1_S196 -> Zak_S691 - cost: 0.86, layer: 58
Fold over basal
Time (min)
Normal
Diabetic
Normal data
1.8
1.6
1.4
1.2
1
0.8
0.6
0.4
0.2
0
-0.2
0
10
20
30
40
50
60

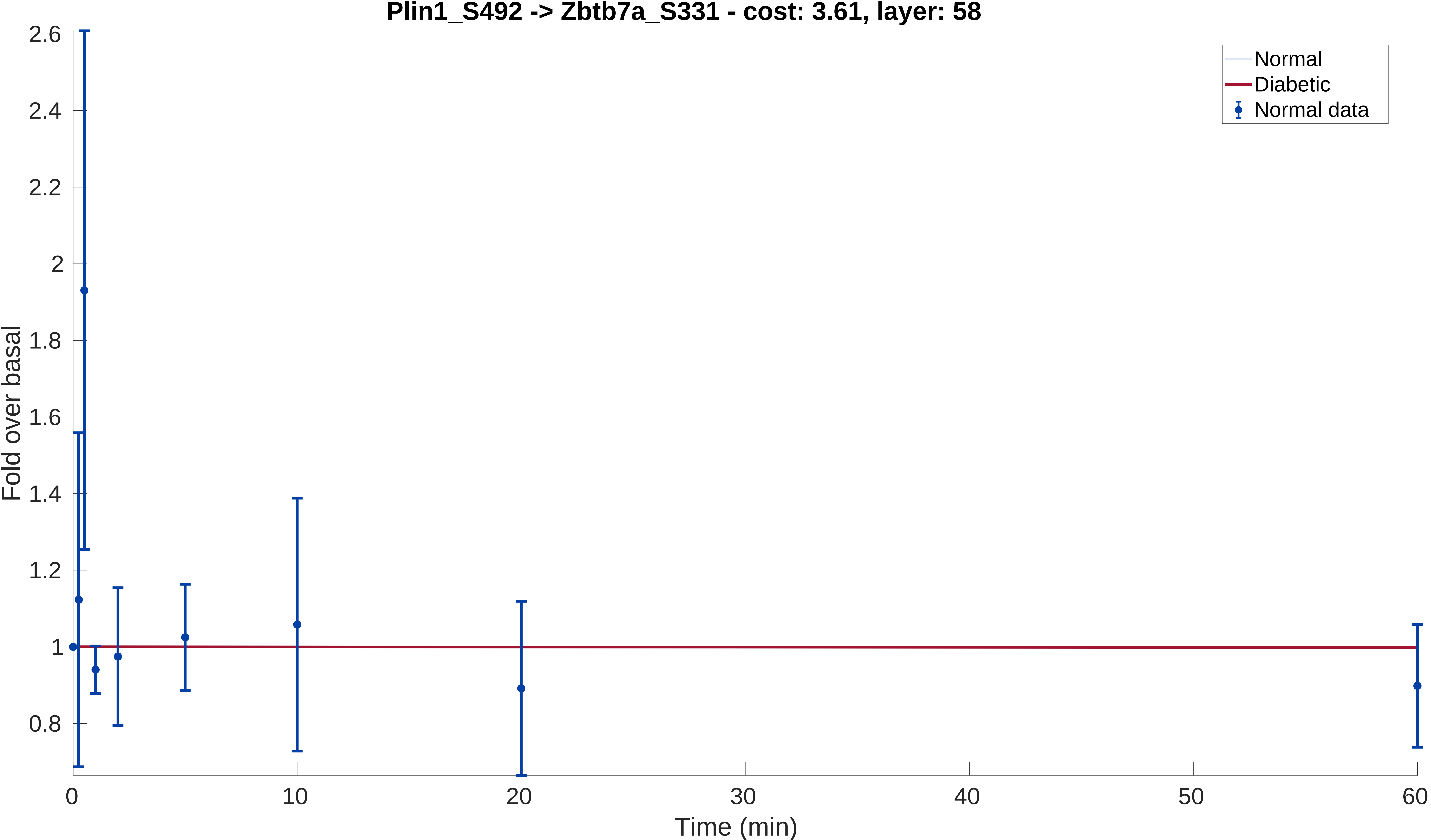

Plin1_S492 -> Zbtb7a_S331 - cost: 3.61, layer: 58
Normal
Diabetic
Normal data
Fold over basal
Time (min)
0.8
1
1.2
1.4
1.6
1.8
2
2.2
2.4
2.6
0
10
20
30
40
50
60

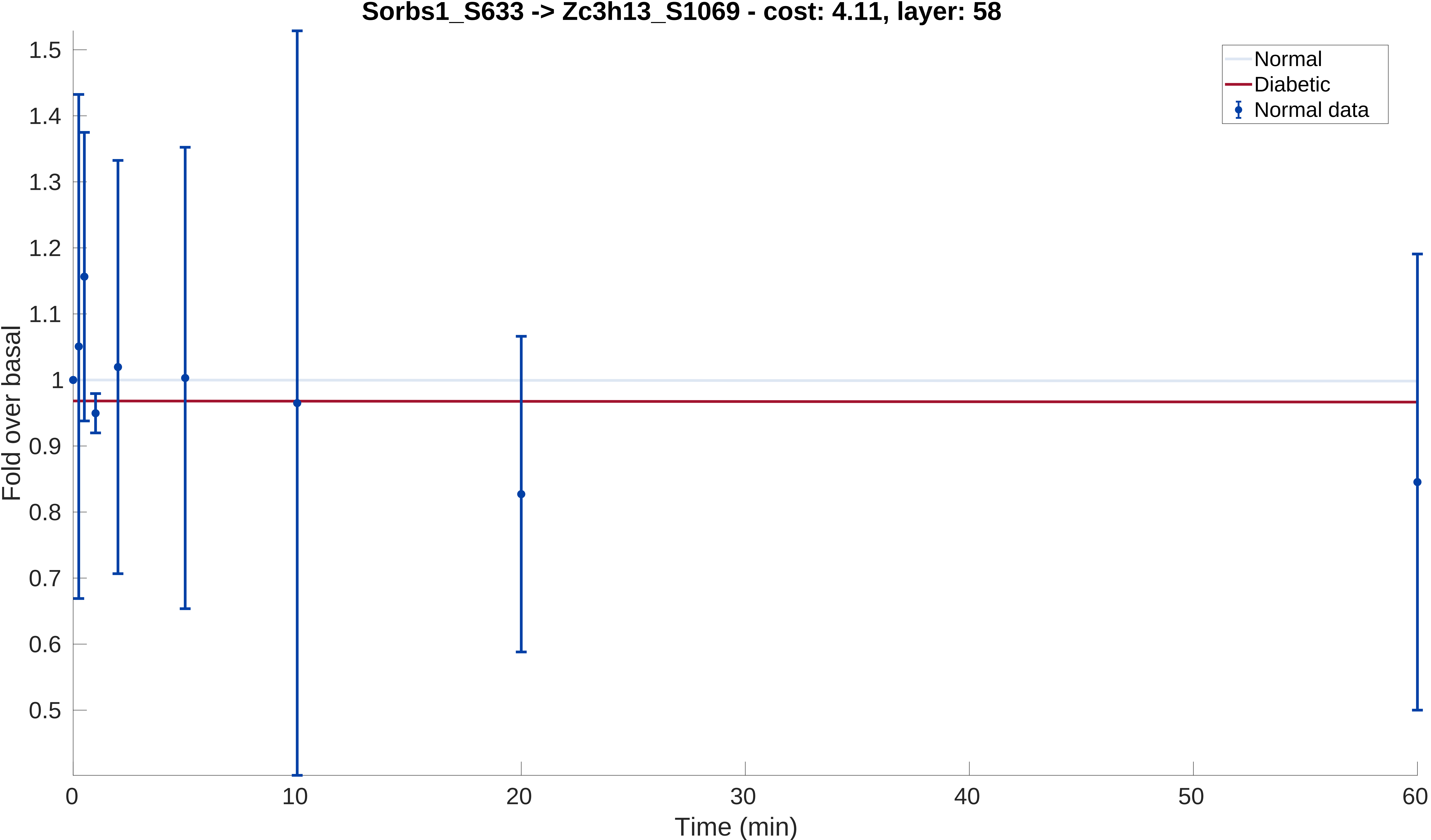
Sorbs1_S633 -> Zc3h13_S1069 - cost: 4.11, layer: 58
Normal
Diabetic
Normal data
Fold over basal
Time (min)
1.5
1.4
1.3
1.2
1.1
1
0.9
0.8
0.7
0.6
0.5
0
10
20
30
40
50
60

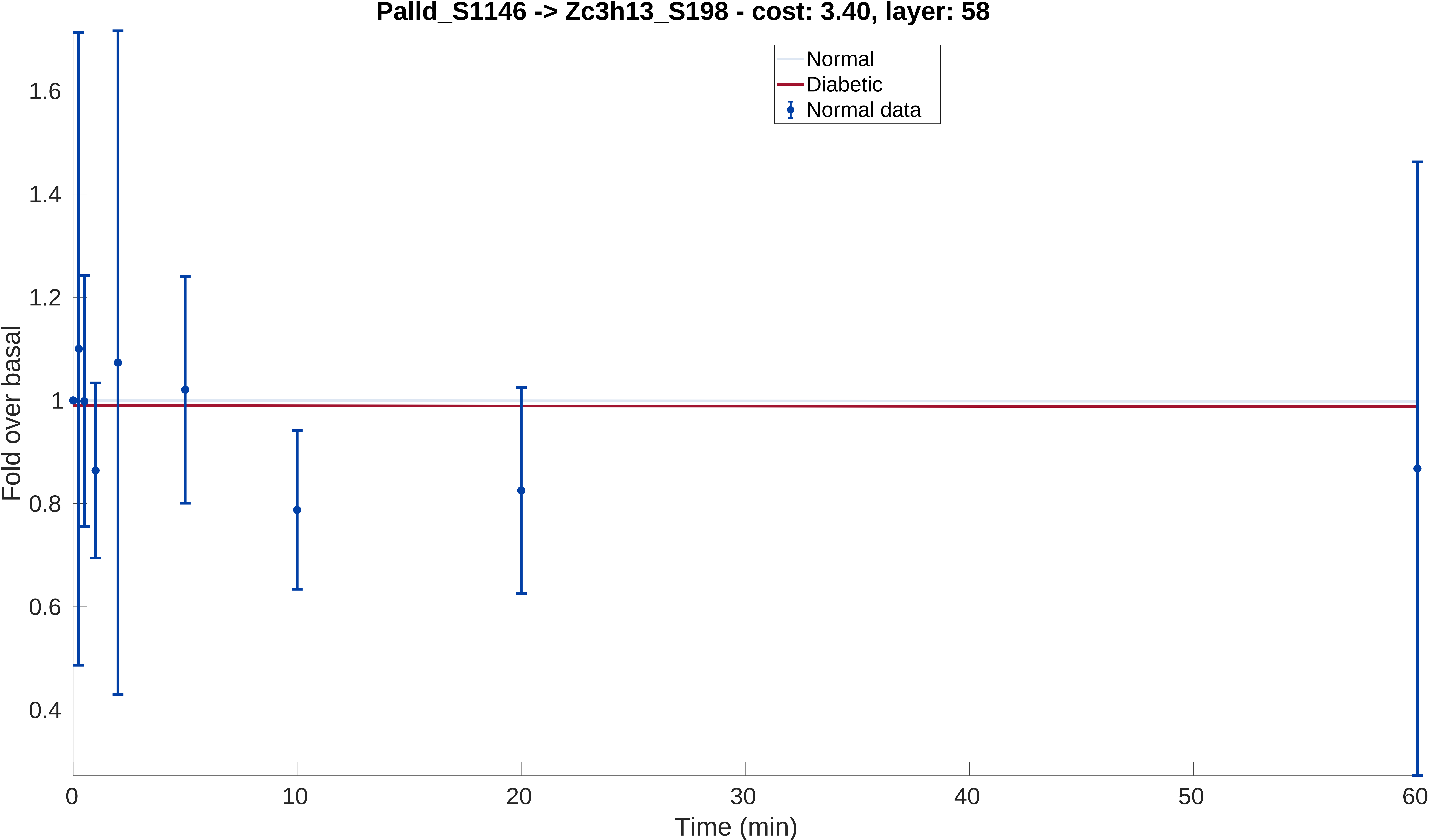

Palld_S1146 -> Zc3h13_S198 - cost: 3.40, layer: 58
Normal
Diabetic
Normal data
Fold over basal
1.6
1.4
1.2
1
0.8
0.6
0.4
0
10
20
30
40
50
60
Time (min)

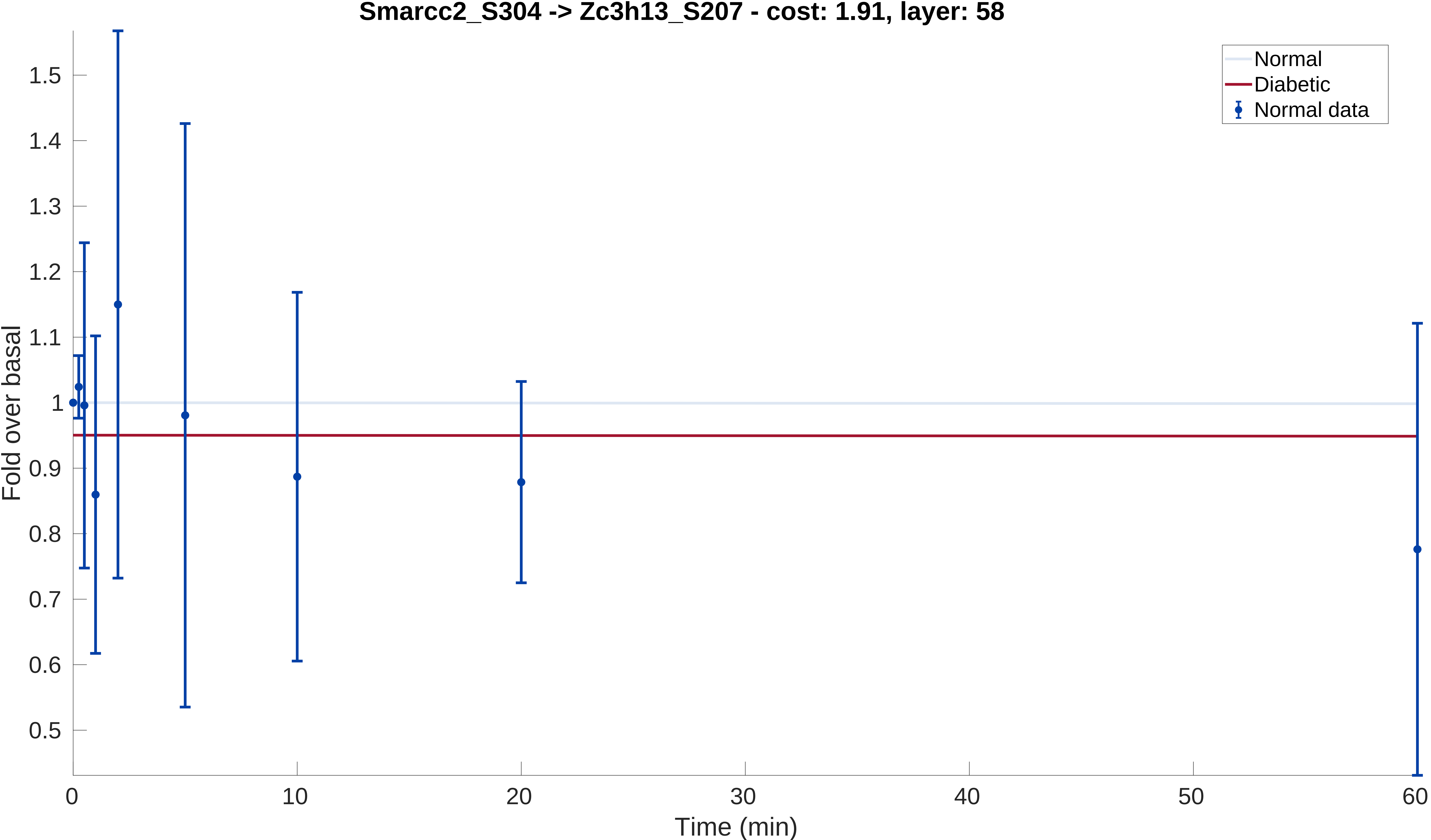

Smarcc2_S304 -> Zc3h13_S207 - cost: 1.91, layer: 58
Normal
Diabetic
Normal data
Fold over basal
1.5
1.4
1.3
1.2
1.1
1
0.9
0.8
0.7
0.6
0.5
0
10
20
30
40
50
60
Time (min)

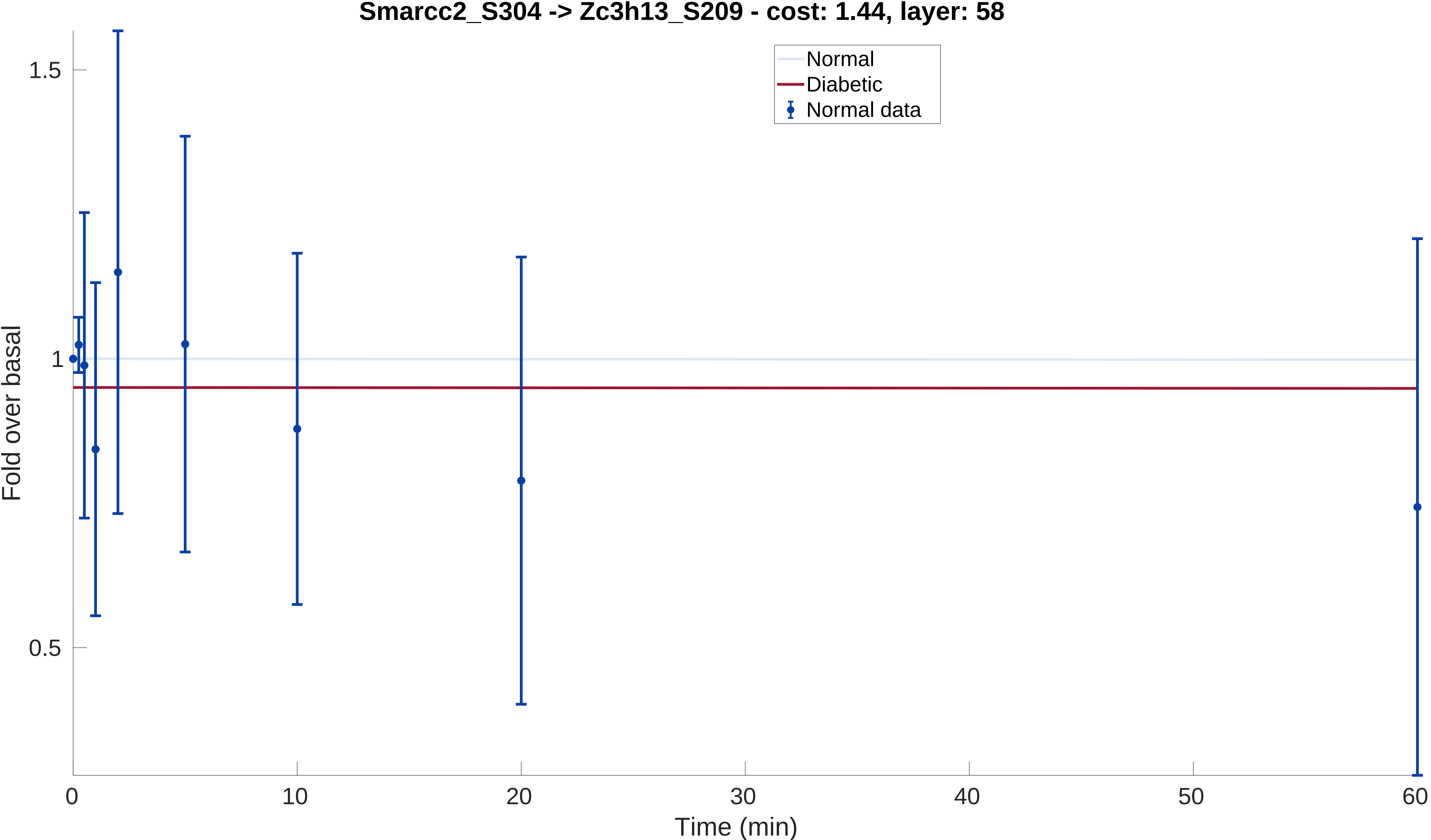

Smarcc2_S304 -> Zc3h13_S209 - cost: 1.44, layer: 58
Normal
Diabetic
Normal data
1.5
1
0.5
Fold over basal
0
10
20
30
40
50
60
Time (min)

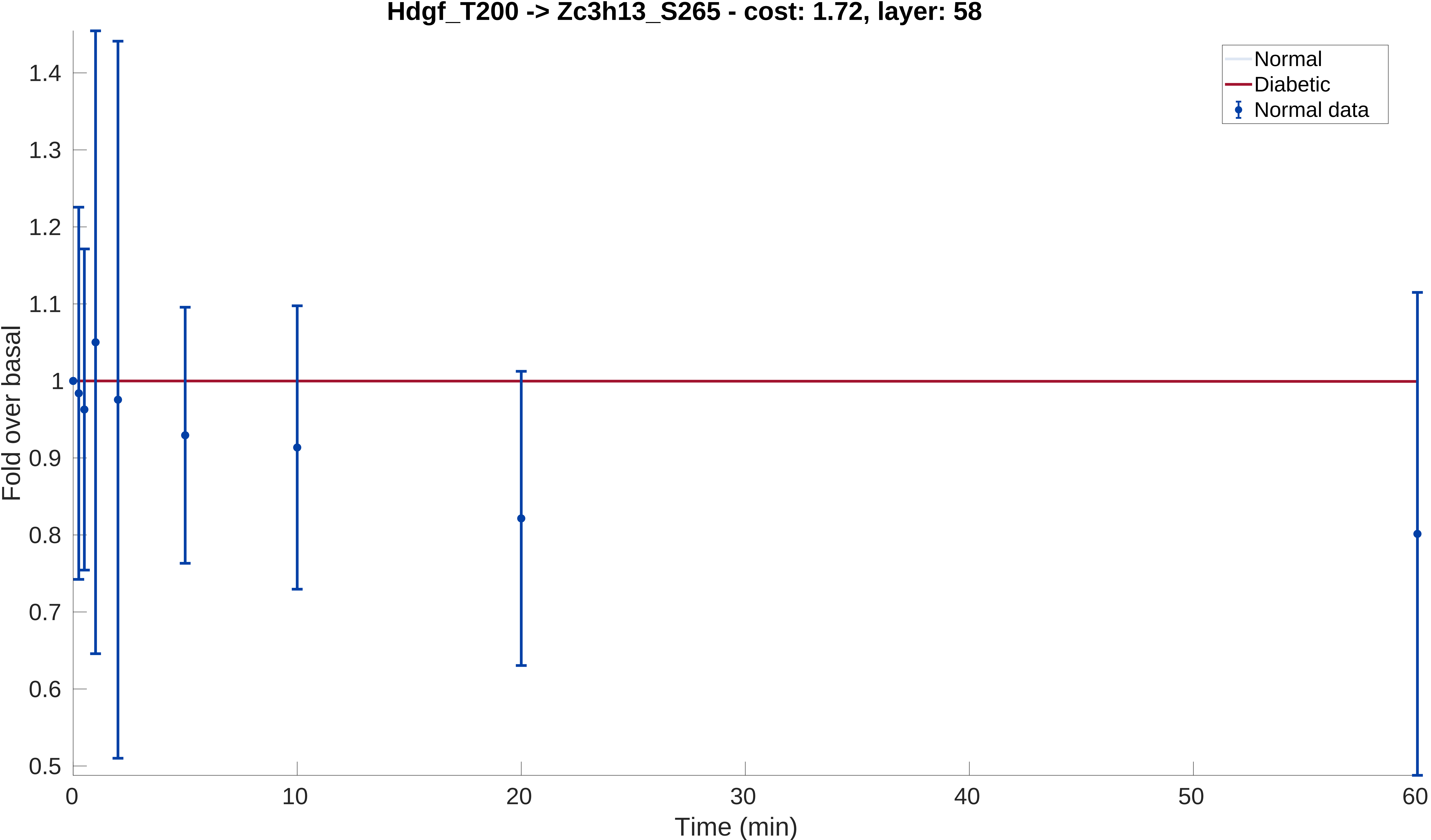
Hdgf_T200 -> Zc3h13_S265 - cost: 1.72, layer: 58
Normal
Diabetic
Normal data
Fold over basal
Time (min)
1.4
1.3
1.2
1.1
1
0.9
0.8
0.7
0.6
0.5
0
10
20
30
40
50
60

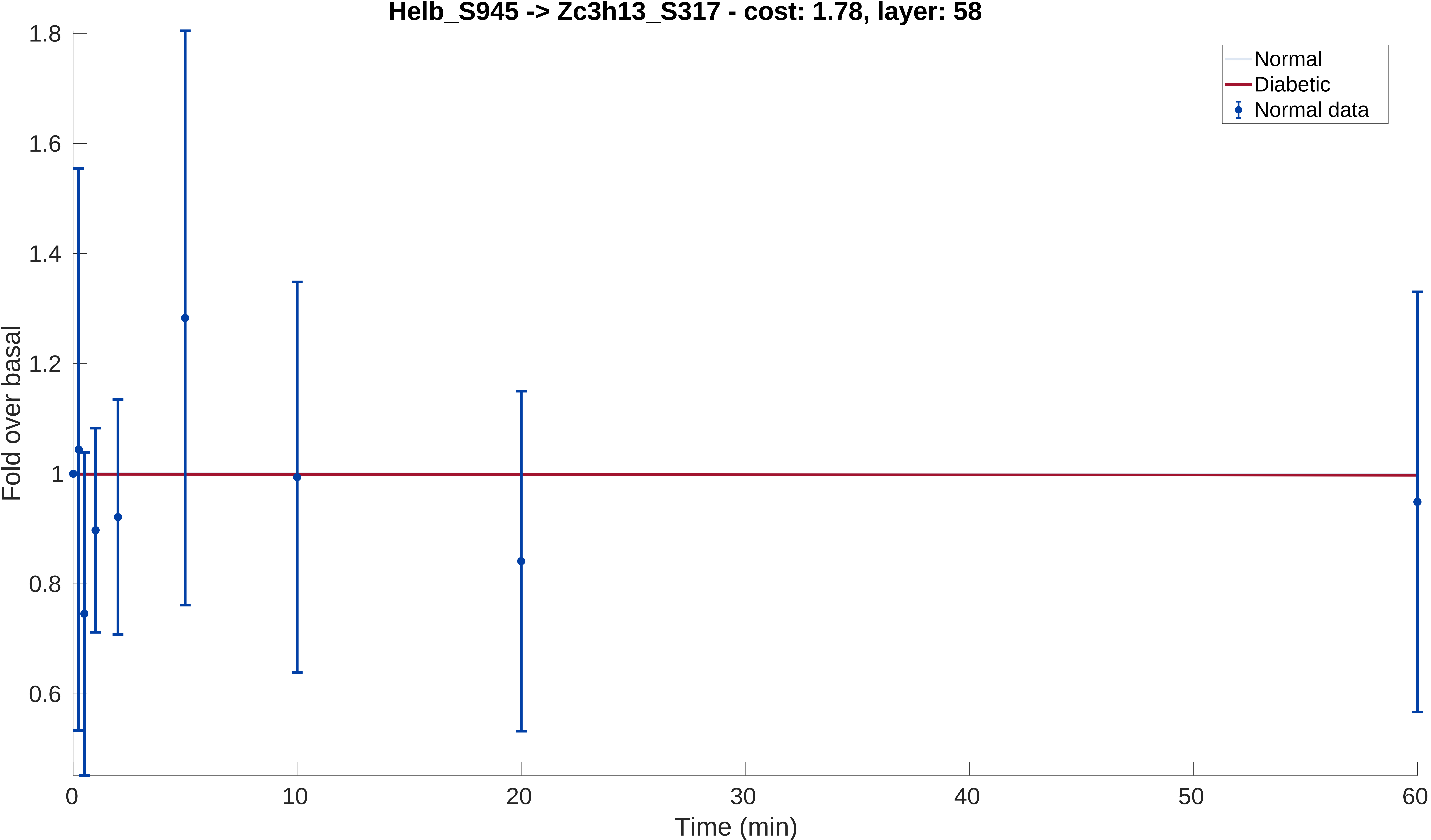

Helb_S945 -> Zc3h13_S317 - cost: 1.78, layer: 58
Normal
Diabetic
Normal data
Fold over basal
1.8
1.6
1.4
1.2
1
0.8
0.6
0
10
20
30
40
50
60
Time (min)

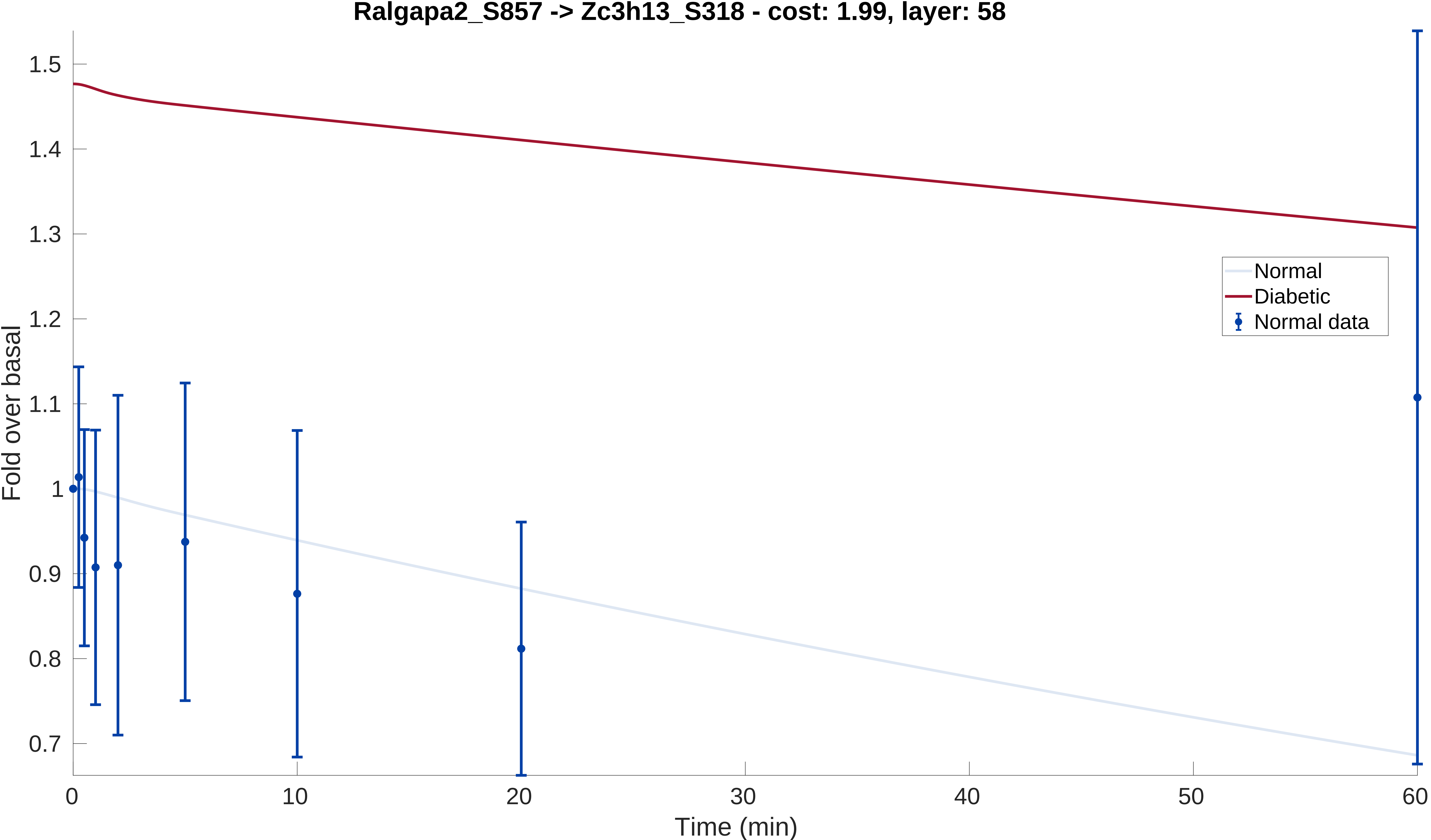

Ralgapa2_S857 -> Zc3h13_S318 - cost: 1.99, layer: 58
Fold over basal
Time (min)
1.5
1.4
1.3
1.2
1.1
1
0.9
0.8
0.7
0
10
20
30
40
50
60
Normal
Diabetic
Normal data

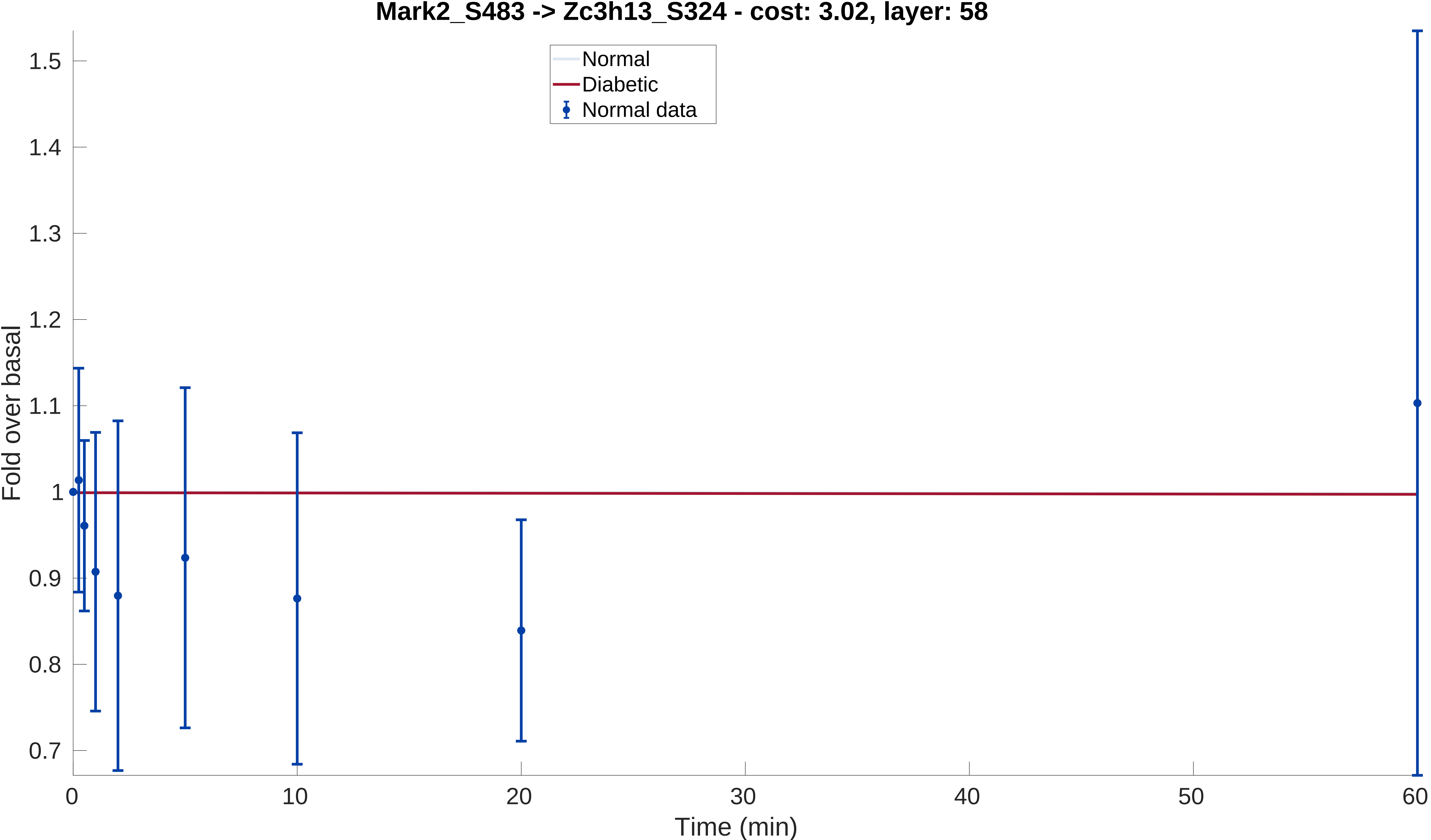

Mark2_S483 -> Zc3h13_S324 - cost: 3.02, layer: 58
Normal
Diabetic
Normal data
Fold over basal
1.5
1.4
1.3
1.2
1.1
1
0.9
0.8
0.7
0
10
20
30
40
50
60
Time (min)

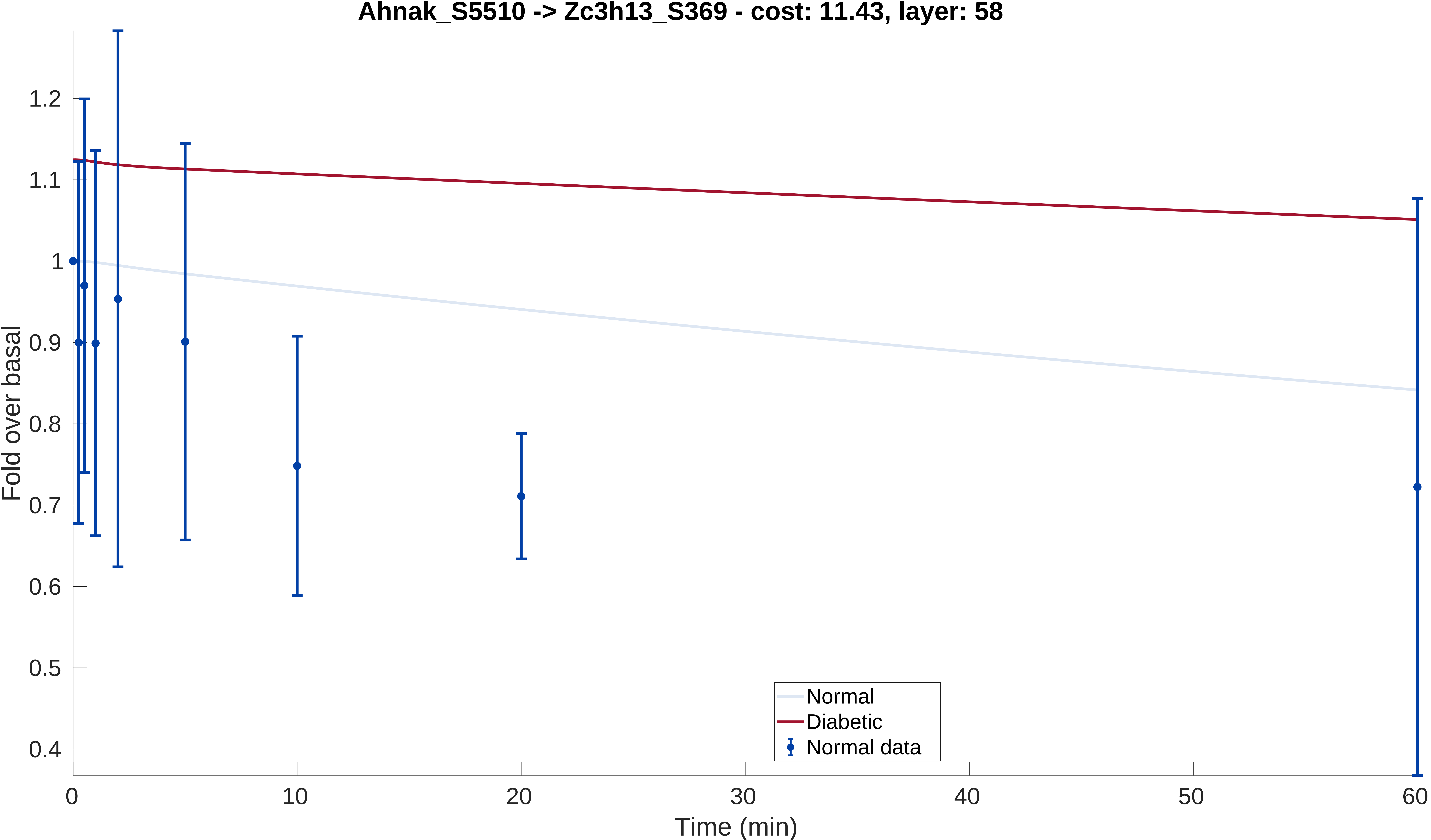

Ahnak_S5510 -> Zc3h13_S369 - cost: 11.43, layer: 58
Fold over basal
Time (min)
1.2
1.1
1
0.9
0.8
0.7
0.6
0.5
0.4
0
10
20
30
40
50
60
Normal
Diabetic
Normal data

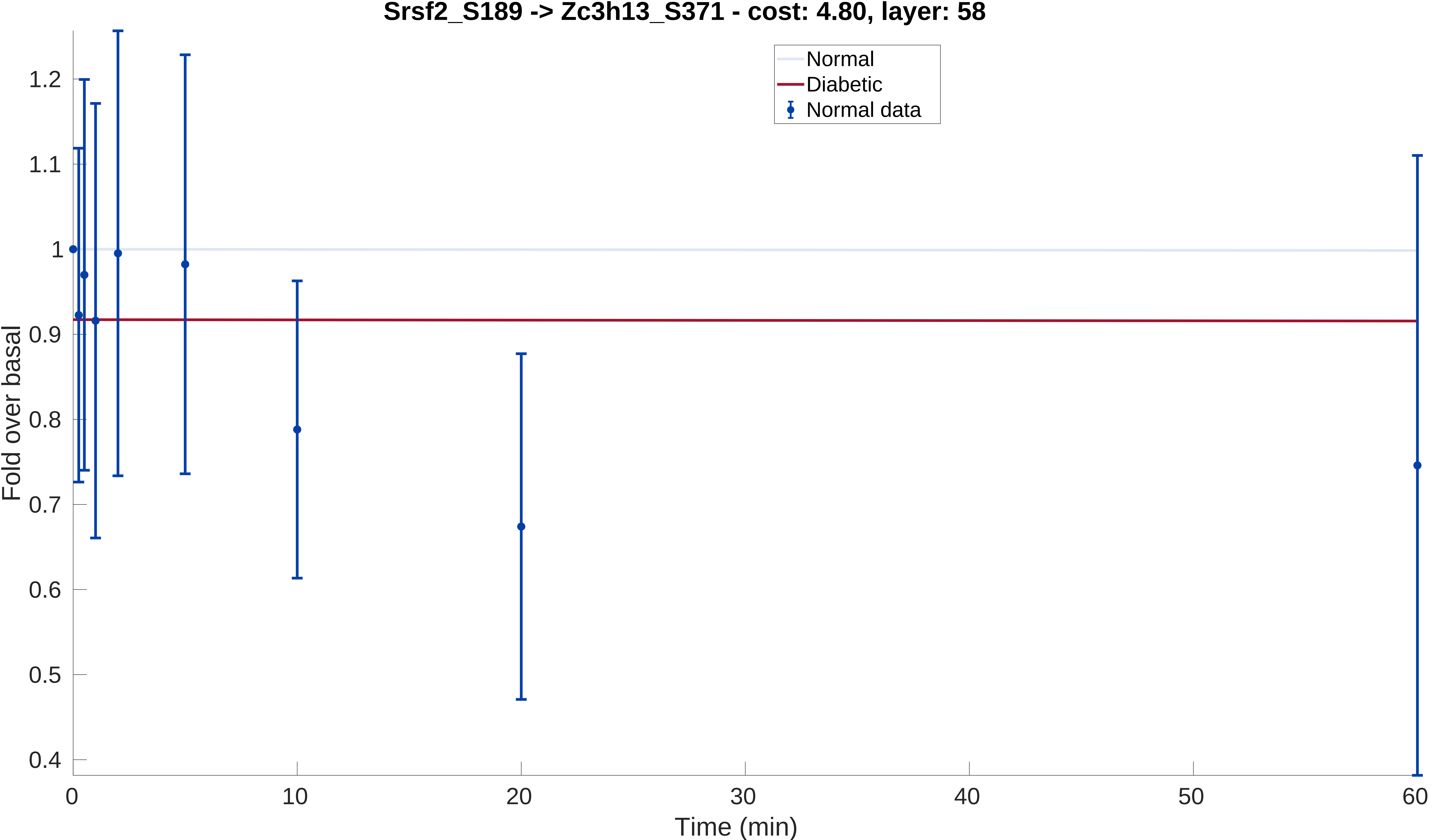
Srsf2_S189 -> Zc3h13_S371 - cost: 4.80, layer: 58
Normal
Diabetic
Normal data
Fold over basal
1.2
1.1
1
0.9
0.8
0.7
0.6
0.5
0.4
0
10
20
30
40
50
60
Time (min)

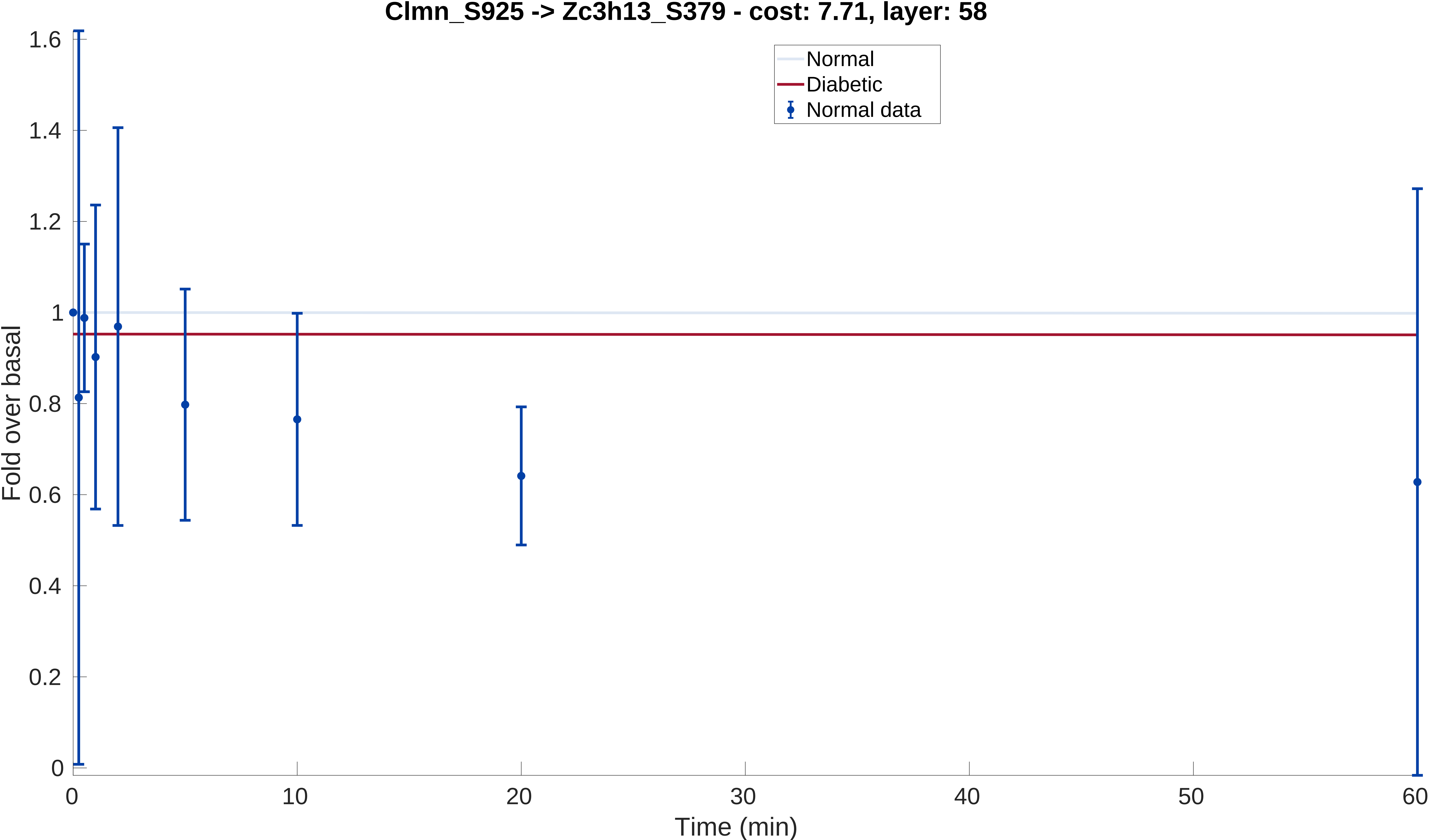
Clmn_S925 -> Zc3h13_S379 - cost: 7.71, layer: 58
Normal
Diabetic
Normal data
Fold over basal
Time (min)
0
0.2
0.4
0.6
0.8
1
1.2
1.4
1.6
0
10
20
30
40
50
60

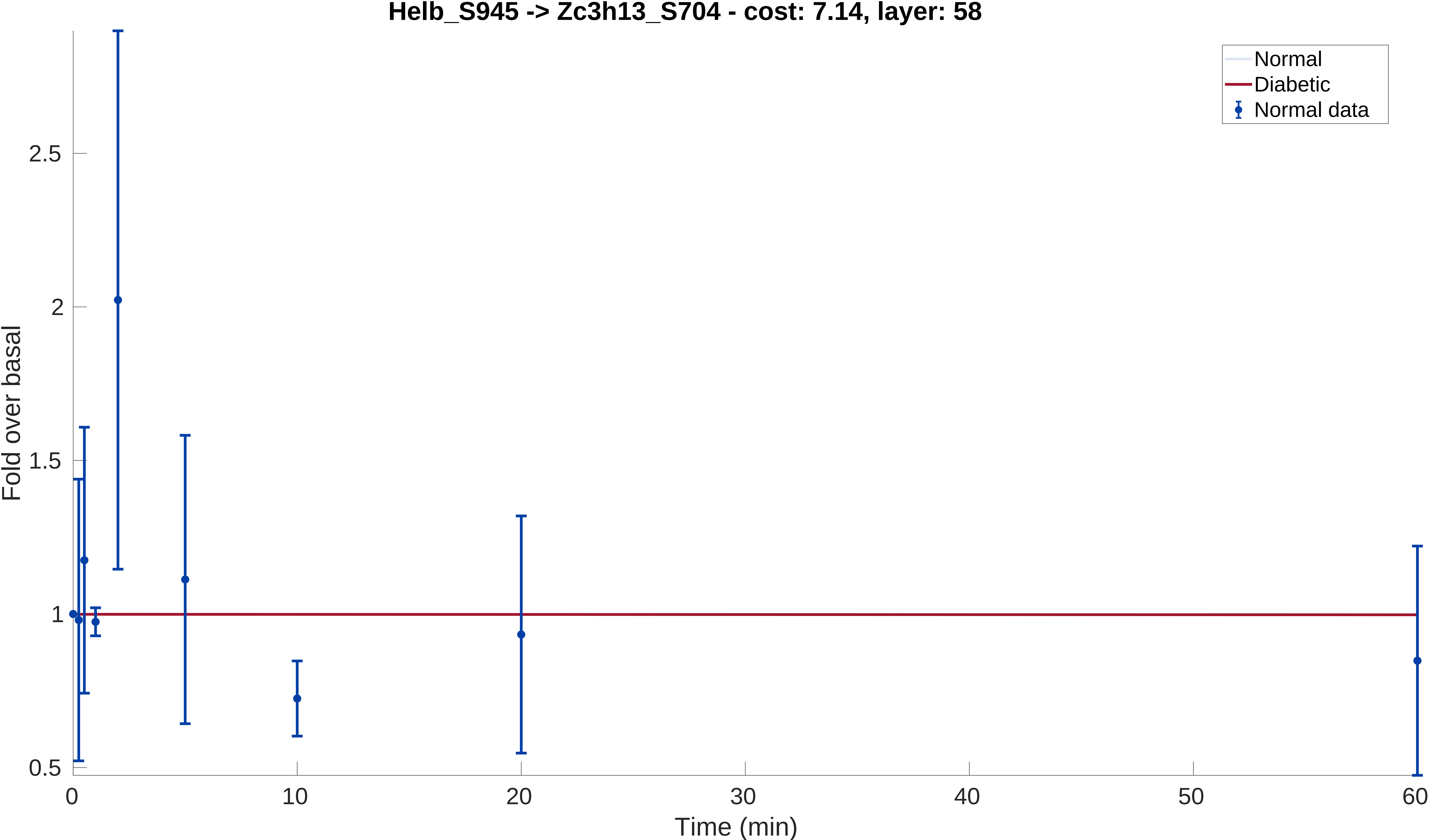
Helb_S945 -> Zc3h13_S704 - cost: 7.14, layer: 58
Normal
Diabetic
Normal data
Fold over basal
2.5
2
1.5
1
0.5
0
10
20
30
40
50
60
Time (min)

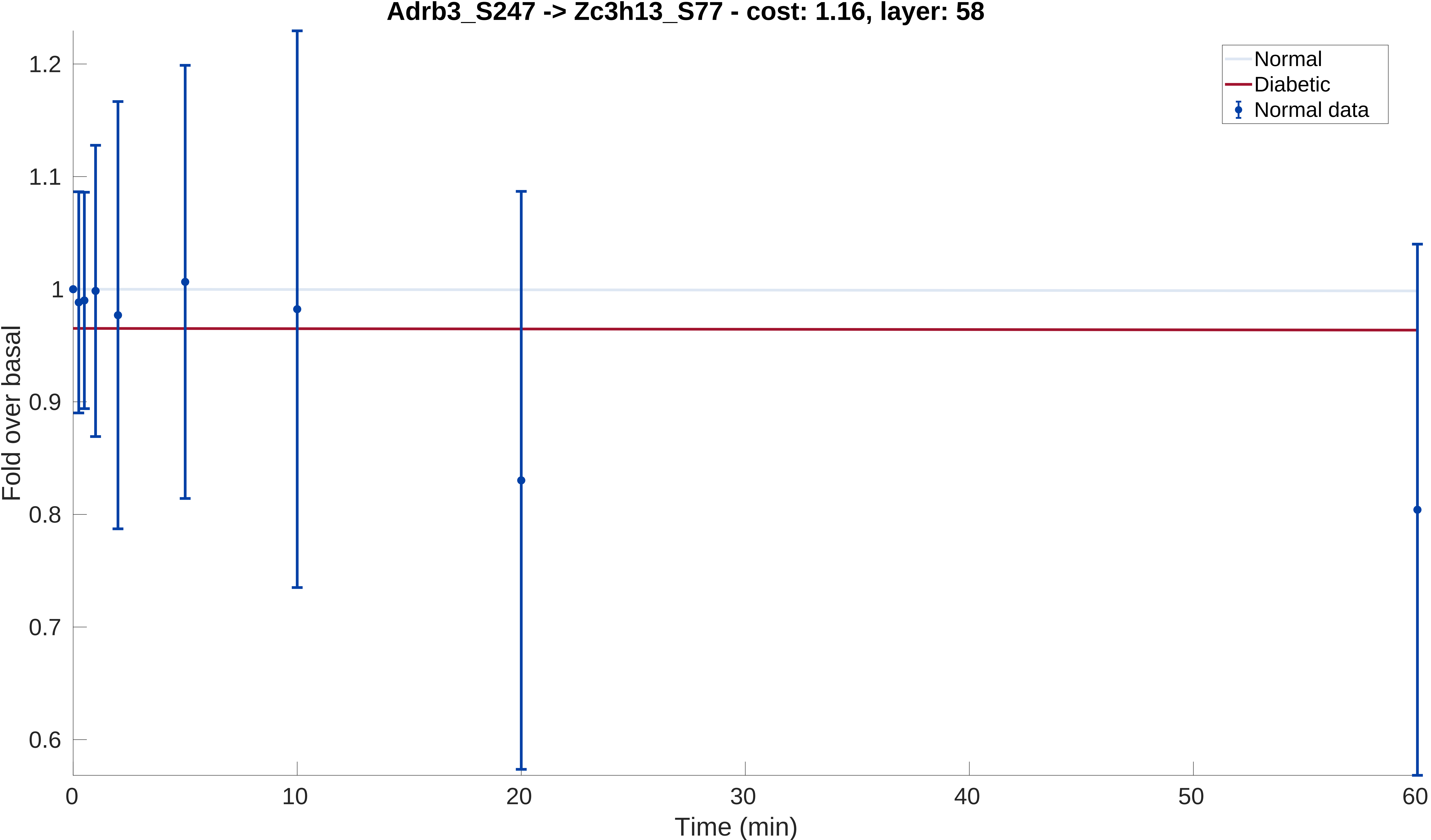

Adrb3_S247 -> Zc3h13_S77 - cost: 1.16, layer: 58
Normal
Diabetic
Normal data
Fold over basal
1.2
1.1
1
0.9
0.8
0.7
0.6
0
10
20
30
40
50
60
Time (min)

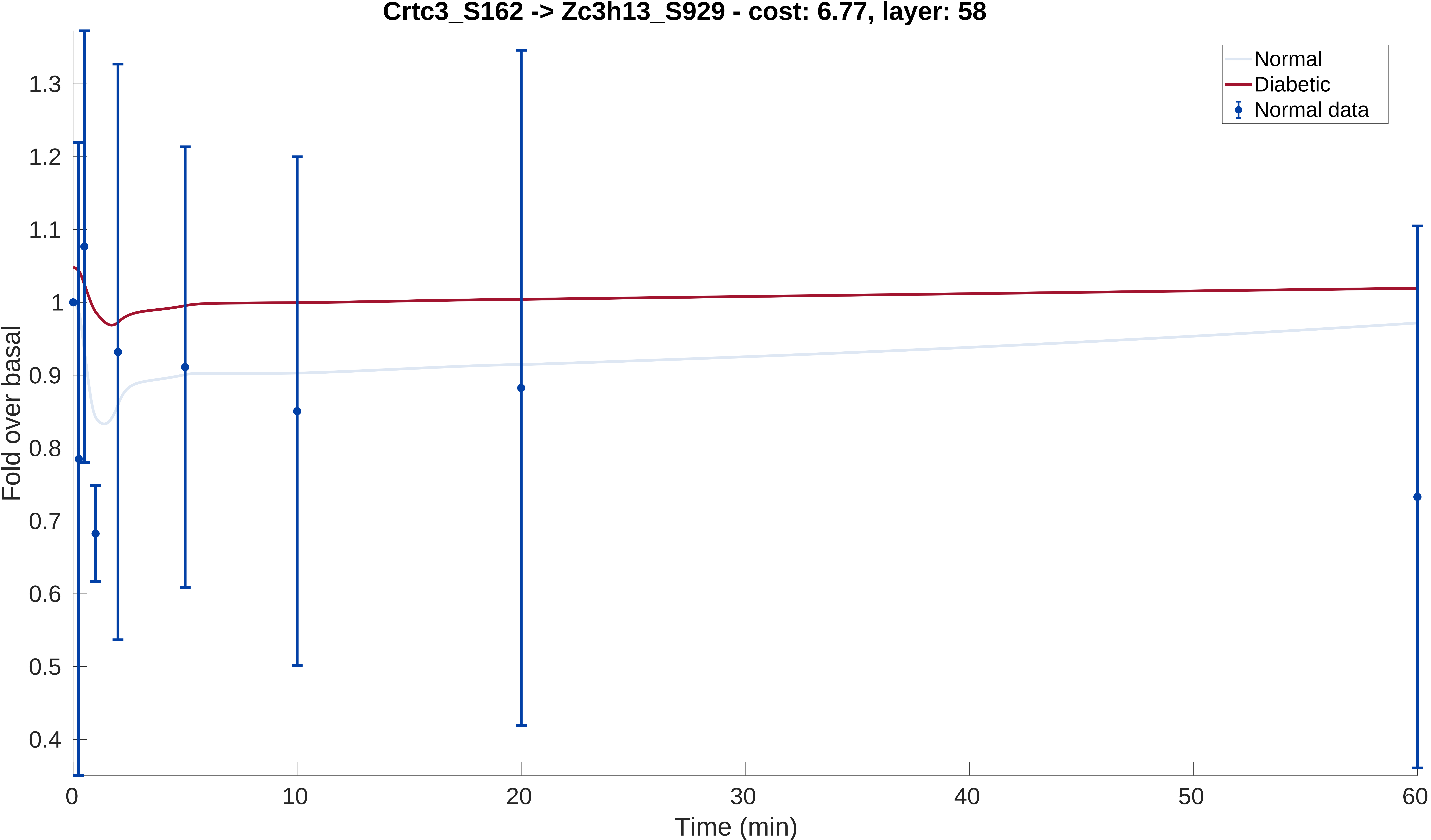
Crtc3_S162 -> Zc3h13_S929 - cost: 6.77, layer: 58
Normal
Diabetic
Normal data
Fold over basal
1.3
1.2
1.1
1
0.9
0.8
0.7
0.6
0.5
0.4
0
10
20
30
40
50
60
Time (min)

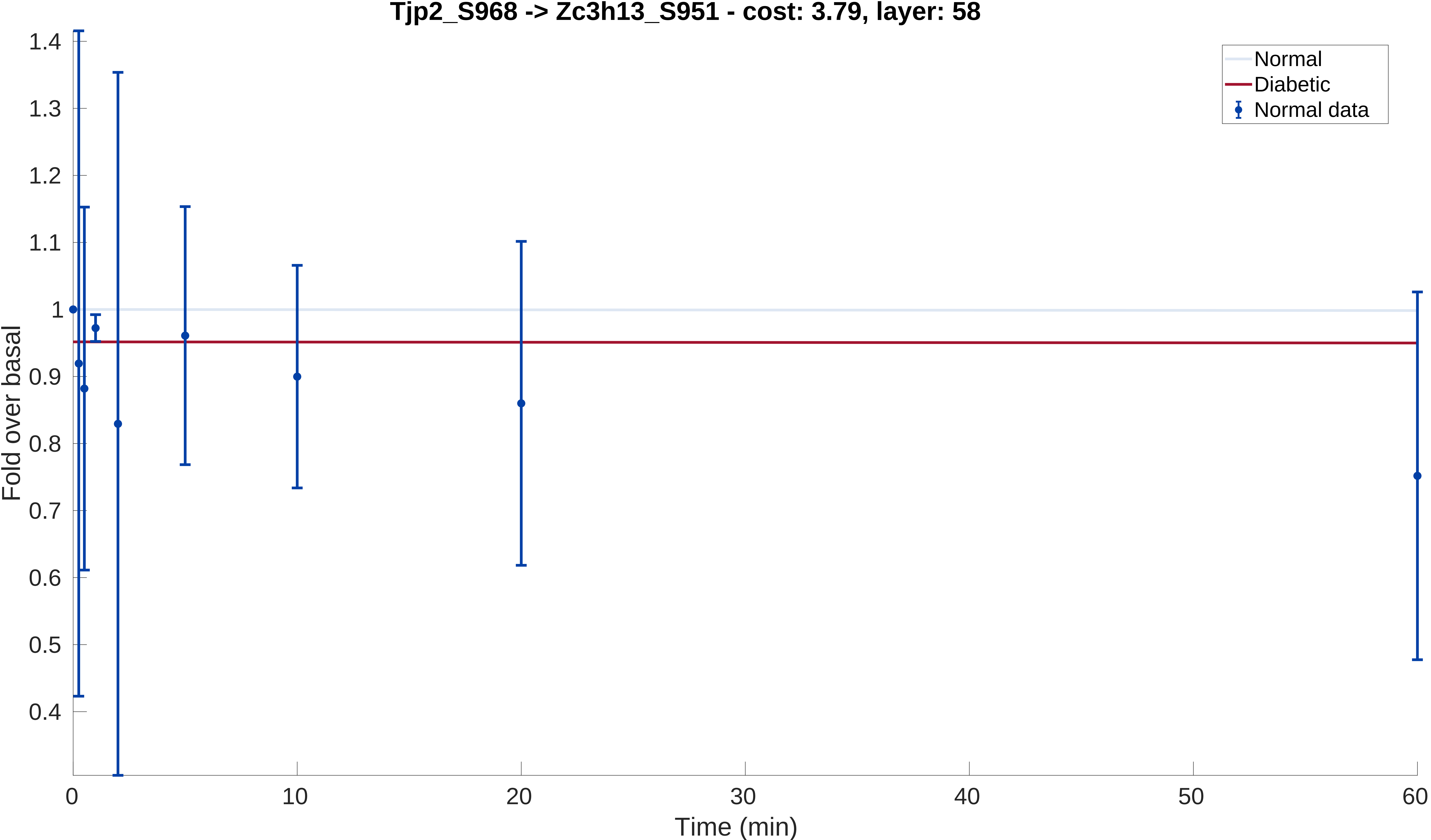
Tjp2_S968 -> Zc3h13_S951 - cost: 3.79, layer: 58
Normal
Diabetic
Normal data
Fold over basal
1.4
1.3
1.2
1.1
1
0.9
0.8
0.7
0.6
0.5
0.4
0
10
20
30
40
50
60
Time (min)

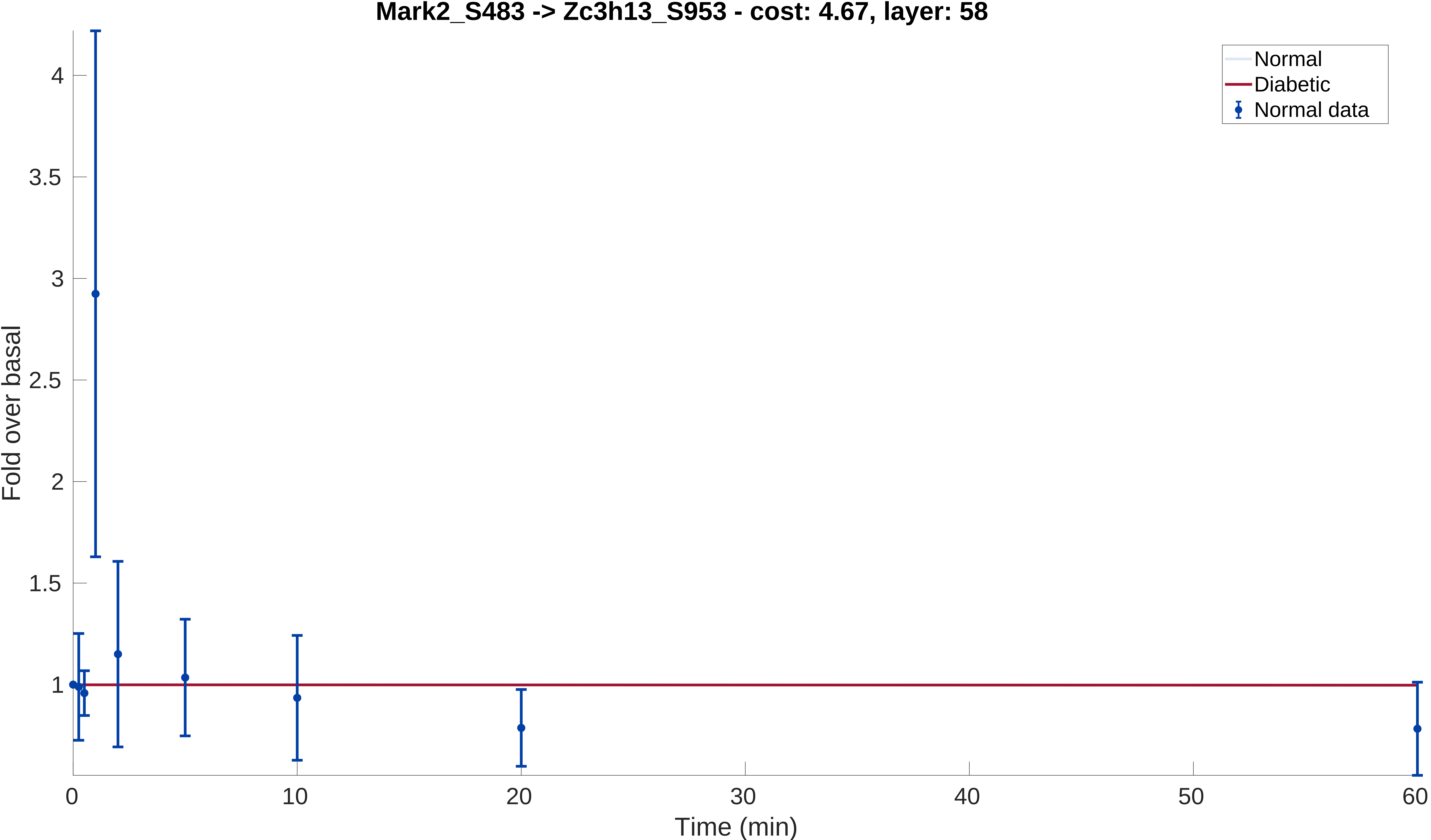

Mark2_S483 -> Zc3h13_S953 - cost: 4.67, layer: 58
Normal
Diabetic
Normal data
Fold over basal
Time (min)
4
3.5
3
2.5
2
1.5
1
0
10
20
30
40
50
60

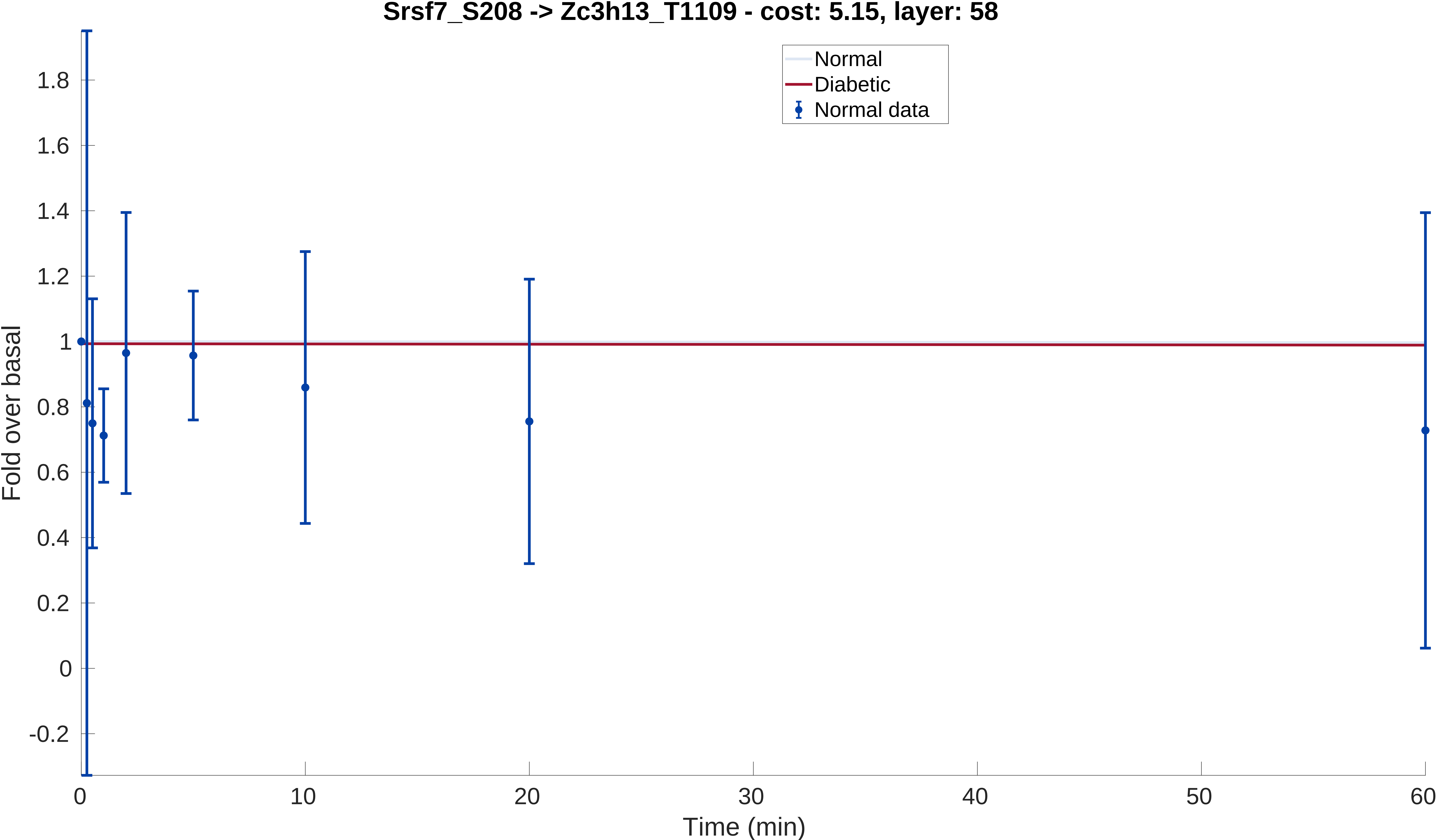
Srsf7_S208 -> Zc3h13_T1109 - cost: 5.15, layer: 58
Normal
Diabetic
Normal data
Fold over basal
Time (min)
-0.2
0
0.2
0.4
0.6
0.8
1
1.2
1.4
1.6
1.8
0
10
20
30
40
50
60

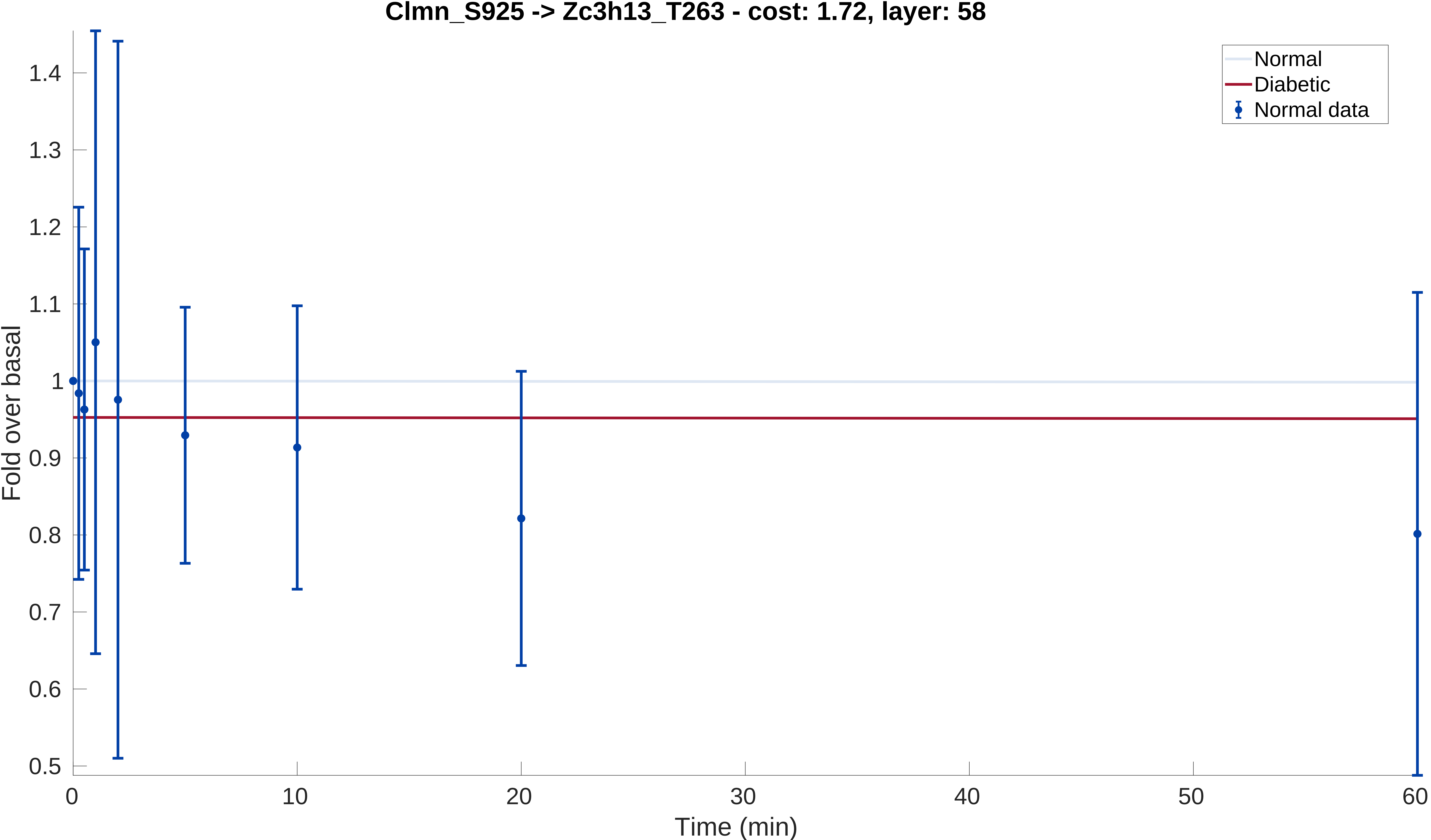

Clmn_S925 -> Zc3h13_T263 - cost: 1.72, layer: 58
Normal
Diabetic
Normal data
Fold over basal
1.4
1.3
1.2
1.1
1
0.9
0.8
0.7
0.6
0.5
0
10
20
30
40
50
60
Time (min)

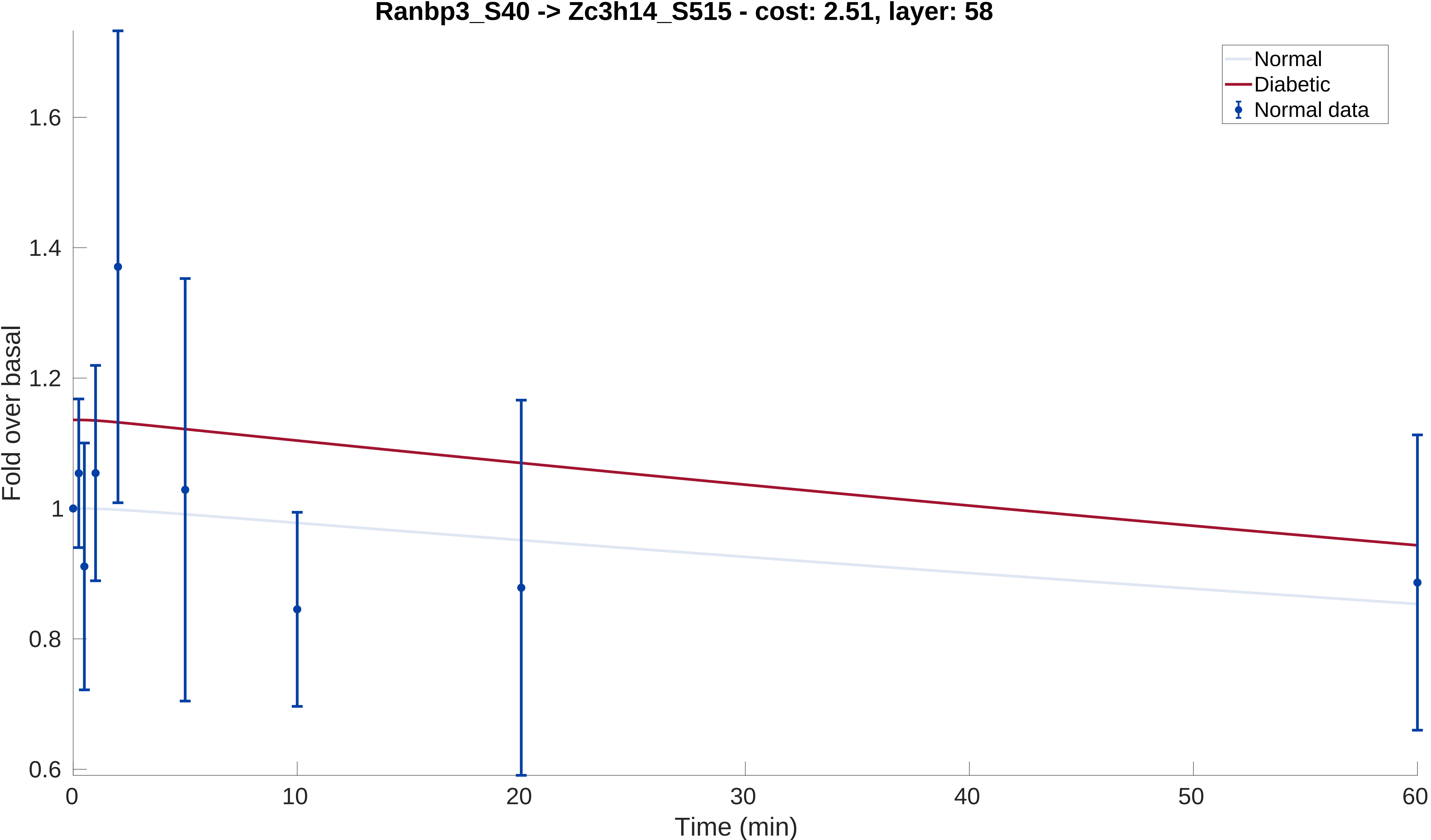
Ranbp3_S40 -> Zc3h14_S515 - cost: 2.51, layer: 58
Normal
Diabetic
Normal data
Fold over basal
Time (min)
0.6
0.8
1
1.2
1.4
1.6
0
10
20
30
40
50
60

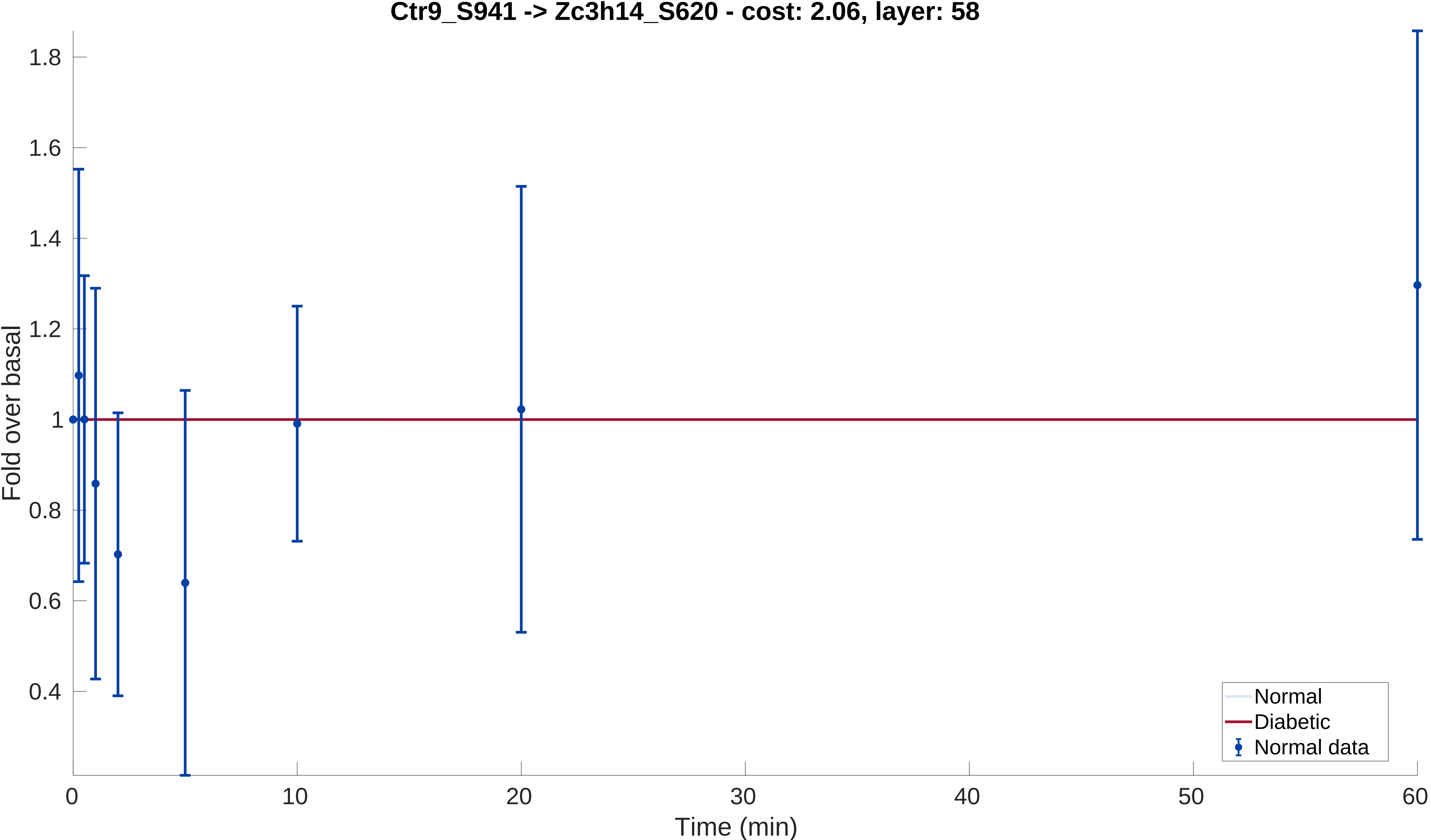
Ctr9_S941 -> Zc3h14_S620 - cost: 2.06, layer: 58
Fold over basal
Time (min)
1.8
1.6
1.4
1.2
1
0.8
0.6
0.4
0
10
20
30
40
50
60
Normal
Diabetic
Normal data

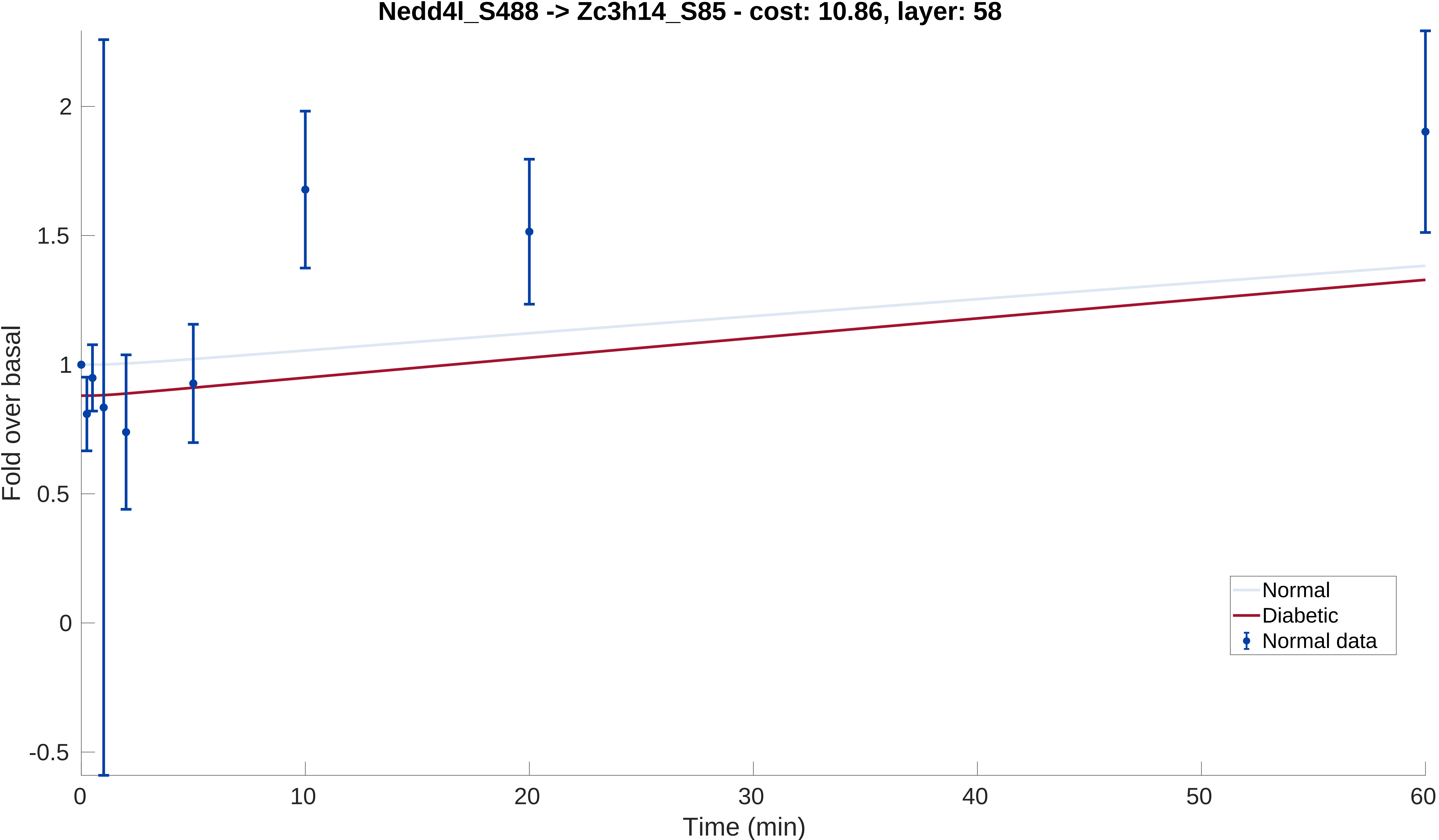

Nedd4l_S488 -> Zc3h14_S85 - cost: 10.86, layer: 58
Fold over basal
2
1.5
1
0.5
0
-0.5
0
10
20
30
40
50
60
Time (min)
Normal
Diabetic
Normal data

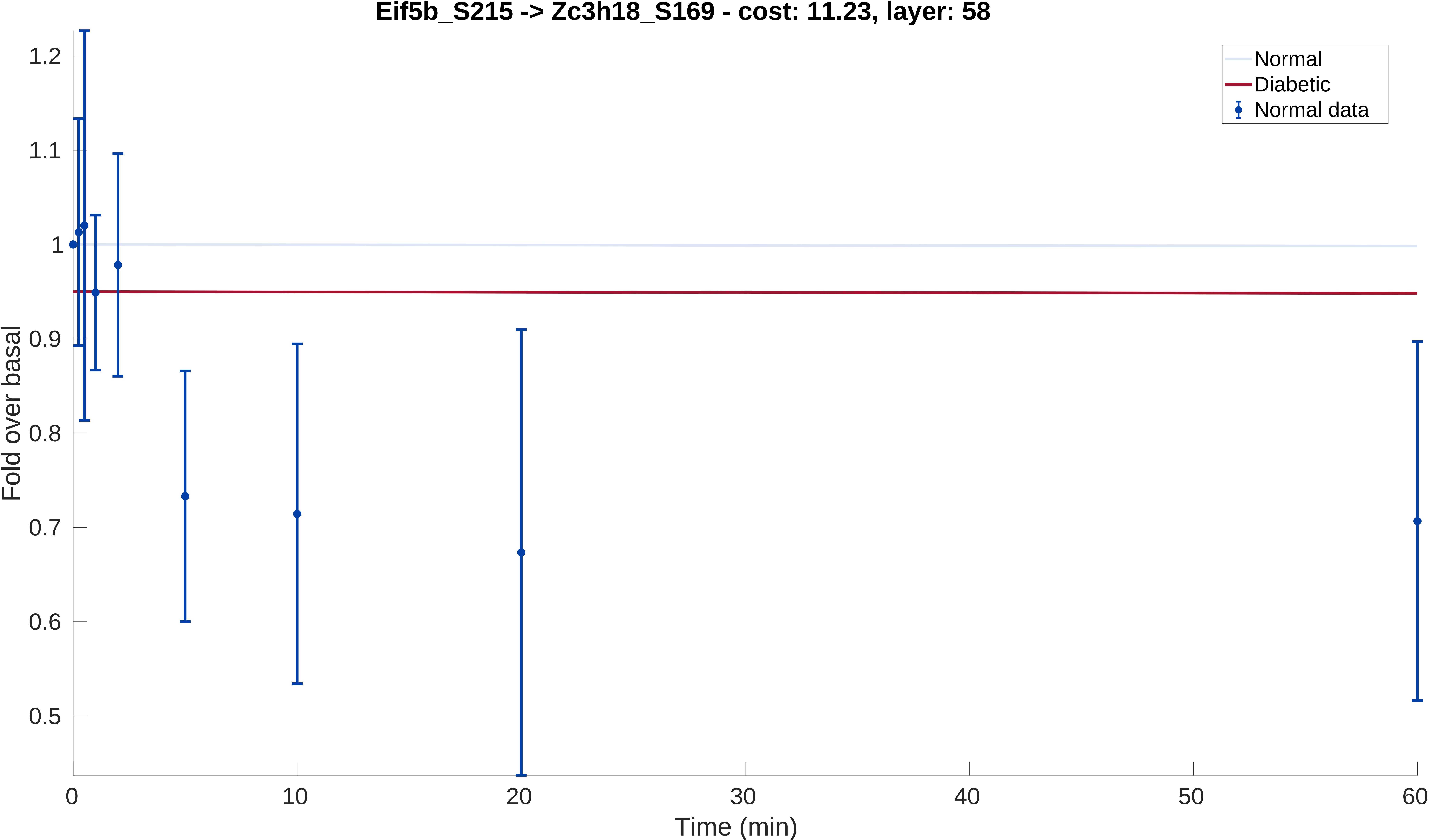
Eif5b_S215 -> Zc3h18_S169 - cost: 11.23, layer: 58
Normal
Diabetic
Normal data
Fold over basal
1.2
1.1
1
0.9
0.8
0.7
0.6
0.5
0
10
20
30
40
50
60
Time (min)

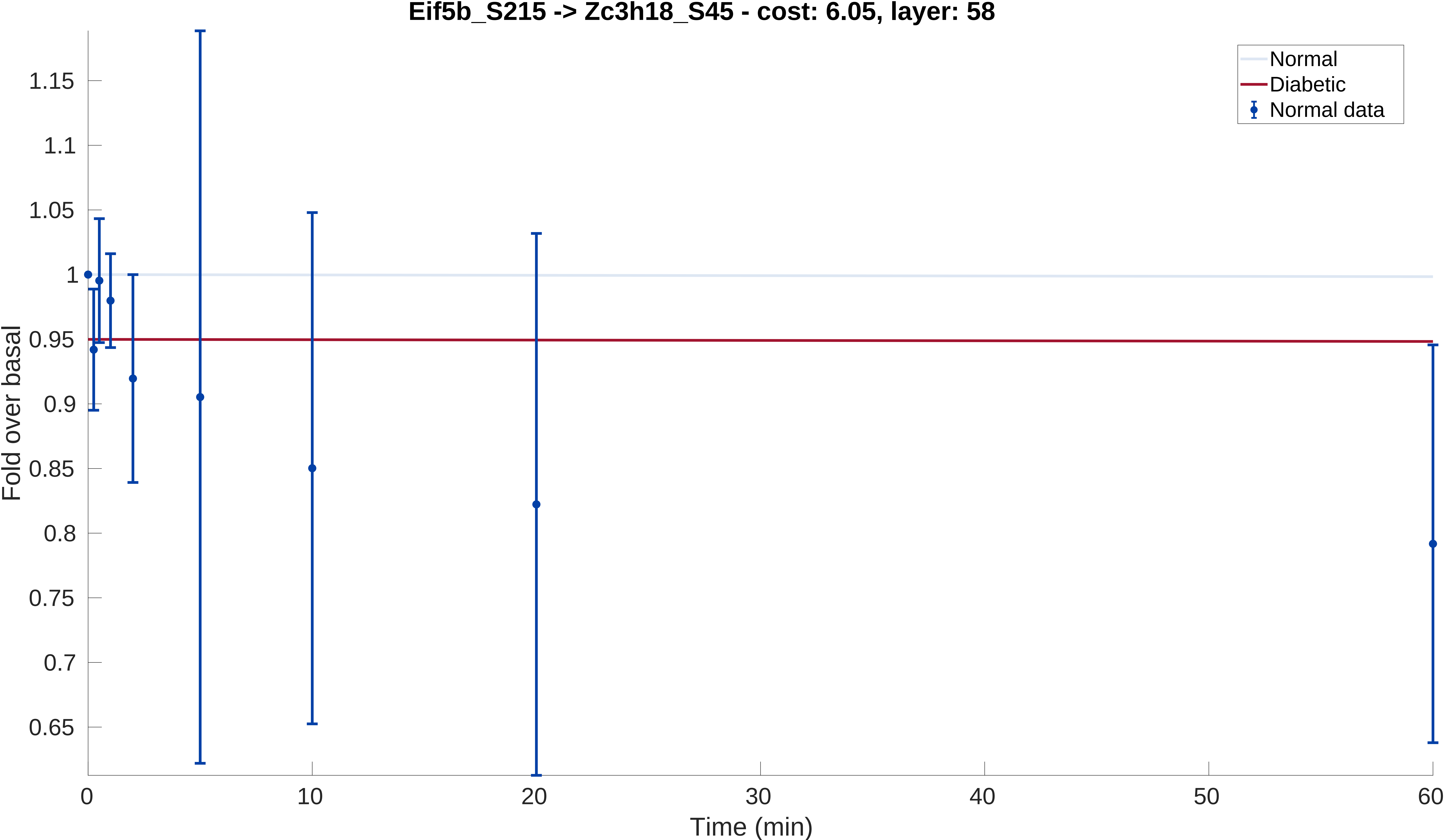

Eif5b_S215 -> Zc3h18_S45 - cost: 6.05, layer: 58
Normal
Diabetic
Normal data
Fold over basal
1.15
1.1
1.05
1
0.95
0.9
0.85
0.8
0.75
0.7
0.65
0
10
20
30
40
50
60
Time (min)

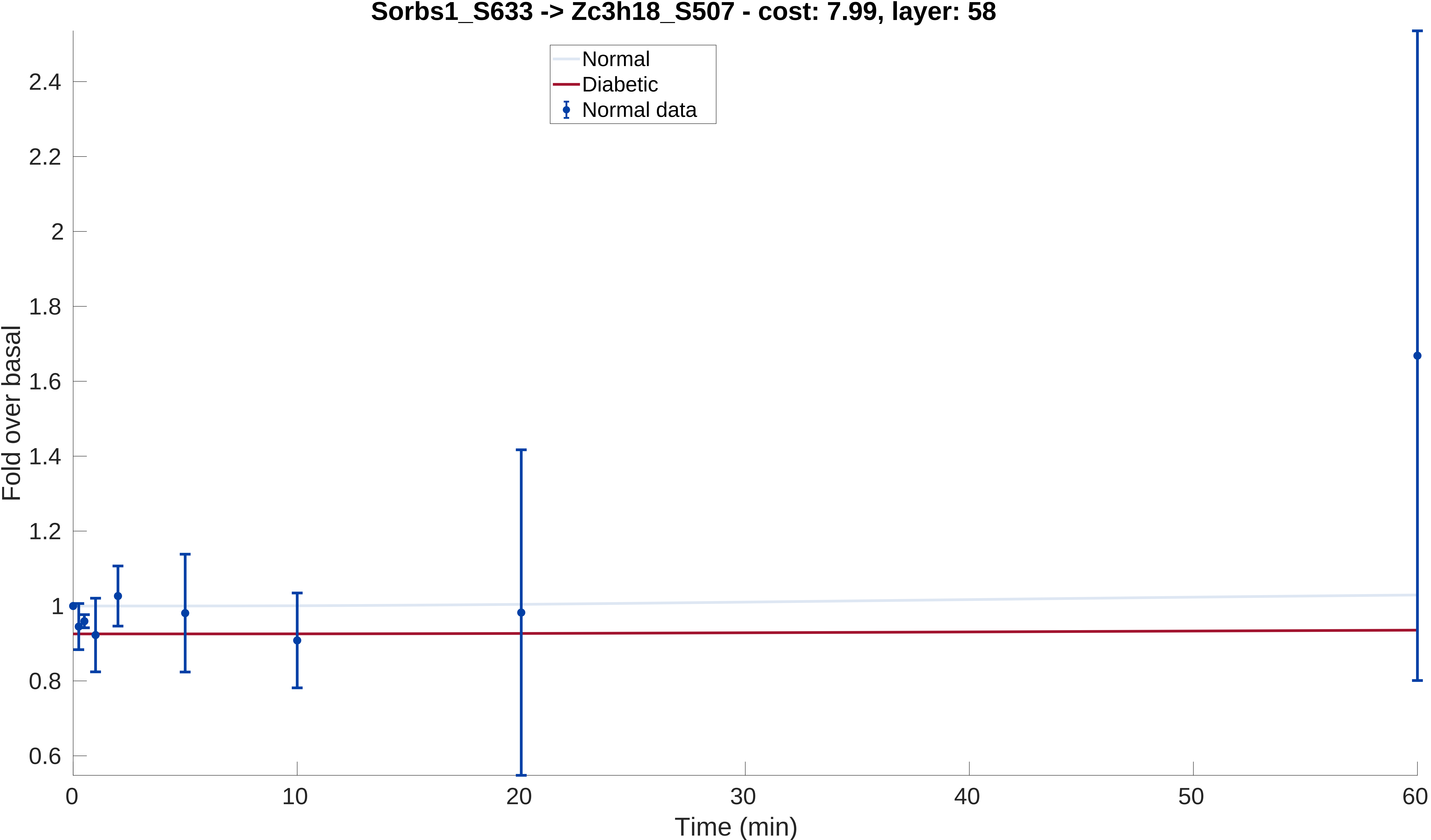

Sorbs1_S633 -> Zc3h18_S507 - cost: 7.99, layer: 58
Normal
Diabetic
Normal data
Fold over basal
2.4
2.2
2
1.8
1.6
1.4
1.2
1
0.8
0.6
0
10
20
30
40
50
60
Time (min)

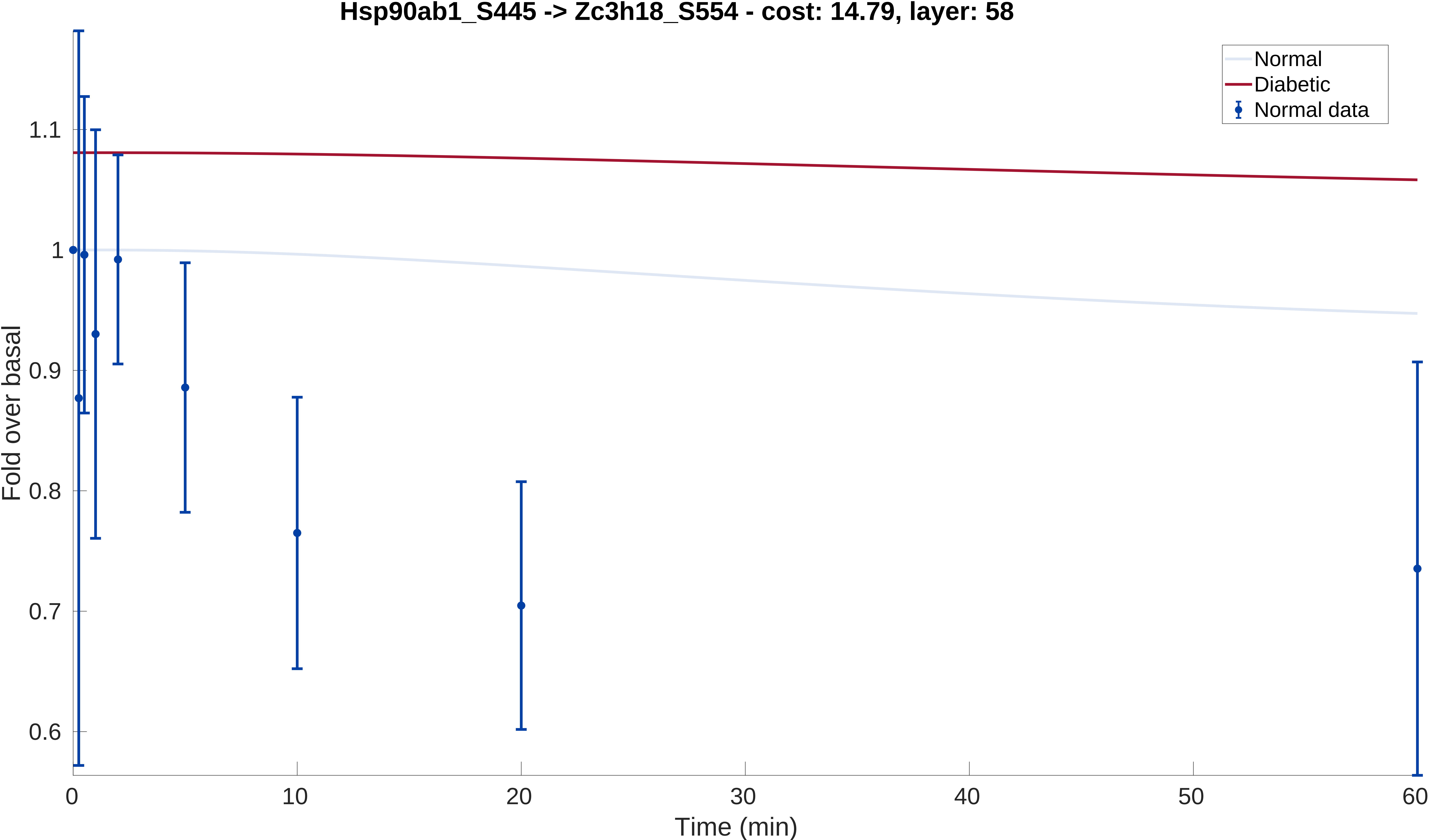
Hsp90ab1_S445 -> Zc3h18_S554 - cost: 14.79, layer: 58
Normal
Diabetic
Normal data
Fold over basal
0.6
0.7
0.8
0.9
1
1.1
Time (min)
0
10
20
30
40
50
60

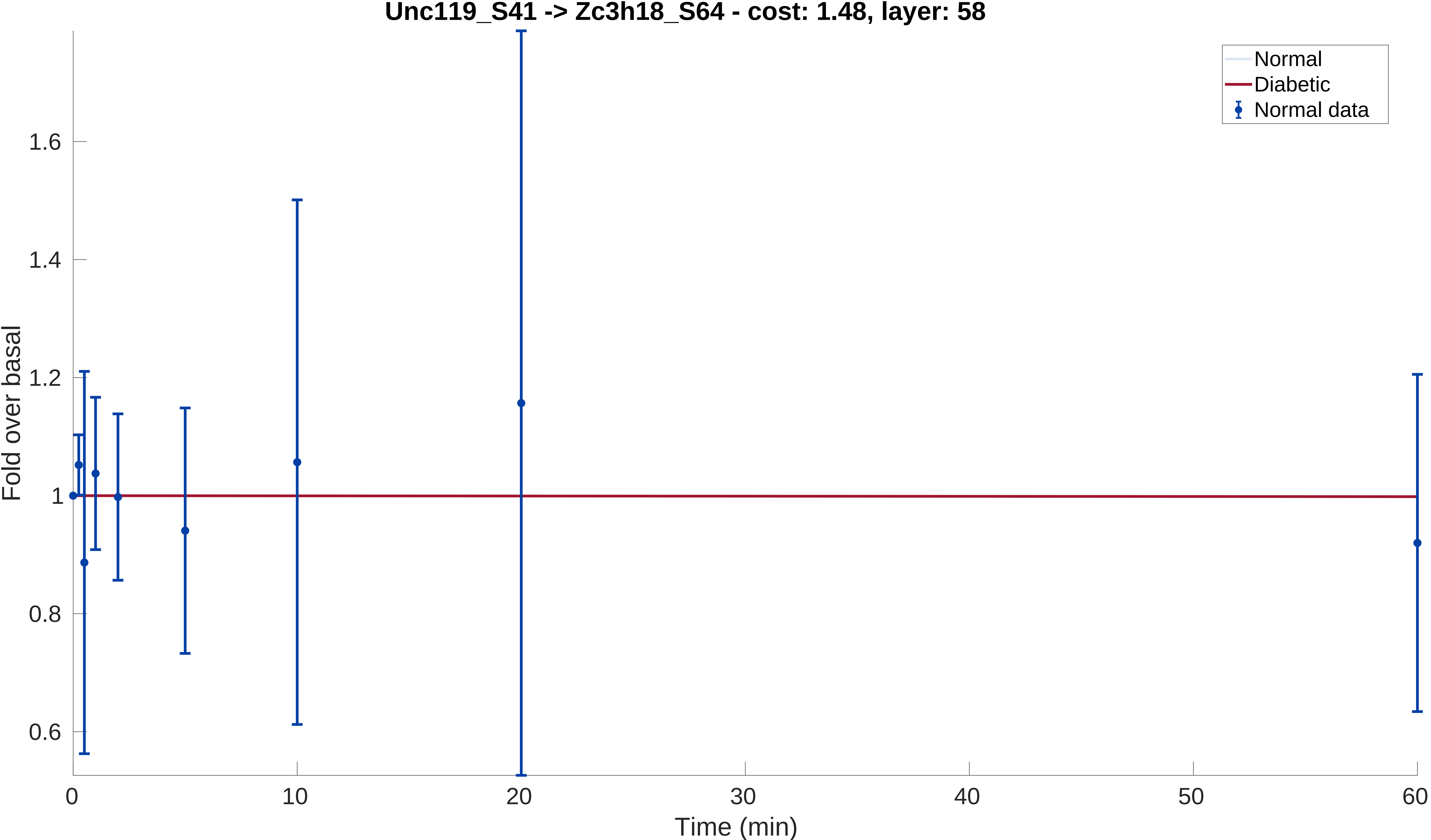
Unc119_S41 -> Zc3h18_S64 - cost: 1.48, layer: 58
Normal
Diabetic
Normal data
Fold over basal
1.6
1.4
1.2
1
0.8
0.6
0
10
20
30
40
50
60
Time (min)

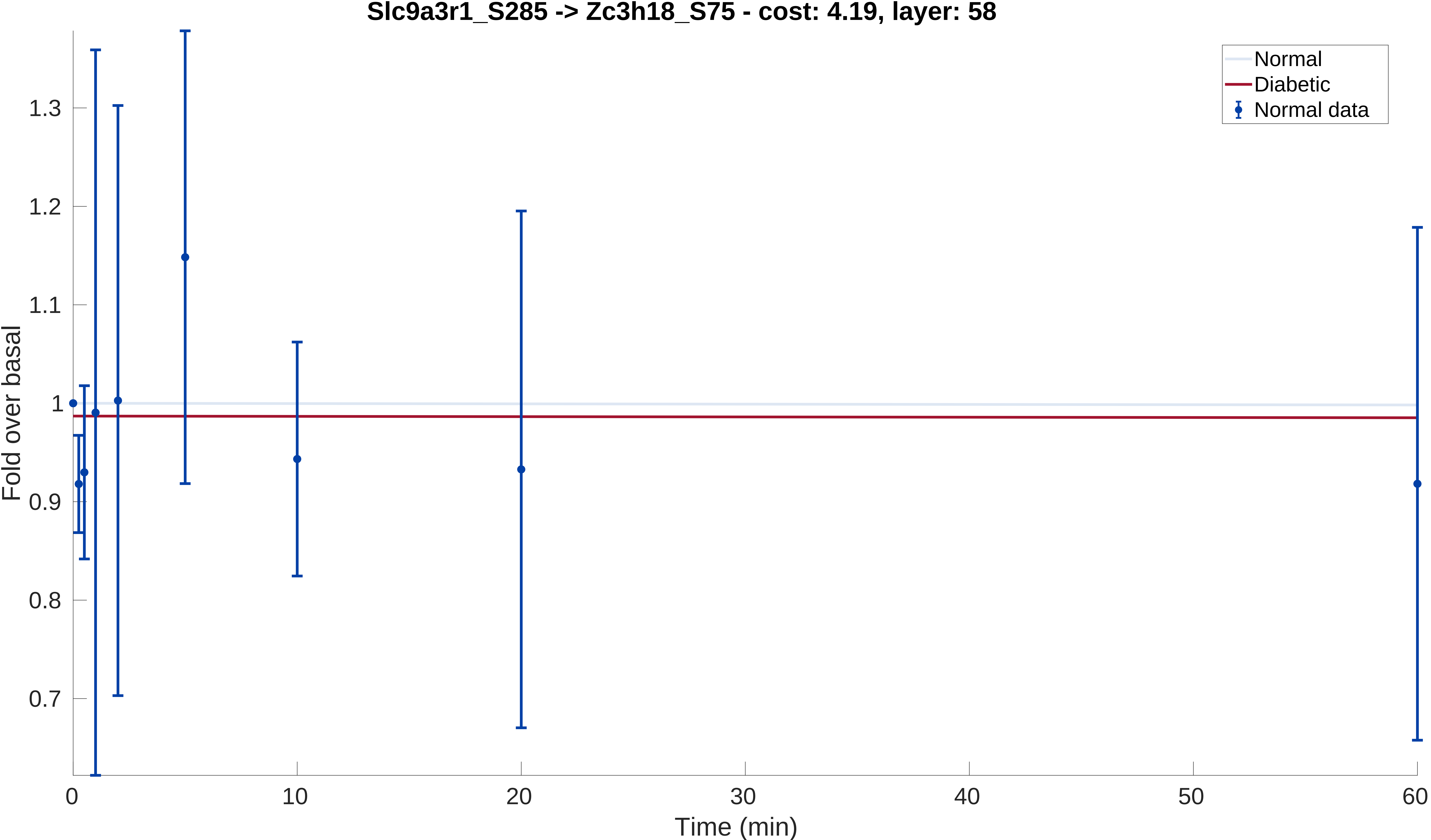

Slc9a3r1_S285 -> Zc3h18_S75 - cost: 4.19, layer: 58
Normal
Diabetic
Normal data
Fold over basal
1.3
1.2
1.1
1
0.9
0.8
0.7
0
10
20
30
40
50
60
Time (min)

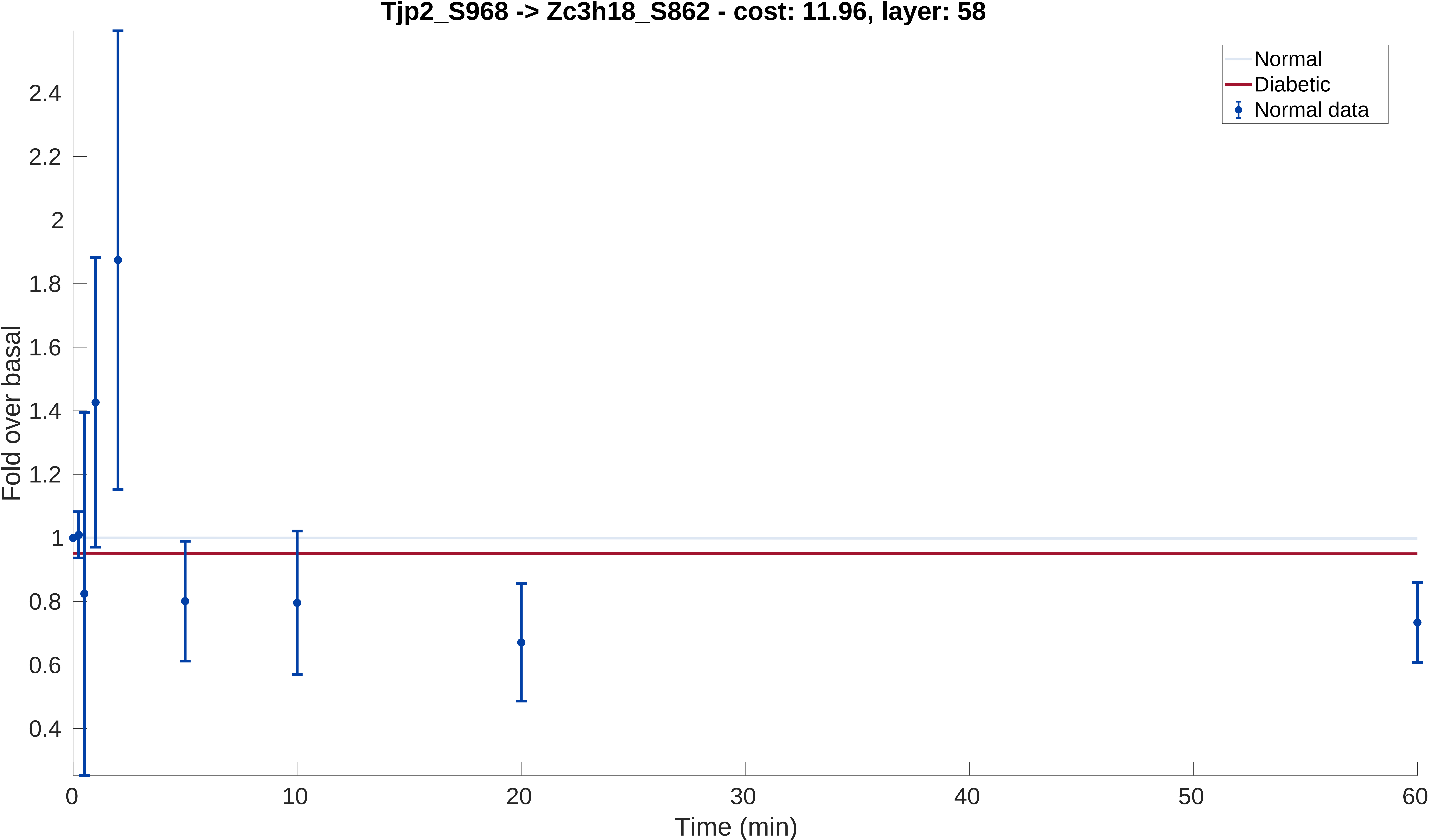

Tjp2_S968 -> Zc3h18_S862 - cost: 11.96, layer: 58
Fold over basal
2.4
2.2
2
1.8
1.6
1.4
1.2
1
0.8
0.6
0.4
0
10
20
30
40
50
60
Time (min)
Normal
Diabetic
Normal data

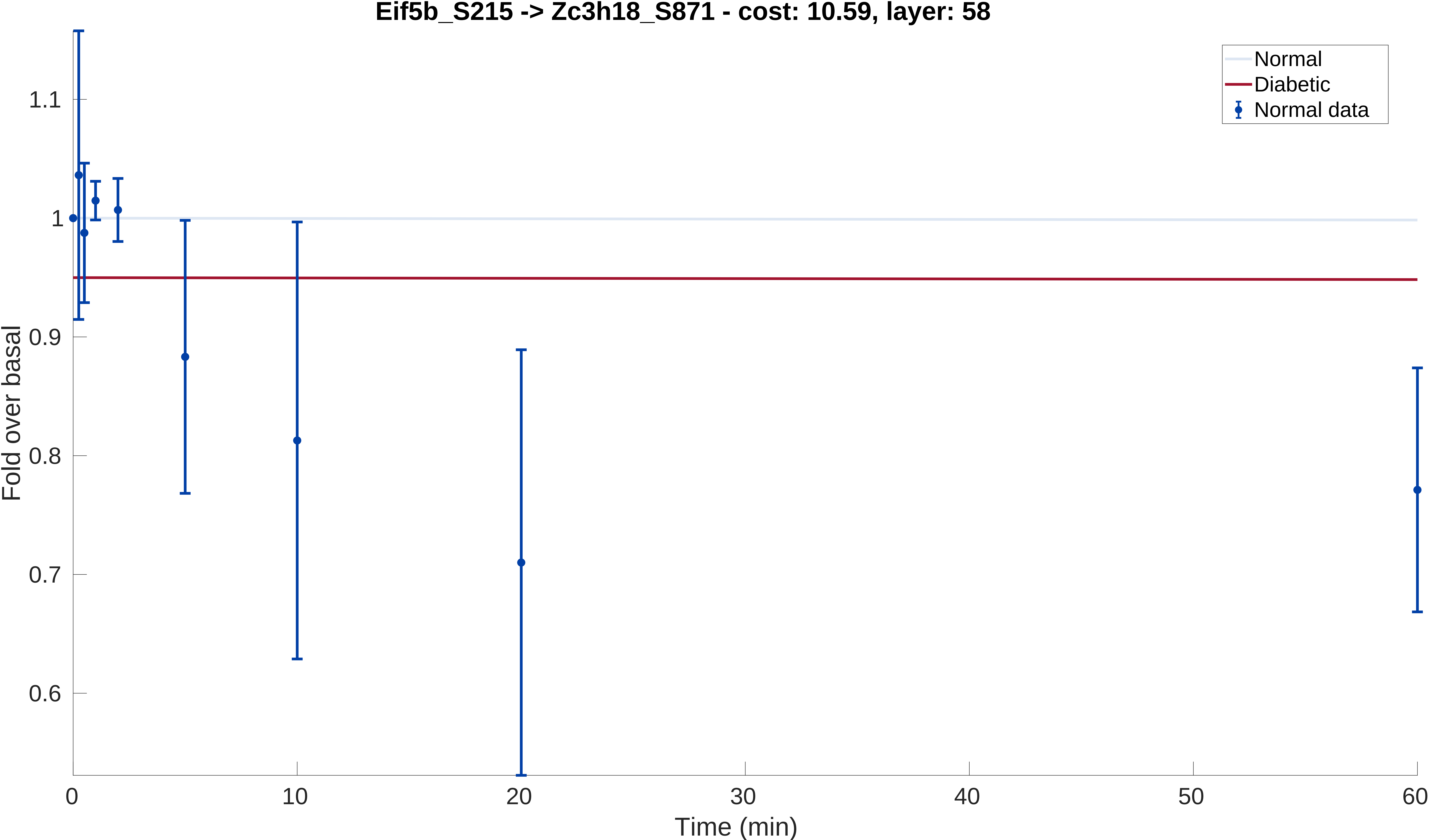

Eif5b_S215 -> Zc3h18_S871 - cost: 10.59, layer: 58
Normal
Diabetic
Normal data
1.1
1
0.9
0.8
0.7
0.6
Fold over basal
0
10
20
30
40
50
60
Time (min)

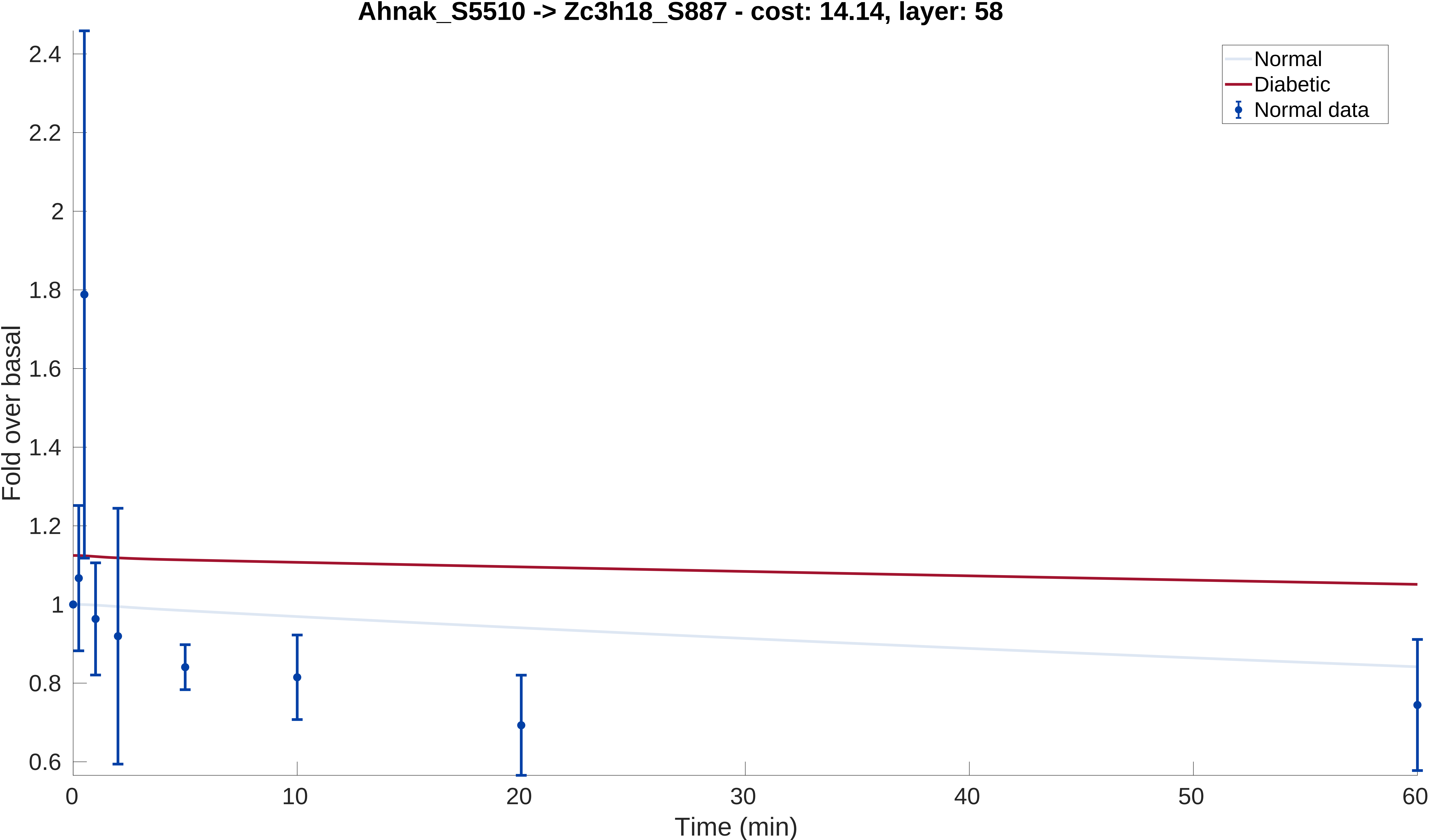

Ahnak_S5510 -> Zc3h18_S887 - cost: 14.14, layer: 58
Normal
Diabetic
Normal data
Fold over basal
2.4
2.2
2
1.8
1.6
1.4
1.2
1
0.8
0.6
0
10
20
30
40
50
60
Time (min)

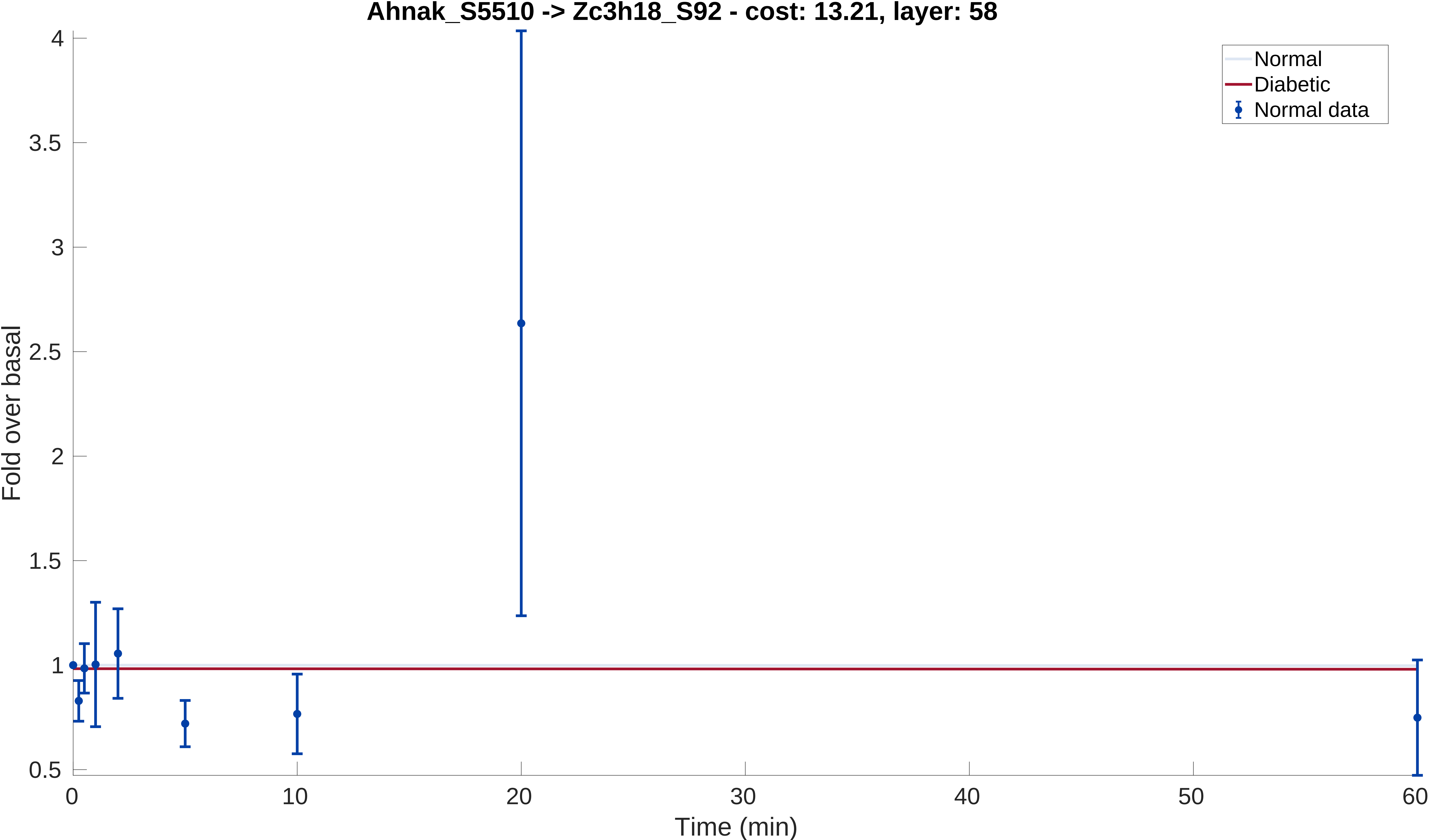

Ahnak_S5510 -> Zc3h18_S92 - cost: 13.21, layer: 58
Normal
Diabetic
Normal data
Fold over basal
Time (min)
4
3.5
3
2.5
2
1.5
1
0.5
0
10
20
30
40
50
60

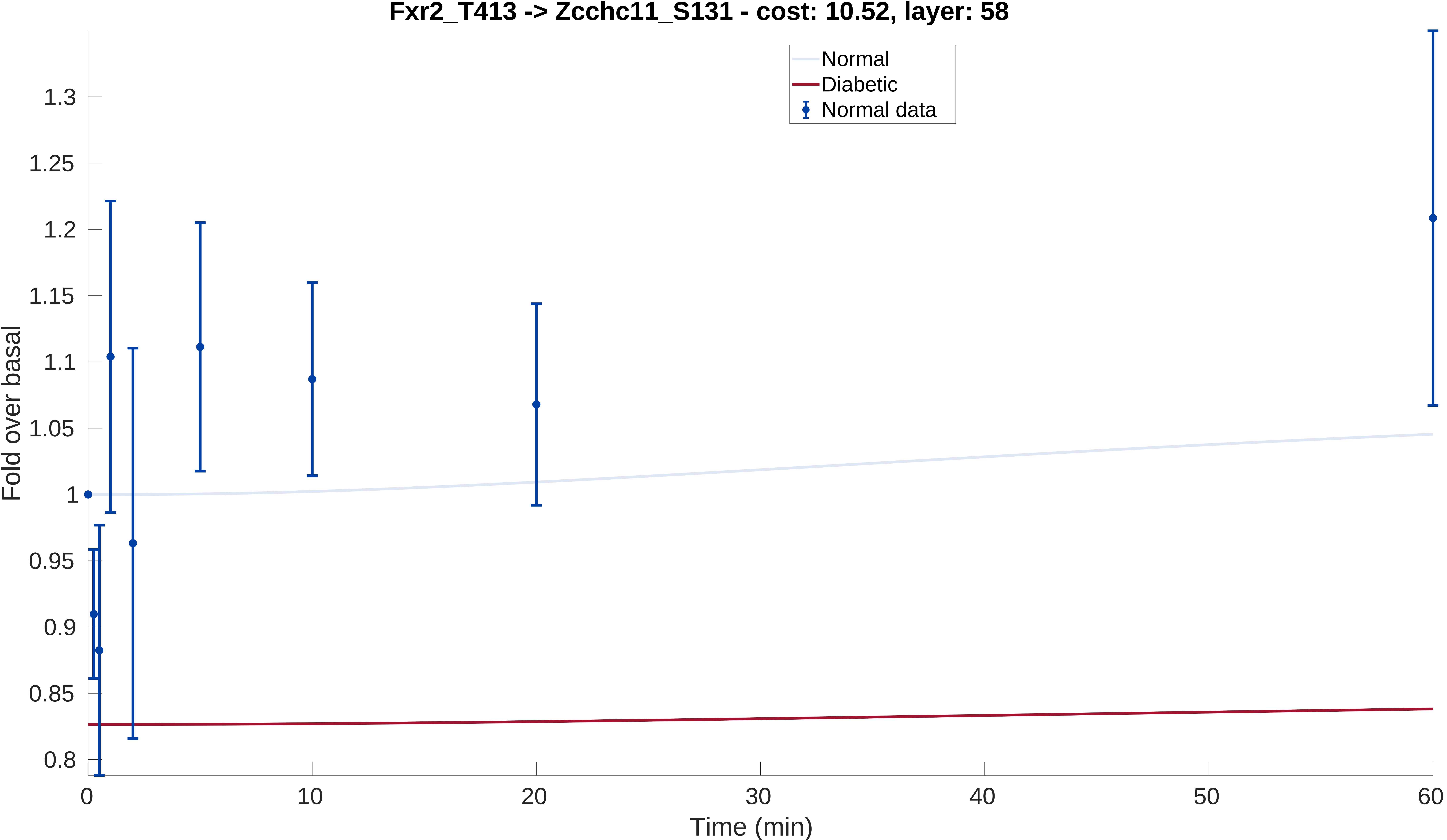

Fxr2_T413 -> Zcchc11_S131 - cost: 10.52, layer: 58
Normal
Diabetic
Normal data
Fold over basal
1.3
1.25
1.2
1.15
1.1
1.05
1
0.95
0.9
0.85
0.8
0
10
20
30
40
50
60
Time (min)

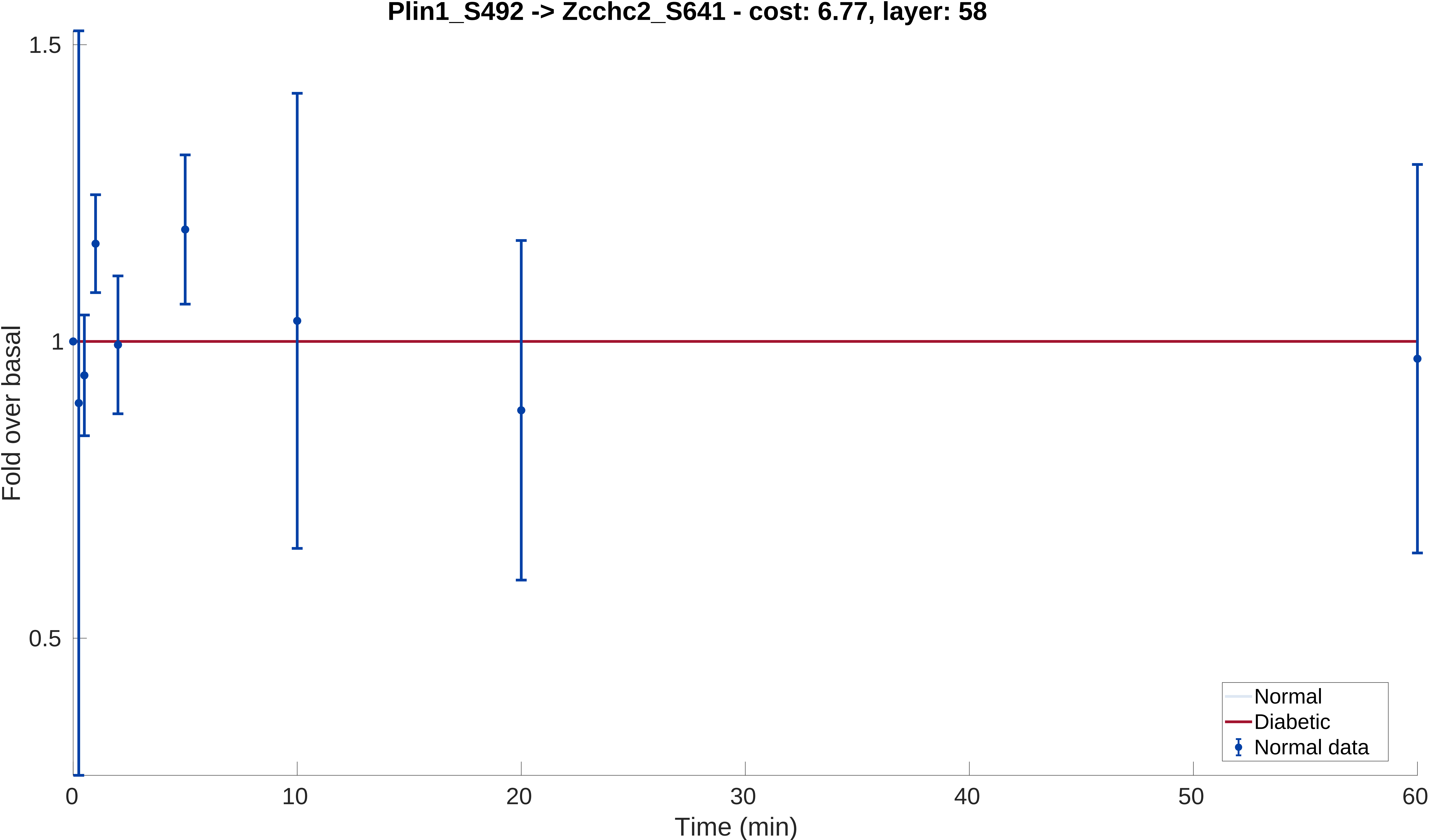

Plin1_S492 -> Zcchc2_S641 - cost: 6.77, layer: 58
Fold over basal
1.5
1
0.5
0
10
20
30
40
50
60
Time (min)
Normal
Diabetic
Normal data

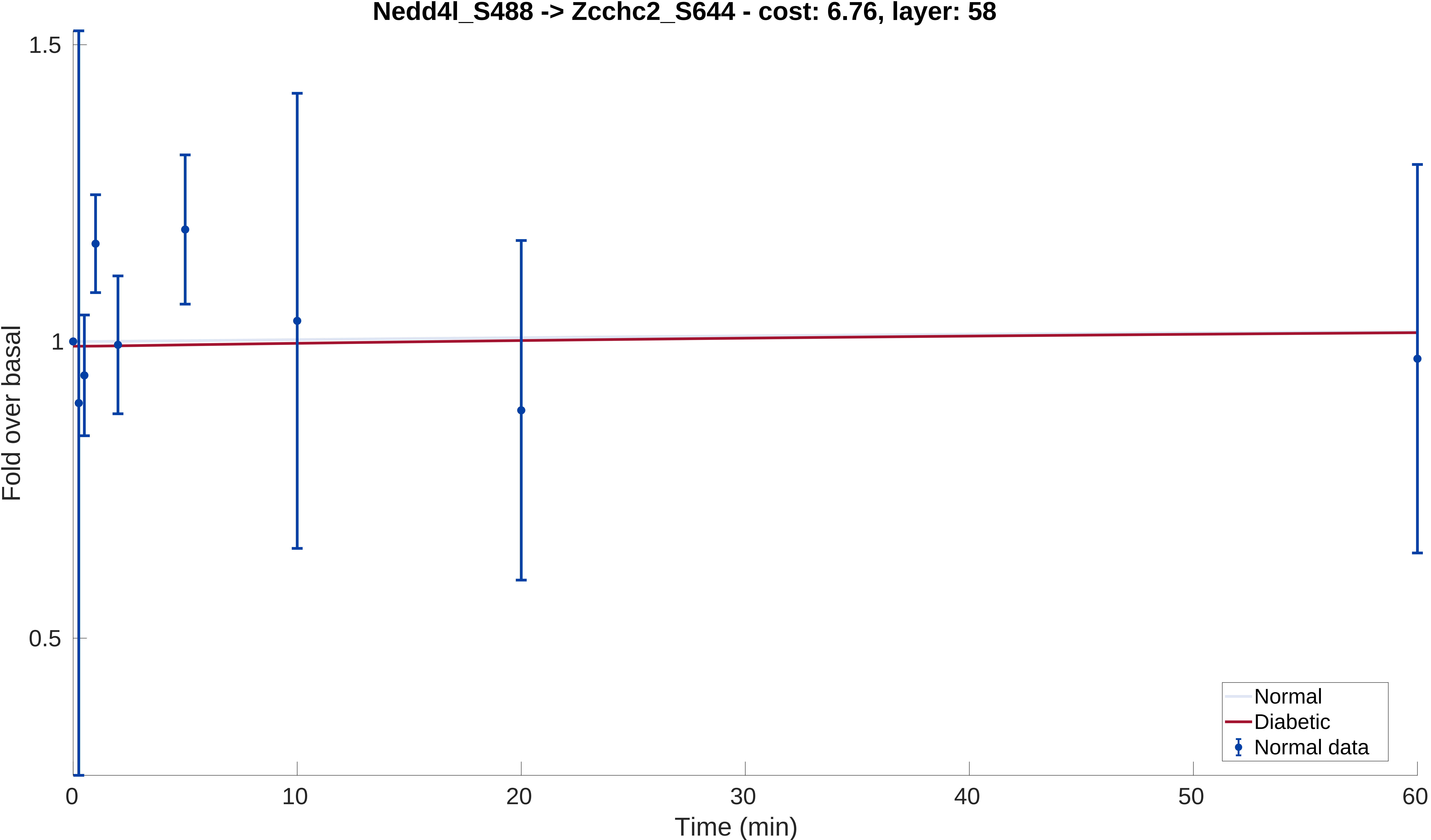

Nedd4l_S488 -> Zcchc2_S644 - cost: 6.76, layer: 58
1.5
1
0.5
Fold over basal
0
10
20
30
40
50
60
Time (min)
Normal
Diabetic
Normal data

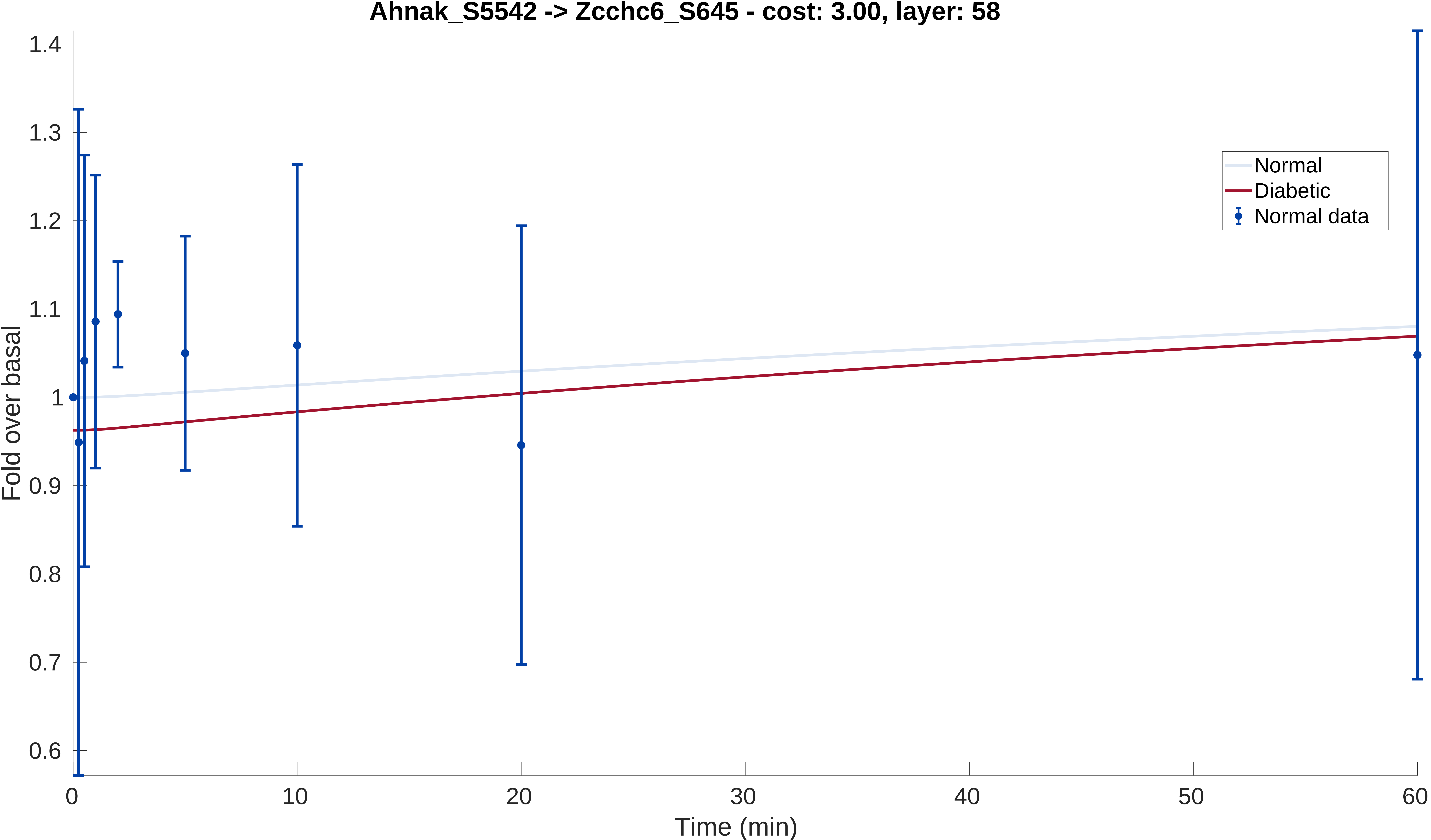

Ahnak_S5542 -> Zcchc6_S645 - cost: 3.00, layer: 58
Fold over basal
Time (min)
Normal
Diabetic
Normal data
1.4
1.3
1.2
1.1
1
0.9
0.8
0.7
0.6
0
10
20
30
40
50
60

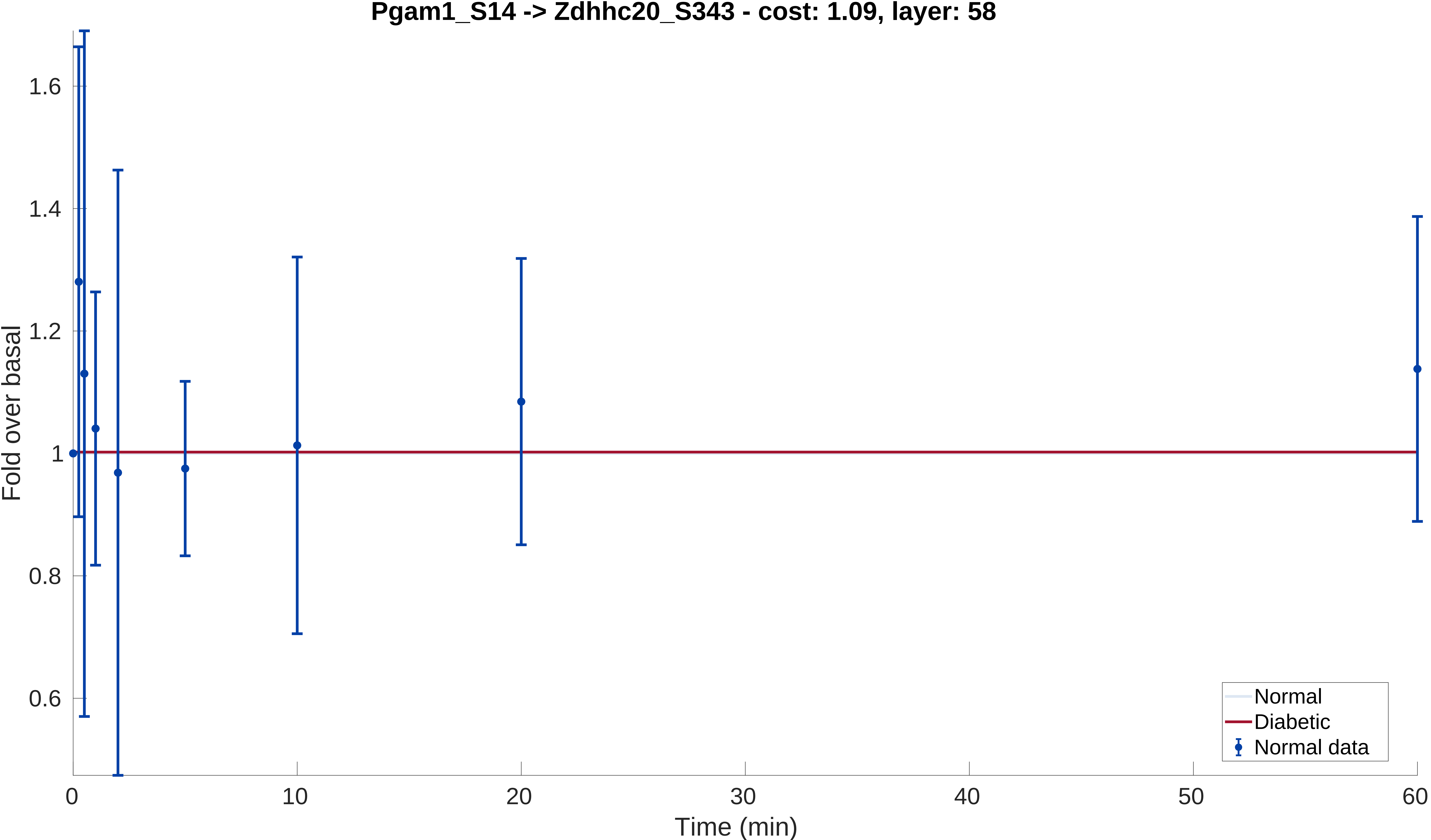

Pgam1_S14 -> Zdhhc20_S343 - cost: 1.09, layer: 58
Fold over basal
1.6
1.4
1.2
1
0.8
0.6
0
10
20
30
40
50
60
Time (min)
Normal
Diabetic
Normal data

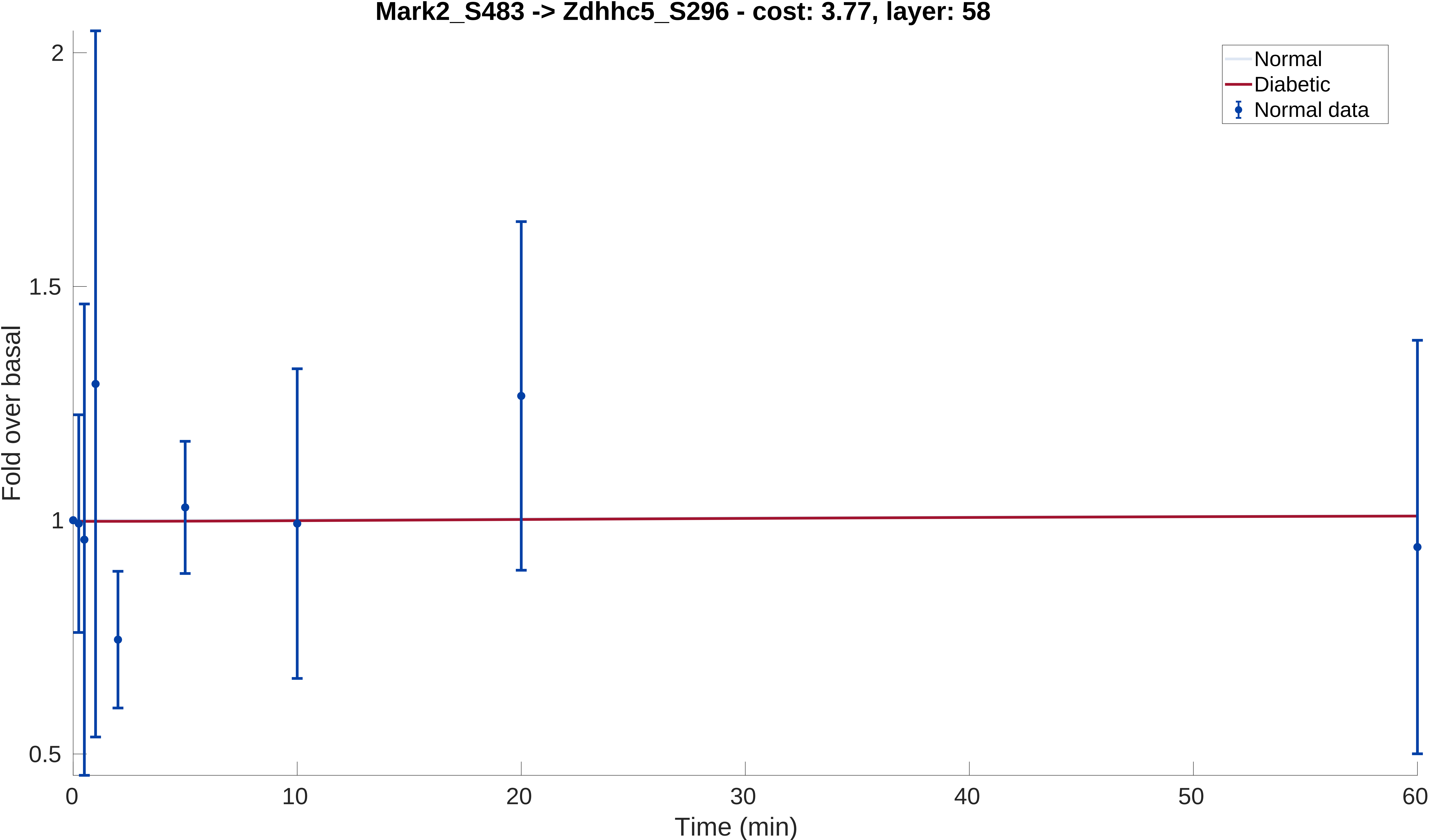

Mark2_S483 -> Zdhhc5_S296 - cost: 3.77, layer: 58
Normal
Diabetic
Normal data
Fold over basal
0.5
1
1.5
2
0
10
20
30
40
50
60
Time (min)

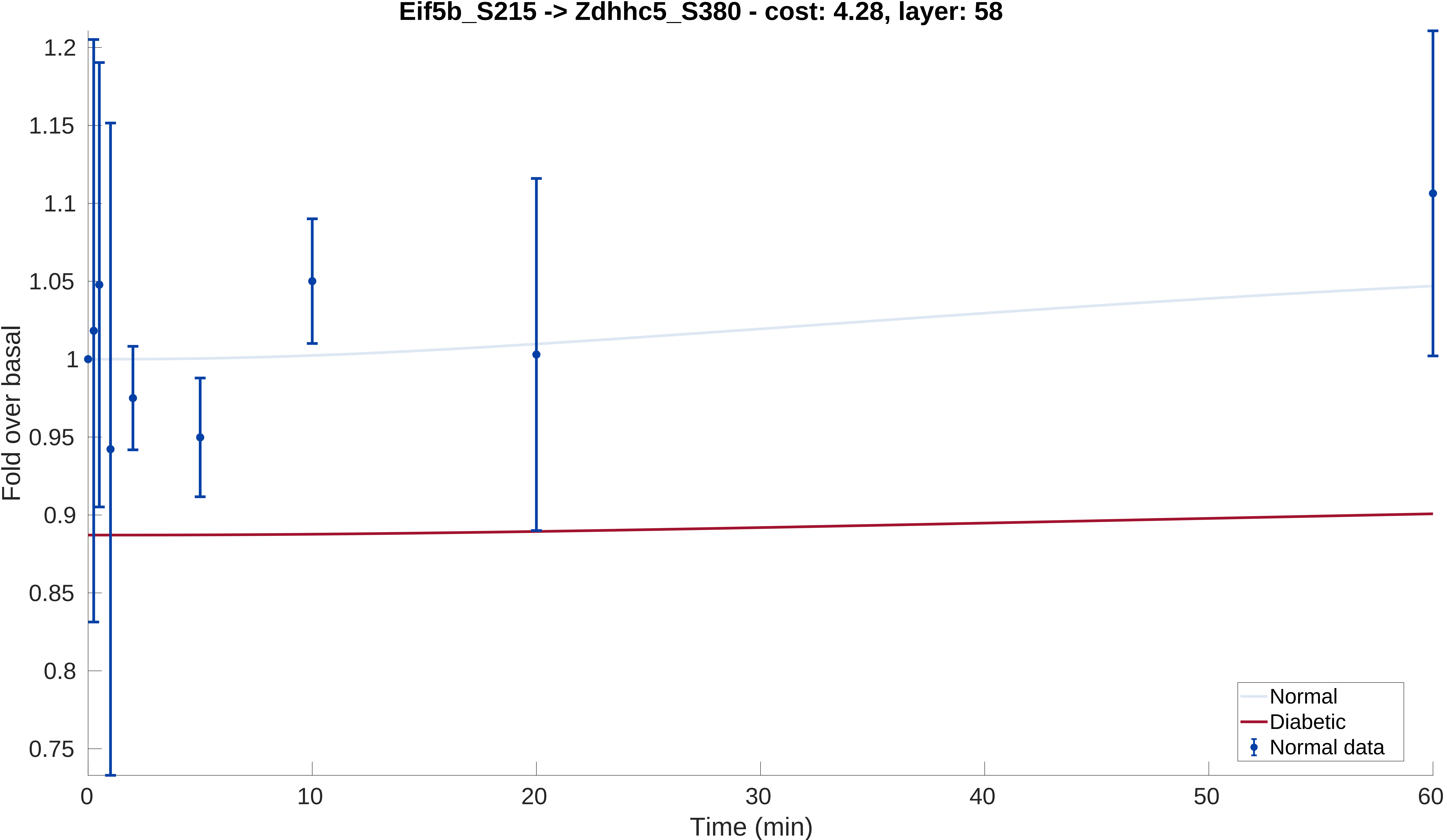

Eif5b_S215 -> Zdhhc5_S380 - cost: 4.28, layer: 58
Fold over basal
1.2
1.15
1.1
1.05
1
0.95
0.9
0.85
0.8
0.75
0
10
20
30
40
50
60
Time (min)
Normal
Diabetic
Normal data

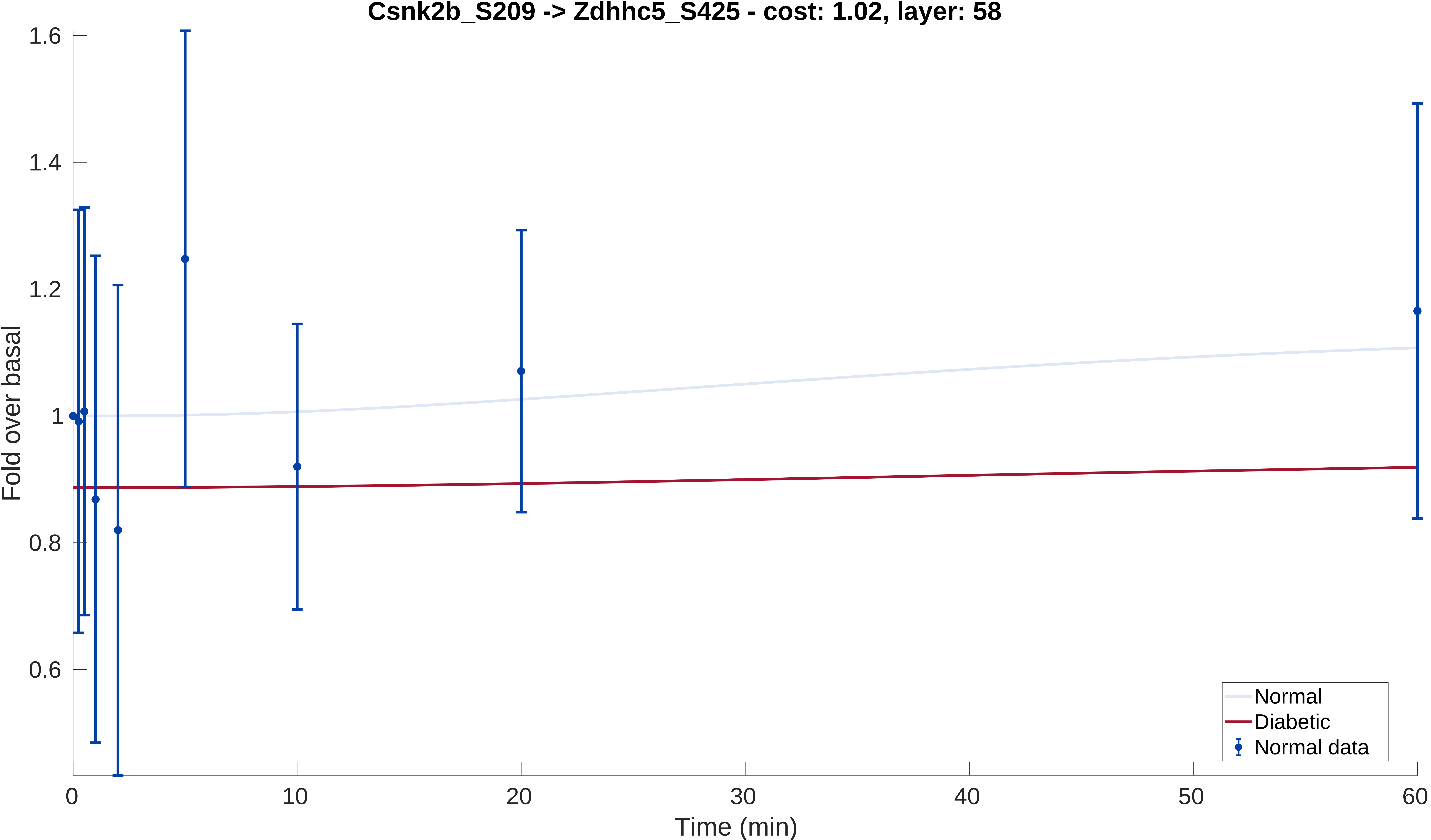

Csnk2b_S209 -> Zdhhc5_S425 - cost: 1.02, layer: 58
1.6
1.4
1.2
1
0.8
0.6
Fold over basal
0
10
20
30
40
50
60
Time (min)
Normal
Diabetic
Normal data

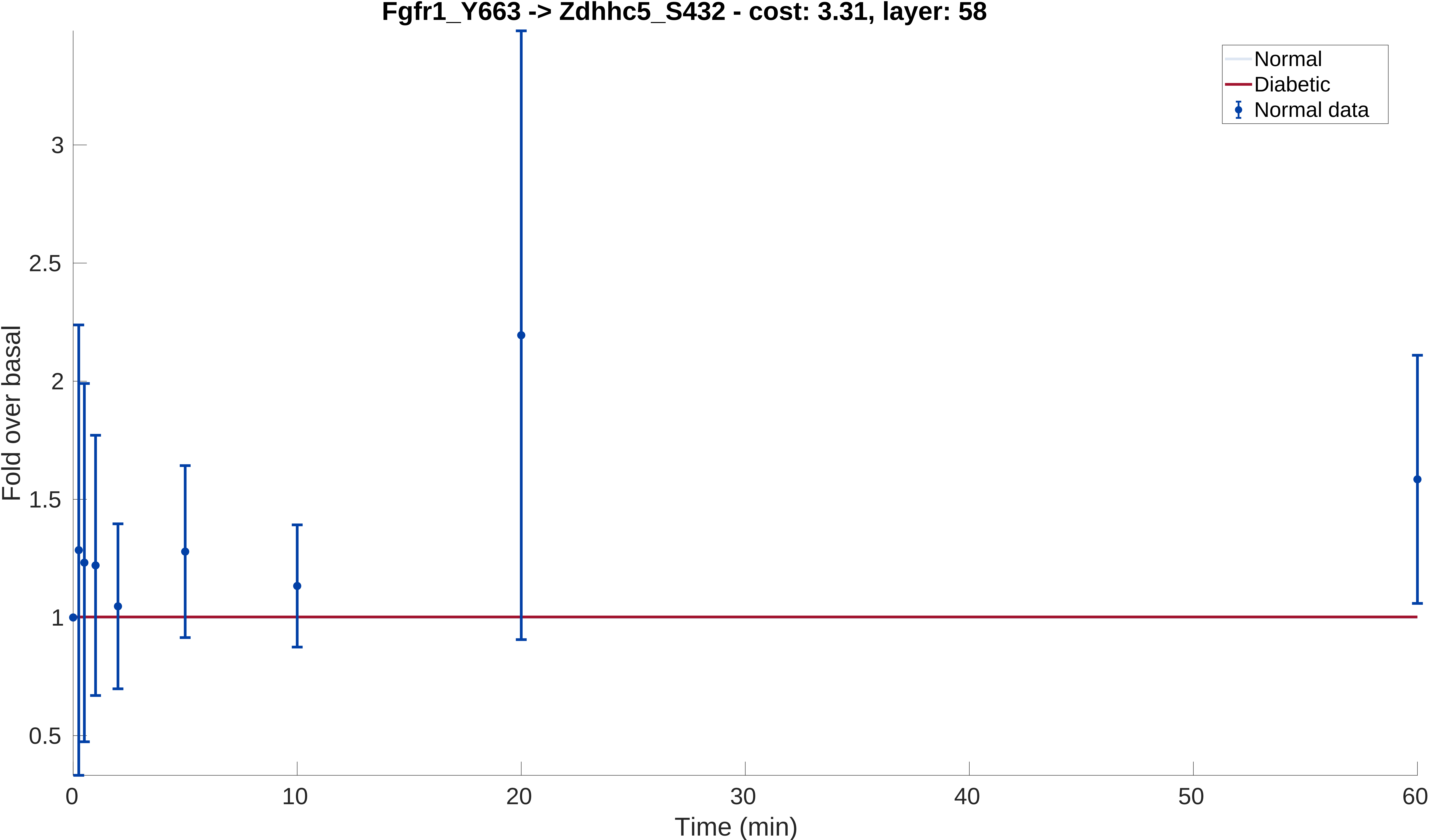

Fgfr1_Y663 -> Zdhhc5_S432 - cost: 3.31, layer: 58
Normal
Diabetic
Normal data
Fold over basal
0.5
1
1.5
2
2.5
3
0
10
20
30
40
50
60
Time (min)

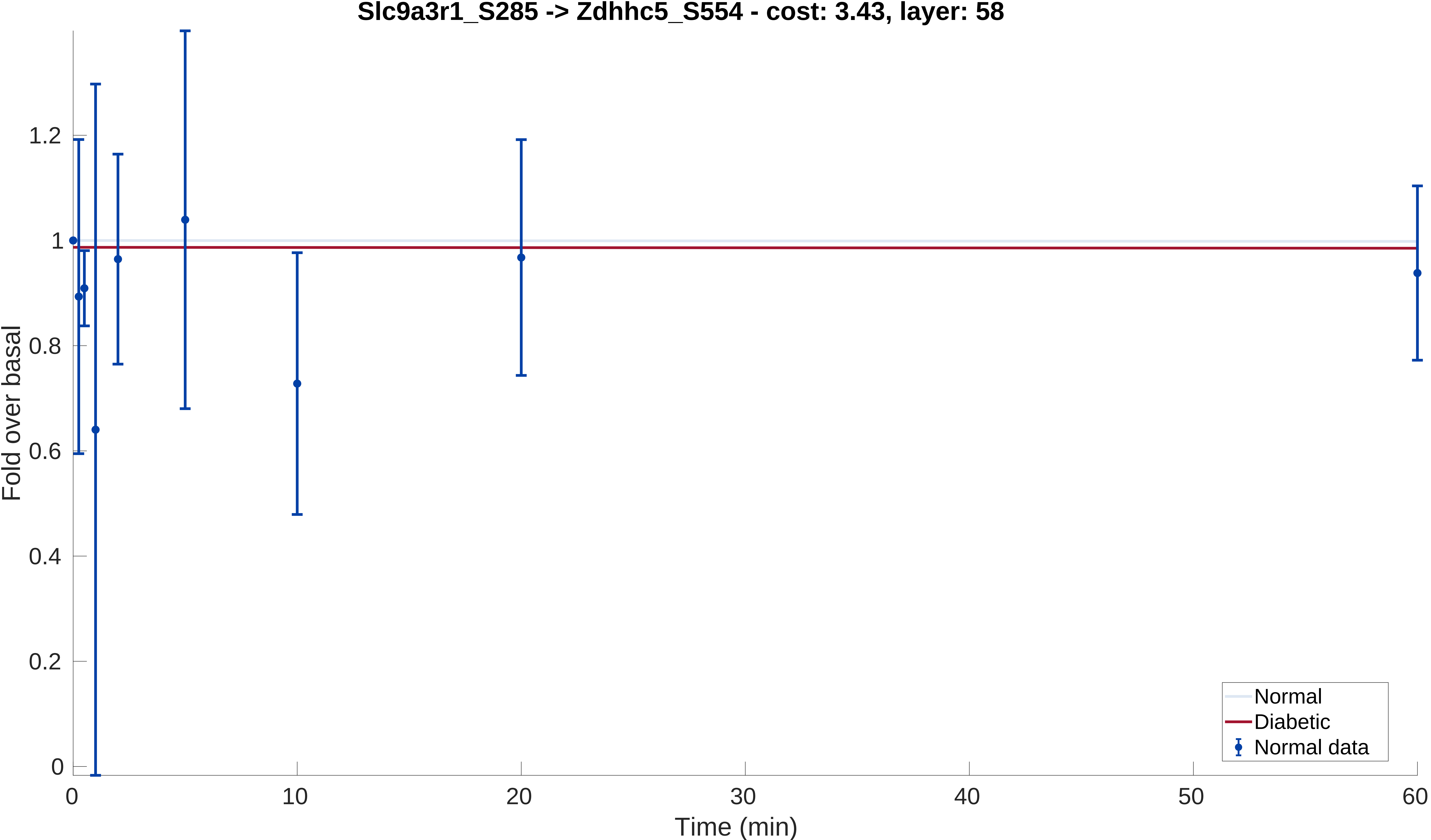

Slc9a3r1_S285 -> Zdhhc5_S554 - cost: 3.43, layer: 58
Fold over basal
0
0.2
0.4
0.6
0.8
1
1.2
0
10
20
30
40
50
60
Time (min)
Normal
Diabetic
Normal data

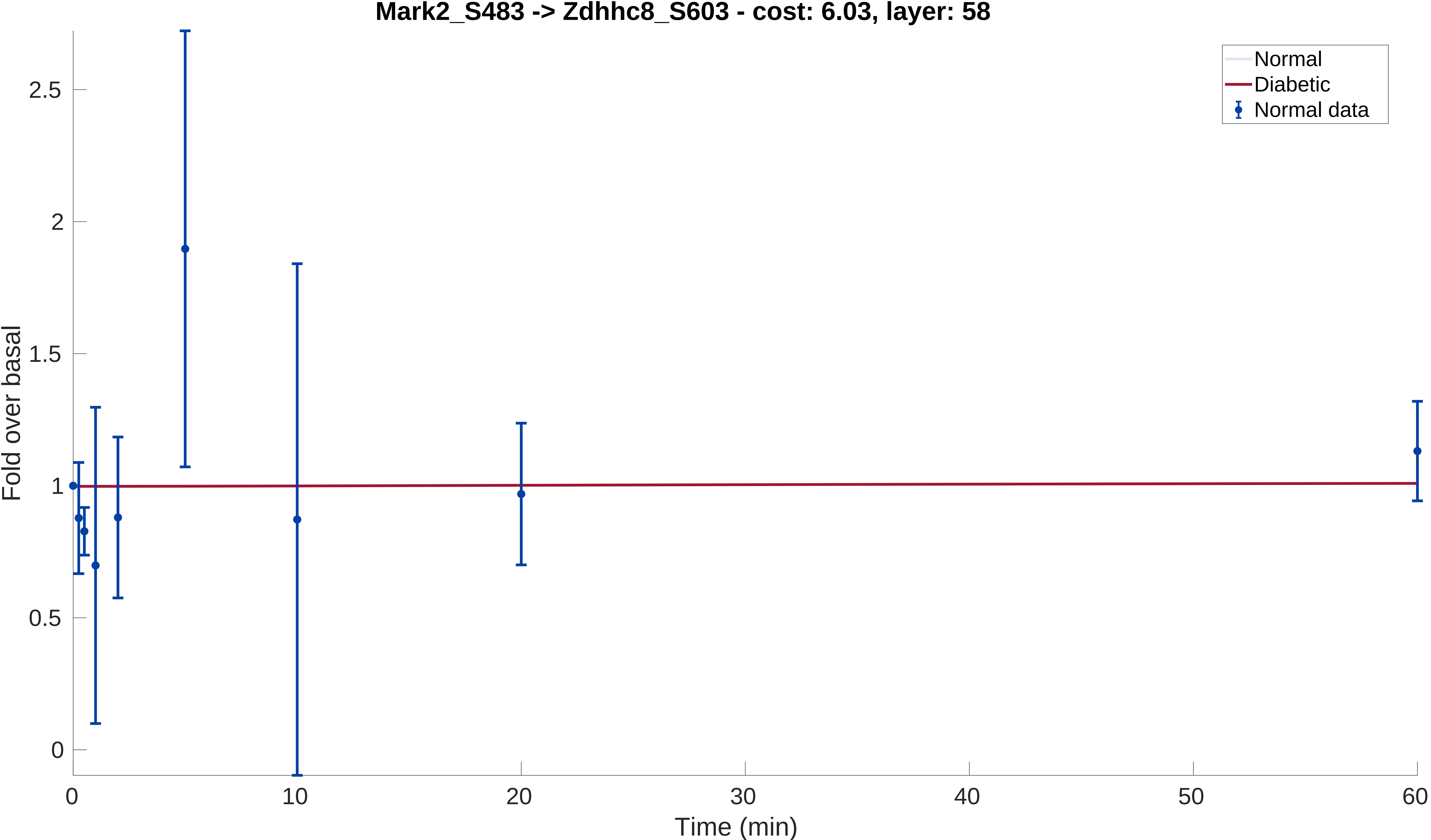
Mark2_S483 -> Zdhhc8_S603 - cost: 6.03, layer: 58
Fold over basal
0
0.5
1
1.5
2
2.5
0
10
20
30
40
50
60
Time (min)
Normal
Diabetic
Normal data

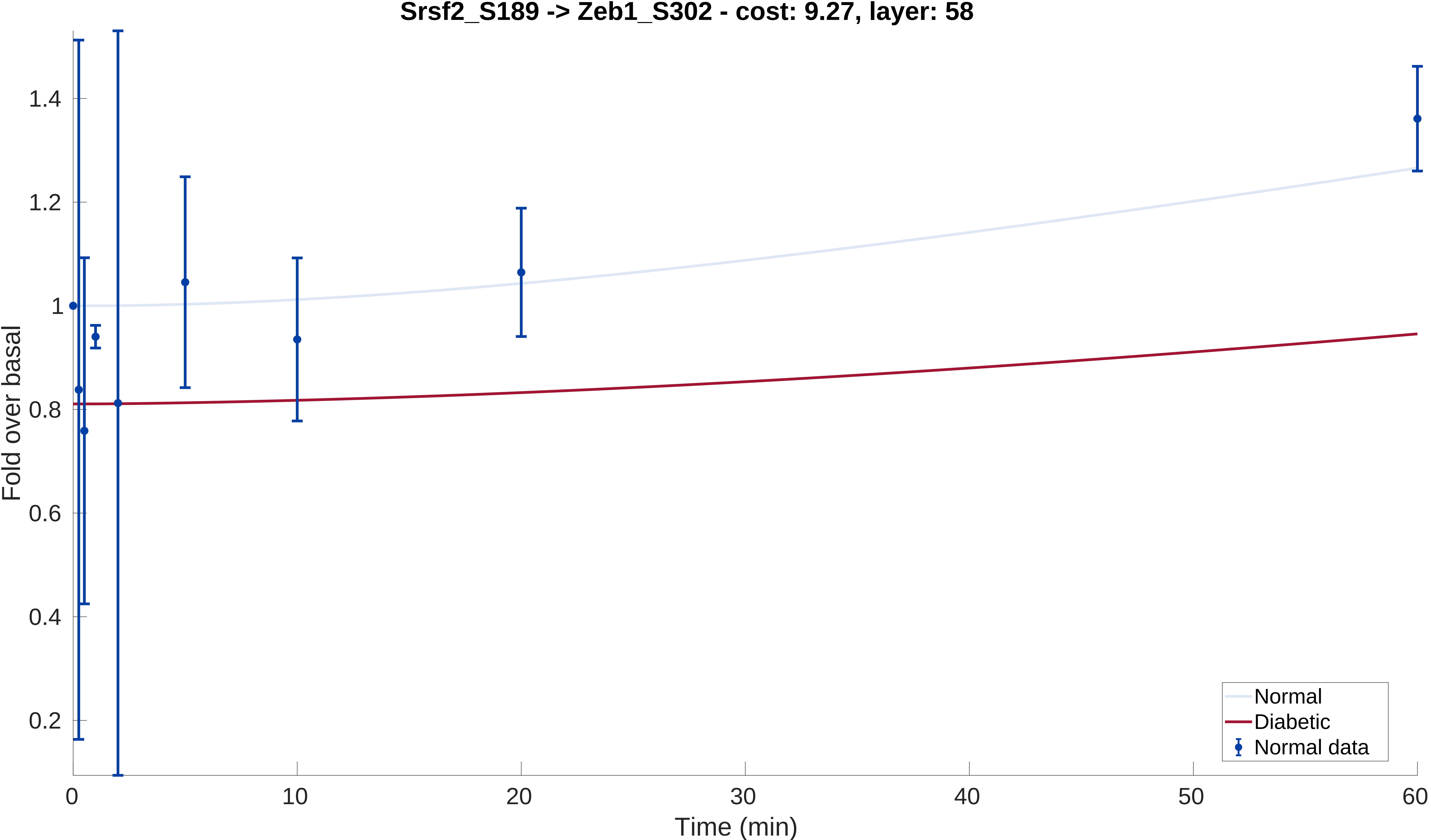

Srsf2_S189 -> Zeb1_S302 - cost: 9.27, layer: 58
Fold over basal
1.4
1.2
1
0.8
0.6
0.4
0.2
0
10
20
30
40
50
60
Time (min)
Normal
Diabetic
Normal data

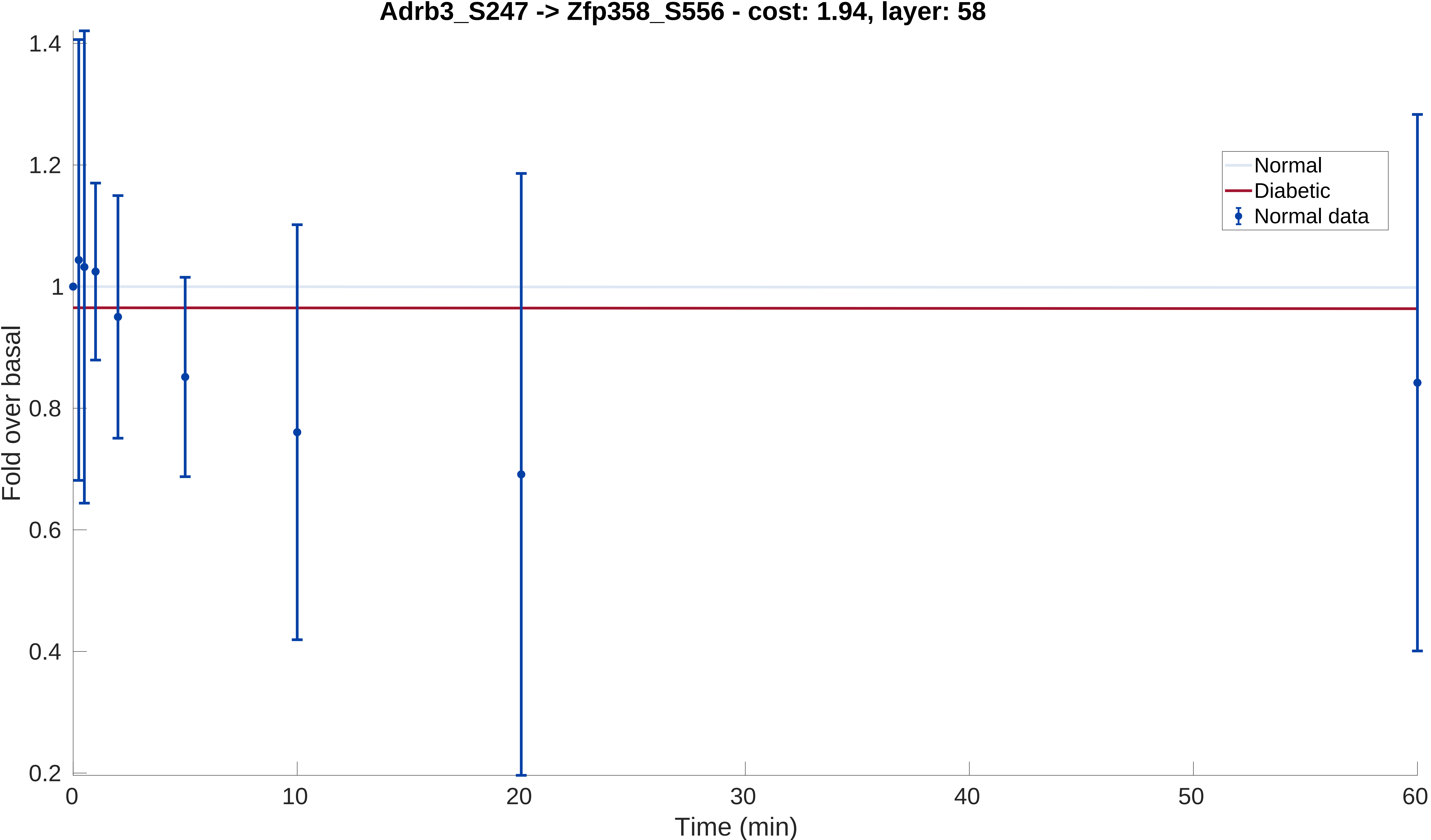
Adrb3_S247 -> Zfp358_S556 - cost: 1.94, layer: 58
Normal
Diabetic
Normal data
Fold over basal
Time (min)
0.2
0.4
0.6
0.8
1
1.2
1.4
0
10
20
30
40
50
60

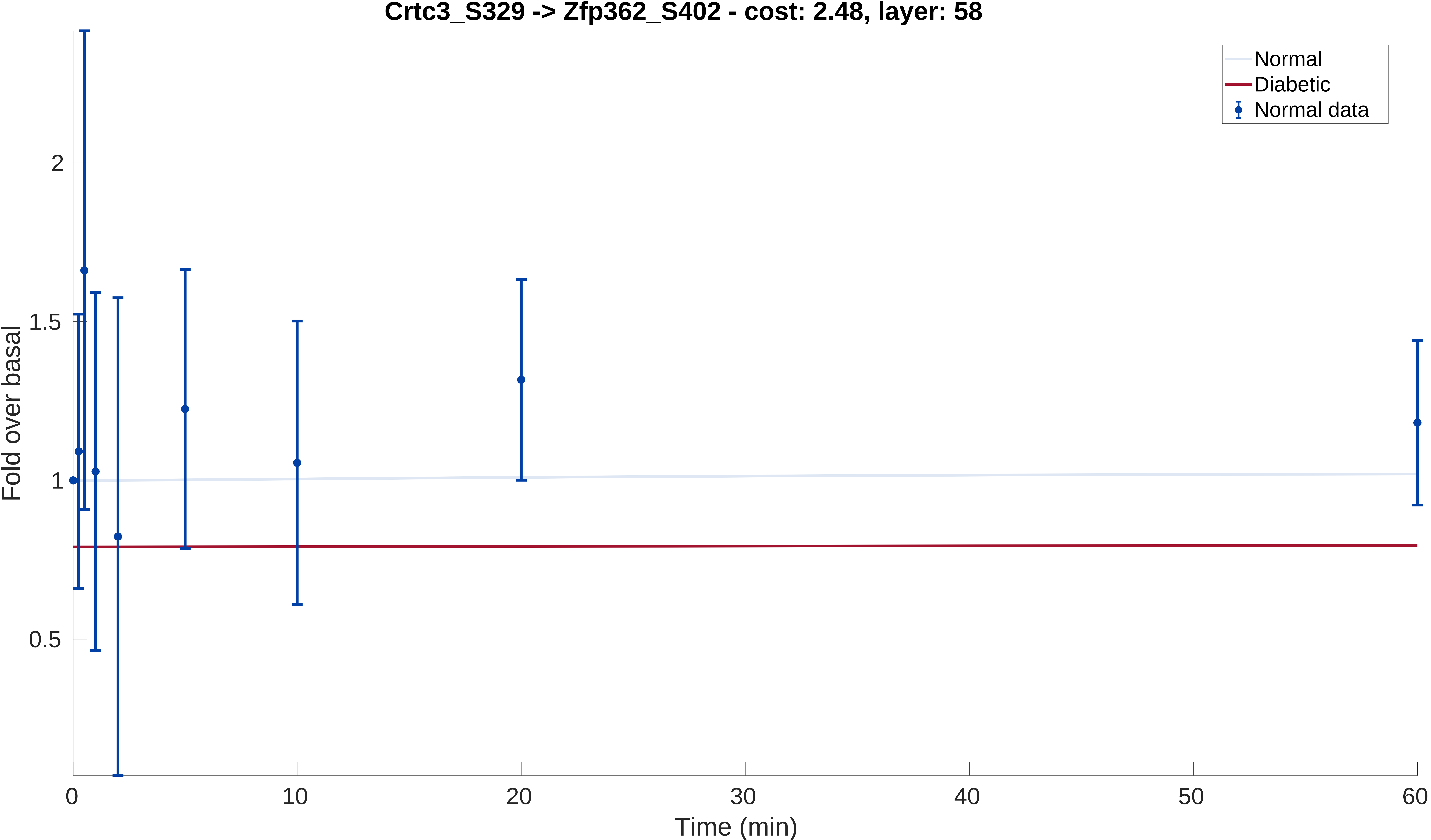

Crtc3_S329 -> Zfp362_S402 - cost: 2.48, layer: 58
Normal
Diabetic
Normal data
Fold over basal
2
1.5
1
0.5
0
10
20
30
40
50
60
Time (min)

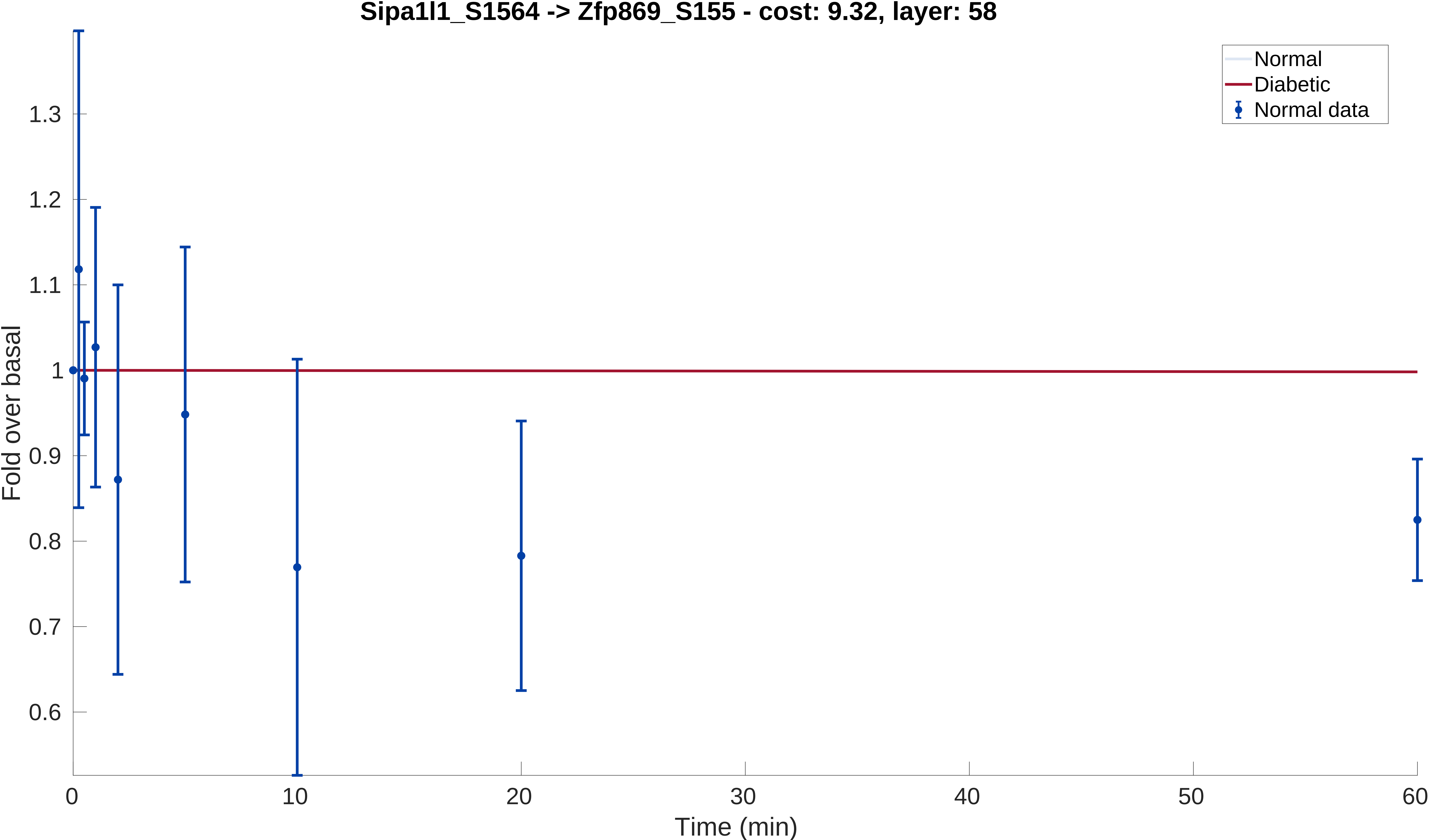

Sipa1l1_S1564 -> Zfp869_S155 - cost: 9.32, layer: 58
Normal
Diabetic
Normal data
Fold over basal
Time (min)
1.3
1.2
1.1
1
0.9
0.8
0.7
0.6
0
10
20
30
40
50
60

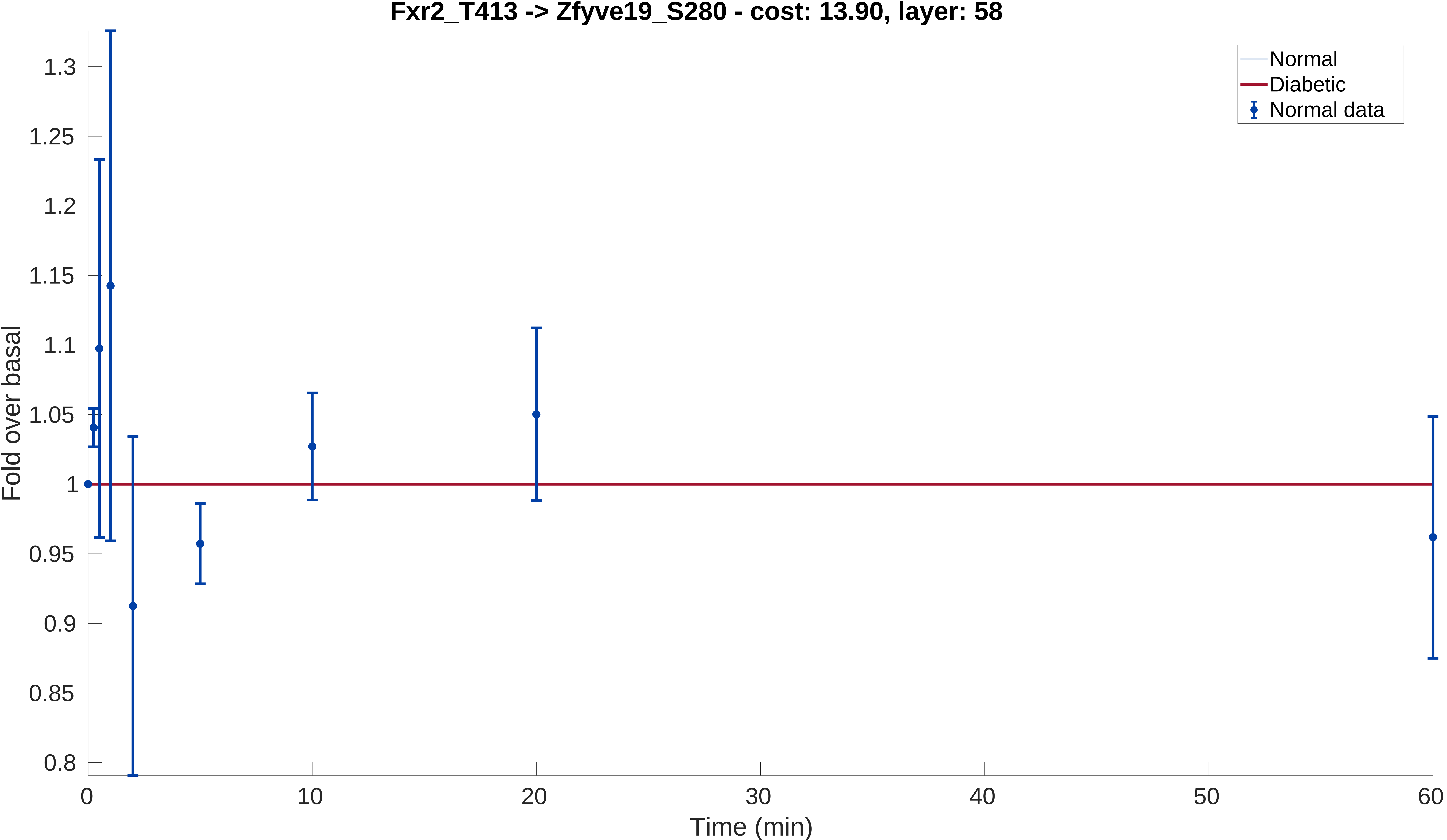

Fxr2_T413 -> Zfyve19_S280 - cost: 13.90, layer: 58
Normal
Diabetic
Normal data
Fold over basal
Time (min)
1.3
1.25
1.2
1.15
1.1
1.05
1
0.95
0.9
0.85
0.8
0
10
20
30
40
50
60

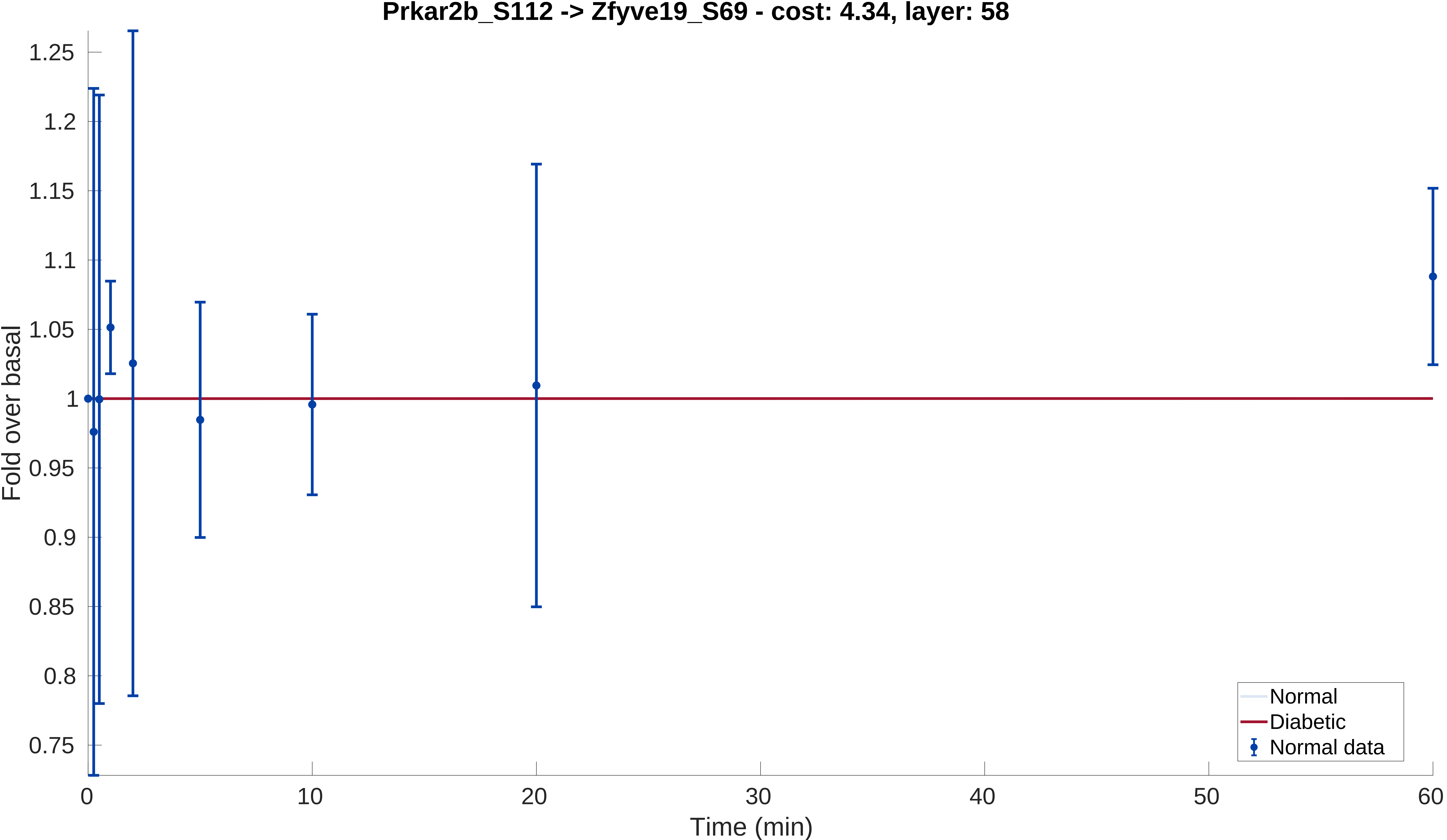
Prkar2b_S112 -> Zfyve19_S69 - cost: 4.34, layer: 58
Fold over basal
1.25
1.2
1.15
1.1
1.05
1
0.95
0.9
0.85
0.8
0.75
0
10
20
30
40
50
60
Time (min)
Normal
Diabetic
Normal data

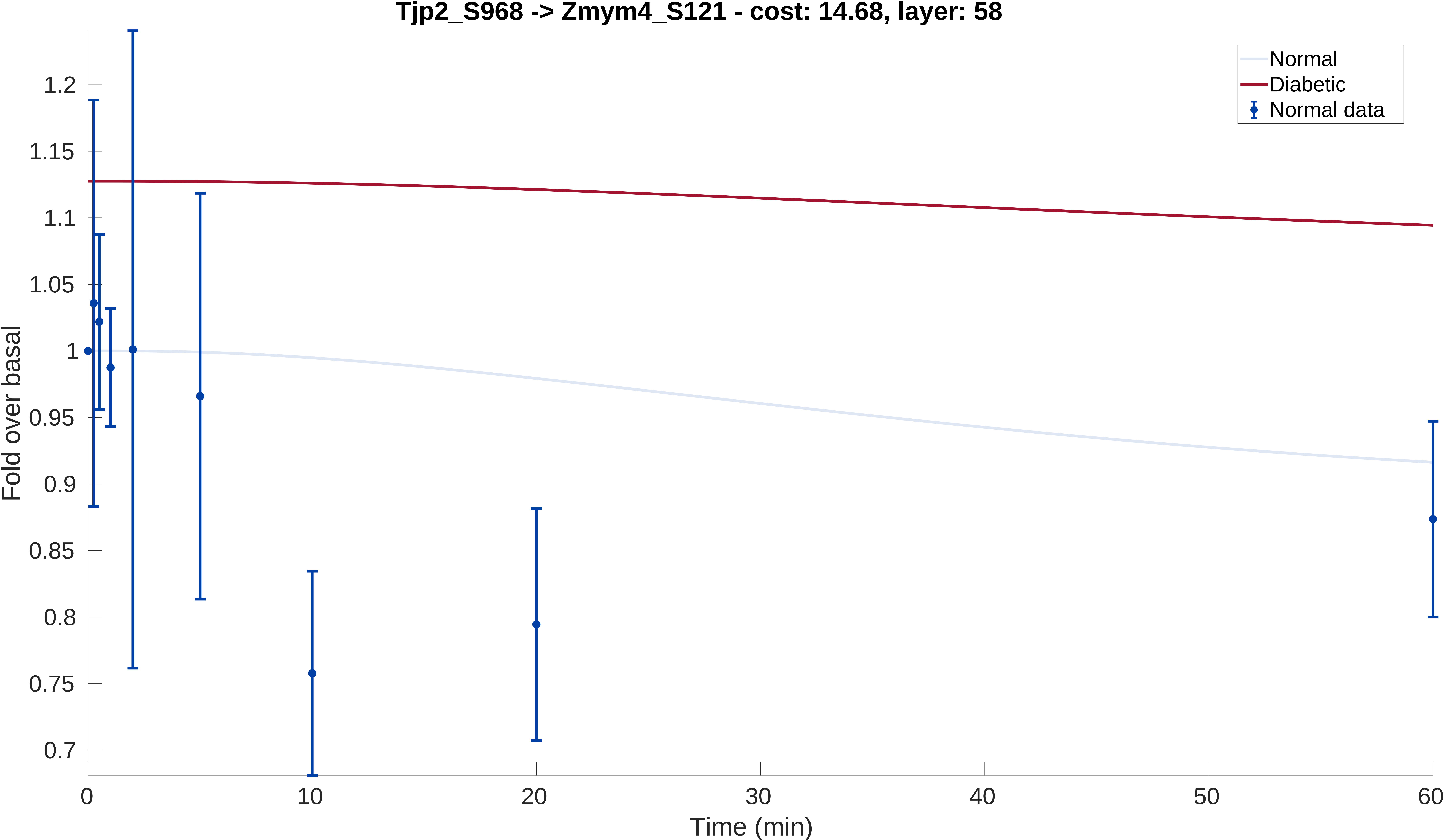

Tjp2_S968 -> Zmym4_S121 - cost: 14.68, layer: 58
Normal
Diabetic
Normal data
Fold over basal
Time (min)
1.2
1.15
1.1
1.05
1
0.95
0.9
0.85
0.8
0.75
0.7
0
10
20
30
40
50
60

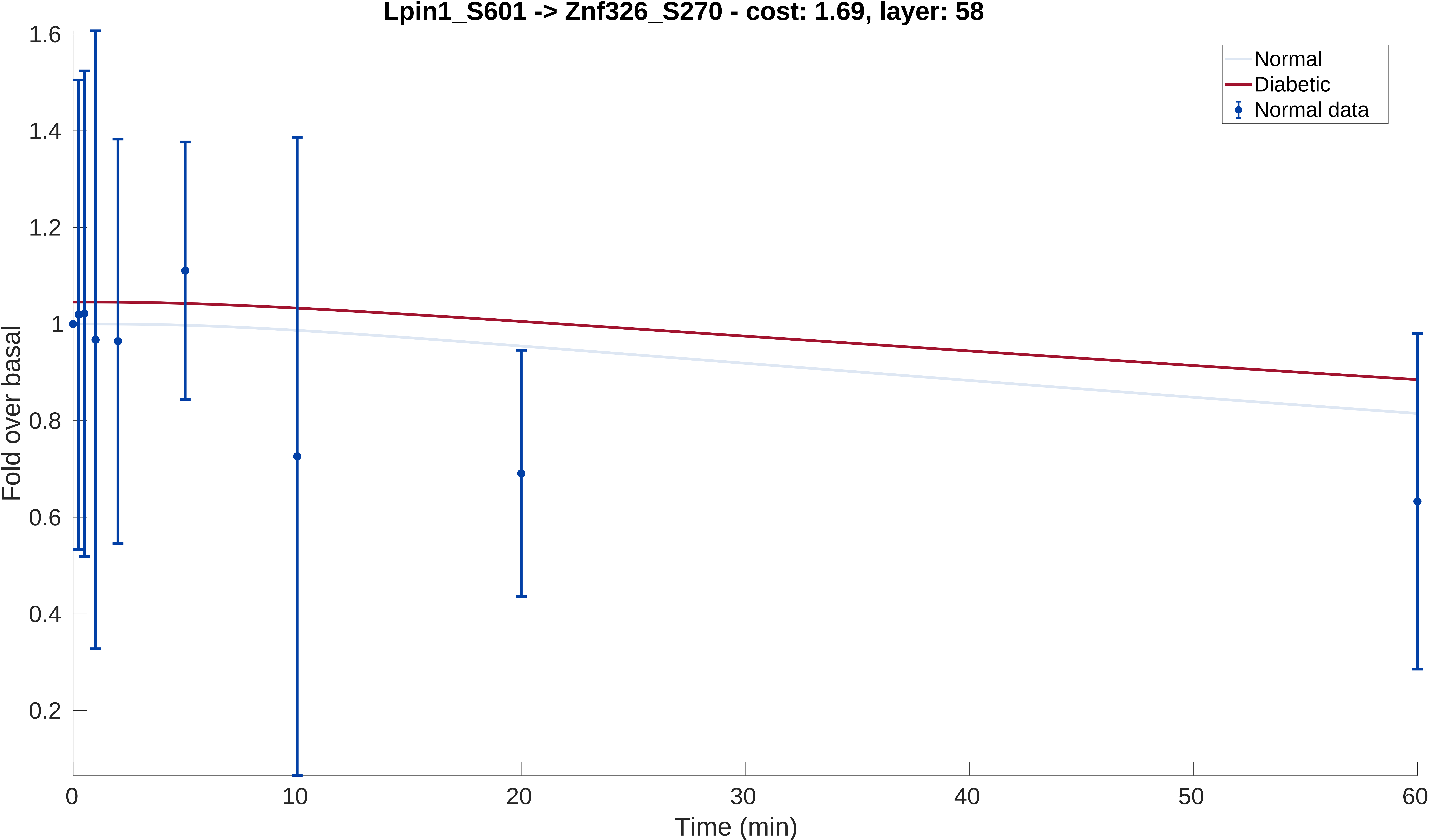

Lpin1_S601 -> Znf326_S270 - cost: 1.69, layer: 58
Normal
Diabetic
Normal data
Fold over basal
Time (min)
1.6
1.4
1.2
1
0.8
0.6
0.4
0.2
0
10
20
30
40
50
60

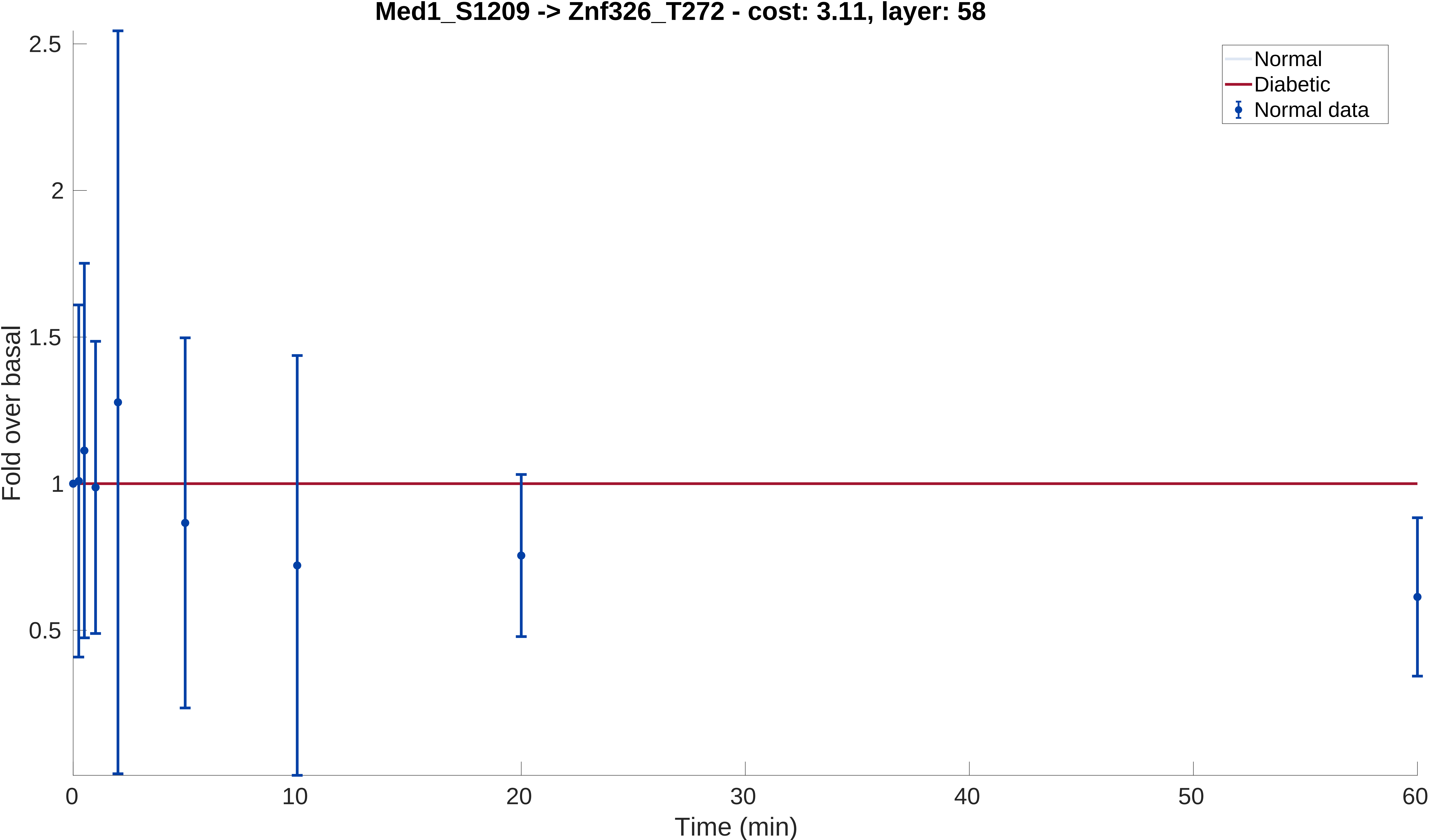
Med1_S1209 -> Znf326_T272 - cost: 3.11, layer: 58
Normal
Diabetic
Normal data
Fold over basal
Time (min)
0.5
1
1.5
2
2.5
0
10
20
30
40
50
60

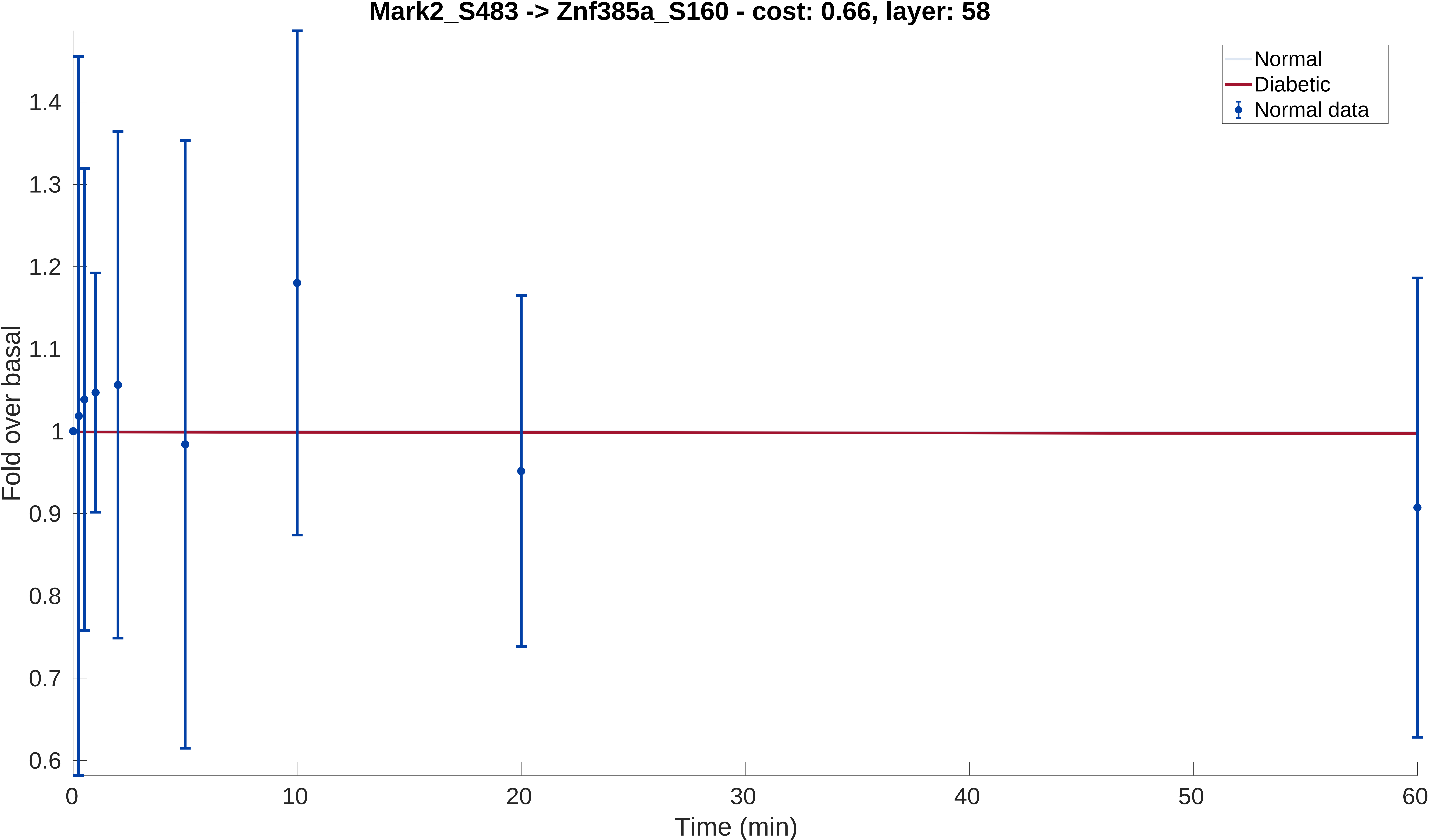

Mark2_S483 -> Znf385a_S160 - cost: 0.66, layer: 58
Normal
Diabetic
Normal data
Fold over basal
Time (min)
0.6
0.7
0.8
0.9
1
1.1
1.2
1.3
1.4
0
10
20
30
40
50
60

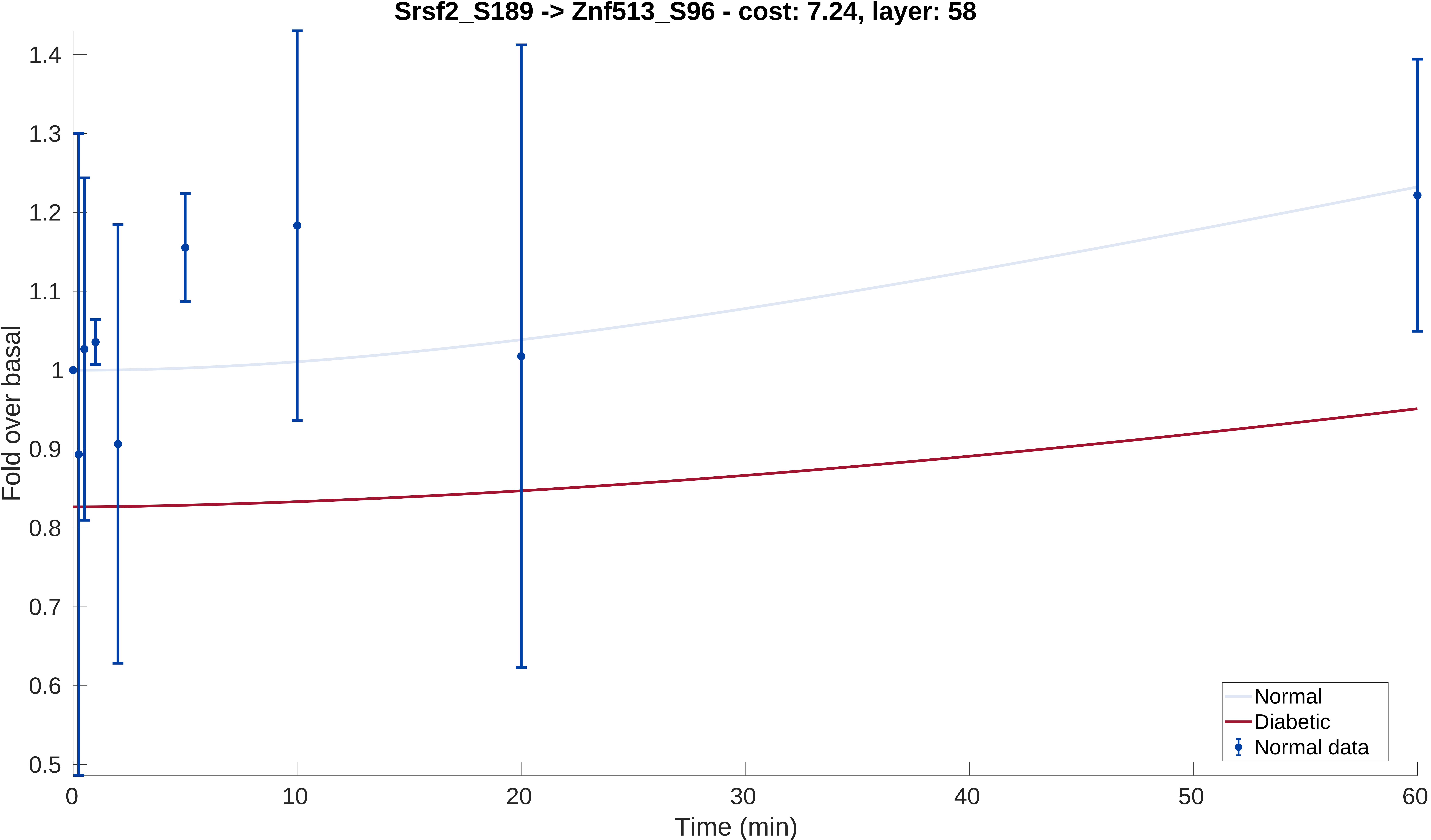
Srsf2_S189 -> Znf513_S96 - cost: 7.24, layer: 58
Fold over basal
1.4
1.3
1.2
1.1
1
0.9
0.8
0.7
0.6
0.5
0
10
20
30
40
50
60
Time (min)
Normal
Diabetic
Normal data

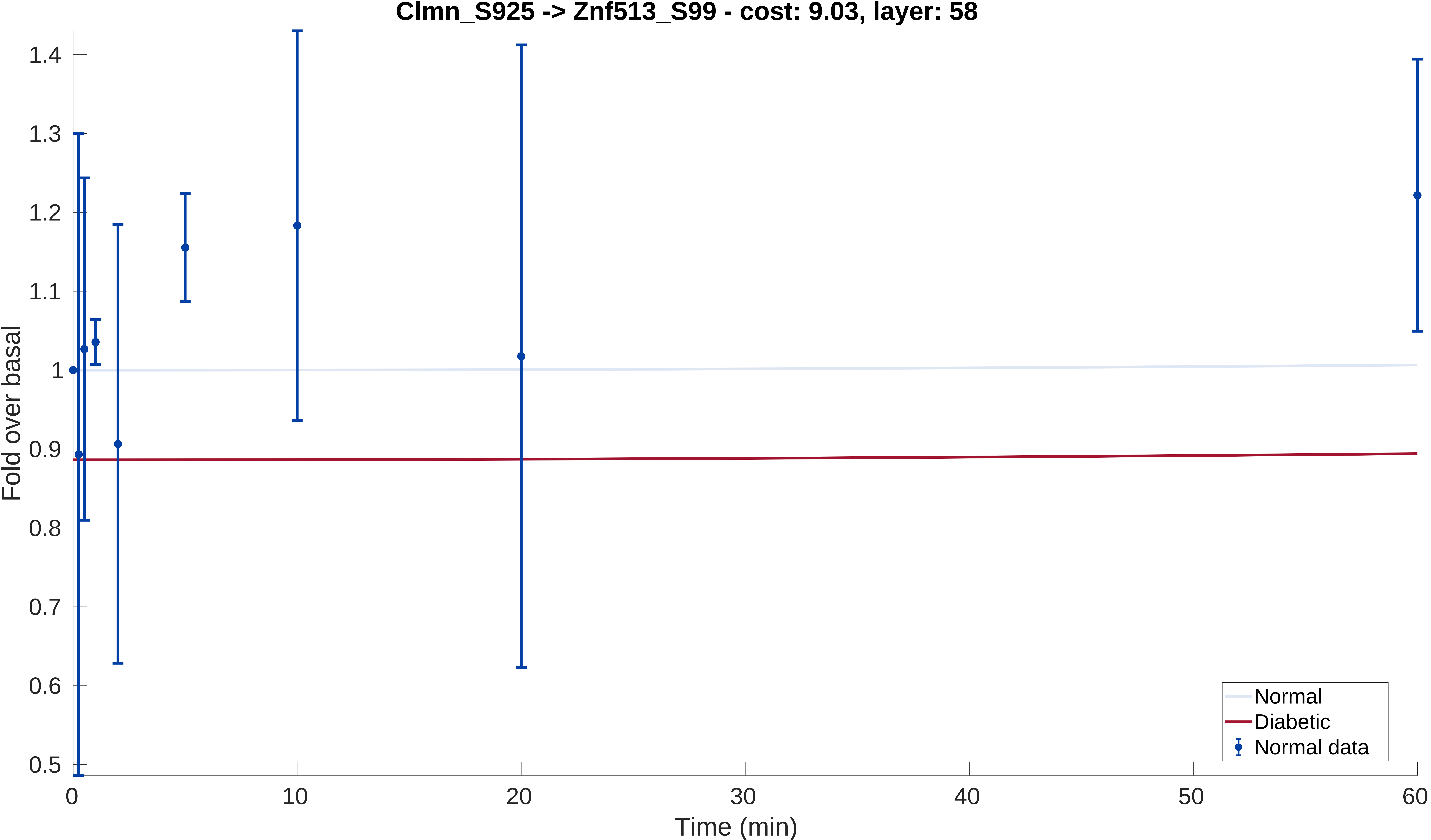

Clmn_S925 -> Znf513_S99 - cost: 9.03, layer: 58
Fold over basal
1.4
1.3
1.2
1.1
1
0.9
0.8
0.7
0.6
0.5
0
10
20
30
40
50
60
Time (min)
Normal
Diabetic
Normal data

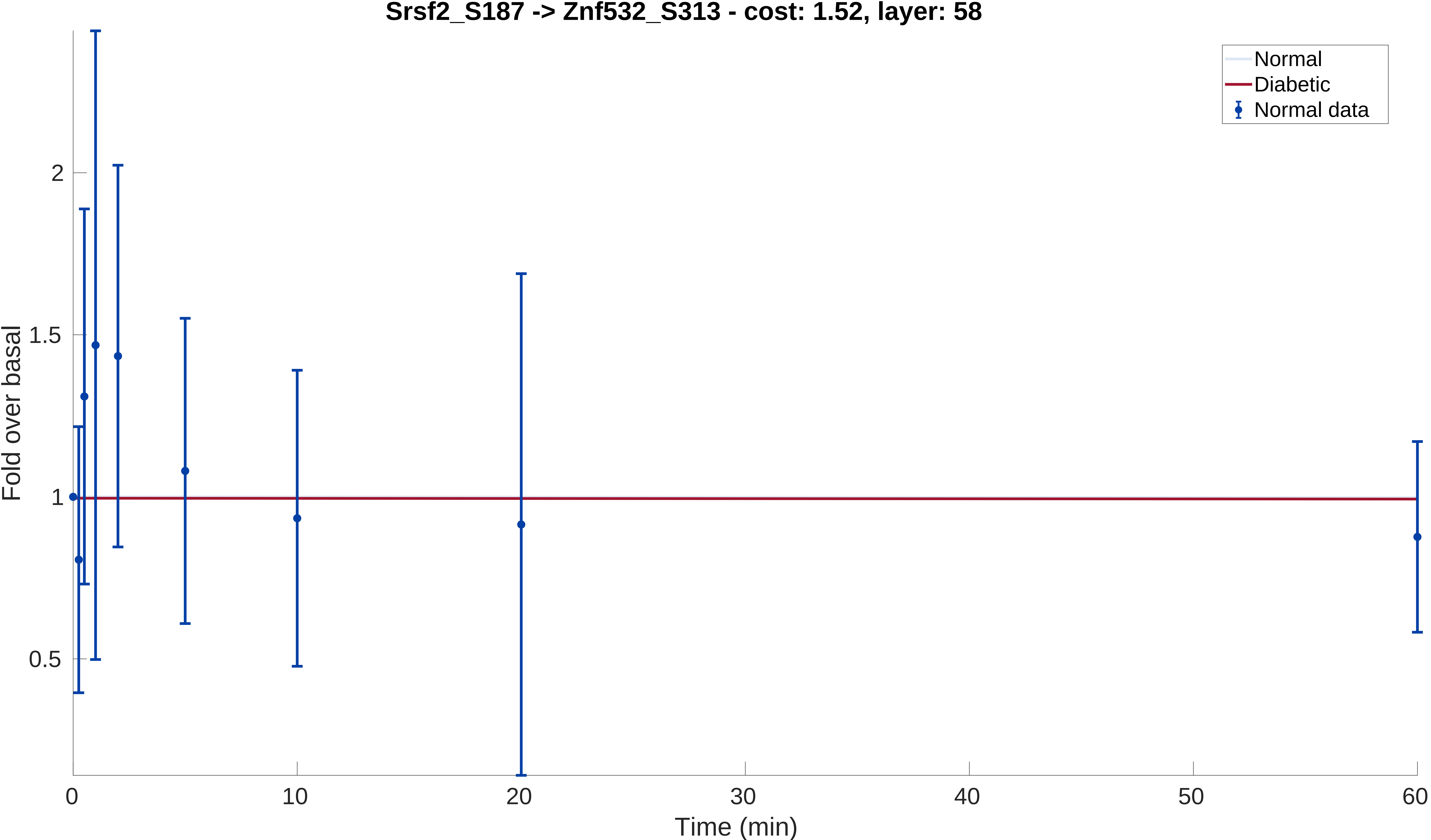

Srsf2_S187 -> Znf532_S313 - cost: 1.52, layer: 58
Normal
Diabetic
Normal data
Fold over basal
Time (min)
0.5
1
1.5
2
0
10
20
30
40
50
60

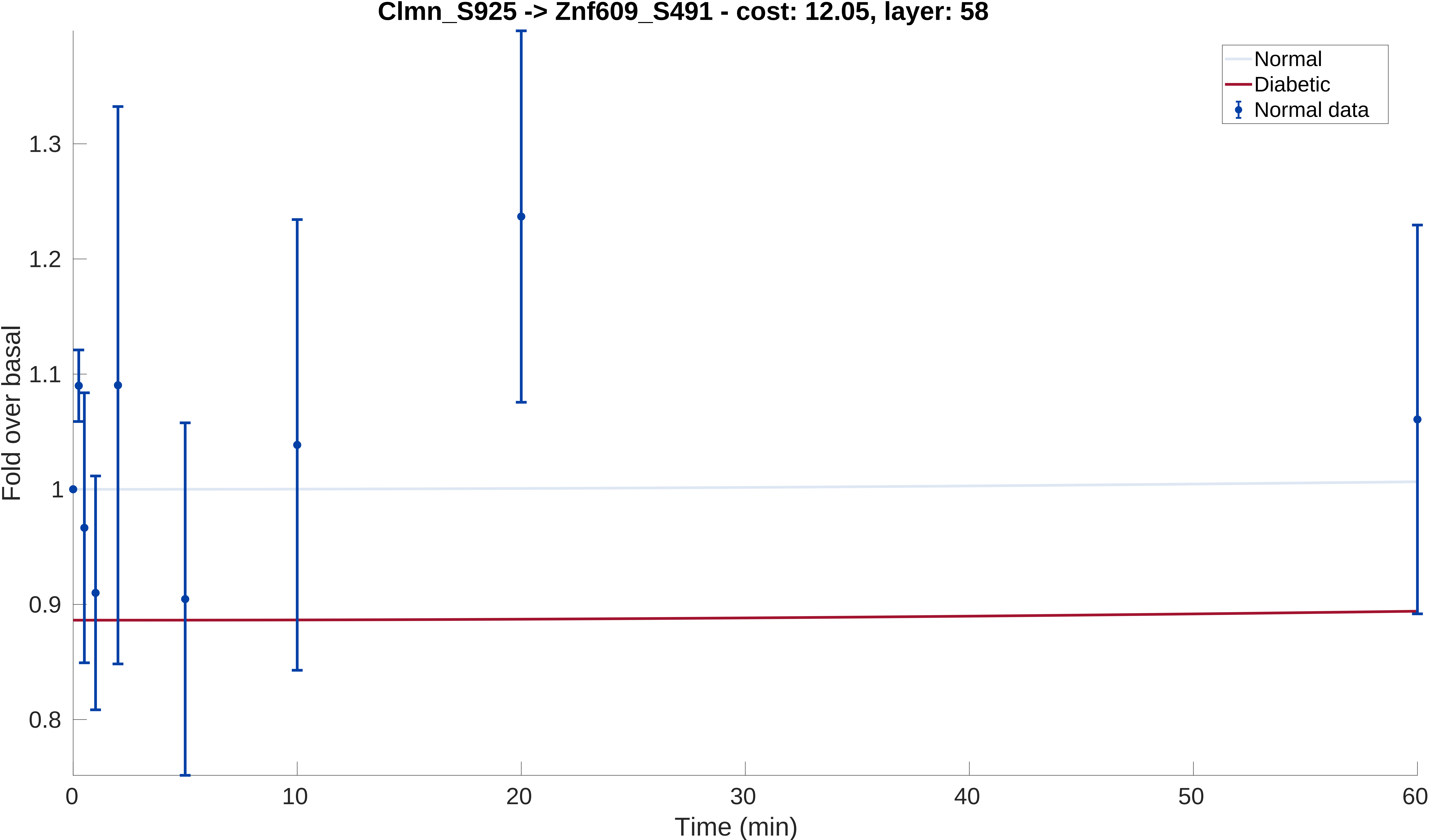
Clmn_S925 -> Znf609_S491 - cost: 12.05, layer: 58
Normal
Diabetic
Normal data
Fold over basal
1.3
1.2
1.1
1
0.9
0.8
0
10
20
30
40
50
60
Time (min)

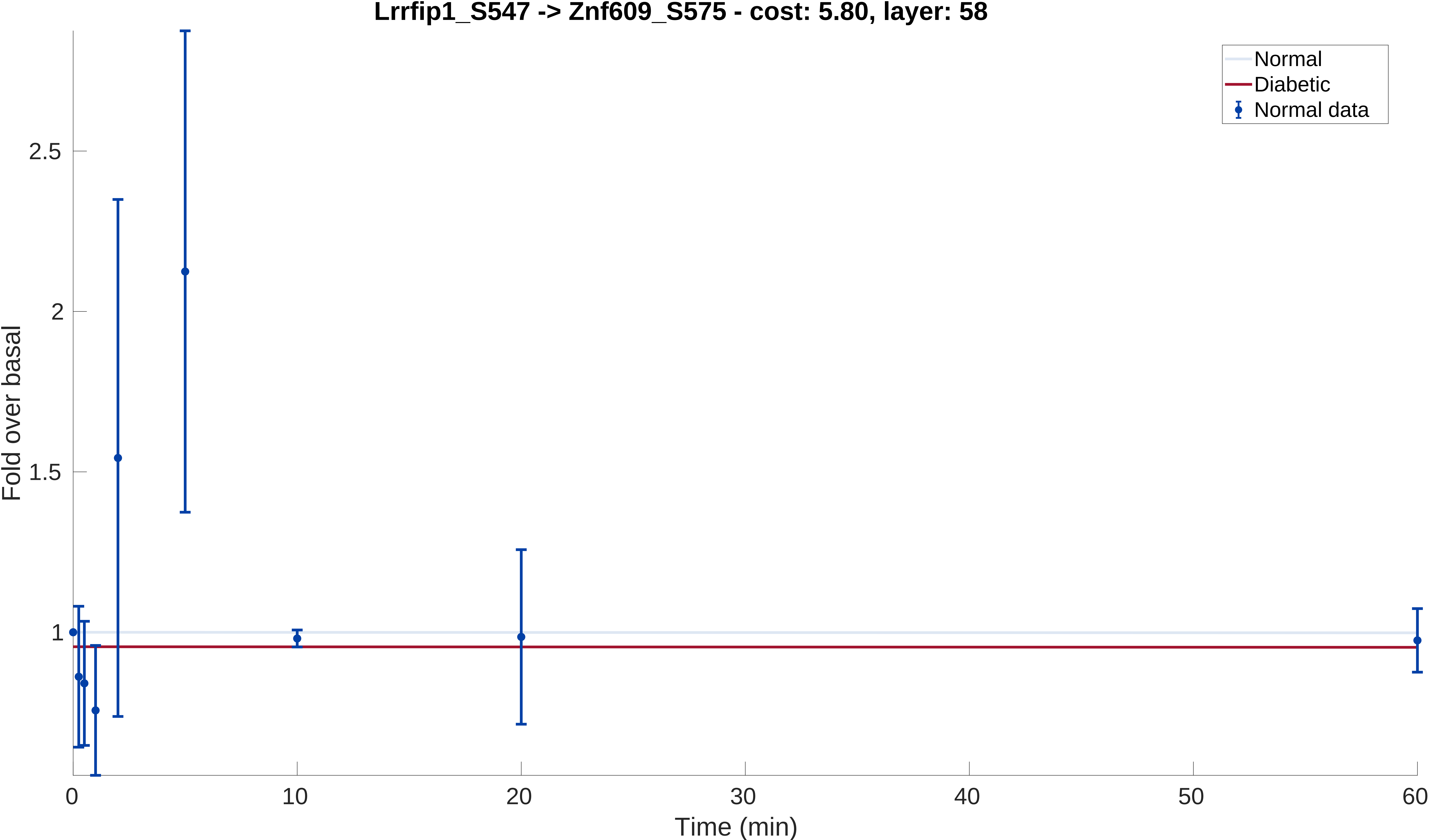
Lrrfip1_S547 -> Znf609_S575 - cost: 5.80, layer: 58
Normal
Diabetic
Normal data
Fold over basal
2.5
2
1.5
1
0
10
20
30
40
50
60
Time (min)

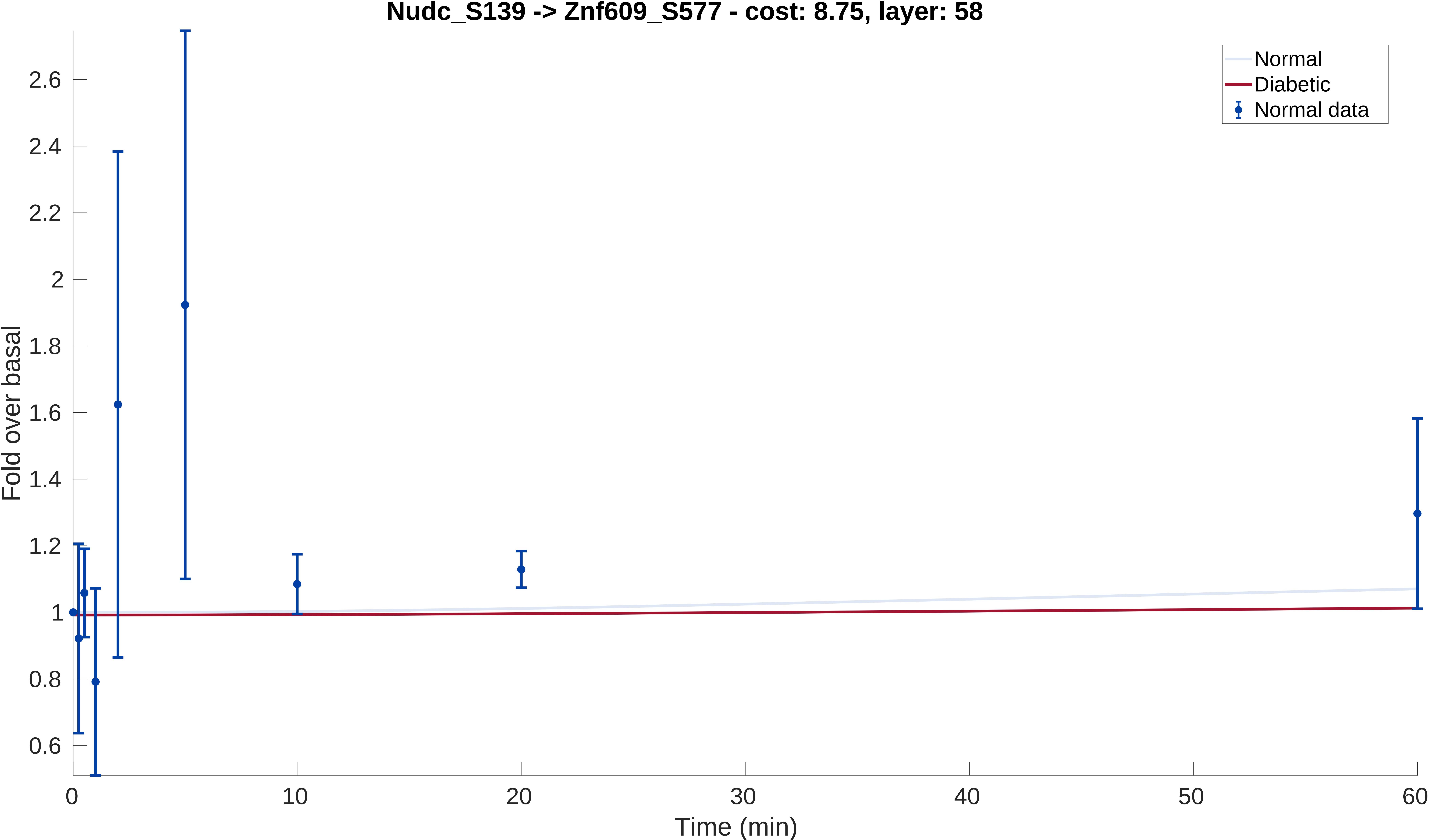

Nudc_S139 -> Znf609_S577 - cost: 8.75, layer: 58
Normal
Diabetic
Normal data
Fold over basal
2.6
2.4
2.2
2
1.8
1.6
1.4
1.2
1
0.8
0.6
0
10
20
30
40
50
60
Time (min)

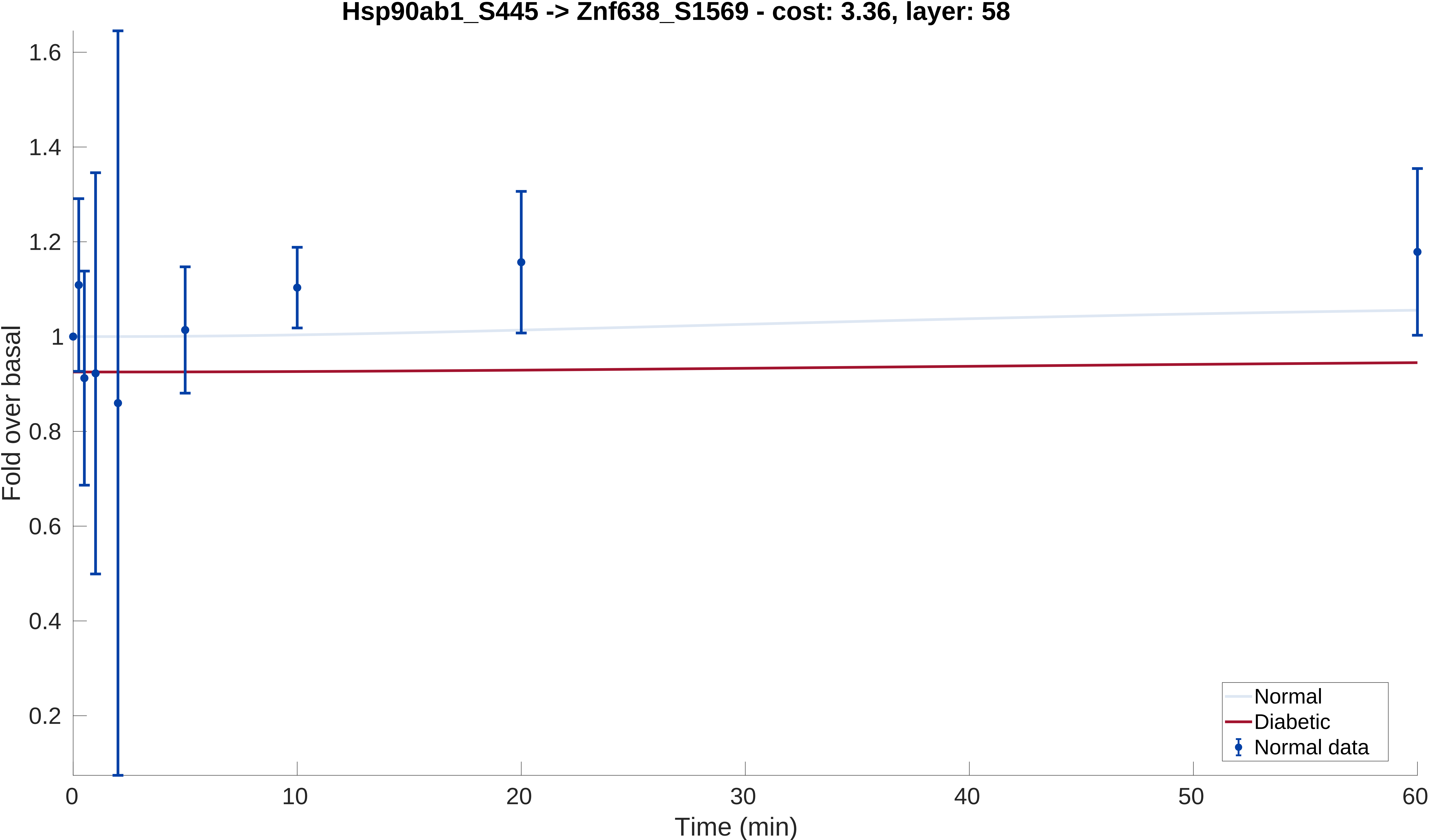

Hsp90ab1_S445 -> Znf638_S1569 - cost: 3.36, layer: 58
Fold over basal
1.6
1.4
1.2
1
0.8
0.6
0.4
0.2
0
10
20
30
40
50
60
Time (min)
Normal
Diabetic
Normal data

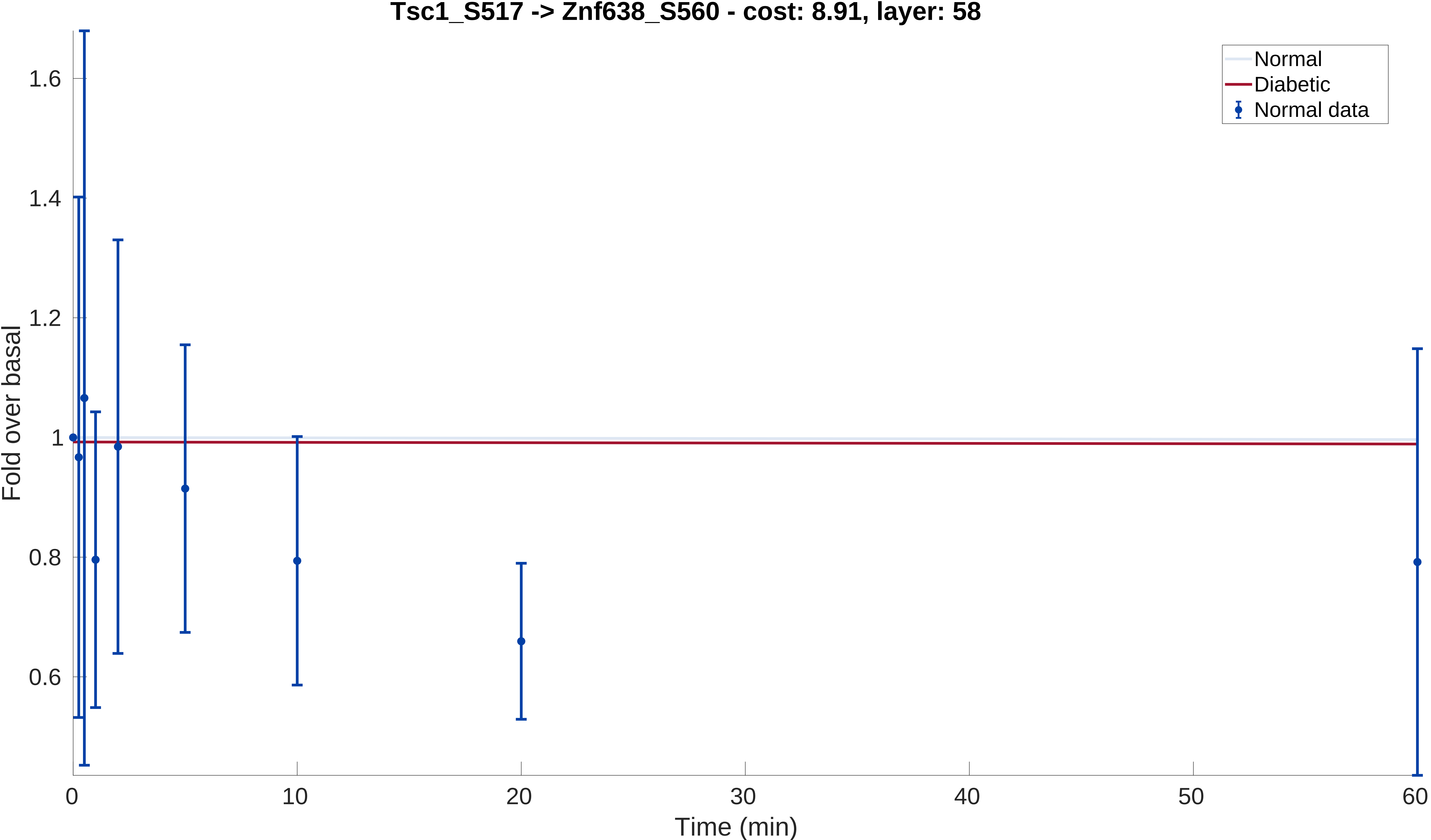

Tsc1_S517 -> Znf638_S560 - cost: 8.91, layer: 58
Normal
Diabetic
Normal data
Fold over basal
1.6
1.4
1.2
1
0.8
0.6
0
10
20
30
40
50
60
Time (min)

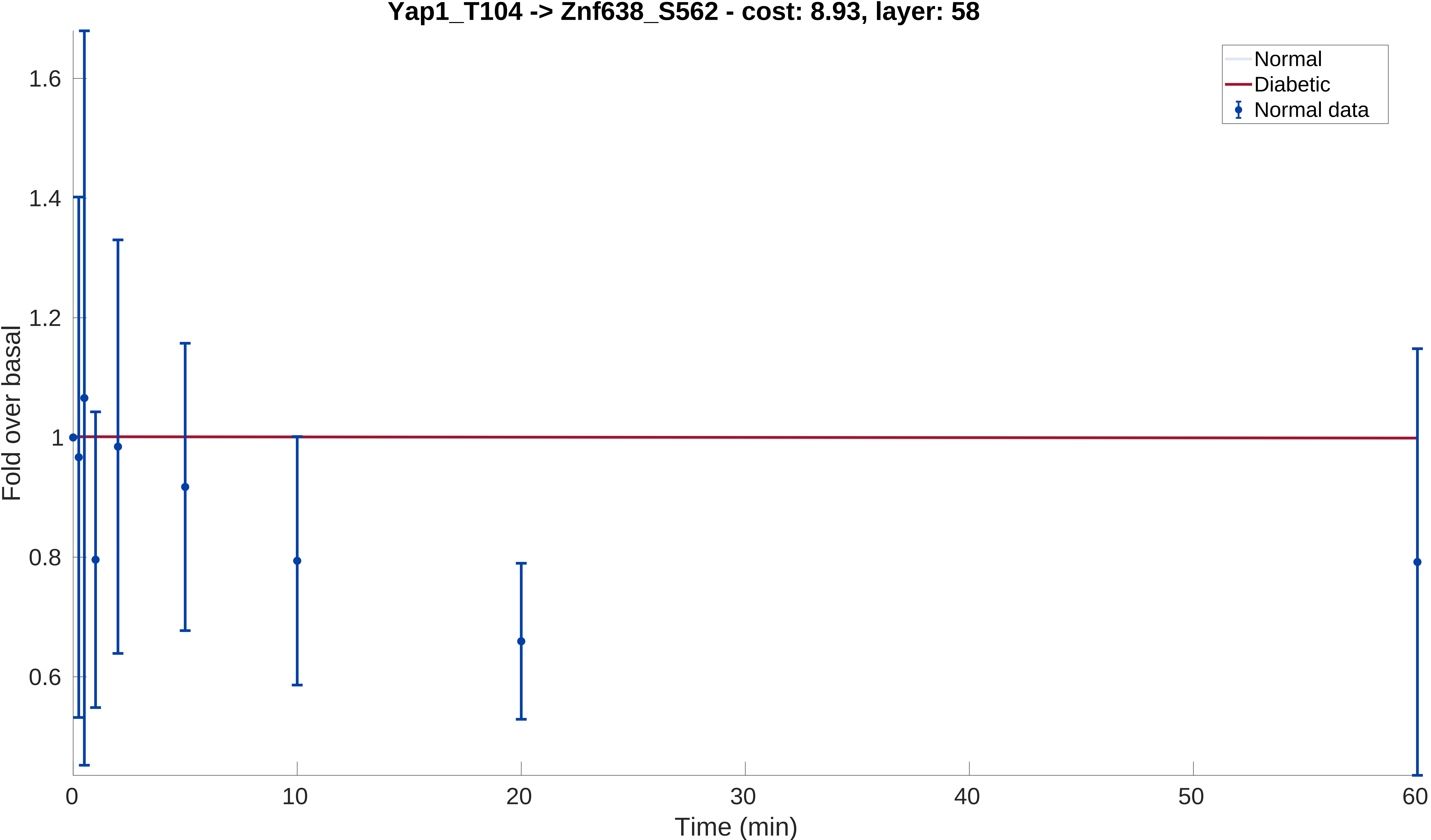
Yap1_T104 -> Znf638_S562 - cost: 8.93, layer: 58
Normal
Diabetic
Normal data
Fold over basal
1.6
1.4
1.2
1
0.8
0.6
0
10
20
30
40
50
60
Time (min)

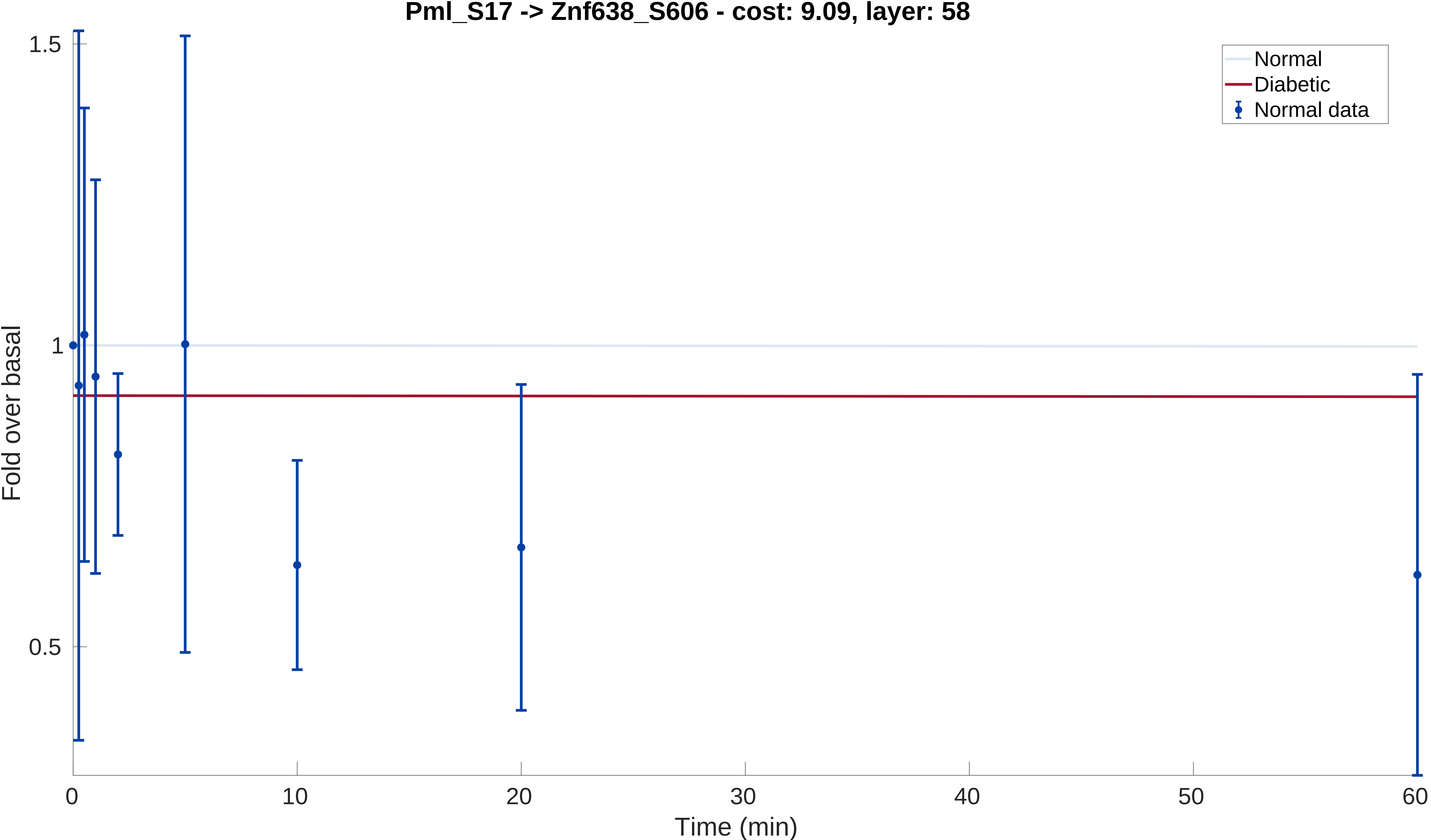

Pml_S17 -> Znf638_S606 - cost: 9.09, layer: 58
Normal
Diabetic
Normal data
Fold over basal
1.5
1
0.5
0
10
20
30
40
50
60
Time (min)

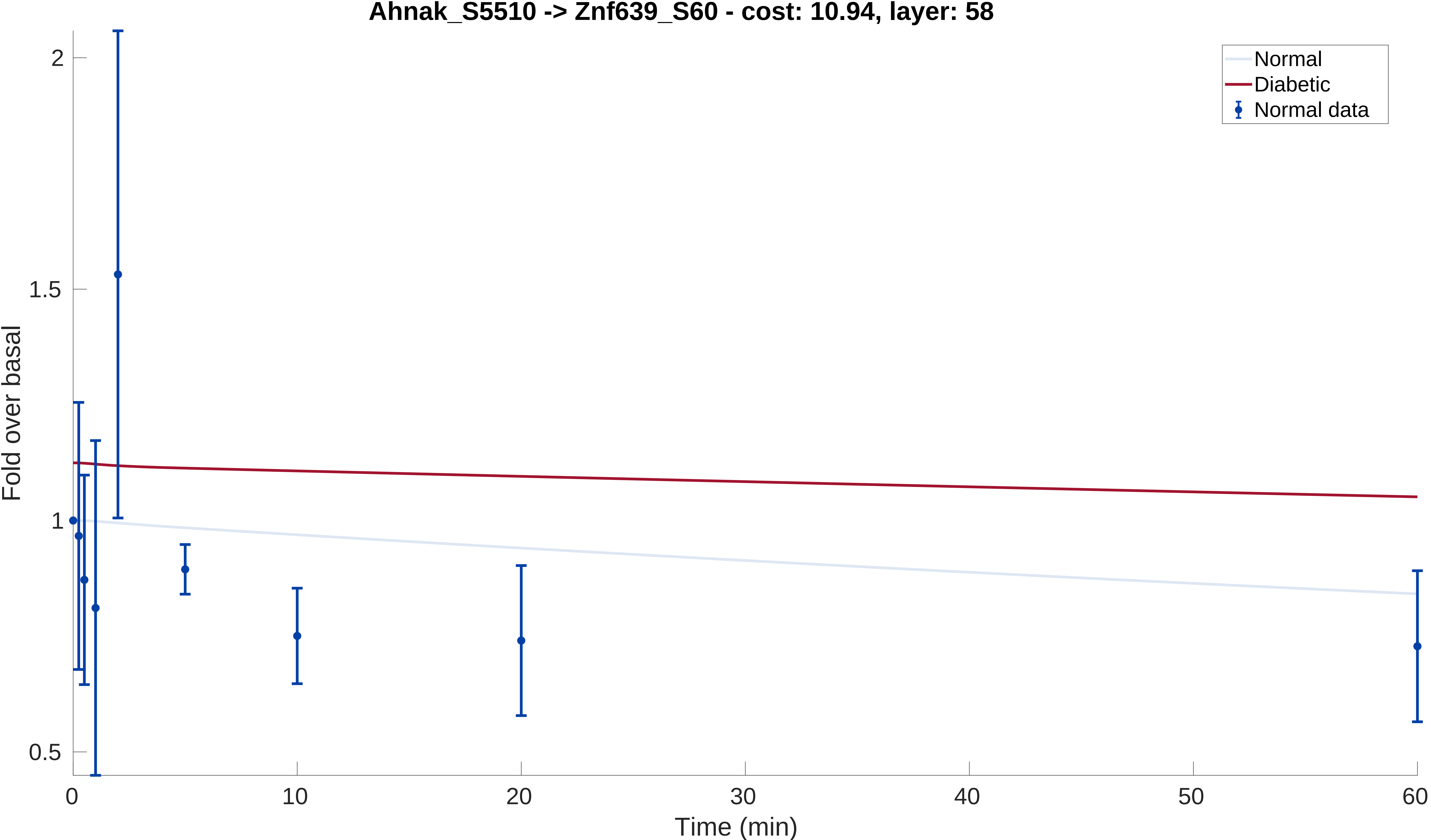
Ahnak_S5510 -> Znf639_S60 - cost: 10.94, layer: 58
Fold over basal
Time (min)
0.5
1
1.5
2
0
10
20
30
40
50
60
Normal
Diabetic
Normal data

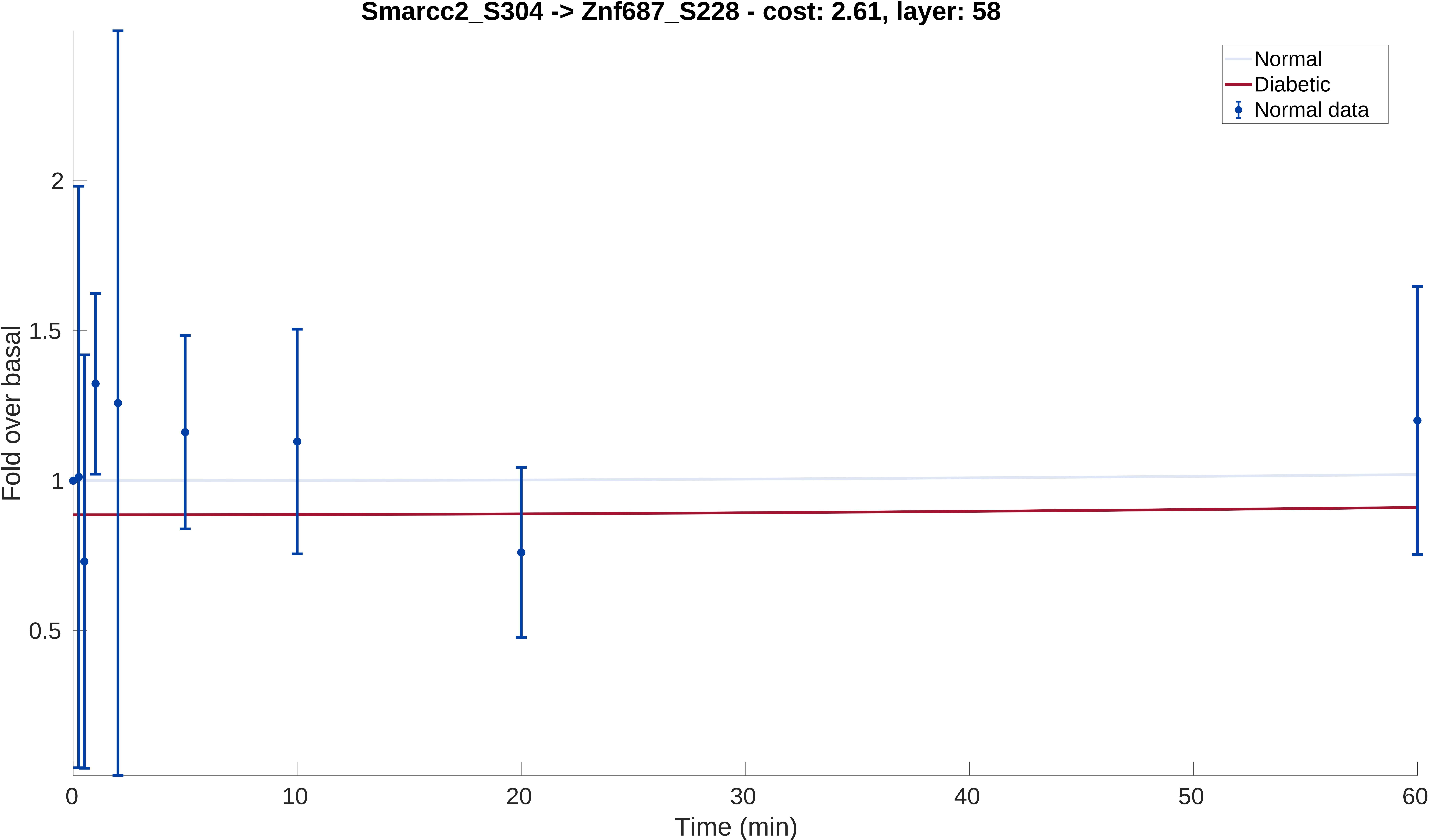

Smarcc2_S304 -> Znf687_S228 - cost: 2.61, layer: 58
Normal
Diabetic
Normal data
Fold over basal
2
1.5
1
0.5
0
10
20
30
40
50
60
Time (min)

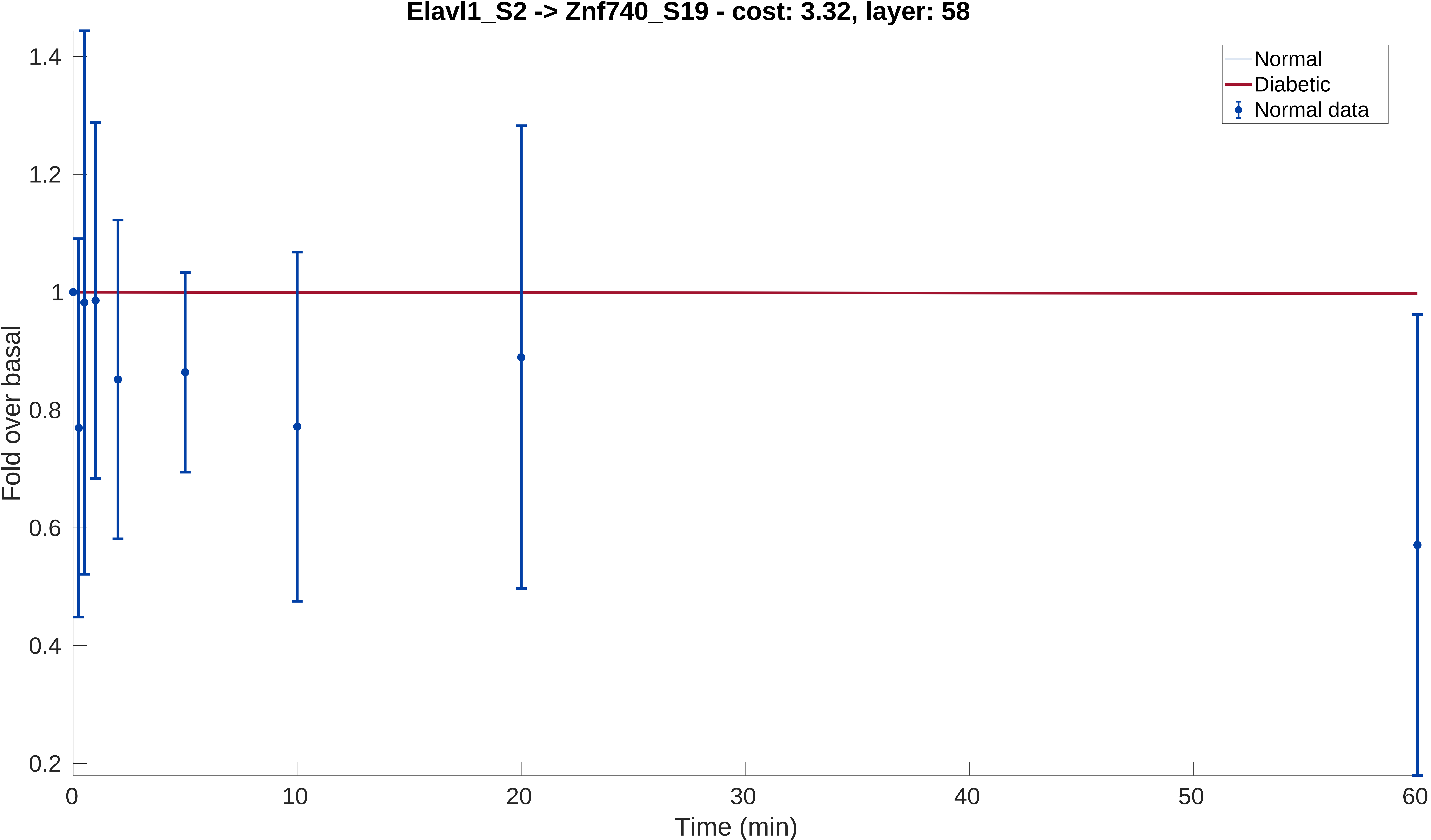
Elavl1_S2 -> Znf740_S19 - cost: 3.32, layer: 58
Normal
Diabetic
Normal data
Fold over basal
Time (min)
1.4
1.2
1
0.8
0.6
0.4
0.2
0
10
20
30
40
50
60

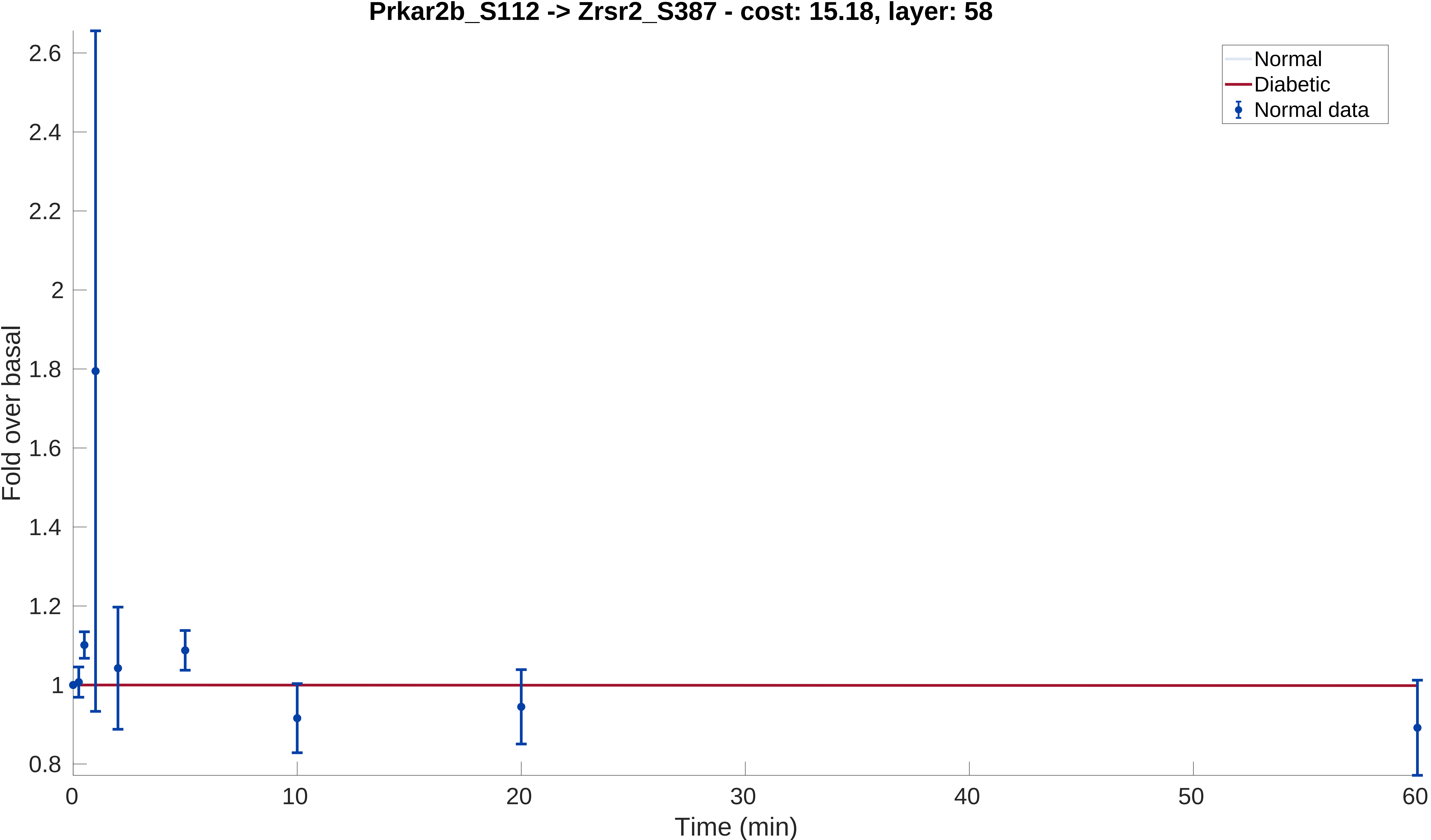

Prkar2b_S112 -> Zrsr2_S387 - cost: 15.18, layer: 58
Normal
Diabetic
Normal data
Fold over basal
2.6
2.4
2.2
2
1.8
1.6
1.4
1.2
1
0.8
0
10
20
30
40
50
60
Time (min)

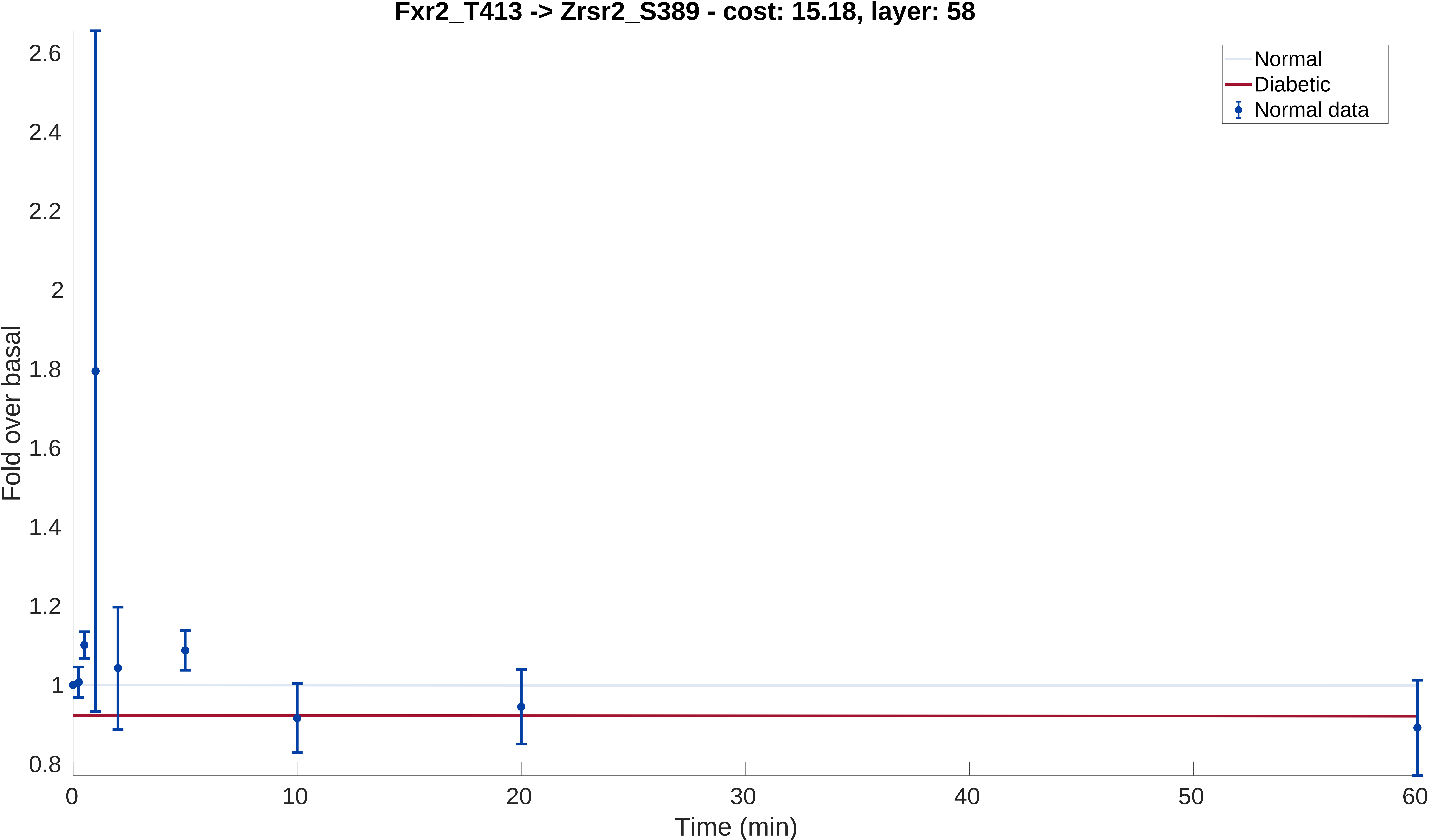

Fxr2_T413 -> Zrsr2_S389 - cost: 15.18, layer: 58
Normal
Diabetic
Normal data
Fold over basal
Time (min)
0.8
1
1.2
1.4
1.6
1.8
2
2.2
2.4
2.6
0
10
20
30
40
50
60

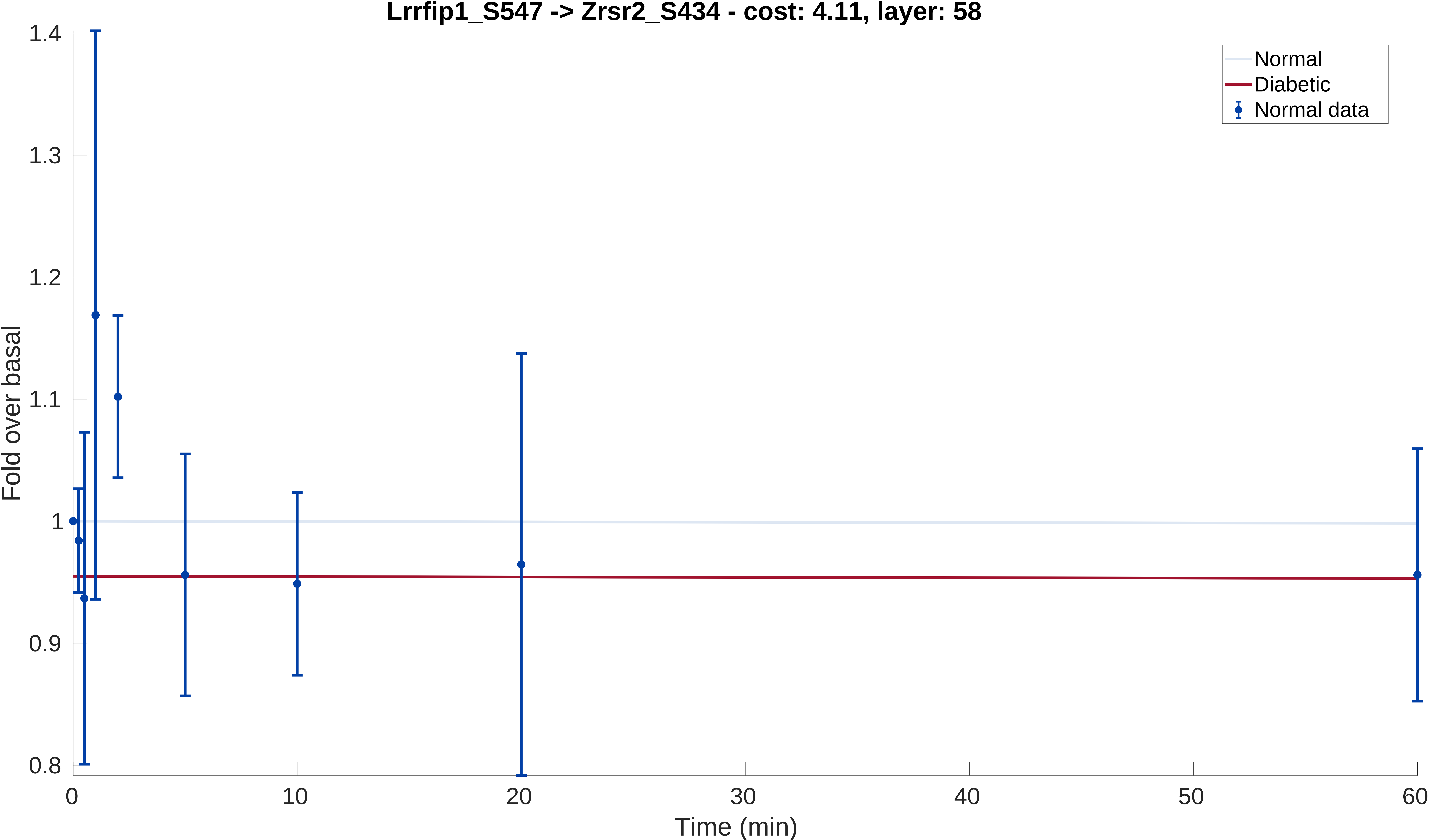

Lrrfip1_S547 -> Zrsr2_S434 - cost: 4.11, layer: 58
Normal
Diabetic
Normal data
Fold over basal
1.4
1.3
1.2
1.1
1
0.9
0.8
0
10
20
30
40
50
60
Time (min)

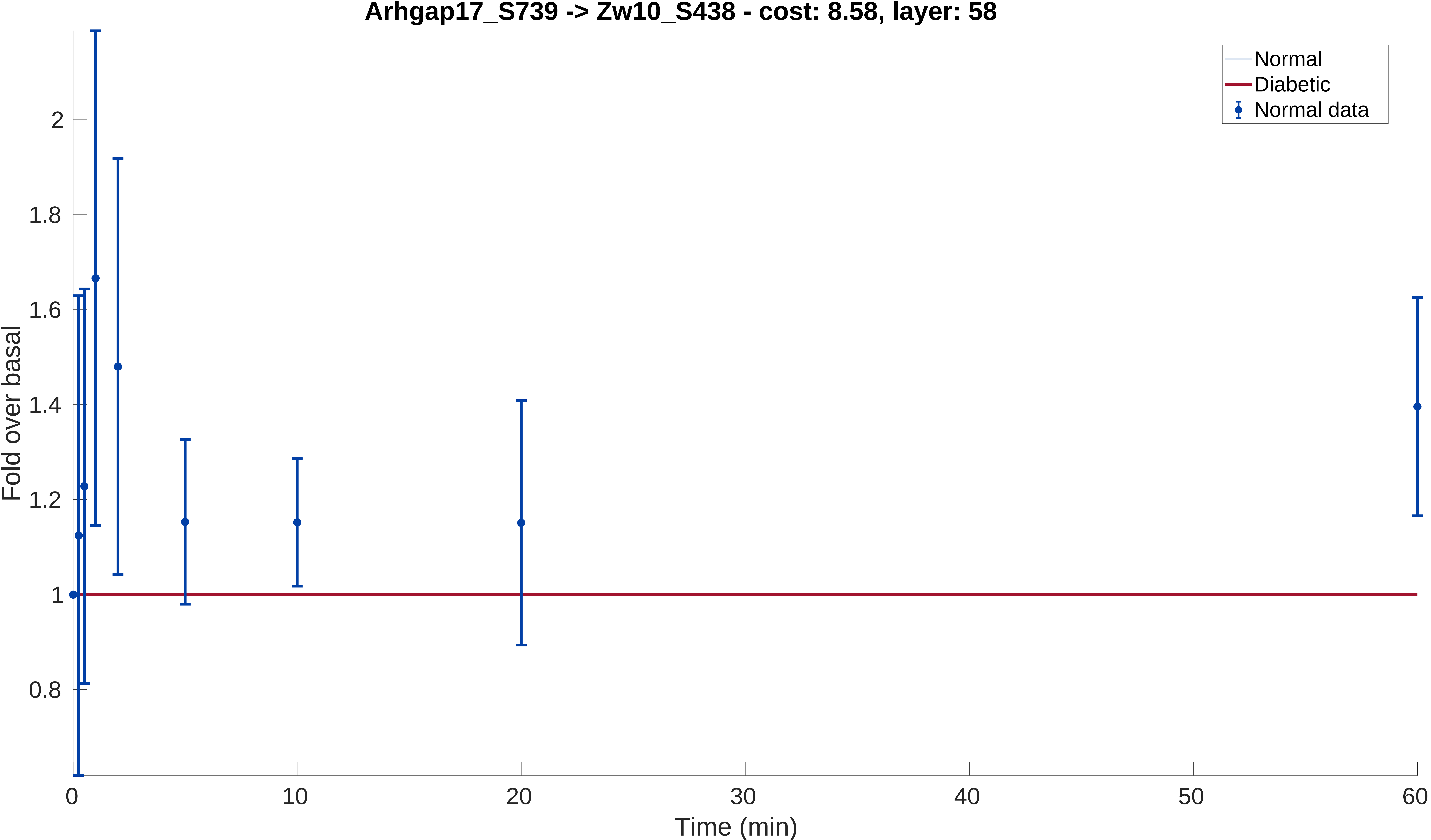

Arhgap17_S739 -> Zw10_S438 - cost: 8.58, layer: 58
Normal
Diabetic
Normal data
Fold over basal
2
1.8
1.6
1.4
1.2
1
0.8
0
10
20
30
40
50
60
Time (min)

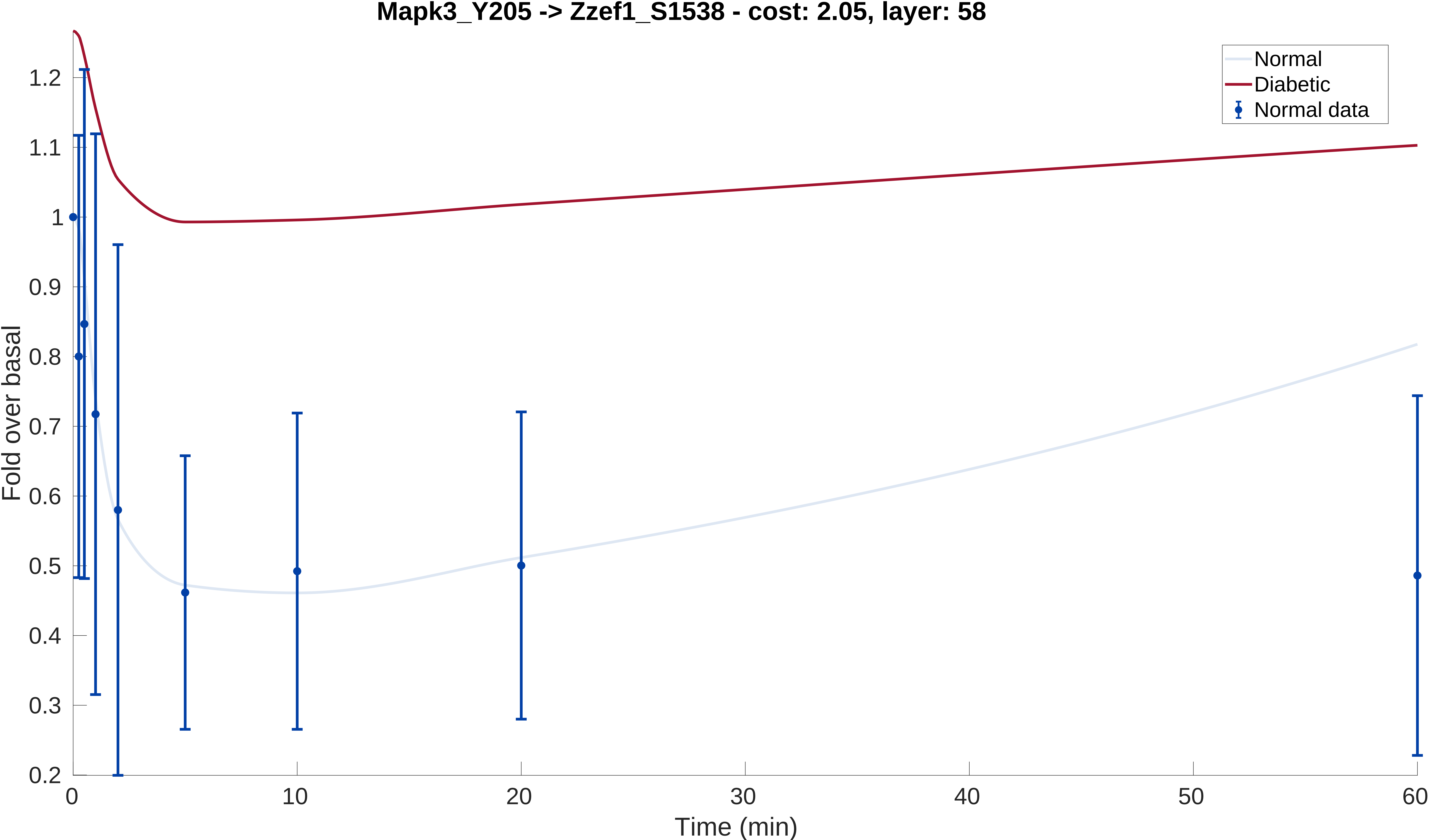
Mapk3_Y205 -> Zzef1_S1538 - cost: 2.05, layer: 58
Normal
Diabetic
Normal data
Fold over basal
1.2
1.1
1
0.9
0.8
0.7
0.6
0.5
0.4
0.3
0.2
0
10
20
30
40
50
60
Time (min)

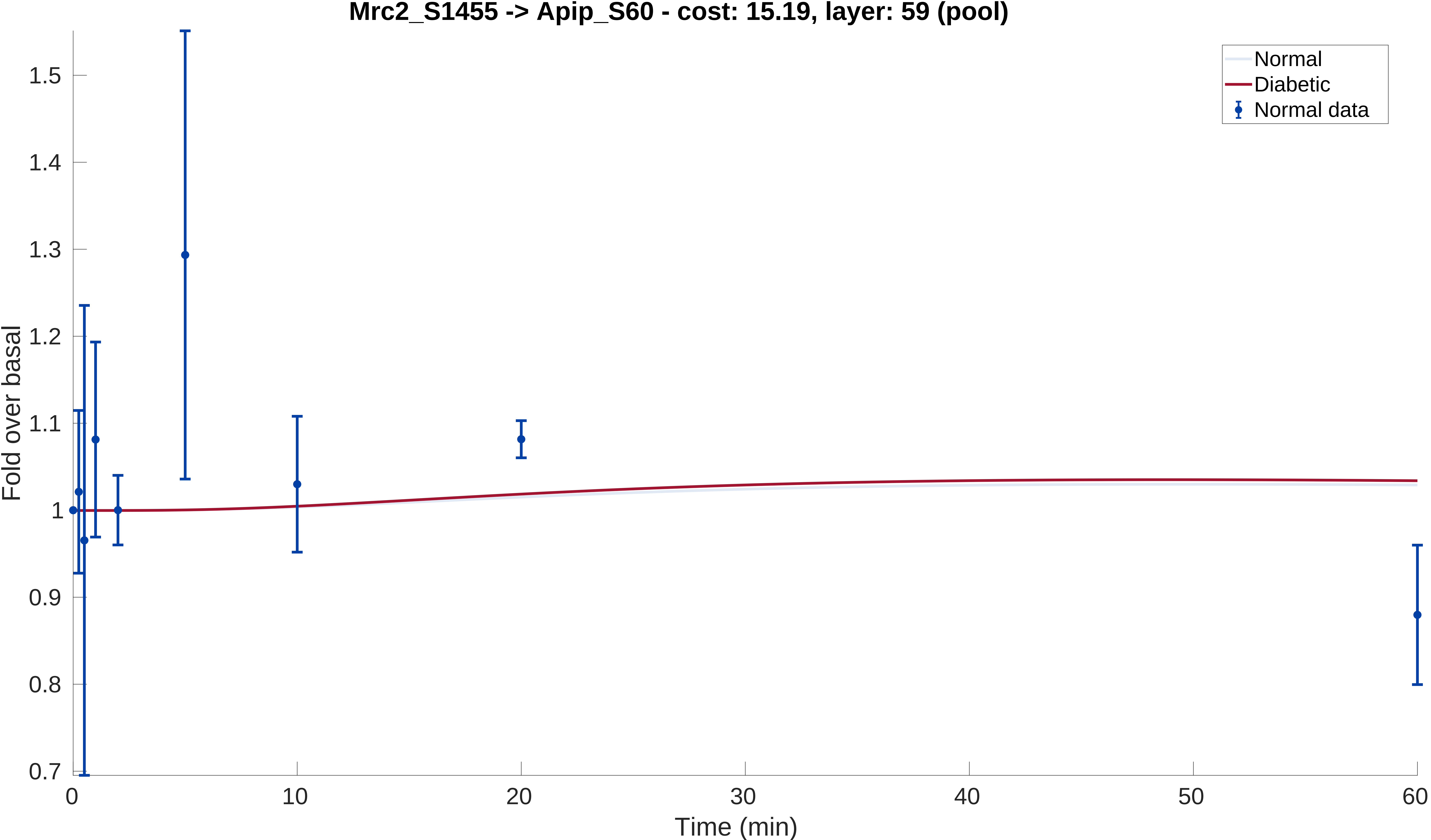
Mrc2_S1455 -> Apip_S60 - cost: 15.19, layer: 59 (pool)
Normal
Diabetic
Normal data
Fold over basal
Time (min)
1.5
1.4
1.3
1.2
1.1
1
0.9
0.8
0.7
0
10
20
30
40
50
60

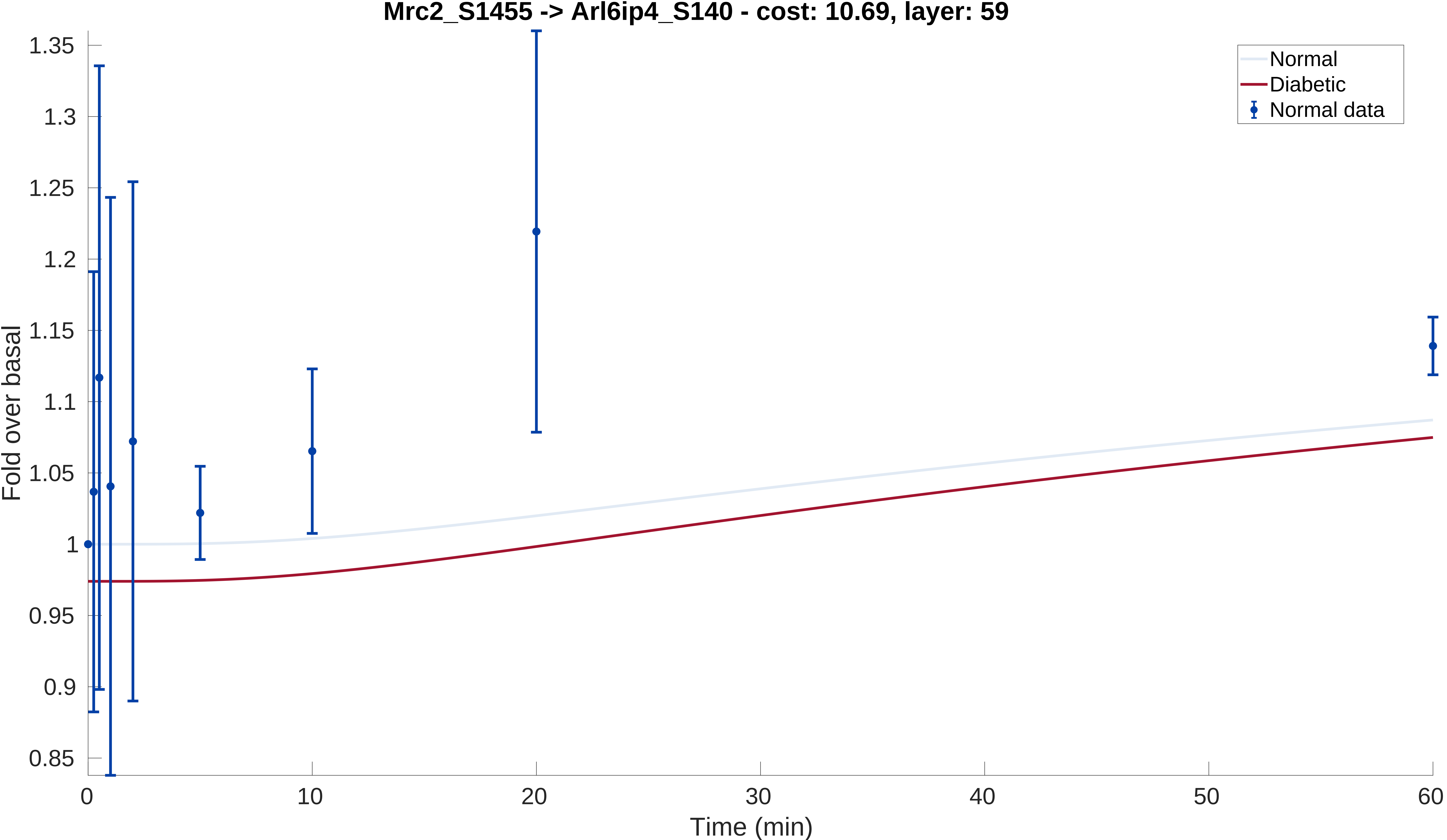
Mrc2_S1455 -> Arl6ip4_S140 - cost: 10.69, layer: 59
Normal
Diabetic
Normal data
Fold over basal
Time (min)
1.35
1.3
1.25
1.2
1.15
1.1
1.05
1
0.95
0.9
0.85
0
10
20
30
40
50
60

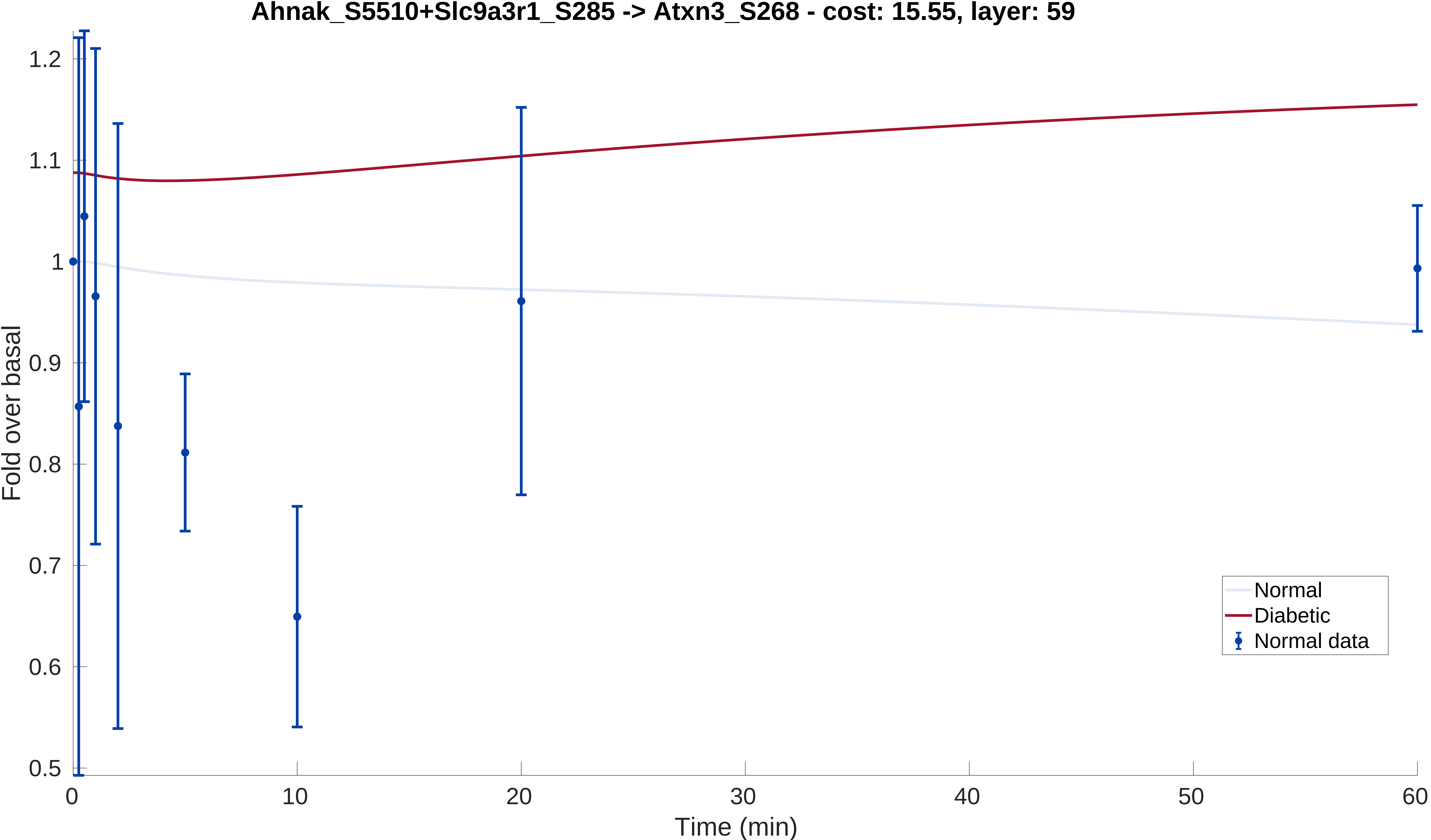

Ahnak_S5510+Slc9a3r1_S285 -> Atxn3_S268 - cost: 15.55, layer: 59
1.2
1.1
1
0.9
0.8
0.7
0.6
0.5
Fold over basal
0
10
20
30
40
50
60
Time (min)
Normal
Diabetic
Normal data

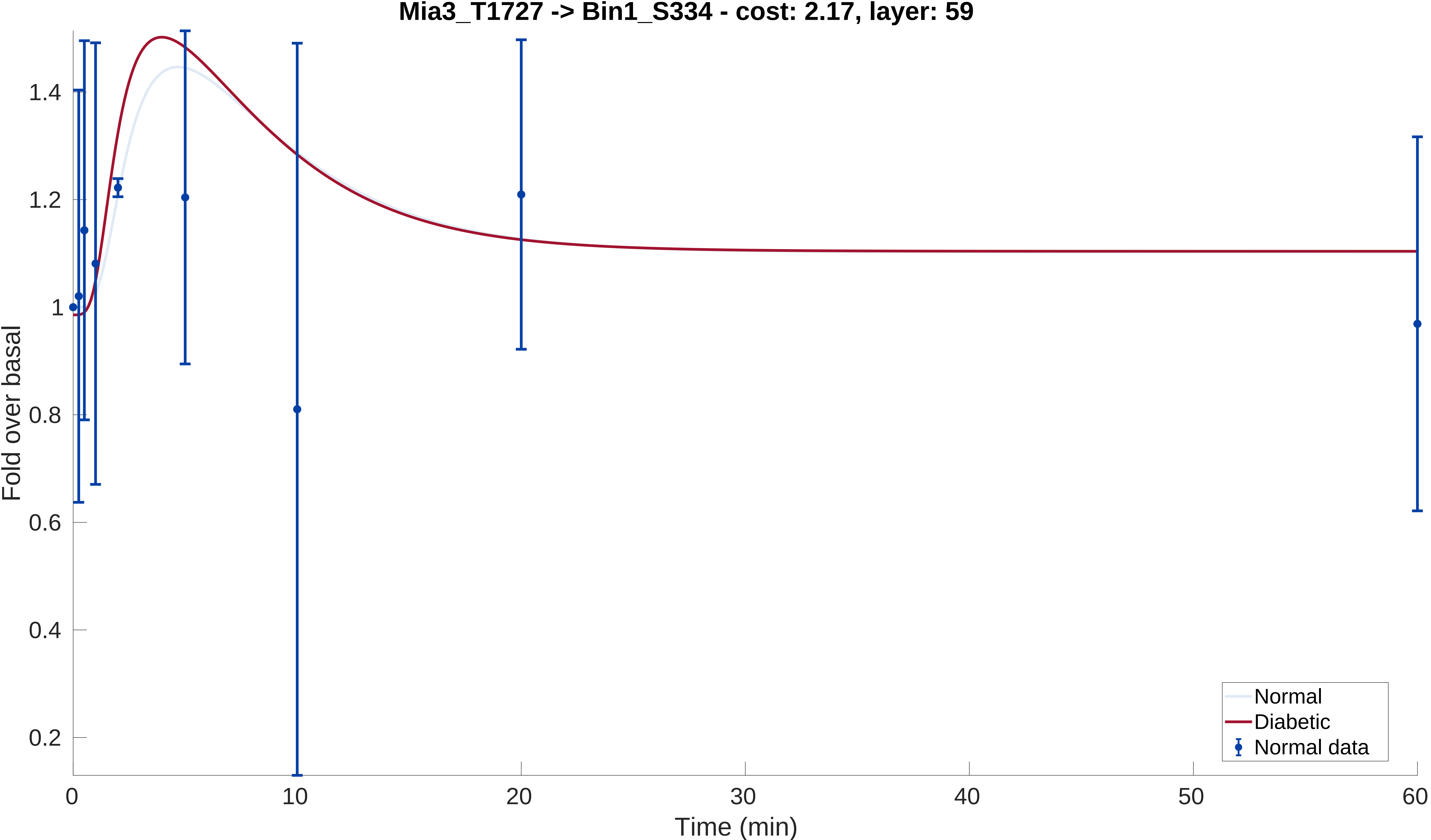

Mia3_T1727 -> Bin1_S334 - cost: 2.17, layer: 59
Fold over basal
1.4
1.2
1
0.8
0.6
0.4
0.2
0
10
20
30
40
50
60
Time (min)
Normal
Diabetic
Normal data

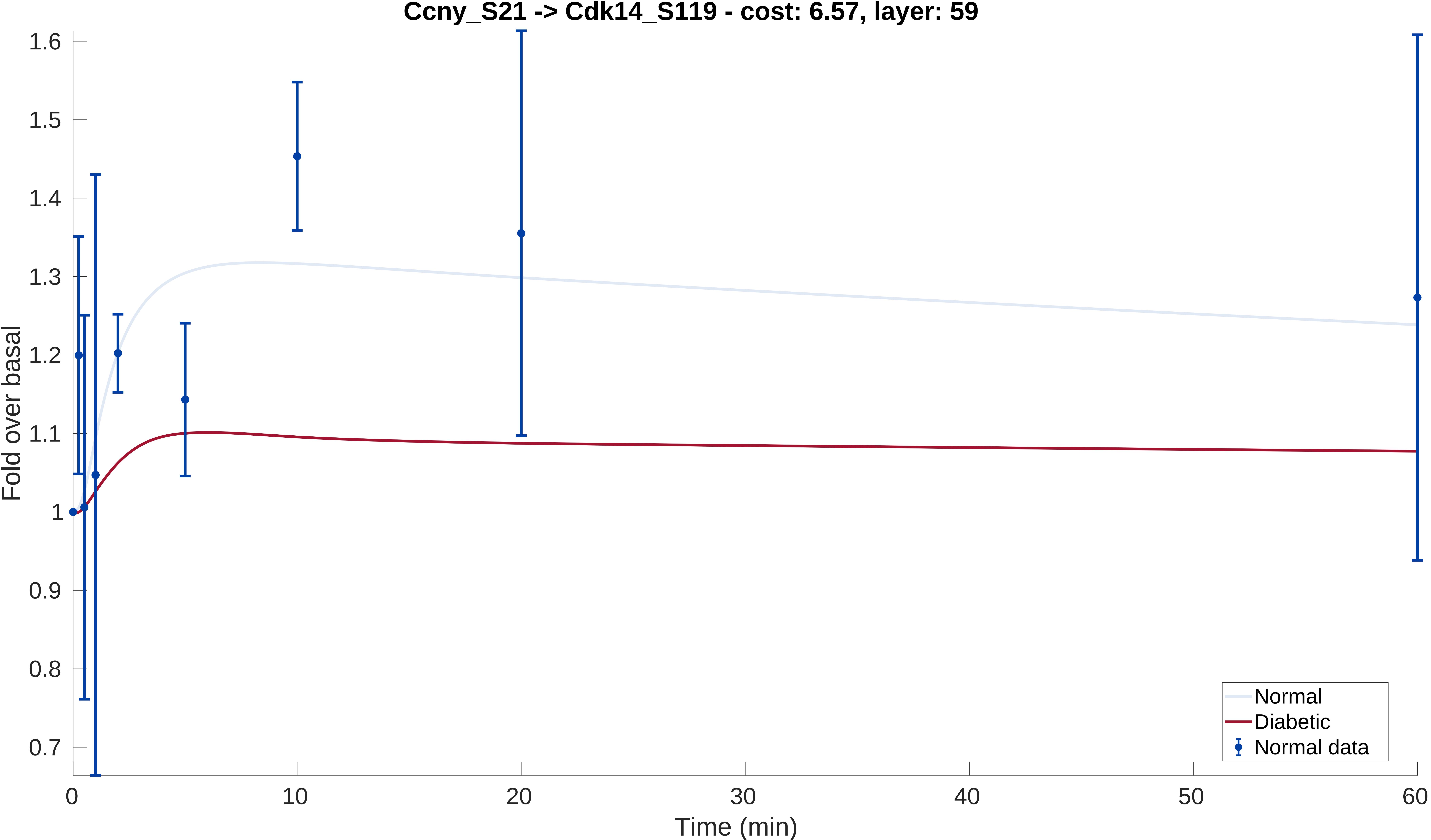
Ccny_S21 -> Cdk14_S119 - cost: 6.57, layer: 59
Fold over basal
1.6
1.5
1.4
1.3
1.2
1.1
1
0.9
0.8
0.7
0
10
20
30
40
50
60
Time (min)
Normal
Diabetic
Normal data

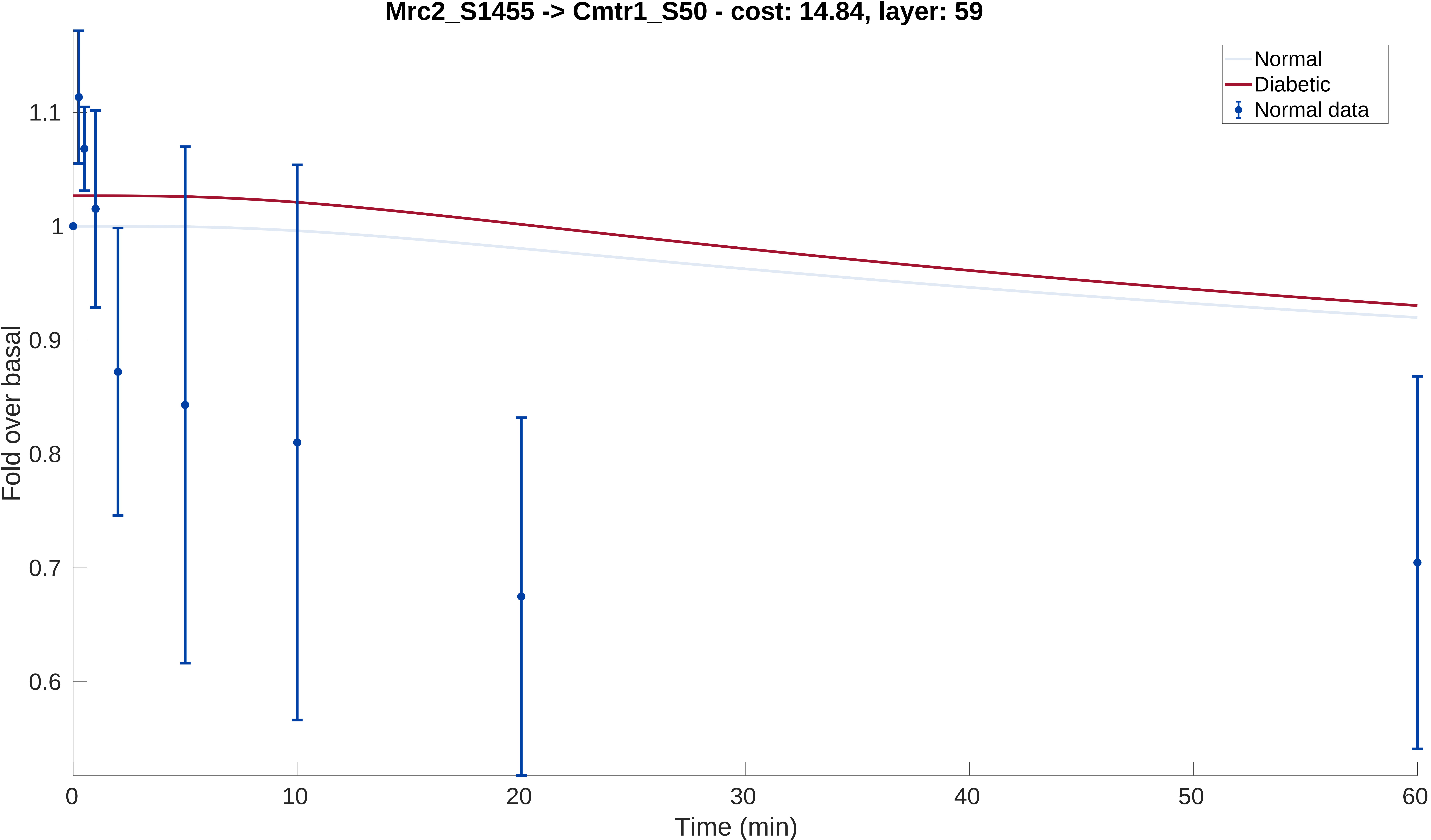

Mrc2_S1455 -> Cmtr1_S50 - cost: 14.84, layer: 59
Normal
Diabetic
Normal data
Fold over basal
1.1
1
0.9
0.8
0.7
0.6
0
10
20
30
40
50
60
Time (min)

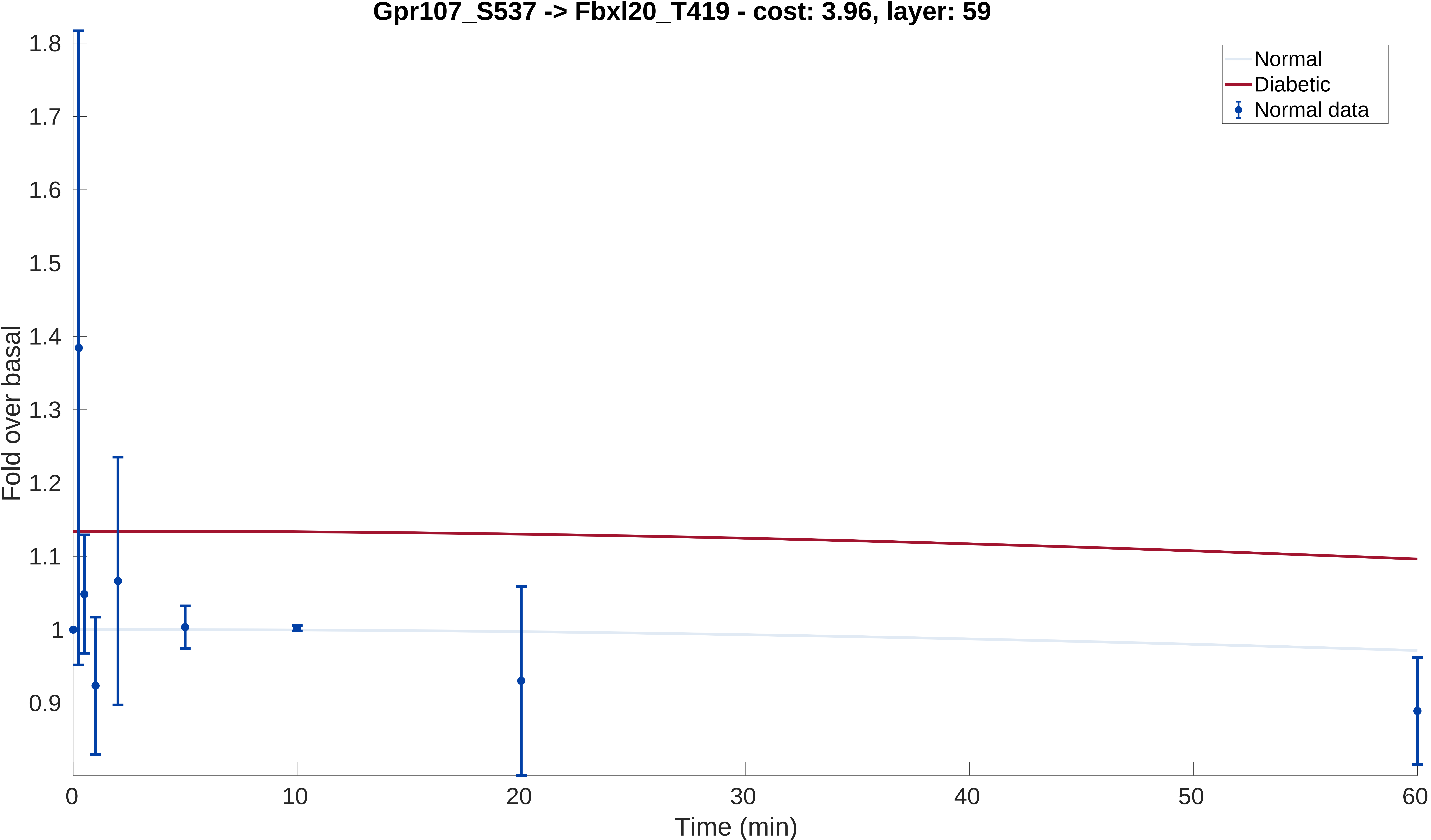

Gpr107_S537 -> Fbxl20_T419 - cost: 3.96, layer: 59
Normal
Diabetic
Normal data
Fold over basal
1.8
1.7
1.6
1.5
1.4
1.3
1.2
1.1
1
0.9
0
10
20
30
40
50
60
Time (min)

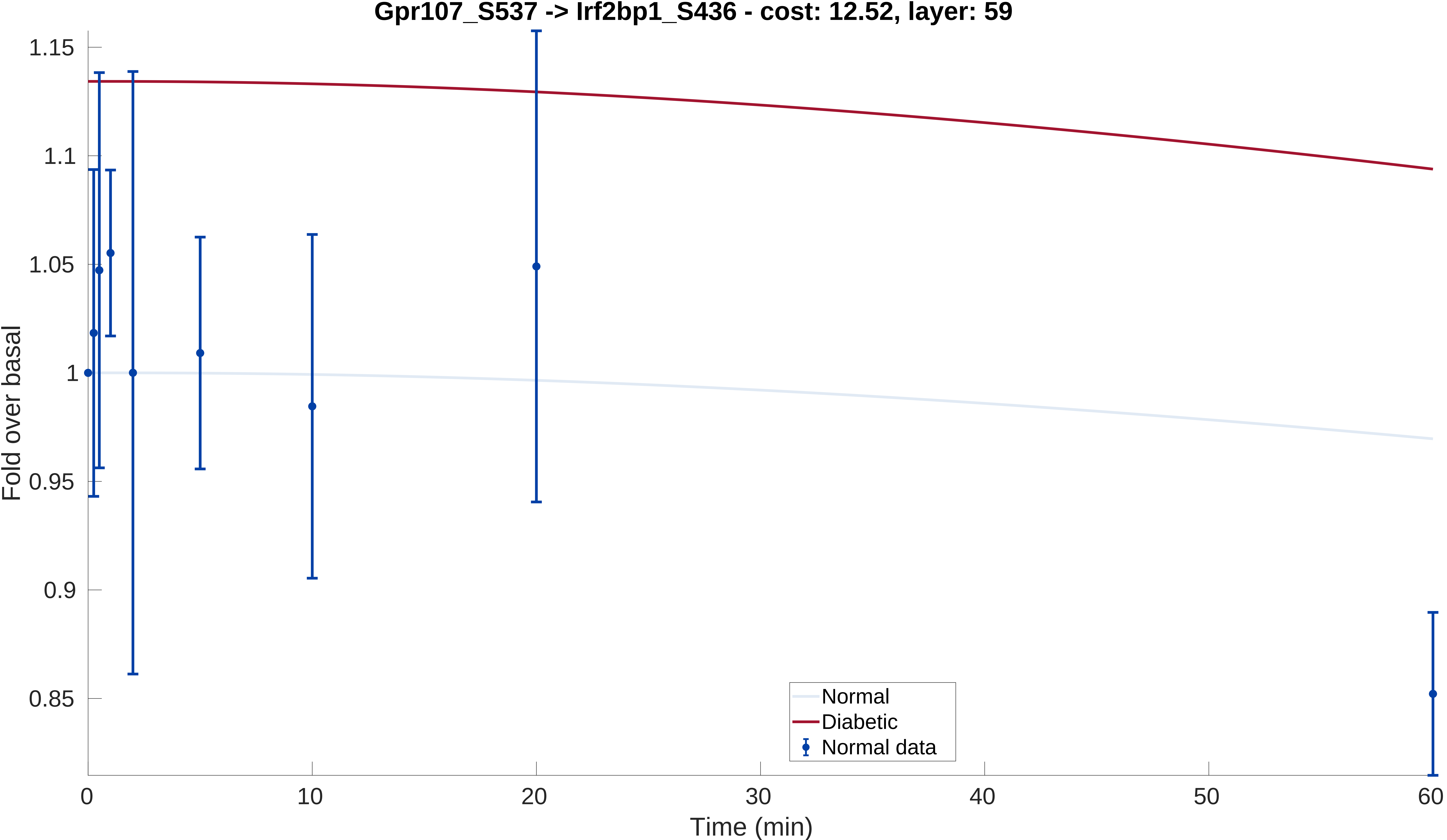

Gpr107_S537 -> Irf2bp1_S436 - cost: 12.52, layer: 59
Fold over basal
1.15
1.1
1.05
1
0.95
0.9
0.85
0
10
20
30
40
50
60
Time (min)
Normal
Diabetic
Normal data

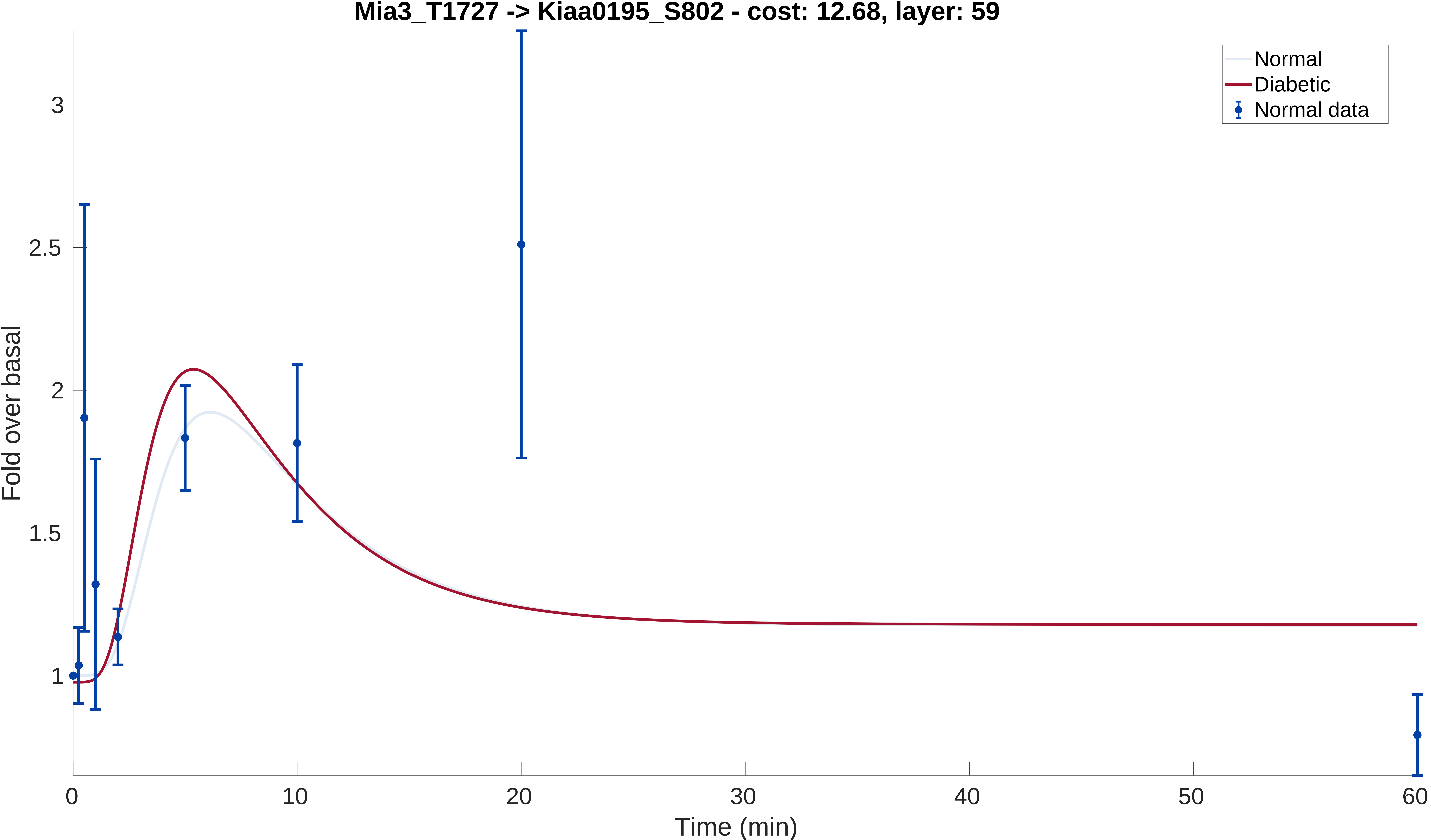

Mia3_T1727 -> Kiaa0195_S802 - cost: 12.68, layer: 59
Normal
Diabetic
Normal data
Fold over basal
3
2.5
2
1.5
1
0
10
20
30
40
50
60
Time (min)

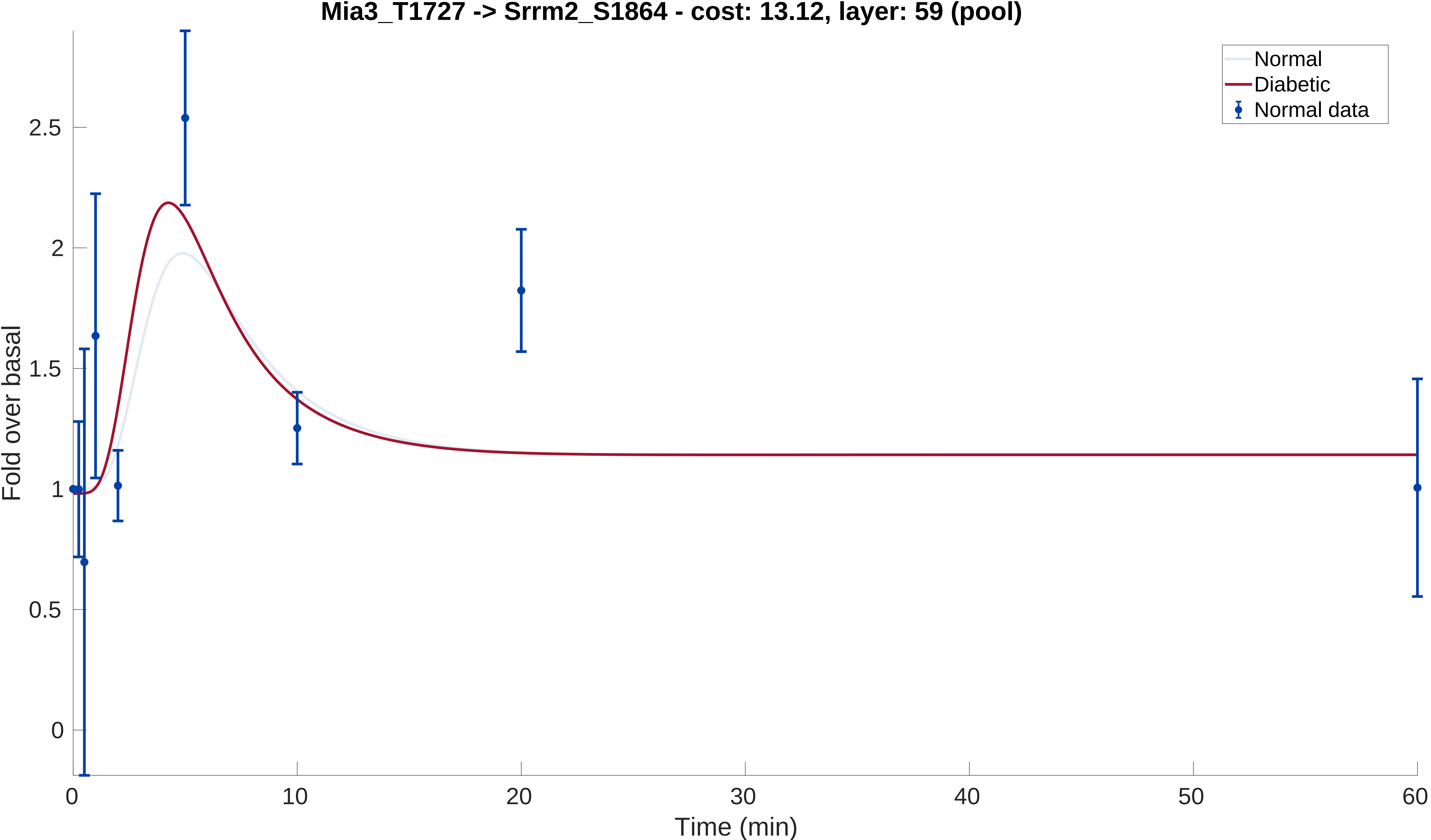

Mia3_T1727 -> Srrm2_S1864 - cost: 13.12, layer: 59 (pool)
Normal
Diabetic
Normal data
Fold over basal
Time (min)
0
0.5
1
1.5
2
2.5
0
10
20
30
40
50
60

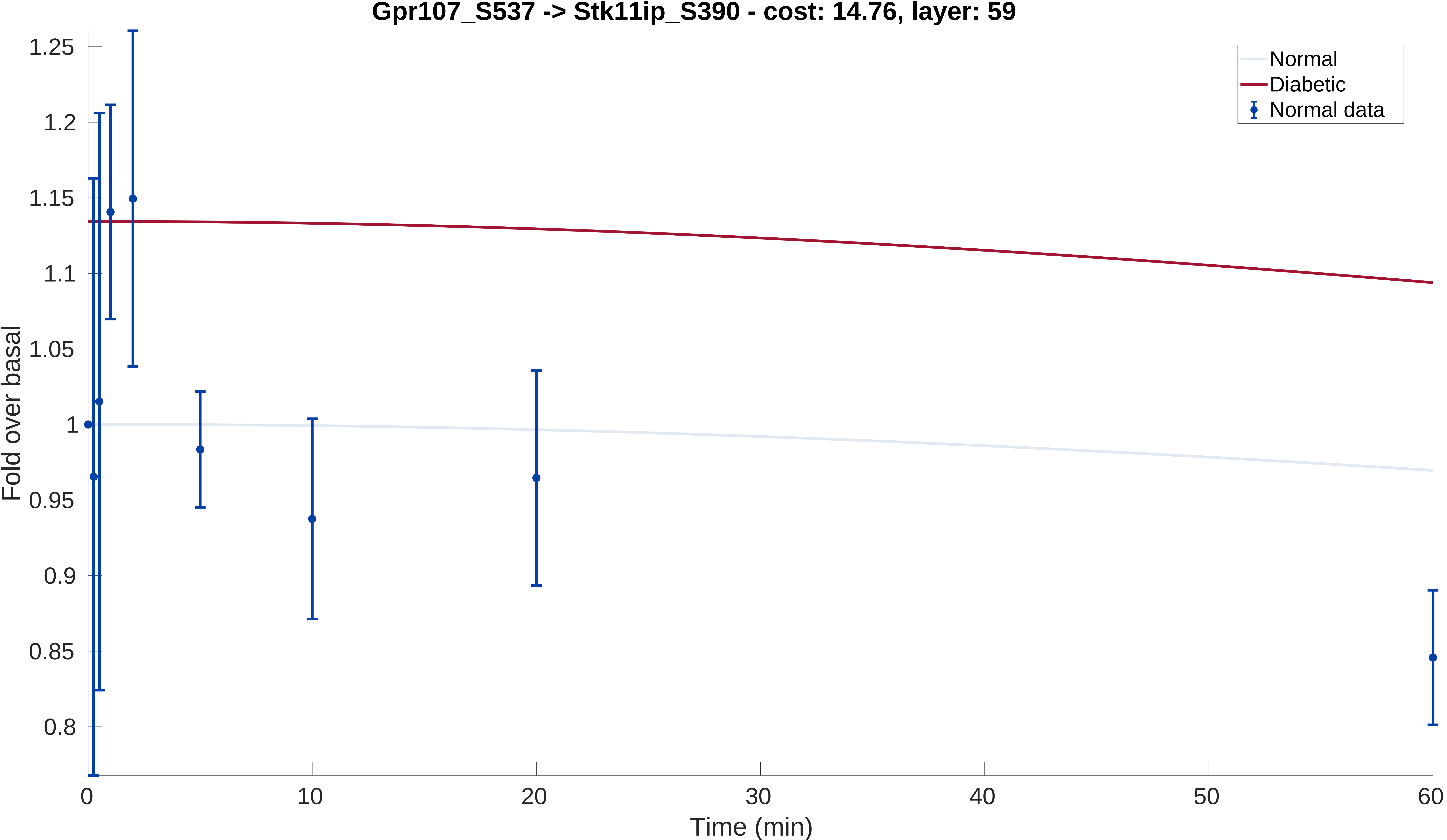

Gpr107_S537 -> Stk11ip_S390 - cost: 14.76, layer: 59
Normal
Diabetic
Normal data
Fold over basal
1.25
1.2
1.15
1.1
1.05
1
0.95
0.9
0.85
0.8
0
10
20
30
40
50
60
Time (min)

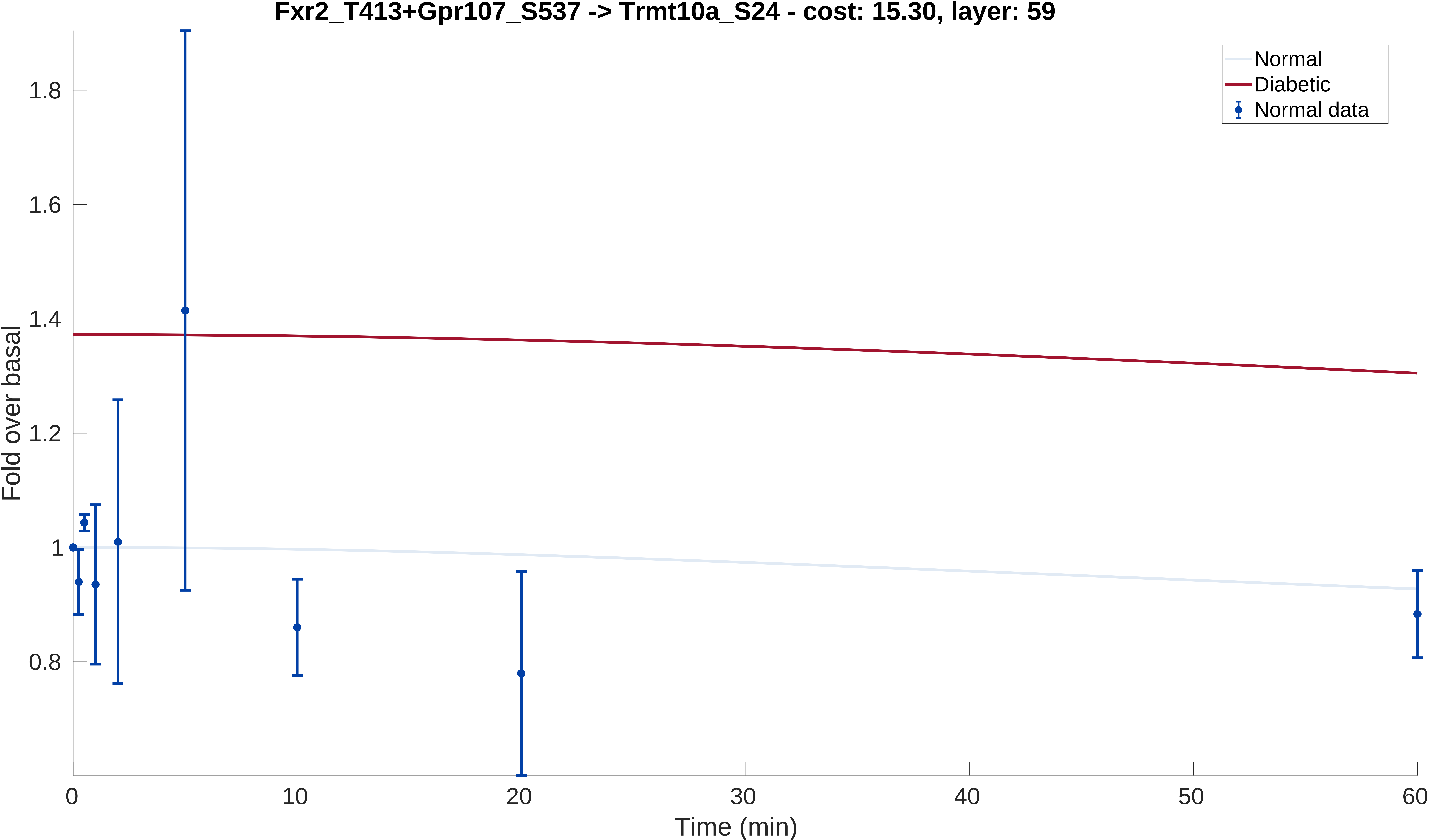

Fxr2_T413+Gpr107_S537 -> Trmt10a_S24 - cost: 15.30, layer: 59
Normal
Diabetic
Normal data
Fold over basal
1.8
1.6
1.4
1.2
1
0.8
0
10
20
30
40
50
60
Time (min)

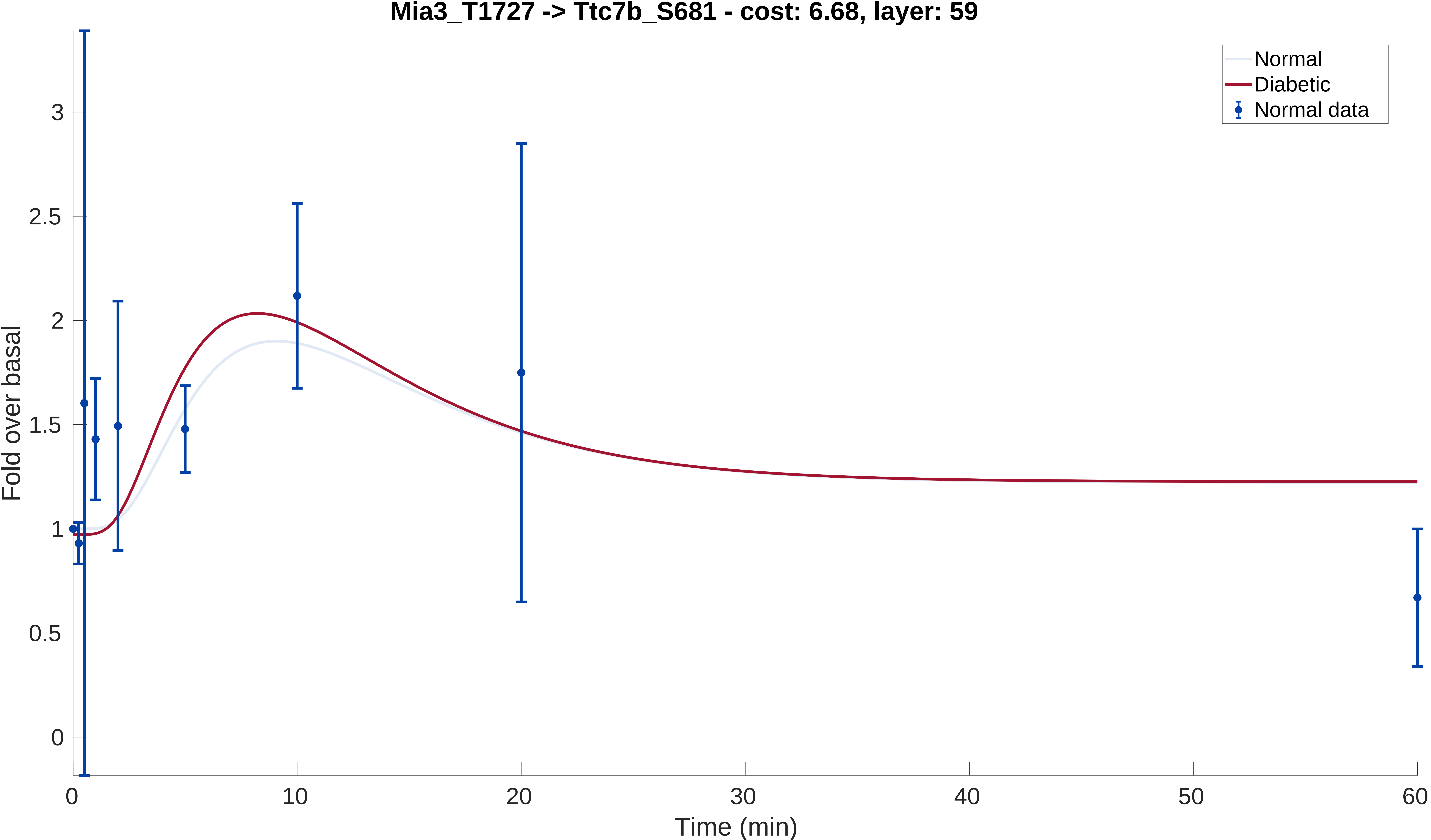

Mia3_T1727 -> Ttc7b_S681 - cost: 6.68, layer: 59
Normal
Diabetic
Normal data
Fold over basal
0
0.5
1
1.5
2
2.5
3
0
10
20
30
40
50
60
Time (min)

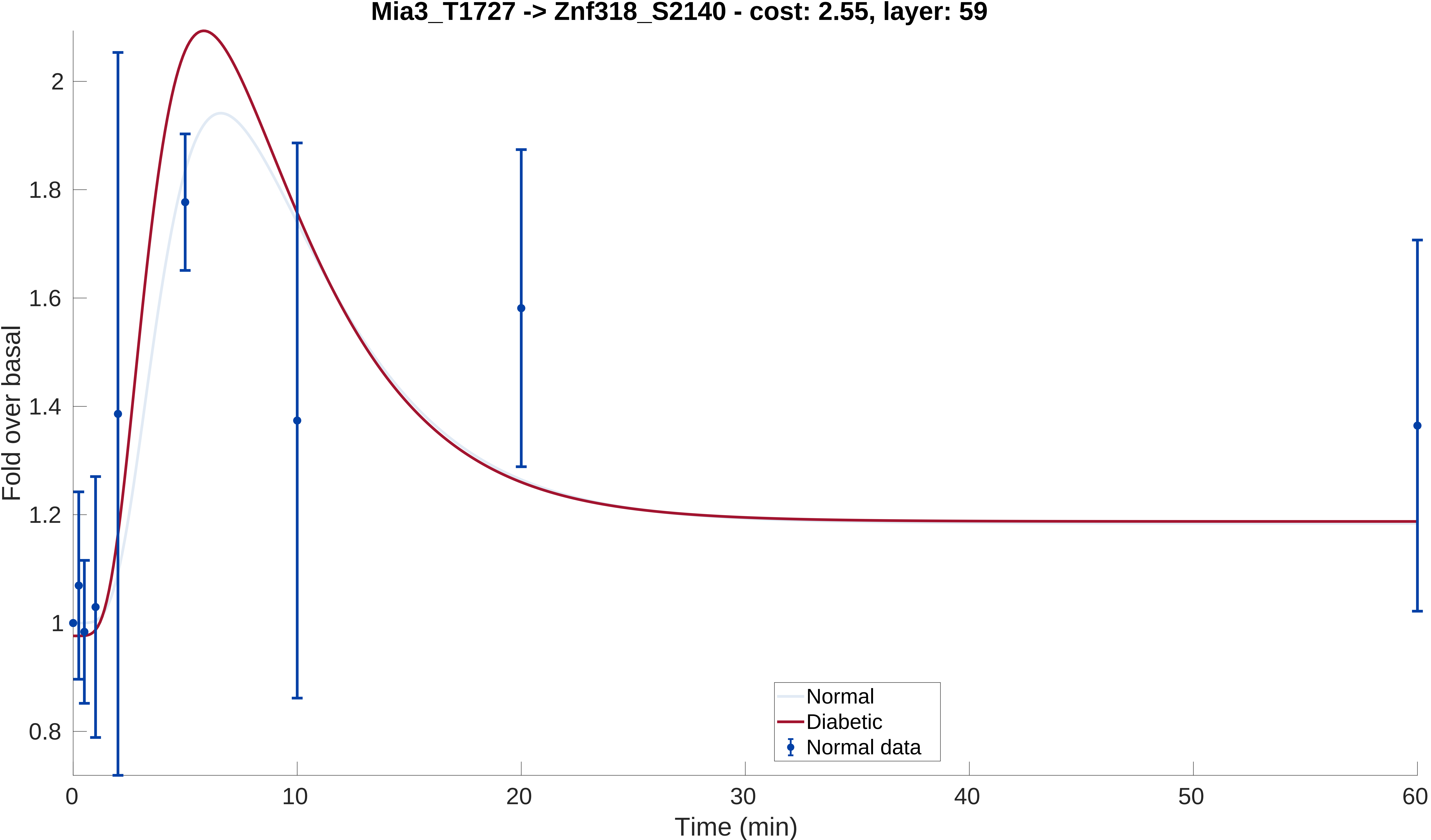

Mia3_T1727 -> Znf318_S2140 - cost: 2.55, layer: 59
Fold over basal
2
1.8
1.6
1.4
1.2
1
0.8
0
10
20
30
40
50
60
Time (min)
Normal
Diabetic
Normal data

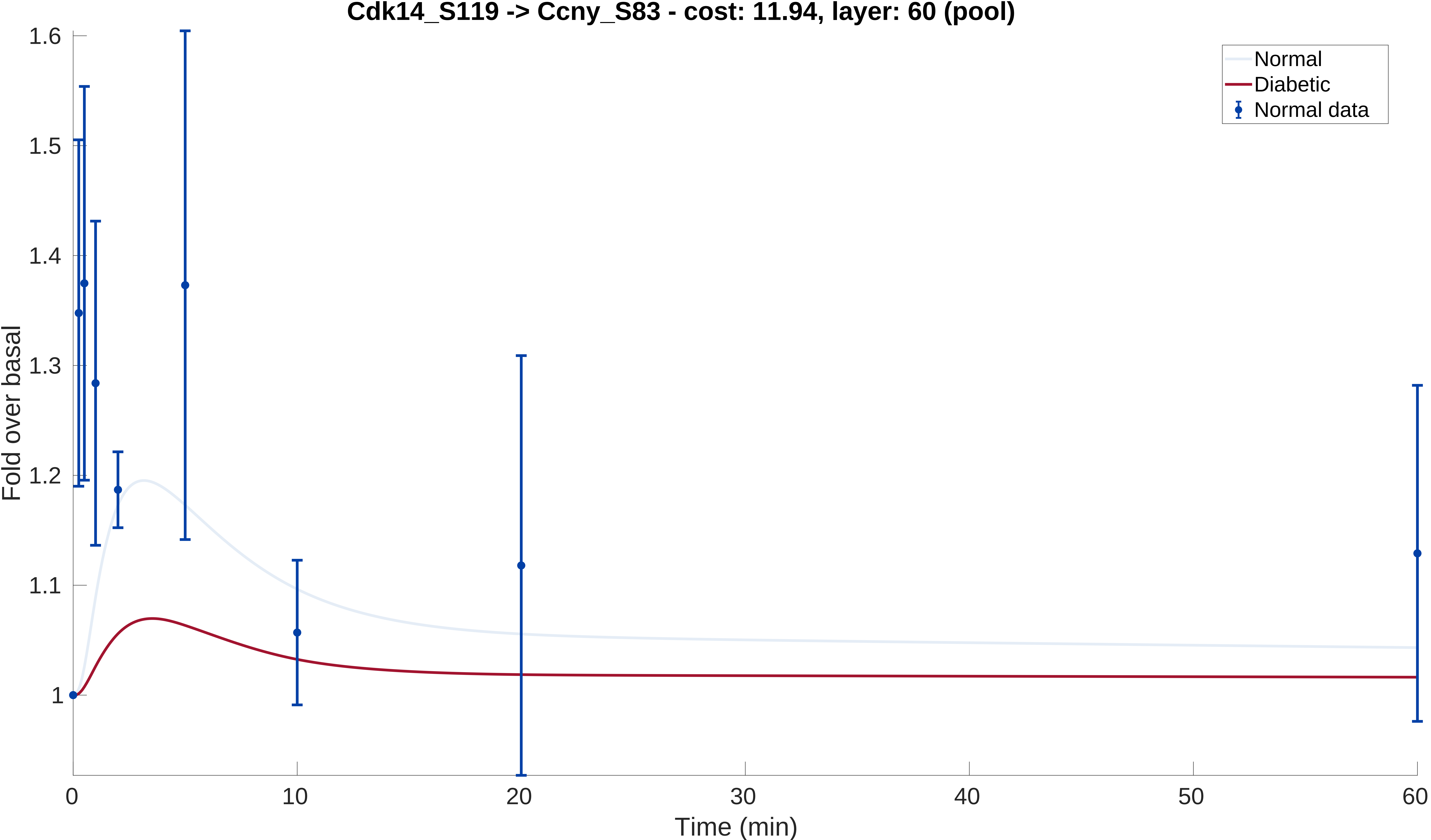

Cdk14_S119 -> Ccny_S83 - cost: 11.94, layer: 60 (pool)
Normal
Diabetic
Normal data
Fold over basal
1.6
1.5
1.4
1.3
1.2
1.1
1
0
10
20
30
40
50
60
Time (min)

#### 2 Insulin stimulation, normal and in type 2 diabetes

The following pages give all simulations of insulin stimulation normally and in type 2 diabetic conditions.

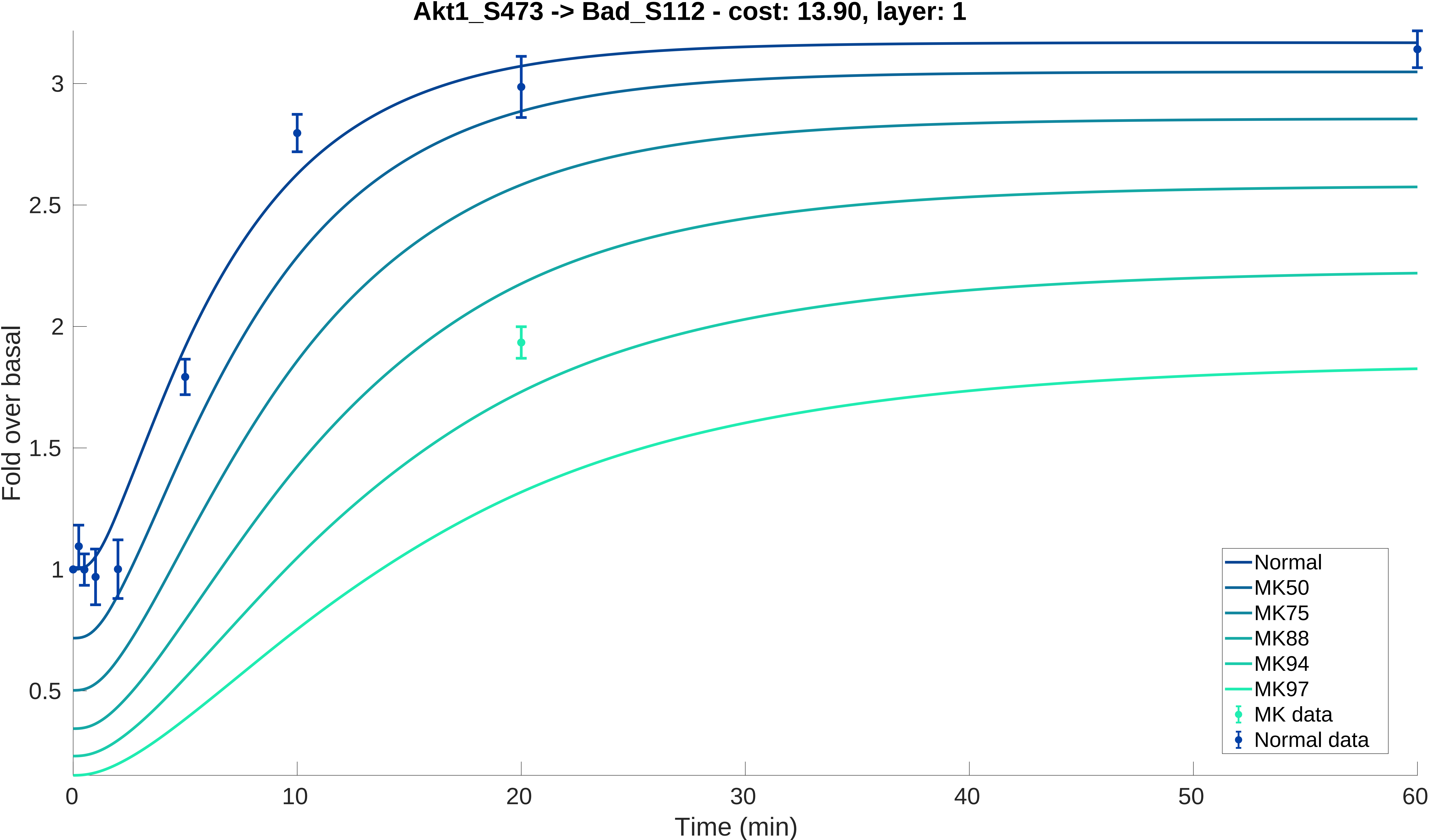

Akt1_S473 -> Bad_S112 - cost: 13.90, layer: 1
Fold over basal
3
2.5
2
1.5
1
0.5
0
10
20
30
40
50
60
Time (min)
Normal
MK50
MK75
MK88
MK94
MK97
MK data
Normal data

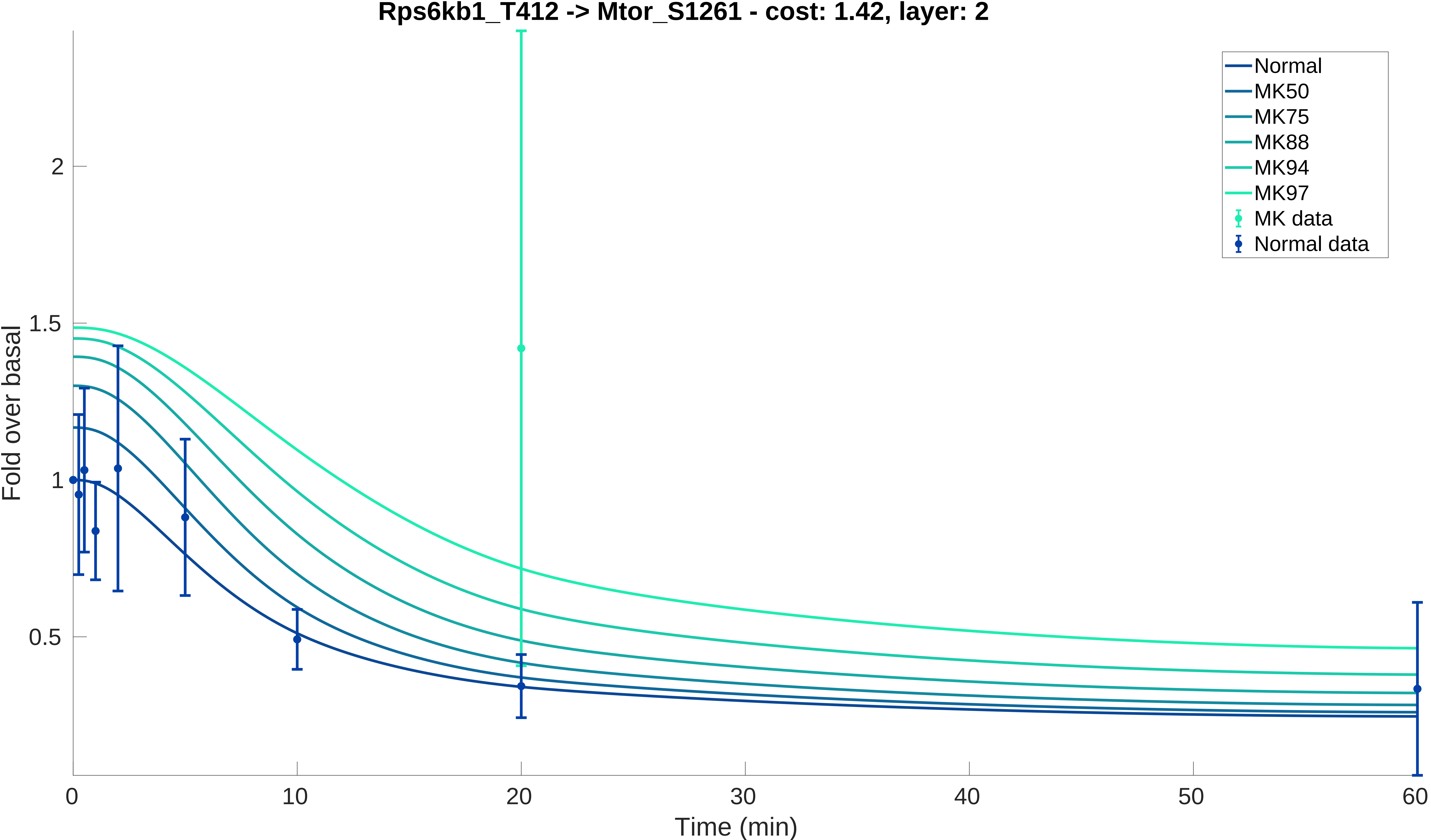

Rps6kb1_T412 -> Mtor_S1261 - cost: 1.42, layer: 2
Normal
MK50
MK75
MK88
MK94
MK97
MK data
Normal data
Fold over basal
Time (min)
0
10
20
30
40
50
60
0.5
1
1.5
2

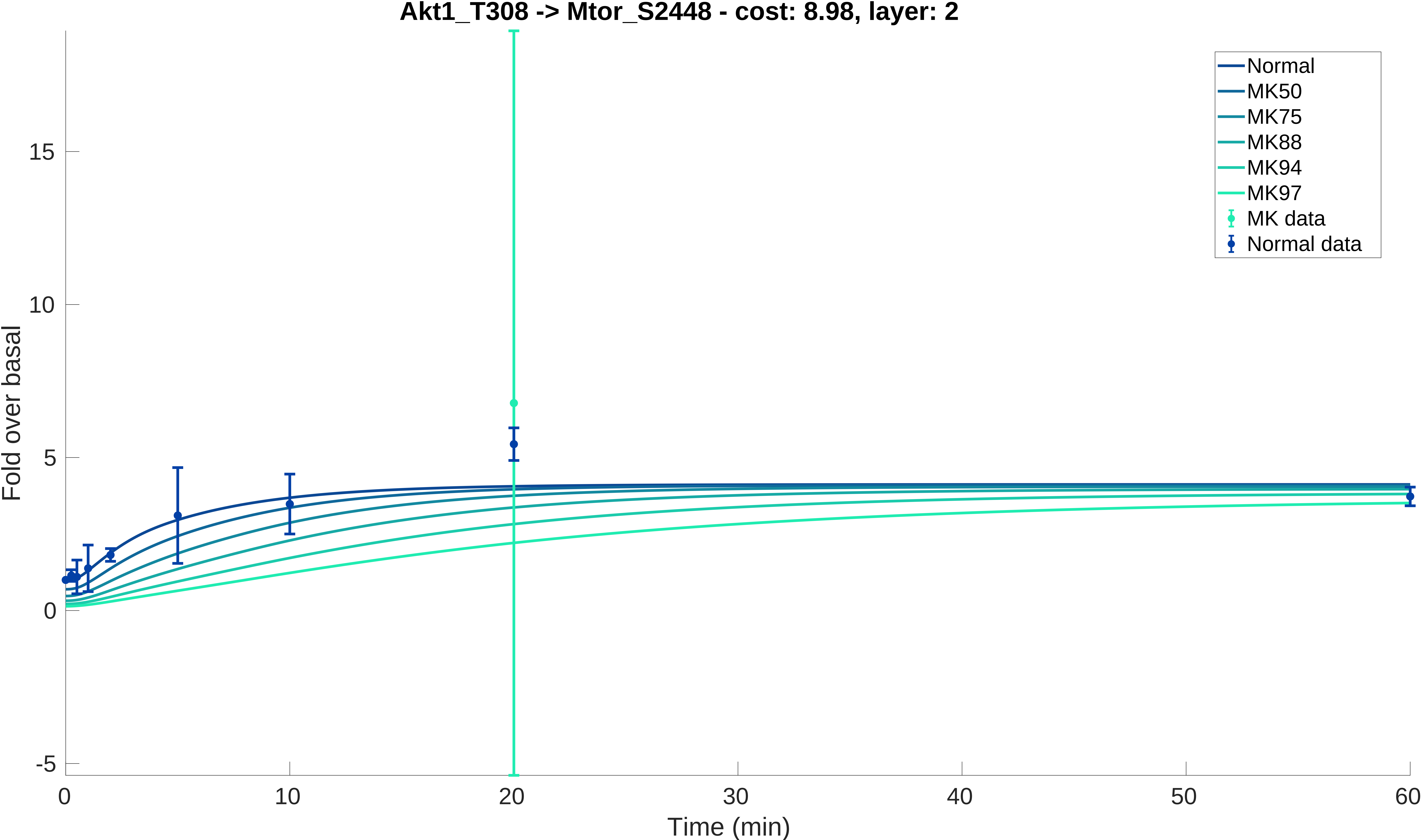

Akt1_T308 -> Mtor_S2448 - cost: 8.98, layer: 2
Fold over basal
Time (min)
15
10
5
0
-5
0
10
20
30
40
50
60
Normal
MK50
MK75
MK88
MK94
MK97
MK data
Normal data

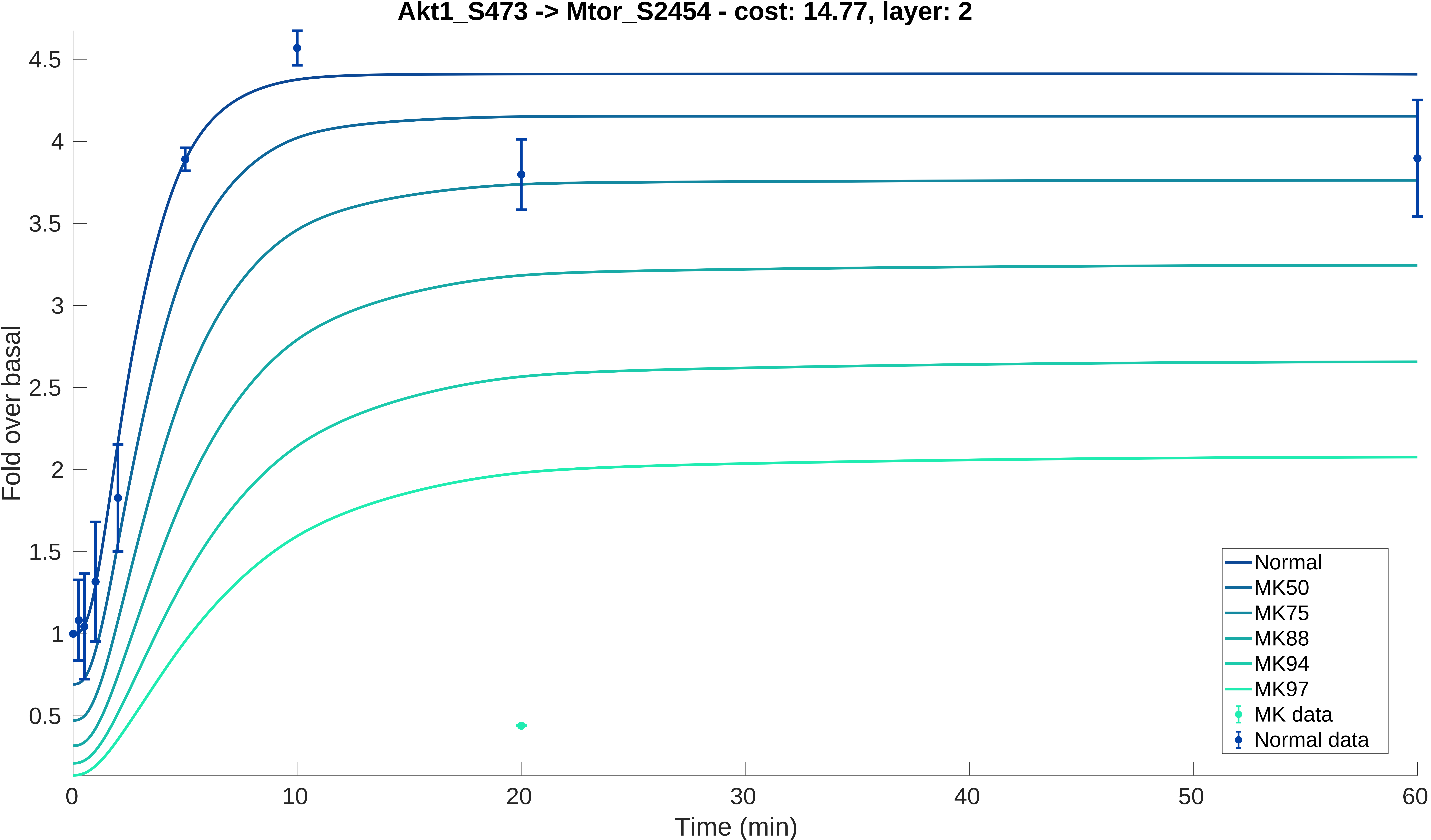

Akt1_S473 -> Mtor_S2454 - cost: 14.77, layer: 2
Fold over basal
Time (min)
4.5
4
3.5
3
2.5
2
1.5
1
0.5
0
10
20
30
40
50
60
Normal
MK50
MK75
MK88
MK94
MK97
MK data
Normal data

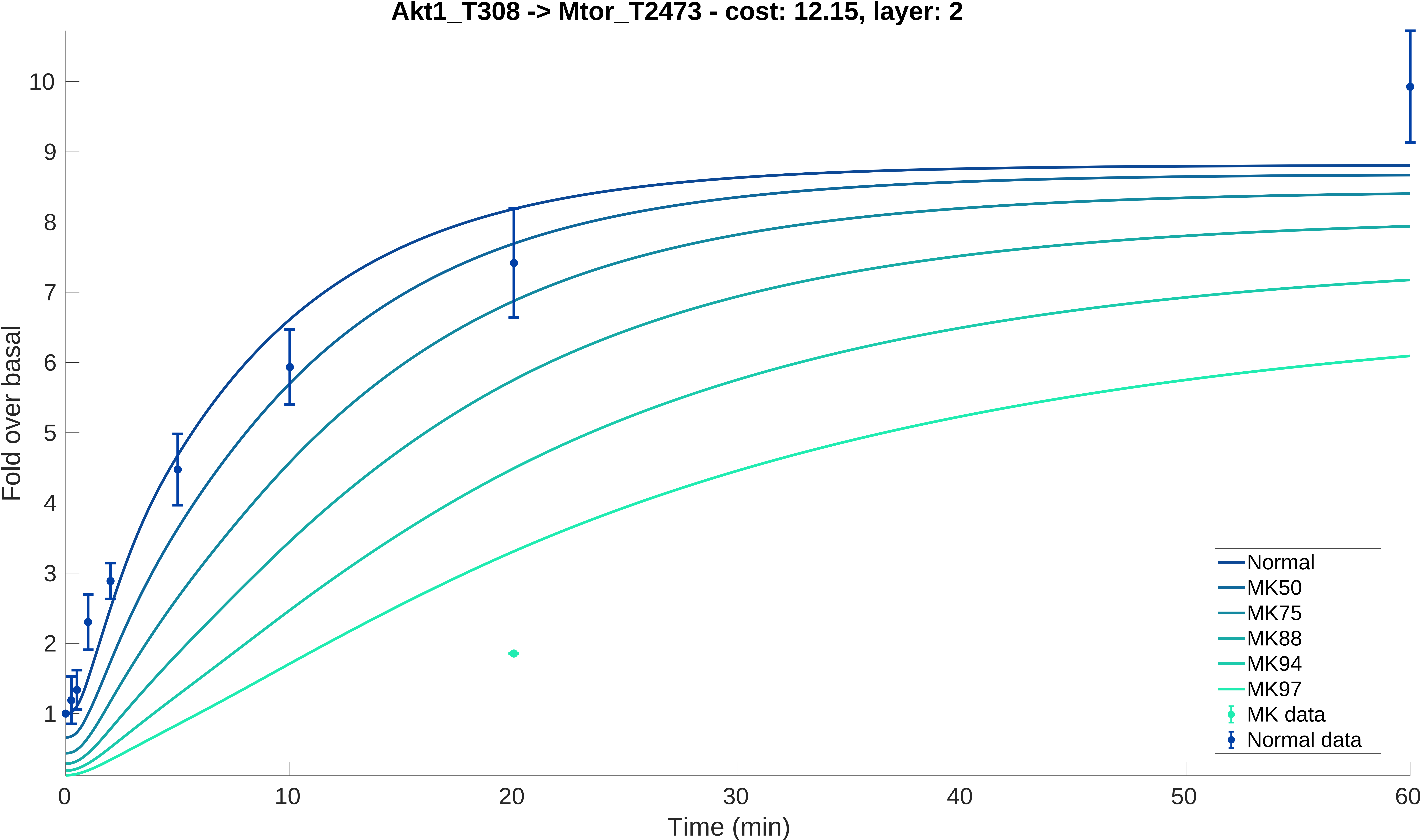

Akt1_T308 -> Mtor_T2473 - cost: 12.15, layer: 2
Fold over basal
Time (min)
0
10
20
30
40
50
60
1
2
3
4
5
6
7
8
9
10
Normal
MK50
MK75
MK88
MK94
MK97
MK data
Normal data

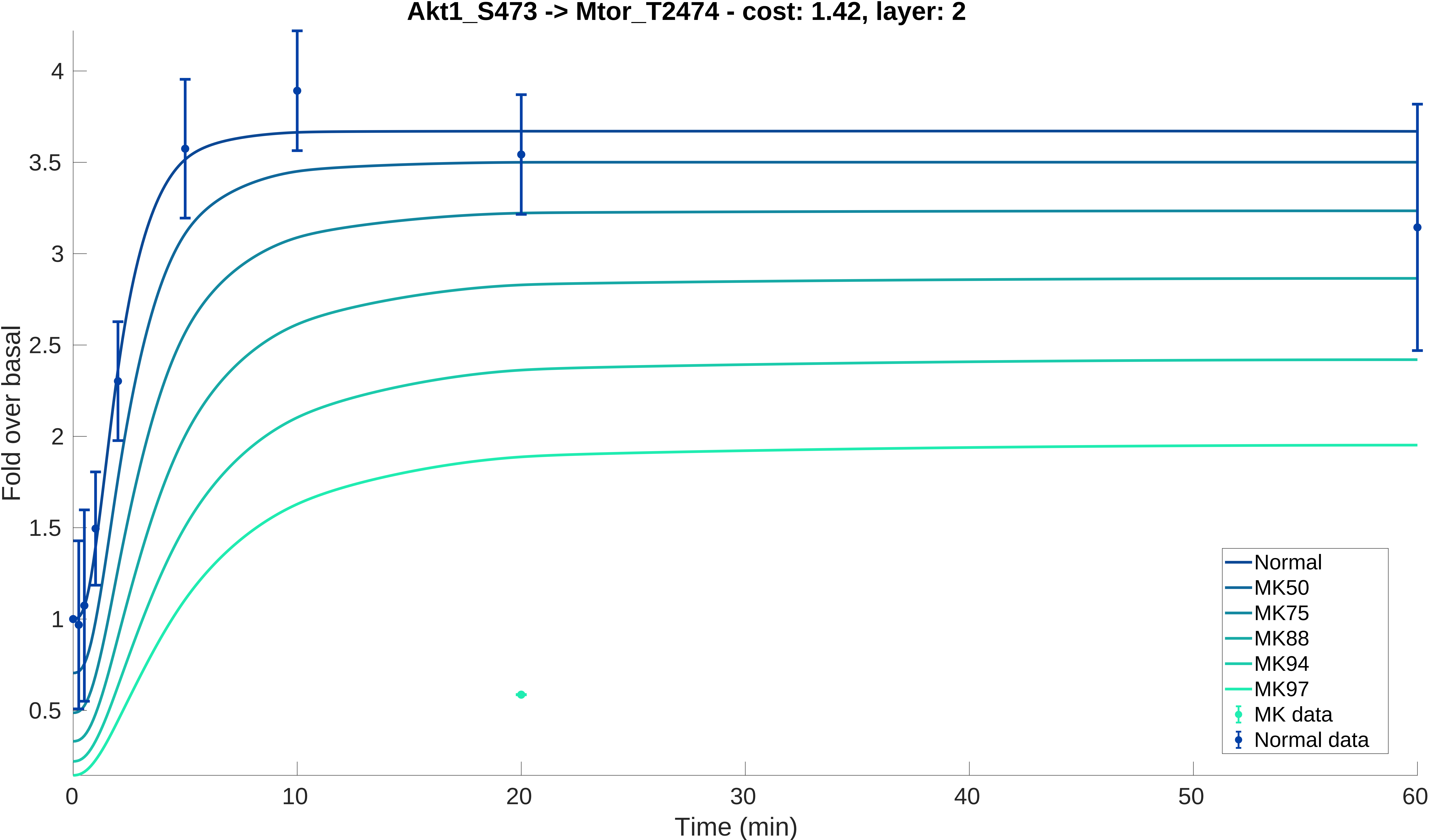

Akt1_S473 -> Mtor_T2474 - cost: 1.42, layer: 2
Fold over basal
Time (min)
0.5
1
1.5
2
2.5
3
3.5
4
0
10
20
30
40
50
60
Normal
MK50
MK75
MK88
MK94
MK97
MK data
Normal data

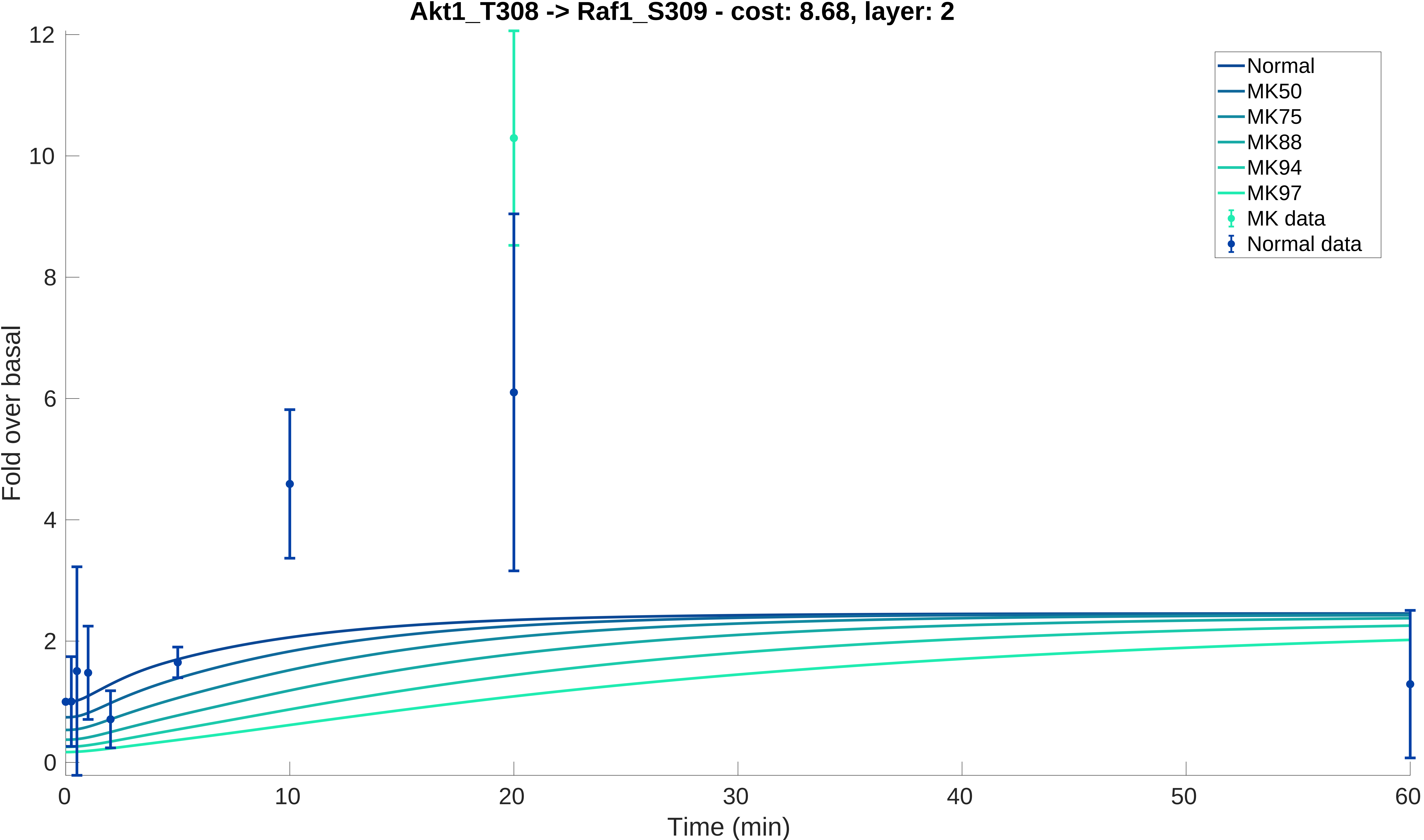

Akt1_T308 -> Raf1_S309 - cost: 8.68, layer: 2
Fold over basal
Time (min)
0
2
4
6
8
10
12
0
10
20
30
40
50
60
Normal
MK50
MK75
MK88
MK94
MK97
MK data
Normal data

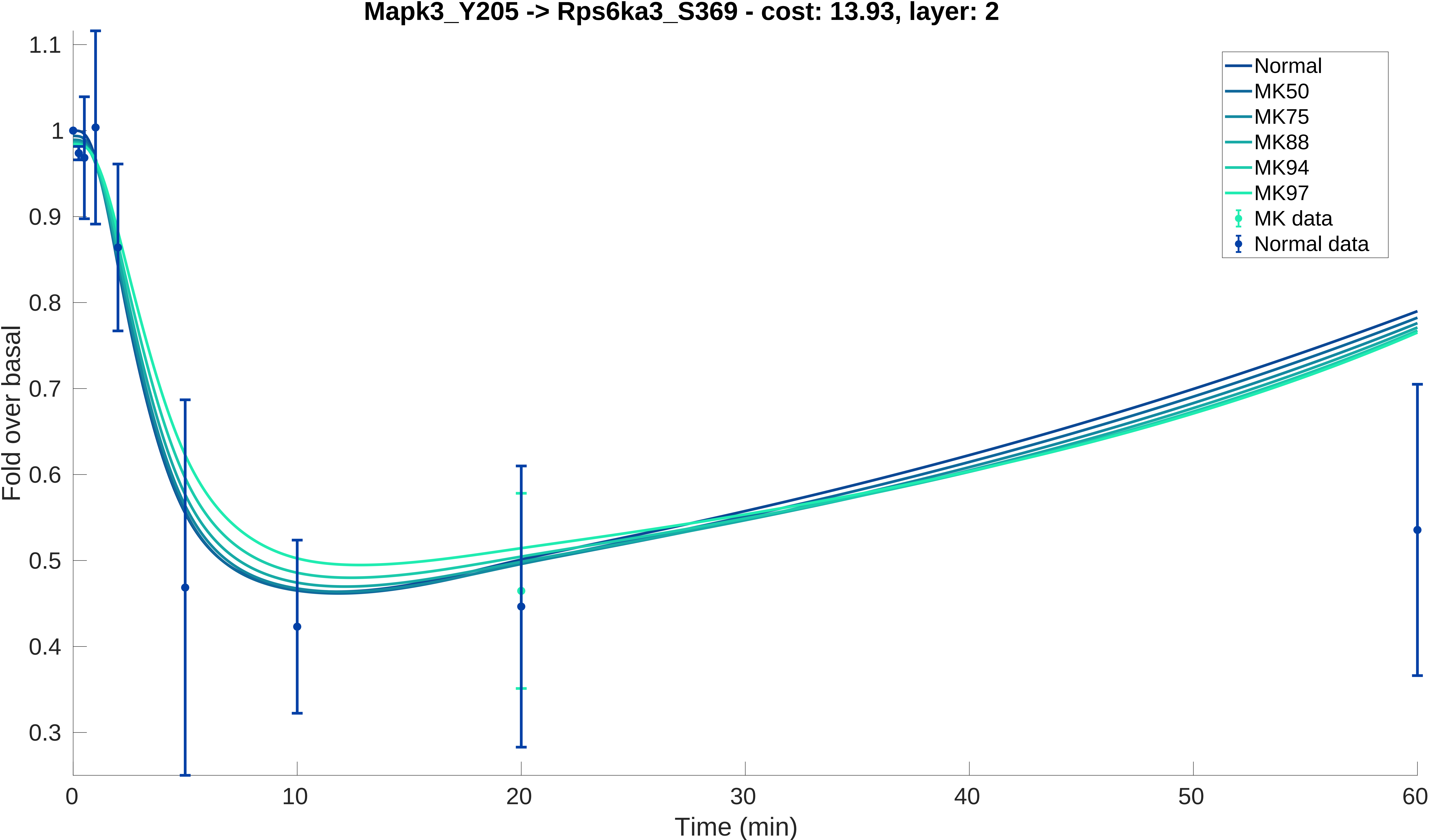
Mapk3_Y205 -> Rps6ka3_S369 - cost: 13.93, layer: 2
Fold over basal
Time (min)
1.1
1
0.9
0.8
0.7
0.6
0.5
0.4
0.3
0
10
20
30
40
50
60
Normal
MK50
MK75
MK88
MK94
MK97
MK data
Normal data

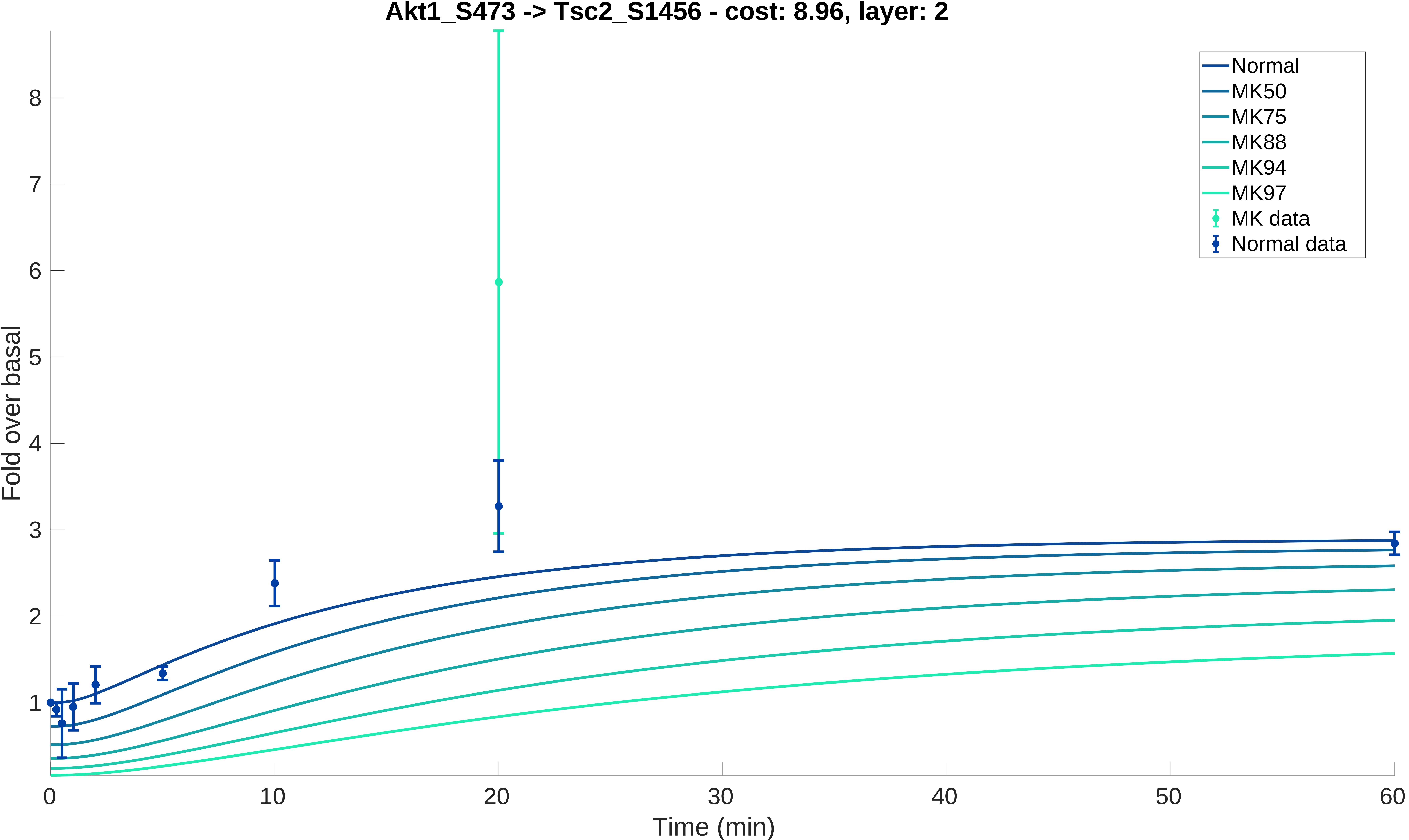
Akt1_S473 -> Tsc2_S1456 - cost: 8.96, layer: 2
Fold over basal
Time (min)
Normal
MK50
MK75
MK88
MK94
MK97
MK data
Normal data
0
10
20
30
40
50
60
1
2
3
4
5
6
7
8

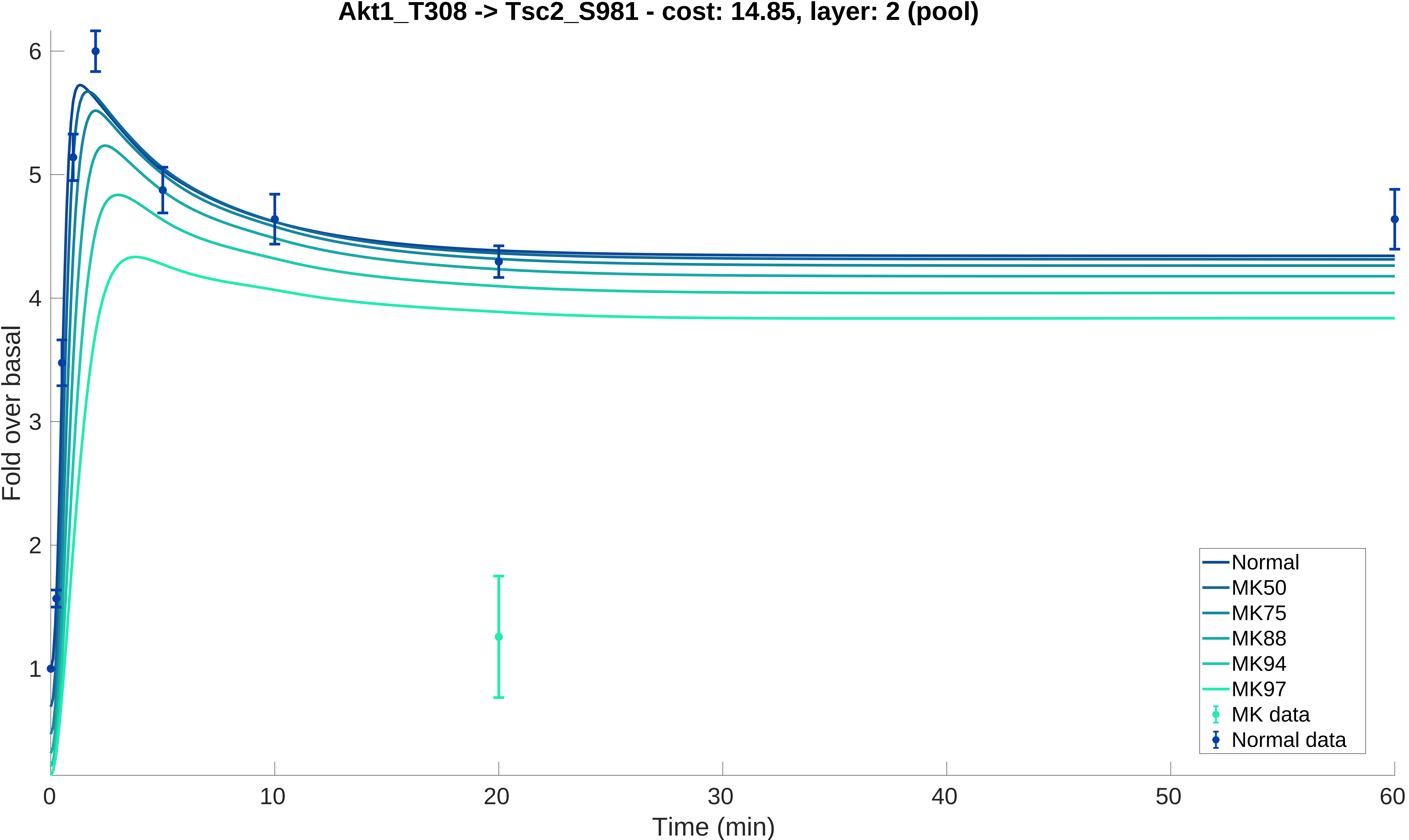

Akt1_T308 -> Tsc2_S981 - cost: 14.85, layer: 2 (pool)
Fold over basal
Time (min)
0
10
20
30
40
50
60
1
2
3
4
5
6
Normal
MK50
MK75
MK88
MK94
MK97
MK data
Normal data

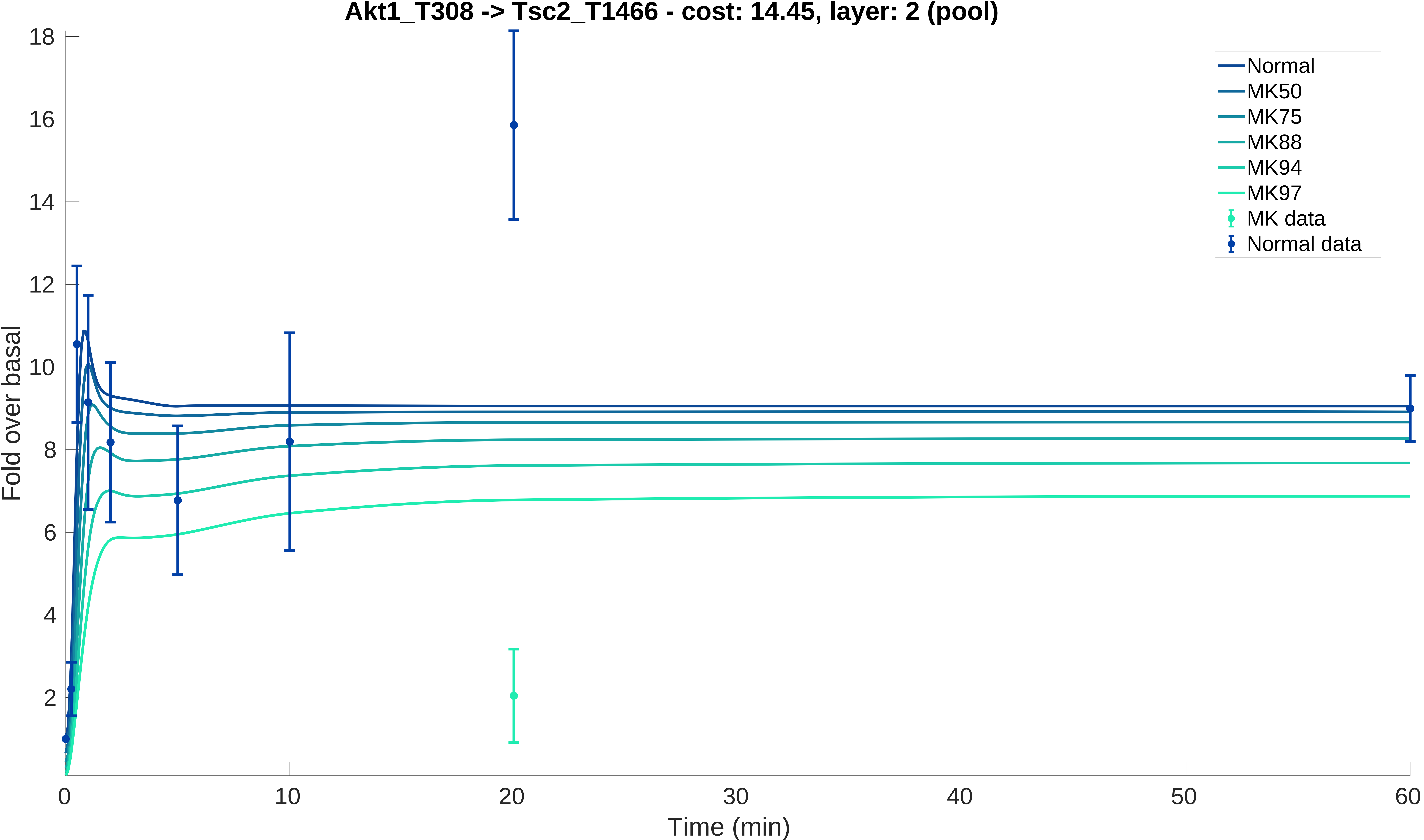
Akt1_T308 -> Tsc2_T1466 - cost: 14.45, layer: 2 (pool)
Fold over basal
Time (min)
0
10
20
30
40
50
60
2
4
6
8
10
12
14
16
18
Normal
MK50
MK75
MK88
MK94
MK97
MK data
Normal data

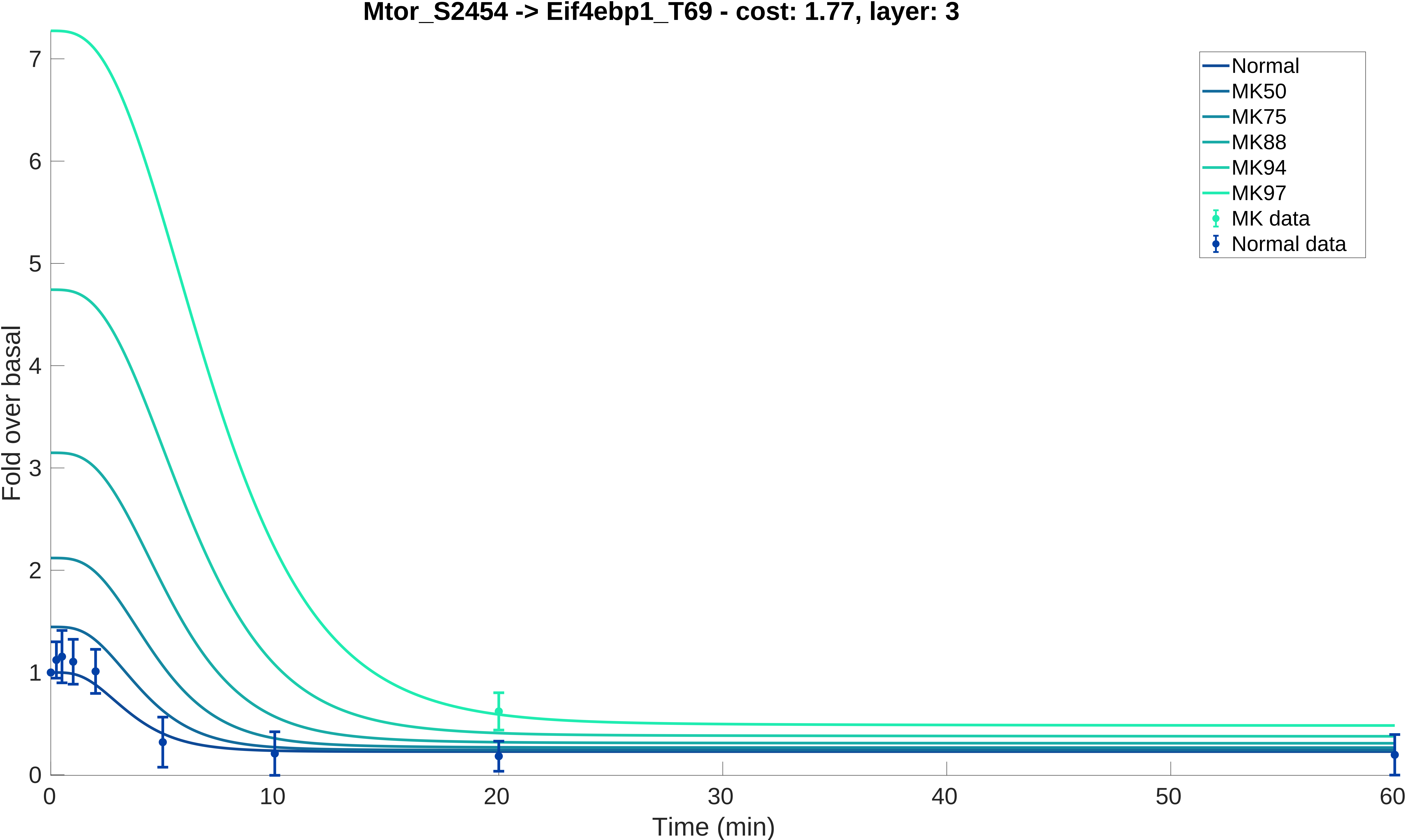
Mtor_S2454 -> Eif4ebp1_T69 - cost: 1.77, layer: 3
Fold over basal
Time (min)
0
1
2
3
4
5
6
7
0
10
20
30
40
50
60
Normal
MK50
MK75
MK88
MK94
MK97
MK data
Normal data

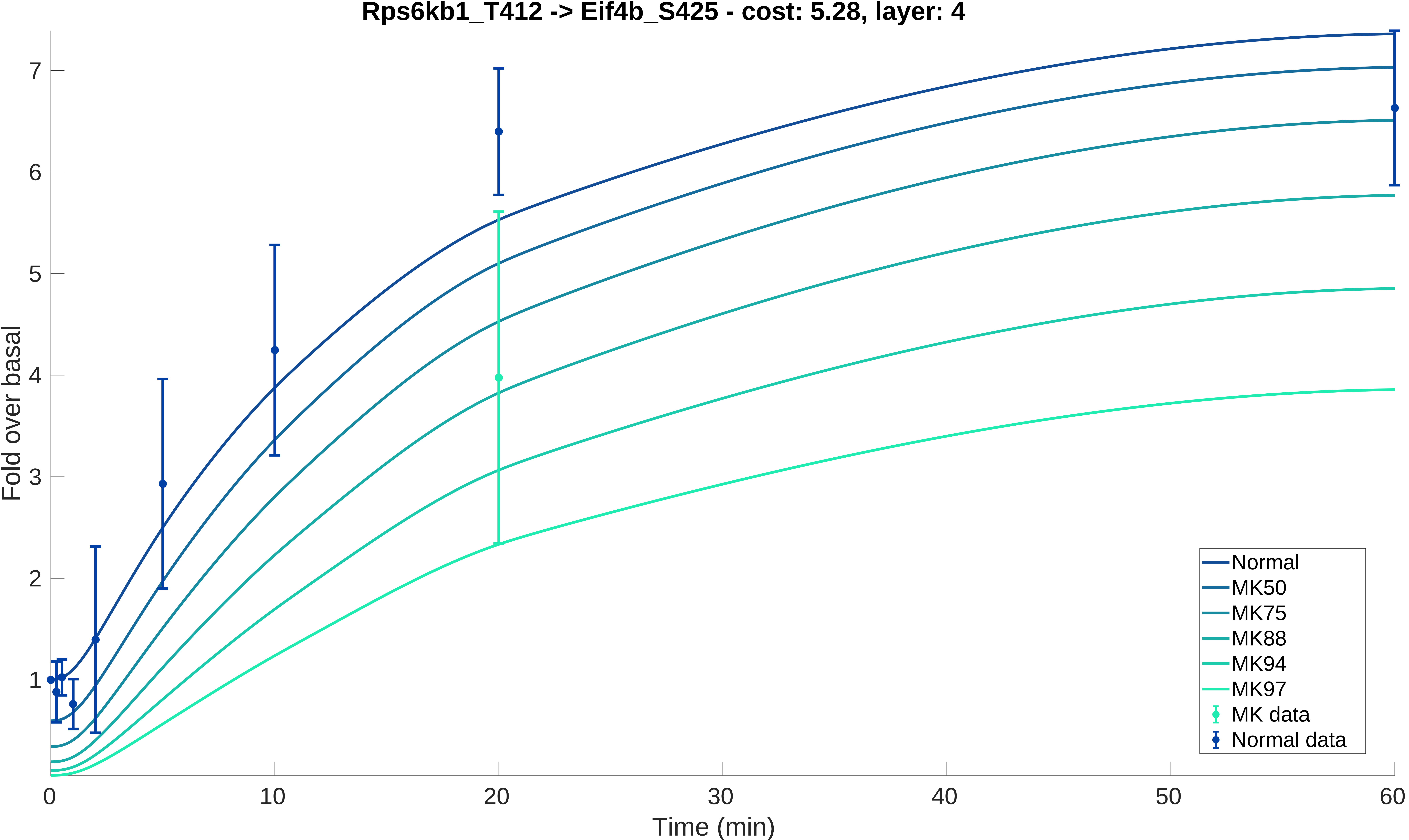

Rps6kb1_T412 -> Eif4b_S425 - cost: 5.28, layer: 4
Fold over basal
Time (min)
0
10
20
30
40
50
60
1
2
3
4
5
6
7
Normal
MK50
MK75
MK88
MK94
MK97
MK data
Normal data

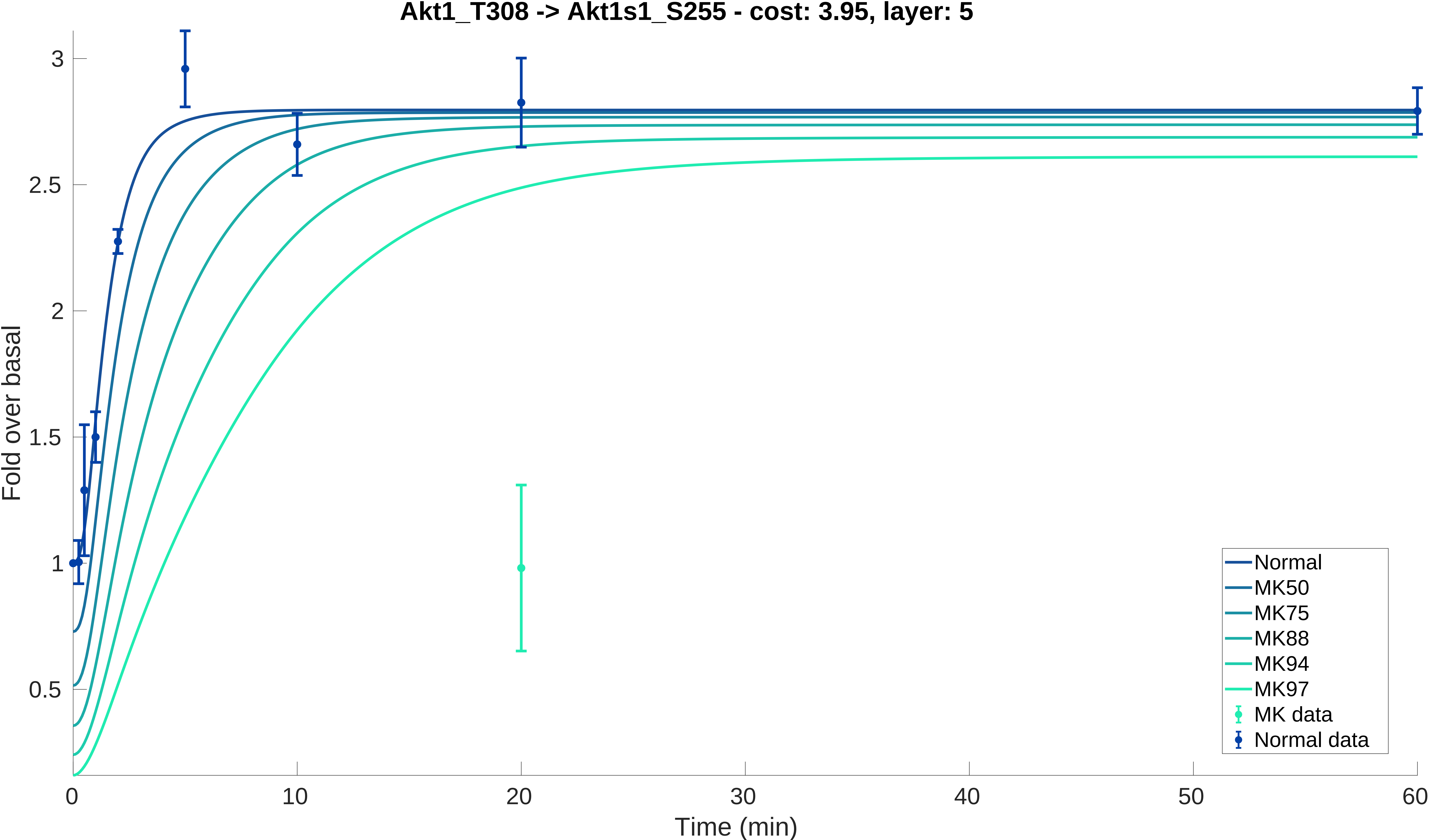
Akt1_T308 -> Akt1s1_S255 - cost: 3.95, layer: 5
Fold over basal
Time (min)
0
10
20
30
40
50
60
0.5
1
1.5
2
2.5
3
Normal
MK50
MK75
MK88
MK94
MK97
MK data
Normal data

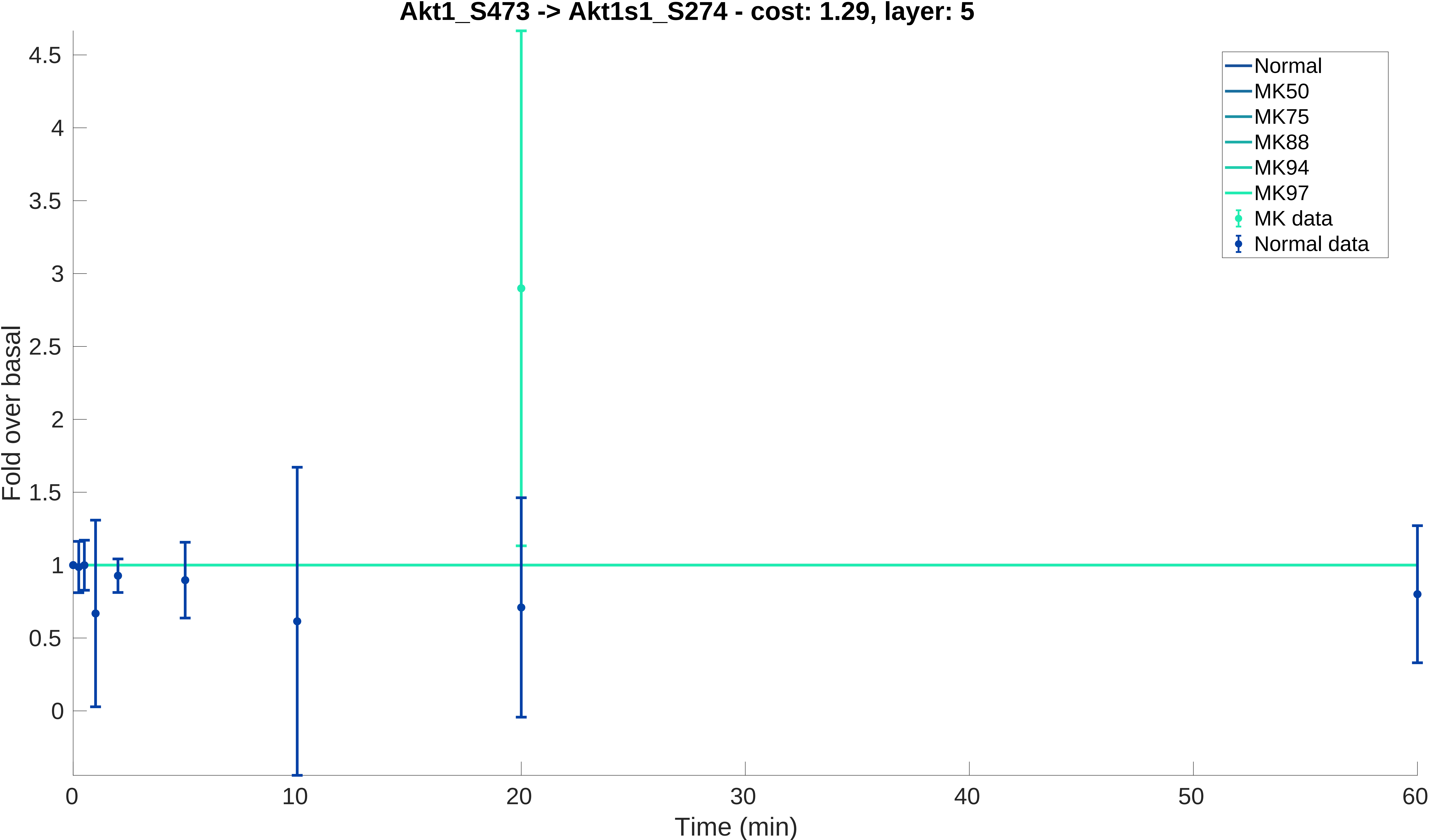

Akt1_S473 -> Akt1s1_S274 - cost: 1.29, layer: 5
Fold over basal
Time (min)
Normal
MK50
MK75
MK88
MK94
MK97
MK data
Normal data
0
0.5
1
1.5
2
2.5
3
3.5
4
4.5
0
10
20
30
40
50
60

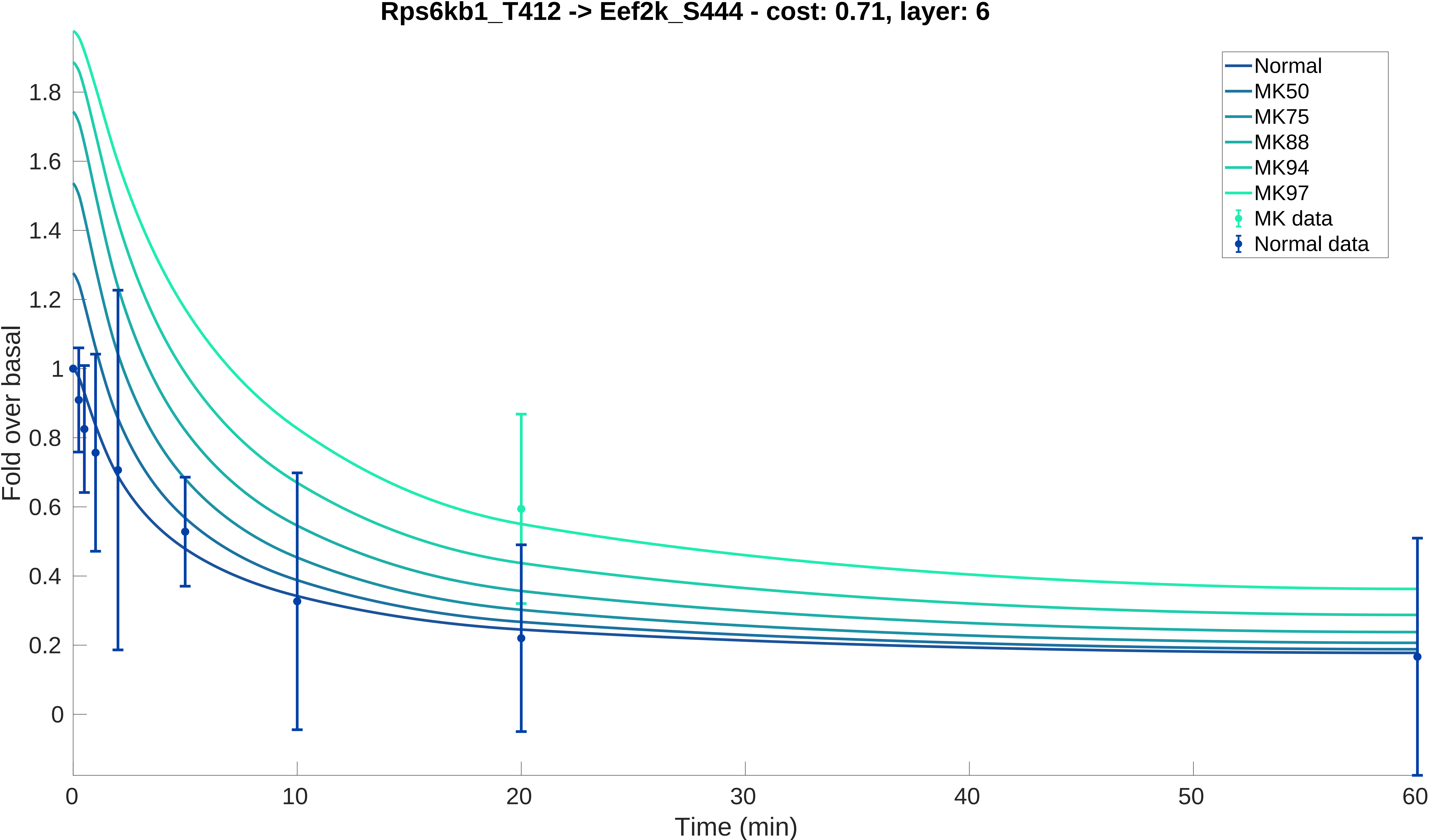

Rps6kb1_T412 -> Eef2k_S444 - cost: 0.71, layer: 6
Normal
MK50
MK75
MK88
MK94
MK97
MK data
Normal data
Fold over basal
1.8
1.6
1.4
1.2
1
0.8
0.6
0.4
0.2
0
0
10
20
30
40
50
60
Time (min)

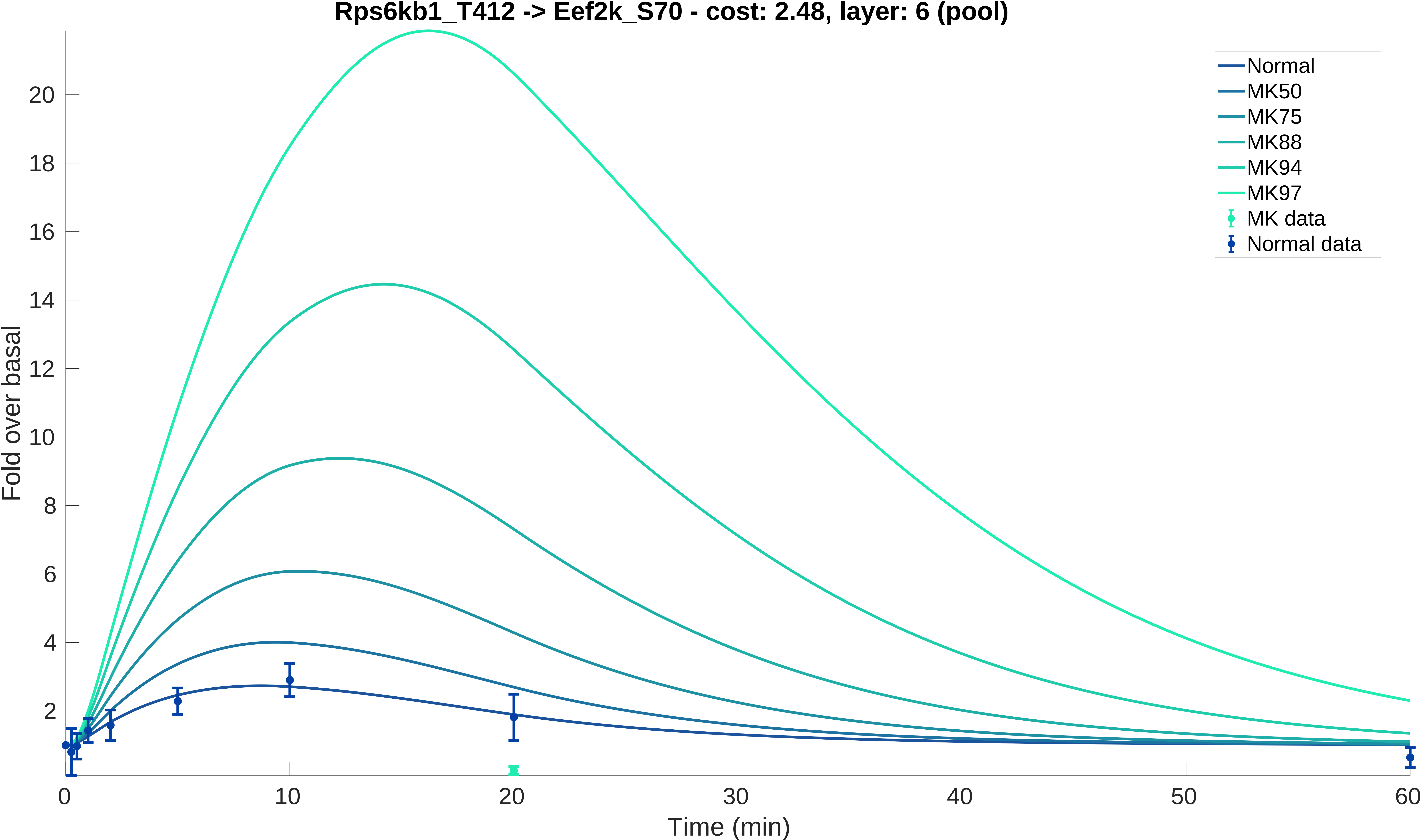

Rps6kb1_T412 -> Eef2k_S70 - cost: 2.48, layer: 6 (pool)
Fold over basal
Time (min)
0
10
20
30
40
50
60
2
4
6
8
10
12
14
16
18
20
Normal
MK50
MK75
MK88
MK94
MK97
MK data
Normal data

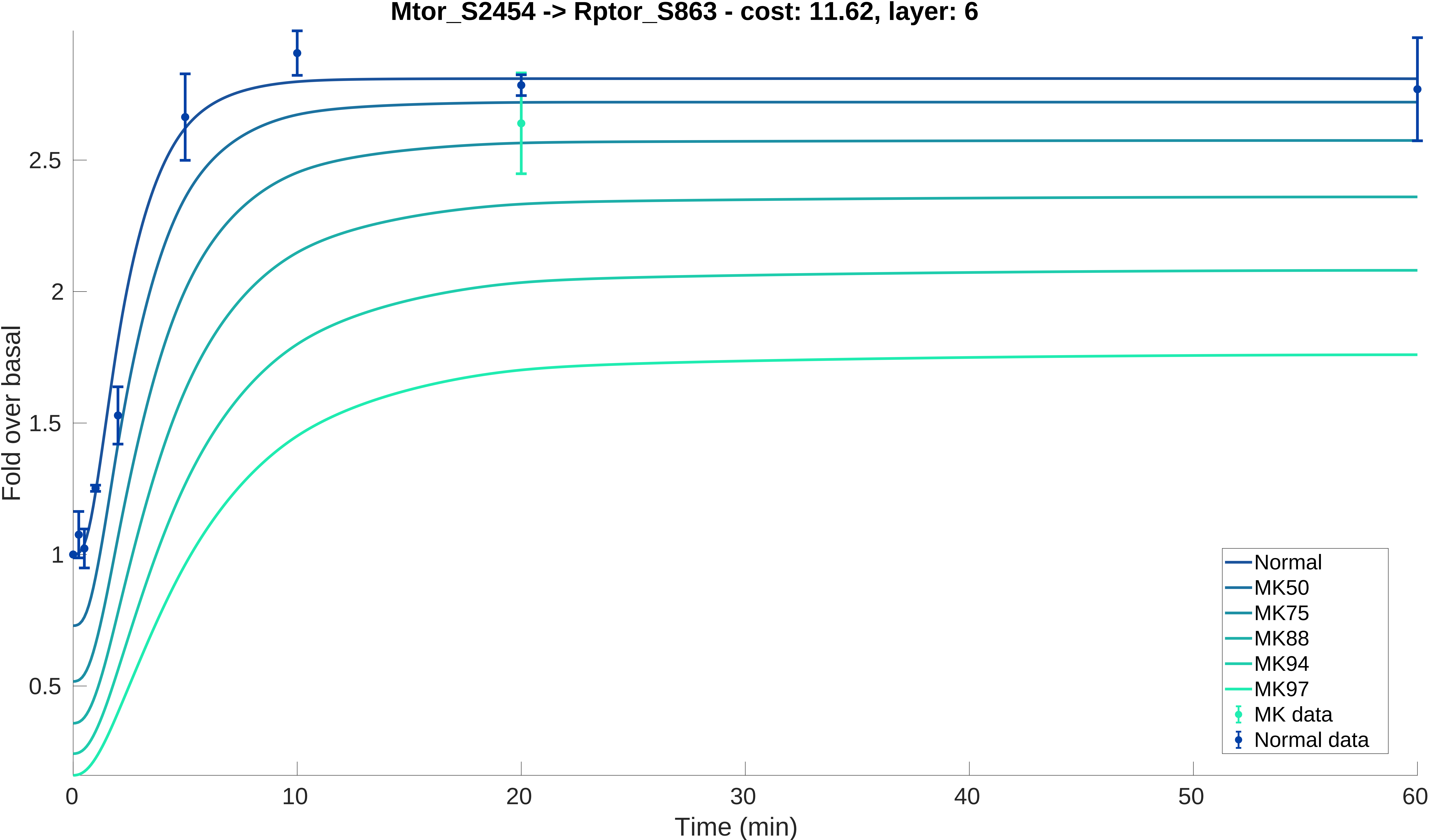

Mtor_S2454 -> Rptor_S863 - cost: 11.62, layer: 6
Fold over basal
0.5
1
1.5
2
2.5
0
10
20
30
40
50
60
Time (min)
Normal
MK50
MK75
MK88
MK94
MK97
MK data
Normal data

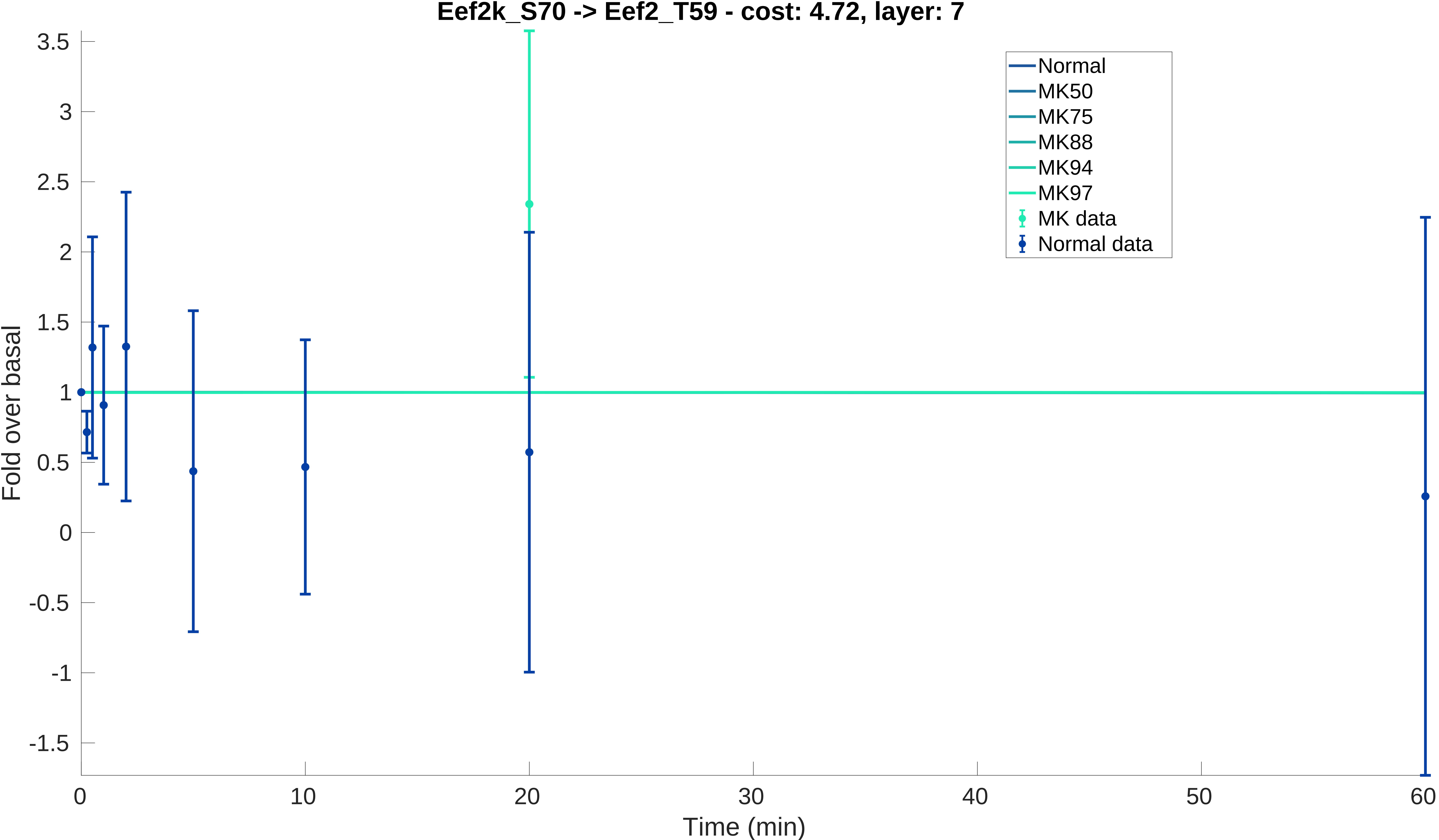

Eef2k_S70 -> Eef2_T59 - cost: 4.72, layer: 7
Fold over basal
Time (min)
3.5
3
2.5
2
1.5
1
0.5
0
-0.5
-1
-1.5
0
10
20
30
40
50
60
Normal
MK50
MK75
MK88
MK94
MK97
MK data
Normal data

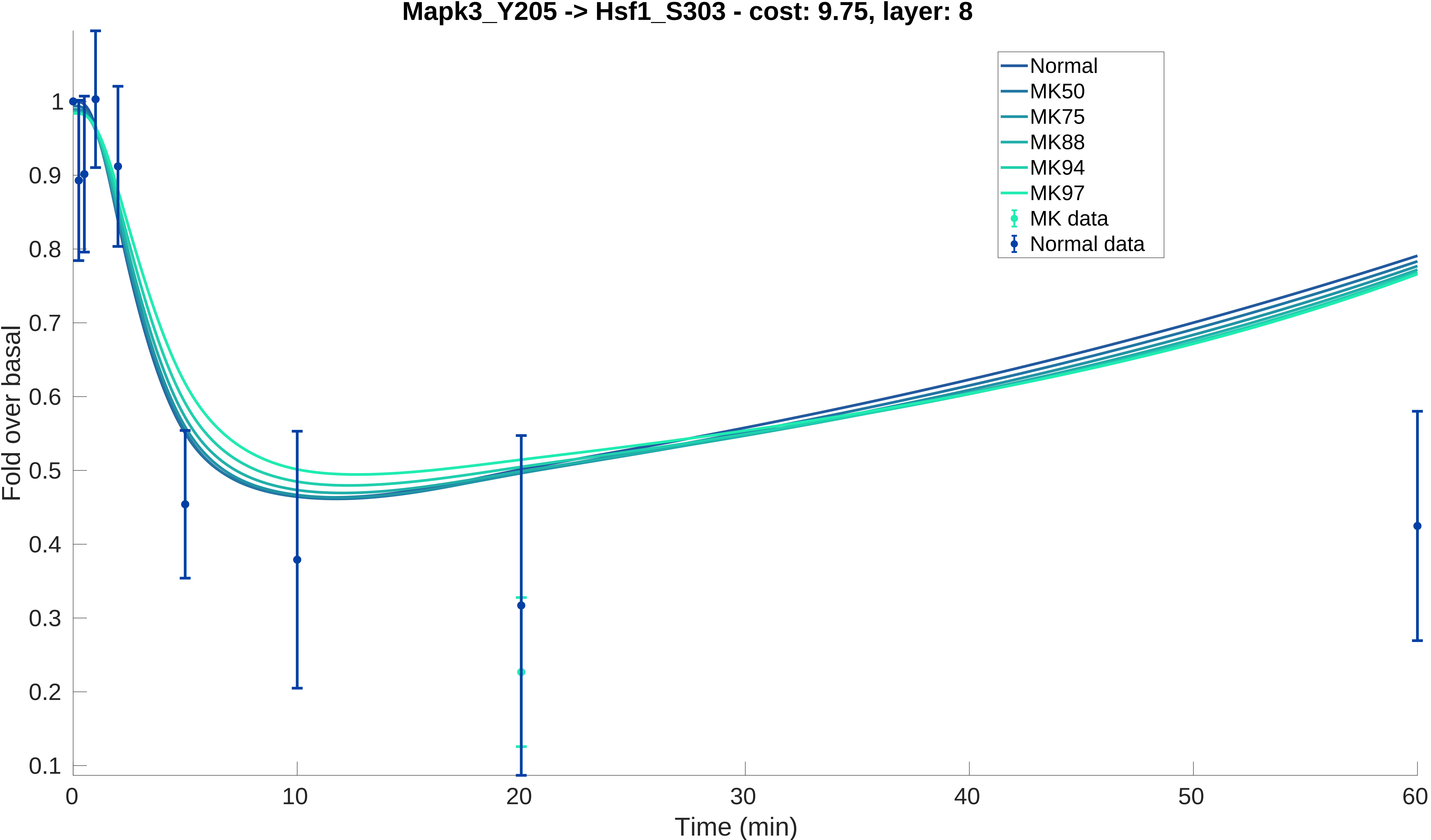

Mapk3_Y205 -> Hsf1_S303 - cost: 9.75, layer: 8
Normal
MK50
MK75
MK88
MK94
MK97
MK data
Normal data
Fold over basal
1
0.9
0.8
0.7
0.6
0.5
0.4
0.3
0.2
0.1
0
10
20
30
40
50
60
Time (min)

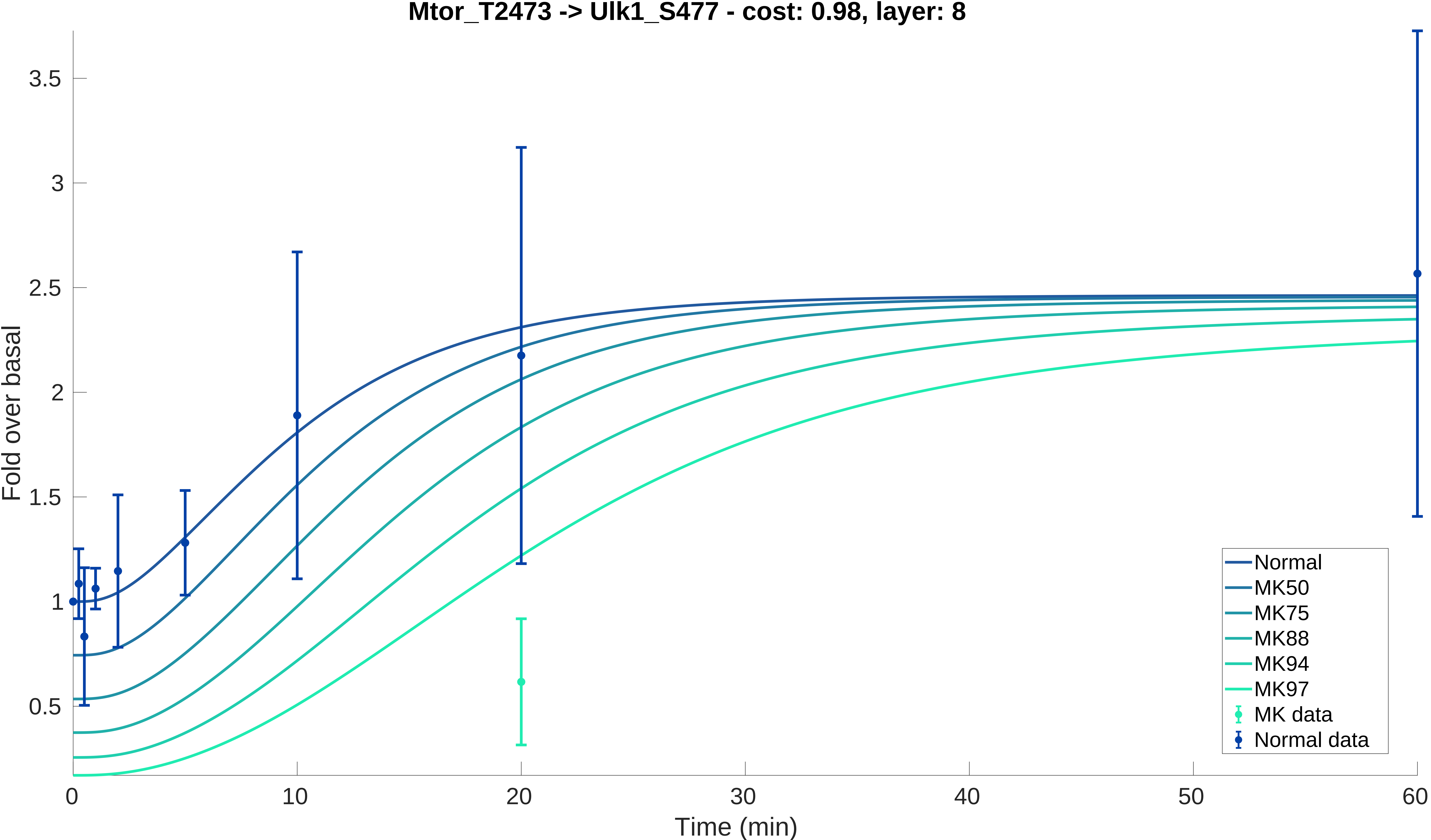

Mtor_T2473 -> Ulk1_S477 - cost: 0.98, layer: 8
Fold over basal
3.5
3
2.5
2
1.5
1
0.5
0
10
20
30
40
50
60
Time (min)
Normal
MK50
MK75
MK88
MK94
MK97
MK data
Normal data

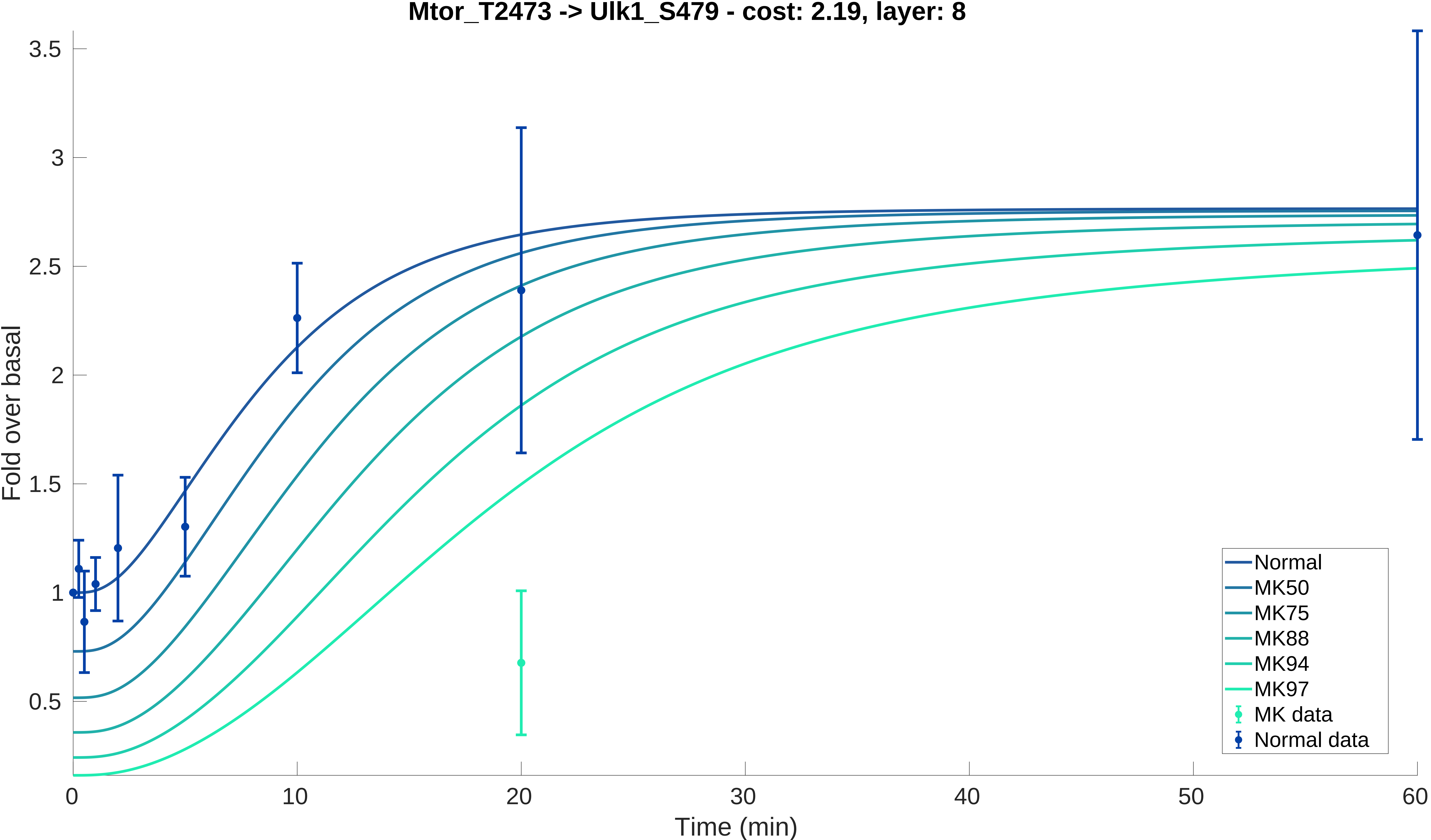
Mtor_T2473 -> Ulk1_S479 - cost: 2.19, layer: 8
Fold over basal
3.5
3
2.5
2
1.5
1
0.5
0
10
20
30
40
50
60
Time (min)
Normal
MK50
MK75
MK88
MK94
MK97
MK data
Normal data

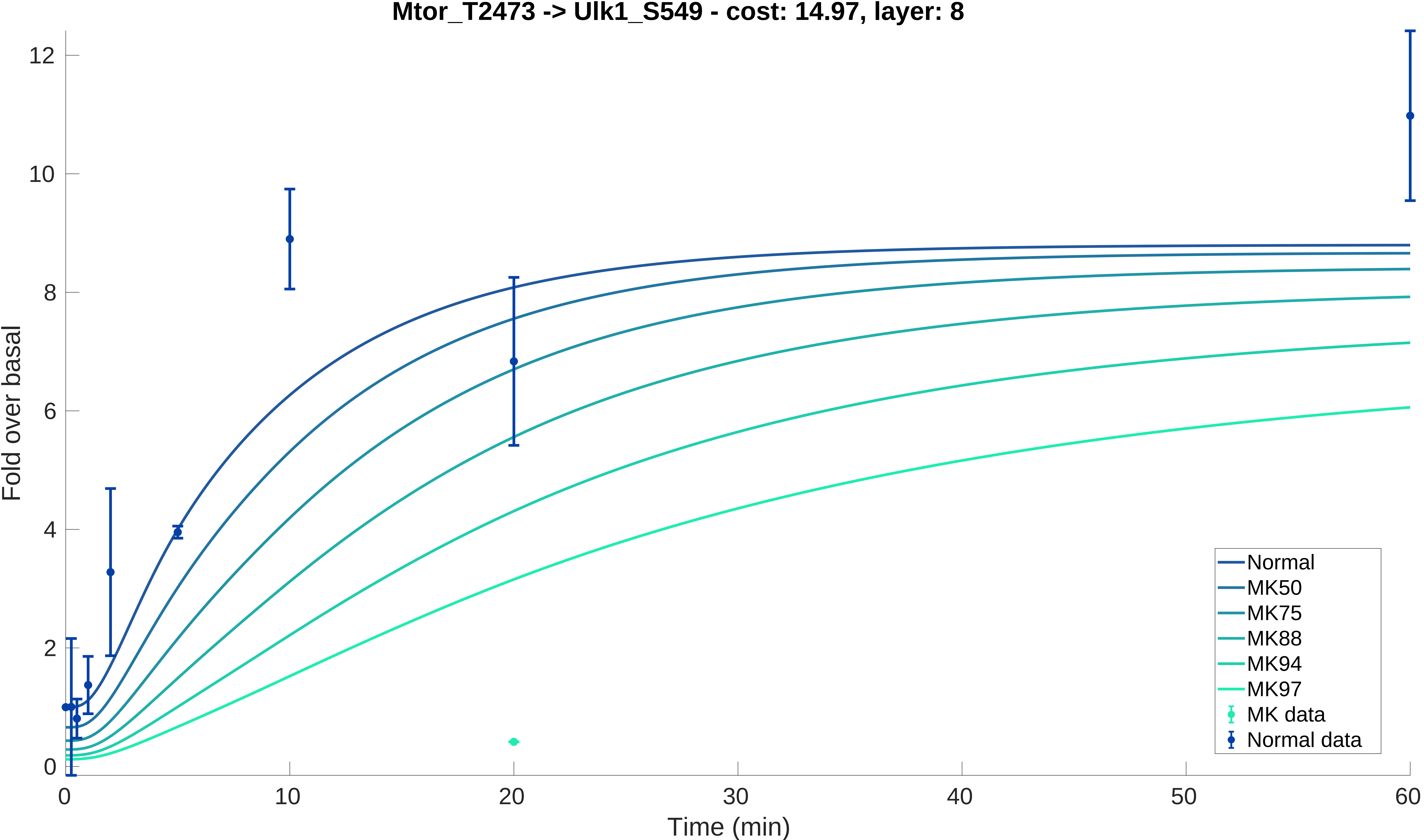

Mtor_T2473 -> Ulk1_S549 - cost: 14.97, layer: 8
Fold over basal
Time (min)
0
2
4
6
8
10
12
0
10
20
30
40
50
60
Normal
MK50
MK75
MK88
MK94
MK97
MK data
Normal data

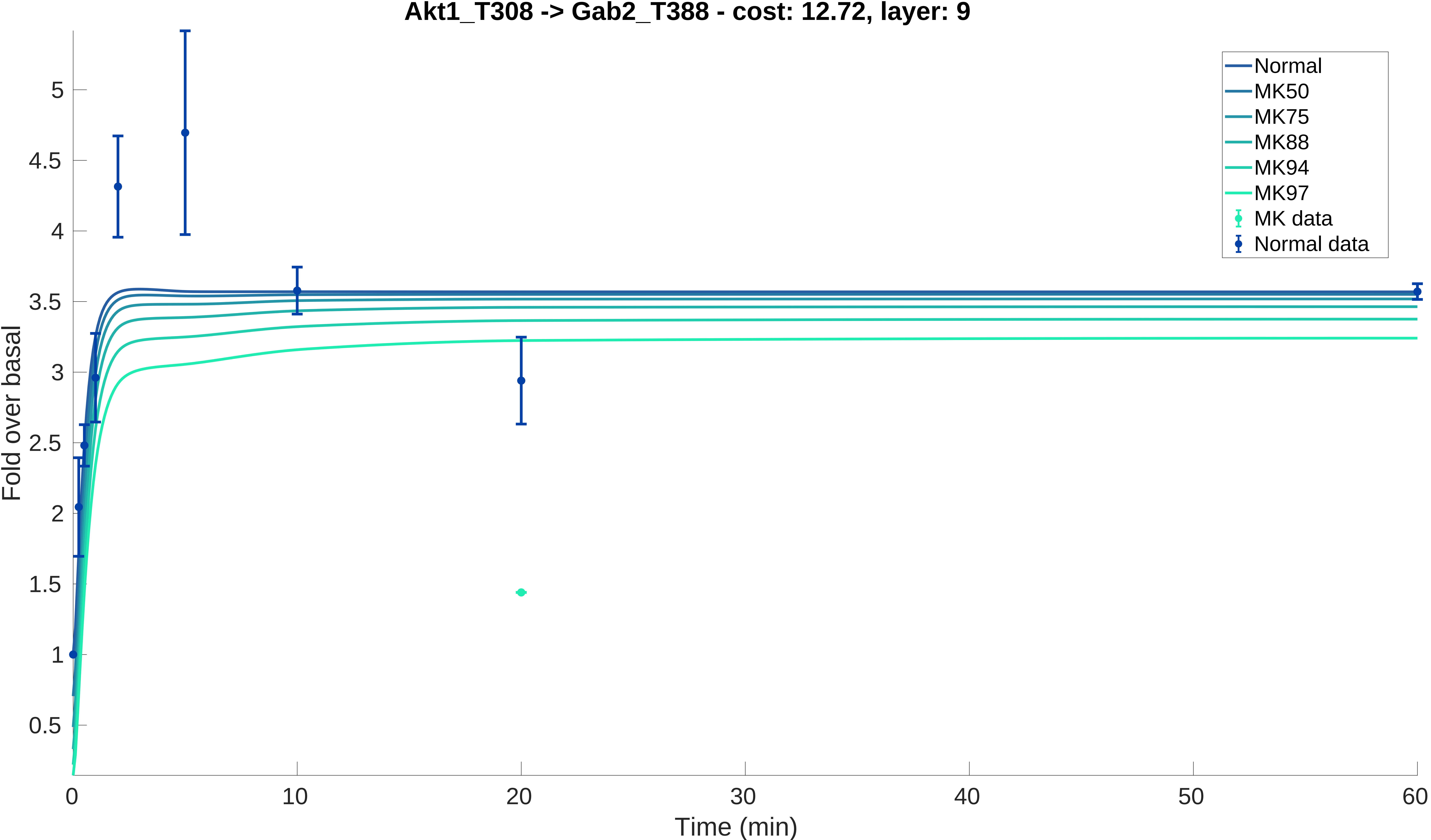

Akt1_T308 -> Gab2_T388 - cost: 12.72, layer: 9
Fold over basal
Time (min)
0
10
20
30
40
50
60
0.5
1
1.5
2
2.5
3
3.5
4
4.5
5
Normal
MK50
MK75
MK88
MK94
MK97
MK data
Normal data

Mapk1_T183 -> Irs1_S343 - cost: 10.06, layer: 9

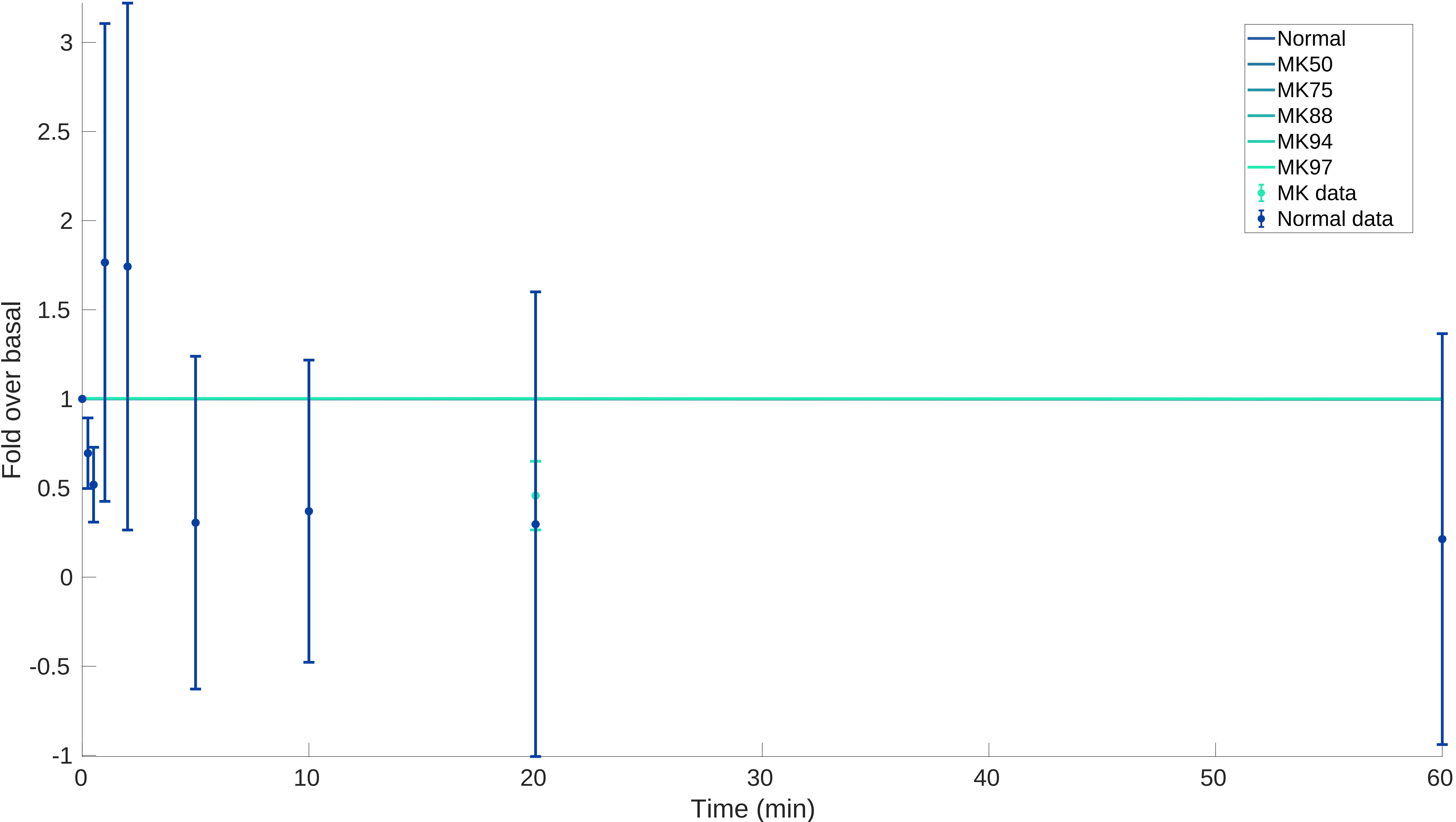

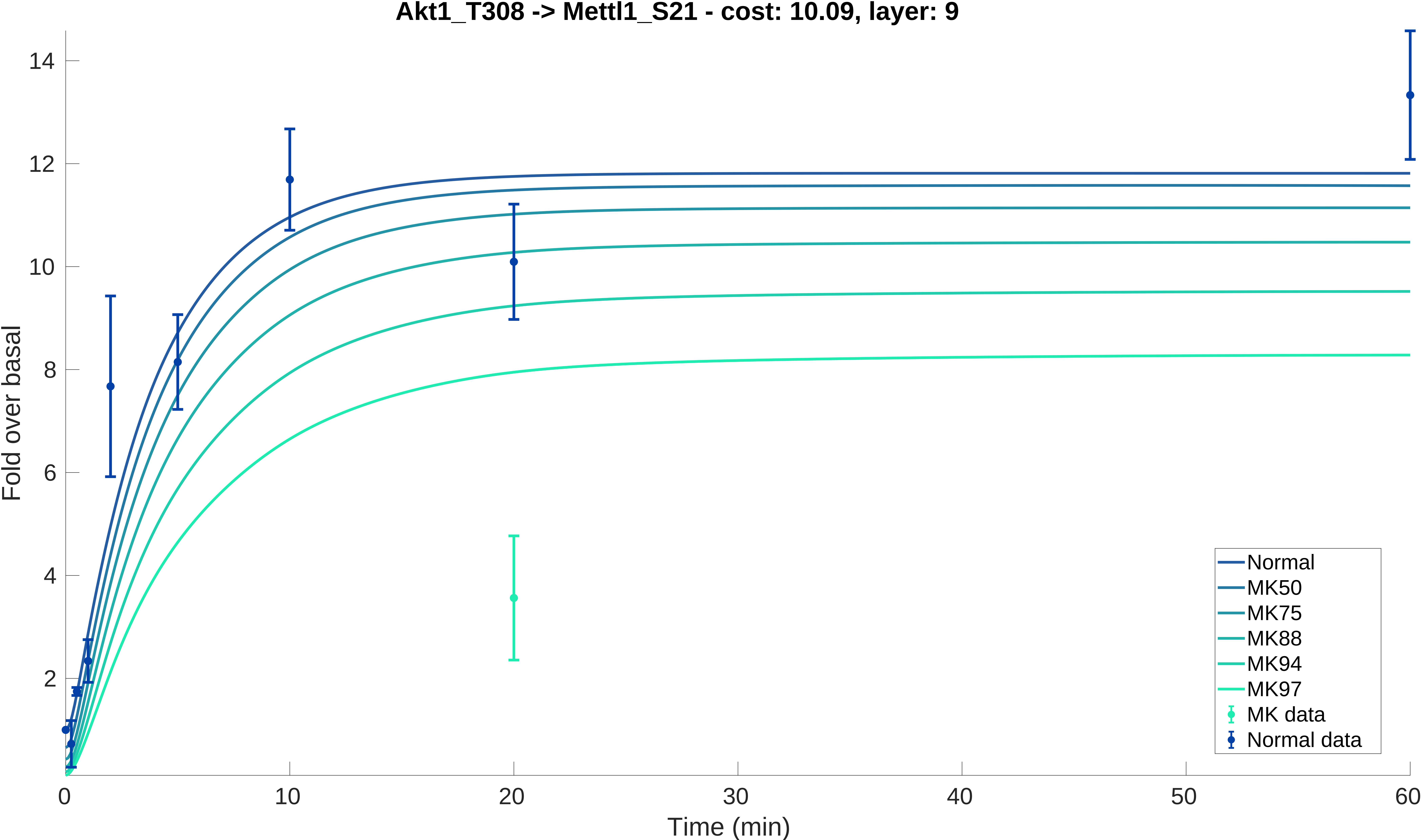
Akt1_T308 -> Mettl1_S21 - cost: 10.09, layer: 9
Fold over basal
Time (min)
0
10
20
30
40
50
60
2
4
6
8
10
12
14
Normal
MK50
MK75
MK88
MK94
MK97
MK data
Normal data

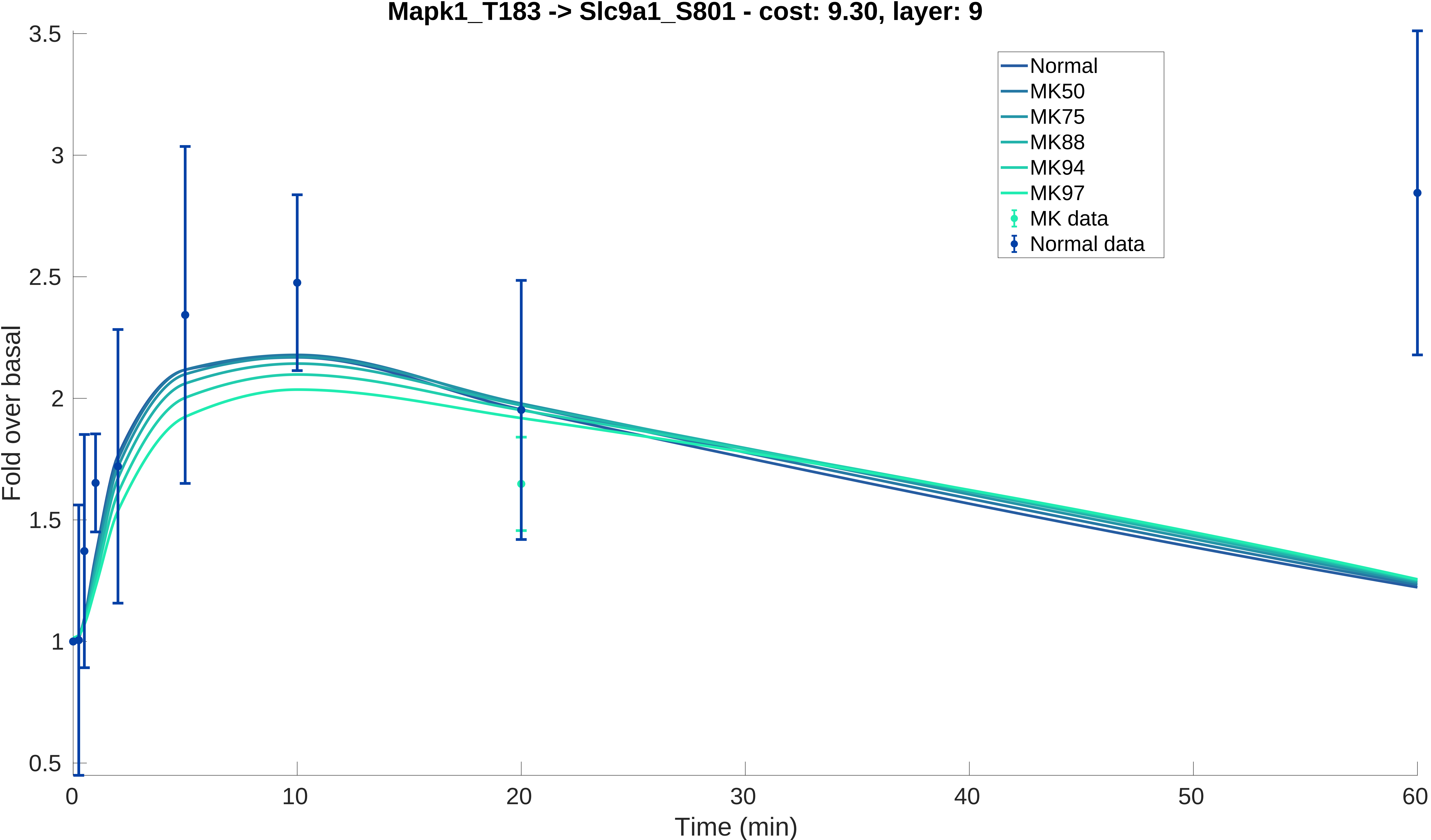

Mapk1_T183 -> Slc9a1_S801 - cost: 9.30, layer: 9
Fold over basal
Time (min)
0.5
1
1.5
2
2.5
3
3.5
0
10
20
30
40
50
60
Normal
MK50
MK75
MK88
MK94
MK97
MK data
Normal data

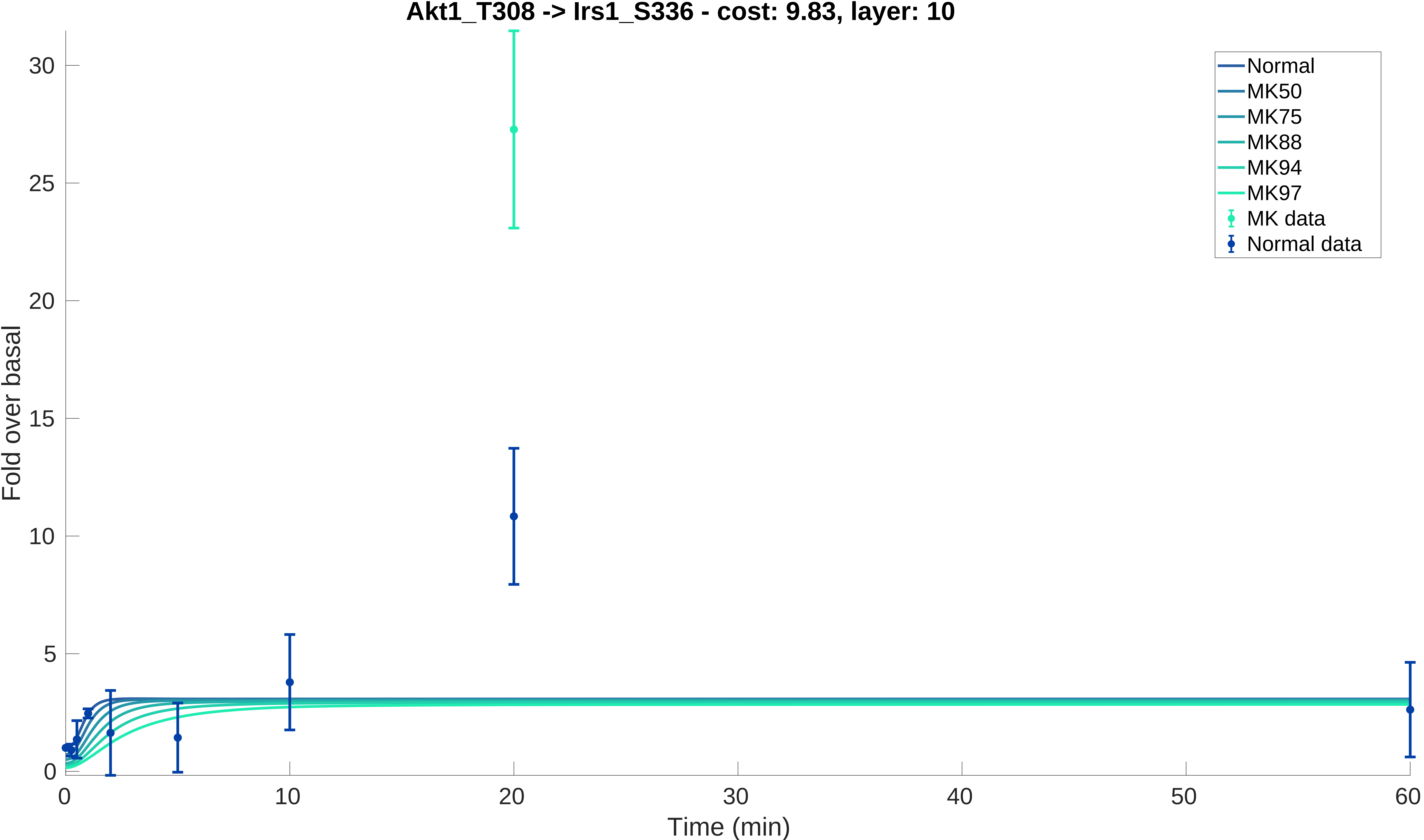
Akt1_T308 -> Irs1_S336 - cost: 9.83, layer: 10
Fold over basal
Time (min)
0
5
10
15
20
25
30
0
10
20
30
40
50
60
Normal
MK50
MK75
MK88
MK94
MK97
MK data
Normal data

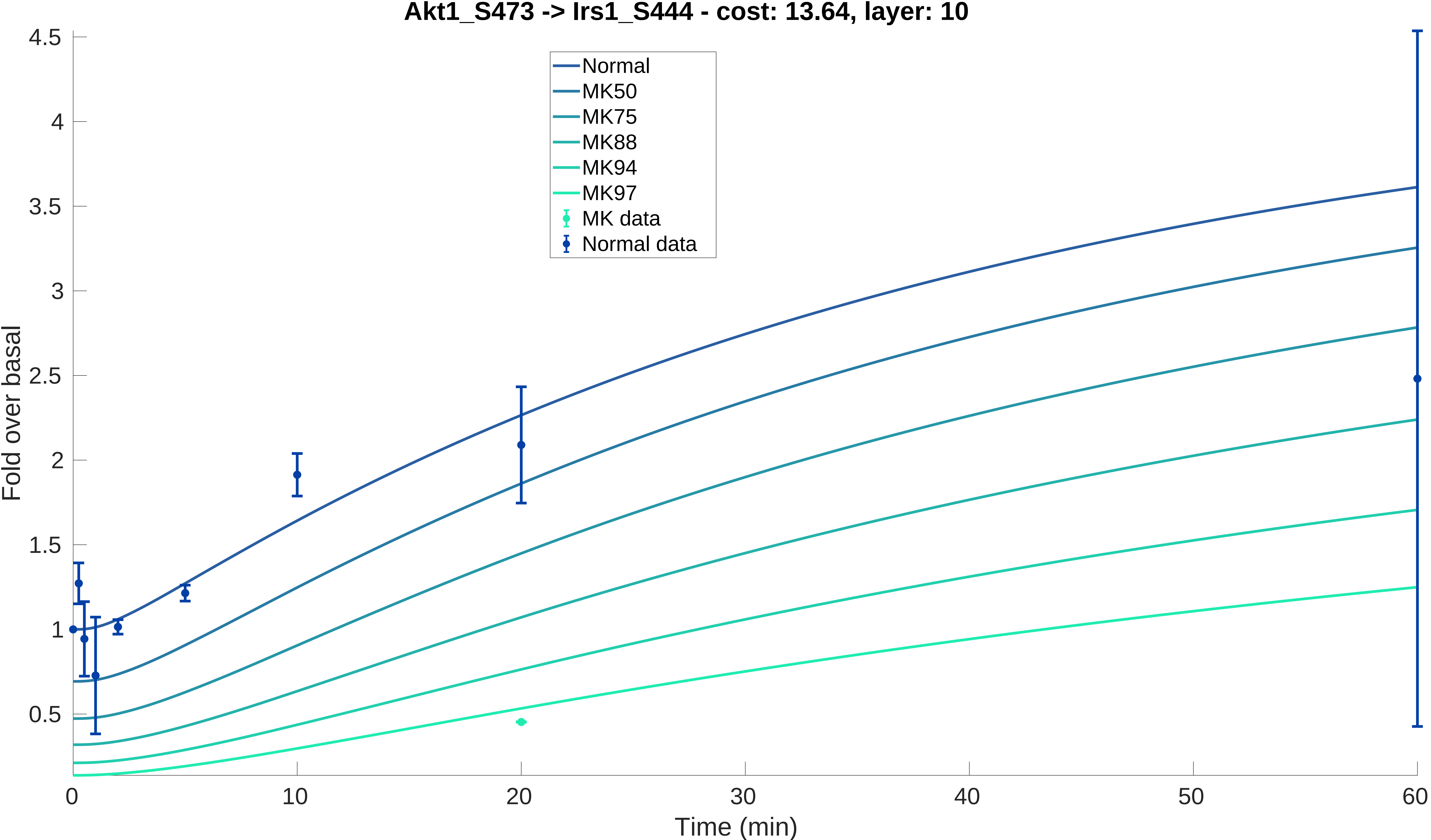
Akt1_S473 -> Irs1_S444 - cost: 13.64, layer: 10
Normal
MK50
MK75
MK88
MK94
MK97
MK data
Normal data
Fold over basal
Time (min)
0.5
1
1.5
2
2.5
3
3.5
4
4.5
0
10
20
30
40
50
60

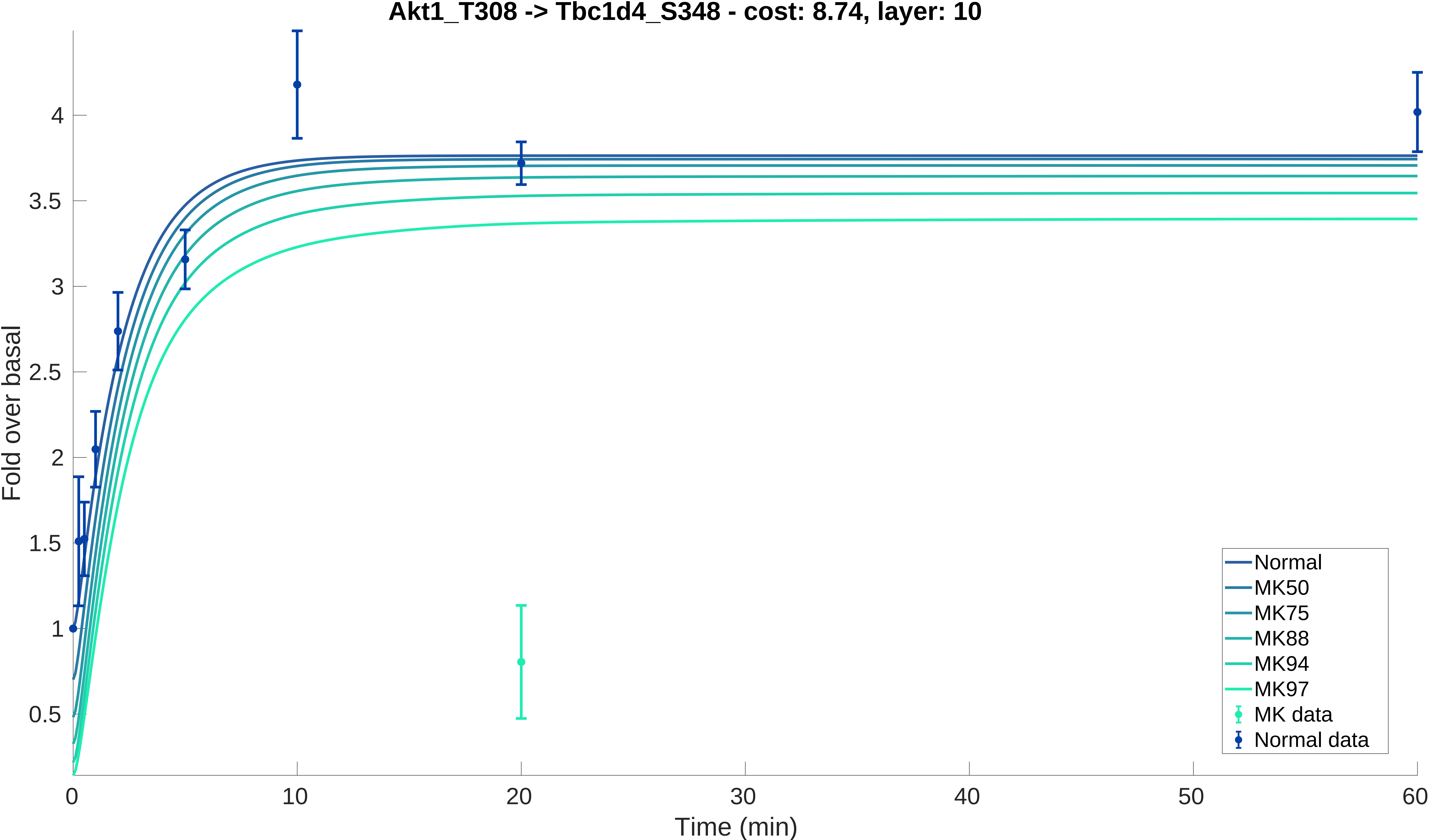

Akt1_T308 -> Tbc1d4_S348 - cost: 8.74, layer: 10
Fold over basal
Time (min)
4
3.5
3
2.5
2
1.5
1
0.5
0
10
20
30
40
50
60
Normal
MK50
MK75
MK88
MK94
MK97
MK data
Normal data

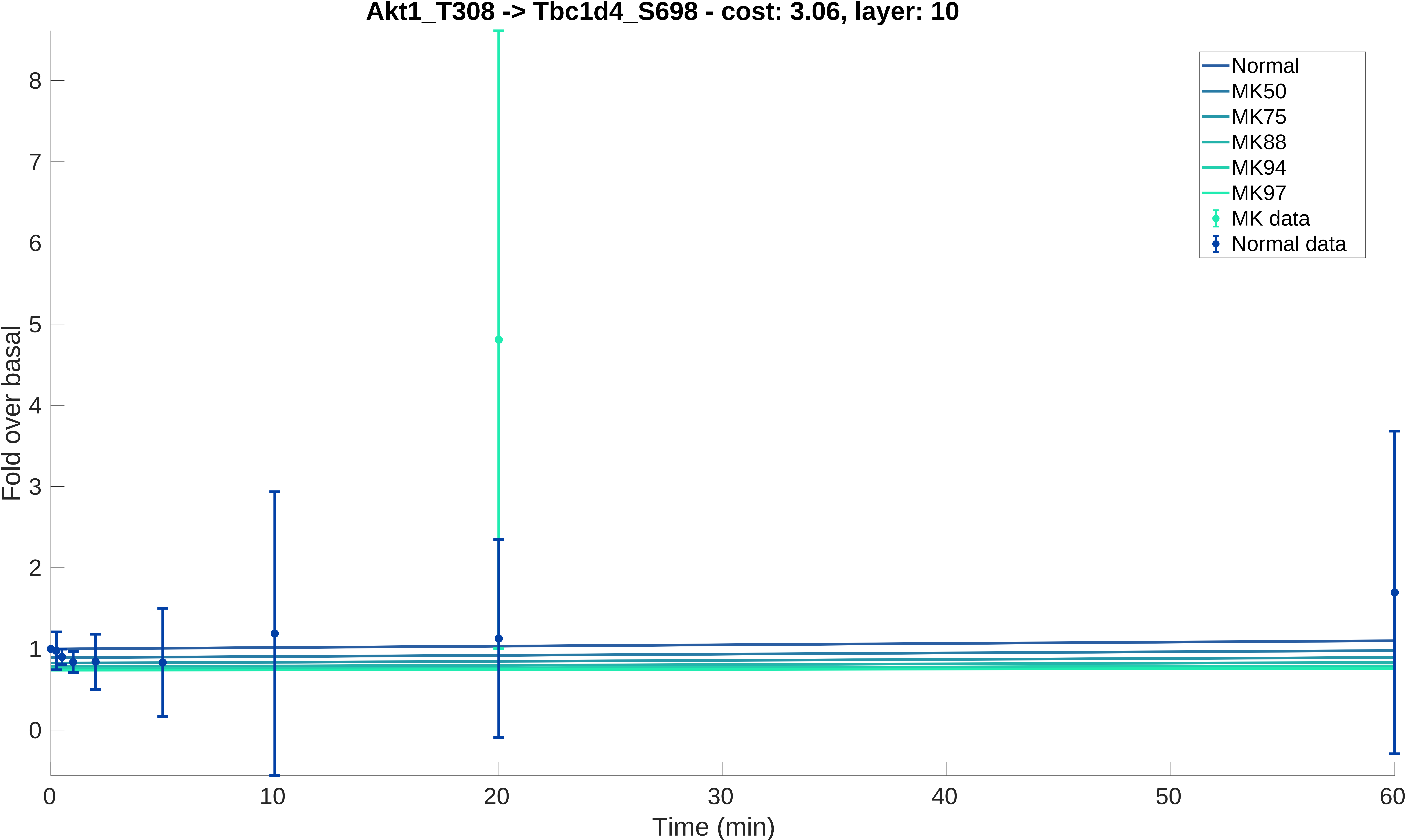
Akt1_T308 -> Tbc1d4_S698 - cost: 3.06, layer: 10
Fold over basal
Time (min)
0
1
2
3
4
5
6
7
8
0
10
20
30
40
50
60
Normal
MK50
MK75
MK88
MK94
MK97
MK data
Normal data

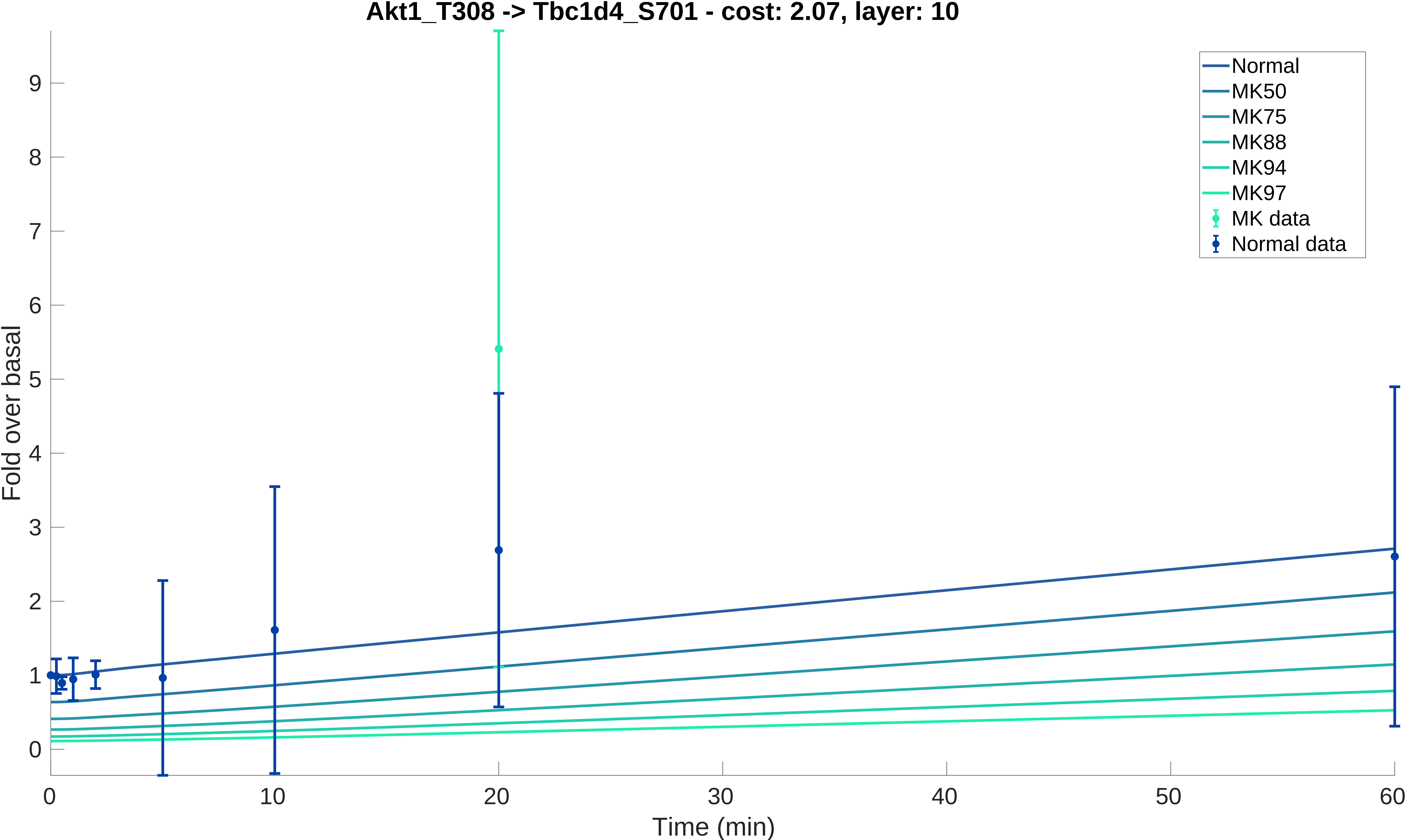
Akt1_T308 -> Tbc1d4_S701 - cost: 2.07, layer: 10
Fold over basal
Time (min)
0
1
2
3
4
5
6
7
8
9
0
10
20
30
40
50
60
Normal
MK50
MK75
MK88
MK94
MK97
MK data
Normal data

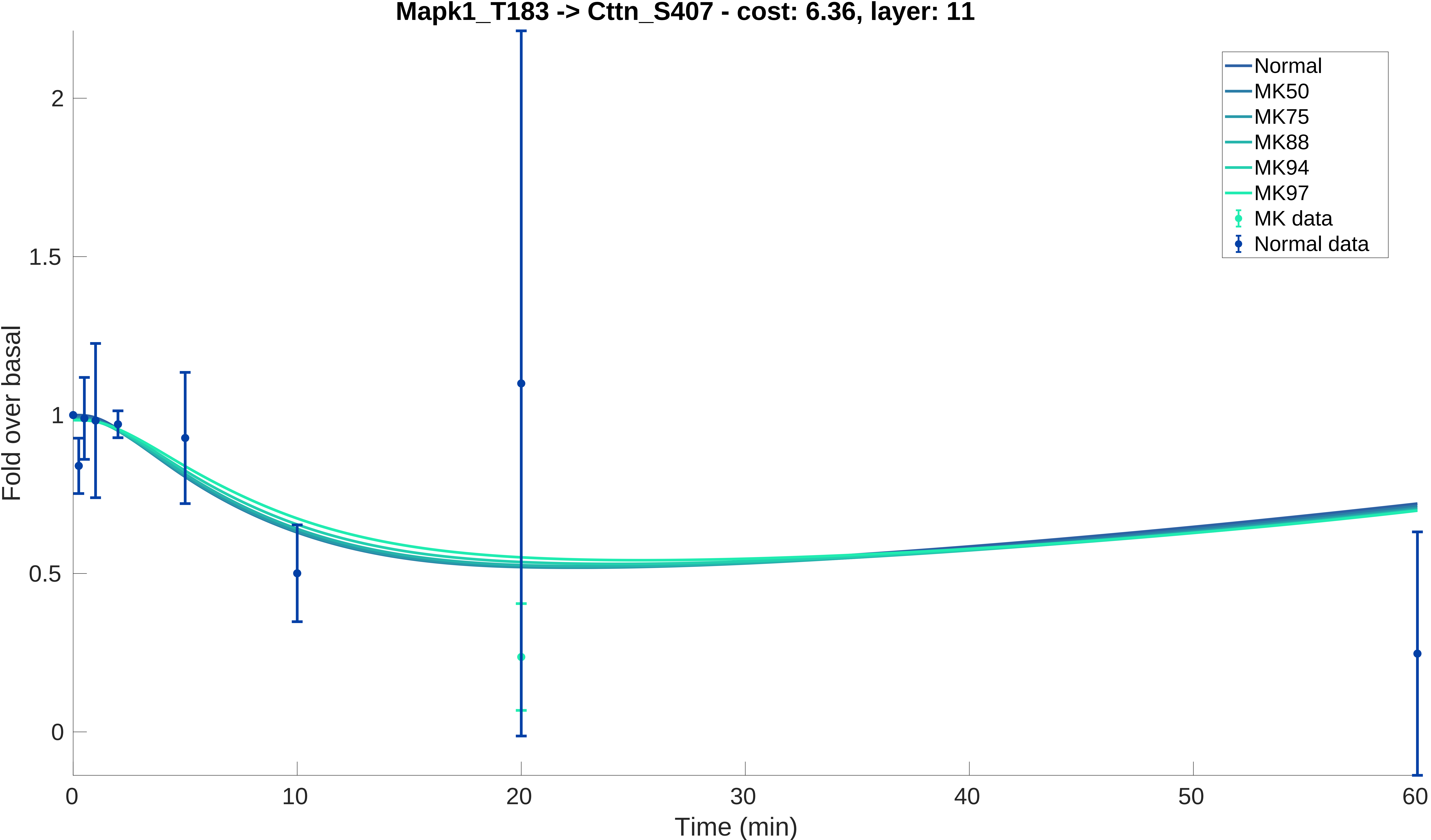

Mapk1_T183 -> Cttn_S407 - cost: 6.36, layer: 11
Normal
MK50
MK75
MK88
MK94
MK97
MK data
Normal data
Fold over basal
Time (min)
0
0.5
1
1.5
2
0
10
20
30
40
50
60

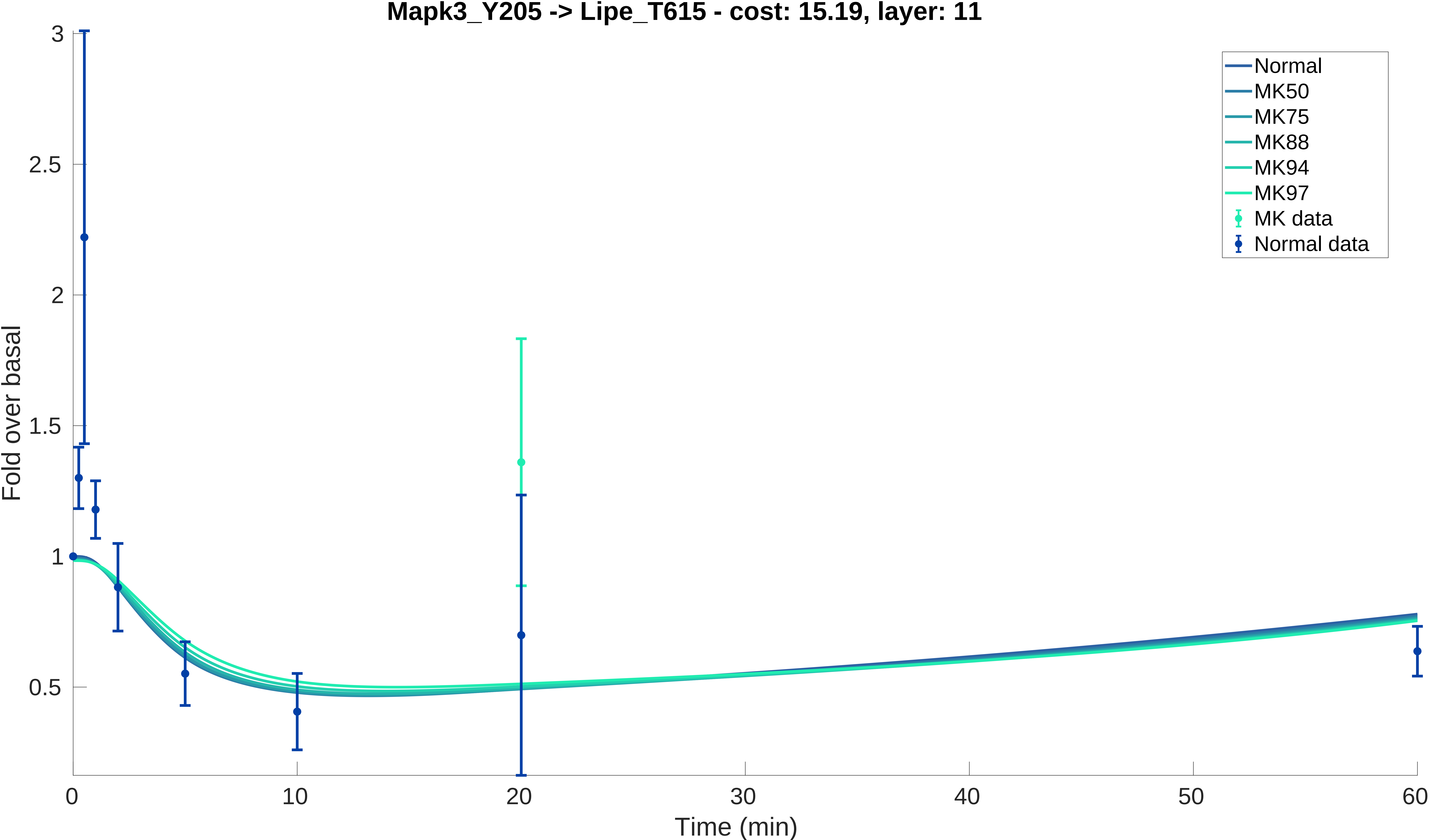

Mapk3_Y205 -> Lipe_T615 - cost: 15.19, layer: 11
Fold over basal
Time (min)
0
10
20
30
40
50
60
0.5
1
1.5
2
2.5
3
Normal
MK50
MK75
MK88
MK94
MK97
MK data
Normal data

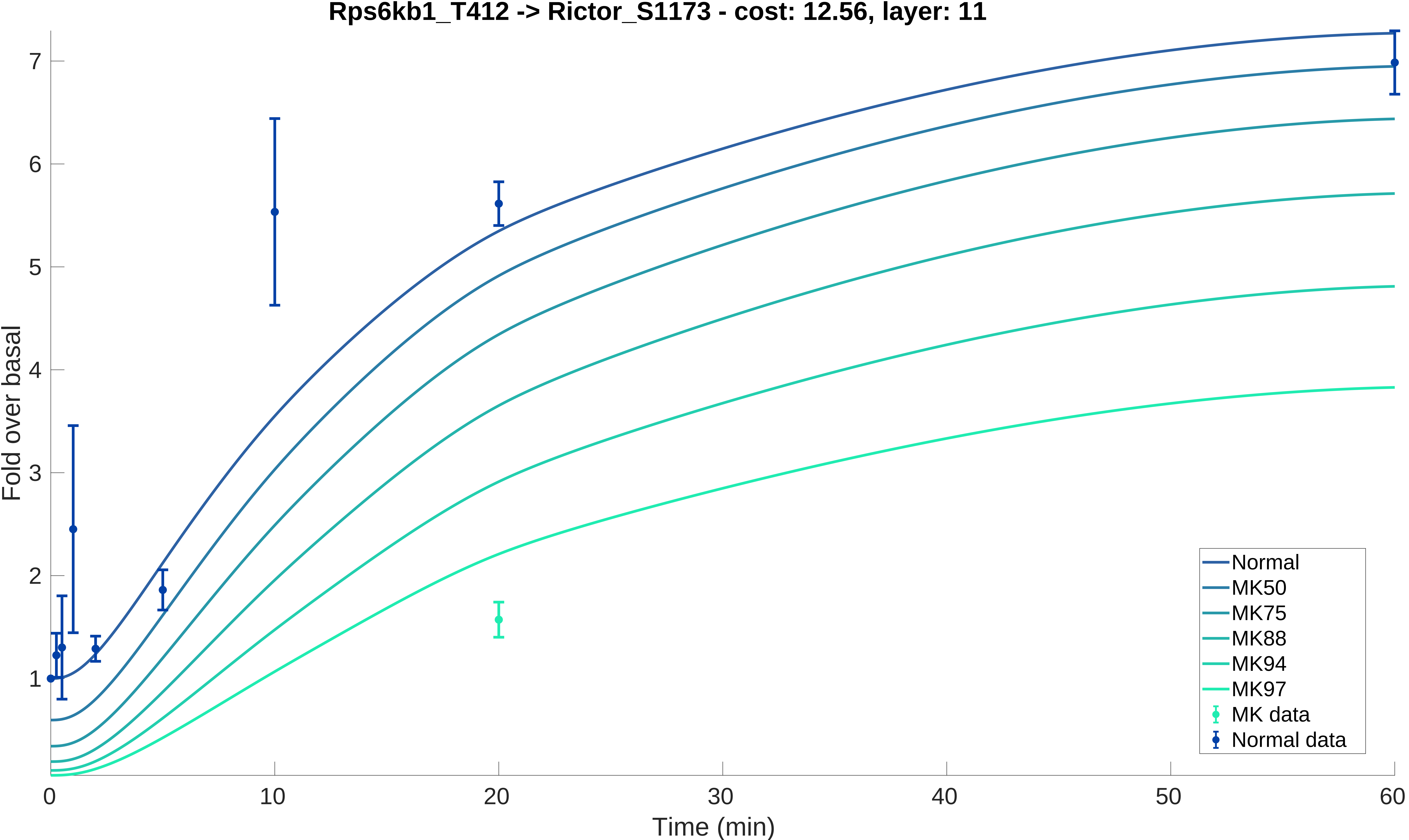

Rps6kb1_T412 -> Rictor_S1173 - cost: 12.56, layer: 11
Fold over basal
Time (min)
0
10
20
30
40
50
60
1
2
3
4
5
6
7
Normal
MK50
MK75
MK88
MK94
MK97
MK data
Normal data

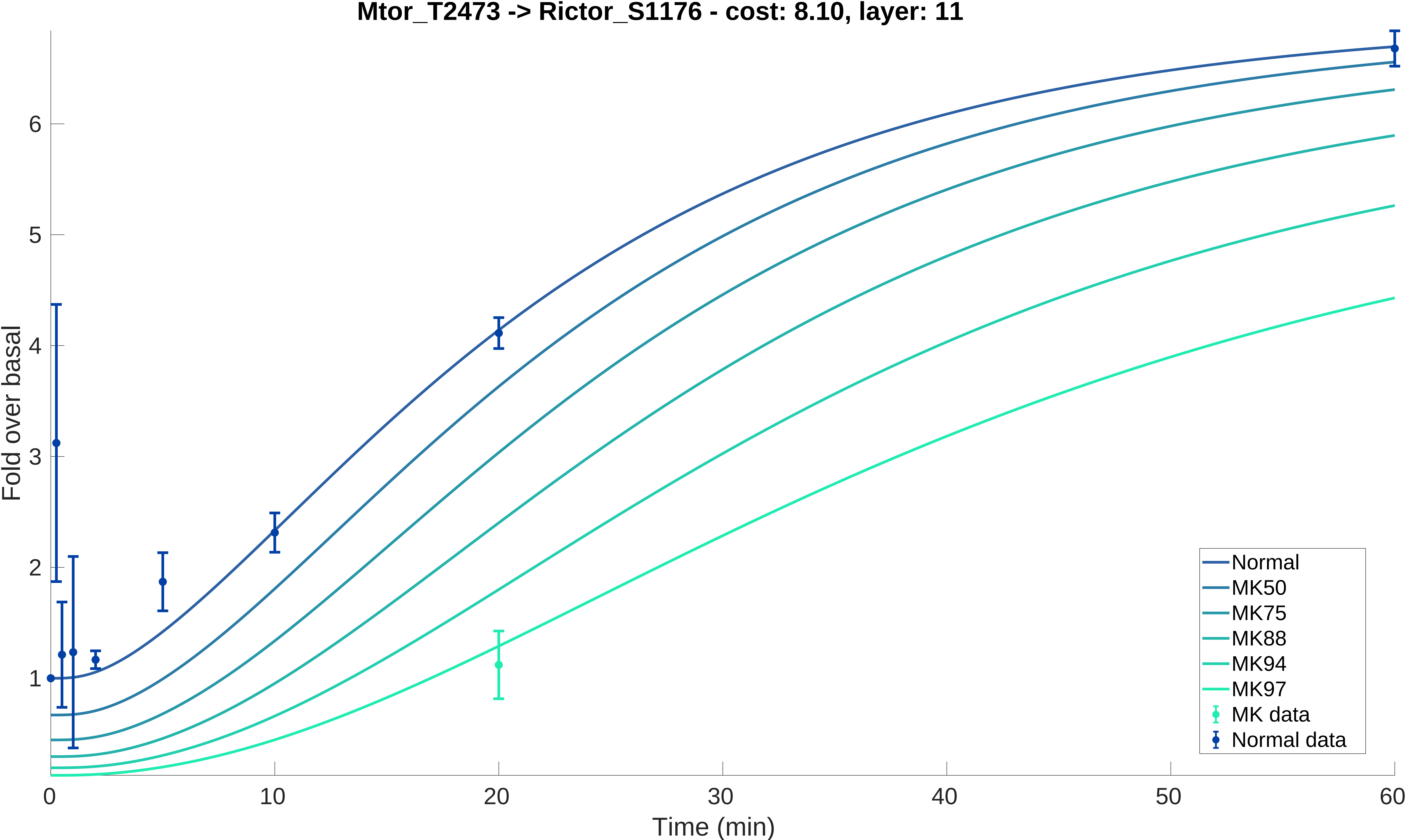

Mtor_T2473 -> Rictor_S1176 - cost: 8.10, layer: 11
Fold over basal
Time (min)
0
10
20
30
40
50
60
1
2
3
4
5
6
Normal
MK50
MK75
MK88
MK94
MK97
MK data
Normal data

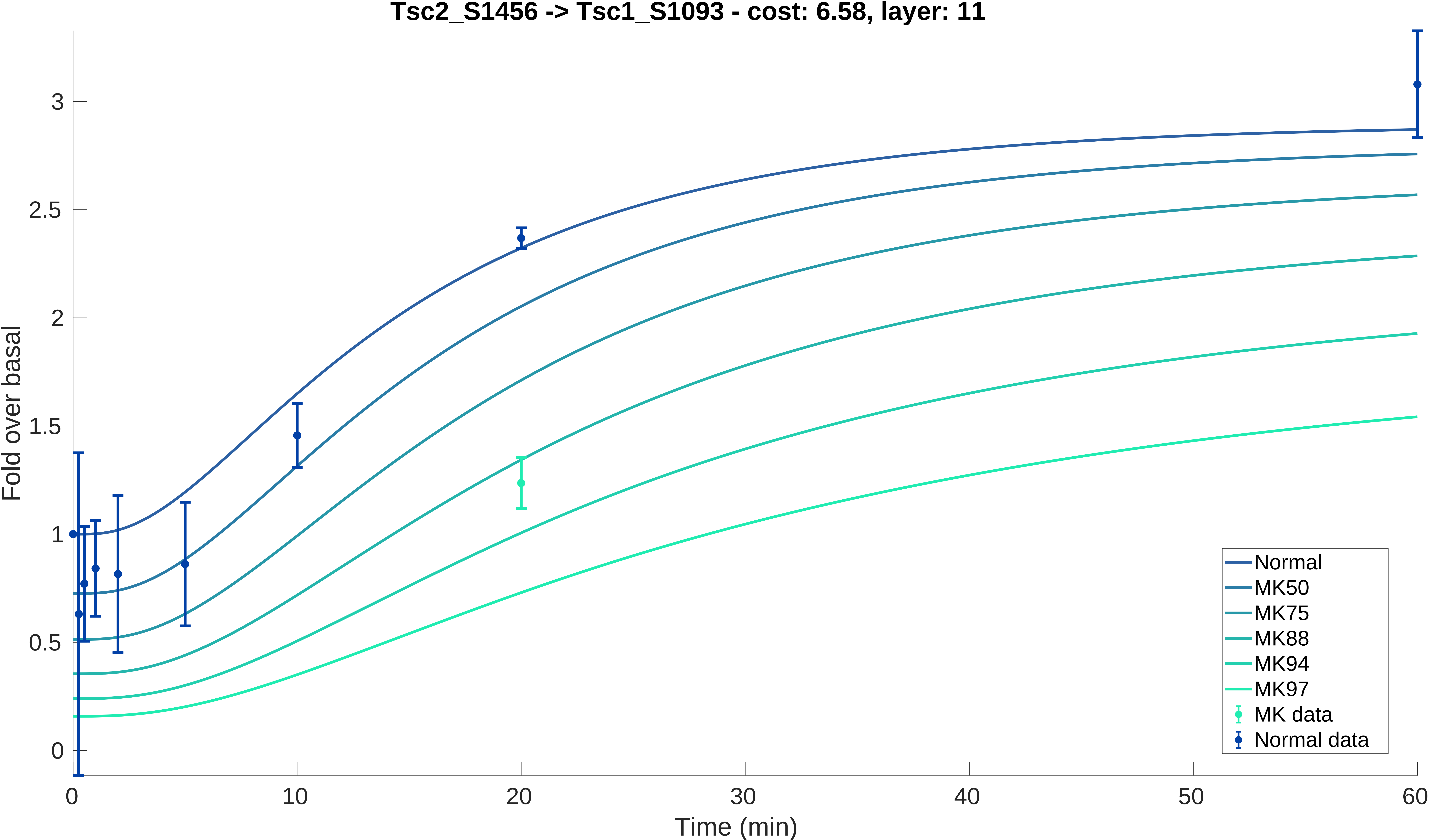

Tsc2_S1456 -> Tsc1_S1093 - cost: 6.58, layer: 11
Fold over basal
Time (min)
0
0.5
1
1.5
2
2.5
3
0
10
20
30
40
50
60
Normal
MK50
MK75
MK88
MK94
MK97
MK data
Normal data

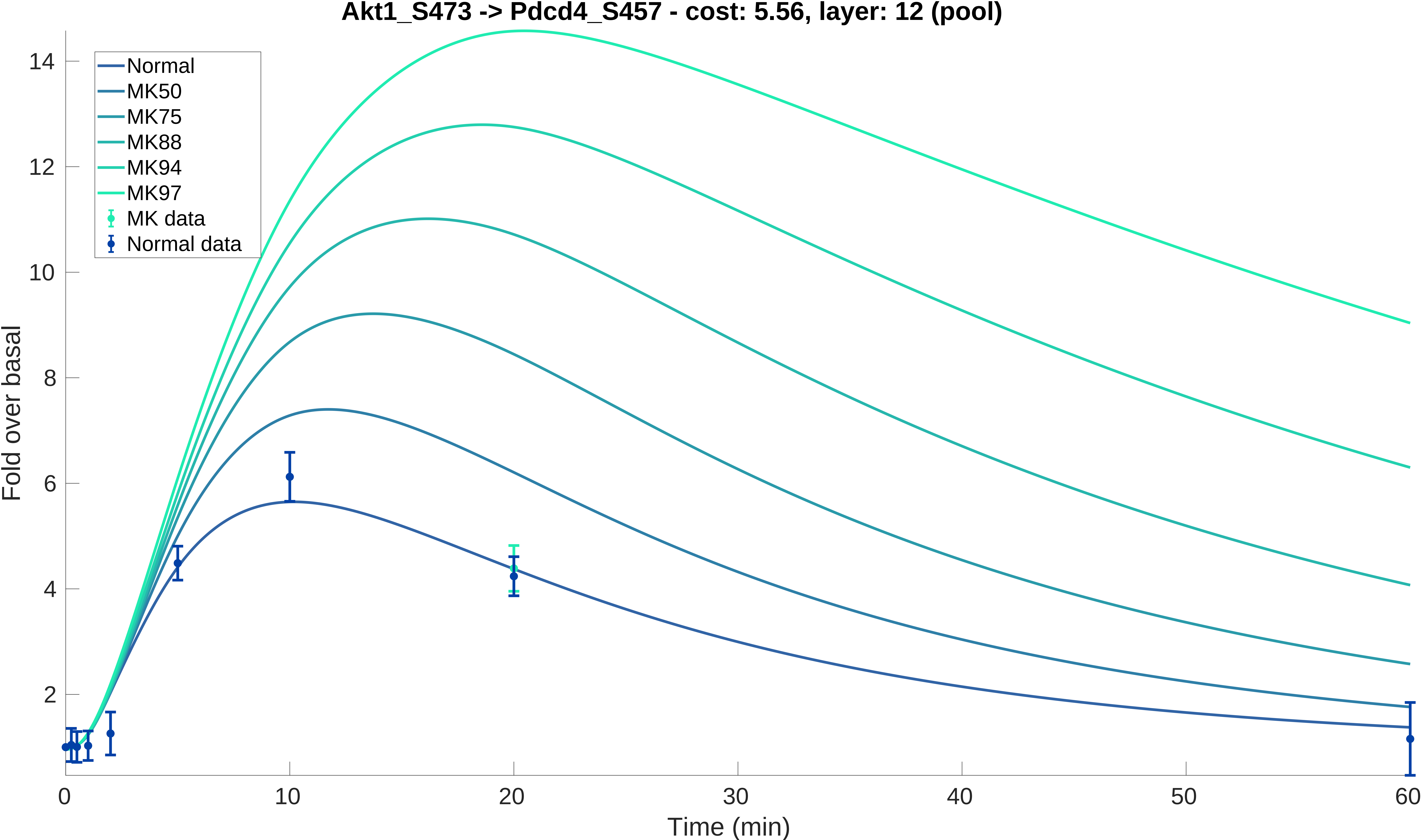

Akt1_S473 -> Pdcd4_S457 - cost: 5.56, layer: 12 (pool)
Normal
MK50
MK75
MK88
MK94
MK97
MK data
Normal data
Fold over basal
Time (min)
0
10
20
30
40
50
60
2
4
6
8
10
12
14

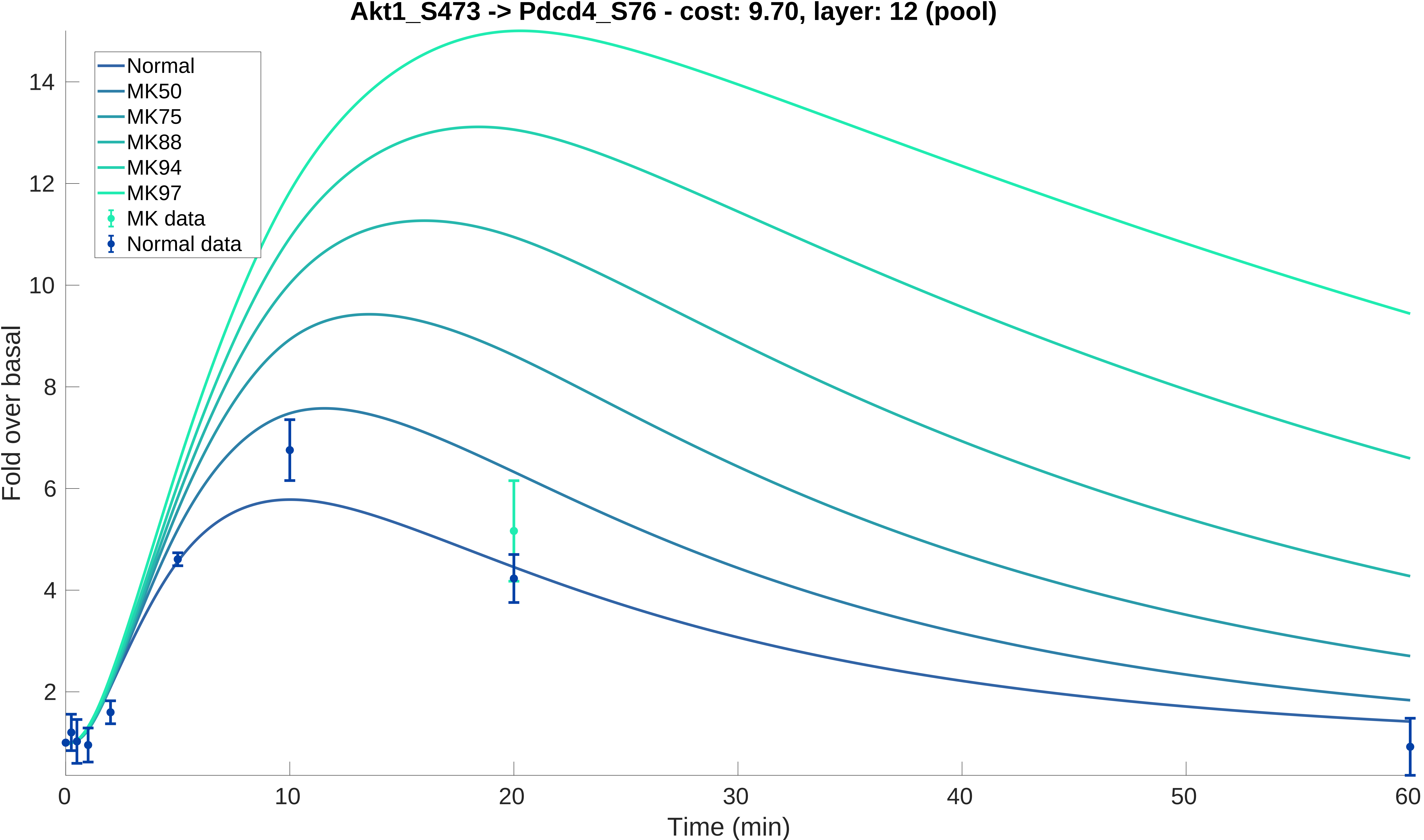

Akt1_S473 -> Pdcd4_S76 - cost: 9.70, layer: 12 (pool)
Normal
MK50
MK75
MK88
MK94
MK97
MK data
Normal data
Fold over basal
Time (min)
14
12
10
8
6
4
2
0
10
20
30
40
50
60

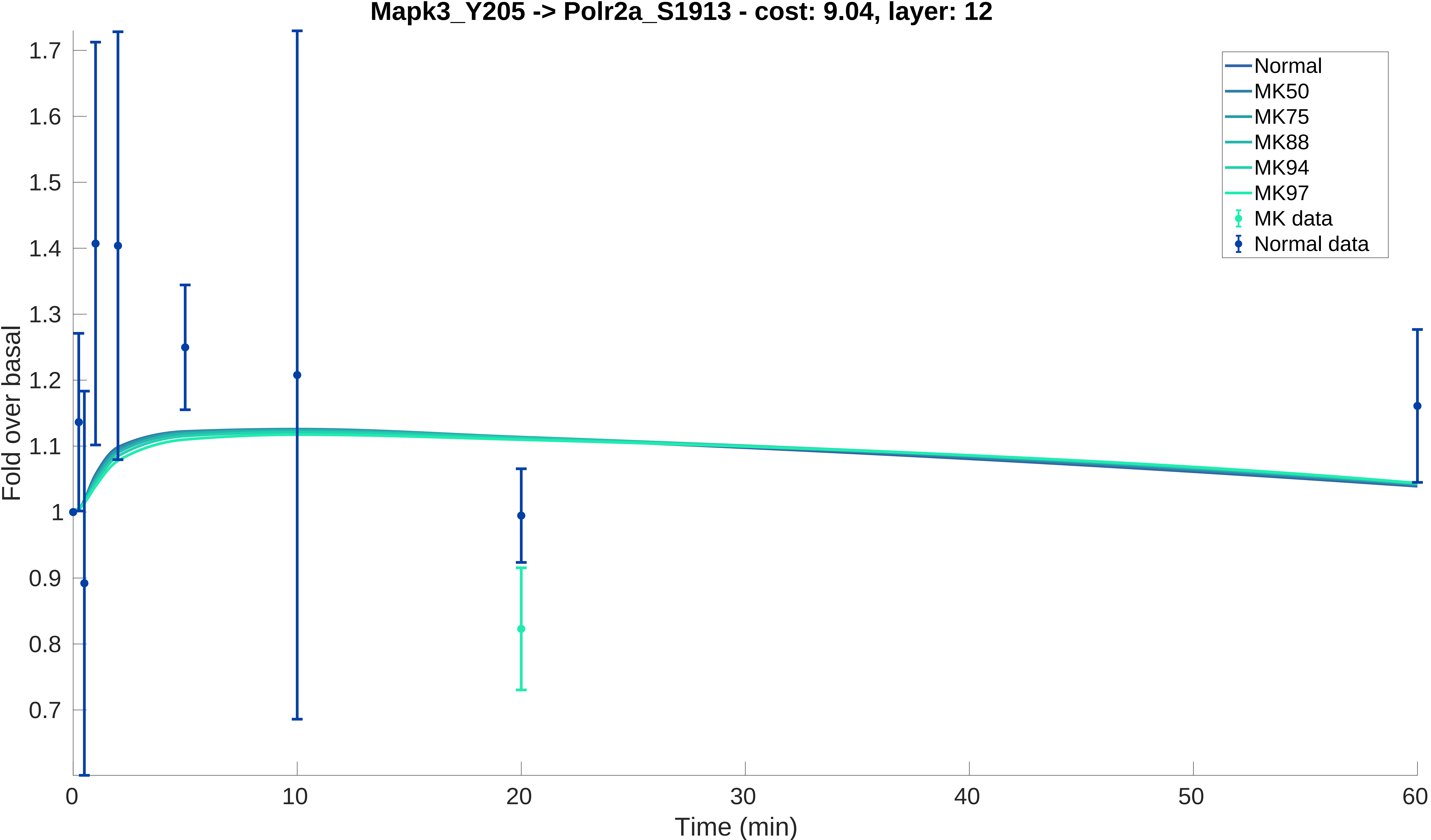

Mapk3_Y205 -> Polr2a_S1913 - cost: 9.04, layer: 12
Normal
MK50
MK75
MK88
MK94
MK97
MK data
Normal data
Fold over basal
Time (min)
1.7
1.6
1.5
1.4
1.3
1.2
1.1
1
0.9
0.8
0.7
0
10
20
30
40
50
60

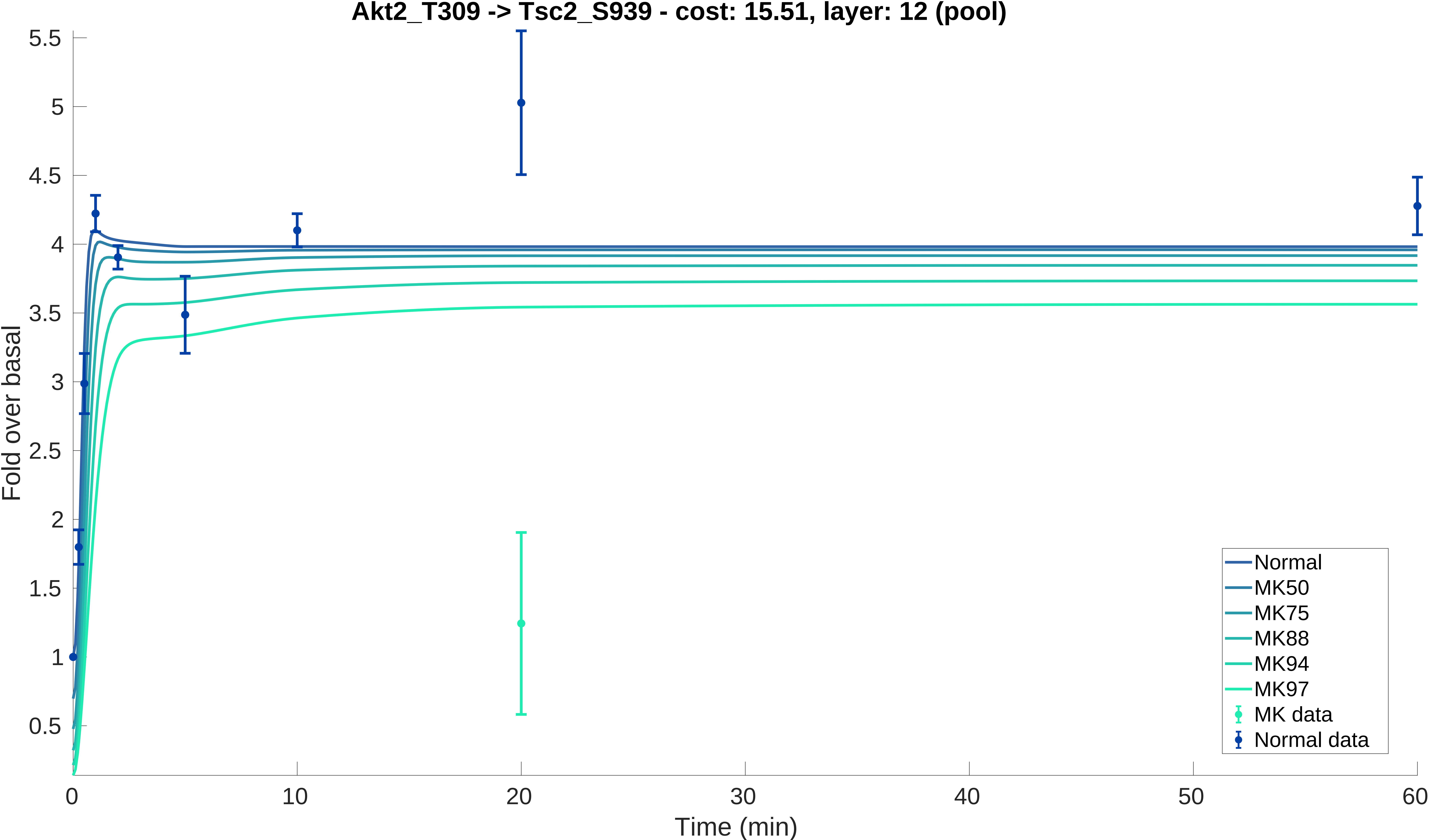
Akt2_T309 -> Tsc2_S939 - cost: 15.51, layer: 12 (pool)
Fold over basal
Time (min)
0.5
1
1.5
2
2.5
3
3.5
4
4.5
5
5.5
0
10
20
30
40
50
60
Normal
MK50
MK75
MK88
MK94
MK97
MK data
Normal data

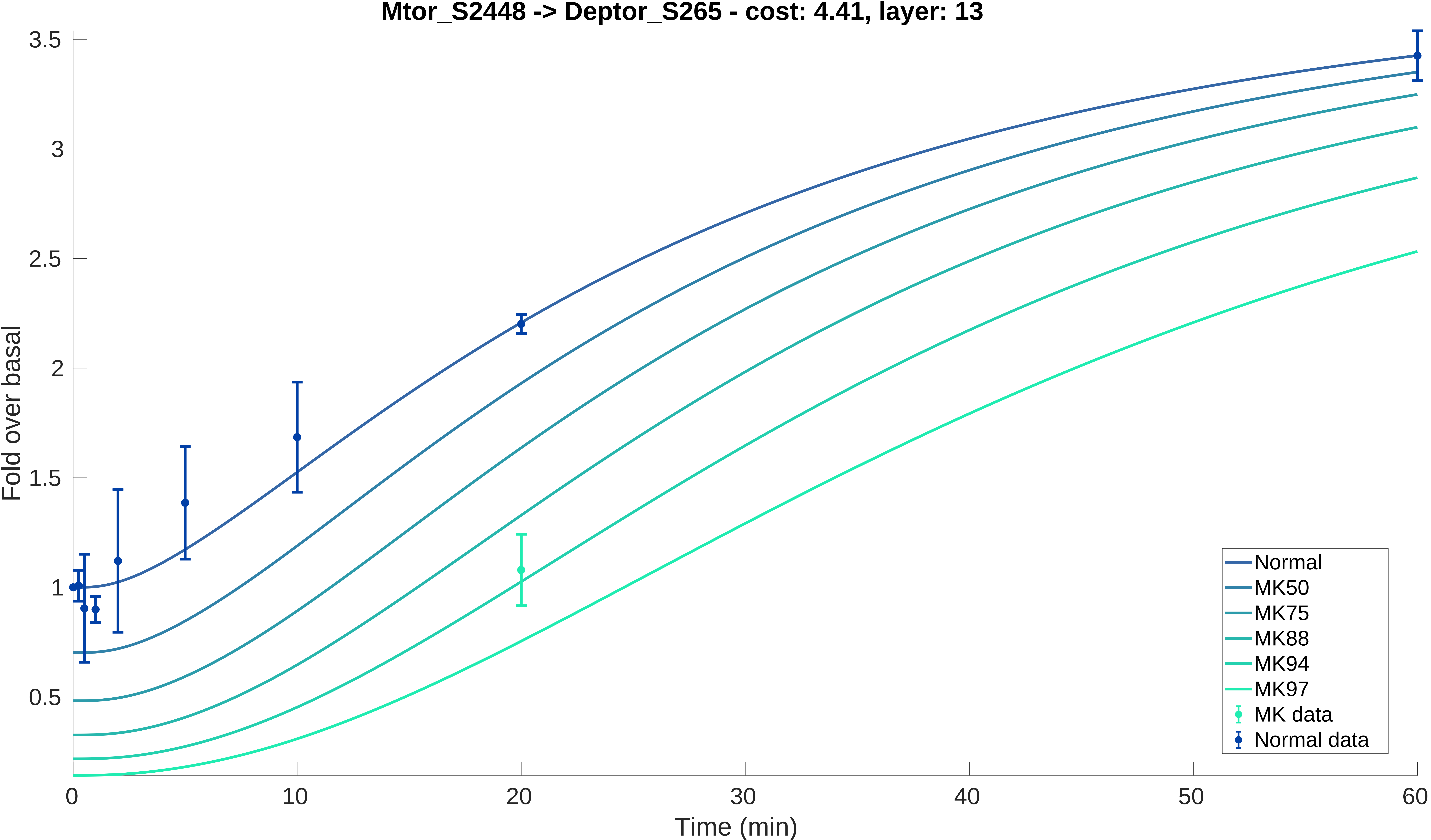
Mtor_S2448 -> Deptor_S265 - cost: 4.41, layer: 13
Fold over basal
Time (min)
0.5
1
1.5
2
2.5
3
3.5
0
10
20
30
40
50
60
Normal
MK50
MK75
MK88
MK94
MK97
MK data
Normal data

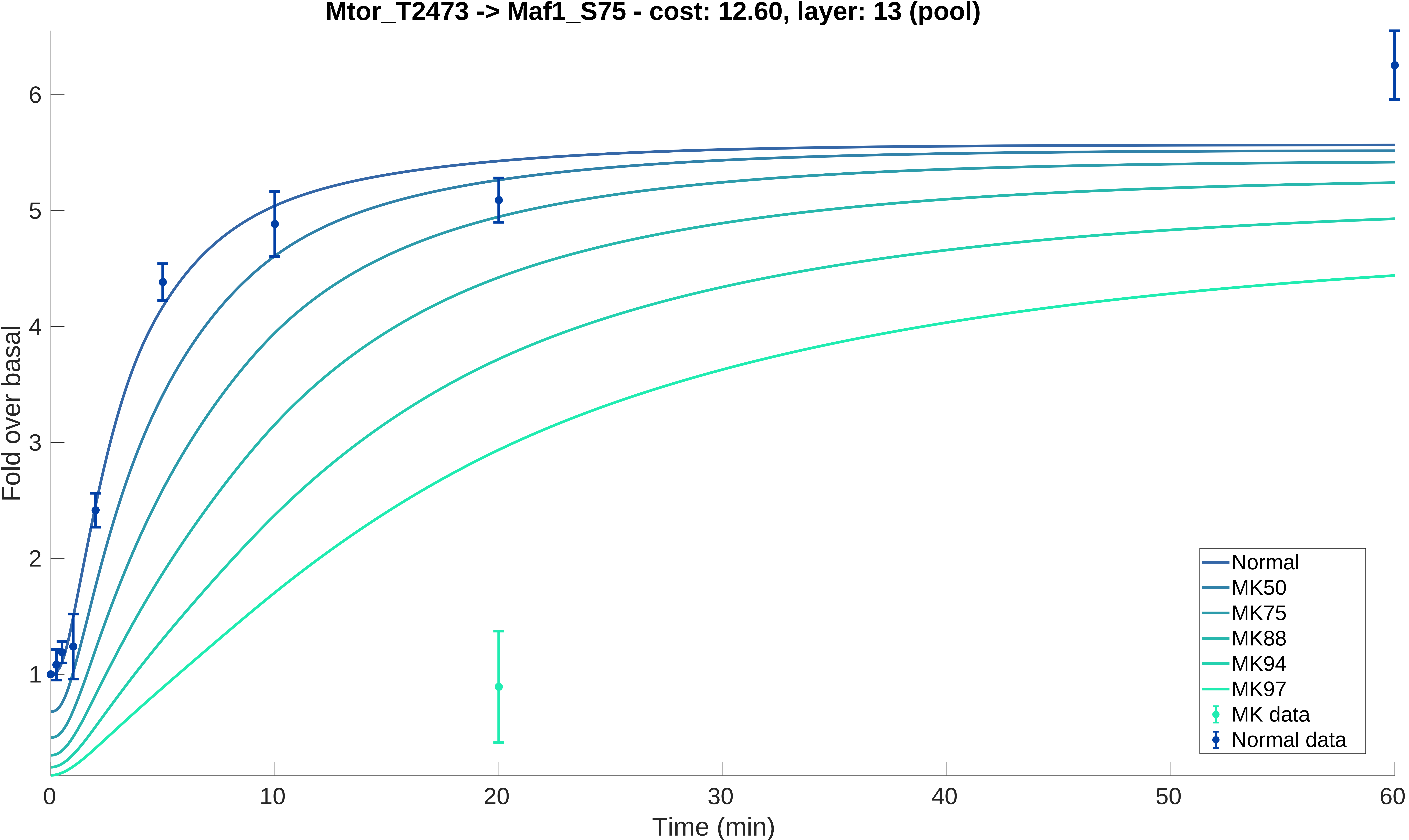

Mtor_T2473 -> Maf1_S75 - cost: 12.60, layer: 13 (pool)
Fold over basal
Time (min)
0
10
20
30
40
50
60
1
2
3
4
5
6
Normal
MK50
MK75
MK88
MK94
MK97
MK data
Normal data

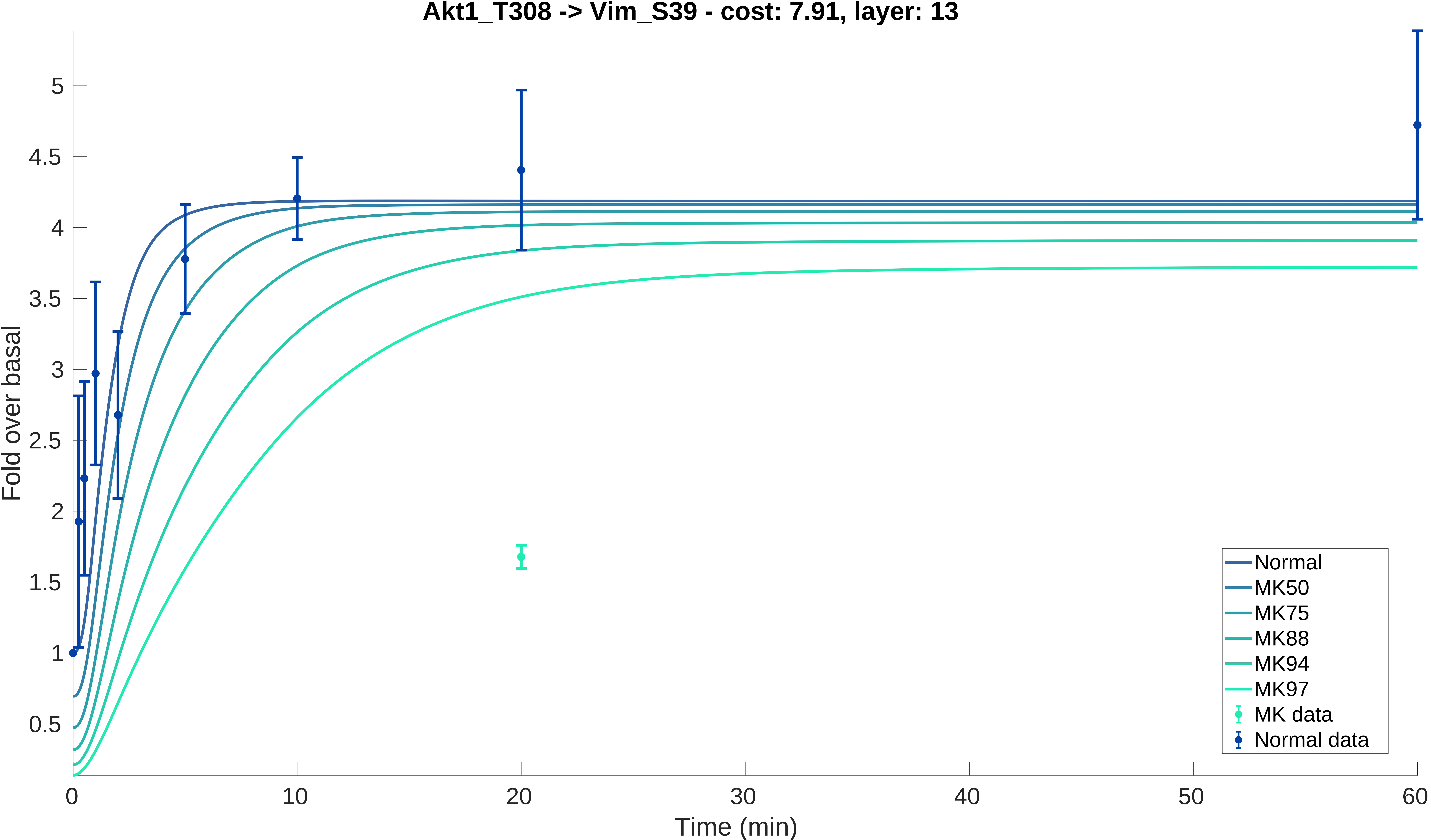
Akt1_T308 -> Vim_S39 - cost: 7.91, layer: 13
Fold over basal
Time (min)
0
10
20
30
40
50
60
0.5
1
1.5
2
2.5
3
3.5
4
4.5
5
Normal
MK50
MK75
MK88
MK94
MK97
MK data
Normal data

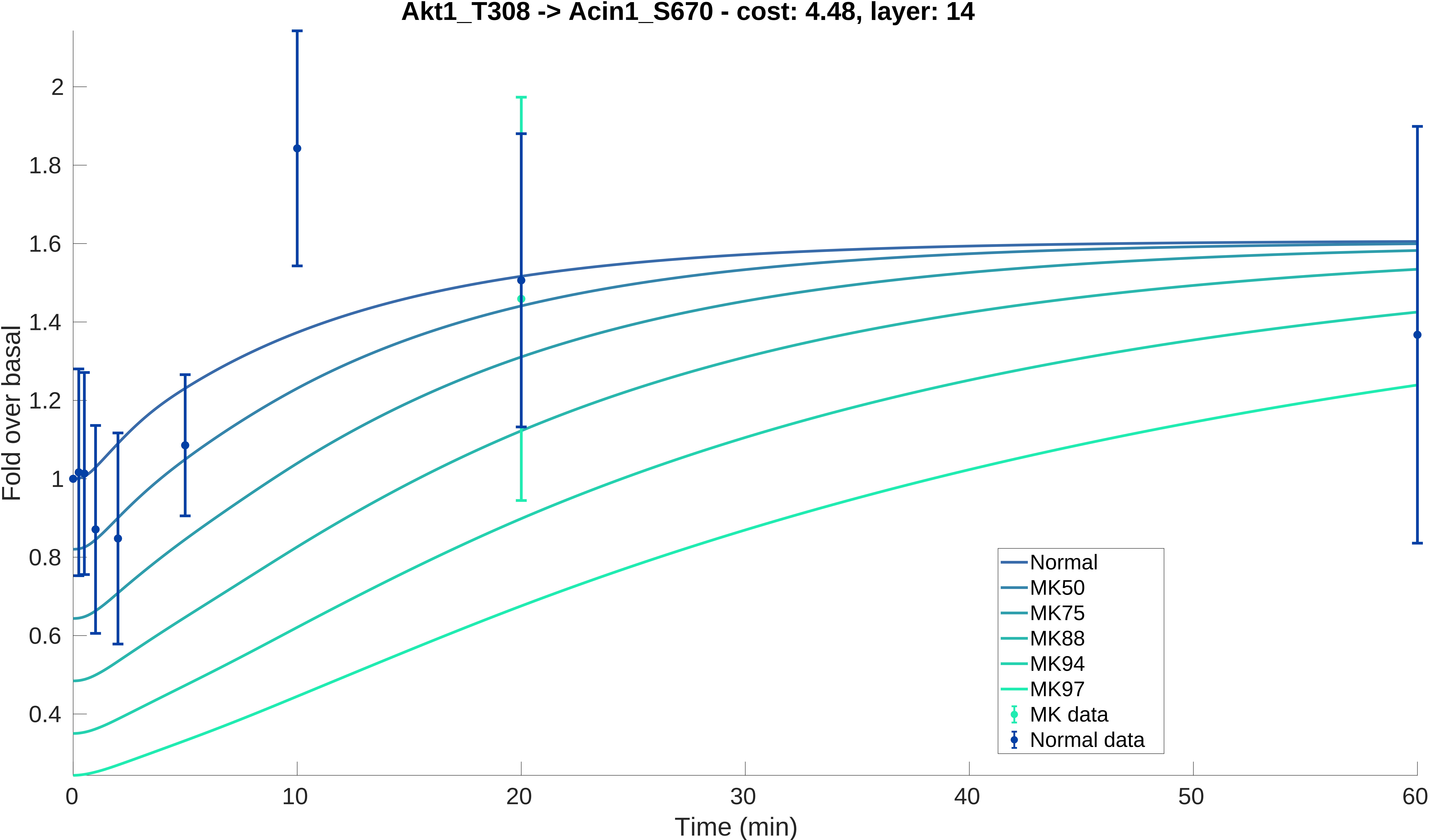

Akt1_T308 -> Acin1_S670 - cost: 4.48, layer: 14
Fold over basal
Time (min)
0
10
20
30
40
50
60
0.4
0.6
0.8
1
1.2
1.4
1.6
1.8
2
Normal
MK50
MK75
MK88
MK94
MK97
MK data
Normal data

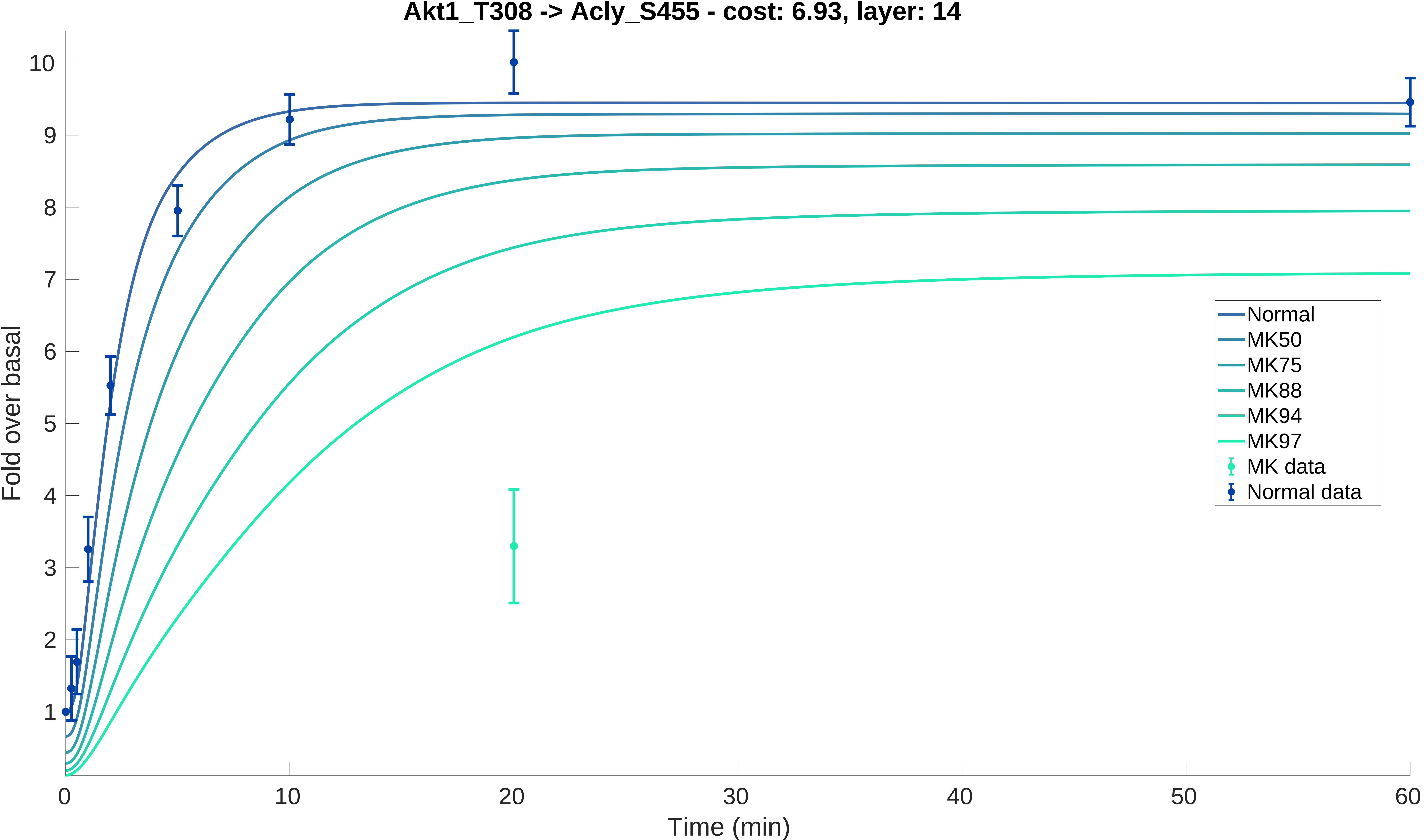

Akt1_T308 -> Acly_S455 - cost: 6.93, layer: 14
Fold over basal
Time (min)
0
10
20
30
40
50
60
1
2
3
4
5
6
7
8
9
10
Normal
MK50
MK75
MK88
MK94
MK97
MK data
Normal data

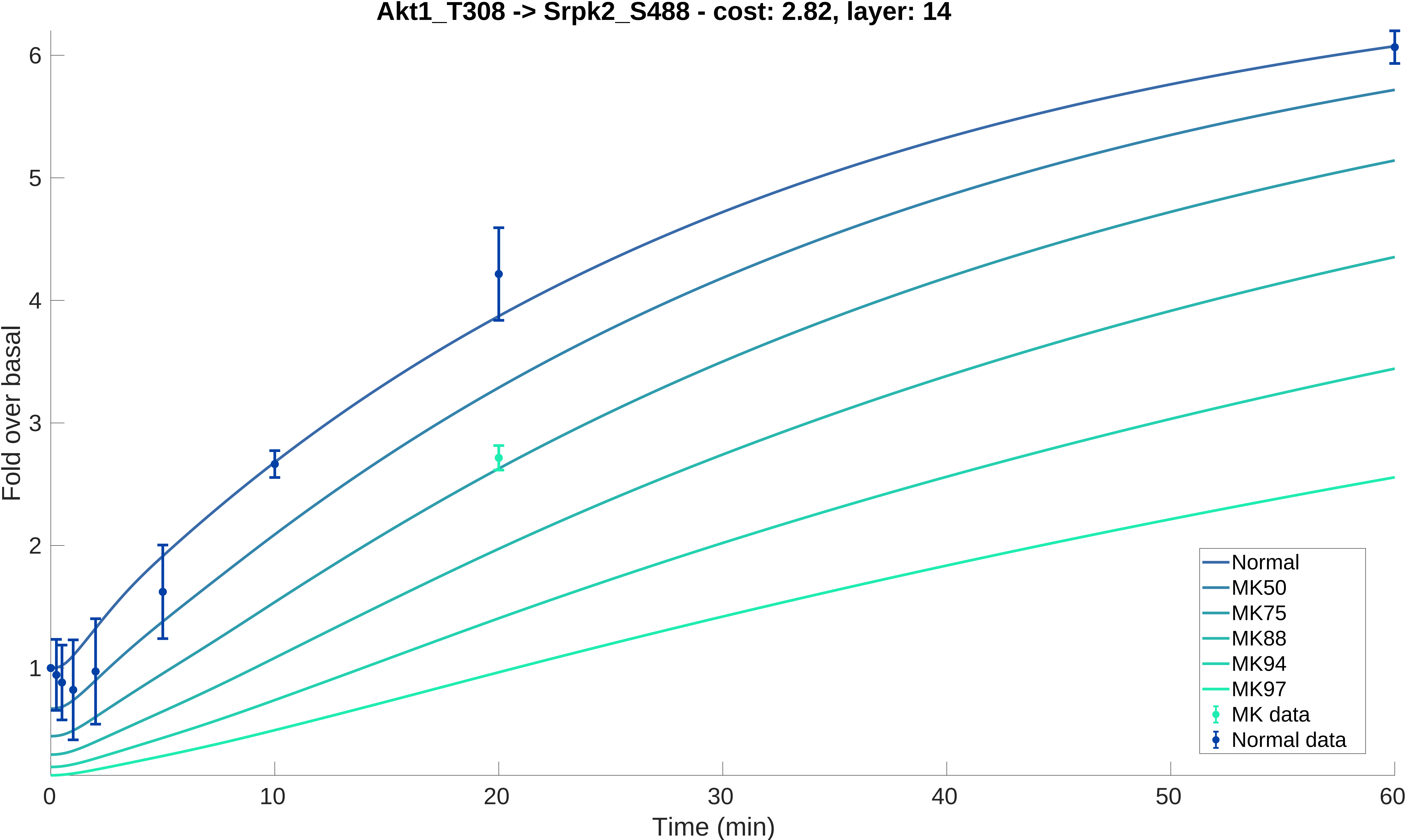
Akt1_T308 -> Srpk2_S488 - cost: 2.82, layer: 14
Fold over basal
Time (min)
0
10
20
30
40
50
60
1
2
3
4
5
6
Normal
MK50
MK75
MK88
MK94
MK97
MK data
Normal data

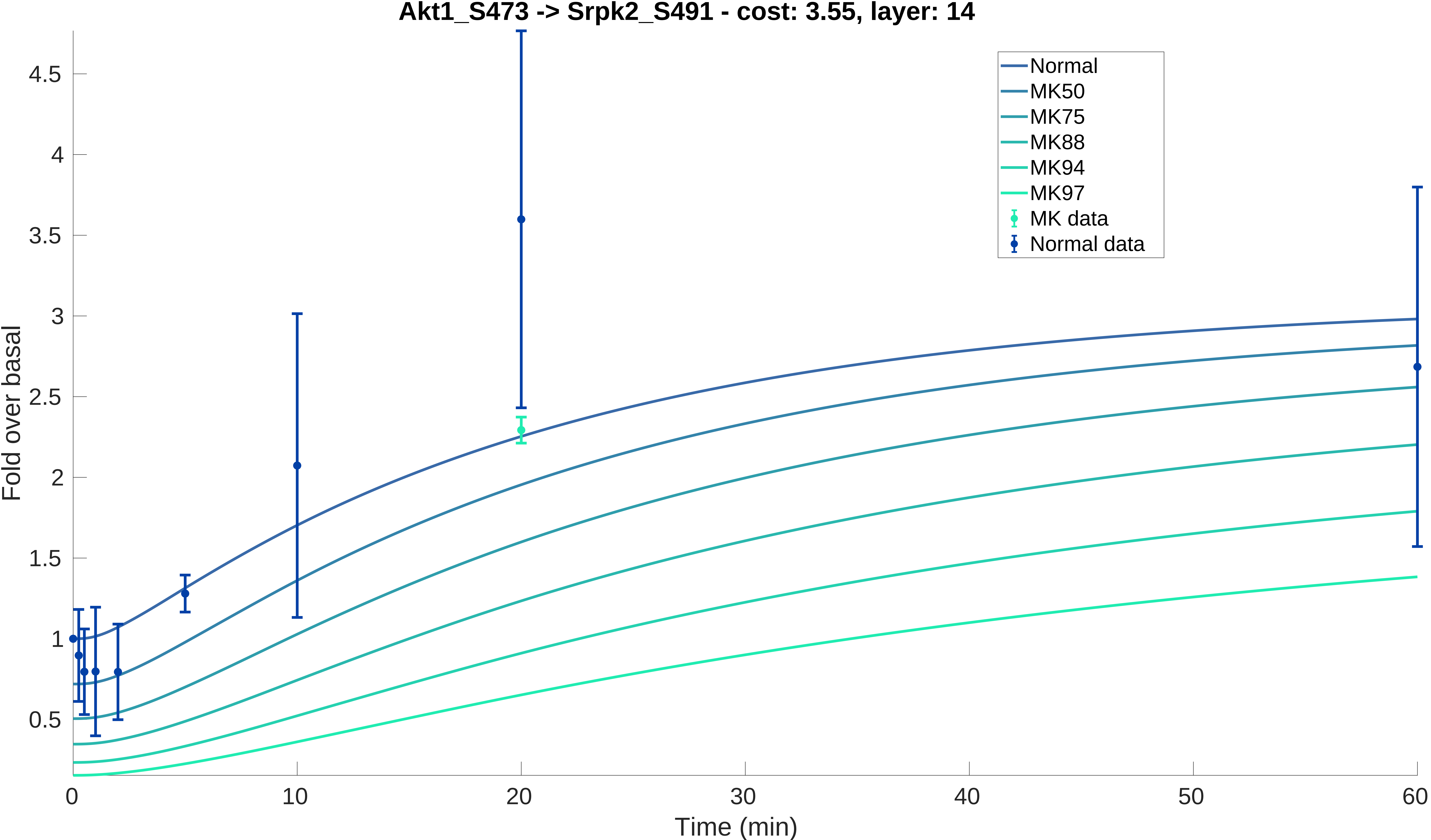

Akt1_S473 -> Srpk2_S491 - cost: 3.55, layer: 14
Normal
MK50
MK75
MK88
MK94
MK97
MK data
Normal data
Fold over basal
0.5
1
1.5
2
2.5
3
3.5
4
4.5
0
10
20
30
40
50
60
Time (min)

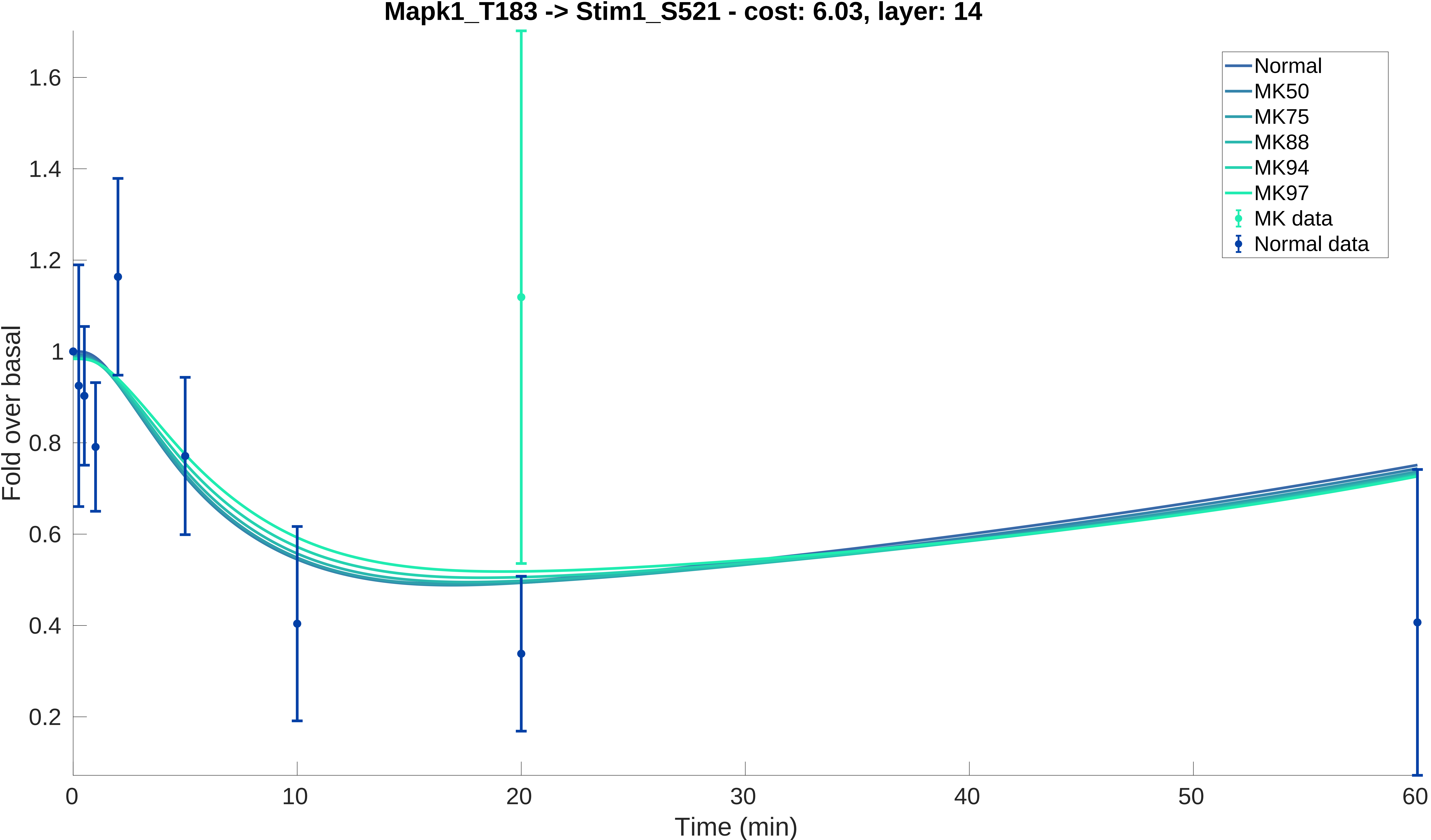
Mapk1_T183 -> Stim1_S521 - cost: 6.03, layer: 14
Fold over basal
Time (min)
0
10
20
30
40
50
60
0.2
0.4
0.6
0.8
1
1.2
1.4
1.6
Normal
MK50
MK75
MK88
MK94
MK97
MK data
Normal data

Mapk1_T183 -> Stim1_S524 - cost: 7.47, layer: 14

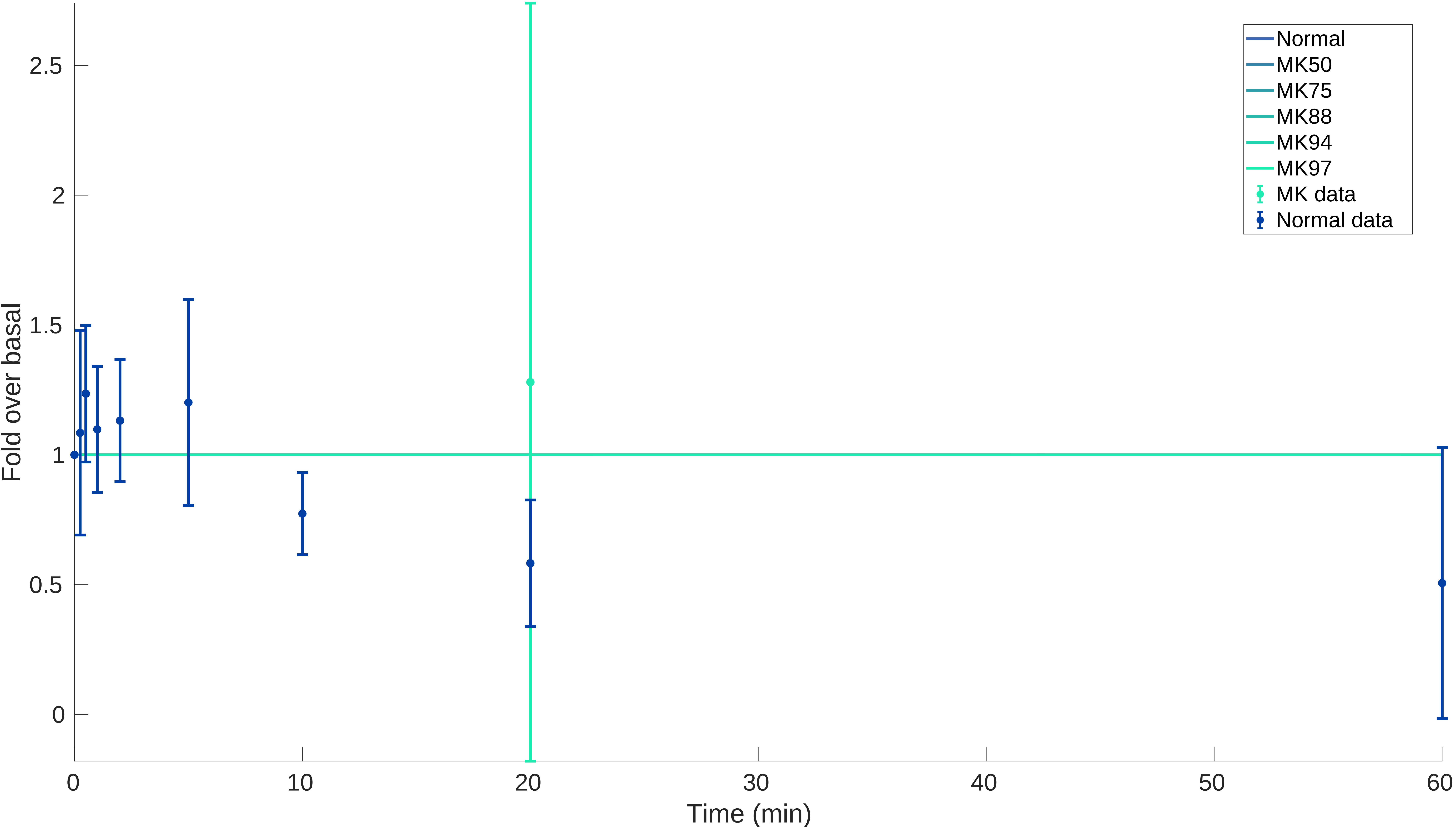

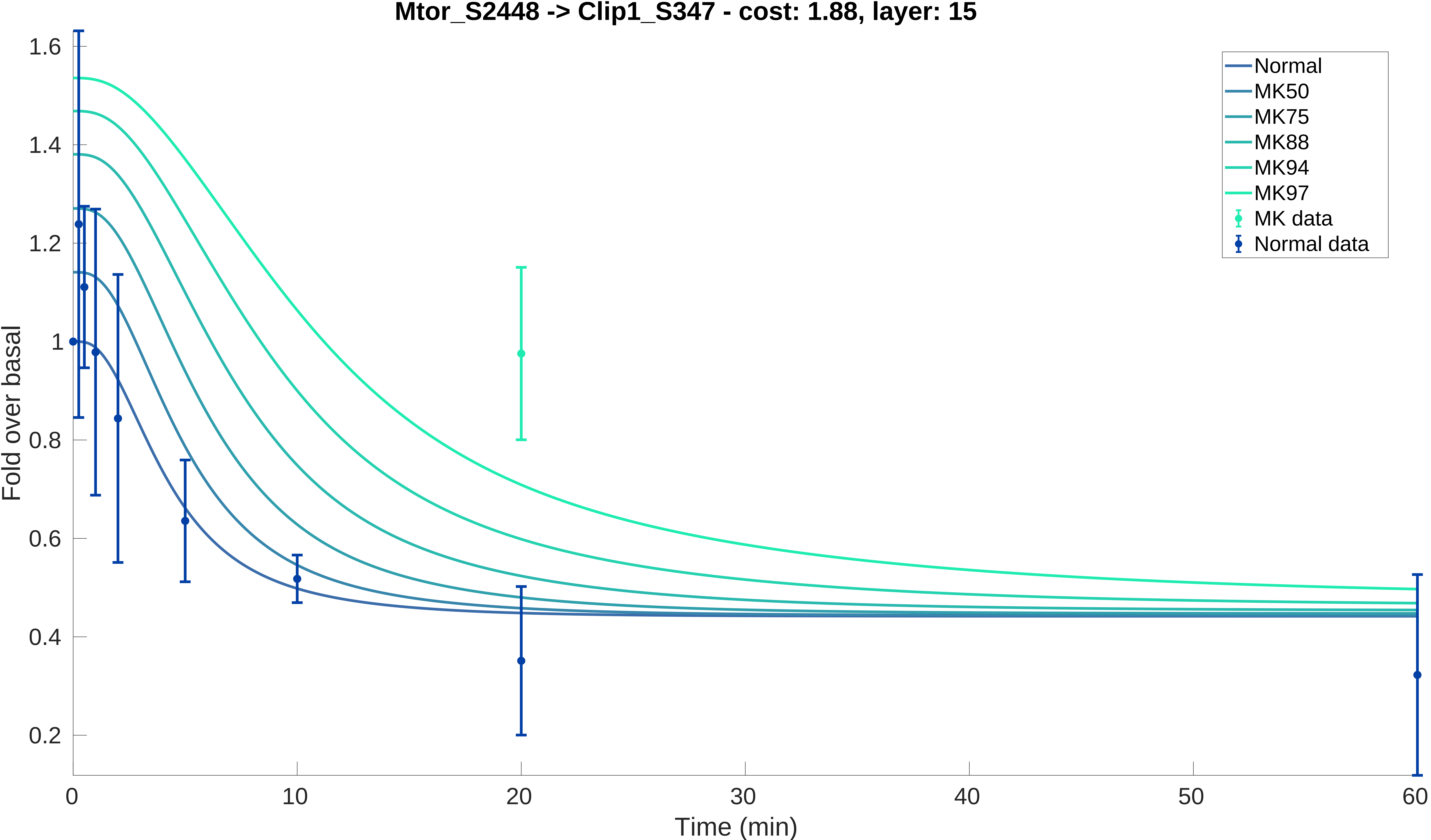

Mtor_S2448 -> Clip1_S347 - cost: 1.88, layer: 15
Normal
MK50
MK75
MK88
MK94
MK97
MK data
Normal data
Fold over basal
Time (min)
0.2
0.4
0.6
0.8
1
1.2
1.4
1.6
0
10
20
30
40
50
60

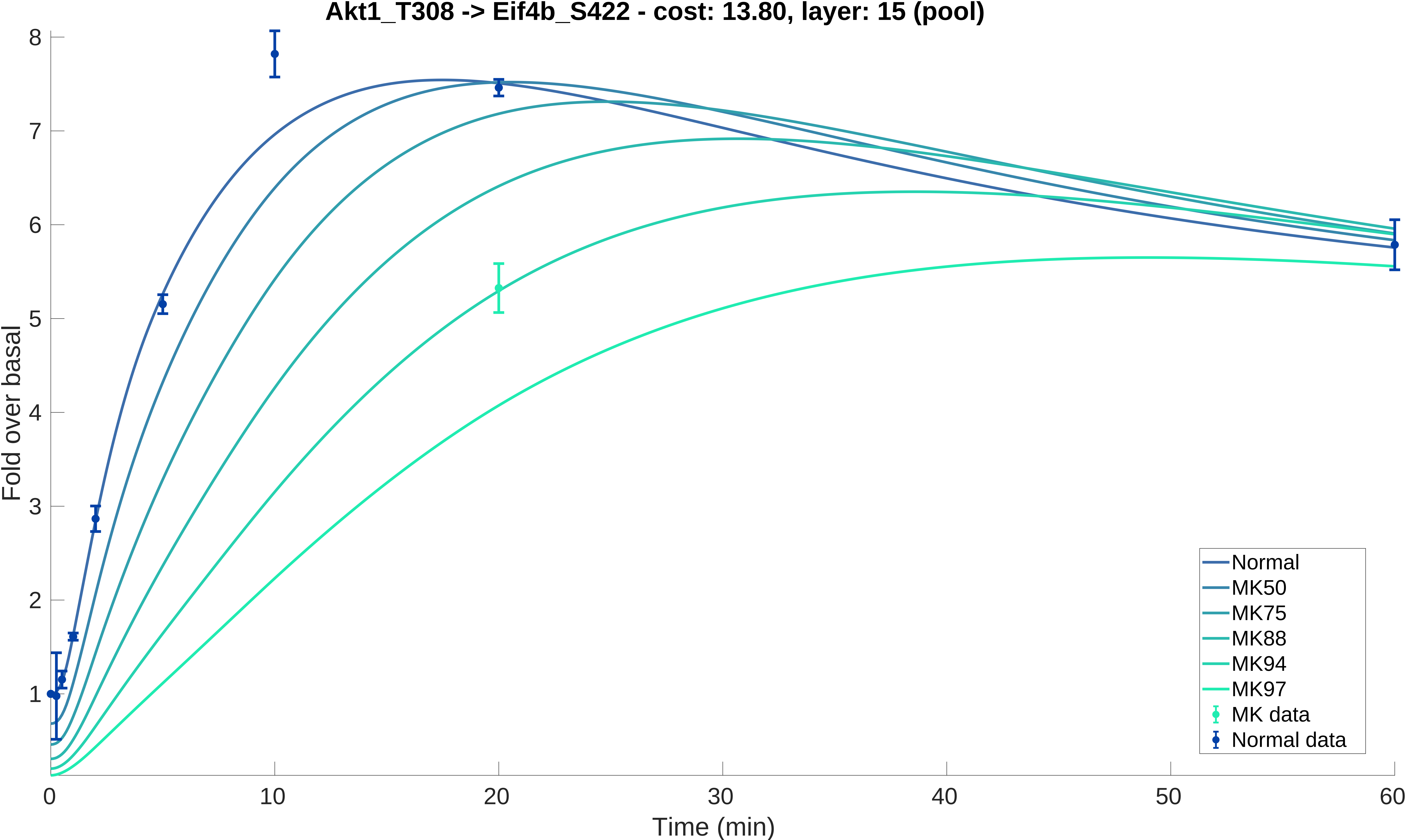
Akt1_T308 -> Eif4b_S422 - cost: 13.80, layer: 15 (pool)
Fold over basal
Time (min)
0
10
20
30
40
50
60
1
2
3
4
5
6
7
8
Normal
MK50
MK75
MK88
MK94
MK97
MK data
Normal data

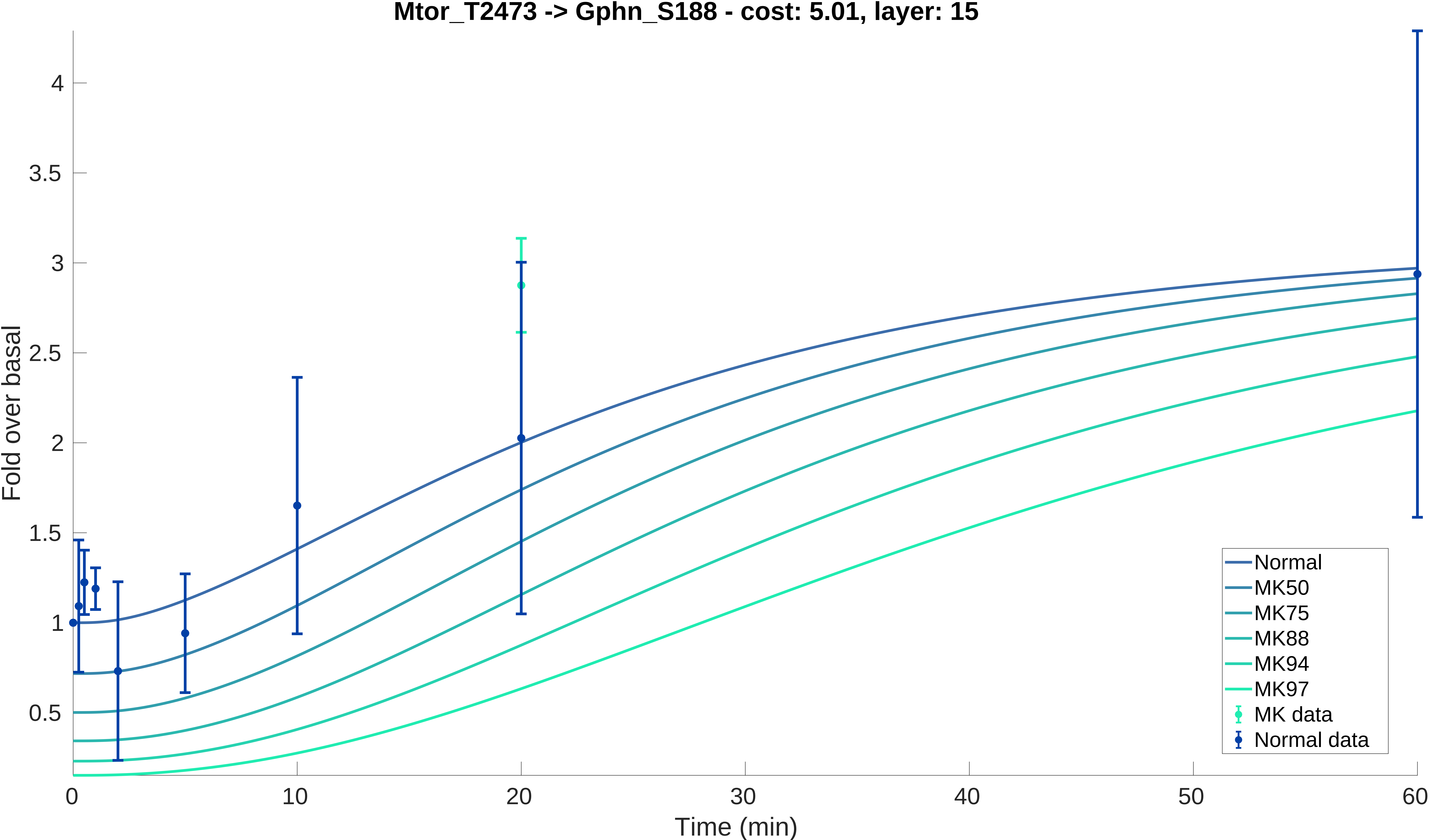
Mtor_T2473 -> Gphn_S188 - cost: 5.01, layer: 15
Fold over basal
Time (min)
0
10
20
30
40
50
60
0.5
1
1.5
2
2.5
3
3.5
4
Normal
MK50
MK75
MK88
MK94
MK97
MK data
Normal data

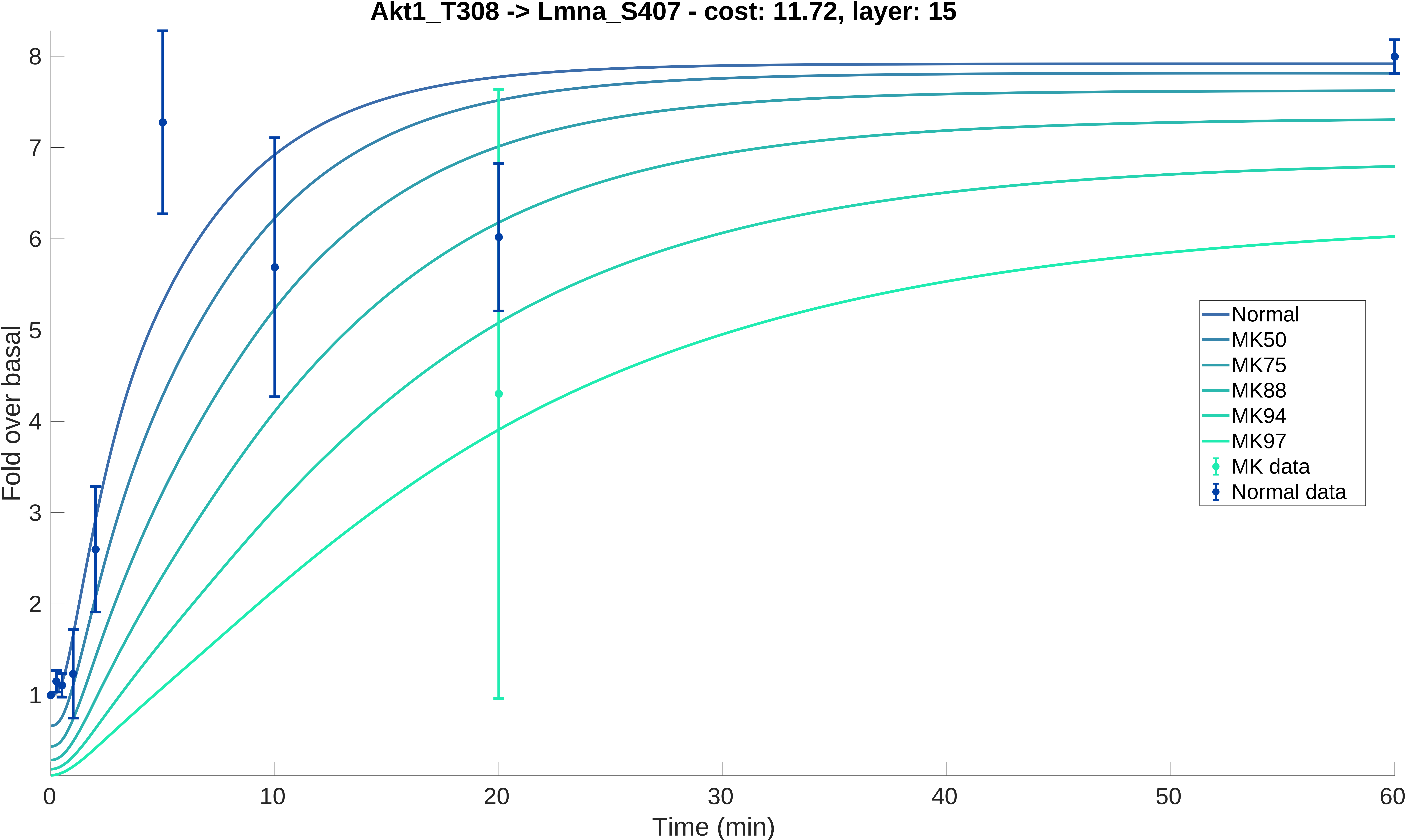

Akt1_T308 -> Lmna_S407 - cost: 11.72, layer: 15
Fold over basal
Time (min)
0
10
20
30
40
50
60
1
2
3
4
5
6
7
8
Normal
MK50
MK75
MK88
MK94
MK97
MK data
Normal data

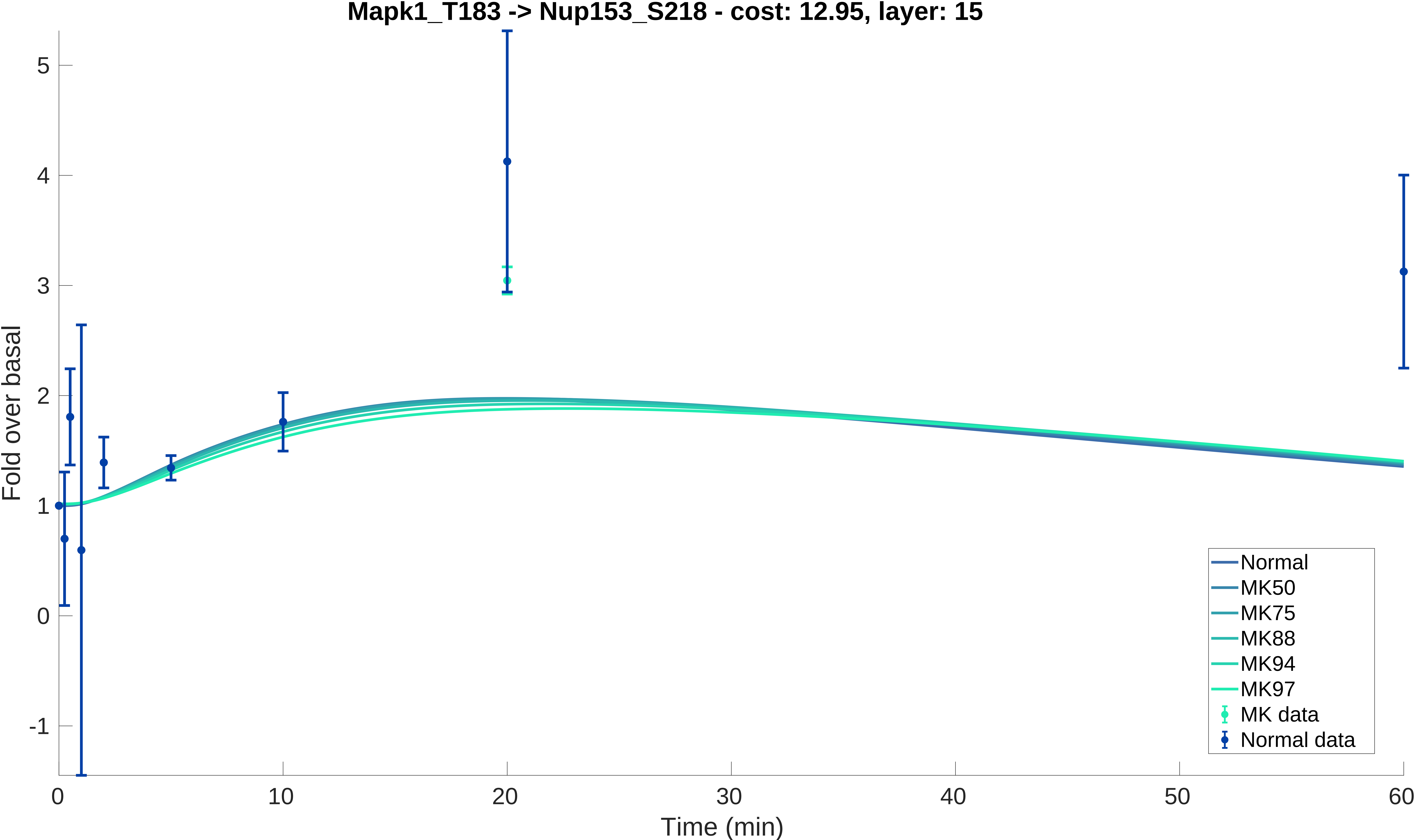

Mapk1_T183 -> Nup153_S218 - cost: 12.95, layer: 15
Fold over basal
Time (min)
5
4
3
2
1
0
-1
0
10
20
30
40
50
60
Normal
MK50
MK75
MK88
MK94
MK97
MK data
Normal data

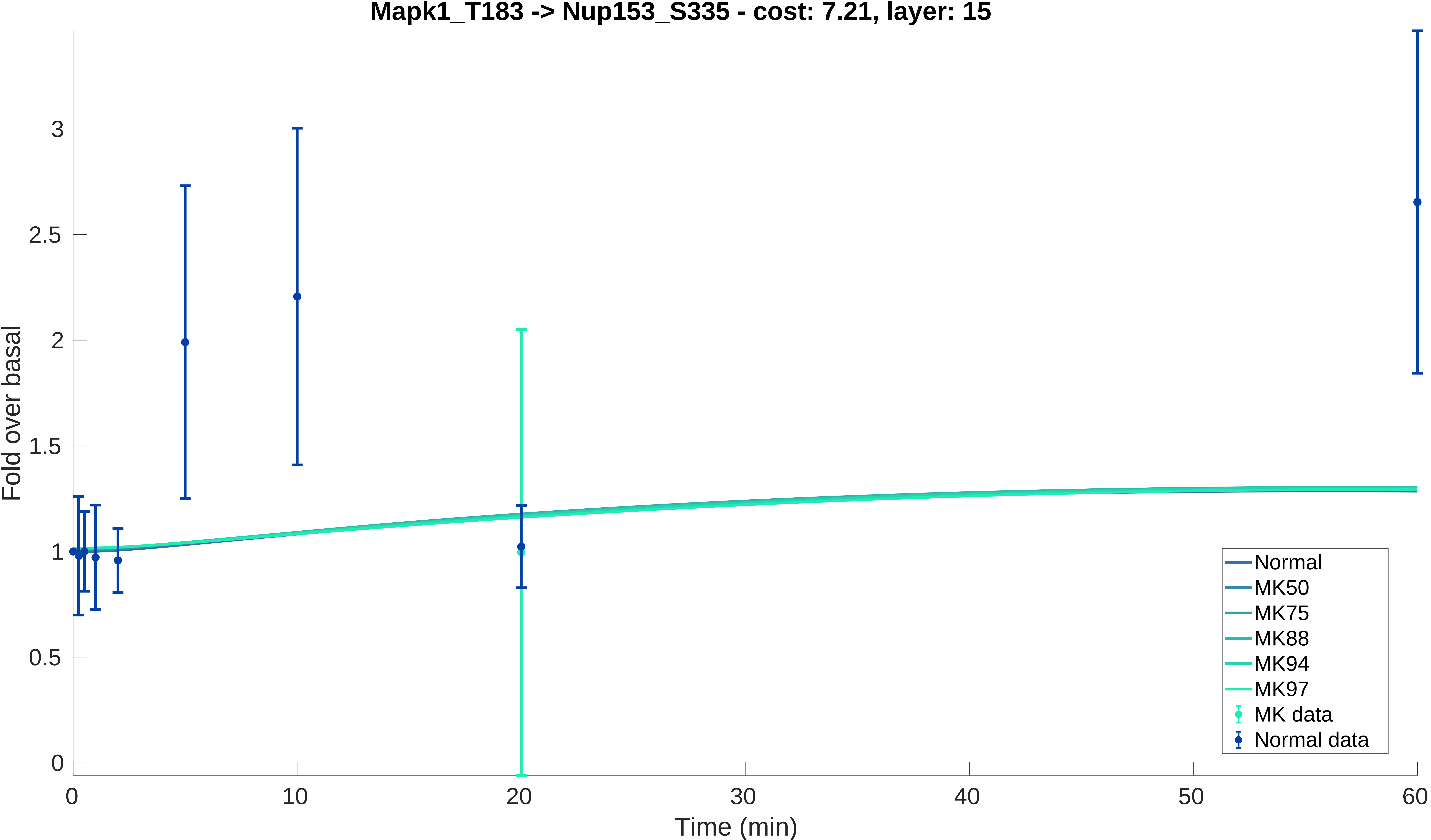

Mapk1_T183 -> Nup153_S335 - cost: 7.21, layer: 15
Fold over basal
Time (min)
0
0.5
1
1.5
2
2.5
3
0
10
20
30
40
50
60
Normal
MK50
MK75
MK88
MK94
MK97
MK data
Normal data

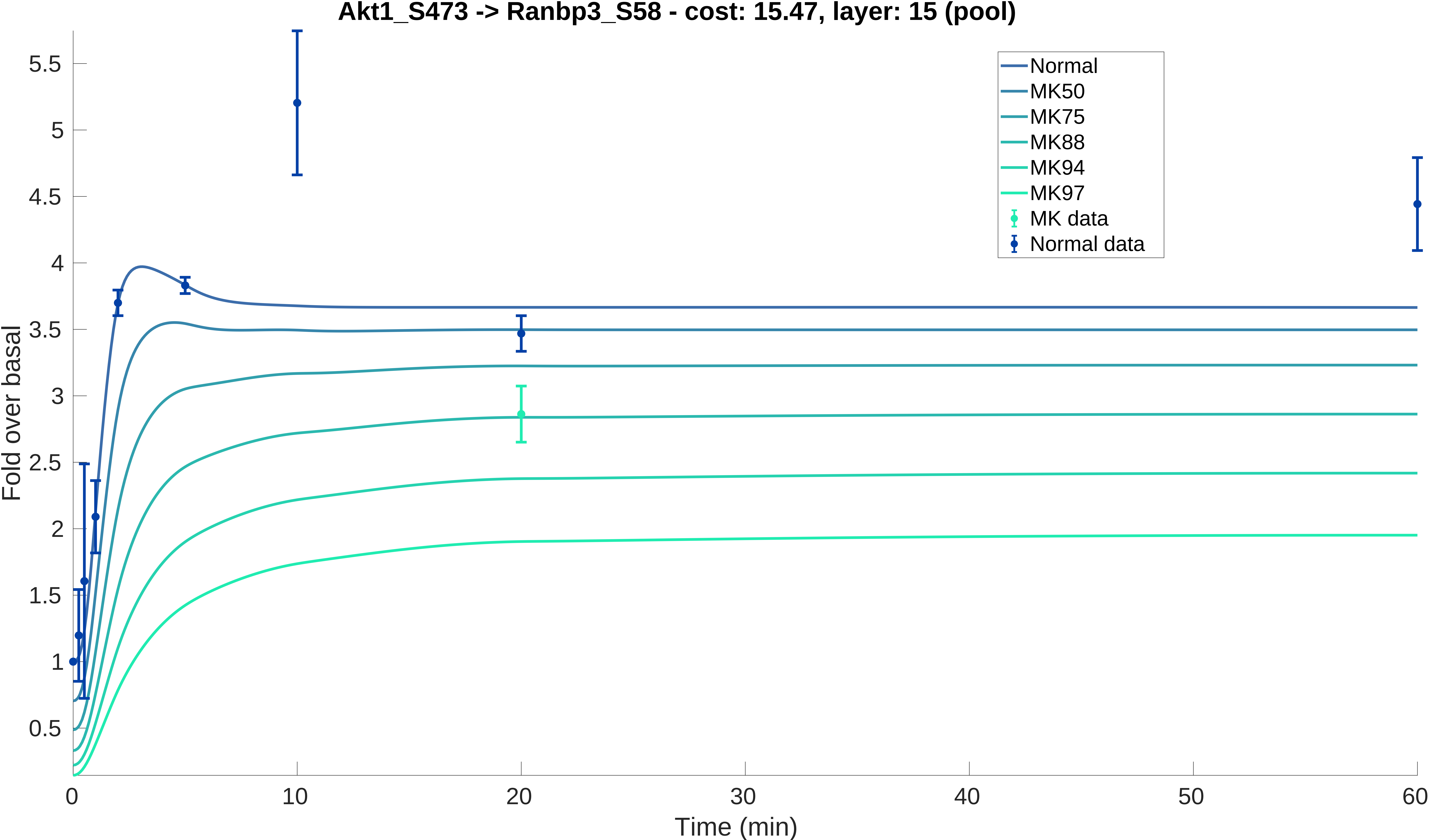

Akt1_S473 -> Ranbp3_S58 - cost: 15.47, layer: 15 (pool)
Normal
MK50
MK75
MK88
MK94
MK97
MK data
Normal data
Fold over basal
Time (min)
0.5
1
1.5
2
2.5
3
3.5
4
4.5
5
5.5
0
10
20
30
40
50
60

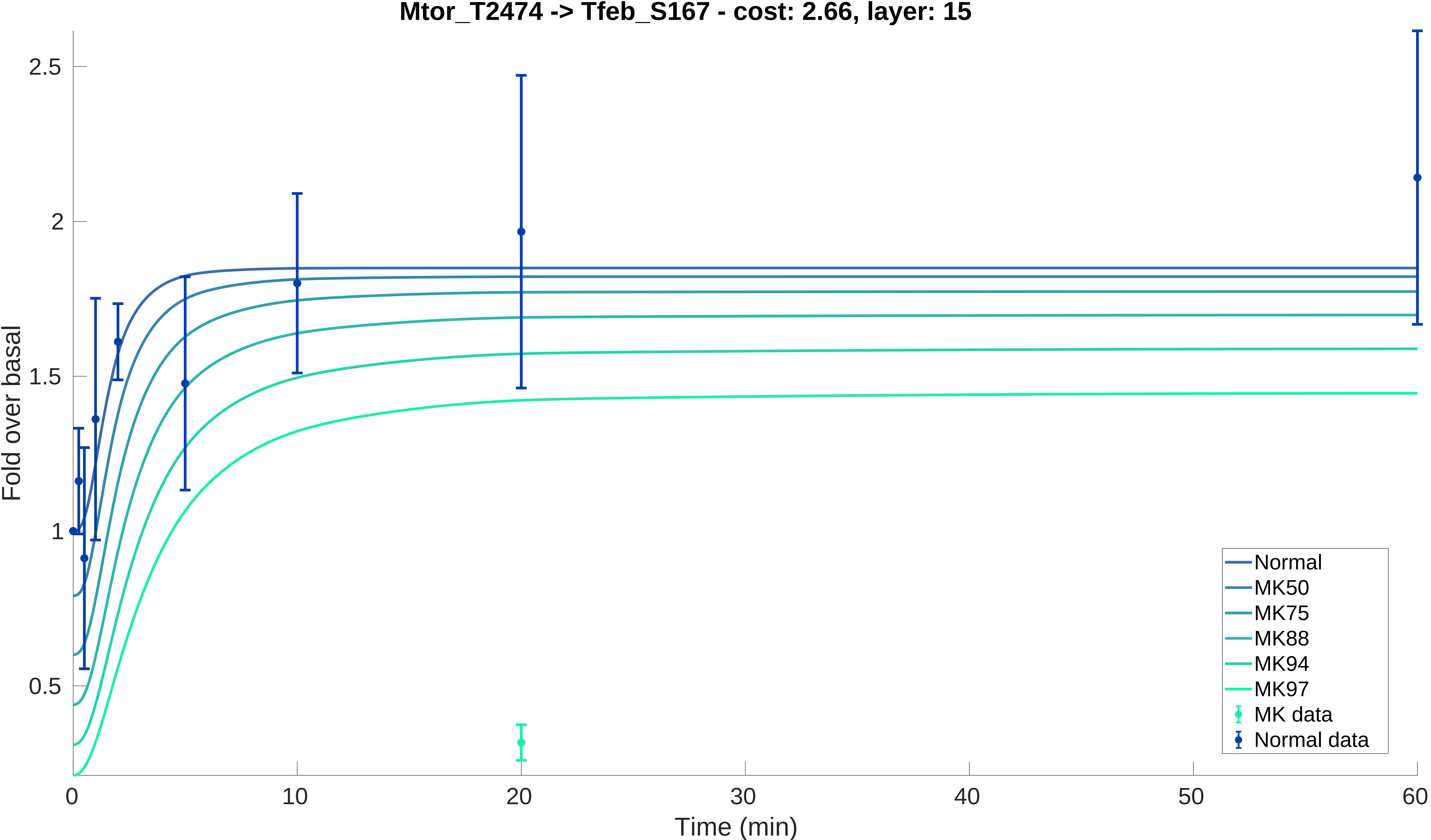

Mtor_T2474 -> Tfeb_S167 - cost: 2.66, layer: 15
Fold over basal
2.5
2
1.5
1
0.5
0
10
20
30
40
50
60
Time (min)
Normal
MK50
MK75
MK88
MK94
MK97
MK data
Normal data

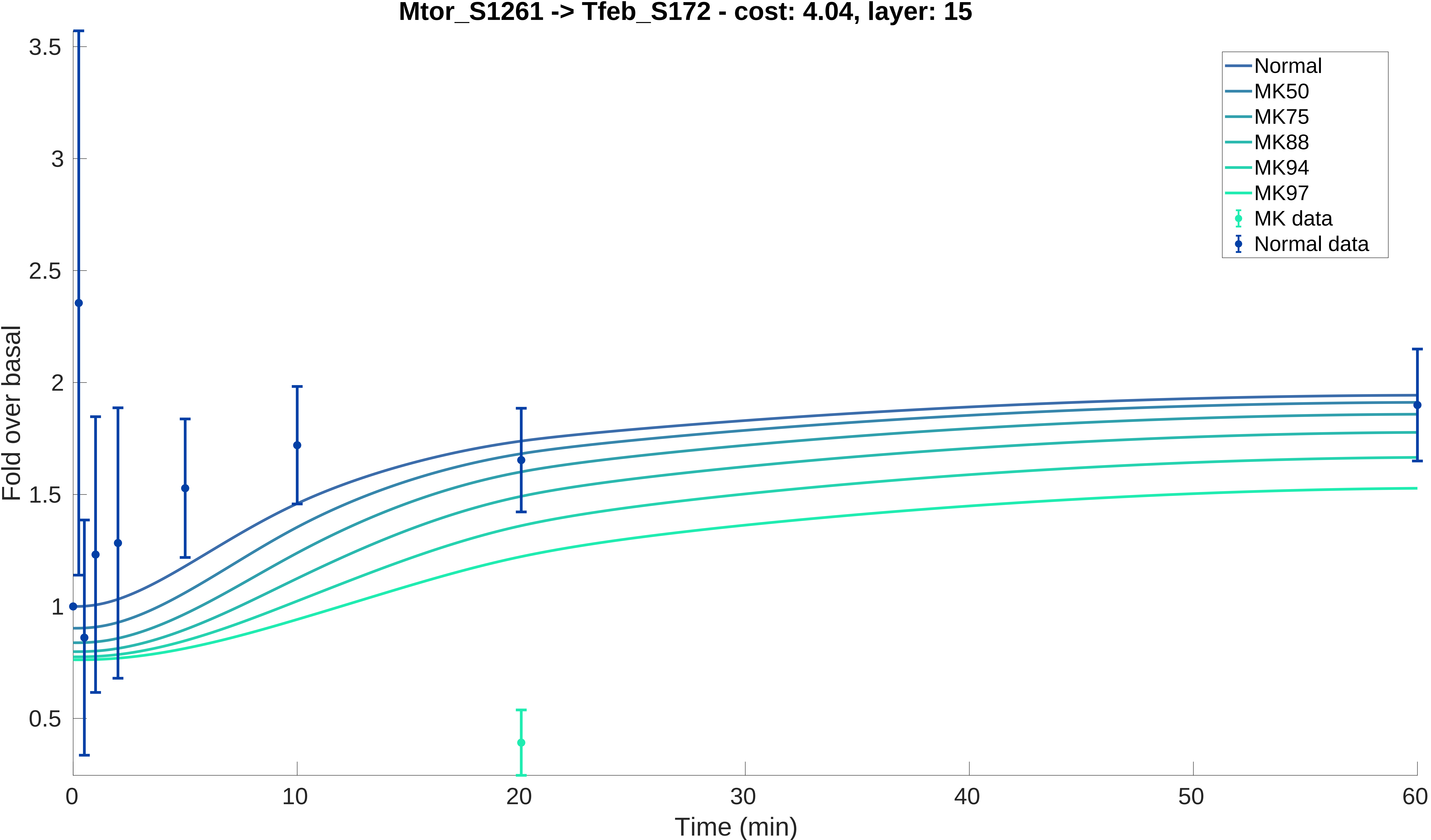

Mtor_S1261 -> Tfeb_S172 - cost: 4.04, layer: 15
Fold over basal
Time (min)
0.5
1
1.5
2
2.5
3
3.5
0
10
20
30
40
50
60
Normal
MK50
MK75
MK88
MK94
MK97
MK data
Normal data

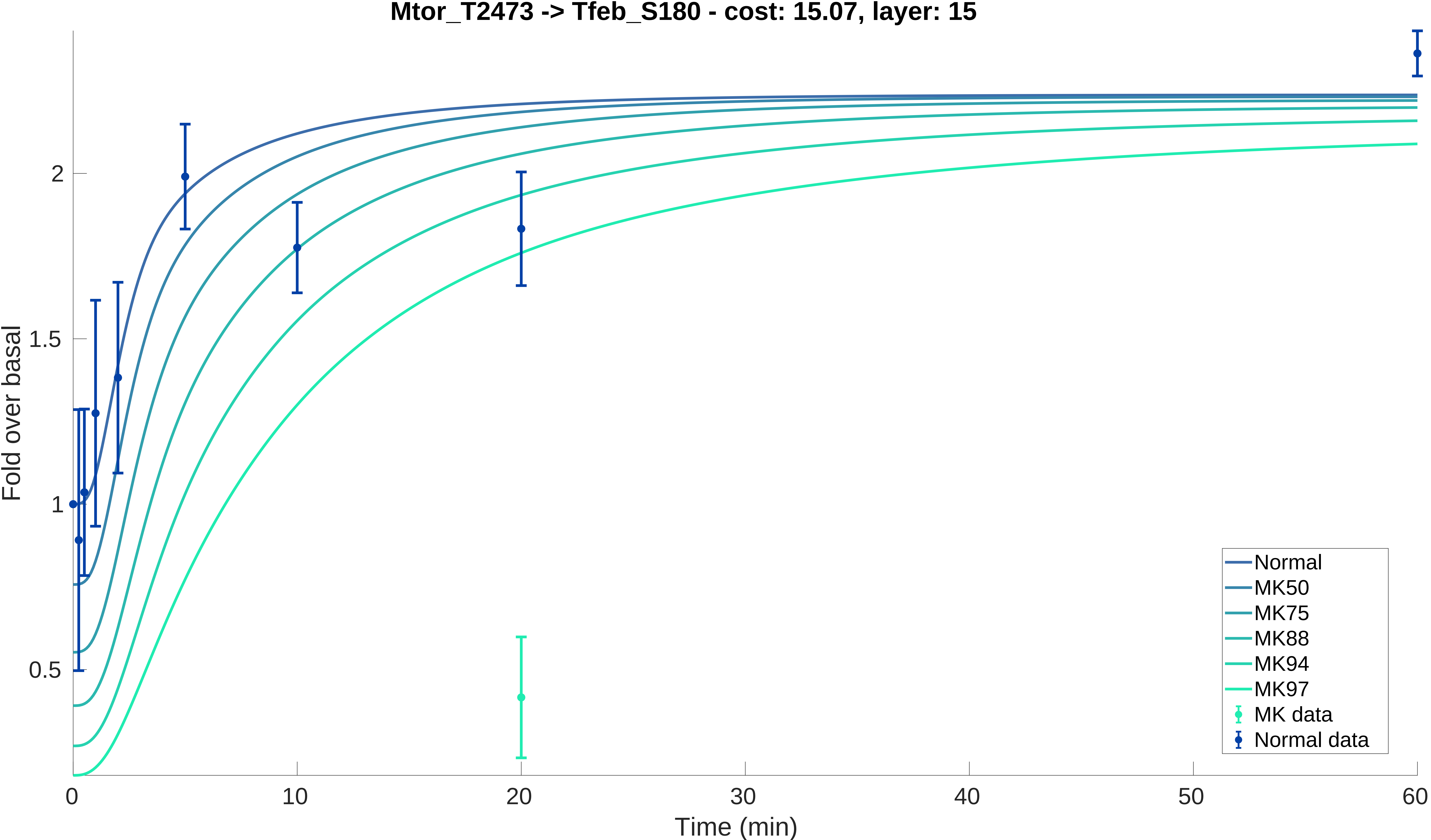

Mtor_T2473 -> Tfeb_S180 - cost: 15.07, layer: 15
Fold over basal
Time (min)
0.5
1
1.5
2
0
10
20
30
40
50
60
Normal
MK50
MK75
MK88
MK94
MK97
MK data
Normal data

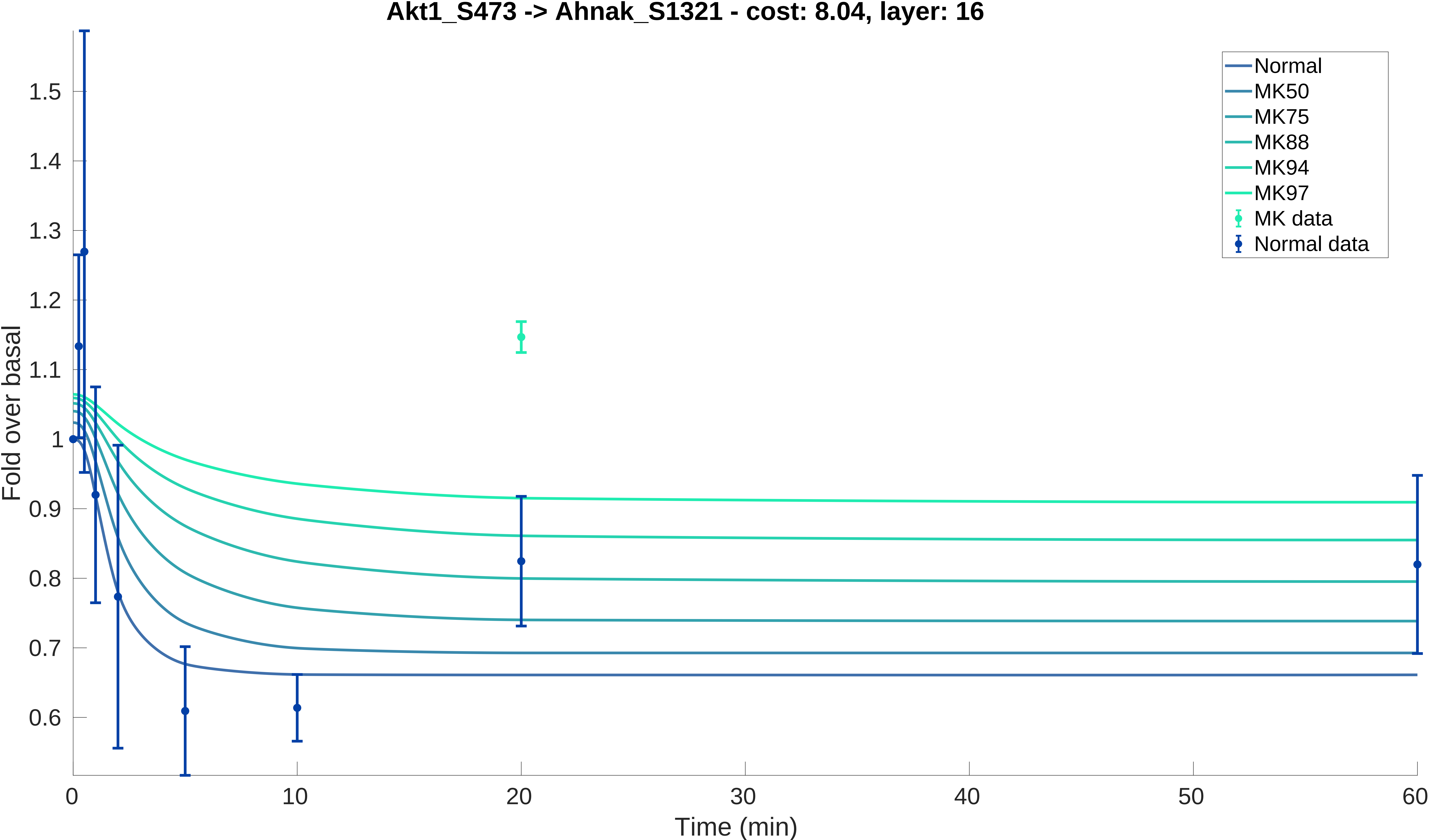

Akt1_S473 -> Ahnak_S1321 - cost: 8.04, layer: 16
Fold over basal
Time (min)
0.6
0.7
0.8
0.9
1
1.1
1.2
1.3
1.4
1.5
0
10
20
30
40
50
60
Normal
MK50
MK75
MK88
MK94
MK97
MK data
Normal data

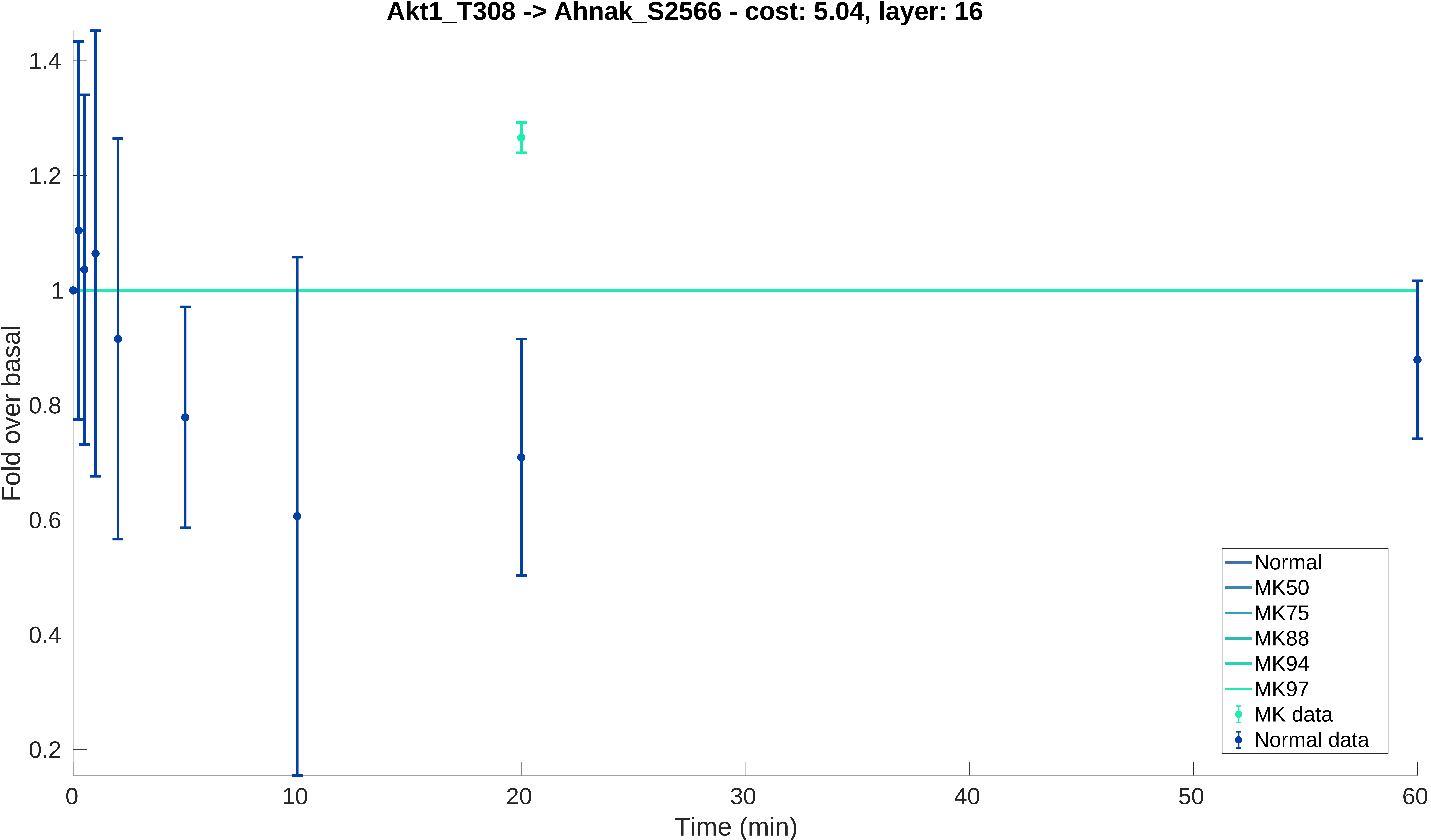
Akt1_T308 -> Ahnak_S2566 - cost: 5.04, layer: 16
Fold over basal
1.4
1.2
1
0.8
0.6
0.4
0.2
0
10
20
30
40
50
60
Time (min)
Normal
MK50
MK75
MK88
MK94
MK97
MK data
Normal data

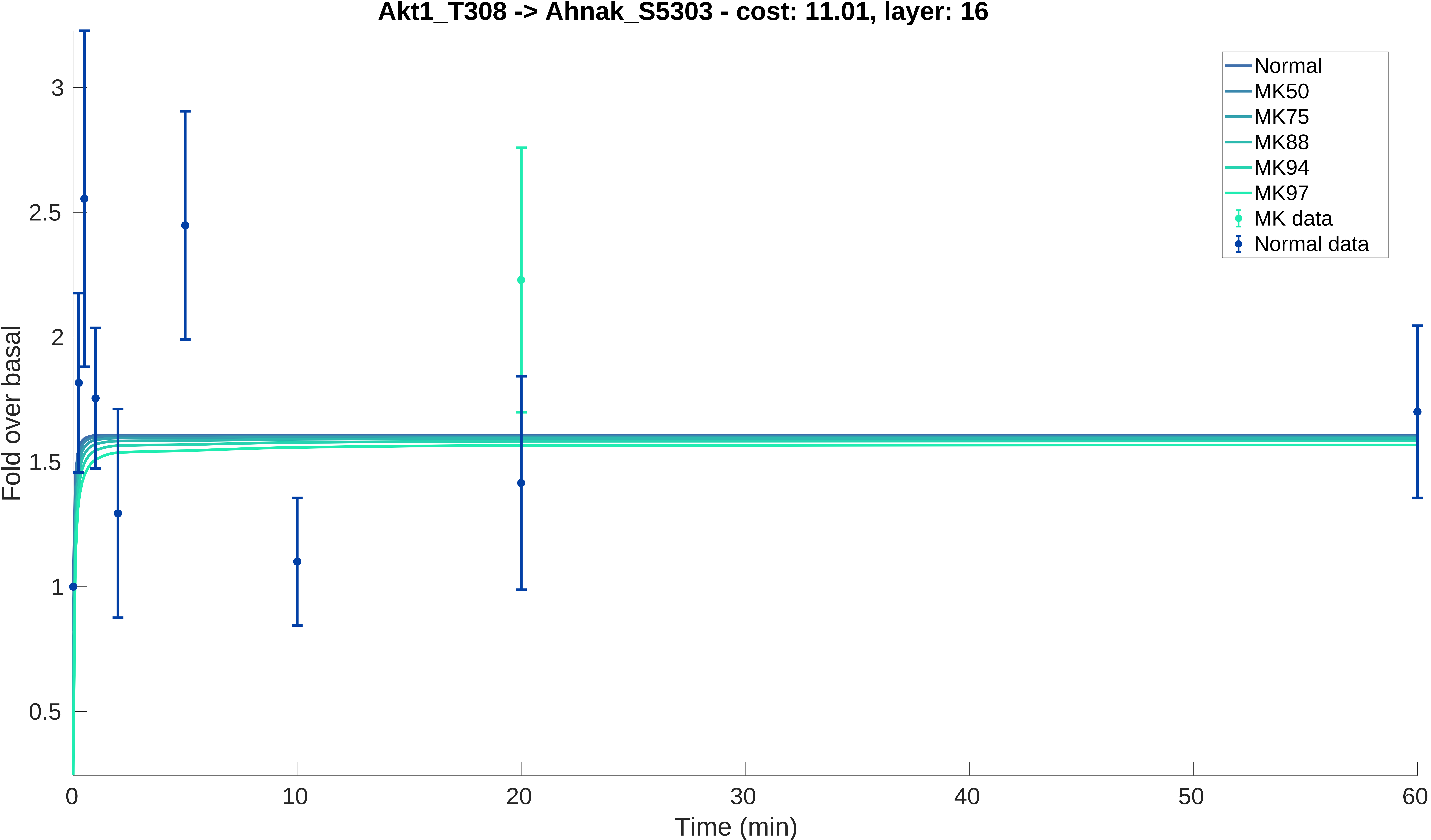

Akt1_T308 -> Ahnak_S5303 - cost: 11.01, layer: 16
Fold over basal
Time (min)
0.5
1
1.5
2
2.5
3
0
10
20
30
40
50
60
Normal
MK50
MK75
MK88
MK94
MK97
MK data
Normal data

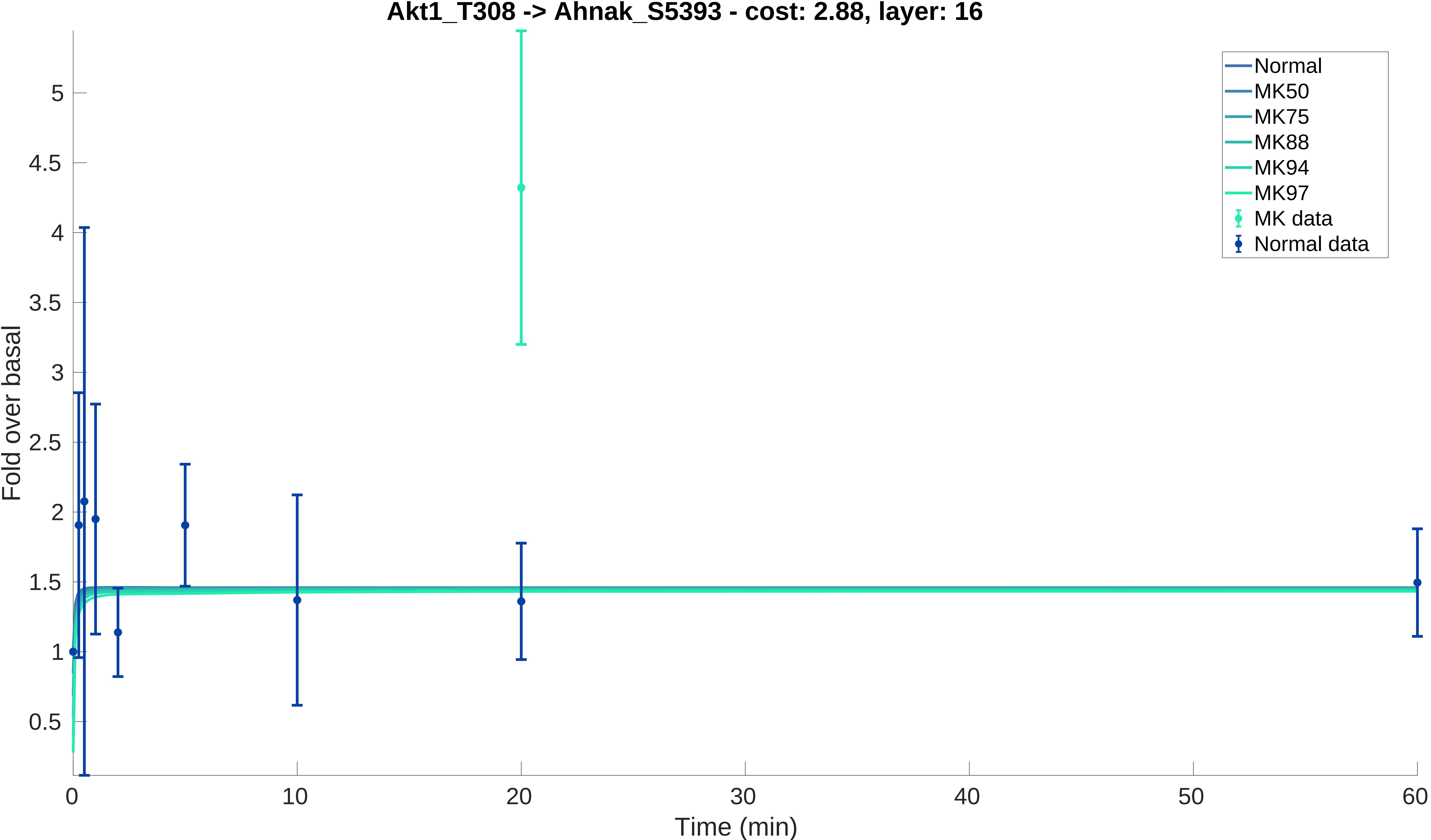
Akt1_T308 -> Ahnak_S5393 - cost: 2.88, layer: 16
Fold over basal
Time (min)
0
10
20
30
40
50
60
0.5
1
1.5
2
2.5
3
3.5
4
4.5
5
Normal
MK50
MK75
MK88
MK94
MK97
MK data
Normal data

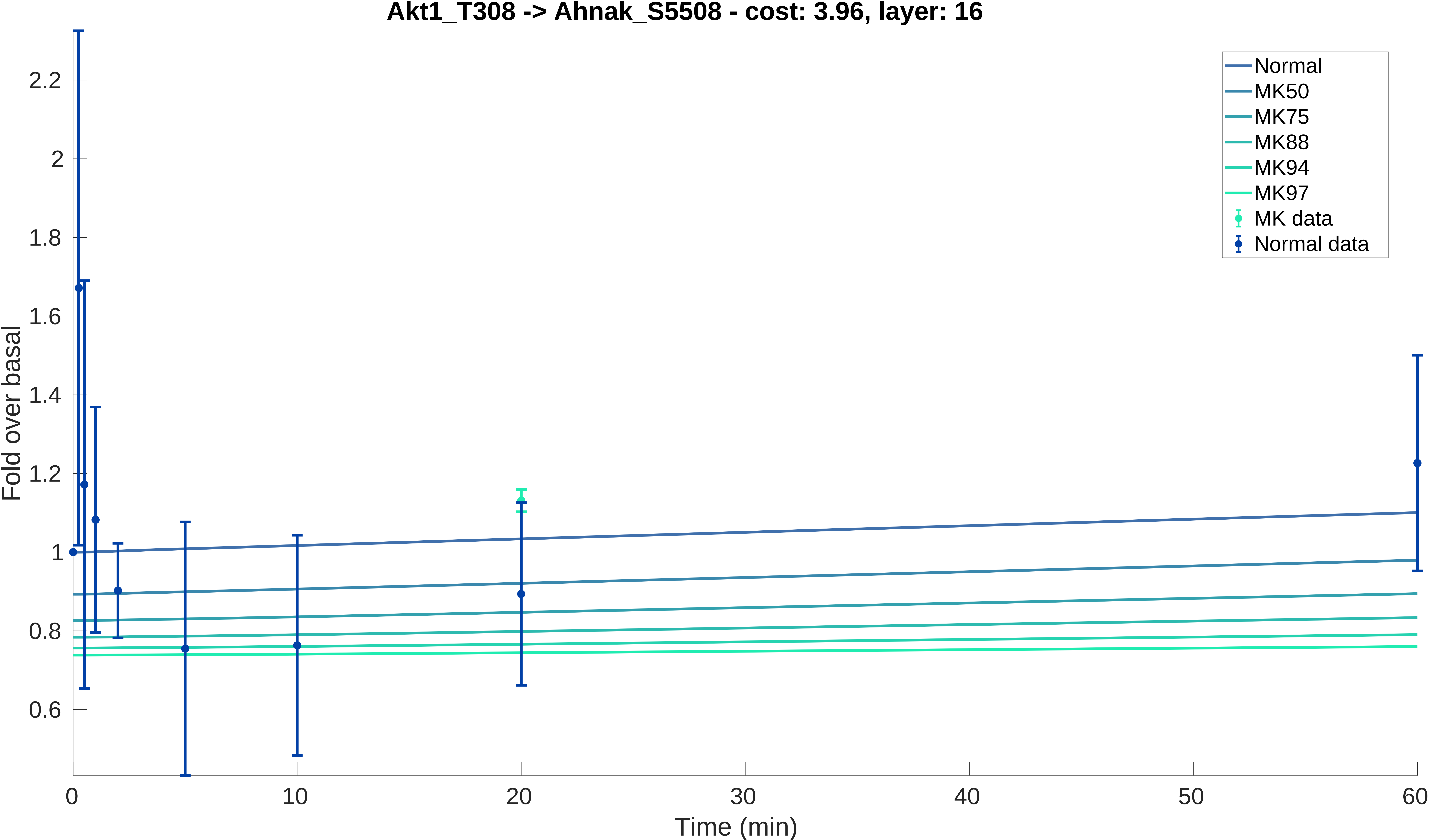
Akt1_T308 -> Ahnak_S5508 - cost: 3.96, layer: 16
Fold over basal
Time (min)
0
10
20
30
40
50
60
0.6
0.8
1
1.2
1.4
1.6
1.8
2
2.2
Normal
MK50
MK75
MK88
MK94
MK97
MK data
Normal data

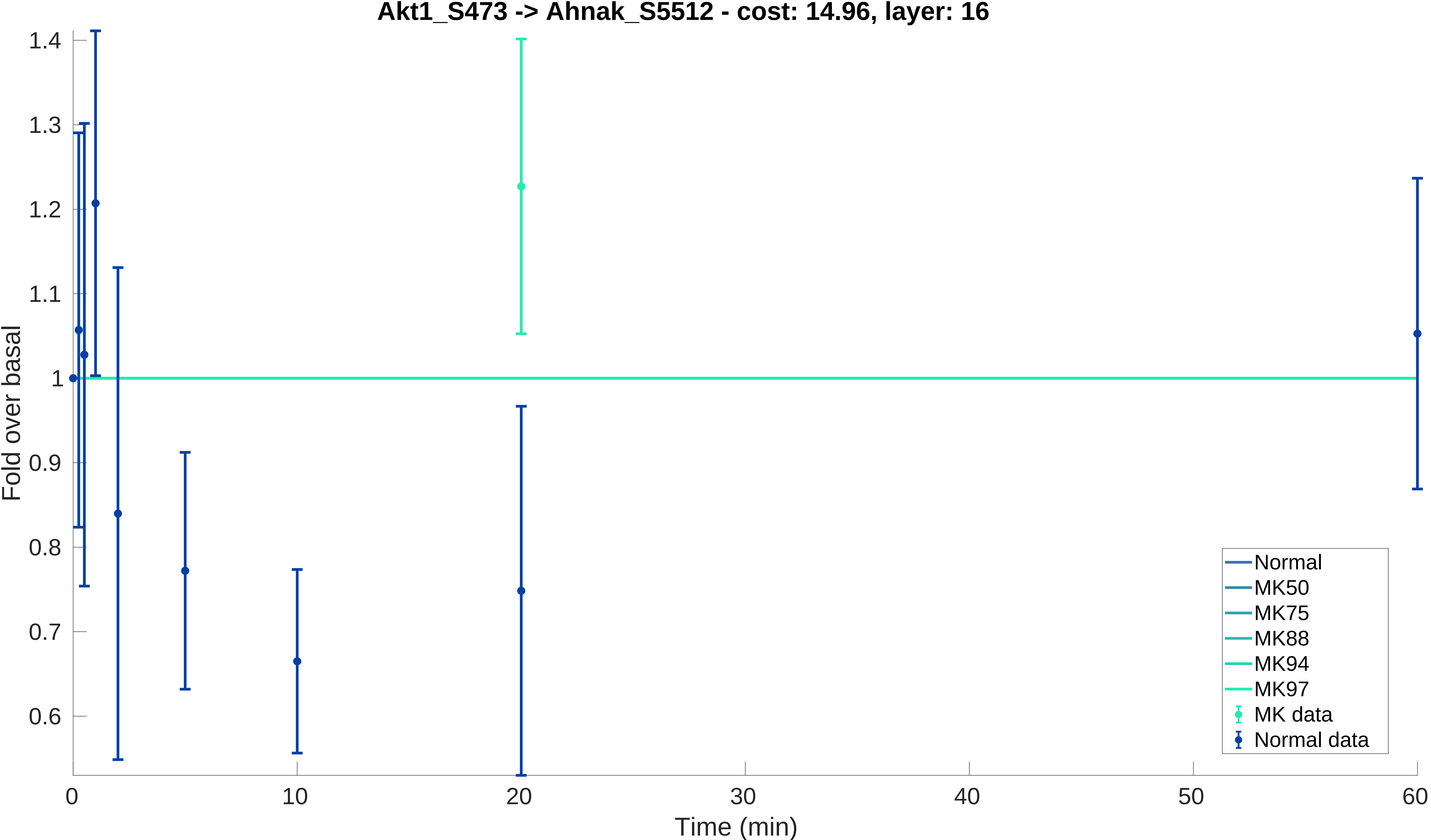

Akt1_S473 -> Ahnak_S5512 - cost: 14.96, layer: 16
Fold over basal
1.4
1.3
1.2
1.1
1
0.9
0.8
0.7
0.6
0
10
20
30
40
50
60
Time (min)
Normal
MK50
MK75
MK88
MK94
MK97
MK data
Normal data

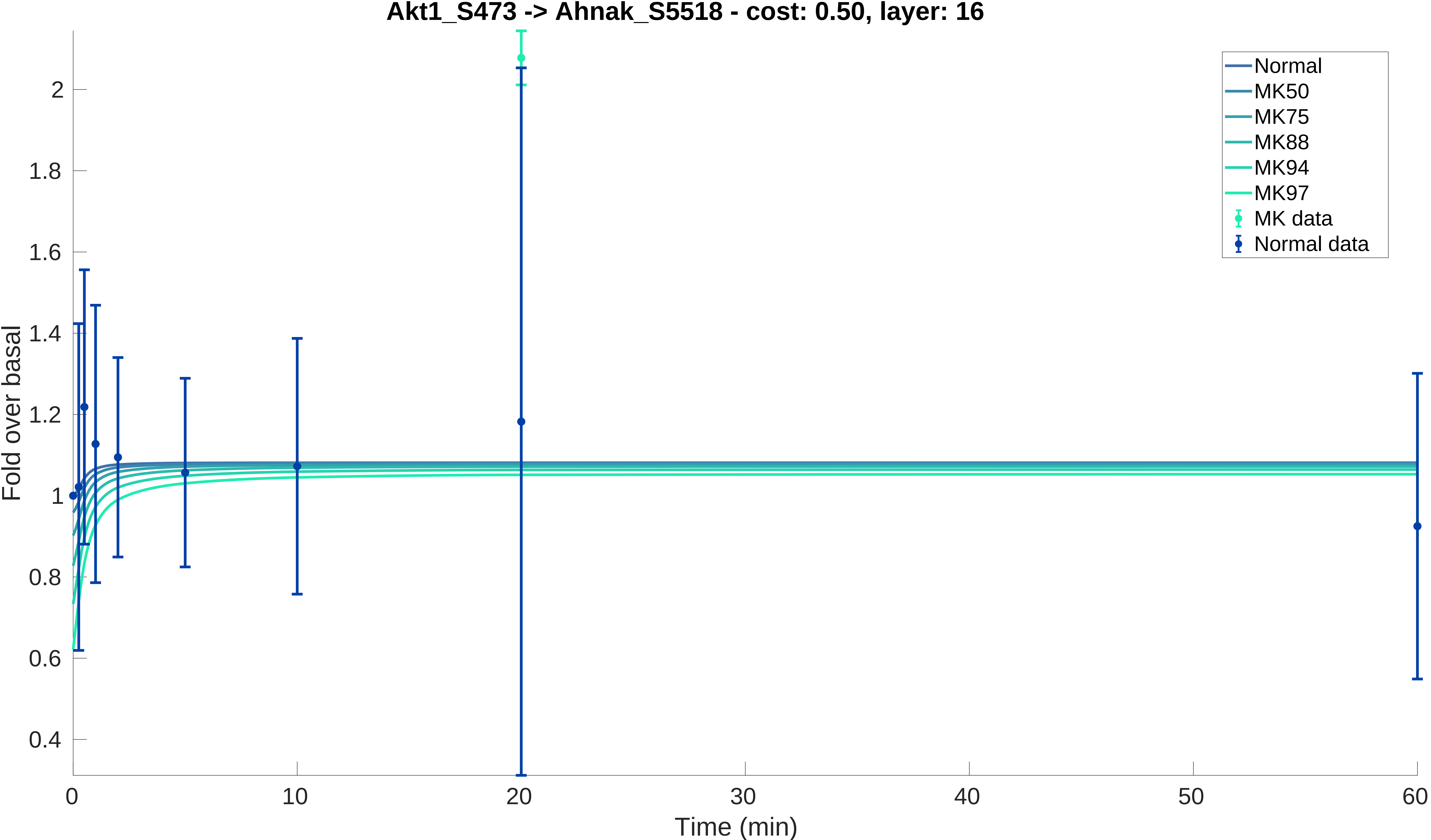
Akt1_S473 -> Ahnak_S5518 - cost: 0.50, layer: 16
Fold over basal
Time (min)
Normal
MK50
MK75
MK88
MK94
MK97
MK data
Normal data
0.4
0.6
0.8
1
1.2
1.4
1.6
1.8
2
0
10
20
30
40
50
60

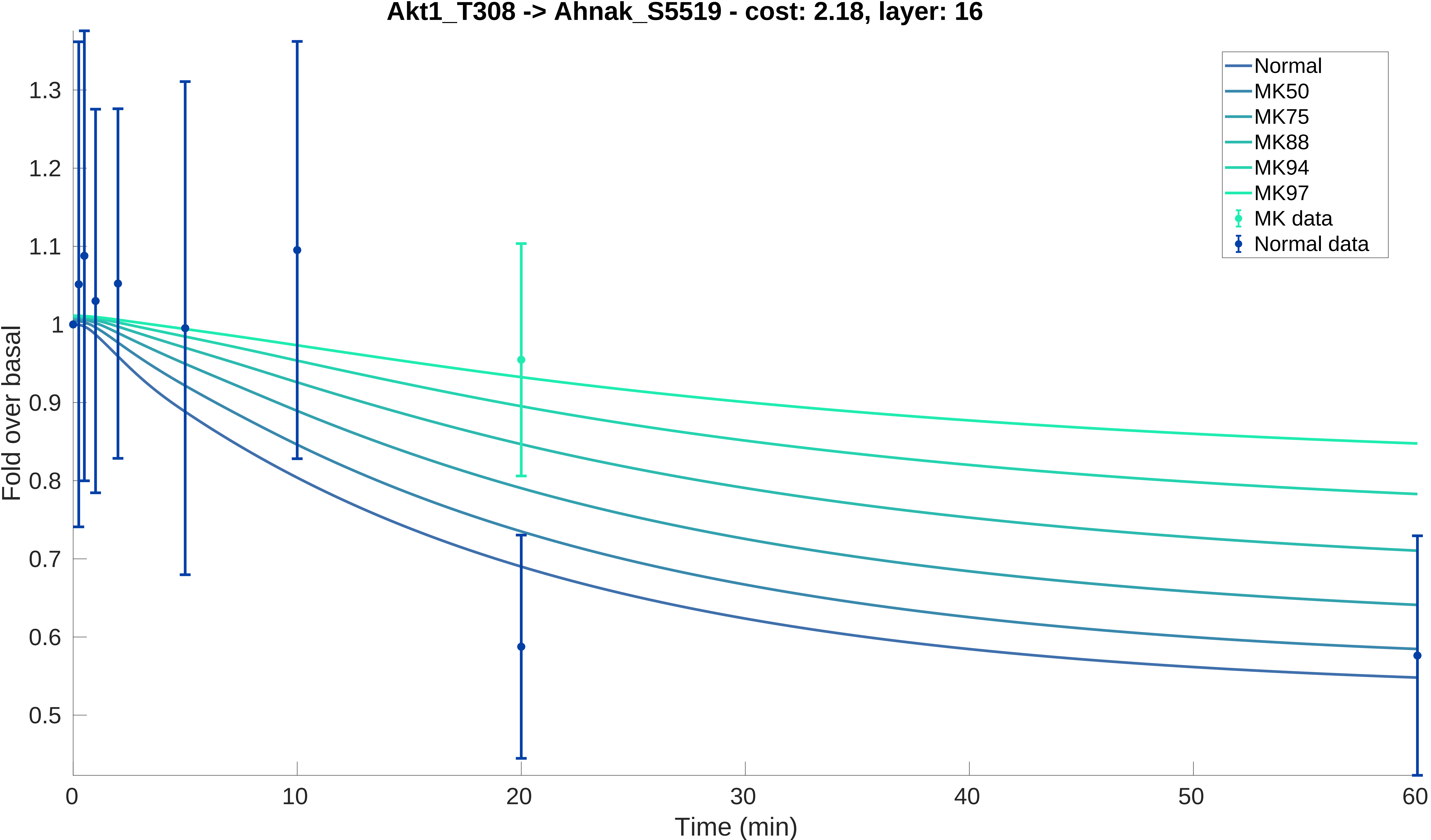

Akt1_T308 -> Ahnak_S5519 - cost: 2.18, layer: 16
Fold over basal
Time (min)
1.3
1.2
1.1
1
0.9
0.8
0.7
0.6
0.5
0
10
20
30
40
50
60
Normal
MK50
MK75
MK88
MK94
MK97
MK data
Normal data

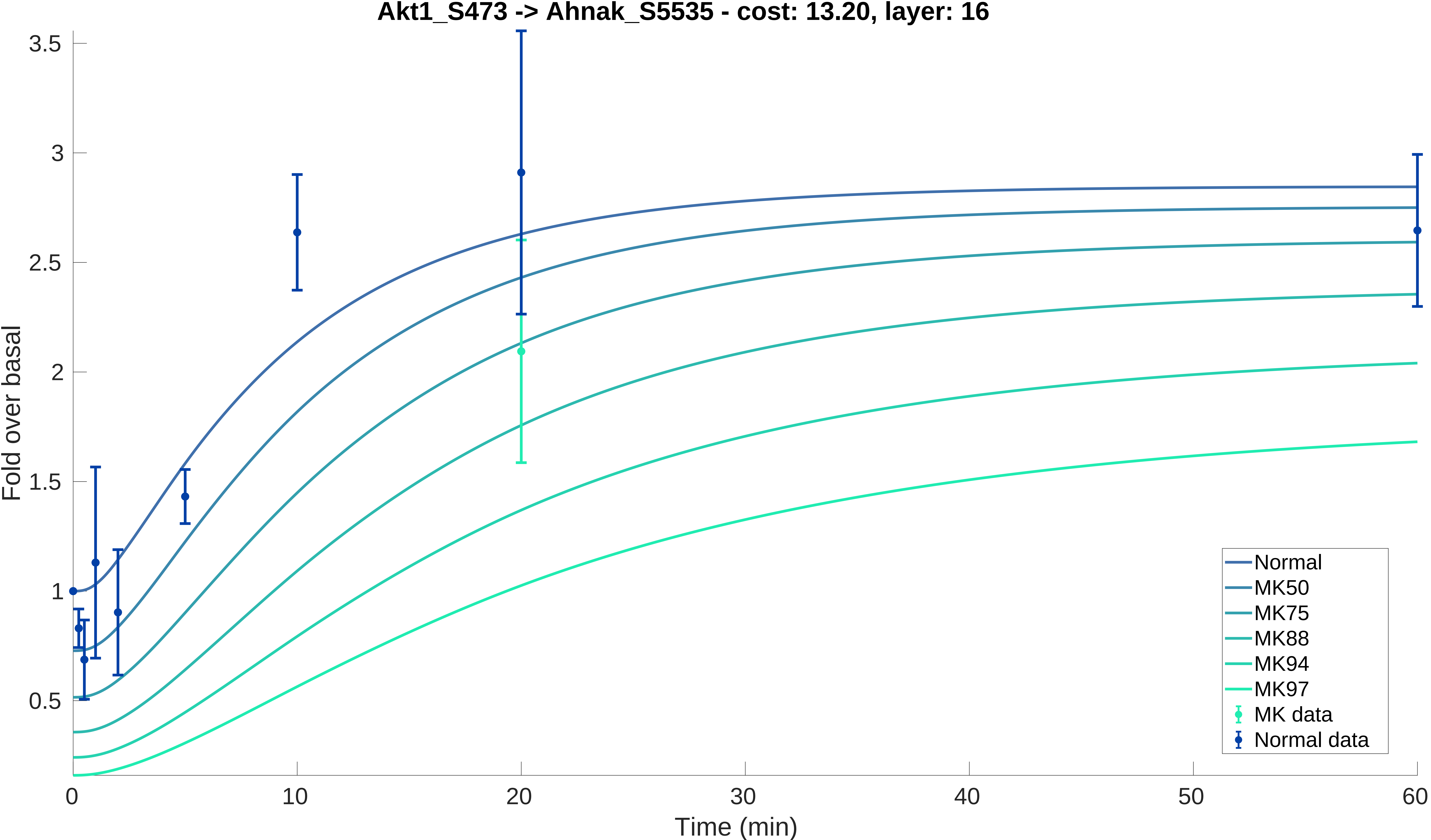
Akt1_S473 -> Ahnak_S5535 - cost: 13.20, layer: 16
Fold over basal
Time (min)
0
10
20
30
40
50
60
0.5
1
1.5
2
2.5
3
3.5
Normal
MK50
MK75
MK88
MK94
MK97
MK data
Normal data

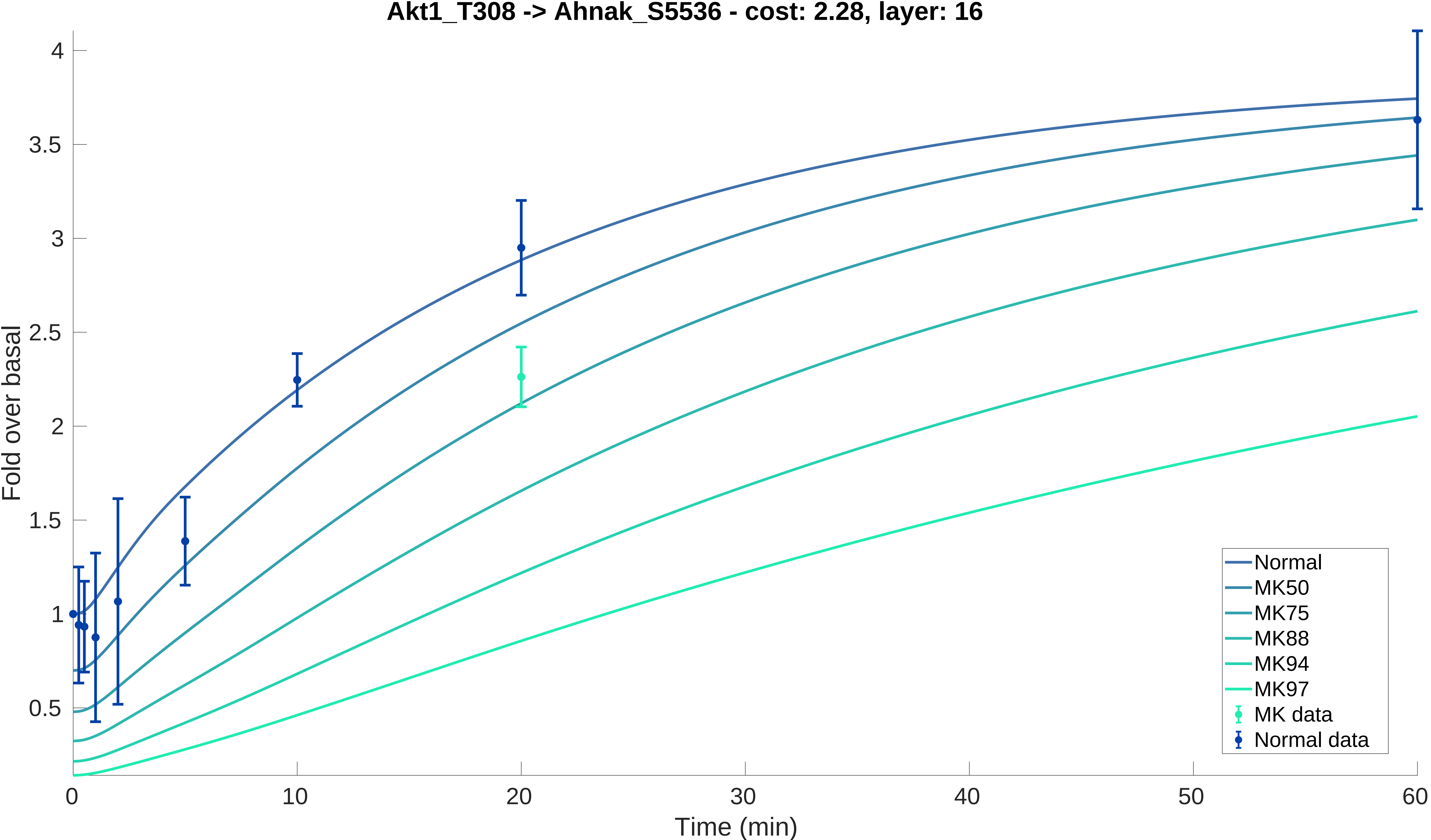

Akt1_T308 -> Ahnak_S5536 - cost: 2.28, layer: 16
Fold over basal
4
3.5
3
2.5
2
1.5
1
0.5
0
10
20
30
40
50
60
Time (min)
Normal
MK50
MK75
MK88
MK94
MK97
MK data
Normal data

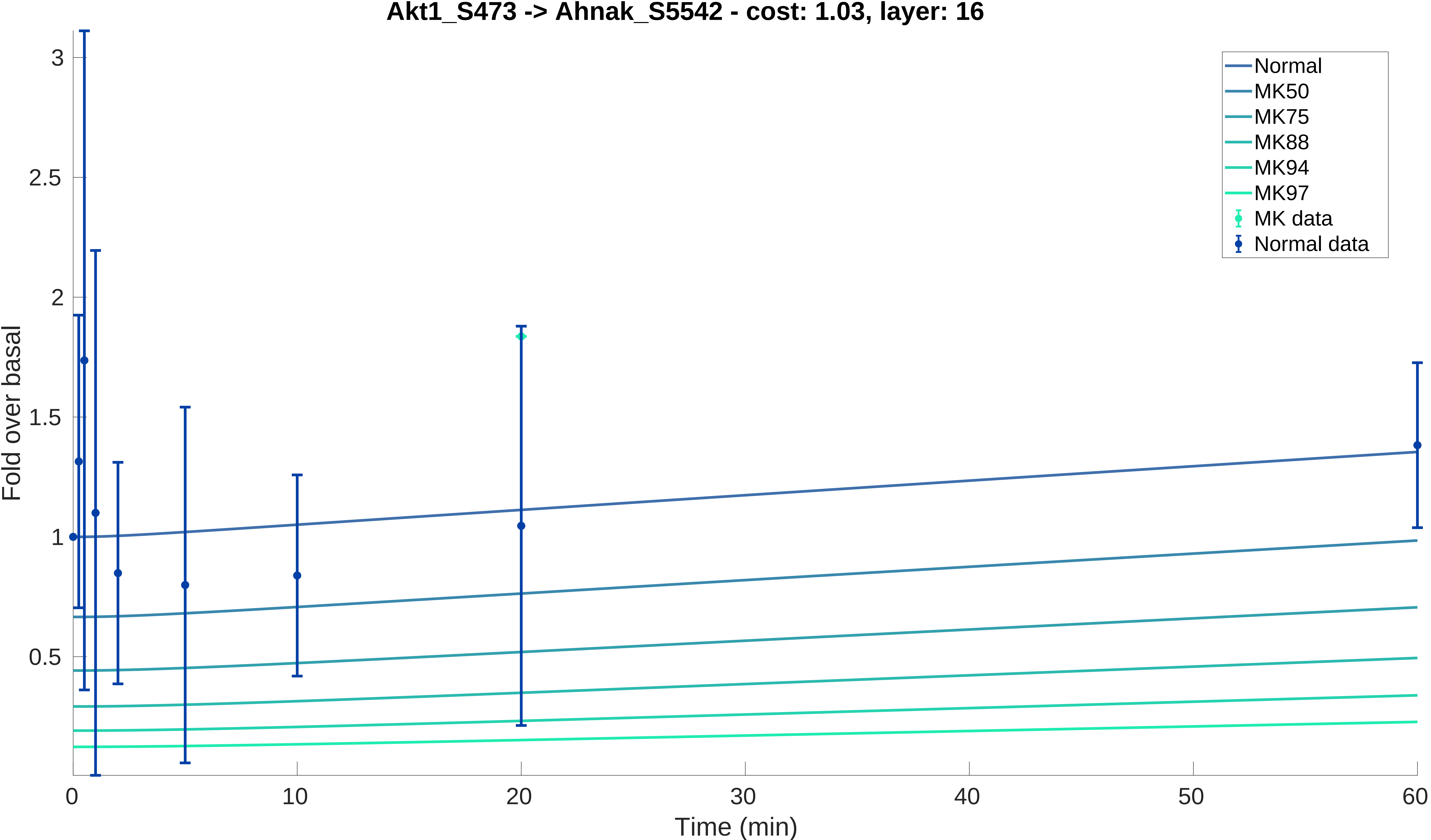

Akt1_S473 -> Ahnak_S5542 - cost: 1.03, layer: 16
Fold over basal
Time (min)
0
0.5
1
1.5
2
2.5
3
0
10
20
30
40
50
60
Normal
MK50
MK75
MK88
MK94
MK97
MK data
Normal data

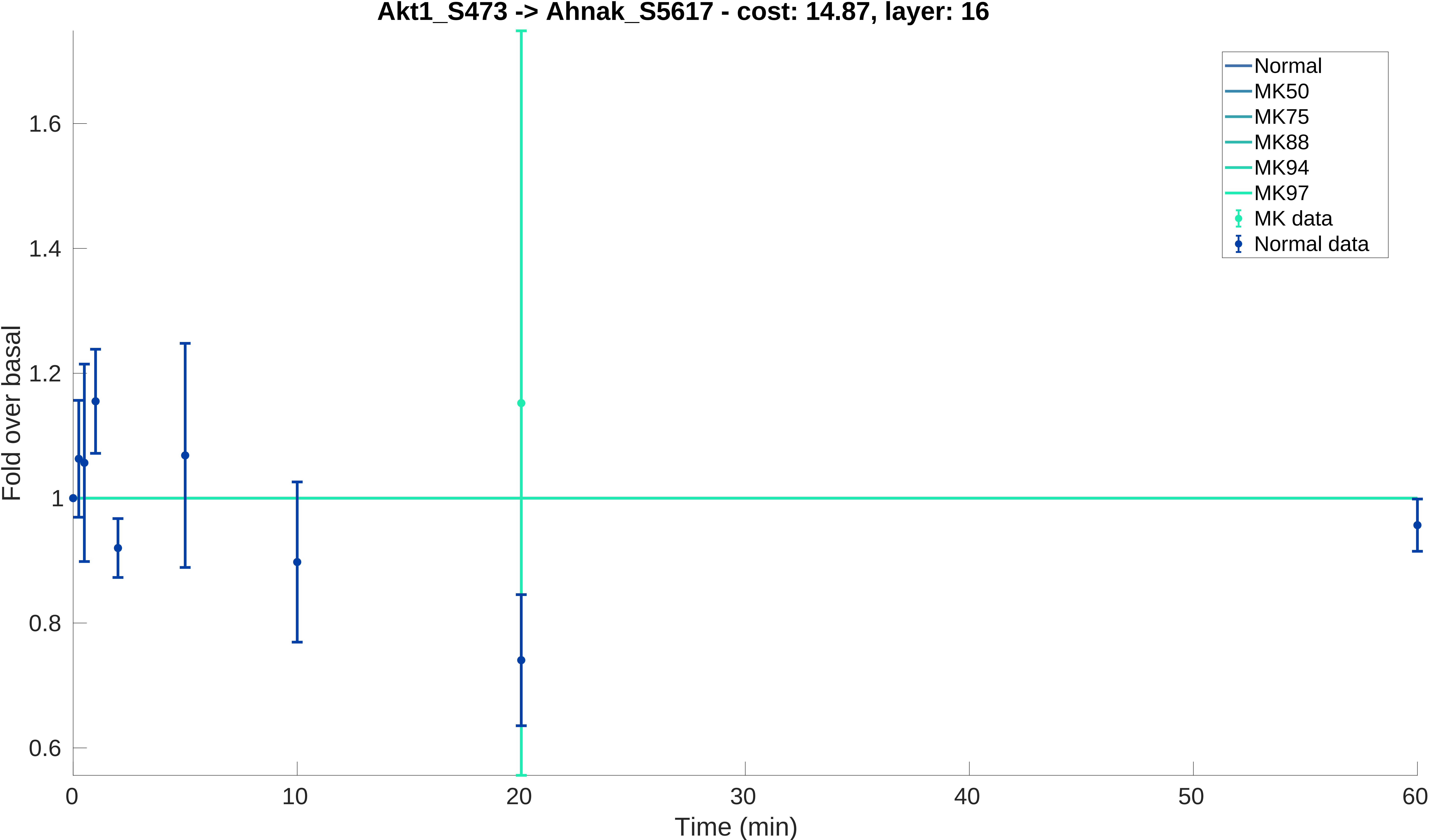
Akt1_S473 -> Ahnak_S5617 - cost: 14.87, layer: 16
Normal
MK50
MK75
MK88
MK94
MK97
MK data
Normal data
Fold over basal
1.6
1.4
1.2
1
0.8
0.6
0
10
20
30
40
50
60
Time (min)

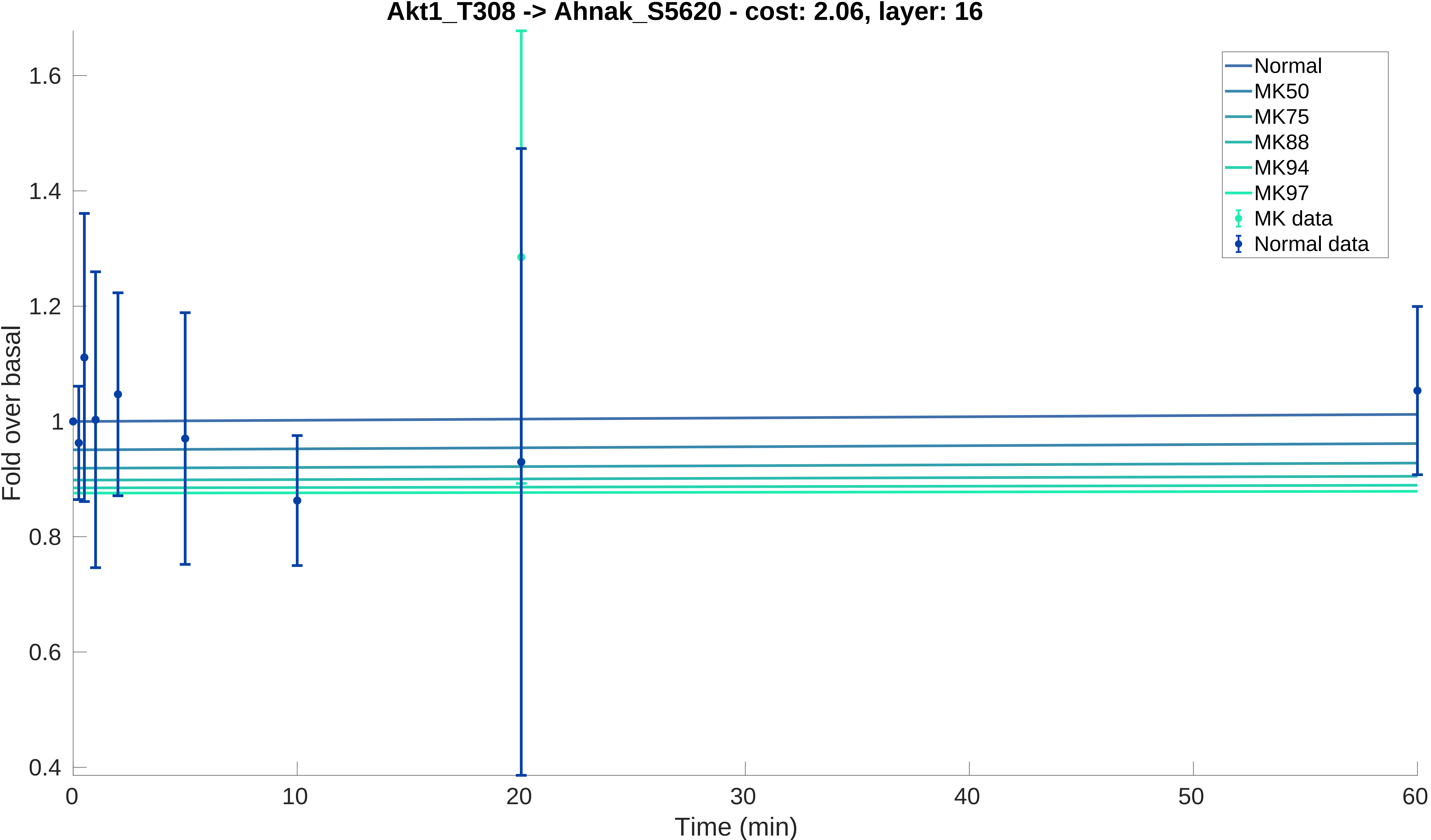

Akt1_T308 -> Ahnak_S5620 - cost: 2.06, layer: 16
Normal
MK50
MK75
MK88
MK94
MK97
MK data
Normal data
Fold over basal
0.4
0.6
0.8
1
1.2
1.4
1.6
0
10
20
30
40
50
60
Time (min)

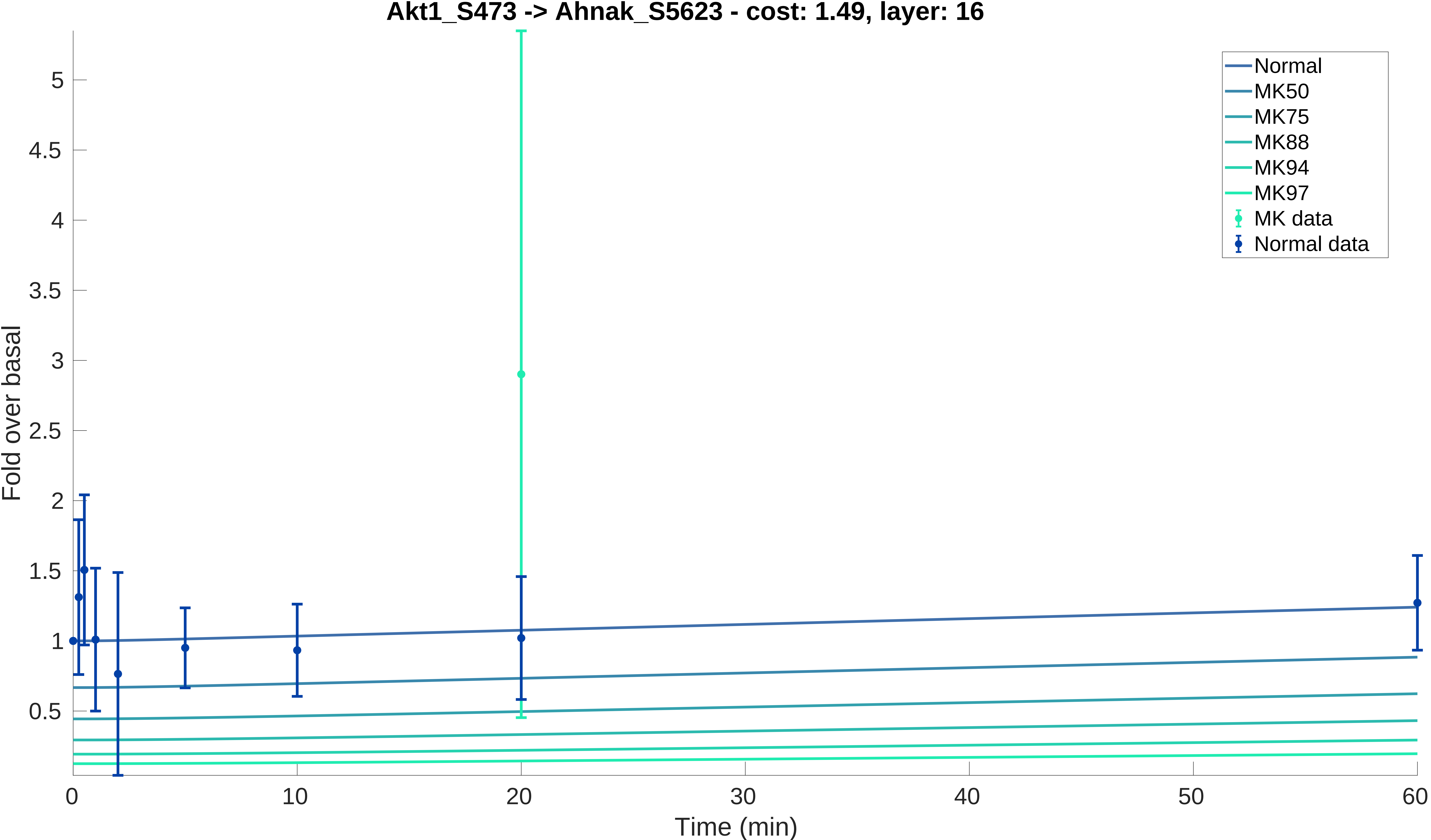
Akt1_S473 -> Ahnak_S5623 - cost: 1.49, layer: 16
Fold over basal
Time (min)
0
10
20
30
40
50
60
0.5
1
1.5
2
2.5
3
3.5
4
4.5
5
Normal
MK50
MK75
MK88
MK94
MK97
MK data
Normal data

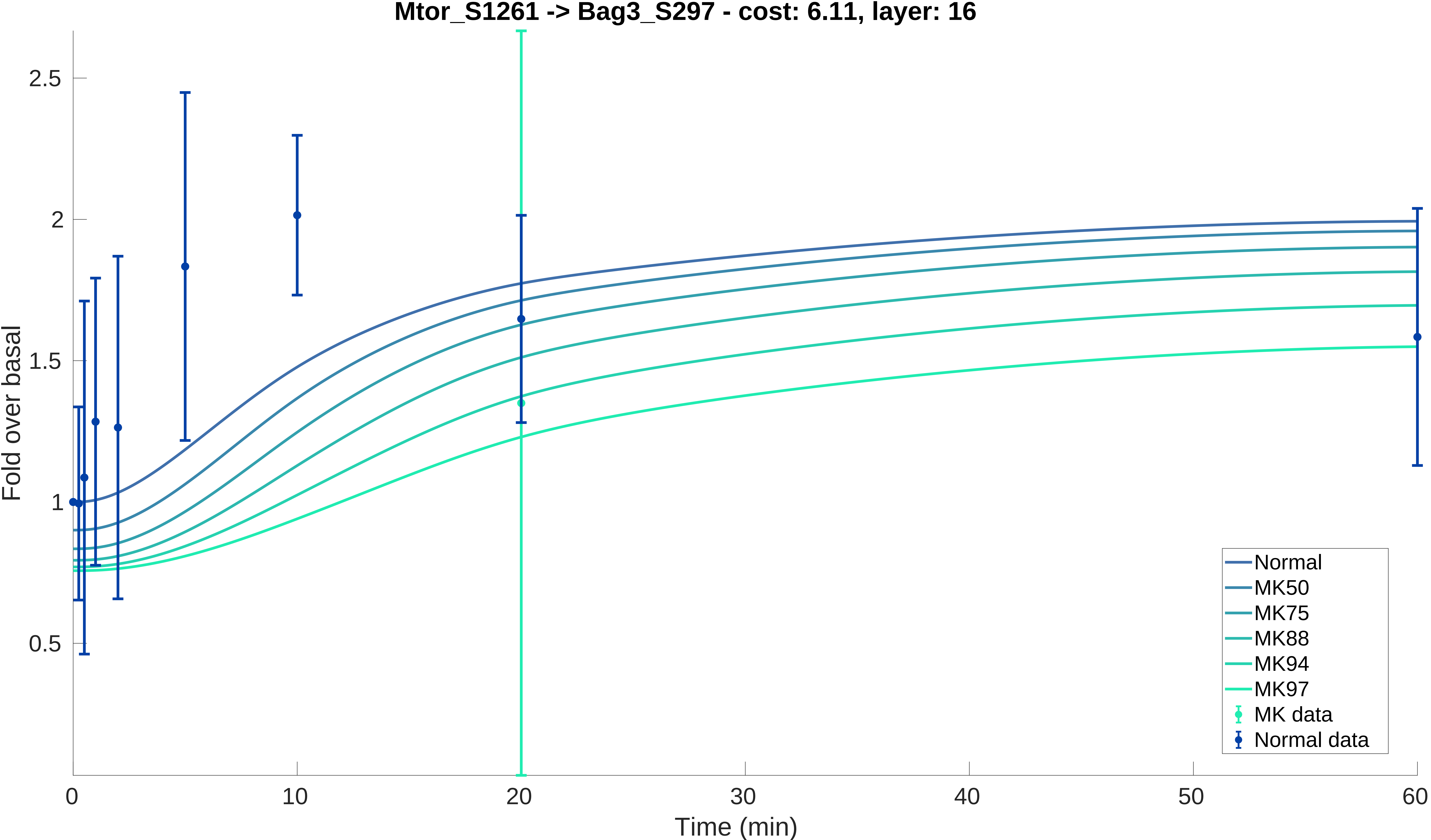

Mtor_S1261 -> Bag3_S297 - cost: 6.11, layer: 16
Fold over basal
2.5
2
1.5
1
0.5
0
10
20
30
40
50
60
Time (min)
Normal
MK50
MK75
MK88
MK94
MK97
MK data
Normal data

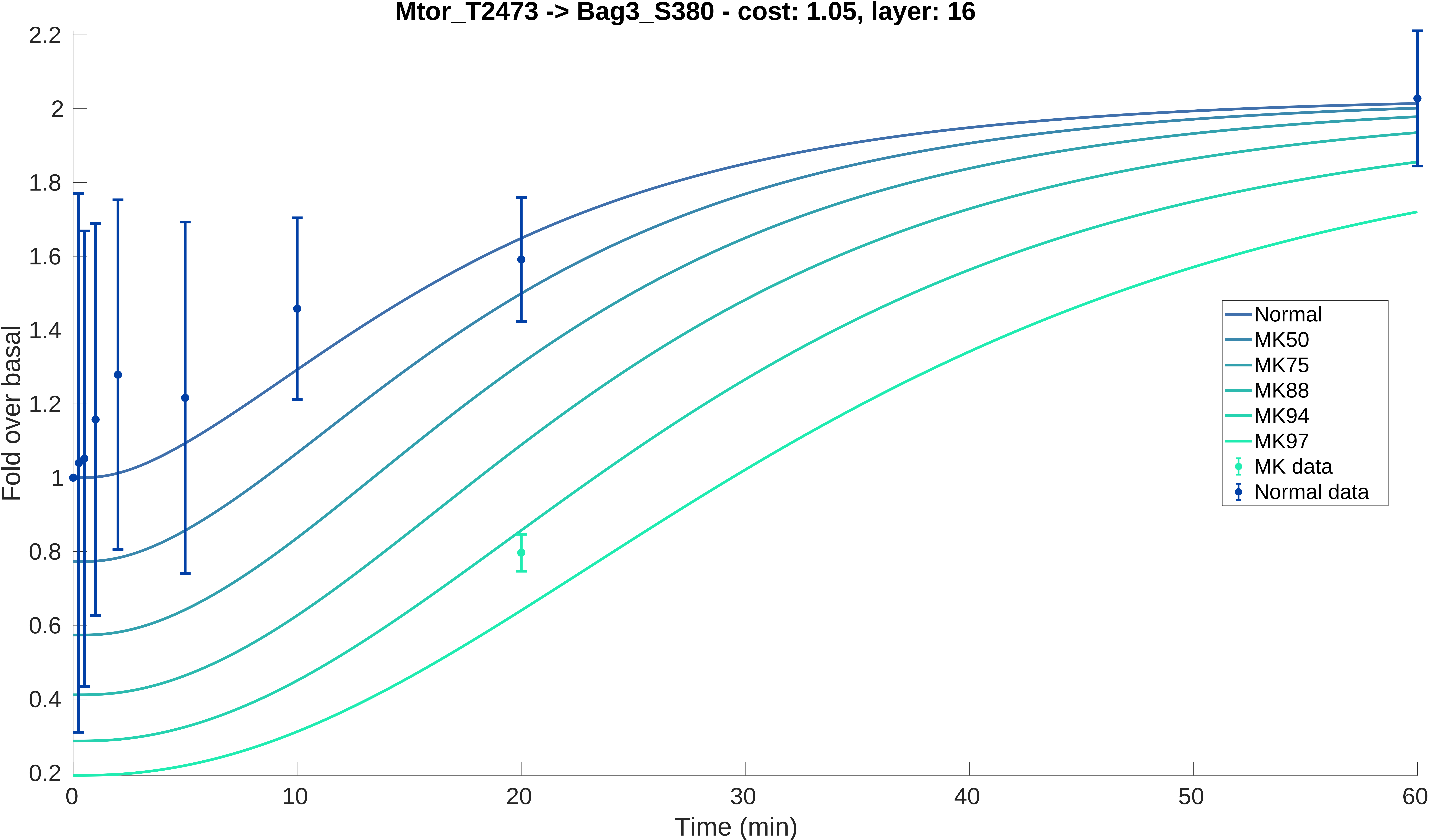

Mtor_T2473 -> Bag3_S380 - cost: 1.05, layer: 16
Fold over basal
Time (min)
0.2
0.4
0.6
0.8
1
1.2
1.4
1.6
1.8
2
2.2
0
10
20
30
40
50
60
Normal
MK50
MK75
MK88
MK94
MK97
MK data
Normal data

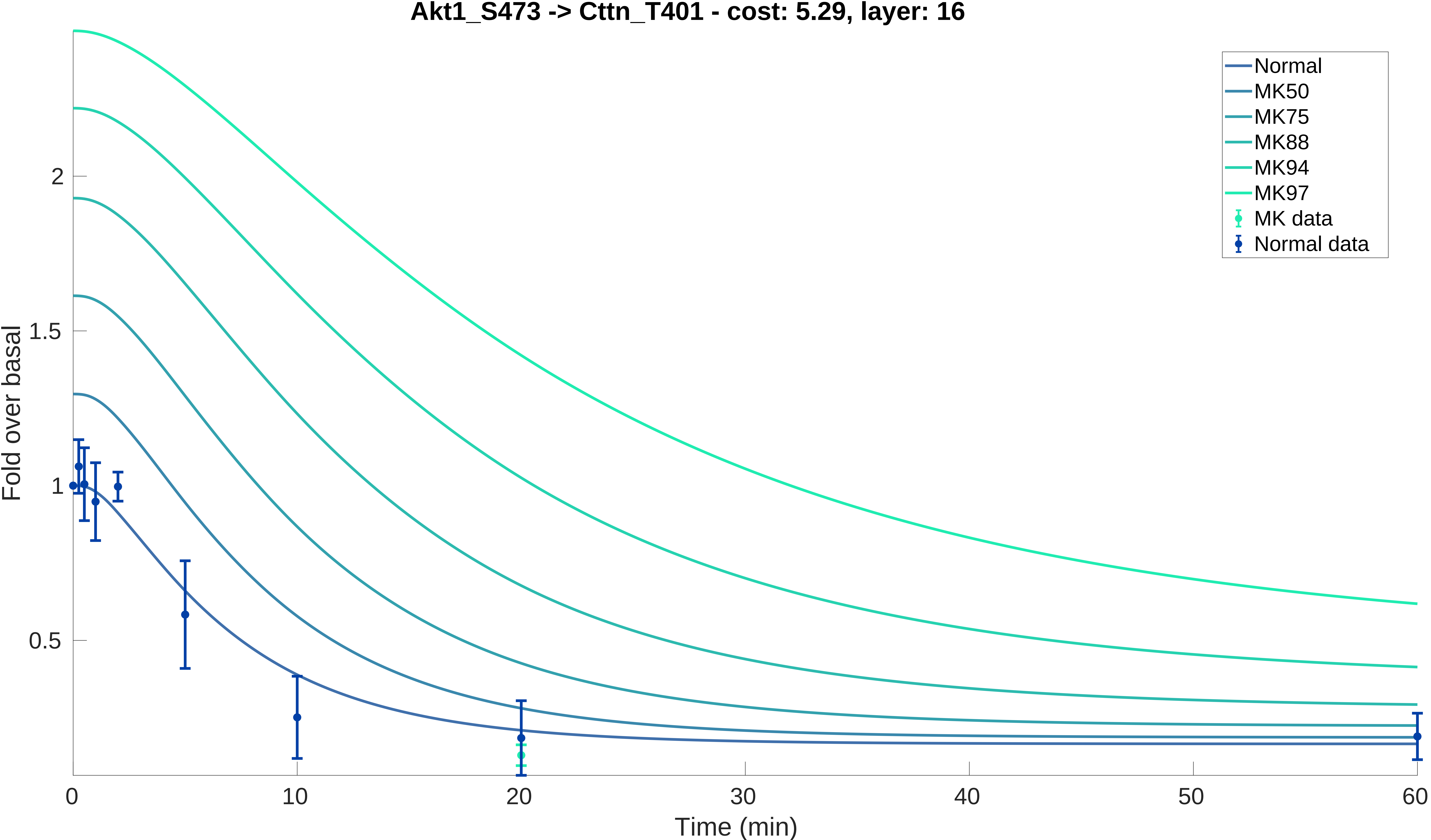

Akt1_S473 -> Cttn_T401 - cost: 5.29, layer: 16
Normal
MK50
MK75
MK88
MK94
MK97
MK data
Normal data
Fold over basal
0.5
1
1.5
2
0
10
20
30
40
50
60
Time (min)

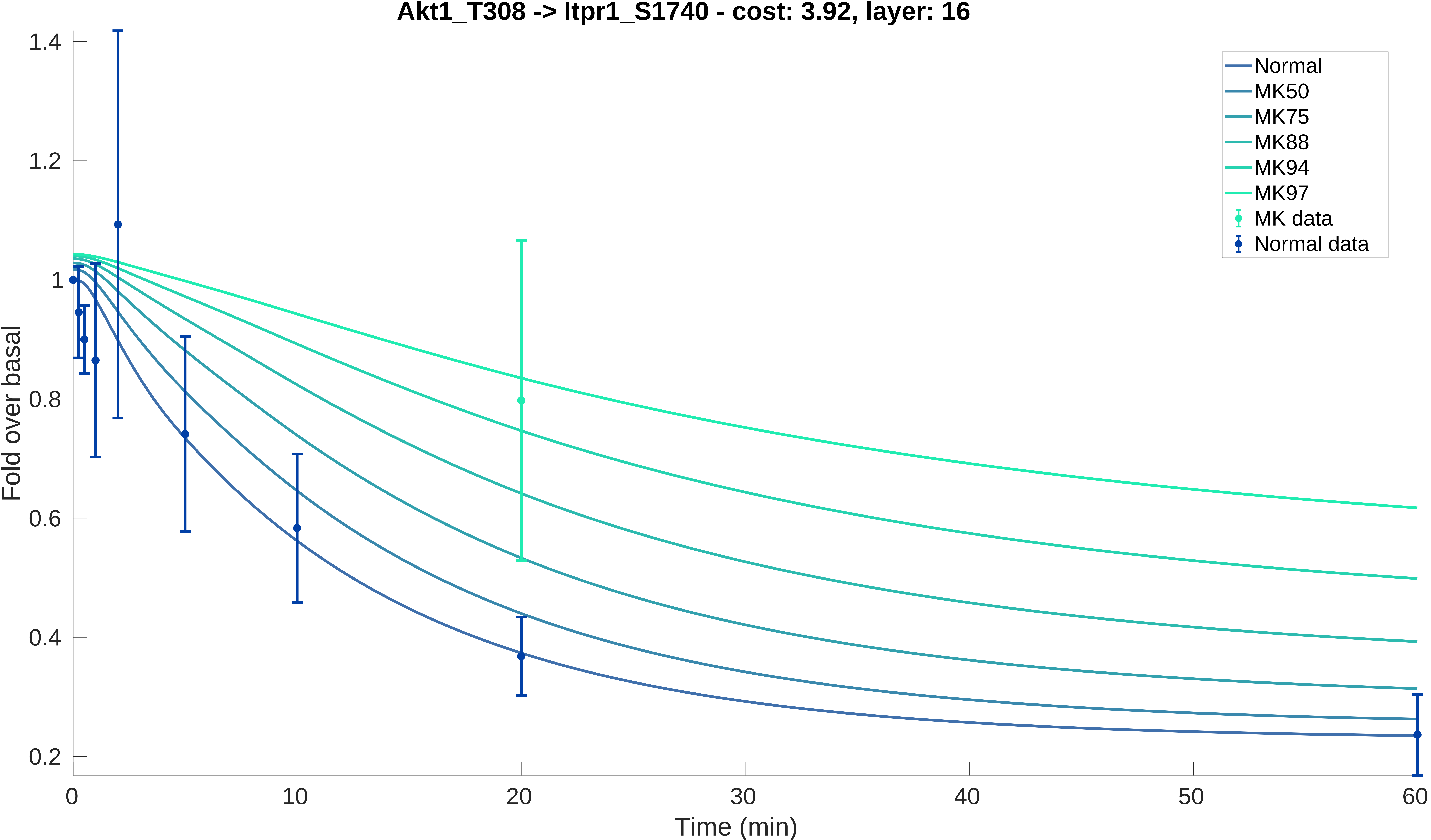

Akt1_T308 -> Itpr1_S1740 - cost: 3.92, layer: 16
1.4
1.2
1
0.8
0.6
0.4
0.2
Fold over basal
0
10
20
30
40
50
60
Time (min)
Normal
MK50
MK75
MK88
MK94
MK97
MK data
Normal data

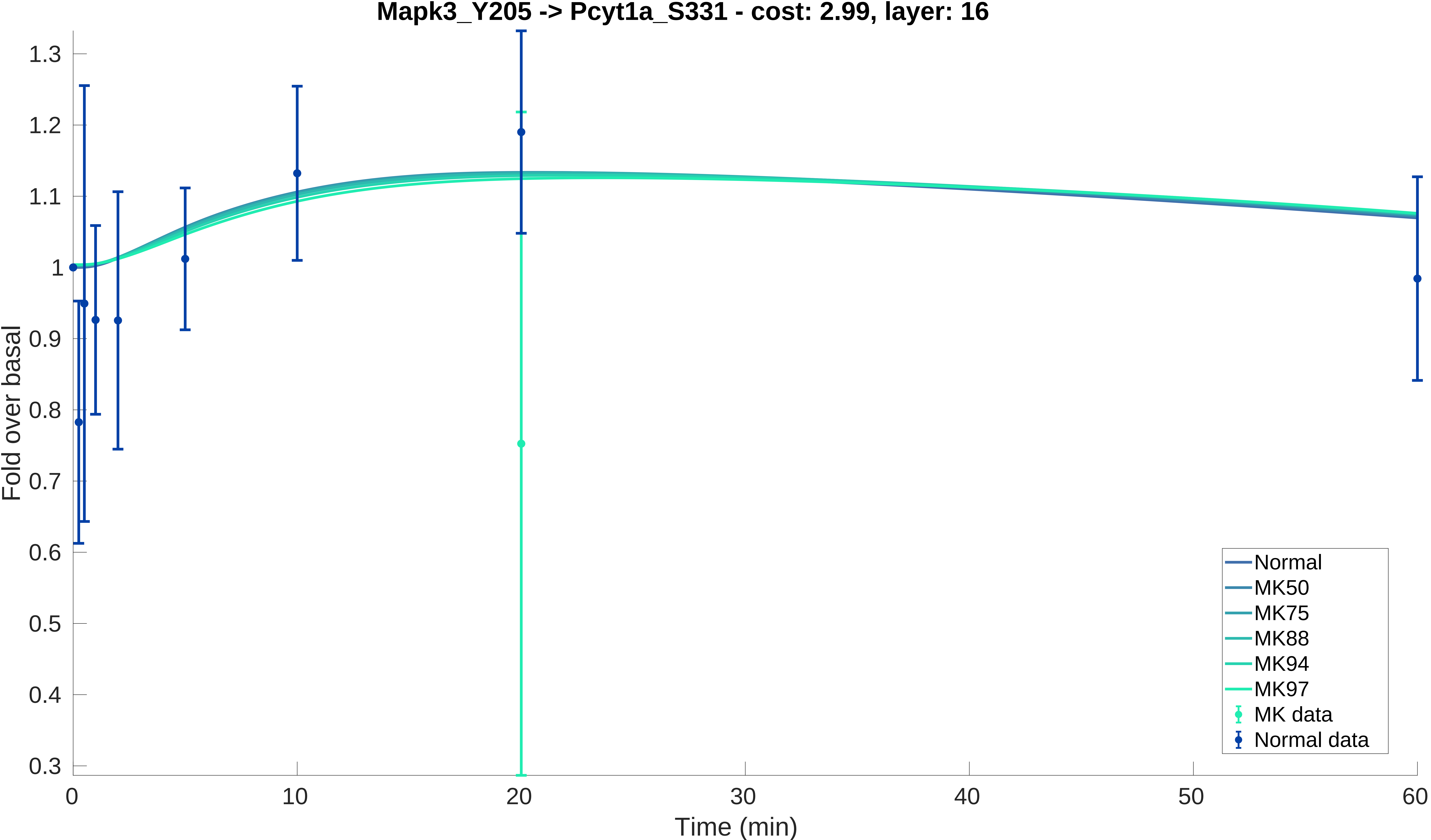
Mapk3_Y205 -> Pcyt1a_S331 - cost: 2.99, layer: 16
Fold over basal
1.3
1.2
1.1
1
0.9
0.8
0.7
0.6
0.5
0.4
0.3
0
10
20
30
40
50
60
Time (min)
Normal
MK50
MK75
MK88
MK94
MK97
MK data
Normal data

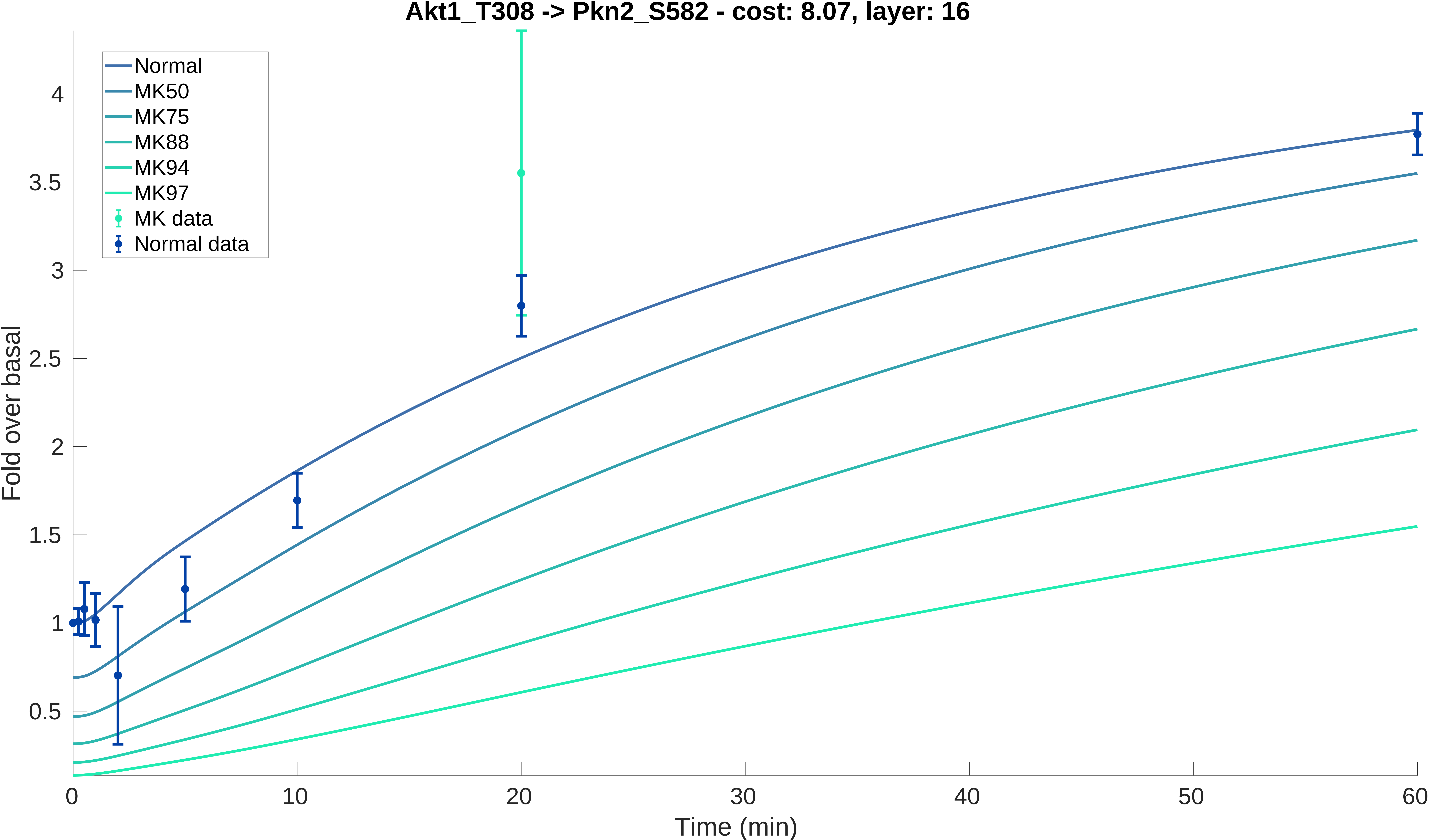
Akt1_T308 -> Pkn2_S582 - cost: 8.07, layer: 16
Normal
MK50
MK75
MK88
MK94
MK97
MK data
Normal data
Fold over basal
Time (min)
0.5
1
1.5
2
2.5
3
3.5
4
0
10
20
30
40
50
60

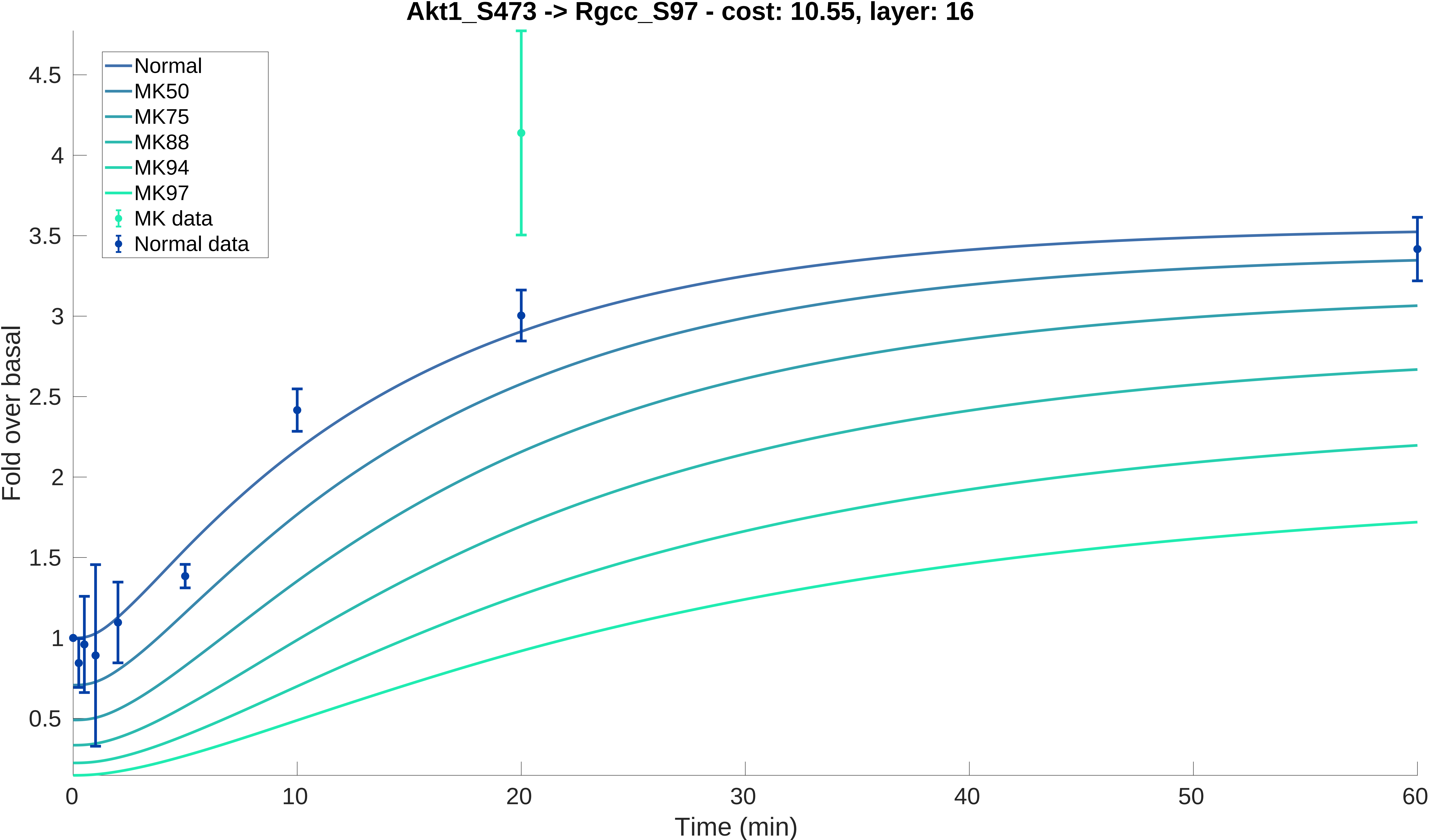

Akt1_S473 -> Rgcc_S97 - cost: 10.55, layer: 16
Normal
MK50
MK75
MK88
MK94
MK97
MK data
Normal data
Fold over basal
4.5
4
3.5
3
2.5
2
1.5
1
0.5
0
10
20
30
40
50
60
Time (min)

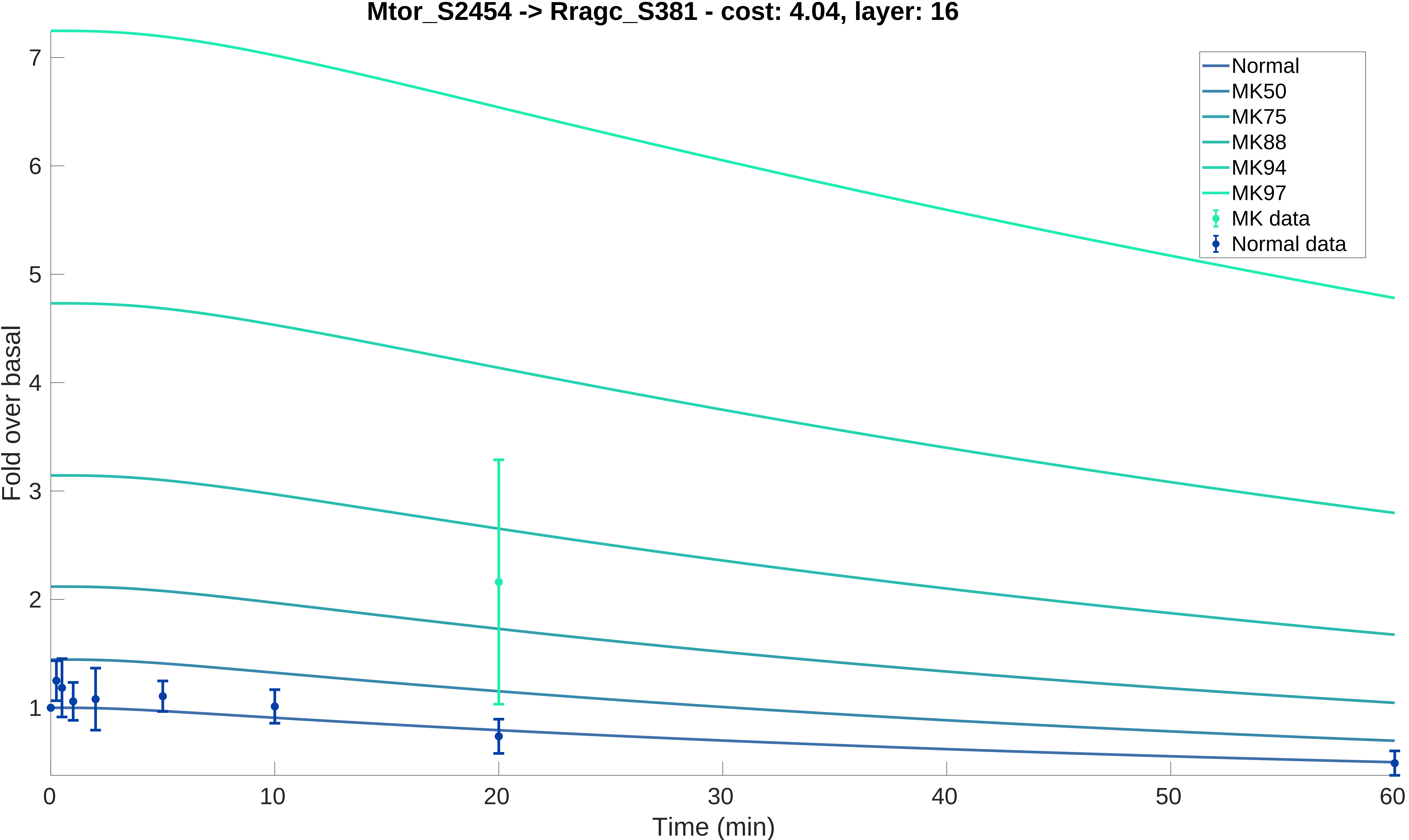

Mtor_S2454 -> Rragc_S381 - cost: 4.04, layer: 16
Fold over basal
Time (min)
0
10
20
30
40
50
60
1
2
3
4
5
6
7
Normal
MK50
MK75
MK88
MK94
MK97
MK data
Normal data

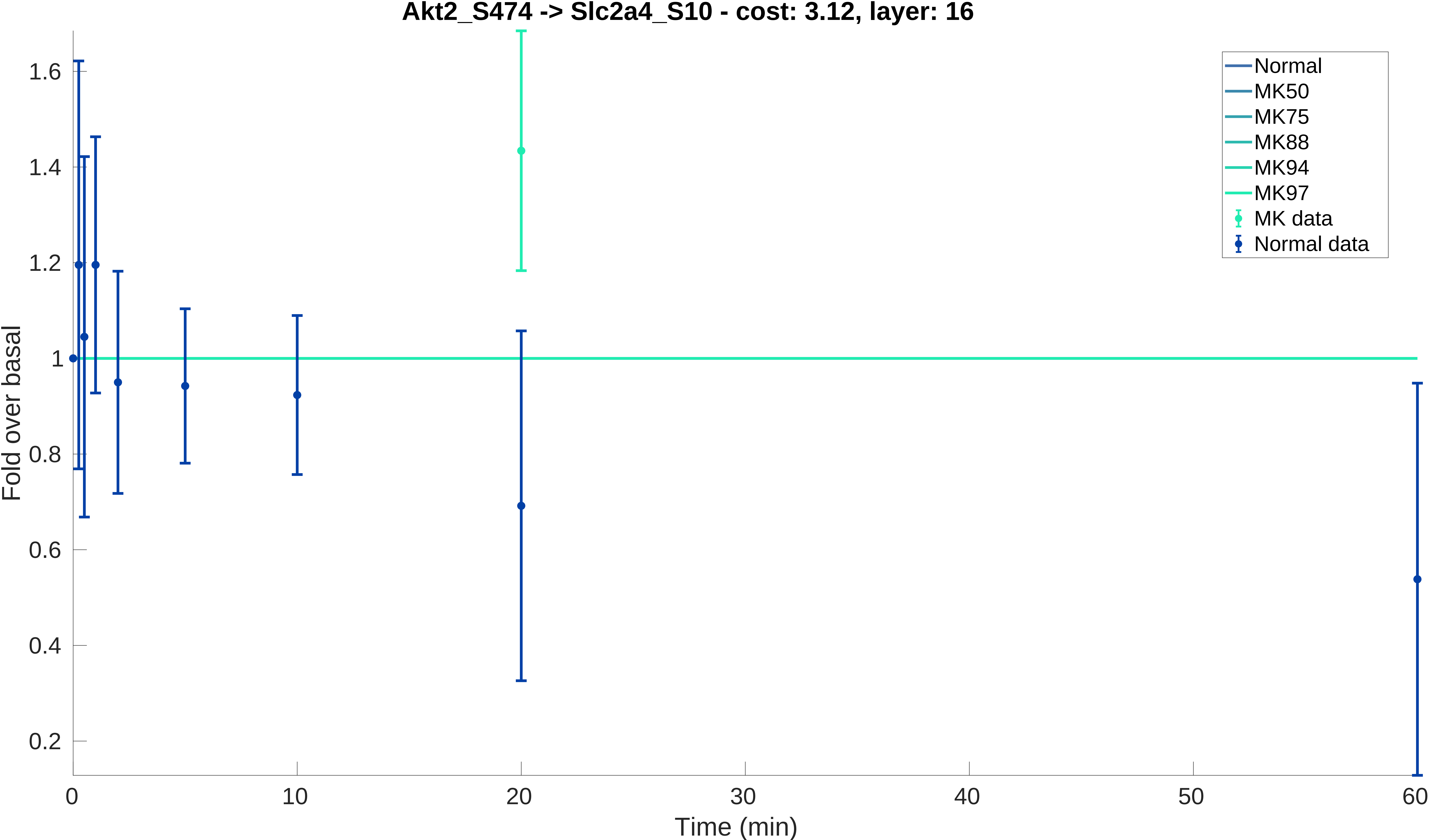

Akt2_S474 -> Slc2a4_S10 - cost: 3.12, layer: 16
Normal
MK50
MK75
MK88
MK94
MK97
MK data
Normal data
Fold over basal
1.6
1.4
1.2
1
0.8
0.6
0.4
0.2
0
10
20
30
40
50
60
Time (min)

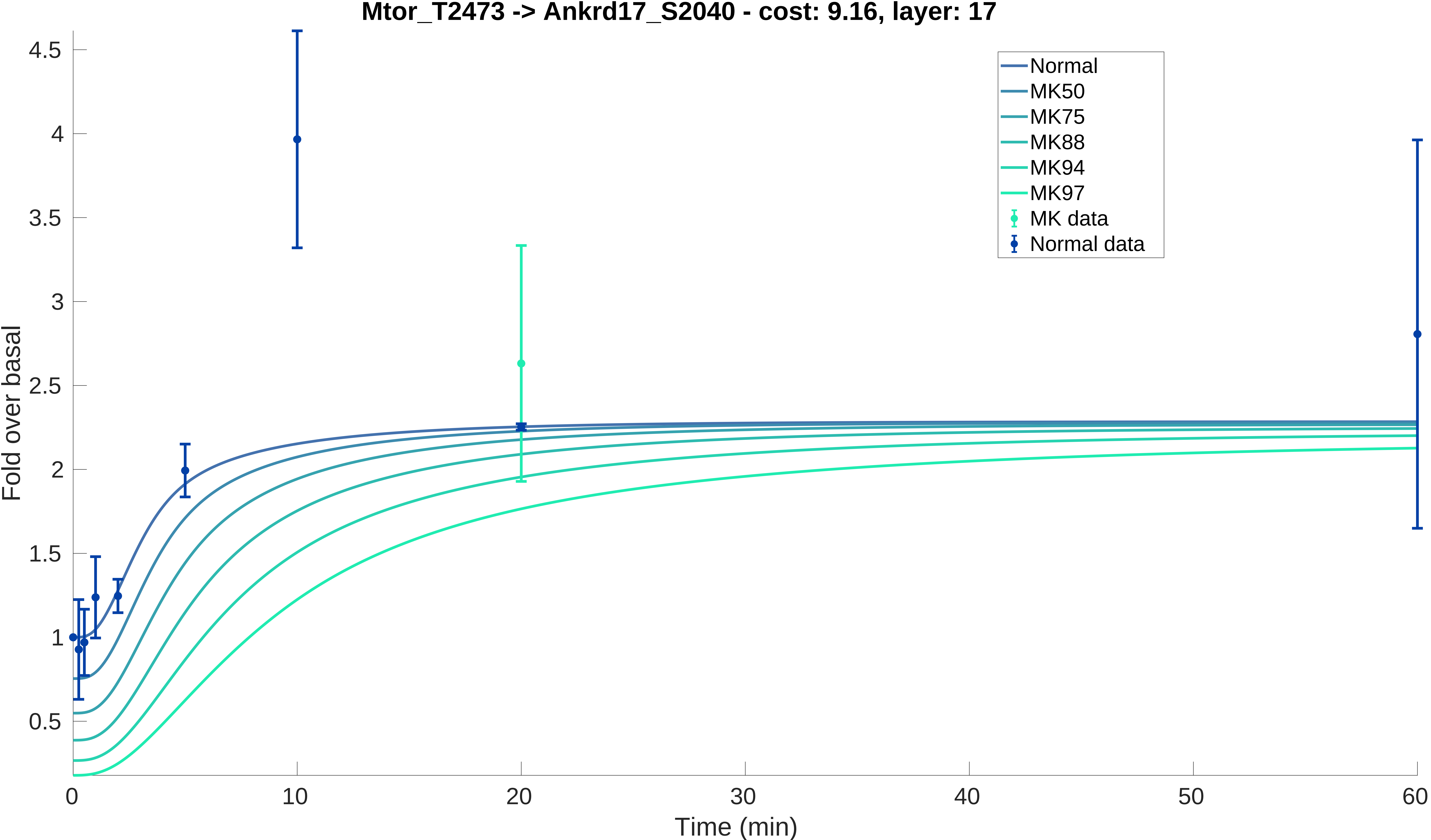

Mtor_T2473 -> Ankrd17_S2040 - cost: 9.16, layer: 17
Normal
MK50
MK75
MK88
MK94
MK97
MK data
Normal data
Fold over basal
Time (min)
0.5
1
1.5
2
2.5
3
3.5
4
4.5
0
10
20
30
40
50
60

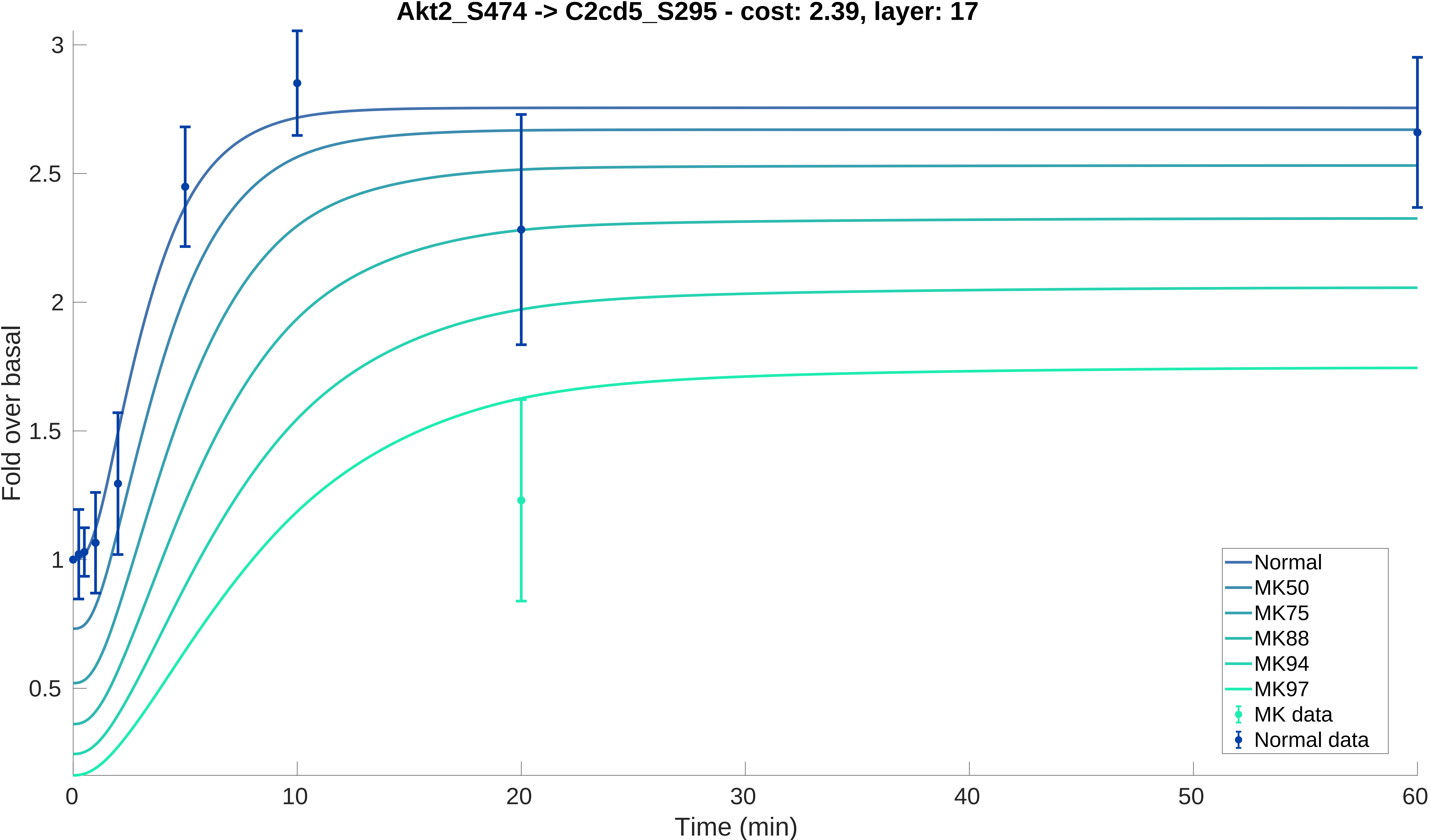

Akt2_S474 -> C2cd5_S295 - cost: 2.39, layer: 17
Fold over basal
Time (min)
0
10
20
30
40
50
60
0.5
1
1.5
2
2.5
3
Normal
MK50
MK75
MK88
MK94
MK97
MK data
Normal data

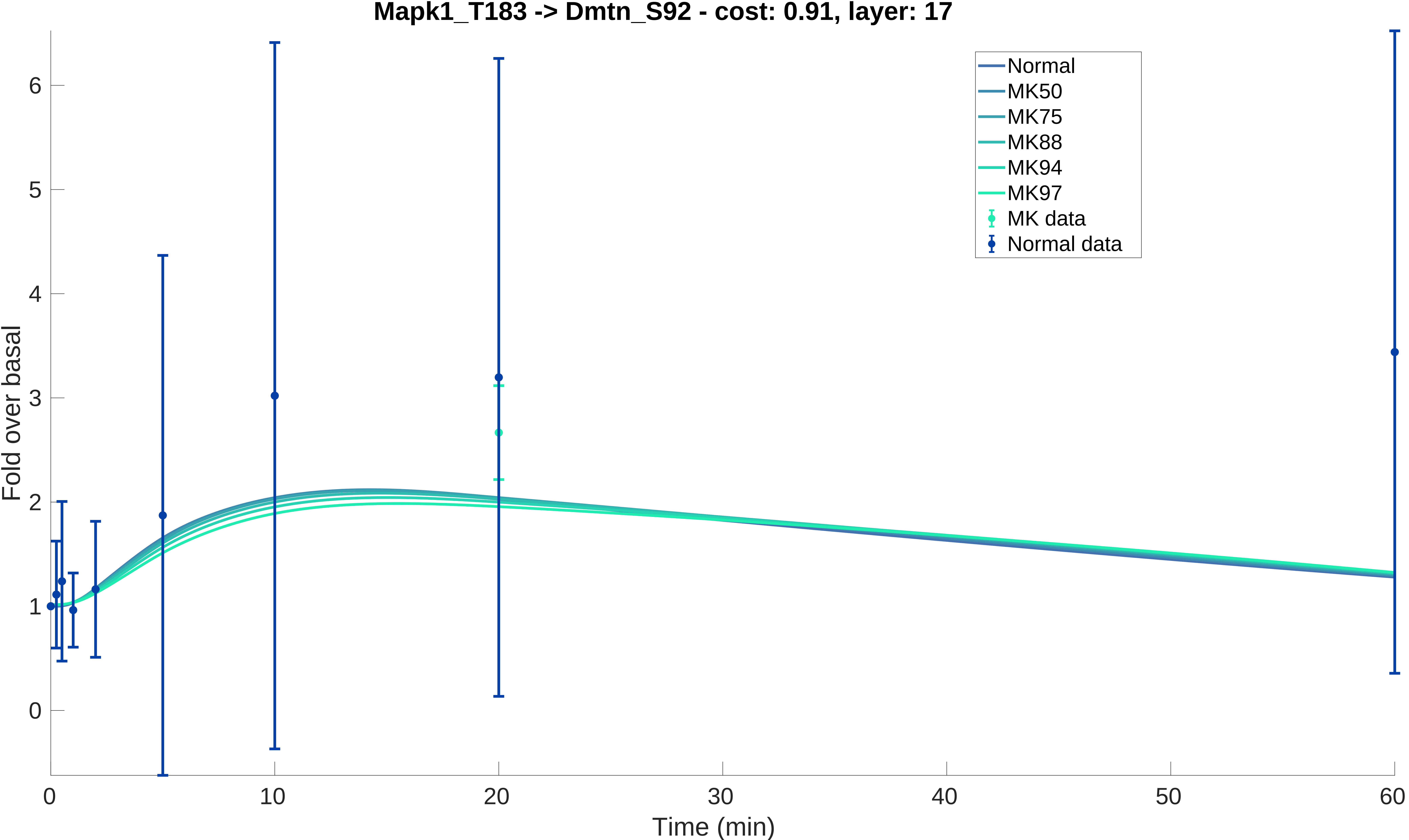

Mapk1_T183 -> Dmtn_S92 - cost: 0.91, layer: 17
Fold over basal
Time (min)
Normal
MK50
MK75
MK88
MK94
MK97
MK data
Normal data
0
1
2
3
4
5
6
0
10
20
30
40
50
60

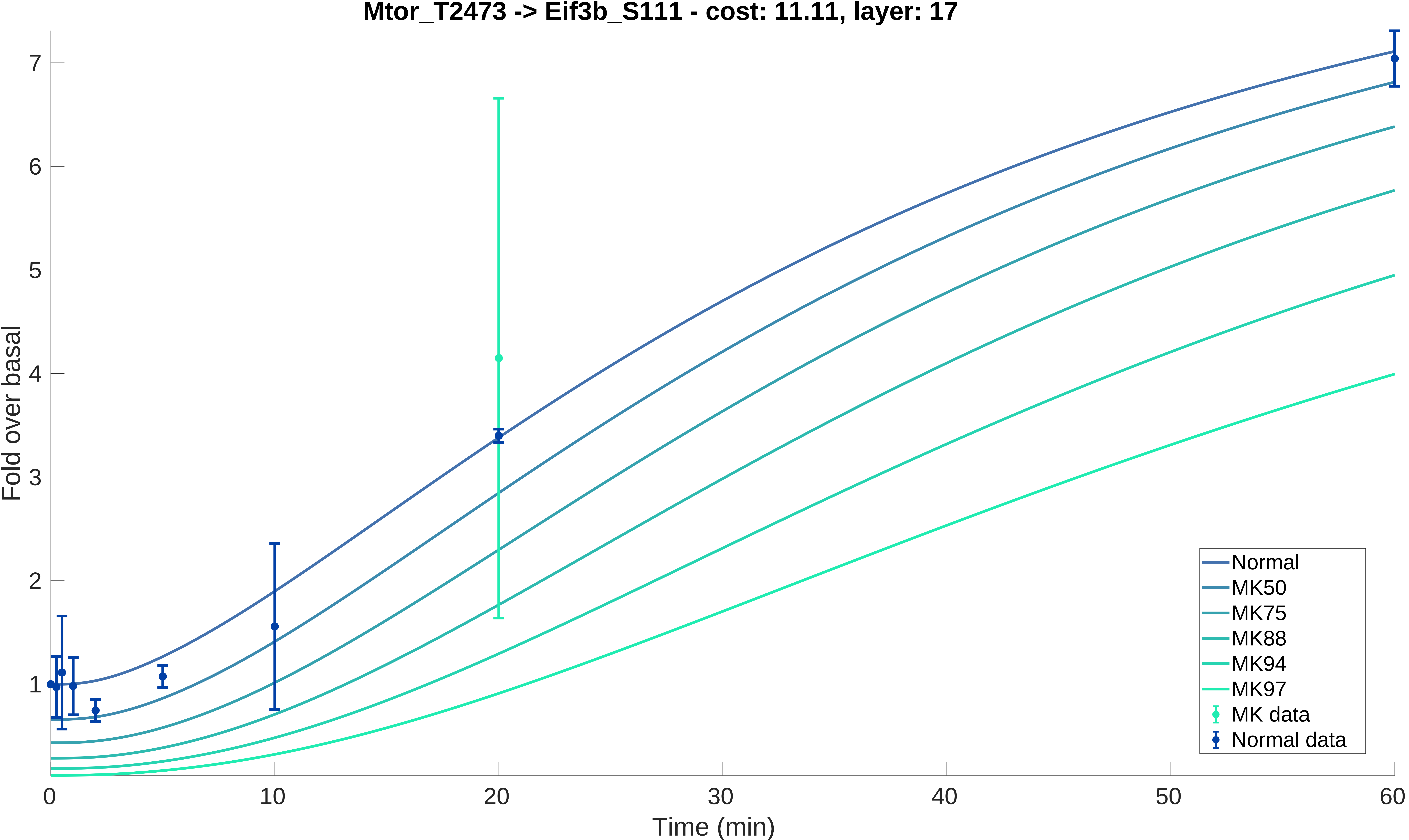

Mtor_T2473 -> Eif3b_S111 - cost: 11.11, layer: 17
Fold over basal
Time (min)
0
10
20
30
40
50
60
1
2
3
4
5
6
7
Normal
MK50
MK75
MK88
MK94
MK97
MK data
Normal data

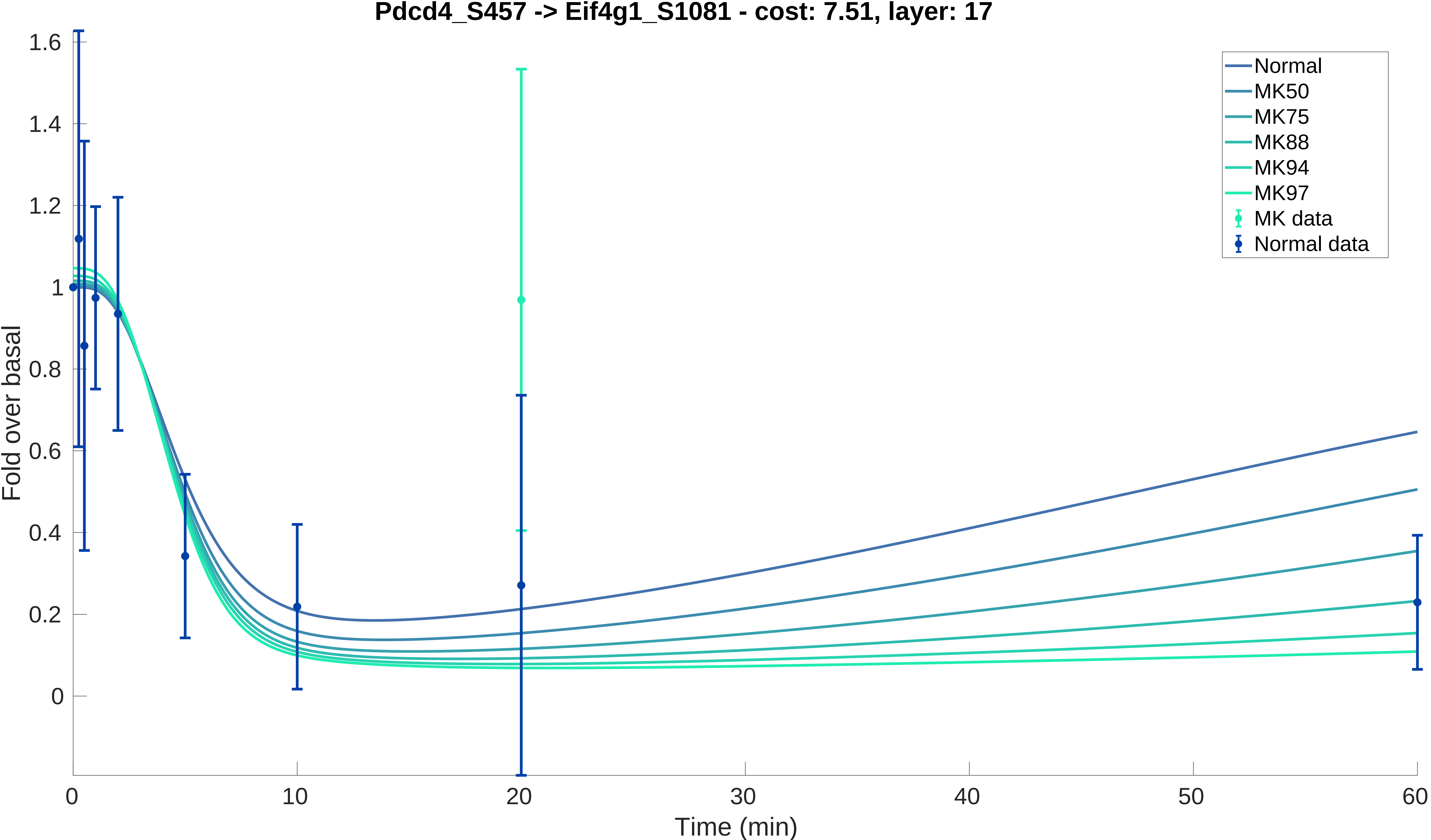

Pdcd4_S457 -> Eif4g1_S1081 - cost: 7.51, layer: 17
1.6
1.4
1.2
1
0.8
0.6
0.4
0.2
0
Fold over basal
0
10
20
30
40
50
60
Time (min)
Normal
MK50
MK75
MK88
MK94
MK97
MK data
Normal data

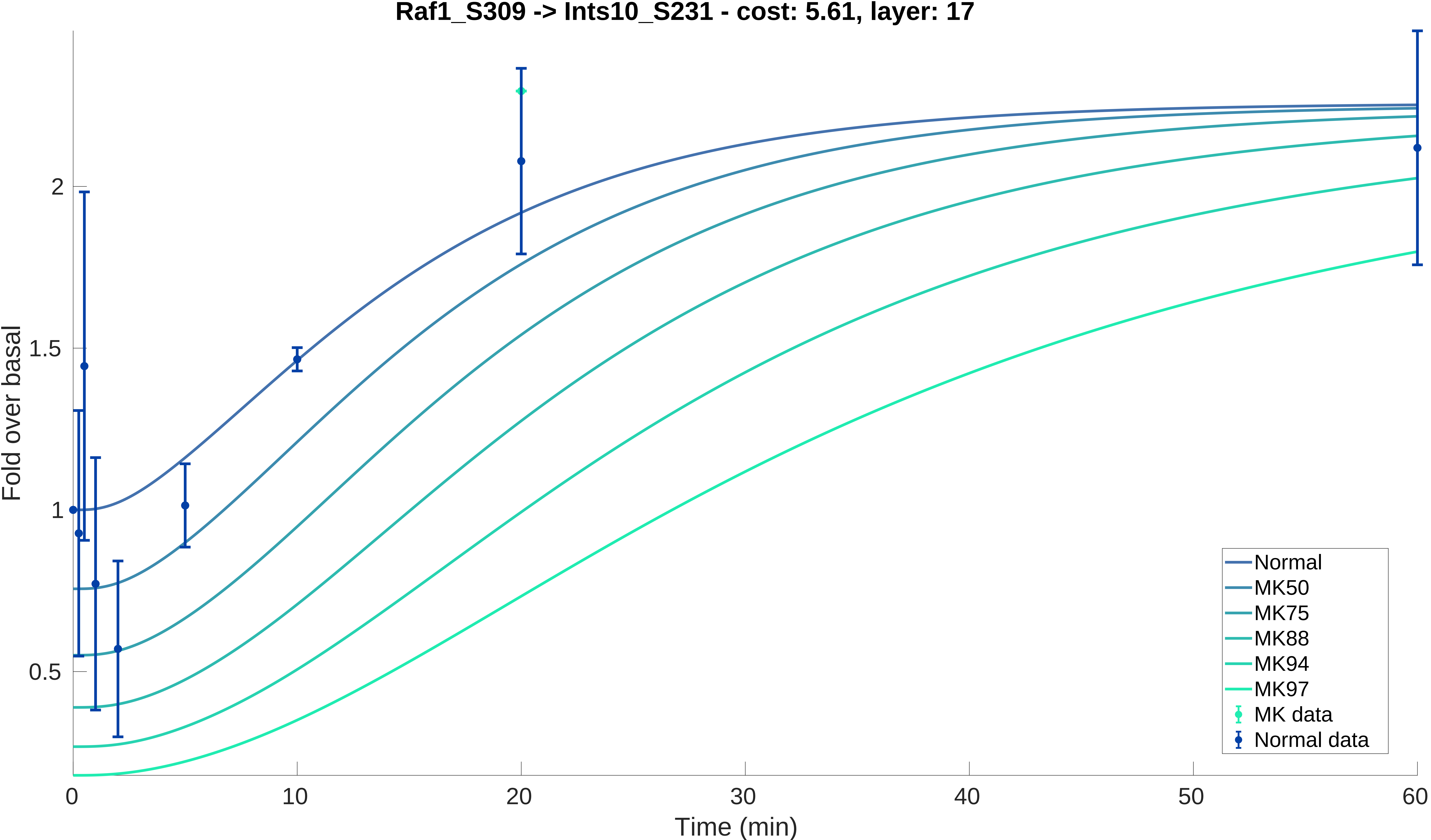
Raf1_S309 -> Ints10_S231 - cost: 5.61, layer: 17
Fold over basal
2
1.5
1
0.5
0
10
20
30
40
50
60
Time (min)
Normal
MK50
MK75
MK88
MK94
MK97
MK data
Normal data

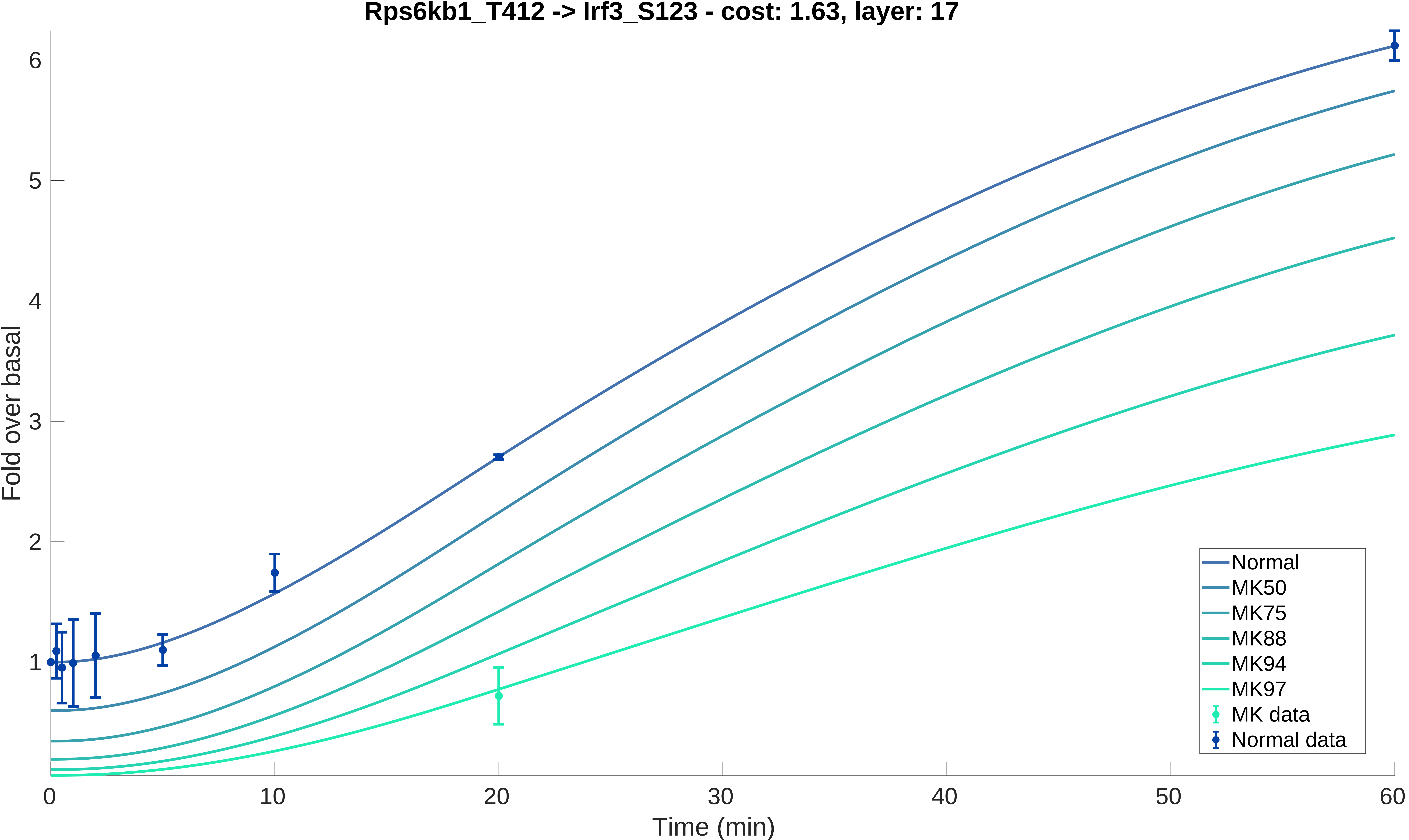
Rps6kb1_T412 -> Irf3_S123 - cost: 1.63, layer: 17
Fold over basal
Time (min)
0
10
20
30
40
50
60
1
2
3
4
5
6
Normal
MK50
MK75
MK88
MK94
MK97
MK data
Normal data

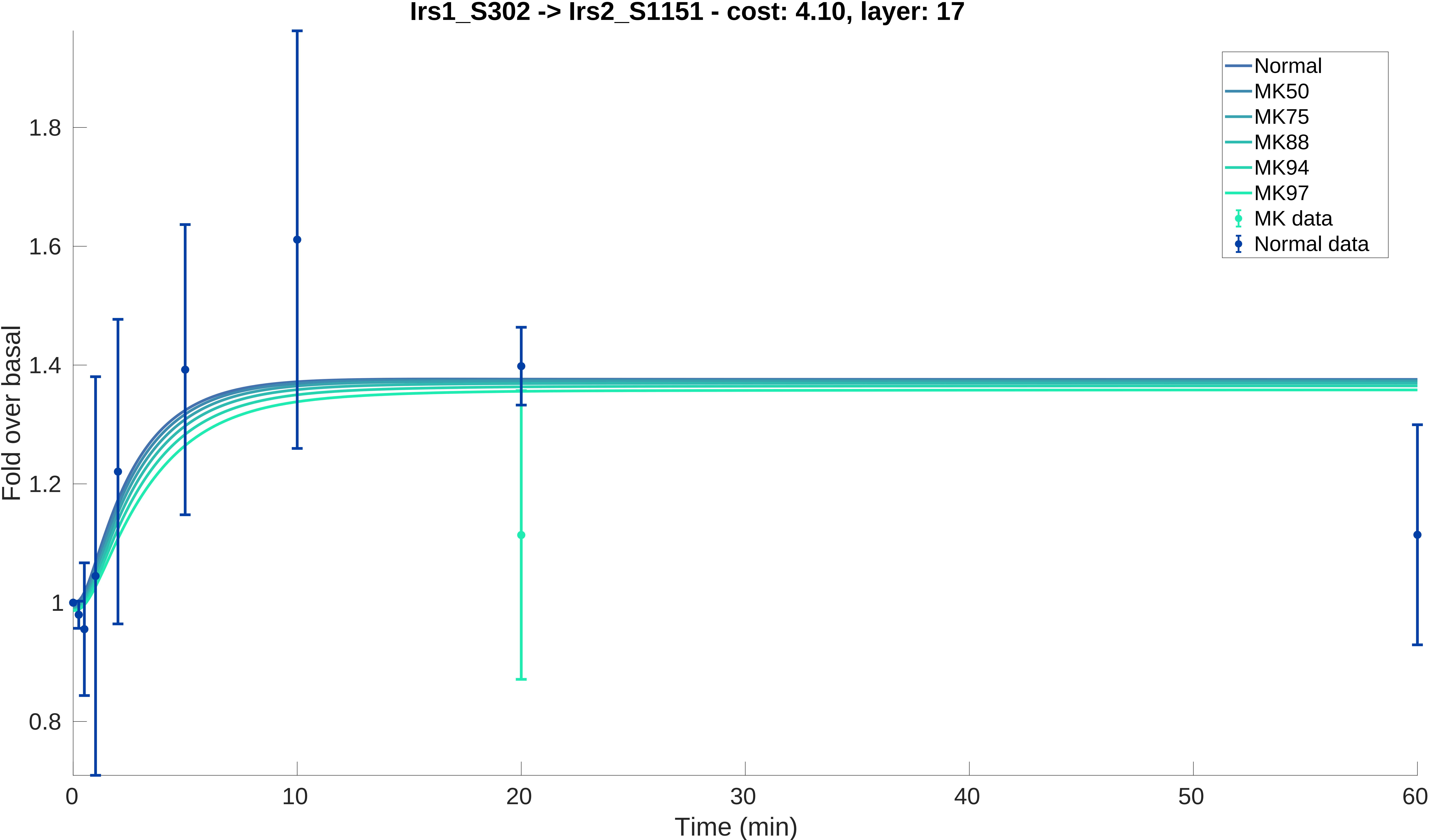
Irs1_S302 -> Irs2_S1151 - cost: 4.10, layer: 17
Fold over basal
Time (min)
0.8
1
1.2
1.4
1.6
1.8
0
10
20
30
40
50
60
Normal
MK50
MK75
MK88
MK94
MK97
MK data
Normal data

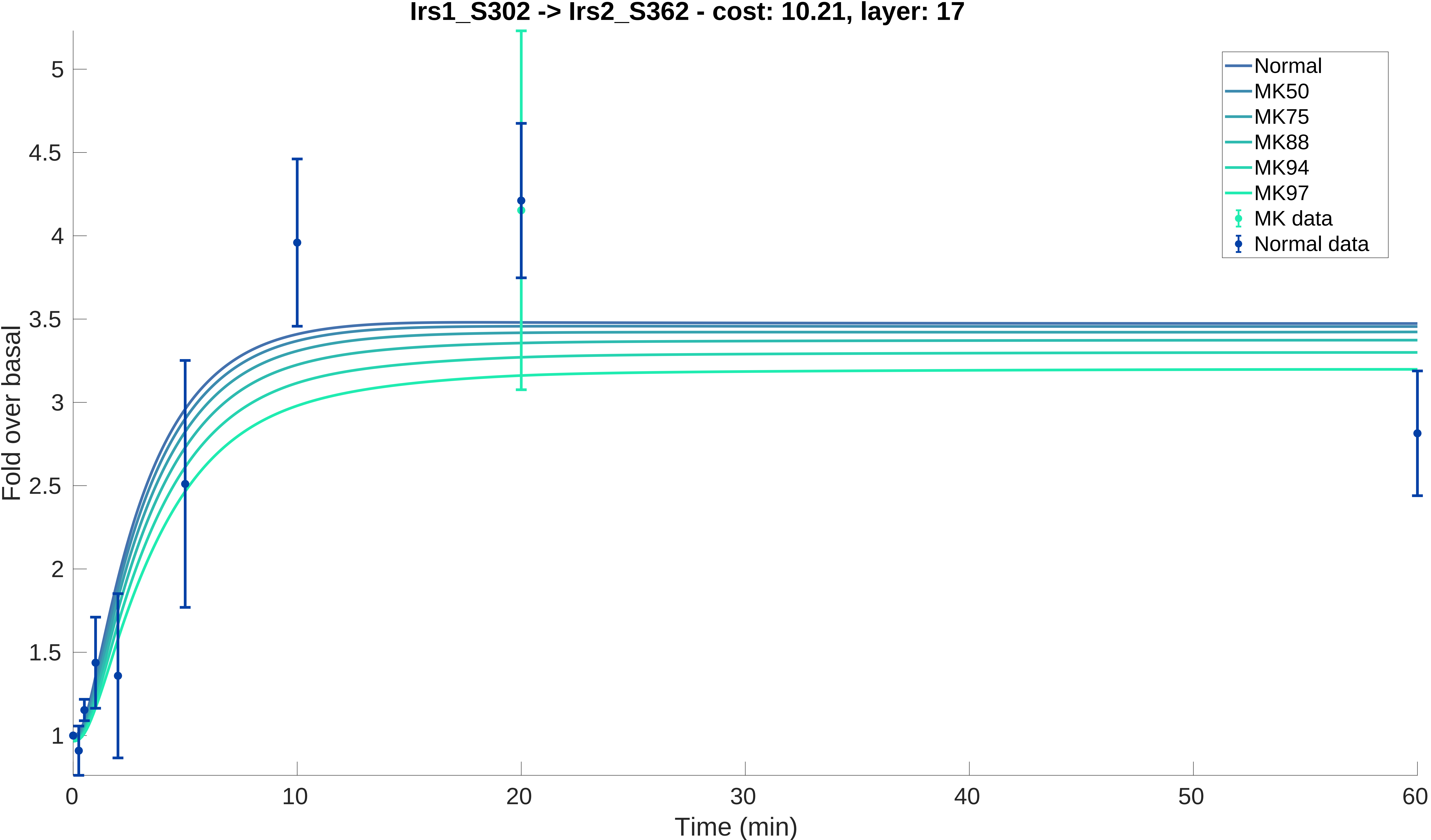

Irs1_S302 -> Irs2_S362 - cost: 10.21, layer: 17
Normal
MK50
MK75
MK88
MK94
MK97
MK data
Normal data
Fold over basal
Time (min)
5
4.5
4
3.5
3
2.5
2
1.5
1
0
10
20
30
40
50
60

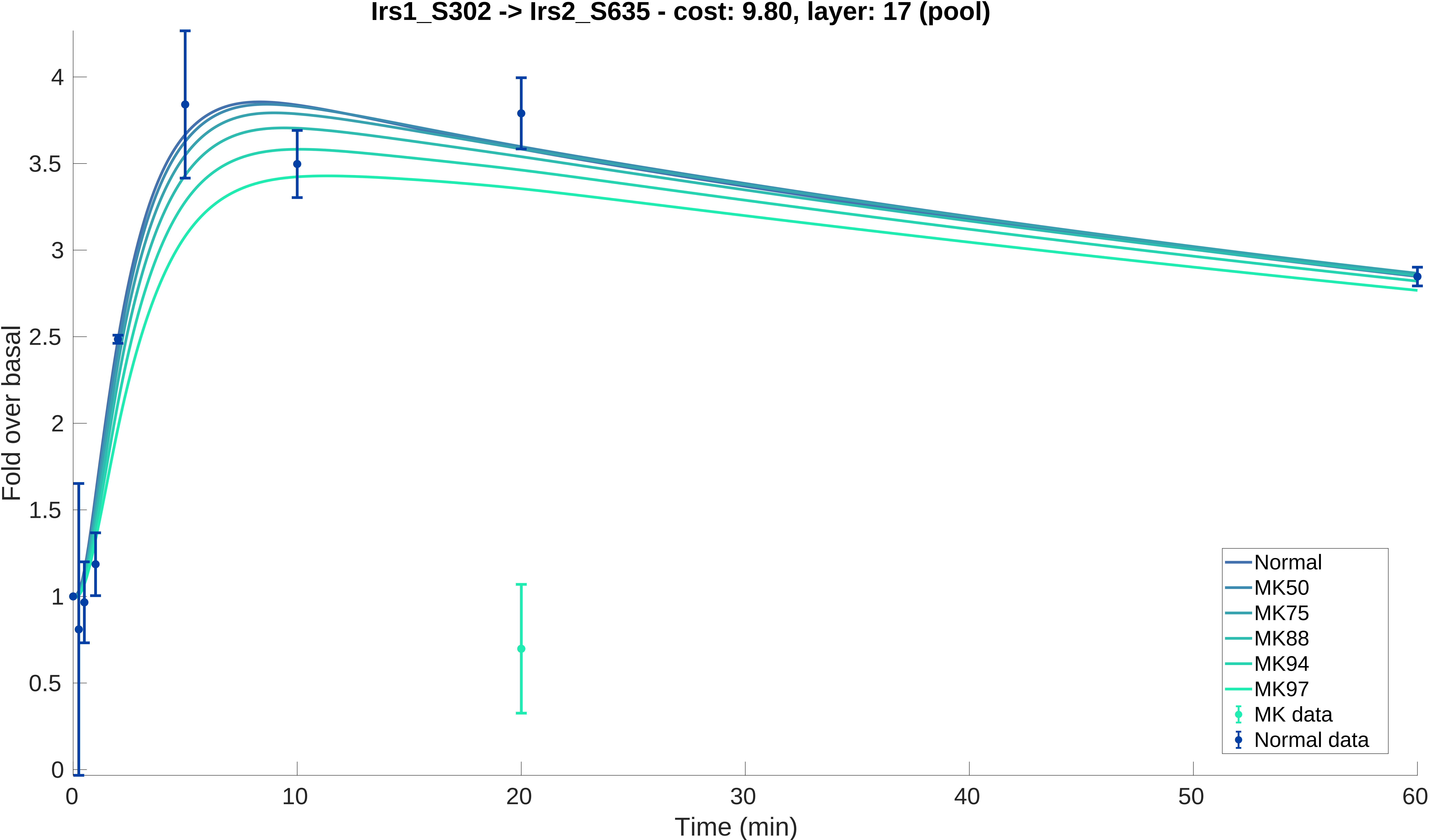
Irs1_S302 -> Irs2_S635 - cost: 9.80, layer: 17 (pool)
Fold over basal
Time (min)
0
0.5
1
1.5
2
2.5
3
3.5
4
0
10
20
30
40
50
60
Normal
MK50
MK75
MK88
MK94
MK97
MK data
Normal data

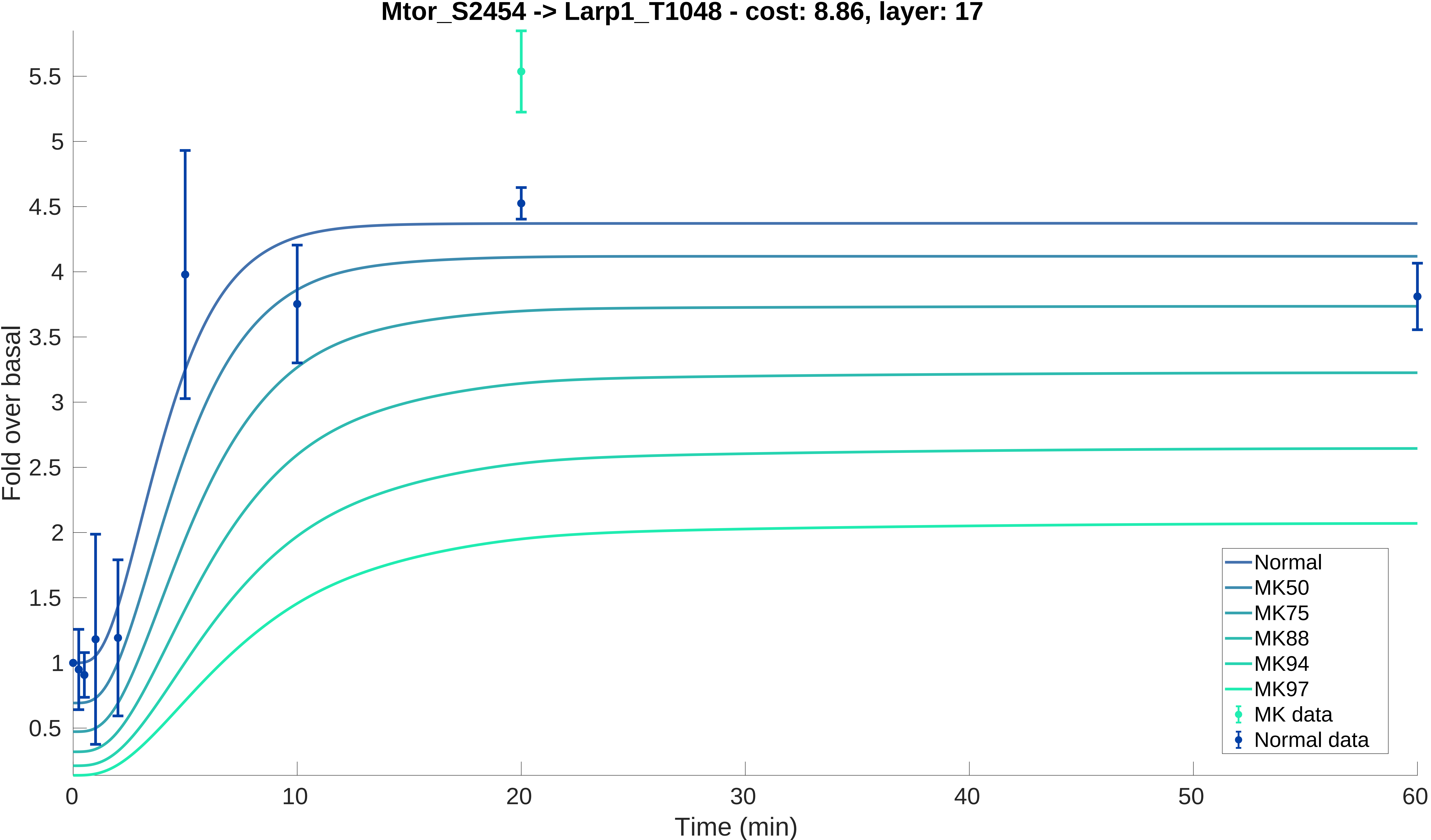
Mtor_S2454 -> Larp1_T1048 - cost: 8.86, layer: 17
Fold over basal
Time (min)
0
10
20
30
40
50
60
0.5
1
1.5
2
2.5
3
3.5
4
4.5
5
5.5
Normal
MK50
MK75
MK88
MK94
MK97
MK data
Normal data

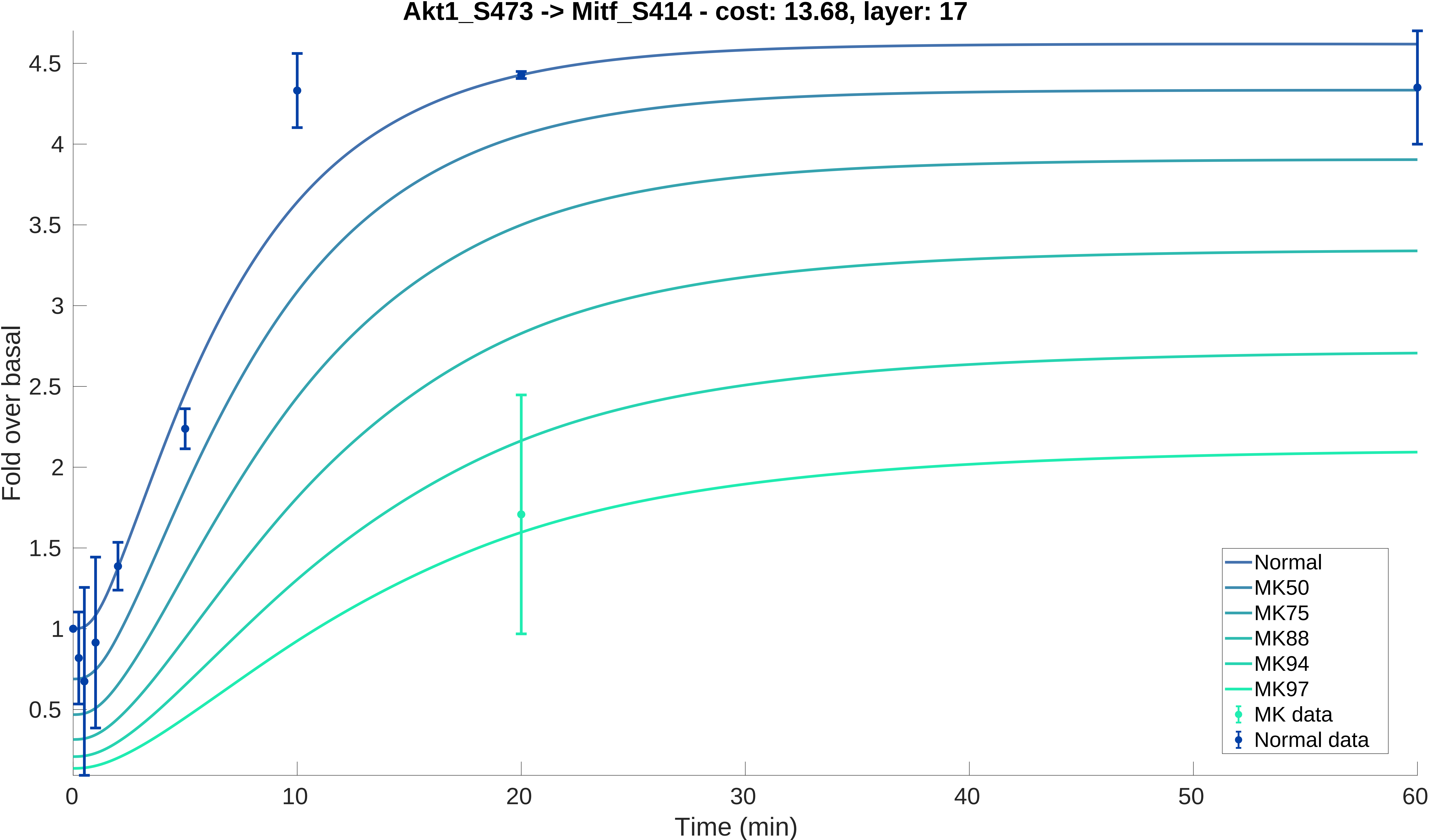
Akt1_S473 -> Mitf_S414 - cost: 13.68, layer: 17
Fold over basal
Time (min)
0.5
1
1.5
2
2.5
3
3.5
4
4.5
0
10
20
30
40
50
60
Normal
MK50
MK75
MK88
MK94
MK97
MK data
Normal data

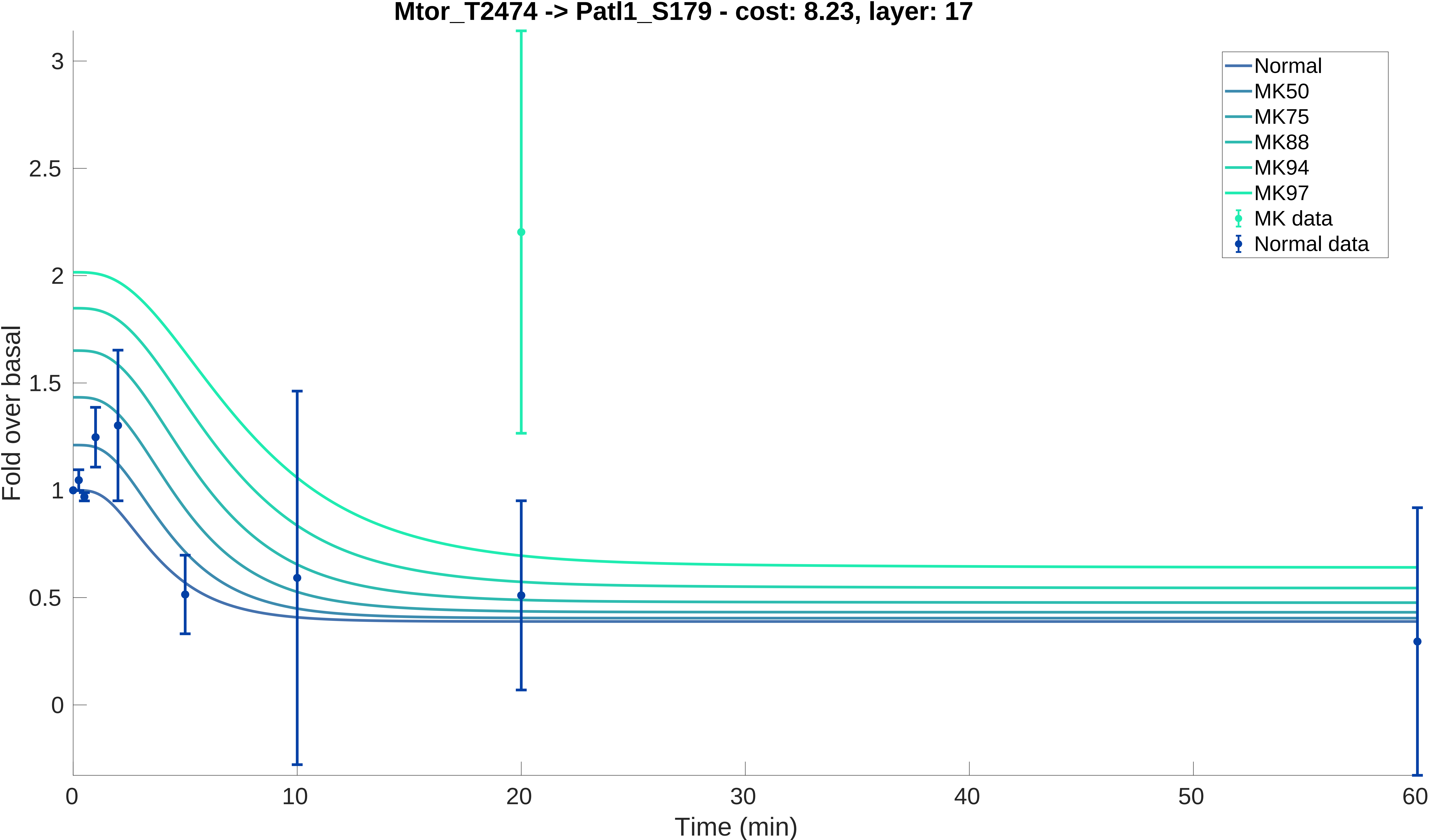

Mtor_T2474 -> Patl1_S179 - cost: 8.23, layer: 17
Fold over basal
Time (min)
0
0.5
1
1.5
2
2.5
3
0
10
20
30
40
50
60
Normal
MK50
MK75
MK88
MK94
MK97
MK data
Normal data

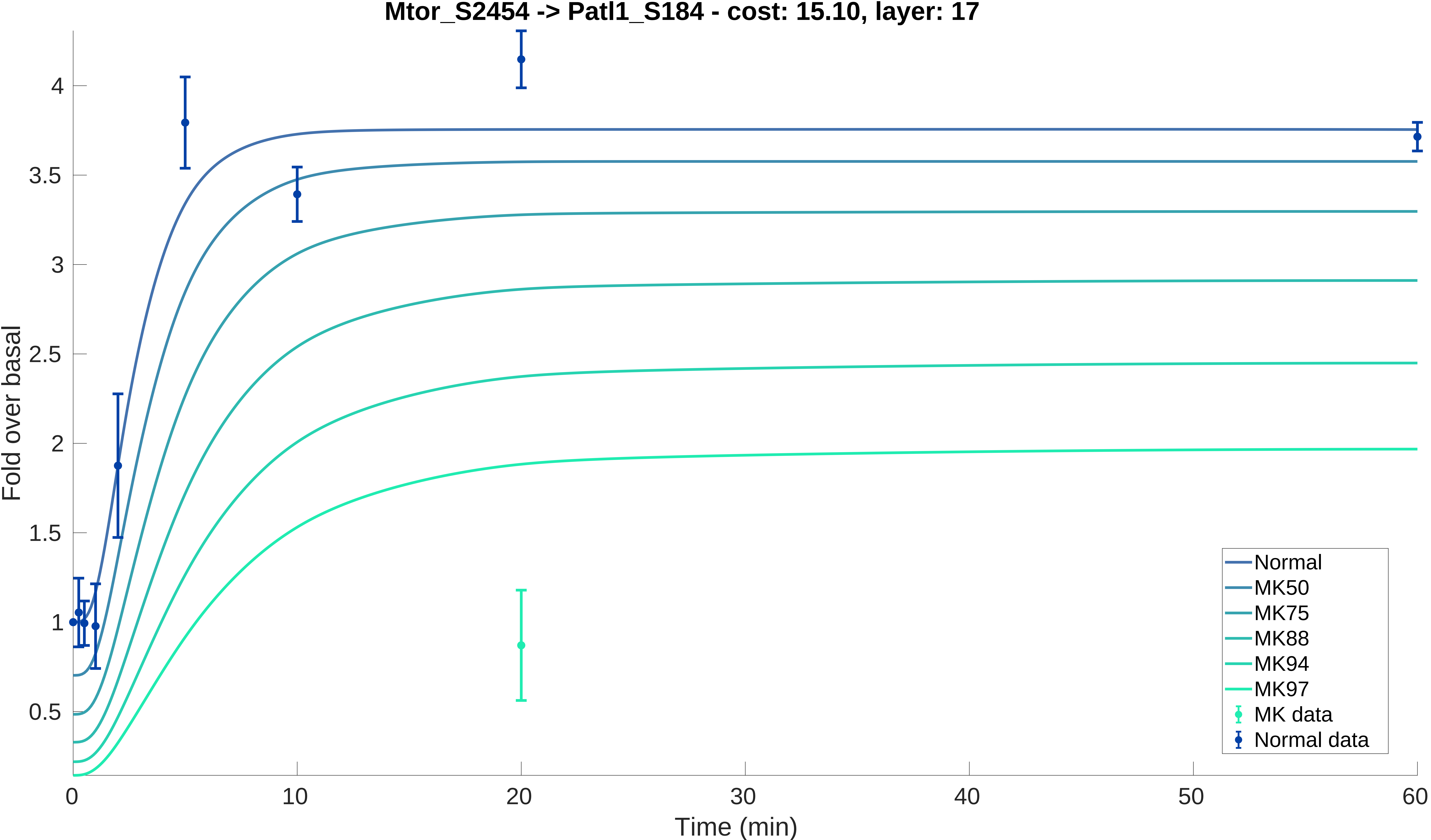
Mtor_S2454 -> Patl1_S184 - cost: 15.10, layer: 17
Fold over basal
4
3.5
3
2.5
2
1.5
1
0.5
0
10
20
30
40
50
60
Time (min)
Normal
MK50
MK75
MK88
MK94
MK97
MK data
Normal data

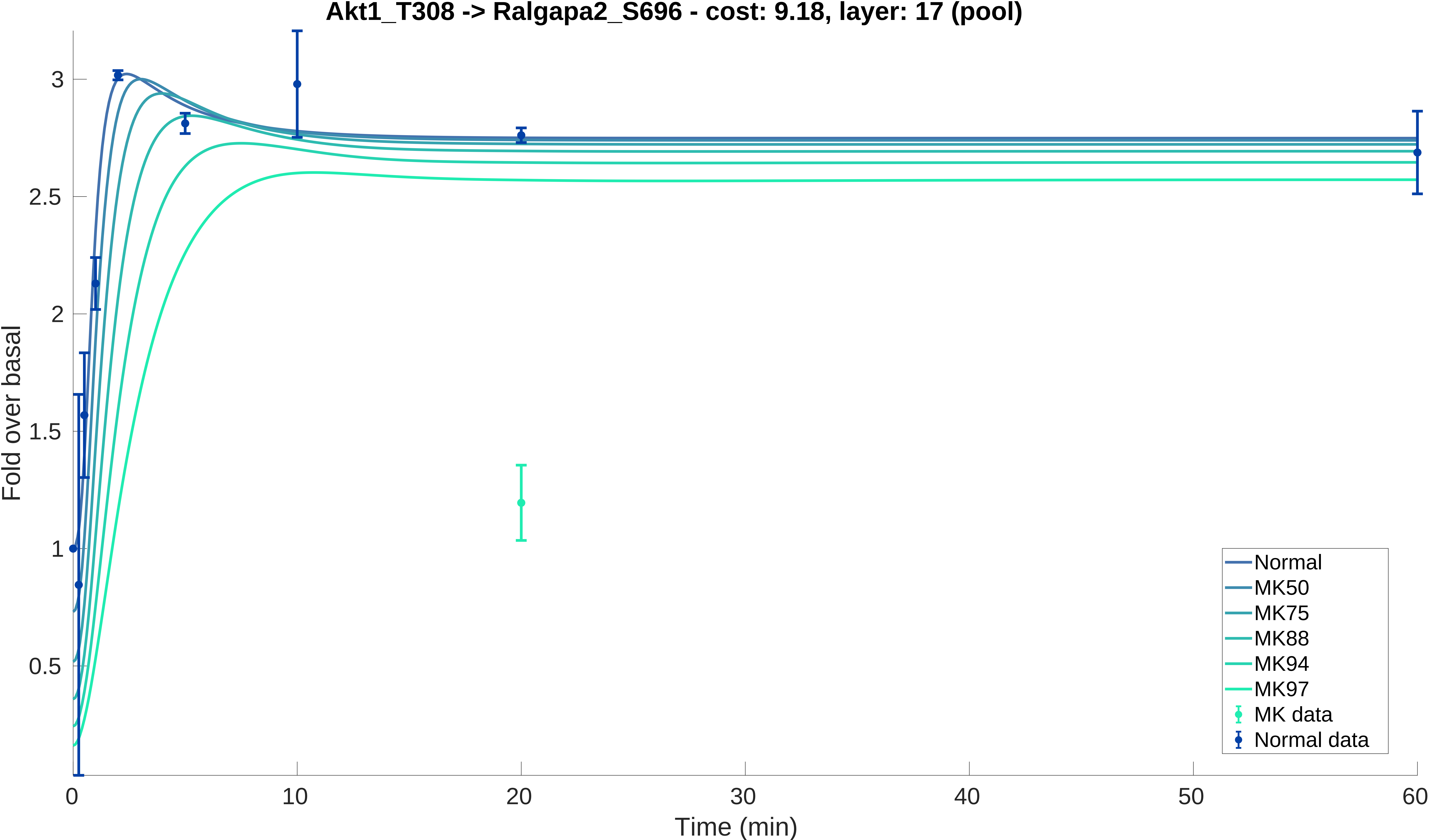

Akt1_T308 -> Ralgapa2_S696 - cost: 9.18, layer: 17 (pool)
Fold over basal
Time (min)
0.5
1
1.5
2
2.5
3
0
10
20
30
40
50
60
Normal
MK50
MK75
MK88
MK94
MK97
MK data
Normal data

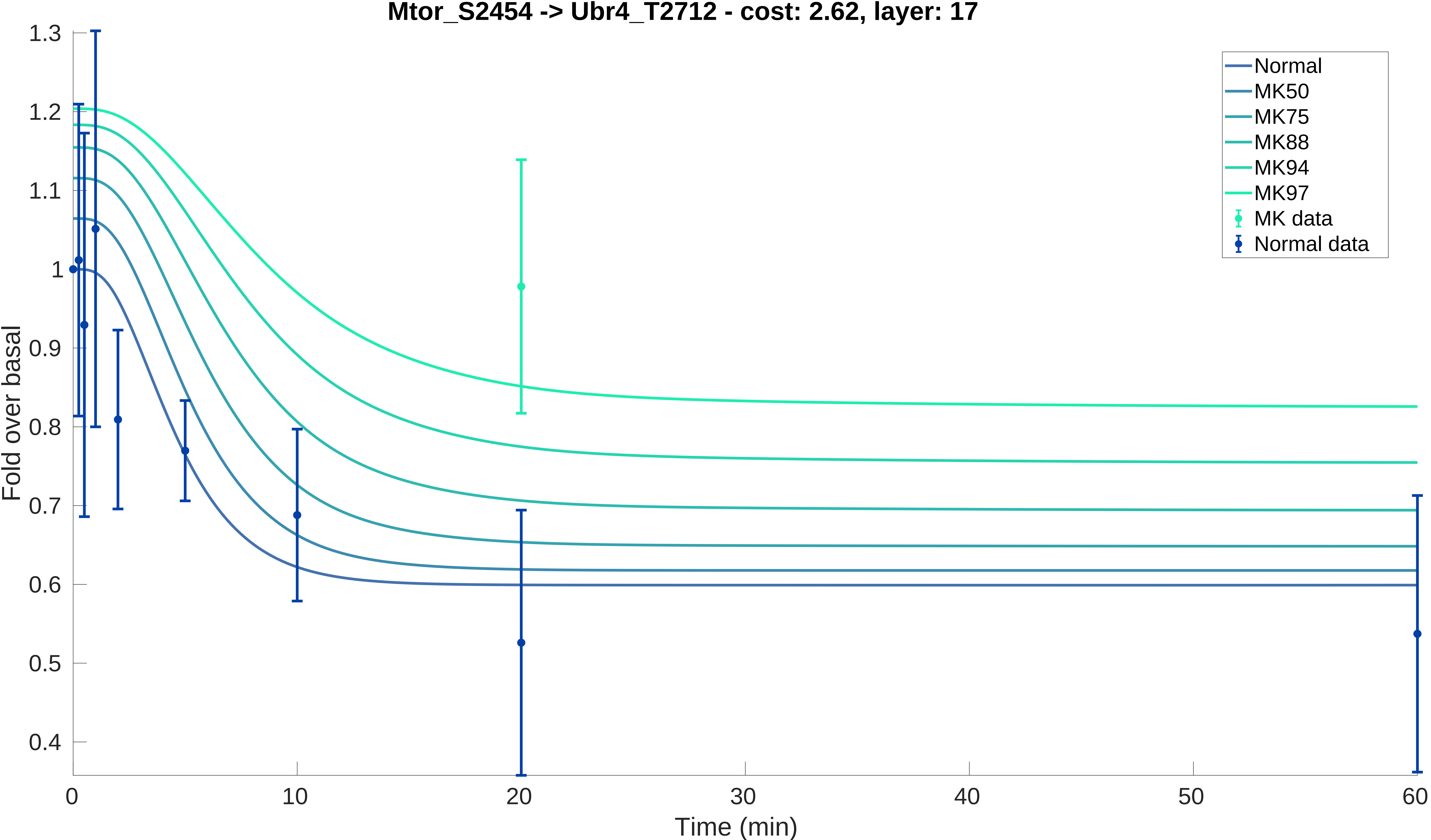

Mtor_S2454 -> Ubr4_T2712 - cost: 2.62, layer: 17
Fold over basal
Time (min)
1.3
1.2
1.1
1
0.9
0.8
0.7
0.6
0.5
0.4
0
10
20
30
40
50
60
Normal
MK50
MK75
MK88
MK94
MK97
MK data
Normal data

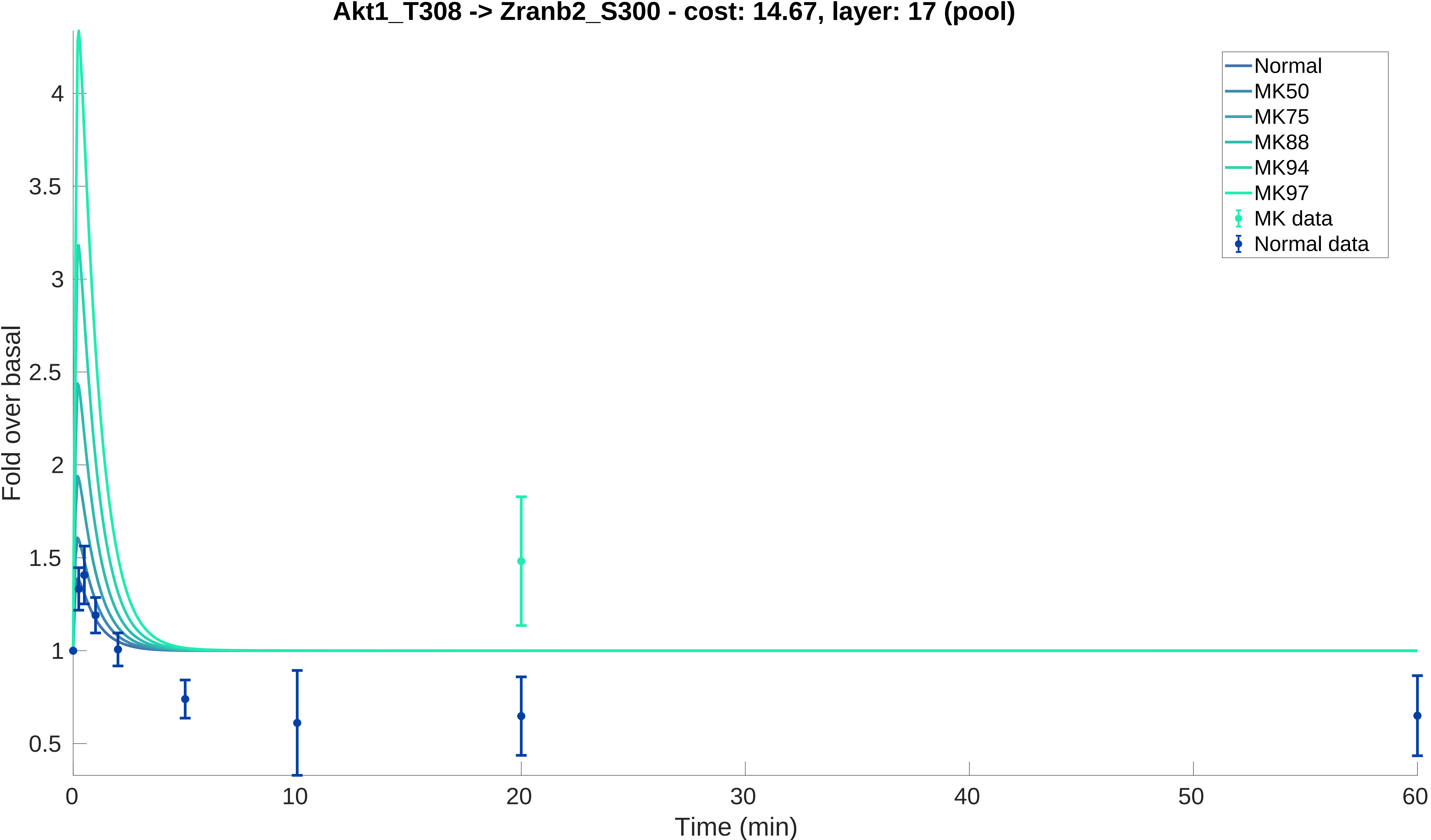
Akt1_T308 -> Zranb2_S300 - cost: 14.67, layer: 17 (pool)
Normal
MK50
MK75
MK88
MK94
MK97
MK data
Normal data
Fold over basal
Time (min)
0.5
1
1.5
2
2.5
3
3.5
4
0
10
20
30
40
50
60

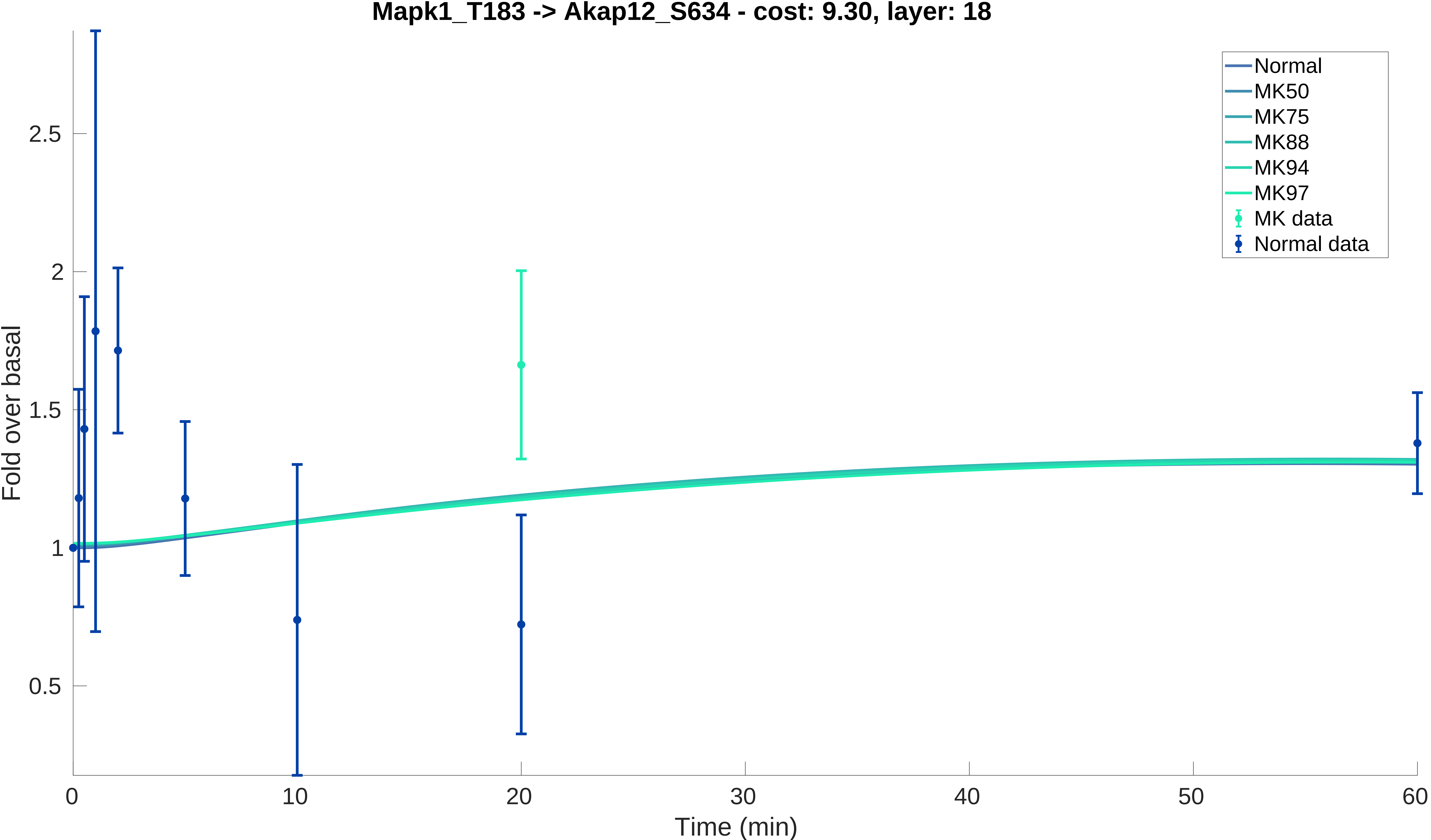

Mapk1_T183 -> Akap12_S634 - cost: 9.30, layer: 18
Normal
MK50
MK75
MK88
MK94
MK97
MK data
Normal data
Fold over basal
2.5
2
1.5
1
0.5
0
10
20
30
40
50
60
Time (min)

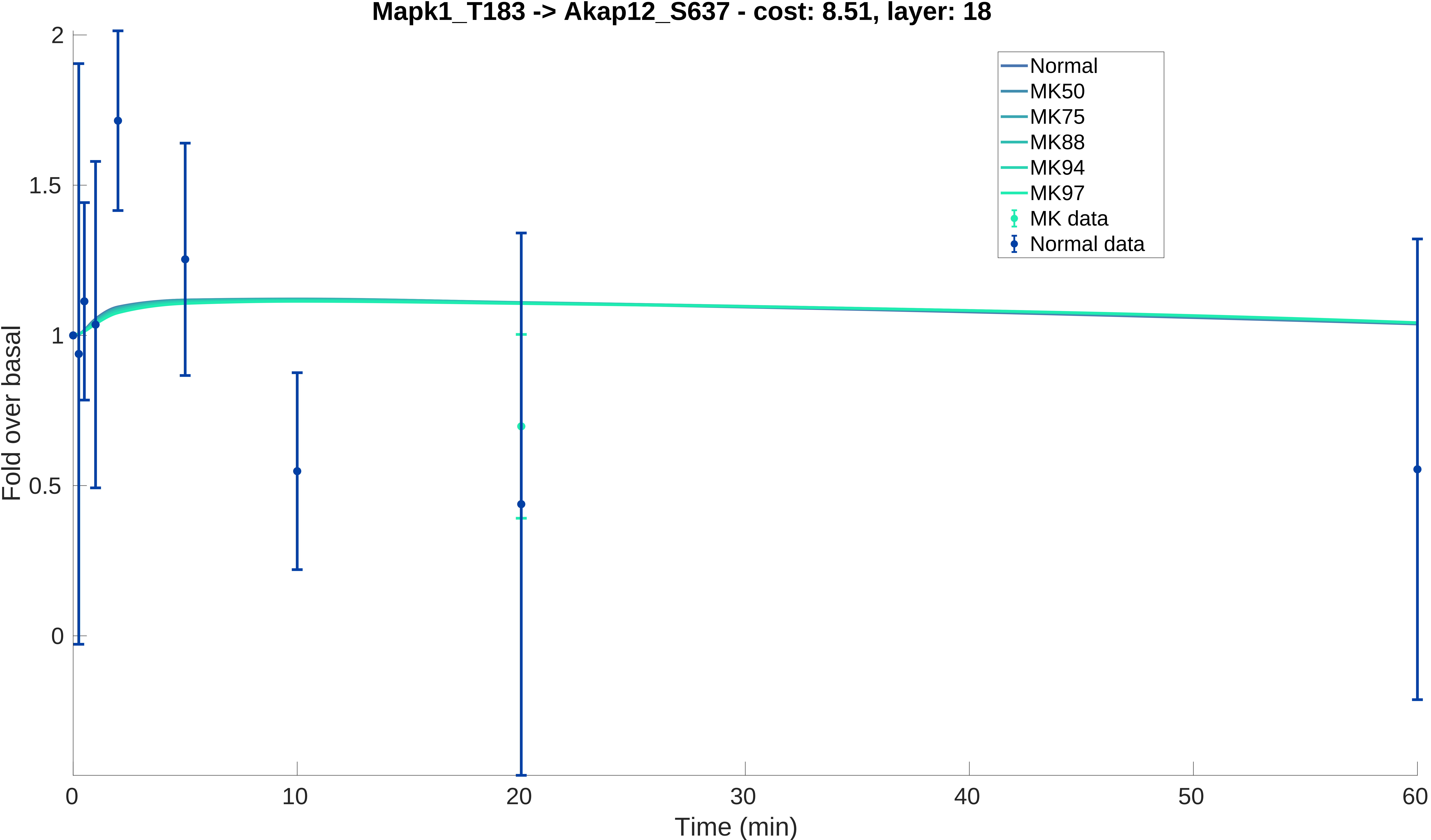

Mapk1_T183 -> Akap12_S637 - cost: 8.51, layer: 18
Fold over basal
Time (min)
2
1.5
1
0.5
0
0
10
20
30
40
50
60
Normal
MK50
MK75
MK88
MK94
MK97
MK data
Normal data

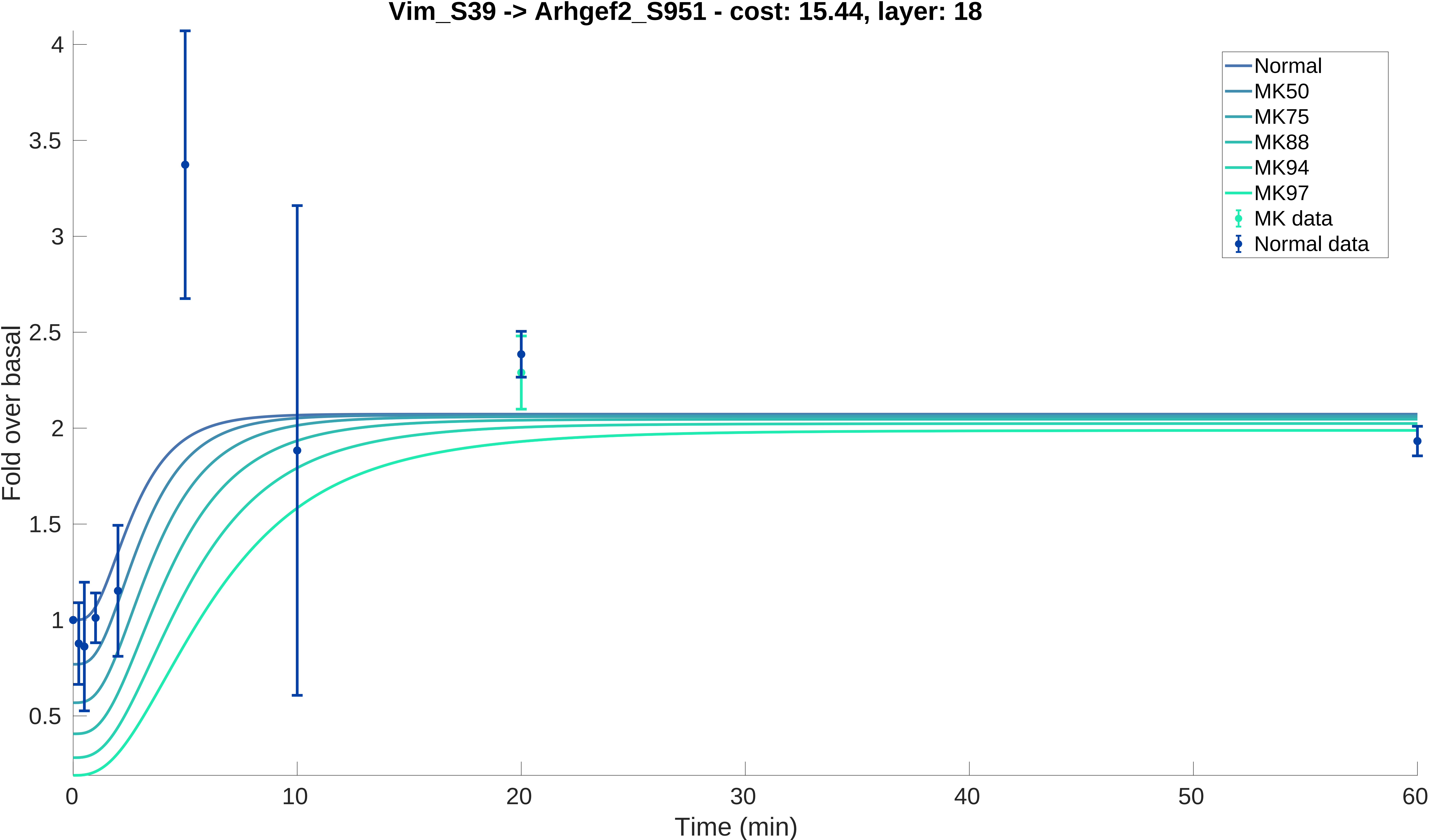

Vim_S39 -> Arhgef2_S951 - cost: 15.44, layer: 18
Fold over basal
Time (min)
0.5
1
1.5
2
2.5
3
3.5
4
0
10
20
30
40
50
60
Normal
MK50
MK75
MK88
MK94
MK97
MK data
Normal data

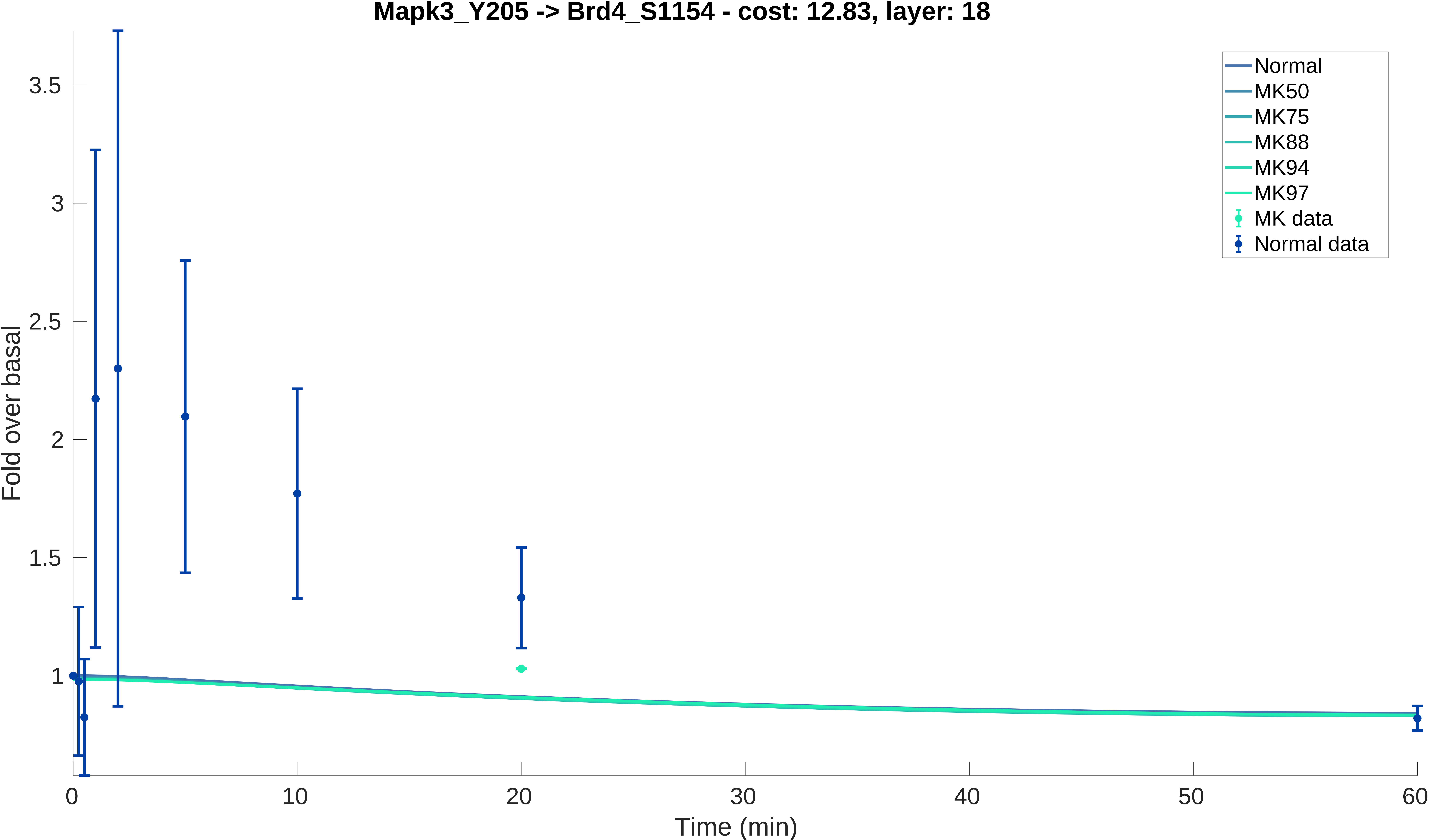

Mapk3_Y205 -> Brd4_S1154 - cost: 12.83, layer: 18
Fold over basal
Time (min)
0
10
20
30
40
50
60
1
1.5
2
2.5
3
3.5
Normal
MK50
MK75
MK88
MK94
MK97
MK data
Normal data

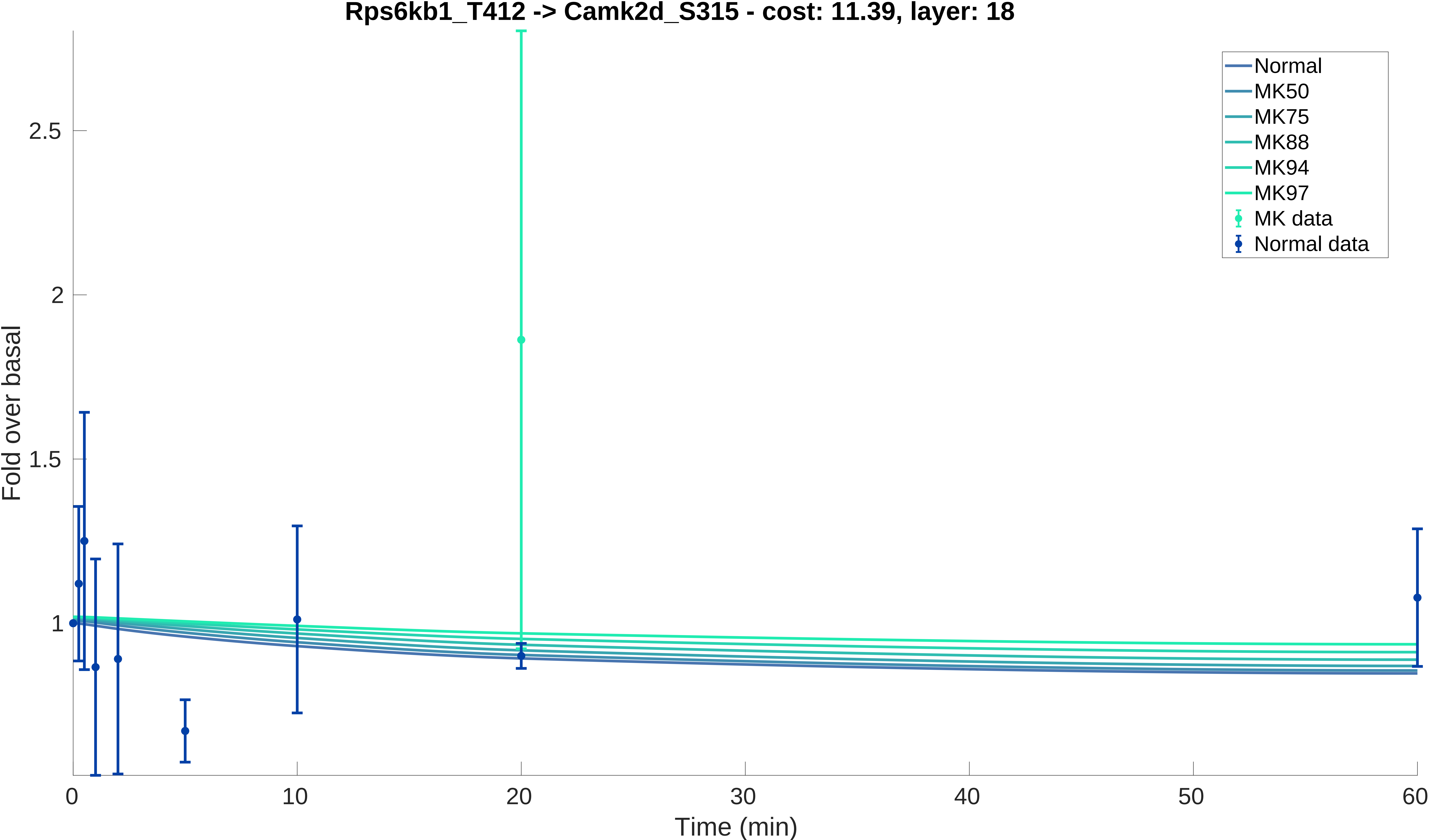
Rps6kb1_T412 -> Camk2d_S315 - cost: 11.39, layer: 18
Fold over basal
2.5
2
1.5
1
Time (min)
0
10
20
30
40
50
60
Normal
MK50
MK75
MK88
MK94
MK97
MK data
Normal data

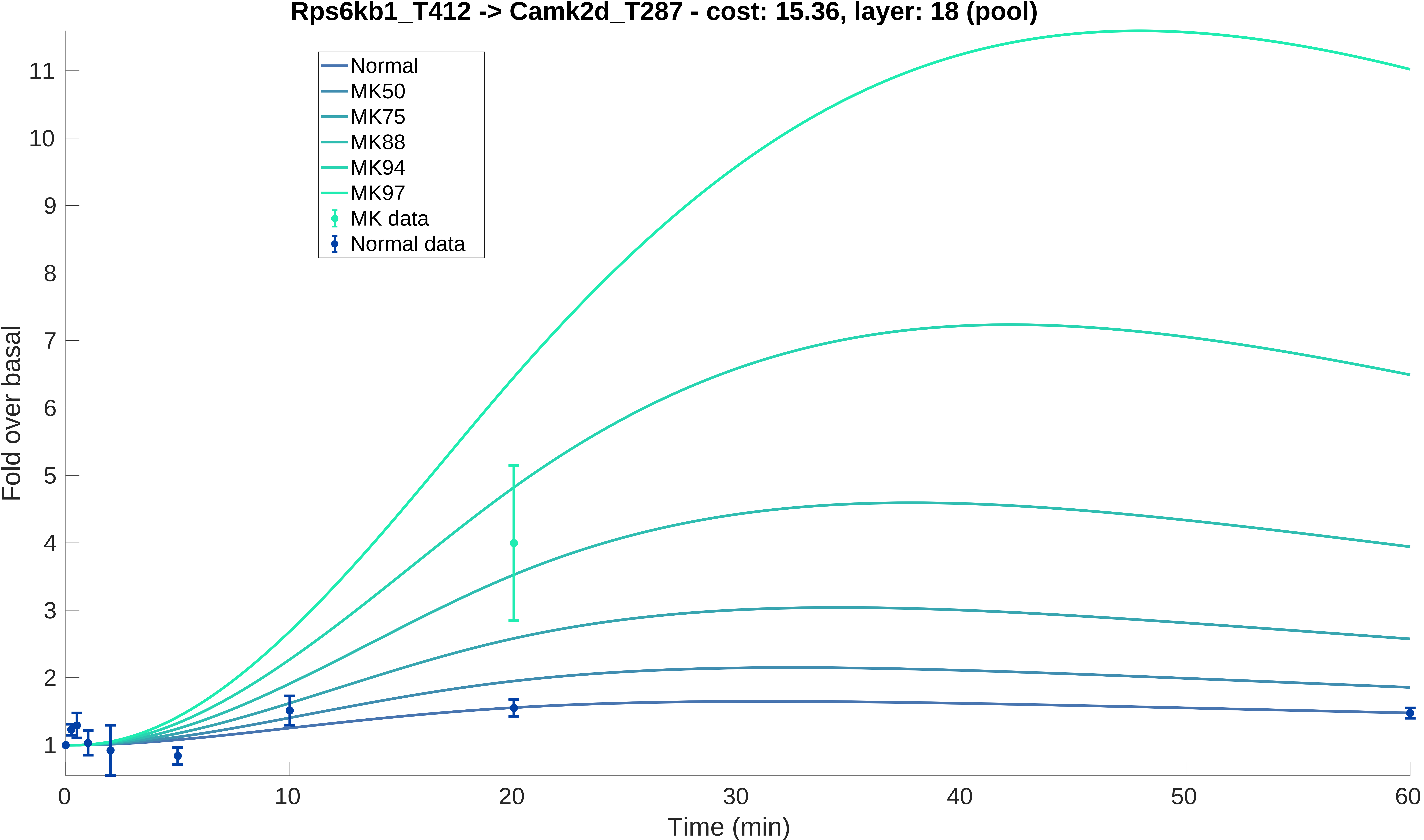

Rps6kb1_T412 -> Camk2d_T287 - cost: 15.36, layer: 18 (pool)
Normal
MK50
MK75
MK88
MK94
MK97
MK data
Normal data
Fold over basal
Time (min)
1
2
3
4
5
6
7
8
9
10
11
0
10
20
30
40
50
60

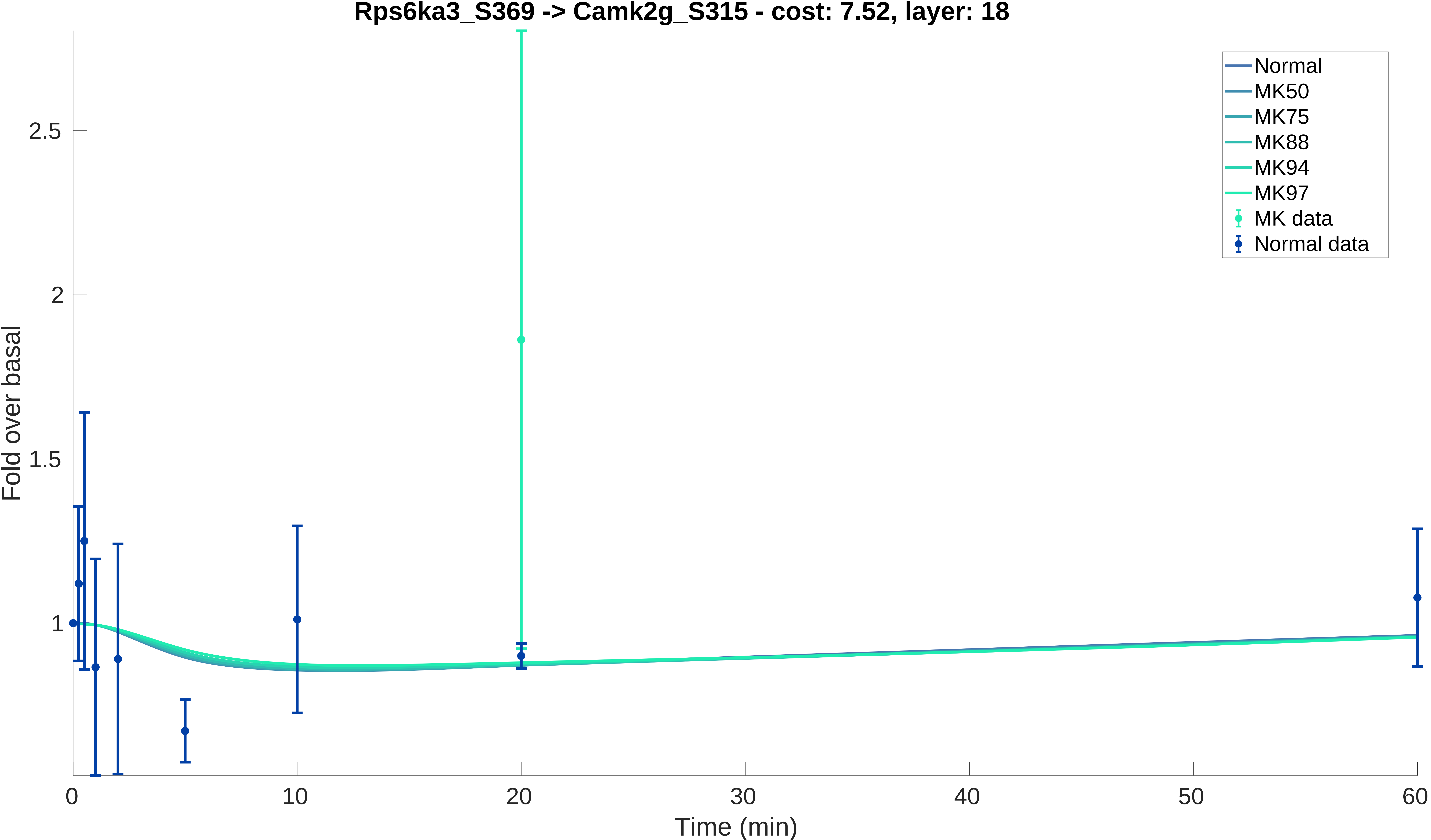

Rps6ka3_S369 -> Camk2g_S315 - cost: 7.52, layer: 18
Fold over basal
Time (min)
0
10
20
30
40
50
60
1
1.5
2
2.5
Normal
MK50
MK75
MK88
MK94
MK97
MK data
Normal data

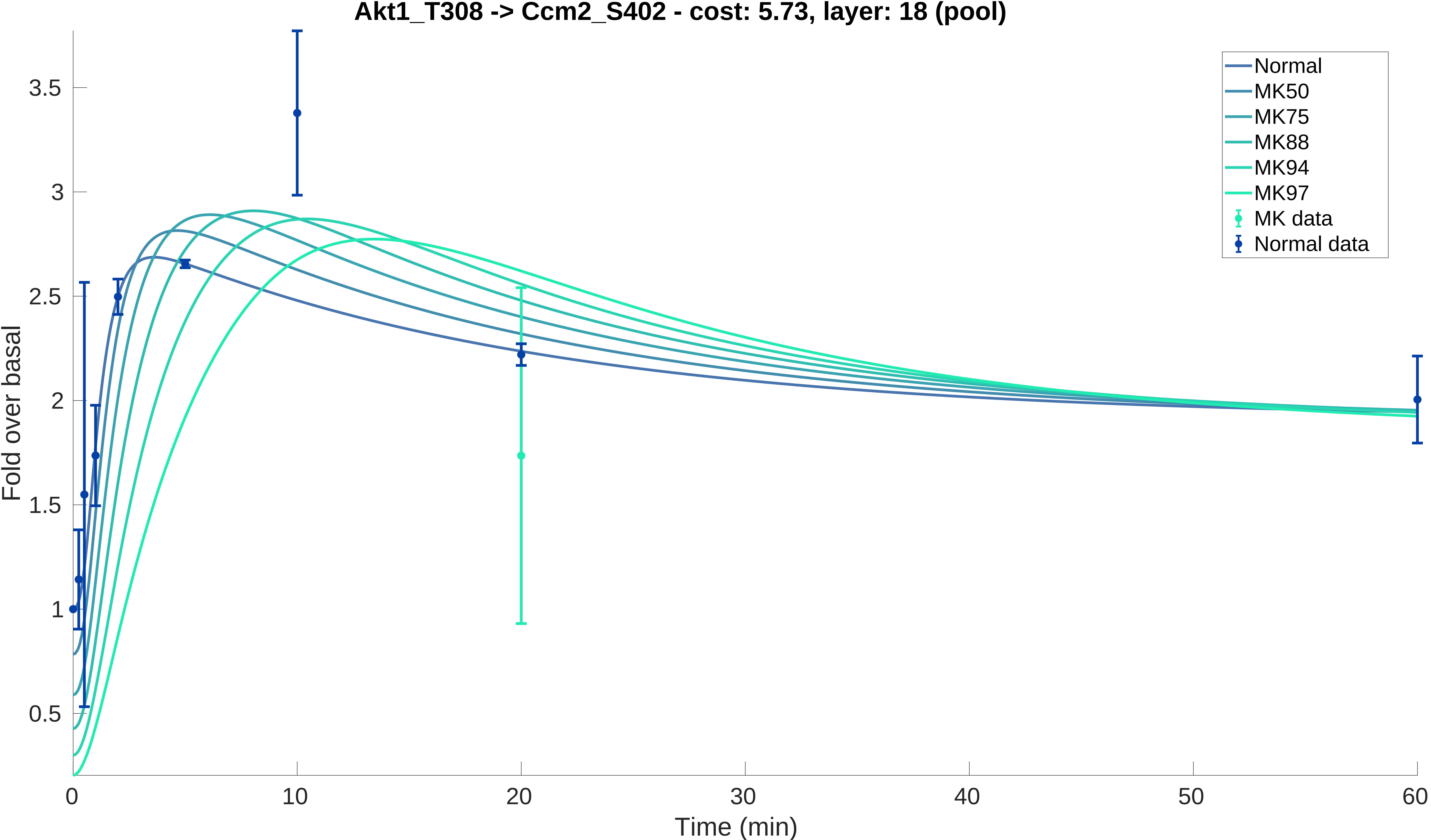

Akt1_T308 -> Ccm2_S402 - cost: 5.73, layer: 18 (pool)
Normal
MK50
MK75
MK88
MK94
MK97
MK data
Normal data
Fold over basal
Time (min)
0.5
1
1.5
2
2.5
3
3.5
0
10
20
30
40
50
60

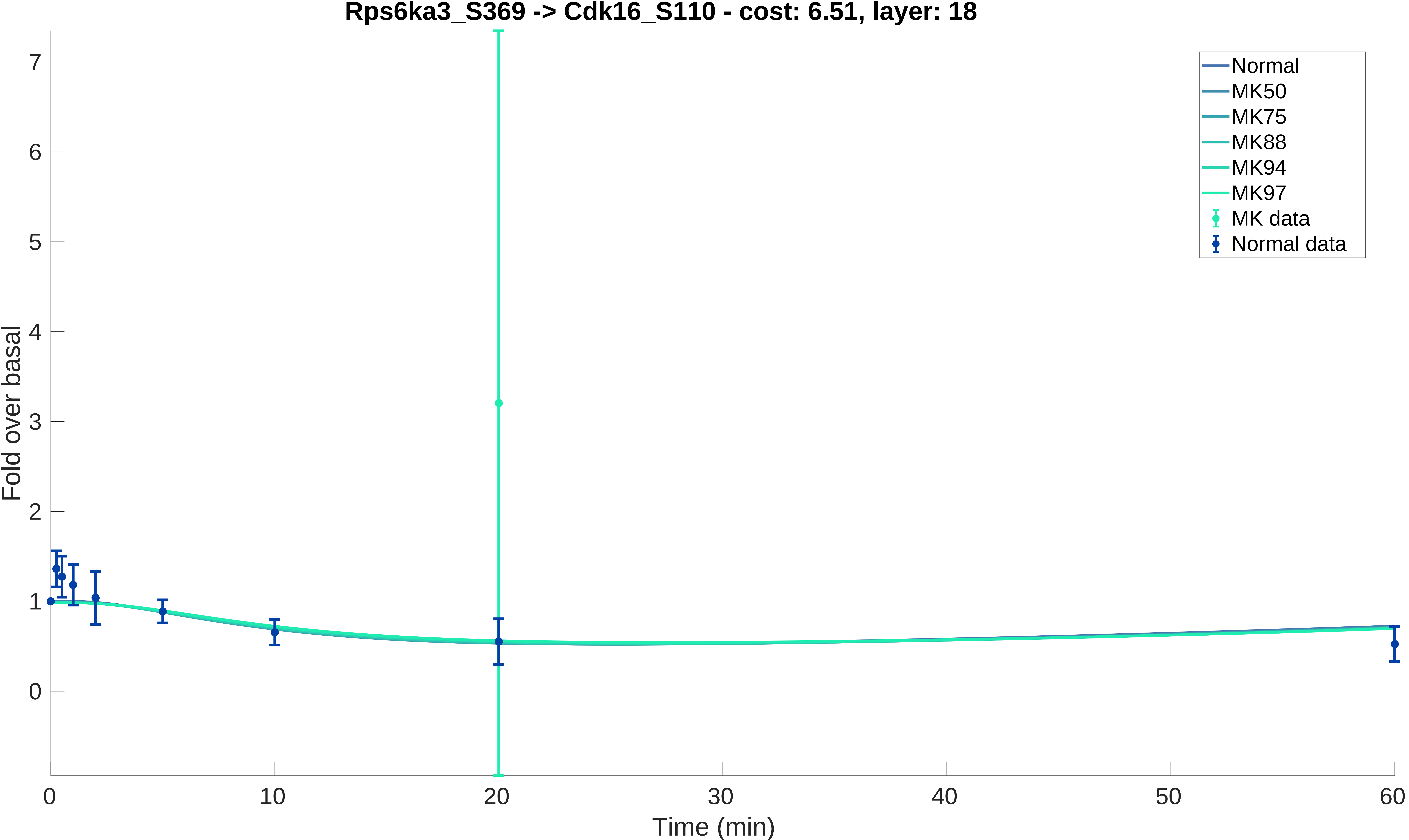

Rps6ka3_S369 -> Cdk16_S110 - cost: 6.51, layer: 18
Fold over basal
Time (min)
0
1
2
3
4
5
6
7
0
10
20
30
40
50
60
Normal
MK50
MK75
MK88
MK94
MK97
MK data
Normal data

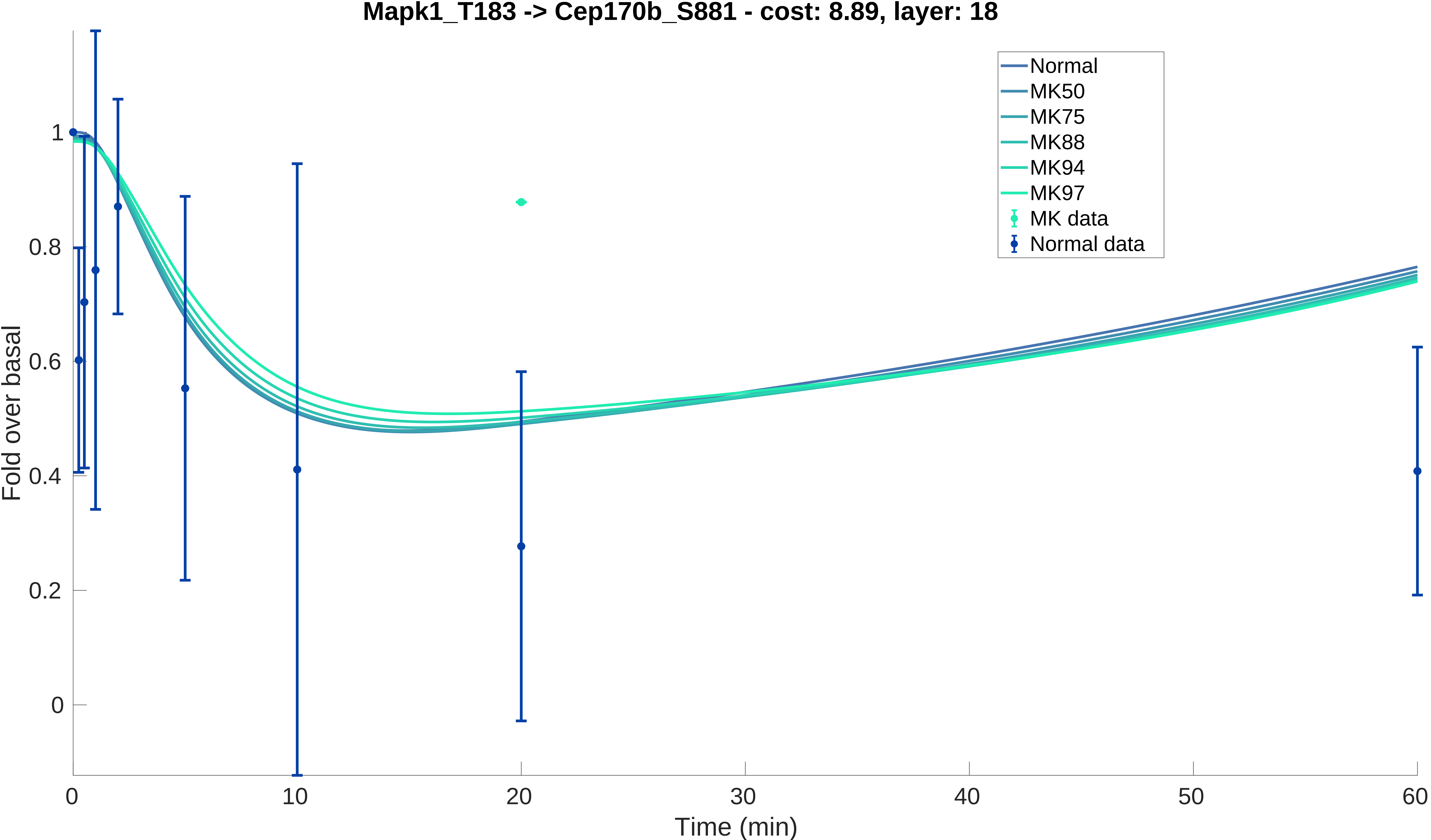

Mapk1_T183 -> Cep170b_S881 - cost: 8.89, layer: 18
Normal
MK50
MK75
MK88
MK94
MK97
MK data
Normal data
Fold over basal
Time (min)
0
0.2
0.4
0.6
0.8
1
0
10
20
30
40
50
60

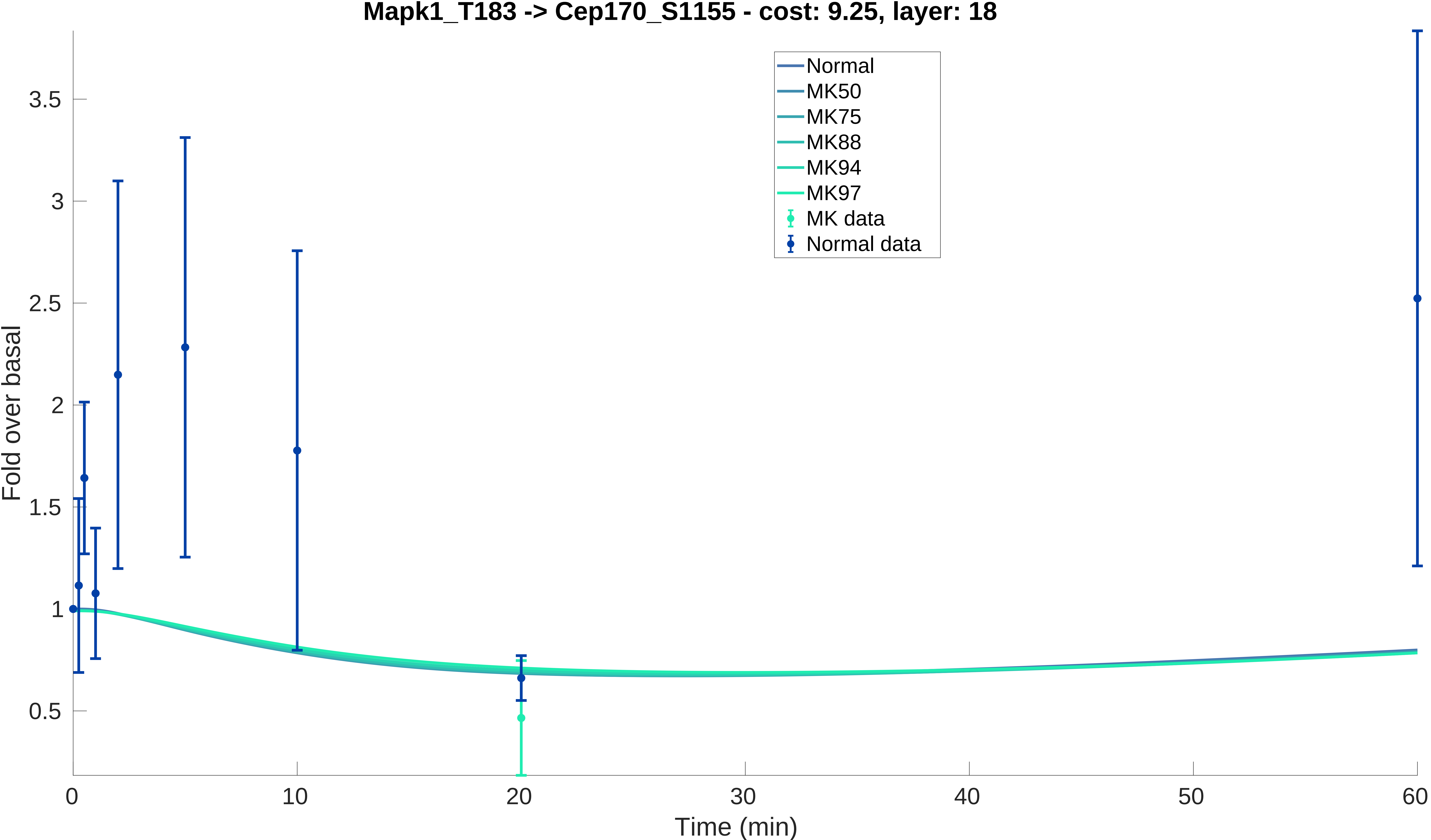
Mapk1_T183 -> Cep170_S1155 - cost: 9.25, layer: 18
Fold over basal
Time (min)
0.5
1
1.5
2
2.5
3
3.5
0
10
20
30
40
50
60
Normal
MK50
MK75
MK88
MK94
MK97
MK data
Normal data

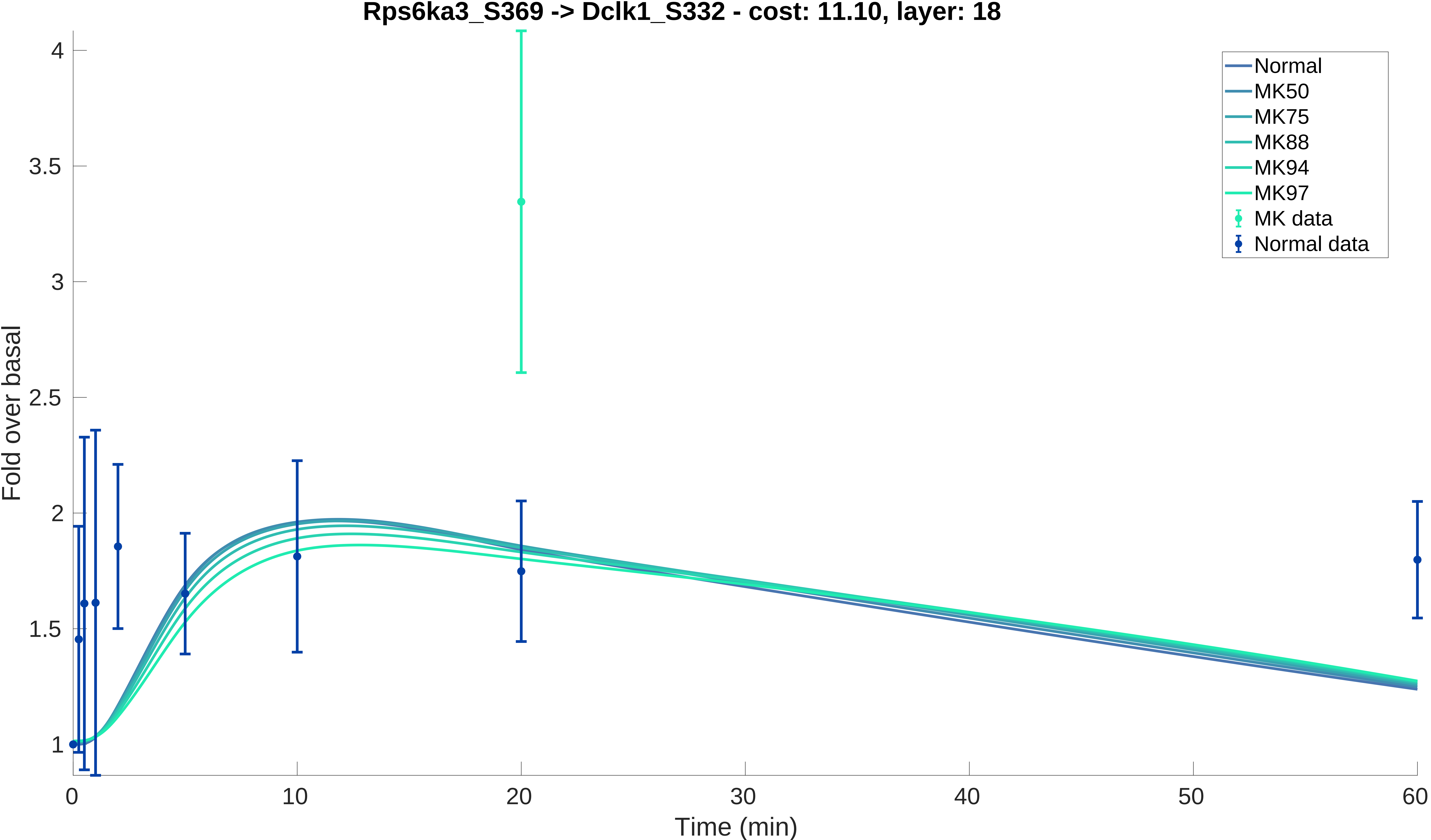

Rps6ka3_S369 -> Dclk1_S332 - cost: 11.10, layer: 18
Fold over basal
Time (min)
4
3.5
3
2.5
2
1.5
1
0
10
20
30
40
50
60
Normal
MK50
MK75
MK88
MK94
MK97
MK data
Normal data

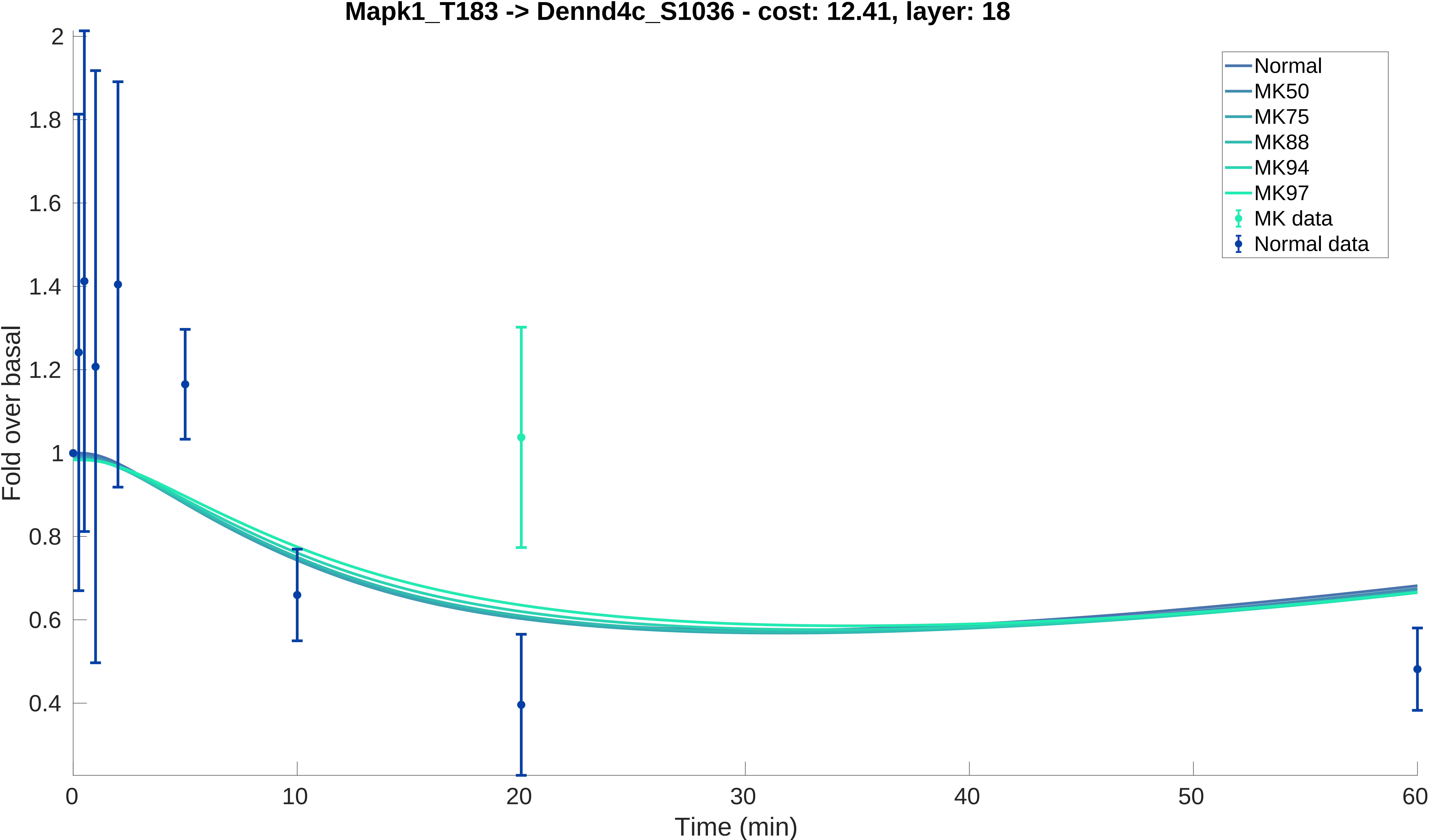
Mapk1_T183 -> Dennd4c_S1036 - cost: 12.41, layer: 18
Fold over basal
Time (min)
0
10
20
30
40
50
60
0.4
0.6
0.8
1
1.2
1.4
1.6
1.8
2
Normal
MK50
MK75
MK88
MK94
MK97
MK data
Normal data

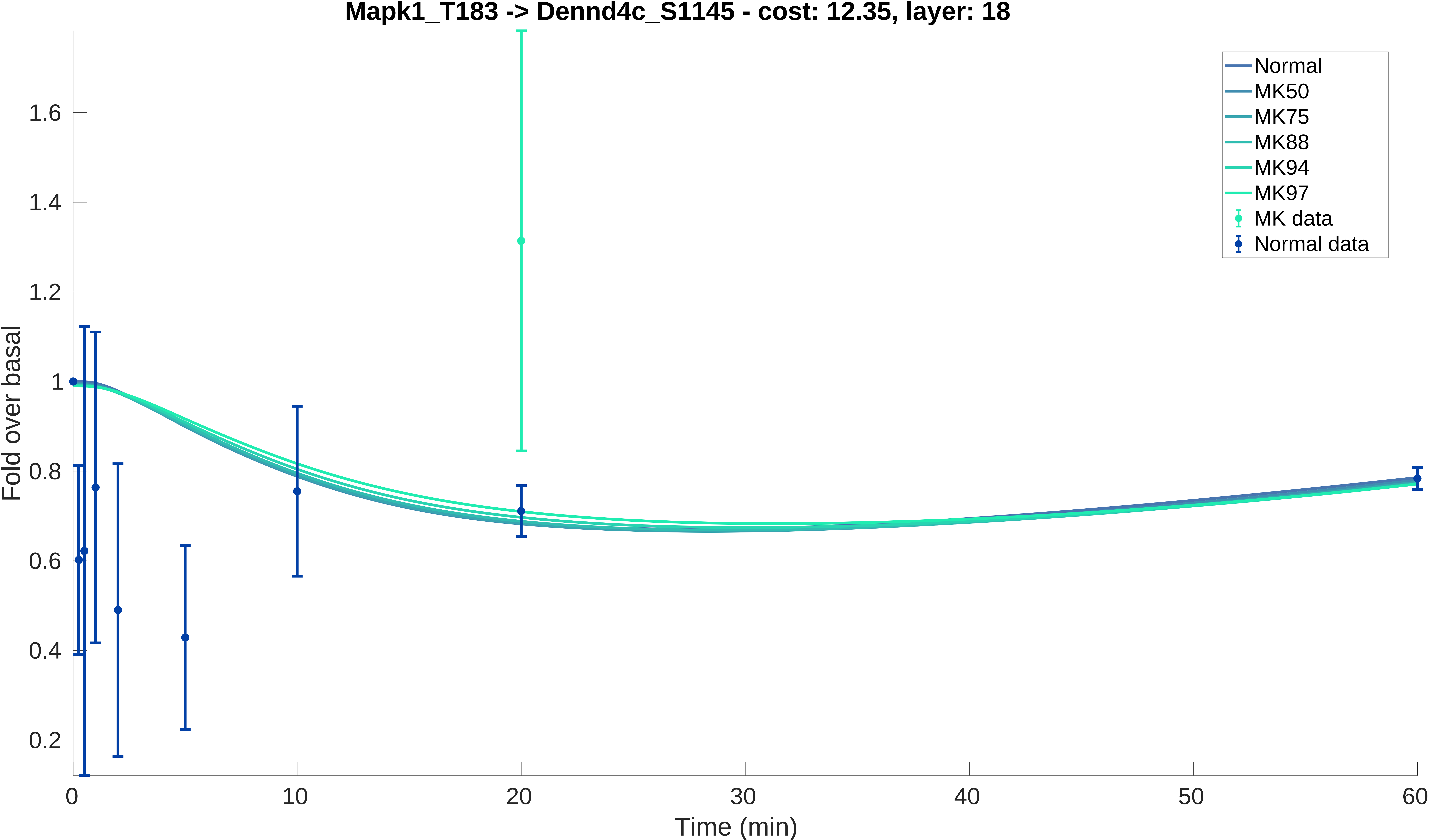

Mapk1_T183 -> Dennd4c_S1145 - cost: 12.35, layer: 18
Fold over basal
Time (min)
0
10
20
30
40
50
60
0.2
0.4
0.6
0.8
1
1.2
1.4
1.6
Normal
MK50
MK75
MK88
MK94
MK97
MK data
Normal data

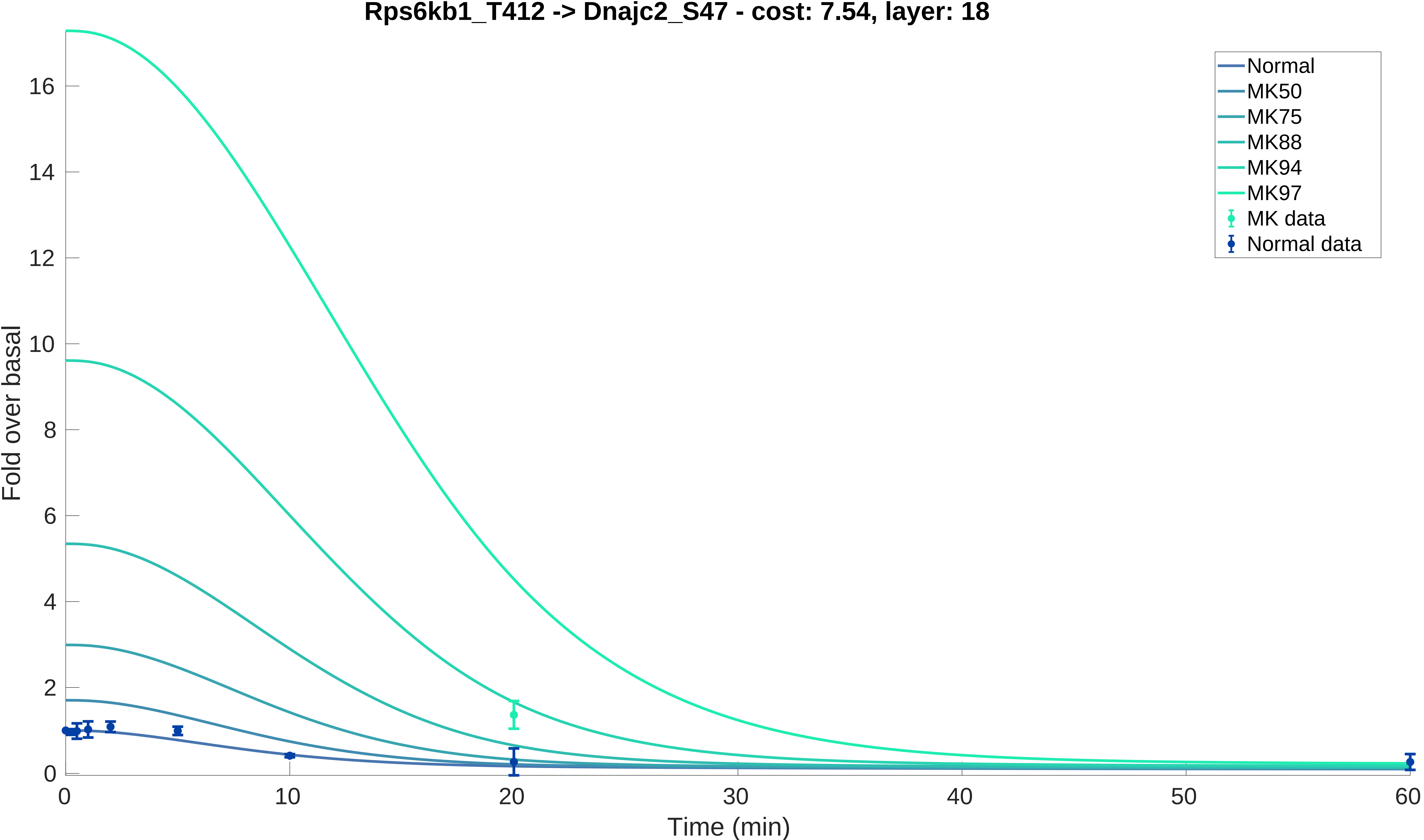
Rps6kb1_T412 -> Dnajc2_S47 - cost: 7.54, layer: 18
Fold over basal
0
2
4
6
8
10
12
14
16
Time (min)
0
10
20
30
40
50
60
Normal
MK50
MK75
MK88
MK94
MK97
MK data
Normal data

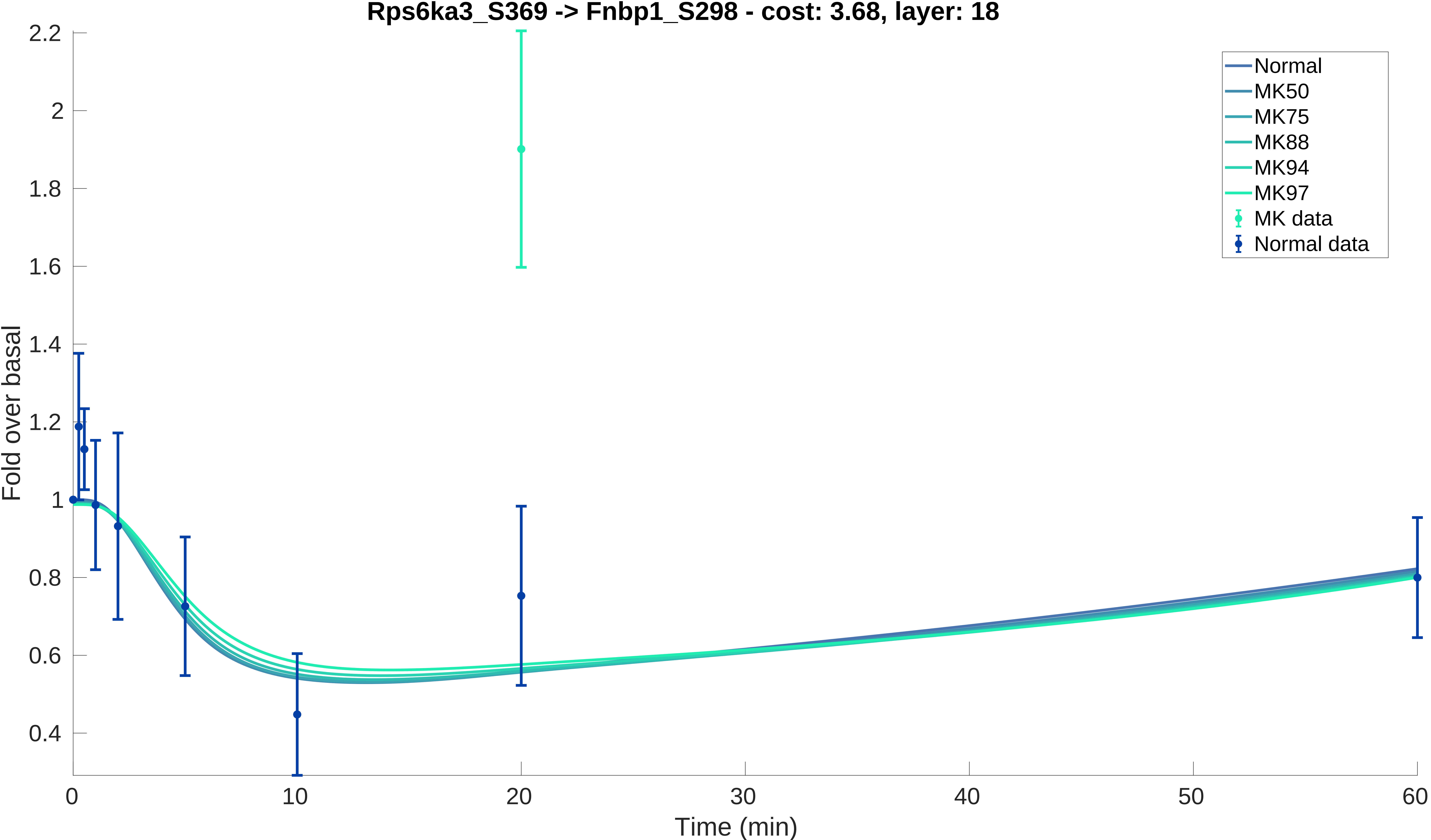

Rps6ka3_S369 -> Fnbp1_S298 - cost: 3.68, layer: 18
Fold over basal
Time (min)
2.2
2
1.8
1.6
1.4
1.2
1
0.8
0.6
0.4
0
10
20
30
40
50
60
Normal
MK50
MK75
MK88
MK94
MK97
MK data
Normal data

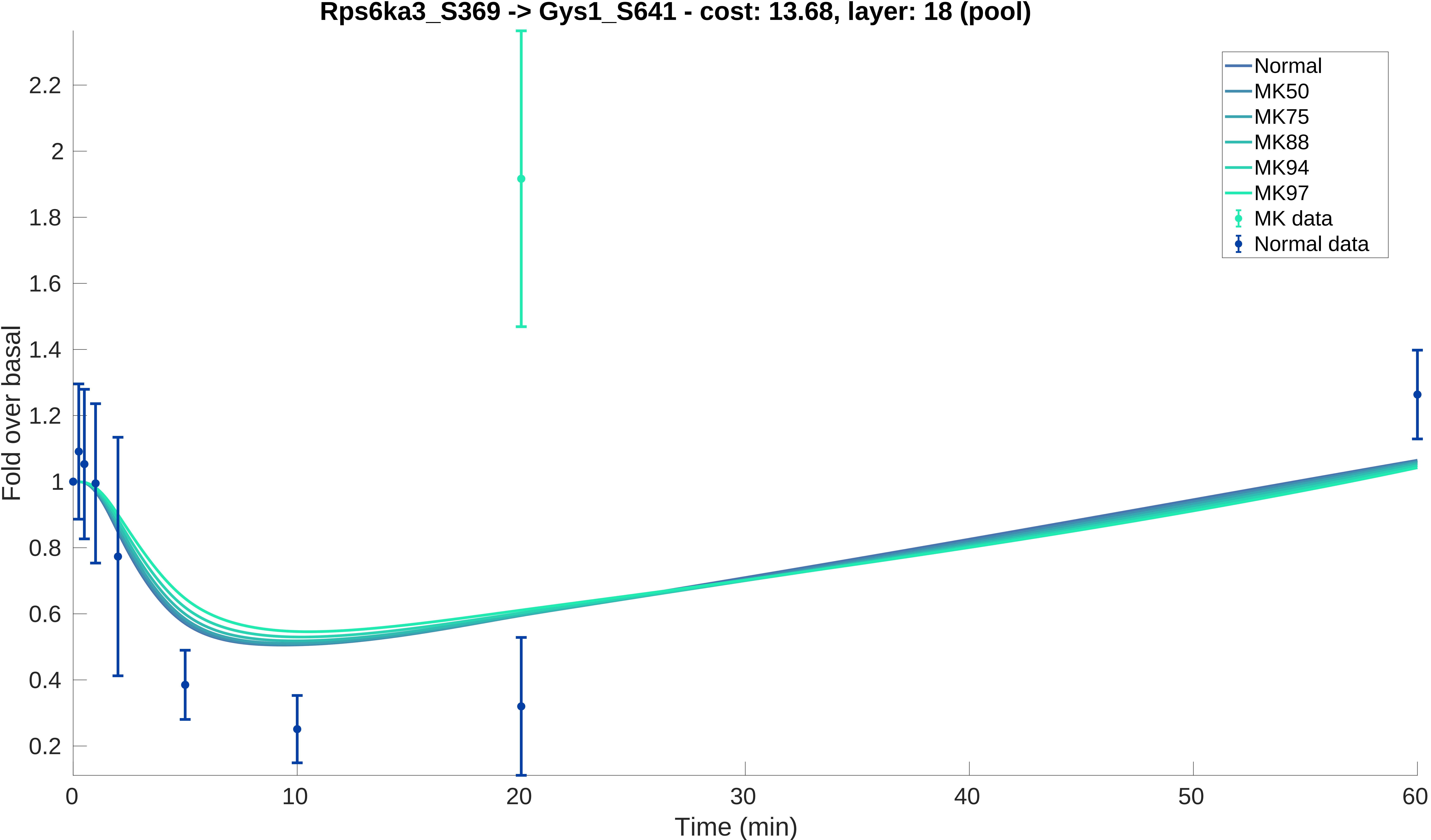

Rps6ka3_S369 -> Gys1_S641 - cost: 13.68, layer: 18 (pool)
Fold over basal
Time (min)
Normal
MK50
MK75
MK88
MK94
MK97
MK data
Normal data
0.2
0.4
0.6
0.8
1
1.2
1.4
1.6
1.8
2
2.2
0
10
20
30
40
50
60

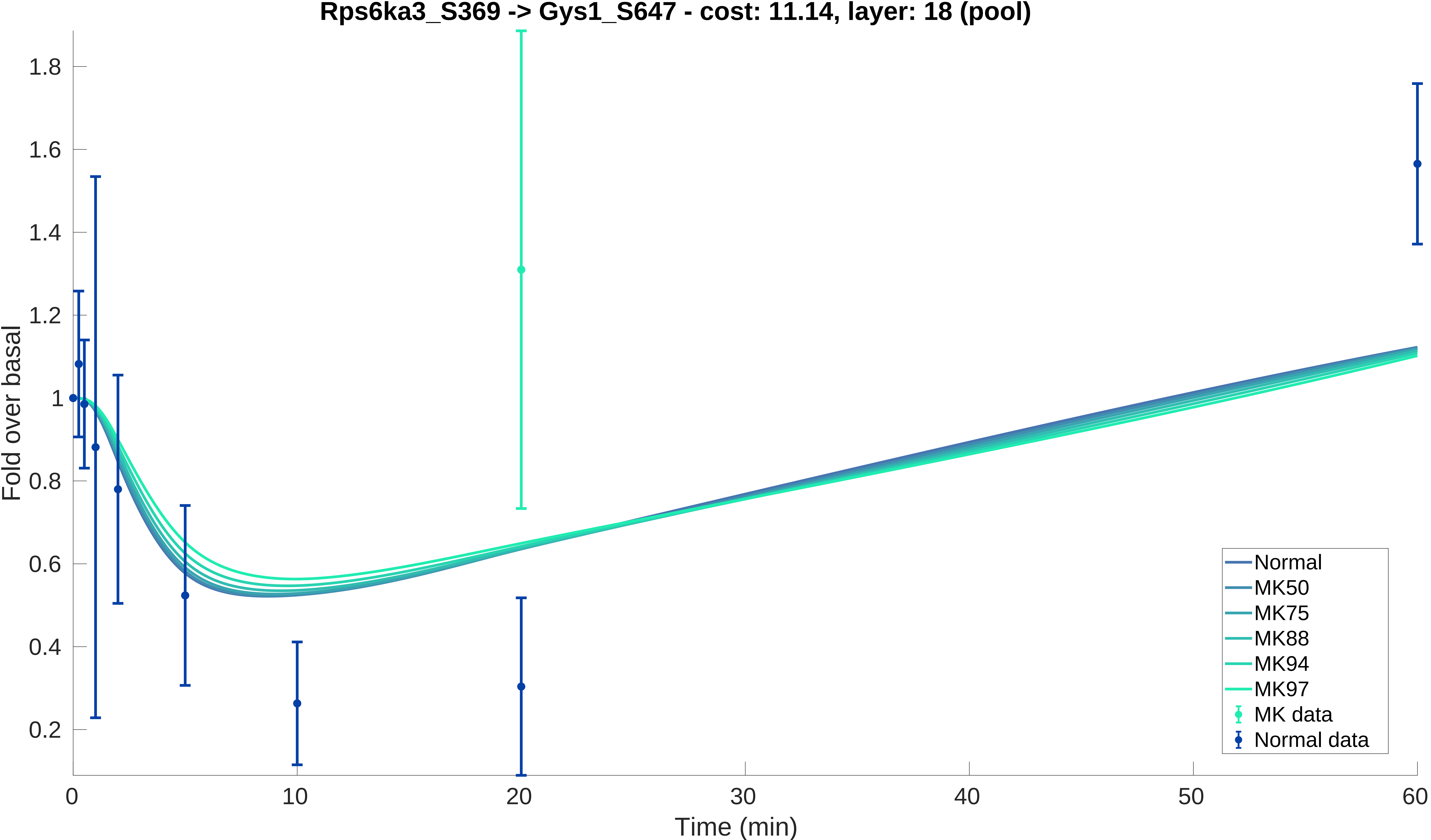
Rps6ka3_S369 -> Gys1_S647 - cost: 11.14, layer: 18 (pool)
Fold over basal
1.8
1.6
1.4
1.2
1
0.8
0.6
0.4
0.2
0
10
20
30
40
50
60
Time (min)
Normal
MK50
MK75
MK88
MK94
MK97
MK data
Normal data

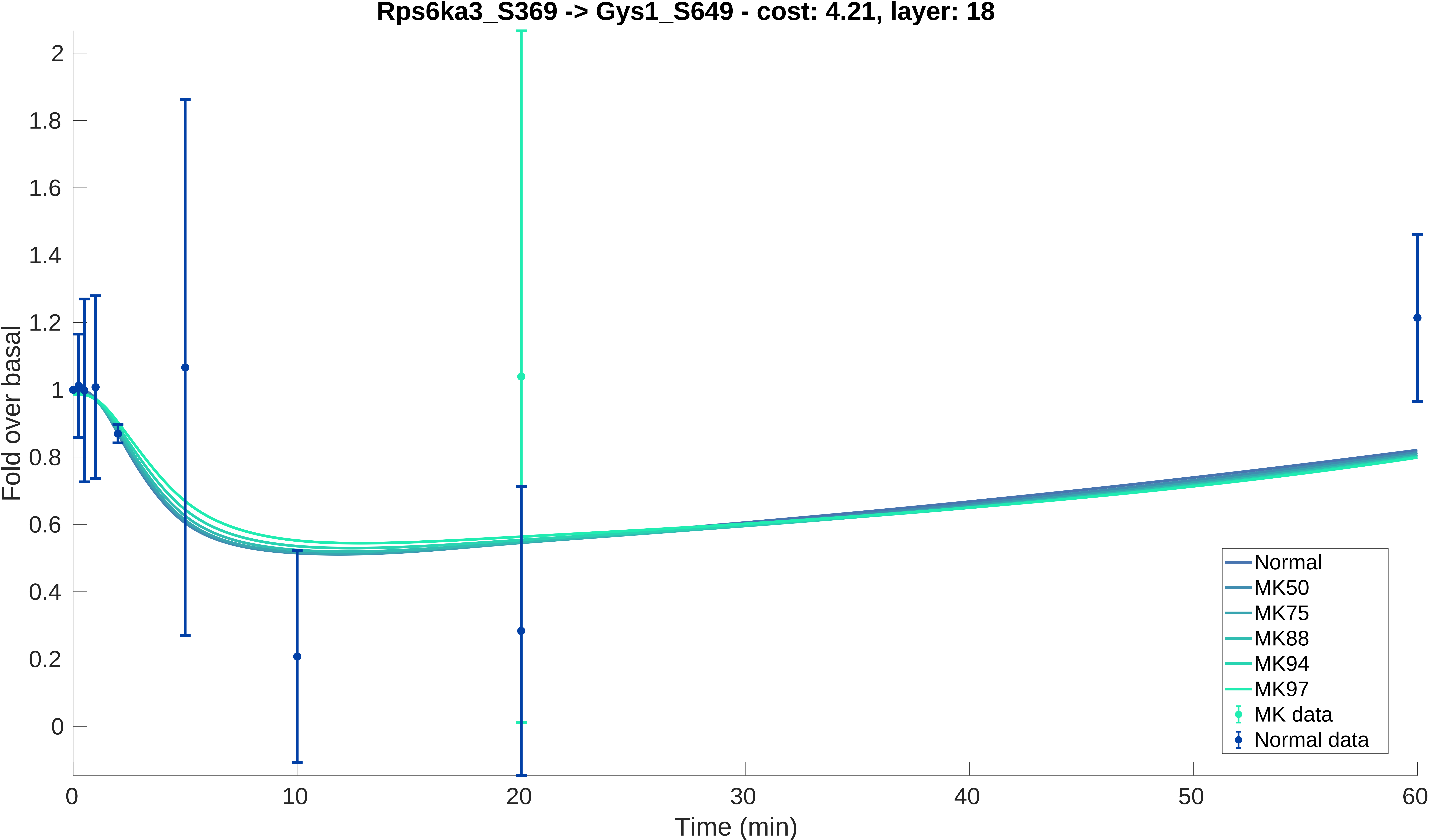

Rps6ka3_S369 -> Gys1_S649 - cost: 4.21, layer: 18
Fold over basal
Time (min)
0
10
20
30
40
50
60
0
0.2
0.4
0.6
0.8
1
1.2
1.4
1.6
1.8
2
Normal
MK50
MK75
MK88
MK94
MK97
MK data
Normal data

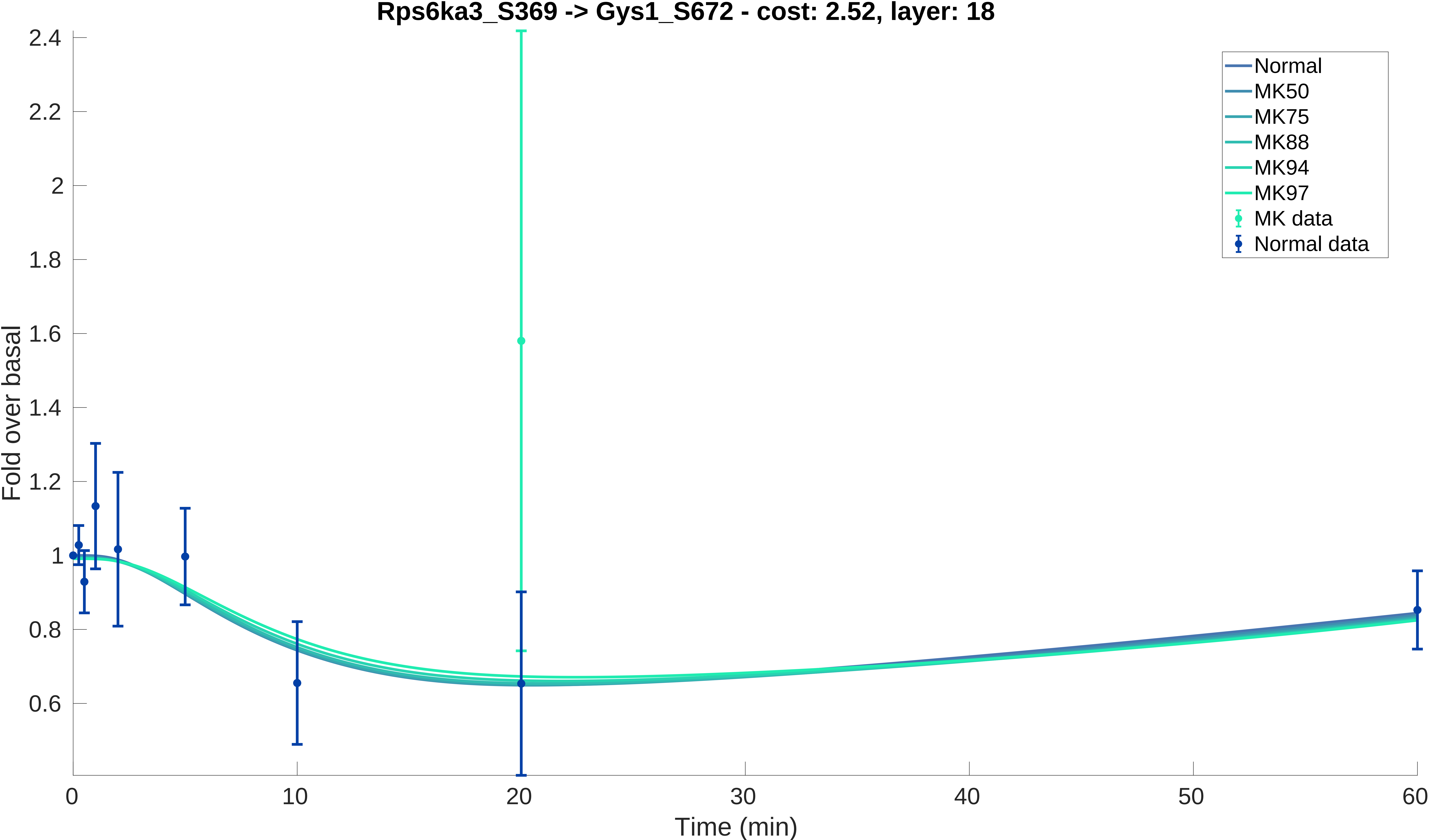

Rps6ka3_S369 -> Gys1_S672 - cost: 2.52, layer: 18
Fold over basal
Time (min)
0.6
0.8
1
1.2
1.4
1.6
1.8
2
2.2
2.4
0
10
20
30
40
50
60
Normal
MK50
MK75
MK88
MK94
MK97
MK data
Normal data

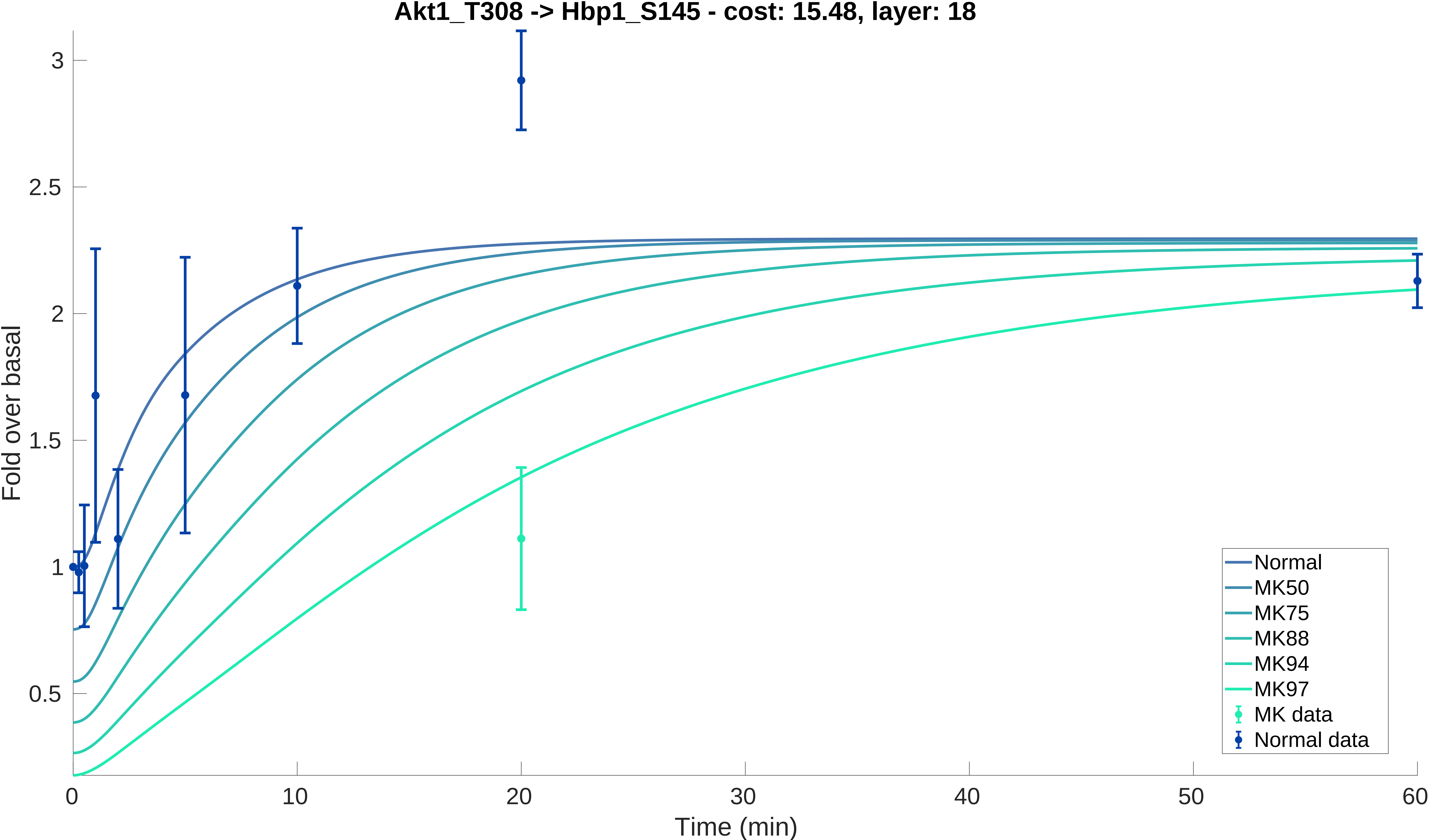

Akt1_T308 -> Hbp1_S145 - cost: 15.48, layer: 18
Fold over basal
3
2.5
2
1.5
1
0.5
0
10
20
30
40
50
60
Time (min)
Normal
MK50
MK75
MK88
MK94
MK97
MK data
Normal data

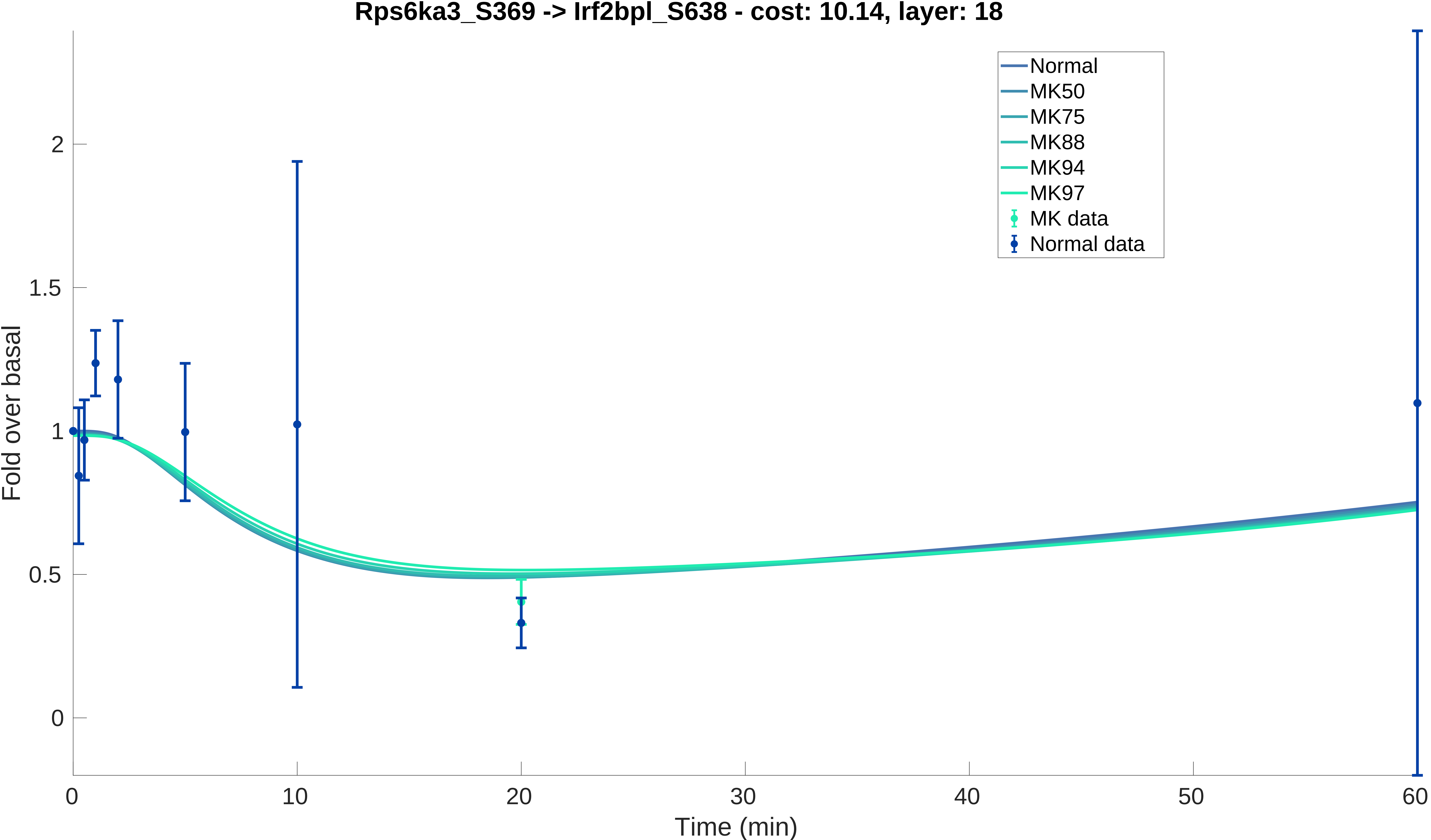
Rps6ka3_S369 -> Irf2bpl_S638 - cost: 10.14, layer: 18
Normal
MK50
MK75
MK88
MK94
MK97
MK data
Normal data
Fold over basal
2
1.5
1
0.5
0
0
10
20
30
40
50
60
Time (min)

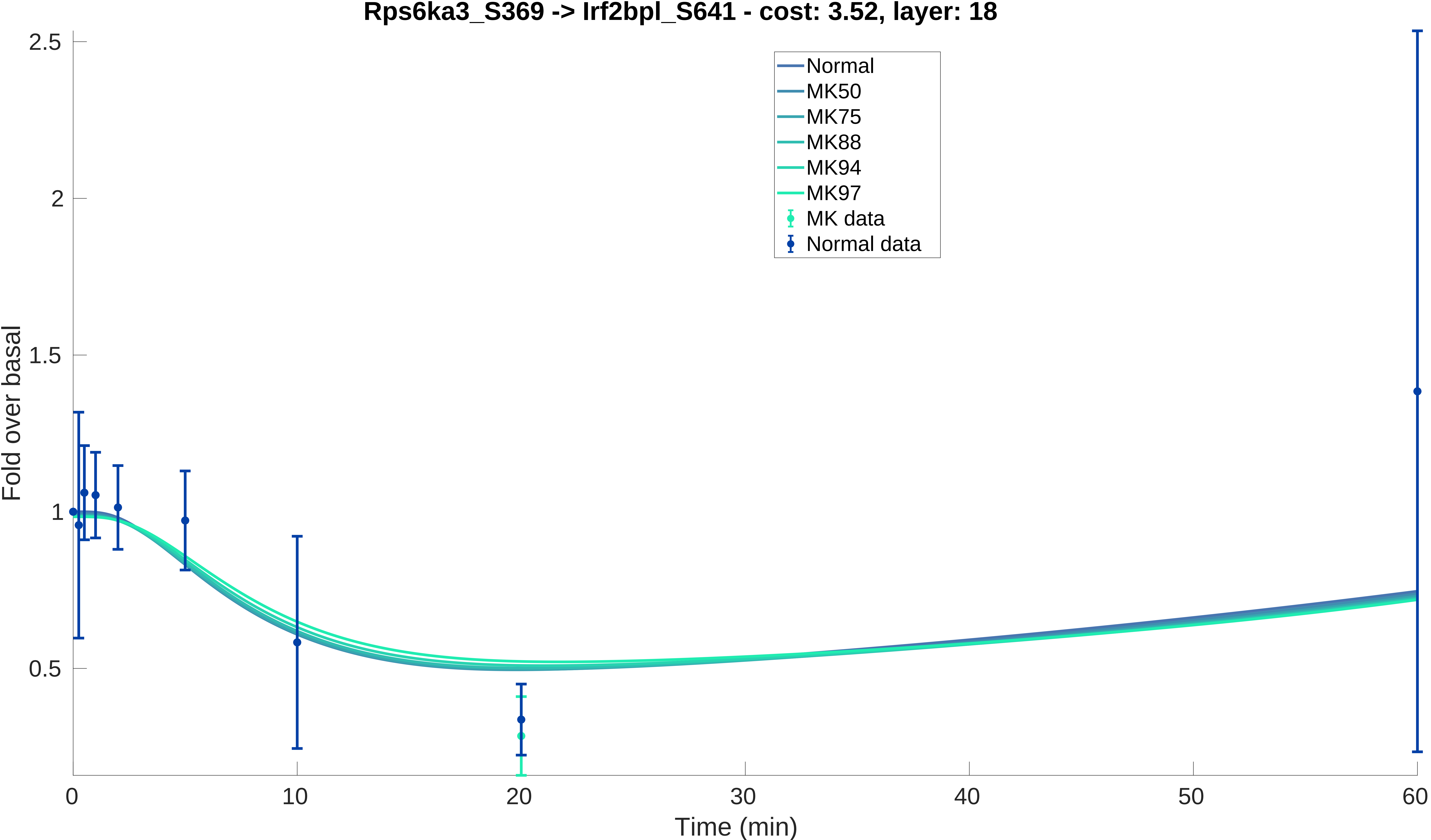
Rps6ka3_S369 -> Irf2bpl_S641 - cost: 3.52, layer: 18
Fold over basal
2.5
2
1.5
1
0.5
0
10
20
30
40
50
60
Time (min)
Normal
MK50
MK75
MK88
MK94
MK97
MK data
Normal data

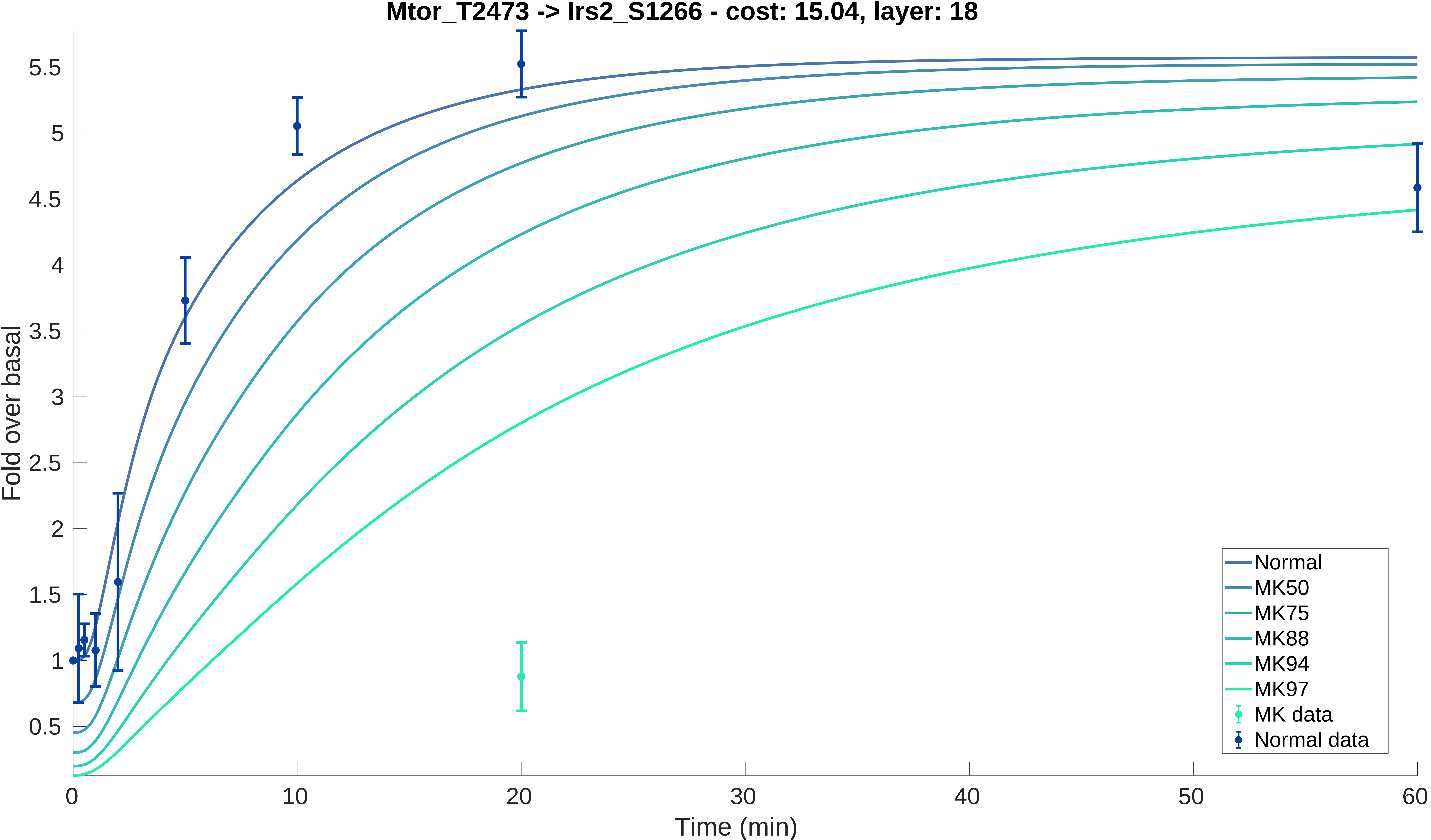

Mtor_T2473 -> Irs2_S1266 - cost: 15.04, layer: 18
Fold over basal
Time (min)
0.5
1
1.5
2
2.5
3
3.5
4
4.5
5
5.5
0
10
20
30
40
50
60
Normal
MK50
MK75
MK88
MK94
MK97
MK data
Normal data

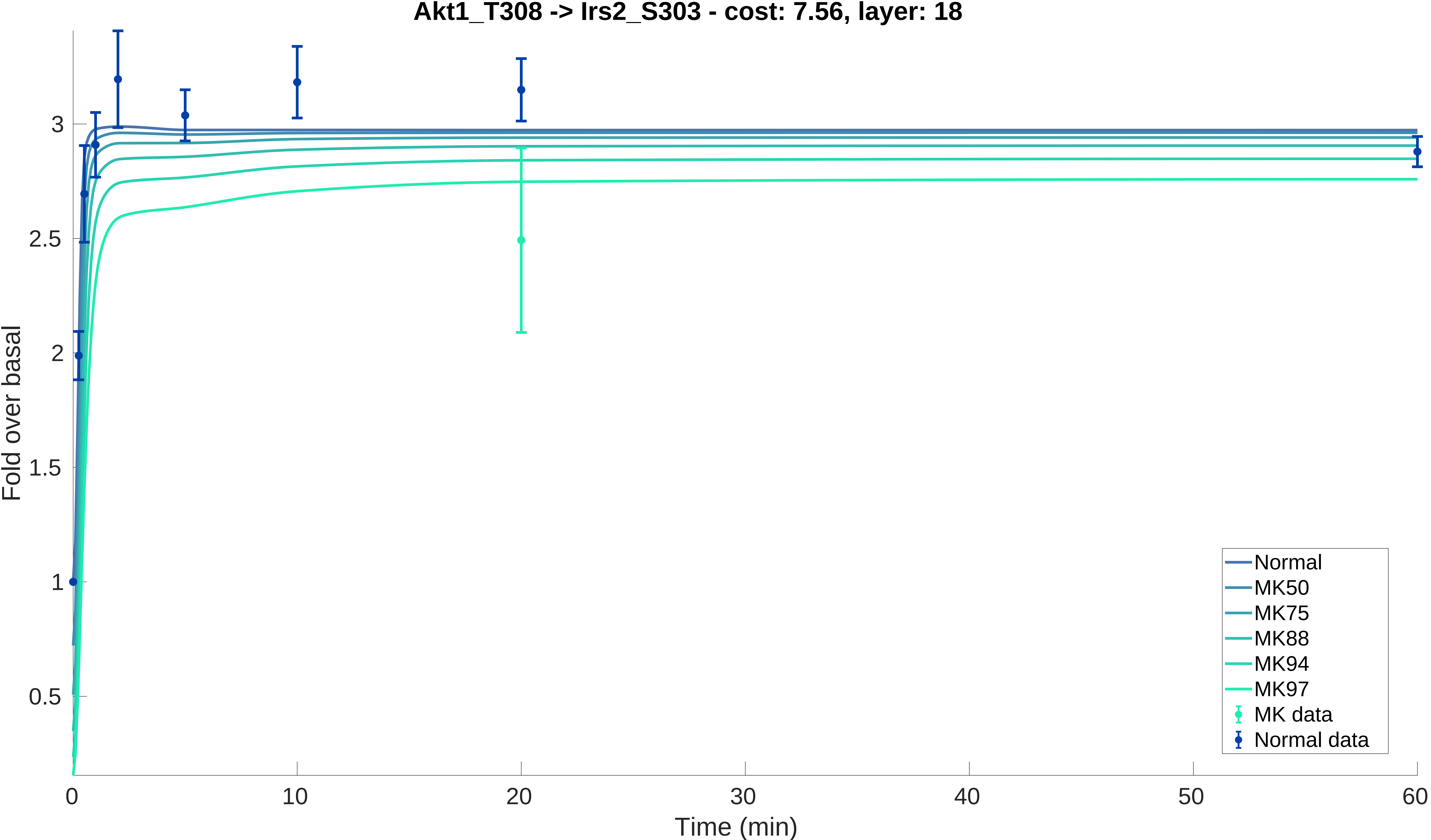

Akt1_T308 -> Irs2_S303 - cost: 7.56, layer: 18
Fold over basal
Time (min)
0.5
1
1.5
2
2.5
3
0
10
20
30
40
50
60
Normal
MK50
MK75
MK88
MK94
MK97
MK data
Normal data

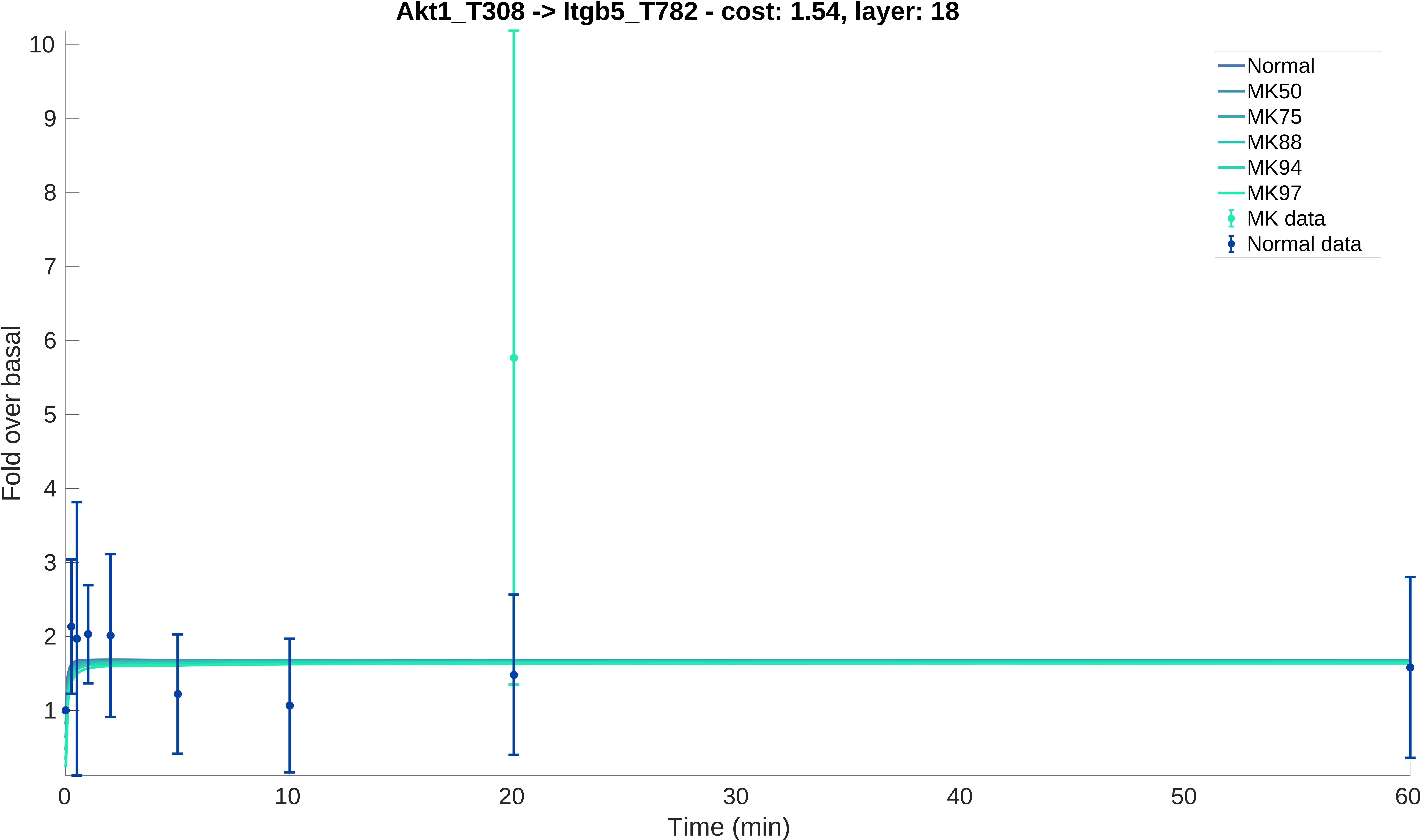

Akt1_T308 -> Itgb5_T782 - cost: 1.54, layer: 18
Fold over basal
Time (min)
Normal
MK50
MK75
MK88
MK94
MK97
MK data
Normal data
10
9
8
7
6
5
4
3
2
1
0
10
20
30
40
50
60

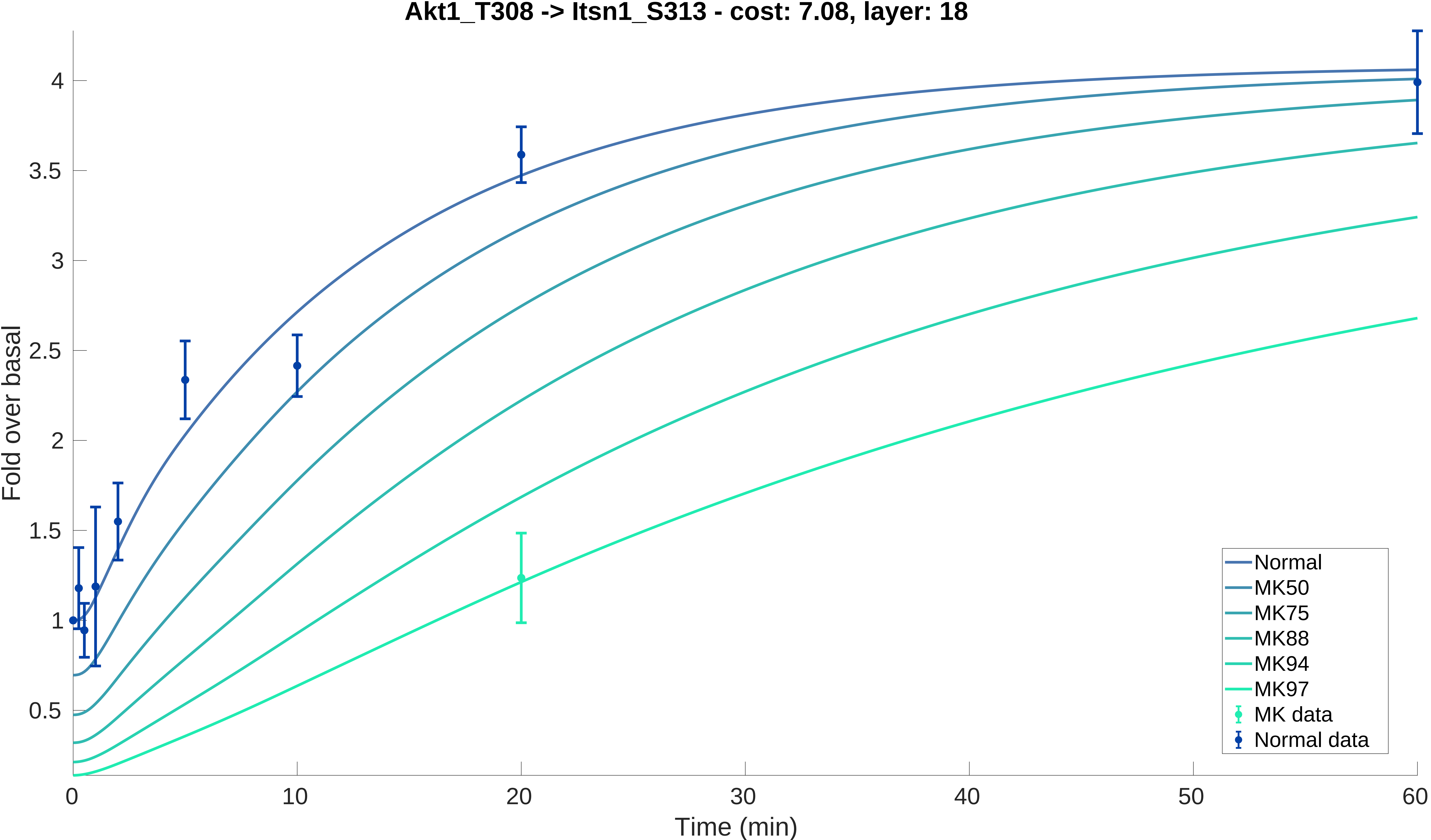
Akt1_T308 -> Itsn1_S313 - cost: 7.08, layer: 18
Fold over basal
Time (min)
0
10
20
30
40
50
60
0.5
1
1.5
2
2.5
3
3.5
4
Normal
MK50
MK75
MK88
MK94
MK97
MK data
Normal data

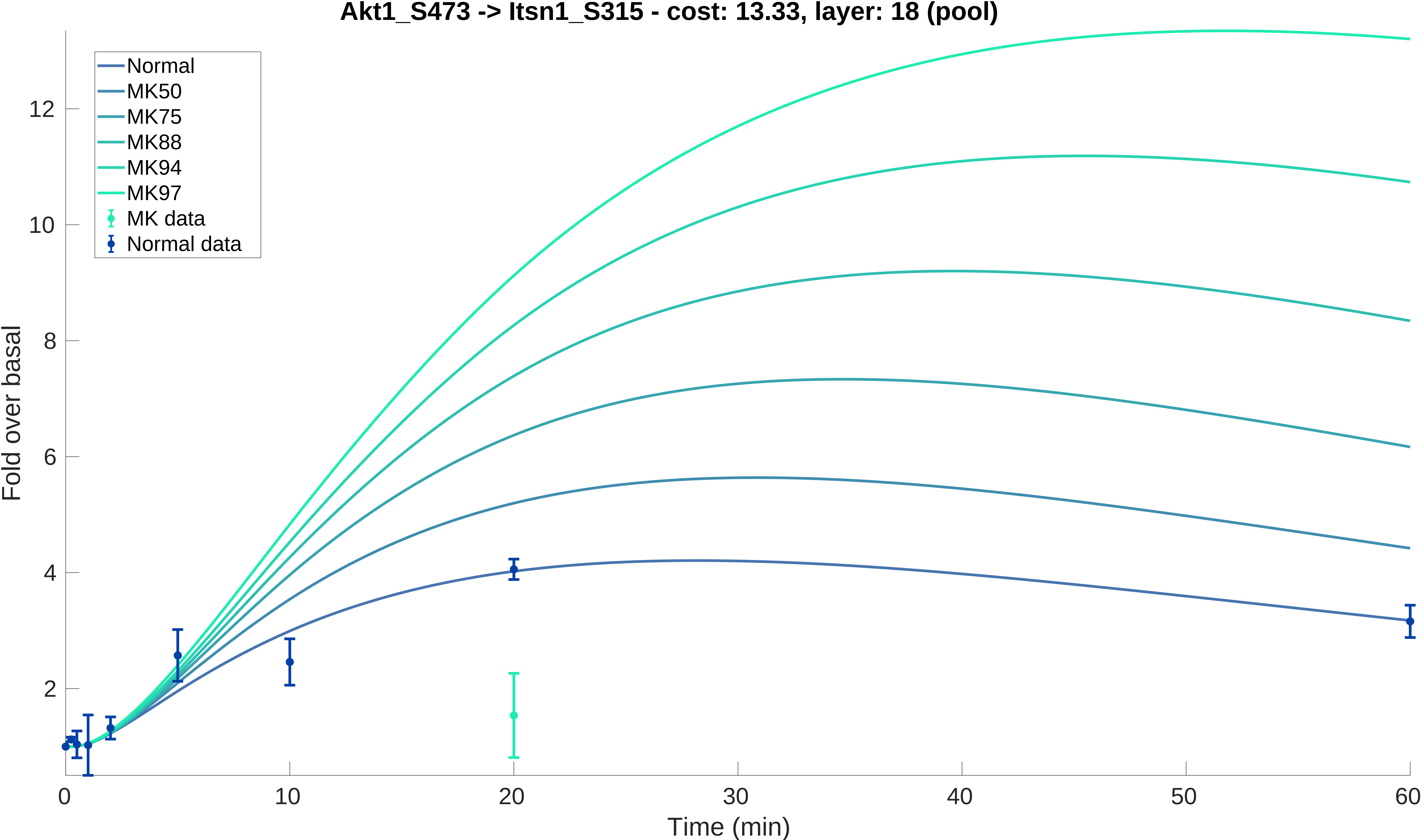
Akt1_S473 -> Itsn1_S315 - cost: 13.33, layer: 18 (pool)
Normal
MK50
MK75
MK88
MK94
MK97
MK data
Normal data
Fold over basal
Time (min)
0
10
20
30
40
50
60
2
4
6
8
10
12

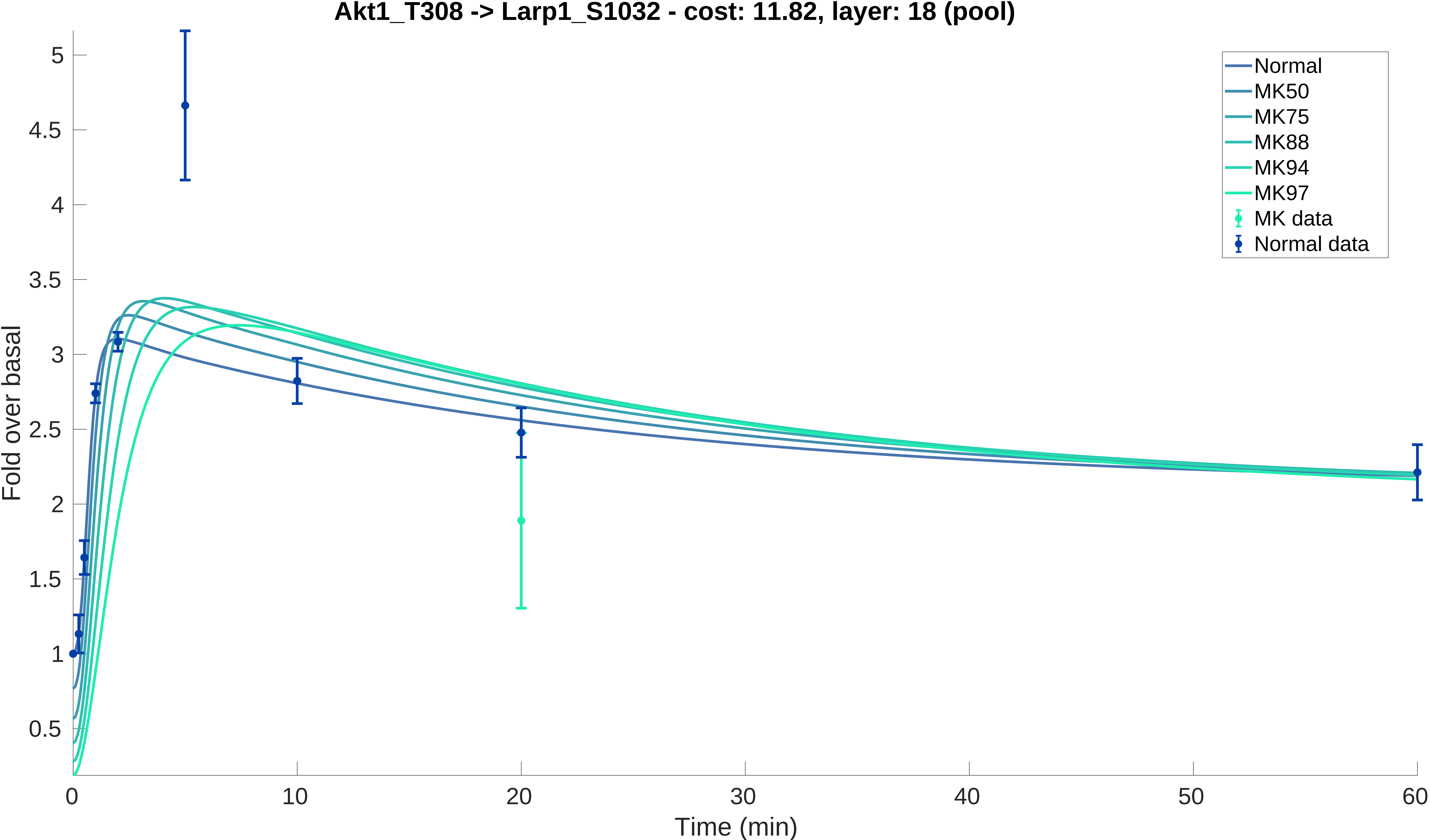
Akt1_T308 -> Larp1_S1032 - cost: 11.82, layer: 18 (pool)
Fold over basal
Time (min)
Normal
MK50
MK75
MK88
MK94
MK97
MK data
Normal data
0.5
1
1.5
2
2.5
3
3.5
4
4.5
5
0
10
20
30
40
50
60

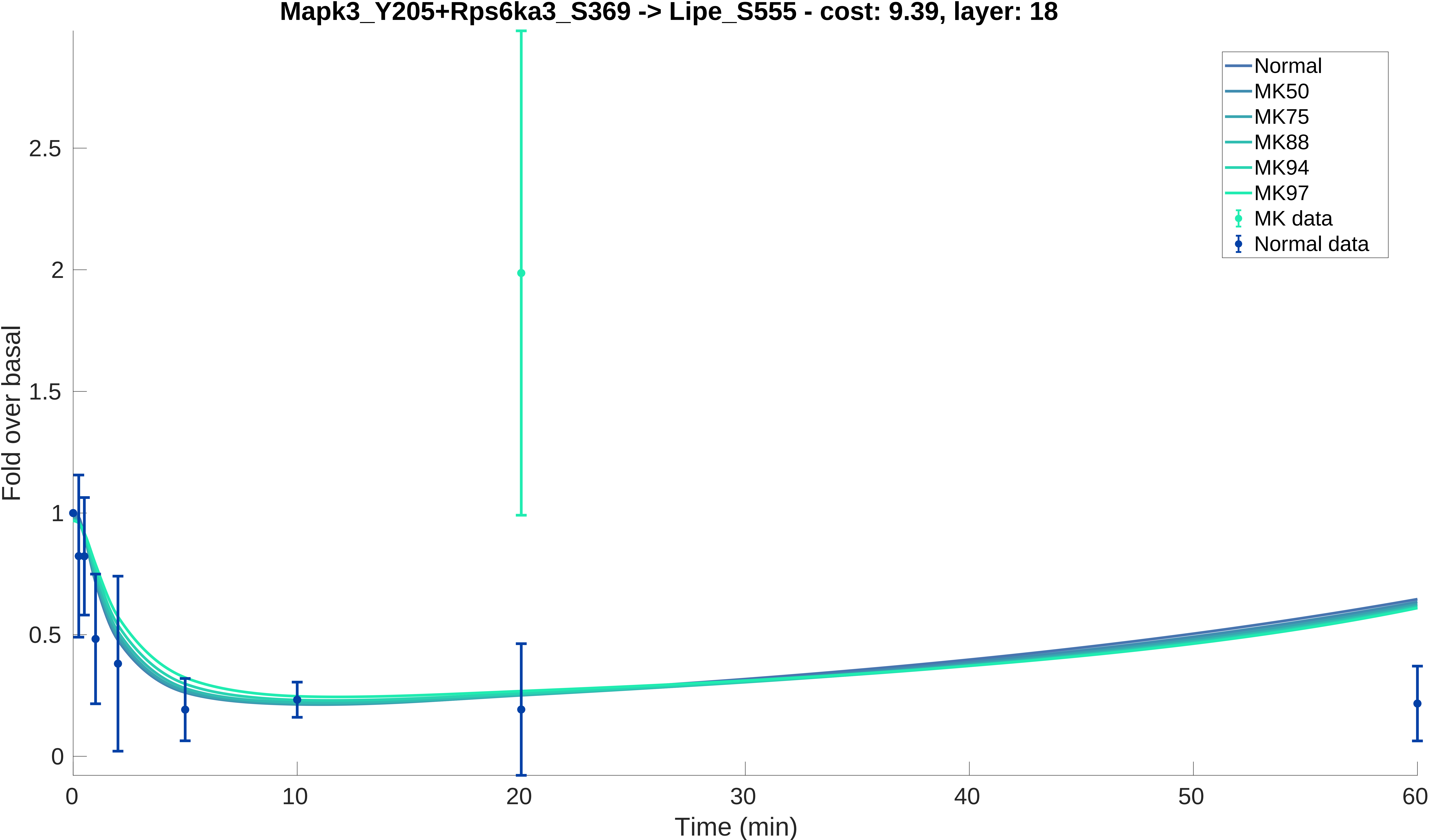
Mapk3_Y205+Rps6ka3_S369 -> Lipe_S555 - cost: 9.39, layer: 18
Fold over basal
Time (min)
0
0.5
1
1.5
2
2.5
0
10
20
30
40
50
60
Normal
MK50
MK75
MK88
MK94
MK97
MK data
Normal data

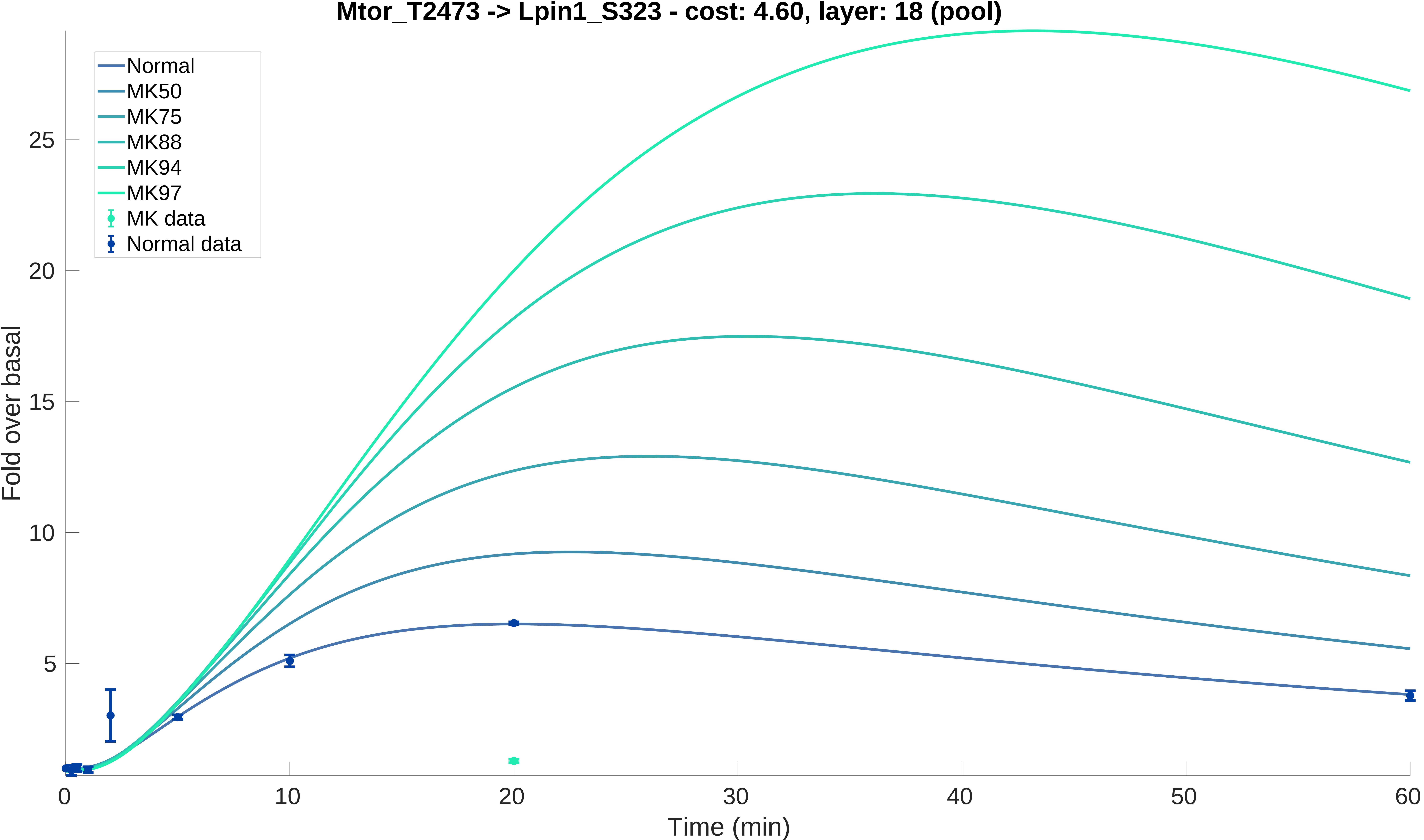

Mtor_T2473 -> Lpin1_S323 - cost: 4.60, layer: 18 (pool)
Normal
MK50
MK75
MK88
MK94
MK97
MK data
Normal data
Fold over basal
Time (min)
0
10
20
30
40
50
60
5
10
15
20
25

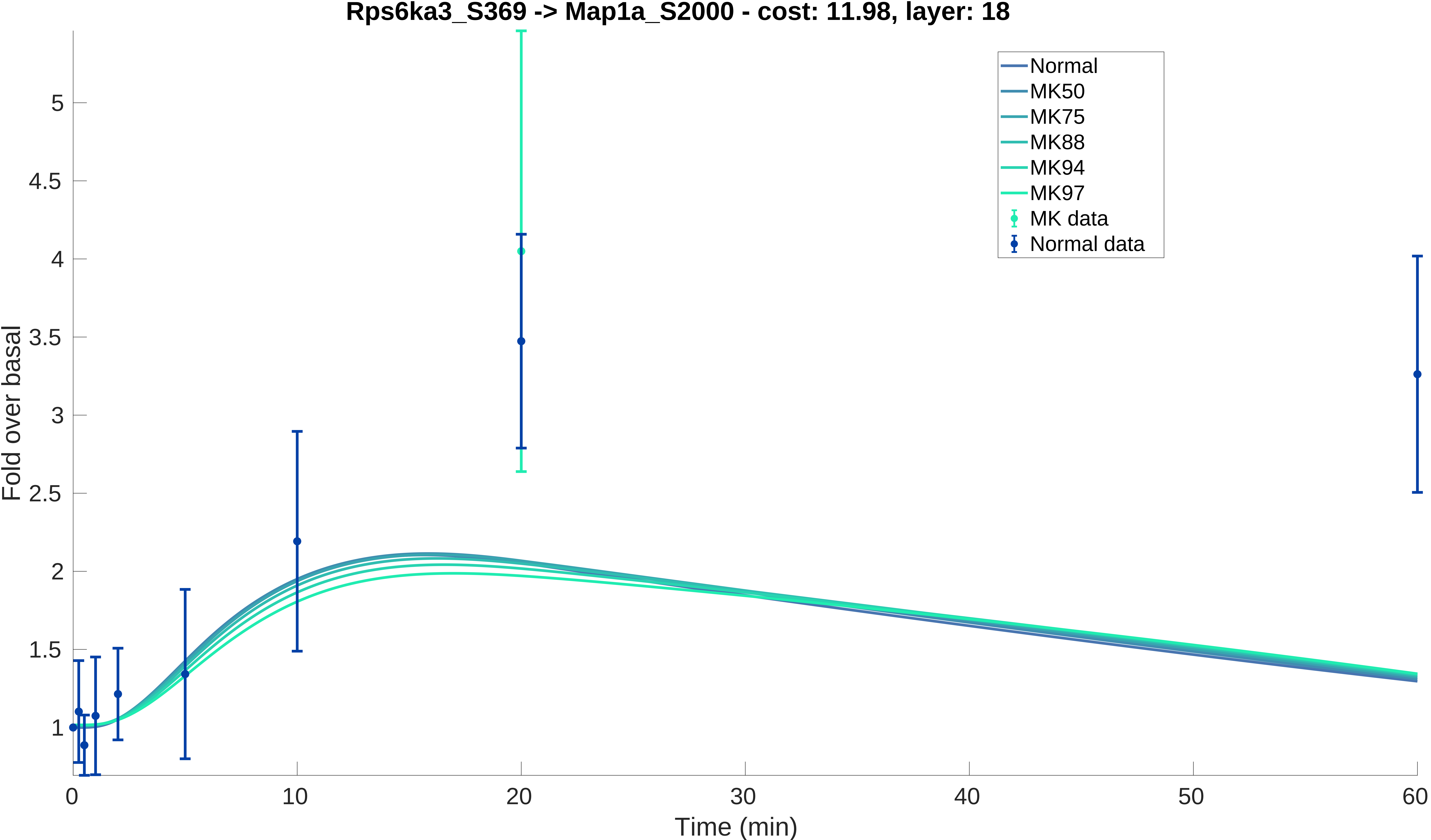

Rps6ka3_S369 -> Map1a_S2000 - cost: 11.98, layer: 18
Fold over basal
Time (min)
Normal
MK50
MK75
MK88
MK94
MK97
MK data
Normal data
5
4.5
4
3.5
3
2.5
2
1.5
1
0
10
20
30
40
50
60

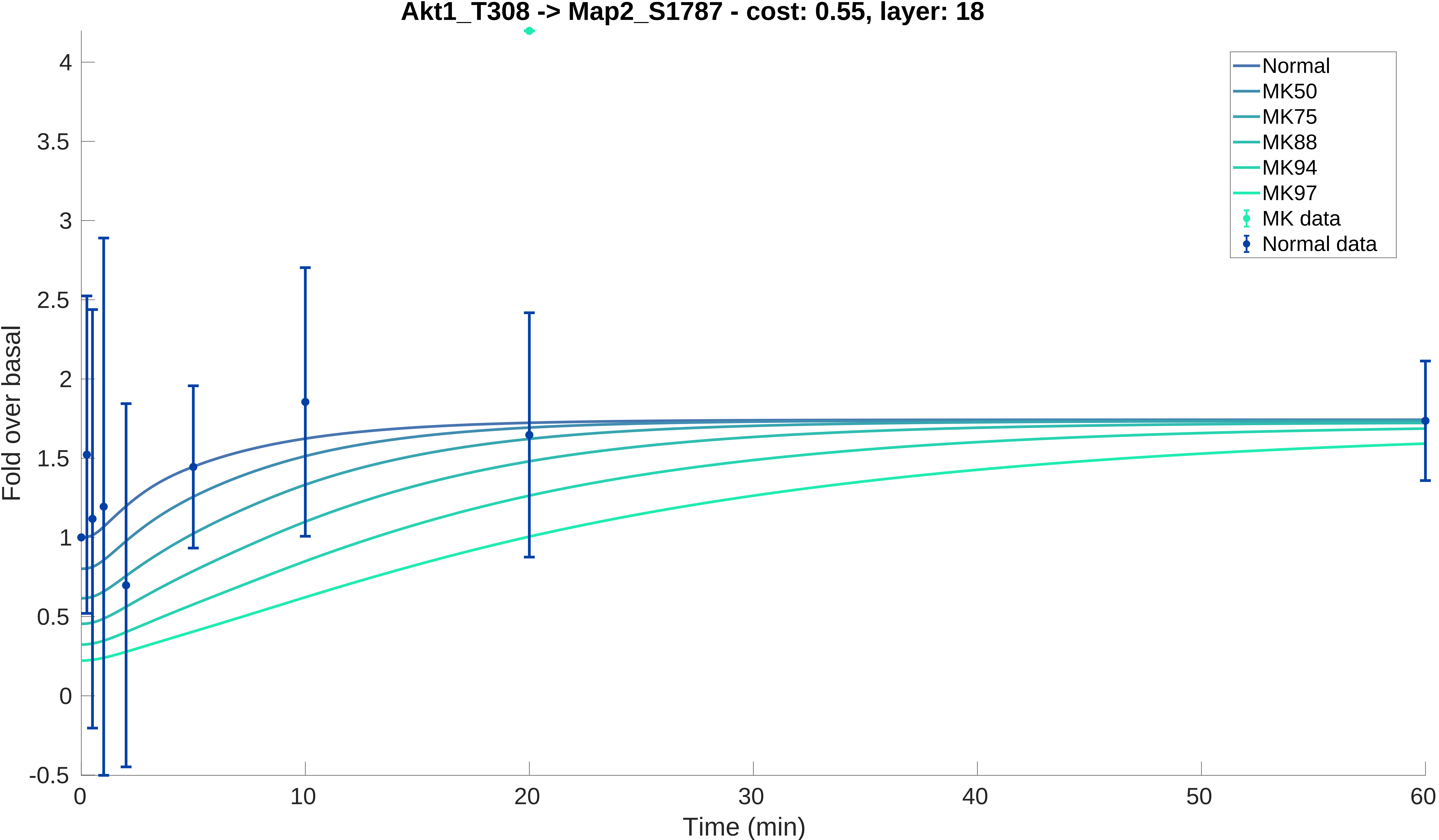

Akt1_T308 -> Map2_S1787 - cost: 0.55, layer: 18
Fold over basal
Time (min)
4
3.5
3
2.5
2
1.5
1
0.5
0
-0.5
0
10
20
30
40
50
60
Normal
MK50
MK75
MK88
MK94
MK97
MK data
Normal data

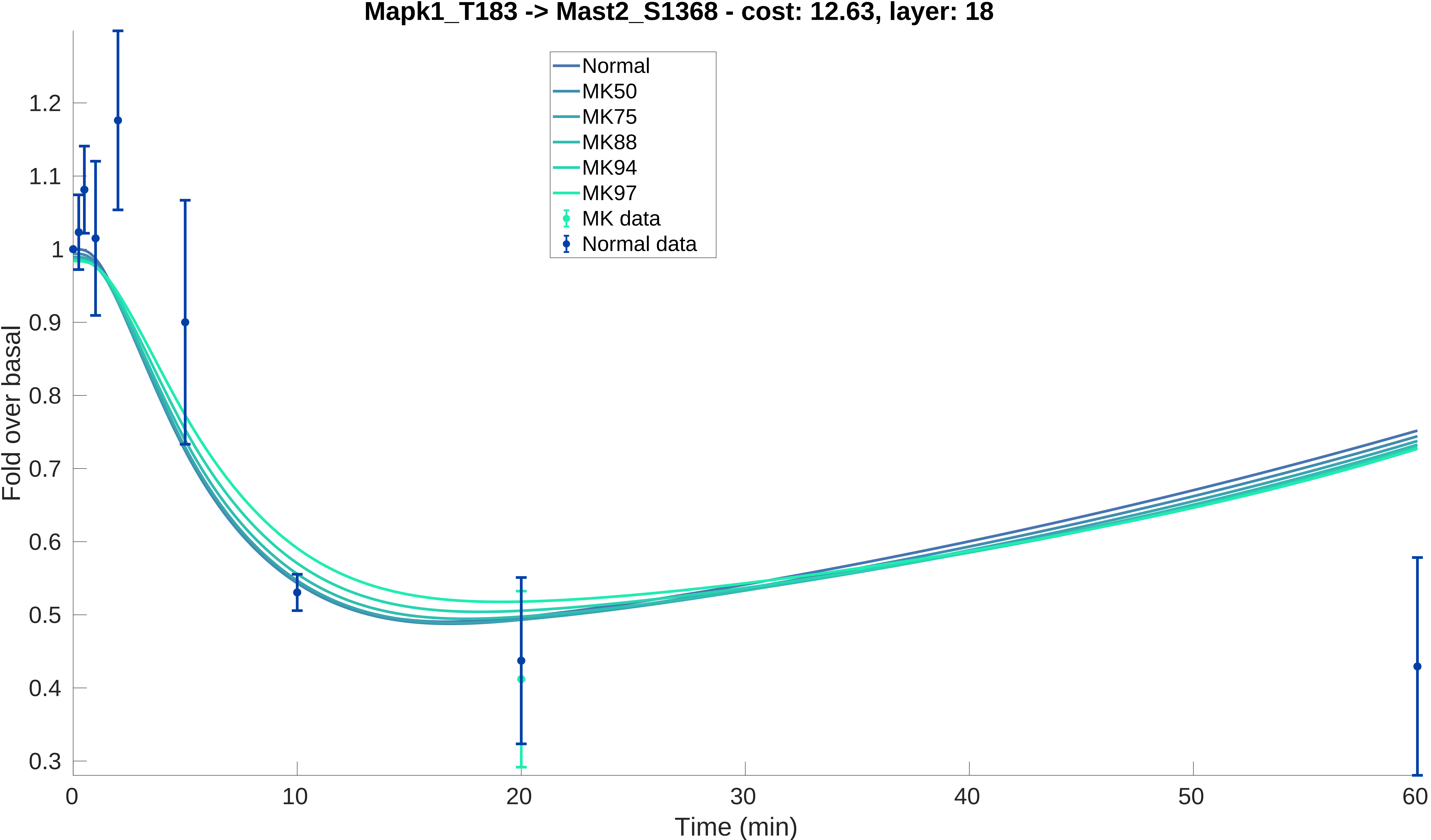

Mapk1_T183 -> Mast2_S1368 - cost: 12.63, layer: 18
Normal
MK50
MK75
MK88
MK94
MK97
MK data
Normal data
Fold over basal
Time (min)
0
10
20
30
40
50
60
0.3
0.4
0.5
0.6
0.7
0.8
0.9
1
1.1
1.2

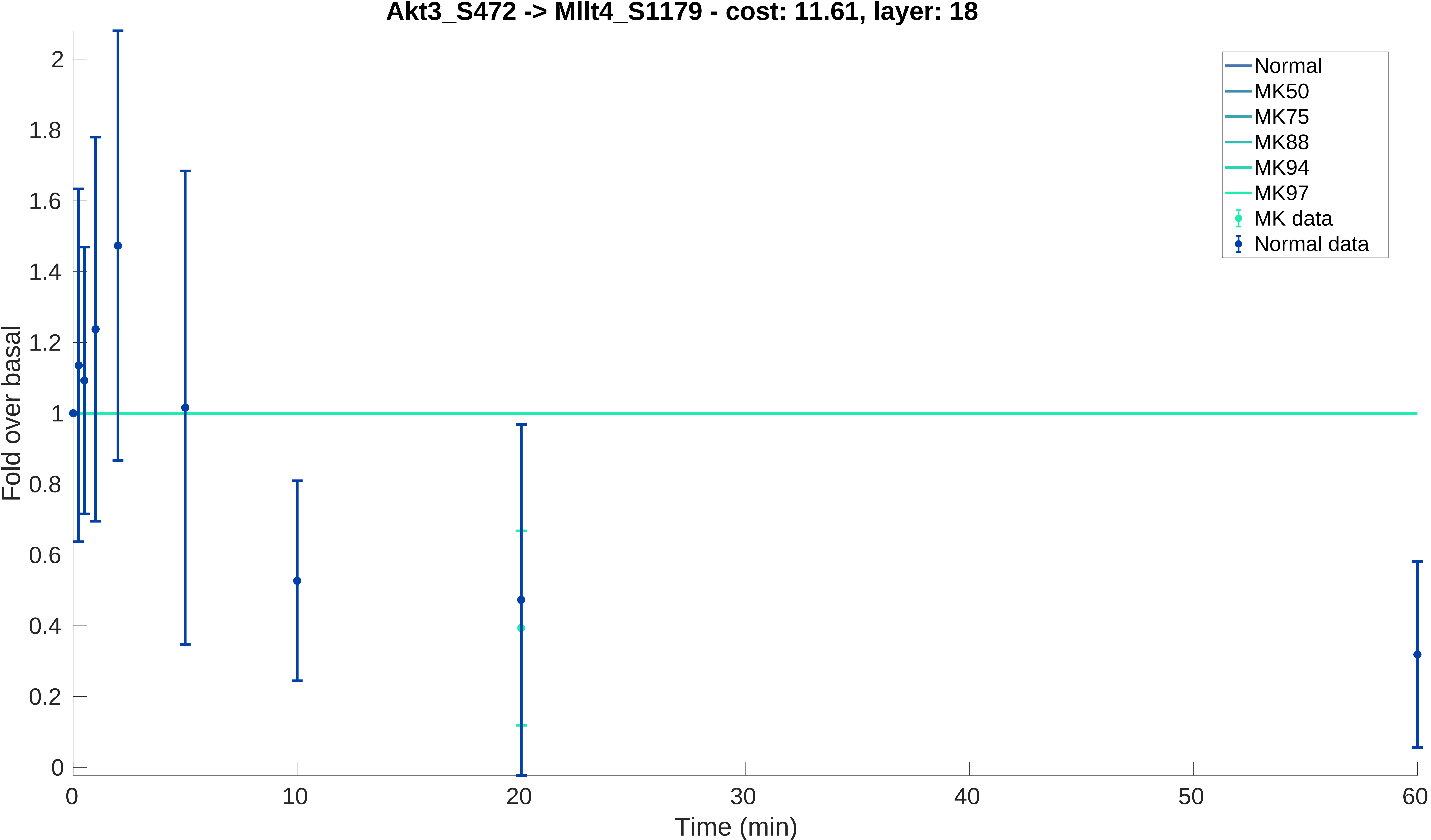

Akt3_S472 -> Mllt4_S1179 - cost: 11.61, layer: 18
Fold over basal
Time (min)
0
0.2
0.4
0.6
0.8
1
1.2
1.4
1.6
1.8
2
0
10
20
30
40
50
60
Normal
MK50
MK75
MK88
MK94
MK97
MK data
Normal data

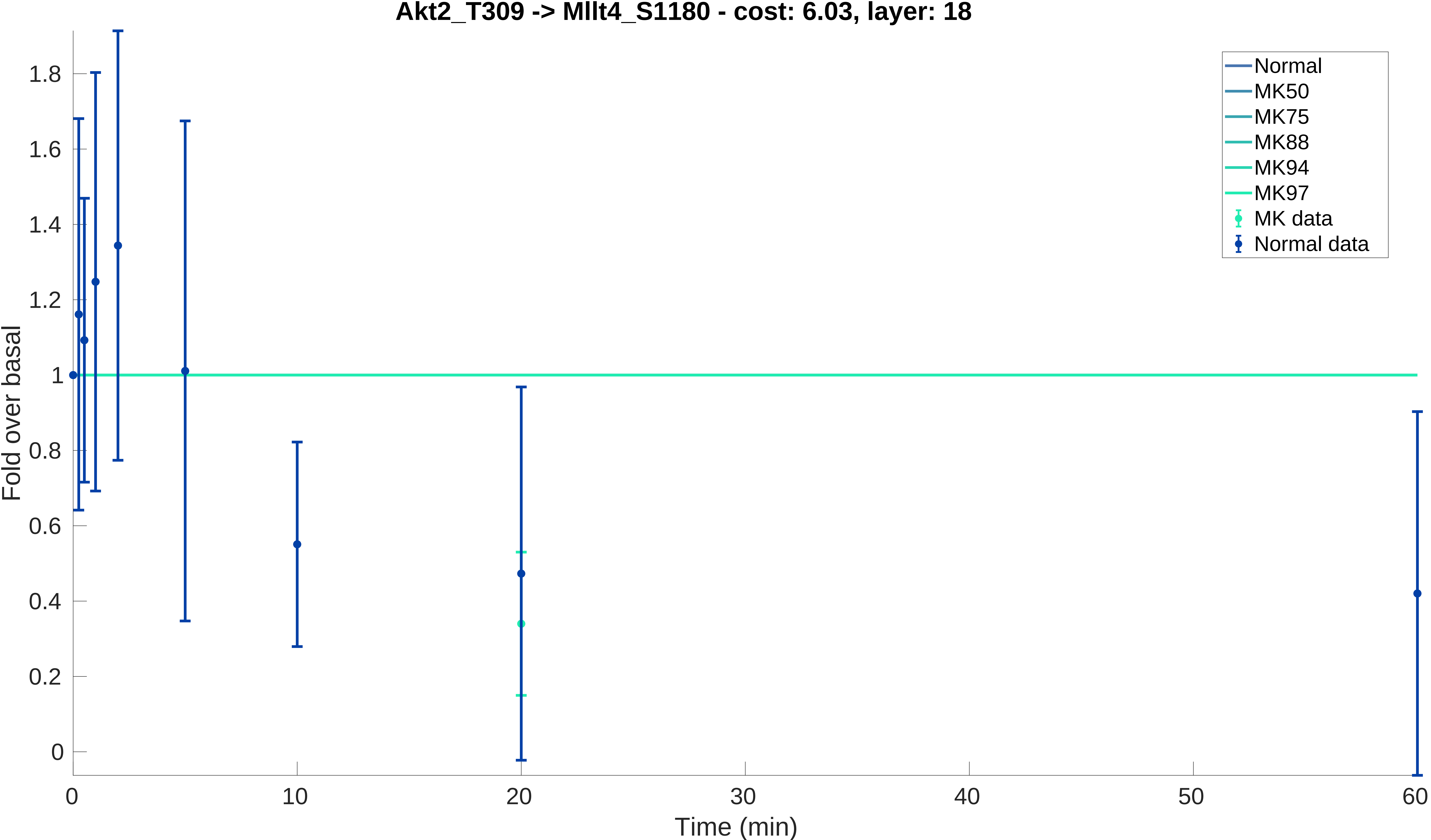

Akt2_T309 -> Mllt4_S1180 - cost: 6.03, layer: 18
Fold over basal
Time (min)
0
0.2
0.4
0.6
0.8
1
1.2
1.4
1.6
1.8
0
10
20
30
40
50
60
Normal
MK50
MK75
MK88
MK94
MK97
MK data
Normal data

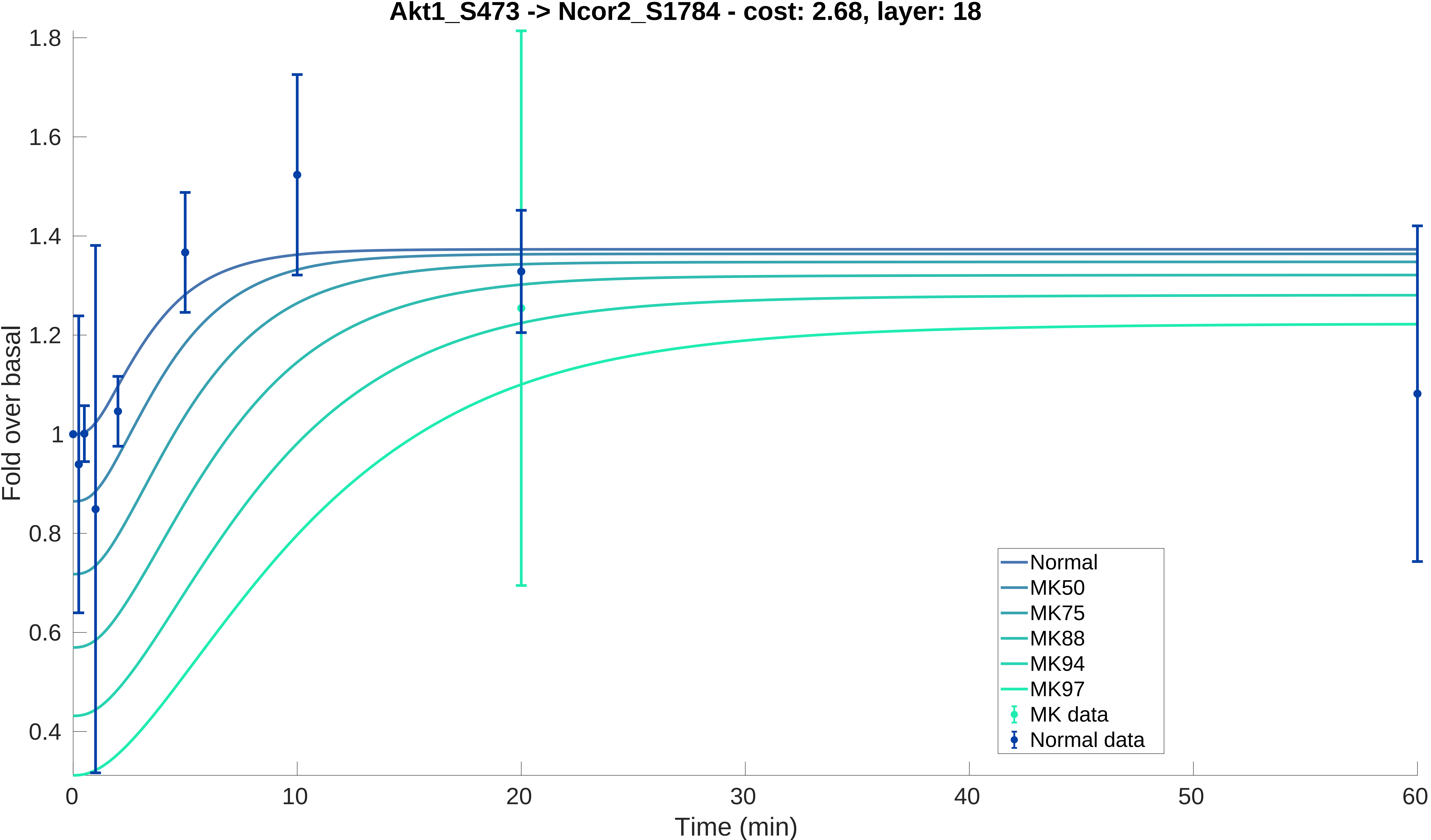

Akt1_S473 -> Ncor2_S1784 - cost: 2.68, layer: 18
Fold over basal
Time (min)
0.4
0.6
0.8
1
1.2
1.4
1.6
1.8
0
10
20
30
40
50
60
Normal
MK50
MK75
MK88
MK94
MK97
MK data
Normal data

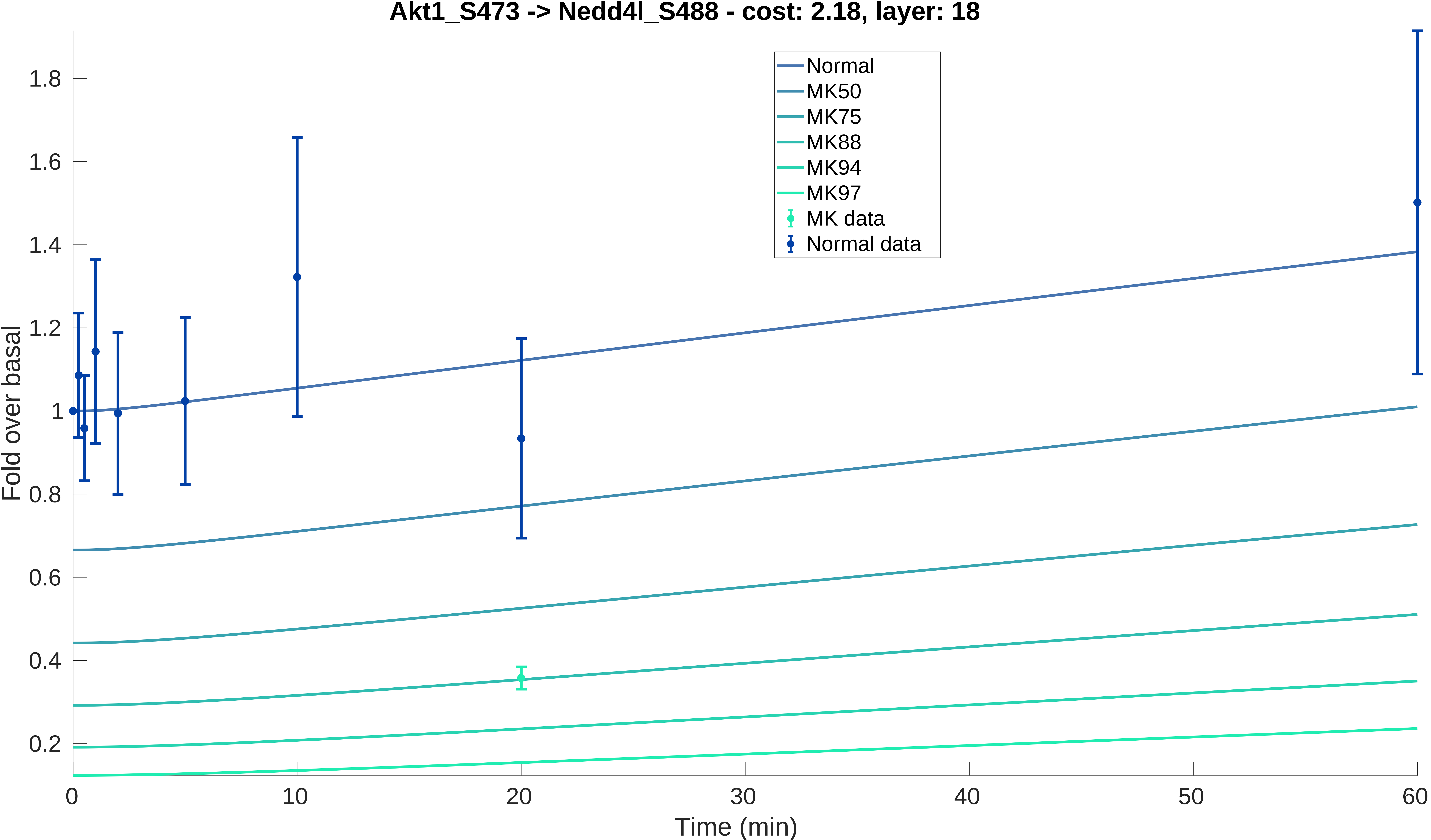

Akt1_S473 -> Nedd4l_S488 - cost: 2.18, layer: 18
Normal
MK50
MK75
MK88
MK94
MK97
MK data
Normal data
Fold over basal
1.8
1.6
1.4
1.2
1
0.8
0.6
0.4
0.2
0
10
20
30
40
50
60
Time (min)

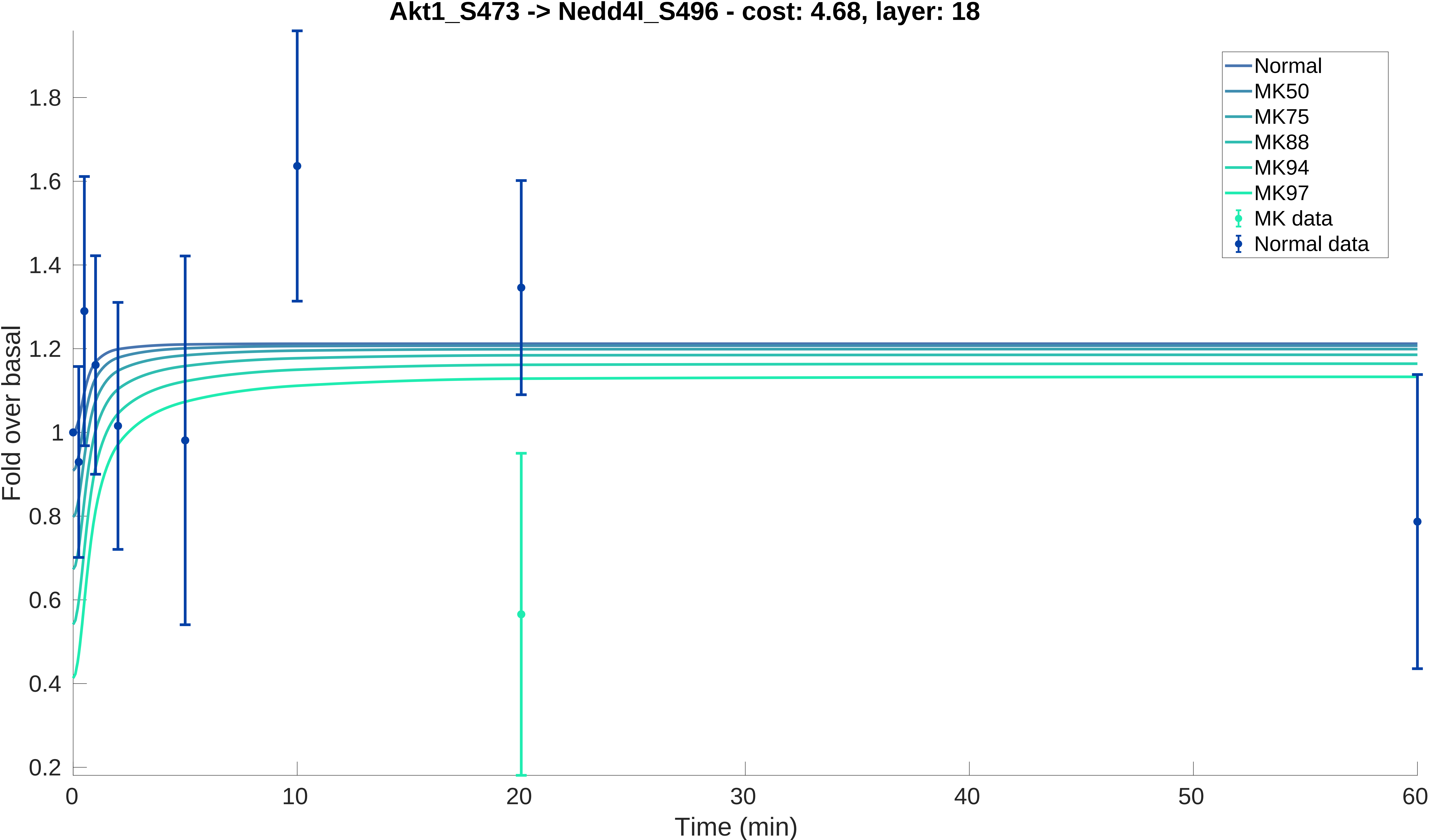

Akt1_S473 -> Nedd4l_S496 - cost: 4.68, layer: 18
Normal
MK50
MK75
MK88
MK94
MK97
MK data
Normal data
Fold over basal
Time (min)
0
10
20
30
40
50
60
0.2
0.4
0.6
0.8
1
1.2
1.4
1.6
1.8

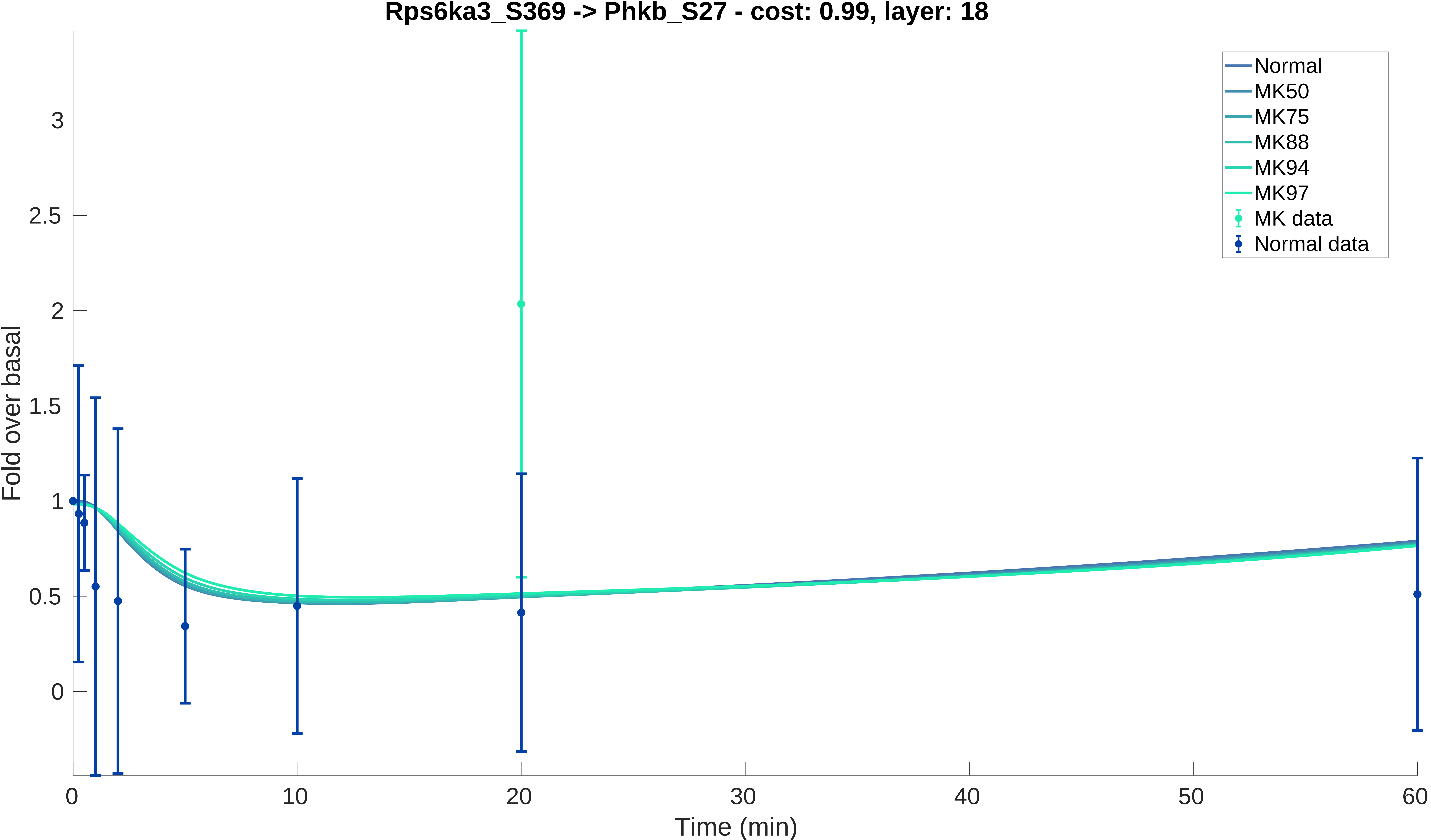
Rps6ka3_S369 -> Phkb_S27 - cost: 0.99, layer: 18
Fold over basal
Time (min)
0
0.5
1
1.5
2
2.5
3
0
10
20
30
40
50
60
Normal
MK50
MK75
MK88
MK94
MK97
MK data
Normal data

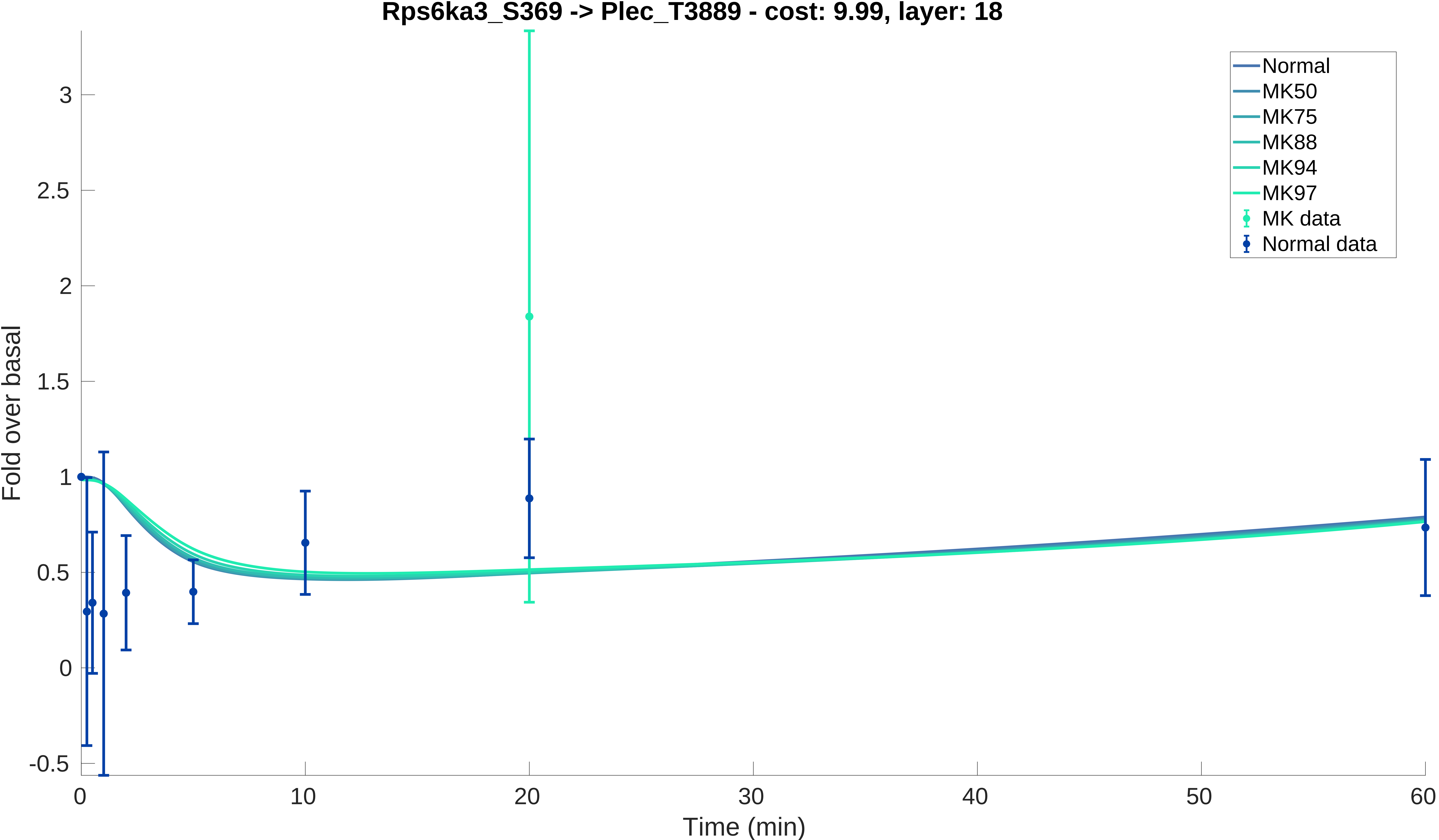
Rps6ka3_S369 -> Plec_T3889 - cost: 9.99, layer: 18
Fold over basal
Time (min)
Normal
MK50
MK75
MK88
MK94
MK97
MK data
Normal data
3
2.5
2
1.5
1
0.5
0
-0.5
0
10
20
30
40
50
60

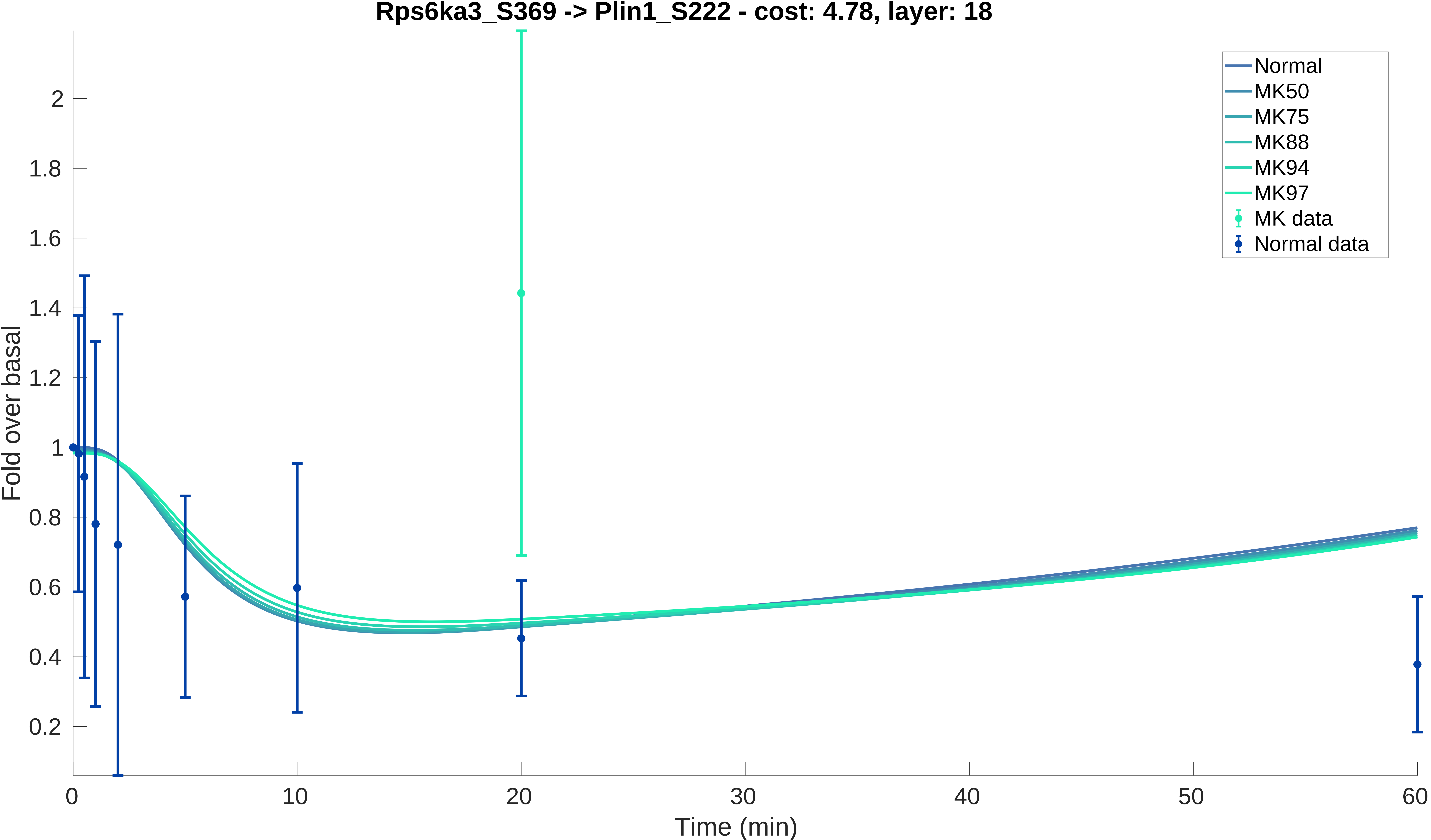

Rps6ka3_S369 -> Plin1_S222 - cost: 4.78, layer: 18
Fold over basal
Time (min)
Normal
MK50
MK75
MK88
MK94
MK97
MK data
Normal data
0
10
20
30
40
50
60
0.2
0.4
0.6
0.8
1
1.2
1.4
1.6
1.8
2

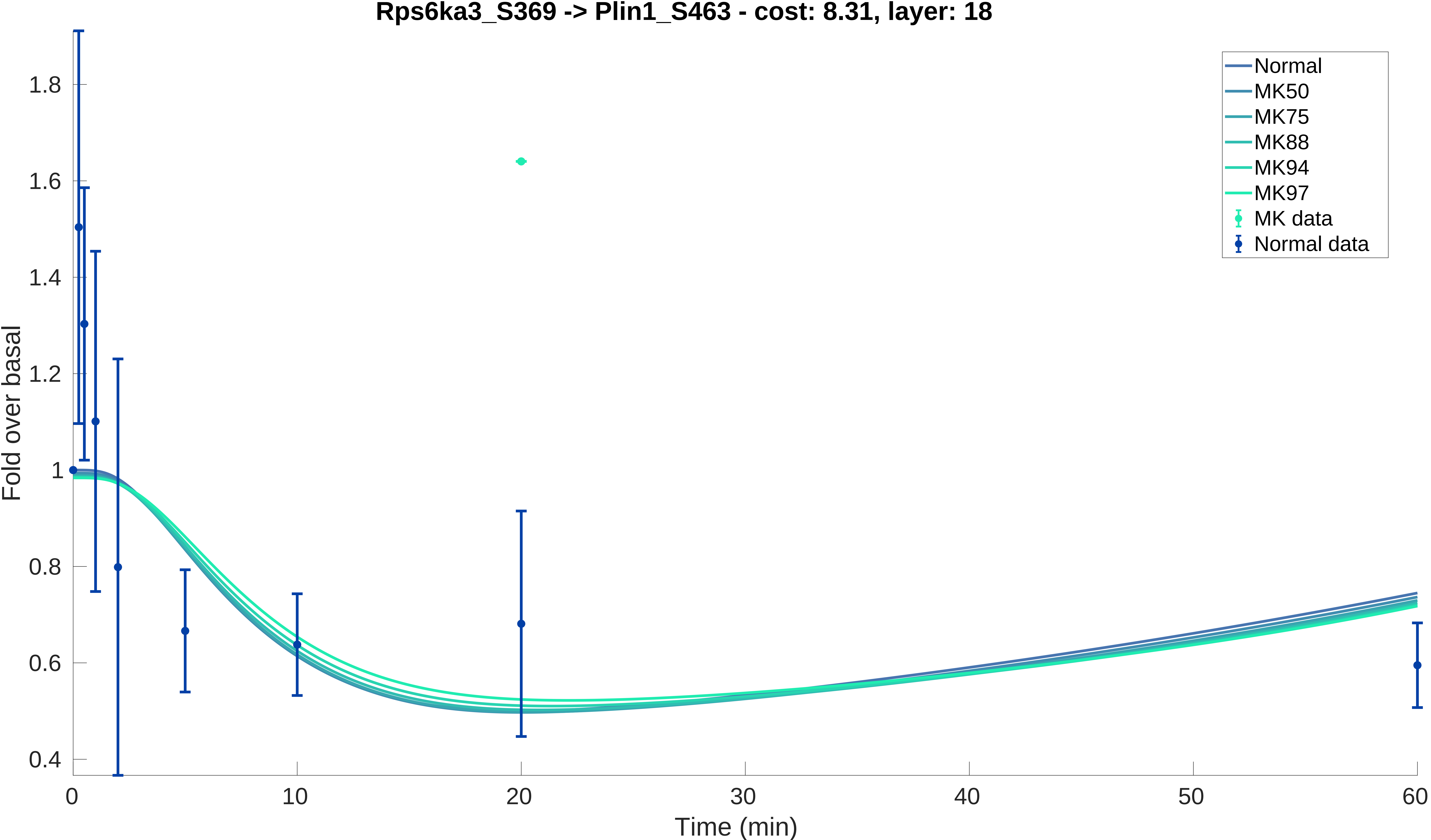

Rps6ka3_S369 -> Plin1_S463 - cost: 8.31, layer: 18
Fold over basal
Time (min)
0.4
0.6
0.8
1
1.2
1.4
1.6
1.8
0
10
20
30
40
50
60
Normal
MK50
MK75
MK88
MK94
MK97
MK data
Normal data

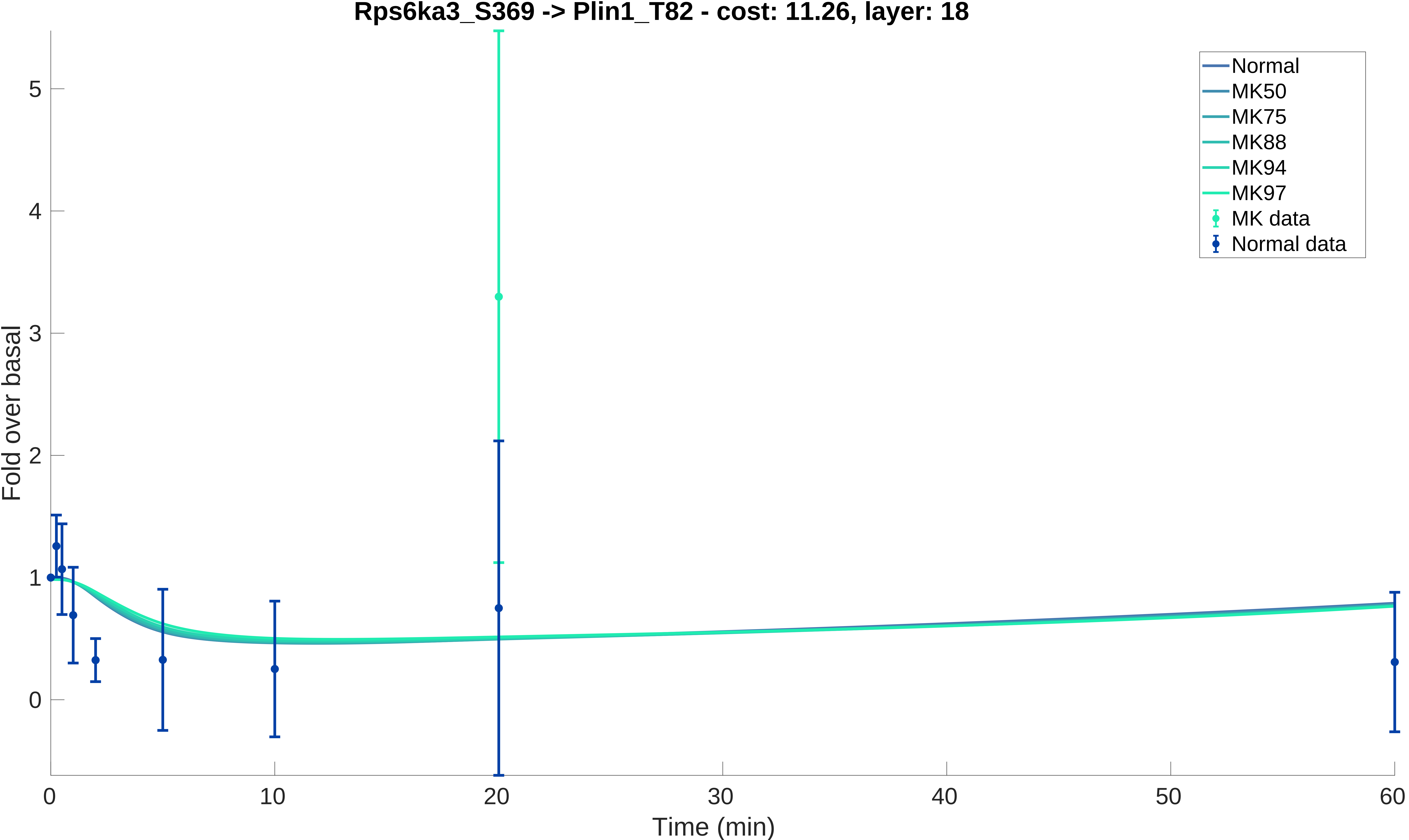
Rps6ka3_S369 -> Plin1_T82 - cost: 11.26, layer: 18
Fold over basal
Time (min)
Normal
MK50
MK75
MK88
MK94
MK97
MK data
Normal data
0
1
2
3
4
5
0
10
20
30
40
50
60

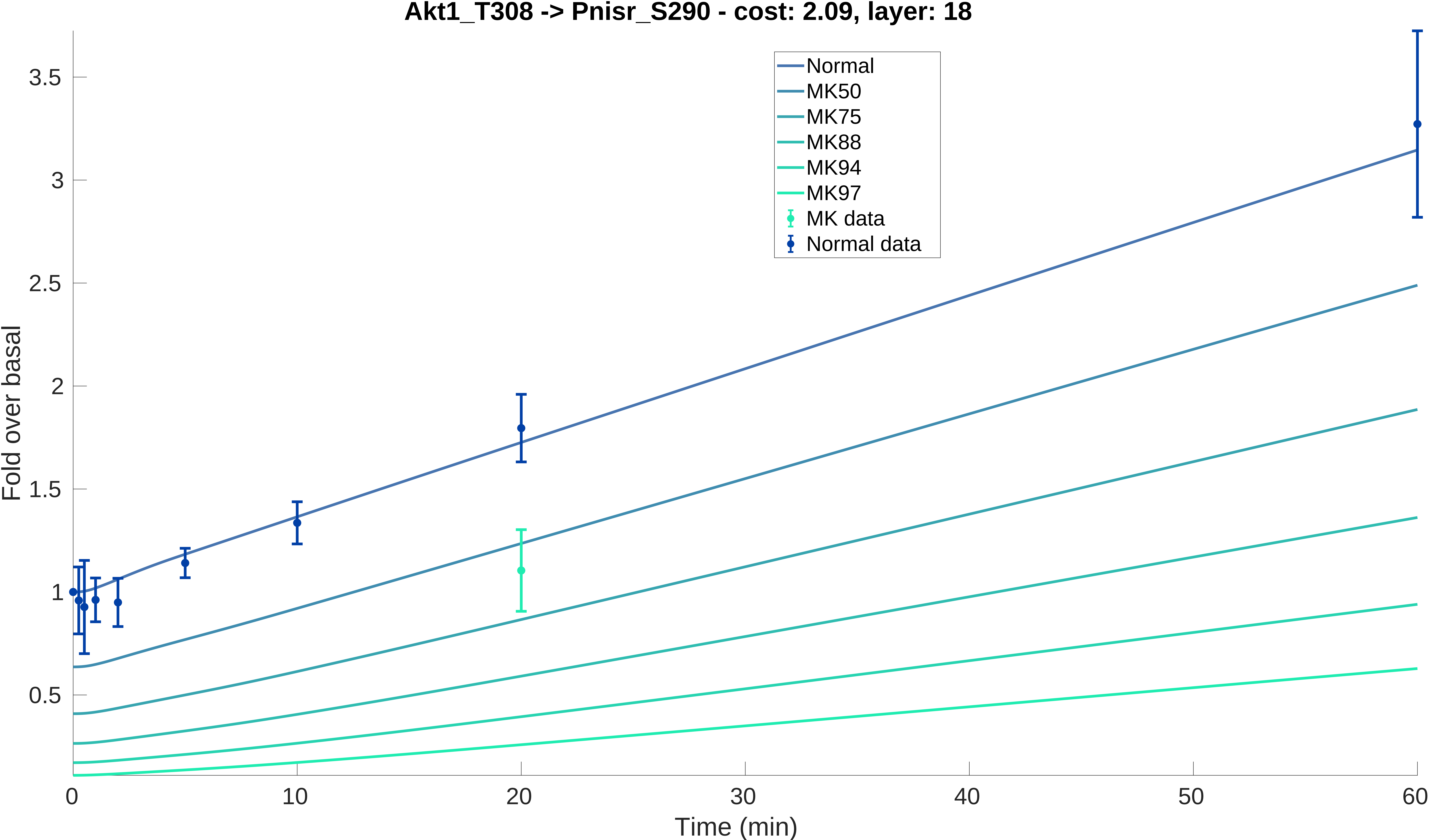
Akt1_T308 -> Pnisr_S290 - cost: 2.09, layer: 18
Fold over basal
Time (min)
Normal
MK50
MK75
MK88
MK94
MK97
MK data
Normal data
0.5
1
1.5
2
2.5
3
3.5
0
10
20
30
40
50
60

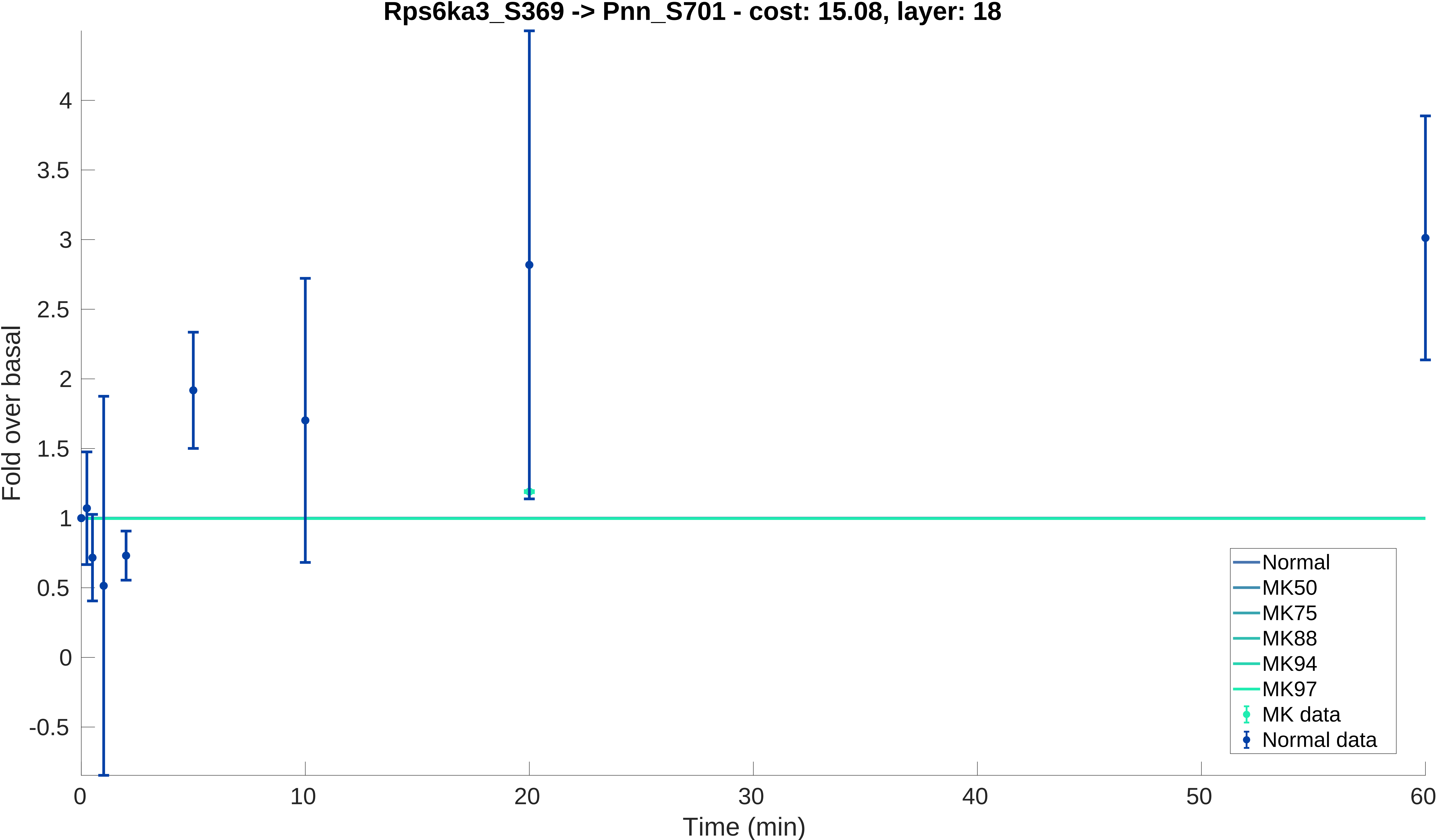

Rps6ka3_S369 -> Pnn_S701 - cost: 15.08, layer: 18
Fold over basal
Time (min)
4
3.5
3
2.5
2
1.5
1
0.5
0
-0.5
0
10
20
30
40
50
60
Normal
MK50
MK75
MK88
MK94
MK97
MK data
Normal data

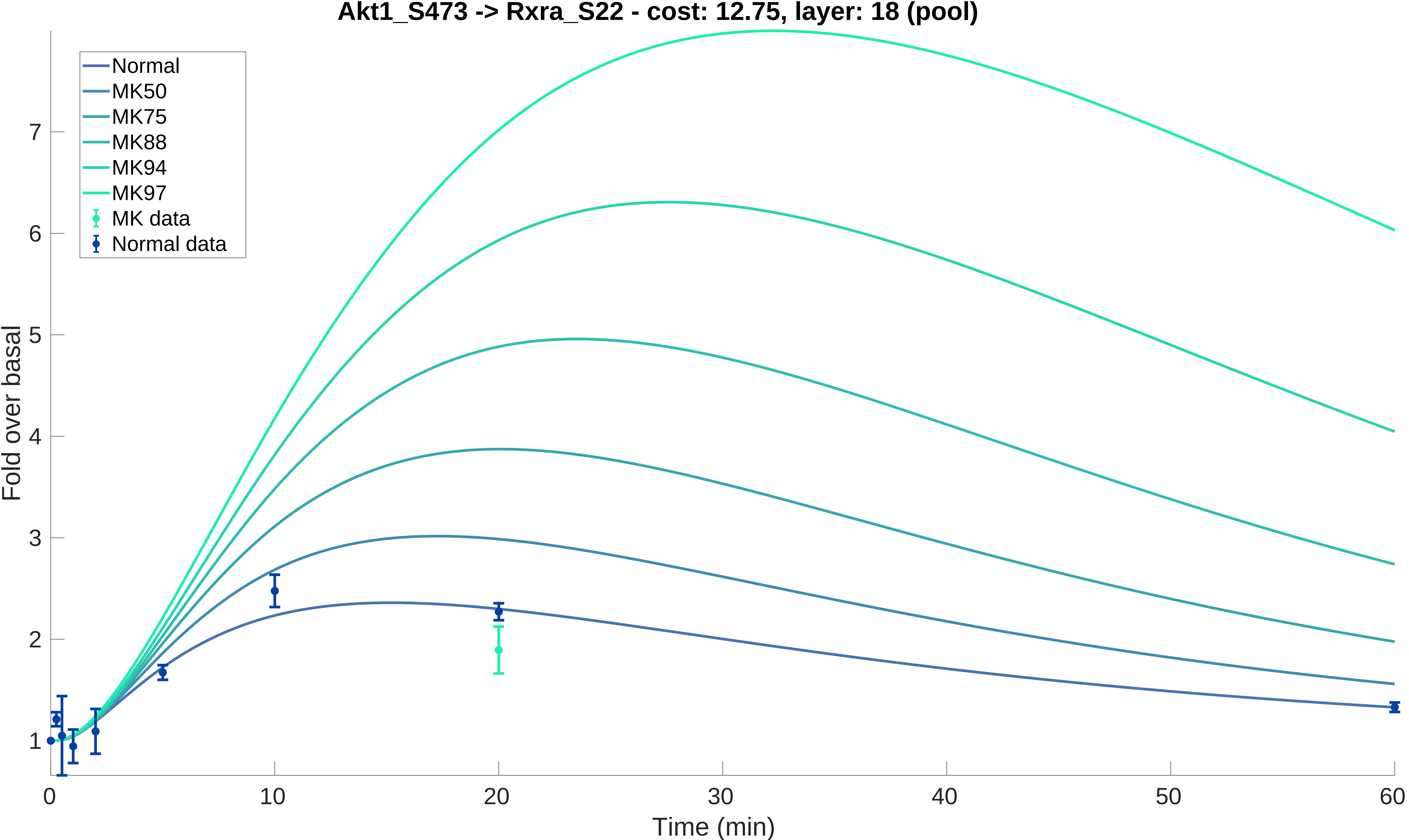

Akt1_S473 -> Rxra_S22 - cost: 12.75, layer: 18 (pool)
Normal
MK50
MK75
MK88
MK94
MK97
MK data
Normal data
Fold over basal
Time (min)
1
2
3
4
5
6
7
0
10
20
30
40
50
60

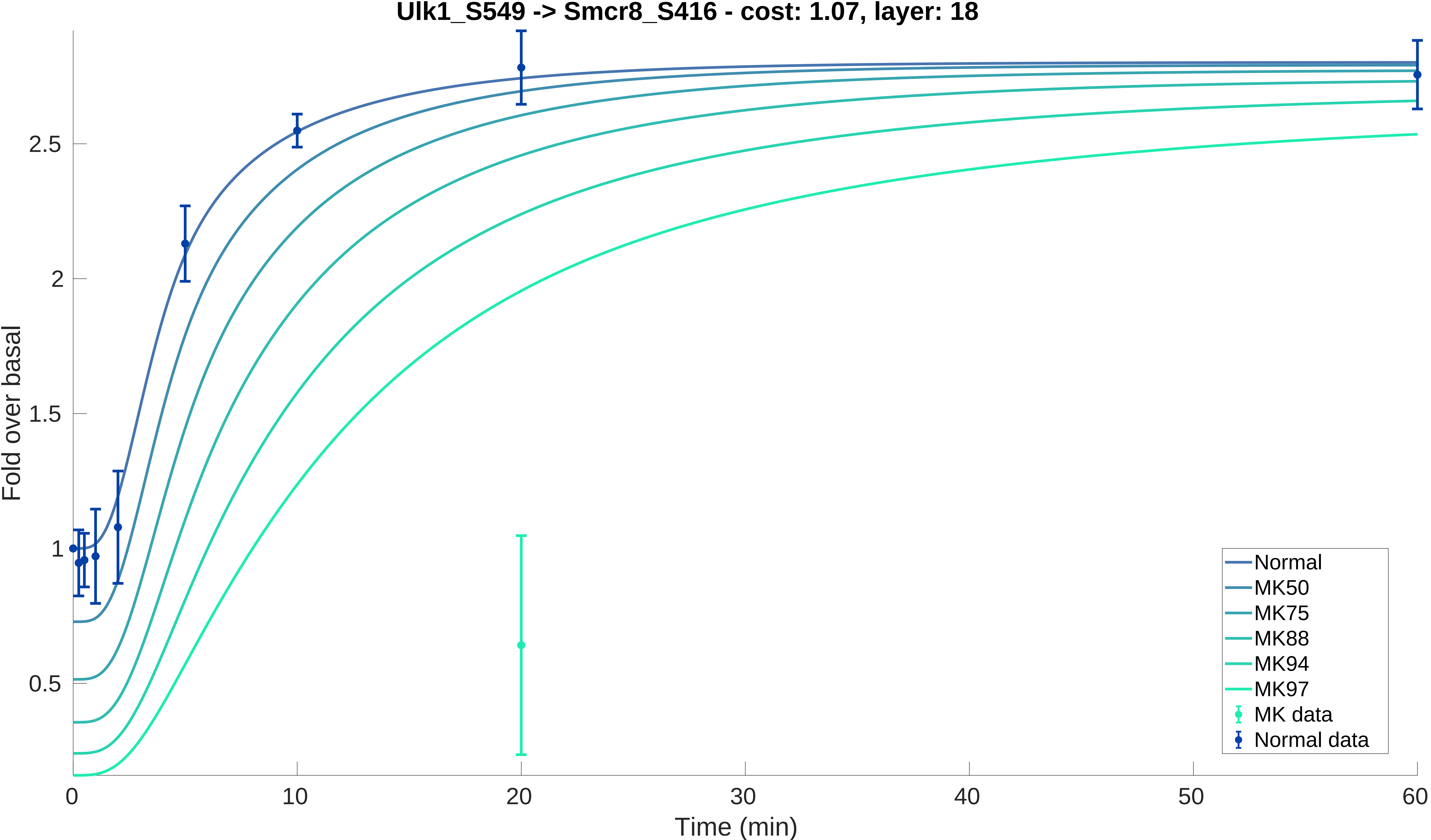

Ulk1_S549 -> Smcr8_S416 - cost: 1.07, layer: 18
Fold over basal
Time (min)
0
10
20
30
40
50
60
0.5
1
1.5
2
2.5
Normal
MK50
MK75
MK88
MK94
MK97
MK data
Normal data

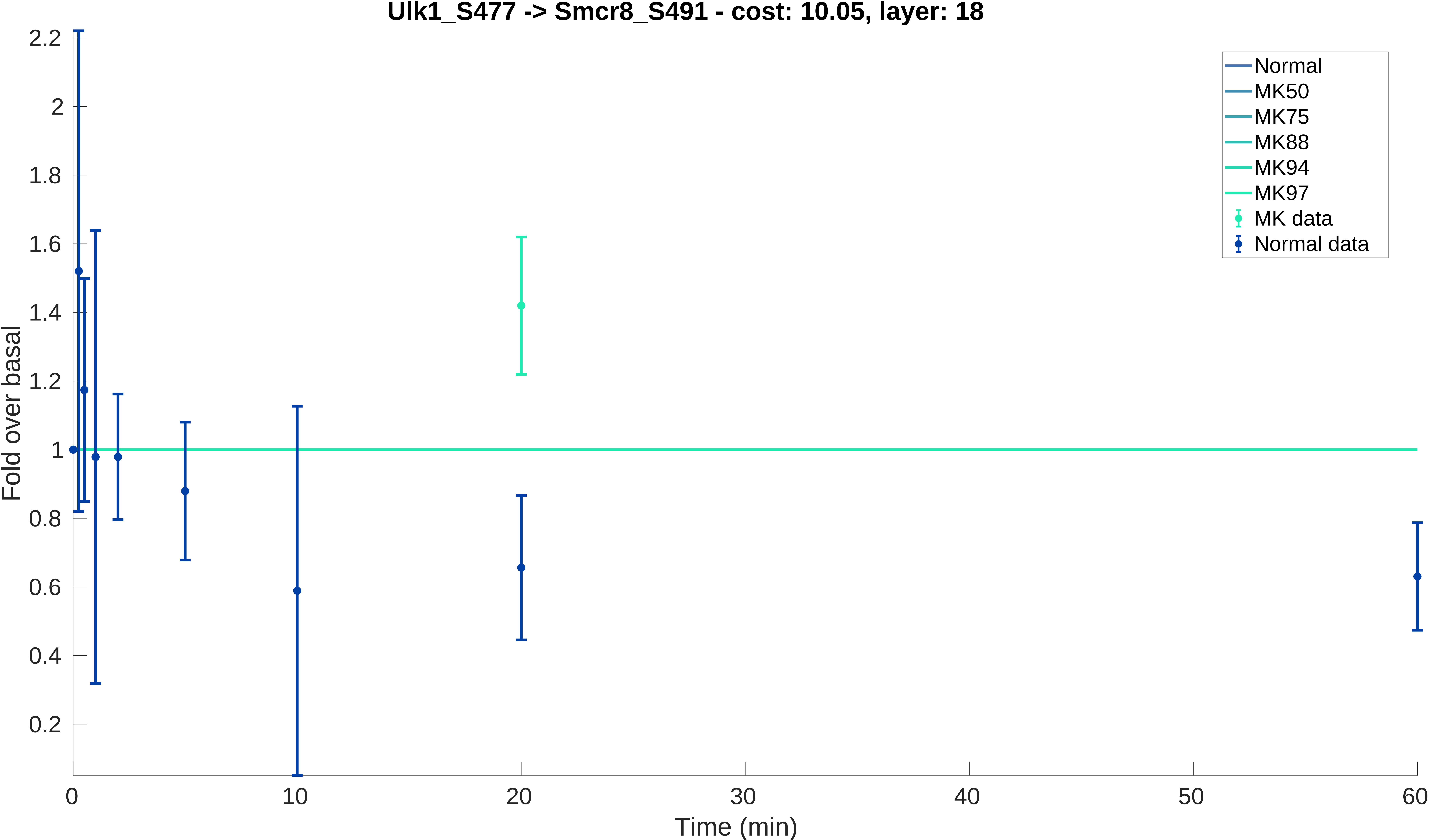

Ulk1_S477 -> Smcr8_S491 - cost: 10.05, layer: 18
Fold over basal
2.2
2
1.8
1.6
1.4
1.2
1
0.8
0.6
0.4
0.2
0
10
20
30
40
50
60
Time (min)
Normal
MK50
MK75
MK88
MK94
MK97
MK data
Normal data

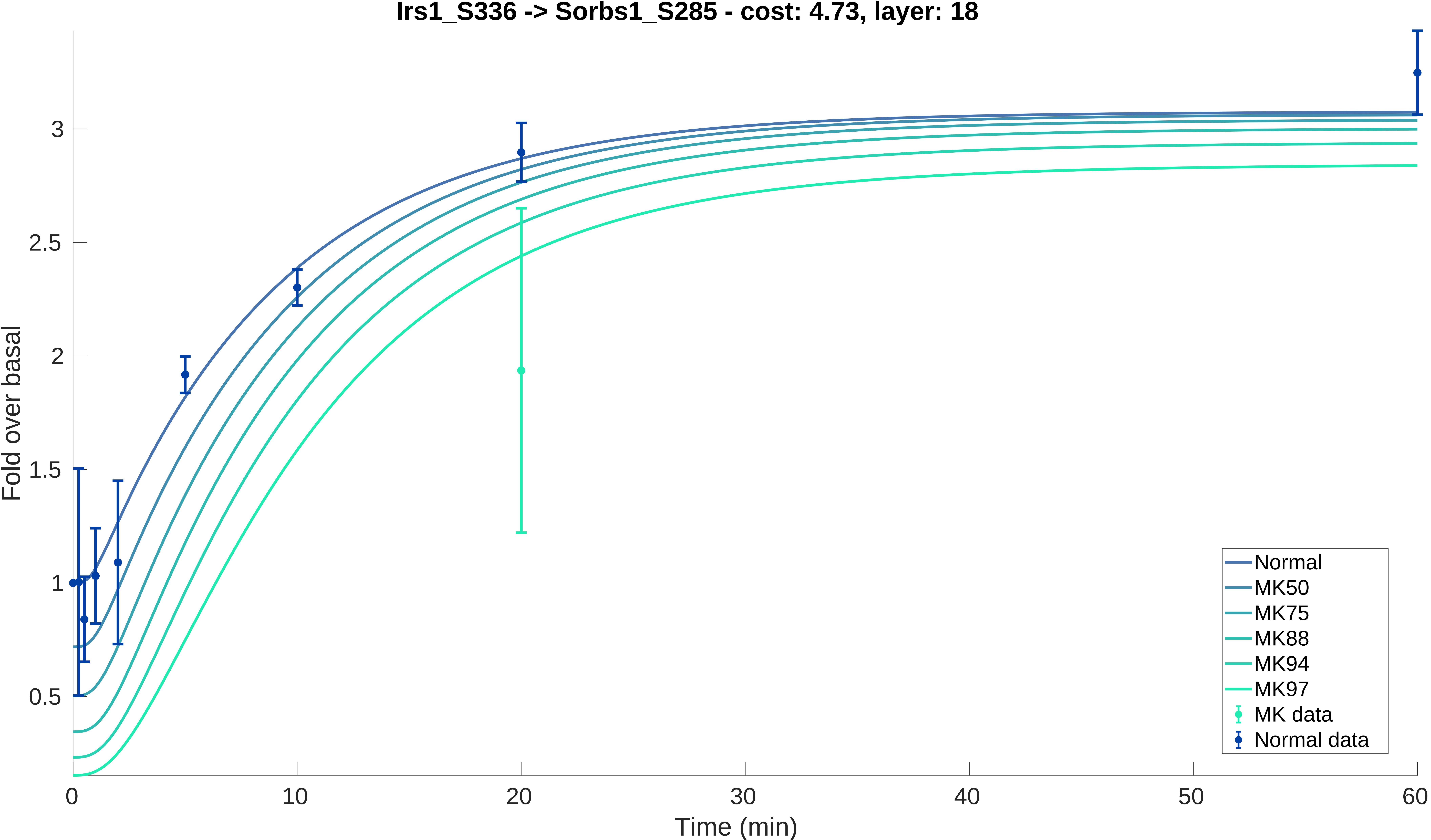
Irs1_S336 -> Sorbs1_S285 - cost: 4.73, layer: 18
Fold over basal
Time (min)
0.5
1
1.5
2
2.5
3
0
10
20
30
40
50
60
Normal
MK50
MK75
MK88
MK94
MK97
MK data
Normal data

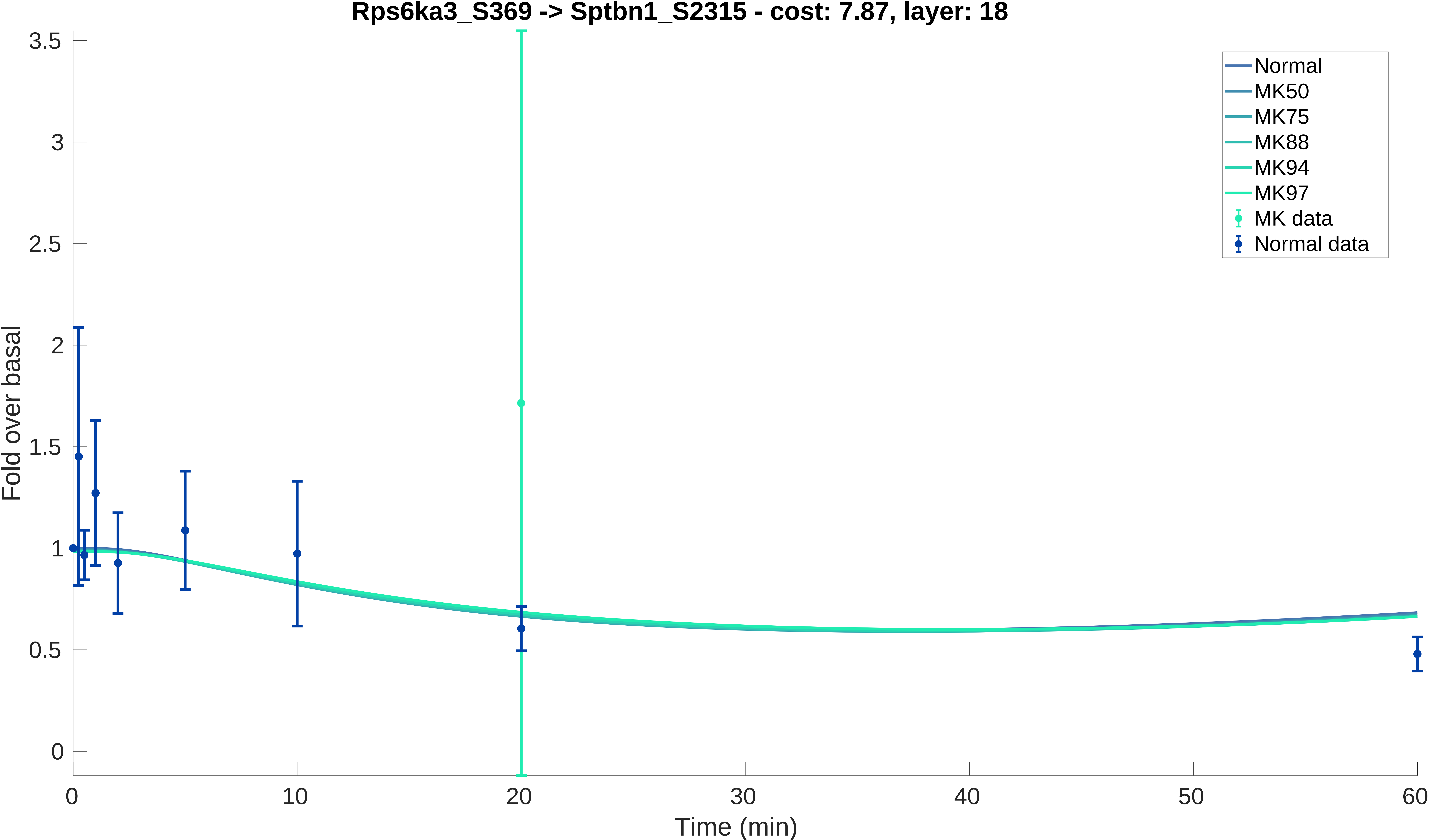

Rps6ka3_S369 -> Sptbn1_S2315 - cost: 7.87, layer: 18
Fold over basal
Time (min)
0
0.5
1
1.5
2
2.5
3
3.5
0
10
20
30
40
50
60
Normal
MK50
MK75
MK88
MK94
MK97
MK data
Normal data

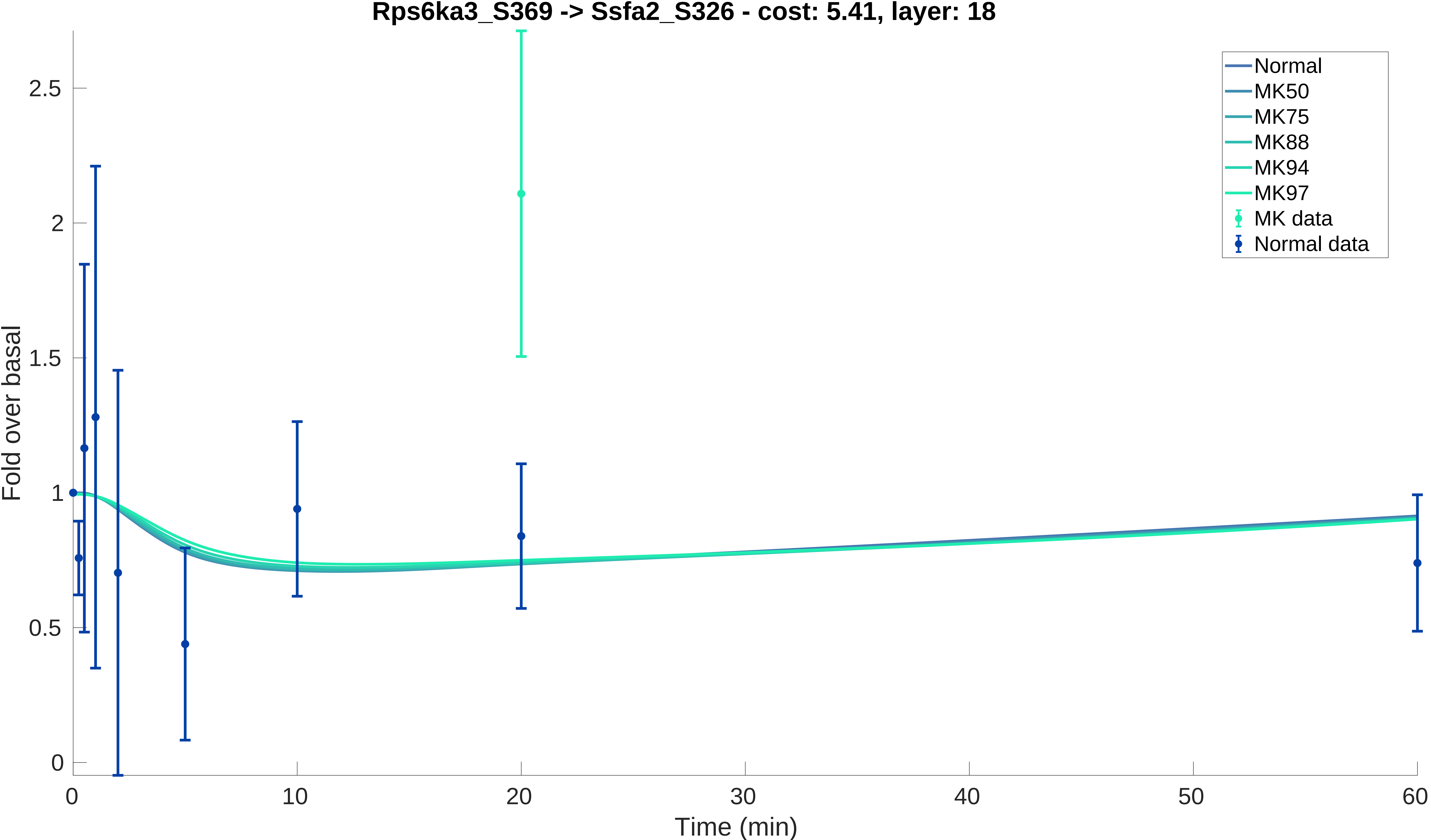

Rps6ka3_S369 -> Ssfa2_S326 - cost: 5.41, layer: 18
Normal
MK50
MK75
MK88
MK94
MK97
MK data
Normal data
Fold over basal
Time (min)
0
0.5
1
1.5
2
2.5
0
10
20
30
40
50
60

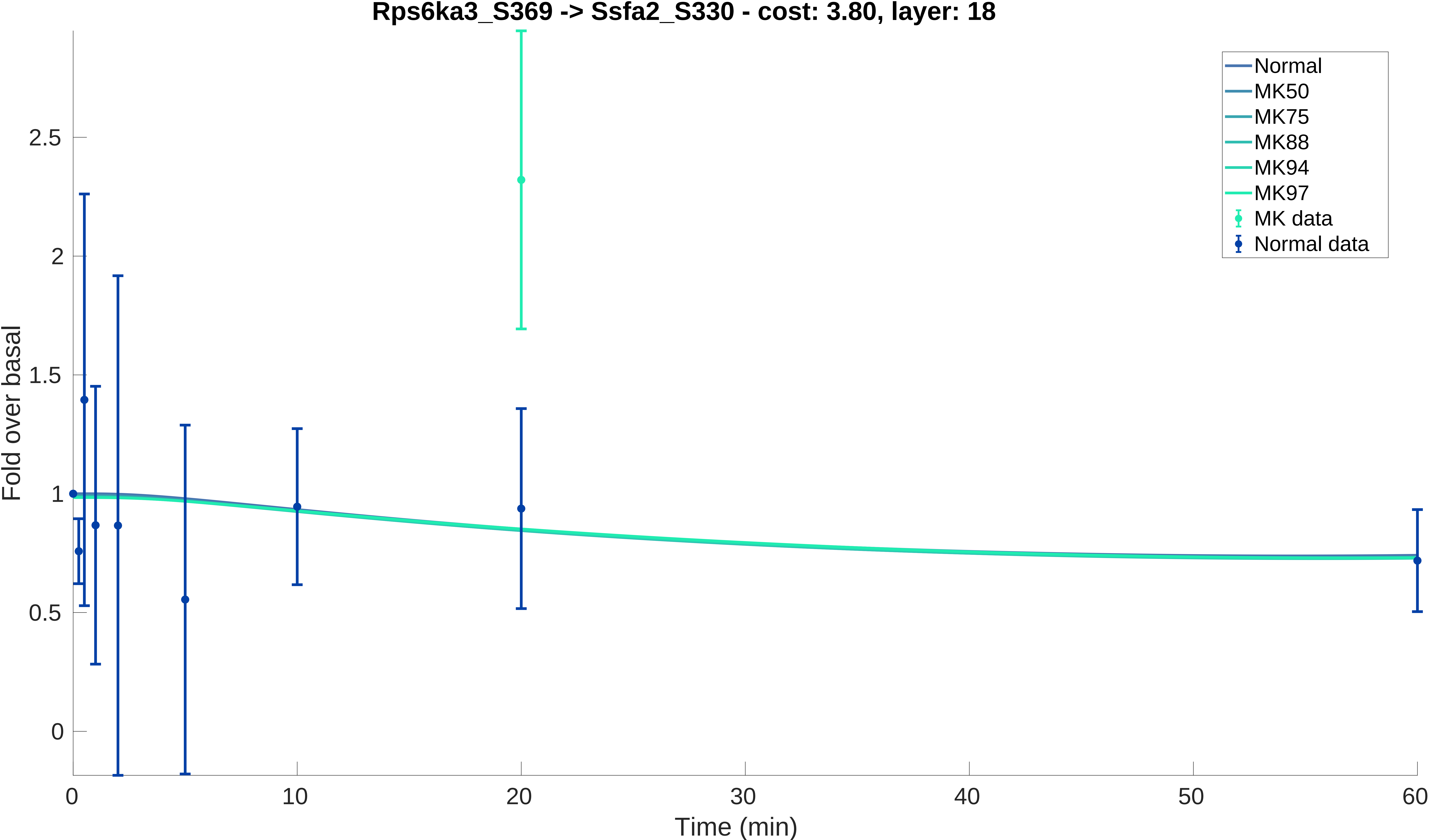

Rps6ka3_S369 -> Ssfa2_S330 - cost: 3.80, layer: 18
Fold over basal
Time (min)
Normal
MK50
MK75
MK88
MK94
MK97
MK data
Normal data
0
0.5
1
1.5
2
2.5
0
10
20
30
40
50
60

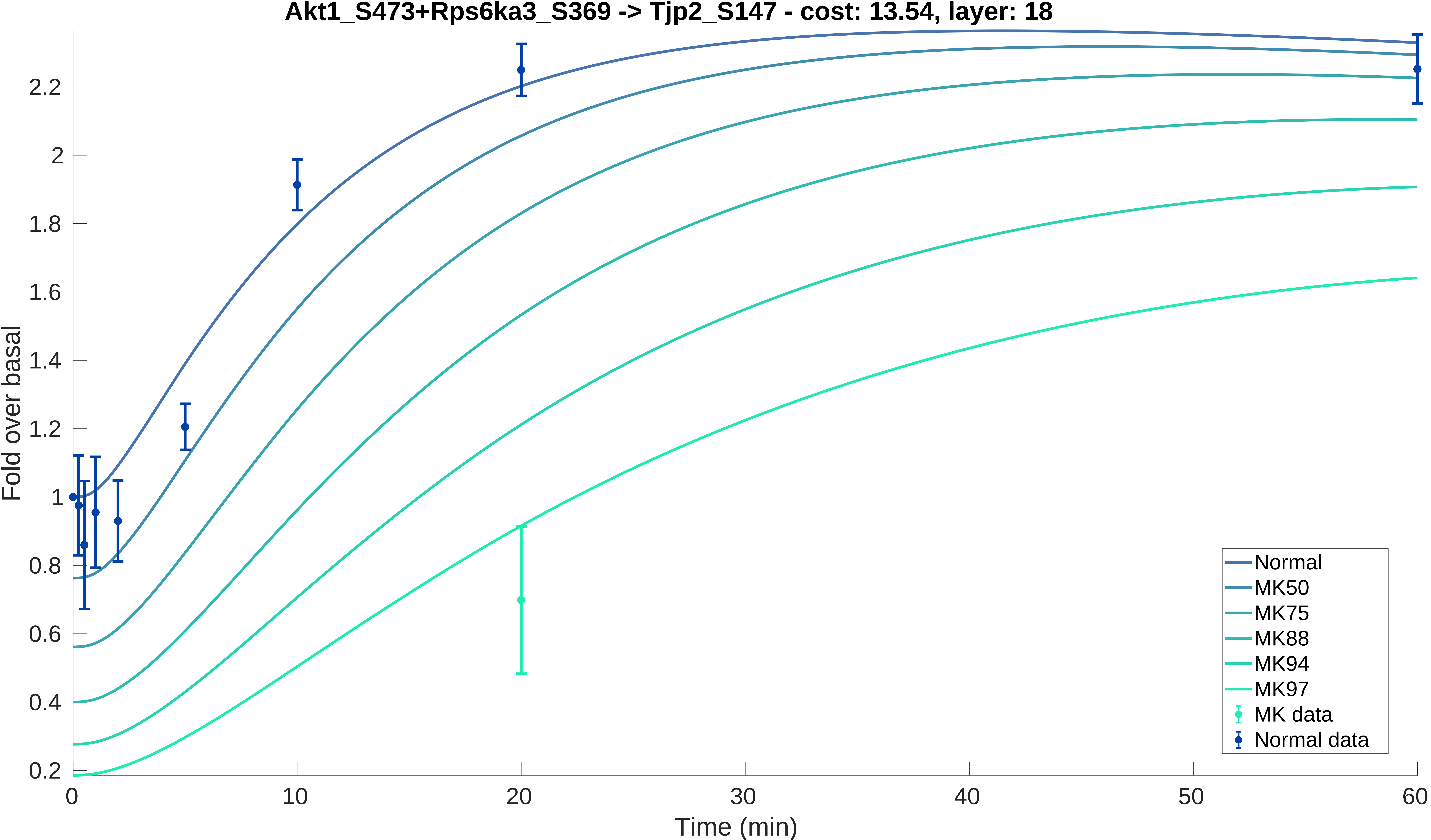

Akt1_S473+Rps6ka3_S369 -> Tjp2_S147 - cost: 13.54, layer: 18
Fold over basal
2.2
2
1.8
1.6
1.4
1.2
1
0.8
0.6
0.4
0.2
0
10
20
30
40
50
60
Time (min)
Normal
MK50
MK75
MK88
MK94
MK97
MK data
Normal data

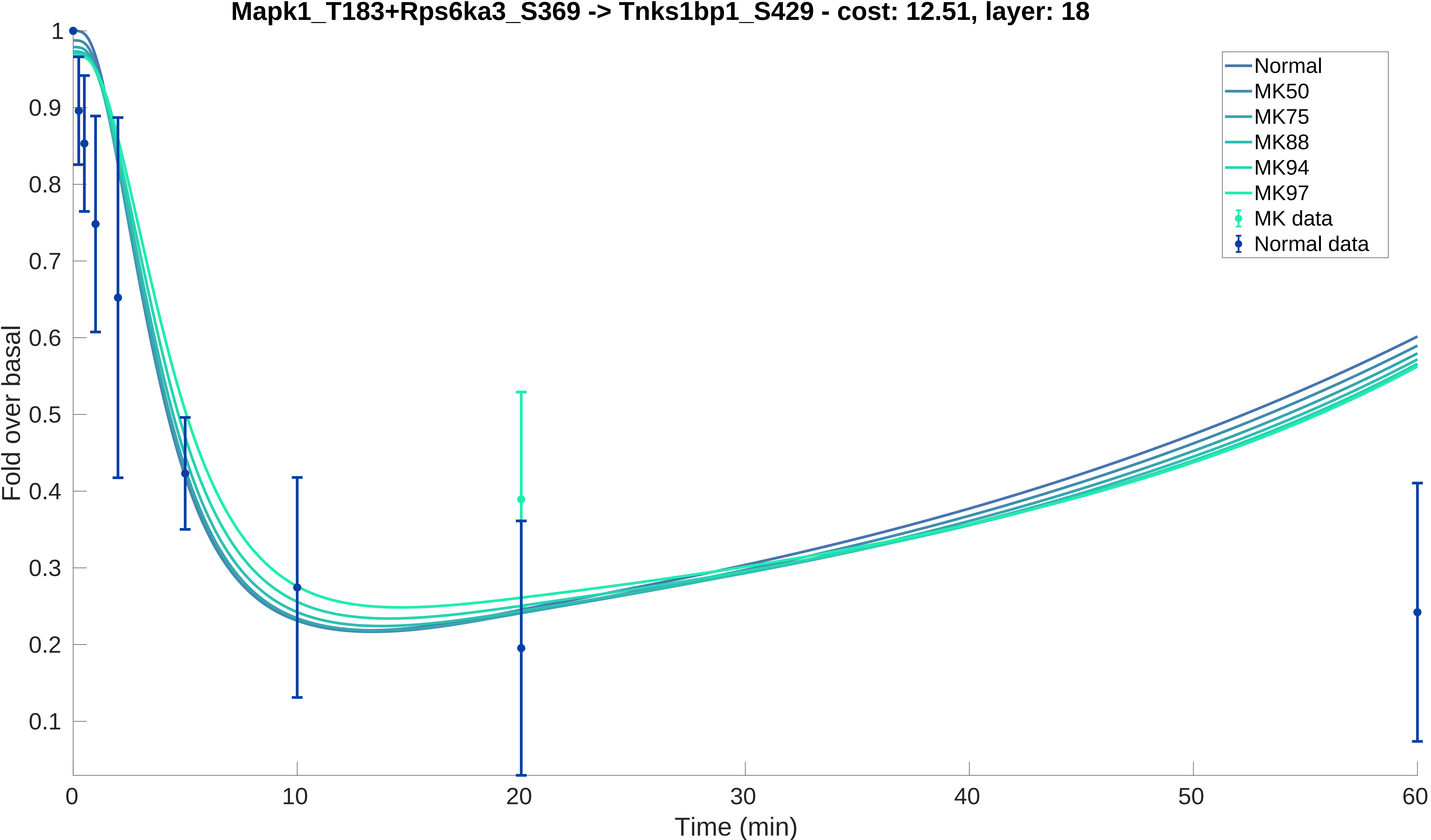
Mapk1_T183+Rps6ka3_S369 -> Tnks1bp1_S429 - cost: 12.51, layer: 18
Fold over basal
Time (min)
Normal
MK50
MK75
MK88
MK94
MK97
MK data
Normal data
0
10
20
30
40
50
60
0.1
0.2
0.3
0.4
0.5
0.6
0.7
0.8
0.9
1

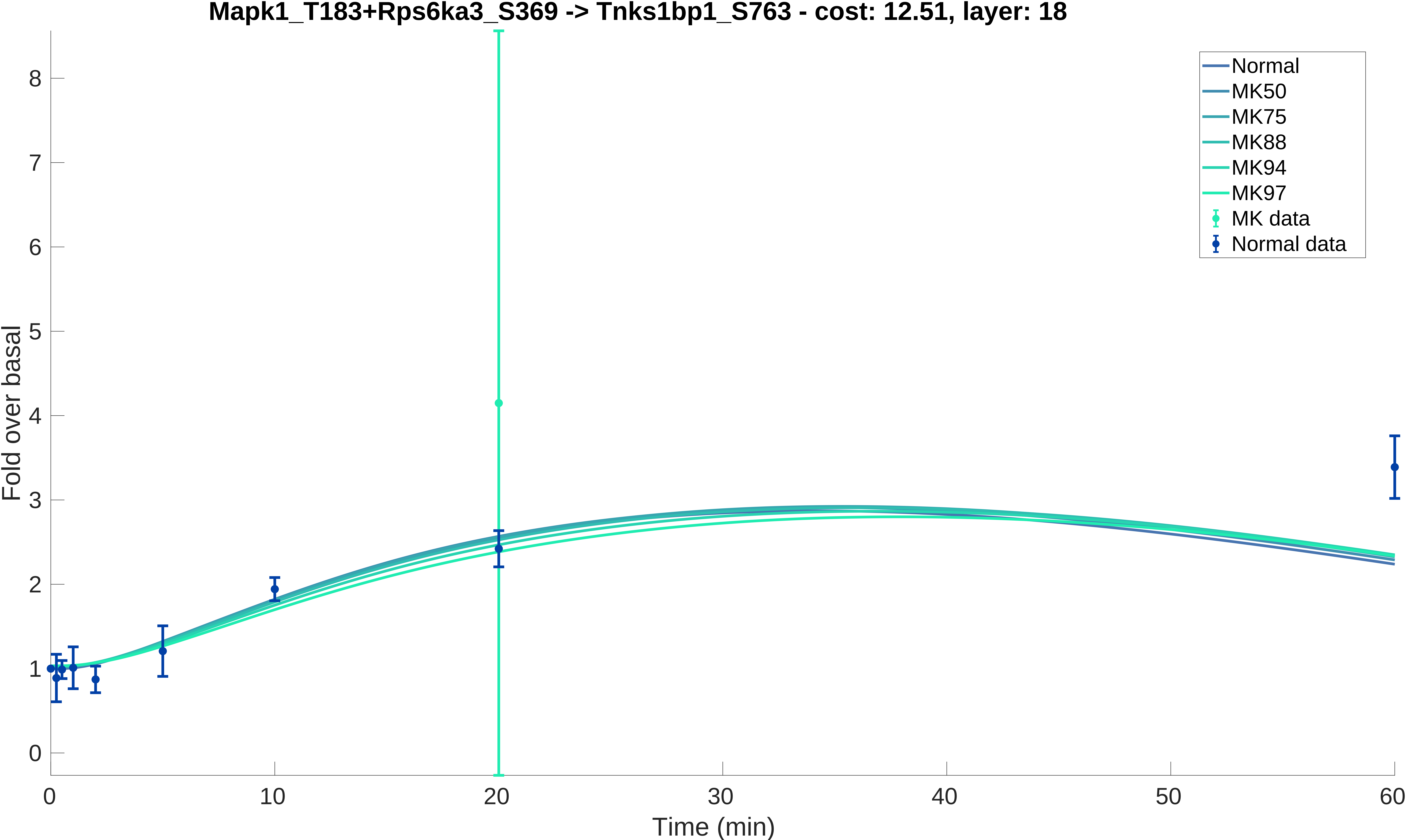

Mapk1_T183+Rps6ka3_S369 -> Tnks1bp1_S763 - cost: 12.51, layer: 18
Fold over basal
Time (min)
0
1
2
3
4
5
6
7
8
0
10
20
30
40
50
60
Normal
MK50
MK75
MK88
MK94
MK97
MK data
Normal data

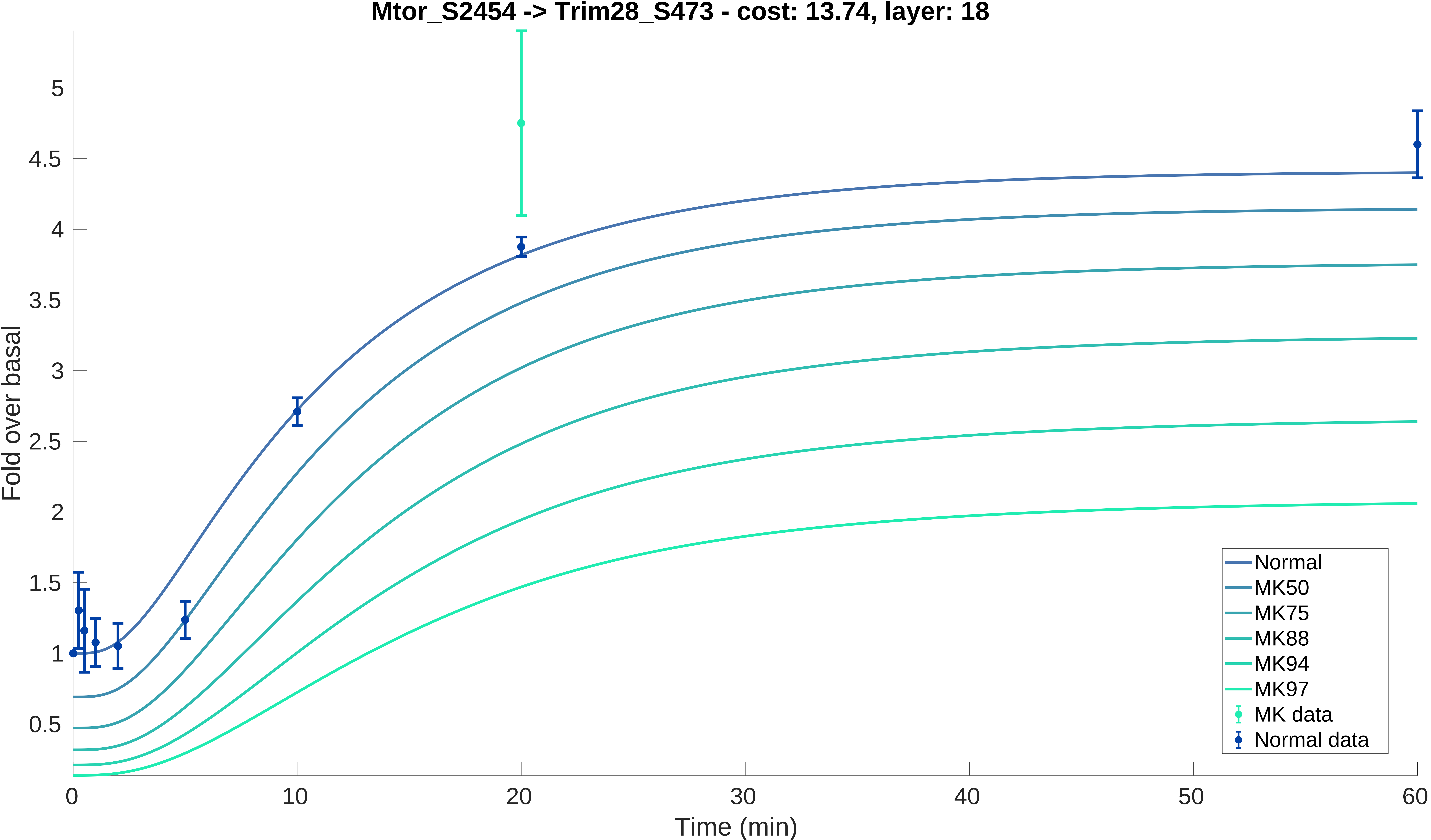

Mtor_S2454 -> Trim28_S473 - cost: 13.74, layer: 18
Fold over basal
Time (min)
0
10
20
30
40
50
60
0.5
1
1.5
2
2.5
3
3.5
4
4.5
5
Normal
MK50
MK75
MK88
MK94
MK97
MK data
Normal data

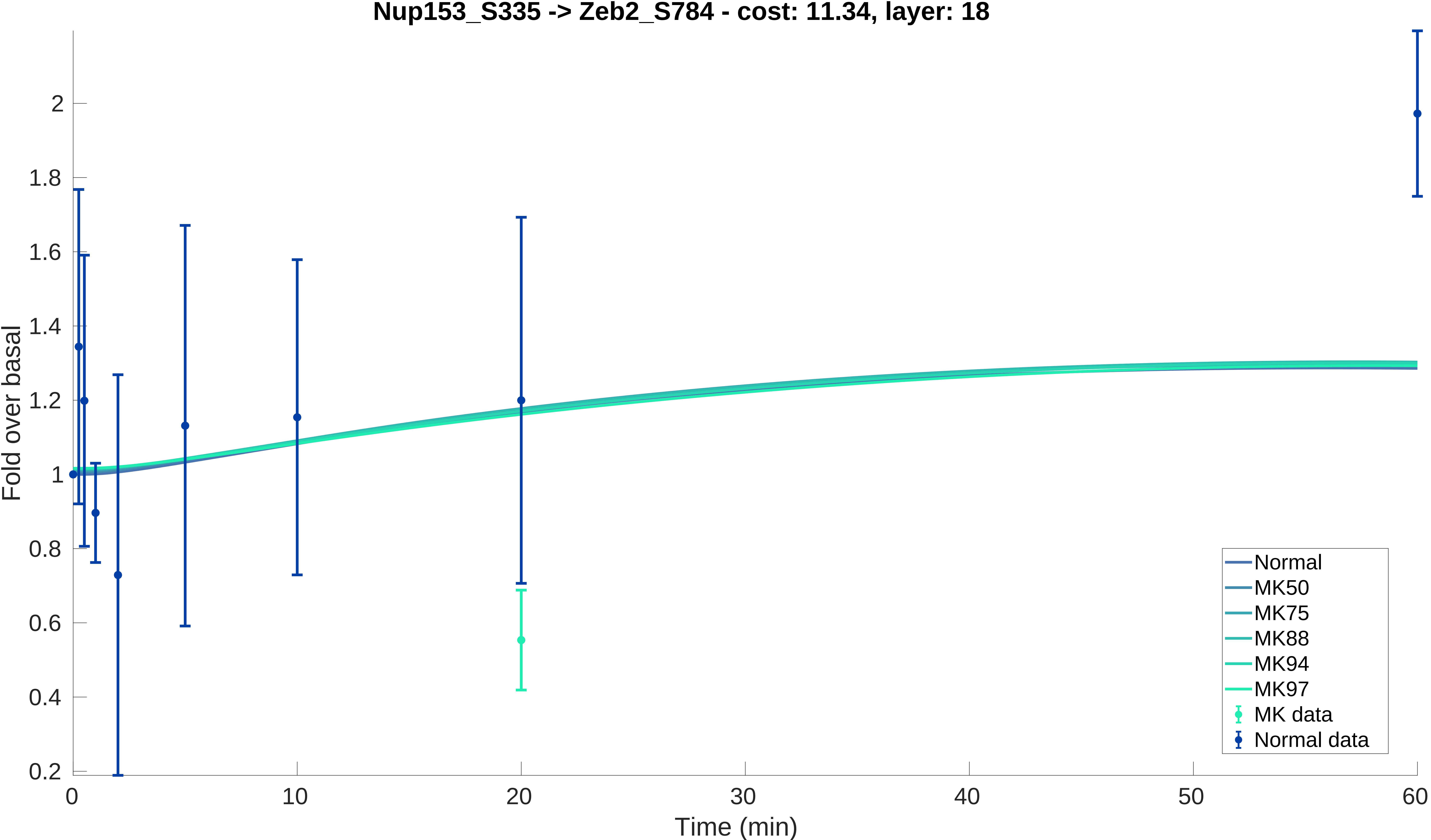
Nup153_S335 -> Zeb2_S784 - cost: 11.34, layer: 18
Fold over basal
Time (min)
0.2
0.4
0.6
0.8
1
1.2
1.4
1.6
1.8
2
0
10
20
30
40
50
60
Normal
MK50
MK75
MK88
MK94
MK97
MK data
Normal data

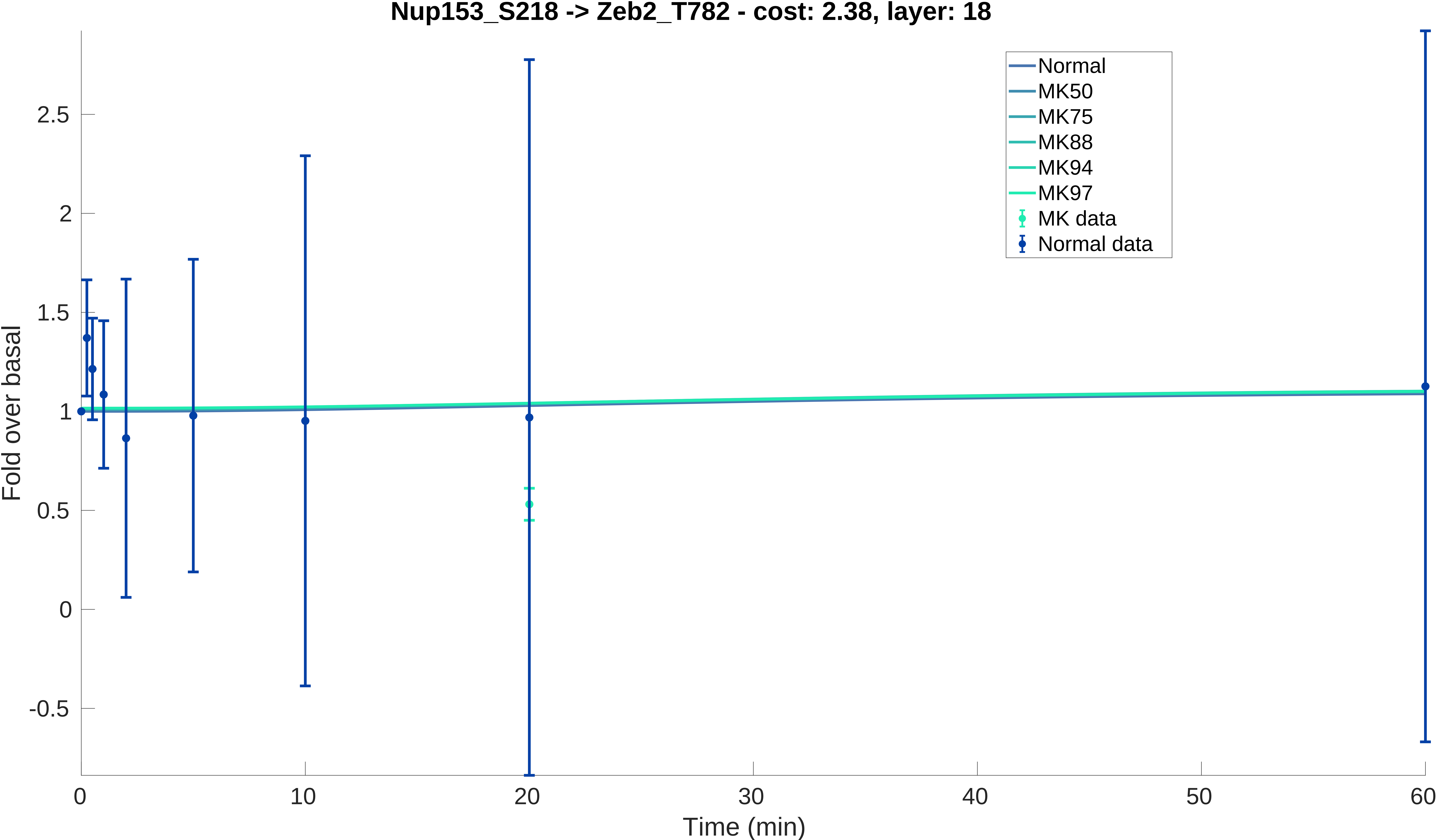
Nup153_S218 -> Zeb2_T782 - cost: 2.38, layer: 18
Normal
MK50
MK75
MK88
MK94
MK97
MK data
Normal data
Fold over basal
Time (min)
2.5
2
1.5
1
0.5
0
-0.5
0
10
20
30
40
50
60

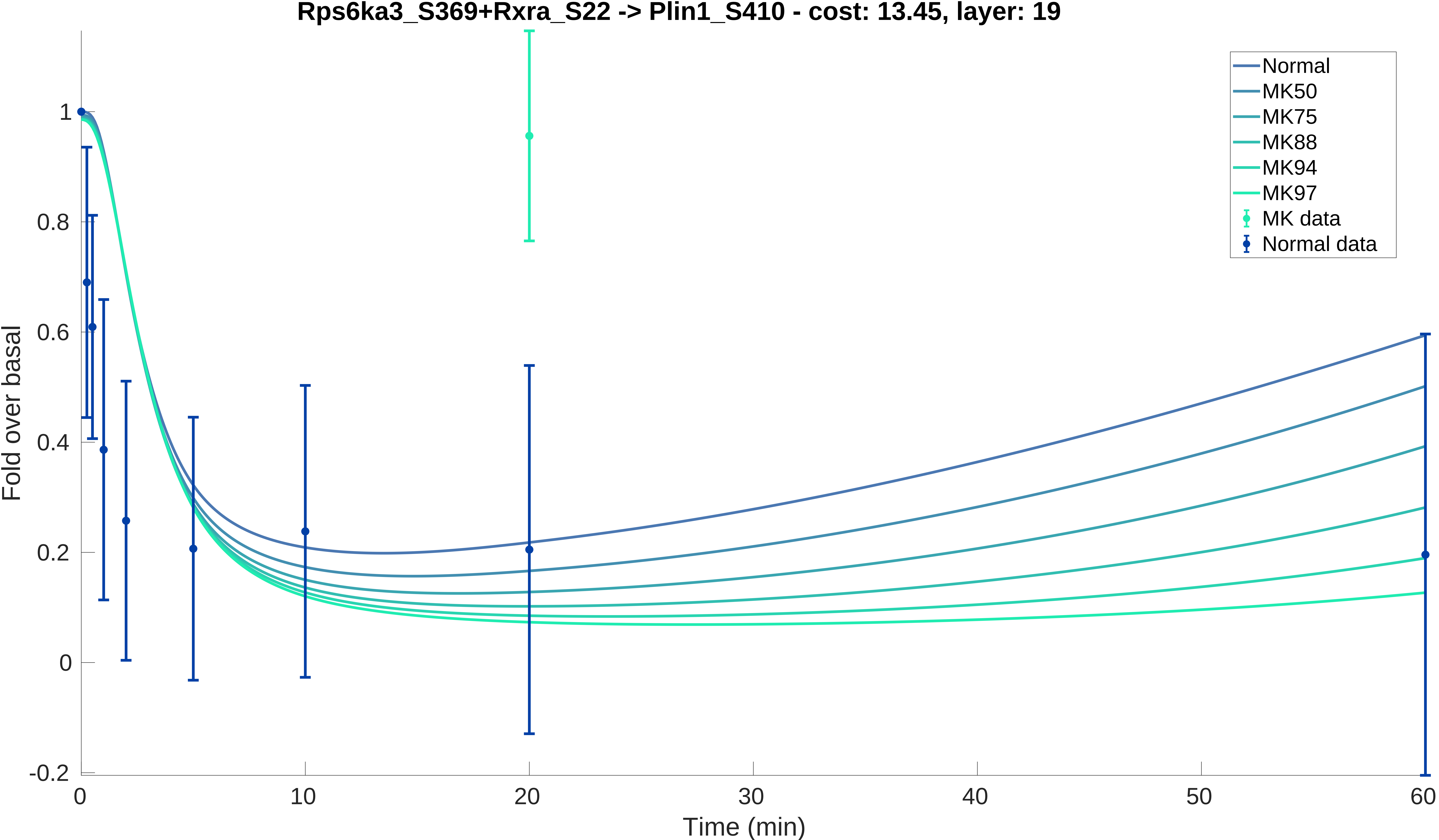

Rps6ka3_S369+Rxra_S22 -> Plin1_S410 - cost: 13.45, layer: 19
Fold over basal
Time (min)
1
0.8
0.6
0.4
0.2
0
-0.2
0
10
20
30
40
50
60
Normal
MK50
MK75
MK88
MK94
MK97
MK data
Normal data

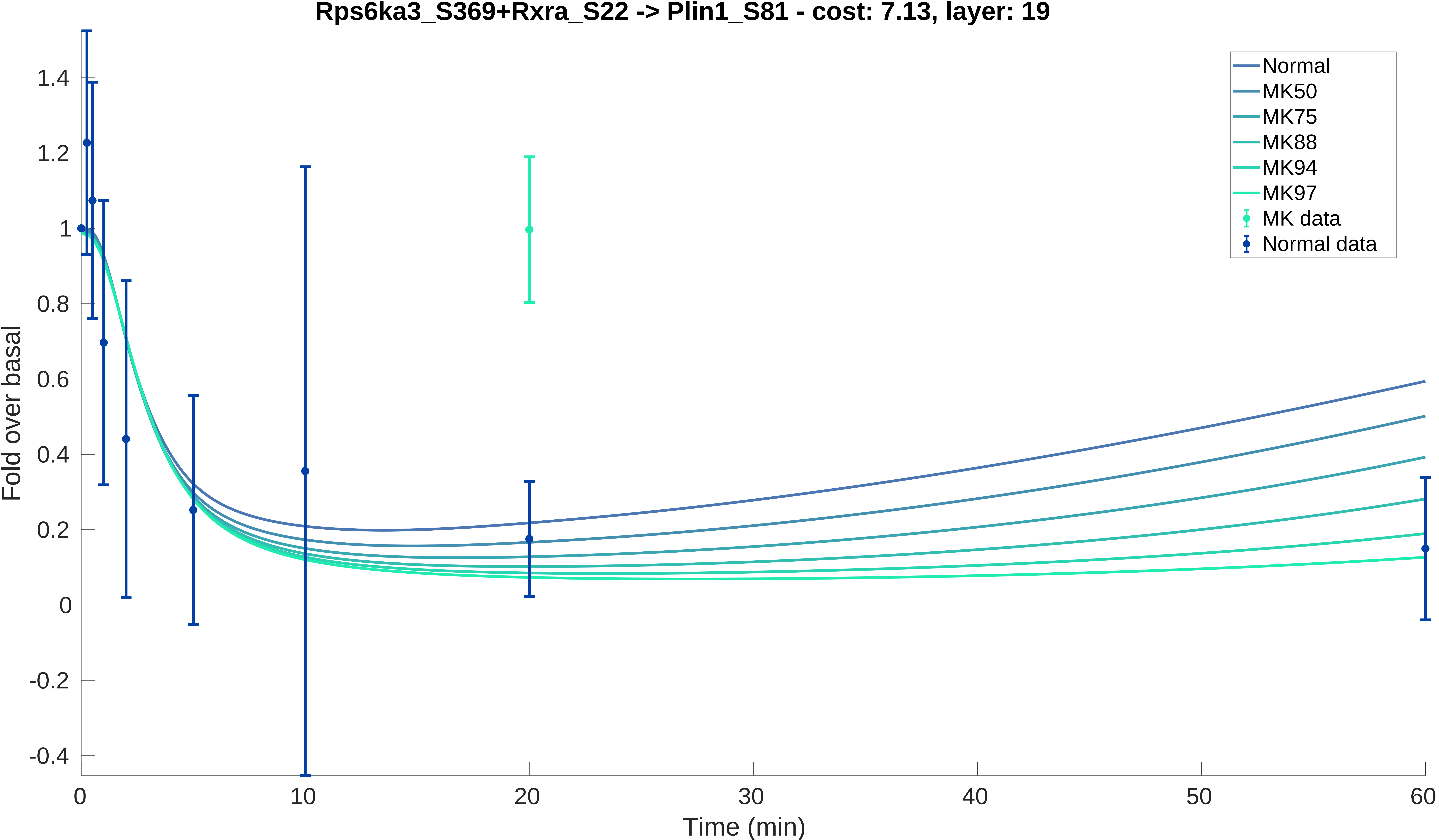

Rps6ka3_S369+Rxra_S22 -> Plin1_S81 - cost: 7.13, layer: 19
Fold over basal
1.4
1.2
1
0.8
0.6
0.4
0.2
0
-0.2
-0.4
0
10
20
30
40
50
60
Time (min)
Normal
MK50
MK75
MK88
MK94
MK97
MK data
Normal data

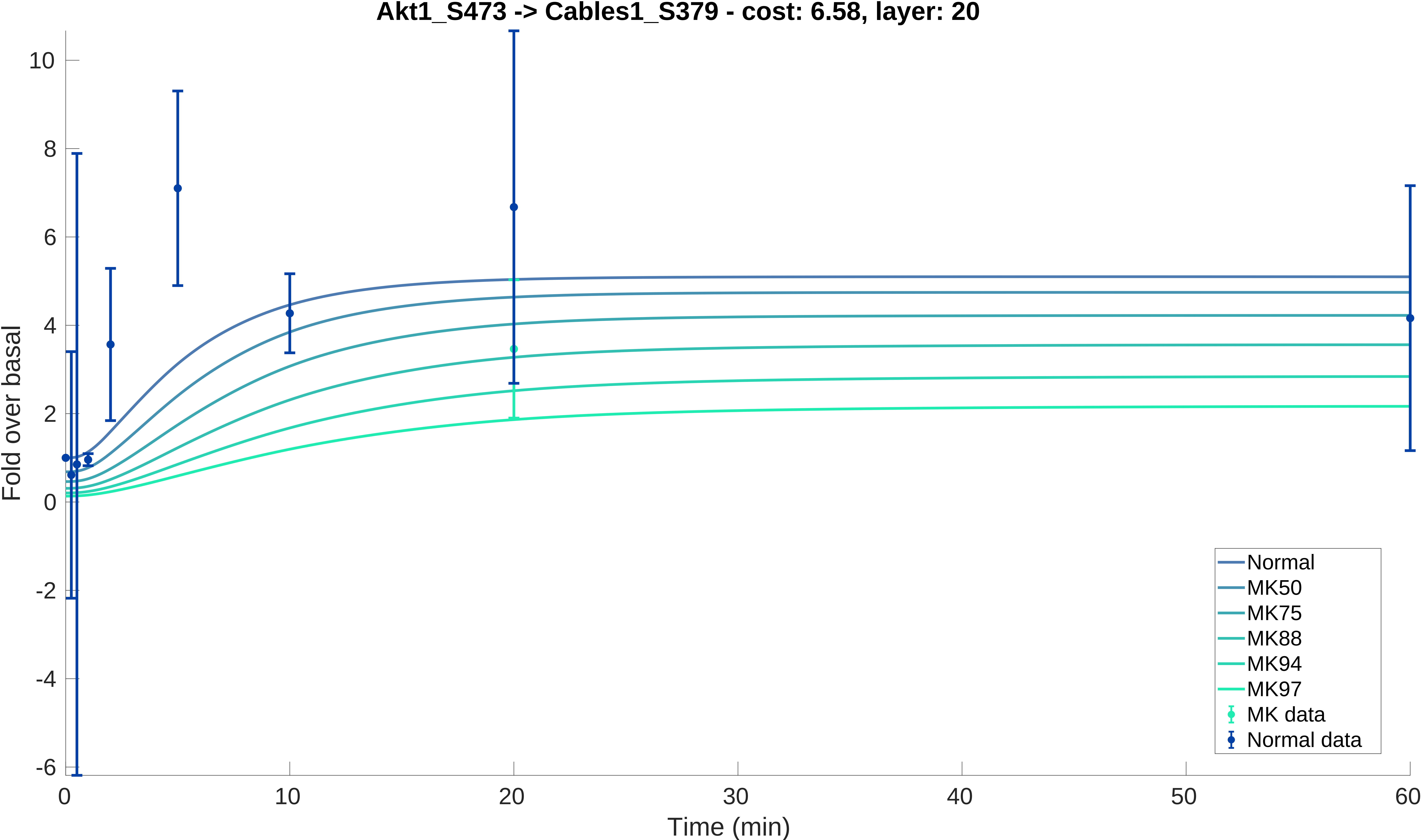

Akt1_S473 -> Cables1_S379 - cost: 6.58, layer: 20
Fold over basal
Time (min)
10
8
6
4
2
0
-2
-4
-6
0
10
20
30
40
50
60
Normal
MK50
MK75
MK88
MK94
MK97
MK data
Normal data

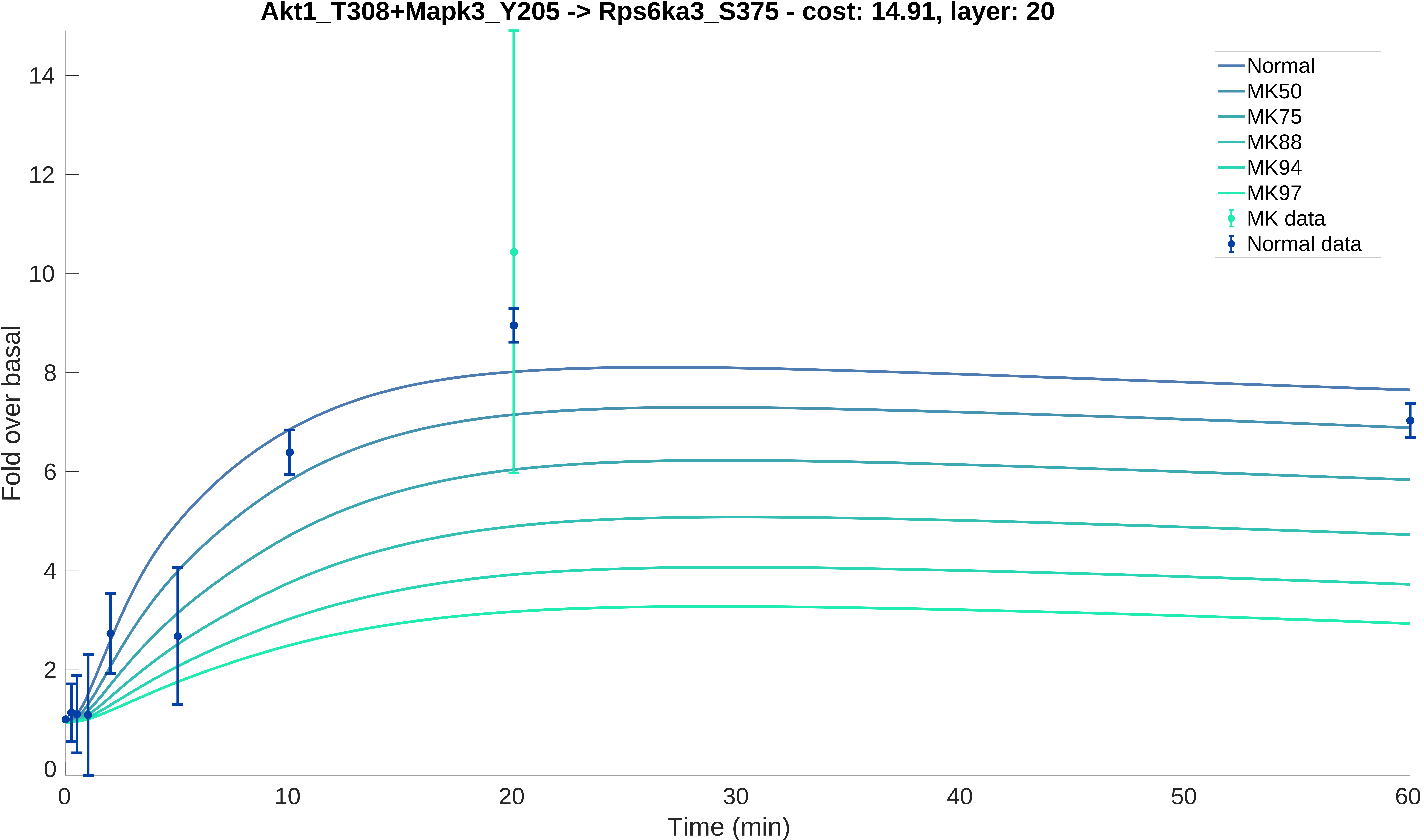

Akt1_T308+Mapk3_Y205 -> Rps6ka3_S375 - cost: 14.91, layer: 20
Fold over basal
Time (min)
0
2
4
6
8
10
12
14
0
10
20
30
40
50
60
Normal
MK50
MK75
MK88
MK94
MK97
MK data
Normal data

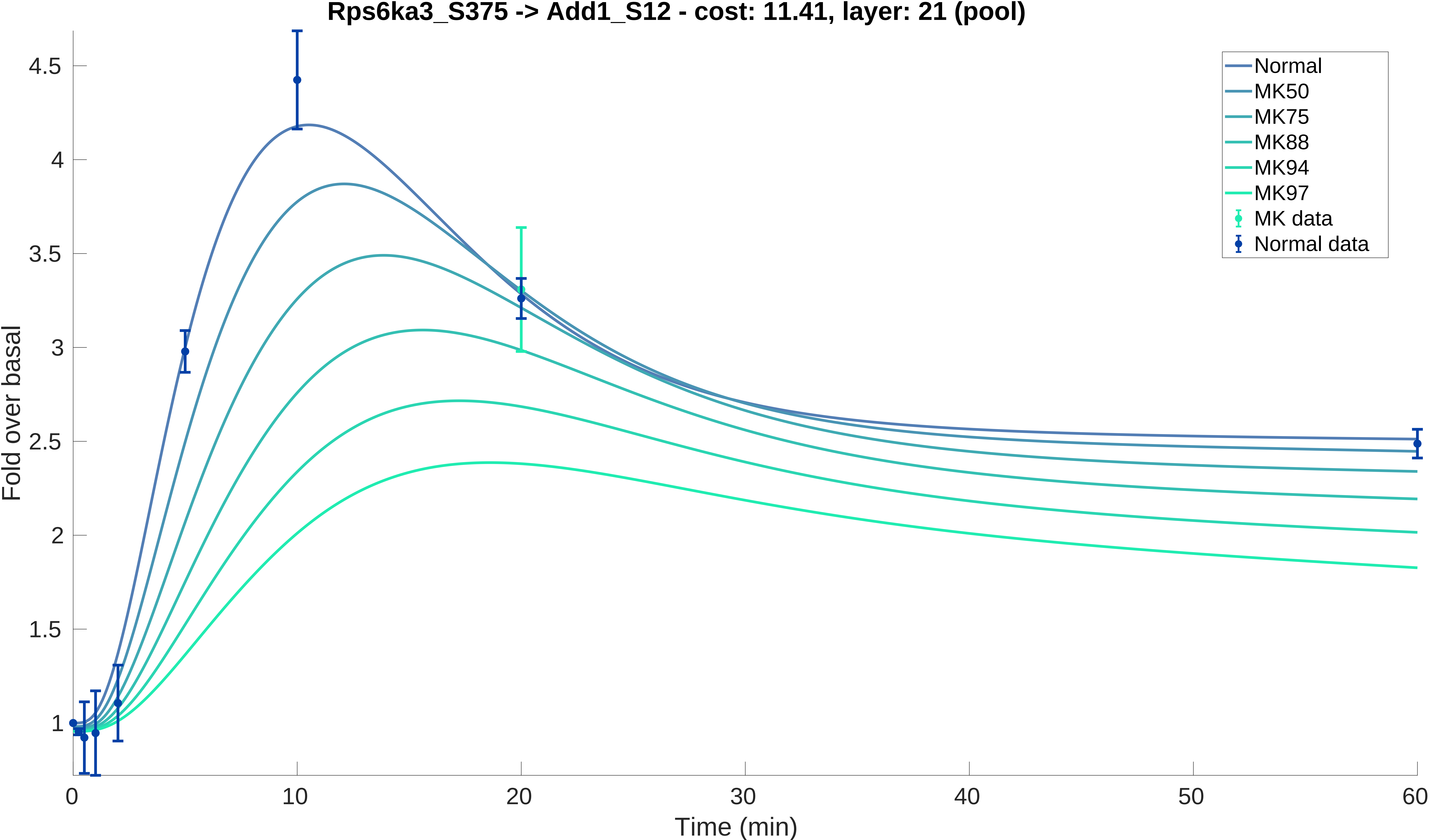

Rps6ka3_S375 -> Add1_S12 - cost: 11.41, layer: 21 (pool)
Fold over basal
Time (min)
4.5
4
3.5
3
2.5
2
1.5
1
0
10
20
30
40
50
60
Normal
MK50
MK75
MK88
MK94
MK97
MK data
Normal data

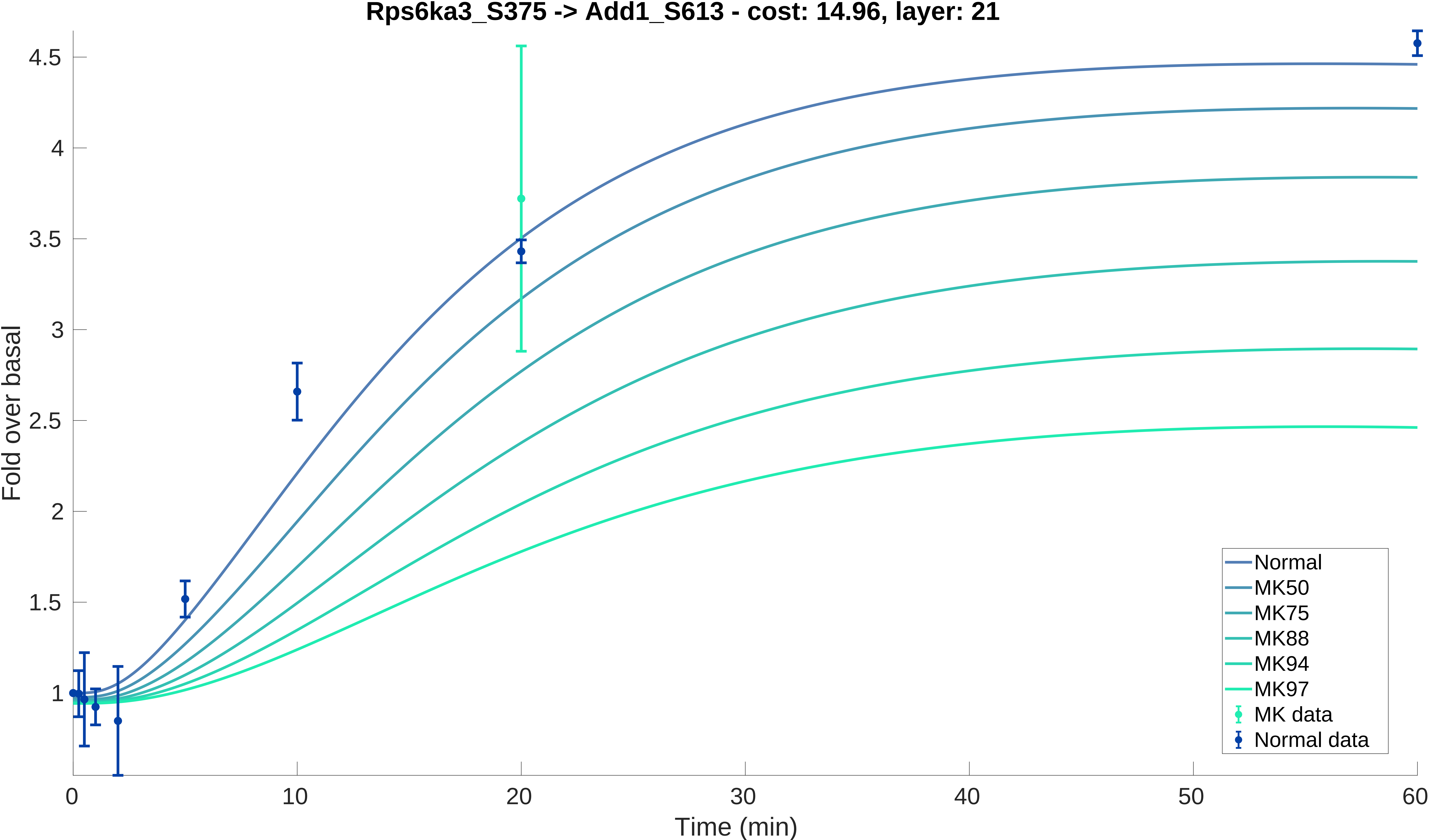
Rps6ka3_S375 -> Add1_S613 - cost: 14.96, layer: 21
Fold over basal
Time (min)
4.5
4
3.5
3
2.5
2
1.5
1
0
10
20
30
40
50
60
Normal
MK50
MK75
MK88
MK94
MK97
MK data
Normal data

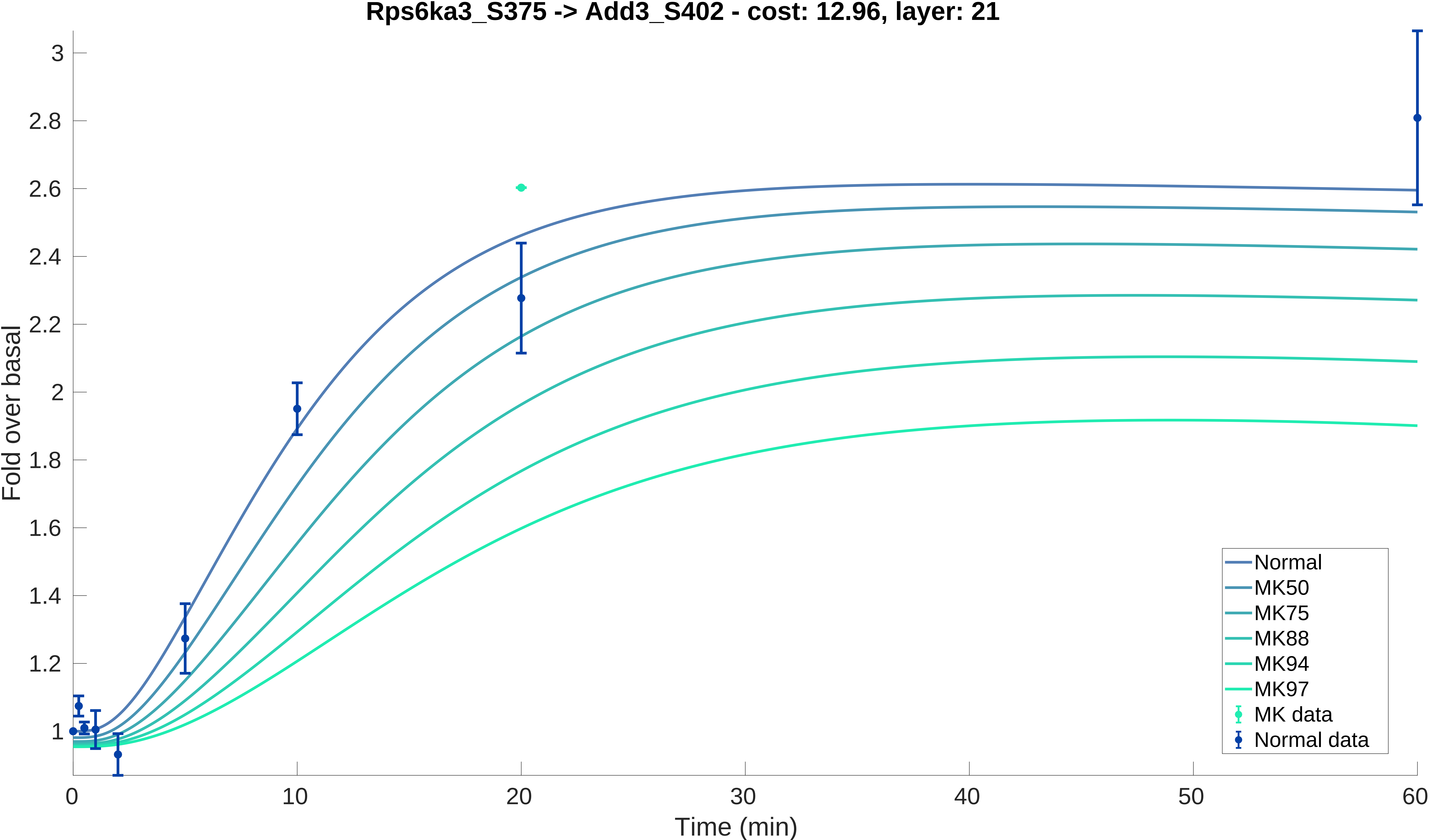

Rps6ka3_S375 -> Add3_S402 - cost: 12.96, layer: 21
Fold over basal
Time (min)
3
2.8
2.6
2.4
2.2
2
1.8
1.6
1.4
1.2
1
0
10
20
30
40
50
60
Normal
MK50
MK75
MK88
MK94
MK97
MK data
Normal data

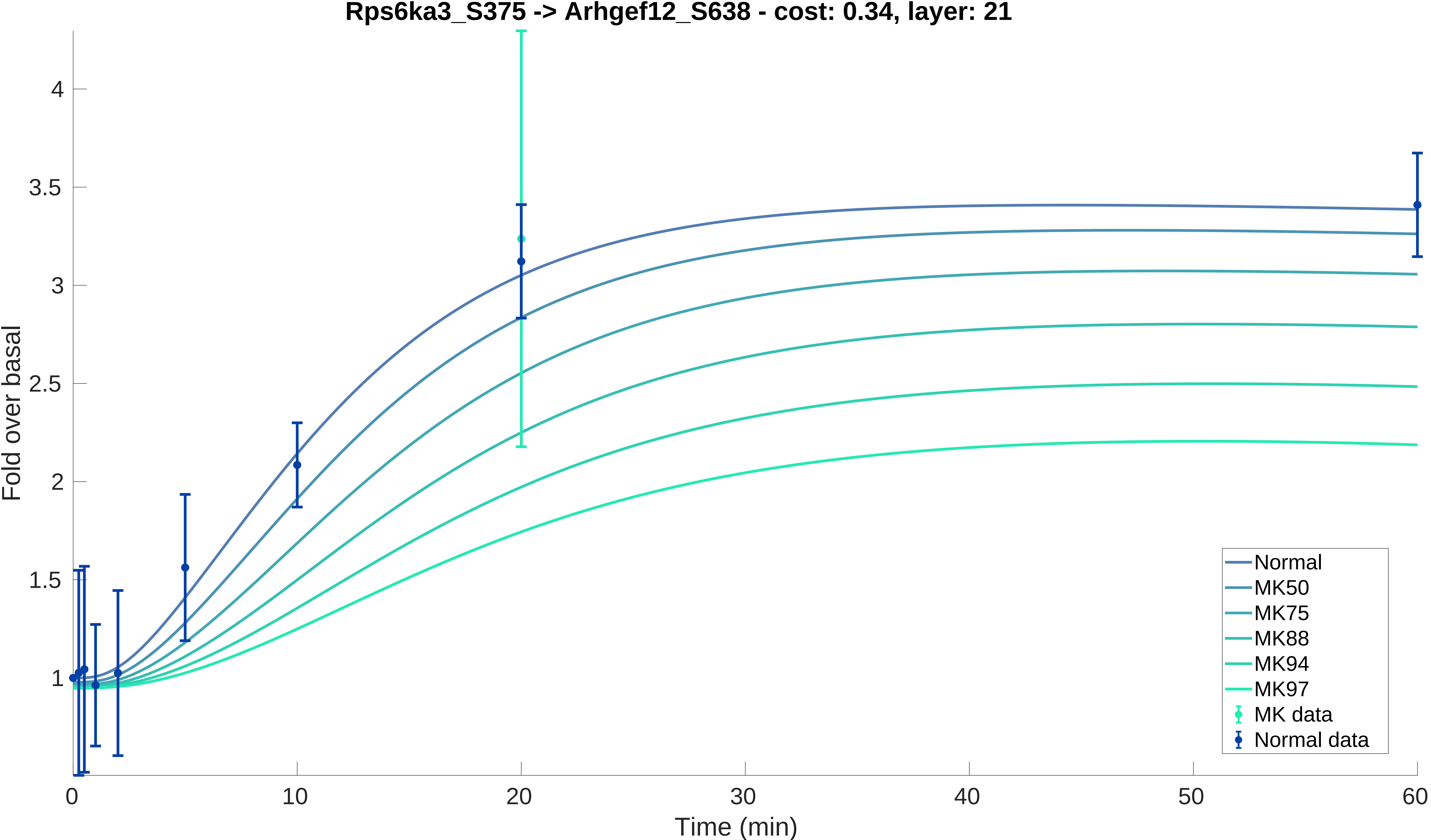
Rps6ka3_S375 -> Arhgef12_S638 - cost: 0.34, layer: 21
Fold over basal
Time (min)
4
3.5
3
2.5
2
1.5
1
0
10
20
30
40
50
60
Normal
MK50
MK75
MK88
MK94
MK97
MK data
Normal data

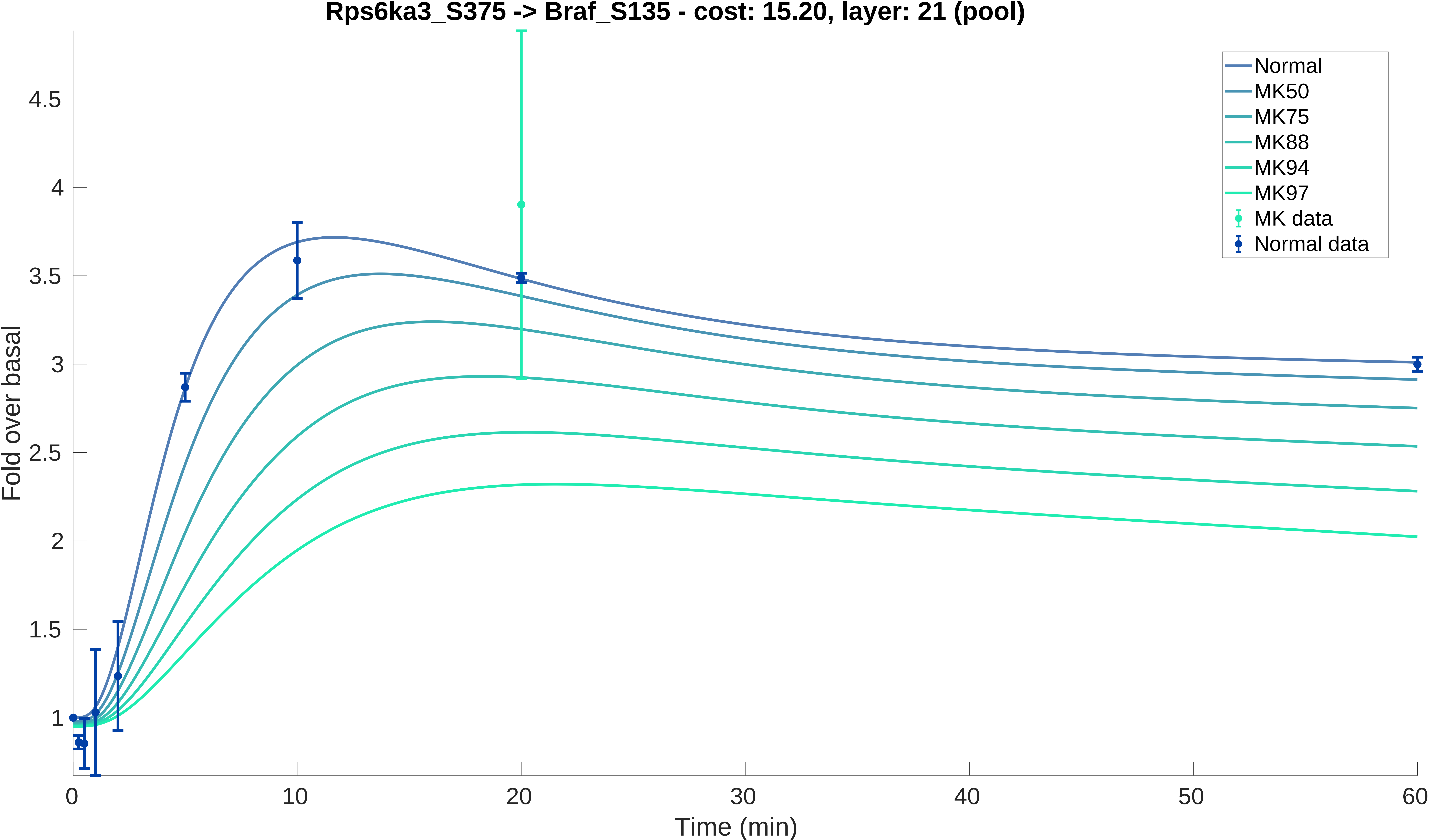
Rps6ka3_S375 -> Braf_S135 - cost: 15.20, layer: 21 (pool)
Normal
MK50
MK75
MK88
MK94
MK97
MK data
Normal data
Fold over basal
Time (min)
4.5
4
3.5
3
2.5
2
1.5
1
0
10
20
30
40
50
60

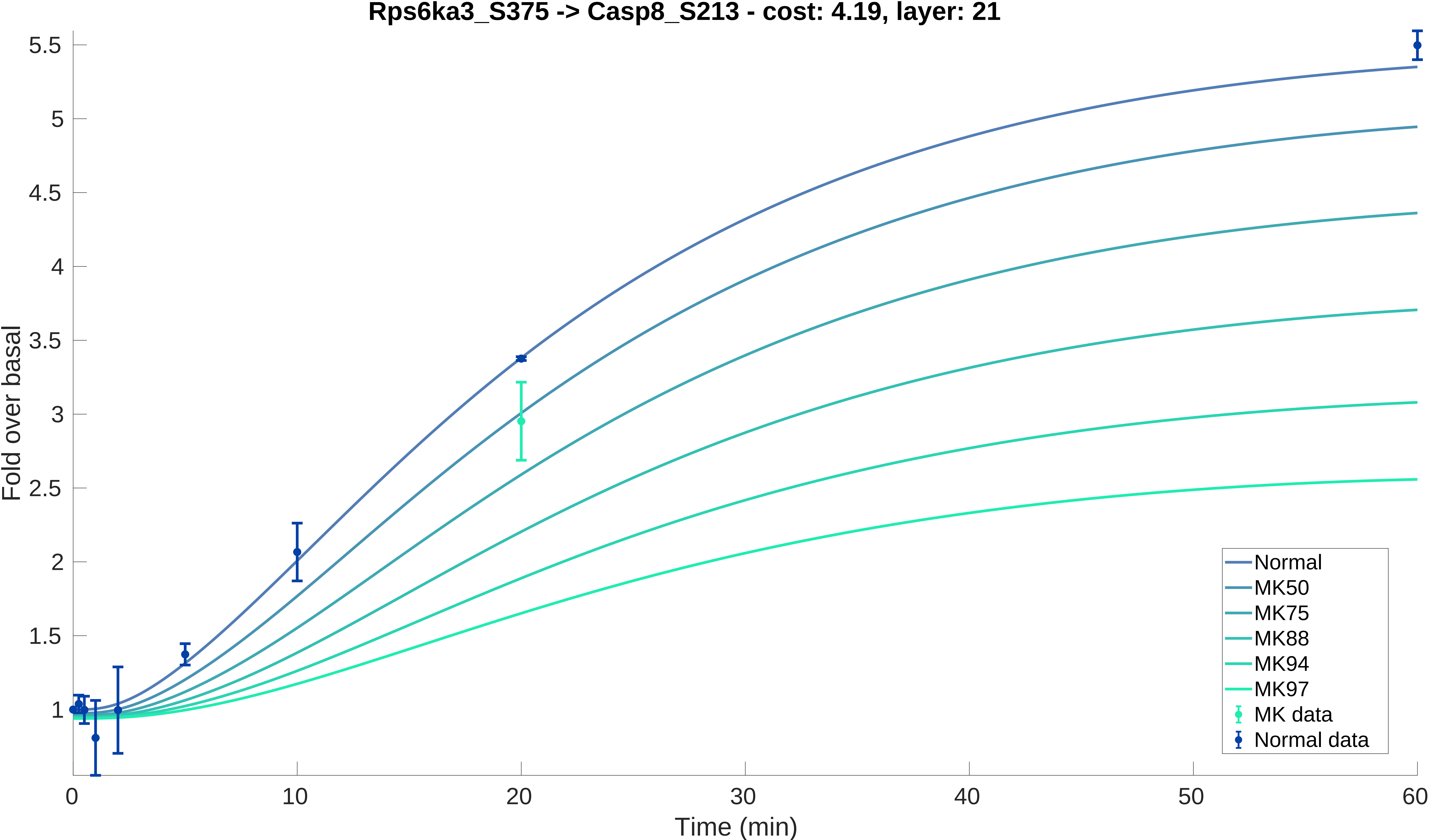
Rps6ka3_S375 -> Casp8_S213 - cost: 4.19, layer: 21
Fold over basal
Time (min)
5.5
5
4.5
4
3.5
3
2.5
2
1.5
1
0
10
20
30
40
50
60
Normal
MK50
MK75
MK88
MK94
MK97
MK data
Normal data

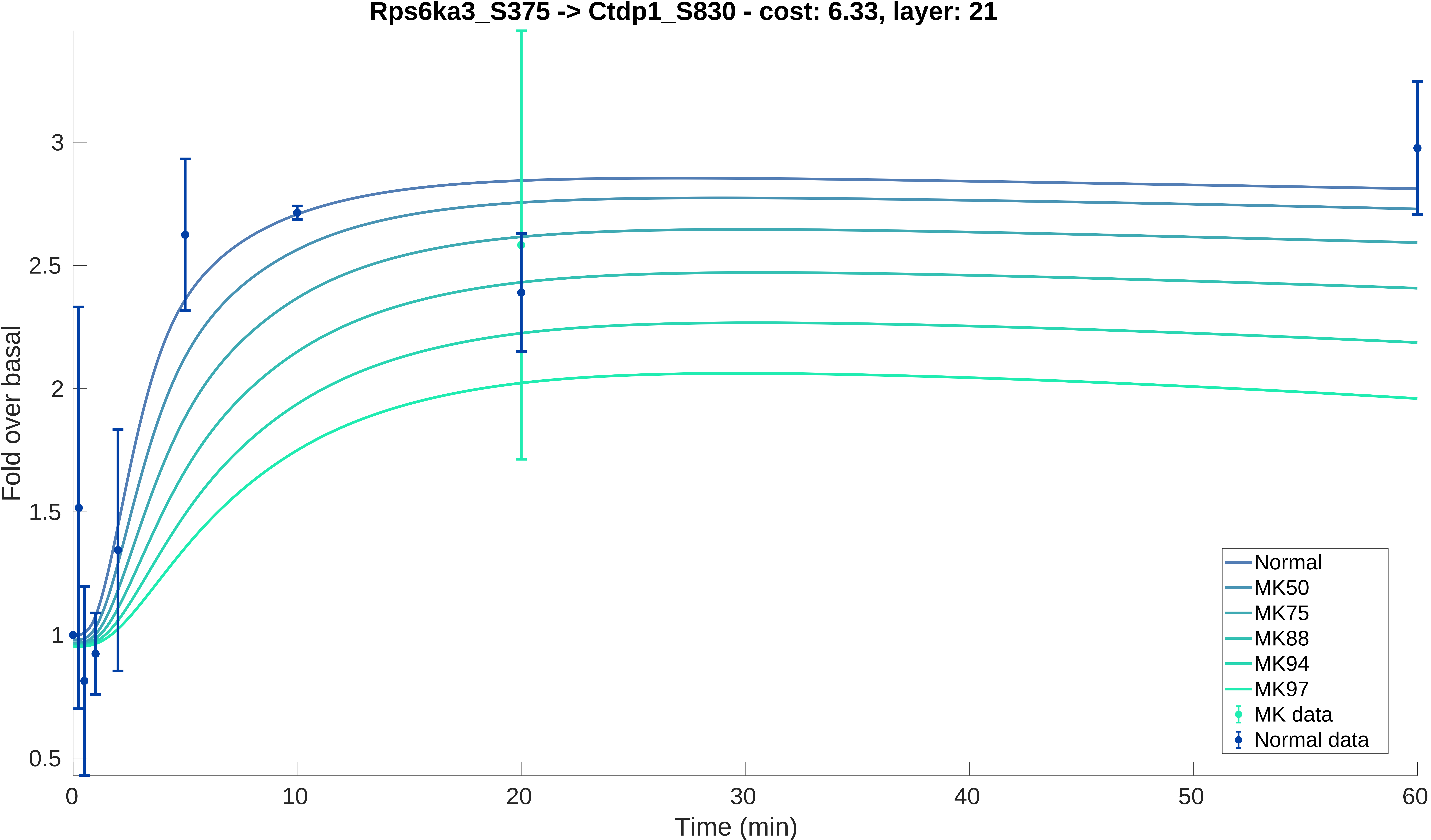

Rps6ka3_S375 -> Ctdp1_S830 - cost: 6.33, layer: 21
Fold over basal
Time (min)
0.5
1
1.5
2
2.5
3
0
10
20
30
40
50
60
Normal
MK50
MK75
MK88
MK94
MK97
MK data
Normal data

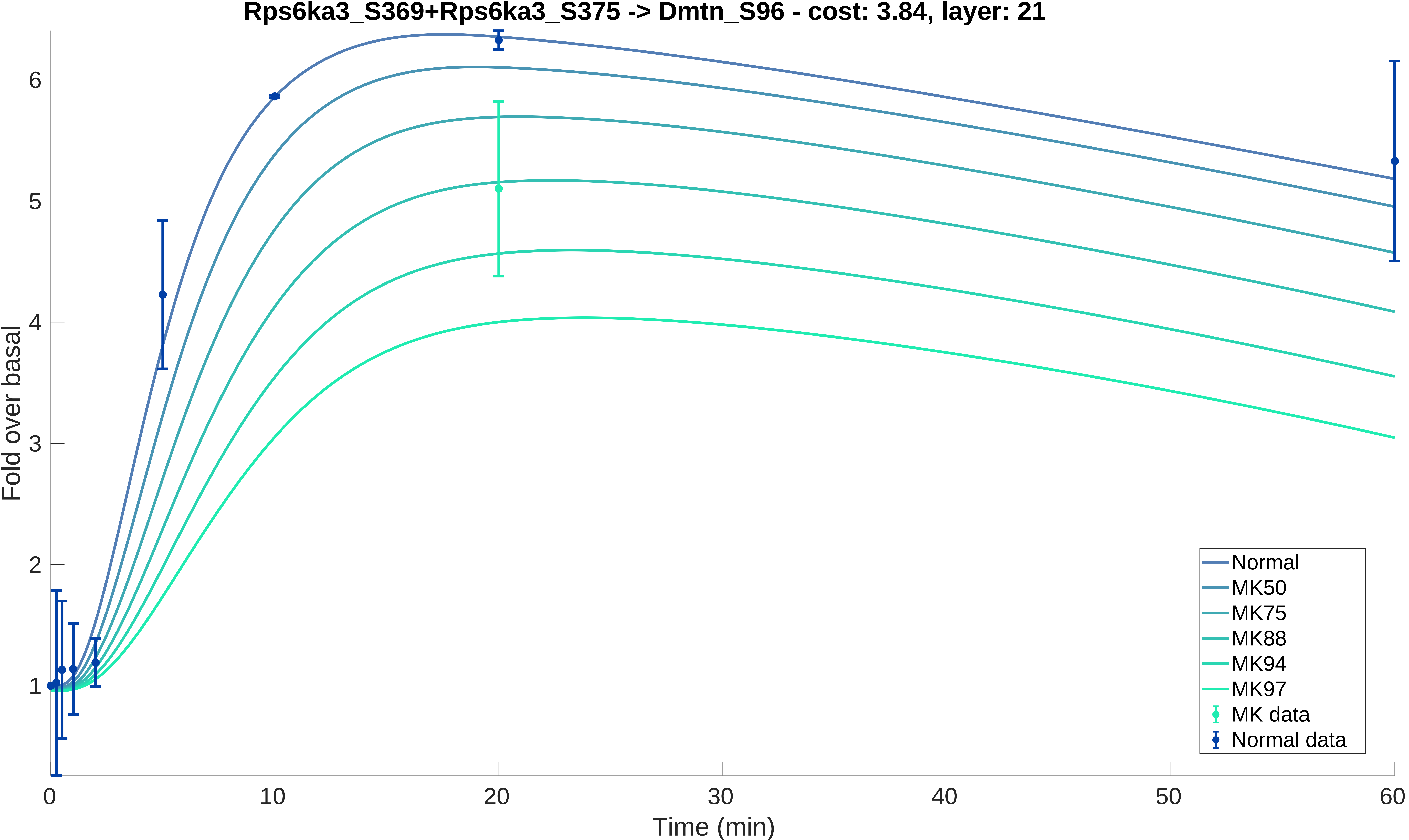

Rps6ka3_S369+Rps6ka3_S375 -> Dmtn_S96 - cost: 3.84, layer: 21
Fold over basal
Time (min)
0
10
20
30
40
50
60
1
2
3
4
5
6
Normal
MK50
MK75
MK88
MK94
MK97
MK data
Normal data

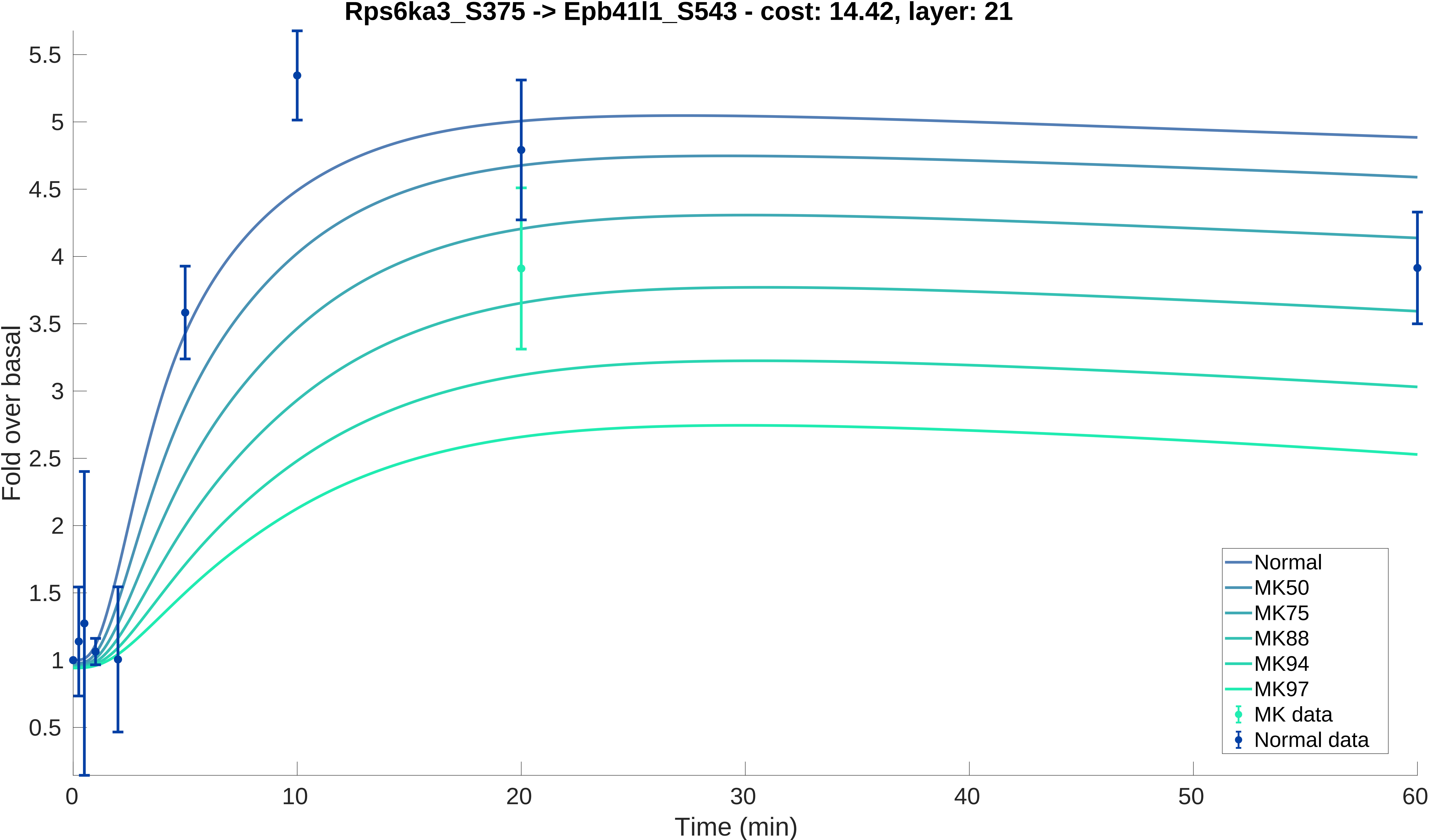
Rps6ka3_S375 -> Epb41l1_S543 - cost: 14.42, layer: 21
Fold over basal
0.5
1
1.5
2
2.5
3
3.5
4
4.5
5
5.5
0
10
20
30
40
50
60
Time (min)
Normal
MK50
MK75
MK88
MK94
MK97
MK data
Normal data

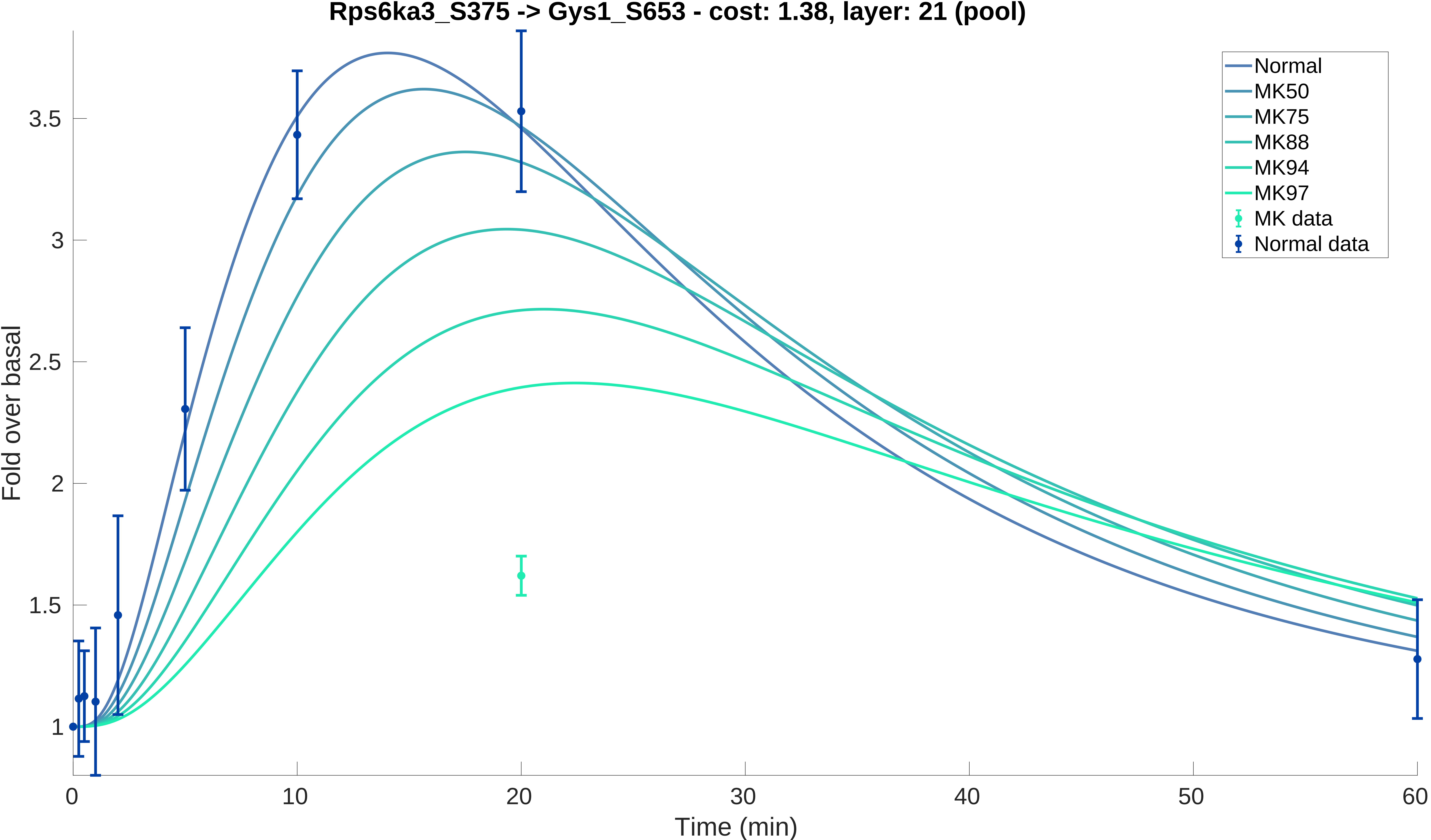

Rps6ka3_S375 -> Gys1_S653 - cost: 1.38, layer: 21 (pool)
Fold over basal
Time (min)
3.5
3
2.5
2
1.5
1
0
10
20
30
40
50
60
Normal
MK50
MK75
MK88
MK94
MK97
MK data
Normal data

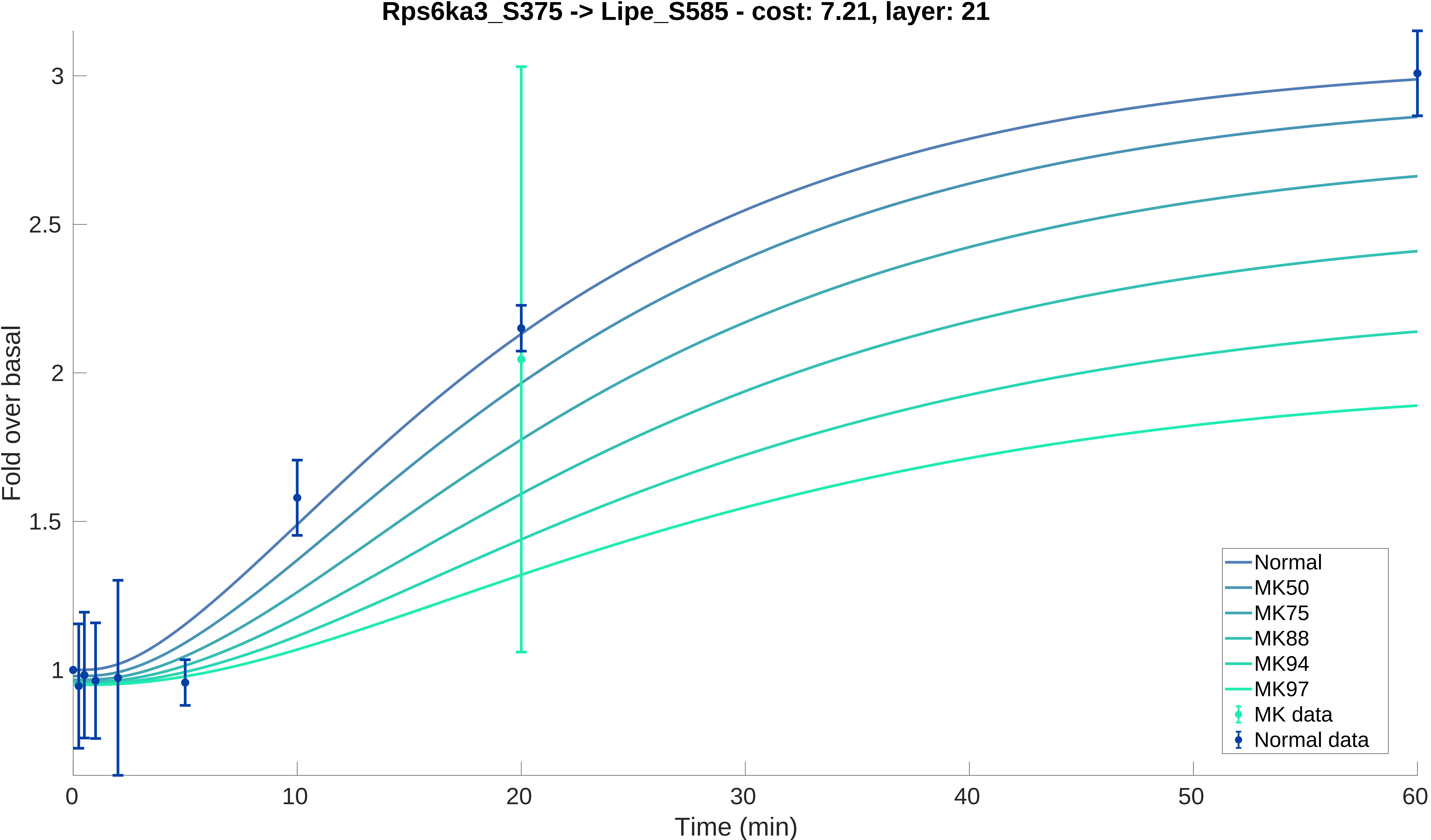
Rps6ka3_S375 -> Lipe_S585 - cost: 7.21, layer: 21
Fold over basal
Time (min)
3
2.5
2
1.5
1
0
10
20
30
40
50
60
Normal
MK50
MK75
MK88
MK94
MK97
MK data
Normal data

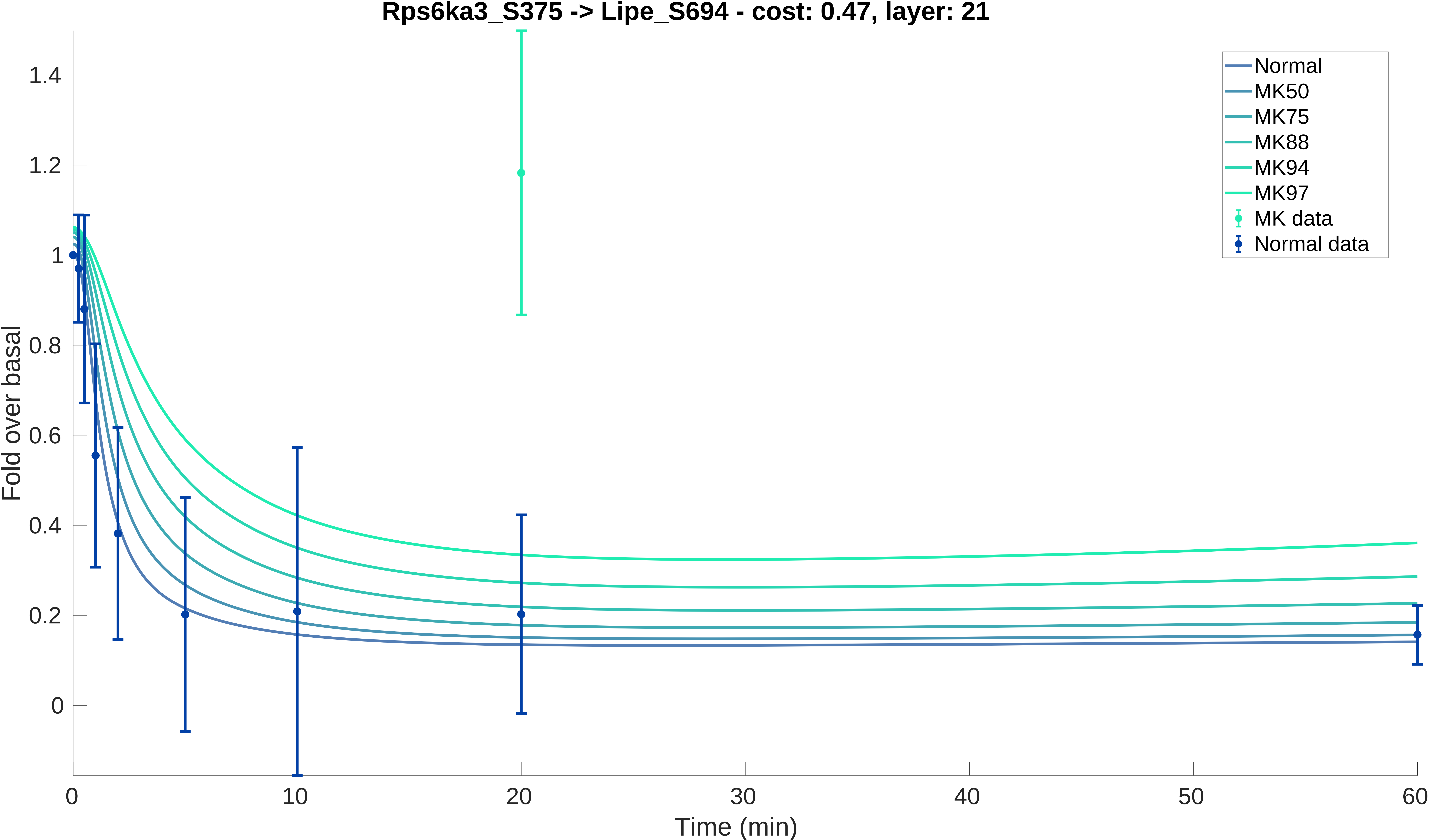
Rps6ka3_S375 -> Lipe_S694 - cost: 0.47, layer: 21
Fold over basal
Time (min)
0
0.2
0.4
0.6
0.8
1
1.2
1.4
0
10
20
30
40
50
60
Normal
MK50
MK75
MK88
MK94
MK97
MK data
Normal data

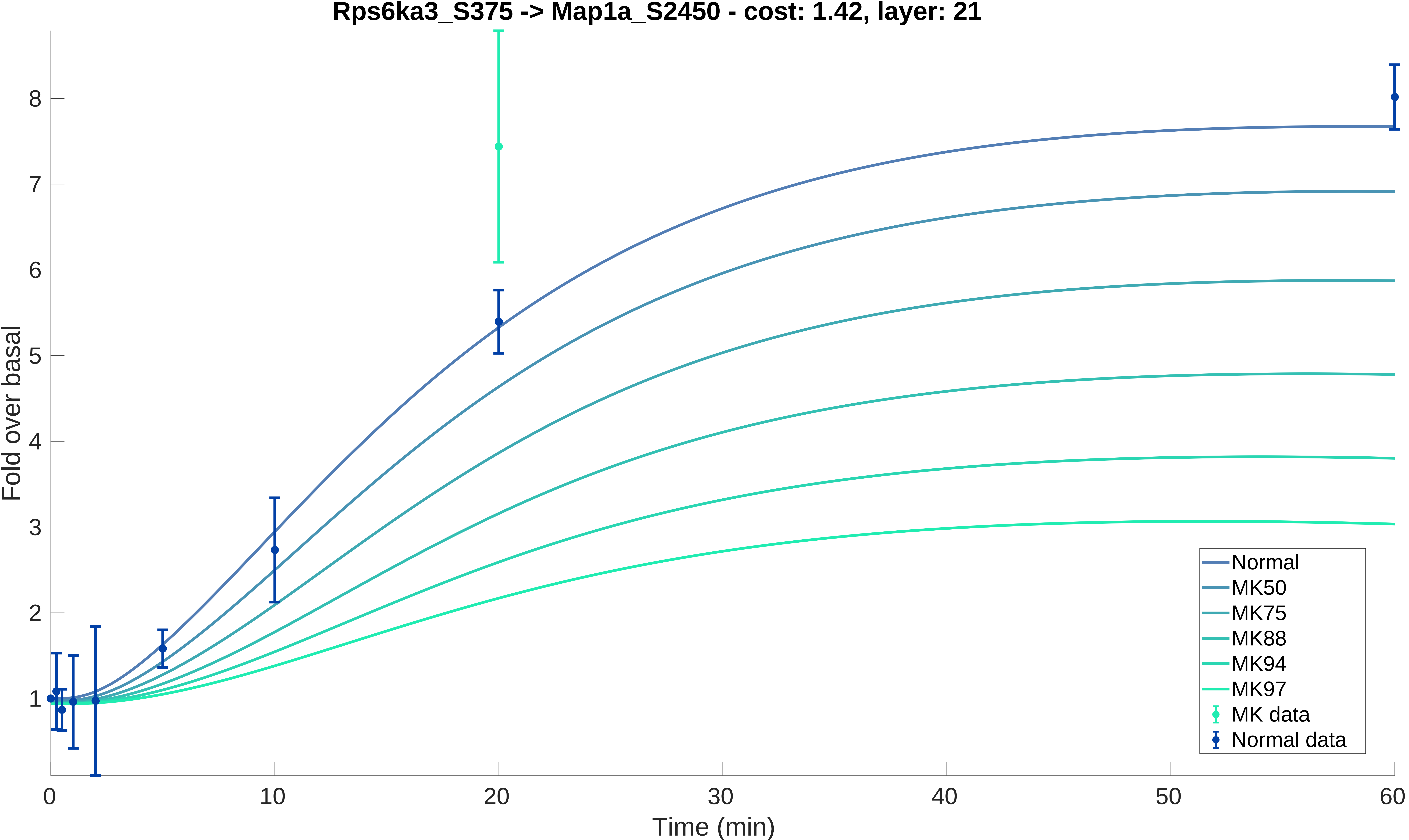

Rps6ka3_S375 -> Map1a_S2450 - cost: 1.42, layer: 21
Fold over basal
Time (min)
0
10
20
30
40
50
60
1
2
3
4
5
6
7
8
Normal
MK50
MK75
MK88
MK94
MK97
MK data
Normal data

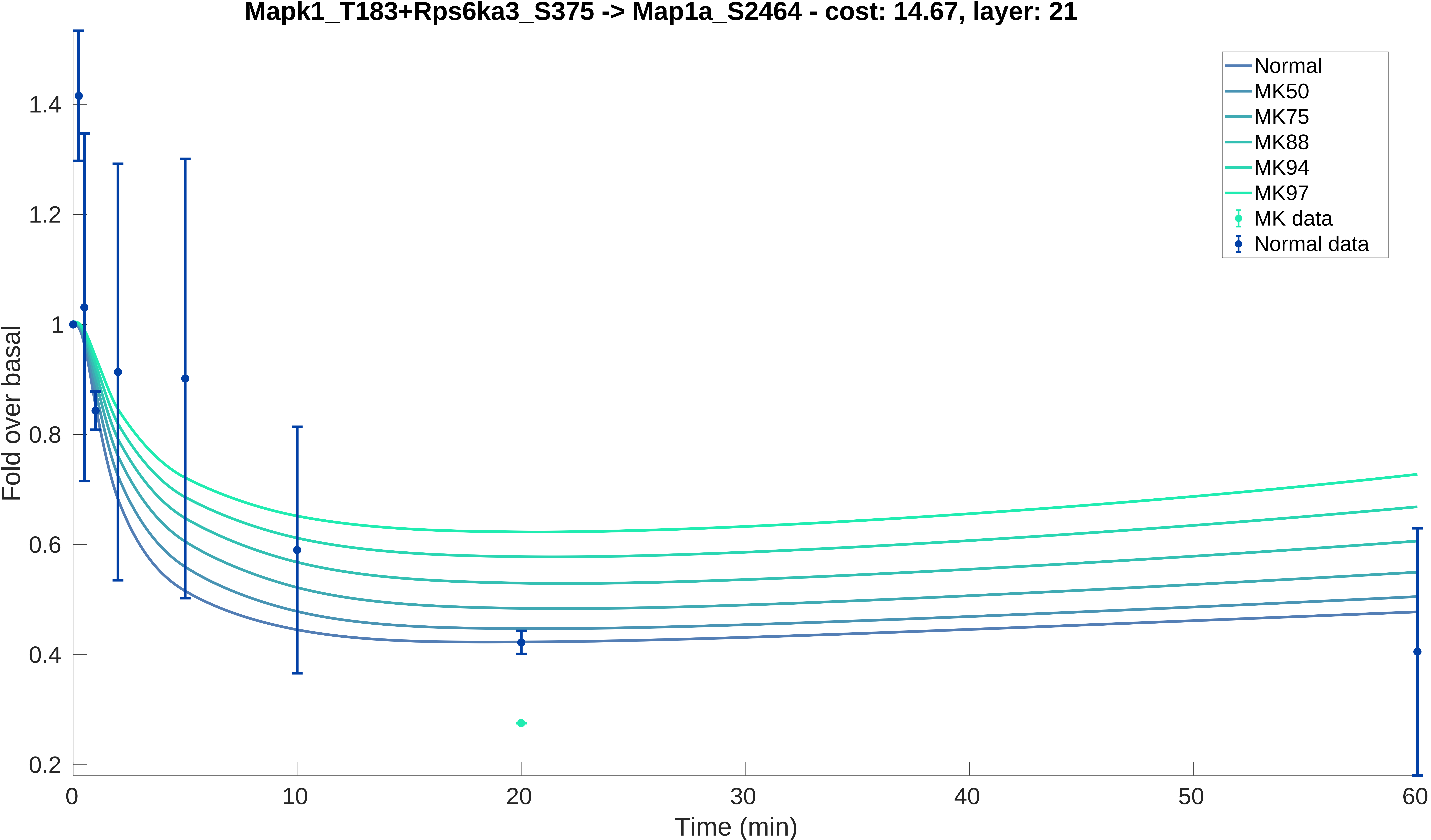

Mapk1_T183+Rps6ka3_S375 -> Map1a_S2464 - cost: 14.67, layer: 21
Normal
MK50
MK75
MK88
MK94
MK97
MK data
Normal data
Fold over basal
1.4
1.2
1
0.8
0.6
0.4
0.2
0
10
20
30
40
50
60
Time (min)

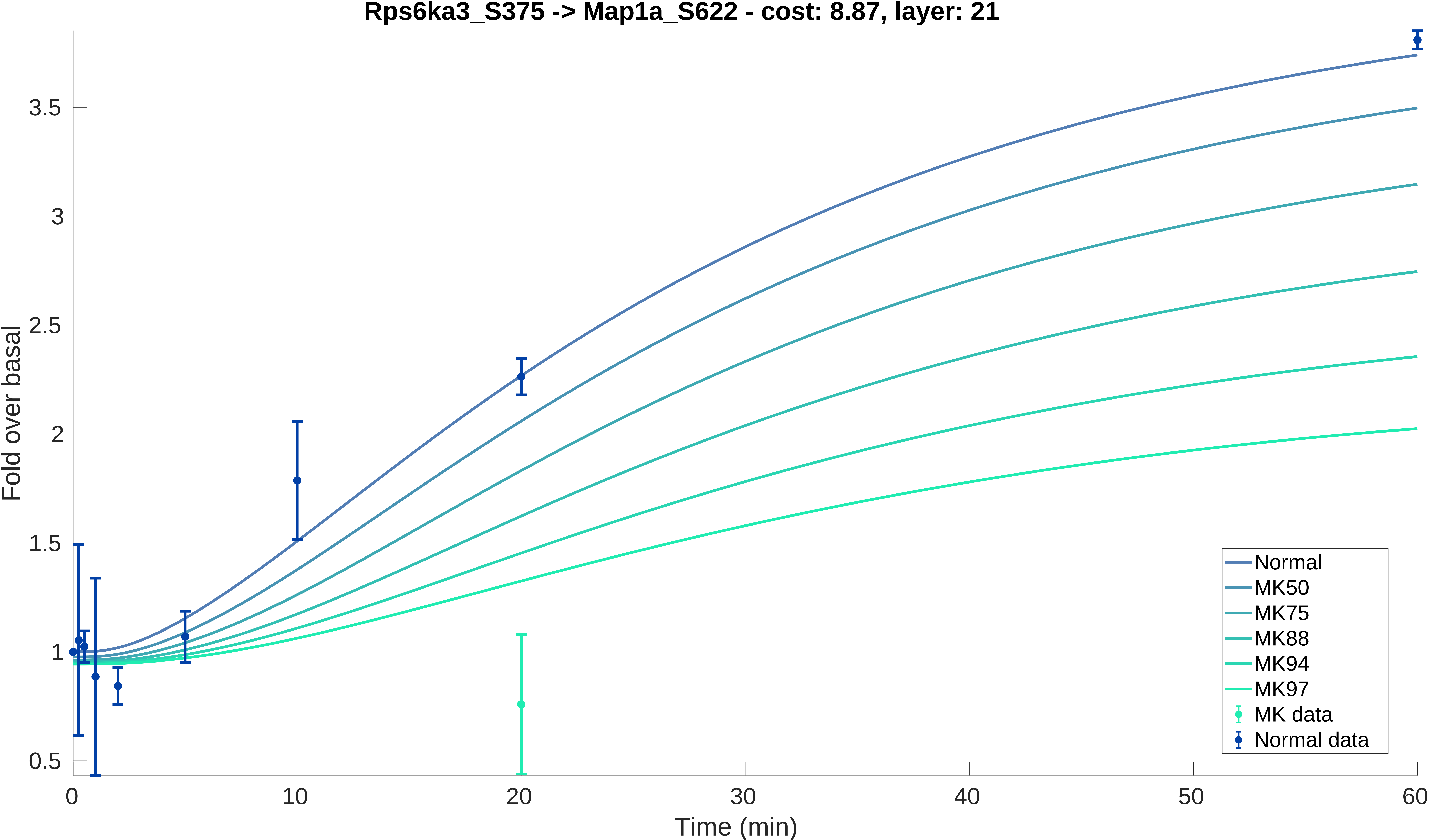

Rps6ka3_S375 -> Map1a_S622 - cost: 8.87, layer: 21
Fold over basal
0.5
1
1.5
2
2.5
3
3.5
0
10
20
30
40
50
60
Time (min)
Normal
MK50
MK75
MK88
MK94
MK97
MK data
Normal data

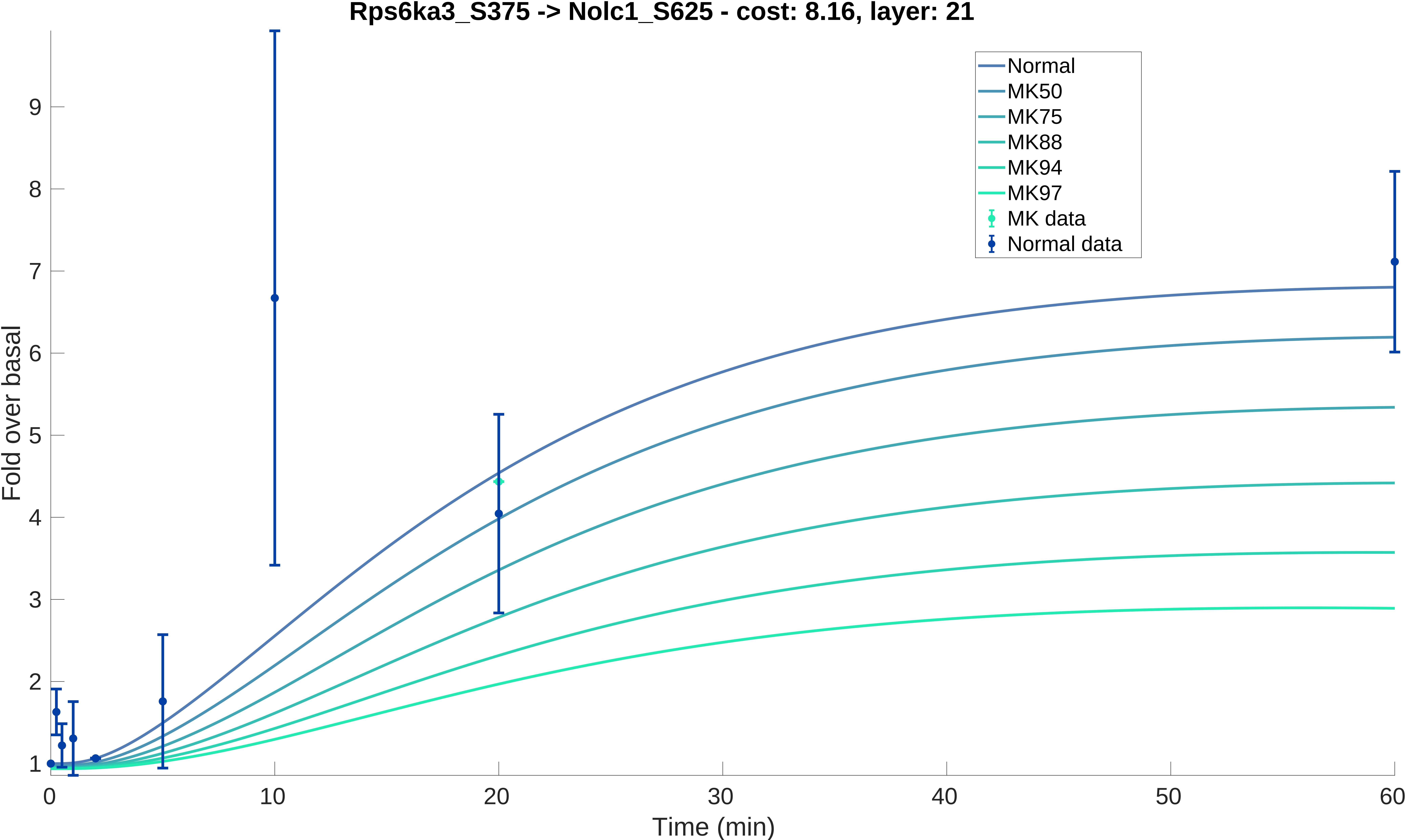
Rps6ka3_S375 -> Nolc1_S625 - cost: 8.16, layer: 21
Normal
MK50
MK75
MK88
MK94
MK97
MK data
Normal data
Fold over basal
Time (min)
1
2
3
4
5
6
7
8
9
0
10
20
30
40
50
60

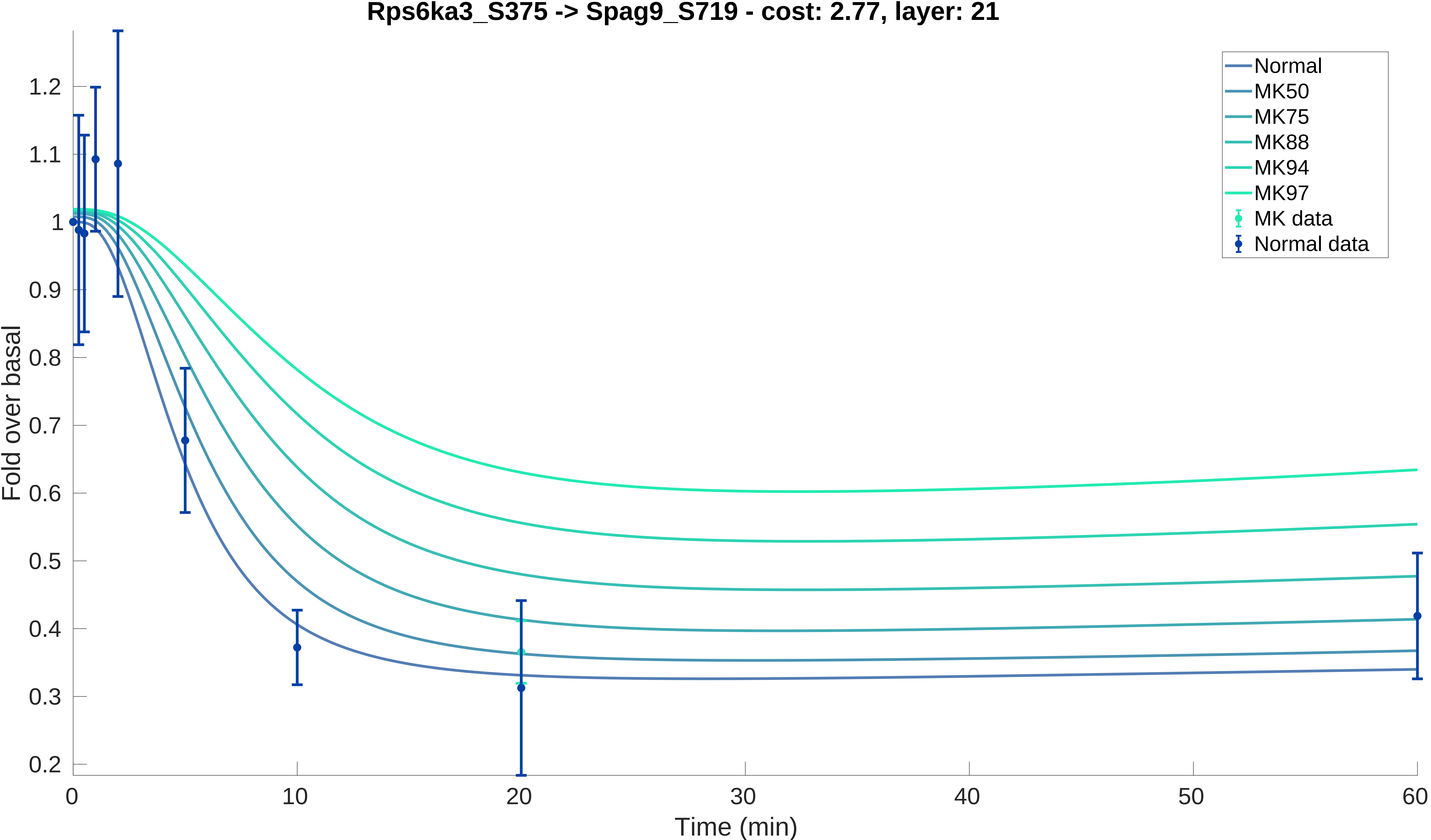

Rps6ka3_S375 -> Spag9_S719 - cost: 2.77, layer: 21
Fold over basal
Time (min)
Normal
MK50
MK75
MK88
MK94
MK97
MK data
Normal data
1.2
1.1
1
0.9
0.8
0.7
0.6
0.5
0.4
0.3
0.2
0
10
20
30
40
50
60

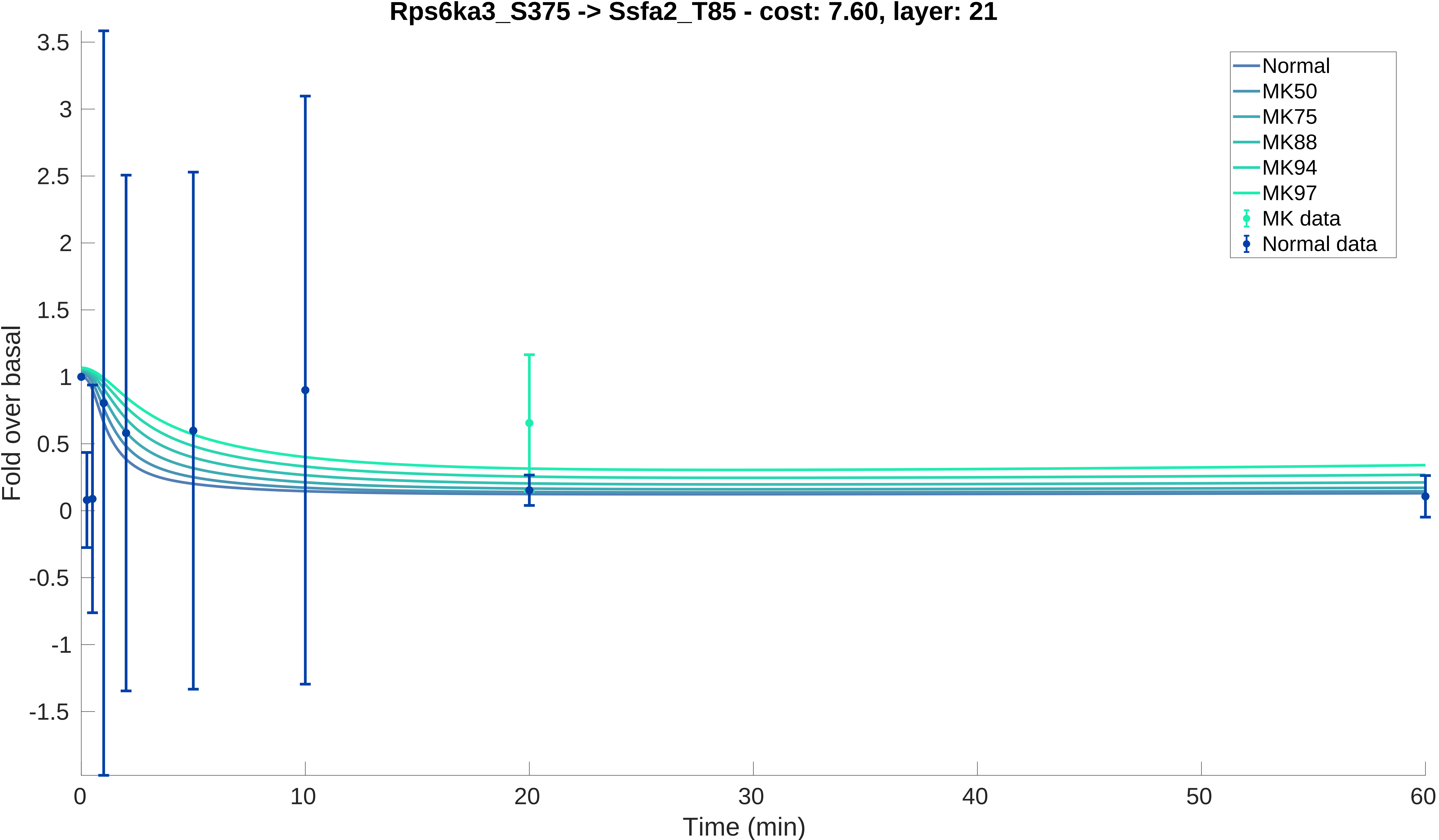

Rps6ka3_S375 -> Ssfa2_T85 - cost: 7.60, layer: 21
Fold over basal
Time (min)
3.5
3
2.5
2
1.5
1
0.5
0
-0.5
-1
-1.5
0
10
20
30
40
50
60
Normal
MK50
MK75
MK88
MK94
MK97
MK data
Normal data

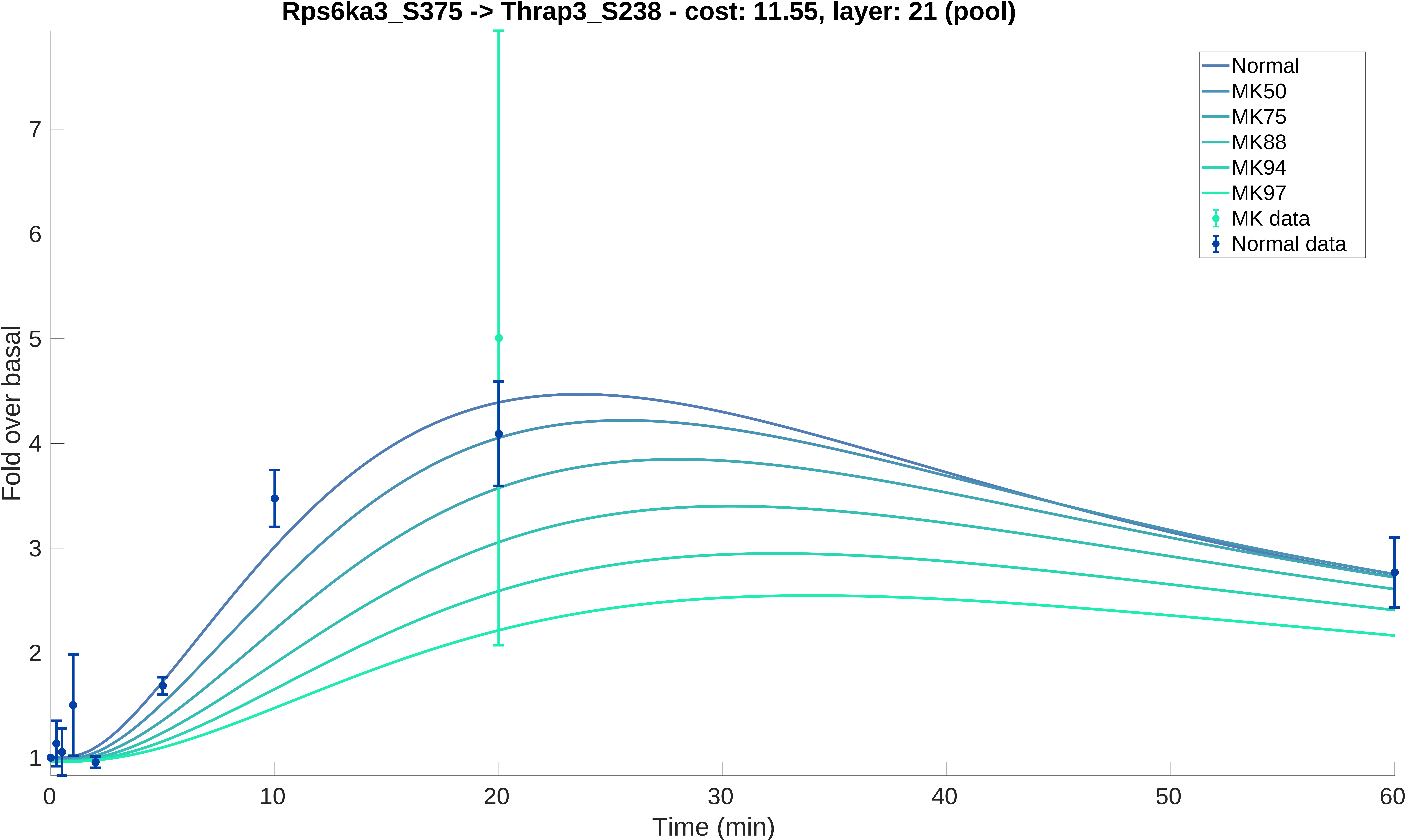

Rps6ka3_S375 -> Thrap3_S238 - cost: 11.55, layer: 21 (pool)
Normal
MK50
MK75
MK88
MK94
MK97
MK data
Normal data
Fold over basal
Time (min)
1
2
3
4
5
6
7
0
10
20
30
40
50
60

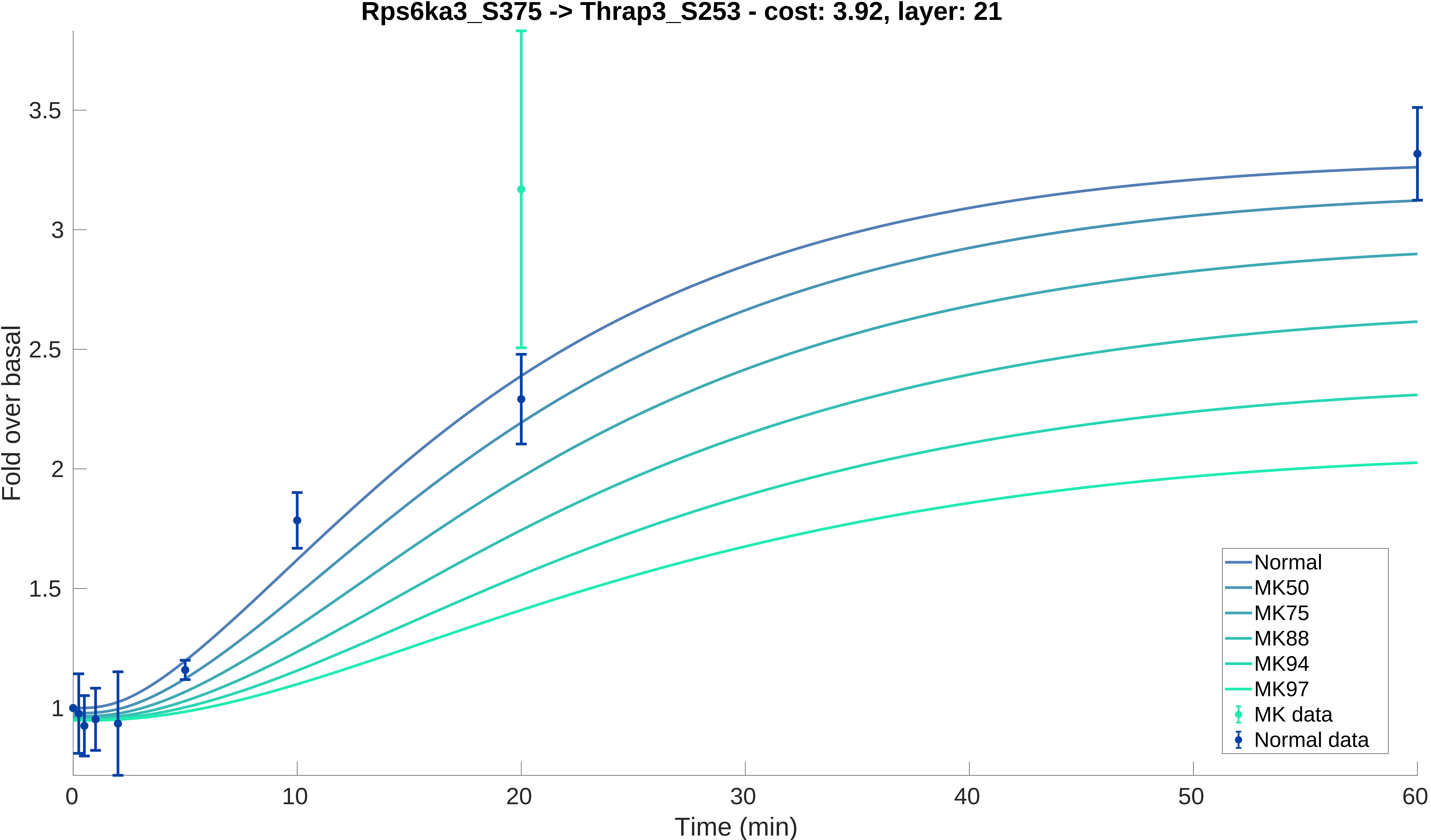

Rps6ka3_S375 -> Thrap3_S253 - cost: 3.92, layer: 21
Fold over basal
Time (min)
3.5
3
2.5
2
1.5
1
0
10
20
30
40
50
60
Normal
MK50
MK75
MK88
MK94
MK97
MK data
Normal data

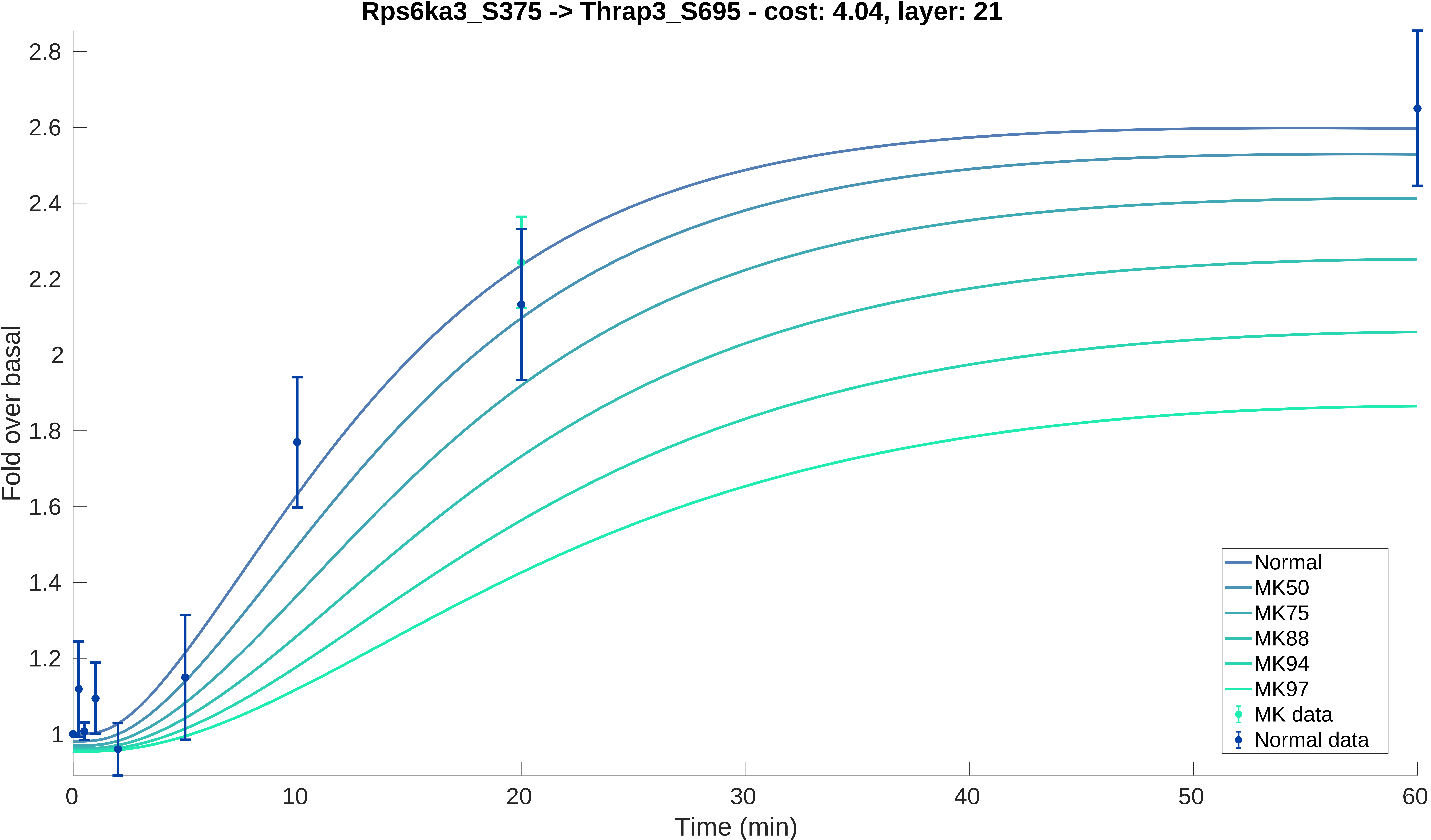

Rps6ka3_S375 -> Thrap3_S695 - cost: 4.04, layer: 21
Fold over basal
2.8
2.6
2.4
2.2
2
1.8
1.6
1.4
1.2
1
0
10
20
30
40
50
60
Time (min)
Normal
MK50
MK75
MK88
MK94
MK97
MK data
Normal data

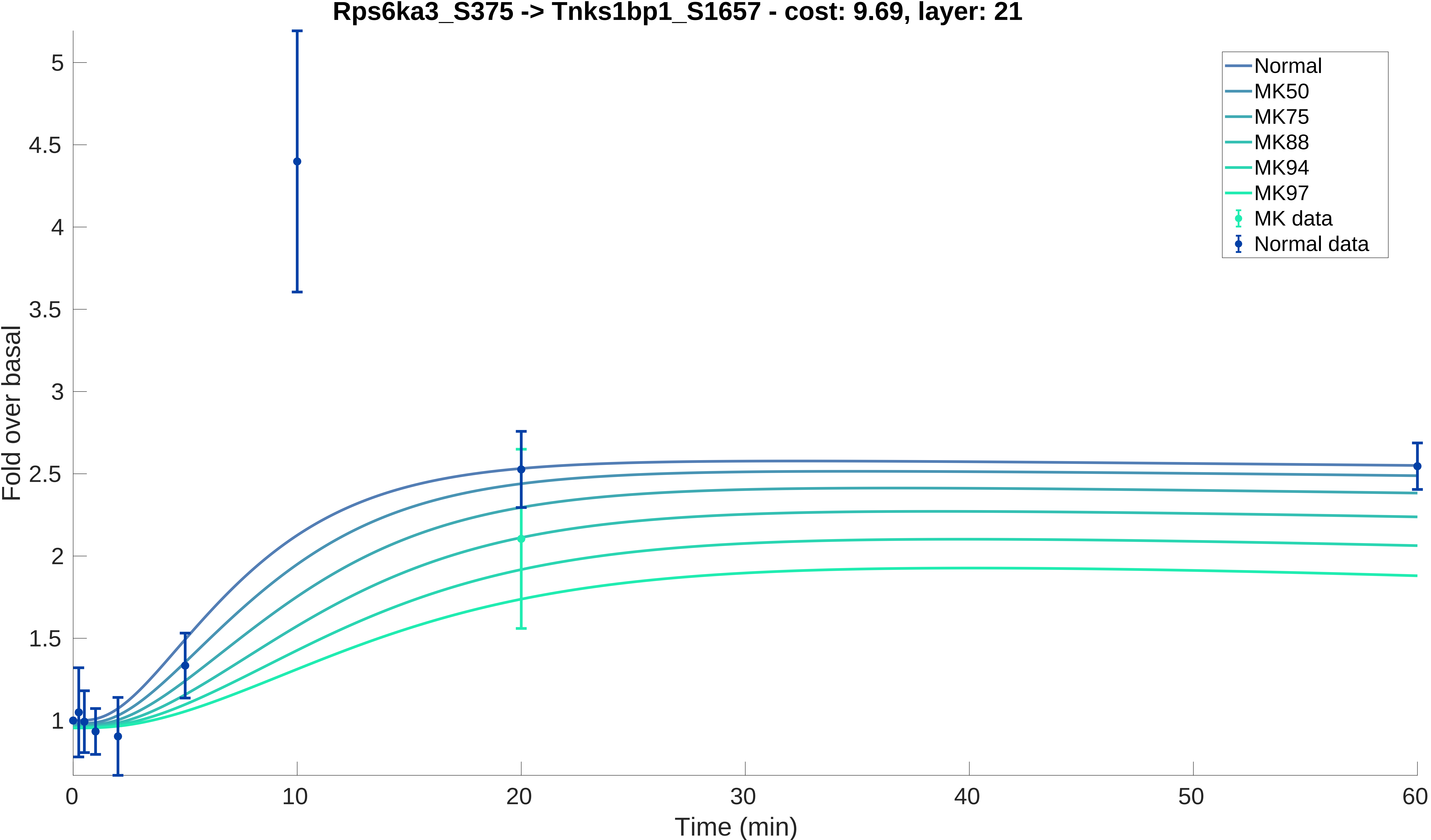

Rps6ka3_S375 -> Tnks1bp1_S1657 - cost: 9.69, layer: 21
Fold over basal
Time (min)
5
4.5
4
3.5
3
2.5
2
1.5
1
0
10
20
30
40
50
60
Normal
MK50
MK75
MK88
MK94
MK97
MK data
Normal data

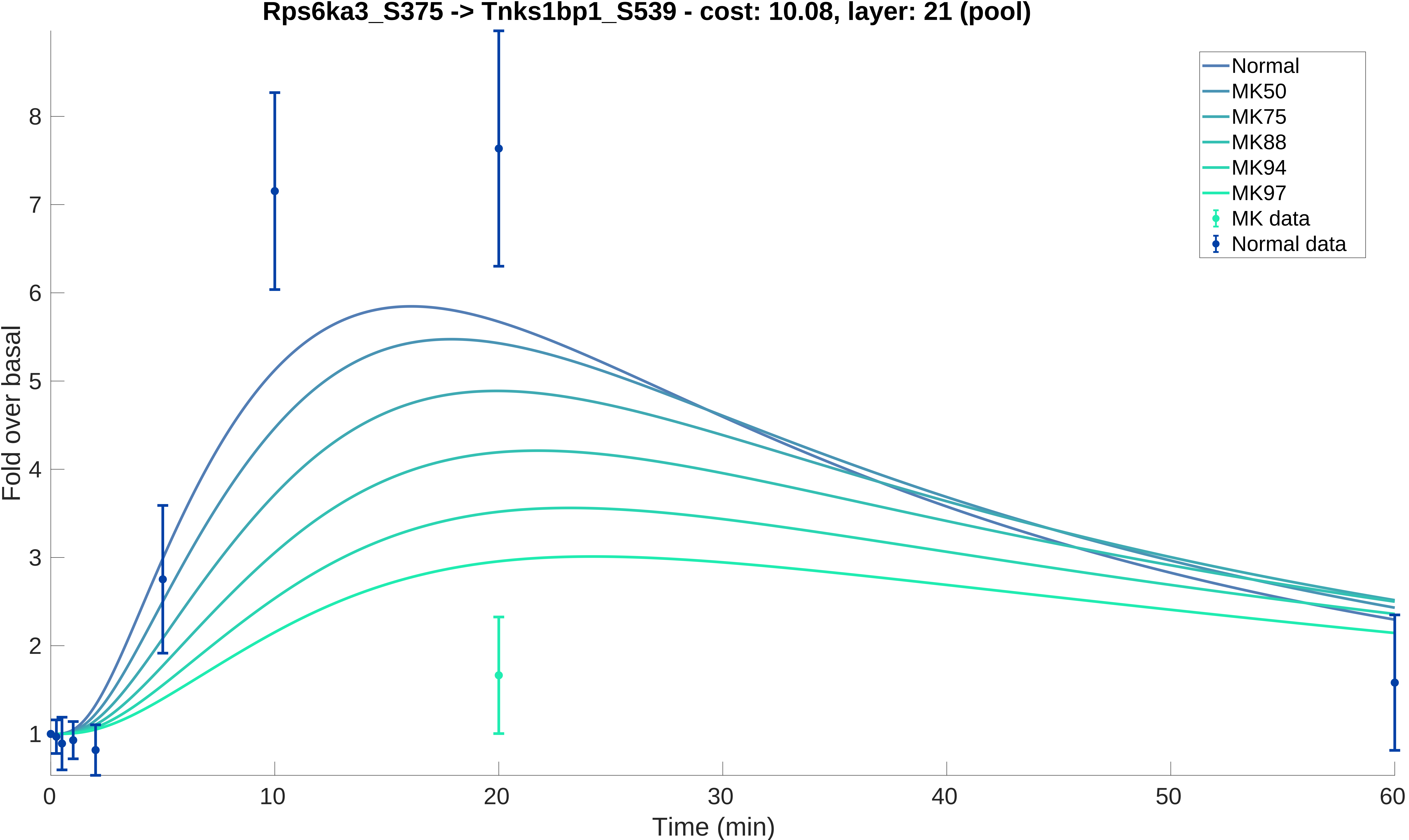

Rps6ka3_S375 -> Tnks1bp1_S539 - cost: 10.08, layer: 21 (pool)
Fold over basal
Time (min)
0
10
20
30
40
50
60
1
2
3
4
5
6
7
8
Normal
MK50
MK75
MK88
MK94
MK97
MK data
Normal data

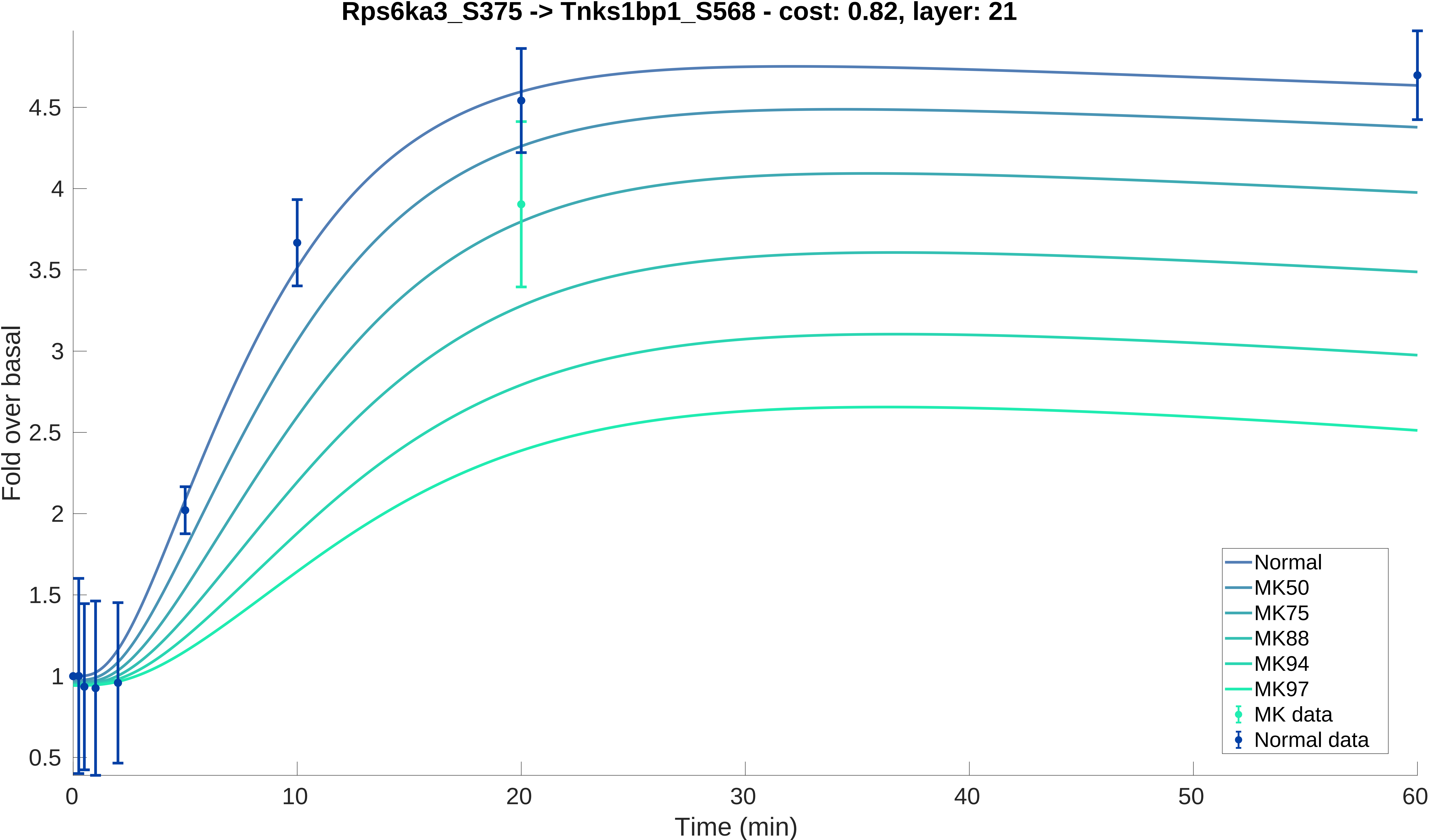

Rps6ka3_S375 -> Tnks1bp1_S568 - cost: 0.82, layer: 21
Fold over basal
0.5
1
1.5
2
2.5
3
3.5
4
4.5
0
10
20
30
40
50
60
Time (min)
Normal
MK50
MK75
MK88
MK94
MK97
MK data
Normal data

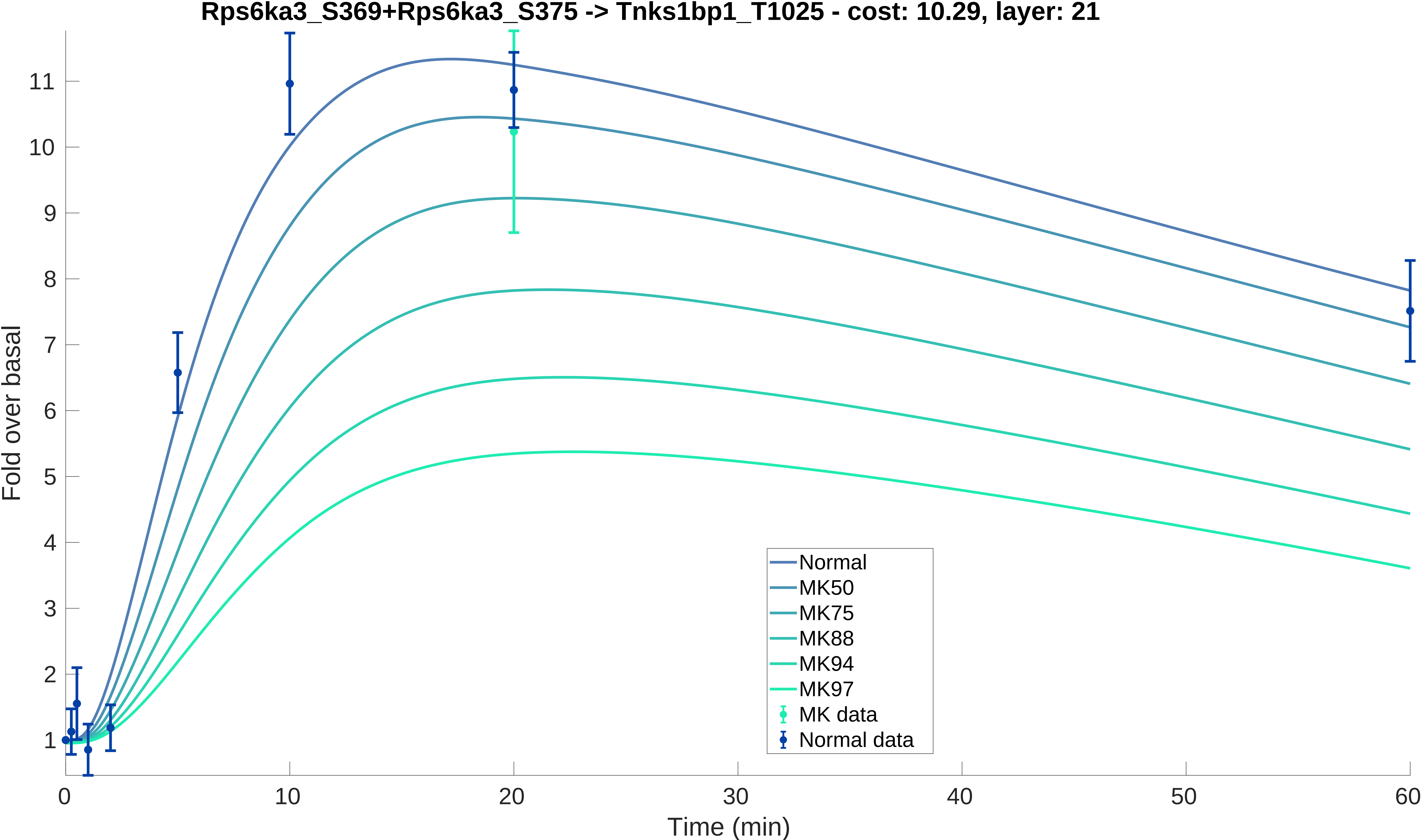

Rps6ka3_S369+Rps6ka3_S375 -> Tnks1bp1_T1025 - cost: 10.29, layer: 21
Fold over basal
Time (min)
0
10
20
30
40
50
60
1
2
3
4
5
6
7
8
9
10
11
Normal
MK50
MK75
MK88
MK94
MK97
MK data
Normal data

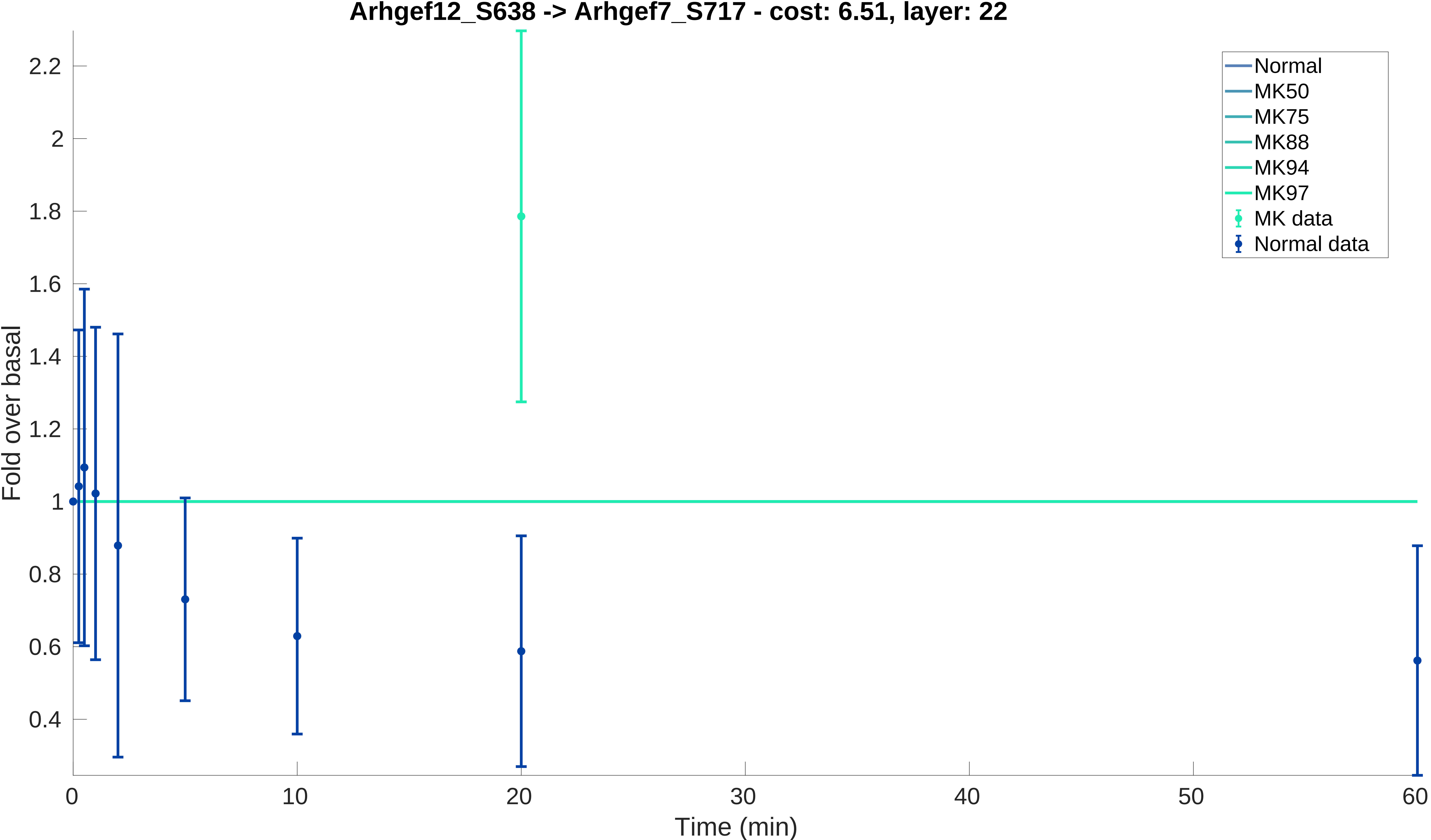
Arhgef12_S638 -> Arhgef7_S717 - cost: 6.51, layer: 22
Fold over basal
Time (min)
0
10
20
30
40
50
60
0.4
0.6
0.8
1
1.2
1.4
1.6
1.8
2
2.2
Normal
MK50
MK75
MK88
MK94
MK97
MK data
Normal data

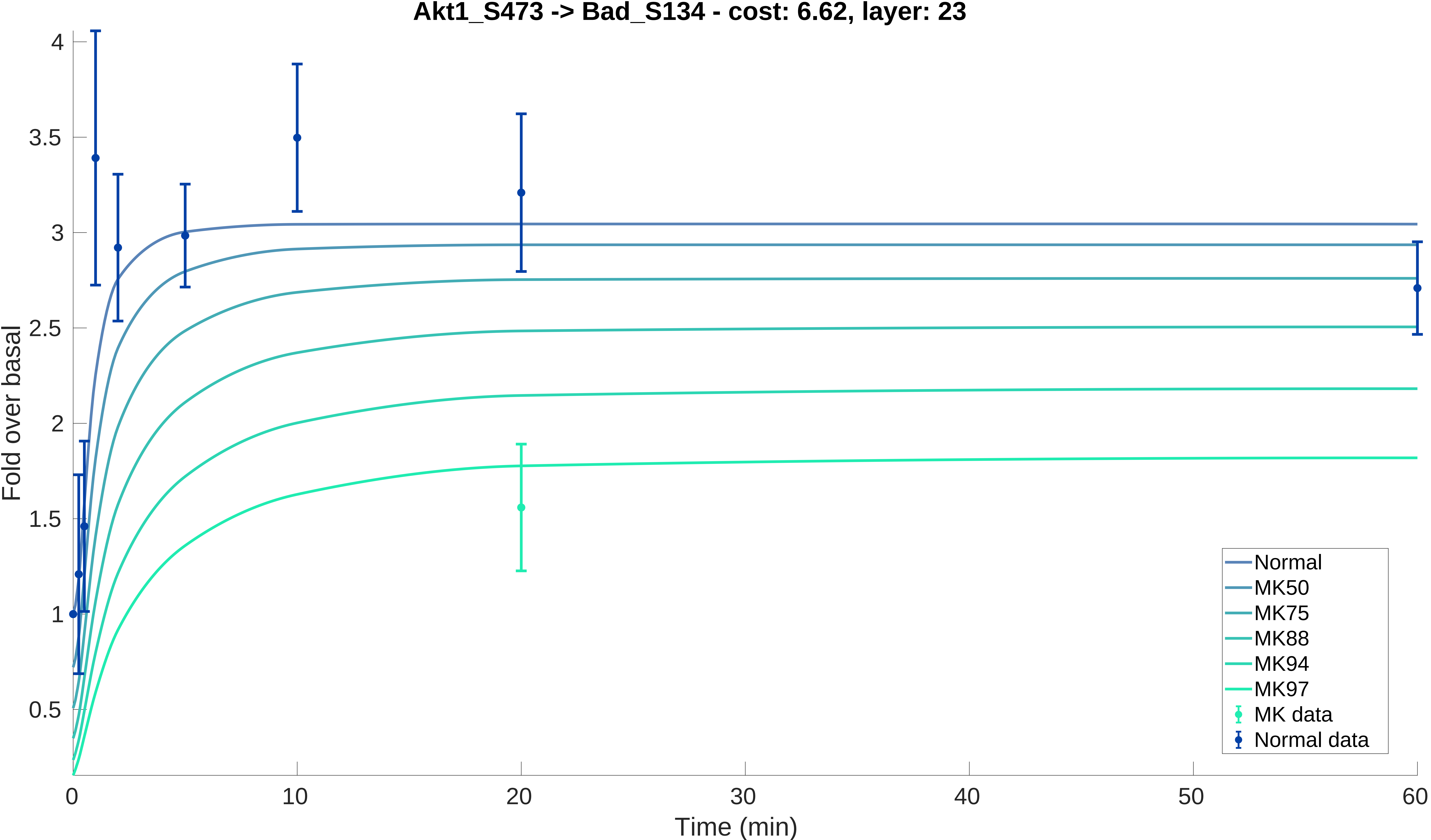

Akt1_S473 -> Bad_S134 - cost: 6.62, layer: 23
Fold over basal
Time (min)
0.5
1
1.5
2
2.5
3
3.5
4
0
10
20
30
40
50
60
Normal
MK50
MK75
MK88
MK94
MK97
MK data
Normal data

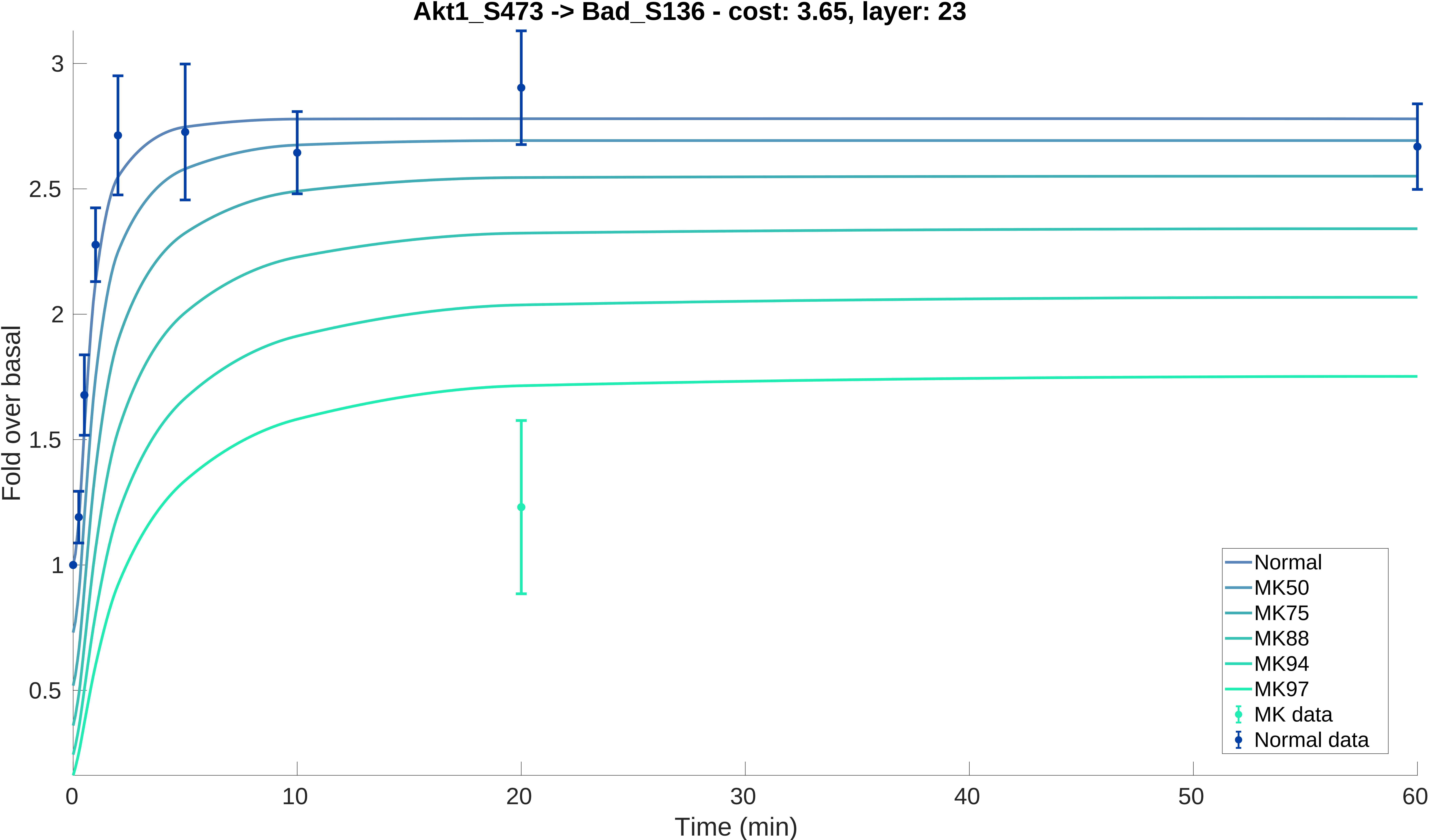

Akt1_S473 -> Bad_S136 - cost: 3.65, layer: 23
Fold over basal
Time (min)
0.5
1
1.5
2
2.5
3
0
10
20
30
40
50
60
Normal
MK50
MK75
MK88
MK94
MK97
MK data
Normal data

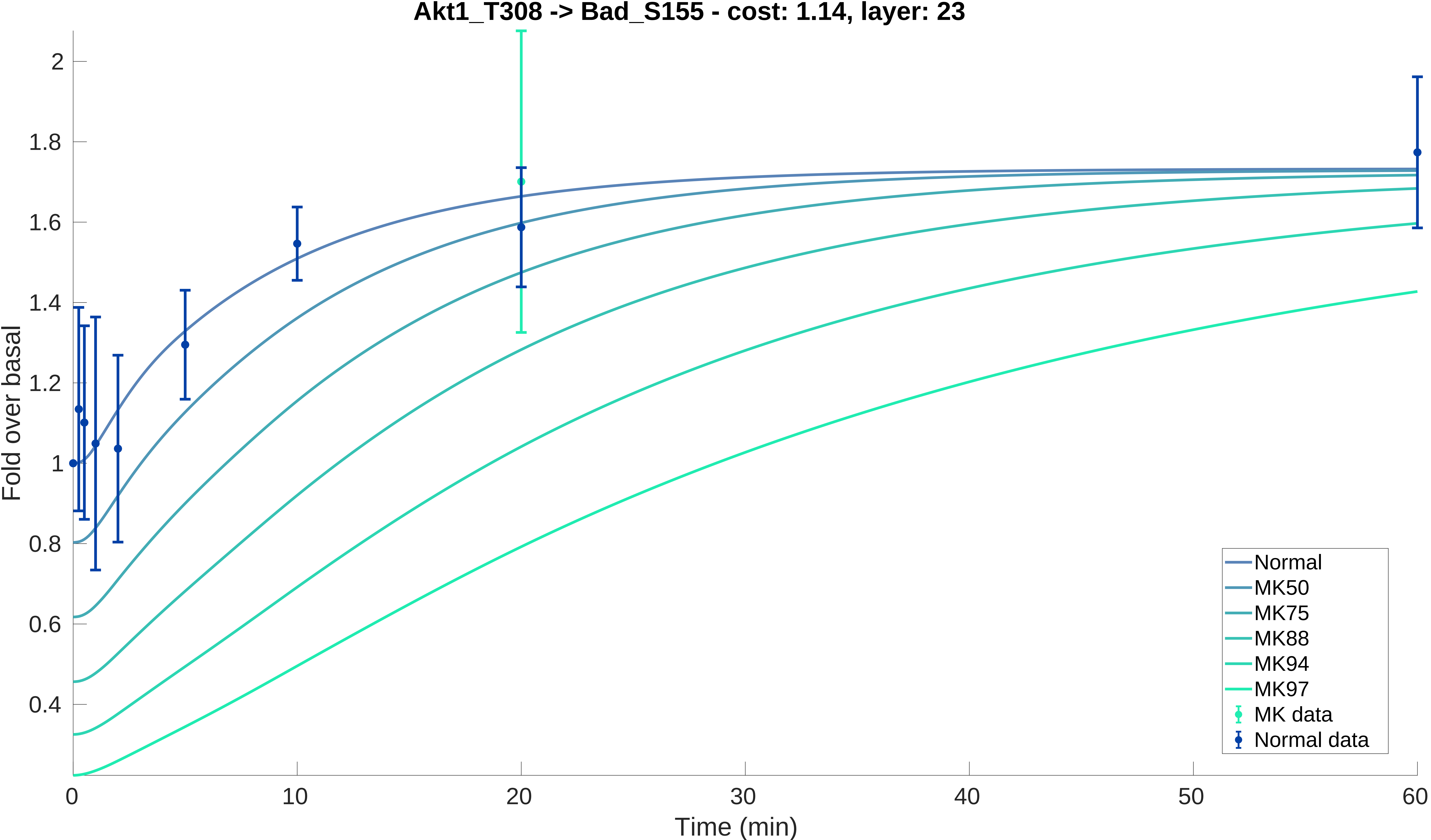

Akt1_T308 -> Bad_S155 - cost: 1.14, layer: 23
Fold over basal
Time (min)
0
10
20
30
40
50
60
0.4
0.6
0.8
1
1.2
1.4
1.6
1.8
2
Normal
MK50
MK75
MK88
MK94
MK97
MK data
Normal data

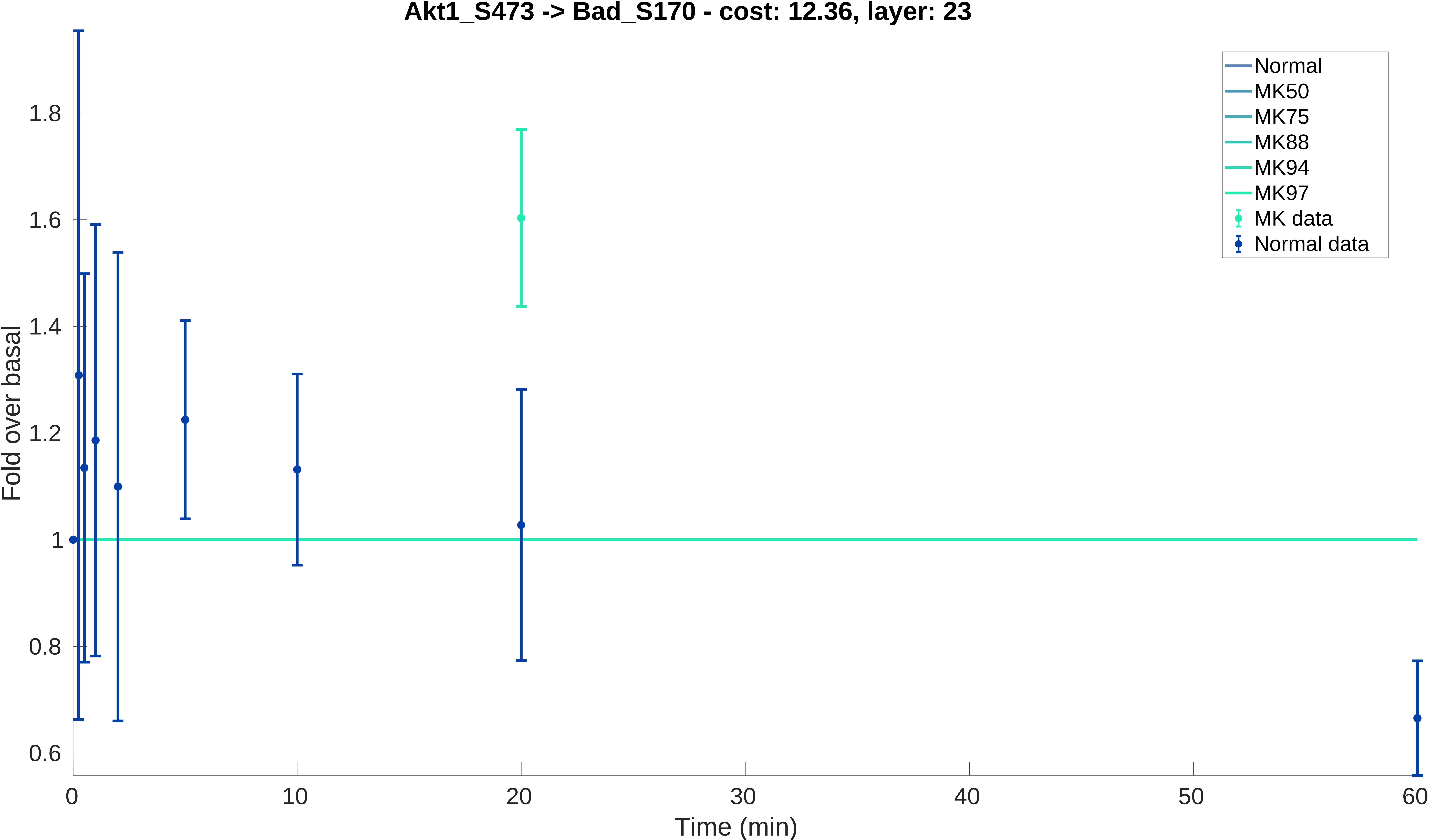
Akt1_S473 -> Bad_S170 - cost: 12.36, layer: 23
Fold over basal
1.8
1.6
1.4
1.2
1
0.8
0.6
0
10
20
30
40
50
60
Time (min)
Normal
MK50
MK75
MK88
MK94
MK97
MK data
Normal data

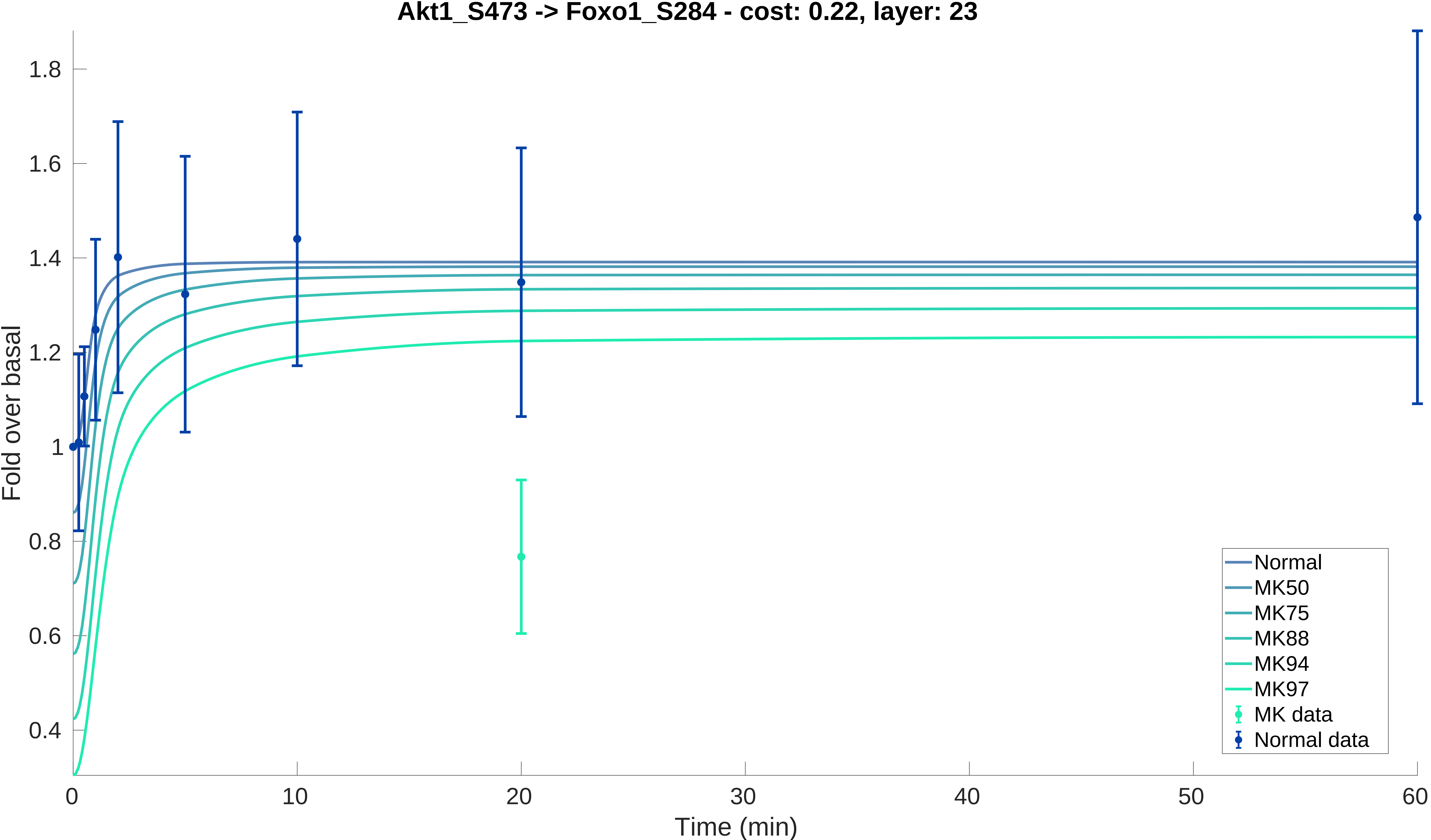

Akt1_S473 -> Foxo1_S284 - cost: 0.22, layer: 23
Fold over basal
1.8
1.6
1.4
1.2
1
0.8
0.6
0.4
0
10
20
30
40
50
60
Time (min)
Normal
MK50
MK75
MK88
MK94
MK97
MK data
Normal data

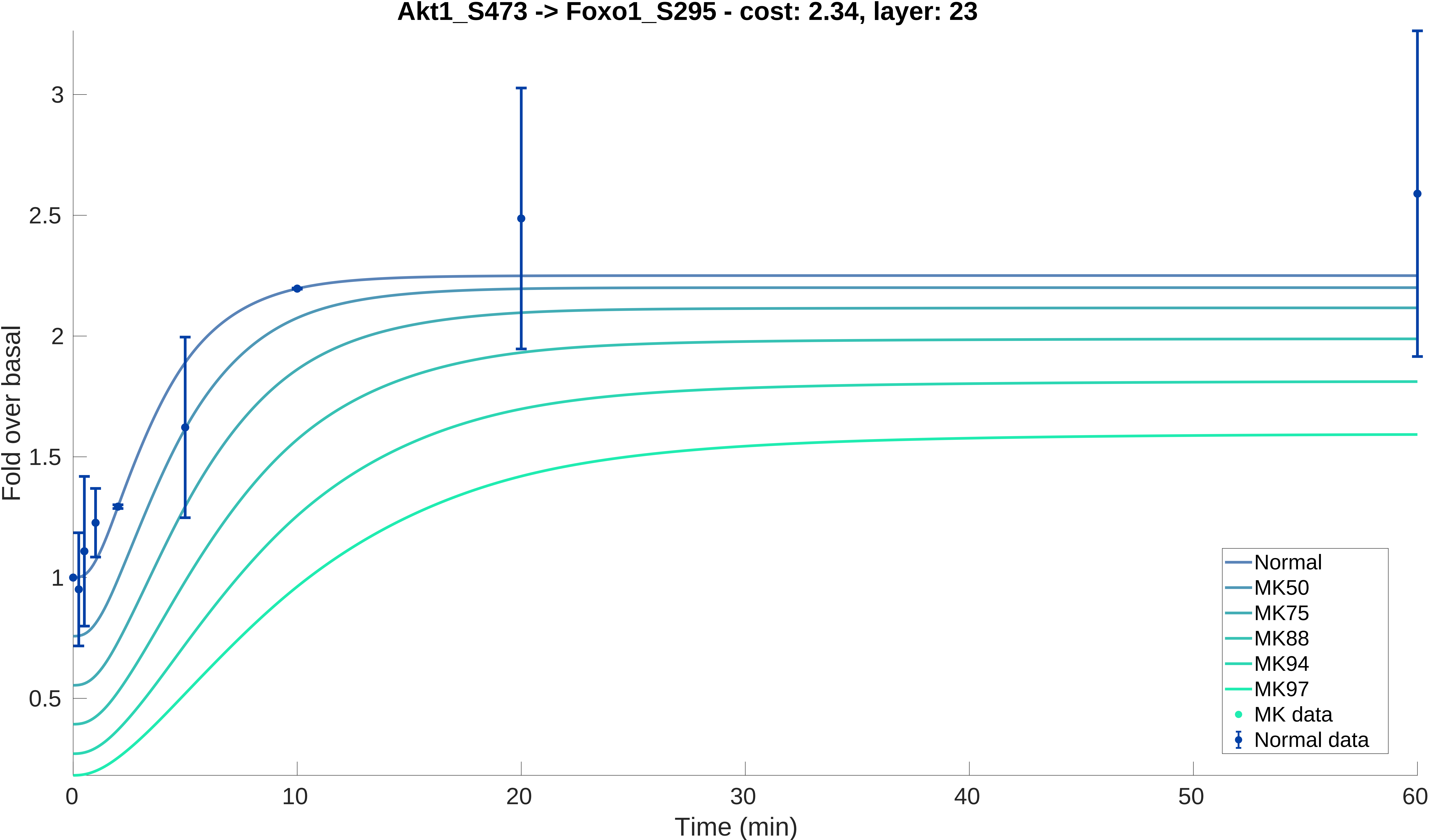
Akt1_S473 -> Foxo1_S295 - cost: 2.34, layer: 23
Fold over basal
Time (min)
0
10
20
30
40
50
60
0.5
1
1.5
2
2.5
3
Normal
MK50
MK75
MK88
MK94
MK97
MK data
Normal data

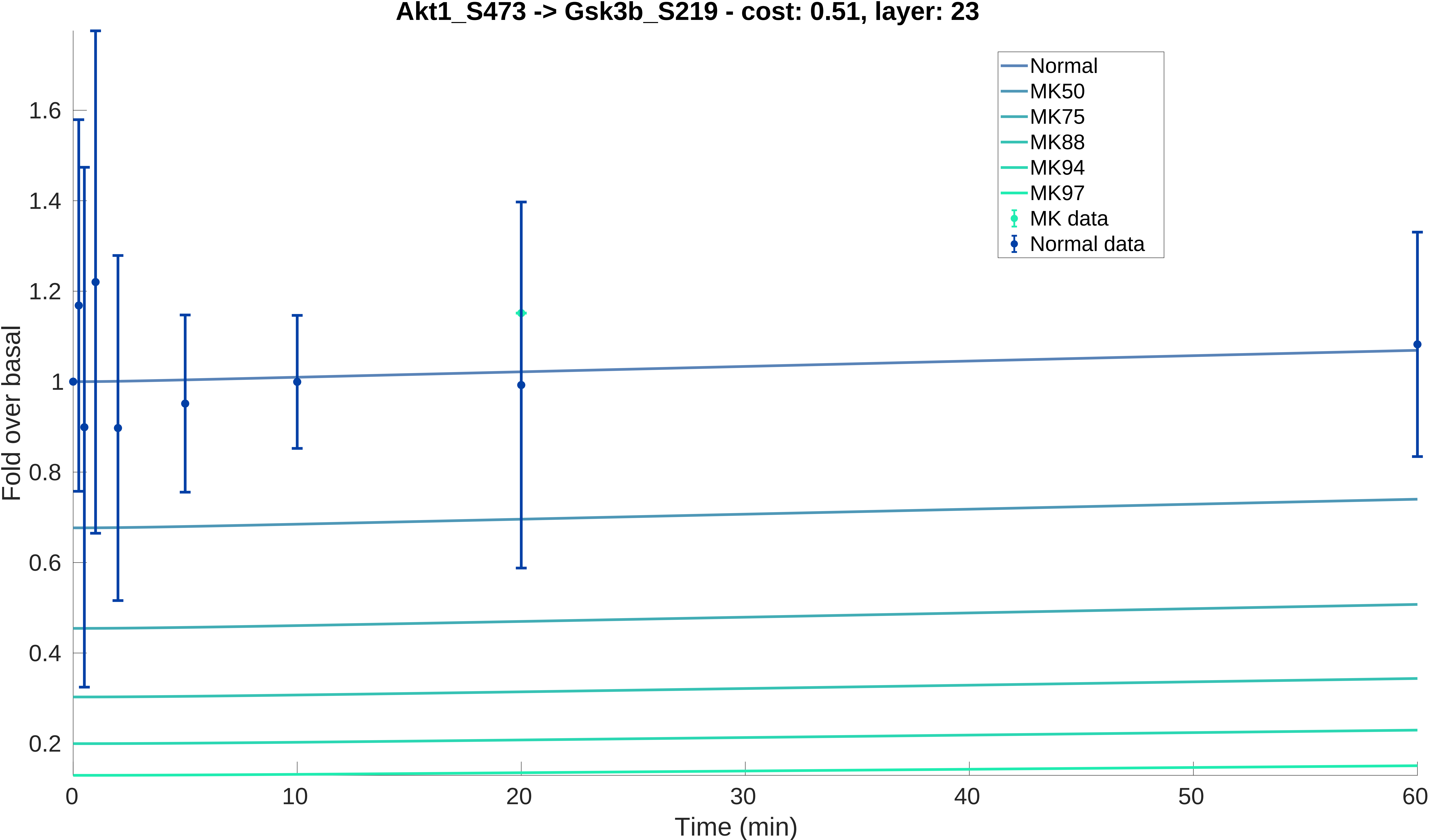

Akt1_S473 -> Gsk3b_S219 - cost: 0.51, layer: 23
Fold over basal
Time (min)
Normal
MK50
MK75
MK88
MK94
MK97
MK data
Normal data
0
10
20
30
40
50
60
0.2
0.4
0.6
0.8
1
1.2
1.4
1.6

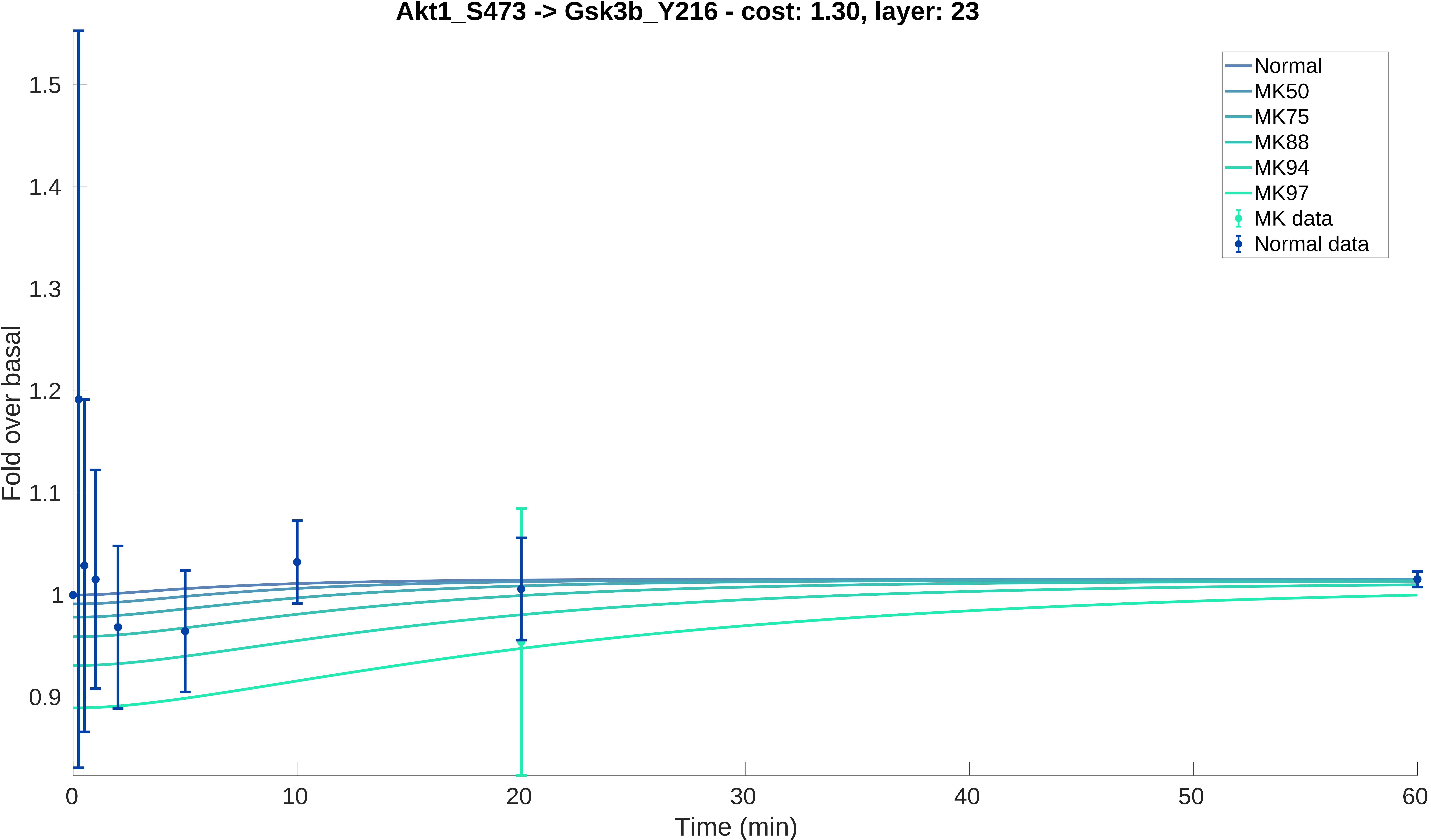

Akt1_S473 -> Gsk3b_Y216 - cost: 1.30, layer: 23
Normal
MK50
MK75
MK88
MK94
MK97
MK data
Normal data
Fold over basal
1.5
1.4
1.3
1.2
1.1
1
0.9
0
10
20
30
40
50
60
Time (min)

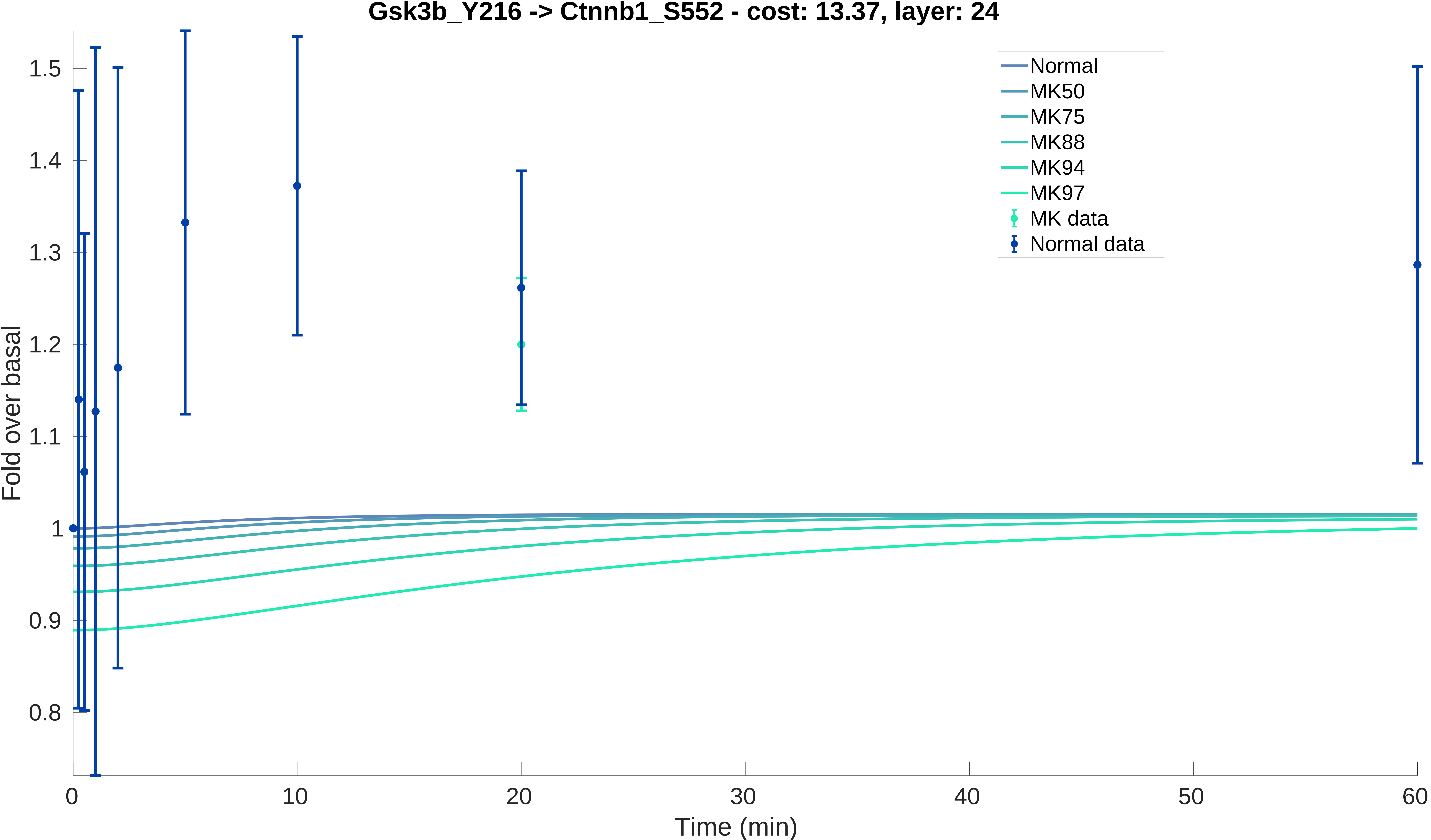
Gsk3b_Y216 -> Ctnnb1_S552 - cost: 13.37, layer: 24
Fold over basal
Time (min)
Normal
MK50
MK75
MK88
MK94
MK97
MK data
Normal data
0.8
0.9
1
1.1
1.2
1.3
1.4
1.5
0
10
20
30
40
50
60

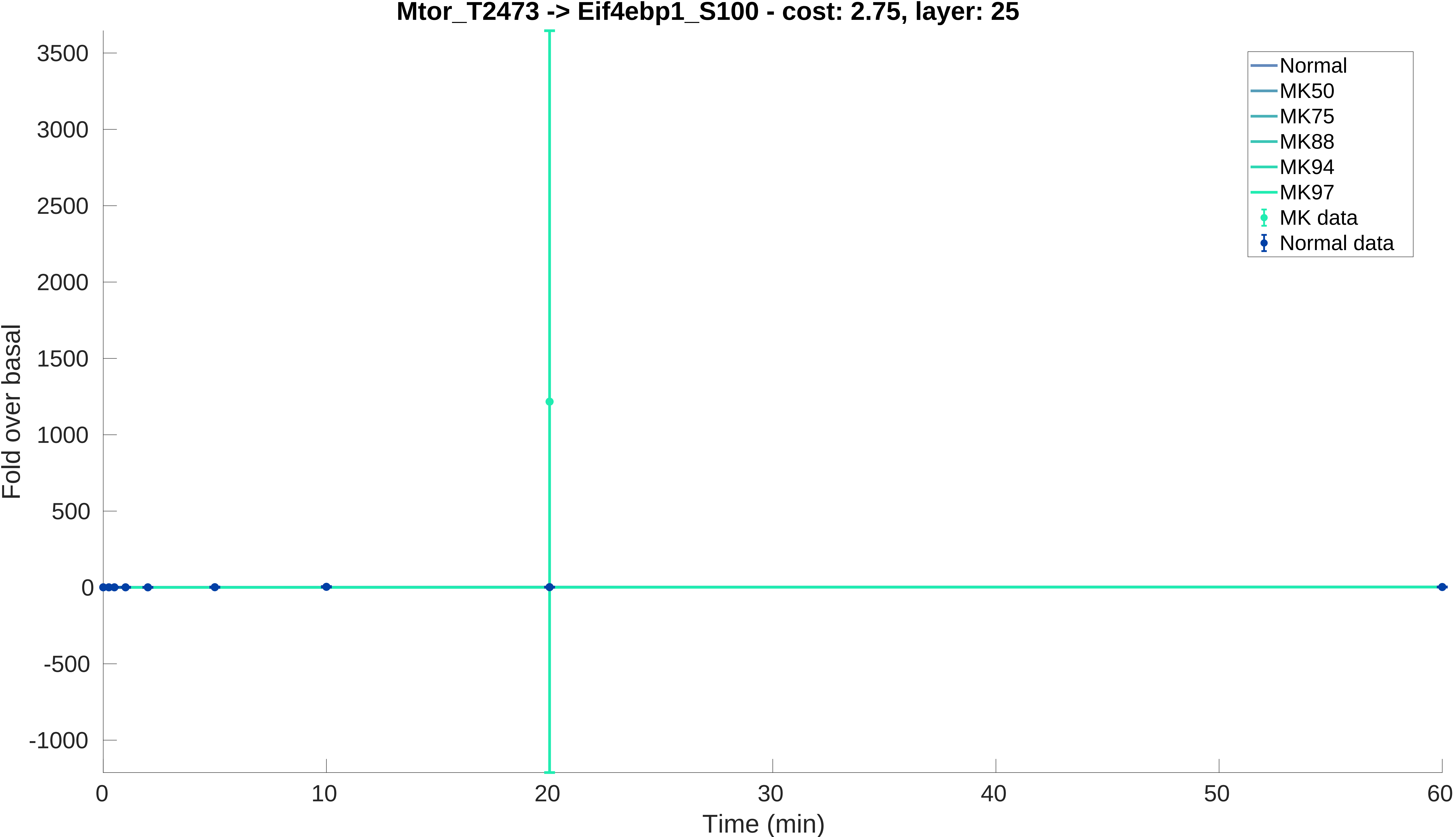

Mtor_T2473 -> Eif4ebp1_S100 - cost: 2.75, layer: 25
Fold over basal
3500
3000
2500
2000
1500
1000
500
0
-500
-1000
0
10
20
30
40
50
60
Time (min)
Normal
MK50
MK75
MK88
MK94
MK97
MK data
Normal data

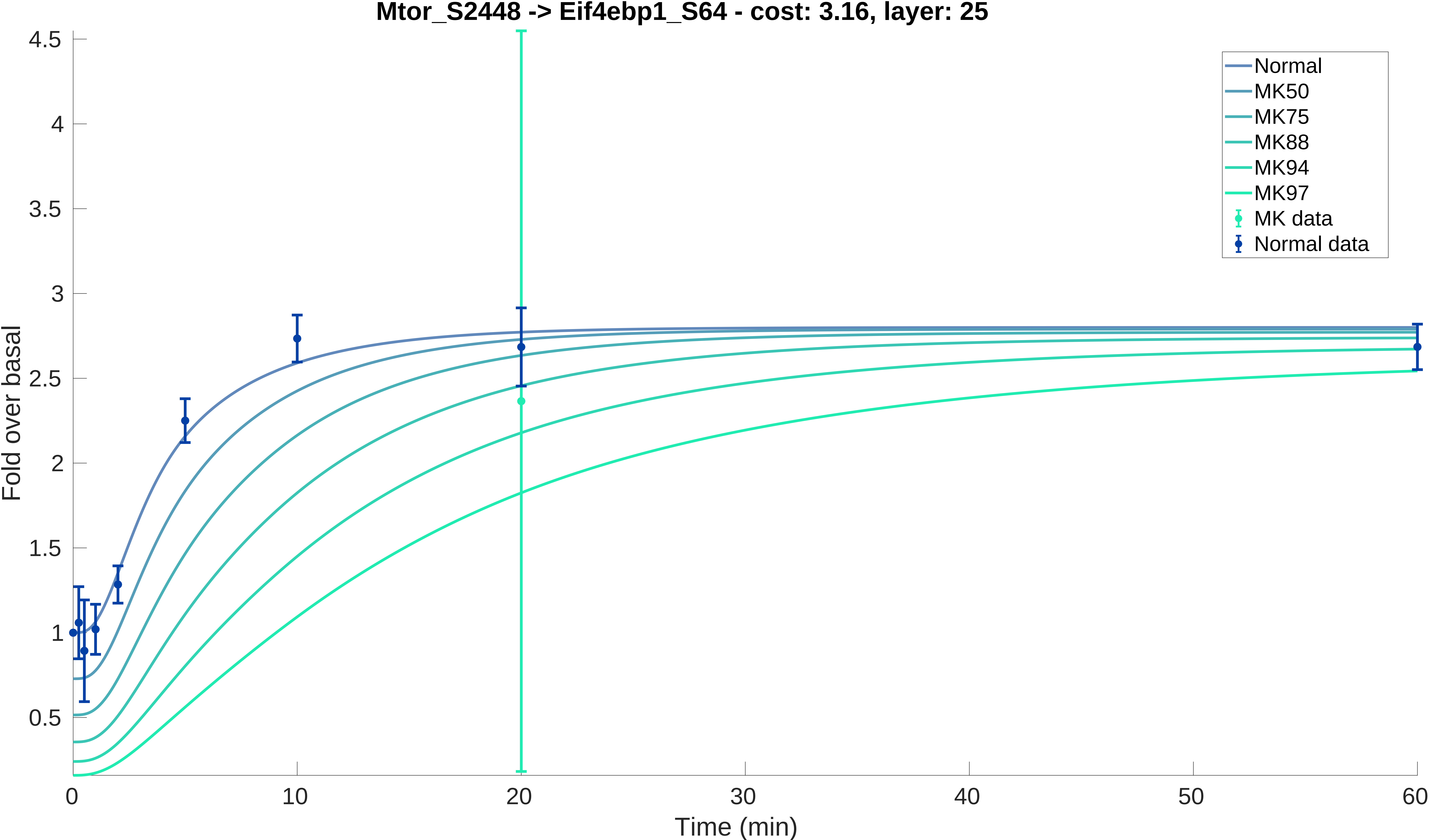

Mtor_S2448 -> Eif4ebp1_S64 - cost: 3.16, layer: 25
Normal
MK50
MK75
MK88
MK94
MK97
MK data
Normal data
Fold over basal
Time (min)
4.5
4
3.5
3
2.5
2
1.5
1
0.5
0
10
20
30
40
50
60

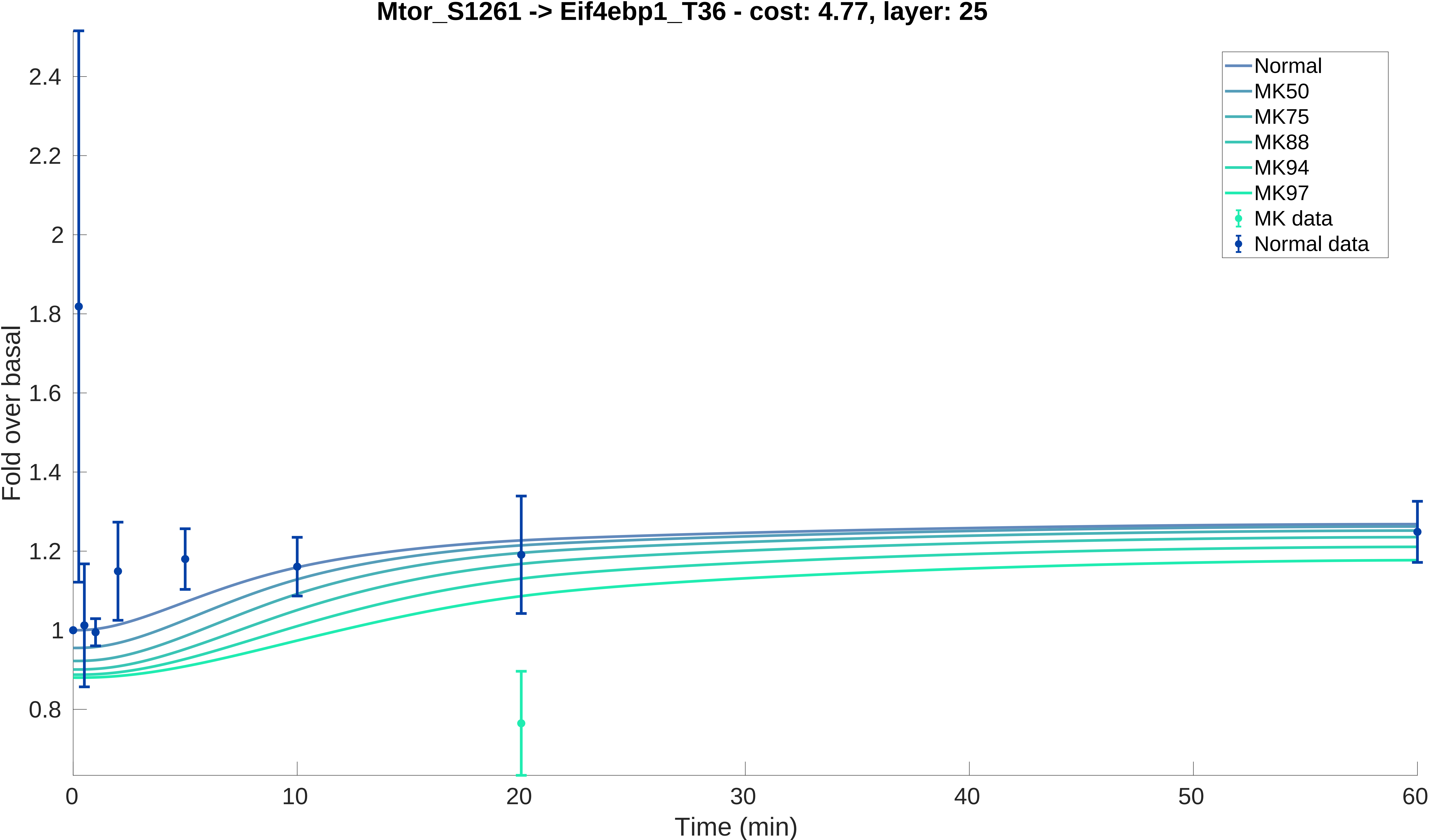
Mtor_S1261 -> Eif4ebp1_T36 - cost: 4.77, layer: 25
Fold over basal
Time (min)
Normal
MK50
MK75
MK88
MK94
MK97
MK data
Normal data
2.4
2.2
2
1.8
1.6
1.4
1.2
1
0.8
0
10
20
30
40
50
60

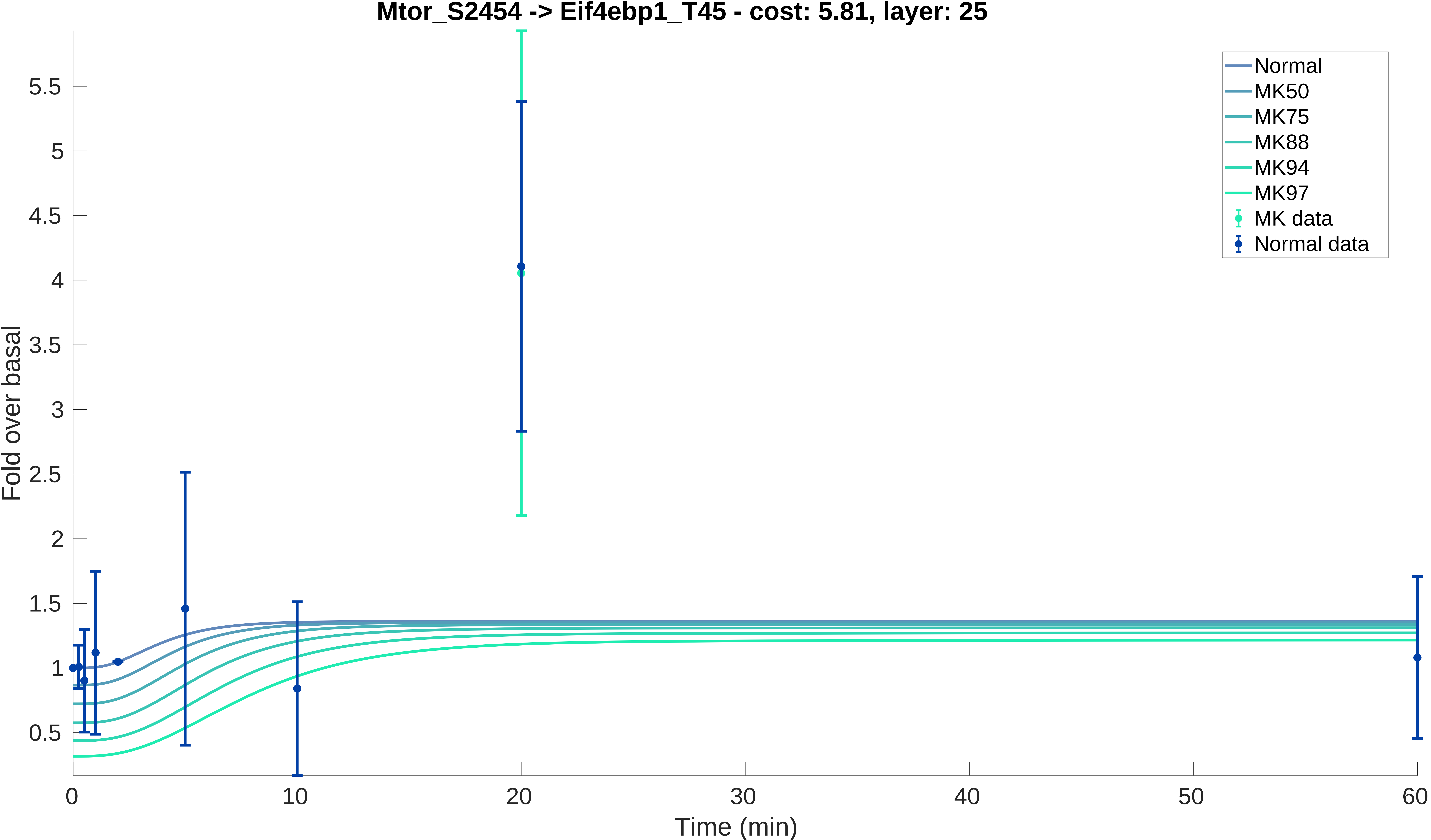

Mtor_S2454 -> Eif4ebp1_T45 - cost: 5.81, layer: 25
Fold over basal
Time (min)
0.5
1
1.5
2
2.5
3
3.5
4
4.5
5
5.5
0
10
20
30
40
50
60
Normal
MK50
MK75
MK88
MK94
MK97
MK data
Normal data

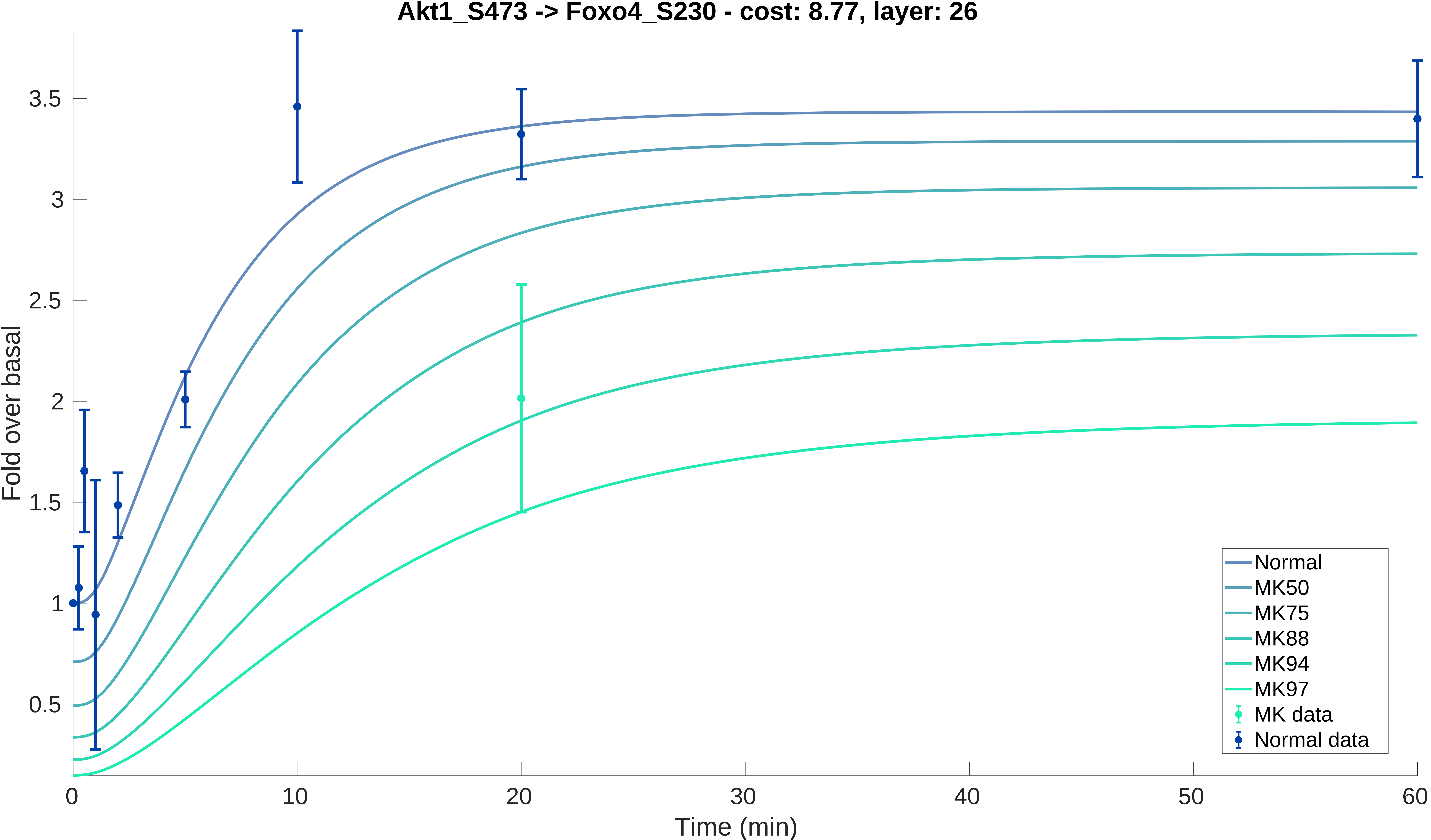

Akt1_S473 -> Foxo4_S230 - cost: 8.77, layer: 26
Fold over basal
3.5
3
2.5
2
1.5
1
0.5
0
10
20
30
40
50
60
Time (min)
Normal
MK50
MK75
MK88
MK94
MK97
MK data
Normal data

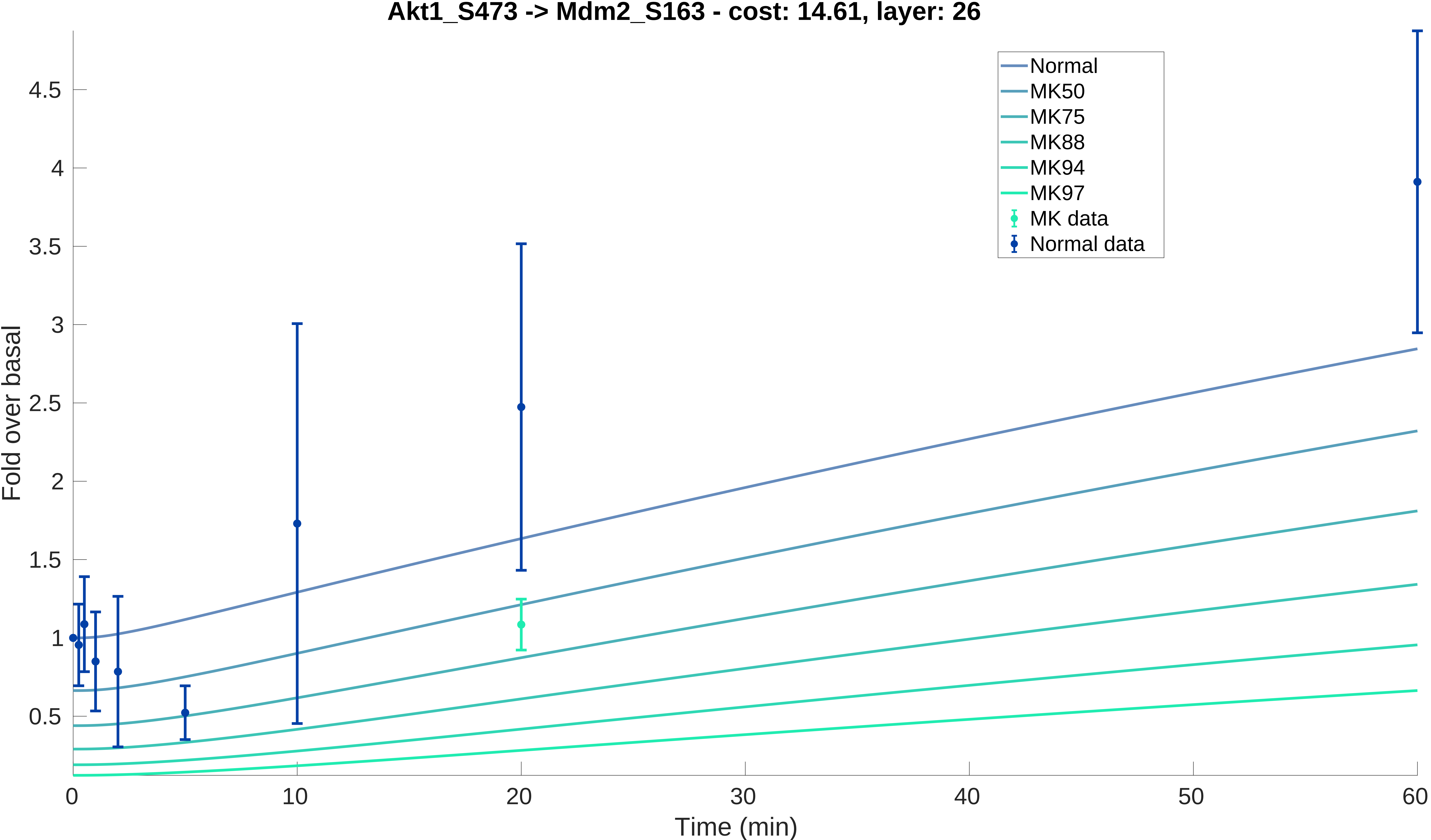

Akt1_S473 -> Mdm2_S163 - cost: 14.61, layer: 26
Normal
MK50
MK75
MK88
MK94
MK97
MK data
Normal data
Fold over basal
Time (min)
0.5
1
1.5
2
2.5
3
3.5
4
4.5
0
10
20
30
40
50
60

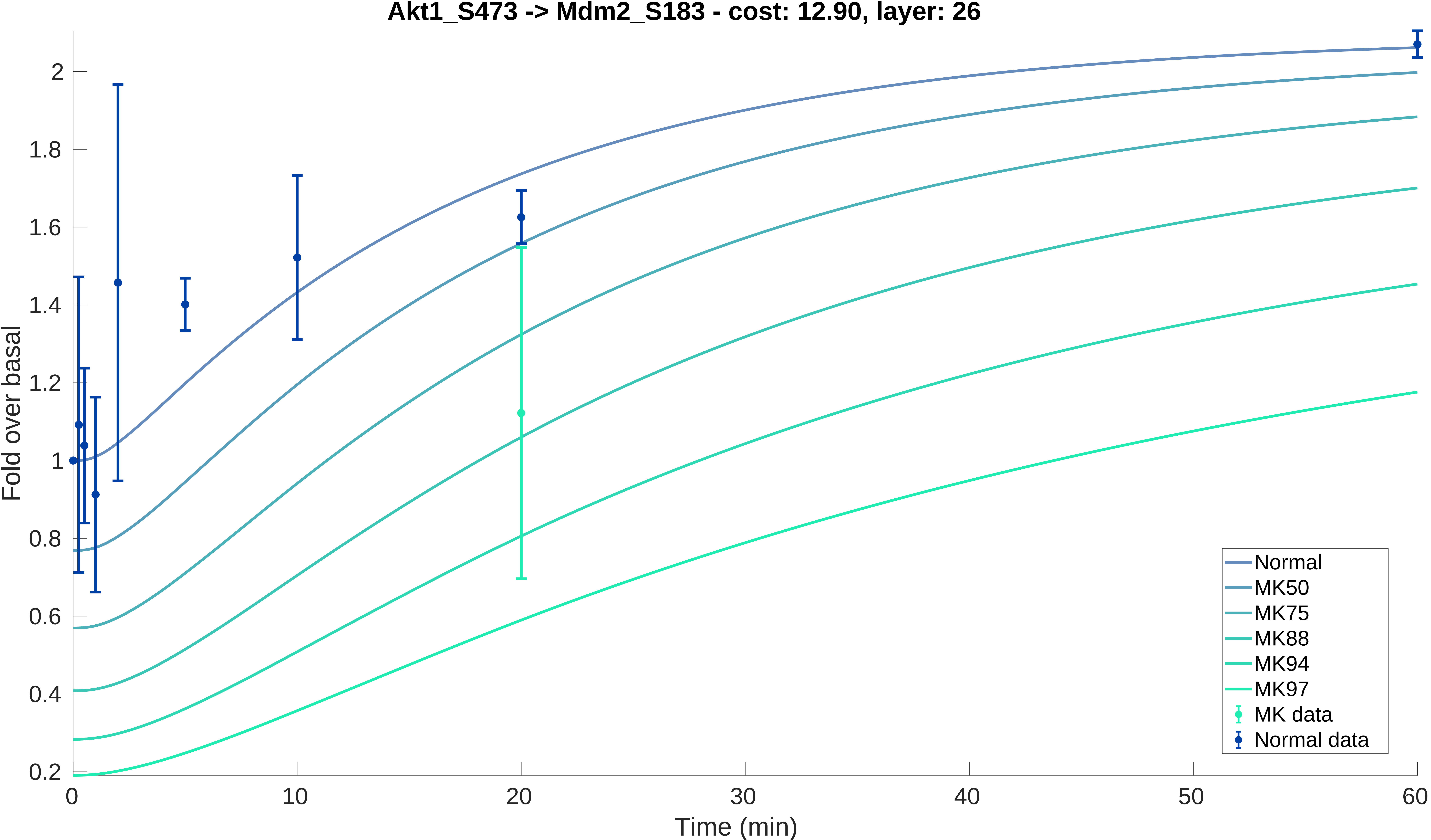

Akt1_S473 -> Mdm2_S183 - cost: 12.90, layer: 26
Fold over basal
Time (min)
0.2
0.4
0.6
0.8
1
1.2
1.4
1.6
1.8
2
0
10
20
30
40
50
60
Normal
MK50
MK75
MK88
MK94
MK97
MK data
Normal data

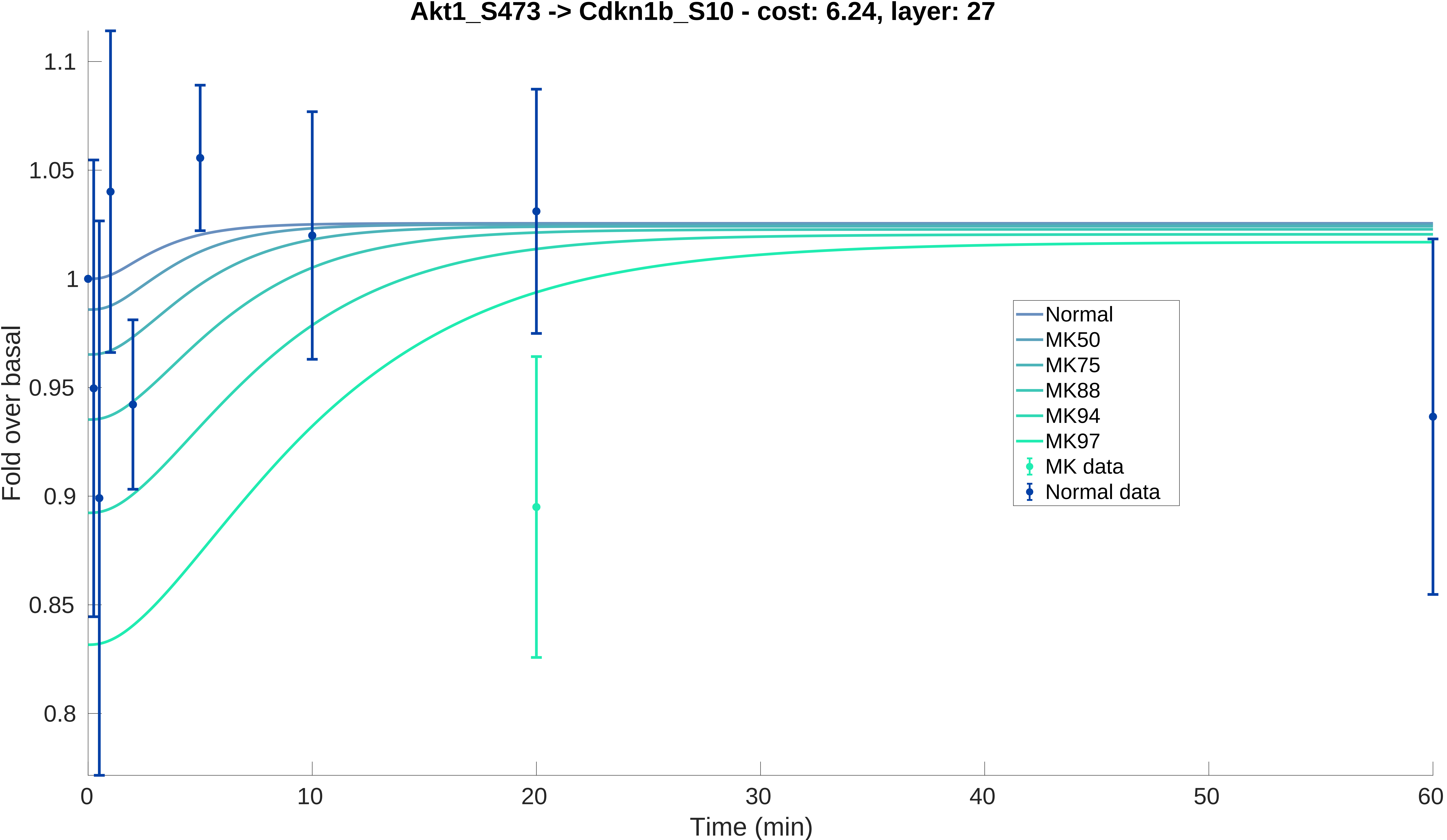

Akt1_S473 -> Cdkn1b_S10 - cost: 6.24, layer: 27
Fold over basal
Time (min)
0.8
0.85
0.9
0.95
1
1.05
1.1
0
10
20
30
40
50
60
Normal
MK50
MK75
MK88
MK94
MK97
MK data
Normal data

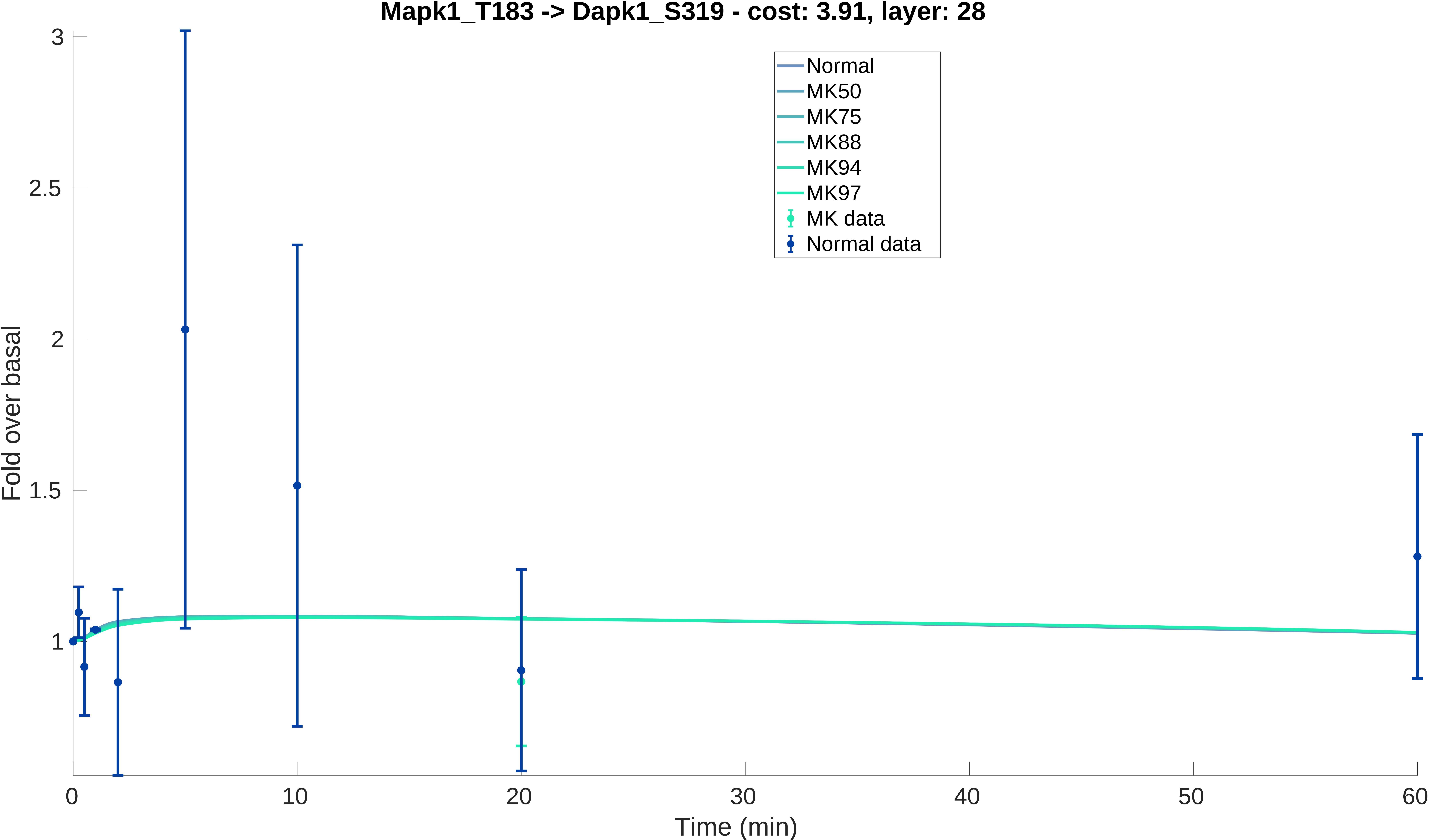
Mapk1_T183 -> Dapk1_S319 - cost: 3.91, layer: 28
Fold over basal
Time (min)
3
2.5
2
1.5
1
0
10
20
30
40
50
60
Normal
MK50
MK75
MK88
MK94
MK97
MK data
Normal data

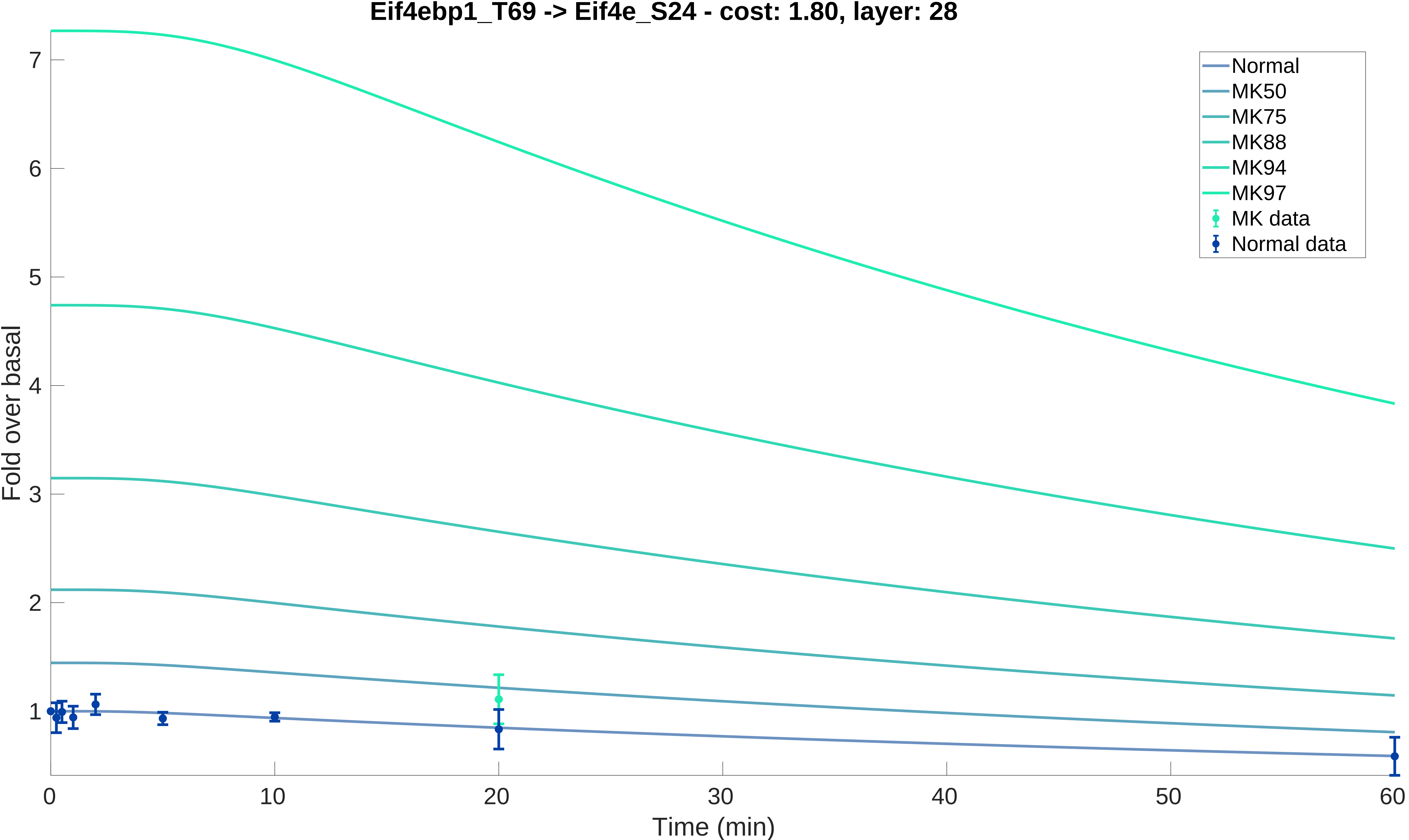

Eif4ebp1_T69 -> Eif4e_S24 - cost: 1.80, layer: 28
Normal
MK50
MK75
MK88
MK94
MK97
MK data
Normal data
Fold over basal
Time (min)
0
10
20
30
40
50
60
1
2
3
4
5
6
7

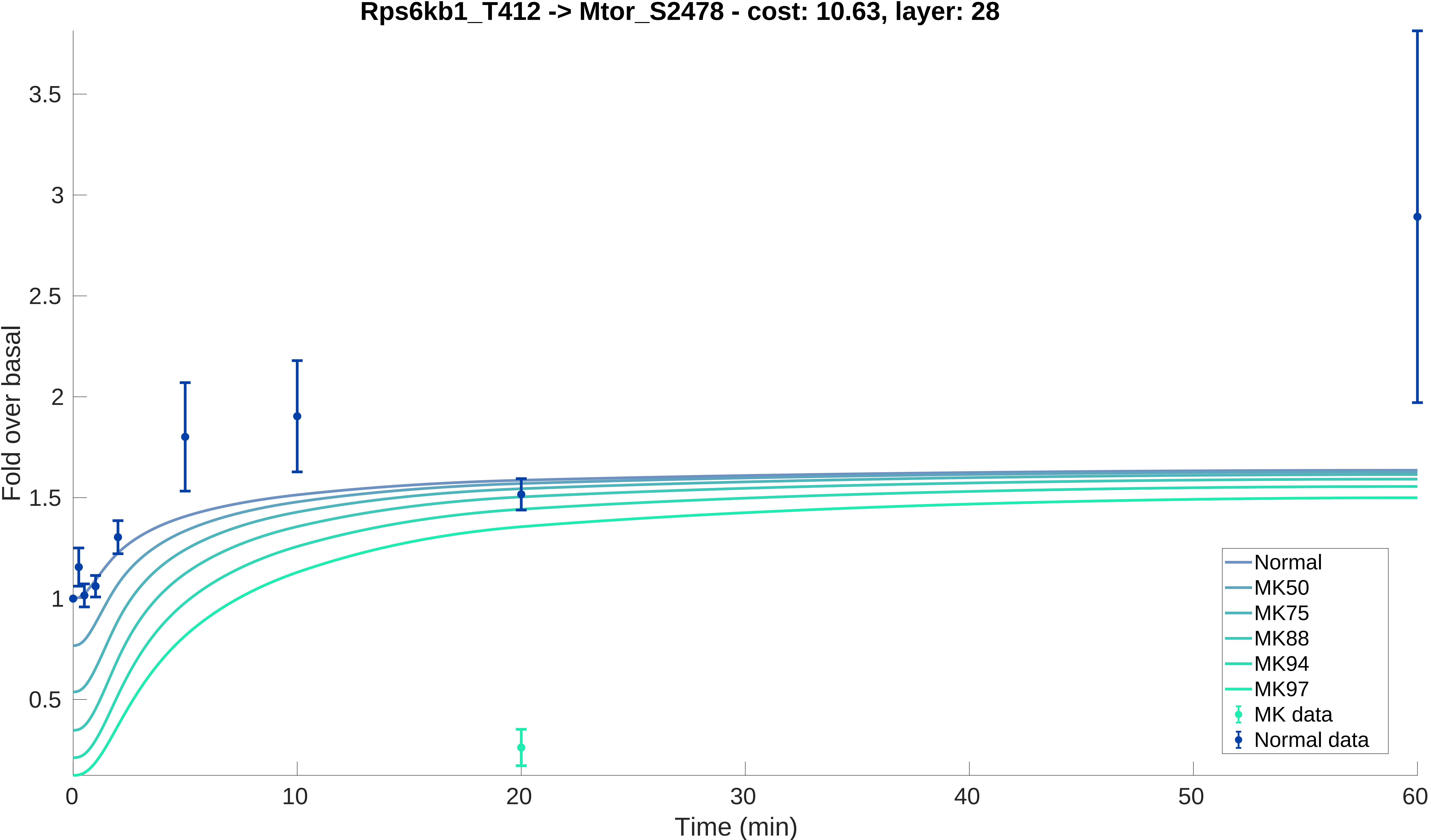

Rps6kb1_T412 -> Mtor_S2478 - cost: 10.63, layer: 28
Fold over basal
Time (min)
0.5
1
1.5
2
2.5
3
3.5
0
10
20
30
40
50
60
Normal
MK50
MK75
MK88
MK94
MK97
MK data
Normal data

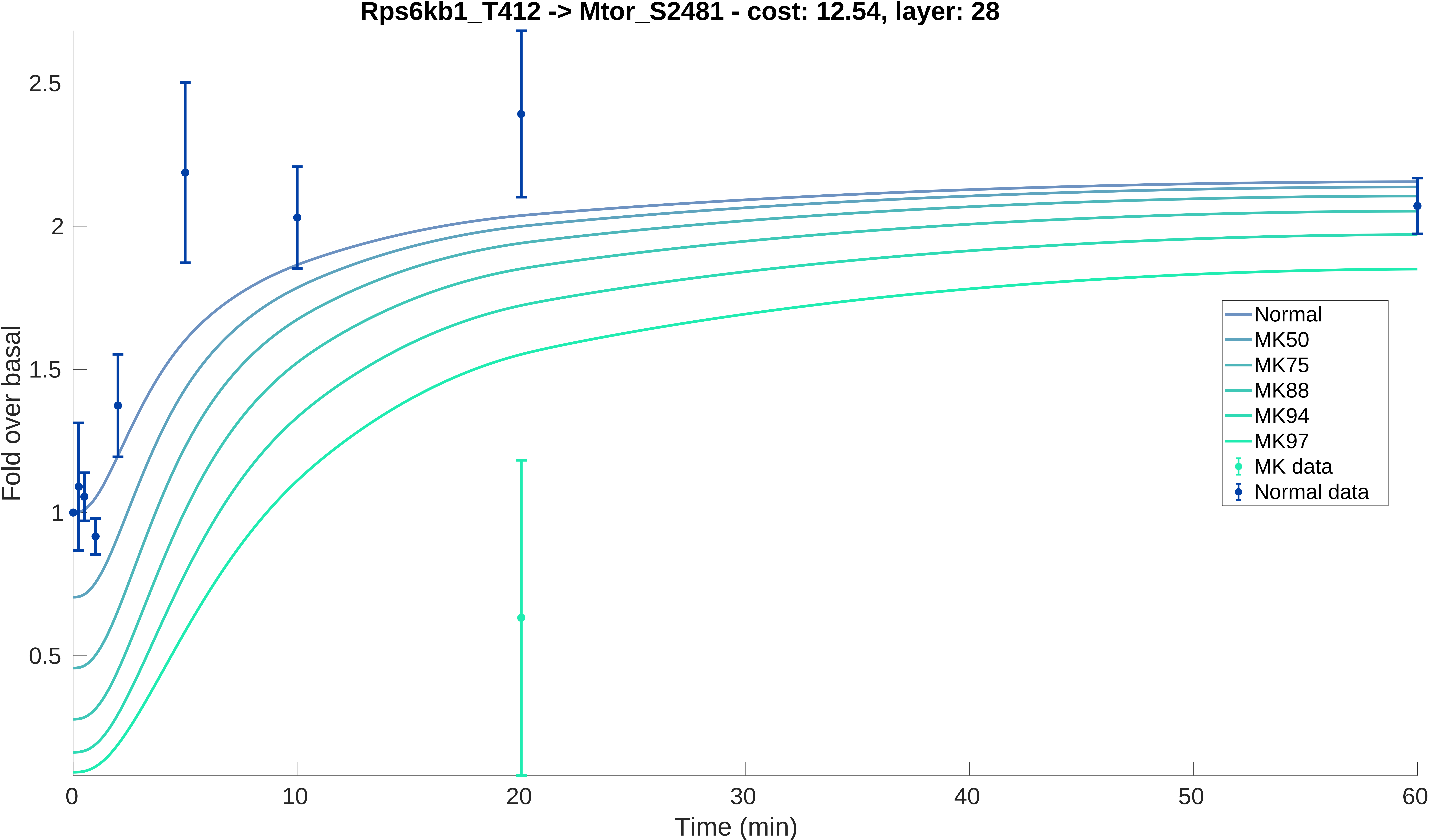
Rps6kb1_T412 -> Mtor_S2481 - cost: 12.54, layer: 28
Fold over basal
Time (min)
0
10
20
30
40
50
60
0.5
1
1.5
2
2.5
Normal
MK50
MK75
MK88
MK94
MK97
MK data
Normal data

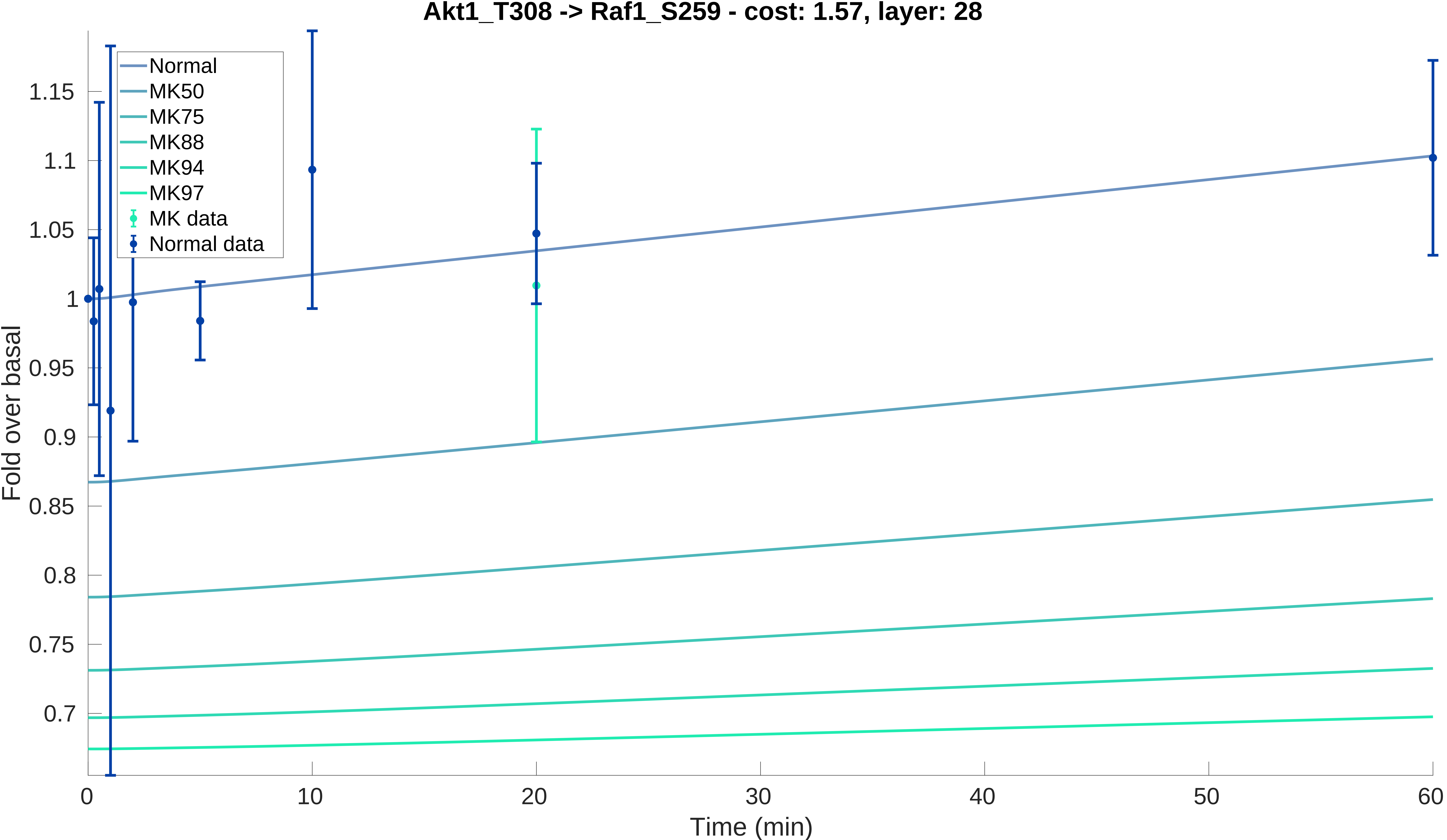

Akt1_T308 -> Raf1_S259 - cost: 1.57, layer: 28
Normal
MK50
MK75
MK88
MK94
MK97
MK data
Normal data
Fold over basal
1.15
1.1
1.05
1
0.95
0.9
0.85
0.8
0.75
0.7
0
10
20
30
40
50
60
Time (min)

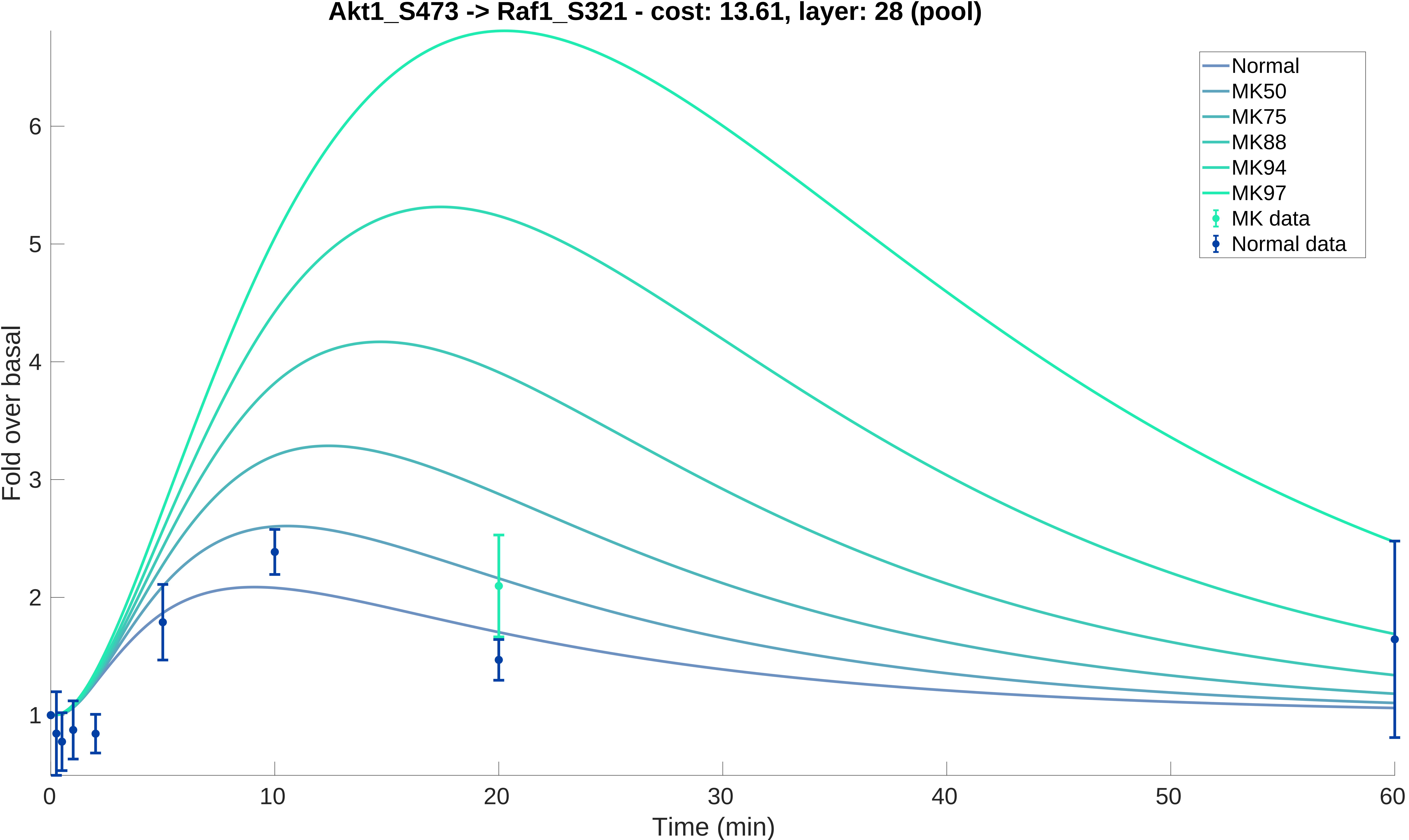

Akt1_S473 -> Raf1_S321 - cost: 13.61, layer: 28 (pool)
Fold over basal
Time (min)
0
10
20
30
40
50
60
1
2
3
4
5
6
Normal
MK50
MK75
MK88
MK94
MK97
MK data
Normal data

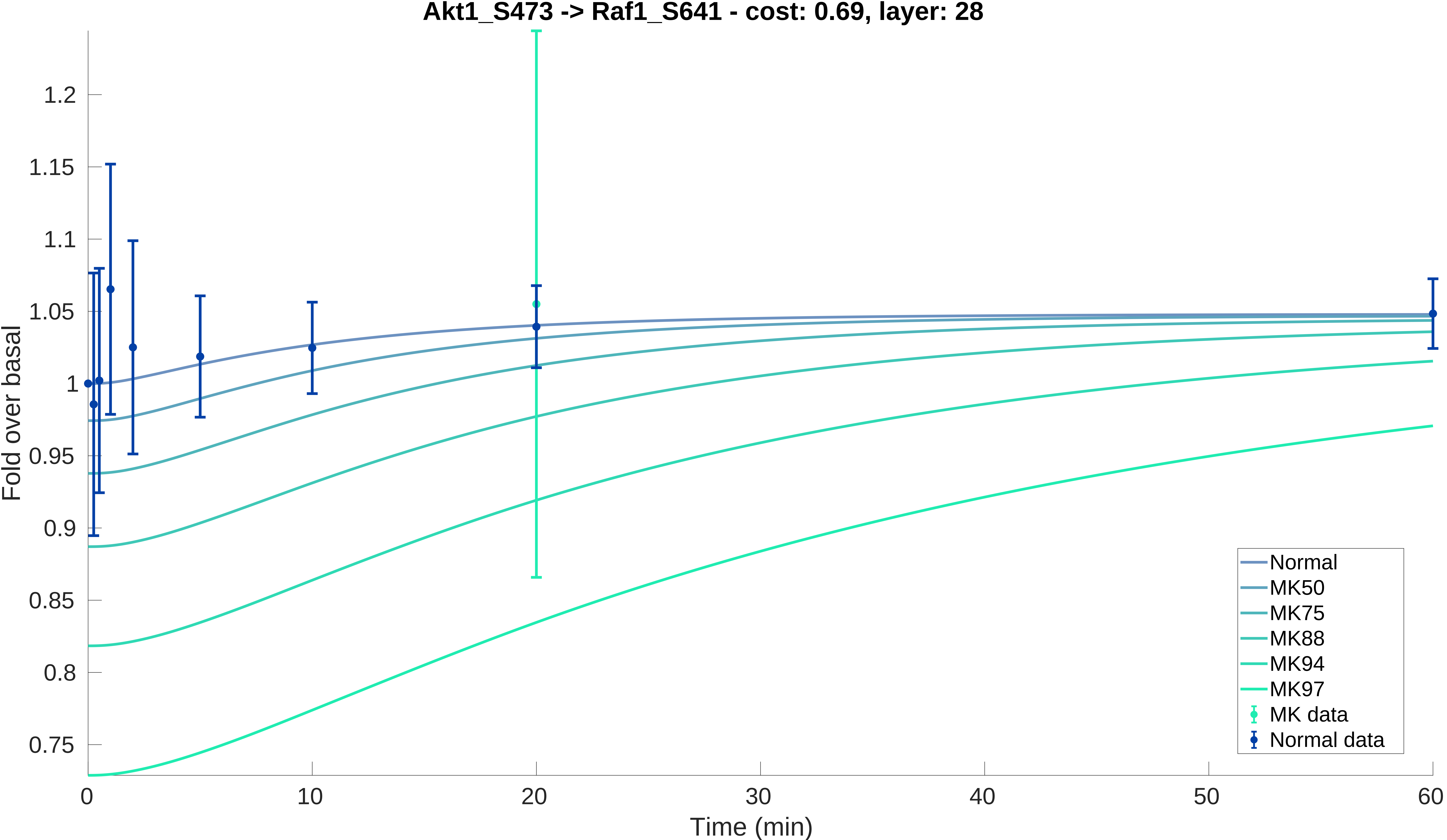

Akt1_S473 -> Raf1_S641 - cost: 0.69, layer: 28
Fold over basal
1.2
1.15
1.1
1.05
1
0.95
0.9
0.85
0.8
0.75
0
10
20
30
40
50
60
Time (min)
Normal
MK50
MK75
MK88
MK94
MK97
MK data
Normal data

#### Mapk3_Y205 -> Rps6ka3_S415 - cost: 2.75, layer: 28

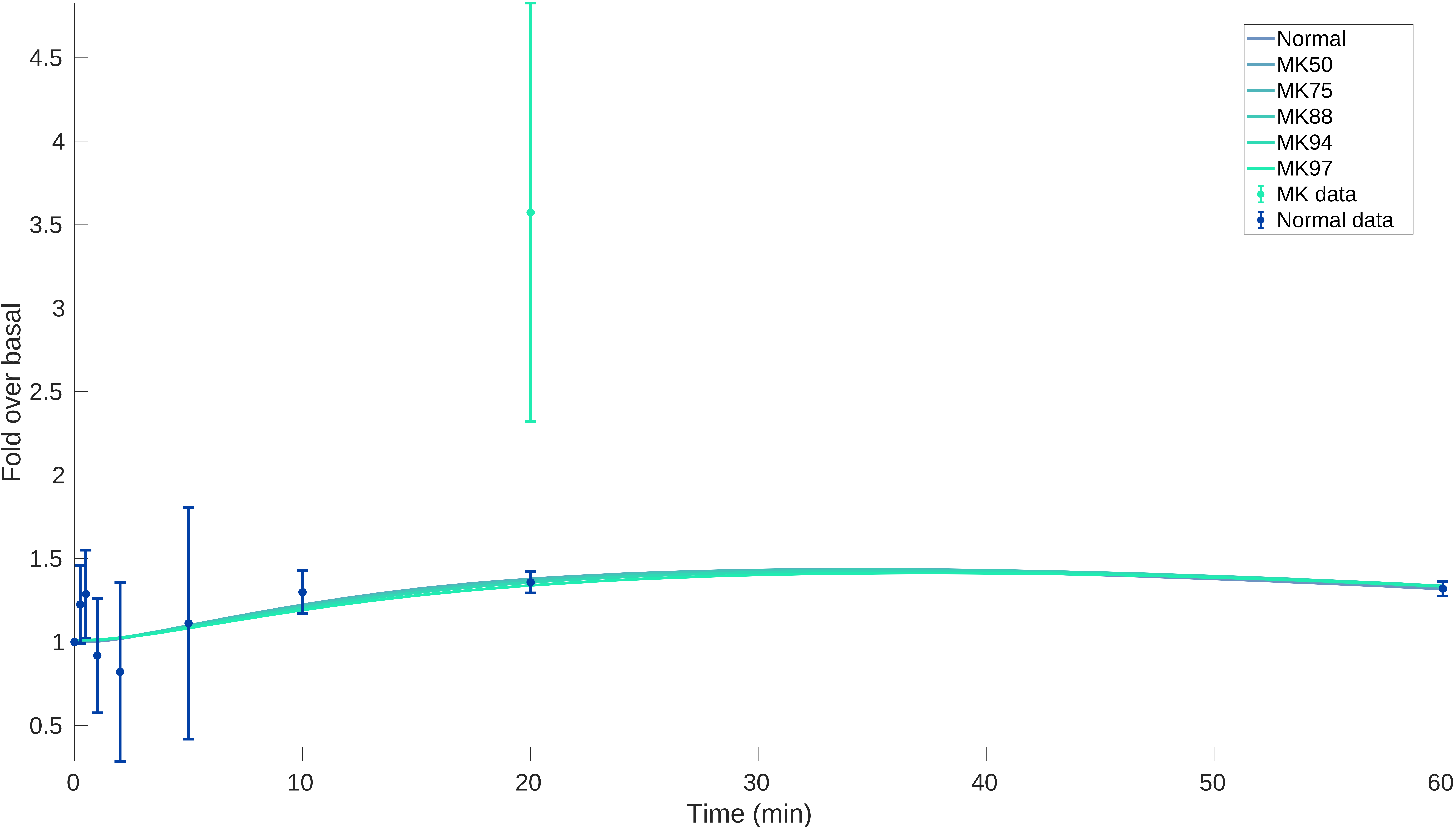

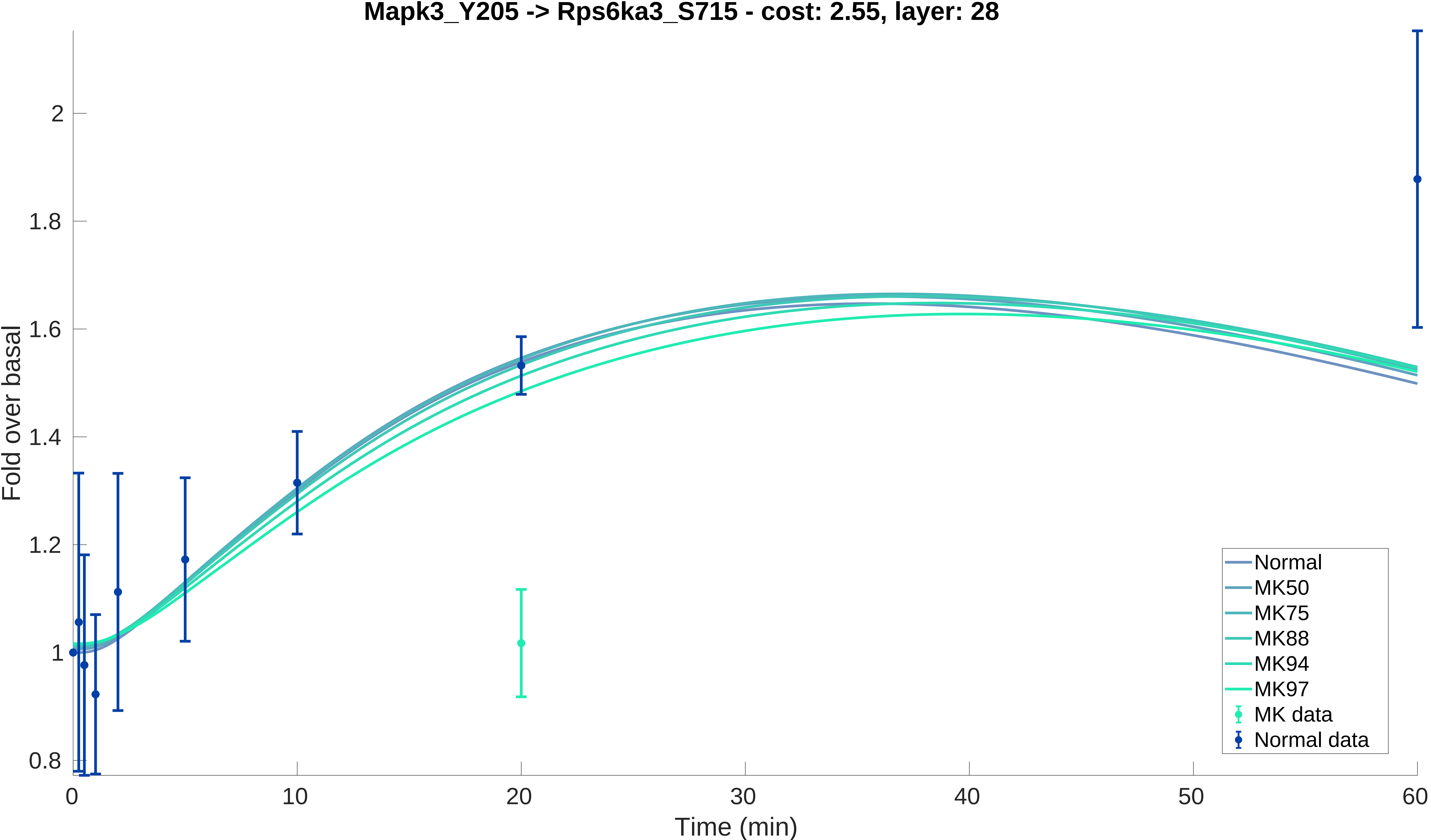

Mapk3_Y205 -> Rps6ka3_S715 - cost: 2.55, layer: 28
Fold over basal
Time (min)
0.8
1
1.2
1.4
1.6
1.8
2
0
10
20
30
40
50
60
Normal
MK50
MK75
MK88
MK94
MK97
MK data
Normal data

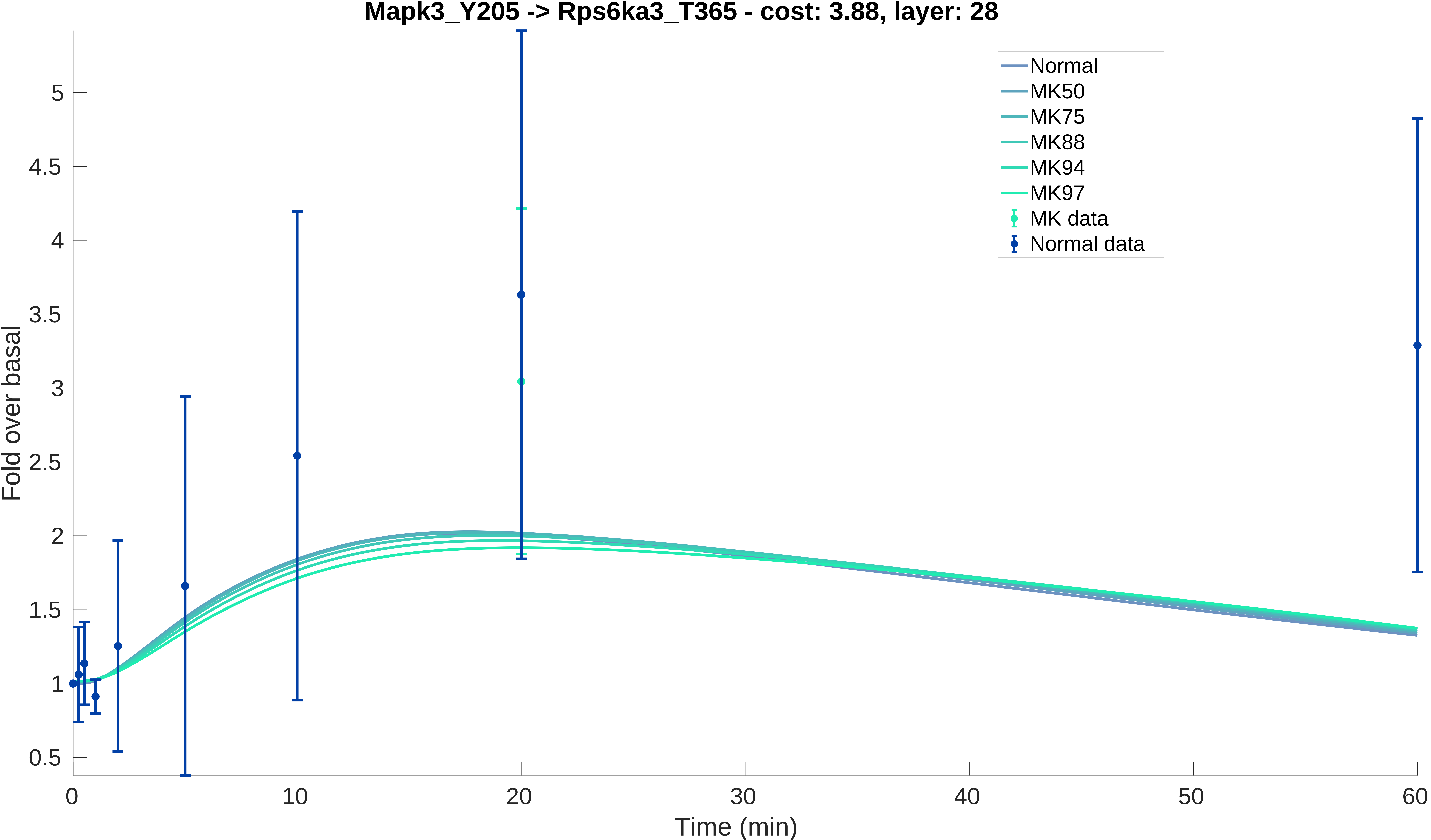

Mapk3_Y205 -> Rps6ka3_T365 - cost: 3.88, layer: 28
Fold over basal
Time (min)
0.5
1
1.5
2
2.5
3
3.5
4
4.5
5
0
10
20
30
40
50
60
Normal
MK50
MK75
MK88
MK94
MK97
MK data
Normal data

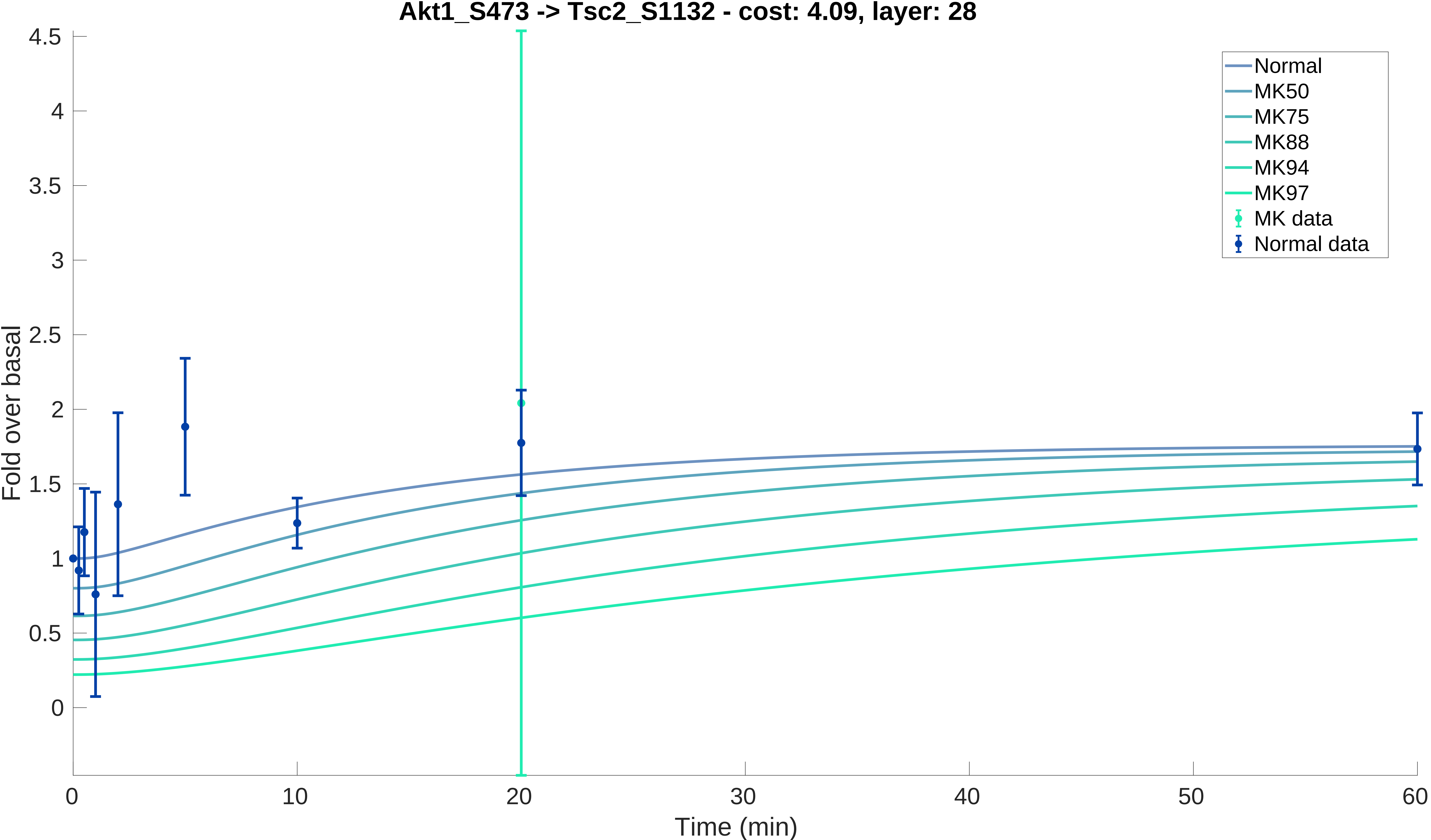

Akt1_S473 -> Tsc2_S1132 - cost: 4.09, layer: 28
Fold over basal
Time (min)
0
0.5
1
1.5
2
2.5
3
3.5
4
4.5
0
10
20
30
40
50
60
Normal
MK50
MK75
MK88
MK94
MK97
MK data
Normal data

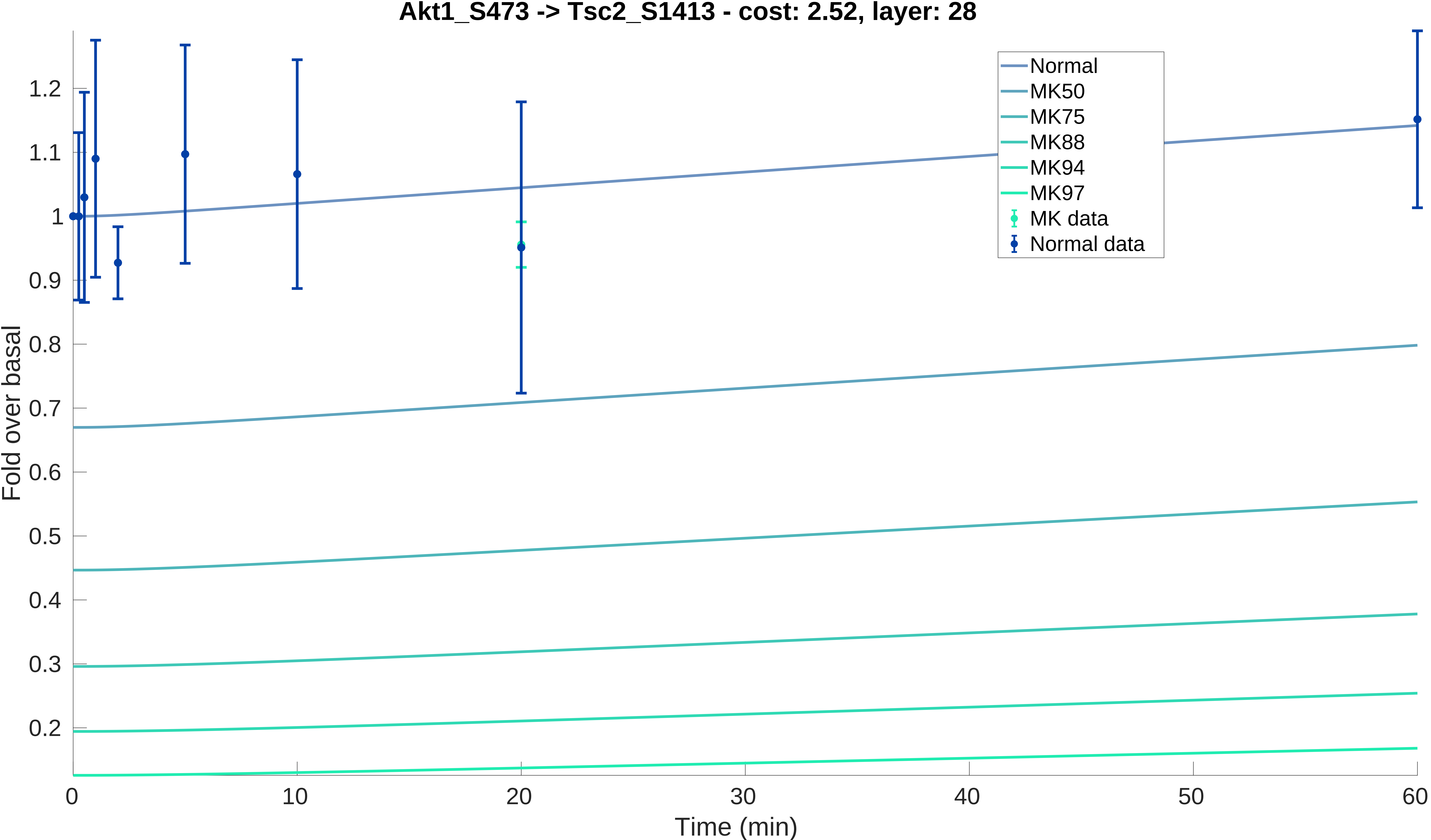

Akt1_S473 -> Tsc2_S1413 - cost: 2.52, layer: 28
Normal
MK50
MK75
MK88
MK94
MK97
MK data
Normal data
Fold over basal
Time (min)
0
10
20
30
40
50
60
0.2
0.3
0.4
0.5
0.6
0.7
0.8
0.9
1
1.1
1.2

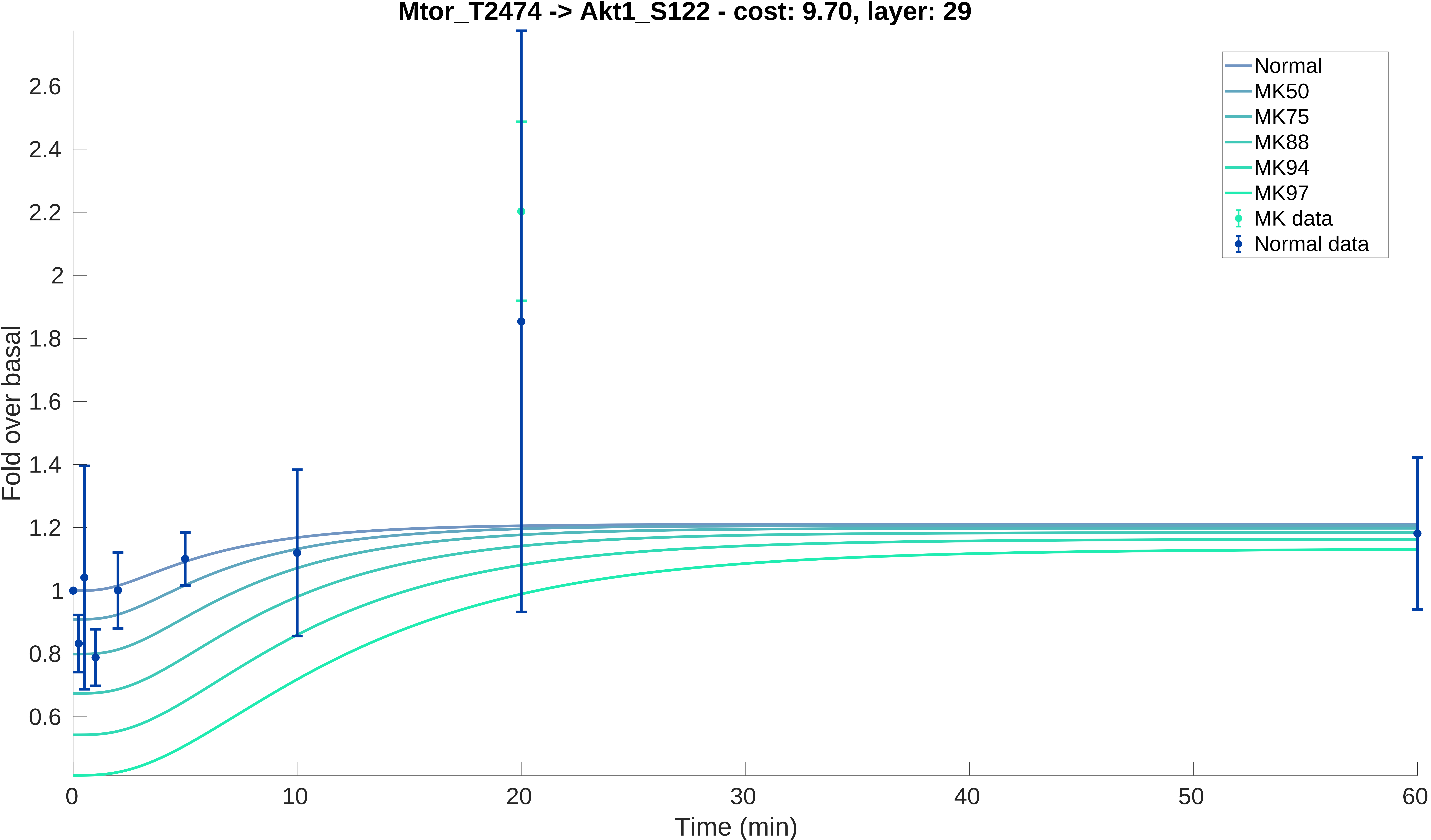

Mtor_T2474 -> Akt1_S122 - cost: 9.70, layer: 29
Fold over basal
Time (min)
Normal
MK50
MK75
MK88
MK94
MK97
MK data
Normal data
2.6
2.4
2.2
2
1.8
1.6
1.4
1.2
1
0.8
0.6
0
10
20
30
40
50
60

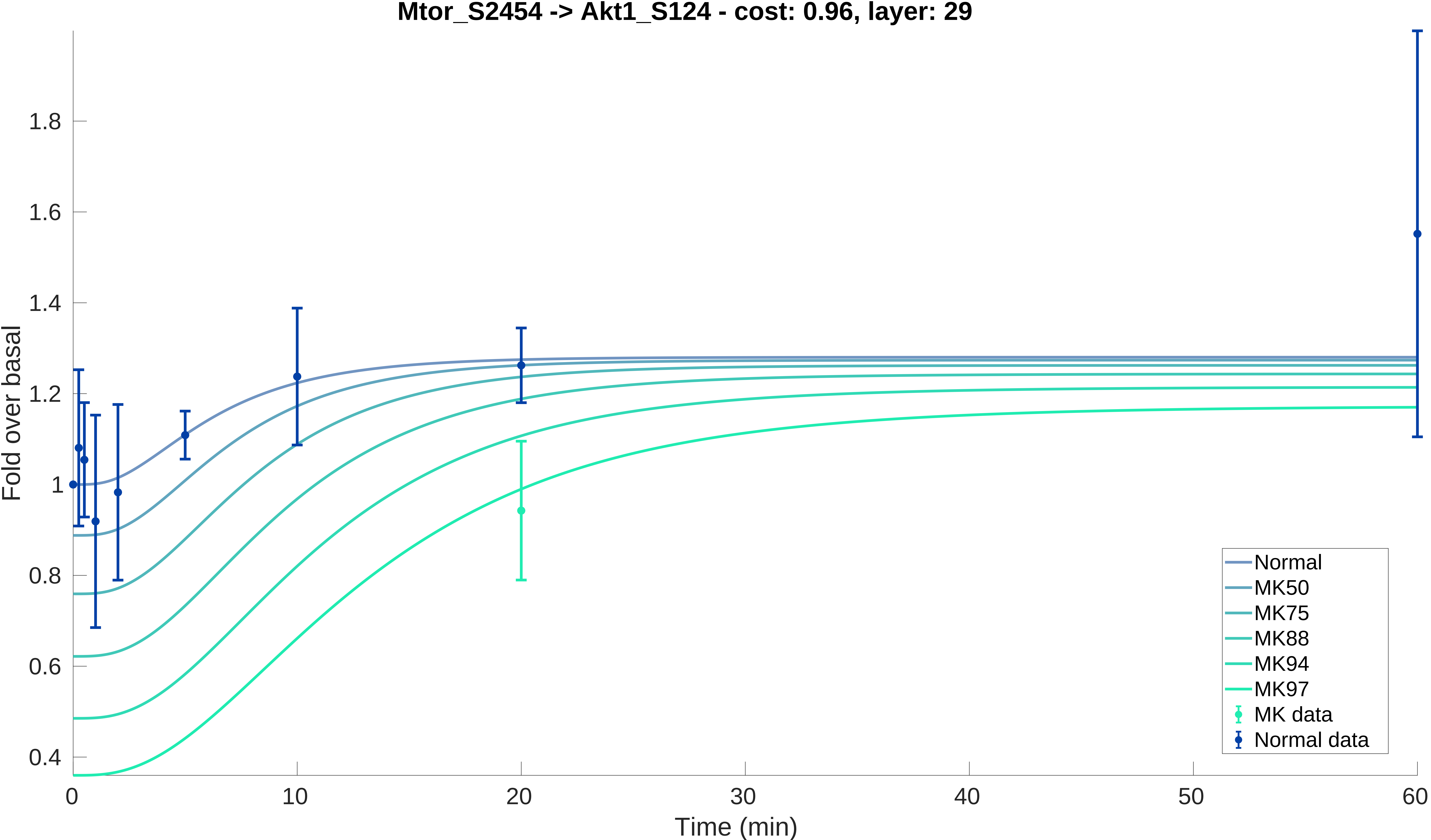

Mtor_S2454 -> Akt1_S124 - cost: 0.96, layer: 29
1.8
1.6
1.4
1.2
1
0.8
0.6
0.4
Fold over basal
0
10
20
30
40
50
60
Time (min)
Normal
MK50
MK75
MK88
MK94
MK97
MK data
Normal data

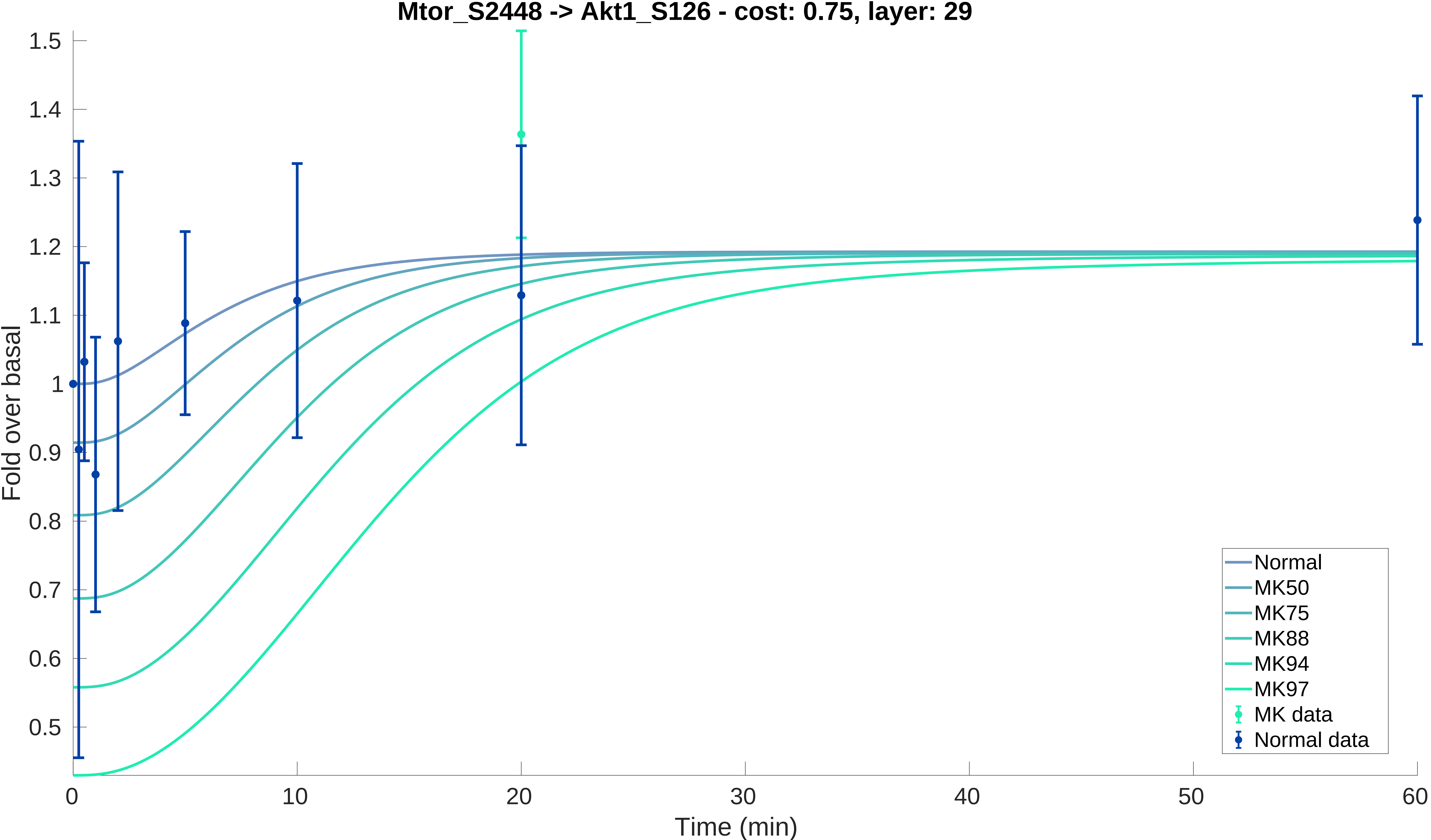

Mtor_S2448 -> Akt1_S126 - cost: 0.75, layer: 29
Fold over basal
1.5
1.4
1.3
1.2
1.1
1
0.9
0.8
0.7
0.6
0.5
0
10
20
30
40
50
60
Time (min)
Normal
MK50
MK75
MK88
MK94
MK97
MK data
Normal data

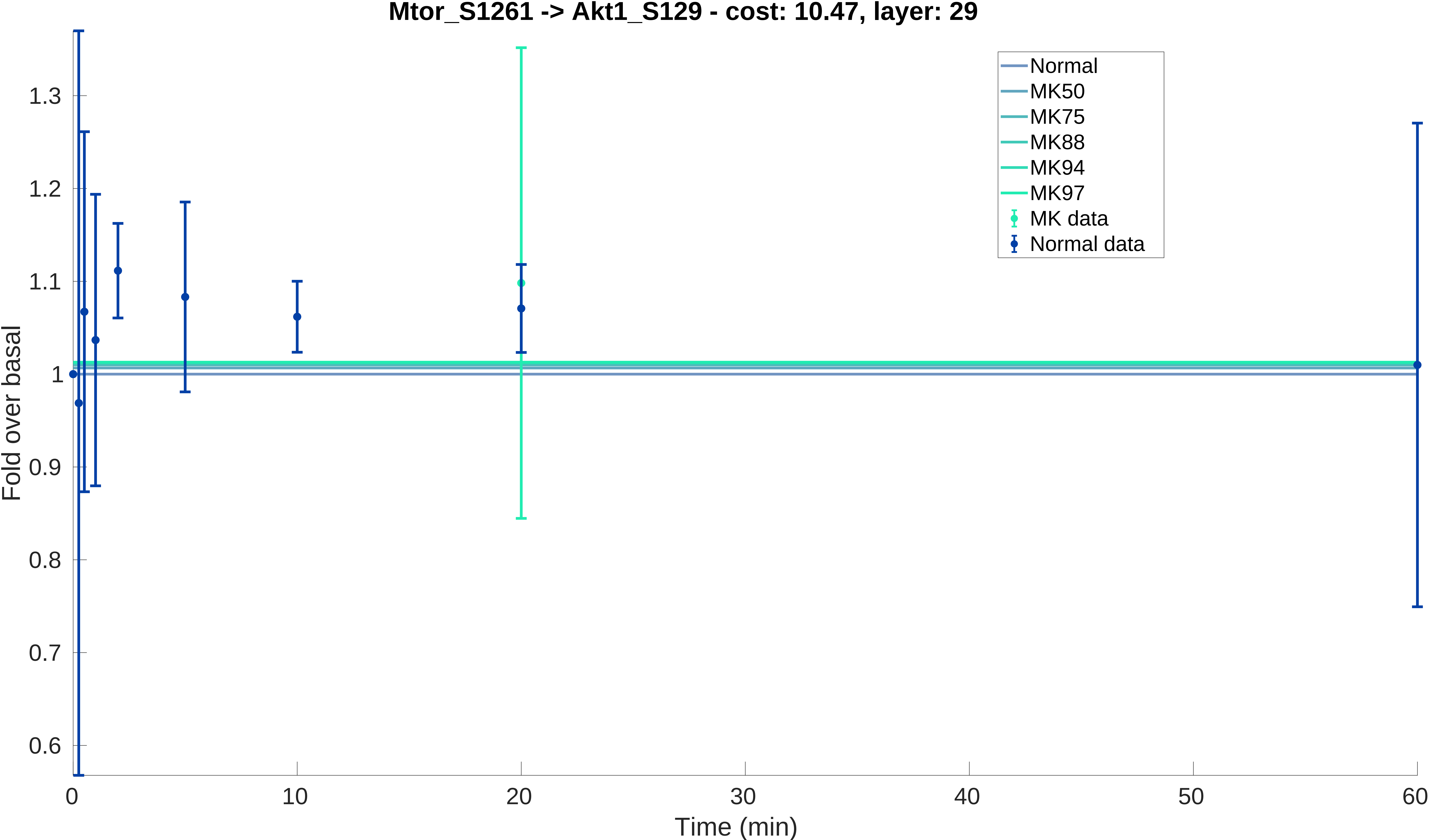

Mtor_S1261 -> Akt1_S129 - cost: 10.47, layer: 29
Normal
MK50
MK75
MK88
MK94
MK97
MK data
Normal data
Fold over basal
1.3
1.2
1.1
1
0.9
0.8
0.7
0.6
0
10
20
30
40
50
60
Time (min)

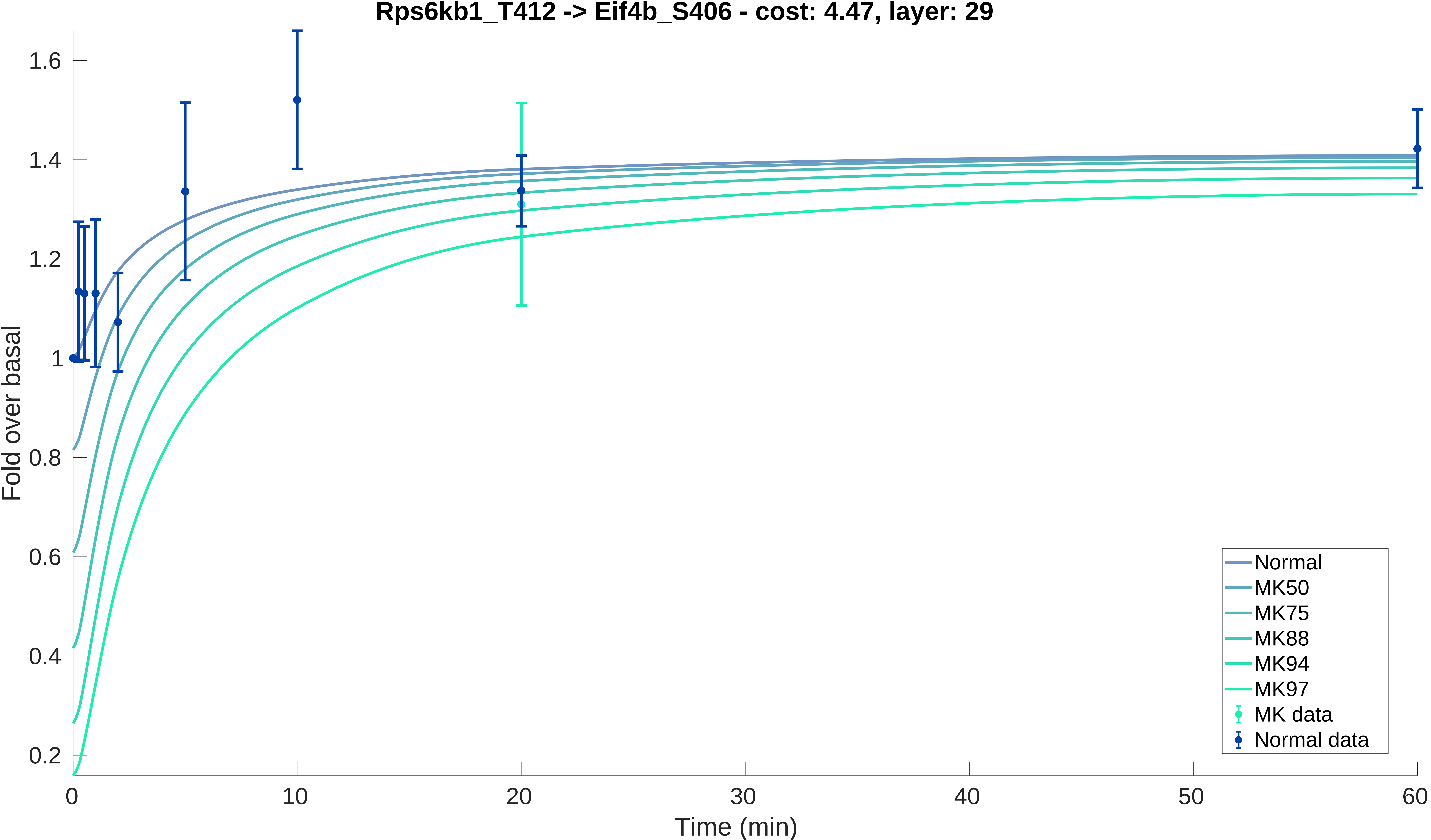
Rps6kb1_T412 -> Eif4b_S406 - cost: 4.47, layer: 29
Fold over basal
1.6
1.4
1.2
1
0.8
0.6
0.4
0.2
0
10
20
30
40
50
60
Time (min)
Normal
MK50
MK75
MK88
MK94
MK97
MK data
Normal data

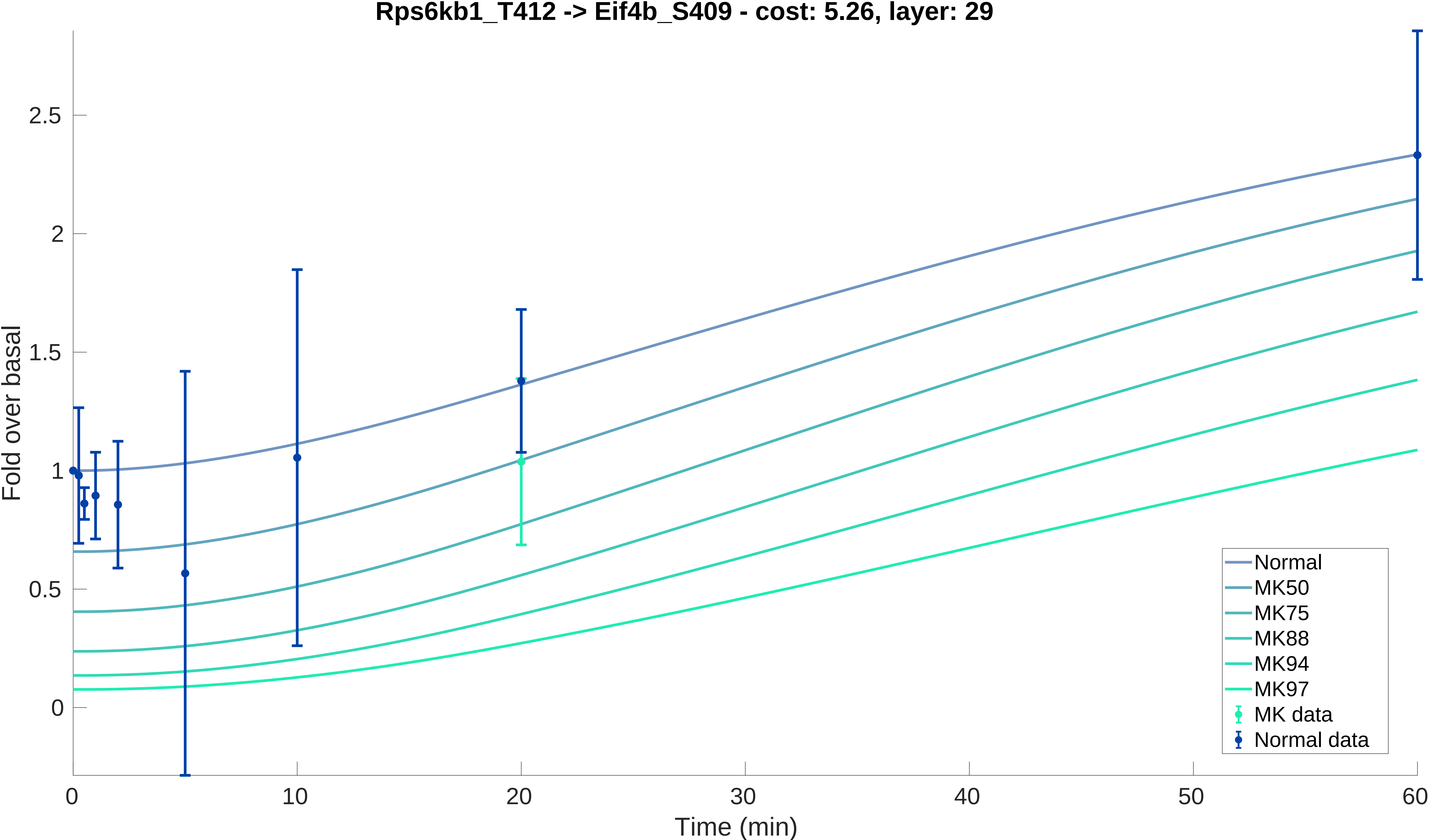

Rps6kb1_T412 -> Eif4b_S409 - cost: 5.26, layer: 29
Fold over basal
Time (min)
2.5
2
1.5
1
0.5
0
0
10
20
30
40
50
60
Normal
MK50
MK75
MK88
MK94
MK97
MK data
Normal data

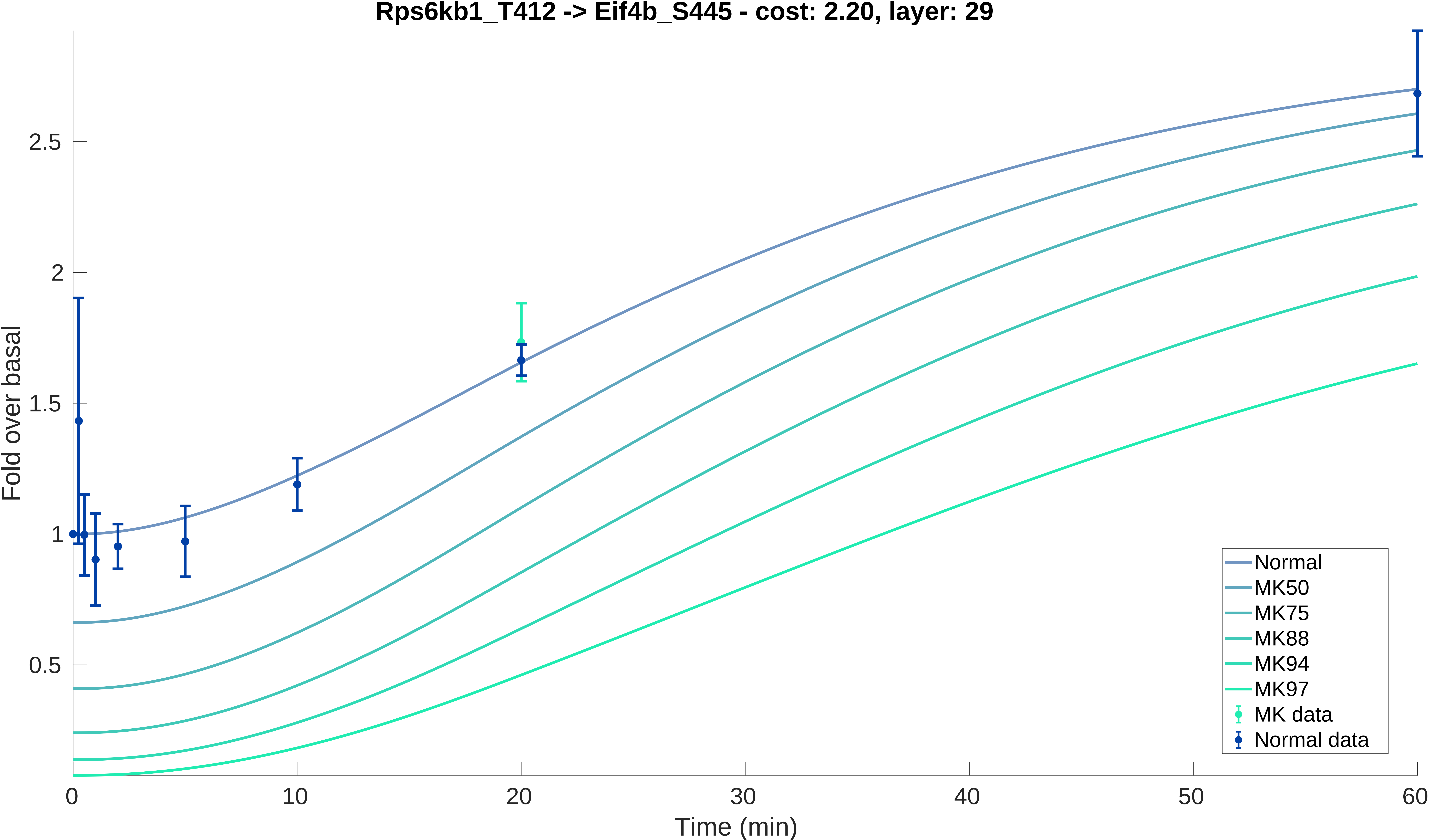

Rps6kb1_T412 -> Eif4b_S445 - cost: 2.20, layer: 29
Fold over basal
2.5
2
1.5
1
0.5
0
10
20
30
40
50
60
Time (min)
Normal
MK50
MK75
MK88
MK94
MK97
MK data
Normal data

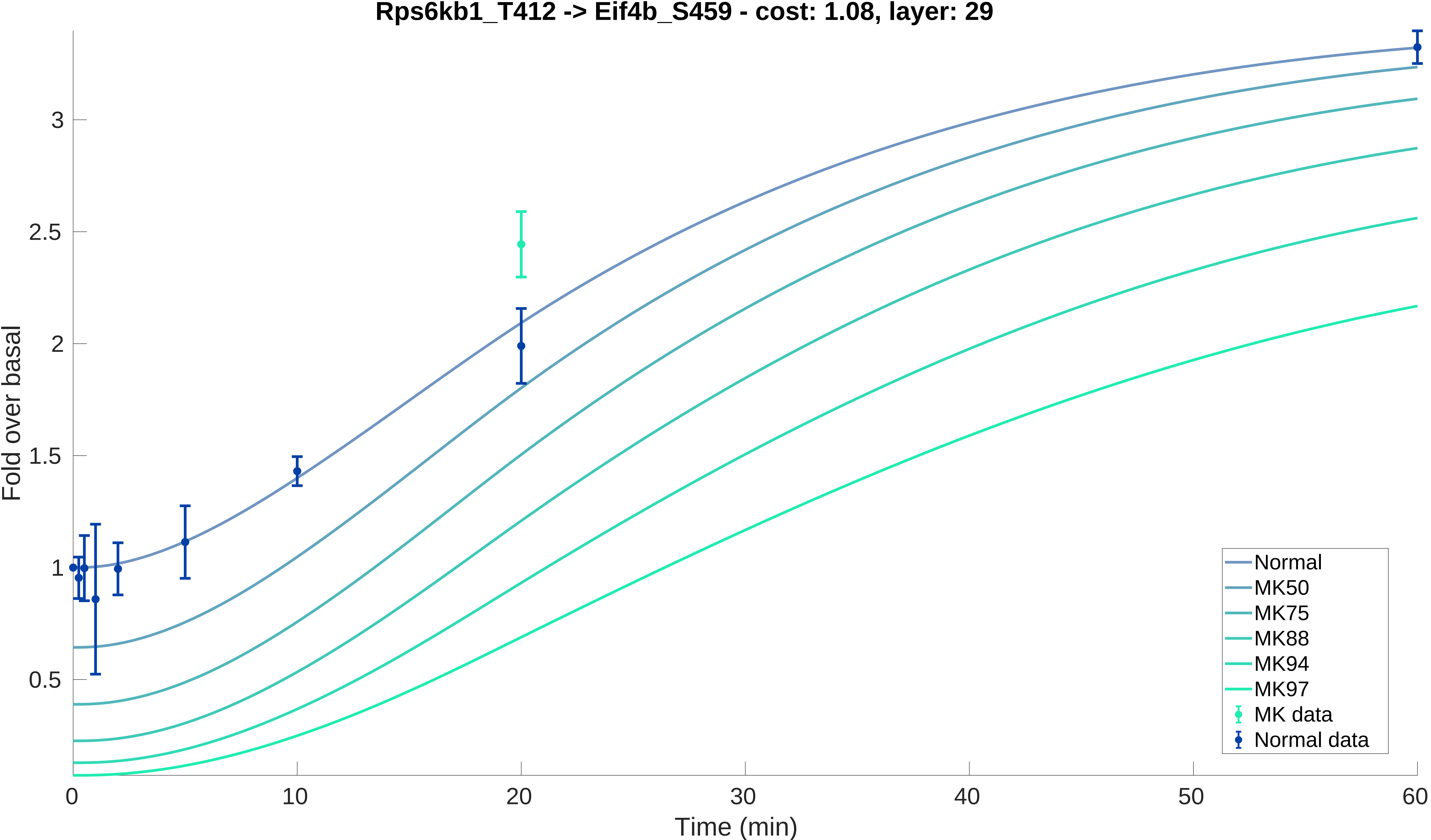

Rps6kb1_T412 -> Eif4b_S459 - cost: 1.08, layer: 29
Fold over basal
Time (min)
0.5
1
1.5
2
2.5
3
0
10
20
30
40
50
60
Normal
MK50
MK75
MK88
MK94
MK97
MK data
Normal data

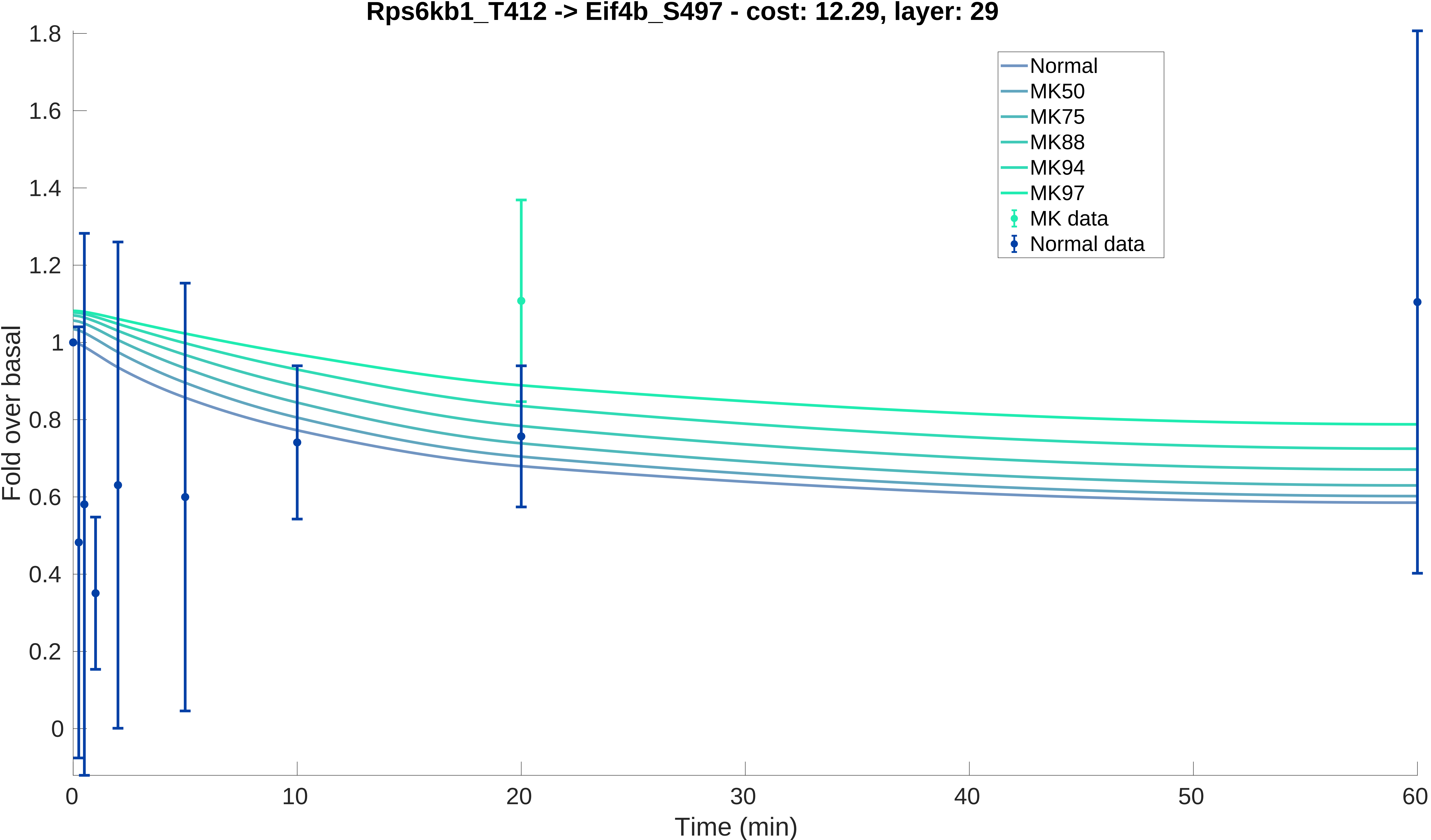
Rps6kb1_T412 -> Eif4b_S497 - cost: 12.29, layer: 29
Fold over basal
Time (min)
Normal
MK50
MK75
MK88
MK94
MK97
MK data
Normal data
0
0.2
0.4
0.6
0.8
1
1.2
1.4
1.6
1.8
0
10
20
30
40
50
60

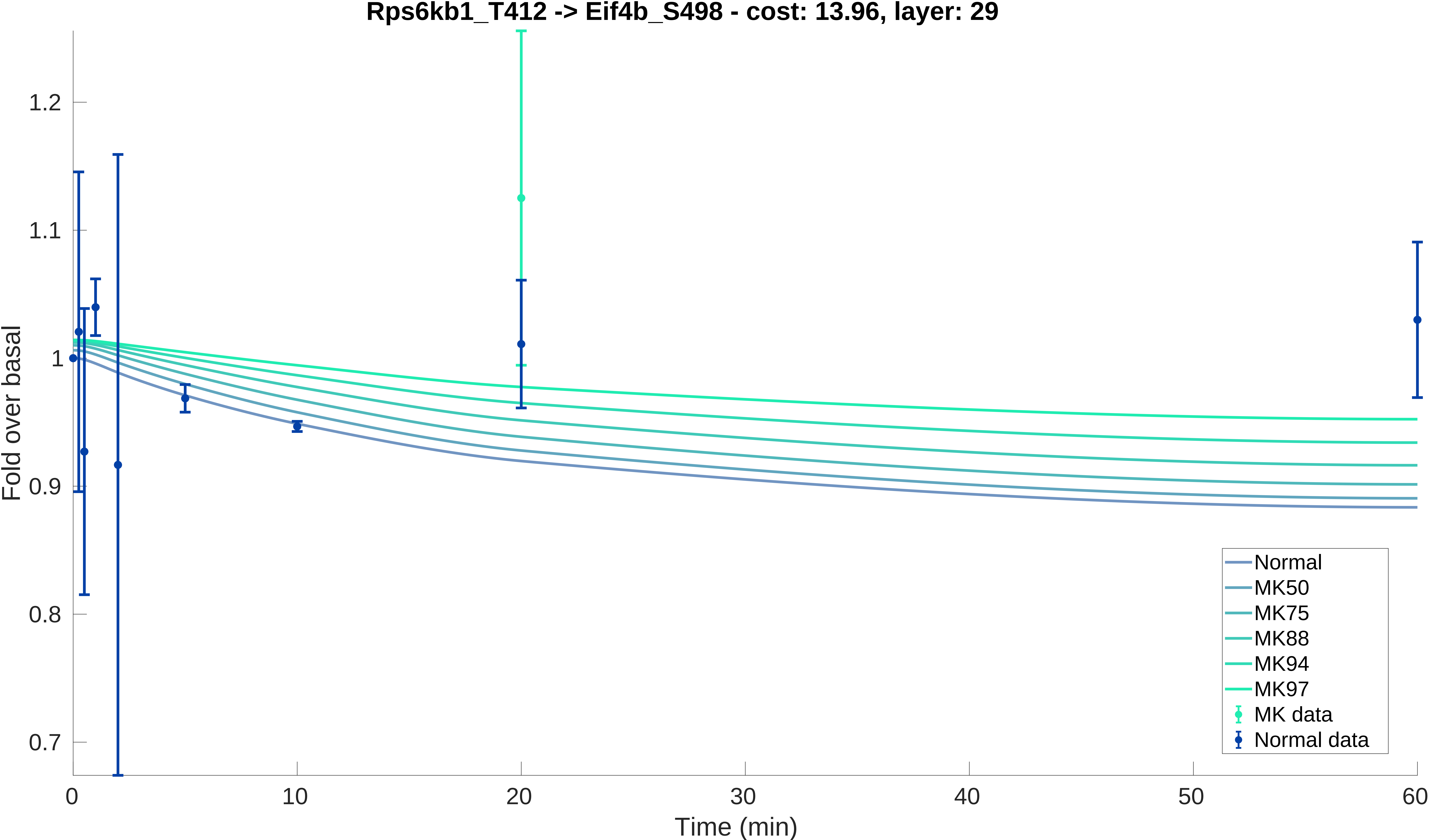

Rps6kb1_T412 -> Eif4b_S498 - cost: 13.96, layer: 29
Fold over basal
1.2
1.1
1
0.9
0.8
0.7
0
10
20
30
40
50
60
Time (min)
Normal
MK50
MK75
MK88
MK94
MK97
MK data
Normal data

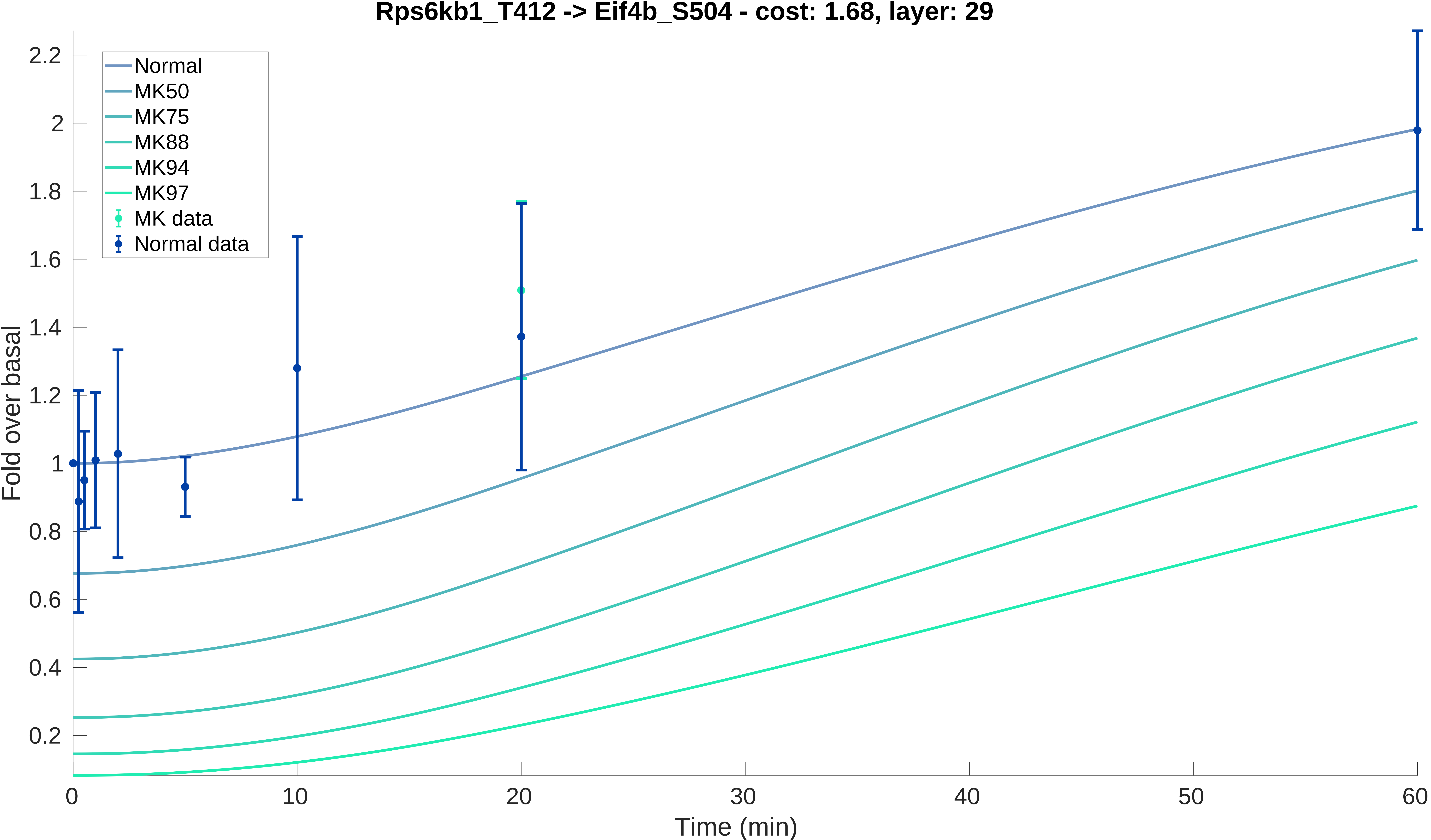

Rps6kb1_T412 -> Eif4b_S504 - cost: 1.68, layer: 29
Normal
MK50
MK75
MK88
MK94
MK97
MK data
Normal data
Fold over basal
Time (min)
0
10
20
30
40
50
60
0.2
0.4
0.6
0.8
1
1.2
1.4
1.6
1.8
2
2.2

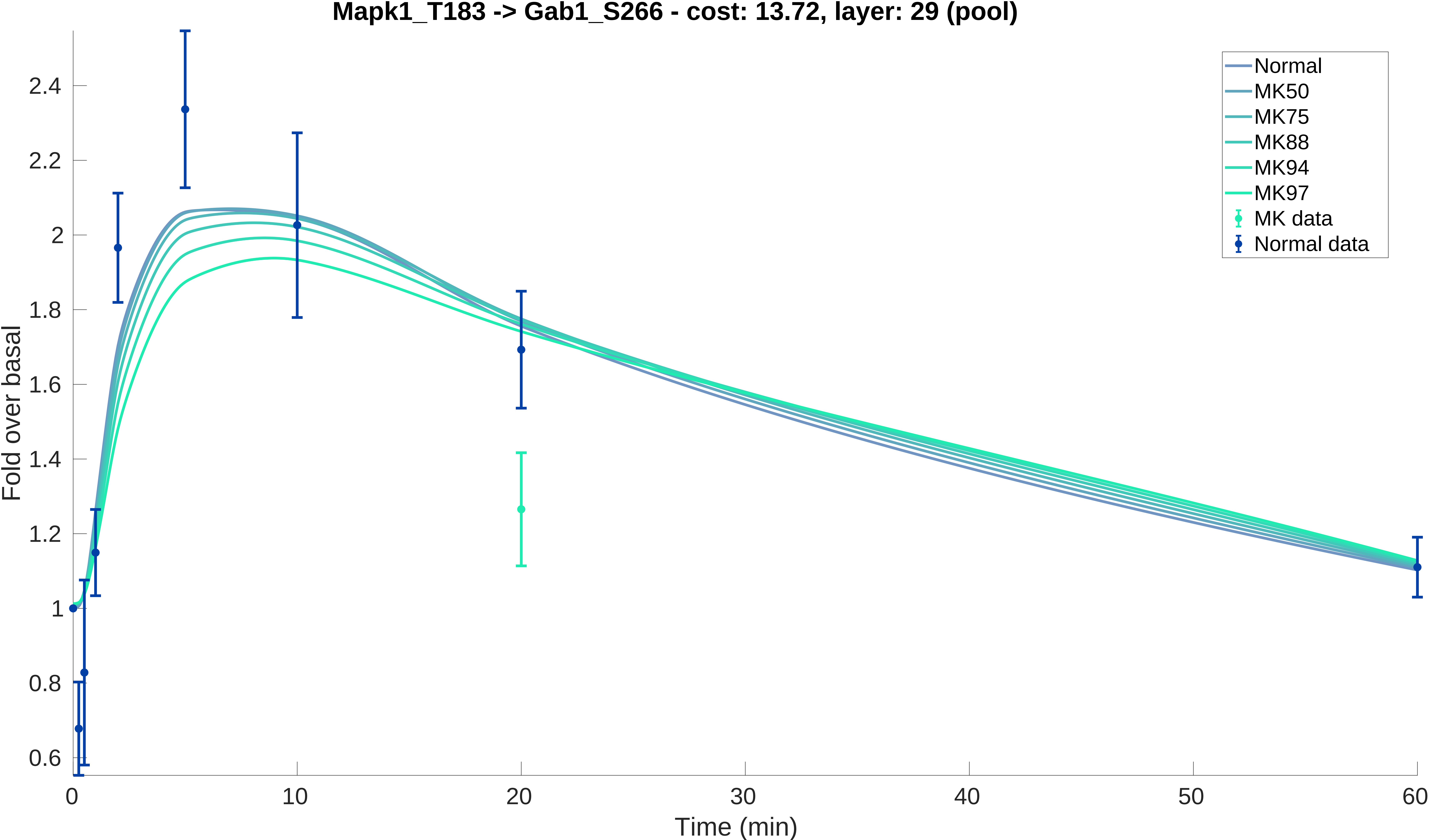
Mapk1_T183 -> Gab1_S266 - cost: 13.72, layer: 29 (pool)
Fold over basal
Time (min)
2.4
2.2
2
1.8
1.6
1.4
1.2
1
0.8
0.6
0
10
20
30
40
50
60
Normal
MK50
MK75
MK88
MK94
MK97
MK data
Normal data

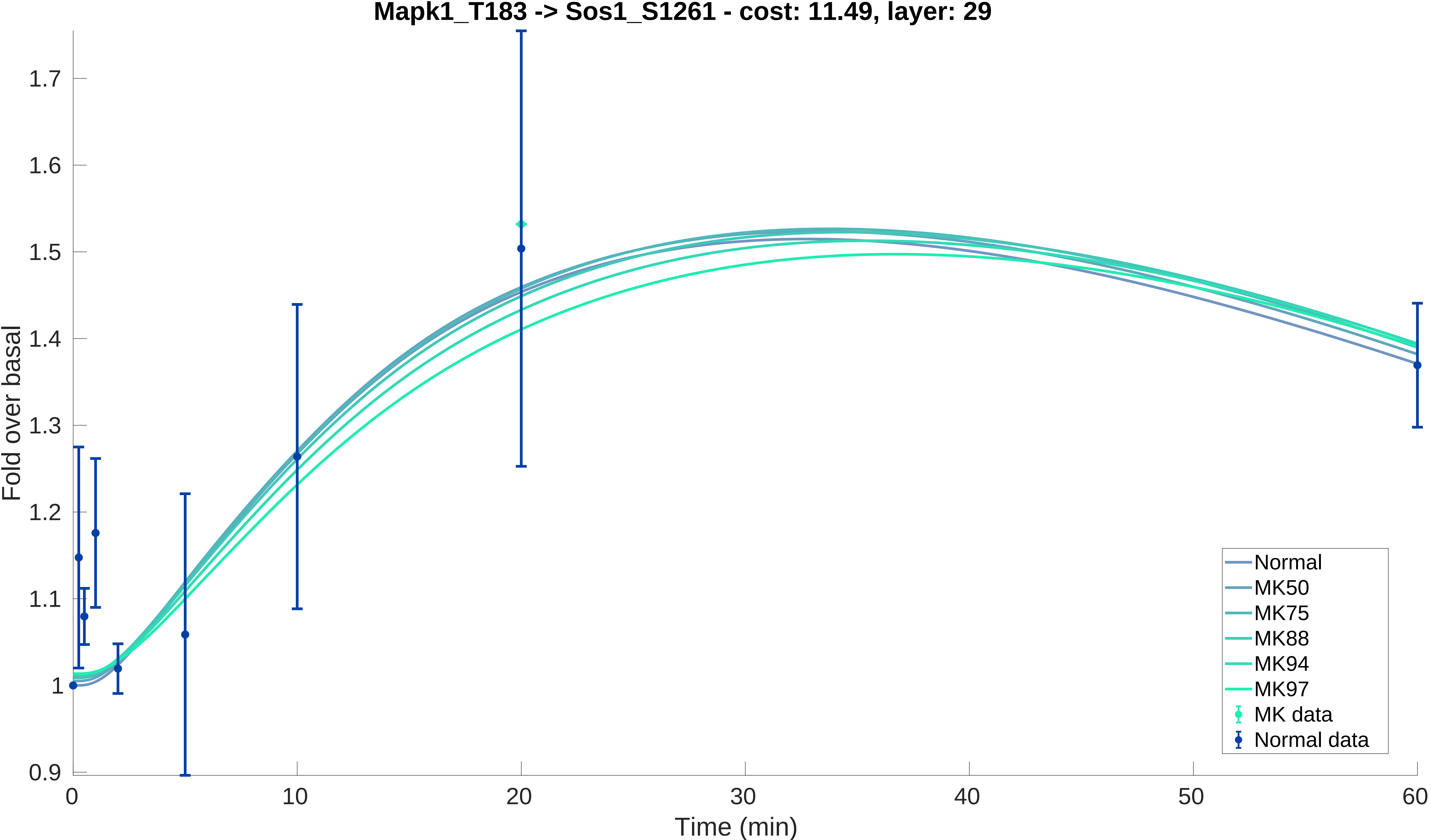

Mapk1_T183 -> Sos1_S1261 - cost: 11.49, layer: 29
Fold over basal
Time (min)
0.9
1
1.1
1.2
1.3
1.4
1.5
1.6
1.7
0
10
20
30
40
50
60
Normal
MK50
MK75
MK88
MK94
MK97
MK data
Normal data

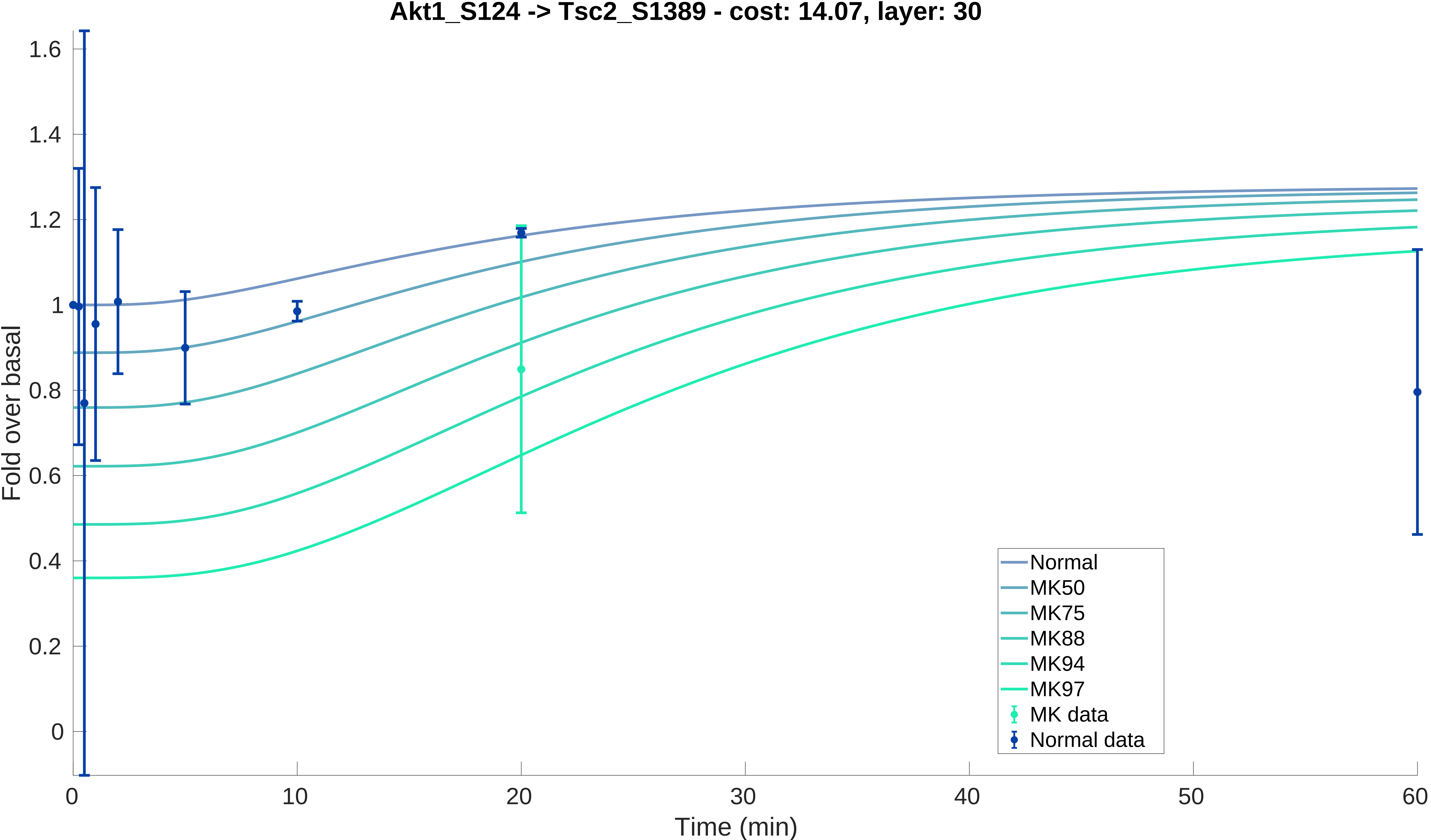
Akt1_S124 -> Tsc2_S1389 - cost: 14.07, layer: 30
Fold over basal
1.6
1.4
1.2
1
0.8
0.6
0.4
0.2
0
0
10
20
30
40
50
60
Time (min)
Normal
MK50
MK75
MK88
MK94
MK97
MK data
Normal data

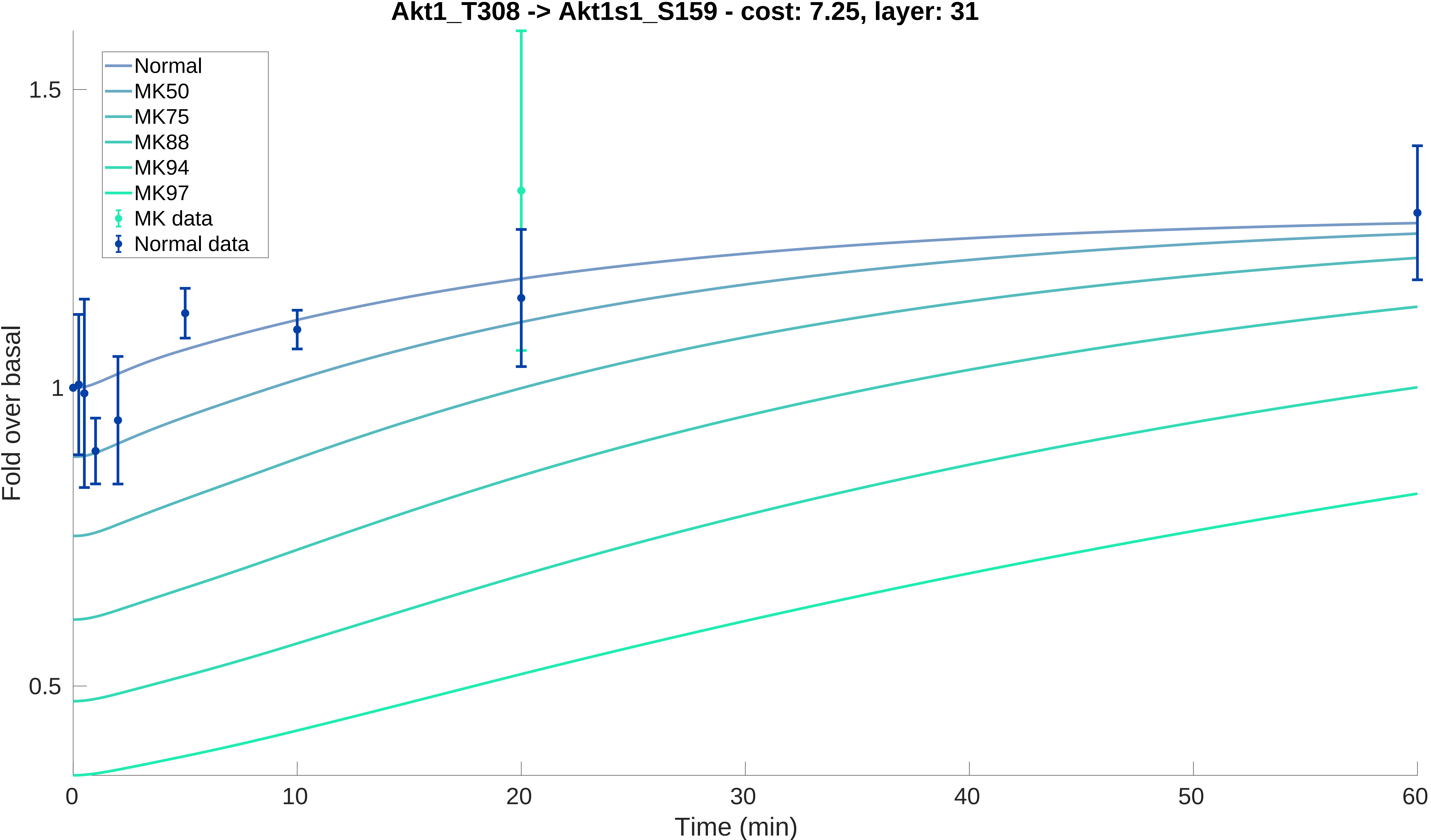
Akt1_T308 -> Akt1s1_S159 - cost: 7.25, layer: 31
Normal
MK50
MK75
MK88
MK94
MK97
MK data
Normal data
Fold over basal
1.5
1
0.5
0
10
20
30
40
50
60
Time (min)

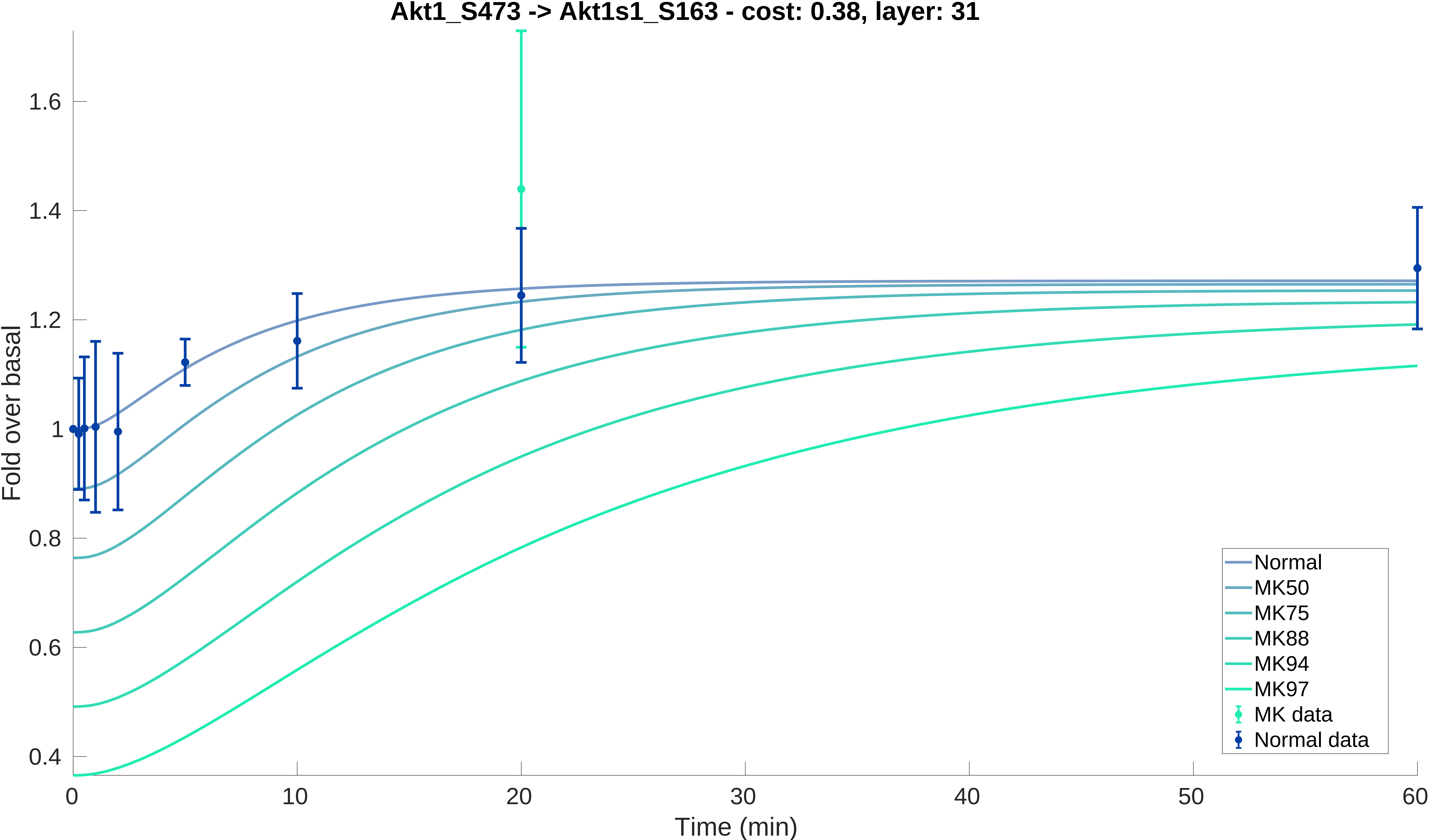

Akt1_S473 -> Akt1s1_S163 - cost: 0.38, layer: 31
Fold over basal
1.6
1.4
1.2
1
0.8
0.6
0.4
0
10
20
30
40
50
60
Time (min)
Normal
MK50
MK75
MK88
MK94
MK97
MK data
Normal data

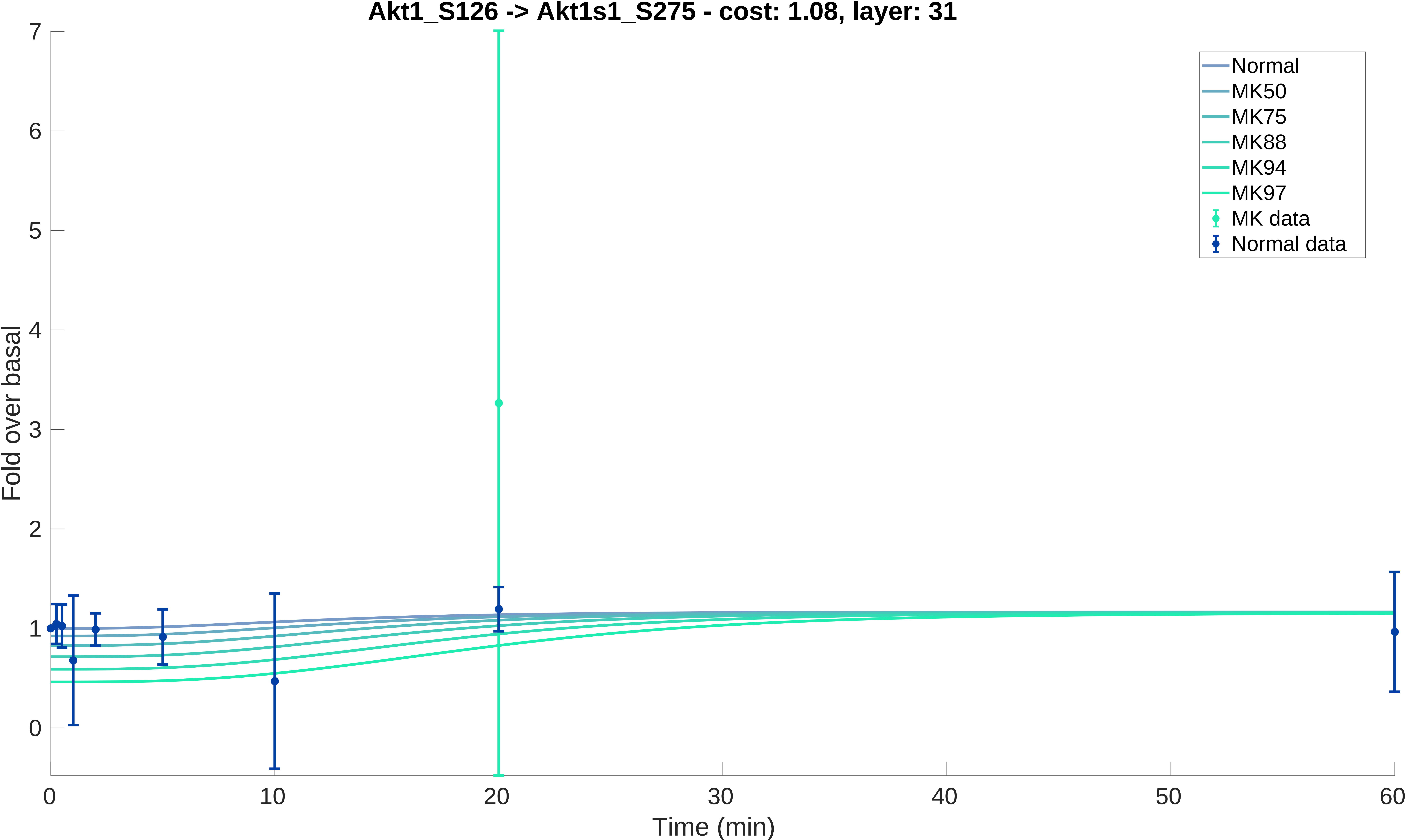

Akt1_S126 -> Akt1s1_S275 - cost: 1.08, layer: 31
Fold over basal
Time (min)
0
1
2
3
4
5
6
7
10
20
30
40
50
60
Normal
MK50
MK75
MK88
MK94
MK97
MK data
Normal data

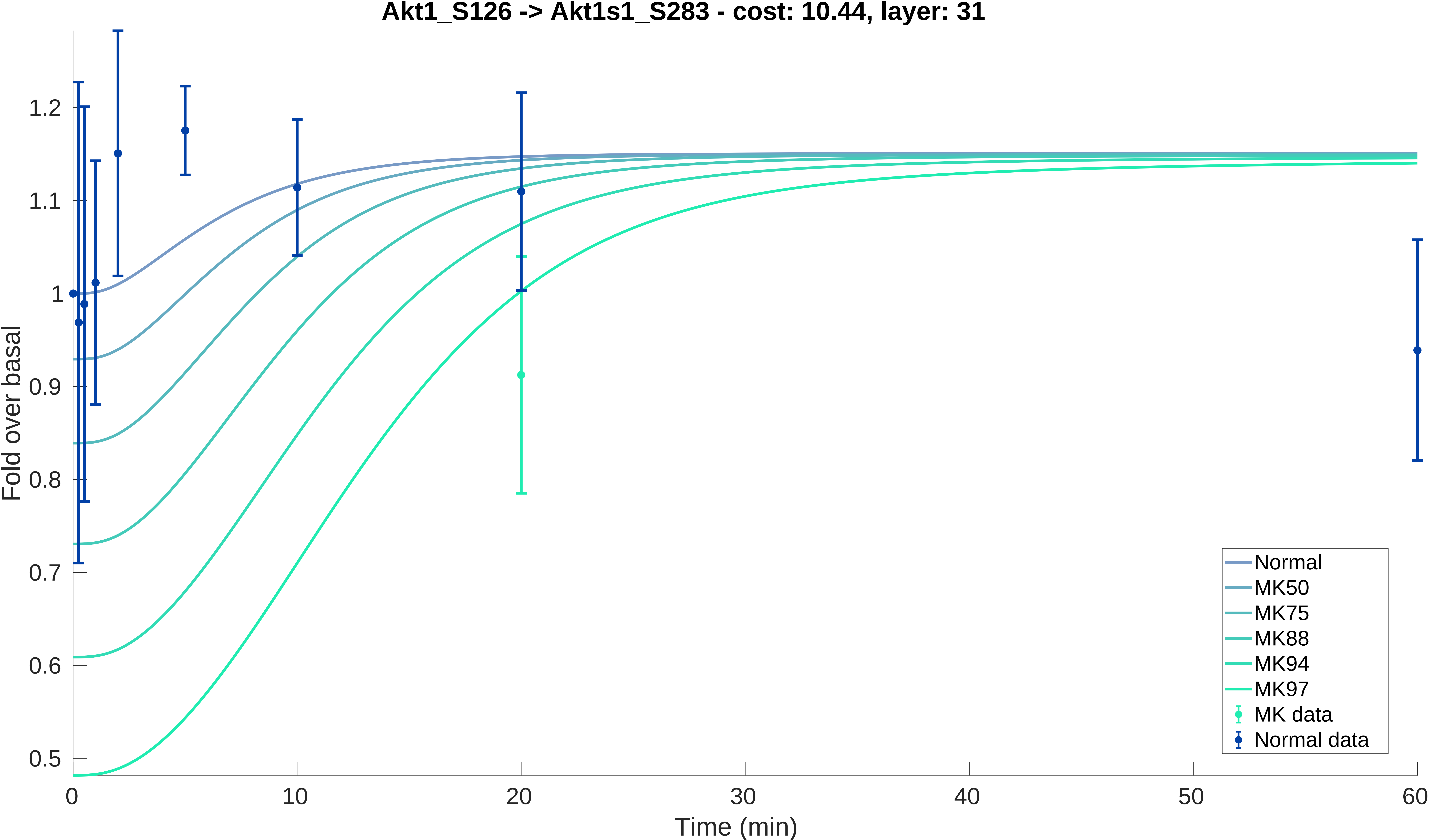

Akt1_S126 -> Akt1s1_S283 - cost: 10.44, layer: 31
Fold over basal
Time (min)
0.5
0.6
0.7
0.8
0.9
1
1.1
1.2
0
10
20
30
40
50
60
Normal
MK50
MK75
MK88
MK94
MK97
MK data
Normal data

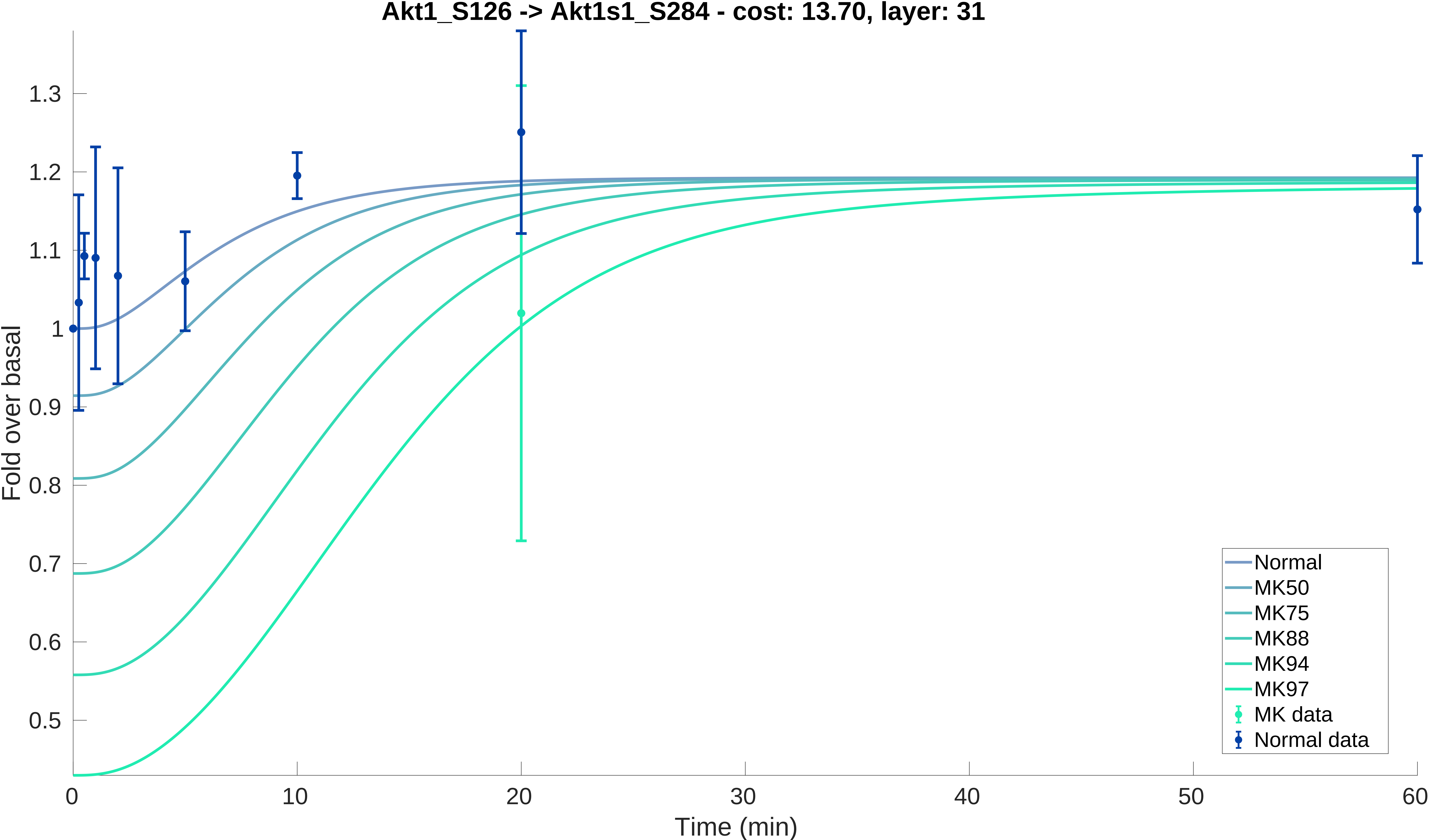
Akt1_S126 -> Akt1s1_S284 - cost: 13.70, layer: 31
Fold over basal
1.3
1.2
1.1
1
0.9
0.8
0.7
0.6
0.5
0
10
20
30
40
50
60
Time (min)
Normal
MK50
MK75
MK88
MK94
MK97
MK data
Normal data

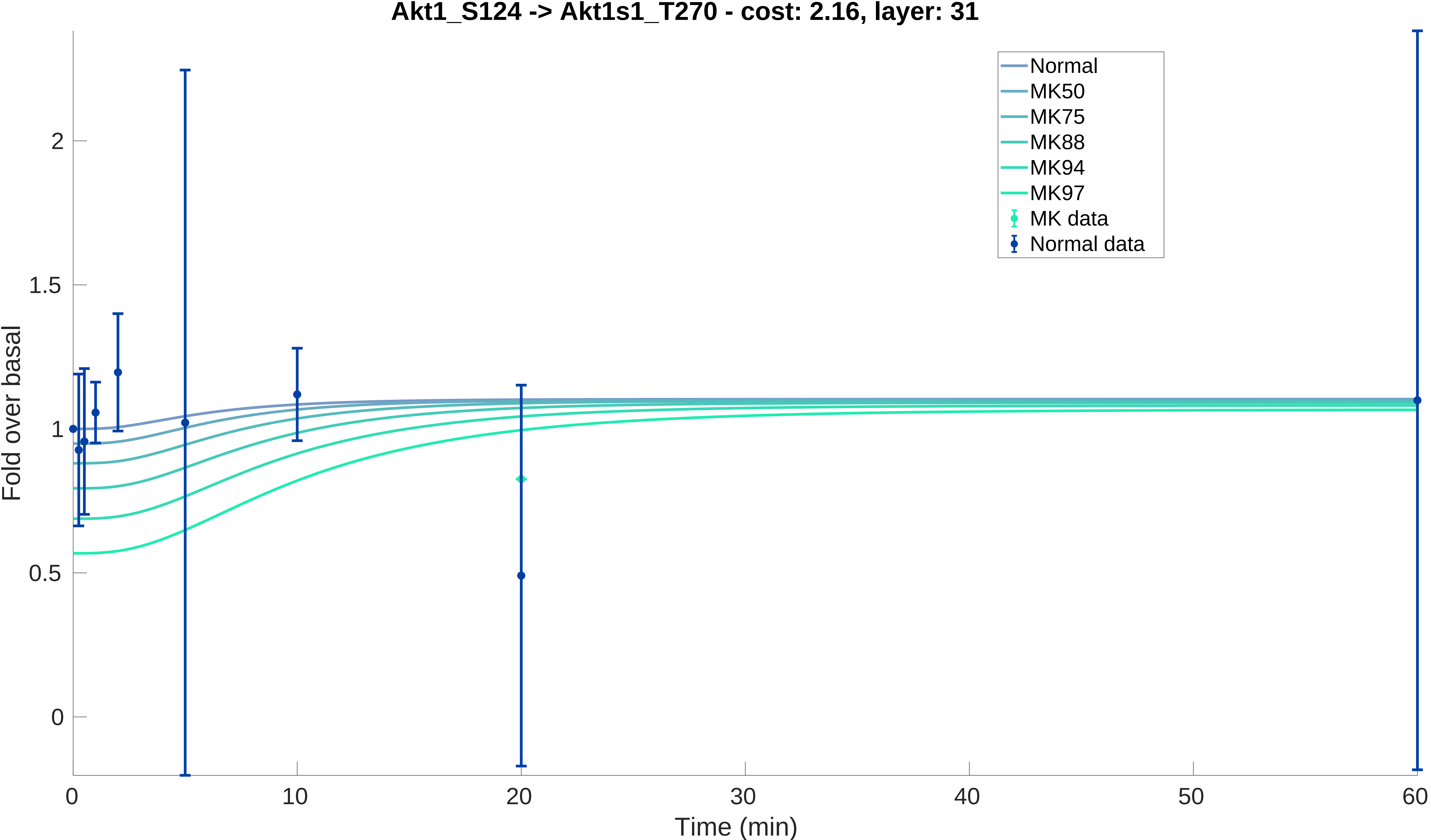

Akt1_S124 -> Akt1s1_T270 - cost: 2.16, layer: 31
Normal
MK50
MK75
MK88
MK94
MK97
MK data
Normal data
Fold over basal
Time (min)
0
0.5
1
1.5
2
0
10
20
30
40
50
60

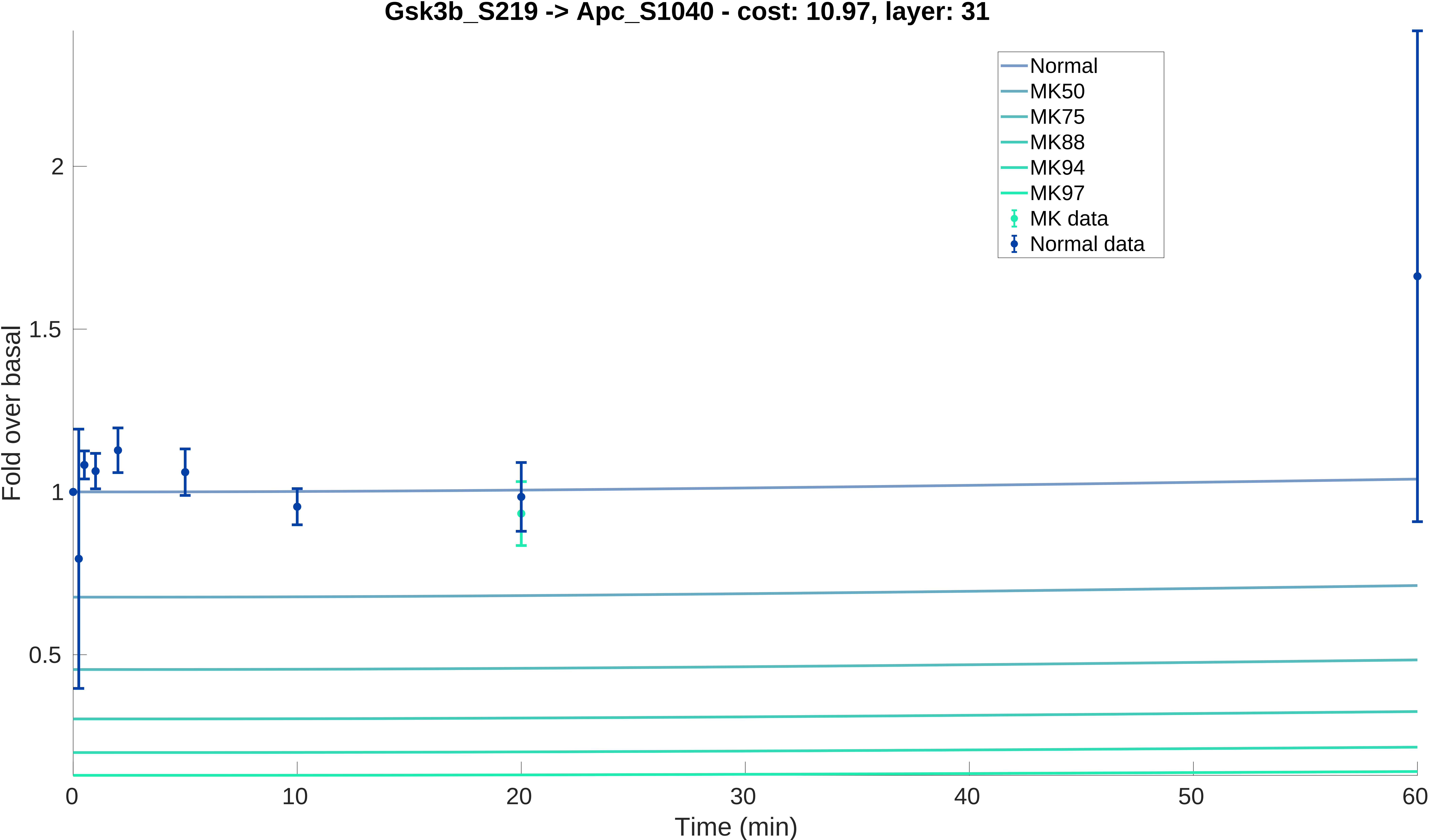

Gsk3b_S219 -> Apc_S1040 - cost: 10.97, layer: 31
Normal
MK50
MK75
MK88
MK94
MK97
MK data
Normal data
Fold over basal
2
1.5
1
0.5
0
10
20
30
40
50
60
Time (min)

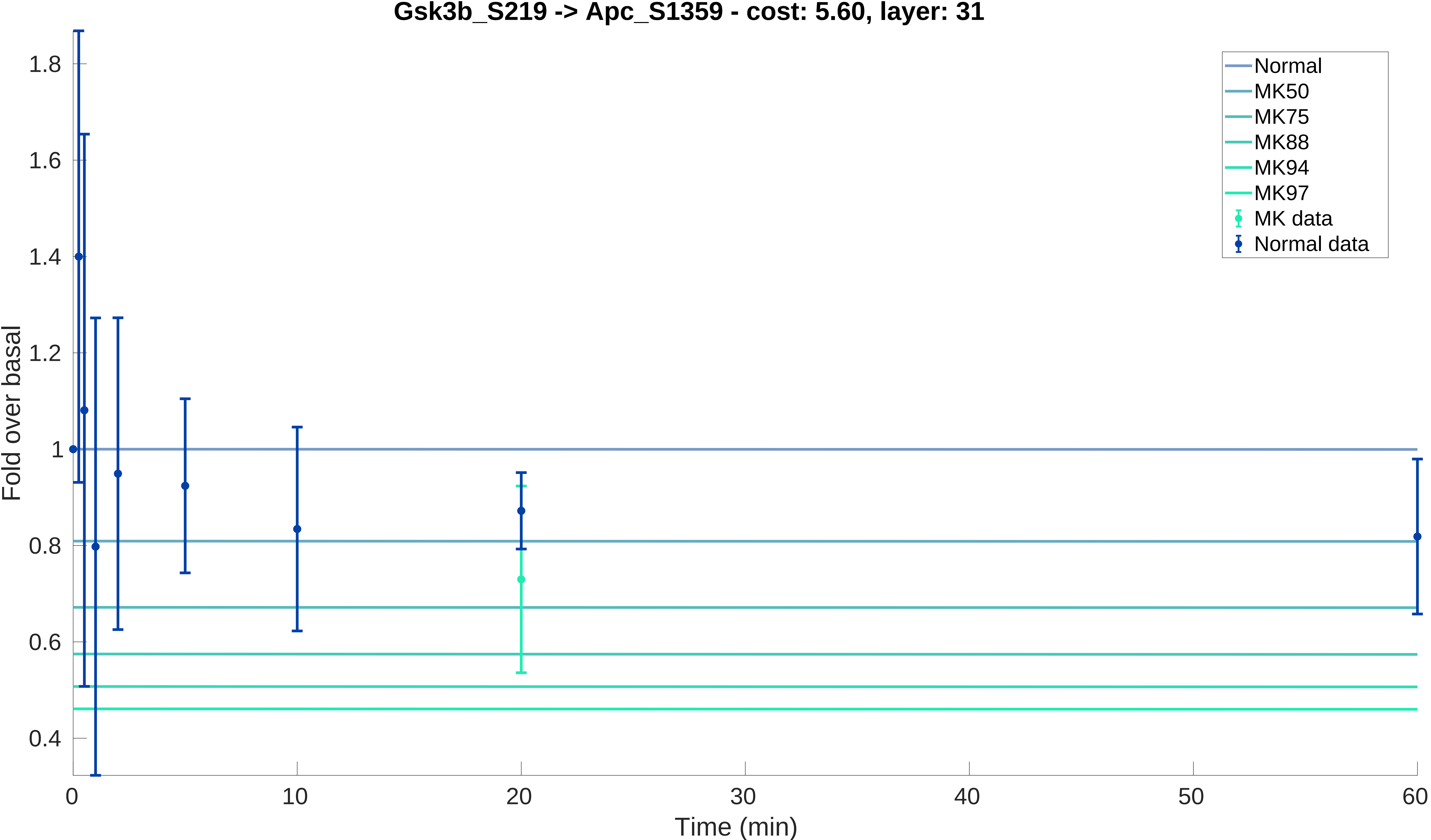

Gsk3b_S219 -> Apc_S1359 - cost: 5.60, layer: 31
Normal
MK50
MK75
MK88
MK94
MK97
MK data
Normal data
Fold over basal
1.8
1.6
1.4
1.2
1
0.8
0.6
0.4
0
10
20
30
40
50
60
Time (min)

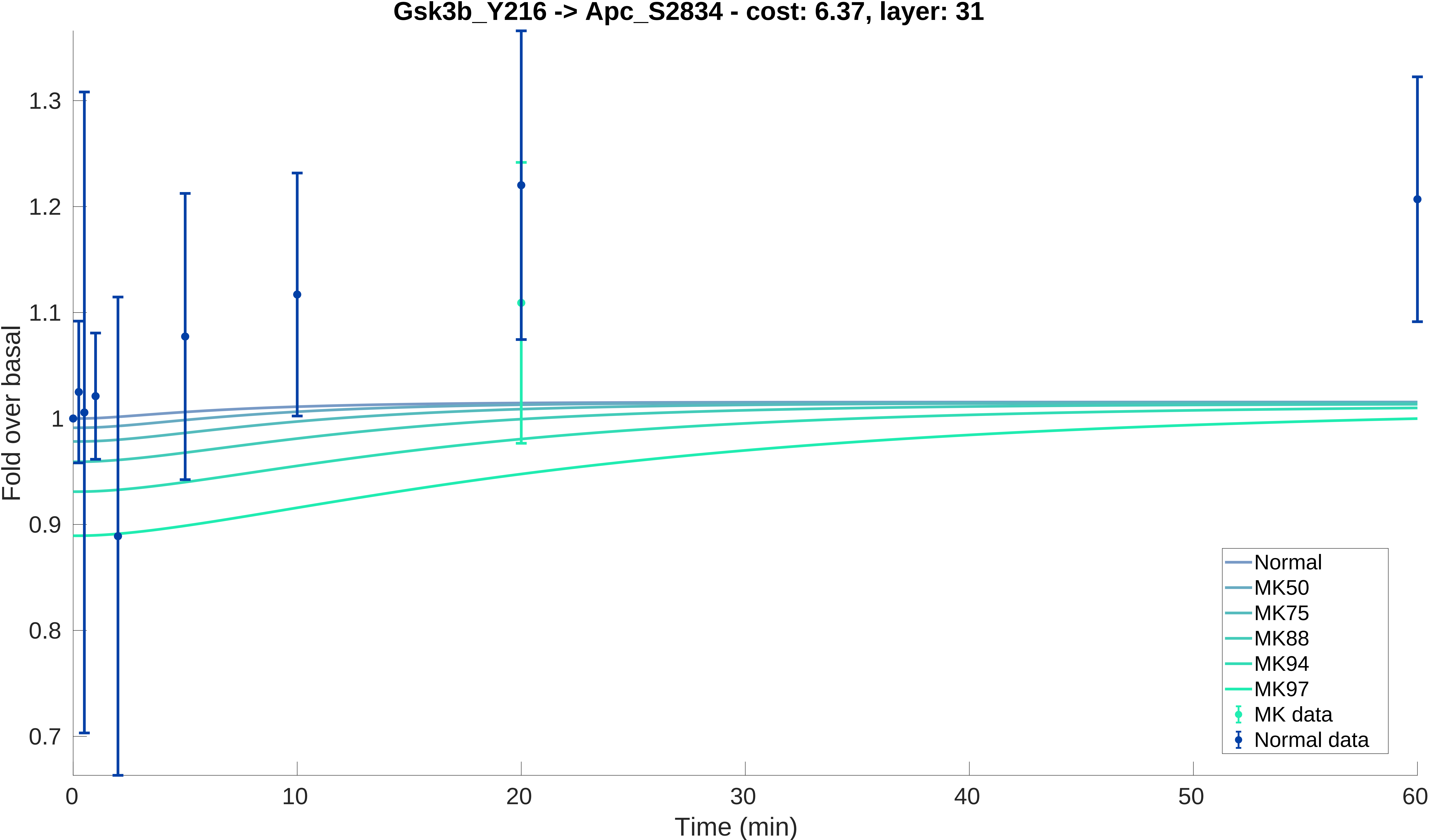

Gsk3b_Y216 -> Apc_S2834 - cost: 6.37, layer: 31
Fold over basal
1.3
1.2
1.1
1
0.9
0.8
0.7
0
10
20
30
40
50
60
Time (min)
Normal
MK50
MK75
MK88
MK94
MK97
MK data
Normal data

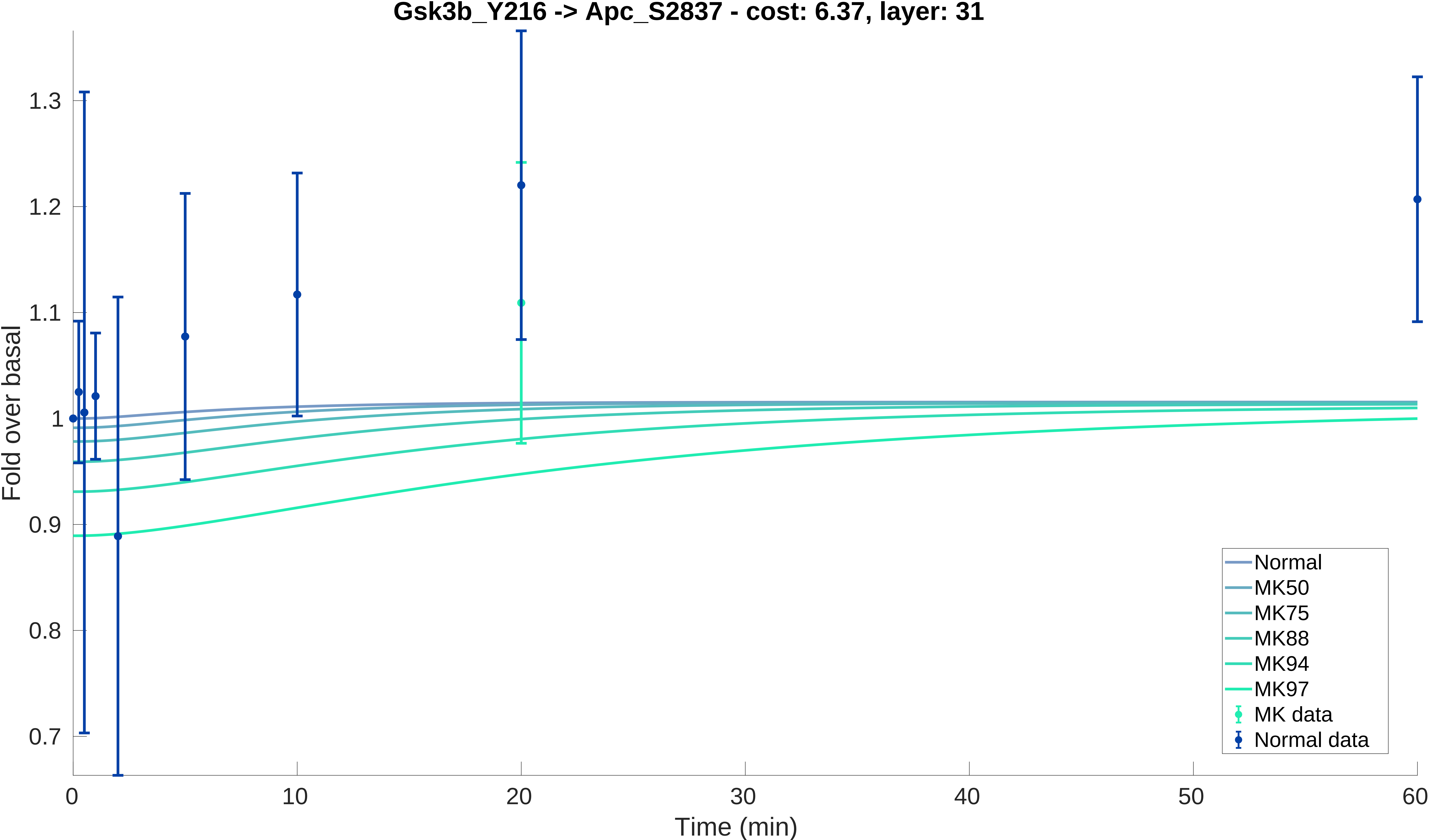

Gsk3b_Y216 -> Apc_S2837 - cost: 6.37, layer: 31
Fold over basal
Time (min)
1.3
1.2
1.1
1
0.9
0.8
0.7
0
10
20
30
40
50
60
Normal
MK50
MK75
MK88
MK94
MK97
MK data
Normal data

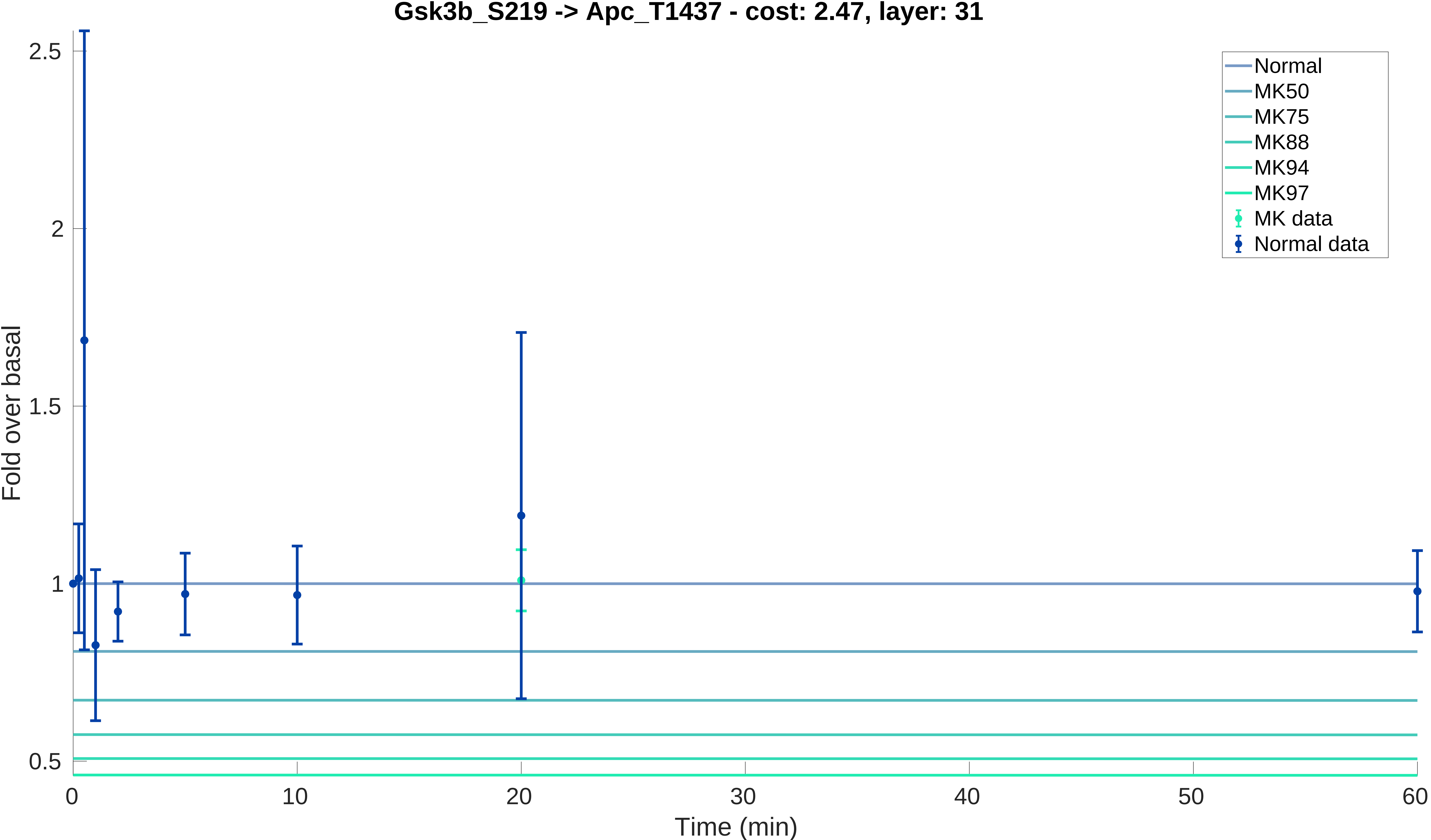
Gsk3b_S219 -> Apc_T1437 - cost: 2.47, layer: 31
Fold over basal
Time (min)
2.5
2
1.5
1
0.5
0
10
20
30
40
50
60
Normal
MK50
MK75
MK88
MK94
MK97
MK data
Normal data

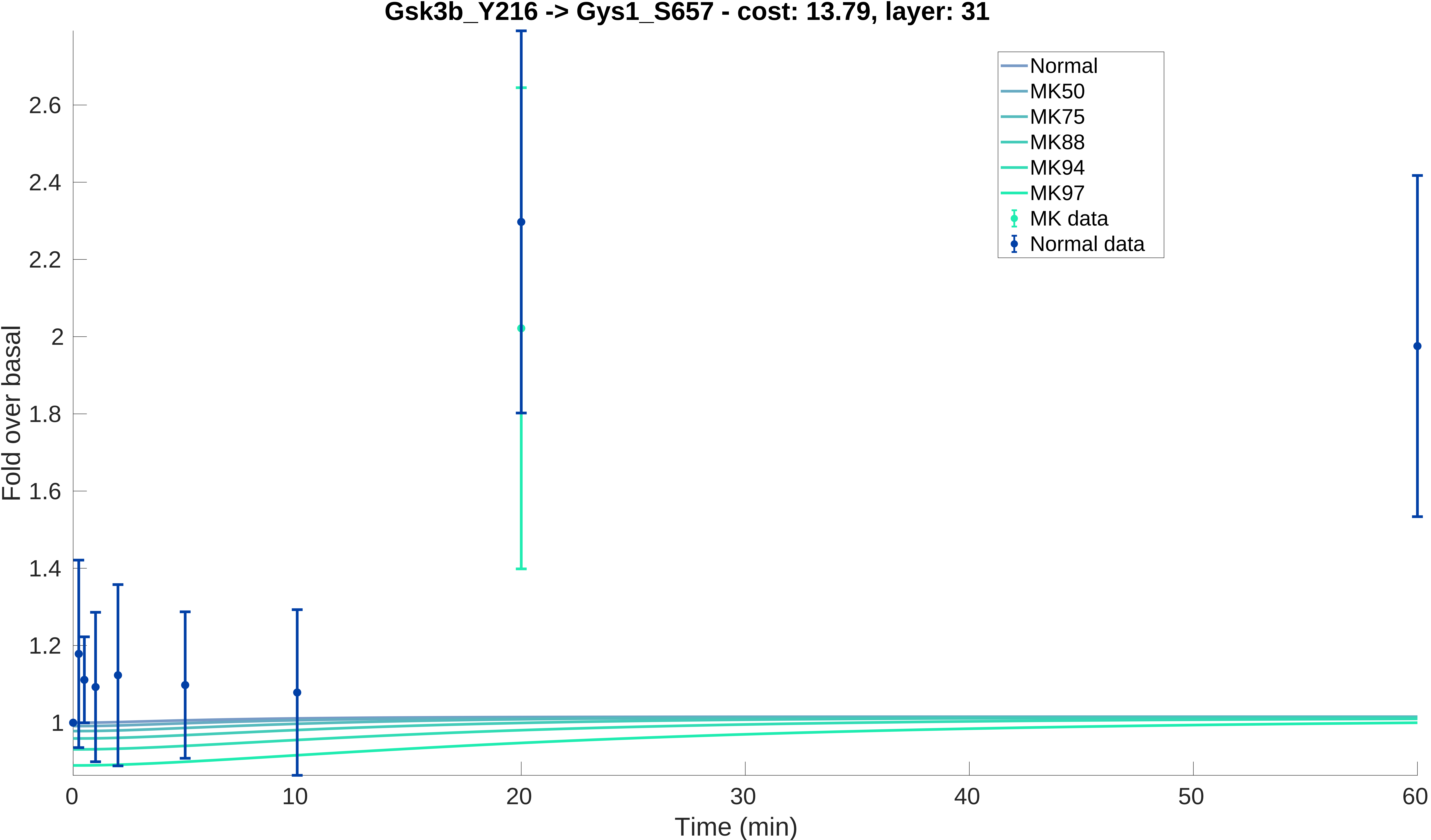
Gsk3b_Y216 -> Gys1_S657 - cost: 13.79, layer: 31
Normal
MK50
MK75
MK88
MK94
MK97
MK data
Normal data
Fold over basal
2.6
2.4
2.2
2
1.8
1.6
1.4
1.2
1
0
10
20
30
40
50
60
Time (min)

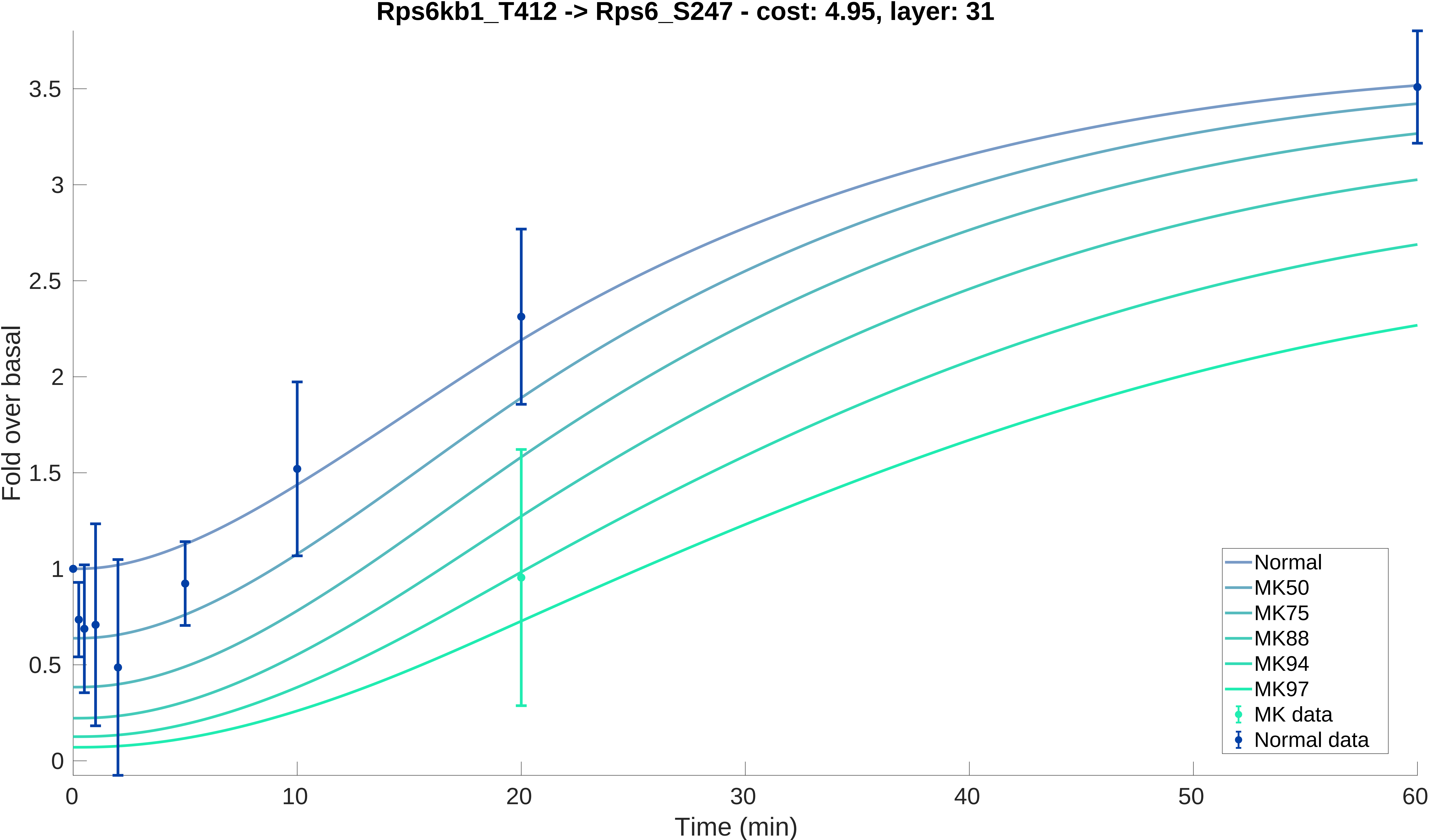

Rps6kb1_T412 -> Rps6_S247 - cost: 4.95, layer: 31
Fold over basal
Time (min)
0
0.5
1
1.5
2
2.5
3
3.5
0
10
20
30
40
50
60
Normal
MK50
MK75
MK88
MK94
MK97
MK data
Normal data

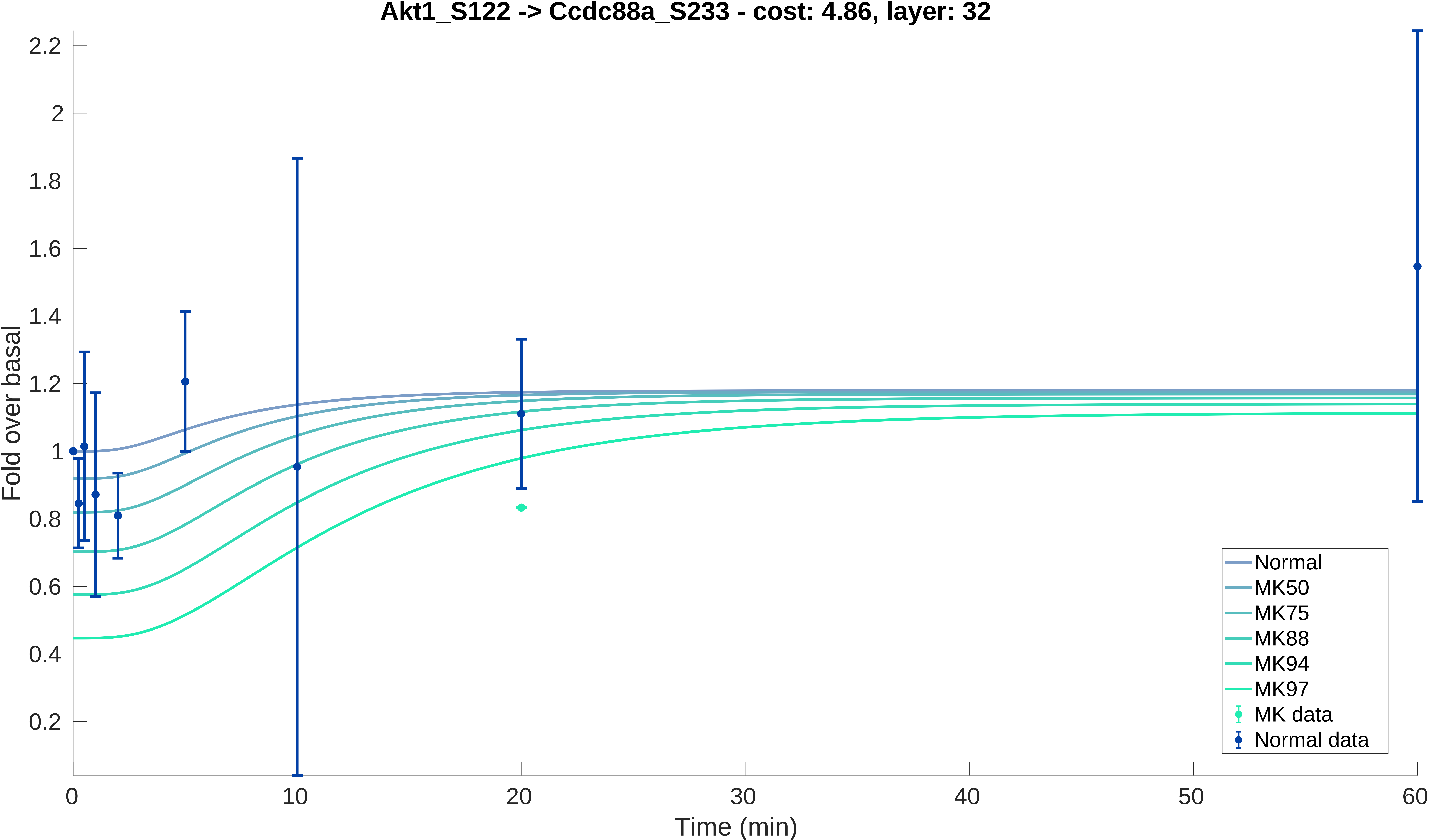
Akt1_S122 -> Ccdc88a_S233 - cost: 4.86, layer: 32
Fold over basal
Time (min)
0.2
0.4
0.6
0.8
1
1.2
1.4
1.6
1.8
2
2.2
0
10
20
30
40
50
60
Normal
MK50
MK75
MK88
MK94
MK97
MK data
Normal data

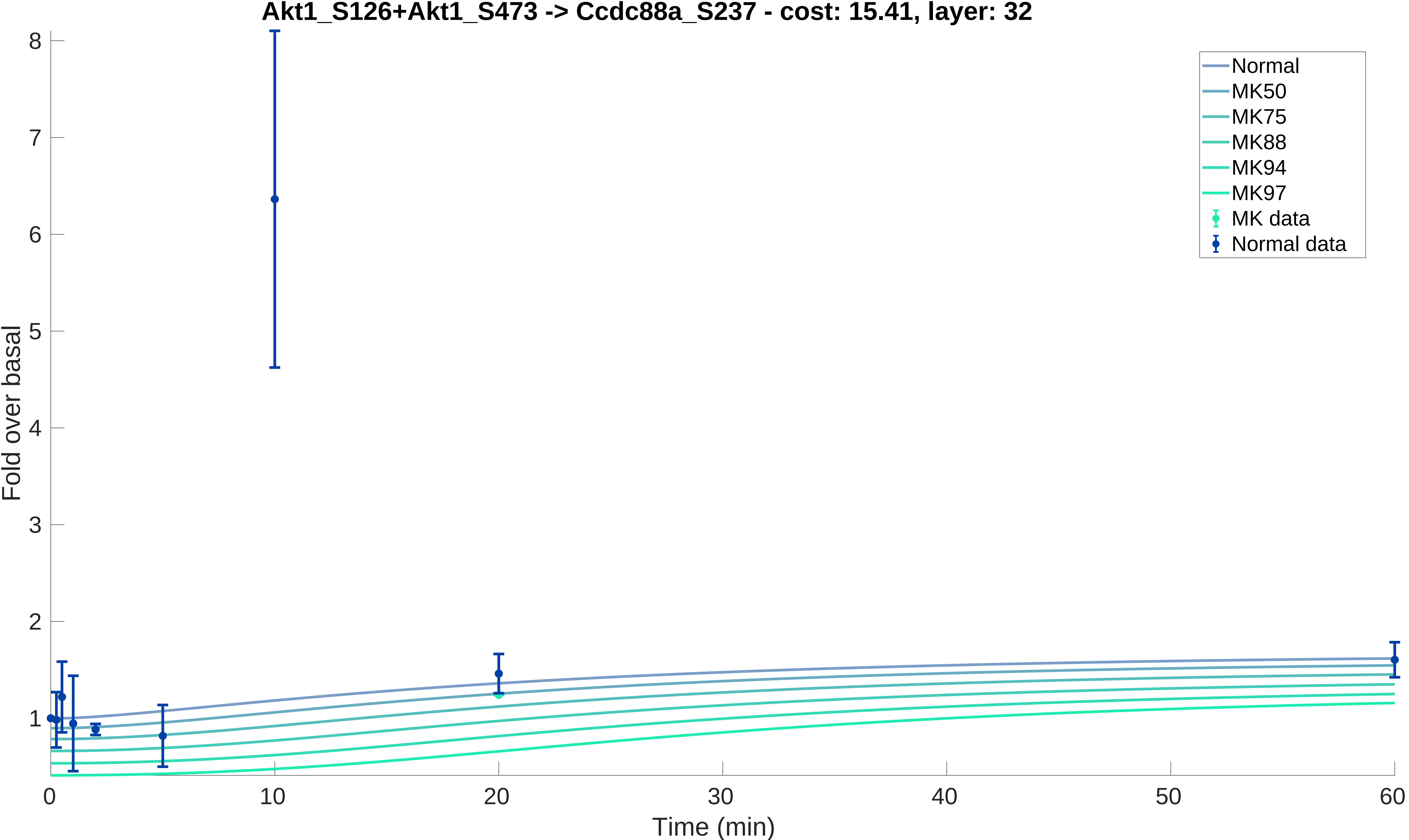
Akt1_S126+Akt1_S473 -> Ccdc88a_S237 - cost: 15.41, layer: 32
Fold over basal
Time (min)
Normal
MK50
MK75
MK88
MK94
MK97
MK data
Normal data
0
10
20
30
40
50
60
1
2
3
4
5
6
7
8

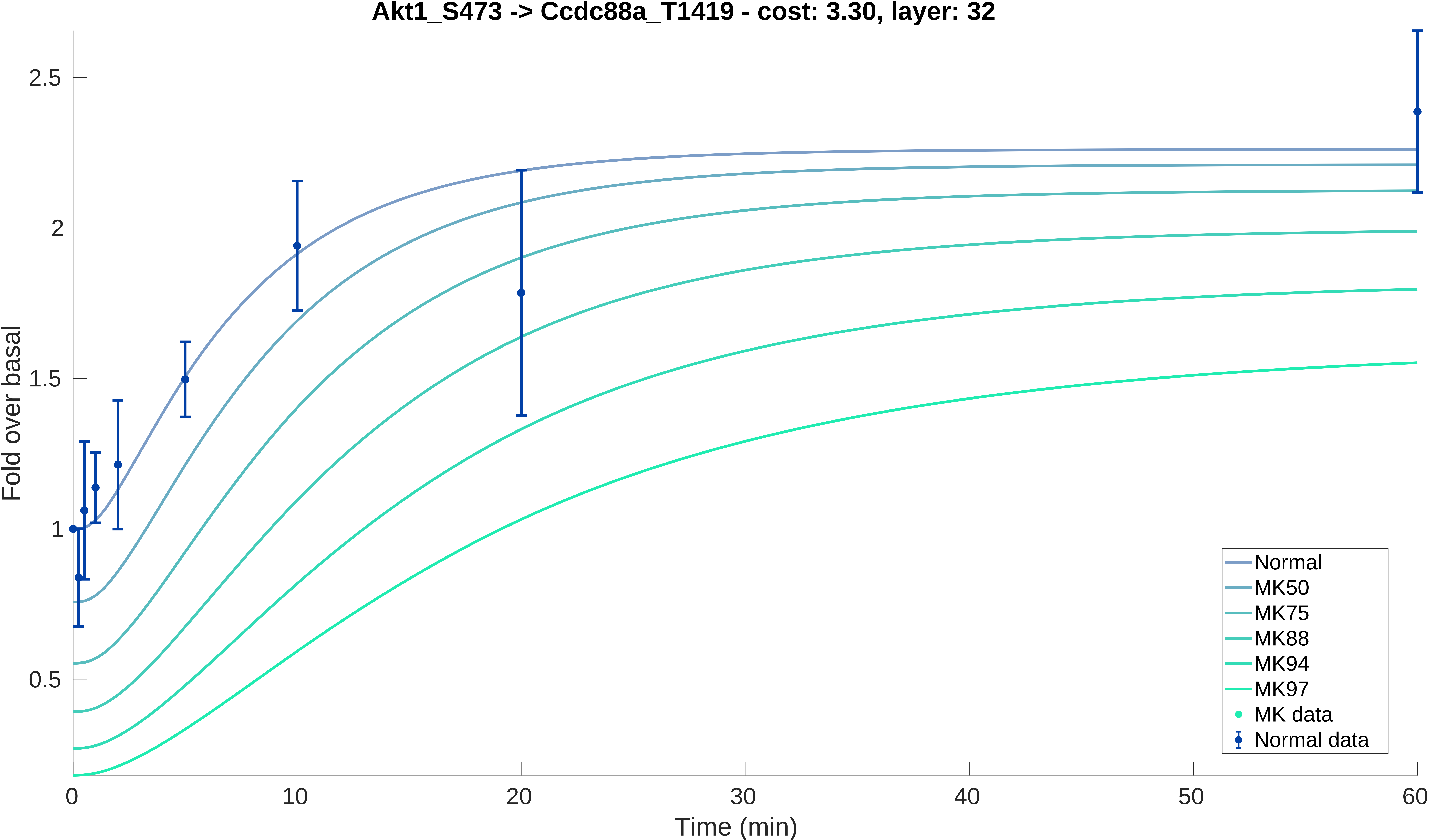

Akt1_S473 -> Ccdc88a_T1419 - cost: 3.30, layer: 32
Fold over basal
2.5
2
1.5
1
0.5
0
10
20
30
40
50
60
Time (min)
Normal
MK50
MK75
MK88
MK94
MK97
MK data
Normal data

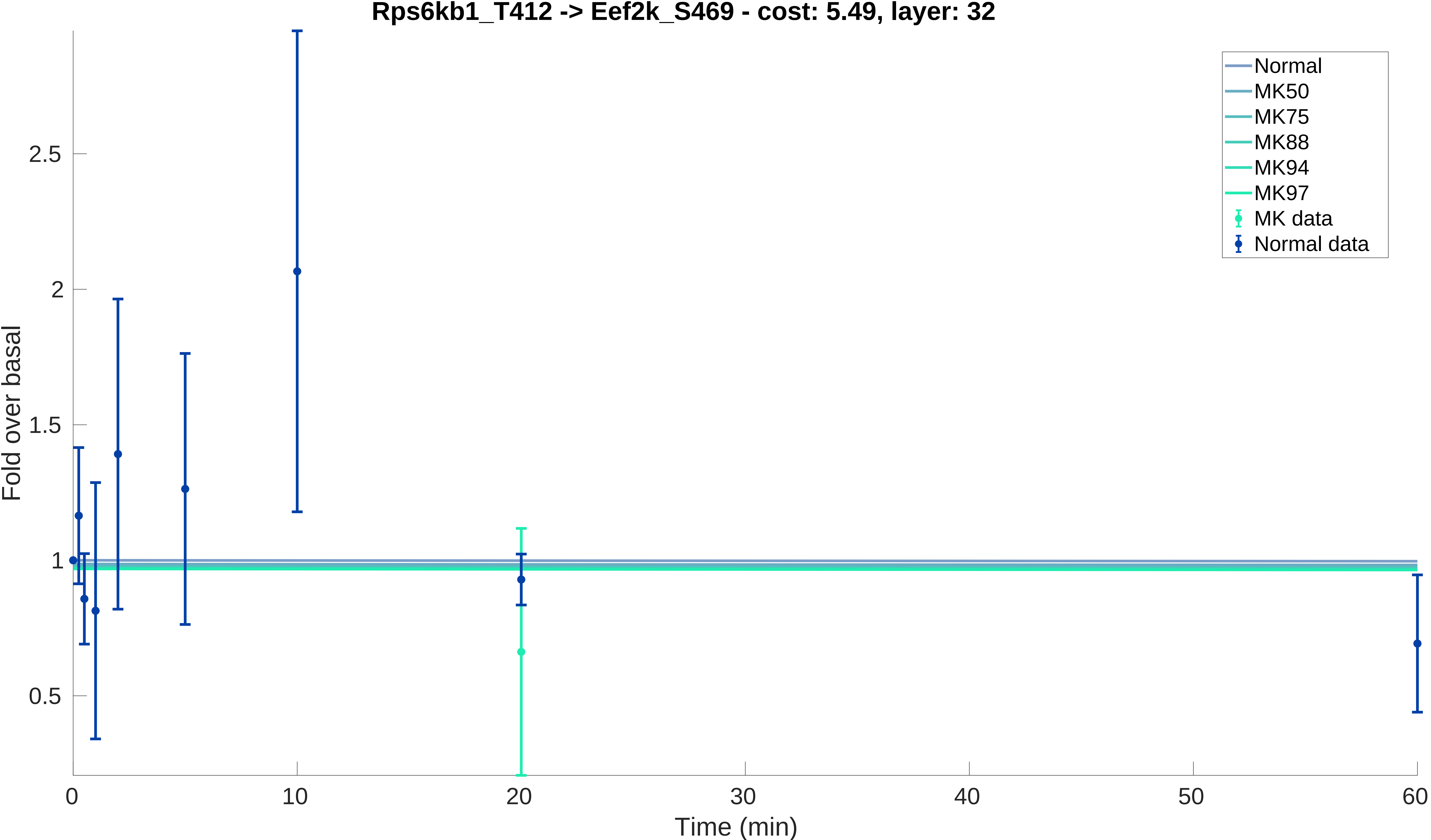

Rps6kb1_T412 -> Eef2k_S469 - cost: 5.49, layer: 32
Normal
MK50
MK75
MK88
MK94
MK97
MK data
Normal data
Fold over basal
0.5
1
1.5
2
2.5
0
10
20
30
40
50
60
Time (min)

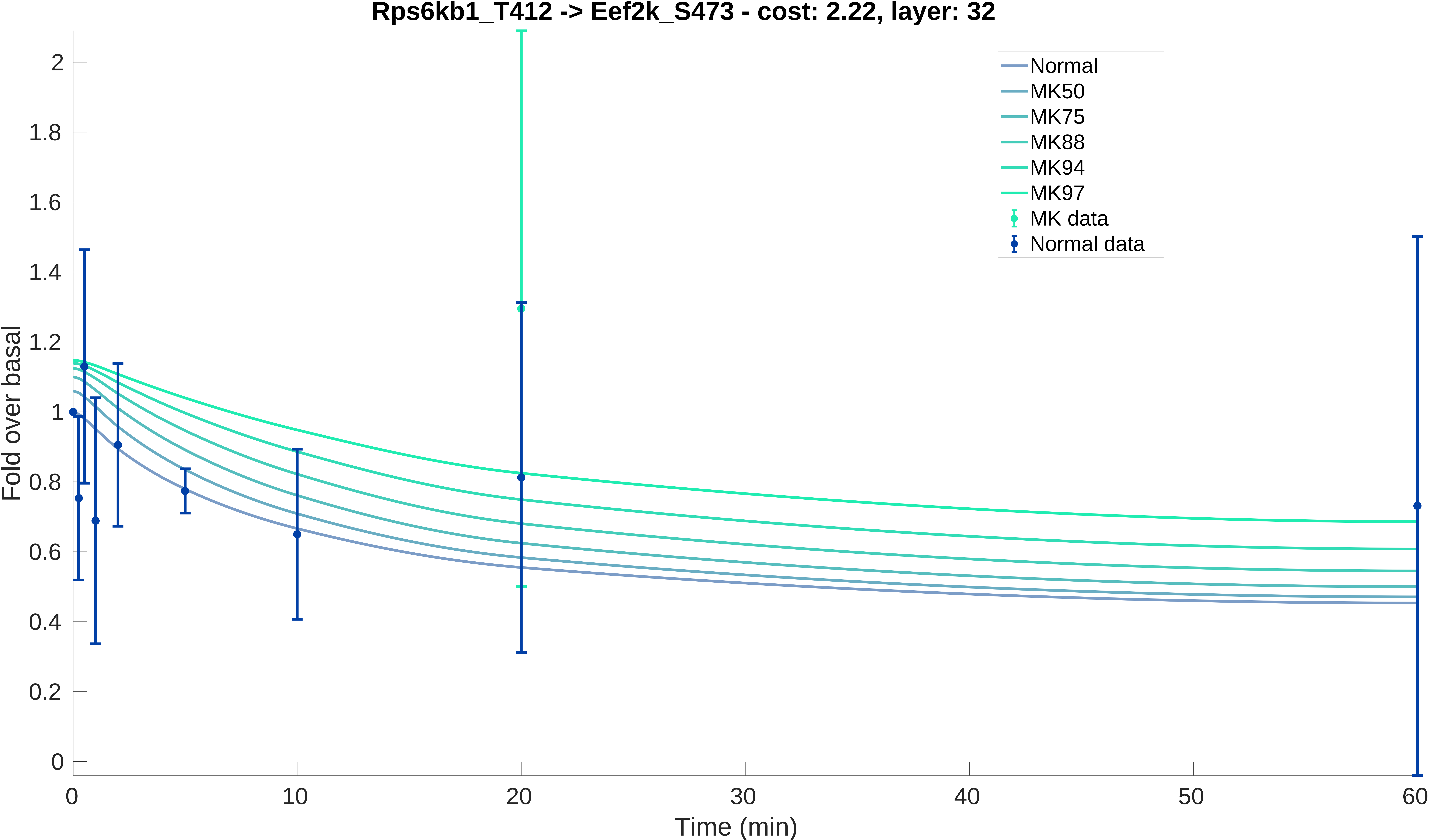

Rps6kb1_T412 -> Eef2k_S473 - cost: 2.22, layer: 32
Normal
MK50
MK75
MK88
MK94
MK97
MK data
Normal data
Fold over basal
Time (min)
0
0.2
0.4
0.6
0.8
1
1.2
1.4
1.6
1.8
2
0
10
20
30
40
50
60

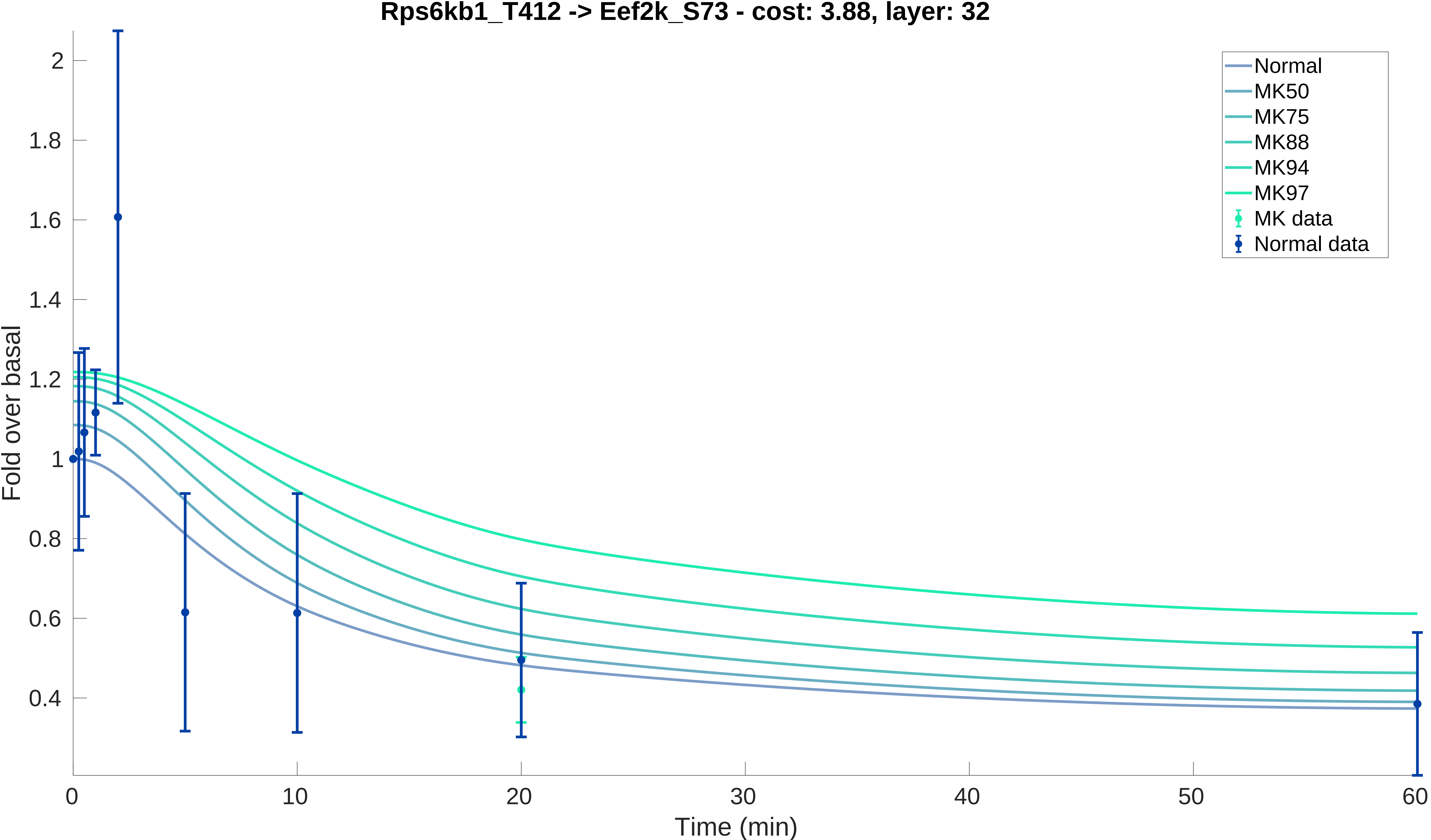

Rps6kb1_T412 -> Eef2k_S73 - cost: 3.88, layer: 32
Fold over basal
Time (min)
Normal
MK50
MK75
MK88
MK94
MK97
MK data
Normal data
0
10
20
30
40
50
60
0.4
0.6
0.8
1
1.2
1.4
1.6
1.8
2

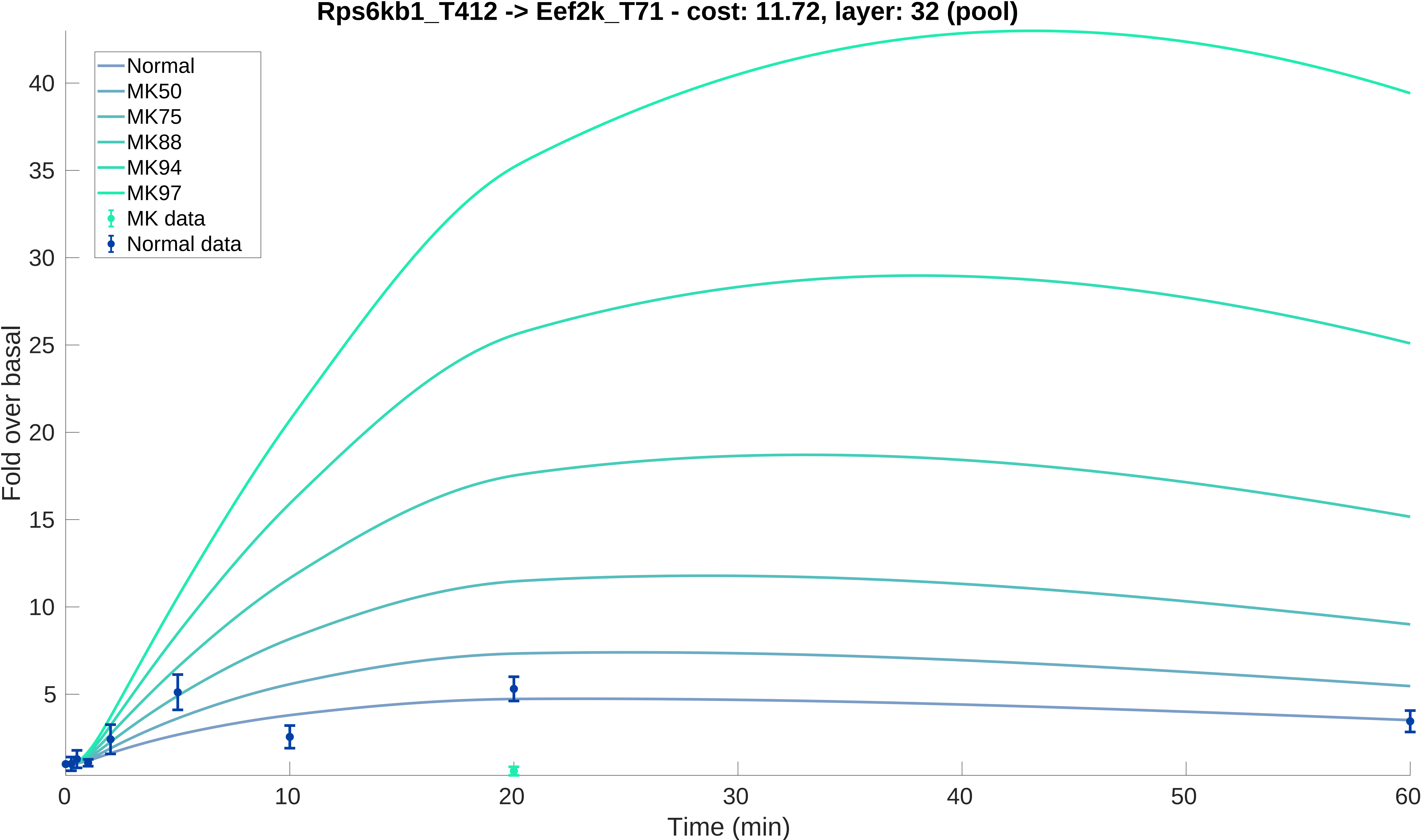

Rps6kb1_T412 -> Eef2k_T71 - cost: 11.72, layer: 32 (pool)
Normal
MK50
MK75
MK88
MK94
MK97
MK data
Normal data
Fold over basal
Time (min)
0
10
20
30
40
50
60
5
15
25
35

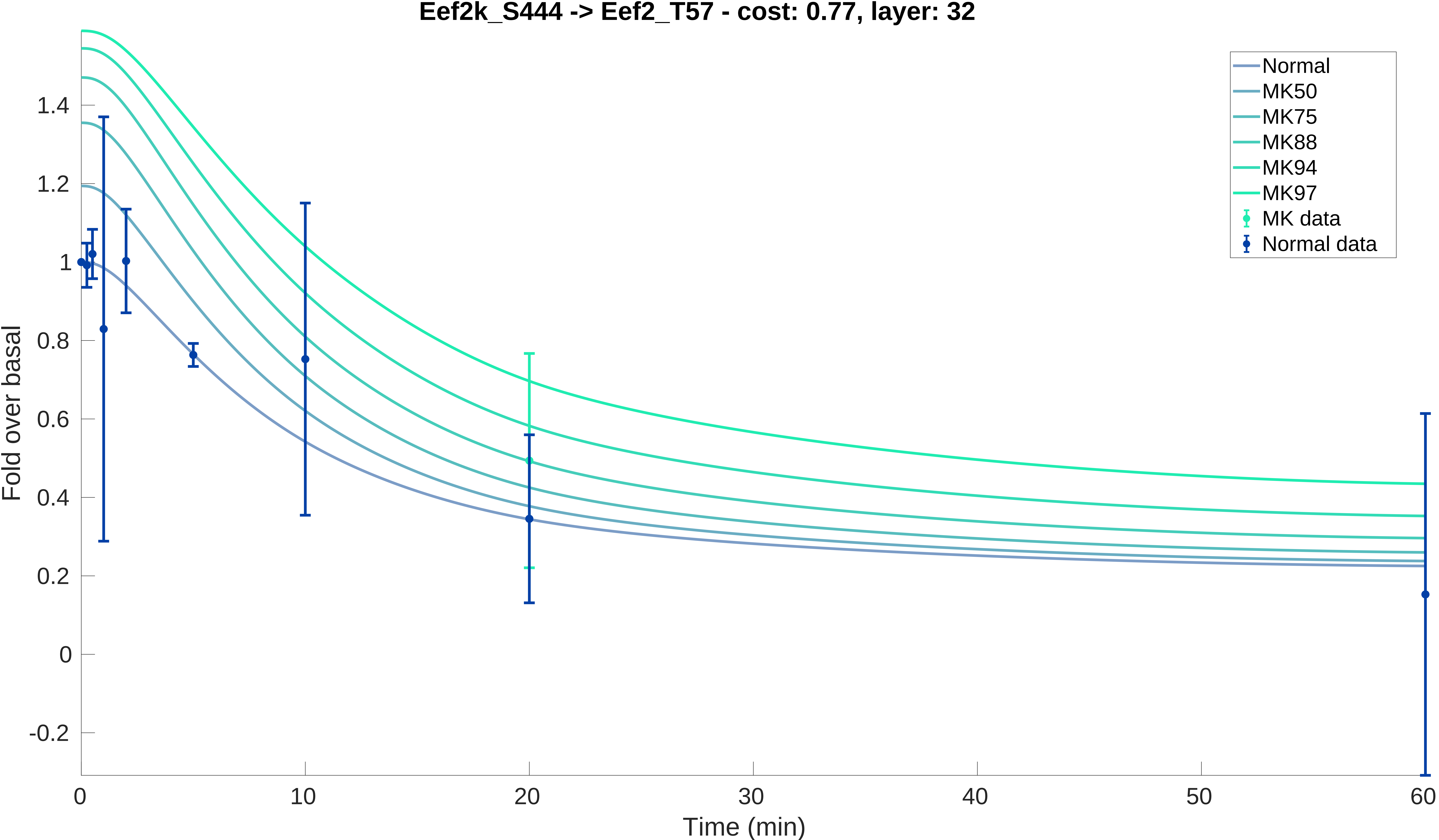

Eef2k_S444 -> Eef2_T57 - cost: 0.77, layer: 32
Fold over basal
Time (min)
1.4
1.2
1
0.8
0.6
0.4
0.2
0
-0.2
0
10
20
30
40
50
60
Normal
MK50
MK75
MK88
MK94
MK97
MK data
Normal data

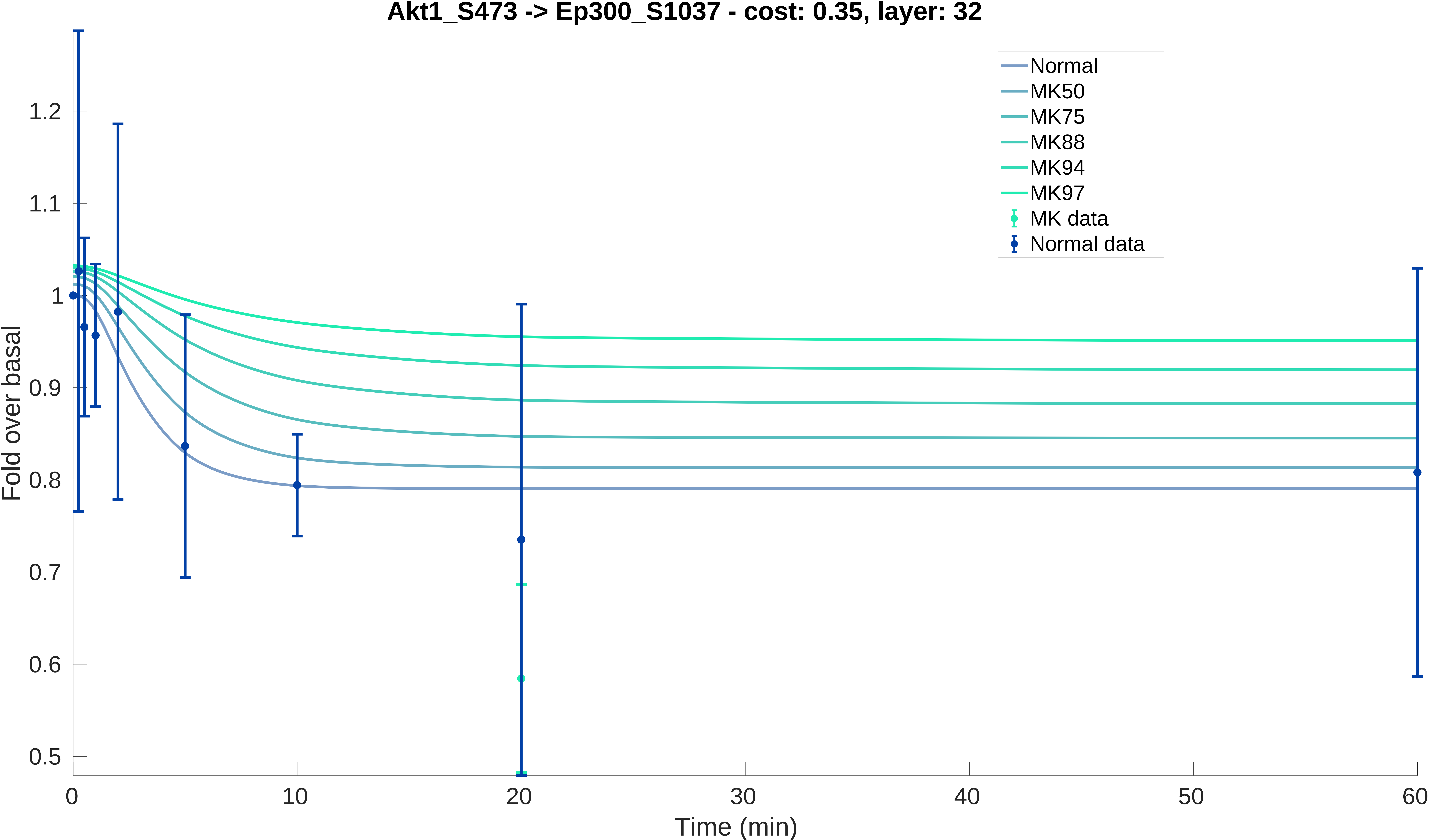

Akt1_S473 -> Ep300_S1037 - cost: 0.35, layer: 32
Normal
MK50
MK75
MK88
MK94
MK97
MK data
Normal data
Fold over basal
1.2
1.1
1
0.9
0.8
0.7
0.6
0.5
0
10
20
30
40
50
60
Time (min)

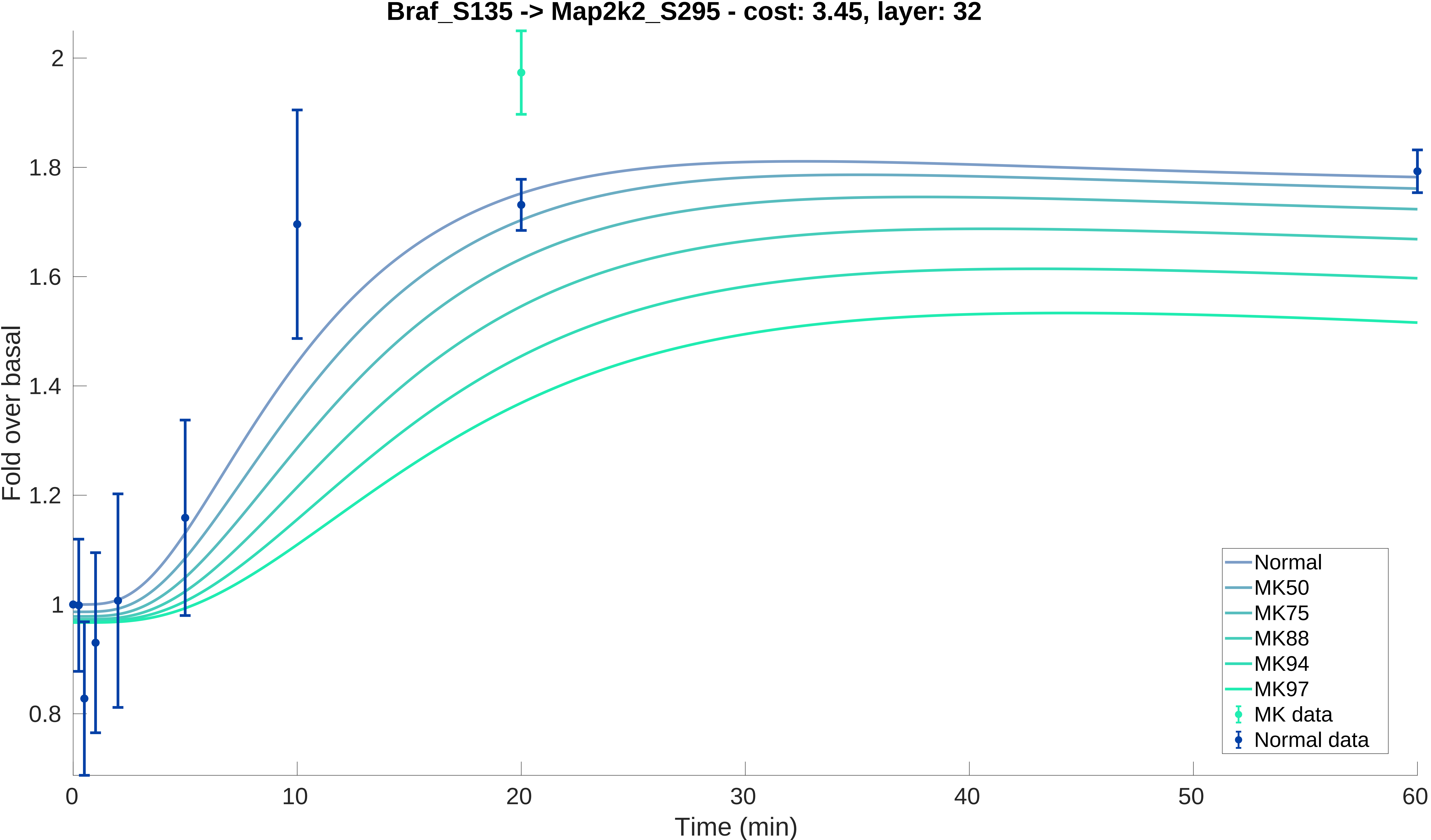

Braf_S135 -> Map2k2_S295 - cost: 3.45, layer: 32
Fold over basal
Time (min)
0.8
1
1.2
1.4
1.6
1.8
2
0
10
20
30
40
50
60
Normal
MK50
MK75
MK88
MK94
MK97
MK data
Normal data

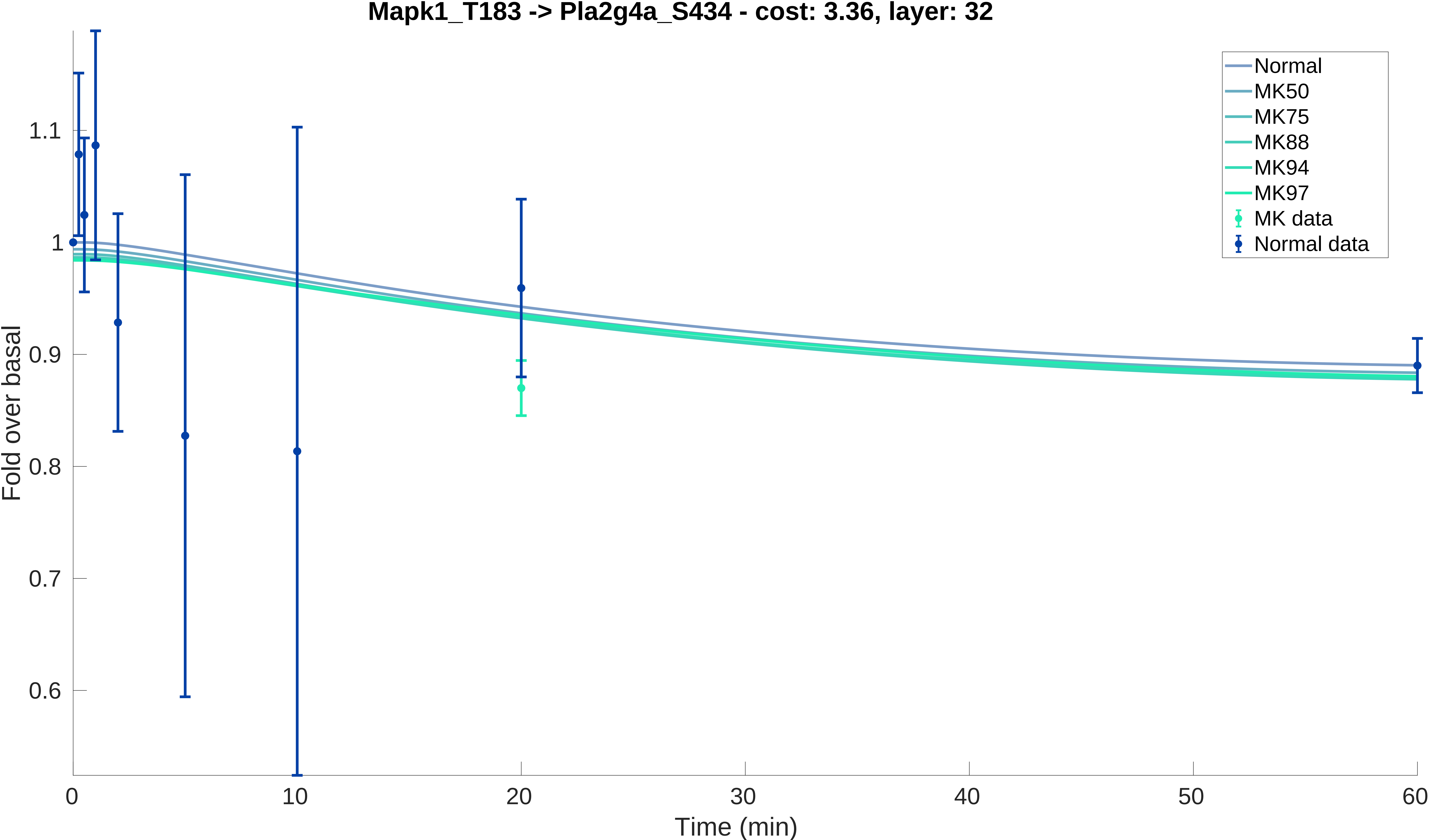

Mapk1_T183 -> Pla2g4a_S434 - cost: 3.36, layer: 32
Normal
MK50
MK75
MK88
MK94
MK97
MK data
Normal data
Fold over basal
1.1
1
0.9
0.8
0.7
0.6
0
10
20
30
40
50
60
Time (min)

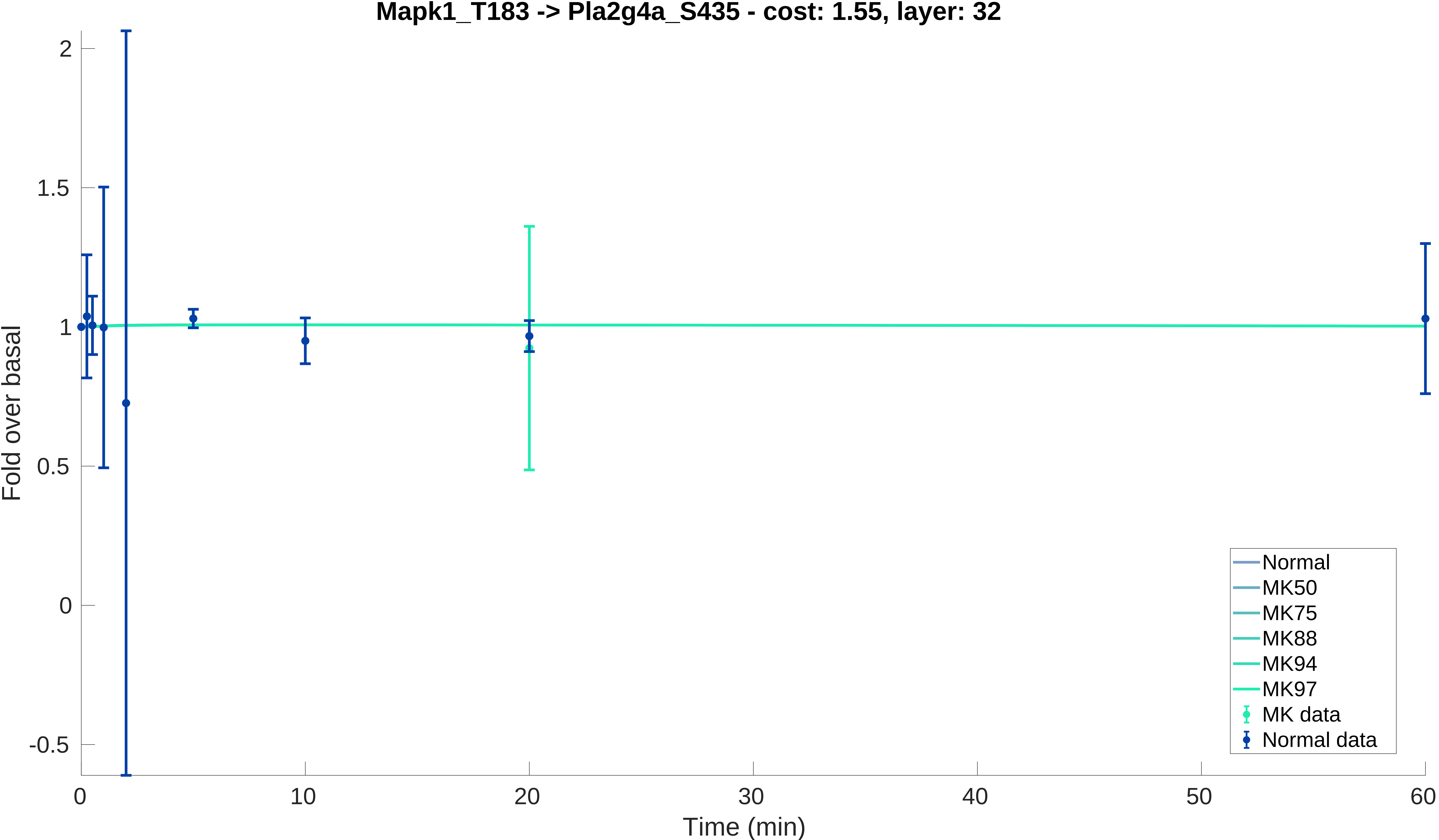

Mapk1_T183 -> Pla2g4a_S435 - cost: 1.55, layer: 32
Fold over basal
Time (min)
2
1.5
1
0.5
0
-0.5
0
10
20
30
40
50
60
Normal
MK50
MK75
MK88
MK94
MK97
MK data
Normal data

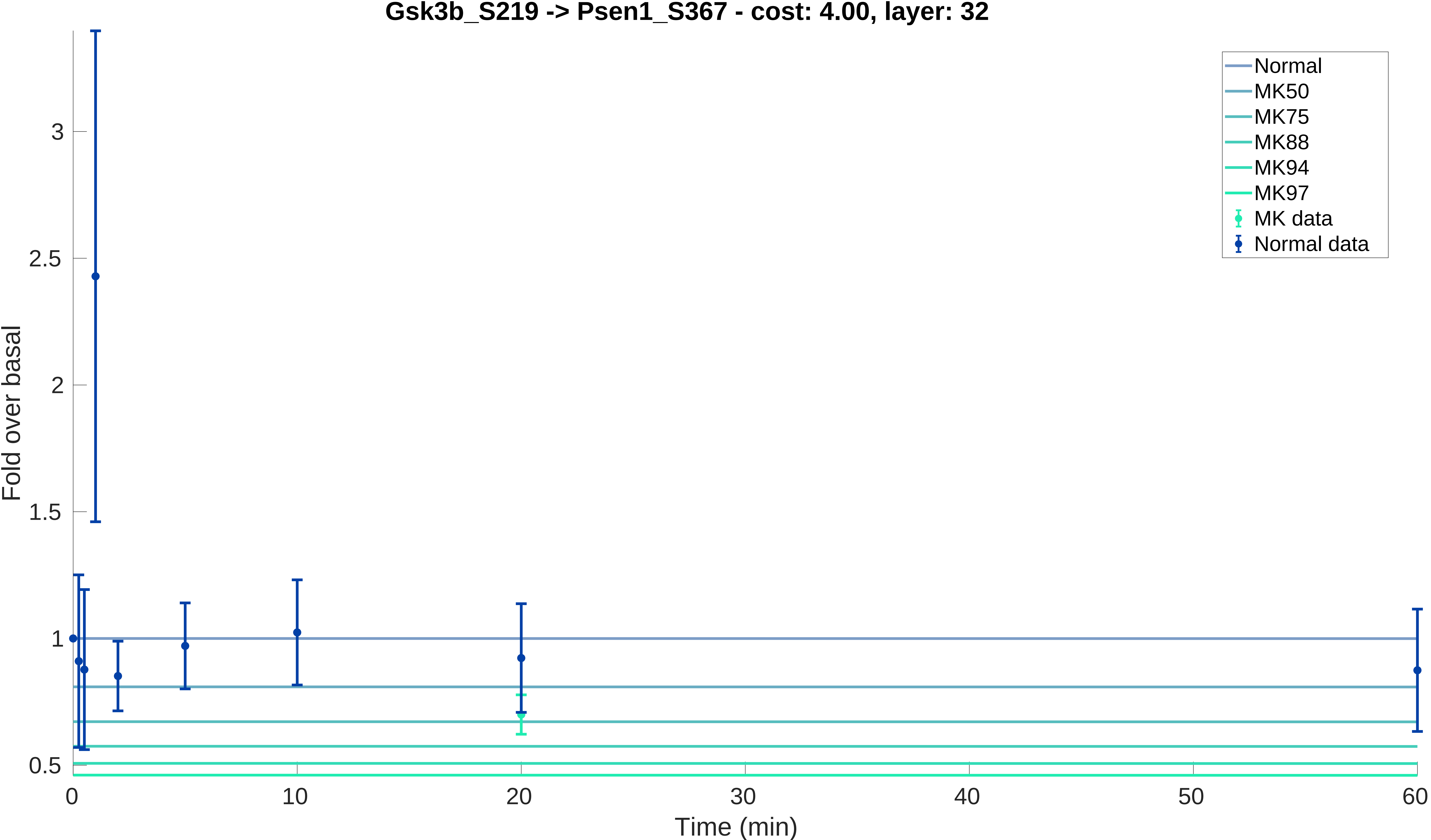

Gsk3b_S219 -> Psen1_S367 - cost: 4.00, layer: 32
Normal
MK50
MK75
MK88
MK94
MK97
MK data
Normal data
Fold over basal
Time (min)
0.5
1
1.5
2
2.5
3
0
10
20
30
40
50
60

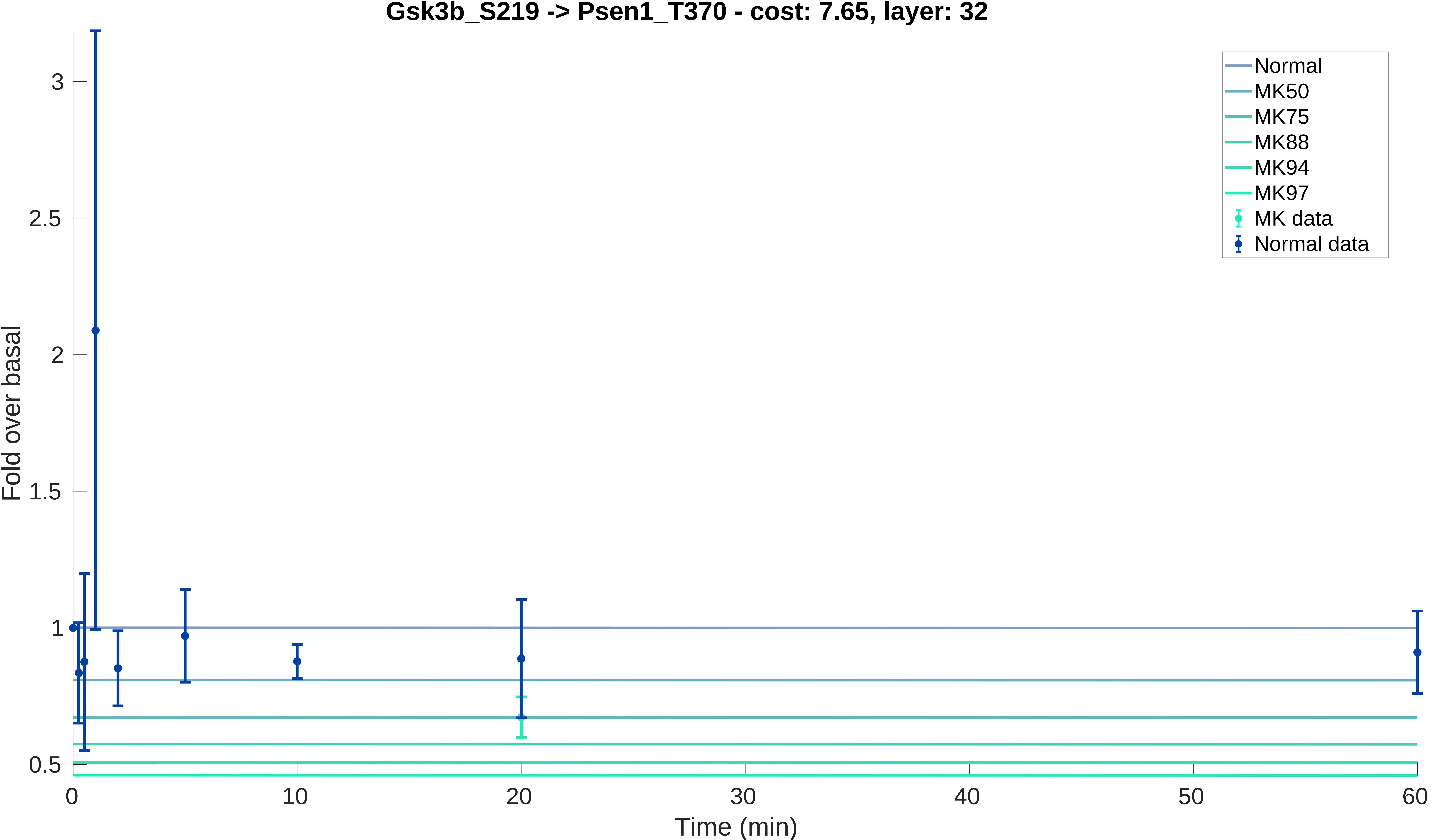

Gsk3b_S219 -> Psen1_T370 - cost: 7.65, layer: 32
Fold over basal
Time (min)
0.5
1
1.5
2
2.5
3
0
10
20
30
40
50
60
Normal
MK50
MK75
MK88
MK94
MK97
MK data
Normal data

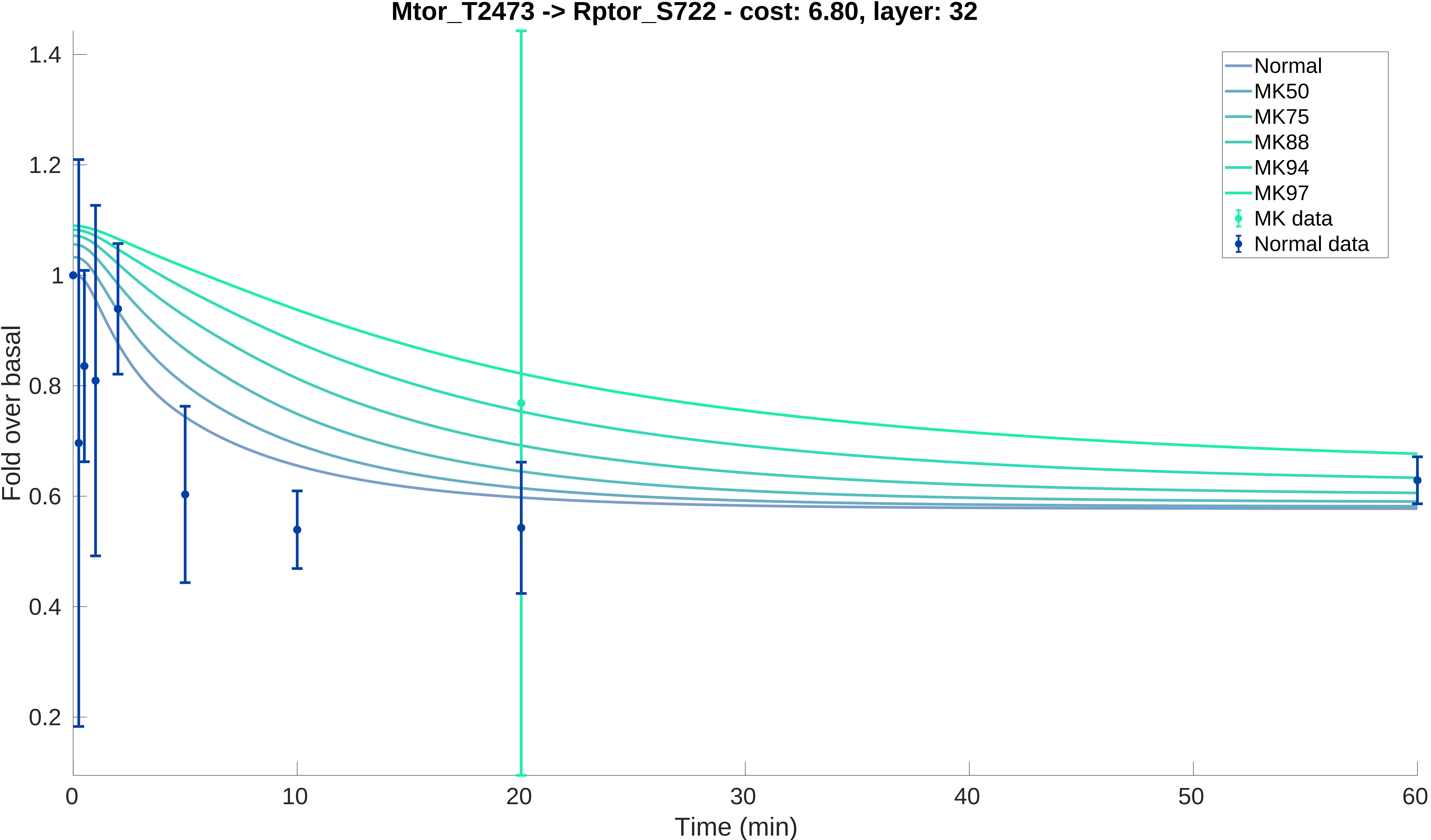

Mtor_T2473 -> Rptor_S722 - cost: 6.80, layer: 32
Normal
MK50
MK75
MK88
MK94
MK97
MK data
Normal data
Fold over basal
Time (min)
0.2
0.4
0.6
0.8
1
1.2
1.4
0
10
20
30
40
50
60

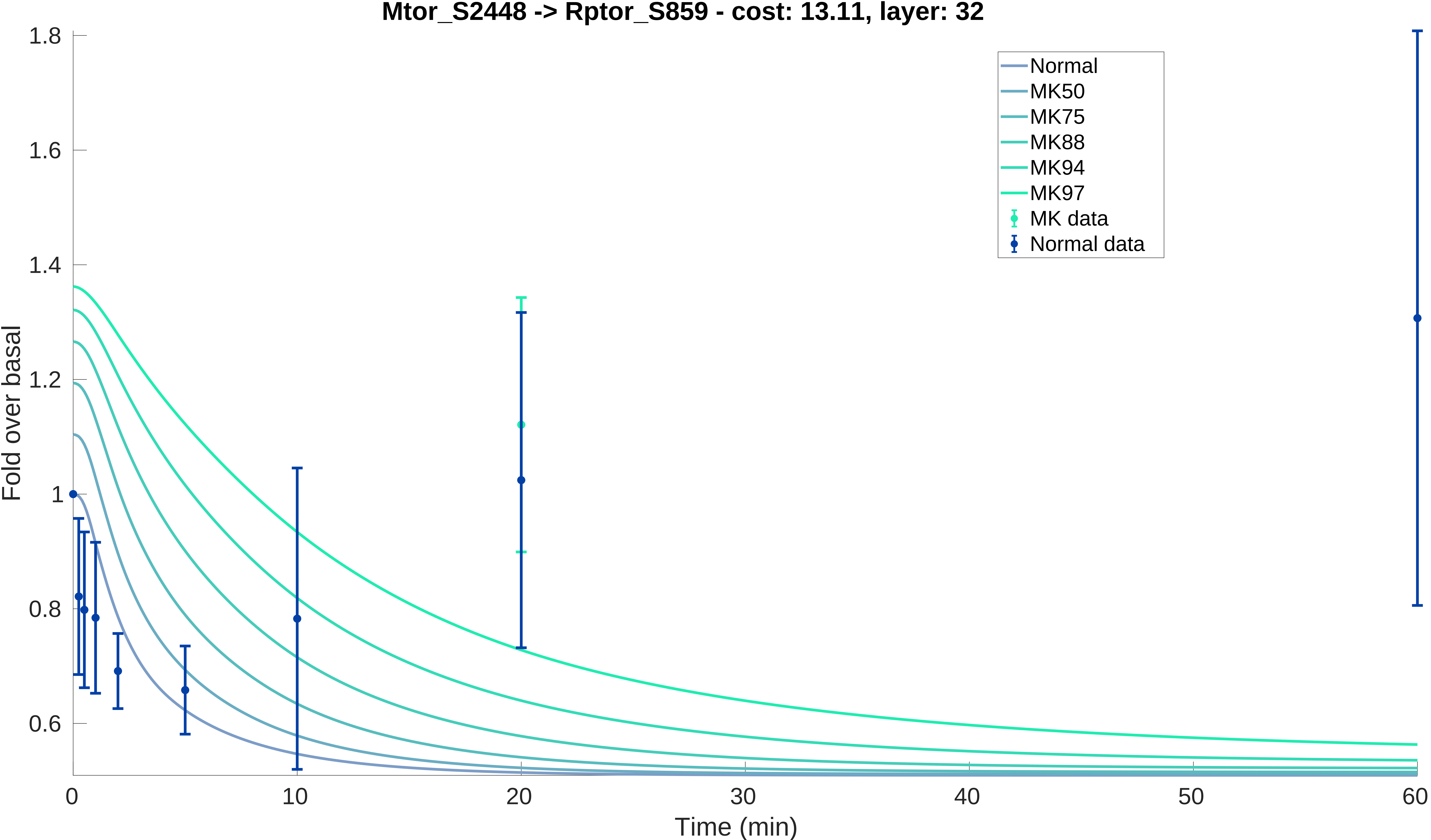

Mtor_S2448 -> Rptor_S859 - cost: 13.11, layer: 32
Normal
MK50
MK75
MK88
MK94
MK97
MK data
Normal data
Fold over basal
Time (min)
1.8
1.6
1.4
1.2
1
0.8
0.6
0
10
20
30
40
50
60

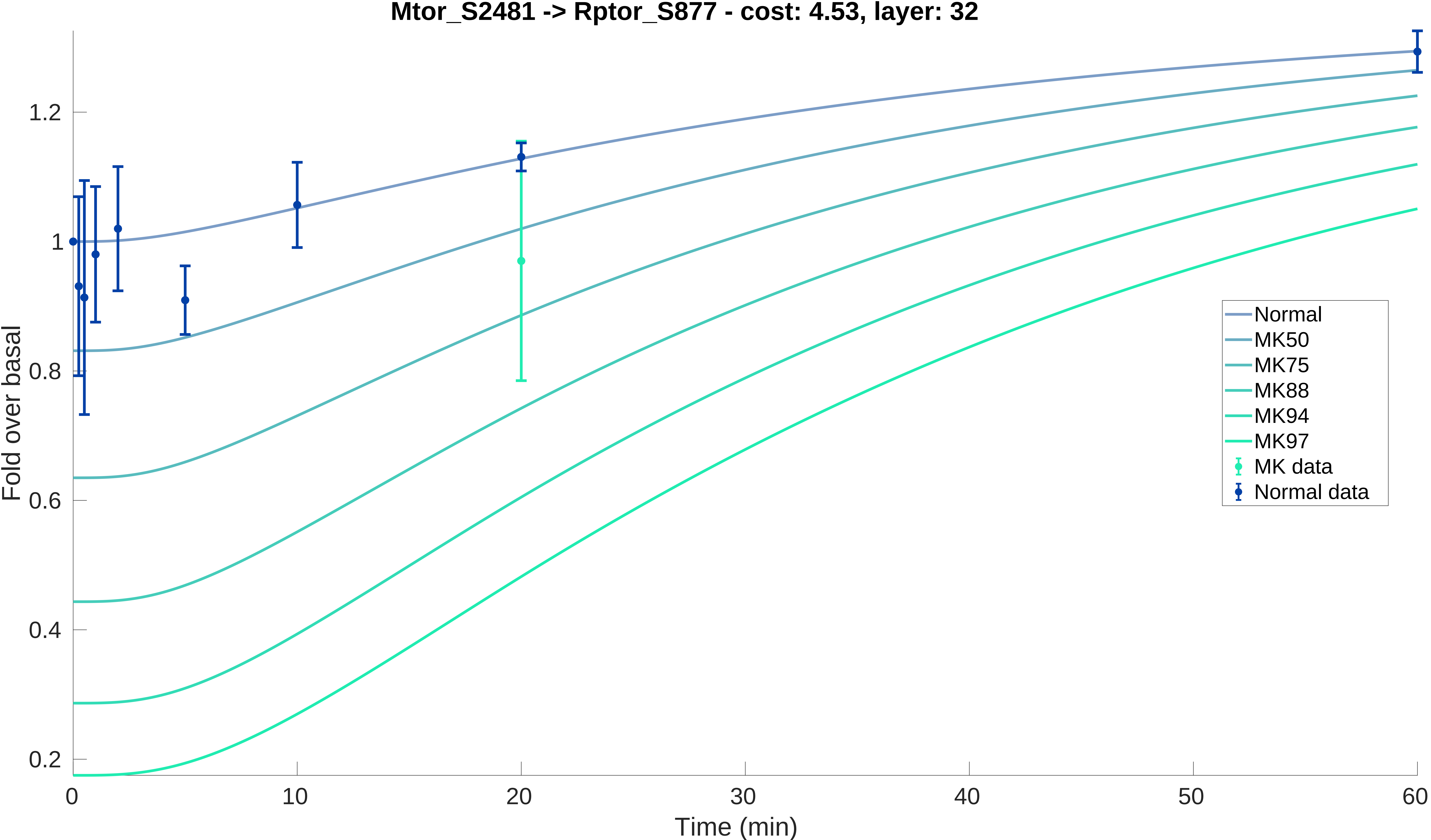

Mtor_S2481 -> Rptor_S877 - cost: 4.53, layer: 32
Fold over basal
1.2
1
0.8
0.6
0.4
0.2
0
10
20
30
40
50
60
Time (min)
Normal
MK50
MK75
MK88
MK94
MK97
MK data
Normal data

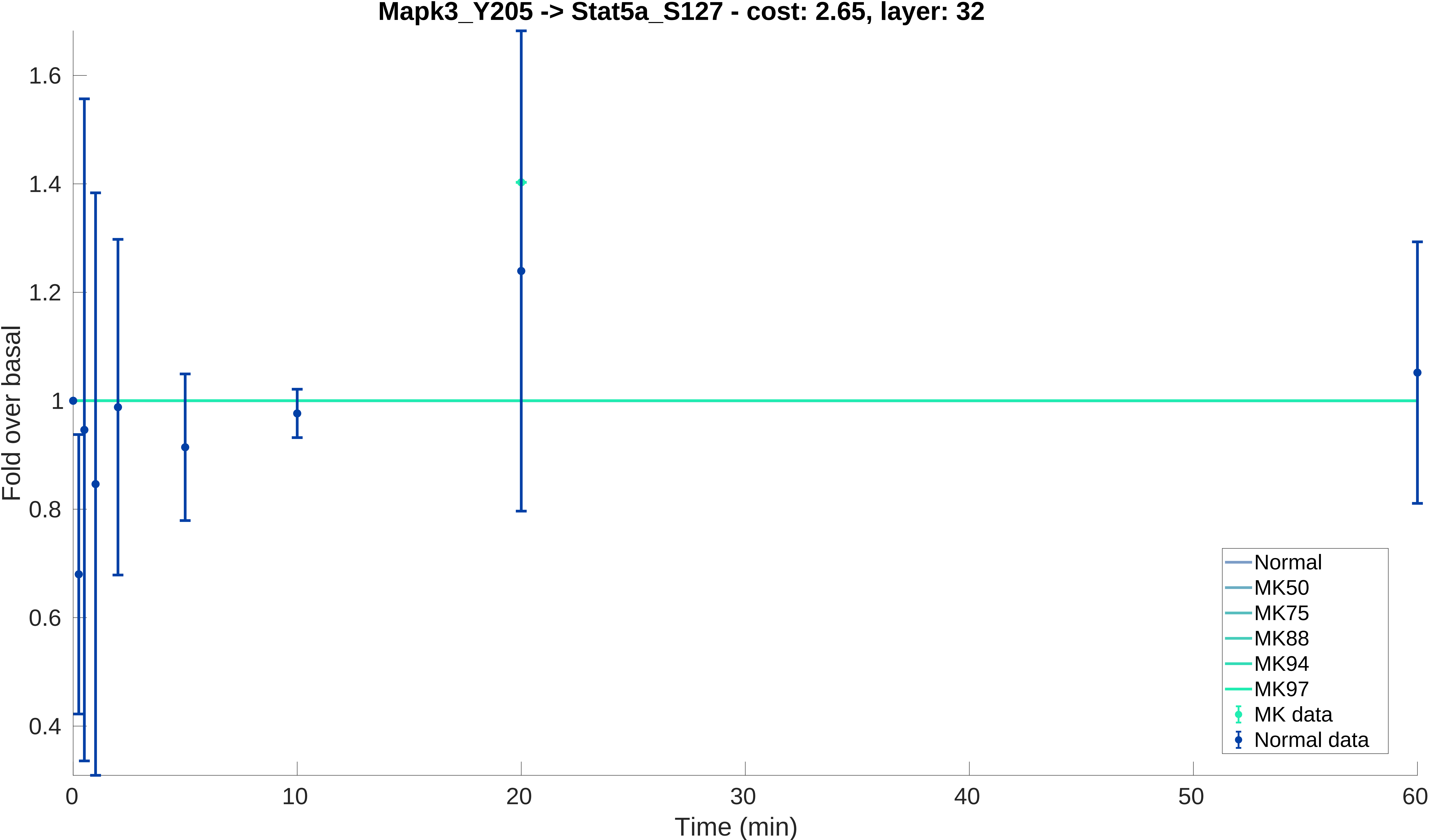

Mapk3_Y205 -> Stat5a_S127 - cost: 2.65, layer: 32
Fold over basal
Time (min)
1.6
1.4
1.2
1
0.8
0.6
0.4
0
10
20
30
40
50
60
Normal
MK50
MK75
MK88
MK94
MK97
MK data
Normal data

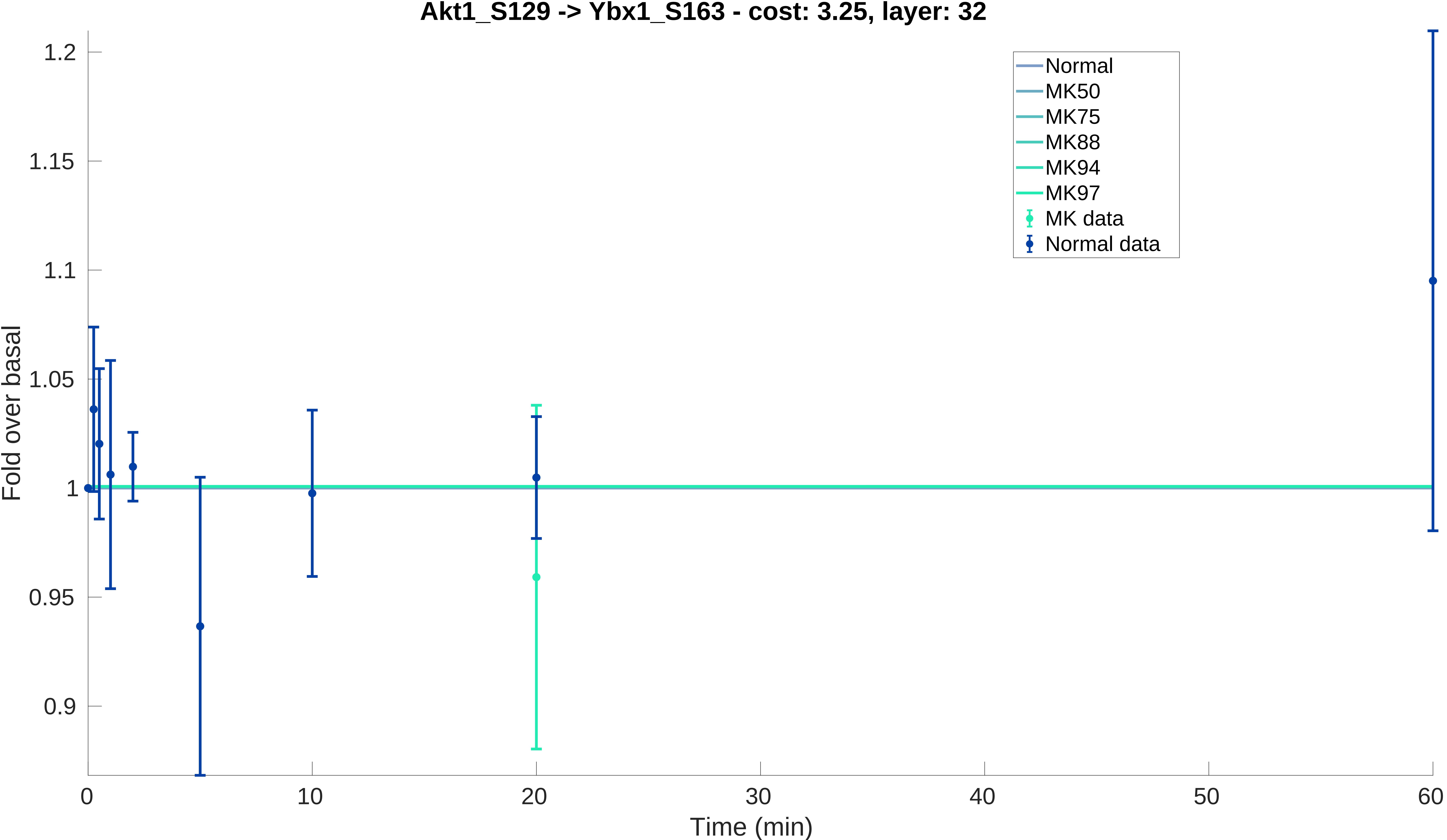

Akt1_S129 -> Ybx1_S163 - cost: 3.25, layer: 32
Fold over basal
1.2
1.15
1.1
1.05
1
0.95
0.9
0
10
20
30
40
50
60
Time (min)
Normal
MK50
MK75
MK88
MK94
MK97
MK data
Normal data

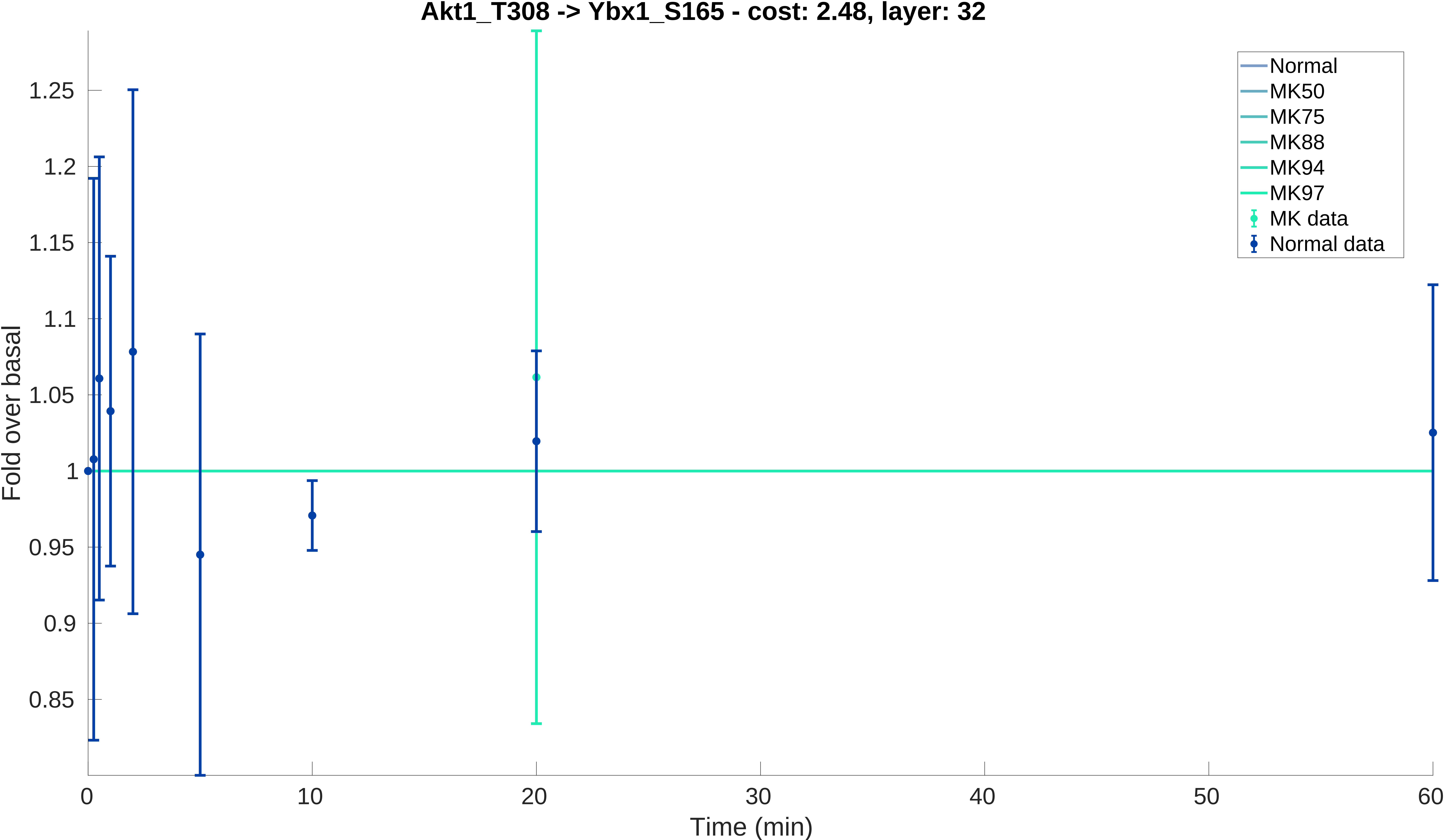

Akt1_T308 -> Ybx1_S165 - cost: 2.48, layer: 32
Normal
MK50
MK75
MK88
MK94
MK97
MK data
Normal data
Fold over basal
1.25
1.2
1.15
1.1
1.05
1
0.95
0.9
0.85
0
10
20
30
40
50
60
Time (min)

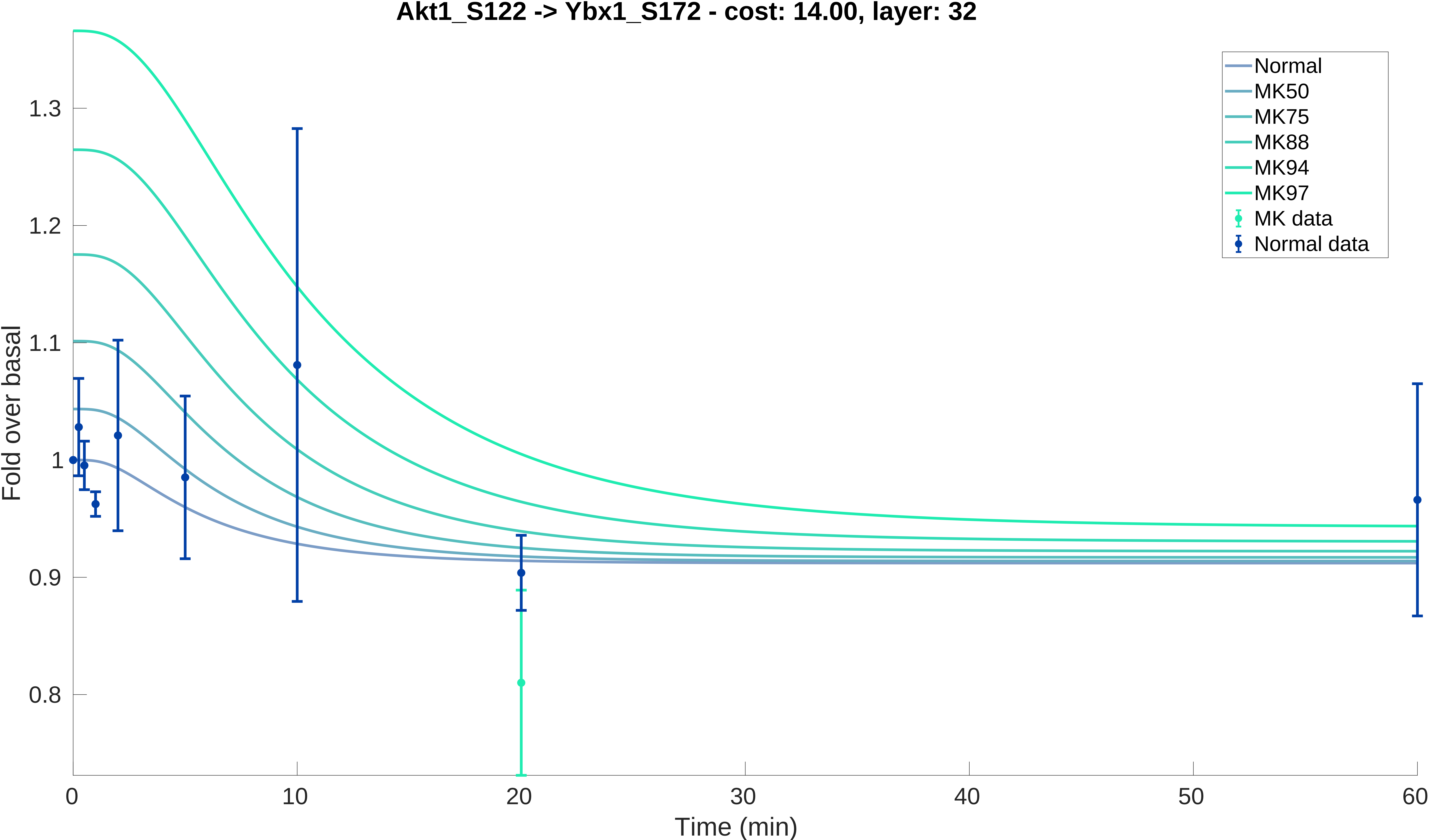
Akt1_S122 -> Ybx1_S172 - cost: 14.00, layer: 32
Normal
MK50
MK75
MK88
MK94
MK97
MK data
Normal data
1.3
1.2
1.1
1
0.9
0.8
Fold over basal
0
10
20
30
40
50
60
Time (min)

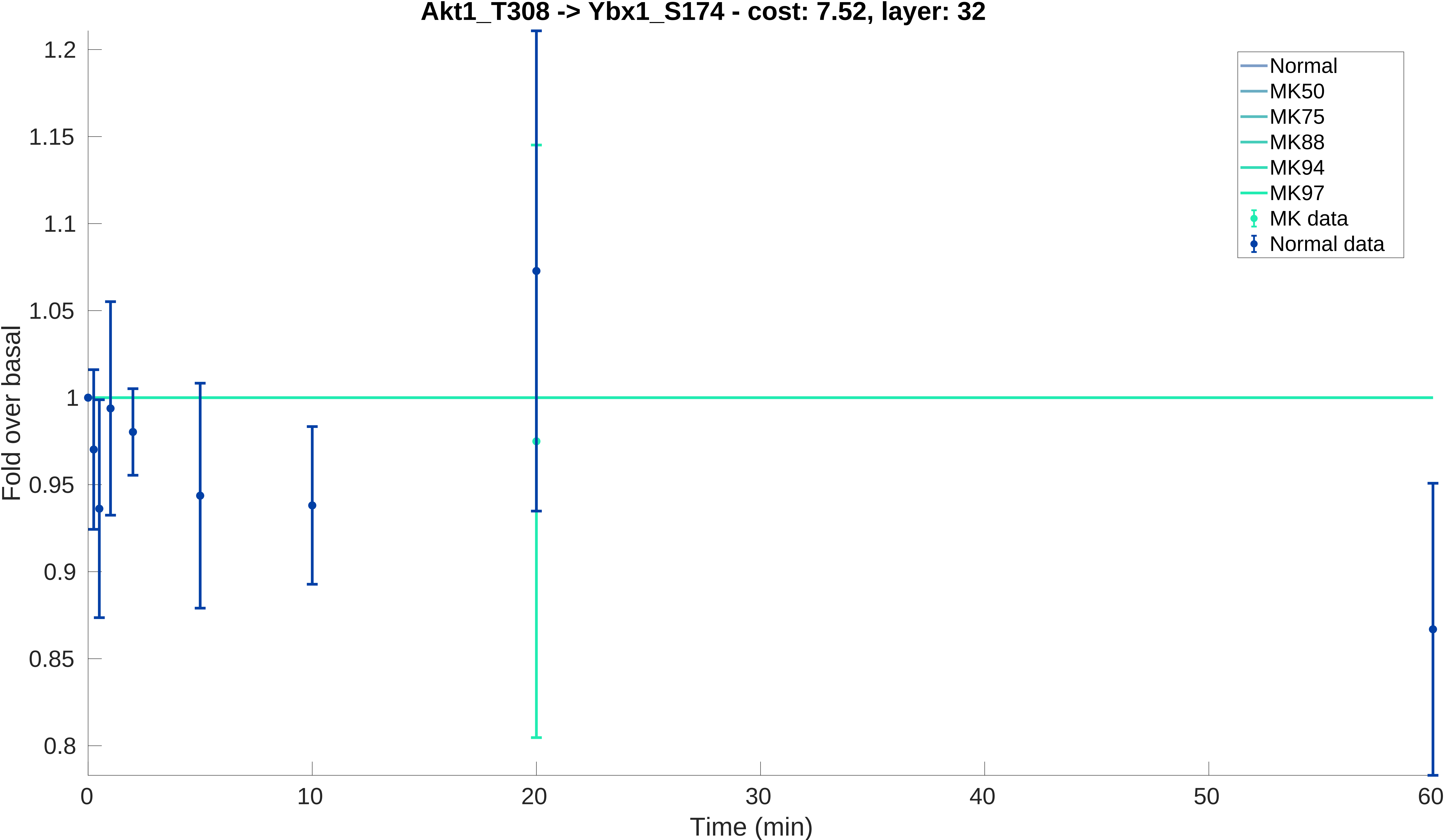
Akt1_T308 -> Ybx1_S174 - cost: 7.52, layer: 32
Fold over basal
1.2
1.15
1.1
1.05
1
0.95
0.9
0.85
0.8
0
10
20
30
40
50
60
Time (min)
Normal
MK50
MK75
MK88
MK94
MK97
MK data
Normal data

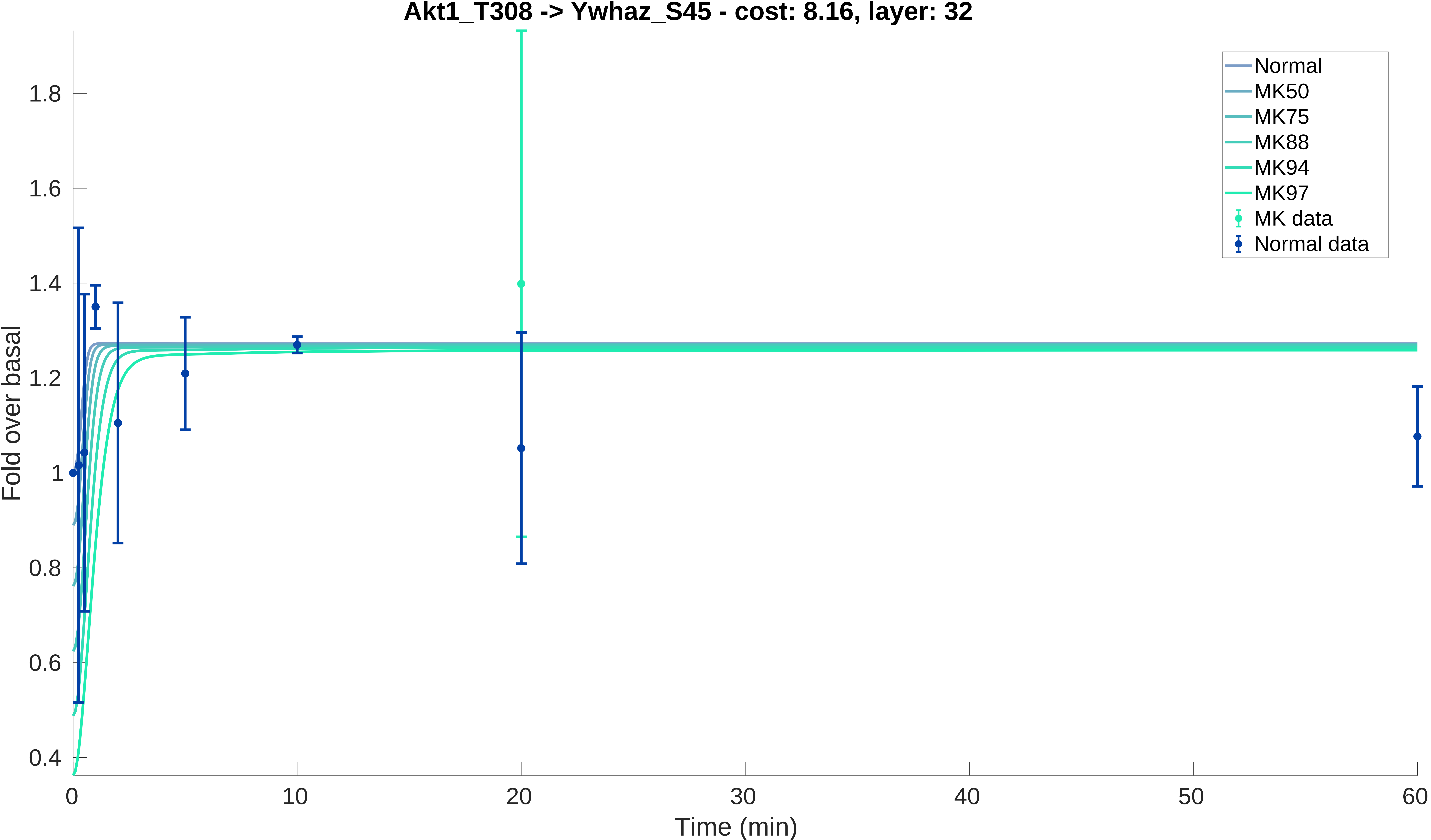

Akt1_T308 -> Ywhaz_S45 - cost: 8.16, layer: 32
Normal
MK50
MK75
MK88
MK94
MK97
MK data
Normal data
Fold over basal
Time (min)
1.8
1.6
1.4
1.2
1
0.8
0.6
0.4
0
10
20
30
40
50
60

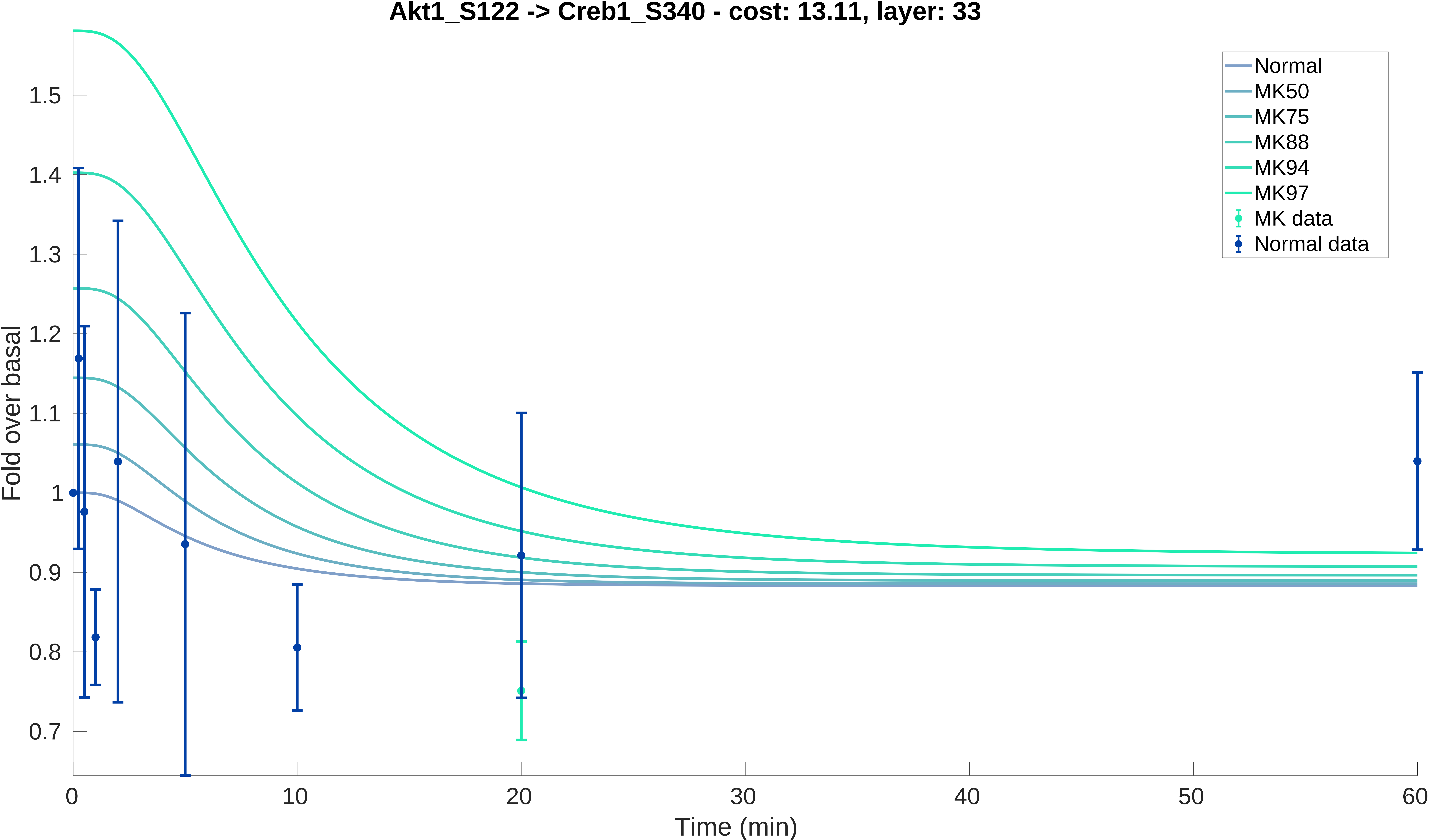

Akt1_S122 -> Creb1_S340 - cost: 13.11, layer: 33
Fold over basal
1.5
1.4
1.3
1.2
1.1
1
0.9
0.8
0.7
Time (min)
0
10
20
30
40
50
60
Normal
MK50
MK75
MK88
MK94
MK97
MK data
Normal data

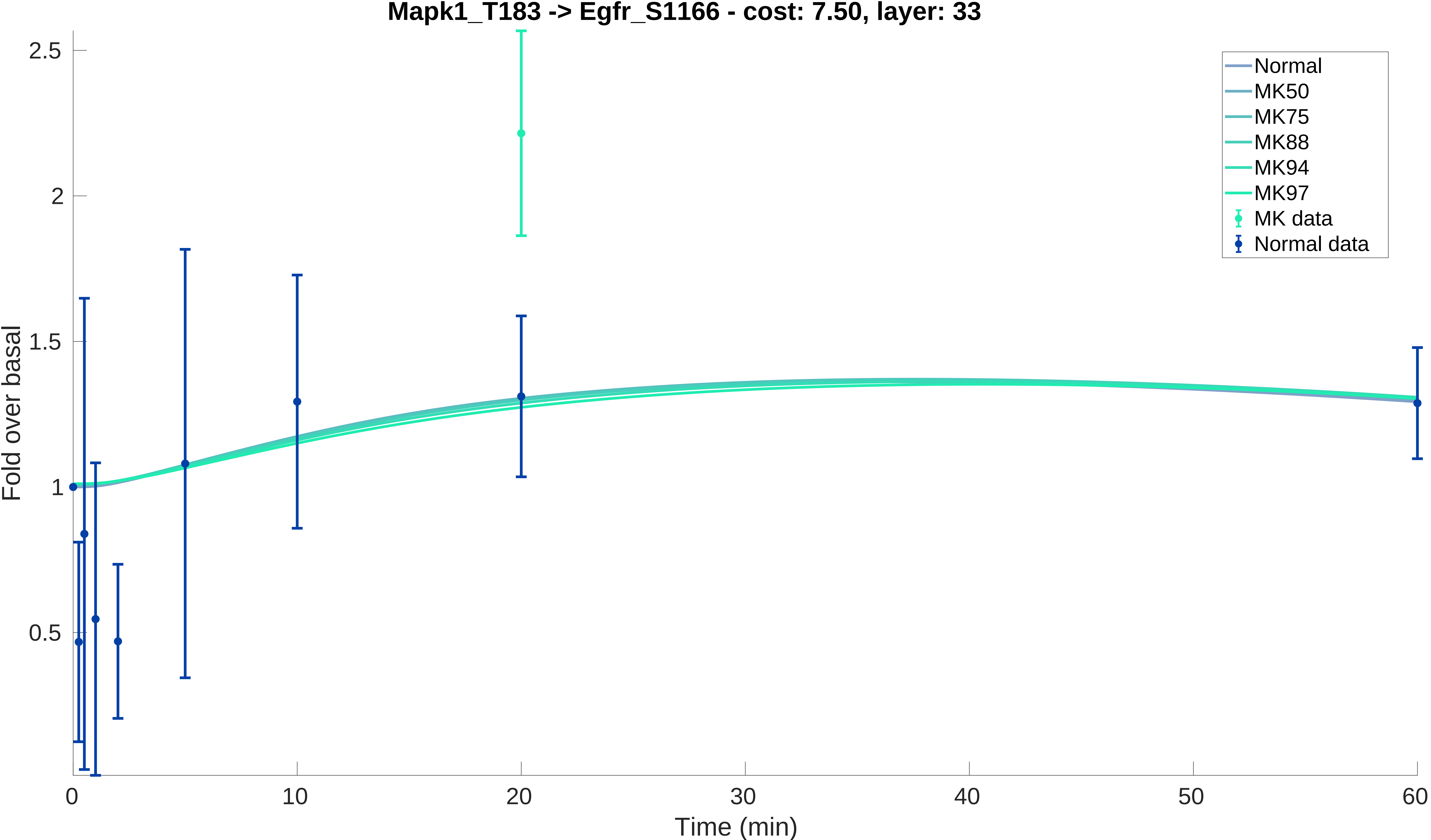

Mapk1_T183 -> Egfr_S1166 - cost: 7.50, layer: 33
Fold over basal
Time (min)
0.5
1
1.5
2
2.5
0
10
20
30
40
50
60
Normal
MK50
MK75
MK88
MK94
MK97
MK data
Normal data

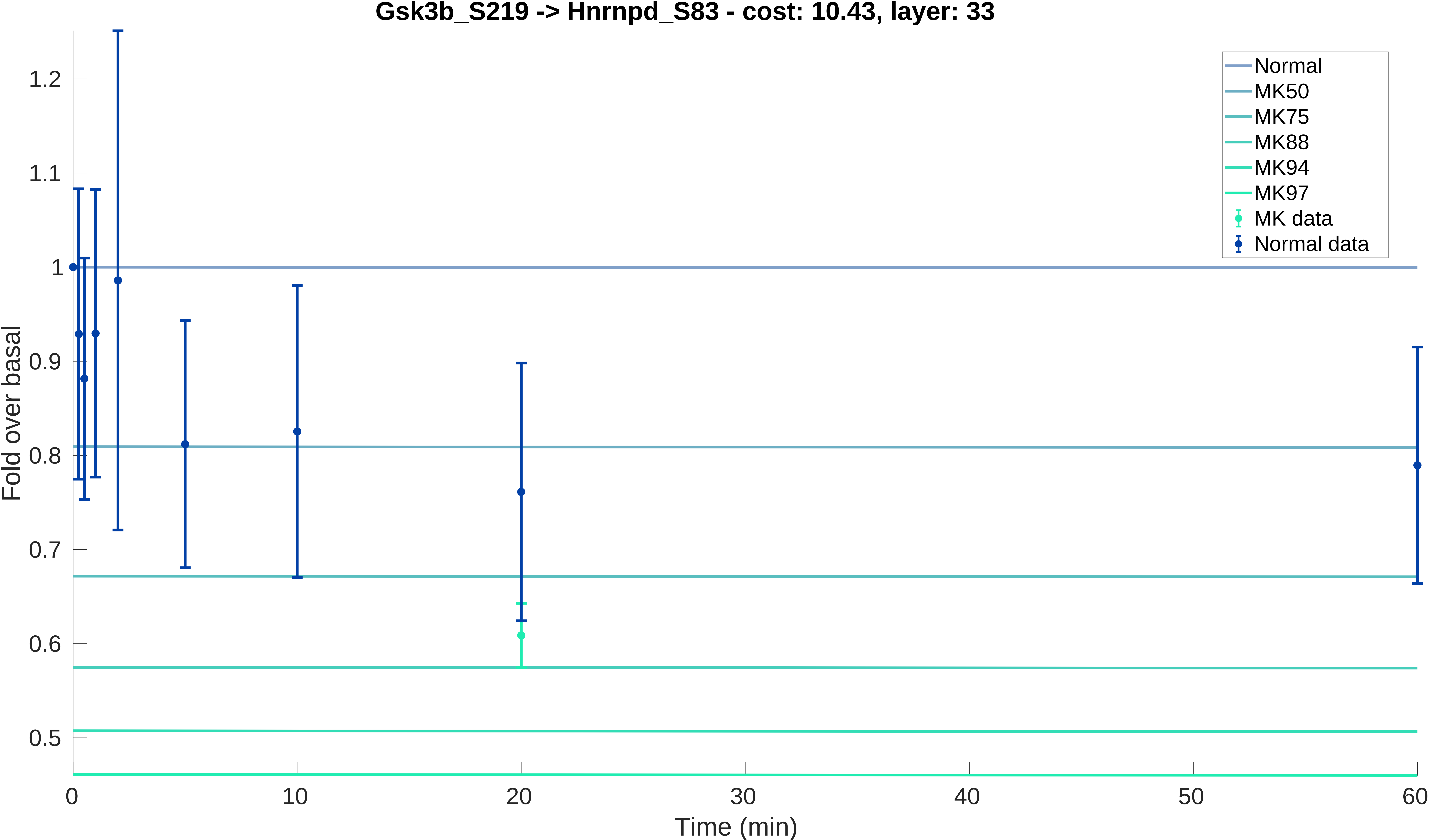

Gsk3b_S219 -> Hnrnpd_S83 - cost: 10.43, layer: 33
Normal
MK50
MK75
MK88
MK94
MK97
MK data
Normal data
Fold over basal
1.2
1.1
1
0.9
0.8
0.7
0.6
0.5
0
10
20
30
40
50
60
Time (min)

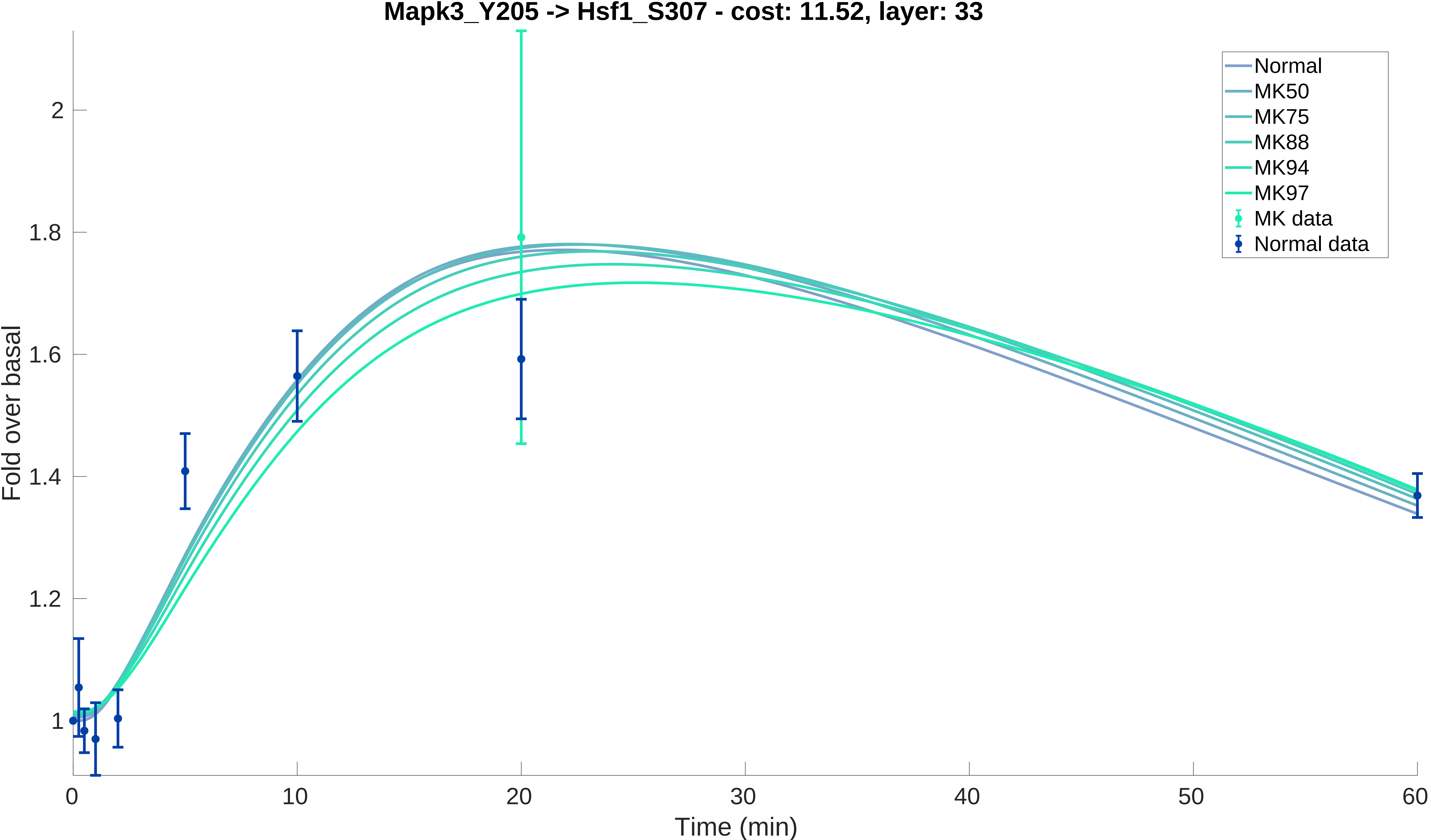

Mapk3_Y205 -> Hsf1_S307 - cost: 11.52, layer: 33
Normal
MK50
MK75
MK88
MK94
MK97
MK data
Normal data
Fold over basal
2
1.8
1.6
1.4
1.2
1
0
10
20
30
40
50
60
Time (min)

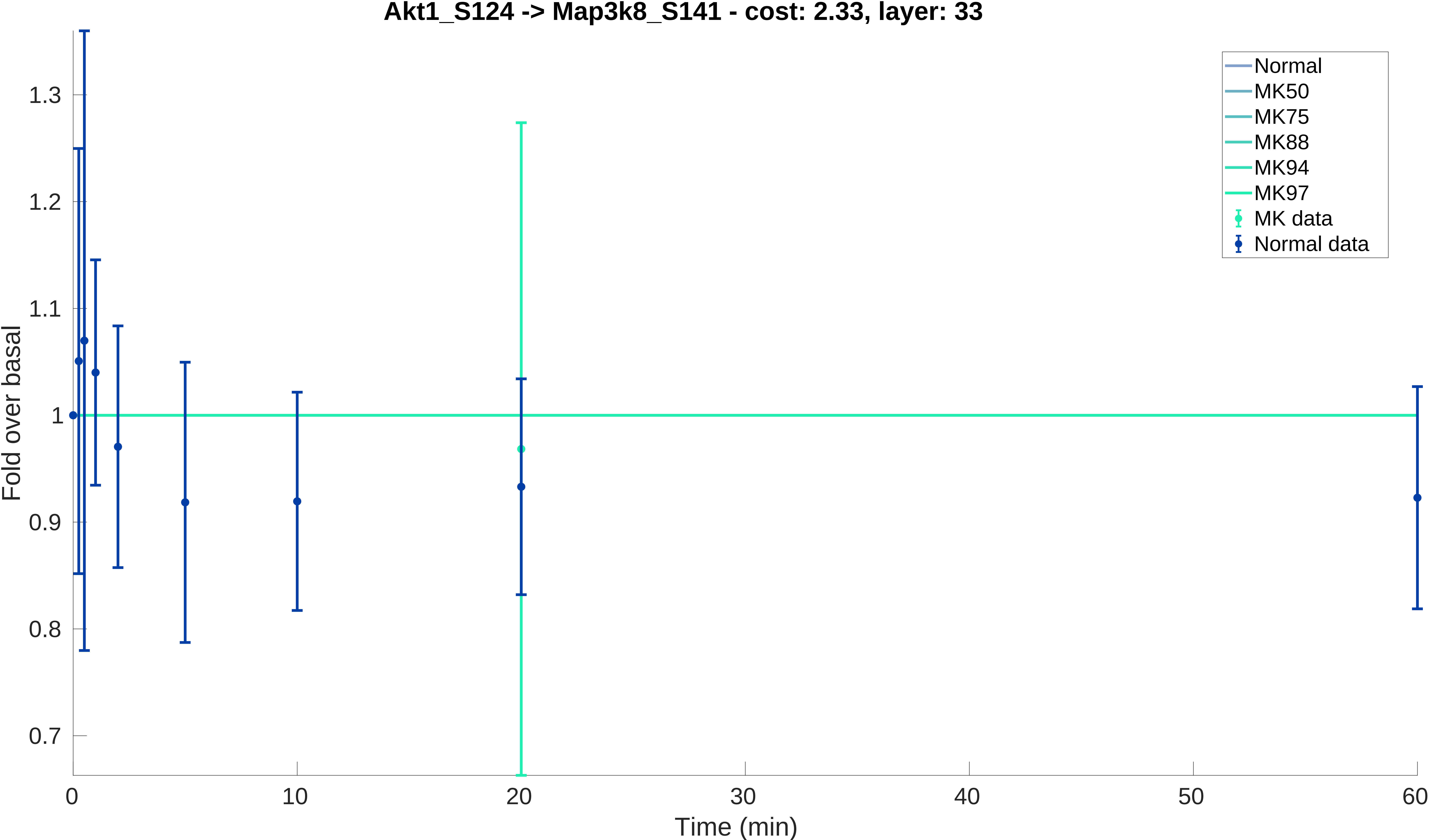

Akt1_S124 -> Map3k8_S141 - cost: 2.33, layer: 33
Fold over basal
Time (min)
1.3
1.2
1.1
1
0.9
0.8
0.7
0
10
20
30
40
50
60
Normal
MK50
MK75
MK88
MK94
MK97
MK data
Normal data

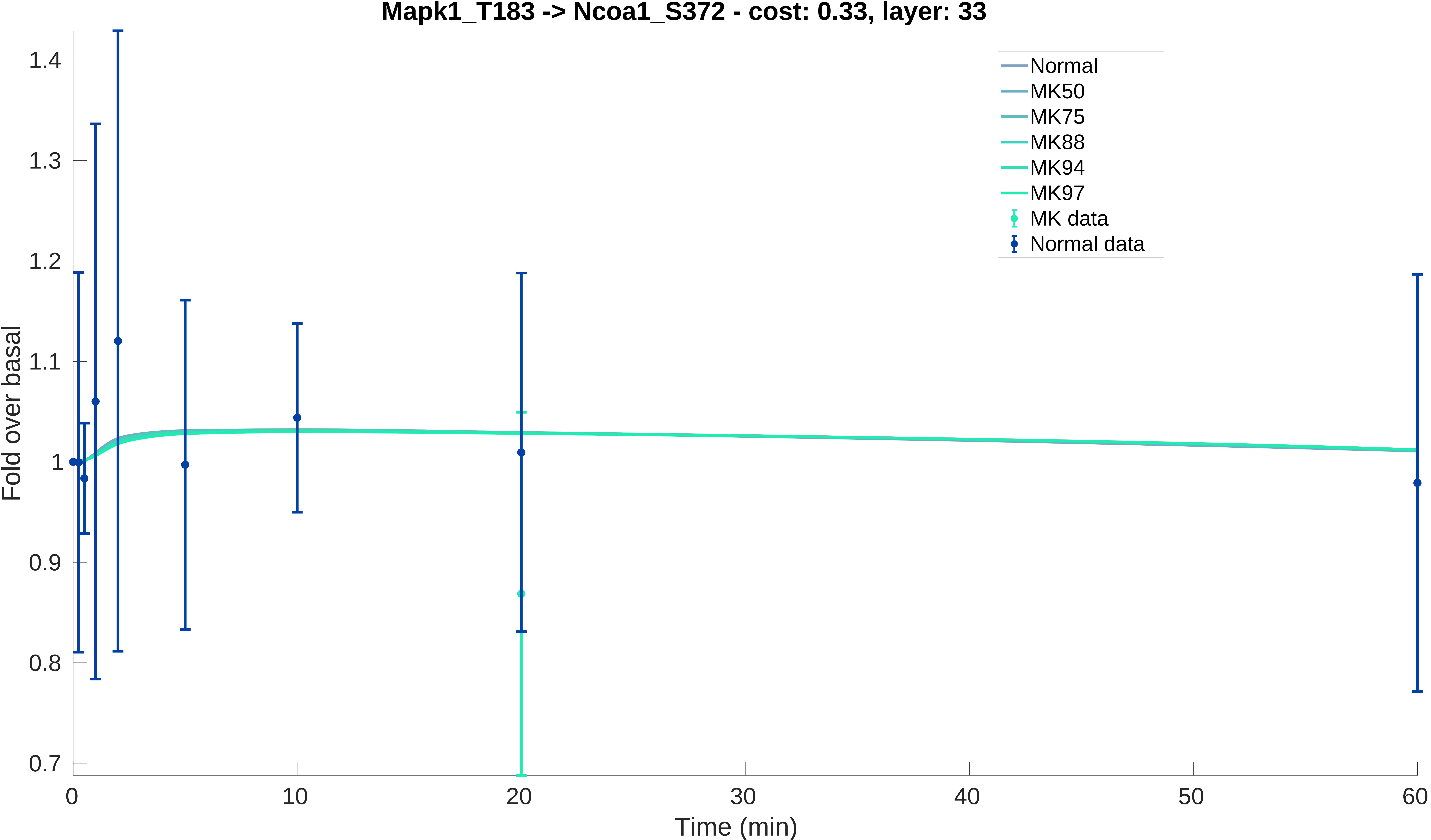

Mapk1_T183 -> Ncoa1_S372 - cost: 0.33, layer: 33
Fold over basal
Time (min)
Normal
MK50
MK75
MK88
MK94
MK97
MK data
Normal data
0.7
0.8
0.9
1
1.1
1.2
1.3
1.4
0
10
20
30
40
50
60

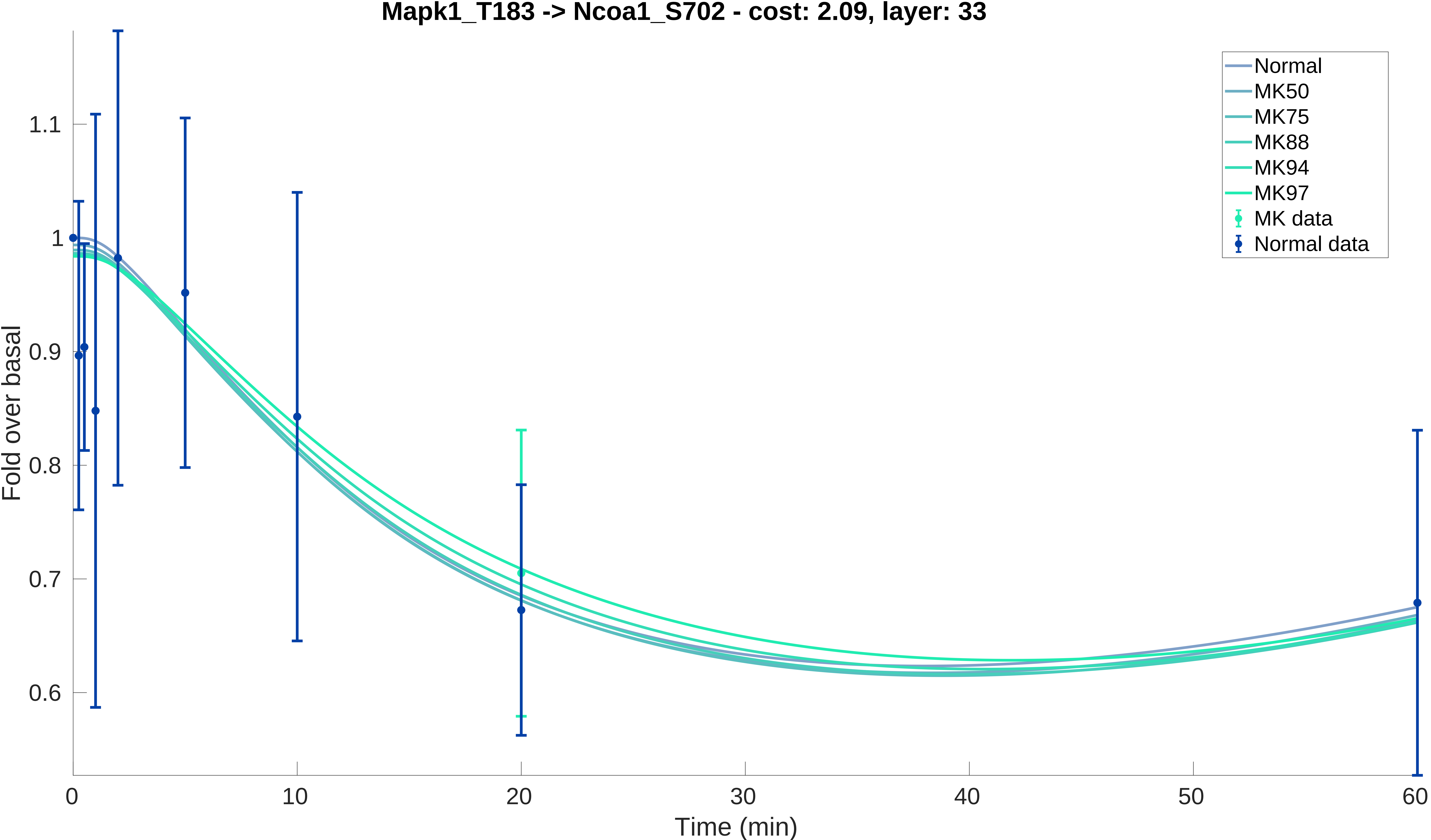
Mapk1_T183 -> Ncoa1_S702 - cost: 2.09, layer: 33
Fold over basal
Time (min)
0.6
0.7
0.8
0.9
1
1.1
0
10
20
30
40
50
60
Normal
MK50
MK75
MK88
MK94
MK97
MK data
Normal data

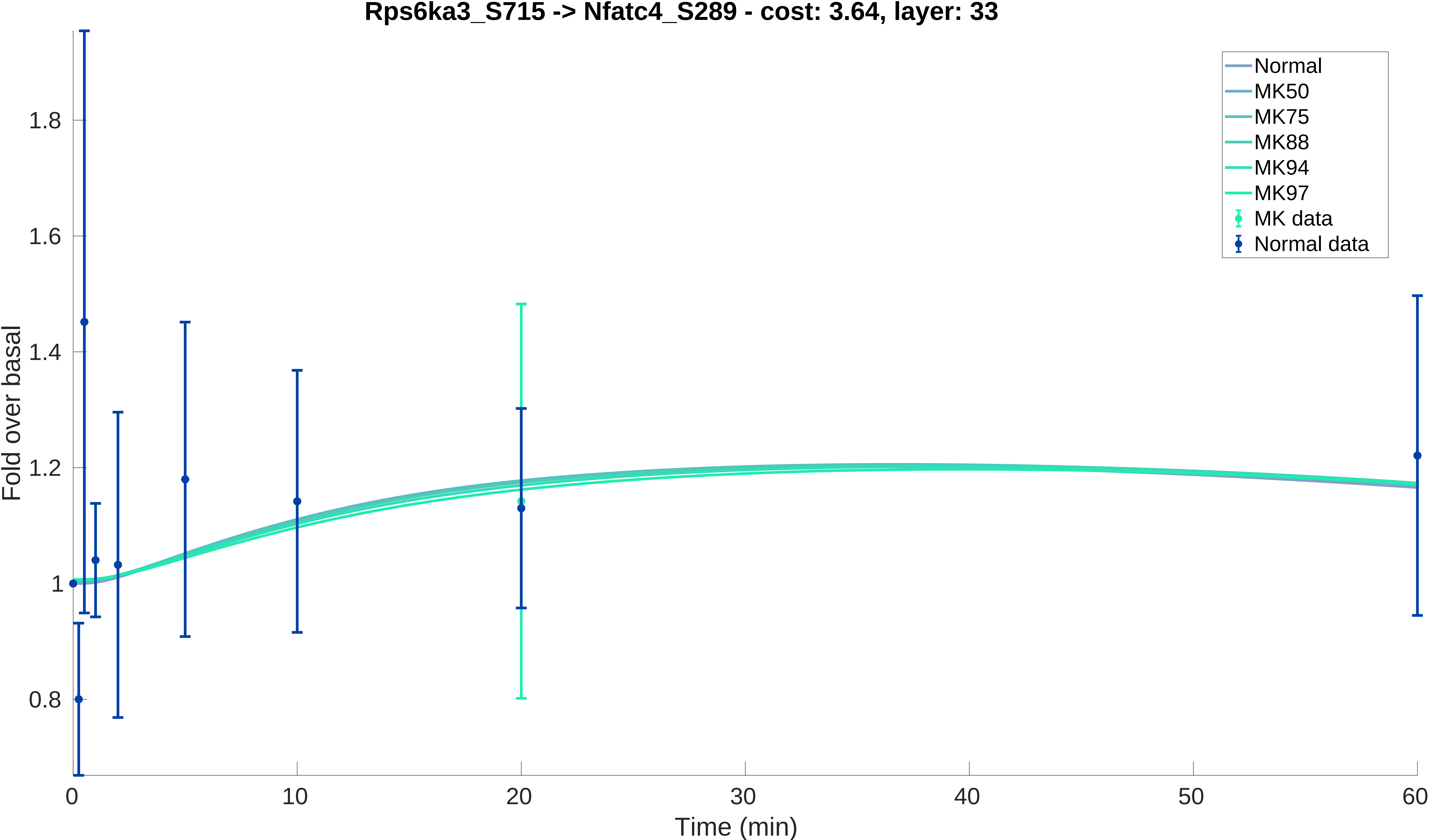

Rps6ka3_S715 -> Nfatc4_S289 - cost: 3.64, layer: 33
Normal
MK50
MK75
MK88
MK94
MK97
MK data
Normal data
Fold over basal
Time (min)
1.8
1.6
1.4
1.2
1
0.8
0
10
20
30
40
50
60

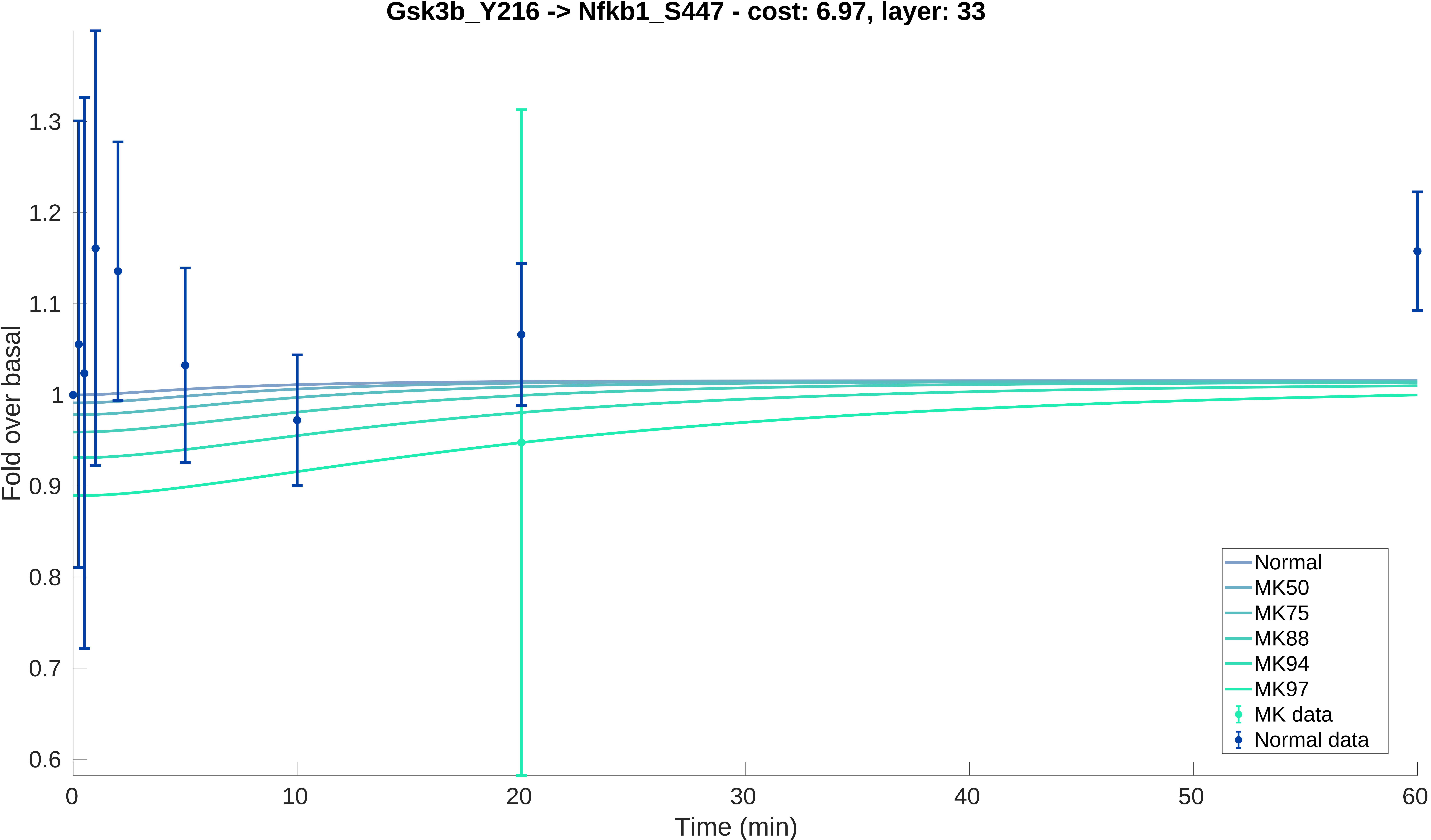

Gsk3b_Y216 -> Nfkb1_S447 - cost: 6.97, layer: 33
Fold over basal
Time (min)
0.6
0.7
0.8
0.9
1
1.1
1.2
1.3
0
10
20
30
40
50
60
Normal
MK50
MK75
MK88
MK94
MK97
MK data
Normal data

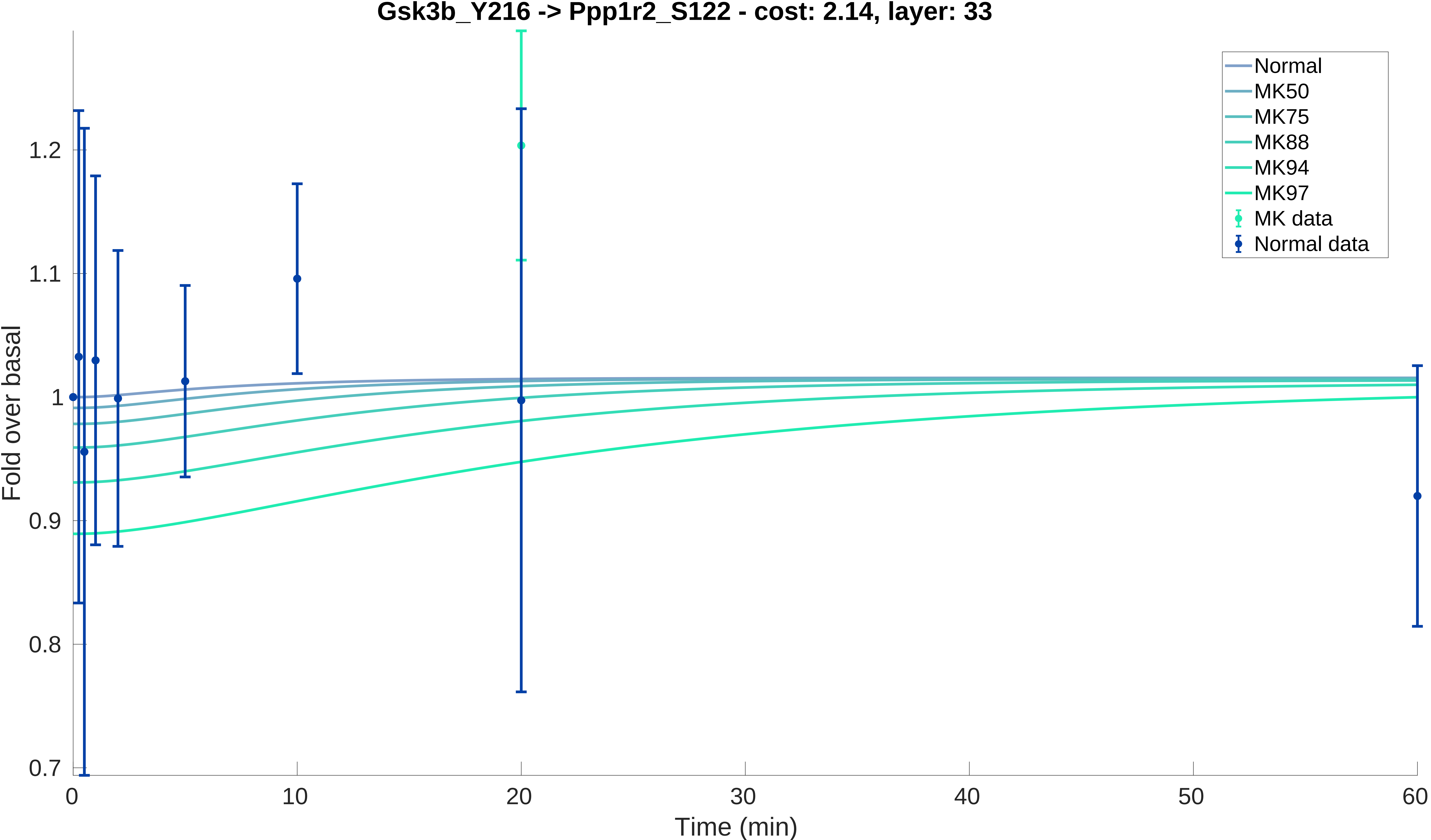

Gsk3b_Y216 -> Ppp1r2_S122 - cost: 2.14, layer: 33
Fold over basal
Time (min)
0.7
0.8
0.9
1
1.1
1.2
0
10
20
30
40
50
60
Normal
MK50
MK75
MK88
MK94
MK97
MK data
Normal data

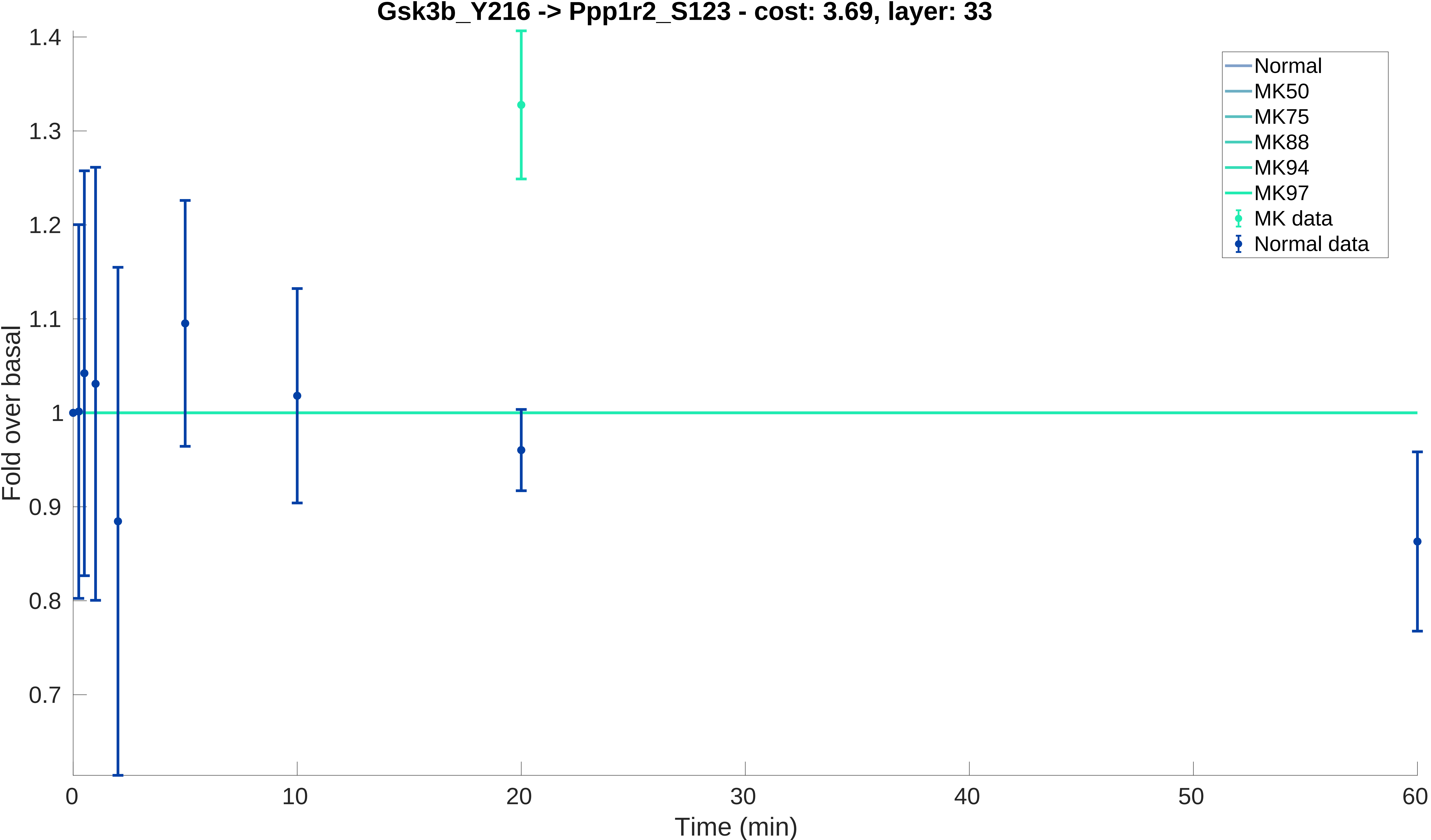
Gsk3b_Y216 -> Ppp1r2_S123 - cost: 3.69, layer: 33
Normal
MK50
MK75
MK88
MK94
MK97
MK data
Normal data
Fold over basal
Time (min)
1.4
1.3
1.2
1.1
1
0.9
0.8
0.7
0
10
20
30
40
50
60

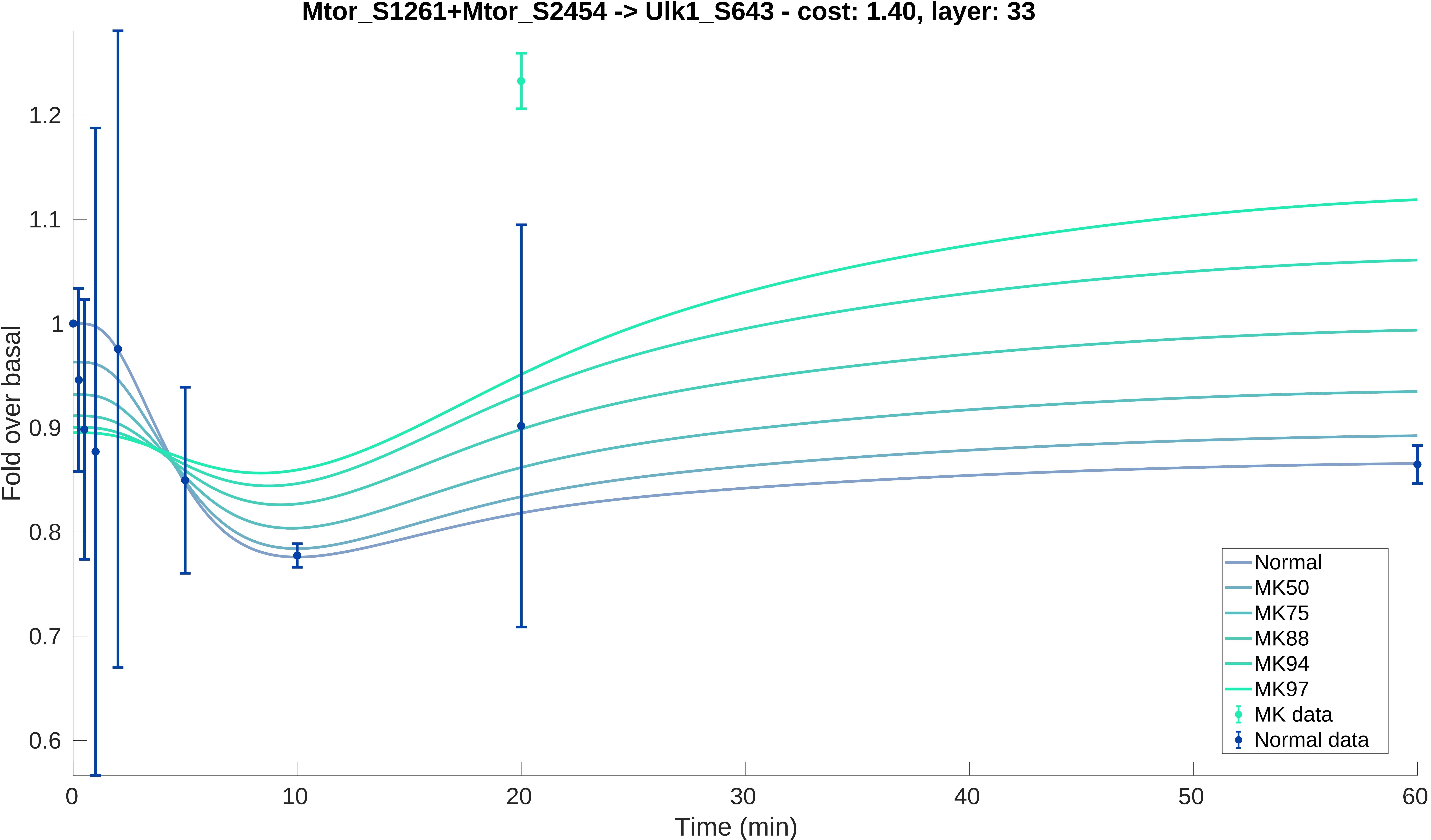

Mtor_S1261+Mtor_S2454 -> Ulk1_S643 - cost: 1.40, layer: 33
Fold over basal
Time (min)
1.2
1.1
1
0.9
0.8
0.7
0.6
0
10
20
30
40
50
60
Normal
MK50
MK75
MK88
MK94
MK97
MK data
Normal data

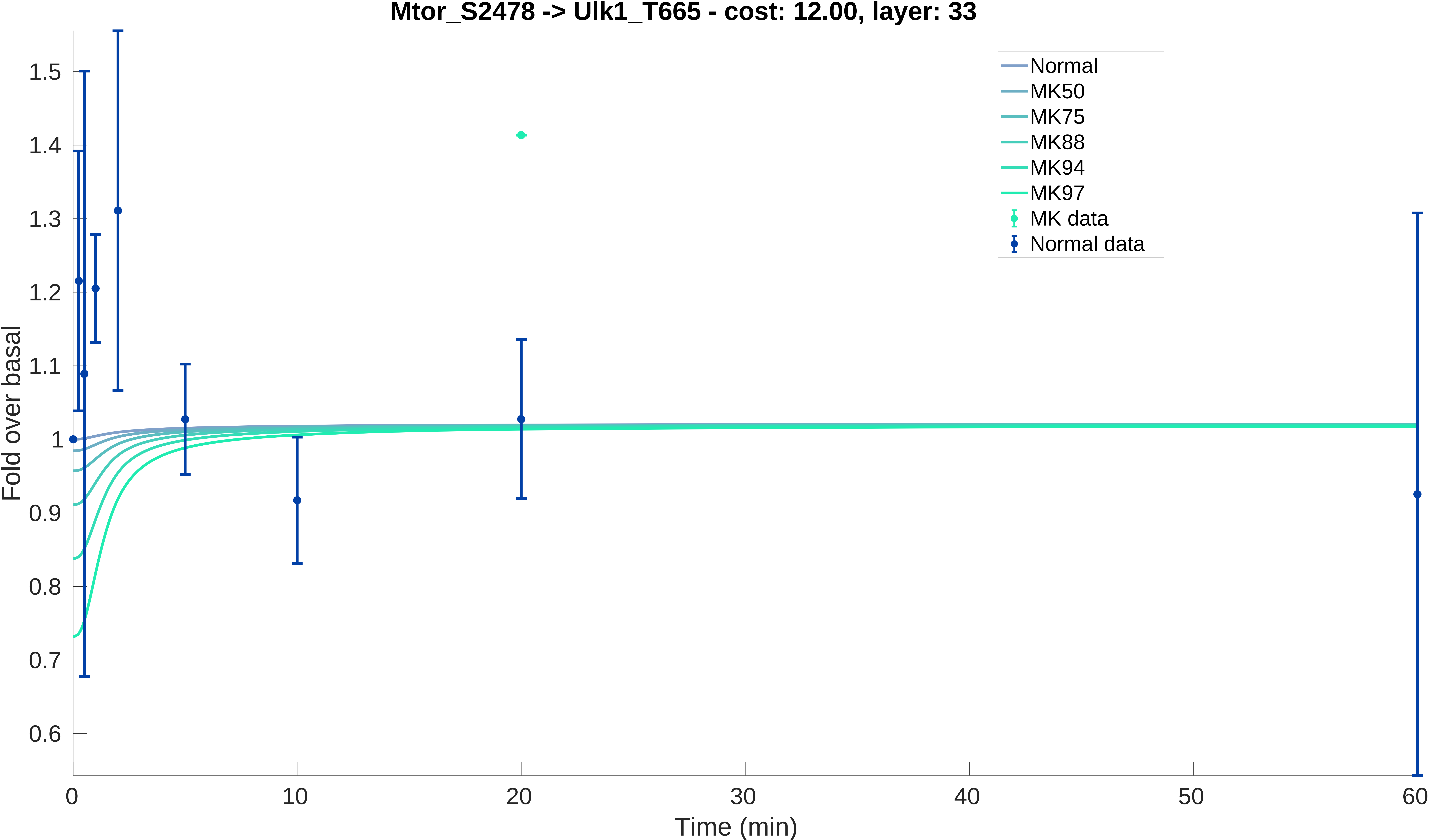

Mtor_S2478 -> Ulk1_T665 - cost: 12.00, layer: 33
Normal
MK50
MK75
MK88
MK94
MK97
MK data
Normal data
Fold over basal
1.5
1.4
1.3
1.2
1.1
1
0.9
0.8
0.7
0.6
0
10
20
30
40
50
60
Time (min)

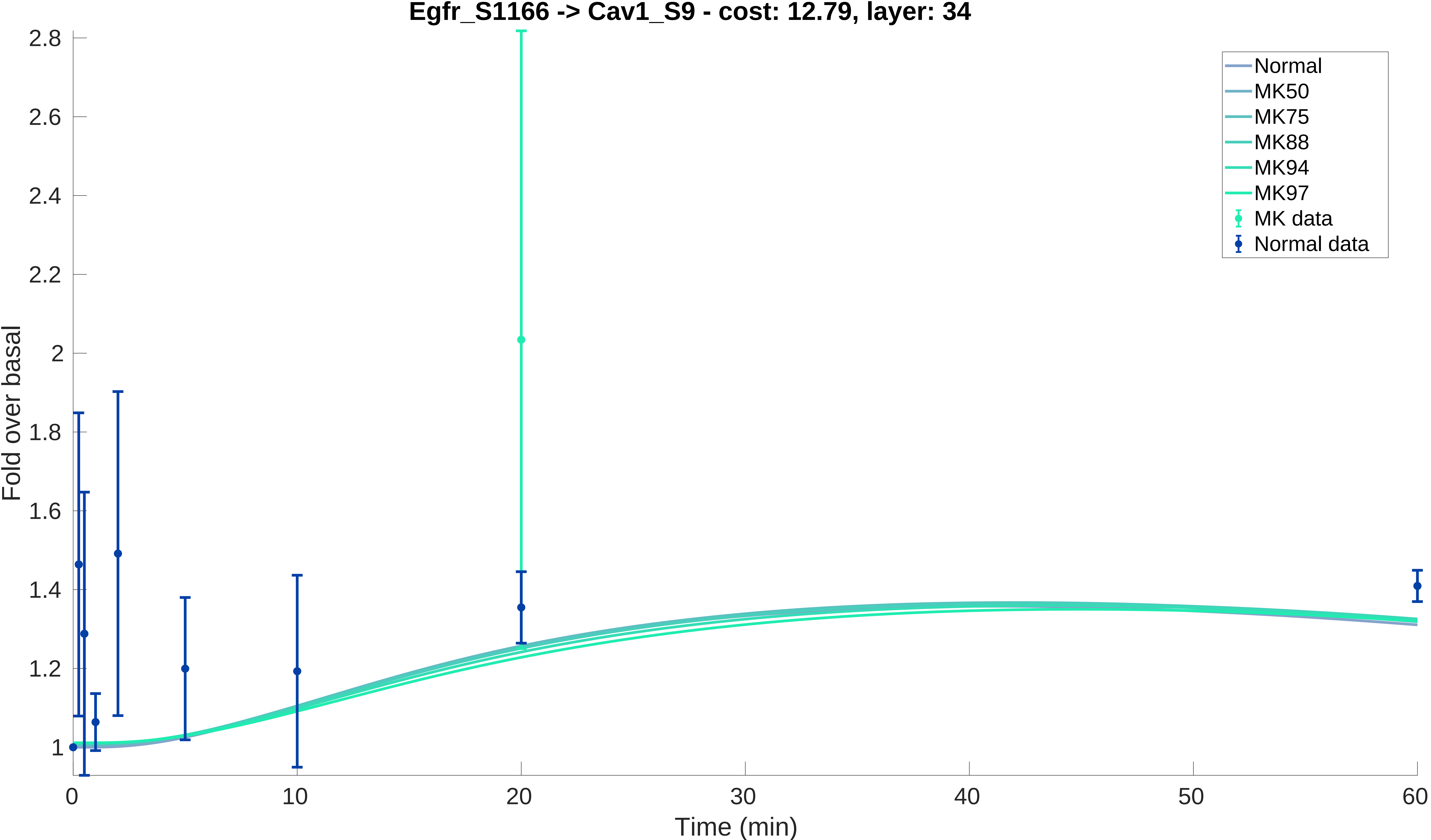

Egfr_S1166 -> Cav1_S9 - cost: 12.79, layer: 34
Fold over basal
2.8
2.6
2.4
2.2
2
1.8
1.6
1.4
1.2
1
0
10
20
30
40
50
60
Time (min)
Normal
MK50
MK75
MK88
MK94
MK97
MK data
Normal data

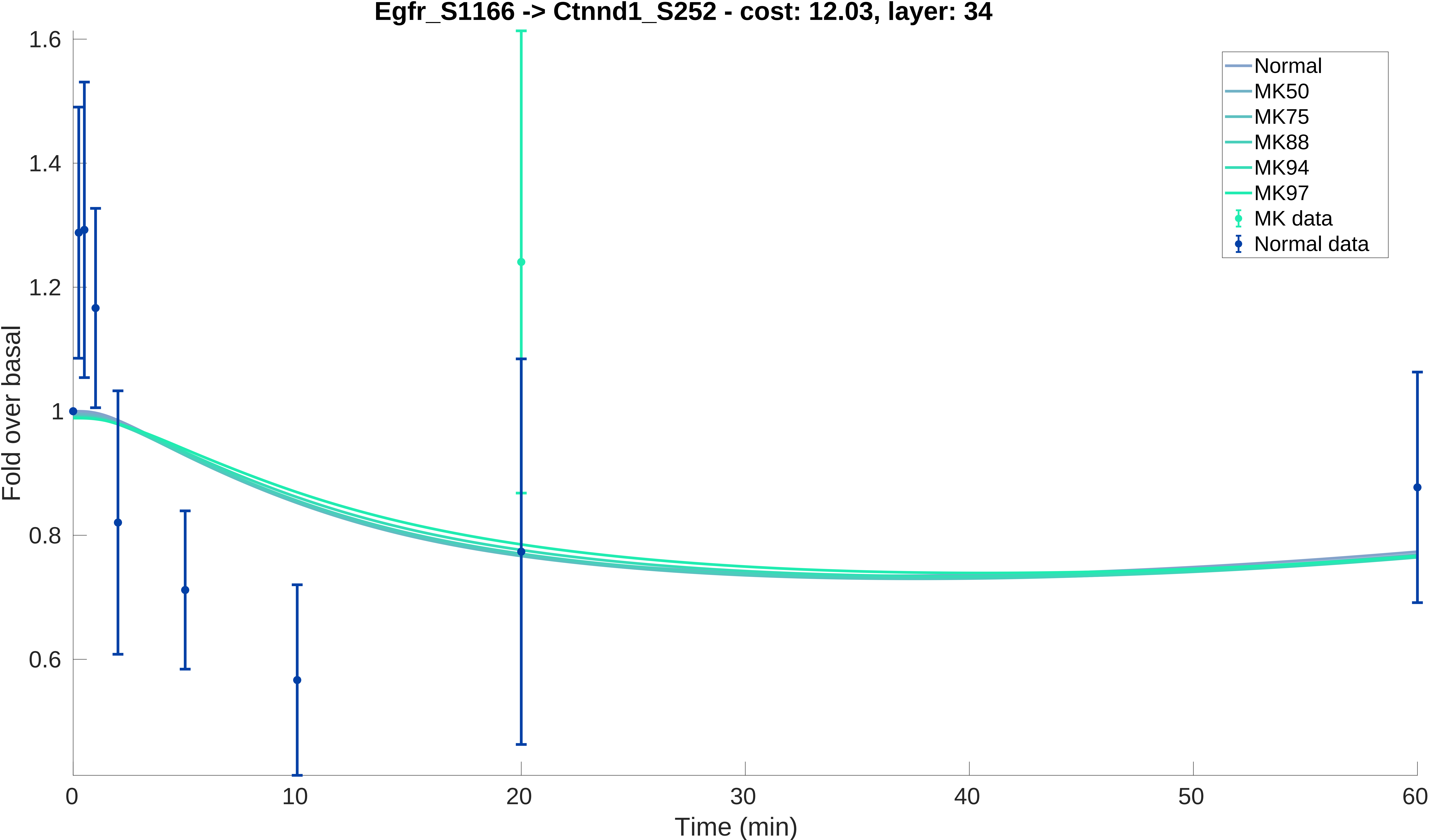

Egfr_S1166 -> Ctnnd1_S252 - cost: 12.03, layer: 34
Fold over basal
1.6
1.4
1.2
1
0.8
0.6
0
10
20
30
40
50
60
Time (min)
Normal
MK50
MK75
MK88
MK94
MK97
MK data
Normal data

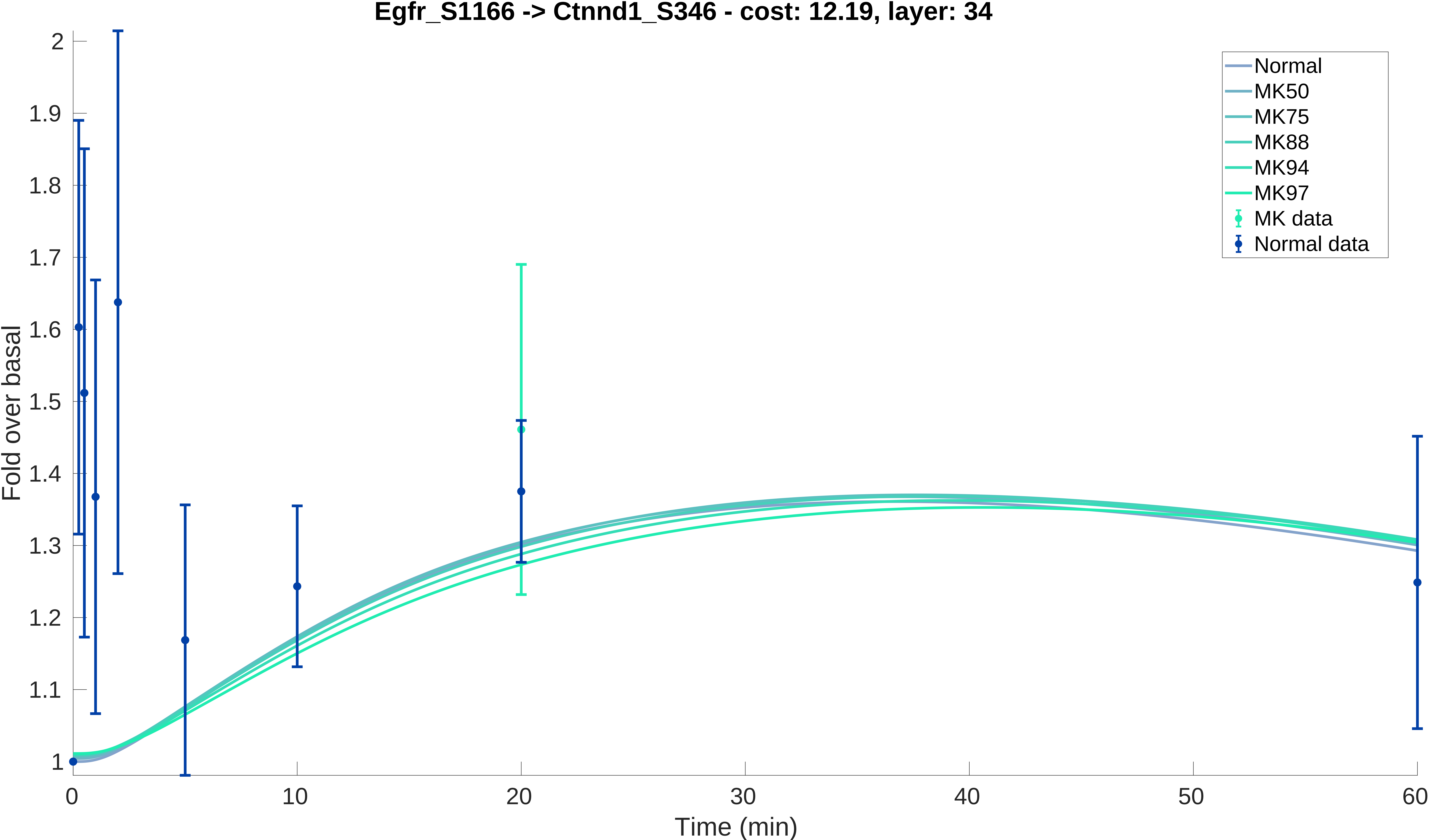

Egfr_S1166 -> Ctnnd1_S346 - cost: 12.19, layer: 34
Fold over basal
Time (min)
Normal
MK50
MK75
MK88
MK94
MK97
MK data
Normal data
2
1.9
1.8
1.7
1.6
1.5
1.4
1.3
1.2
1.1
1
0
10
20
30
40
50
60

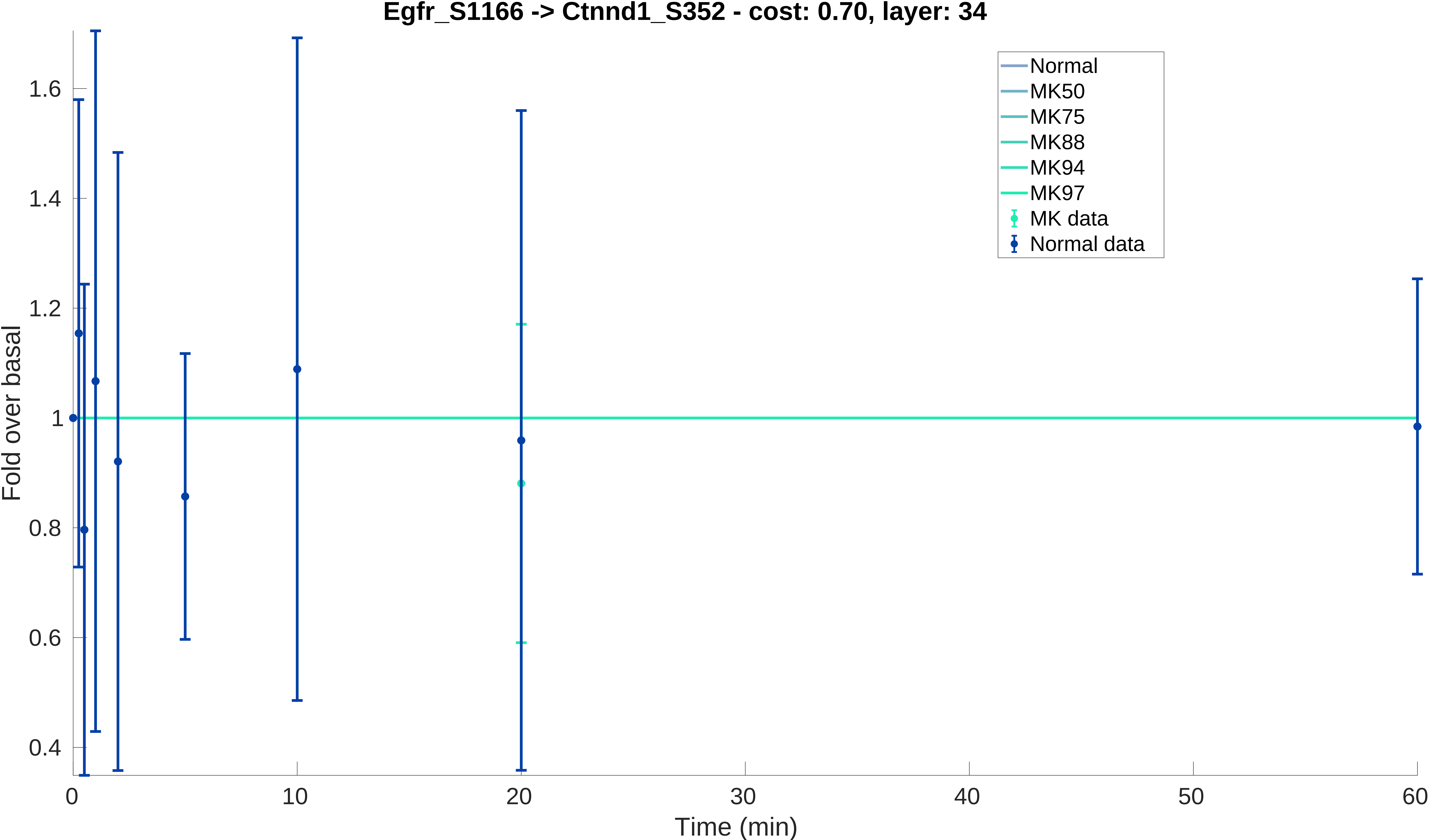

Egfr_S1166 -> Ctnnd1_S352 - cost: 0.70, layer: 34
Fold over basal
Time (min)
0
10
20
30
40
50
60
0.4
0.6
0.8
1
1.2
1.4
1.6
Normal
MK50
MK75
MK88
MK94
MK97
MK data
Normal data

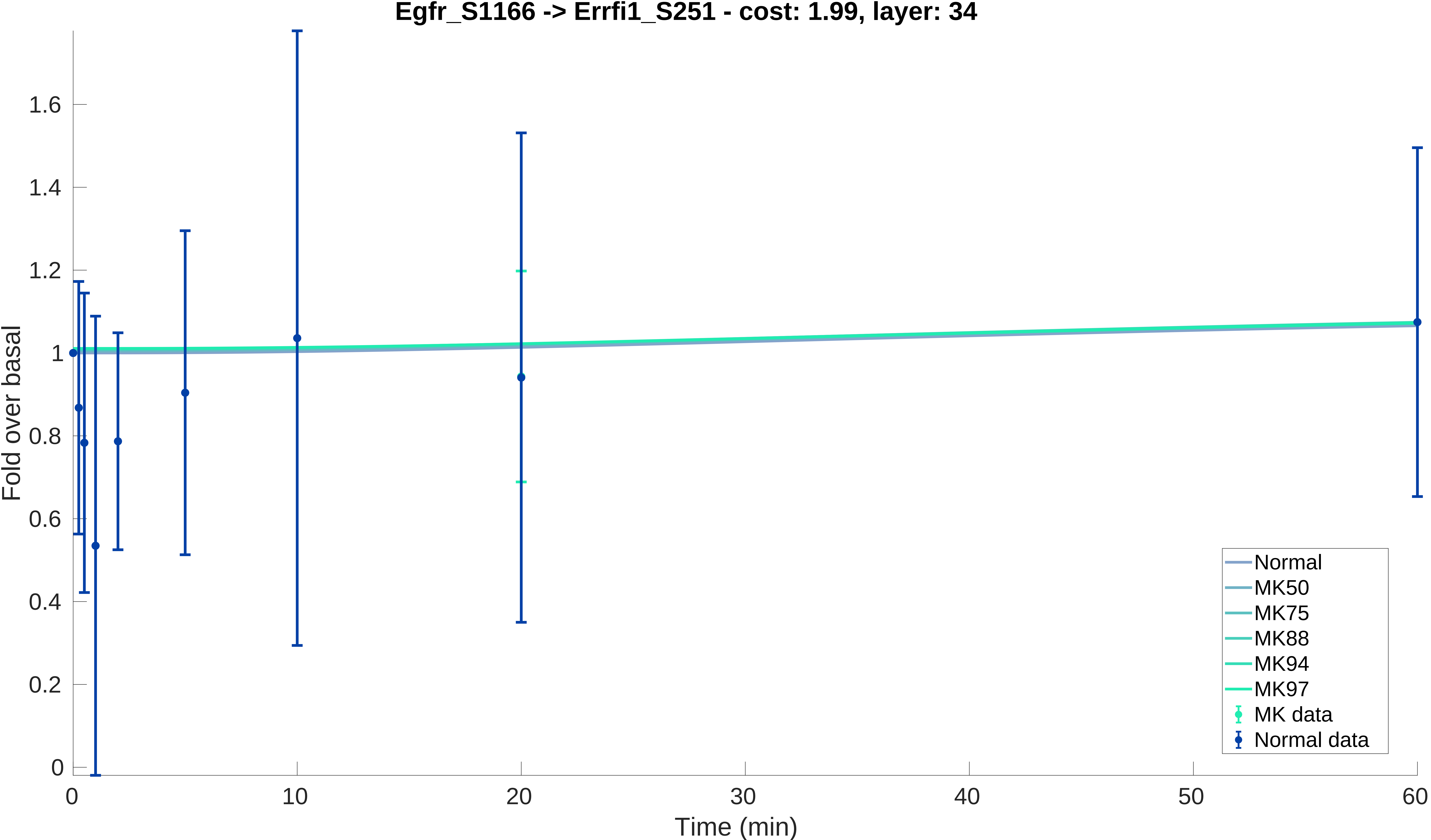

Egfr_S1166 -> Errfi1_S251 - cost: 1.99, layer: 34
Fold over basal
0
0.2
0.4
0.6
0.8
1
1.2
1.4
1.6
Time (min)
0
10
20
30
40
50
60
Normal
MK50
MK75
MK88
MK94
MK97
MK data
Normal data

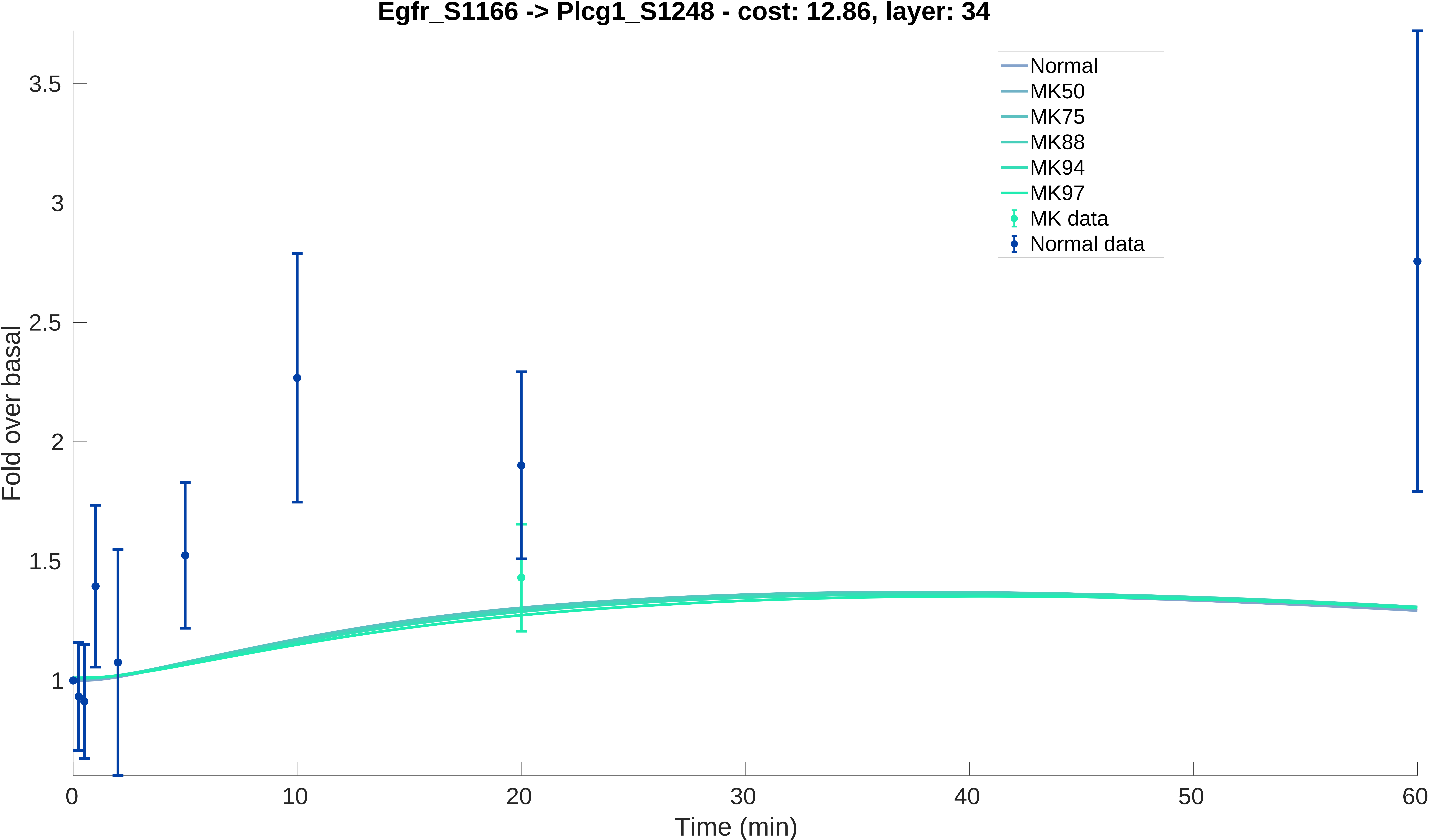

Egfr_S1166 -> Plcg1_S1248 - cost: 12.86, layer: 34
Fold over basal
Time (min)
0
10
20
30
40
50
60
1
1.5
2
2.5
3
3.5
Normal
MK50
MK75
MK88
MK94
MK97
MK data
Normal data

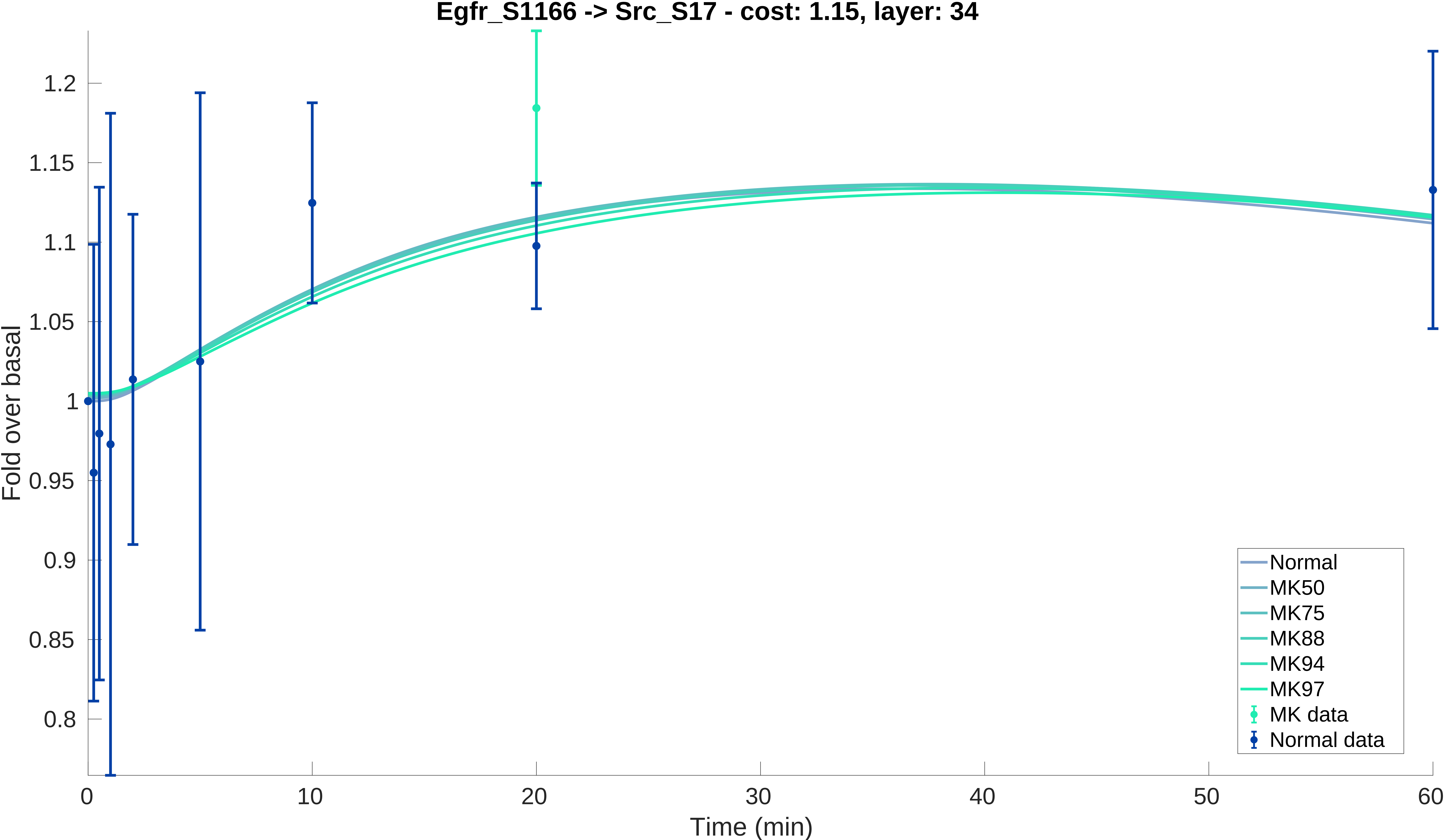

Egfr_S1166 -> Src_S17 - cost: 1.15, layer: 34
Fold over basal
Time (min)
0.8
0.85
0.9
0.95
1
1.05
1.1
1.15
1.2
0
10
20
30
40
50
60
Normal
MK50
MK75
MK88
MK94
MK97
MK data
Normal data

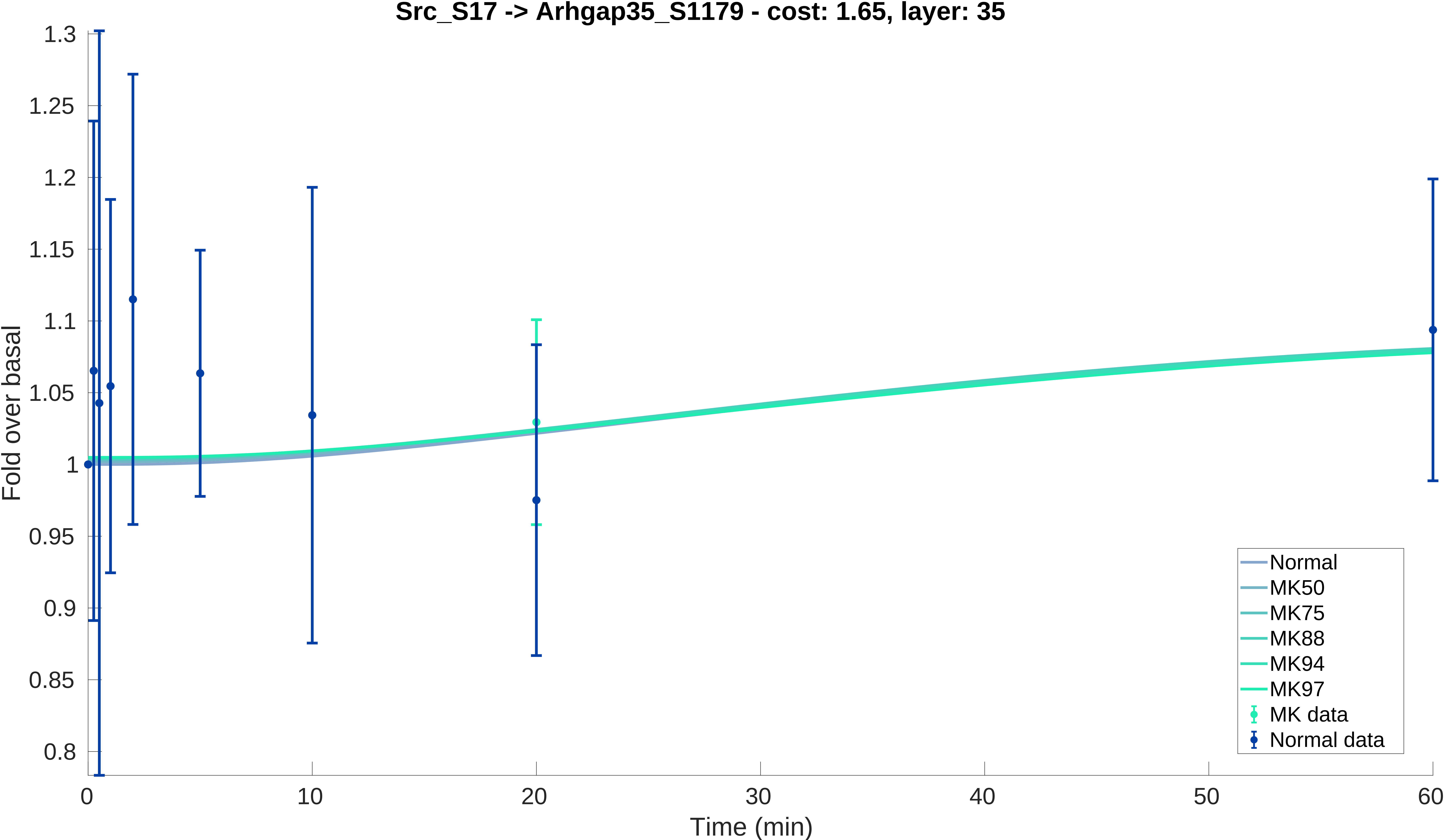

Src_S17 -> Arhgap35_S1179 - cost: 1.65, layer: 35
Fold over basal
Time (min)
1.3
1.25
1.2
1.15
1.1
1.05
1
0.95
0.9
0.85
0.8
0
10
20
30
40
50
60
Normal
MK50
MK75
MK88
MK94
MK97
MK data
Normal data

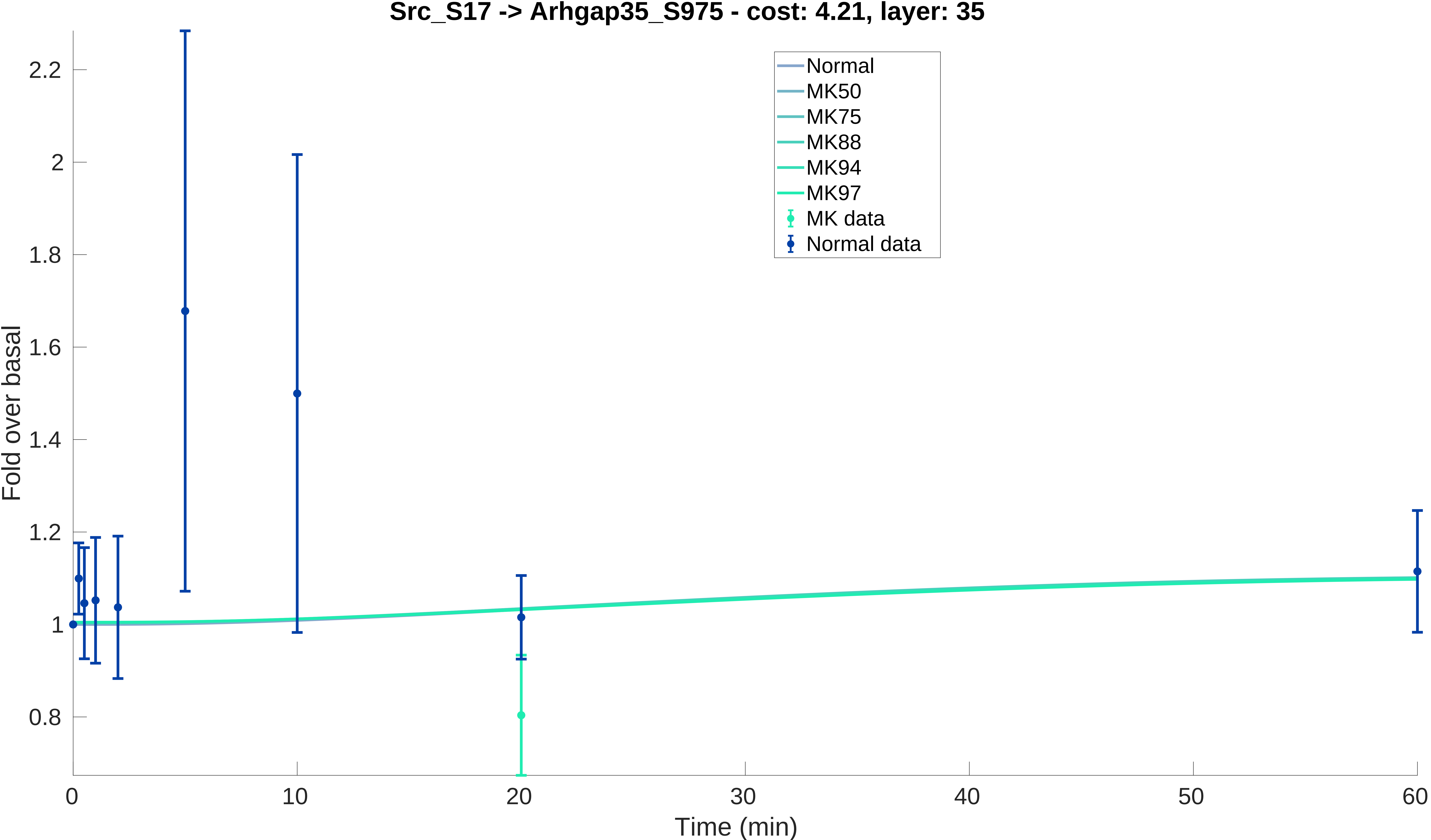

Src_S17 -> Arhgap35_S975 - cost: 4.21, layer: 35
Normal
MK50
MK75
MK88
MK94
MK97
MK data
Normal data
Fold over basal
Time (min)
0.8
1
1.2
1.4
1.6
1.8
2
2.2
0
10
20
30
40
50
60

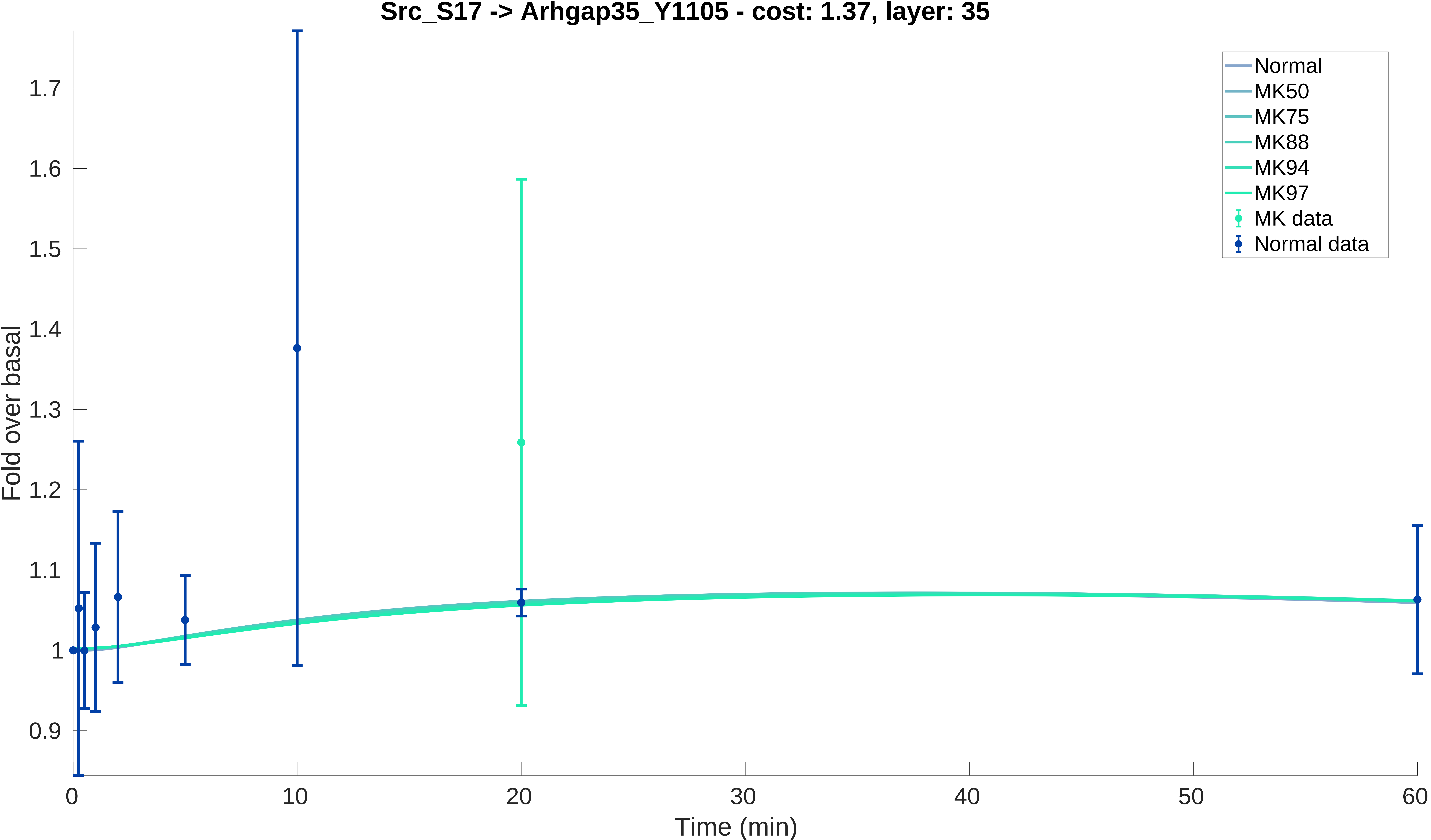
Src_S17 -> Arhgap35_Y1105 - cost: 1.37, layer: 35
Fold over basal
Time (min)
0.9
1
1.1
1.2
1.3
1.4
1.5
1.6
1.7
0
10
20
30
40
50
60
Normal
MK50
MK75
MK88
MK94
MK97
MK data
Normal data

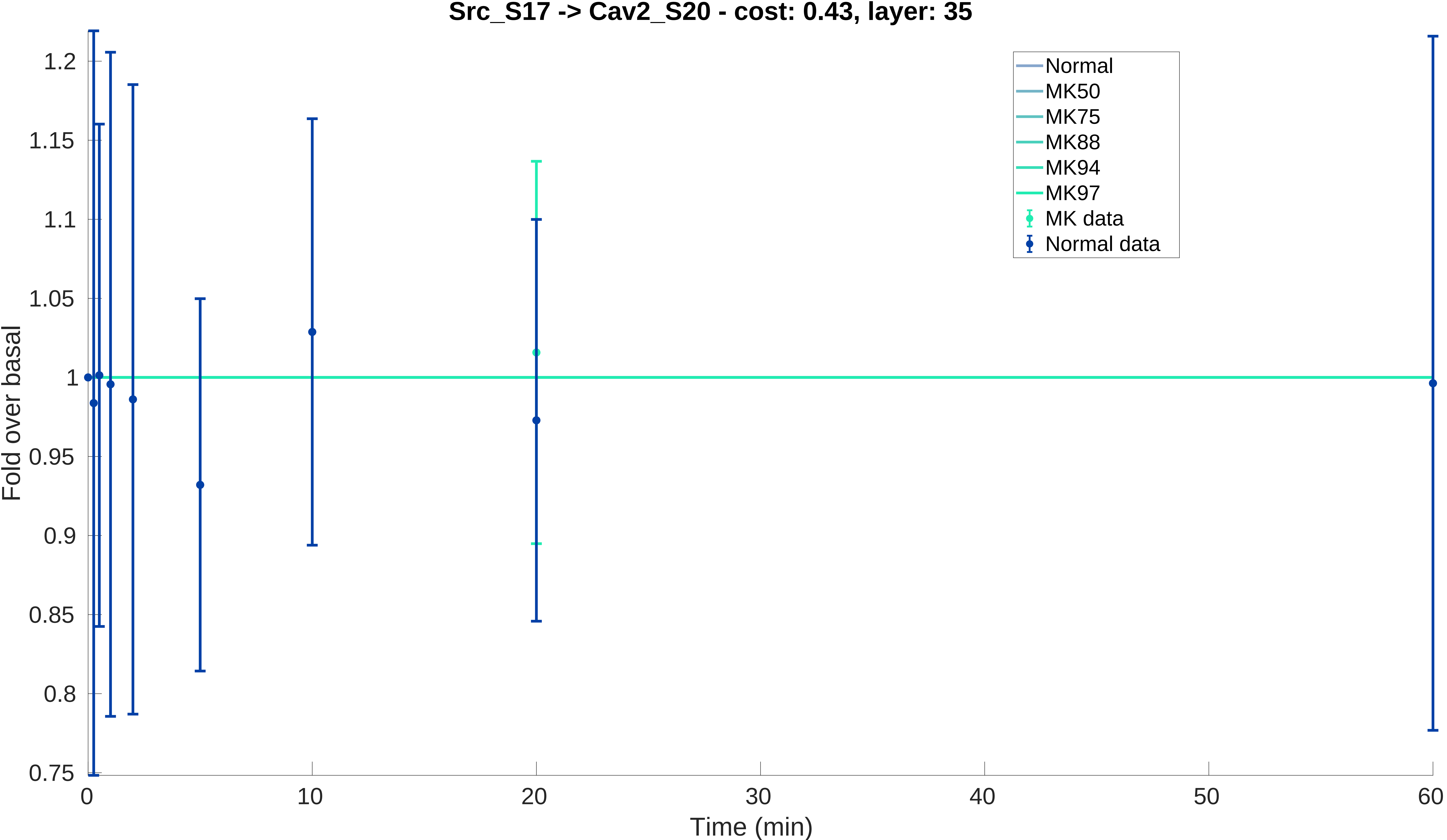

Src_S17 -> Cav2_S20 - cost: 0.43, layer: 35
Normal
MK50
MK75
MK88
MK94
MK97
MK data
Normal data
Fold over basal
Time (min)
1.2
1.15
1.1
1.05
1
0.95
0.9
0.85
0.8
0.75
0
10
20
30
40
50
60

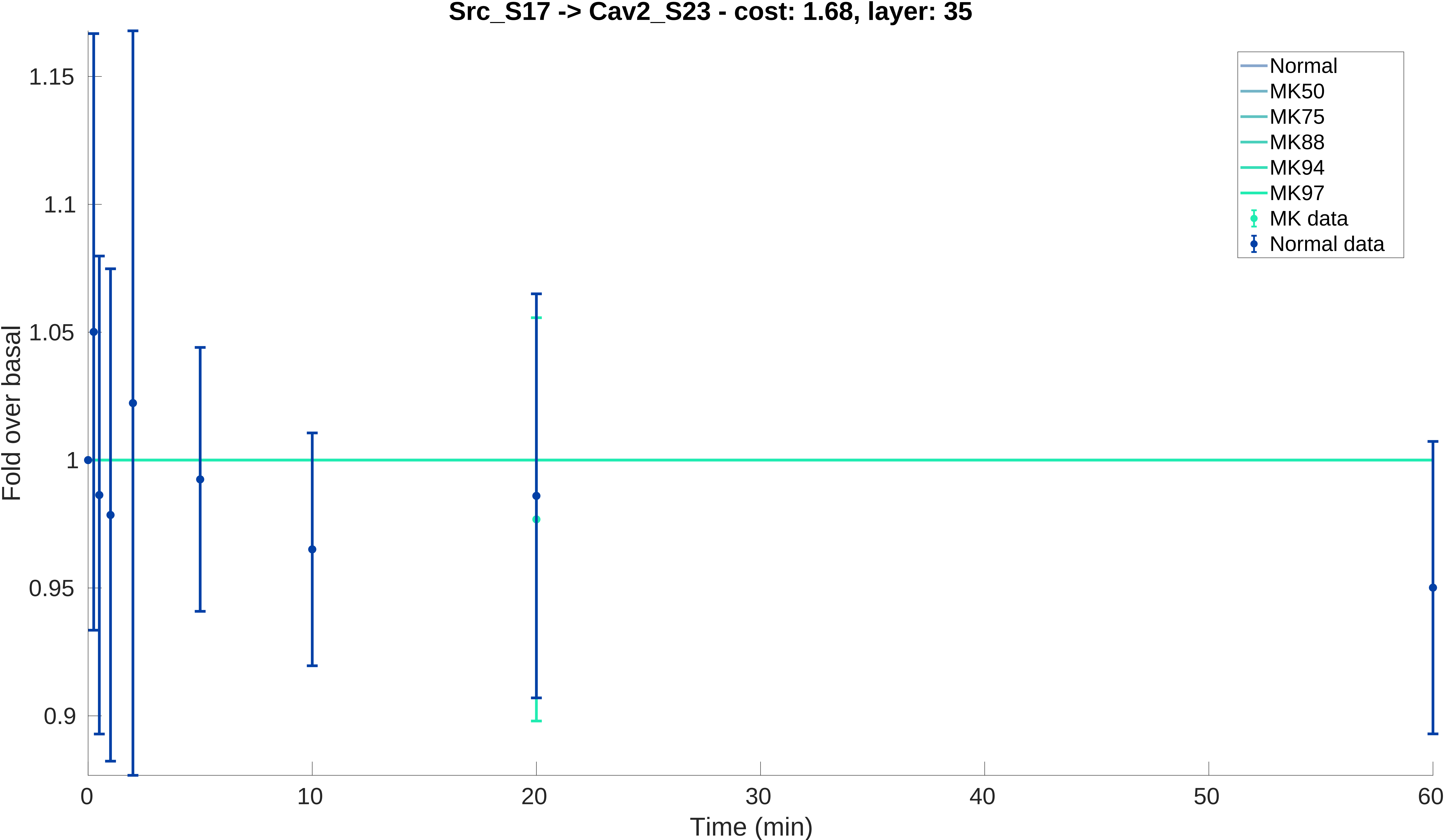

Src_S17 -> Cav2_S23 - cost: 1.68, layer: 35
Normal
MK50
MK75
MK88
MK94
MK97
MK data
Normal data
Fold over basal
1.15
1.1
1.05
1
0.95
0.9
0
10
20
30
40
50
60
Time (min)

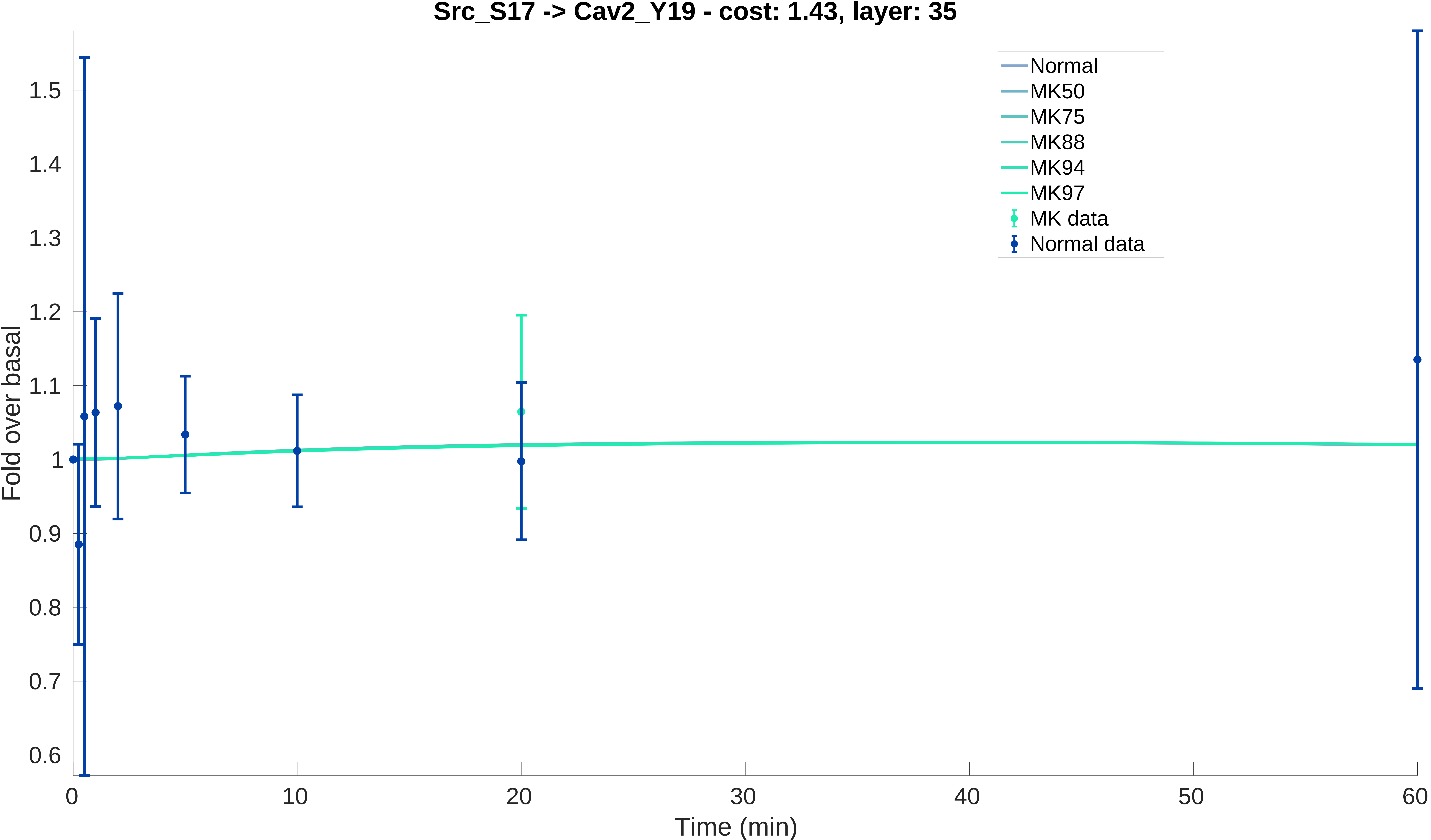

Src_S17 -> Cav2_Y19 - cost: 1.43, layer: 35
Fold over basal
Time (min)
Normal
MK50
MK75
MK88
MK94
MK97
MK data
Normal data
0.6
0.7
0.8
0.9
1
1.1
1.2
1.3
1.4
1.5
0
10
20
30
40
50
60

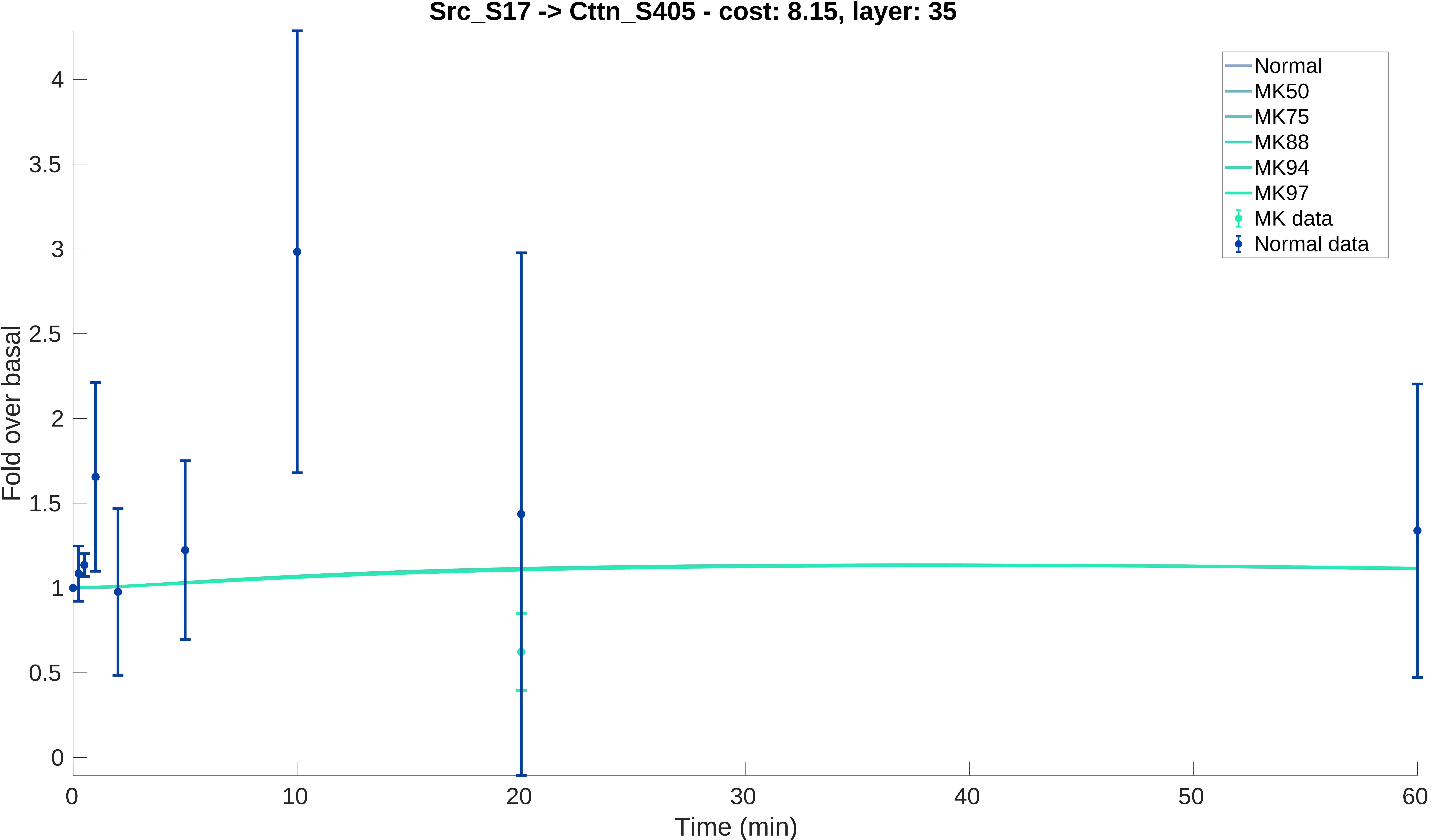

Src_S17 -> Cttn_S405 - cost: 8.15, layer: 35
Normal
MK50
MK75
MK88
MK94
MK97
MK data
Normal data
Fold over basal
0
0.5
1
1.5
2
2.5
3
3.5
4
0
10
20
30
40
50
60
Time (min)

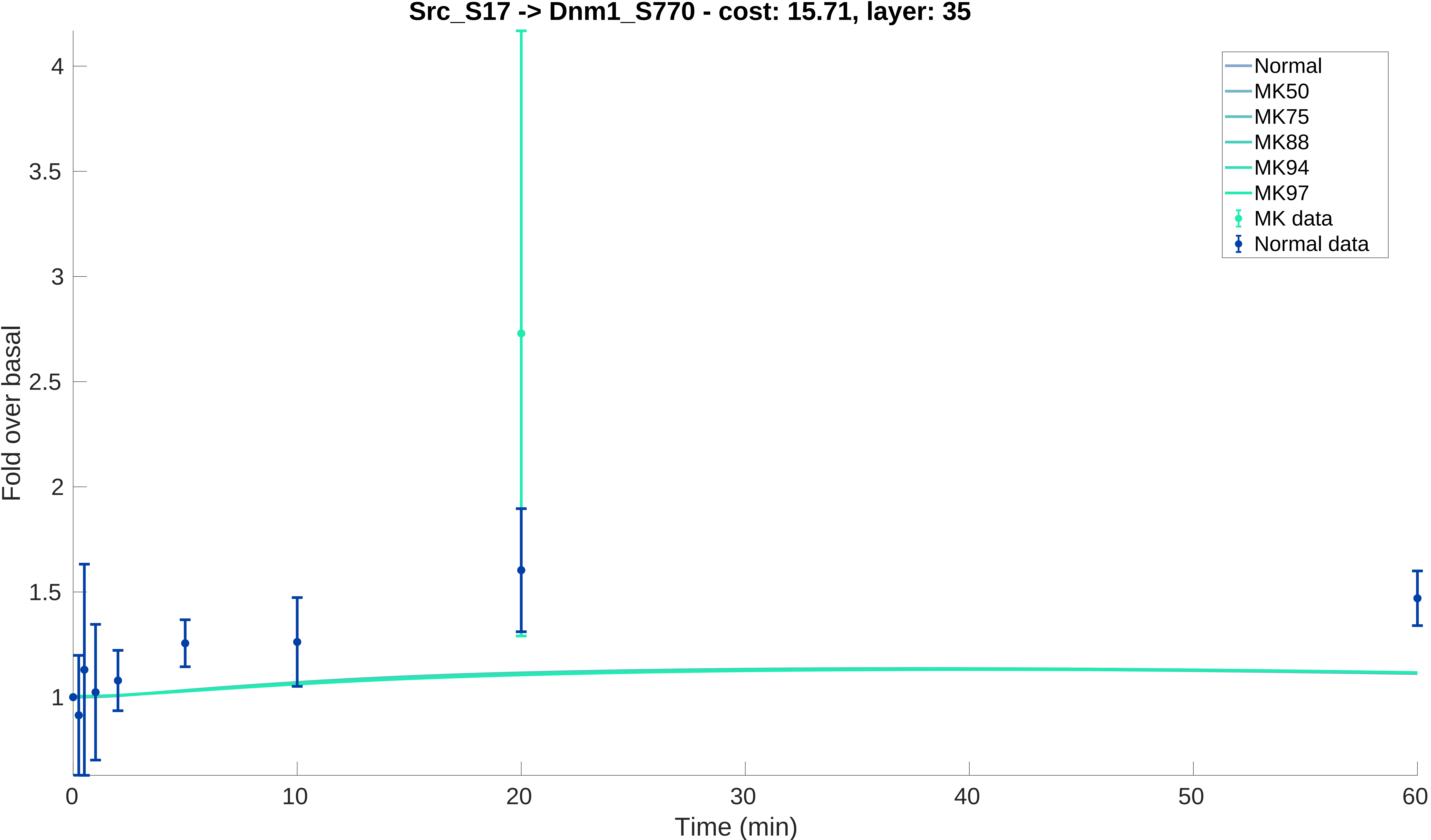

Src_S17 -> Dnm1_S770 - cost: 15.71, layer: 35
Fold over basal
Time (min)
4
3.5
3
2.5
2
1.5
1
0
10
20
30
40
50
60
Normal
MK50
MK75
MK88
MK94
MK97
MK data
Normal data

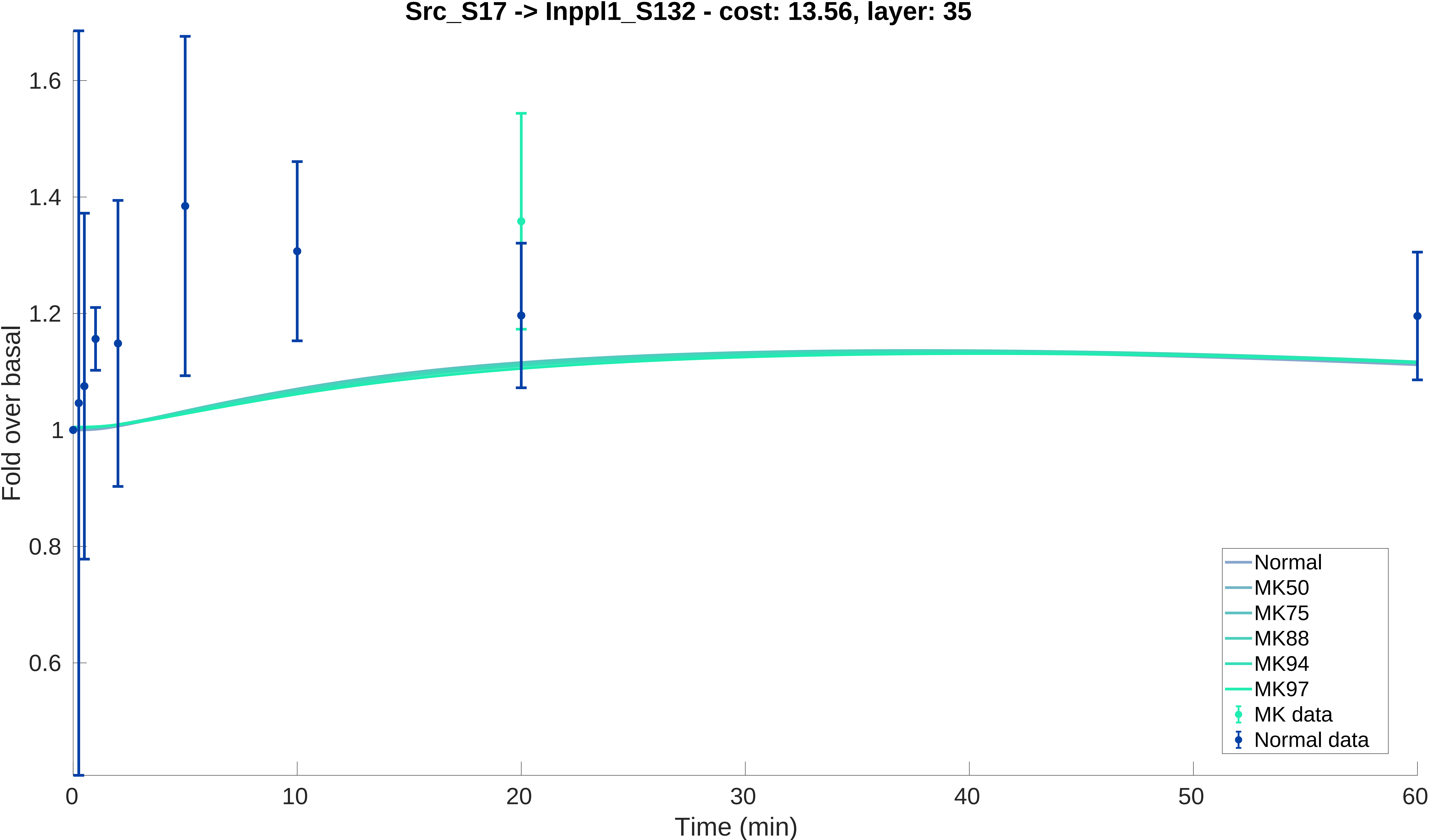

Src_S17 -> Inppl1_S132 - cost: 13.56, layer: 35
Fold over basal
1.6
1.4
1.2
1
0.8
0.6
0
10
20
30
40
50
60
Time (min)
Normal
MK50
MK75
MK88
MK94
MK97
MK data
Normal data

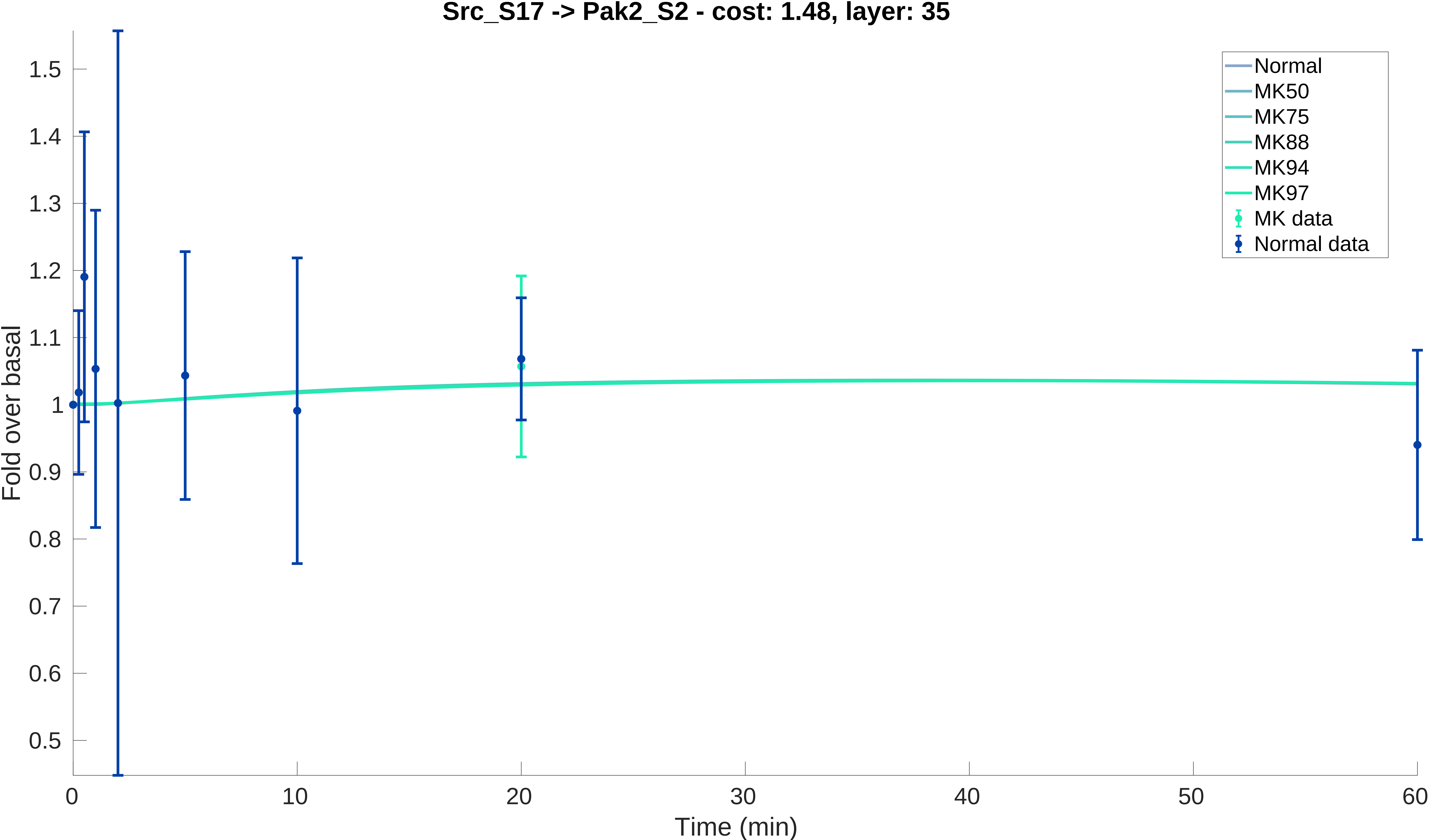

Src_S17 -> Pak2_S2 - cost: 1.48, layer: 35
Fold over basal
Time (min)
0.5
0.6
0.7
0.8
0.9
1
1.1
1.2
1.3
1.4
1.5
0
10
20
30
40
50
60
Normal
MK50
MK75
MK88
MK94
MK97
MK data
Normal data

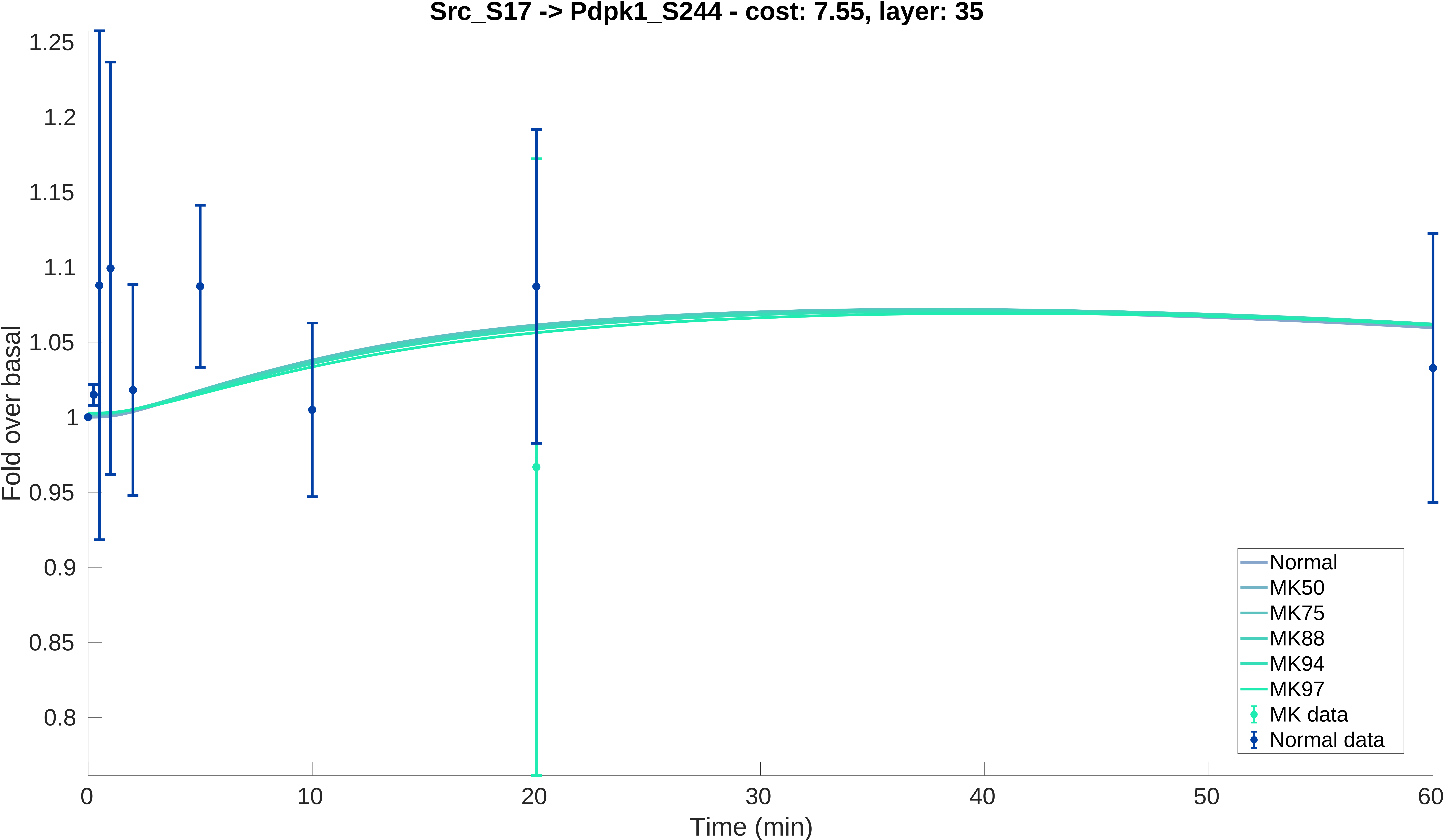

Src_S17 -> Pdpk1_S244 - cost: 7.55, layer: 35
Fold over basal
Time (min)
1.25
1.2
1.15
1.1
1.05
1
0.95
0.9
0.85
0.8
0
10
20
30
40
50
60
Normal
MK50
MK75
MK88
MK94
MK97
MK data
Normal data

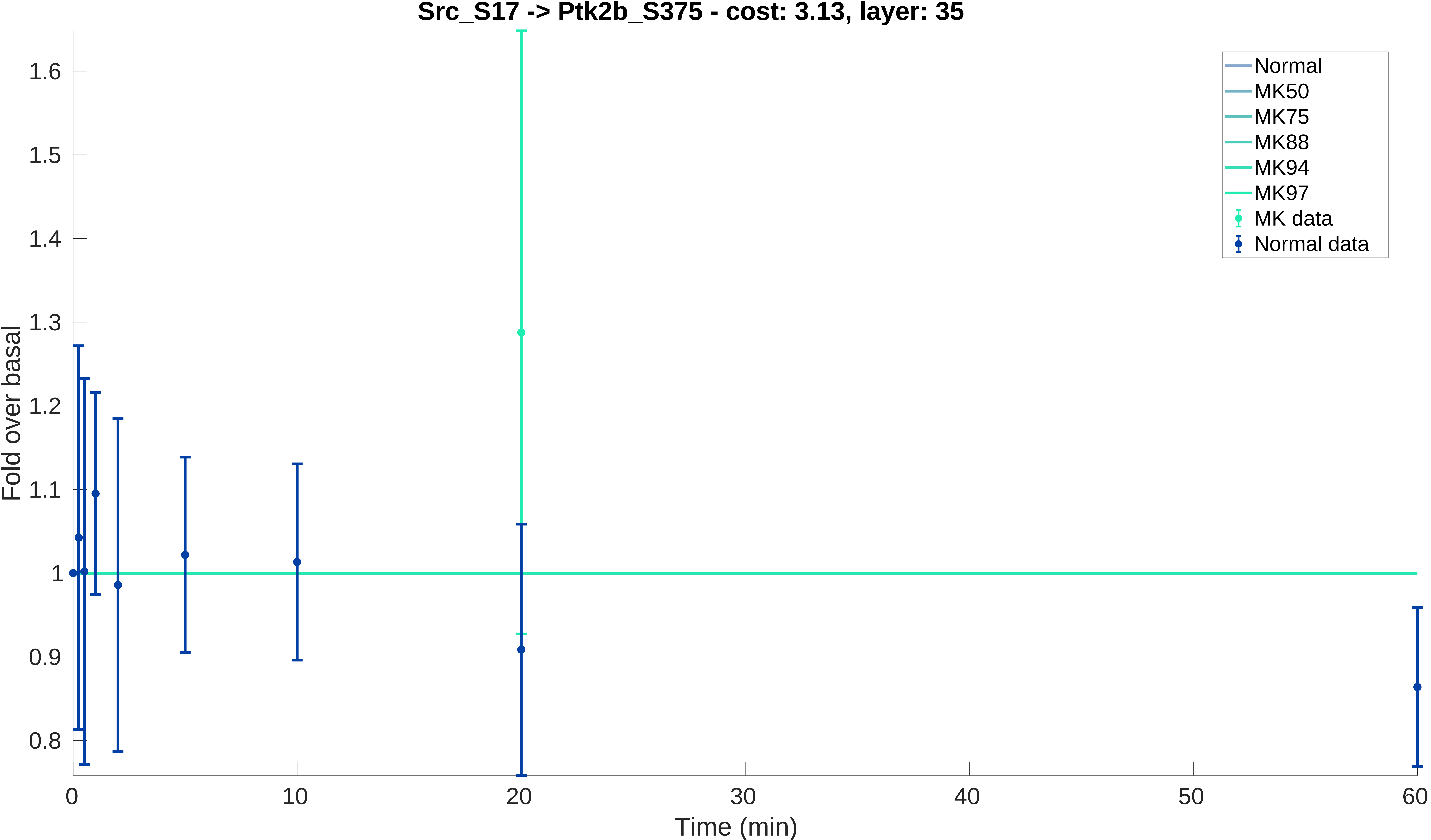

Src_S17 -> Ptk2b_S375 - cost: 3.13, layer: 35
Normal
MK50
MK75
MK88
MK94
MK97
MK data
Normal data
Fold over basal
1.6
1.5
1.4
1.3
1.2
1.1
1
0.9
0.8
0
10
20
30
40
50
60
Time (min)

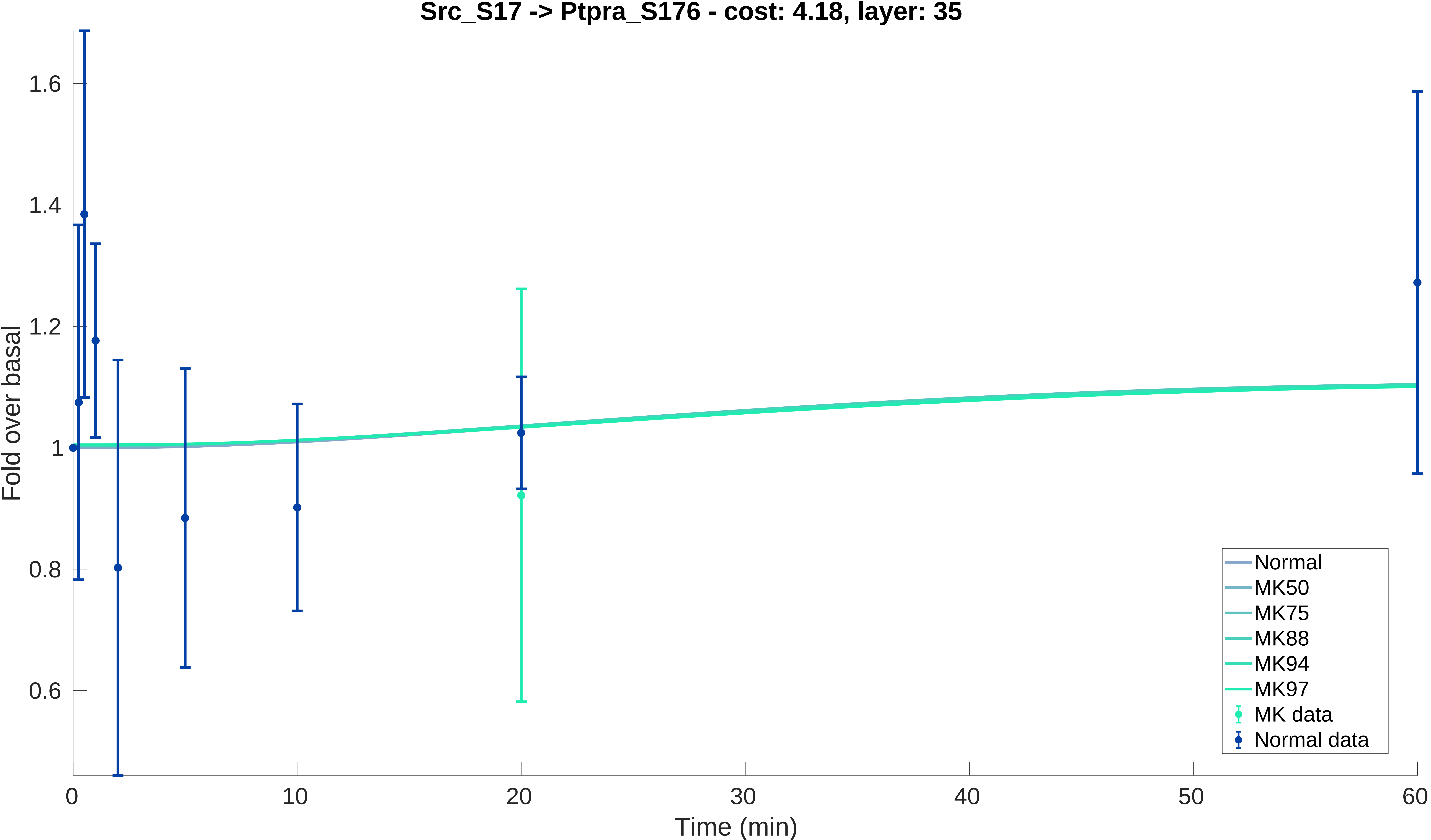
Src_S17 -> Ptpra_S176 - cost: 4.18, layer: 35
Fold over basal
1.6
1.4
1.2
1
0.8
0.6
0
10
20
30
40
50
60
Time (min)
Normal
MK50
MK75
MK88
MK94
MK97
MK data
Normal data

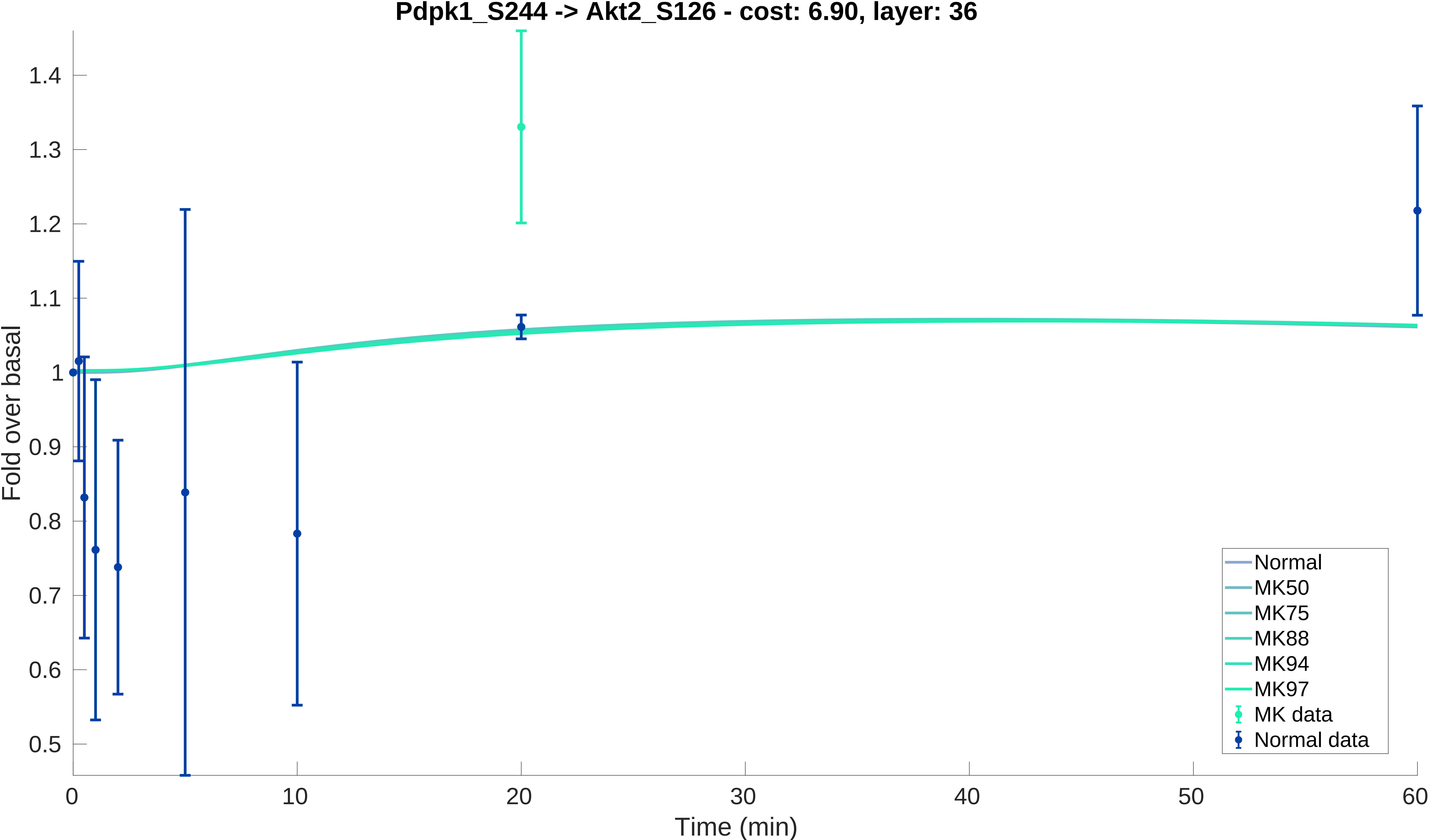
Pdpk1_S244 -> Akt2_S126 - cost: 6.90, layer: 36
Fold over basal
1.4
1.3
1.2
1.1
1
0.9
0.8
0.7
0.6
0.5
0
10
20
30
40
50
60
Time (min)
Normal
MK50
MK75
MK88
MK94
MK97
MK data
Normal data

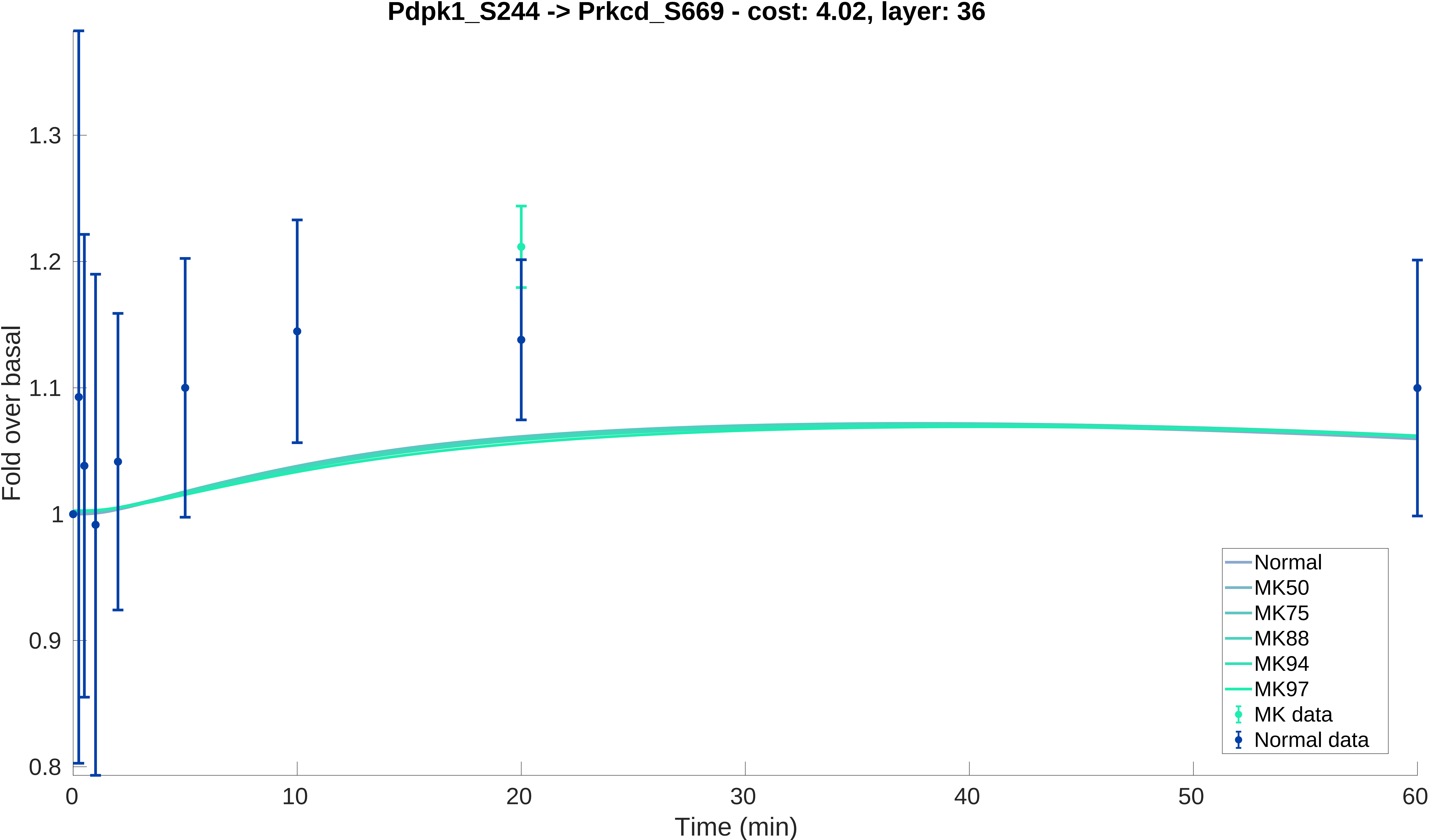

Pdpk1_S244 -> Prkcd_S669 - cost: 4.02, layer: 36
Fold over basal
1.3
1.2
1.1
1
0.9
0.8
0
10
20
30
40
50
60
Time (min)
Normal
MK50
MK75
MK88
MK94
MK97
MK data
Normal data

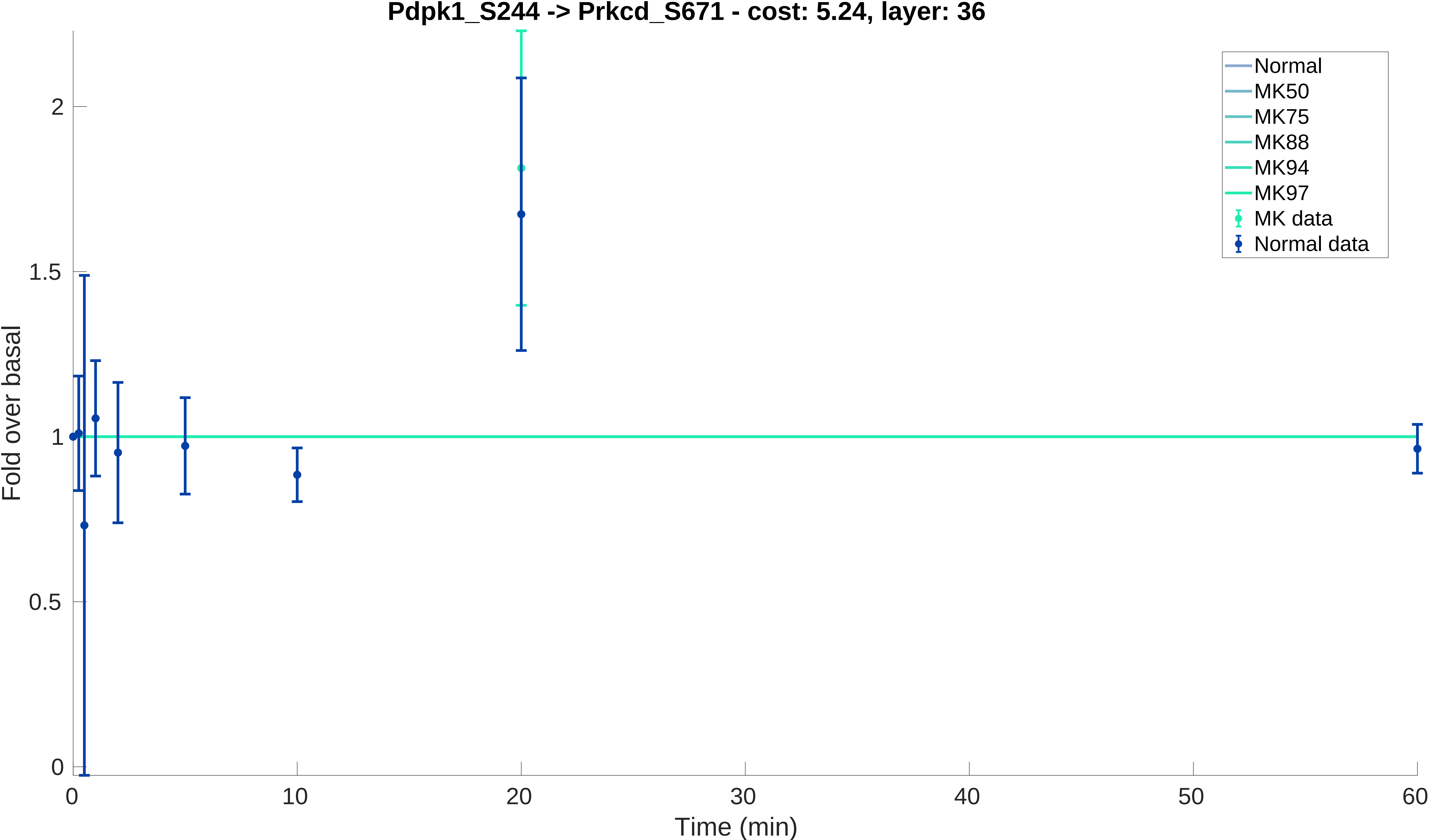
Pdpk1_S244 -> Prkcd_S671 - cost: 5.24, layer: 36
Fold over basal
Time (min)
0
0.5
1
1.5
2
0
10
20
30
40
50
60
Normal
MK50
MK75
MK88
MK94
MK97
MK data
Normal data

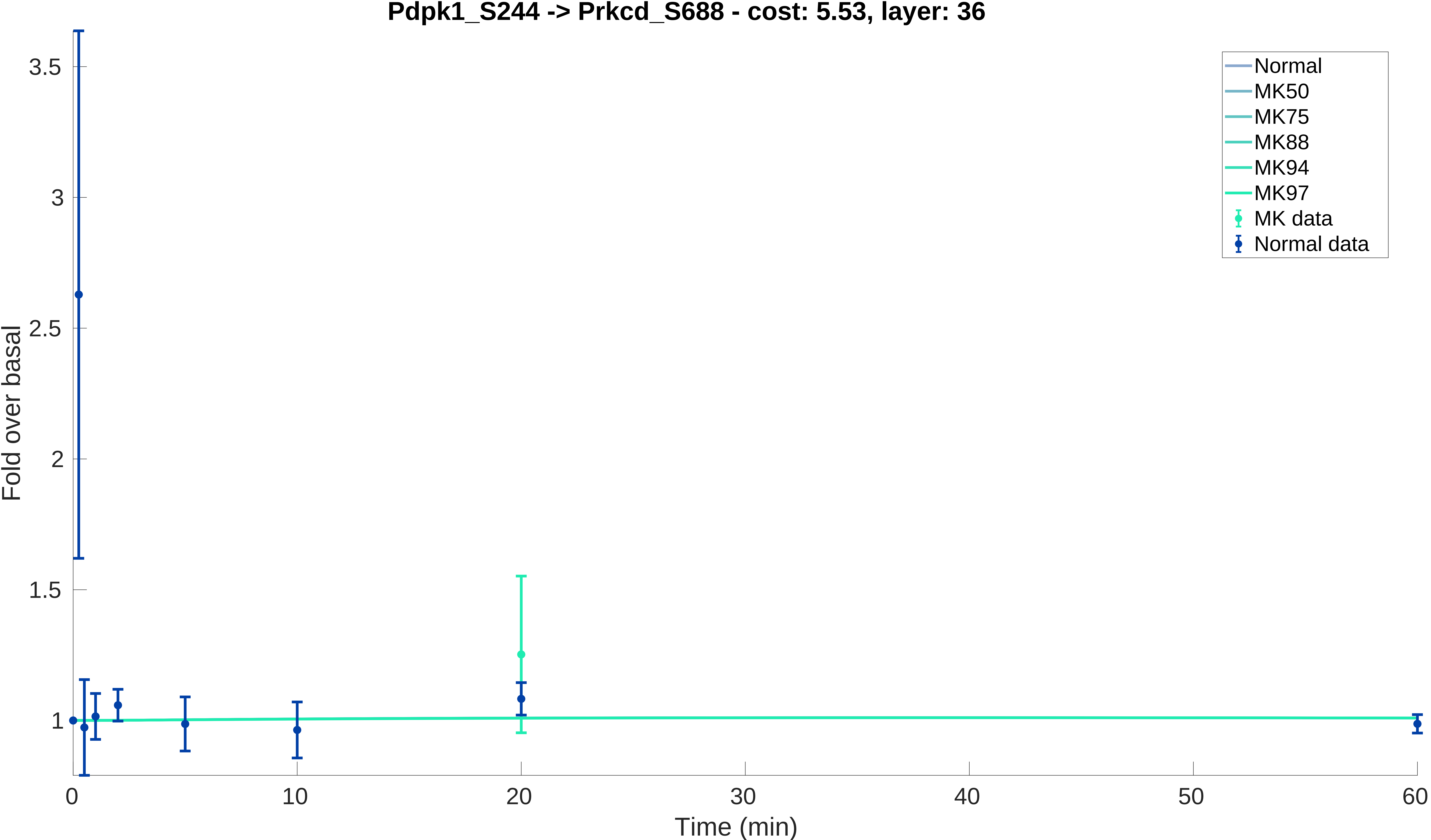

Pdpk1_S244 -> Prkcd_S688 - cost: 5.53, layer: 36
Fold over basal
Time (min)
Normal
MK50
MK75
MK88
MK94
MK97
MK data
Normal data
3.5
3
2.5
2
1.5
1
0
10
20
30
40
50
60

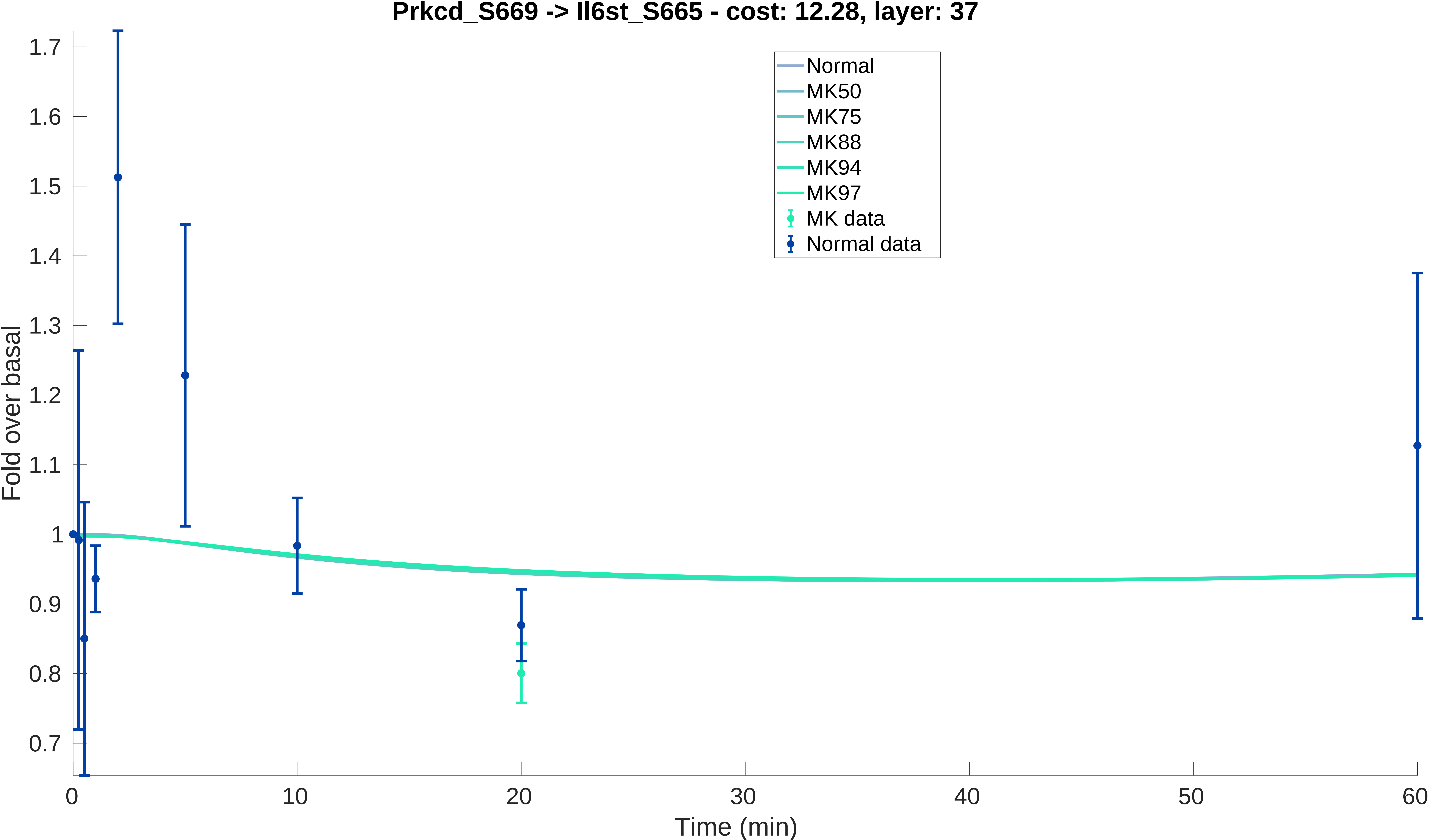

Prkcd_S669 -> Il6st_S665 - cost: 12.28, layer: 37
Fold over basal
Time (min)
Normal
MK50
MK75
MK88
MK94
MK97
MK data
Normal data
1.7
1.6
1.5
1.4
1.3
1.2
1.1
1
0.9
0.8
0.7
0
10
20
30
40
50
60

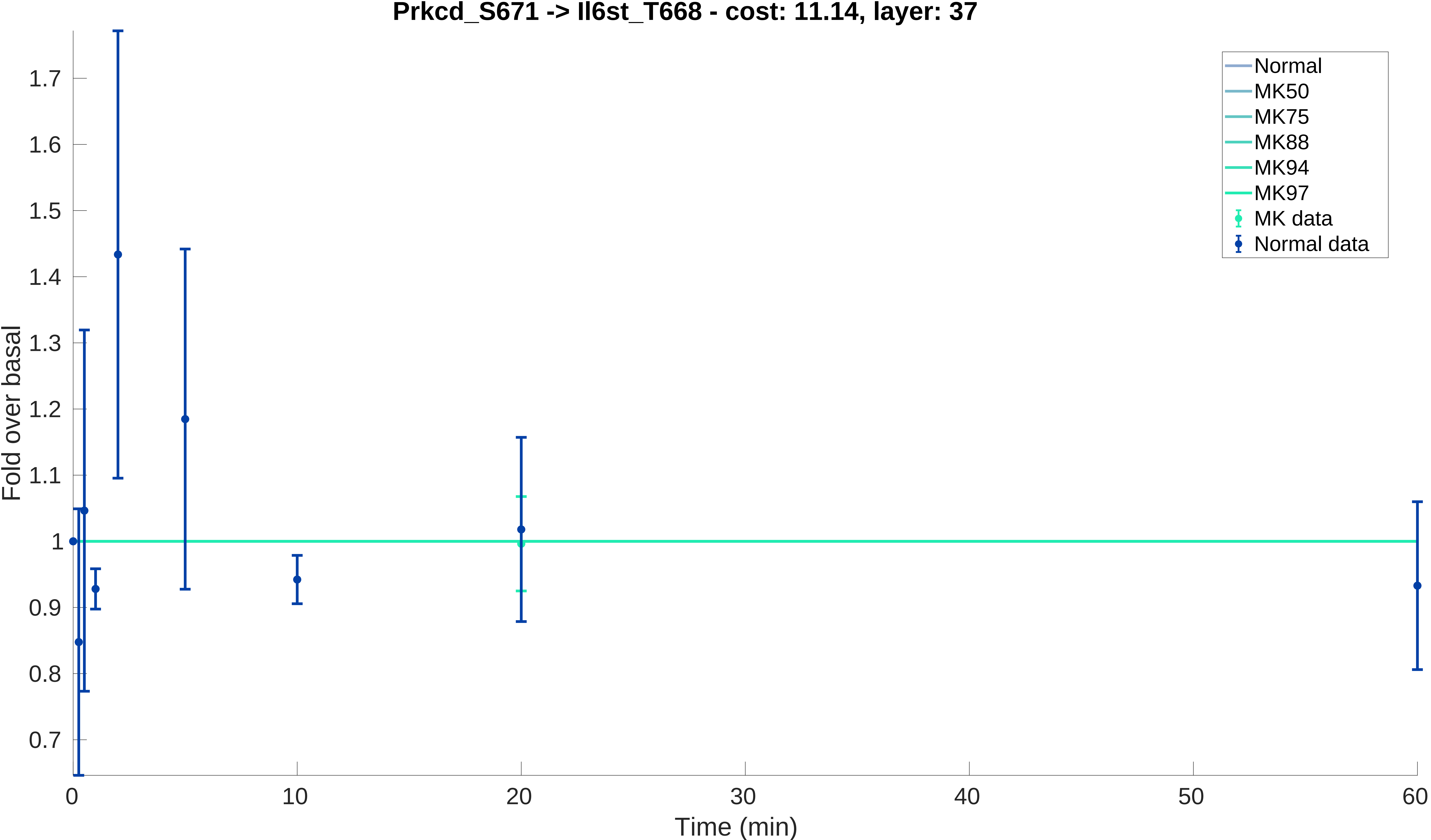

Prkcd_S671 -> Il6st_T668 - cost: 11.14, layer: 37
Fold over basal
Time (min)
1.7
1.6
1.5
1.4
1.3
1.2
1.1
1
0.9
0.8
0.7
0
10
20
30
40
50
60
Normal
MK50
MK75
MK88
MK94
MK97
MK data
Normal data

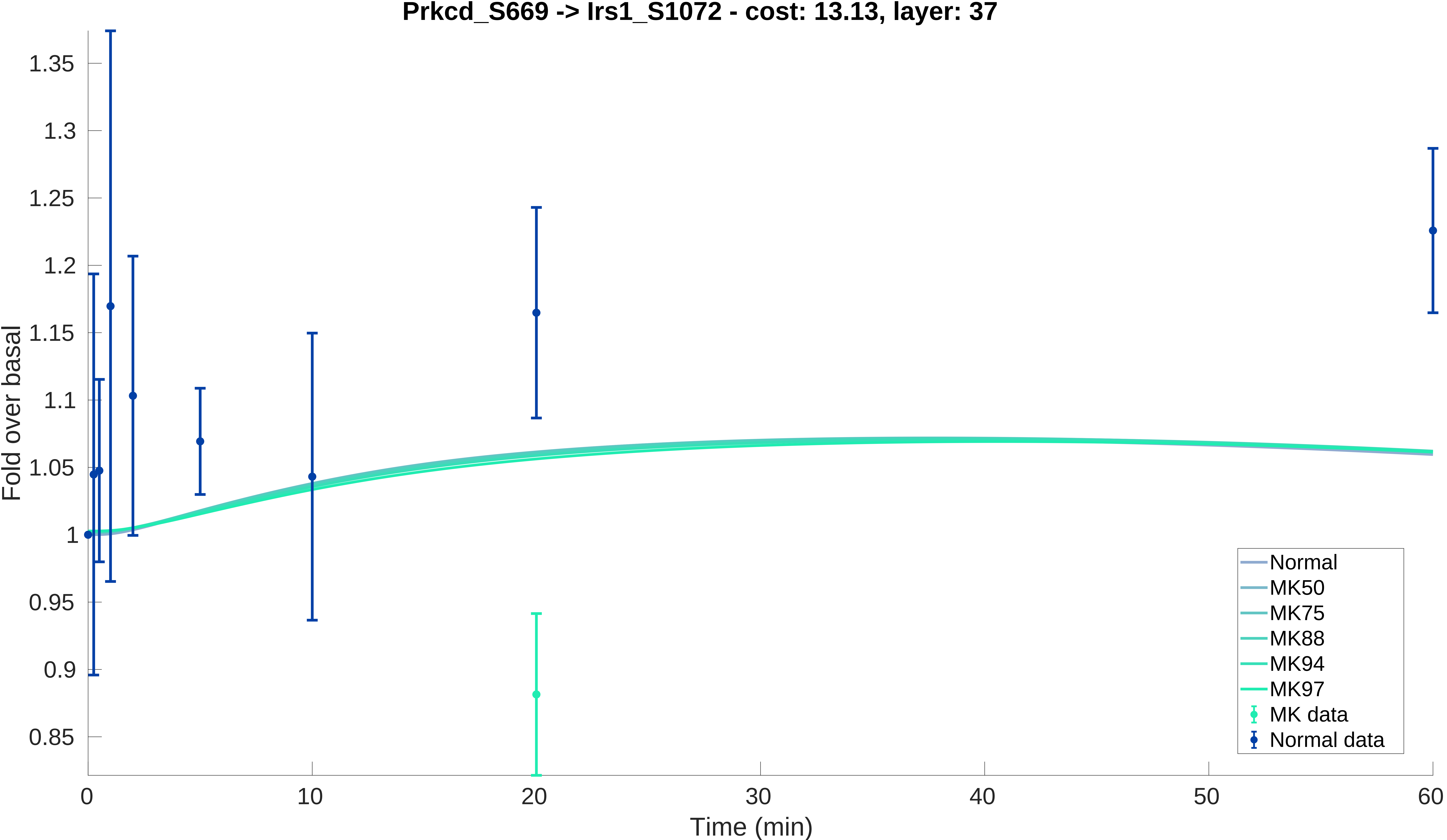

Prkcd_S669 -> Irs1_S1072 - cost: 13.13, layer: 37
Fold over basal
1.35
1.3
1.25
1.2
1.15
1.1
1.05
1
0.95
0.9
0.85
0
10
20
30
40
50
60
Time (min)
Normal
MK50
MK75
MK88
MK94
MK97
MK data
Normal data

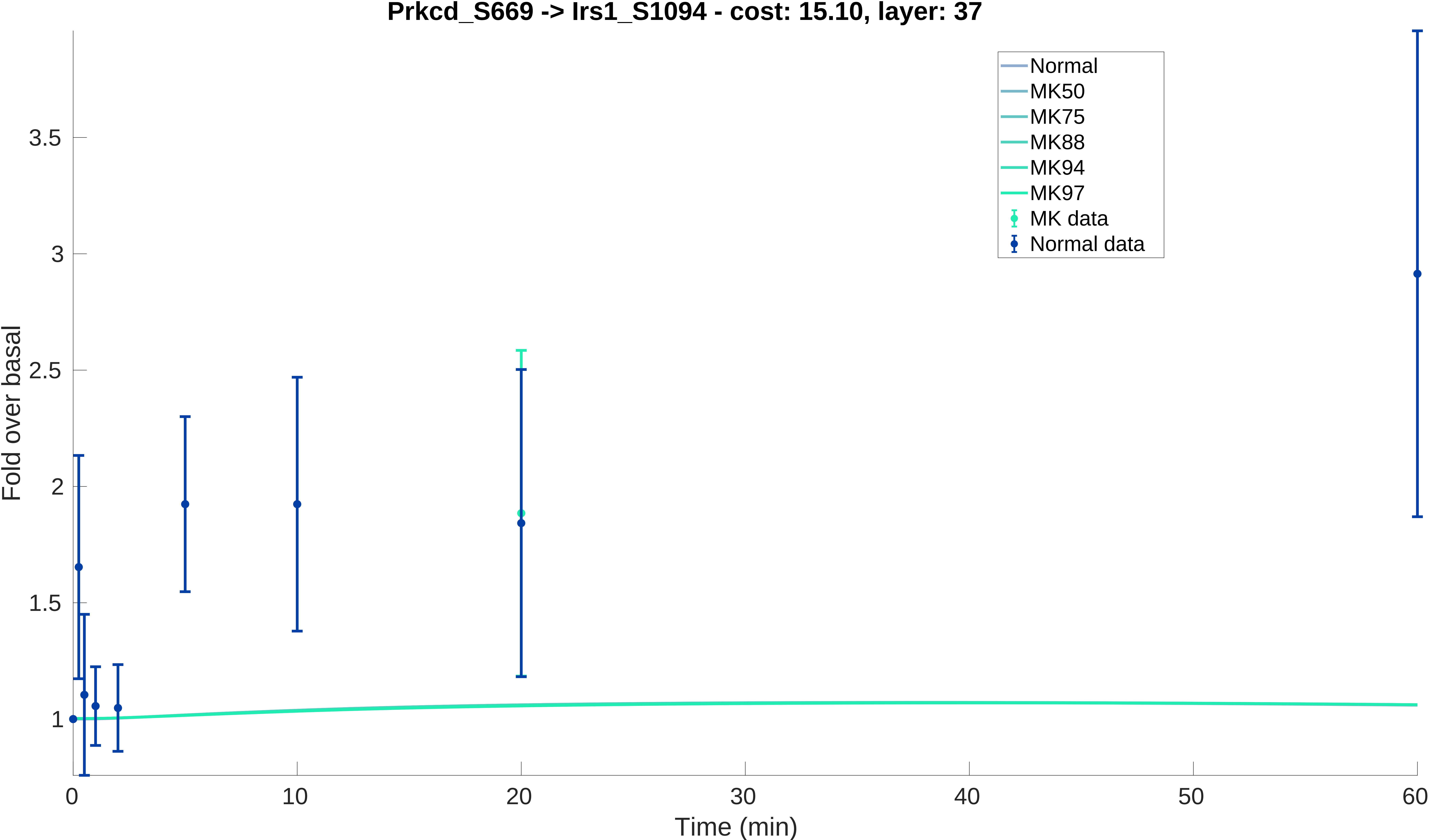

Prkcd_S669 -> Irs1_S1094 - cost: 15.10, layer: 37
Normal
MK50
MK75
MK88
MK94
MK97
MK data
Normal data
Fold over basal
3.5
3
2.5
2
1.5
1
0
10
20
30
40
50
60
Time (min)

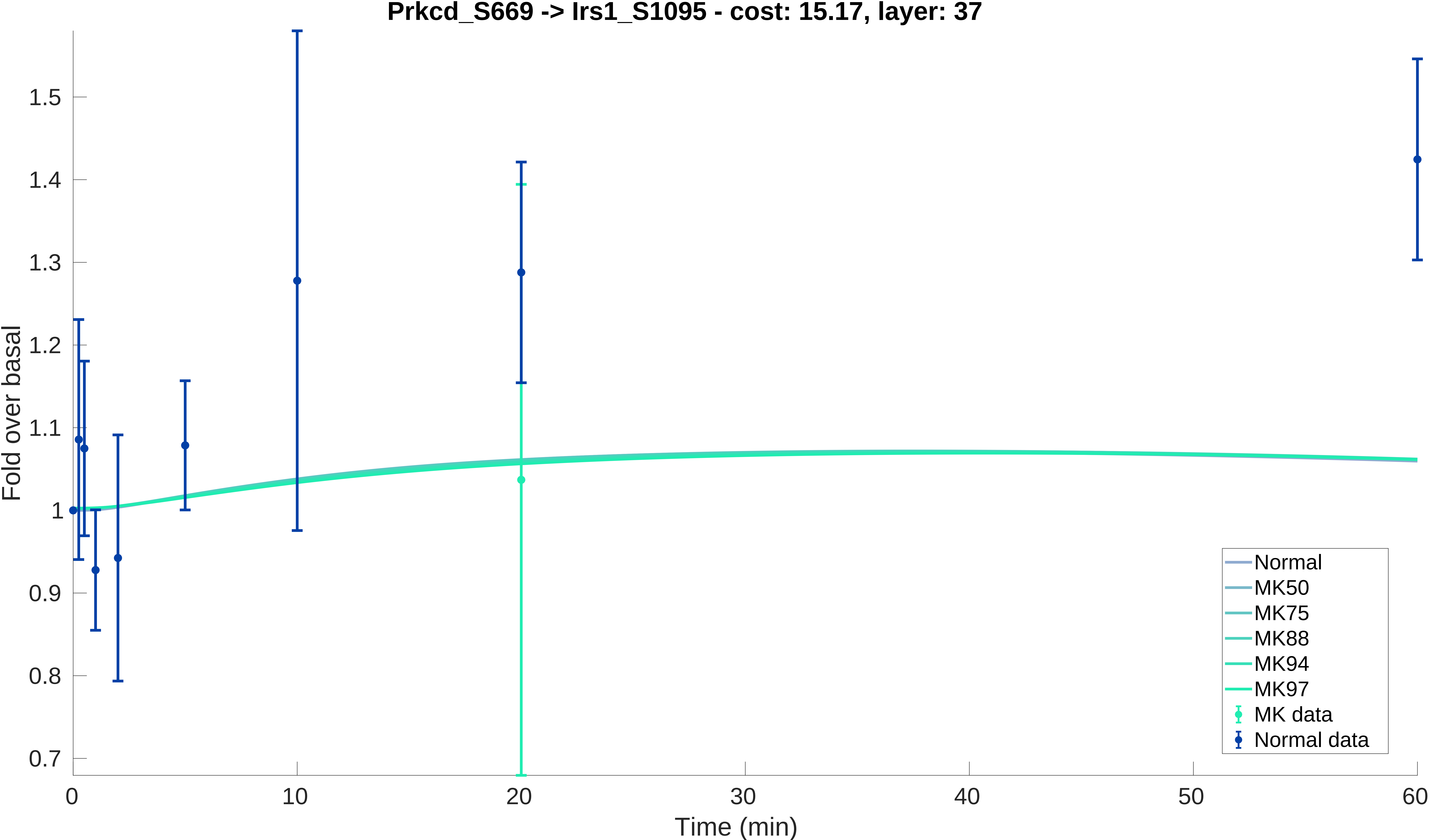

Prkcd_S669 -> Irs1_S1095 - cost: 15.17, layer: 37
Fold over basal
Time (min)
0.7
0.8
0.9
1
1.1
1.2
1.3
1.4
1.5
0
10
20
30
40
50
60
Normal
MK50
MK75
MK88
MK94
MK97
MK data
Normal data

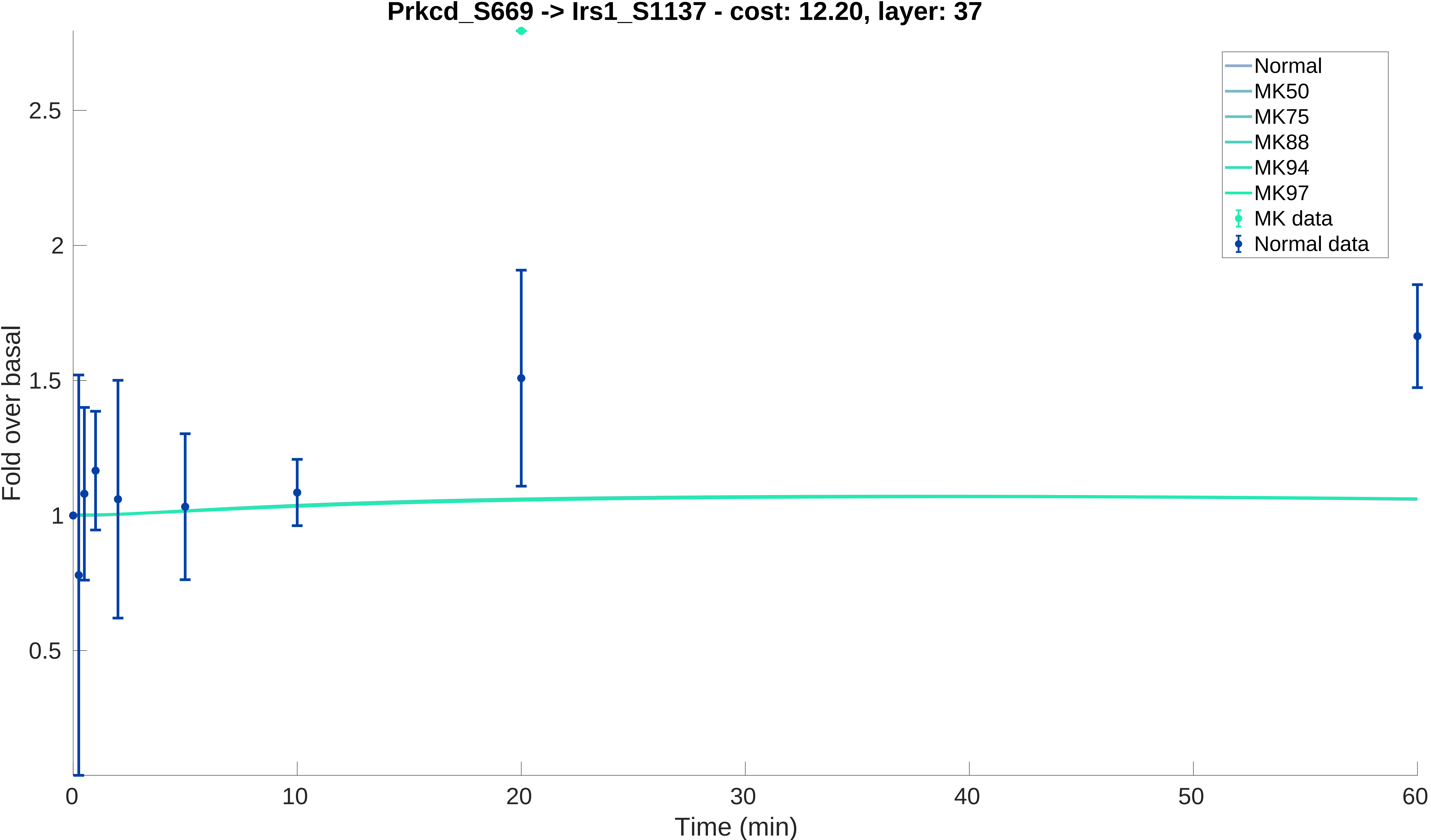

Prkcd_S669 -> Irs1_S1137 - cost: 12.20, layer: 37
Fold over basal
Time (min)
0.5
1
1.5
2
2.5
0
10
20
30
40
50
60
Normal
MK50
MK75
MK88
MK94
MK97
MK data
Normal data

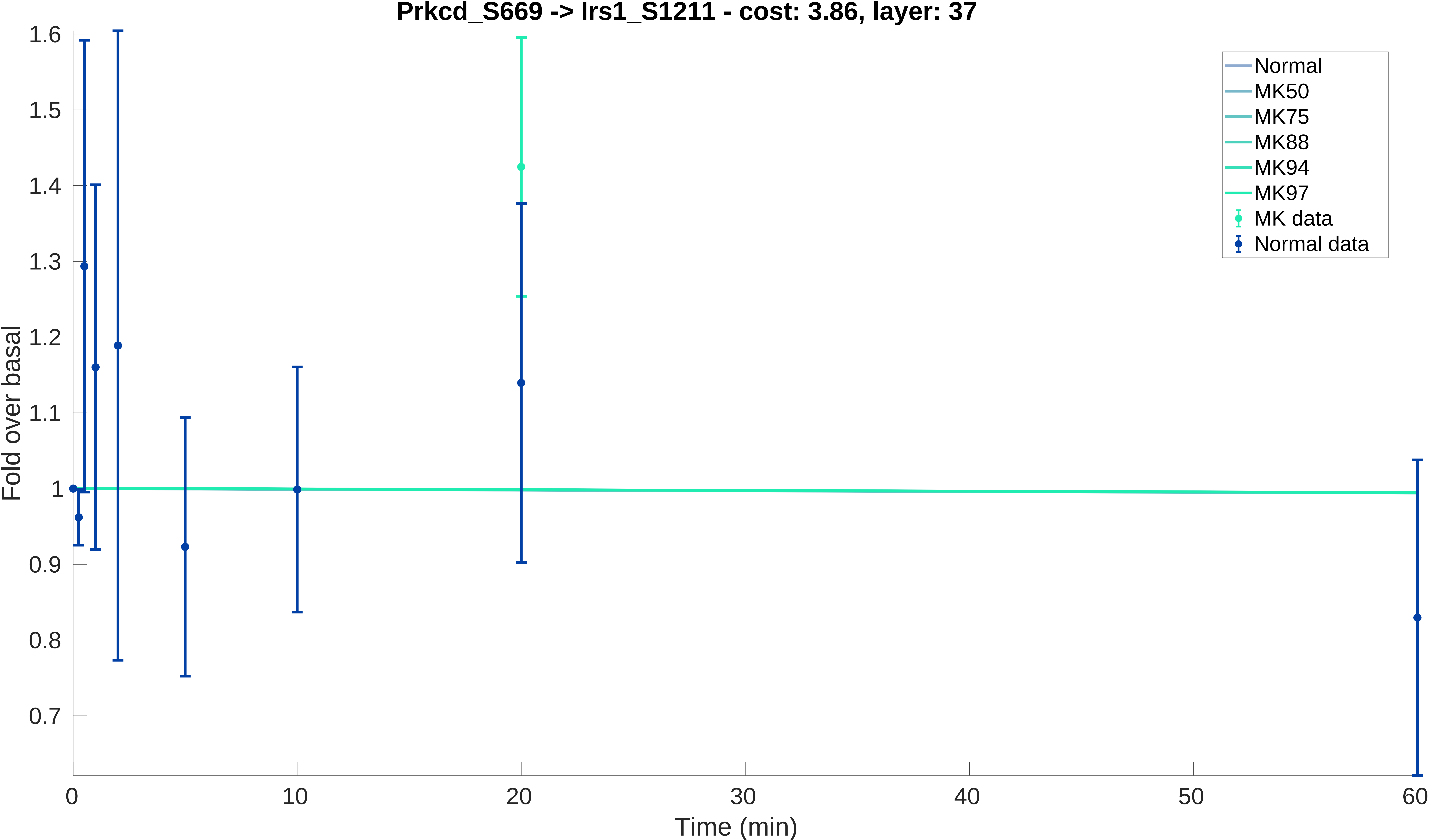

Prkcd_S669 -> Irs1_S1211 - cost: 3.86, layer: 37
Fold over basal
Time (min)
1.6
1.5
1.4
1.3
1.2
1.1
1
0.9
0.8
0.7
0
10
20
30
40
50
60
Normal
MK50
MK75
MK88
MK94
MK97
MK data
Normal data

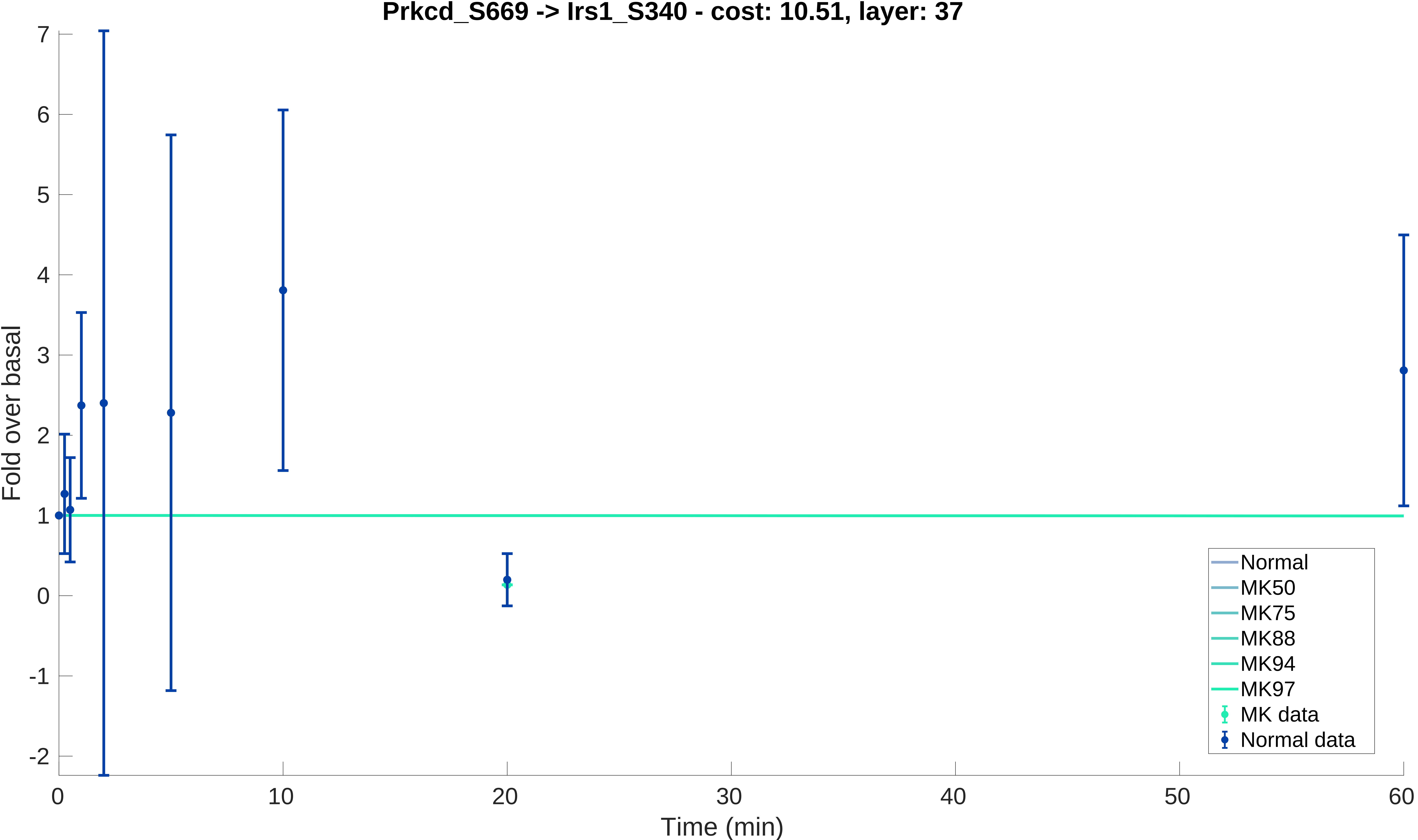
Prkcd_S669 -> Irs1_S340 - cost: 10.51, layer: 37
Fold over basal
Time (min)
7
6
5
4
3
2
1
0
-1
-2
0
10
20
30
40
50
60
Normal
MK50
MK75
MK88
MK94
MK97
MK data
Normal data

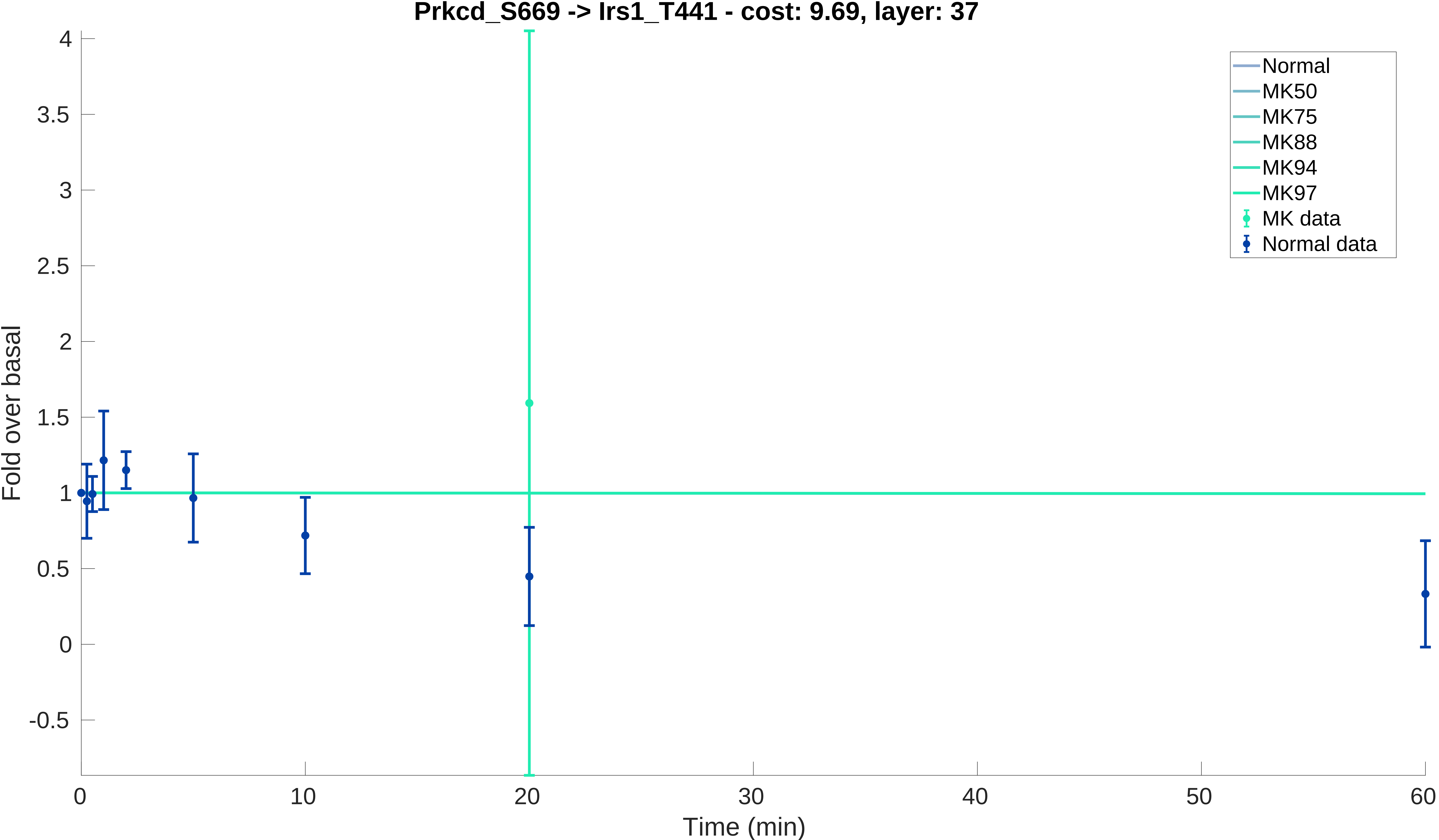

Prkcd_S669 -> Irs1_T441 - cost: 9.69, layer: 37
Fold over basal
Time (min)
4
3.5
3
2.5
2
1.5
1
0.5
0
-0.5
0
10
20
30
40
50
60
Normal
MK50
MK75
MK88
MK94
MK97
MK data
Normal data

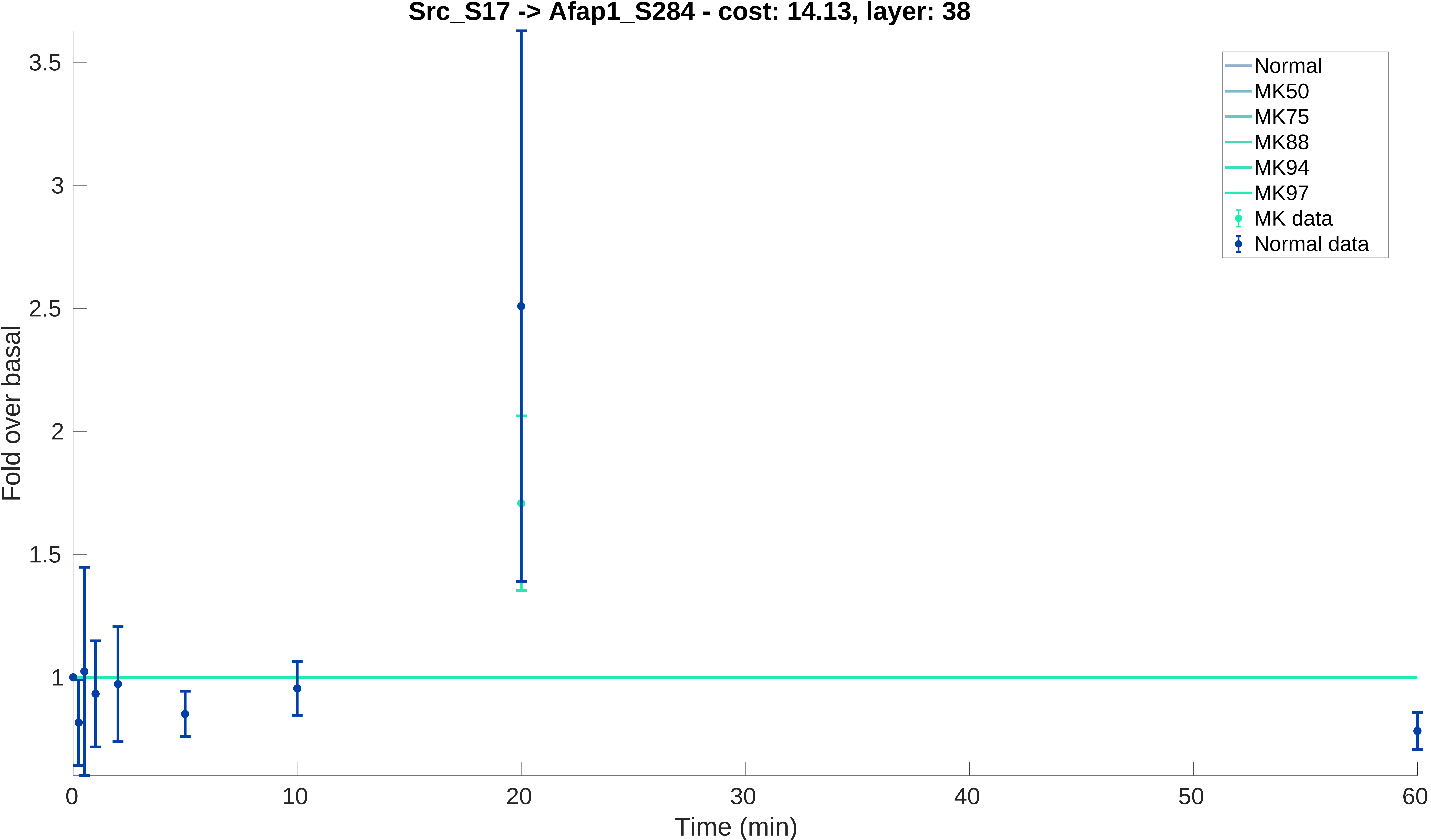

Src_S17 -> Afap1_S284 - cost: 14.13, layer: 38
Fold over basal
Time (min)
3.5
3
2.5
2
1.5
1
0
10
20
30
40
50
60
Normal
MK50
MK75
MK88
MK94
MK97
MK data
Normal data

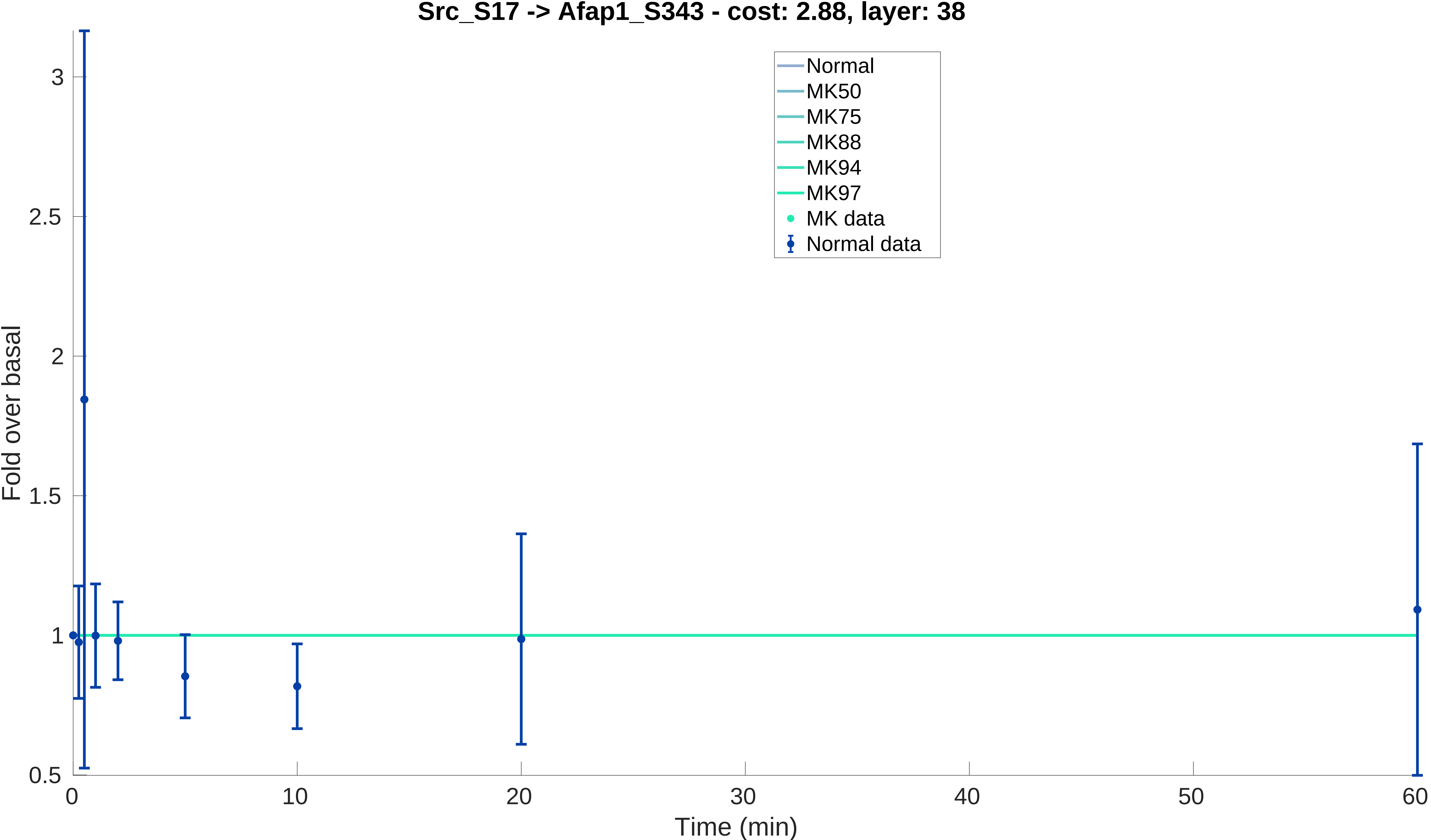
Src_S17 -> Afap1_S343 - cost: 2.88, layer: 38
Normal
MK50
MK75
MK88
MK94
MK97
MK data
Normal data
Fold over basal
Time (min)
0.5
1
1.5
2
2.5
3
0
10
20
30
40
50
60

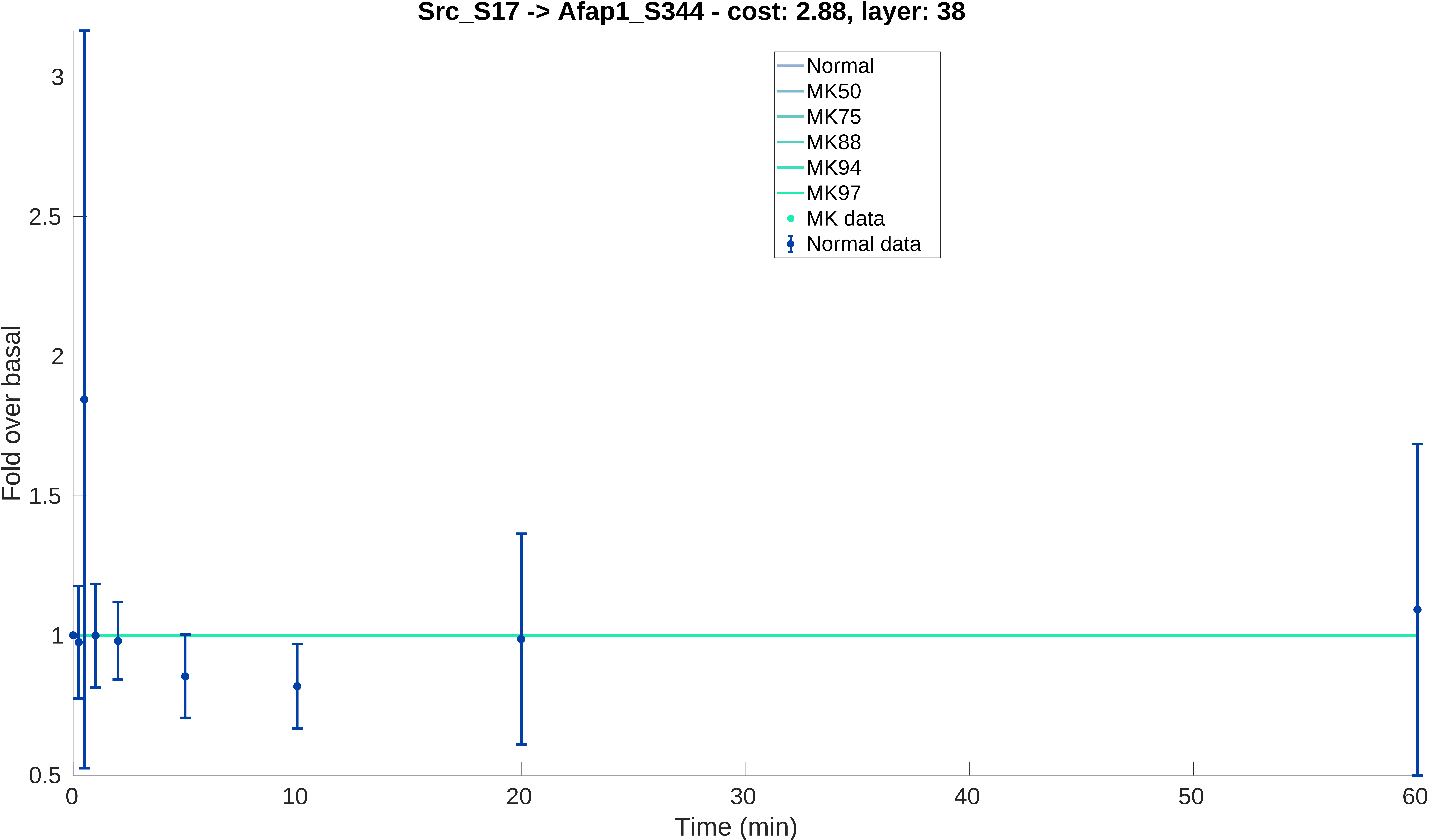

Src_S17 -> Afap1_S344 - cost: 2.88, layer: 38
Normal
MK50
MK75
MK88
MK94
MK97
MK data
Normal data
Fold over basal
Time (min)
0.5
1
1.5
2
2.5
3
0
10
20
30
40
50
60

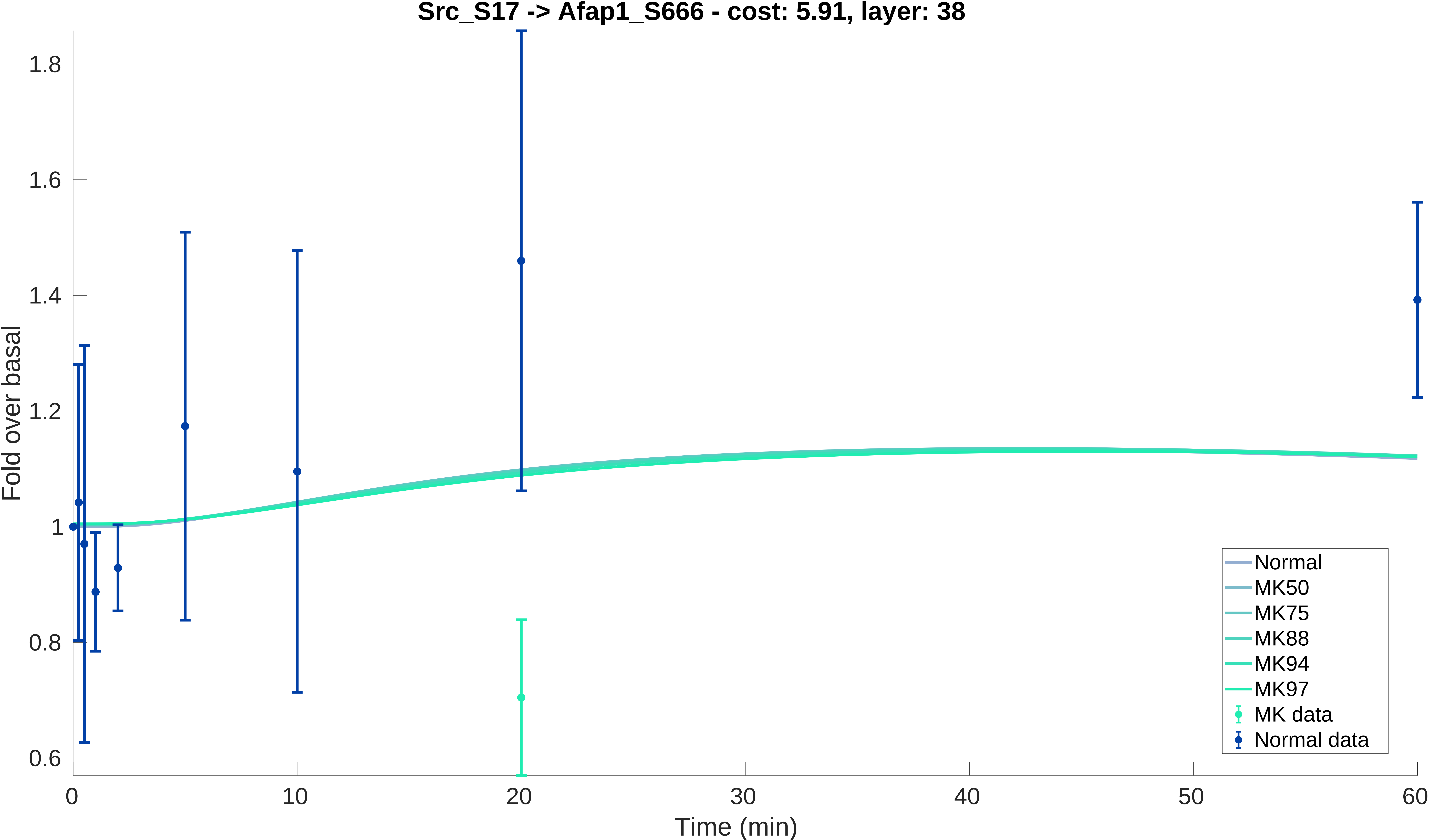

Src_S17 -> Afap1_S666 - cost: 5.91, layer: 38
Fold over basal
Time (min)
0
10
20
30
40
50
60
0.6
0.8
1
1.2
1.4
1.6
1.8
Normal
MK50
MK75
MK88
MK94
MK97
MK data
Normal data

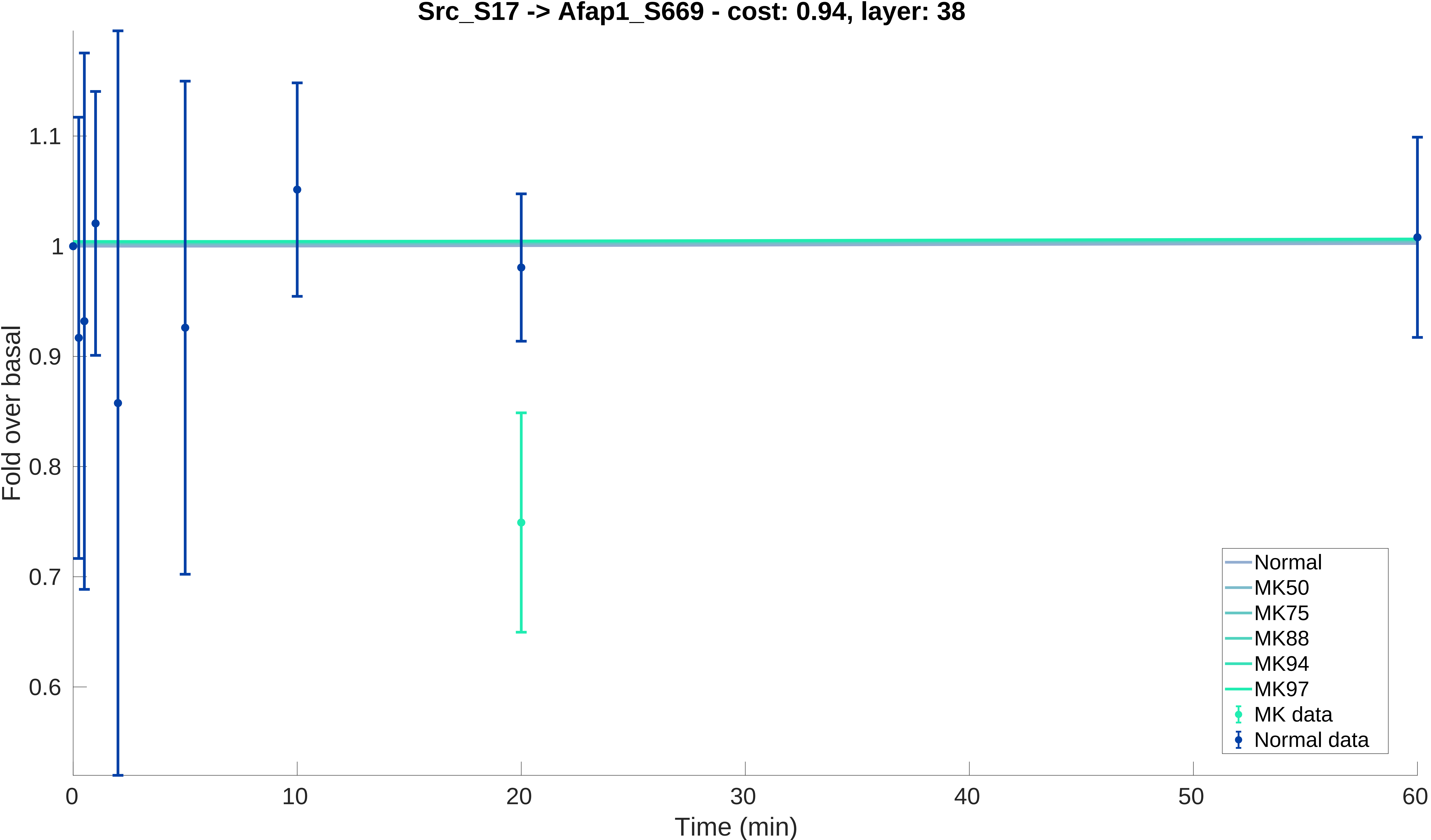

Src_S17 -> Afap1_S669 - cost: 0.94, layer: 38
Fold over basal
Time (min)
0.6
0.7
0.8
0.9
1
1.1
0
10
20
30
40
50
60
Normal
MK50
MK75
MK88
MK94
MK97
MK data
Normal data

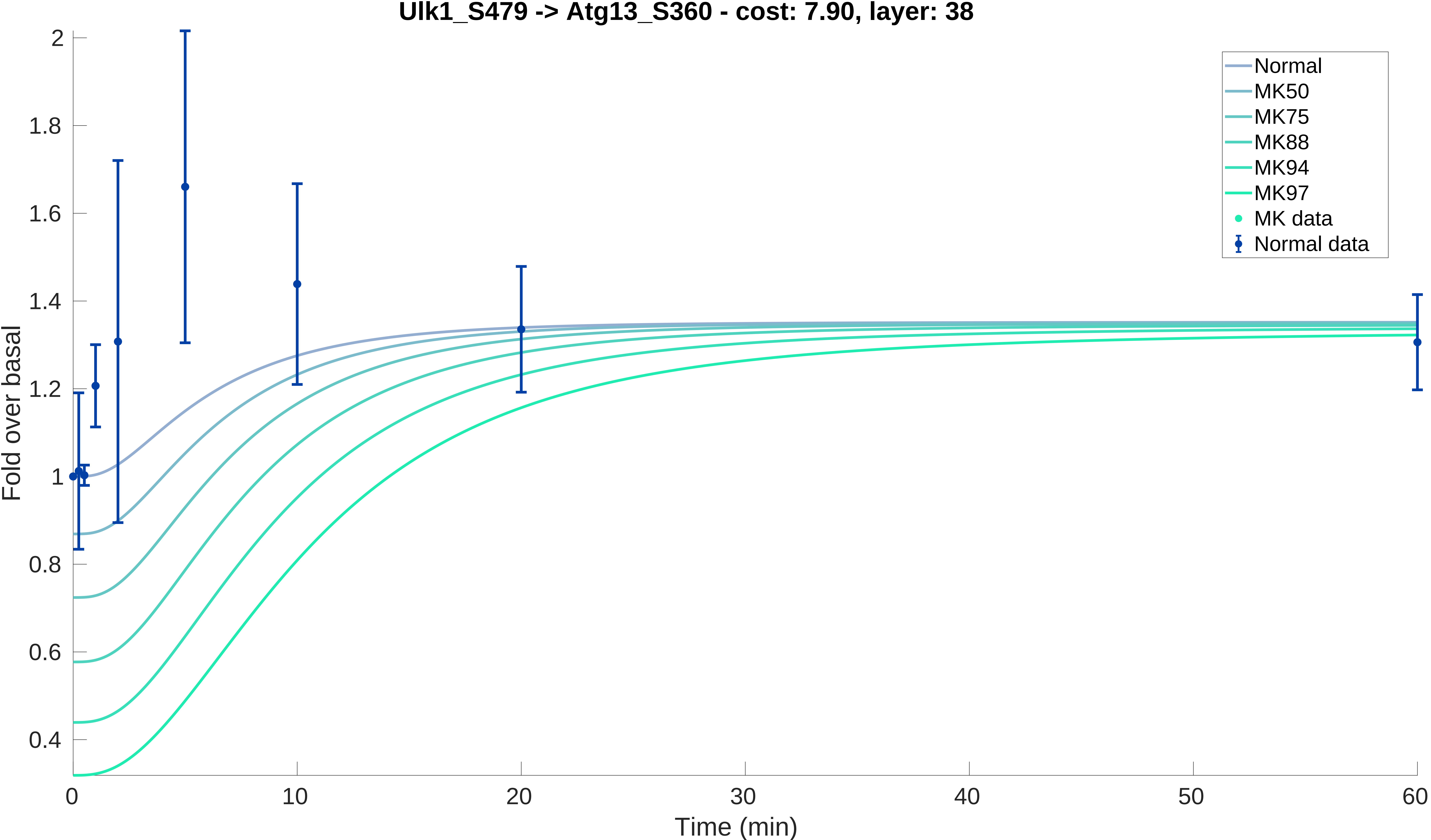

Ulk1_S479 -> Atg13_S360 - cost: 7.90, layer: 38
Fold over basal
Time (min)
Normal
MK50
MK75
MK88
MK94
MK97
MK data
Normal data
0.4
0.6
0.8
1
1.2
1.4
1.6
1.8
2
0
10
20
30
40
50
60

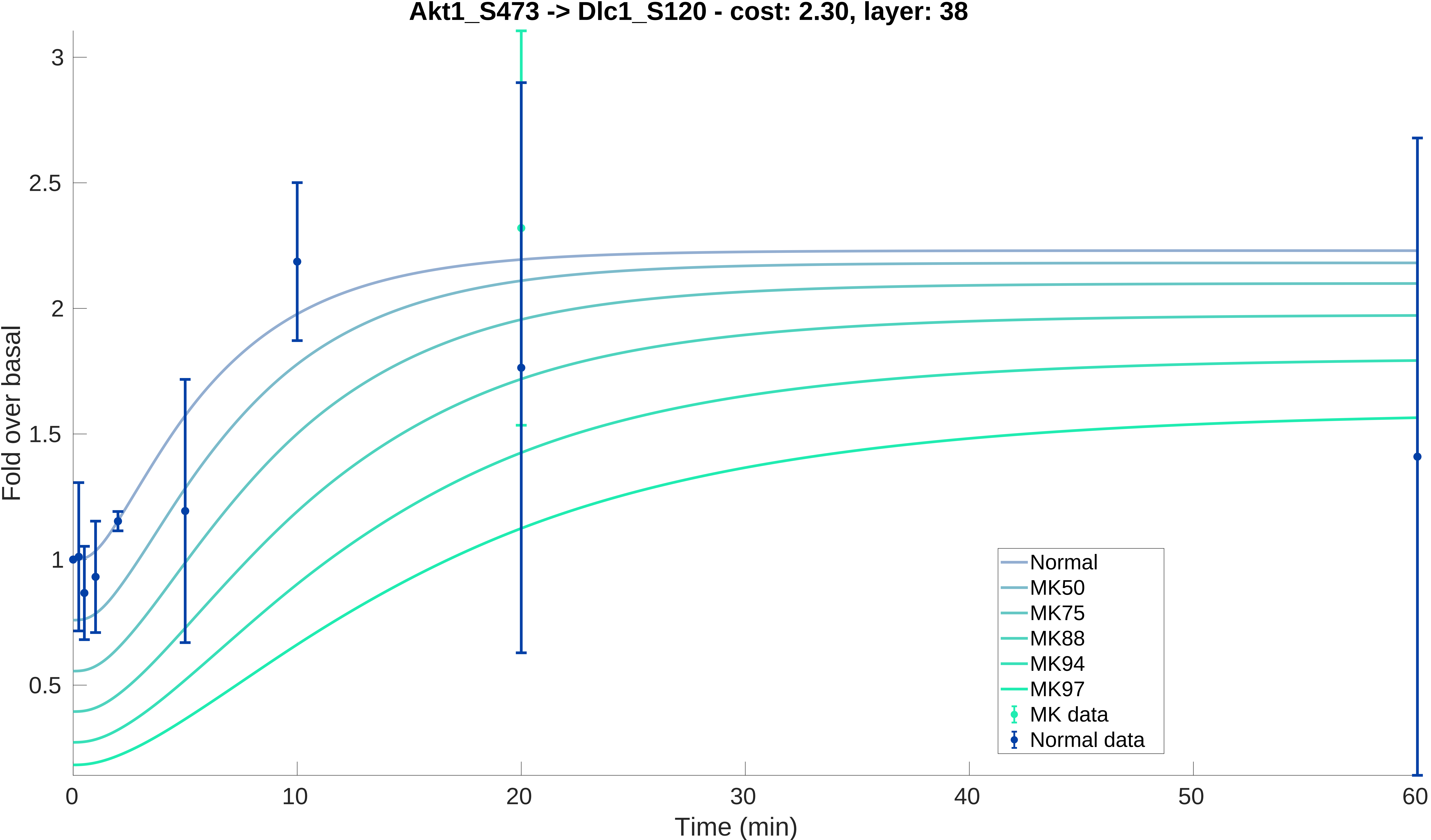
Akt1_S473 -> Dlc1_S120 - cost: 2.30, layer: 38
Fold over basal
3
2.5
2
1.5
1
0.5
0
10
20
30
40
50
60
Time (min)
Normal
MK50
MK75
MK88
MK94
MK97
MK data
Normal data

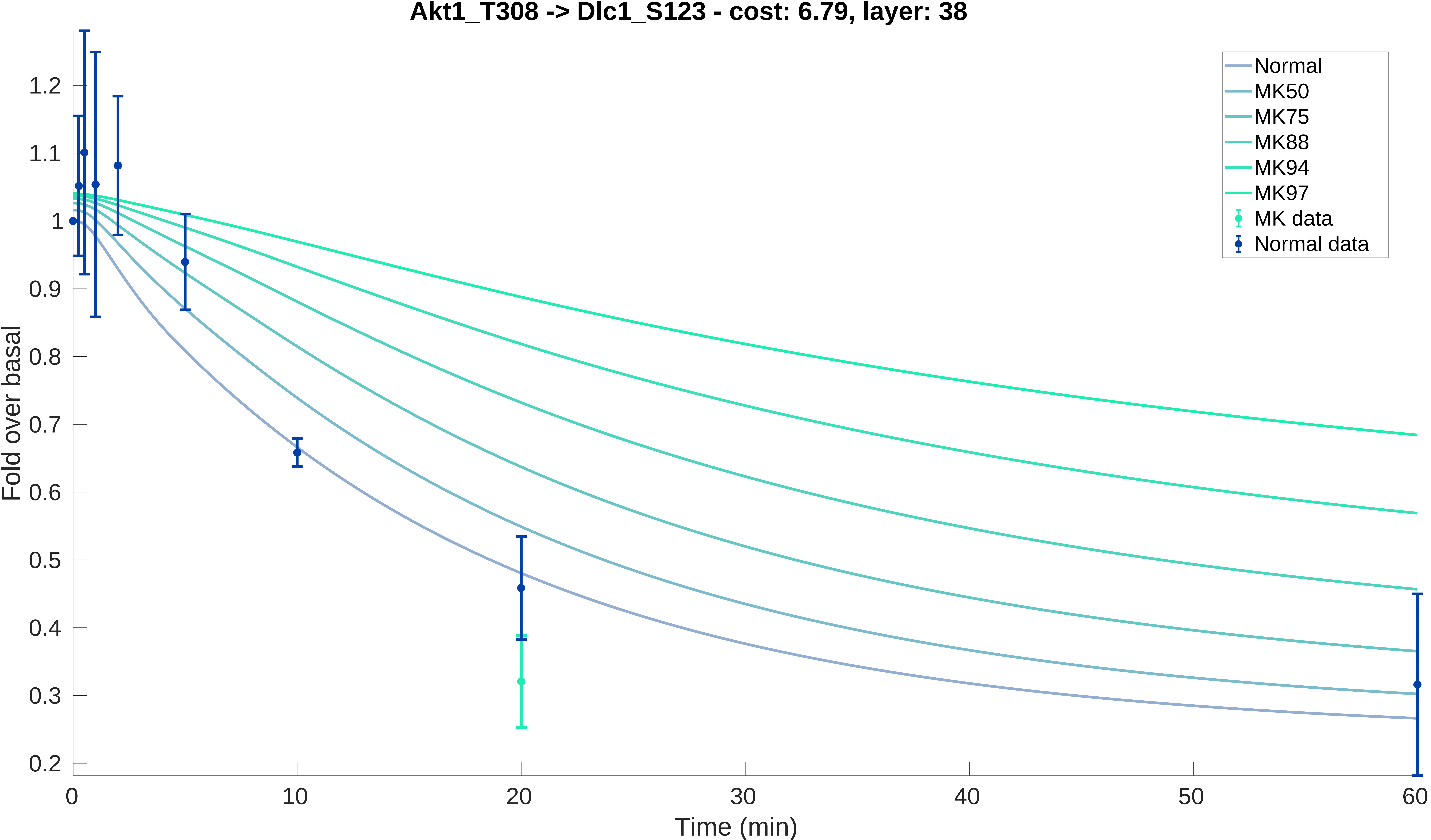

Akt1_T308 -> Dlc1_S123 - cost: 6.79, layer: 38
Fold over basal
Time (min)
0
10
20
30
40
50
60
0.2
0.3
0.4
0.5
0.6
0.7
0.8
0.9
1
1.1
1.2
Normal
MK50
MK75
MK88
MK94
MK97
MK data
Normal data

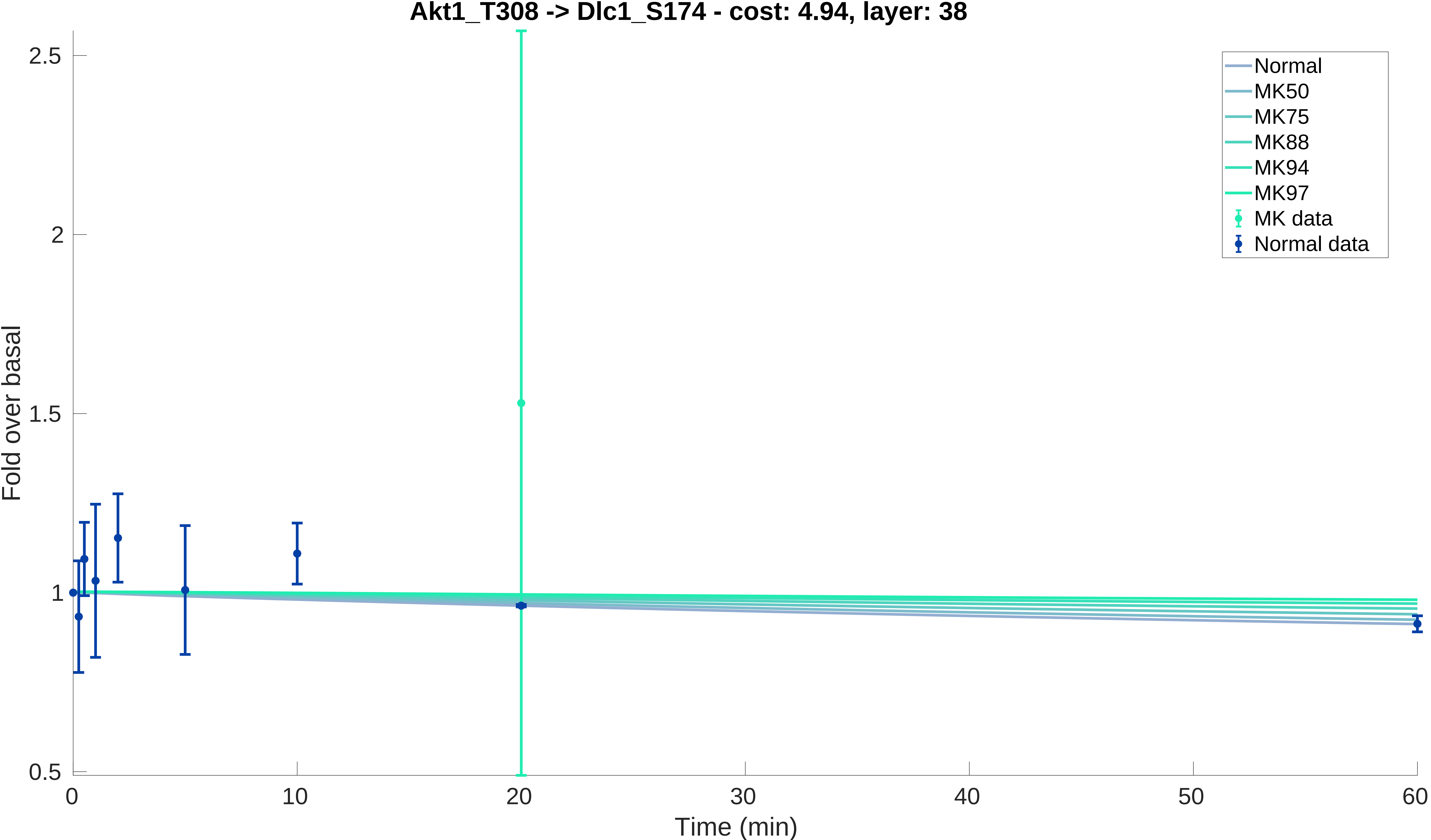

Akt1_T308 -> Dlc1_S174 - cost: 4.94, layer: 38
Fold over basal
Time (min)
0.5
1
1.5
2
2.5
0
10
20
30
40
50
60
Normal
MK50
MK75
MK88
MK94
MK97
MK data
Normal data

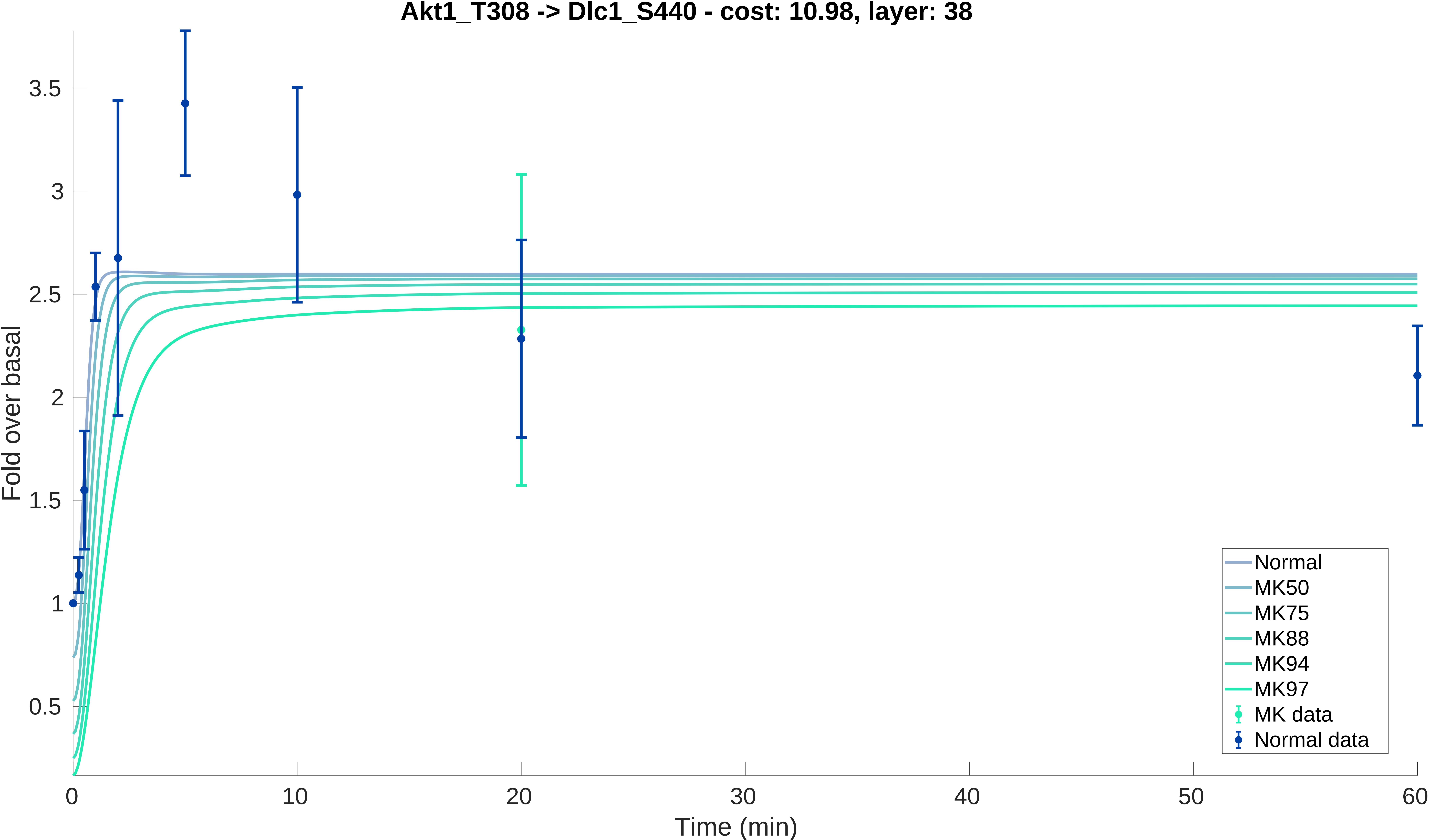

Akt1_T308 -> Dlc1_S440 - cost: 10.98, layer: 38
Fold over basal
3.5
3
2.5
2
1.5
1
0.5
0
10
20
30
40
50
60
Time (min)
Normal
MK50
MK75
MK88
MK94
MK97
MK data
Normal data

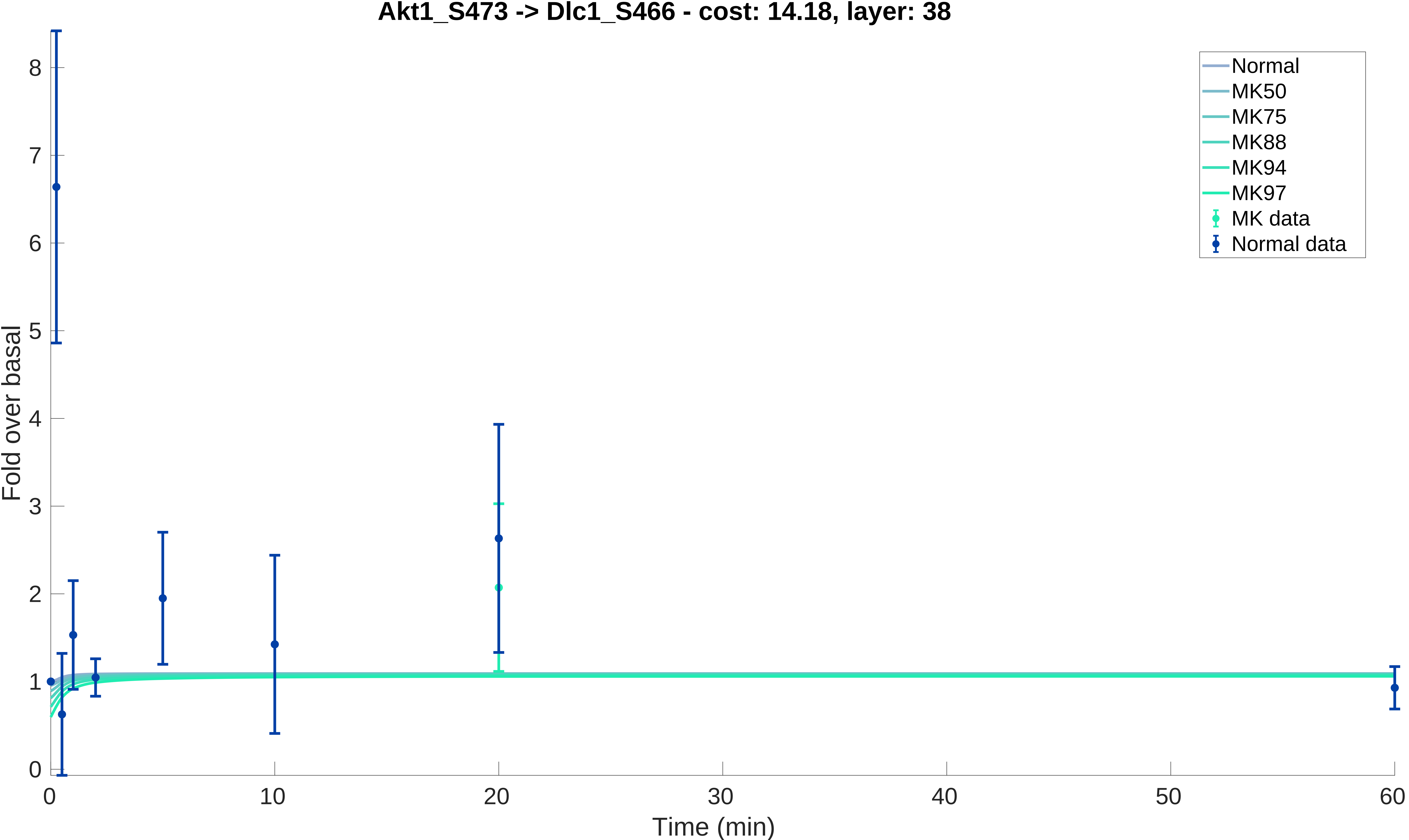

Akt1_S473 -> Dlc1_S466 - cost: 14.18, layer: 38
Fold over basal
Time (min)
0
1
2
3
4
5
6
7
8
0
10
20
30
40
50
60
Normal
MK50
MK75
MK88
MK94
MK97
MK data
Normal data

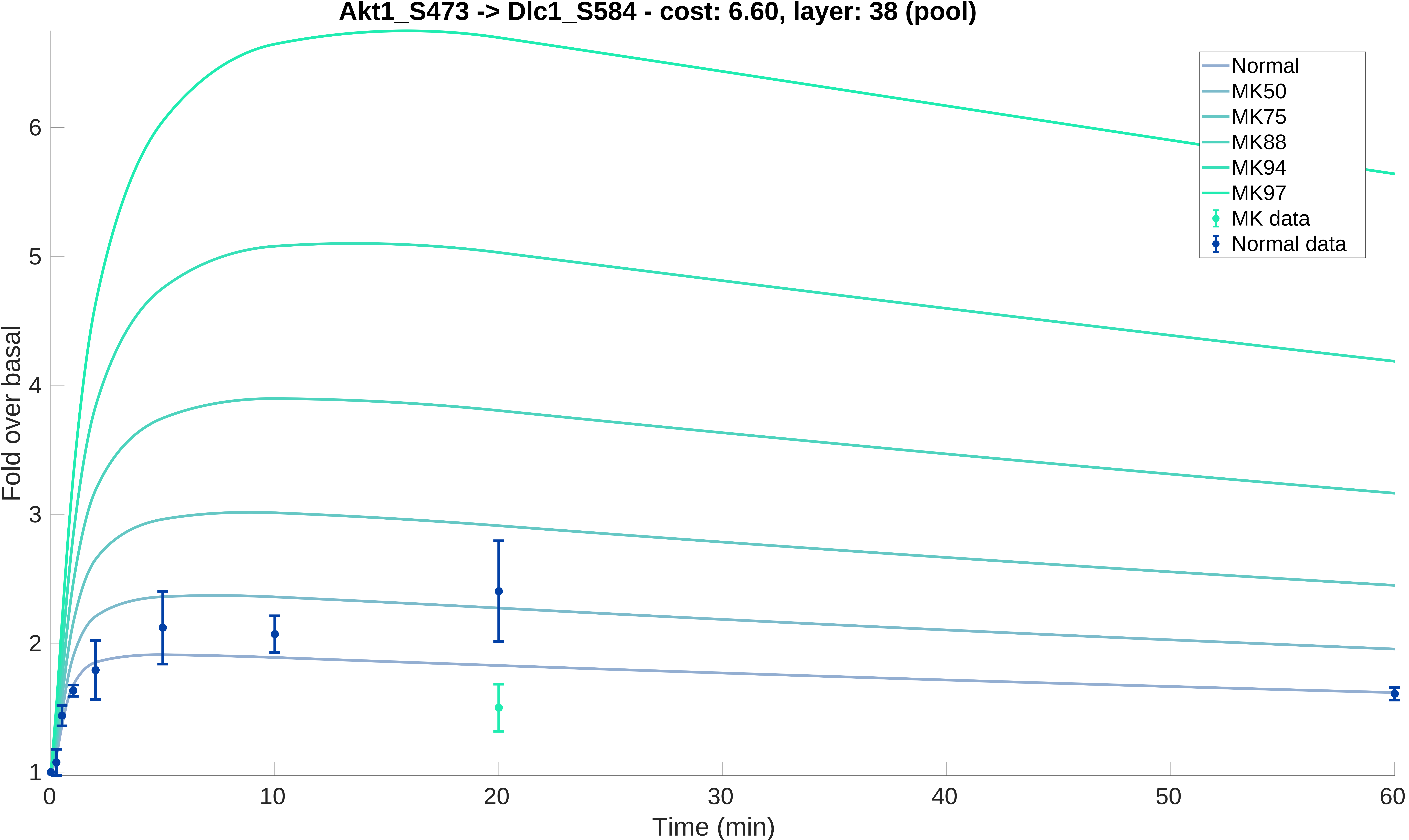

Akt1_S473 -> Dlc1_S584 - cost: 6.60, layer: 38 (pool)
Fold over basal
Time (min)
1
2
3
4
5
6
0
10
20
30
40
50
60
Normal
MK50
MK75
MK88
MK94
MK97
MK data
Normal data

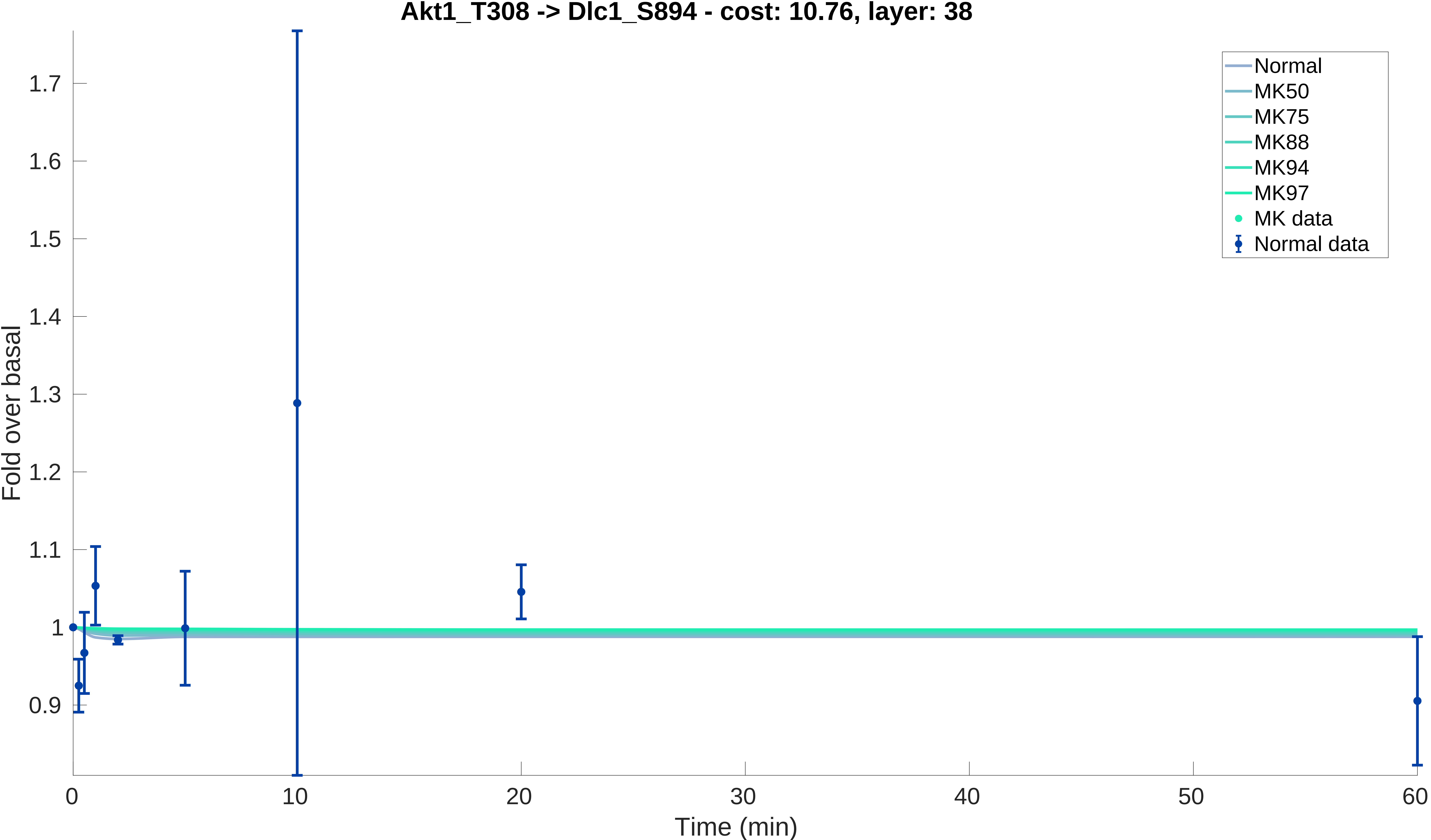

Akt1_T308 -> Dlc1_S894 - cost: 10.76, layer: 38
Fold over basal
Time (min)
0
10
20
30
40
50
60
0.9
1
1.1
1.2
1.3
1.4
1.5
1.6
1.7
Normal
MK50
MK75
MK88
MK94
MK97
MK data
Normal data

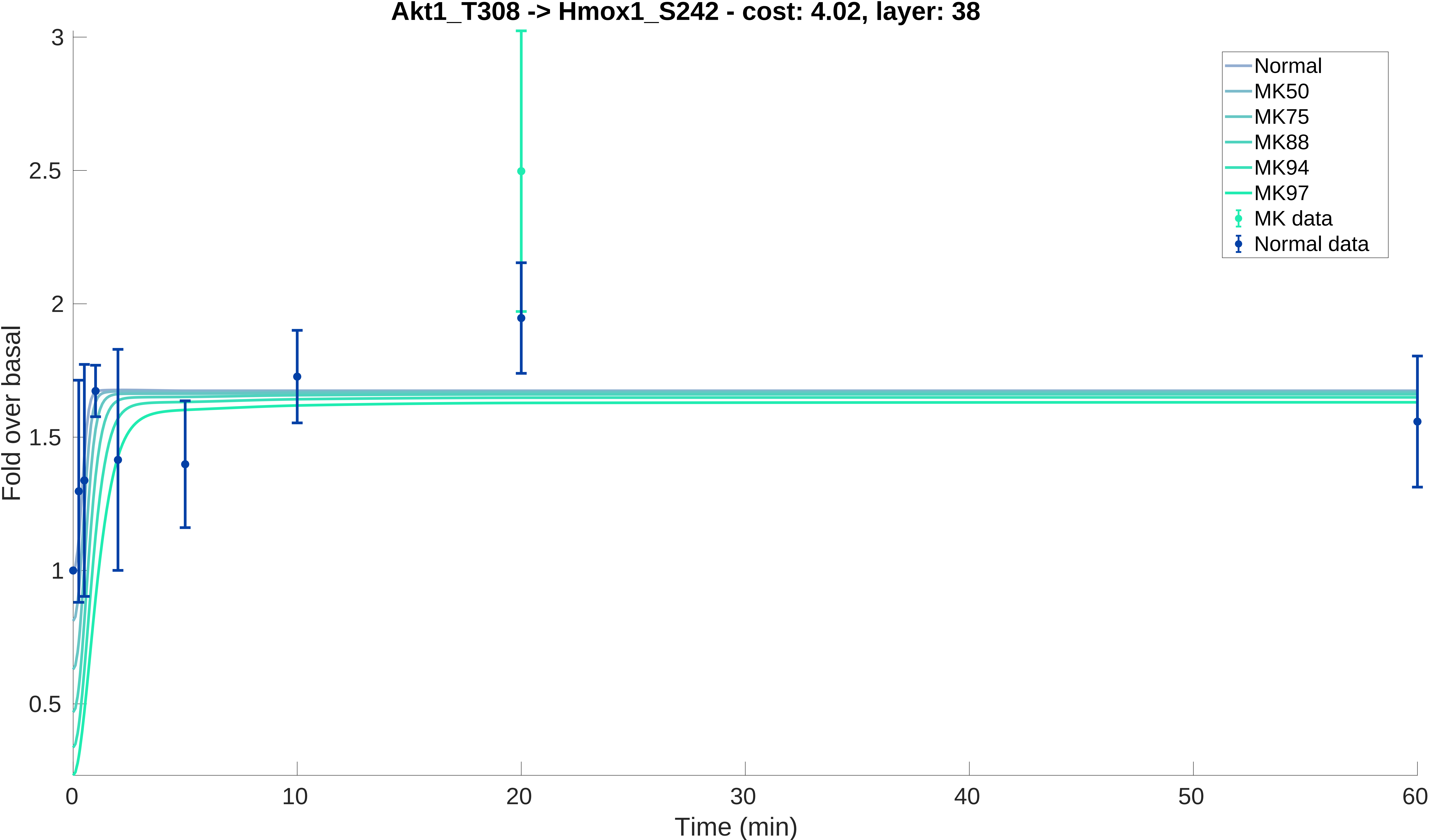

Akt1_T308 -> Hmox1_S242 - cost: 4.02, layer: 38
Normal
MK50
MK75
MK88
MK94
MK97
MK data
Normal data
Fold over basal
Time (min)
0.5
1
1.5
2
2.5
3
0
10
20
30
40
50
60

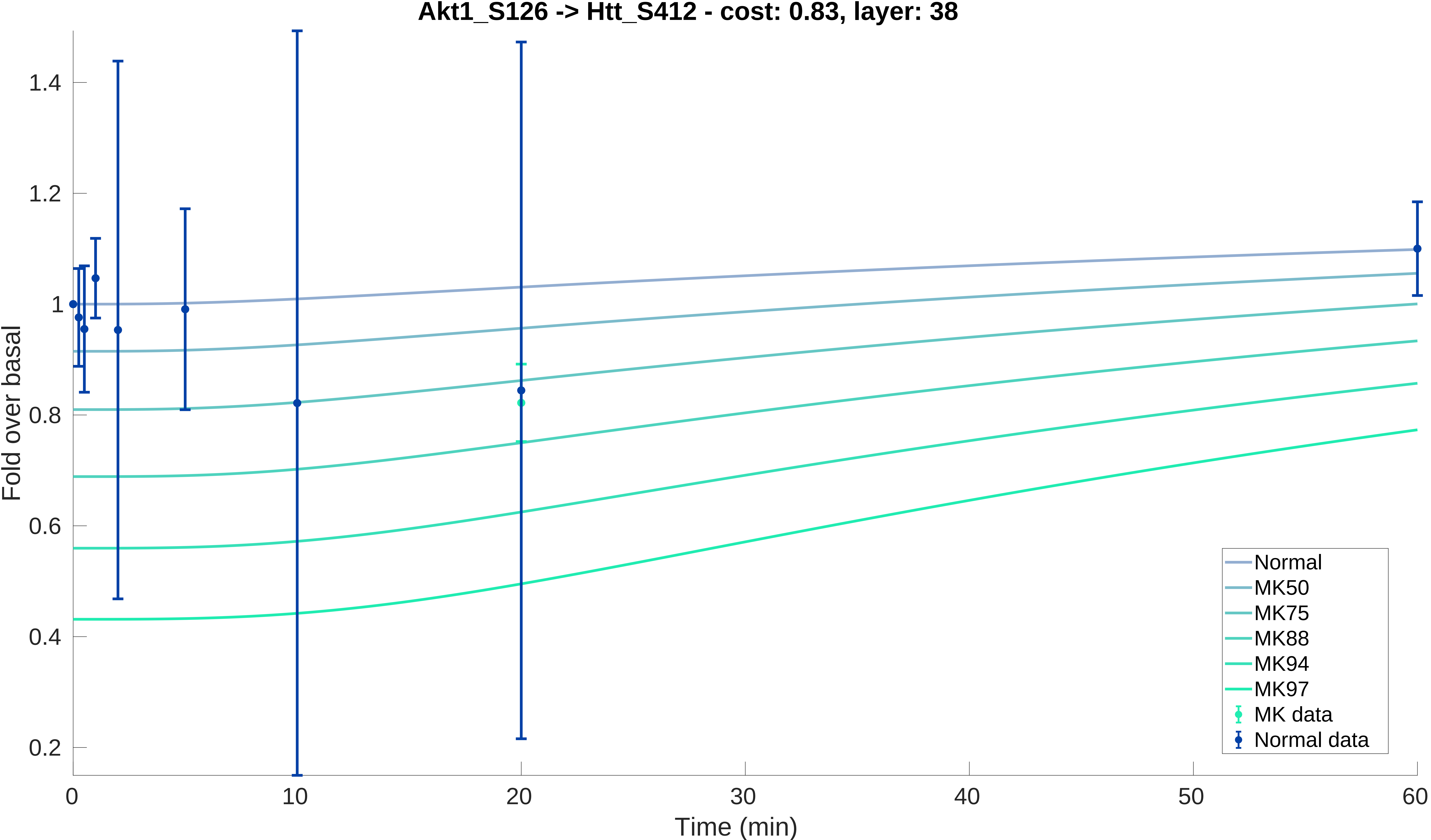
Akt1_S126 -> Htt_S412 - cost: 0.83, layer: 38
Fold over basal
Time (min)
0.2
0.4
0.6
0.8
1
1.2
1.4
0
10
20
30
40
50
60
Normal
MK50
MK75
MK88
MK94
MK97
MK data
Normal data

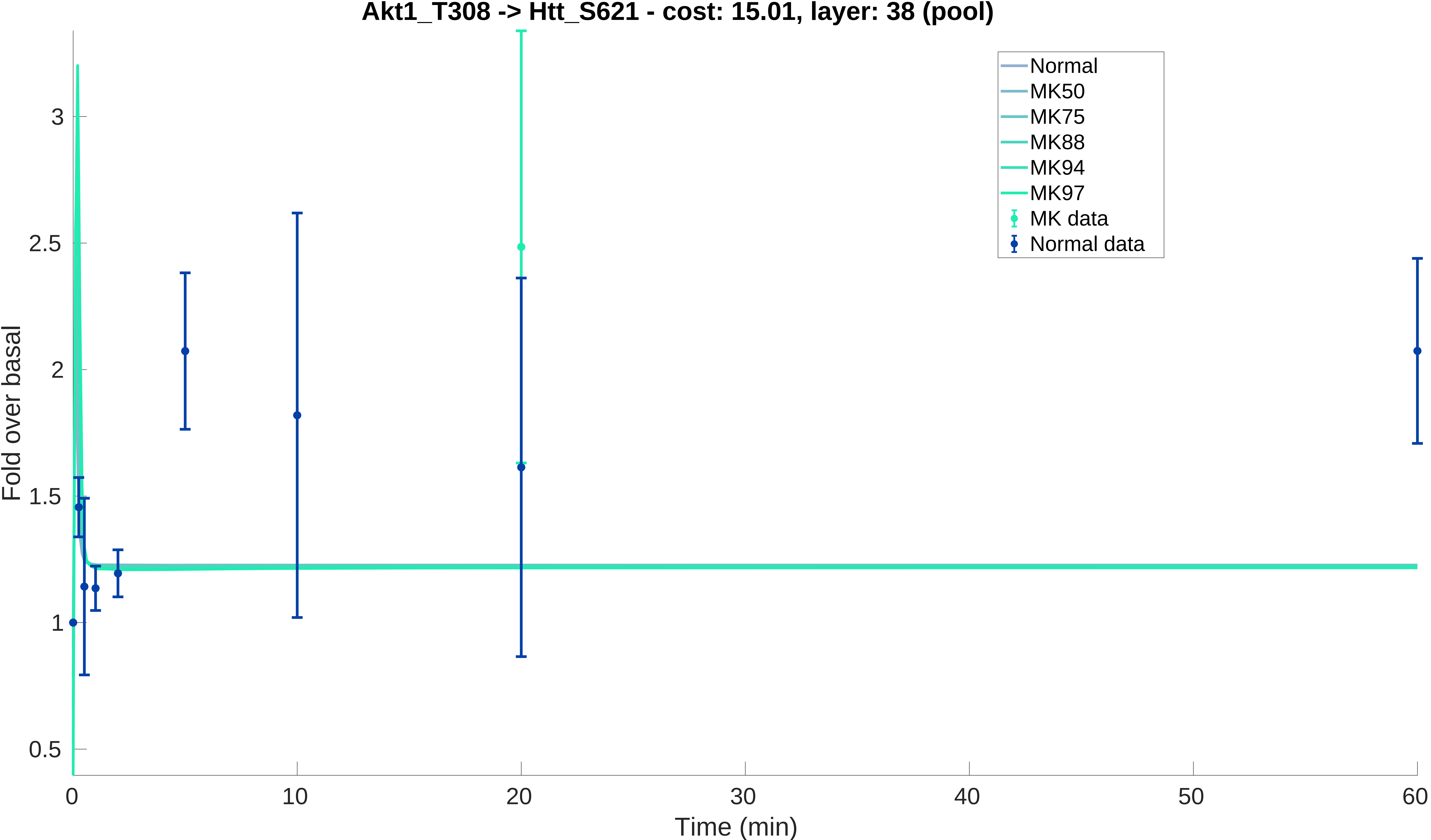

Akt1_T308 -> Htt_S621 - cost: 15.01, layer: 38 (pool)
Normal
MK50
MK75
MK88
MK94
MK97
MK data
Normal data
Fold over basal
Time (min)
0.5
1
1.5
2
2.5
3
0
10
20
30
40
50
60

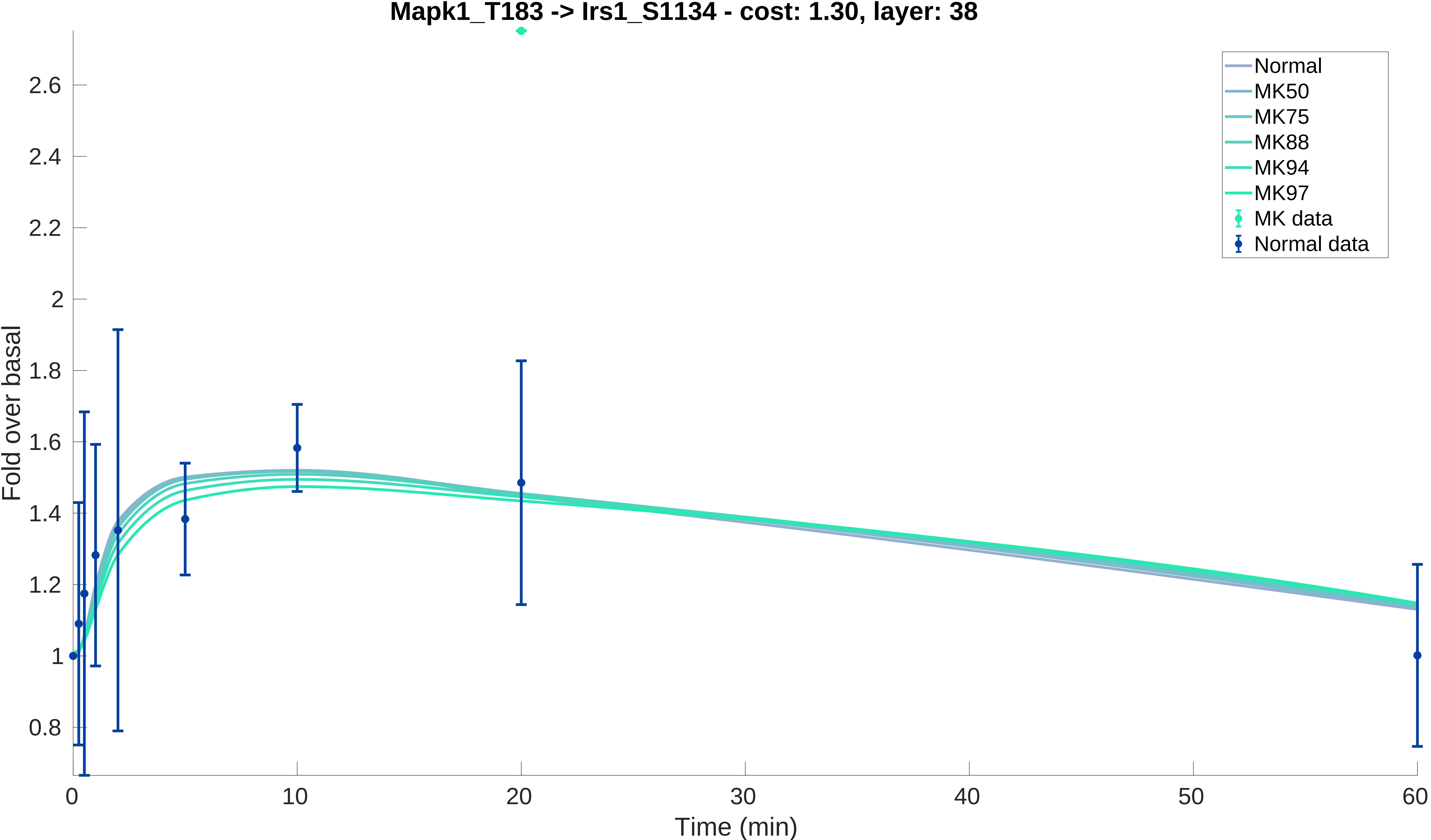

Mapk1_T183 -> Irs1_S1134 - cost: 1.30, layer: 38
Fold over basal
Time (min)
Normal
MK50
MK75
MK88
MK94
MK97
MK data
Normal data
0.8
1
1.2
1.4
1.6
1.8
2
2.2
2.4
2.6
0
10
20
30
40
50
60

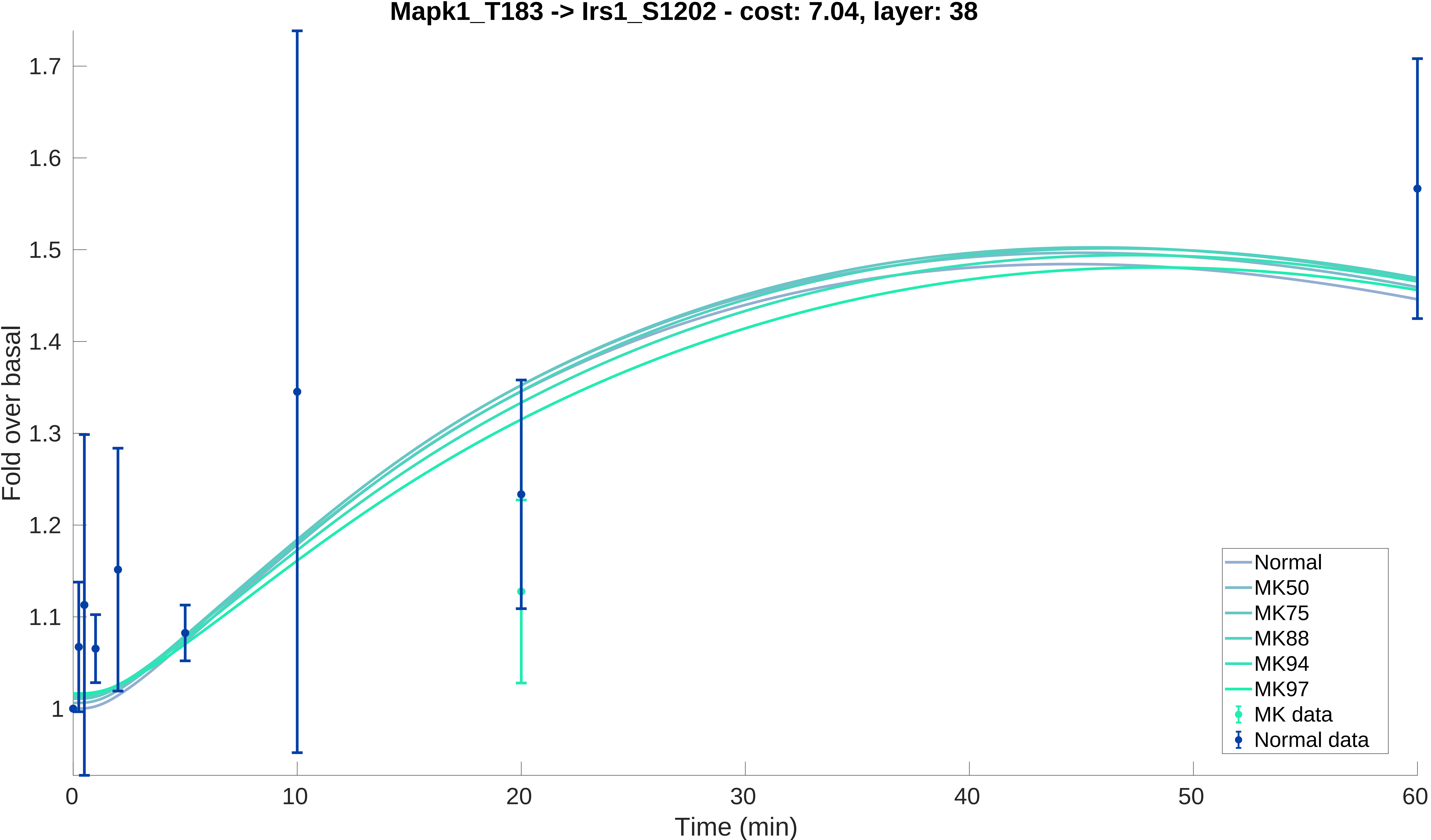
Mapk1_T183 -> Irs1_S1202 - cost: 7.04, layer: 38
Fold over basal
Time (min)
1
1.1
1.2
1.3
1.4
1.5
1.6
1.7
0
10
20
30
40
50
60
Normal
MK50
MK75
MK88
MK94
MK97
MK data
Normal data

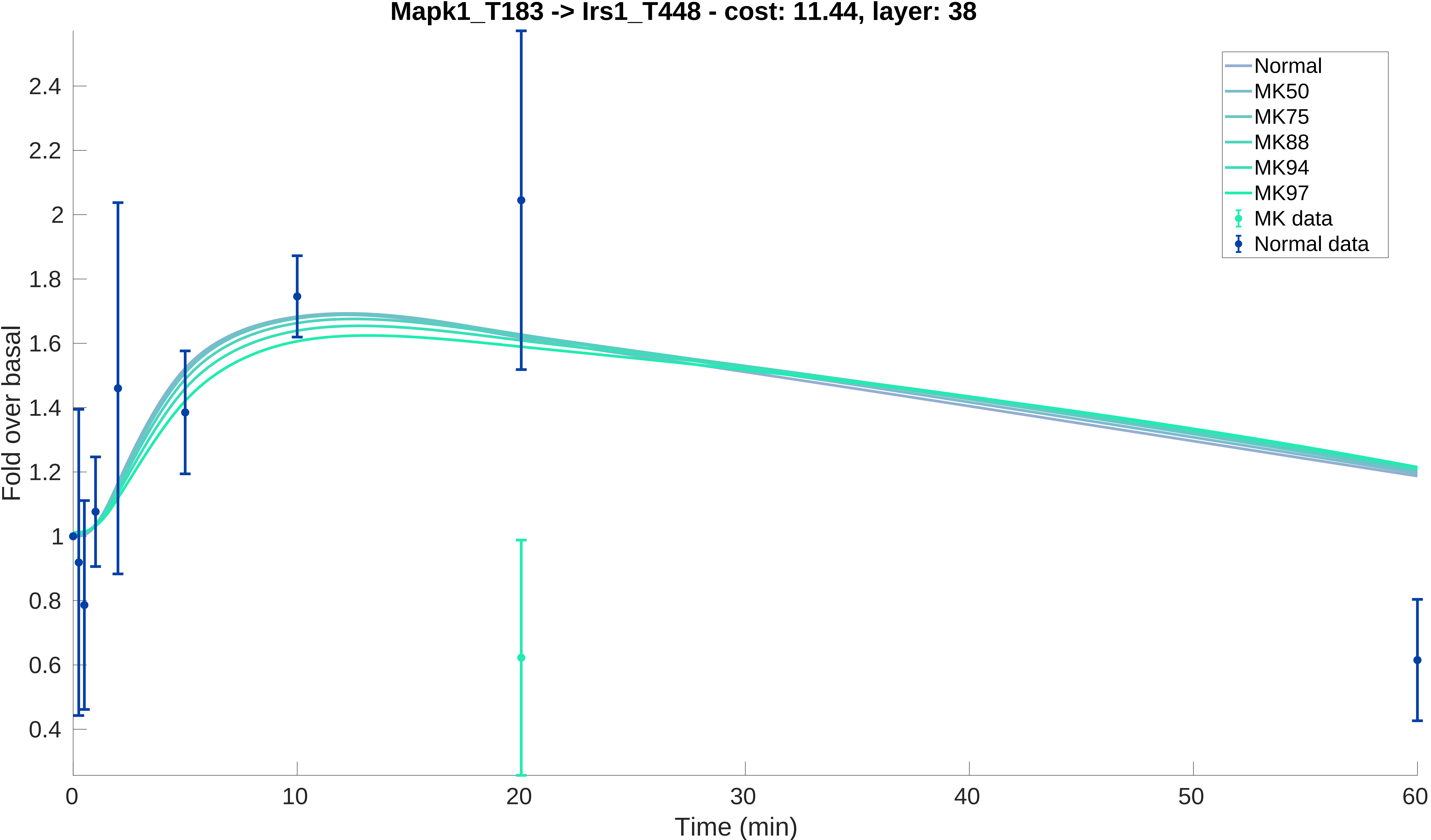

Mapk1_T183 -> Irs1_T448 - cost: 11.44, layer: 38
Fold over basal
Time (min)
0
10
20
30
40
50
60
0.4
0.6
0.8
1
1.2
1.4
1.6
1.8
2
2.2
2.4
Normal
MK50
MK75
MK88
MK94
MK97
MK data
Normal data

Akt1_S129 -> Pea15_S104 - cost: 7.12, layer: 38

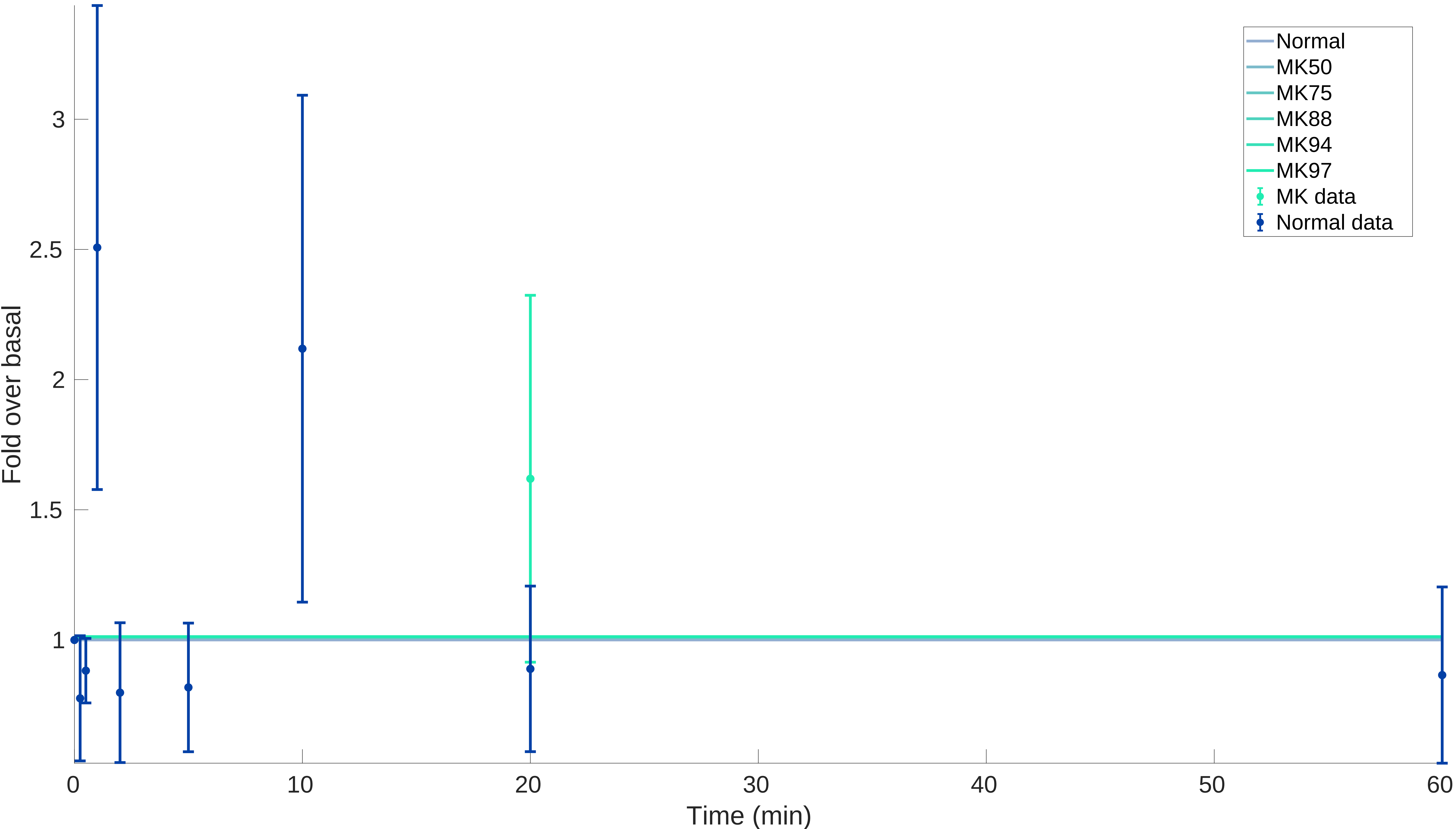

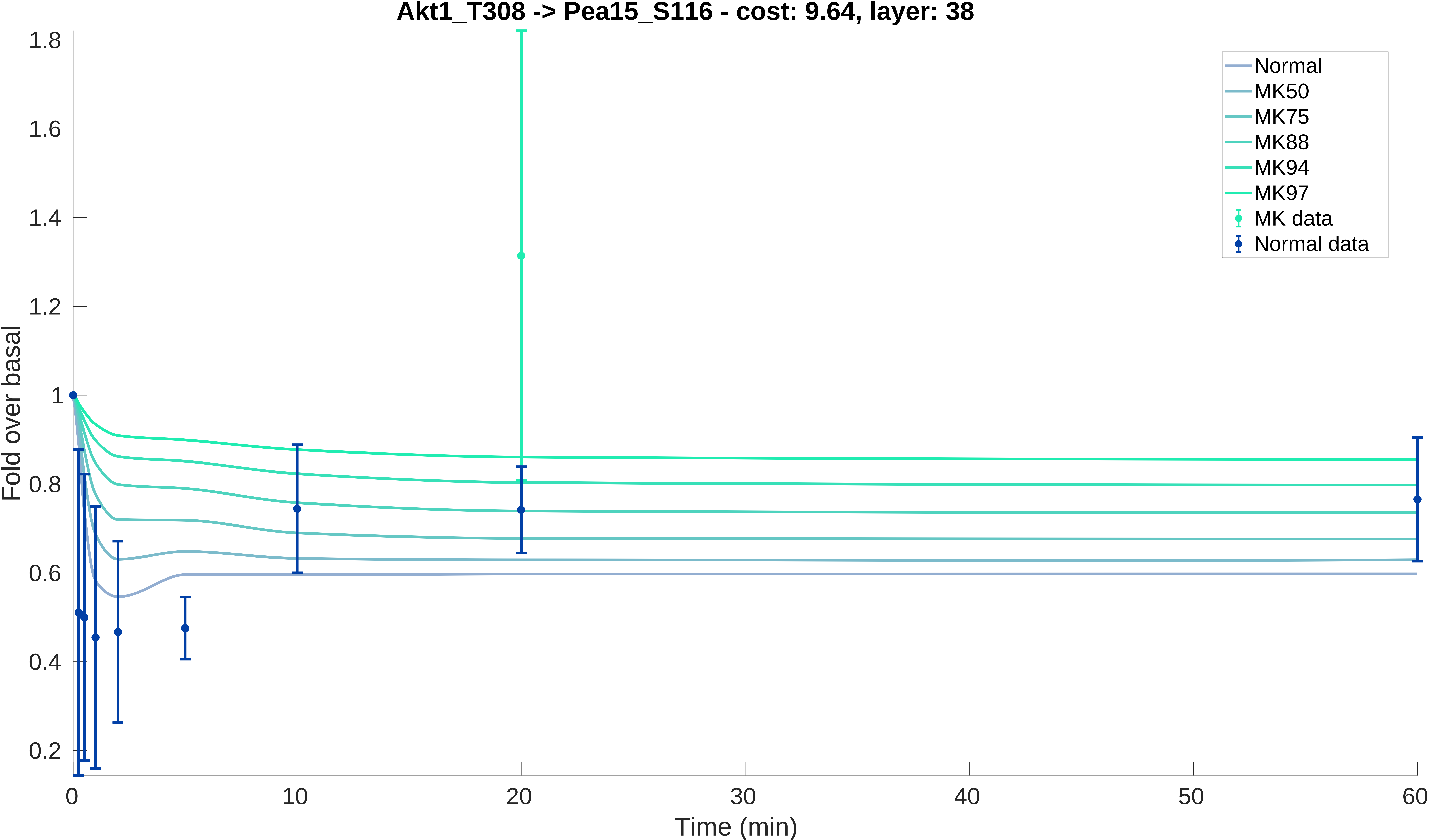

Akt1_T308 -> Pea15_S116 - cost: 9.64, layer: 38
Fold over basal
1.8
1.6
1.4
1.2
1
0.8
0.6
0.4
0.2
0
10
20
30
40
50
60
Time (min)
Normal
MK50
MK75
MK88
MK94
MK97
MK data
Normal data

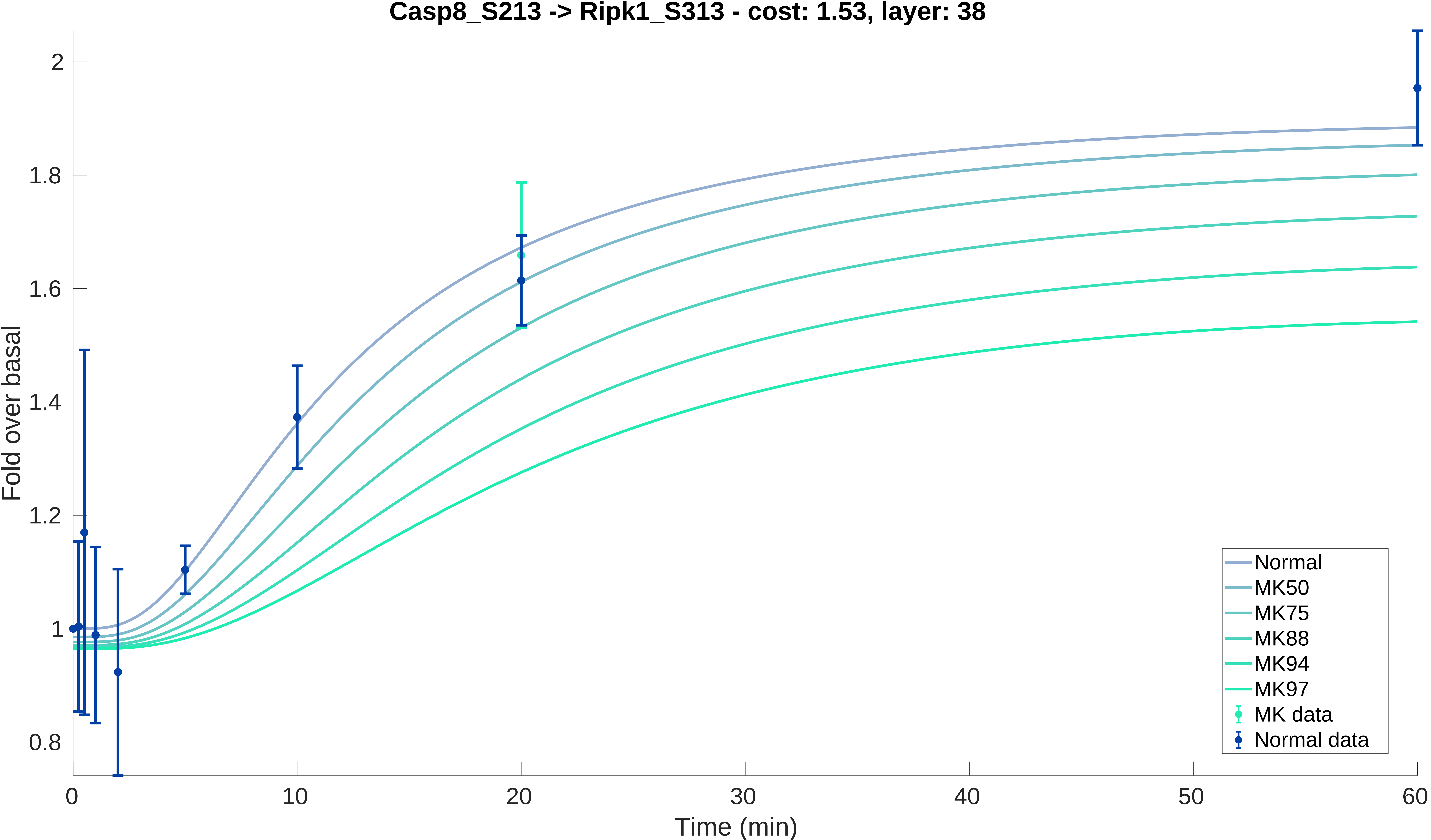

Casp8_S213 -> Ripk1_S313 - cost: 1.53, layer: 38
Fold over basal
Time (min)
0
10
20
30
40
50
60
0.8
1
1.2
1.4
1.6
1.8
2
Normal
MK50
MK75
MK88
MK94
MK97
MK data
Normal data

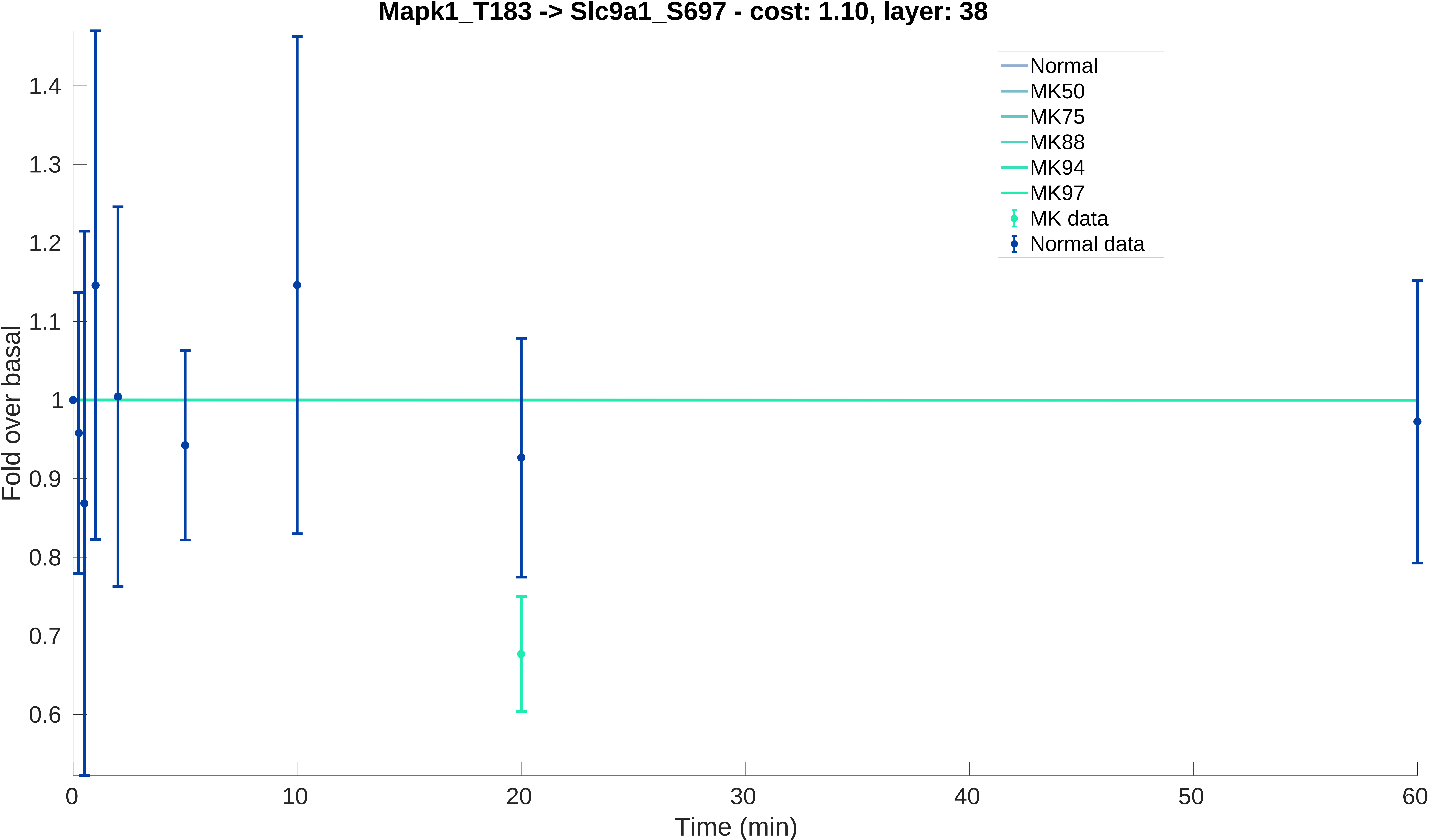

Mapk1_T183 -> Slc9a1_S697 - cost: 1.10, layer: 38
Normal
MK50
MK75
MK88
MK94
MK97
MK data
Normal data
Fold over basal
1.4
1.3
1.2
1.1
1
0.9
0.8
0.7
0.6
0
10
20
30
40
50
60
Time (min)

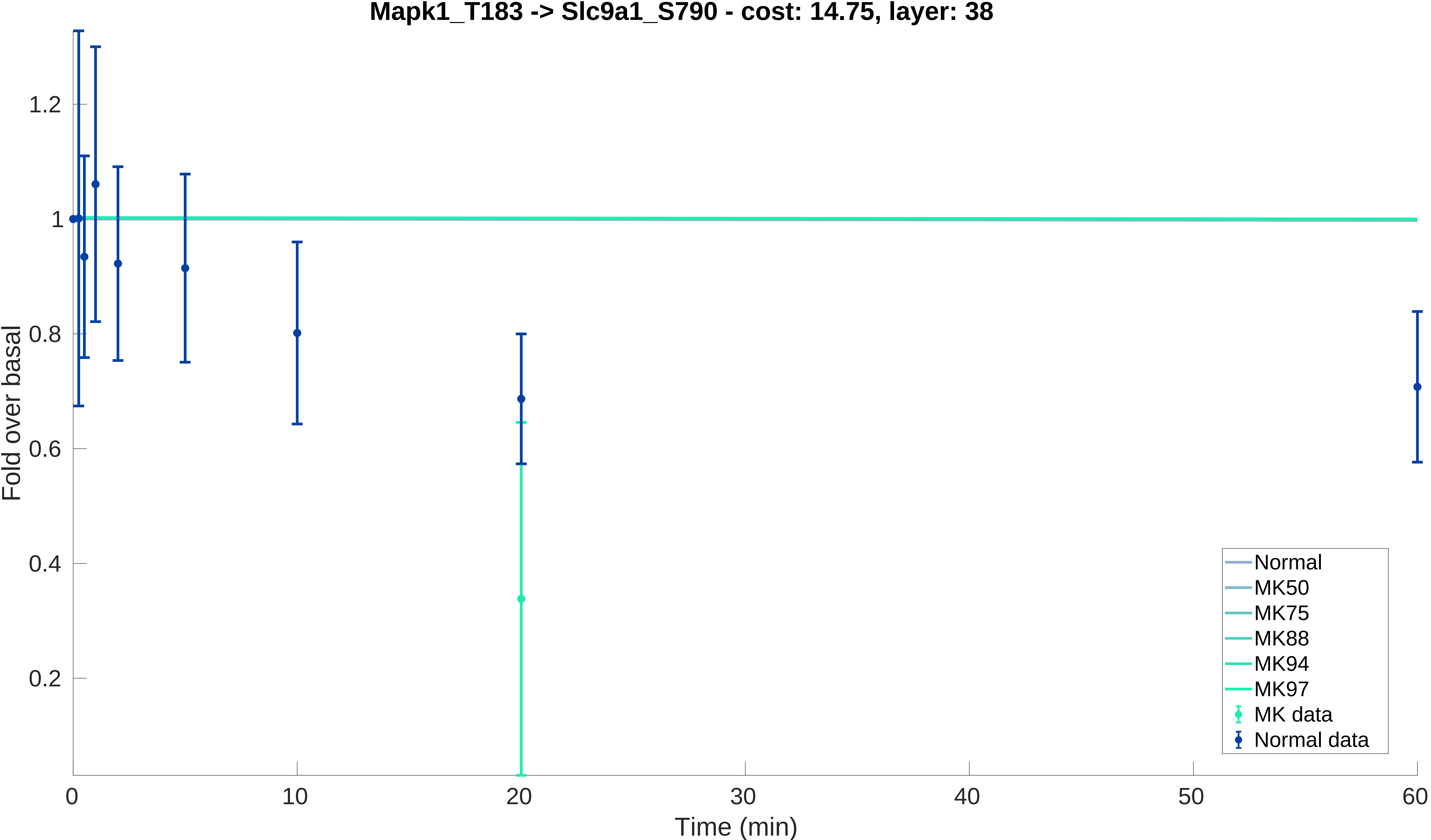
Mapk1_T183 -> Slc9a1_S790 - cost: 14.75, layer: 38
Fold over basal
1.2
1
0.8
0.6
0.4
0.2
0
10
20
30
40
50
60
Time (min)
Normal
MK50
MK75
MK88
MK94
MK97
MK data
Normal data

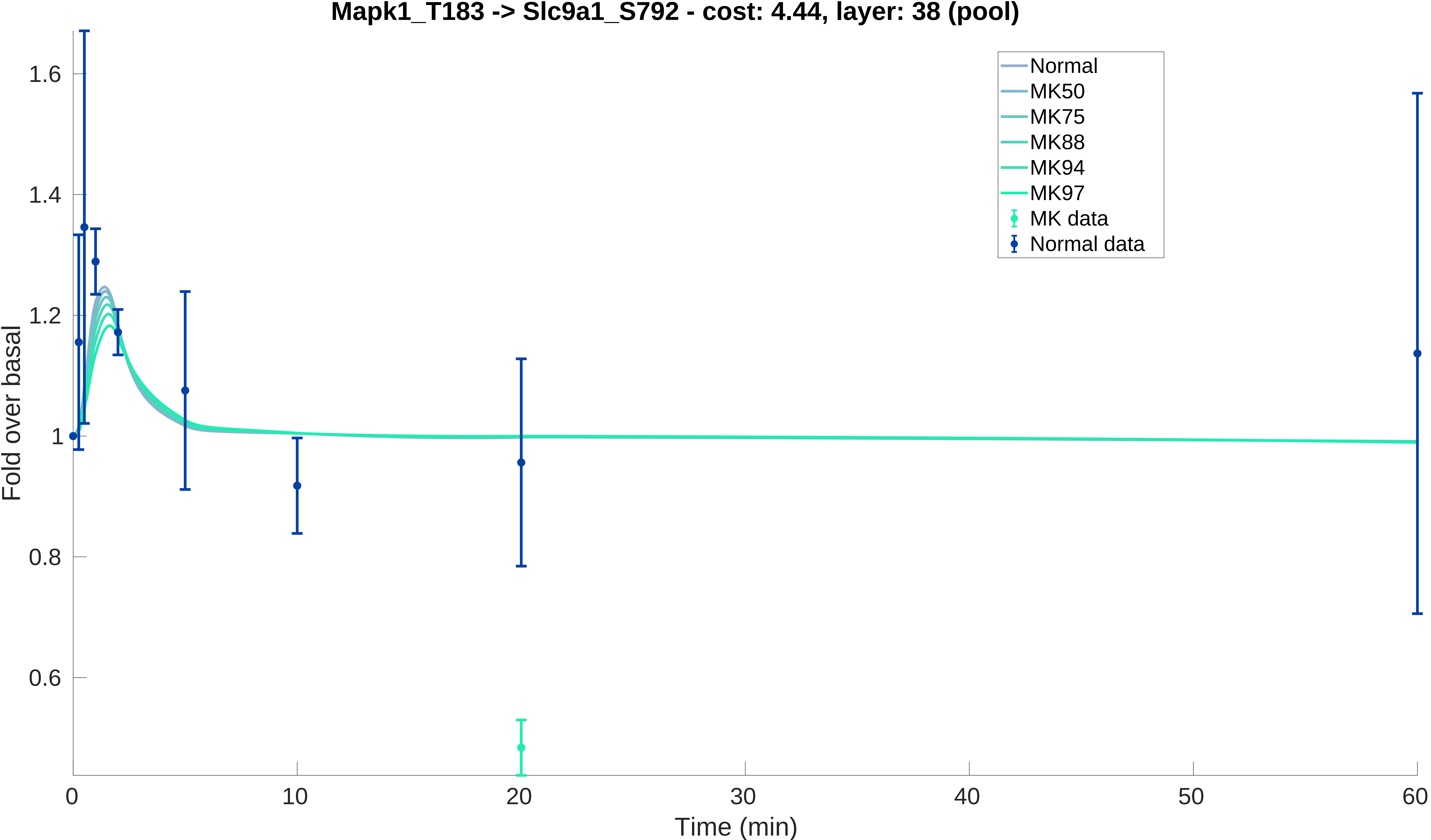

Mapk1_T183 -> Slc9a1_S792 - cost: 4.44, layer: 38 (pool)
Normal
MK50
MK75
MK88
MK94
MK97
MK data
Normal data
Fold over basal
1.6
1.4
1.2
1
0.8
0.6
0
10
20
30
40
50
60
Time (min)

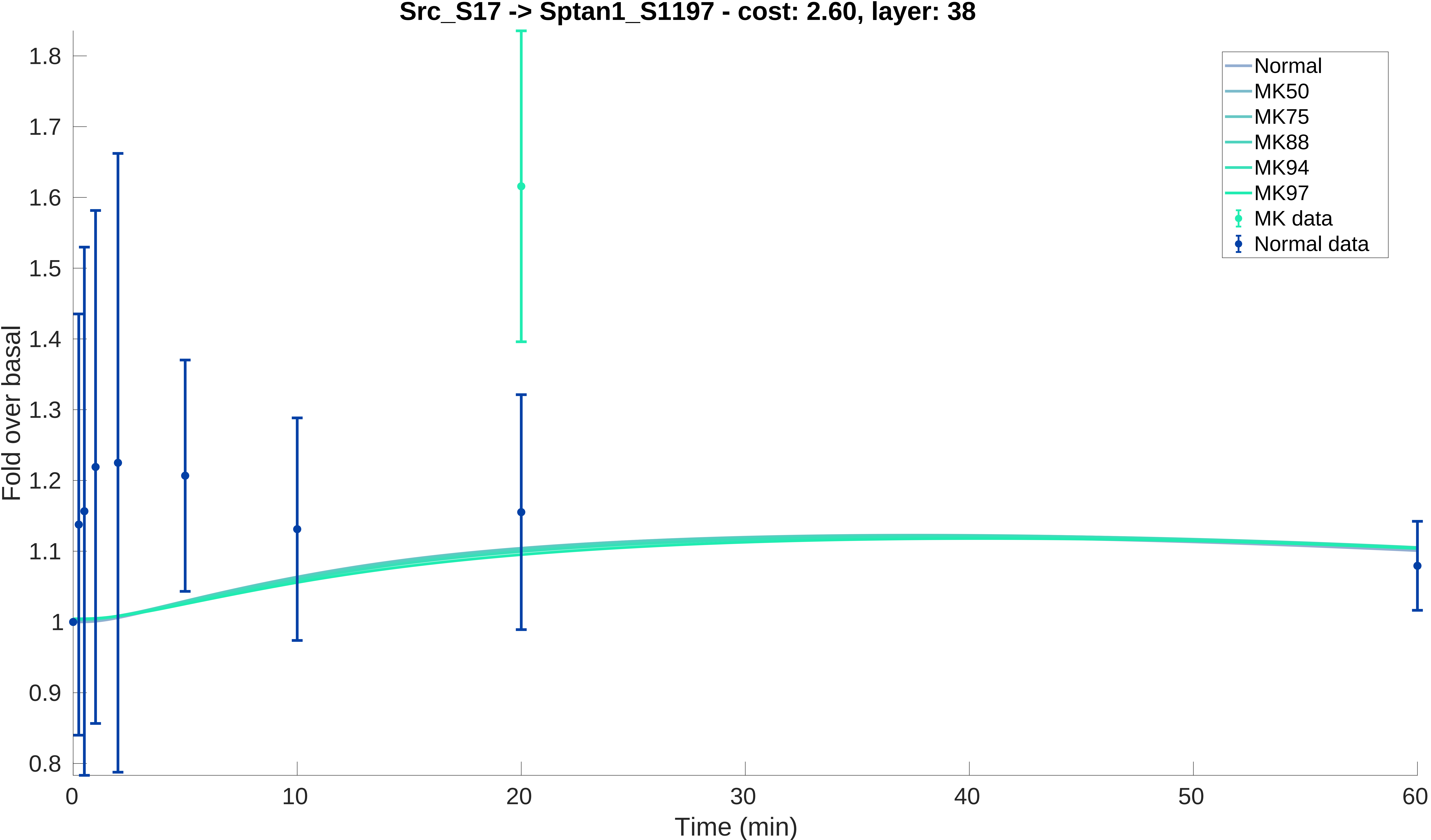

Src_S17 -> Sptan1_S1197 - cost: 2.60, layer: 38
Fold over basal
Time (min)
Normal
MK50
MK75
MK88
MK94
MK97
MK data
Normal data
0.8
0.9
1
1.1
1.2
1.3
1.4
1.5
1.6
1.7
1.8
0
10
20
30
40
50
60

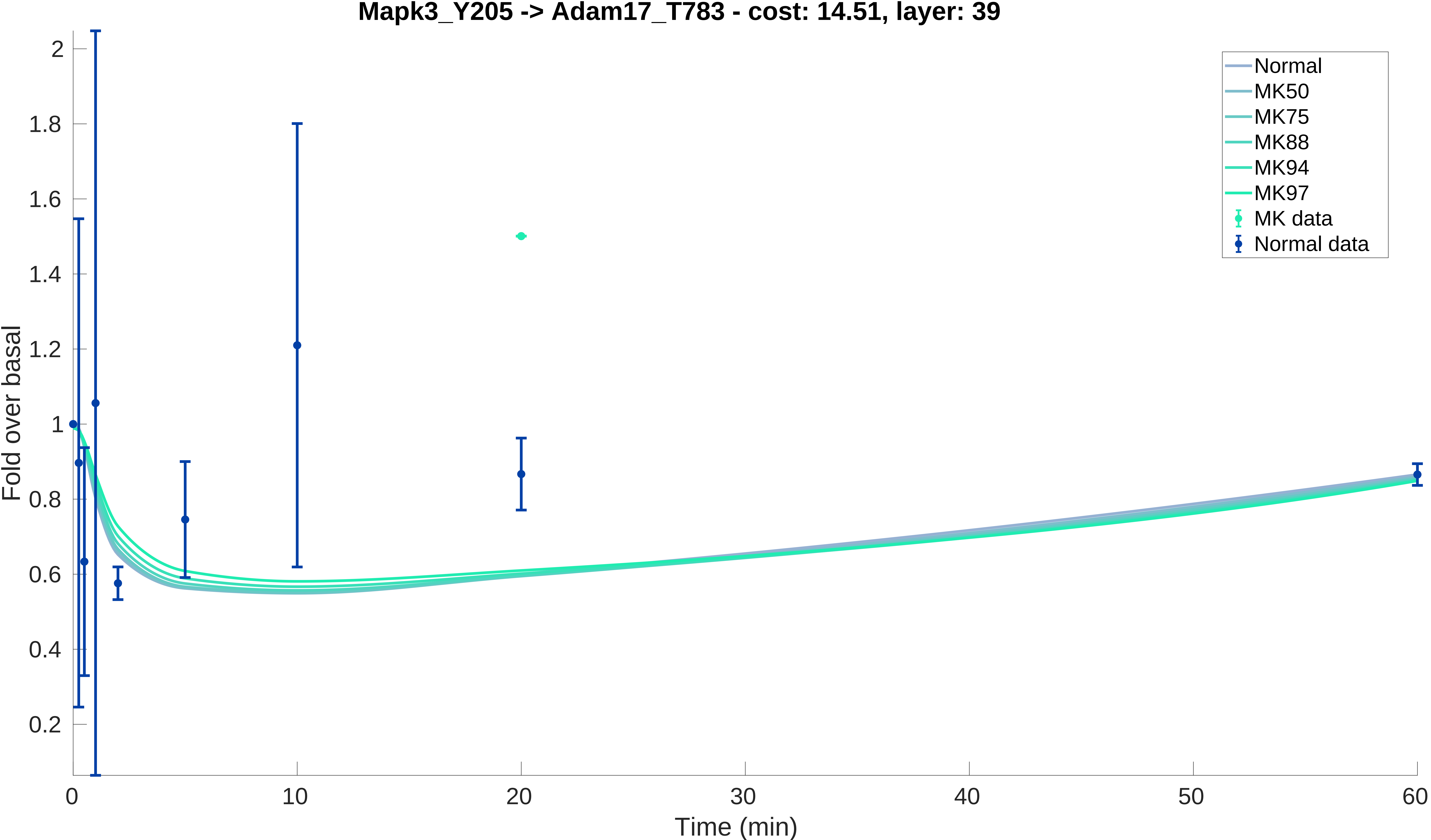

Mapk3_Y205 -> Adam17_T783 - cost: 14.51, layer: 39
Fold over basal
Time (min)
0
10
20
30
40
50
60
0.2
0.4
0.6
0.8
1
1.2
1.4
1.6
1.8
2
Normal
MK50
MK75
MK88
MK94
MK97
MK data
Normal data

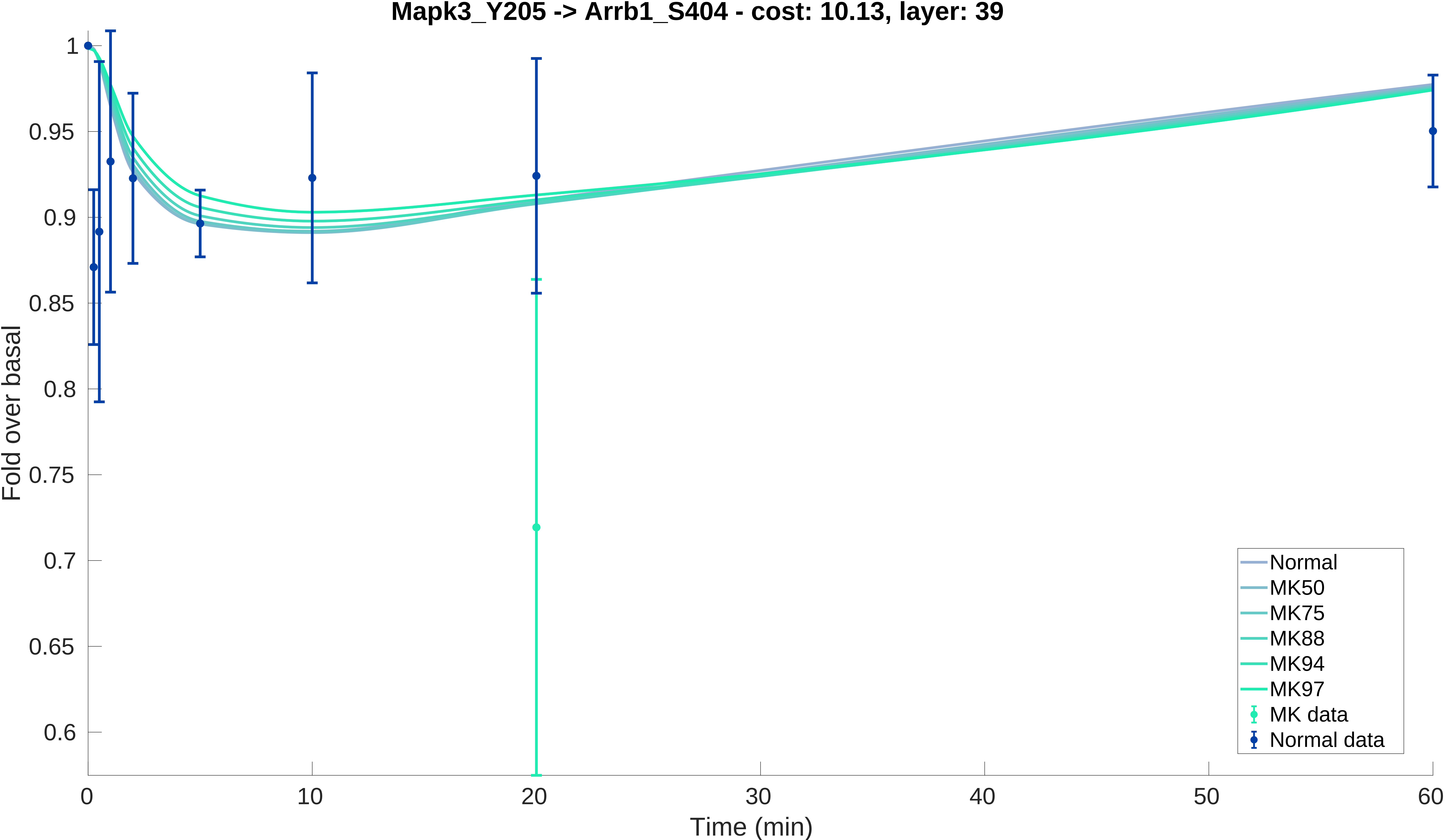
Mapk3_Y205 -> Arrb1_S404 - cost: 10.13, layer: 39
Fold over basal
Time (min)
1
0.95
0.9
0.85
0.8
0.75
0.7
0.65
0.6
0
10
20
30
40
50
60
Normal
MK50
MK75
MK88
MK94
MK97
MK data
Normal data

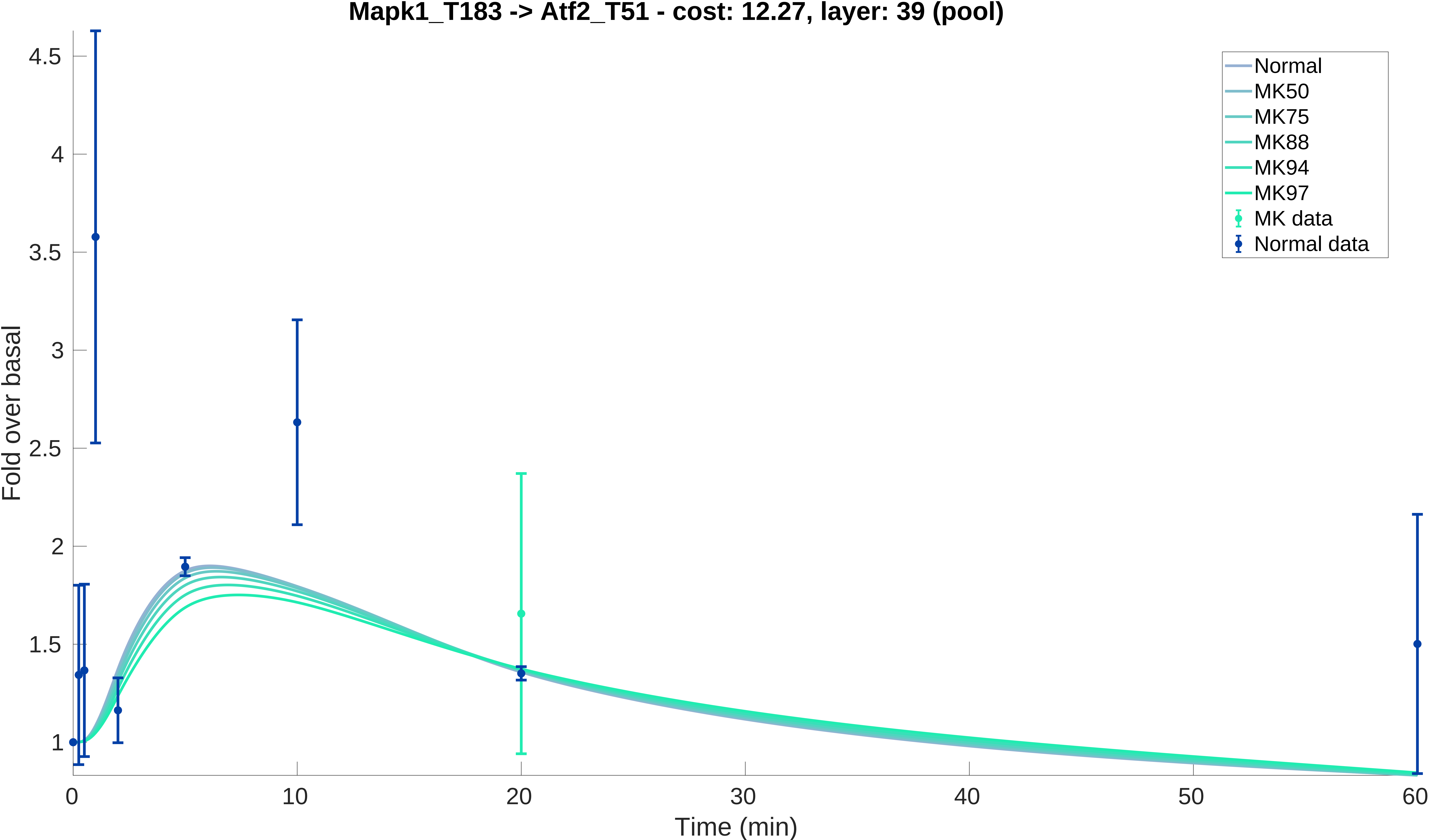

Mapk1_T183 -> Atf2_T51 - cost: 12.27, layer: 39 (pool)
Fold over basal
4.5
4
3.5
3
2.5
2
1.5
1
0
10
20
30
40
50
60
Time (min)
Normal
MK50
MK75
MK88
MK94
MK97
MK data
Normal data

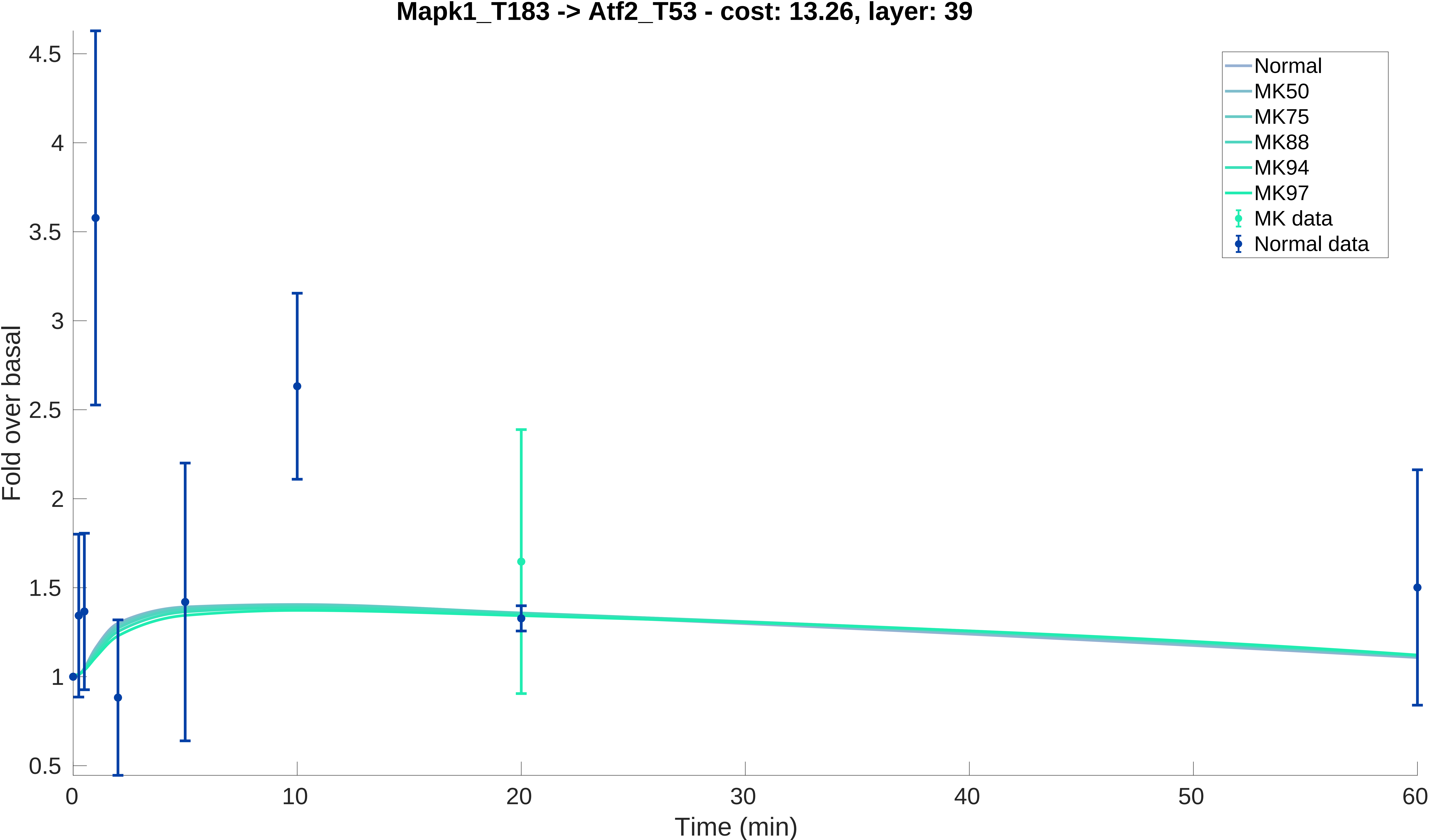
Mapk1_T183 -> Atf2_T53 - cost: 13.26, layer: 39
Fold over basal
Time (min)
4.5
4
3.5
3
2.5
2
1.5
1
0.5
0
10
20
30
40
50
60
Normal
MK50
MK75
MK88
MK94
MK97
MK data
Normal data

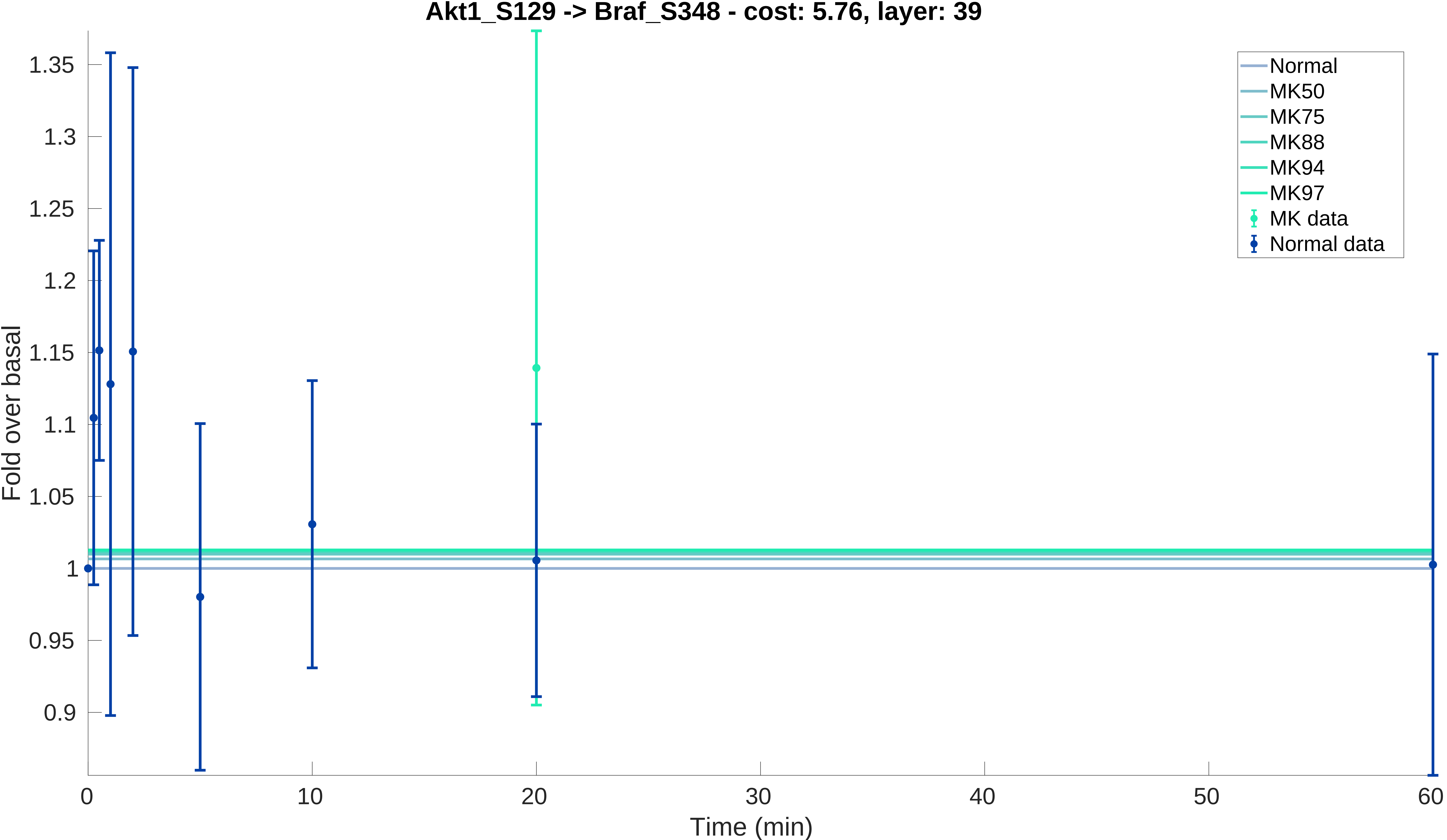

Akt1_S129 -> Braf_S348 - cost: 5.76, layer: 39
Normal
MK50
MK75
MK88
MK94
MK97
MK data
Normal data
Fold over basal
Time (min)
1.35
1.3
1.25
1.2
1.15
1.1
1.05
1
0.95
0.9
0
10
20
30
40
50
60

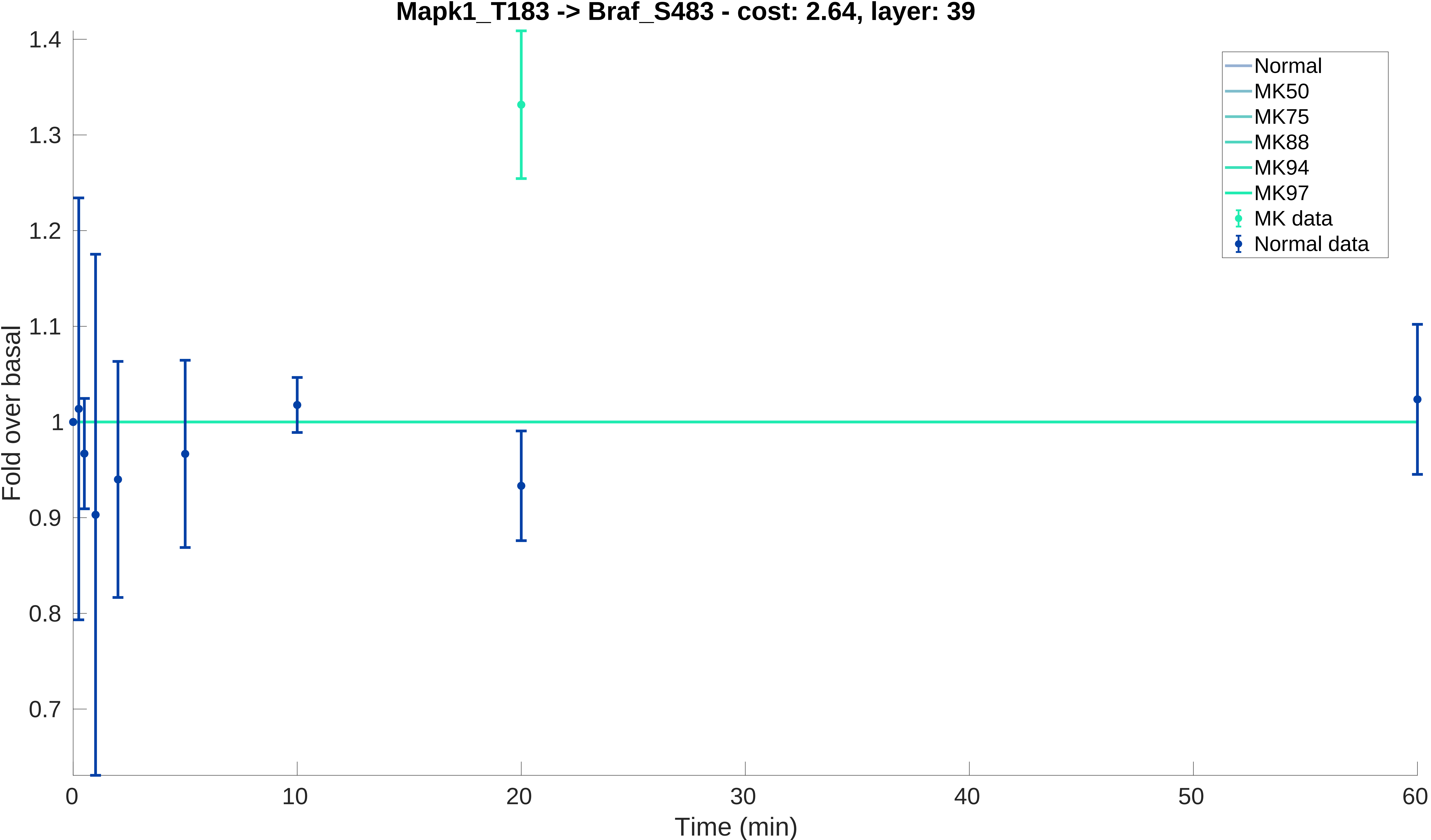

Mapk1_T183 -> Braf_S483 - cost: 2.64, layer: 39
1.4
1.3
1.2
1.1
1
0.9
0.8
0.7
Fold over basal
0
10
20
30
40
50
60
Time (min)
Normal
MK50
MK75
MK88
MK94
MK97
MK data
Normal data

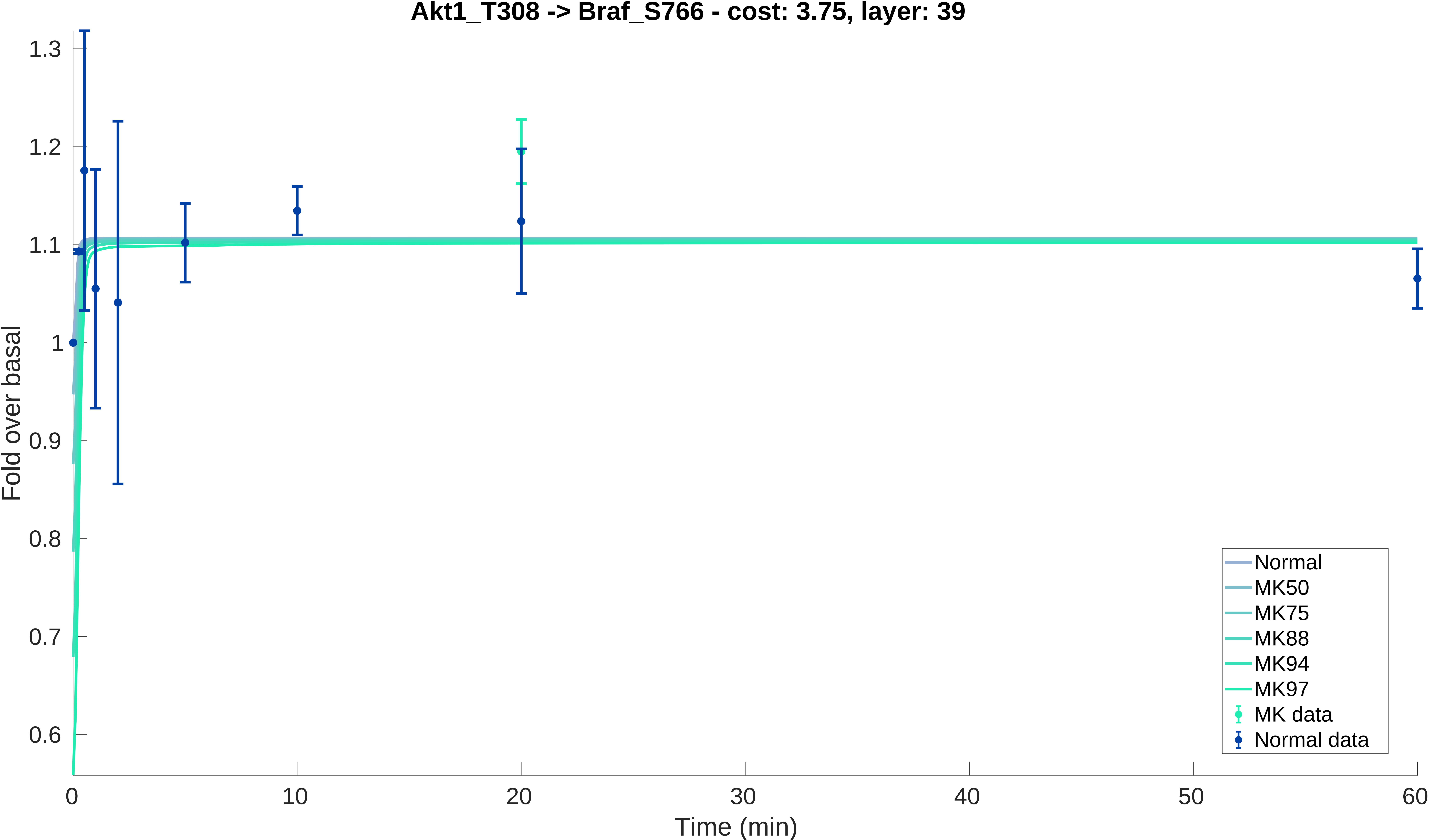

Akt1_T308 -> Braf_S766 - cost: 3.75, layer: 39
Fold over basal
Time (min)
1.3
1.2
1.1
1
0.9
0.8
0.7
0.6
0
10
20
30
40
50
60
Normal
MK50
MK75
MK88
MK94
MK97
MK data
Normal data

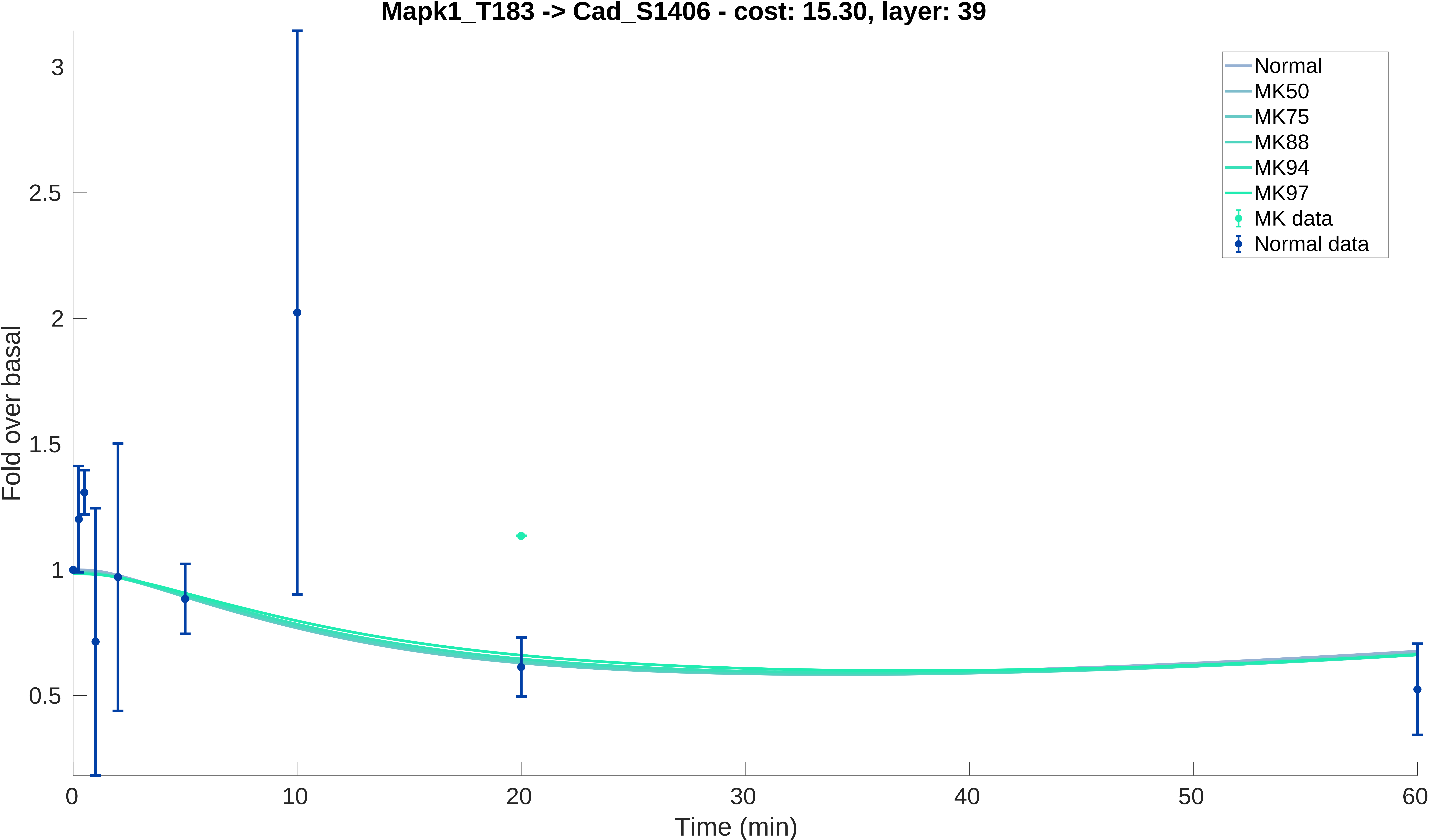
Mapk1_T183 -> Cad_S1406 - cost: 15.30, layer: 39
Fold over basal
Time (min)
0.5
1
1.5
2
2.5
3
0
10
20
30
40
50
60
Normal
MK50
MK75
MK88
MK94
MK97
MK data
Normal data

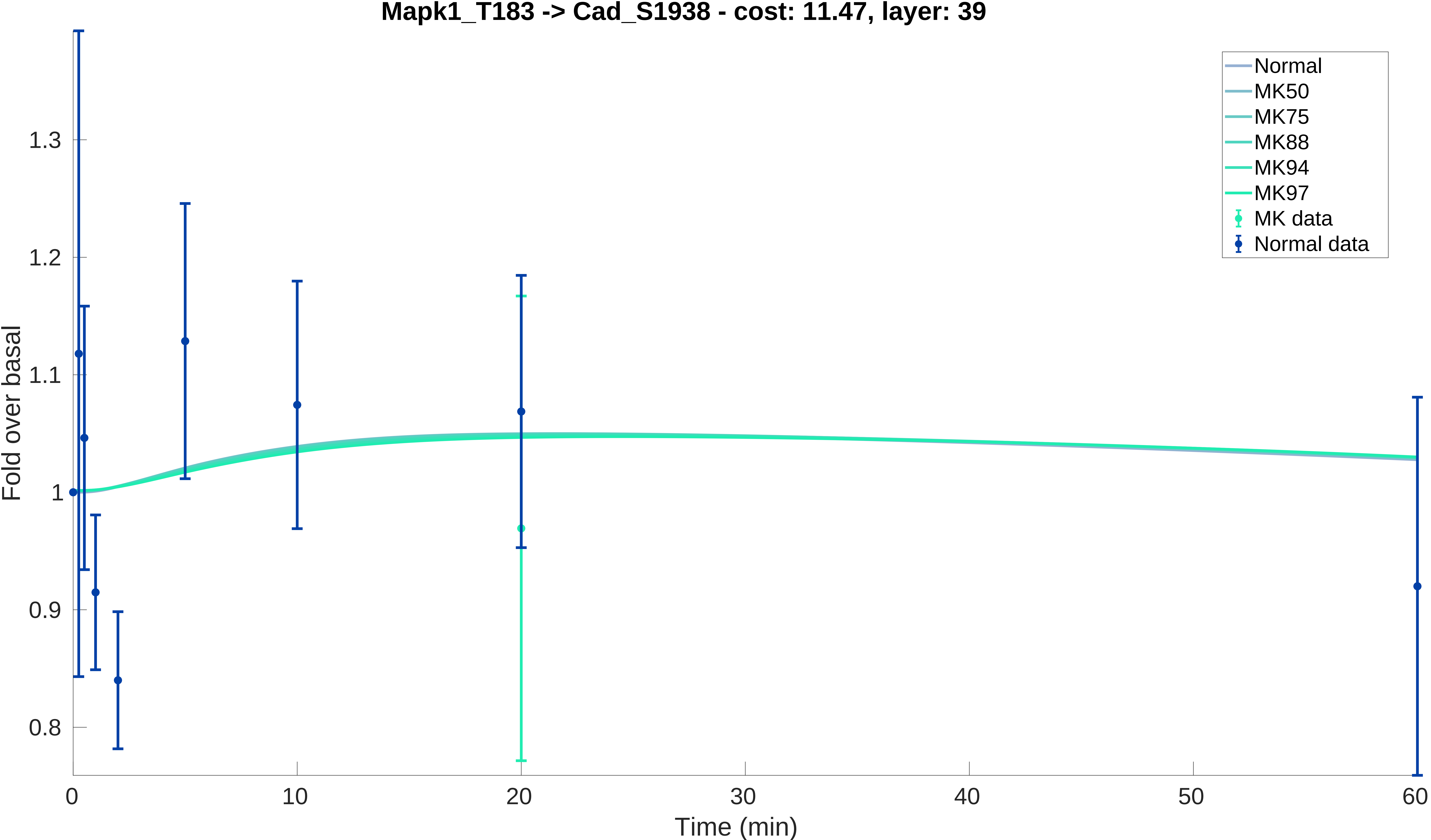
Mapk1_T183 -> Cad_S1938 - cost: 11.47, layer: 39
Fold over basal
Time (min)
0
10
20
30
40
50
60
0.8
0.9
1
1.1
1.2
1.3
Normal
MK50
MK75
MK88
MK94
MK97
MK data
Normal data

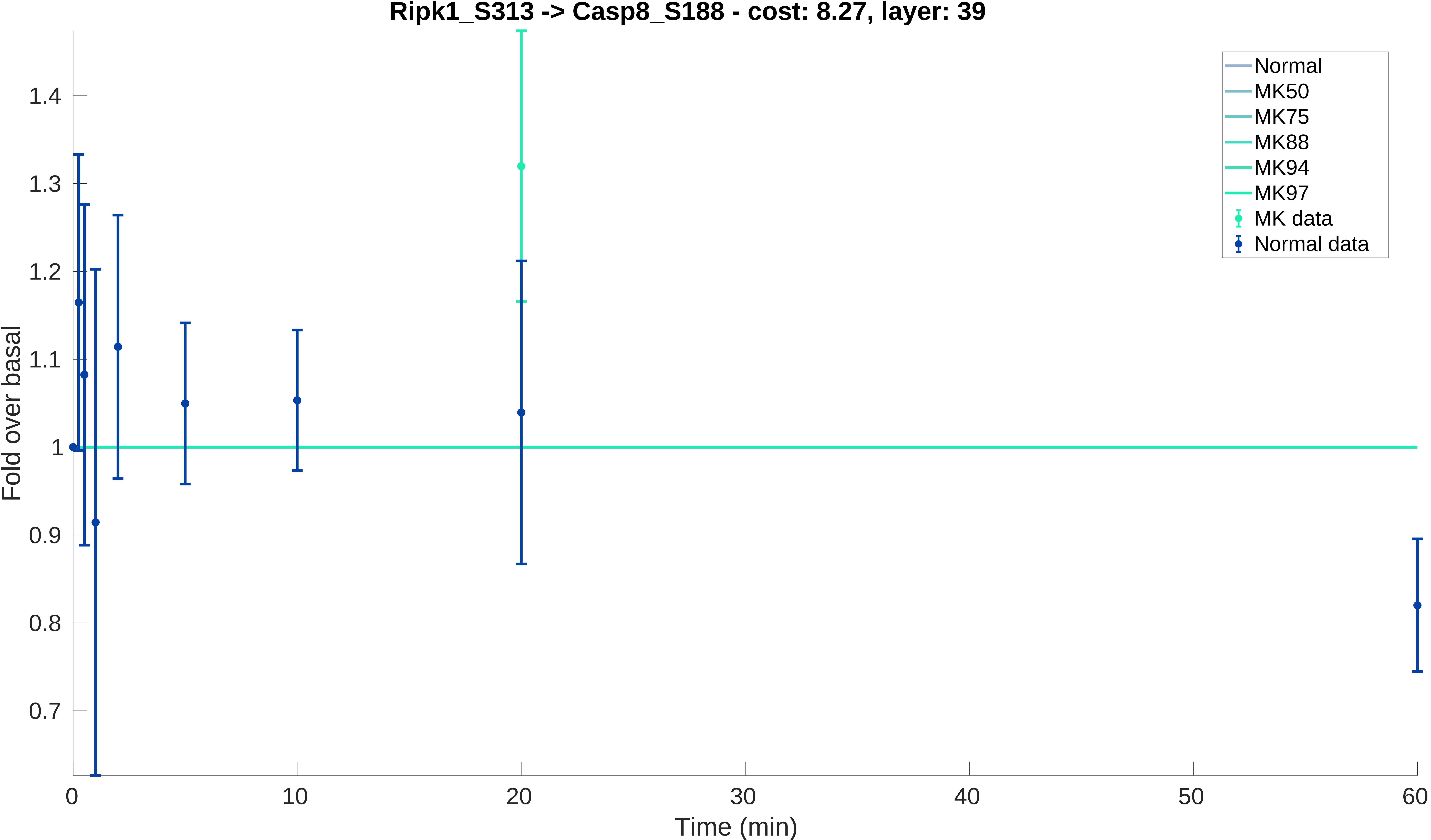
Ripk1_S313 -> Casp8_S188 - cost: 8.27, layer: 39
Fold over basal
Time (min)
1.4
1.3
1.2
1.1
1
0.9
0.8
0.7
0
10
20
30
40
50
60
Normal
MK50
MK75
MK88
MK94
MK97
MK data
Normal data

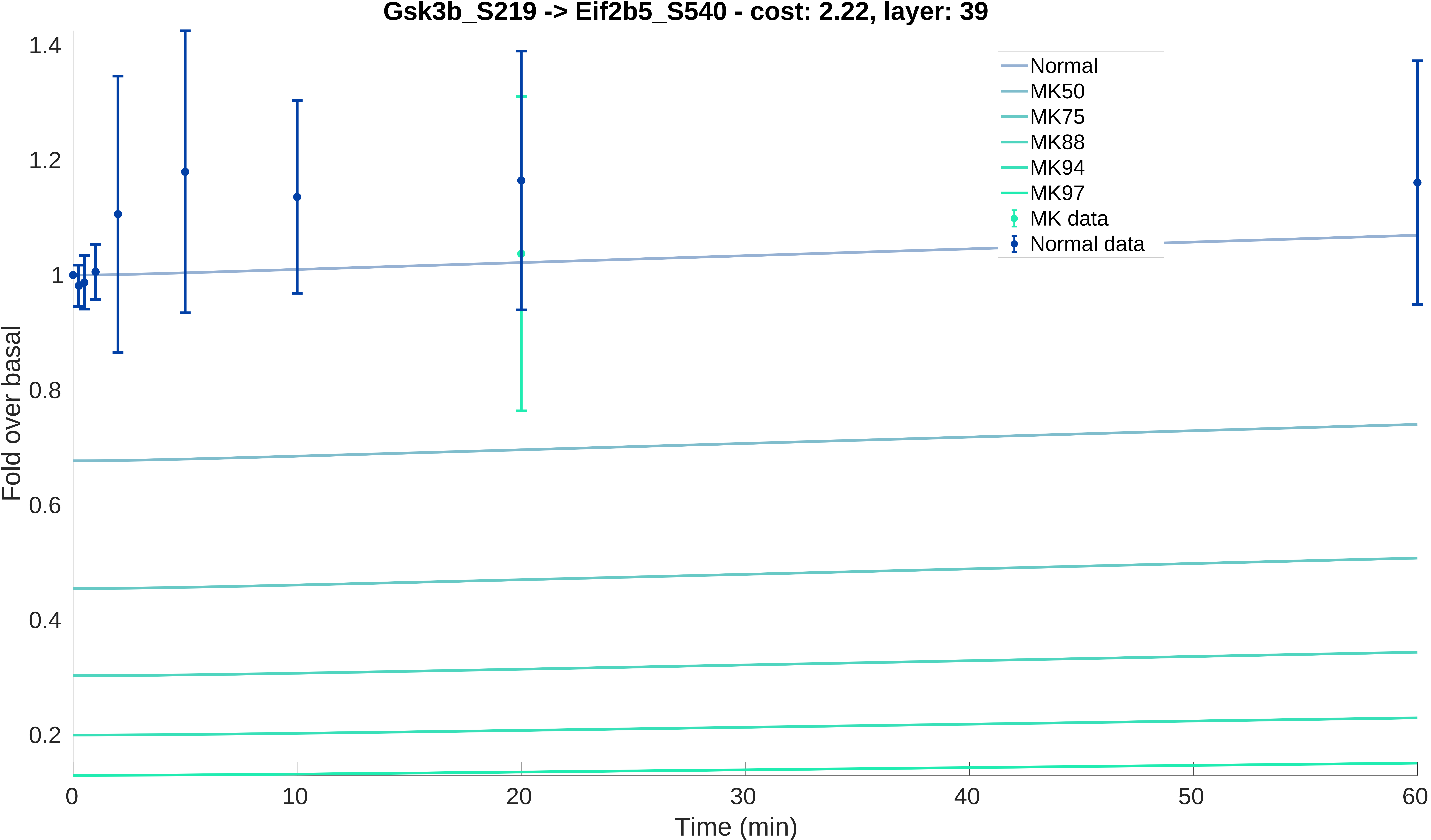

Gsk3b_S219 -> Eif2b5_S540 - cost: 2.22, layer: 39
Fold over basal
Time (min)
0
10
20
30
40
50
60
0.2
0.4
0.6
0.8
1
1.2
1.4
Normal
MK50
MK75
MK88
MK94
MK97
MK data
Normal data

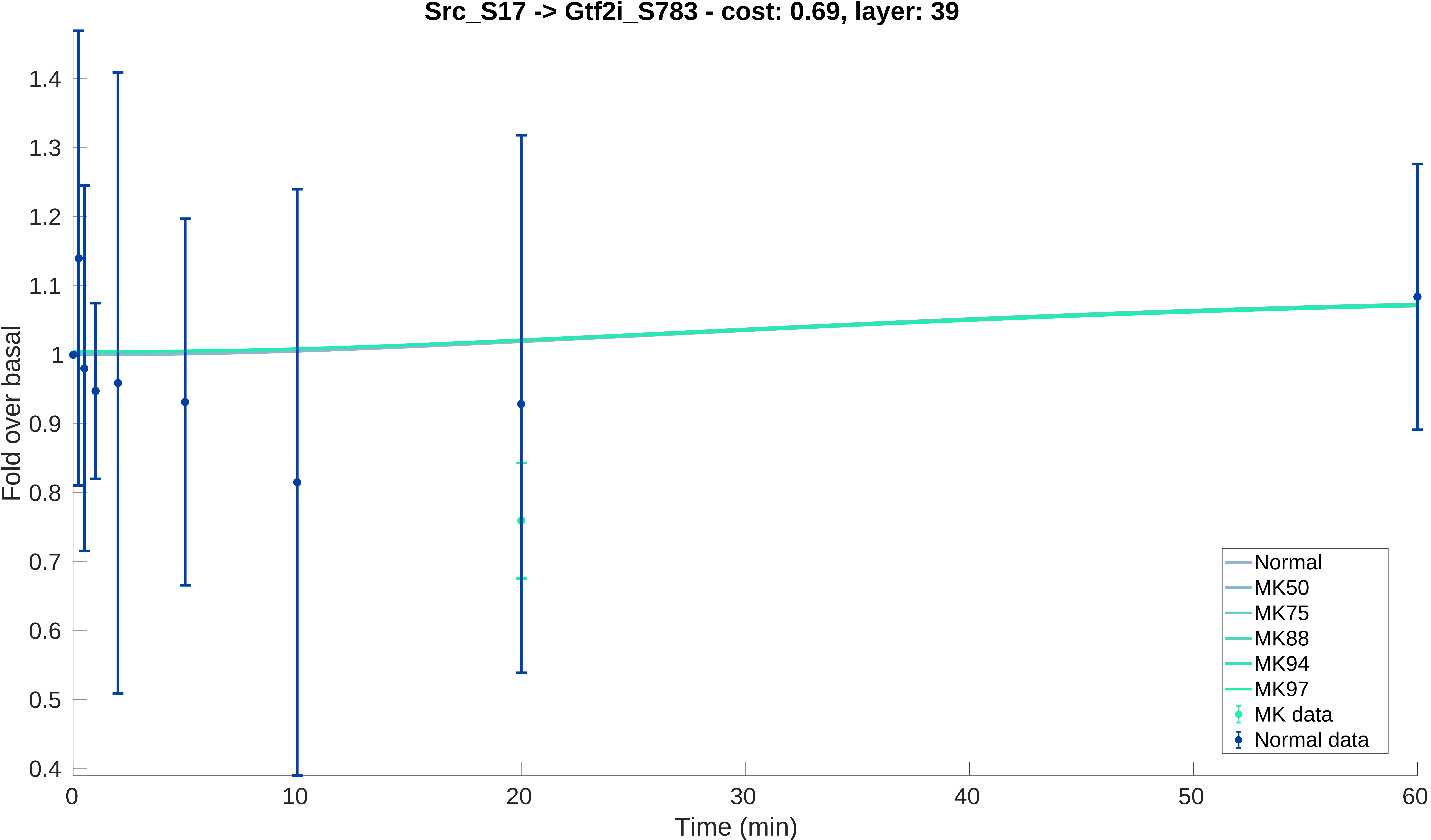

Src_S17 -> Gtf2i_S783 - cost: 0.69, layer: 39
Fold over basal
1.4
1.3
1.2
1.1
1
0.9
0.8
0.7
0.6
0.5
0.4
0
10
20
30
40
50
60
Time (min)
Normal
MK50
MK75
MK88
MK94
MK97
MK data
Normal data

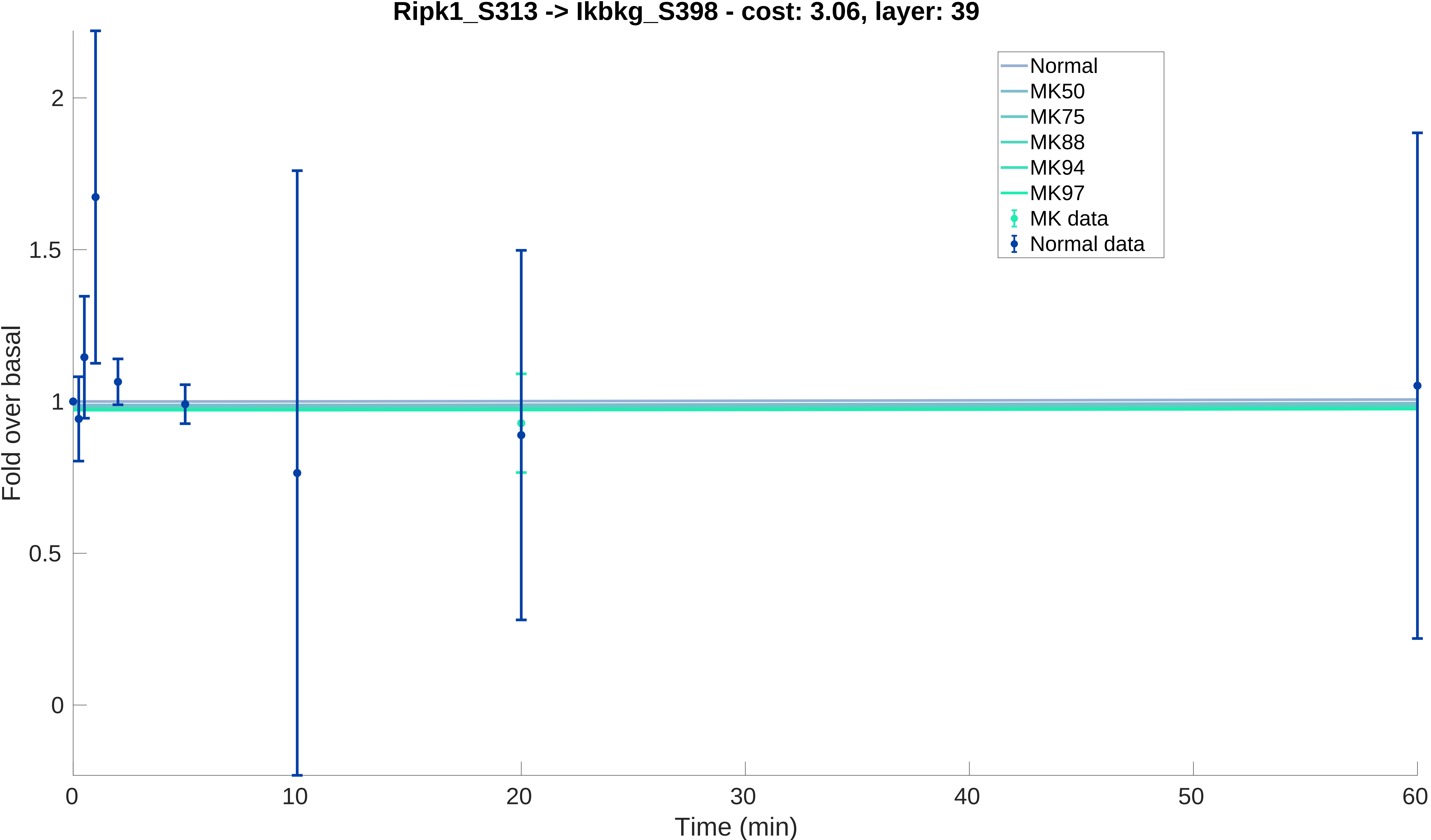

Ripk1_S313 -> Ikbkg_S398 - cost: 3.06, layer: 39
Normal
MK50
MK75
MK88
MK94
MK97
MK data
Normal data
Fold over basal
2
1.5
1
0.5
0
0
10
20
30
40
50
60
Time (min)

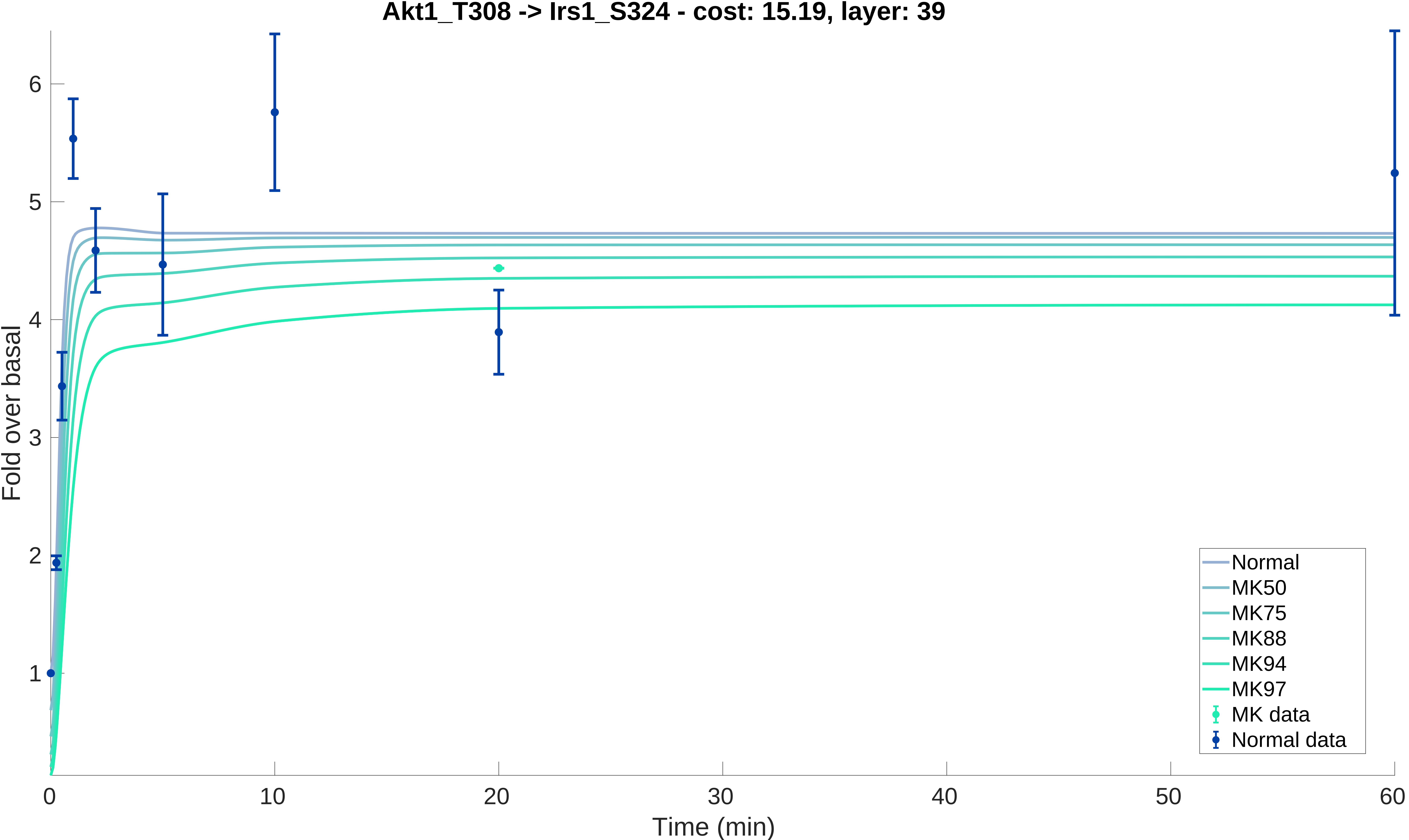

Akt1_T308 -> Irs1_S324 - cost: 15.19, layer: 39
Fold over basal
Time (min)
0
10
20
30
40
50
60
1
2
3
4
5
6
Normal
MK50
MK75
MK88
MK94
MK97
MK data
Normal data

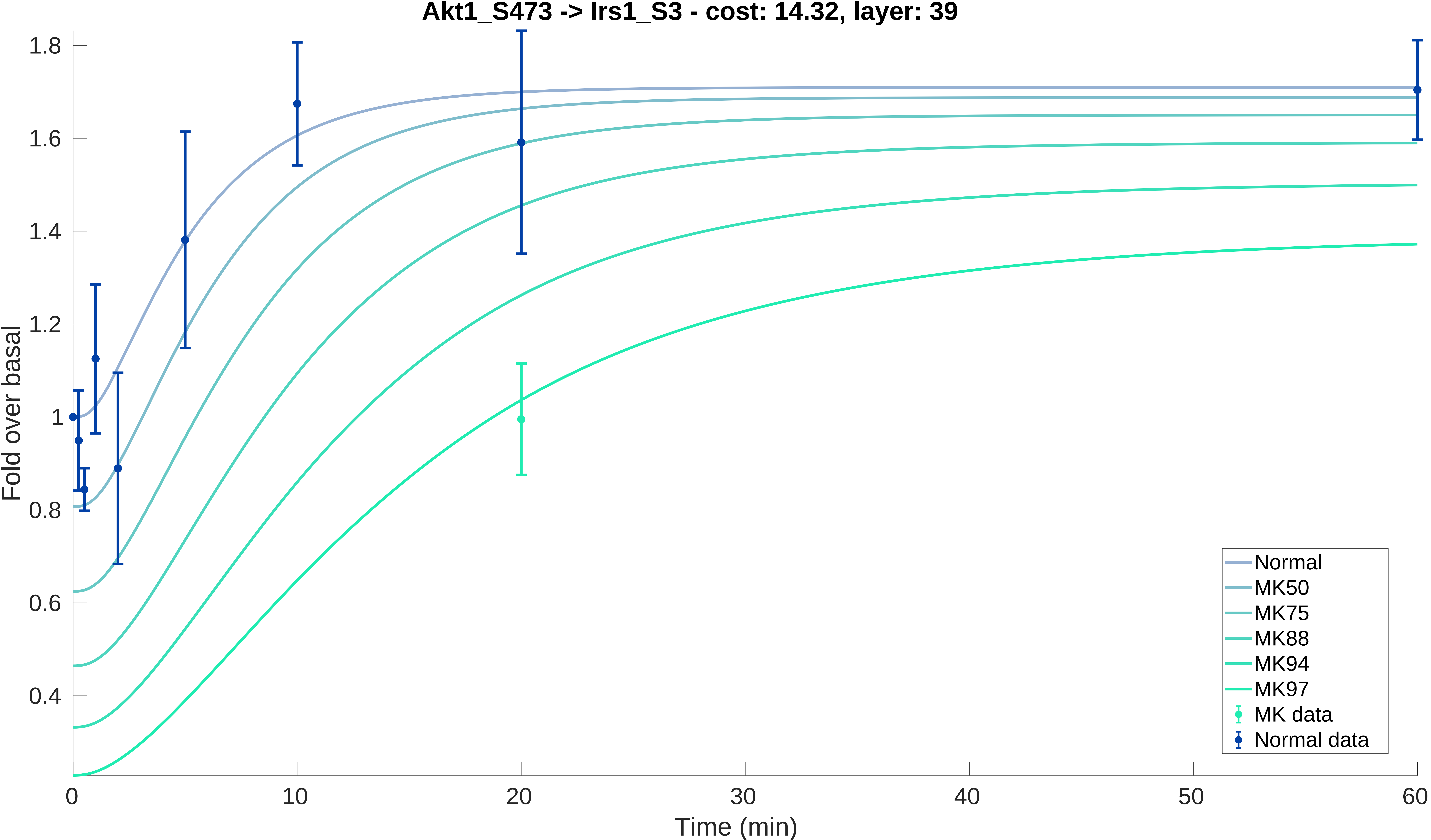
Akt1_S473 -> Irs1_S3 - cost: 14.32, layer: 39
Fold over basal
Time (min)
0
10
20
30
40
50
60
0.4
0.6
0.8
1
1.2
1.4
1.6
1.8
Normal
MK50
MK75
MK88
MK94
MK97
MK data
Normal data

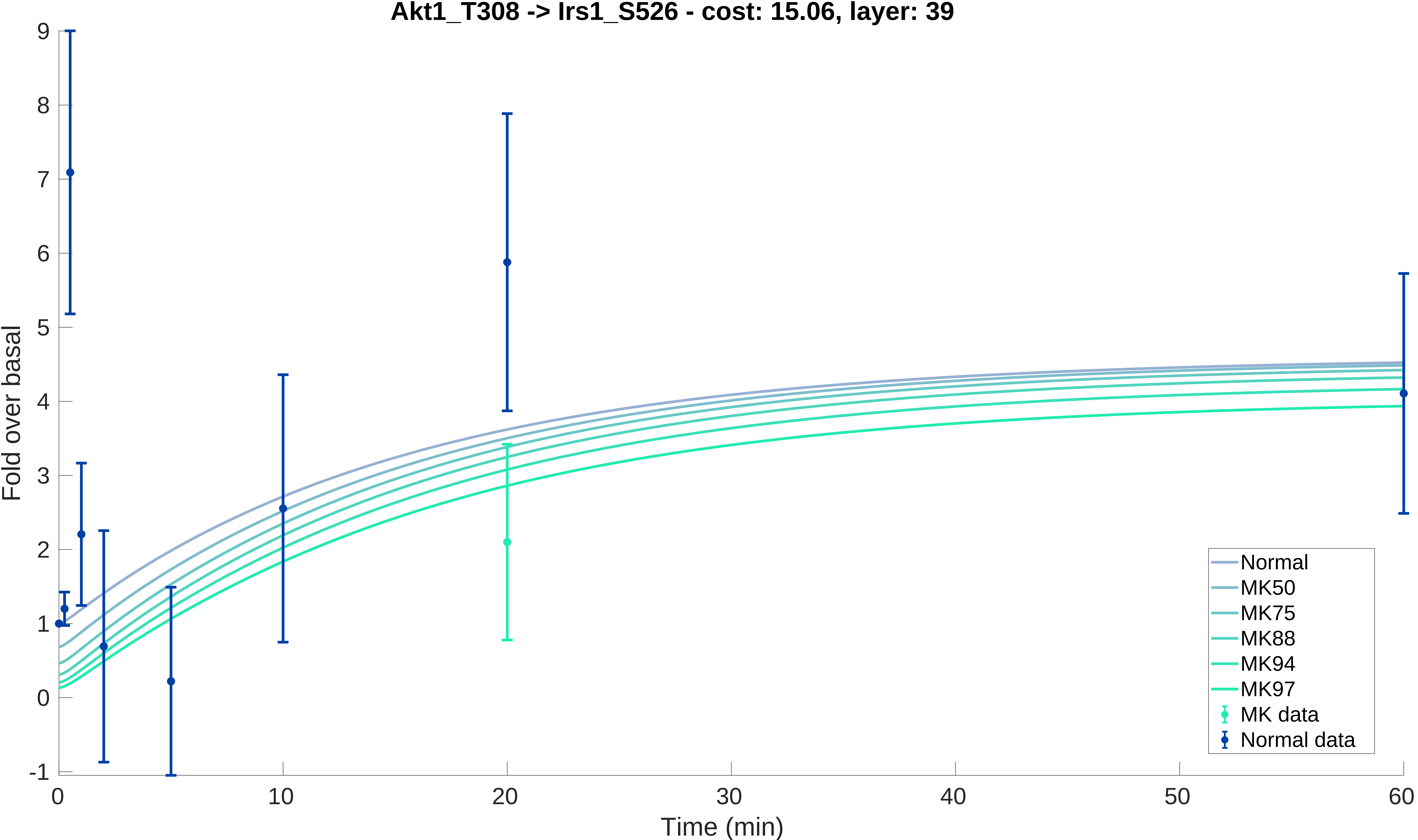
Akt1_T308 -> Irs1_S526 - cost: 15.06, layer: 39
Fold over basal
Time (min)
9
8
7
6
5
4
3
2
1
0
-1
0
10
20
30
40
50
60
Normal
MK50
MK75
MK88
MK94
MK97
MK data
Normal data

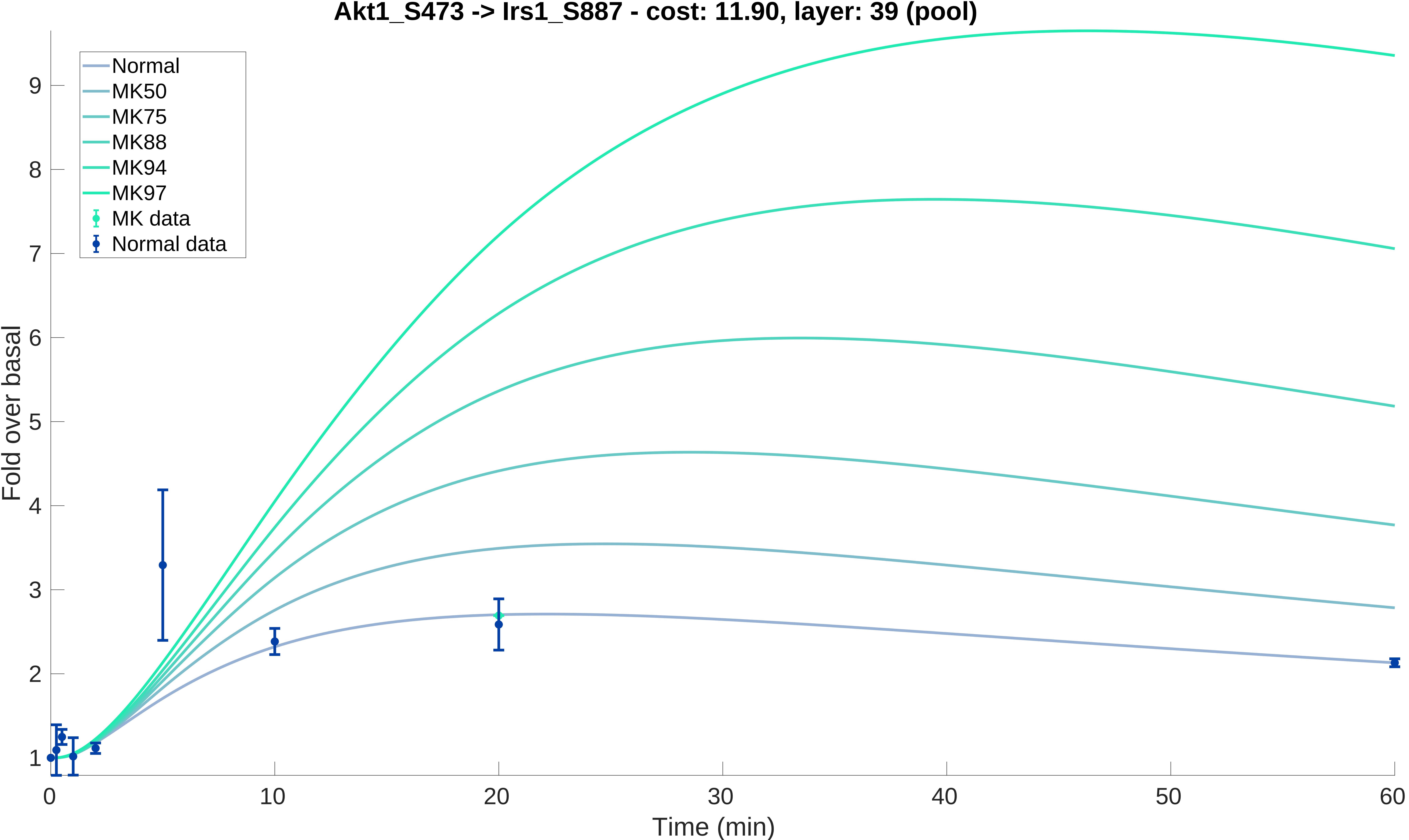
Akt1_S473 -> Irs1_S887 - cost: 11.90, layer: 39 (pool)
Normal
MK50
MK75
MK88
MK94
MK97
MK data
Normal data
Fold over basal
Time (min)
1
2
3
4
5
6
7
8
9
0
10
20
30
40
50
60

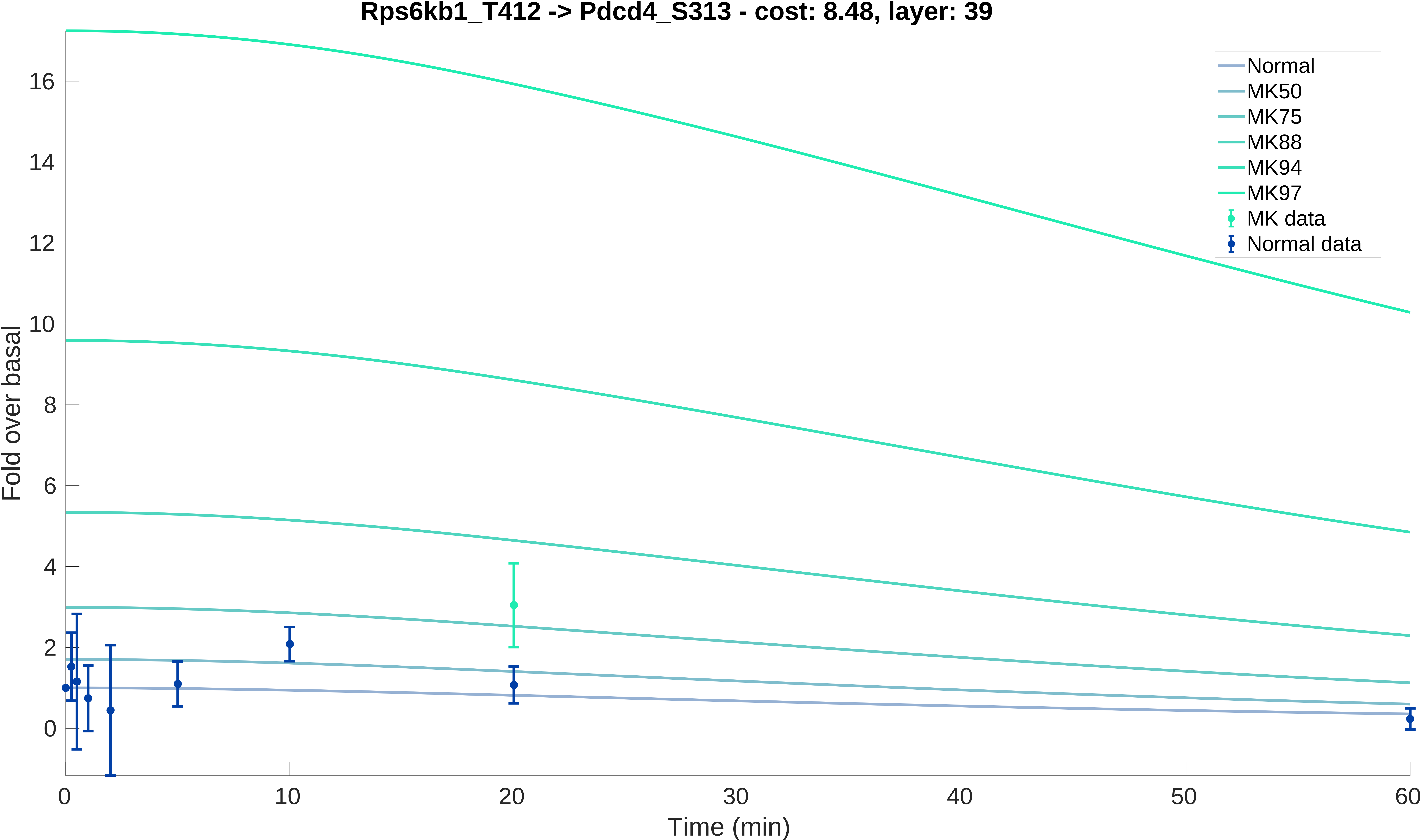

Rps6kb1_T412 -> Pdcd4_S313 - cost: 8.48, layer: 39
Fold over basal
Time (min)
0
2
4
6
8
10
12
14
16
0
10
20
30
40
50
60
Normal
MK50
MK75
MK88
MK94
MK97
MK data
Normal data

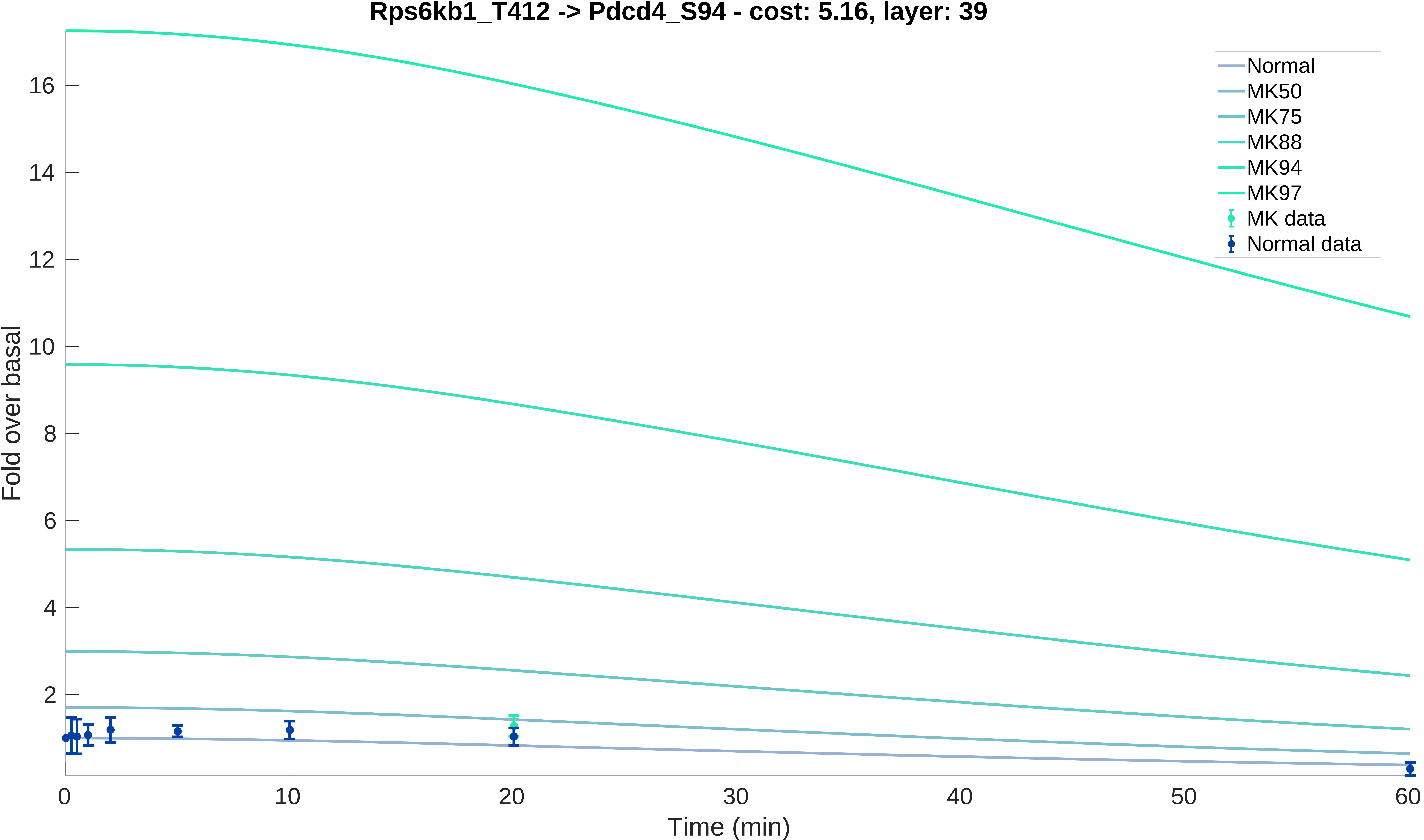

Rps6kb1_T412 -> Pdcd4_S94 - cost: 5.16, layer: 39
Normal
MK50
MK75
MK88
MK94
MK97
MK data
Normal data
Fold over basal
16
14
12
10
8
6
4
2
0
10
20
30
40
50
60
Time (min)

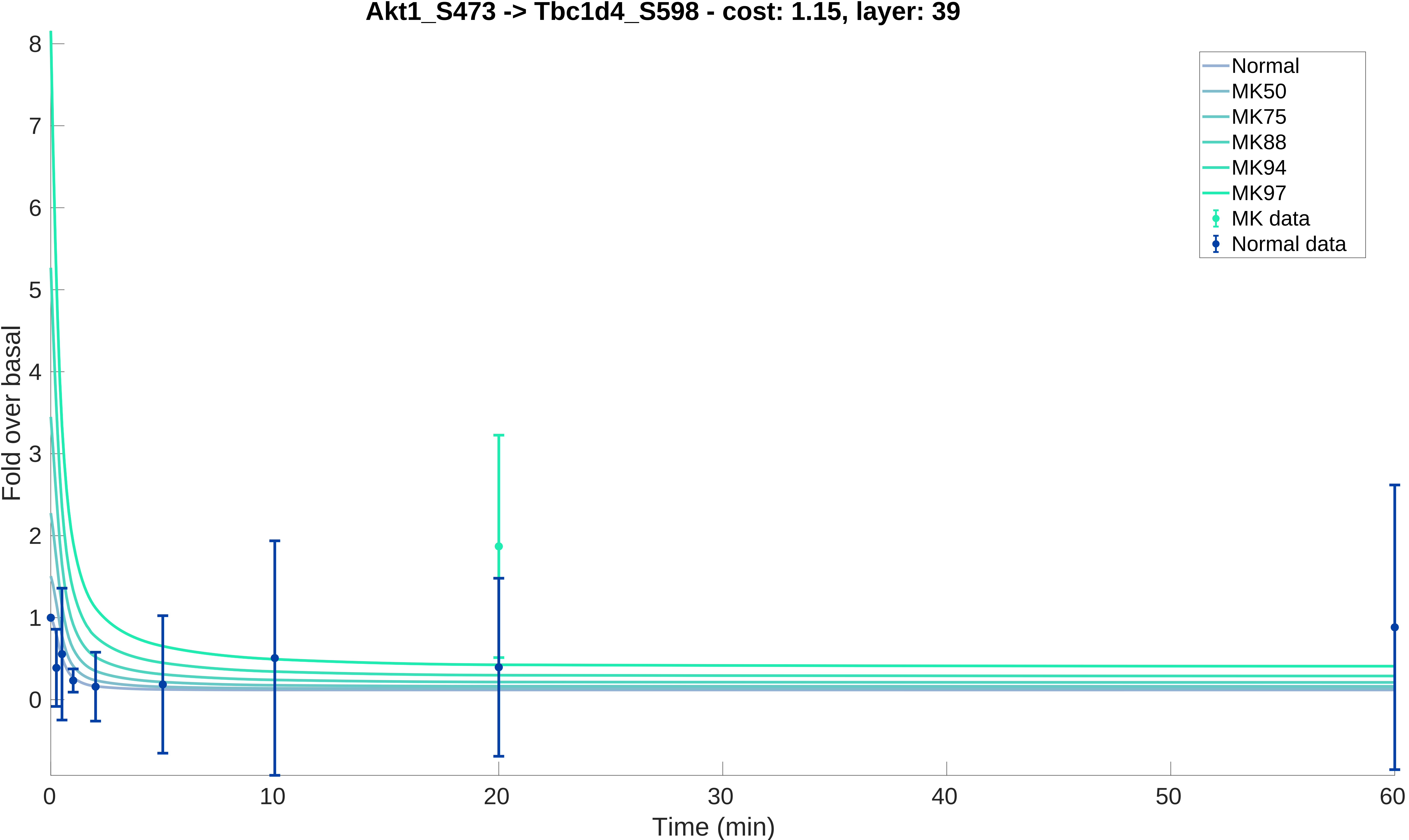

Akt1_S473 -> Tbc1d4_S598 - cost: 1.15, layer: 39
Fold over basal
Time (min)
0
1
2
3
4
5
6
7
8
10
20
30
40
50
60
Normal
MK50
MK75
MK88
MK94
MK97
MK data
Normal data

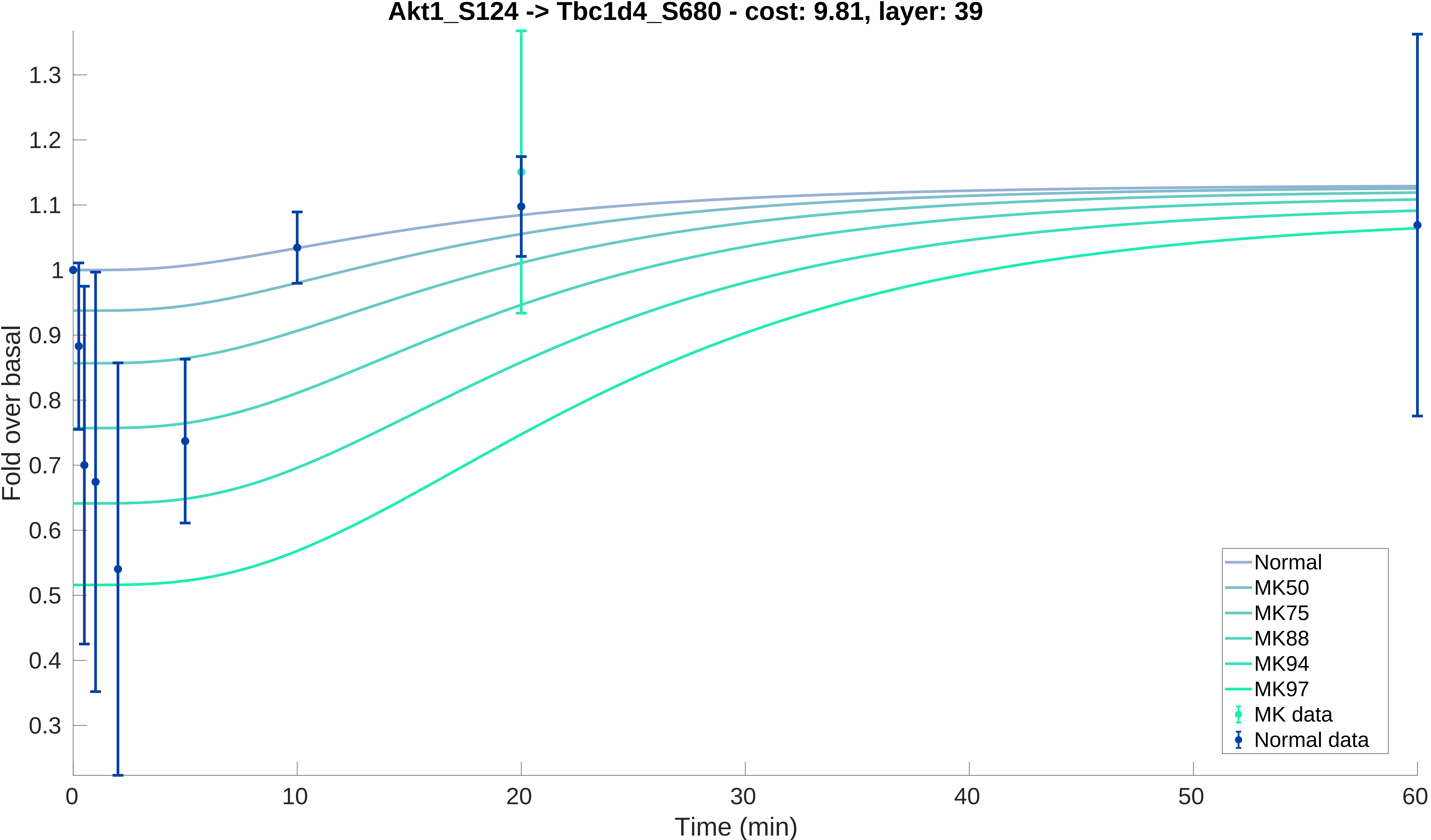

Akt1_S124 -> Tbc1d4_S680 - cost: 9.81, layer: 39
Fold over basal
Time (min)
1.3
1.2
1.1
1
0.9
0.8
0.7
0.6
0.5
0.4
0.3
0
10
20
30
40
50
60
Normal
MK50
MK75
MK88
MK94
MK97
MK data
Normal data

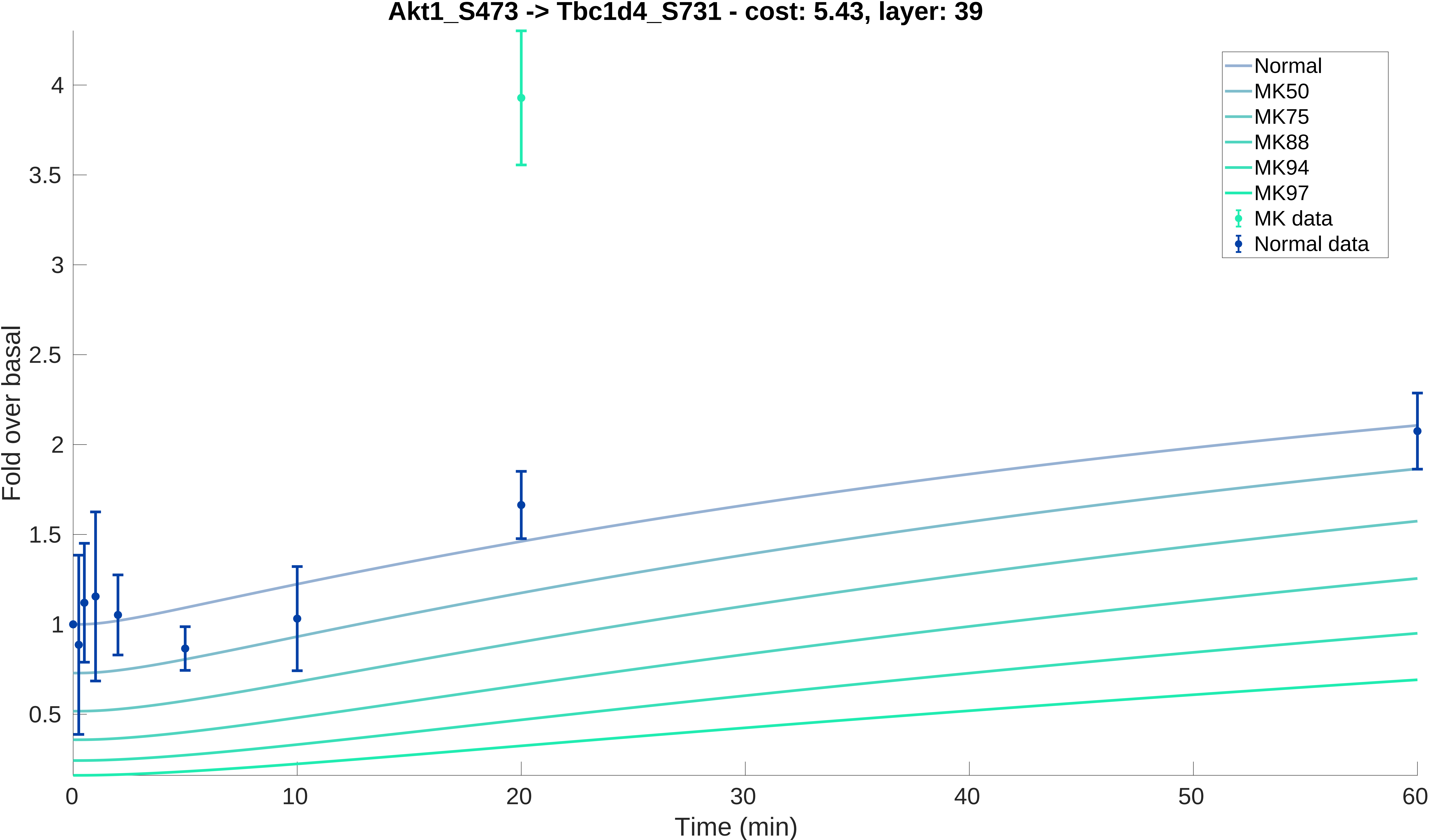

Akt1_S473 -> Tbc1d4_S731 - cost: 5.43, layer: 39
Fold over basal
Time (min)
0.5
1
1.5
2
2.5
3
3.5
4
0
10
20
30
40
50
60
Normal
MK50
MK75
MK88
MK94
MK97
MK data
Normal data

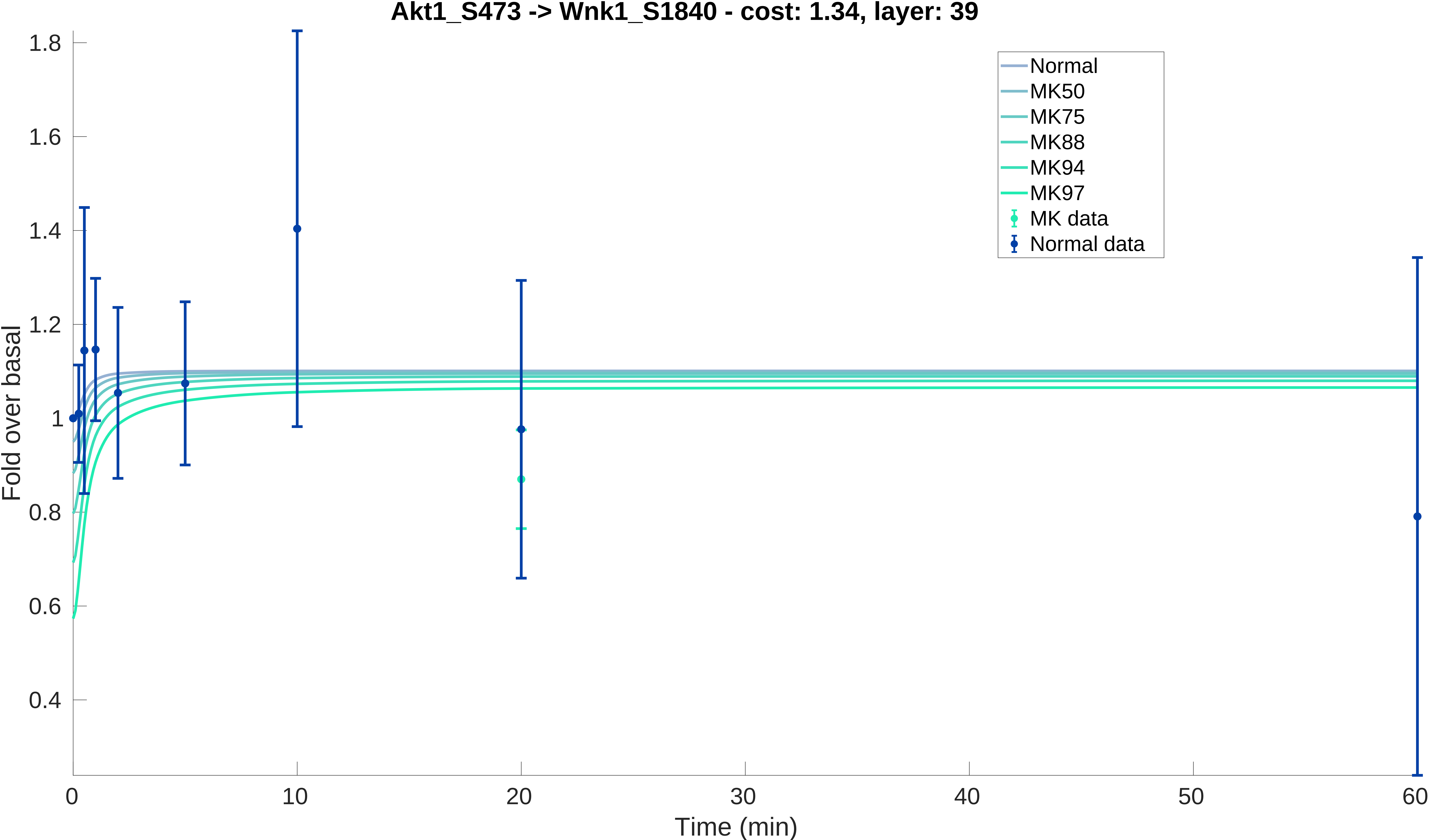

Akt1_S473 -> Wnk1_S1840 - cost: 1.34, layer: 39
Fold over basal
Time (min)
1.8
1.6
1.4
1.2
1
0.8
0.6
0.4
0
10
20
30
40
50
60
Normal
MK50
MK75
MK88
MK94
MK97
MK data
Normal data

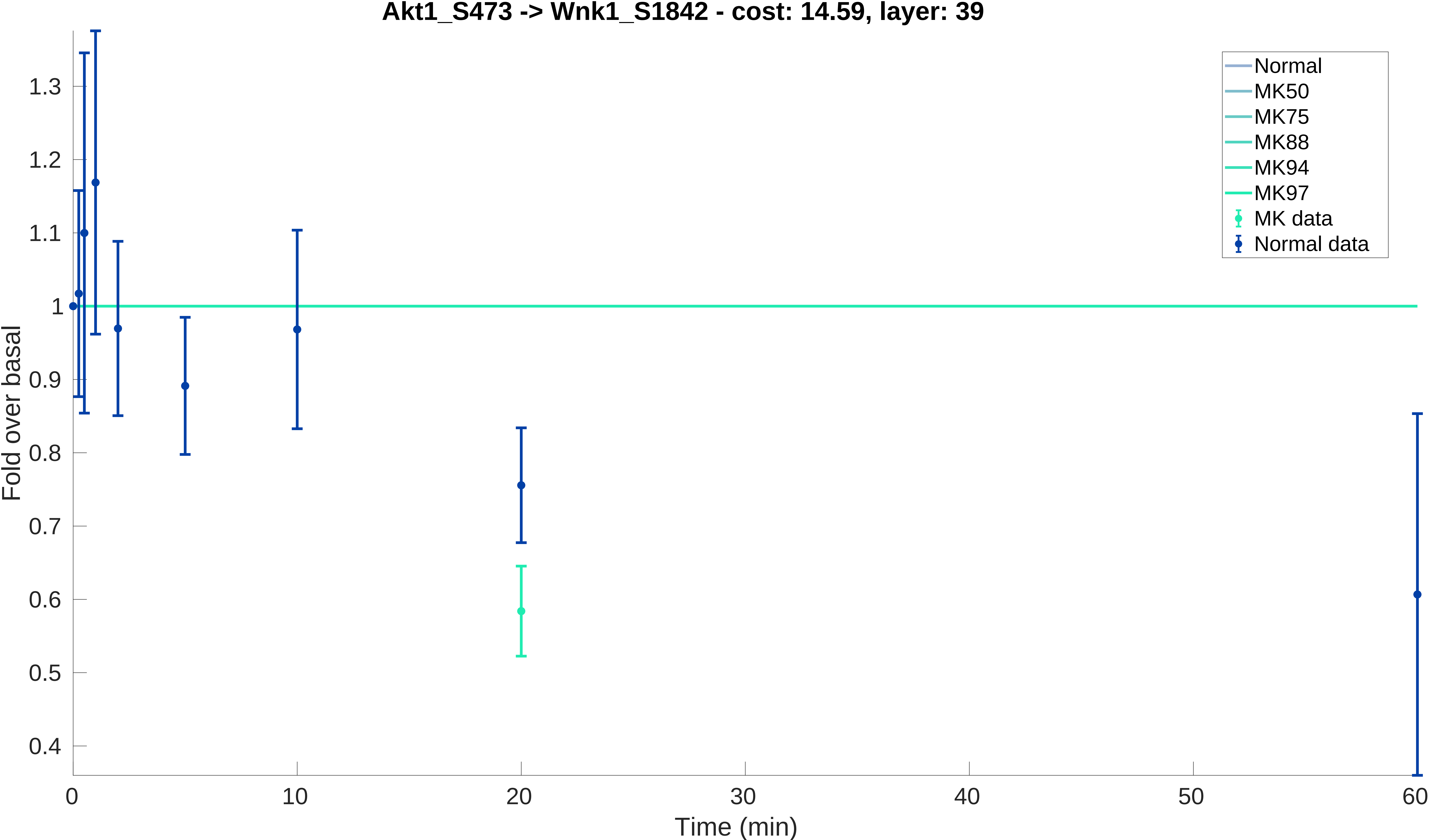
Akt1_S473 -> Wnk1_S1842 - cost: 14.59, layer: 39
Fold over basal
Time (min)
0.4
0.5
0.6
0.7
0.8
0.9
1
1.1
1.2
1.3
0
10
20
30
40
50
60
Normal
MK50
MK75
MK88
MK94
MK97
MK data
Normal data

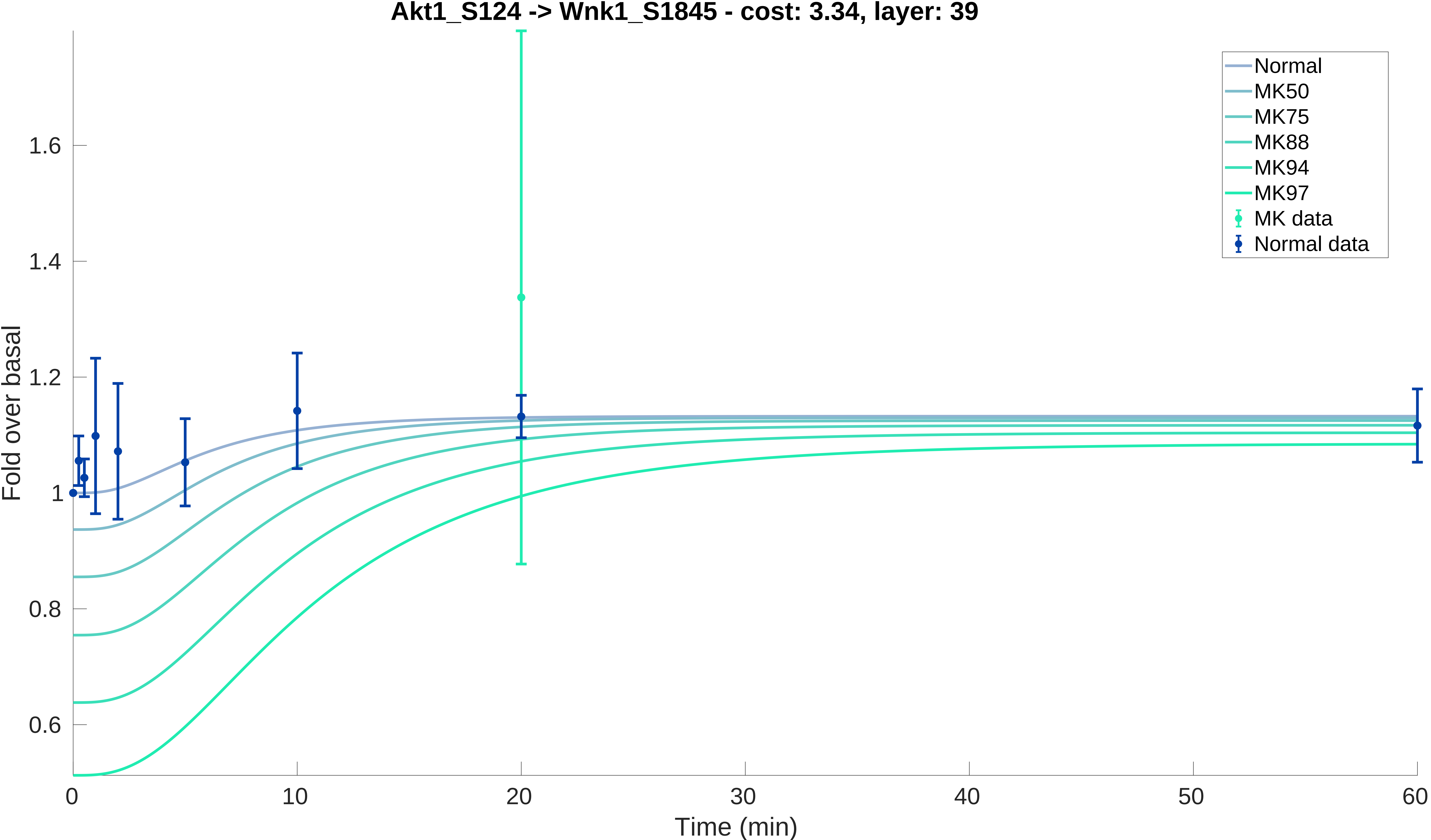

Akt1_S124 -> Wnk1_S1845 - cost: 3.34, layer: 39
Fold over basal
Time (min)
0
10
20
30
40
50
60
0.6
0.8
1
1.2
1.4
1.6
Normal
MK50
MK75
MK88
MK94
MK97
MK data
Normal data

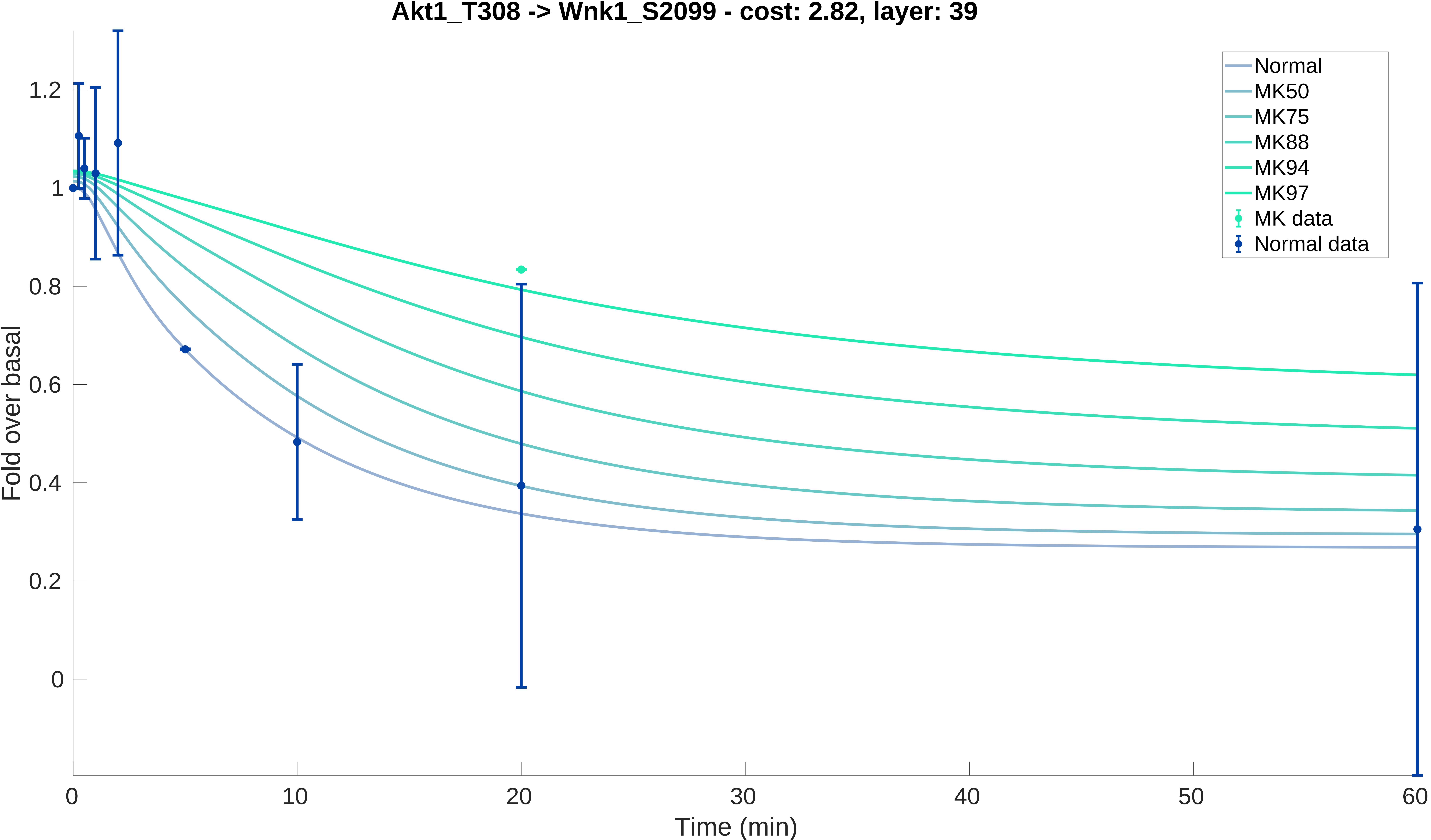

Akt1_T308 -> Wnk1_S2099 - cost: 2.82, layer: 39
Fold over basal
Time (min)
Normal
MK50
MK75
MK88
MK94
MK97
MK data
Normal data
1.2
1
0.8
0.6
0.4
0.2
0
0
10
20
30
40
50
60

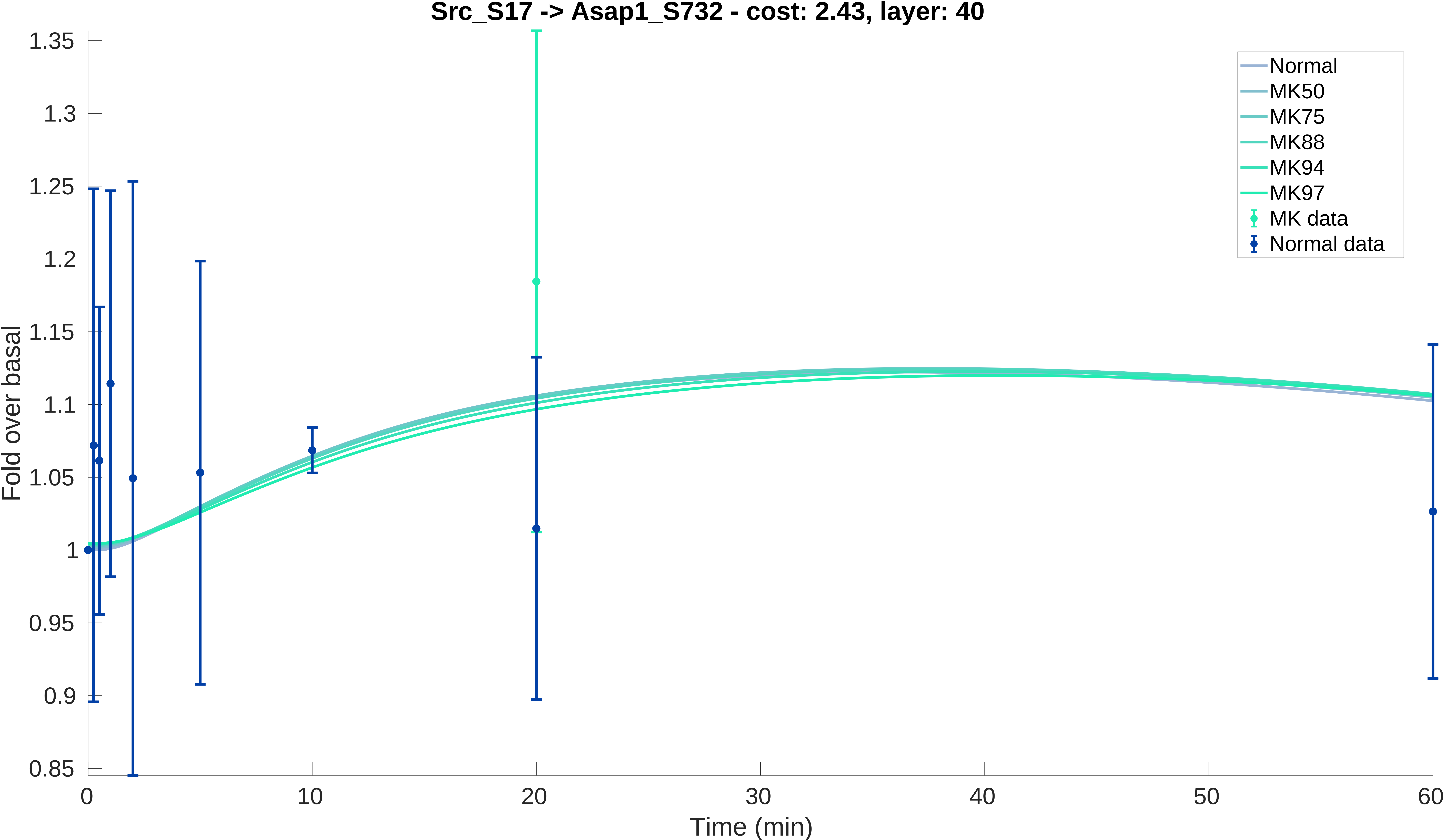

Src_S17 -> Asap1_S732 - cost: 2.43, layer: 40
Normal
MK50
MK75
MK88
MK94
MK97
MK data
Normal data
Fold over basal
Time (min)
1.35
1.3
1.25
1.2
1.15
1.1
1.05
1
0.95
0.9
0.85
0
10
20
30
40
50
60

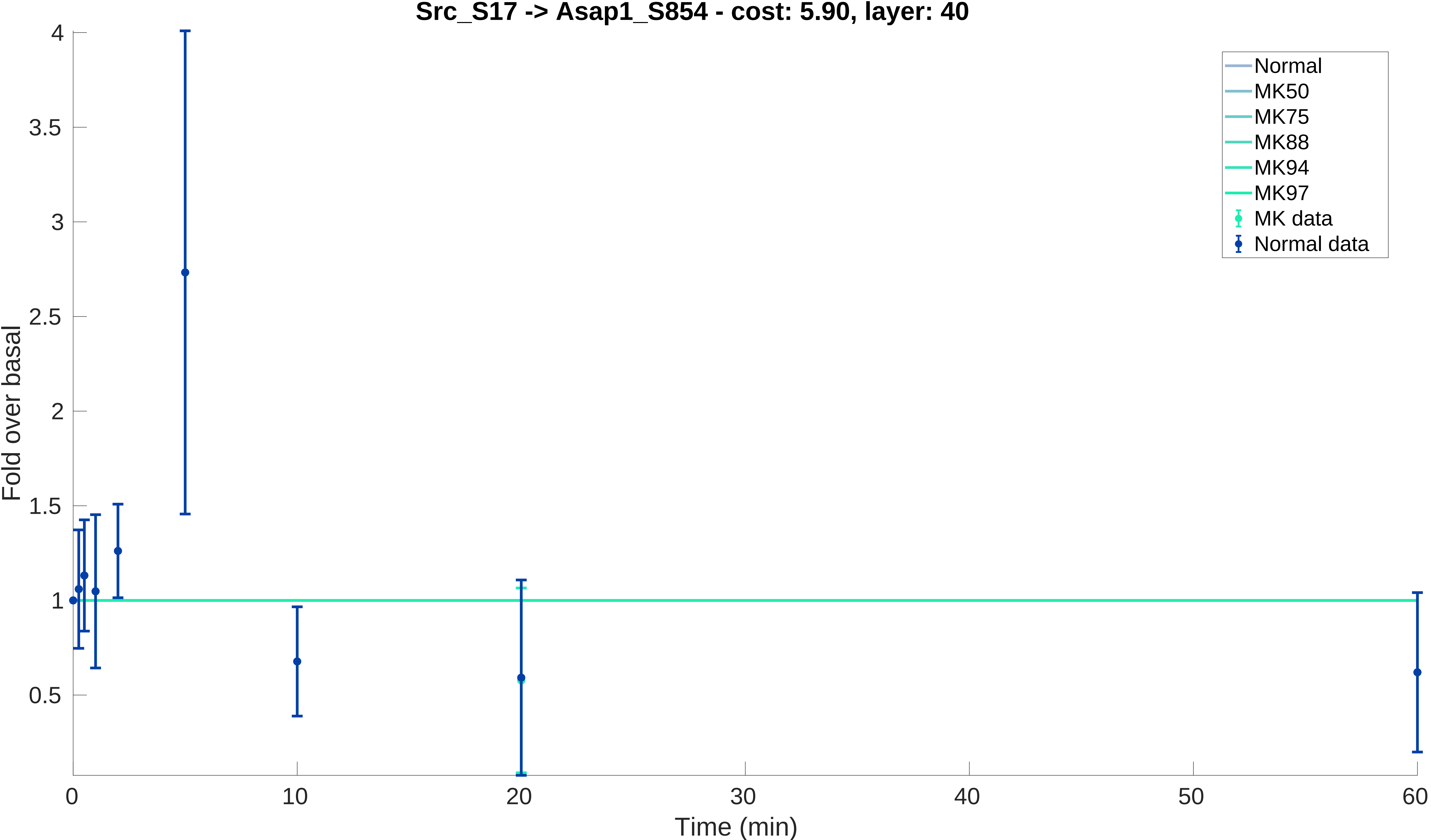

Src_S17 -> Asap1_S854 - cost: 5.90, layer: 40
Fold over basal
4
3.5
3
2.5
2
1.5
1
0.5
0
10
20
30
40
50
60
Time (min)
Normal
MK50
MK75
MK88
MK94
MK97
MK data
Normal data

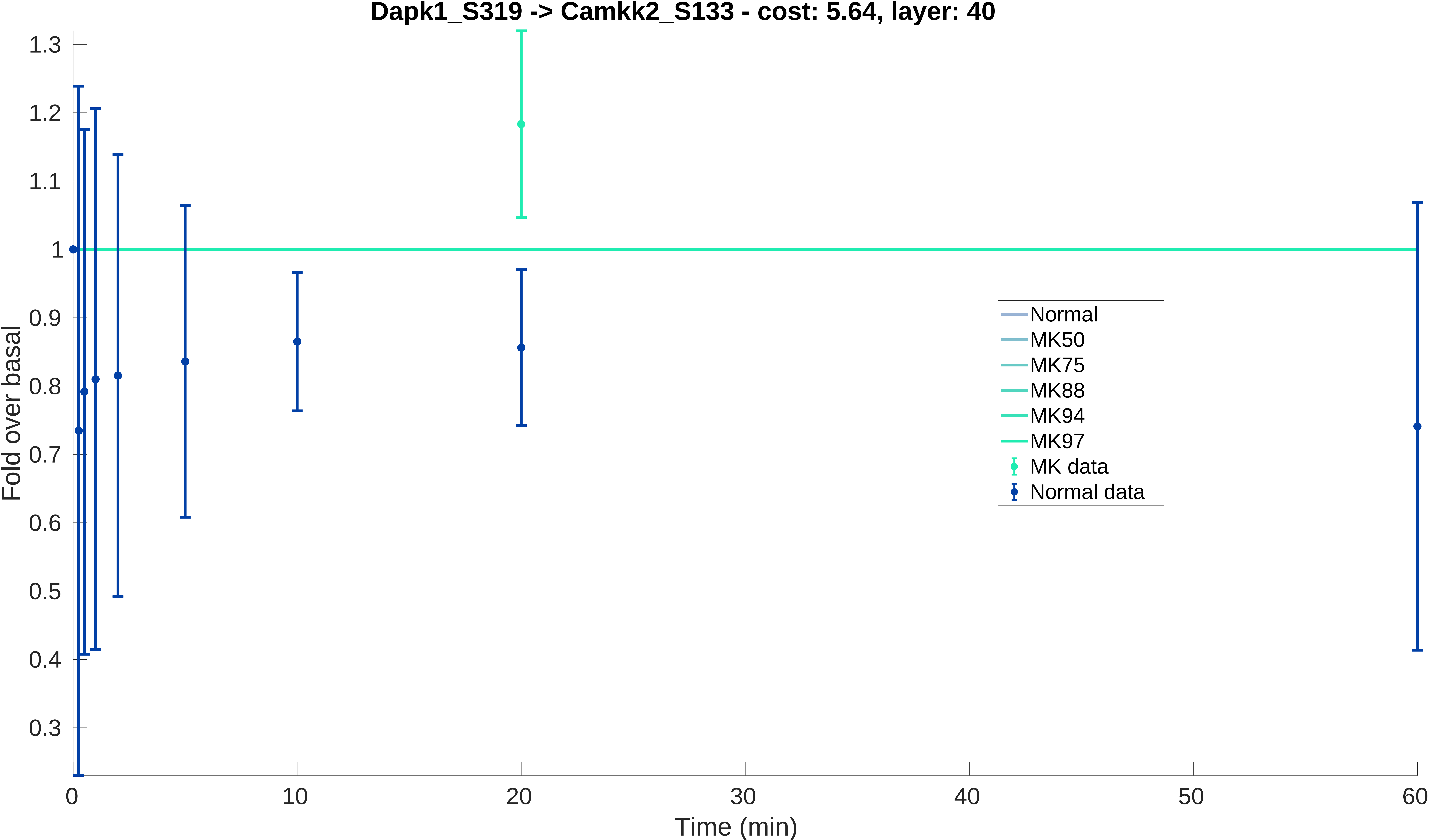

Dapk1_S319 -> Camkk2_S133 - cost: 5.64, layer: 40
Fold over basal
Time (min)
1.3
1.2
1.1
1
0.9
0.8
0.7
0.6
0.5
0.4
0.3
0
10
20
30
40
50
60
Normal
MK50
MK75
MK88
MK94
MK97
MK data
Normal data

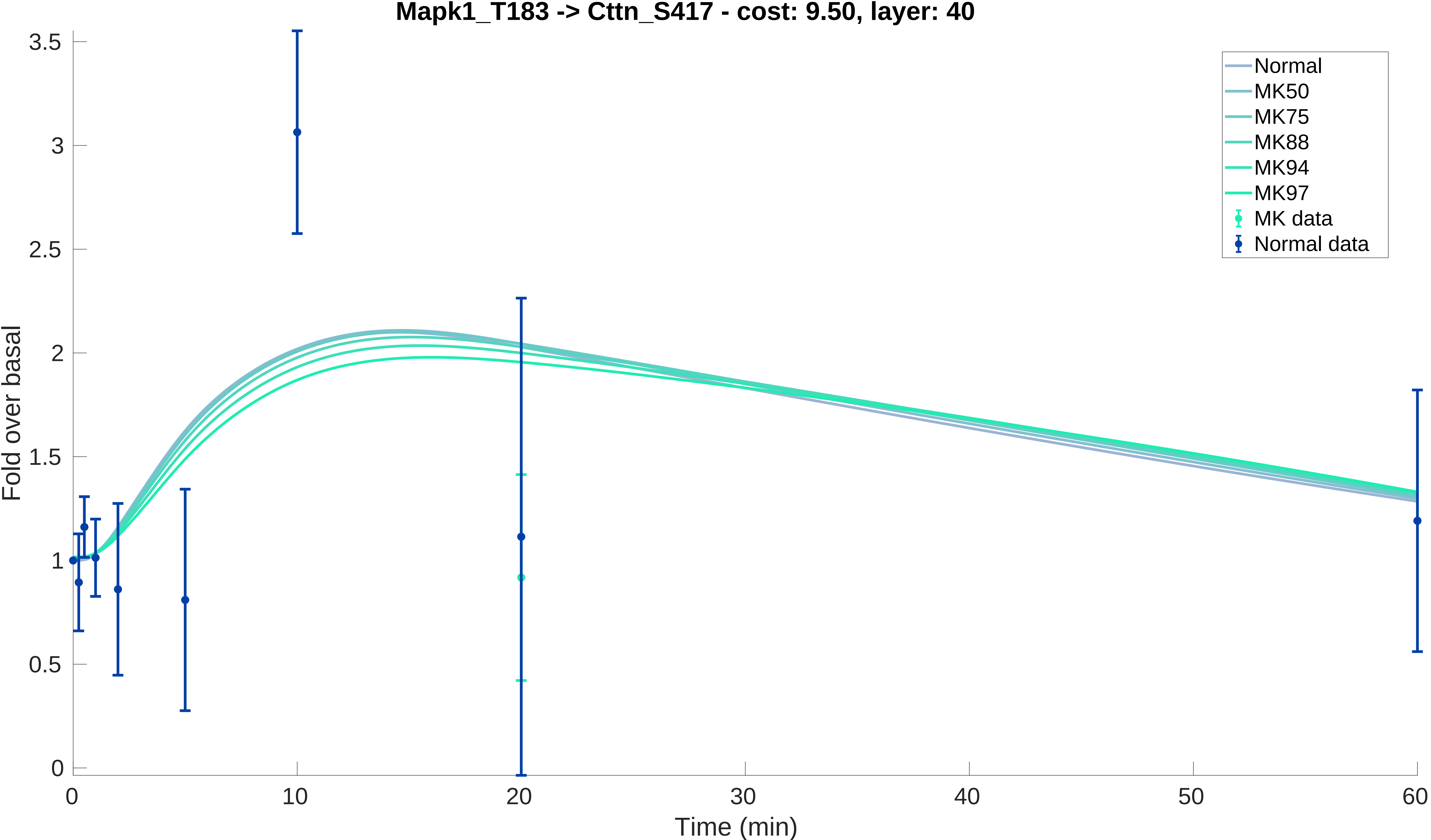
Mapk1_T183 -> Cttn_S417 - cost: 9.50, layer: 40
Fold over basal
Time (min)
0
0.5
1
1.5
2
2.5
3
3.5
0
10
20
30
40
50
60
Normal
MK50
MK75
MK88
MK94
MK97
MK data
Normal data

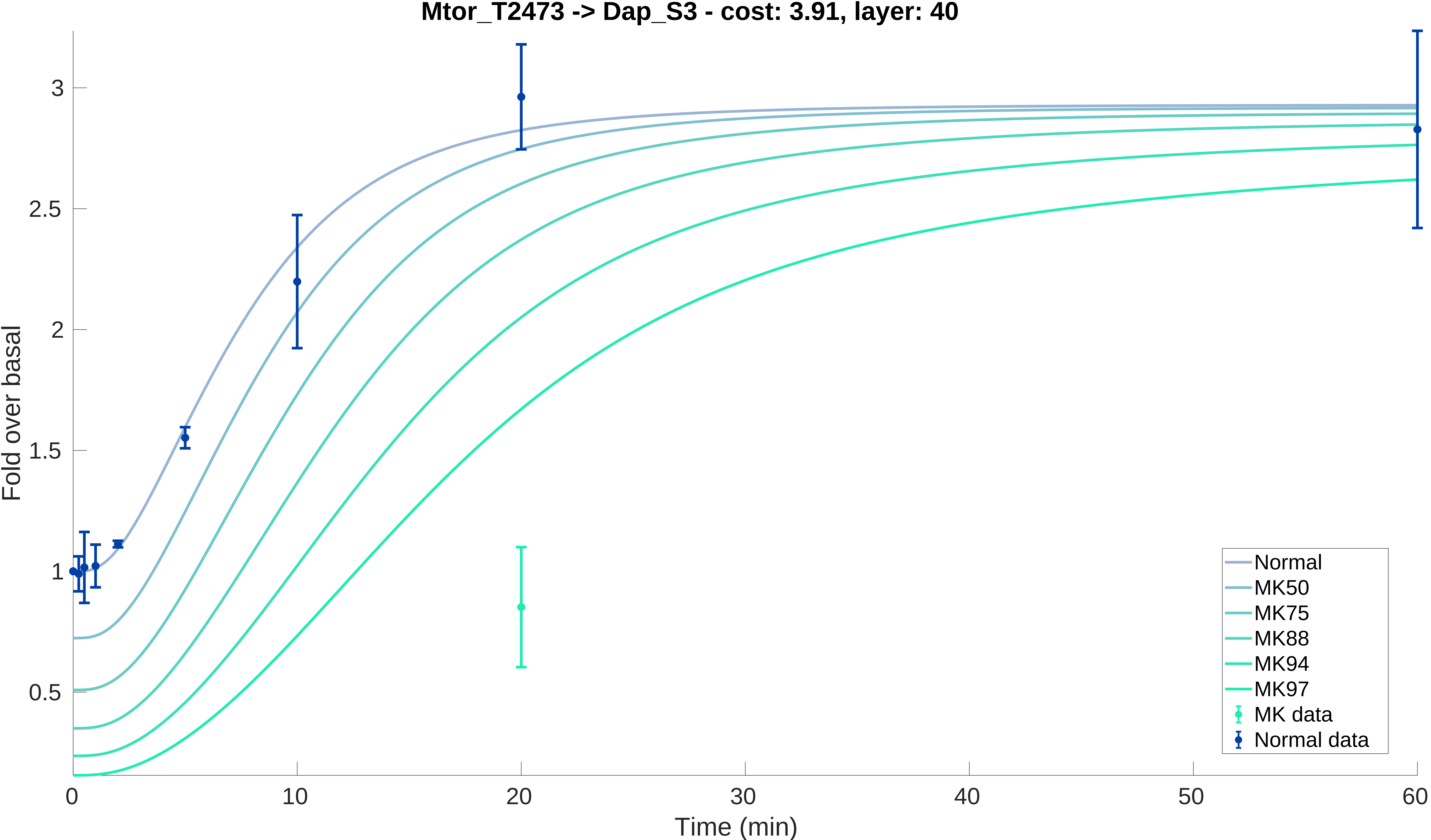
Mtor_T2473 -> Dap_S3 - cost: 3.91, layer: 40
Fold over basal
Time (min)
0
10
20
30
40
50
60
0.5
1
1.5
2
2.5
3
Normal
MK50
MK75
MK88
MK94
MK97
MK data
Normal data

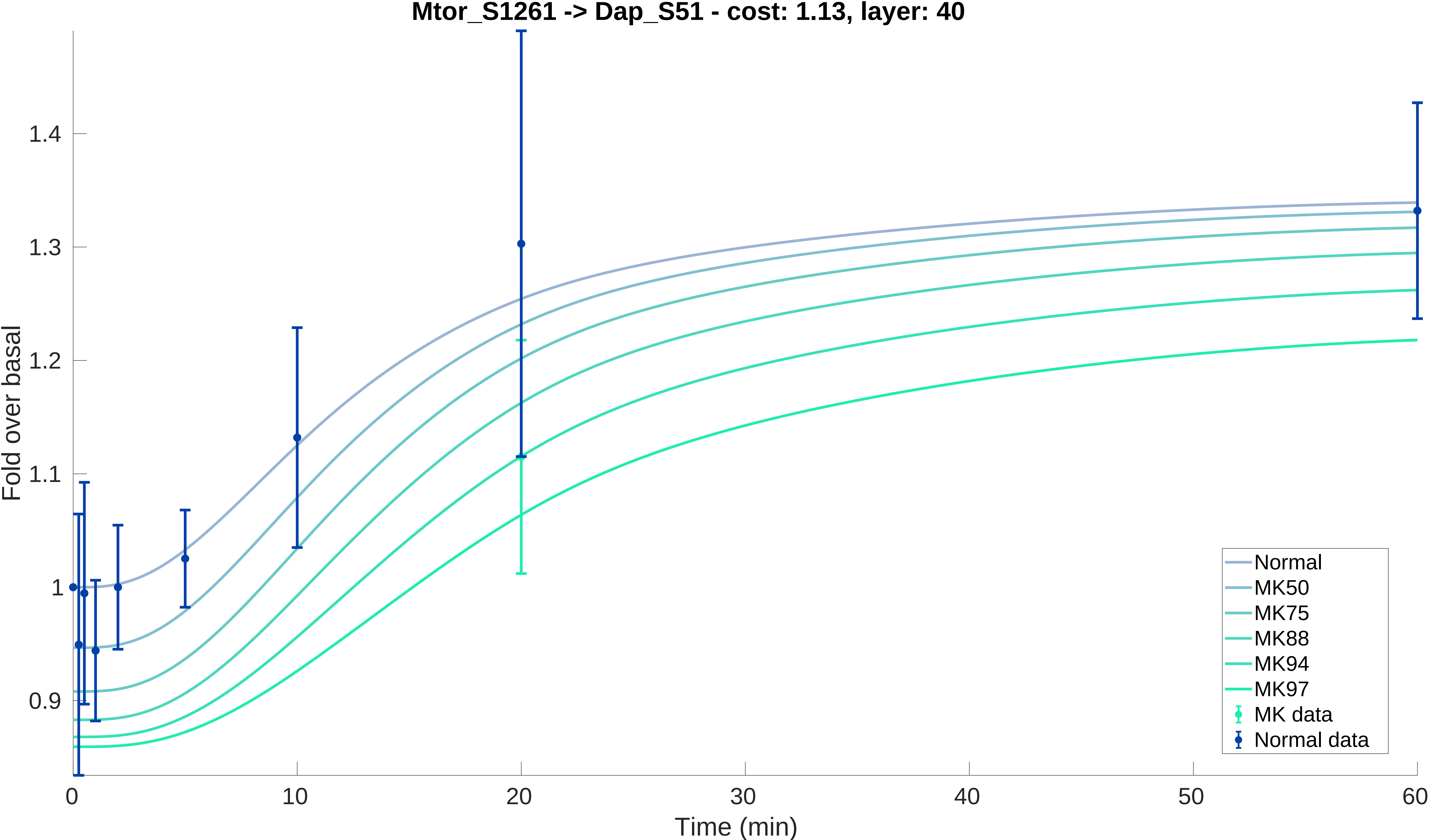

Mtor_S1261 -> Dap_S51 - cost: 1.13, layer: 40
Fold over basal
1.4
1.3
1.2
1.1
1
0.9
0
10
20
30
40
50
60
Time (min)
Normal
MK50
MK75
MK88
MK94
MK97
MK data
Normal data

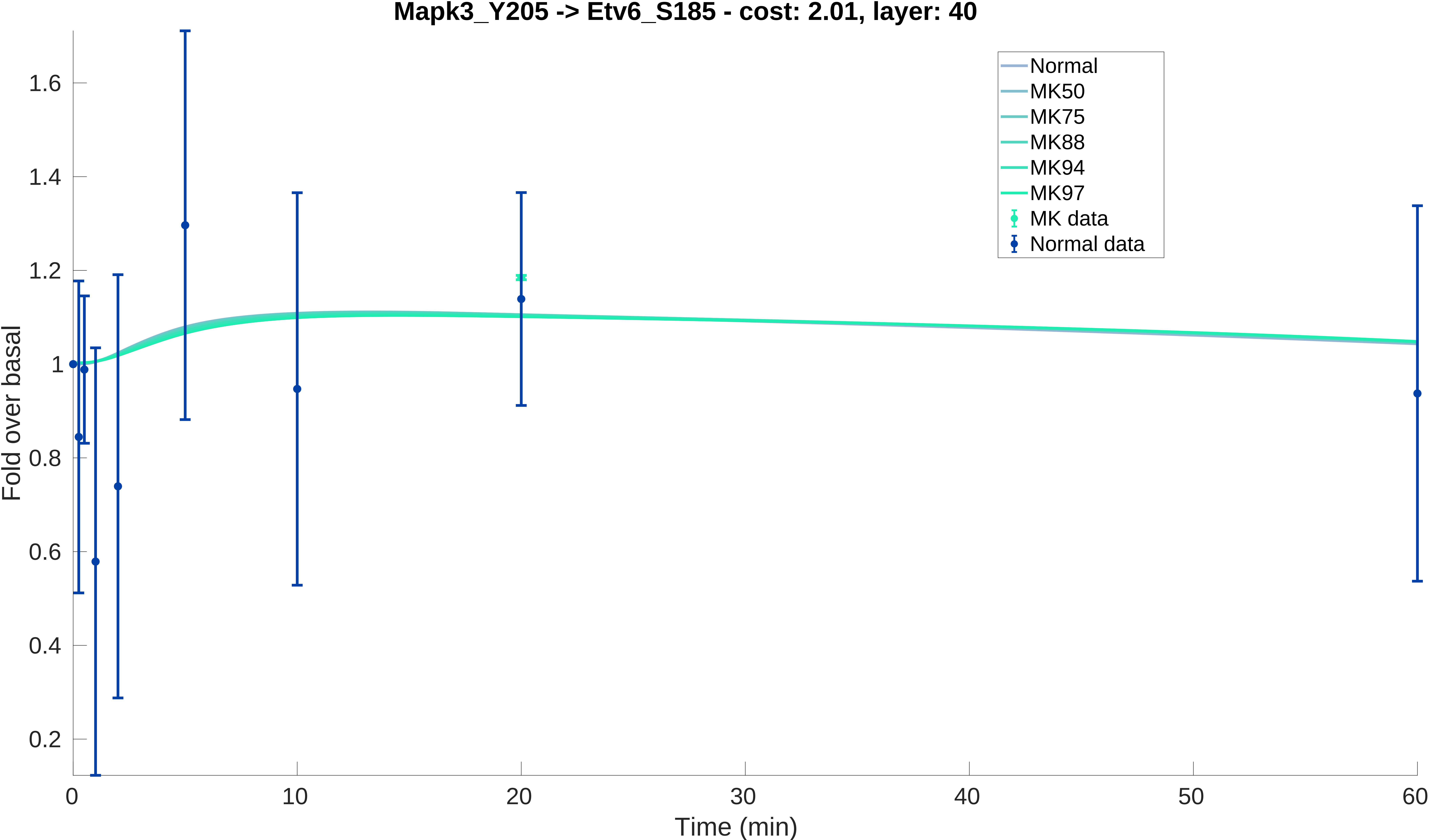

Mapk3_Y205 -> Etv6_S185 - cost: 2.01, layer: 40
Normal
MK50
MK75
MK88
MK94
MK97
MK data
Normal data
Fold over basal
1.6
1.4
1.2
1
0.8
0.6
0.4
0.2
0
10
20
30
40
50
60
Time (min)

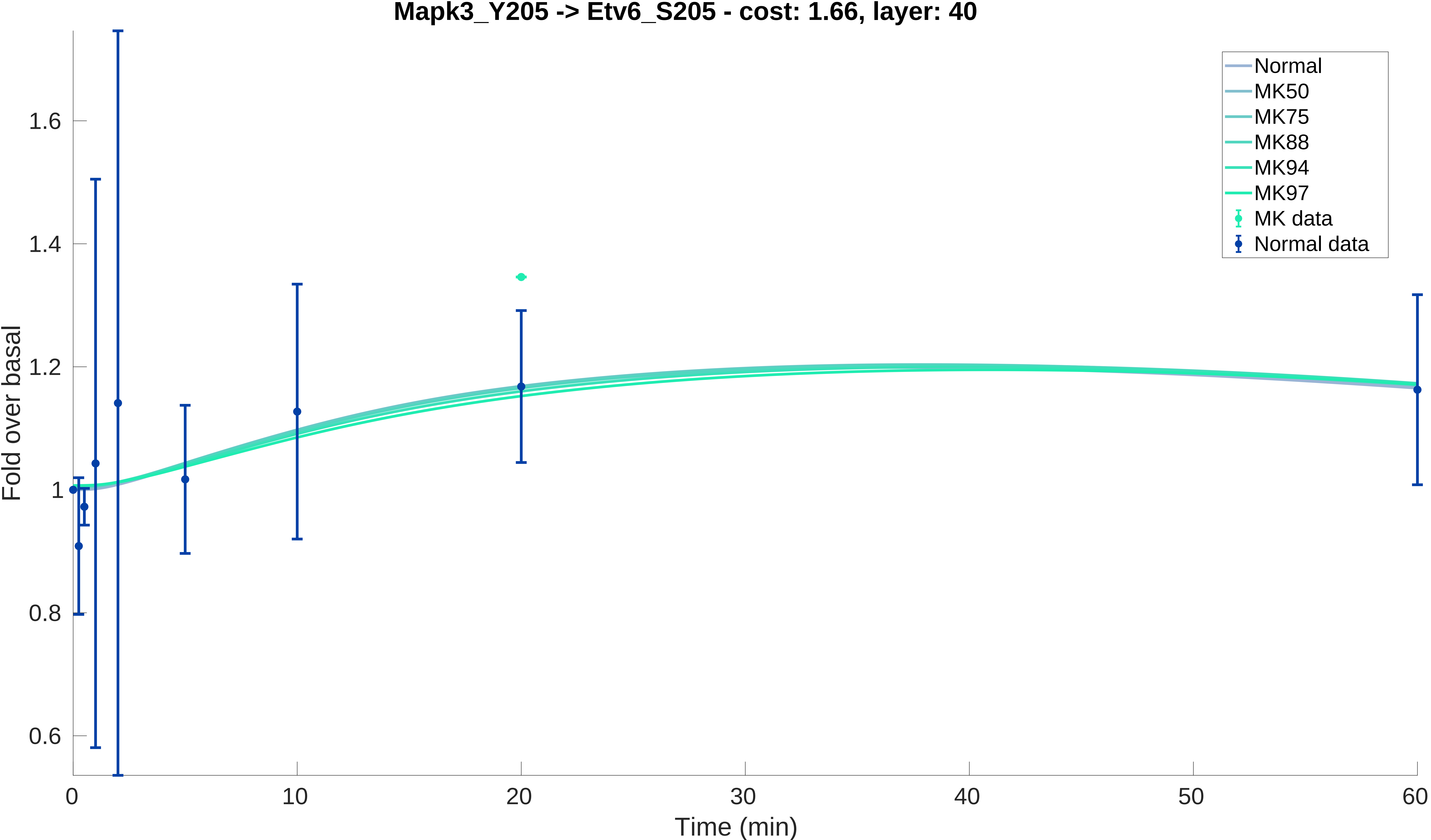
Mapk3_Y205 -> Etv6_S205 - cost: 1.66, layer: 40
Fold over basal
Time (min)
0.6
0.8
1
1.2
1.4
1.6
0
10
20
30
40
50
60
Normal
MK50
MK75
MK88
MK94
MK97
MK data
Normal data

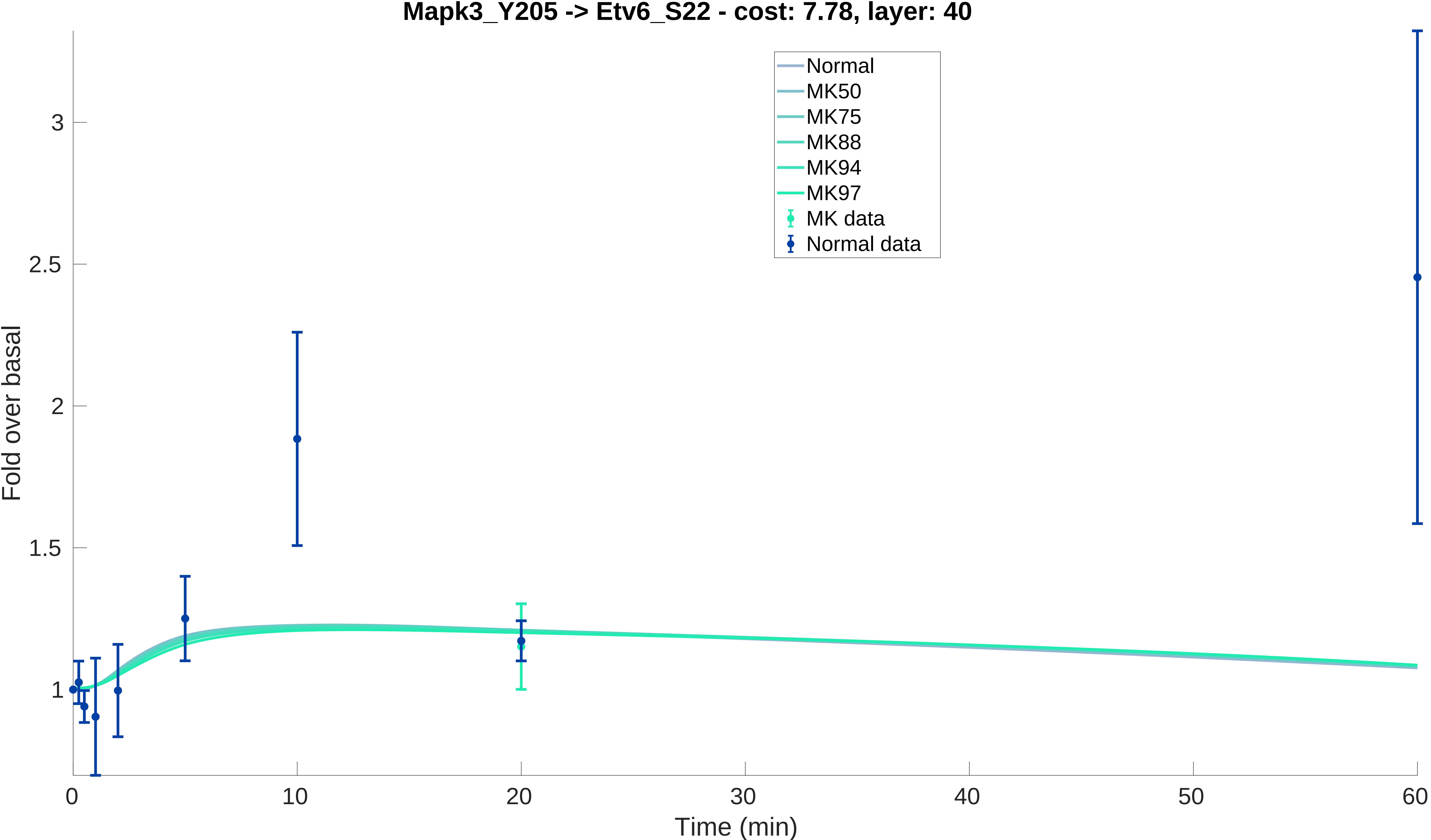
Mapk3_Y205 -> Etv6_S22 - cost: 7.78, layer: 40
Normal
MK50
MK75
MK88
MK94
MK97
MK data
Normal data
Fold over basal
3
2.5
2
1.5
1
0
10
20
30
40
50
60
Time (min)

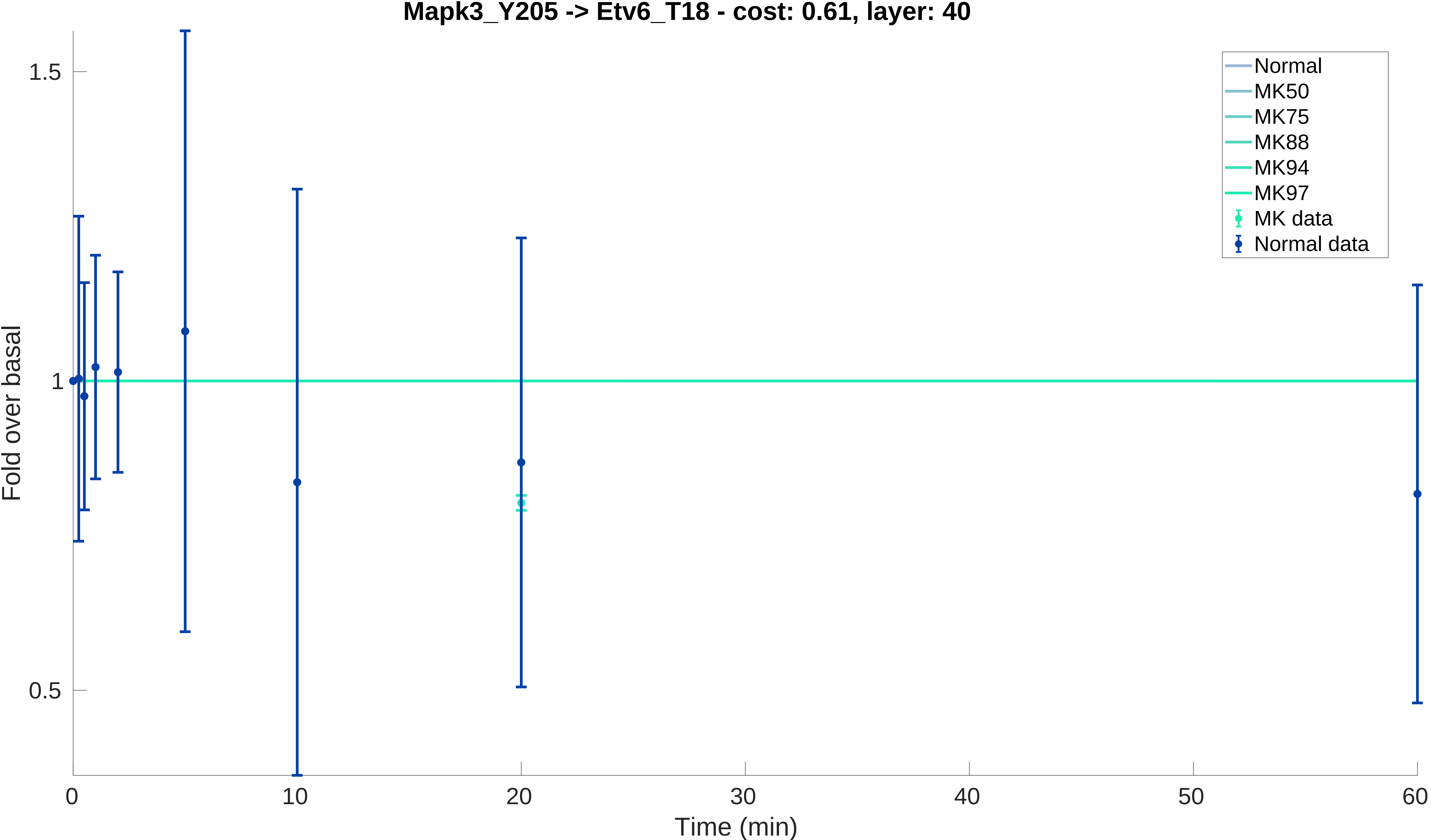

Mapk3_Y205 -> Etv6_T18 - cost: 0.61, layer: 40
Normal
MK50
MK75
MK88
MK94
MK97
MK data
Normal data
Fold over basal
1.5
1
0.5
0
10
20
30
40
50
60
Time (min)

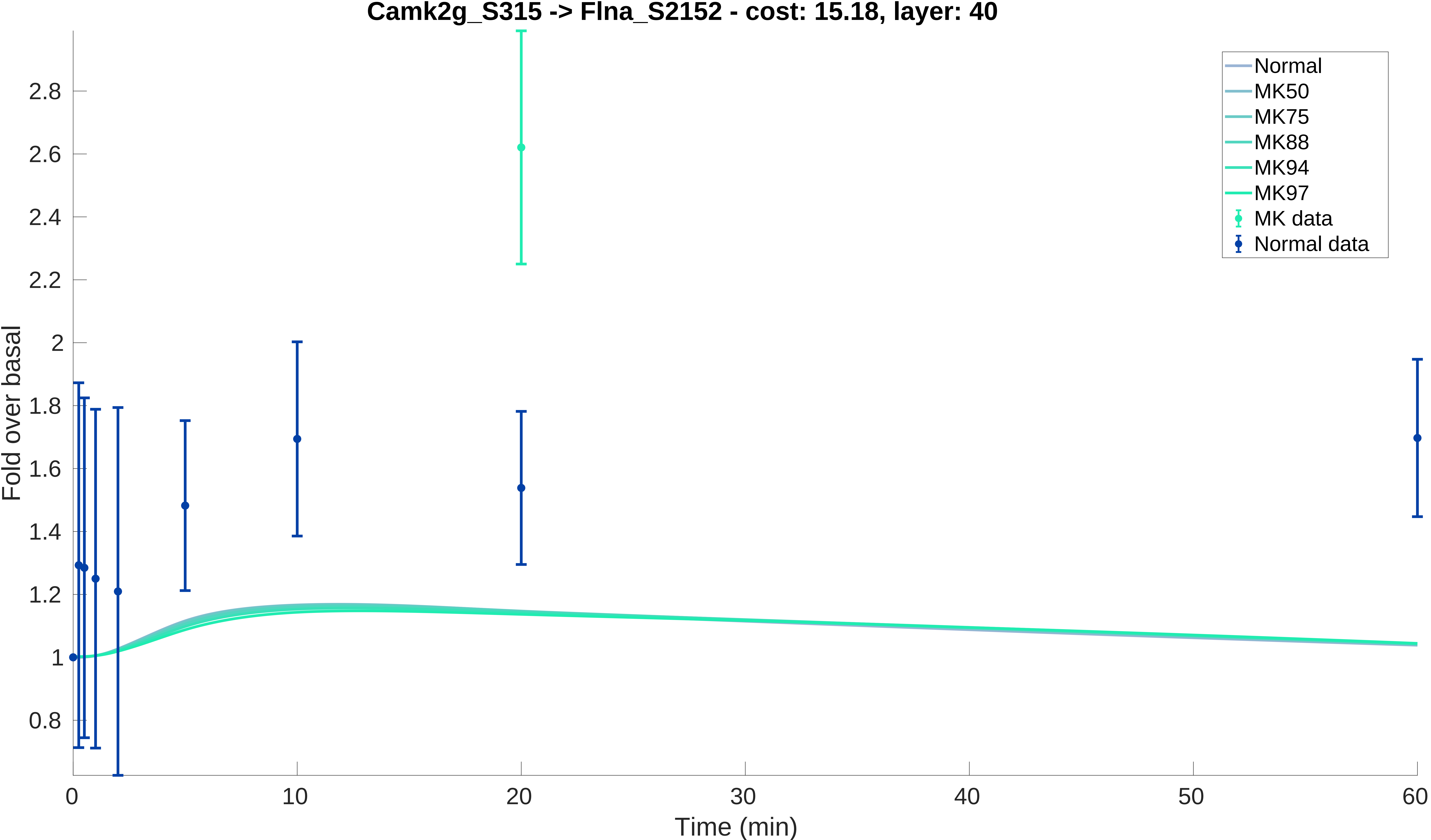

Camk2g_S315 -> Flna_S2152 - cost: 15.18, layer: 40
Normal
MK50
MK75
MK88
MK94
MK97
MK data
Normal data
Fold over basal
Time (min)
0.8
1
1.2
1.4
1.6
1.8
2
2.2
2.4
2.6
2.8
0
10
20
30
40
50
60

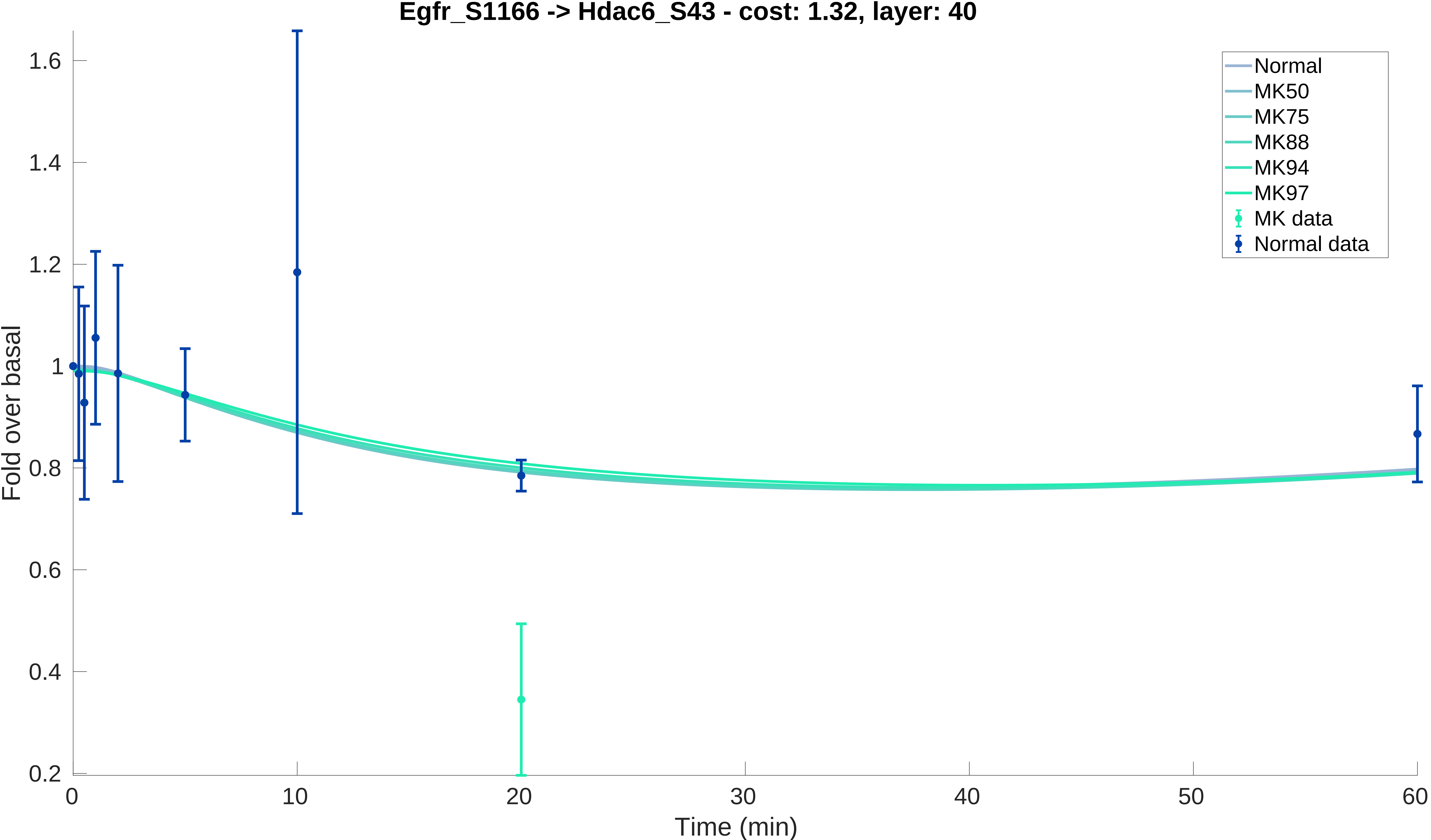
Egfr_S1166 -> Hdac6_S43 - cost: 1.32, layer: 40
Fold over basal
1.6
1.4
1.2
1
0.8
0.6
0.4
0.2
0
10
20
30
40
50
60
Time (min)
Normal
MK50
MK75
MK88
MK94
MK97
MK data
Normal data

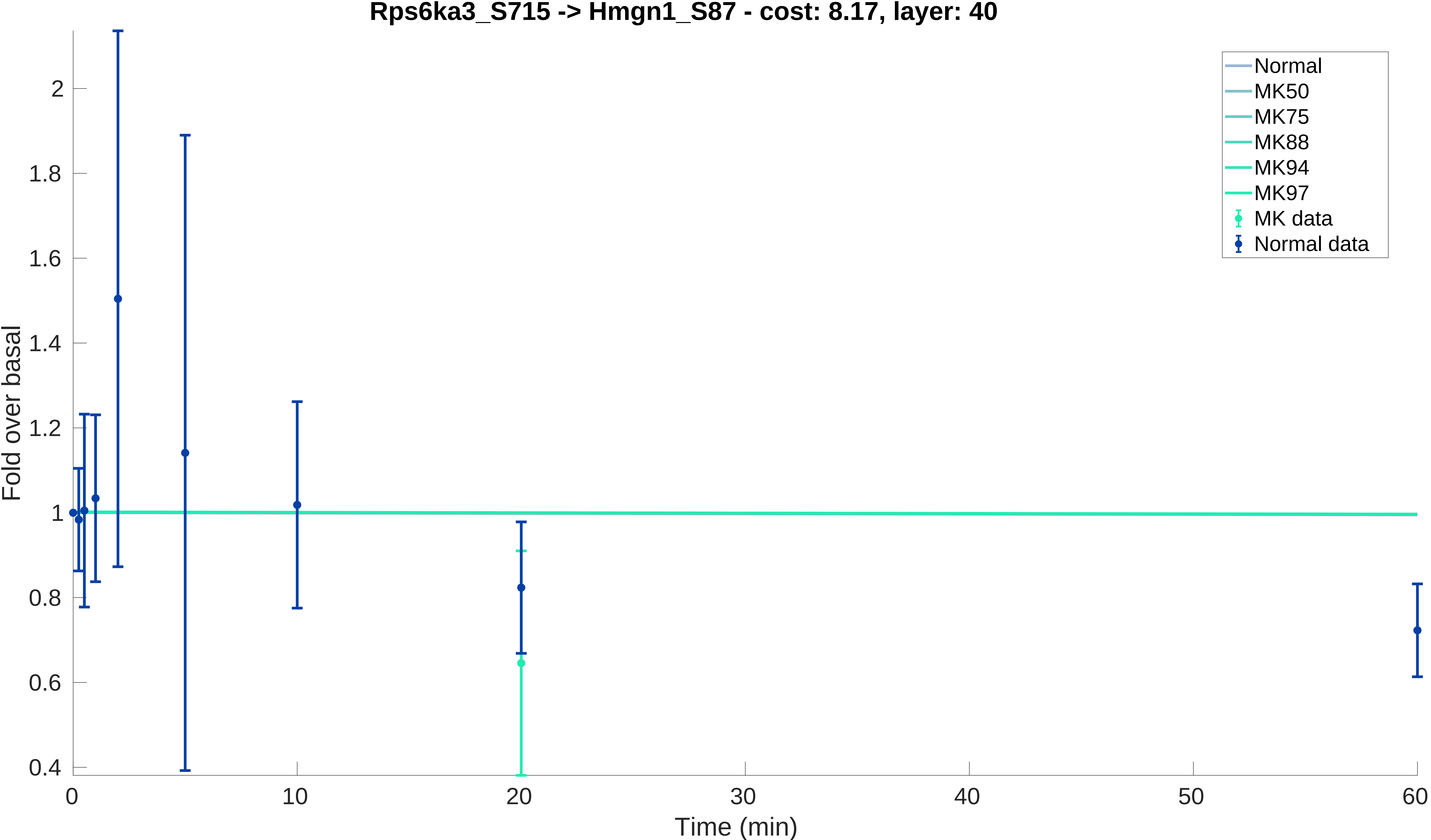
Rps6ka3_S715 -> Hmgn1_S87 - cost: 8.17, layer: 40
Fold over basal
Time (min)
0
10
20
30
40
50
60
0.4
0.6
0.8
1
1.2
1.4
1.6
1.8
2
Normal
MK50
MK75
MK88
MK94
MK97
MK data
Normal data

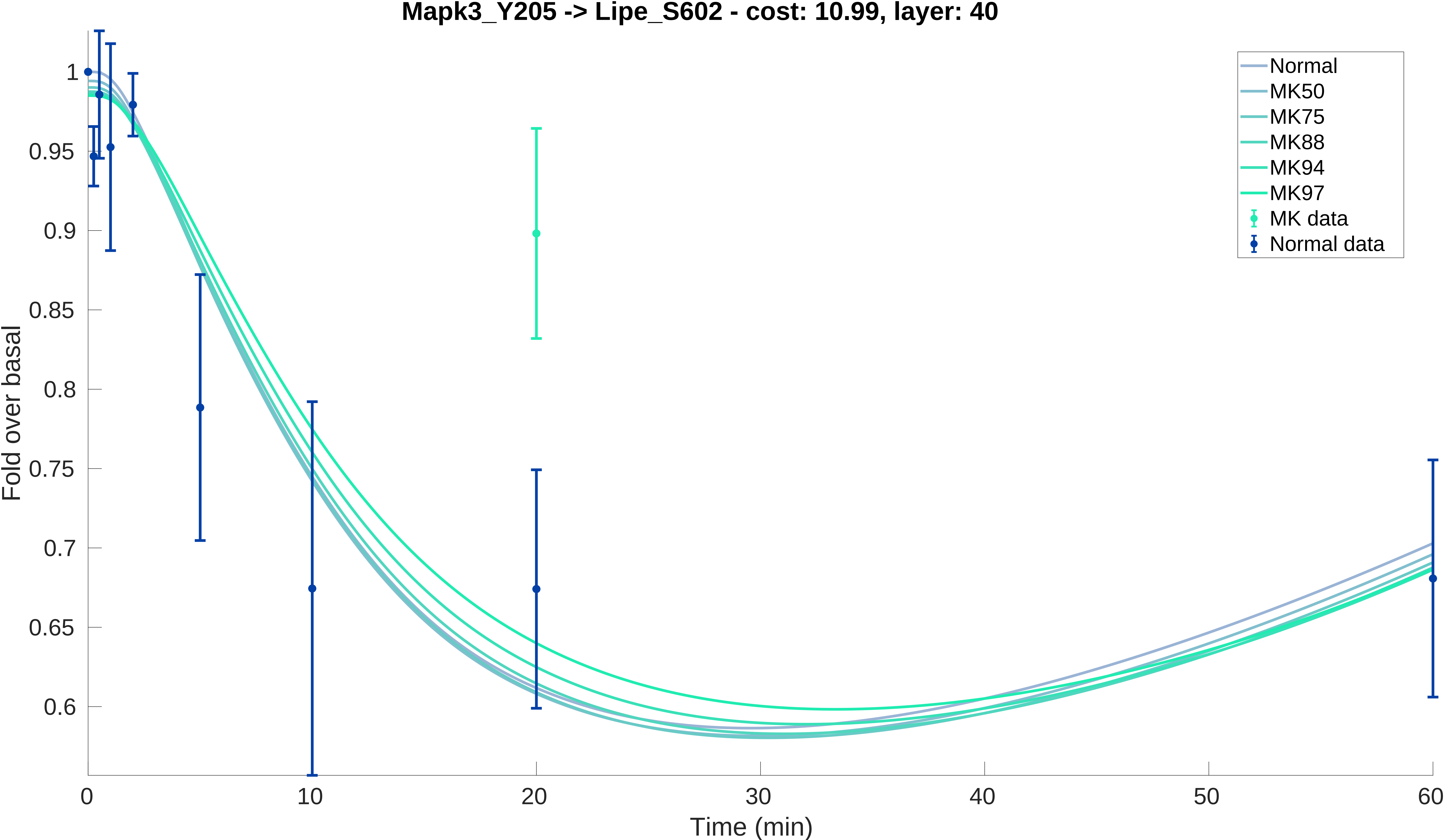

Mapk3_Y205 -> Lipe_S602 - cost: 10.99, layer: 40
Fold over basal
1
0.95
0.9
0.85
0.8
0.75
0.7
0.65
0.6
0
10
20
30
40
50
60
Time (min)
Normal
MK50
MK75
MK88
MK94
MK97
MK data
Normal data

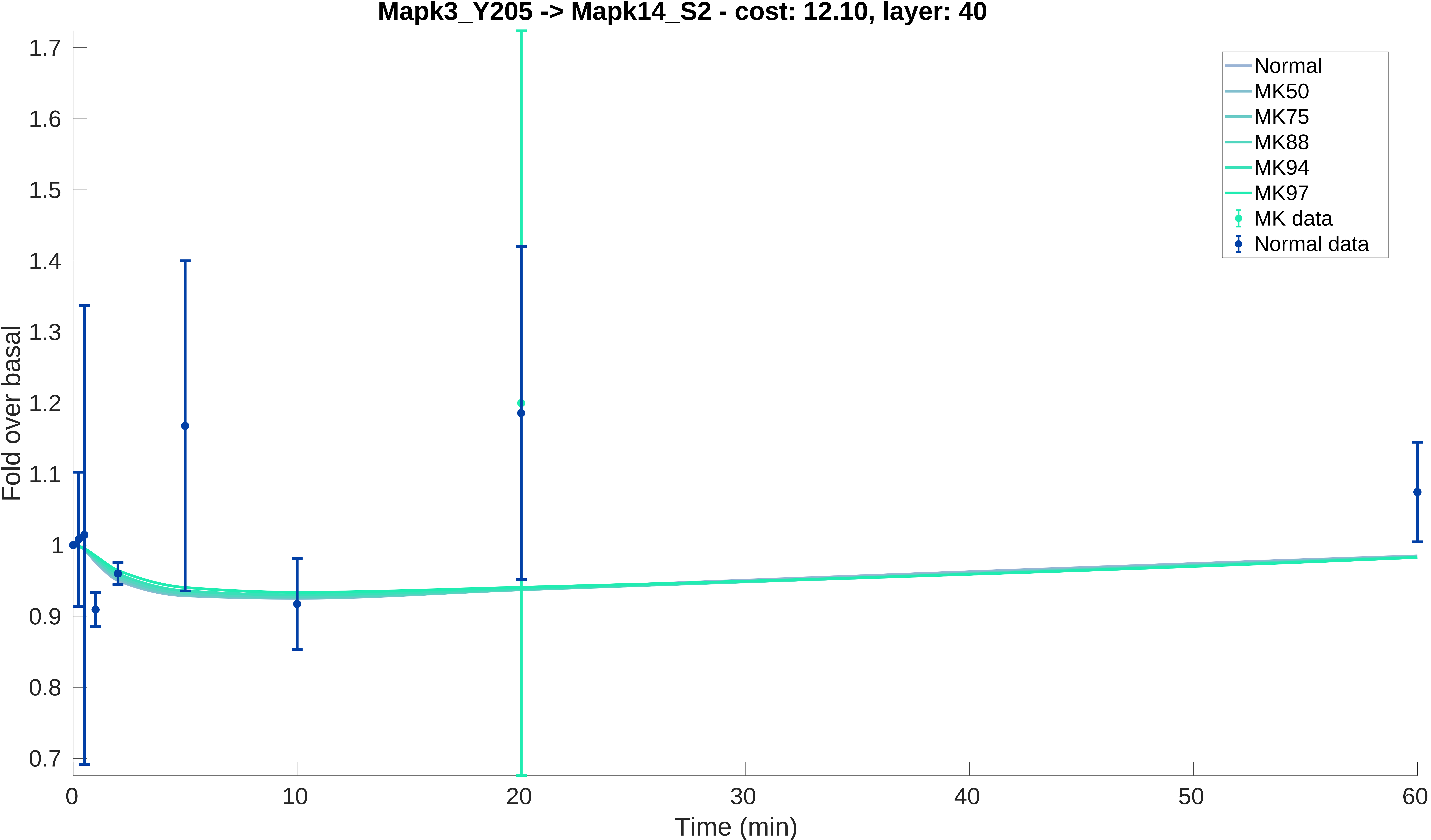

Mapk3_Y205 -> Mapk14_S2 - cost: 12.10, layer: 40
Fold over basal
Time (min)
1.7
1.6
1.5
1.4
1.3
1.2
1.1
1
0.9
0.8
0.7
0
10
20
30
40
50
60
Normal
MK50
MK75
MK88
MK94
MK97
MK data
Normal data

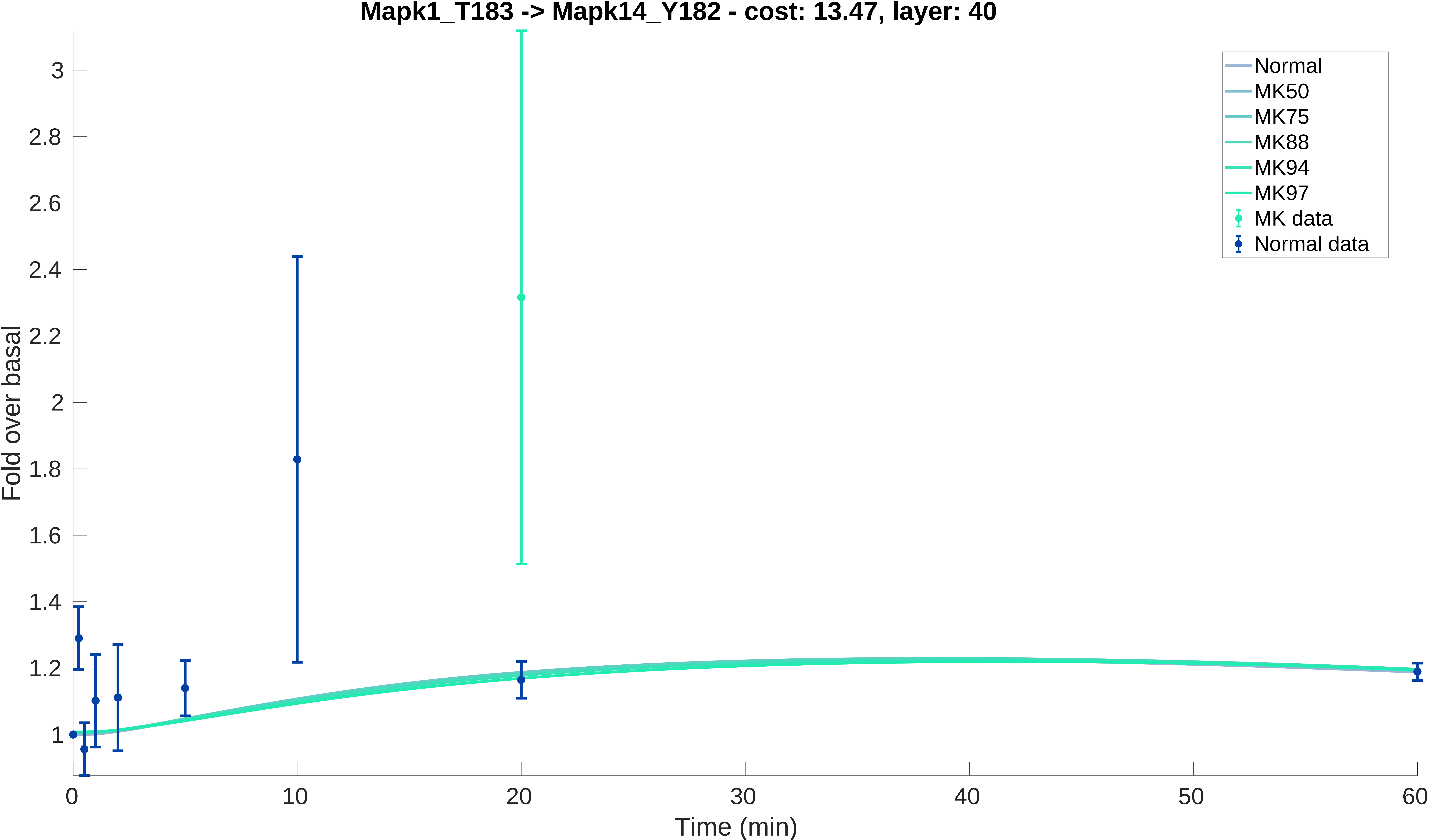

Mapk1_T183 -> Mapk14_Y182 - cost: 13.47, layer: 40
Fold over basal
Time (min)
0
10
20
30
40
50
60
1
1.2
1.4
1.6
1.8
2
2.2
2.4
2.6
2.8
3
Normal
MK50
MK75
MK88
MK94
MK97
MK data
Normal data

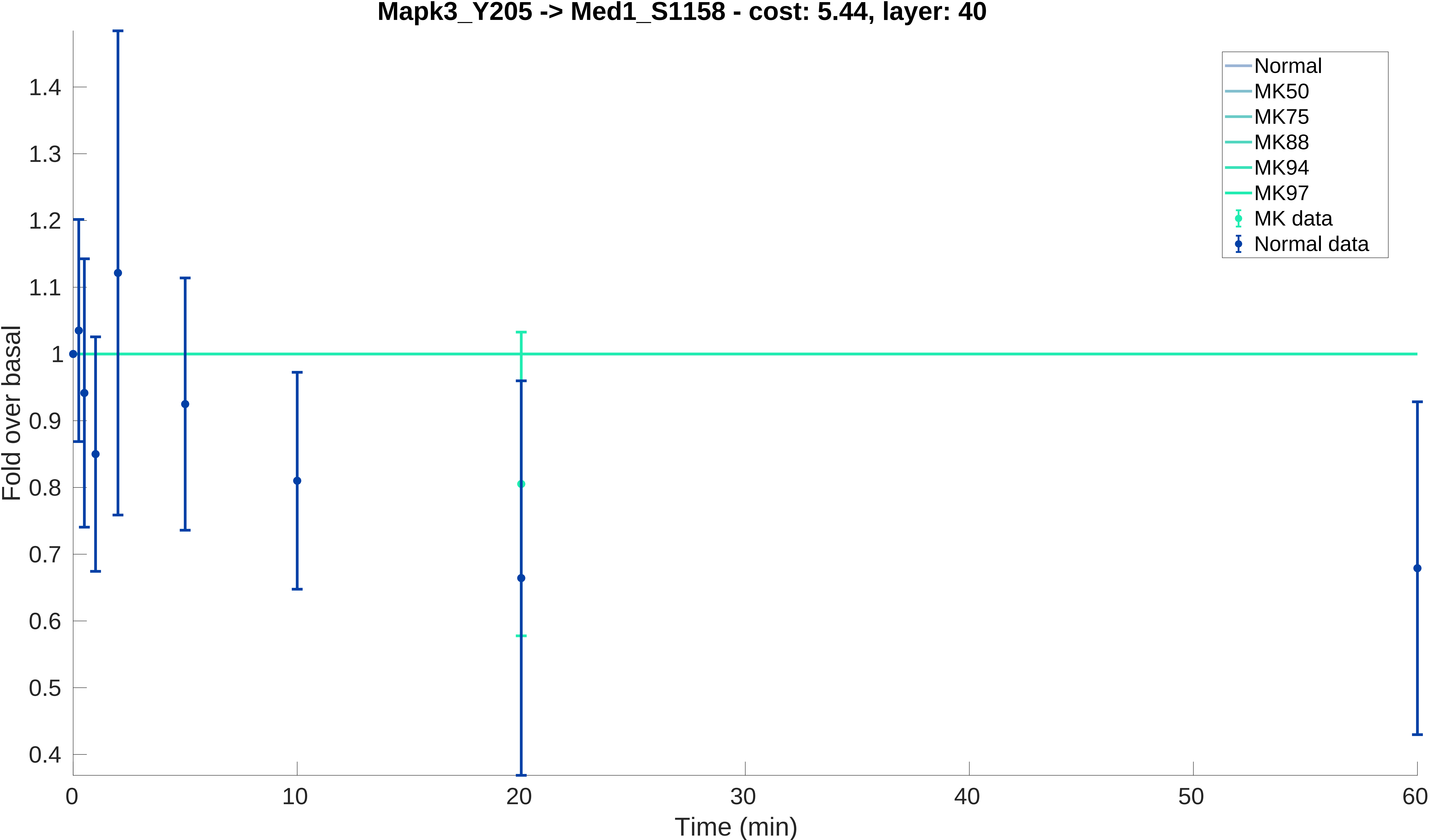
Mapk3_Y205 -> Med1_S1158 - cost: 5.44, layer: 40
Fold over basal
Time (min)
1.4
1.3
1.2
1.1
1
0.9
0.8
0.7
0.6
0.5
0.4
0
10
20
30
40
50
60
Normal
MK50
MK75
MK88
MK94
MK97
MK data
Normal data

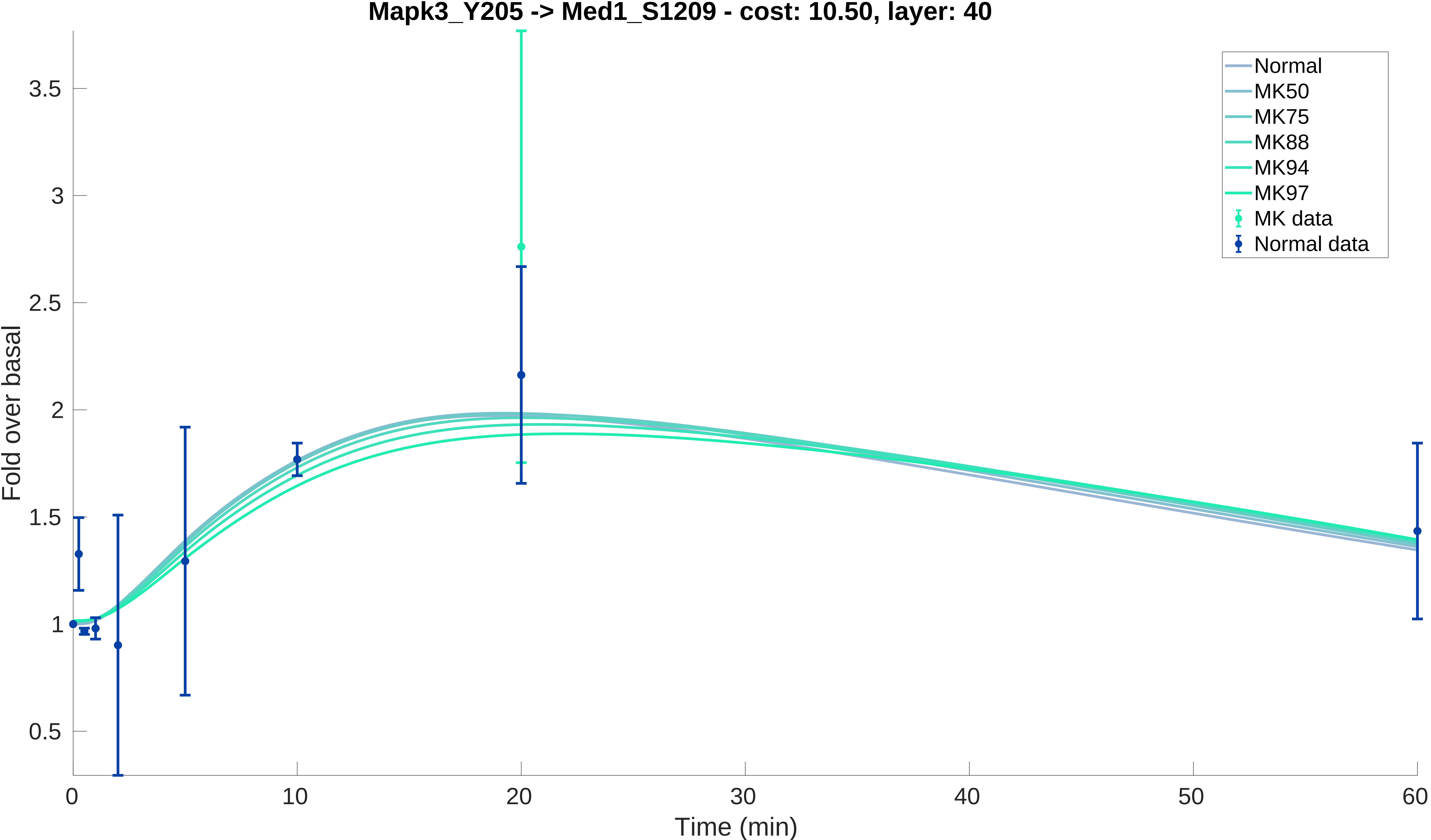
Mapk3_Y205 -> Med1_S1209 - cost: 10.50, layer: 40
Fold over basal
Time (min)
0.5
1
1.5
2
2.5
3
3.5
0
10
20
30
40
50
60
Normal
MK50
MK75
MK88
MK94
MK97
MK data
Normal data

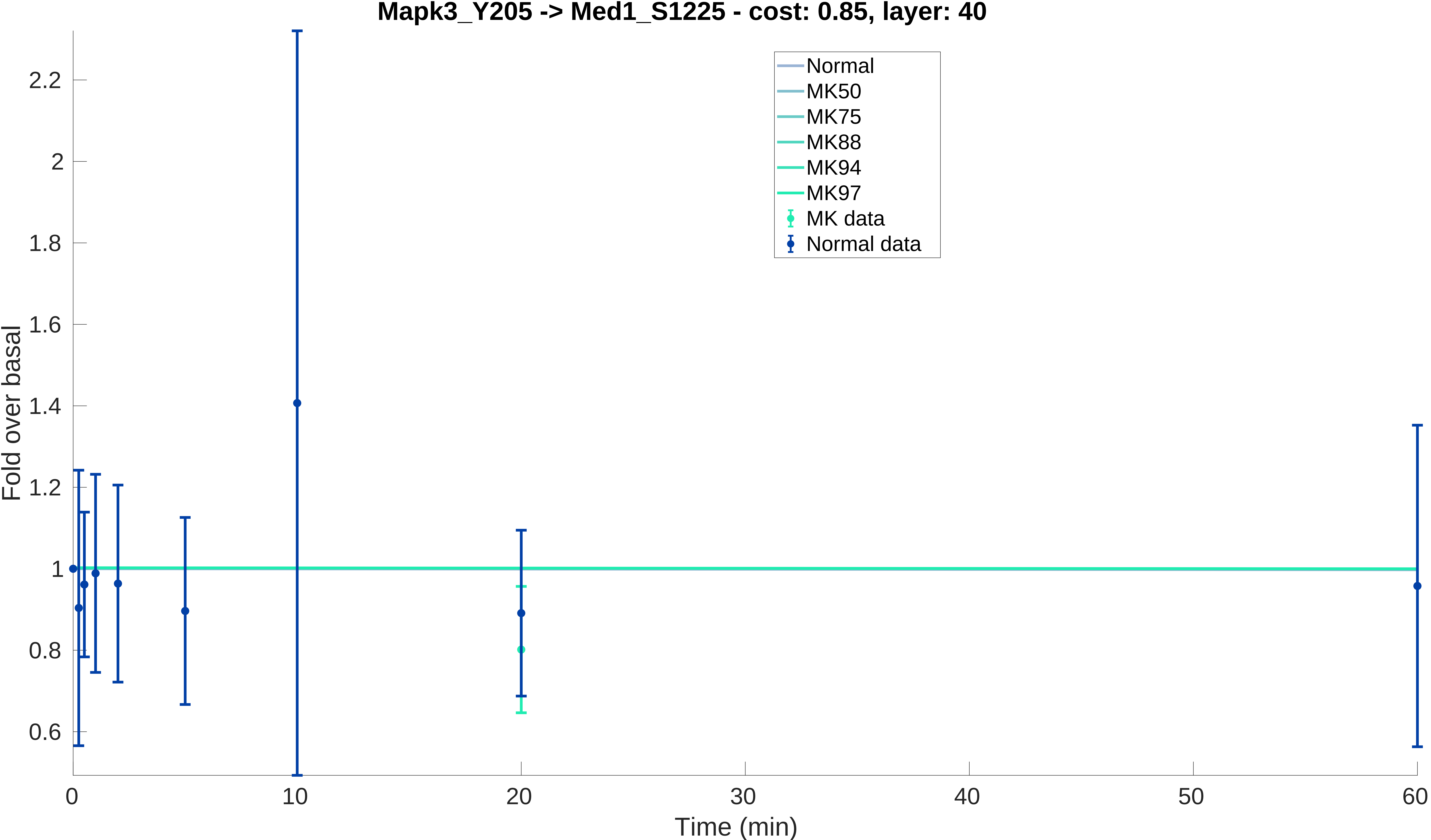

Mapk3_Y205 -> Med1_S1225 - cost: 0.85, layer: 40
Normal
MK50
MK75
MK88
MK94
MK97
MK data
Normal data
Fold over basal
2.2
2
1.8
1.6
1.4
1.2
1
0.8
0.6
0
10
20
30
40
50
60
Time (min)

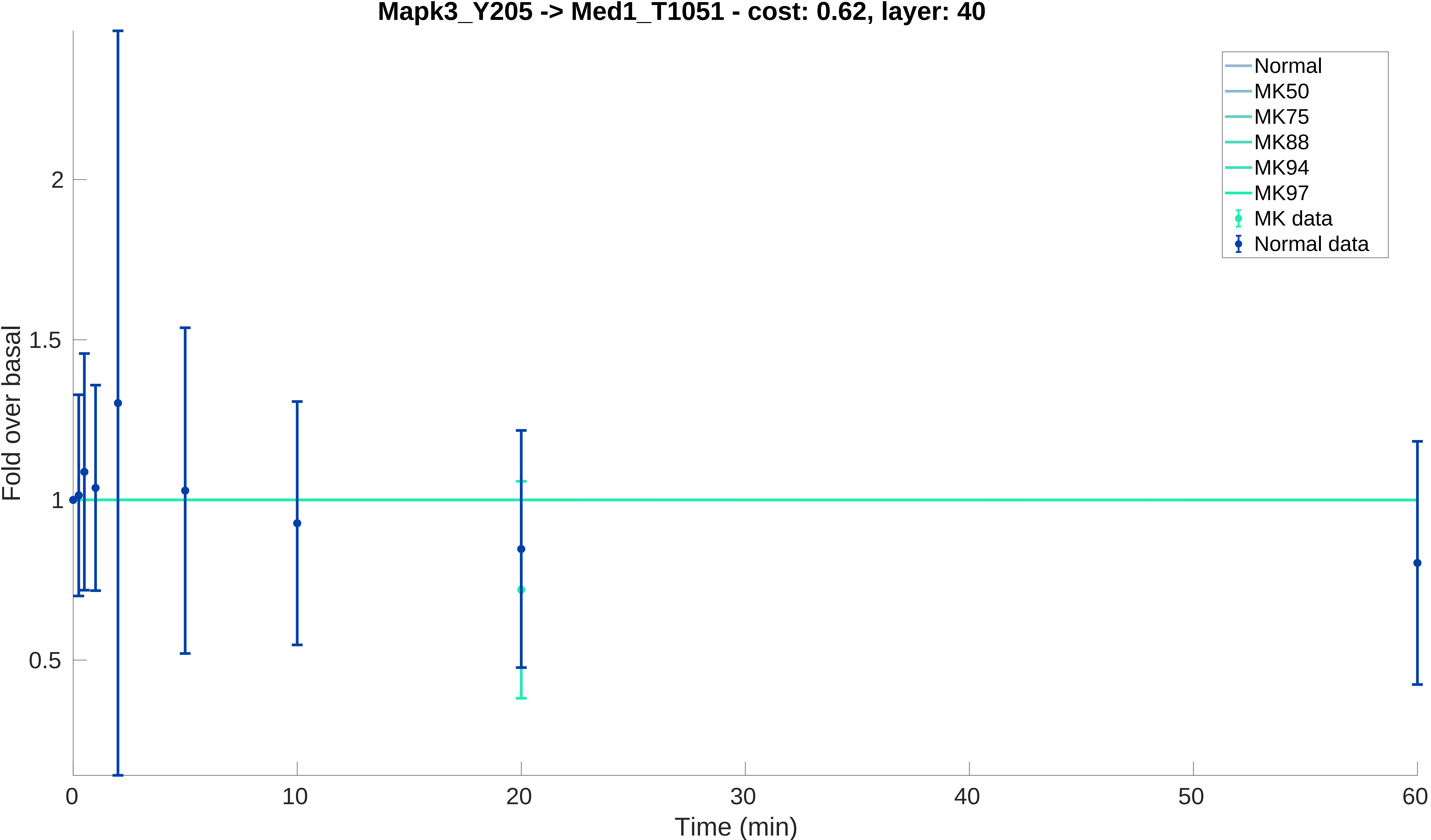

Mapk3_Y205 -> Med1_T1051 - cost: 0.62, layer: 40
Fold over basal
0.5
1
1.5
2
0
10
20
30
40
50
60
Time (min)
Normal
MK50
MK75
MK88
MK94
MK97
MK data
Normal data

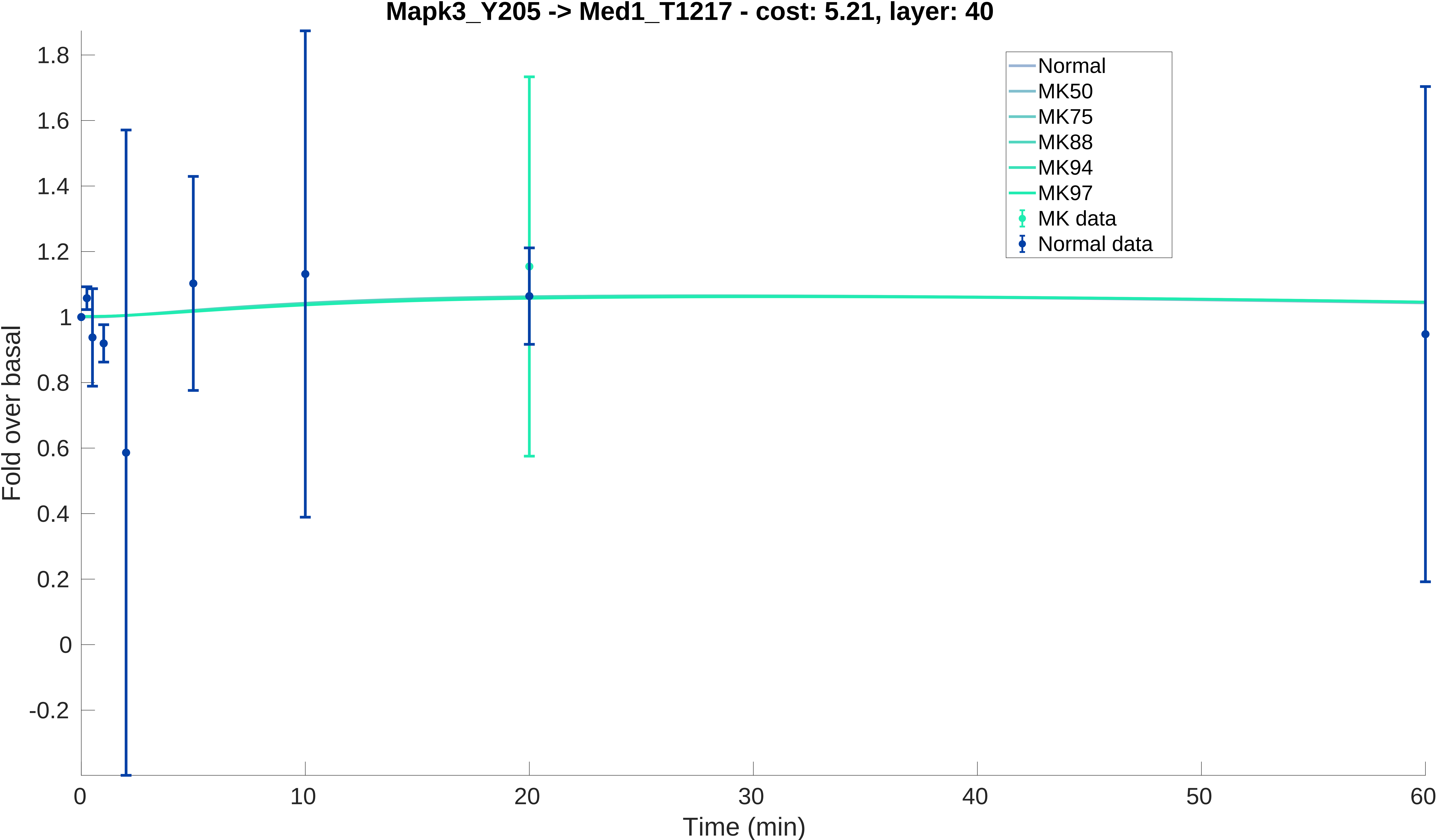

Mapk3_Y205 -> Med1_T1217 - cost: 5.21, layer: 40
Fold over basal
Time (min)
1.8
1.6
1.4
1.2
1
0.8
0.6
0.4
0.2
0
-0.2
0
10
20
30
40
50
60
Normal
MK50
MK75
MK88
MK94
MK97
MK data
Normal data

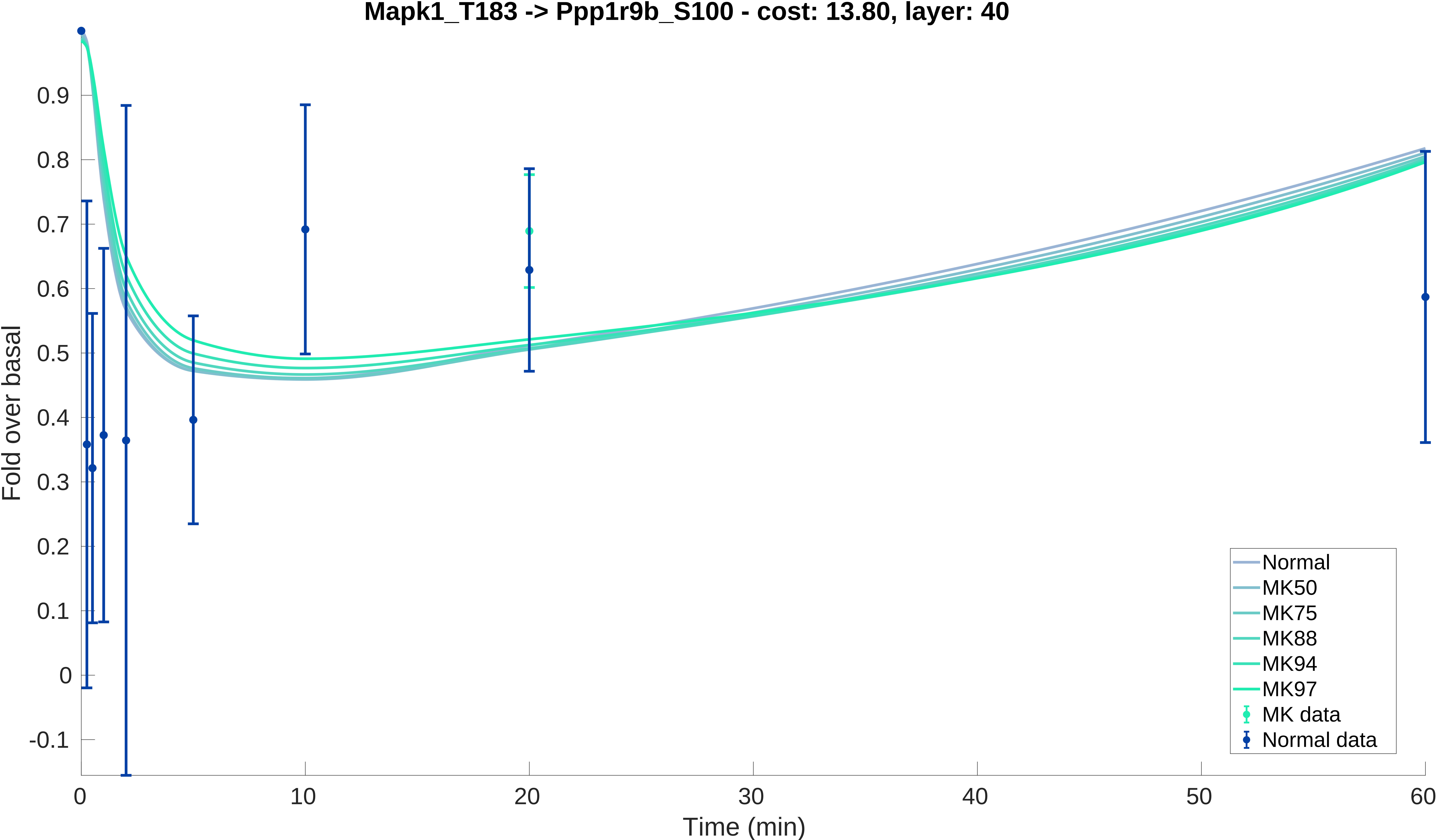

Mapk1_T183 -> Ppp1r9b_S100 - cost: 13.80, layer: 40
Fold over basal
Time (min)
0.9
0.8
0.7
0.6
0.5
0.4
0.3
0.2
0.1
0
-0.1
0
10
20
30
40
50
60
Normal
MK50
MK75
MK88
MK94
MK97
MK data
Normal data

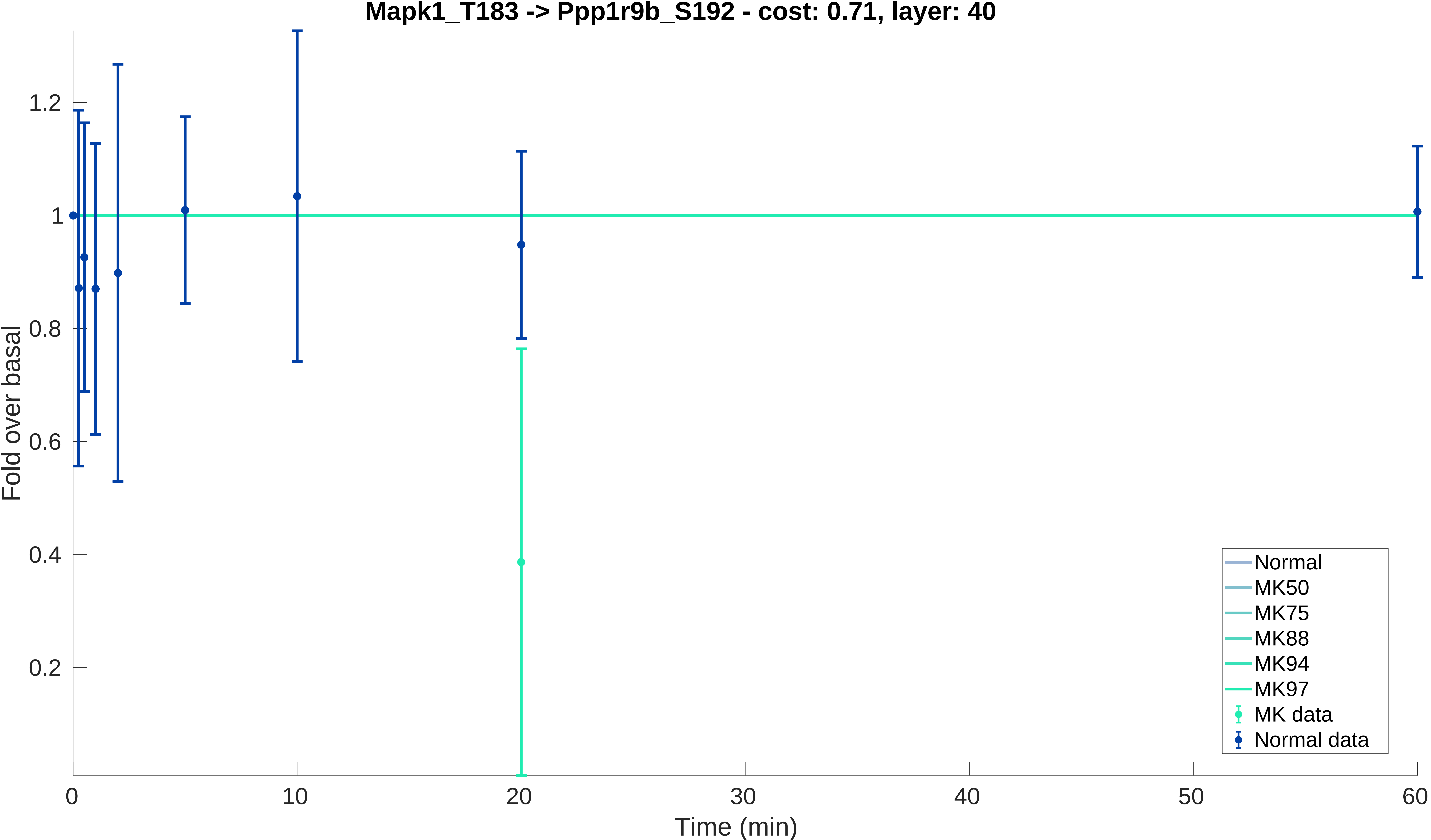

Mapk1_T183 -> Ppp1r9b_S192 - cost: 0.71, layer: 40
Fold over basal
1.2
1
0.8
0.6
0.4
0.2
0
10
20
30
40
50
60
Time (min)
Normal
MK50
MK75
MK88
MK94
MK97
MK data
Normal data

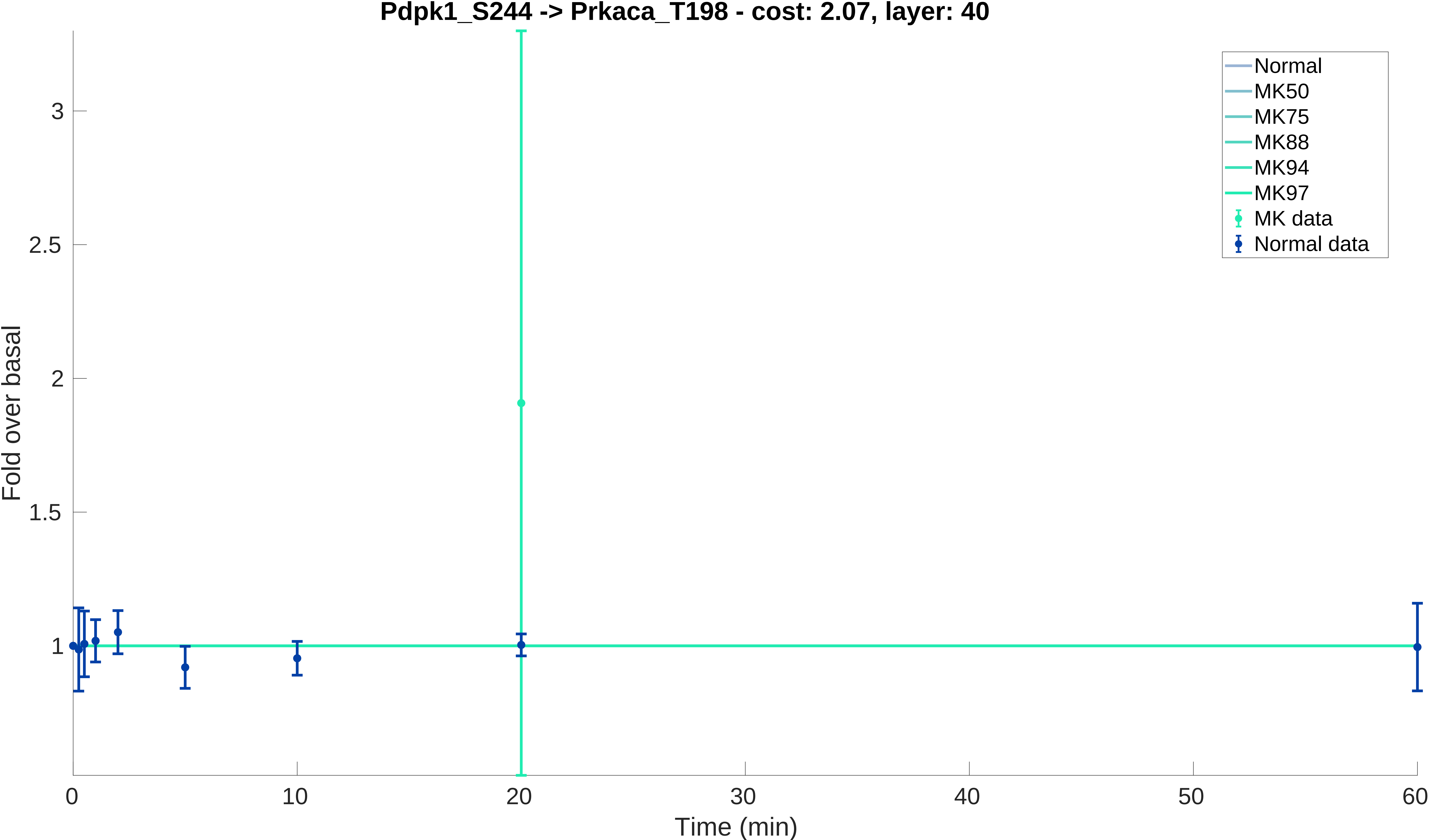
Pdpk1_S244 -> Prkaca_T198 - cost: 2.07, layer: 40
Fold over basal
Time (min)
0
10
20
30
40
50
60
1
1.5
2
2.5
3
Normal
MK50
MK75
MK88
MK94
MK97
MK data
Normal data

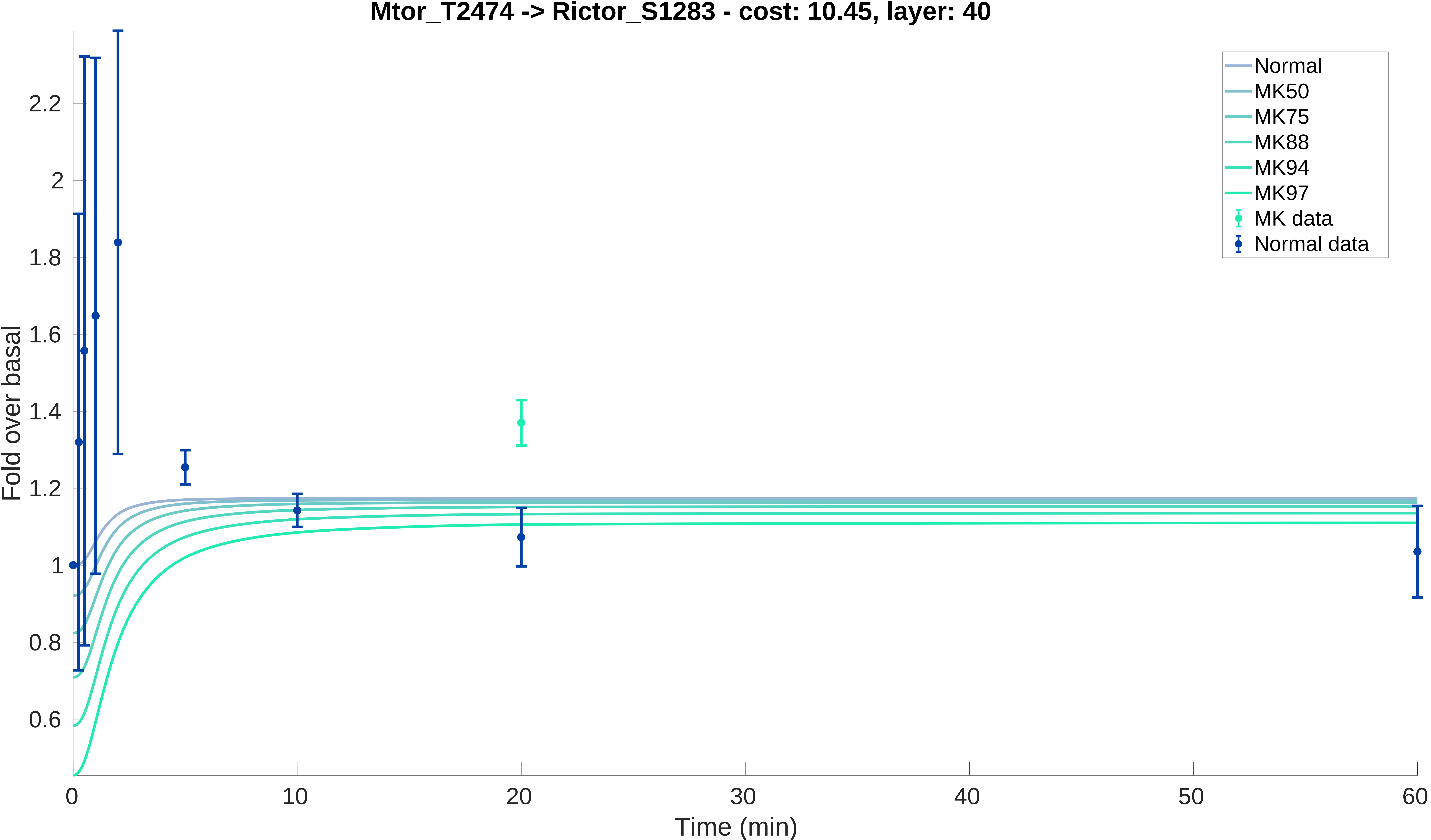

Mtor_T2474 -> Rictor_S1283 - cost: 10.45, layer: 40
Normal
MK50
MK75
MK88
MK94
MK97
MK data
Normal data
Fold over basal
0.6
0.8
1
1.2
1.4
1.6
1.8
2
2.2
0
10
20
30
40
50
60
Time (min)

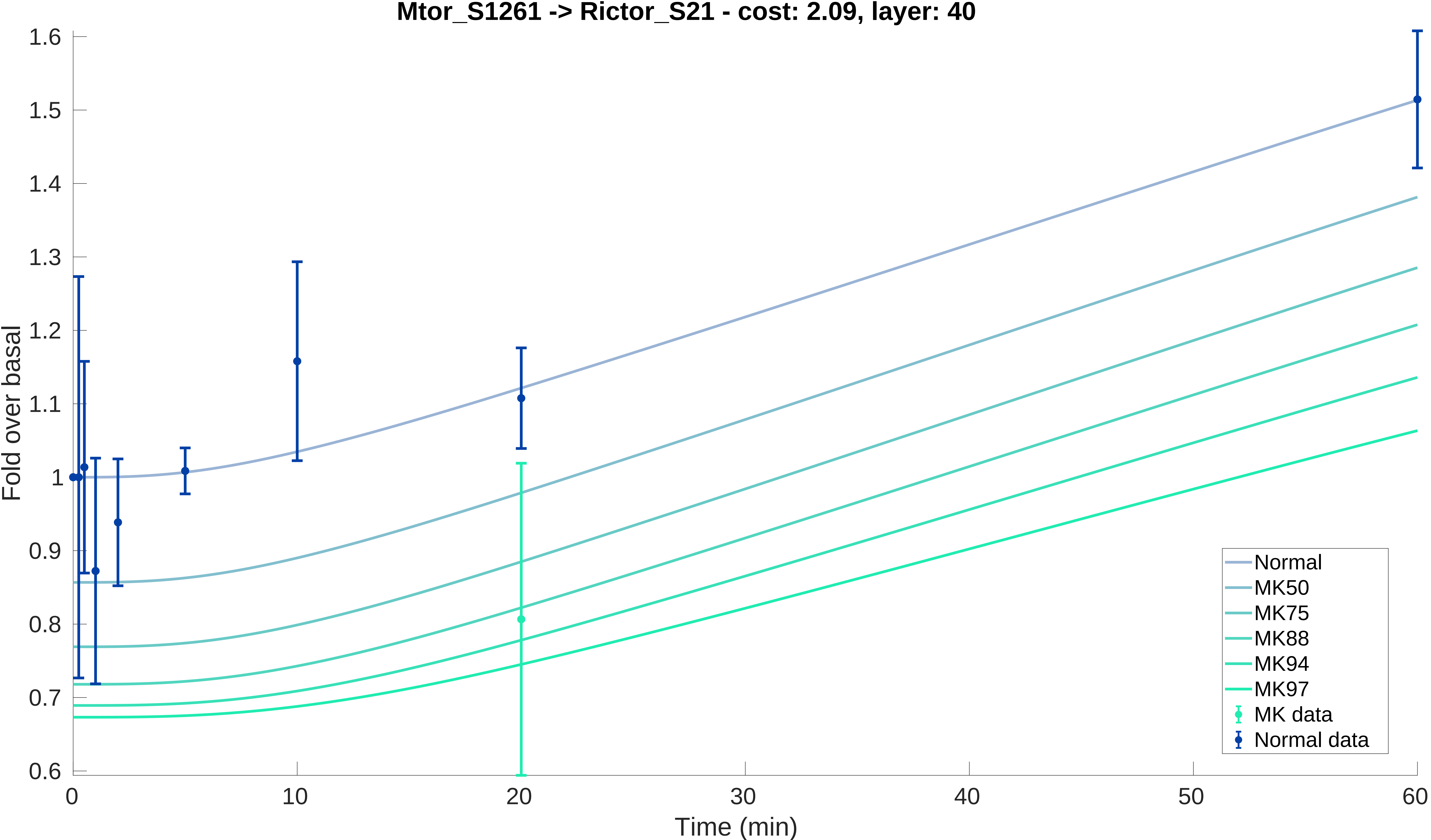
Mtor_S1261 -> Rictor_S21 - cost: 2.09, layer: 40
Fold over basal
Time (min)
0.6
0.7
0.8
0.9
1
1.1
1.2
1.3
1.4
1.5
1.6
0
10
20
30
40
50
60
Normal
MK50
MK75
MK88
MK94
MK97
MK data
Normal data

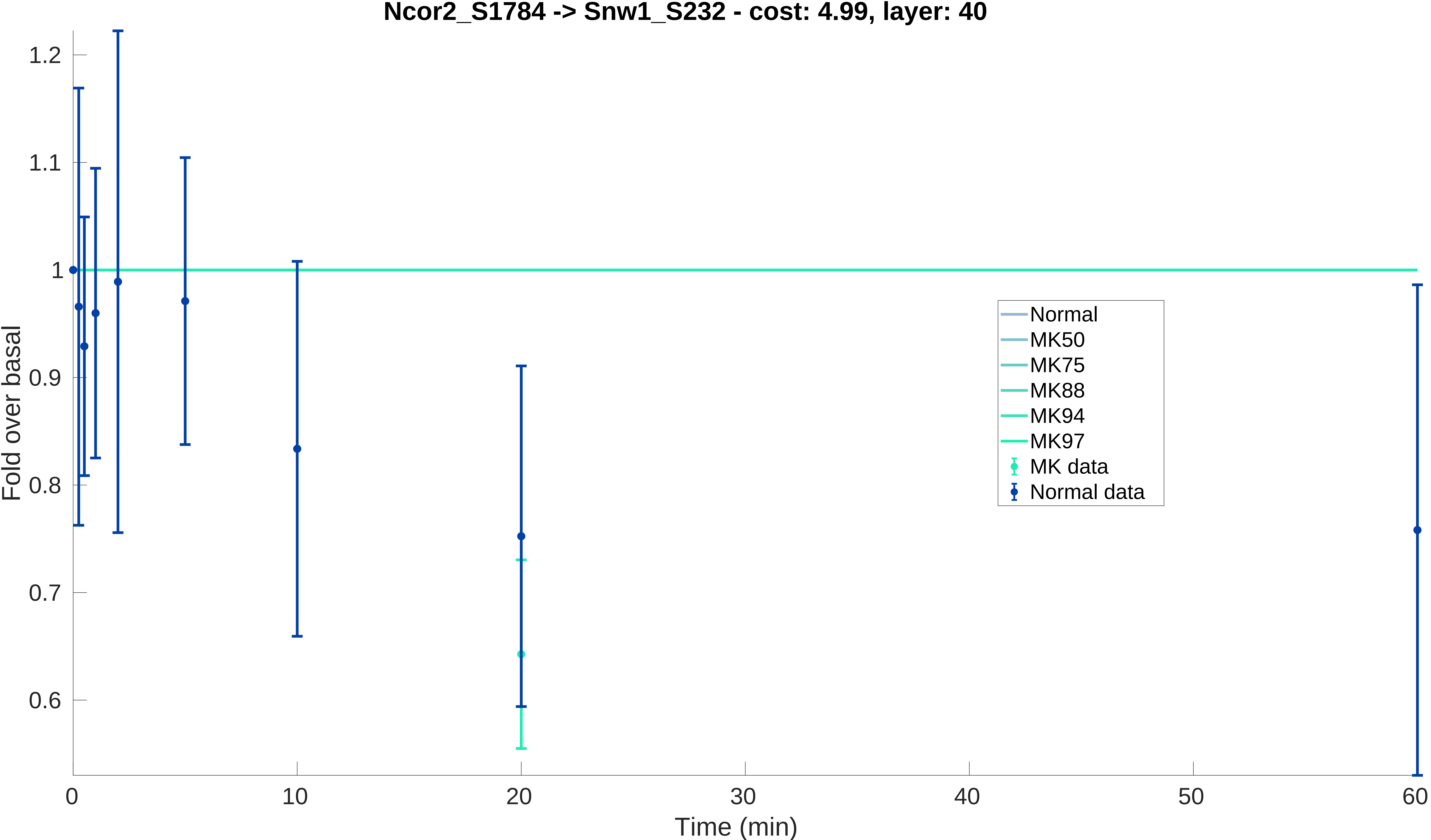

Ncor2_S1784 -> Snw1_S232 - cost: 4.99, layer: 40
Fold over basal
Time (min)
0.6
0.7
0.8
0.9
1
1.1
1.2
0
10
20
30
40
50
60
Normal
MK50
MK75
MK88
MK94
MK97
MK data
Normal data

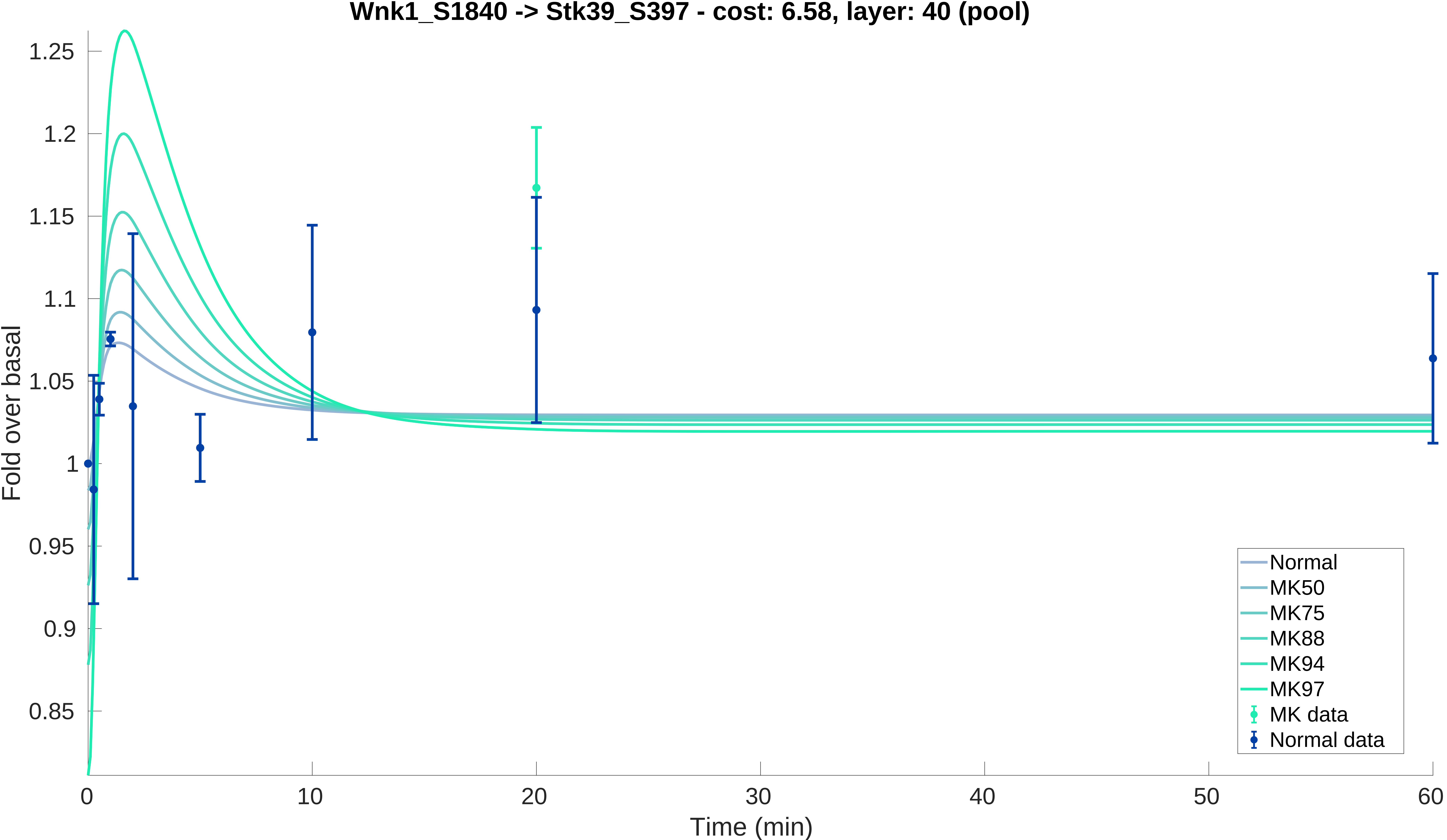
Wnk1_S1840 -> Stk39_S397 - cost: 6.58, layer: 40 (pool)
Fold over basal
Time (min)
1.25
1.2
1.15
1.1
1.05
1
0.95
0.9
0.85
0
10
20
30
40
50
60
Normal
MK50
MK75
MK88
MK94
MK97
MK data
Normal data

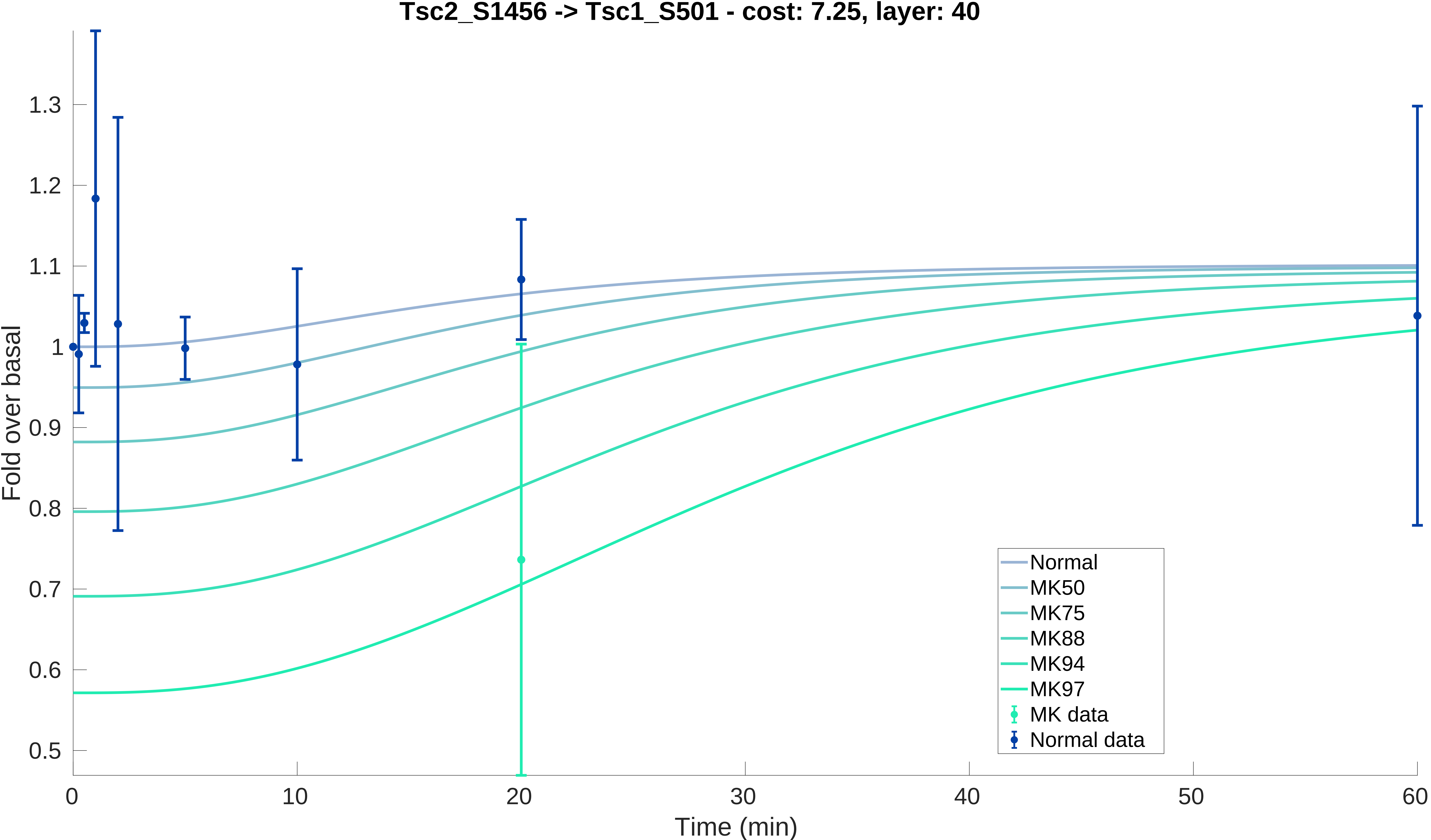
Tsc2_S1456 -> Tsc1_S501 - cost: 7.25, layer: 40
Fold over basal
Time (min)
0.5
0.6
0.7
0.8
0.9
1
1.1
1.2
1.3
0
10
20
30
40
50
60
Normal
MK50
MK75
MK88
MK94
MK97
MK data
Normal data

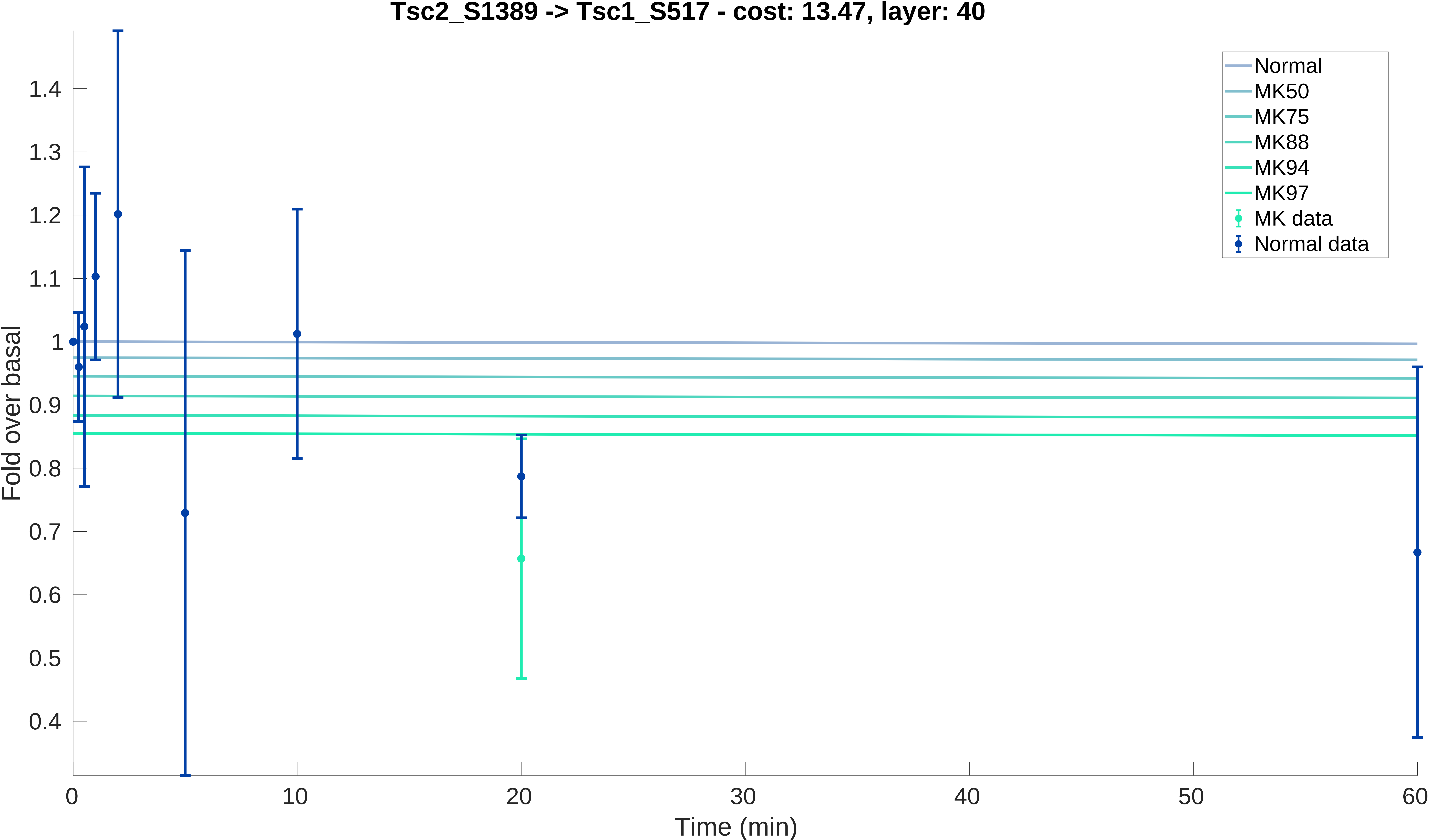
Tsc2_S1389 -> Tsc1_S517 - cost: 13.47, layer: 40
Fold over basal
Time (min)
1.4
1.3
1.2
1.1
1
0.9
0.8
0.7
0.6
0.5
0.4
0
10
20
30
40
50
60
Normal
MK50
MK75
MK88
MK94
MK97
MK data
Normal data

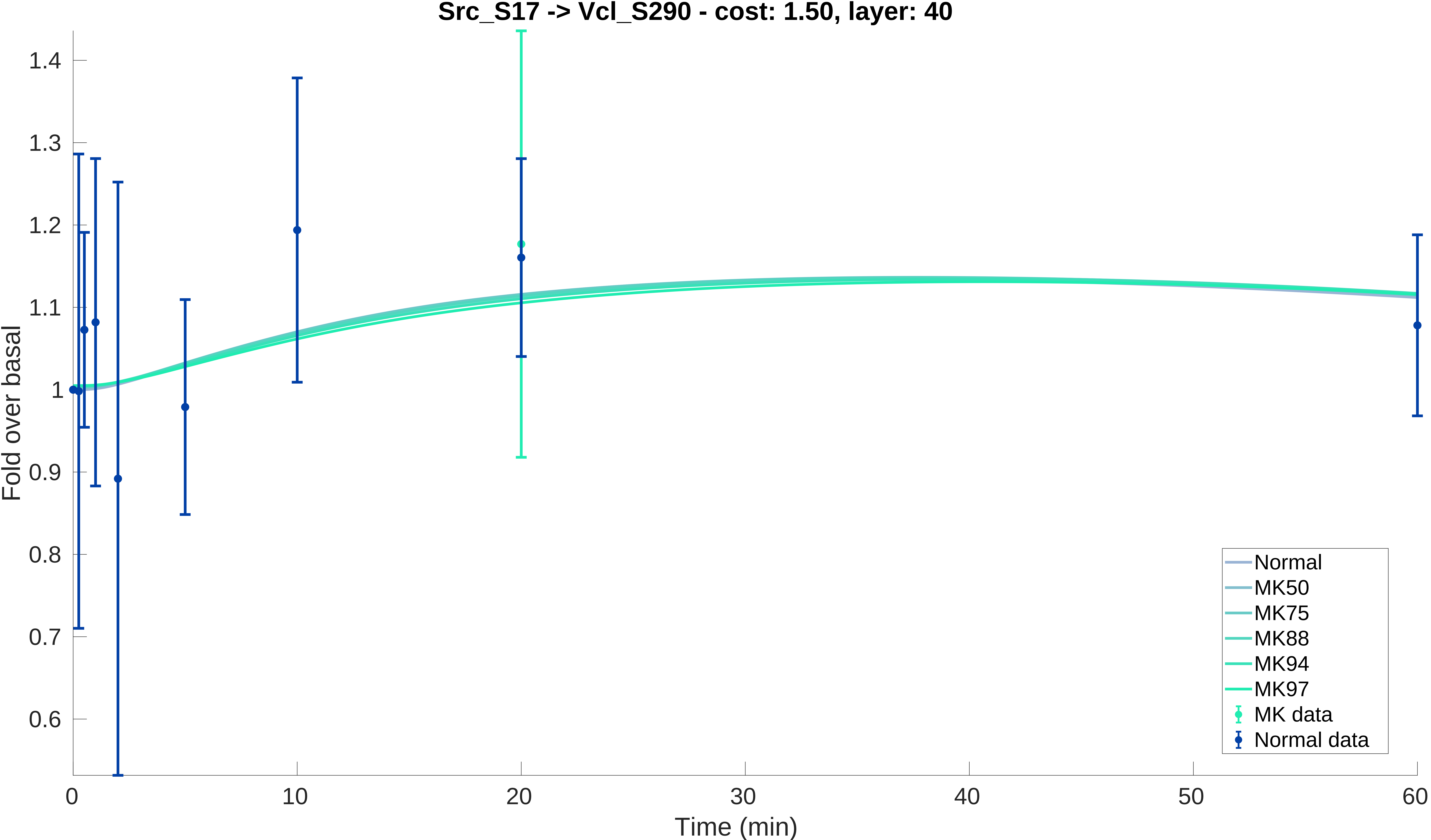

Src_S17 -> Vcl_S290 - cost: 1.50, layer: 40
Fold over basal
1.4
1.3
1.2
1.1
1
0.9
0.8
0.7
0.6
0
10
20
30
40
50
60
Time (min)
Normal
MK50
MK75
MK88
MK94
MK97
MK data
Normal data

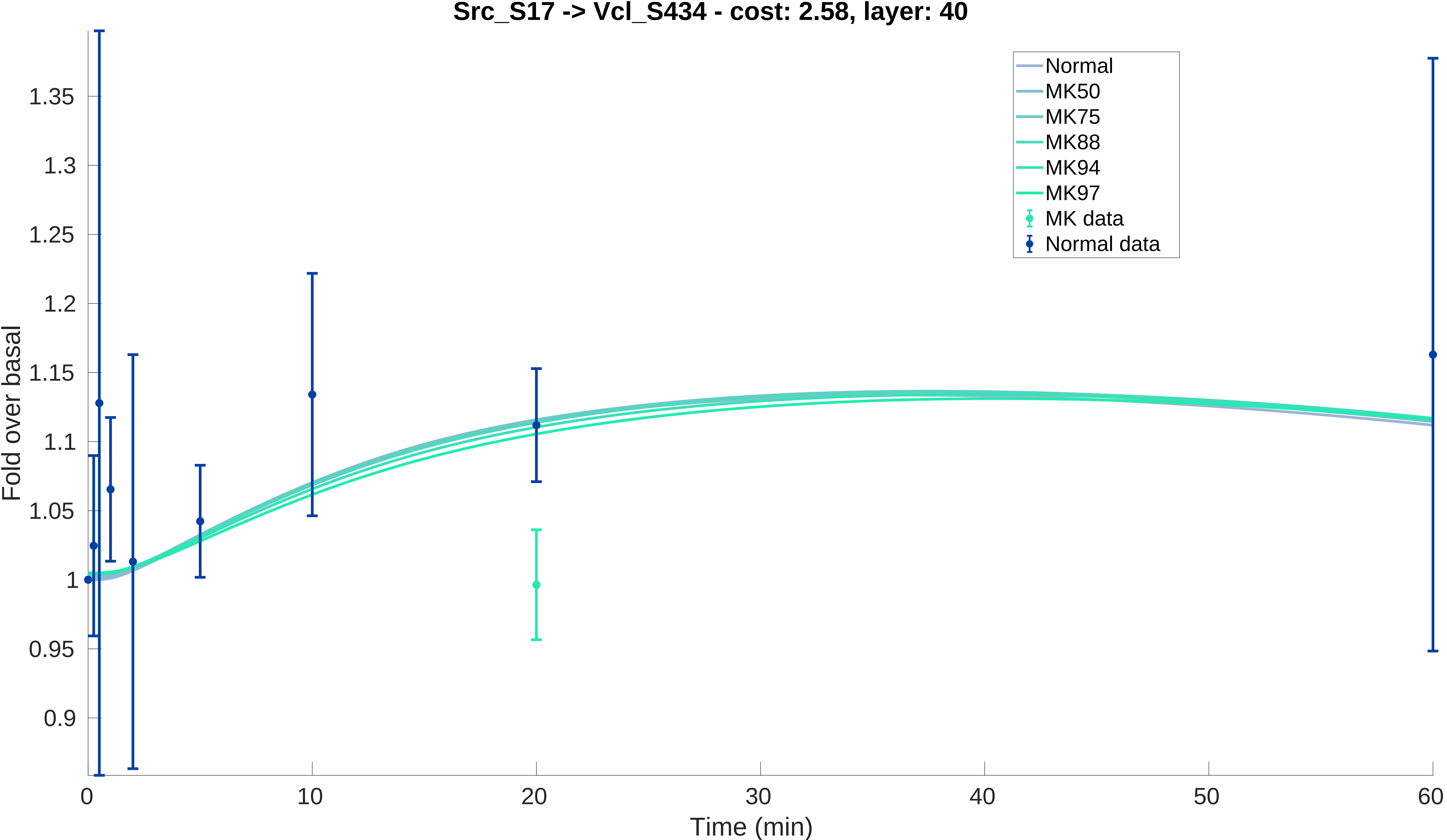

Src_S17 -> Vcl_S434 - cost: 2.58, layer: 40
Normal
MK50
MK75
MK88
MK94
MK97
MK data
Normal data
Fold over basal
1.35
1.3
1.25
1.2
1.15
1.1
1.05
1
0.95
0.9
0
10
20
30
40
50
60
Time (min)

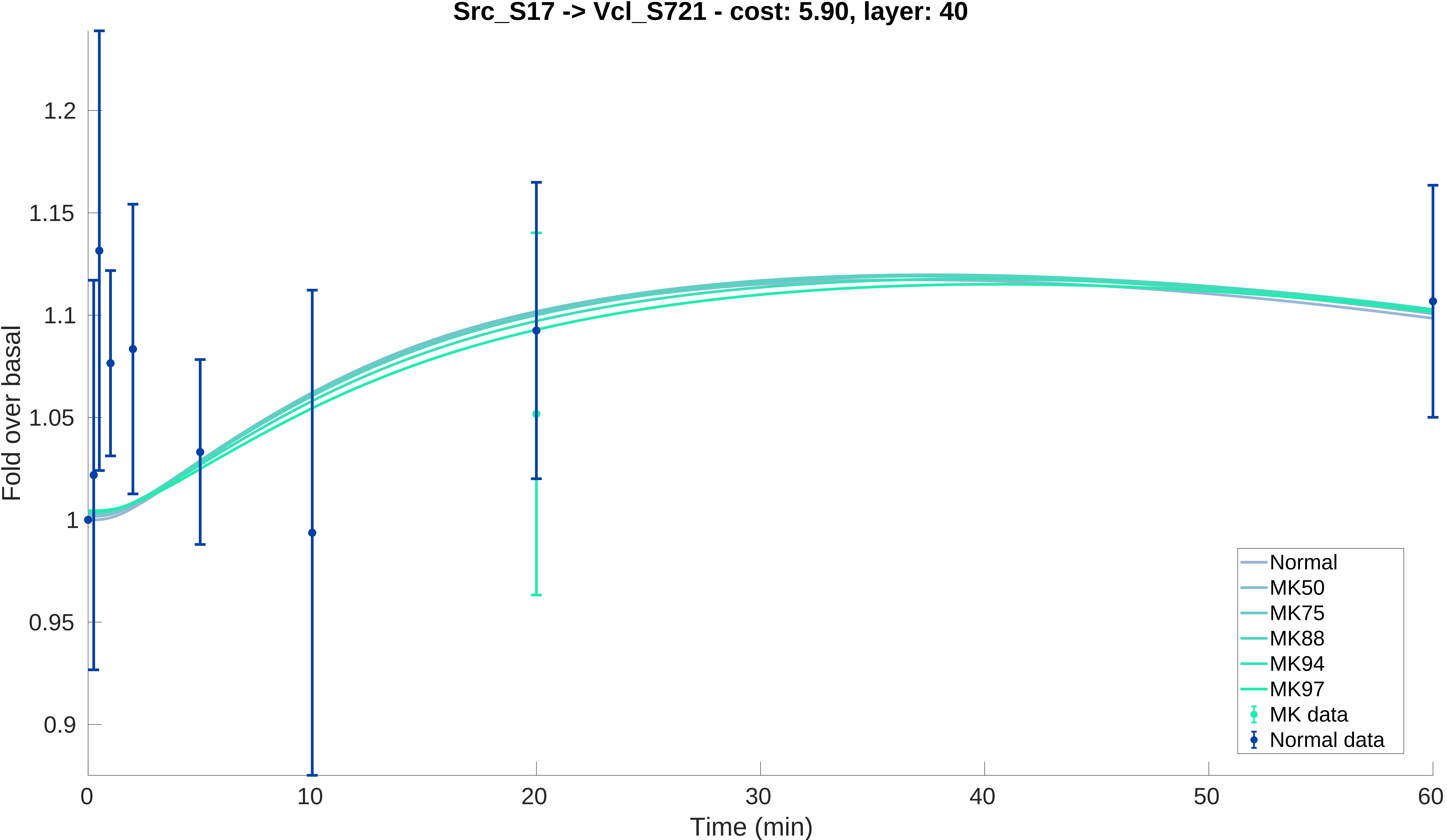

Src_S17 -> Vcl_S721 - cost: 5.90, layer: 40
Fold over basal
1.2
1.15
1.1
1.05
1
0.95
0.9
0
10
20
30
40
50
60
Time (min)
Normal
MK50
MK75
MK88
MK94
MK97
MK data
Normal data

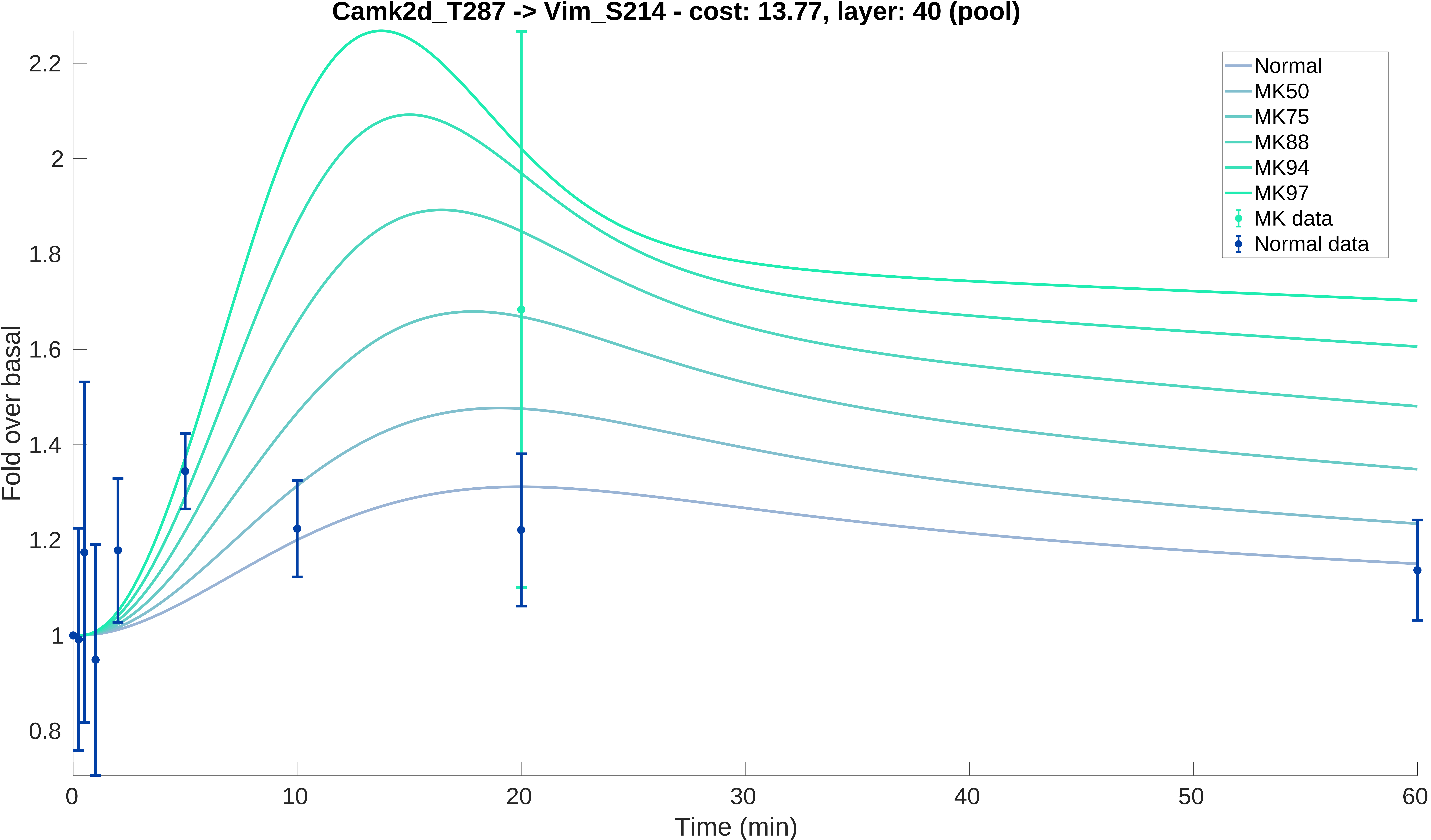
Camk2d_T287 -> Vim_S214 - cost: 13.77, layer: 40 (pool)
Fold over basal
Time (min)
0.8
1
1.2
1.4
1.6
1.8
2
2.2
0
10
20
30
40
50
60
Normal
MK50
MK75
MK88
MK94
MK97
MK data
Normal data

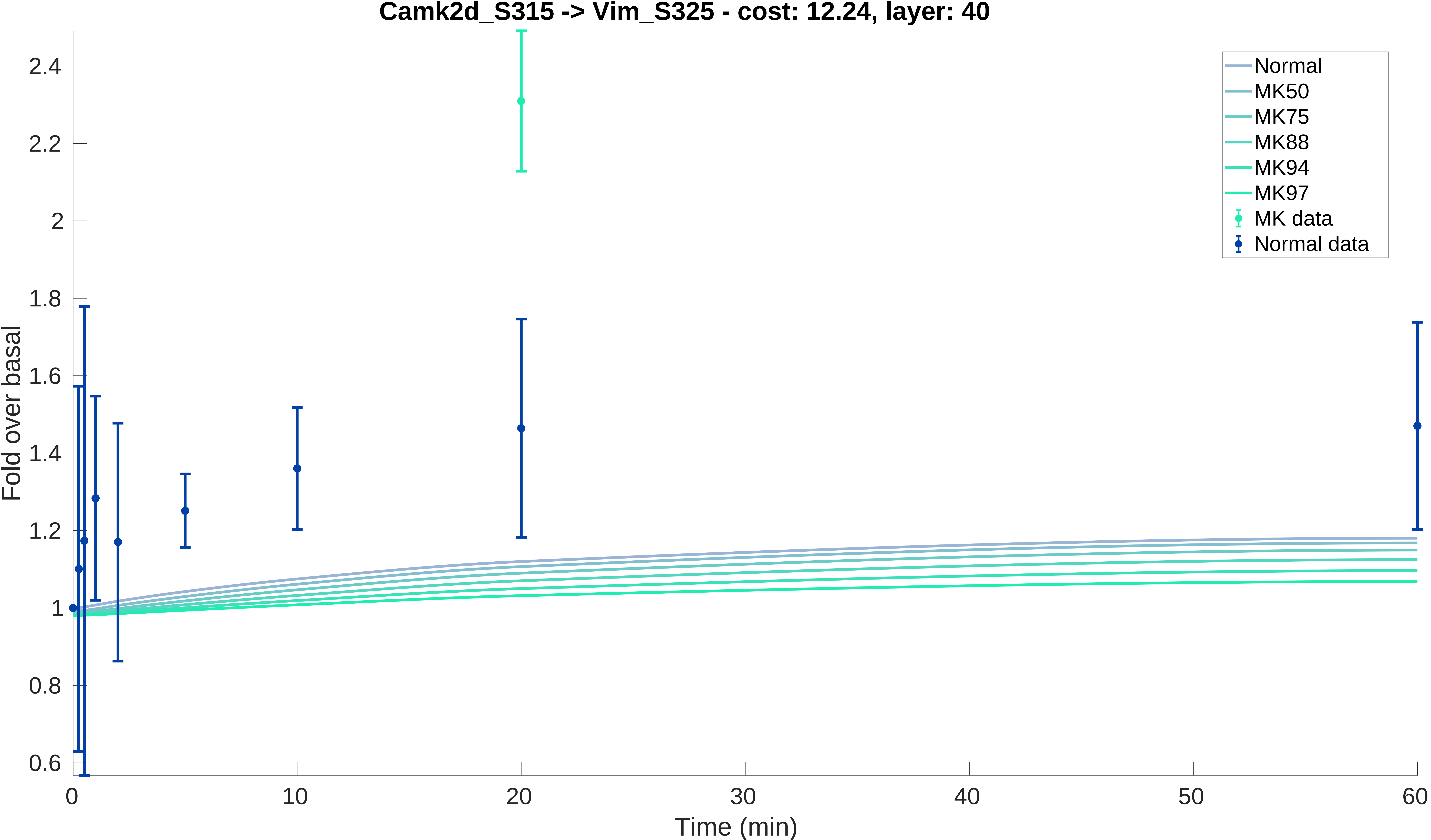

Camk2d_S315 -> Vim_S325 - cost: 12.24, layer: 40
Fold over basal
Time (min)
0.6
0.8
1
1.2
1.4
1.6
1.8
2
2.2
2.4
0
10
20
30
40
50
60
Normal
MK50
MK75
MK88
MK94
MK97
MK data
Normal data

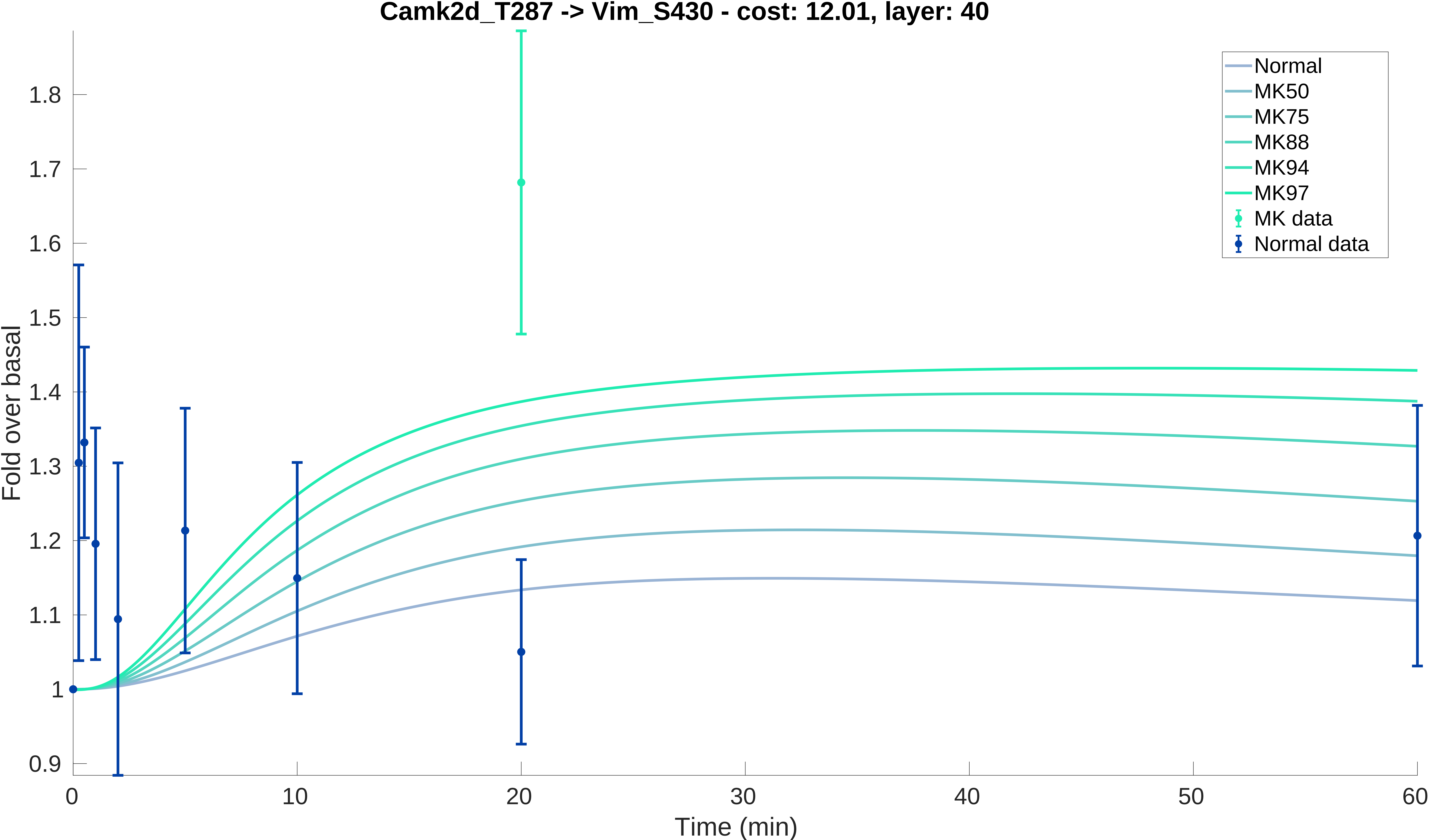

Camk2d_T287 -> Vim_S430 - cost: 12.01, layer: 40
Fold over basal
Time (min)
0.9
1
1.1
1.2
1.3
1.4
1.5
1.6
1.7
1.8
0
10
20
30
40
50
60
Normal
MK50
MK75
MK88
MK94
MK97
MK data
Normal data

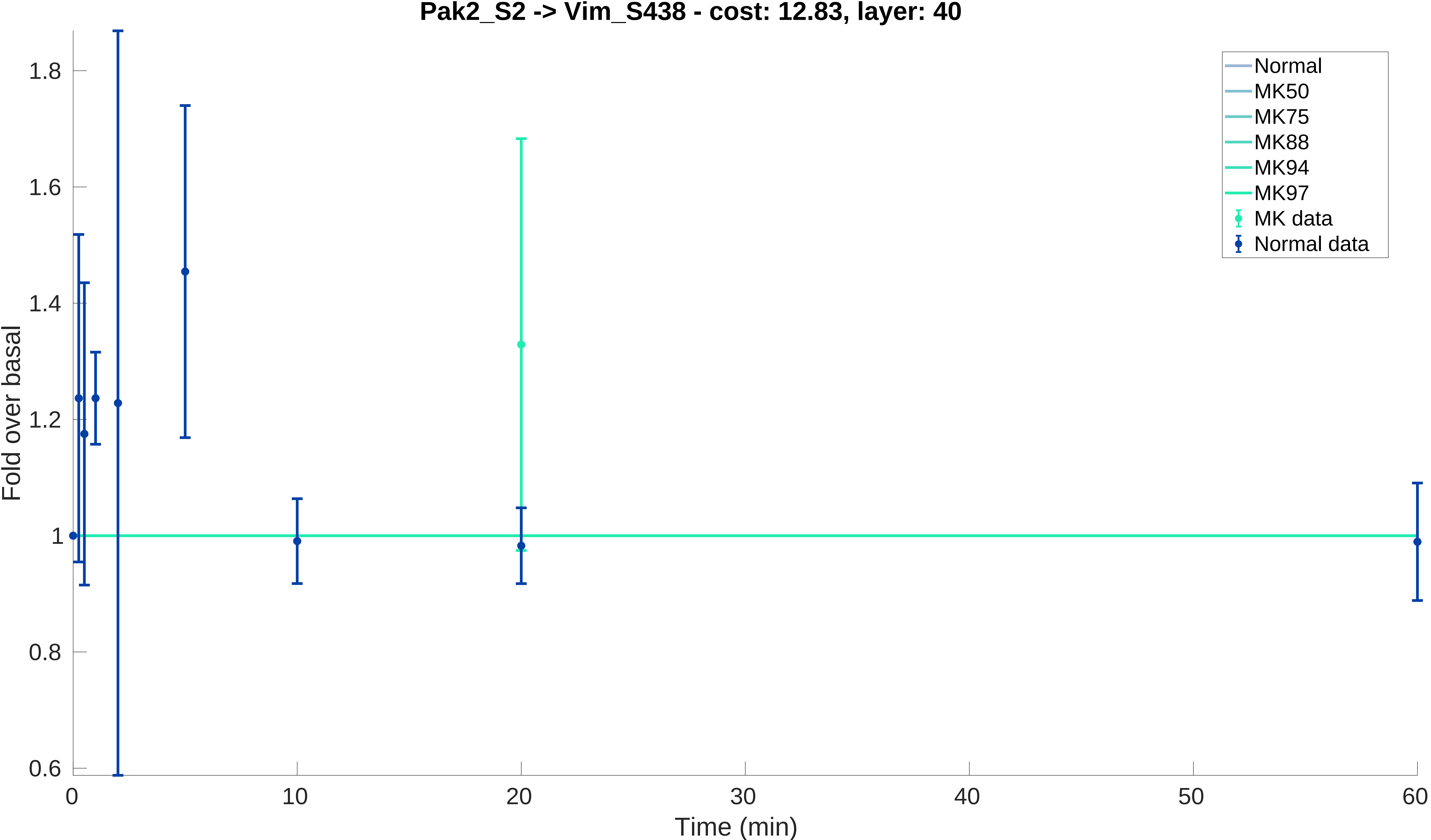

Pak2_S2 -> Vim_S438 - cost: 12.83, layer: 40
Fold over basal
Time (min)
0.6
0.8
1
1.2
1.4
1.6
1.8
0
10
20
30
40
50
60
Normal
MK50
MK75
MK88
MK94
MK97
MK data
Normal data

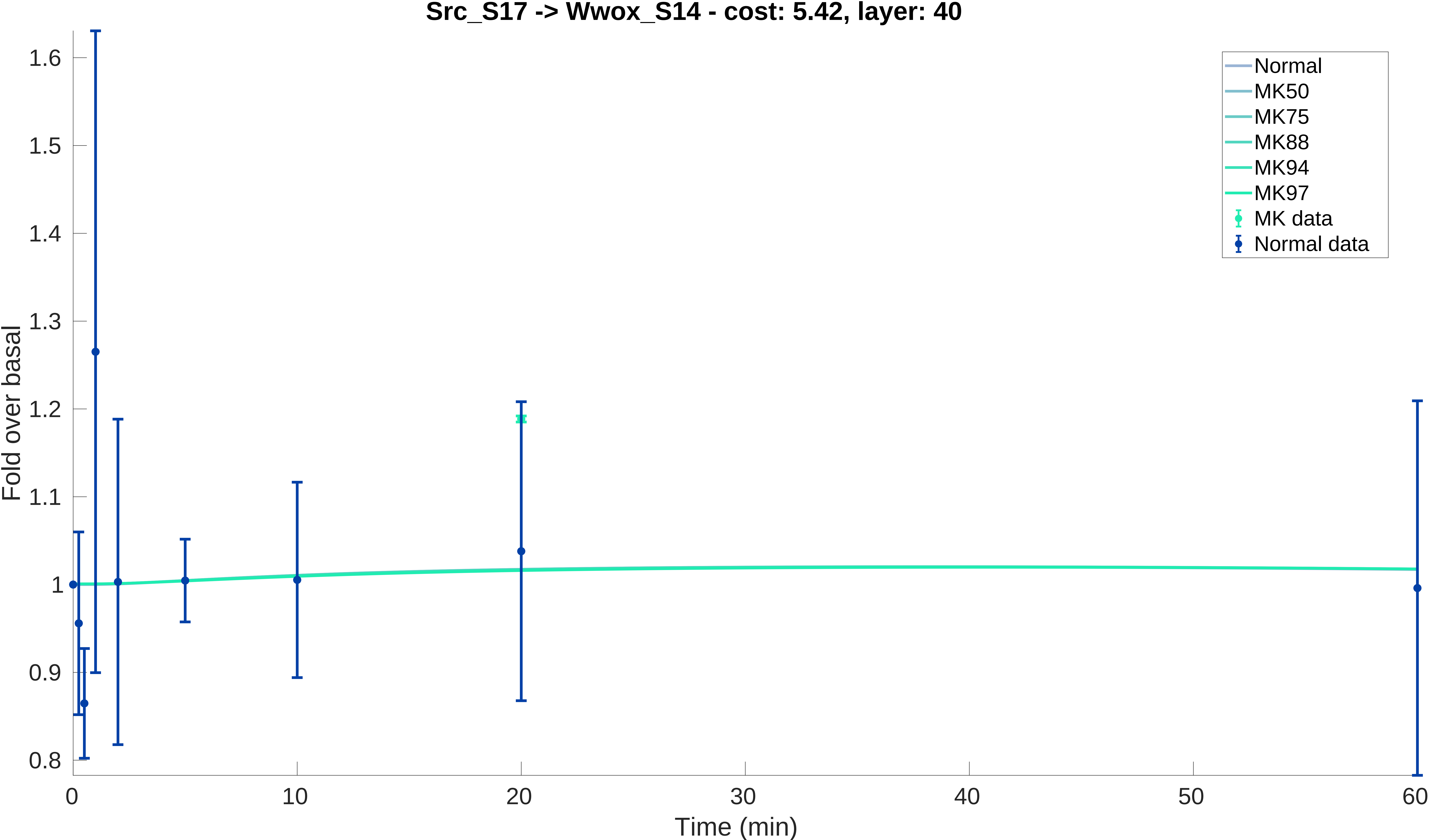

Src_S17 -> Wwox_S14 - cost: 5.42, layer: 40
Normal
MK50
MK75
MK88
MK94
MK97
MK data
Normal data
Fold over basal
Time (min)
0.8
0.9
1
1.1
1.2
1.3
1.4
1.5
1.6
0
10
20
30
40
50
60

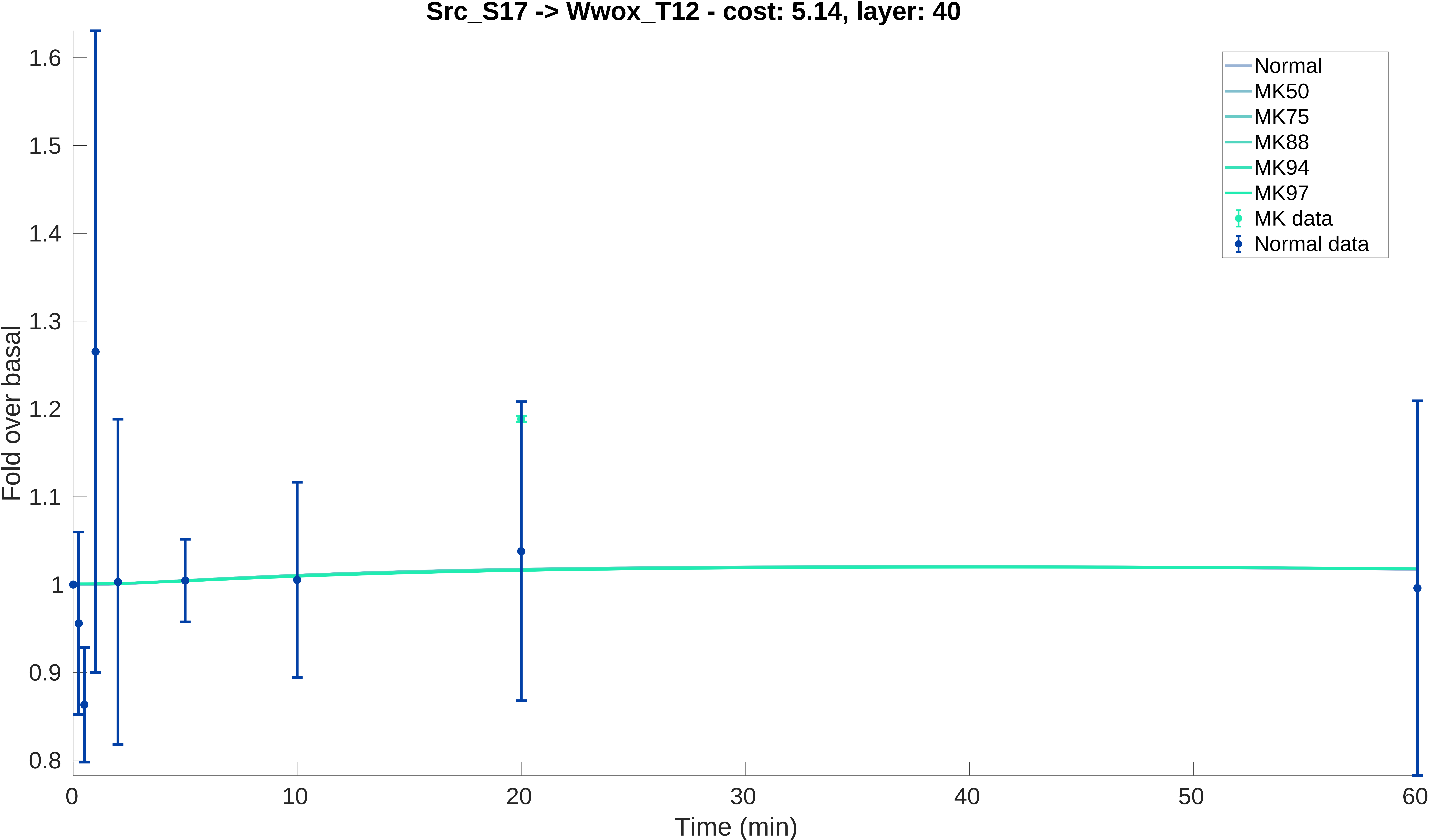

Src_S17 -> Wwox_T12 - cost: 5.14, layer: 40
Fold over basal
Time (min)
0.8
0.9
1
1.1
1.2
1.3
1.4
1.5
1.6
0
10
20
30
40
50
60
Normal
MK50
MK75
MK88
MK94
MK97
MK data
Normal data

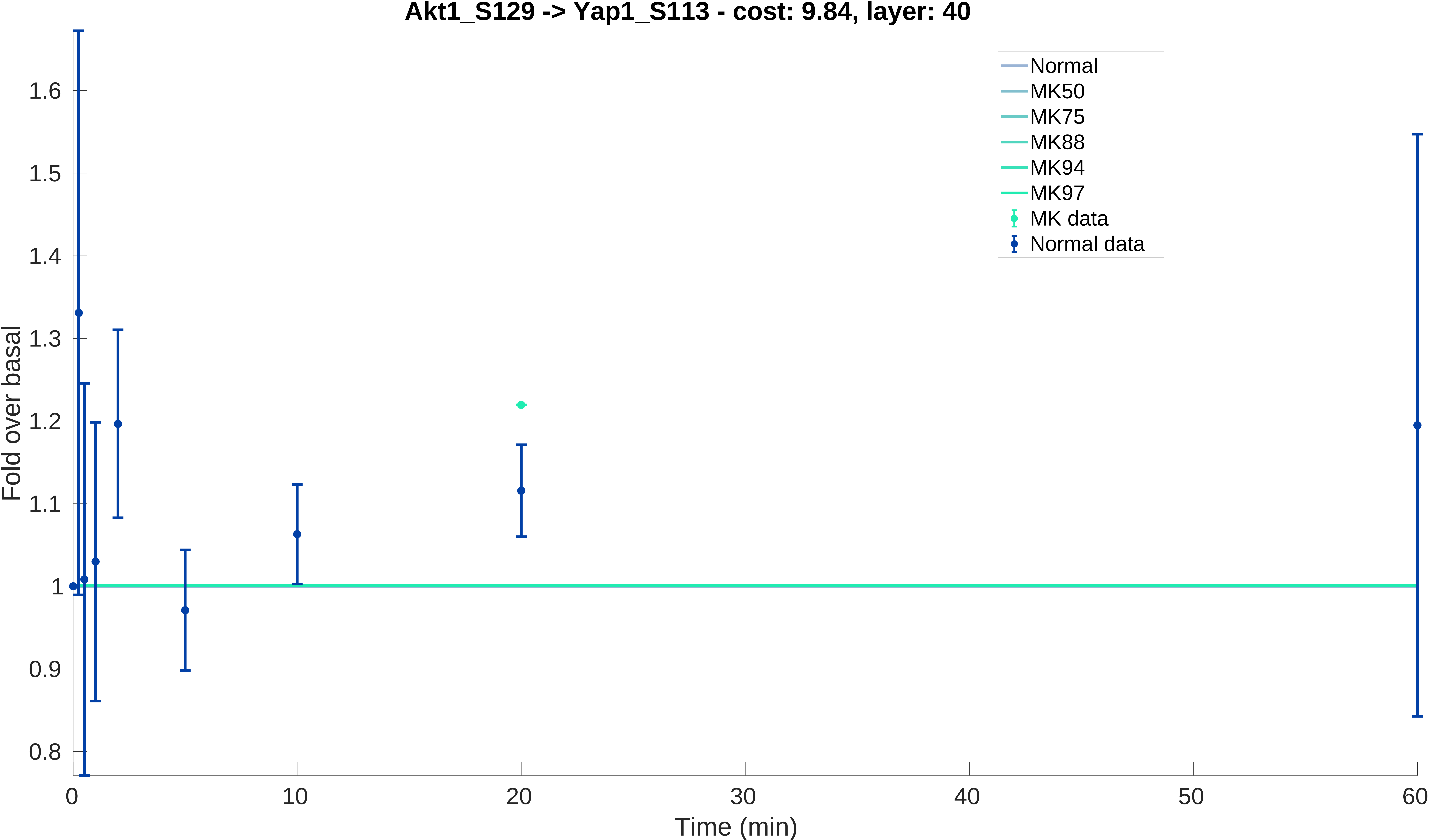

Akt1_S129 -> Yap1_S113 - cost: 9.84, layer: 40
Fold over basal
Time (min)
1.6
1.5
1.4
1.3
1.2
1.1
1
0.9
0.8
0
10
20
30
40
50
60
Normal
MK50
MK75
MK88
MK94
MK97
MK data
Normal data

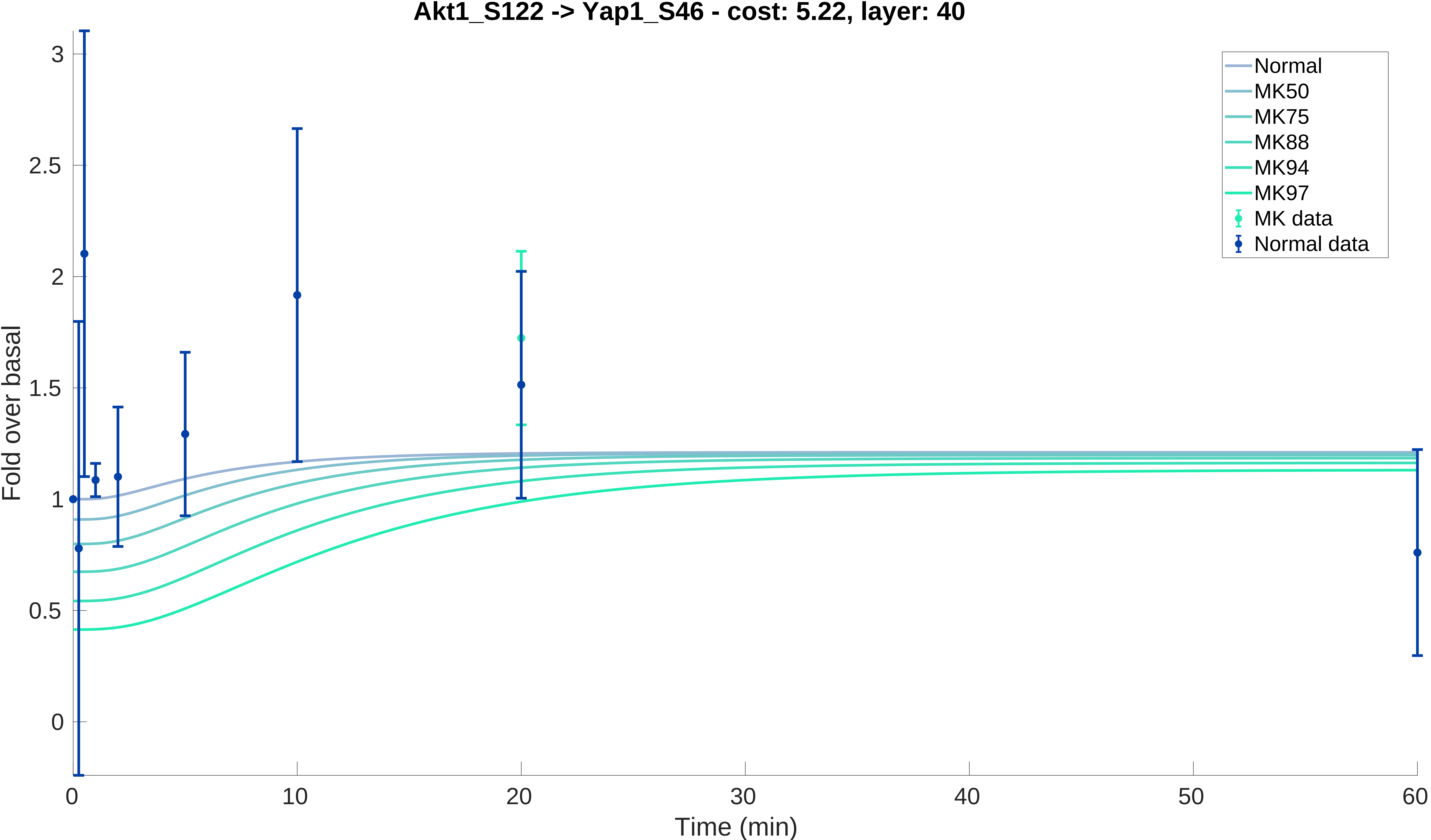
Akt1_S122 -> Yap1_S46 - cost: 5.22, layer: 40
Fold over basal
Time (min)
0
0.5
1
1.5
2
2.5
3
0
10
20
30
40
50
60
Normal
MK50
MK75
MK88
MK94
MK97
MK data
Normal data

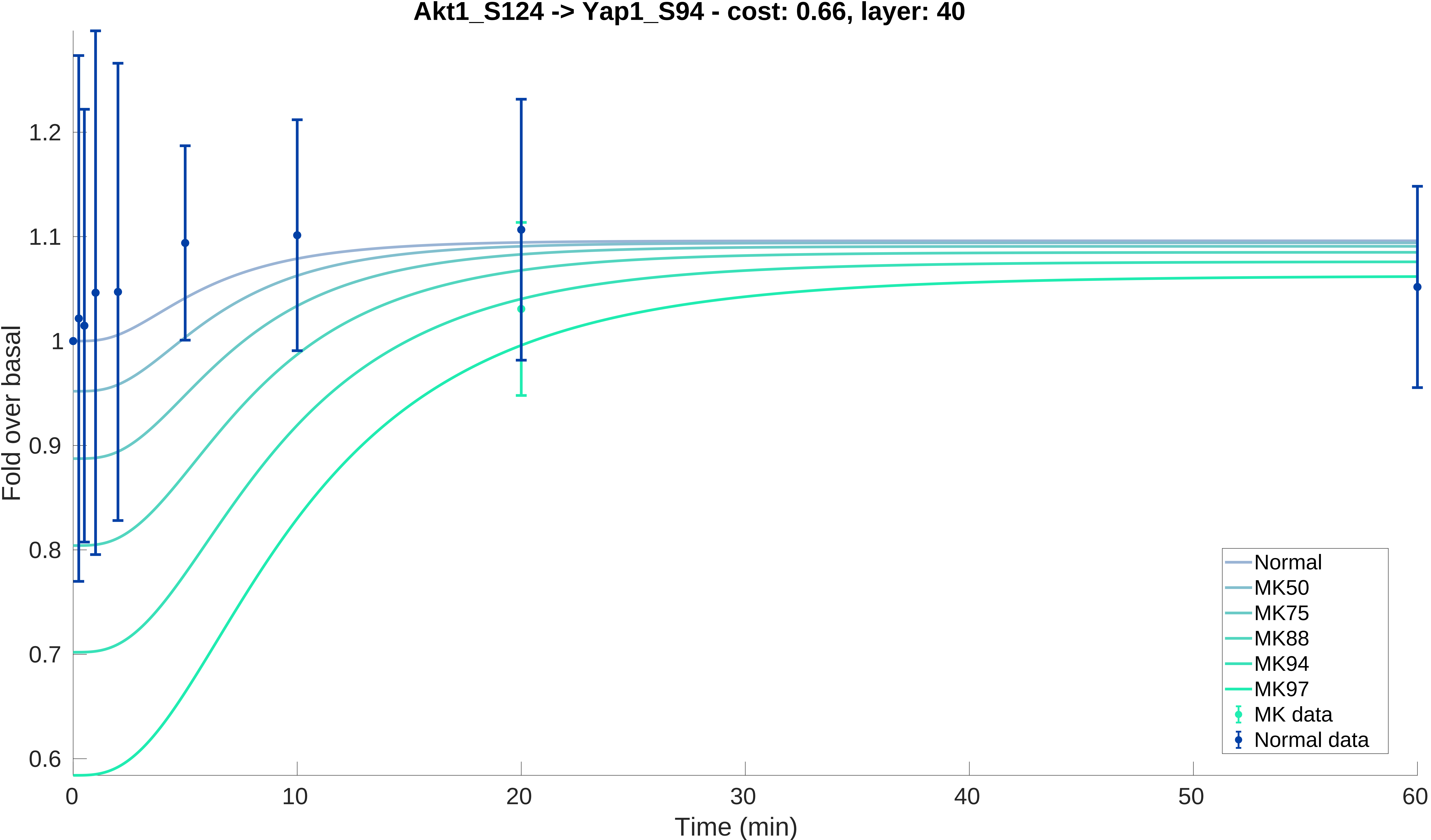

Akt1_S124 -> Yap1_S94 - cost: 0.66, layer: 40
Fold over basal
Time (min)
0.6
0.7
0.8
0.9
1
1.1
1.2
0
10
20
30
40
50
60
Normal
MK50
MK75
MK88
MK94
MK97
MK data
Normal data

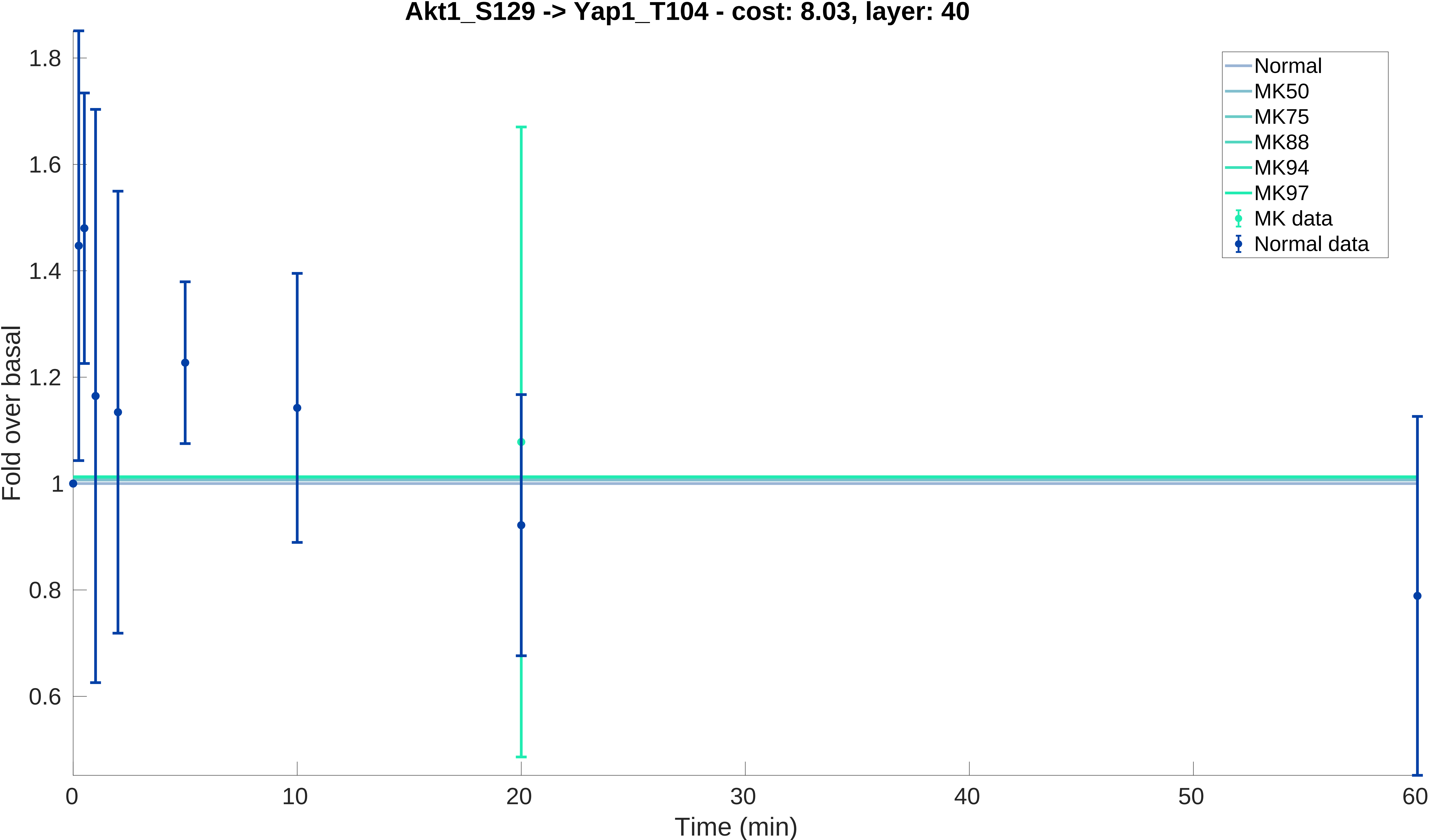

Akt1_S129 -> Yap1_T104 - cost: 8.03, layer: 40
Normal
MK50
MK75
MK88
MK94
MK97
MK data
Normal data
Fold over basal
1.8
1.6
1.4
1.2
1
0.8
0.6
0
10
20
30
40
50
60
Time (min)

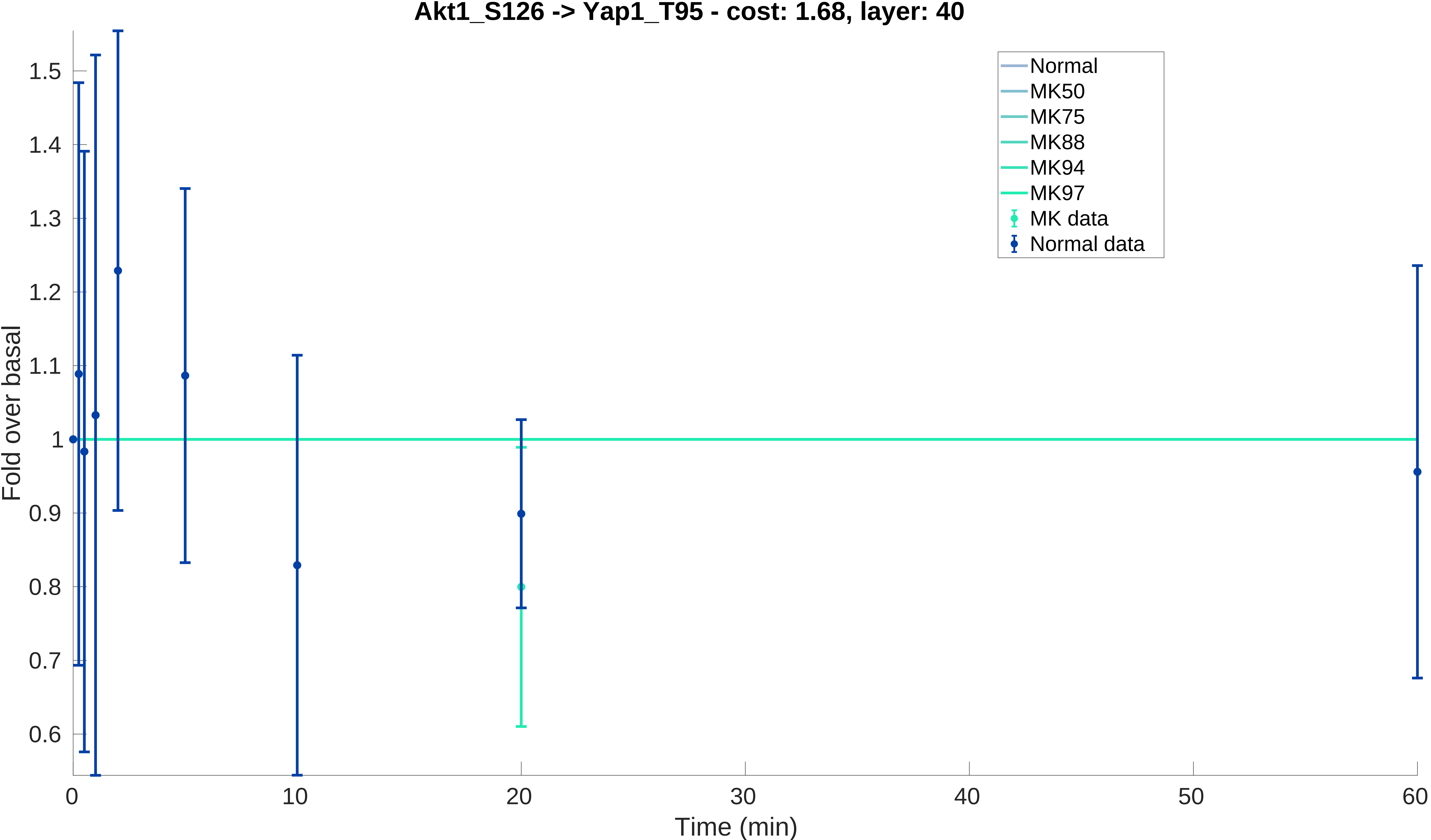

Akt1_S126 -> Yap1_T95 - cost: 1.68, layer: 40
Fold over basal
Time (min)
0.6
0.7
0.8
0.9
1
1.1
1.2
1.3
1.4
1.5
0
10
20
30
40
50
60
Normal
MK50
MK75
MK88
MK94
MK97
MK data
Normal data

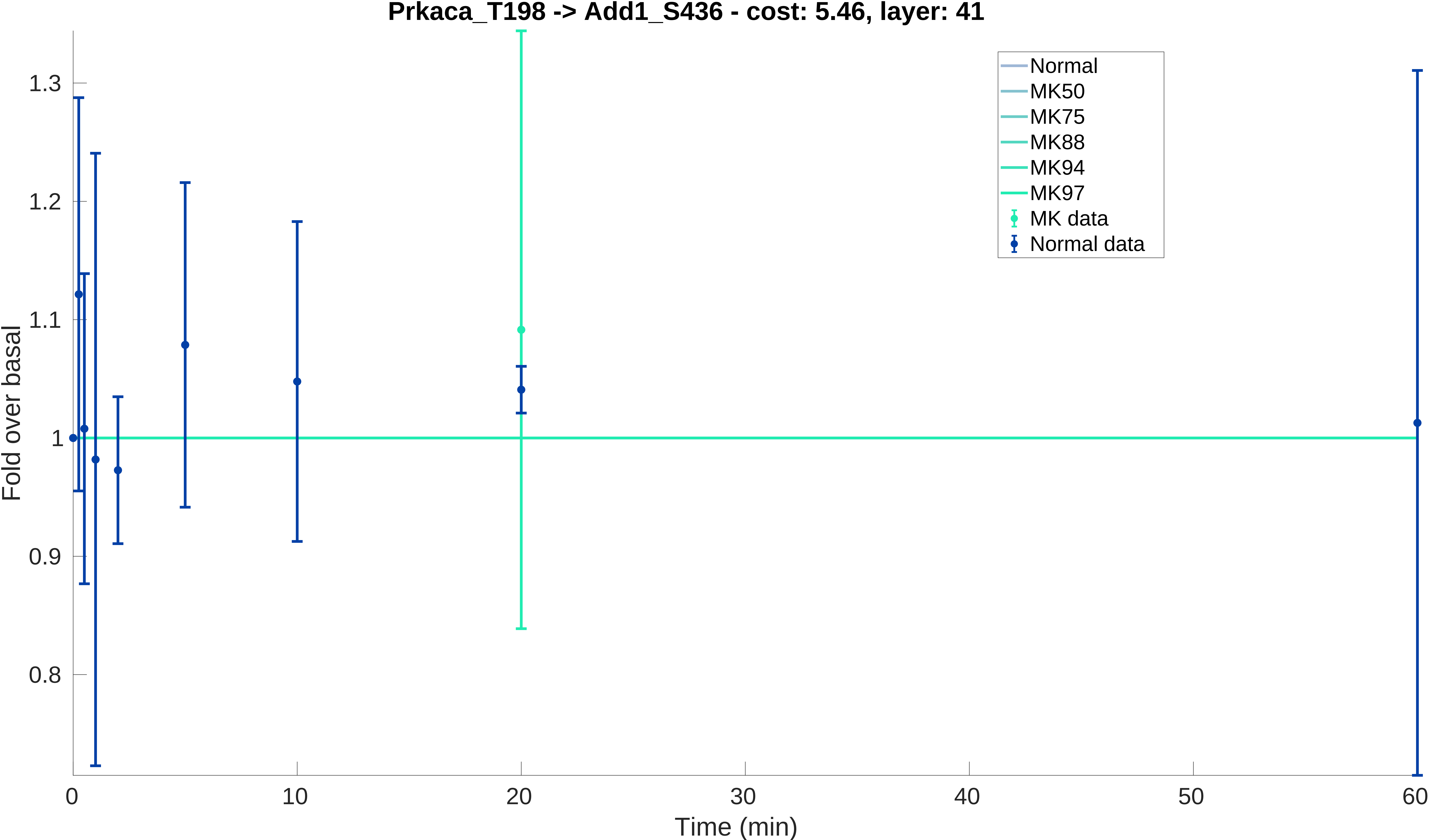

Prkaca_T198 -> Add1_S436 - cost: 5.46, layer: 41
Normal
MK50
MK75
MK88
MK94
MK97
MK data
Normal data
Fold over basal
Time (min)
0
10
20
30
40
50
60
0.8
0.9
1
1.1
1.2
1.3

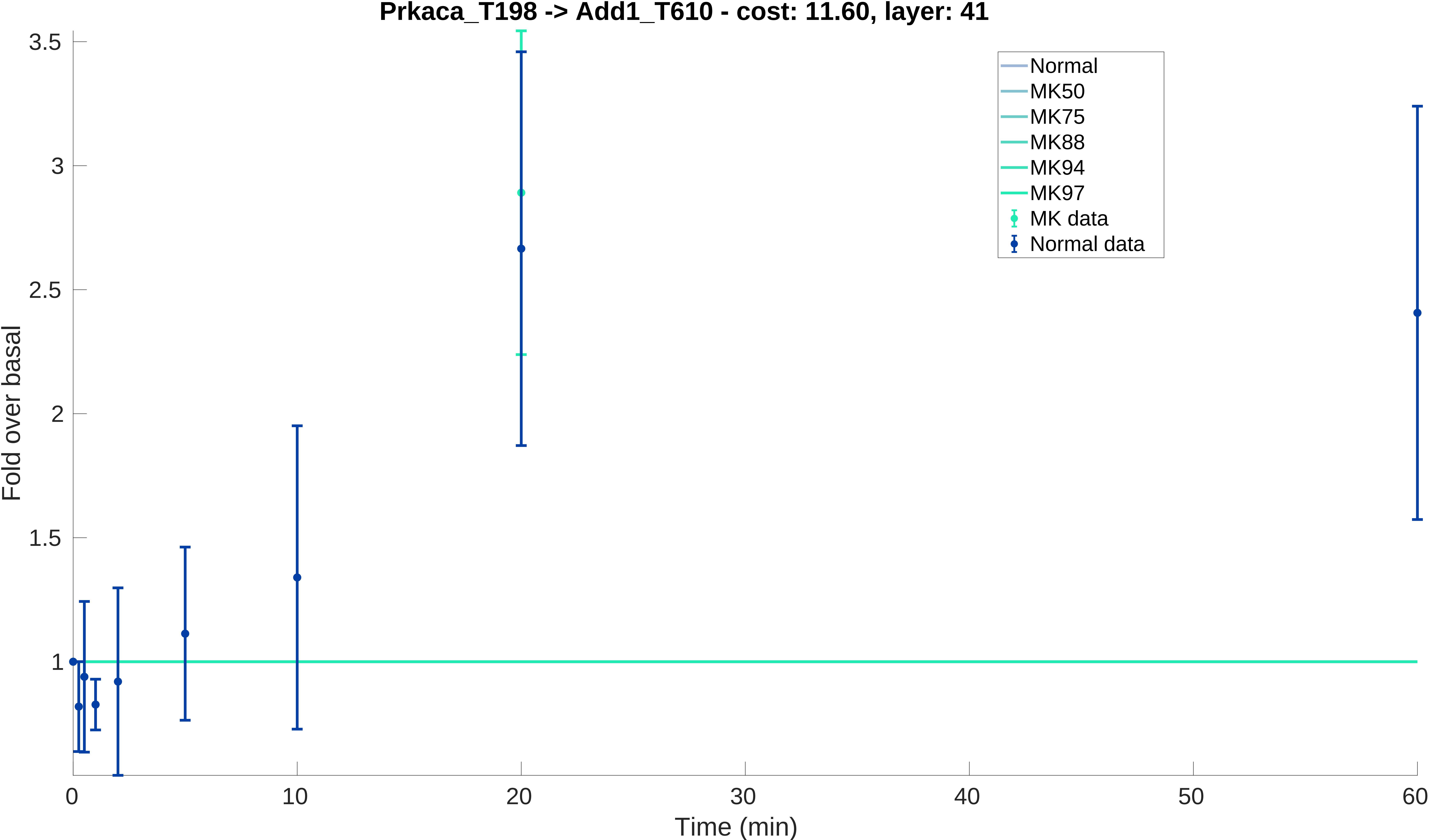
Prkaca_T198 -> Add1_T610 - cost: 11.60, layer: 41
Fold over basal
3.5
3
2.5
2
1.5
1
0
10
20
30
40
50
60
Time (min)
Normal
MK50
MK75
MK88
MK94
MK97
MK data
Normal data

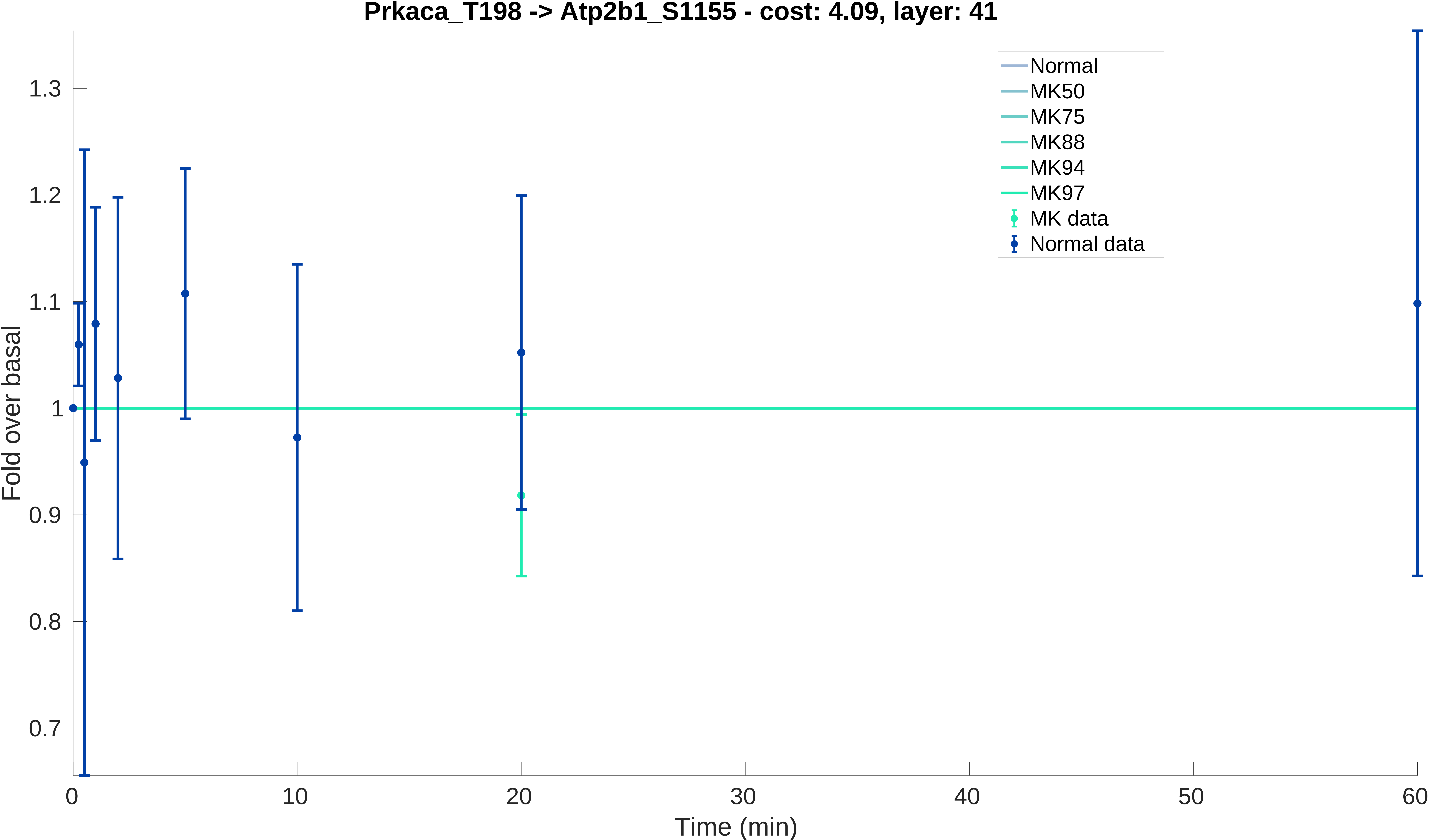
Prkaca_T198 -> Atp2b1_S1155 - cost: 4.09, layer: 41
Normal
MK50
MK75
MK88
MK94
MK97
MK data
Normal data
Fold over basal
1.3
1.2
1.1
1
0.9
0.8
0.7
0
10
20
30
40
50
60
Time (min)

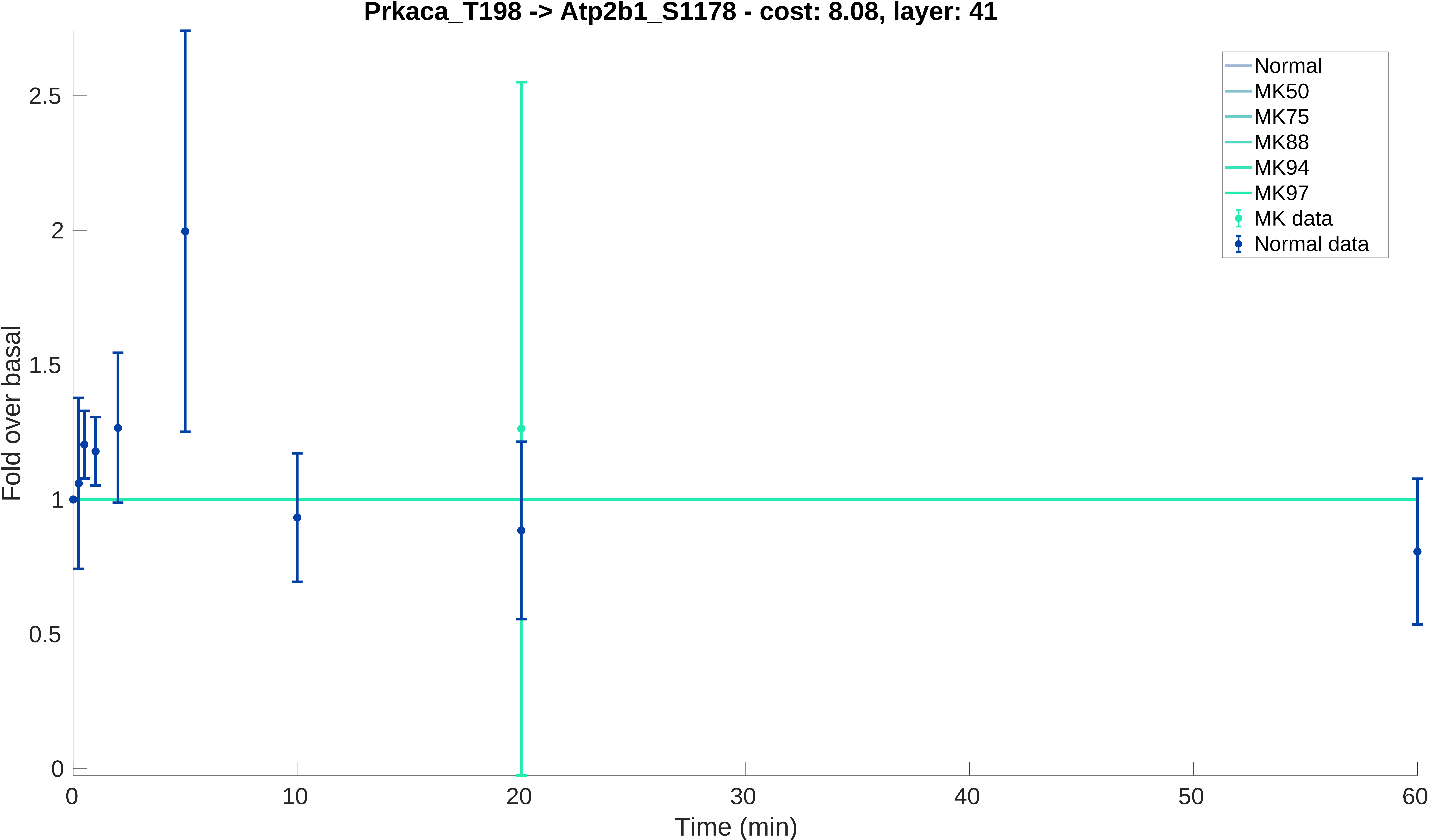

Prkaca_T198 -> Atp2b1_S1178 - cost: 8.08, layer: 41
Fold over basal
Time (min)
0
0.5
1
1.5
2
2.5
0
10
20
30
40
50
60
Normal
MK50
MK75
MK88
MK94
MK97
MK data
Normal data

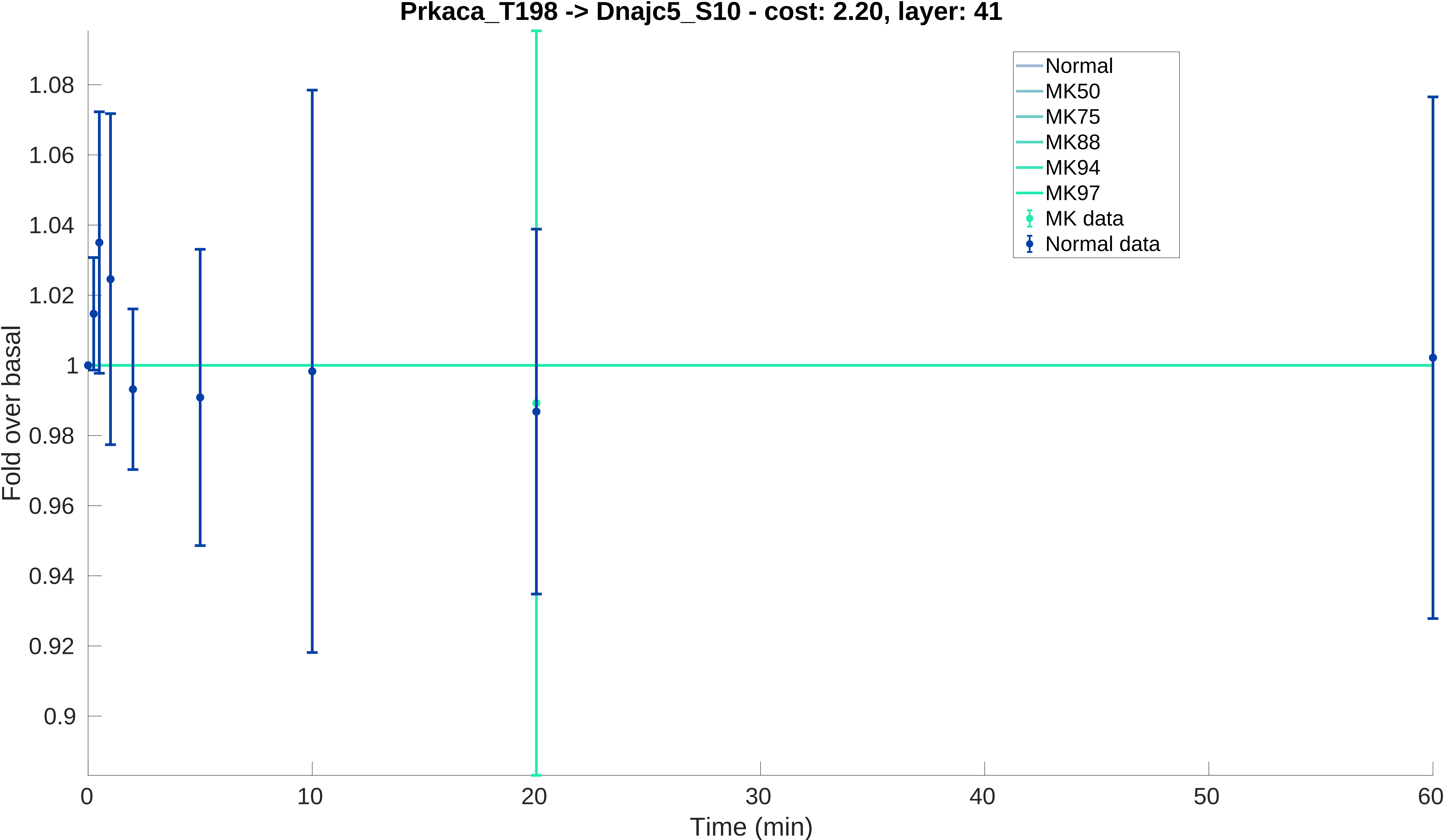

Prkaca_T198 -> Dnajc5_S10 - cost: 2.20, layer: 41
Normal
MK50
MK75
MK88
MK94
MK97
MK data
Normal data
Fold over basal
Time (min)
1.08
1.06
1.04
1.02
1
0.98
0.96
0.94
0.92
0.9
0
10
20
30
40
50
60

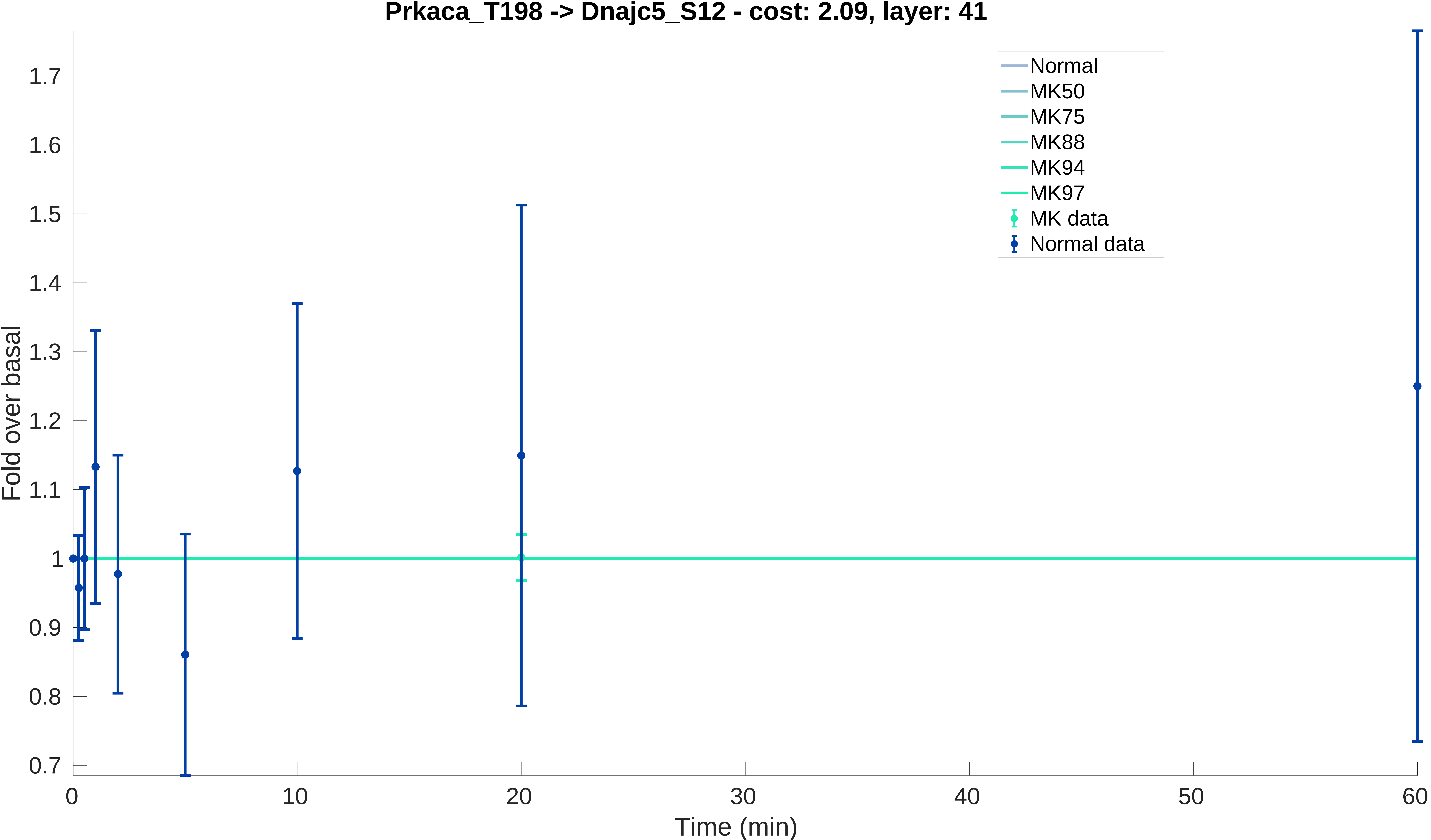
Prkaca_T198 -> Dnajc5_S12 - cost: 2.09, layer: 41
Fold over basal
Time (min)
0.7
0.8
0.9
1
1.1
1.2
1.3
1.4
1.5
1.6
1.7
0
10
20
30
40
50
60
Normal
MK50
MK75
MK88
MK94
MK97
MK data
Normal data

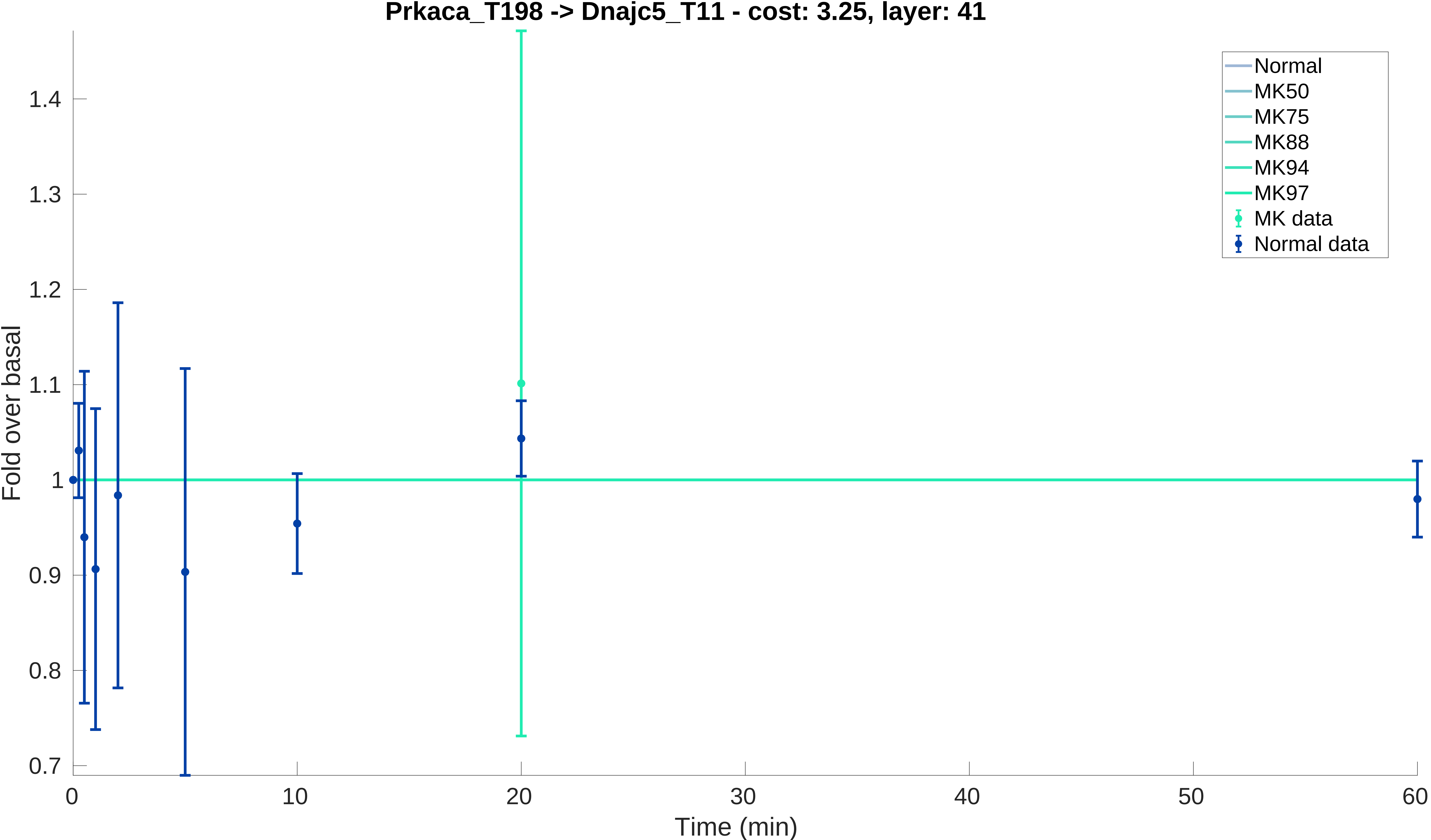

Prkaca_T198 -> Dnajc5_T11 - cost: 3.25, layer: 41
Fold over basal
Time (min)
1.4
1.3
1.2
1.1
1
0.9
0.8
0.7
0
10
20
30
40
50
60
Normal
MK50
MK75
MK88
MK94
MK97
MK data
Normal data

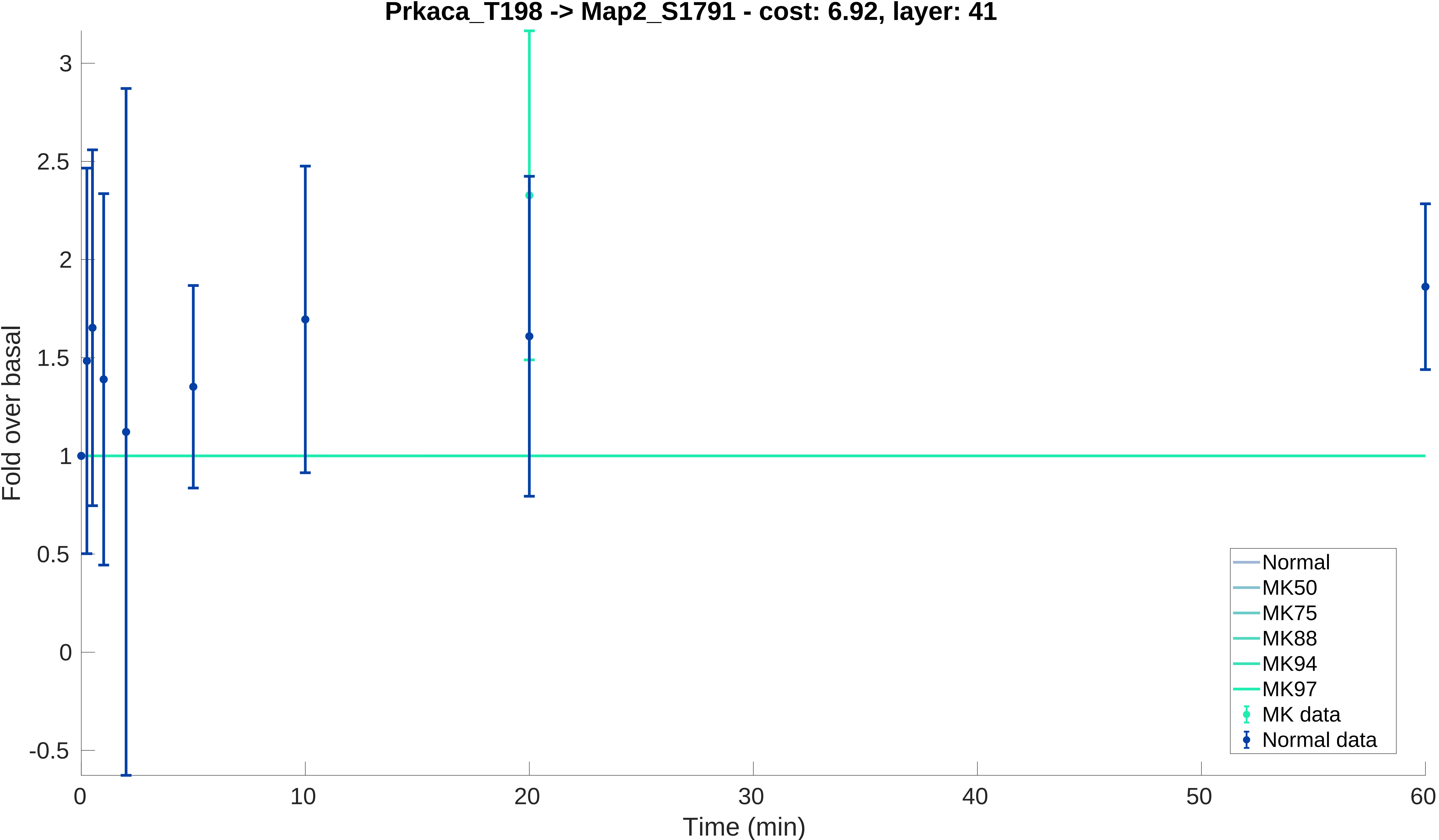

Prkaca_T198 -> Map2_S1791 - cost: 6.92, layer: 41
Fold over basal
Time (min)
3
2.5
2
1.5
1
0.5
0
-0.5
0
10
20
30
40
50
60
Normal
MK50
MK75
MK88
MK94
MK97
MK data
Normal data

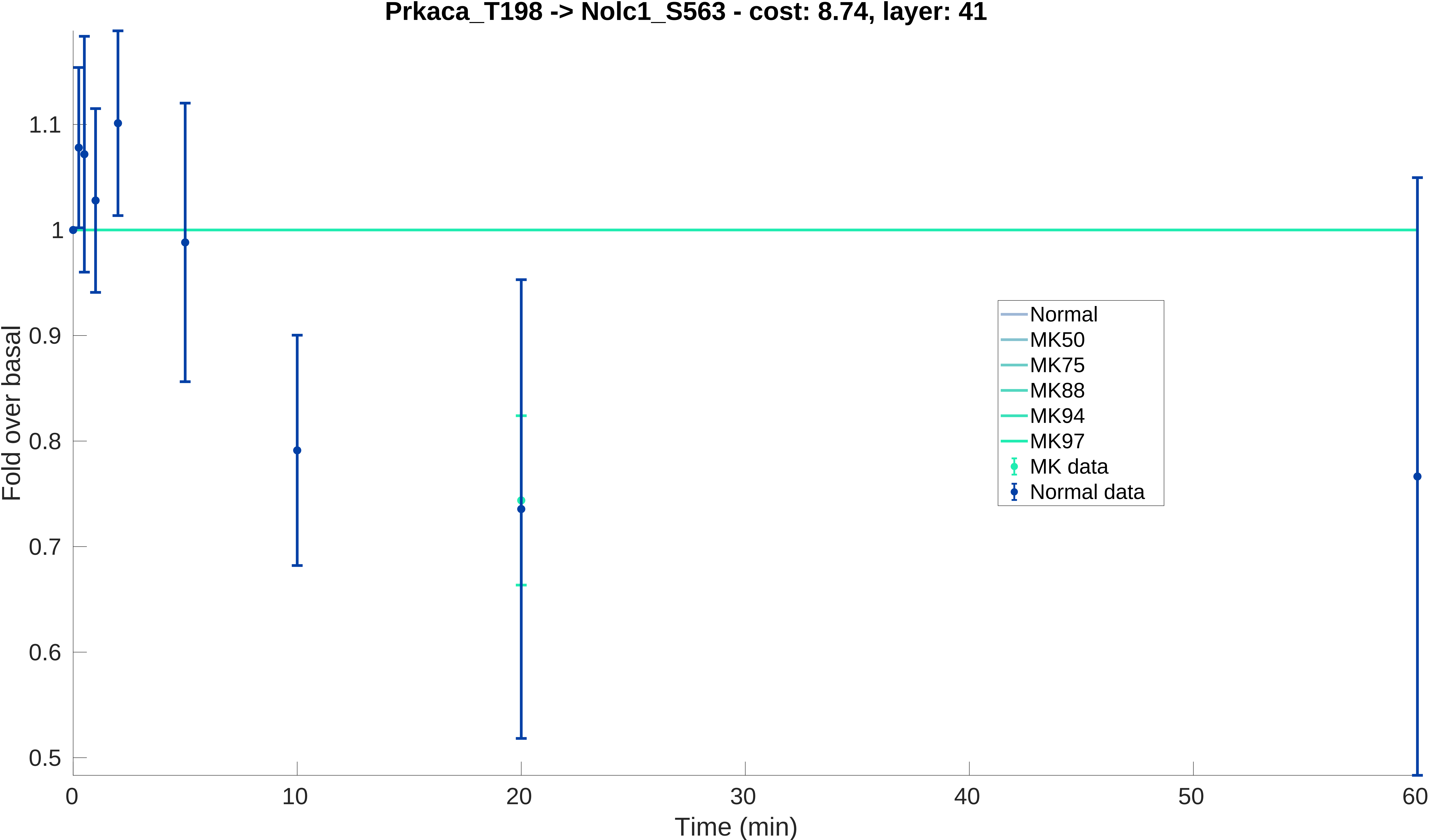

Prkaca_T198 -> Nolc1_S563 - cost: 8.74, layer: 41
Fold over basal
1.1
1
0.9
0.8
0.7
0.6
0.5
0
10
20
30
40
50
60
Time (min)
Normal
MK50
MK75
MK88
MK94
MK97
MK data
Normal data

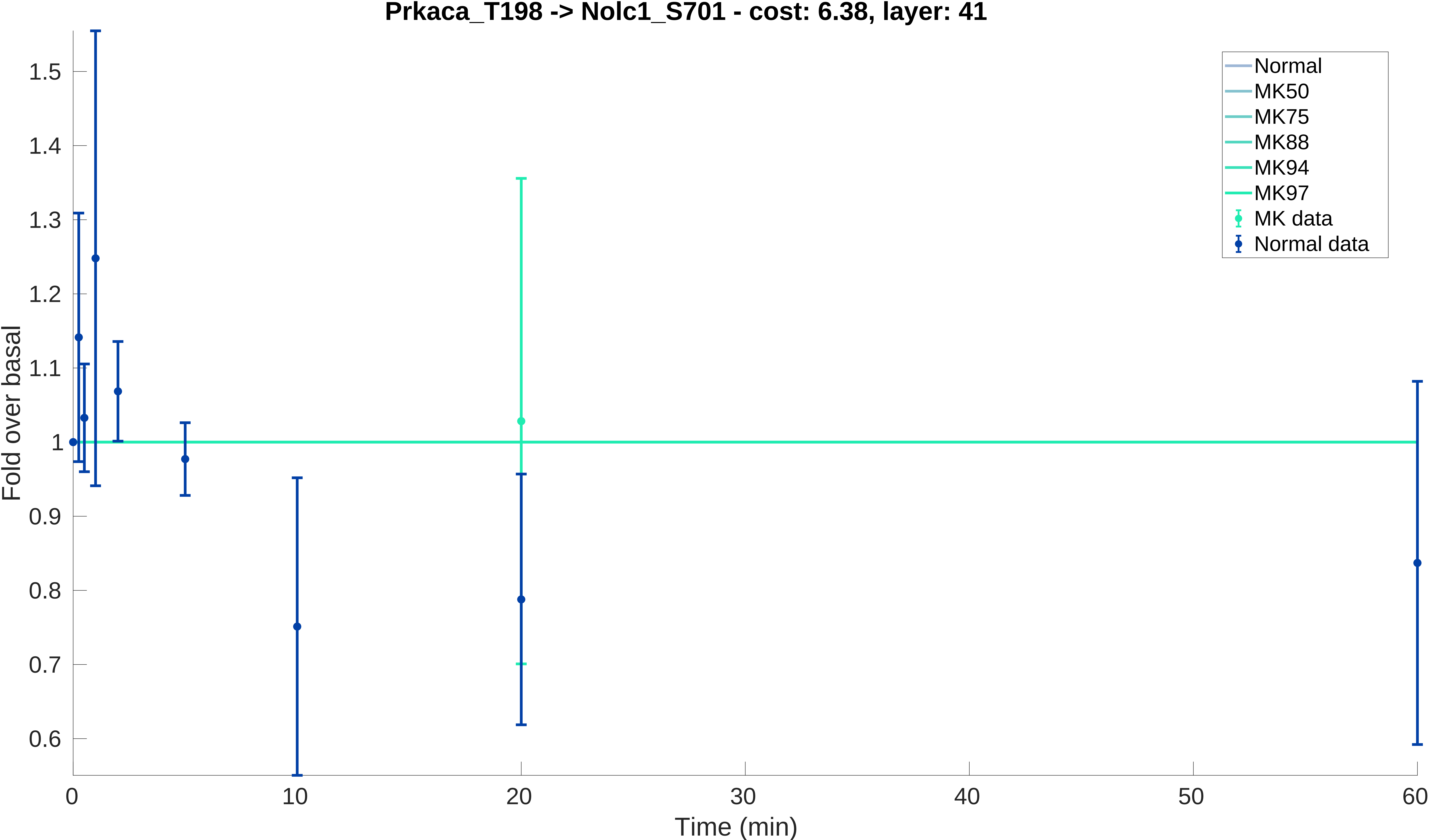

Prkaca_T198 -> Nolc1_S701 - cost: 6.38, layer: 41
Fold over basal
Time (min)
0.6
0.7
0.8
0.9
1
1.1
1.2
1.3
1.4
1.5
0
10
20
30
40
50
60
Normal
MK50
MK75
MK88
MK94
MK97
MK data
Normal data

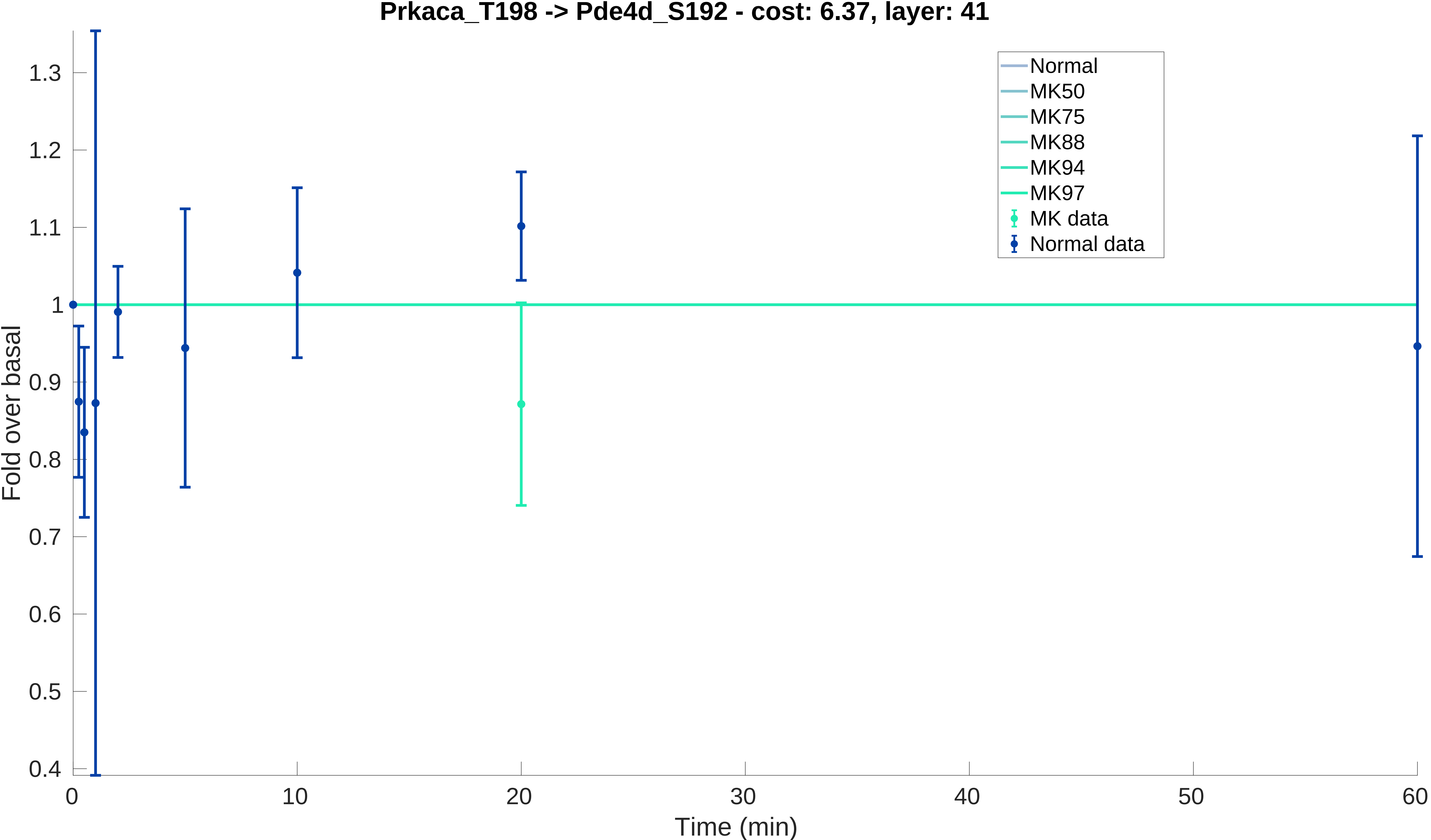
Prkaca_T198 -> Pde4d_S192 - cost: 6.37, layer: 41
Fold over basal
Time (min)
0.4
0.5
0.6
0.7
0.8
0.9
1
1.1
1.2
1.3
0
10
20
30
40
50
60
Normal
MK50
MK75
MK88
MK94
MK97
MK data
Normal data

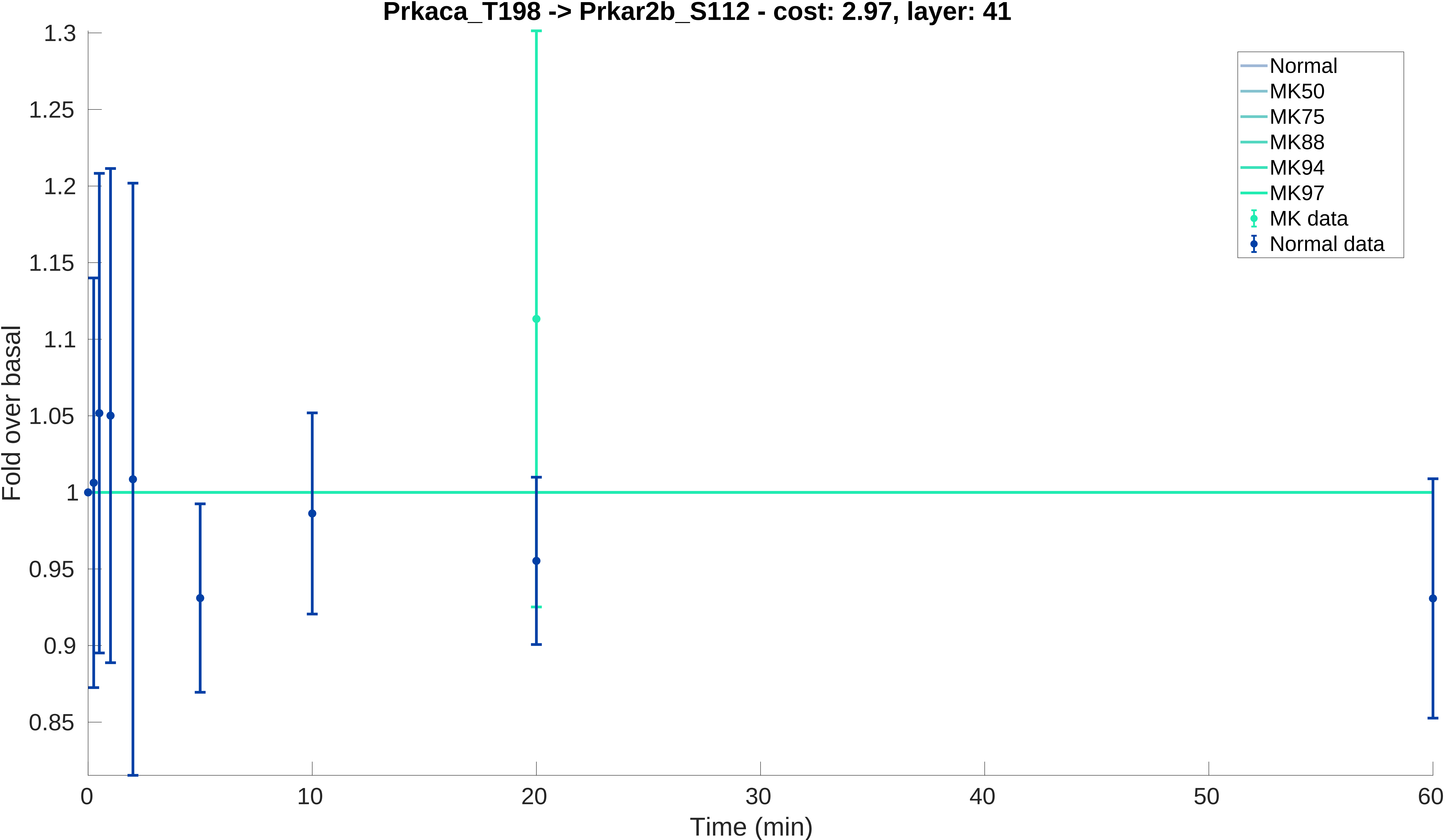

Prkaca_T198 -> Prkar2b_S112 - cost: 2.97, layer: 41
Fold over basal
Time (min)
1.3
1.25
1.2
1.15
1.1
1.05
1
0.95
0.9
0.85
0
10
20
30
40
50
60
Normal
MK50
MK75
MK88
MK94
MK97
MK data
Normal data

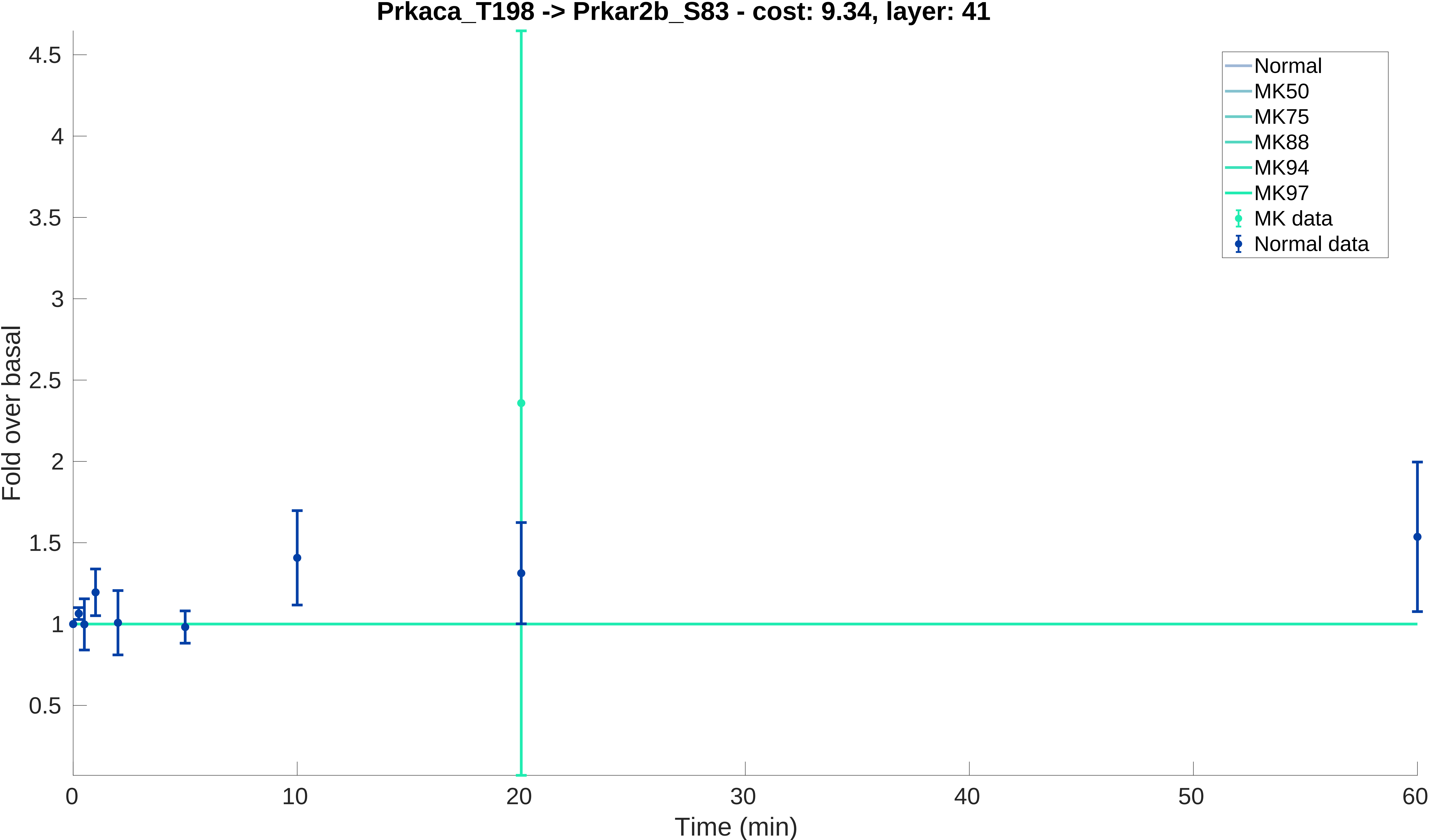
Prkaca_T198 -> Prkar2b_S83 - cost: 9.34, layer: 41
Fold over basal
Time (min)
0.5
1
1.5
2
2.5
3
3.5
4
4.5
0
10
20
30
40
50
60
Normal
MK50
MK75
MK88
MK94
MK97
MK data
Normal data

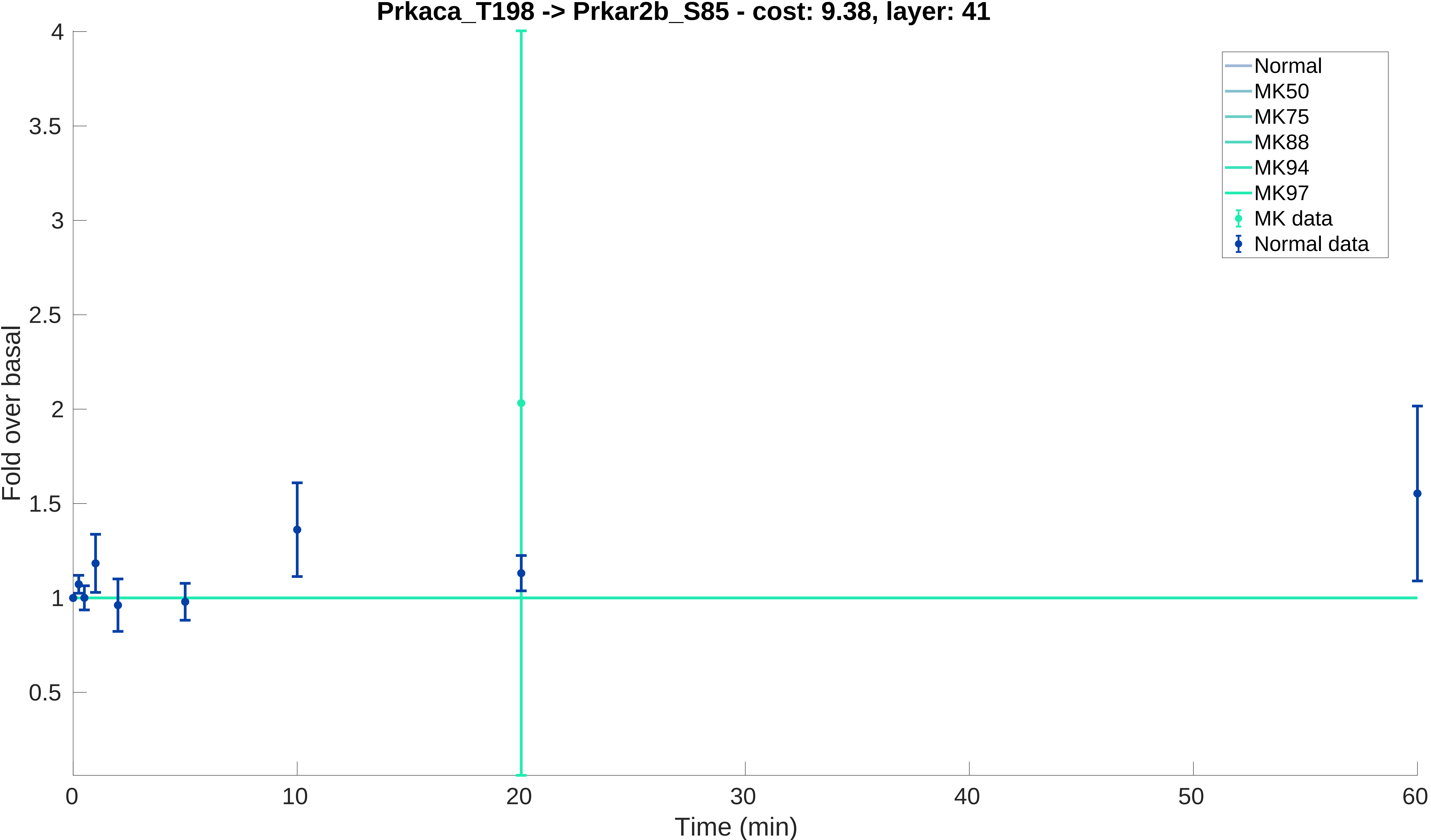

Prkaca_T198 -> Prkar2b_S85 - cost: 9.38, layer: 41
Fold over basal
Time (min)
0.5
1
1.5
2
2.5
3
3.5
4
0
10
20
30
40
50
60
Normal
MK50
MK75
MK88
MK94
MK97
MK data
Normal data

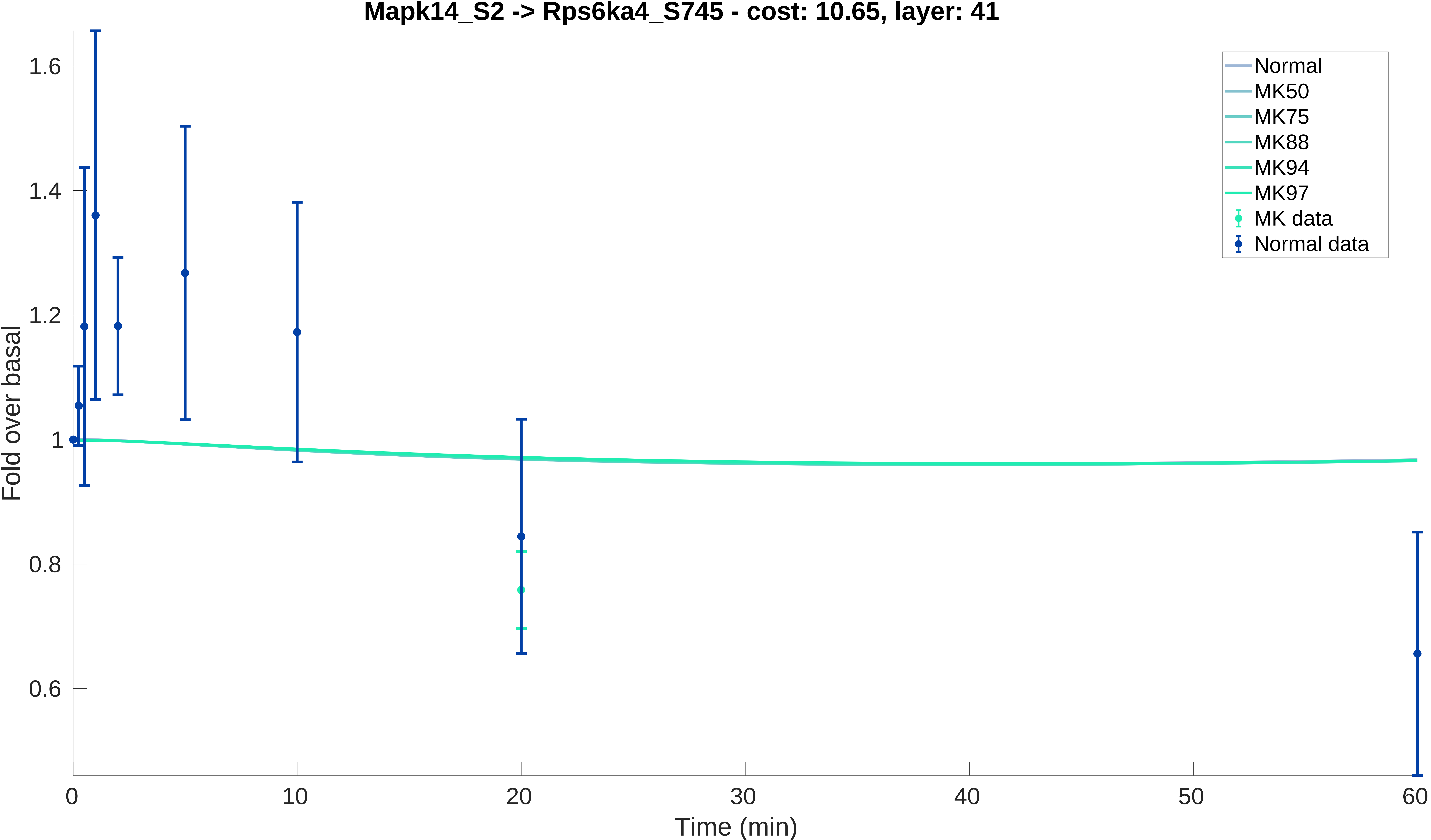
Mapk14_S2 -> Rps6ka4_S745 - cost: 10.65, layer: 41
Fold over basal
1.6
1.4
1.2
1
0.8
0.6
0
10
20
30
40
50
60
Time (min)
Normal
MK50
MK75
MK88
MK94
MK97
MK data
Normal data

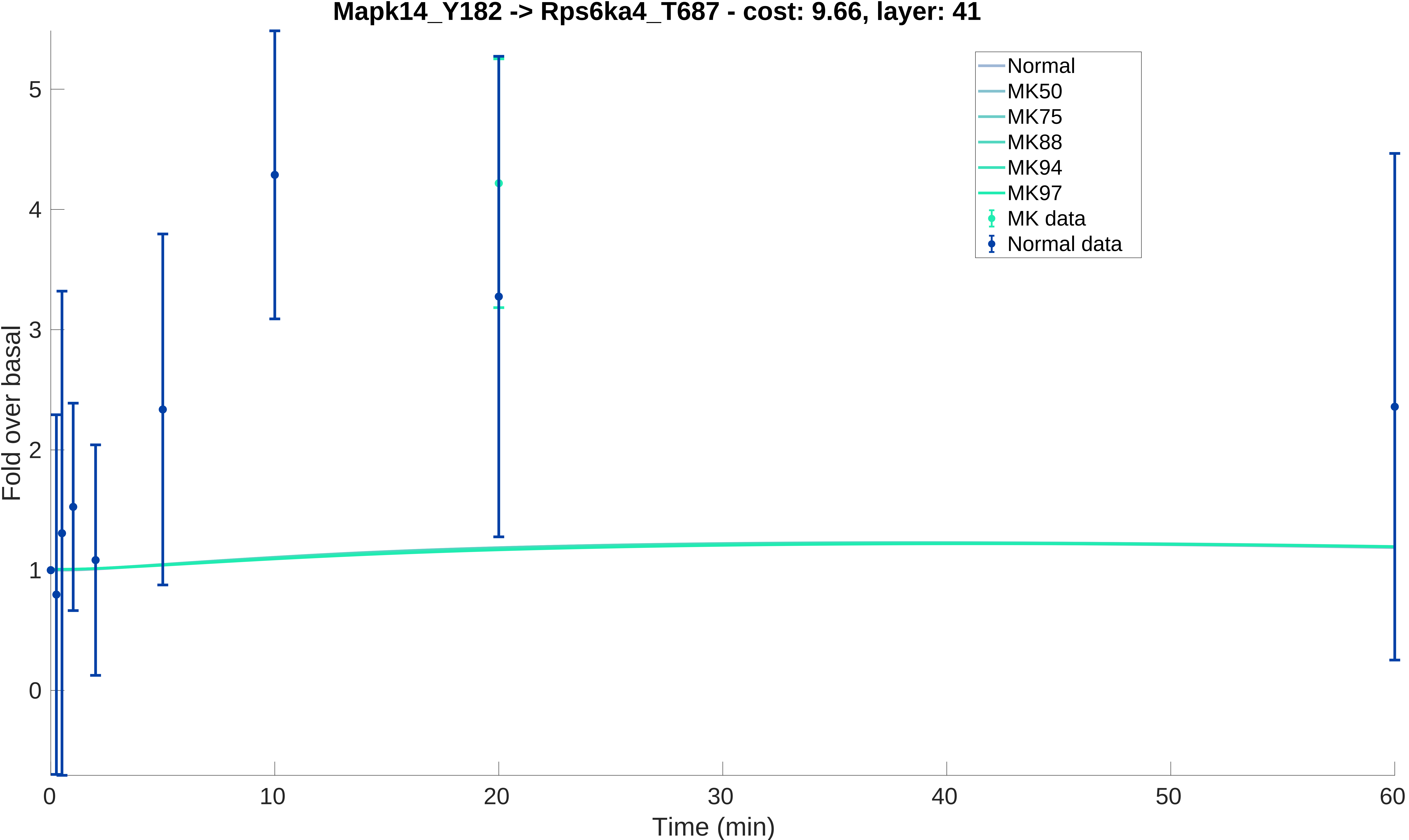

Mapk14_Y182 -> Rps6ka4_T687 - cost: 9.66, layer: 41
Fold over basal
Time (min)
0
1
2
3
4
5
0
10
20
30
40
50
60
Normal
MK50
MK75
MK88
MK94
MK97
MK data
Normal data

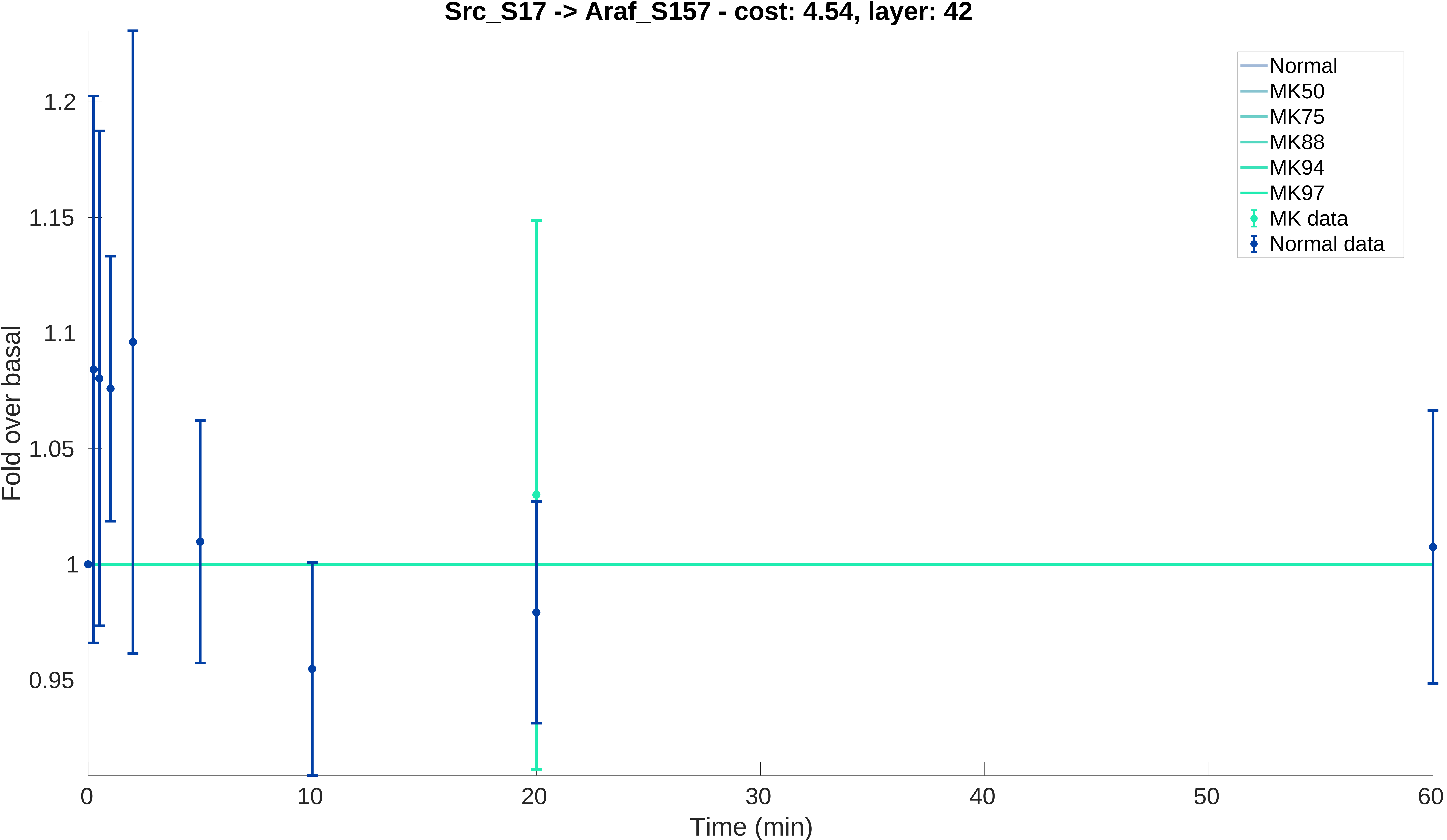

Src_S17 -> Araf_S157 - cost: 4.54, layer: 42
Normal
MK50
MK75
MK88
MK94
MK97
MK data
Normal data
Fold over basal
Time (min)
1.2
1.15
1.1
1.05
1
0.95
0
10
20
30
40
50
60

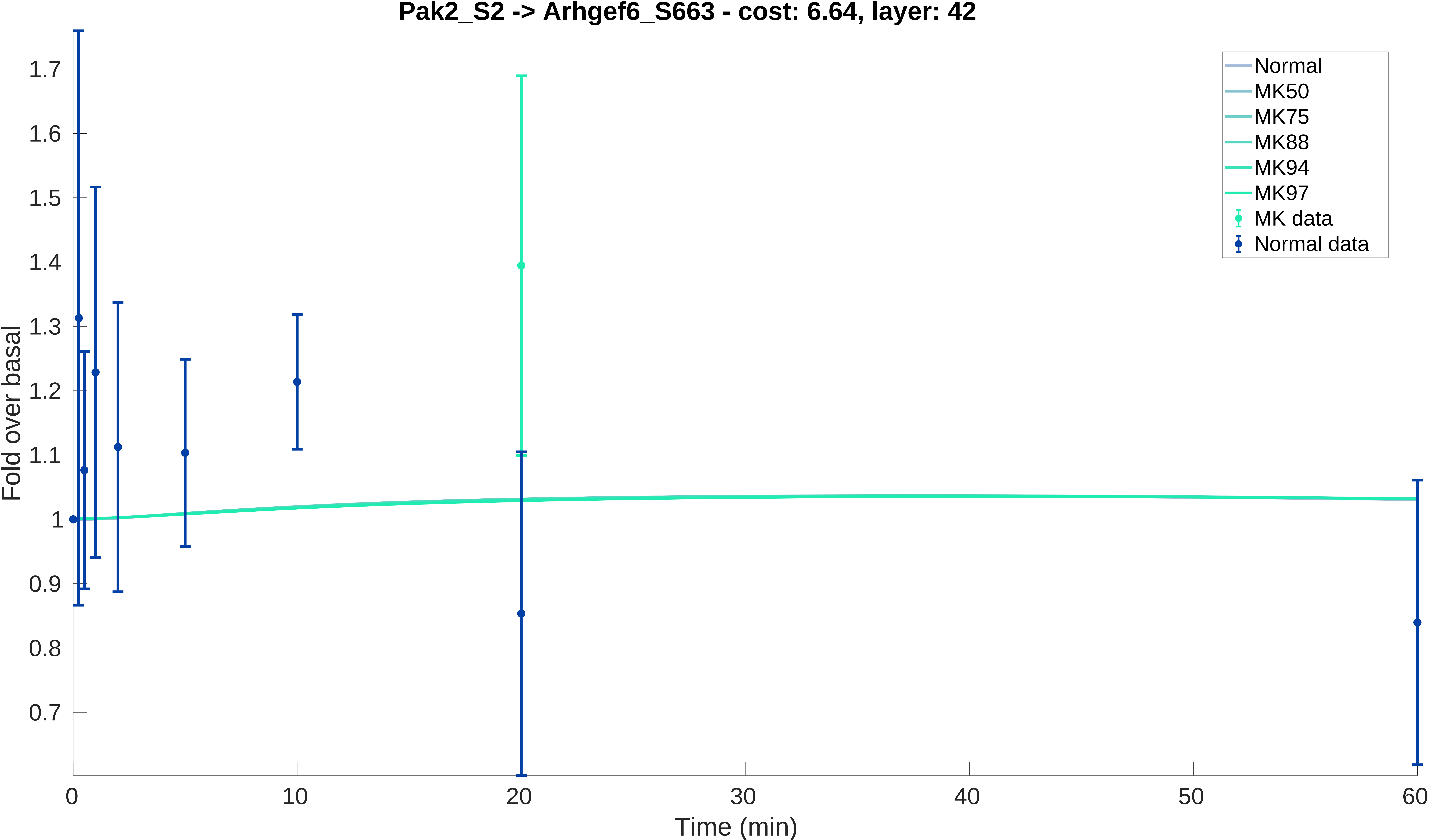

Pak2_S2 -> Arhgef6_S663 - cost: 6.64, layer: 42
Fold over basal
Time (min)
0.7
0.8
0.9
1
1.1
1.2
1.3
1.4
1.5
1.6
1.7
0
10
20
30
40
50
60
Normal
MK50
MK75
MK88
MK94
MK97
MK data
Normal data

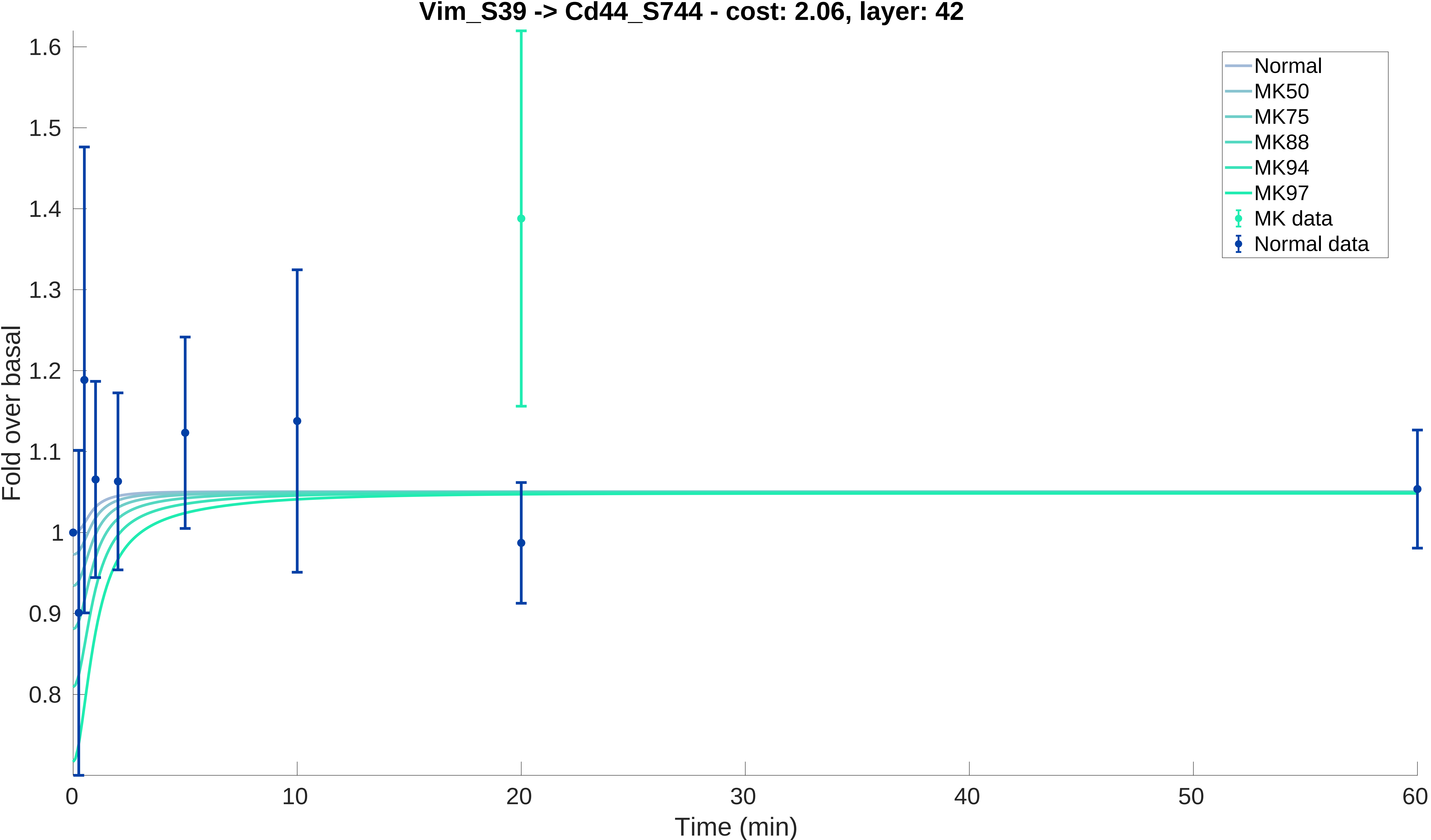

Vim_S39 -> Cd44_S744 - cost: 2.06, layer: 42
Fold over basal
Time (min)
0
10
20
30
40
50
60
0.8
0.9
1
1.1
1.2
1.3
1.4
1.5
1.6
Normal
MK50
MK75
MK88
MK94
MK97
MK data
Normal data

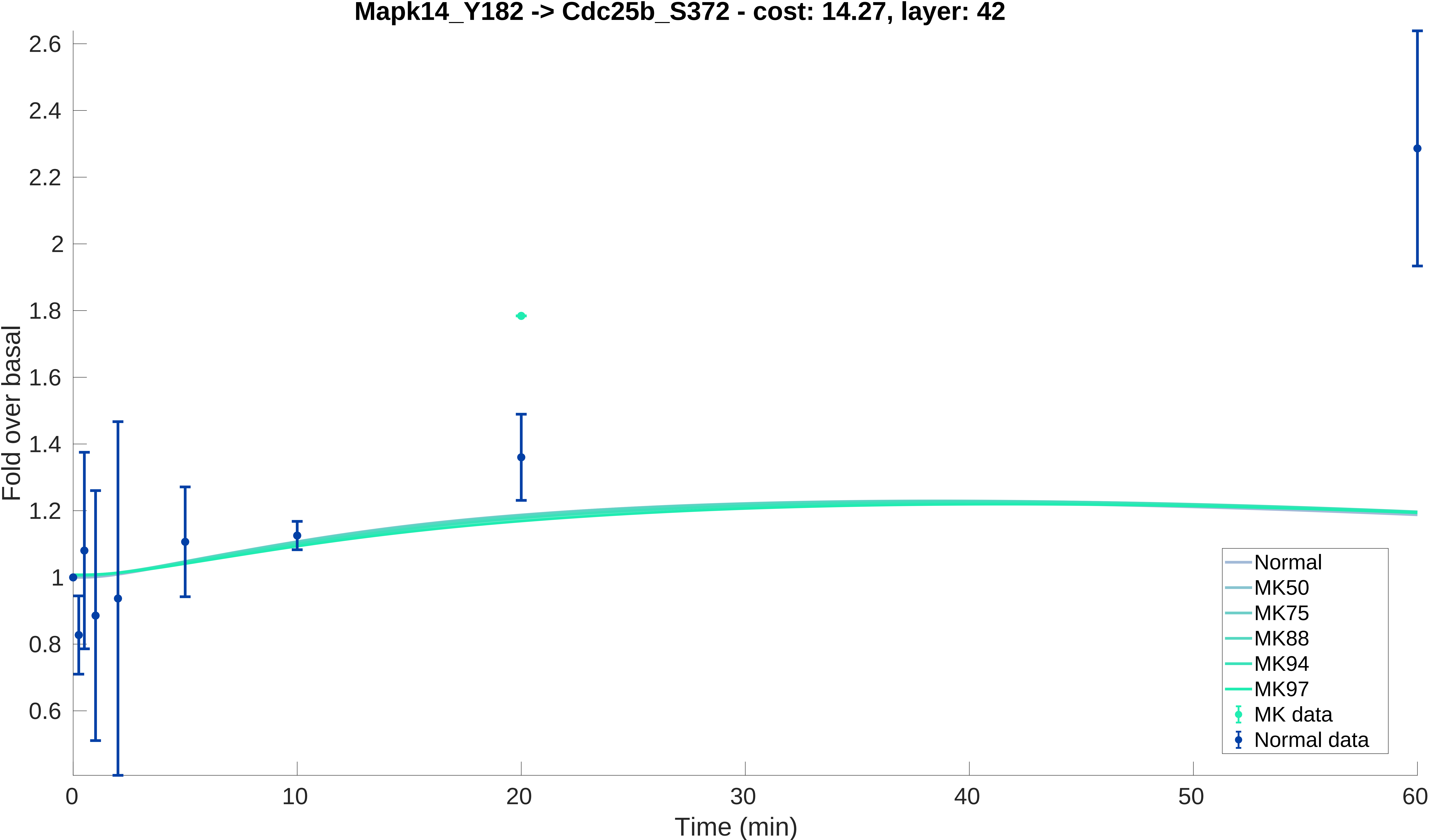
Mapk14_Y182 -> Cdc25b_S372 - cost: 14.27, layer: 42
Fold over basal
2.6
2.4
2.2
2
1.8
1.6
1.4
1.2
1
0.8
0.6
0
10
20
30
40
50
60
Time (min)
Normal
MK50
MK75
MK88
MK94
MK97
MK data
Normal data

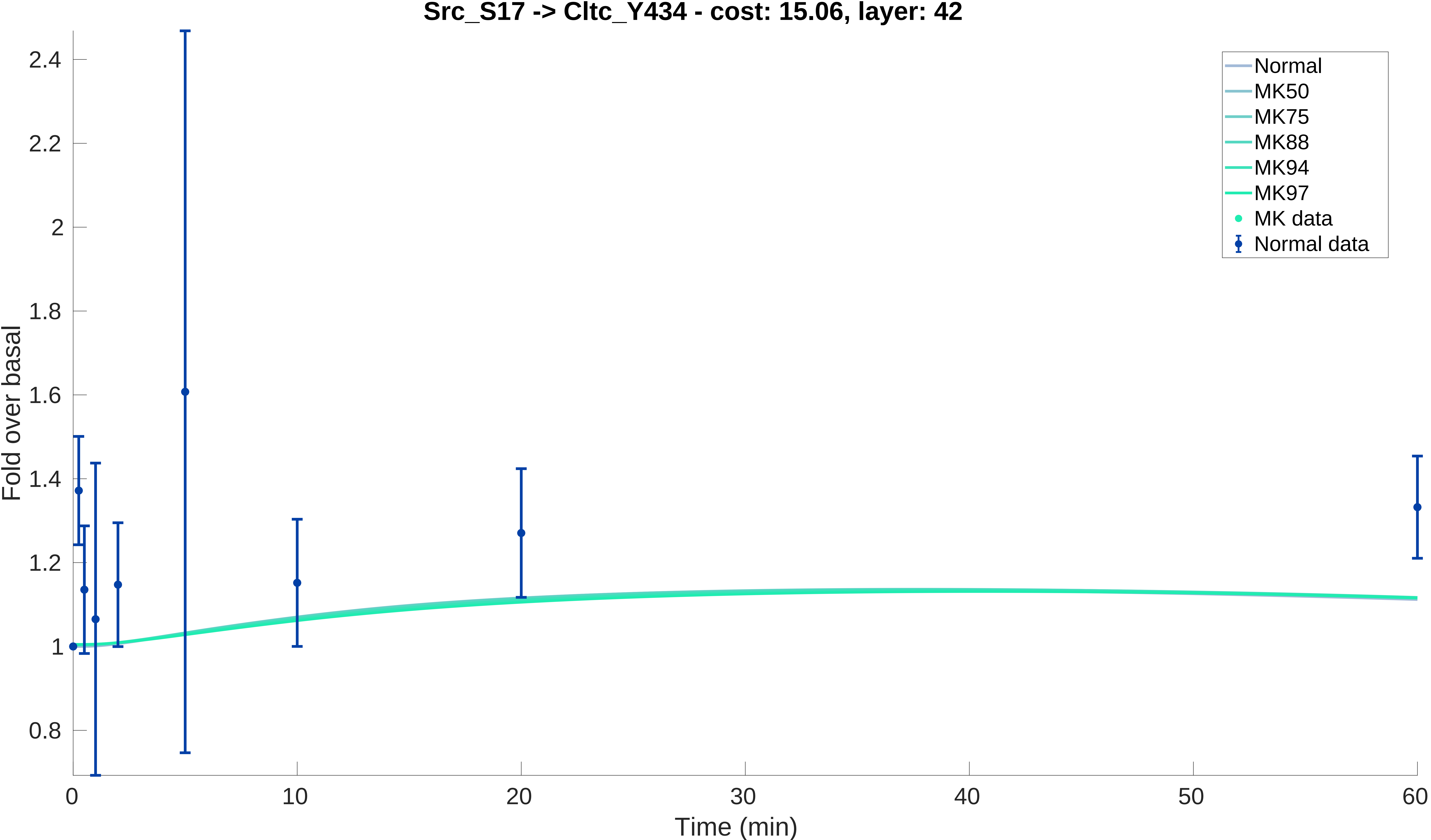

Src_S17 -> Cltc_Y434 - cost: 15.06, layer: 42
Fold over basal
Time (min)
0.8
1
1.2
1.4
1.6
1.8
2
2.2
2.4
0
10
20
30
40
50
60
Normal
MK50
MK75
MK88
MK94
MK97
MK data
Normal data

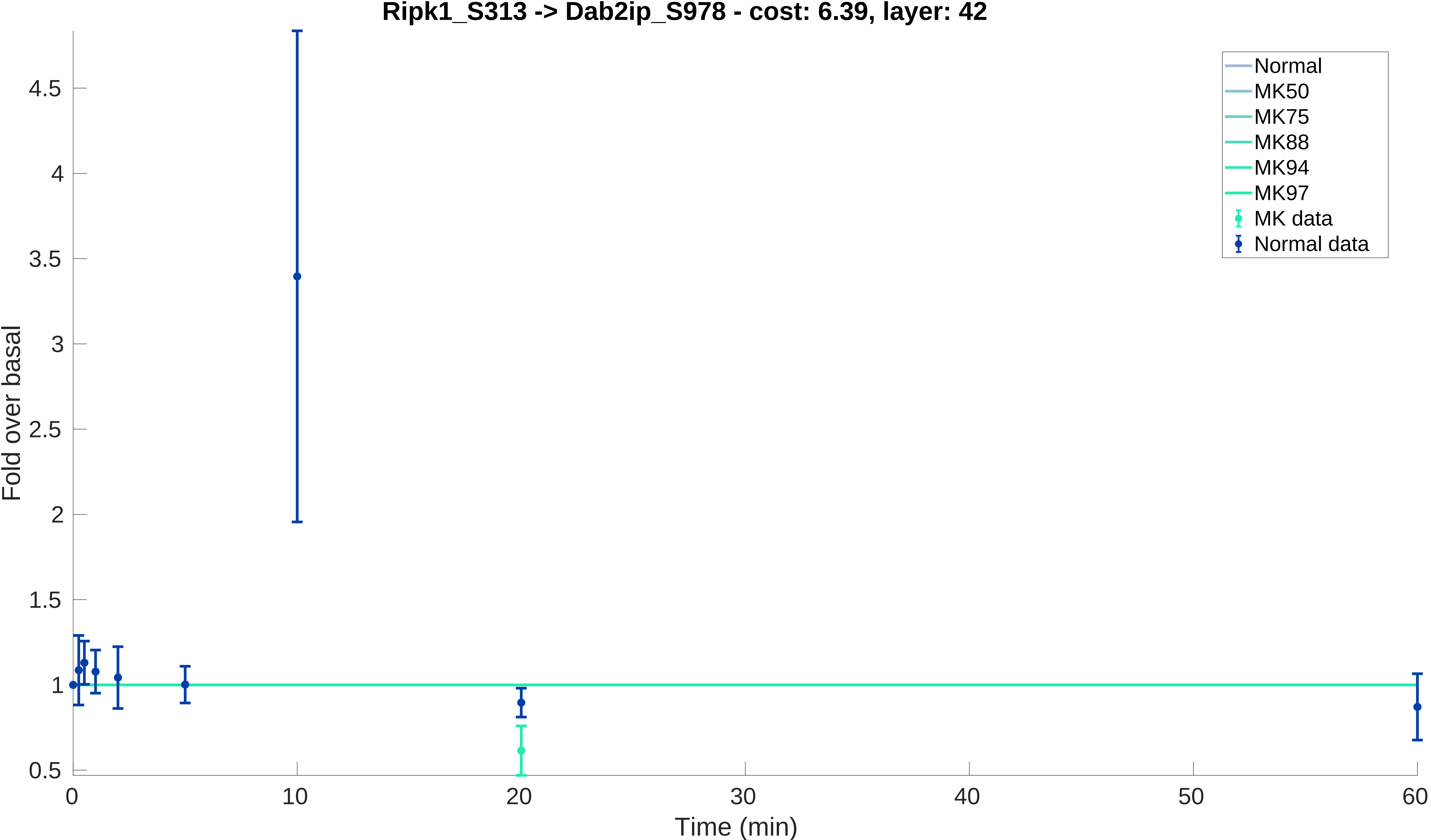

Ripk1_S313 -> Dab2ip_S978 - cost: 6.39, layer: 42
Fold over basal
Time (min)
0.5
1
1.5
2
2.5
3
3.5
4
4.5
0
10
20
30
40
50
60
Normal
MK50
MK75
MK88
MK94
MK97
MK data
Normal data

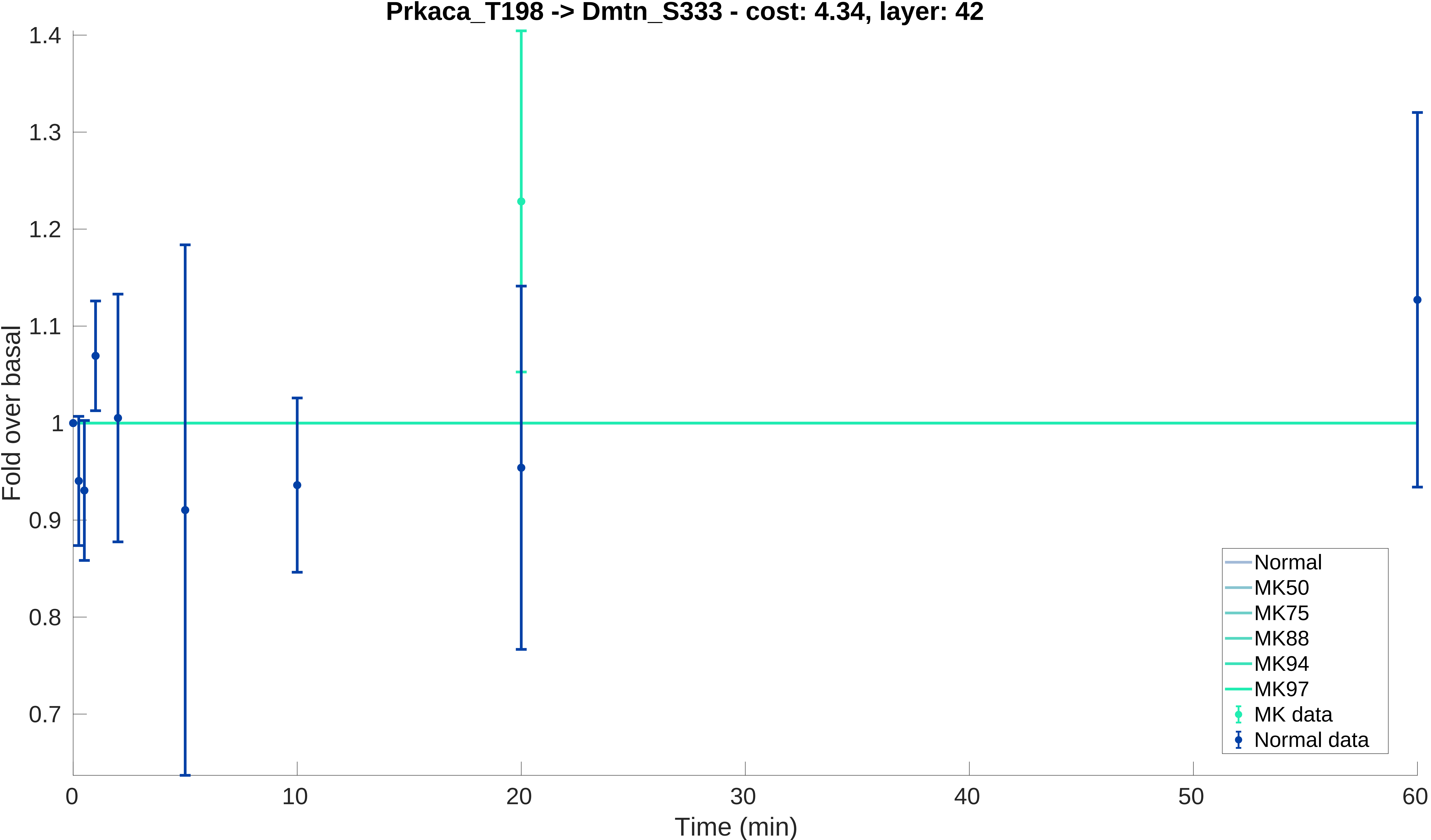
Prkaca_T198 -> Dmtn_S333 - cost: 4.34, layer: 42
Fold over basal
Time (min)
1.4
1.3
1.2
1.1
1
0.9
0.8
0.7
0
10
20
30
40
50
60
Normal
MK50
MK75
MK88
MK94
MK97
MK data
Normal data

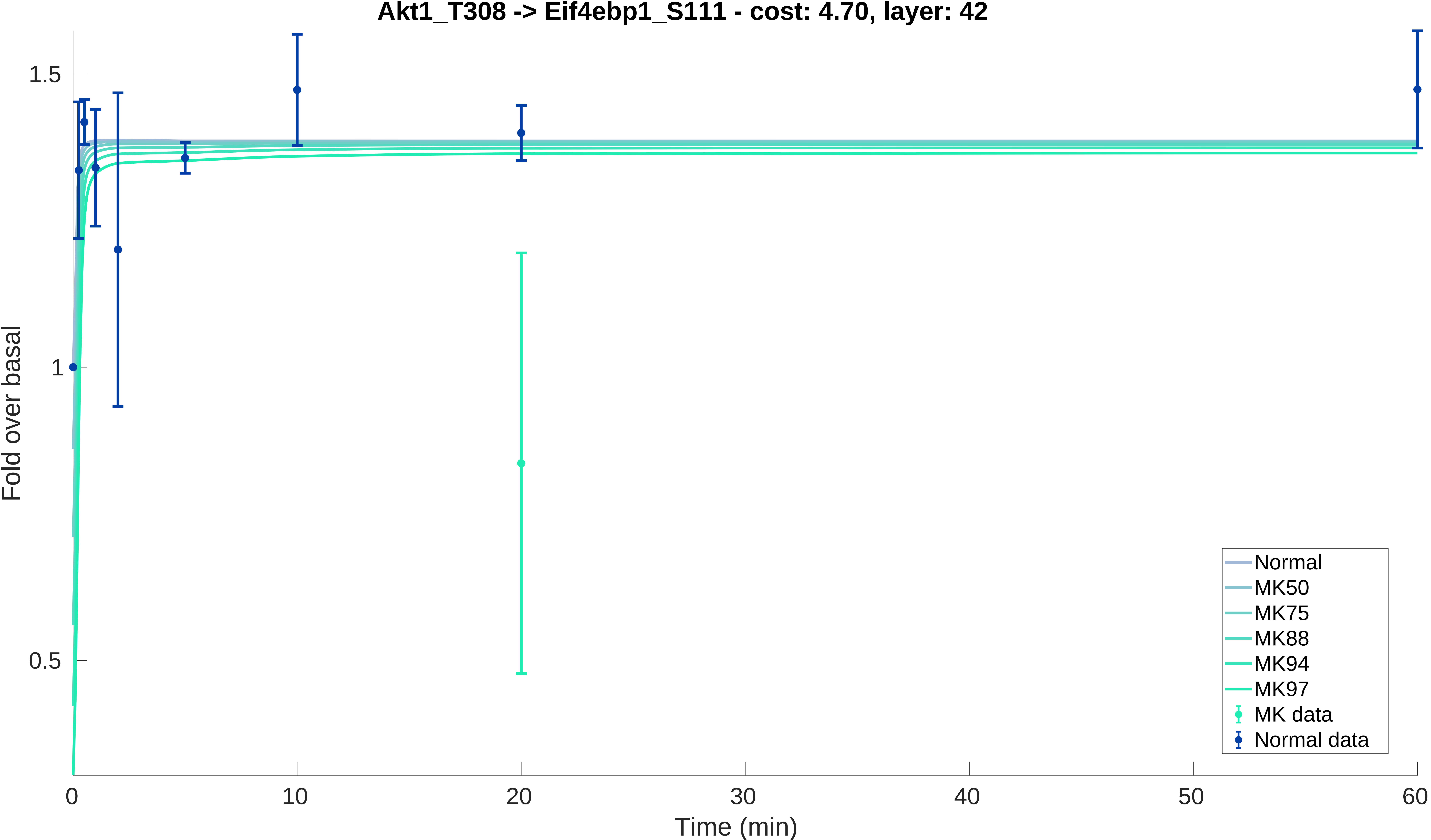

Akt1_T308 -> Eif4ebp1_S111 - cost: 4.70, layer: 42
Fold over basal
Time (min)
1.5
1
0.5
0
10
20
30
40
50
60
Normal
MK50
MK75
MK88
MK94
MK97
MK data
Normal data

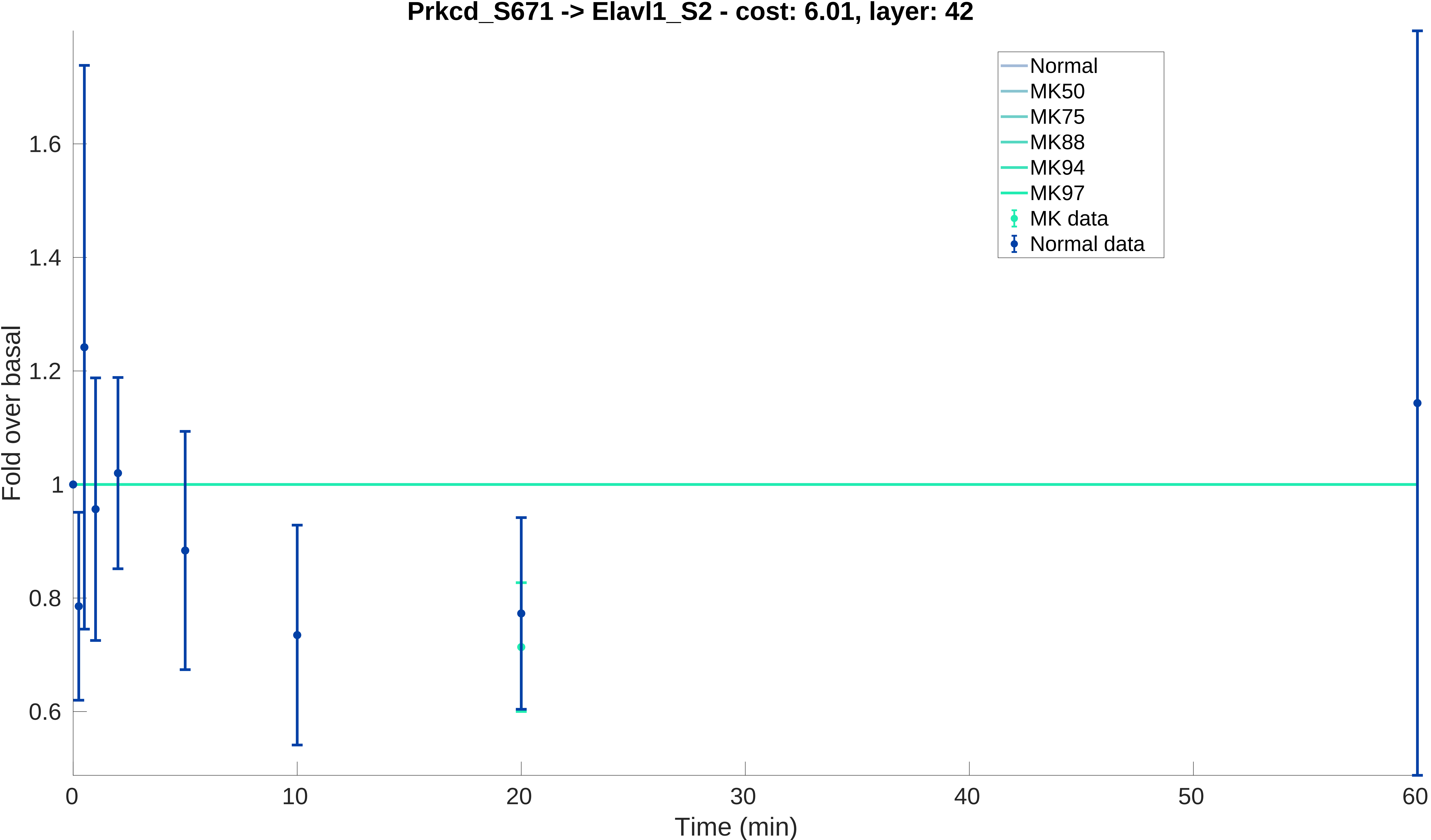
Prrkcd_S671 -> Elavl1_S2 - cost: 6.01, layer: 42
Fold over basal
Time (min)
0.6
0.8
1
1.2
1.4
1.6
0
10
20
30
40
50
60
Normal
MK50
MK75
MK88
MK94
MK97
MK data
Normal data

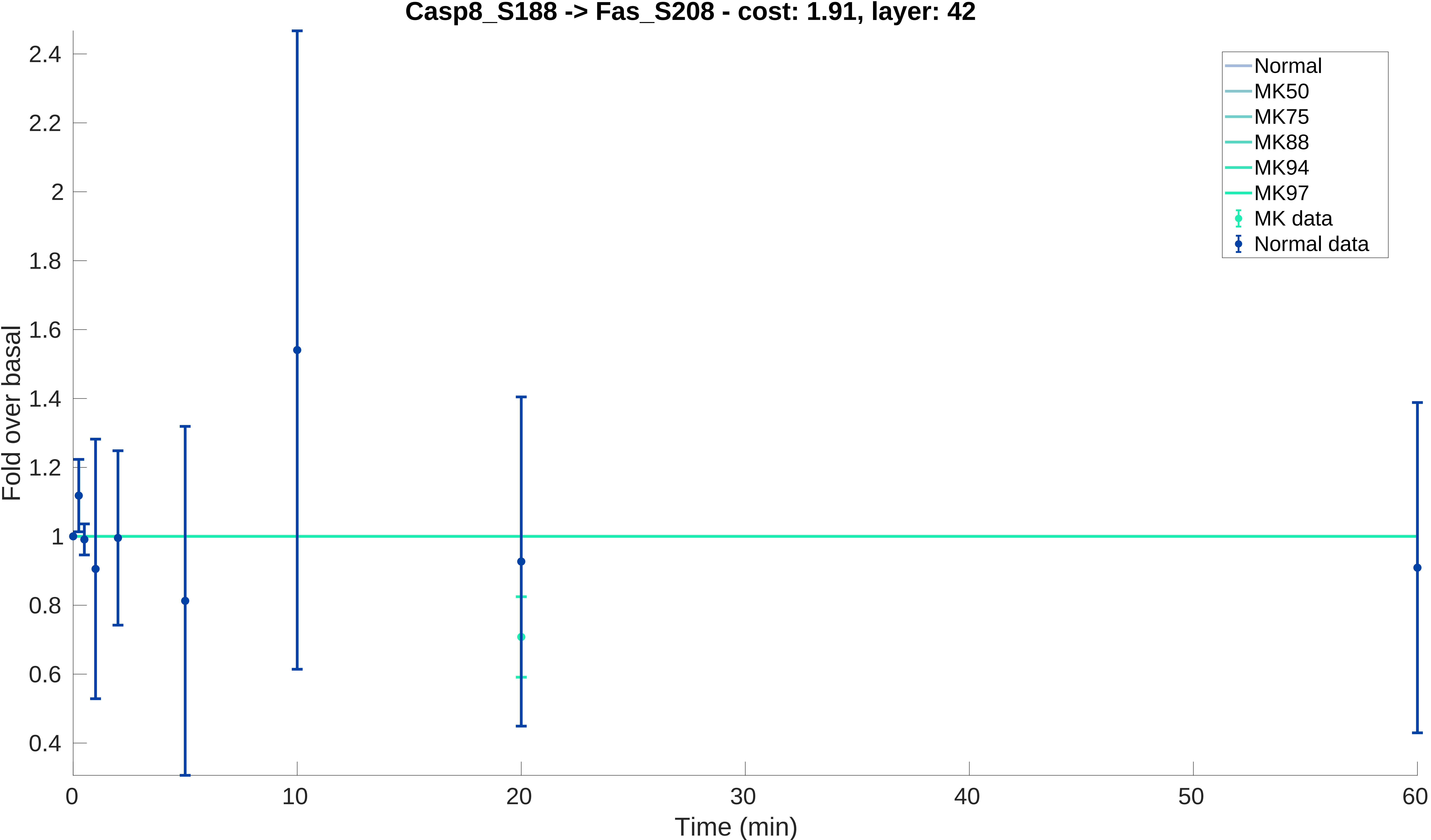
Casp8_S188 -> Fas_S208 - cost: 1.91, layer: 42
Fold over basal
Time (min)
0
10
20
30
40
50
60
0.4
0.6
0.8
1
1.2
1.4
1.6
1.8
2
2.2
2.4
Normal
MK50
MK75
MK88
MK94
MK97
MK data
Normal data

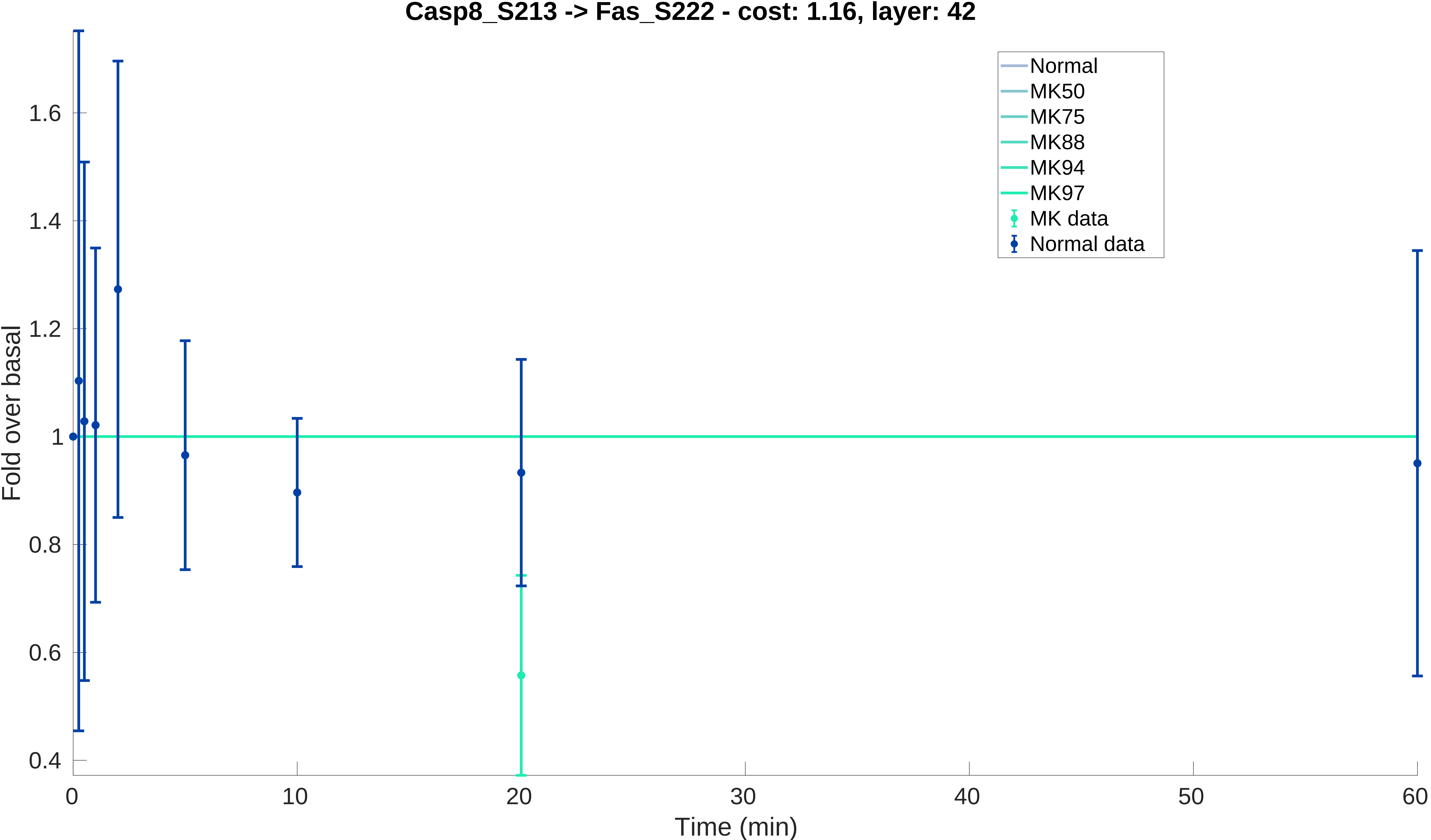

Casp8_S213 -> Fas_S222 - cost: 1.16, layer: 42
Normal
MK50
MK75
MK88
MK94
MK97
MK data
Normal data
Fold over basal
1.6
1.4
1.2
1
0.8
0.6
0.4
0
10
20
30
40
50
60
Time (min)

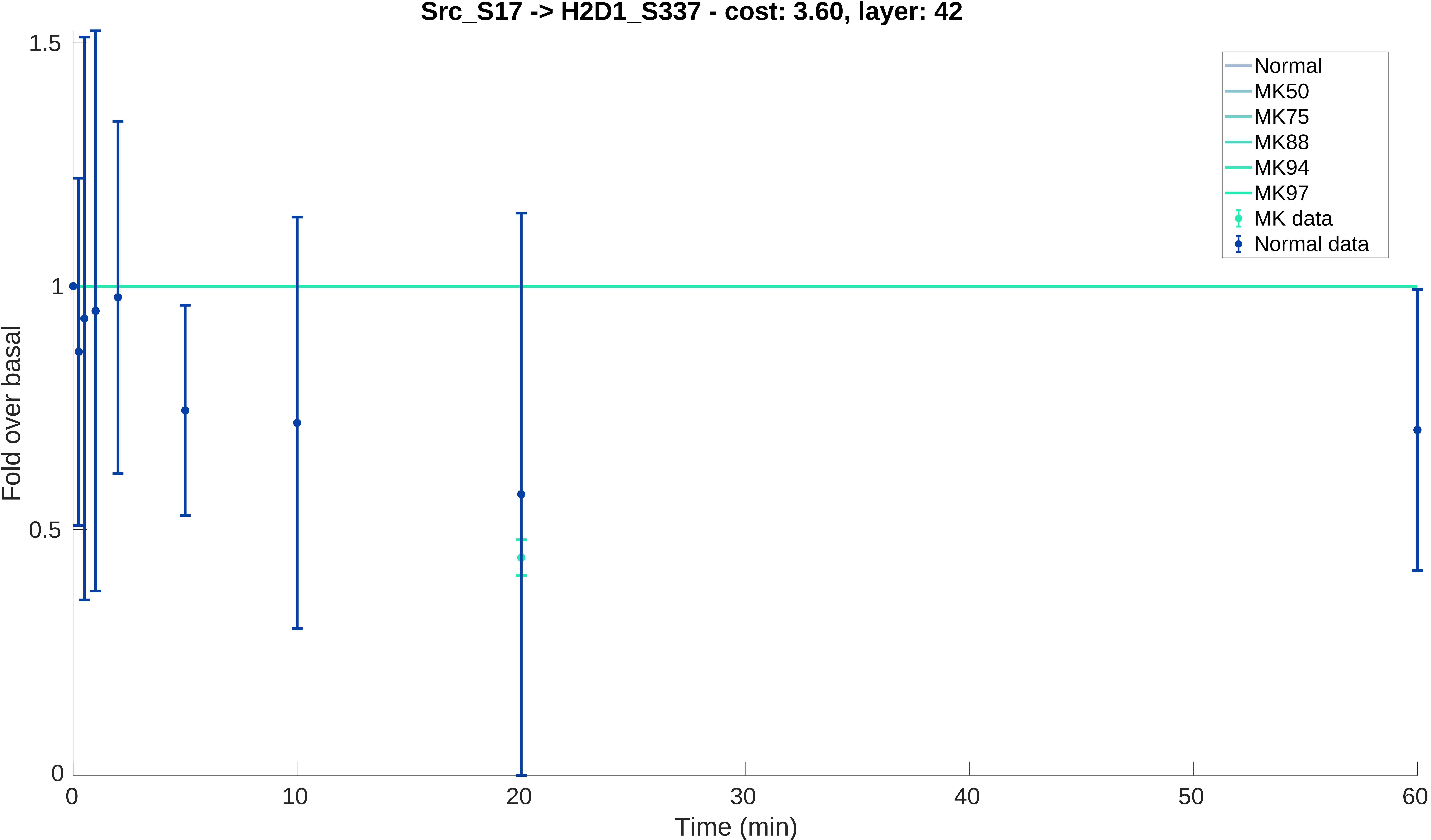

Src_S17 -> H2D1_S337 - cost: 3.60, layer: 42
Fold over basal
1.5
1
0.5
0
Time (min)
0
10
20
30
40
50
60
Normal
MK50
MK75
MK88
MK94
MK97
MK data
Normal data

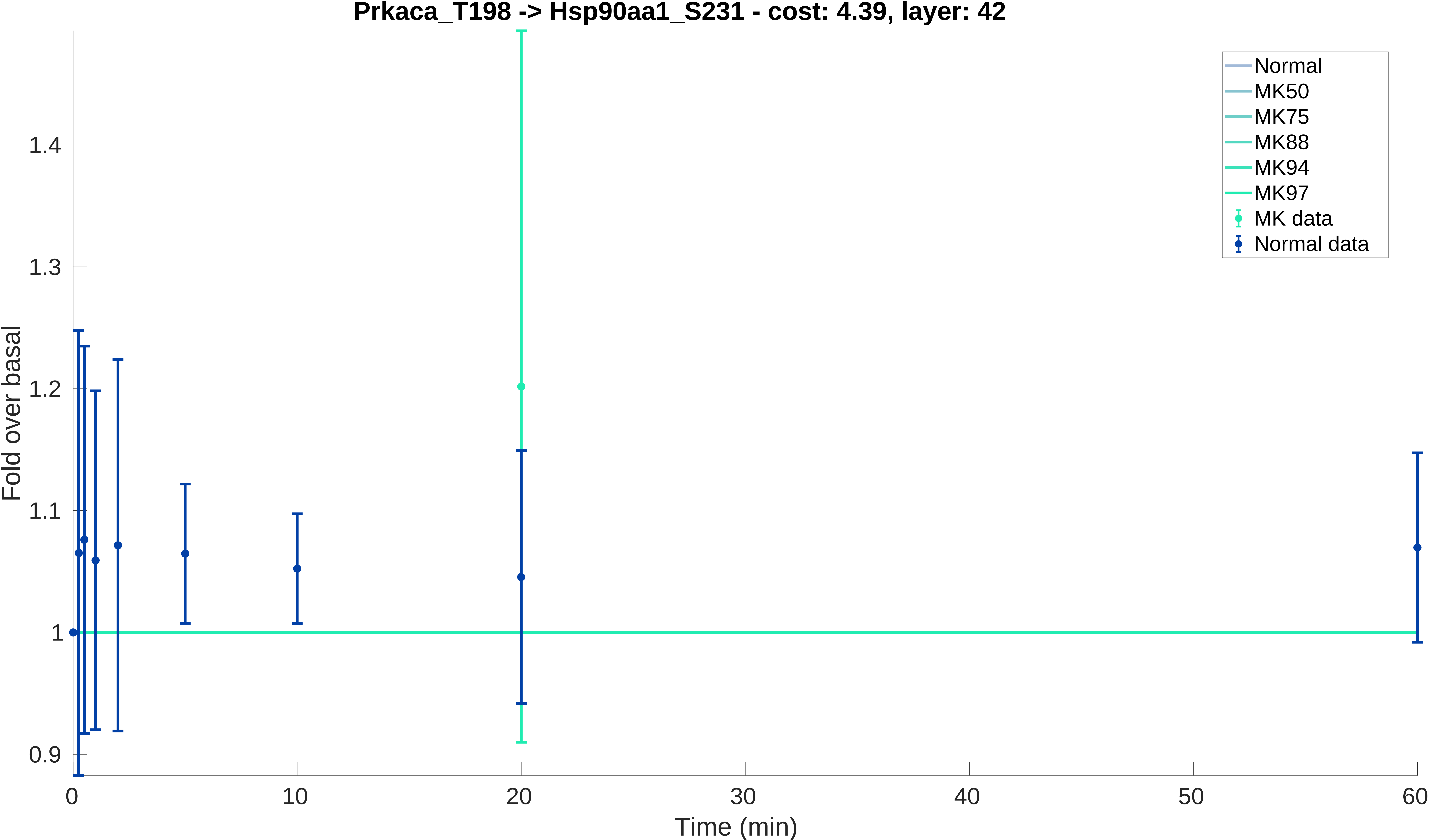
Prkaca_T198 -> Hsp90aa1_S231 - cost: 4.39, layer: 42
Fold over basal
Time (min)
0.9
1
1.1
1.2
1.3
1.4
0
10
20
30
40
50
60
Normal
MK50
MK75
MK88
MK94
MK97
MK data
Normal data

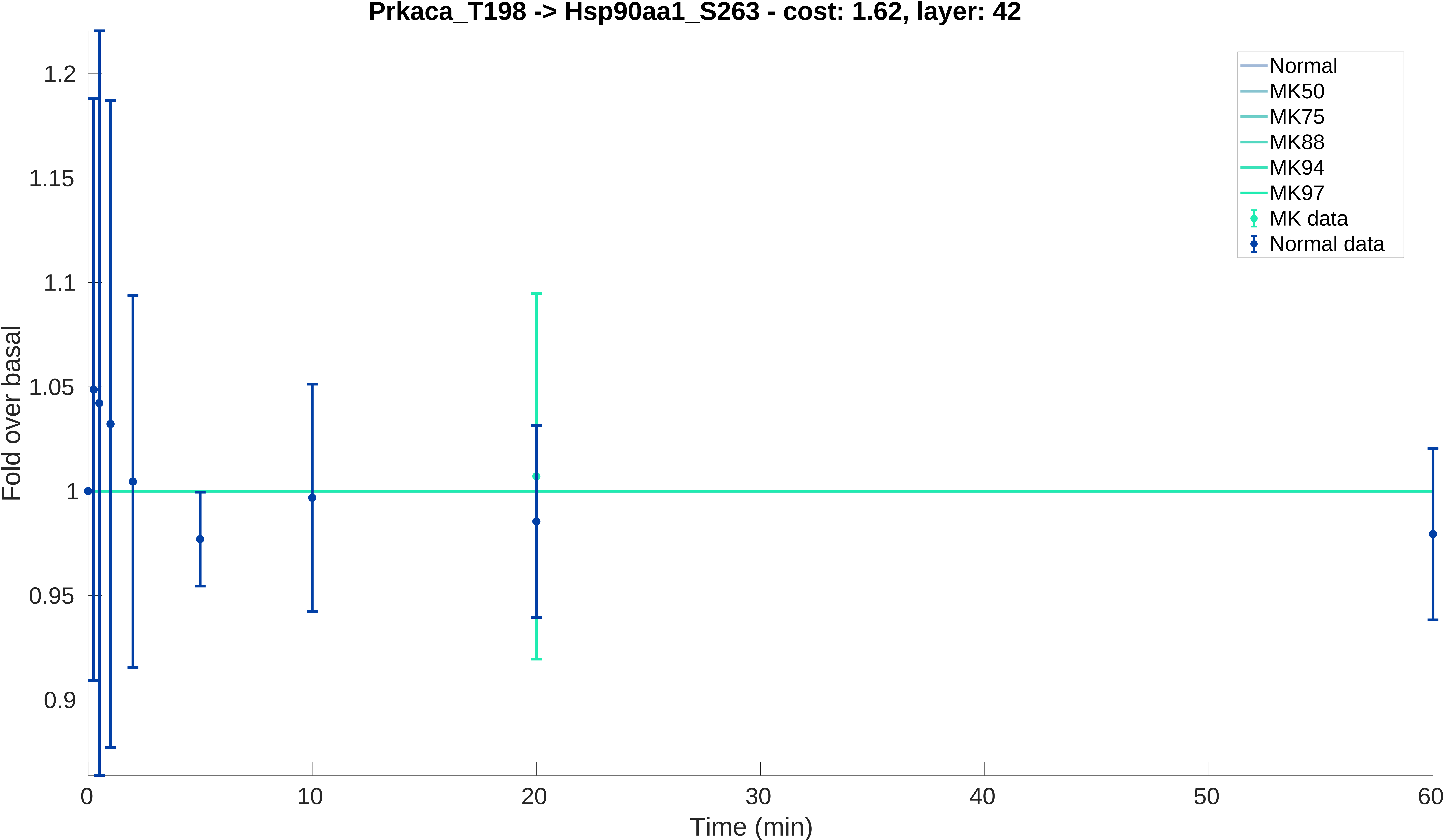

Prrkaca_T198 -> Hsp90aa1_S263 - cost: 1.62, layer: 42
Normal
MK50
MK75
MK88
MK94
MK97
MK data
Normal data
Fold over basal
1.2
1.15
1.1
1.05
1
0.95
0.9
0
10
20
30
40
50
60
Time (min)

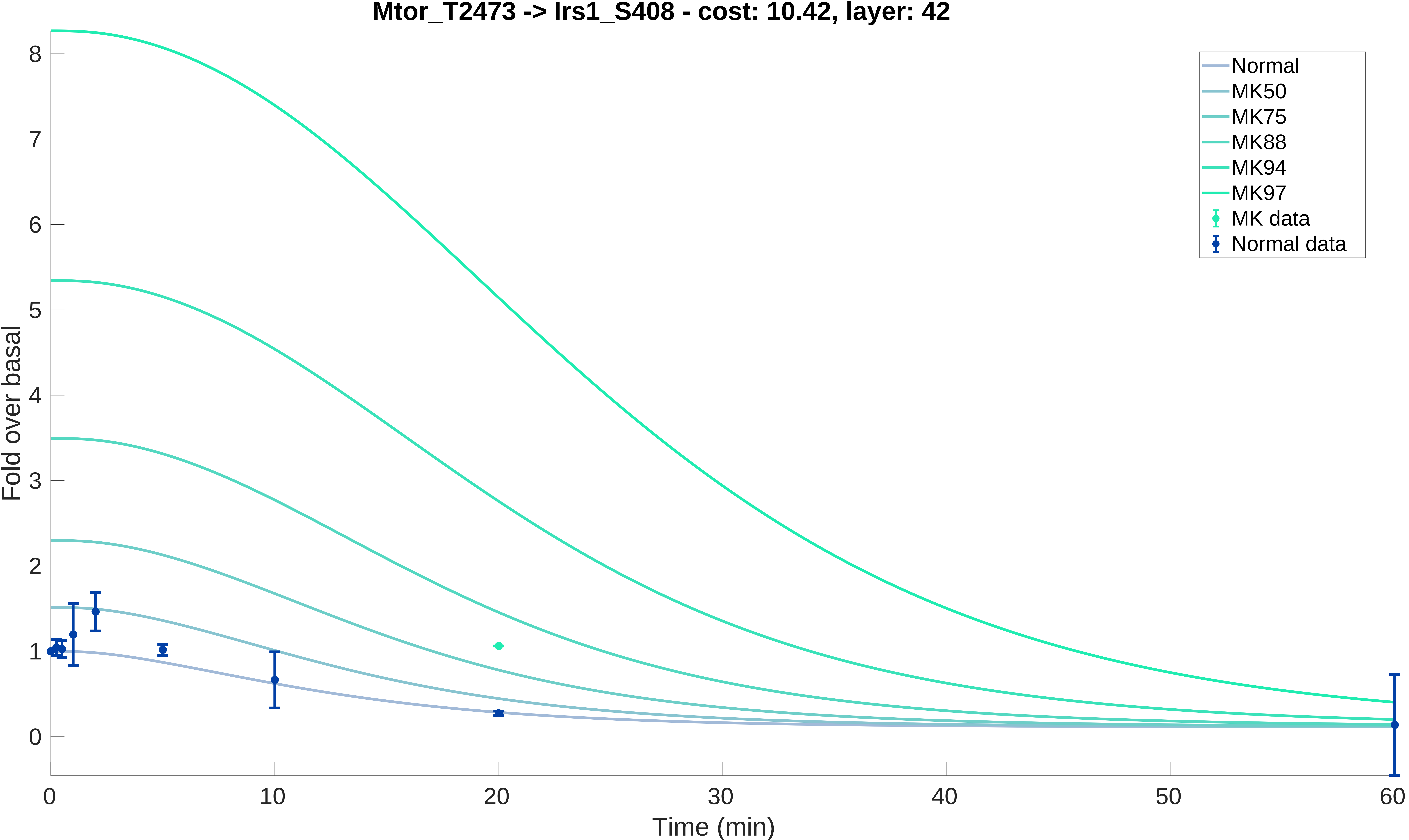

Mtor_T2473 -> Irs1_S408 - cost: 10.42, layer: 42
Fold over basal
Time (min)
0
1
2
3
4
5
6
7
8
0
10
20
30
40
50
60
Normal
MK50
MK75
MK88
MK94
MK97
MK data
Normal data

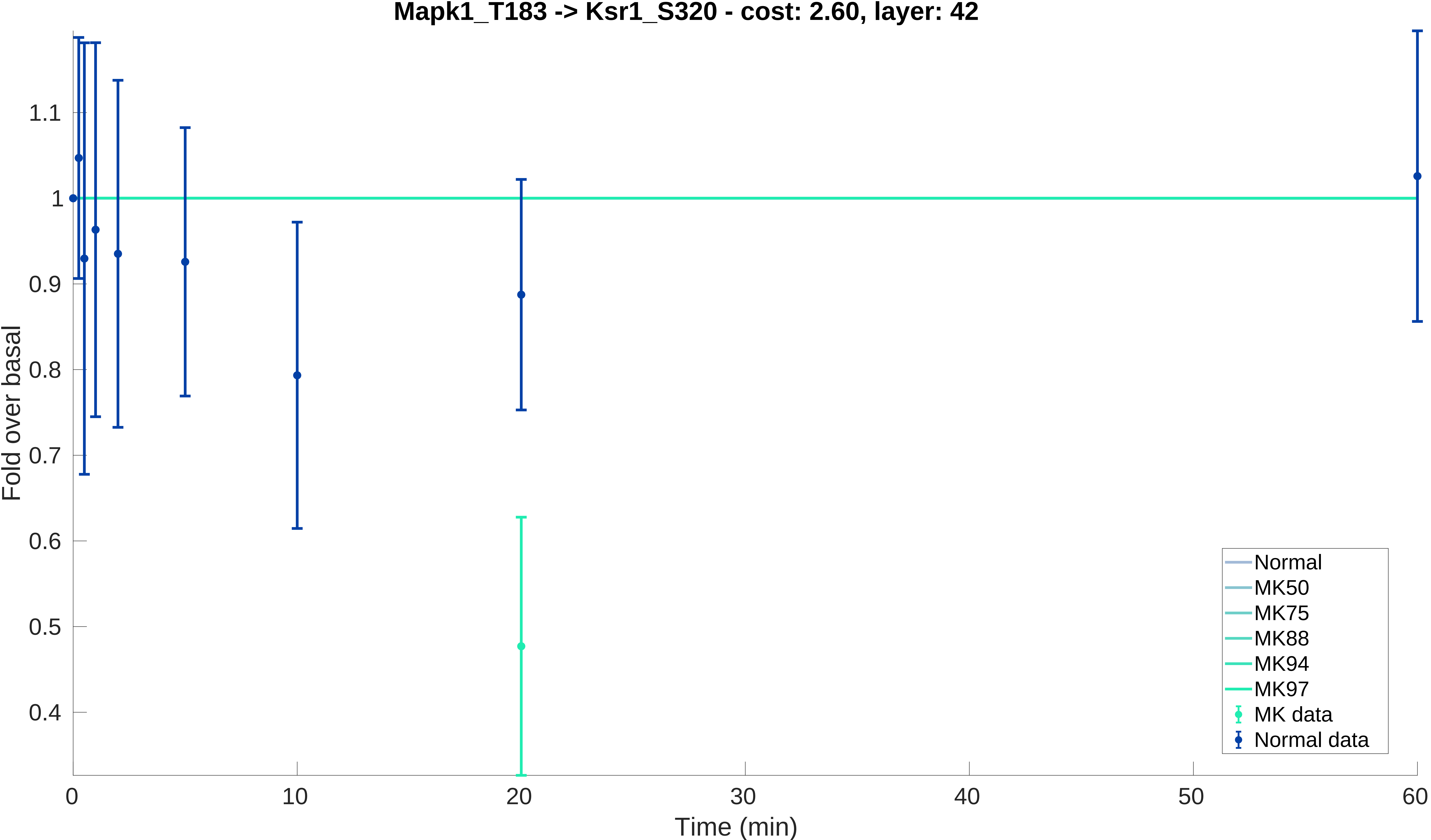

Mapk1_T183 -> Ksr1_S320 - cost: 2.60, layer: 42
Fold over basal
1.1
1
0.9
0.8
0.7
0.6
0.5
0.4
0
10
20
30
40
50
60
Time (min)
Normal
MK50
MK75
MK88
MK94
MK97
MK data
Normal data

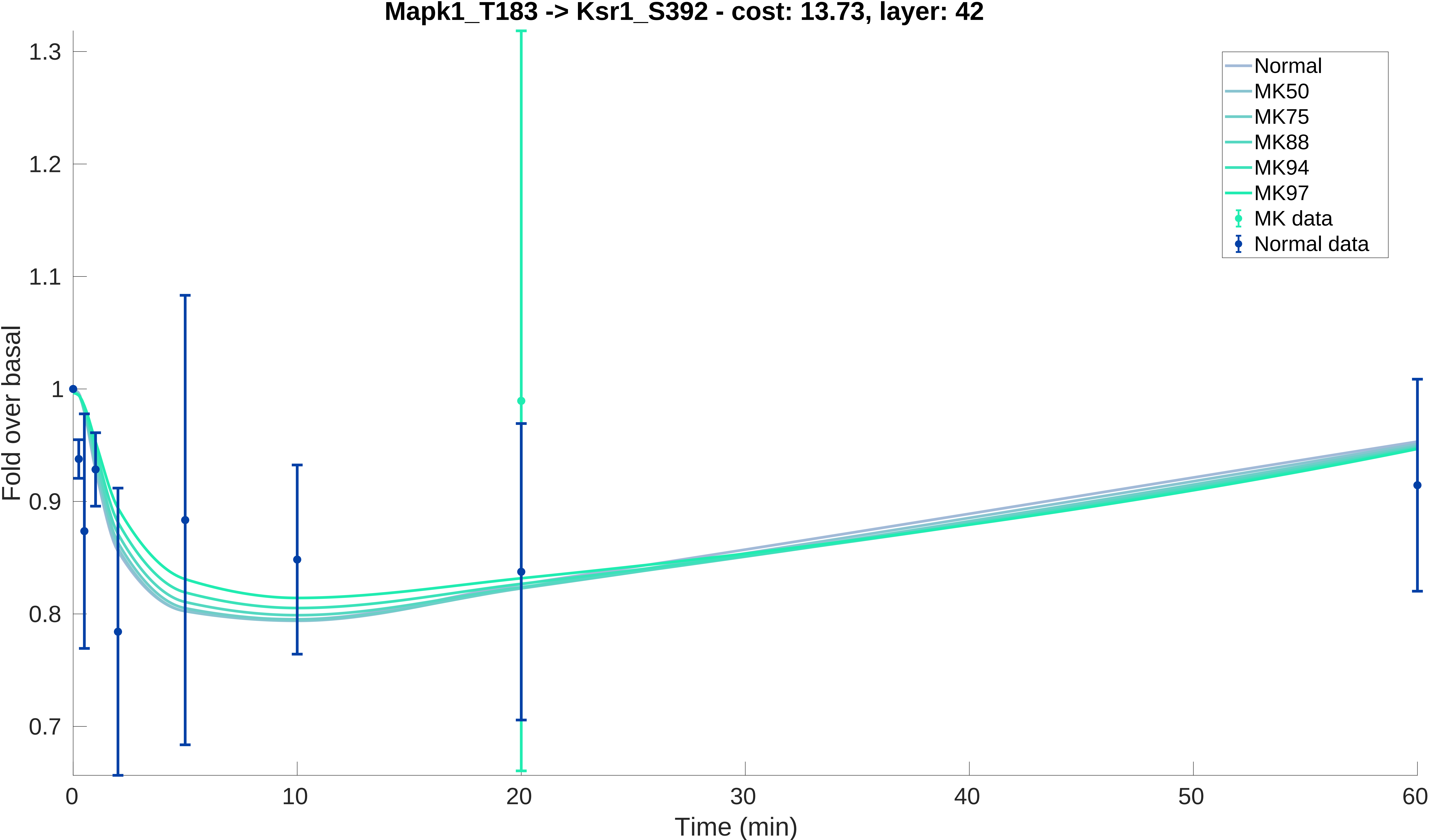
Mapk1_T183 -> Ksr1_S392 - cost: 13.73, layer: 42
Fold over basal
Time (min)
1.3
1.2
1.1
1
0.9
0.8
0.7
0
10
20
30
40
50
60
Normal
MK50
MK75
MK88
MK94
MK97
MK data
Normal data

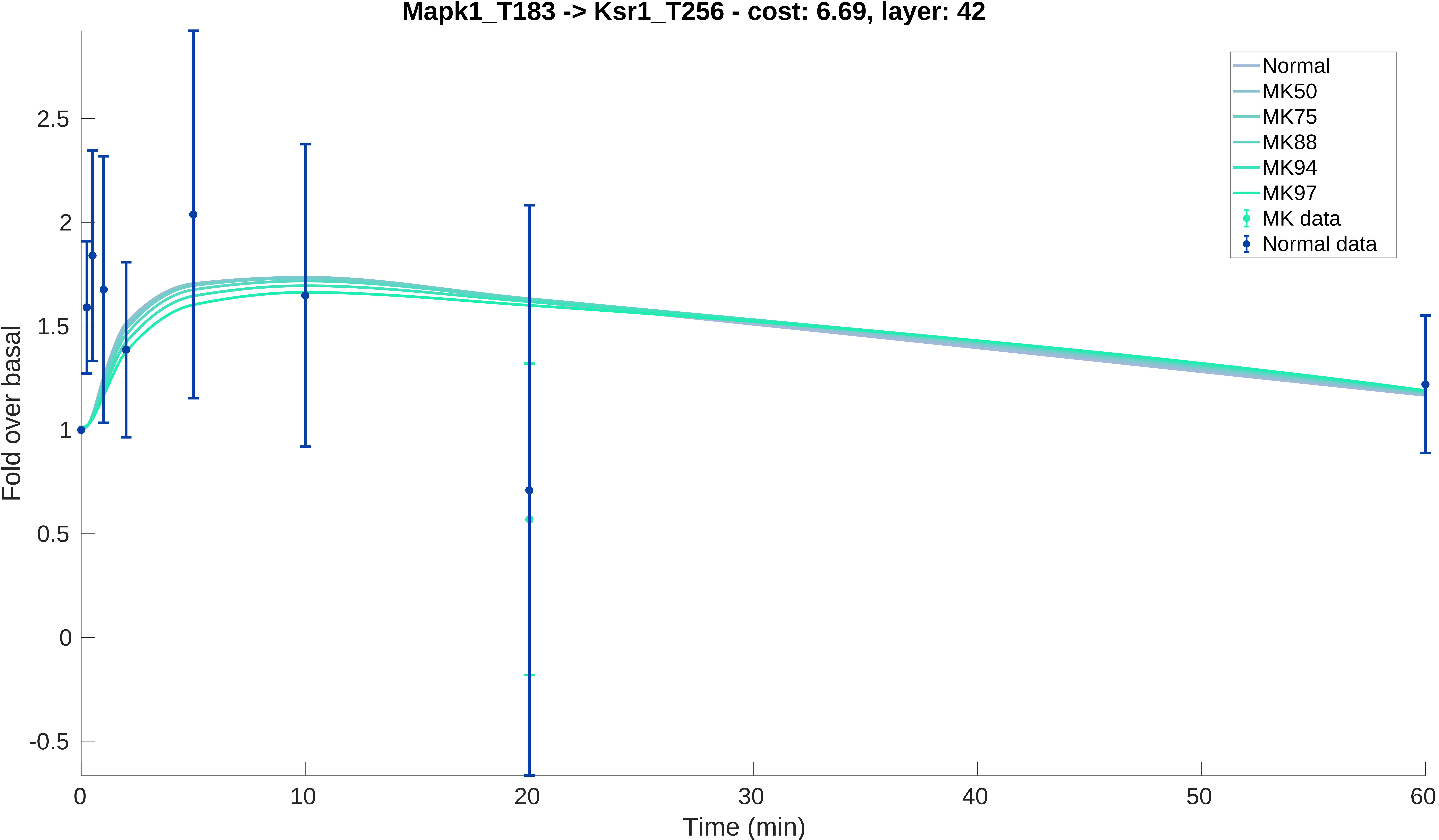

Mapk1_T183 -> Ksr1_T256 - cost: 6.69, layer: 42
Fold over basal
Time (min)
2.5
2
1.5
1
0.5
0
-0.5
0
10
20
30
40
50
60
Normal
MK50
MK75
MK88
MK94
MK97
MK data
Normal data

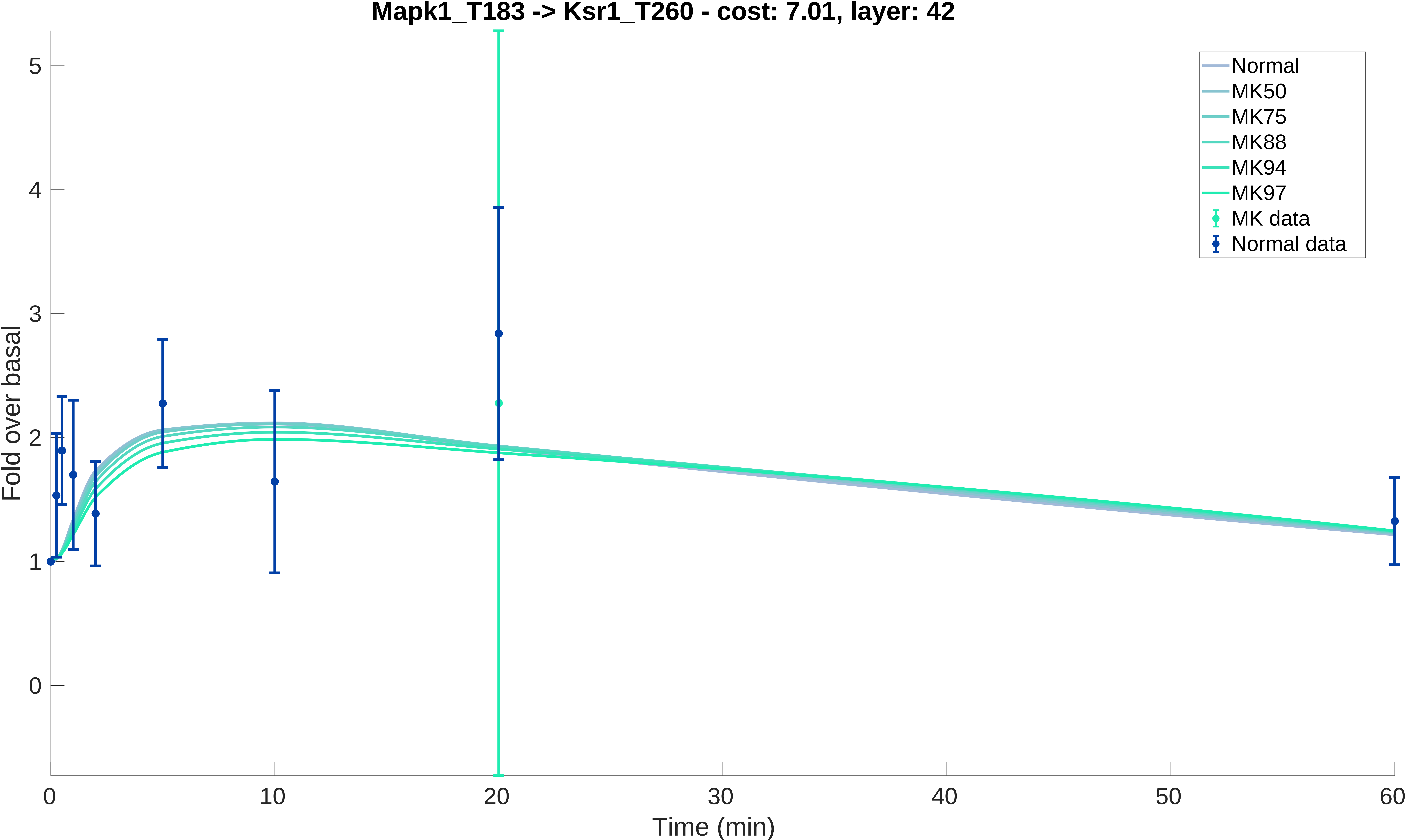

Mapk1_T183 -> Ksr1_T260 - cost: 7.01, layer: 42
Fold over basal
Time (min)
0
1
2
3
4
5
0
10
20
30
40
50
60
Normal
MK50
MK75
MK88
MK94
MK97
MK data
Normal data

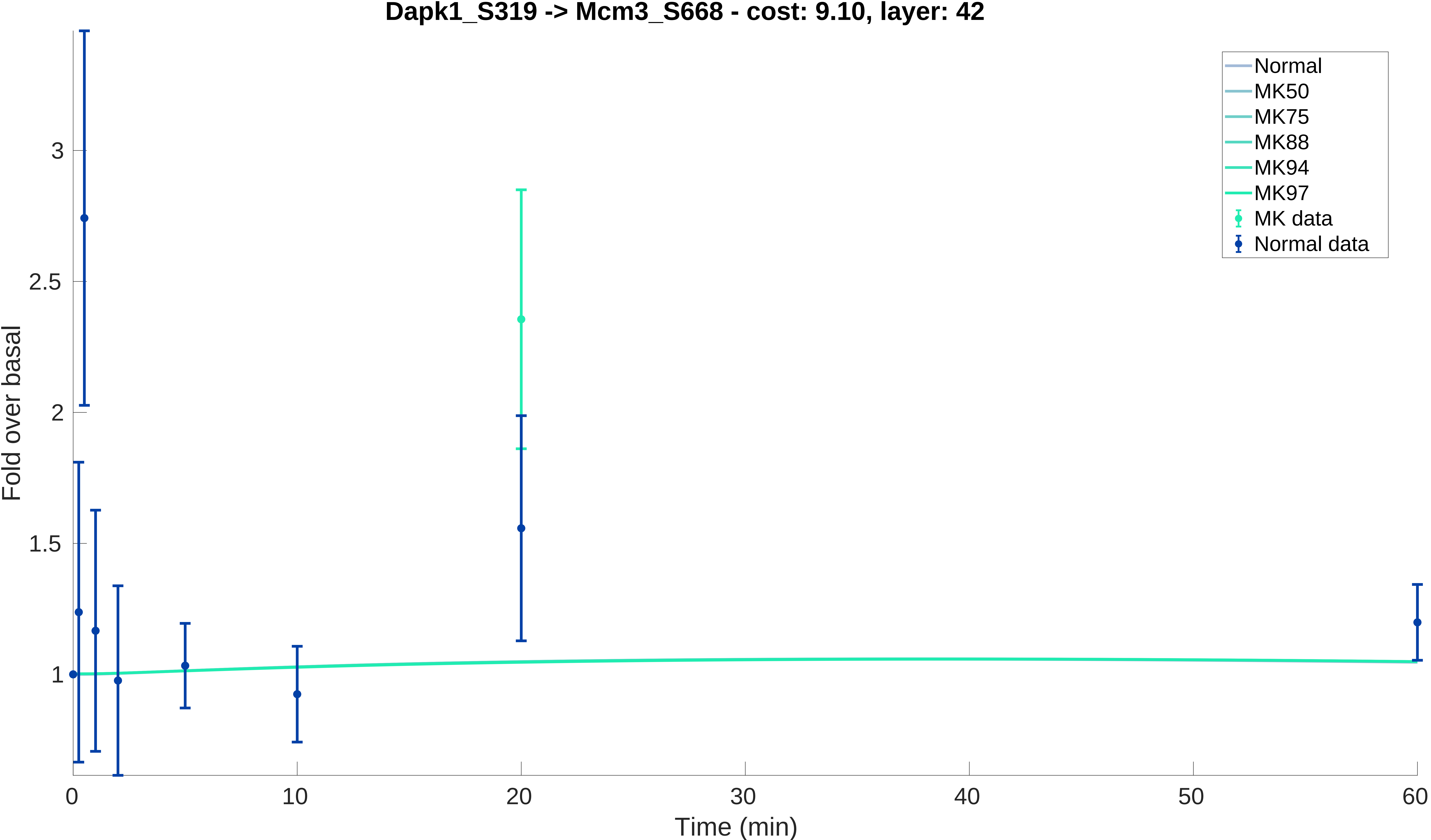

Dapk1_S319 -> Mcm3_S668 - cost: 9.10, layer: 42
Normal
MK50
MK75
MK88
MK94
MK97
MK data
Normal data
Fold over basal
3
2.5
2
1.5
1
0
10
20
30
40
50
60
Time (min)

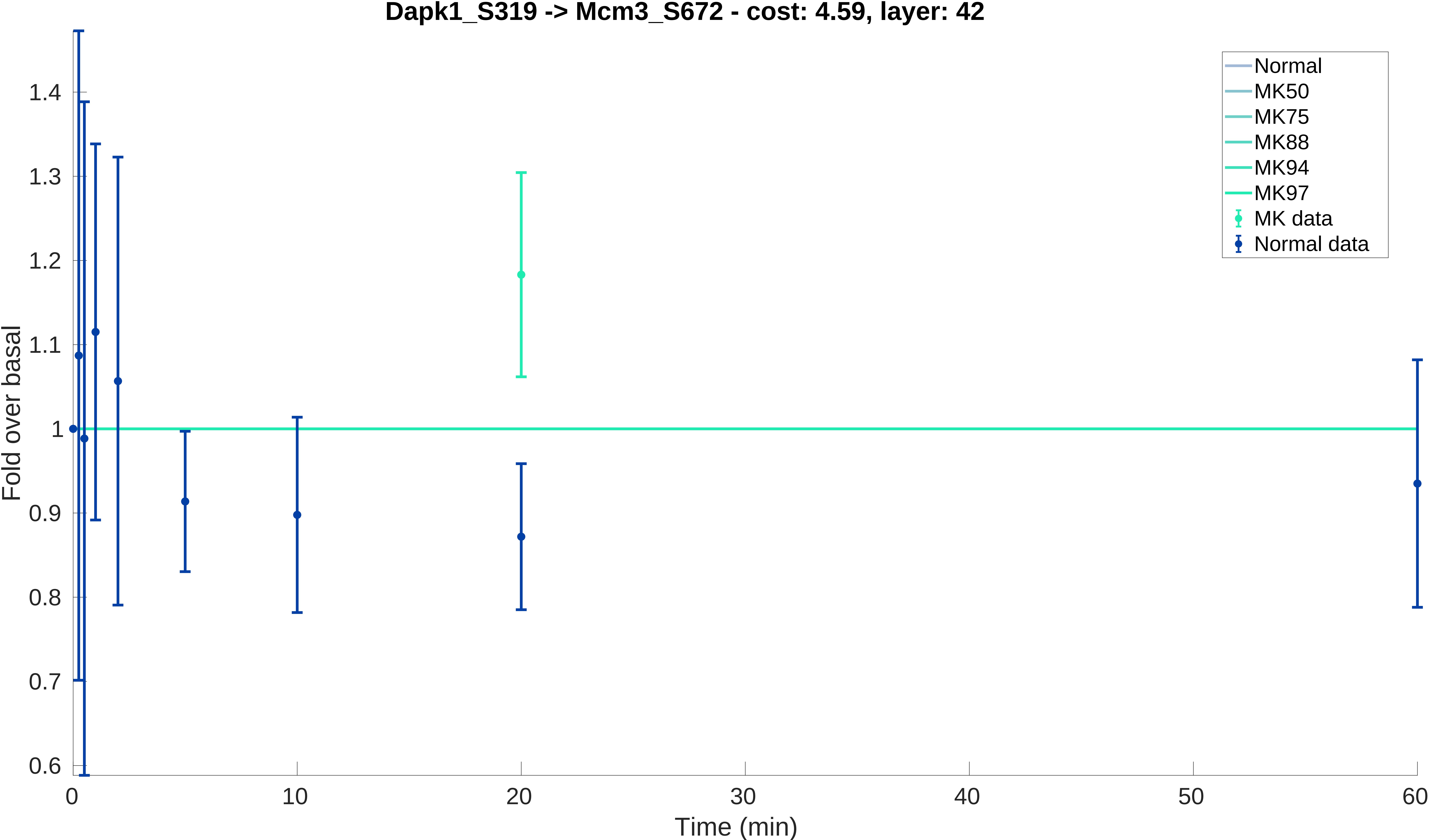

Dapk1_S319 -> Mcm3_S672 - cost: 4.59, layer: 42
Normal
MK50
MK75
MK88
MK94
MK97
MK data
Normal data
Fold over basal
1.4
1.3
1.2
1.1
1
0.9
0.8
0.7
0.6
0
10
20
30
40
50
60
Time (min)

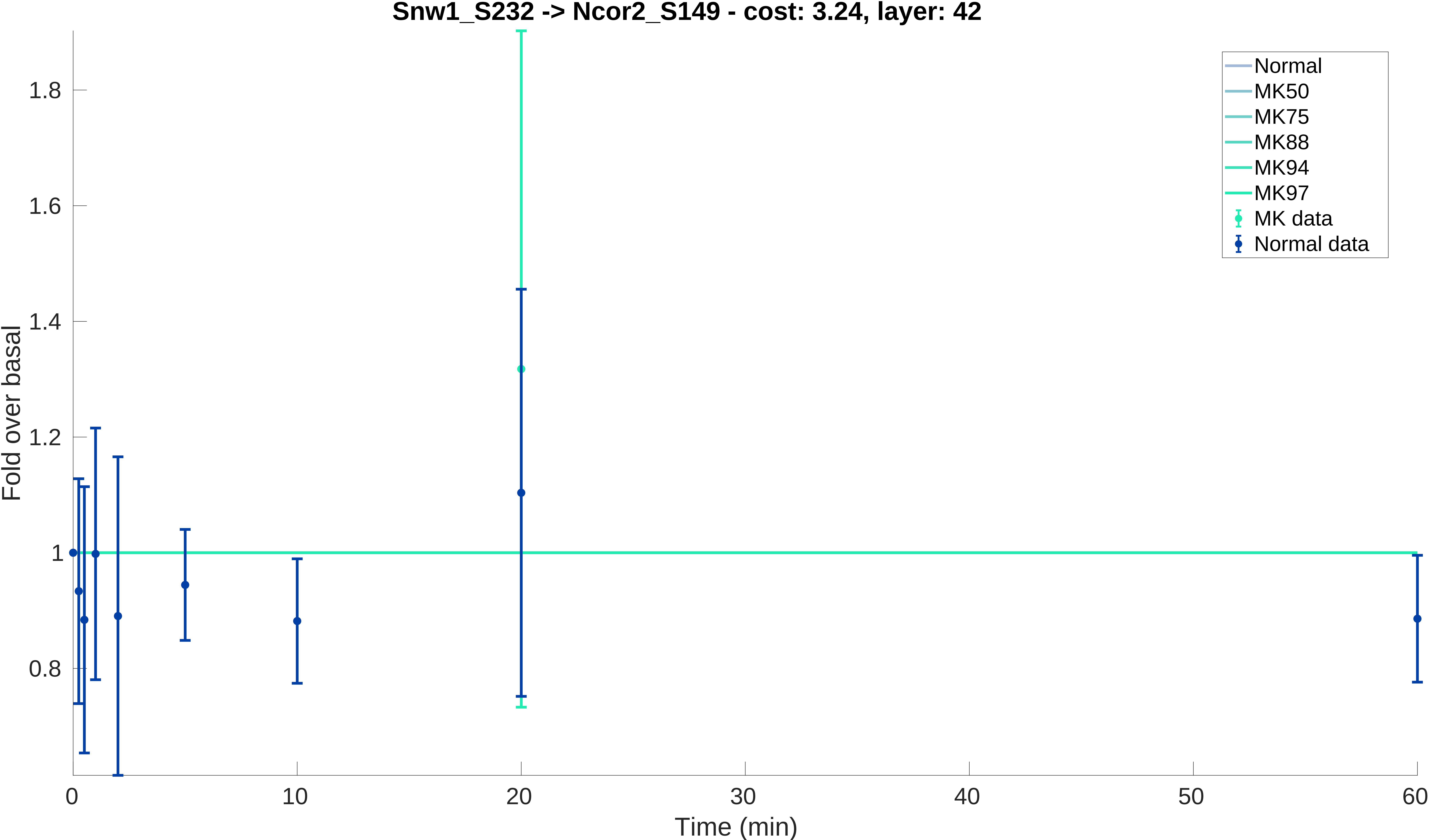

Snw1_S232 -> Ncor2_S149 - cost: 3.24, layer: 42
Fold over basal
1.8
1.6
1.4
1.2
1
0.8
0
10
20
30
40
50
60
Time (min)
Normal
MK50
MK75
MK88
MK94
MK97
MK data
Normal data

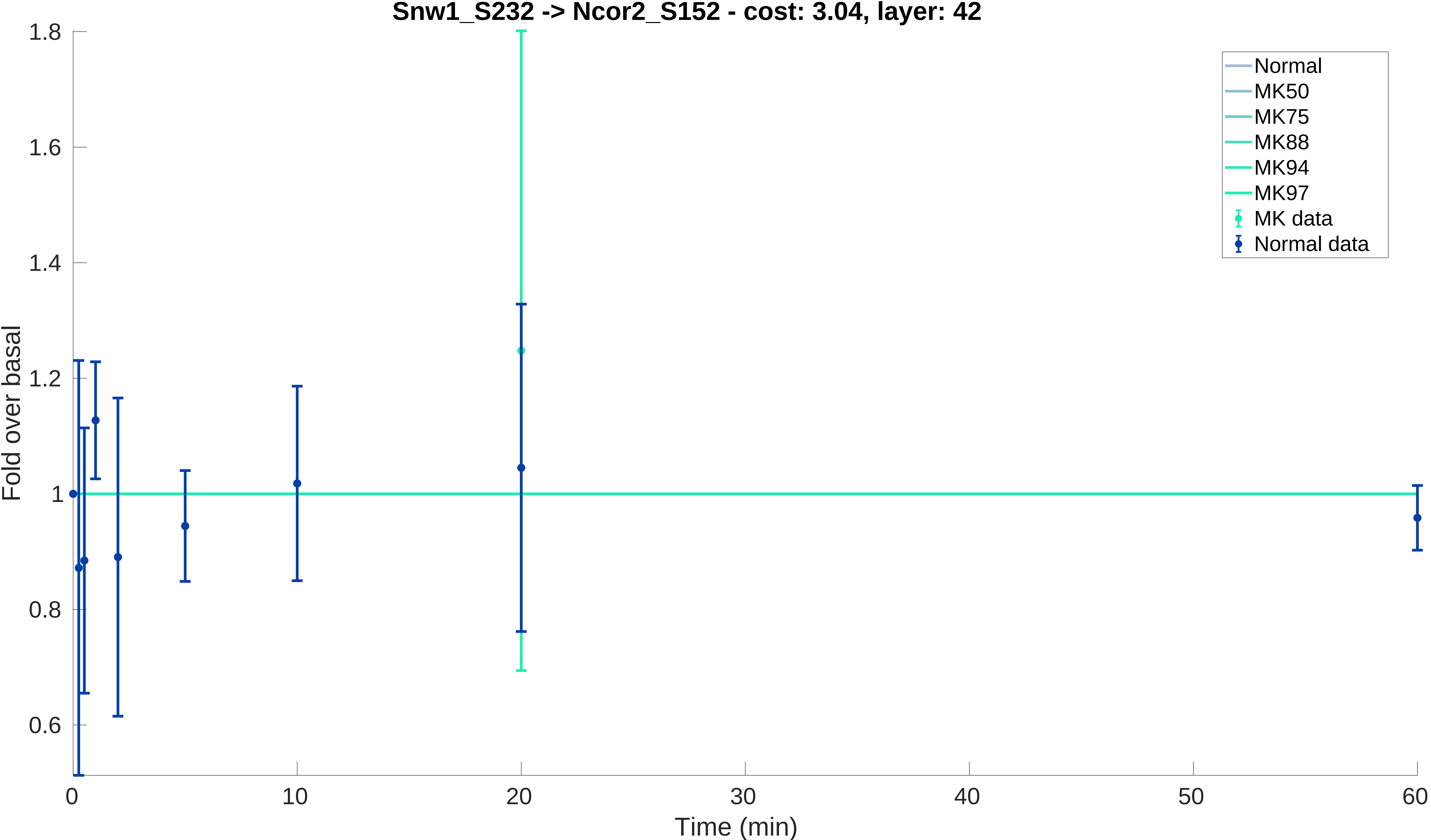

Snw1_S232 -> Ncor2_S152 - cost: 3.04, layer: 42
Fold over basal
Time (min)
1.8
1.6
1.4
1.2
1
0.8
0.6
0
10
20
30
40
50
60
Normal
MK50
MK75
MK88
MK94
MK97
MK data
Normal data

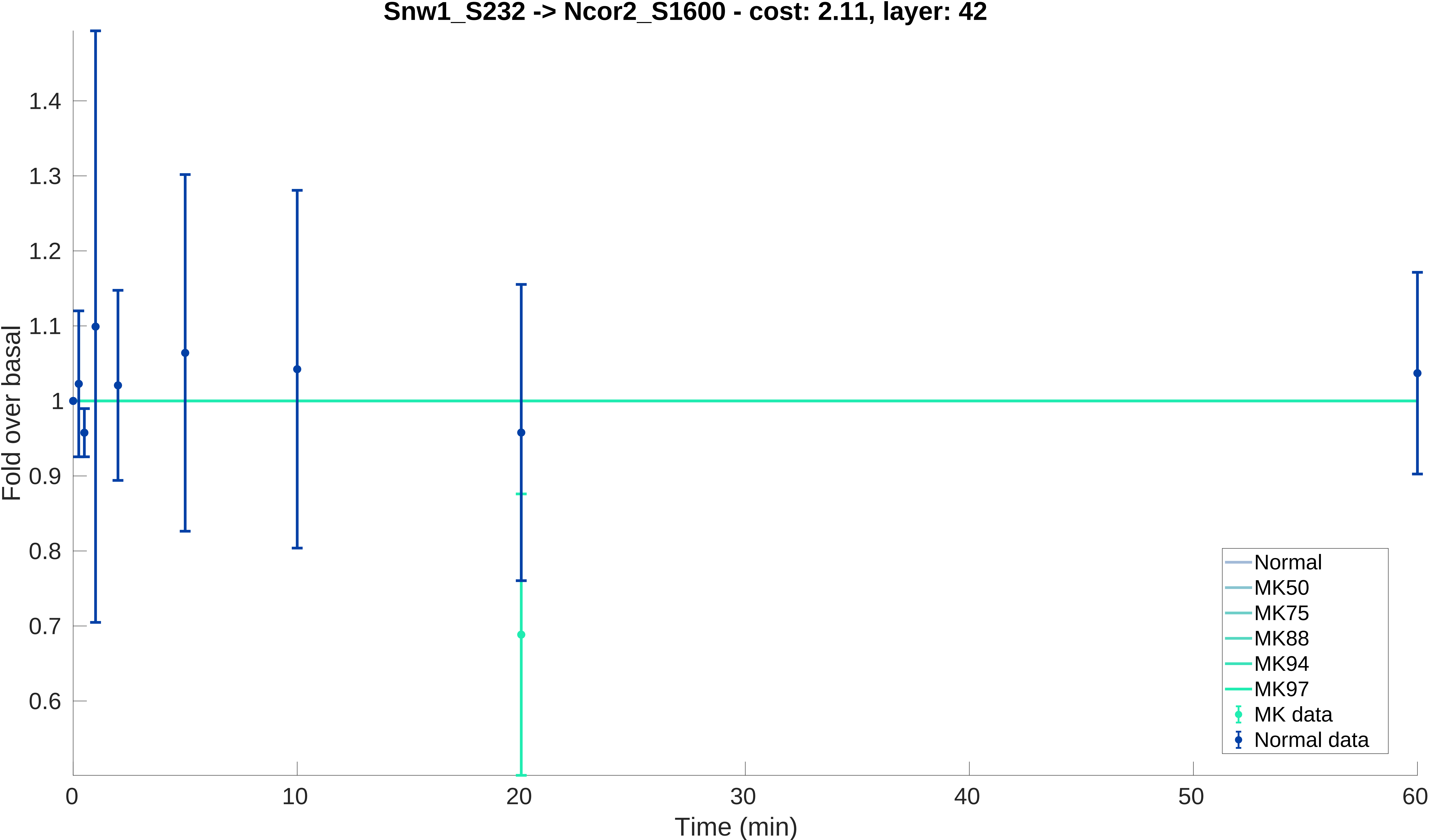

Snw1_S232 -> Ncor2_S1600 - cost: 2.11, layer: 42
Fold over basal
Time (min)
1.4
1.3
1.2
1.1
1
0.9
0.8
0.7
0.6
0
10
20
30
40
50
60
Normal
MK50
MK75
MK88
MK94
MK97
MK data
Normal data

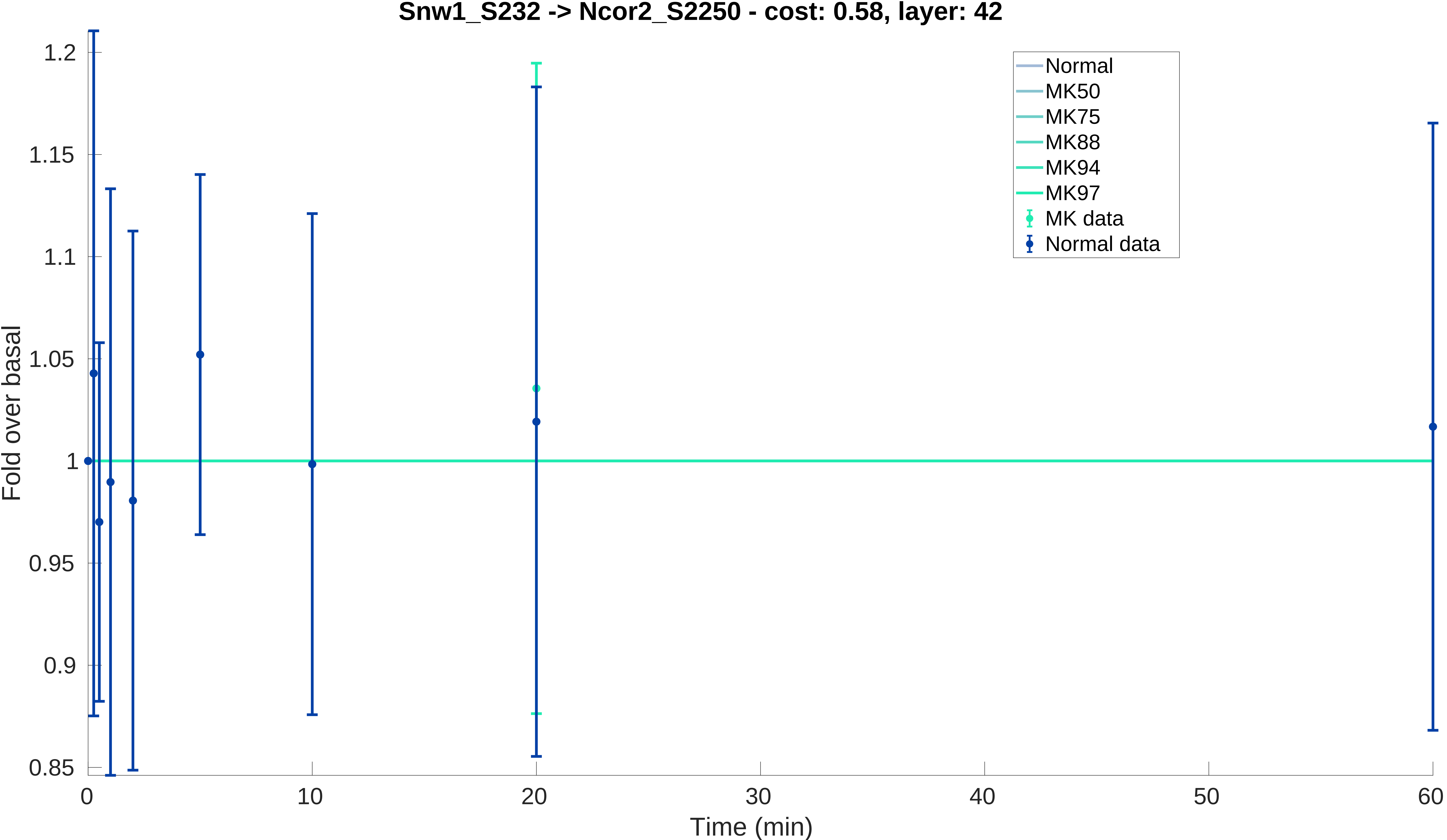

Snw1_S232 -> Ncor2_S2250 - cost: 0.58, layer: 42
Normal
MK50
MK75
MK88
MK94
MK97
MK data
Normal data
Fold over basal
1.2
1.15
1.1
1.05
1
0.95
0.9
0.85
Time (min)
0
10
20
30
40
50
60

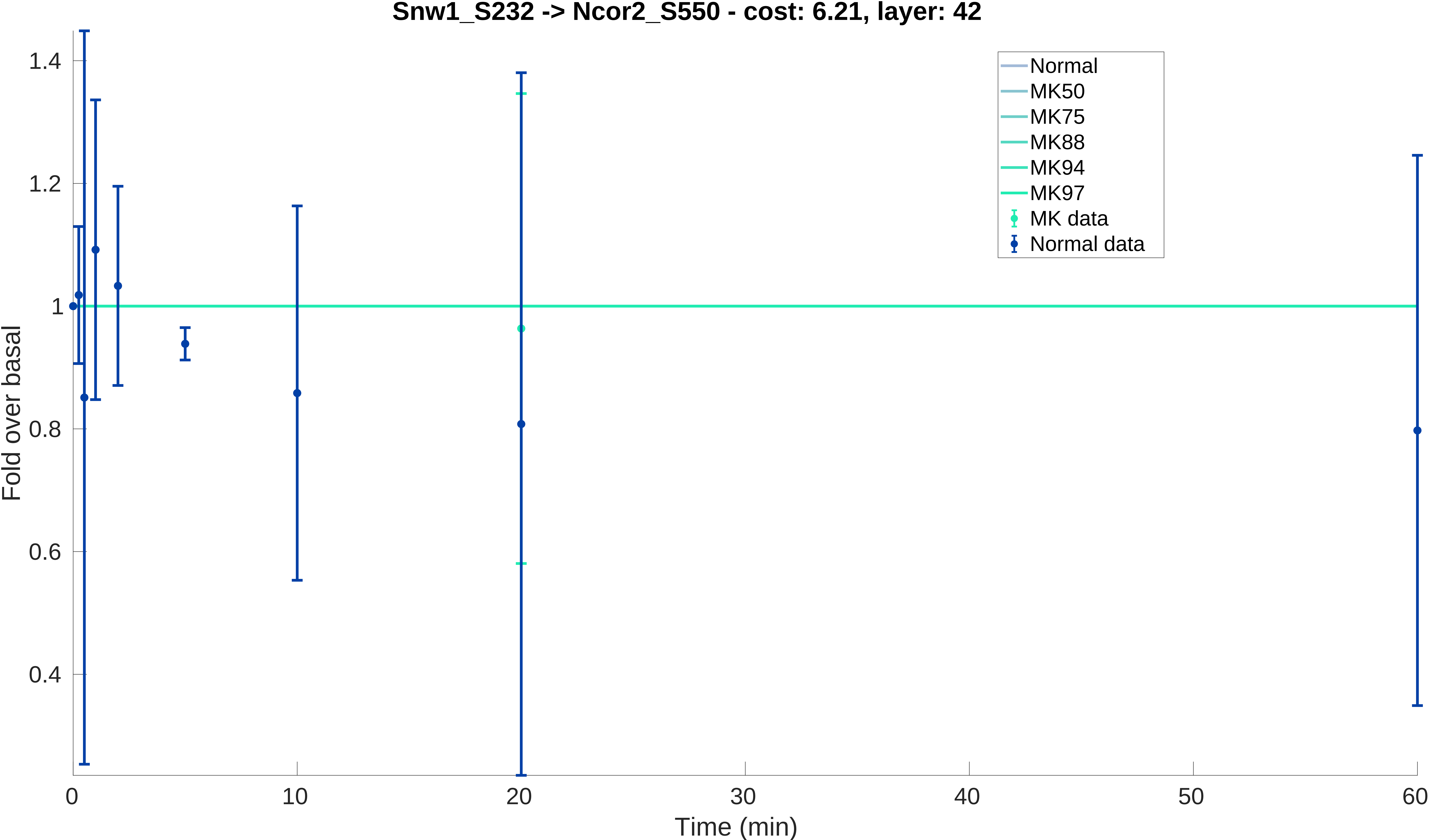
Snw1_S232 -> Ncor2_S550 - cost: 6.21, layer: 42
Fold over basal
Time (min)
0
10
20
30
40
50
60
0.4
0.6
0.8
1
1.2
1.4
Normal
MK50
MK75
MK88
MK94
MK97
MK data
Normal data

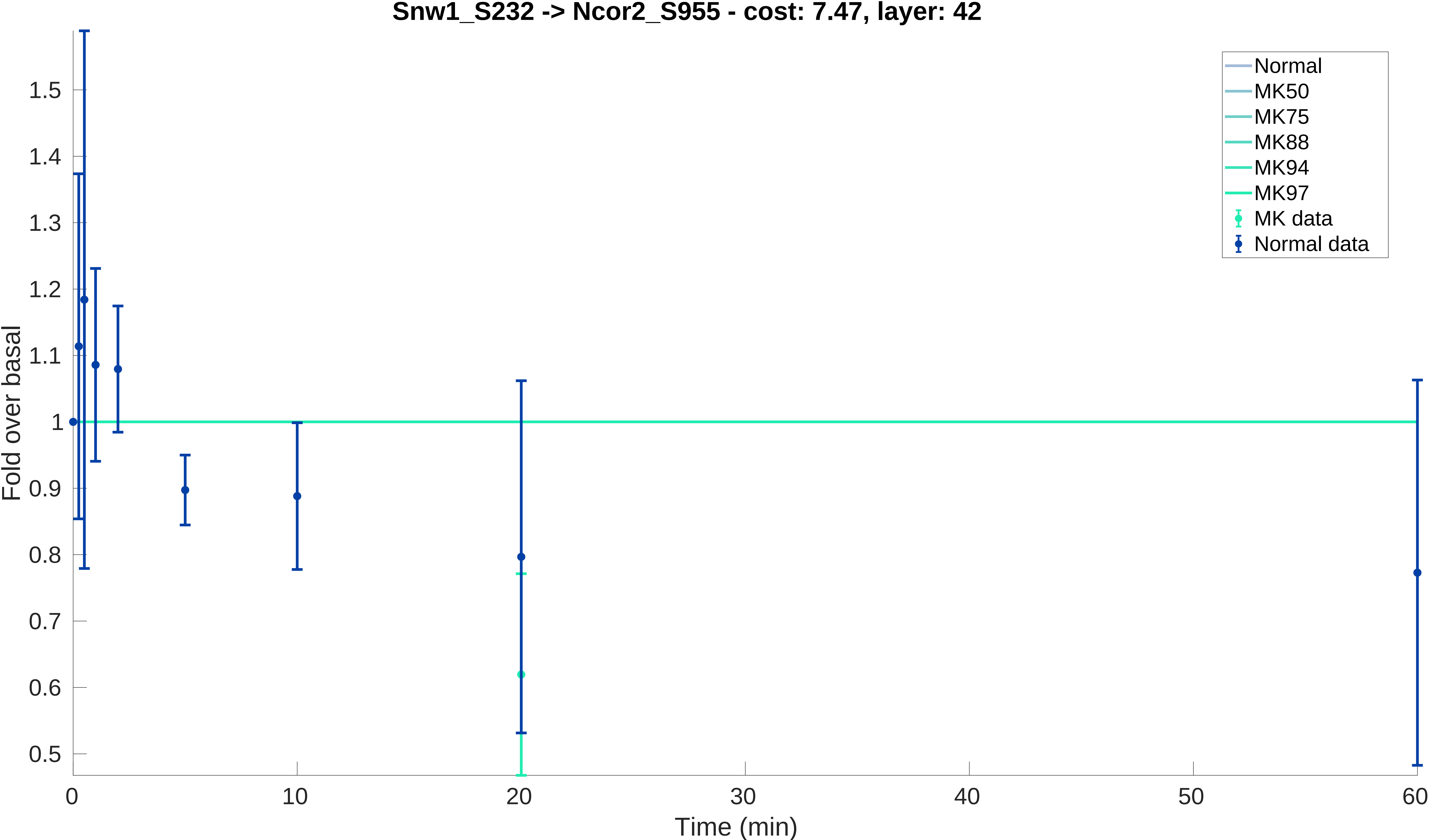

Snw1_S232 -> Ncor2_S955 - cost: 7.47, layer: 42
Fold over basal
Time (min)
0.5
0.6
0.7
0.8
0.9
1
1.1
1.2
1.3
1.4
1.5
0
10
20
30
40
50
60
Normal
MK50
MK75
MK88
MK94
MK97
MK data
Normal data

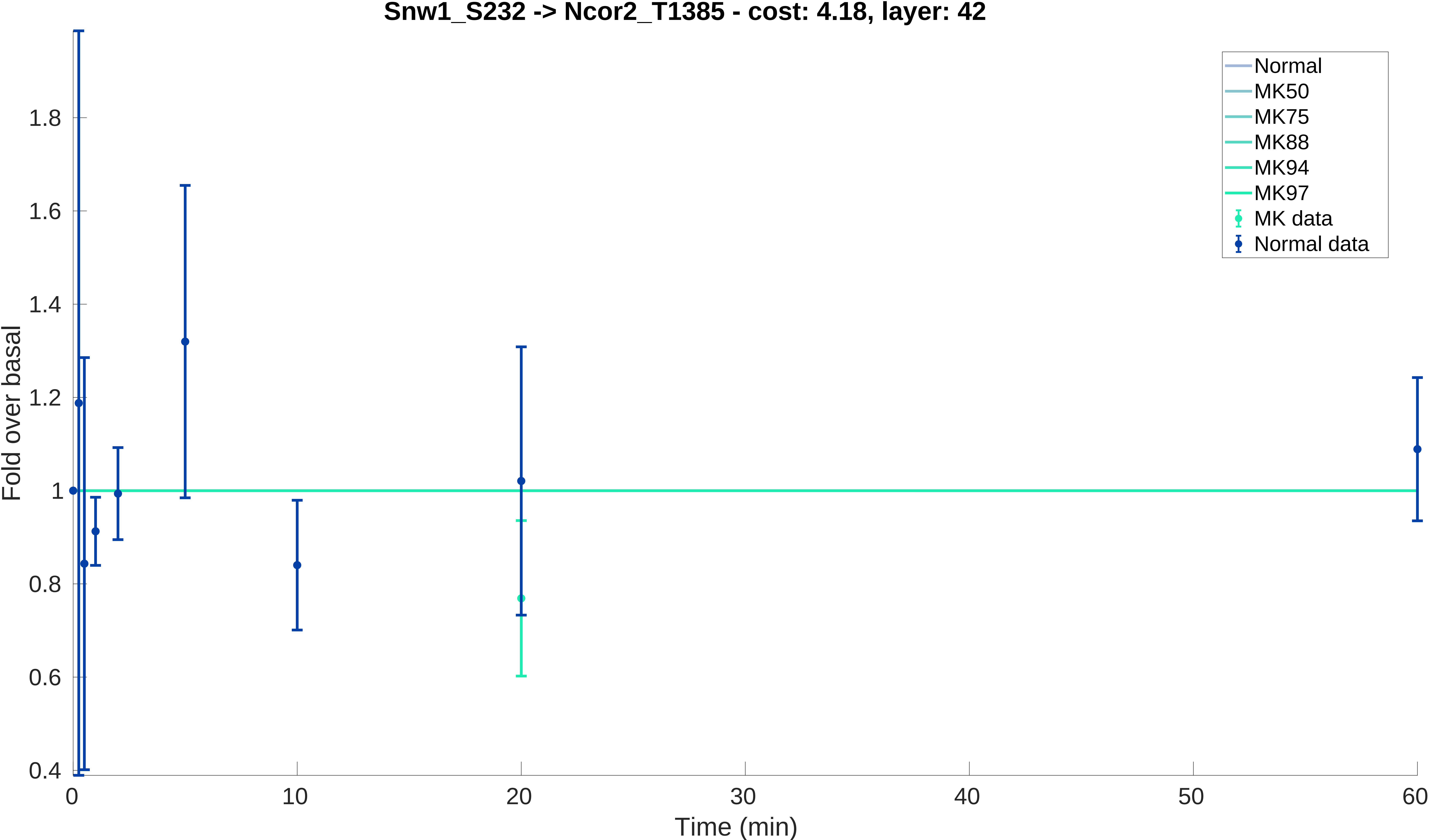

Snw1_S232 -> Ncor2_T1385 - cost: 4.18, layer: 42
Normal
MK50
MK75
MK88
MK94
MK97
MK data
Normal data
Fold over basal
Time (min)
0.4
0.6
0.8
1
1.2
1.4
1.6
1.8
0
10
20
30
40
50
60

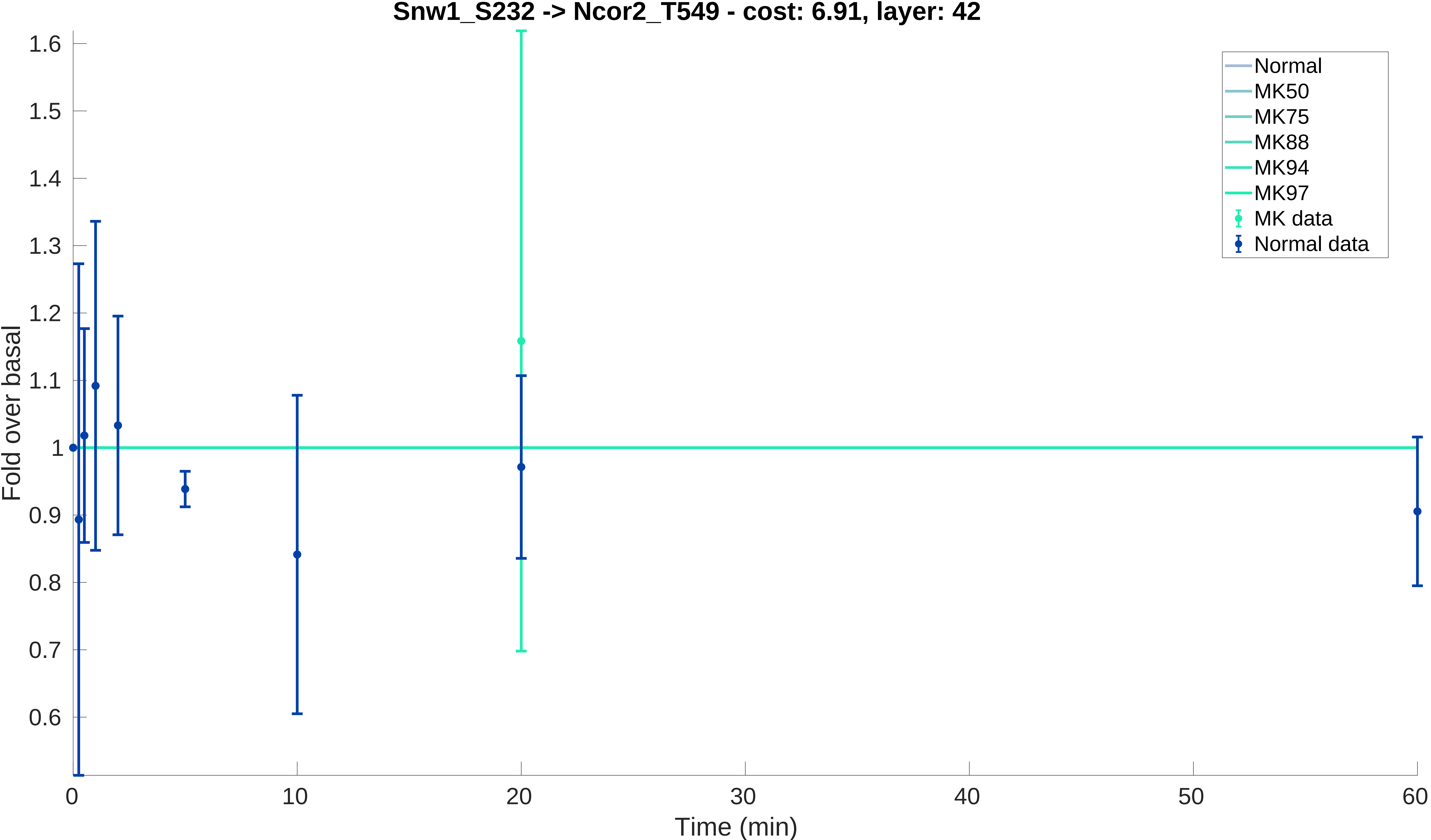

Snw1_S232 -> Ncor2_T549 - cost: 6.91, layer: 42
Fold over basal
Time (min)
0
10
20
30
40
50
60
0.6
0.7
0.8
0.9
1
1.1
1.2
1.3
1.4
1.5
1.6
Normal
MK50
MK75
MK88
MK94
MK97
MK data
Normal data

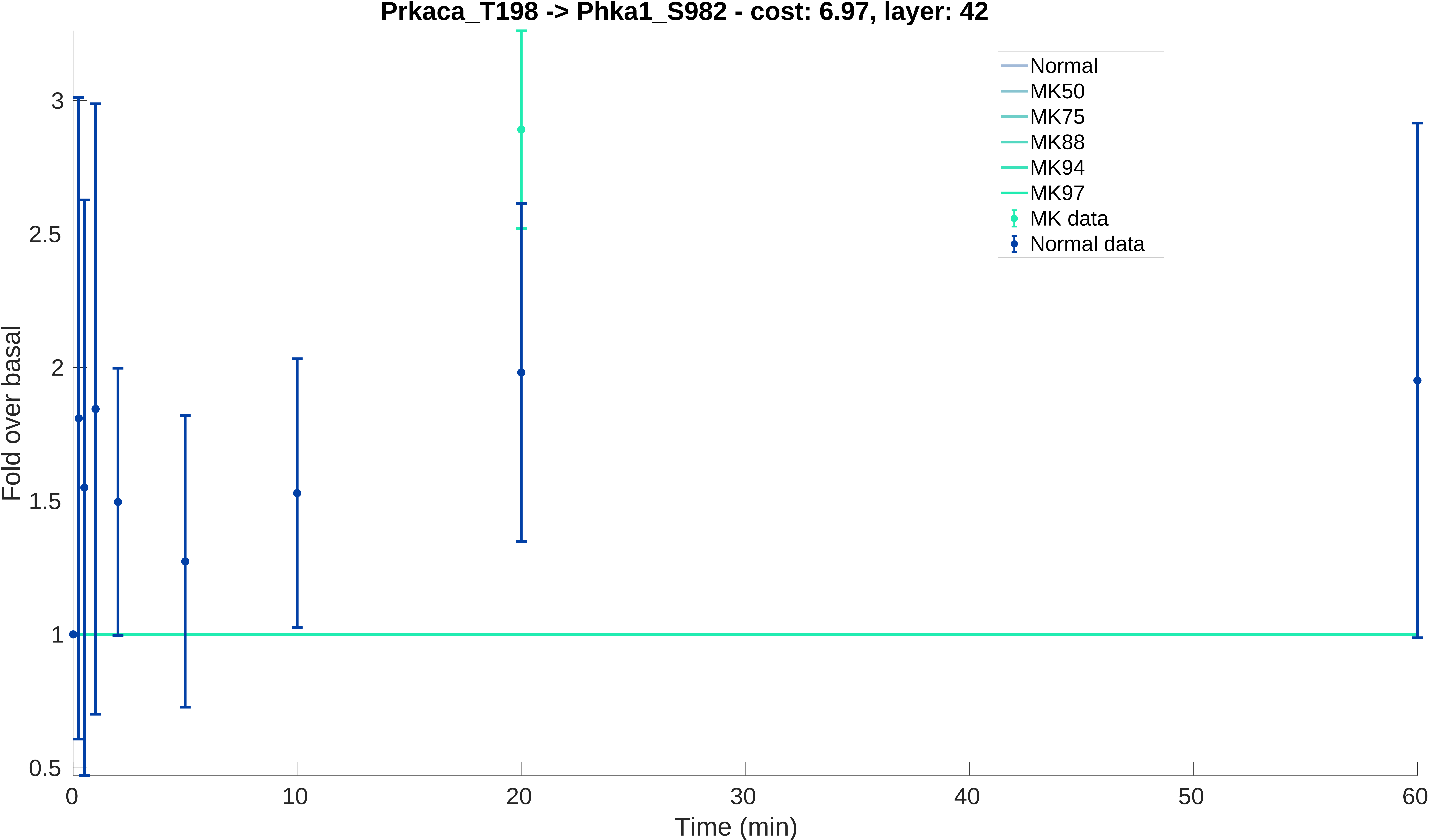

Prkaca_T198 -> Phka1_S982 - cost: 6.97, layer: 42
Normal
MK50
MK75
MK88
MK94
MK97
MK data
Normal data
Fold over basal
Time (min)
0.5
1
1.5
2
2.5
3
0
10
20
30
40
50
60

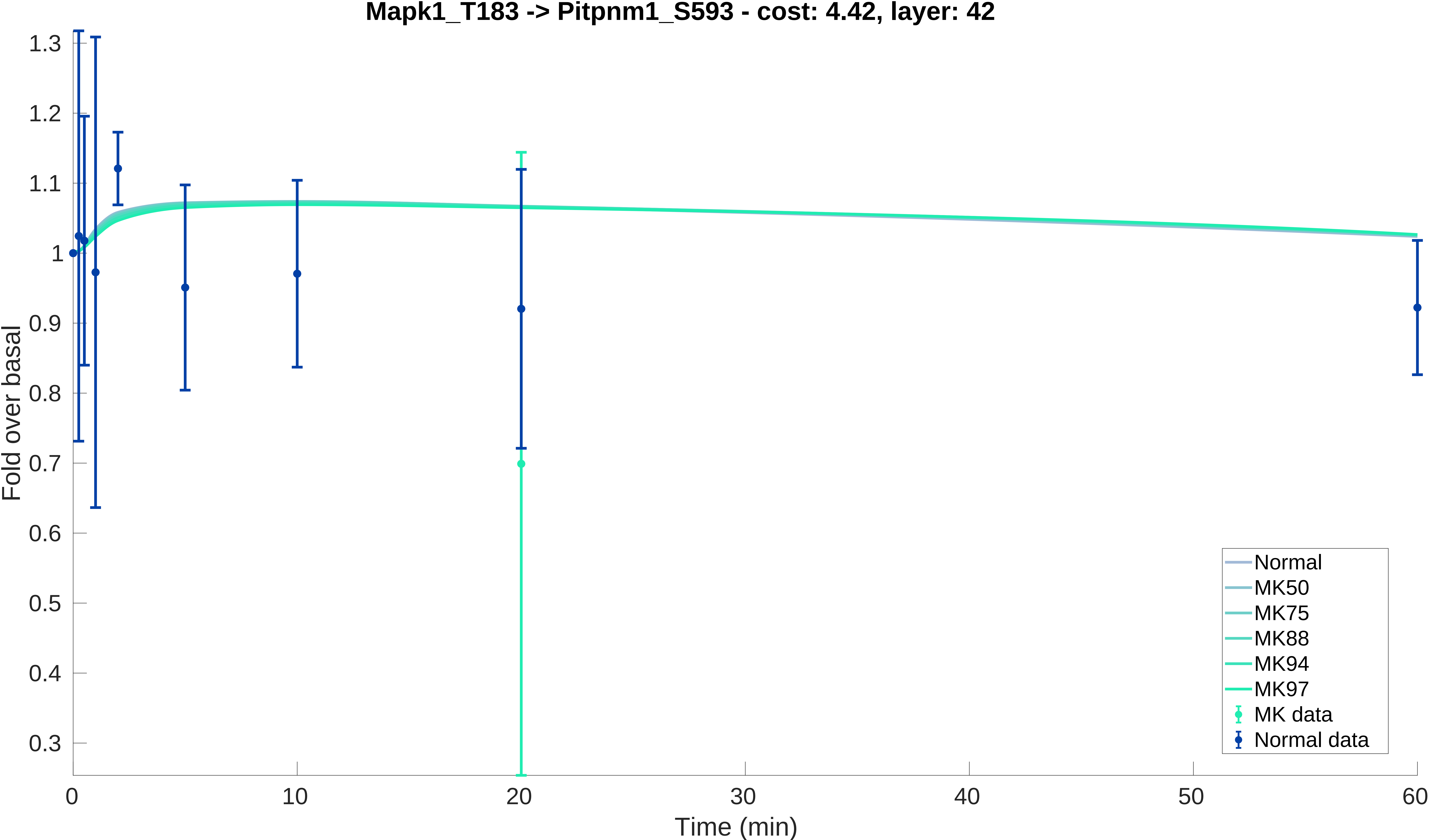

Mapk1_T183 -> Pitpnm1_S593 - cost: 4.42, layer: 42
Fold over basal
1.3
1.2
1.1
1
0.9
0.8
0.7
0.6
0.5
0.4
0.3
0
10
20
30
40
50
60
Time (min)
Normal
MK50
MK75
MK88
MK94
MK97
MK data
Normal data

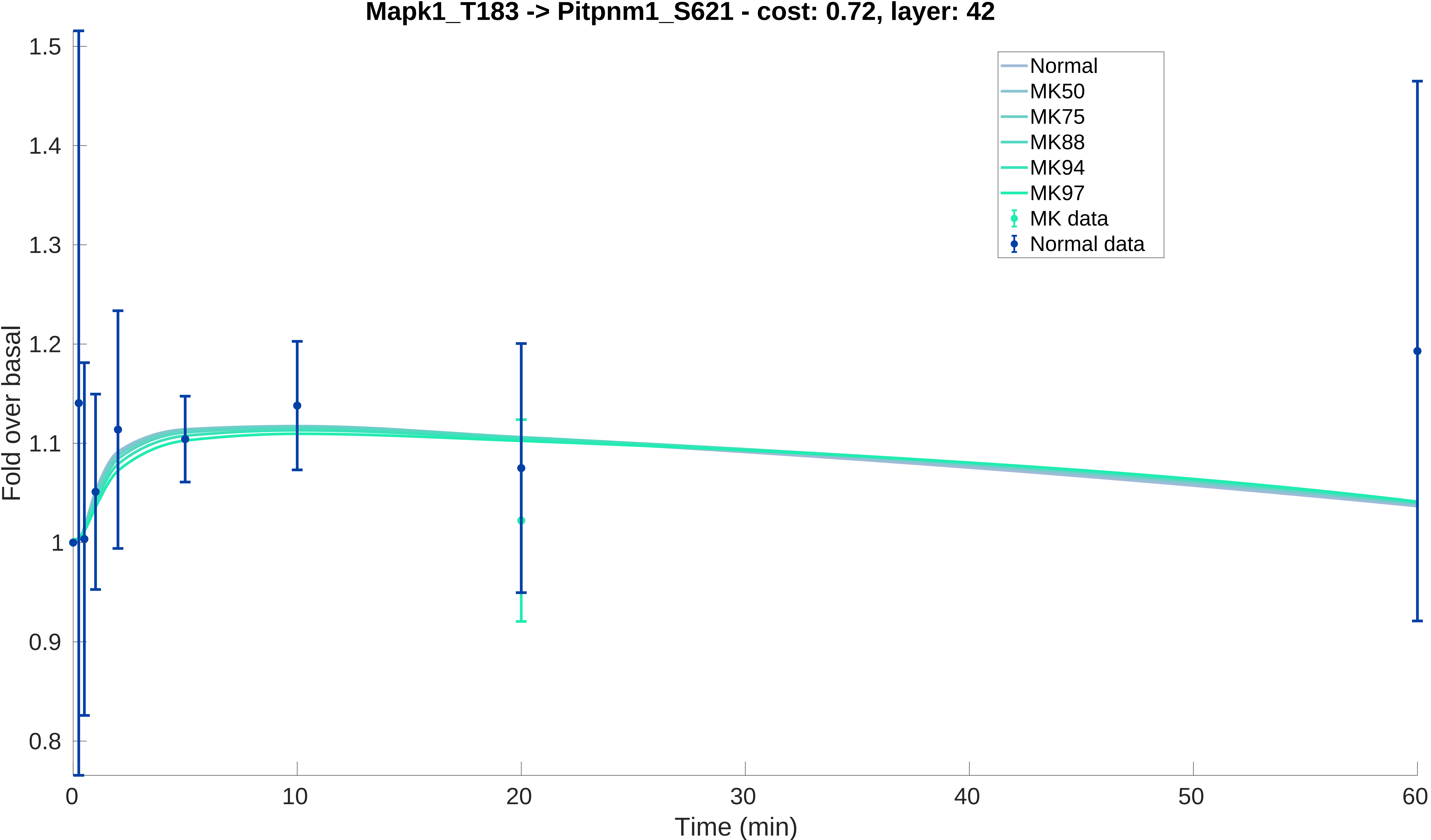

Mapk1_T183 -> Pitpnm1_S621 - cost: 0.72, layer: 42
Fold over basal
Time (min)
1.5
1.4
1.3
1.2
1.1
1
0.9
0.8
0
10
20
30
40
50
60
Normal
MK50
MK75
MK88
MK94
MK97
MK data
Normal data

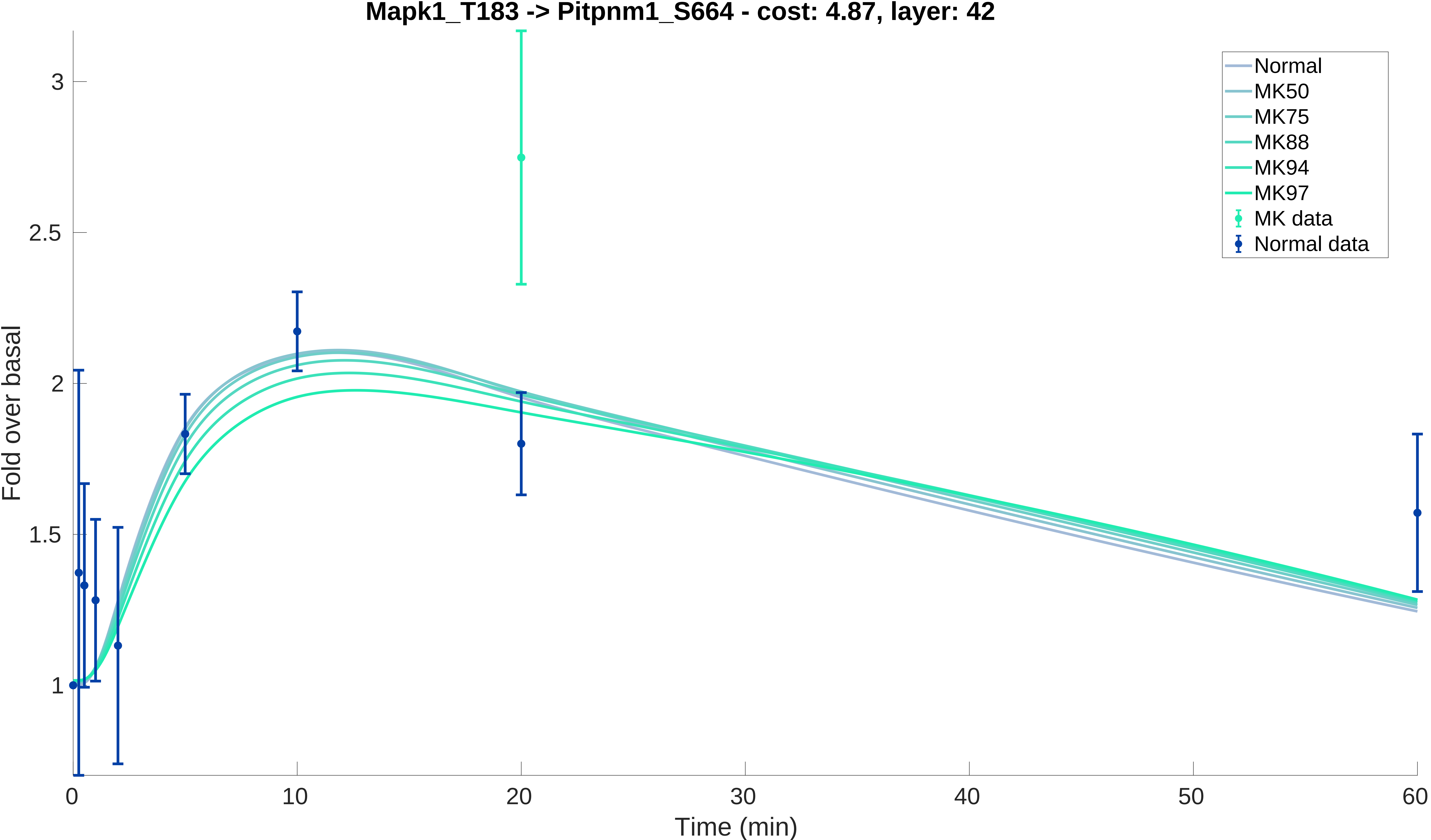

Mapk1_T183 -> Pitpnm1_S664 - cost: 4.87, layer: 42
Fold over basal
Time (min)
Normal
MK50
MK75
MK88
MK94
MK97
MK data
Normal data
3
2.5
2
1.5
1
0
10
20
30
40
50
60

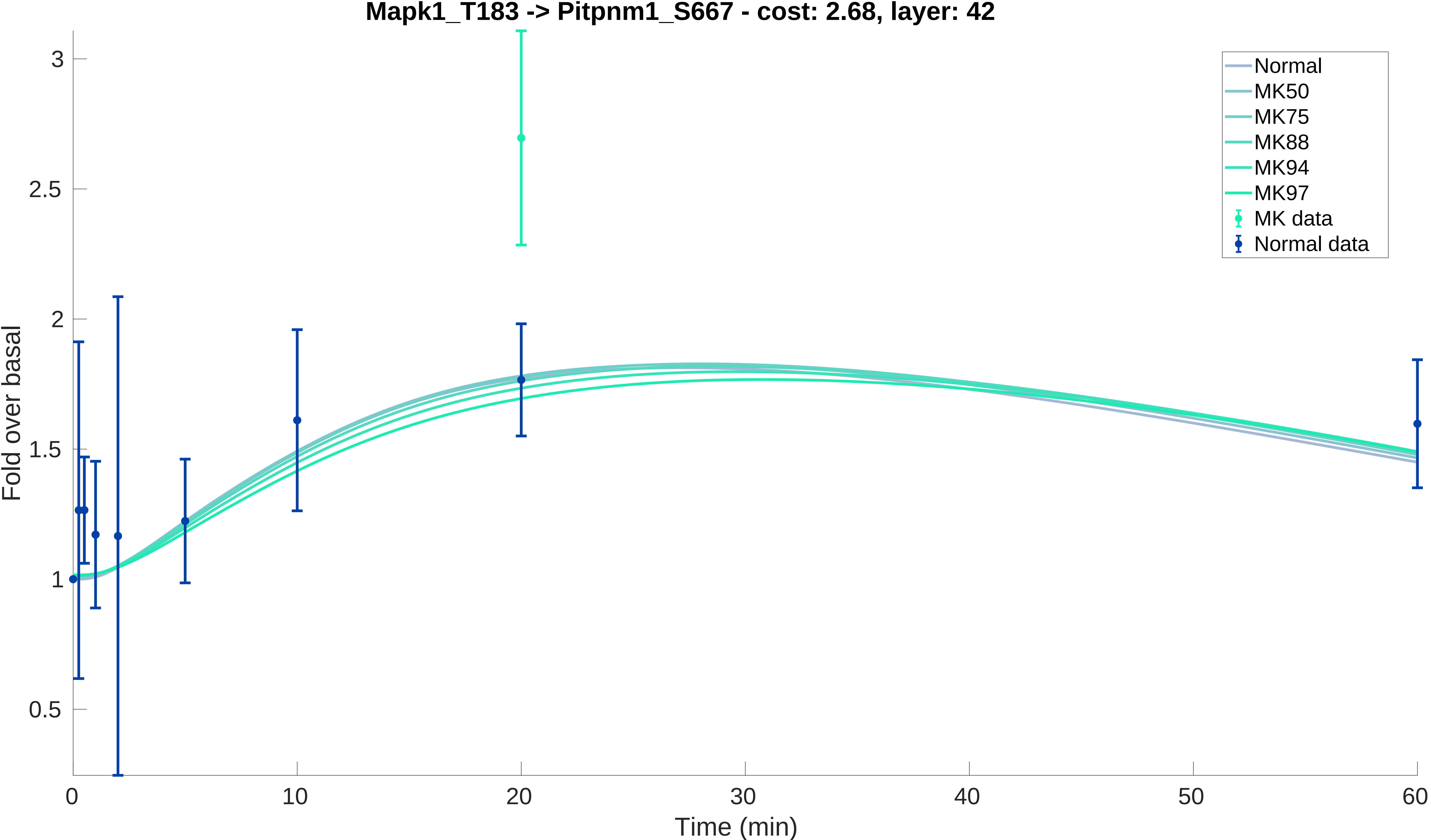

Mapk1_T183 -> Pitpnm1_S667 - cost: 2.68, layer: 42
Normal
MK50
MK75
MK88
MK94
MK97
MK data
Normal data
Fold over basal
Time (min)
3
2.5
2
1.5
1
0.5
0
10
20
30
40
50
60

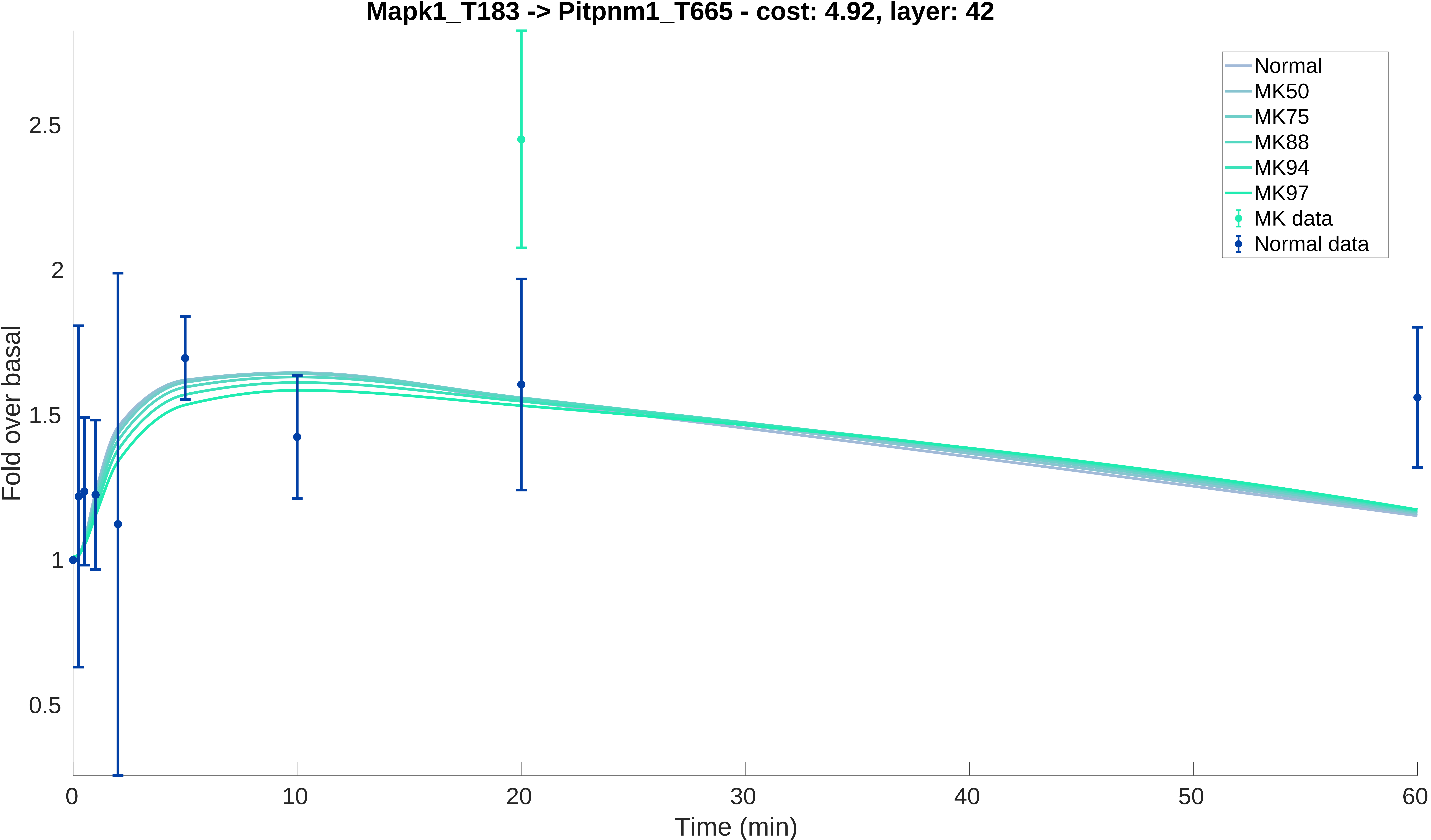
Mapk1_T183 -> Pitpnm1_T665 - cost: 4.92, layer: 42
Fold over basal
Time (min)
0
10
20
30
40
50
60
0.5
1
1.5
2
2.5
Normal
MK50
MK75
MK88
MK94
MK97
MK data
Normal data

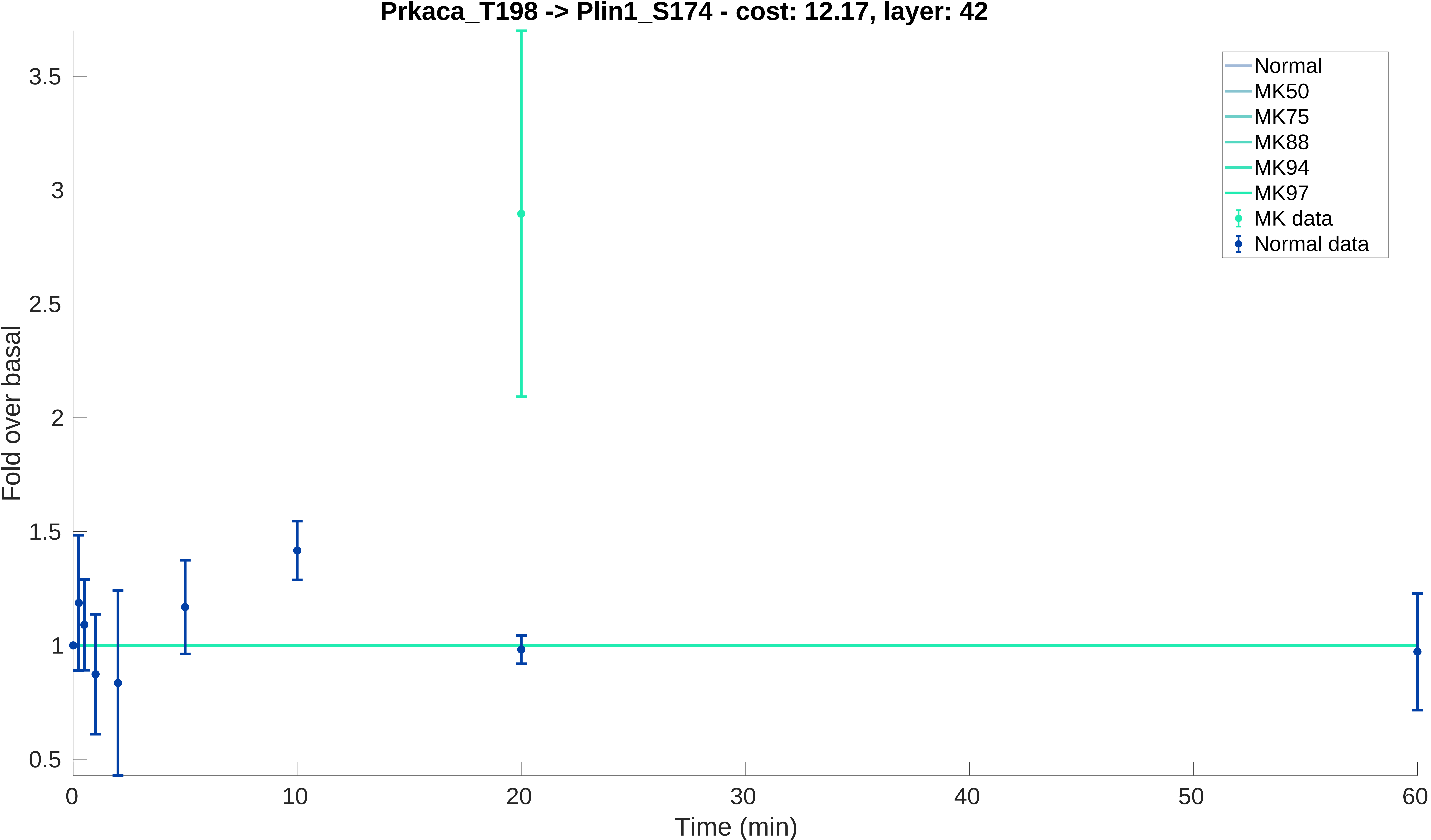

Prrkaca_T198 -> Plin1_S174 - cost: 12.17, layer: 42
Fold over basal
Time (min)
3.5
3
2.5
2
1.5
1
0.5
0
10
20
30
40
50
60
Normal
MK50
MK75
MK88
MK94
MK97
MK data
Normal data

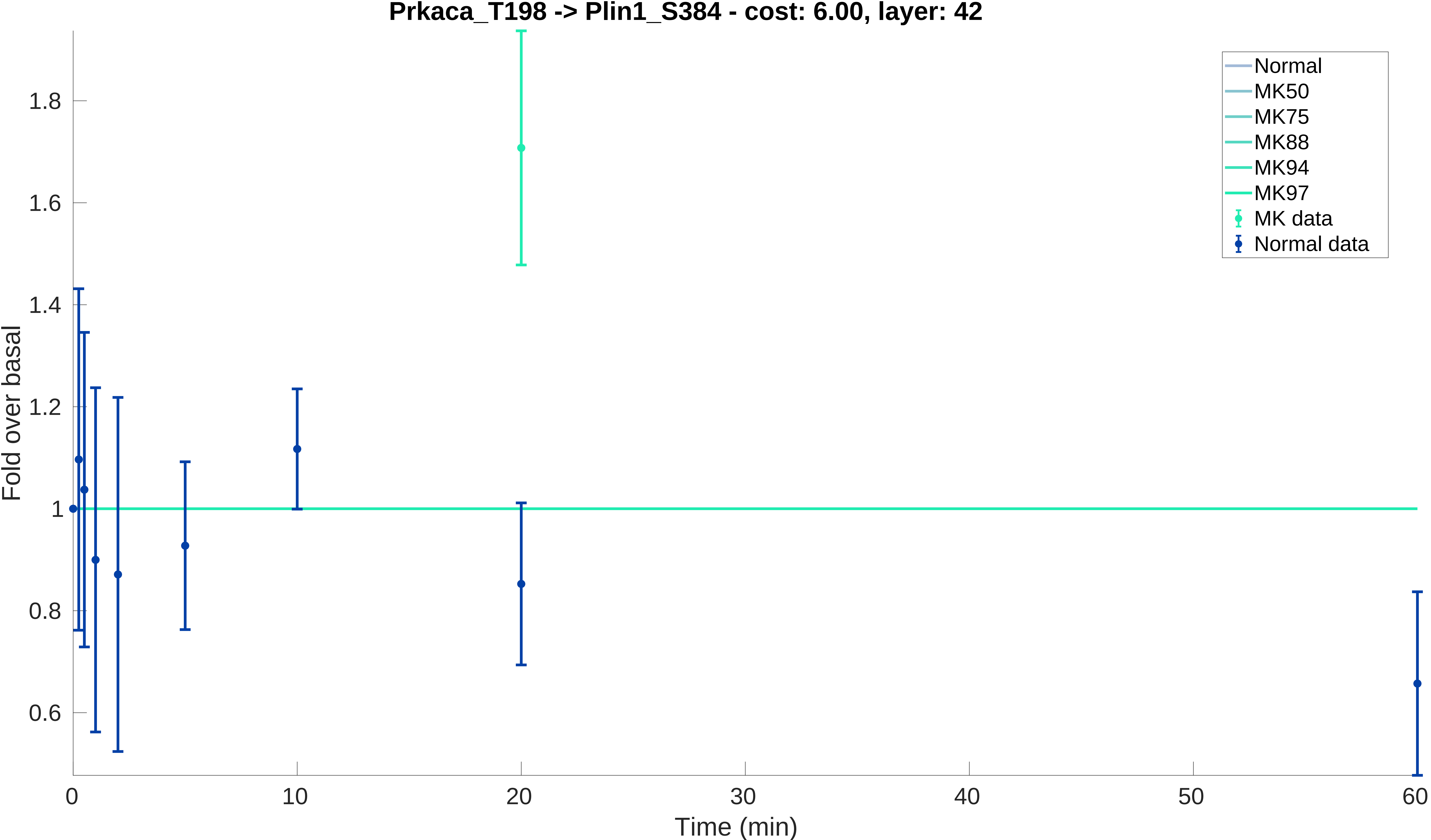

Prkaca_T198 -> Plin1_S384 - cost: 6.00, layer: 42
Normal
MK50
MK75
MK88
MK94
MK97
MK data
Normal data
Fold over basal
1.8
1.6
1.4
1.2
1
0.8
0.6
0
10
20
30
40
50
60
Time (min)

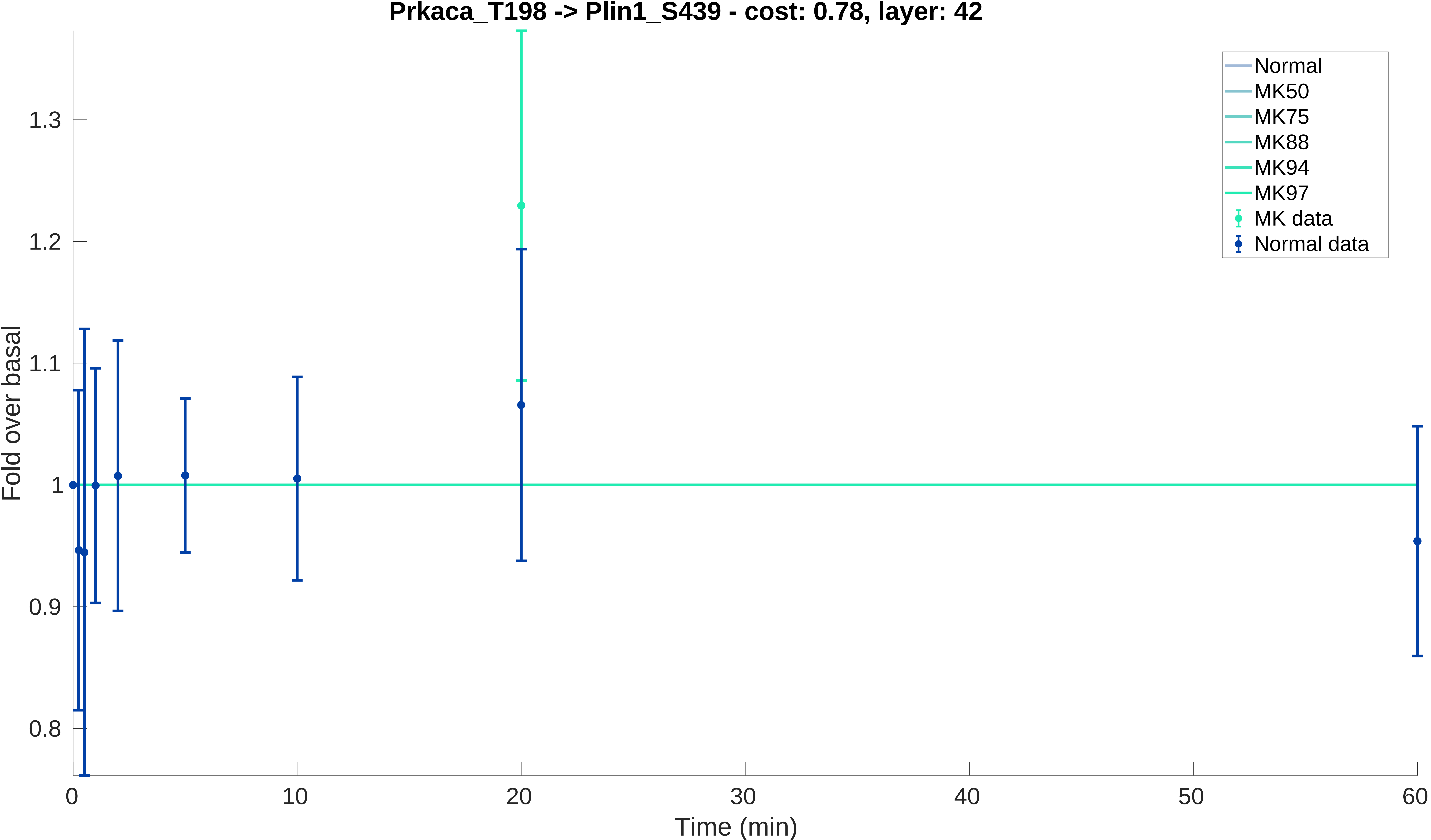

Prkaca_T198 -> Plin1_S439 - cost: 0.78, layer: 42
Fold over basal
Time (min)
0.8
0.9
1
1.1
1.2
1.3
0
10
20
30
40
50
60
Normal
MK50
MK75
MK88
MK94
MK97
MK data
Normal data

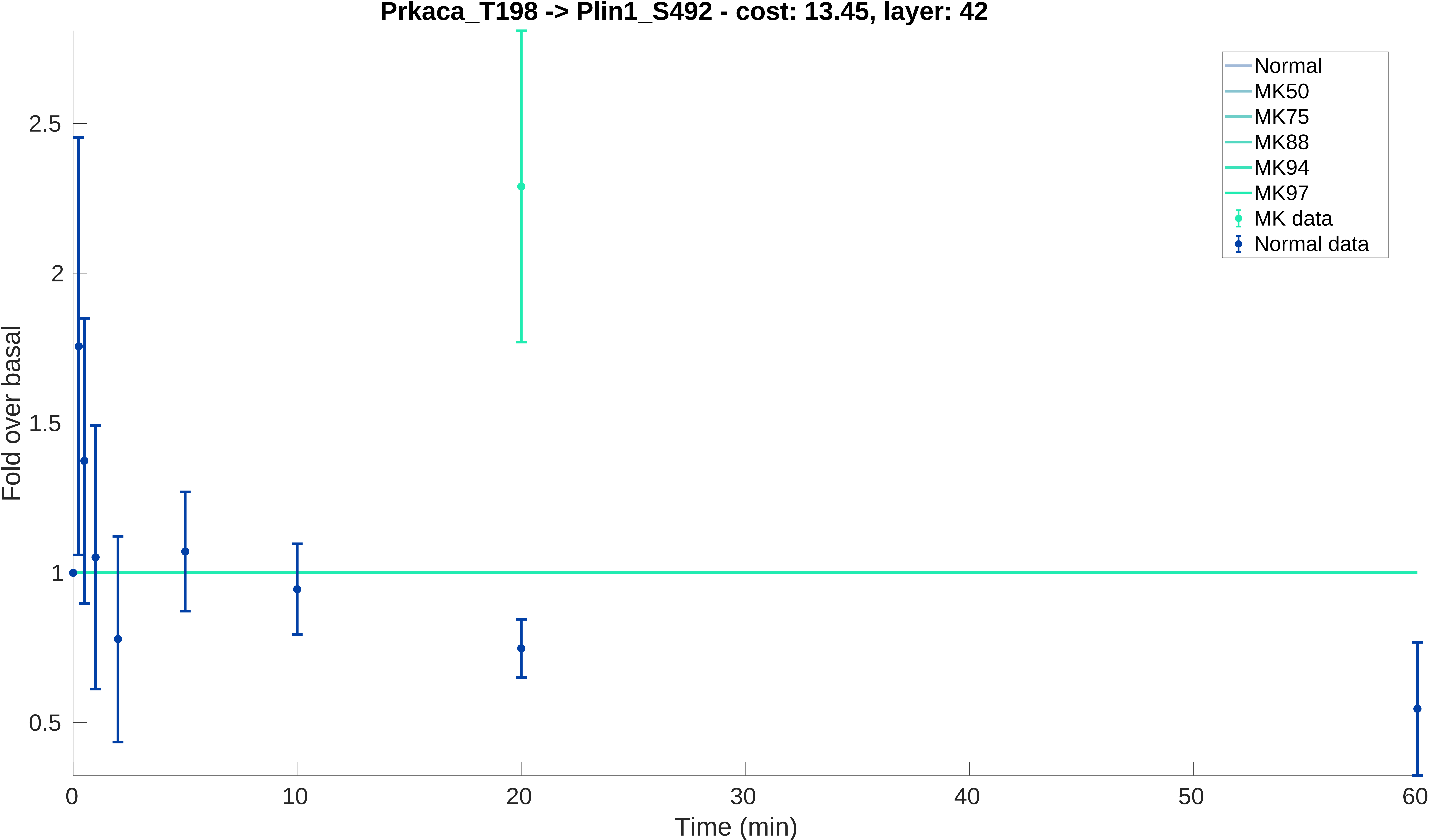

Prkaca_T198 -> Plin1_S492 - cost: 13.45, layer: 42
Normal
MK50
MK75
MK88
MK94
MK97
MK data
Normal data
Fold over basal
2.5
2
1.5
1
0.5
0
10
20
30
40
50
60
Time (min)

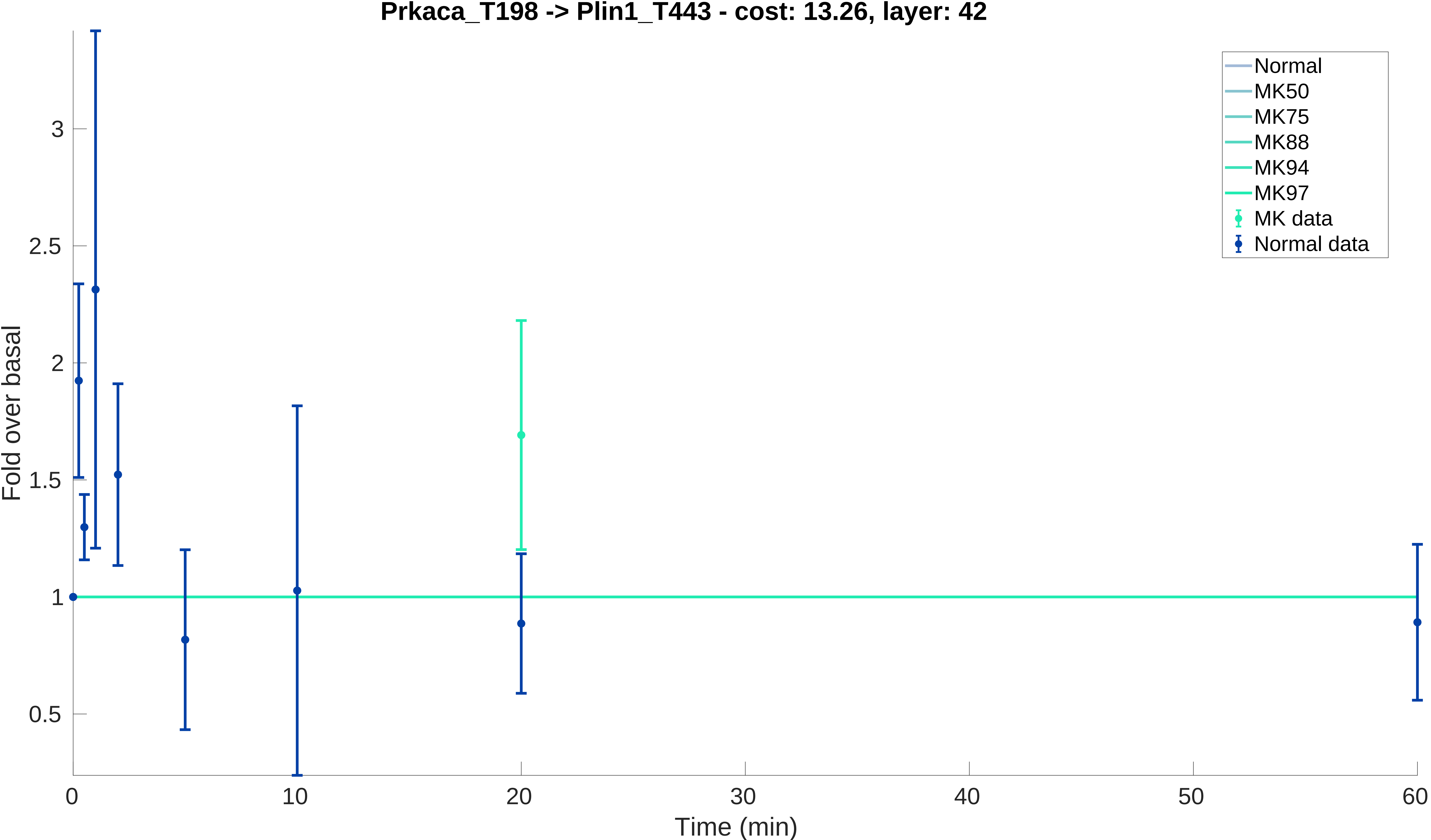

Prkaca_T198 -> Plin1_T443 - cost: 13.26, layer: 42
Fold over basal
Time (min)
0.5
1
1.5
2
2.5
3
0
10
20
30
40
50
60
Normal
MK50
MK75
MK88
MK94
MK97
MK data
Normal data

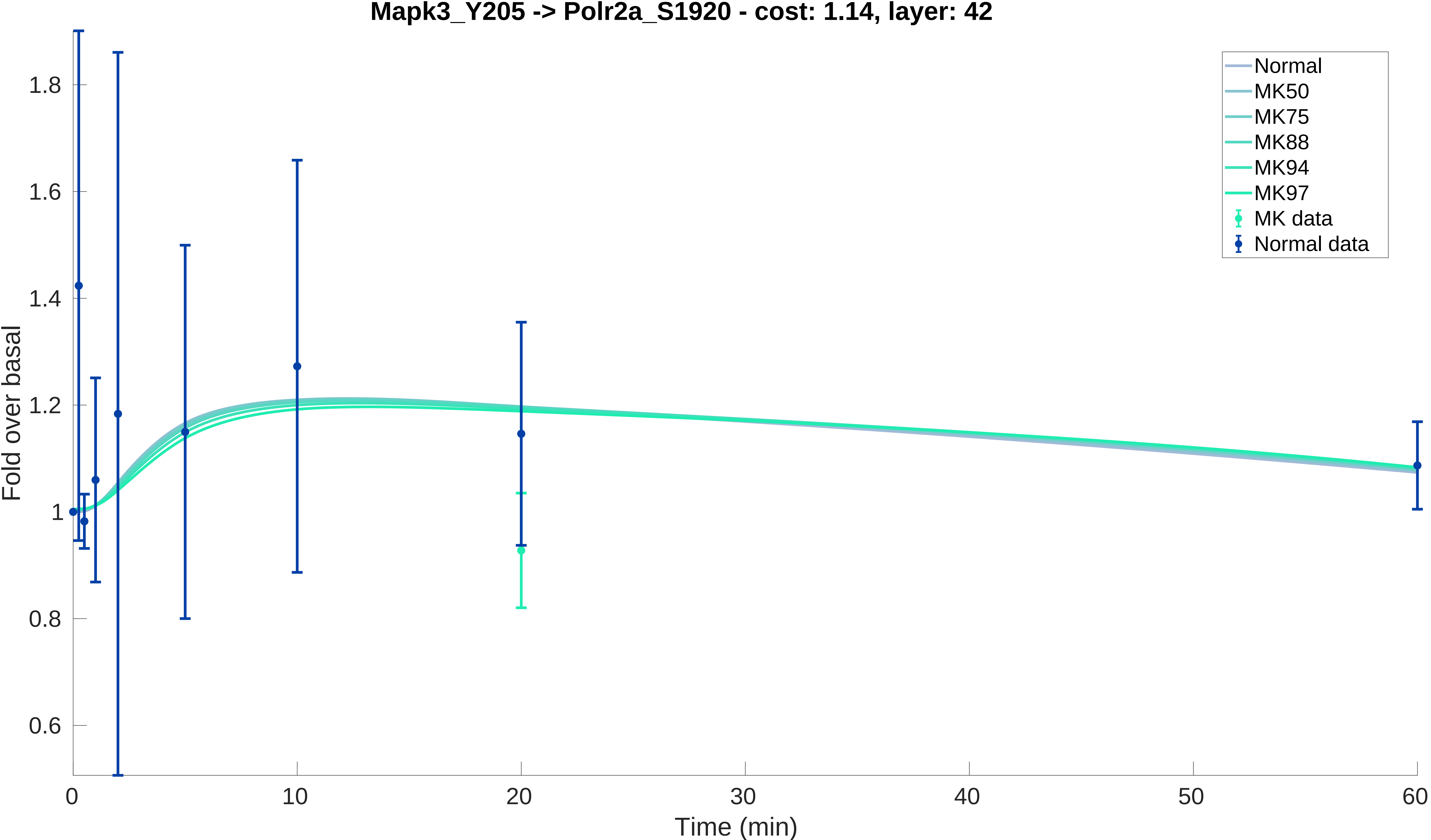

Mapk3_Y205 -> Polr2a_S1920 - cost: 1.14, layer: 42
Normal
MK50
MK75
MK88
MK94
MK97
MK data
Normal data
Fold over basal
Time (min)
0.6
0.8
1
1.2
1.4
1.6
1.8
0
10
20
30
40
50
60

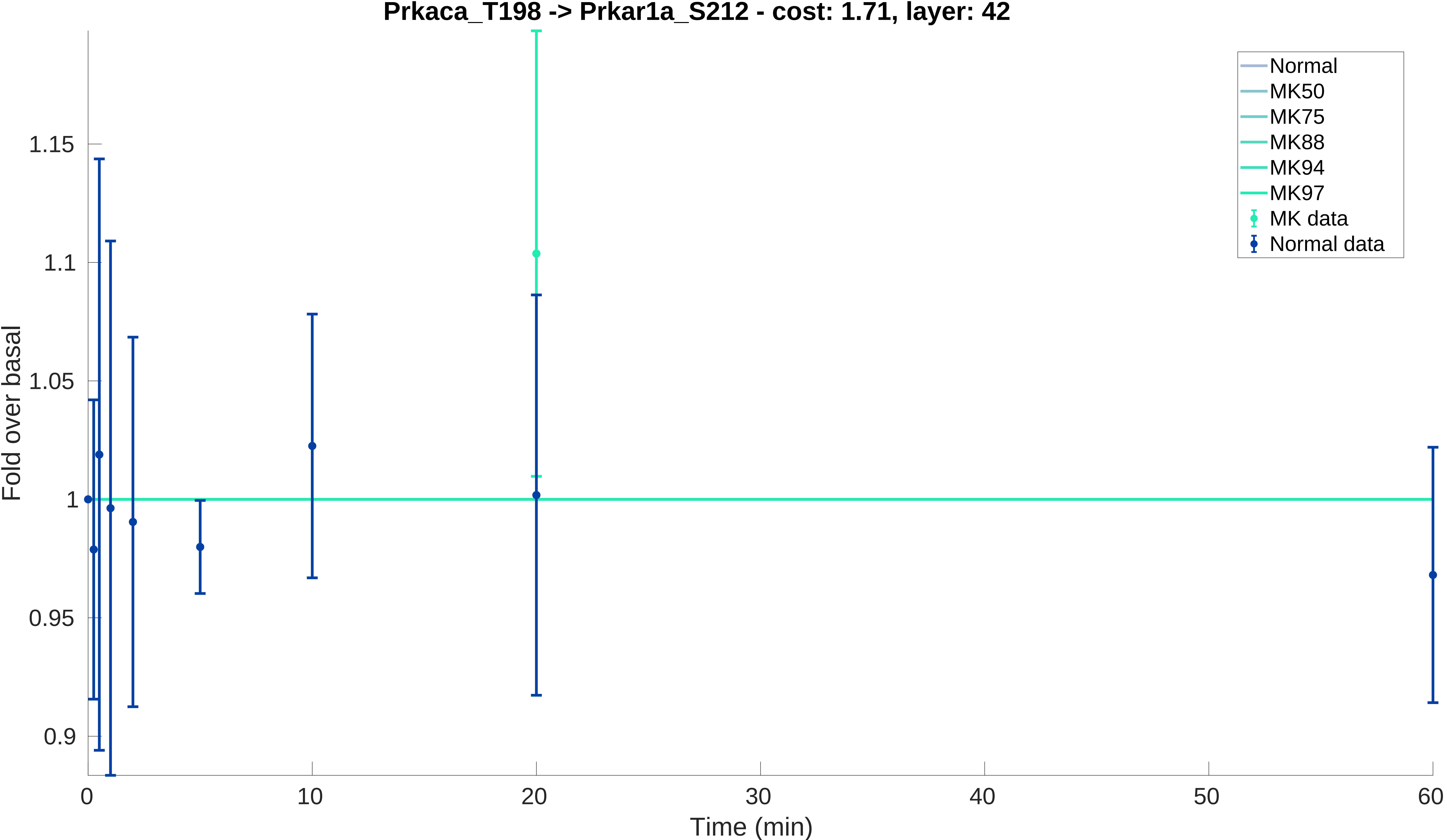
Prkaca_T198 -> Prkar1a_S212 - cost: 1.71, layer: 42
Fold over basal
1.15
1.1
1.05
1
0.95
0.9
0
10
20
30
40
50
60
Time (min)
Normal
MK50
MK75
MK88
MK94
MK97
MK data
Normal data

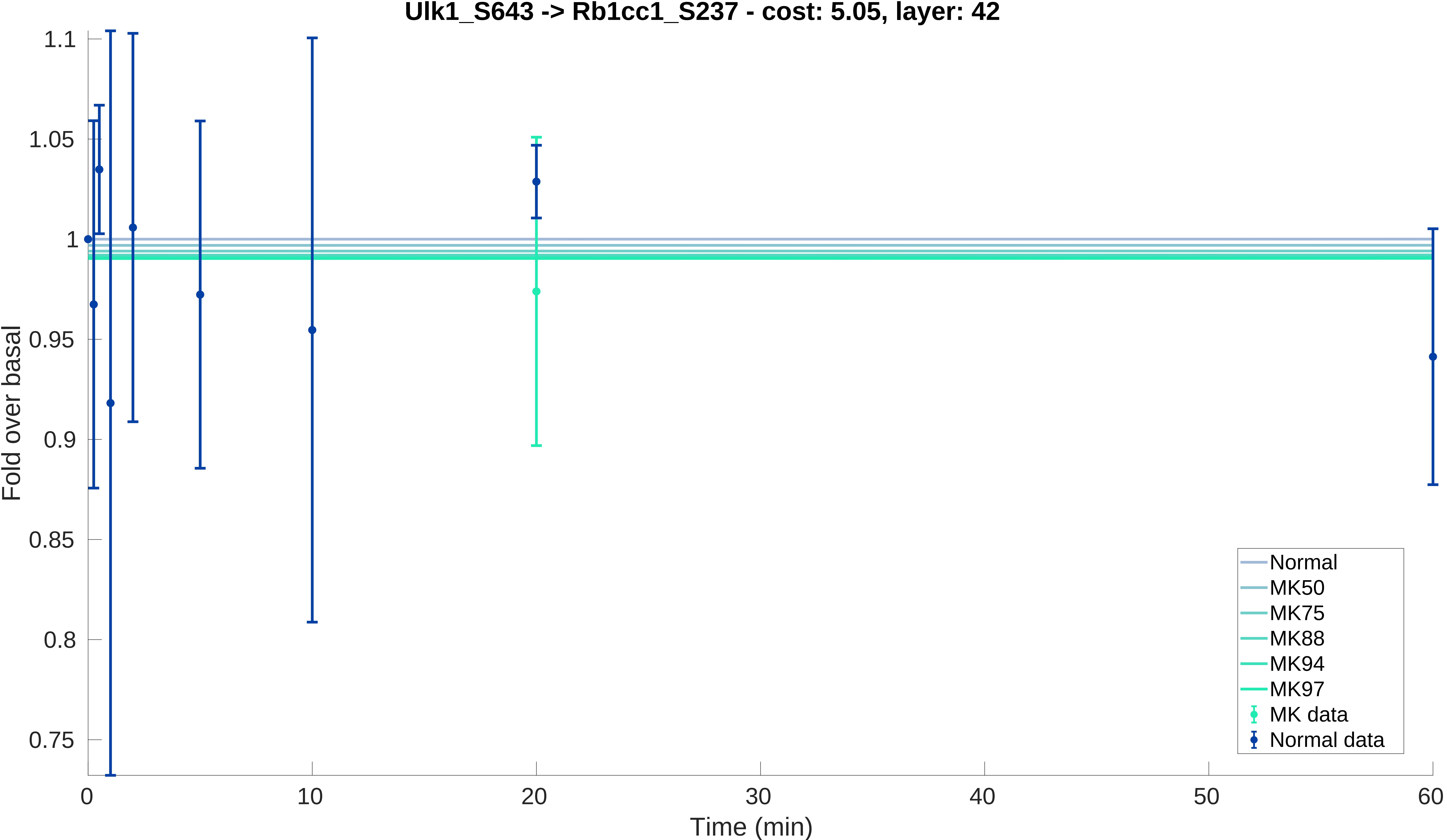

Ulk1_S643 -> Rb1cc1_S237 - cost: 5.05, layer: 42
1.1
1.05
1
0.95
0.9
0.85
0.8
0.75
Fold over basal
0
10
20
30
40
50
60
Time (min)
Normal
MK50
MK75
MK88
MK94
MK97
MK data
Normal data

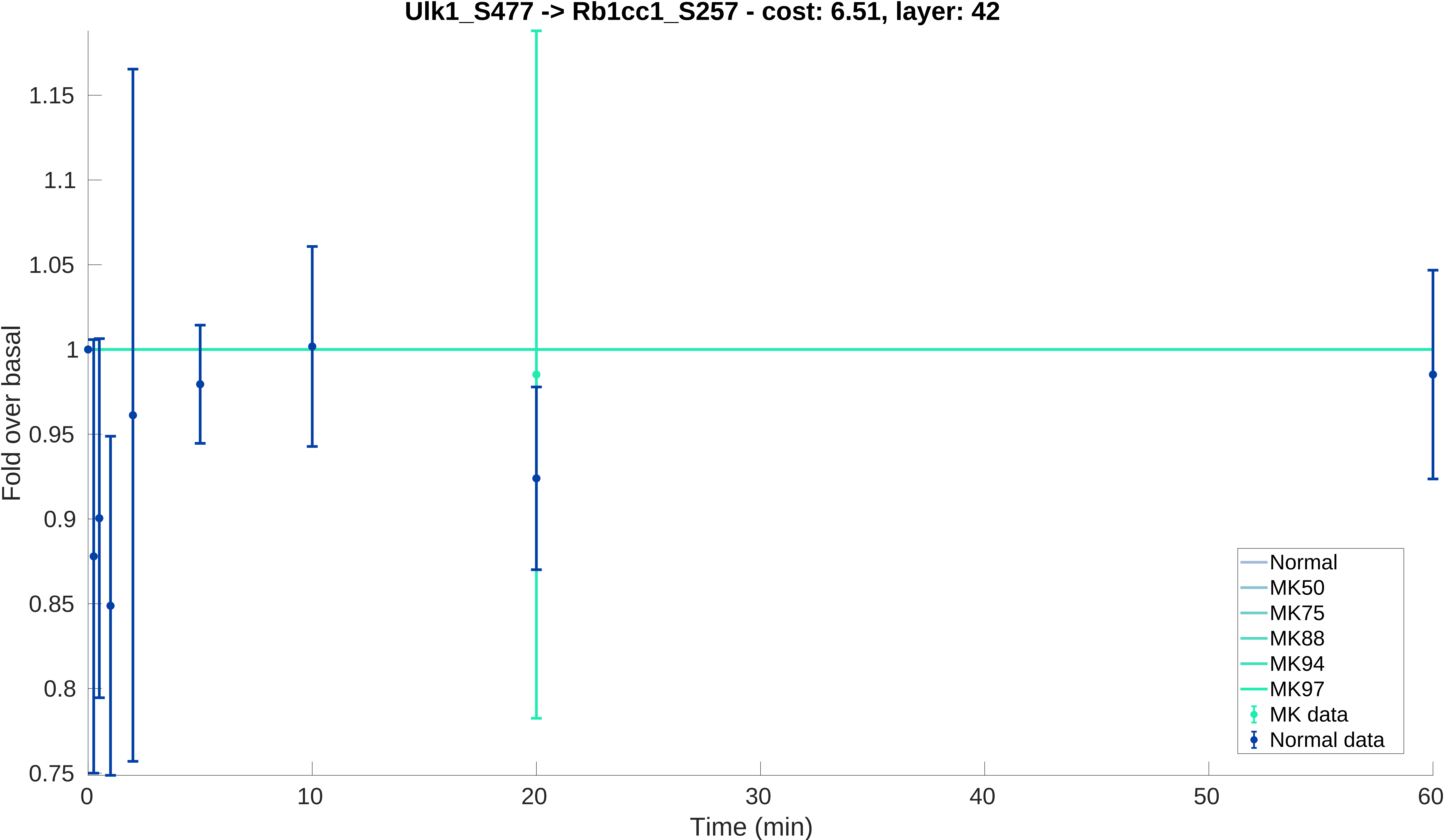

Ulk1_S477 -> Rb1cc1_S257 - cost: 6.51, layer: 42
Fold over basal
1.15
1.1
1.05
1
0.95
0.9
0.85
0.8
0.75
0
10
20
30
40
50
60
Time (min)
Normal
MK50
MK75
MK88
MK94
MK97
MK data
Normal data

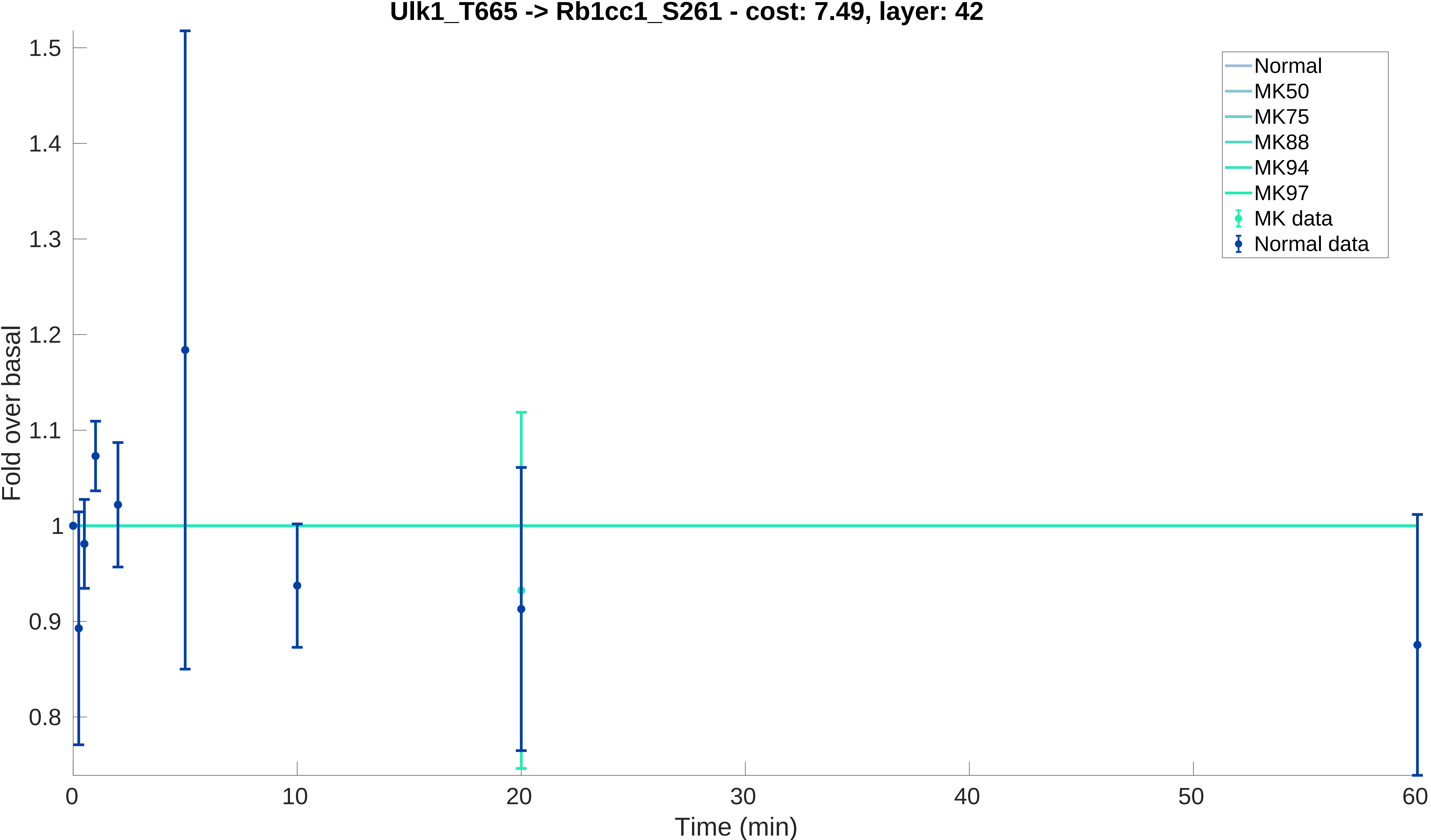

Ulk1_T665 -> Rb1cc1_S261 - cost: 7.49, layer: 42
Fold over basal
Time (min)
1.5
1.4
1.3
1.2
1.1
1
0.9
0.8
0
10
20
30
40
50
60
Normal
MK50
MK75
MK88
MK94
MK97
MK data
Normal data

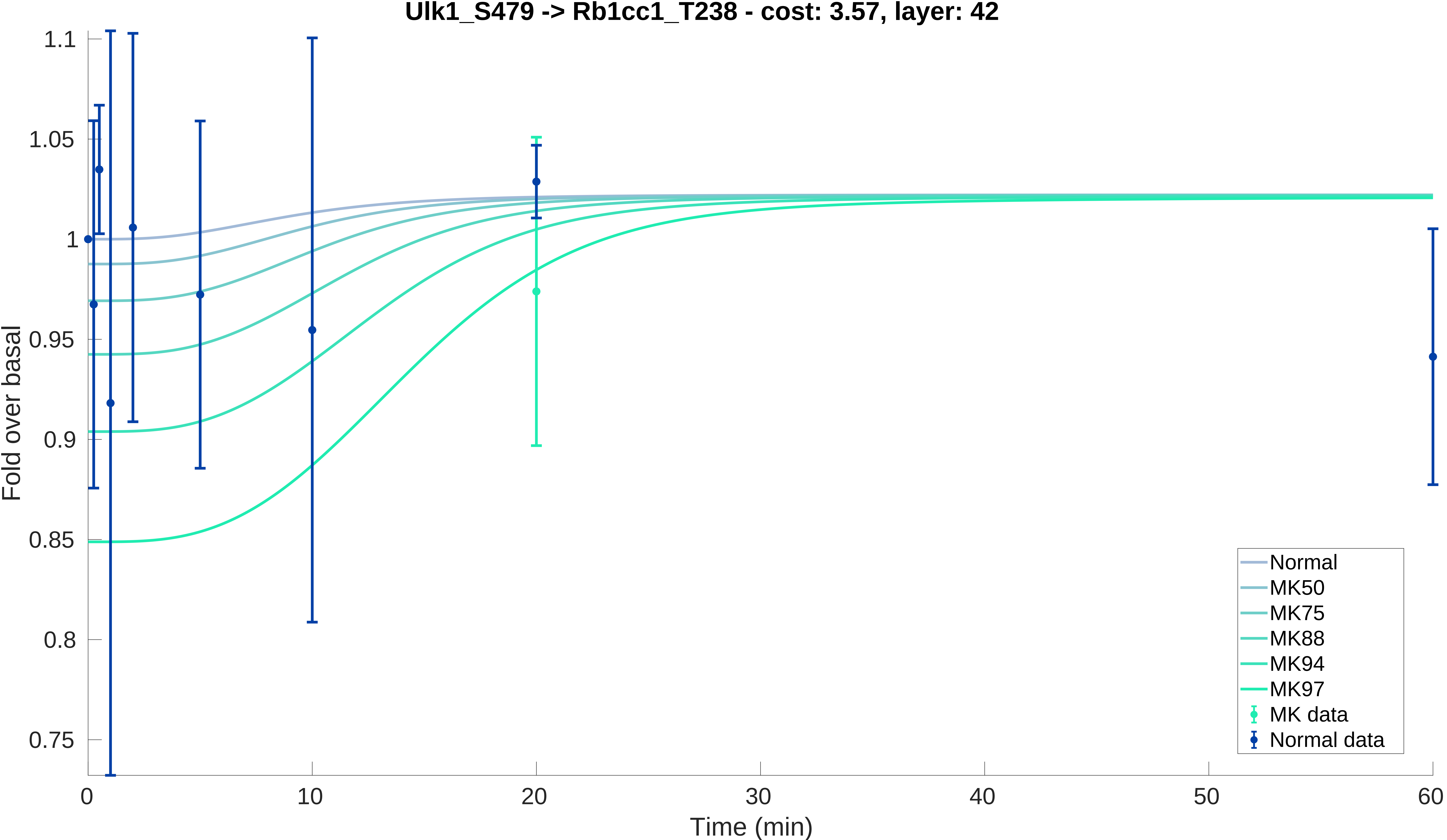

Ulk1_S479 -> Rb1cc1_T238 - cost: 3.57, layer: 42
Fold over basal
1.1
1.05
1
0.95
0.9
0.85
0.8
0.75
0
10
20
30
40
50
60
Time (min)
Normal
MK50
MK75
MK88
MK94
MK97
MK data
Normal data

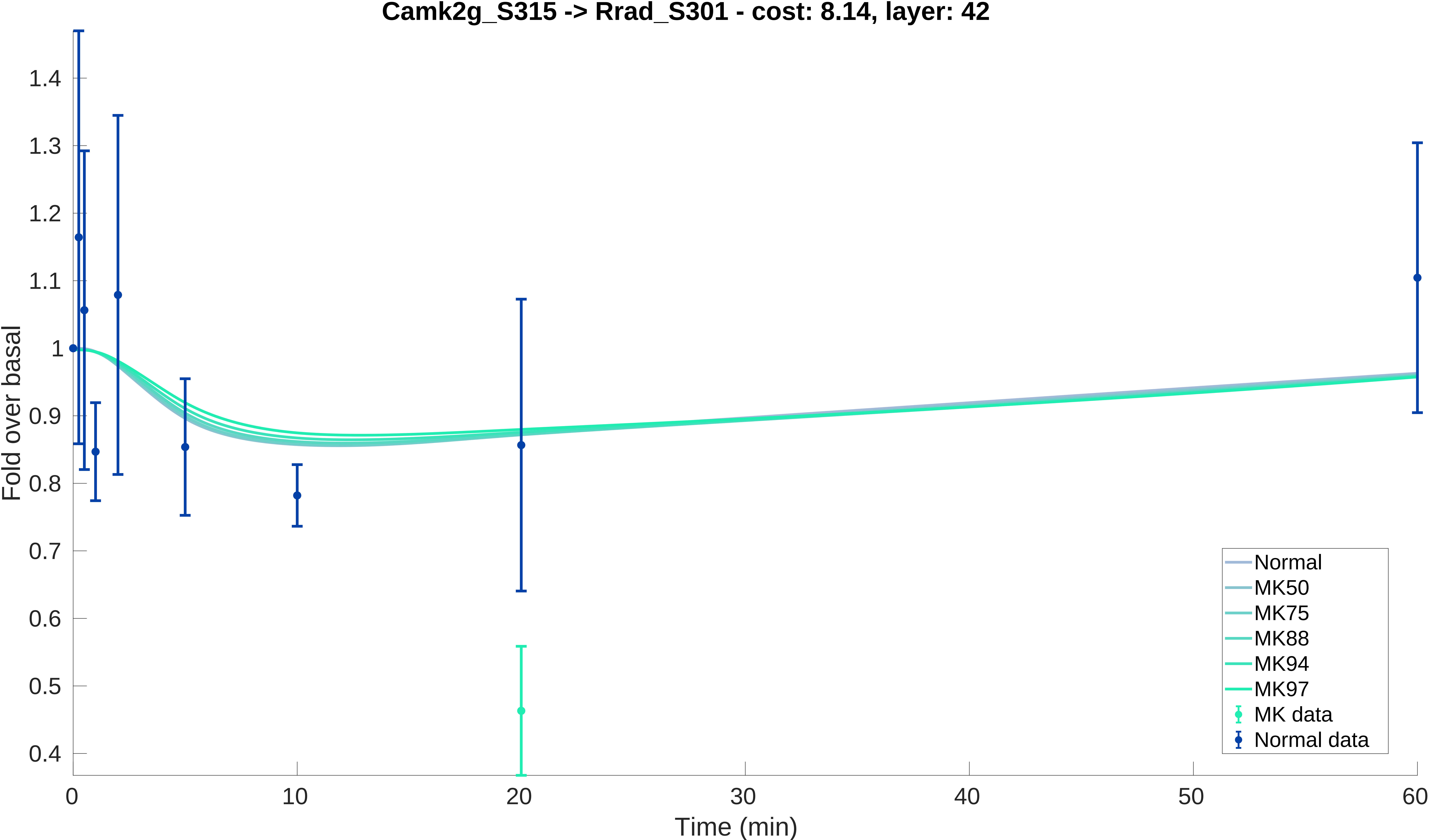

Camk2g_S315 -> Rrad_S301 - cost: 8.14, layer: 42
Fold over basal
Time (min)
0
10
20
30
40
50
60
0.4
0.5
0.6
0.7
0.8
0.9
1
1.1
1.2
1.3
1.4
Normal
MK50
MK75
MK88
MK94
MK97
MK data
Normal data

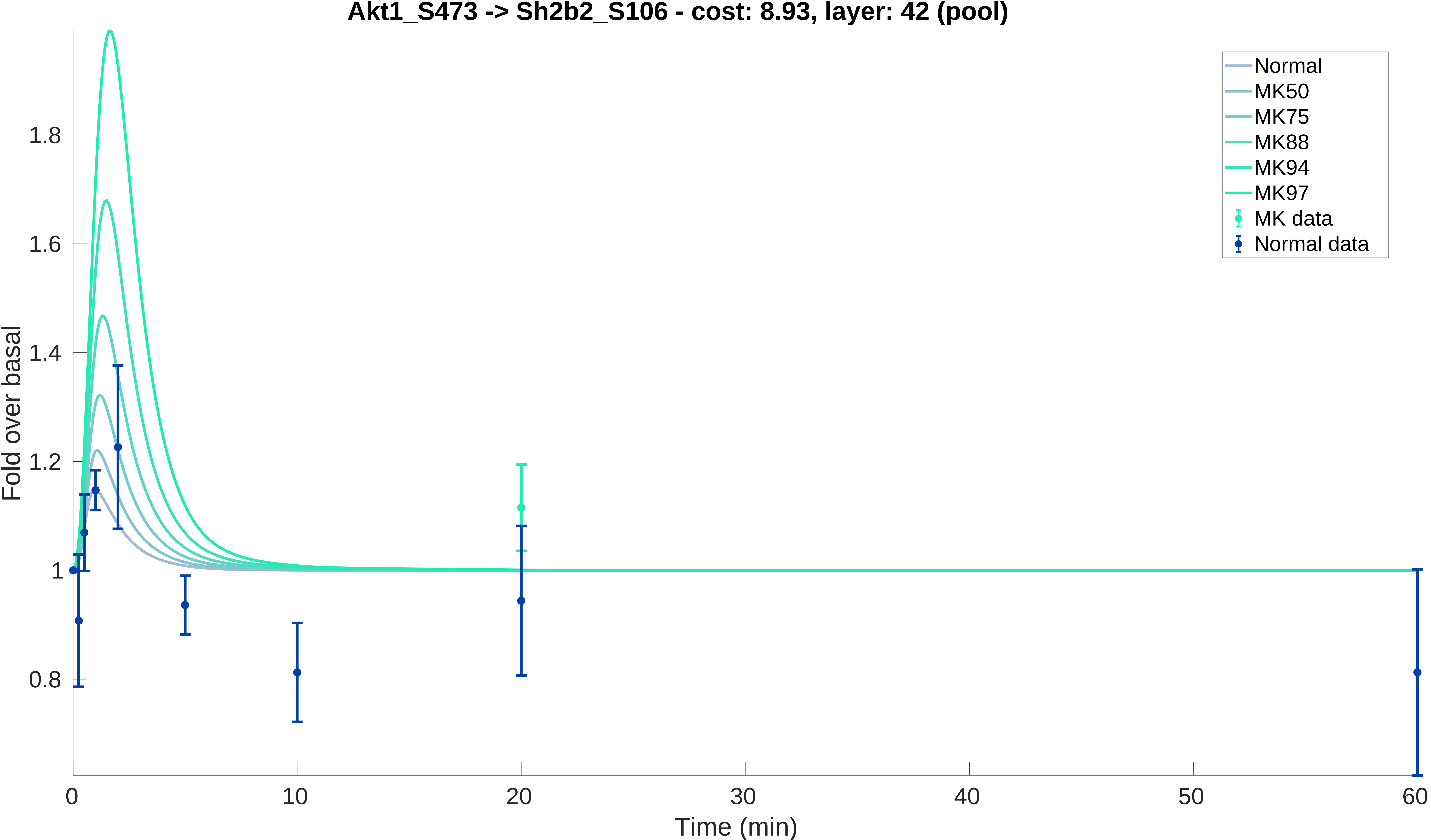

Akt1_S473 -> Sh2b2_S106 - cost: 8.93, layer: 42 (pool)
Fold over basal
1.8
1.6
1.4
1.2
1
0.8
0
10
20
30
40
50
60
Time (min)
Normal
MK50
MK75
MK88
MK94
MK97
MK data
Normal data

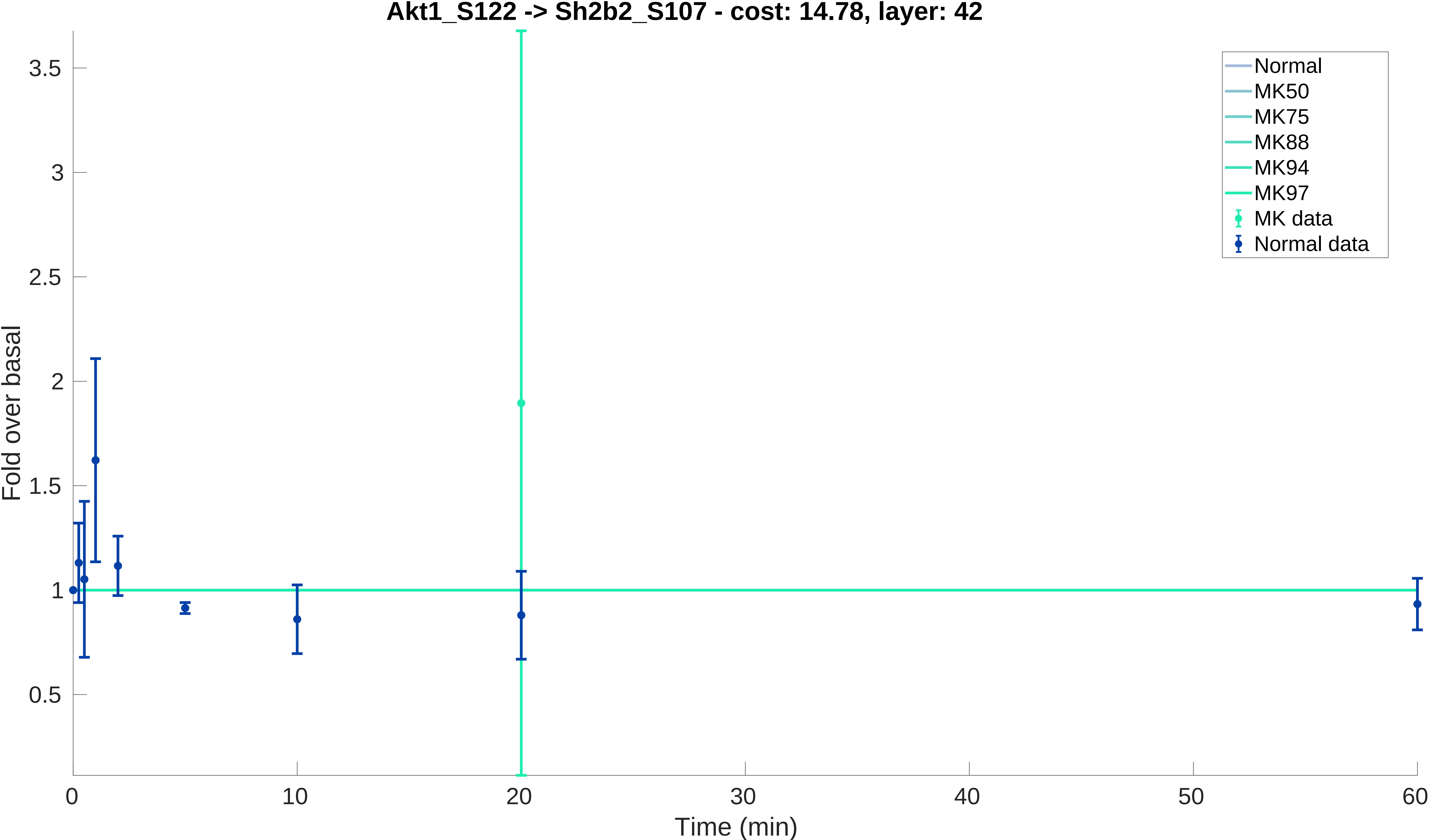
Akt1_S122 -> Sh2b2_S107 - cost: 14.78, layer: 42
Fold over basal
Time (min)
0
10
20
30
40
50
60
0.5
1
1.5
2
2.5
3
3.5
Normal
MK50
MK75
MK88
MK94
MK97
MK data
Normal data

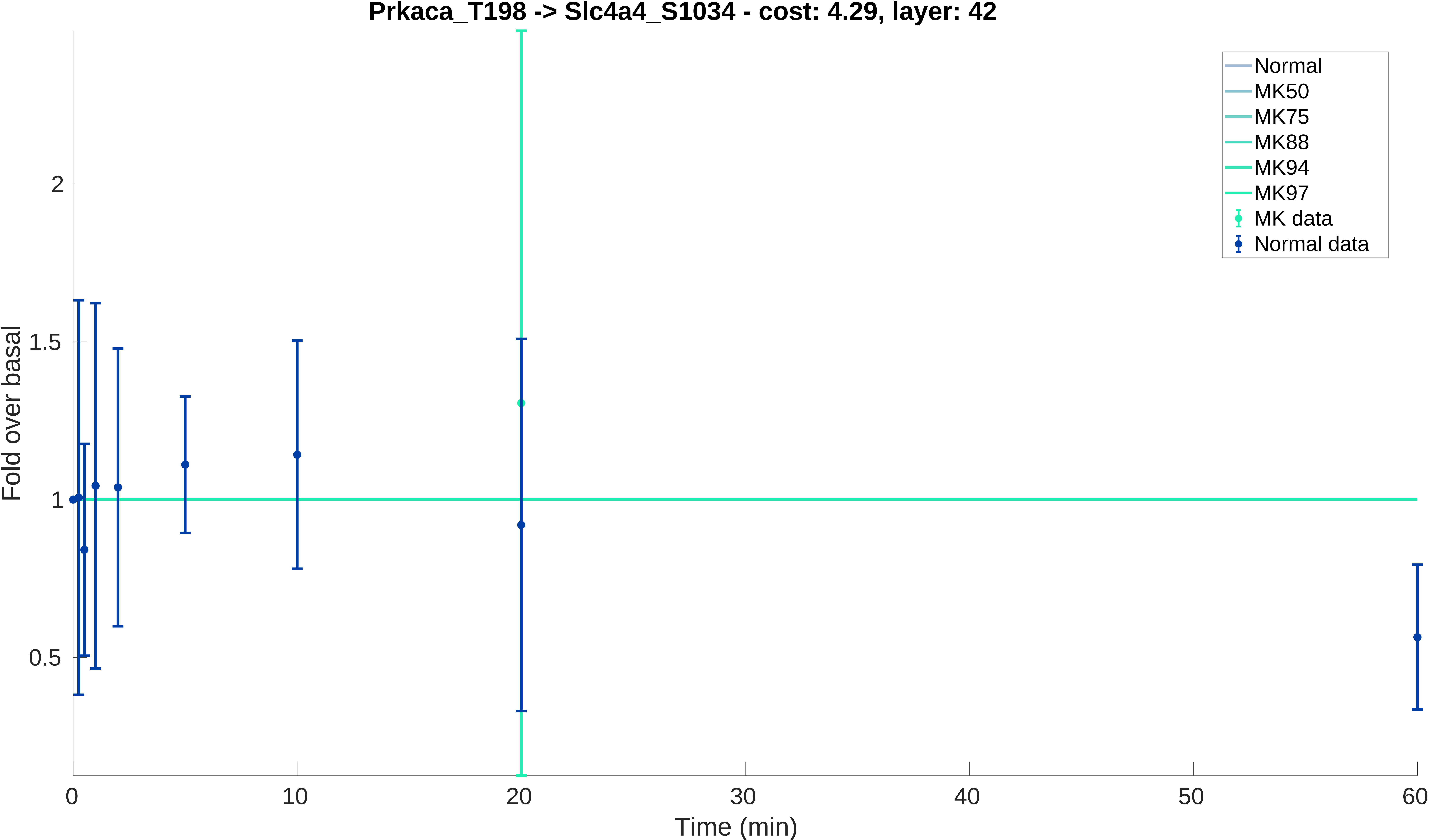
Prrkaca_T198 -> Slc4a4_S1034 - cost: 4.29, layer: 42
Fold over basal
Time (min)
0
10
20
30
40
50
60
0.5
1
1.5
2
Normal
MK50
MK75
MK88
MK94
MK97
MK data
Normal data

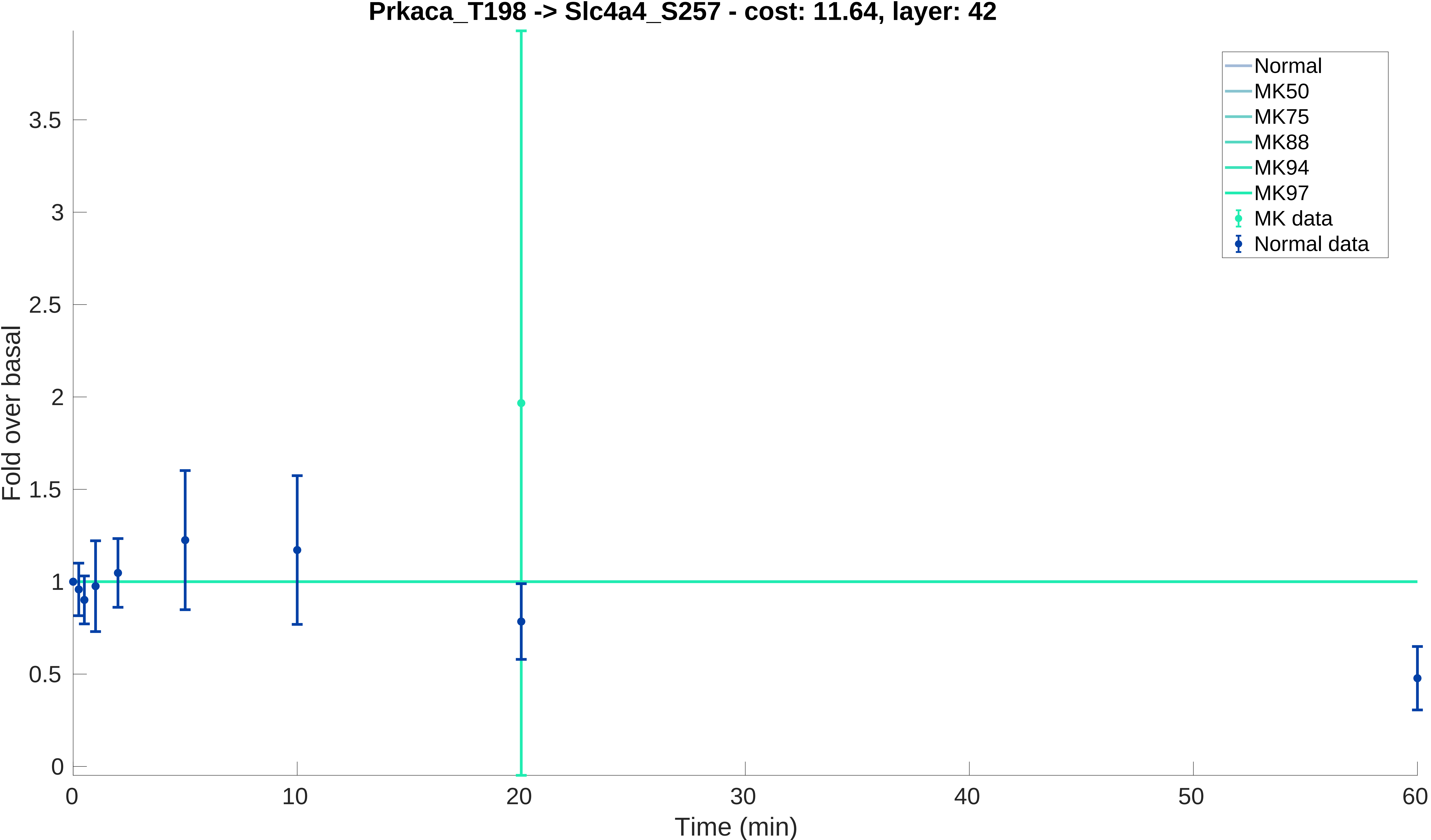

Prkaca_T198 -> Slc4a4_S257 - cost: 11.64, layer: 42
Fold over basal
Time (min)
0
0.5
1
1.5
2
2.5
3
3.5
0
10
20
30
40
50
60
Normal
MK50
MK75
MK88
MK94
MK97
MK data
Normal data

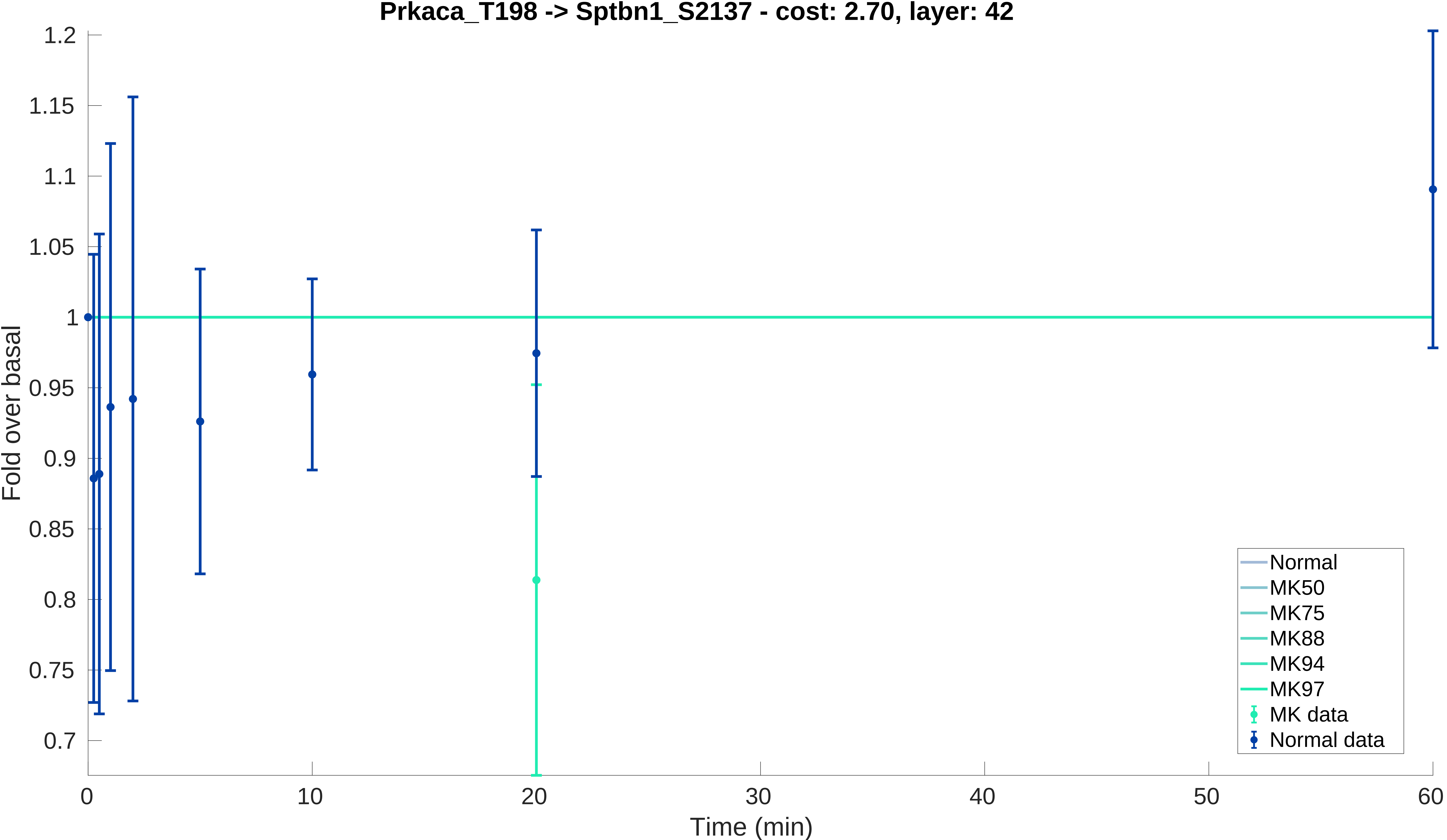

Prkaca_T198 -> Sptbn1_S2137 - cost: 2.70, layer: 42
Fold over basal
Time (min)
0
10
20
30
40
50
60
0.7
0.75
0.8
0.85
0.9
0.95
1
1.05
1.1
1.15
1.2
Normal
MK50
MK75
MK88
MK94
MK97
MK data
Normal data

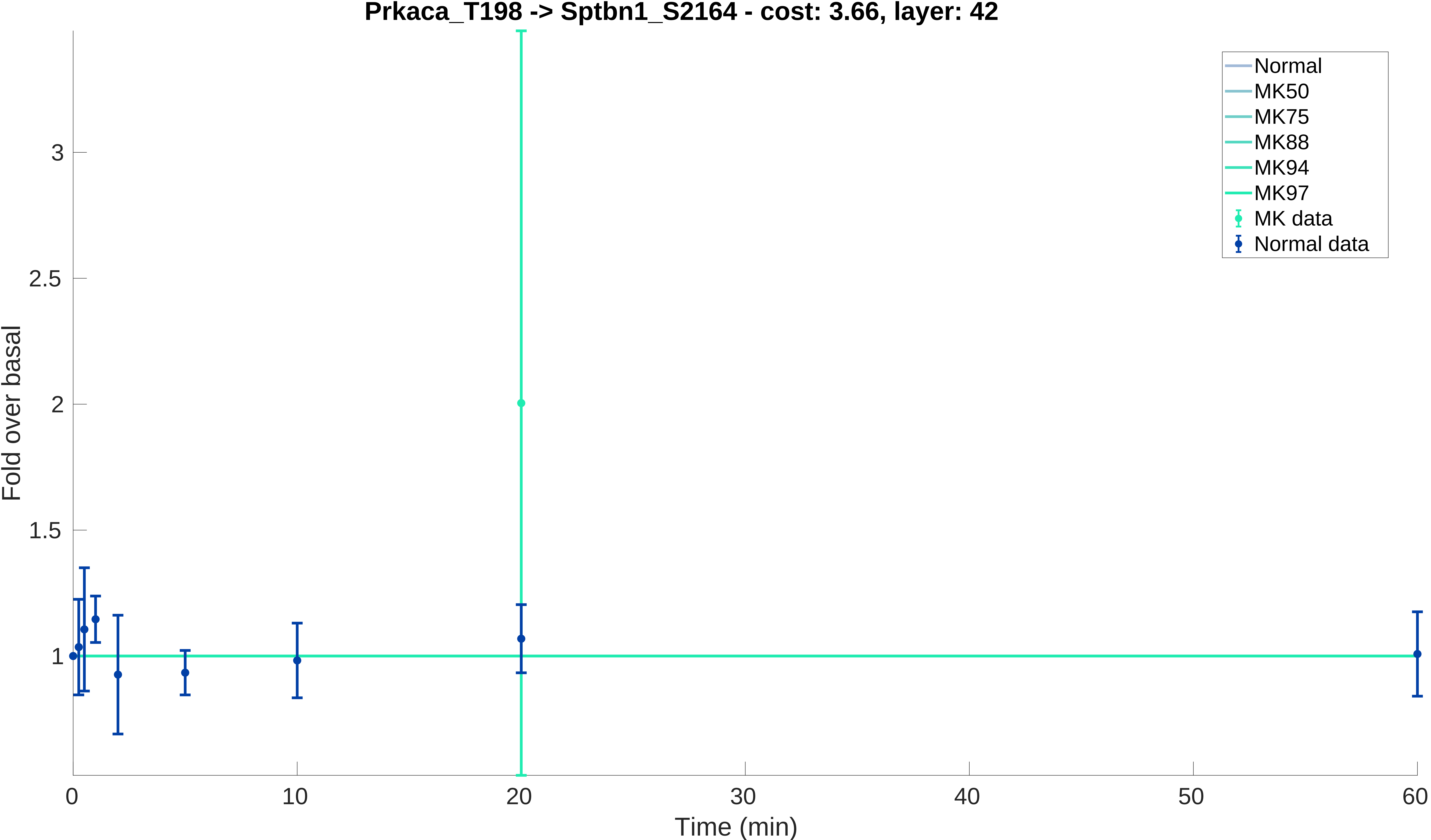

Prkaca_T198 -> Sptbn1_S2164 - cost: 3.66, layer: 42
Fold over basal
3
2.5
2
1.5
1
0
10
20
30
40
50
60
Time (min)
Normal
MK50
MK75
MK88
MK94
MK97
MK data
Normal data

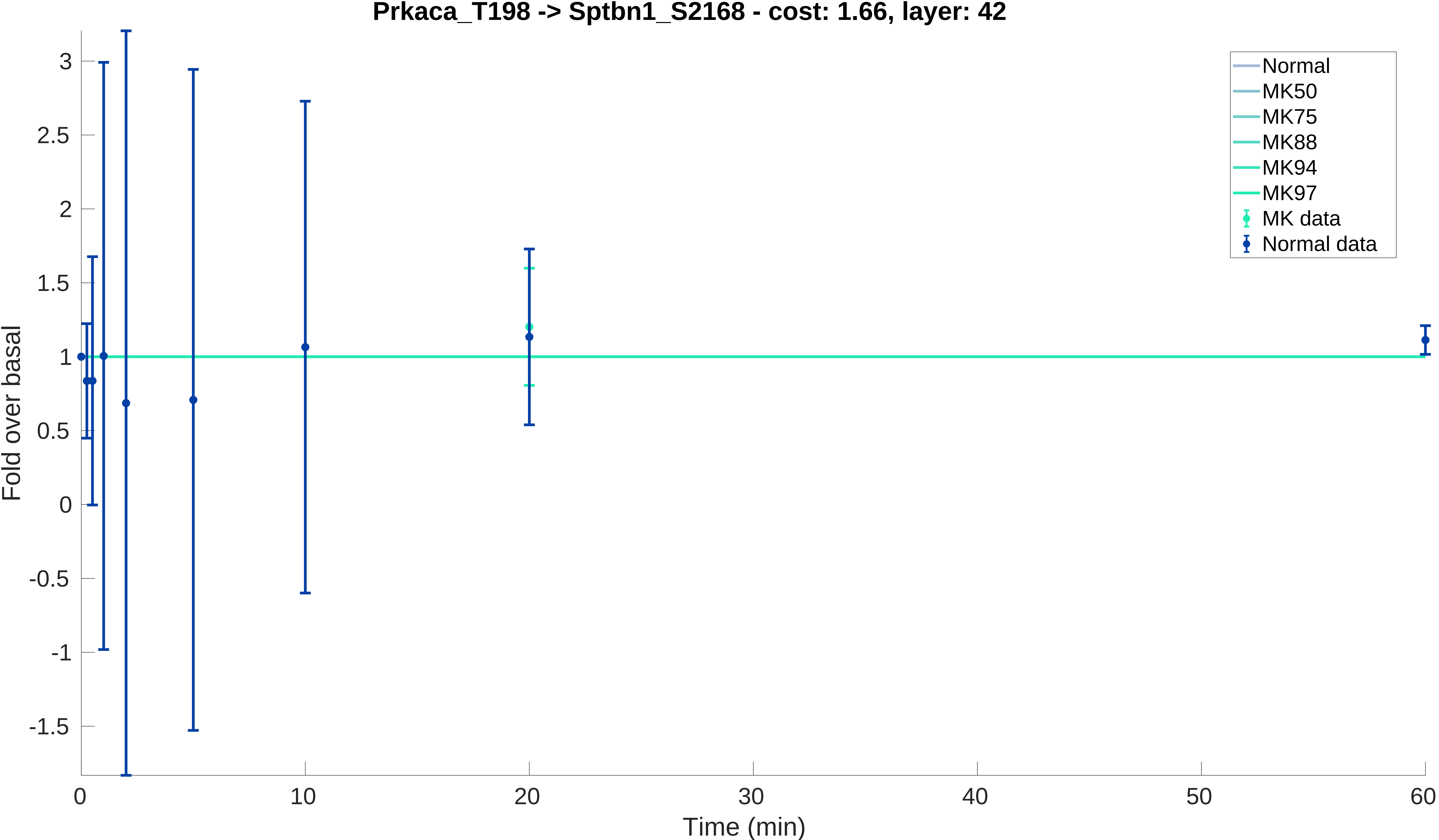

Prkaca_T198 -> Sptbn1_S2168 - cost: 1.66, layer: 42
Fold over basal
Time (min)
3
2.5
2
1.5
1
0.5
0
-0.5
-1
-1.5
0
10
20
30
40
50
60
Normal
MK50
MK75
MK88
MK94
MK97
MK data
Normal data

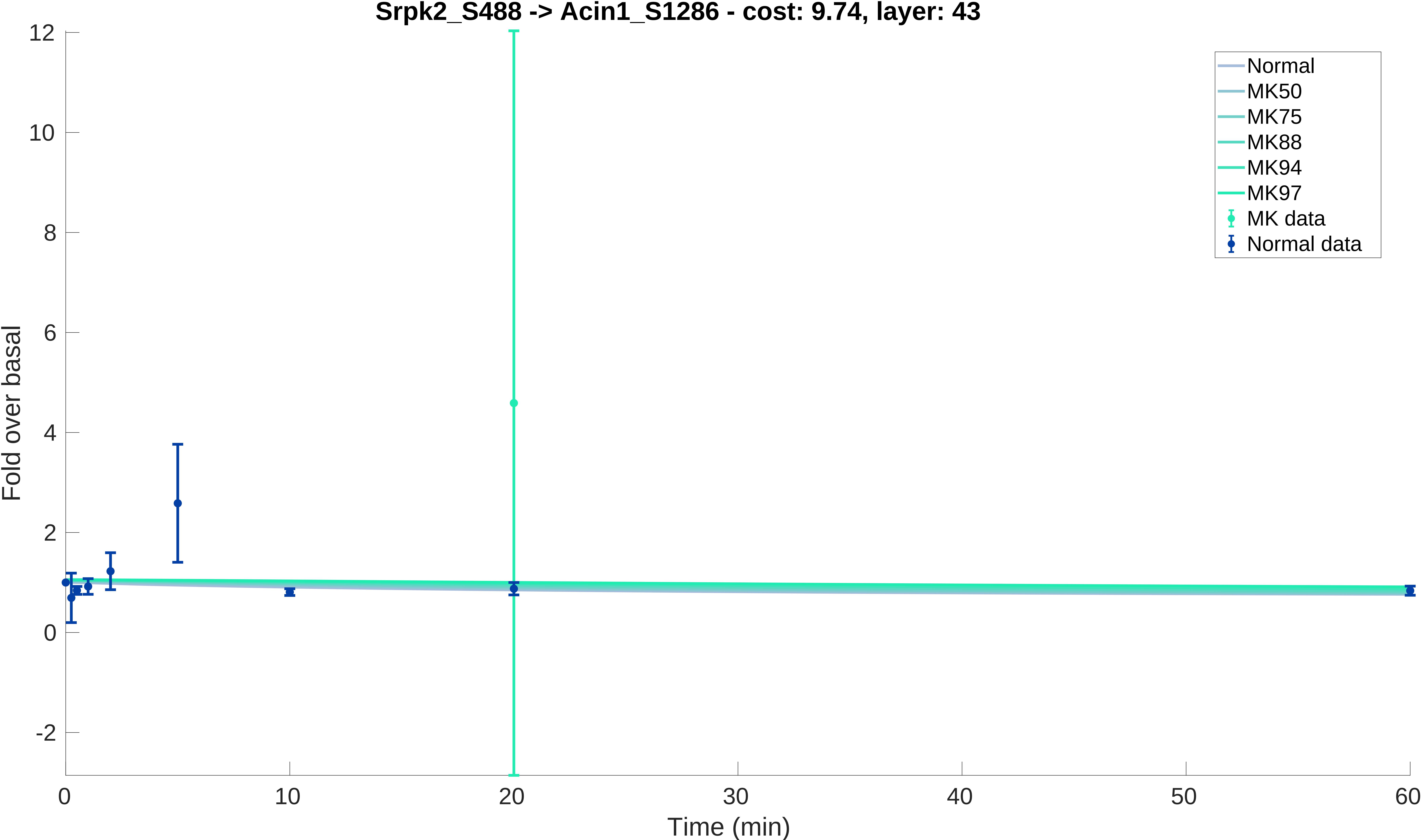
Srpk2_S488 -> Acin1_S1286 - cost: 9.74, layer: 43
Fold over basal
Time (min)
12
10
8
6
4
2
0
-2
0
10
20
30
40
50
60
Normal
MK50
MK75
MK88
MK94
MK97
MK data
Normal data

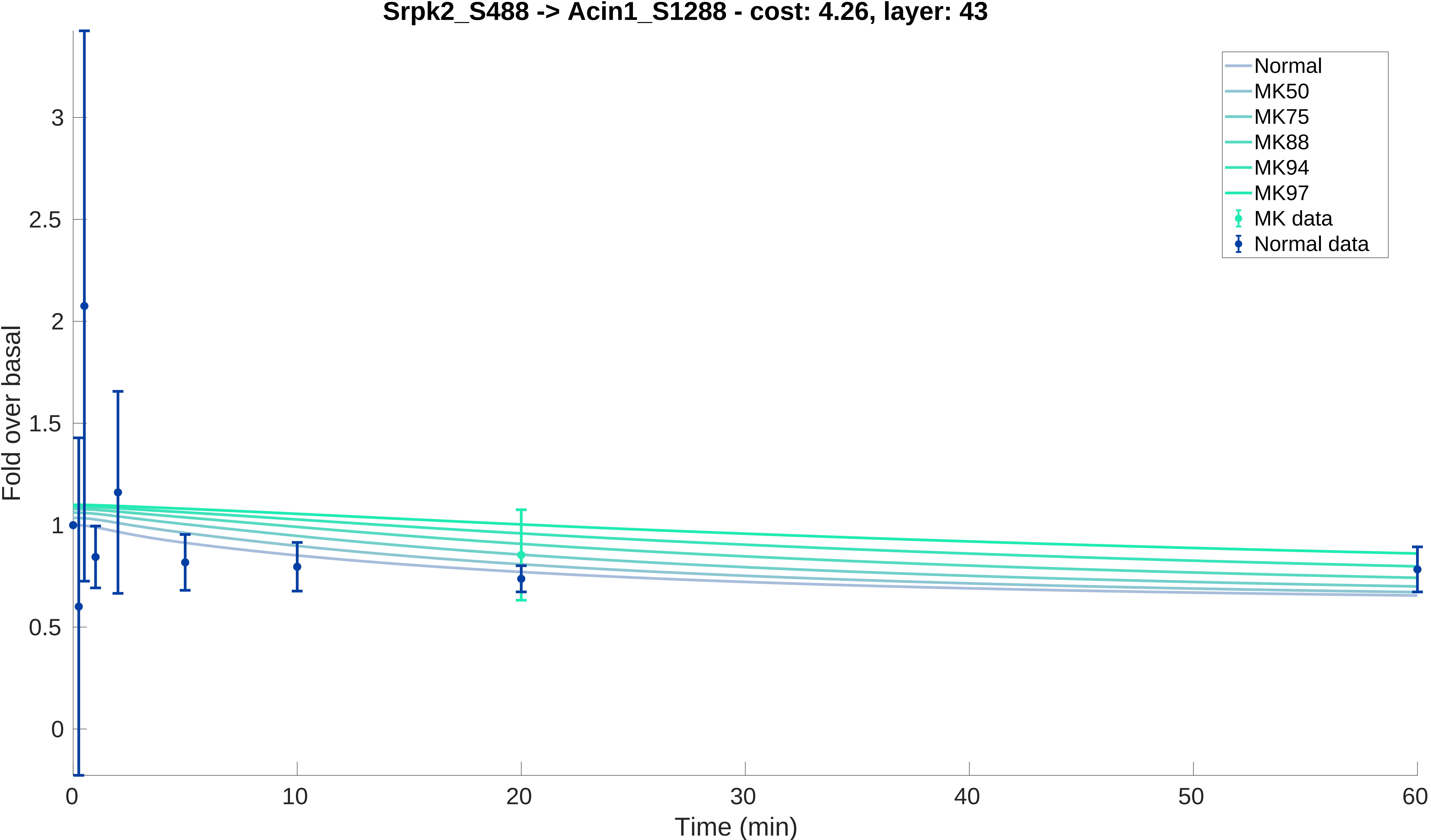

Srpk2_S488 -> Acin1_S1288 - cost: 4.26, layer: 43
Fold over basal
Time (min)
0
0.5
1
1.5
2
2.5
3
0
10
20
30
40
50
60
Normal
MK50
MK75
MK88
MK94
MK97
MK data
Normal data

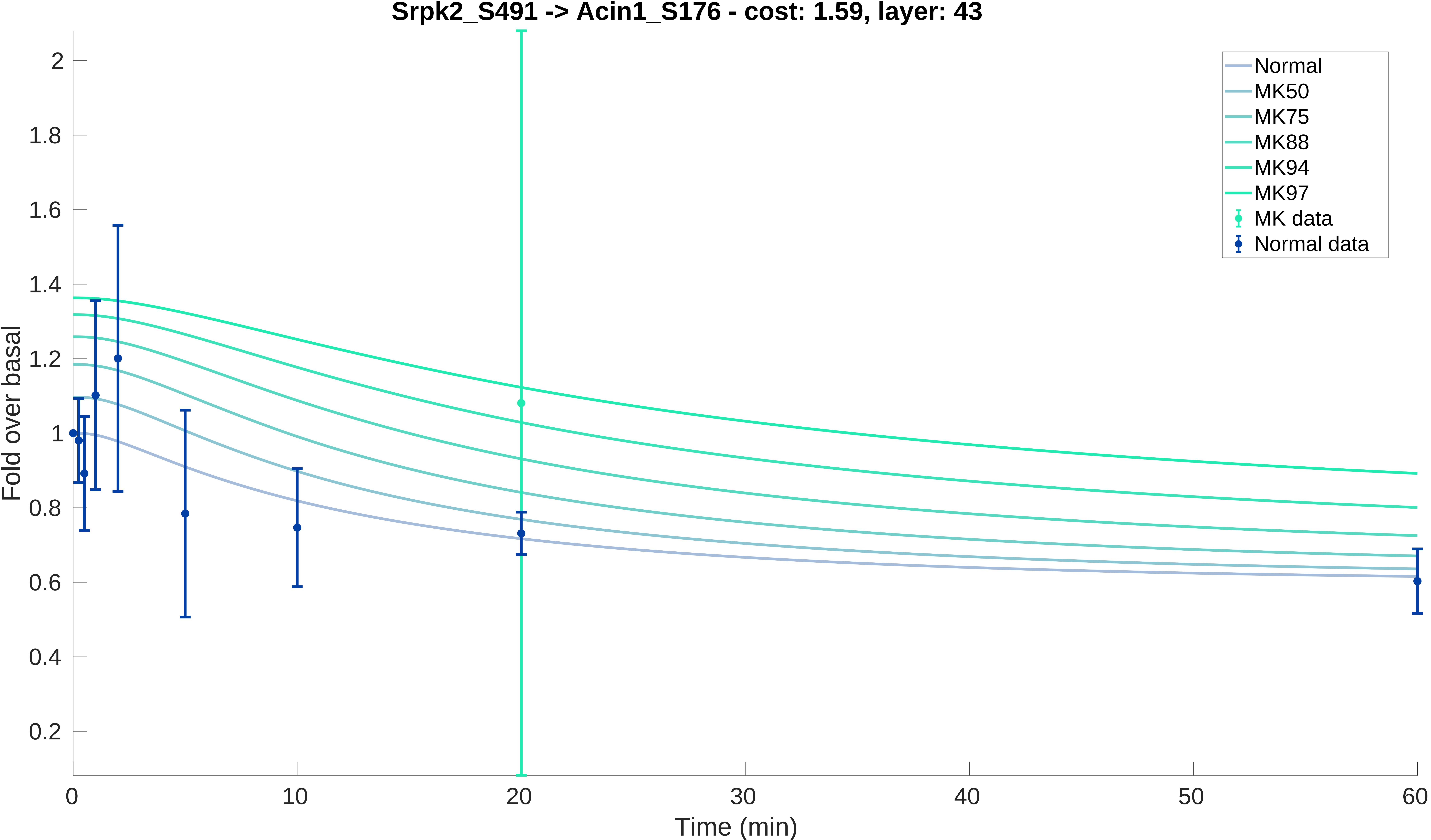

Srpk2_S491 -> Acin1_S176 - cost: 1.59, layer: 43
Fold over basal
Time (min)
0.2
0.4
0.6
0.8
1
1.2
1.4
1.6
1.8
2
0
10
20
30
40
50
60
Normal
MK50
MK75
MK88
MK94
MK97
MK data
Normal data

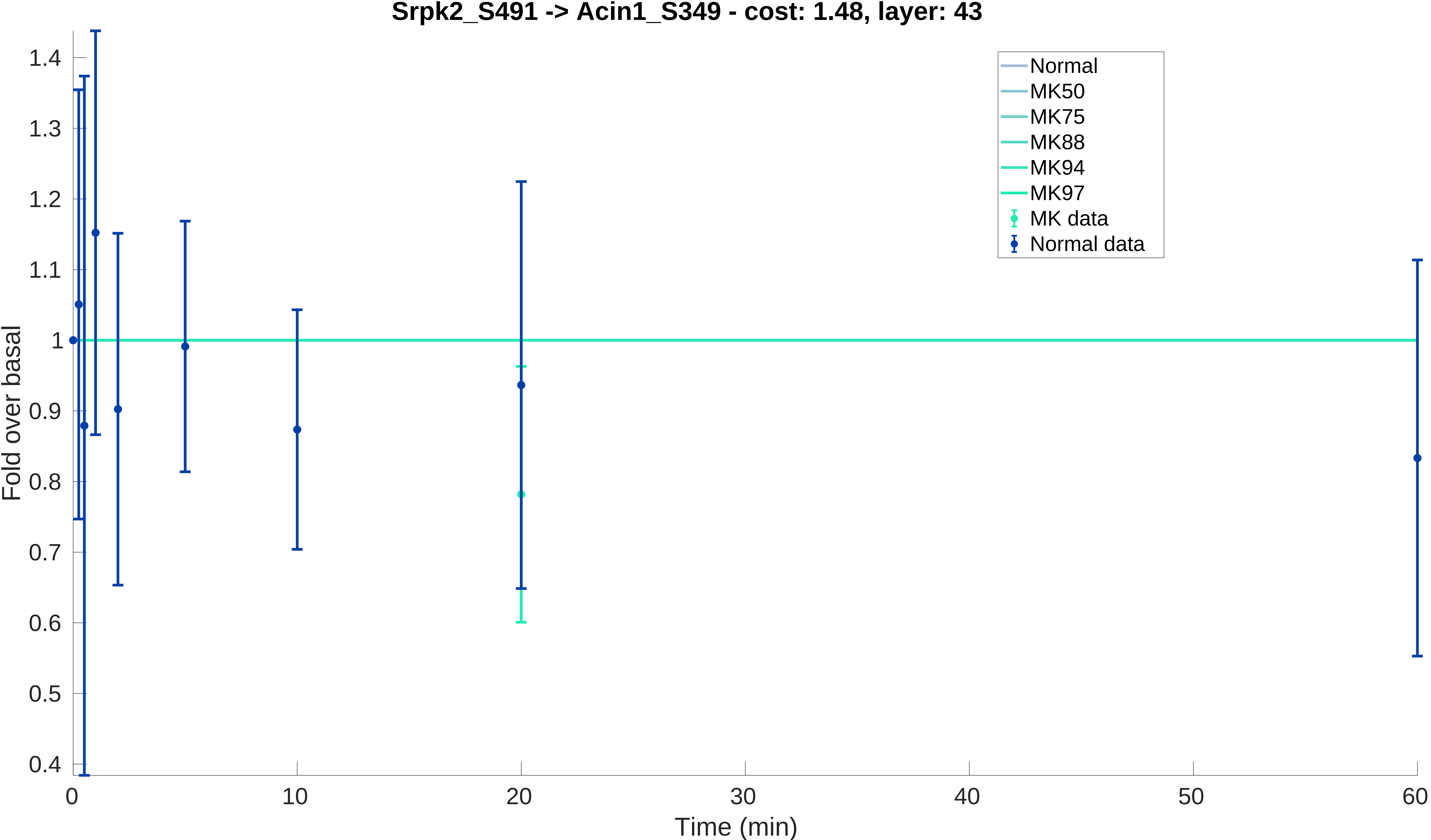

Srpk2_S491 -> Acin1_S349 - cost: 1.48, layer: 43
Fold over basal
Time (min)
0.4
0.5
0.6
0.7
0.8
0.9
1
1.1
1.2
1.3
1.4
0
10
20
30
40
50
60
Normal
MK50
MK75
MK88
MK94
MK97
MK data
Normal data

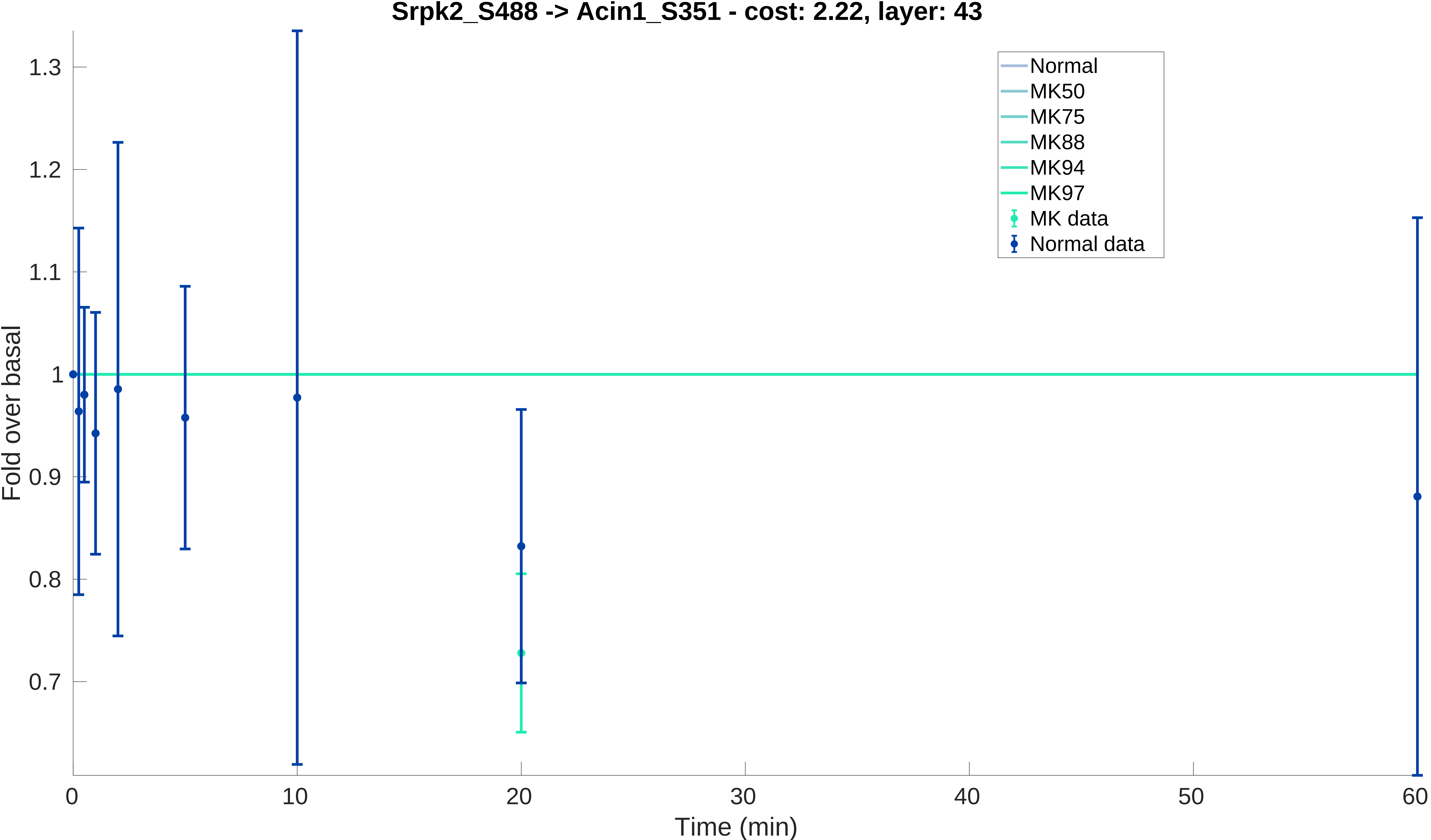

Srpk2_S488 -> Acin1_S351 - cost: 2.22, layer: 43
Fold over basal
Time (min)
Normal
MK50
MK75
MK88
MK94
MK97
MK data
Normal data
1.3
1.2
1.1
1
0.9
0.8
0.7
0
10
20
30
40
50
60

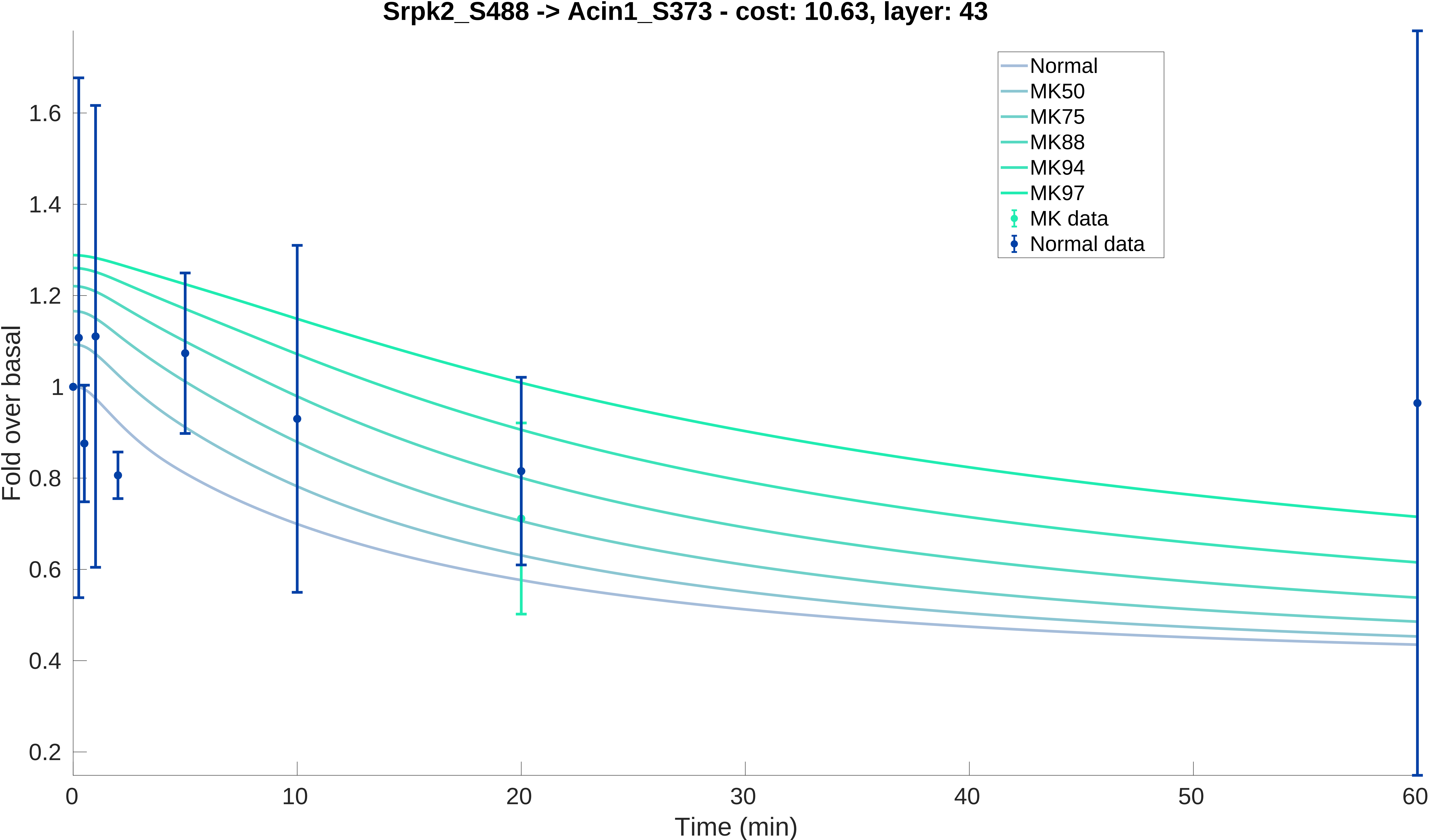

Srpk2_S488 -> Acin1_S373 - cost: 10.63, layer: 43
Normal
MK50
MK75
MK88
MK94
MK97
MK data
Normal data
Fold over basal
1.6
1.4
1.2
1
0.8
0.6
0.4
0.2
0
10
20
30
40
50
60
Time (min)

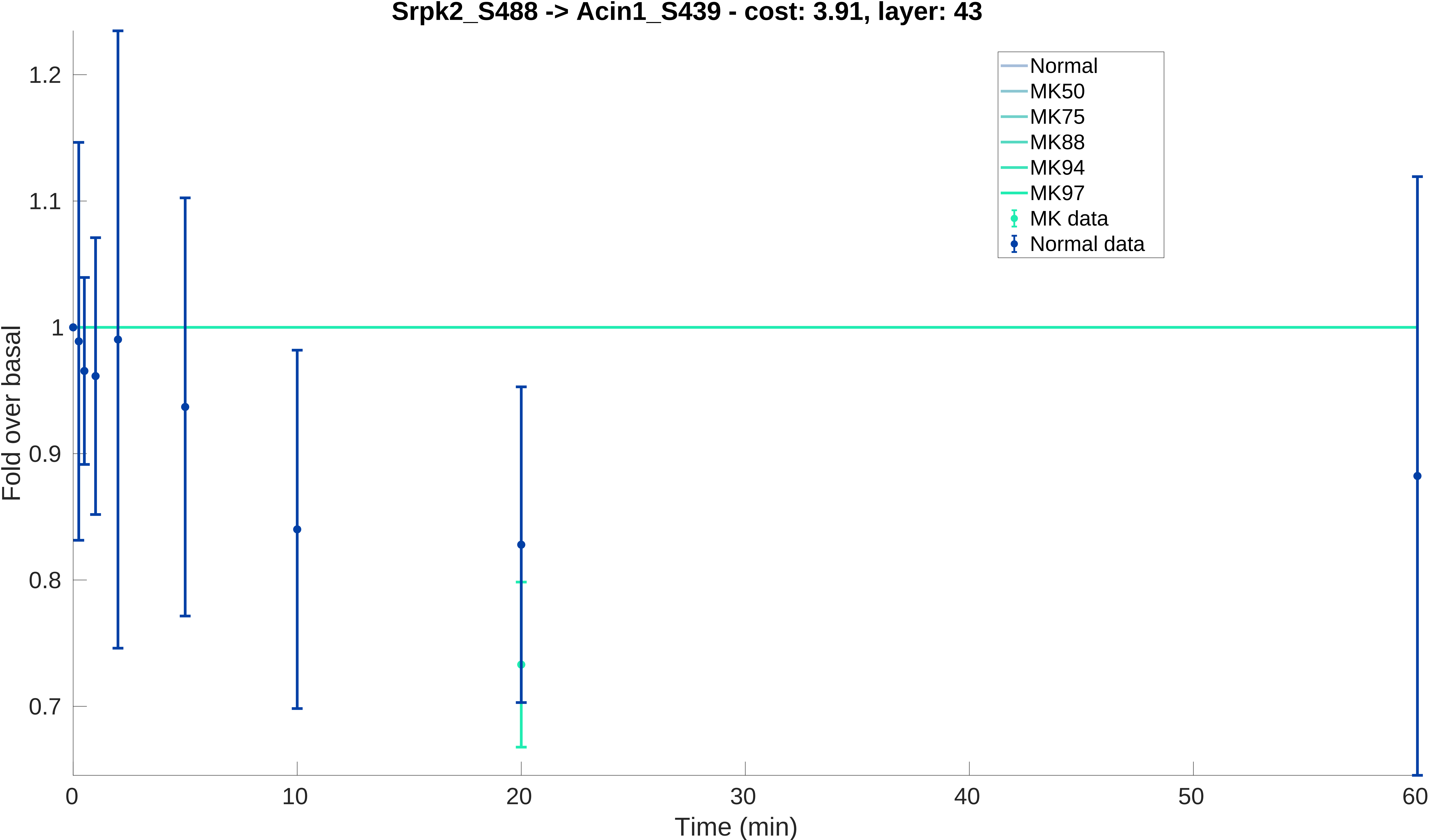
Srpk2_S488 -> Acin1_S439 - cost: 3.91, layer: 43
Fold over basal
Time (min)
Normal
MK50
MK75
MK88
MK94
MK97
MK data
Normal data
1.2
1.1
1
0.9
0.8
0.7
0
10
20
30
40
50
60

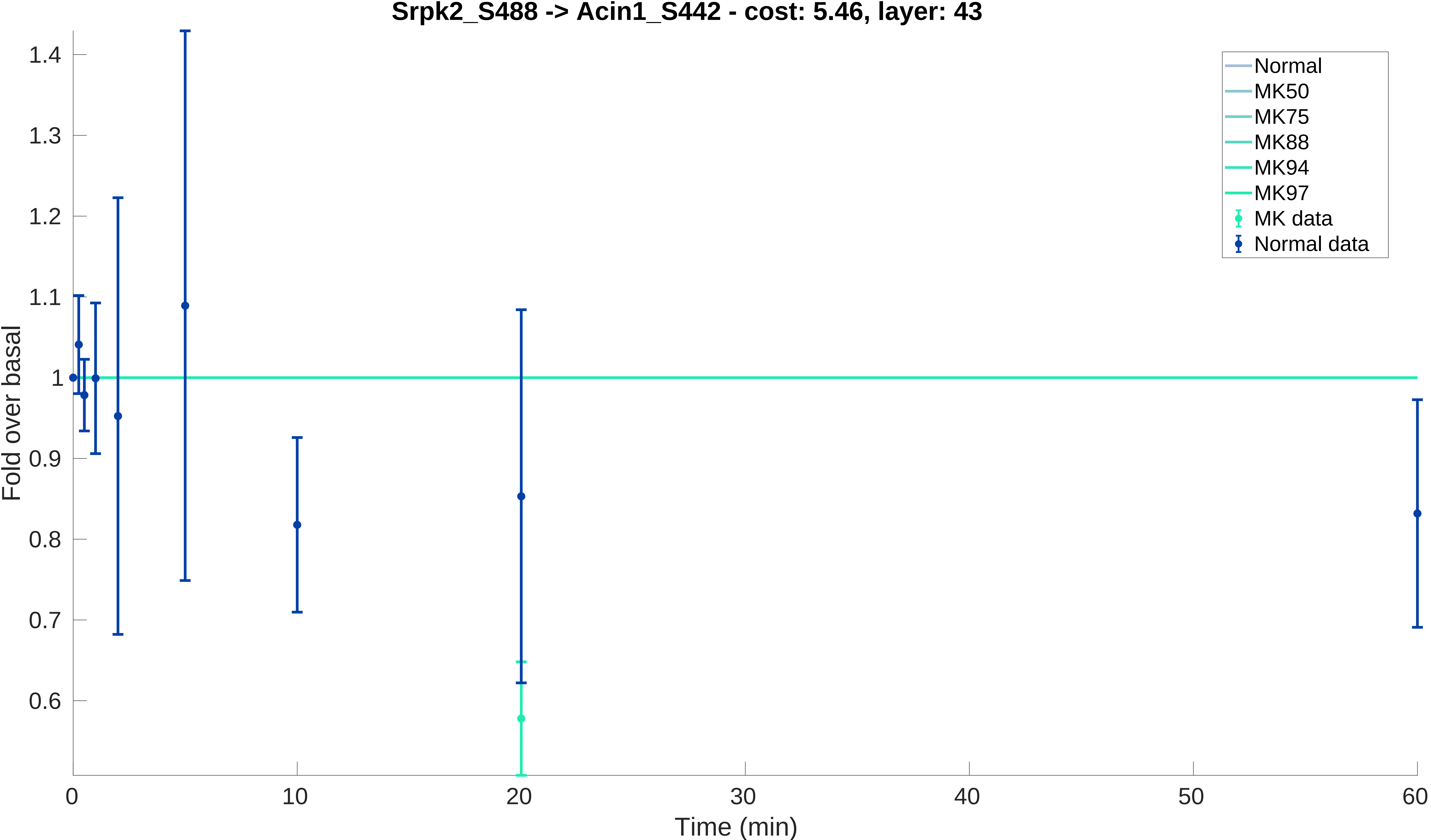

Srpk2_S488 -> Acin1_S442 - cost: 5.46, layer: 43
Normal
MK50
MK75
MK88
MK94
MK97
MK data
Normal data
Fold over basal
Time (min)
1.4
1.3
1.2
1.1
1
0.9
0.8
0.7
0.6
0
10
20
30
40
50
60

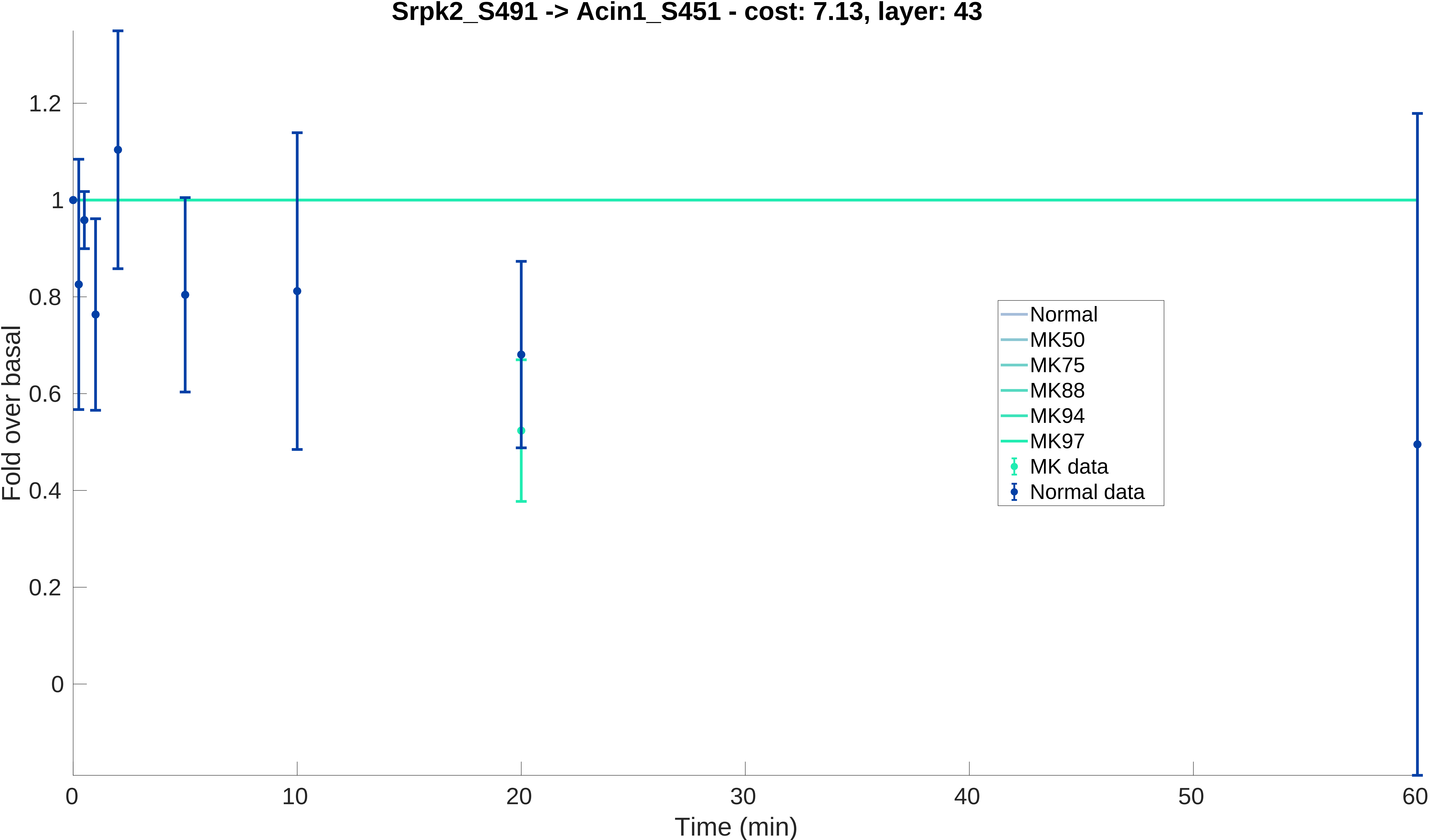

Srpk2_S491 -> Acin1_S451 - cost: 7.13, layer: 43
Fold over basal
Time (min)
1.2
1
0.8
0.6
0.4
0.2
0
0
10
20
30
40
50
60
Normal
MK50
MK75
MK88
MK94
MK97
MK data
Normal data

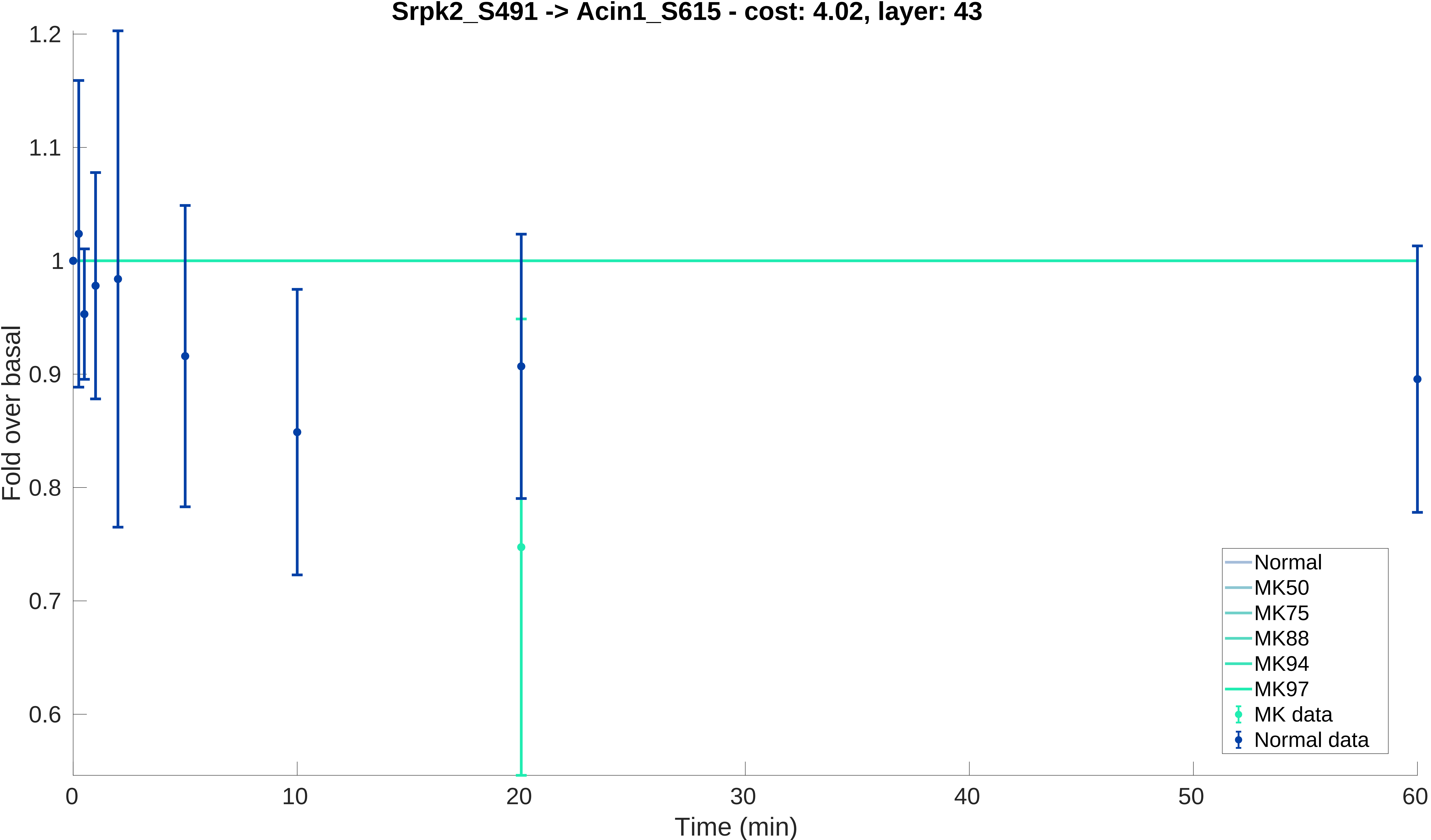
Srpk2_S491 -> Acin1_S615 - cost: 4.02, layer: 43
Fold over basal
1.2
1.1
1
0.9
0.8
0.7
0.6
0
10
20
30
40
50
60
Time (min)
Normal
MK50
MK75
MK88
MK94
MK97
MK data
Normal data

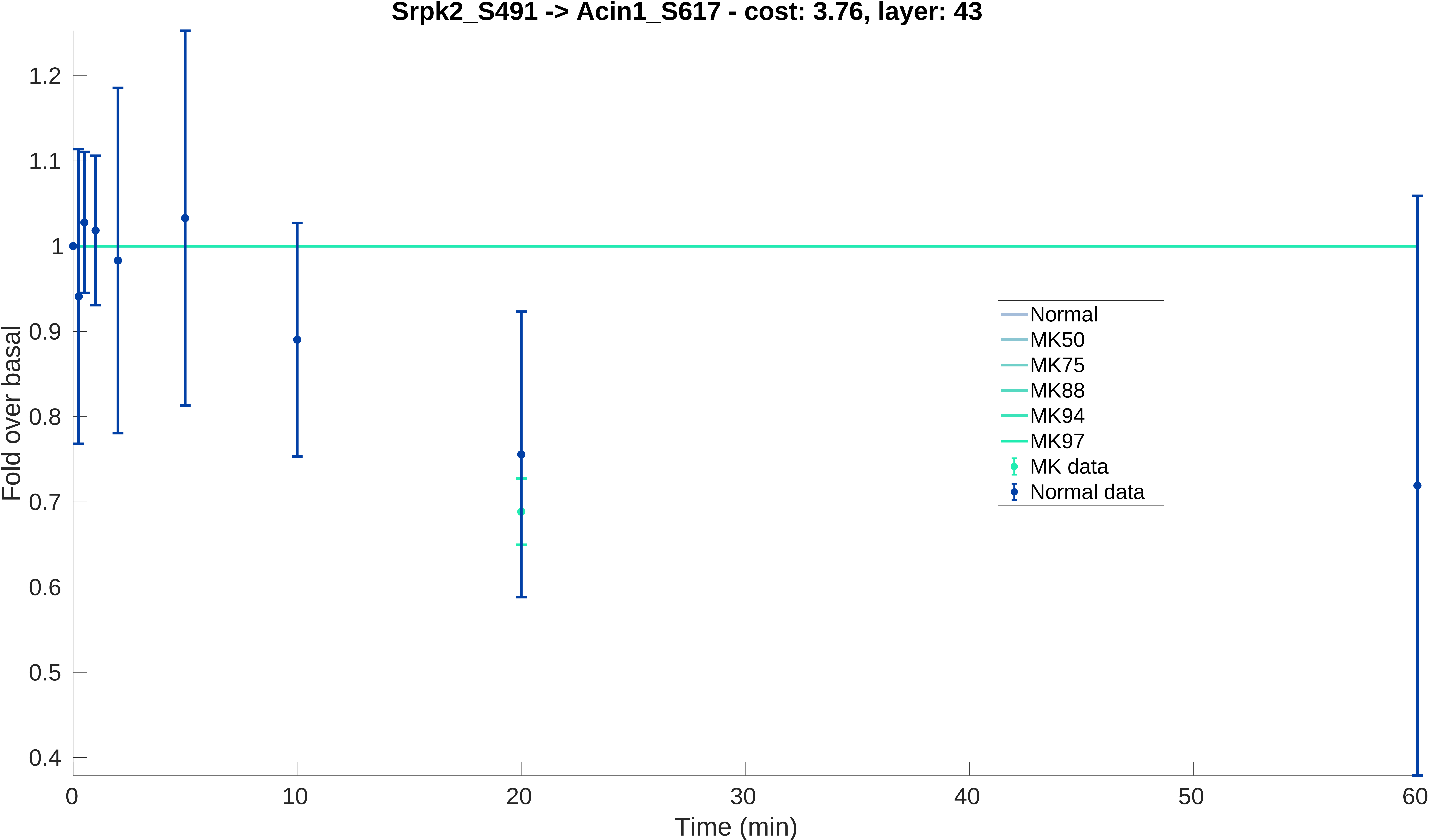

Srpk2_S491 -> Acin1_S617 - cost: 3.76, layer: 43
Fold over basal
Time (min)
0.4
0.5
0.6
0.7
0.8
0.9
1
1.1
1.2
0
10
20
30
40
50
60
Normal
MK50
MK75
MK88
MK94
MK97
MK data
Normal data

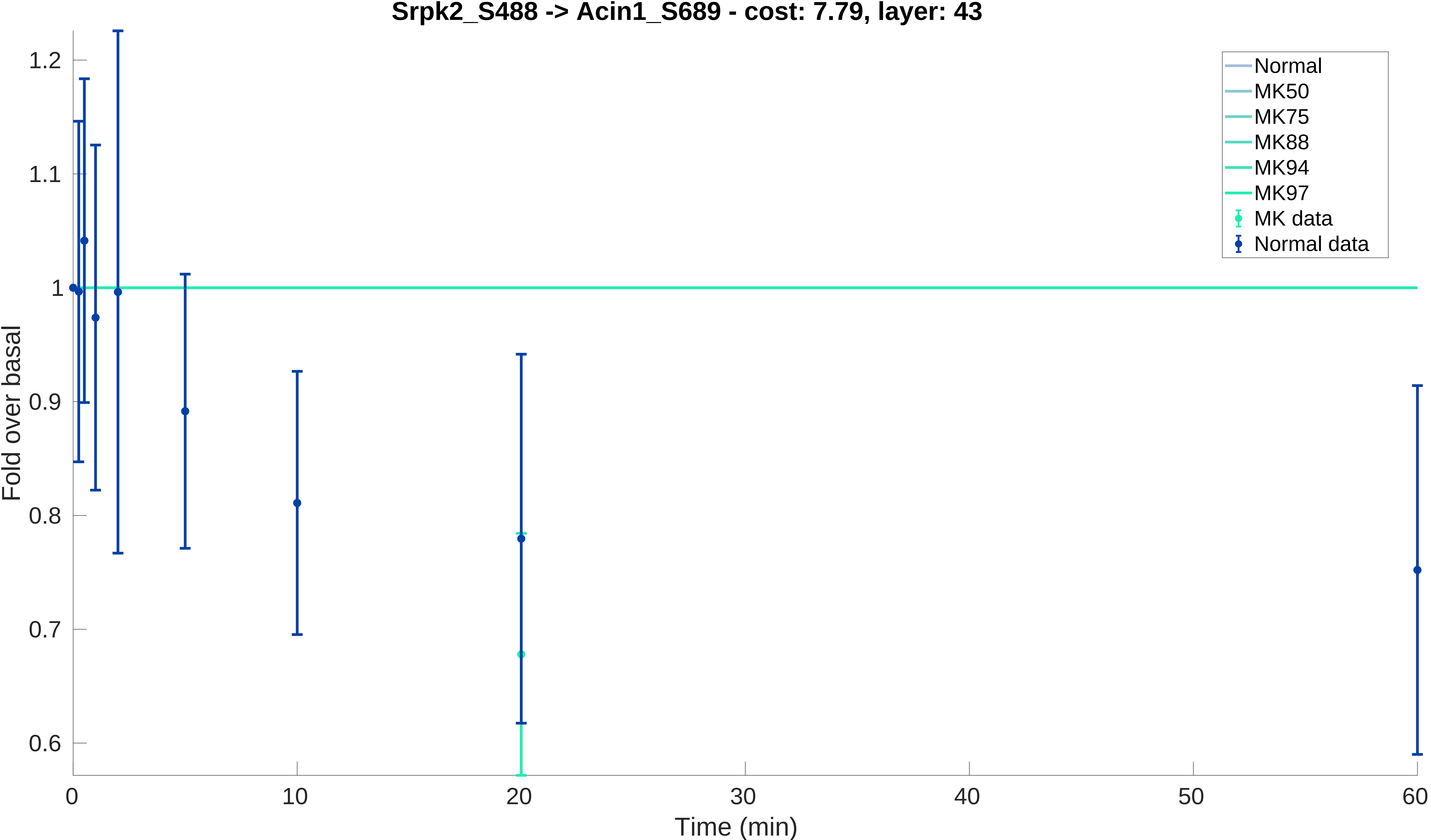

Srpk2_S488 -> Acin1_S689 - cost: 7.79, layer: 43
Normal
MK50
MK75
MK88
MK94
MK97
MK data
Normal data
Fold over basal
Time (min)
1.2
1.1
1
0.9
0.8
0.7
0.6
0
10
20
30
40
50
60

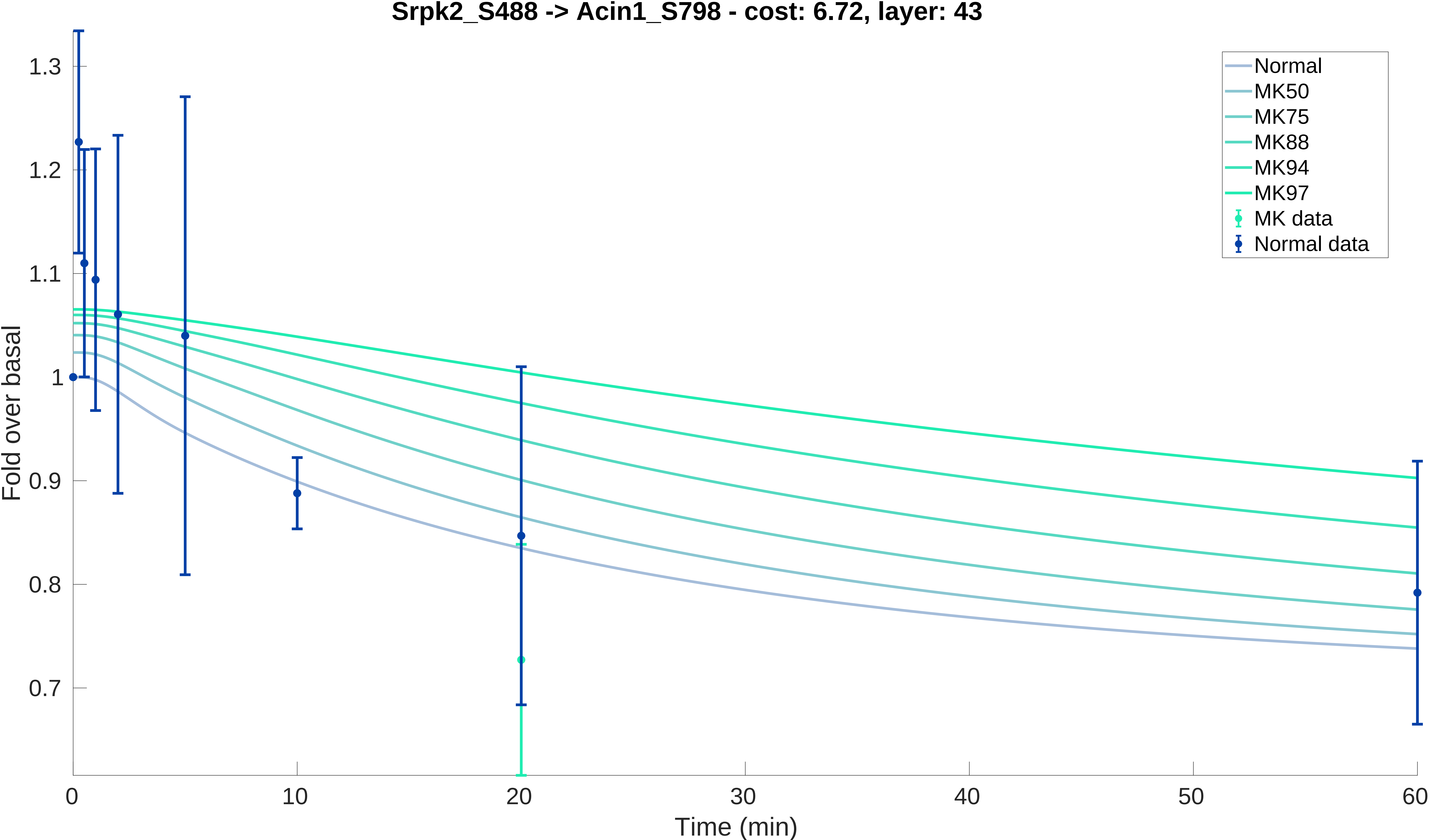

Srpk2_S488 -> Acin1_S798 - cost: 6.72, layer: 43
Normal
MK50
MK75
MK88
MK94
MK97
MK data
Normal data
1.3
1.2
1.1
1
0.9
0.8
0.7
Fold over basal
0
10
20
30
40
50
60
Time (min)

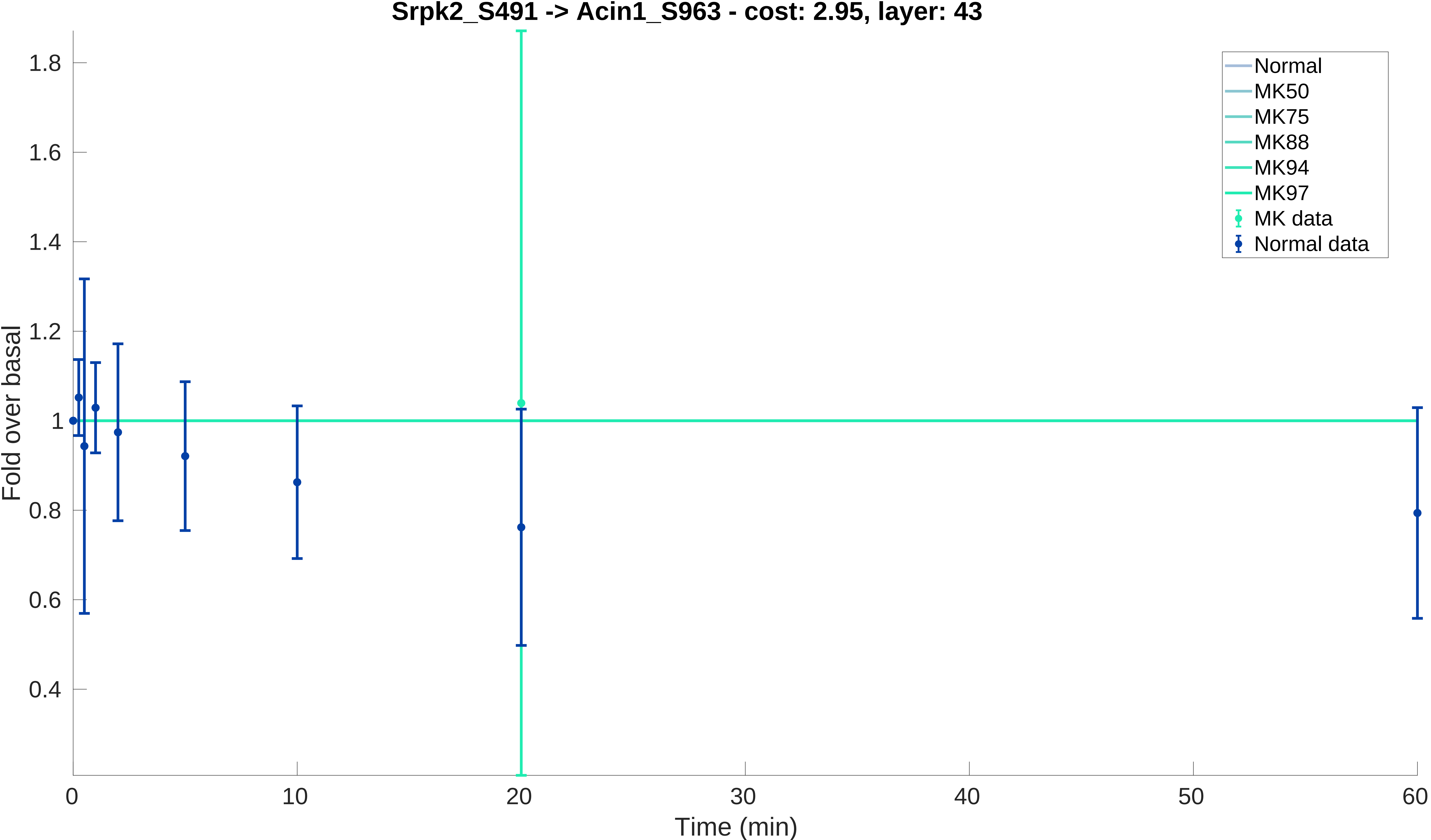

Srpk2_S491 -> Acin1_S963 - cost: 2.95, layer: 43
Fold over basal
1.8
1.6
1.4
1.2
1
0.8
0.6
0.4
0
10
20
30
40
50
60
Time (min)
Normal
MK50
MK75
MK88
MK94
MK97
MK data
Normal data

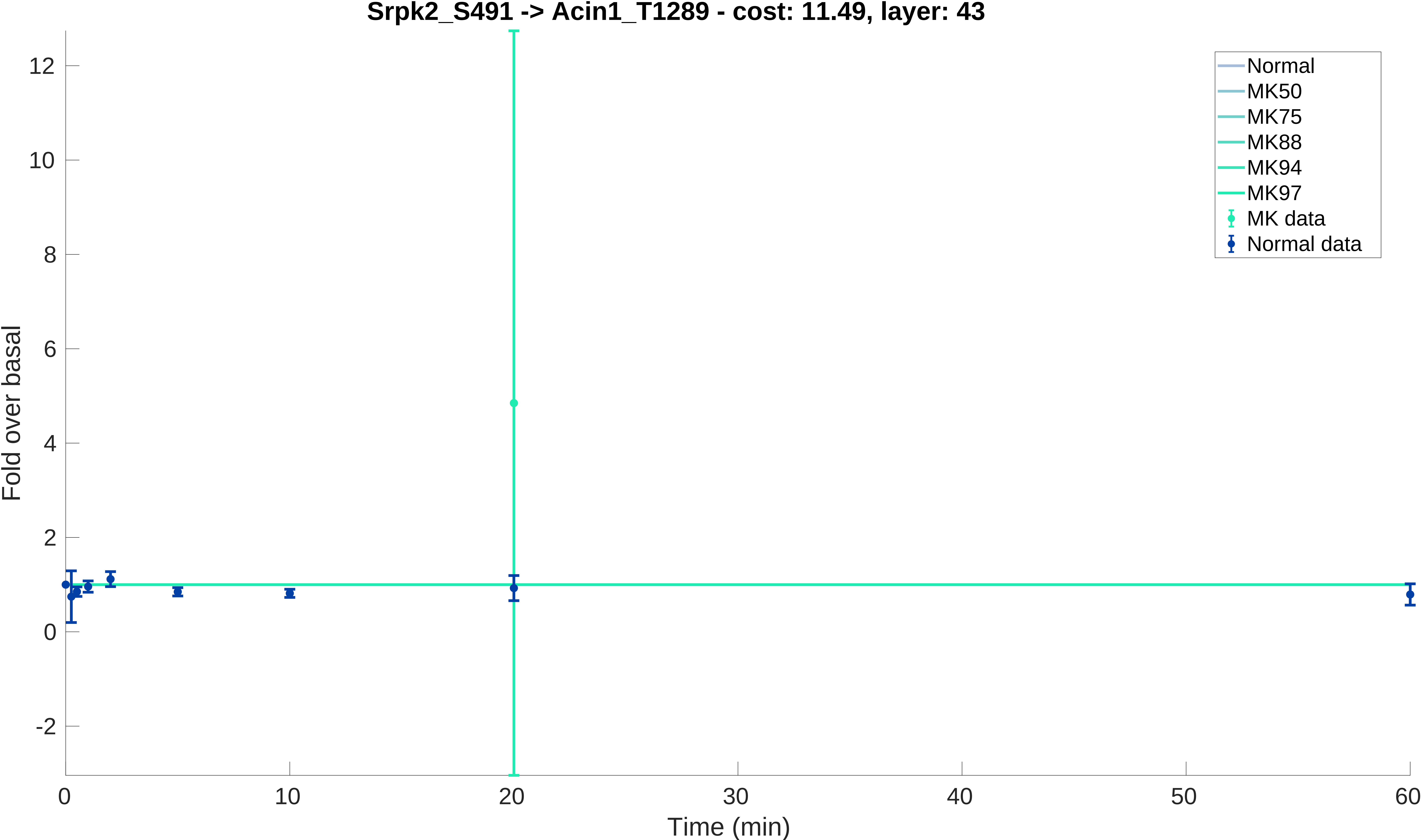

Srpk2_S491 -> Acin1_T1289 - cost: 11.49, layer: 43
Fold over basal
Time (min)
12
10
8
6
4
2
0
-2
0
10
20
30
40
50
60
Normal
MK50
MK75
MK88
MK94
MK97
MK data
Normal data

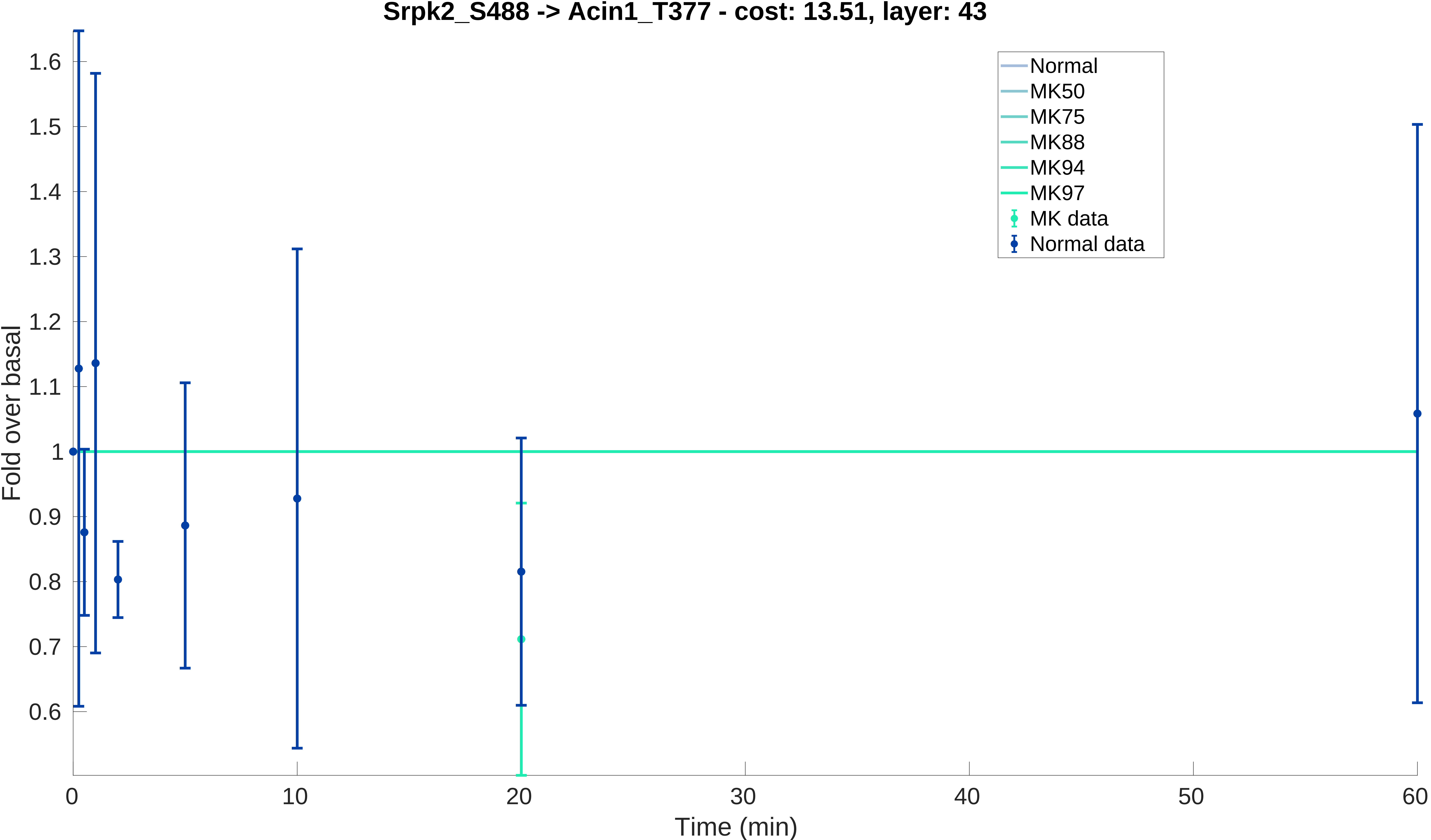

Srpk2_S488 -> Acin1_T377 - cost: 13.51, layer: 43
Normal
MK50
MK75
MK88
MK94
MK97
MK data
Normal data
Fold over basal
Time (min)
1.6
1.5
1.4
1.3
1.2
1.1
1
0.9
0.8
0.7
0.6
0
10
20
30
40
50
60

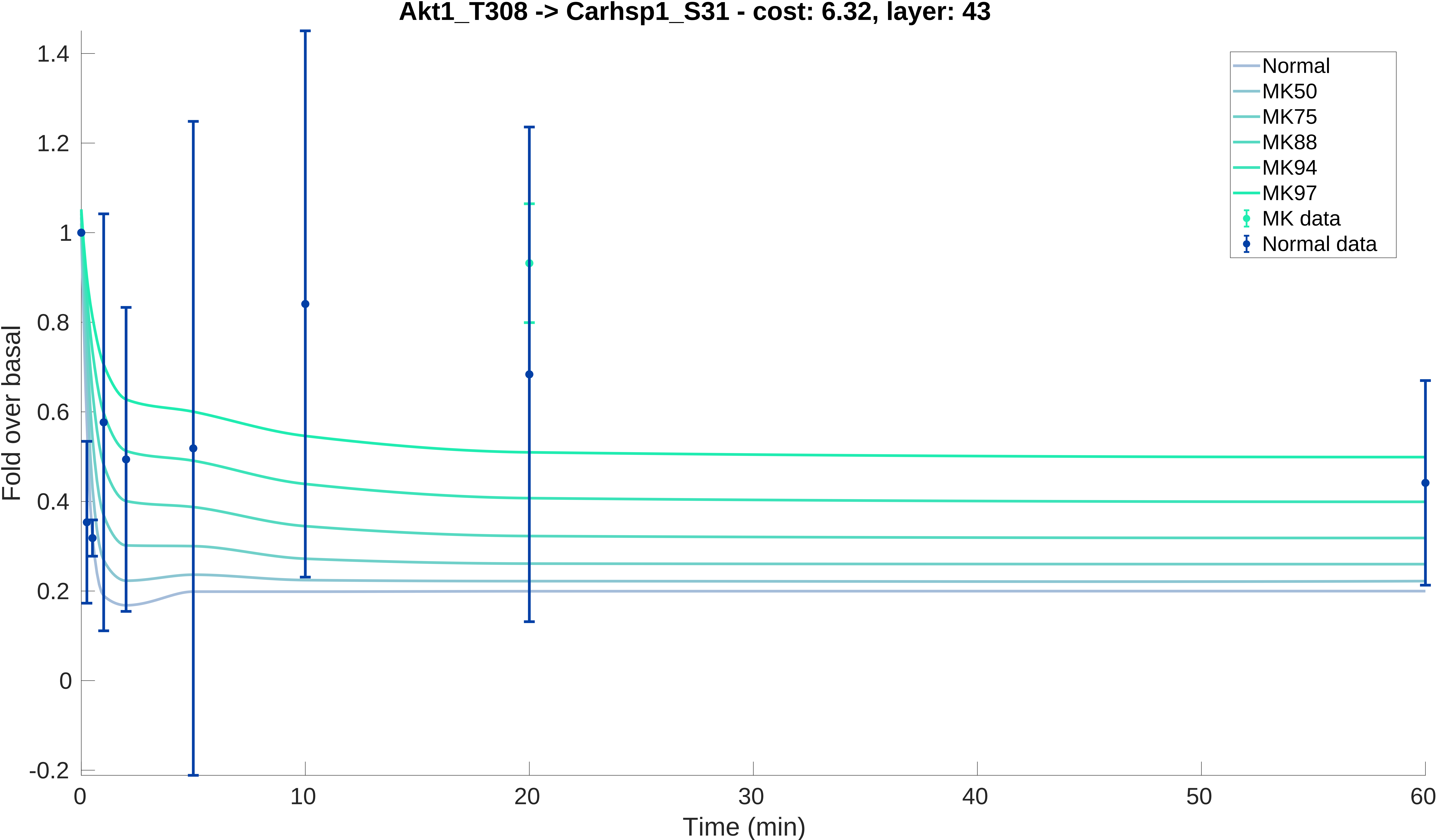

Akt1_T308 -> Carhsp1_S31 - cost: 6.32, layer: 43
Fold over basal
Time (min)
Normal
MK50
MK75
MK88
MK94
MK97
MK data
Normal data
1.4
1.2
1
0.8
0.6
0.4
0.2
0
-0.2
0
10
20
30
40
50
60

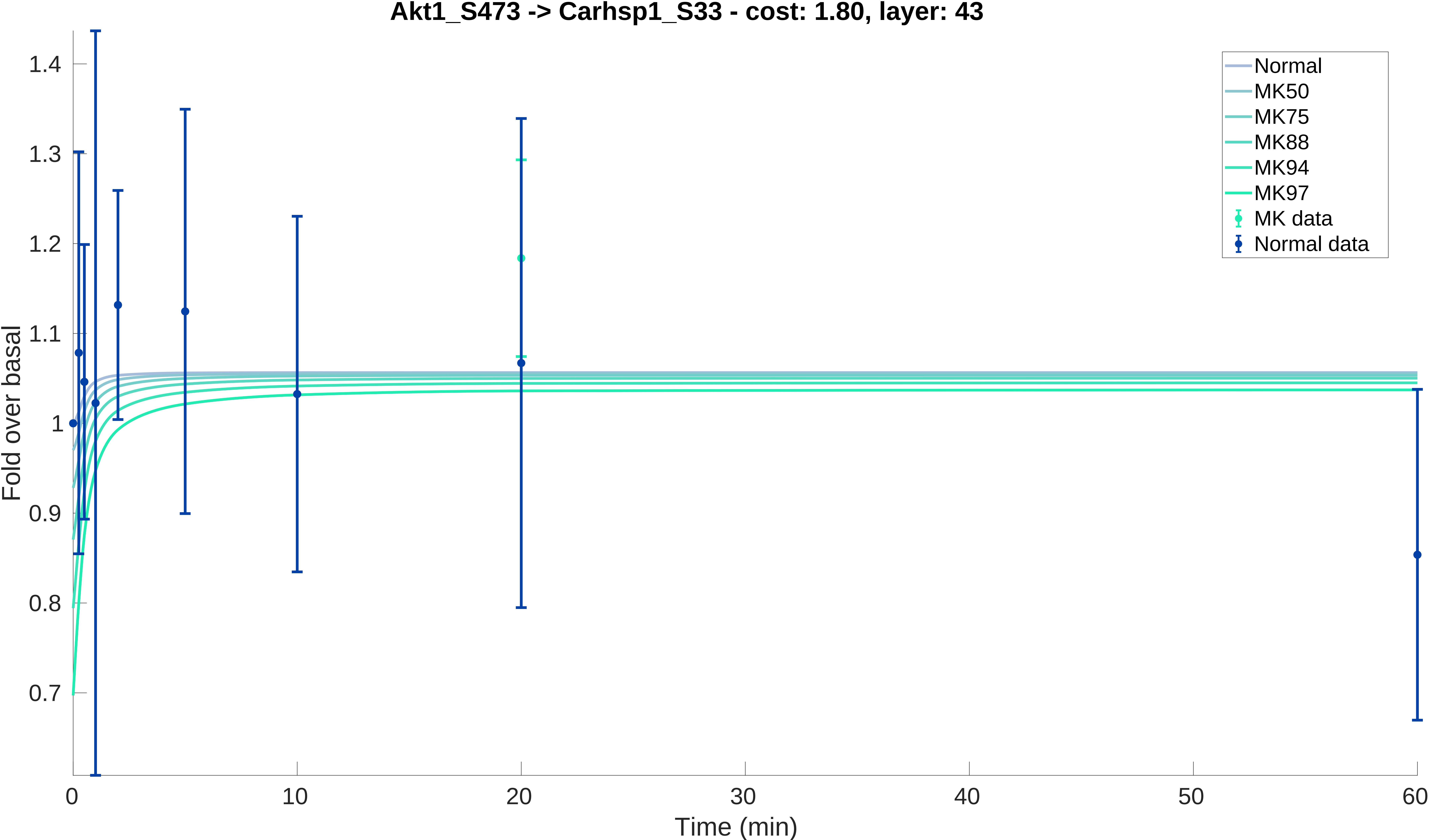

Akt1_S473 -> Carhsp1_S33 - cost: 1.80, layer: 43
Fold over basal
Time (min)
1.4
1.3
1.2
1.1
1
0.9
0.8
0.7
0
10
20
30
40
50
60
Normal
MK50
MK75
MK88
MK94
MK97
MK data
Normal data

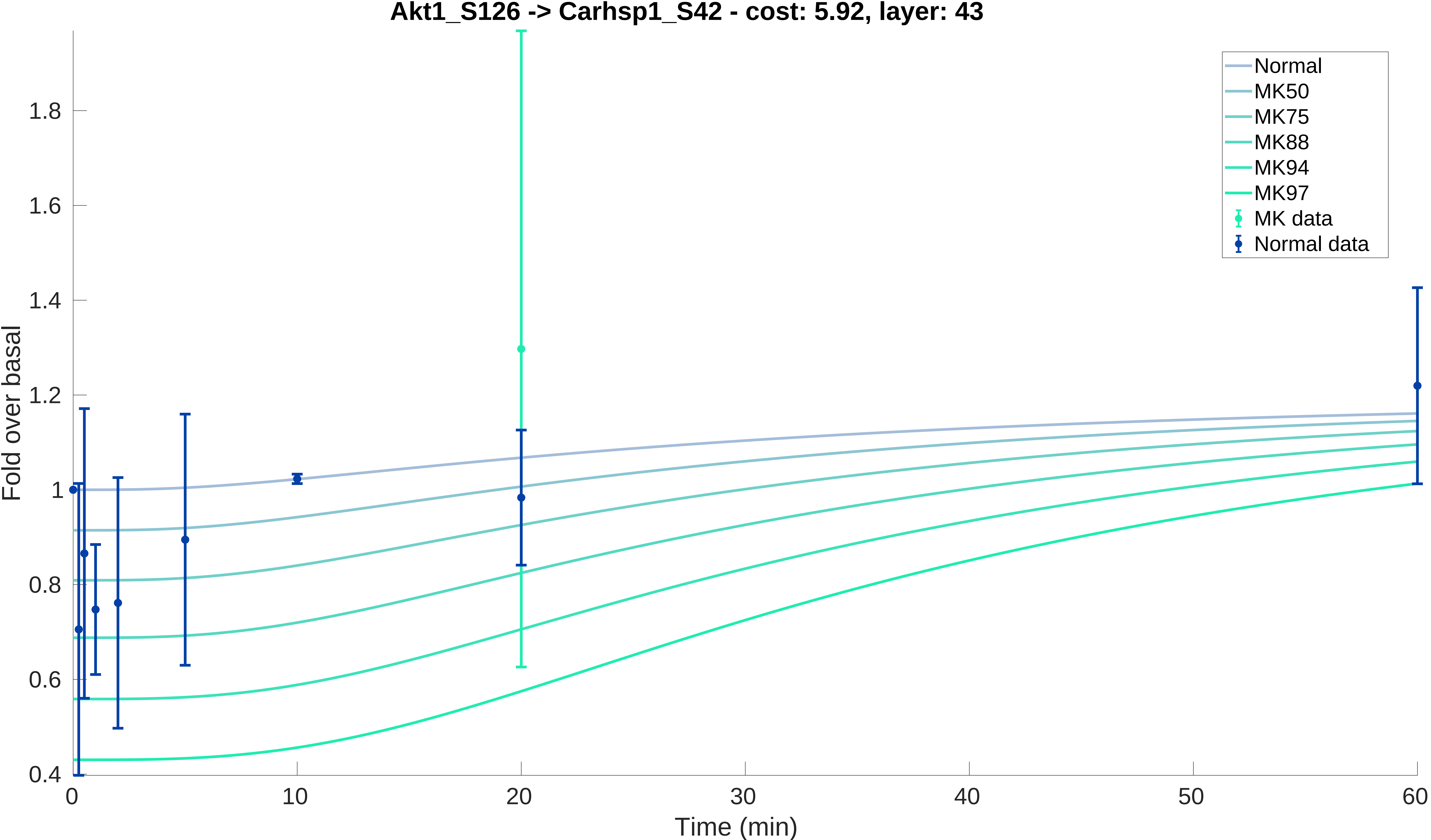

Akt1_S126 -> Carhsp1_S42 - cost: 5.92, layer: 43
Normal
MK50
MK75
MK88
MK94
MK97
MK data
Normal data
Fold over basal
Time (min)
0.4
0.6
0.8
1
1.2
1.4
1.6
1.8
0
10
20
30
40
50
60

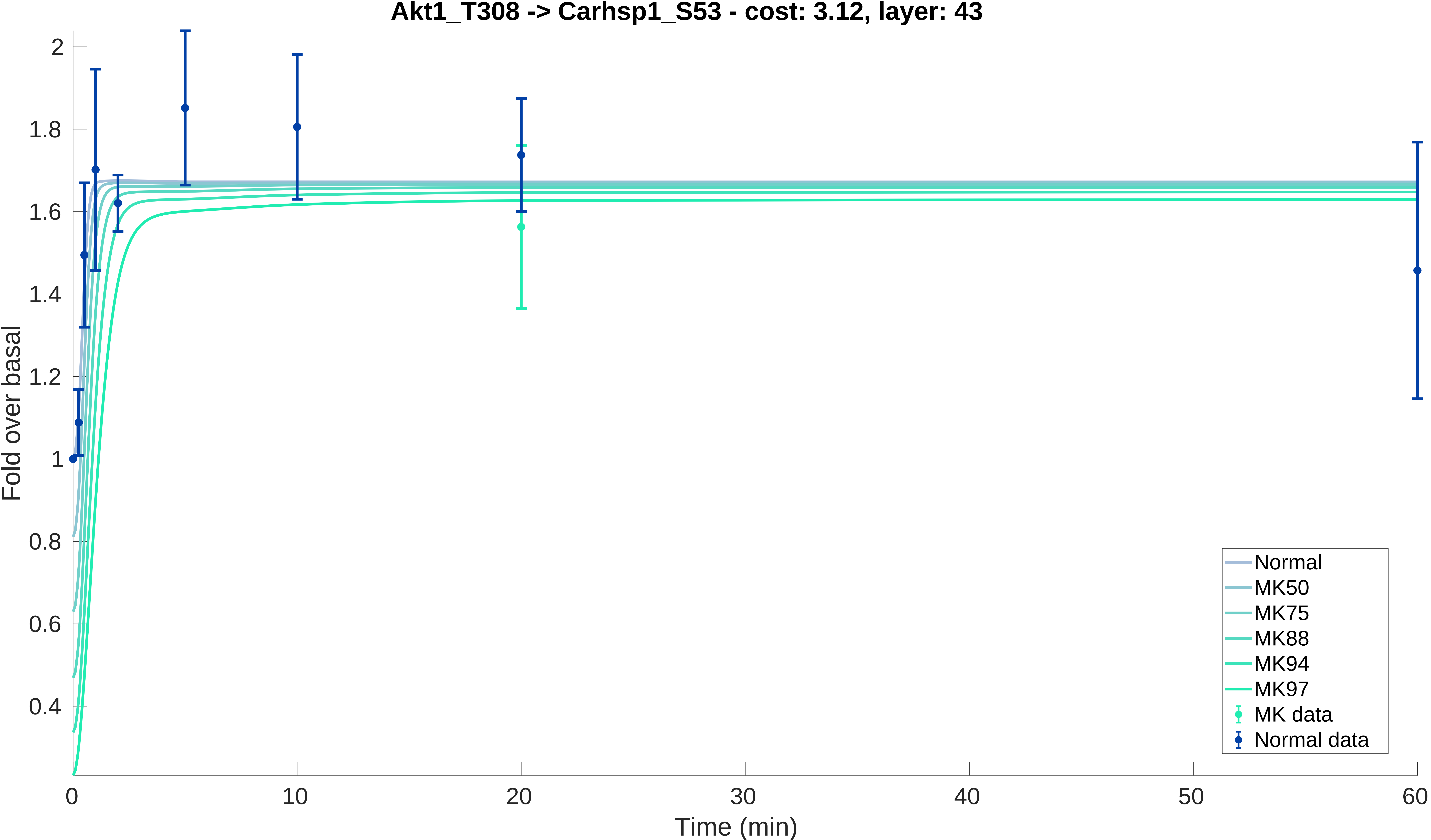

Akt1_T308 -> Carhsp1_S53 - cost: 3.12, layer: 43
Fold over basal
Time (min)
2
1.8
1.6
1.4
1.2
1
0.8
0.6
0.4
0
10
20
30
40
50
60
Normal
MK50
MK75
MK88
MK94
MK97
MK data
Normal data

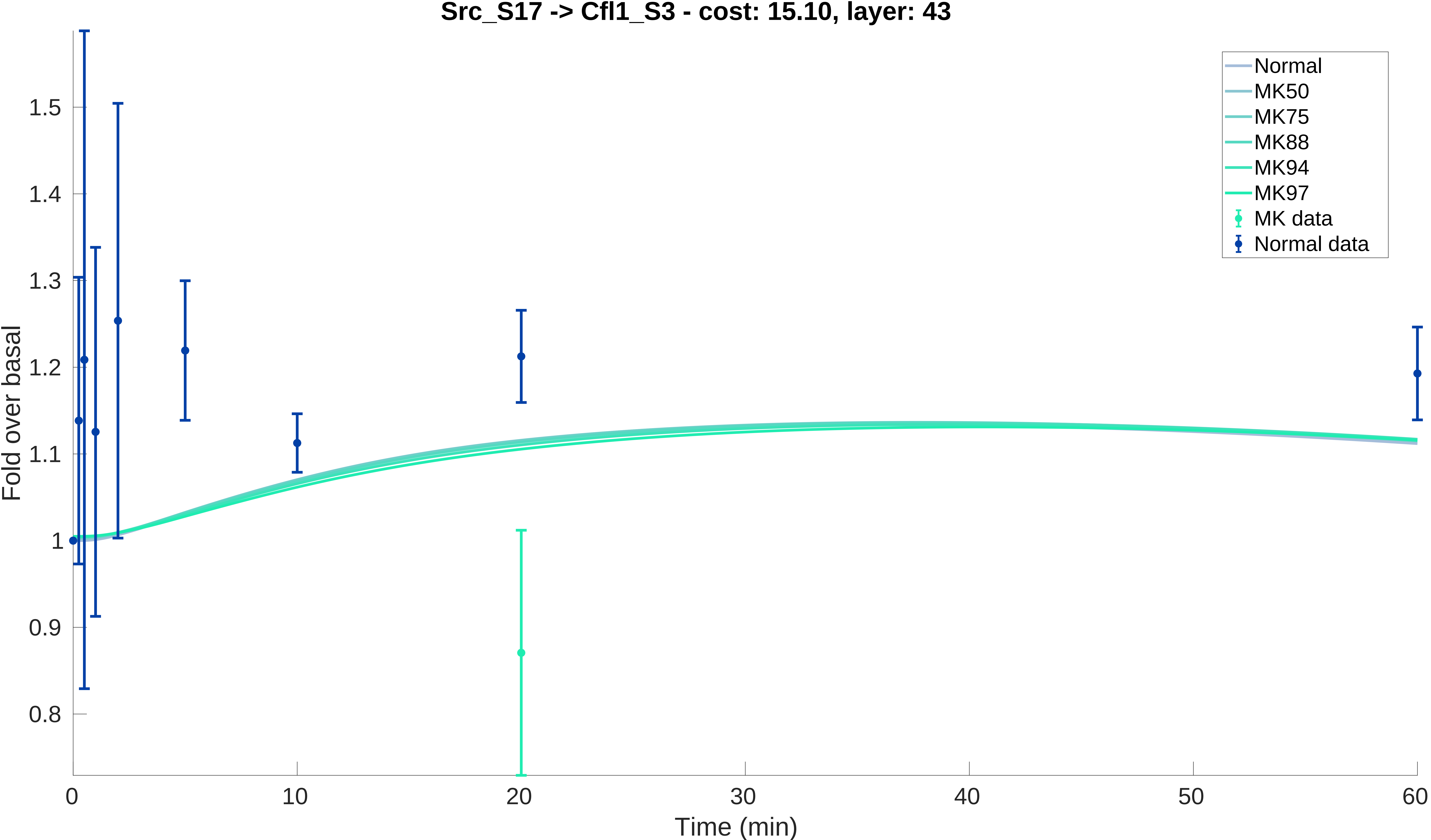

Src_S17 -> Cfl1_S3 - cost: 15.10, layer: 43
Fold over basal
1.5
1.4
1.3
1.2
1.1
1
0.9
0.8
0
10
20
30
40
50
60
Time (min)
Normal
MK50
MK75
MK88
MK94
MK97
MK data
Normal data

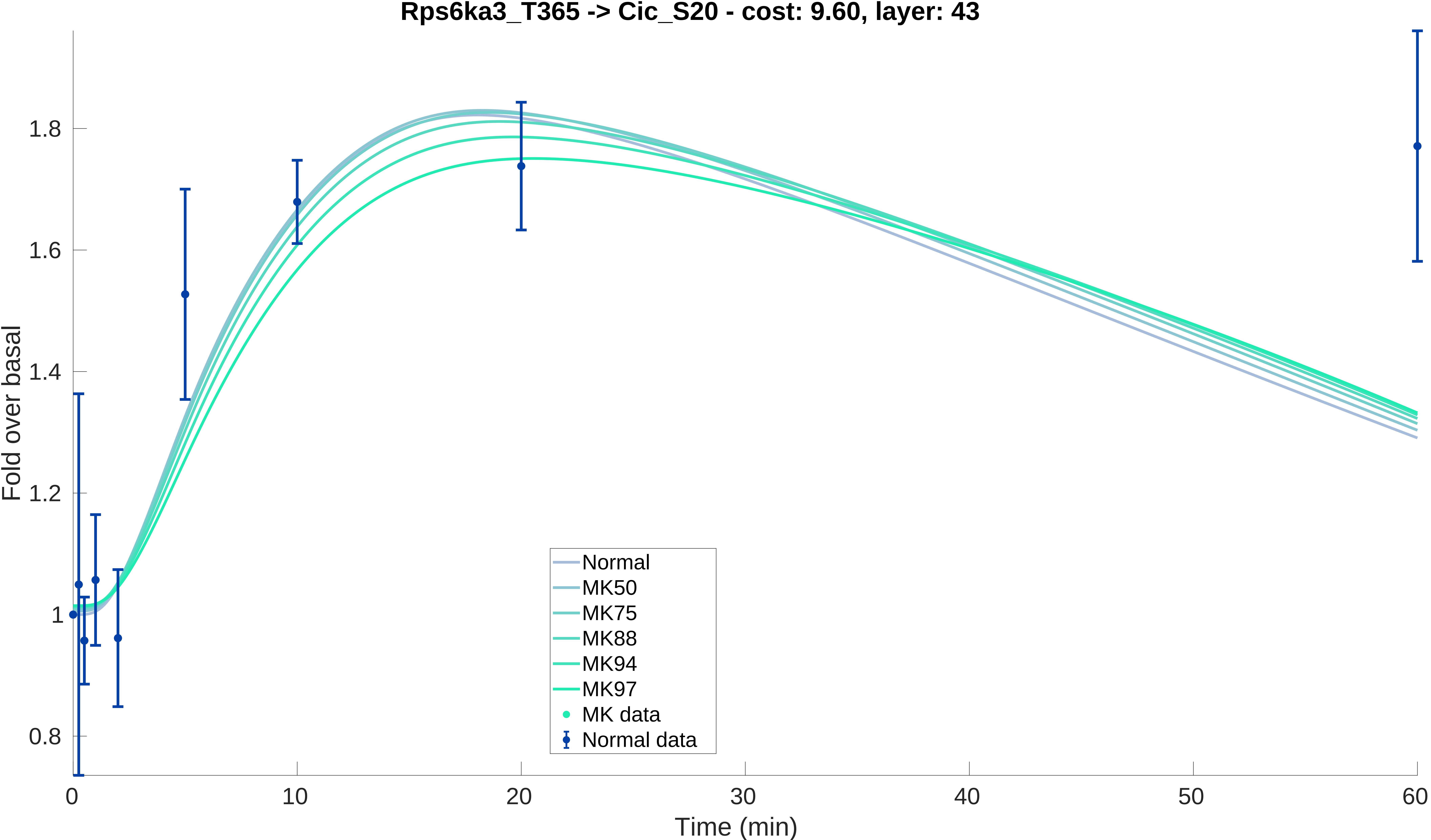
Rps6ka3_T365 -> Cic_S20 - cost: 9.60, layer: 43
Fold over basal
1.8
1.6
1.4
1.2
1
0.8
0
10
20
30
40
50
60
Time (min)
Normal
MK50
MK75
MK88
MK94
MK97
MK data
Normal data

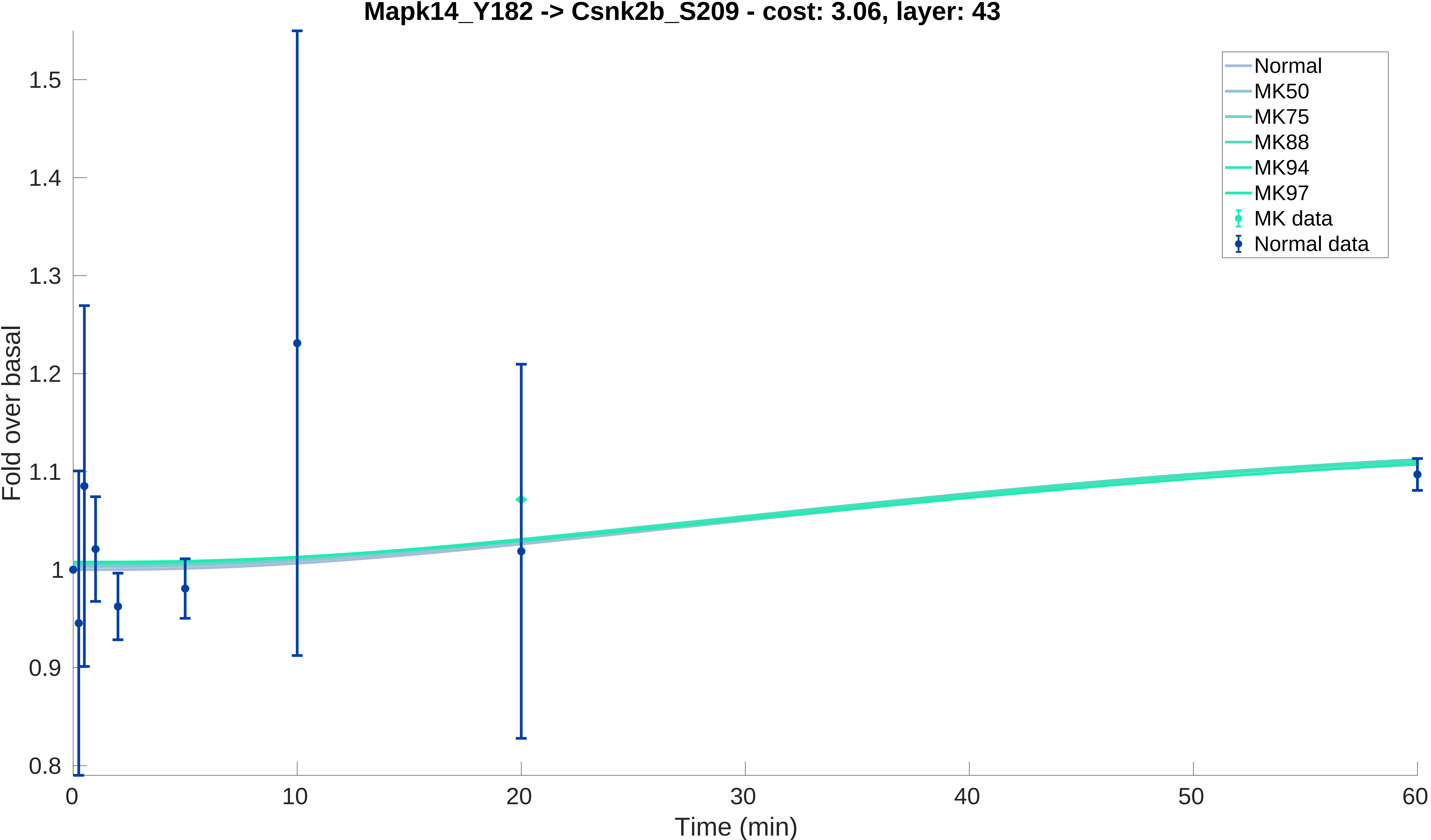
Mapk14_Y182 -> Csnk2b_S209 - cost: 3.06, layer: 43
Fold over basal
Time (min)
0.8
0.9
1
1.1
1.2
1.3
1.4
1.5
0
10
20
30
40
50
60
Normal
MK50
MK75
MK88
MK94
MK97
MK data
Normal data

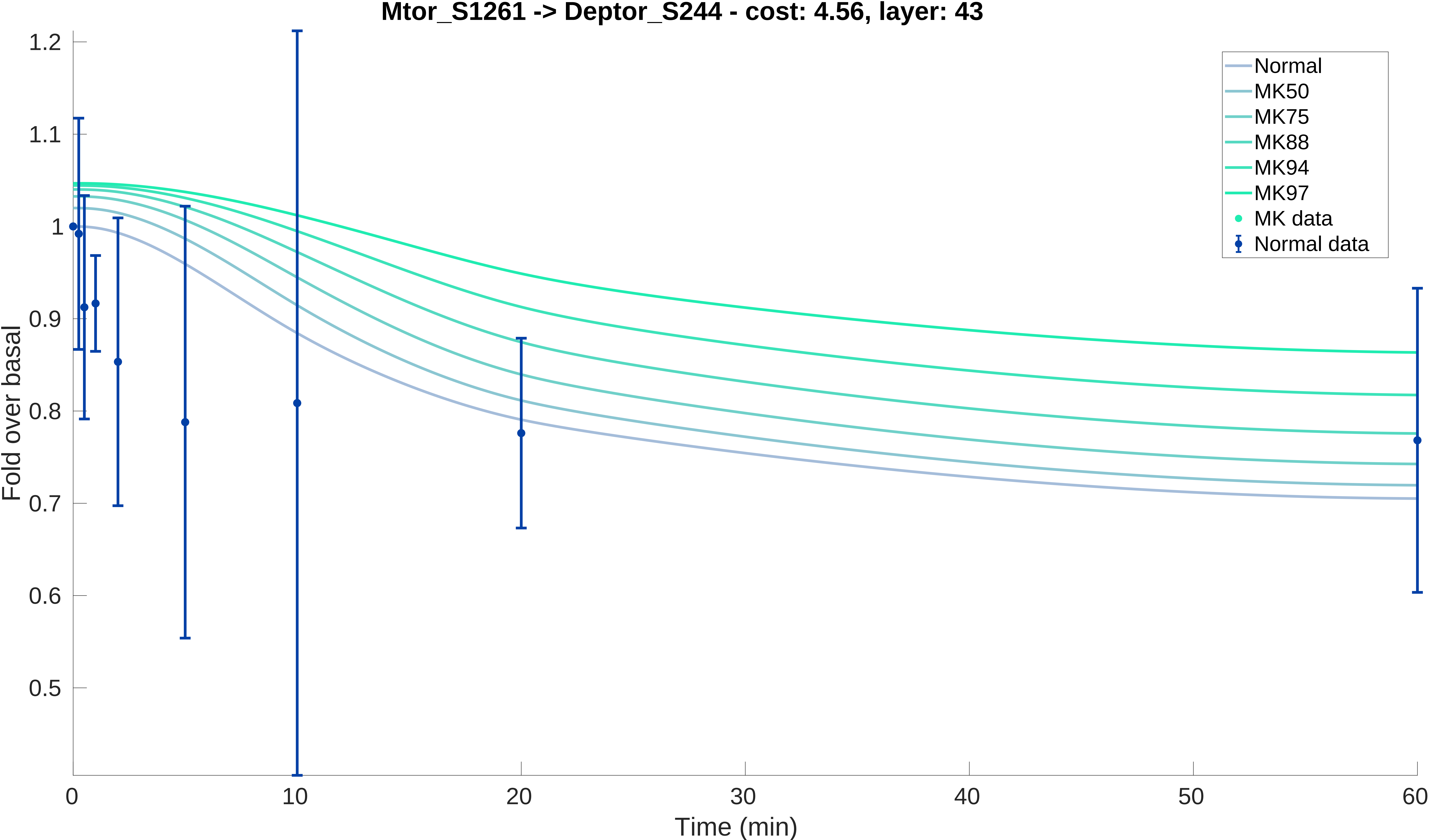

Mtor_S1261 -> Deptor_S244 - cost: 4.56, layer: 43
Normal
MK50
MK75
MK88
MK94
MK97
MK data
Normal data
Fold over basal
Time (min)
1.2
1.1
1
0.9
0.8
0.7
0.6
0.5
0
10
20
30
40
50
60

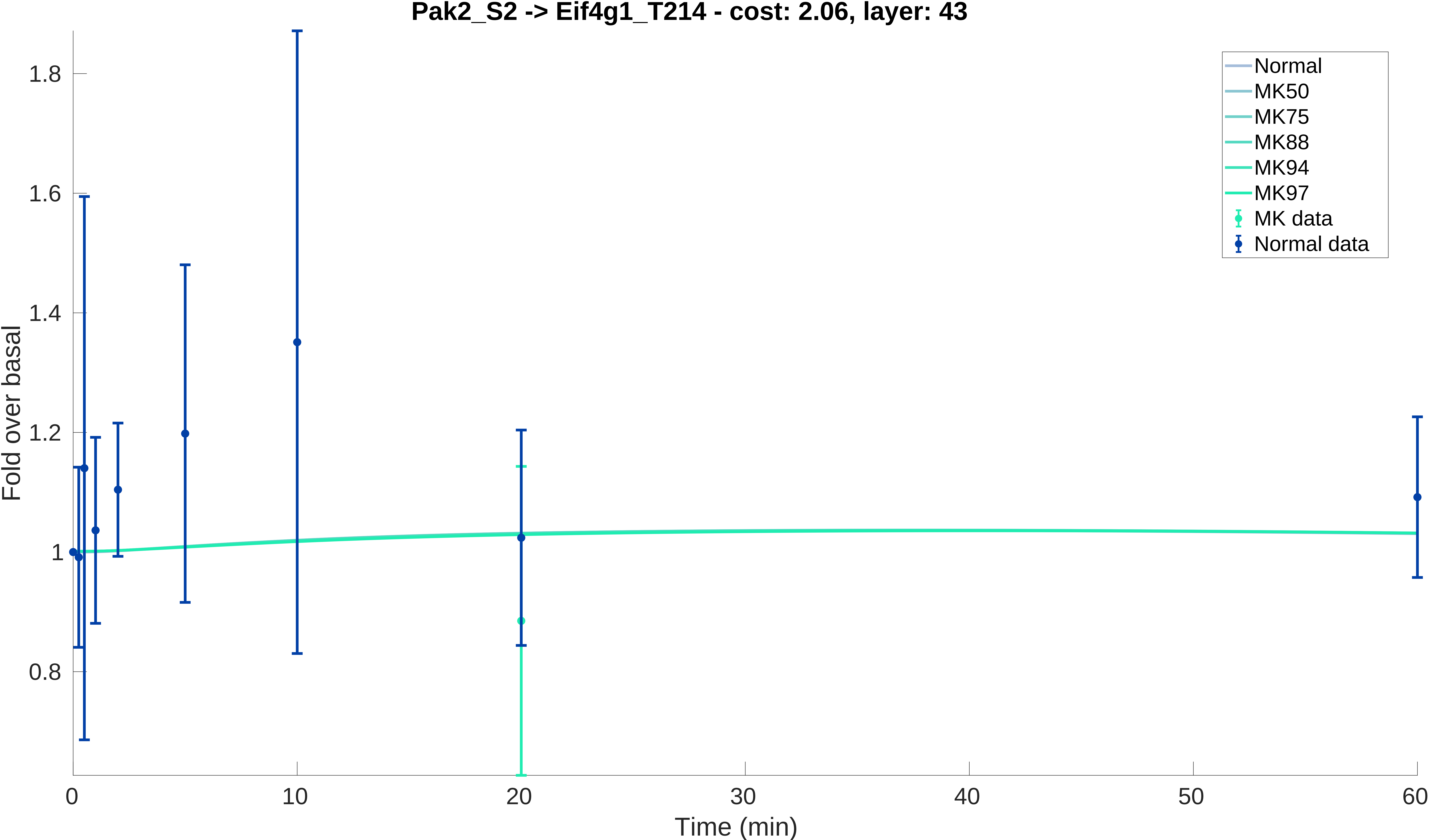

Pak2_S2 -> Eif4g1_T214 - cost: 2.06, layer: 43
Fold over basal
1.8
1.6
1.4
1.2
1
0.8
0
10
20
30
40
50
60
Time (min)
Normal
MK50
MK75
MK88
MK94
MK97
MK data
Normal data

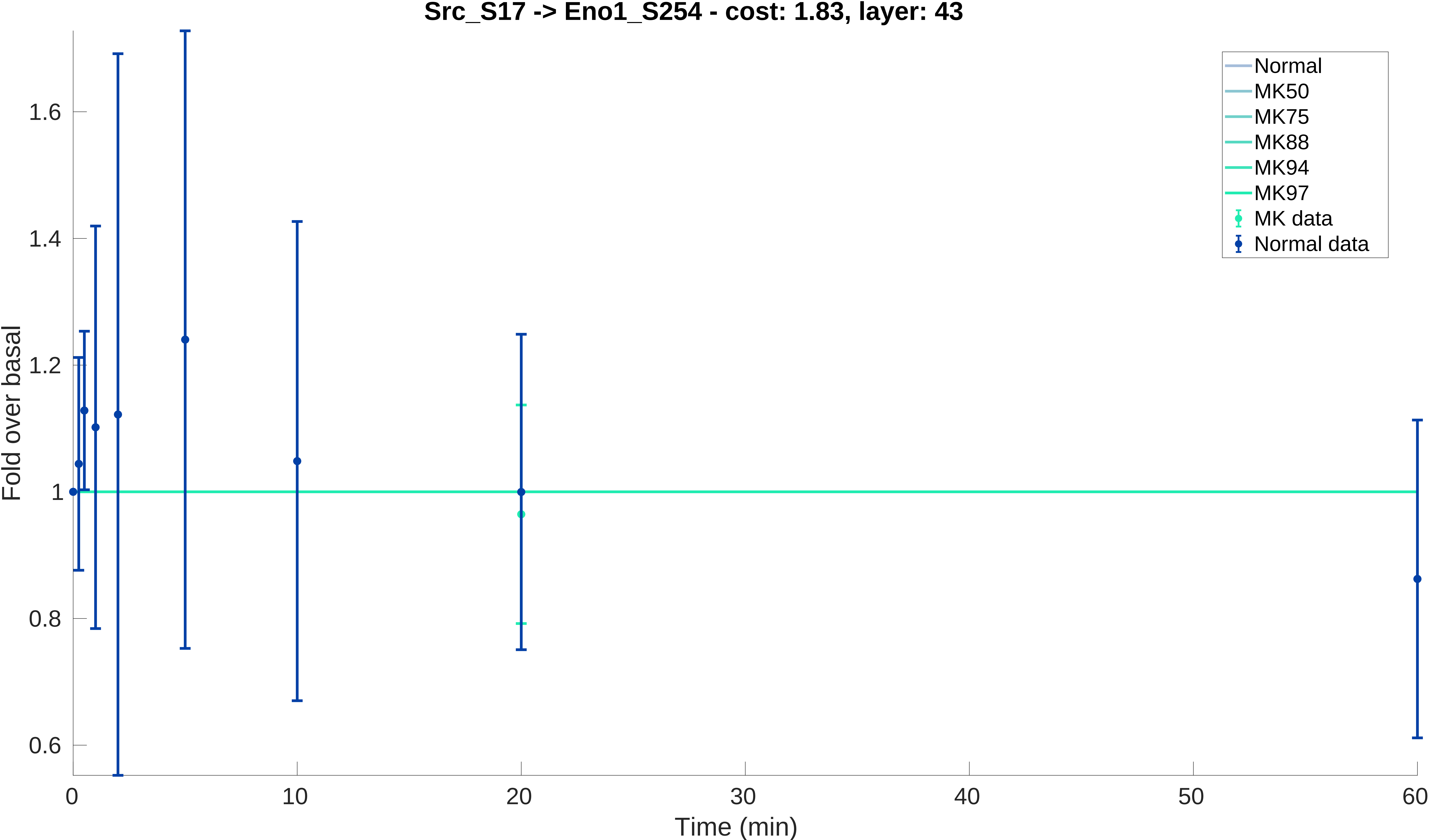

Src_S17 -> Eno1_S254 - cost: 1.83, layer: 43
Fold over basal
1.6
1.4
1.2
1
0.8
0.6
0
10
20
30
40
50
60
Time (min)
Normal
MK50
MK75
MK88
MK94
MK97
MK data
Normal data

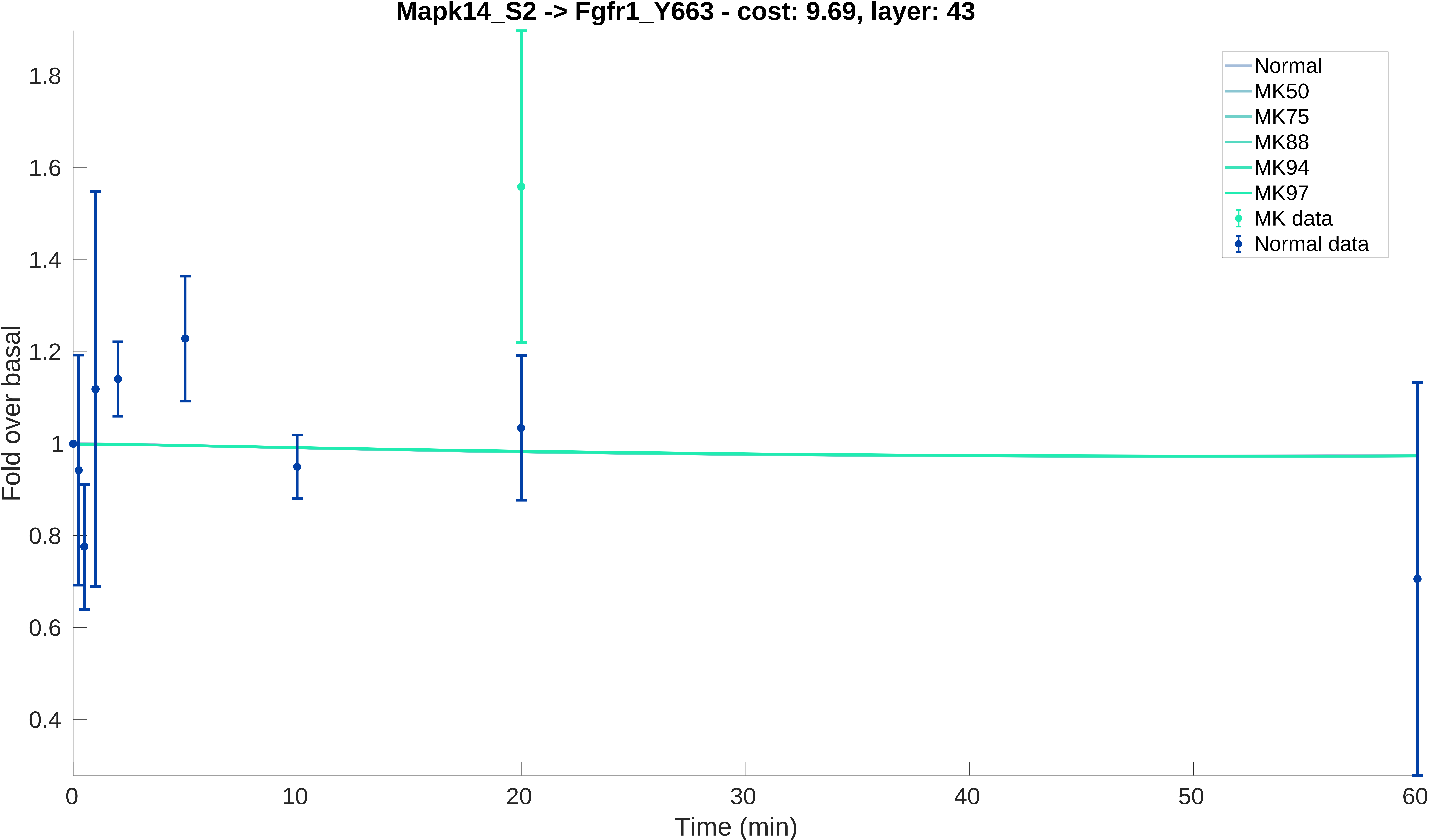

Mapk14_S2 -> Fgfr1_Y663 - cost: 9.69, layer: 43
Fold over basal
Time (min)
0.4
0.6
0.8
1
1.2
1.4
1.6
1.8
0
10
20
30
40
50
60
Normal
MK50
MK75
MK88
MK94
MK97
MK data
Normal data

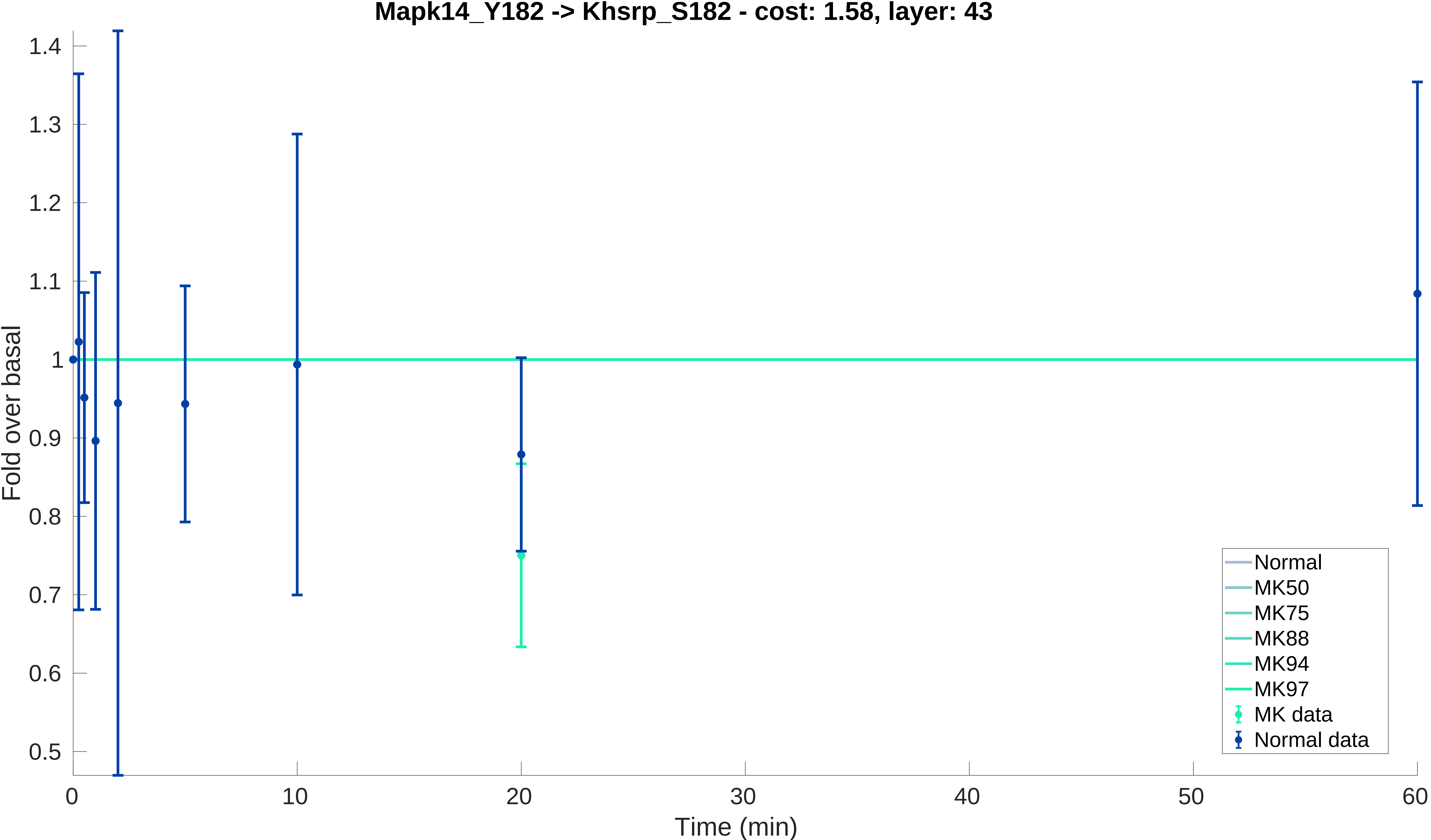

Mapk14_Y182 -> Khsrp_S182 - cost: 1.58, layer: 43
Fold over basal
Time (min)
1.4
1.3
1.2
1.1
1
0.9
0.8
0.7
0.6
0.5
0
10
20
30
40
50
60
Normal
MK50
MK75
MK88
MK94
MK97
MK data
Normal data

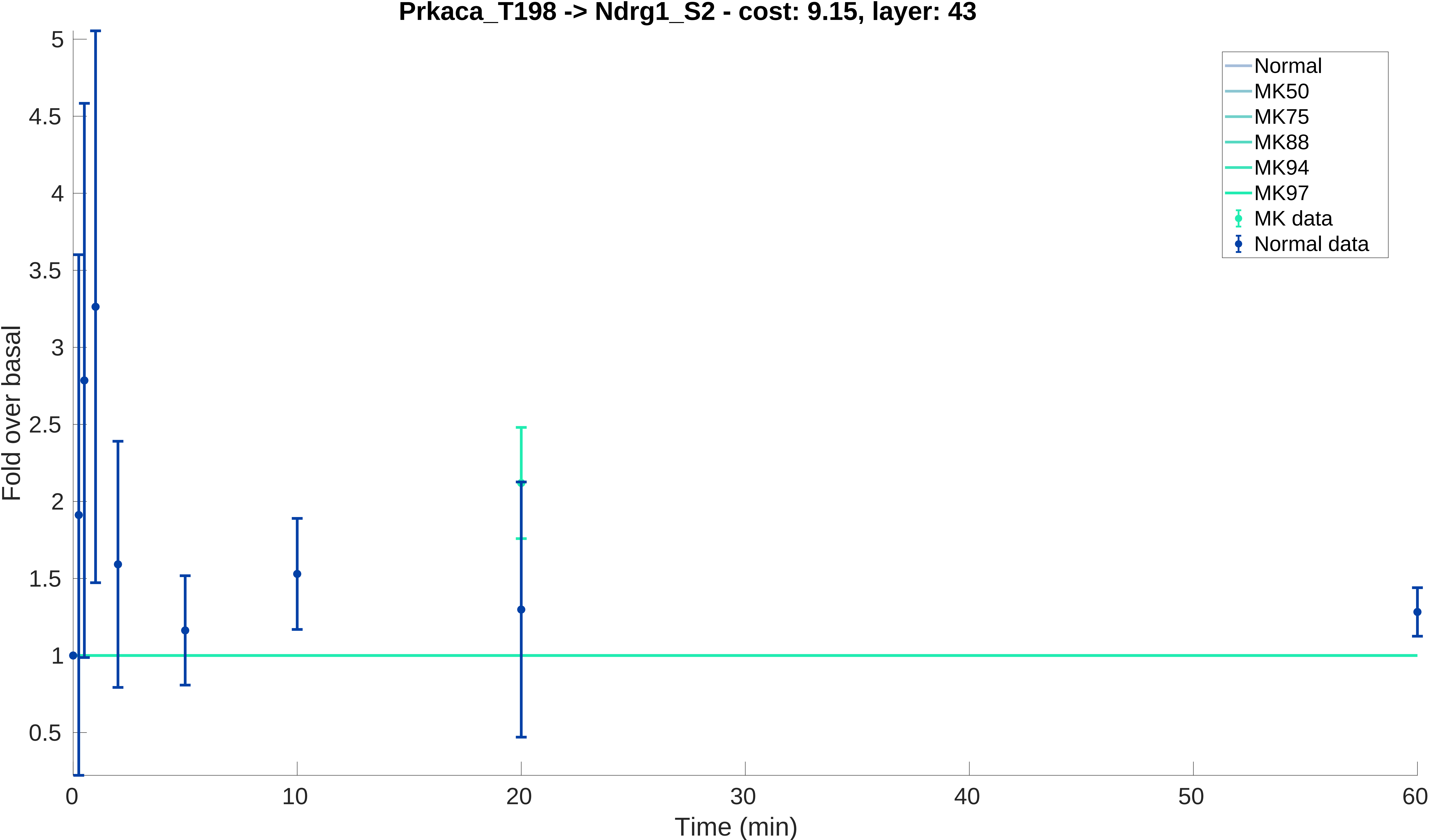
Prrkaca_T198 -> Ndrg1_S2 - cost: 9.15, layer: 43
Fold over basal
Time (min)
0
10
20
30
40
50
60
0.5
1
1.5
2
2.5
3
3.5
4
4.5
5
Normal
MK50
MK75
MK88
MK94
MK97
MK data
Normal data

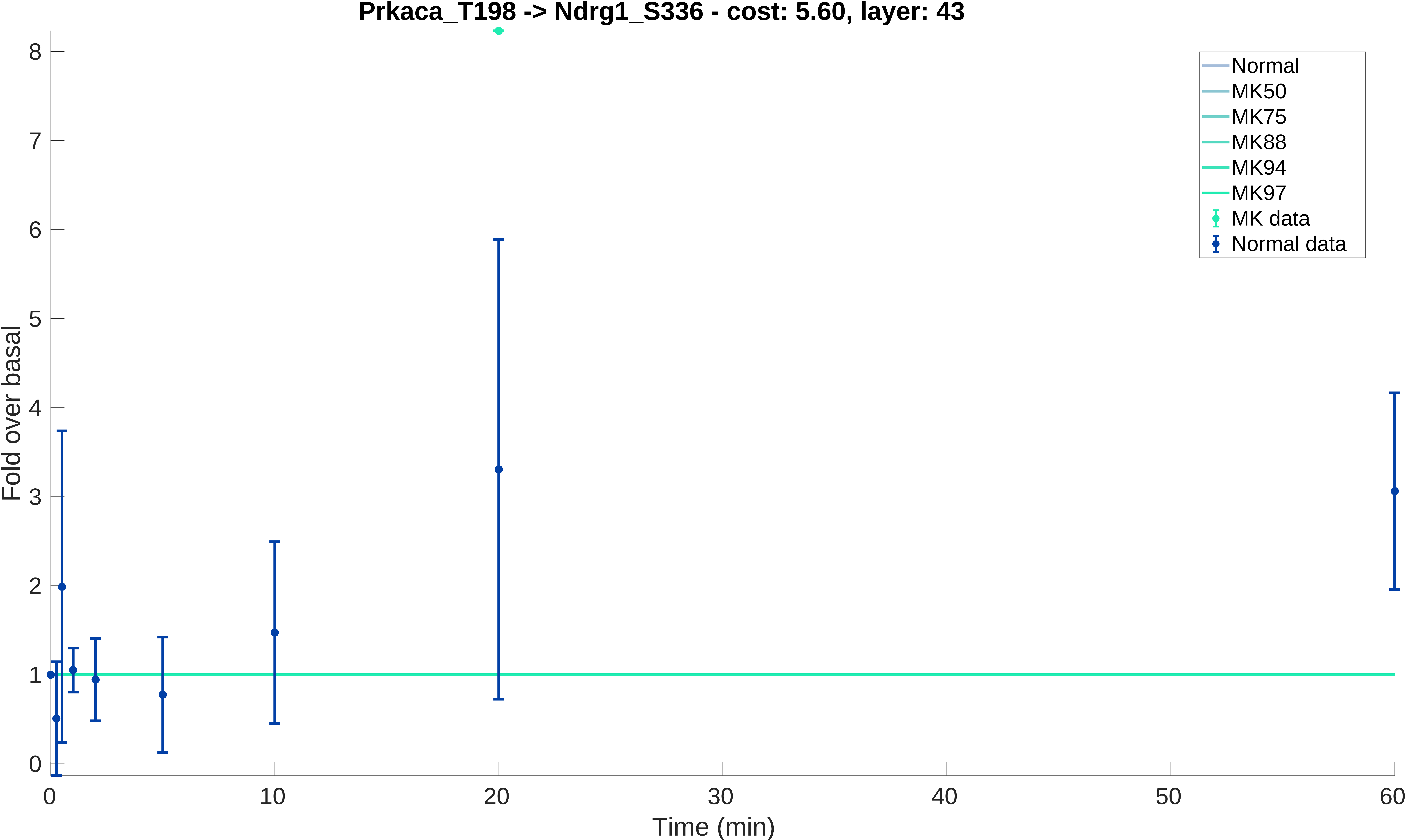

Prkaca_T198 -> Ndrg1_S336 - cost: 5.60, layer: 43
Fold over basal
Time (min)
0
1
2
3
4
5
6
7
8
0
10
20
30
40
50
60
Normal
MK50
MK75
MK88
MK94
MK97
MK data
Normal data

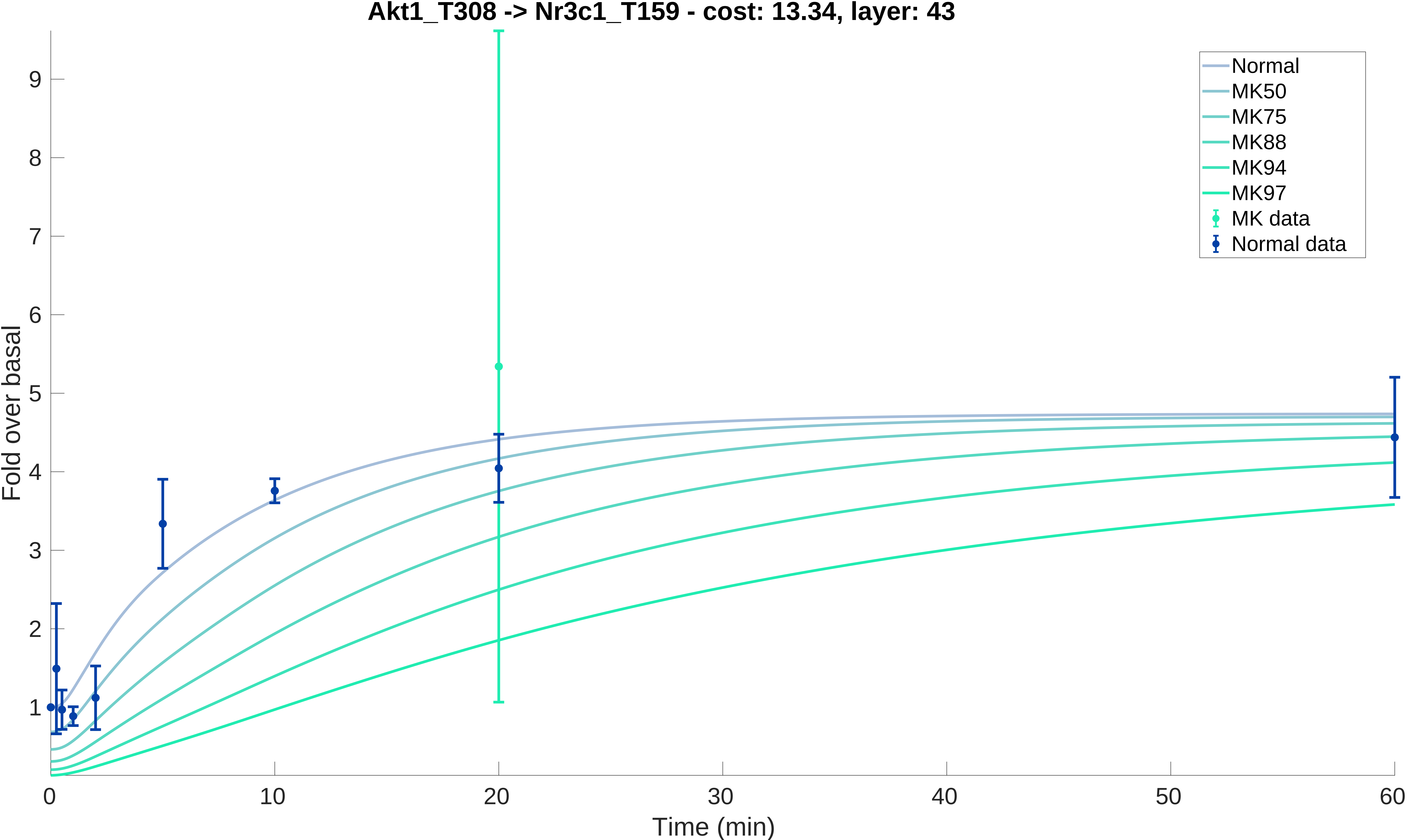

Akt1_T308 -> Nr3c1_T159 - cost: 13.34, layer: 43
Fold over basal
Time (min)
0
10
20
30
40
50
60
1
2
3
4
5
6
7
8
9
Normal
MK50
MK75
MK88
MK94
MK97
MK data
Normal data

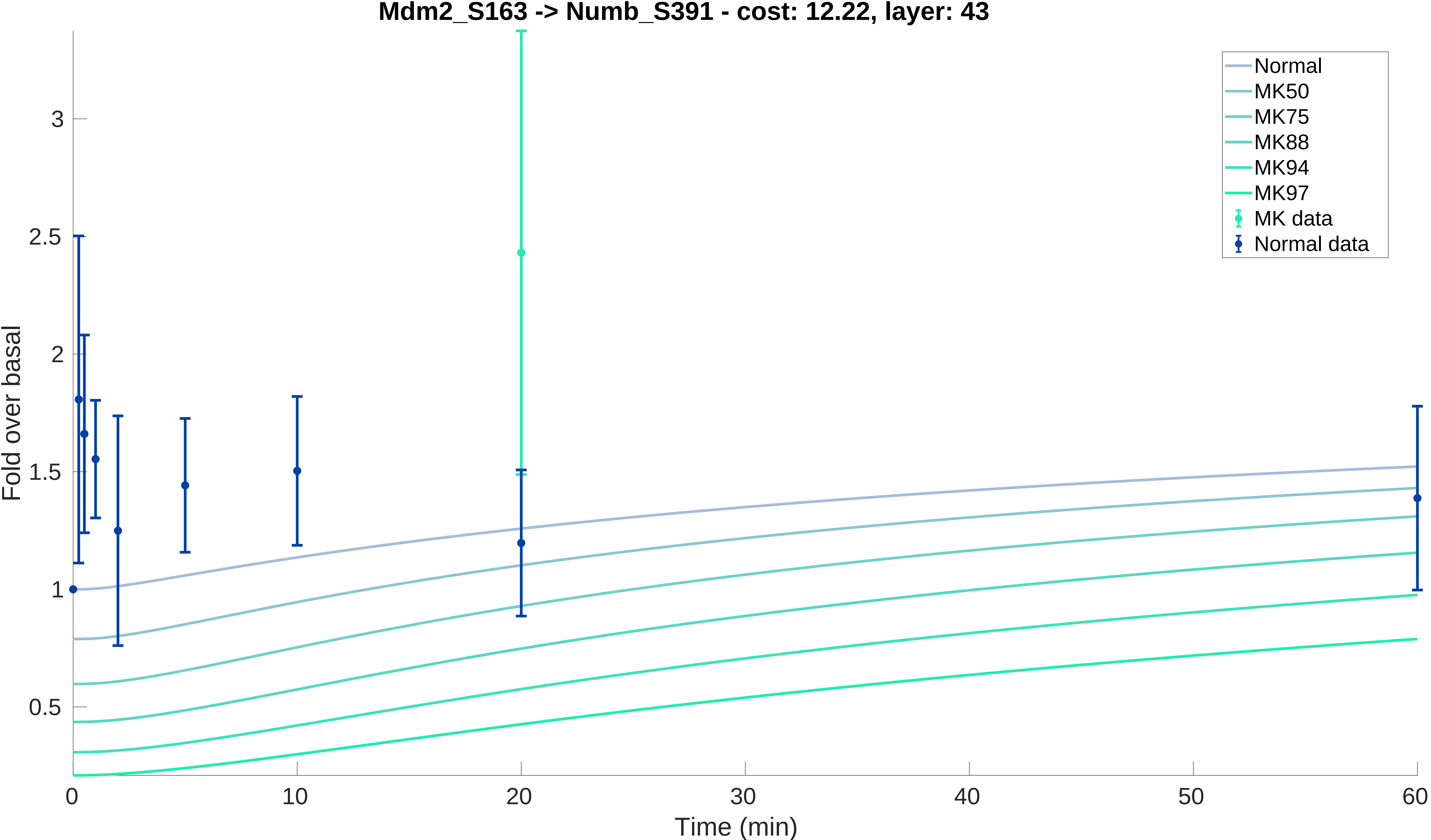

Mdm2_S163 -> Numb_S391 - cost: 12.22, layer: 43
Fold over basal
Time (min)
0
10
20
30
40
50
60
0.5
1
1.5
2
2.5
3
Normal
MK50
MK75
MK88
MK94
MK97
MK data
Normal data

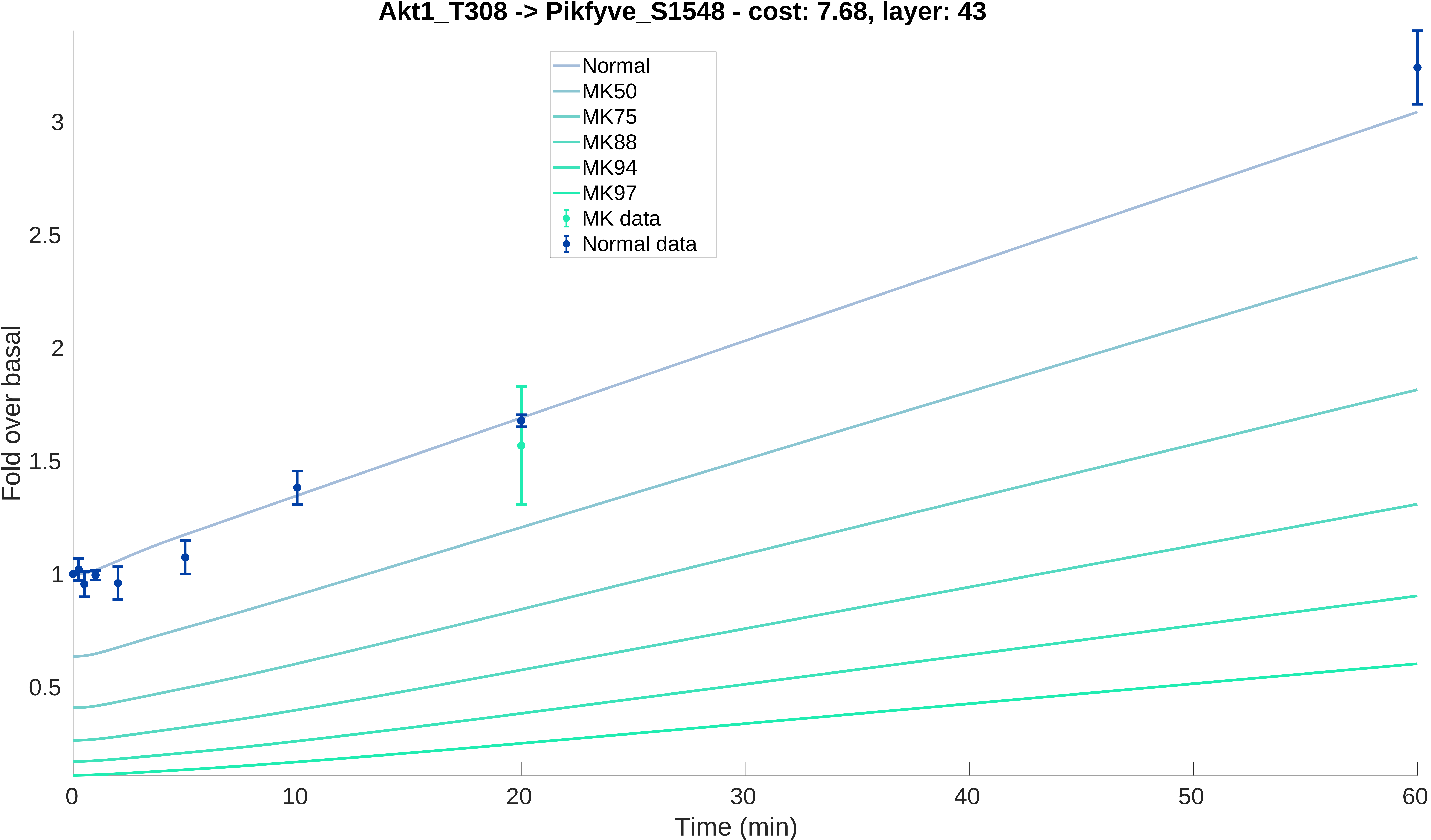

Akt1_T308 -> Pikfyve_S1548 - cost: 7.68, layer: 43
Normal
MK50
MK75
MK88
MK94
MK97
MK data
Normal data
Fold over basal
0.5
1
1.5
2
2.5
3
0
10
20
30
40
50
60
Time (min)

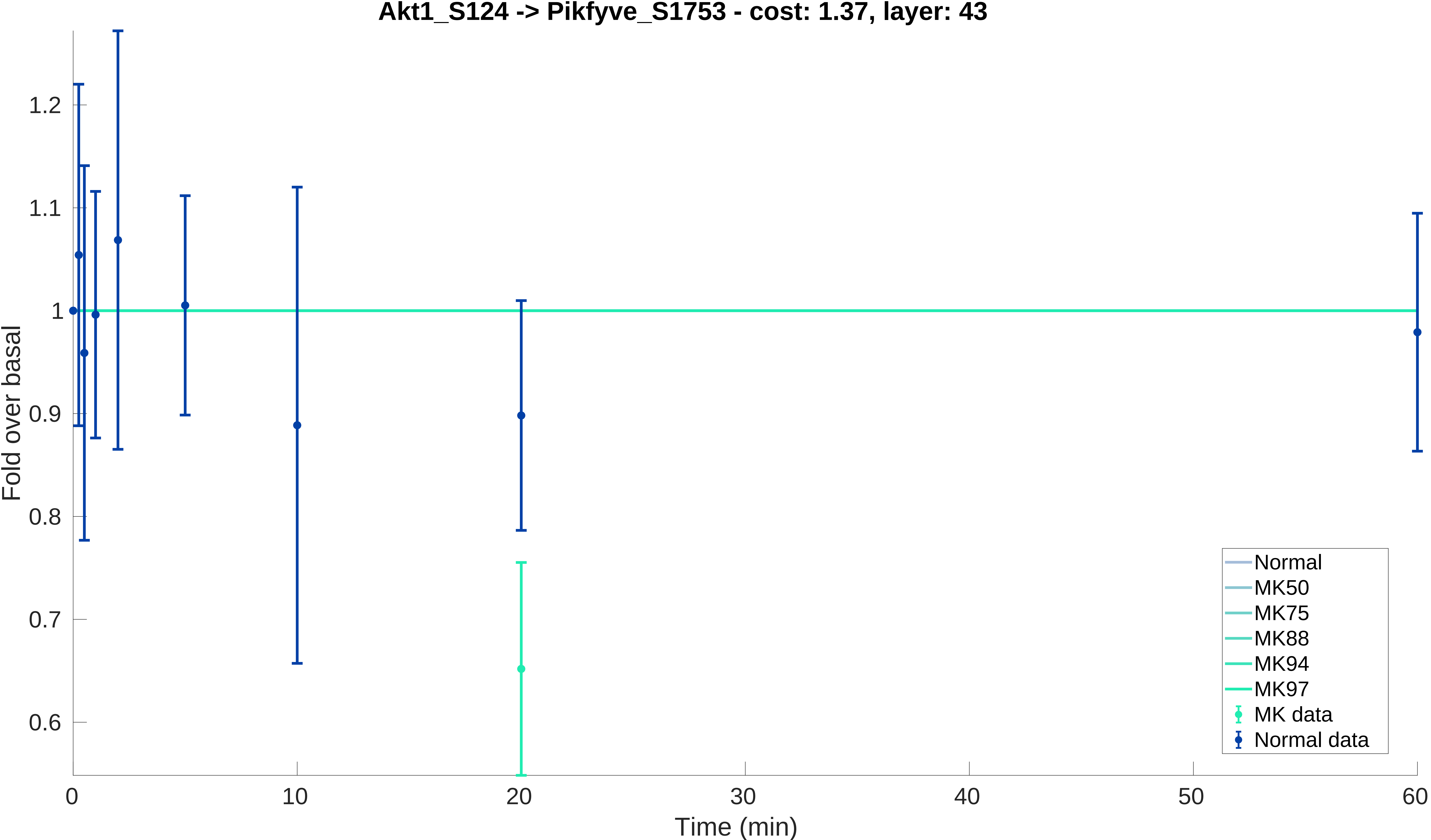

Akt1_S124 -> Pikfyve_S1753 - cost: 1.37, layer: 43
Fold over basal
1.2
1.1
1
0.9
0.8
0.7
0.6
0
10
20
30
40
50
60
Time (min)
Normal
MK50
MK75
MK88
MK94
MK97
MK data
Normal data

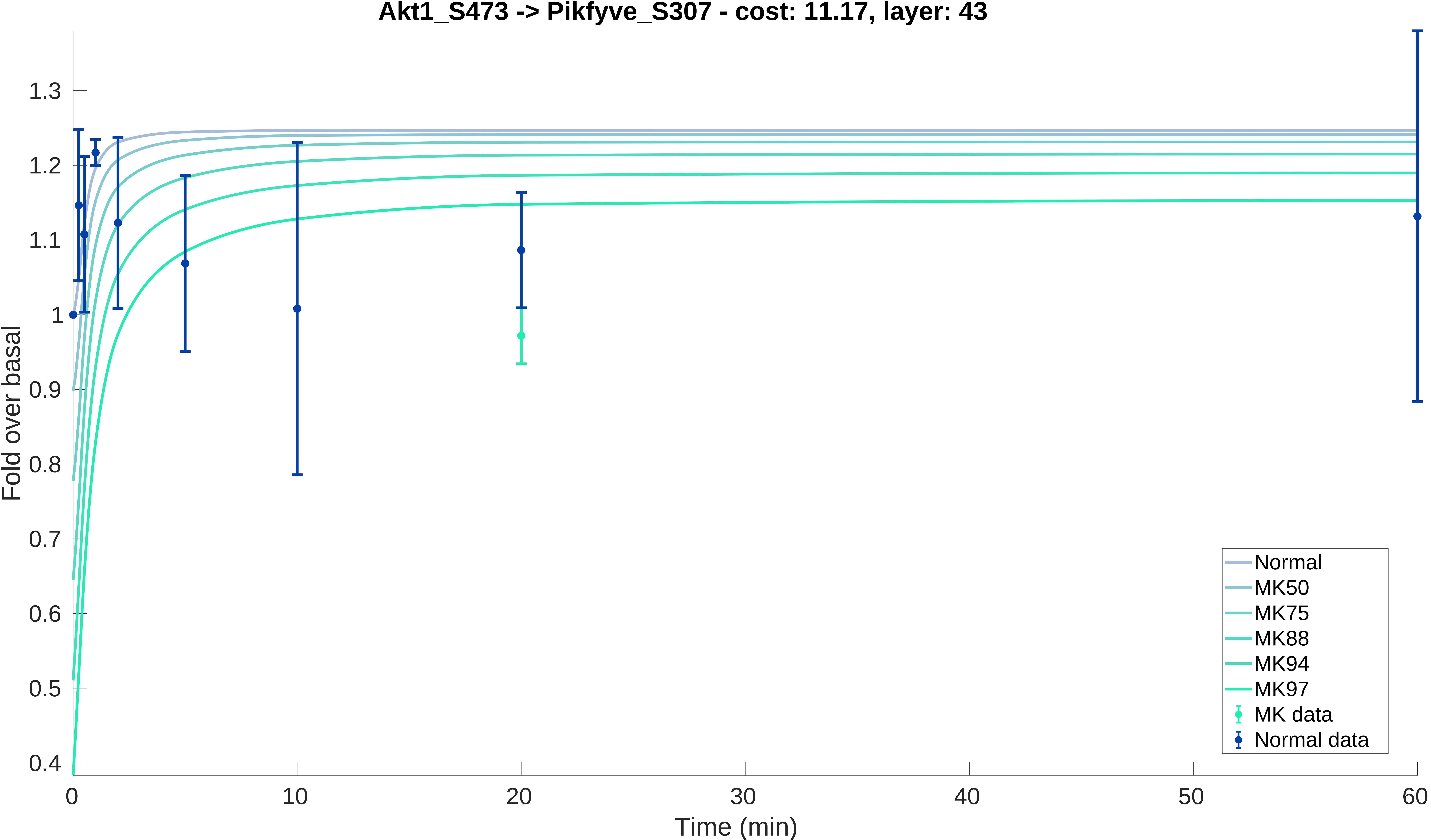
Akt1_S473 -> Pikfyve_S307 - cost: 11.17, layer: 43
Fold over basal
Time (min)
1.3
1.2
1.1
1
0.9
0.8
0.7
0.6
0.5
0.4
0
10
20
30
40
50
60
Normal
MK50
MK75
MK88
MK94
MK97
MK data
Normal data

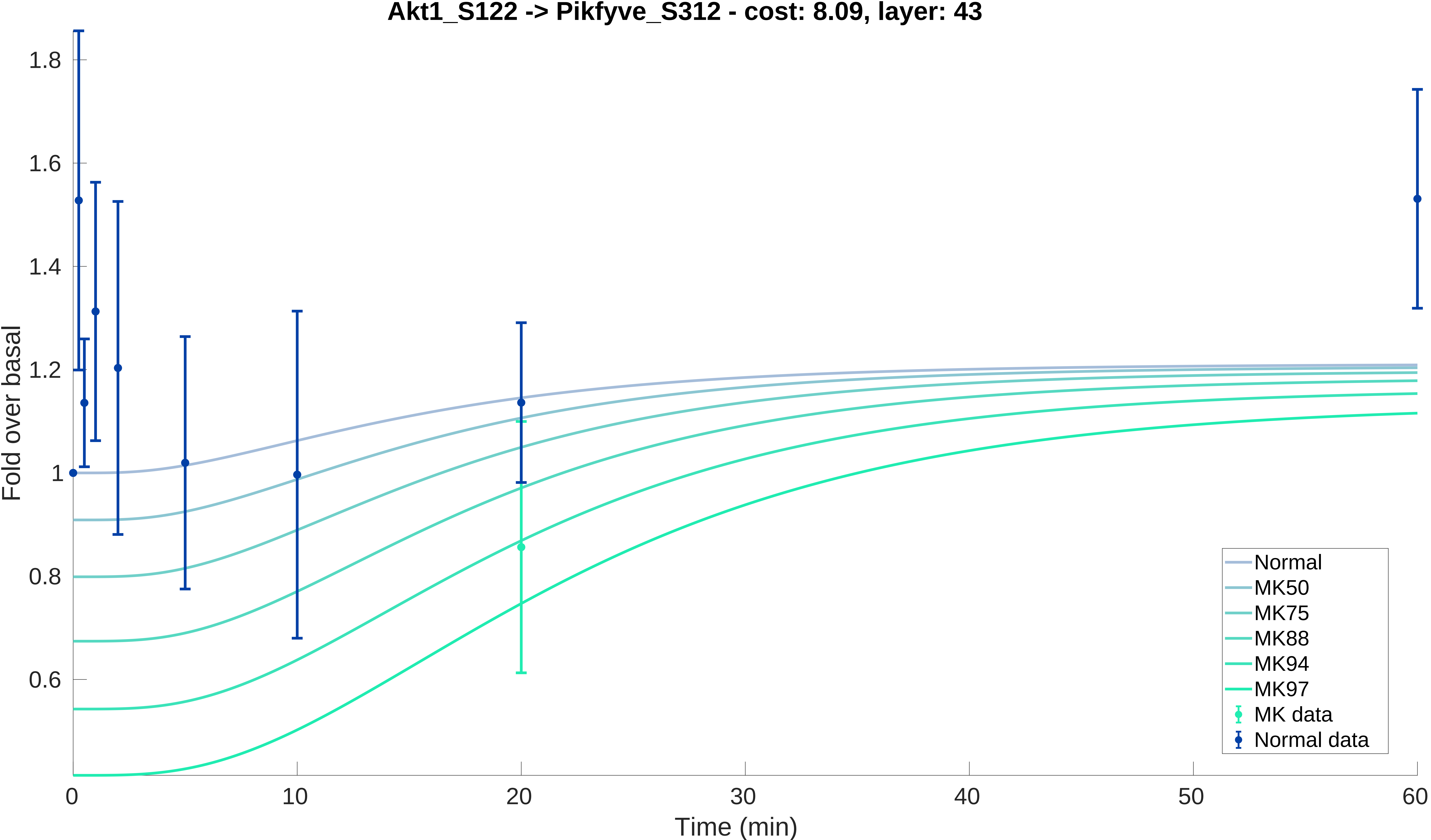

Akt1_S122 -> Pikfyve_S312 - cost: 8.09, layer: 43
Fold over basal
1.8
1.6
1.4
1.2
1
0.8
0.6
0
10
20
30
40
50
60
Time (min)
Normal
MK50
MK75
MK88
MK94
MK97
MK data
Normal data

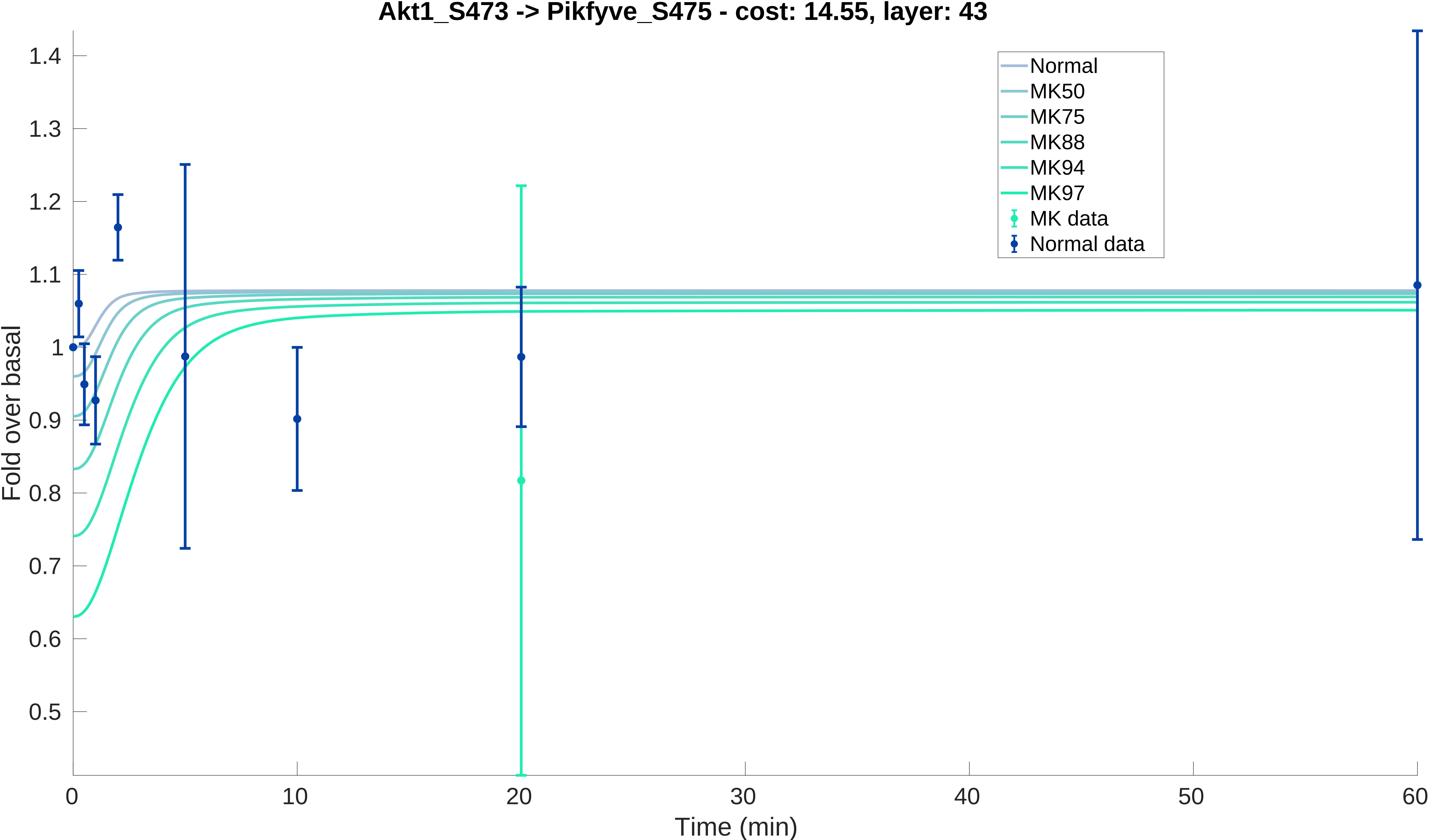

Akt1_S473 -> Pikfyve_S475 - cost: 14.55, layer: 43
Fold over basal
Time (min)
Normal
MK50
MK75
MK88
MK94
MK97
MK data
Normal data
0.5
0.6
0.7
0.8
0.9
1
1.1
1.2
1.3
1.4
0
10
20
30
40
50
60

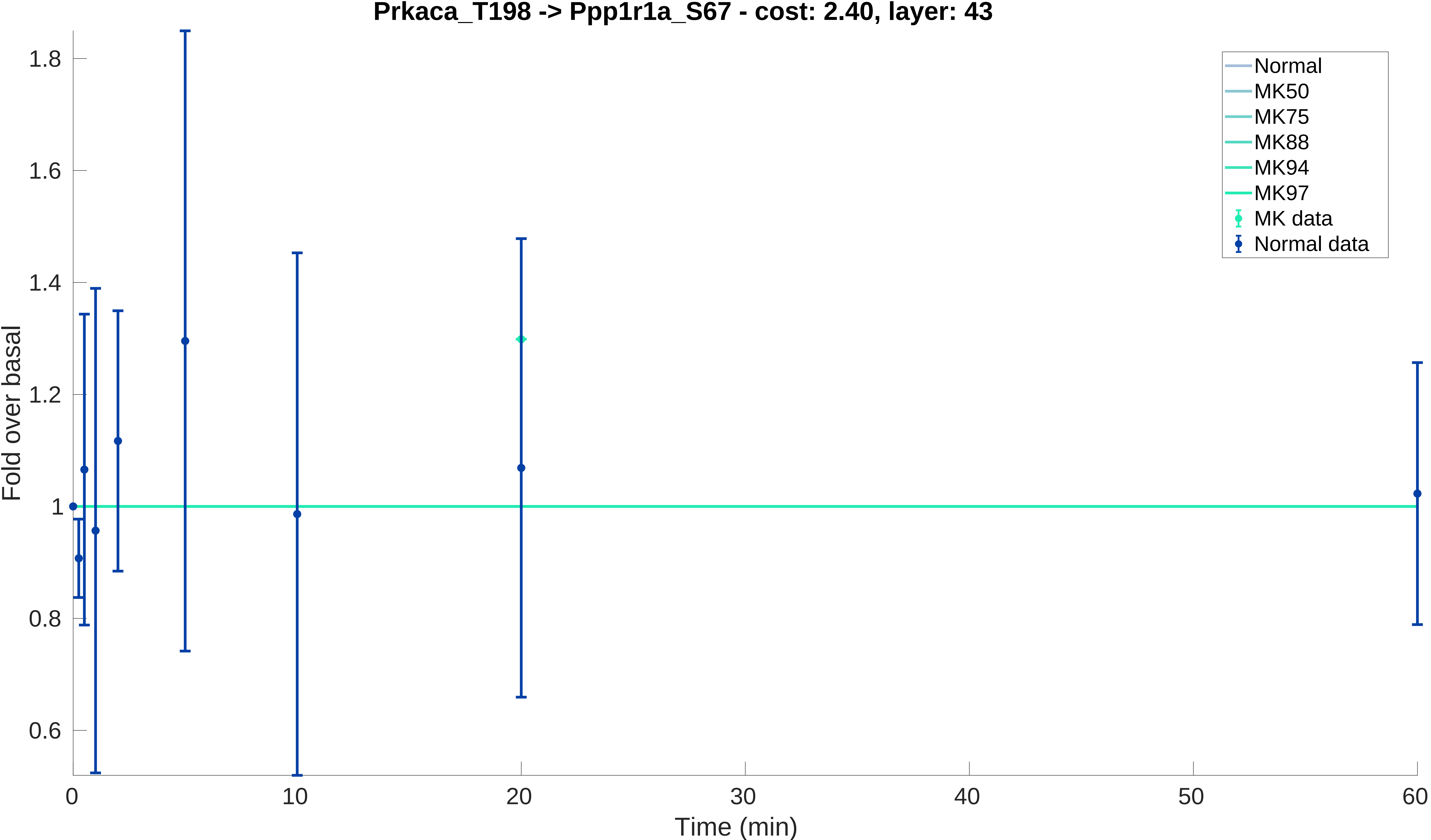

Prkaca_T198 -> Ppp1r1a_S67 - cost: 2.40, layer: 43
Fold over basal
1.8
1.6
1.4
1.2
1
0.8
0.6
0
10
20
30
40
50
60
Time (min)
Normal
MK50
MK75
MK88
MK94
MK97
MK data
Normal data

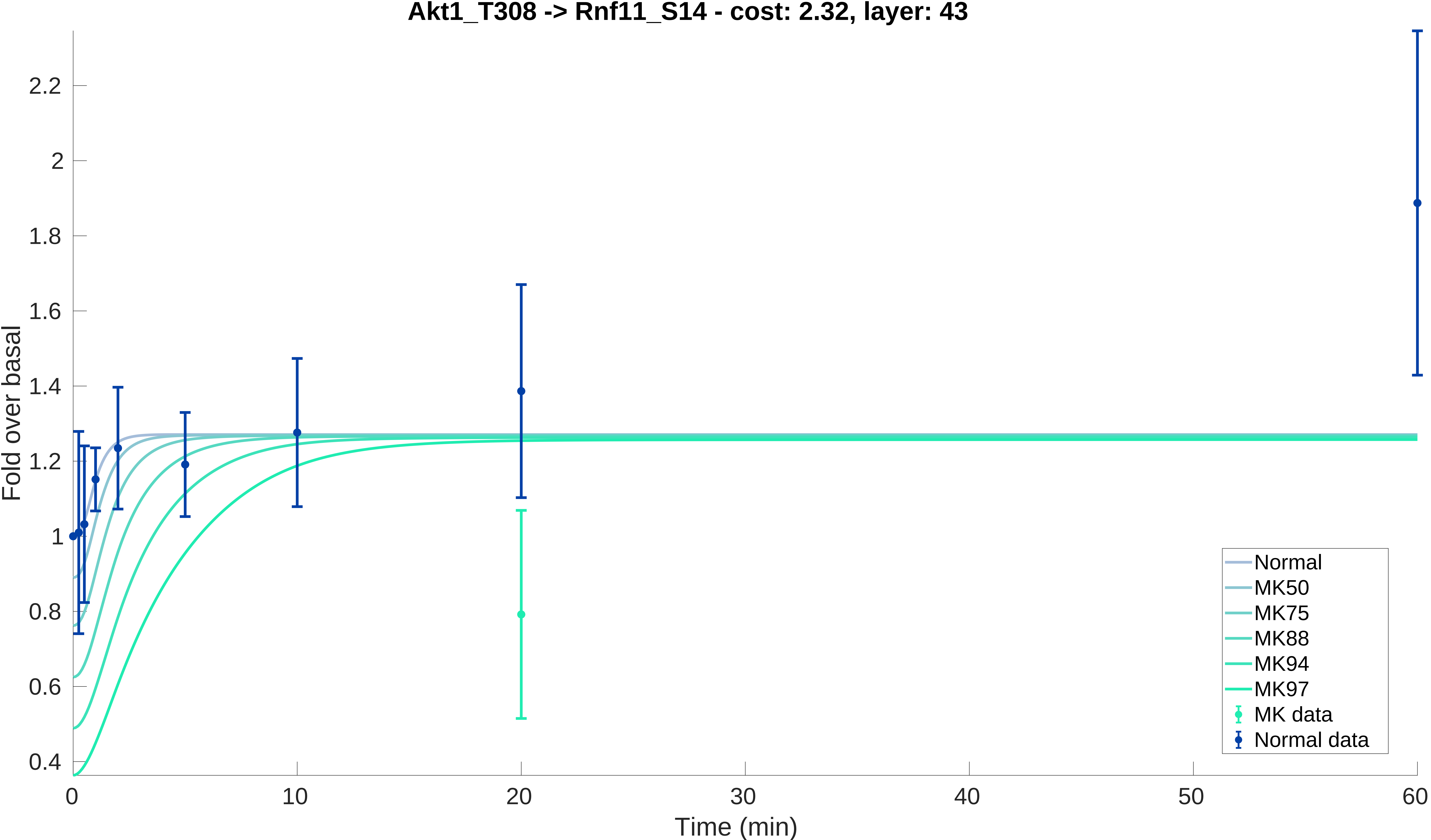
Akt1_T308 -> Rnf11_S14 - cost: 2.32, layer: 43
Fold over basal
Time (min)
0.4
0.6
0.8
1
1.2
1.4
1.6
1.8
2
2.2
0
10
20
30
40
50
60
Normal
MK50
MK75
MK88
MK94
MK97
MK data
Normal data

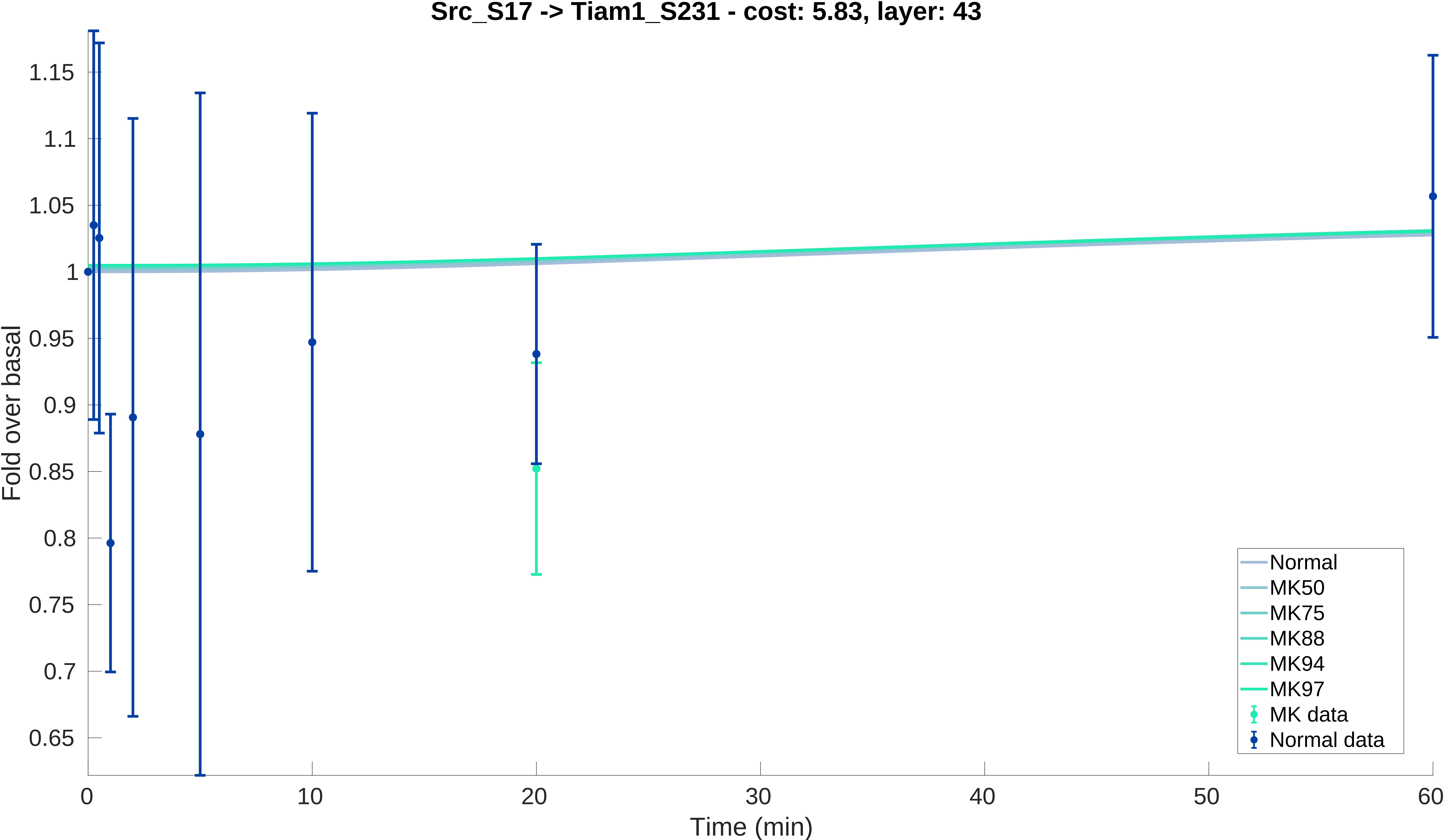

Src_S17 -> Tiam1_S231 - cost: 5.83, layer: 43
Fold over basal
Time (min)
0
10
20
30
40
50
60
1.15
1.1
1.05
1
0.95
0.9
0.85
0.8
0.75
0.7
0.65
Normal
MK50
MK75
MK88
MK94
MK97
MK data
Normal data

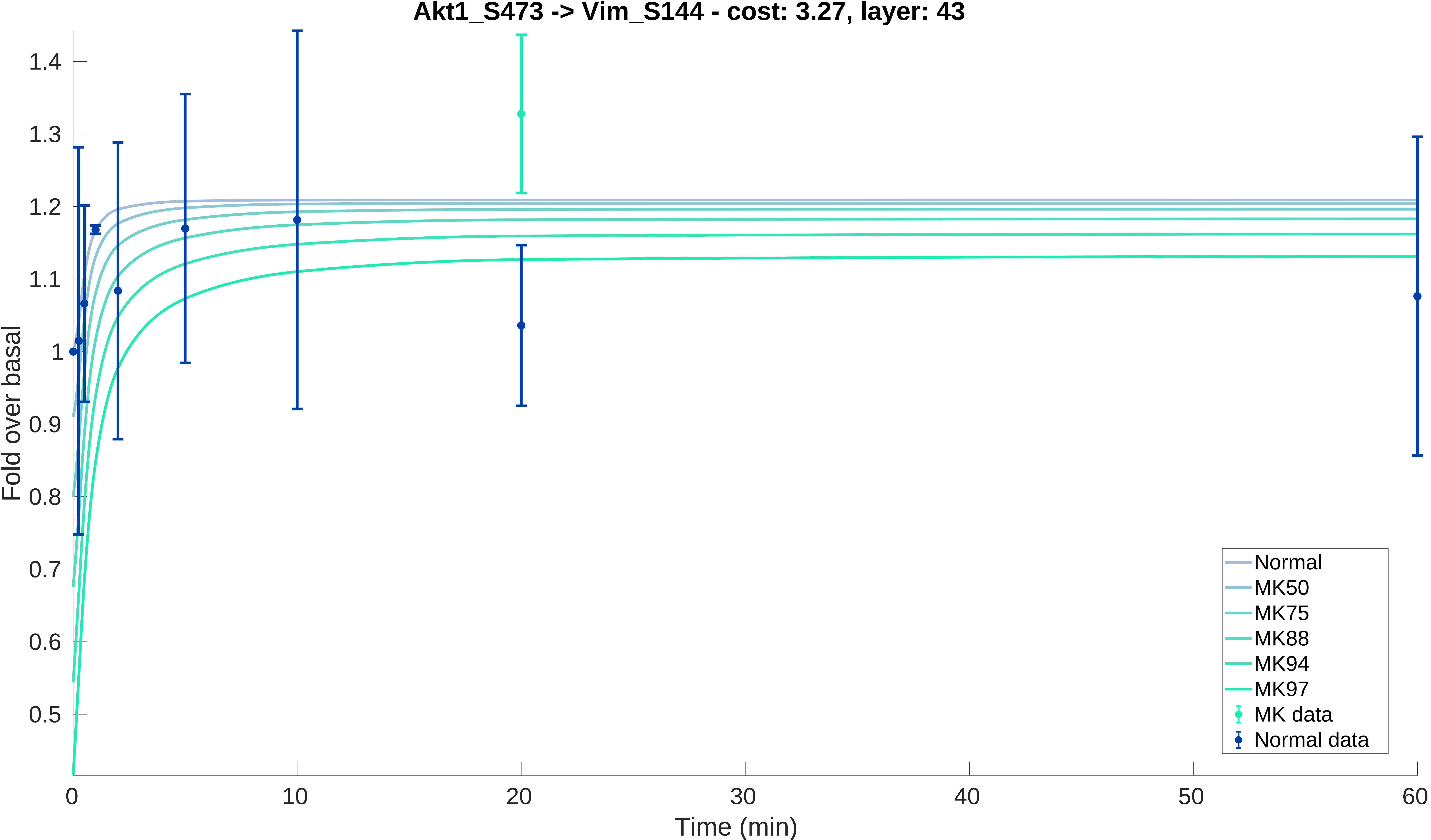
Akt1_S473 -> Vim_S144 - cost: 3.27, layer: 43
Fold over basal
Time (min)
0
10
20
30
40
50
60
0.5
0.6
0.7
0.8
0.9
1
1.1
1.2
1.3
1.4
Normal
MK50
MK75
MK88
MK94
MK97
MK data
Normal data

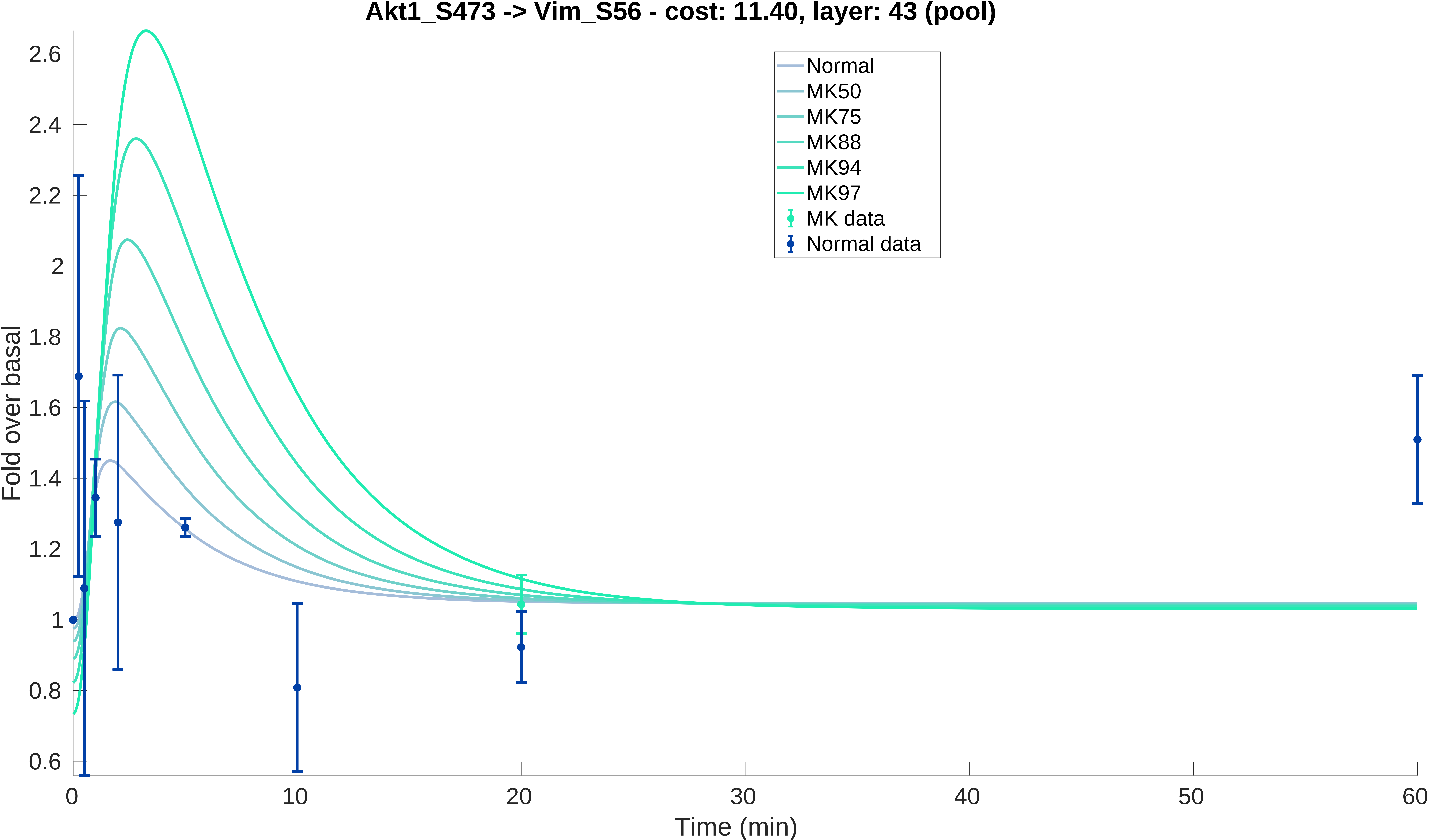
Akt1_S473 -> Vim_S56 - cost: 11.40, layer: 43 (pool)
Fold over basal
Time (min)
0.6
0.8
1
1.2
1.4
1.6
1.8
2
2.2
2.4
2.6
0
10
20
30
40
50
60
Normal
MK50
MK75
MK88
MK94
MK97
MK data
Normal data

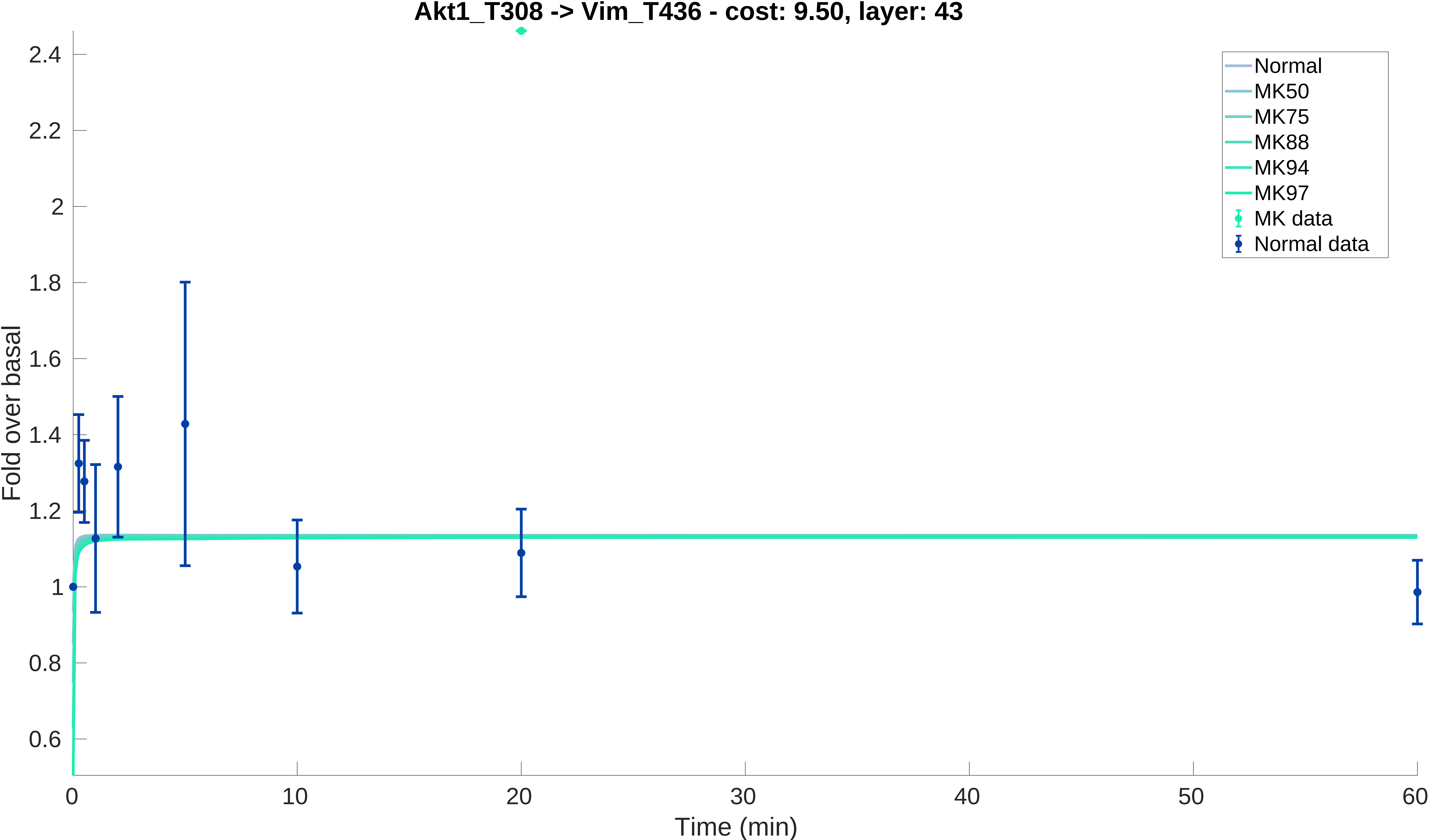

Akt1_T308 -> Vim_T436 - cost: 9.50, layer: 43
Fold over basal
Time (min)
Normal
MK50
MK75
MK88
MK94
MK97
MK data
Normal data
2.4
2.2
2
1.8
1.6
1.4
1.2
1
0.8
0.6
0
10
20
30
40
50
60

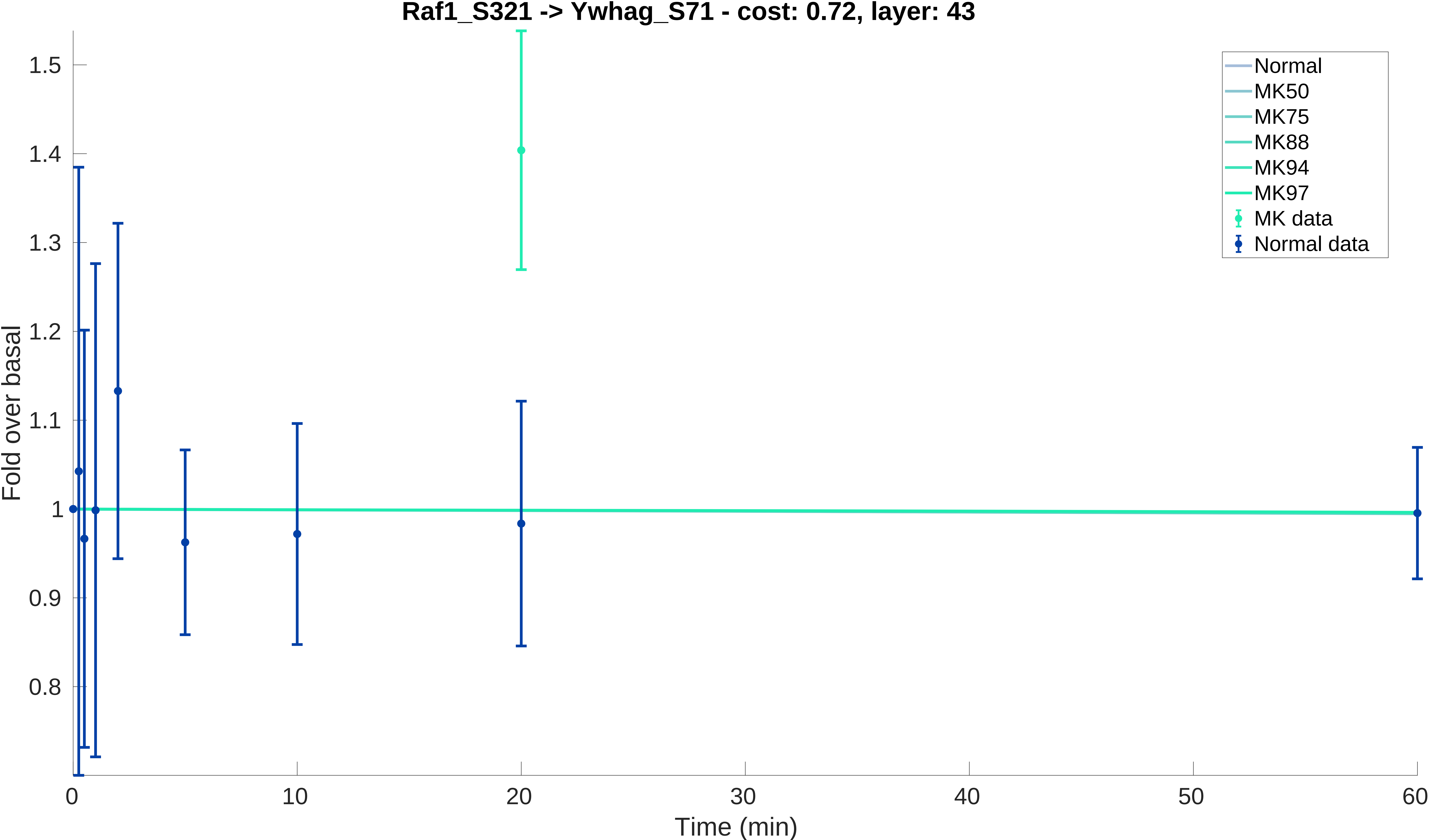

Raf1_S321 -> Ywhag_S71 - cost: 0.72, layer: 43
Fold over basal
Time (min)
1.5
1.4
1.3
1.2
1.1
1
0.9
0.8
0
10
20
30
40
50
60
Normal
MK50
MK75
MK88
MK94
MK97
MK data
Normal data

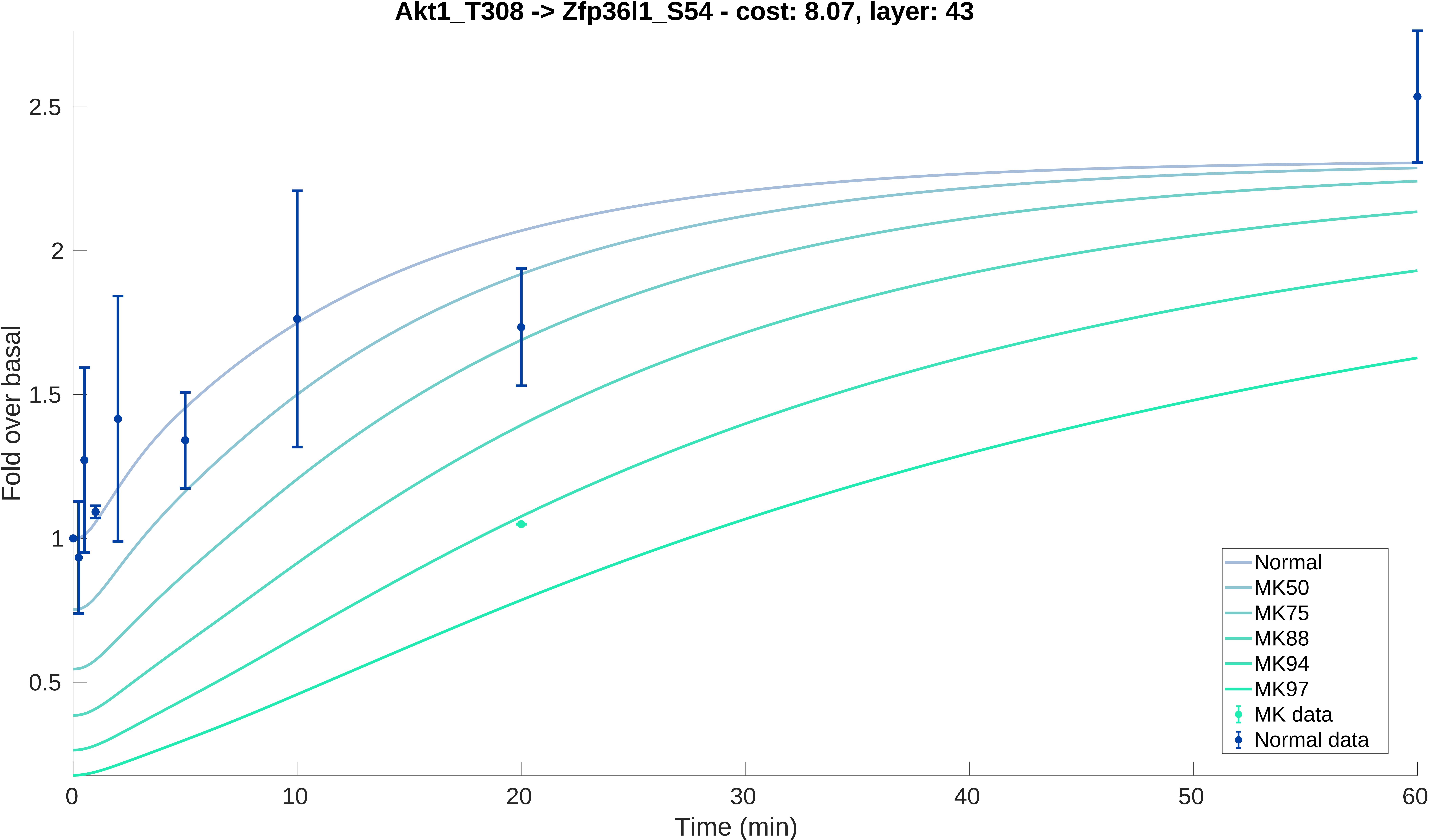

Akt1_T308 -> Zfp36l1_S54 - cost: 8.07, layer: 43
Fold over basal
Time (min)
0.5
1
1.5
2
2.5
0
10
20
30
40
50
60
Normal
MK50
MK75
MK88
MK94
MK97
MK data
Normal data

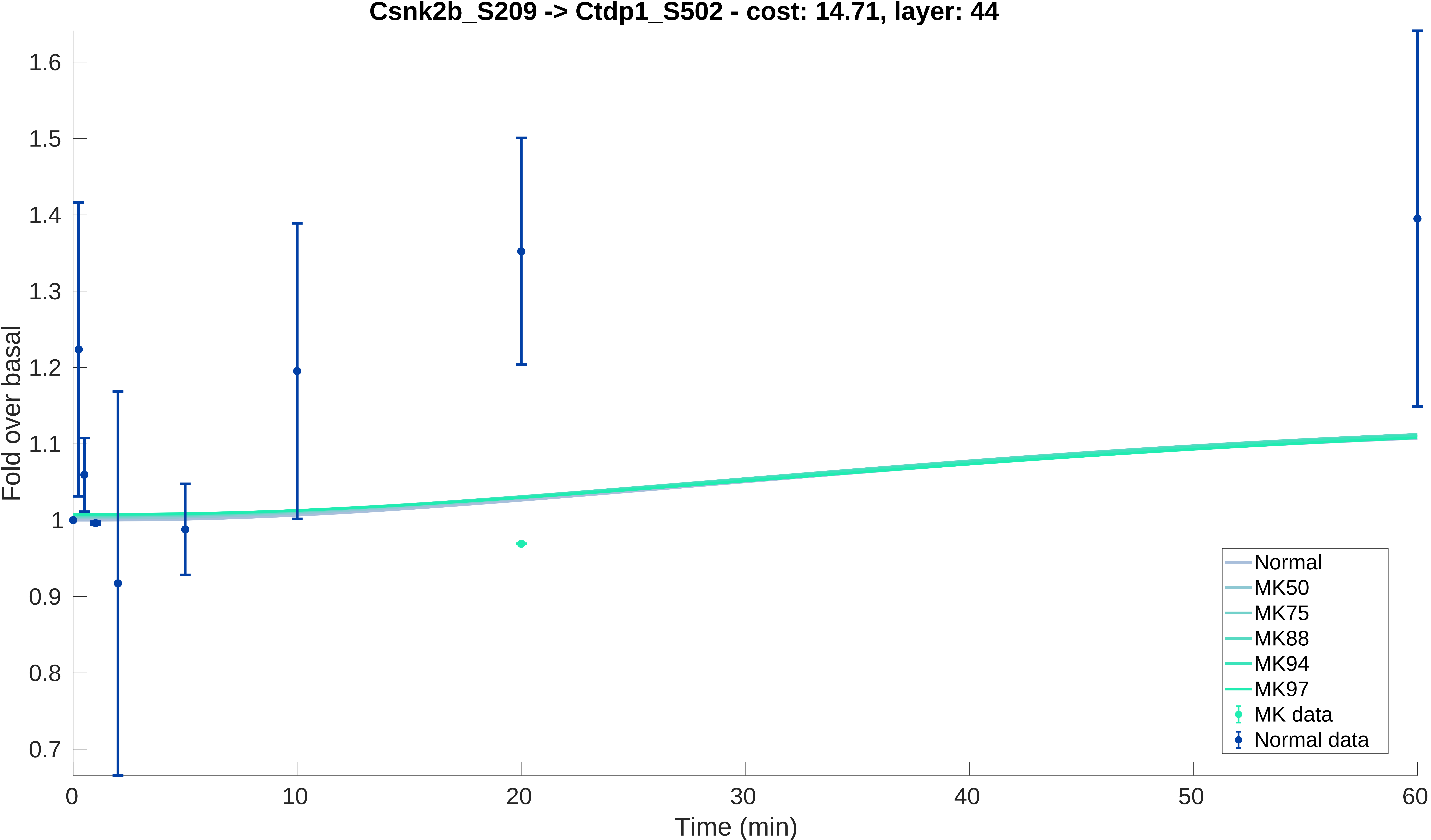

Csnk2b_S209 -> Ctdp1_S502 - cost: 14.71, layer: 44
Fold over basal
1.6
1.5
1.4
1.3
1.2
1.1
1
0.9
0.8
0.7
0
10
20
30
40
50
60
Time (min)
Normal
MK50
MK75
MK88
MK94
MK97
MK data
Normal data

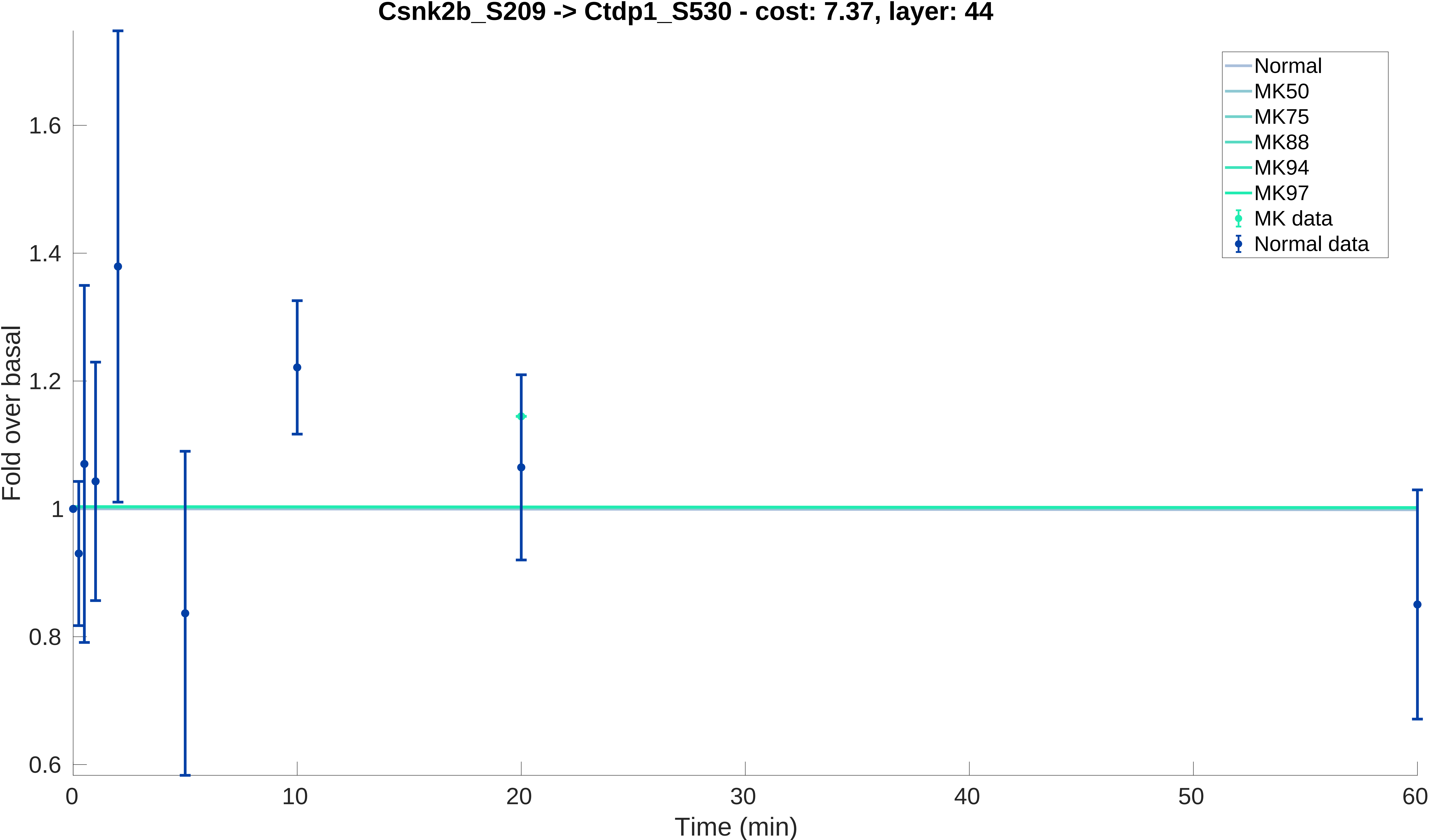

Csnk2b_S209 -> Ctdp1_S530 - cost: 7.37, layer: 44
Fold over basal
Time (min)
1.6
1.4
1.2
1
0.8
0.6
0
10
20
30
40
50
60
Normal
MK50
MK75
MK88
MK94
MK97
MK data
Normal data

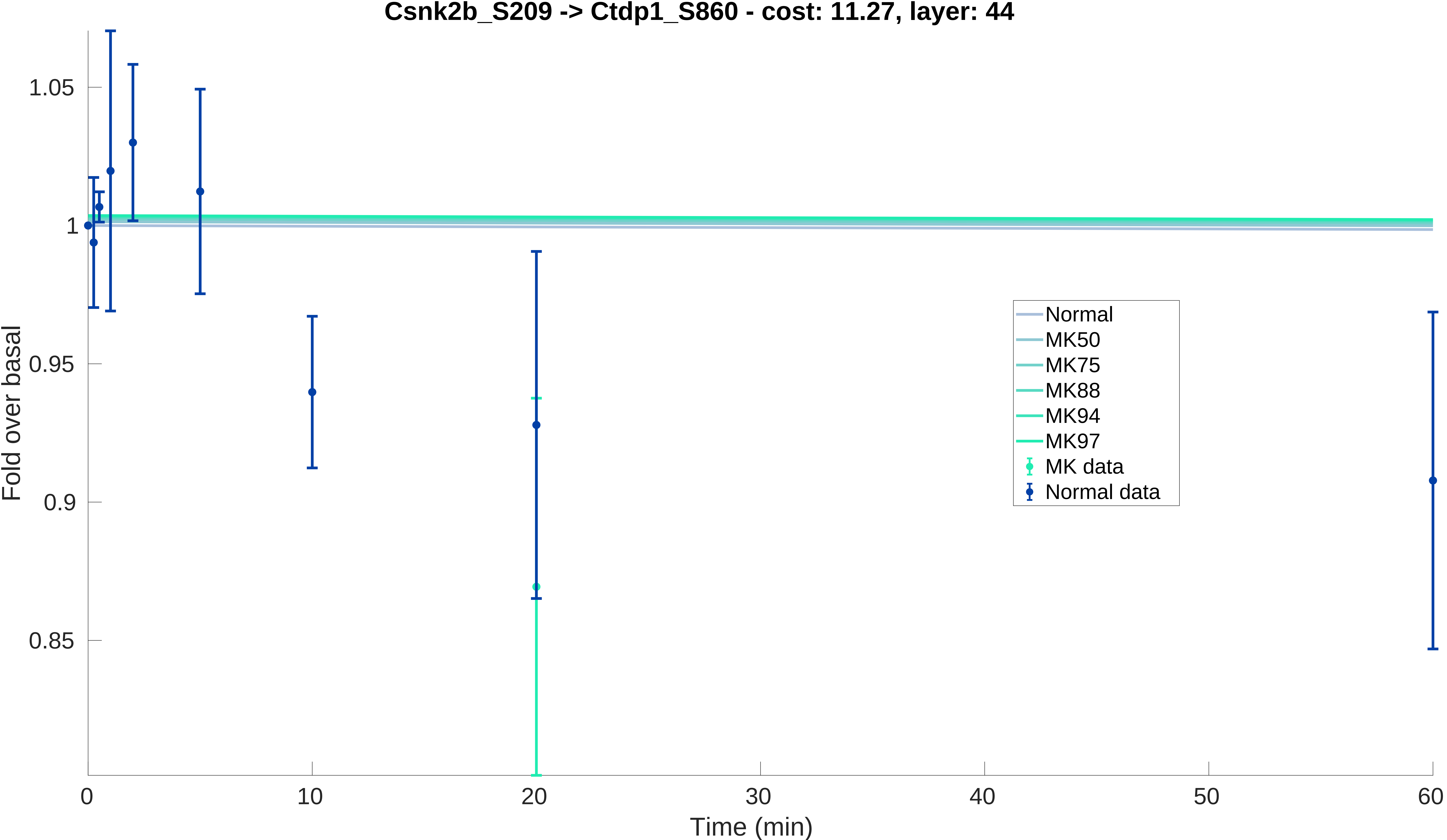

Csnk2b_S209 -> Ctdp1_S860 - cost: 11.27, layer: 44
Fold over basal
1.05
1
0.95
0.9
0.85
0
10
20
30
40
50
60
Time (min)
Normal
MK50
MK75
MK88
MK94
MK97
MK data
Normal data

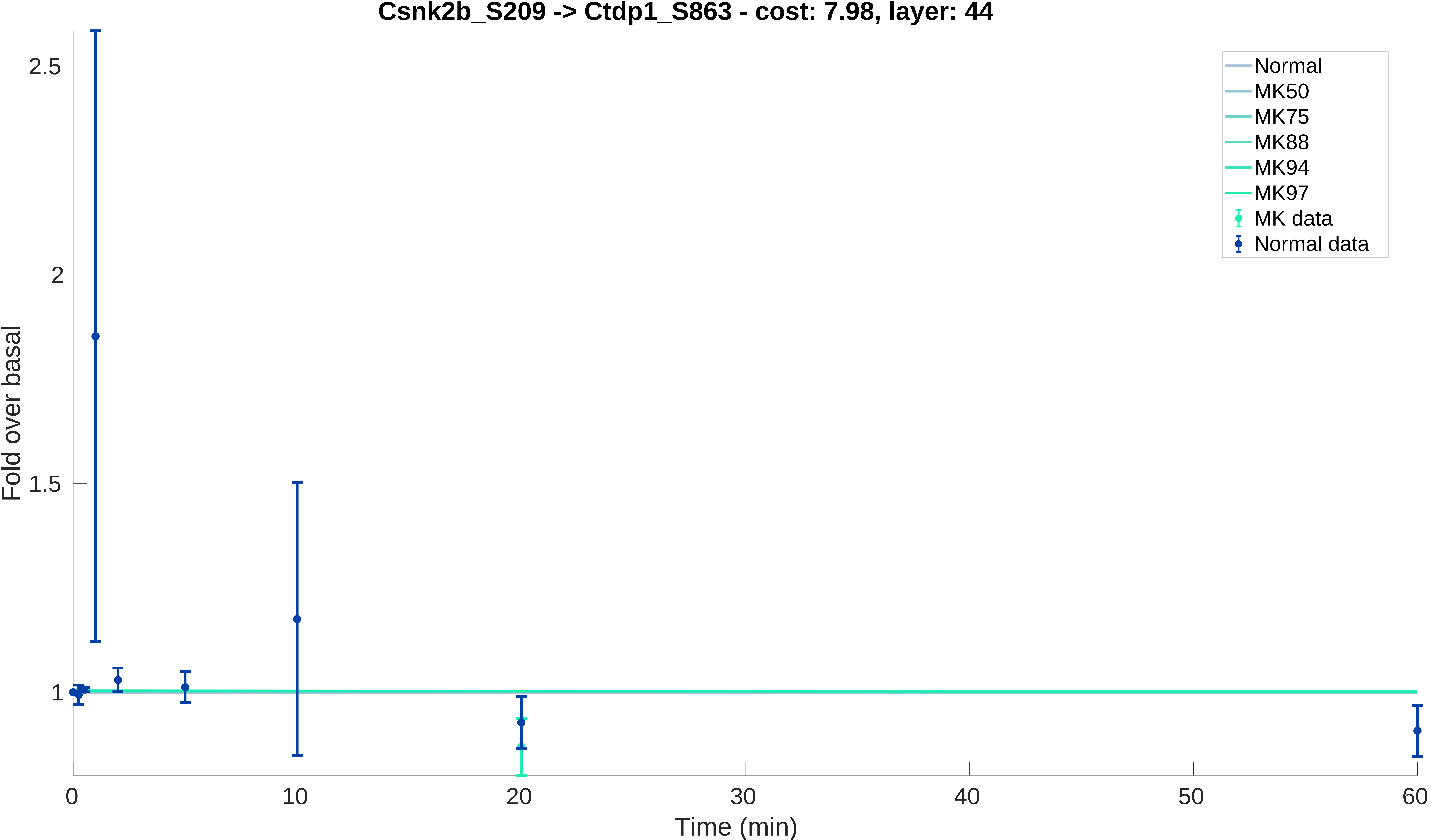

Csnk2b_S209 -> Ctdp1_S863 - cost: 7.98, layer: 44
Normal
MK50
MK75
MK88
MK94
MK97
MK data
Normal data
Fold over basal
2.5
2
1.5
1
0
10
20
30
40
50
60
Time (min)

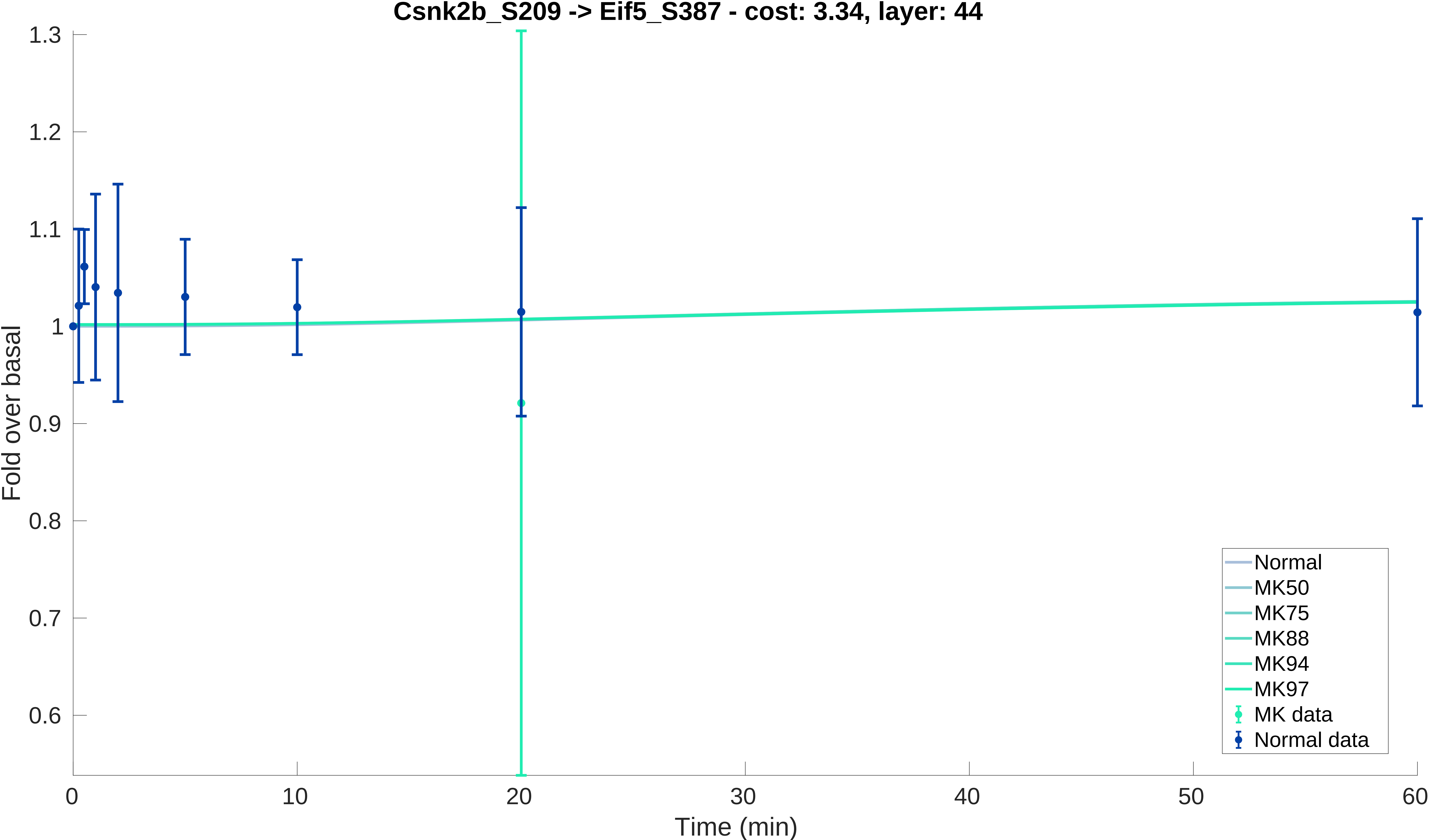

Csnk2b_S209 -> Eif5_S387 - cost: 3.34, layer: 44
Fold over basal
Time (min)
1.3
1.2
1.1
1
0.9
0.8
0.7
0.6
0
10
20
30
40
50
60
Normal
MK50
MK75
MK88
MK94
MK97
MK data
Normal data

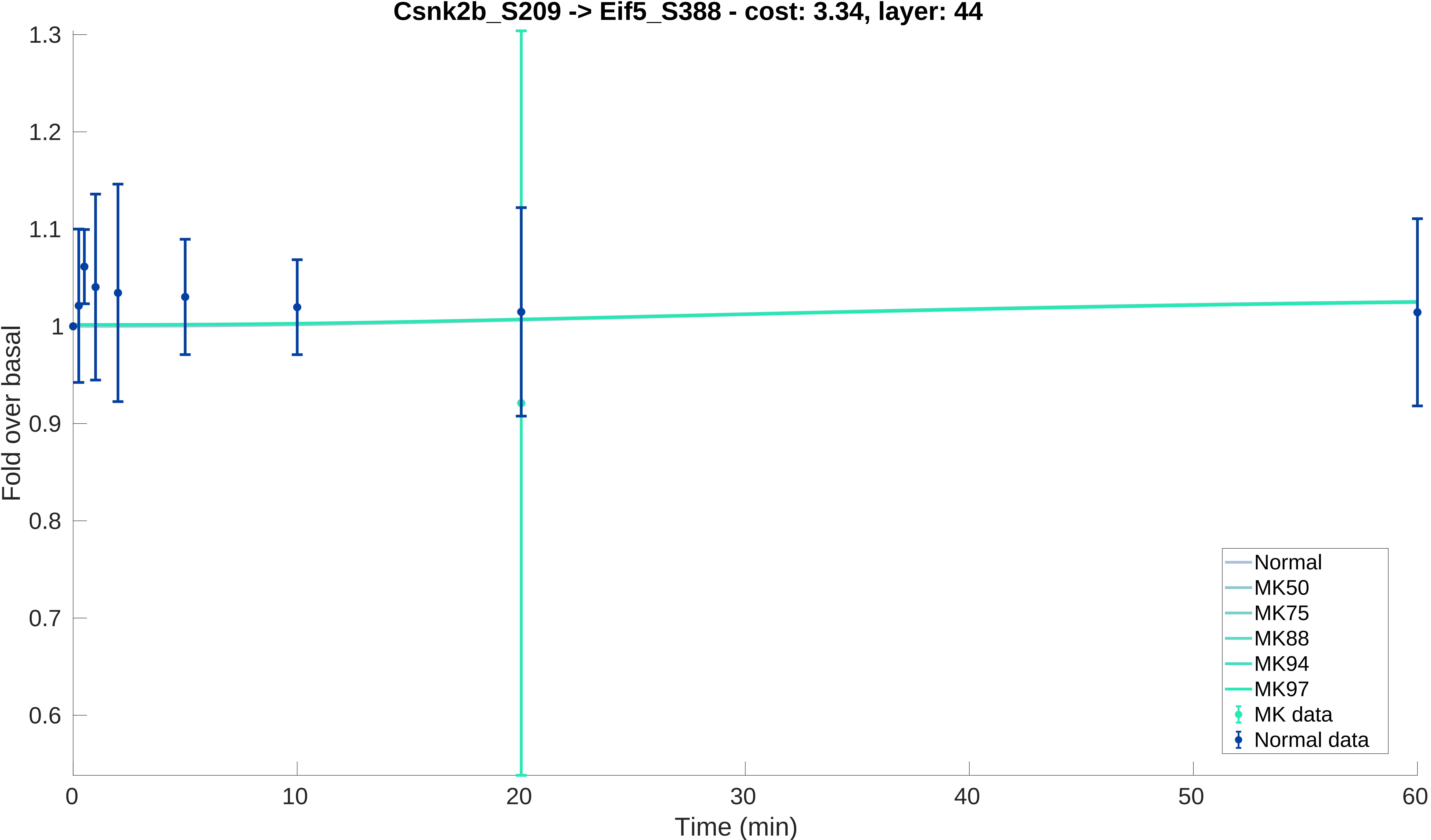

Csnk2b_S209 -> Eif5_S388 - cost: 3.34, layer: 44
Fold over basal
1.3
1.2
1.1
1
0.9
0.8
0.7
0.6
0
10
20
30
40
50
60
Time (min)
Normal
MK50
MK75
MK88
MK94
MK97
MK data
Normal data

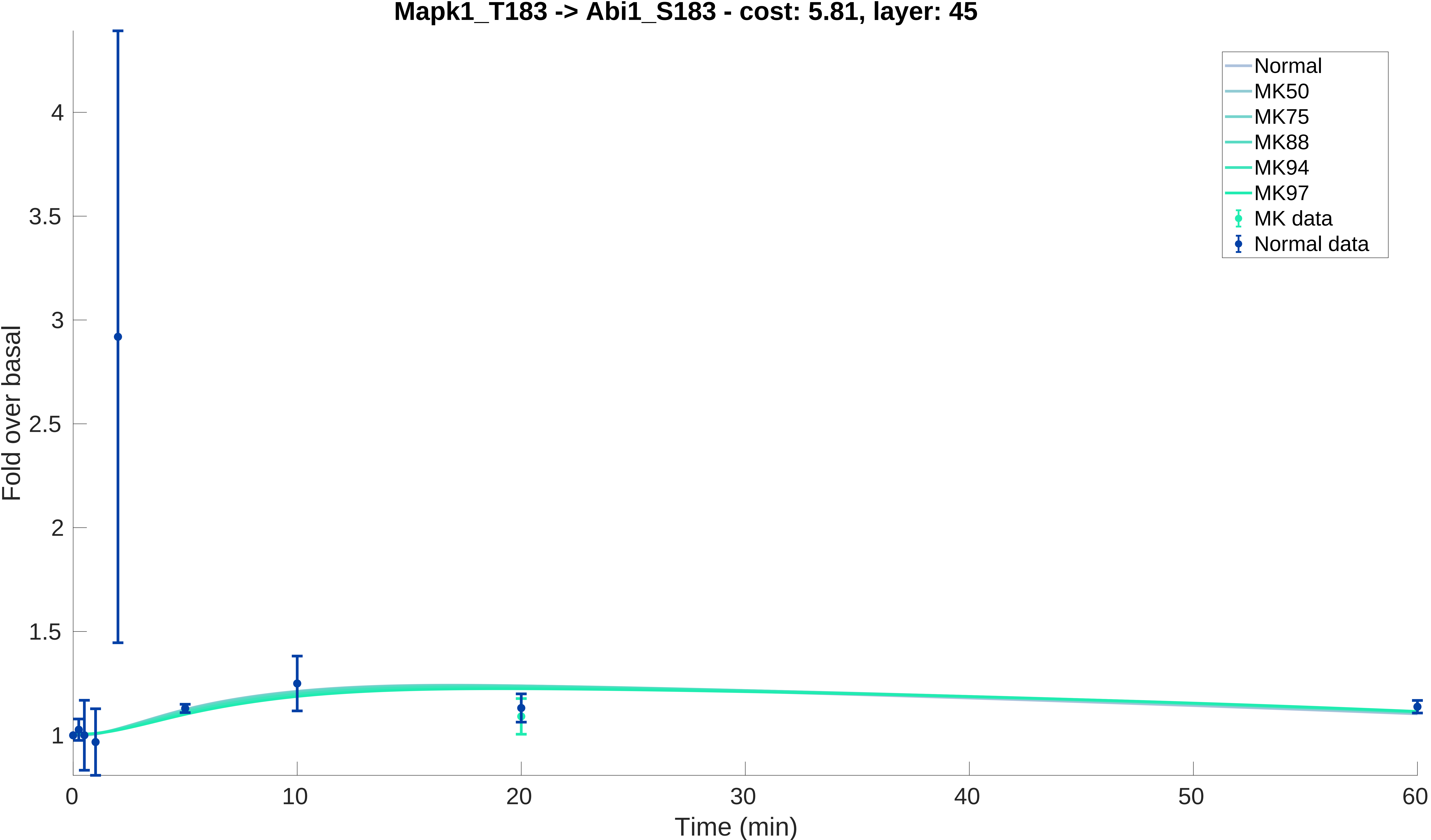

Mapk1_T183 -> Abi1_S183 - cost: 5.81, layer: 45
Fold over basal
Time (min)
Normal
MK50
MK75
MK88
MK94
MK97
MK data
Normal data
1
1.5
2
2.5
3
3.5
4
0
10
20
30
40
50
60

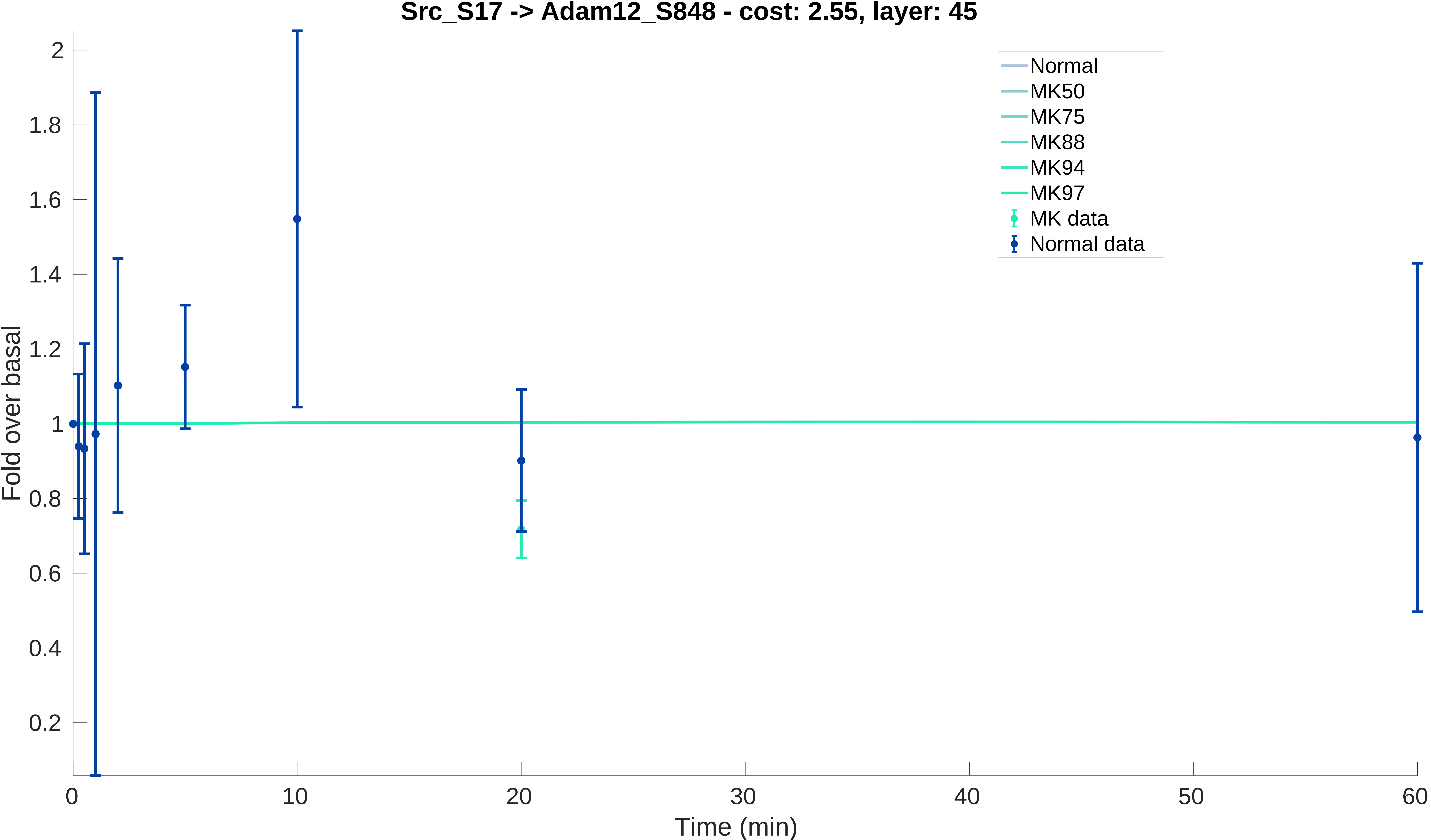

Src_S17 -> Adam12_S848 - cost: 2.55, layer: 45
Fold over basal
Time (min)
Normal
MK50
MK75
MK88
MK94
MK97
MK data
Normal data
2
1.8
1.6
1.4
1.2
1
0.8
0.6
0.4
0.2
0
10
20
30
40
50
60

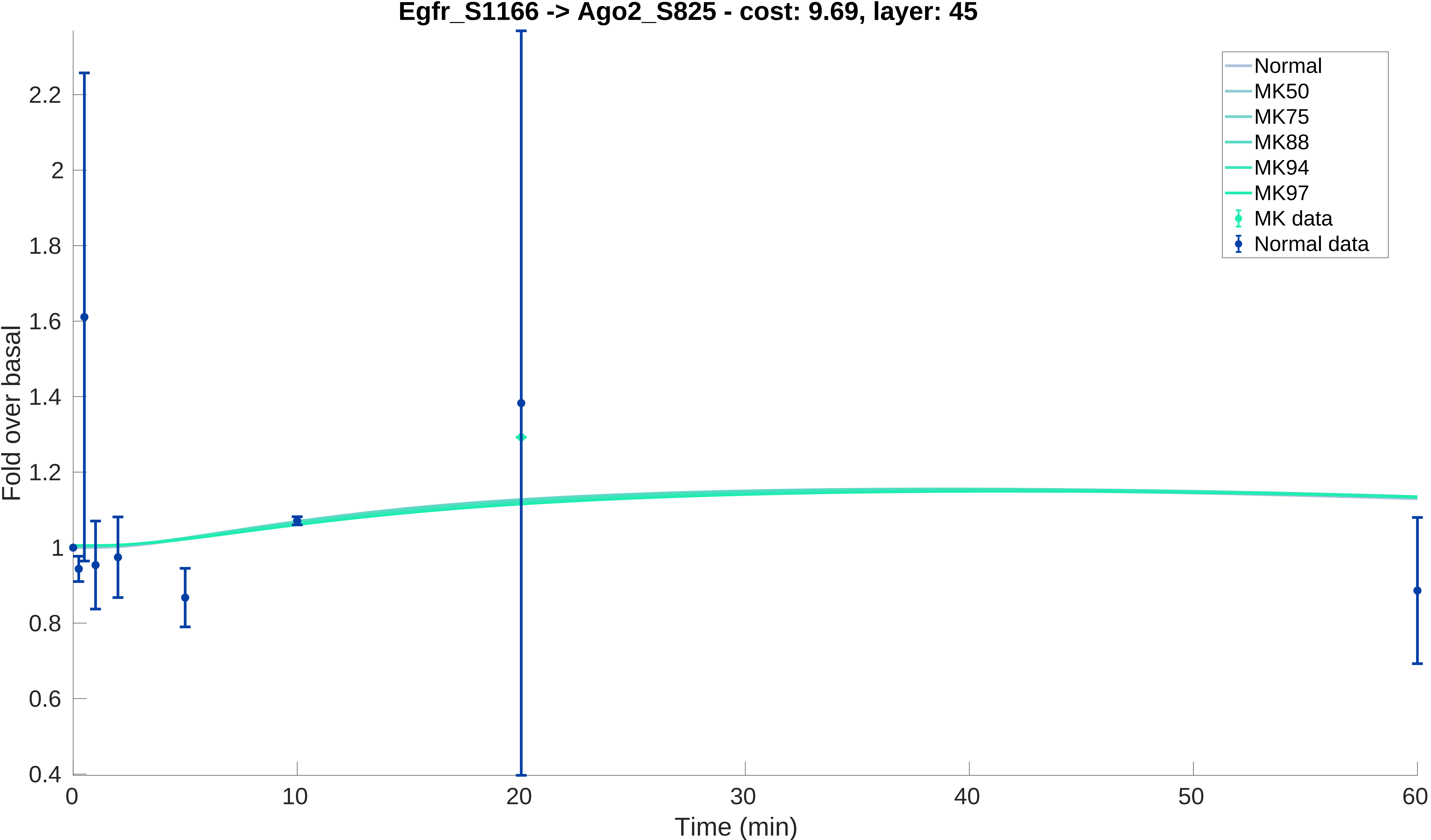
Egfr_S1166 -> Ago2_S825 - cost: 9.69, layer: 45
Fold over basal
Time (min)
0.4
0.6
0.8
1
1.2
1.4
1.6
1.8
2
2.2
0
10
20
30
40
50
60
Normal
MK50
MK75
MK88
MK94
MK97
MK data
Normal data

Prkcd_S671 -> Bag3_S138 - cost: 12.79, layer: 45

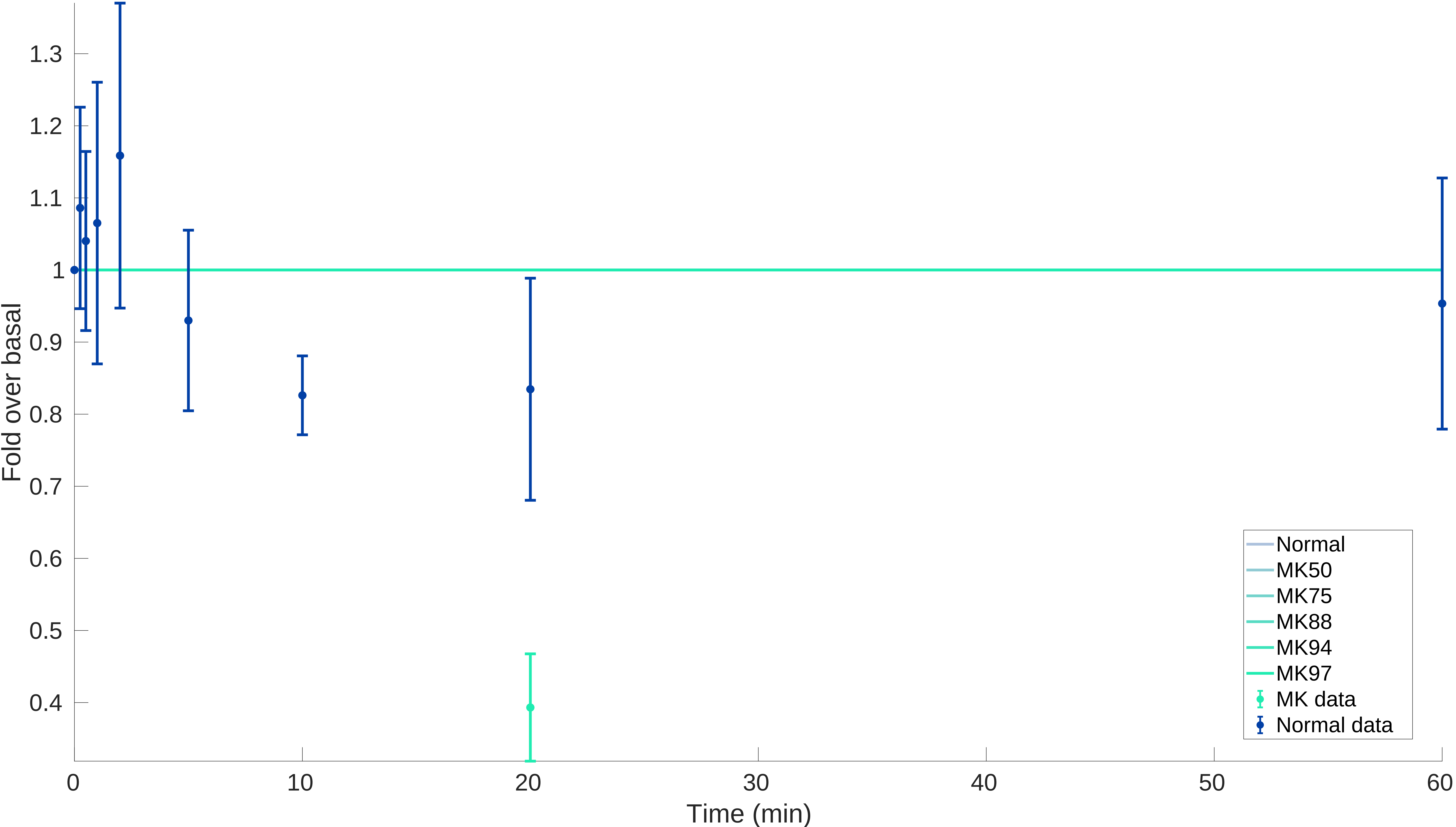

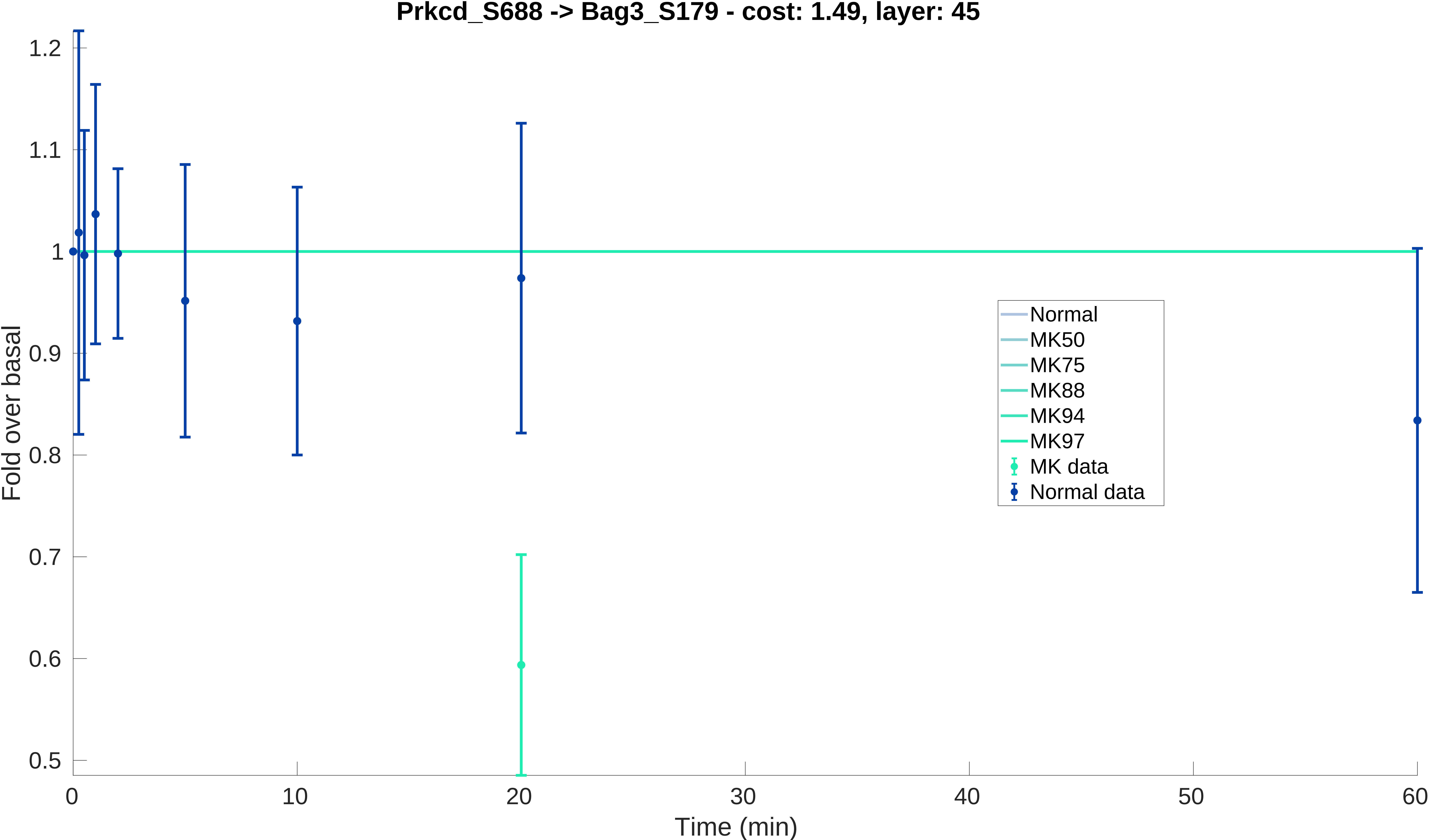

Prkcd_S688 -> Bag3_S179 - cost: 1.49, layer: 45
Fold over basal
1.2
1.1
1
0.9
0.8
0.7
0.6
0.5
0
10
20
30
40
50
60
Time (min)
Normal
MK50
MK75
MK88
MK94
MK97
MK data
Normal data

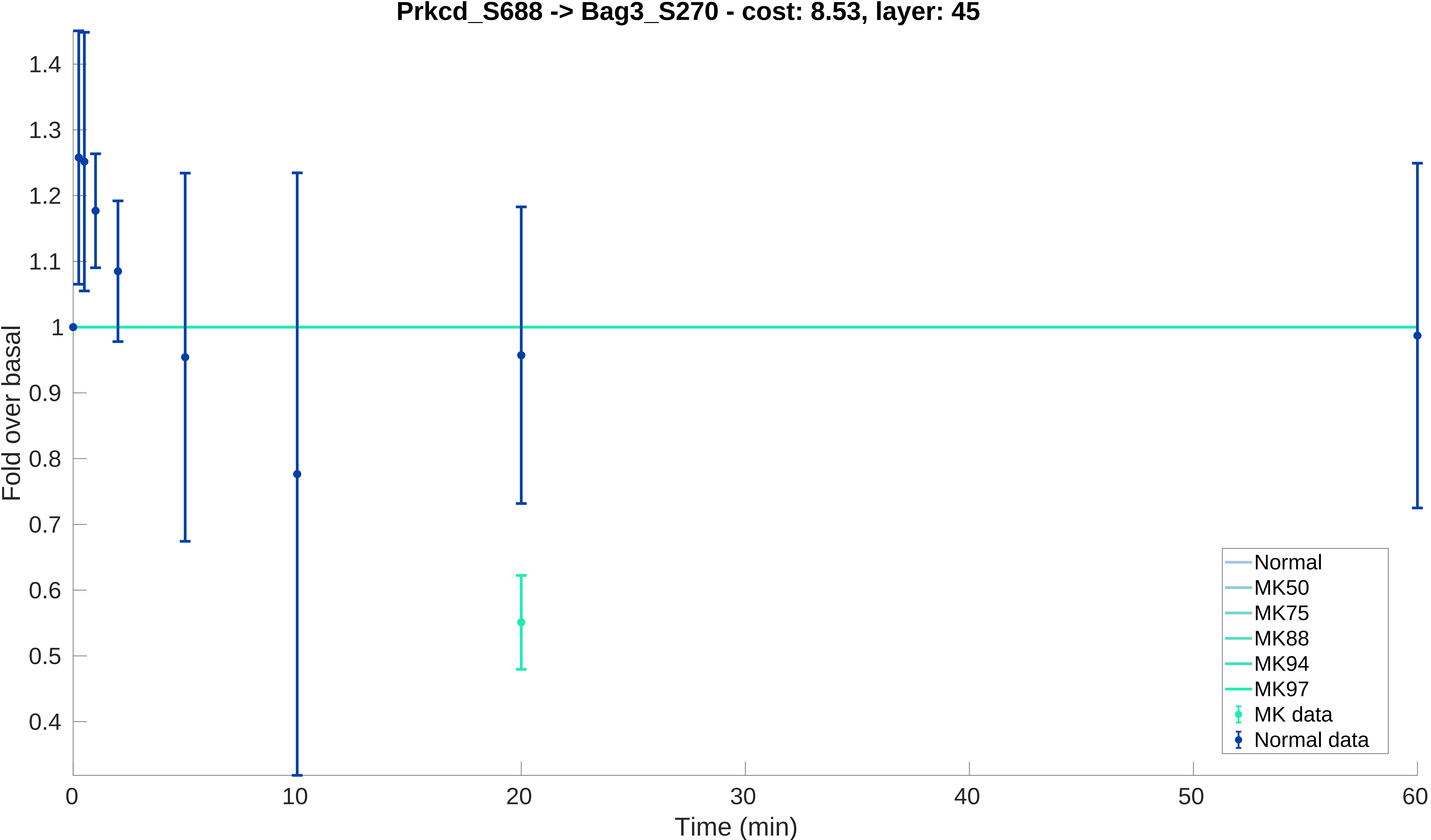

Prkcd_S688 -> Bag3_S270 - cost: 8.53, layer: 45
Fold over basal
Time (min)
1.4
1.3
1.2
1.1
1
0.9
0.8
0.7
0.6
0.5
0.4
0
10
20
30
40
50
60
Normal
MK50
MK75
MK88
MK94
MK97
MK data
Normal data

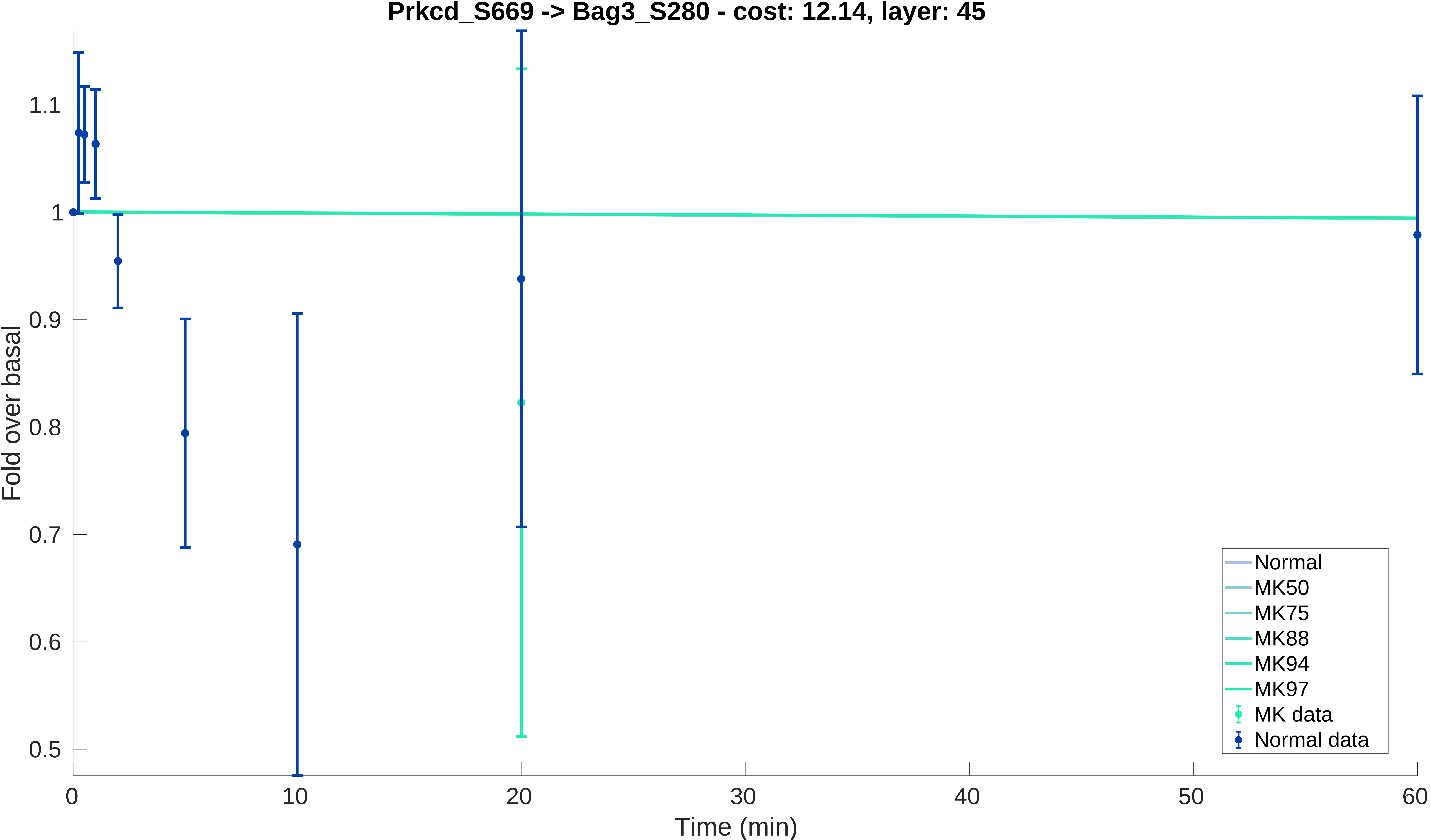

Prkcd_S669 -> Bag3_S280 - cost: 12.14, layer: 45
Fold over basal
Time (min)
0
10
20
30
40
50
60
0.5
0.6
0.7
0.8
0.9
1
1.1
Normal
MK50
MK75
MK88
MK94
MK97
MK data
Normal data

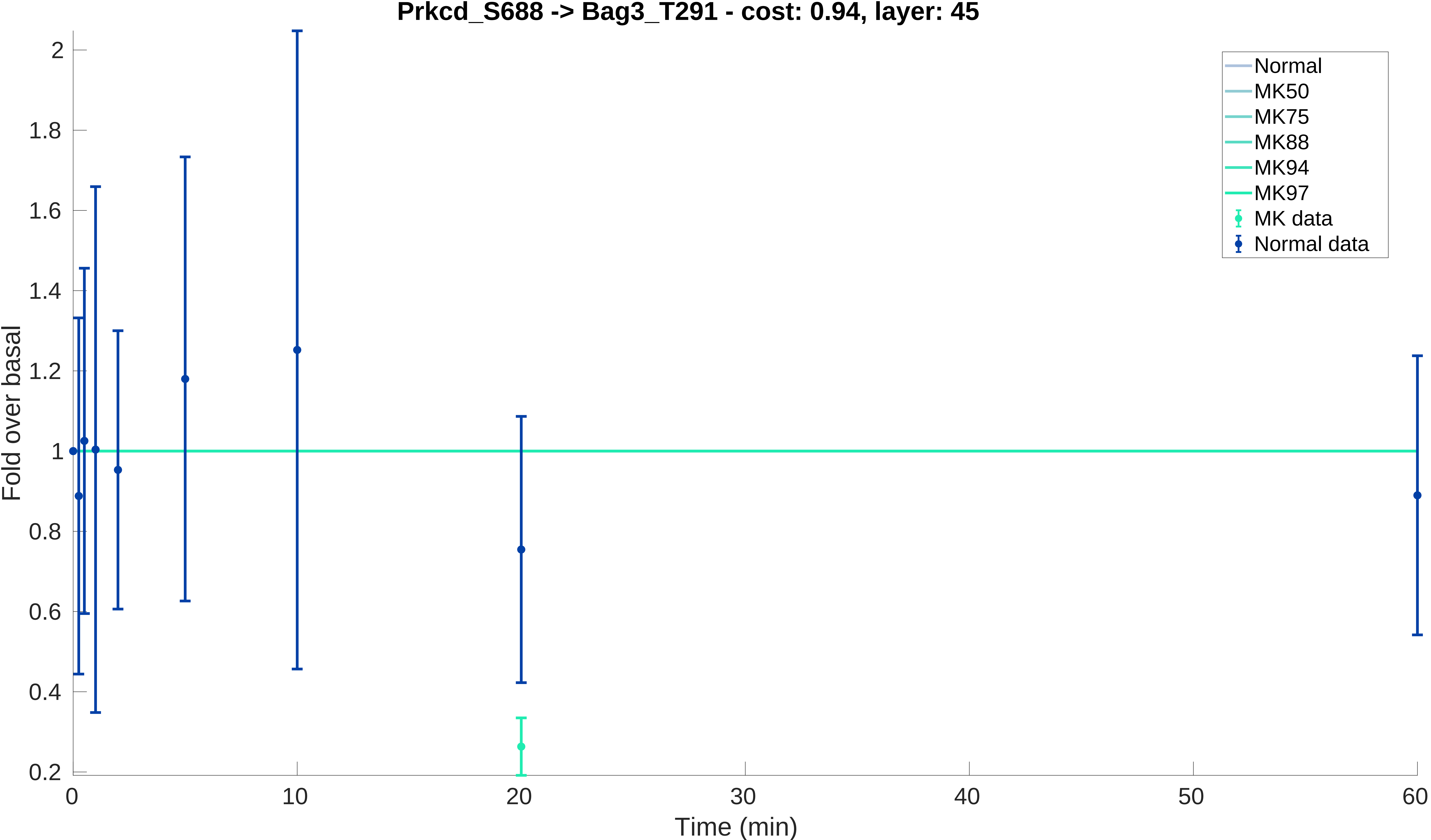

Prkcd_S688 -> Bag3_T291 - cost: 0.94, layer: 45
Fold over basal
Time (min)
0.2
0.4
0.6
0.8
1
1.2
1.4
1.6
1.8
2
0
10
20
30
40
50
60
Normal
MK50
MK75
MK88
MK94
MK97
MK data
Normal data

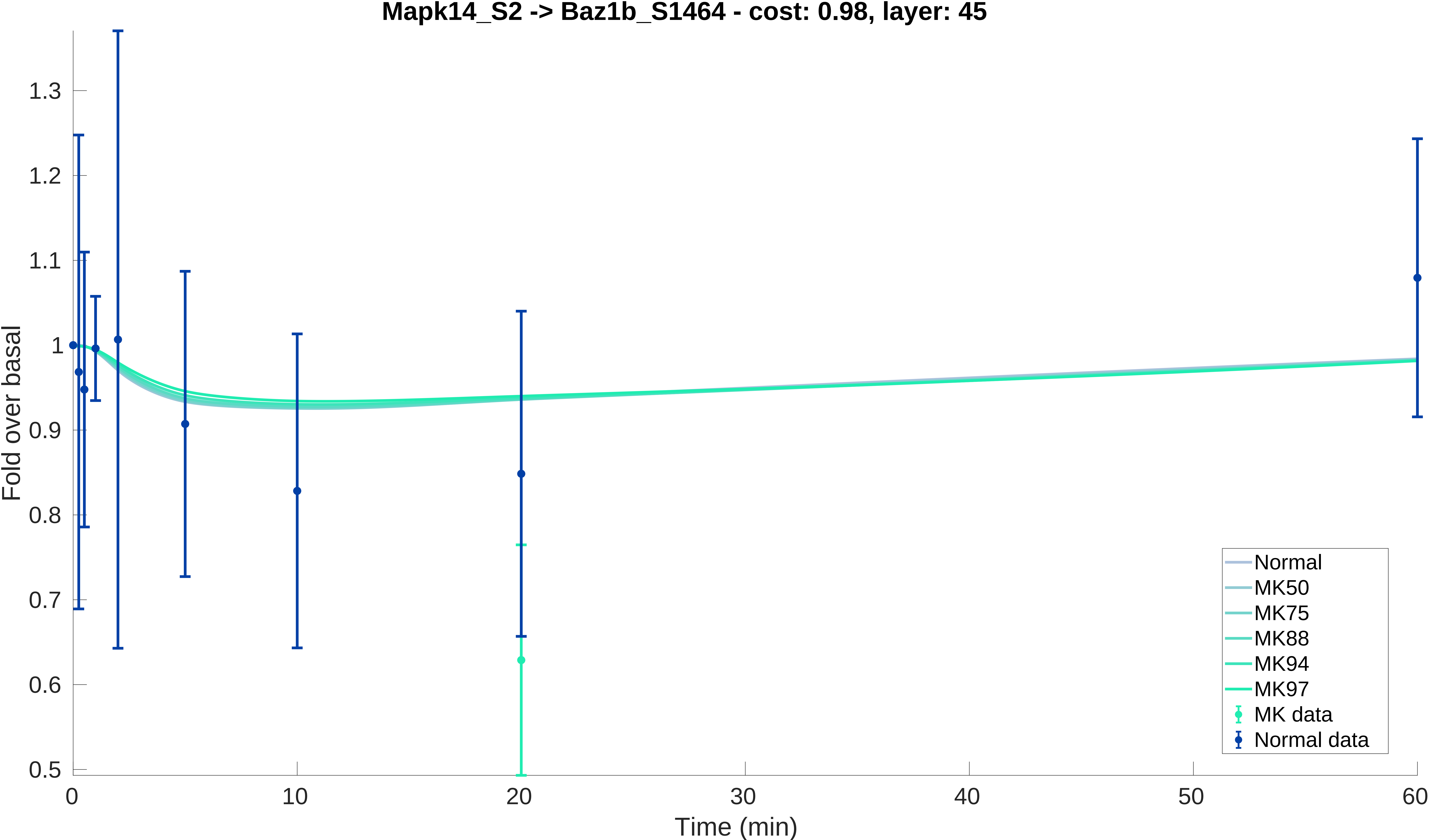

Mapk14_S2 -> Baz1b_S1464 - cost: 0.98, layer: 45
Fold over basal
Time (min)
1.3
1.2
1.1
1
0.9
0.8
0.7
0.6
0.5
0
10
20
30
40
50
60
Normal
MK50
MK75
MK88
MK94
MK97
MK data
Normal data

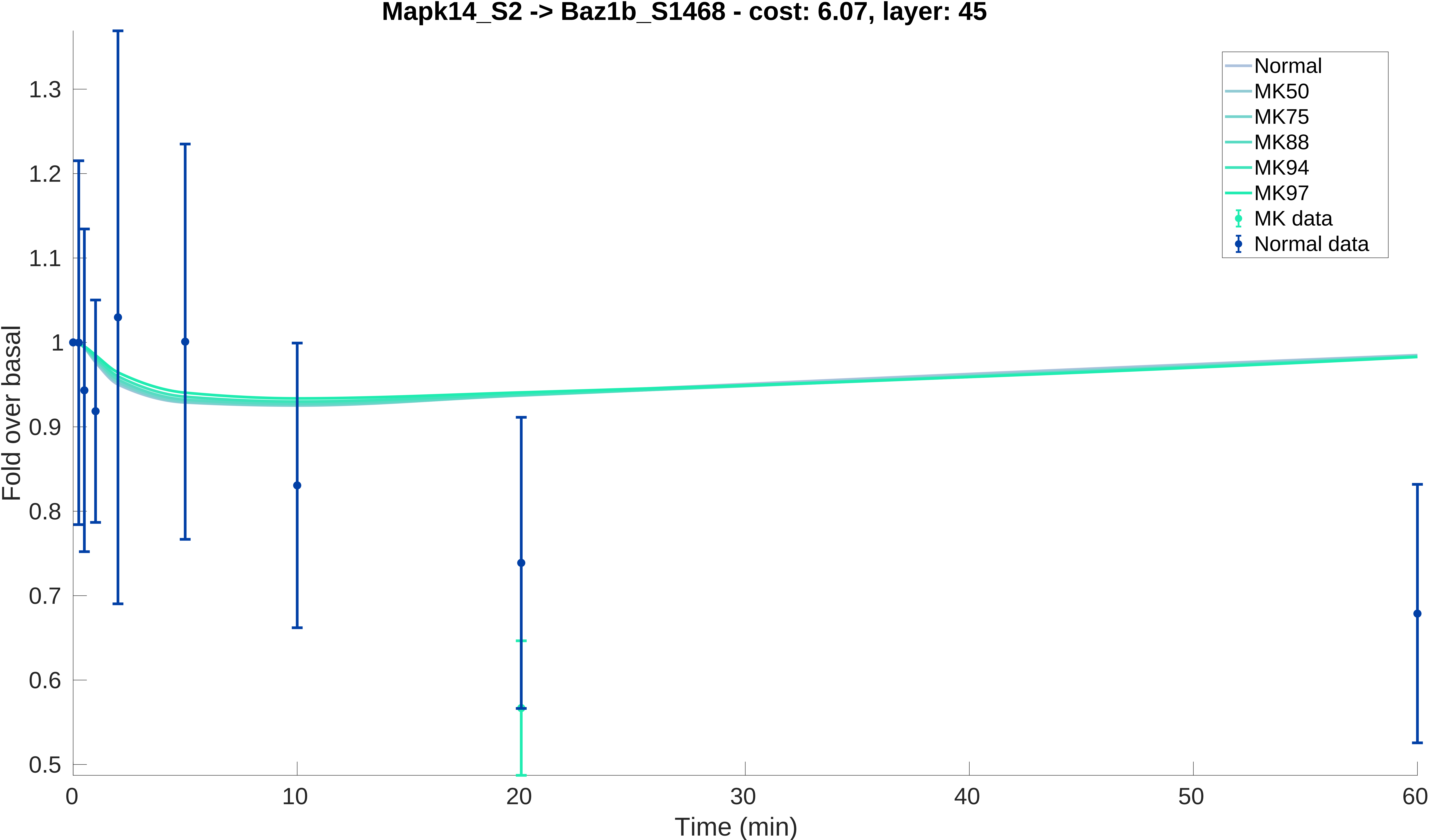

Mapk14_S2 -> Baz1b_S1468 - cost: 6.07, layer: 45
Normal
MK50
MK75
MK88
MK94
MK97
MK data
Normal data
Fold over basal
1.3
1.2
1.1
1
0.9
0.8
0.7
0.6
0.5
0
10
20
30
40
50
60
Time (min)

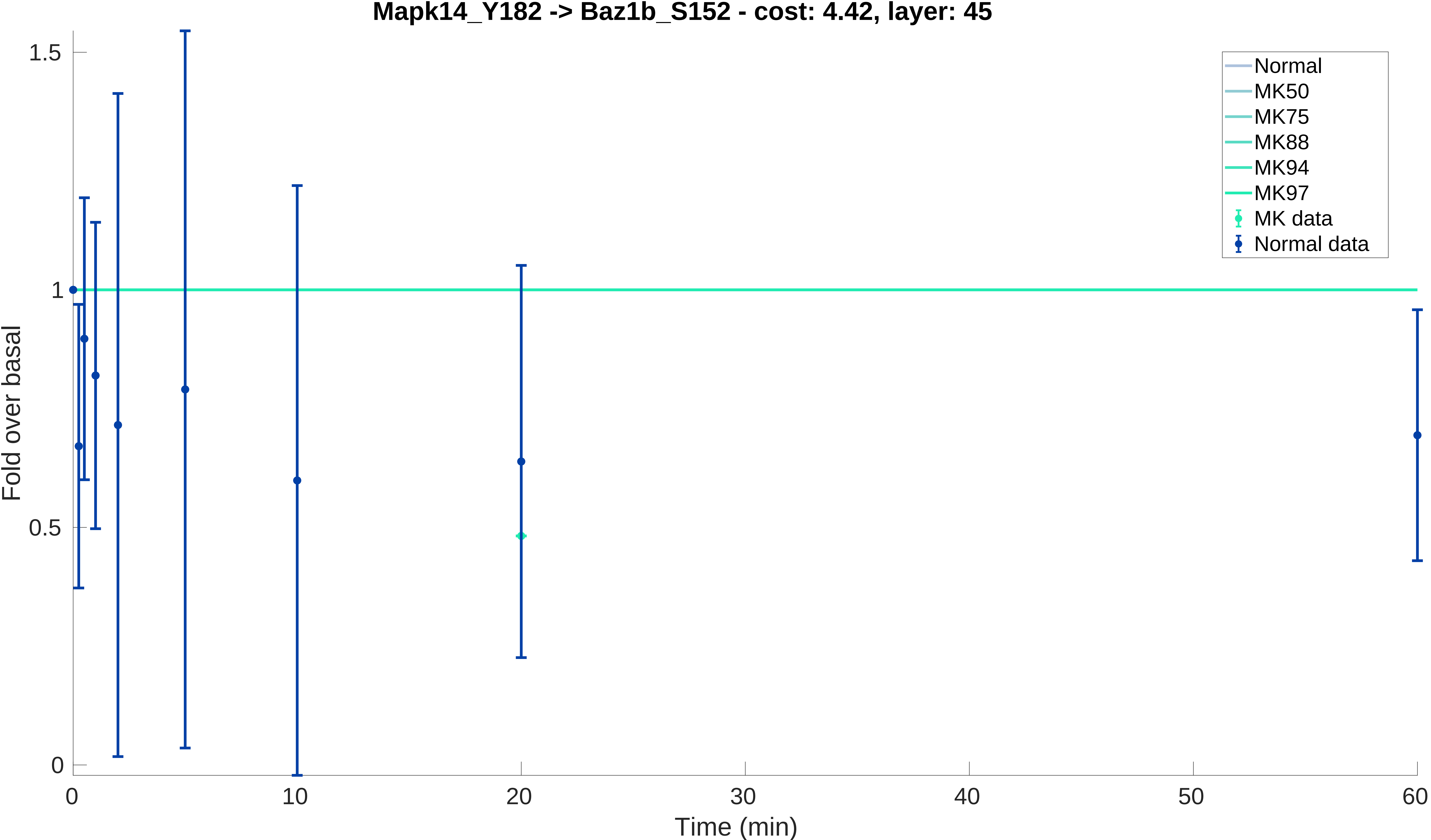

Mapk14_Y182 -> Baz1b_S152 - cost: 4.42, layer: 45
Fold over basal
1.5
1
0.5
0
Time (min)
0
10
20
30
40
50
60
Normal
MK50
MK75
MK88
MK94
MK97
MK data
Normal data

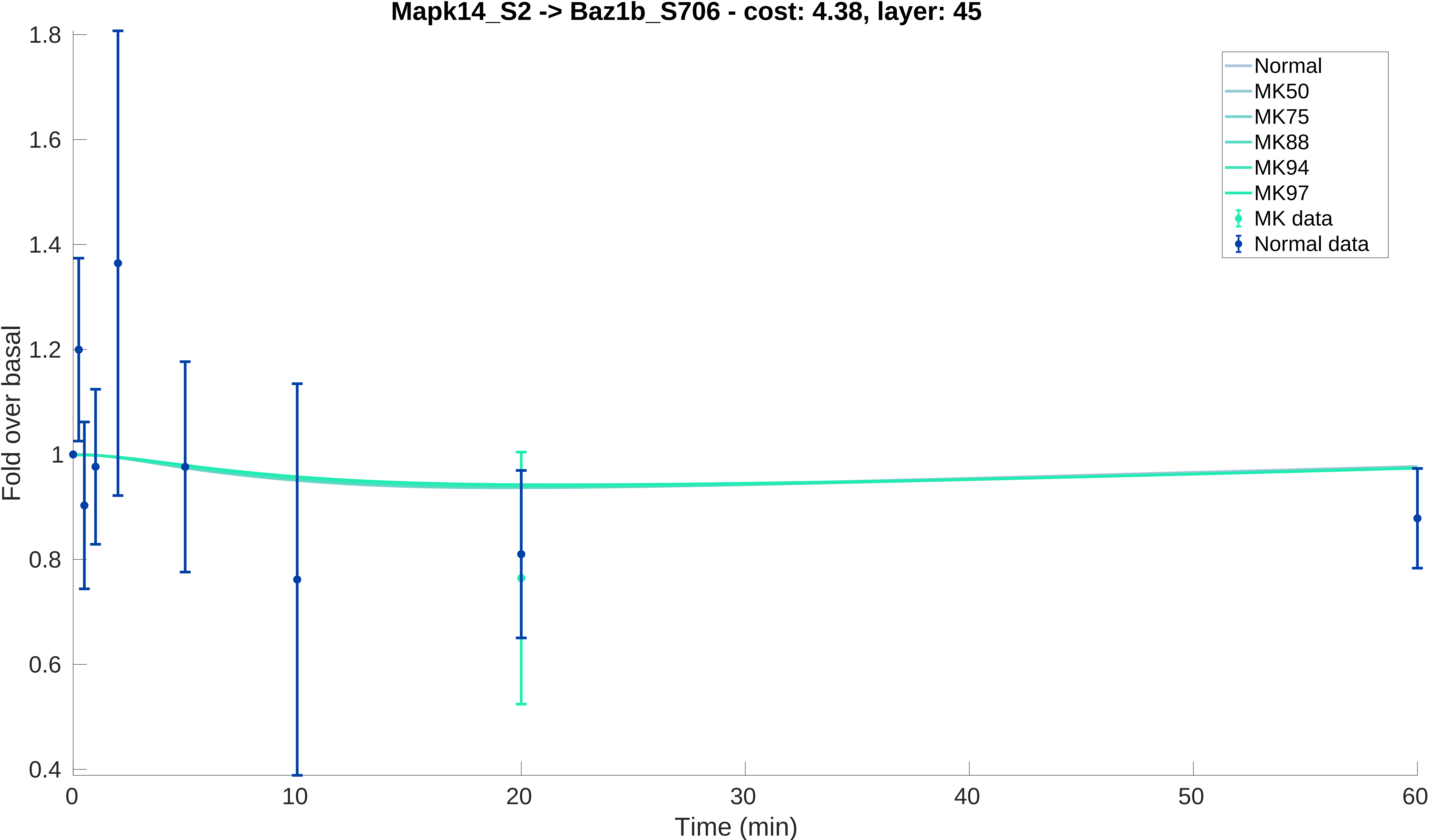
Mapk14_S2 -> Baz1b_S706 - cost: 4.38, layer: 45
Fold over basal
1.8
1.6
1.4
1.2
1
0.8
0.6
0.4
0
10
20
30
40
50
60
Time (min)
Normal
MK50
MK75
MK88
MK94
MK97
MK data
Normal data

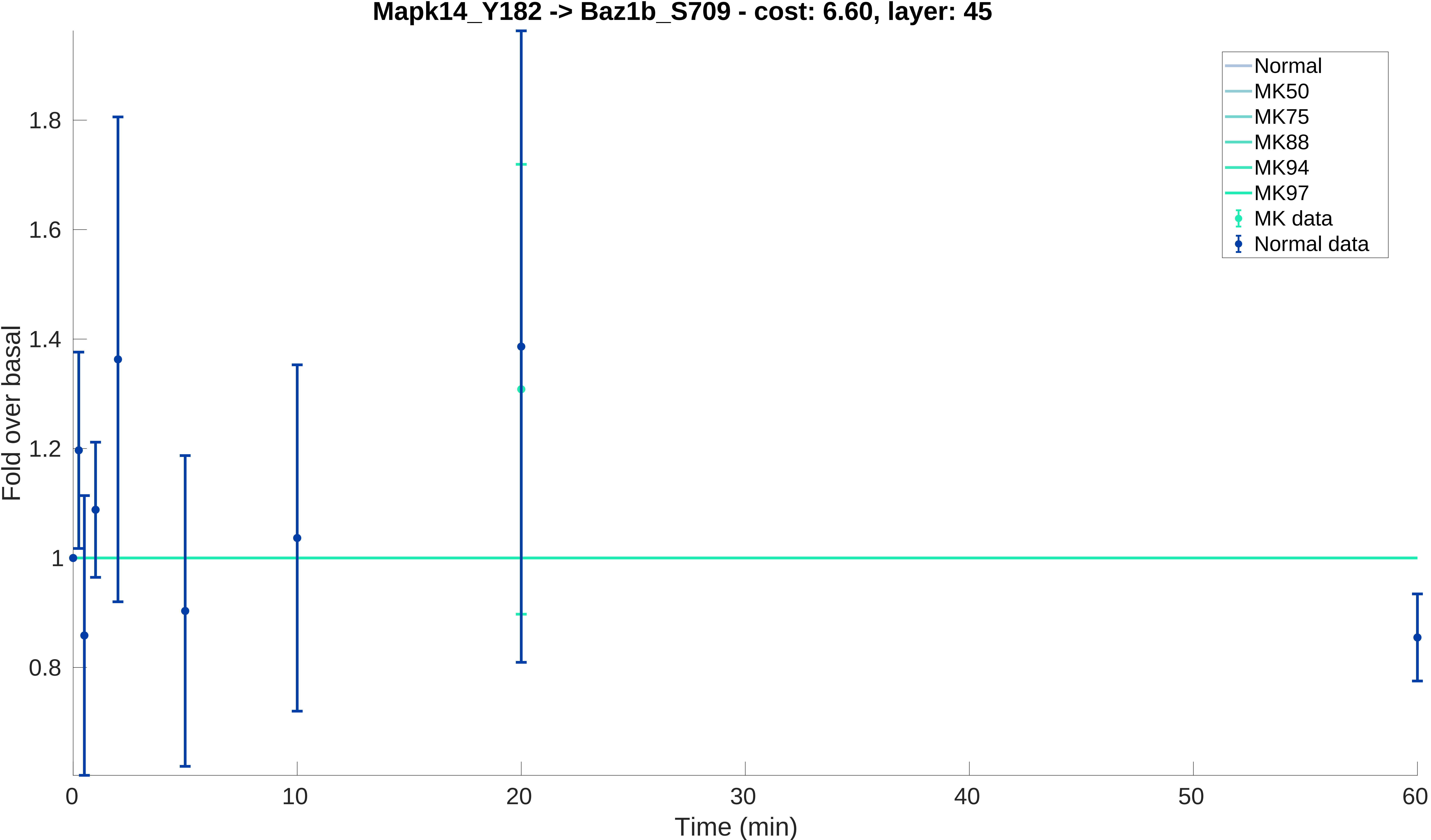

Mapk14_Y182 -> Baz1b_S709 - cost: 6.60, layer: 45
Normal
MK50
MK75
MK88
MK94
MK97
MK data
Normal data
Fold over basal
Time (min)
0.8
1
1.2
1.4
1.6
1.8
0
10
20
30
40
50
60

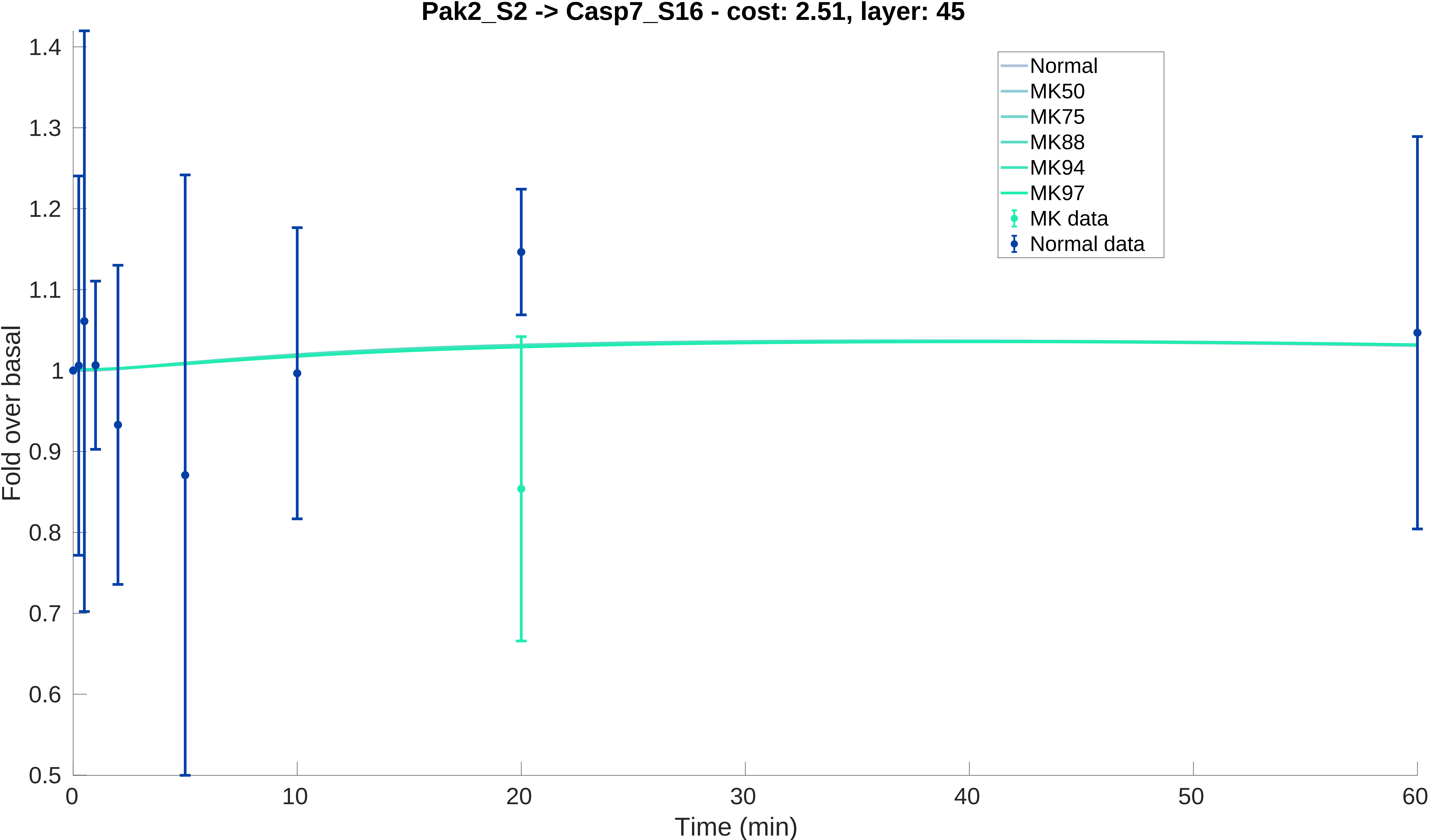
Pak2_S2 -> Casp7_S16 - cost: 2.51, layer: 45
Normal
MK50
MK75
MK88
MK94
MK97
MK data
Normal data
Fold over basal
1.4
1.3
1.2
1.1
1
0.9
0.8
0.7
0.6
0.5
0
10
20
30
40
50
60
Time (min)

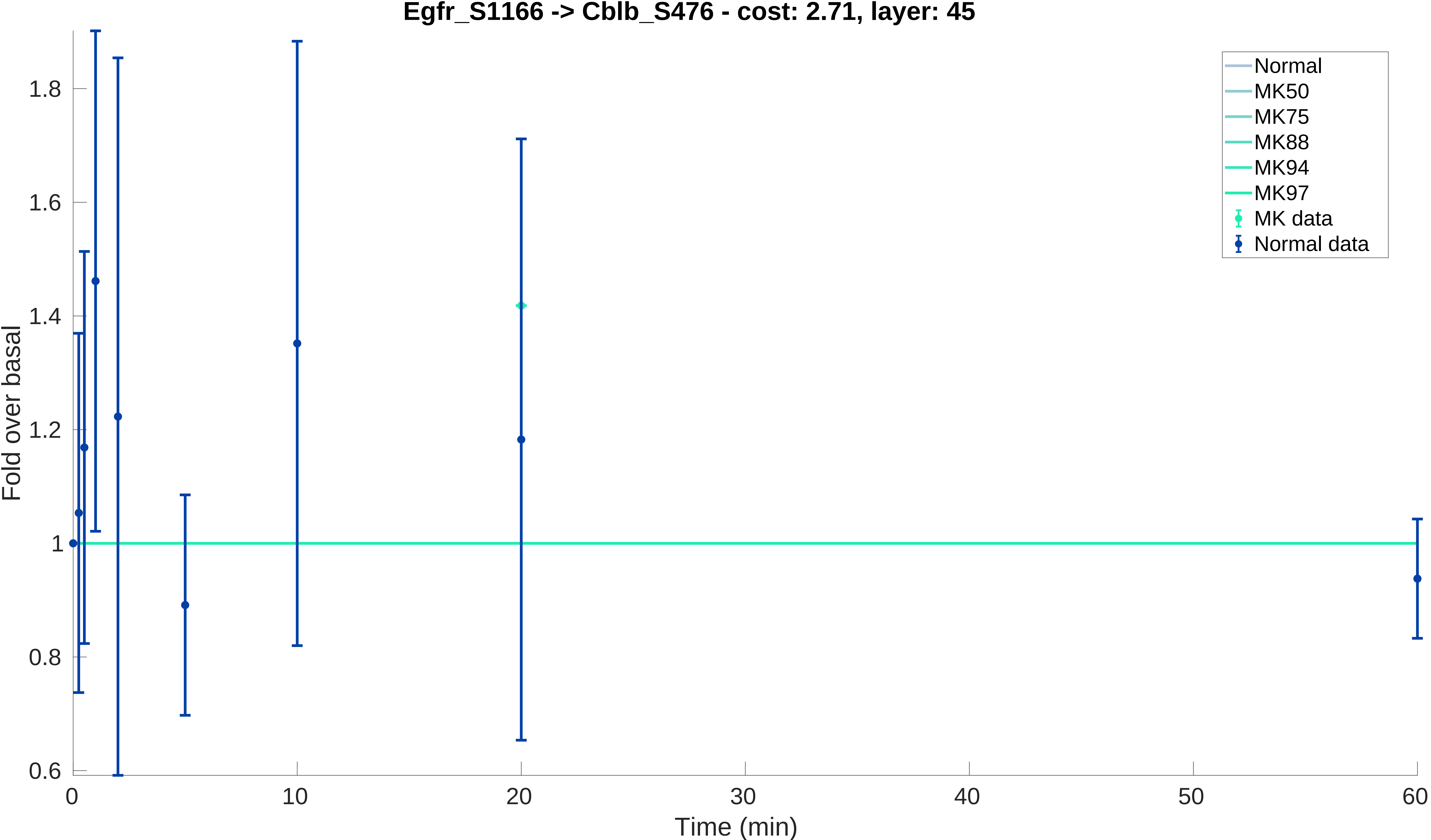

Egfr_S1166 -> Cblb_S476 - cost: 2.71, layer: 45
Fold over basal
Time (min)
0.6
0.8
1
1.2
1.4
1.6
1.8
0
10
20
30
40
50
60
Normal
MK50
MK75
MK88
MK94
MK97
MK data
Normal data

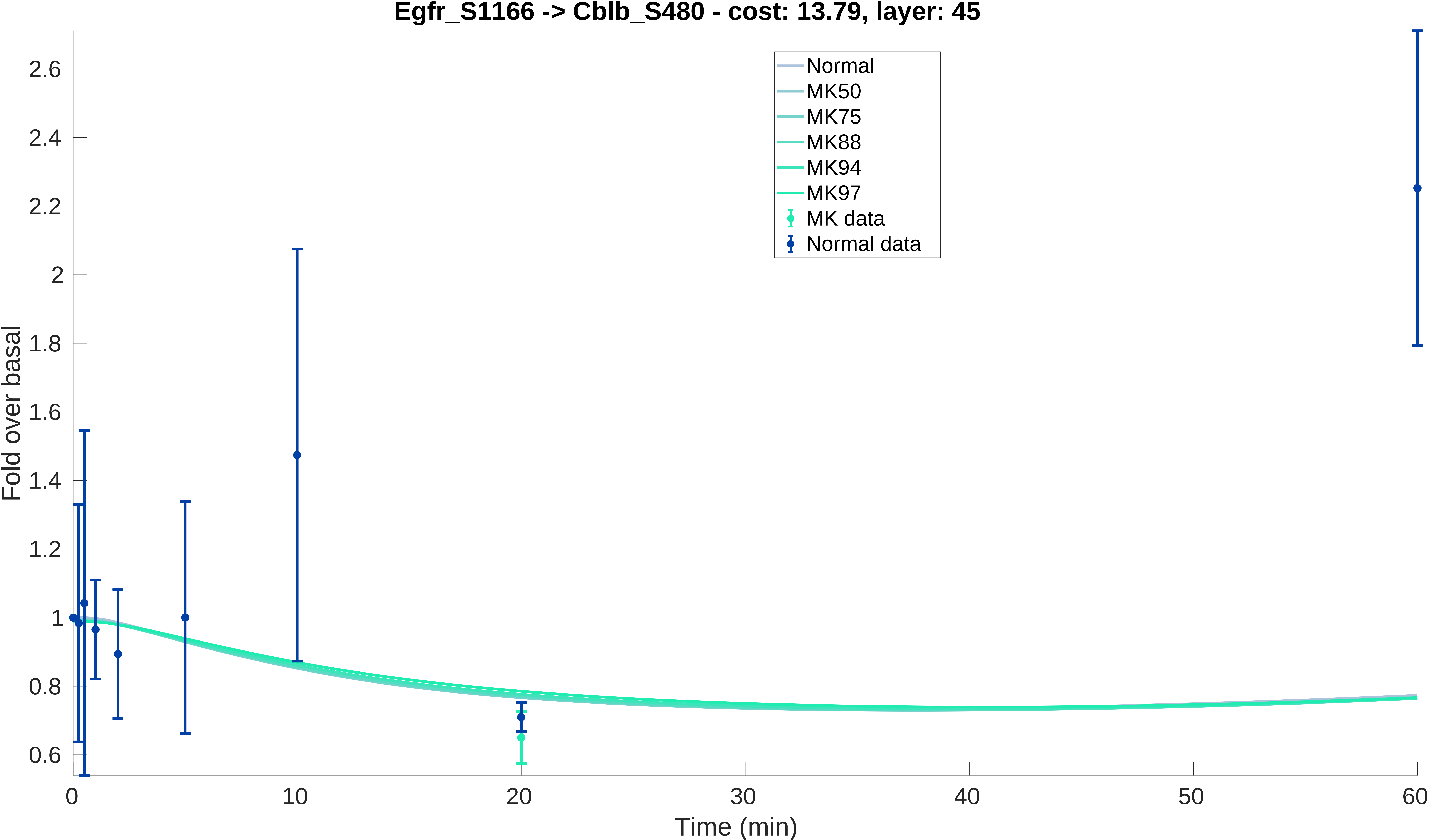
Egfr_S1166 -> Cblb_S480 - cost: 13.79, layer: 45
Normal
MK50
MK75
MK88
MK94
MK97
MK data
Normal data
Fold over basal
Time (min)
2.6
2.4
2.2
2
1.8
1.6
1.4
1.2
1
0.8
0.6
0
10
20
30
40
50
60

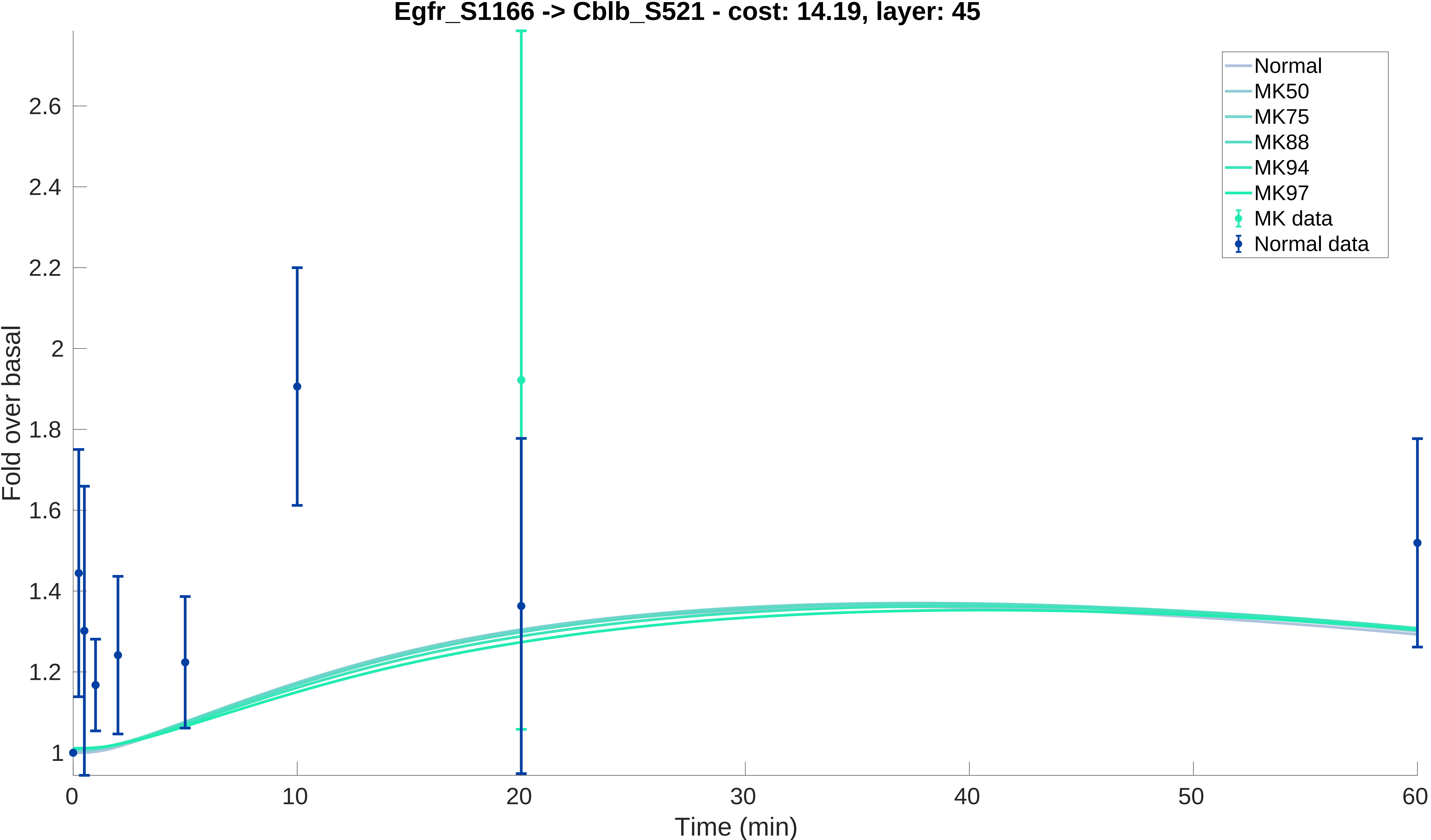
Egfr_S1166 -> Cblb_S521 - cost: 14.19, layer: 45
Fold over basal
Time (min)
Normal
MK50
MK75
MK88
MK94
MK97
MK data
Normal data
2.6
2.4
2.2
2
1.8
1.6
1.4
1.2
1
0
10
20
30
40
50
60

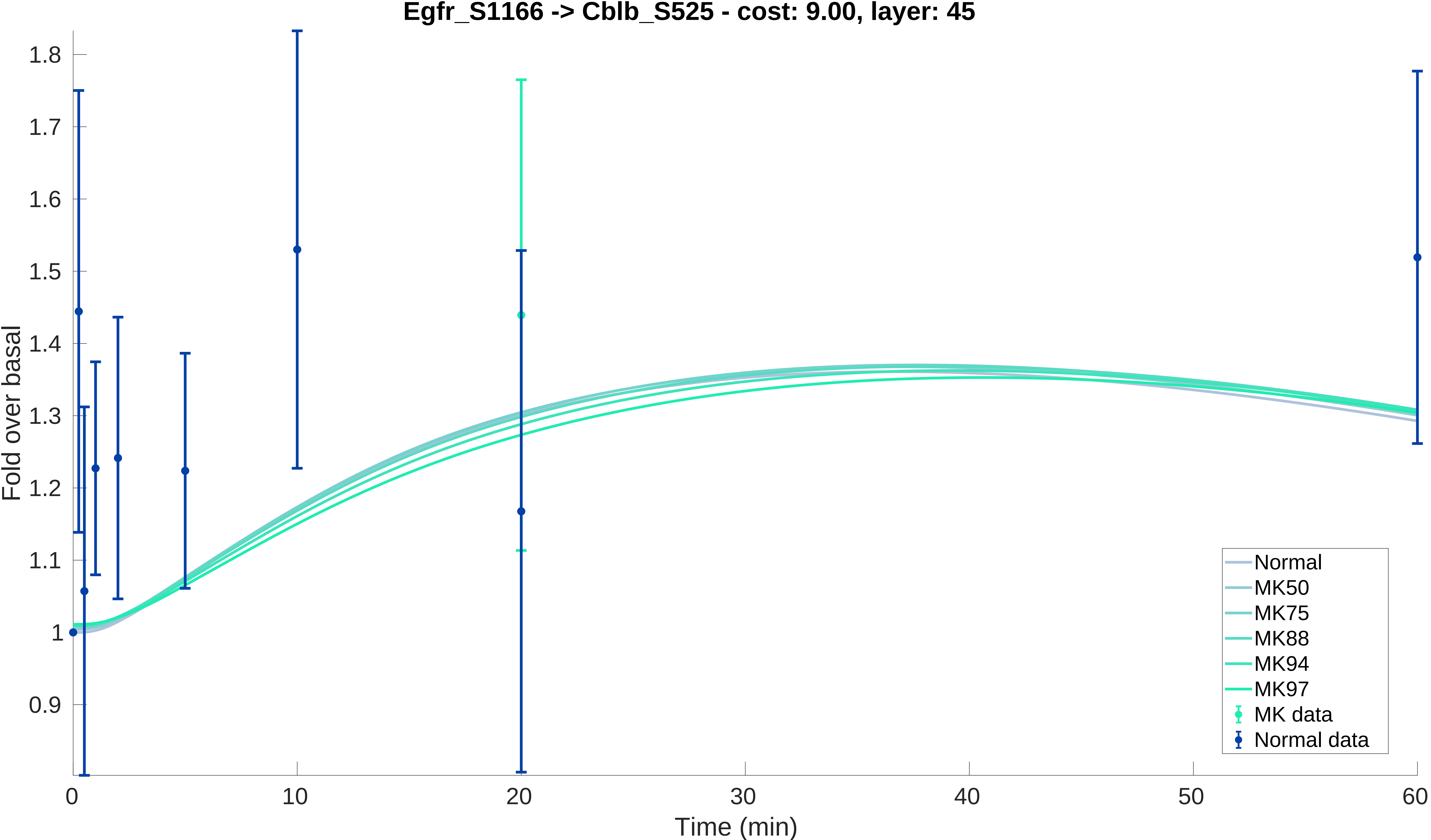

Egfr_S1166 -> Cblb_S525 - cost: 9.00, layer: 45
Fold over basal
1.8
1.7
1.6
1.5
1.4
1.3
1.2
1.1
1
0.9
0
10
20
30
40
50
60
Time (min)
Normal
MK50
MK75
MK88
MK94
MK97
MK data
Normal data

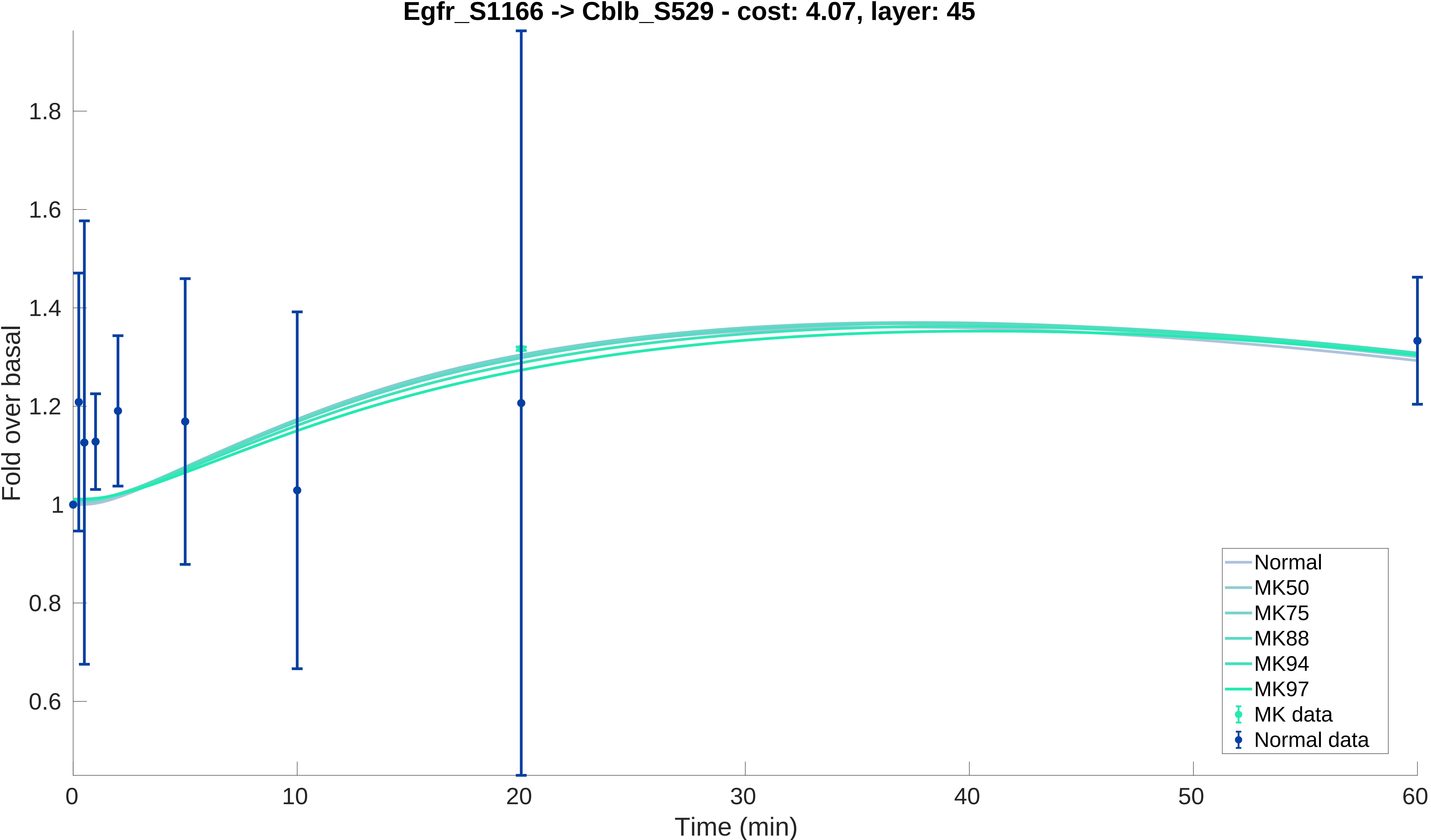

Egfr_S1166 -> Cblb_S529 - cost: 4.07, layer: 45
Fold over basal
1.8
1.6
1.4
1.2
1
0.8
0.6
0
10
20
30
40
50
60
Time (min)
Normal
MK50
MK75
MK88
MK94
MK97
MK data
Normal data

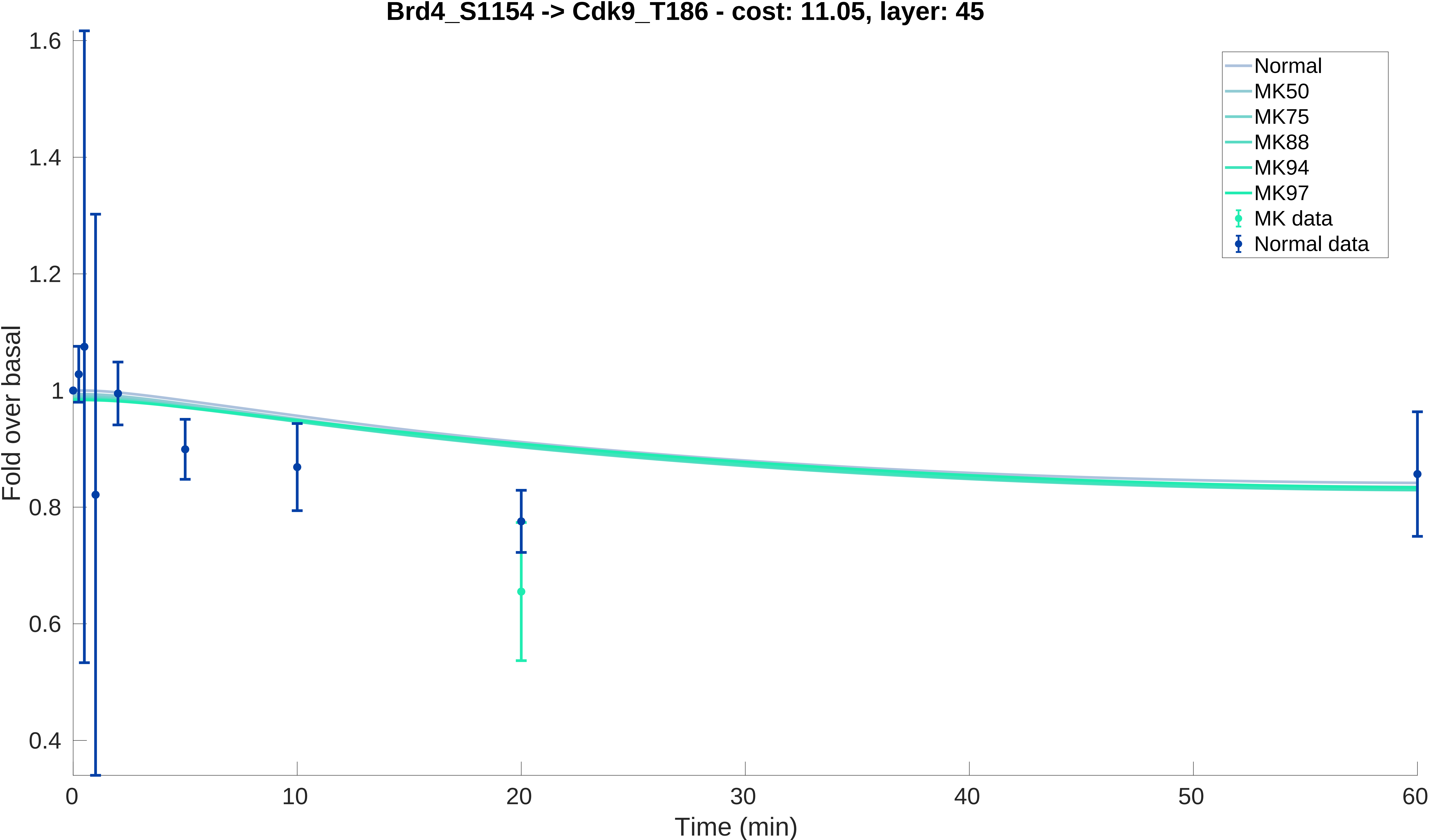
Brd4_S1154 -> Cdk9_T186 - cost: 11.05, layer: 45
Fold over basal
Time (min)
0.4
0.6
0.8
1
1.2
1.4
1.6
0
10
20
30
40
50
60
Normal
MK50
MK75
MK88
MK94
MK97
MK data
Normal data

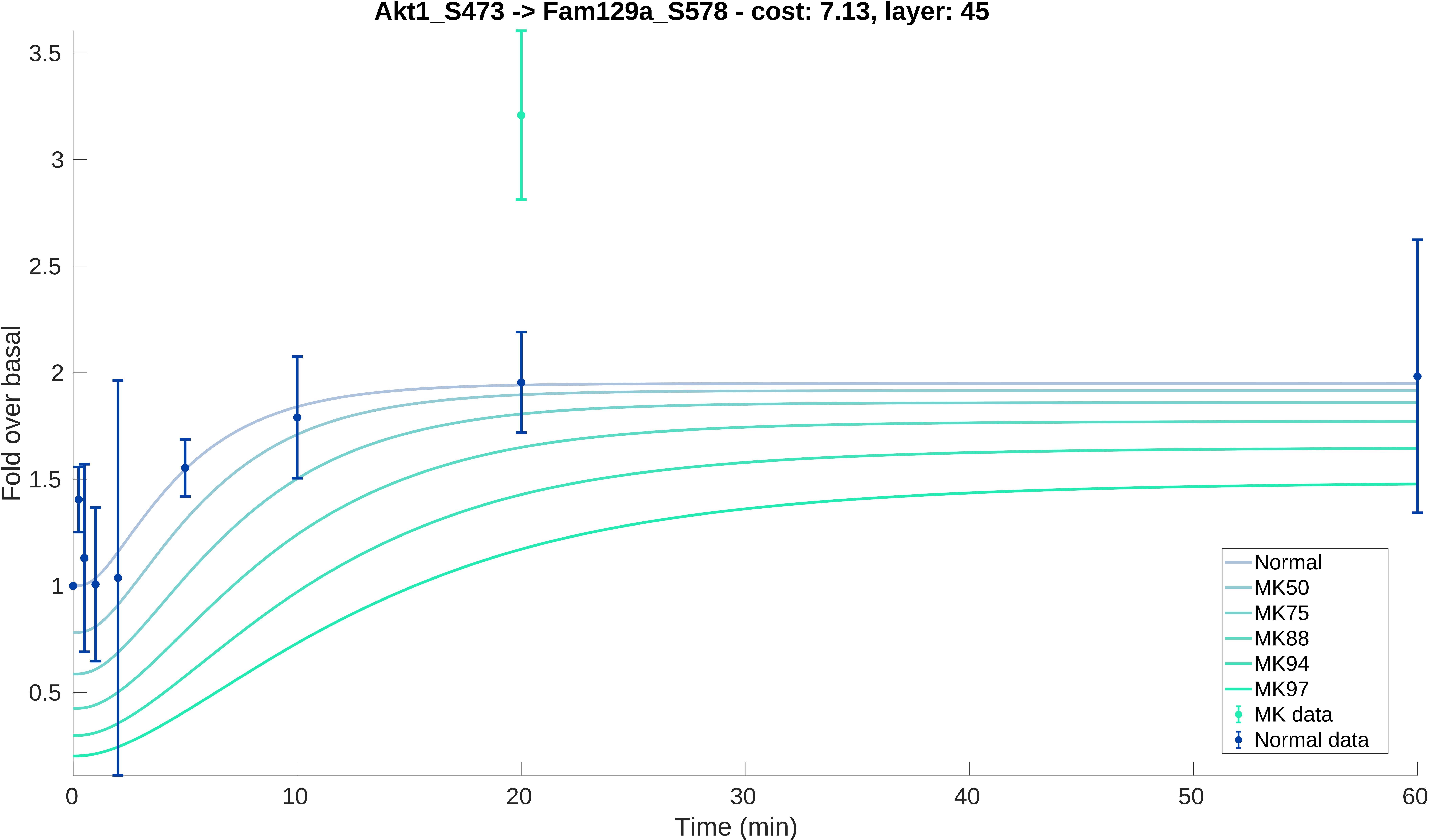

Akt1_S473 -> Fam129a_S578 - cost: 7.13, layer: 45
Fold over basal
Time (min)
3.5
3
2.5
2
1.5
1
0.5
0
10
20
30
40
50
60
Normal
MK50
MK75
MK88
MK94
MK97
MK data
Normal data

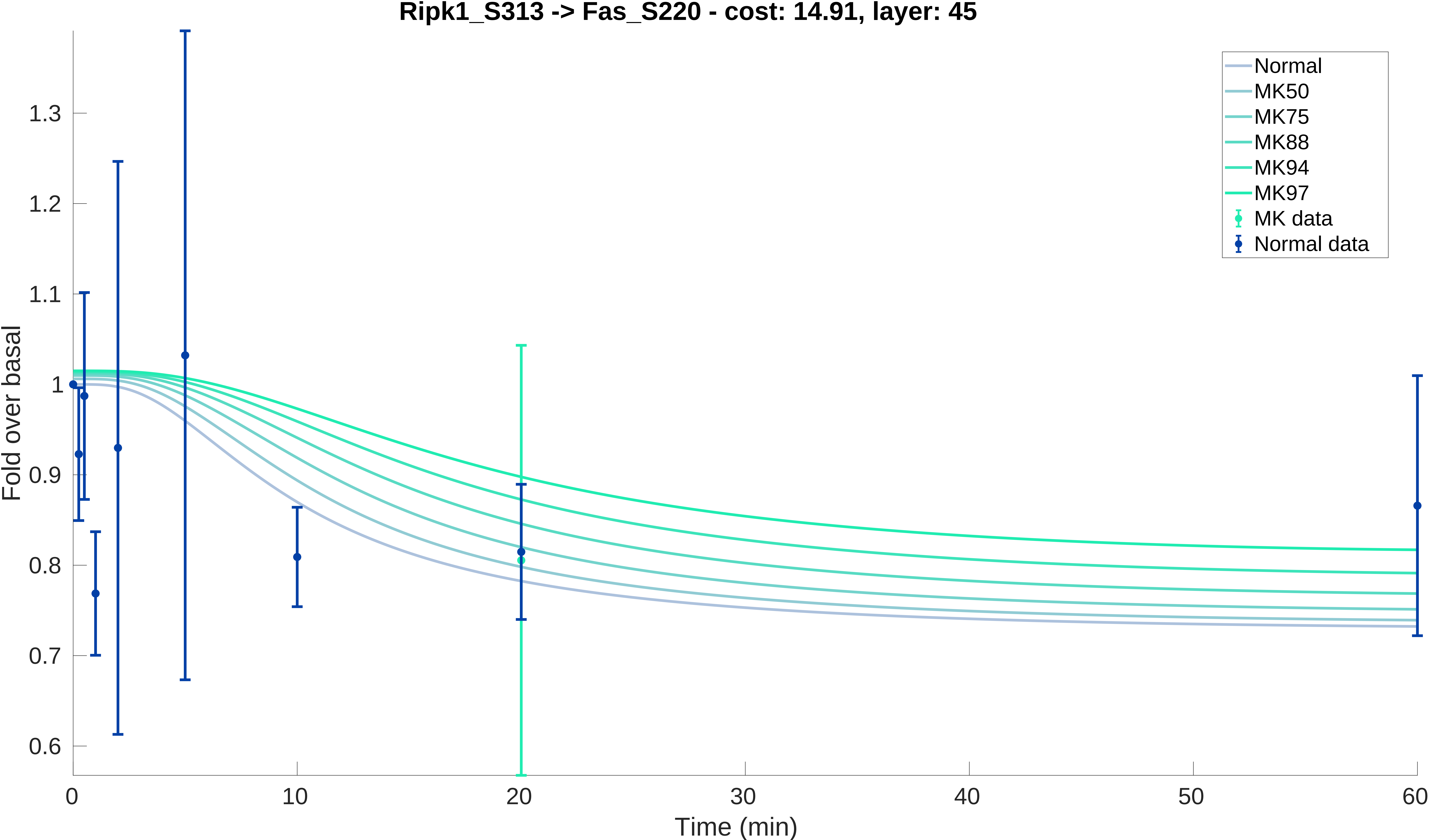

Ripk1_S313 -> Fas_S220 - cost: 14.91, layer: 45
Fold over basal
Time (min)
1.3
1.2
1.1
1
0.9
0.8
0.7
0.6
0
10
20
30
40
50
60
Normal
MK50
MK75
MK88
MK94
MK97
MK data
Normal data

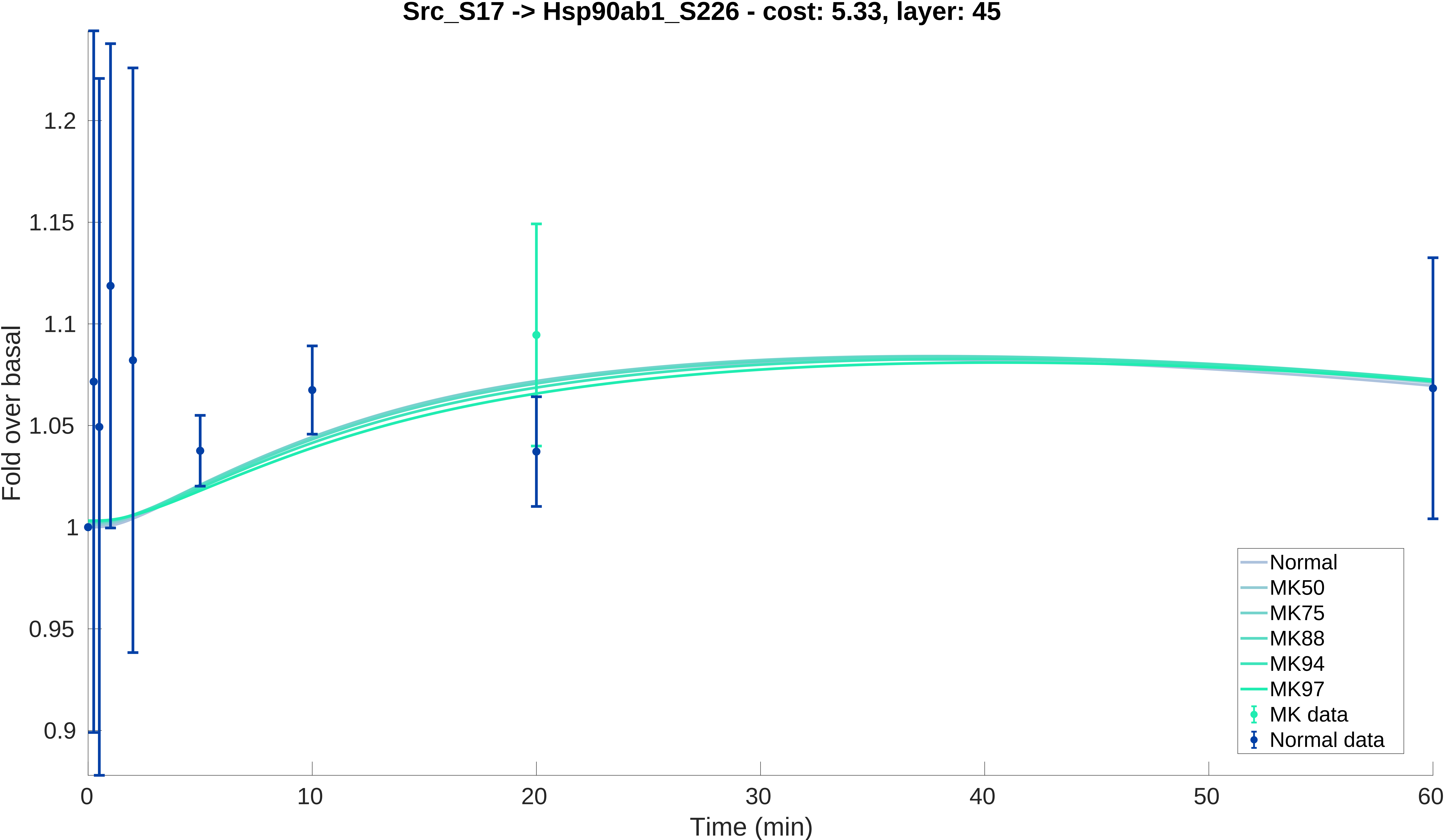

Src_S17 -> Hsp90ab1_S226 - cost: 5.33, layer: 45
Fold over basal
1.2
1.15
1.1
1.05
1
0.95
0.9
0
10
20
30
40
50
60
Time (min)
Normal
MK50
MK75
MK88
MK94
MK97
MK data
Normal data

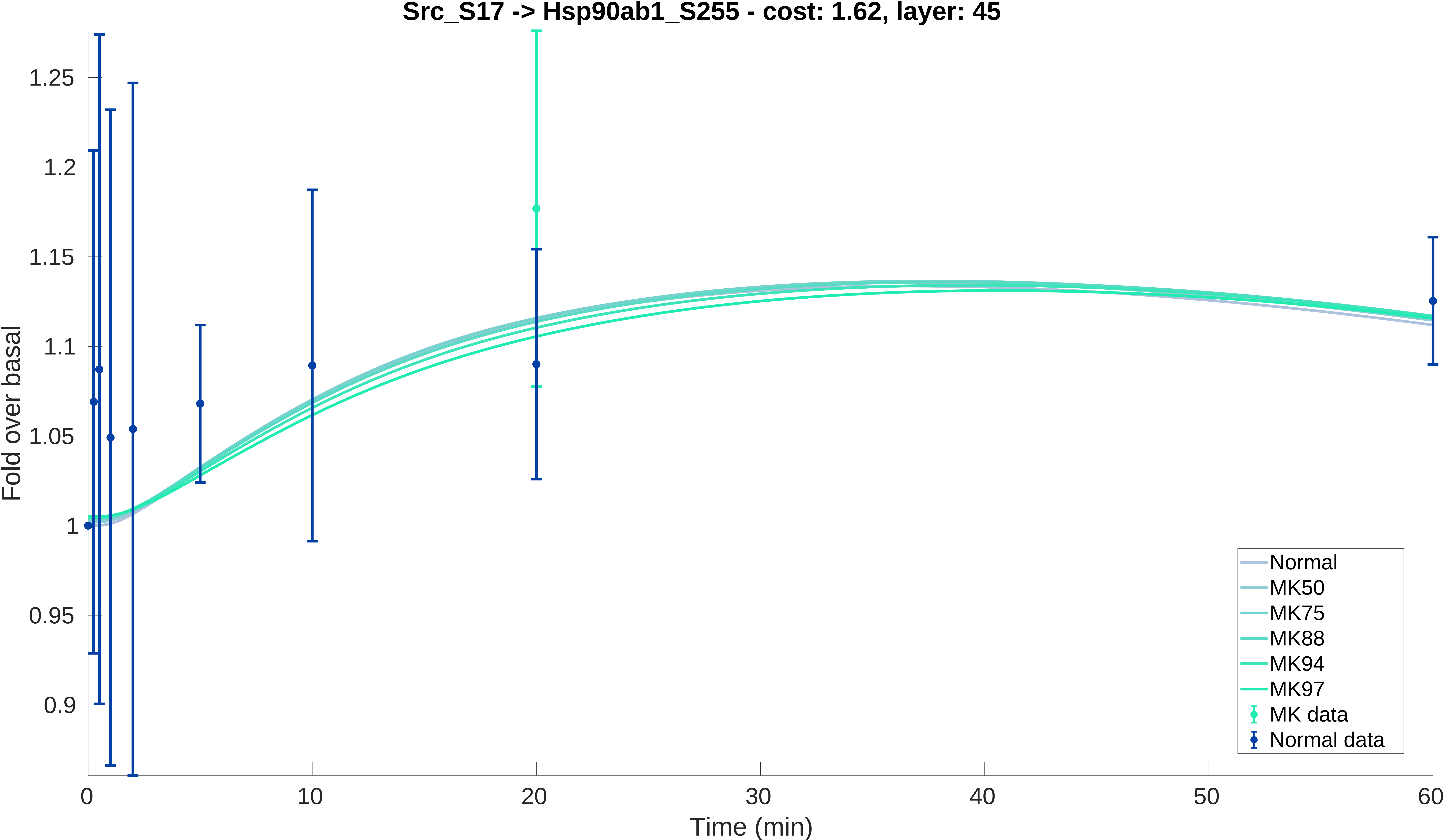

Src_S17 -> Hsp90ab1_S255 - cost: 1.62, layer: 45
Fold over basal
1.25
1.2
1.15
1.1
1.05
1
0.95
0.9
0
10
20
30
40
50
60
Time (min)
Normal
MK50
MK75
MK88
MK94
MK97
MK data
Normal data

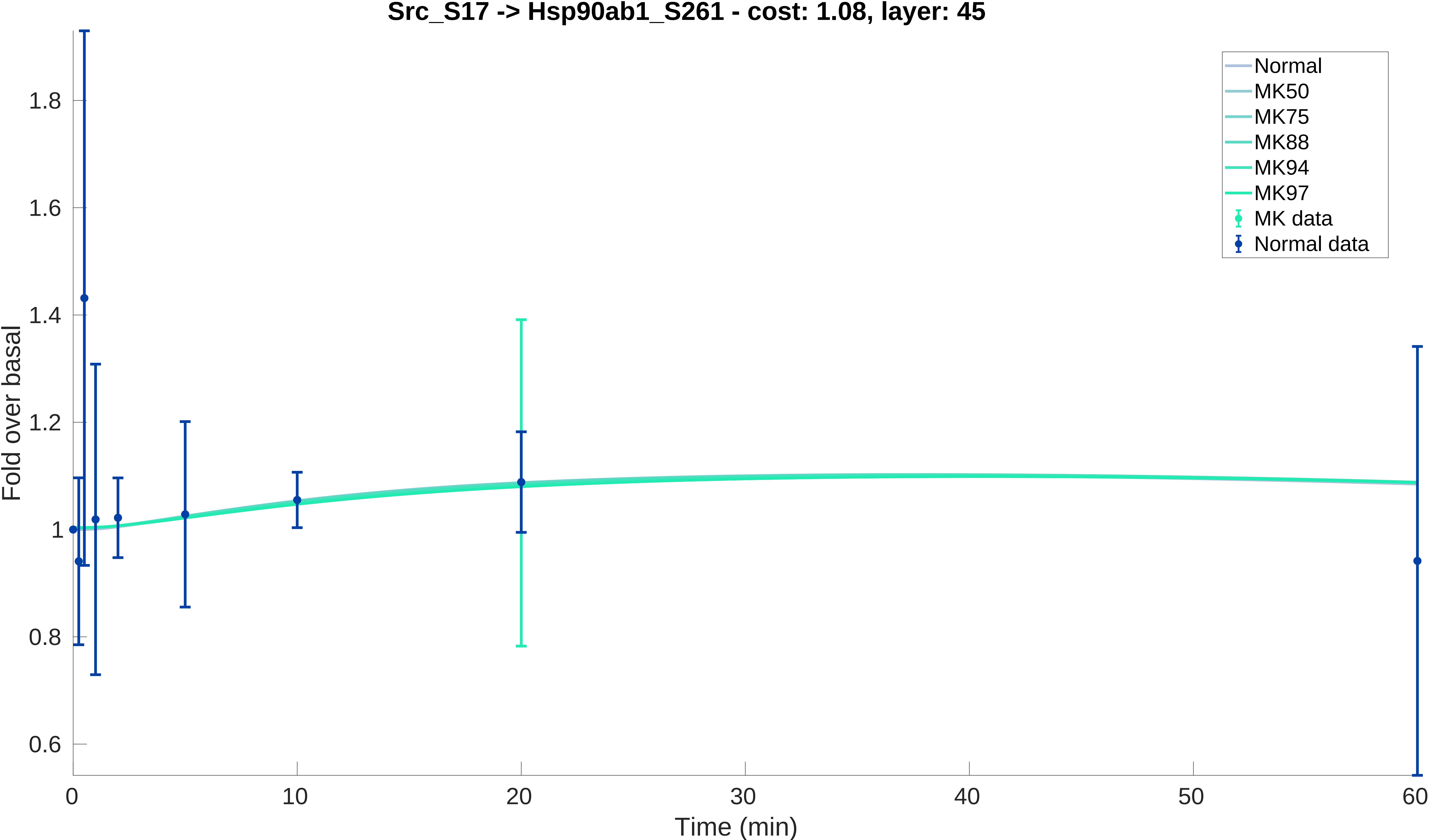
Src_S17 -> Hsp90ab1_S261 - cost: 1.08, layer: 45
Fold over basal
Time (min)
0.6
0.8
1
1.2
1.4
1.6
1.8
0
10
20
30
40
50
60
Normal
MK50
MK75
MK88
MK94
MK97
MK data
Normal data

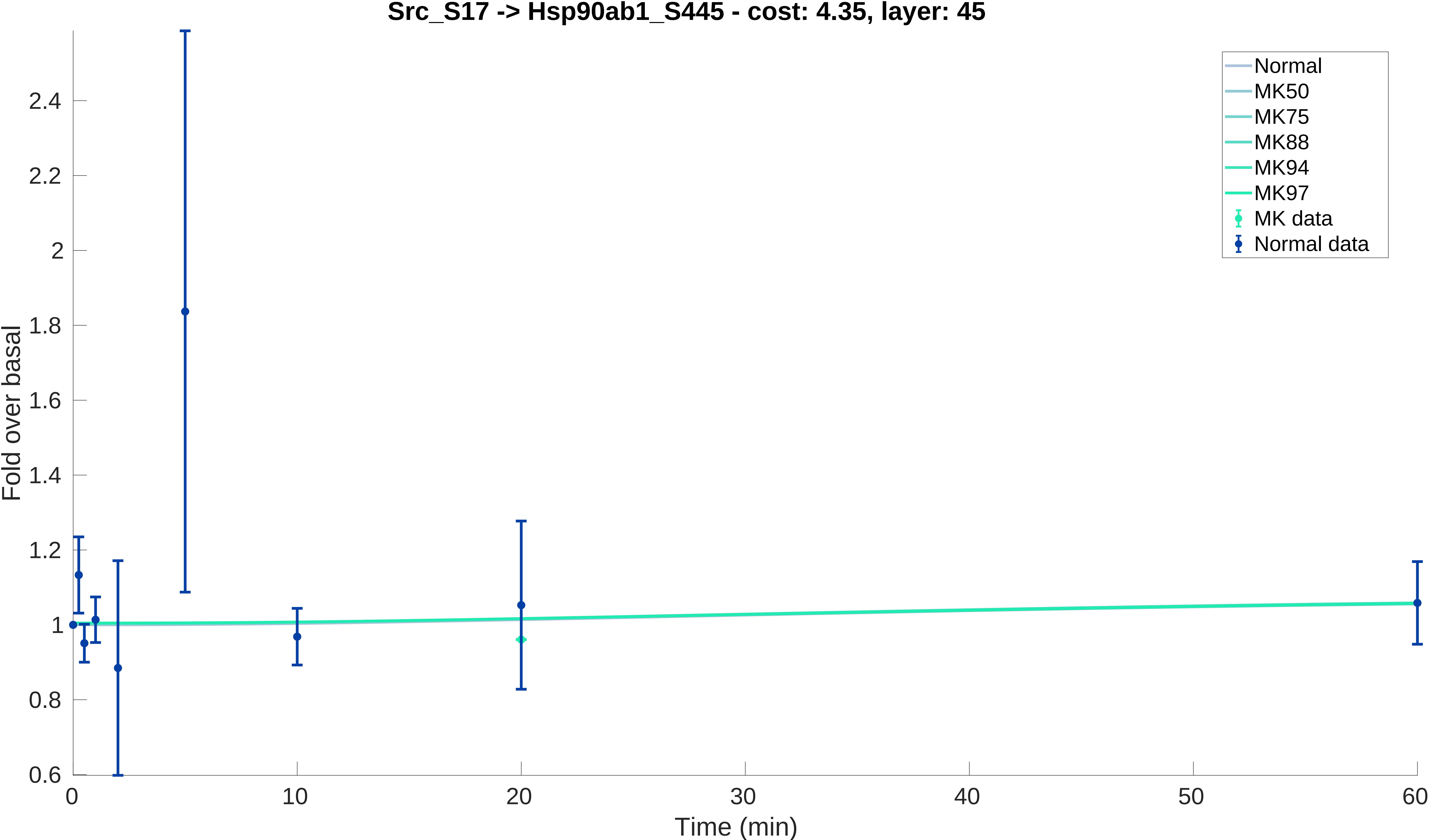
Src_S17 -> Hsp90ab1_S445 - cost: 4.35, layer: 45
Fold over basal
Time (min)
0.6
0.8
1
1.2
1.4
1.6
1.8
2
2.2
2.4
0
10
20
30
40
50
60
Normal
MK50
MK75
MK88
MK94
MK97
MK data
Normal data

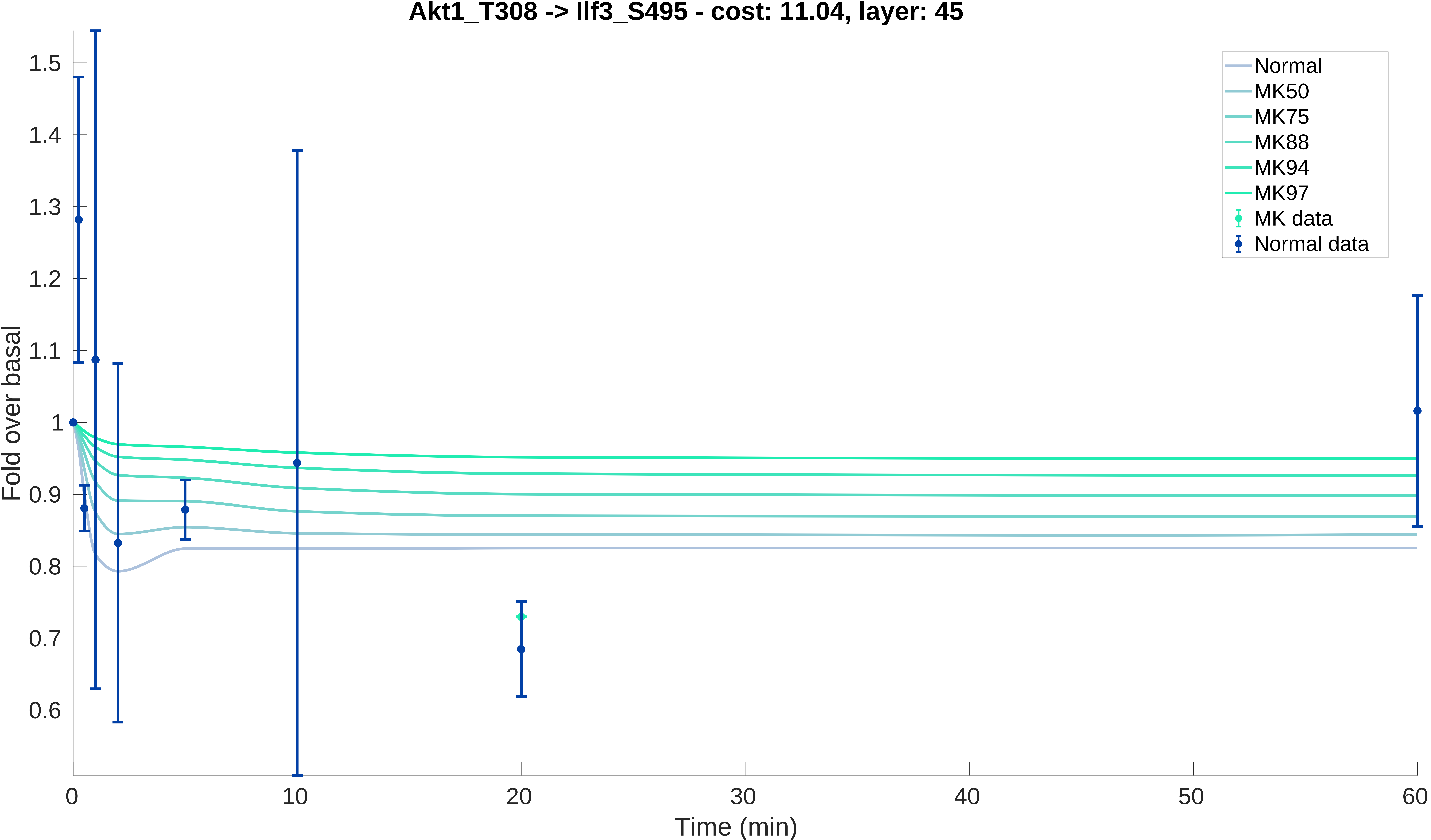

Akt1_T308 -> Ilf3_S495 - cost: 11.04, layer: 45
Fold over basal
1.5
1.4
1.3
1.2
1.1
1
0.9
0.8
0.7
0.6
0
10
20
30
40
50
60
Time (min)
Normal
MK50
MK75
MK88
MK94
MK97
MK data
Normal data

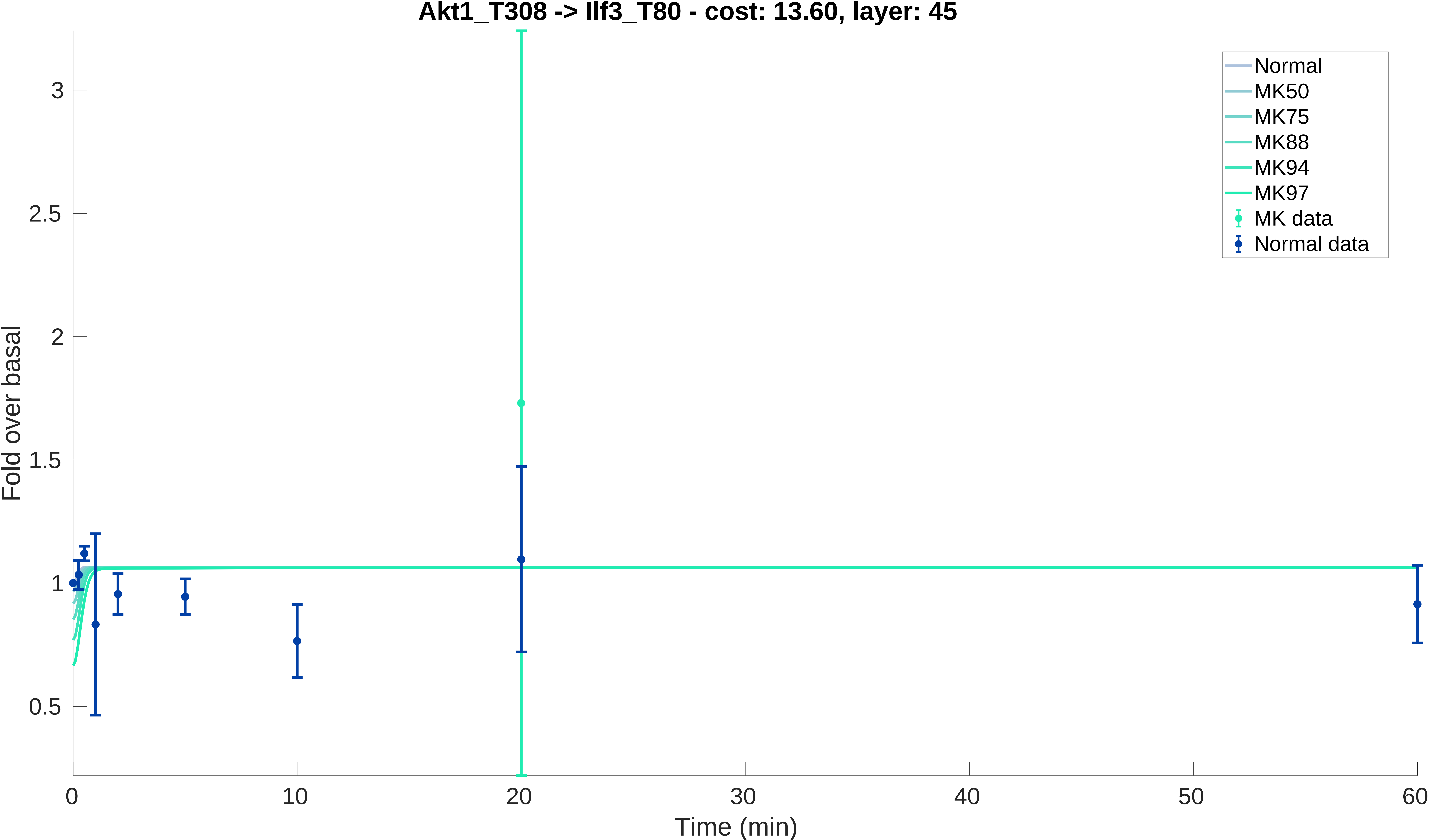

Akt1_T308 -> Ilf3_T80 - cost: 13.60, layer: 45
Normal
MK50
MK75
MK88
MK94
MK97
MK data
Normal data
Fold over basal
3
2.5
2
1.5
1
0.5
0
10
20
30
40
50
60
Time (min)

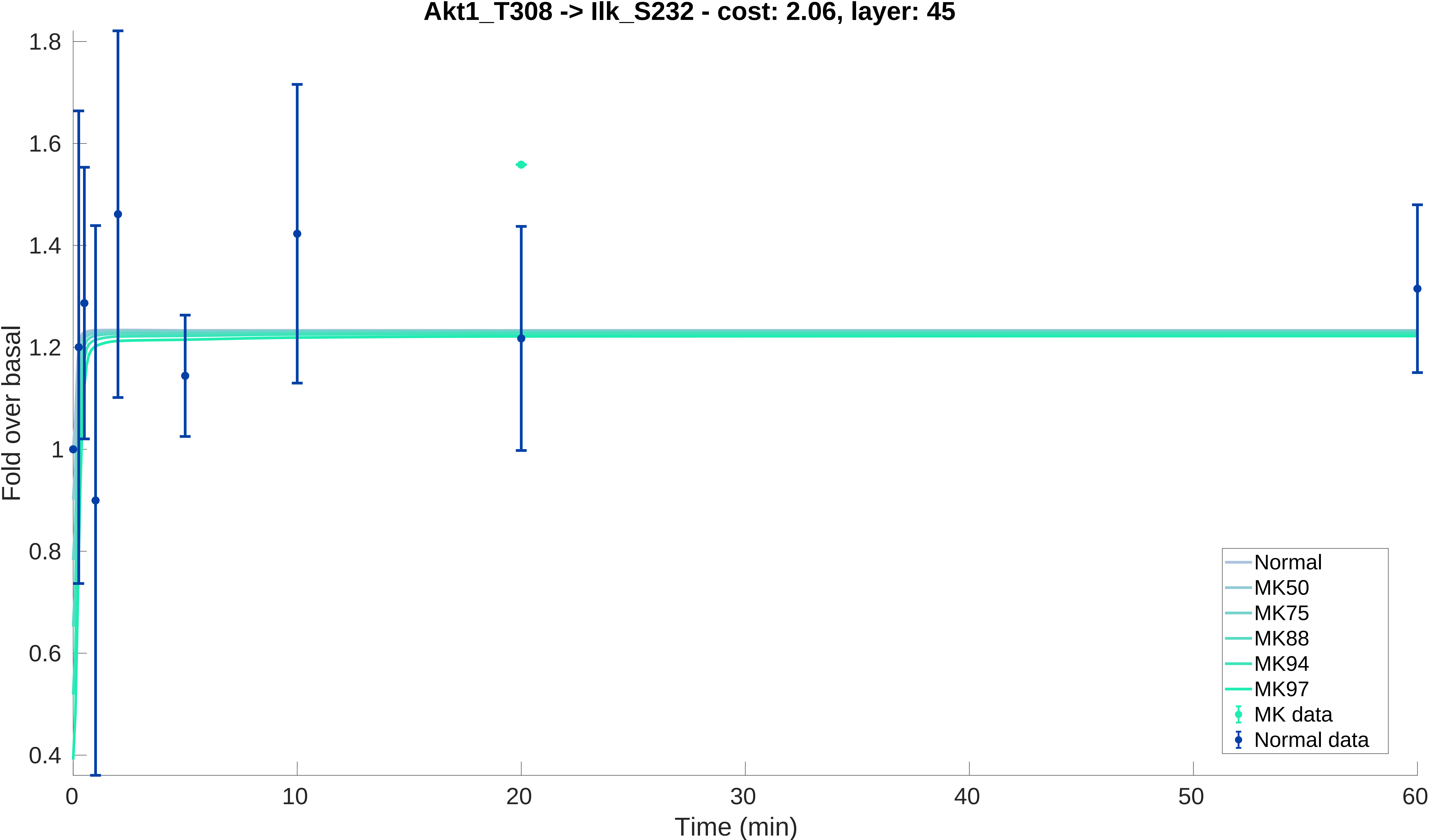

Akt1_T308 -> Ilk_S232 - cost: 2.06, layer: 45
Fold over basal
Time (min)
0.4
0.6
0.8
1
1.2
1.4
1.6
1.8
0
10
20
30
40
50
60
Normal
MK50
MK75
MK88
MK94
MK97
MK data
Normal data

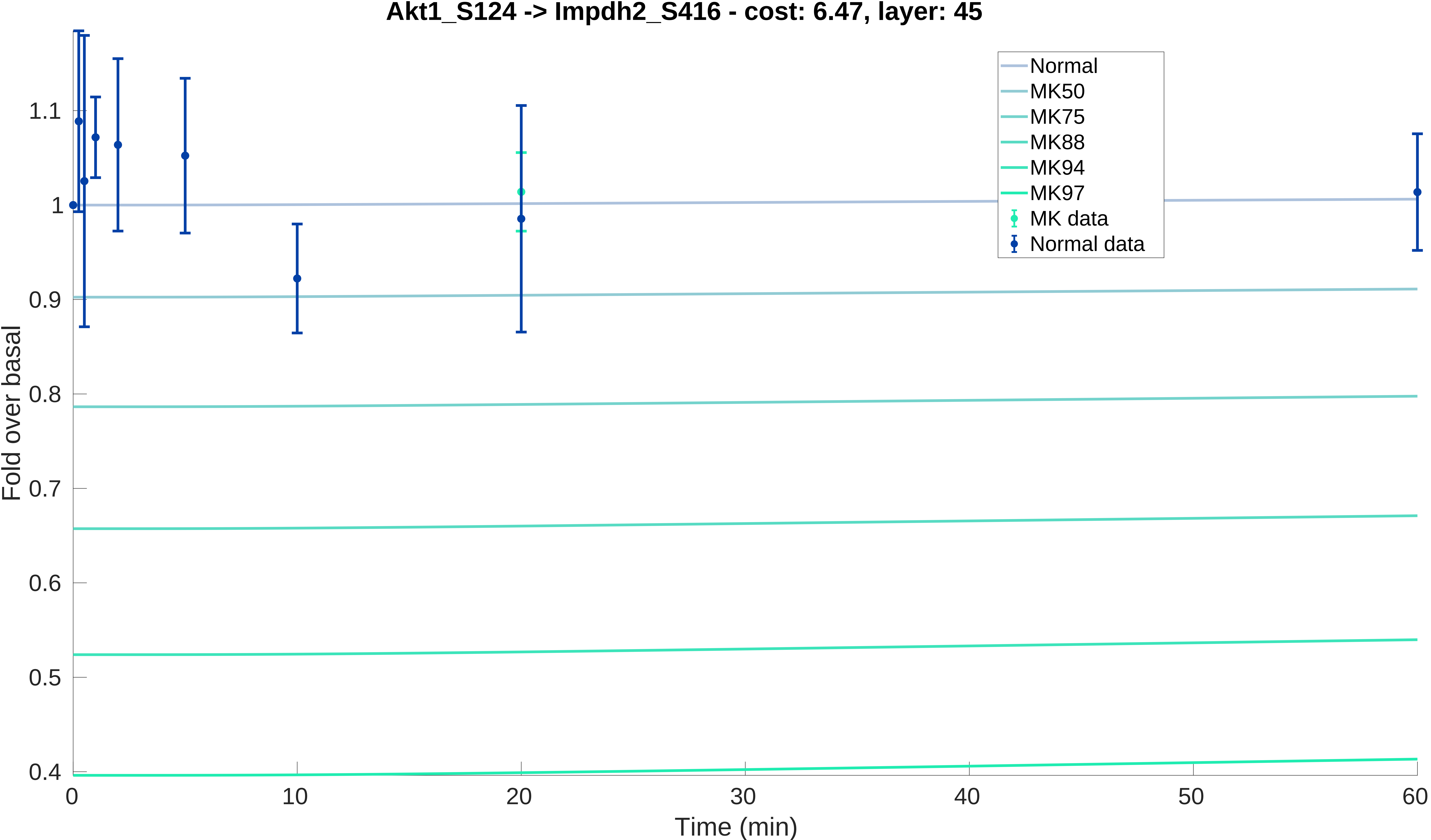

Akt1_S124 -> Impdh2_S416 - cost: 6.47, layer: 45
Normal
MK50
MK75
MK88
MK94
MK97
MK data
Normal data
Fold over basal
Time (min)
1.1
1
0.9
0.8
0.7
0.6
0.5
0.4
0
10
20
30
40
50
60

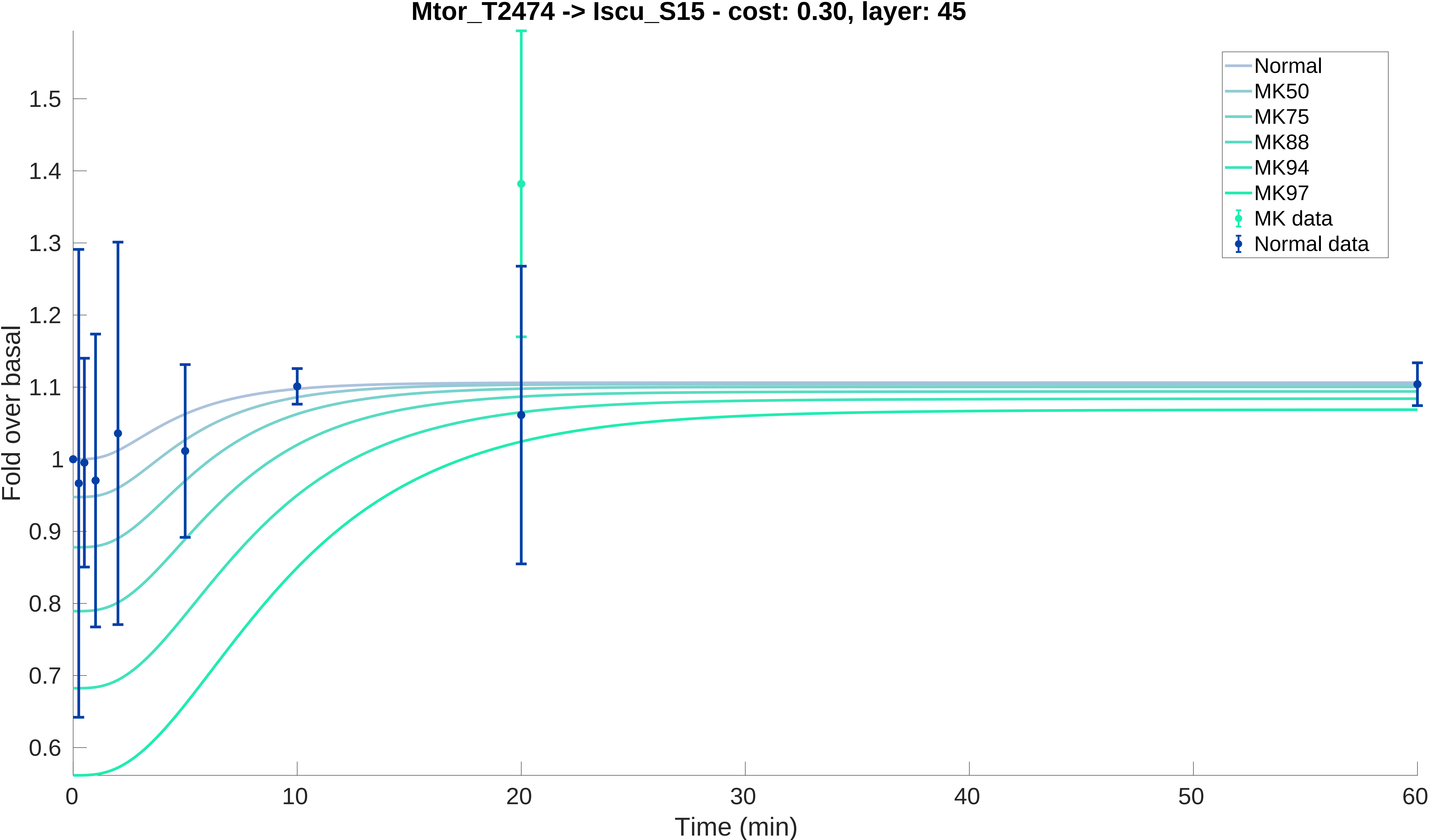

Mtor_T2474 -> Iscu_S15 - cost: 0.30, layer: 45
Normal
MK50
MK75
MK88
MK94
MK97
MK data
Normal data
Fold over basal
1.5
1.4
1.3
1.2
1.1
1
0.9
0.8
0.7
0.6
0
10
20
30
40
50
60
Time (min)

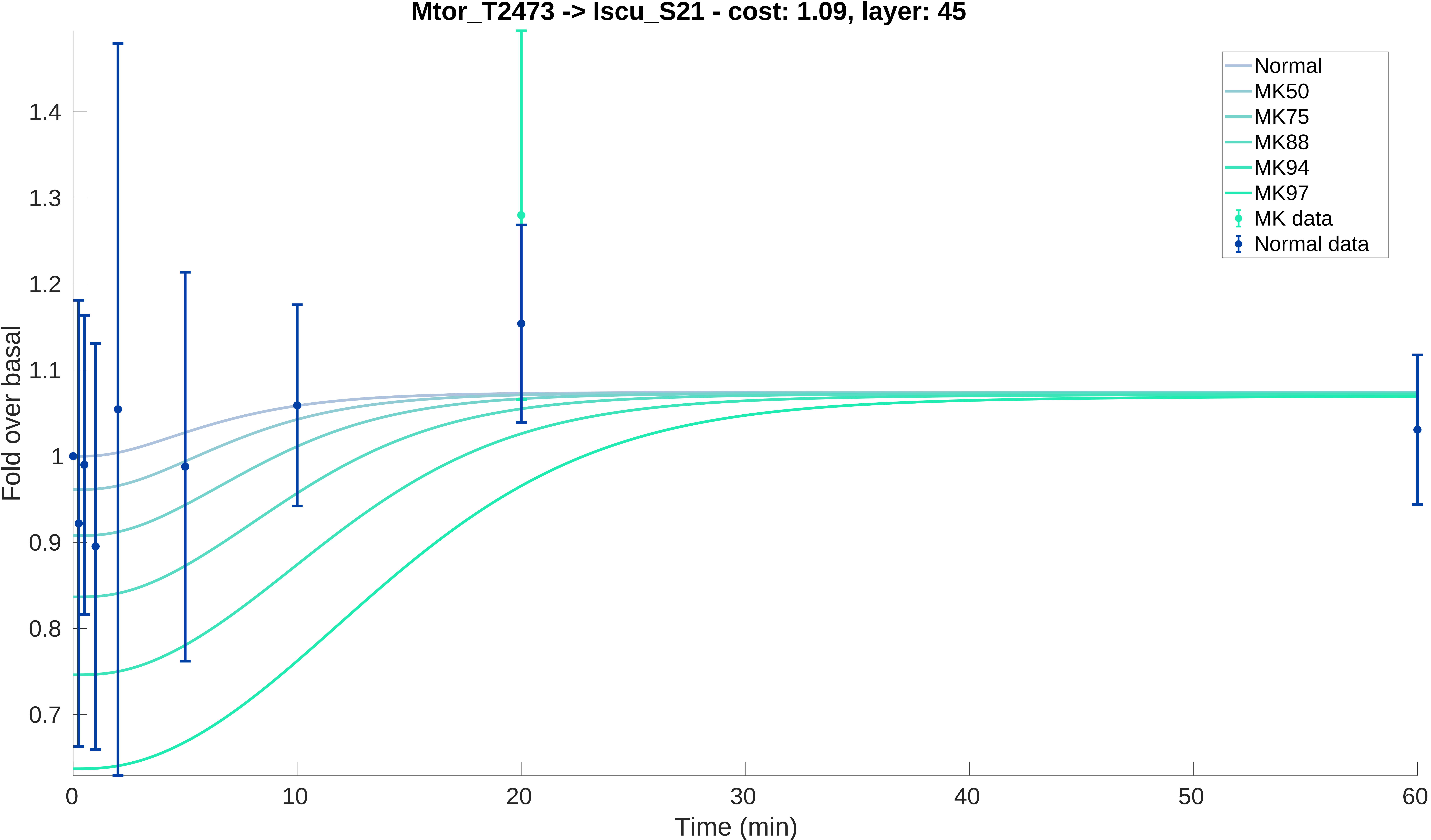

Mtor_T2473 -> Iscu_S21 - cost: 1.09, layer: 45
Fold over basal
Time (min)
1.4
1.3
1.2
1.1
1
0.9
0.8
0.7
0
10
20
30
40
50
60
Normal
MK50
MK75
MK88
MK94
MK97
MK data
Normal data

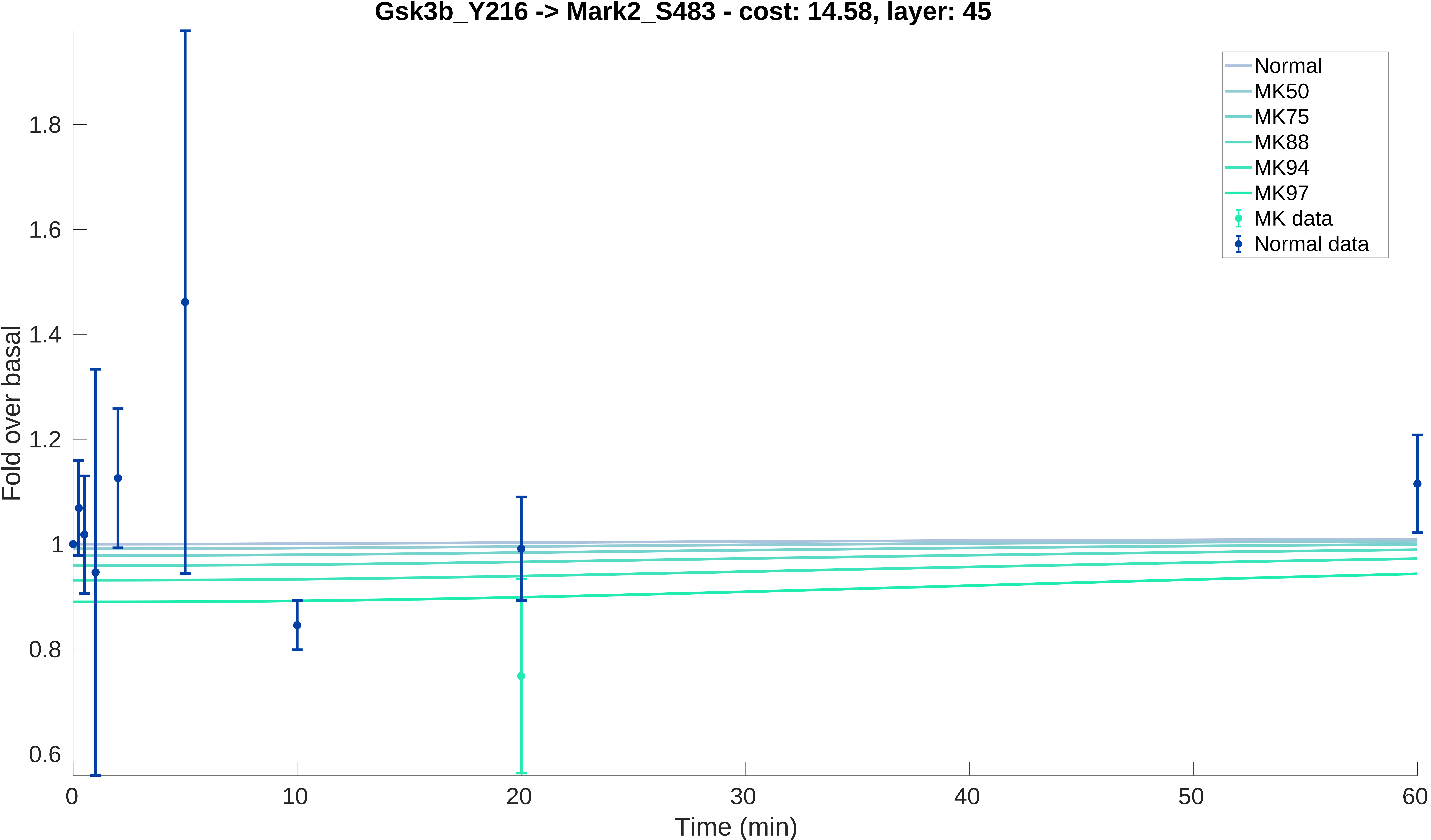
Gsk3b_Y216 -> Mark2_S483 - cost: 14.58, layer: 45
Fold over basal
Time (min)
Normal
MK50
MK75
MK88
MK94
MK97
MK data
Normal data
0.6
0.8
1
1.2
1.4
1.6
1.8
0
10
20
30
40
50
60

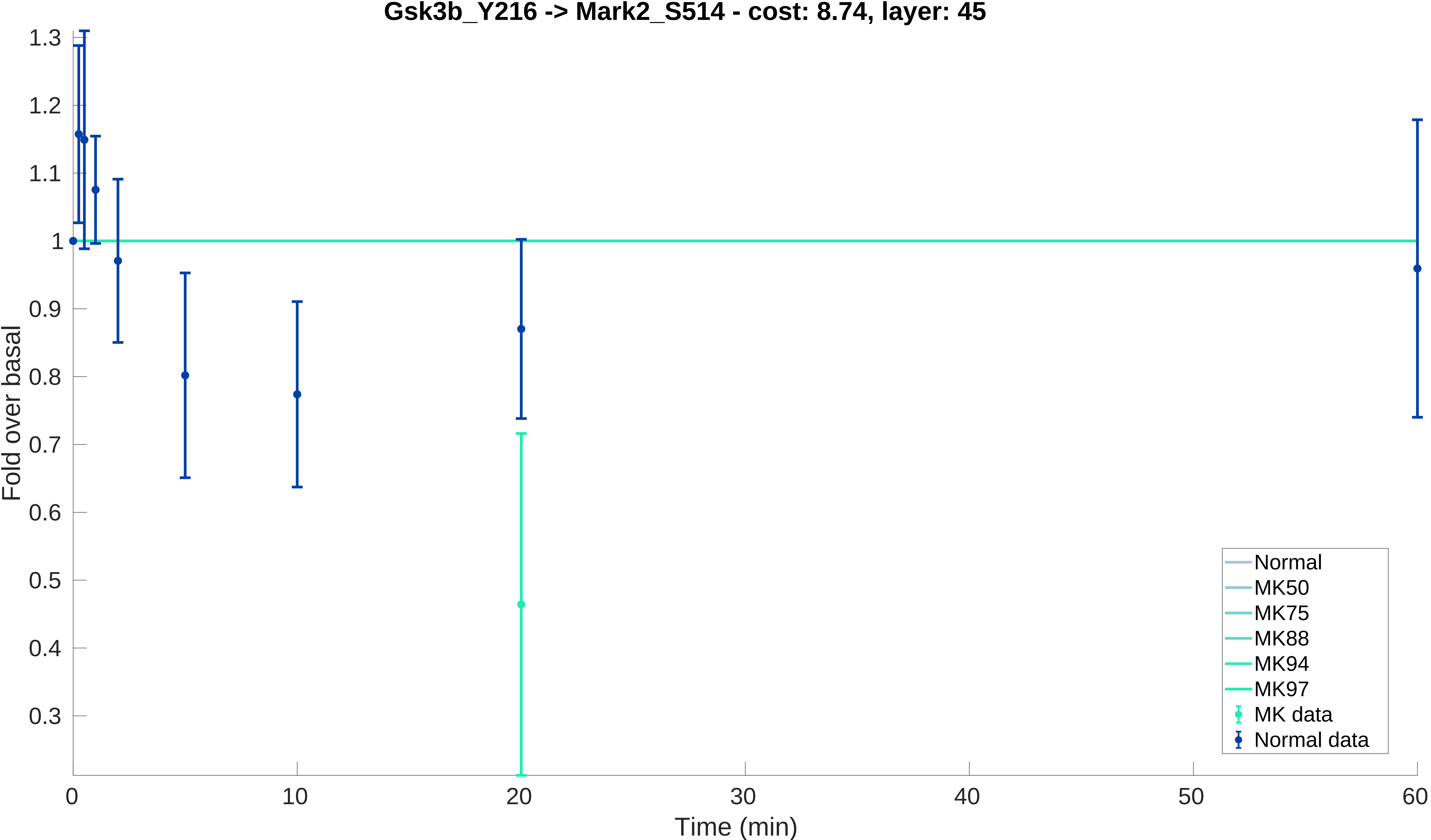

Gsk3b_Y216 -> Mark2_S514 - cost: 8.74, layer: 45
Fold over basal
Time (min)
1.3
1.2
1.1
1
0.9
0.8
0.7
0.6
0.5
0.4
0.3
0
10
20
30
40
50
60
Normal
MK50
MK75
MK88
MK94
MK97
MK data
Normal data

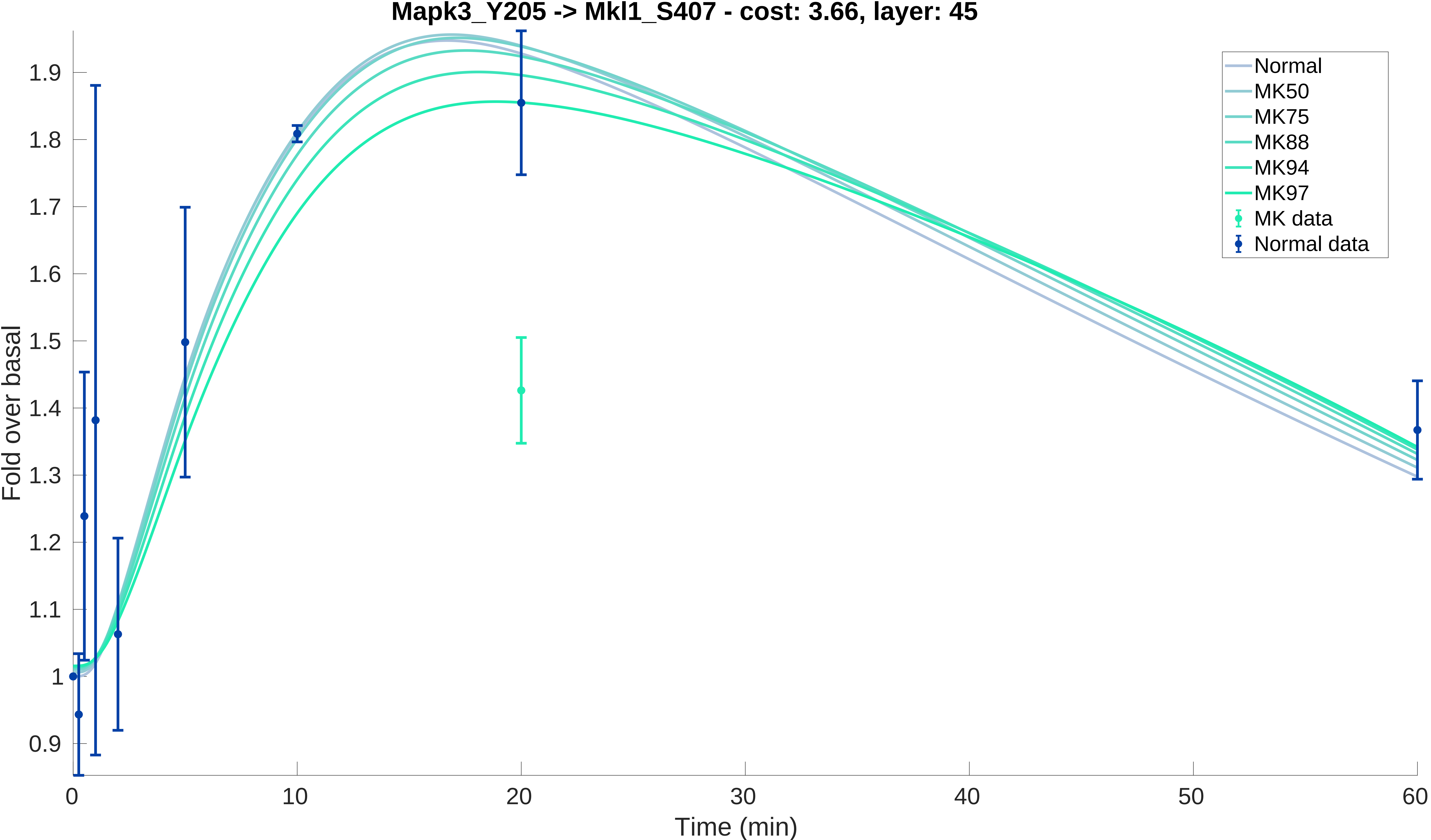
Mapk3_Y205 -> Mkl1_S407 - cost: 3.66, layer: 45
Fold over basal
0.9
1
1.1
1.2
1.3
1.4
1.5
1.6
1.7
1.8
1.9
0
10
20
30
40
50
60
Time (min)
Normal
MK50
MK75
MK88
MK94
MK97
MK data
Normal data

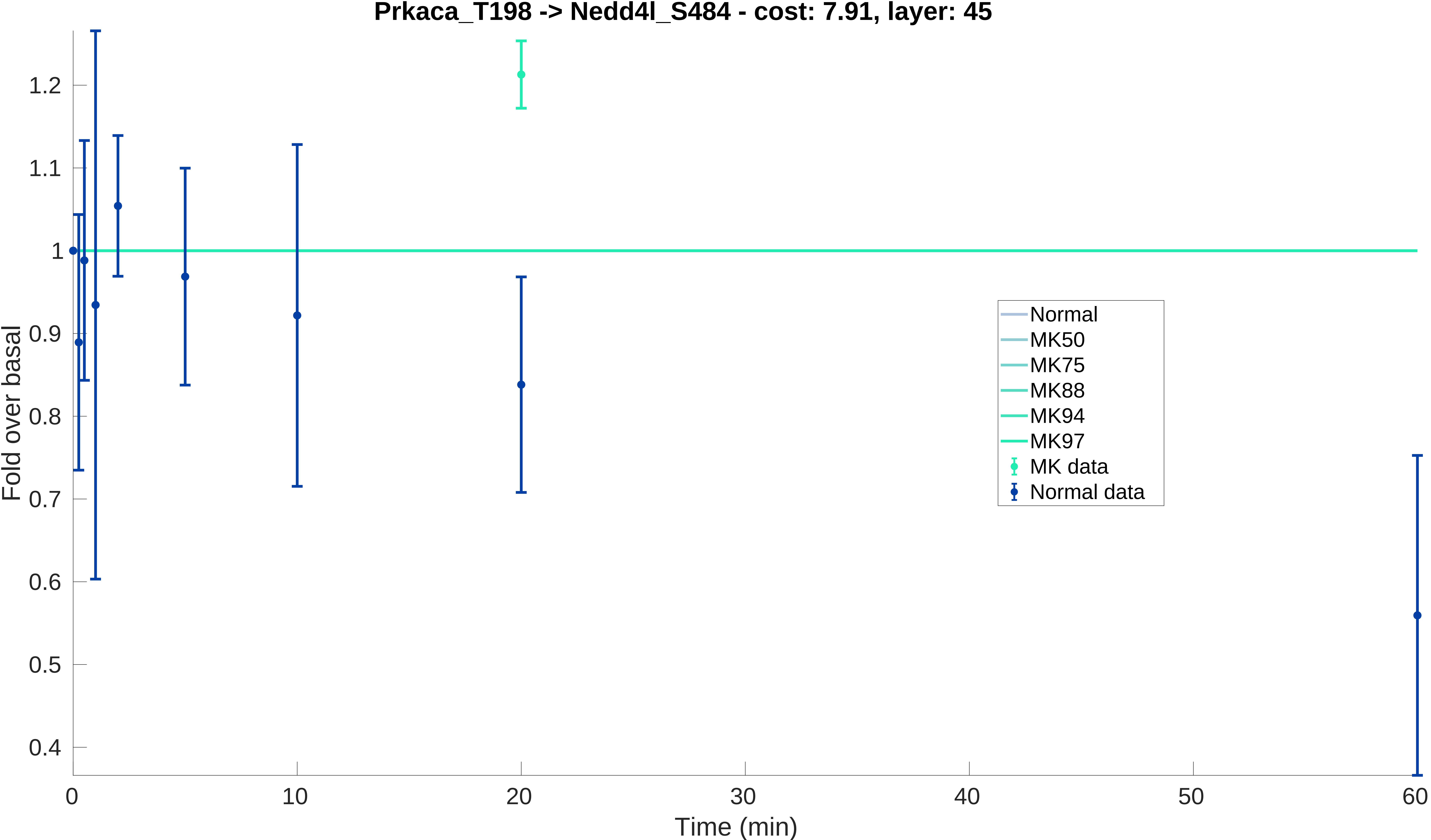

Prkaca_T198 -> Nedd4l_S484 - cost: 7.91, layer: 45
Fold over basal
1.2
1.1
1
0.9
0.8
0.7
0.6
0.5
0.4
0
10
20
30
40
50
60
Time (min)
Normal
MK50
MK75
MK88
MK94
MK97
MK data
Normal data

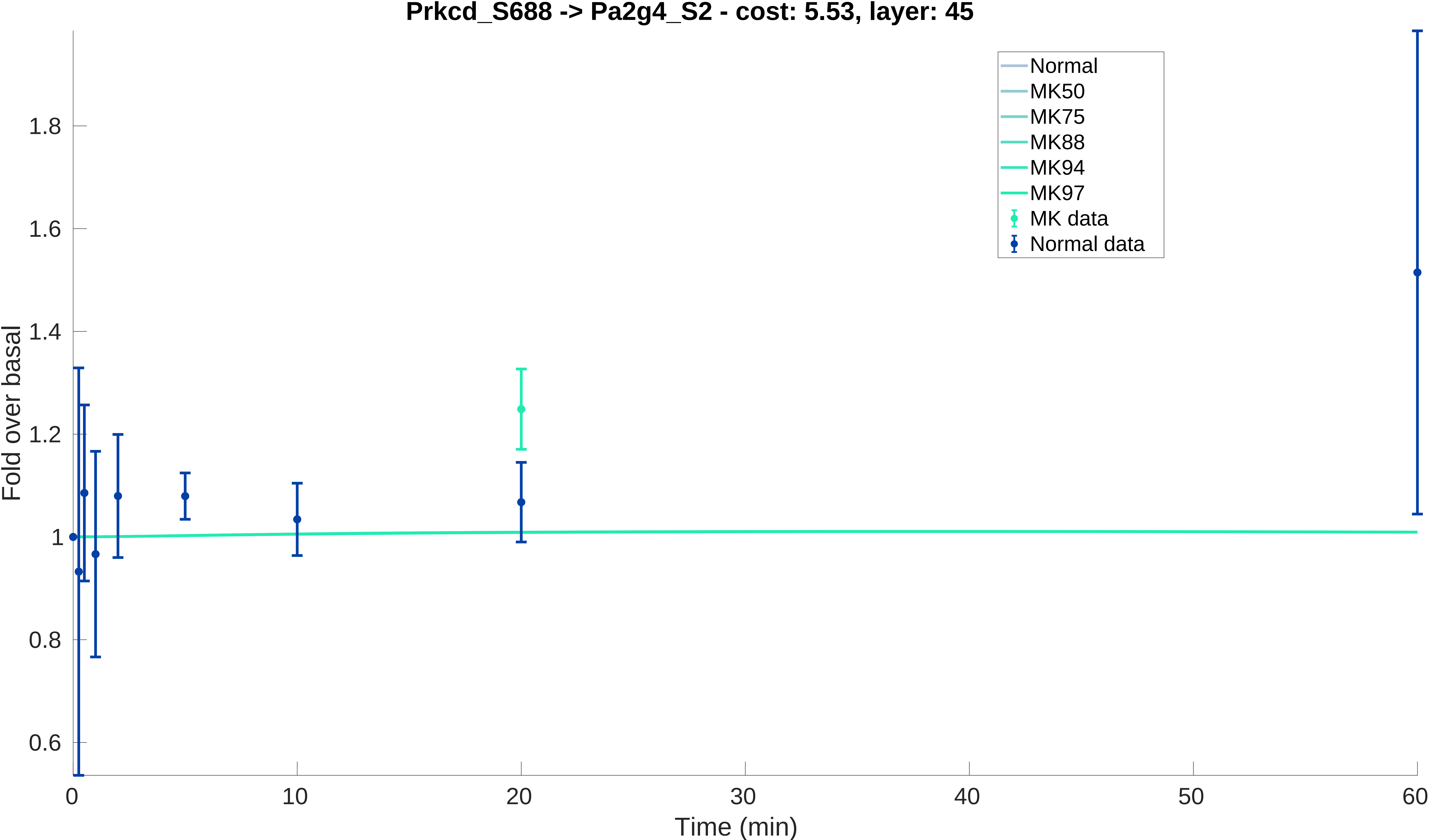

Prkcd_S688 -> Pa2g4_S2 - cost: 5.53, layer: 45
Fold over basal
Time (min)
Normal
MK50
MK75
MK88
MK94
MK97
MK data
Normal data
0.6
0.8
1
1.2
1.4
1.6
1.8
0
10
20
30
40
50
60

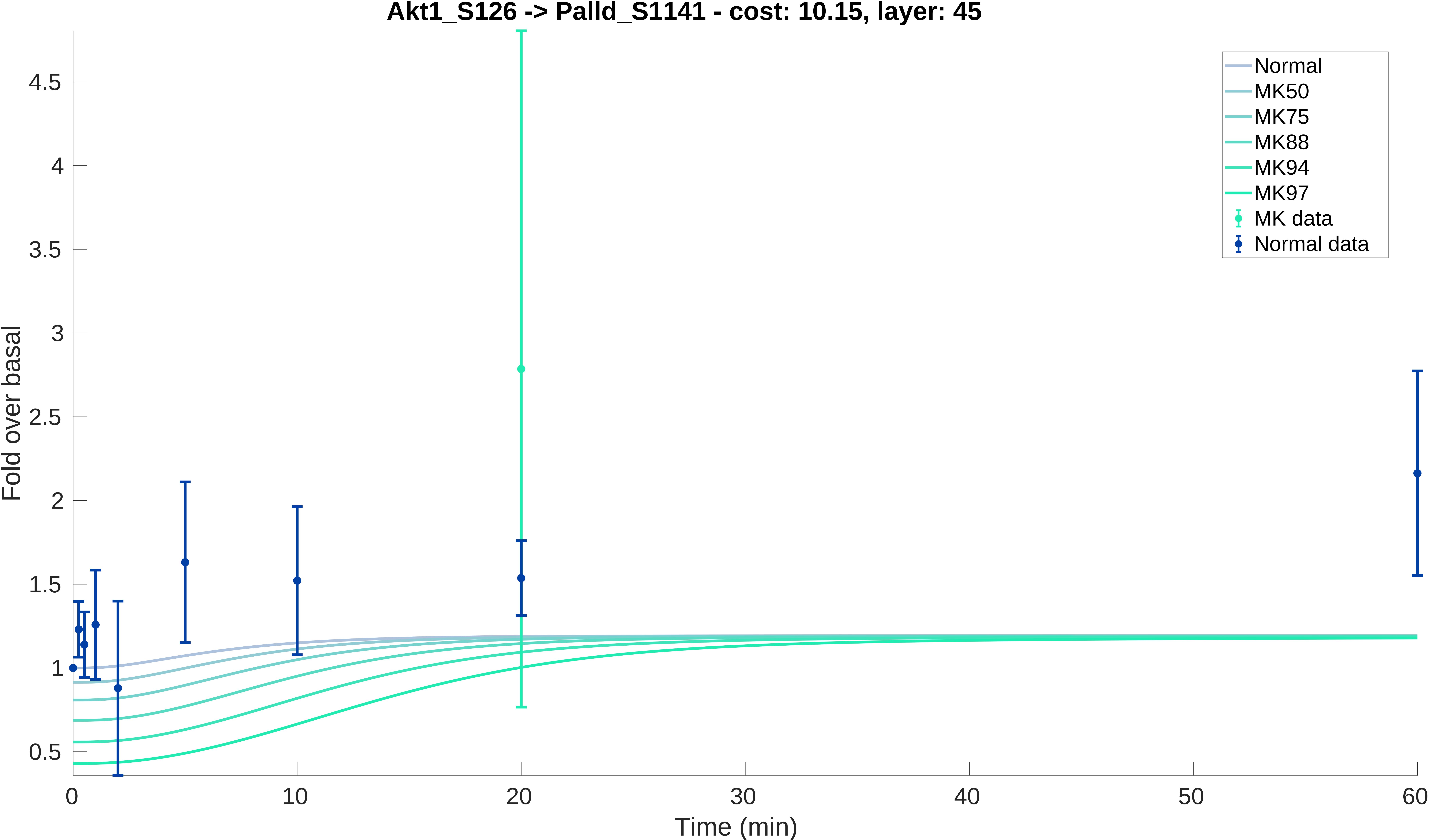
Akt1_S126 -> Palld_S1141 - cost: 10.15, layer: 45
Fold over basal
Time (min)
0.5
1
1.5
2
2.5
3
3.5
4
4.5
0
10
20
30
40
50
60
Normal
MK50
MK75
MK88
MK94
MK97
MK data
Normal data

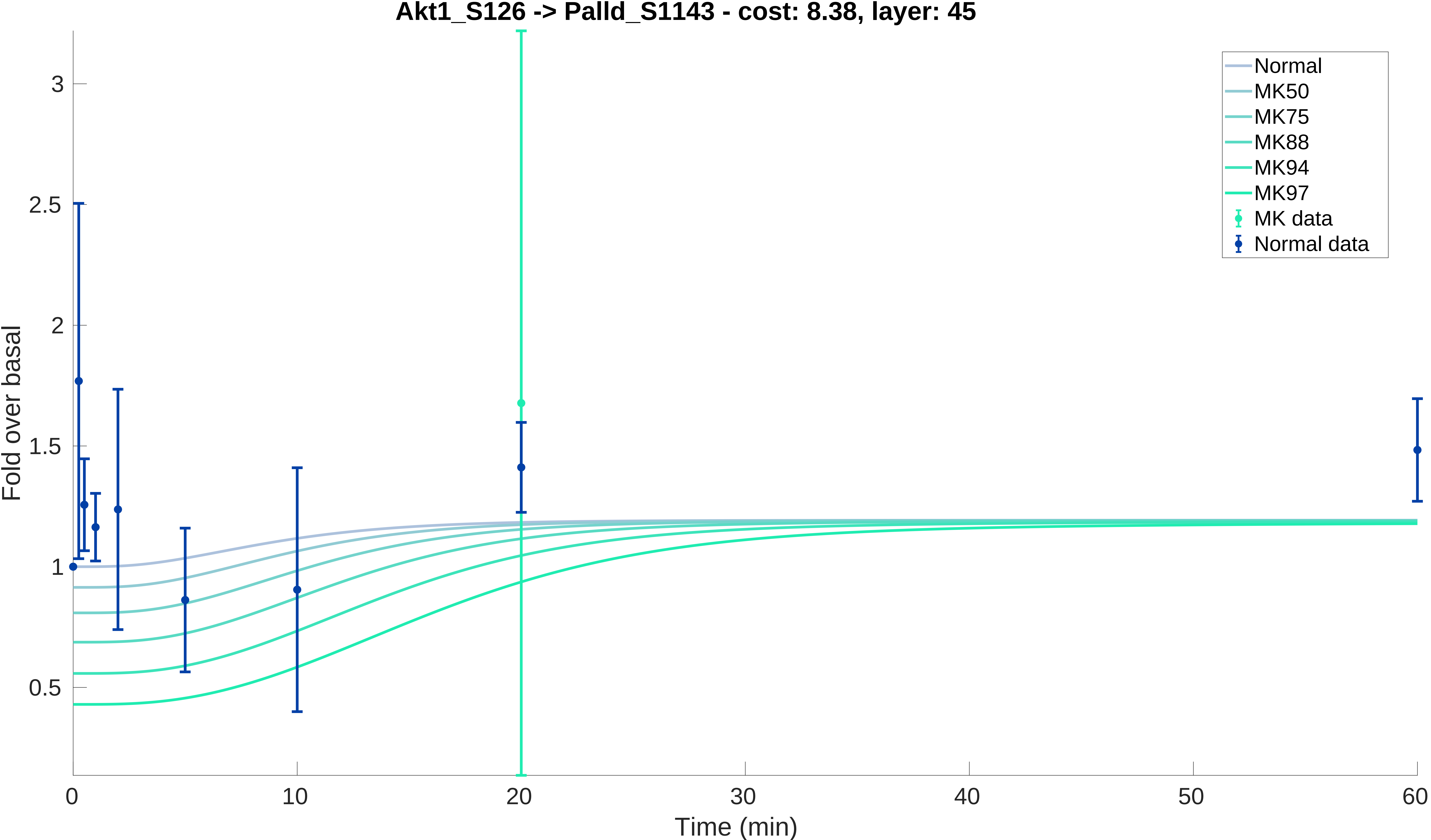

Akt1_S126 -> Palld_S1143 - cost: 8.38, layer: 45
Fold over basal
Time (min)
0.5
1
1.5
2
2.5
3
0
10
20
30
40
50
60
Normal
MK50
MK75
MK88
MK94
MK97
MK data
Normal data

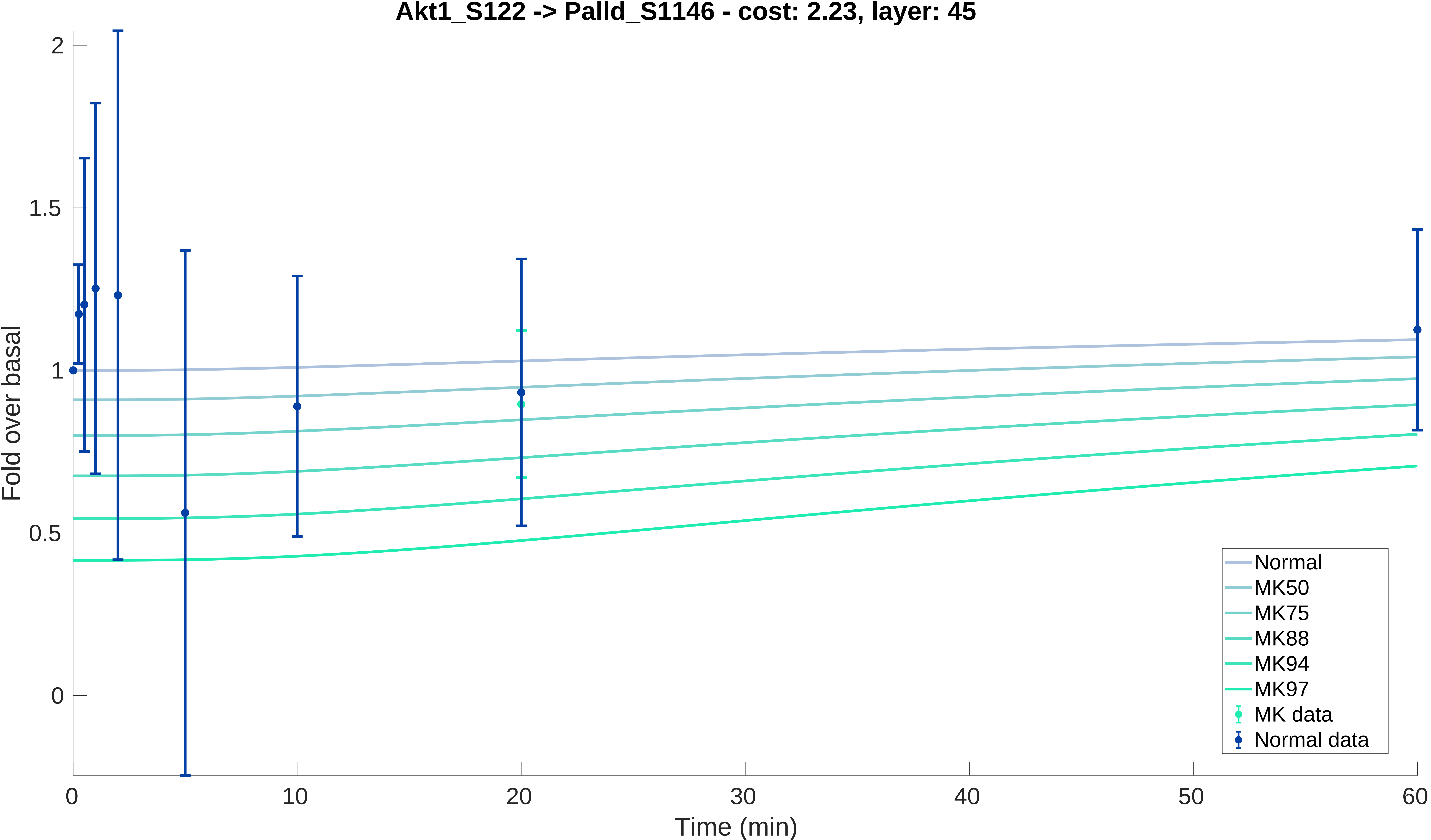
Akt1_S122 -> Palld_S1146 - cost: 2.23, layer: 45
Fold over basal
Time (min)
0
0.5
1
1.5
2
0
10
20
30
40
50
60
Normal
MK50
MK75
MK88
MK94
MK97
MK data
Normal data

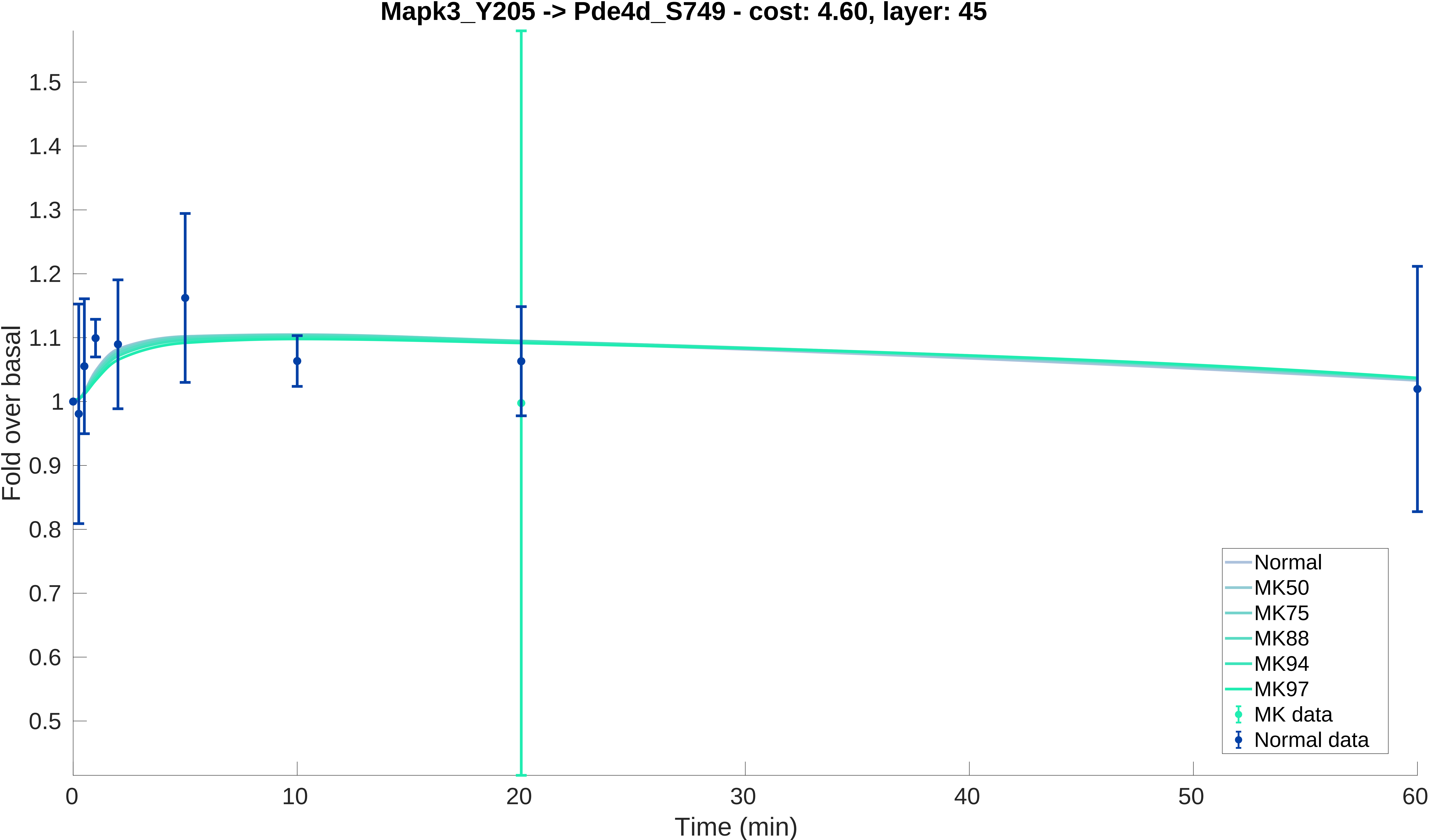

Mapk3_Y205 -> Pde4d_S749 - cost: 4.60, layer: 45
Fold over basal
1.5
1.4
1.3
1.2
1.1
1
0.9
0.8
0.7
0.6
0.5
0
10
20
30
40
50
60
Time (min)
Normal
MK50
MK75
MK88
MK94
MK97
MK data
Normal data

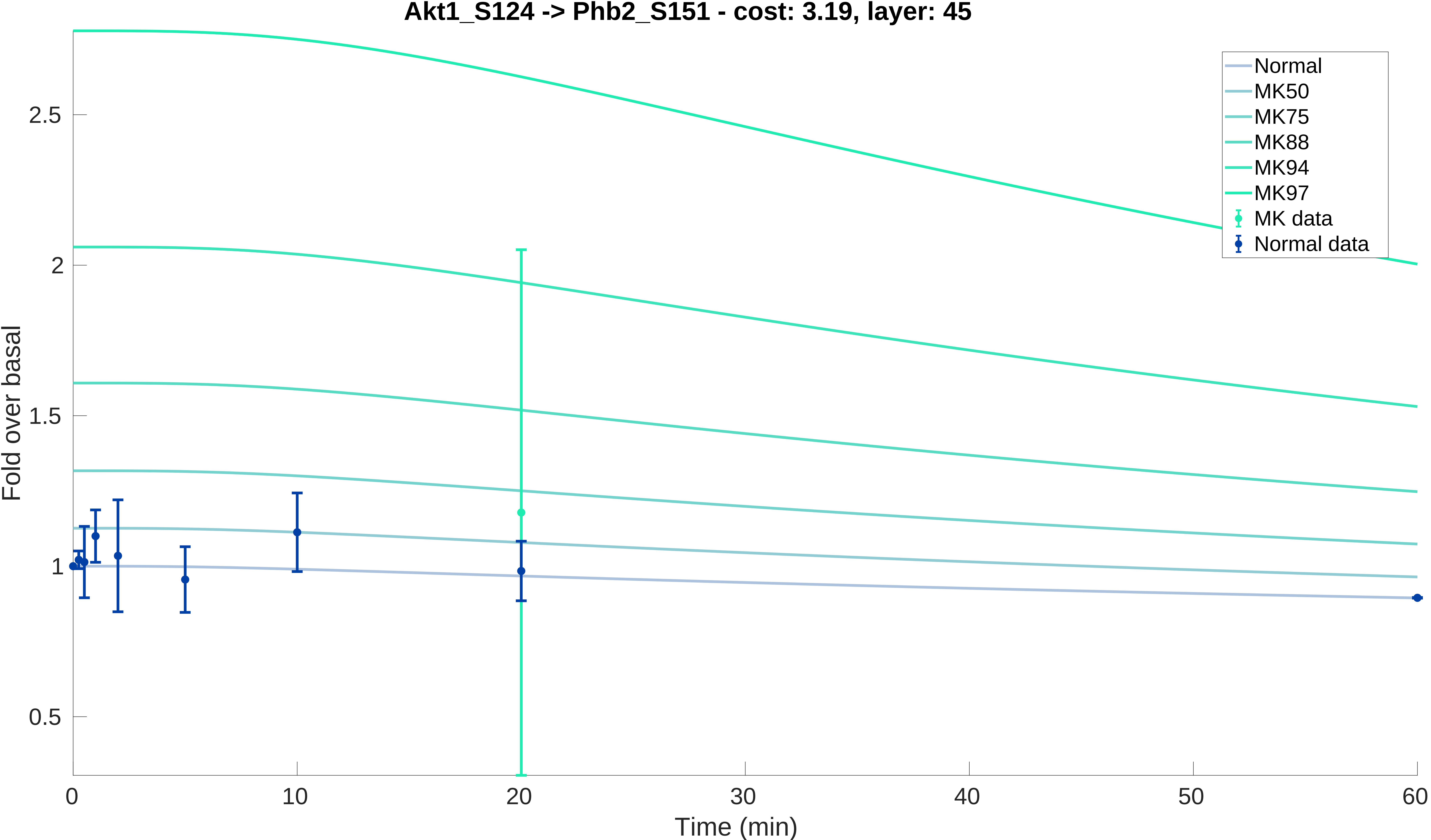

Akt1_S124 -> Phb2_S151 - cost: 3.19, layer: 45
Normal
MK50
MK75
MK88
MK94
MK97
MK data
Normal data
Fold over basal
2.5
2
1.5
1
0.5
0
10
20
30
40
50
60
Time (min)

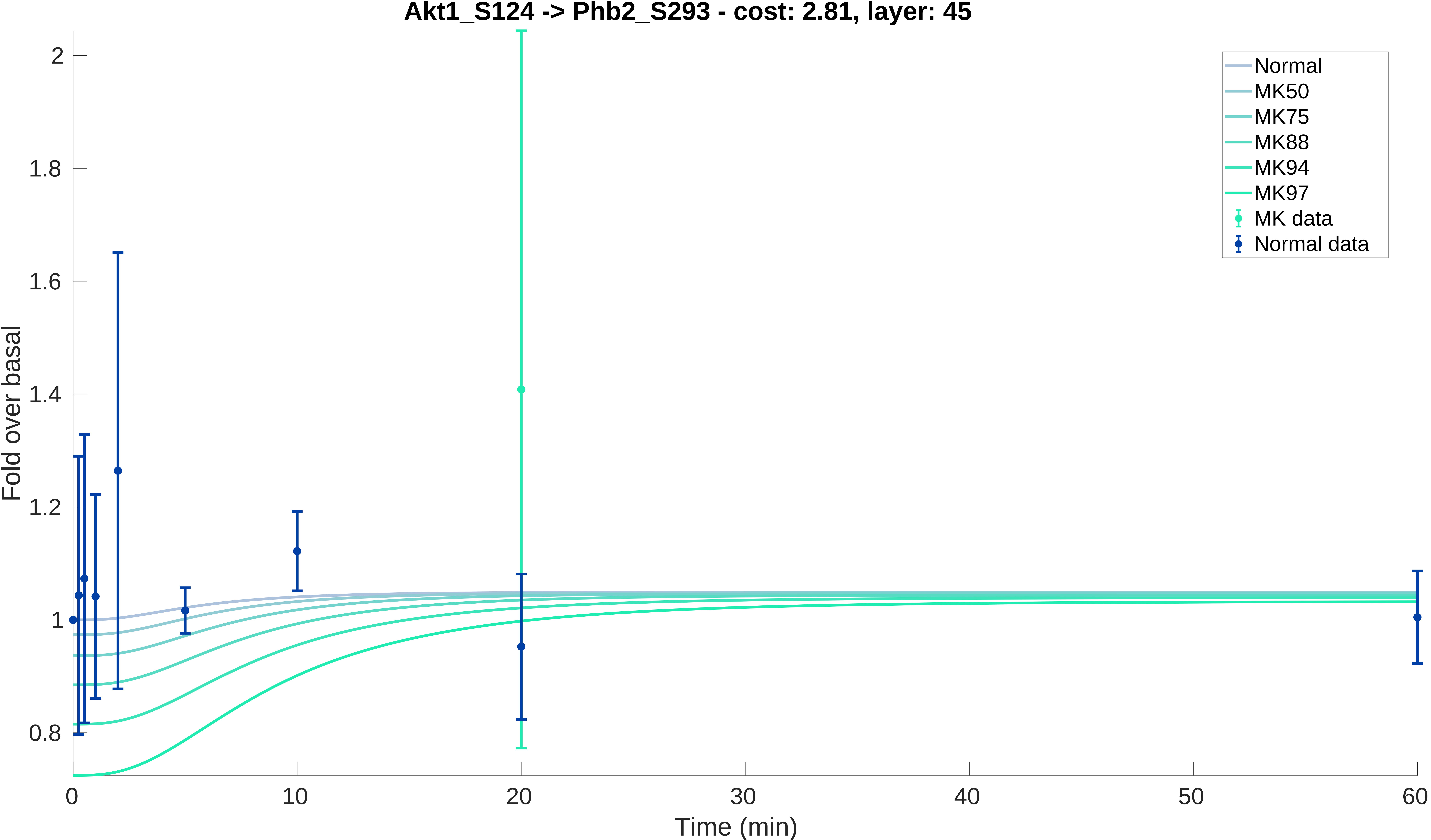

Akt1_S124 -> Phb2_S293 - cost: 2.81, layer: 45
Normal
MK50
MK75
MK88
MK94
MK97
MK data
Normal data
Fold over basal
Time (min)
0.8
1
1.2
1.4
1.6
1.8
2
0
10
20
30
40
50
60

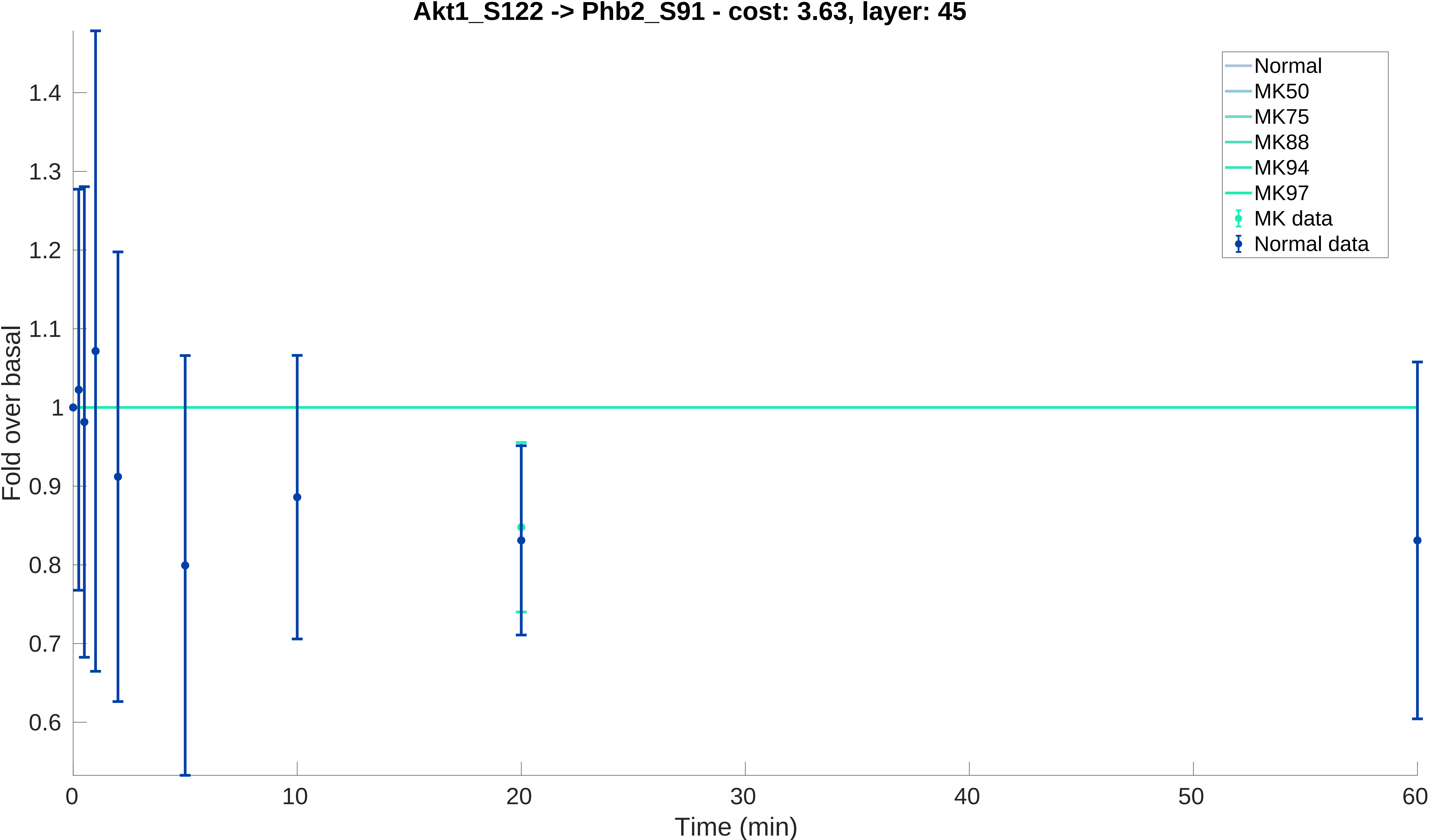
Akt1_S122 -> Phb2_S91 - cost: 3.63, layer: 45
Fold over basal
Time (min)
1.4
1.3
1.2
1.1
1
0.9
0.8
0.7
0.6
0
10
20
30
40
50
60
Normal
MK50
MK75
MK88
MK94
MK97
MK data
Normal data

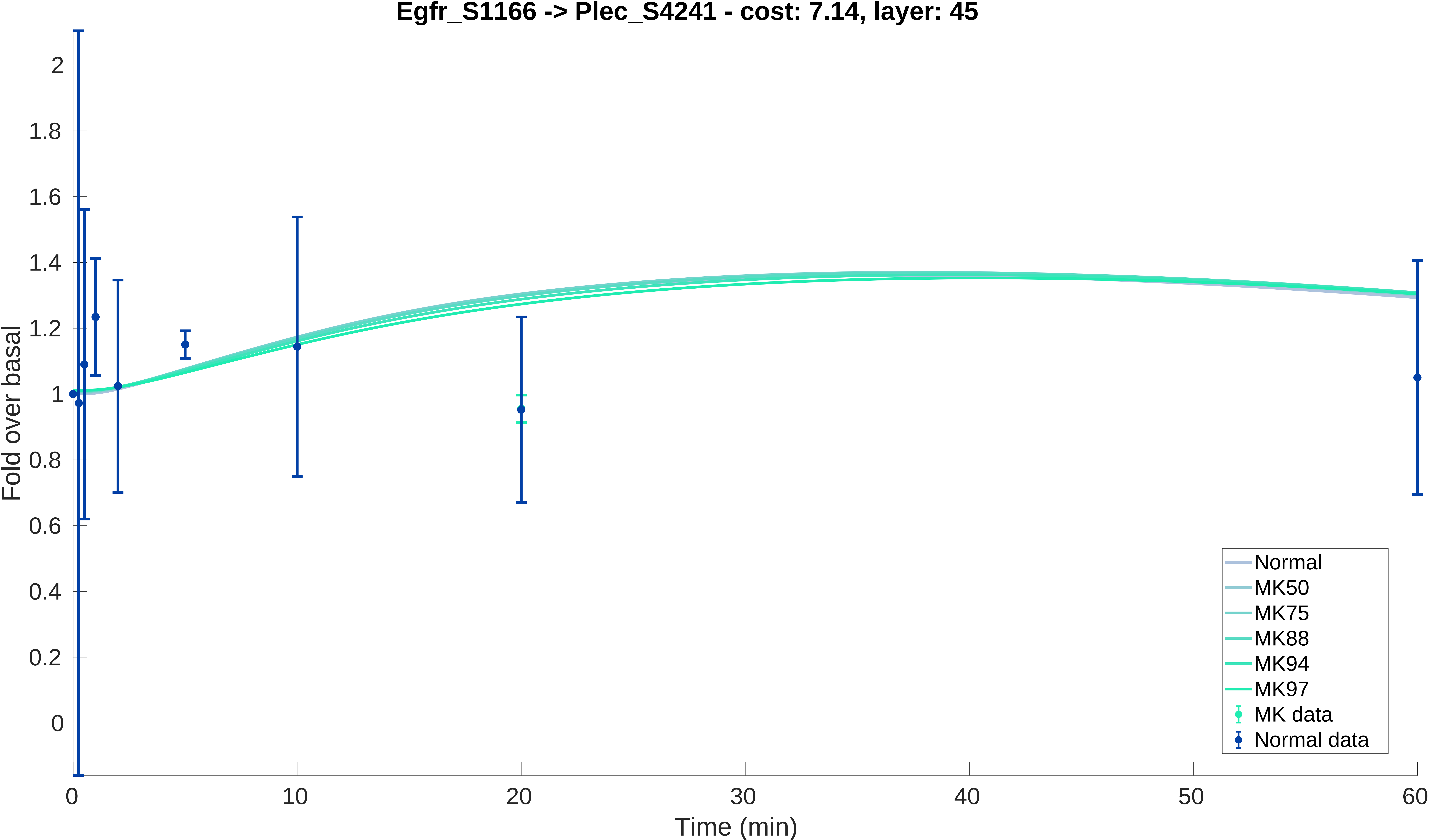

Egfr_S1166 -> Plec_S4241 - cost: 7.14, layer: 45
Fold over basal
Time (min)
2
1.8
1.6
1.4
1.2
1
0.8
0.6
0.4
0.2
0
10
20
30
40
50
60
Normal
MK50
MK75
MK88
MK94
MK97
MK data
Normal data

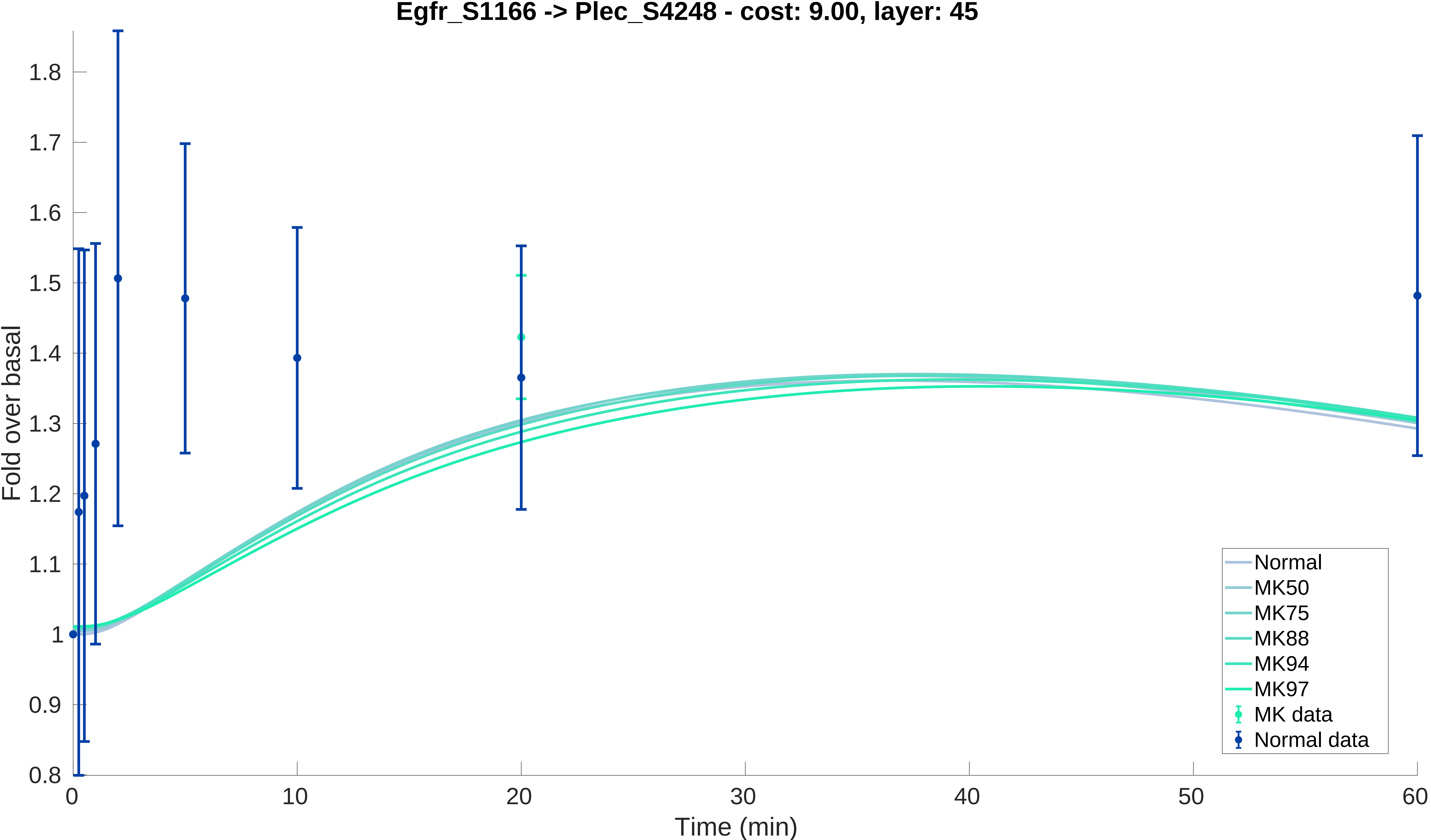

Egfr_S1166 -> Plec_S4248 - cost: 9.00, layer: 45
Fold over basal
1.8
1.7
1.6
1.5
1.4
1.3
1.2
1.1
1
0.9
0.8
0
10
20
30
40
50
60
Time (min)
Normal
MK50
MK75
MK88
MK94
MK97
MK data
Normal data

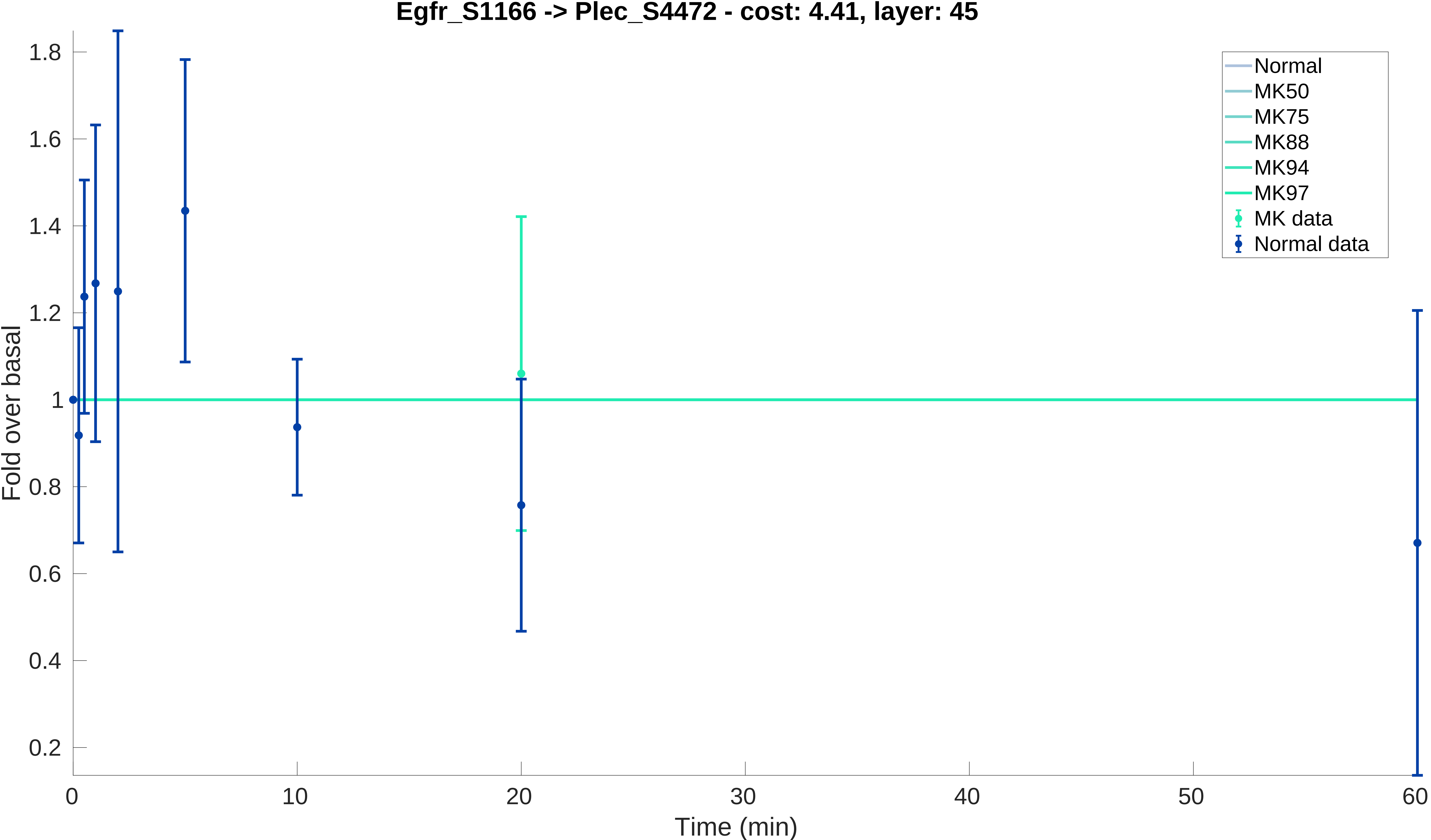

Egfr_S1166 -> Plec_S4472 - cost: 4.41, layer: 45
Fold over basal
Time (min)
1.8
1.6
1.4
1.2
1
0.8
0.6
0.4
0.2
0
10
20
30
40
50
60
Normal
MK50
MK75
MK88
MK94
MK97
MK data
Normal data

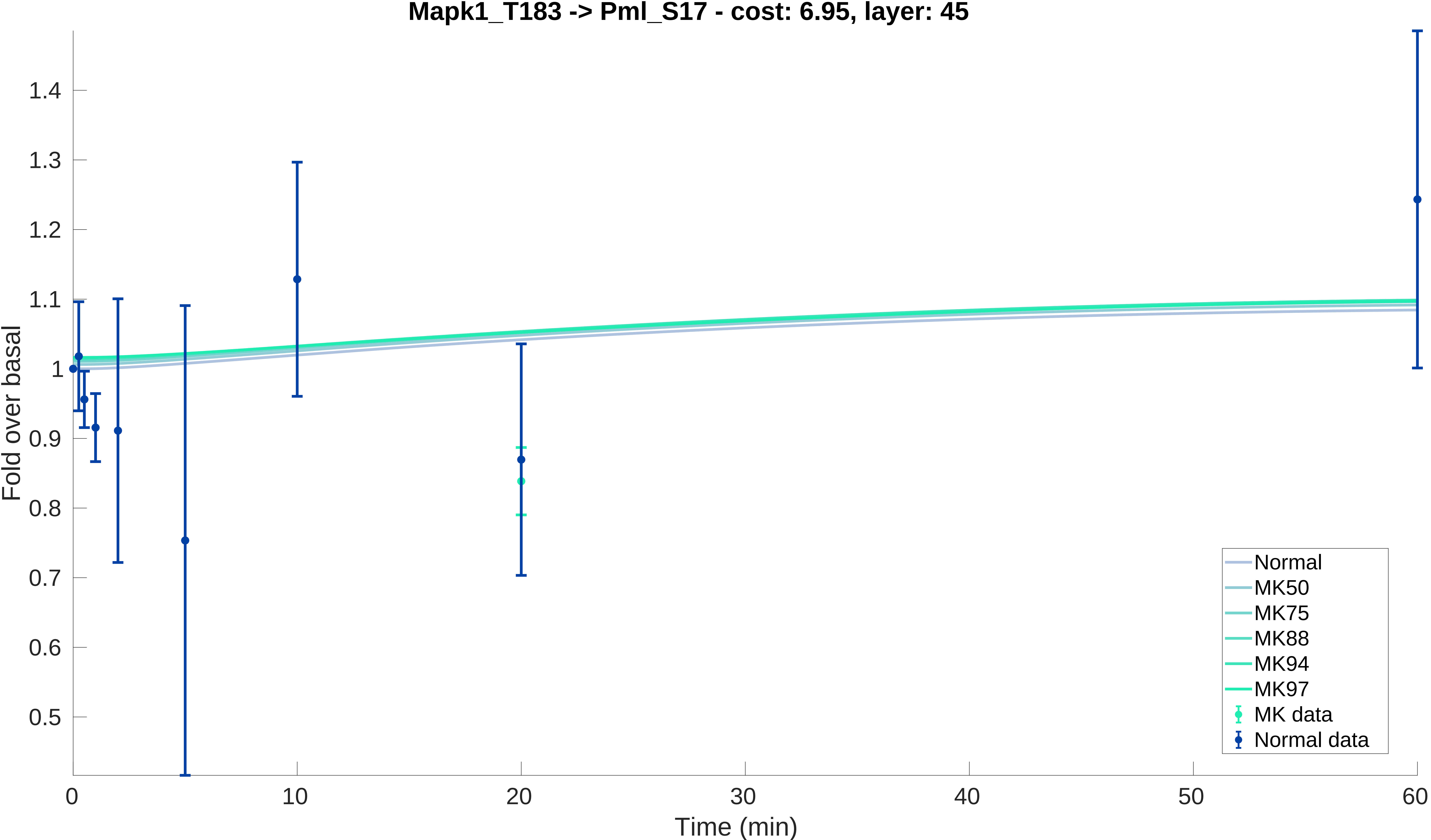

Mapk1_T183 -> Pml_S17 - cost: 6.95, layer: 45
Fold over basal
Time (min)
0.5
0.6
0.7
0.8
0.9
1
1.1
1.2
1.3
1.4
0
10
20
30
40
50
60
Normal
MK50
MK75
MK88
MK94
MK97
MK data
Normal data

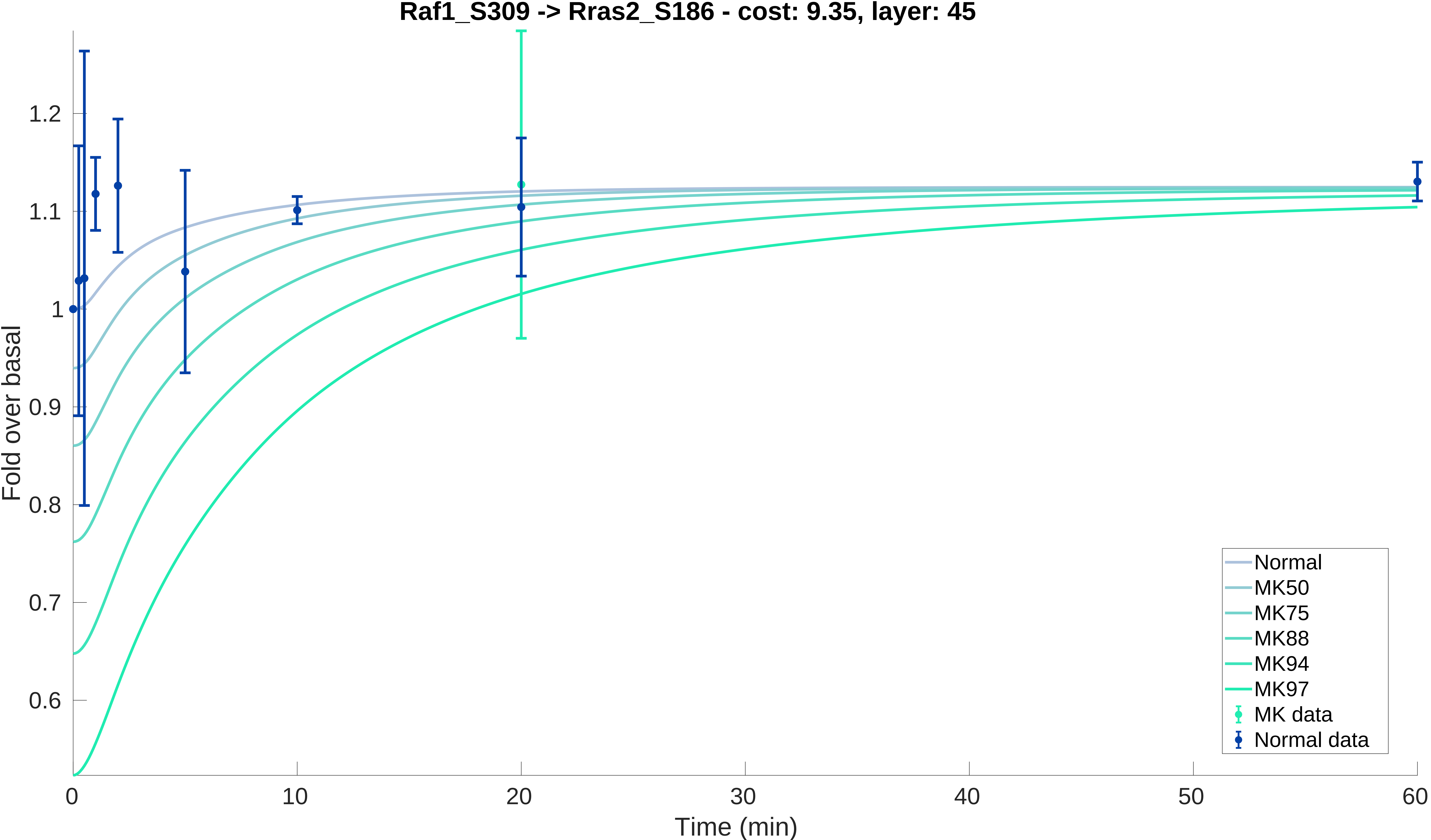

Raf1_S309 -> Rras2_S186 - cost: 9.35, layer: 45
Fold over basal
Time (min)
0.6
0.7
0.8
0.9
1
1.1
1.2
0
10
20
30
40
50
60
Normal
MK50
MK75
MK88
MK94
MK97
MK data
Normal data

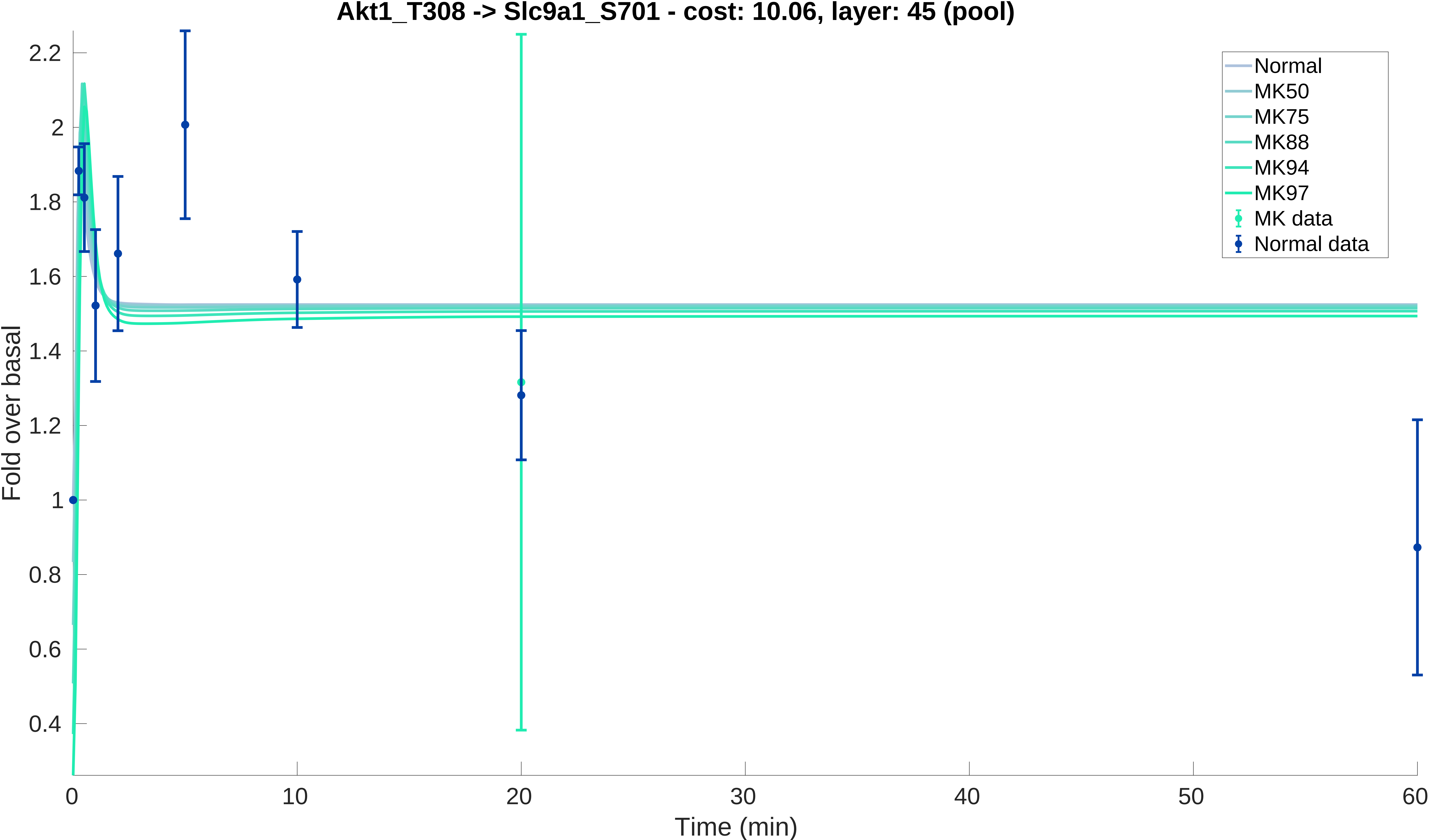

Akt1_T308 -> Slc9a1_S701 - cost: 10.06, layer: 45 (pool)
Fold over basal
Time (min)
0.4
0.6
0.8
1
1.2
1.4
1.6
1.8
2
2.2
0
10
20
30
40
50
60
Normal
MK50
MK75
MK88
MK94
MK97
MK data
Normal data

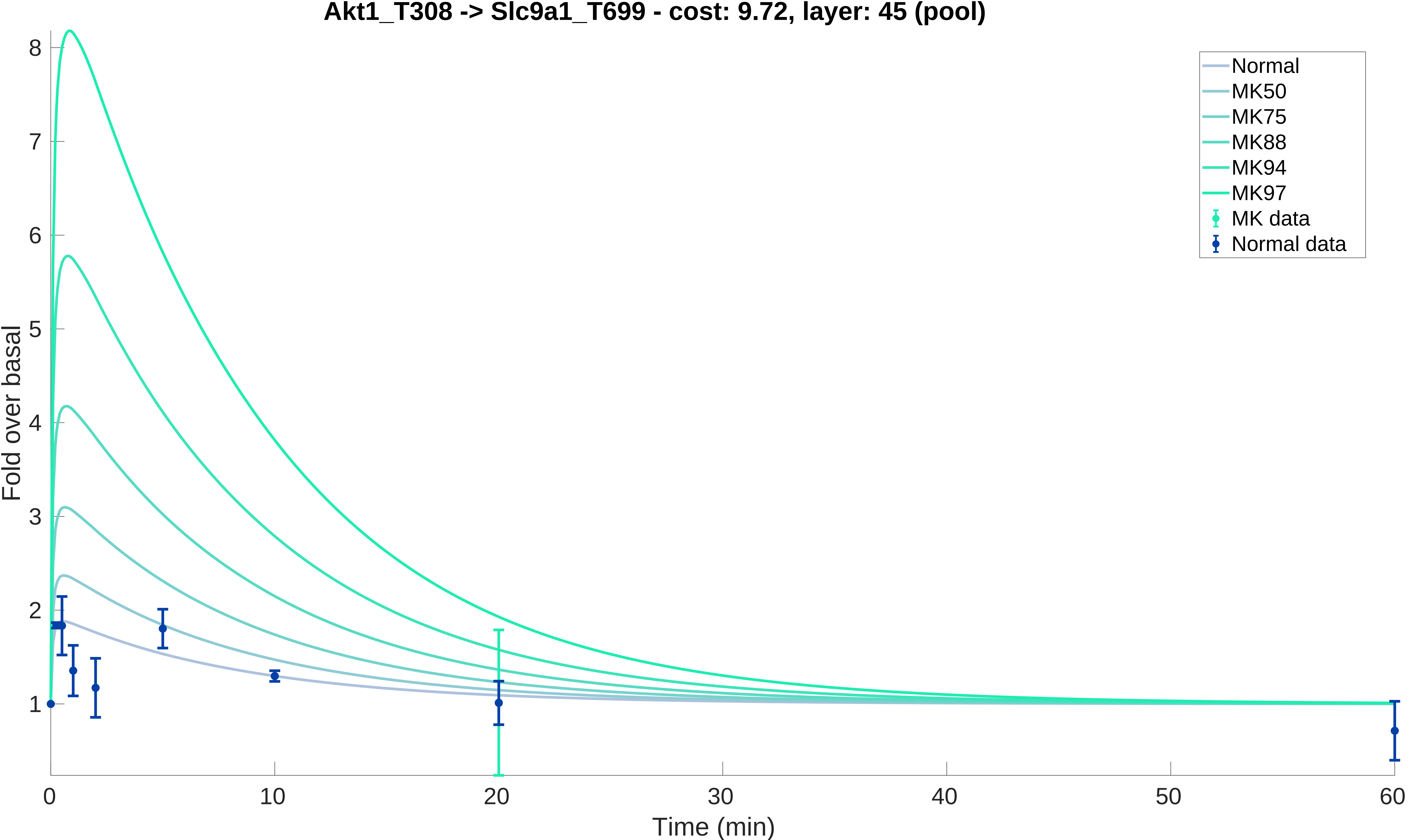
Akt1_T308 -> Slc9a1_T699 - cost: 9.72, layer: 45 (pool)
Fold over basal
Time (min)
Normal
MK50
MK75
MK88
MK94
MK97
MK data
Normal data
0
10
20
30
40
50
60
1
2
3
4
5
6
7
8

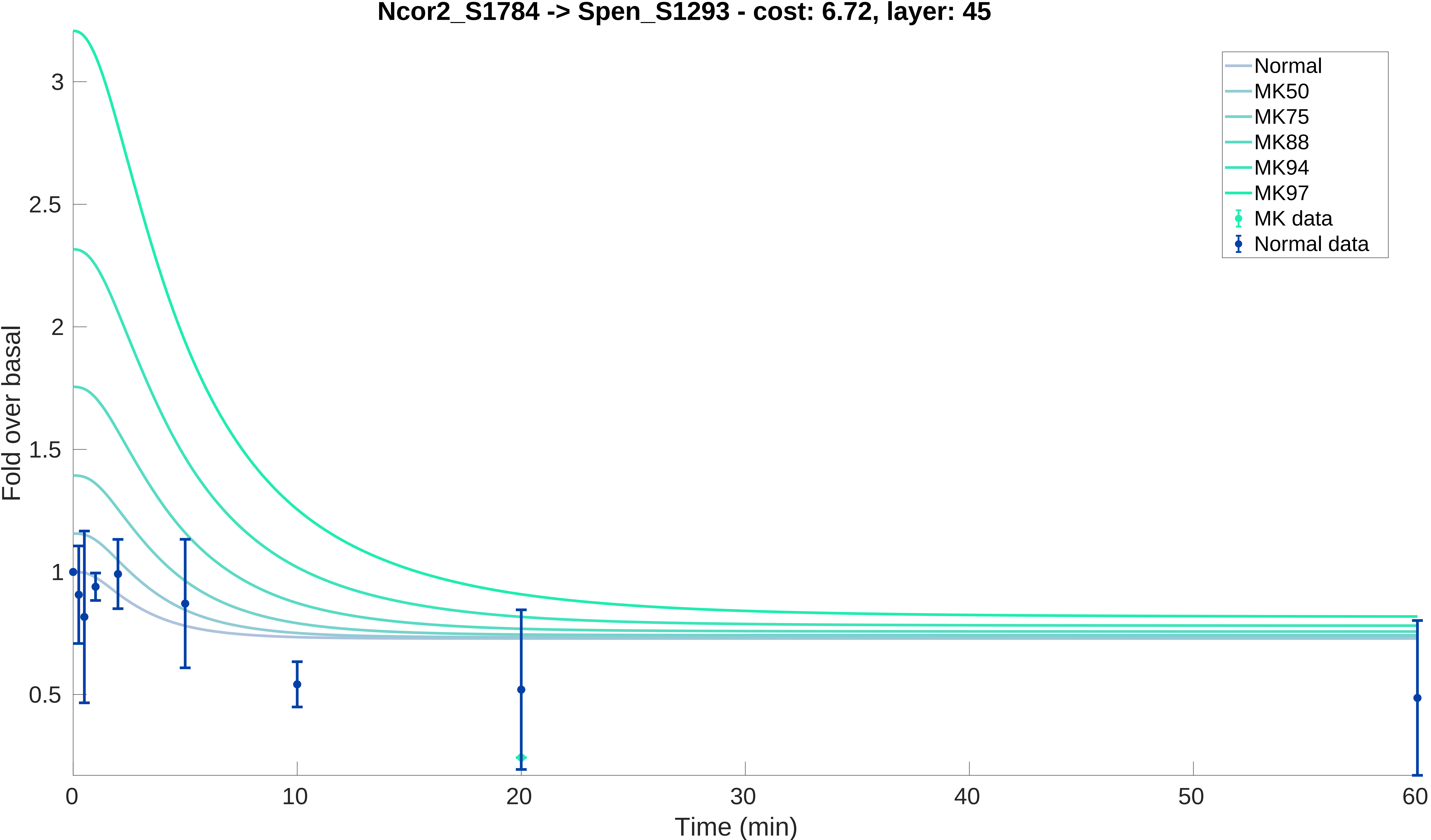

Ncor2_S1784 -> Spen_S1293 - cost: 6.72, layer: 45
Fold over basal
Time (min)
0.5
1
1.5
2
2.5
3
0
10
20
30
40
50
60
Normal
MK50
MK75
MK88
MK94
MK97
MK data
Normal data

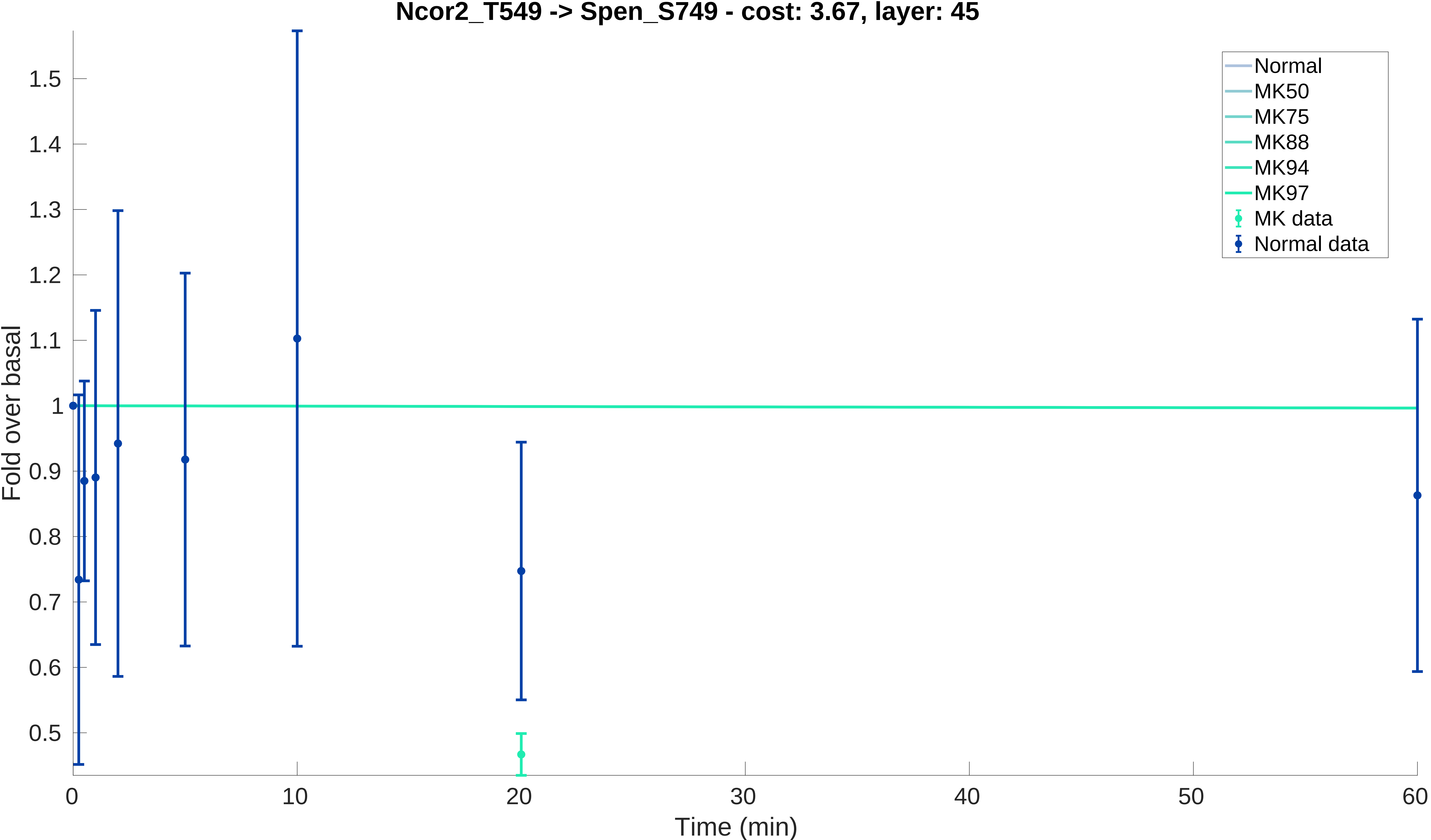

Ncor2_T549 -> Spen_S749 - cost: 3.67, layer: 45
Fold over basal
Time (min)
1.5
1.4
1.3
1.2
1.1
1
0.9
0.8
0.7
0.6
0.5
0
10
20
30
40
50
60
Normal
MK50
MK75
MK88
MK94
MK97
MK data
Normal data

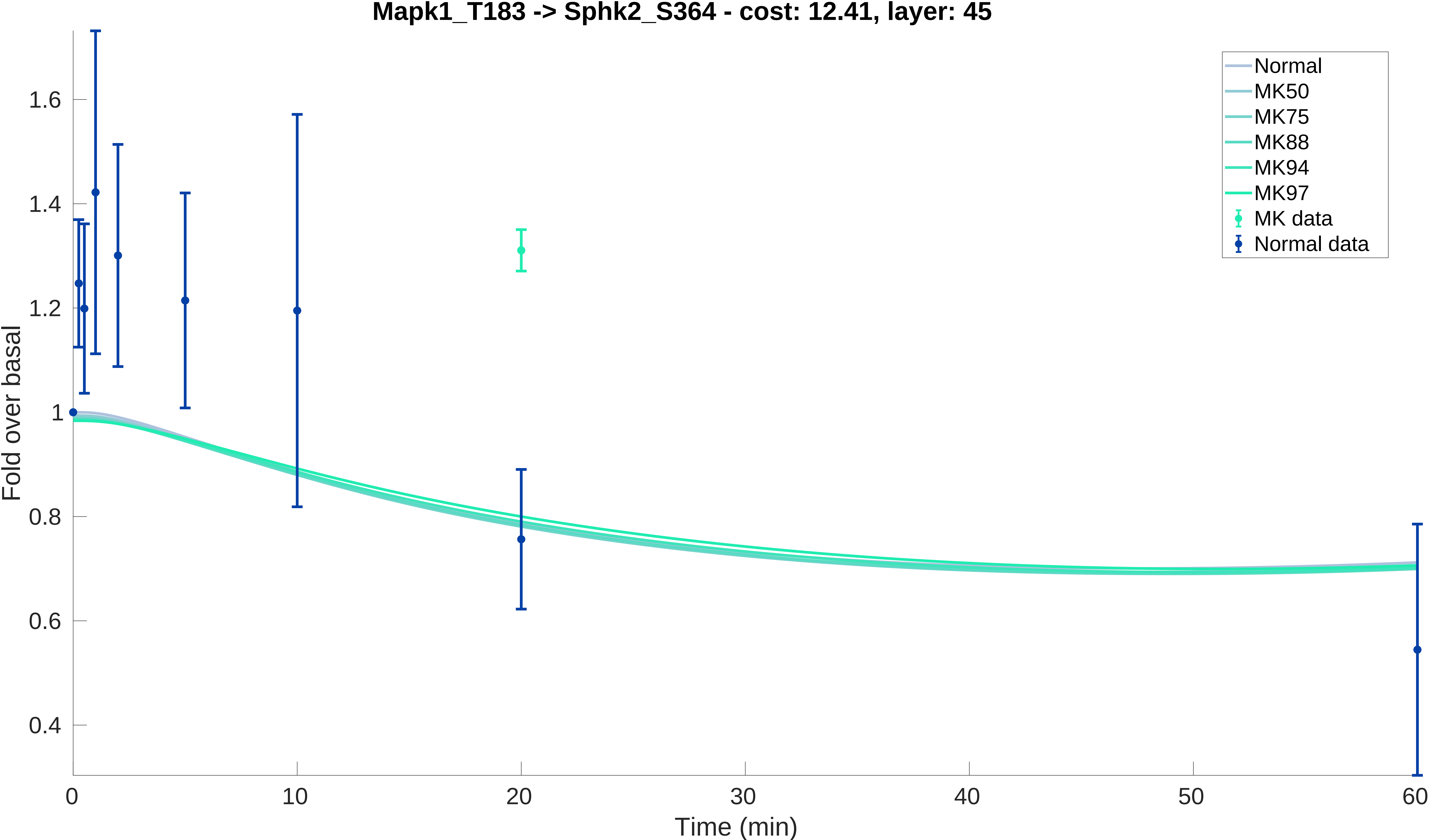
Mapk1_T183 -> Sphk2_S364 - cost: 12.41, layer: 45
Fold over basal
Time (min)
0.4
0.6
0.8
1
1.2
1.4
1.6
0
10
20
30
40
50
60
Normal
MK50
MK75
MK88
MK94
MK97
MK data
Normal data

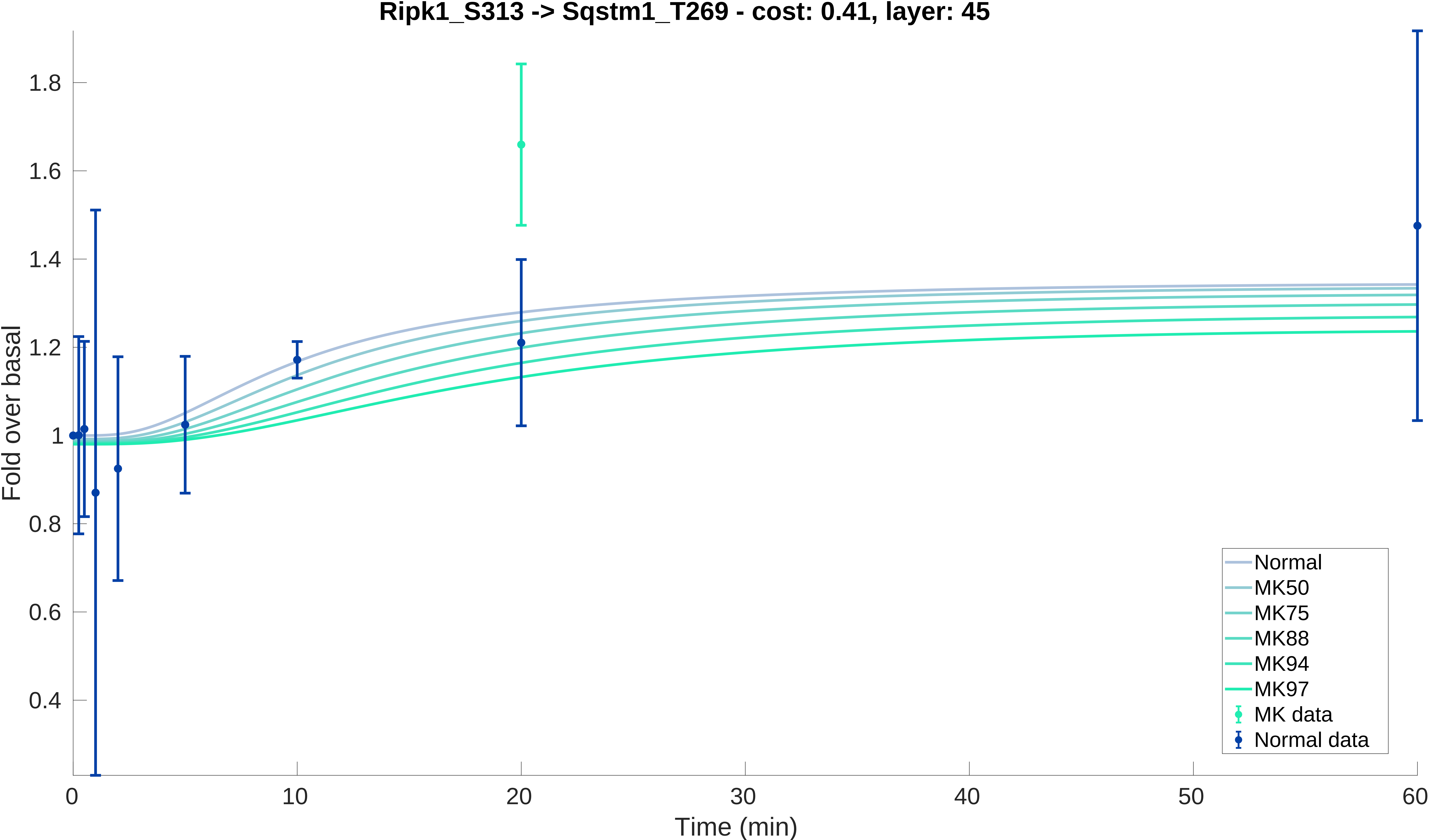
Ripk1_S313 -> Sqstm1_T269 - cost: 0.41, layer: 45
Fold over basal
1.8
1.6
1.4
1.2
1
0.8
0.6
0.4
0
10
20
30
40
50
60
Time (min)
Normal
MK50
MK75
MK88
MK94
MK97
MK data
Normal data

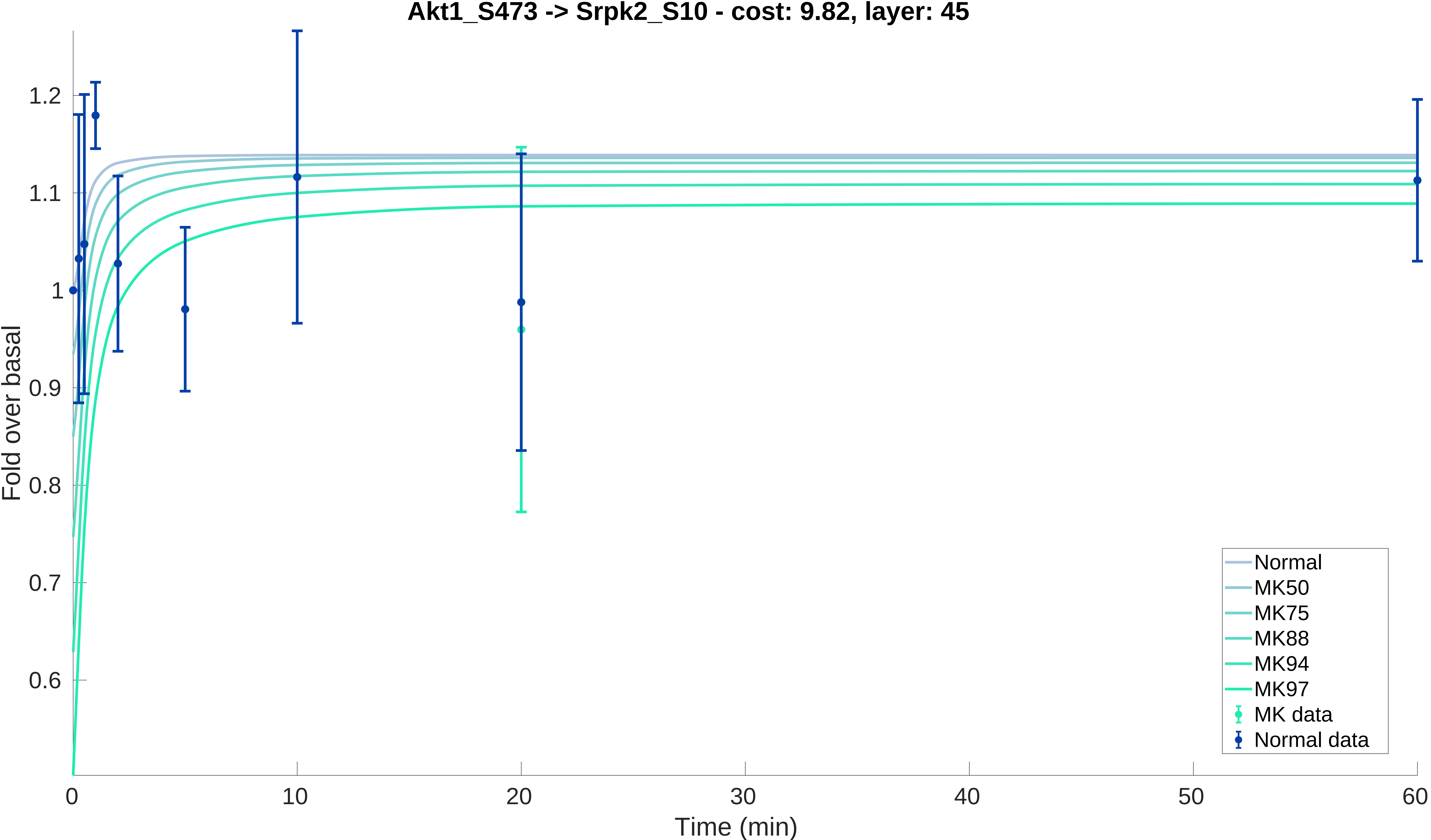
Akt1_S473 -> Srpk2_S10 - cost: 9.82, layer: 45
Fold over basal
Time (min)
0.6
0.7
0.8
0.9
1
1.1
1.2
0
10
20
30
40
50
60
Normal
MK50
MK75
MK88
MK94
MK97
MK data
Normal data

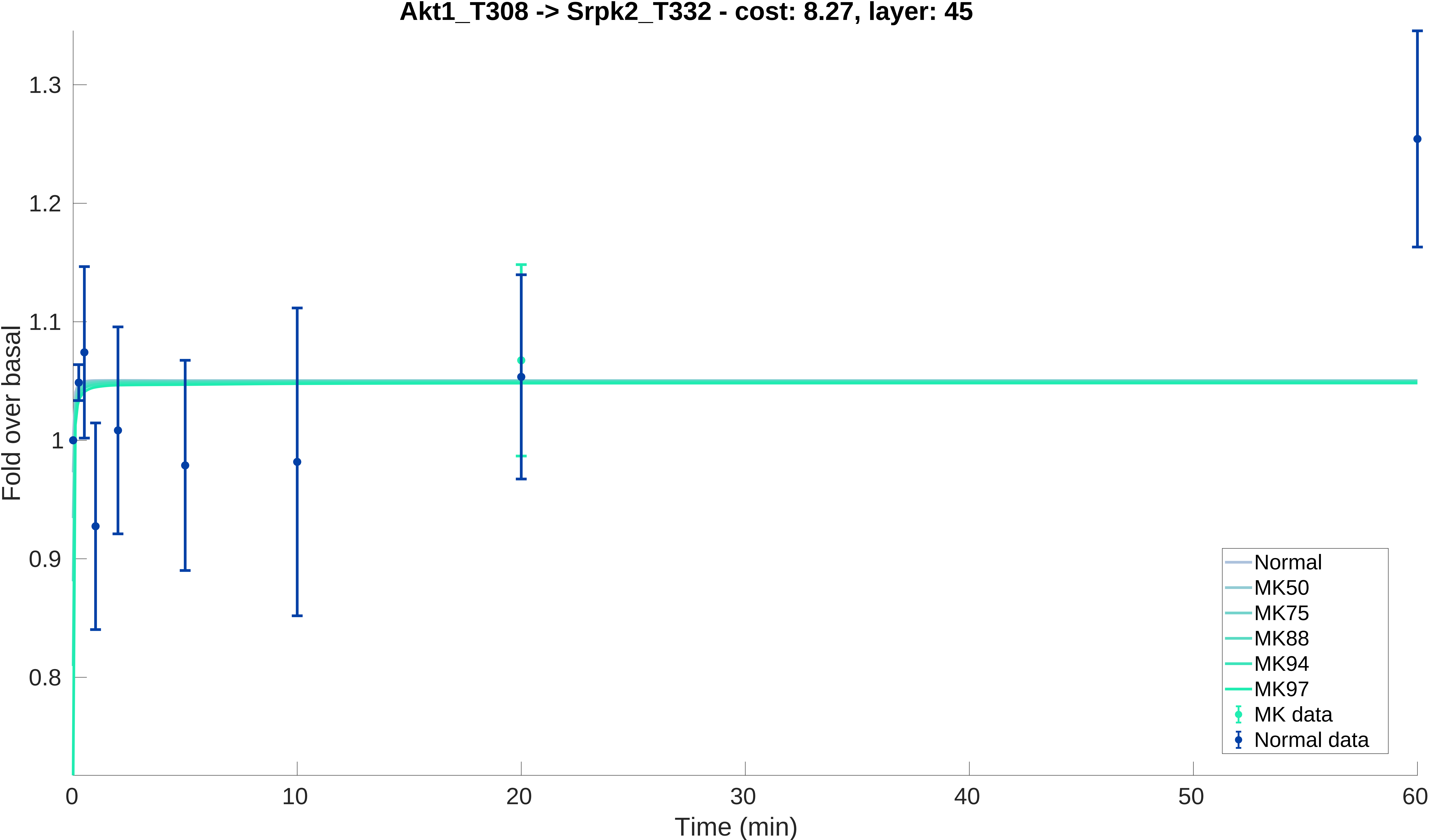

Akt1_T308 -> Srpk2_T332 - cost: 8.27, layer: 45
Fold over basal
Time (min)
0.8
0.9
1
1.1
1.2
1.3
0
10
20
30
40
50
60
Normal
MK50
MK75
MK88
MK94
MK97
MK data
Normal data

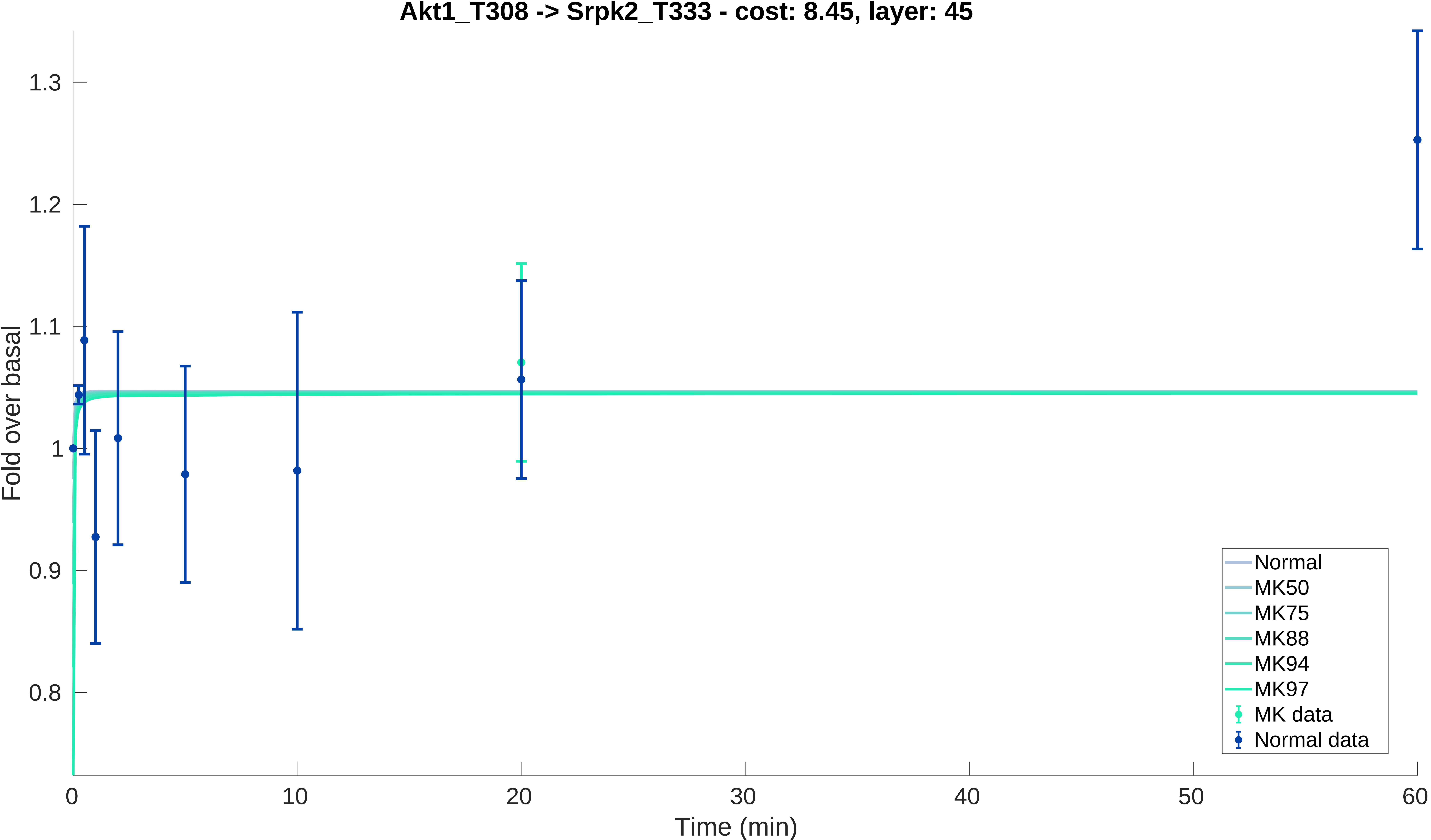
Akt1_T308 -> Srpk2_T333 - cost: 8.45, layer: 45
Fold over basal
Time (min)
1.3
1.2
1.1
1
0.9
0.8
0
10
20
30
40
50
60
Normal
MK50
MK75
MK88
MK94
MK97
MK data
Normal data

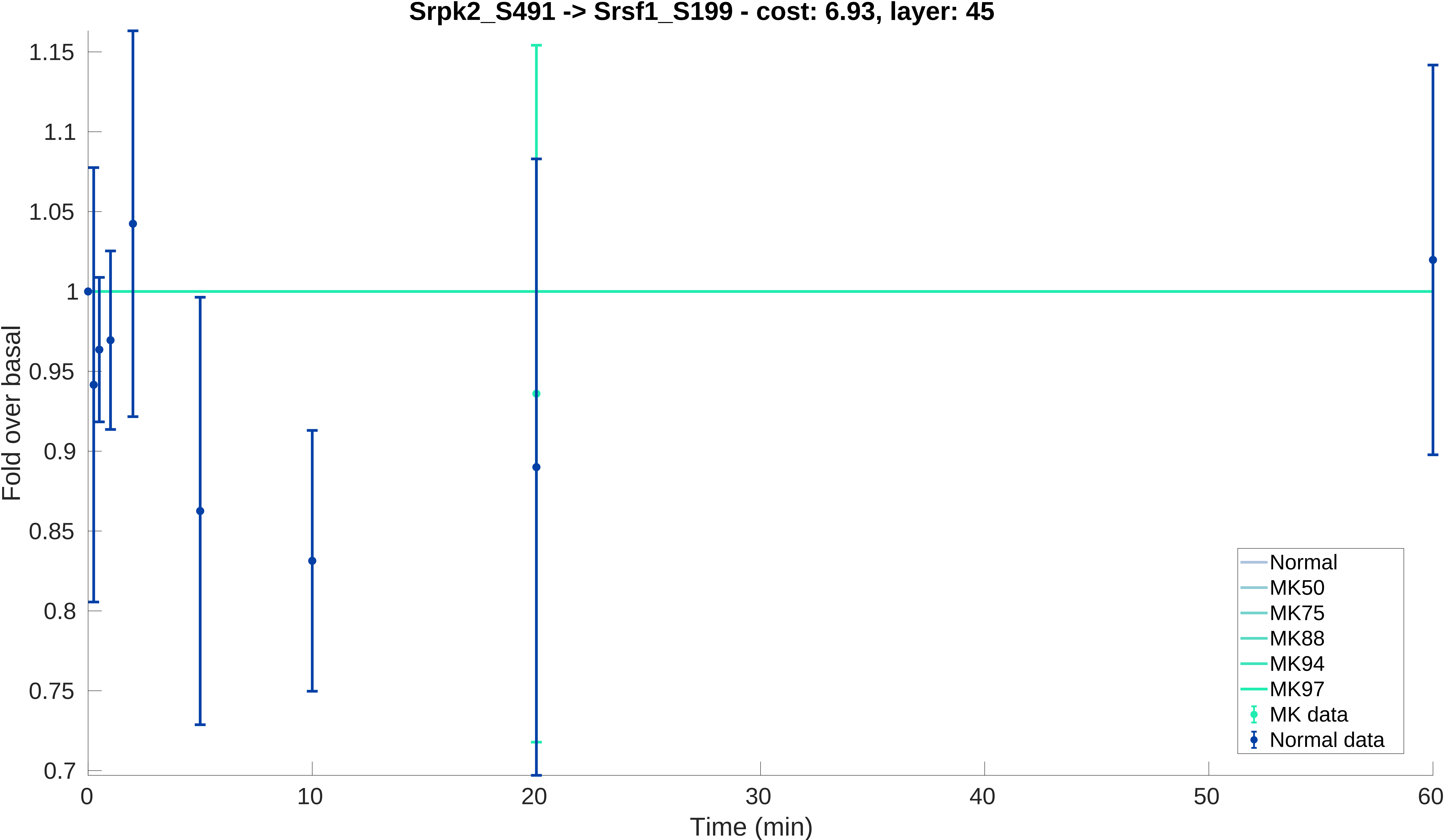
Srpk2_S491 -> Srsf1_S199 - cost: 6.93, layer: 45
Fold over basal
Time (min)
1.15
1.1
1.05
1
0.95
0.9
0.85
0.8
0.75
0.7
0
10
20
30
40
50
60
Normal
MK50
MK75
MK88
MK94
MK97
MK data
Normal data

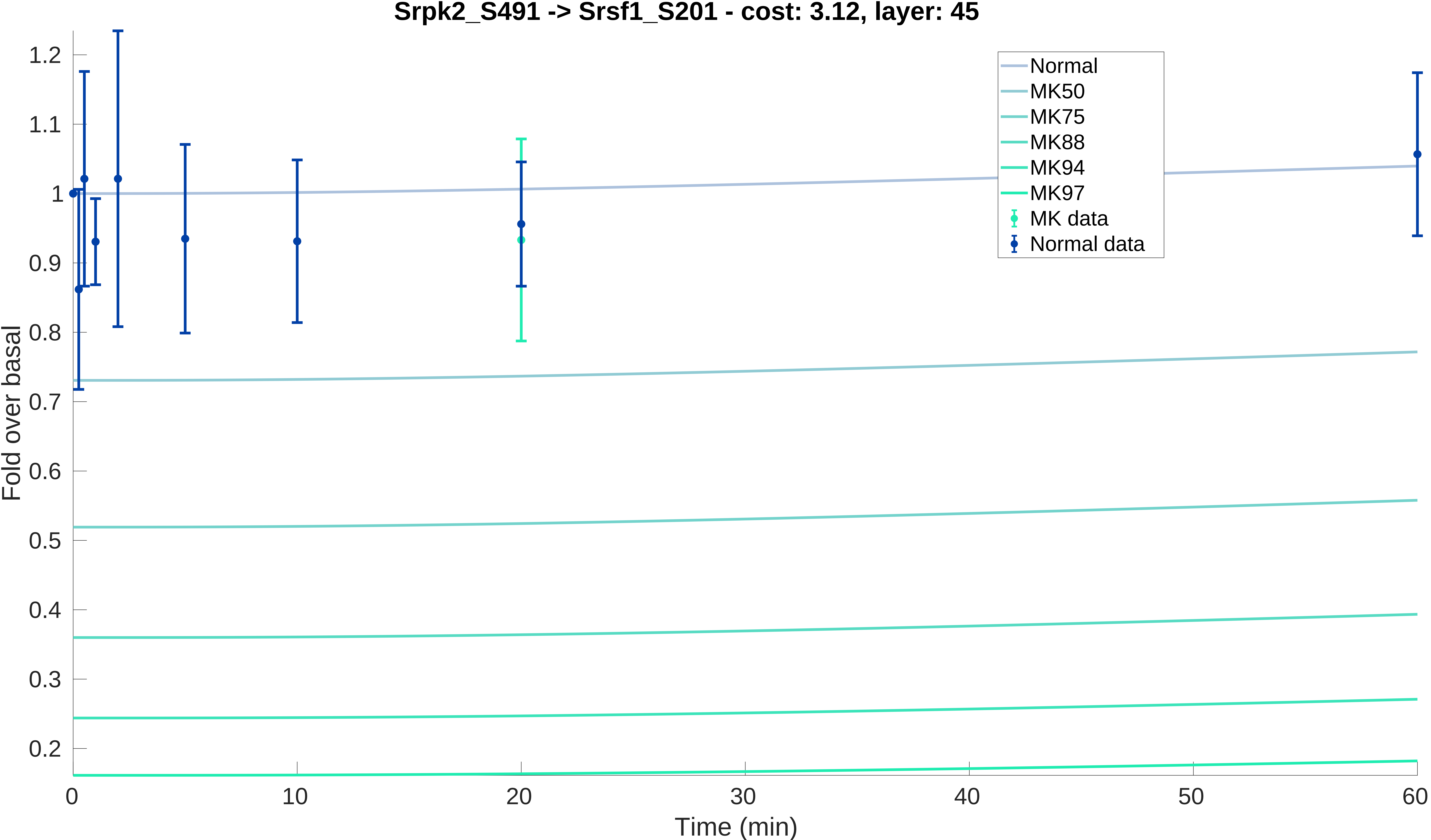
Srpk2_S491 -> Srsf1_S201 - cost: 3.12, layer: 45
Fold over basal
Time (min)
1.2
1.1
1
0.9
0.8
0.7
0.6
0.5
0.4
0.3
0.2
0
10
20
30
40
50
60
Normal
MK50
MK75
MK88
MK94
MK97
MK data
Normal data

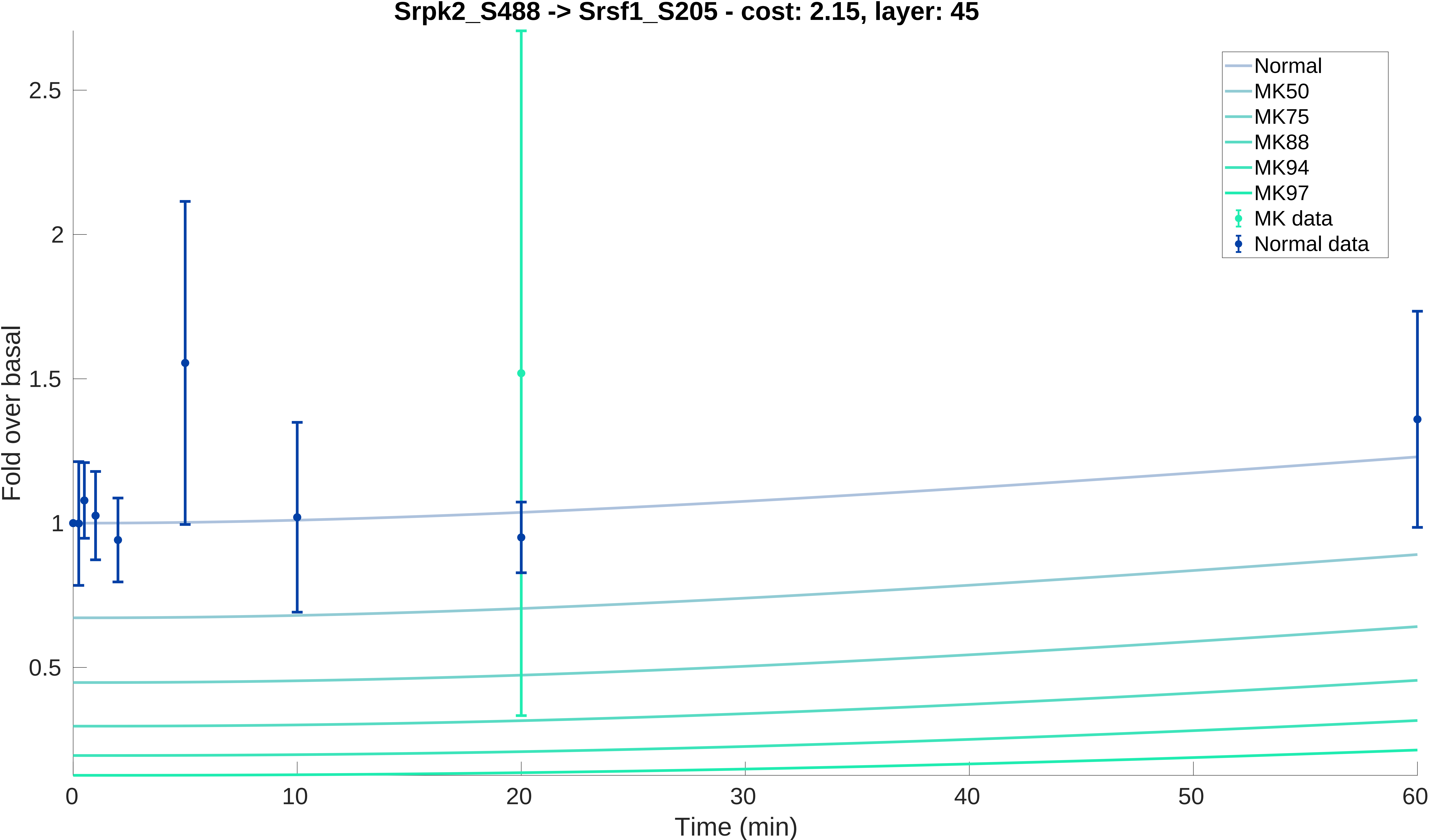
Srpk2_S488 -> Srsf1_S205 - cost: 2.15, layer: 45
Fold over basal
Time (min)
0.5
1
1.5
2
2.5
0
10
20
30
40
50
60
Normal
MK50
MK75
MK88
MK94
MK97
MK data
Normal data

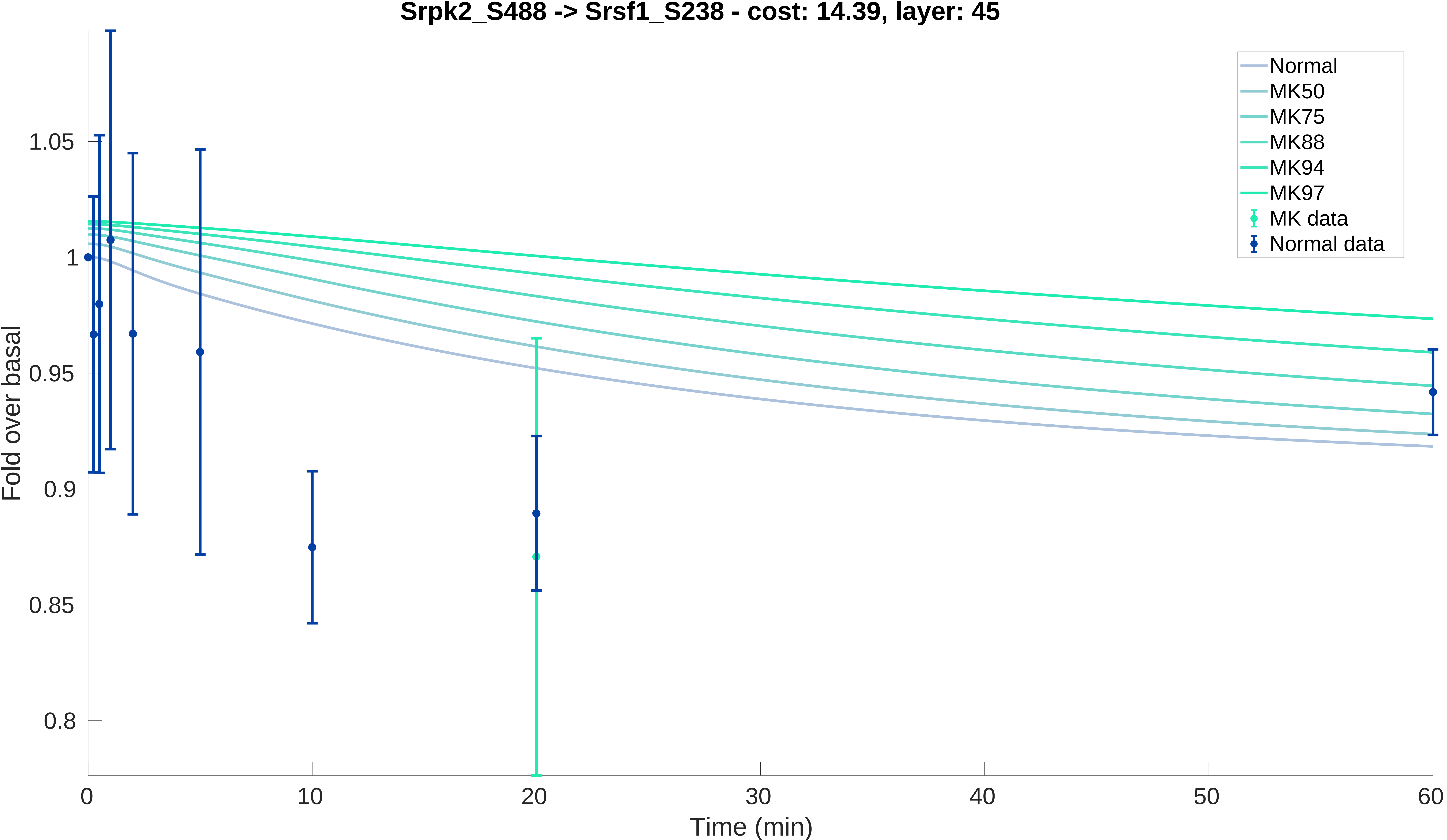

Srpk2_S488 -> Srsf1_S238 - cost: 14.39, layer: 45
Normal
MK50
MK75
MK88
MK94
MK97
MK data
Normal data
Fold over basal
1.05
1
0.95
0.9
0.85
0.8
Time (min)
0
10
20
30
40
50
60

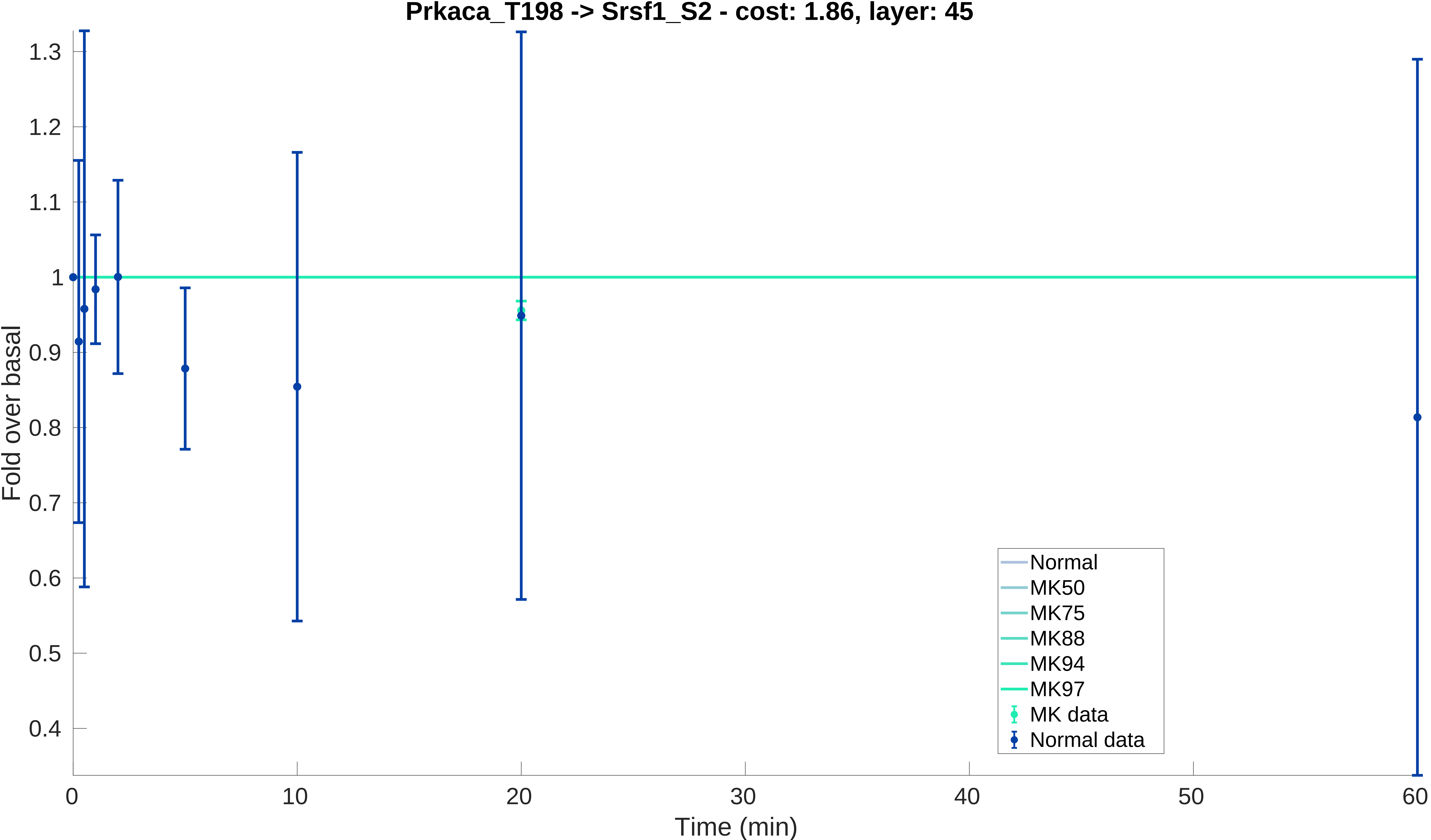
Prkaca_T198 -> Srsf1_S2 - cost: 1.86, layer: 45
1.3
1.2
1.1
1
0.9
0.8
0.7
0.6
0.5
0.4
Fold over basal
0
10
20
30
40
50
60
Time (min)
Normal
MK50
MK75
MK88
MK94
MK97
MK data
Normal data

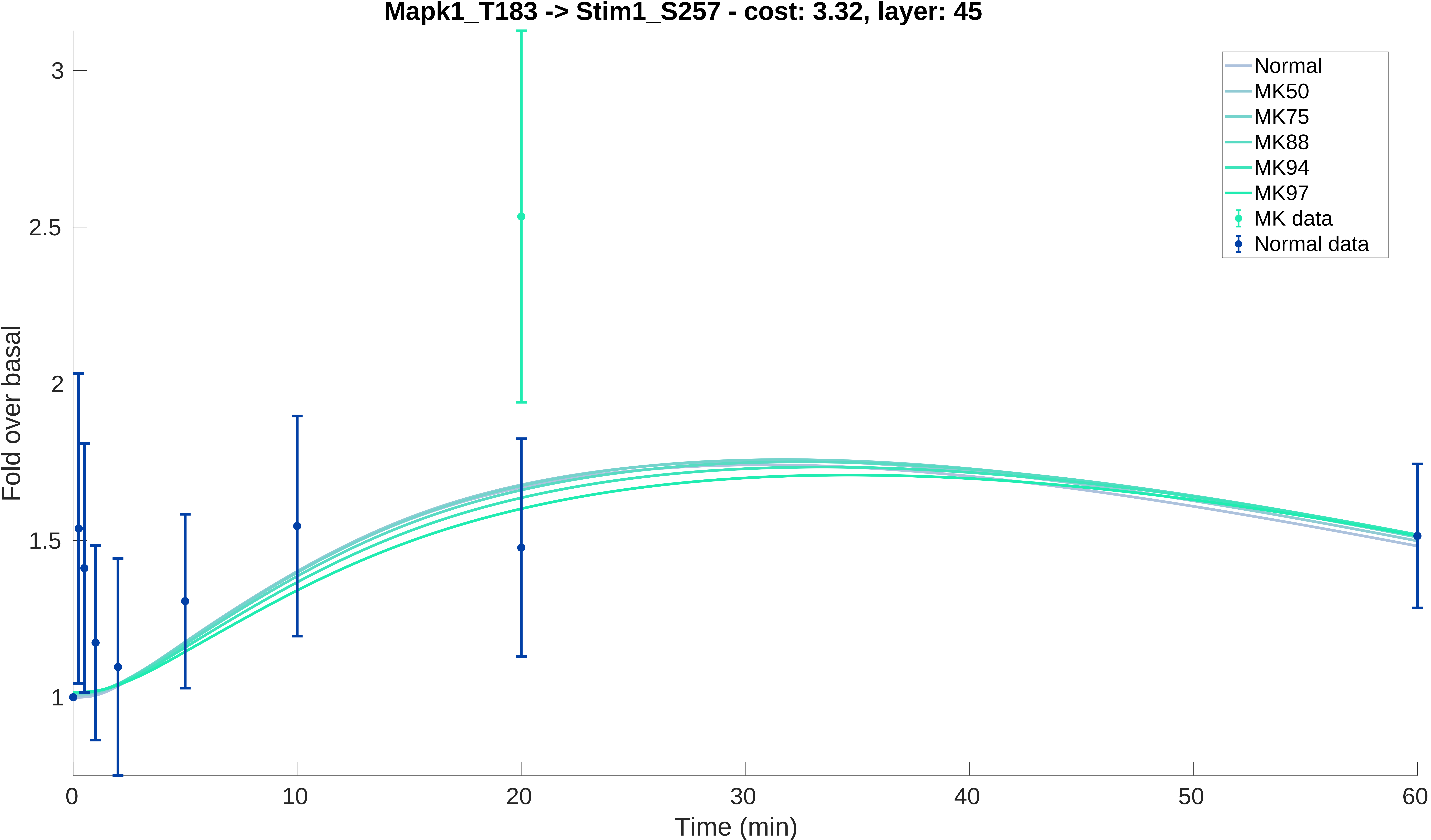
Mapk1_T183 -> Stim1_S257 - cost: 3.32, layer: 45
Normal
MK50
MK75
MK88
MK94
MK97
MK data
Normal data
Fold over basal
3
2.5
2
1.5
1
0
10
20
30
40
50
60
Time (min)

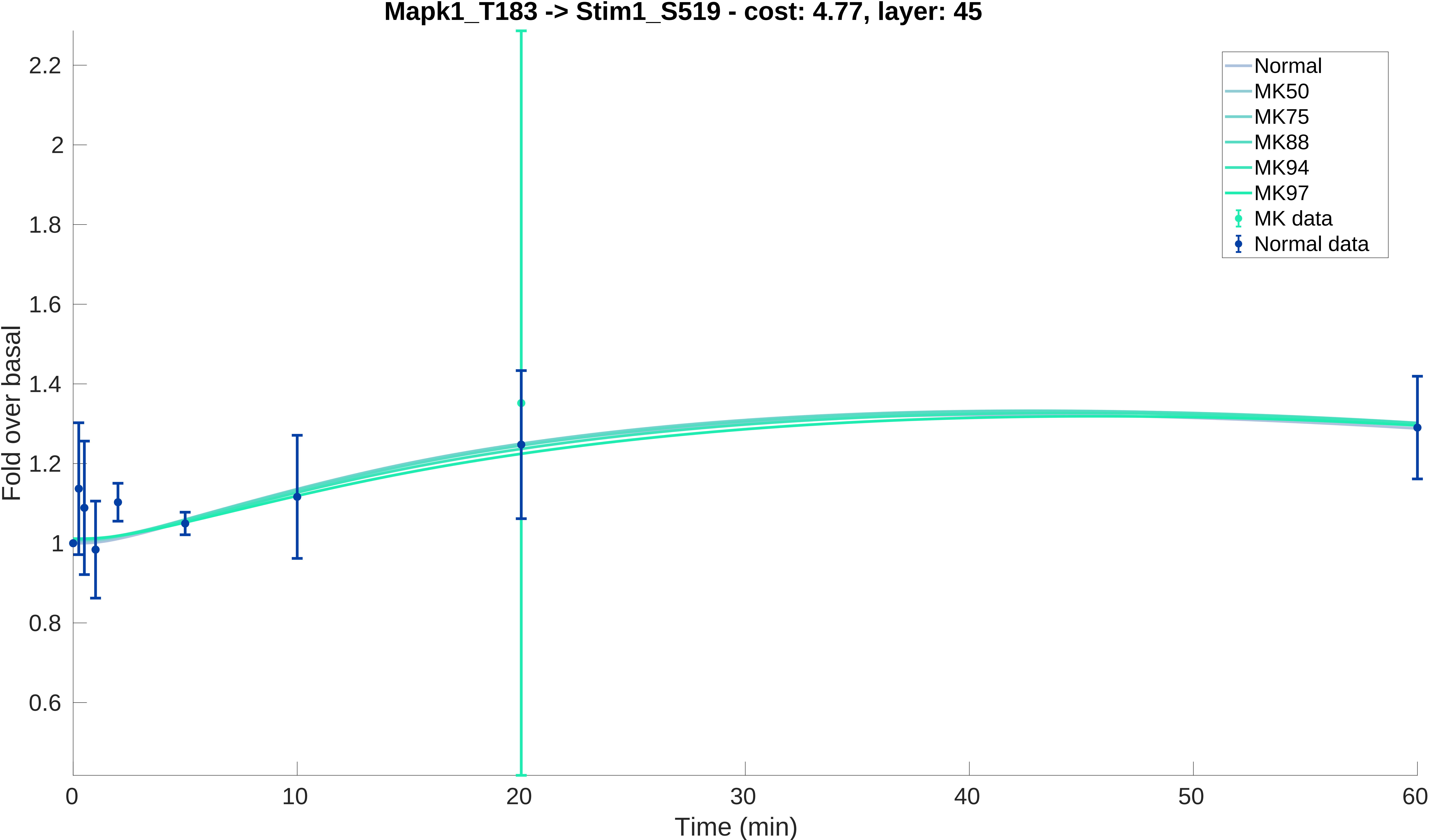

Mapk1_T183 -> Stim1_S519 - cost: 4.77, layer: 45
Normal
MK50
MK75
MK88
MK94
MK97
MK data
Normal data
Fold over basal
2.2
2
1.8
1.6
1.4
1.2
1
0.8
0.6
0
10
20
30
40
50
60
Time (min)

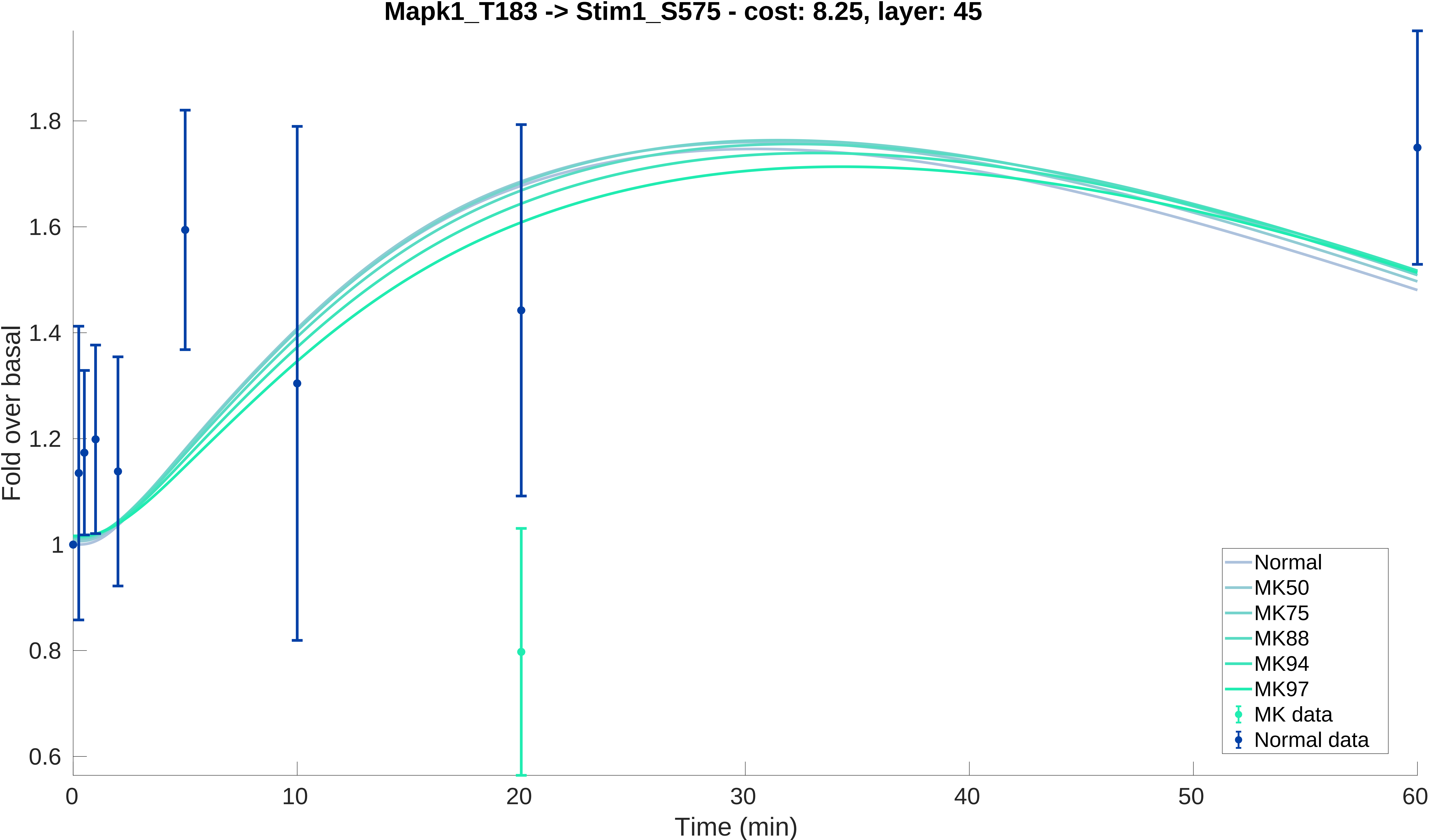

Mapk1_T183 -> Stim1_S575 - cost: 8.25, layer: 45
Fold over basal
1.8
1.6
1.4
1.2
1
0.8
0.6
0
10
20
30
40
50
60
Time (min)
Normal
MK50
MK75
MK88
MK94
MK97
MK data
Normal data

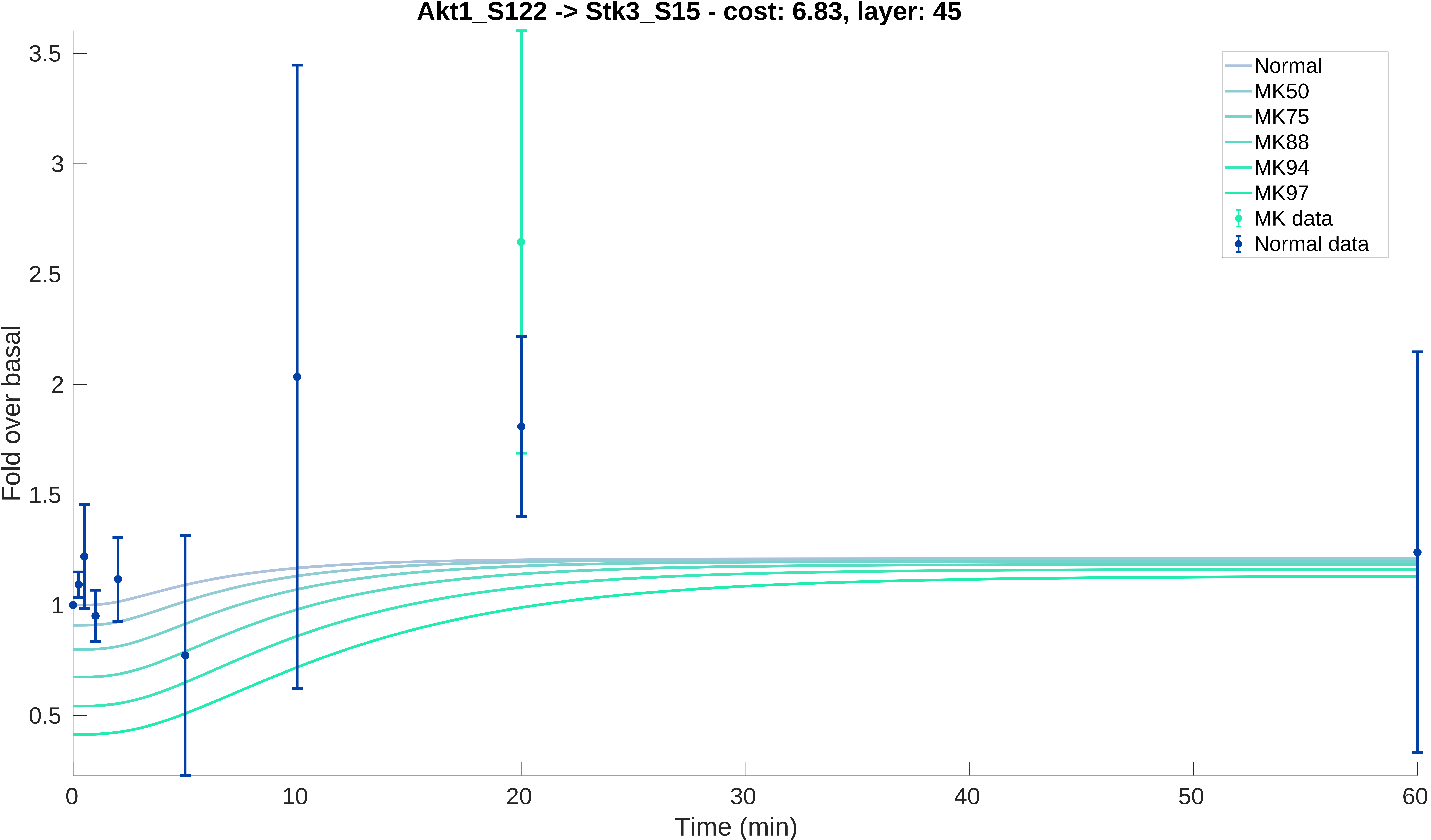

Akt1_S122 -> Stk3_S15 - cost: 6.83, layer: 45
Fold over basal
Time (min)
0.5
1
1.5
2
2.5
3
3.5
0
10
20
30
40
50
60
Normal
MK50
MK75
MK88
MK94
MK97
MK data
Normal data

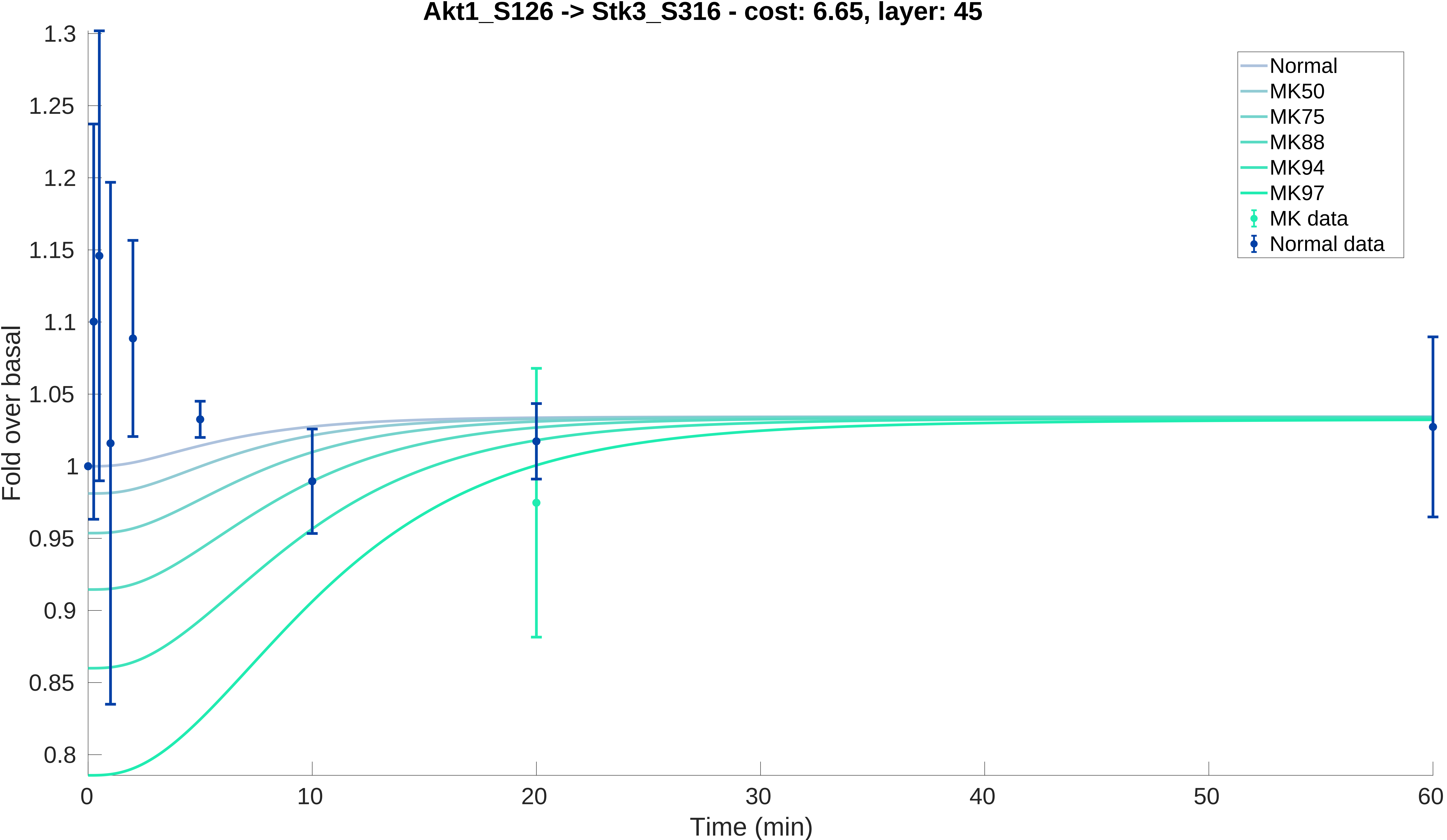

Akt1_S126 -> Stk3_S316 - cost: 6.65, layer: 45
Normal
MK50
MK75
MK88
MK94
MK97
MK data
Normal data
Fold over basal
1.3
1.25
1.2
1.15
1.1
1.05
1
0.95
0.9
0.85
0.8
0
10
20
30
40
50
60
Time (min)

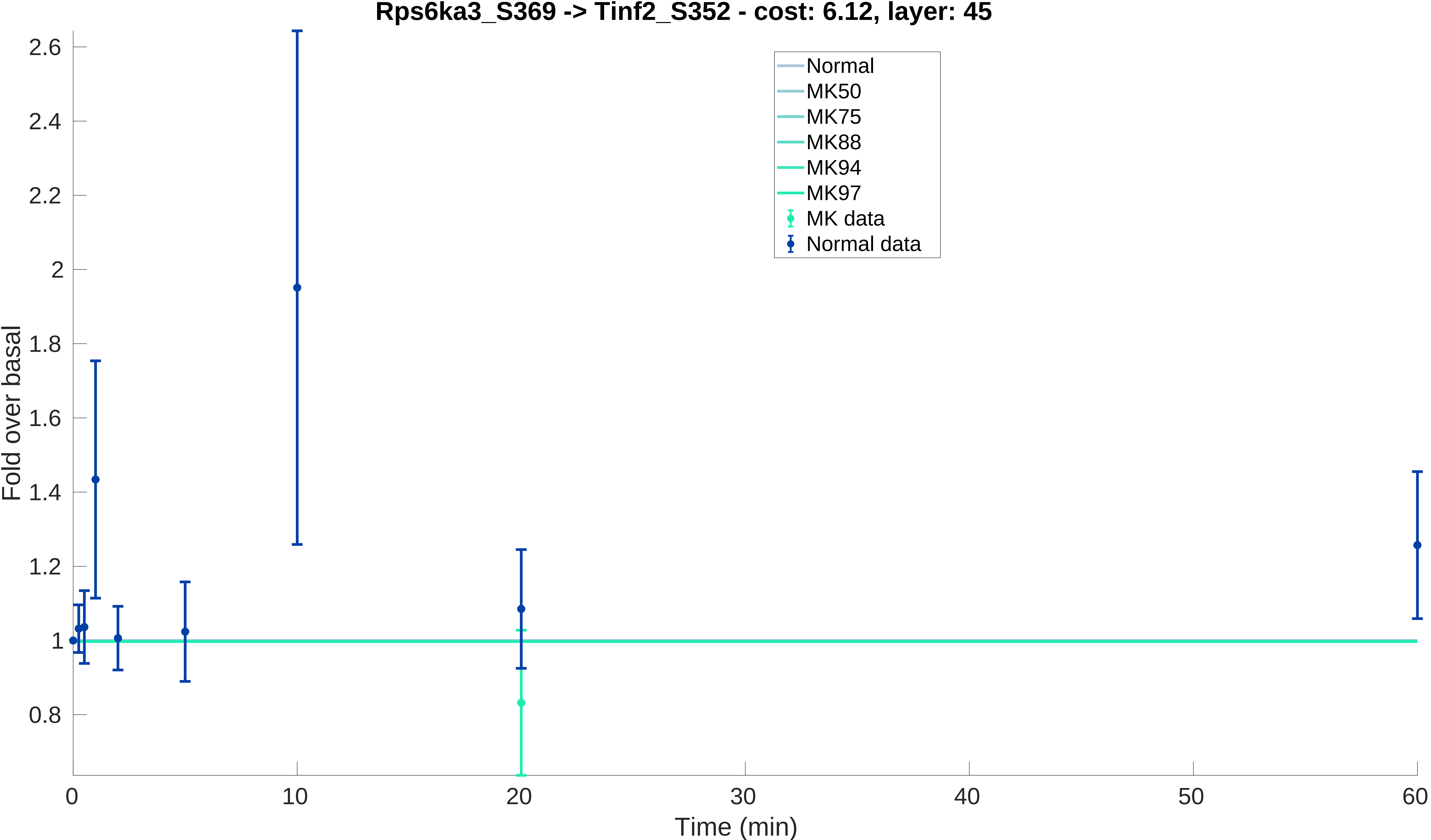

Rps6ka3_S369 -> Tinf2_S352 - cost: 6.12, layer: 45
Normal
MK50
MK75
MK88
MK94
MK97
MK data
Normal data
Fold over basal
2.6
2.4
2.2
2
1.8
1.6
1.4
1.2
1
0.8
0
10
20
30
40
50
60
Time (min)

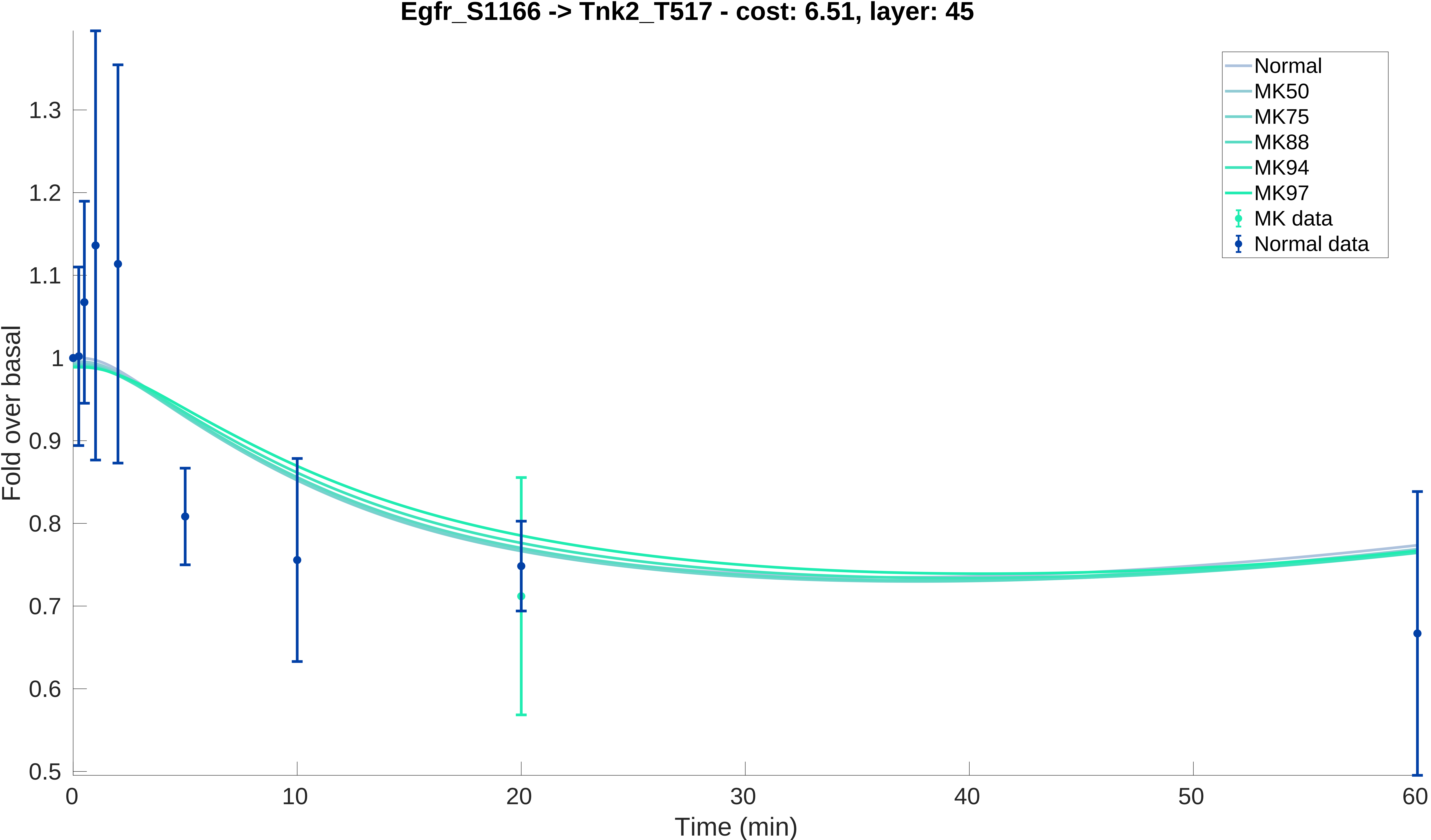

Egfr_S1166 -> Tnk2_T517 - cost: 6.51, layer: 45
Normal
MK50
MK75
MK88
MK94
MK97
MK data
Normal data
Fold over basal
Time (min)
1.3
1.2
1.1
1
0.9
0.8
0.7
0.6
0.5
0
10
20
30
40
50
60

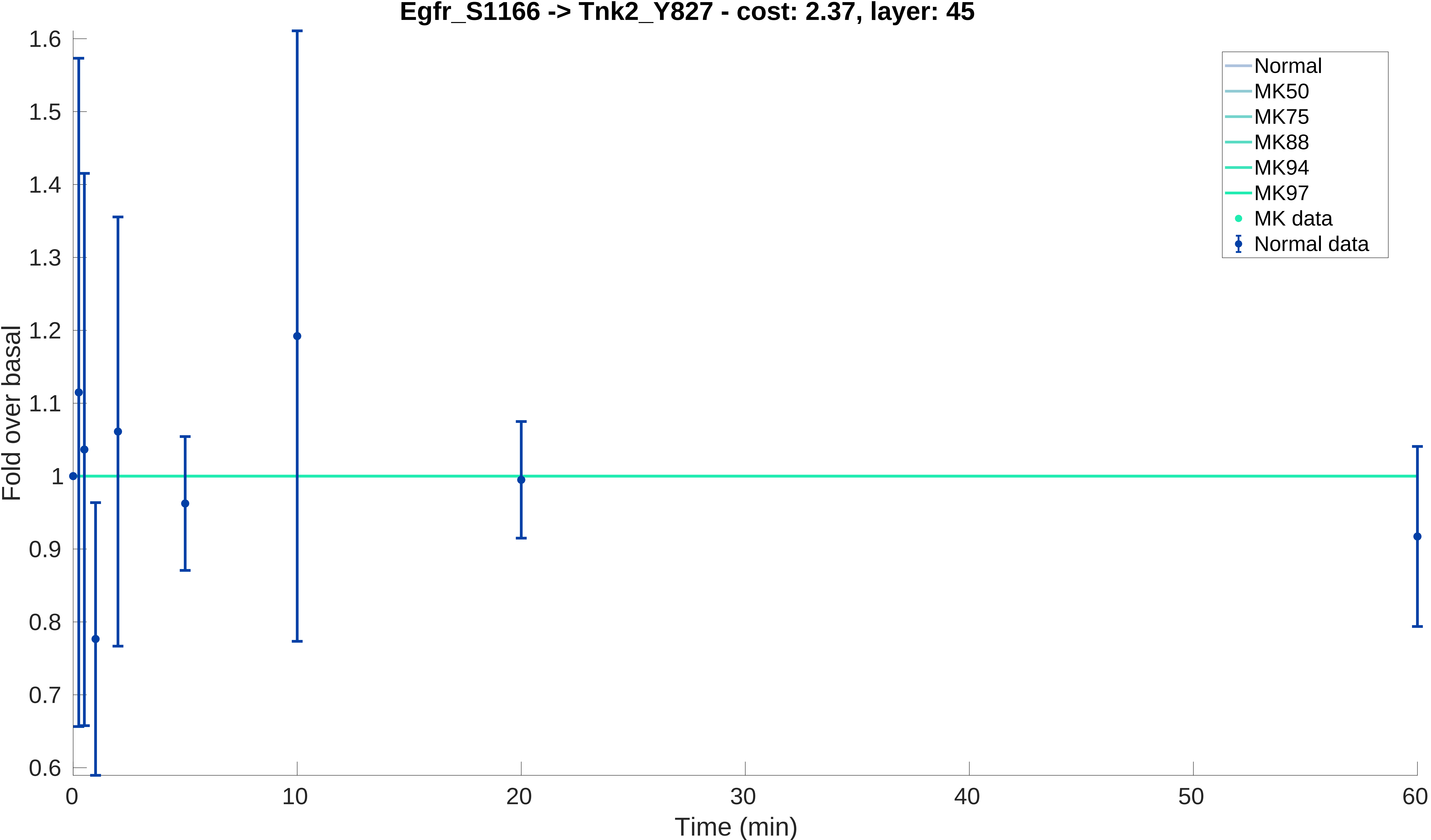

Egfr_S1166 -> Tnk2_Y827 - cost: 2.37, layer: 45
Normal
MK50
MK75
MK88
MK94
MK97
MK data
Normal data
Fold over basal
Time (min)
0.6
0.7
0.8
0.9
1
1.1
1.2
1.3
1.4
1.5
1.6
0
10
20
30
40
50
60

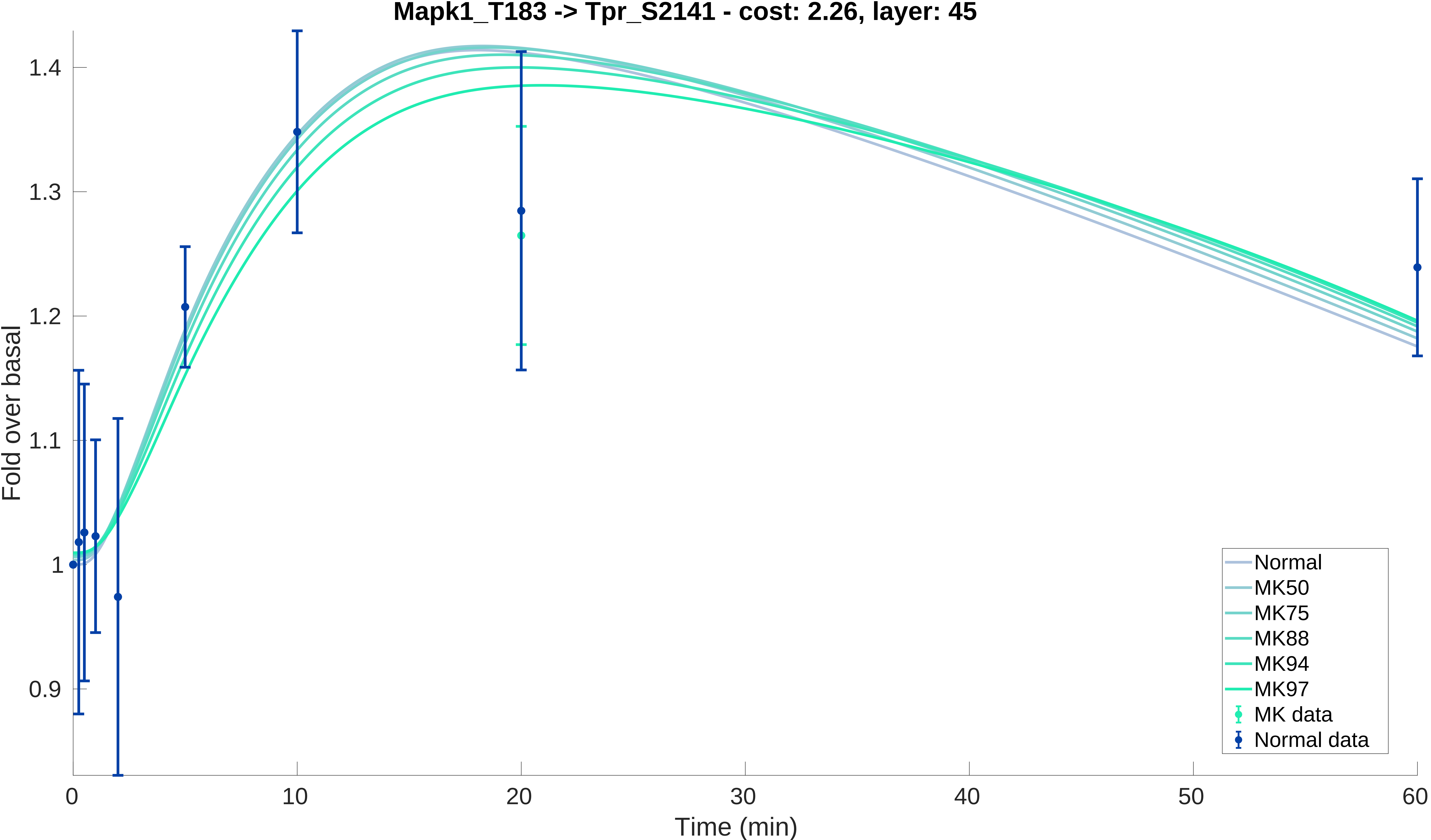

Mapk1_T183 -> Tpr_S2141 - cost: 2.26, layer: 45
Fold over basal
0.9
1
1.1
1.2
1.3
1.4
0
10
20
30
40
50
60
Time (min)
Normal
MK50
MK75
MK88
MK94
MK97
MK data
Normal data

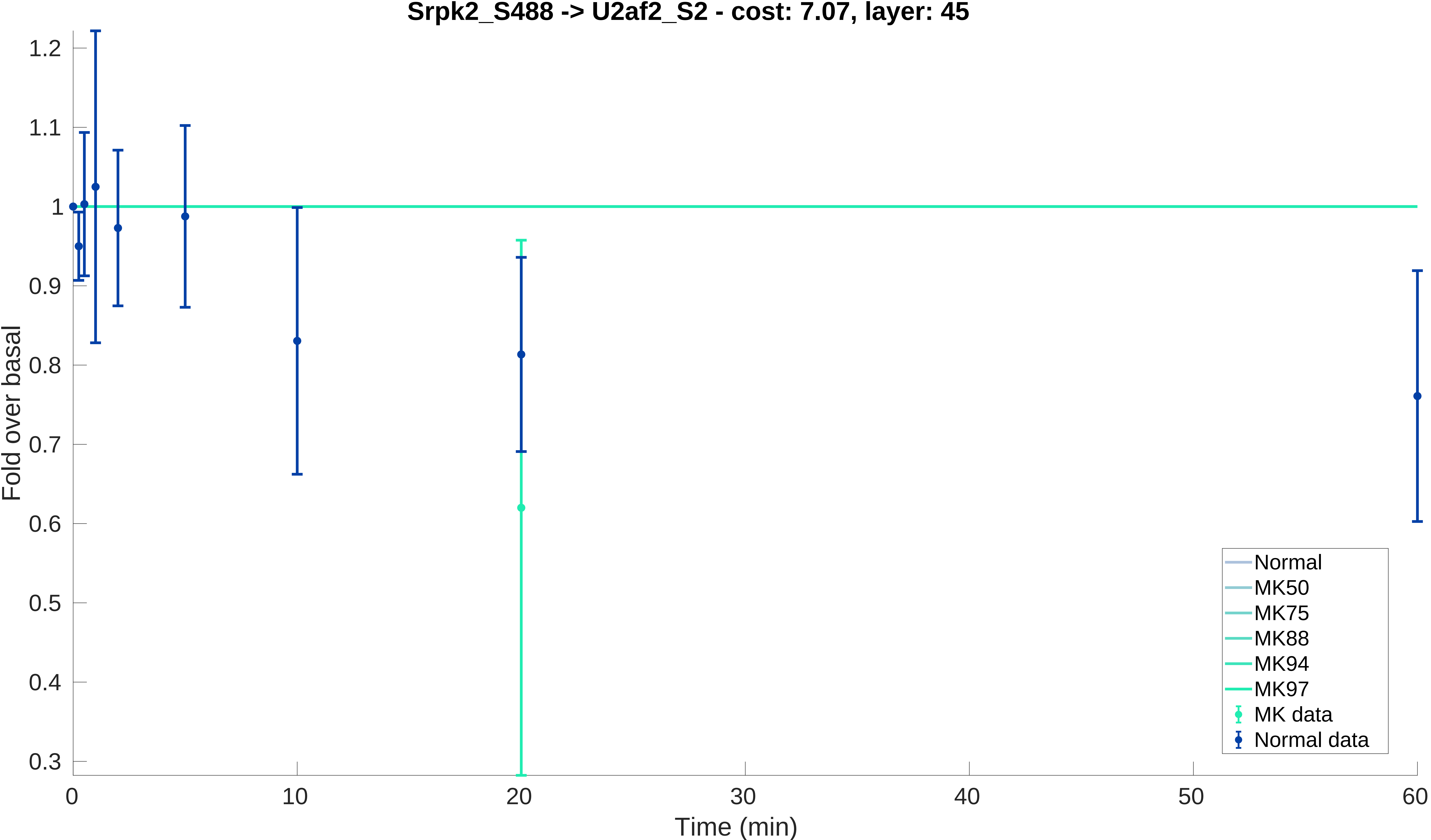

Srpk2_S488 -> U2af2_S2 - cost: 7.07, layer: 45
Fold over basal
1.2
1.1
1
0.9
0.8
0.7
0.6
0.5
0.4
0.3
0
10
20
30
40
50
60
Time (min)
Normal
MK50
MK75
MK88
MK94
MK97
MK data
Normal data

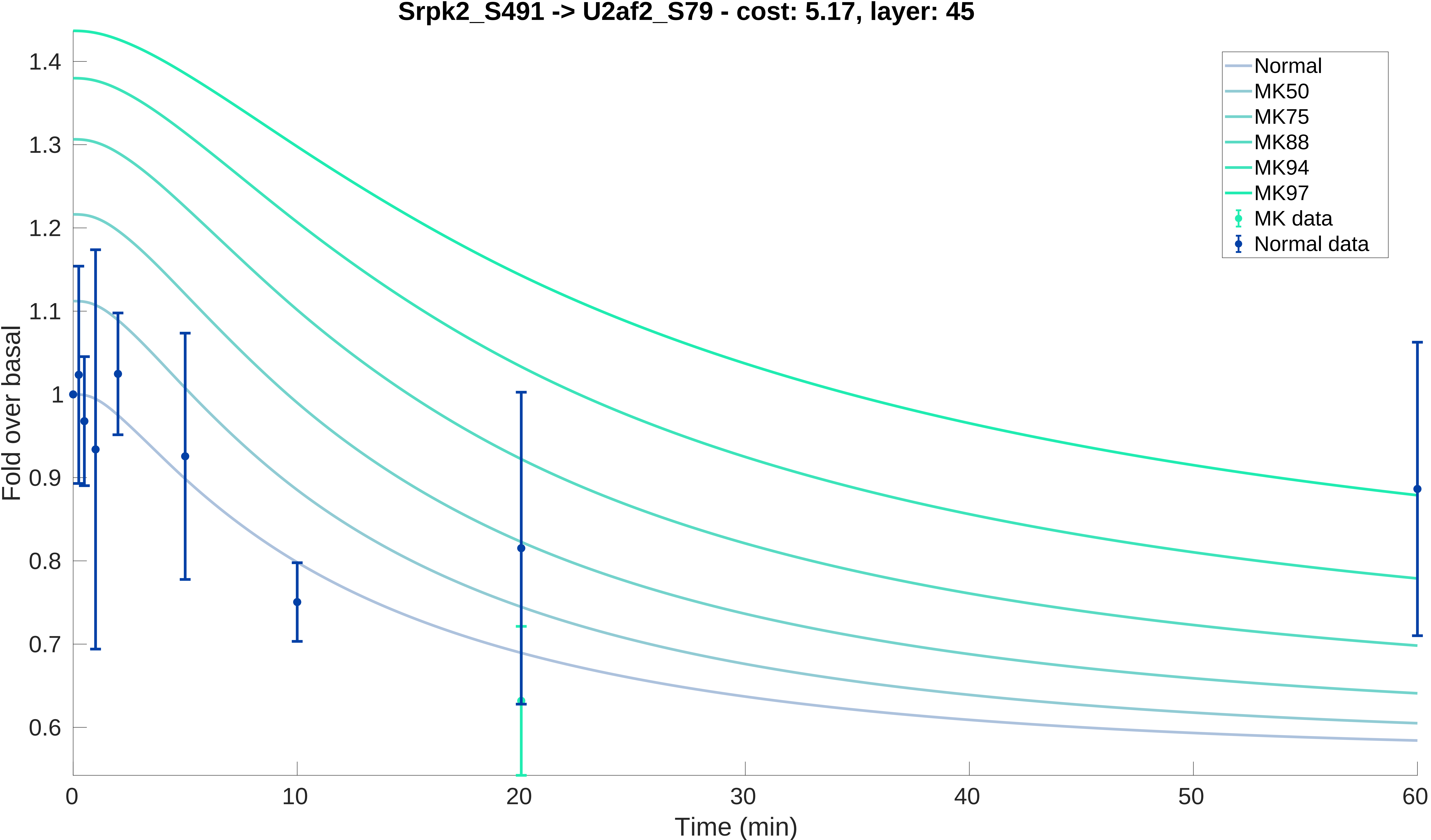

Srpk2_S491 -> U2af2_S79 - cost: 5.17, layer: 45
Fold over basal
Time (min)
0.6
0.7
0.8
0.9
1
1.1
1.2
1.3
1.4
0
10
20
30
40
50
60
Normal
MK50
MK75
MK88
MK94
MK97
MK data
Normal data

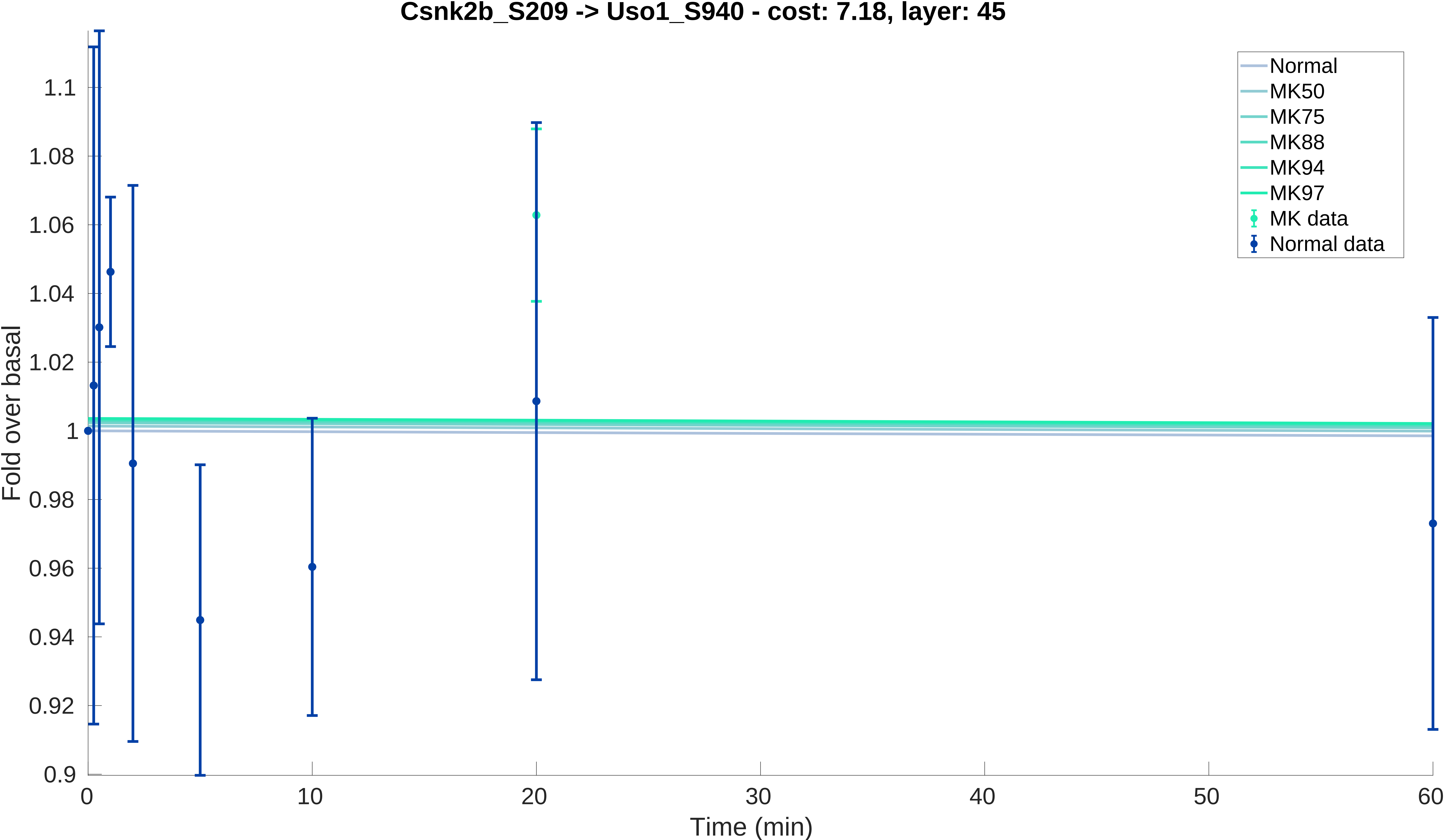

Csnk2b_S209 -> Uso1_S940 - cost: 7.18, layer: 45
Fold over basal
Time (min)
Normal
MK50
MK75
MK88
MK94
MK97
MK data
Normal data
1.1
1.08
1.06
1.04
1.02
1
0.98
0.96
0.94
0.92
0.9
0
10
20
30
40
50
60

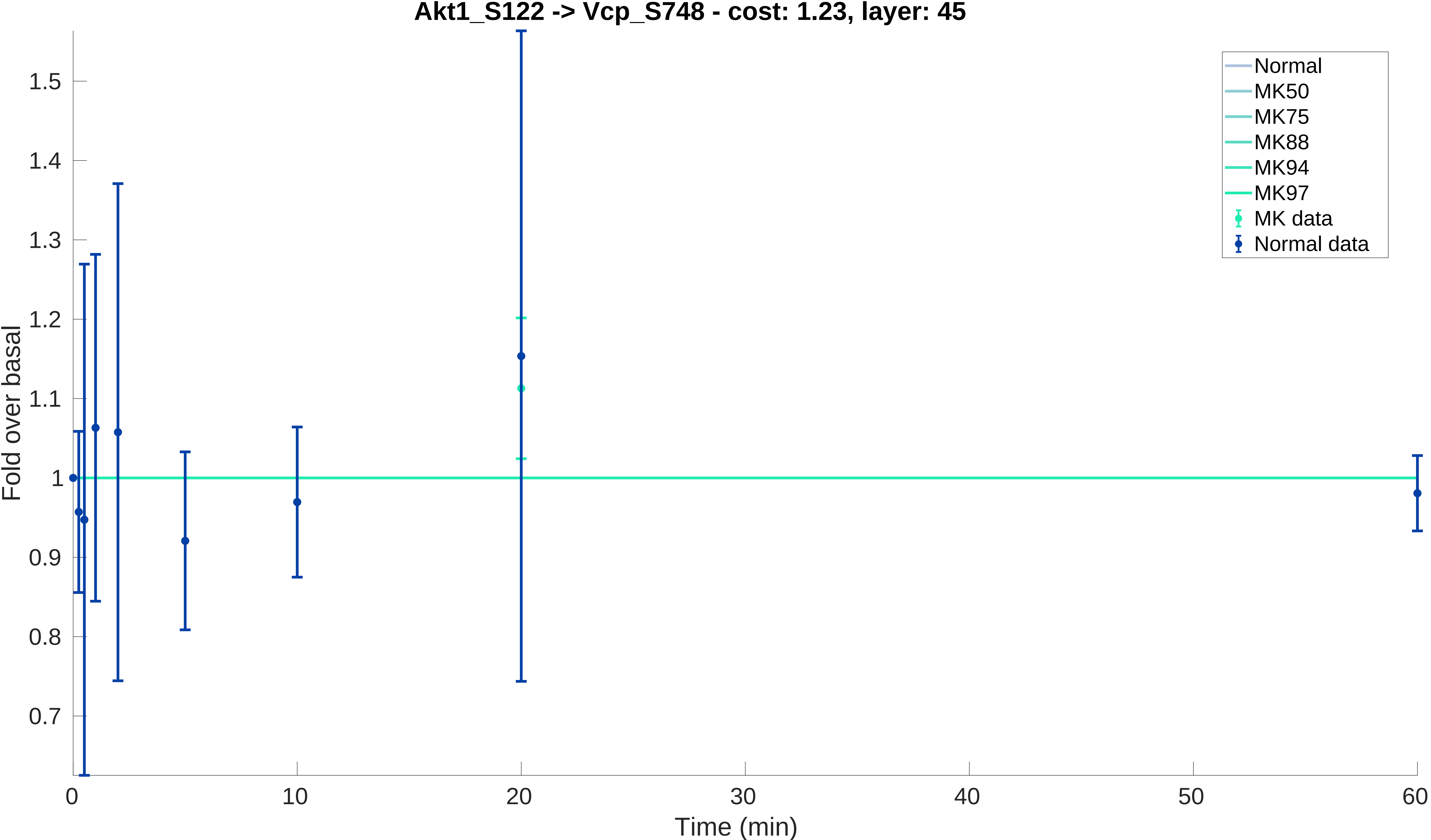

Akt1_S122 -> Vcp_S748 - cost: 1.23, layer: 45
Fold over basal
Time (min)
1.5
1.4
1.3
1.2
1.1
1
0.9
0.8
0.7
0
10
20
30
40
50
60
Normal
MK50
MK75
MK88
MK94
MK97
MK data
Normal data

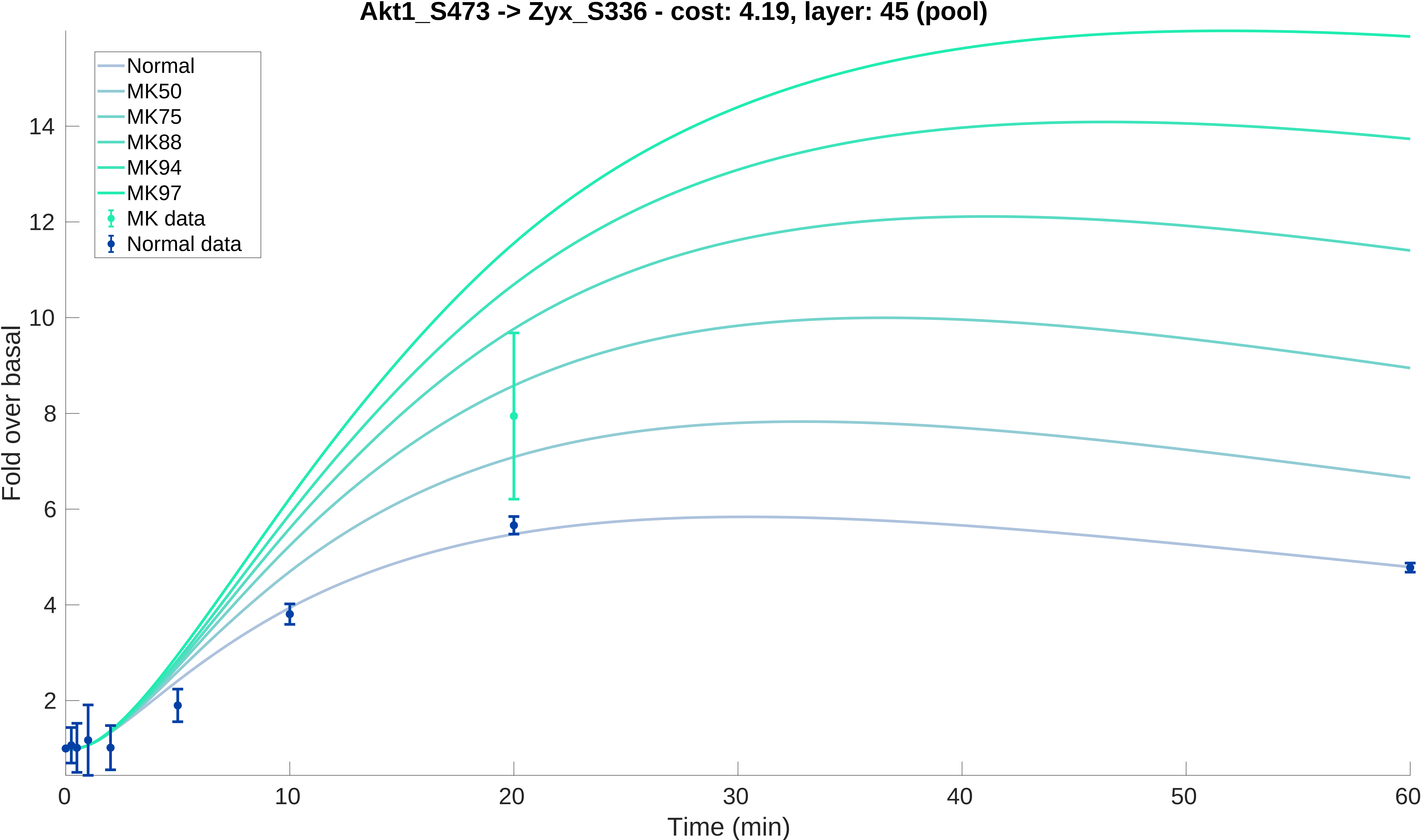

Akt1_S473 -> Zyx_S336 - cost: 4.19, layer: 45 (pool)
Normal
MK50
MK75
MK88
MK94
MK97
MK data
Normal data
Fold over basal
Time (min)
0
10
20
30
40
50
60
2
4
6
8
10
12
14

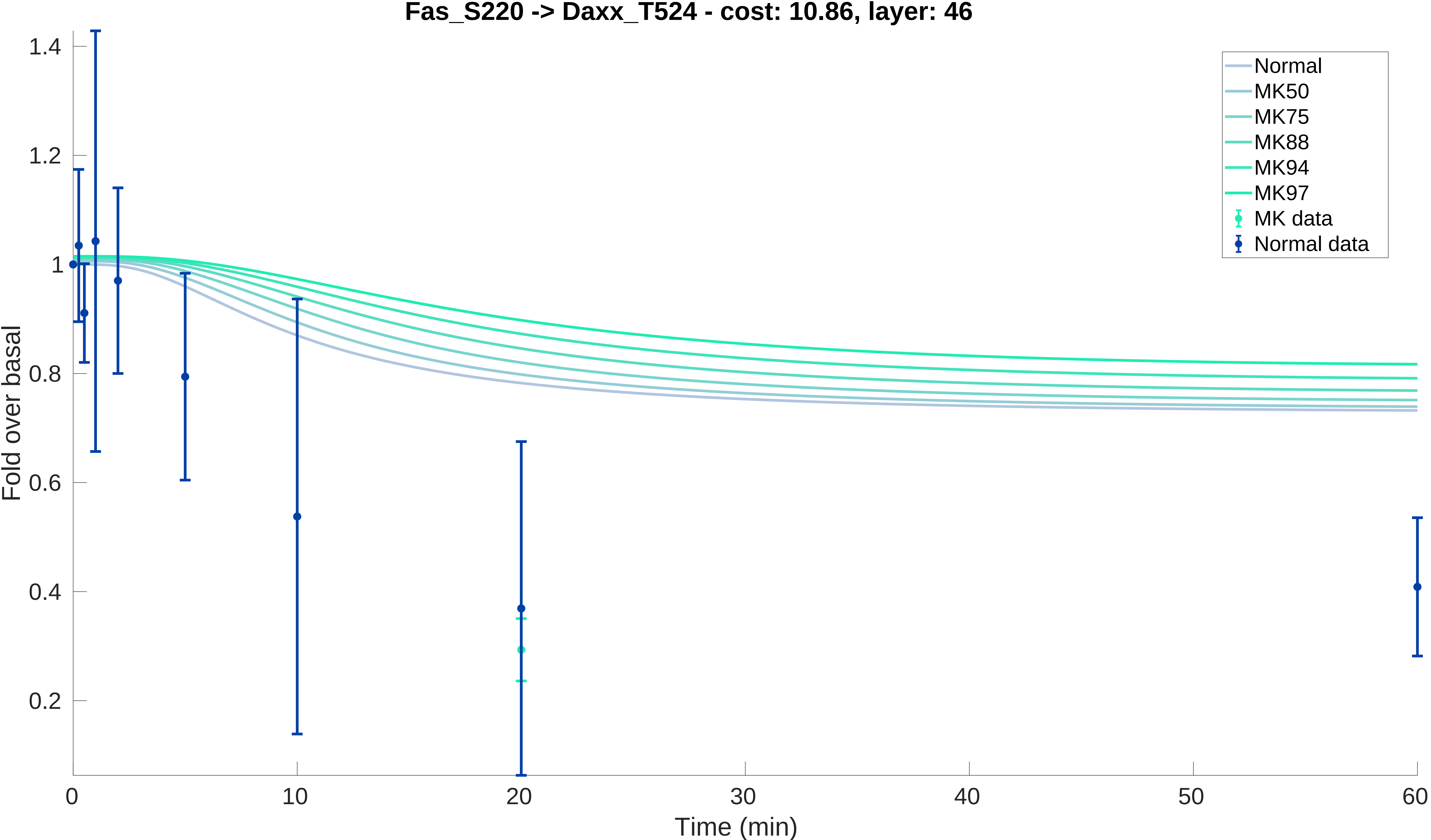

Fas_S220 -> Daxx_T524 - cost: 10.86, layer: 46
Normal
MK50
MK75
MK88
MK94
MK97
MK data
Normal data
Fold over basal
Time (min)
1.4
1.2
1
0.8
0.6
0.4
0.2
0
10
20
30
40
50
60

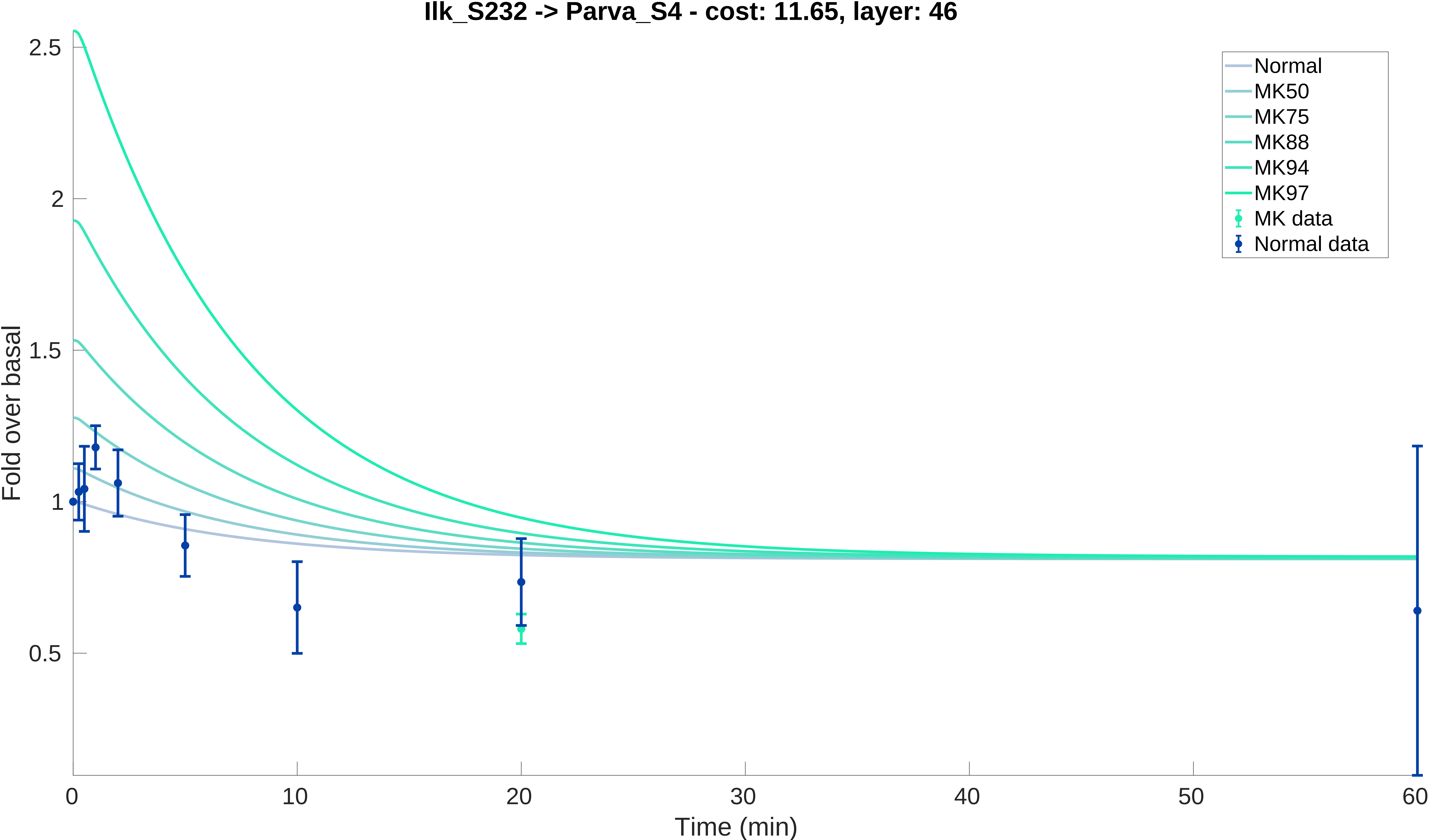
Ilk_S232 -> Parva_S4 - cost: 11.65, layer: 46
Fold over basal
Time (min)
0
10
20
30
40
50
60
0.5
1
1.5
2
2.5
Normal
MK50
MK75
MK88
MK94
MK97
MK data
Normal data

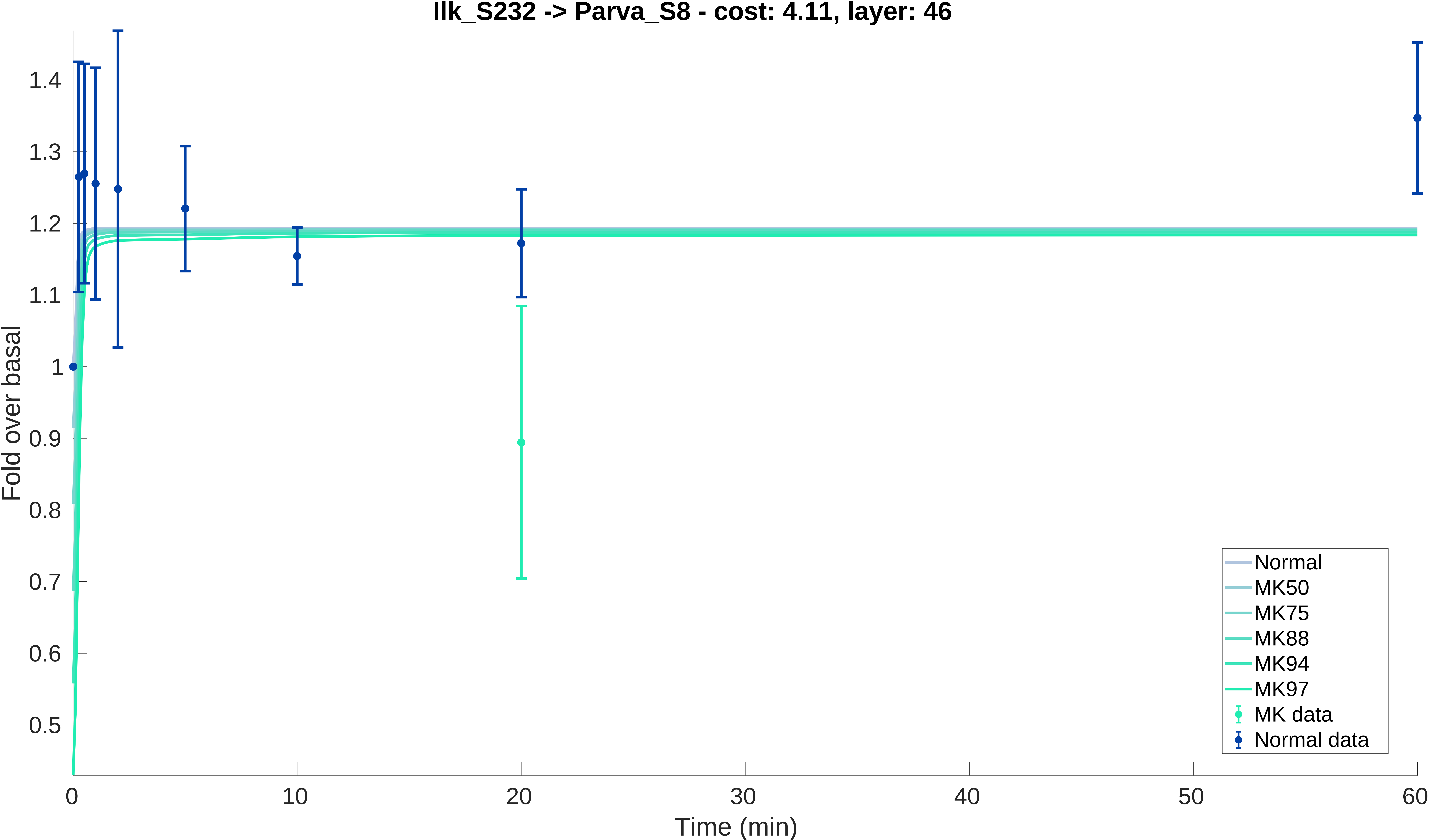

Ilk_S232 -> Parva_S8 - cost: 4.11, layer: 46
Fold over basal
Time (min)
0
10
20
30
40
50
60
0.5
0.6
0.7
0.8
0.9
1
1.1
1.2
1.3
1.4
Normal
MK50
MK75
MK88
MK94
MK97
MK data
Normal data

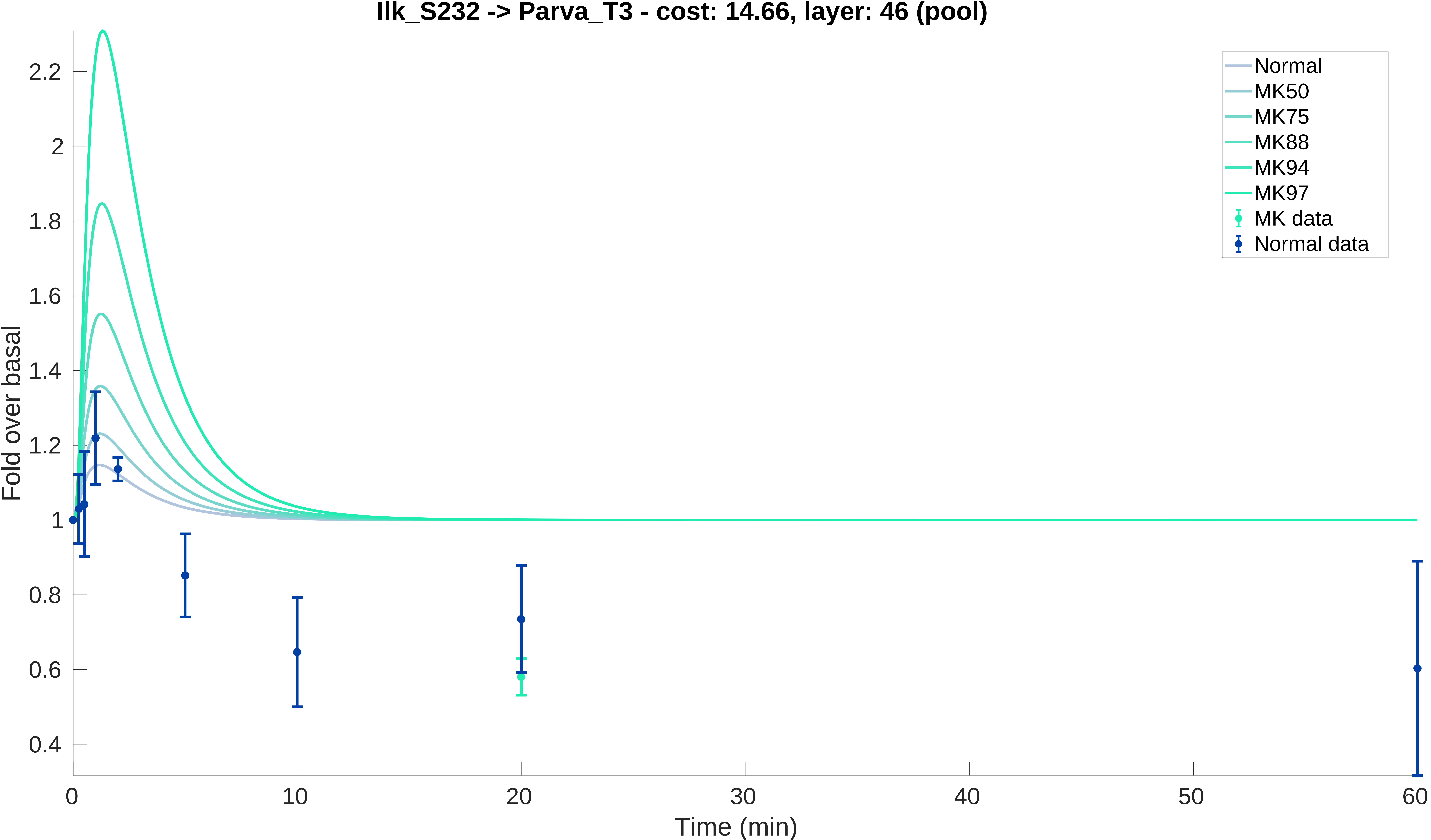

Ilk_S232 -> Parva_T3 - cost: 14.66, layer: 46 (pool)
Fold over basal
Time (min)
0.4
0.6
0.8
1
1.2
1.4
1.6
1.8
2
2.2
0
10
20
30
40
50
60
Normal
MK50
MK75
MK88
MK94
MK97
MK data
Normal data

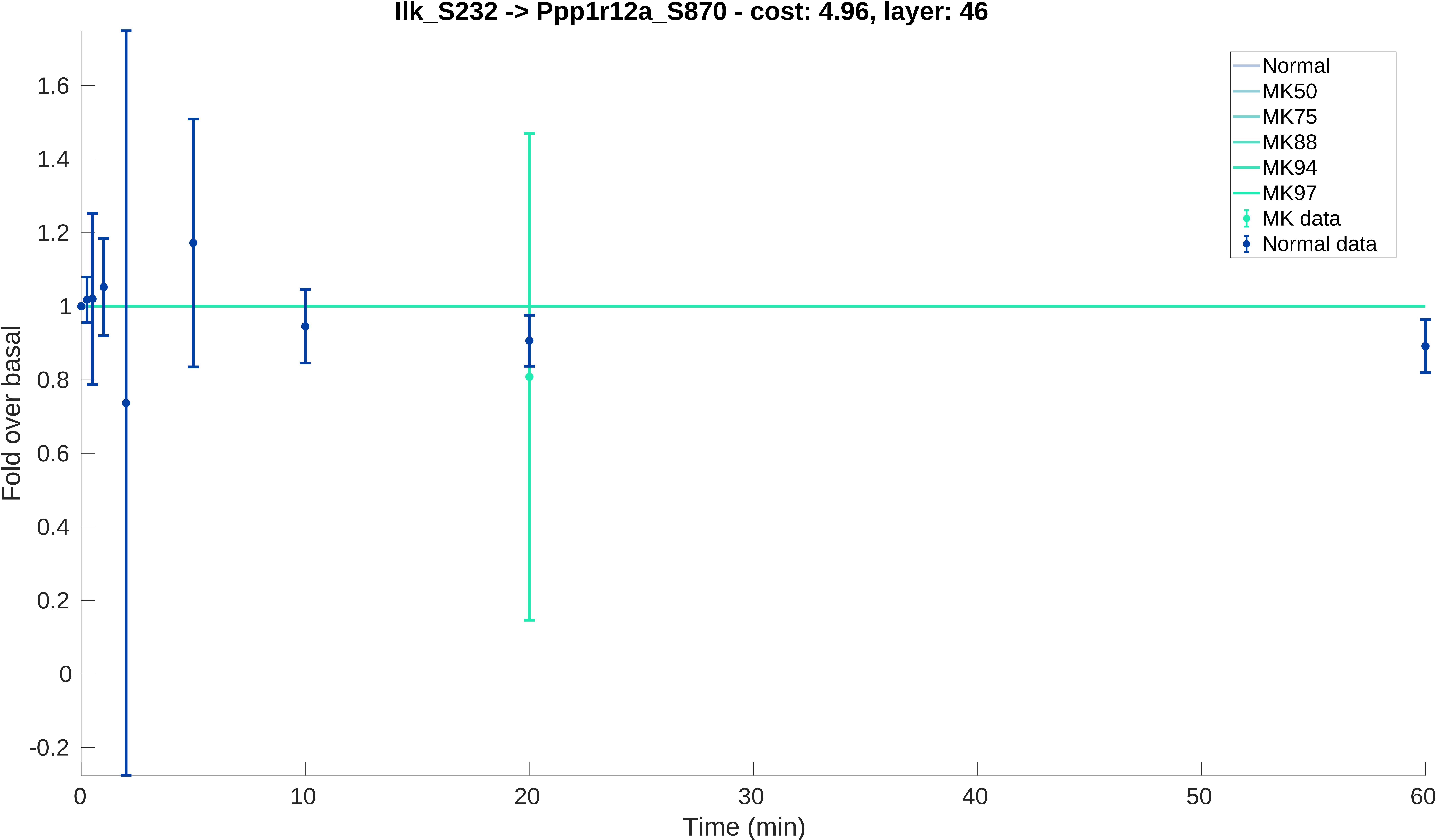

Ilk_S232 -> Ppp1r12a_S870 - cost: 4.96, layer: 46
Normal
MK50
MK75
MK88
MK94
MK97
MK data
Normal data
Fold over basal
1.6
1.4
1.2
1
0.8
0.6
0.4
0.2
0
-0.2
0
10
20
30
40
50
60
Time (min)

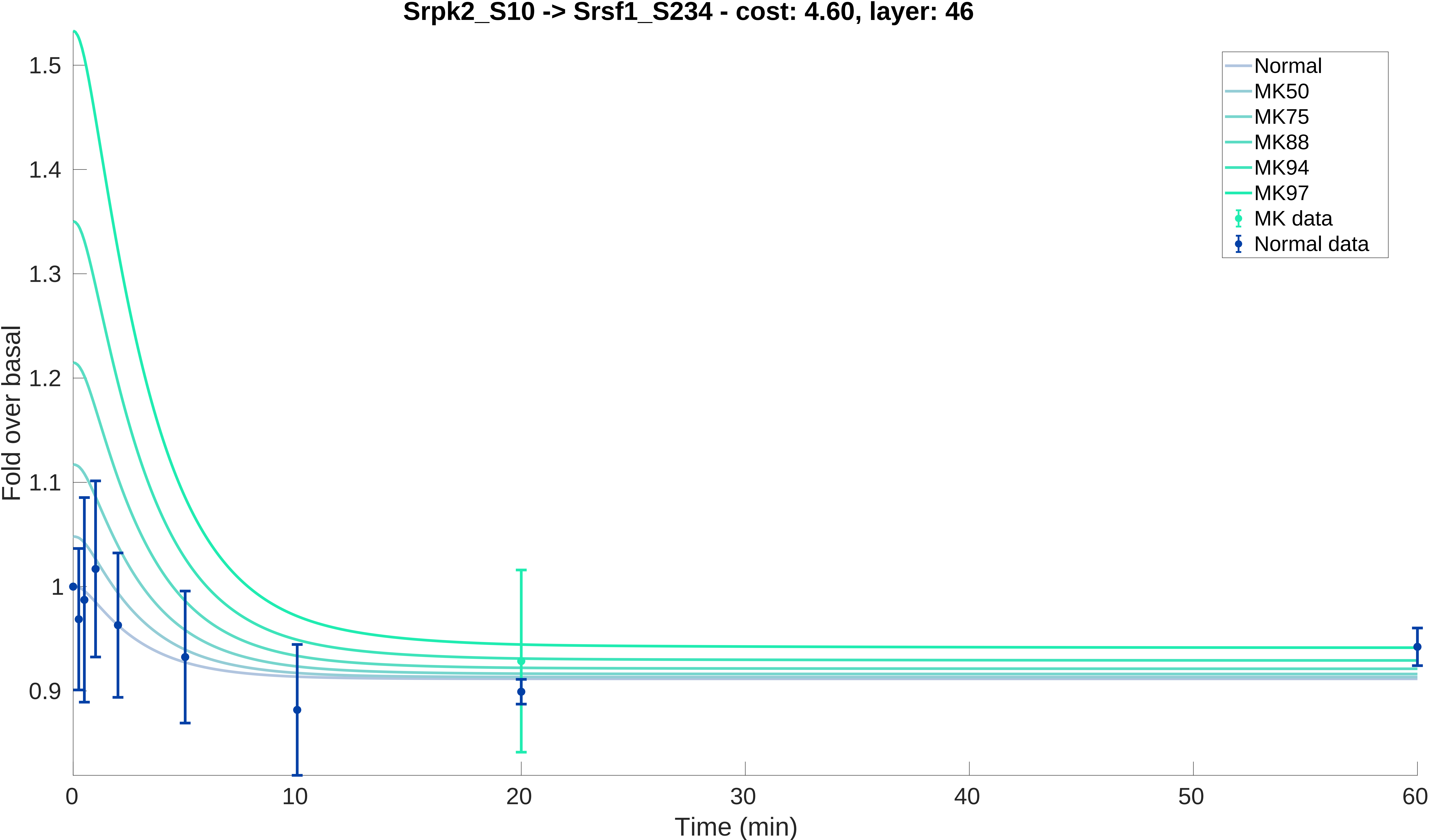

Srpk2_S10 -> Srsf1_S234 - cost: 4.60, layer: 46
Normal
MK50
MK75
MK88
MK94
MK97
MK data
Normal data
Fold over basal
1.5
1.4
1.3
1.2
1.1
1
0.9
0
10
20
30
40
50
60
Time (min)

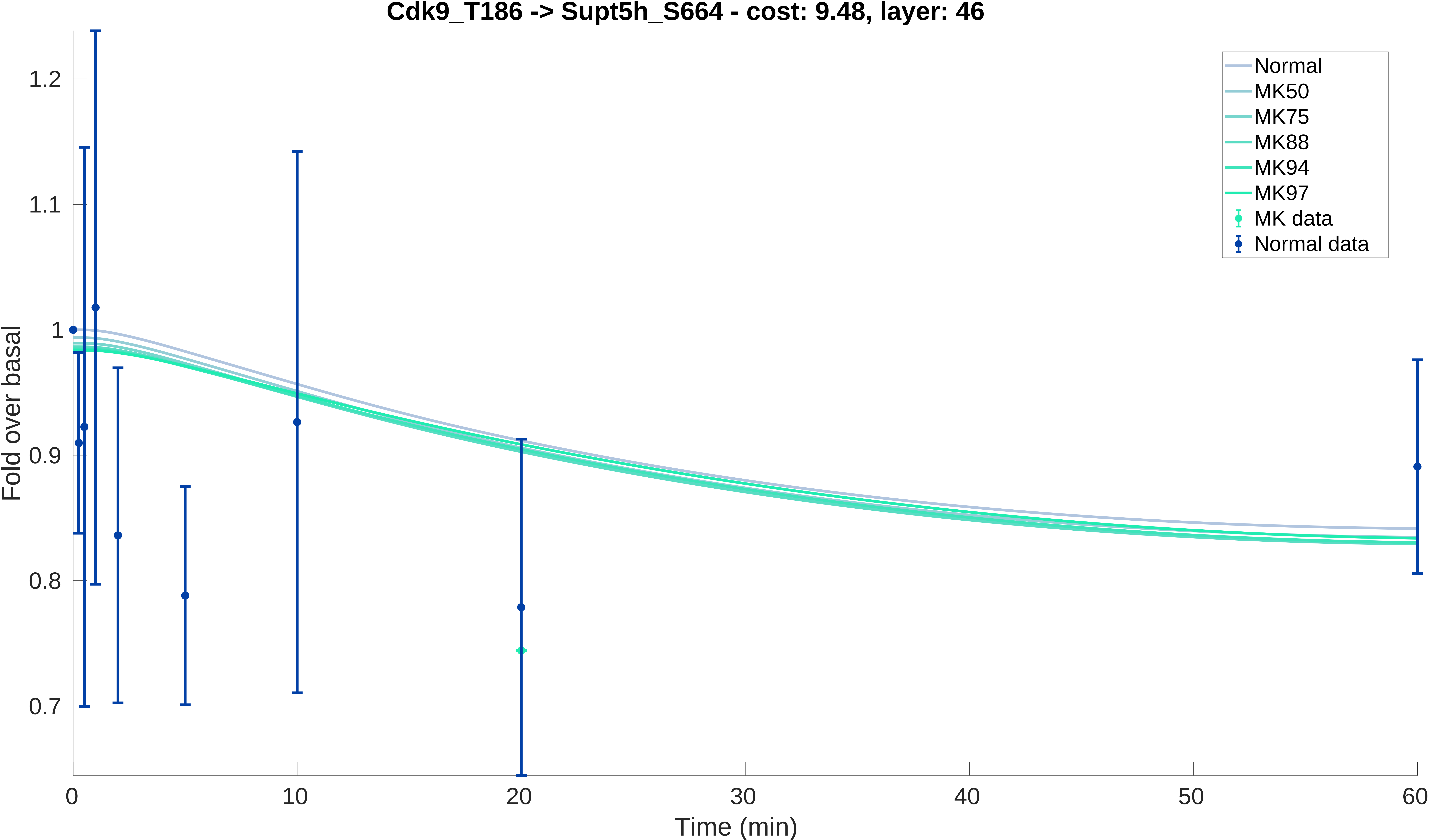

Cdk9_T186 -> Supt5h_S664 - cost: 9.48, layer: 46
Normal
MK50
MK75
MK88
MK94
MK97
MK data
Normal data
Fold over basal
1.2
1.1
1
0.9
0.8
0.7
0
10
20
30
40
50
60
Time (min)

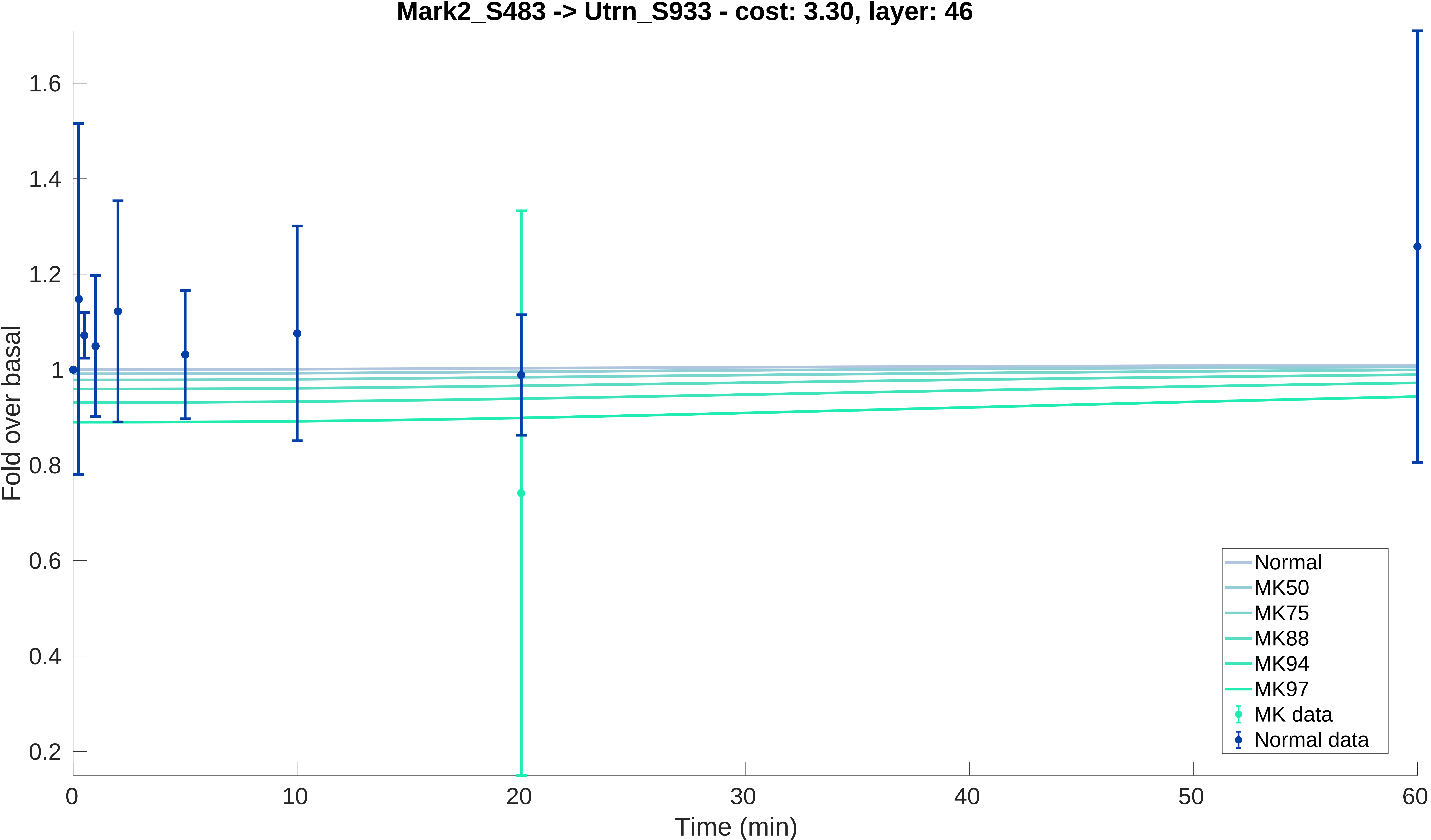
Mark2_S483 -> Utrn_S933 - cost: 3.30, layer: 46
Fold over basal
1.6
1.4
1.2
1
0.8
0.6
0.4
0.2
0
10
20
30
40
50
60
Time (min)
Normal
MK50
MK75
MK88
MK94
MK97
MK data
Normal data

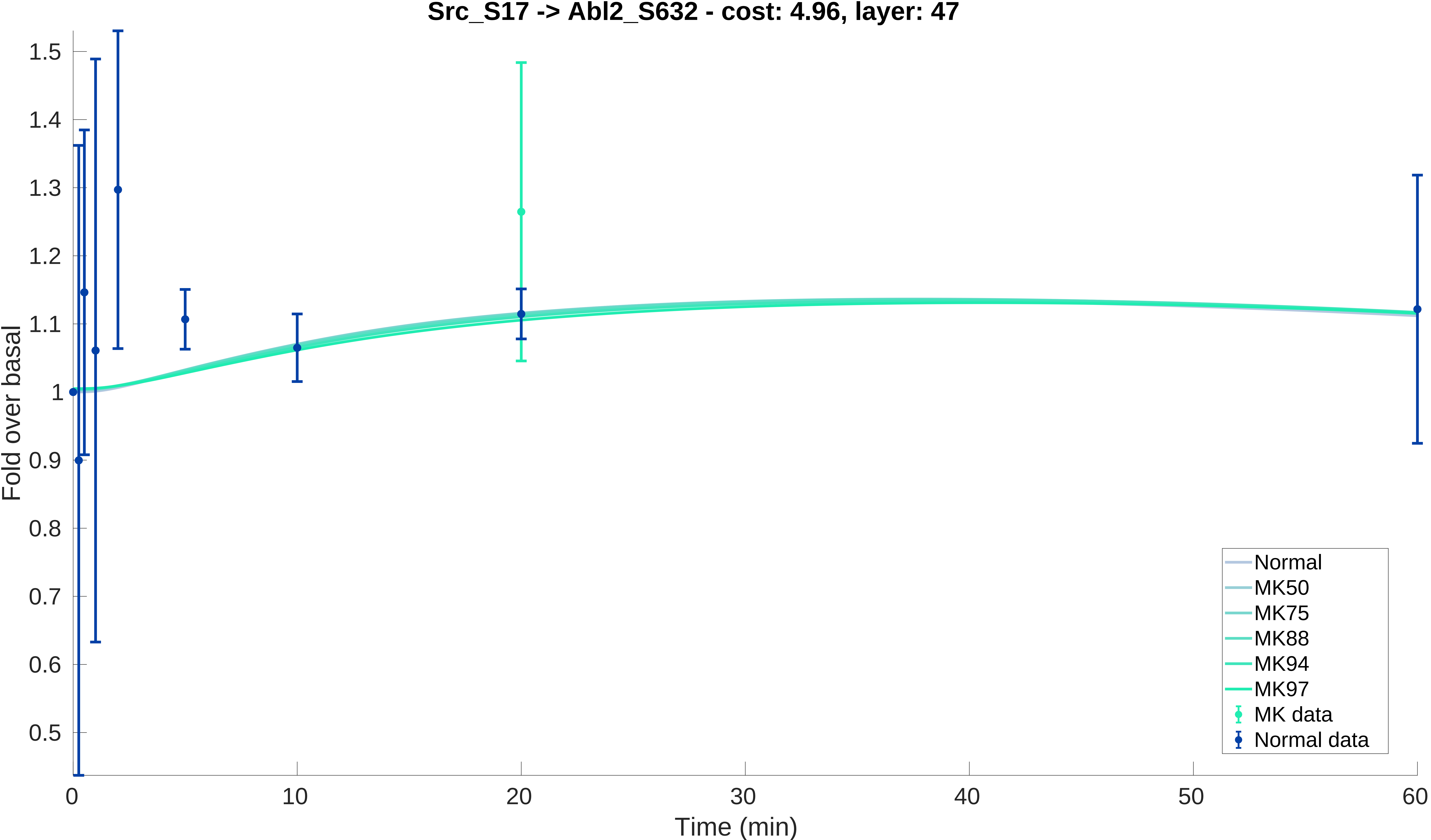

Src_S17 -> Abl2_S632 - cost: 4.96, layer: 47
Fold over basal
1.5
1.4
1.3
1.2
1.1
1
0.9
0.8
0.7
0.6
0.5
0
10
20
30
40
50
60
Time (min)
Normal
MK50
MK75
MK88
MK94
MK97
MK data
Normal data

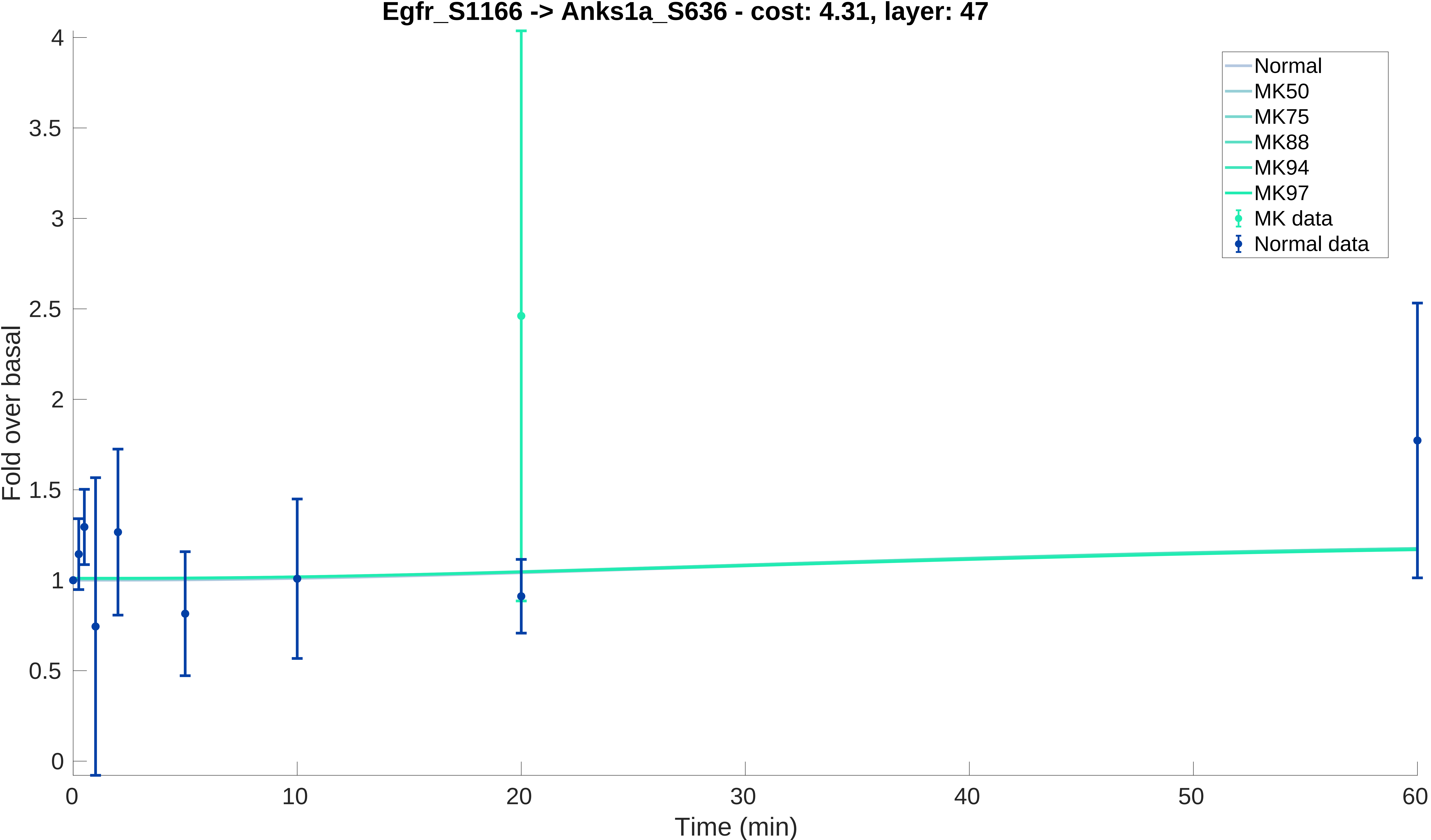
Egfr_S1166 -> Anks1a_S636 - cost: 4.31, layer: 47
Fold over basal
Time (min)
0
0.5
1
1.5
2
2.5
3
3.5
4
0
10
20
30
40
50
60
Normal
MK50
MK75
MK88
MK94
MK97
MK data
Normal data

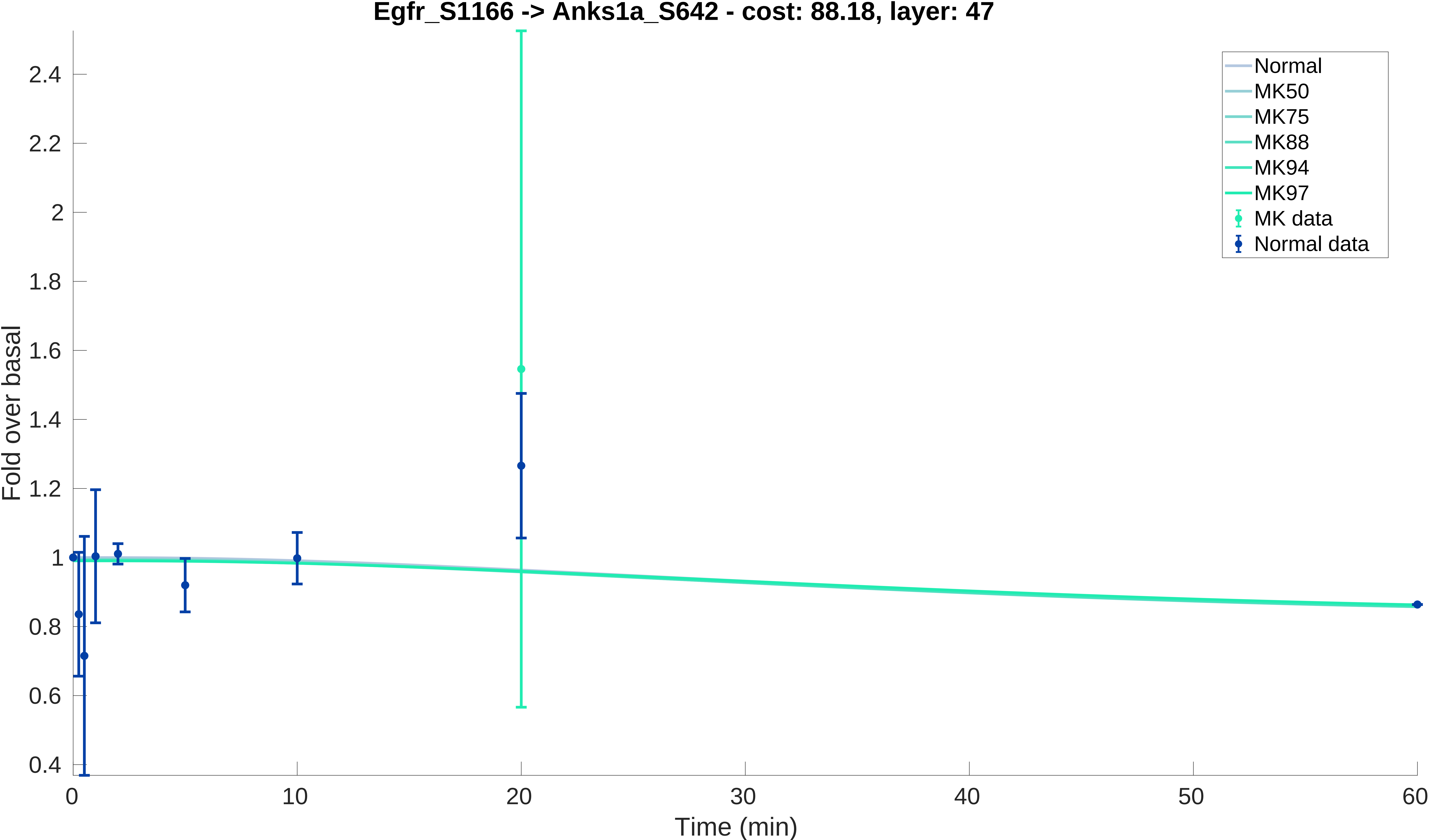

Egfr_S1166 -> Anks1a_S642 - cost: 88.18, layer: 47
Fold over basal
Time (min)
0.4
0.6
0.8
1
1.2
1.4
1.6
1.8
2
2.2
2.4
0
10
20
30
40
50
60
Normal
MK50
MK75
MK88
MK94
MK97
MK data
Normal data

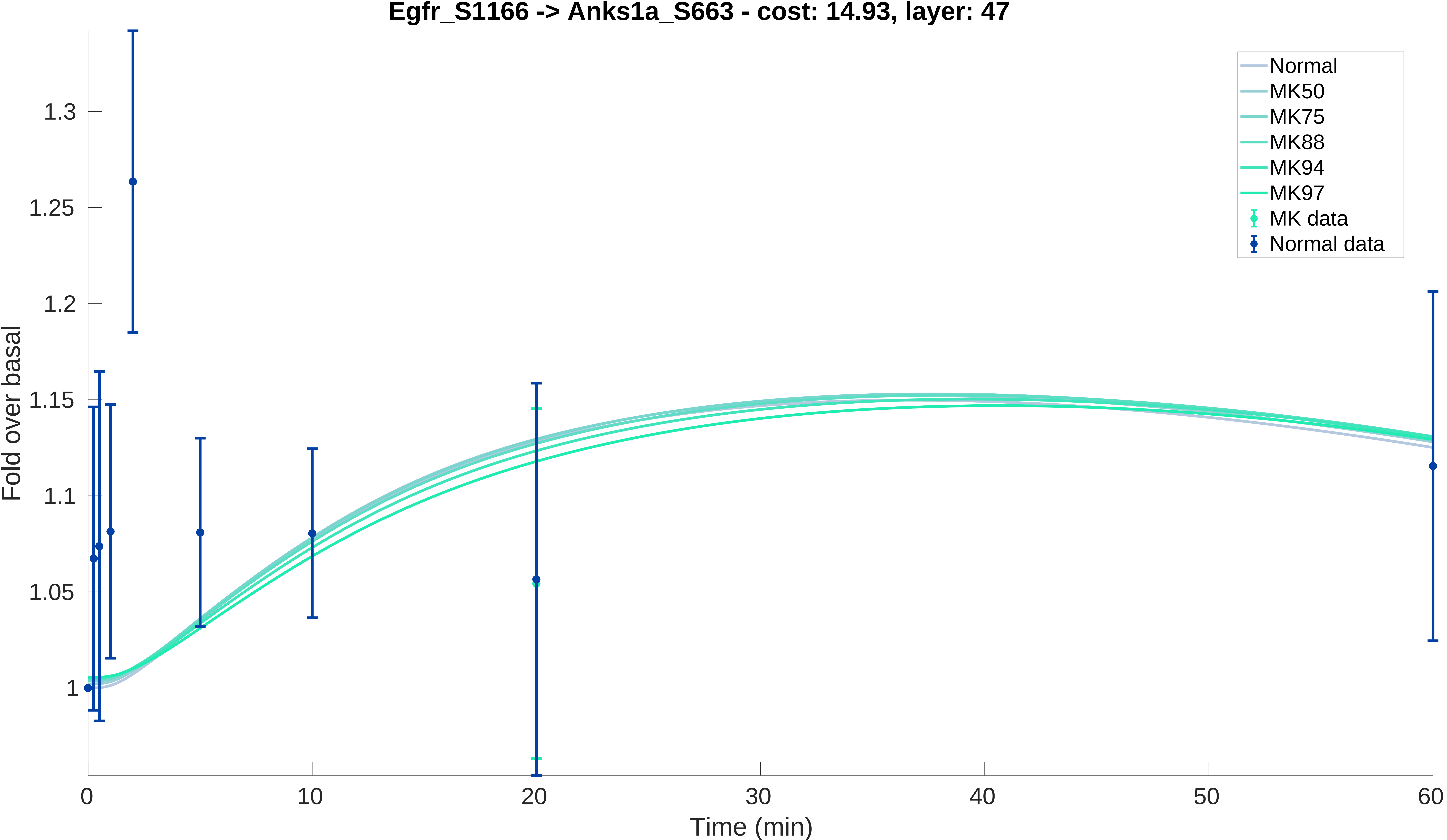
Egfr_S1166 -> Anks1a_S663 - cost: 14.93, layer: 47
Fold over basal
Time (min)
1.3
1.25
1.2
1.15
1.1
1.05
1
0
10
20
30
40
50
60
Normal
MK50
MK75
MK88
MK94
MK97
MK data
Normal data

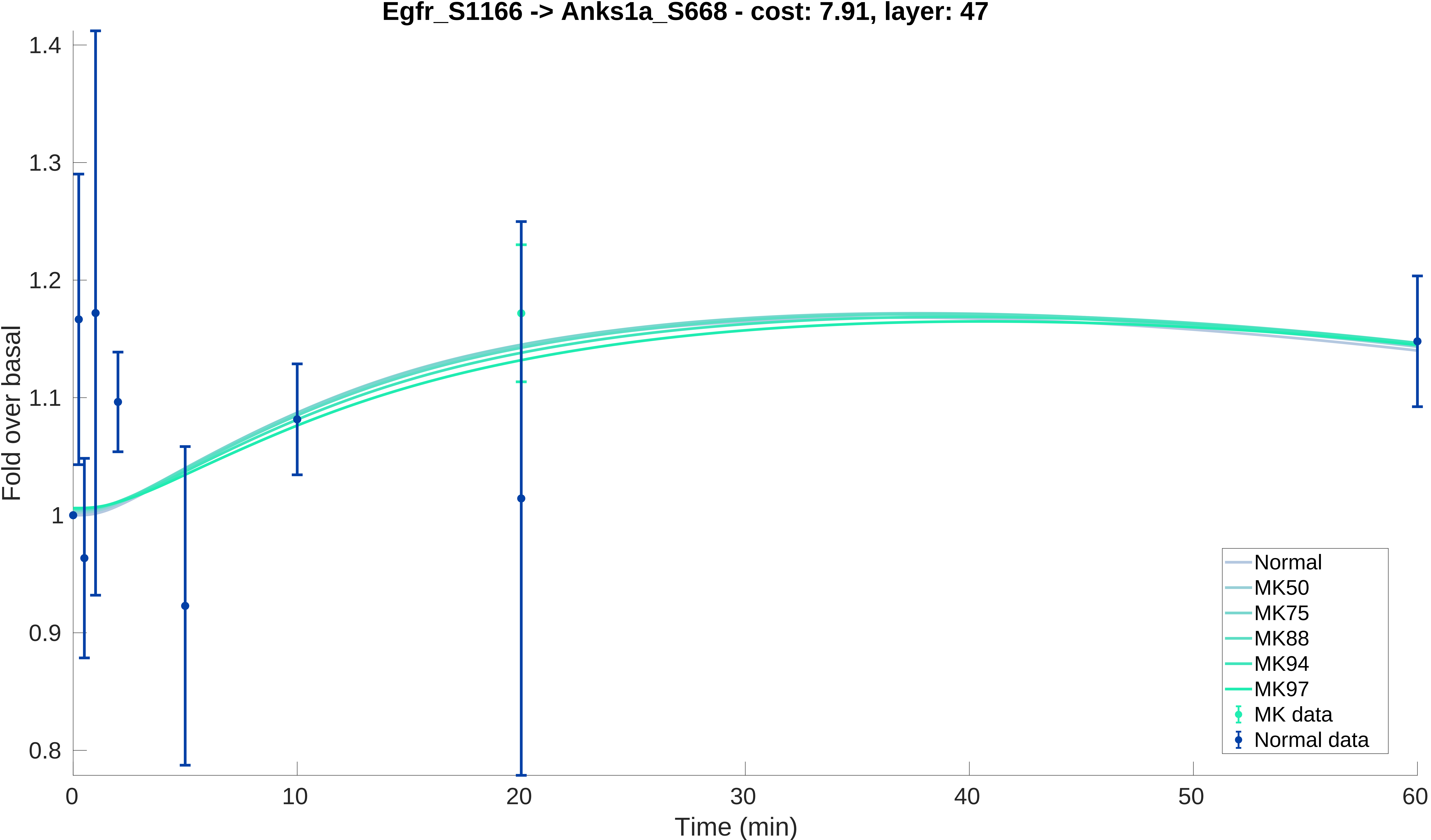

Egfr_S1166 -> Anks1a_S668 - cost: 7.91, layer: 47
Fold over basal
Time (min)
1.4
1.3
1.2
1.1
1
0.9
0.8
0
10
20
30
40
50
60
Normal
MK50
MK75
MK88
MK94
MK97
MK data
Normal data

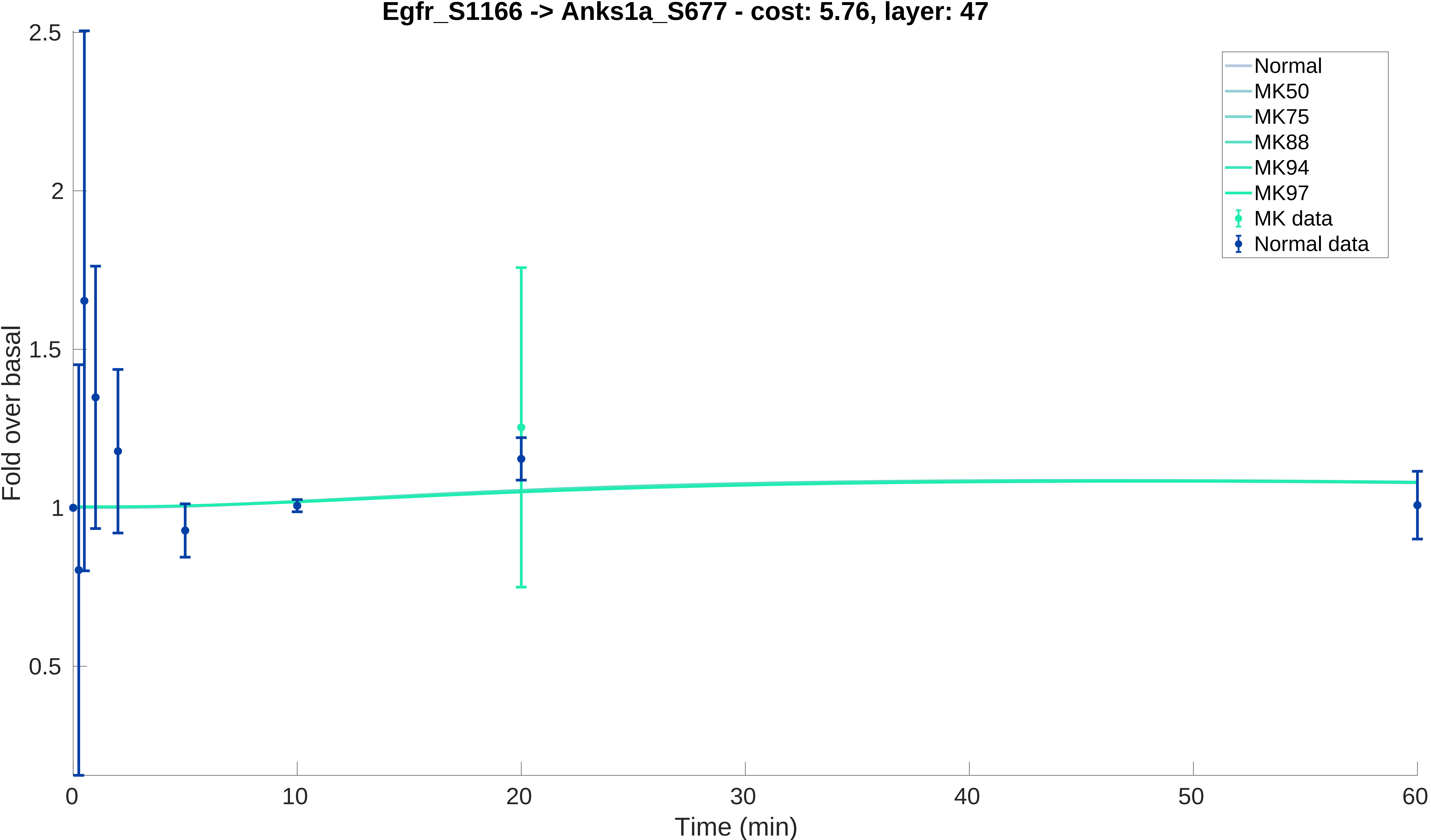

Egfr_S1166 -> Anks1a_S677 - cost: 5.76, layer: 47
Fold over basal
2.5
2
1.5
1
0.5
0
10
20
30
40
50
60
Time (min)
Normal
MK50
MK75
MK88
MK94
MK97
MK data
Normal data

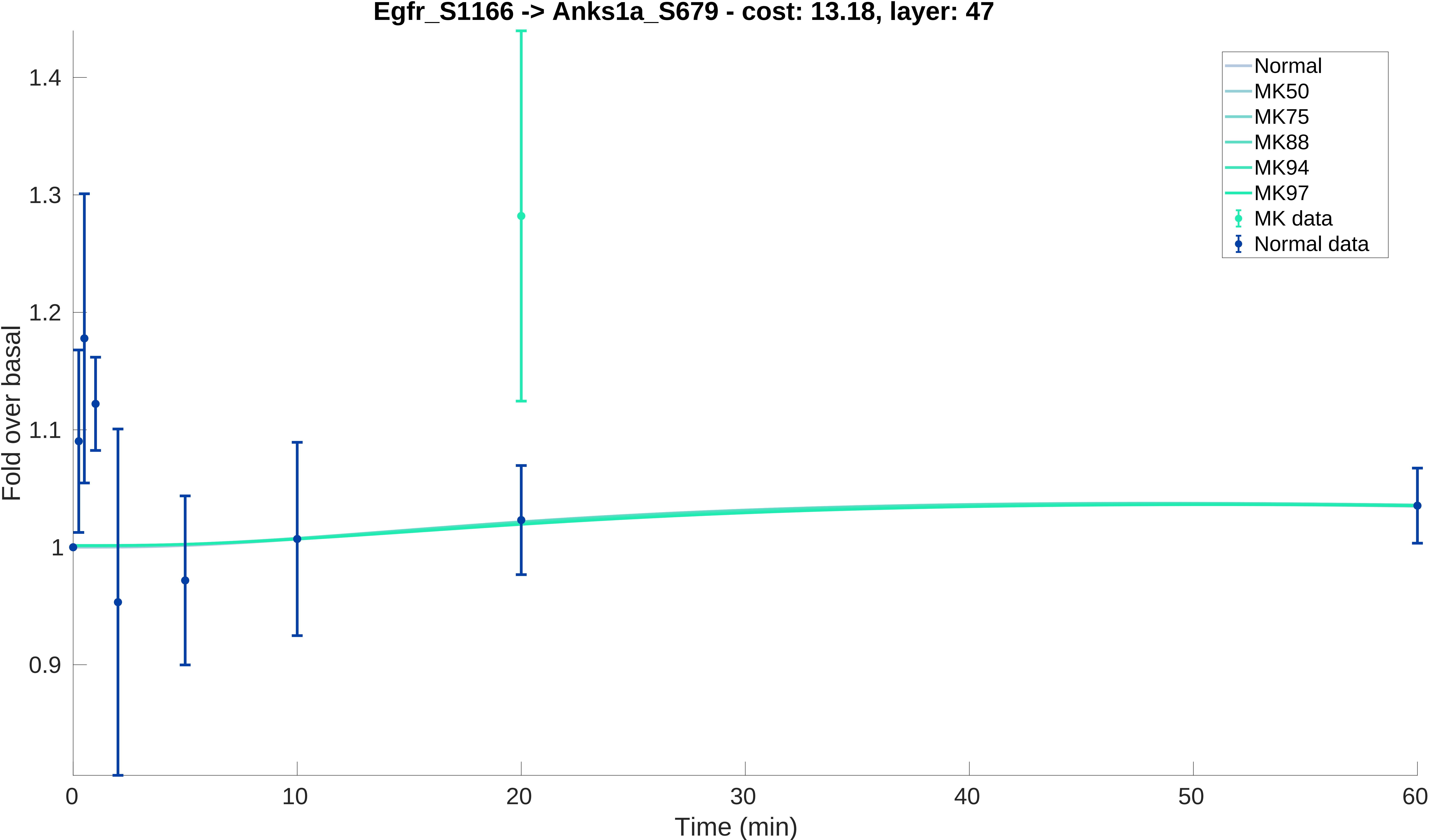

Egfr_S1166 -> Anks1a_S679 - cost: 13.18, layer: 47
Fold over basal
Time (min)
1.4
1.3
1.2
1.1
1
0.9
0
10
20
30
40
50
60
Normal
MK50
MK75
MK88
MK94
MK97
MK data
Normal data

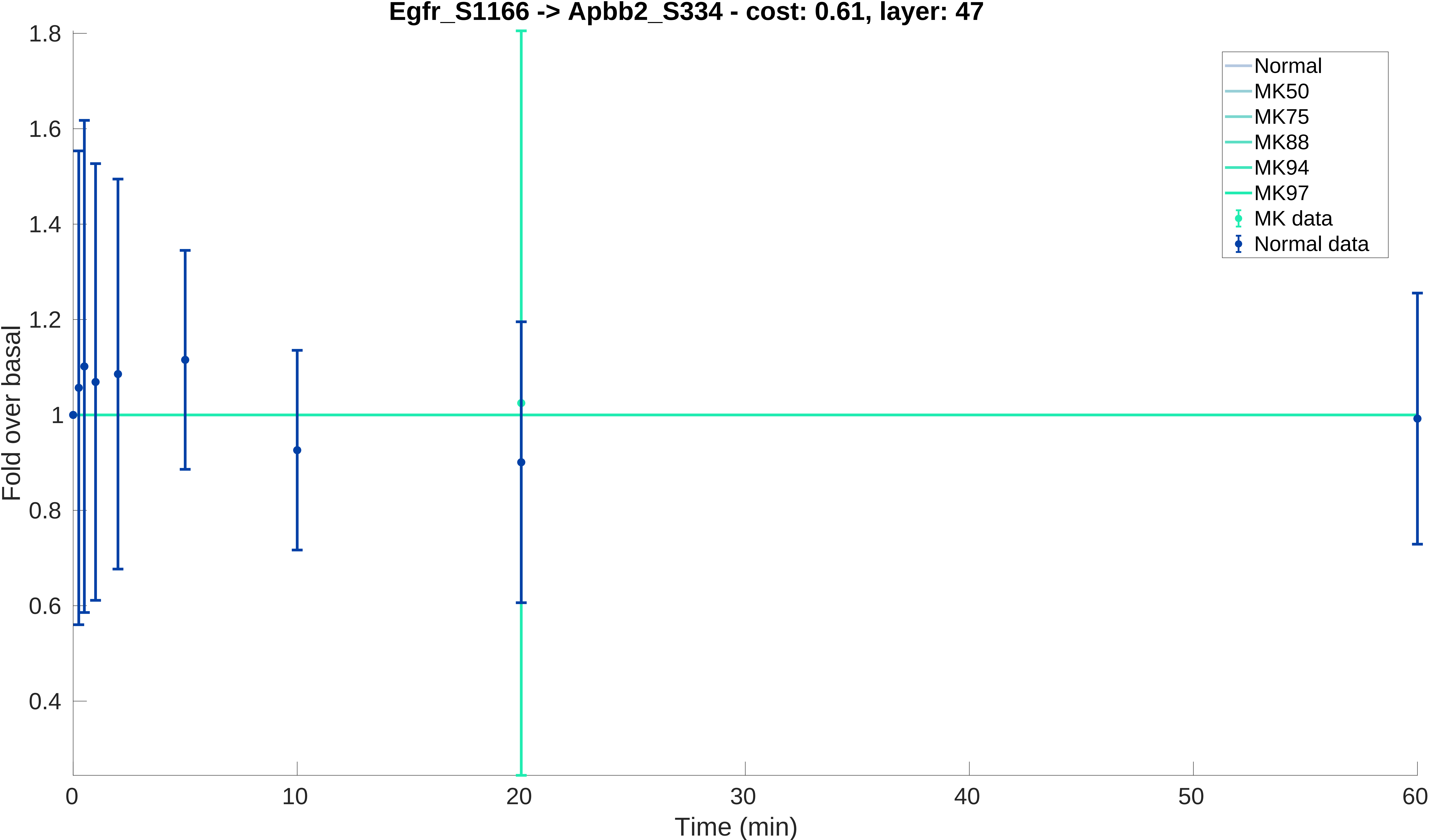

Egfr_S1166 -> Apbb2_S334 - cost: 0.61, layer: 47
Fold over basal
Time (min)
1.8
1.6
1.4
1.2
1
0.8
0.6
0.4
0
10
20
30
40
50
60
Normal
MK50
MK75
MK88
MK94
MK97
MK data
Normal data

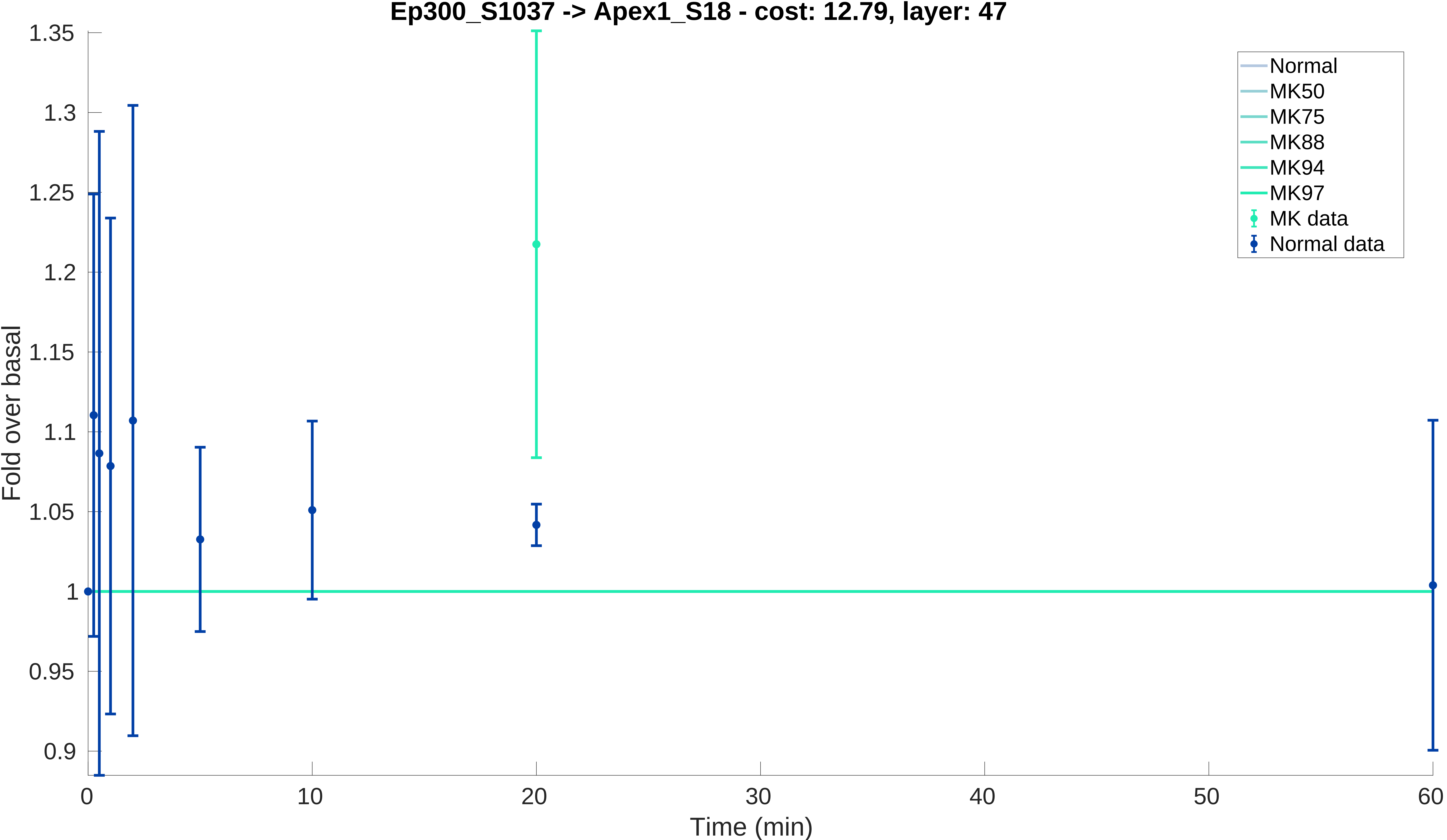
Ep300_S1037 -> Apex1_S18 - cost: 12.79, layer: 47
Fold over basal
Time (min)
1.35
1.3
1.25
1.2
1.15
1.1
1.05
1
0.95
0.9
0
10
20
30
40
50
60
Normal
MK50
MK75
MK88
MK94
MK97
MK data
Normal data

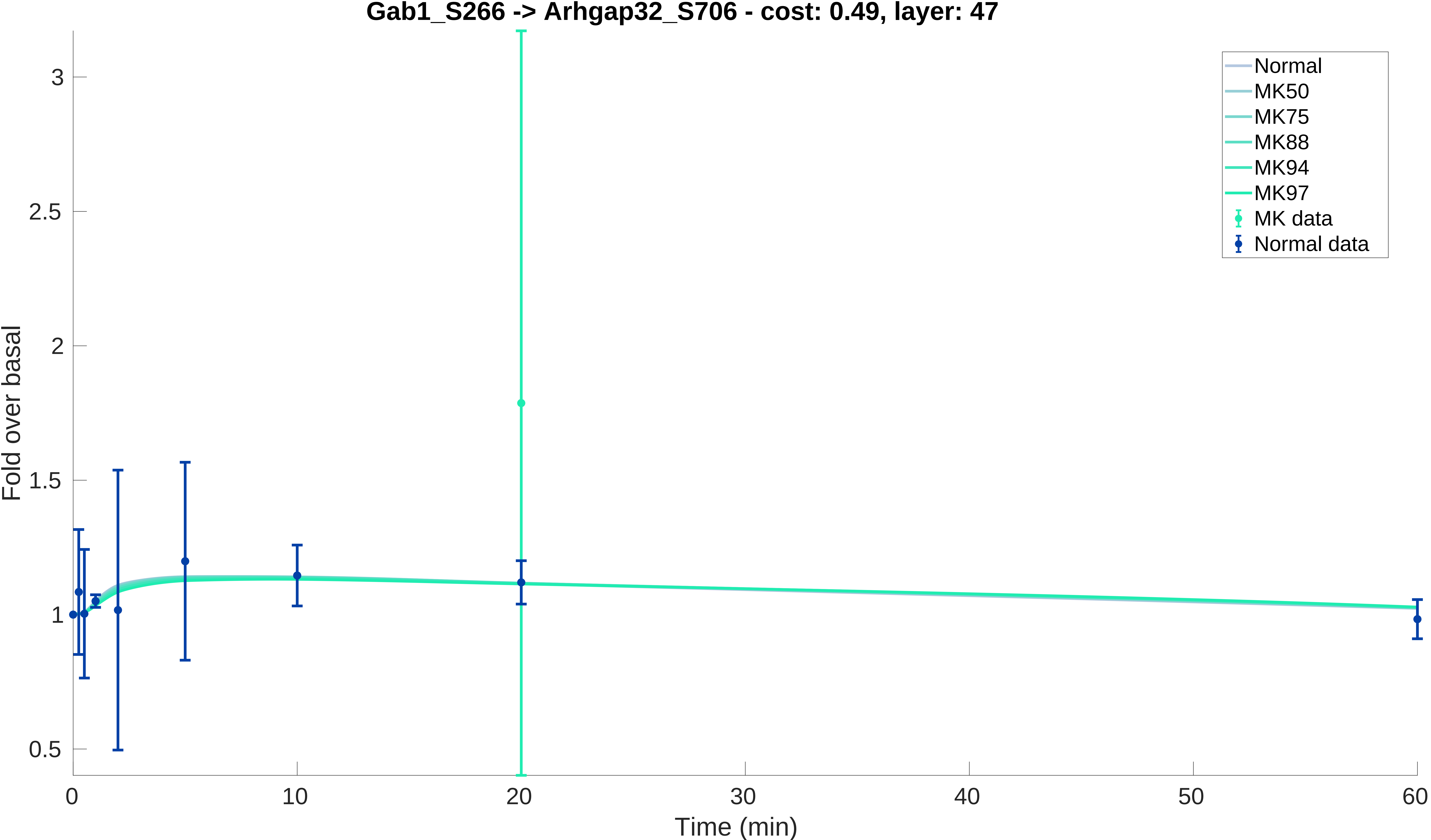
Gab1_S266 -> Arhgap32_S706 - cost: 0.49, layer: 47
Fold over basal
Time (min)
0.5
1
1.5
2
2.5
3
0
10
20
30
40
50
60
Normal
MK50
MK75
MK88
MK94
MK97
MK data
Normal data

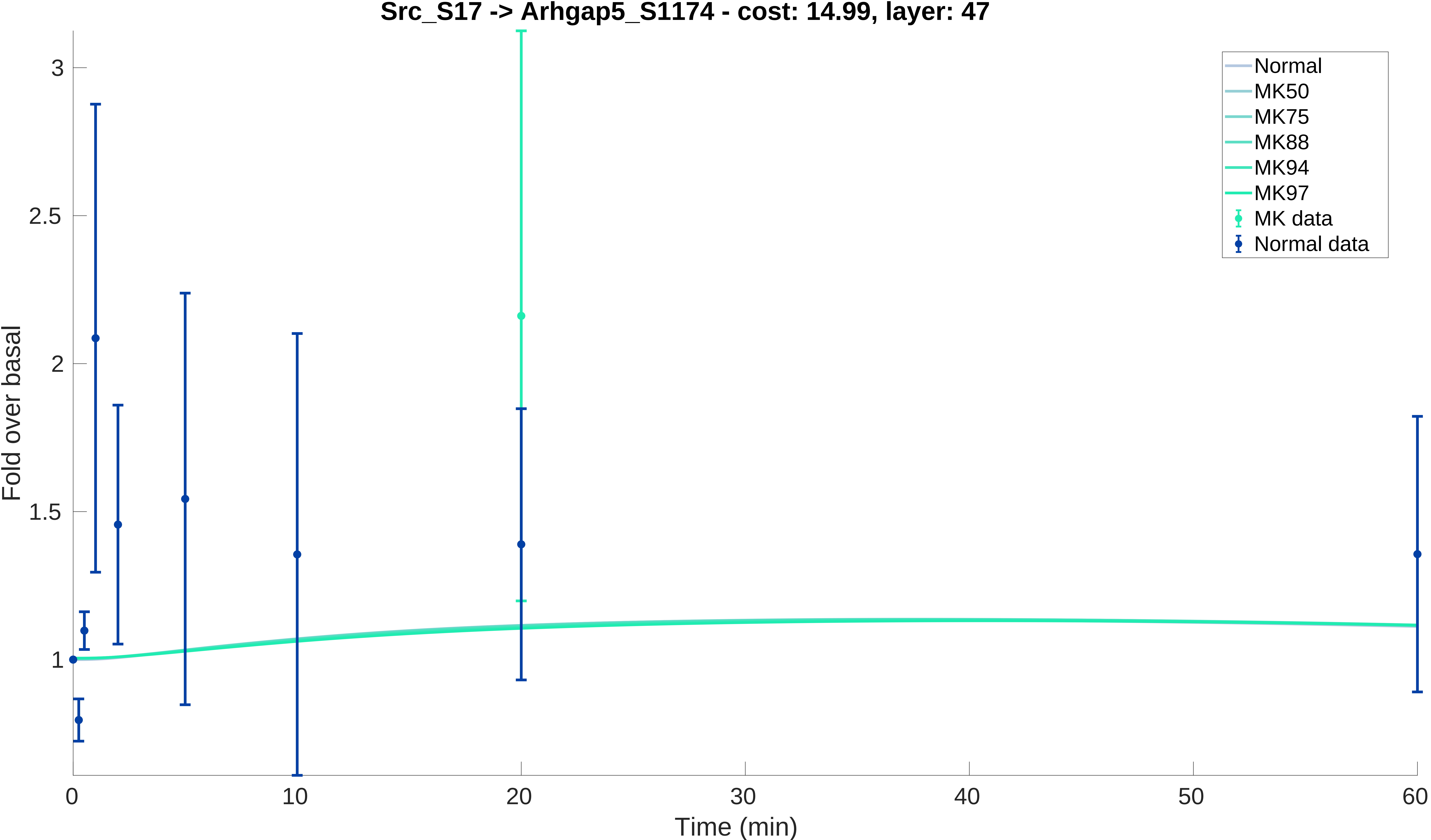

Src_S17 -> Arhgap5_S1174 - cost: 14.99, layer: 47
Fold over basal
Time (min)
3
2.5
2
1.5
1
0
10
20
30
40
50
60
Normal
MK50
MK75
MK88
MK94
MK97
MK data
Normal data

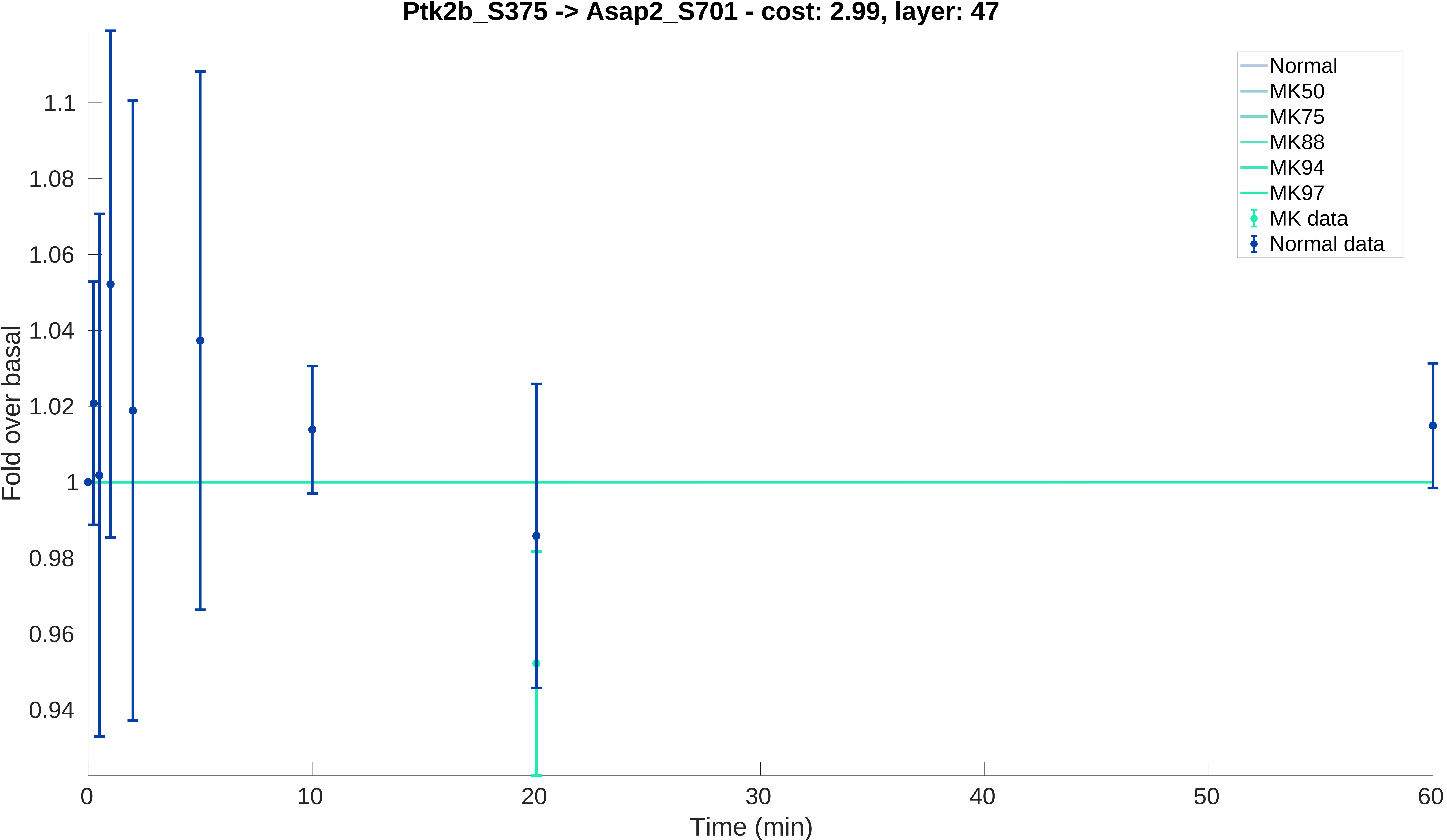

Ptk2b_S375 -> Asap2_S701 - cost: 2.99, layer: 47
Fold over basal
1.1
1.08
1.06
1.04
1.02
1
0.98
0.96
0.94
0
10
20
30
40
50
60
Time (min)
Normal
MK50
MK75
MK88
MK94
MK97
MK data
Normal data

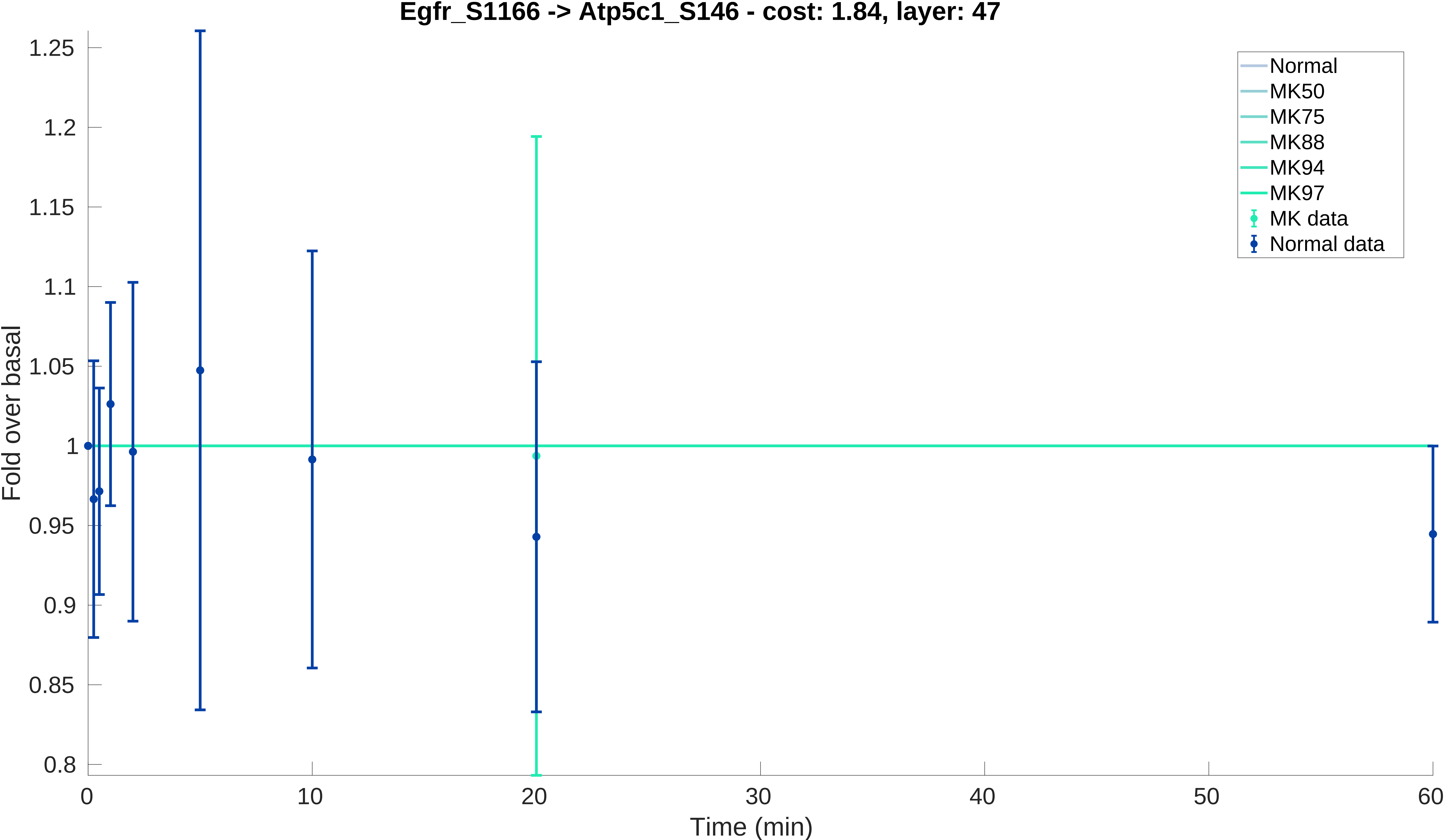

Egfr_S1166 -> Atp5c1_S146 - cost: 1.84, layer: 47
Fold over basal
Time (min)
0
10
20
30
40
50
60
0.8
0.85
0.9
0.95
1
1.05
1.1
1.15
1.2
1.25
Normal
MK50
MK75
MK88
MK94
MK97
MK data
Normal data

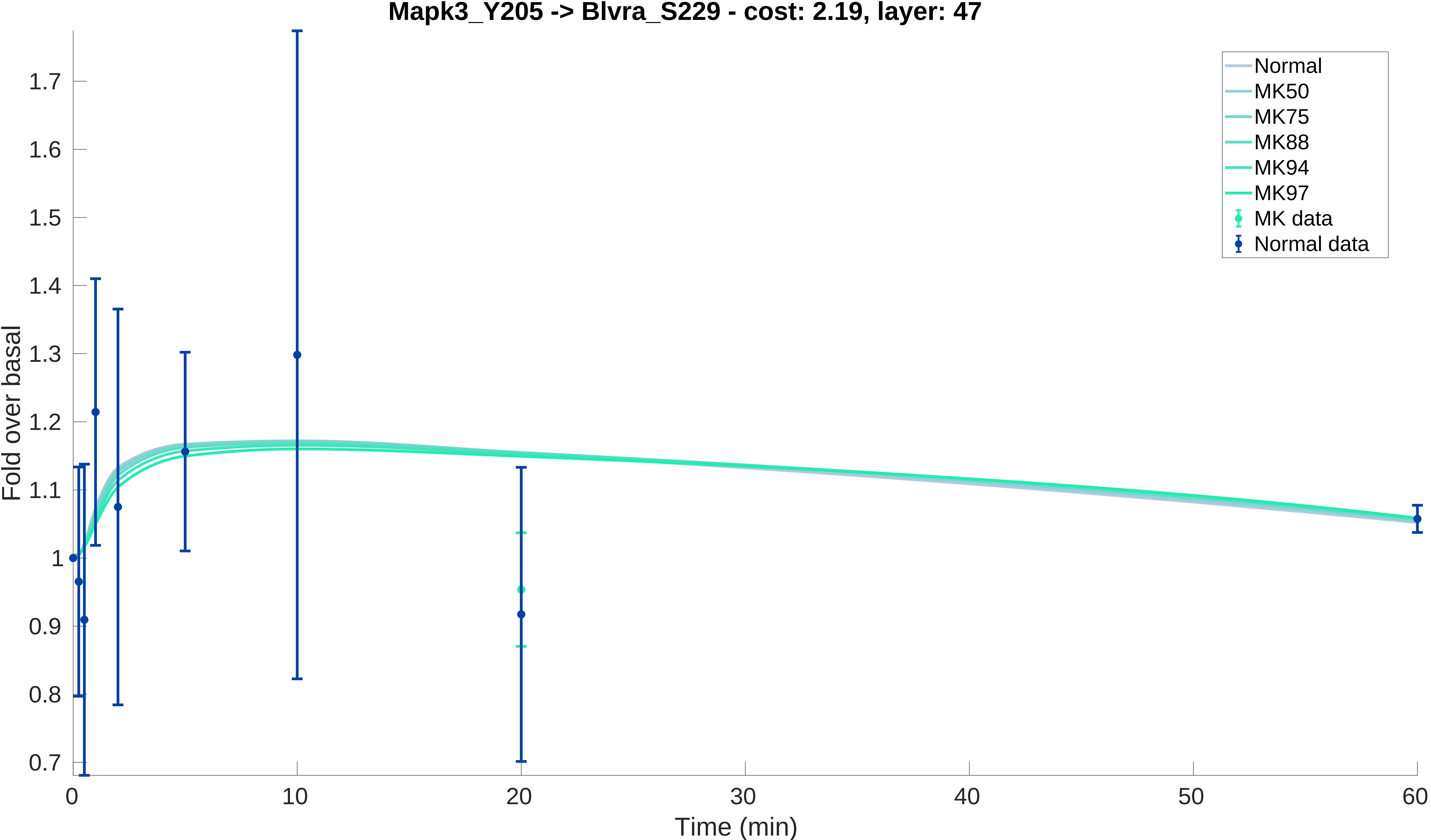
Mapk3_Y205 -> Blvra_S229 - cost: 2.19, layer: 47
Fold over basal
Time (min)
Normal
MK50
MK75
MK88
MK94
MK97
MK data
Normal data
0.7
0.8
0.9
1
1.1
1.2
1.3
1.4
1.5
1.6
1.7
0
10
20
30
40
50
60

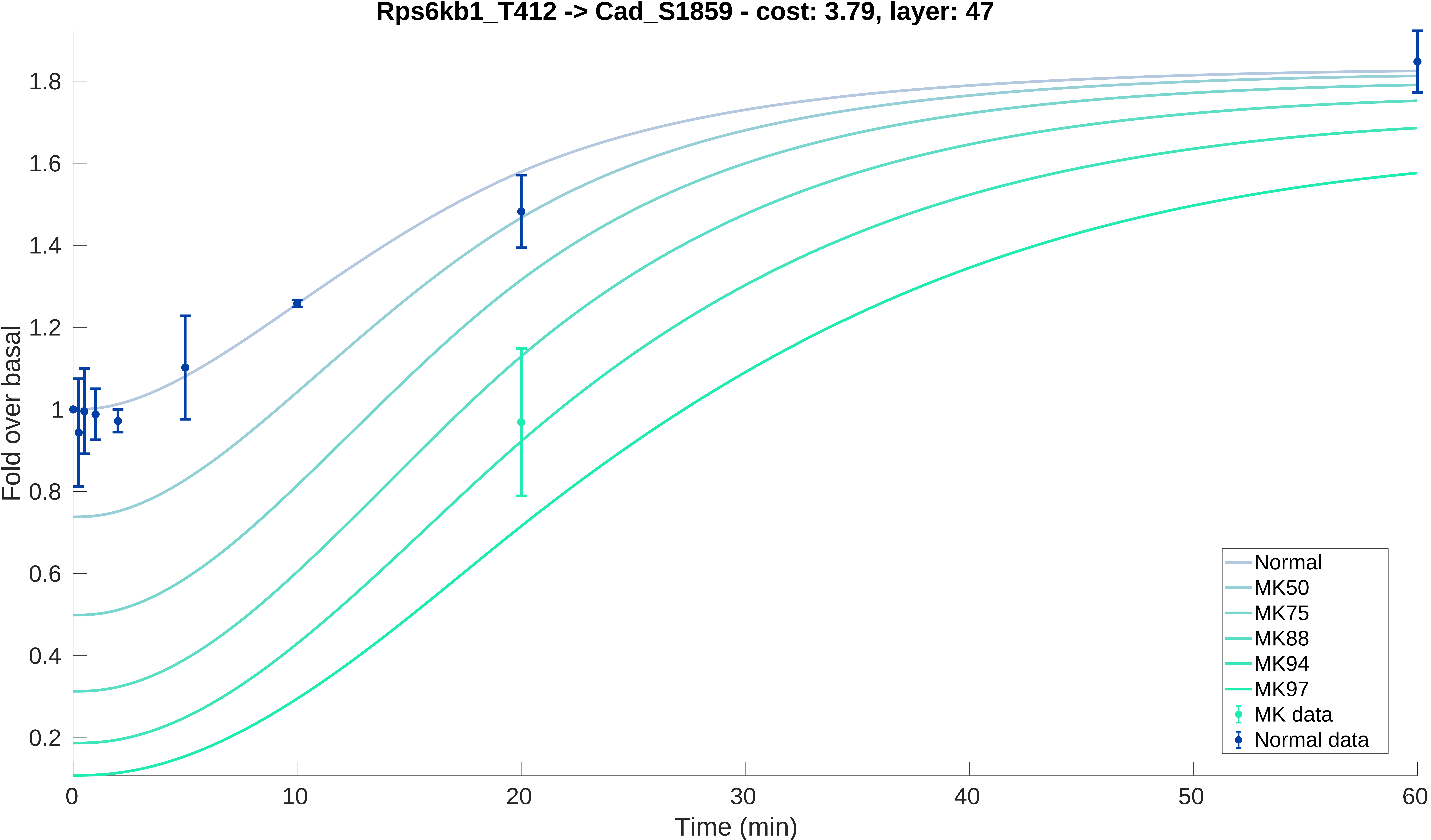
Rps6kb1_T412 -> Cad_S1859 - cost: 3.79, layer: 47
Fold over basal
Time (min)
0
10
20
30
40
50
60
0.2
0.4
0.6
0.8
1
1.2
1.4
1.6
1.8
Normal
MK50
MK75
MK88
MK94
MK97
MK data
Normal data

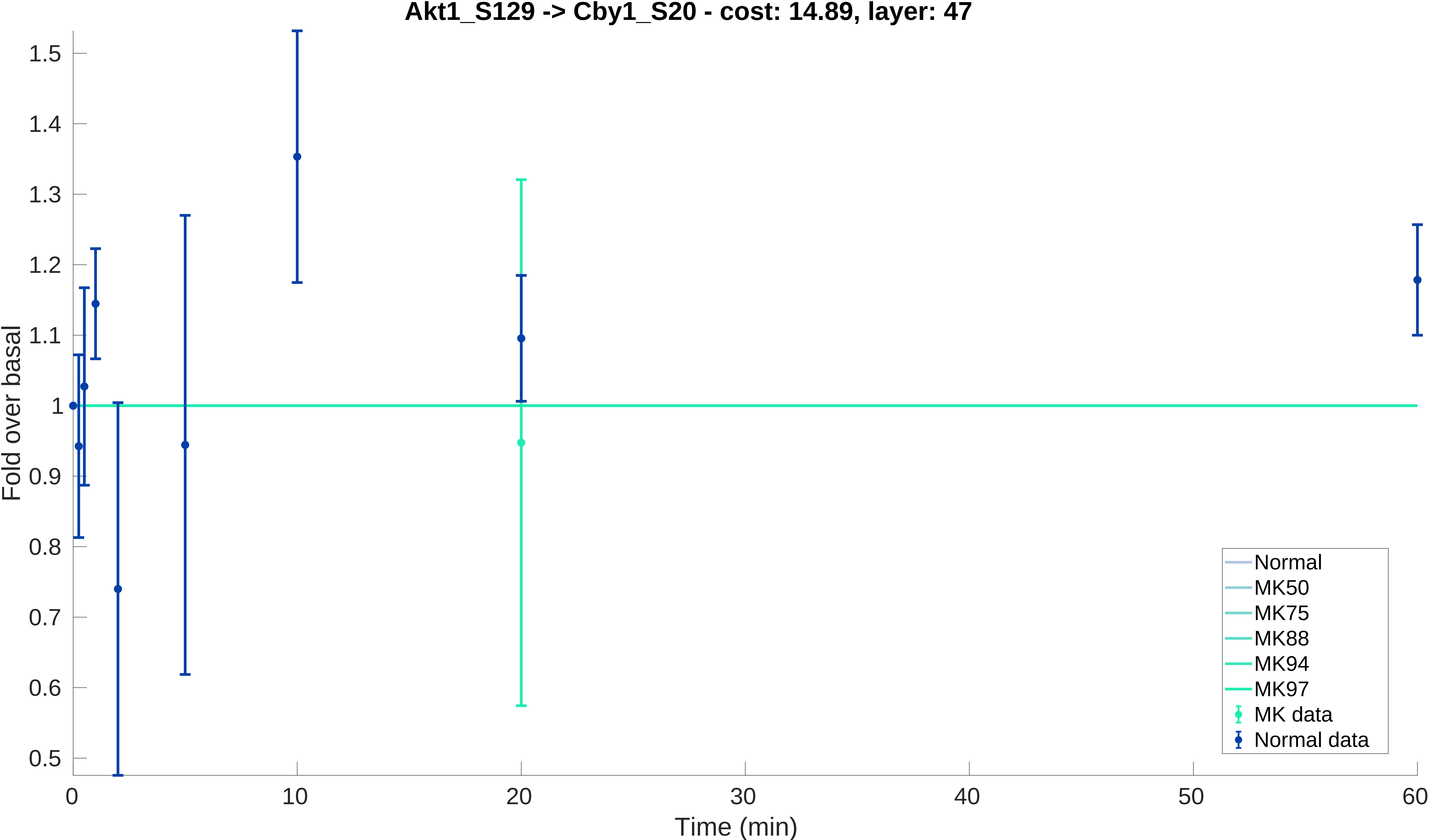

Akt1_S129 -> Cby1_S20 - cost: 14.89, layer: 47
Fold over basal
1.5
1.4
1.3
1.2
1.1
1
0.9
0.8
0.7
0.6
0.5
0
10
20
30
40
50
60
Time (min)
Normal
MK50
MK75
MK88
MK94
MK97
MK data
Normal data

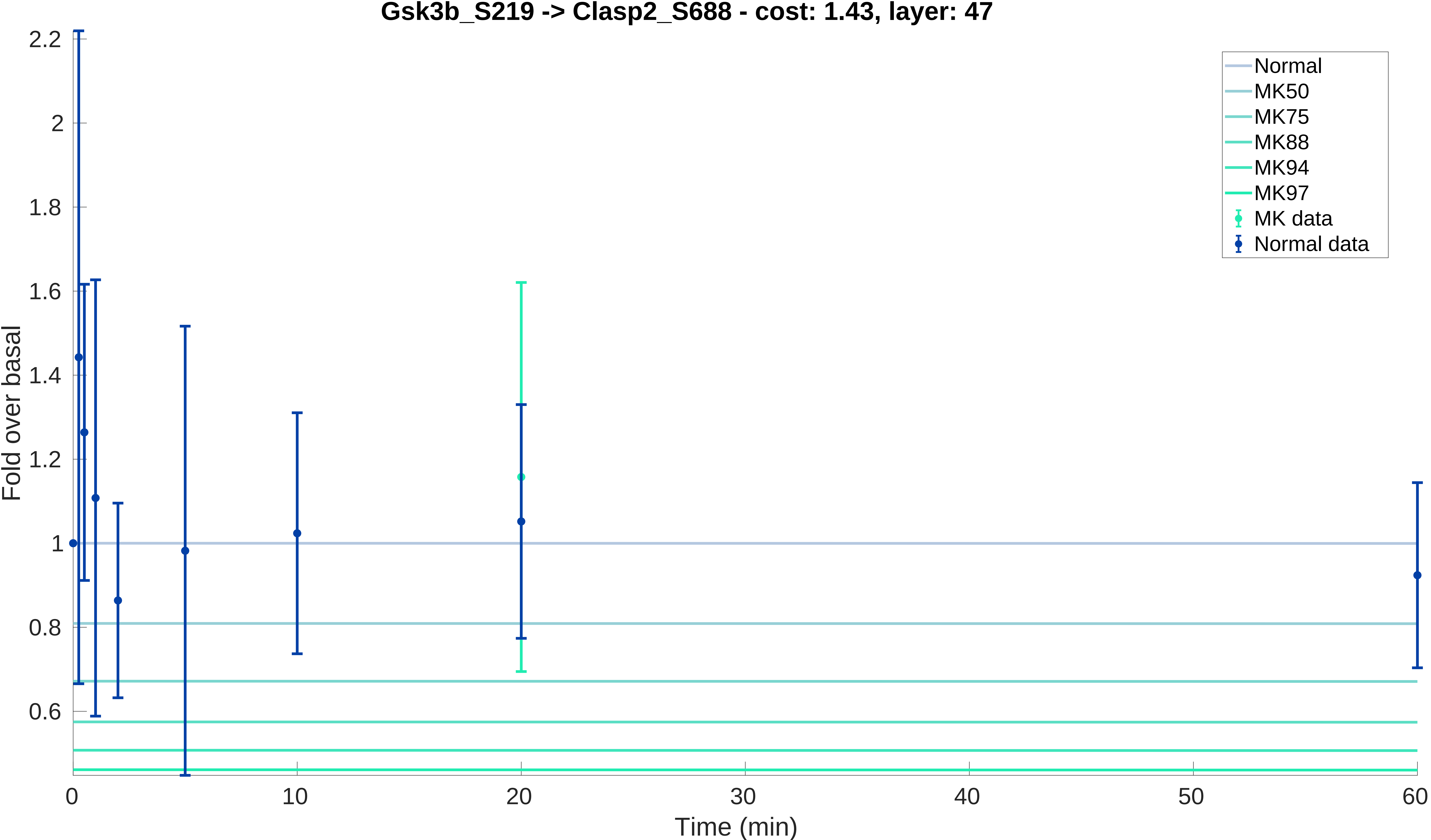

Gsk3b_S219 -> Clasp2_S688 - cost: 1.43, layer: 47
Fold over basal
2.2
2
1.8
1.6
1.4
1.2
1
0.8
0.6
0
10
20
30
40
50
60
Time (min)
Normal
MK50
MK75
MK88
MK94
MK97
MK data
Normal data

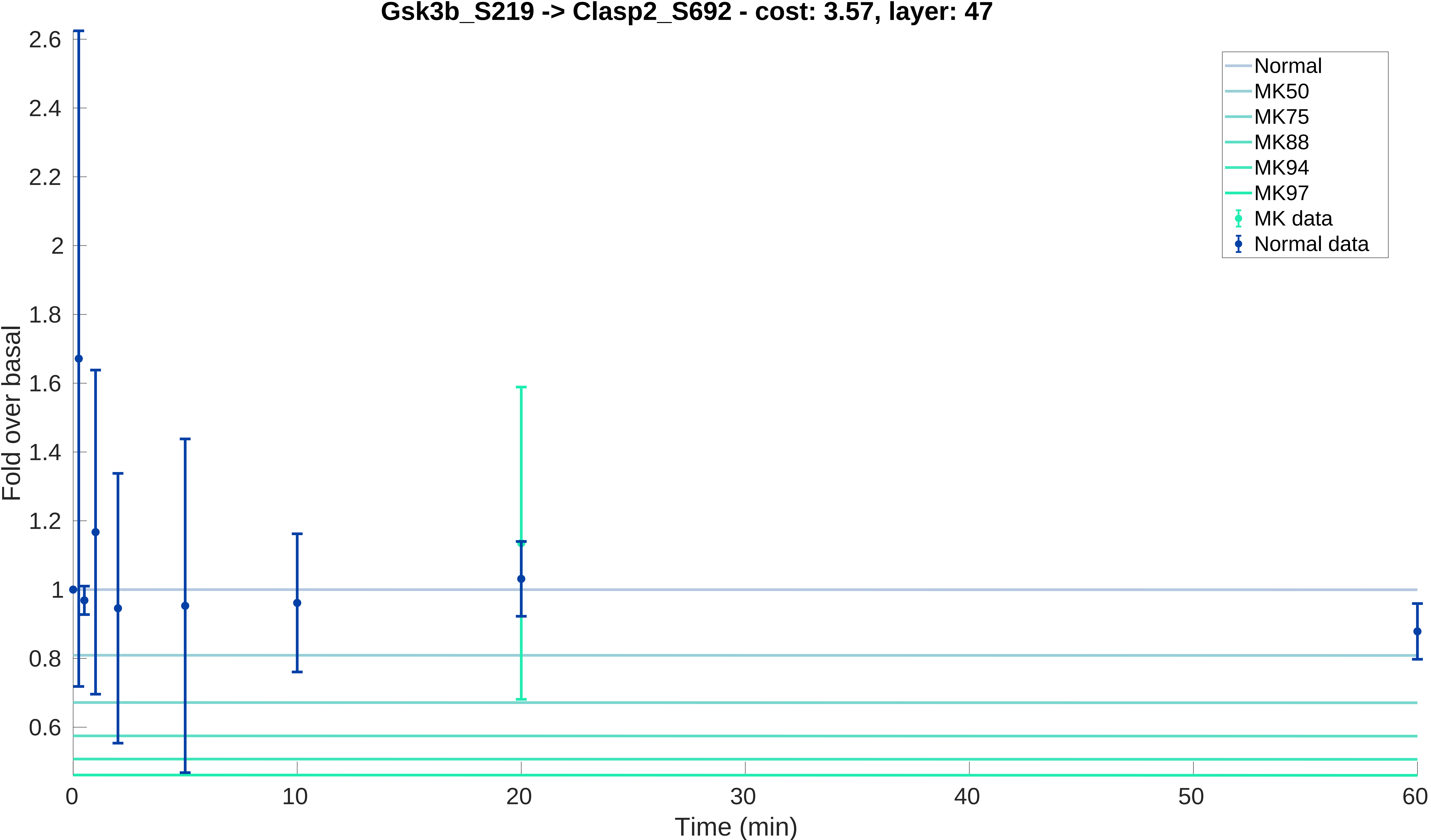
Gsk3b_S219 -> Clasp2_S692 - cost: 3.57, layer: 47
Fold over basal
2.6
2.4
2.2
2
1.8
1.6
1.4
1.2
1
0.8
0.6
0
10
20
30
40
50
60
Time (min)
Normal
MK50
MK75
MK88
MK94
MK97
MK data
Normal data

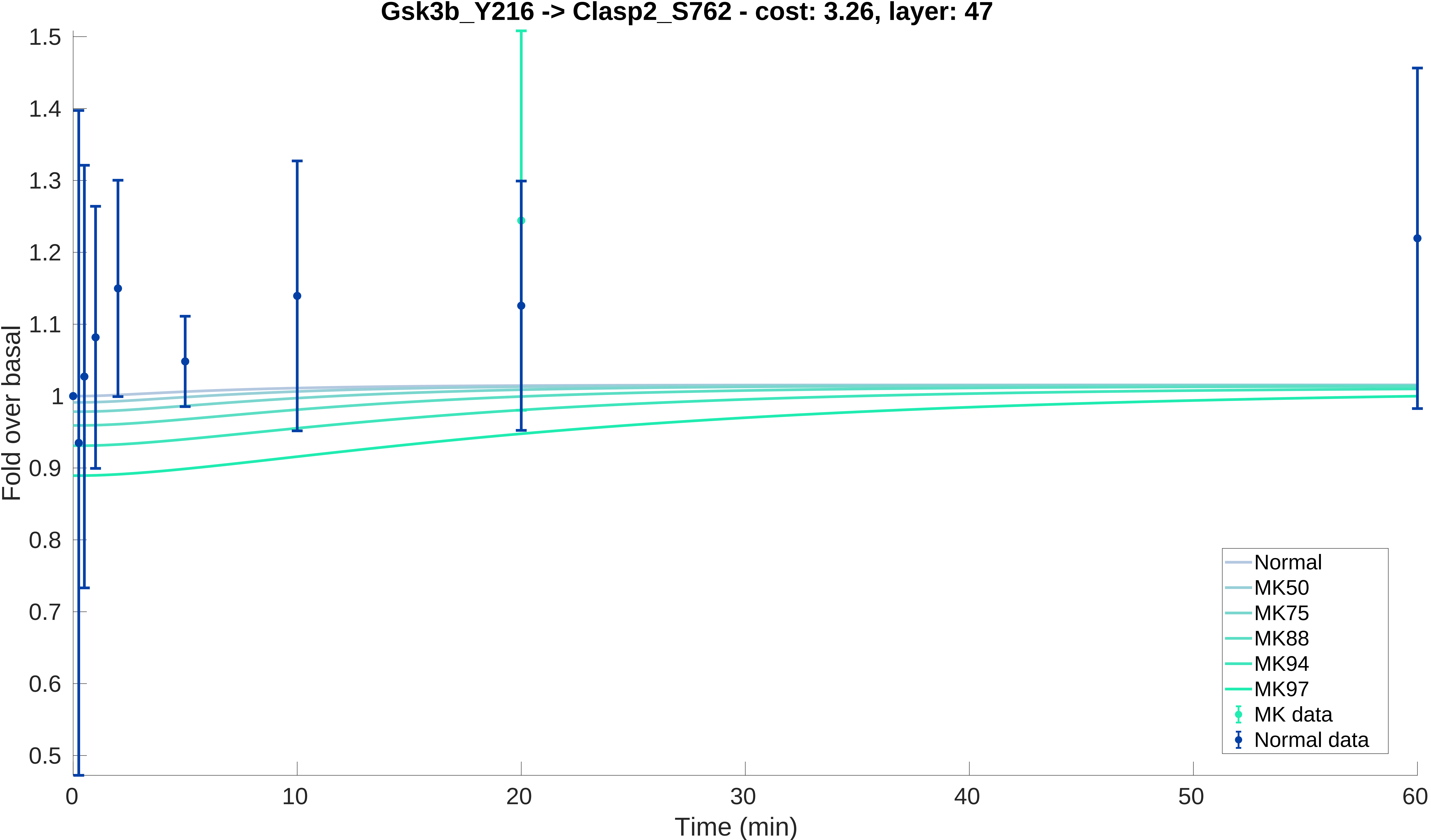

Gsk3b_Y216 -> Clasp2_S762 - cost: 3.26, layer: 47
Fold over basal
Time (min)
1.5
1.4
1.3
1.2
1.1
1
0.9
0.8
0.7
0.6
0.5
0
10
20
30
40
50
60
Normal
MK50
MK75
MK88
MK94
MK97
MK data
Normal data

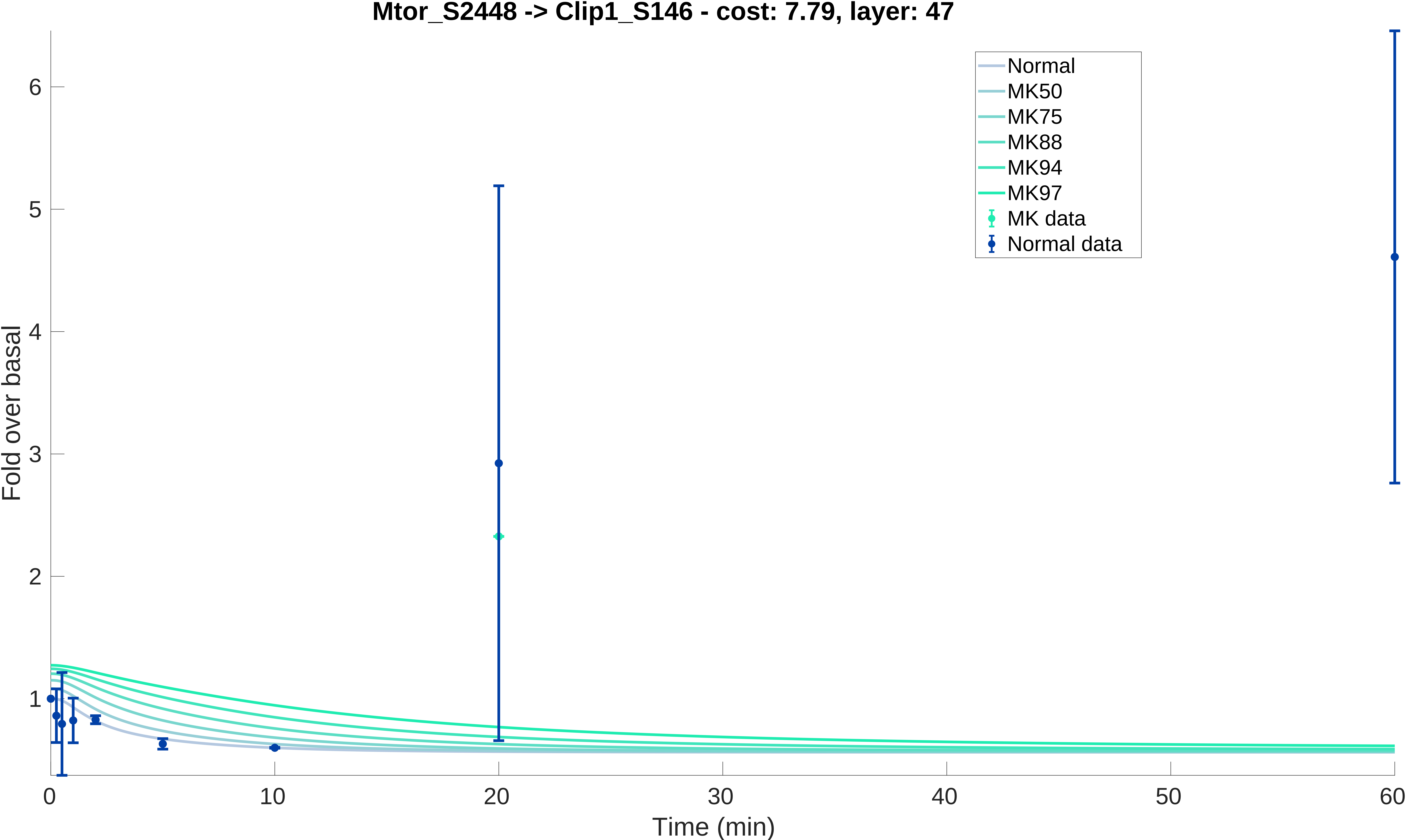
Mtor_S2448 -> Clip1_S146 - cost: 7.79, layer: 47
Fold over basal
Time (min)
0
10
20
30
40
50
60
1
2
3
4
5
6
Normal
MK50
MK75
MK88
MK94
MK97
MK data
Normal data

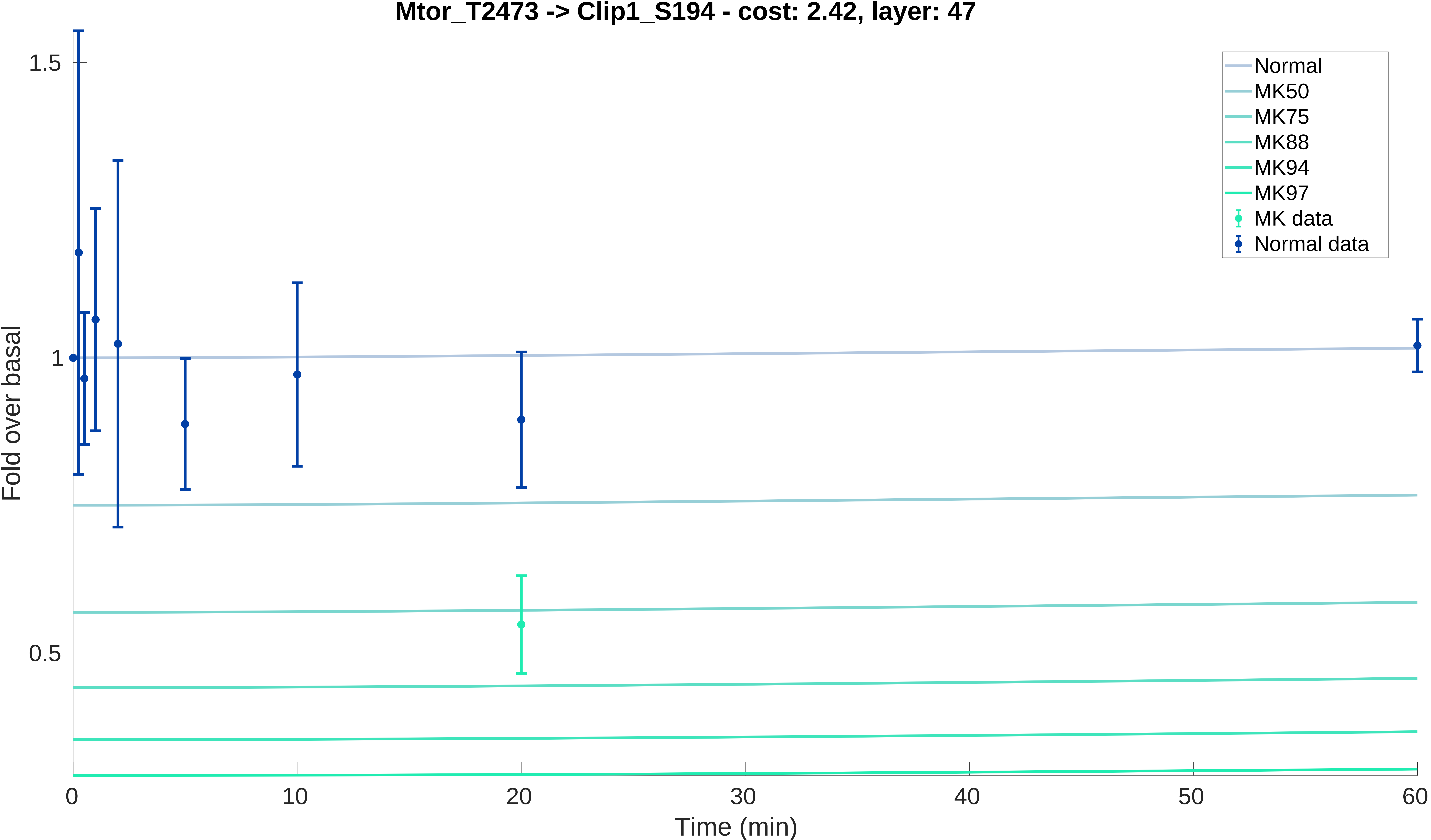

Mtor_T2473 -> Clip1_S194 - cost: 2.42, layer: 47
Normal
MK50
MK75
MK88
MK94
MK97
MK data
Normal data
Fold over basal
1.5
1
0.5
0
10
20
30
40
50
60
Time (min)

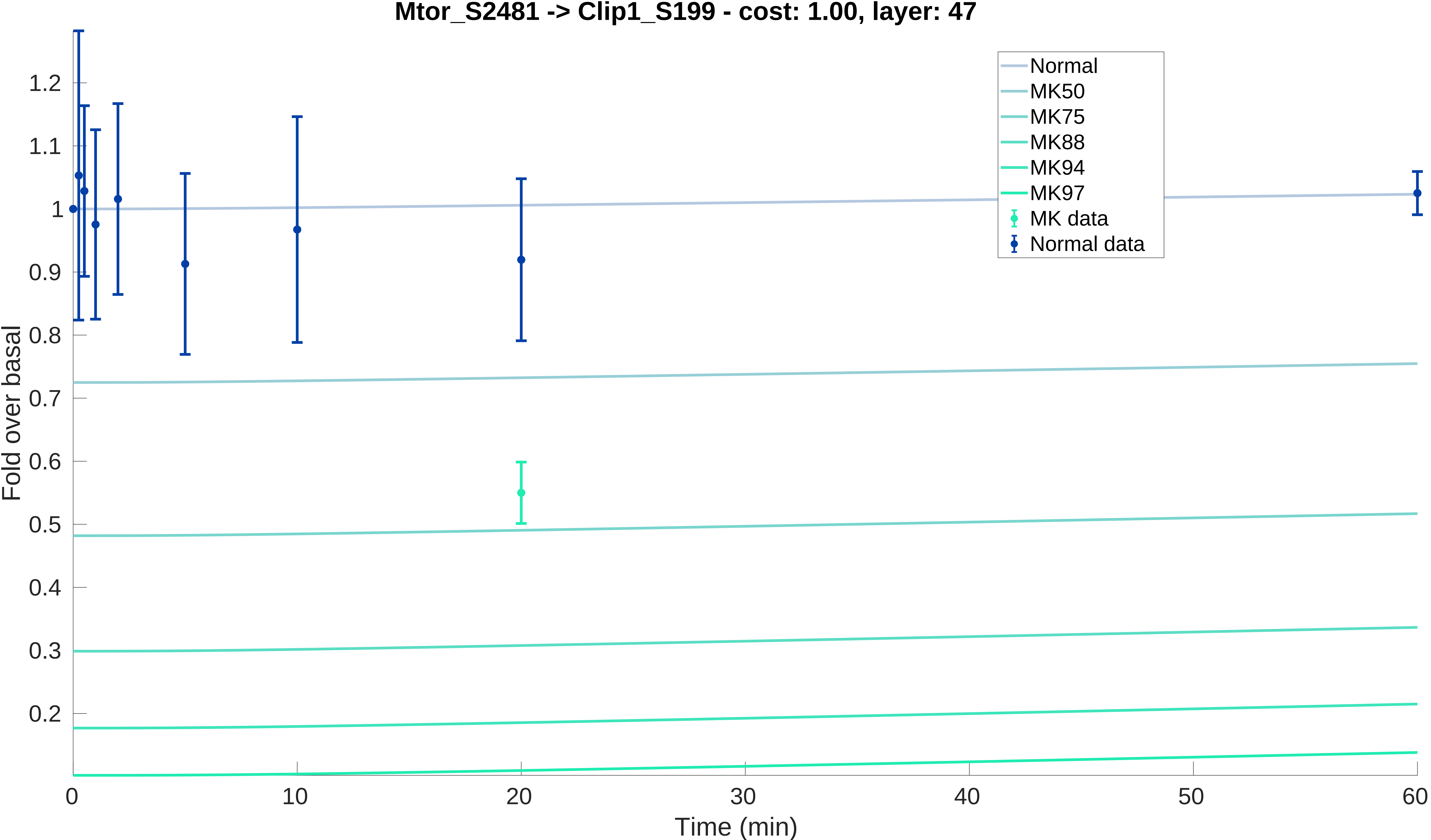

Mtor_S2481 -> Clip1_S199 - cost: 1.00, layer: 47
Normal
MK50
MK75
MK88
MK94
MK97
MK data
Normal data
Fold over basal
Time (min)
0
10
20
30
40
50
60
0.2
0.3
0.4
0.5
0.6
0.7
0.8
0.9
1
1.1
1.2

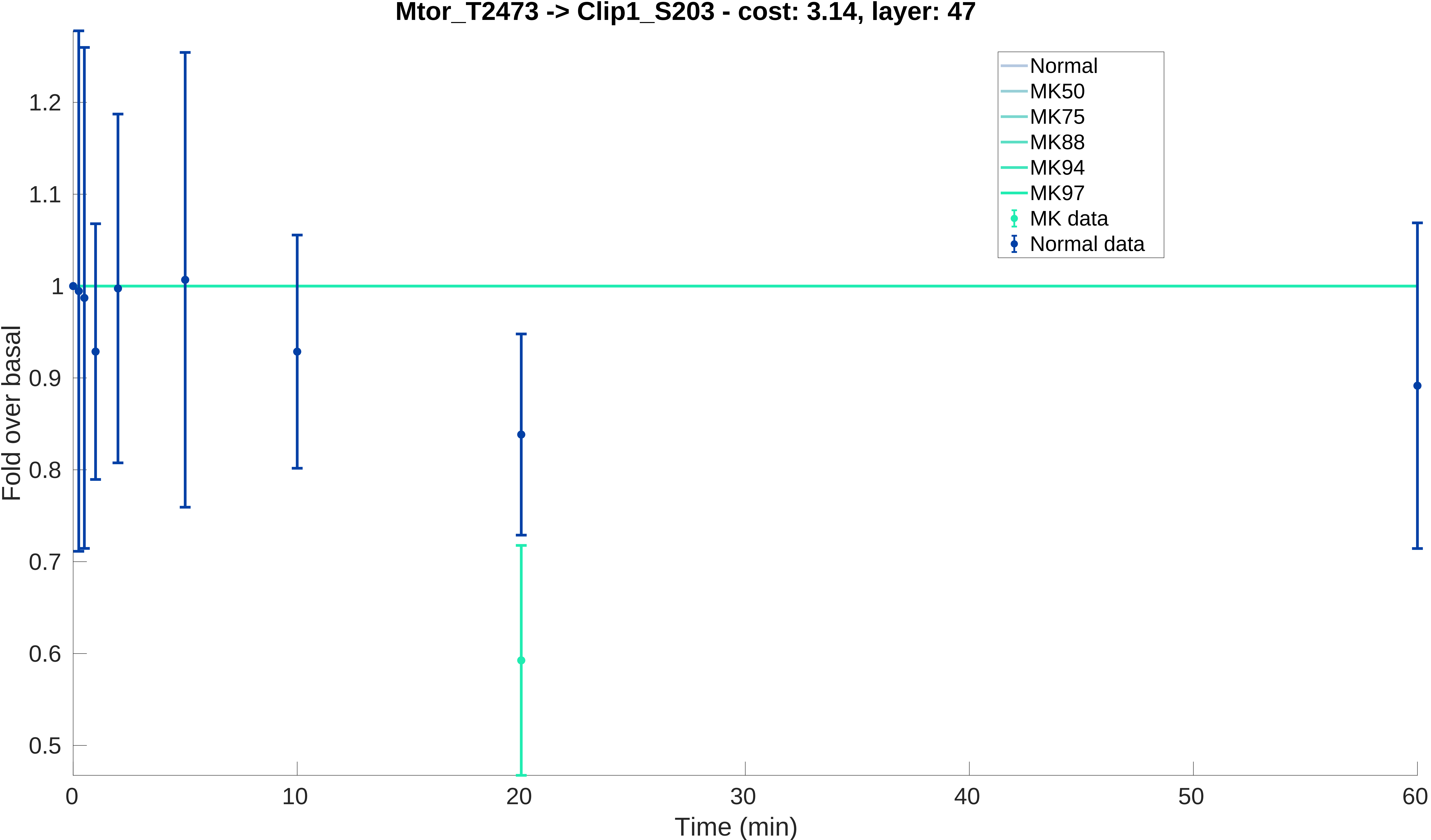

Mtor_T2473 -> Clip1_S203 - cost: 3.14, layer: 47
Normal
MK50
MK75
MK88
MK94
MK97
MK data
Normal data
Fold over basal
1.2
1.1
1
0.9
0.8
0.7
0.6
0.5
0
10
20
30
40
50
60
Time (min)

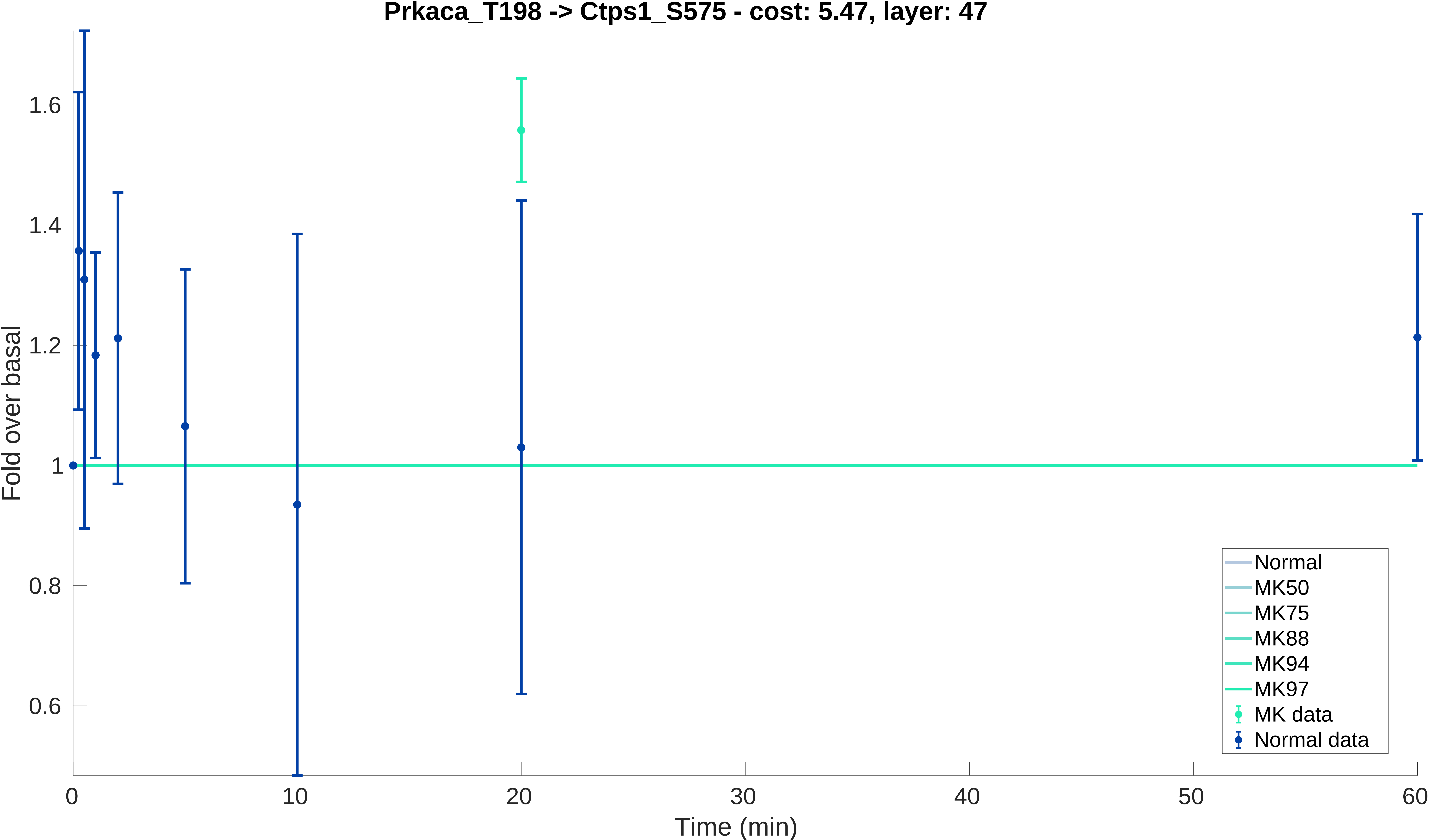
Prkaca_T198 -> Ctps1_S575 - cost: 5.47, layer: 47
Fold over basal
1.6
1.4
1.2
1
0.8
0.6
0
10
20
30
40
50
60
Time (min)
Normal
MK50
MK75
MK88
MK94
MK97
MK data
Normal data

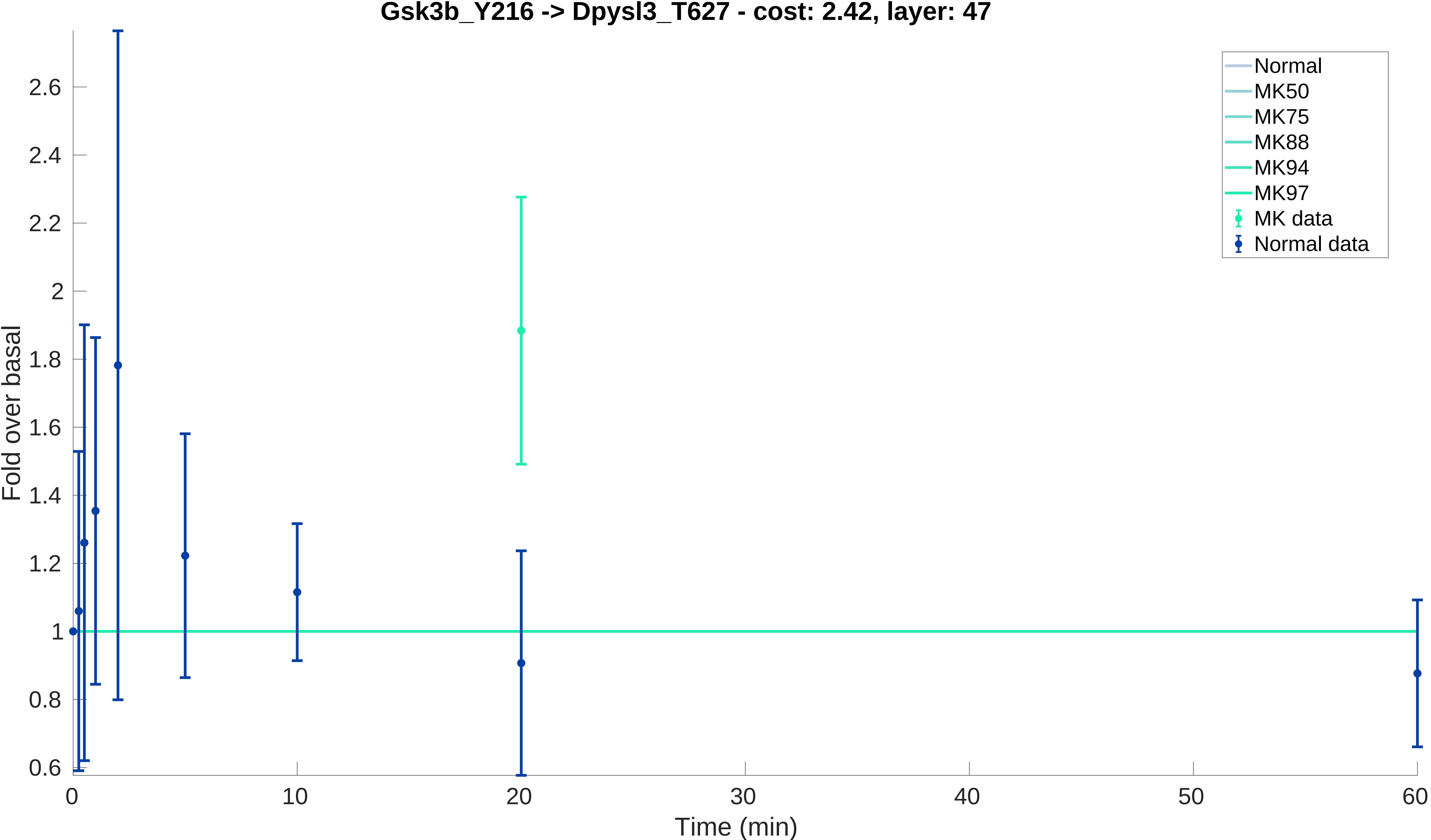

Gsk3b_Y216 -> Dpysl3_T627 - cost: 2.42, layer: 47
Fold over basal
Time (min)
0.6
0.8
1
1.2
1.4
1.6
1.8
2
2.2
2.4
2.6
0
10
20
30
40
50
60
Normal
MK50
MK75
MK88
MK94
MK97
MK data
Normal data

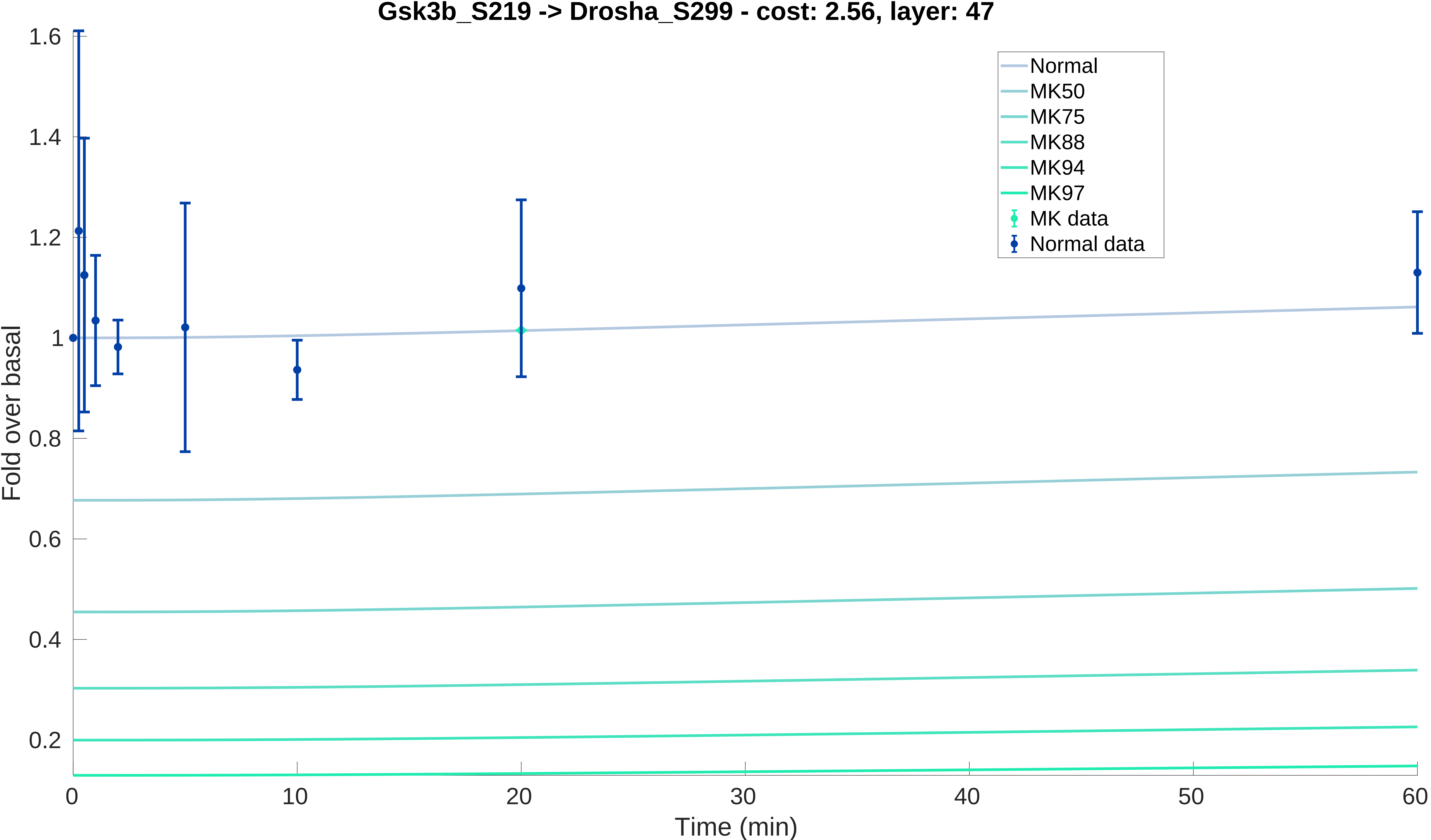

Gsk3b_S219 -> Drosha_S299 - cost: 2.56, layer: 47
Fold over basal
Time (min)
0
10
20
30
40
50
60
0.2
0.4
0.6
0.8
1
1.2
1.4
1.6
Normal
MK50
MK75
MK88
MK94
MK97
MK data
Normal data

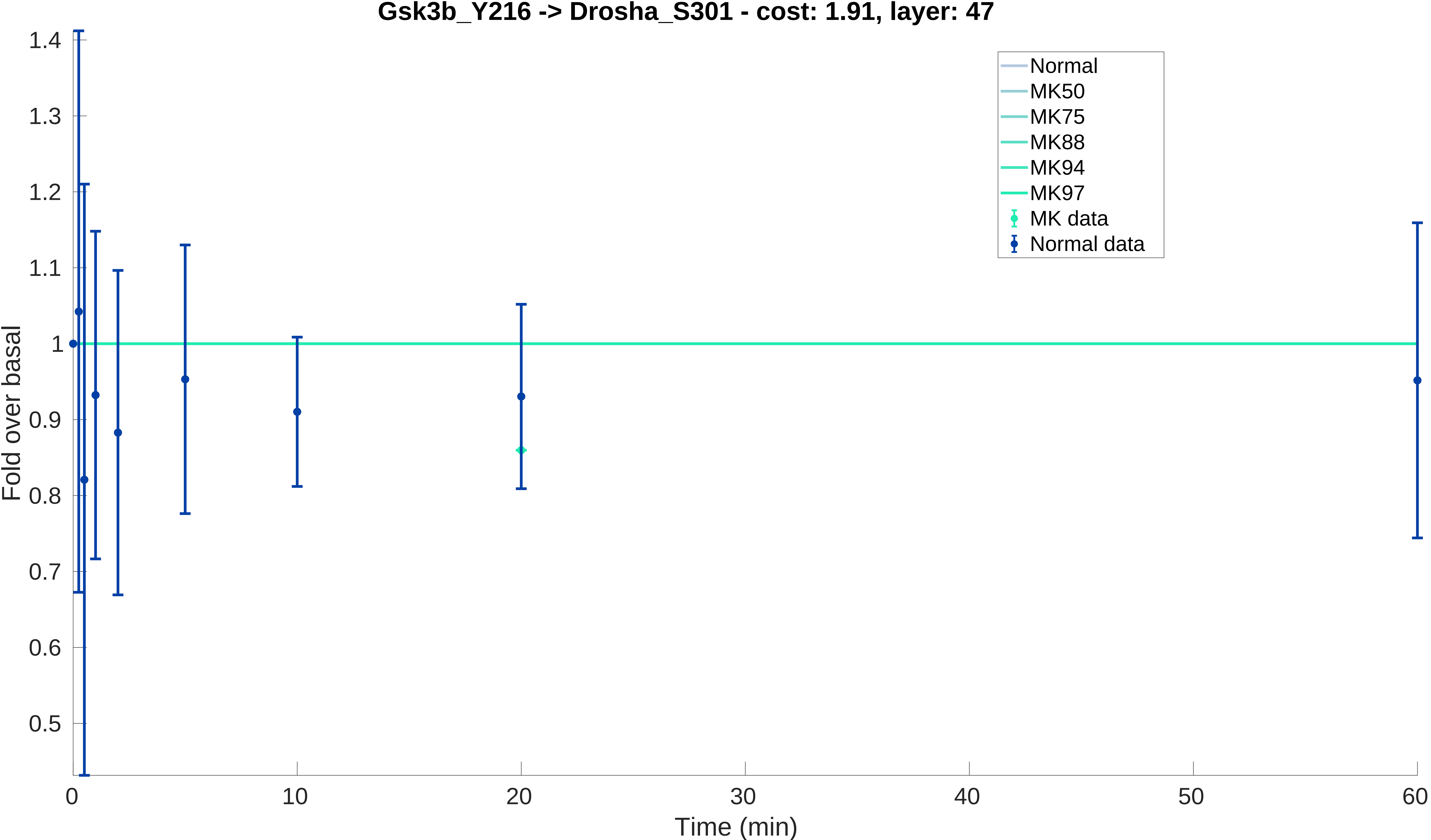

Gsk3b_Y216 -> Drosha_S301 - cost: 1.91, layer: 47
Fold over basal
Time (min)
Normal
MK50
MK75
MK88
MK94
MK97
MK data
Normal data
1.4
1.3
1.2
1.1
1
0.9
0.8
0.7
0.6
0.5
0
10
20
30
40
50
60

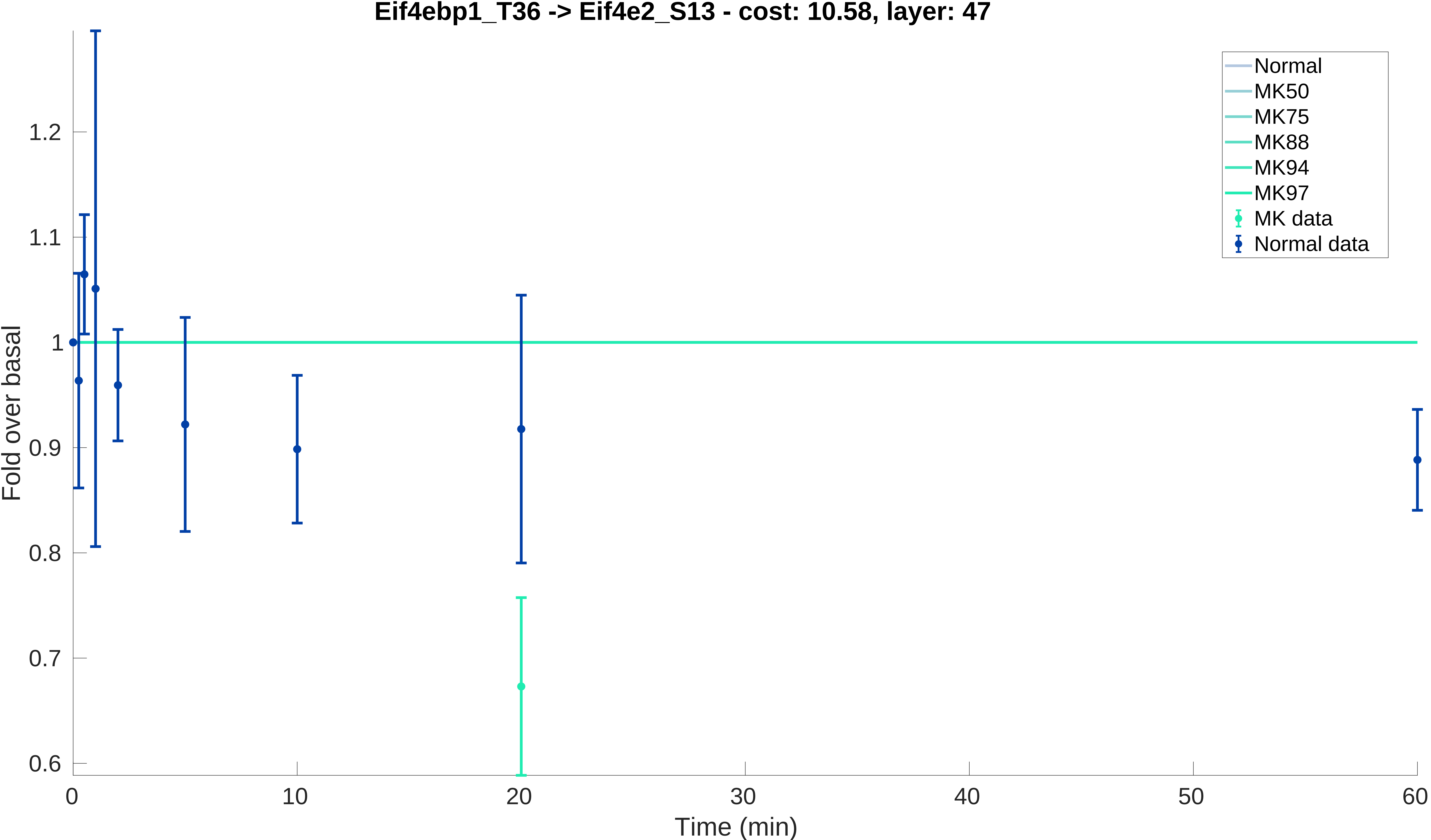
Eif4ebp1_T36 -> Eif4e2_S13 - cost: 10.58, layer: 47
Fold over basal
1.2
1.1
1
0.9
0.8
0.7
0.6
0
10
20
30
40
50
60
Time (min)
Normal
MK50
MK75
MK88
MK94
MK97
MK data
Normal data

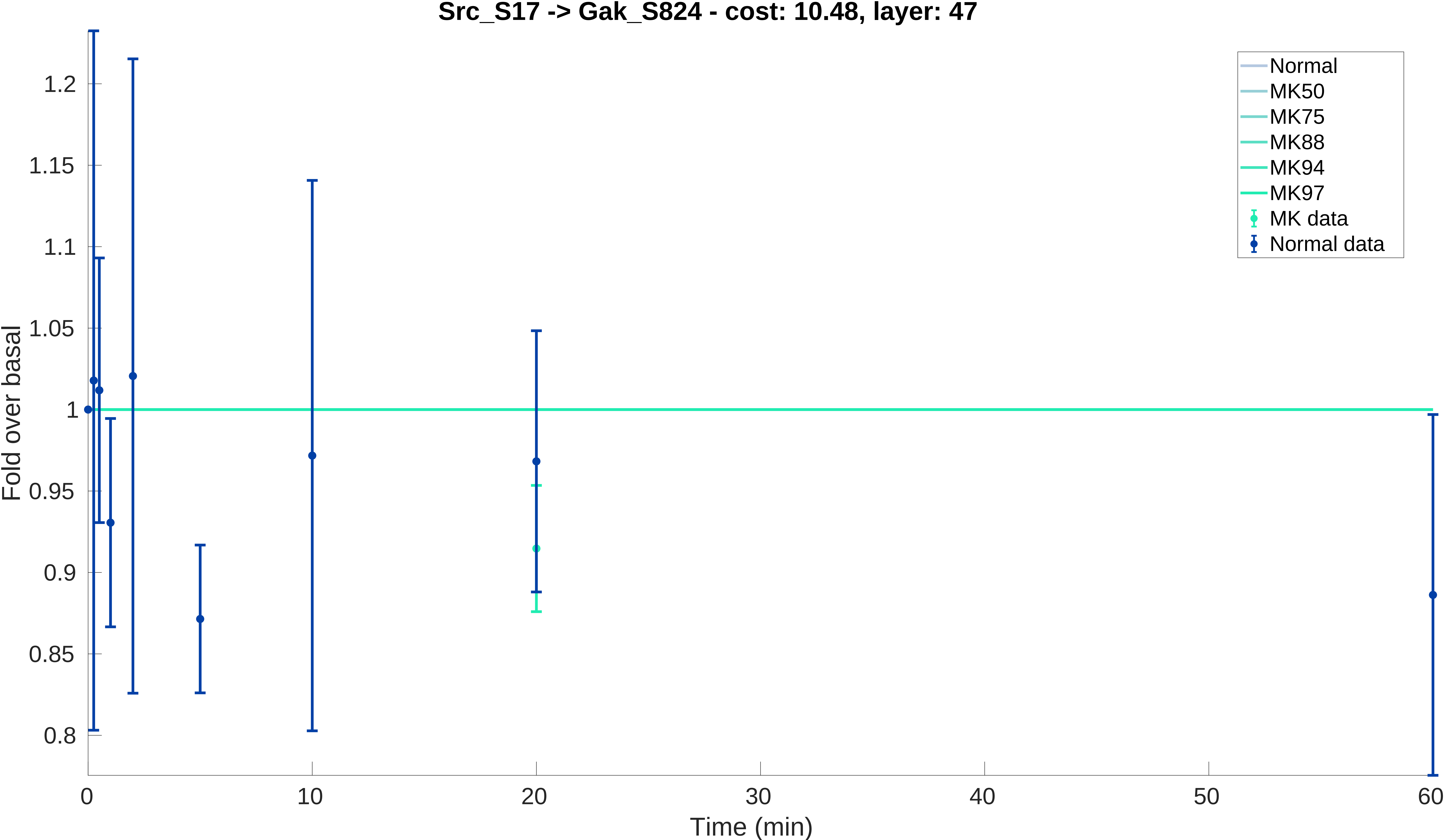

Src_S17 -> Gak_S824 - cost: 10.48, layer: 47
Normal
MK50
MK75
MK88
MK94
MK97
MK data
Normal data
1.2
1.15
1.1
1.05
1
0.95
0.9
0.85
0.8
Fold over basal
0
10
20
30
40
50
60
Time (min)

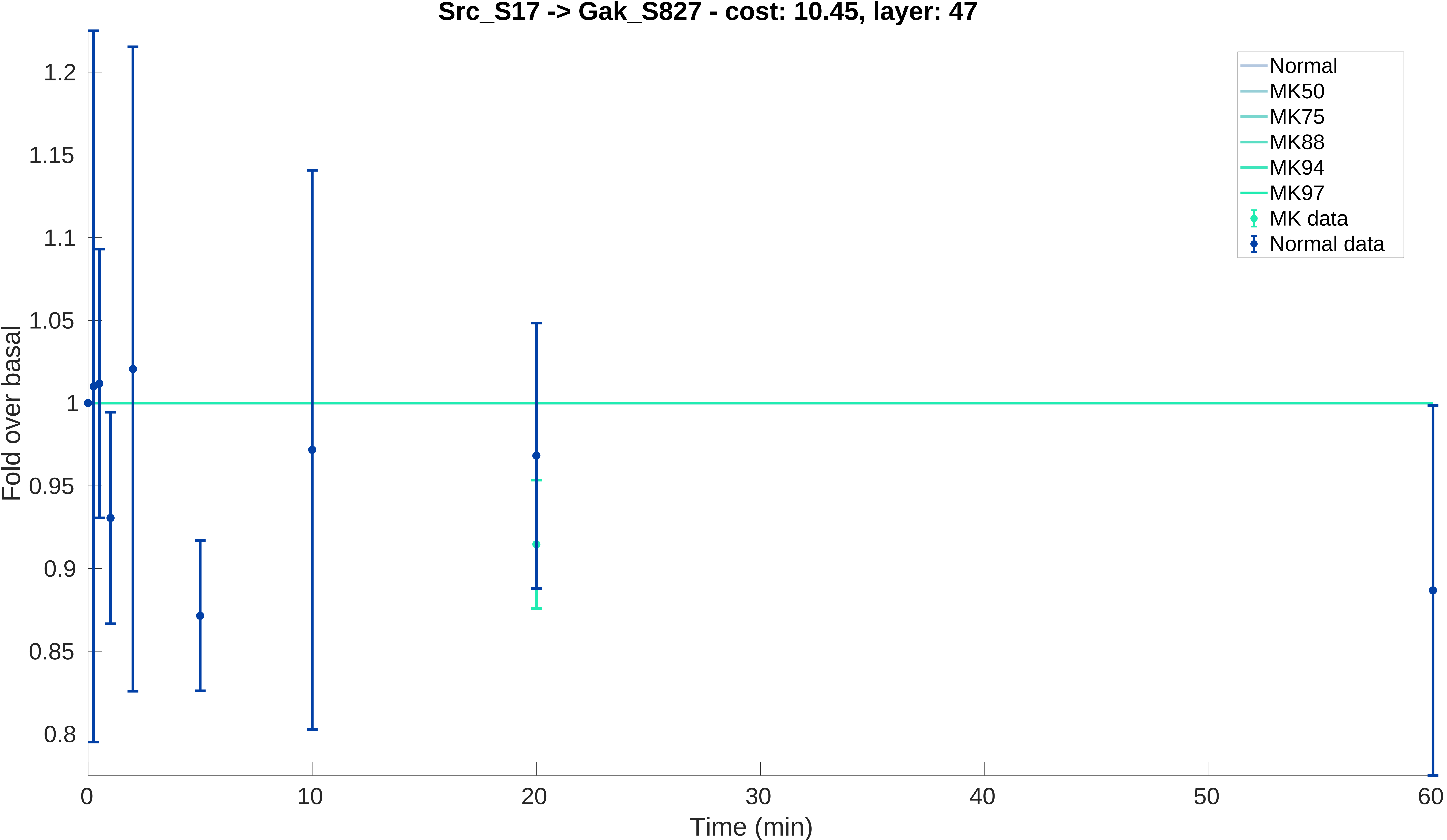

Src_S17 -> Gak_S827 - cost: 10.45, layer: 47
Fold over basal
1.2
1.15
1.1
1.05
1
0.95
0.9
0.85
0.8
0
10
20
30
40
50
60
Time (min)
Normal
MK50
MK75
MK88
MK94
MK97
MK data
Normal data

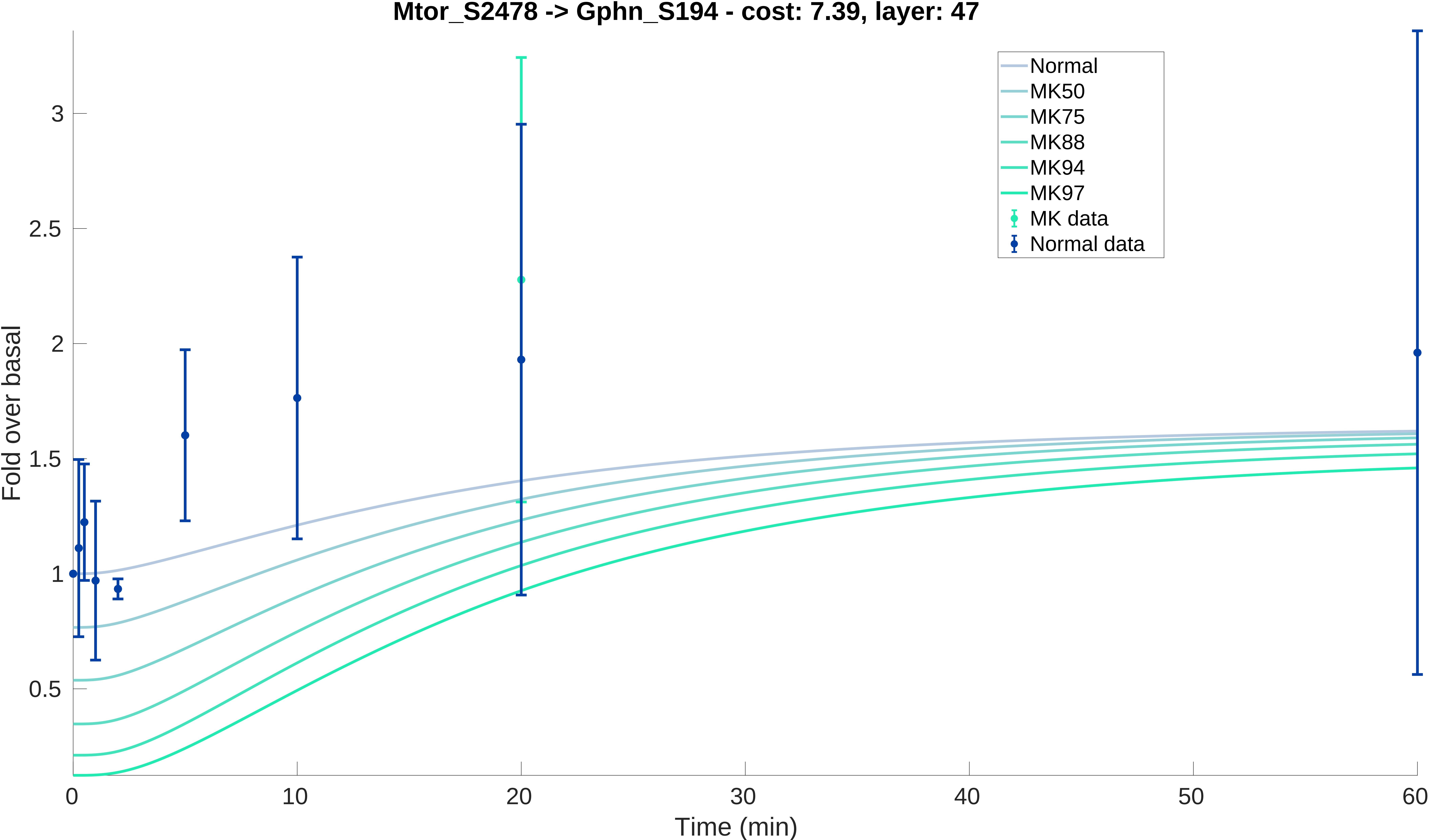

Mtor_S2478 -> Gphn_S194 - cost: 7.39, layer: 47
Fold over basal
Time (min)
Normal
MK50
MK75
MK88
MK94
MK97
MK data
Normal data
0.5
1
1.5
2
2.5
3
0
10
20
30
40
50
60

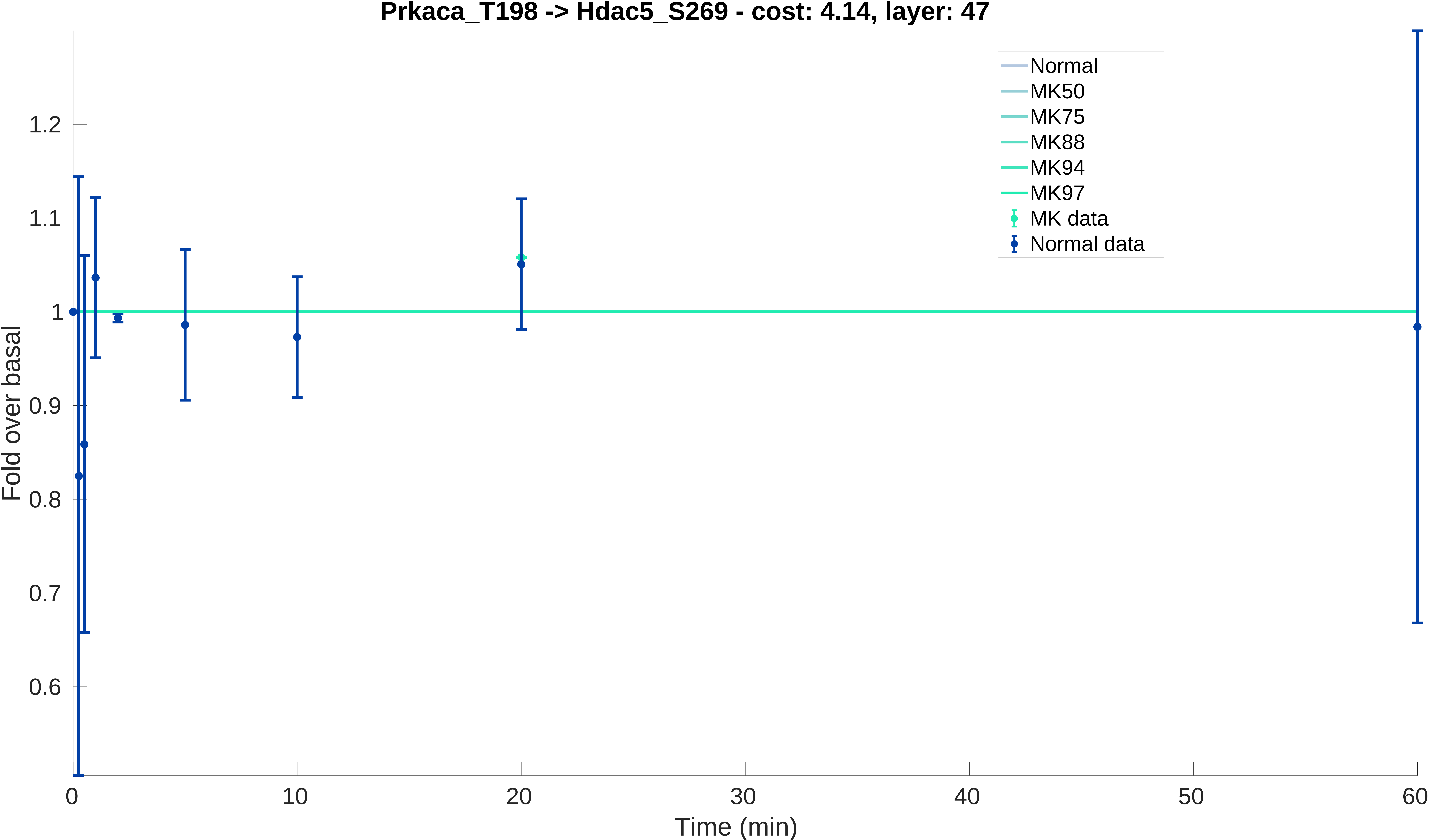

Prkaca_T198 -> Hdac5_S269 - cost: 4.14, layer: 47
Normal
MK50
MK75
MK88
MK94
MK97
MK data
Normal data
Fold over basal
1.2
1.1
1
0.9
0.8
0.7
0.6
0
10
20
30
40
50
60
Time (min)

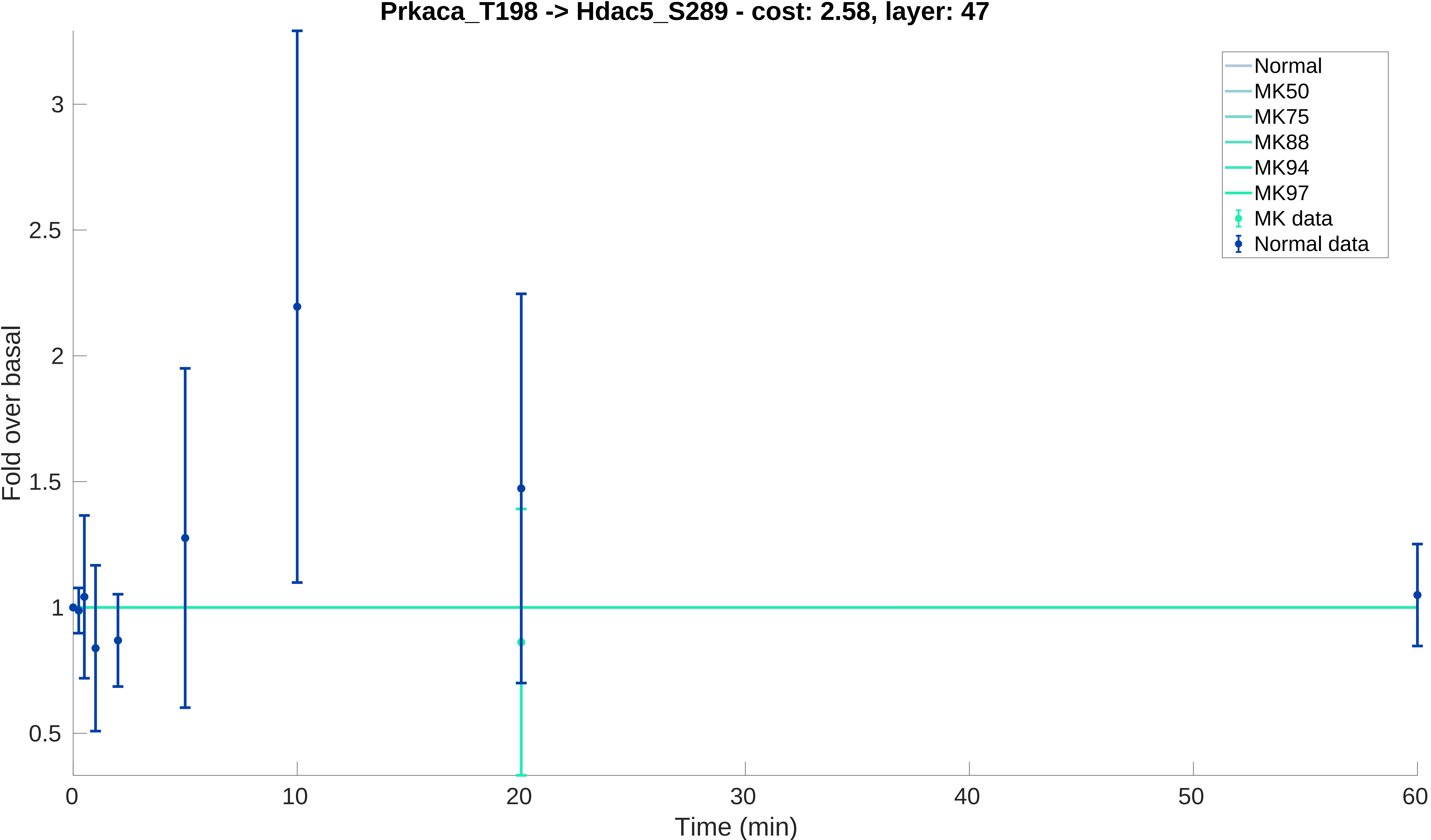

Prkaca_T198 -> Hdac5_S289 - cost: 2.58, layer: 47
Normal
MK50
MK75
MK88
MK94
MK97
MK data
Normal data
Fold over basal
0.5
1
1.5
2
2.5
3
0
10
20
30
40
50
60
Time (min)

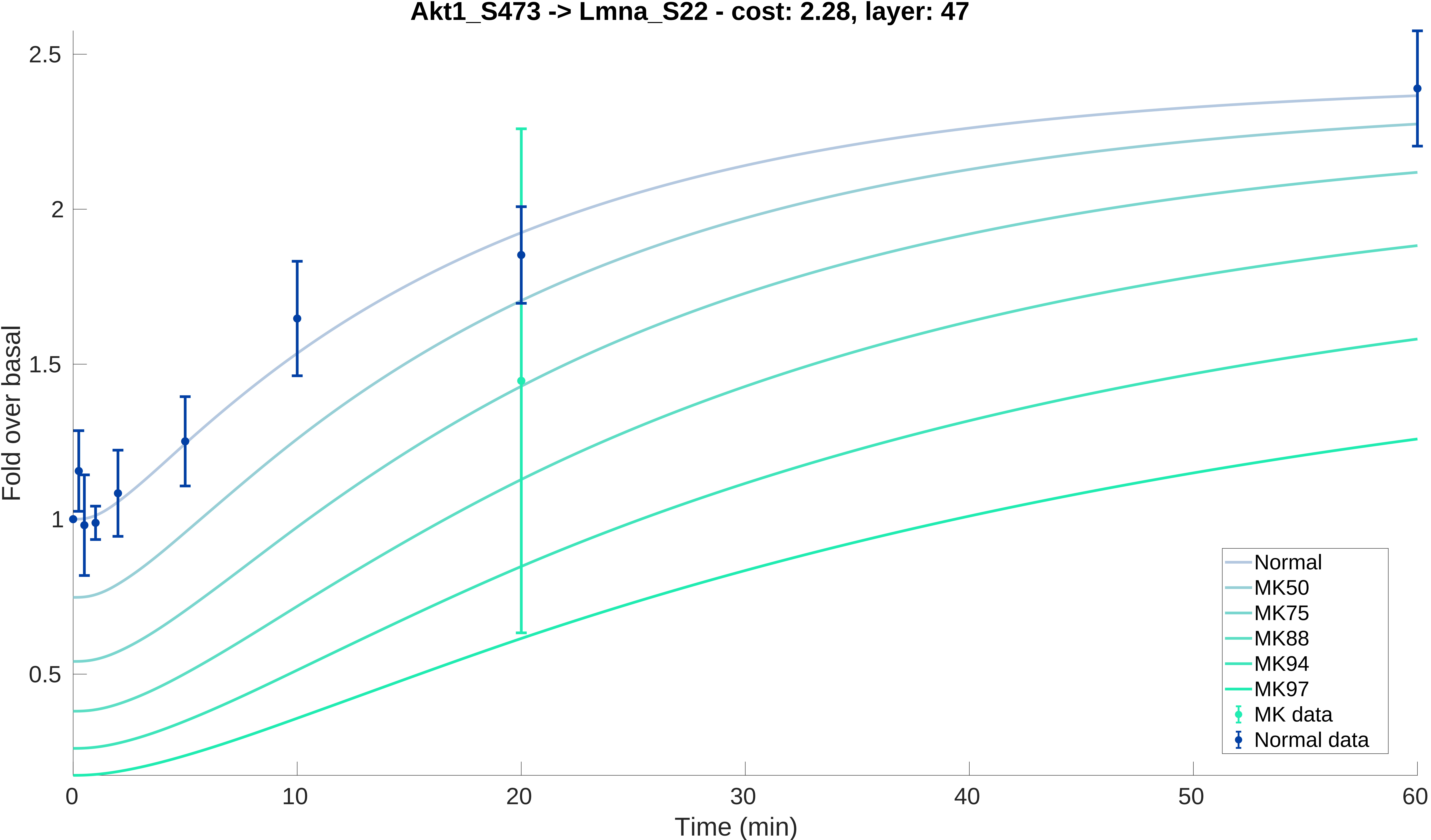
Akt1_S473 -> Lmna_S22 - cost: 2.28, layer: 47
Fold over basal
2.5
2
1.5
1
0.5
0
10
20
30
40
50
60
Time (min)
Normal
MK50
MK75
MK88
MK94
MK97
MK data
Normal data

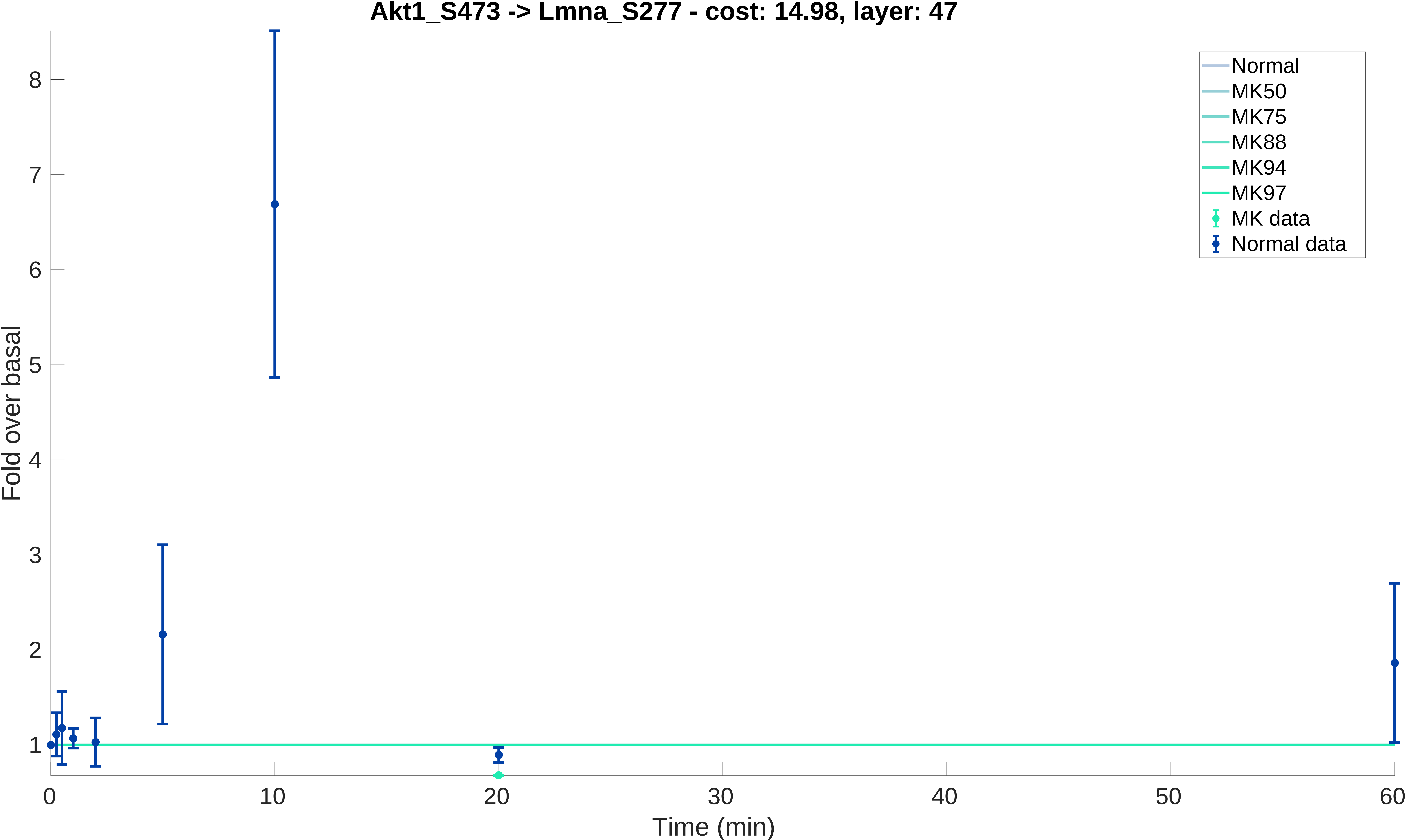
Akt1_S473 -> Lmna_S277 - cost: 14.98, layer: 47
Fold over basal
Time (min)
0
10
20
30
40
50
60
1
2
3
4
5
6
7
8
Normal
MK50
MK75
MK88
MK94
MK97
MK data
Normal data

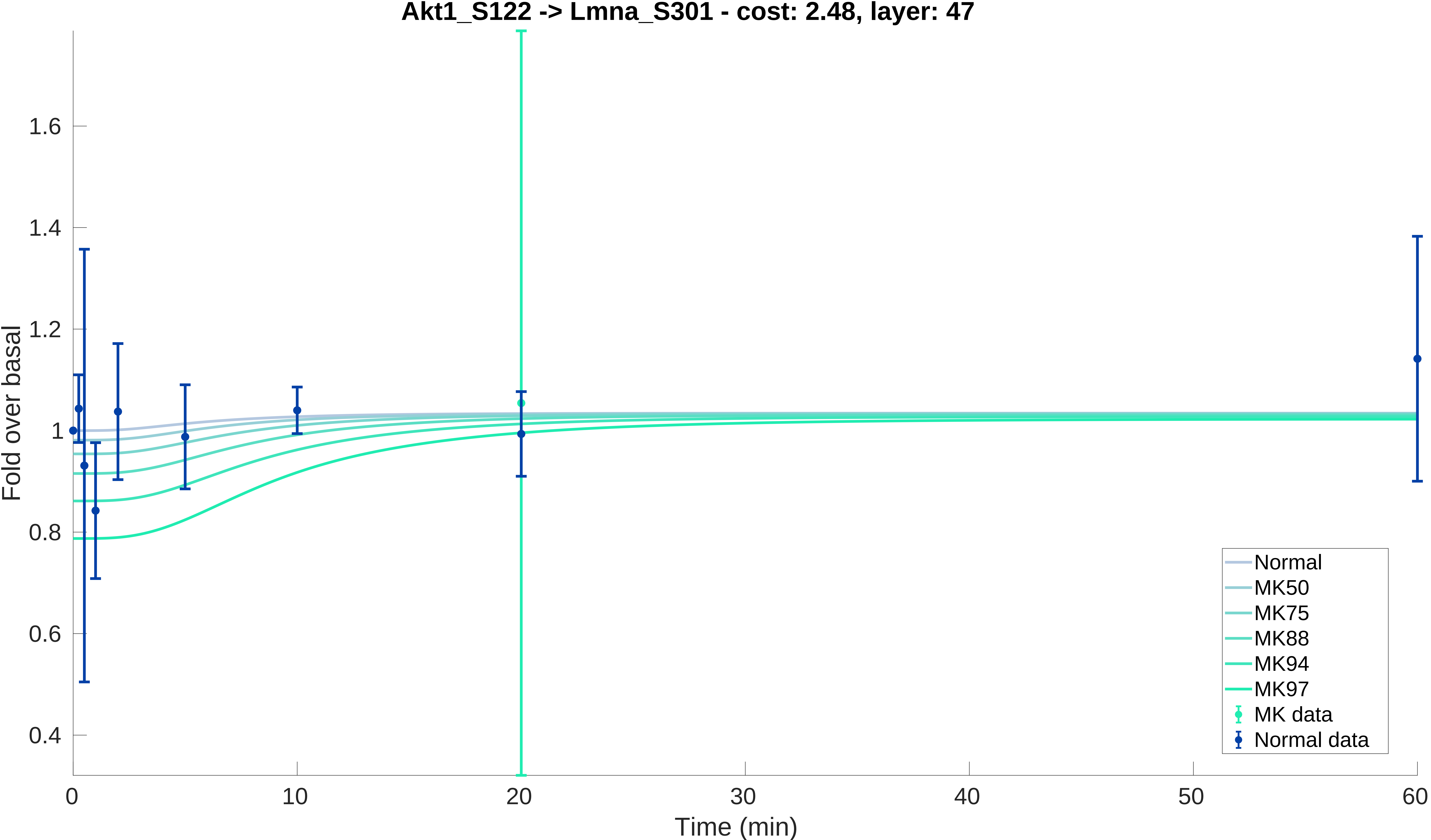

Akt1_S122 -> Lmna_S301 - cost: 2.48, layer: 47
Fold over basal
Time (min)
1.6
1.4
1.2
1
0.8
0.6
0.4
0
10
20
30
40
50
60
Normal
MK50
MK75
MK88
MK94
MK97
MK data
Normal data

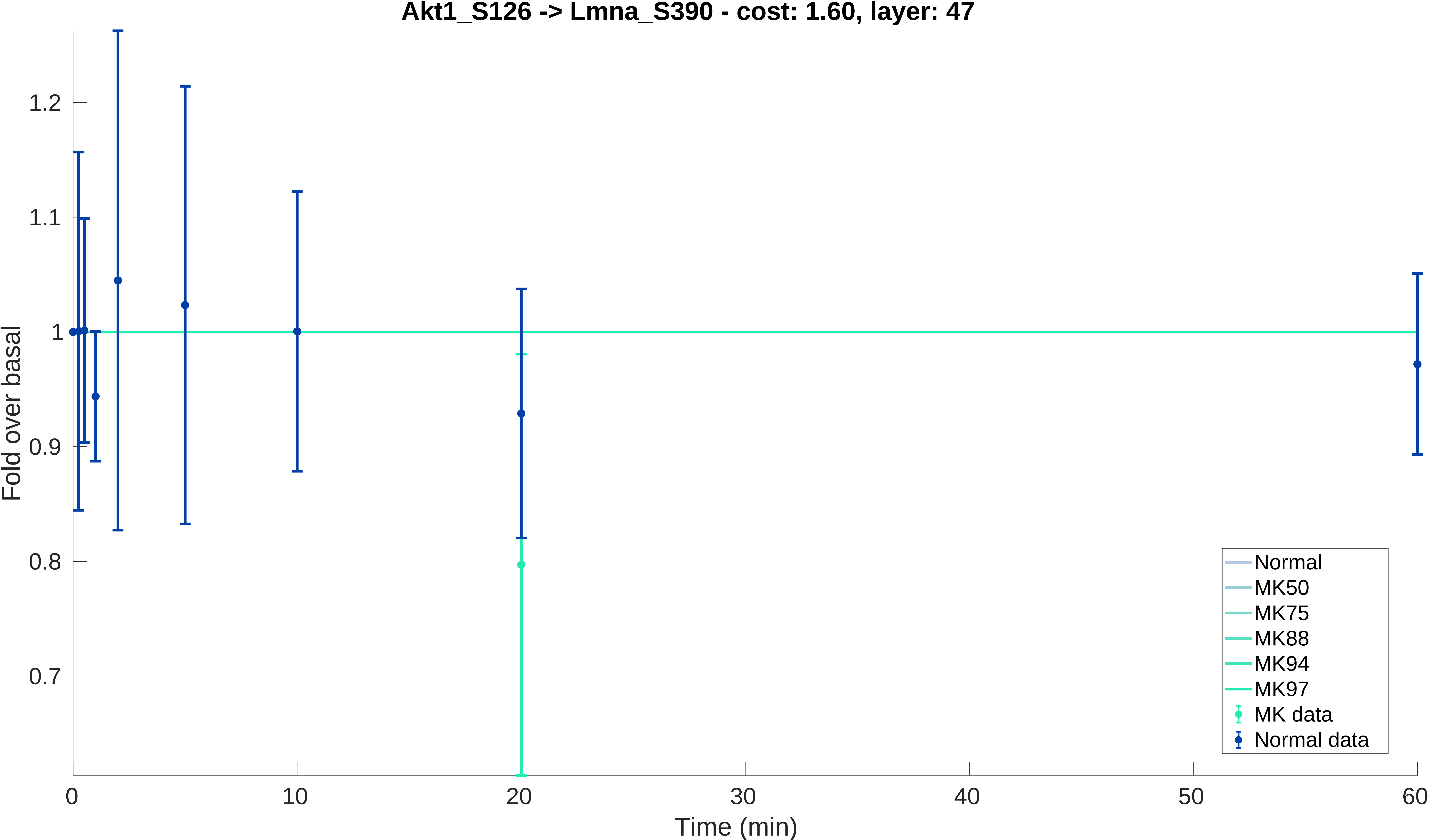
Akt1_S126 -> Lmna_S390 - cost: 1.60, layer: 47
Fold over basal
Time (min)
1.2
1.1
1
0.9
0.8
0.7
0
10
20
30
40
50
60
Normal
MK50
MK75
MK88
MK94
MK97
MK data
Normal data

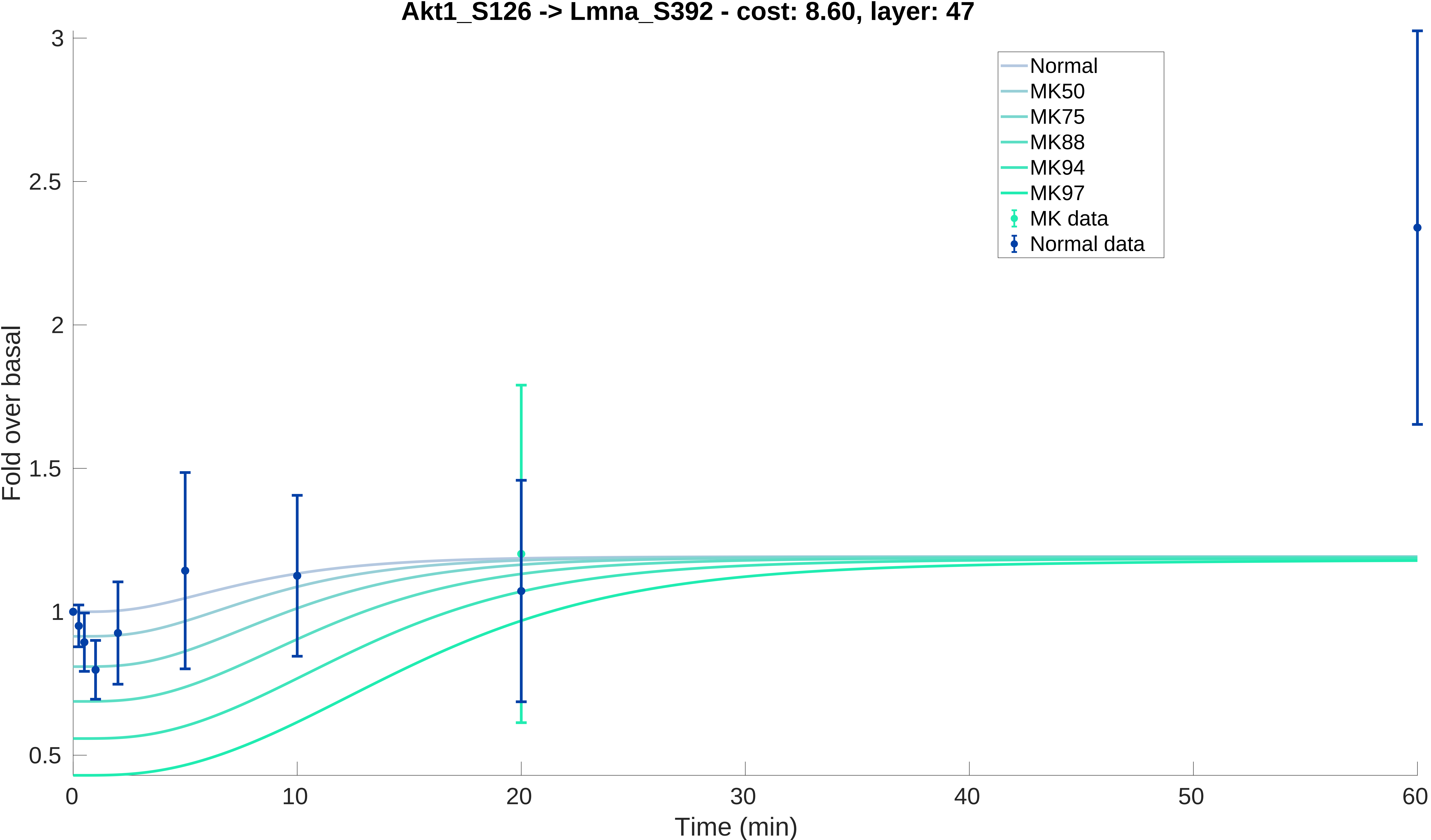
Akt1_S126 -> Lmna_S392 - cost: 8.60, layer: 47
Normal
MK50
MK75
MK88
MK94
MK97
MK data
Normal data
Fold over basal
Time (min)
0.5
1
1.5
2
2.5
3
0
10
20
30
40
50
60

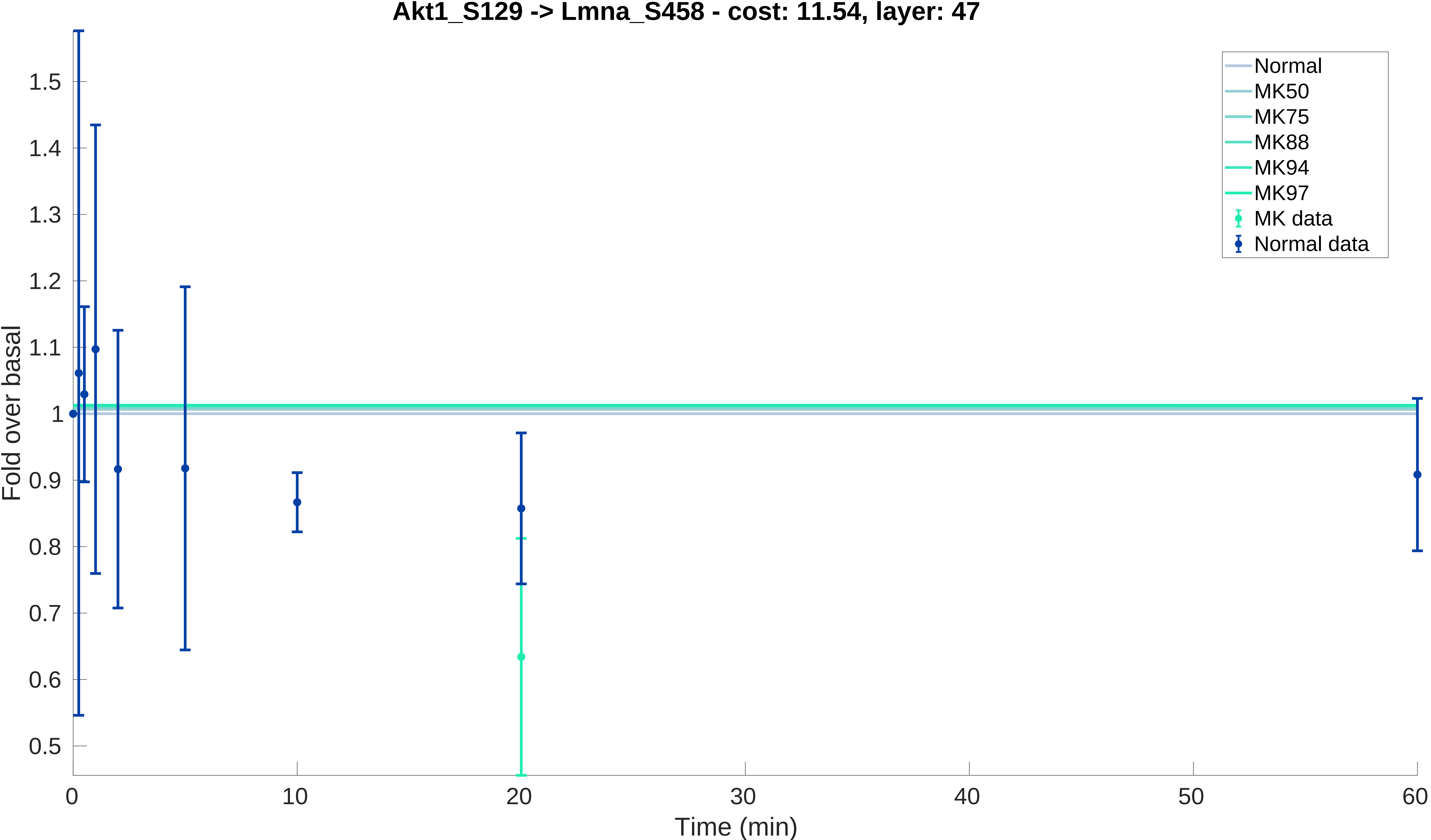

Akt1_S129 -> Lmna_S458 - cost: 11.54, layer: 47
Normal
MK50
MK75
MK88
MK94
MK97
MK data
Normal data
Fold over basal
1.5
1.4
1.3
1.2
1.1
1
0.9
0.8
0.7
0.6
0.5
0
10
20
30
40
50
60
Time (min)

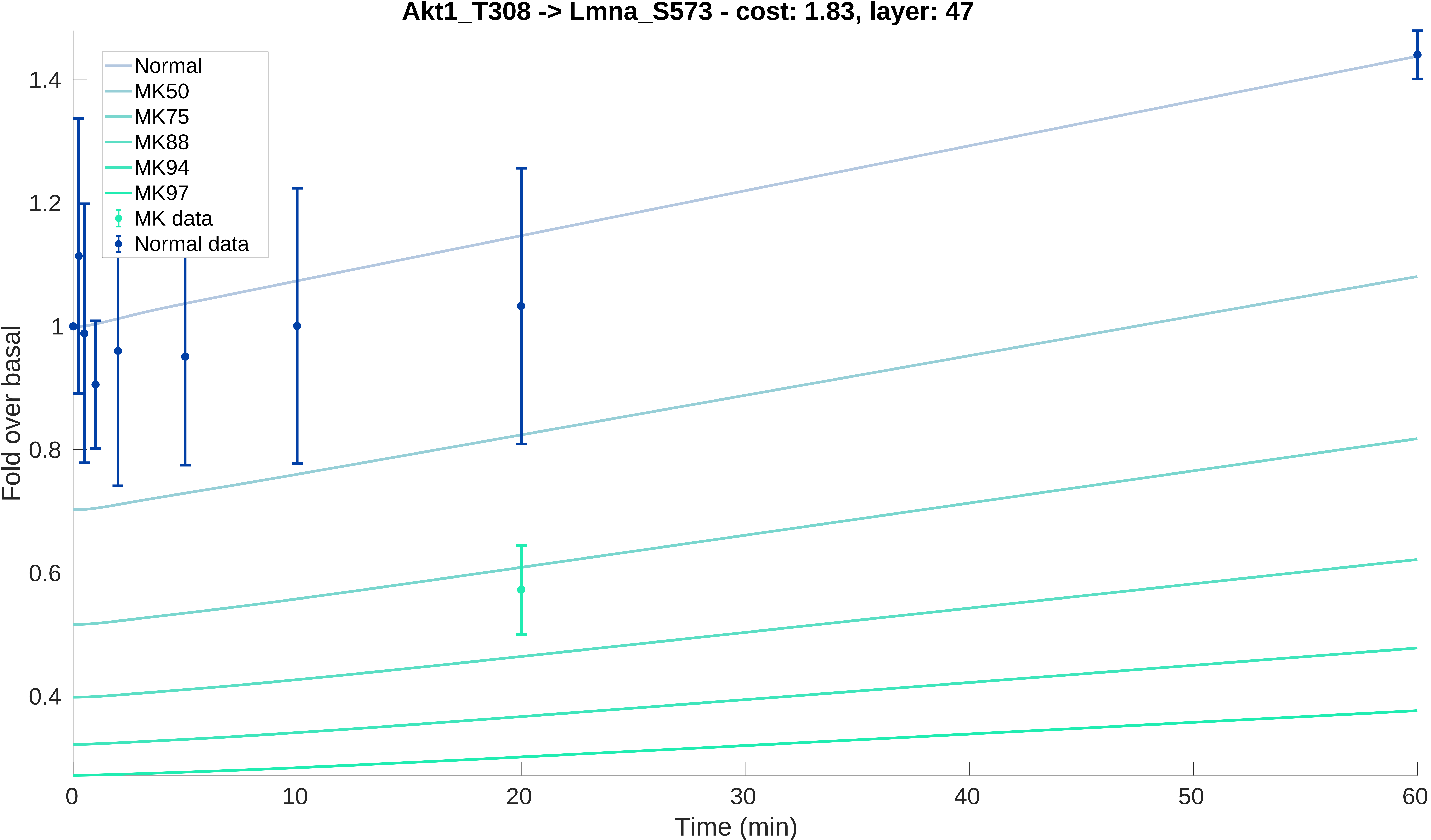

Akt1_T308 -> Lmna_S573 - cost: 1.83, layer: 47
Normal
MK50
MK75
MK88
MK94
MK97
MK data
Normal data
Fold over basal
Time (min)
0.4
0.6
0.8
1
1.2
1.4
0
10
20
30
40
50
60

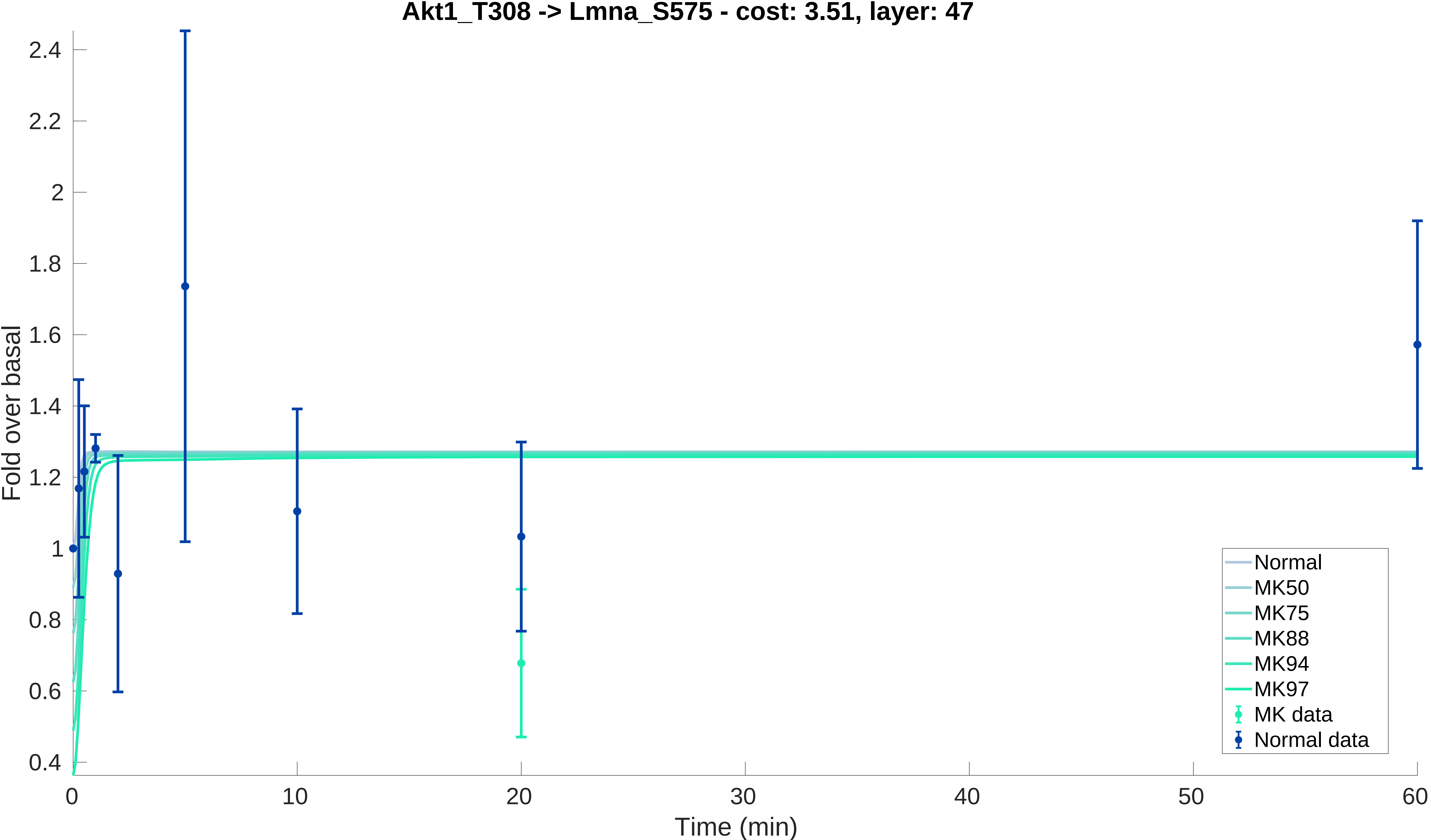

Akt1_T308 -> Lmna_S575 - cost: 3.51, layer: 47
Fold over basal
2.4
2.2
2
1.8
1.6
1.4
1.2
1
0.8
0.6
0.4
0
10
20
30
40
50
60
Time (min)
Normal
MK50
MK75
MK88
MK94
MK97
MK data
Normal data

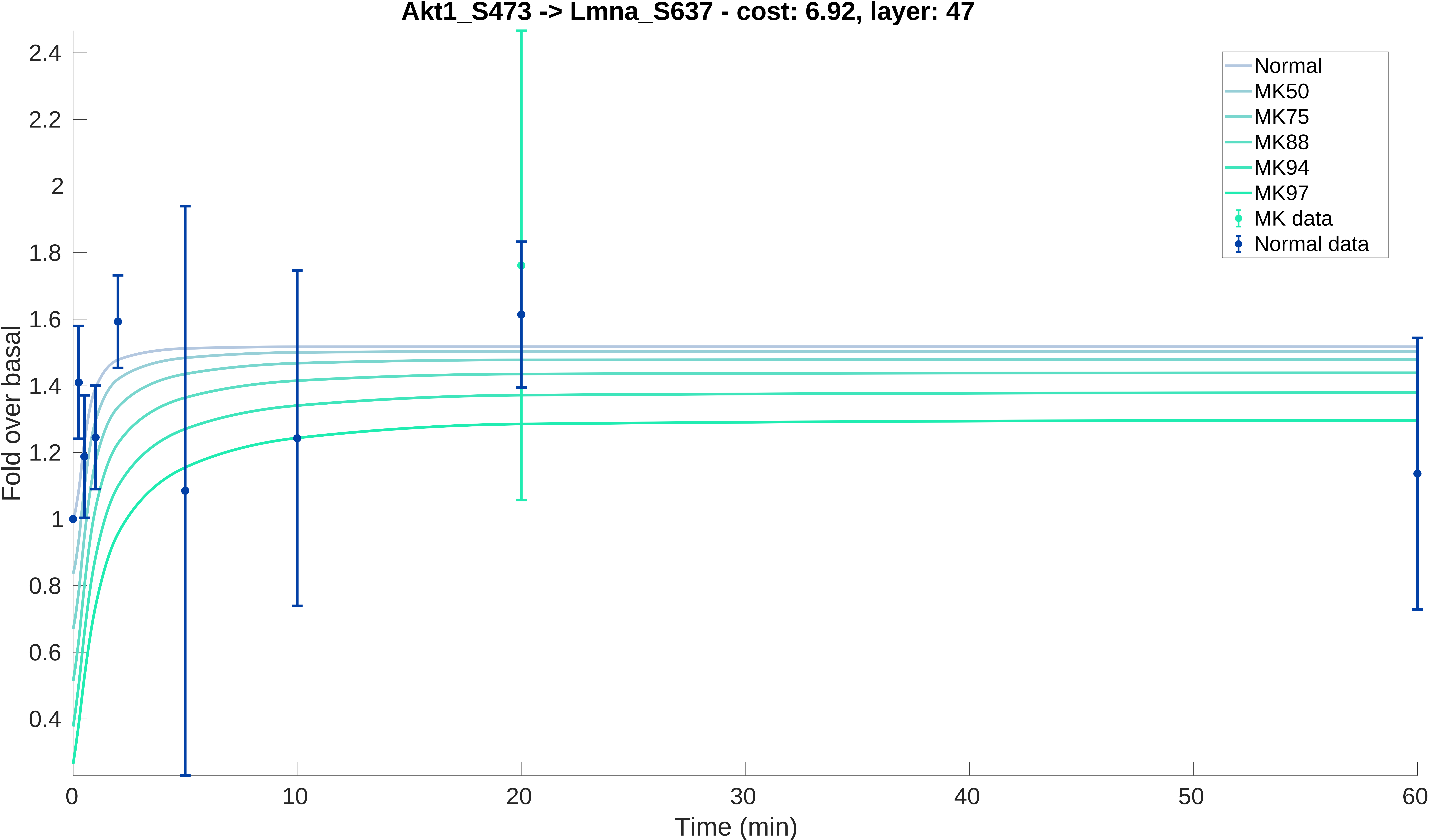

Akt1_S473 -> Lmna_S637 - cost: 6.92, layer: 47
Fold over basal
Time (min)
0.4
0.6
0.8
1
1.2
1.4
1.6
1.8
2
2.2
2.4
0
10
20
30
40
50
60
Normal
MK50
MK75
MK88
MK94
MK97
MK data
Normal data

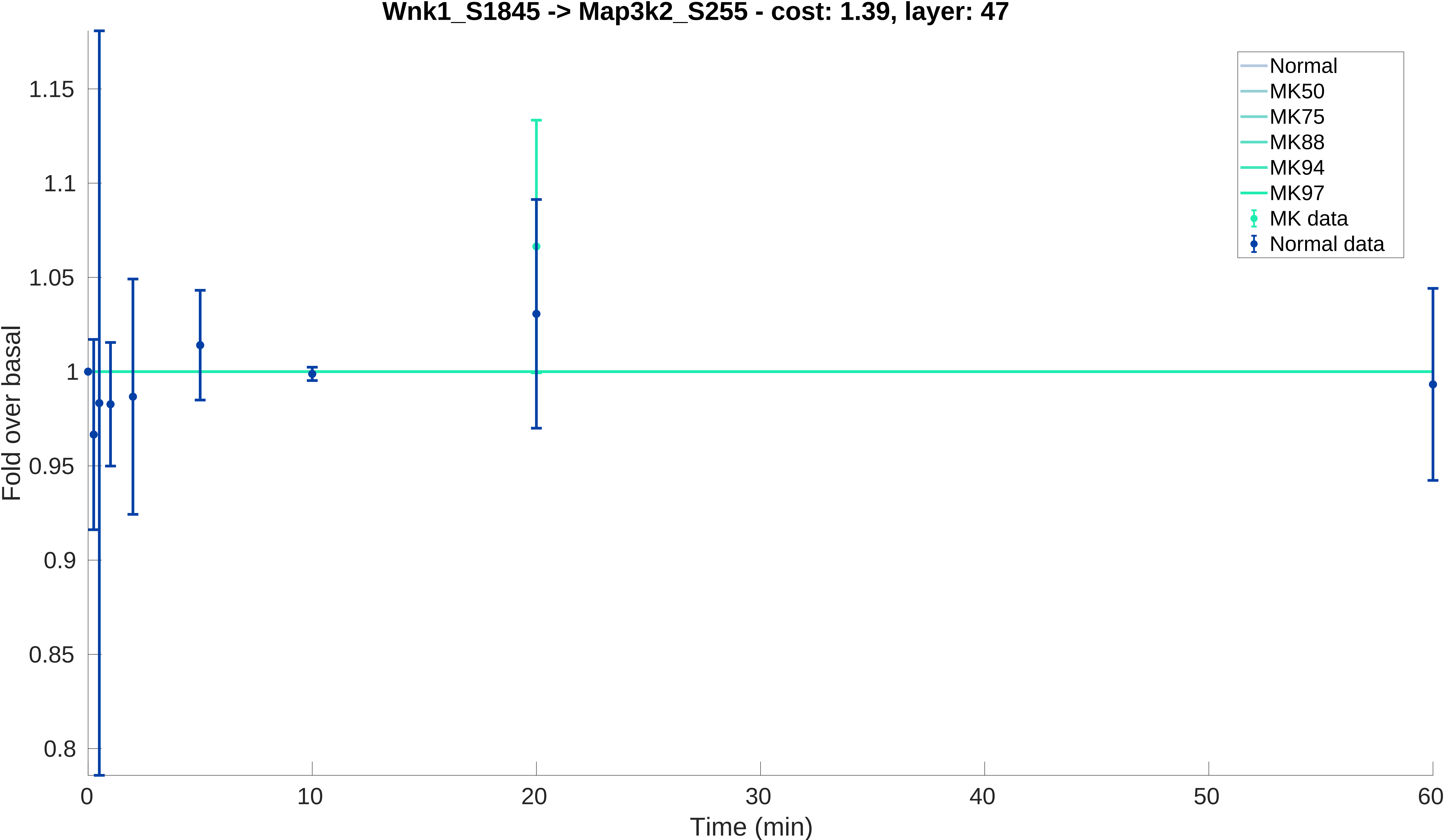

Wnk1_S1845 -> Map3k2_S255 - cost: 1.39, layer: 47
Normal
MK50
MK75
MK88
MK94
MK97
MK data
Normal data
Fold over basal
1.15
1.1
1.05
1
0.95
0.9
0.85
0.8
0
10
20
30
40
50
60
Time (min)

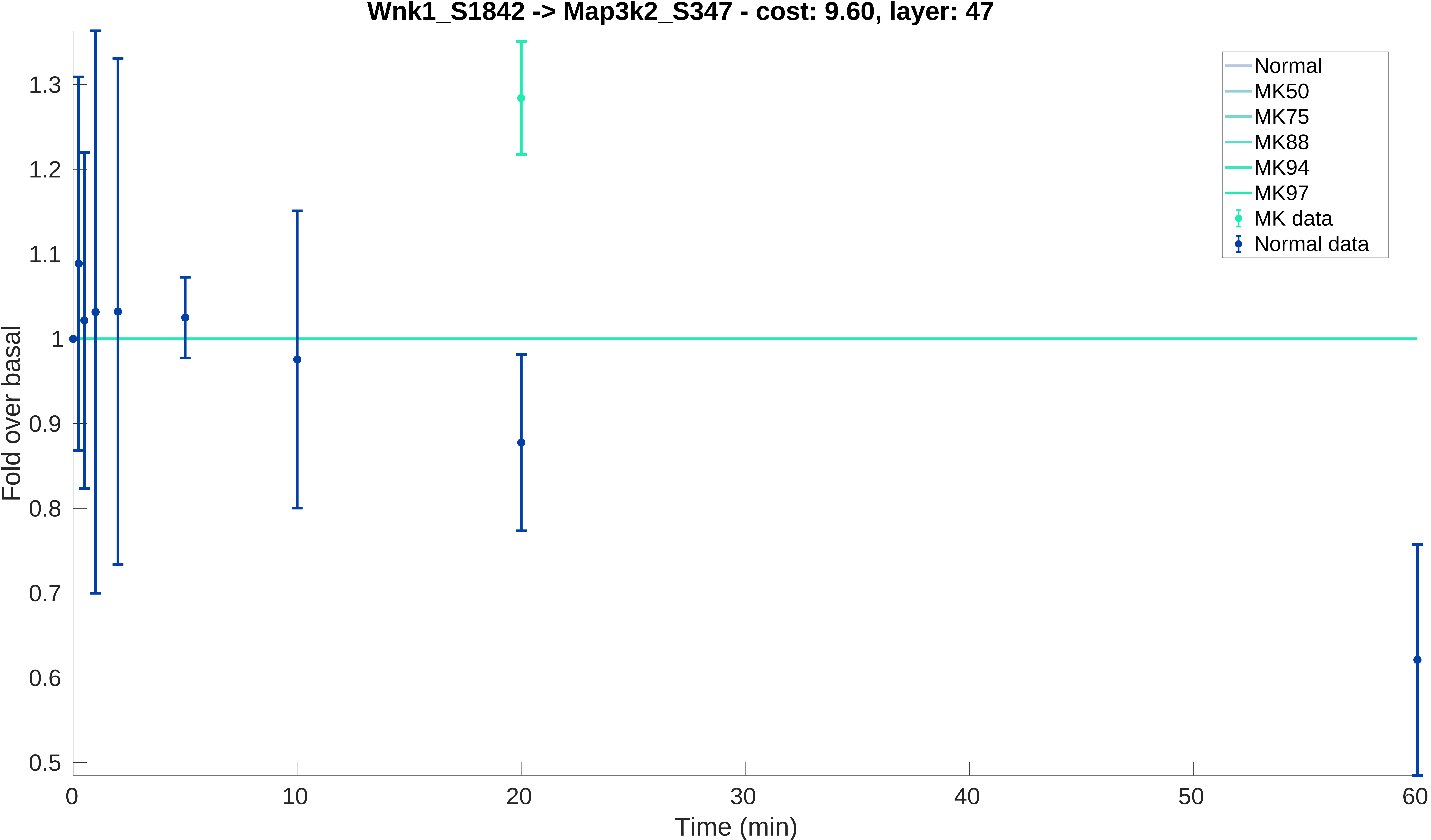

Wnk1_S1842 -> Map3k2_S347 - cost: 9.60, layer: 47
Fold over basal
Time (min)
1.3
1.2
1.1
1
0.9
0.8
0.7
0.6
0.5
0
10
20
30
40
50
60
Normal
MK50
MK75
MK88
MK94
MK97
MK data
Normal data

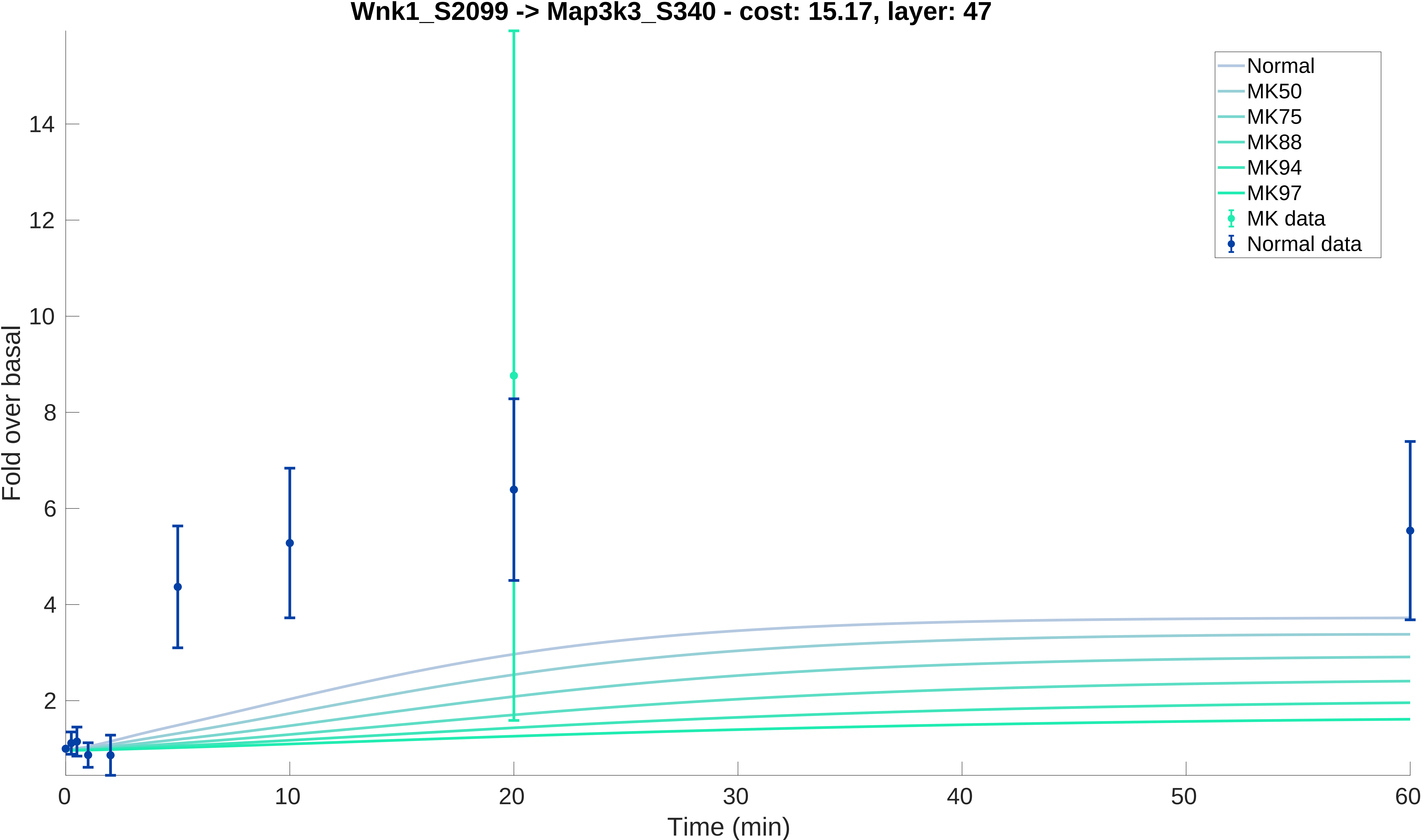

Wnk1_S2099 -> Map3k3_S340 - cost: 15.17, layer: 47
Fold over basal
Time (min)
0
10
20
30
40
50
60
2
4
6
8
10
12
14
Normal
MK50
MK75
MK88
MK94
MK97
MK data
Normal data

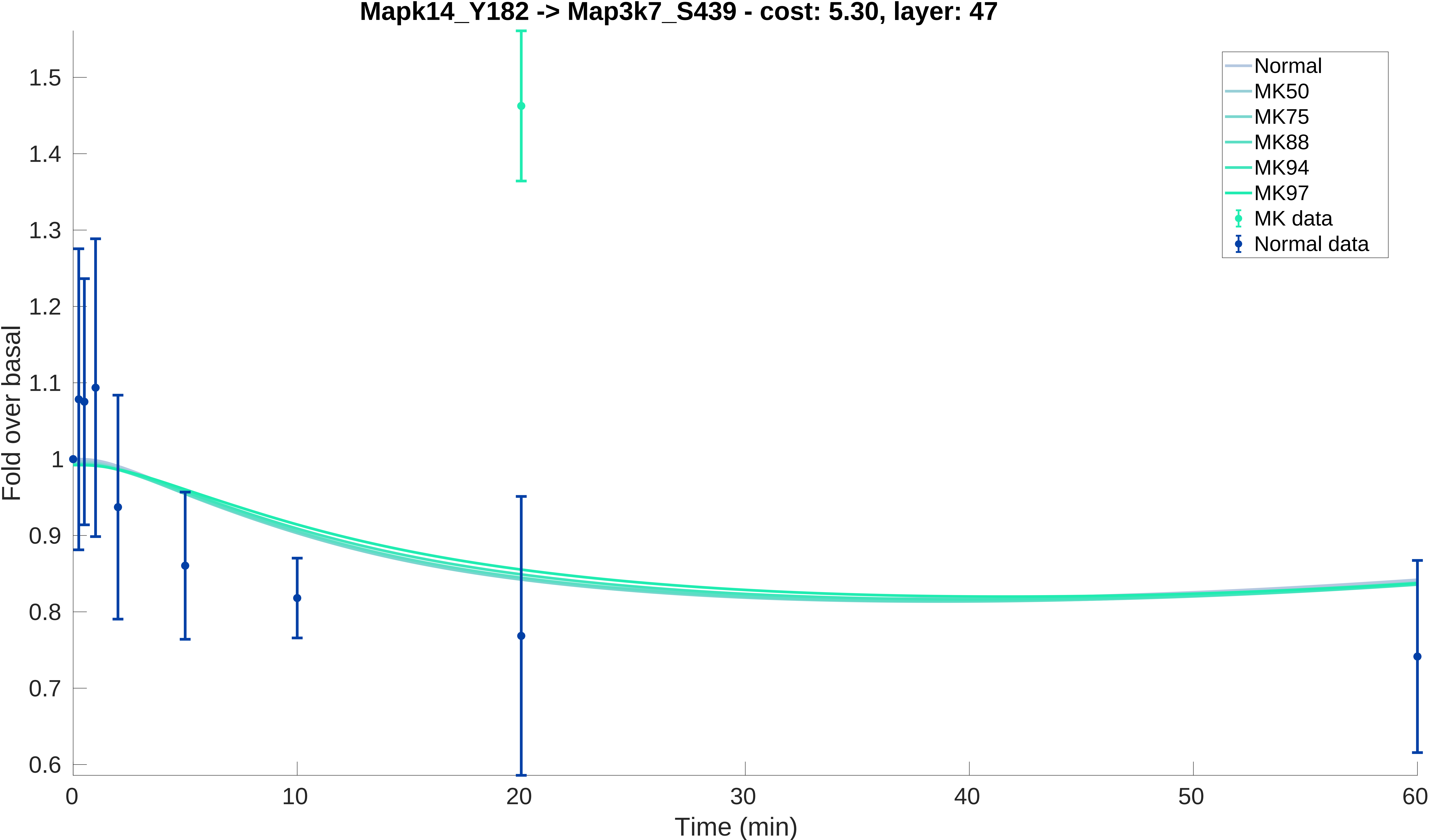

Mapk14_Y182 -> Map3k7_S439 - cost: 5.30, layer: 47
Fold over basal
Time (min)
Normal
MK50
MK75
MK88
MK94
MK97
MK data
Normal data
1.5
1.4
1.3
1.2
1.1
1
0.9
0.8
0.7
0.6
0
10
20
30
40
50
60

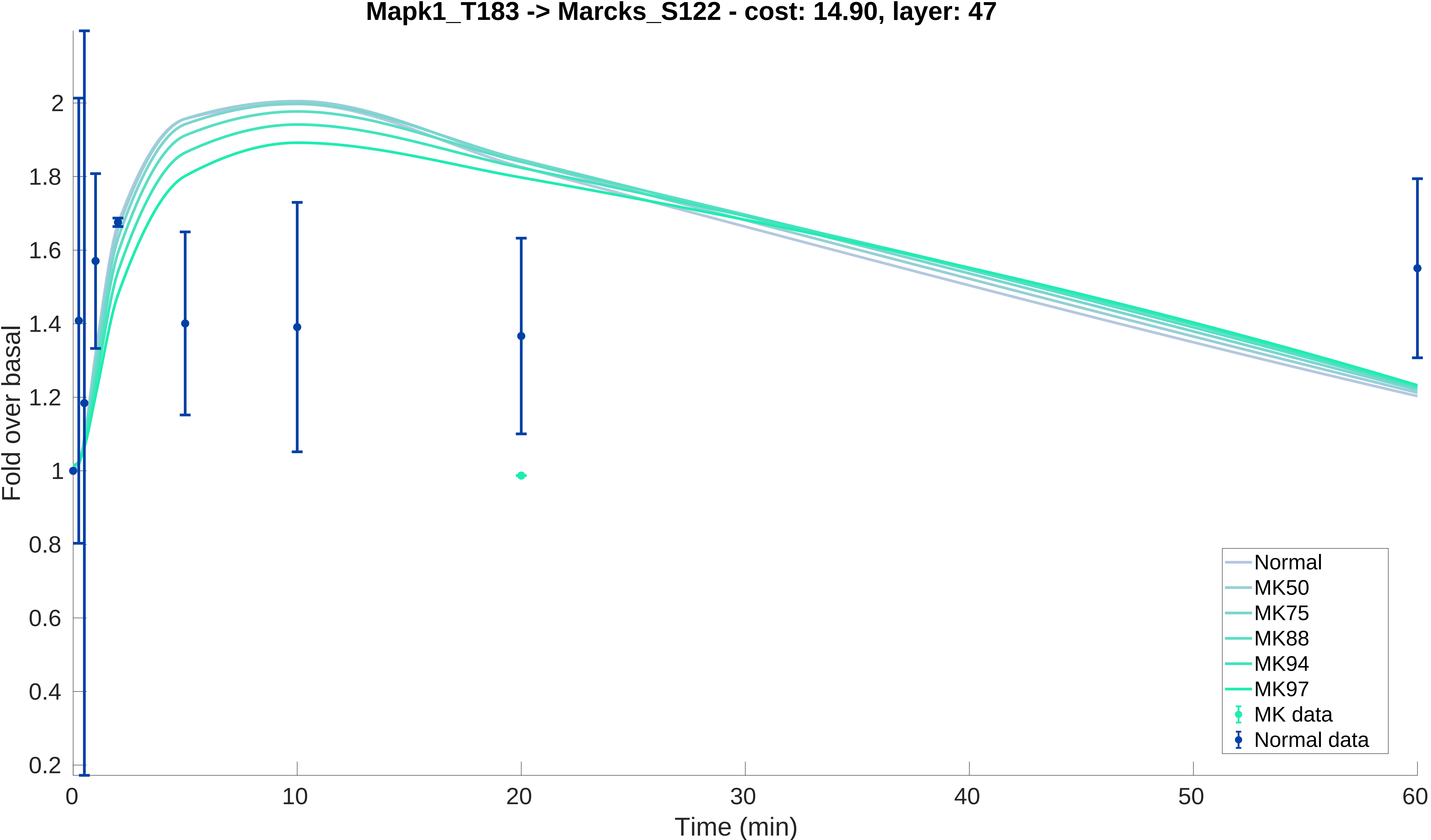

Mapk1_T183 -> Marcks_S122 - cost: 14.90, layer: 47
Fold over basal
Time (min)
0
10
20
30
40
50
60
0.2
0.4
0.6
0.8
1
1.2
1.4
1.6
1.8
2
Normal
MK50
MK75
MK88
MK94
MK97
MK data
Normal data

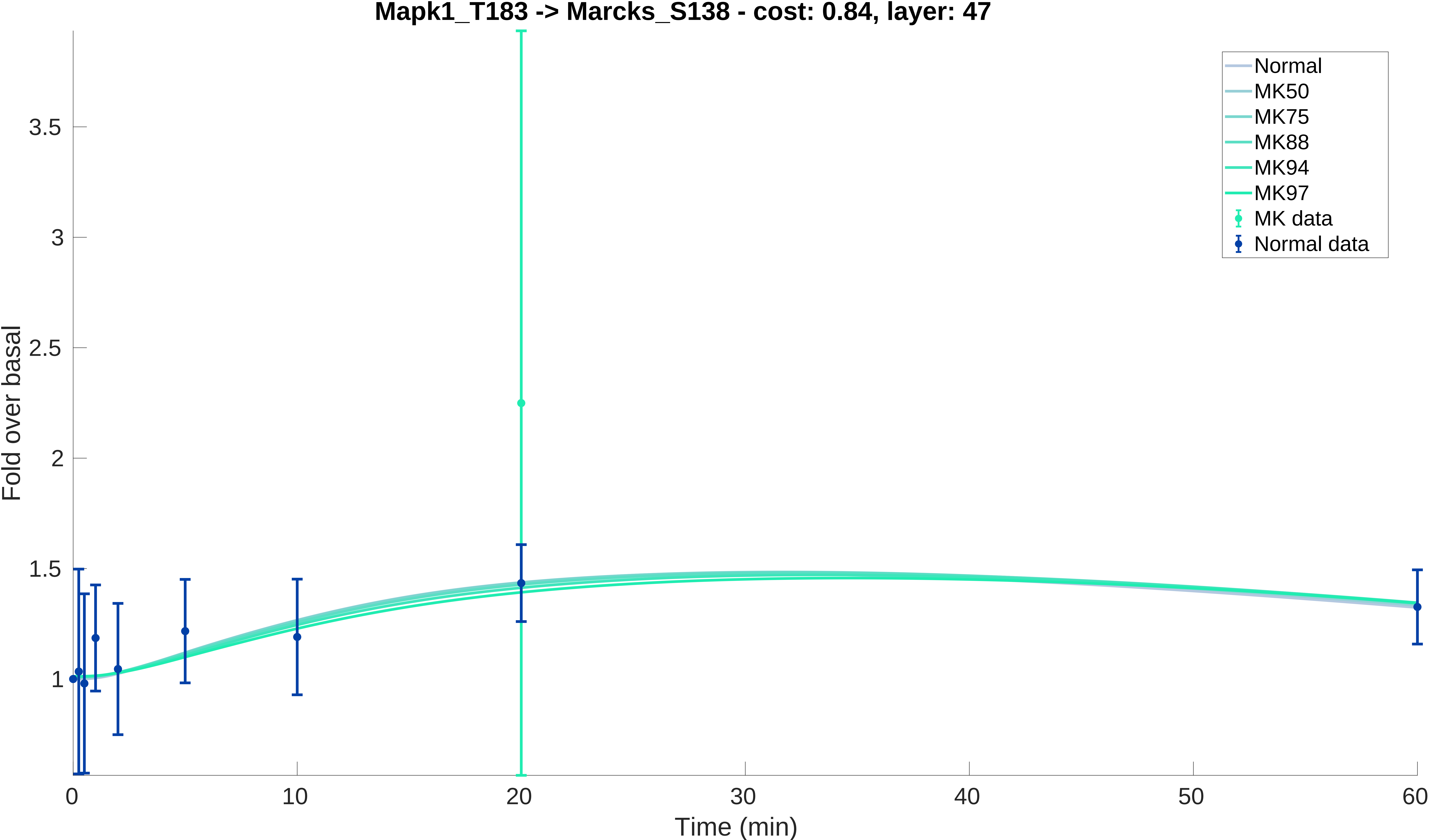

Mapk1_T183 -> Marcks_S138 - cost: 0.84, layer: 47
Fold over basal
Time (min)
3.5
3
2.5
2
1.5
1
0
10
20
30
40
50
60
Normal
MK50
MK75
MK88
MK94
MK97
MK data
Normal data

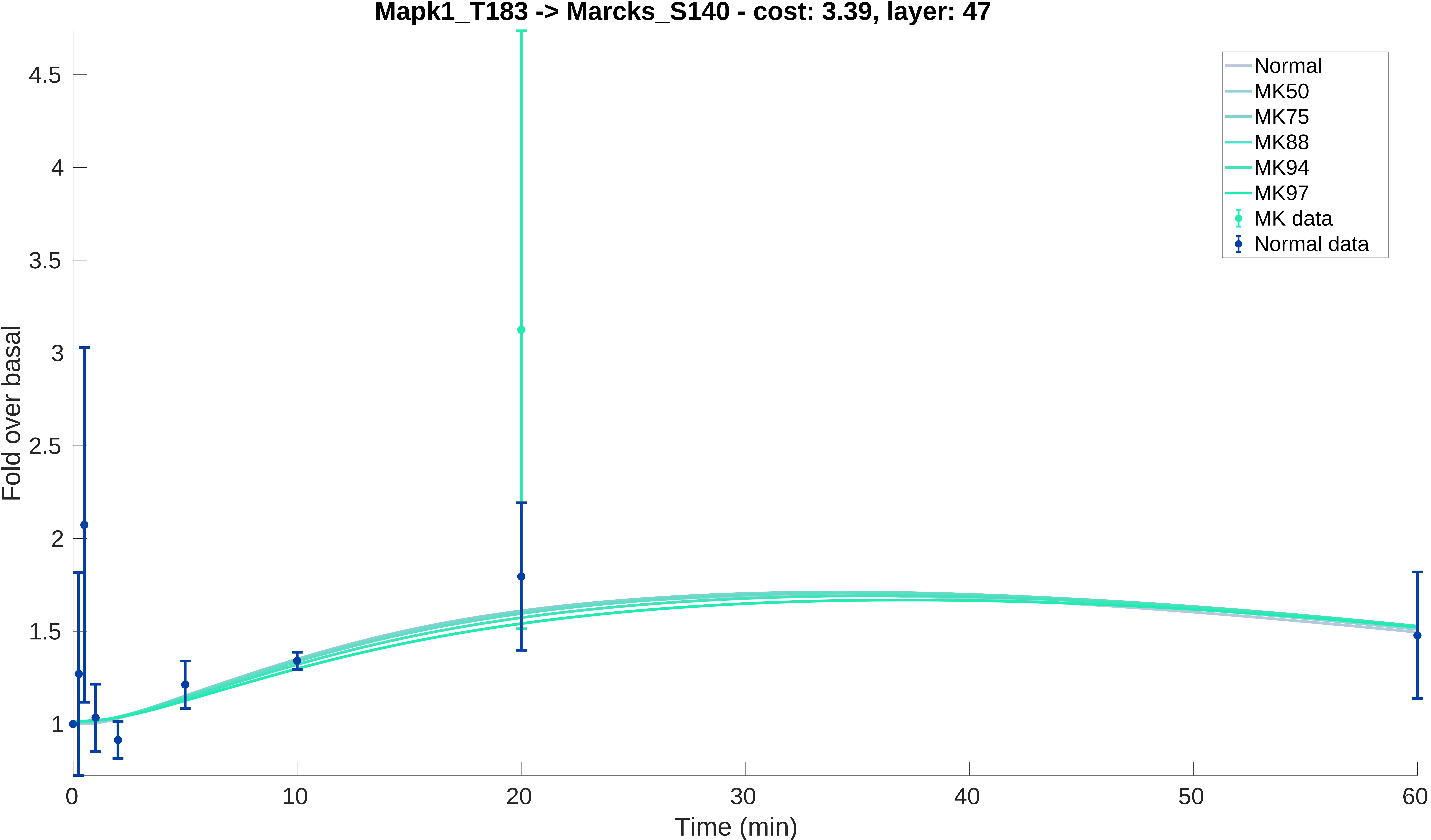

Mapk1_T183 -> Marcks_S140 - cost: 3.39, layer: 47
Fold over basal
Time (min)
Normal
MK50
MK75
MK88
MK94
MK97
MK data
Normal data
4.5
4
3.5
3
2.5
2
1.5
1
0
10
20
30
40
50
60

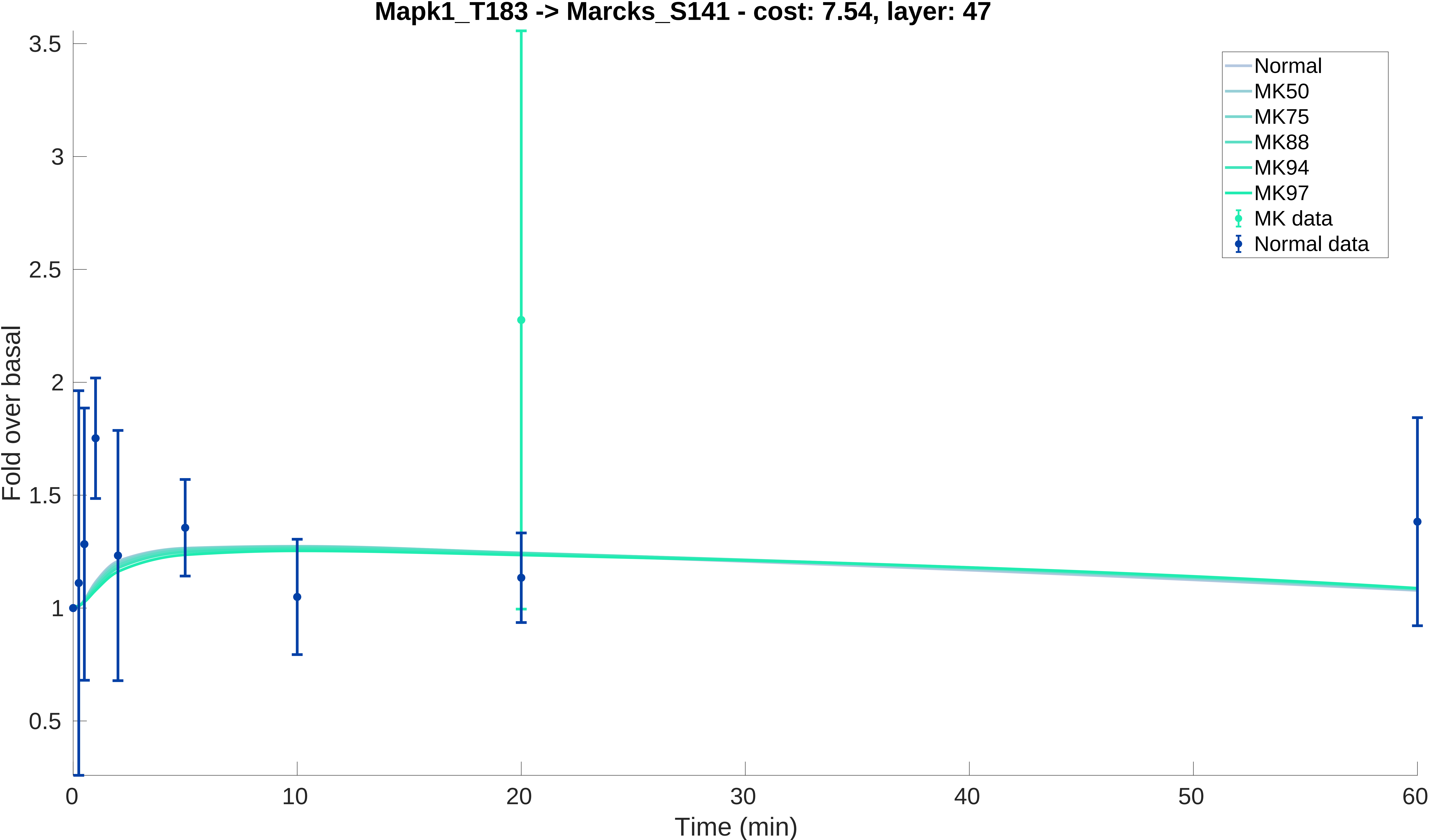

Mapk1_T183 -> Marcks_S141 - cost: 7.54, layer: 47
Fold over basal
Time (min)
3.5
3
2.5
2
1.5
1
0.5
0
10
20
30
40
50
60
Normal
MK50
MK75
MK88
MK94
MK97
MK data
Normal data

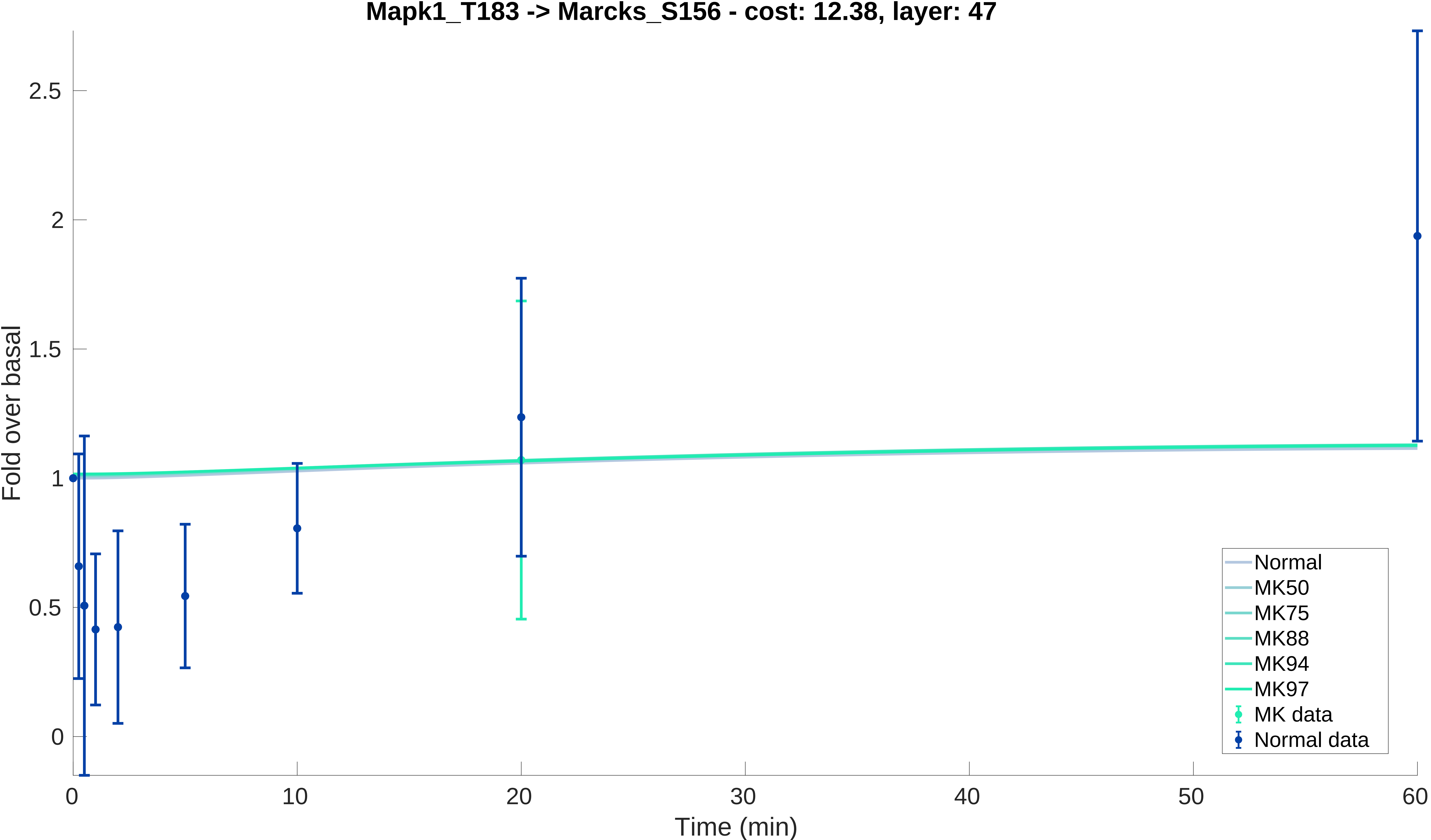

Mapk1_T183 -> Marcks_S156 - cost: 12.38, layer: 47
Fold over basal
Time (min)
0
0.5
1
1.5
2
2.5
0
10
20
30
40
50
60
Normal
MK50
MK75
MK88
MK94
MK97
MK data
Normal data

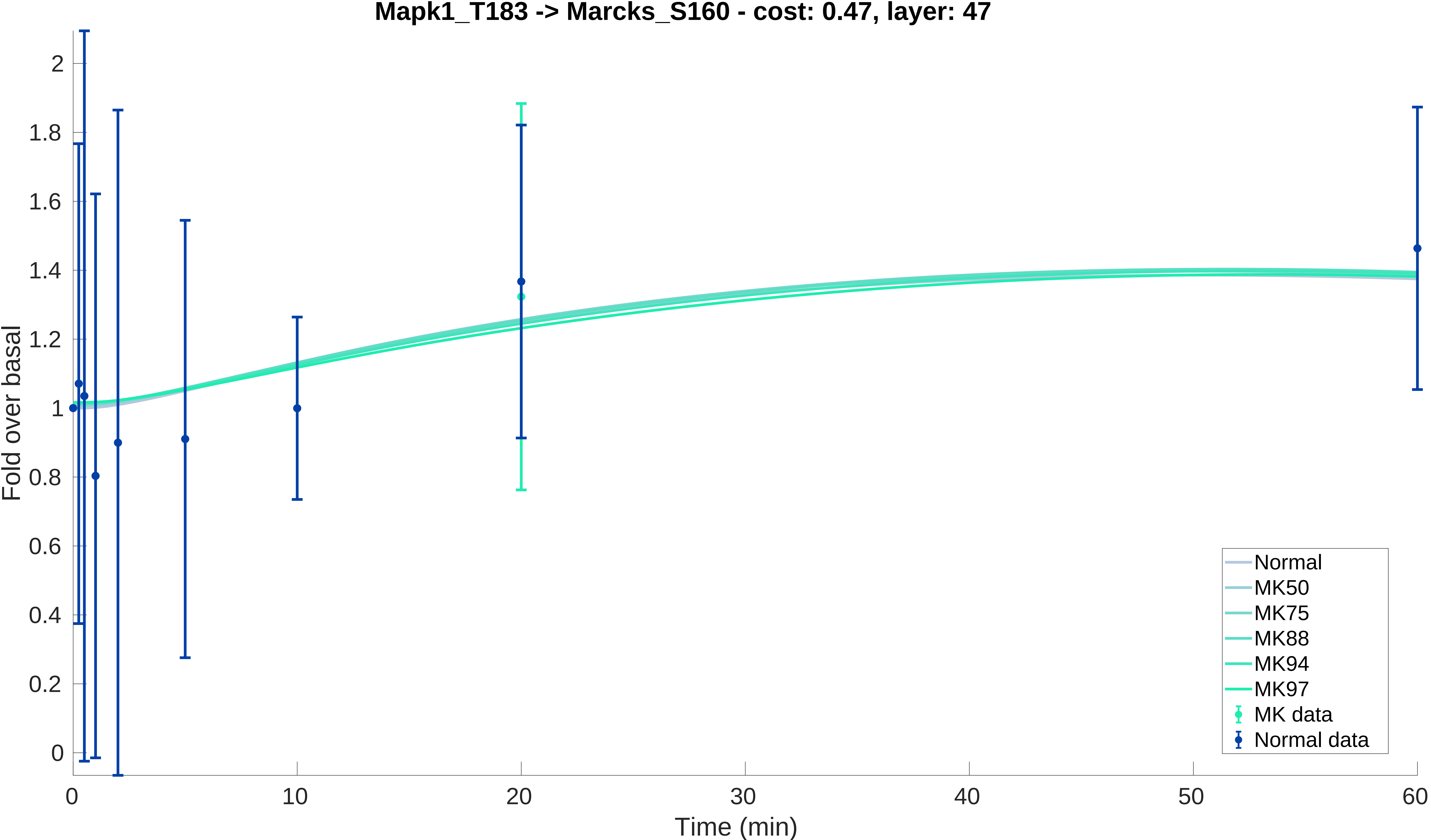

Mapk1_T183 -> Marcks_S160 - cost: 0.47, layer: 47
Fold over basal
Time (min)
0
0.2
0.4
0.6
0.8
1
1.2
1.4
1.6
1.8
2
0
10
20
30
40
50
60
Normal
MK50
MK75
MK88
MK94
MK97
MK data
Normal data

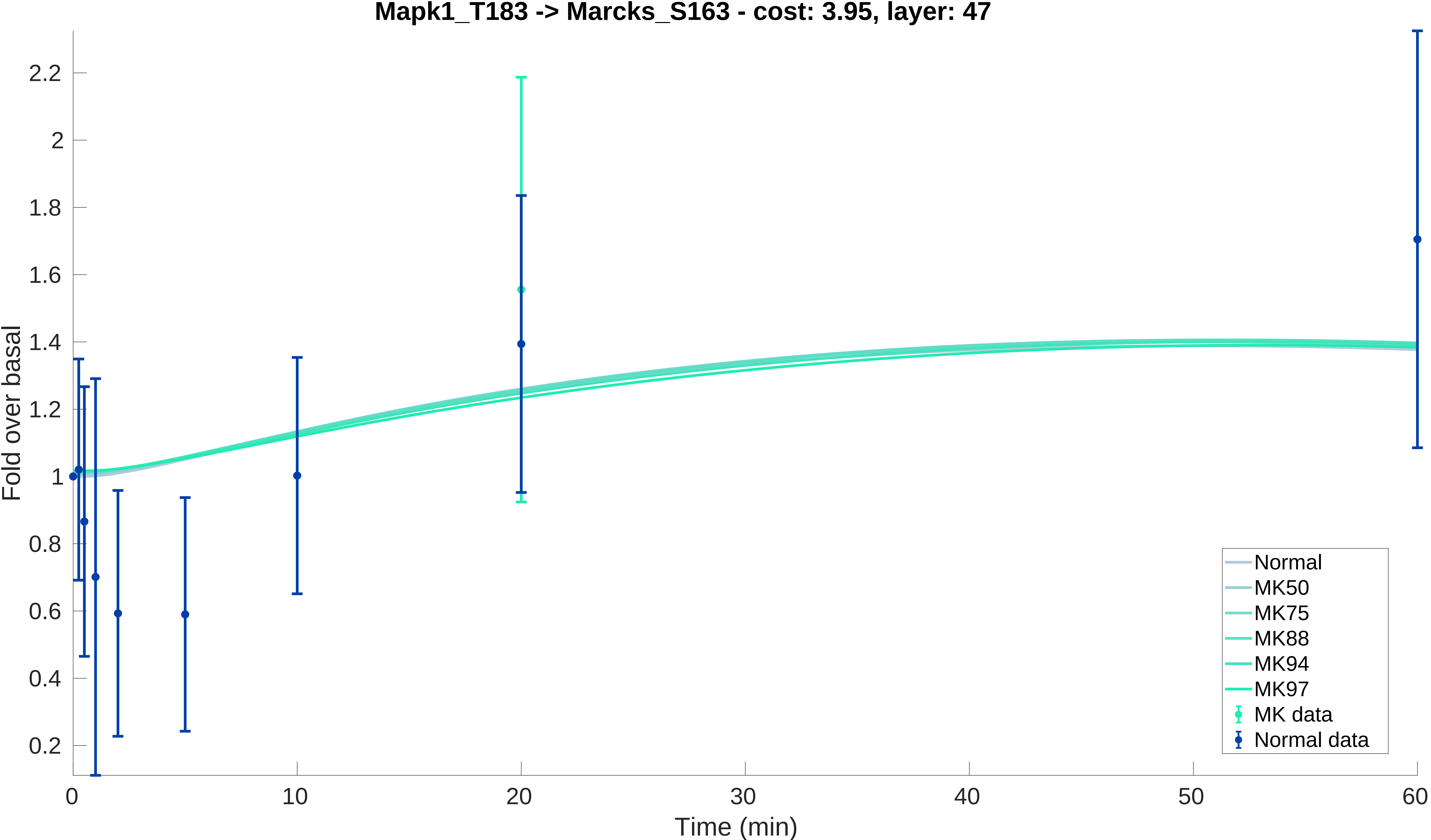

Mapk1_T183 -> Marcks_S163 - cost: 3.95, layer: 47
Fold over basal
Time (min)
0.2
0.4
0.6
0.8
1
1.2
1.4
1.6
1.8
2
2.2
0
10
20
30
40
50
60
Normal
MK50
MK75
MK88
MK94
MK97
MK data
Normal data

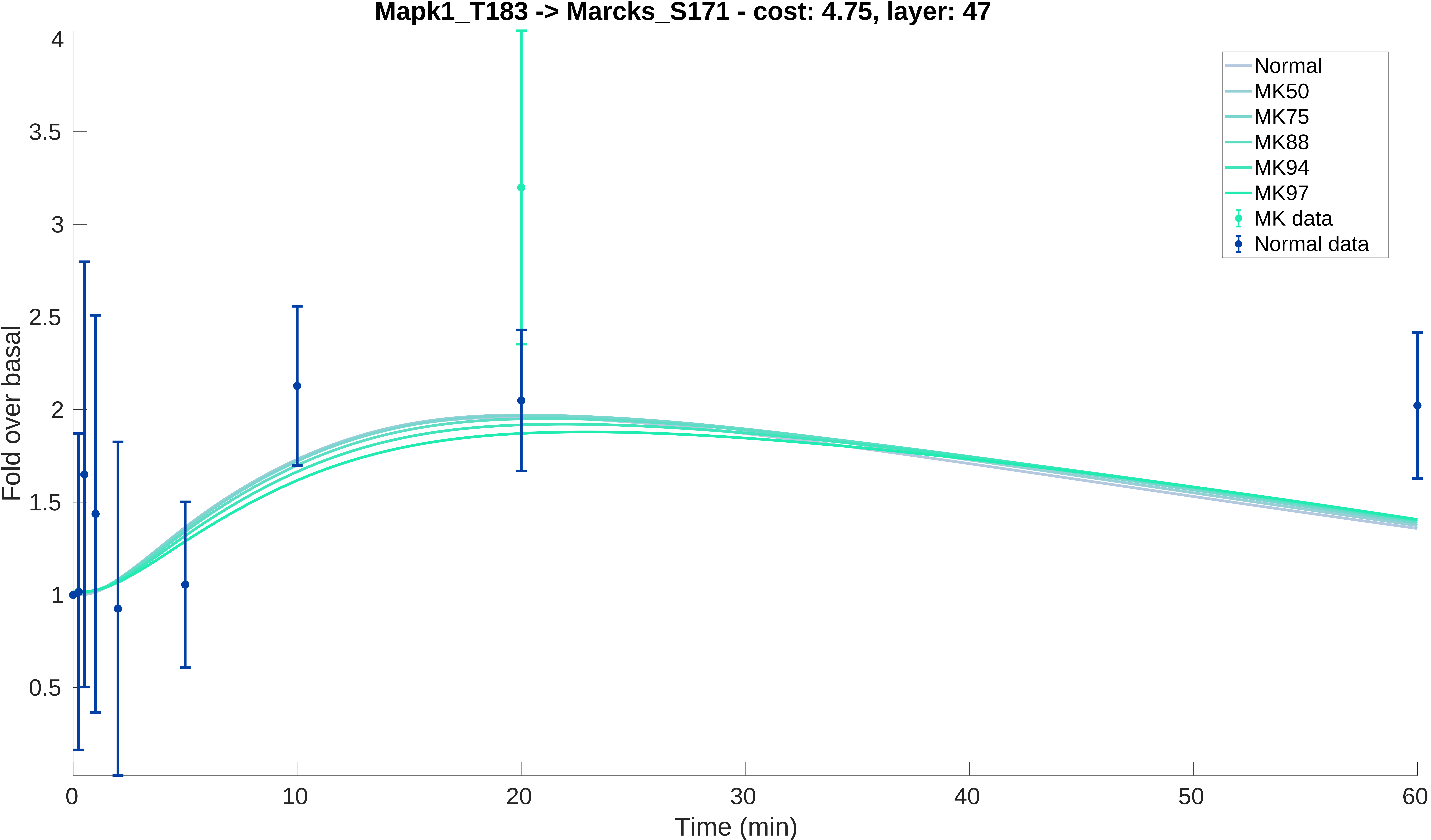

Mapk1_T183 -> Marcks_S171 - cost: 4.75, layer: 47
Fold over basal
Time (min)
0.5
1
1.5
2
2.5
3
3.5
4
0
10
20
30
40
50
60
Normal
MK50
MK75
MK88
MK94
MK97
MK data
Normal data

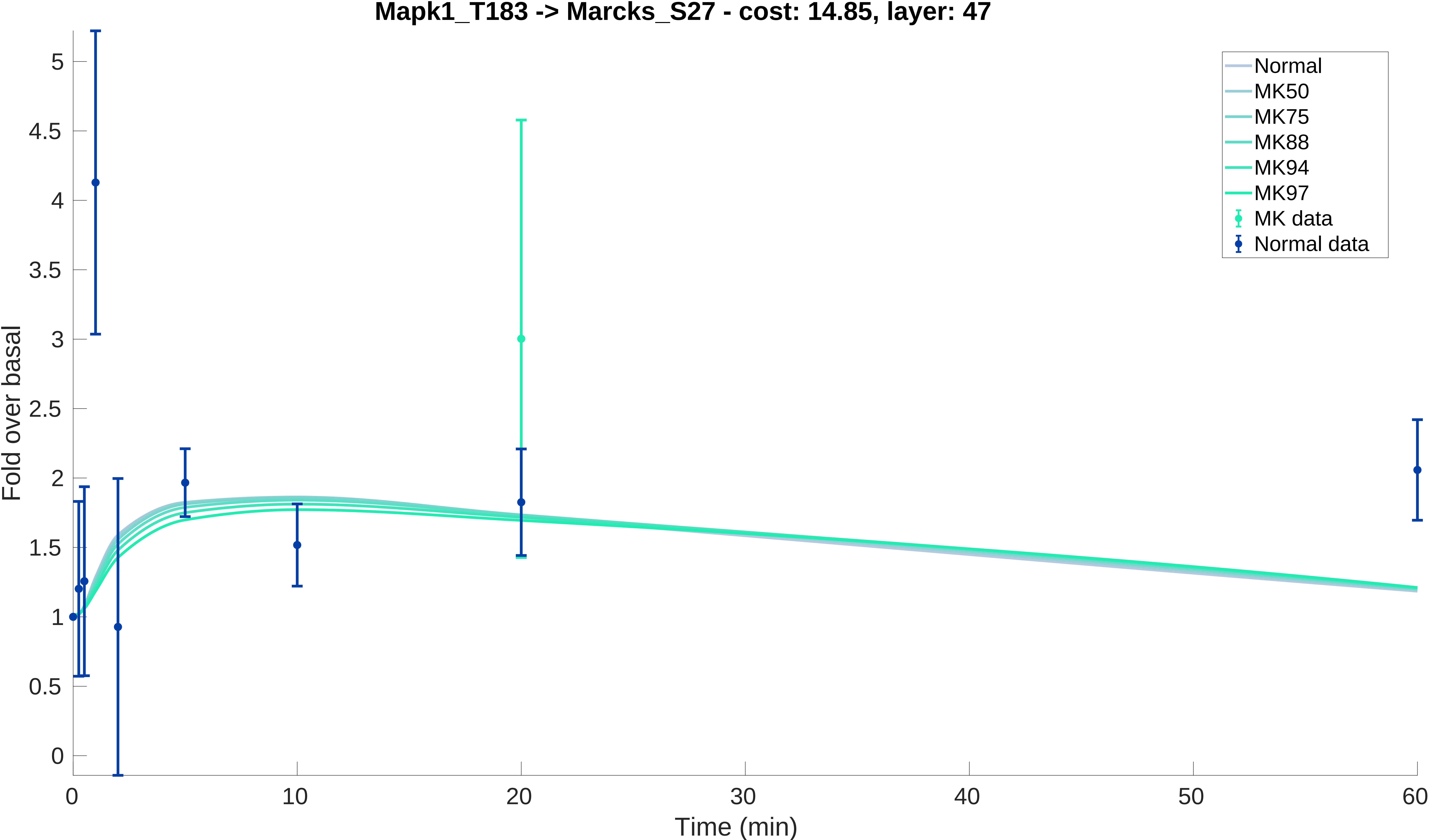

Mapk1_T183 -> Marcks_S27 - cost: 14.85, layer: 47
Fold over basal
Time (min)
0
0.5
1
1.5
2
2.5
3
3.5
4
4.5
5
0
10
20
30
40
50
60
Normal
MK50
MK75
MK88
MK94
MK97
MK data
Normal data

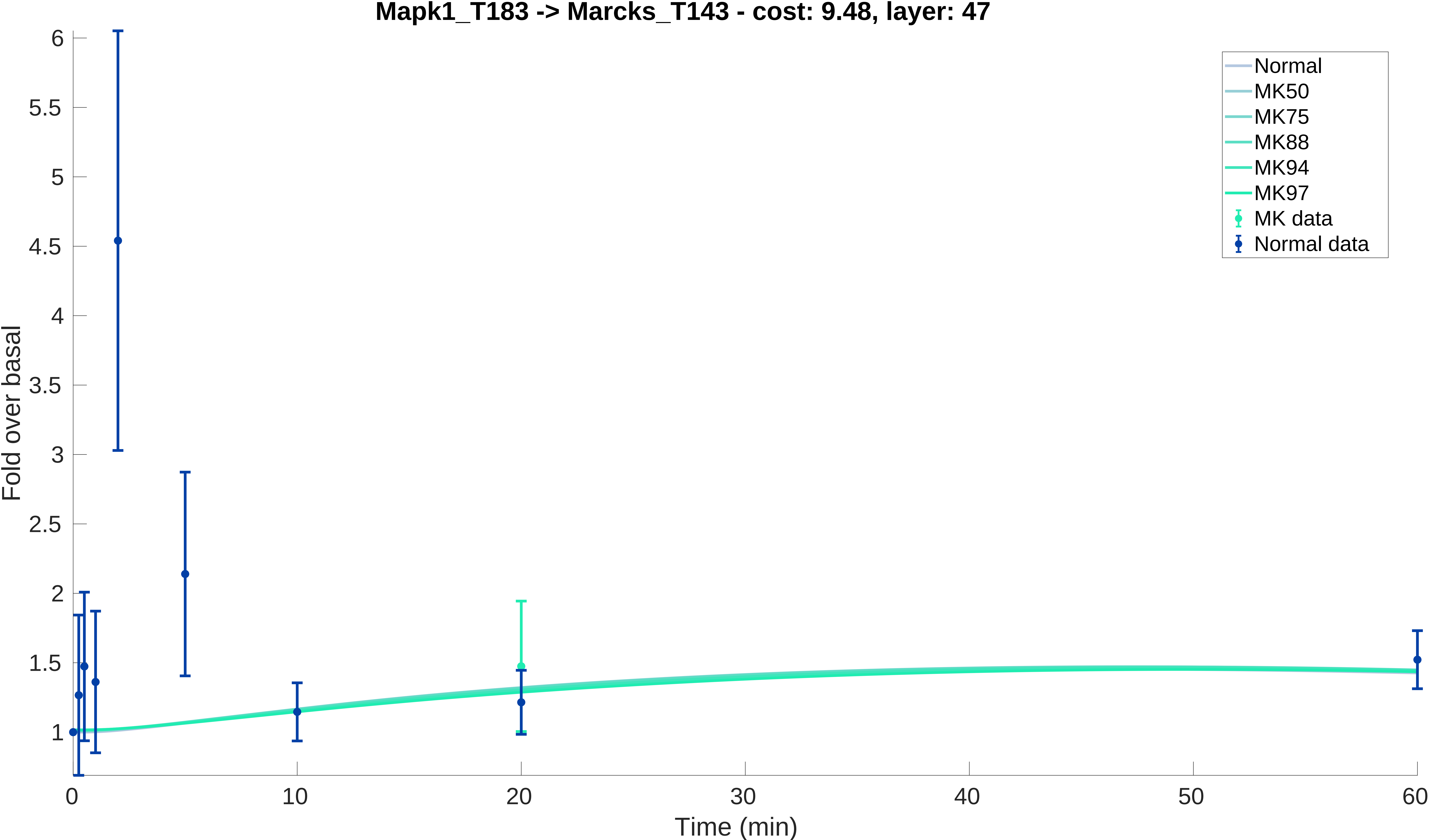

Mapk1_T183 -> Marcks_T143 - cost: 9.48, layer: 47
Fold over basal
Time (min)
6
5.5
5
4.5
4
3.5
3
2.5
2
1.5
1
0
10
20
30
40
50
60
Normal
MK50
MK75
MK88
MK94
MK97
MK data
Normal data

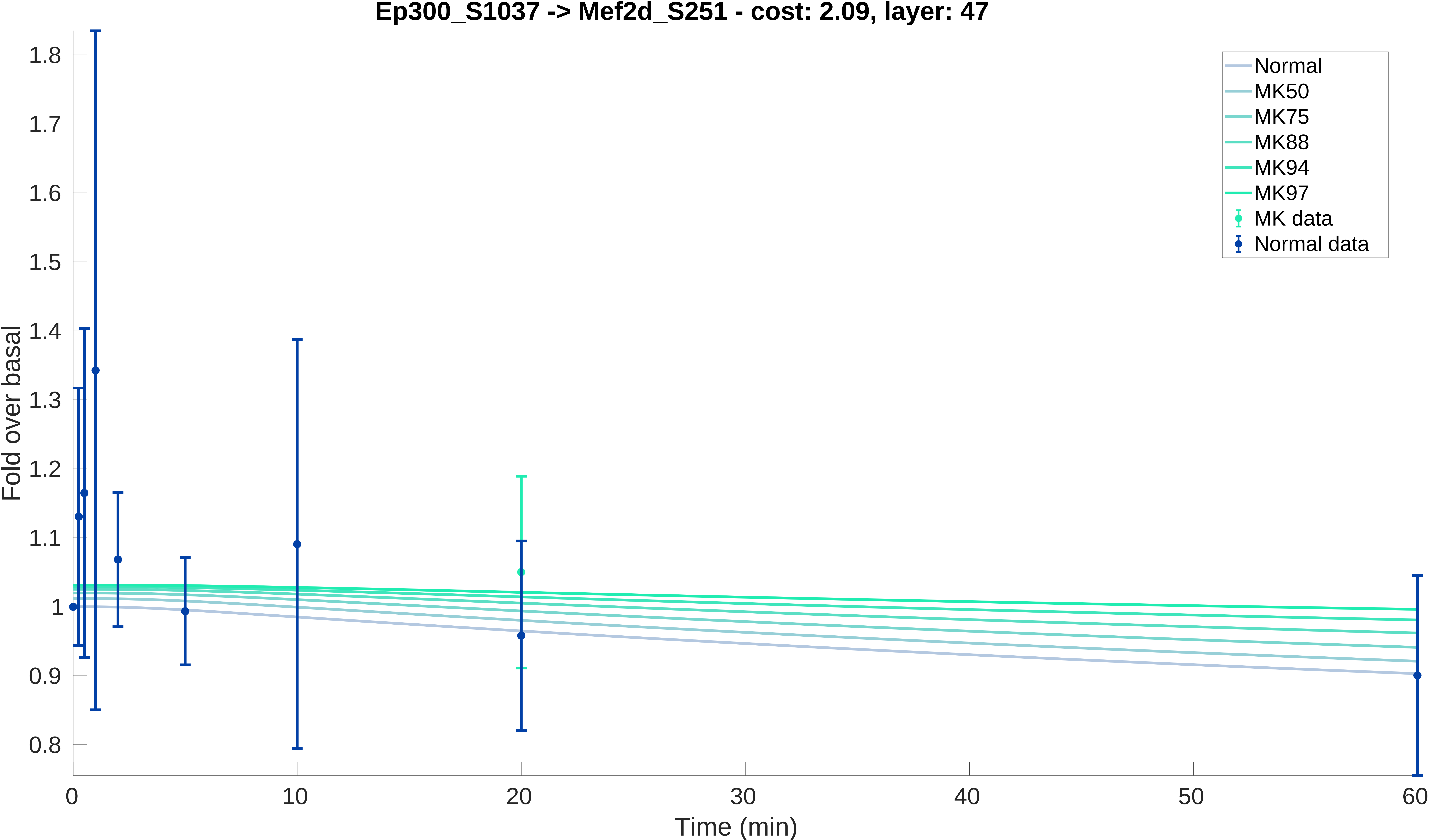

Ep300_S1037 -> Mef2d_S251 - cost: 2.09, layer: 47
Fold over basal
Time (min)
0.8
0.9
1
1.1
1.2
1.3
1.4
1.5
1.6
1.7
1.8
0
10
20
30
40
50
60
Normal
MK50
MK75
MK88
MK94
MK97
MK data
Normal data

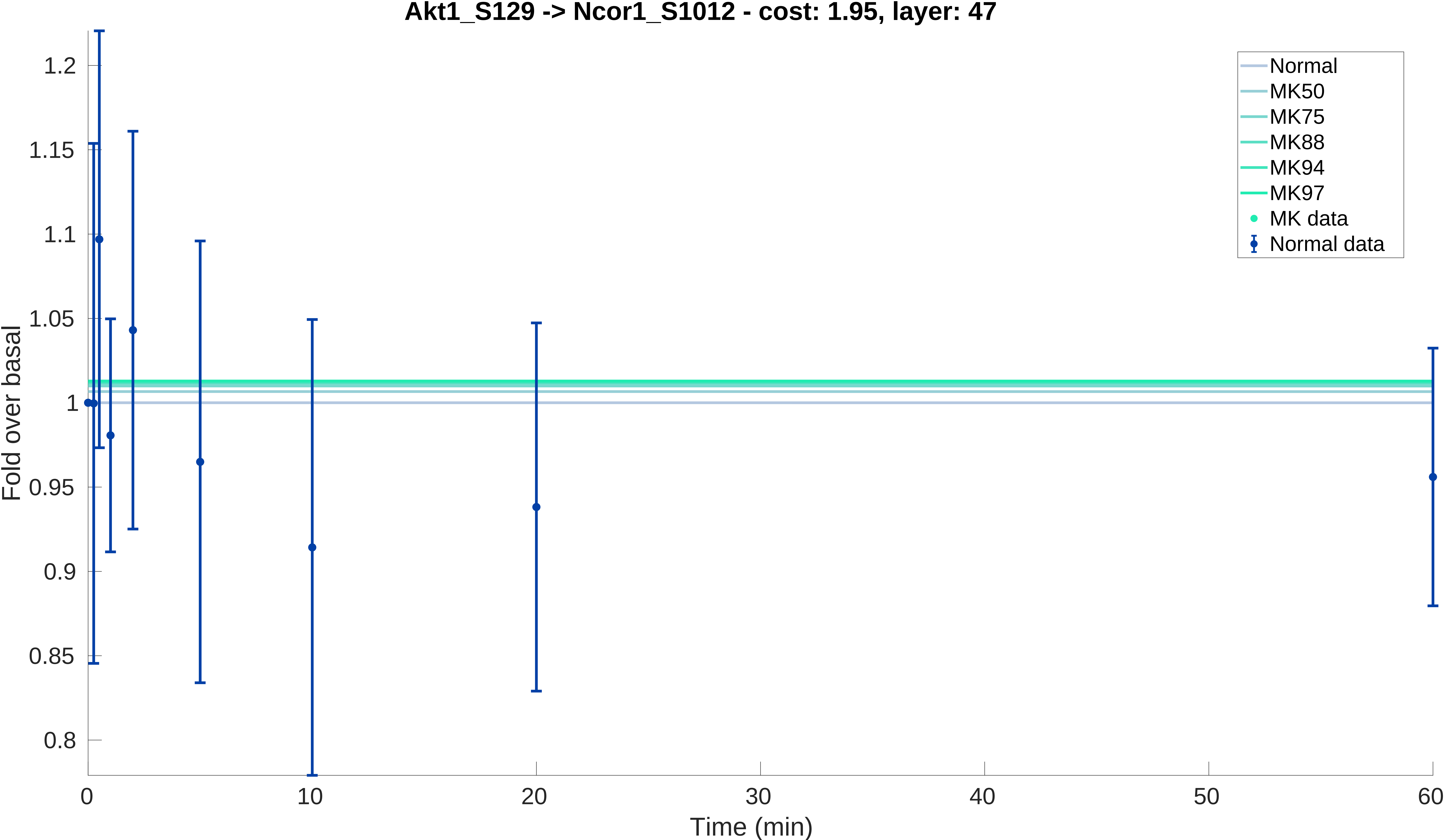

Akt1_S129 -> Ncor1_S1012 - cost: 1.95, layer: 47
1.2
1.15
1.1
1.05
1
0.95
0.9
0.85
0.8
Fold over basal
0
10
20
30
40
50
60
Time (min)
Normal
MK50
MK75
MK88
MK94
MK97
MK data
Normal data

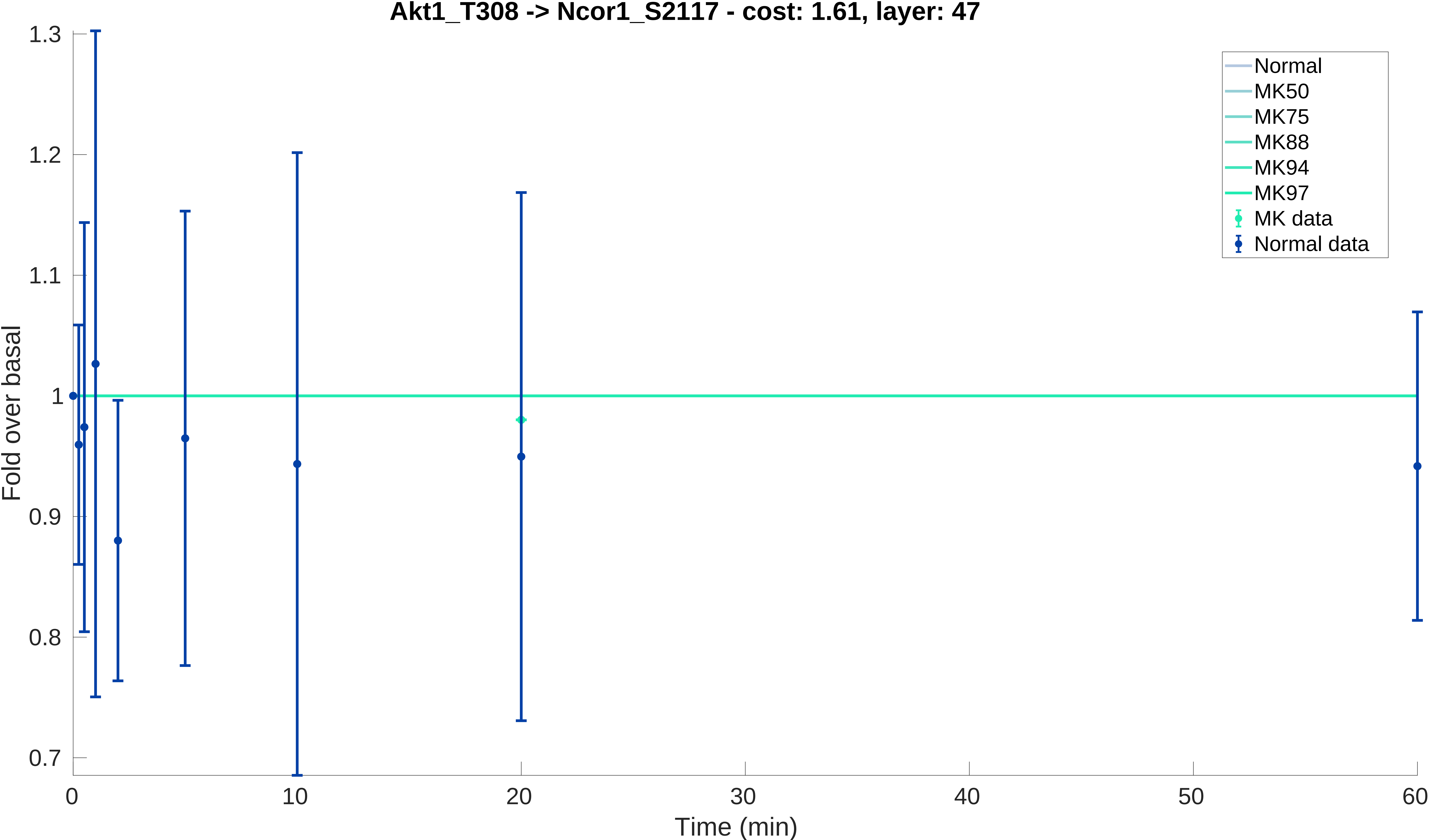

Akt1_T308 -> Ncor1_S2117 - cost: 1.61, layer: 47
Fold over basal
Time (min)
1.3
1.2
1.1
1
0.9
0.8
0.7
0
10
20
30
40
50
60
Normal
MK50
MK75
MK88
MK94
MK97
MK data
Normal data

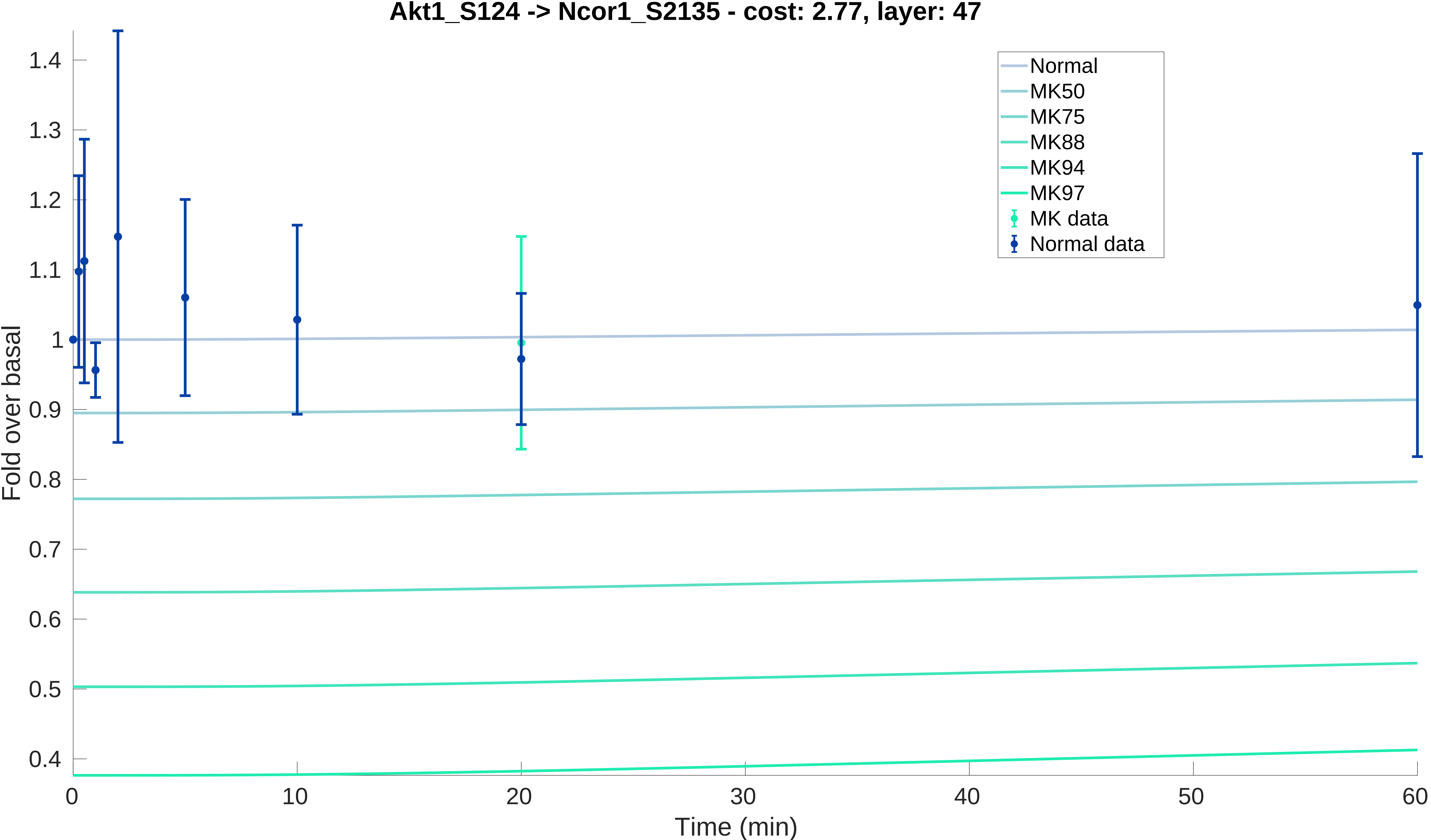

Akt1_S124 -> Ncor1_S2135 - cost: 2.77, layer: 47
Normal
MK50
MK75
MK88
MK94
MK97
MK data
Normal data
Fold over basal
Time (min)
1.4
1.3
1.2
1.1
1
0.9
0.8
0.7
0.6
0.5
0.4
0
10
20
30
40
50
60

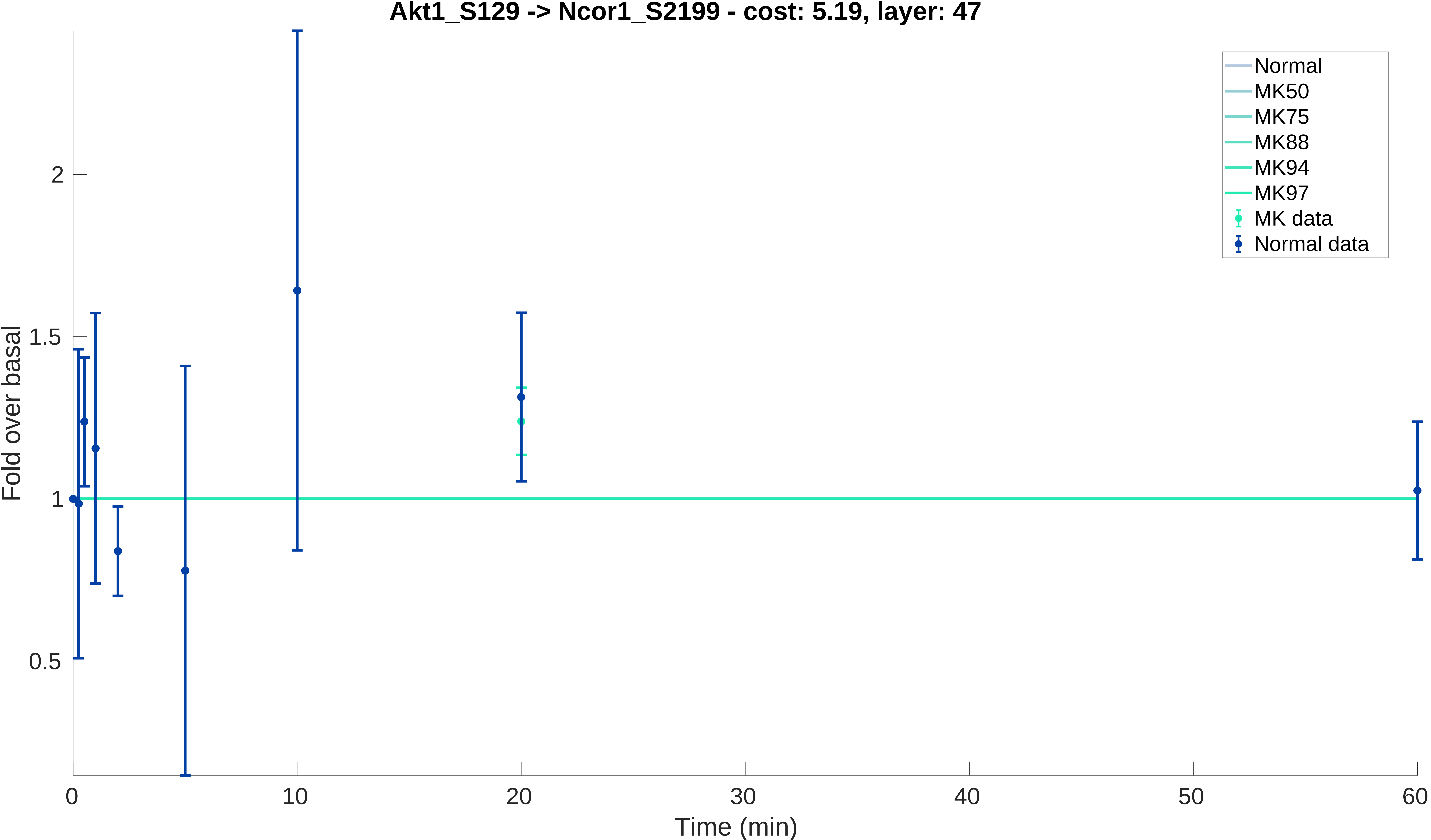
Akt1_S129 -> Ncor1_S2199 - cost: 5.19, layer: 47
Fold over basal
Time (min)
0.5
1
1.5
2
0
10
20
30
40
50
60
Normal
MK50
MK75
MK88
MK94
MK97
MK data
Normal data

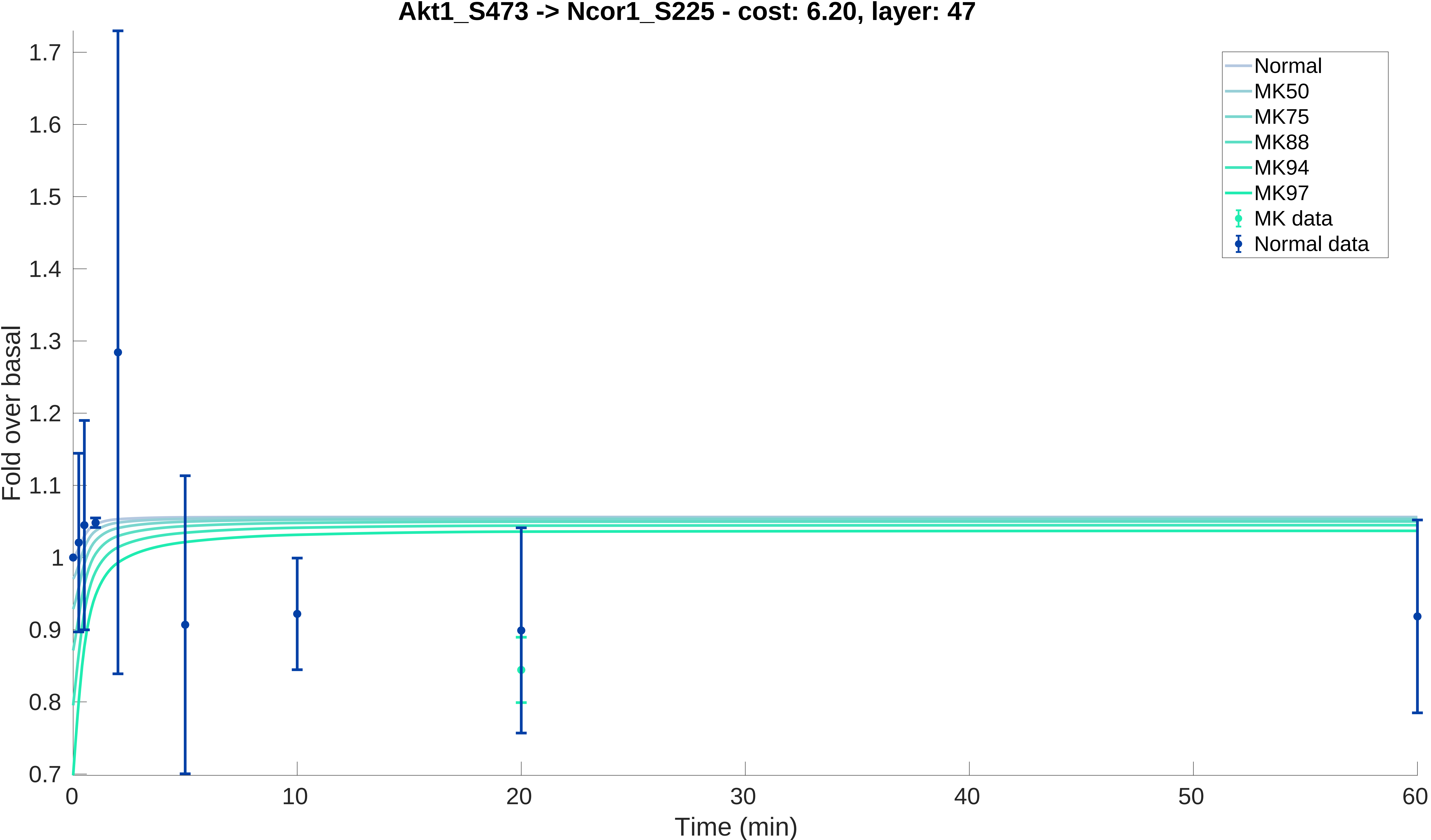
Akt1_S473 -> Ncor1_S225 - cost: 6.20, layer: 47
Fold over basal
Time (min)
0.7
0.8
0.9
1
1.1
1.2
1.3
1.4
1.5
1.6
1.7
0
10
20
30
40
50
60
Normal
MK50
MK75
MK88
MK94
MK97
MK data
Normal data

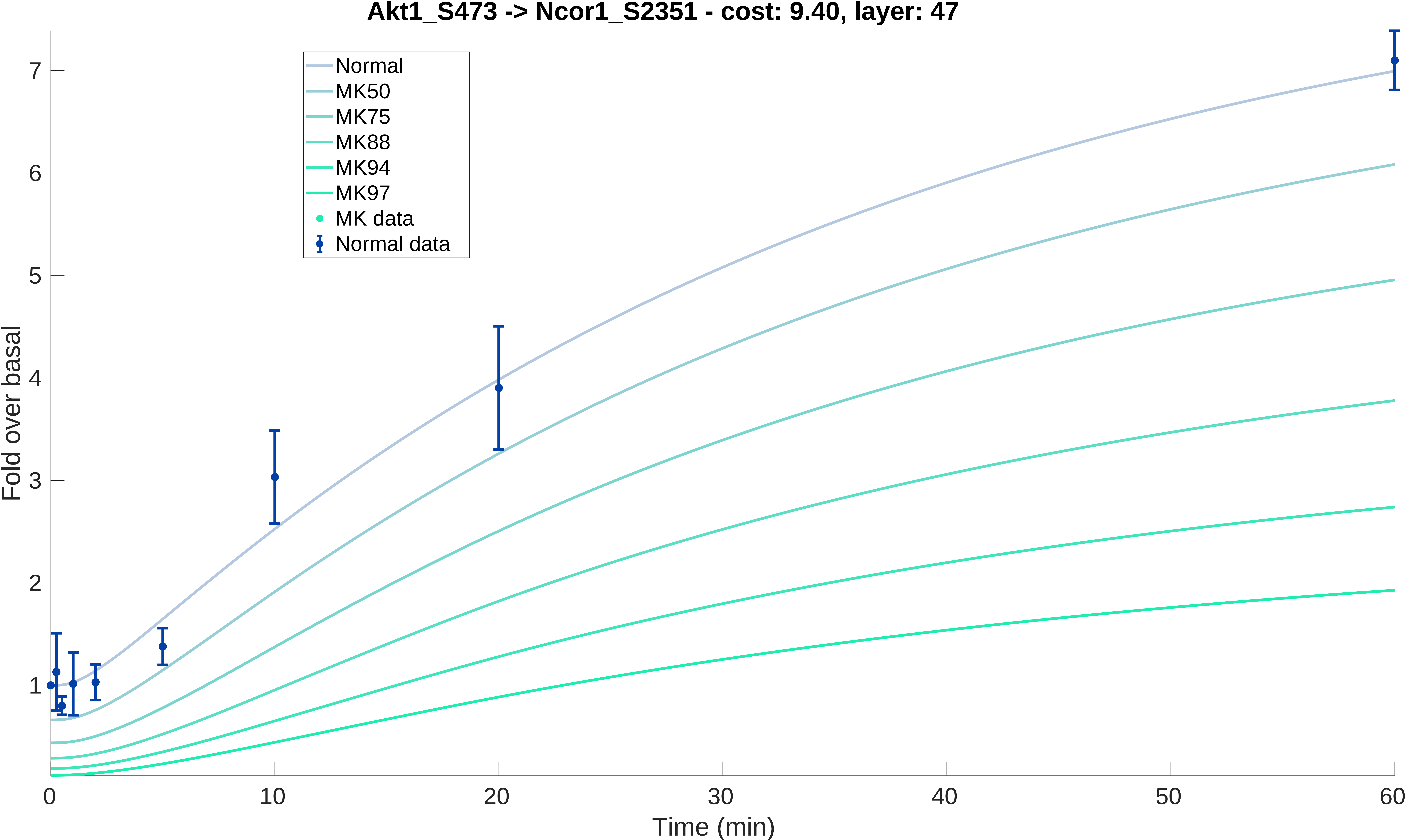
Akt1_S473 -> Ncor1_S2351 - cost: 9.40, layer: 47
Normal
MK50
MK75
MK88
MK94
MK97
MK data
Normal data
Fold over basal
Time (min)
0
10
20
30
40
50
60
1
2
3
4
5
6
7

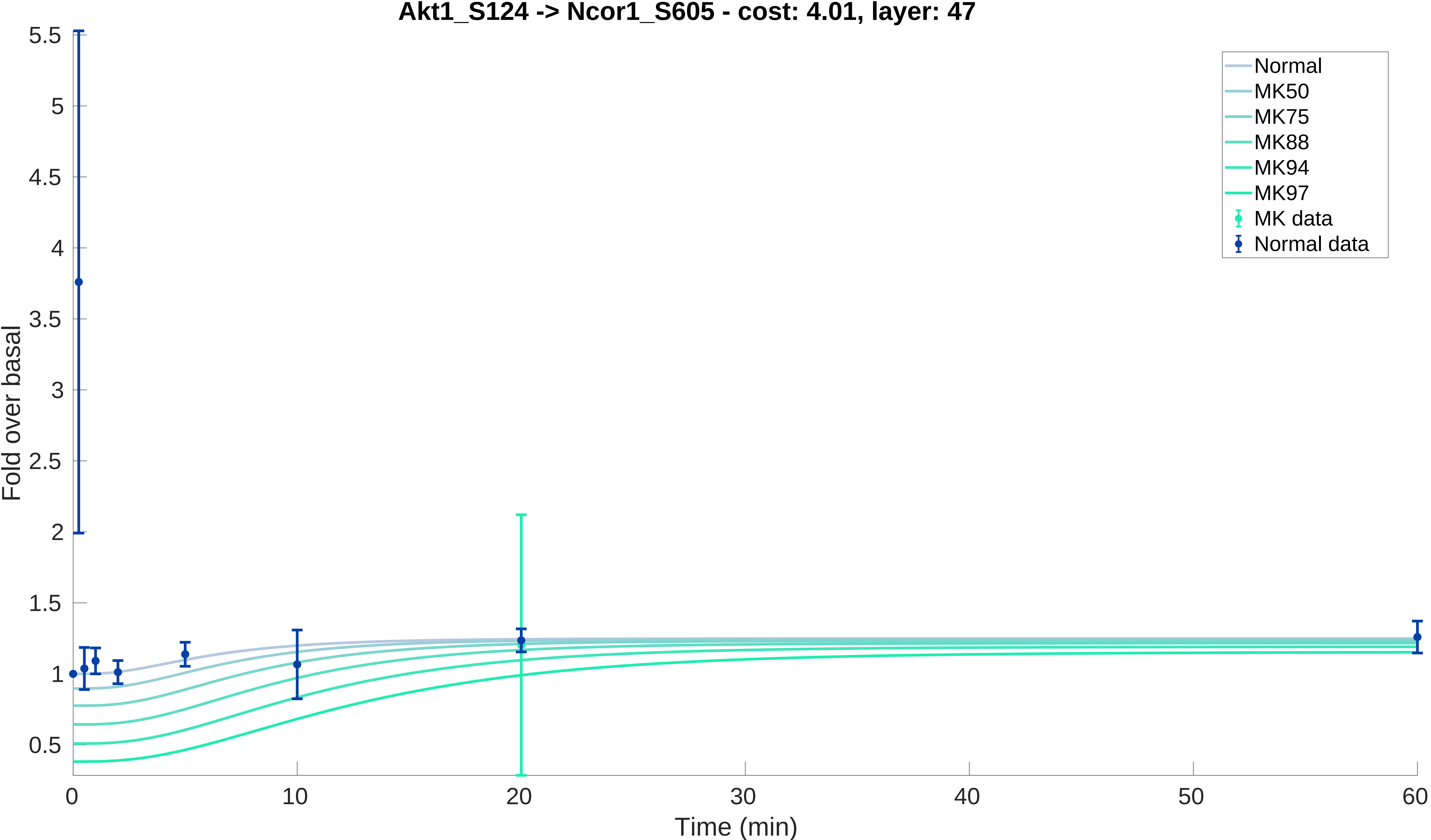

Akt1_S124 -> Ncor1_S605 - cost: 4.01, layer: 47
Fold over basal
Time (min)
0.5
1
1.5
2
2.5
3
3.5
4
4.5
5
5.5
0
10
20
30
40
50
60
Normal
MK50
MK75
MK88
MK94
MK97
MK data
Normal data

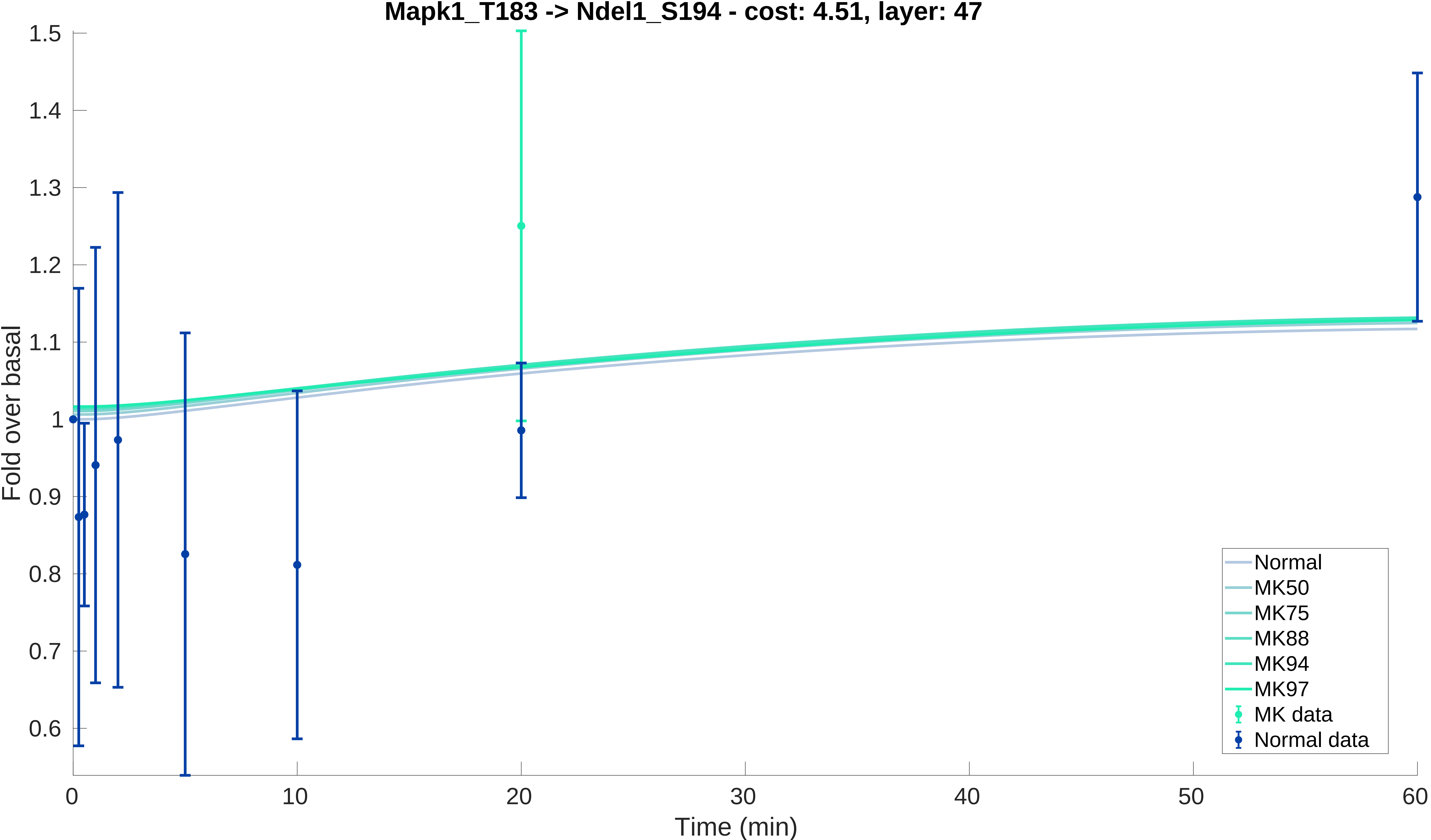

Mapk1_T183 -> Ndel1_S194 - cost: 4.51, layer: 47
Fold over basal
Time (min)
1.5
1.4
1.3
1.2
1.1
1
0.9
0.8
0.7
0.6
0
10
20
30
40
50
60
Normal
MK50
MK75
MK88
MK94
MK97
MK data
Normal data

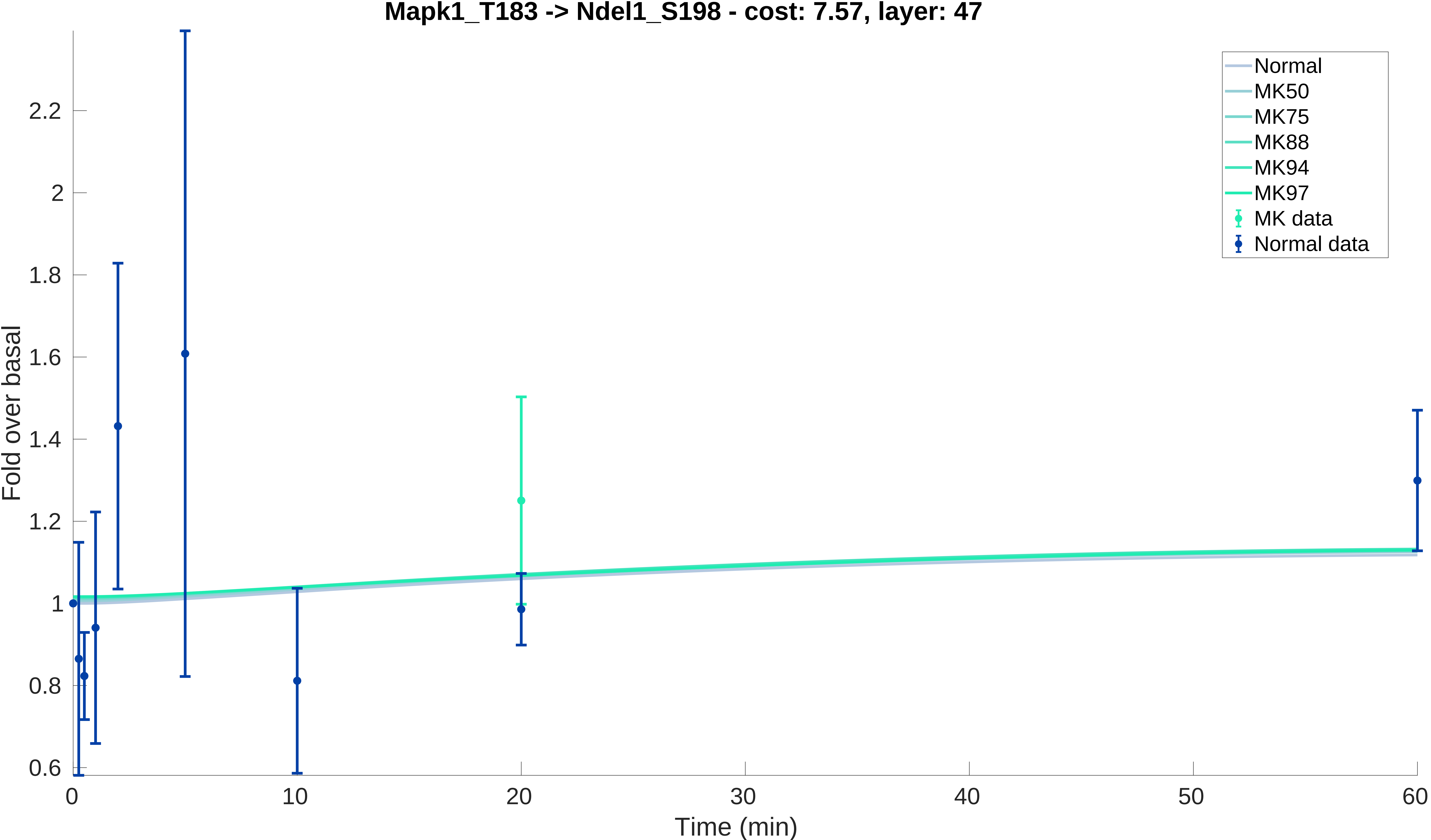

Mapk1_T183 -> Ndel1_S198 - cost: 7.57, layer: 47
Fold over basal
Time (min)
0.6
0.8
1
1.2
1.4
1.6
1.8
2
2.2
0
10
20
30
40
50
60
Normal
MK50
MK75
MK88
MK94
MK97
MK data
Normal data

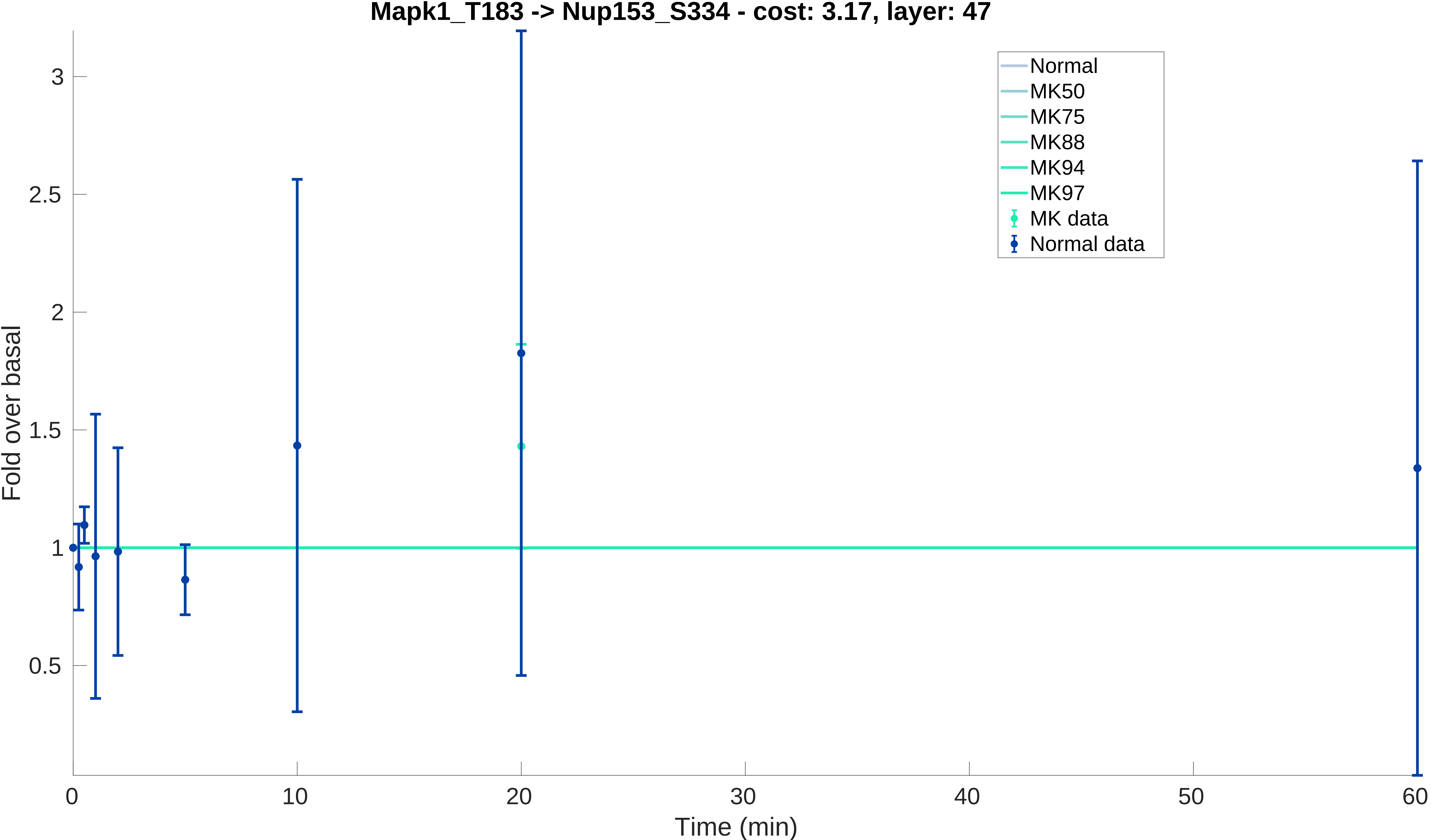

Mapk1_T183 -> Nup153_S334 - cost: 3.17, layer: 47
Fold over basal
3
2.5
2
1.5
1
0.5
0
10
20
30
40
50
60
Time (min)
Normal
MK50
MK75
MK88
MK94
MK97
MK data
Normal data

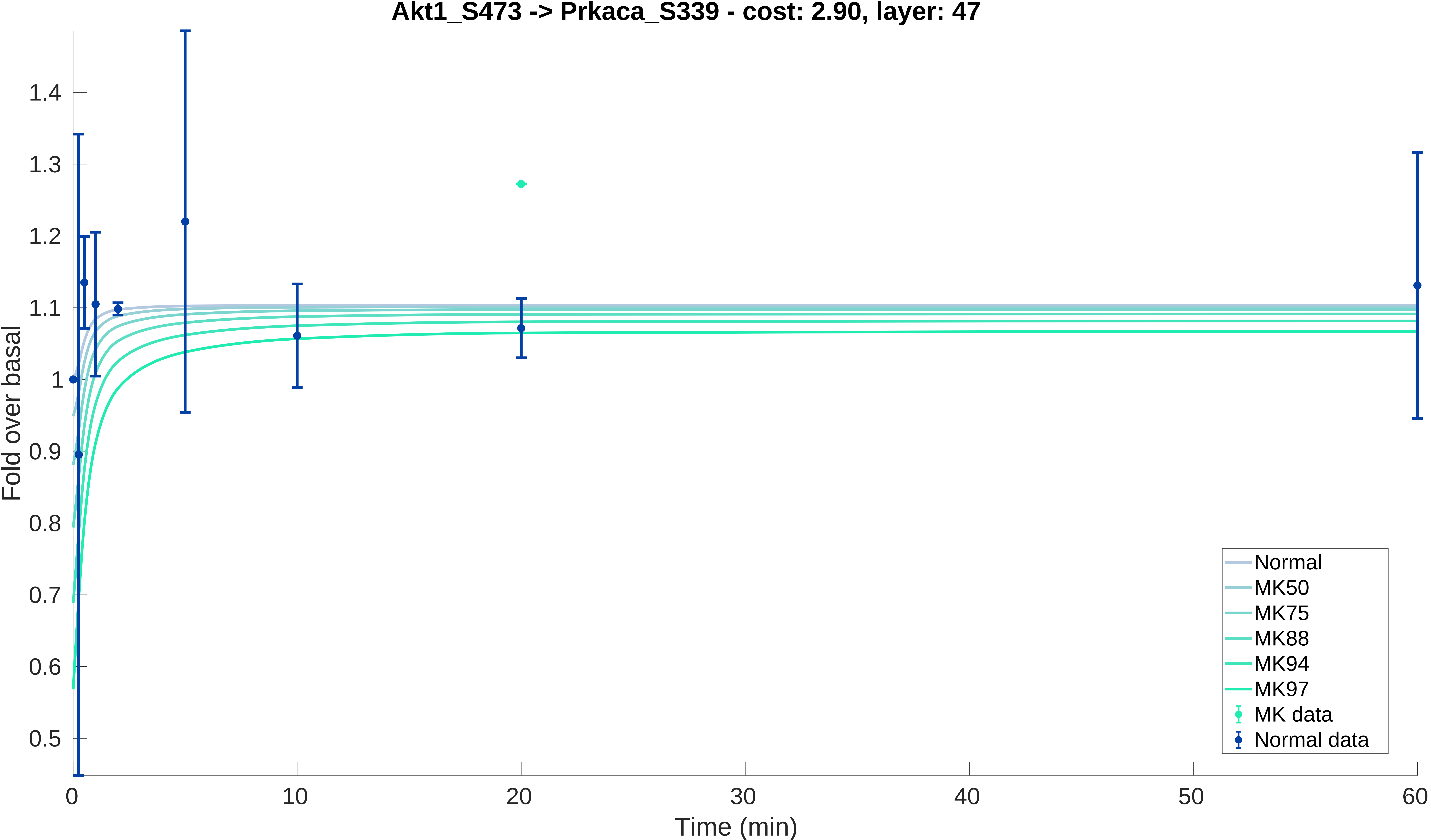
Akt1_S473 -> Prkaca_S339 - cost: 2.90, layer: 47
Fold over basal
Time (min)
1.4
1.3
1.2
1.1
1
0.9
0.8
0.7
0.6
0.5
0
10
20
30
40
50
60
Normal
MK50
MK75
MK88
MK94
MK97
MK data
Normal data

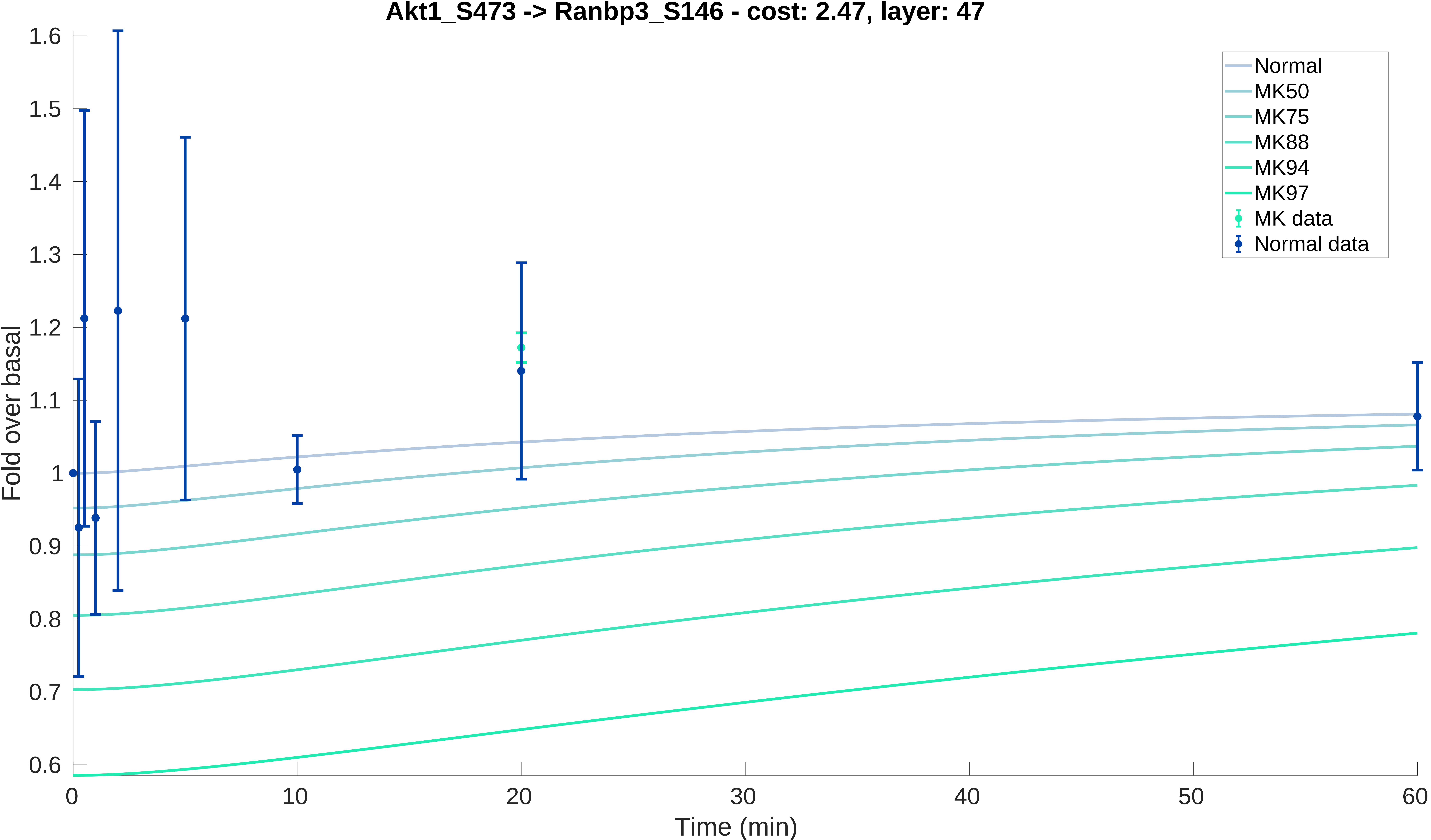
Akt1_S473 -> Ranbp3_S146 - cost: 2.47, layer: 47
Fold over basal
Time (min)
1.6
1.5
1.4
1.3
1.2
1.1
1
0.9
0.8
0.7
0.6
0
10
20
30
40
50
60
Normal
MK50
MK75
MK88
MK94
MK97
MK data
Normal data

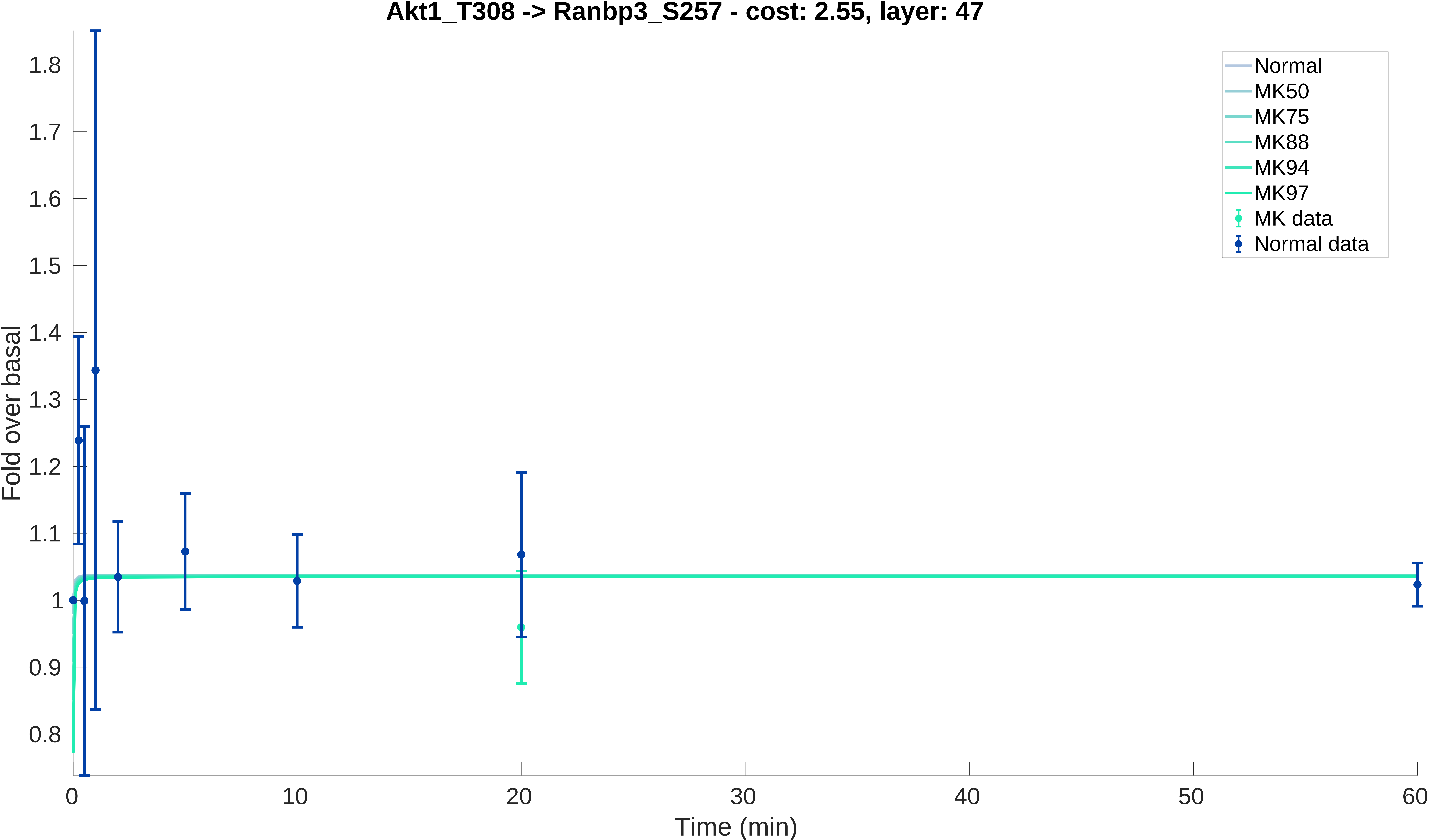

Akt1_T308 -> Ranbp3_S257 - cost: 2.55, layer: 47
Fold over basal
1.8
1.7
1.6
1.5
1.4
1.3
1.2
1.1
1
0.9
0.8
0
10
20
30
40
50
60
Time (min)
Normal
MK50
MK75
MK88
MK94
MK97
MK data
Normal data

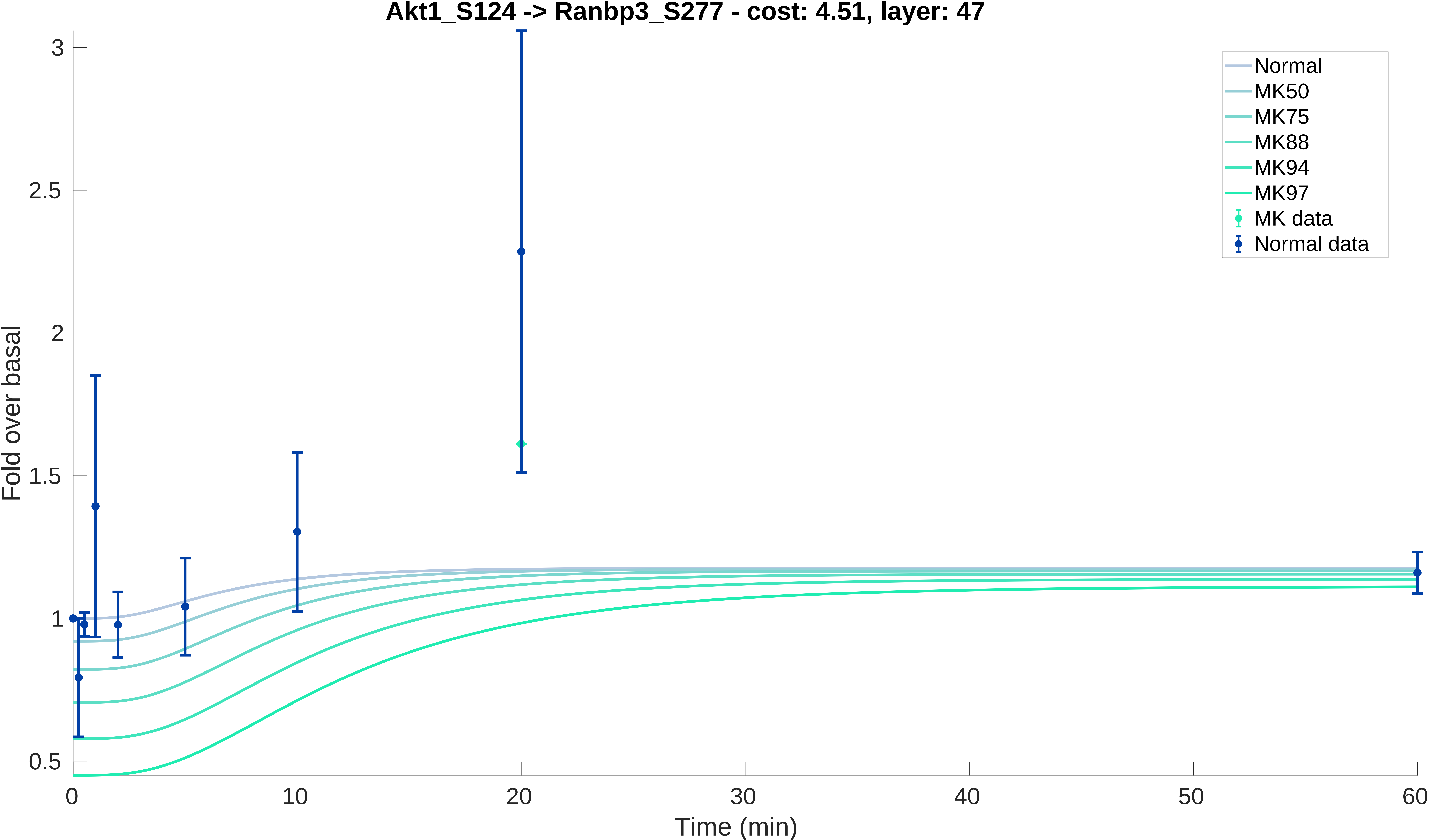

Akt1_S124 -> Ranbp3_S277 - cost: 4.51, layer: 47
Fold over basal
Time (min)
0.5
1
1.5
2
2.5
3
0
10
20
30
40
50
60
Normal
MK50
MK75
MK88
MK94
MK97
MK data
Normal data

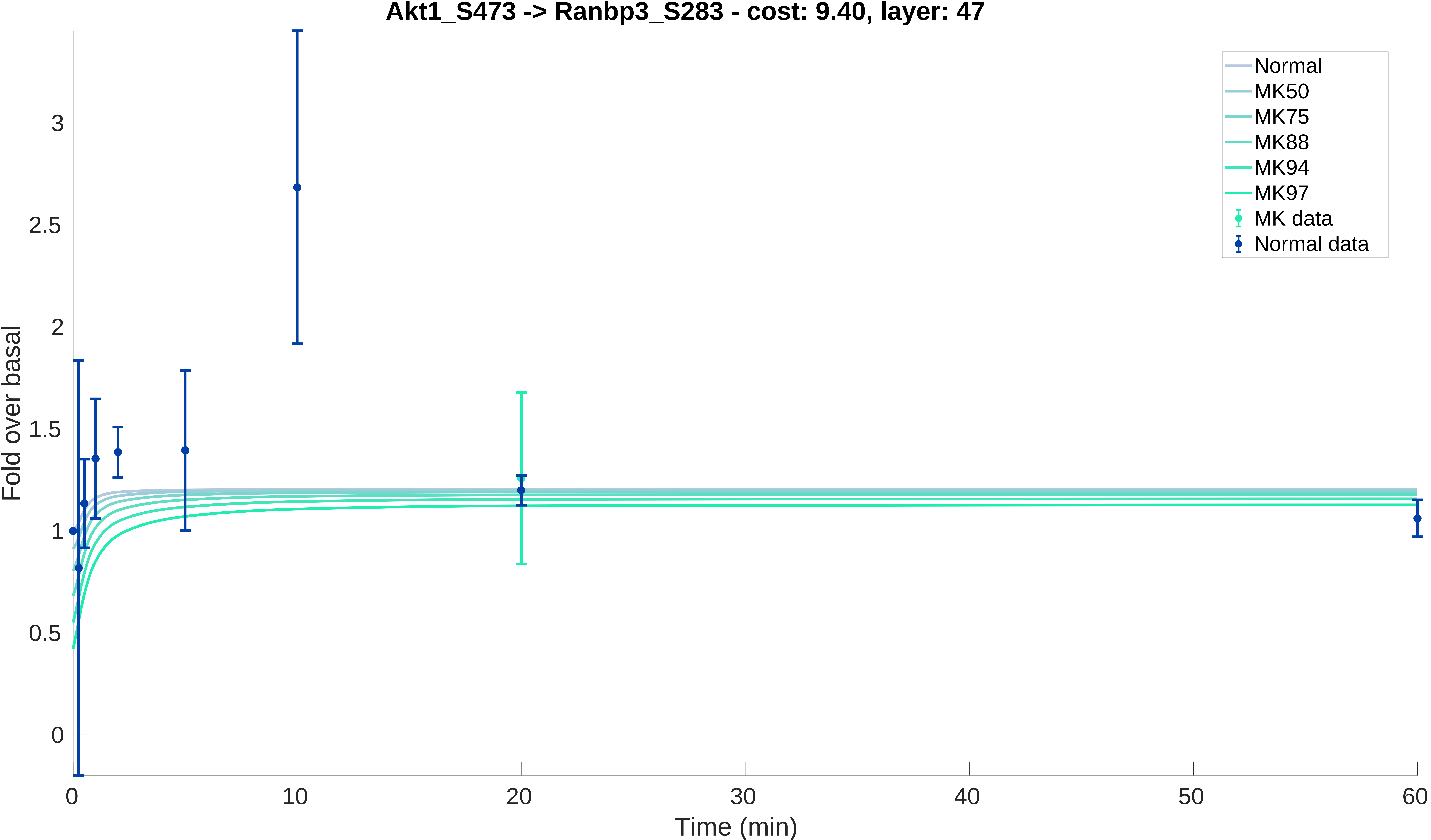

Akt1_S473 -> Ranbp3_S283 - cost: 9.40, layer: 47
Fold over basal
Time (min)
0
0.5
1
1.5
2
2.5
3
0
10
20
30
40
50
60
Normal
MK50
MK75
MK88
MK94
MK97
MK data
Normal data

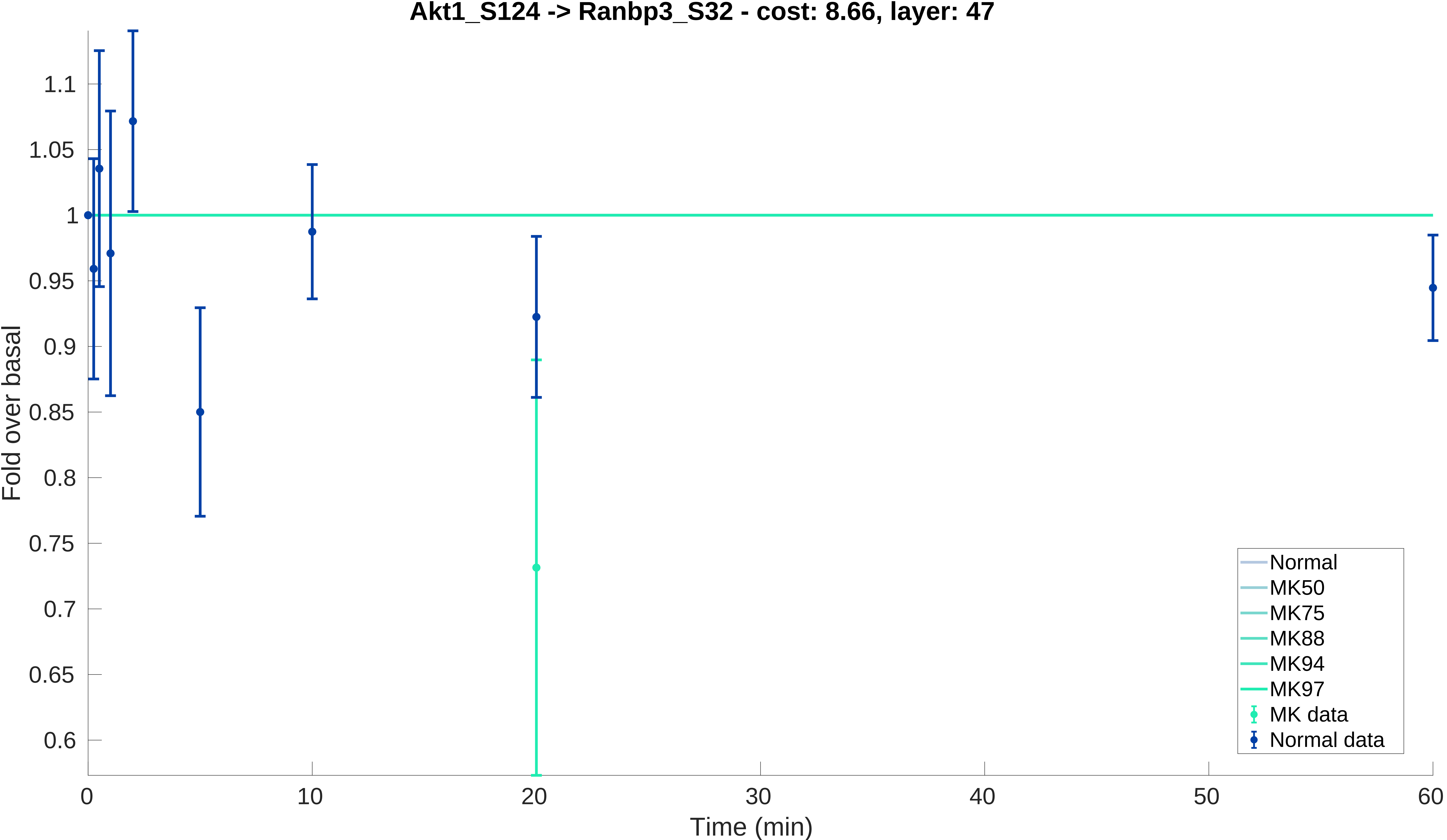

Akt1_S124 -> Ranbp3_S32 - cost: 8.66, layer: 47
Fold over basal
1.1
1.05
1
0.95
0.9
0.85
0.8
0.75
0.7
0.65
0.6
0
10
20
30
40
50
60
Time (min)
Normal
MK50
MK75
MK88
MK94
MK97
MK data
Normal data

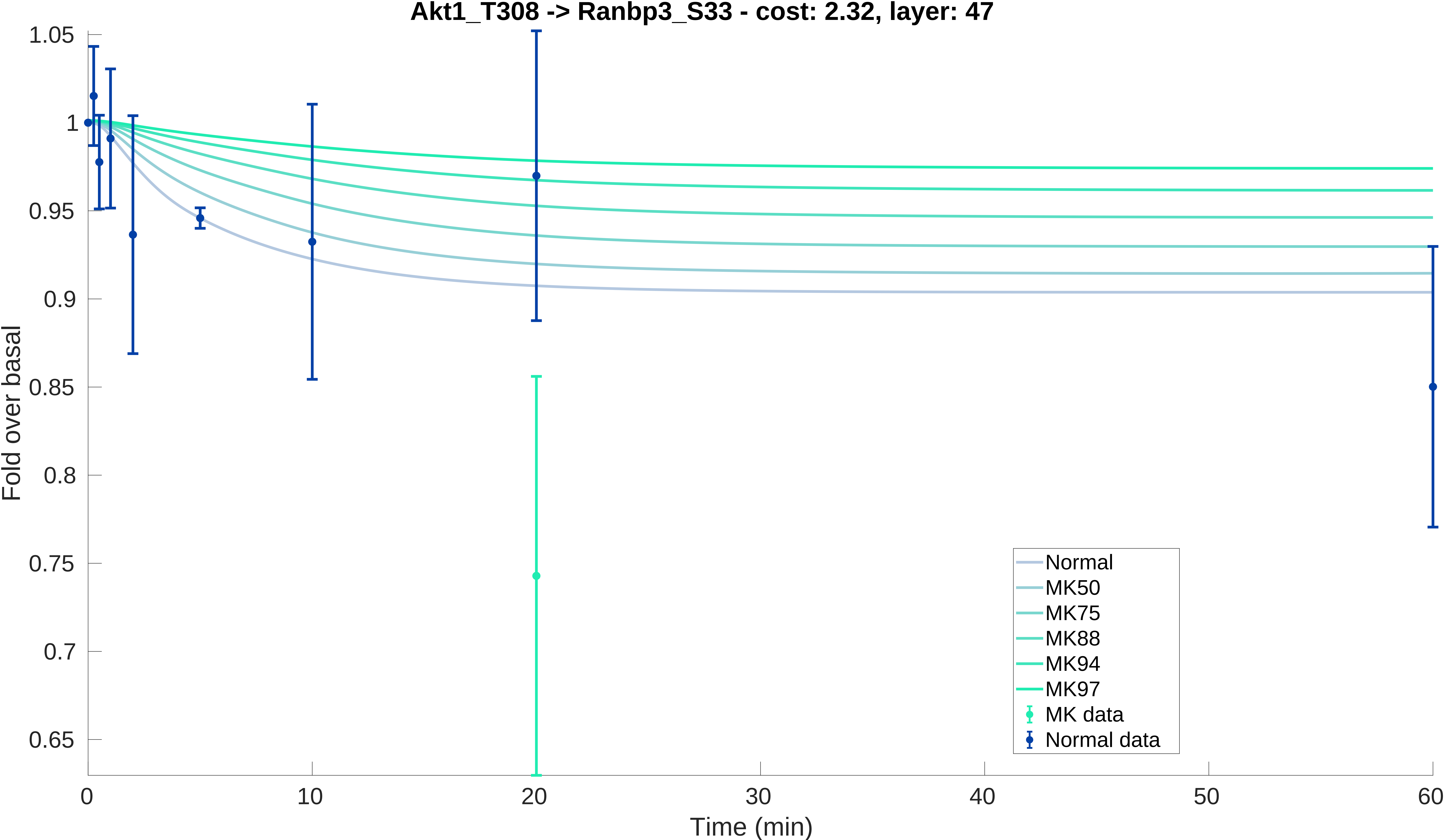

Akt1_T308 -> Ranbp3_S33 - cost: 2.32, layer: 47
Fold over basal
1.05
1
0.95
0.9
0.85
0.8
0.75
0.7
0.65
0
10
20
30
40
50
60
Time (min)
Normal
MK50
MK75
MK88
MK94
MK97
MK data
Normal data

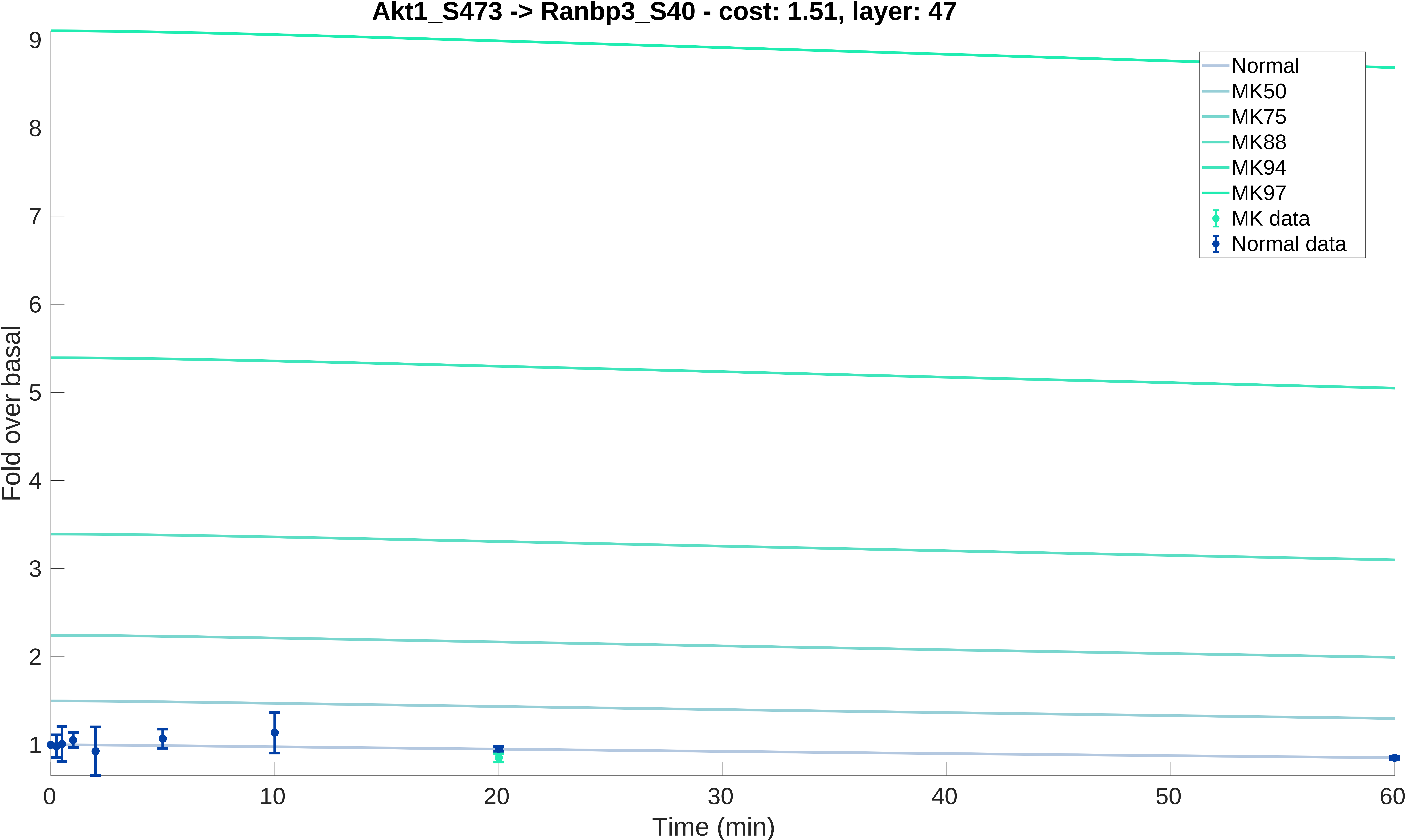

Akt1_S473 -> Ranbp3_S40 - cost: 1.51, layer: 47
Fold over basal
Time (min)
0
10
20
30
40
50
60
1
2
3
4
5
6
7
8
9
Normal
MK50
MK75
MK88
MK94
MK97
MK data
Normal data

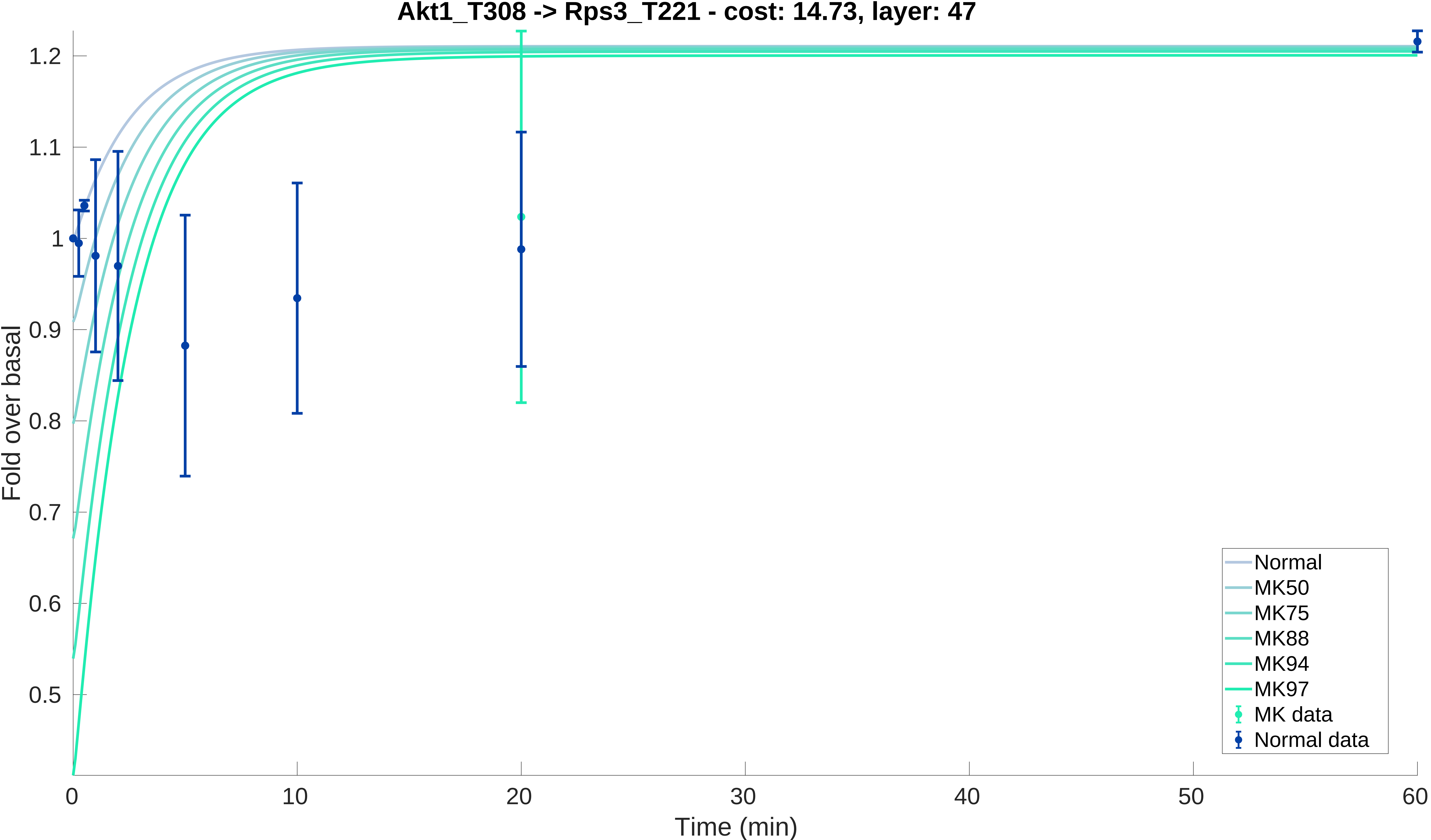
Akt1_T308 -> Rps3_T221 - cost: 14.73, layer: 47
Fold over basal
Time (min)
1.2
1.1
1
0.9
0.8
0.7
0.6
0.5
0
10
20
30
40
50
60
Normal
MK50
MK75
MK88
MK94
MK97
MK data
Normal data

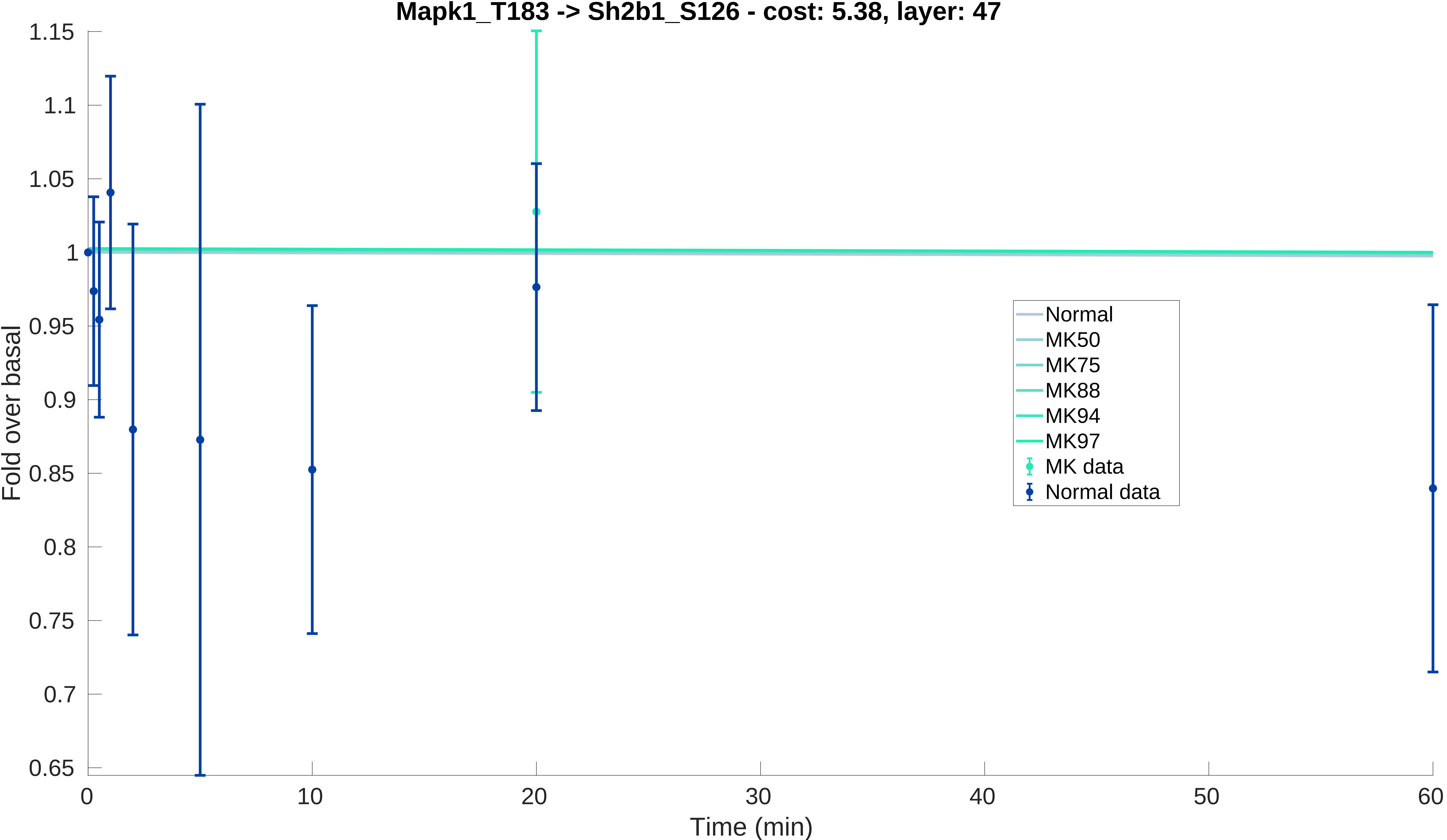

Mapk1_T183 -> Sh2b1_S126 - cost: 5.38, layer: 47
Fold over basal
1.15
1.1
1.05
1
0.95
0.9
0.85
0.8
0.75
0.7
0.65
0
10
20
30
40
50
60
Time (min)
Normal
MK50
MK75
MK88
MK94
MK97
MK data
Normal data

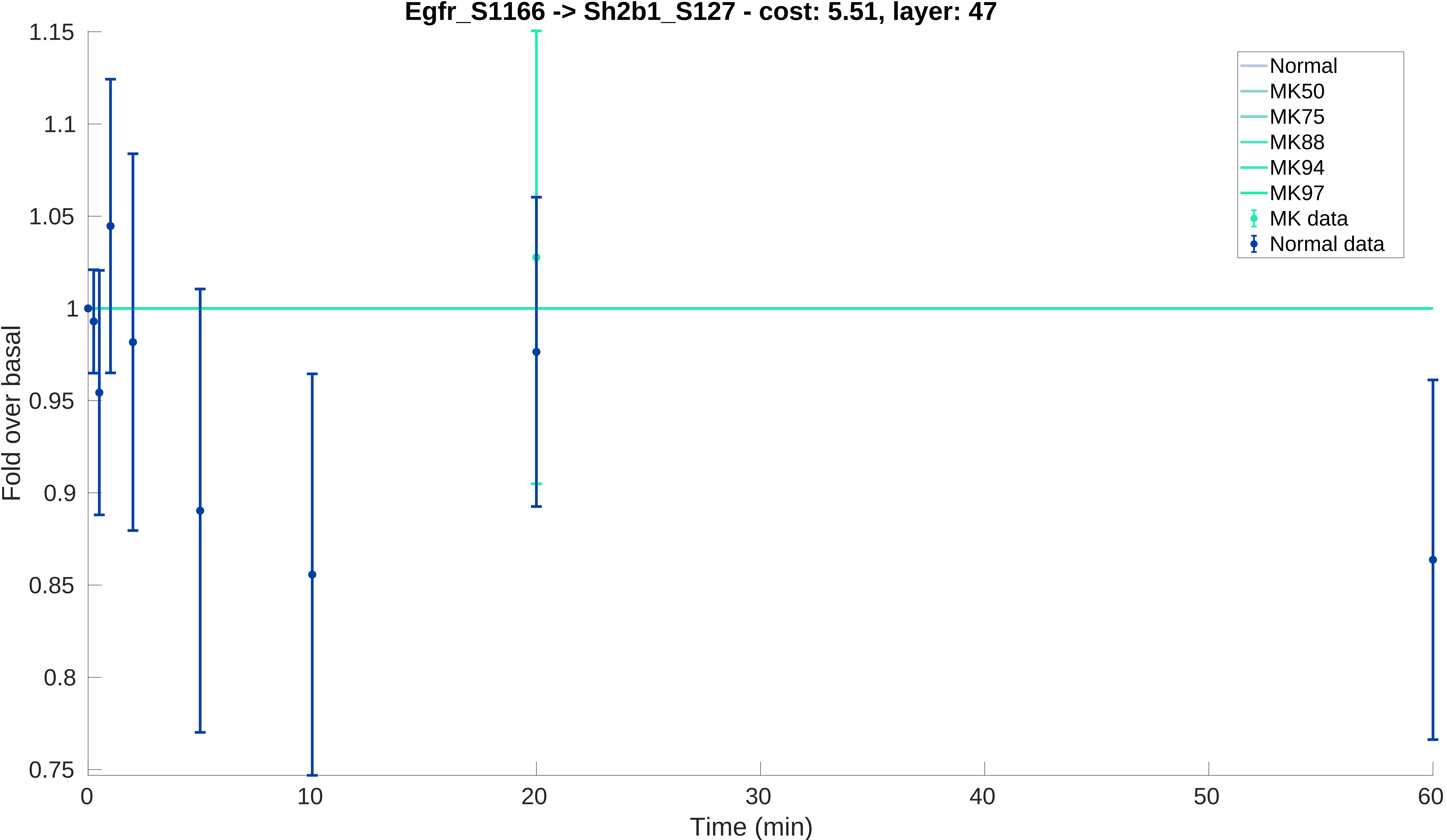

Egfr_S1166 -> Sh2b1_S127 - cost: 5.51, layer: 47
Fold over basal
Time (min)
1.15
1.1
1.05
1
0.95
0.9
0.85
0.8
0.75
0
10
20
30
40
50
60
Normal
MK50
MK75
MK88
MK94
MK97
MK data
Normal data

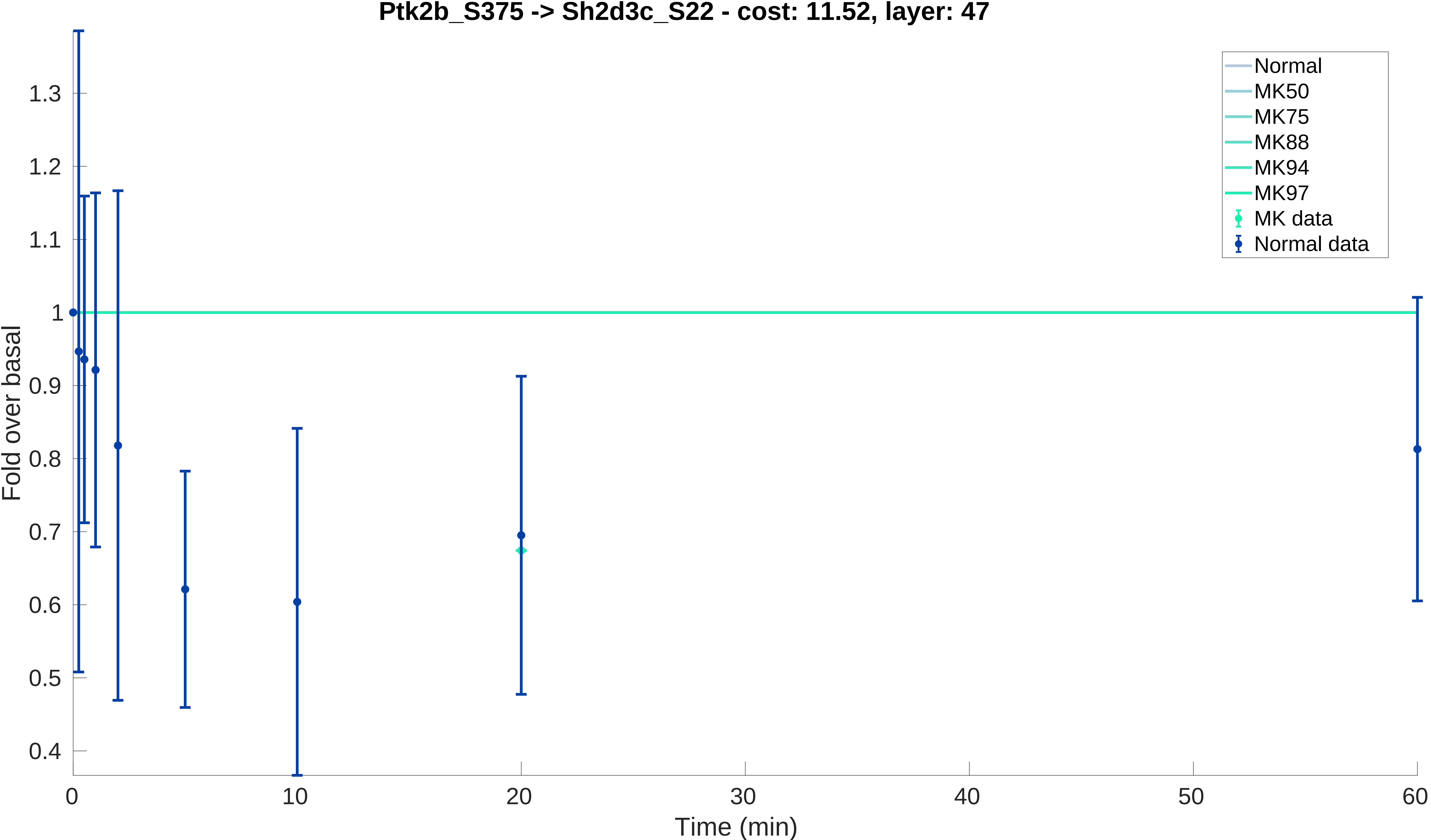

Ptk2b_S375 -> Sh2d3c_S22 - cost: 11.52, layer: 47
Normal
MK50
MK75
MK88
MK94
MK97
MK data
Normal data
Fold over basal
1.3
1.2
1.1
1
0.9
0.8
0.7
0.6
0.5
0.4
0
10
20
30
40
50
60
Time (min)

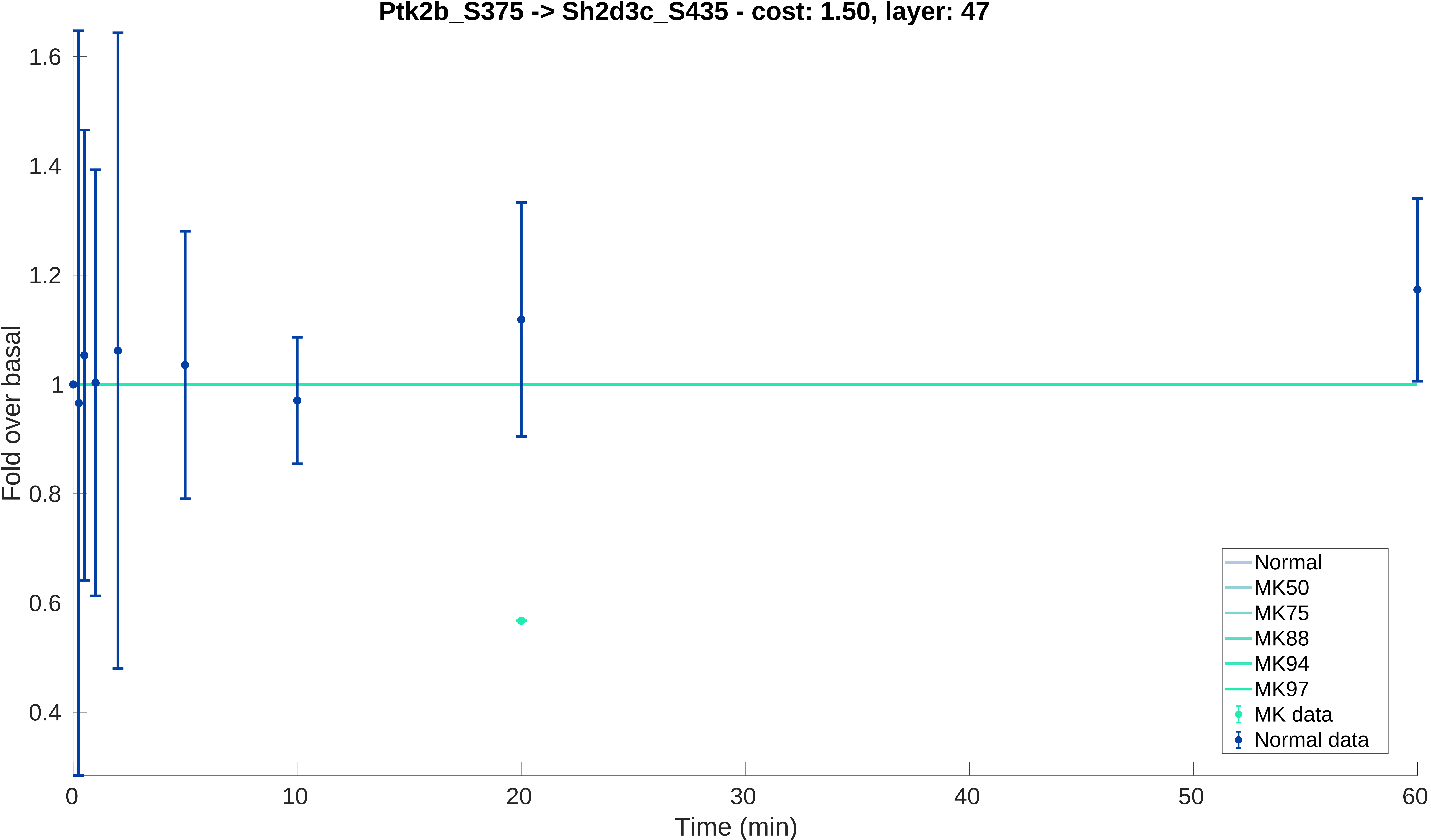
Ptk2b_S375 -> Sh2d3c_S435 - cost: 1.50, layer: 47
1.6
1.4
1.2
1
0.8
0.6
0.4
Fold over basal
0
10
20
30
40
50
60
Time (min)
Normal
MK50
MK75
MK88
MK94
MK97
MK data
Normal data

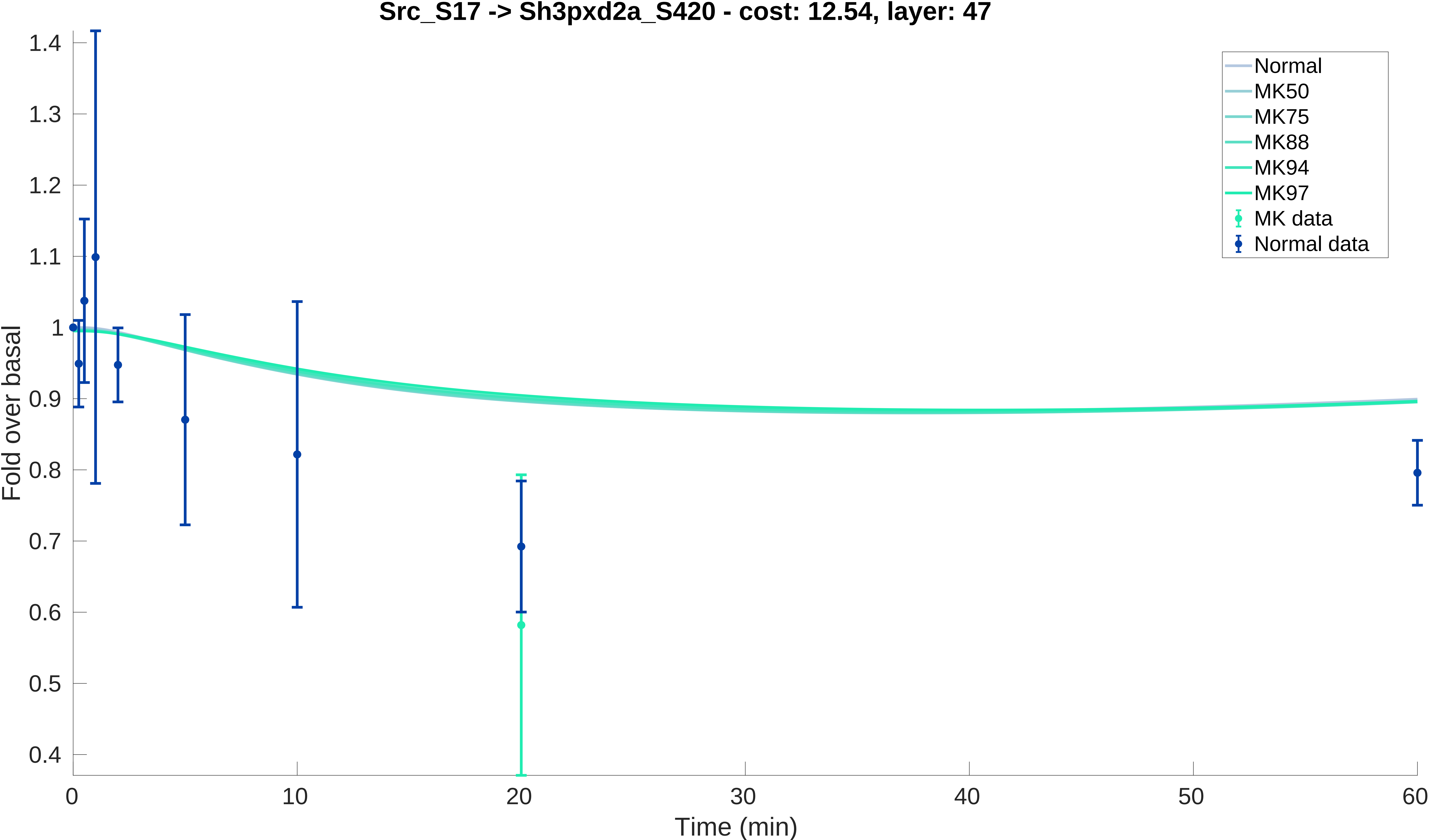

Src_S17 -> Sh3pxd2a_S420 - cost: 12.54, layer: 47
Fold over basal
Time (min)
1.4
1.3
1.2
1.1
1
0.9
0.8
0.7
0.6
0.5
0.4
0
10
20
30
40
50
60
Normal
MK50
MK75
MK88
MK94
MK97
MK data
Normal data

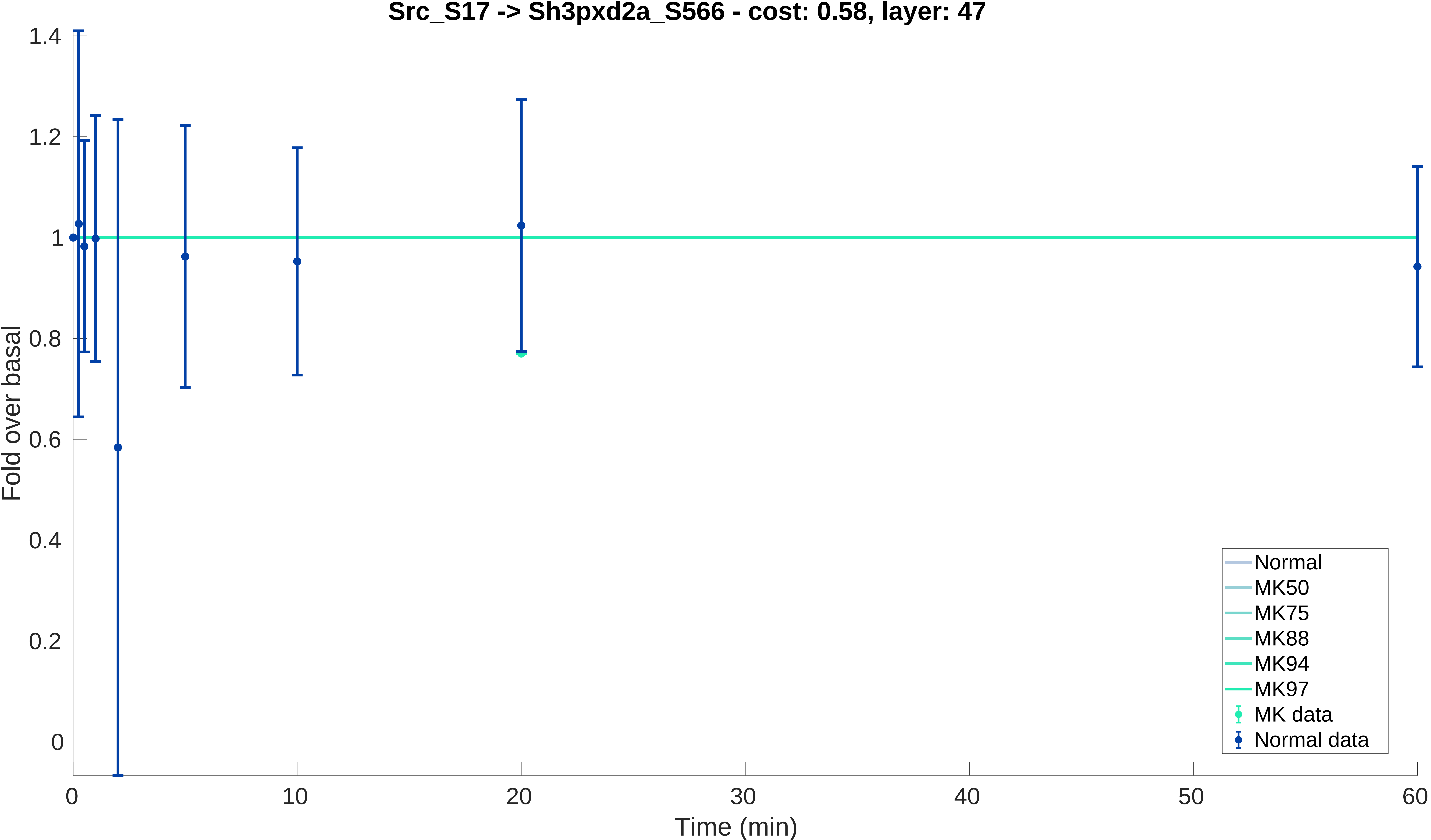

Src_S17 -> Sh3pxd2a_S566 - cost: 0.58, layer: 47
Fold over basal
1.4
1.2
1
0.8
0.6
0.4
0.2
0
0
10
20
30
40
50
60
Time (min)
Normal
MK50
MK75
MK88
MK94
MK97
MK data
Normal data

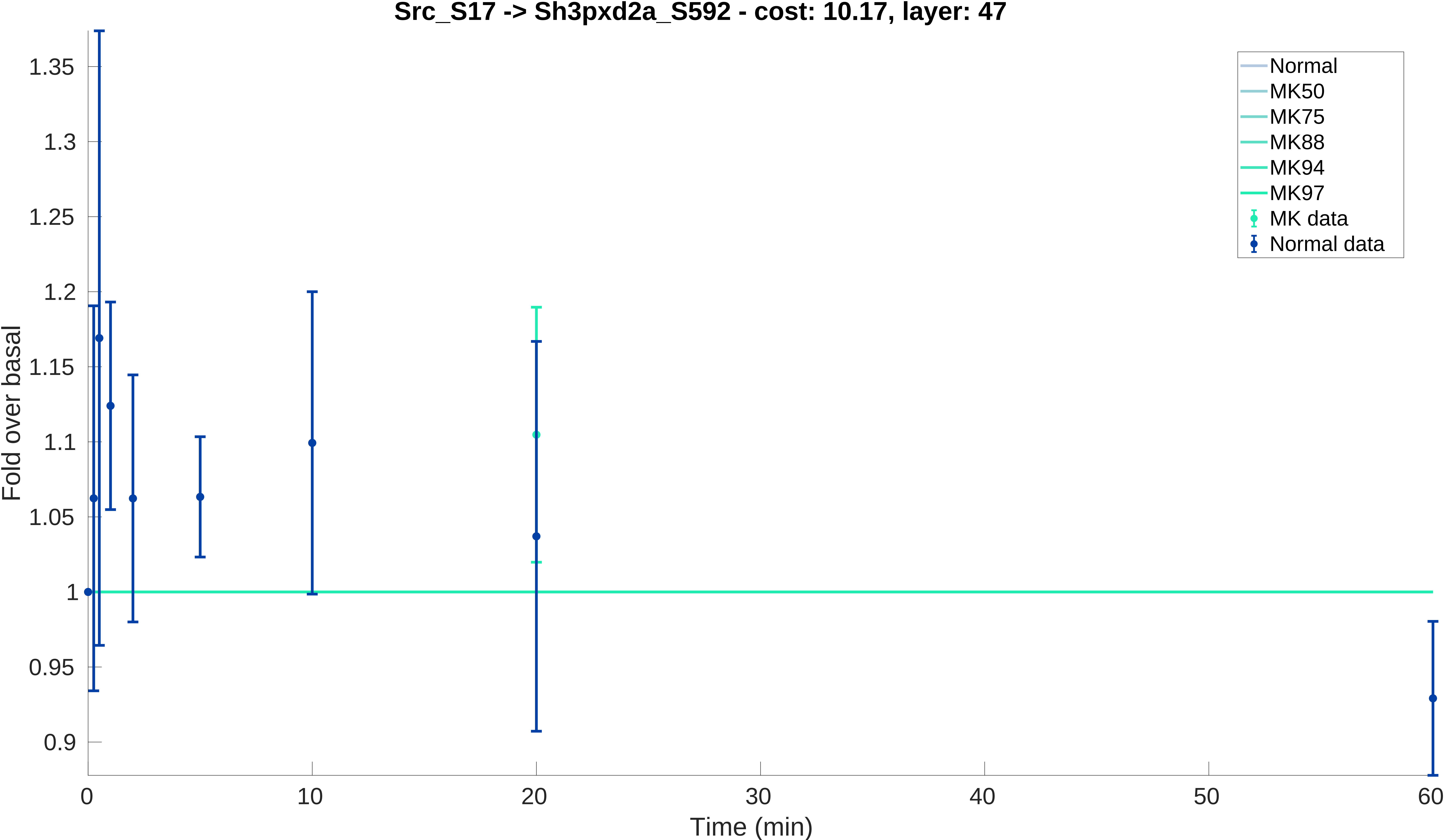

Src_S17 -> Sh3pxd2a_S592 - cost: 10.17, layer: 47
Fold over basal
Time (min)
0.9
0.95
1
1.05
1.1
1.15
1.2
1.25
1.3
1.35
0
10
20
30
40
50
60
Normal
MK50
MK75
MK88
MK94
MK97
MK data
Normal data

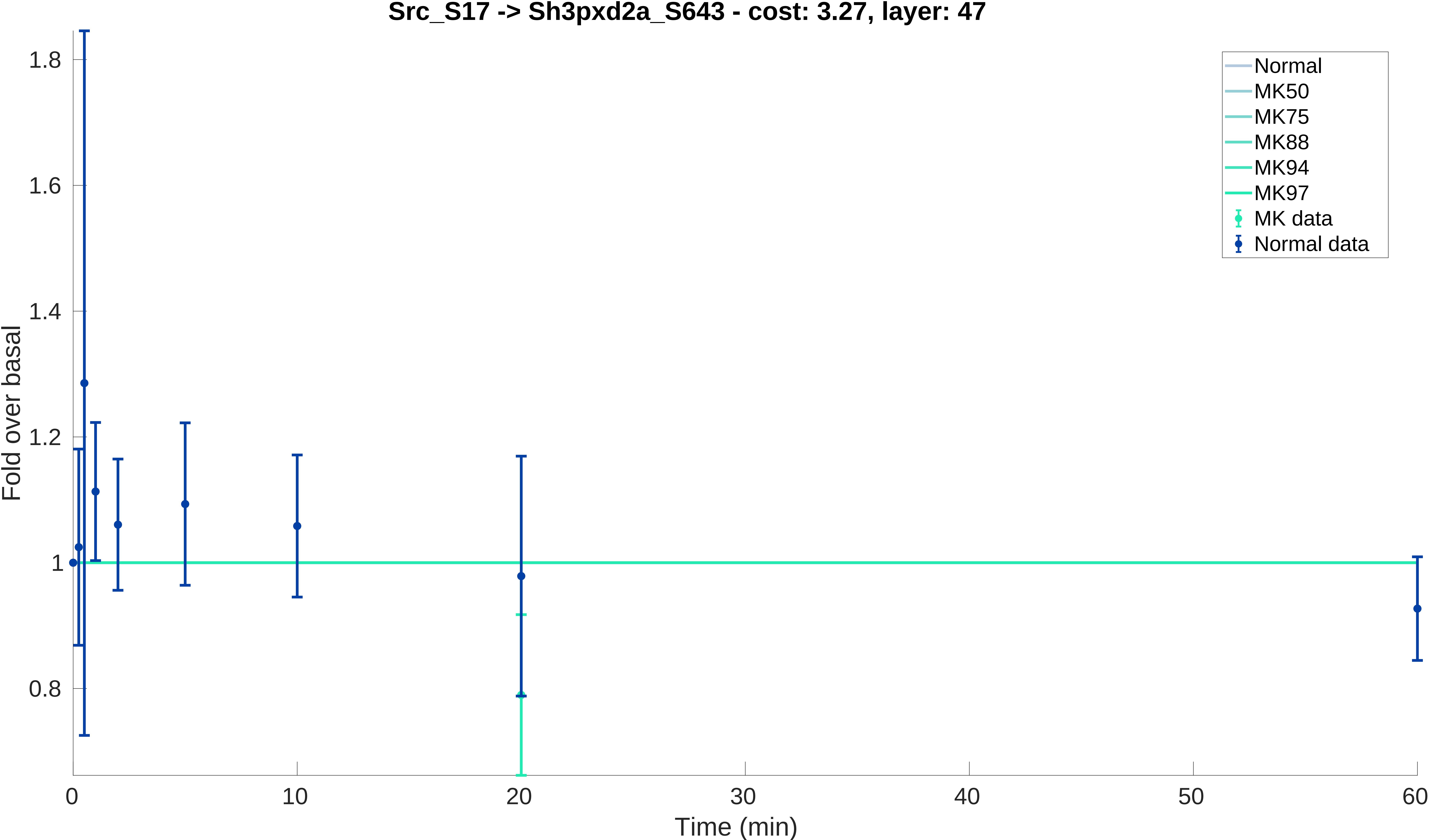

Src_S17 -> Sh3pxd2a_S643 - cost: 3.27, layer: 47
Fold over basal
Time (min)
0.8
1
1.2
1.4
1.6
1.8
0
10
20
30
40
50
60
Normal
MK50
MK75
MK88
MK94
MK97
MK data
Normal data

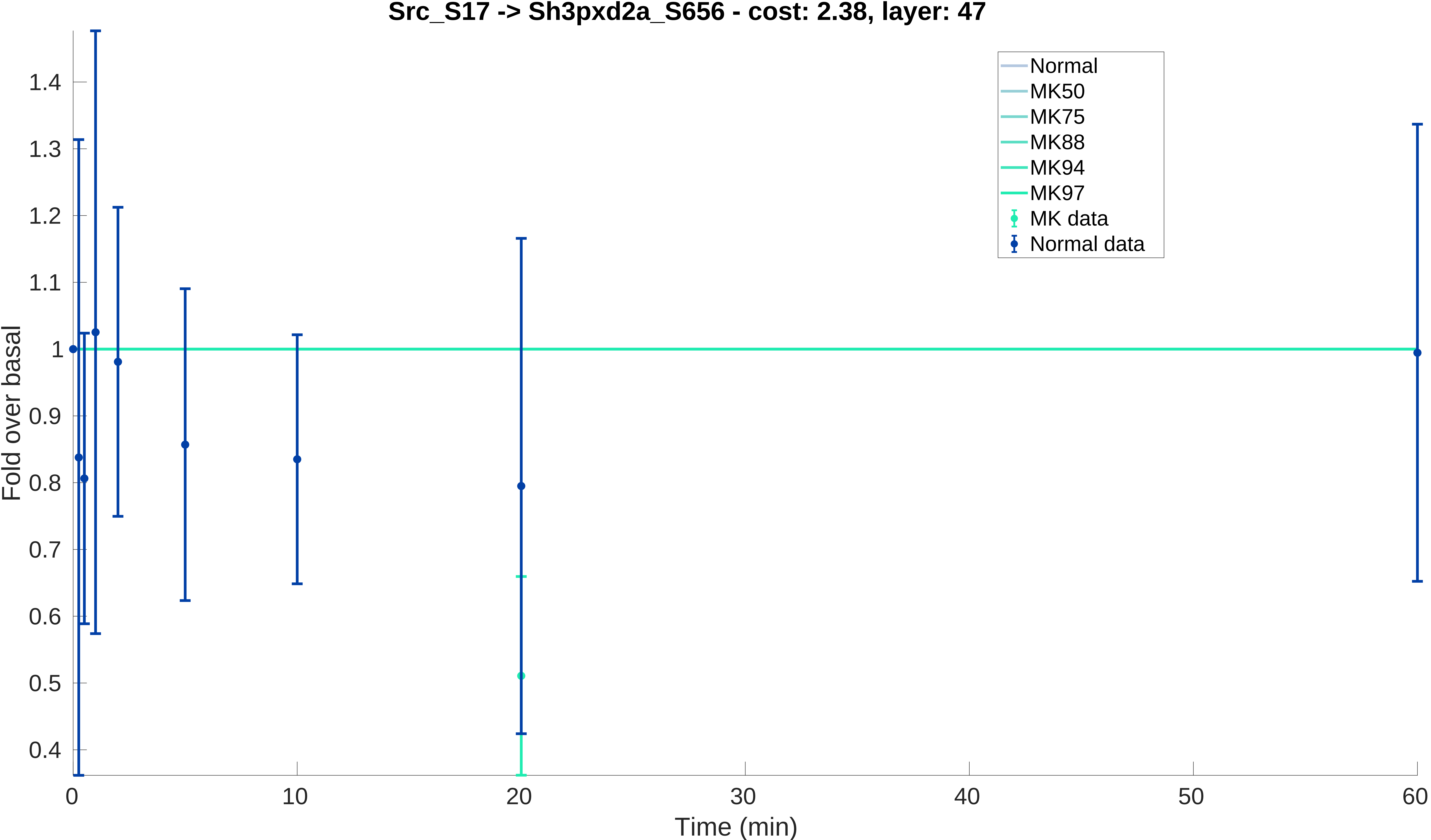
Src_S17 -> Sh3pxd2a_S656 - cost: 2.38, layer: 47
Normal
MK50
MK75
MK88
MK94
MK97
MK data
Normal data
Fold over basal
Time (min)
0
10
20
30
40
50
60
0.4
0.5
0.6
0.7
0.8
0.9
1
1.1
1.2
1.3
1.4

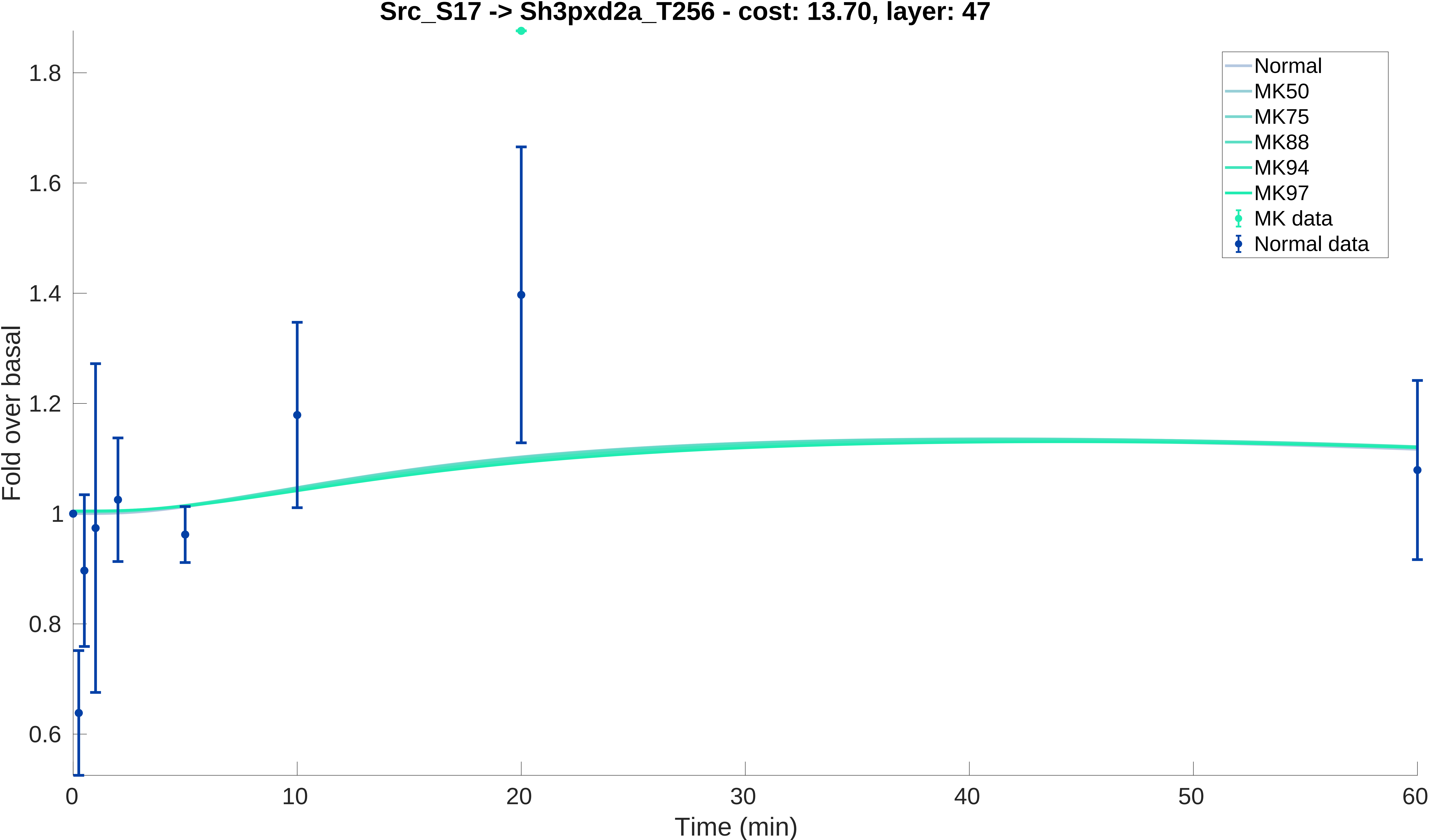

Src_S17 -> Sh3pxd2a_T256 - cost: 13.70, layer: 47
Fold over basal
Time (min)
0.6
0.8
1
1.2
1.4
1.6
1.8
0
10
20
30
40
50
60
Normal
MK50
MK75
MK88
MK94
MK97
MK data
Normal data

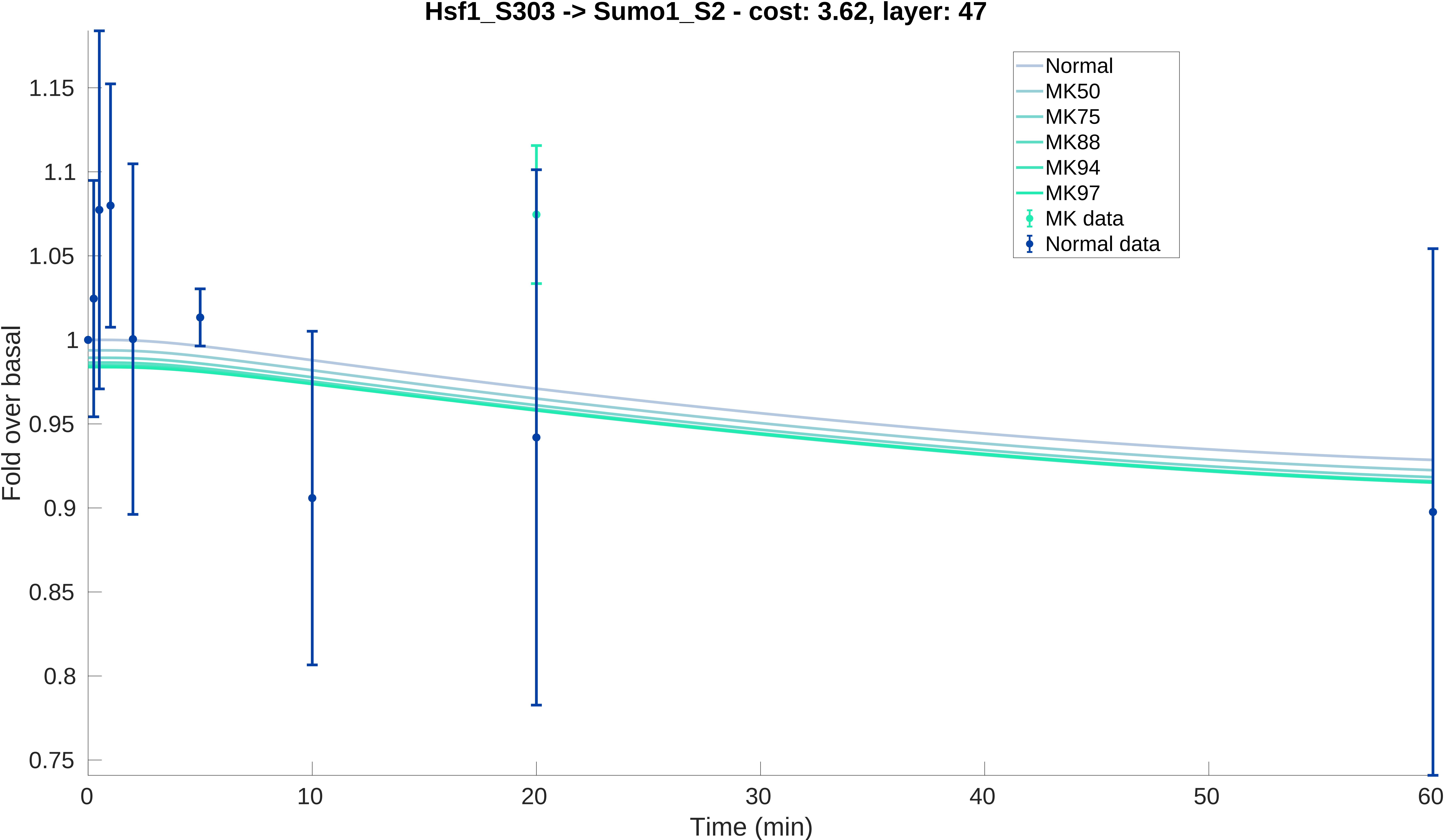

Hsf1_S303 -> Sumo1_S2 - cost: 3.62, layer: 47
Normal
MK50
MK75
MK88
MK94
MK97
MK data
Normal data
Fold over basal
Time (min)
1.15
1.1
1.05
1
0.95
0.9
0.85
0.8
0.75
0
10
20
30
40
50
60

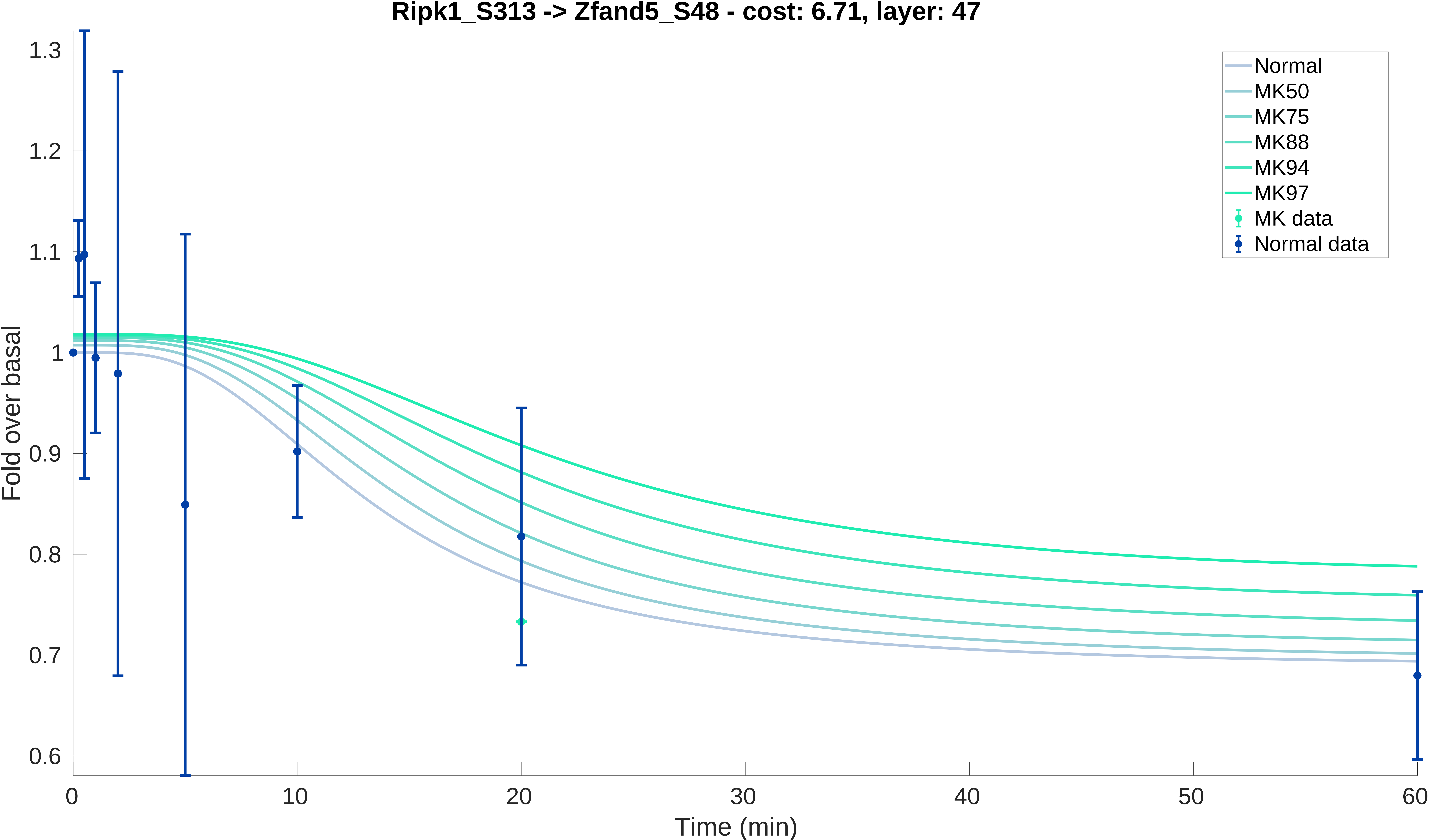

Ripk1_S313 -> Zfand5_S48 - cost: 6.71, layer: 47
Fold over basal
Time (min)
1.3
1.2
1.1
1
0.9
0.8
0.7
0.6
0
10
20
30
40
50
60
Normal
MK50
MK75
MK88
MK94
MK97
MK data
Normal data

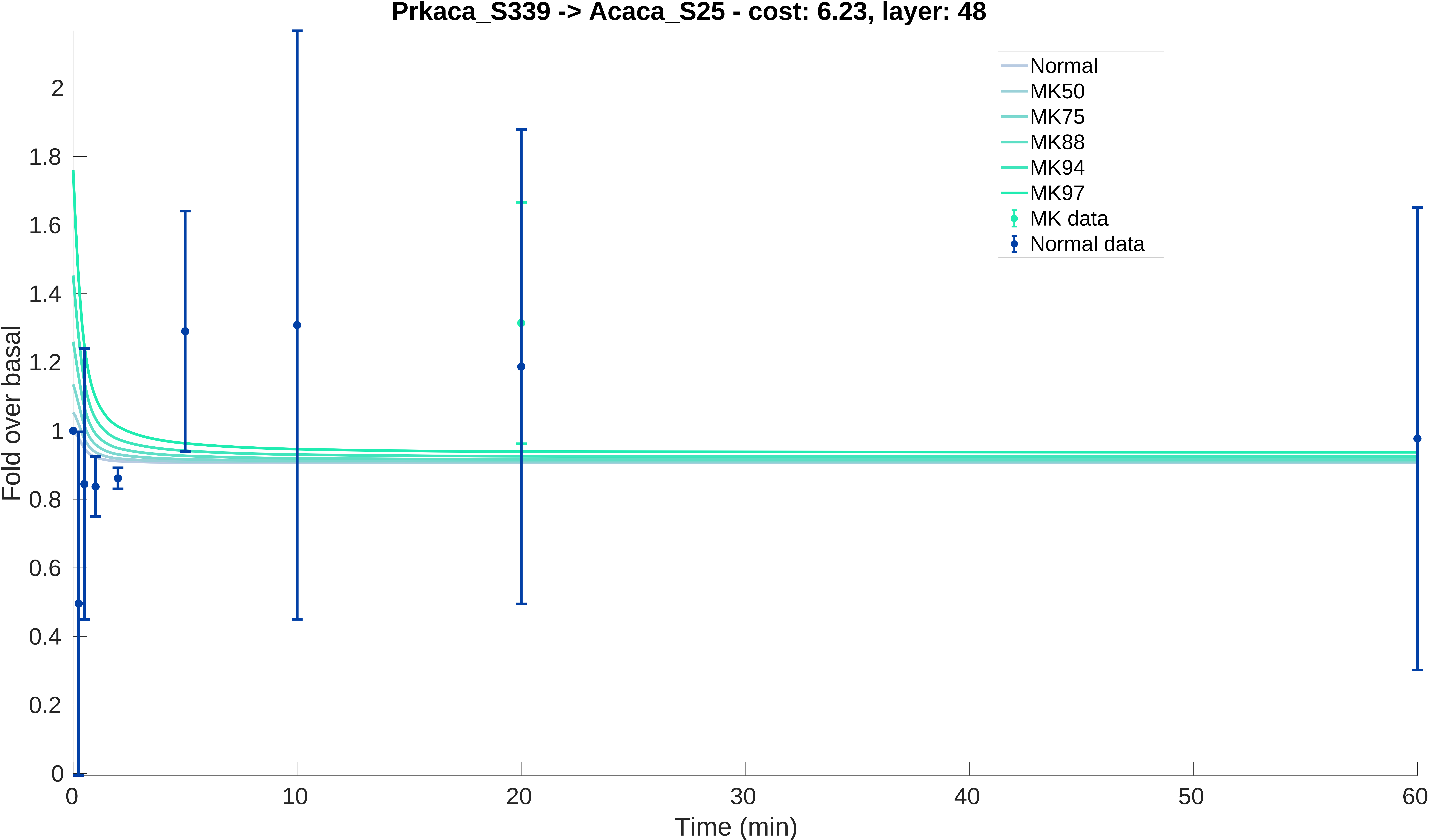

Prkaca_S339 -> Acaca_S25 - cost: 6.23, layer: 48
Normal
MK50
MK75
MK88
MK94
MK97
MK data
Normal data
Fold over basal
Time (min)
0
0.2
0.4
0.6
0.8
1
1.2
1.4
1.6
1.8
2
0
10
20
30
40
50
60

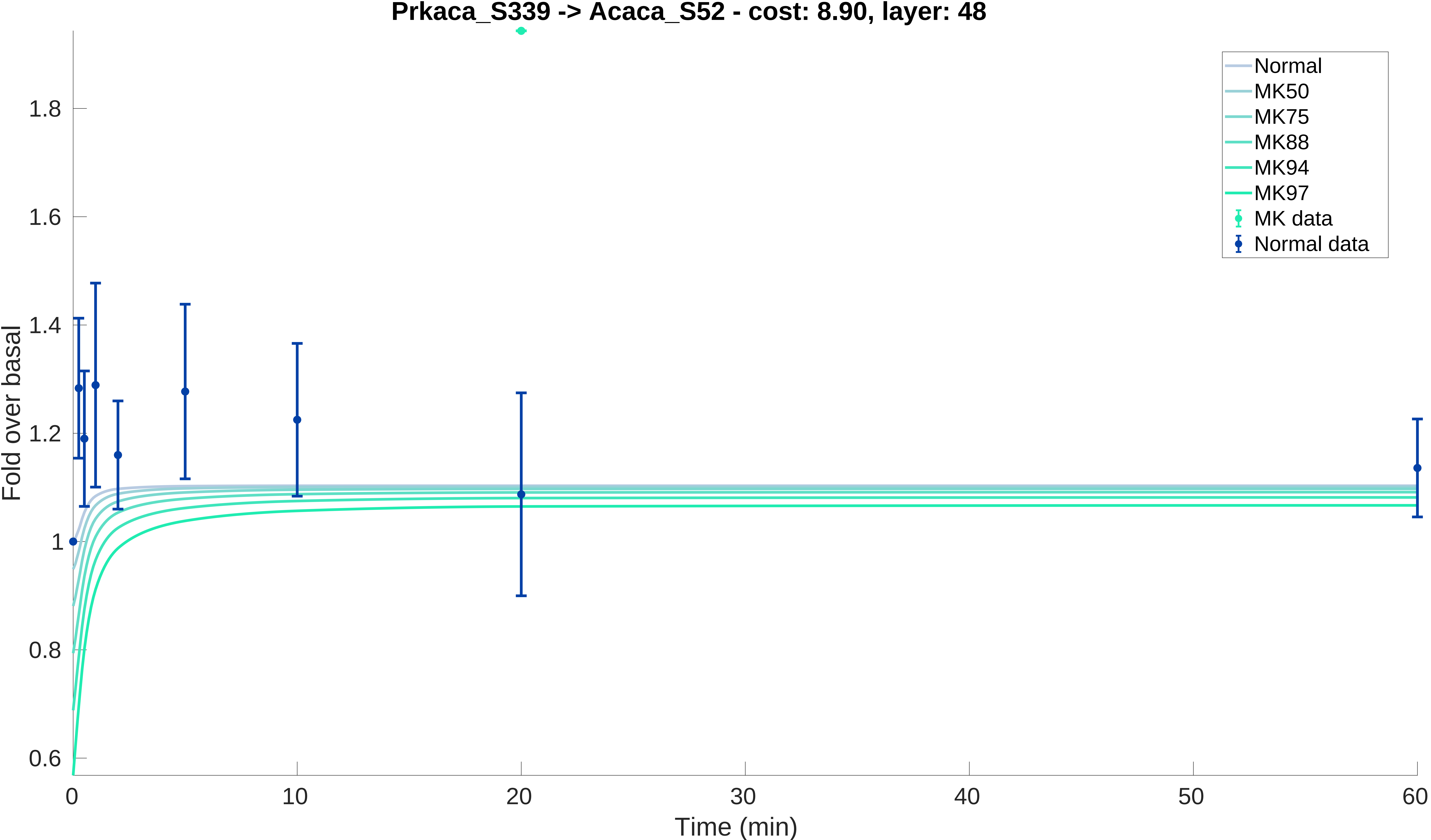

Prkaca_S339 -> Acaca_S52 - cost: 8.90, layer: 48
Normal
MK50
MK75
MK88
MK94
MK97
MK data
Normal data
Fold over basal
Time (min)
0.6
0.8
1
1.2
1.4
1.6
1.8
0
10
20
30
40
50
60

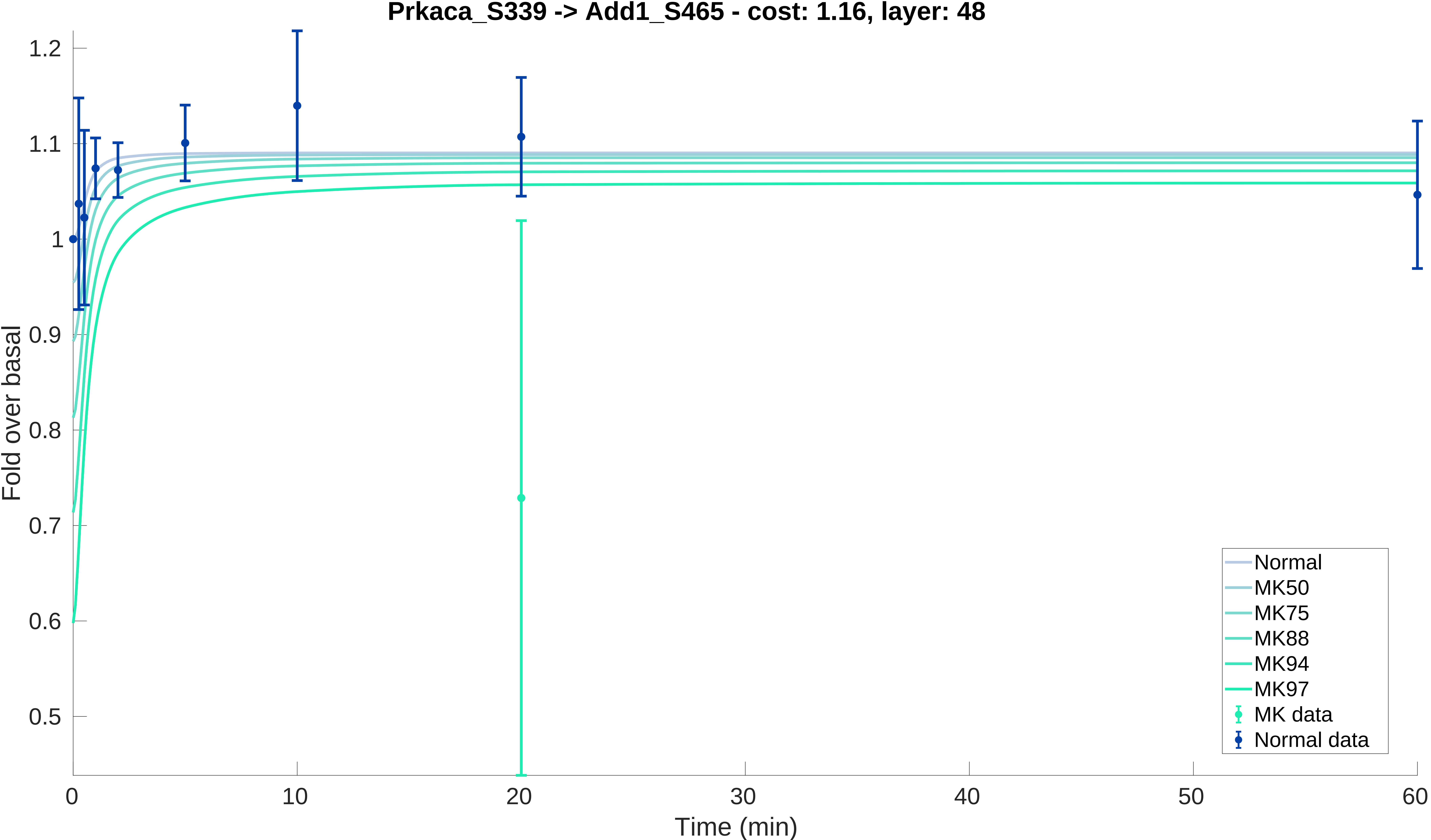
Prkaca_S339 -> Add1_S465 - cost: 1.16, layer: 48
1.2
1.1
1
0.9
0.8
0.7
0.6
0.5
Fold over basal
0
10
20
30
40
50
60
Time (min)
Normal
MK50
MK75
MK88
MK94
MK97
MK data
Normal data

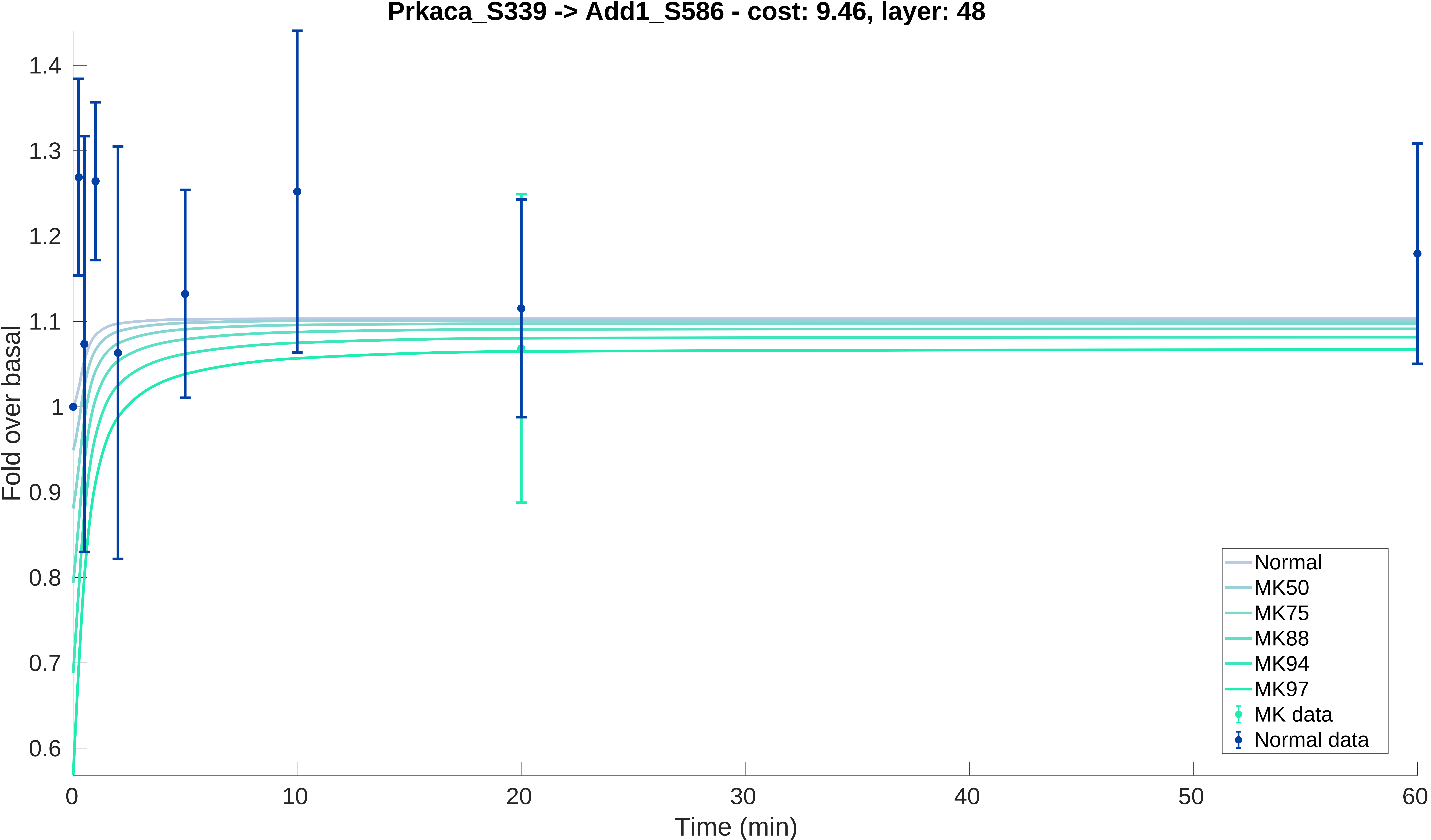
Prkaca_S339 -> Add1_S586 - cost: 9.46, layer: 48
Fold over basal
Time (min)
1.4
1.3
1.2
1.1
1
0.9
0.8
0.7
0.6
0
10
20
30
40
50
60
Normal
MK50
MK75
MK88
MK94
MK97
MK data
Normal data

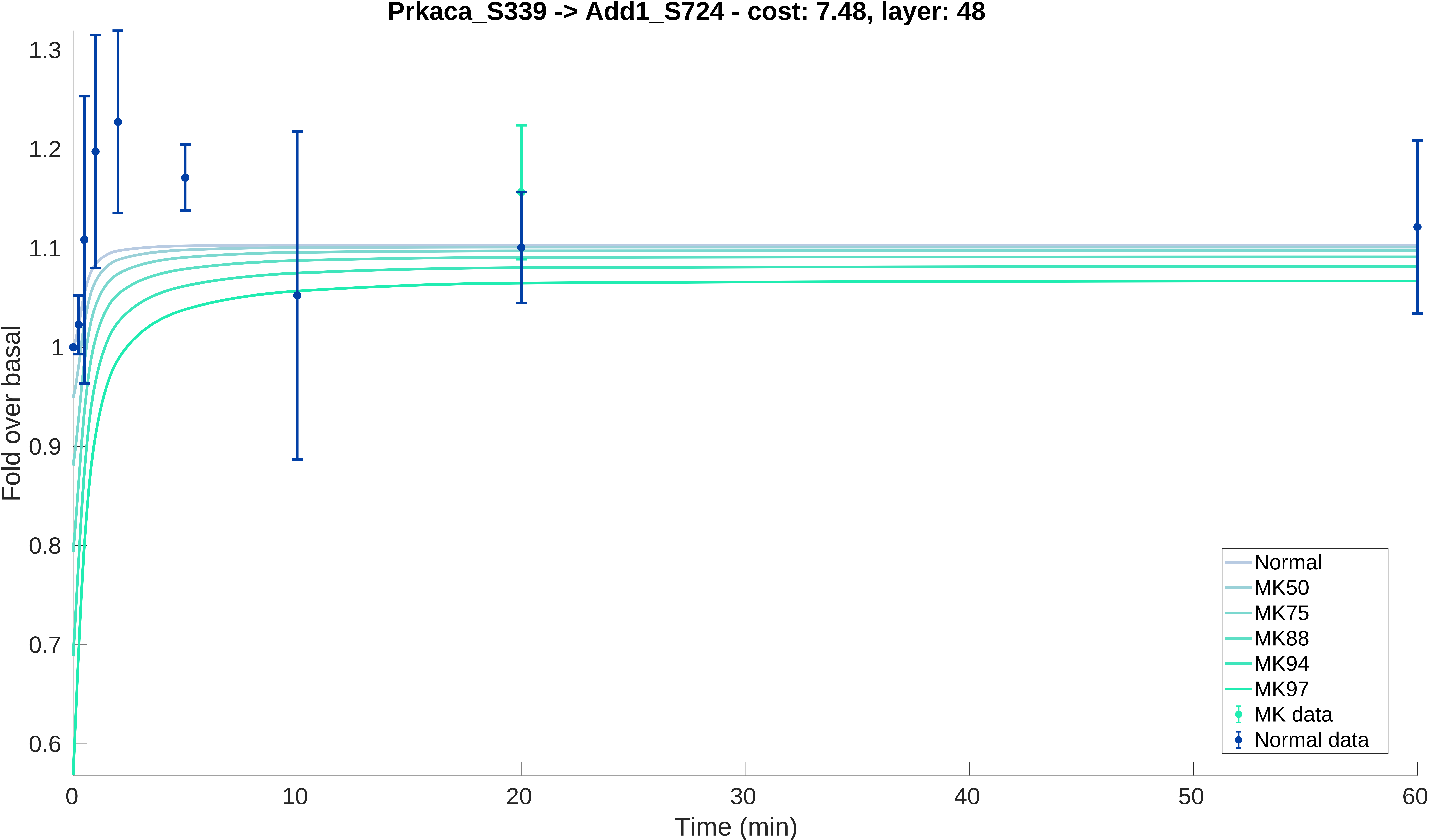

Prkaca_S339 -> Add1_S724 - cost: 7.48, layer: 48
Fold over basal
1.3
1.2
1.1
1
0.9
0.8
0.7
0.6
0
10
20
30
40
50
60
Time (min)
Normal
MK50
MK75
MK88
MK94
MK97
MK data
Normal data

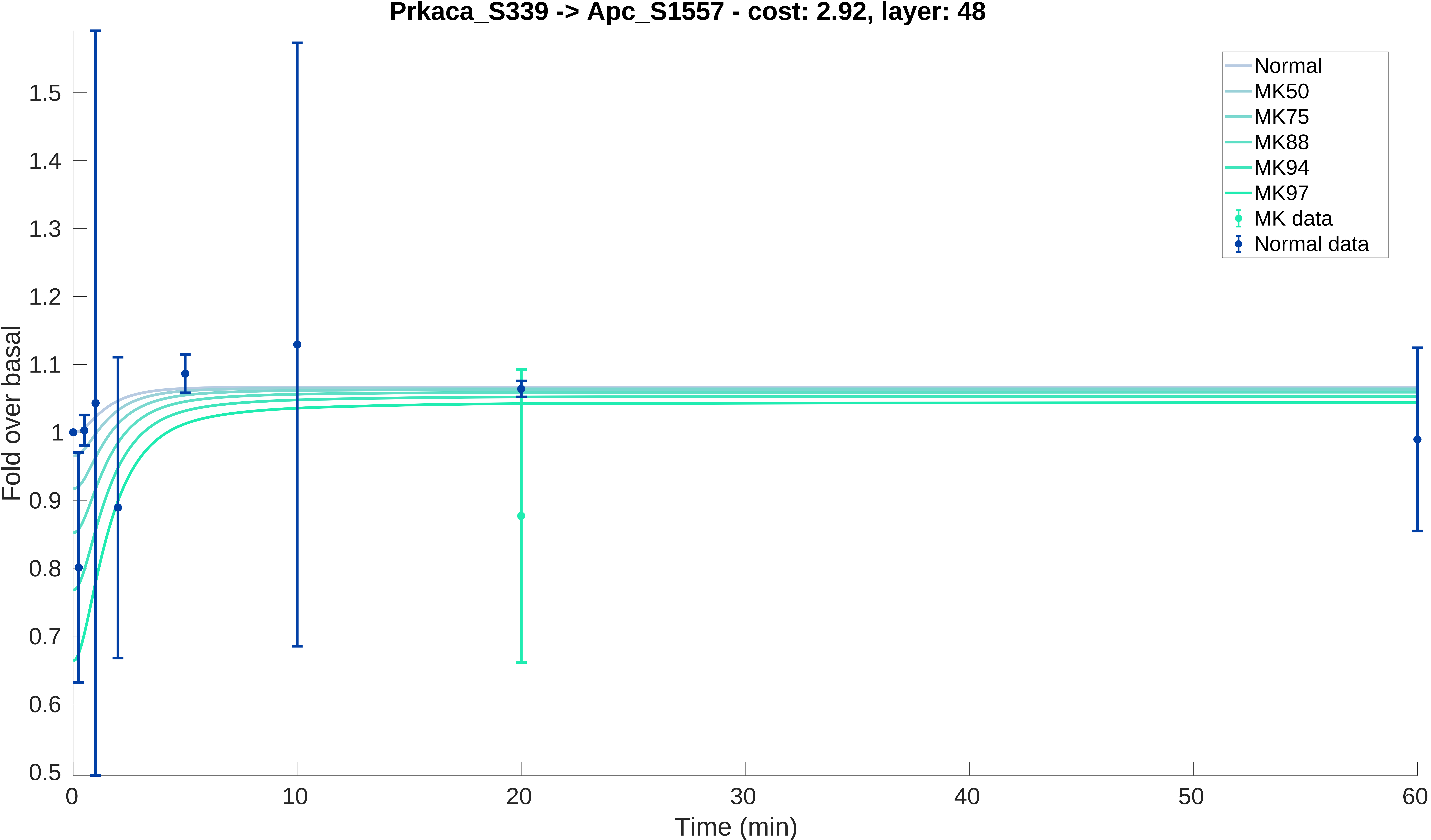

Prkaca_S339 -> Apc_S1557 - cost: 2.92, layer: 48
Fold over basal
Time (min)
0.5
0.6
0.7
0.8
0.9
1
1.1
1.2
1.3
1.4
1.5
0
10
20
30
40
50
60
Normal
MK50
MK75
MK88
MK94
MK97
MK data
Normal data

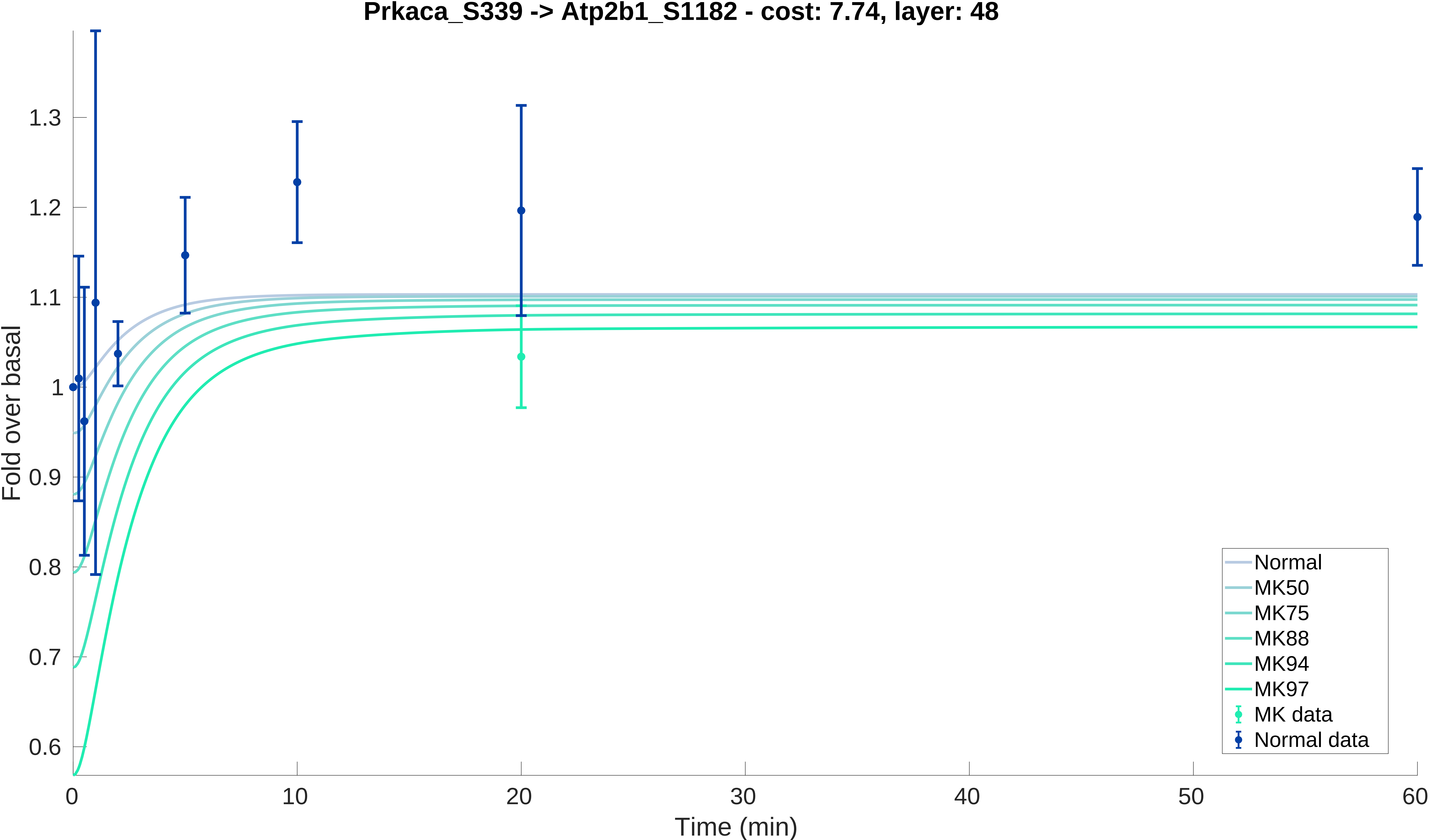
Prkaca_S339 -> Atp2b1_S1182 - cost: 7.74, layer: 48
Fold over basal
Time (min)
0.6
0.7
0.8
0.9
1
1.1
1.2
1.3
0
10
20
30
40
50
60
Normal
MK50
MK75
MK88
MK94
MK97
MK data
Normal data

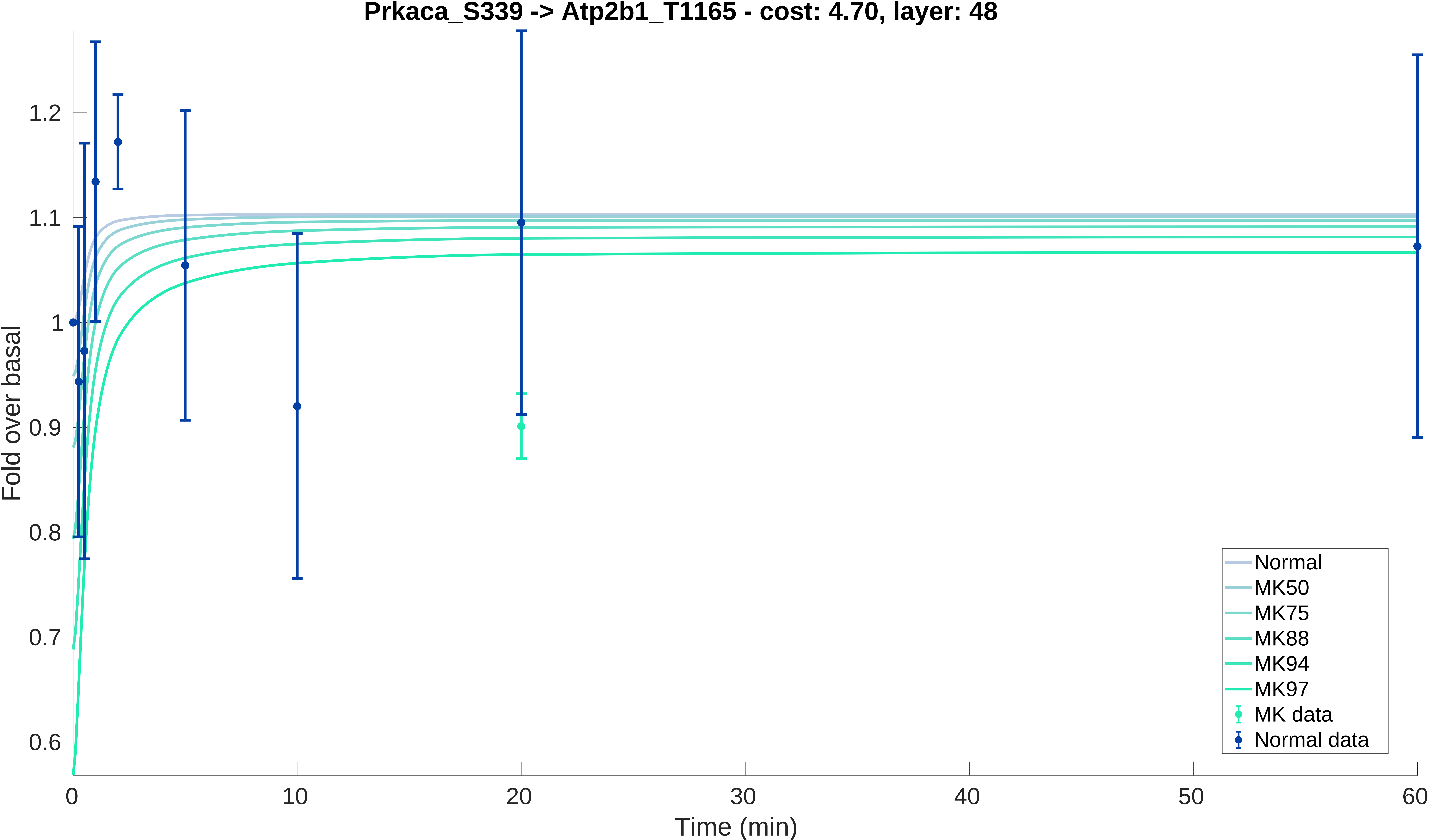

Prkaca_S339 -> Atp2b1_T1165 - cost: 4.70, layer: 48
Fold over basal
Time (min)
0.6
0.7
0.8
0.9
1
1.1
1.2
0
10
20
30
40
50
60
Normal
MK50
MK75
MK88
MK94
MK97
MK data
Normal data

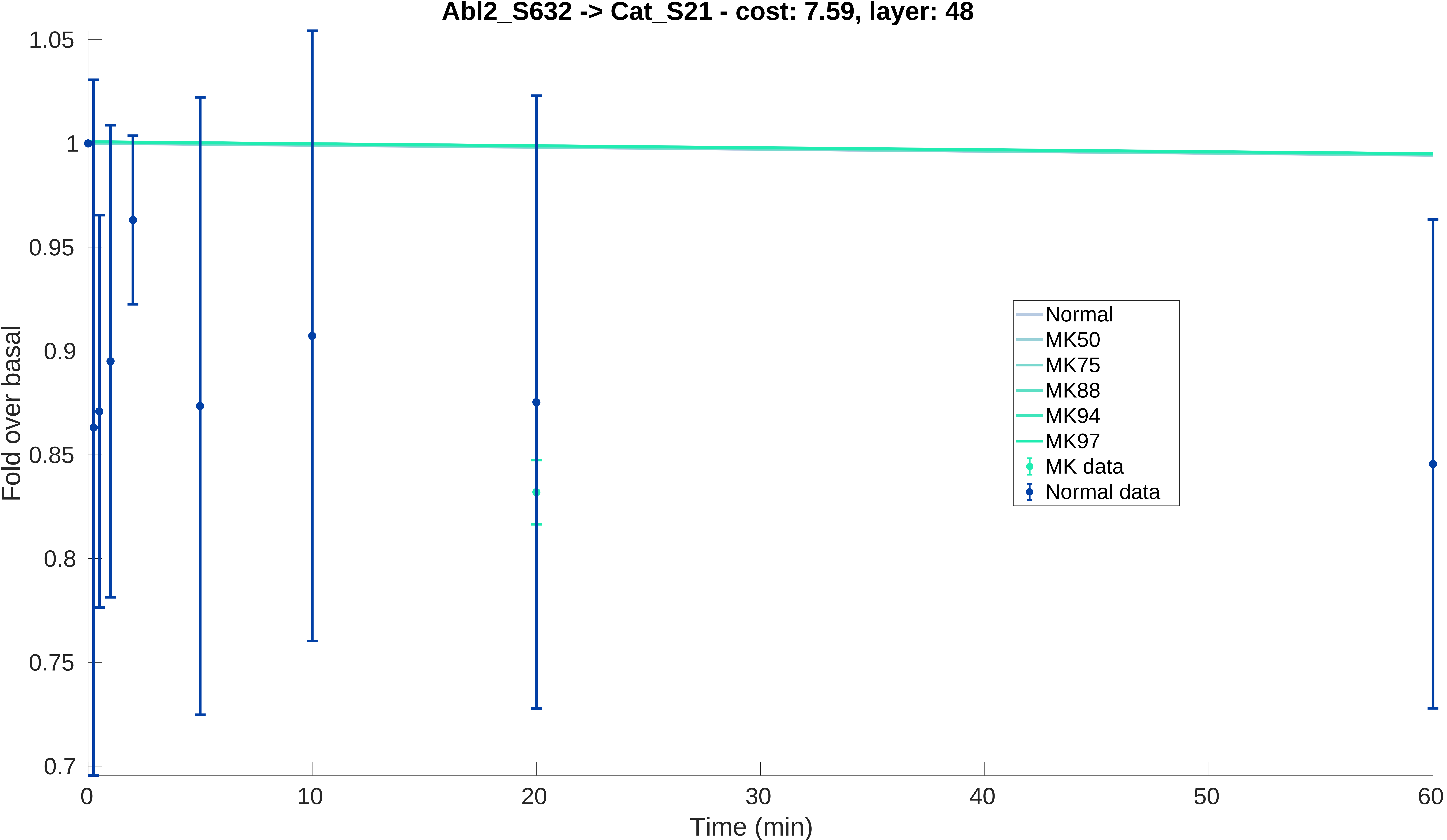

Abl2_S632 -> Cat_S21 - cost: 7.59, layer: 48
Fold over basal
Time (min)
1.05
1
0.95
0.9
0.85
0.8
0.75
0.7
0
10
20
30
40
50
60
Normal
MK50
MK75
MK88
MK94
MK97
MK data
Normal data

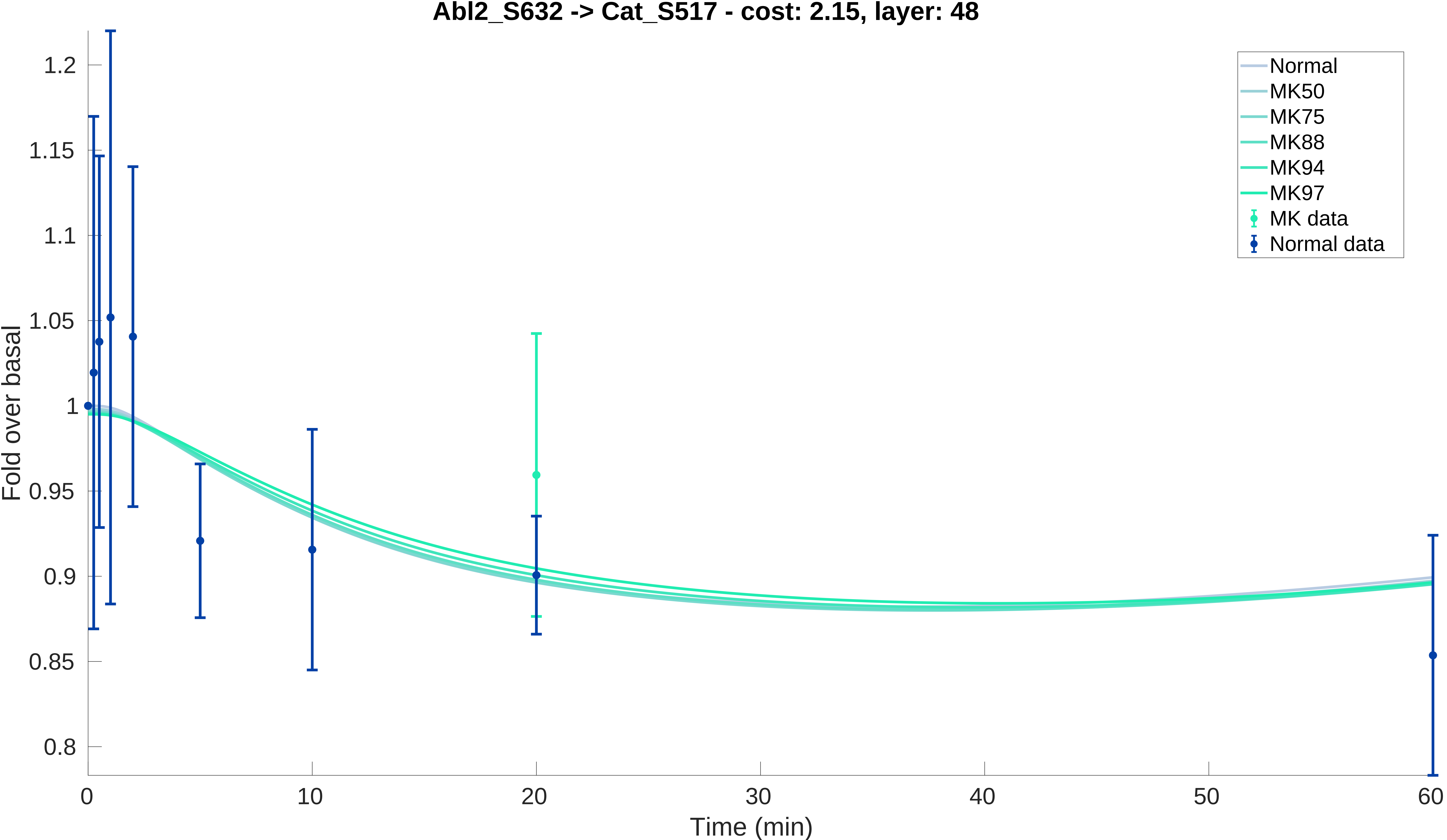

Abl2_S632 -> Cat_S517 - cost: 2.15, layer: 48
Fold over basal
Time (min)
1.2
1.15
1.1
1.05
1
0.95
0.9
0.85
0.8
0
10
20
30
40
50
60
Normal
MK50
MK75
MK88
MK94
MK97
MK data
Normal data

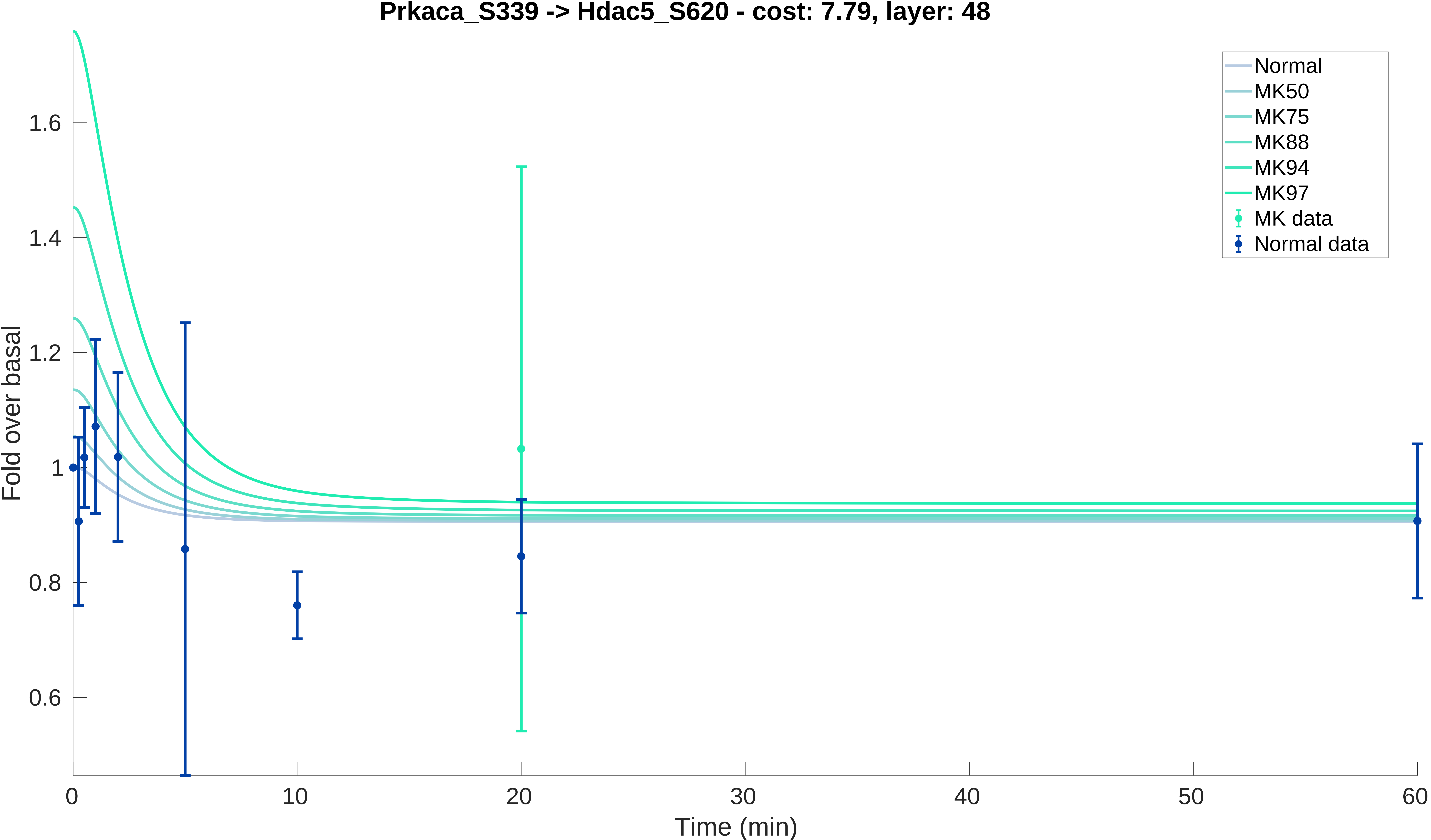

Prkaca_S339 -> Hdac5_S620 - cost: 7.79, layer: 48
Normal
MK50
MK75
MK88
MK94
MK97
MK data
Normal data
Fold over basal
1.6
1.4
1.2
1
0.8
0.6
0
10
20
30
40
50
60
Time (min)

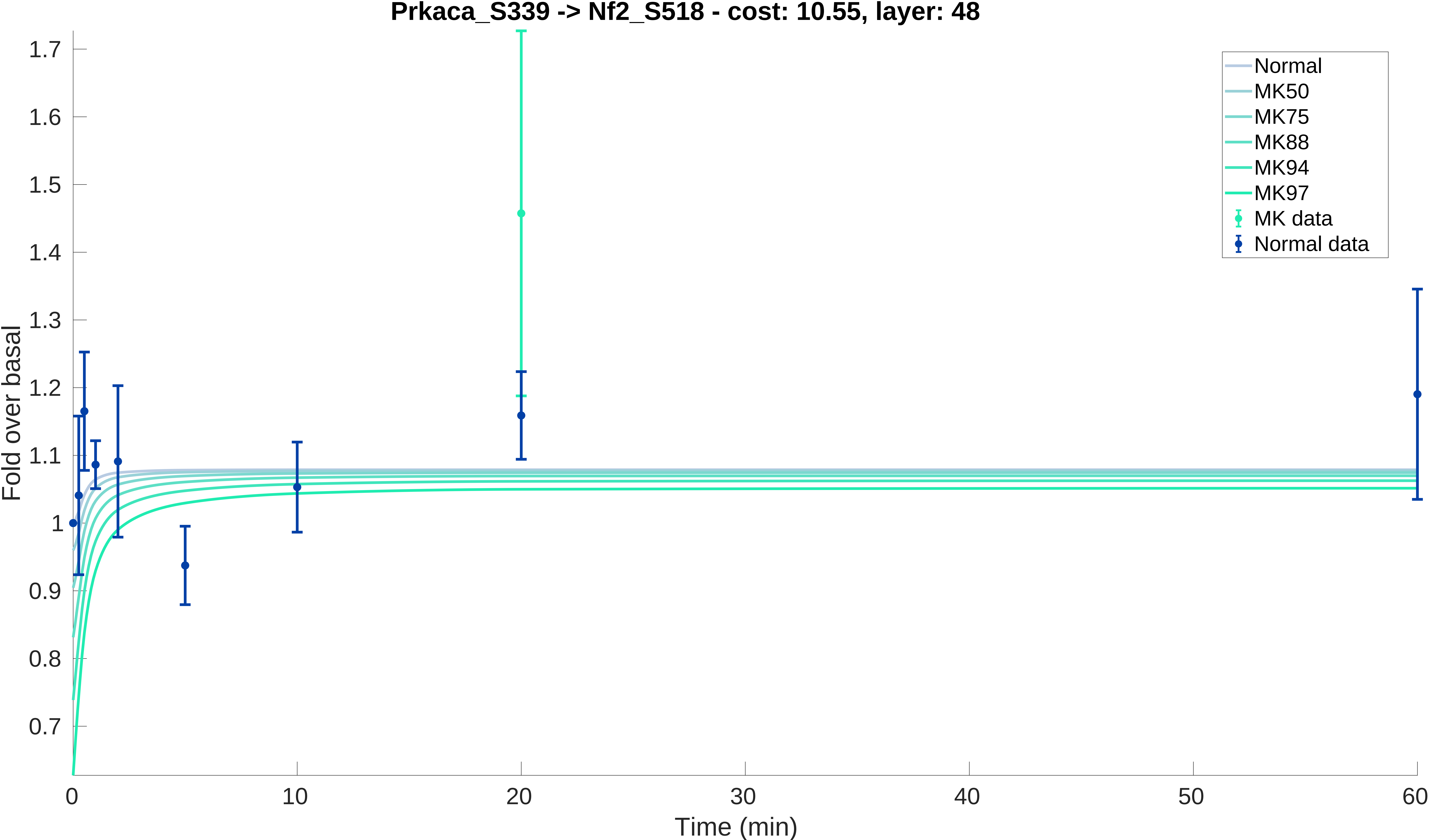

Prkaca_S339 -> Nf2_S518 - cost: 10.55, layer: 48
Fold over basal
Time (min)
1.7
1.6
1.5
1.4
1.3
1.2
1.1
1
0.9
0.8
0.7
0
10
20
30
40
50
60
Normal
MK50
MK75
MK88
MK94
MK97
MK data
Normal data

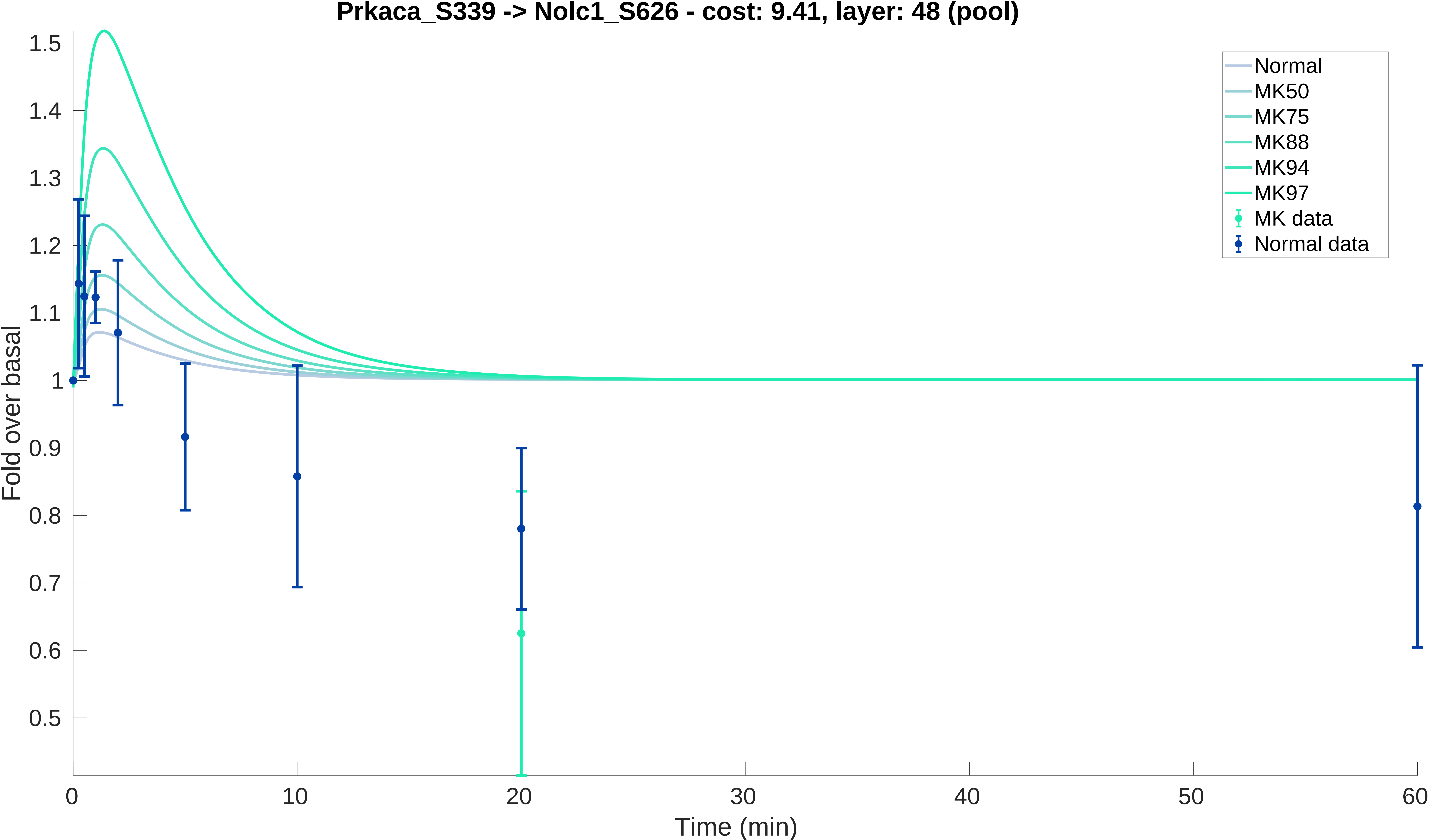

Prkaca_S339 -> Nolc1_S626 - cost: 9.41, layer: 48 (pool)
Fold over basal
Time (min)
0.5
0.6
0.7
0.8
0.9
1
1.1
1.2
1.3
1.4
1.5
0
10
20
30
40
50
60
Normal
MK50
MK75
MK88
MK94
MK97
MK data
Normal data

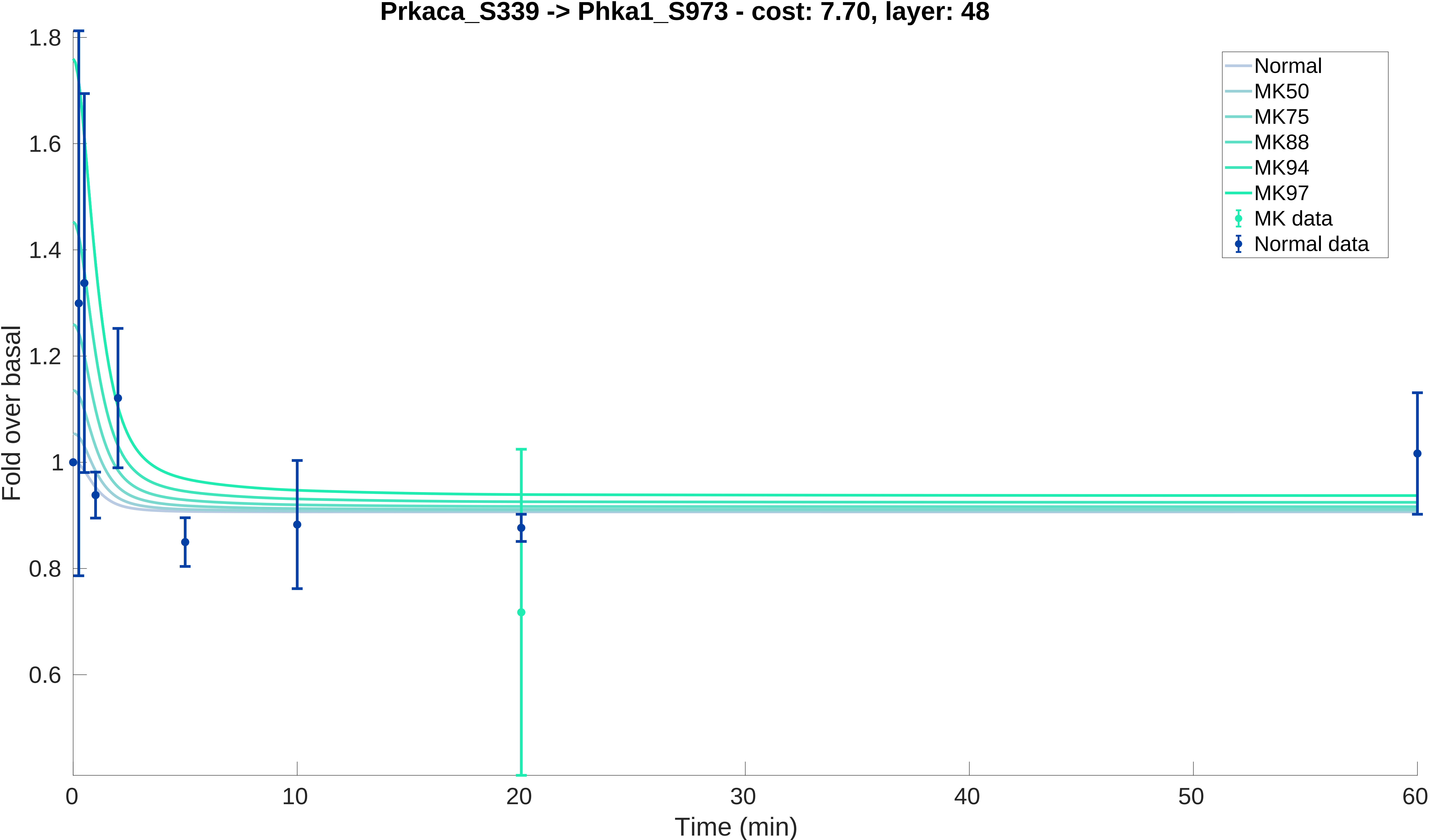

Prrkaca_S339 -> Phka1_S973 - cost: 7.70, layer: 48
Fold over basal
Time (min)
0.6
0.8
1
1.2
1.4
1.6
1.8
0
10
20
30
40
50
60
Normal
MK50
MK75
MK88
MK94
MK97
MK data
Normal data

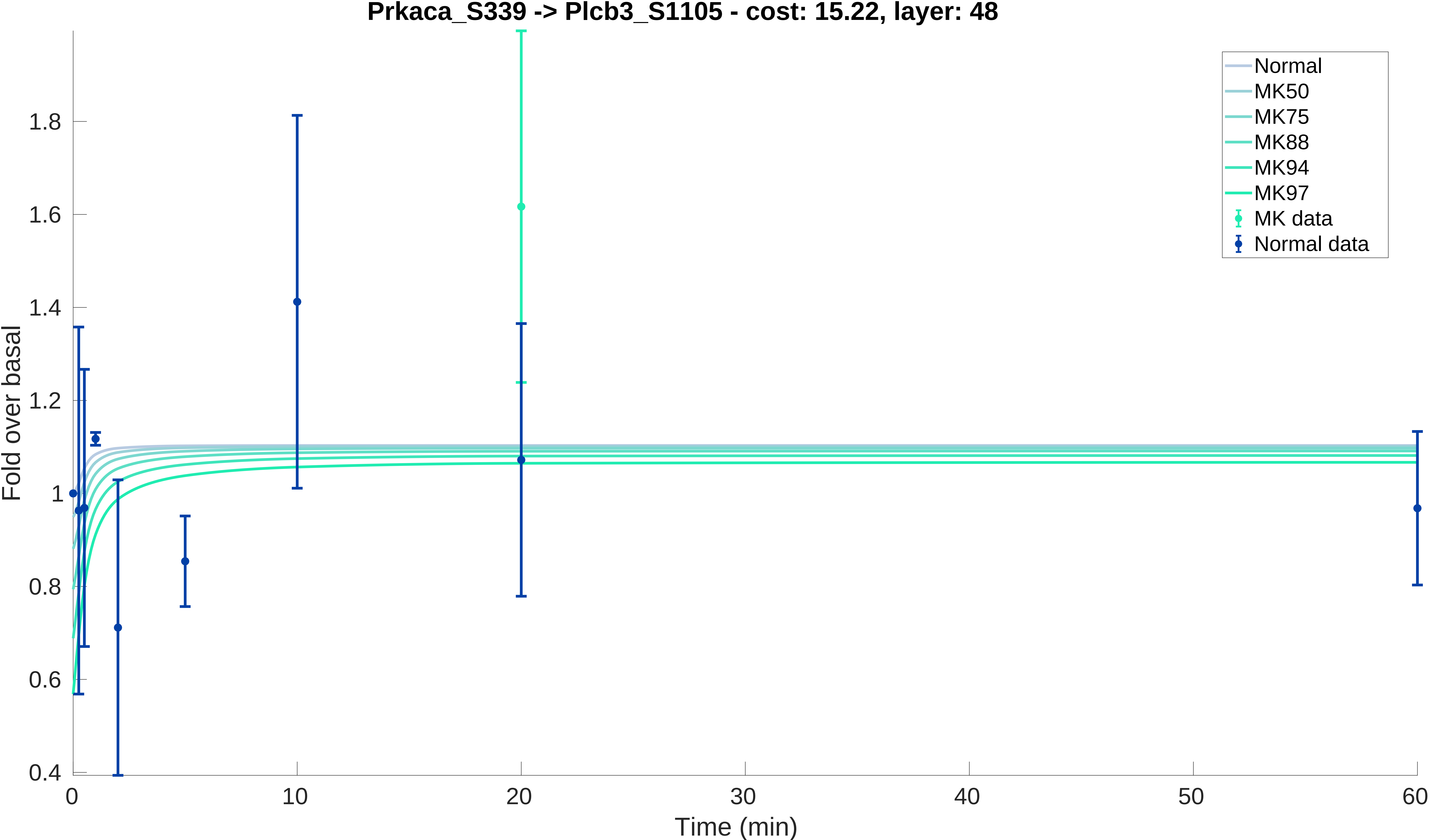

Prkaca_S339 -> Plcb3_S1105 - cost: 15.22, layer: 48
Fold over basal
Time (min)
0.4
0.6
0.8
1
1.2
1.4
1.6
1.8
0
10
20
30
40
50
60
Normal
MK50
MK75
MK88
MK94
MK97
MK data
Normal data

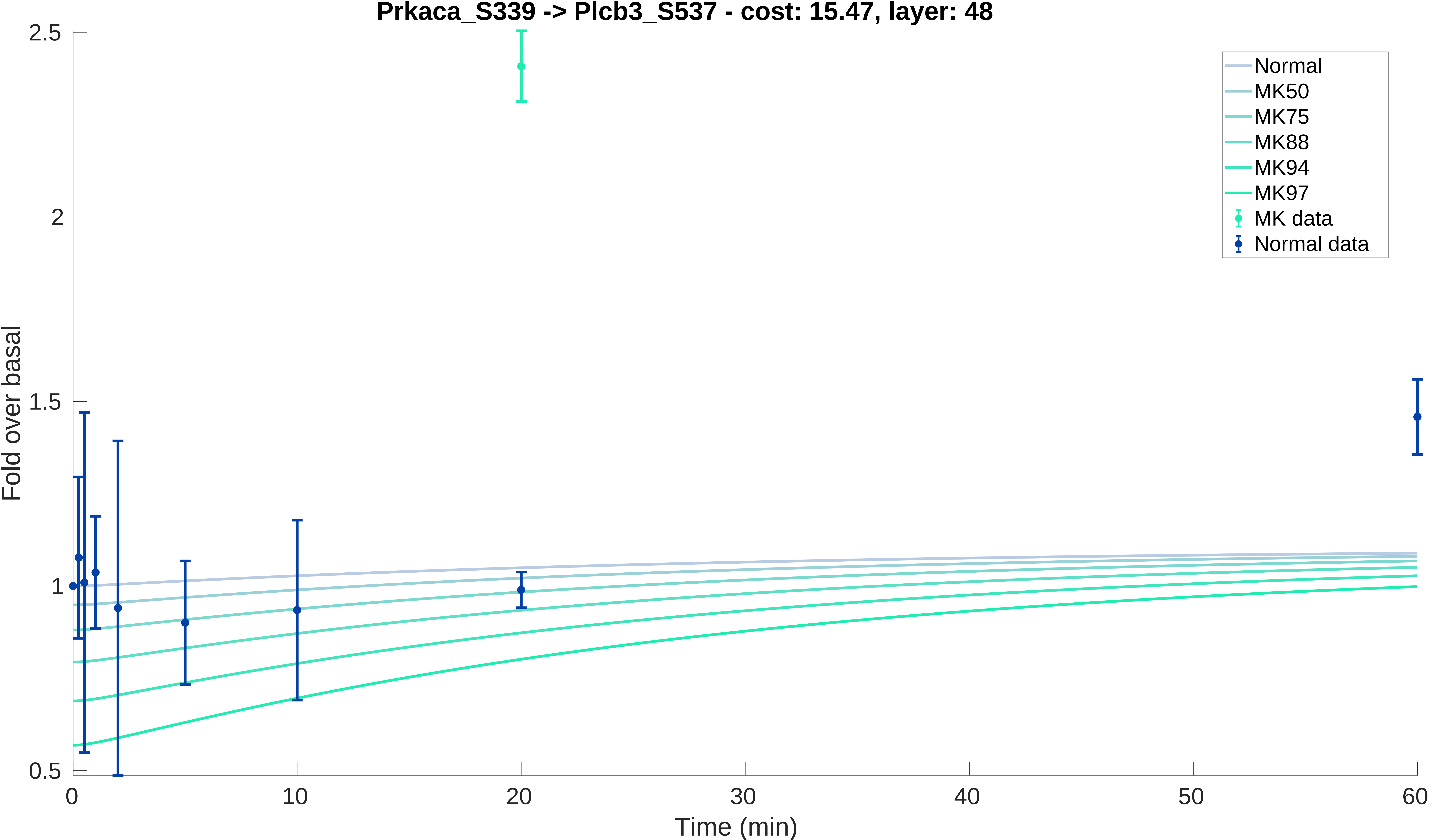

Prkaca_S339 -> Plcb3_S537 - cost: 15.47, layer: 48
Fold over basal
Time (min)
0.5
1
1.5
2
2.5
0
10
20
30
40
50
60
Normal
MK50
MK75
MK88
MK94
MK97
MK data
Normal data

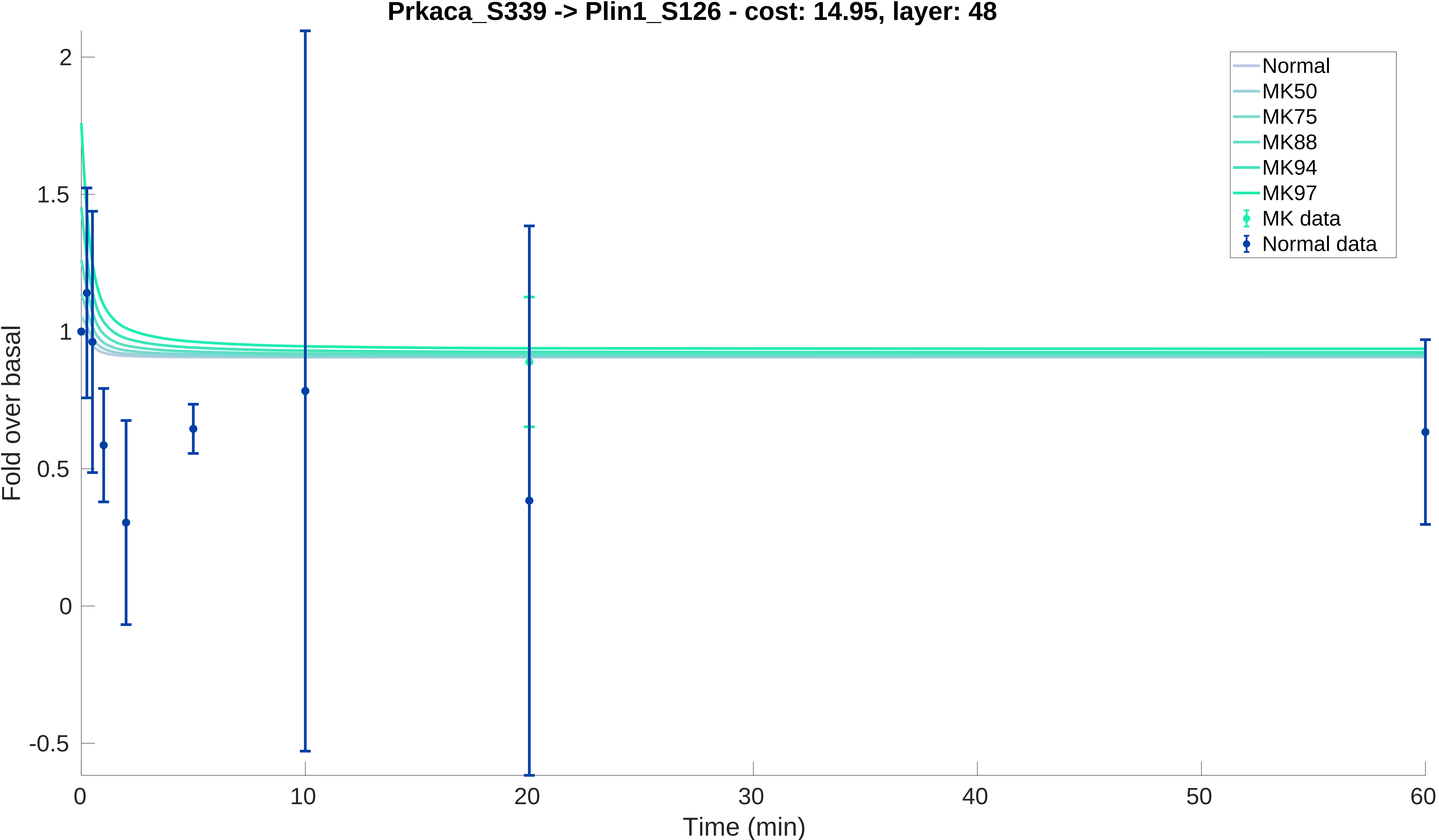

Prkaca_S339 -> Plin1_S126 - cost: 14.95, layer: 48
Normal
MK50
MK75
MK88
MK94
MK97
MK data
Normal data
Fold over basal
2
1.5
1
0.5
0
-0.5
0
10
20
30
40
50
60
Time (min)

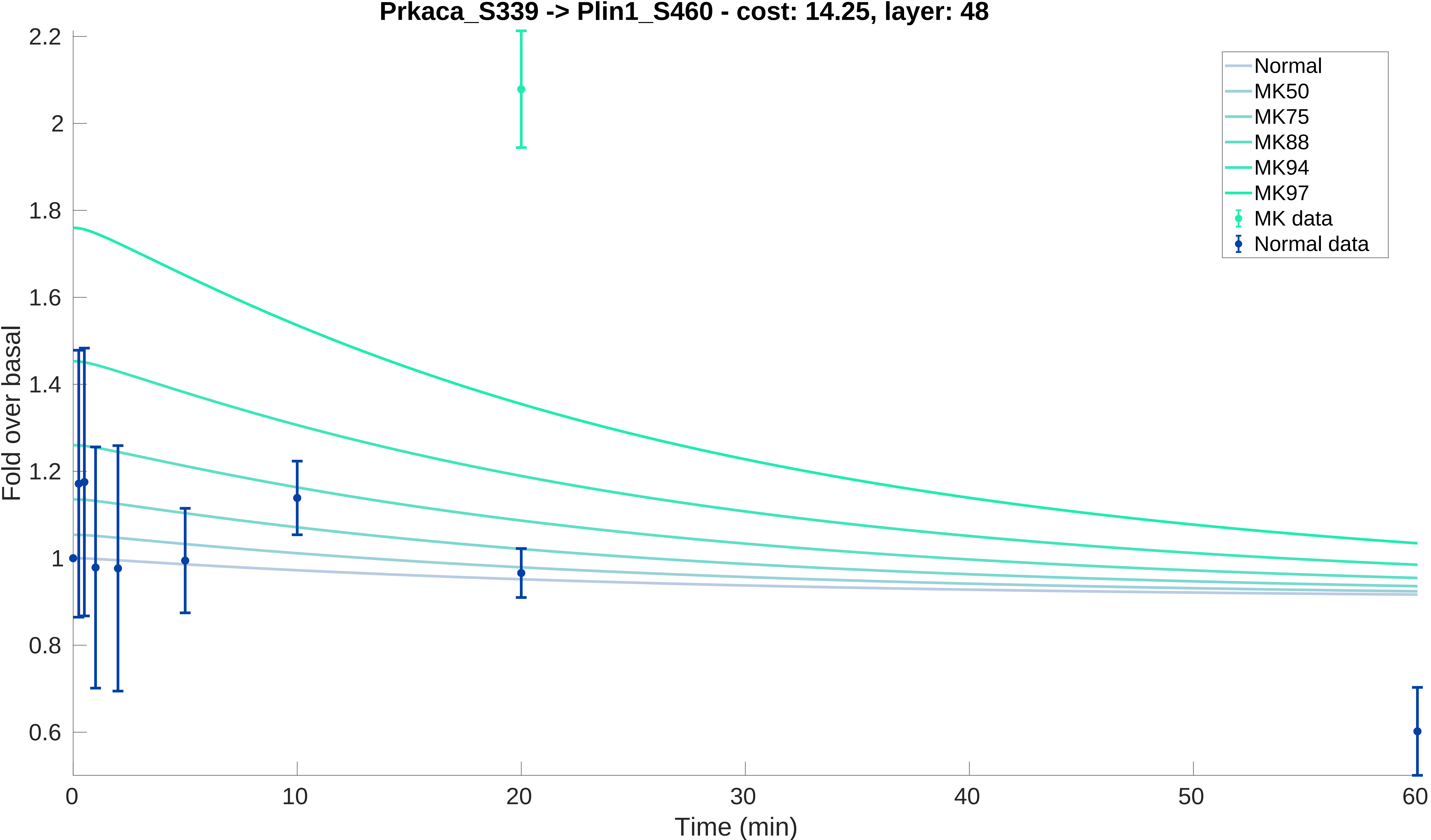
Prrkaca_S339 -> Plin1_S460 - cost: 14.25, layer: 48
Fold over basal
2.2
2
1.8
1.6
1.4
1.2
1
0.8
0.6
0
10
20
30
40
50
60
Time (min)
Normal
MK50
MK75
MK88
MK94
MK97
MK data
Normal data

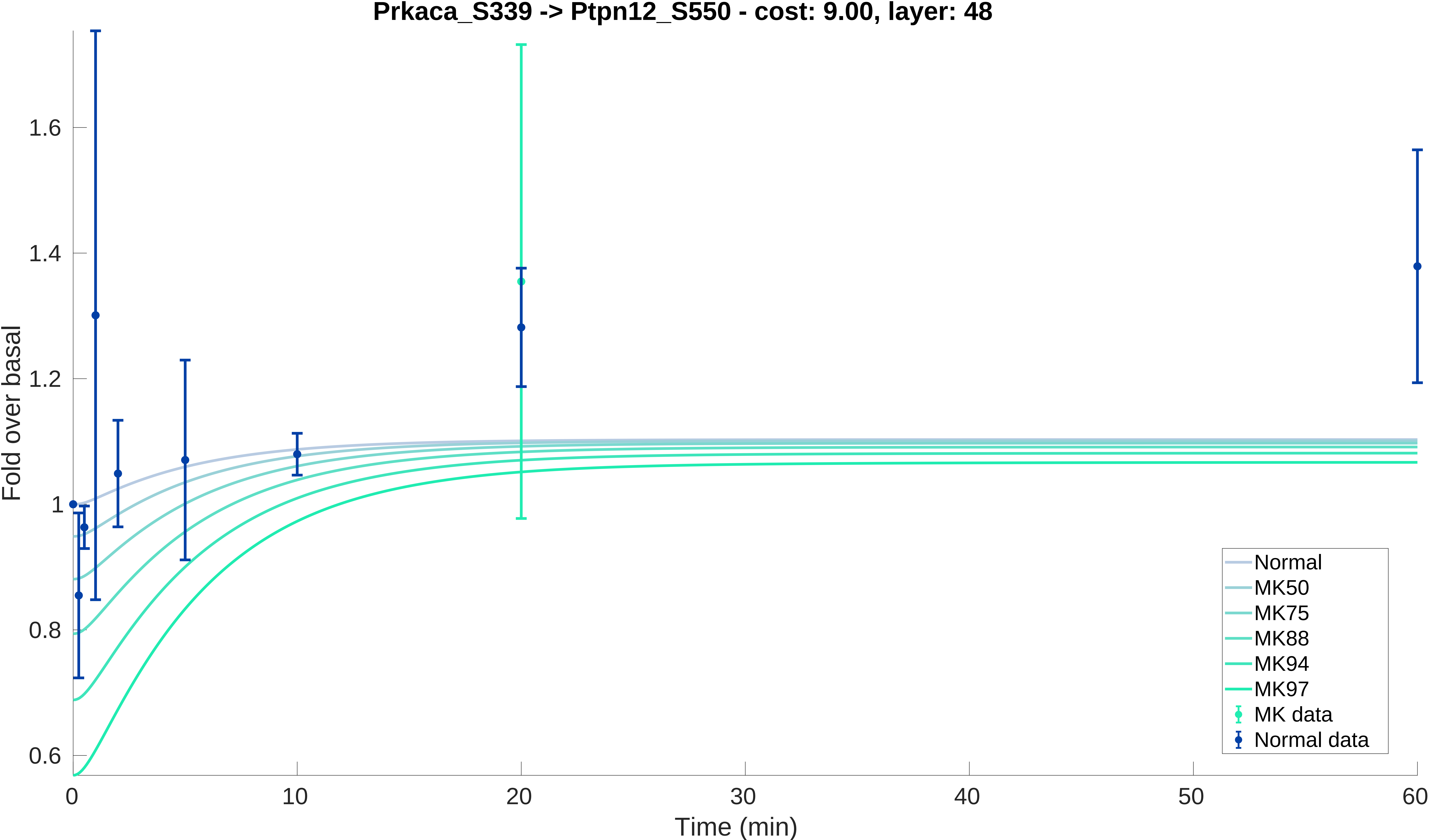

Prkaca_S339 -> Ptpn12_S550 - cost: 9.00, layer: 48
Fold over basal
1.6
1.4
1.2
1
0.8
0.6
0
10
20
30
40
50
60
Time (min)
Normal
MK50
MK75
MK88
MK94
MK97
MK data
Normal data

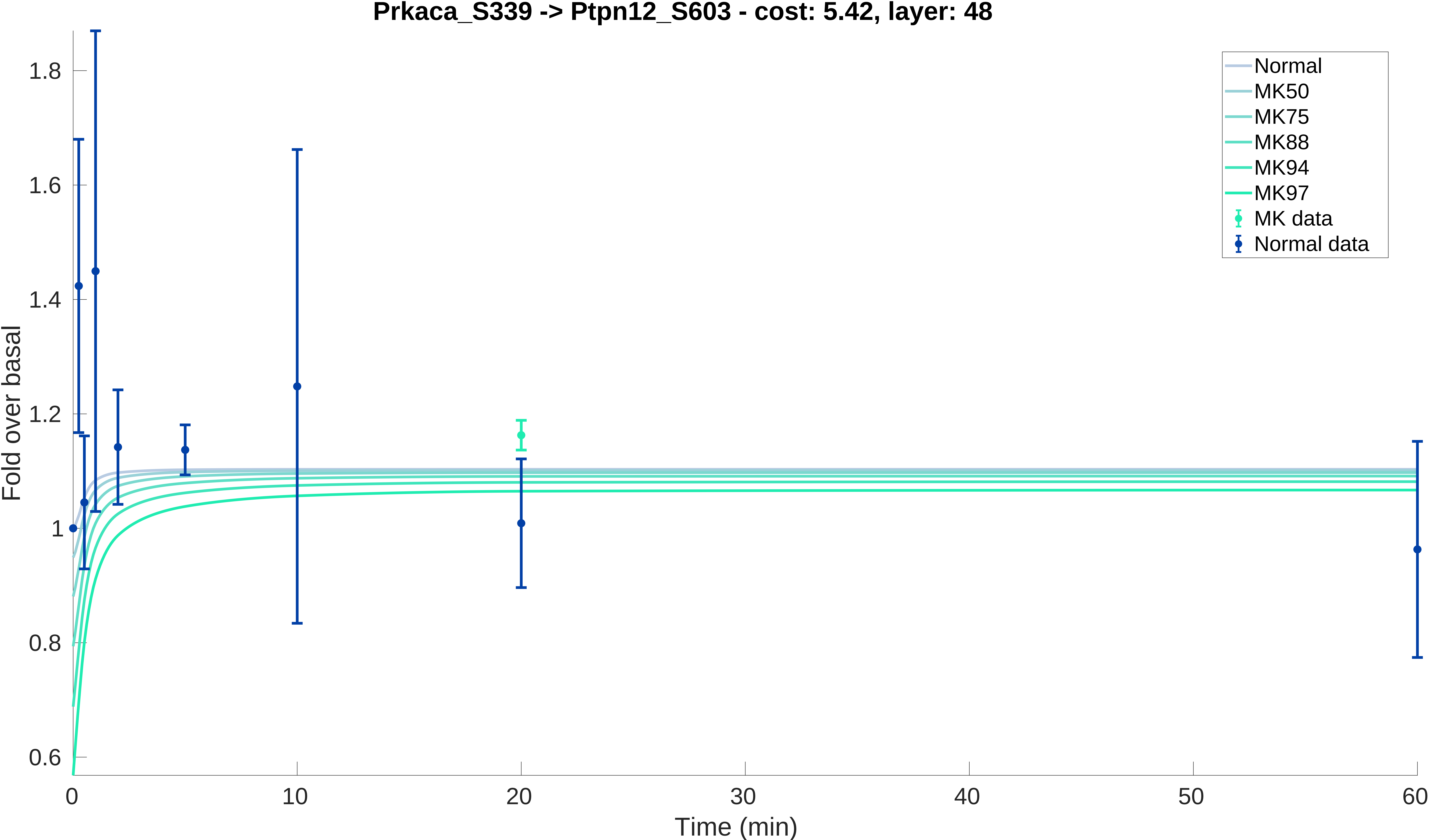

Prrkaca_S339 -> Ptpn12_S603 - cost: 5.42, layer: 48
Fold over basal
Time (min)
0.6
0.8
1
1.2
1.4
1.6
1.8
0
10
20
30
40
50
60
Normal
MK50
MK75
MK88
MK94
MK97
MK data
Normal data

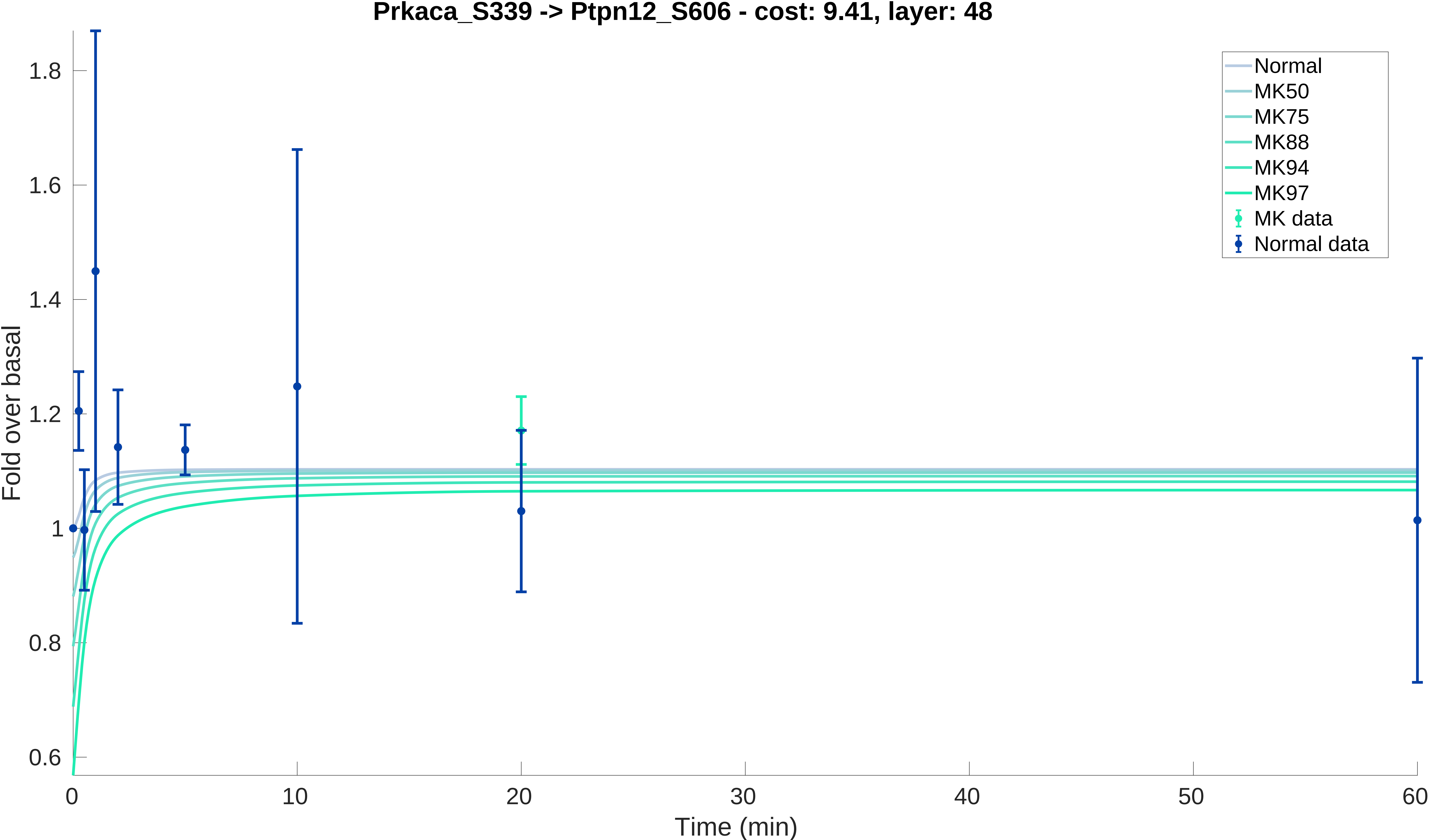

Prrkaca_S339 -> Ptpn12_S606 - cost: 9.41, layer: 48
Fold over basal
Time (min)
Normal
MK50
MK75
MK88
MK94
MK97
MK data
Normal data
1.8
1.6
1.4
1.2
1
0.8
0.6
0
10
20
30
40
50
60

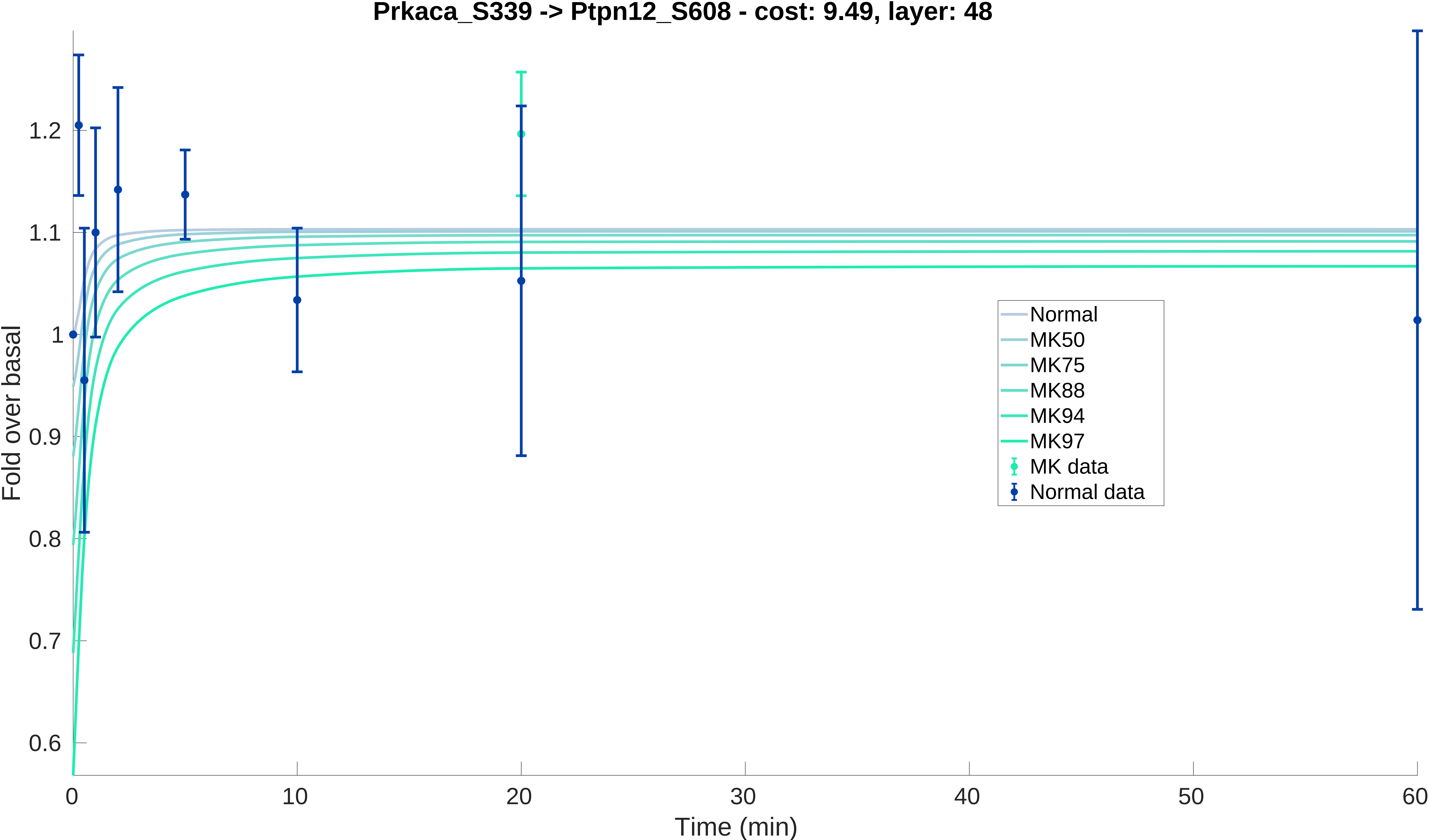

Prkaca_S339 -> Ptpn12_S608 - cost: 9.49, layer: 48
Fold over basal
1.2
1.1
1
0.9
0.8
0.7
0.6
0
10
20
30
40
50
60
Time (min)
Normal
MK50
MK75
MK88
MK94
MK97
MK data
Normal data

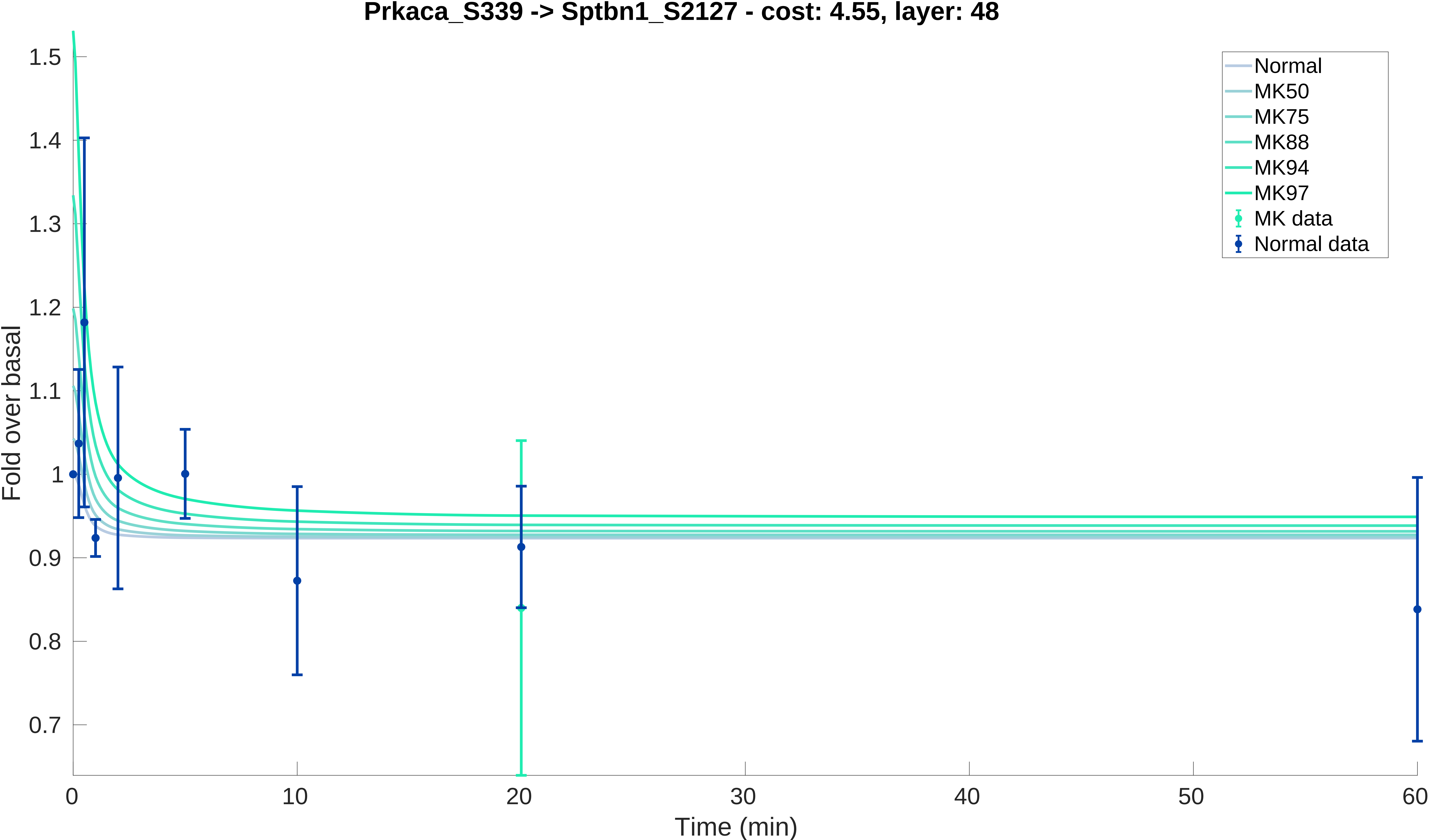
Prkaca_S339 -> Sptbn1_S2127 - cost: 4.55, layer: 48
Fold over basal
Time (min)
0.7
0.8
0.9
1
1.1
1.2
1.3
1.4
1.5
0
10
20
30
40
50
60
Normal
MK50
MK75
MK88
MK94
MK97
MK data
Normal data

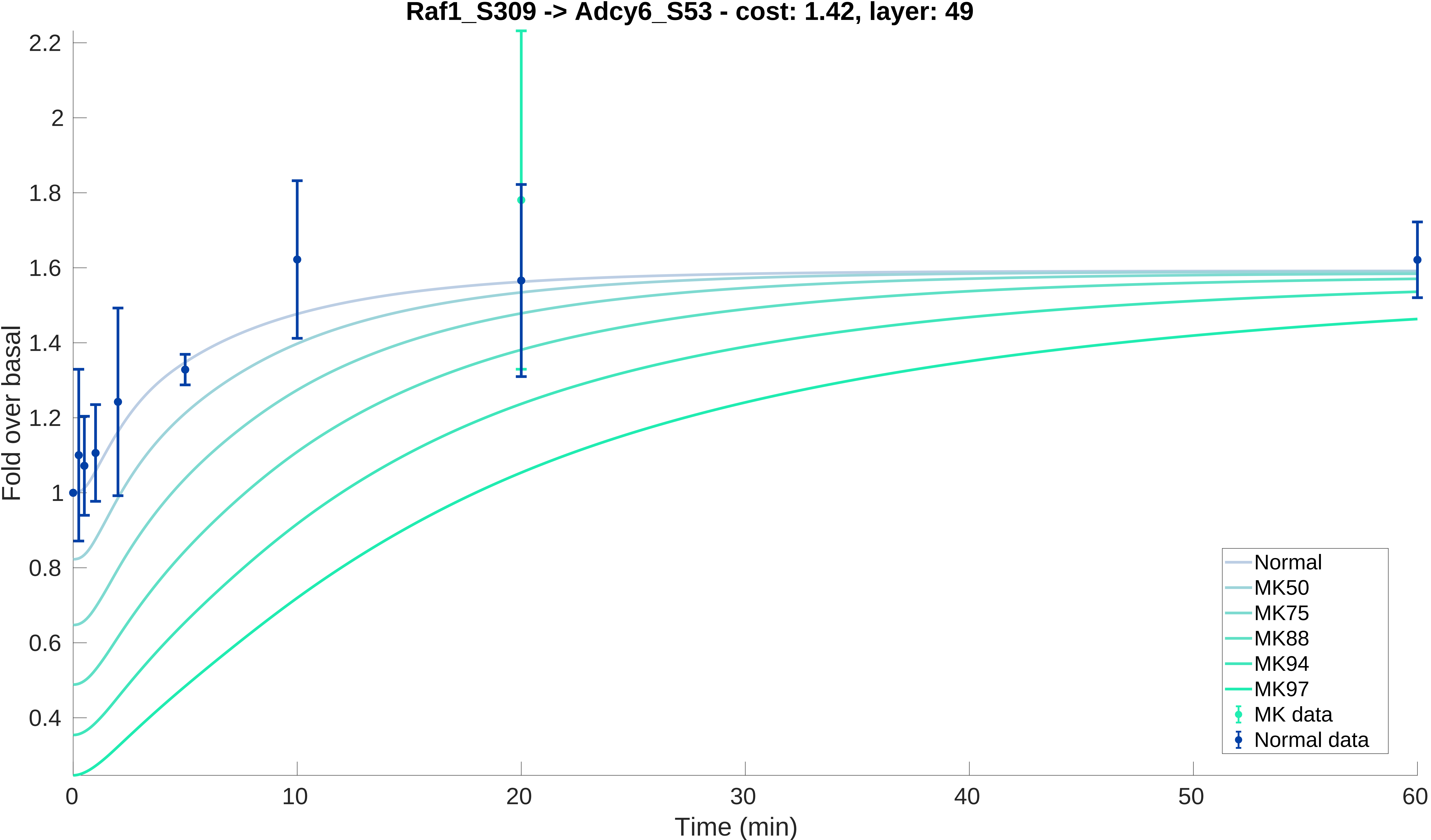

Raf1_S309 -> Adcy6_S53 - cost: 1.42, layer: 49
Fold over basal
Time (min)
0.4
0.6
0.8
1
1.2
1.4
1.6
1.8
2
2.2
0
10
20
30
40
50
60
Normal
MK50
MK75
MK88
MK94
MK97
MK data
Normal data

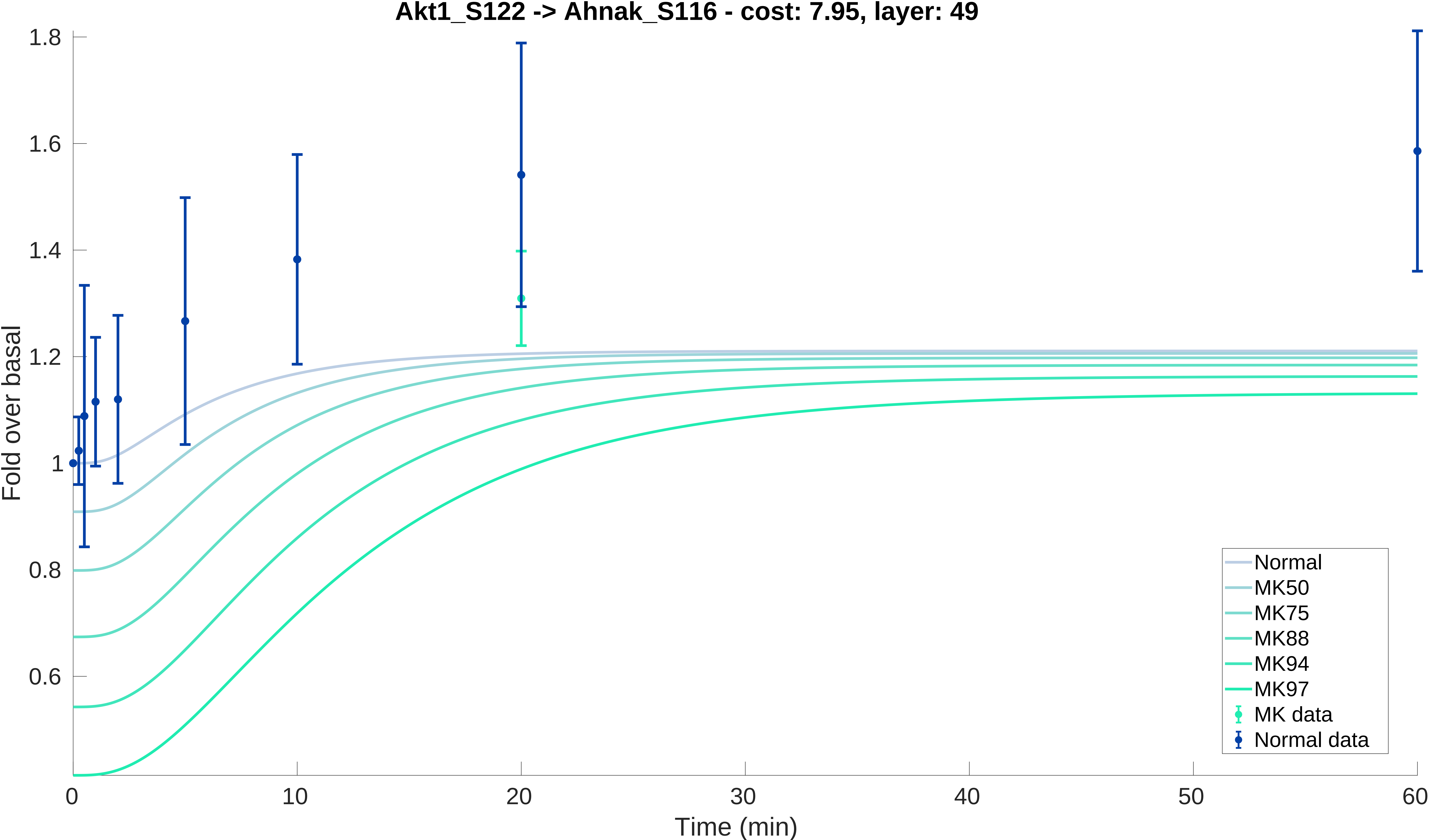

Akt1_S122 -> Ahnak_S116 - cost: 7.95, layer: 49
Fold over basal
Time (min)
1.8
1.6
1.4
1.2
1
0.8
0.6
0
10
20
30
40
50
60
Normal
MK50
MK75
MK88
MK94
MK97
MK data
Normal data

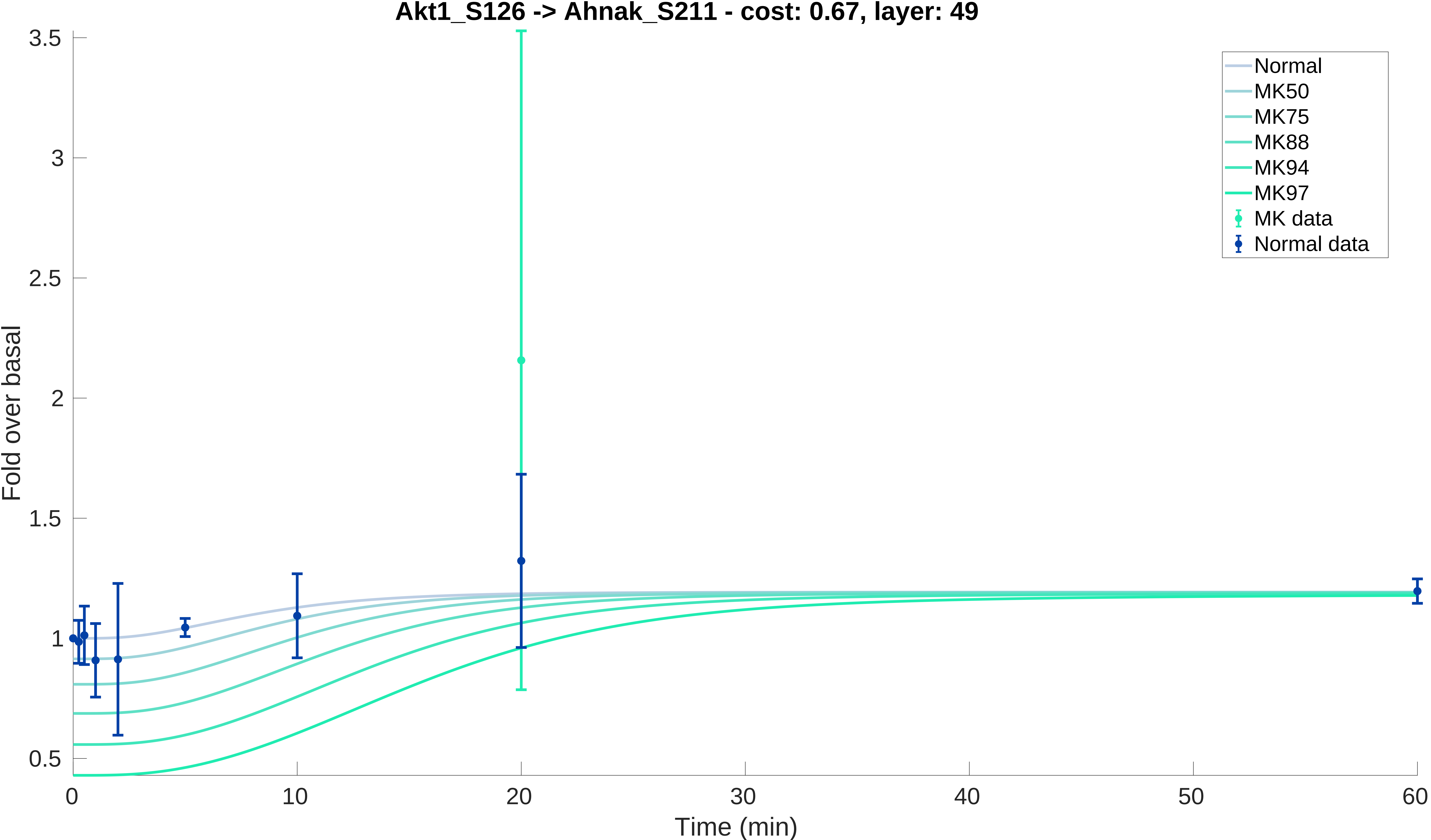

Akt1_S126 -> Ahnak_S211 - cost: 0.67, layer: 49
Fold over basal
Time (min)
3.5
3
2.5
2
1.5
1
0.5
0
10
20
30
40
50
60
Normal
MK50
MK75
MK88
MK94
MK97
MK data
Normal data

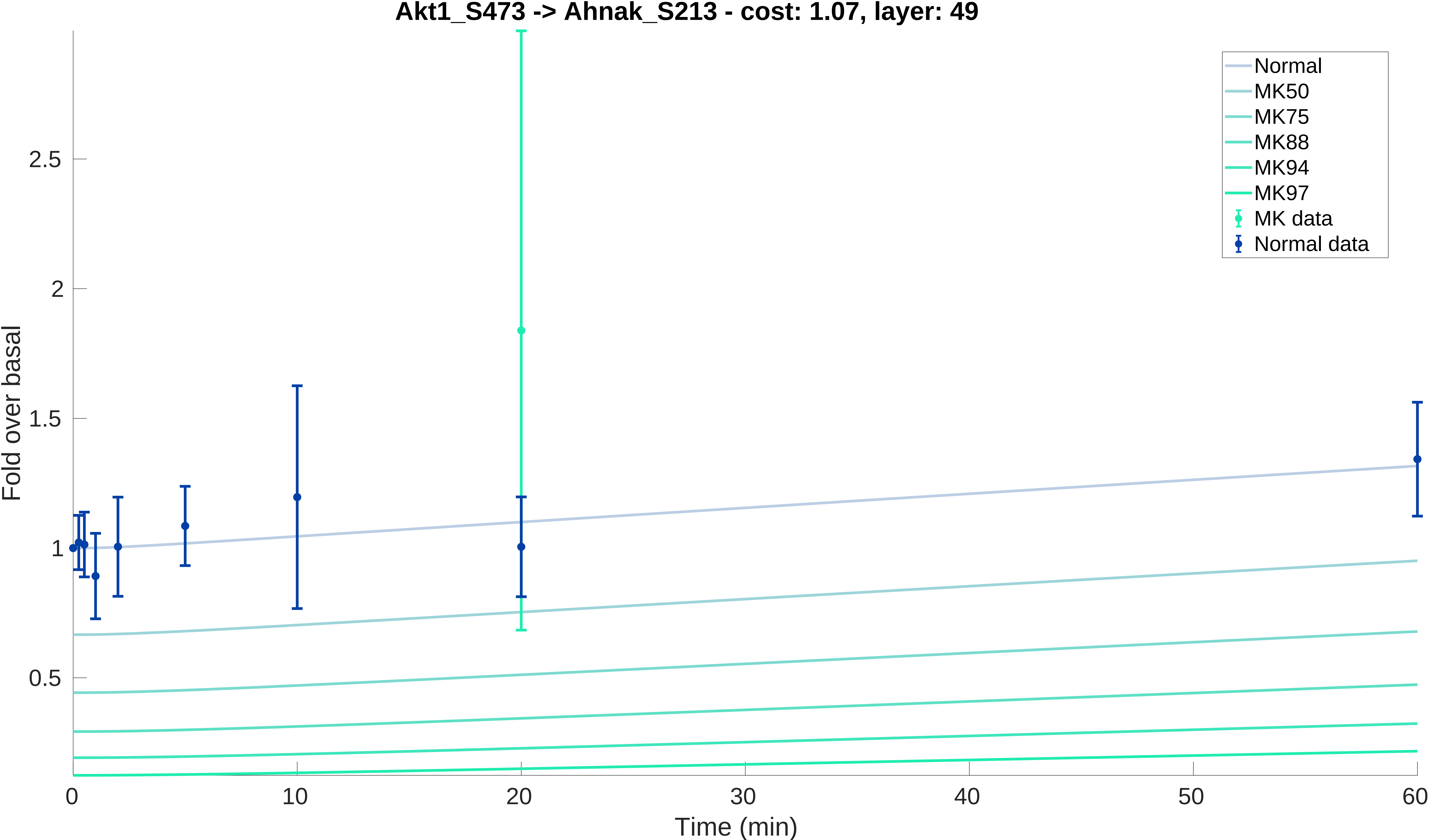

Akt1_S473 -> Ahnak_S213 - cost: 1.07, layer: 49
Normal
MK50
MK75
MK88
MK94
MK97
MK data
Normal data
Fold over basal
2.5
2
1.5
1
0.5
0
10
20
30
40
50
60
Time (min)

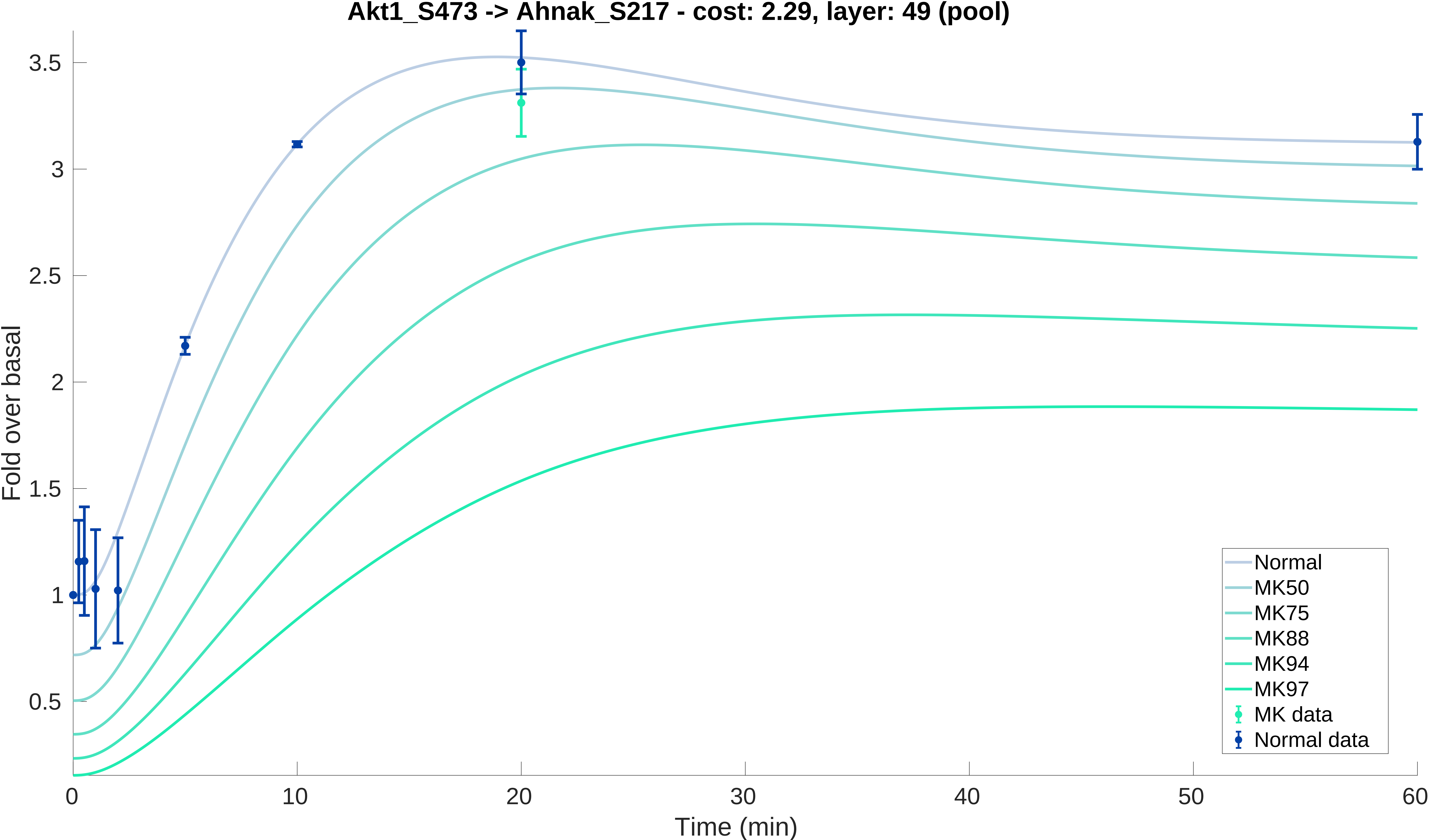

Akt1_S473 -> Ahnak_S217 - cost: 2.29, layer: 49 (pool)
3.5
3
2.5
2
1.5
1
0.5
Fold over basal
0
10
20
30
40
50
60
Time (min)
Normal
MK50
MK75
MK88
MK94
MK97
MK data
Normal data

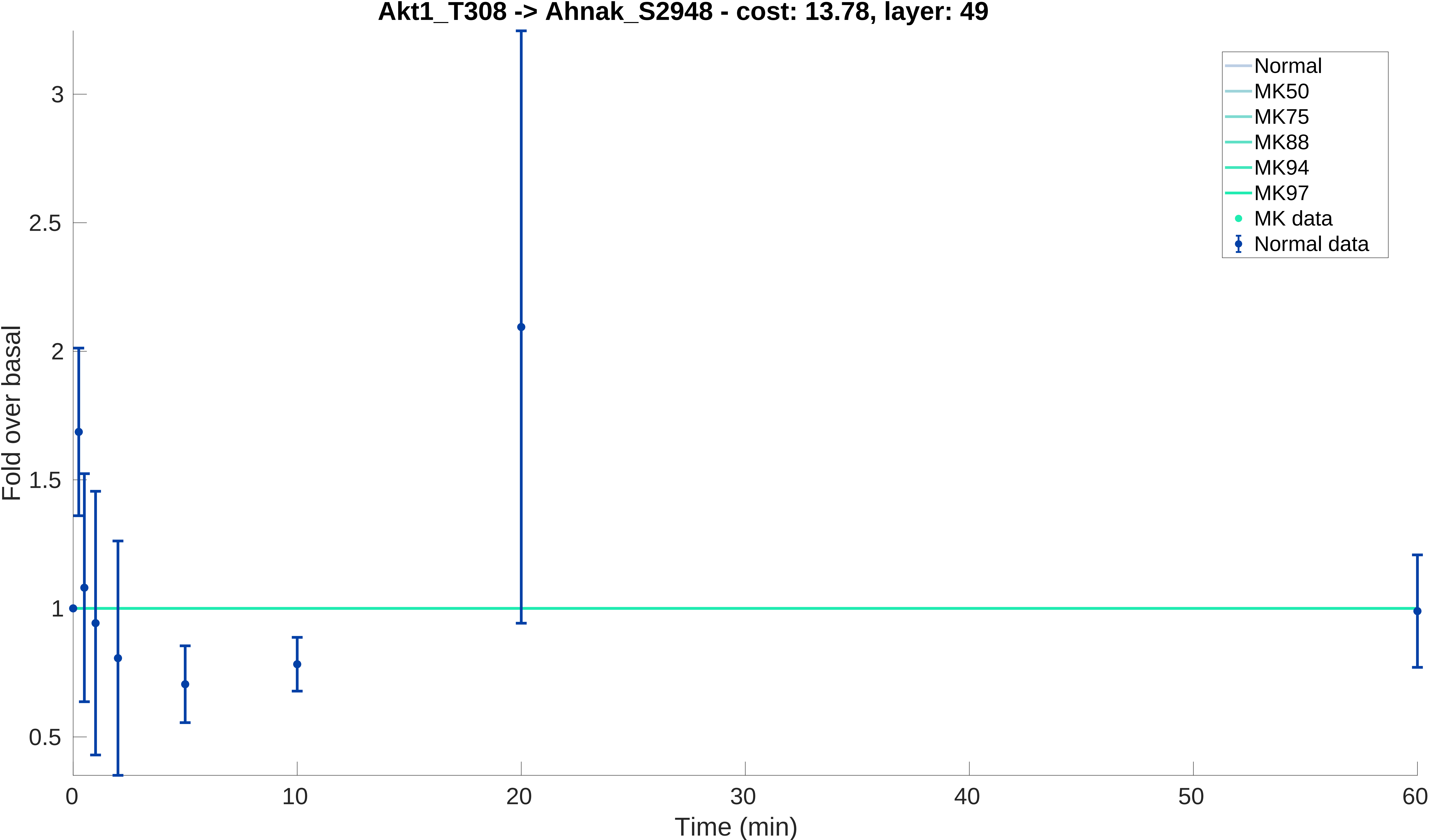

Akt1_T308 -> Ahnak_S2948 - cost: 13.78, layer: 49
Normal
MK50
MK75
MK88
MK94
MK97
MK data
Normal data
Fold over basal
Time (min)
0.5
1
1.5
2
2.5
3
0
10
20
30
40
50
60

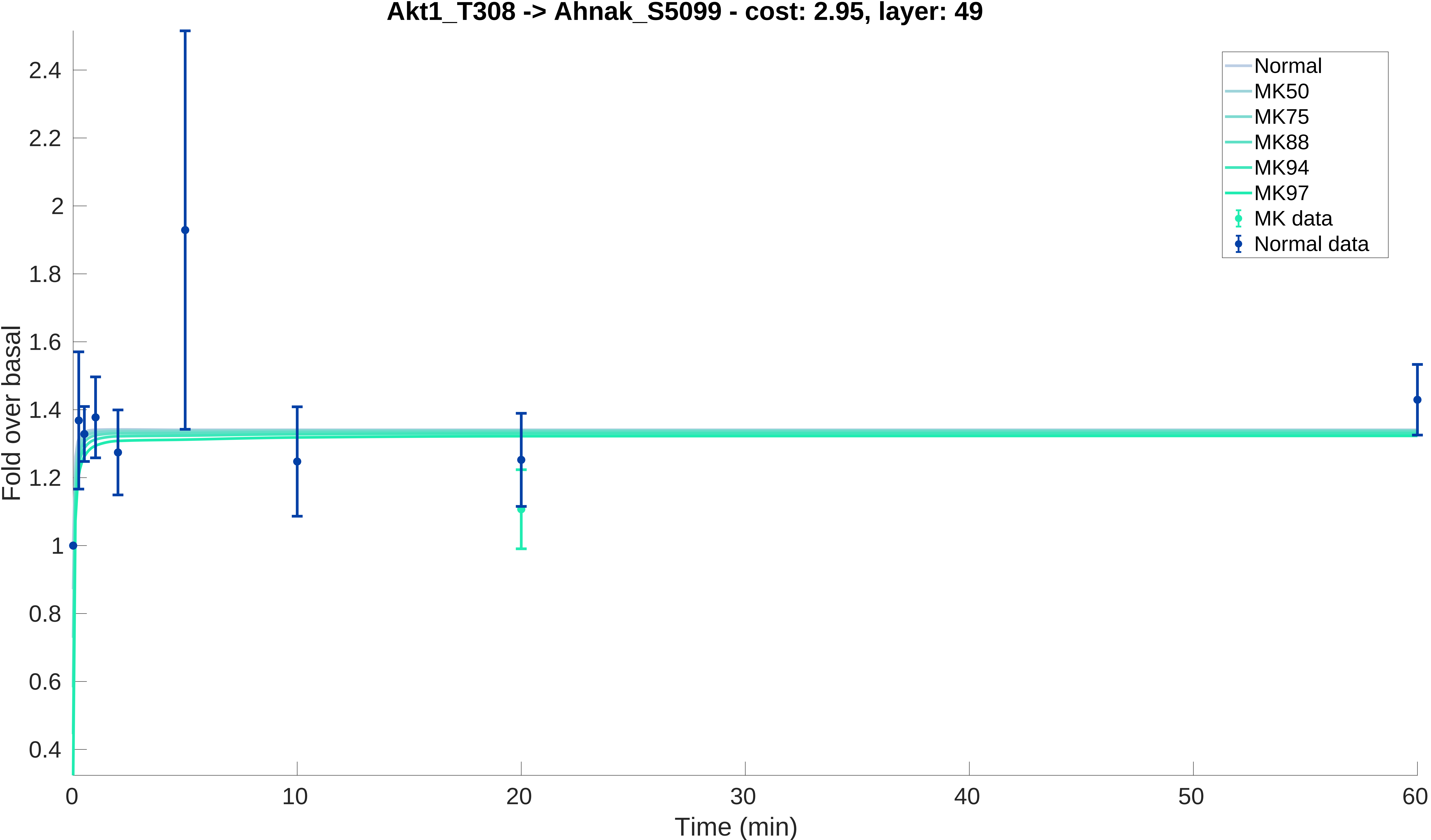

Akt1_T308 -> Ahnak_S5099 - cost: 2.95, layer: 49
Fold over basal
Time (min)
Normal
MK50
MK75
MK88
MK94
MK97
MK data
Normal data
2.4
2.2
2
1.8
1.6
1.4
1.2
1
0.8
0.6
0.4
0
10
20
30
40
50
60

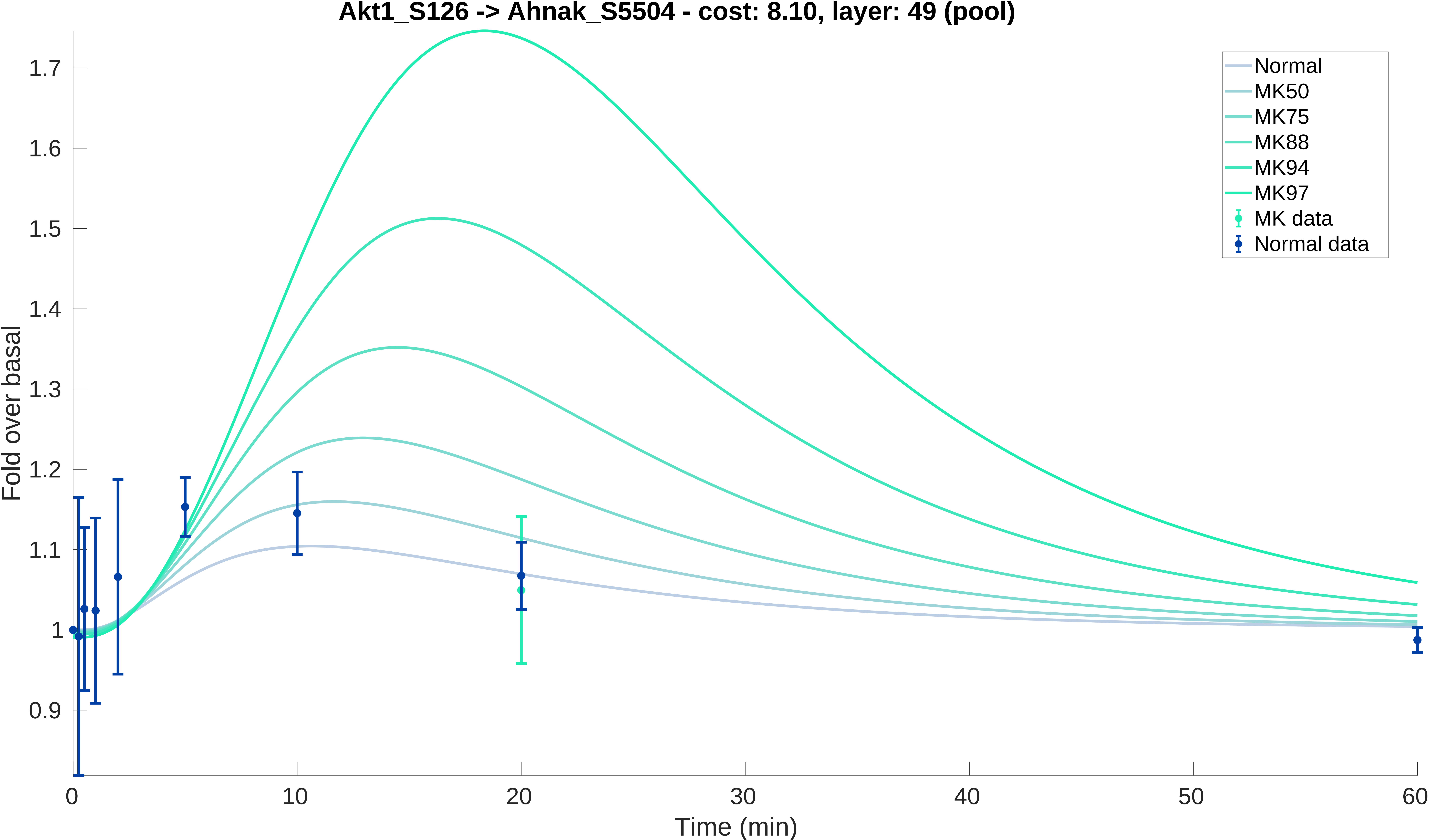

Akt1_S126 -> Ahnak_S5504 - cost: 8.10, layer: 49 (pool)
Normal
MK50
MK75
MK88
MK94
MK97
MK data
Normal data
Fold over basal
Time (min)
0.9
1
1.1
1.2
1.3
1.4
1.5
1.6
1.7
0
10
20
30
40
50
60

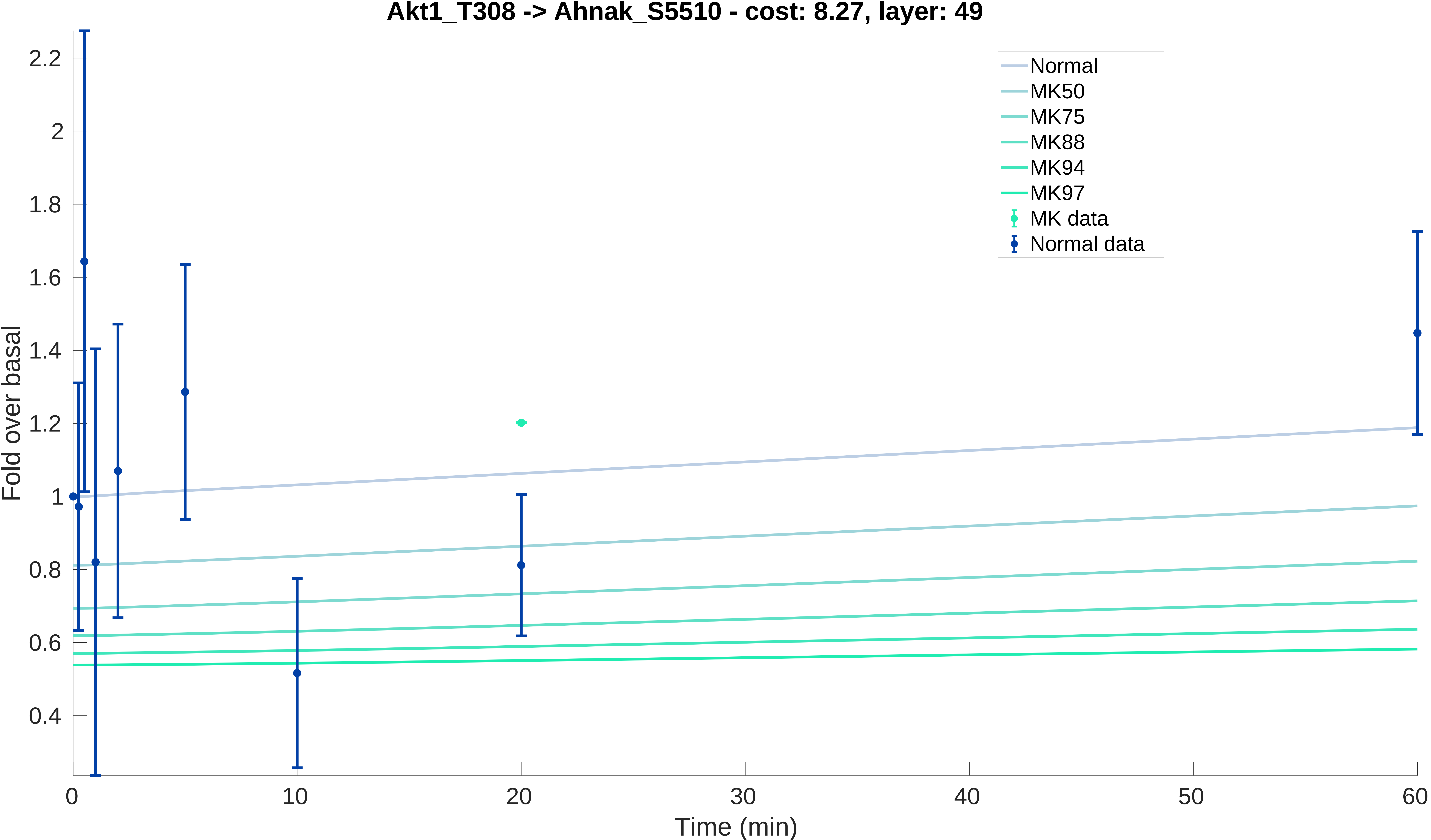
Akt1_T308 -> Ahnak_S5510 - cost: 8.27, layer: 49
Fold over basal
2.2
2
1.8
1.6
1.4
1.2
1
0.8
0.6
0.4
0
10
20
30
40
50
60
Time (min)
Normal
MK50
MK75
MK88
MK94
MK97
MK data
Normal data

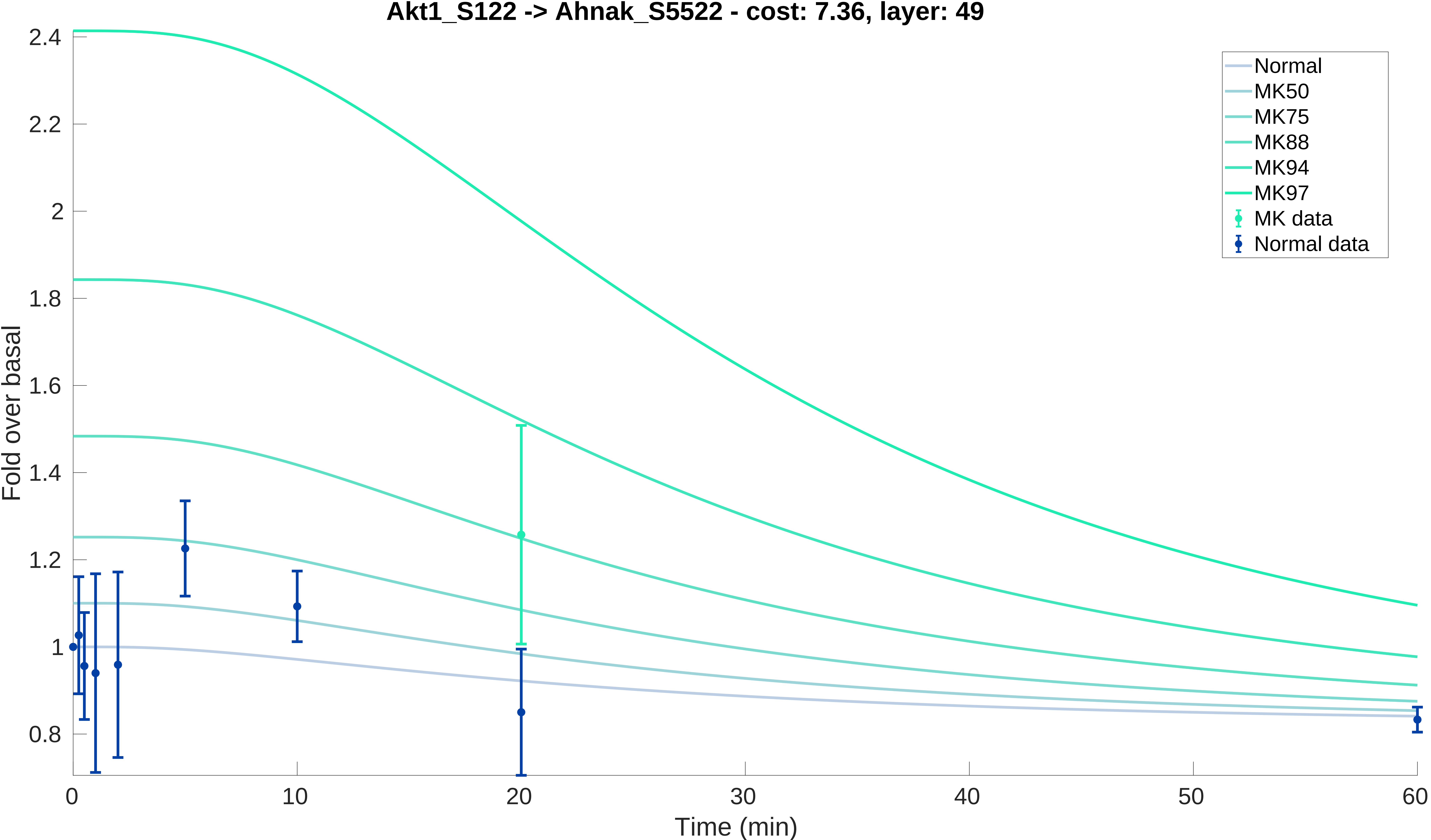

Akt1_S122 -> Ahnak_S5522 - cost: 7.36, layer: 49
Fold over basal
Time (min)
2.4
2.2
2
1.8
1.6
1.4
1.2
1
0.8
0
10
20
30
40
50
60
Normal
MK50
MK75
MK88
MK94
MK97
MK data
Normal data

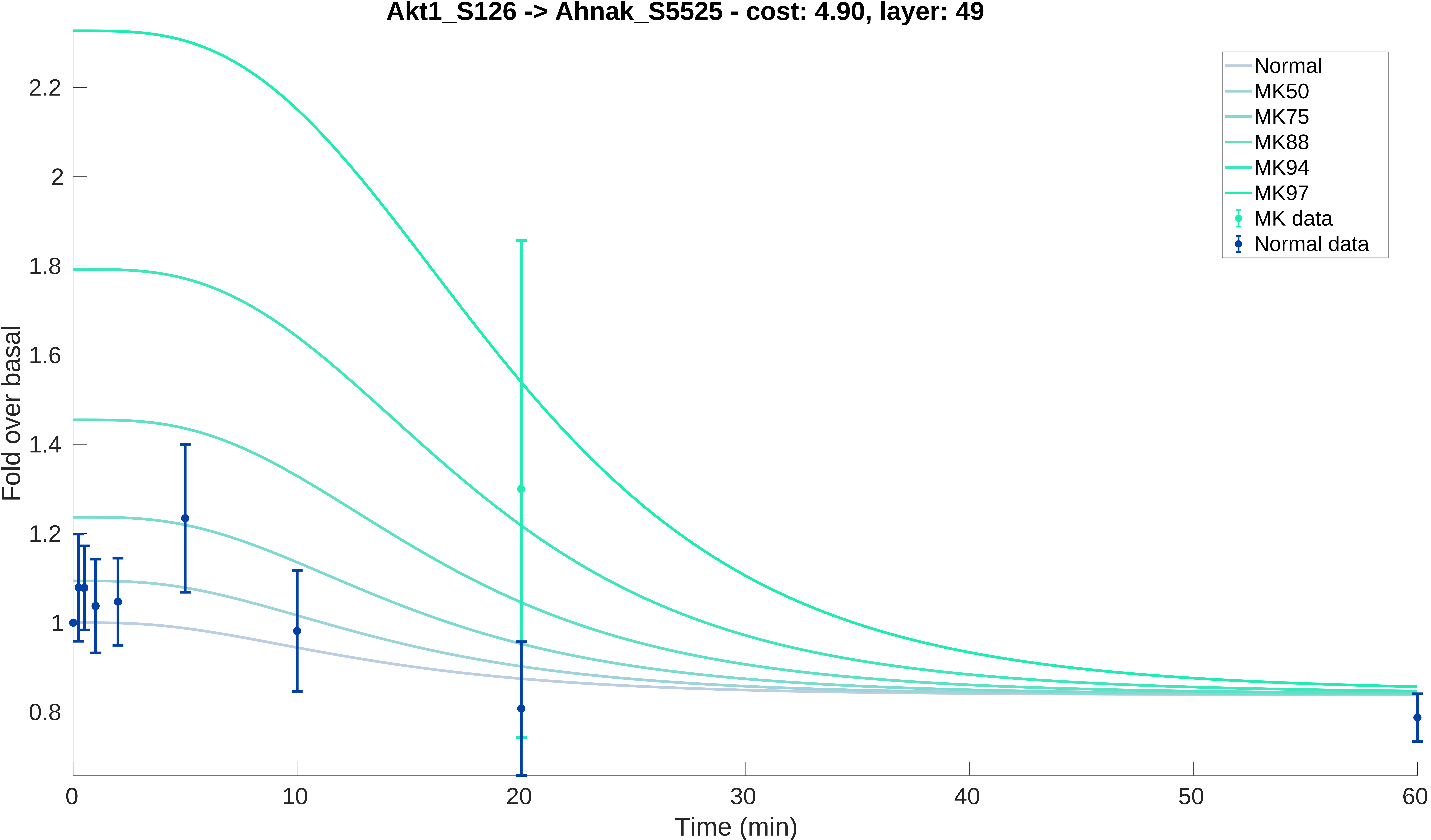

Akt1_S126 -> Ahnak_S5525 - cost: 4.90, layer: 49
Fold over basal
Time (min)
0.8
1
1.2
1.4
1.6
1.8
2
2.2
0
10
20
30
40
50
60
Normal
MK50
MK75
MK88
MK94
MK97
MK data
Normal data

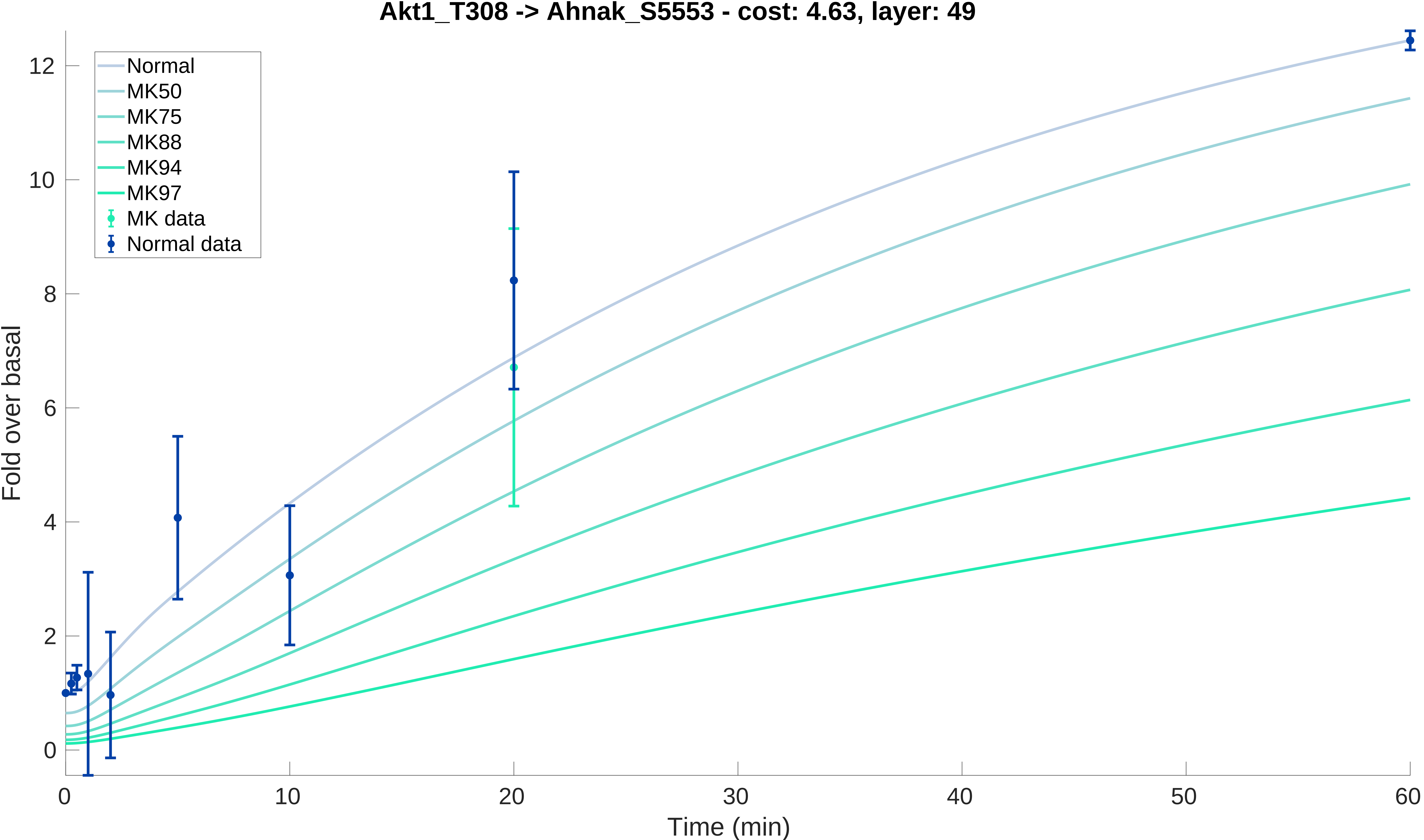
Akt1_T308 -> Ahnak_S5553 - cost: 4.63, layer: 49
Normal
MK50
MK75
MK88
MK94
MK97
MK data
Normal data
Fold over basal
Time (min)
0
2
4
6
8
10
12
0
10
20
30
40
50
60

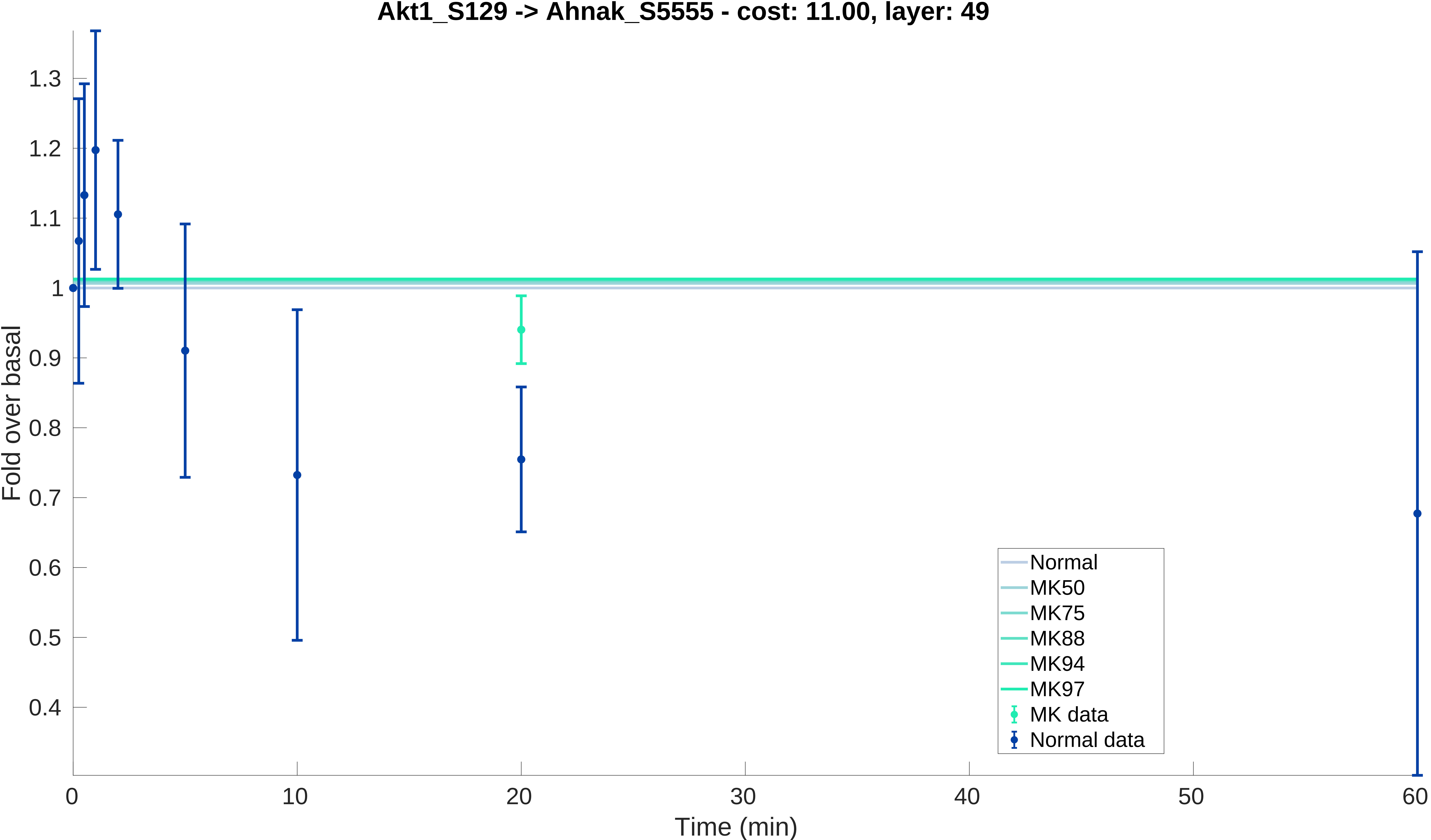

Akt1_S129 -> Ahnak_S5555 - cost: 11.00, layer: 49
Fold over basal
1.3
1.2
1.1
1
0.9
0.8
0.7
0.6
0.5
0.4
0
10
20
30
40
50
60
Time (min)
Normal
MK50
MK75
MK88
MK94
MK97
MK data
Normal data

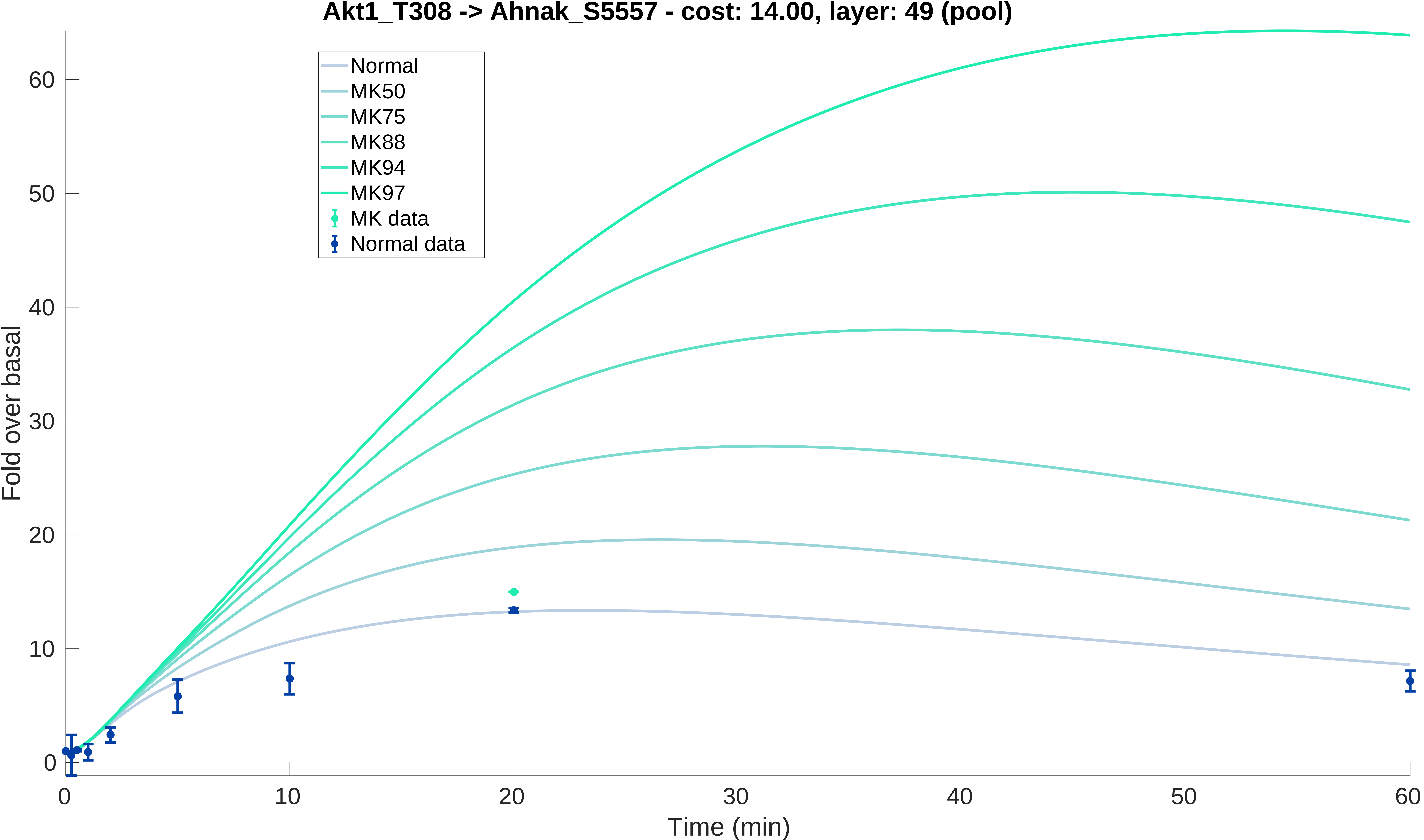

Akt1_T308 -> Ahnak_S5557 - cost: 14.00, layer: 49 (pool)
Normal
MK50
MK75
MK88
MK94
MK97
MK data
Normal data
Fold over basal
0
10
20
30
40
50
60
Time (min)
0
10
20
30
40
50
60

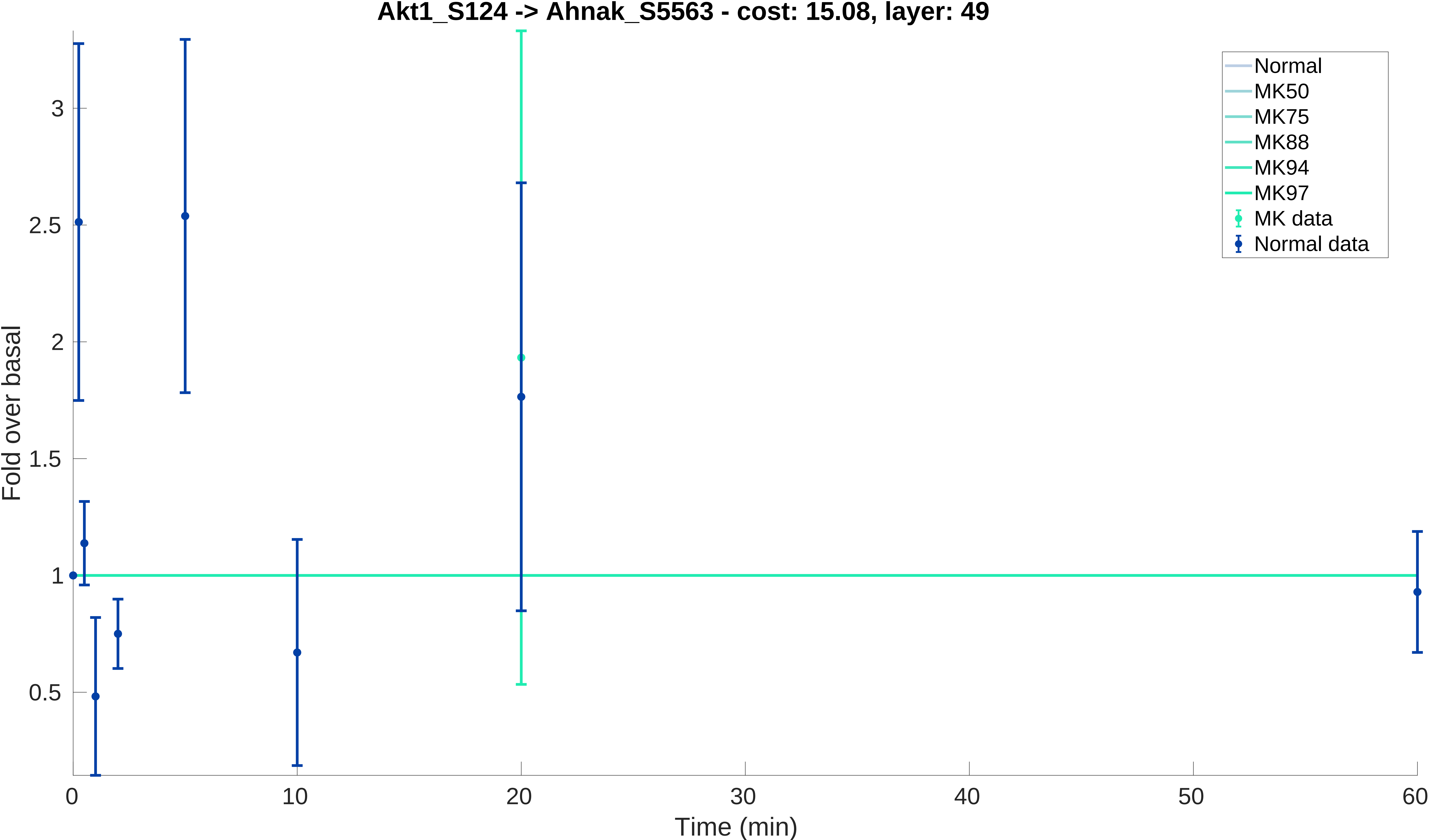

Akt1_S124 -> Ahnak_S5563 - cost: 15.08, layer: 49
Fold over basal
Time (min)
0.5
1
1.5
2
2.5
3
0
10
20
30
40
50
60
Normal
MK50
MK75
MK88
MK94
MK97
MK data
Normal data

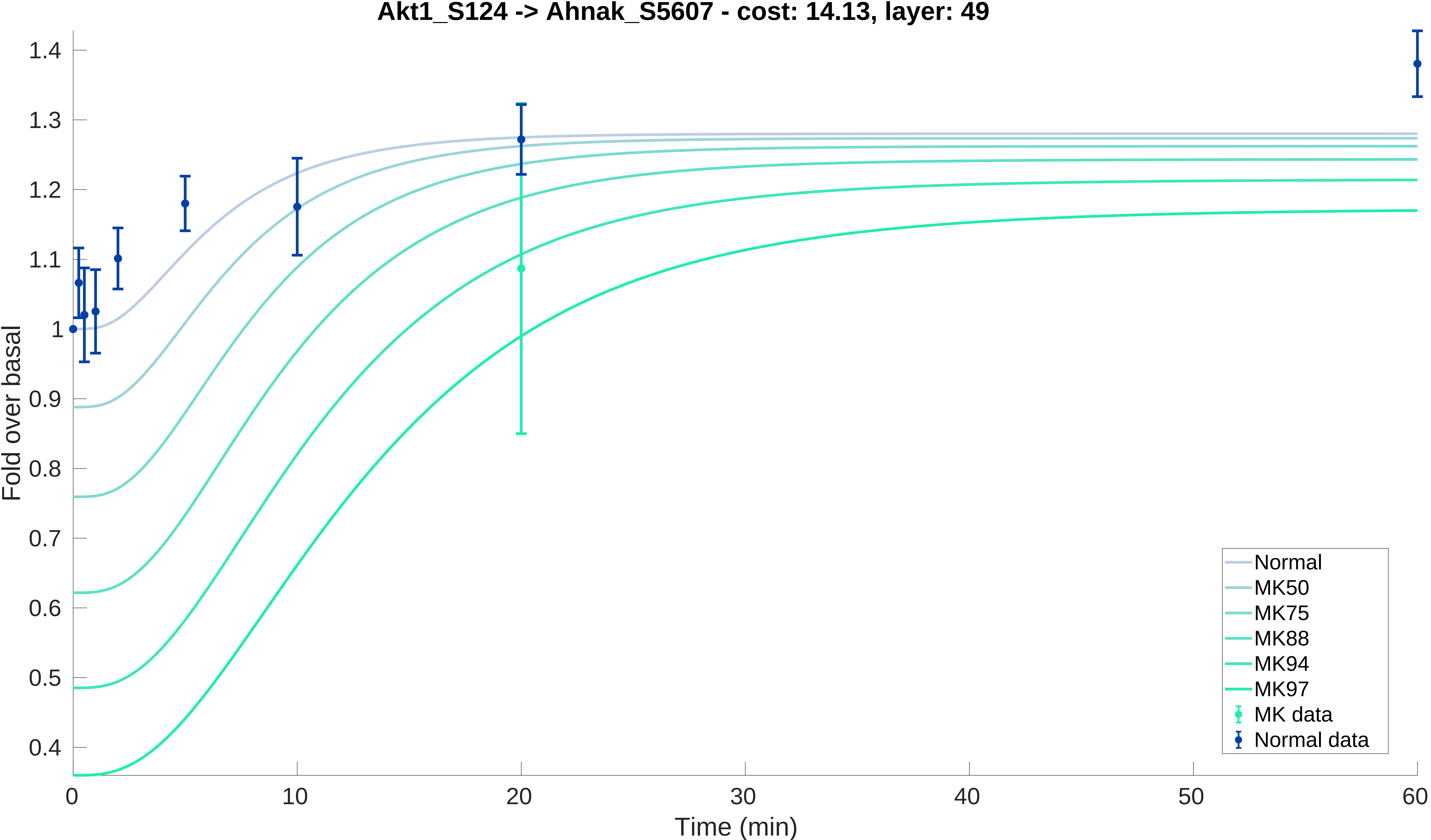

Akt1_S124 -> Ahnak_S5607 - cost: 14.13, layer: 49
Fold over basal
Time (min)
1.4
1.3
1.2
1.1
1
0.9
0.8
0.7
0.6
0.5
0.4
0
10
20
30
40
50
60
Normal
MK50
MK75
MK88
MK94
MK97
MK data
Normal data

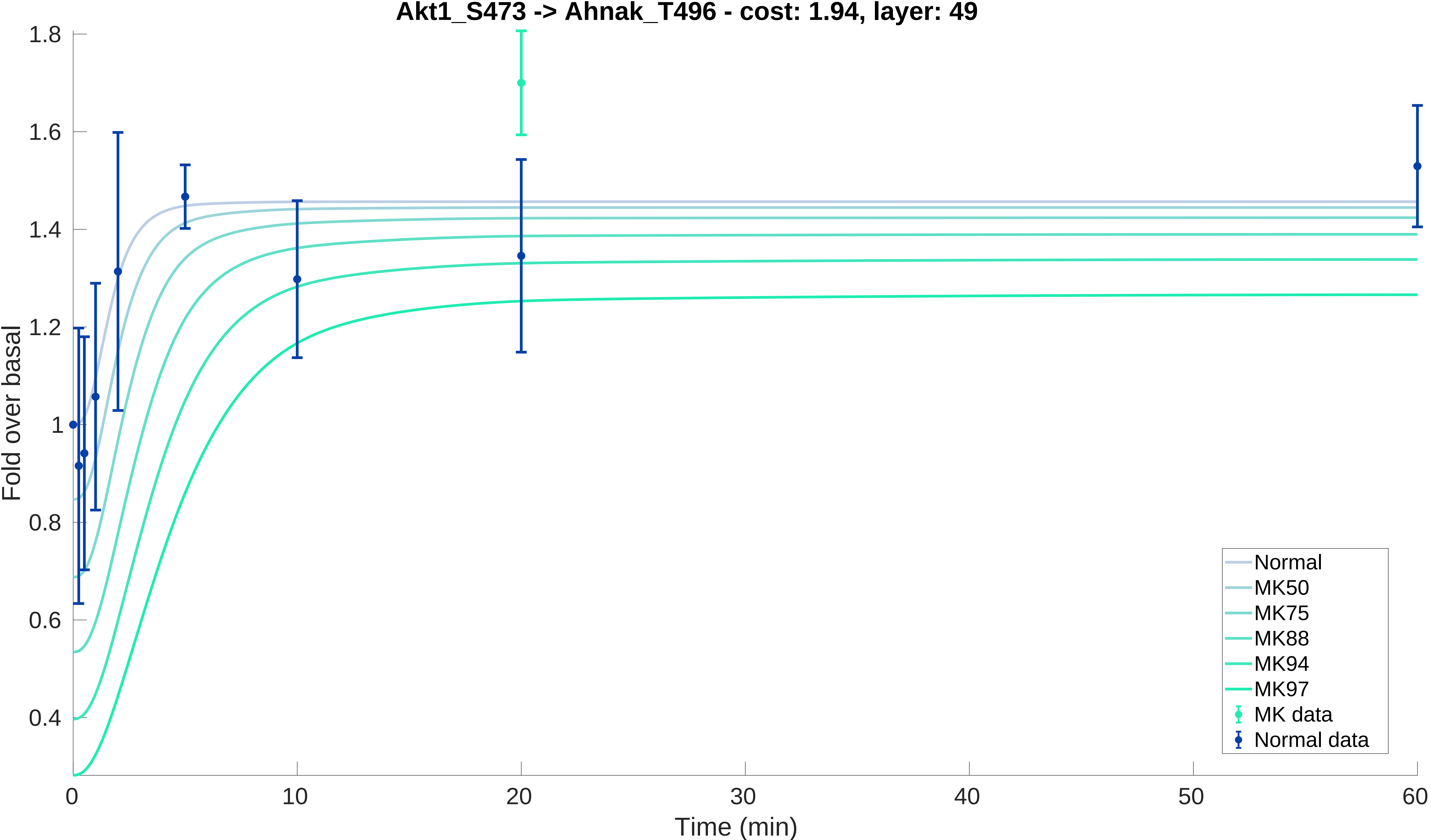

Akt1_S473 -> Ahnak_T496 - cost: 1.94, layer: 49
Fold over basal
1.8
1.6
1.4
1.2
1
0.8
0.6
0.4
0
10
20
30
40
50
60
Time (min)
Normal
MK50
MK75
MK88
MK94
MK97
MK data
Normal data

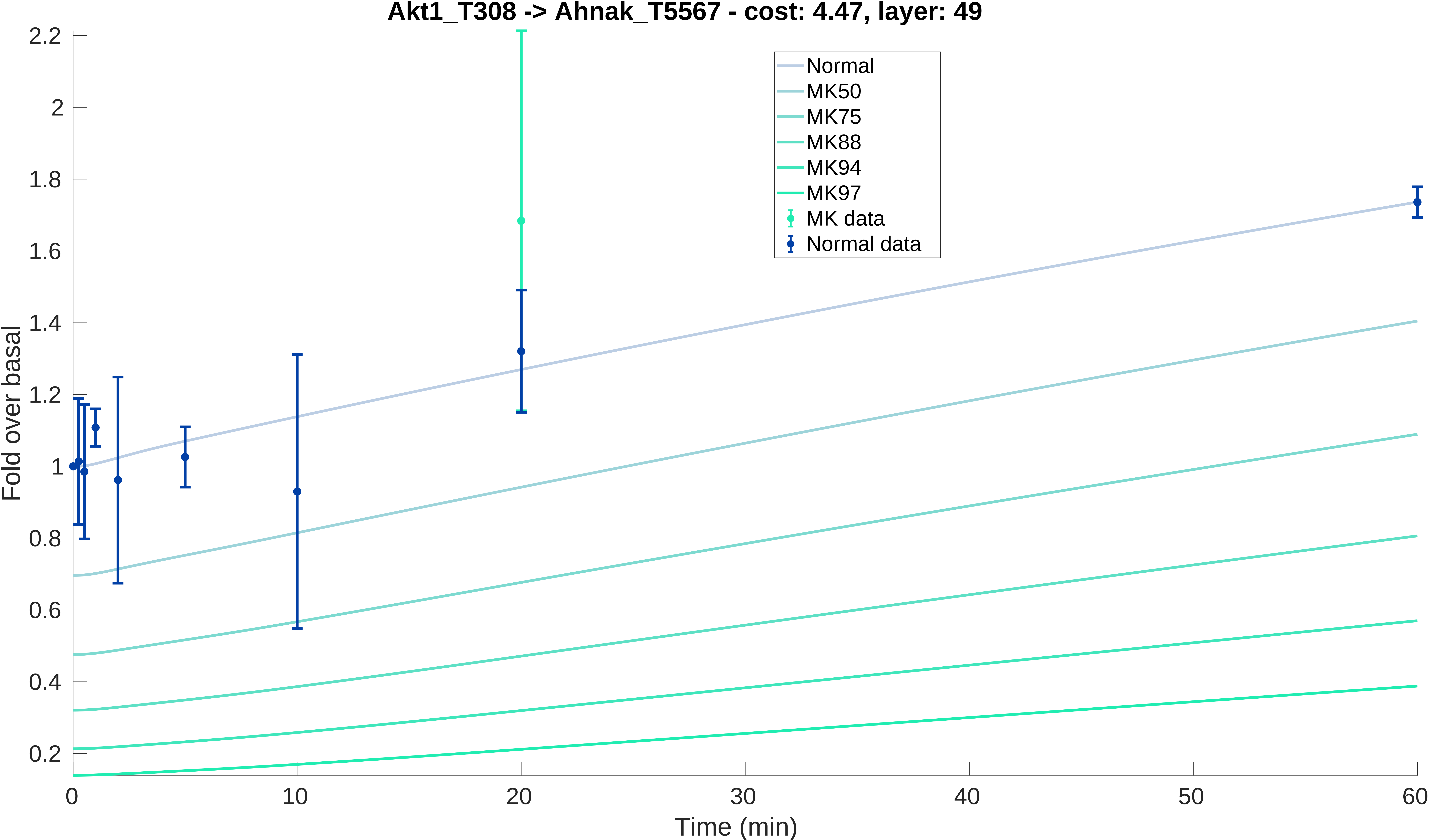

Akt1_T308 -> Ahnak_T5567 - cost: 4.47, layer: 49
Normal
MK50
MK75
MK88
MK94
MK97
MK data
Normal data
Fold over basal
Time (min)
2.2
2
1.8
1.6
1.4
1.2
1
0.8
0.6
0.4
0.2
0
10
20
30
40
50
60

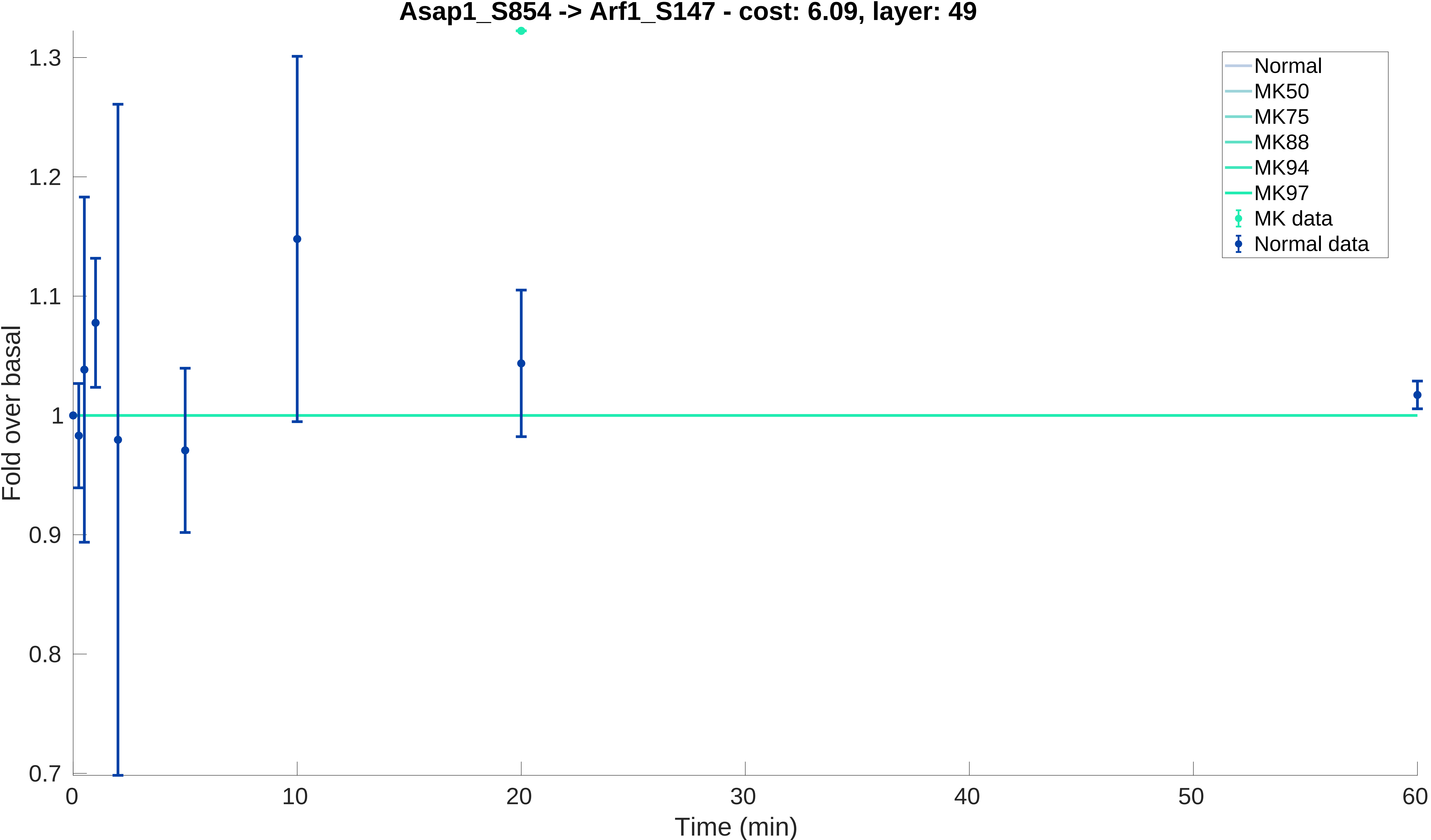
Asap1_S854 -> Arf1_S147 - cost: 6.09, layer: 49
Fold over basal
Time (min)
1.3
1.2
1.1
1
0.9
0.8
0.7
0
10
20
30
40
50
60
Normal
MK50
MK75
MK88
MK94
MK97
MK data
Normal data

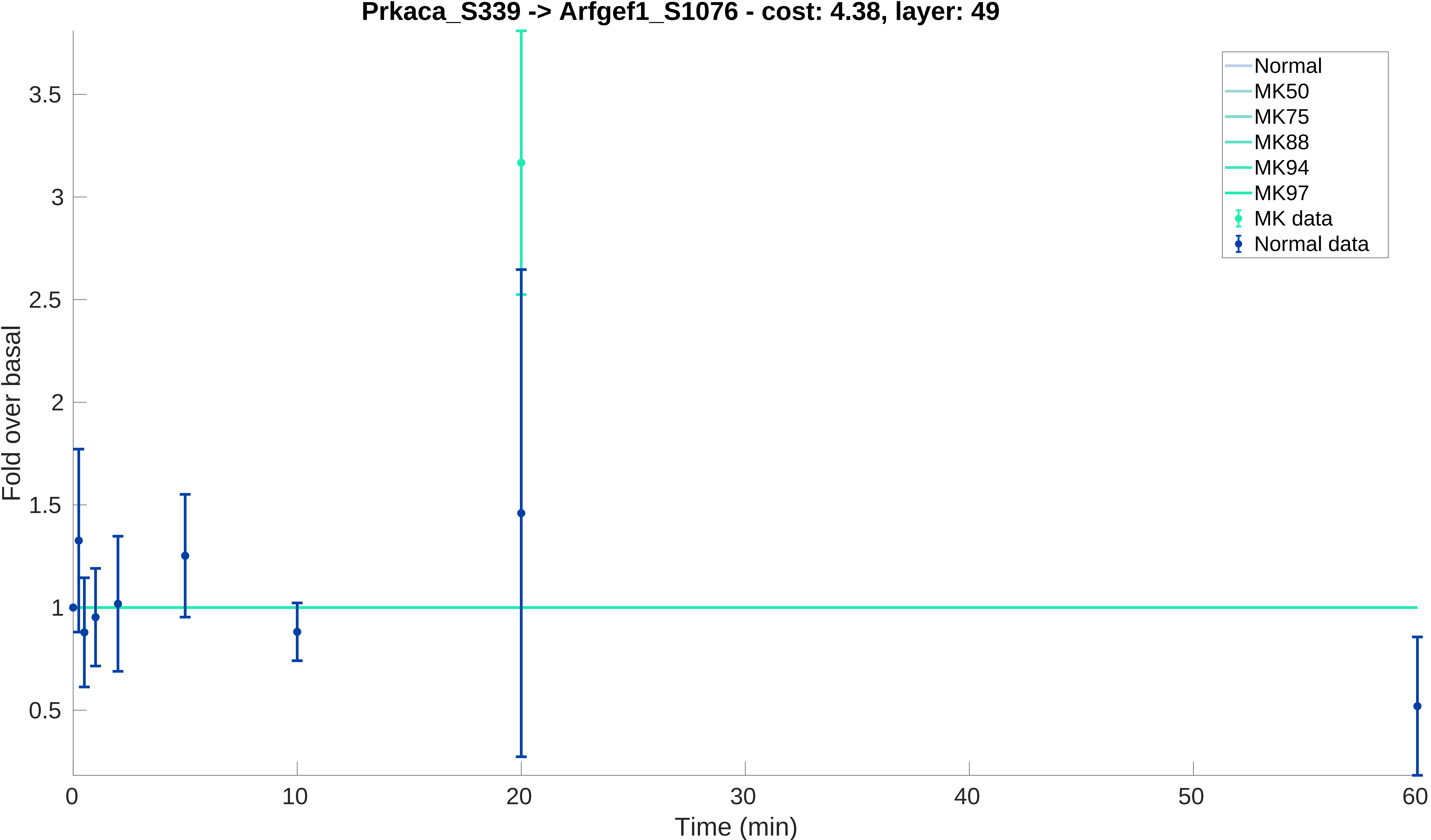

Prkaca_S339 -> Arfgef1_S1076 - cost: 4.38, layer: 49
Fold over basal
Time (min)
0.5
1
1.5
2
2.5
3
3.5
0
10
20
30
40
50
60
Normal
MK50
MK75
MK88
MK94
MK97
MK data
Normal data

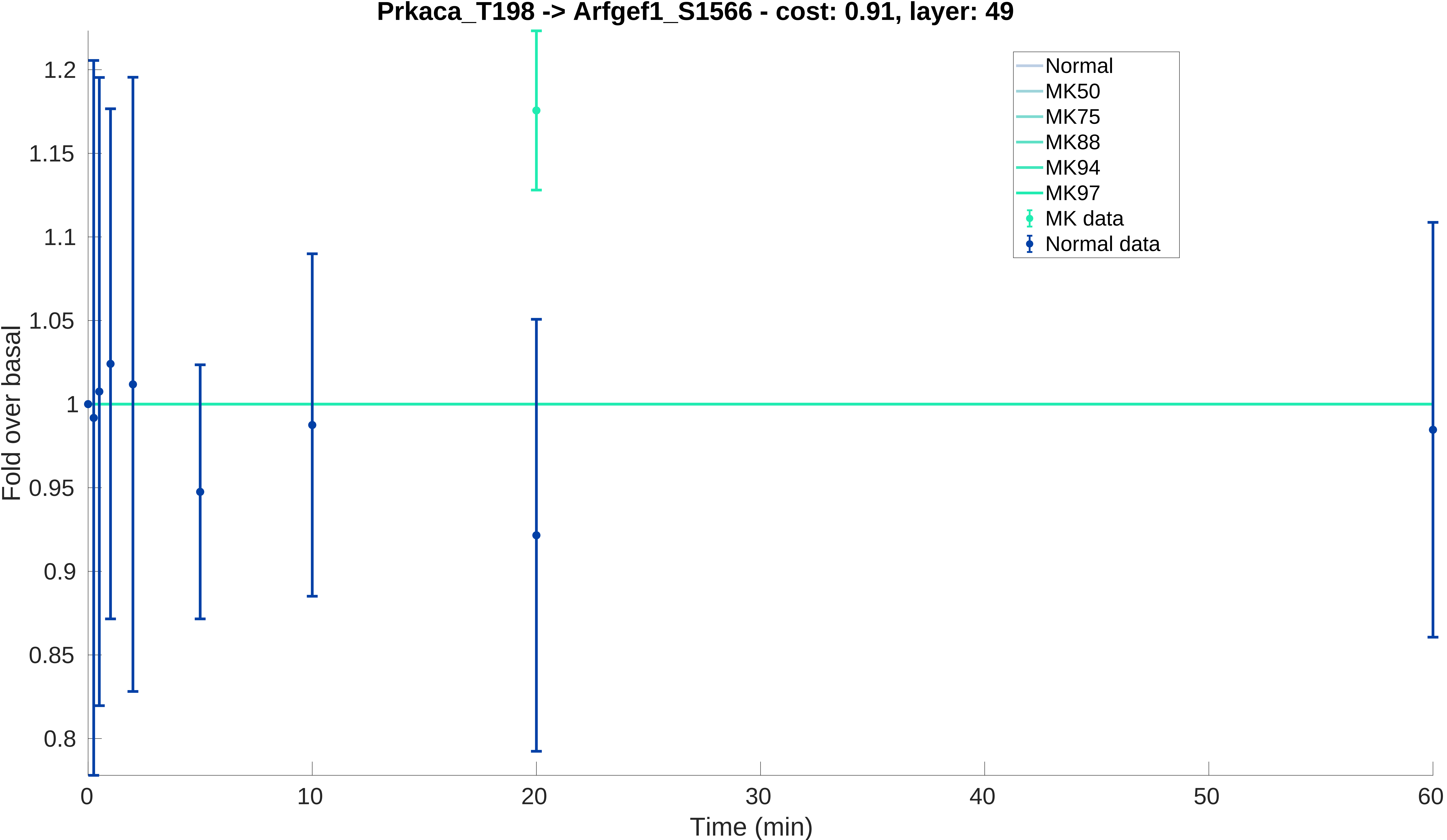

Prkaca_T198 -> Arfgef1_S1566 - cost: 0.91, layer: 49
Normal
MK50
MK75
MK88
MK94
MK97
MK data
Normal data
Fold over basal
1.2
1.15
1.1
1.05
1
0.95
0.9
0.85
0.8
0
10
20
30
40
50
60
Time (min)

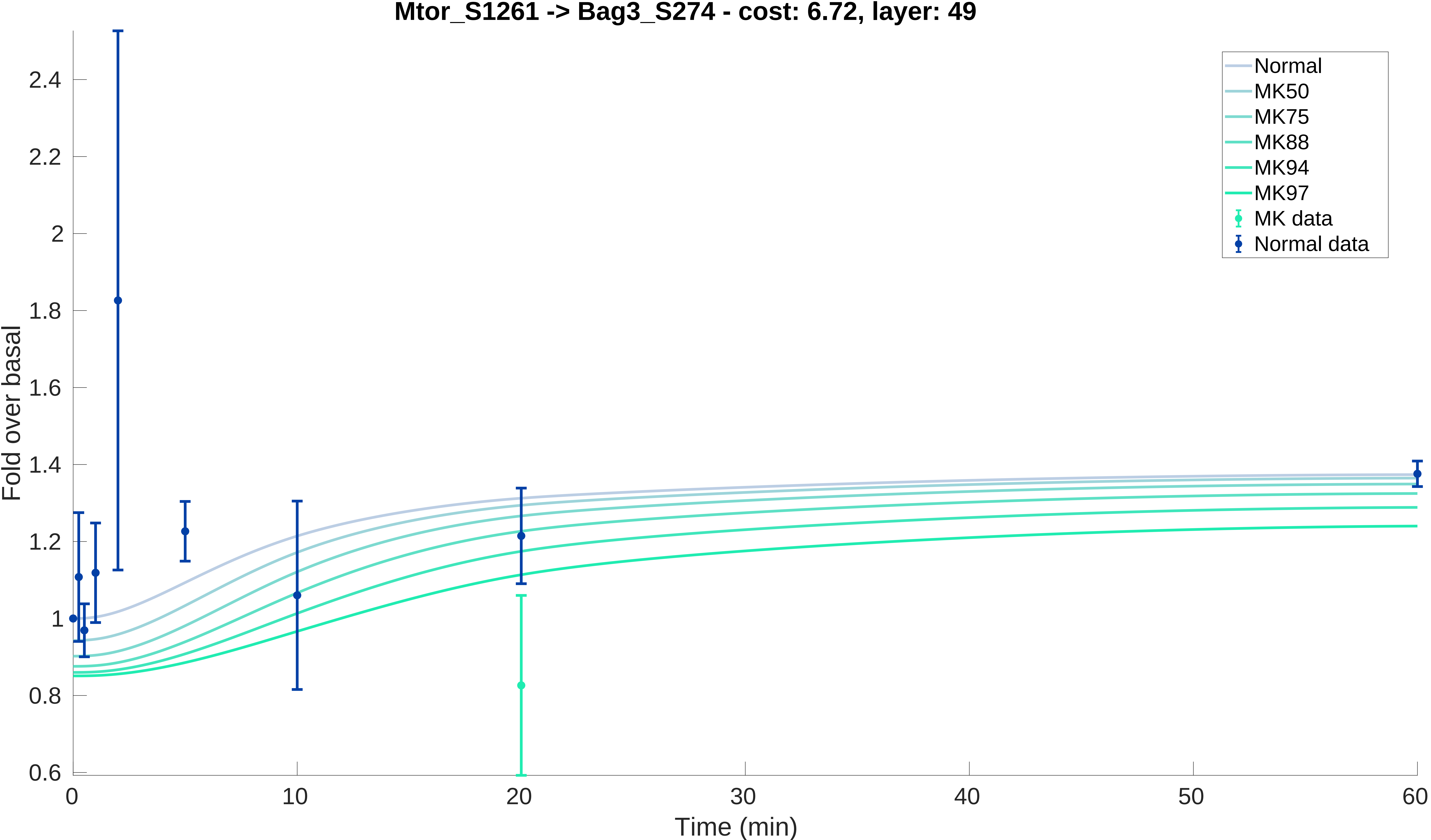

Mtor_S1261 -> Bag3_S274 - cost: 6.72, layer: 49
Fold over basal
Time (min)
0.6
0.8
1
1.2
1.4
1.6
1.8
2
2.2
2.4
0
10
20
30
40
50
60
Normal
MK50
MK75
MK88
MK94
MK97
MK data
Normal data

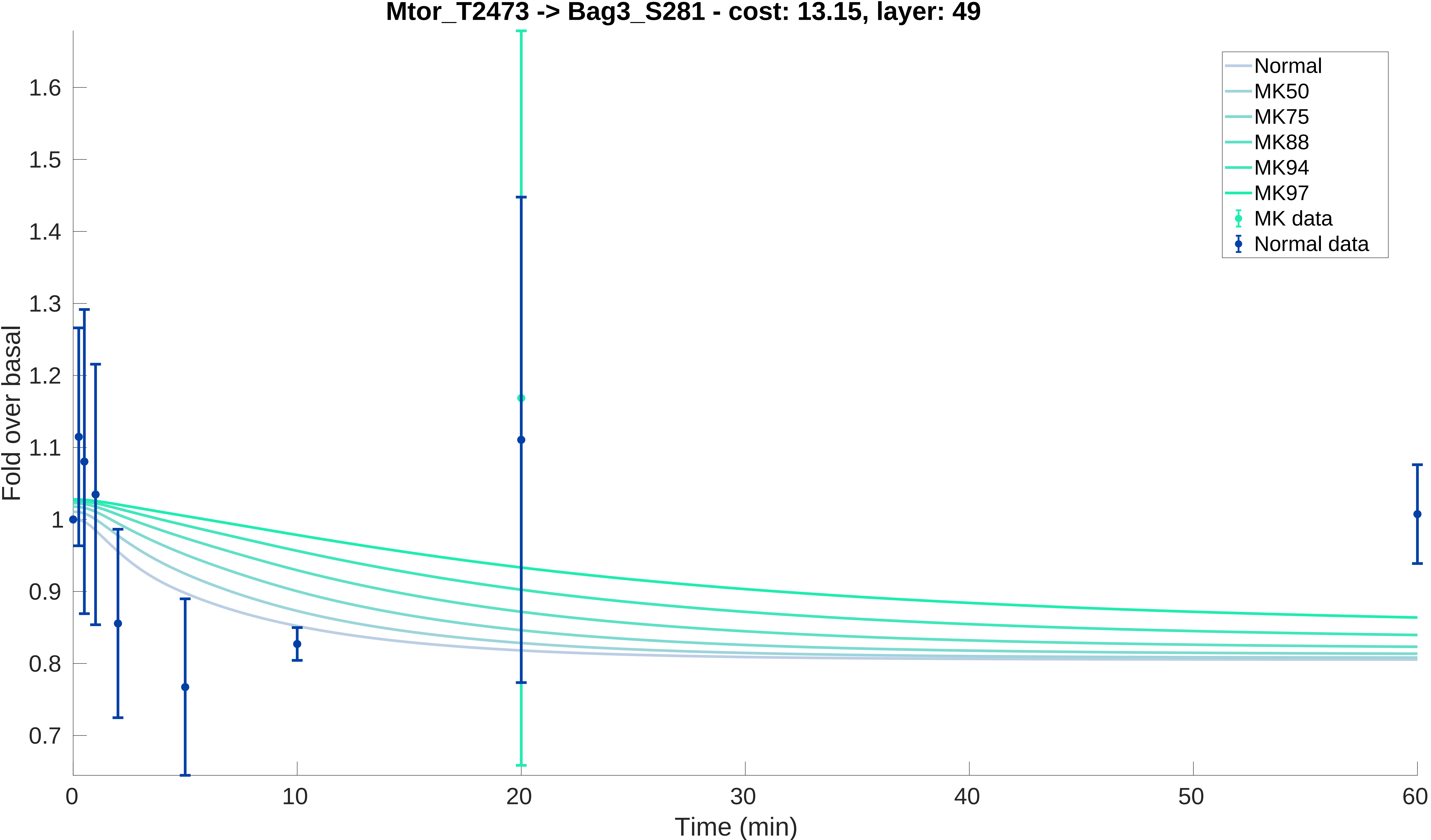

Mtor_T2473 -> Bag3_S281 - cost: 13.15, layer: 49
Fold over basal
Time (min)
Normal
MK50
MK75
MK88
MK94
MK97
MK data
Normal data
0.7
0.8
0.9
1
1.1
1.2
1.3
1.4
1.5
1.6
0
10
20
30
40
50
60

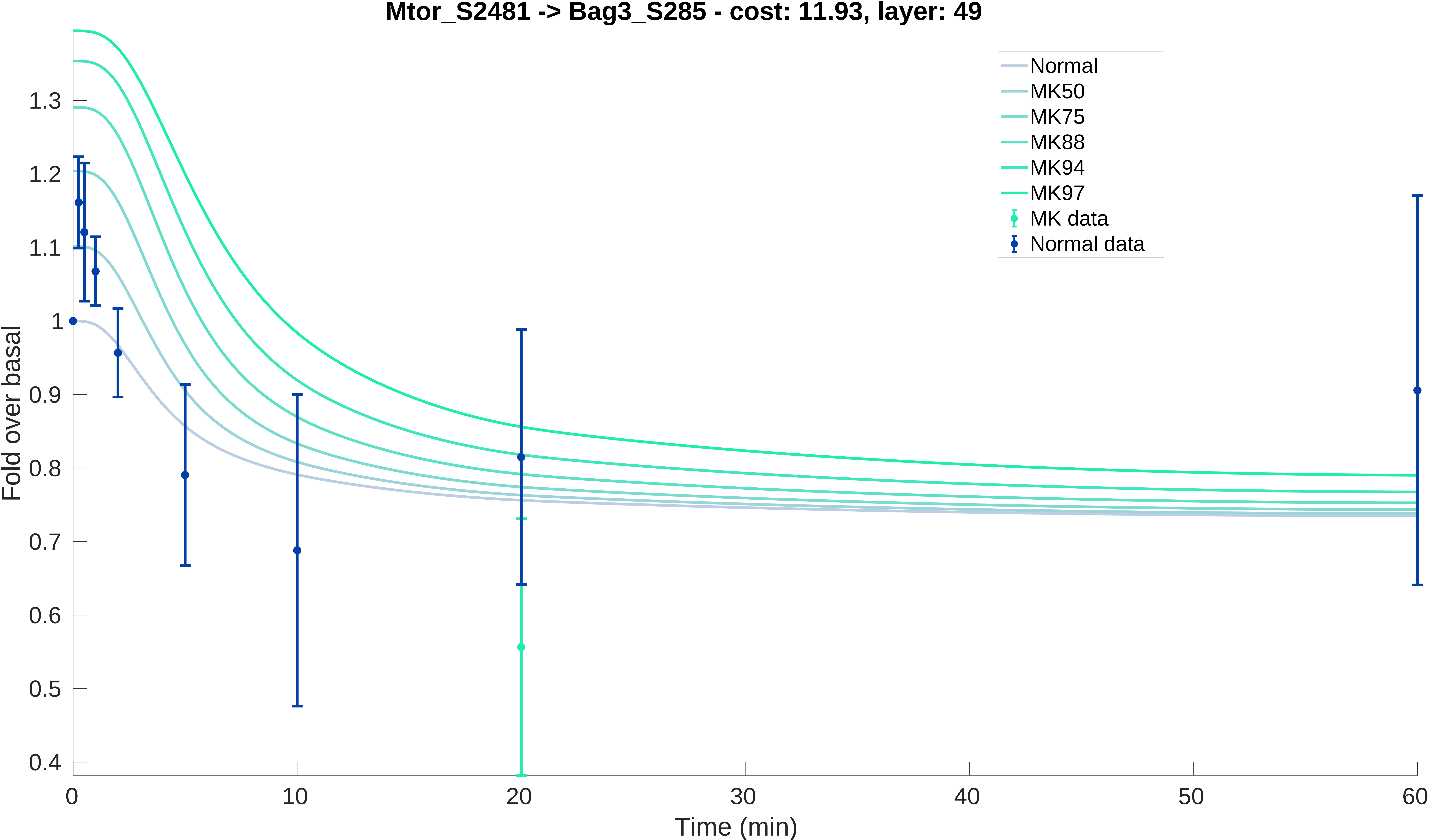

Mtor_S2481 -> Bag3_S285 - cost: 11.93, layer: 49
Normal
MK50
MK75
MK88
MK94
MK97
MK data
Normal data
Fold over basal
1.3
1.2
1.1
1
0.9
0.8
0.7
0.6
0.5
0.4
0
10
20
30
40
50
60
Time (min)

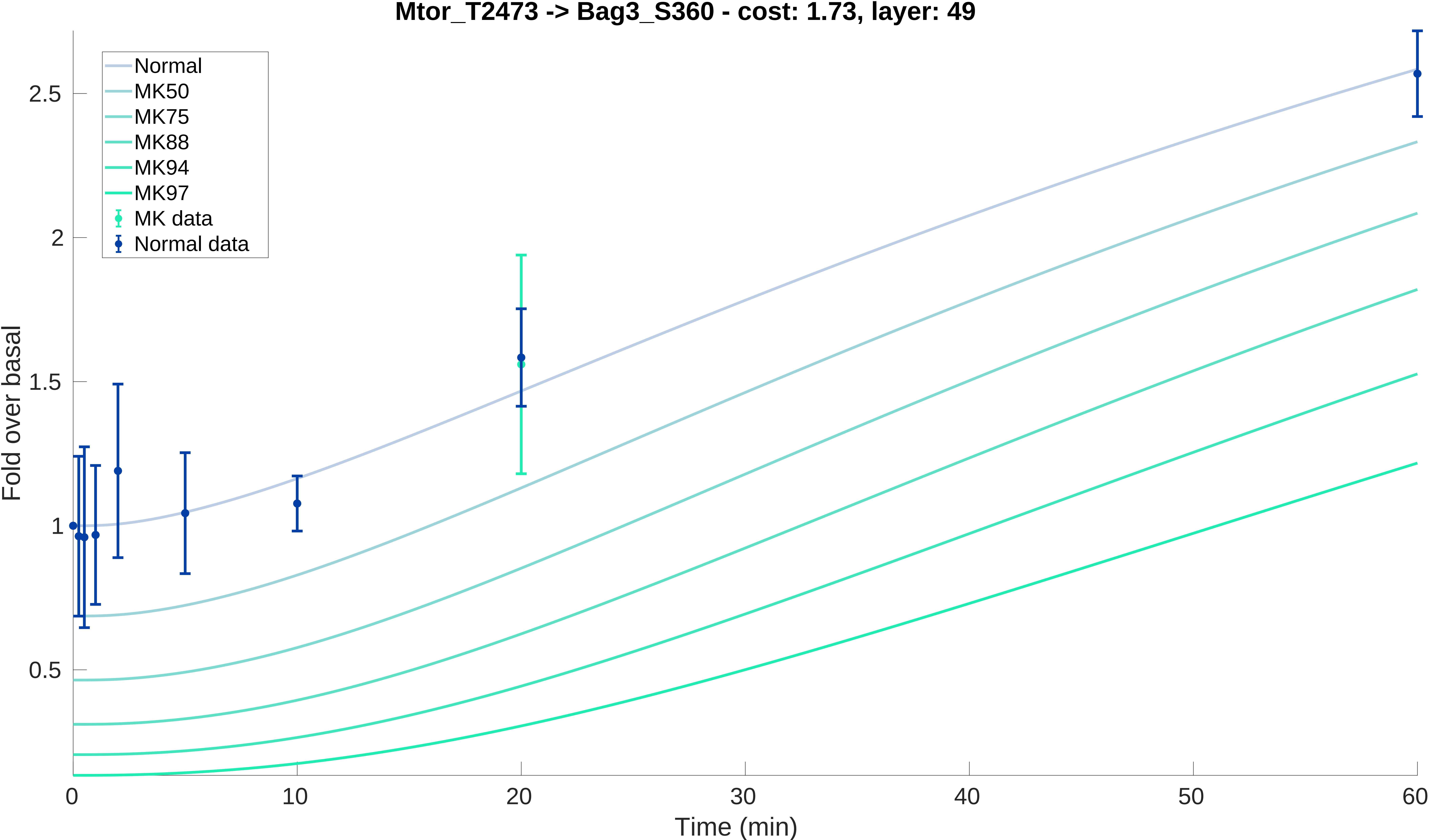
Mtor_T2473 -> Bag3_S360 - cost: 1.73, layer: 49
Normal
MK50
MK75
MK88
MK94
MK97
MK data
Normal data
Fold over basal
2.5
2
1.5
1
0.5
0
10
20
30
40
50
60
Time (min)

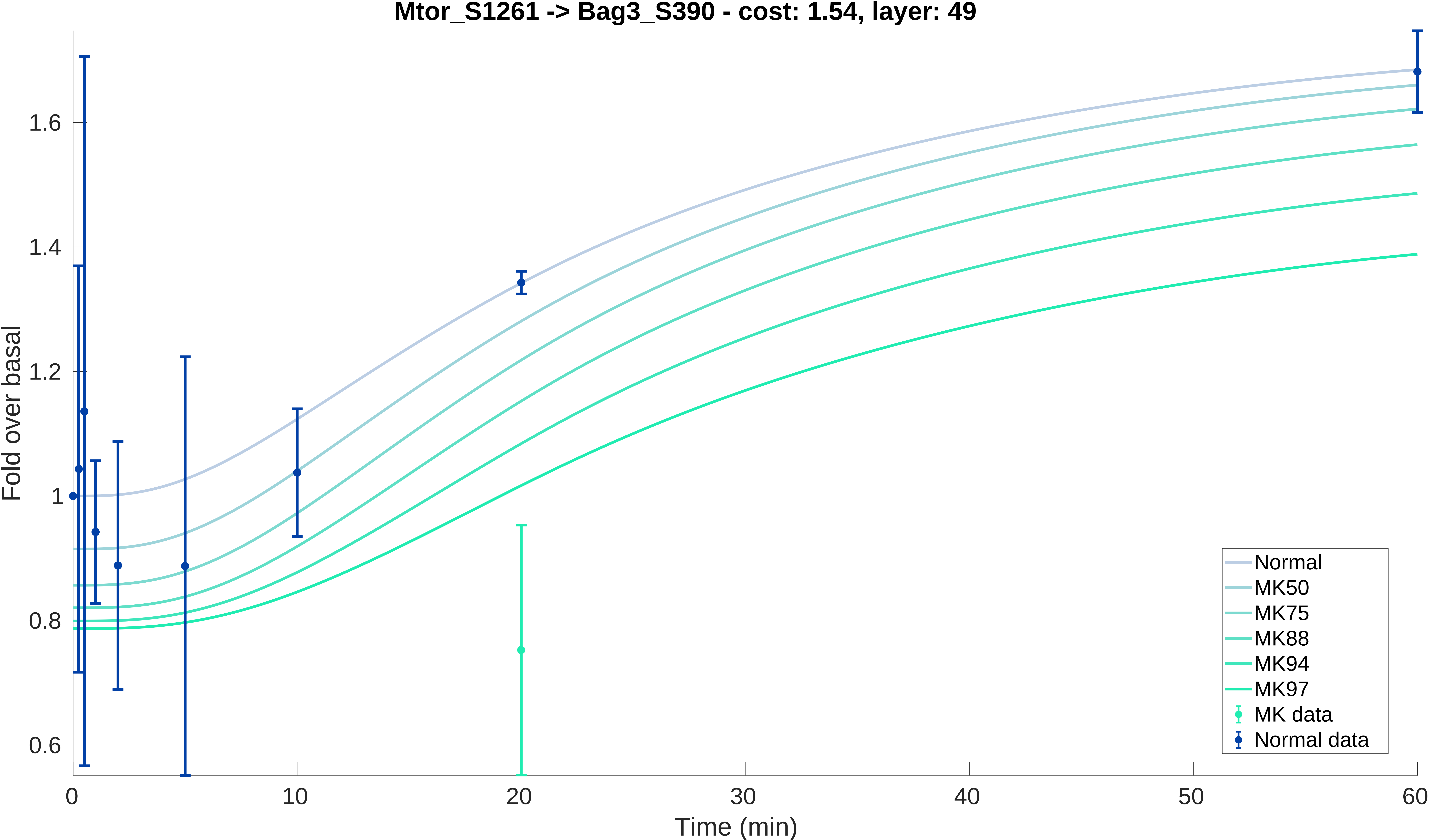

Mtor_S1261 -> Bag3_S390 - cost: 1.54, layer: 49
Fold over basal
Time (min)
0.6
0.8
1
1.2
1.4
1.6
0
10
20
30
40
50
60
Normal
MK50
MK75
MK88
MK94
MK97
MK data
Normal data

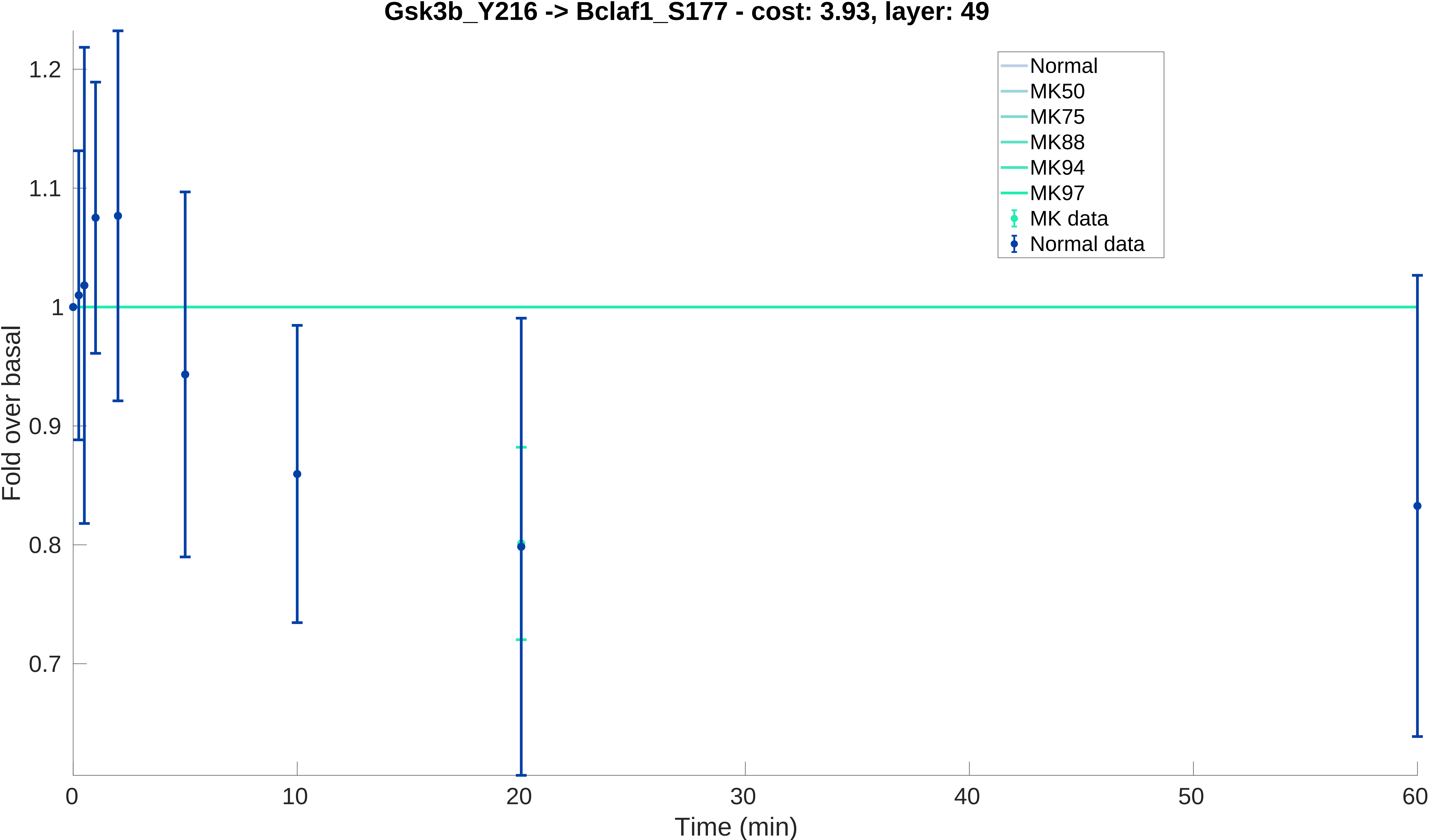

Gsk3b_Y216 -> Bclaf1_S177 - cost: 3.93, layer: 49
Fold over basal
Time (min)
1.2
1.1
1
0.9
0.8
0.7
0
10
20
30
40
50
60
Normal
MK50
MK75
MK88
MK94
MK97
MK data
Normal data

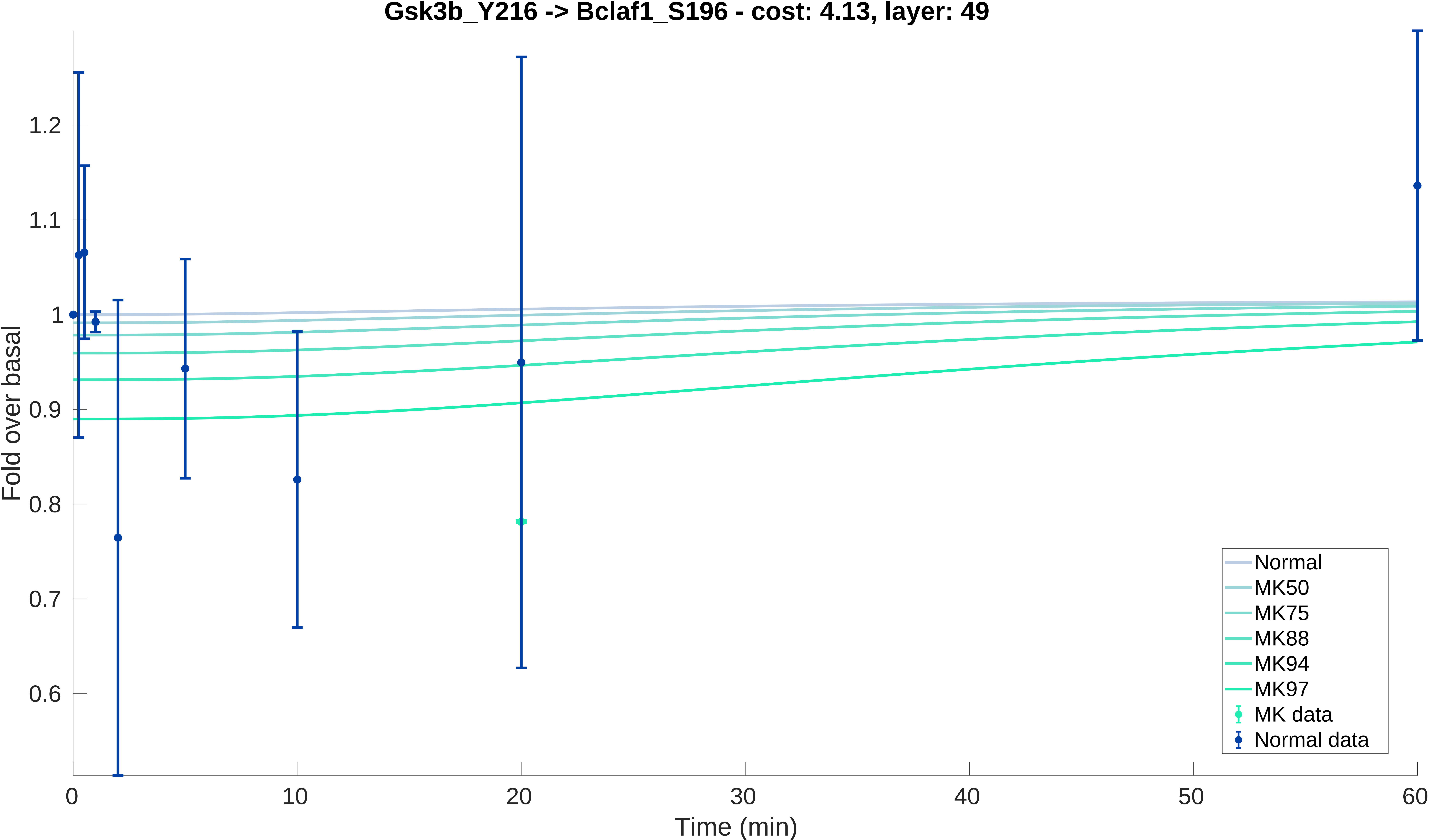

Gsk3b_Y216 -> Bclaf1_S196 - cost: 4.13, layer: 49
Fold over basal
1.2
1.1
1
0.9
0.8
0.7
0.6
0
10
20
30
40
50
60
Time (min)
Normal
MK50
MK75
MK88
MK94
MK97
MK data
Normal data

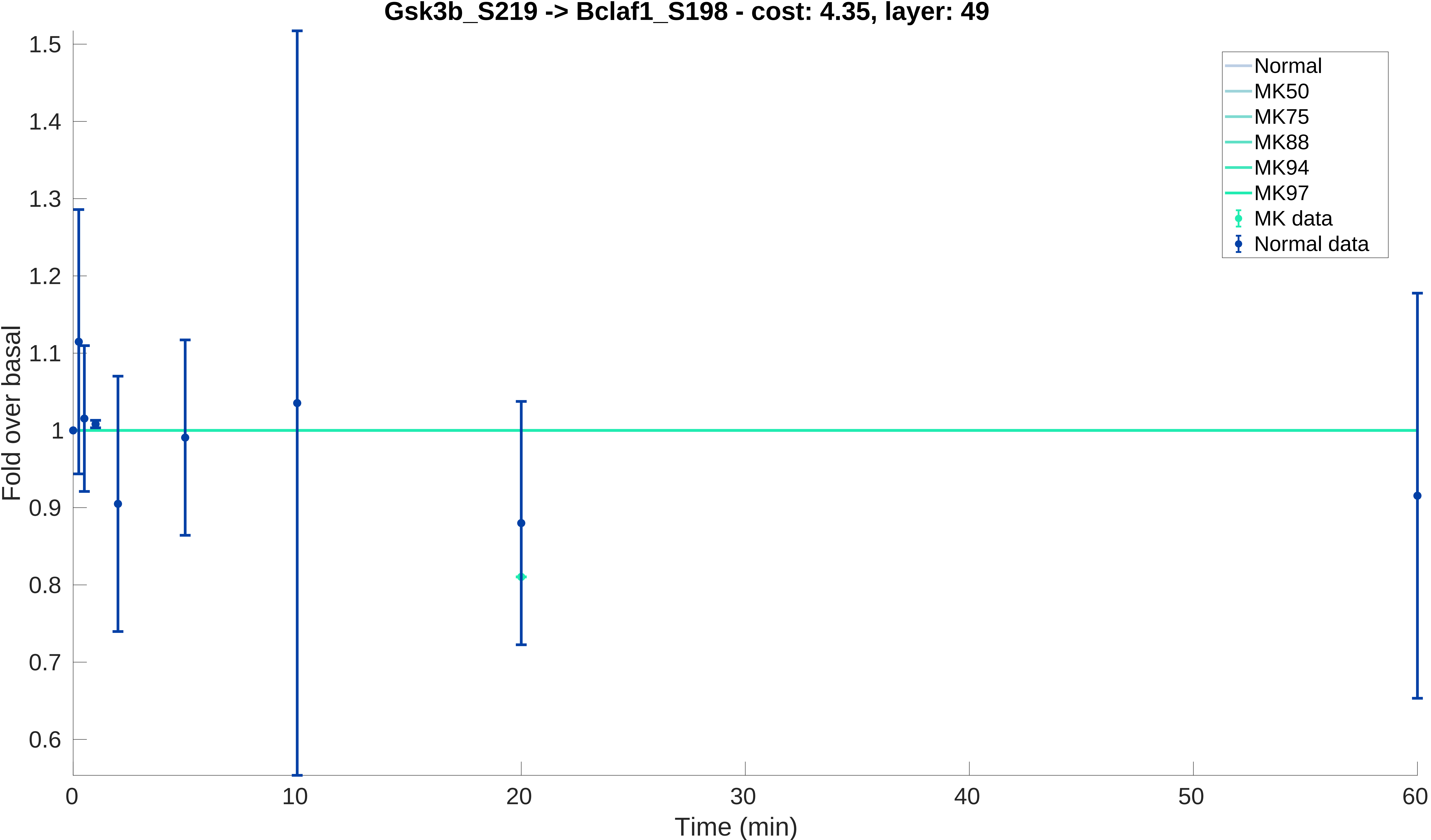

Gsk3b_S219 -> Bclaf1_S198 - cost: 4.35, layer: 49
Fold over basal
Time (min)
Normal
MK50
MK75
MK88
MK94
MK97
MK data
Normal data
0.6
0.7
0.8
0.9
1
1.1
1.2
1.3
1.4
1.5
0
10
20
30
40
50
60

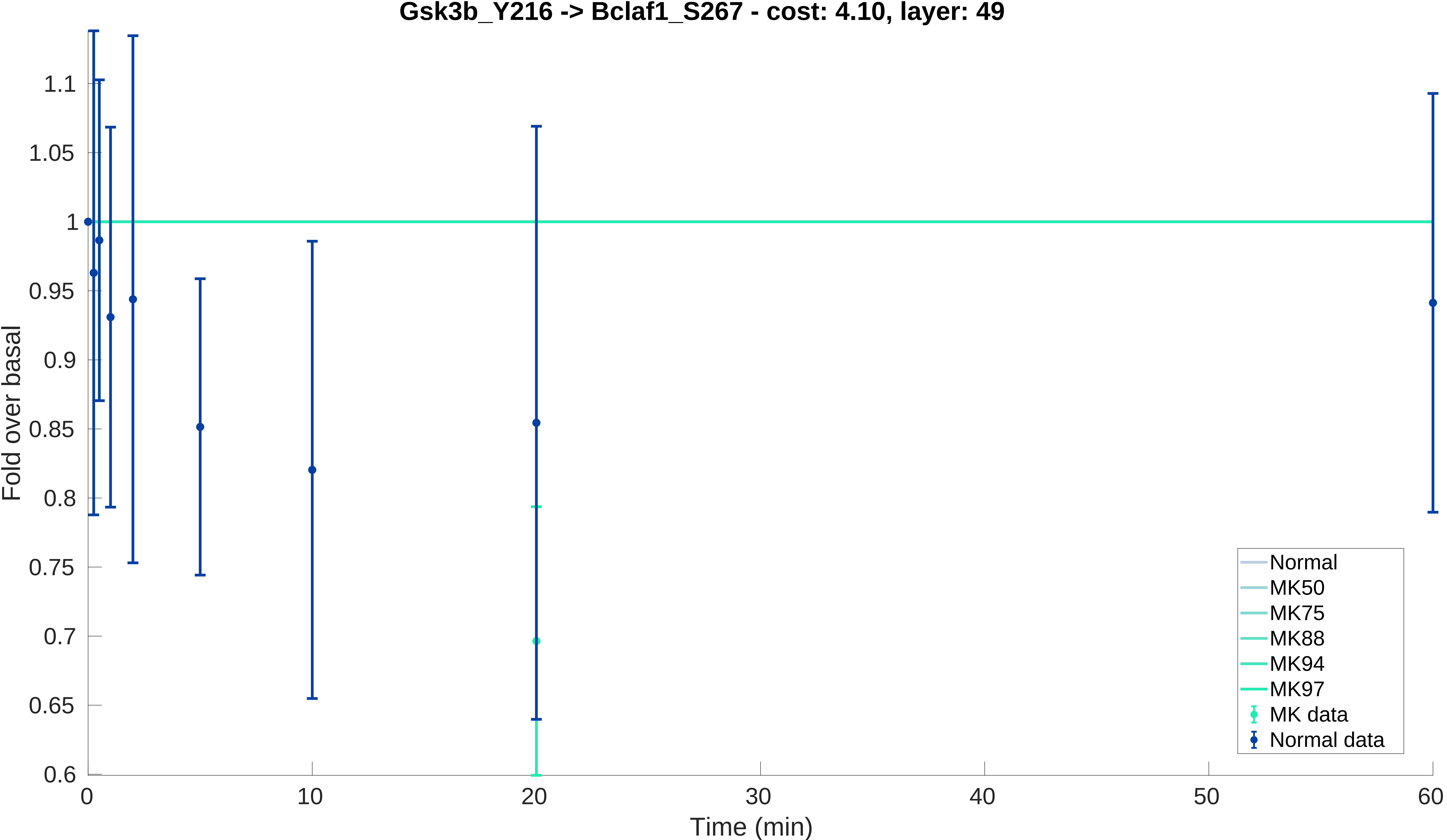

Gsk3b_Y216 -> Bclaf1_S267 - cost: 4.10, layer: 49
Fold over basal
Time (min)
1.1
1.05
1
0.95
0.9
0.85
0.8
0.75
0.7
0.65
0.6
0
10
20
30
40
50
60
Normal
MK50
MK75
MK88
MK94
MK97
MK data
Normal data

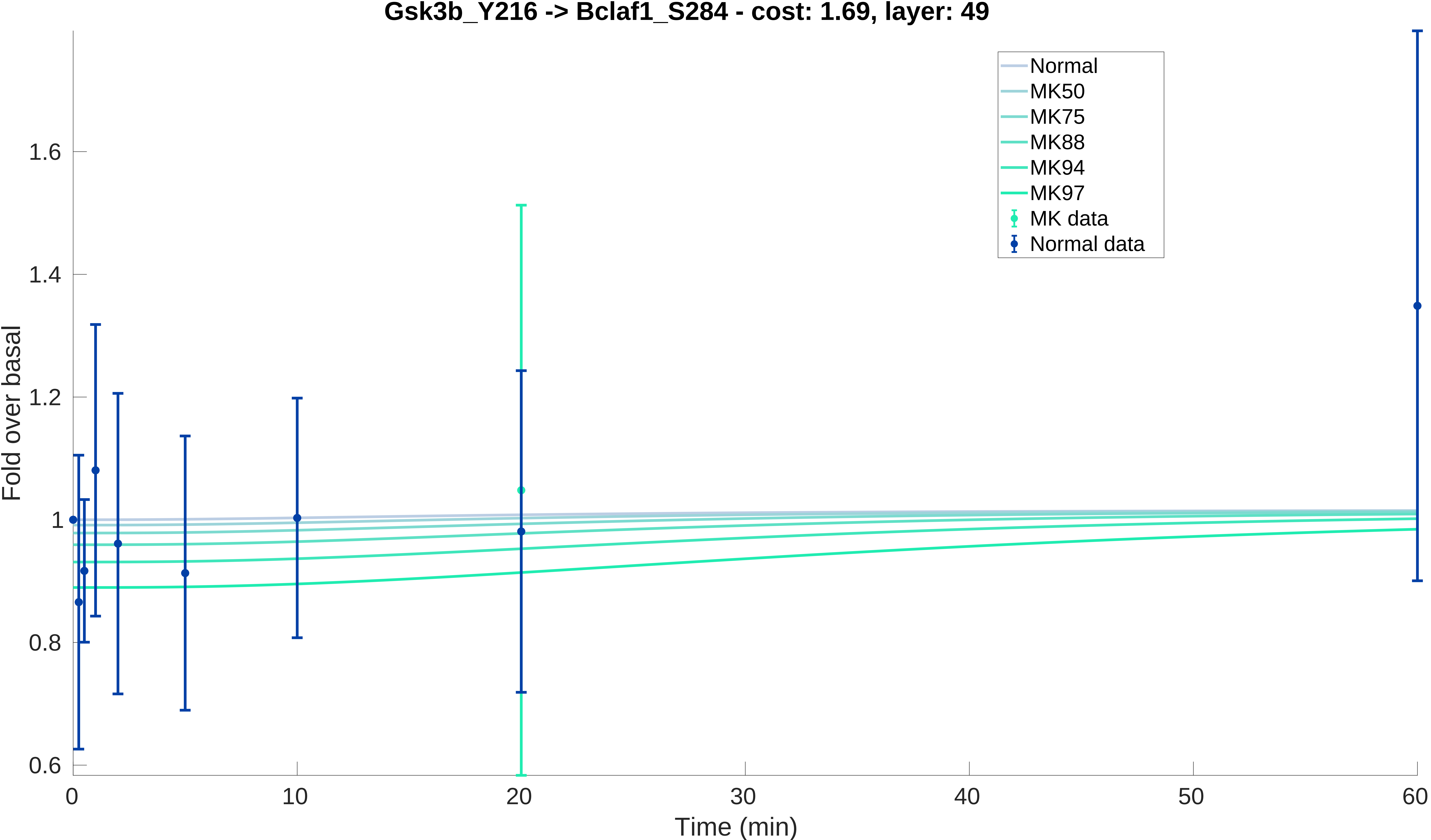

Gsk3b_Y216 -> Bclaf1_S284 - cost: 1.69, layer: 49
Fold over basal
Time (min)
0.6
0.8
1
1.2
1.4
1.6
0
10
20
30
40
50
60
Normal
MK50
MK75
MK88
MK94
MK97
MK data
Normal data

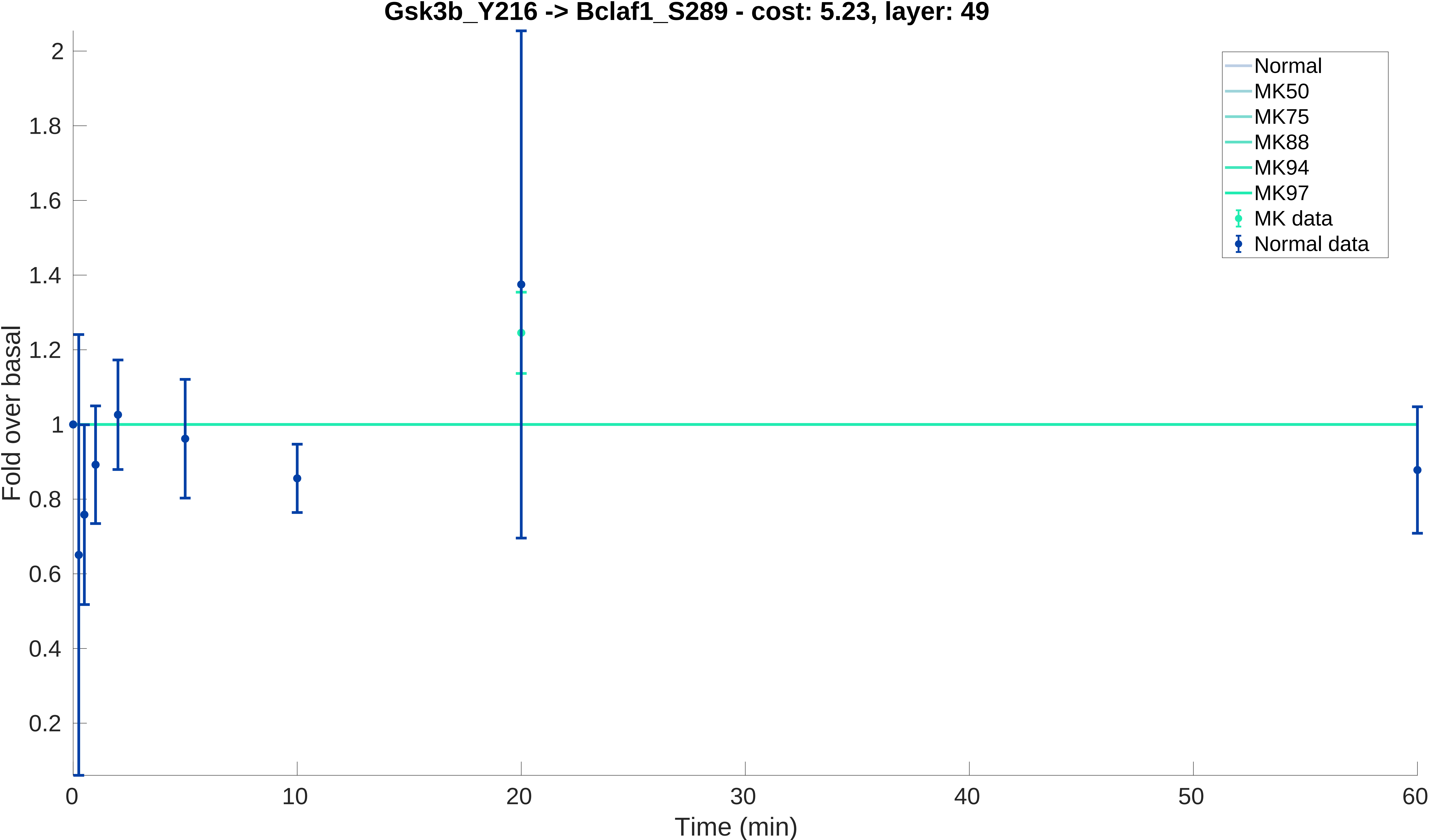

Gsk3b_Y216 -> Bclaf1_S289 - cost: 5.23, layer: 49
Fold over basal
Time (min)
0
10
20
30
40
50
60
0.2
0.4
0.6
0.8
1
1.2
1.4
1.6
1.8
2
Normal
MK50
MK75
MK88
MK94
MK97
MK data
Normal data

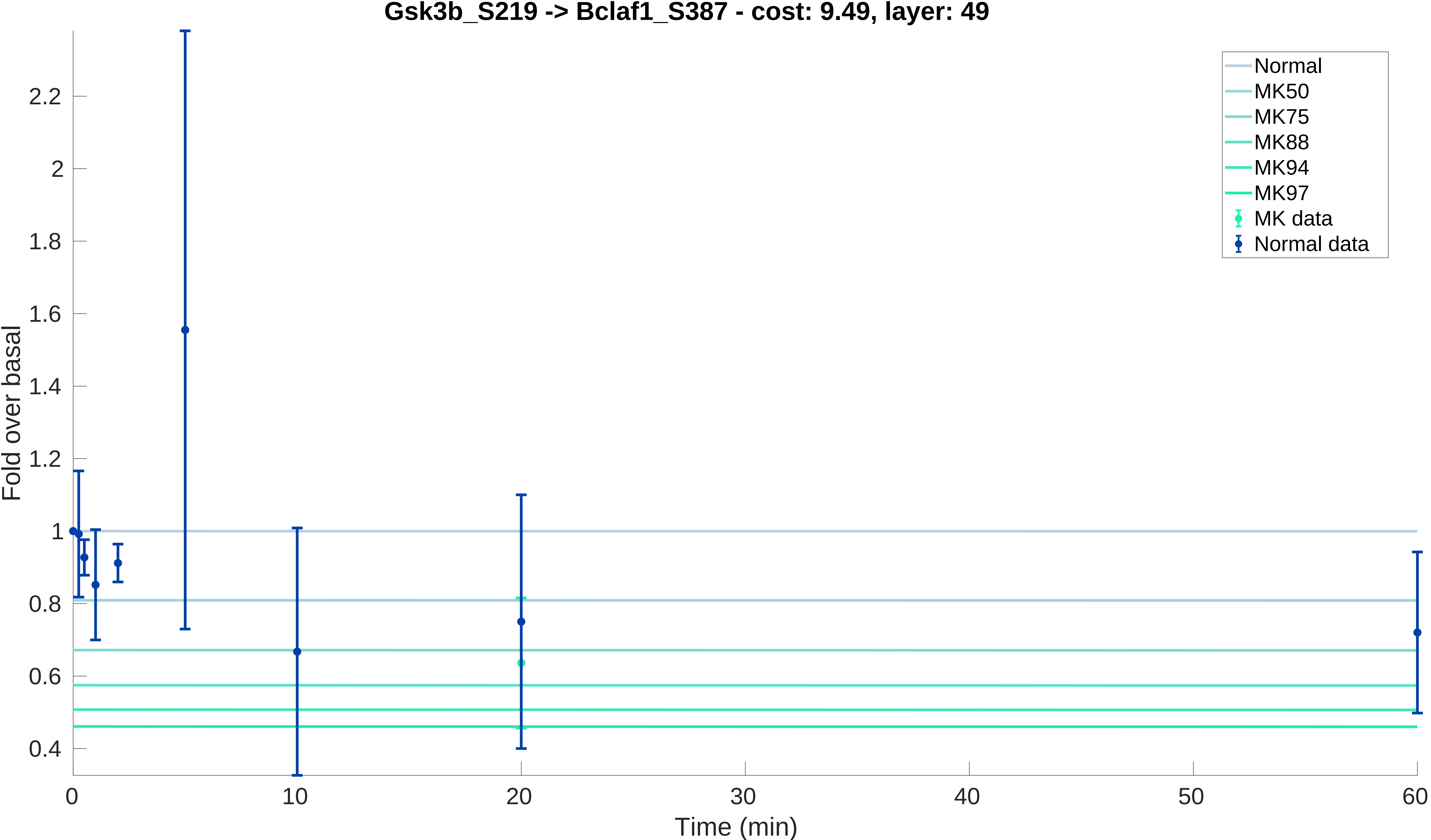

Gsk3b_S219 -> Bclaf1_S387 - cost: 9.49, layer: 49
Fold over basal
2.2
2
1.8
1.6
1.4
1.2
1
0.8
0.6
0.4
0
10
20
30
40
50
60
Time (min)
Normal
MK50
MK75
MK88
MK94
MK97
MK data
Normal data

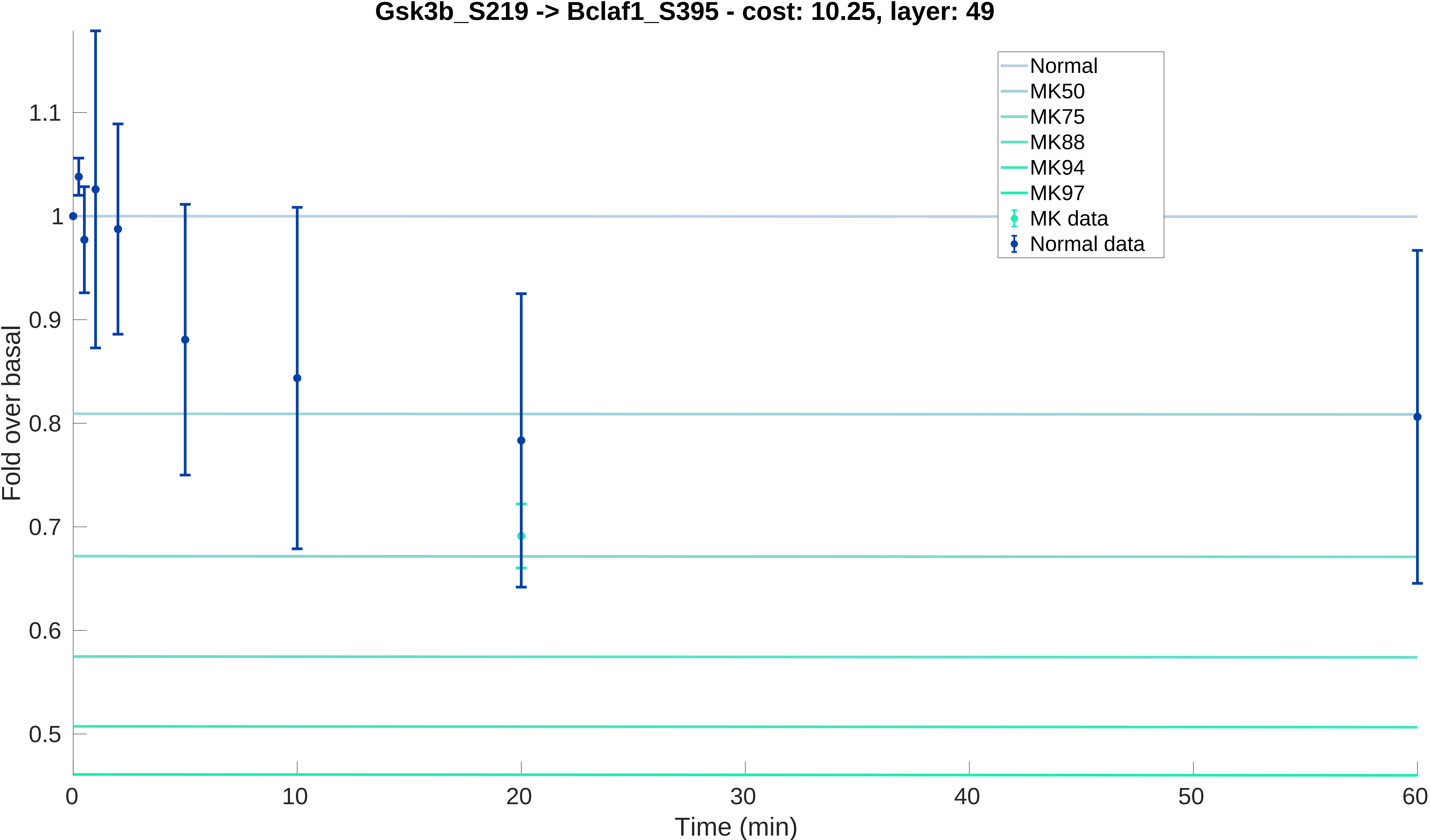

Gsk3b_S219 -> Bclaf1_S395 - cost: 10.25, layer: 49
Normal
MK50
MK75
MK88
MK94
MK97
MK data
Normal data
Fold over basal
1.1
1
0.9
0.8
0.7
0.6
0.5
0
10
20
30
40
50
60
Time (min)

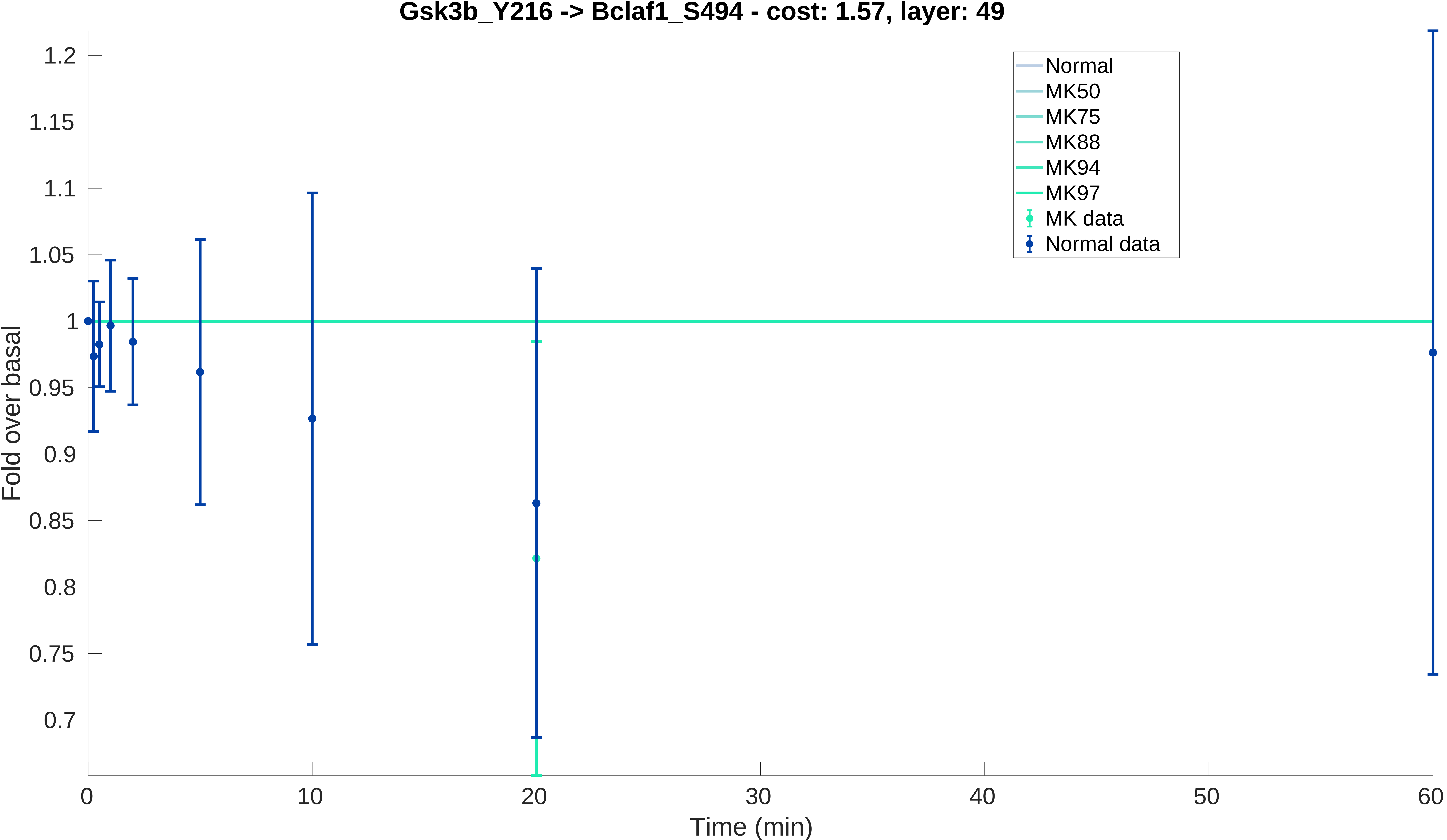
Gsk3b_Y216 -> Bclaf1_S494 - cost: 1.57, layer: 49
Normal
MK50
MK75
MK88
MK94
MK97
MK data
Normal data
Fold over basal
1.2
1.15
1.1
1.05
1
0.95
0.9
0.85
0.8
0.75
0.7
0
10
20
30
40
50
60
Time (min)

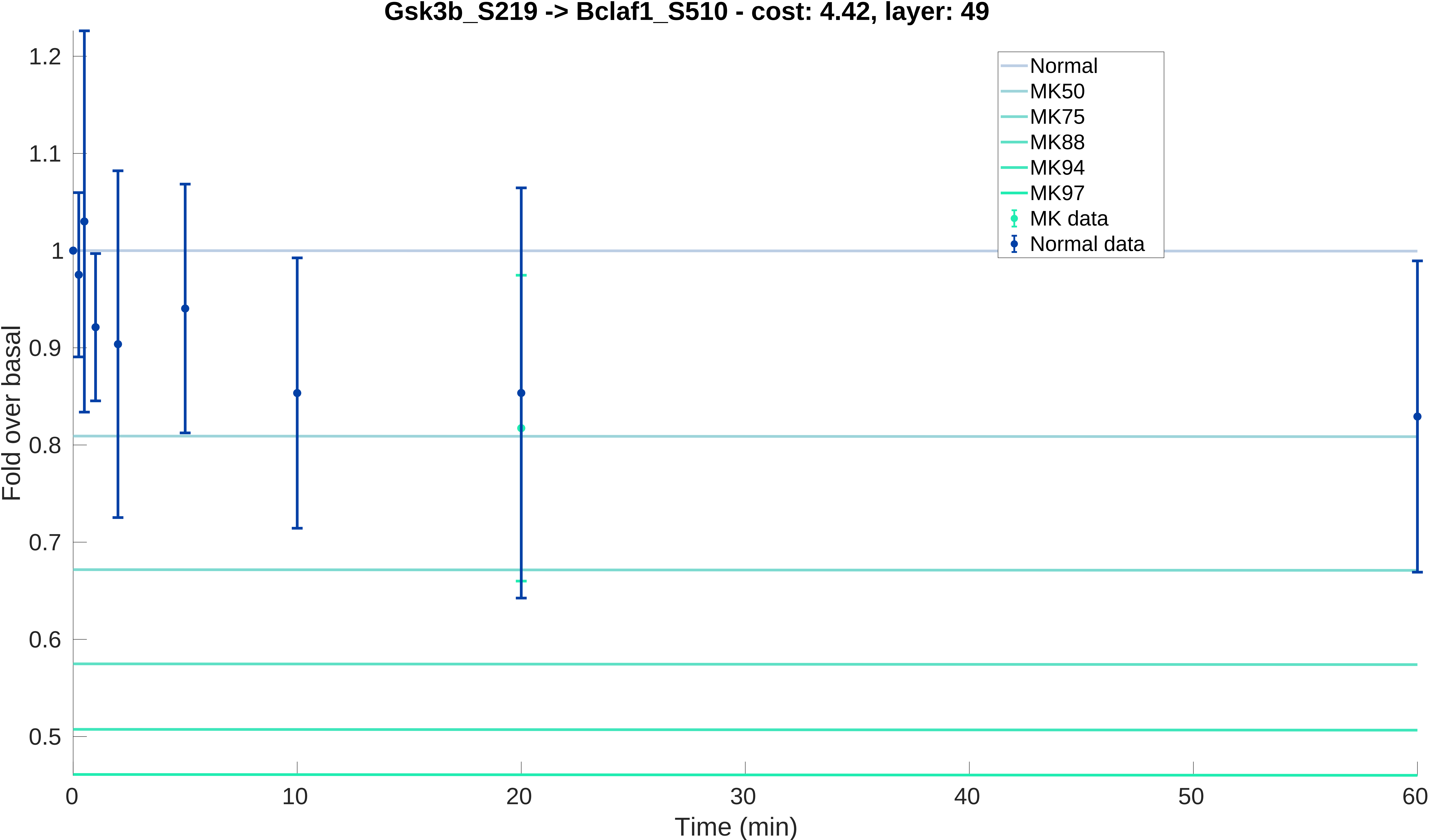

Gsk3b_S219 -> Bclaf1_S510 - cost: 4.42, layer: 49
Normal
MK50
MK75
MK88
MK94
MK97
MK data
Normal data
Fold over basal
1.2
1.1
1
0.9
0.8
0.7
0.6
0.5
0
10
20
30
40
50
60
Time (min)

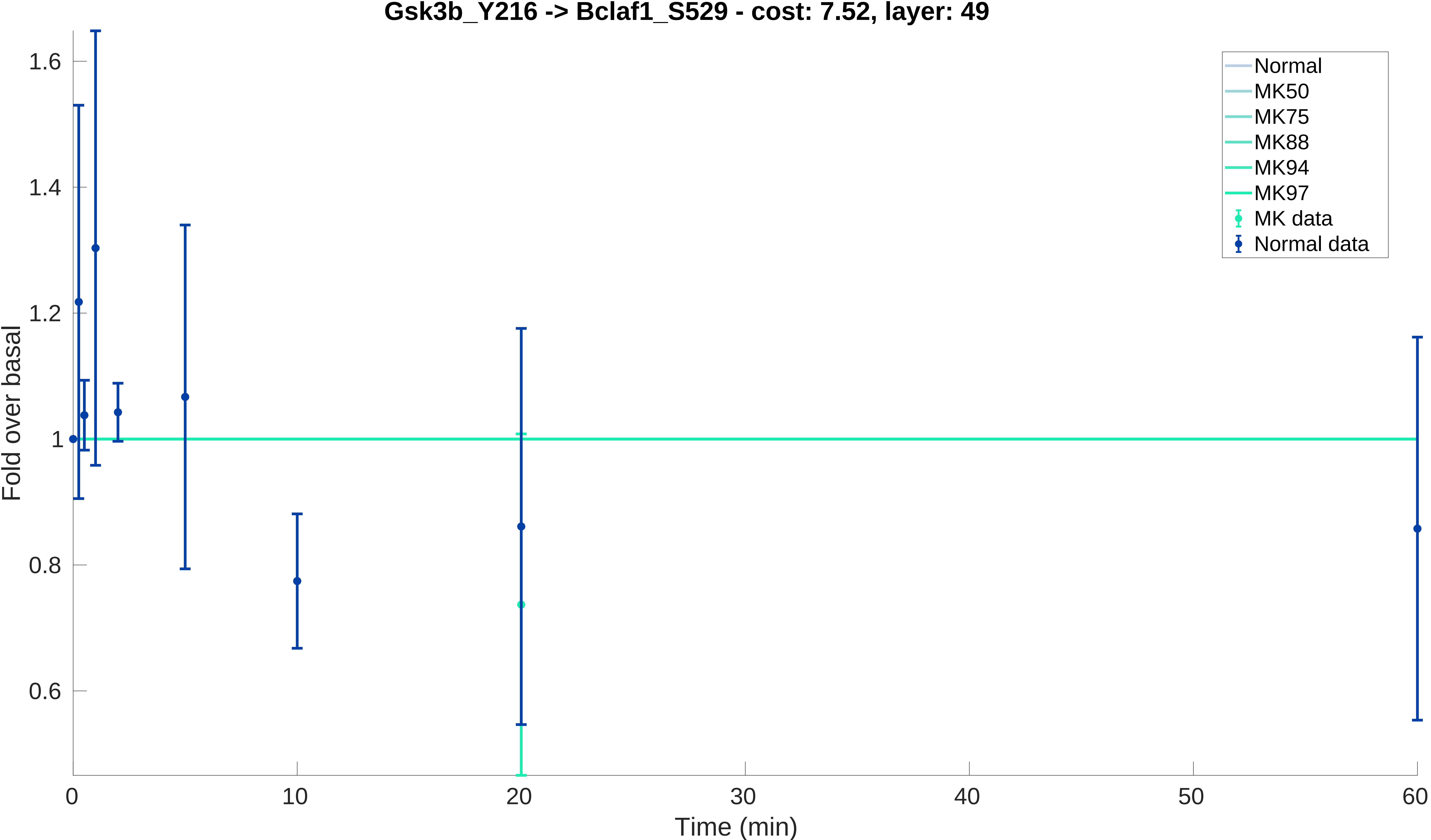

Gsk3b_Y216 -> Bclaf1_S529 - cost: 7.52, layer: 49
Fold over basal
Time (min)
1.6
1.4
1.2
1
0.8
0.6
0
10
20
30
40
50
60
Normal
MK50
MK75
MK88
MK94
MK97
MK data
Normal data

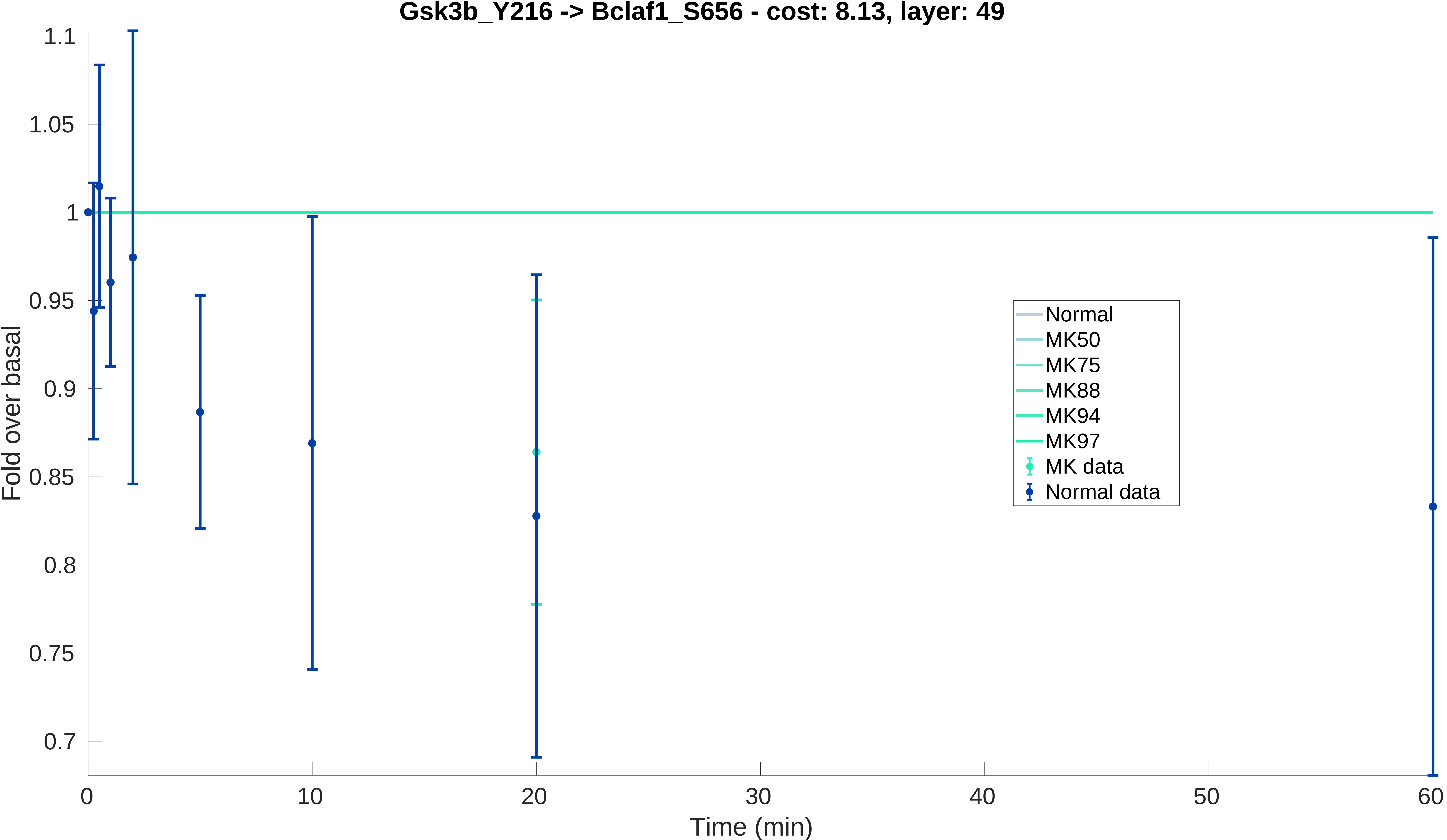

Gsk3b_Y216 -> Bclaf1_S656 - cost: 8.13, layer: 49
Fold over basal
1.1
1.05
1
0.95
0.9
0.85
0.8
0.75
0.7
0
10
20
30
40
50
60
Time (min)
Normal
MK50
MK75
MK88
MK94
MK97
MK data
Normal data

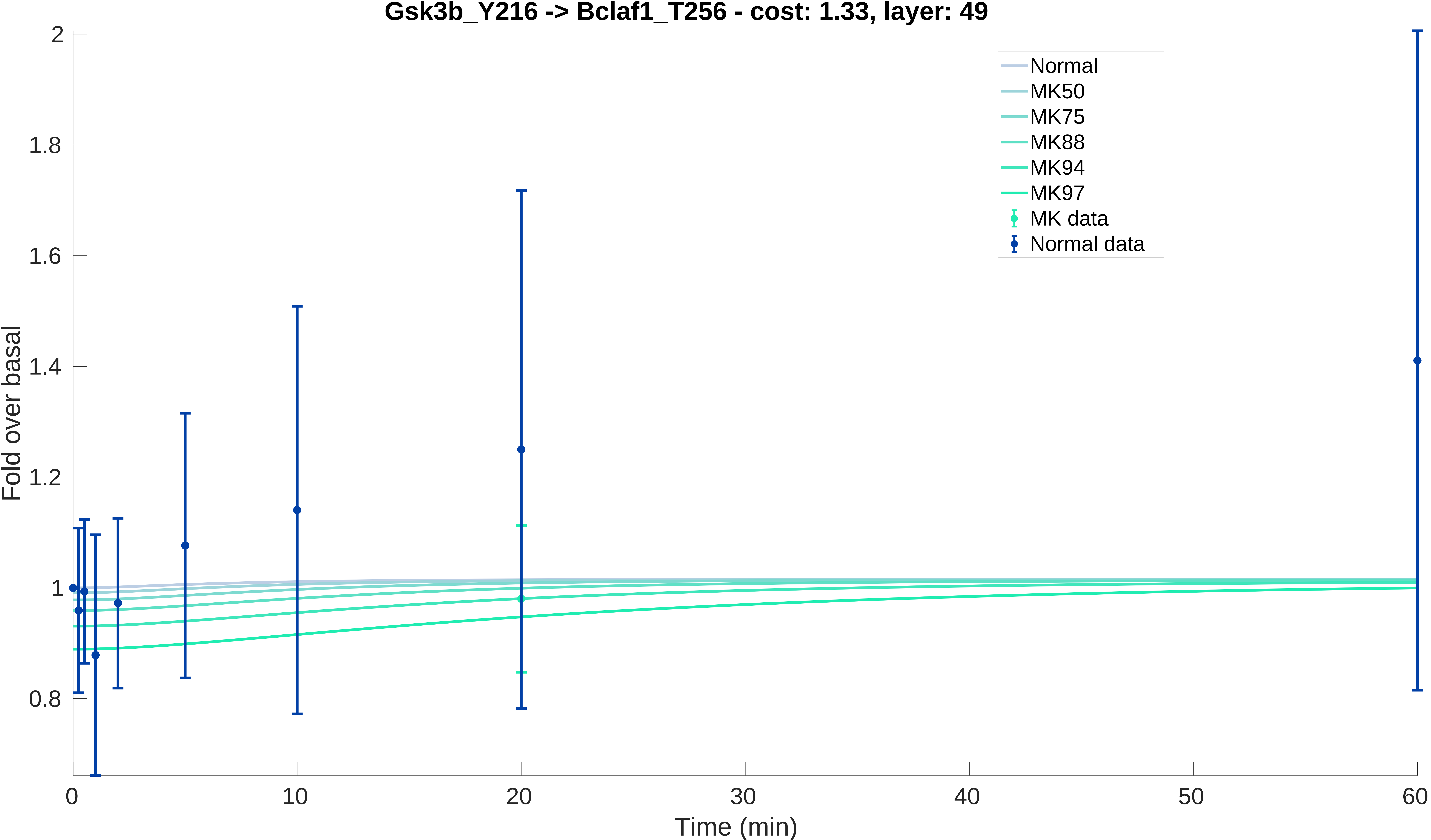

Gsk3b_Y216 -> Bclaf1_T256 - cost: 1.33, layer: 49
Fold over basal
Time (min)
0.8
1
1.2
1.4
1.6
1.8
2
0
10
20
30
40
50
60
Normal
MK50
MK75
MK88
MK94
MK97
MK data
Normal data

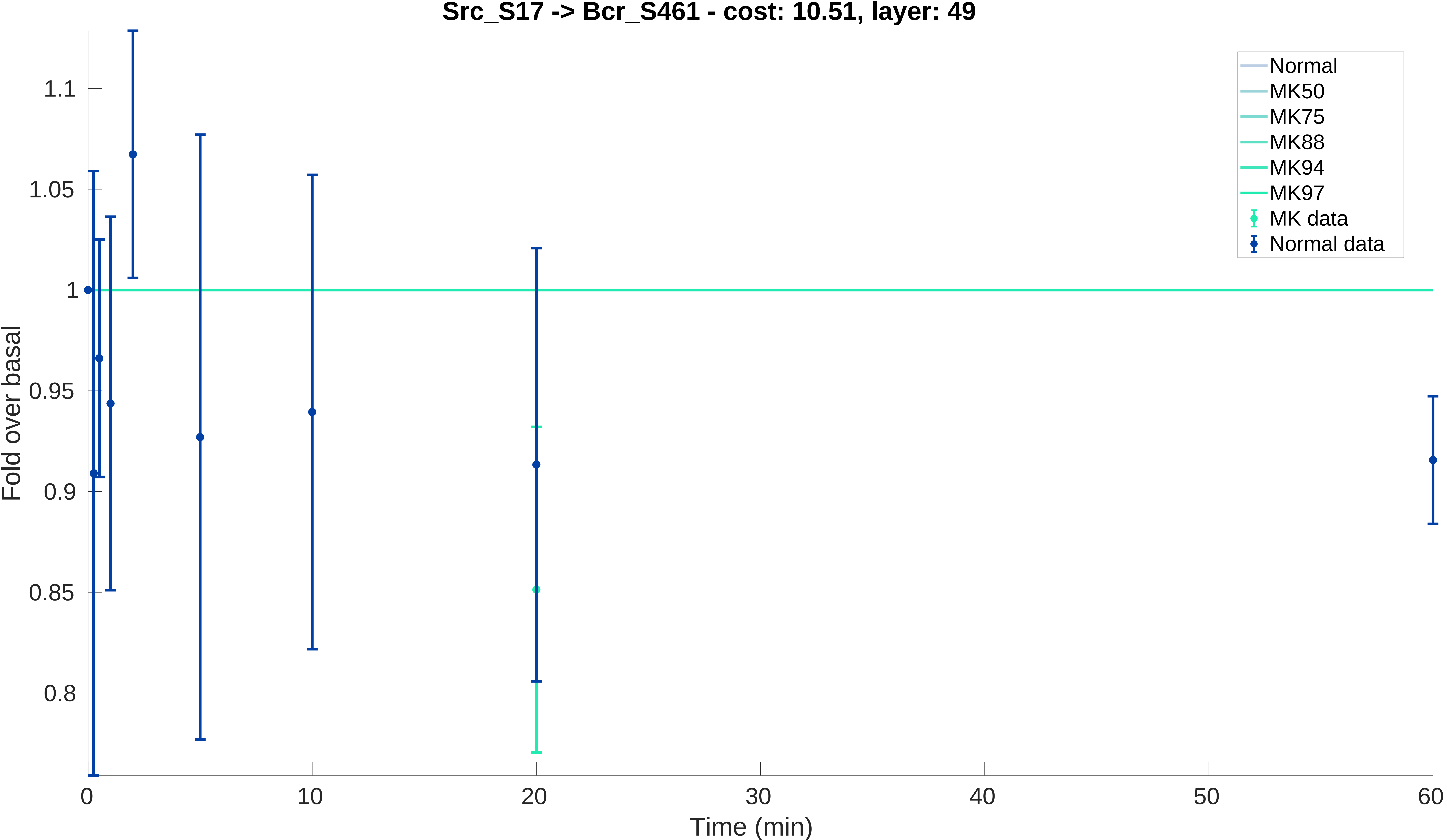

Src_S17 -> Bcr_S461 - cost: 10.51, layer: 49
Fold over basal
1.1
1.05
1
0.95
0.9
0.85
0.8
0
10
20
30
40
50
60
Time (min)
Normal
MK50
MK75
MK88
MK94
MK97
MK data
Normal data

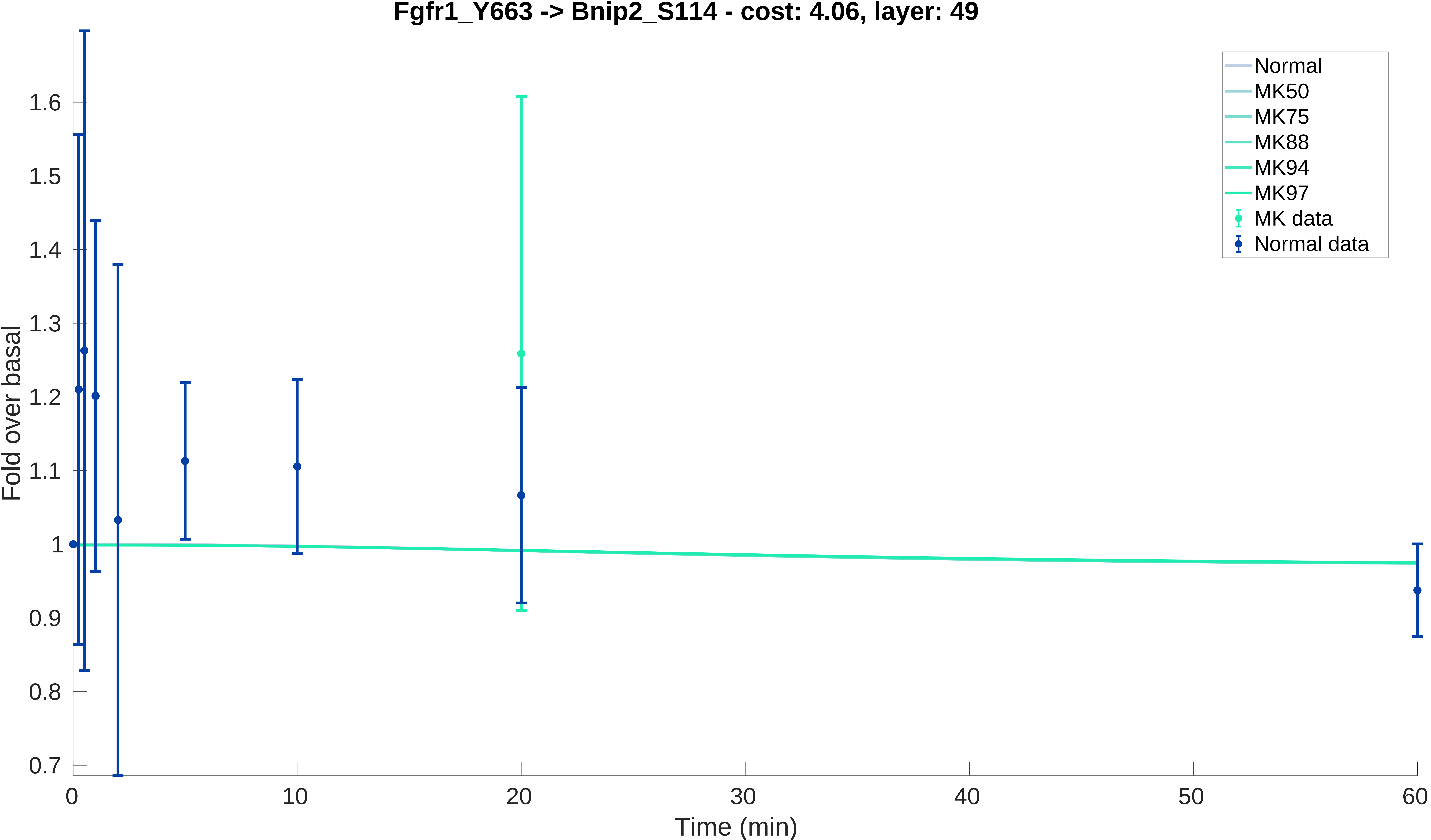

Fgfr1_Y663 -> Bnip2_S114 - cost: 4.06, layer: 49
Fold over basal
Time (min)
0.7
0.8
0.9
1
1.1
1.2
1.3
1.4
1.5
1.6
0
10
20
30
40
50
60
Normal
MK50
MK75
MK88
MK94
MK97
MK data
Normal data

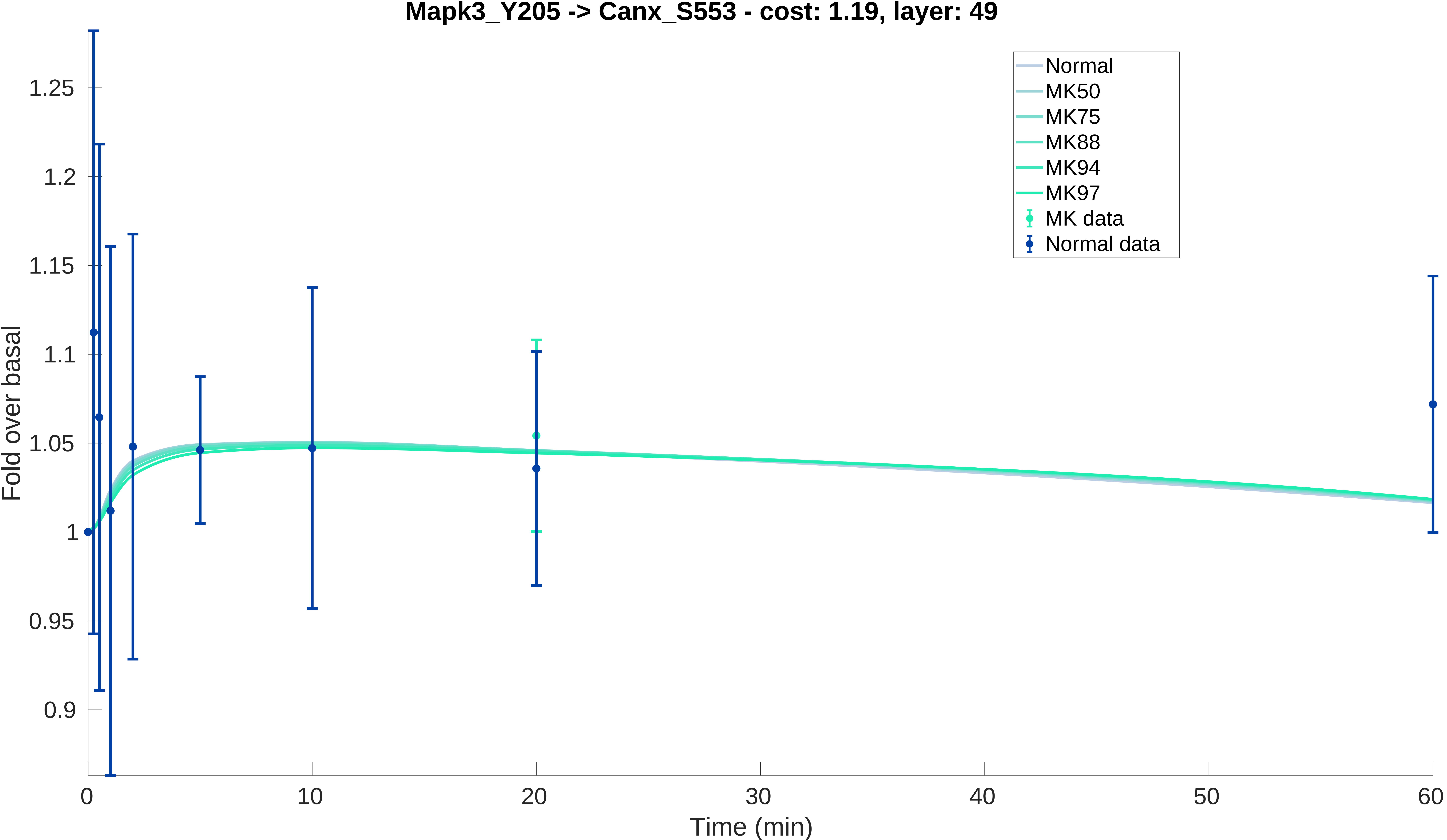

Mapk3_Y205 -> Canx_S553 - cost: 1.19, layer: 49
Normal
MK50
MK75
MK88
MK94
MK97
MK data
Normal data
Fold over basal
1.25
1.2
1.15
1.1
1.05
1
0.95
0.9
0
10
20
30
40
50
60
Time (min)

Mapk3_Y205 -> Canx_S563 - cost: 10.55, layer: 49

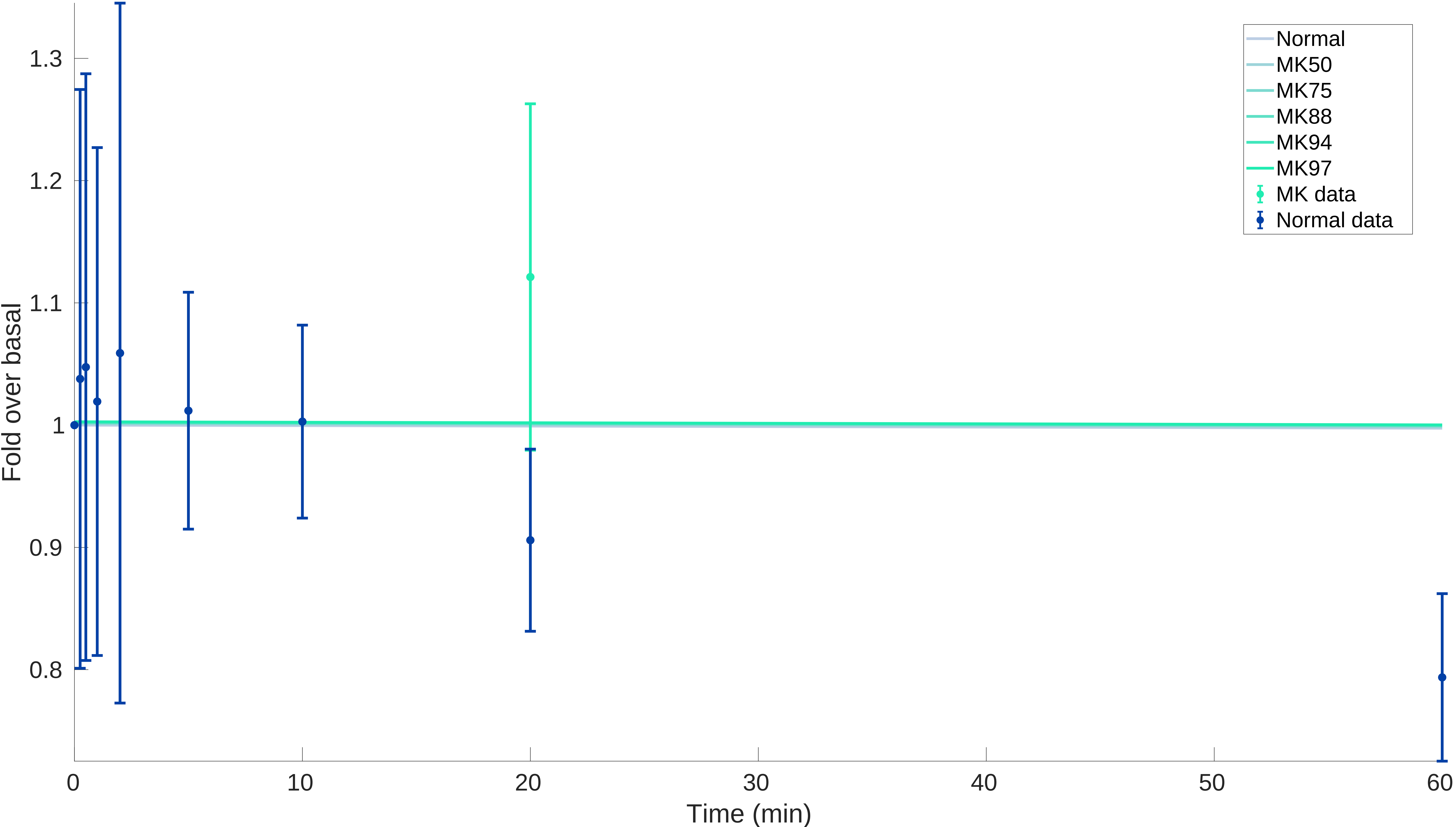

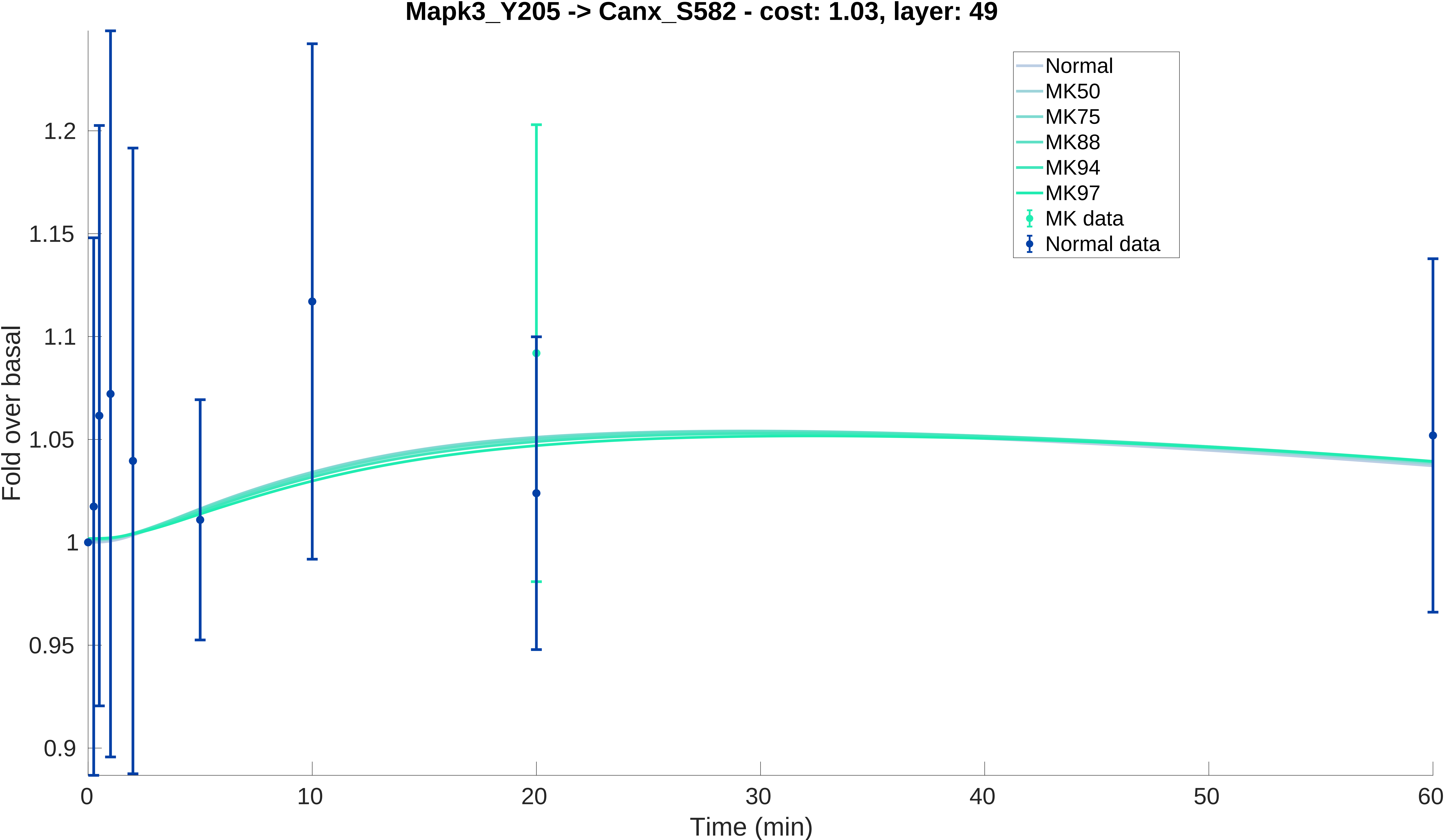

Mapk3_Y205 -> Canx_S582 - cost: 1.03, layer: 49
Normal
MK50
MK75
MK88
MK94
MK97
MK data
Normal data
Fold over basal
1.2
1.15
1.1
1.05
1
0.95
0.9
0
10
20
30
40
50
60
Time (min)

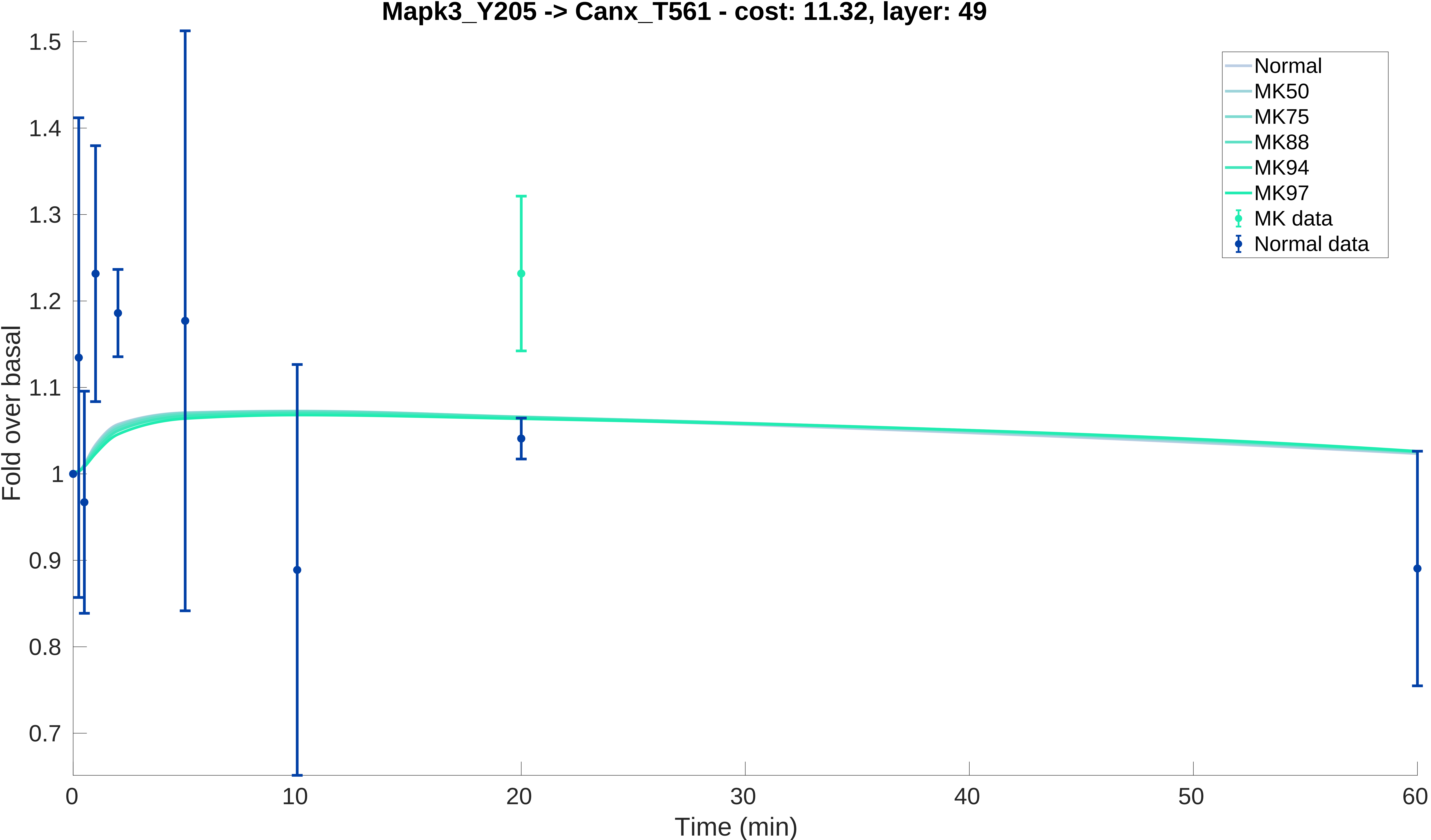

Mapk3_Y205 -> Canx_T561 - cost: 11.32, layer: 49
Fold over basal
Time (min)
0
10
20
30
40
50
60
0.7
0.8
0.9
1
1.1
1.2
1.3
1.4
1.5
Normal
MK50
MK75
MK88
MK94
MK97
MK data
Normal data

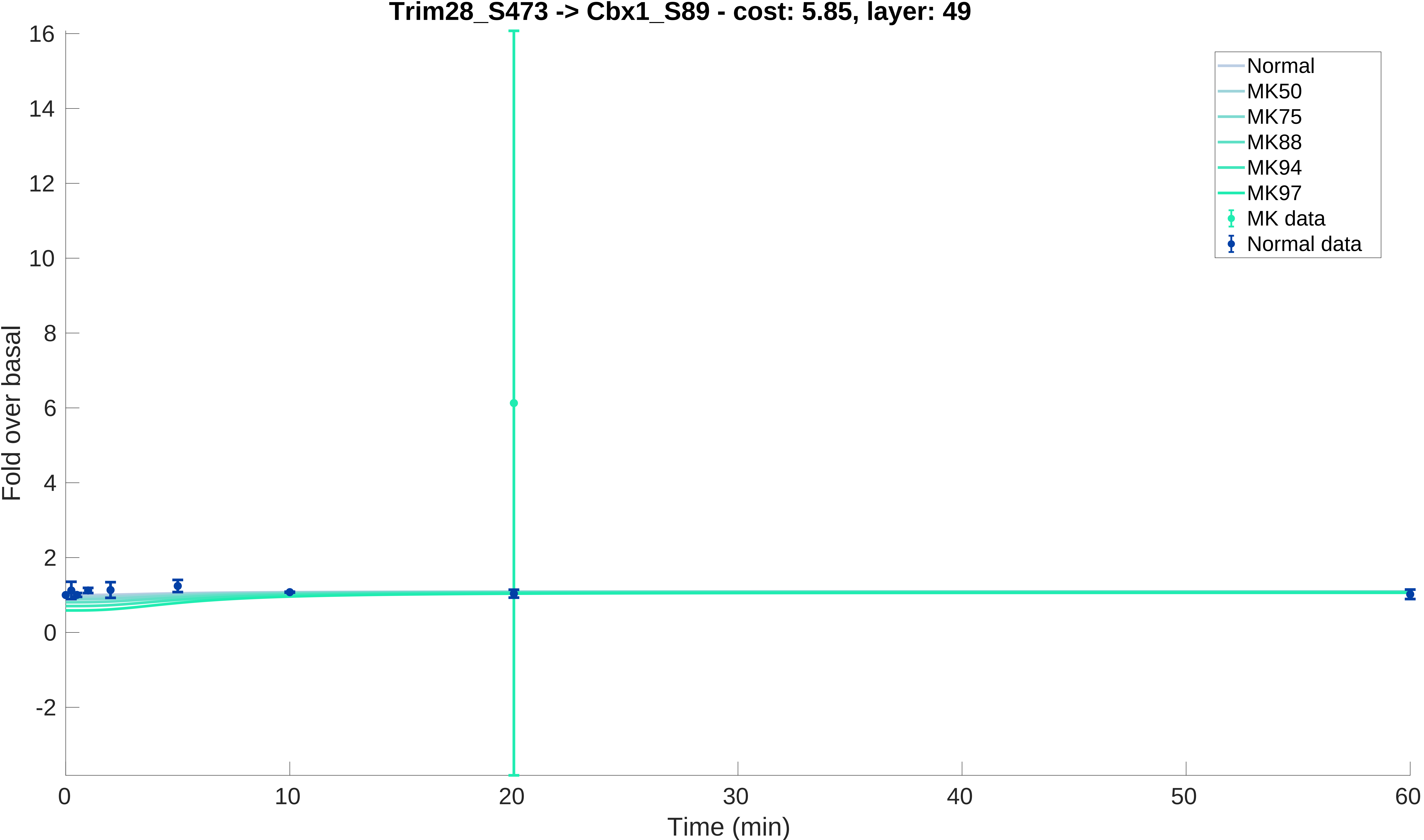

Trim28_S473 -> Cbx1_S89 - cost: 5.85, layer: 49
Fold over basal
Time (min)
Normal
MK50
MK75
MK88
MK94
MK97
MK data
Normal data
16
14
12
10
8
6
4
2
0
-2
0
10
20
30
40
50
60

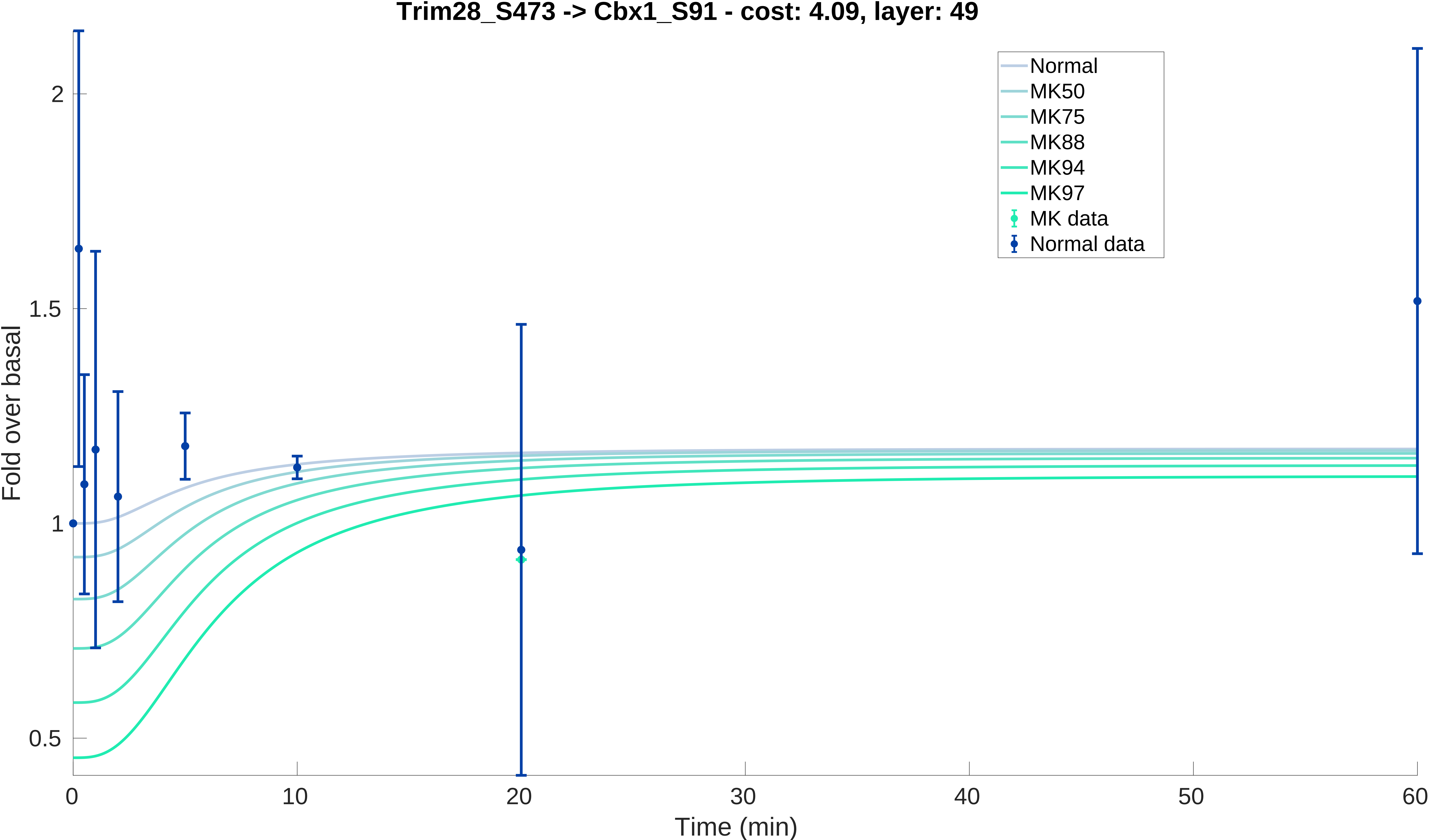

Trim28_S473 -> Cbx1_S91 - cost: 4.09, layer: 49
Normal
MK50
MK75
MK88
MK94
MK97
MK data
Normal data
Fold over basal
Time (min)
0.5
1
1.5
2
0
10
20
30
40
50
60

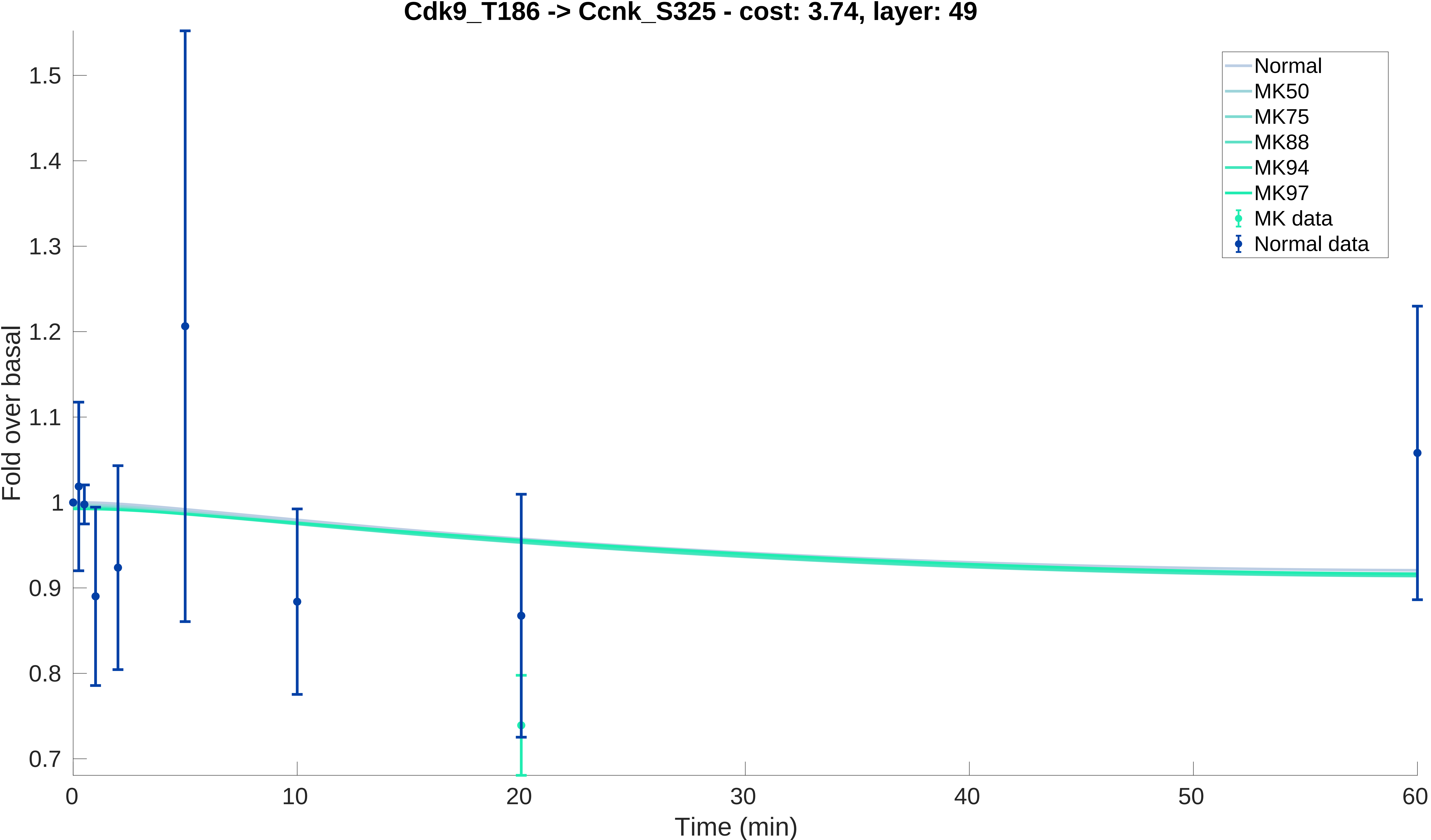
Cdk9_T186 -> Ccnk_S325 - cost: 3.74, layer: 49
Fold over basal
Time (min)
0.7
0.8
0.9
1
1.1
1.2
1.3
1.4
1.5
0
10
20
30
40
50
60
Normal
MK50
MK75
MK88
MK94
MK97
MK data
Normal data

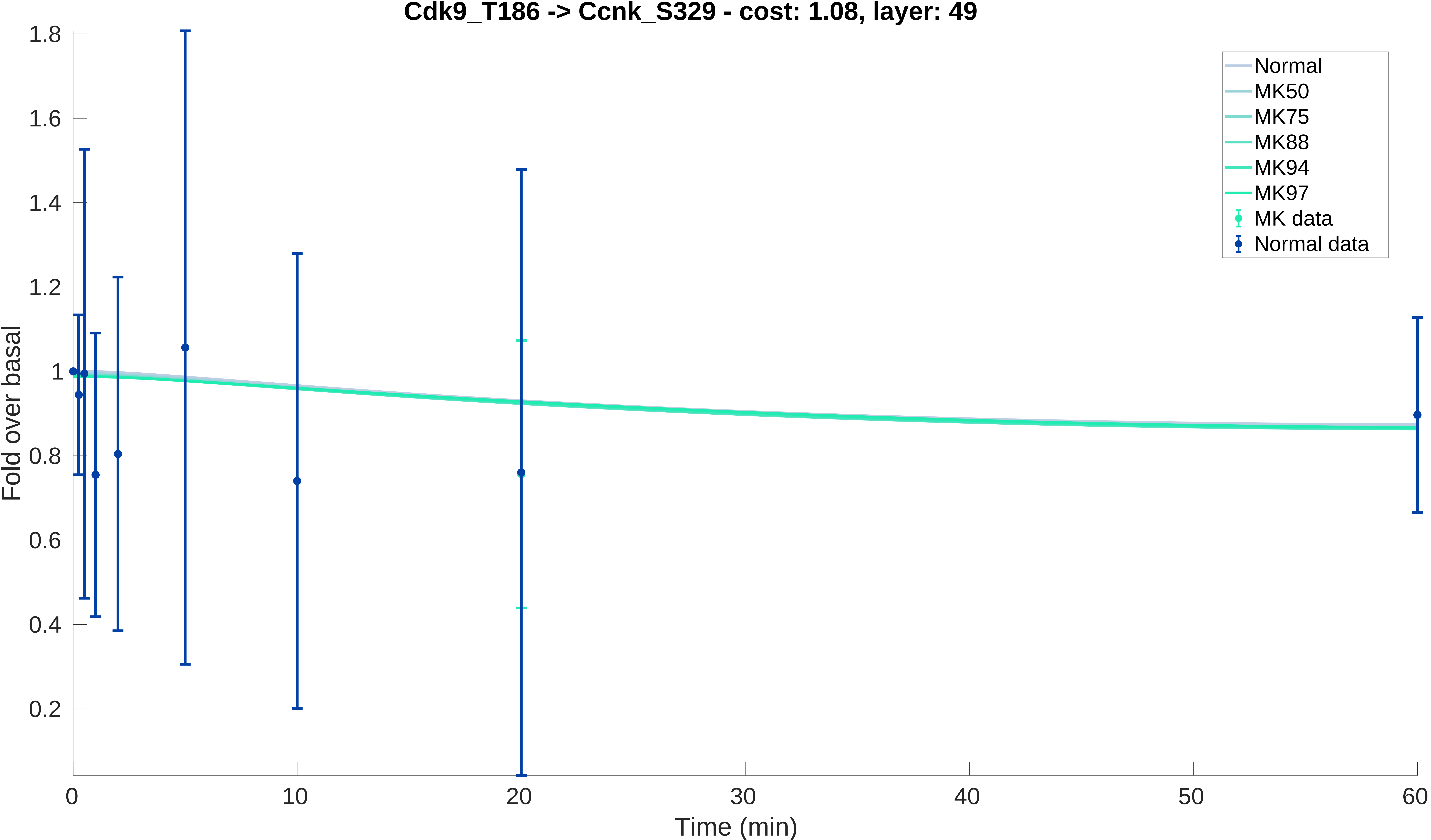

Cdk9_T186 -> Ccnk_S329 - cost: 1.08, layer: 49
Fold over basal
Time (min)
1.8
1.6
1.4
1.2
1
0.8
0.6
0.4
0.2
0
10
20
30
40
50
60
Normal
MK50
MK75
MK88
MK94
MK97
MK data
Normal data

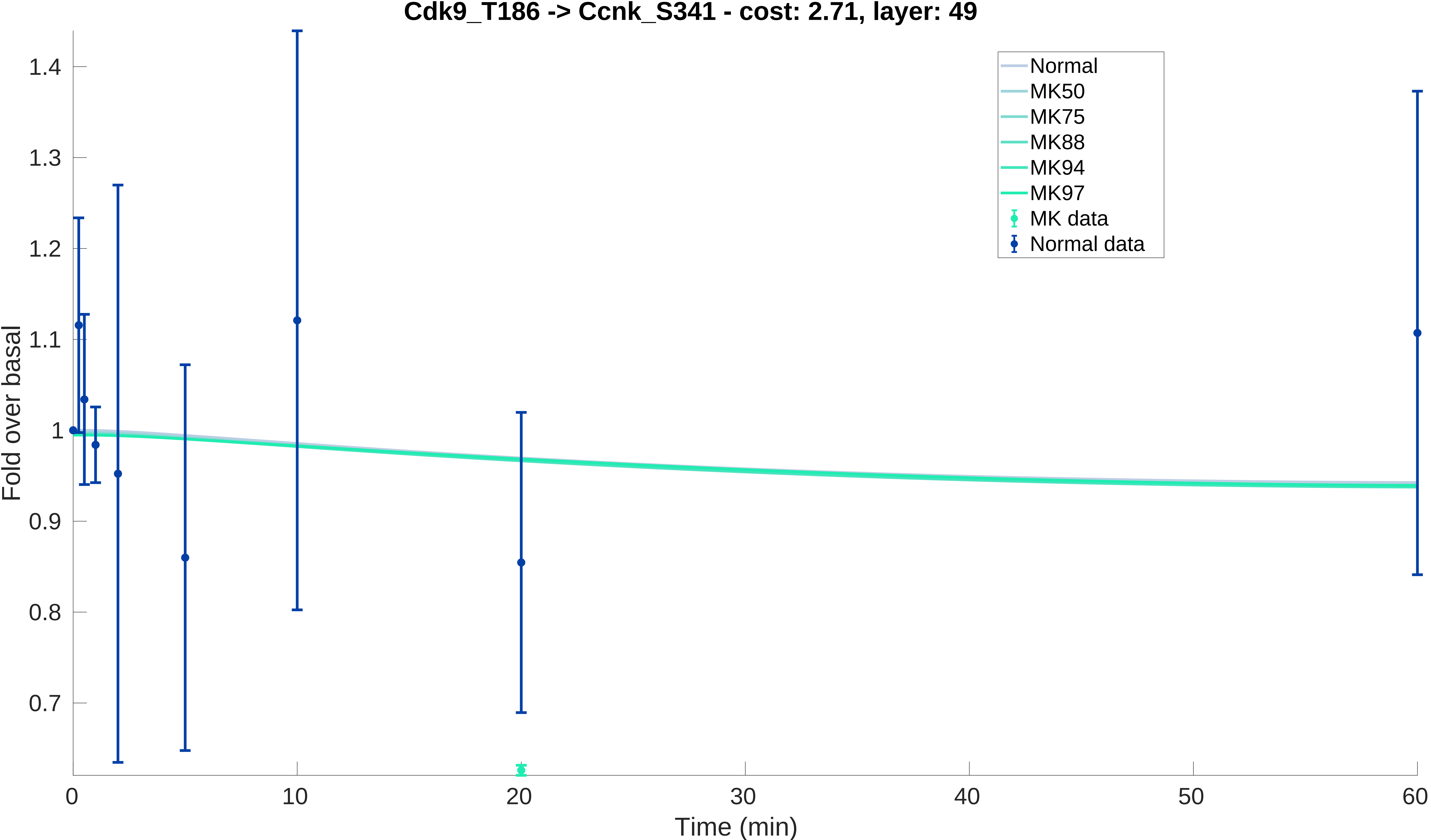
Cdk9_T186 -> Ccnk_S341 - cost: 2.71, layer: 49
Fold over basal
Time (min)
0.7
0.8
0.9
1
1.1
1.2
1.3
1.4
0
10
20
30
40
50
60
Normal
MK50
MK75
MK88
MK94
MK97
MK data
Normal data

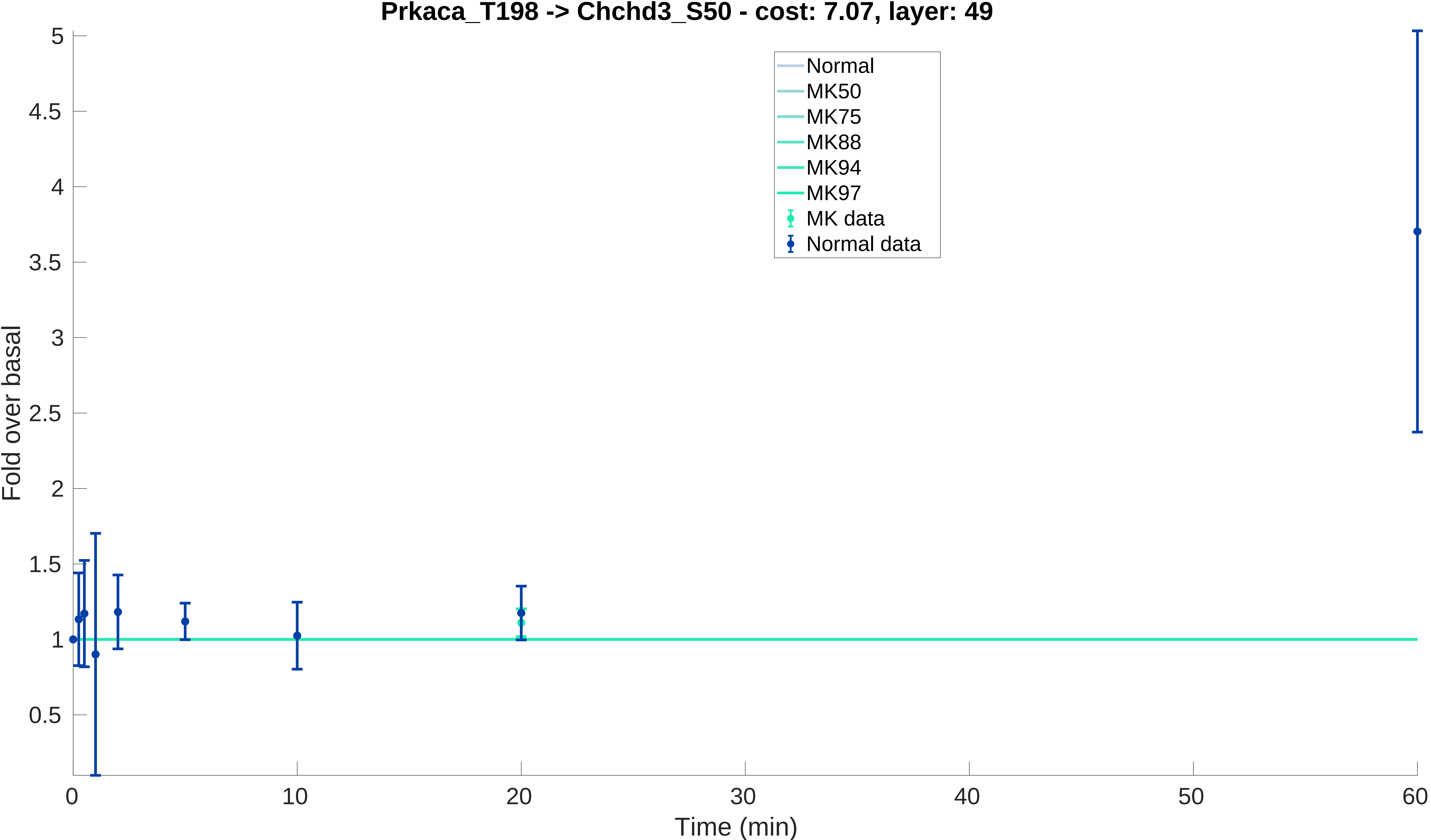
Prkaca_T198 -> Chchd3_S50 - cost: 7.07, layer: 49
Fold over basal
Time (min)
0
10
20
30
40
50
60
0.5
1
1.5
2
2.5
3
3.5
4
4.5
5
Normal
MK50
MK75
MK88
MK94
MK97
MK data
Normal data

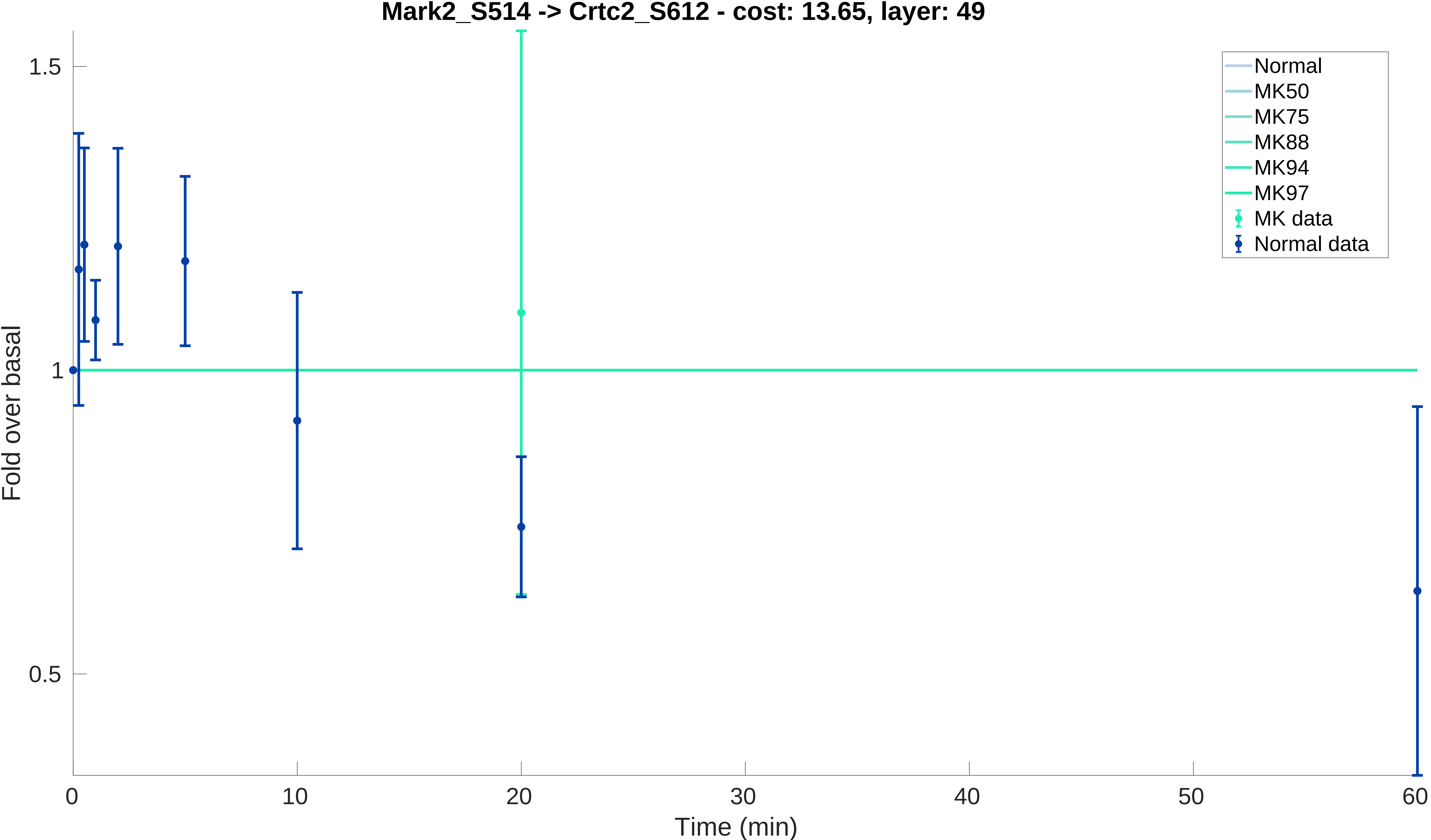

Mark2_S514 -> Crtc2_S612 - cost: 13.65, layer: 49
1.5
1
0.5
Fold over basal
0
10
20
30
40
50
60
Time (min)
Normal
MK50
MK75
MK88
MK94
MK97
MK data
Normal data

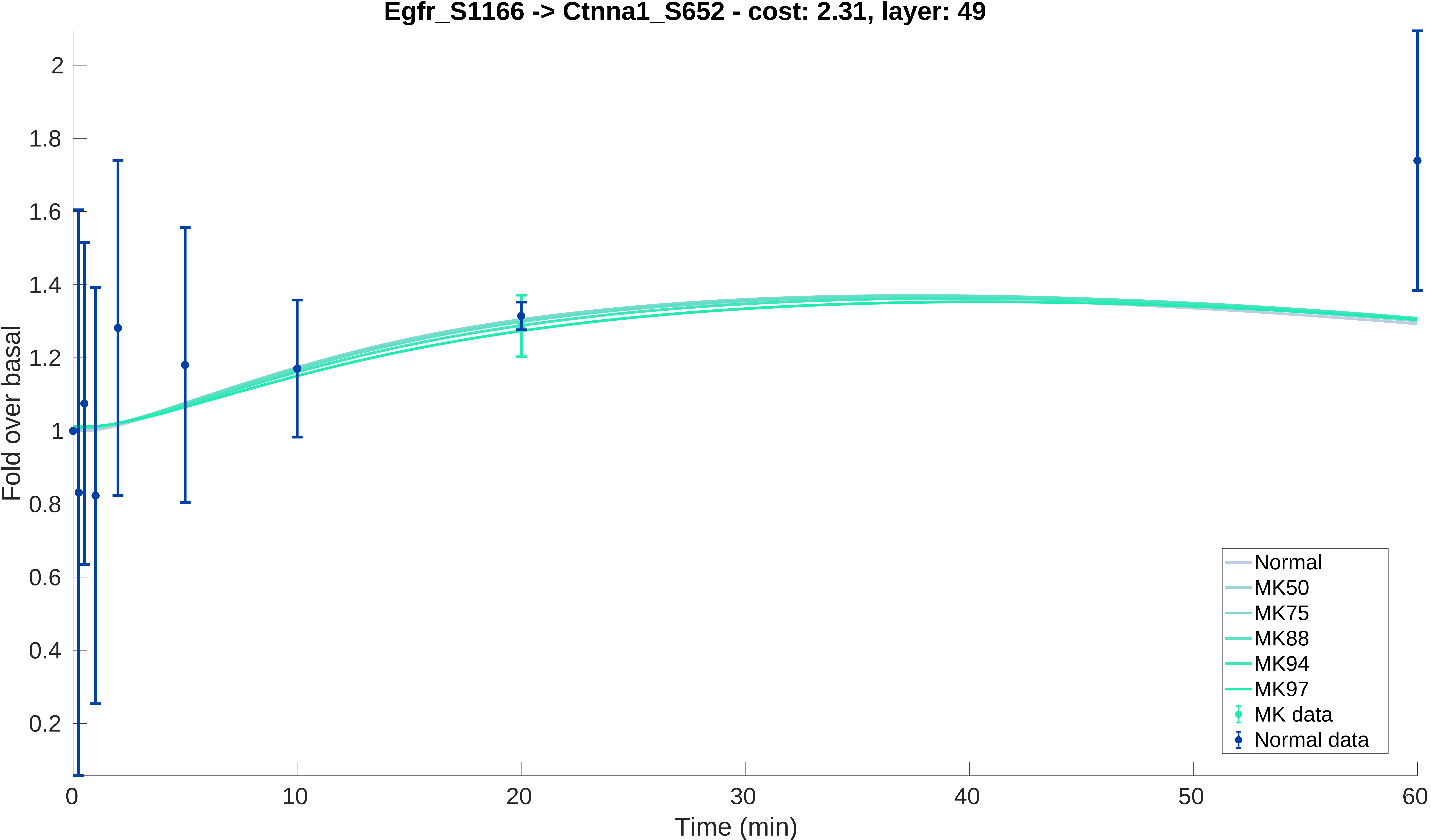
Egfr_S1166 -> Ctnna1_S652 - cost: 2.31, layer: 49
Fold over basal
Time (min)
0
10
20
30
40
50
60
0.2
0.4
0.6
0.8
1
1.2
1.4
1.6
1.8
2
Normal
MK50
MK75
MK88
MK94
MK97
MK data
Normal data

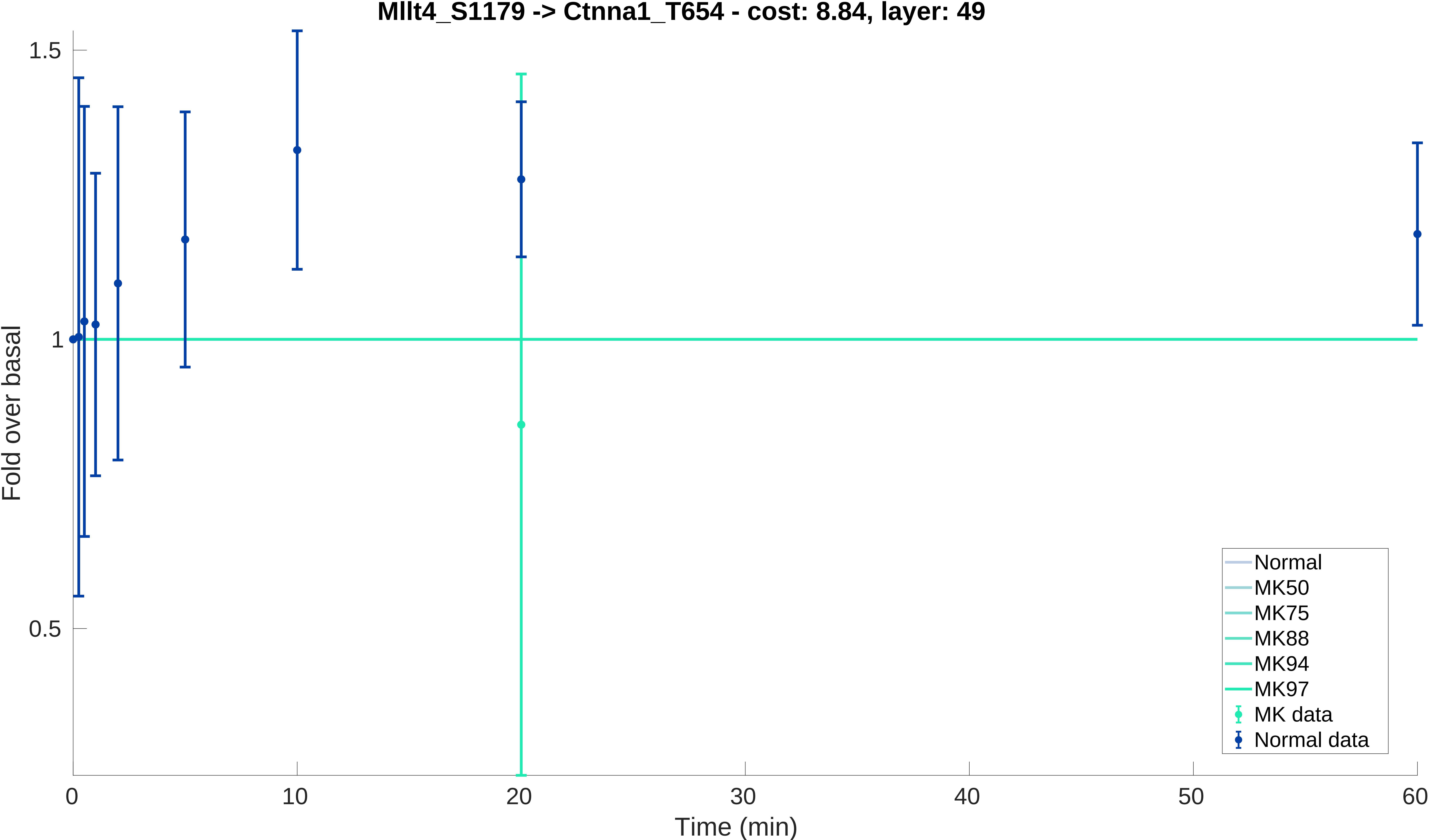

Mllt4_S1179 -> Ctnna1_T654 - cost: 8.84, layer: 49
1.5
1
0.5
Fold over basal
0
10
20
30
40
50
60
Time (min)
Normal
MK50
MK75
MK88
MK94
MK97
MK data
Normal data

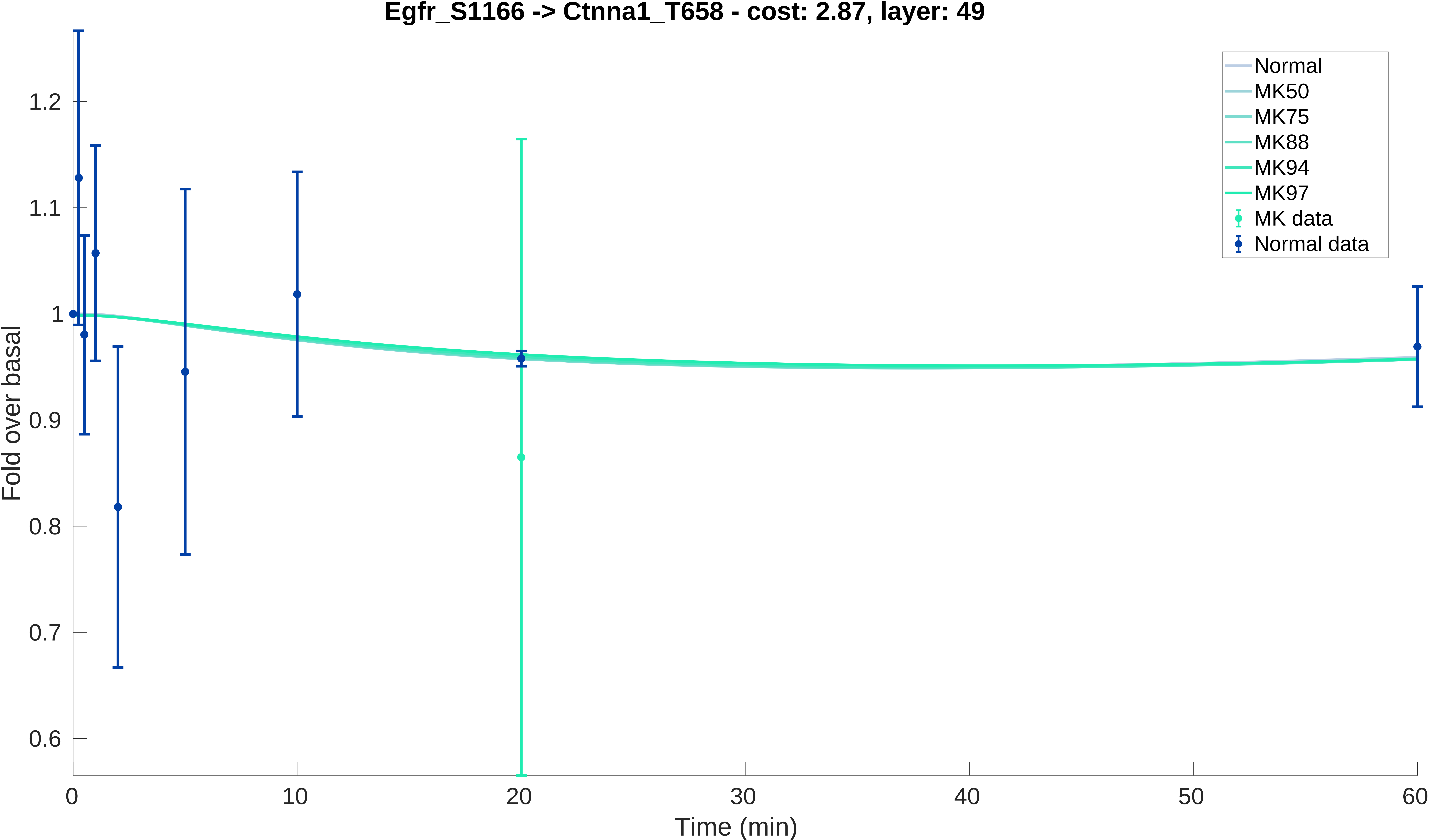

Egfr_S1166 -> Ctnna1_T658 - cost: 2.87, layer: 49
Normal
MK50
MK75
MK88
MK94
MK97
MK data
Normal data
Fold over basal
1.2
1.1
1
0.9
0.8
0.7
0.6
0
10
20
30
40
50
60
Time (min)

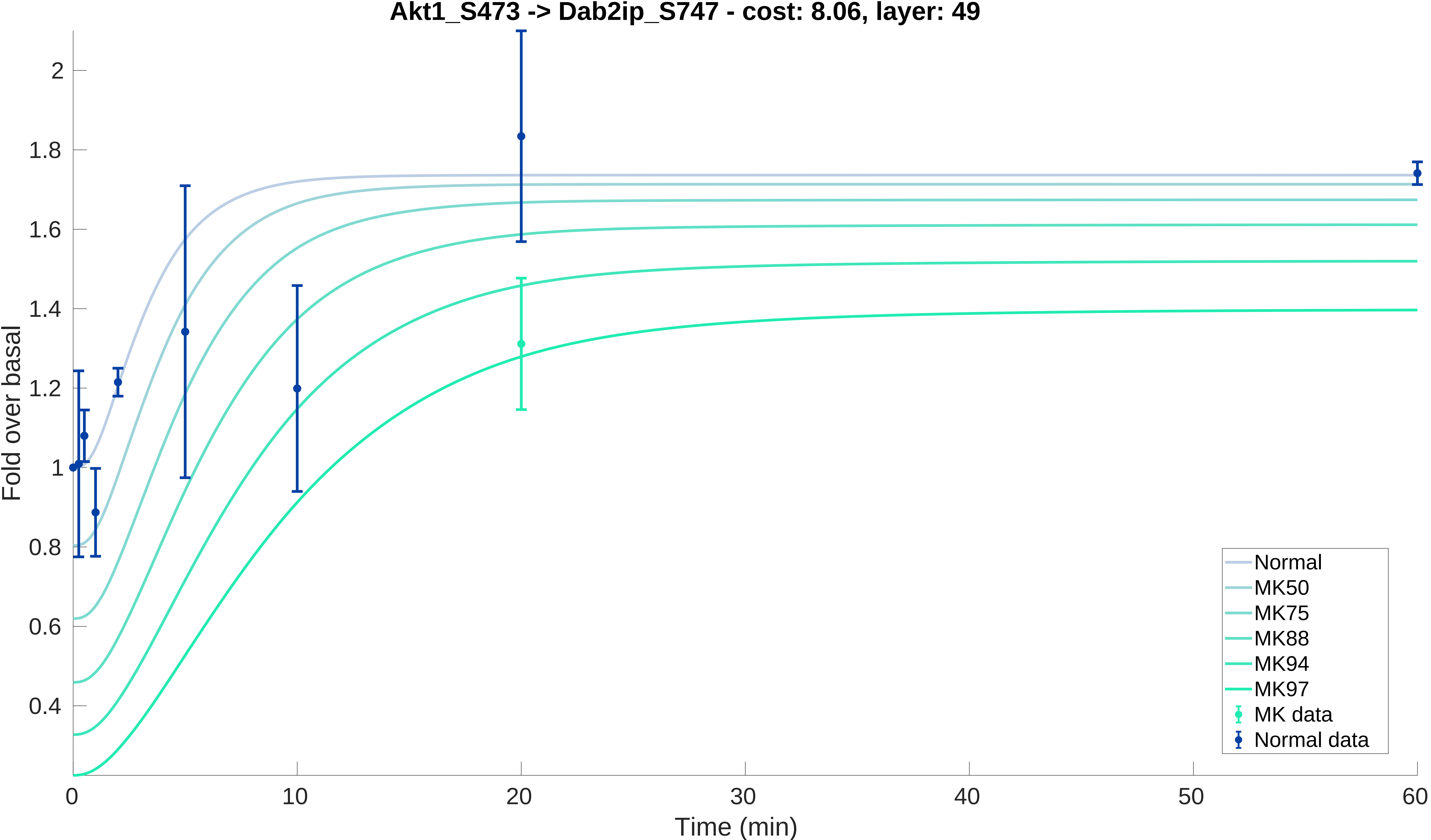

Akt1_S473 -> Dab2ip_S747 - cost: 8.06, layer: 49
Fold over basal
Time (min)
0.4
0.6
0.8
1
1.2
1.4
1.6
1.8
2
0
10
20
30
40
50
60
Normal
MK50
MK75
MK88
MK94
MK97
MK data
Normal data

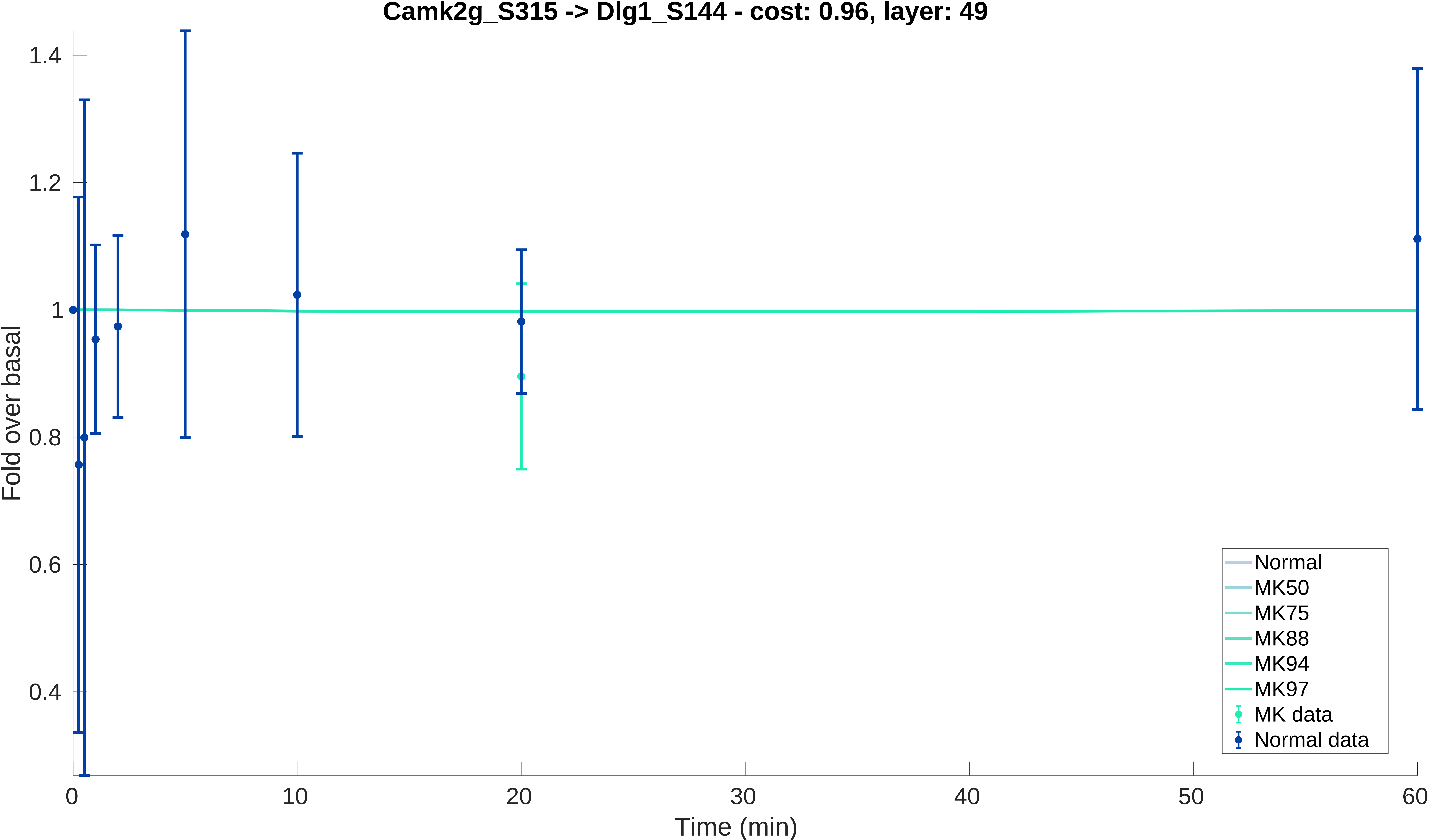

Camk2g_S315 -> Dlg1_S144 - cost: 0.96, layer: 49
1.4
1.2
1
0.8
0.6
0.4
Fold over basal
0
10
20
30
40
50
60
Time (min)
Normal
MK50
MK75
MK88
MK94
MK97
MK data
Normal data

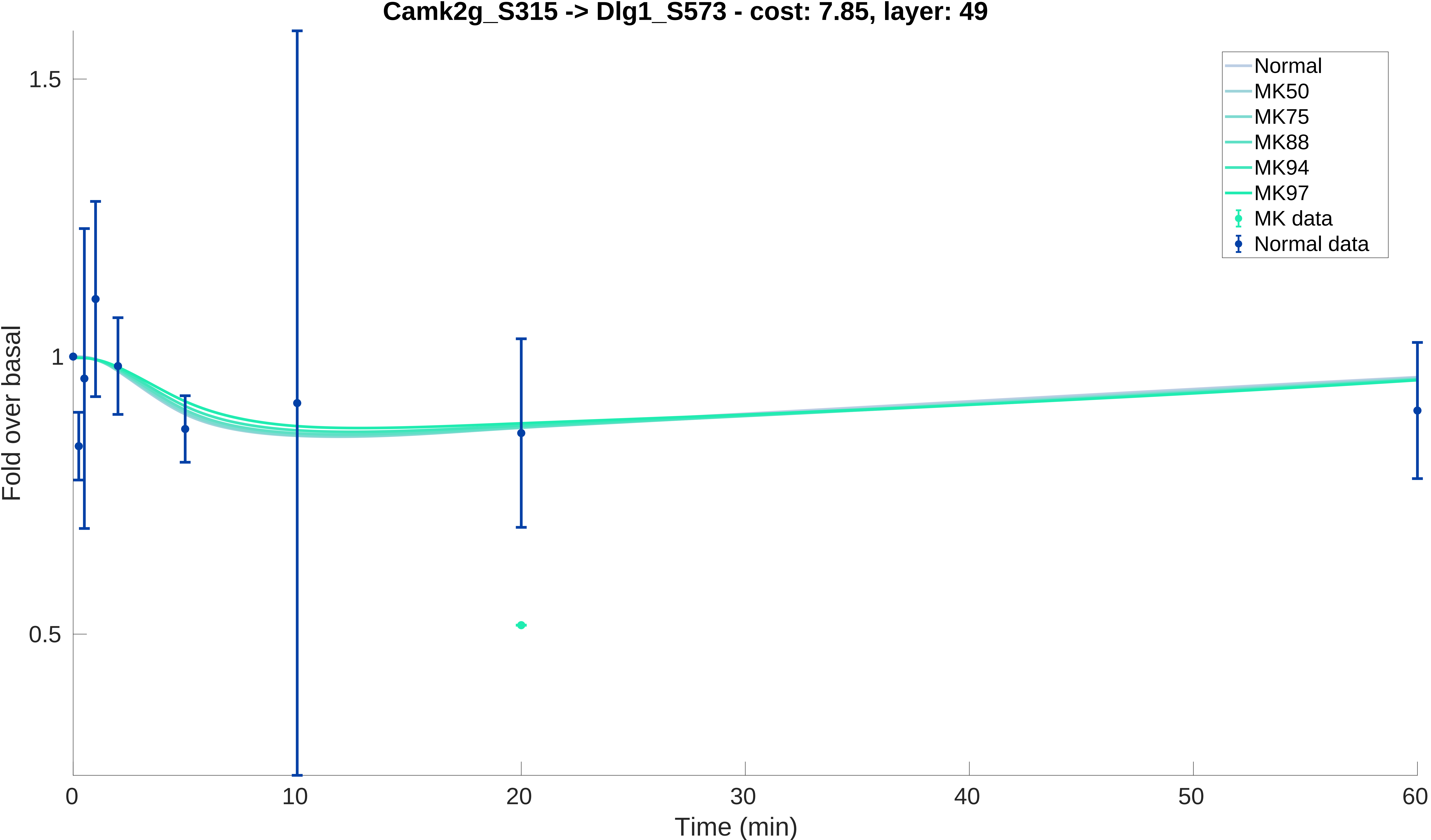

Camk2g_S315 -> Dlg1_S573 - cost: 7.85, layer: 49
Normal
MK50
MK75
MK88
MK94
MK97
MK data
Normal data
Fold over basal
1.5
1
0.5
0
10
20
30
40
50
60
Time (min)

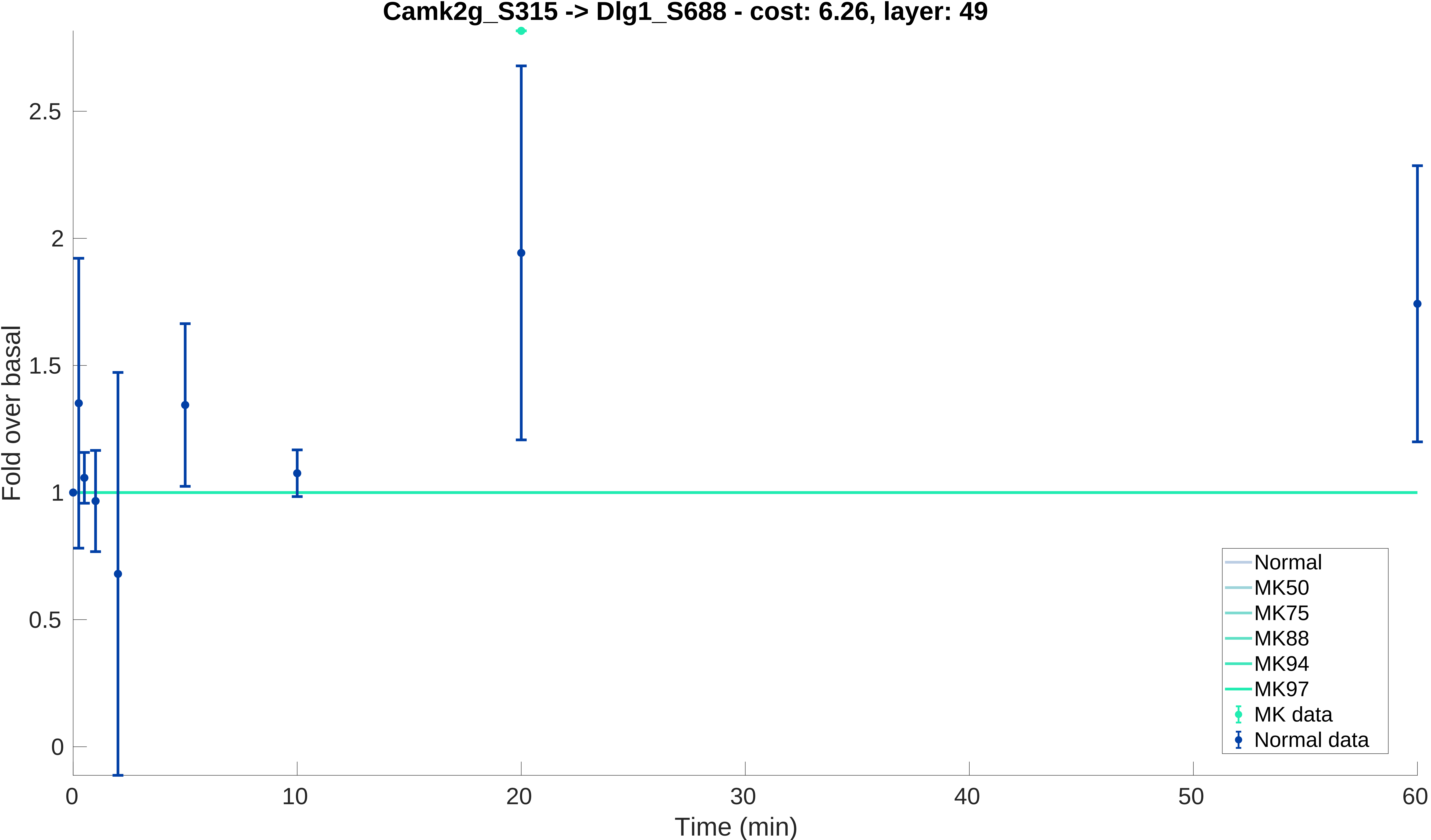
Camk2g_S315 -> Dlg1_S688 - cost: 6.26, layer: 49
Fold over basal
Time (min)
0
0.5
1
1.5
2
2.5
0
10
20
30
40
50
60
Normal
MK50
MK75
MK88
MK94
MK97
MK data
Normal data

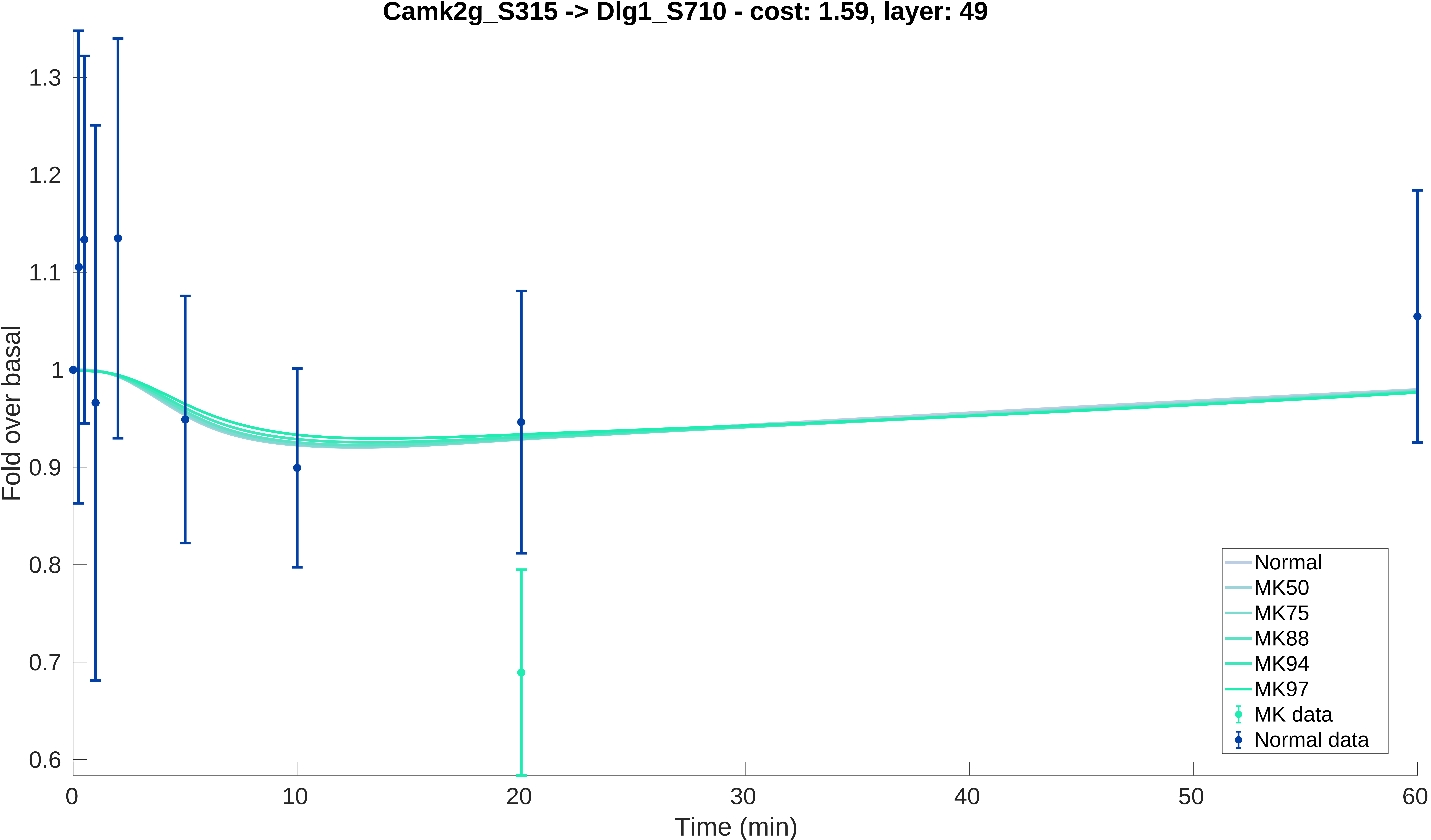

Camk2g_S315 -> Dlg1_S710 - cost: 1.59, layer: 49
Fold over basal
1.3
1.2
1.1
1
0.9
0.8
0.7
0.6
0
10
20
30
40
50
60
Time (min)
Normal
MK50
MK75
MK88
MK94
MK97
MK data
Normal data

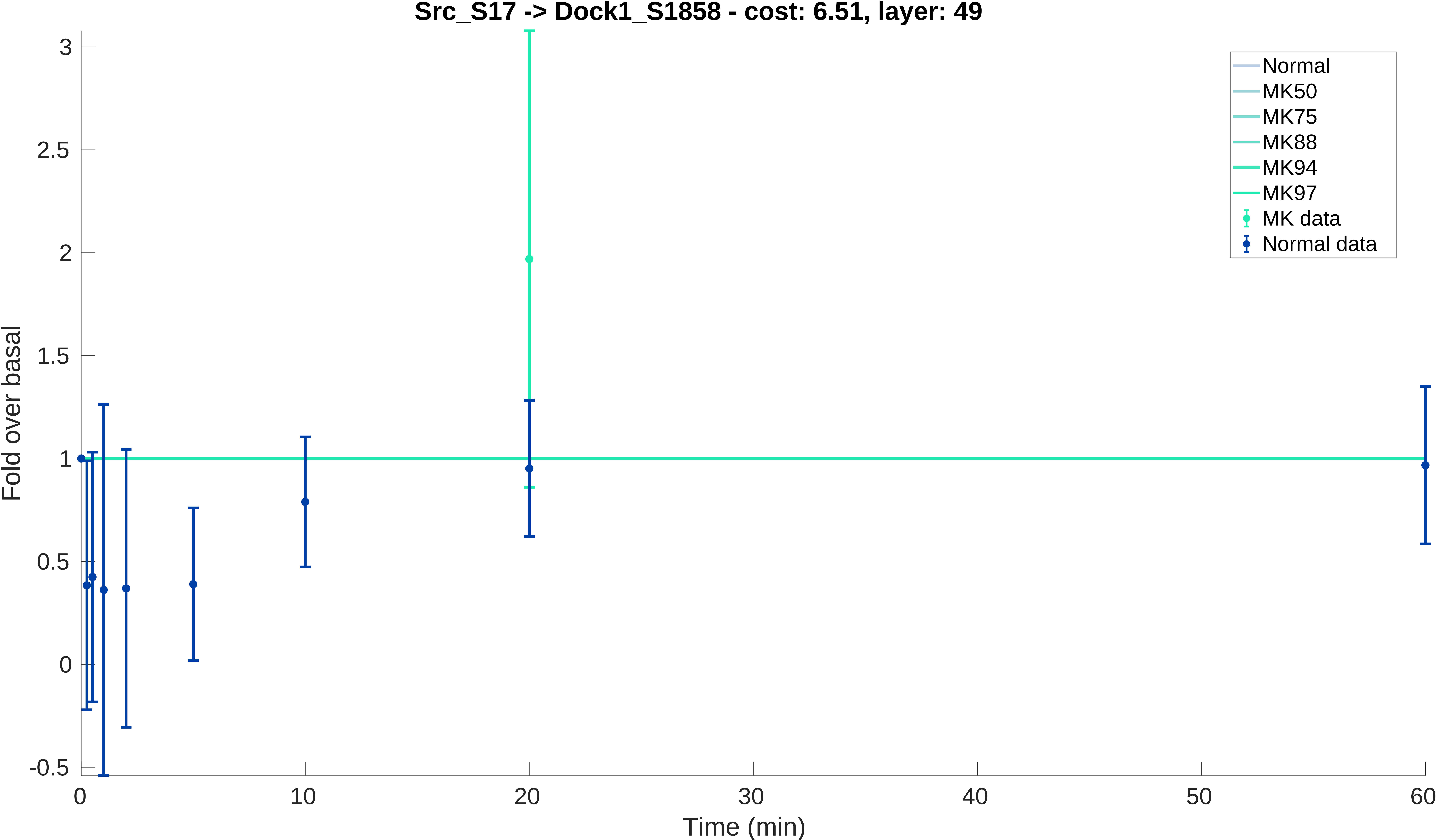

Src_S17 -> Dock1_S1858 - cost: 6.51, layer: 49
Fold over basal
Time (min)
3
2.5
2
1.5
1
0.5
0
-0.5
0
10
20
30
40
50
60
Normal
MK50
MK75
MK88
MK94
MK97
MK data
Normal data

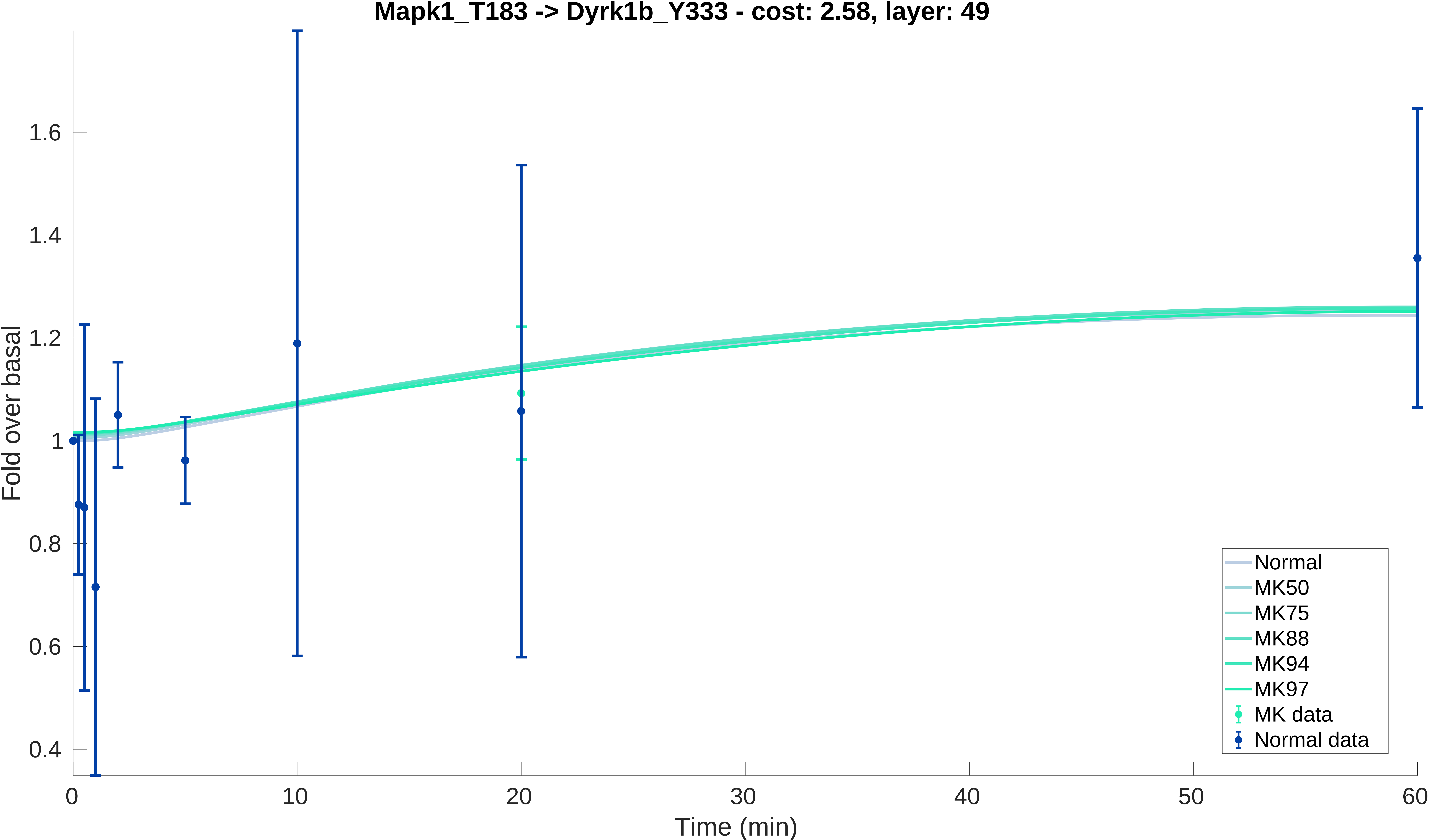

Mapk1_T183 -> Dyrk1b_Y333 - cost: 2.58, layer: 49
Fold over basal
1.6
1.4
1.2
1
0.8
0.6
0.4
0
10
20
30
40
50
60
Time (min)
Normal
MK50
MK75
MK88
MK94
MK97
MK data
Normal data

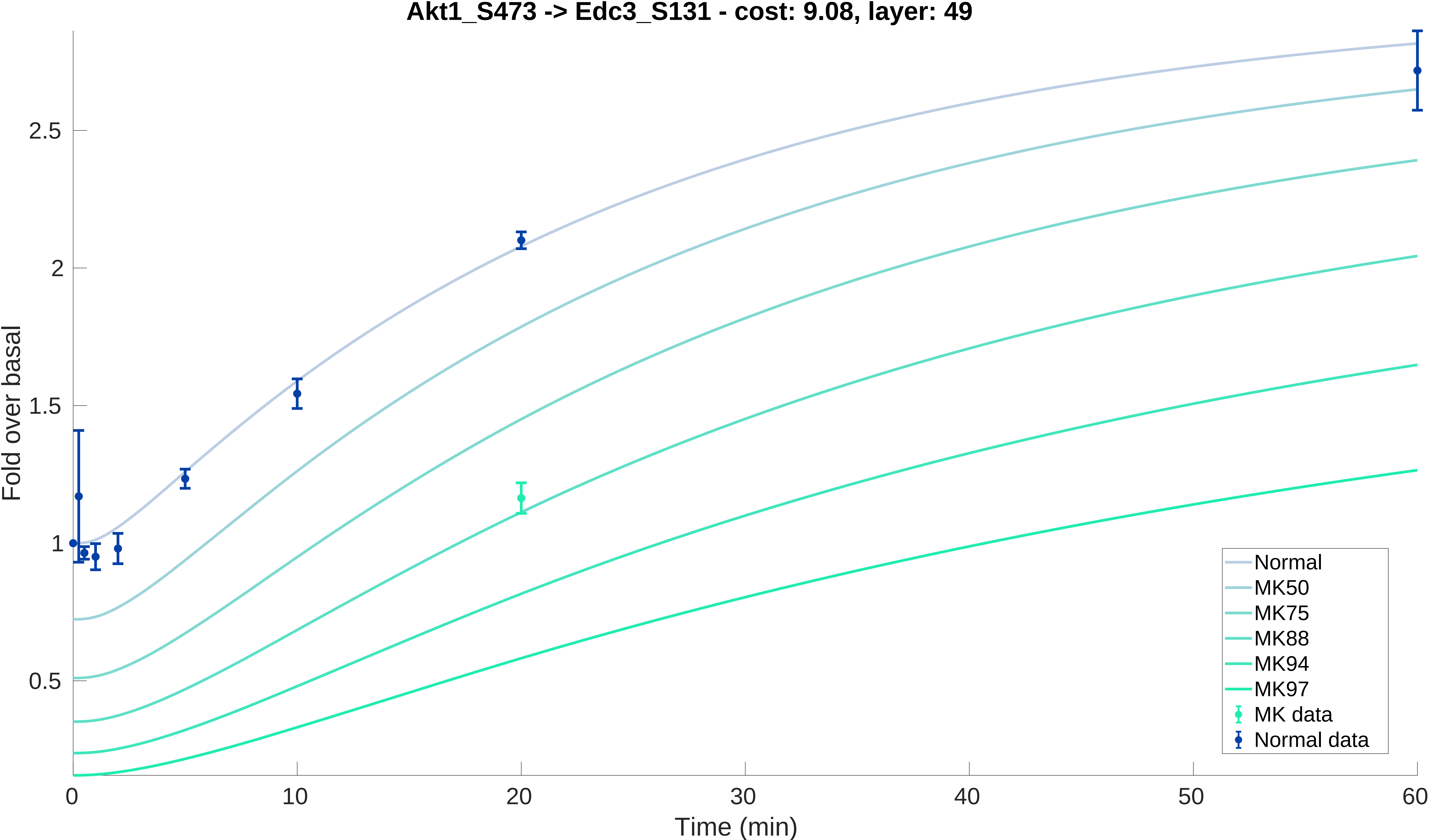
Akt1_S473 -> Edc3_S131 - cost: 9.08, layer: 49
Fold over basal
2.5
2
1.5
1
0.5
0
10
20
30
40
50
60
Time (min)
Normal
MK50
MK75
MK88
MK94
MK97
MK data
Normal data

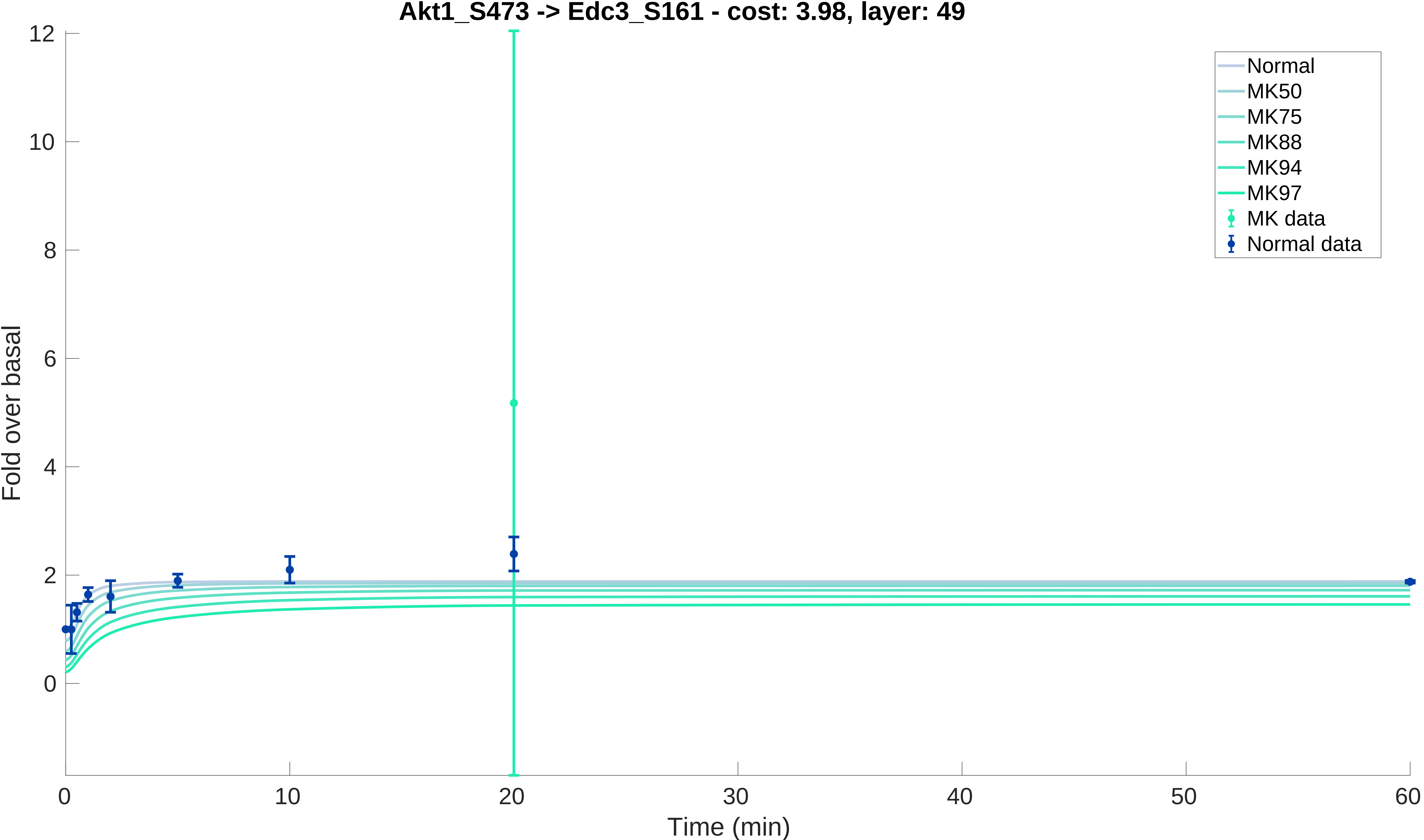

Akt1_S473 -> Edc3_S161 - cost: 3.98, layer: 49
Fold over basal
Time (min)
12
10
8
6
4
2
0
0
10
20
30
40
50
60
Normal
MK50
MK75
MK88
MK94
MK97
MK data
Normal data

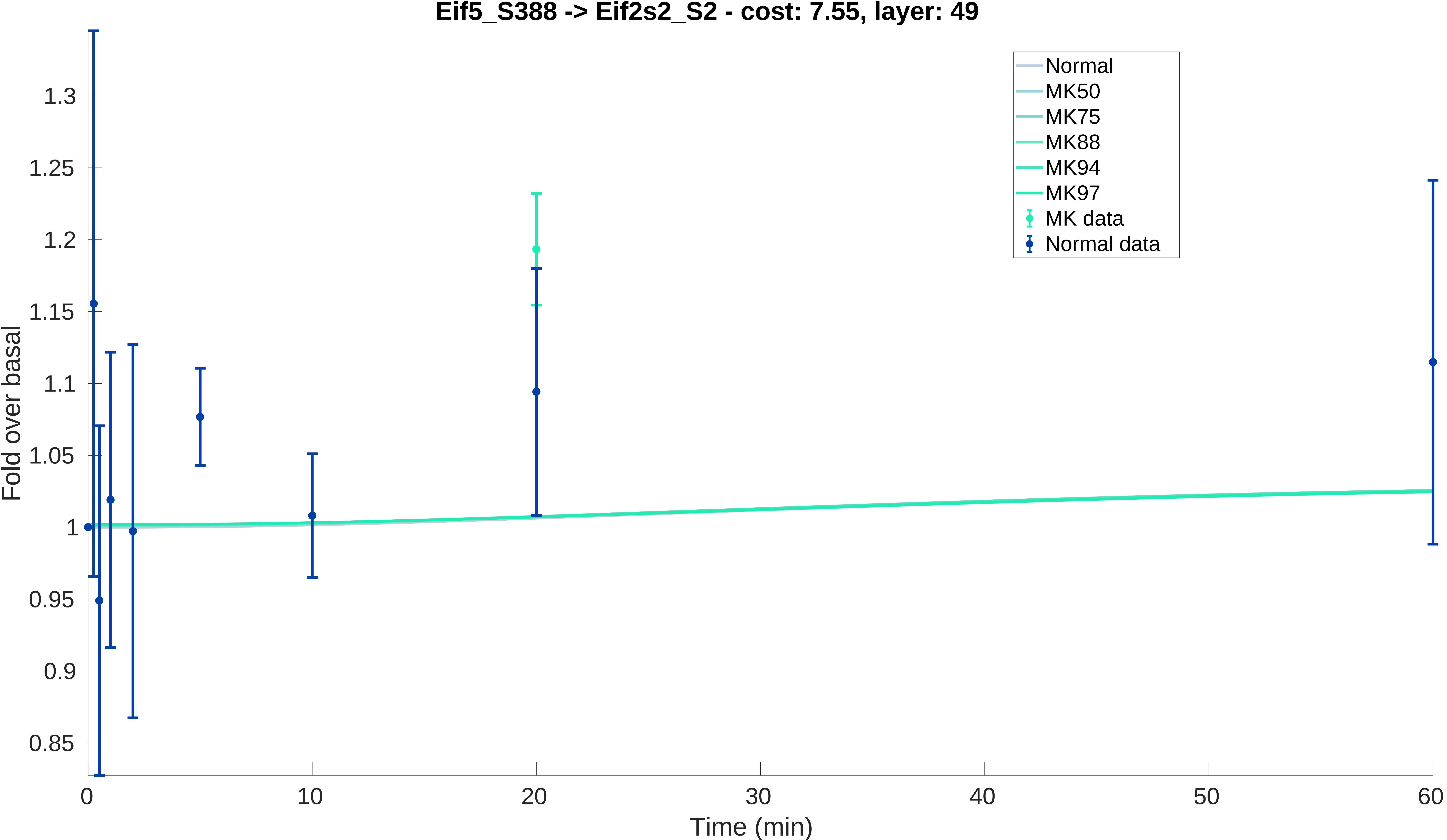

Eif5_S388 -> Eif2s2_S2 - cost: 7.55, layer: 49
Fold over basal
1.3
1.25
1.2
1.15
1.1
1.05
1
0.95
0.9
0.85
0
10
20
30
40
50
60
Time (min)
Normal
MK50
MK75
MK88
MK94
MK97
MK data
Normal data

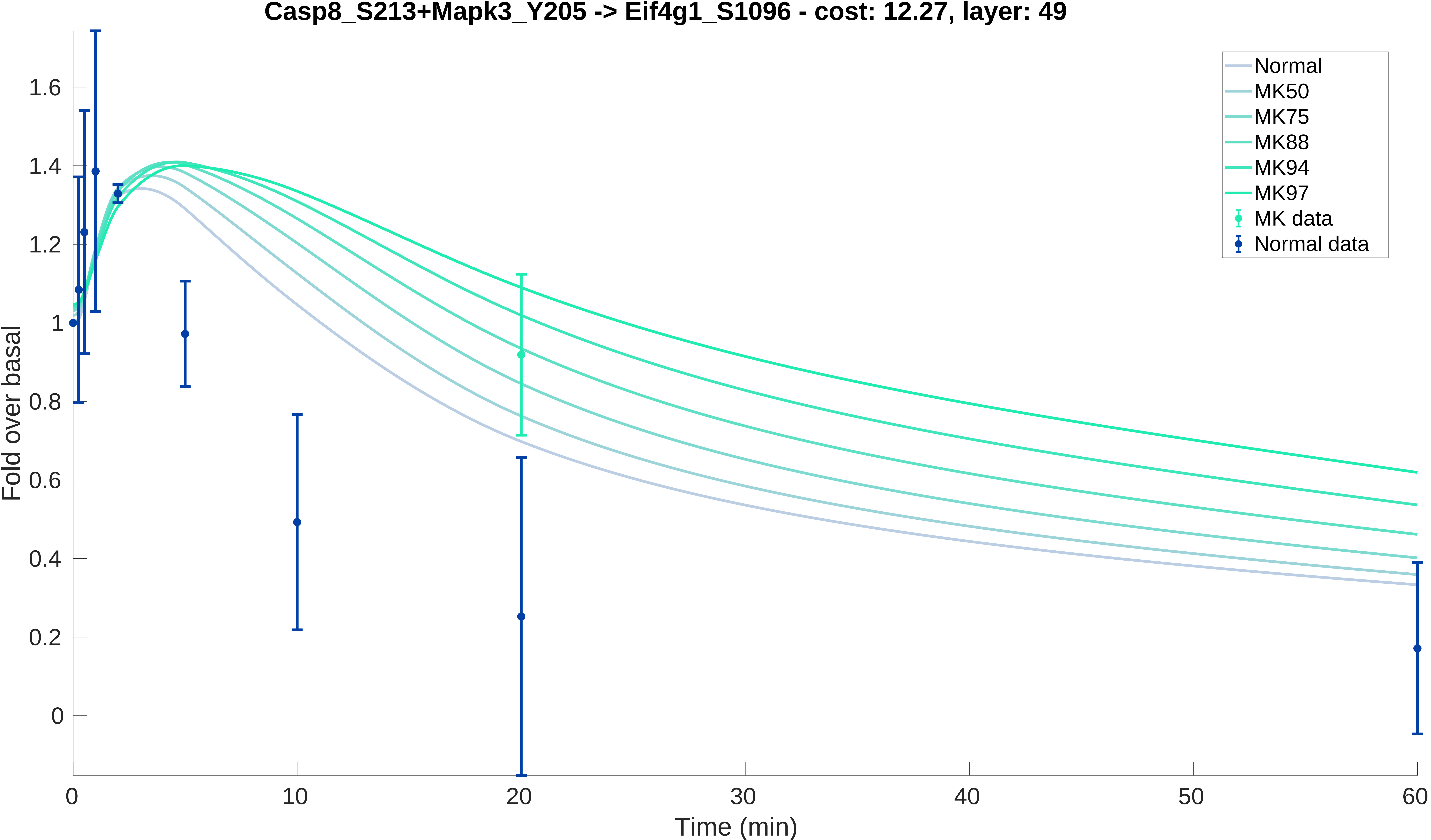

Casp8_S213+Mapk3_Y205 -> Eif4g1_S1096 - cost: 12.27, layer: 49
Fold over basal
Time (min)
0
0.2
0.4
0.6
0.8
1
1.2
1.4
1.6
0
10
20
30
40
50
60
Normal
MK50
MK75
MK88
MK94
MK97
MK data
Normal data

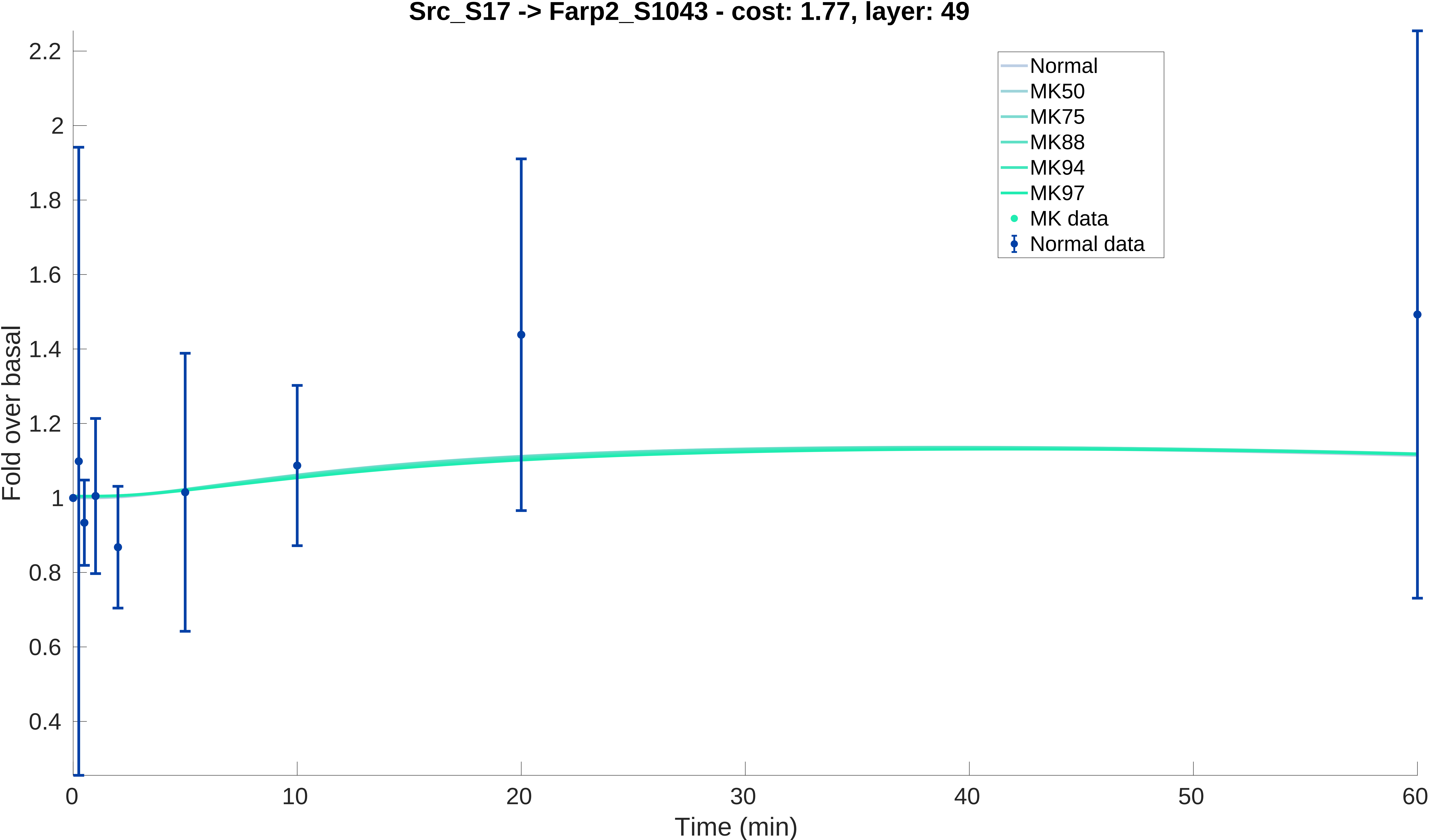
Src_S17 -> Farp2_S1043 - cost: 1.77, layer: 49
Fold over basal
Time (min)
Normal
MK50
MK75
MK88
MK94
MK97
MK data
Normal data
0.4
0.6
0.8
1
1.2
1.4
1.6
1.8
2
2.2
0
10
20
30
40
50
60

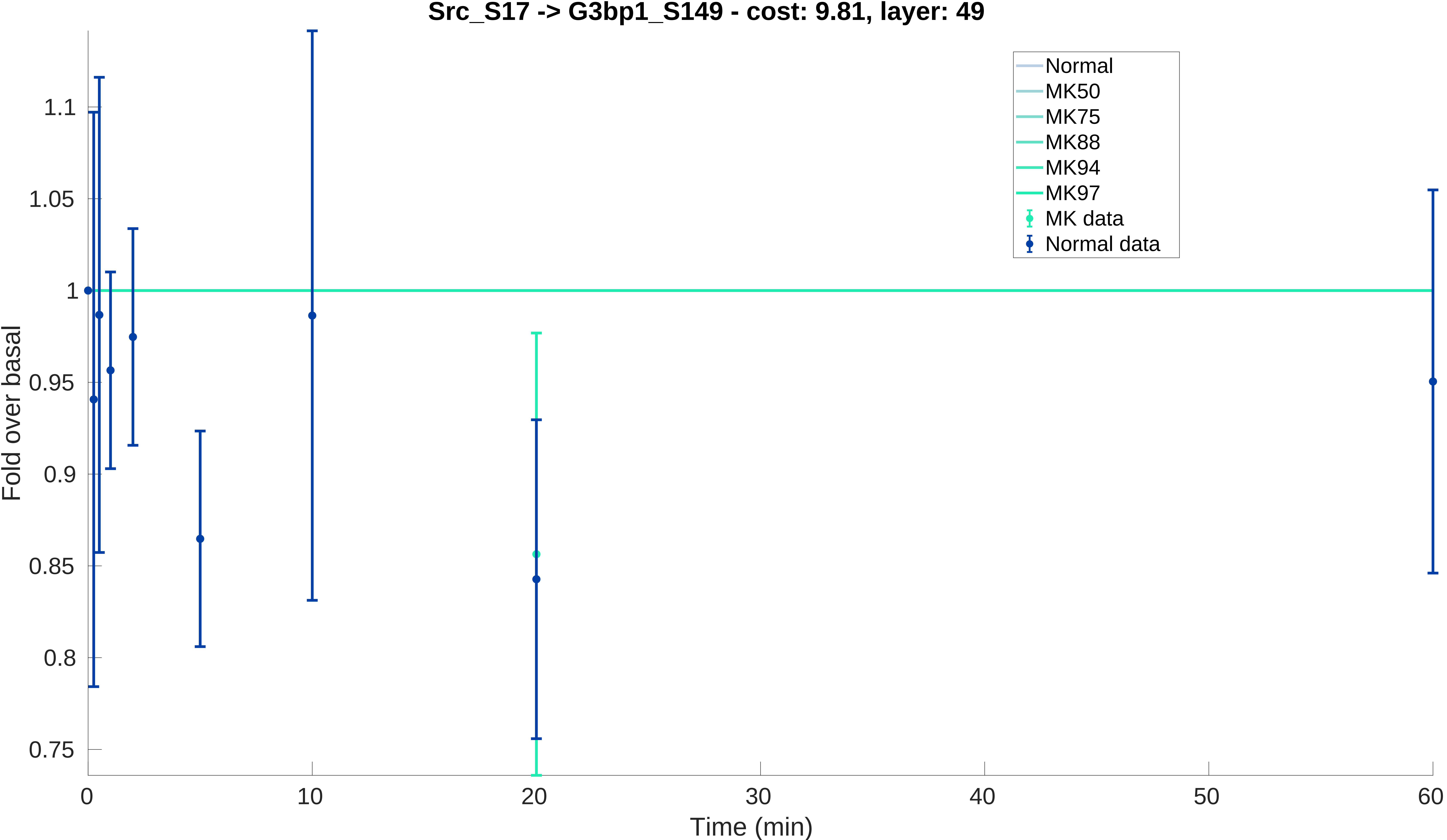
Src_S17 -> G3bp1_S149 - cost: 9.81, layer: 49
Normal
MK50
MK75
MK88
MK94
MK97
MK data
Normal data
Fold over basal
1.1
1.05
1
0.95
0.9
0.85
0.8
0.75
0
10
20
30
40
50
60
Time (min)

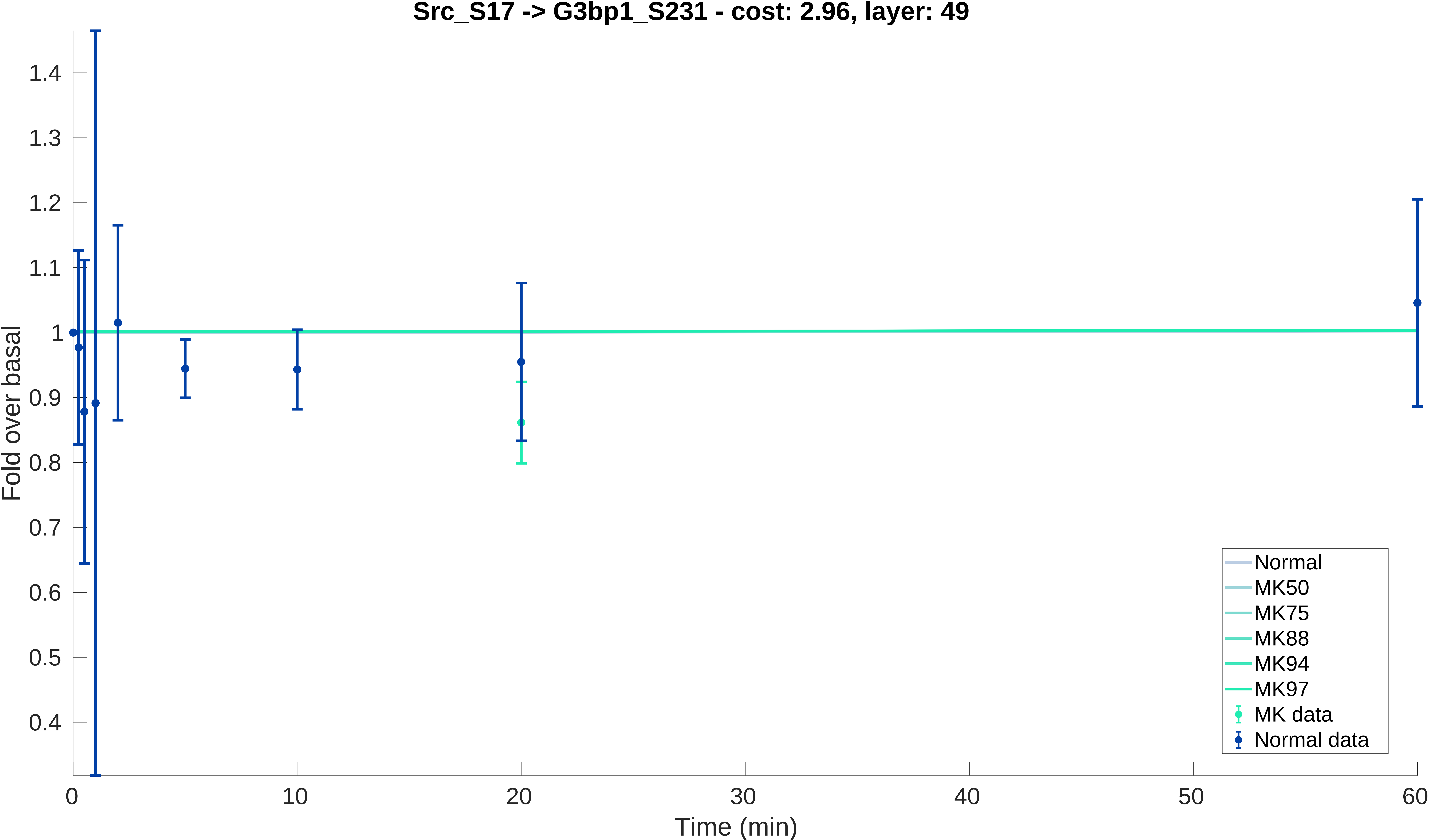
Src_S17 -> G3bp1_S231 - cost: 2.96, layer: 49
Fold over basal
1.4
1.3
1.2
1.1
1
0.9
0.8
0.7
0.6
0.5
0.4
0
10
20
30
40
50
60
Time (min)
Normal
MK50
MK75
MK88
MK94
MK97
MK data
Normal data

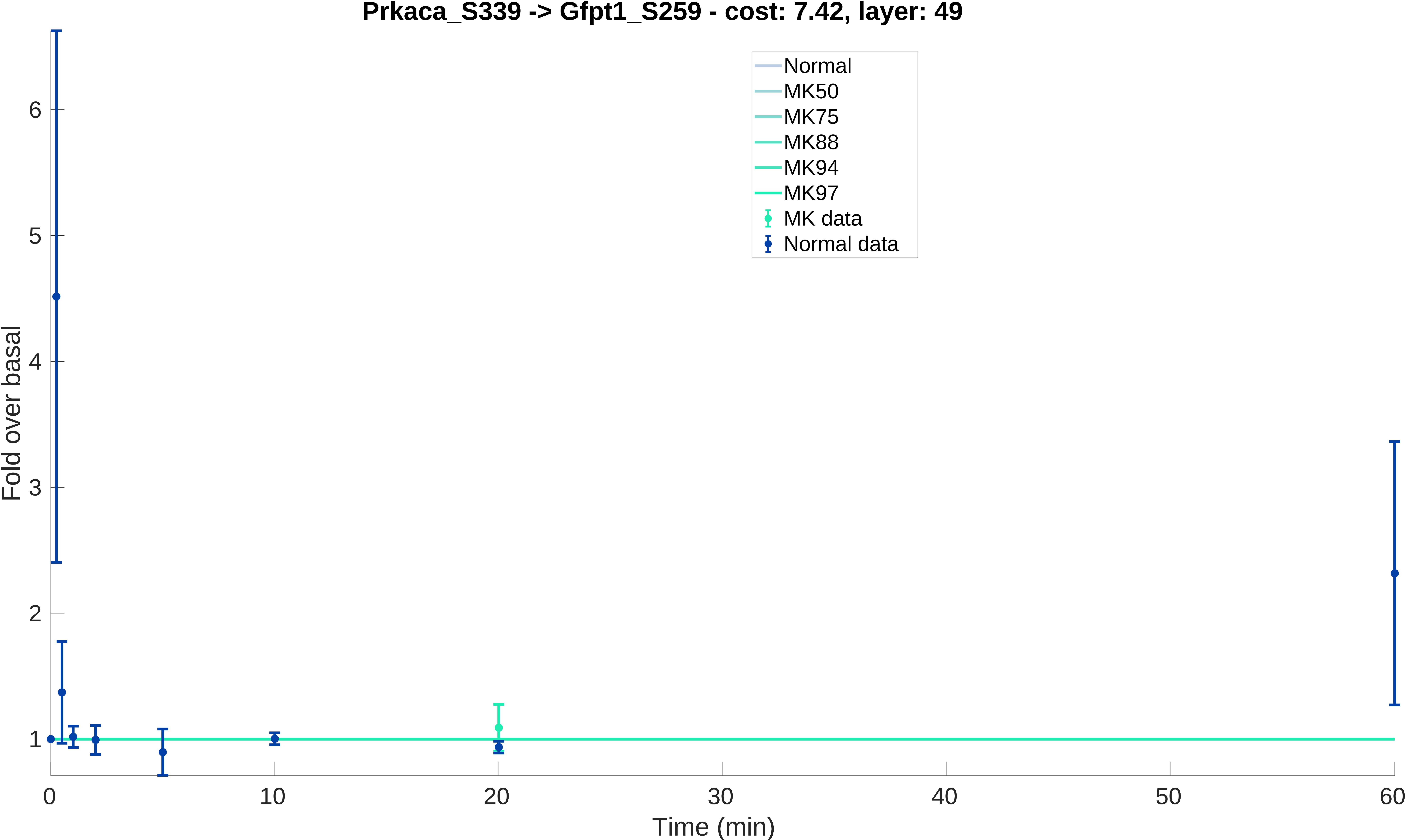

Prkaca_S339 -> Gfpt1_S259 - cost: 7.42, layer: 49
Normal
MK50
MK75
MK88
MK94
MK97
MK data
Normal data
Fold over basal
1
2
3
4
5
6
0
10
20
30
40
50
60
Time (min)

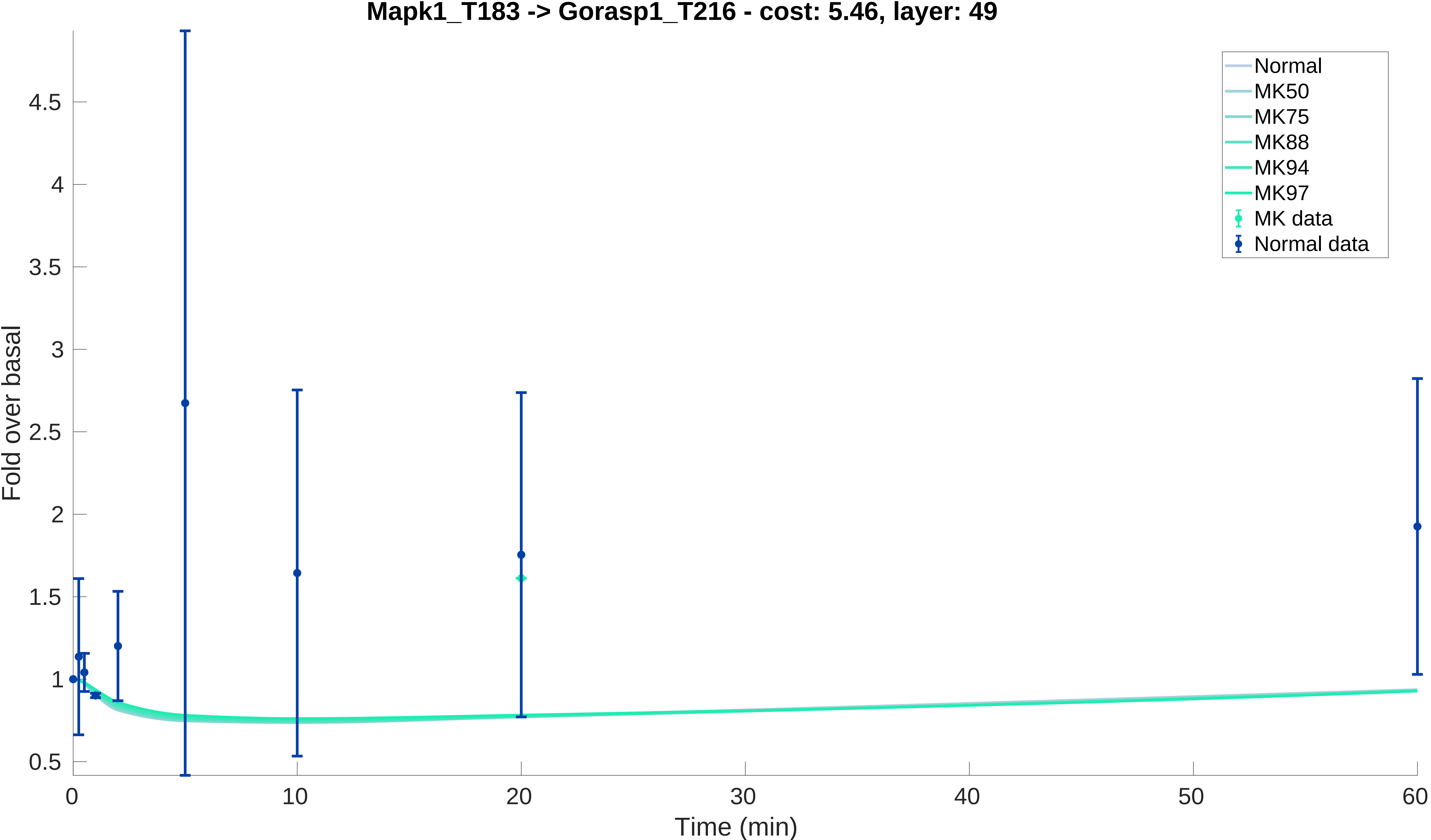

Mapk1_T183 -> Gorasp1_T216 - cost: 5.46, layer: 49
Normal
MK50
MK75
MK88
MK94
MK97
MK data
Normal data
Fold over basal
Time (min)
0
10
20
30
40
50
60
0.5
1
1.5
2
2.5
3
3.5
4
4.5

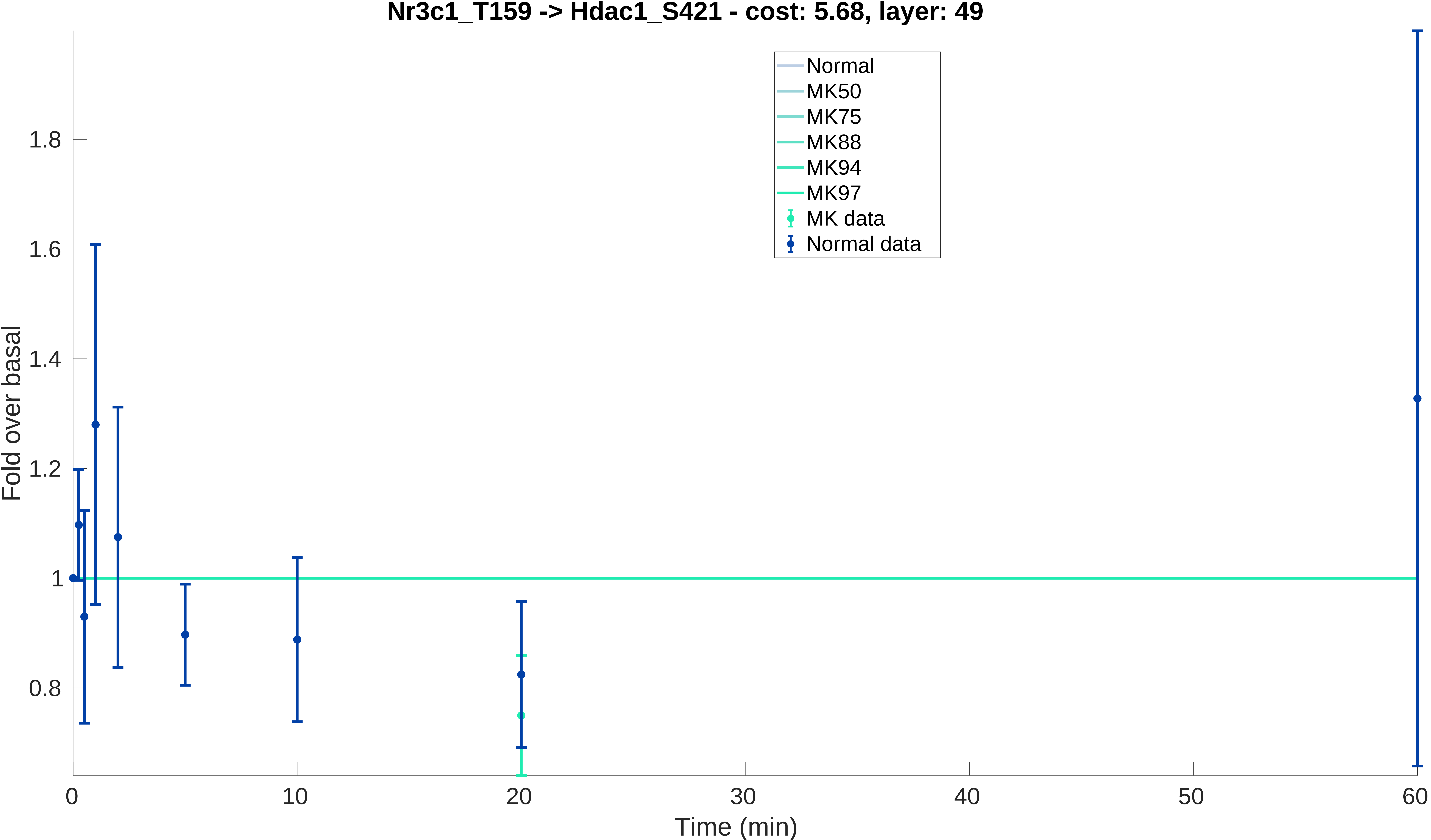
Nr3c1_T159 -> Hdac1_S421 - cost: 5.68, layer: 49
Normal
MK50
MK75
MK88
MK94
MK97
MK data
Normal data
Fold over basal
1.8
1.6
1.4
1.2
1
0.8
0
10
20
30
40
50
60
Time (min)

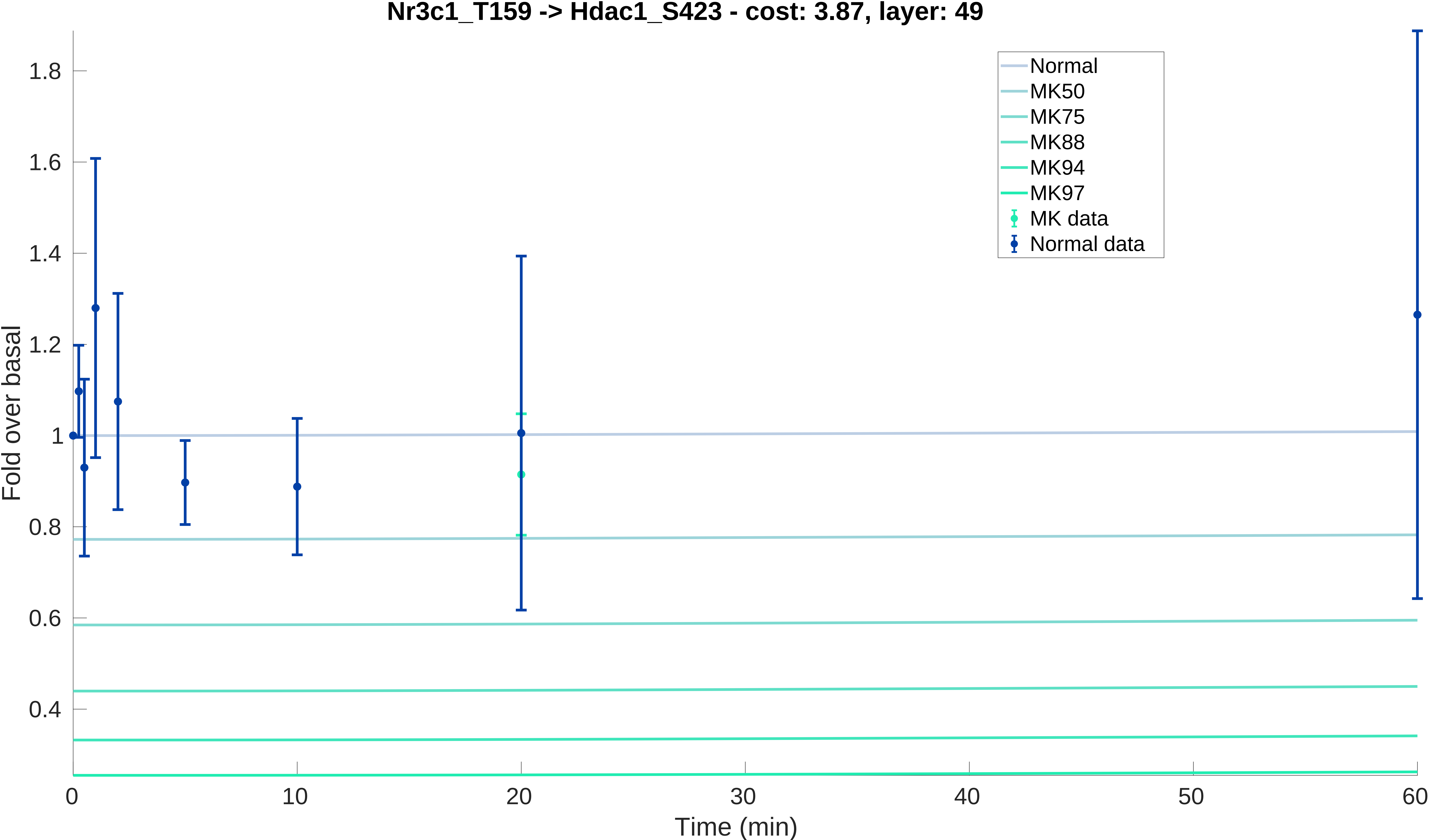

Nr3c1_T159 -> Hdac1_S423 - cost: 3.87, layer: 49
Fold over basal
1.8
1.6
1.4
1.2
1
0.8
0.6
0.4
Time (min)
0
10
20
30
40
50
60
Normal
MK50
MK75
MK88
MK94
MK97
MK data
Normal data

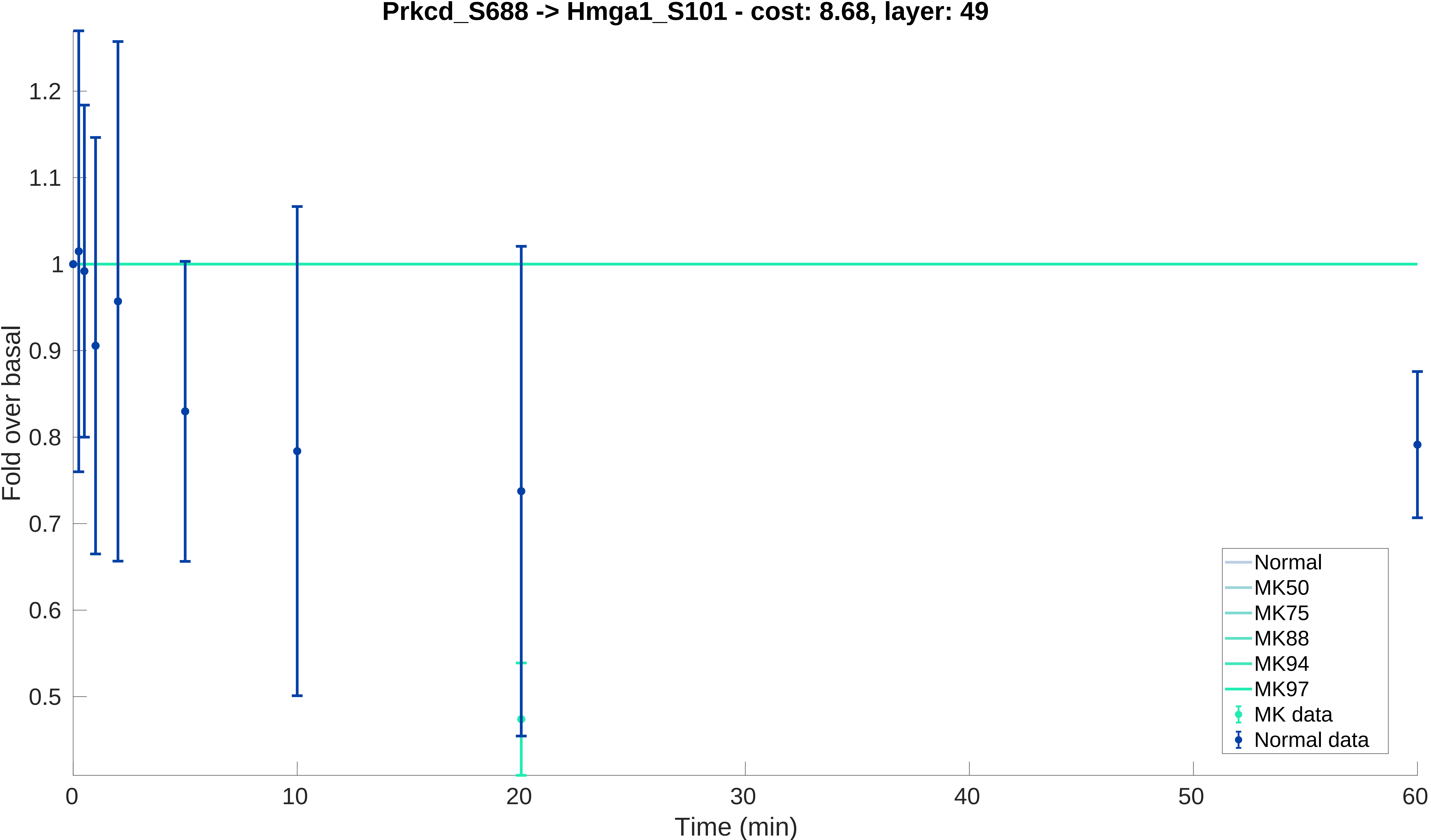

Prkcd_S688 -> Hmga1_S101 - cost: 8.68, layer: 49
Fold over basal
1.2
1.1
1
0.9
0.8
0.7
0.6
0.5
0
10
20
30
40
50
60
Time (min)
Normal
MK50
MK75
MK88
MK94
MK97
MK data
Normal data

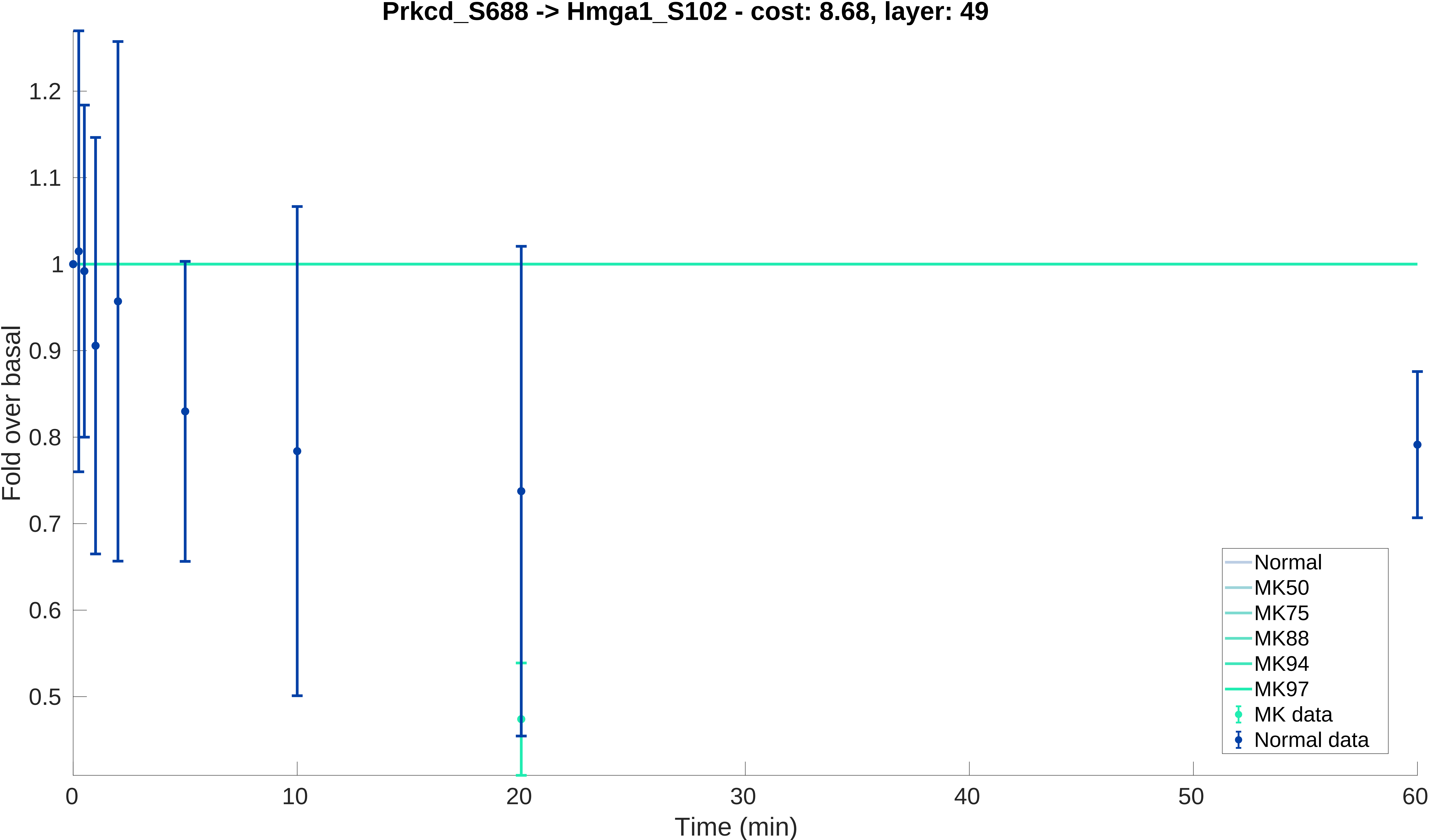

Prkcd_S688 -> Hmga1_S102 - cost: 8.68, layer: 49
Fold over basal
Time (min)
1.2
1.1
1
0.9
0.8
0.7
0.6
0.5
0
10
20
30
40
50
60
Normal
MK50
MK75
MK88
MK94
MK97
MK data
Normal data

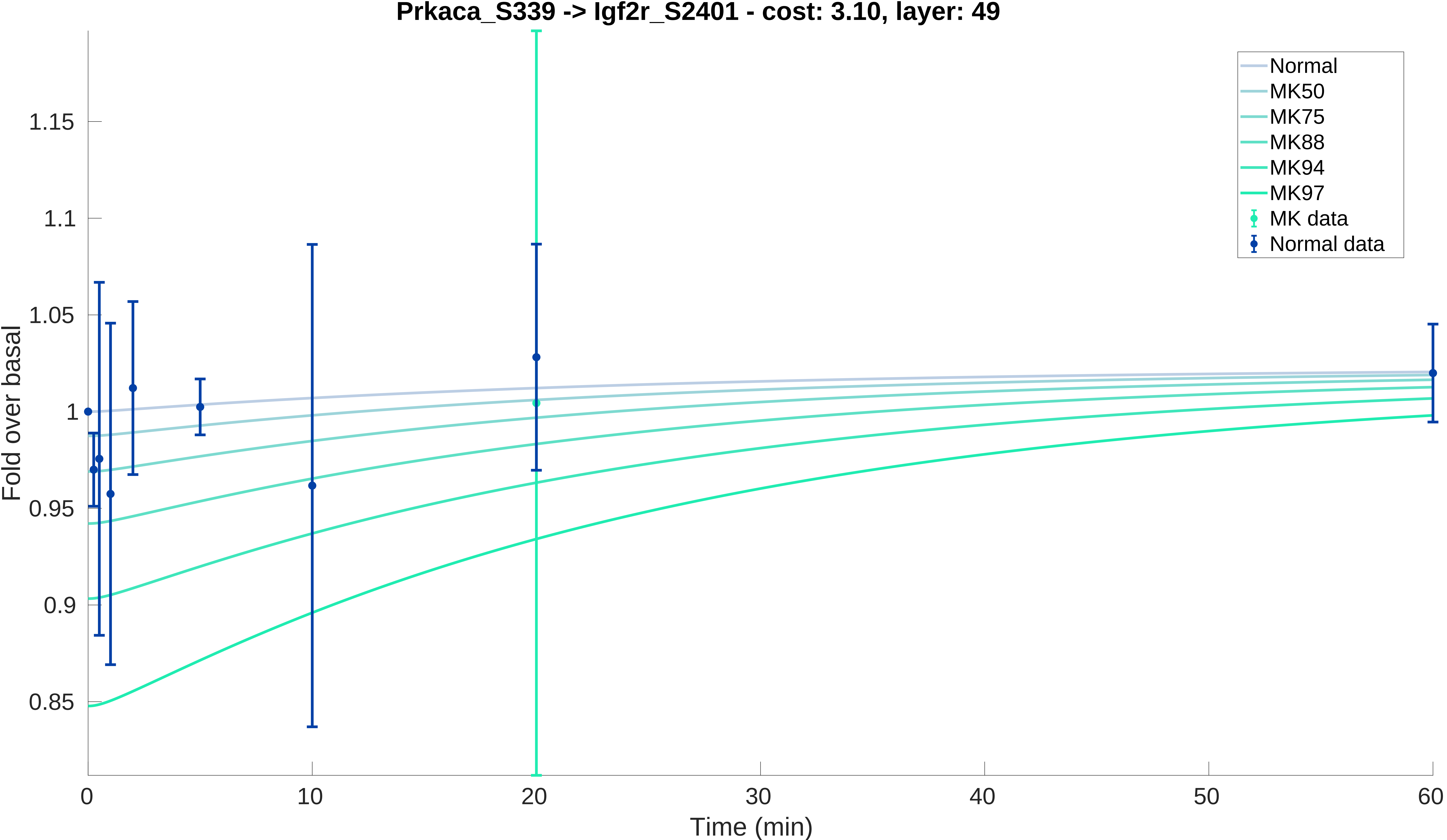
Prkaca_S339 -> Igf2r_S2401 - cost: 3.10, layer: 49
Fold over basal
1.15
1.1
1.05
1
0.95
0.9
0.85
0
10
20
30
40
50
60
Time (min)
Normal
MK50
MK75
MK88
MK94
MK97
MK data
Normal data

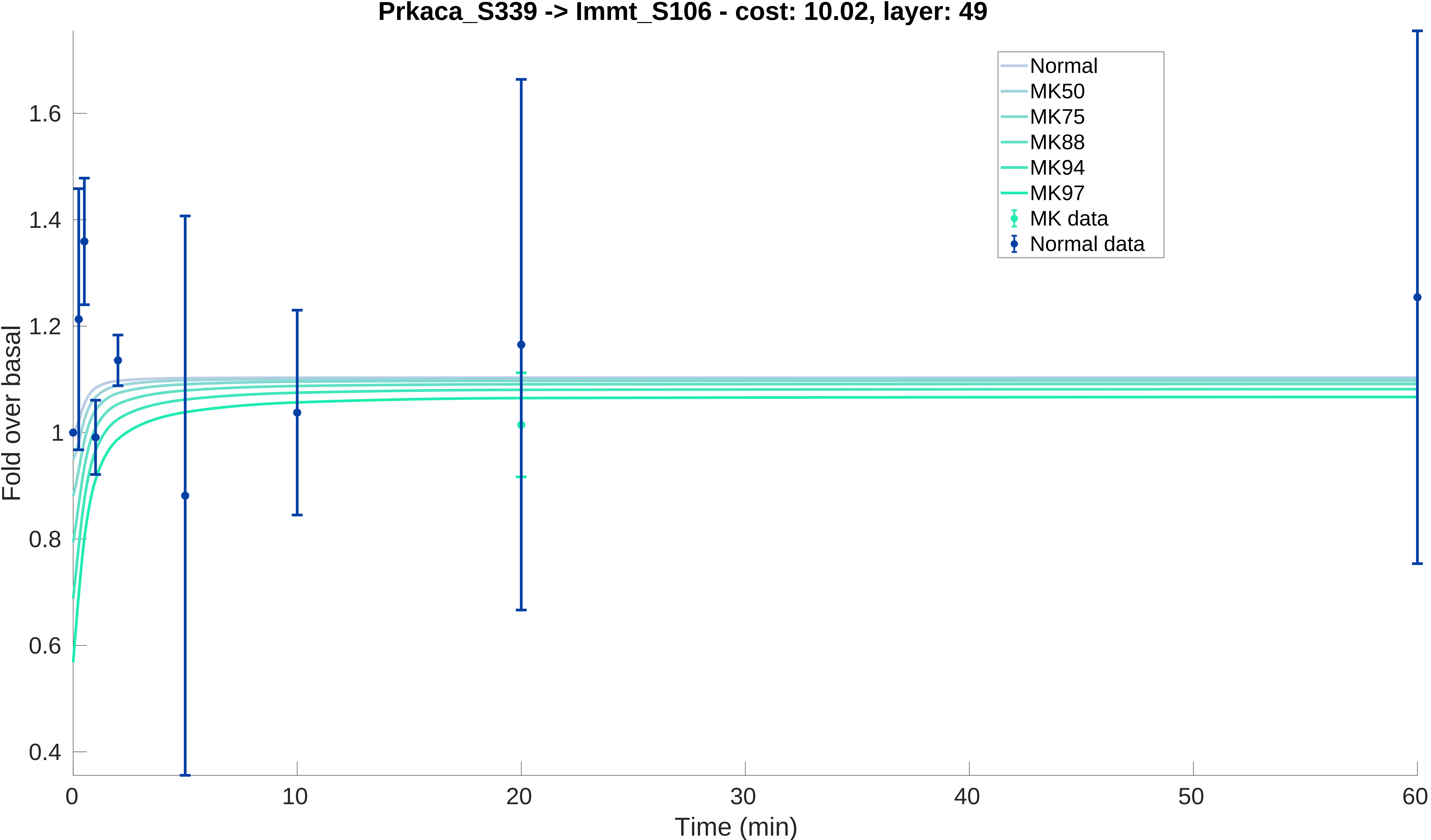

Prkaca_S339 -> Immt_S106 - cost: 10.02, layer: 49
Normal
MK50
MK75
MK88
MK94
MK97
MK data
Normal data
Fold over basal
1.6
1.4
1.2
1
0.8
0.6
0.4
0
10
20
30
40
50
60
Time (min)

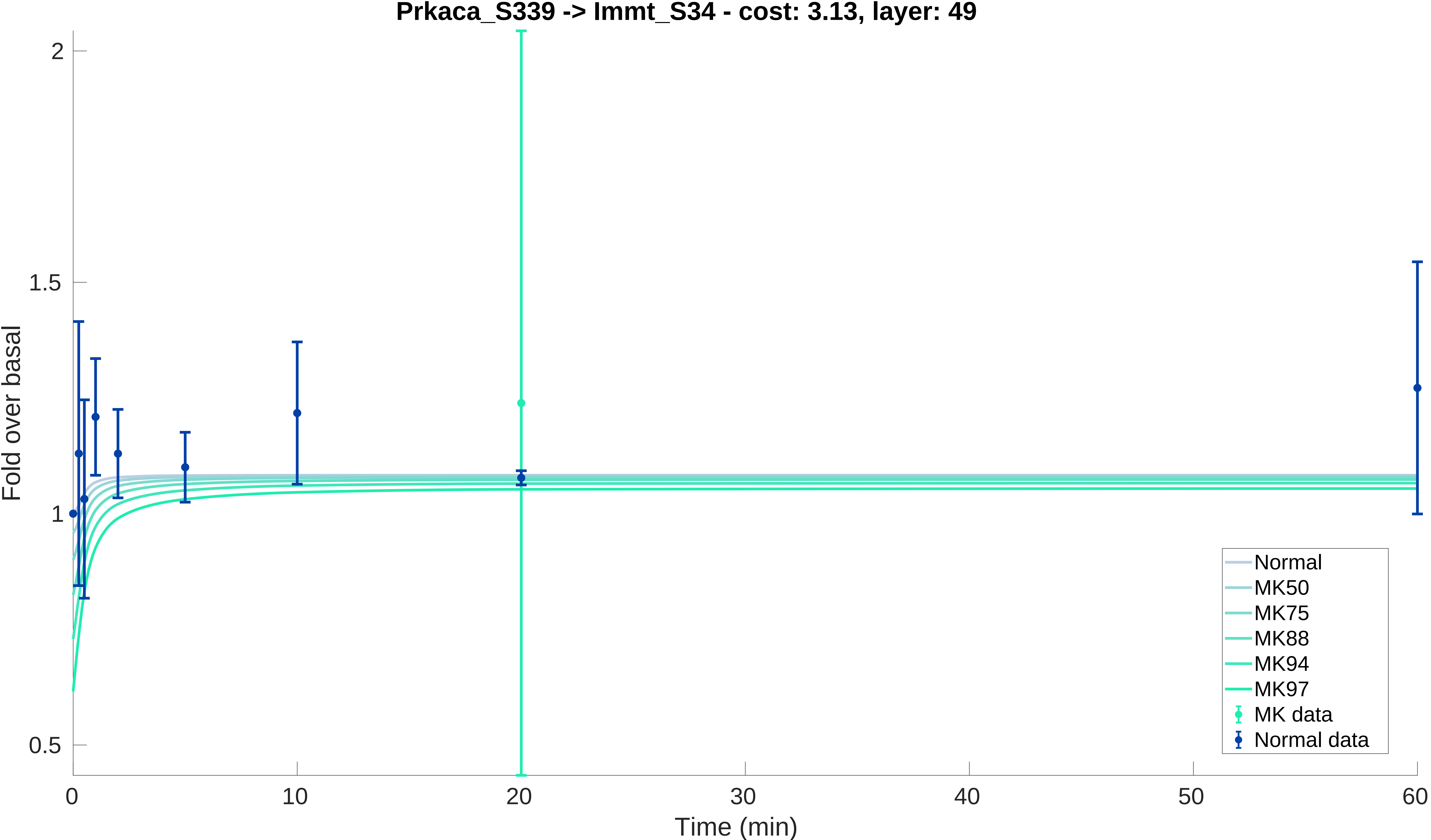

Prkaca_S339 -> Immt_S34 - cost: 3.13, layer: 49
Fold over basal
0.5
1
1.5
2
0
10
20
30
40
50
60
Time (min)
Normal
MK50
MK75
MK88
MK94
MK97
MK data
Normal data

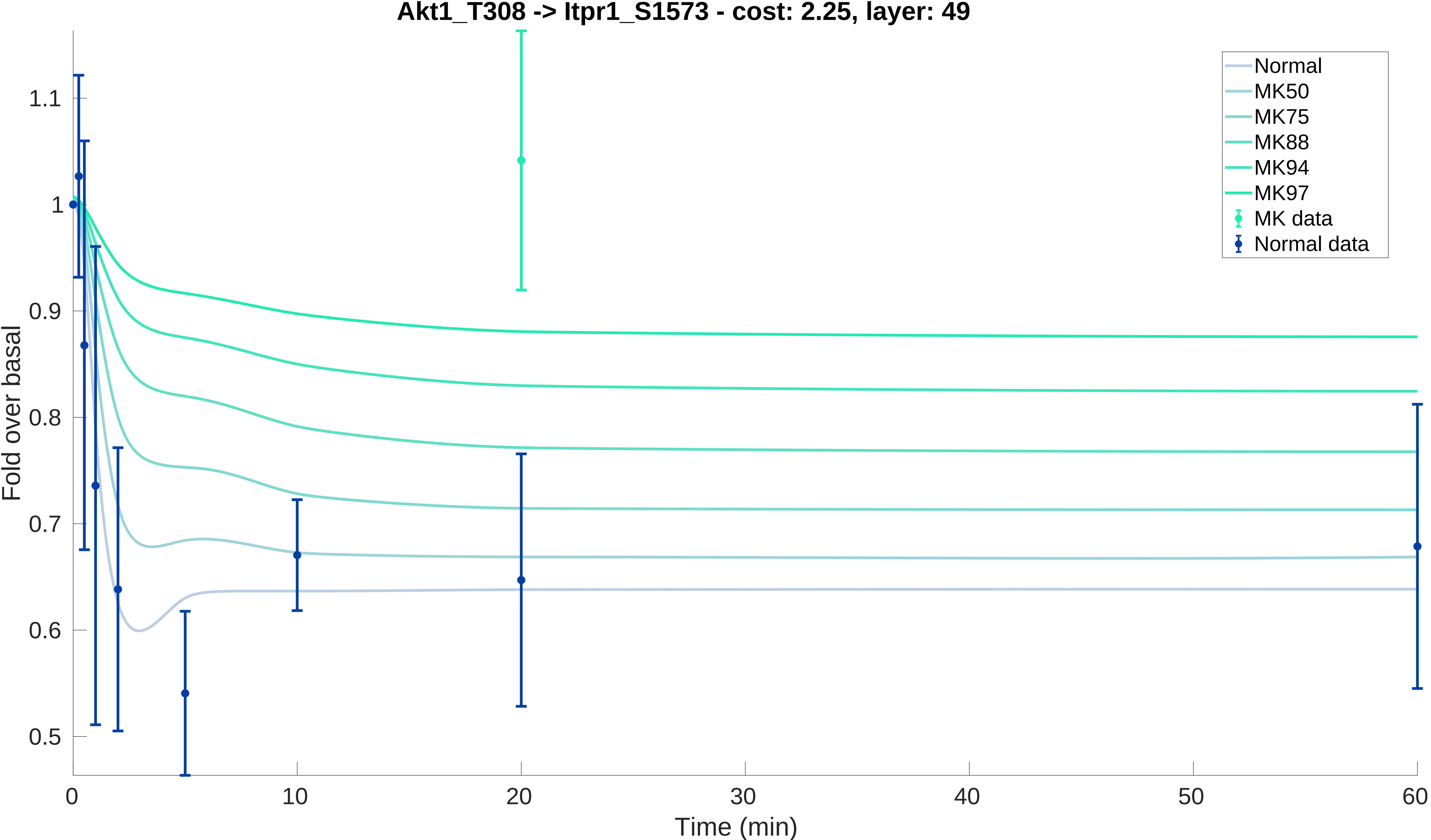
Akt1_T308 -> Itpr1_S1573 - cost: 2.25, layer: 49
Fold over basal
Time (min)
0.5
0.6
0.7
0.8
0.9
1
1.1
0
10
20
30
40
50
60
Normal
MK50
MK75
MK88
MK94
MK97
MK data
Normal data

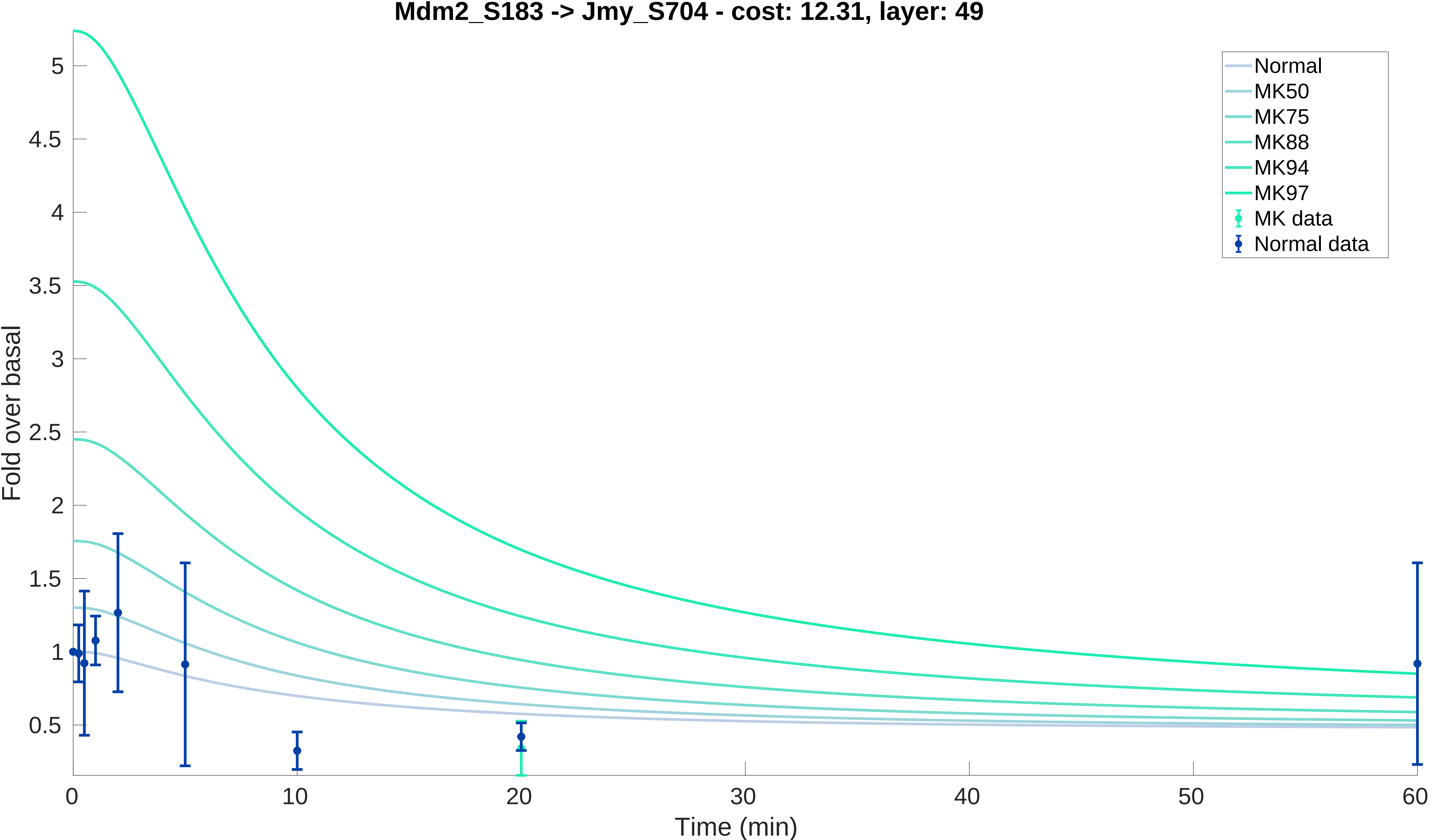

Mdm2_S183 -> Jmy_S704 - cost: 12.31, layer: 49
Fold over basal
Time (min)
0
10
20
30
40
50
60
0.5
1
1.5
2
2.5
3
3.5
4
4.5
5
Normal
MK50
MK75
MK88
MK94
MK97
MK data
Normal data

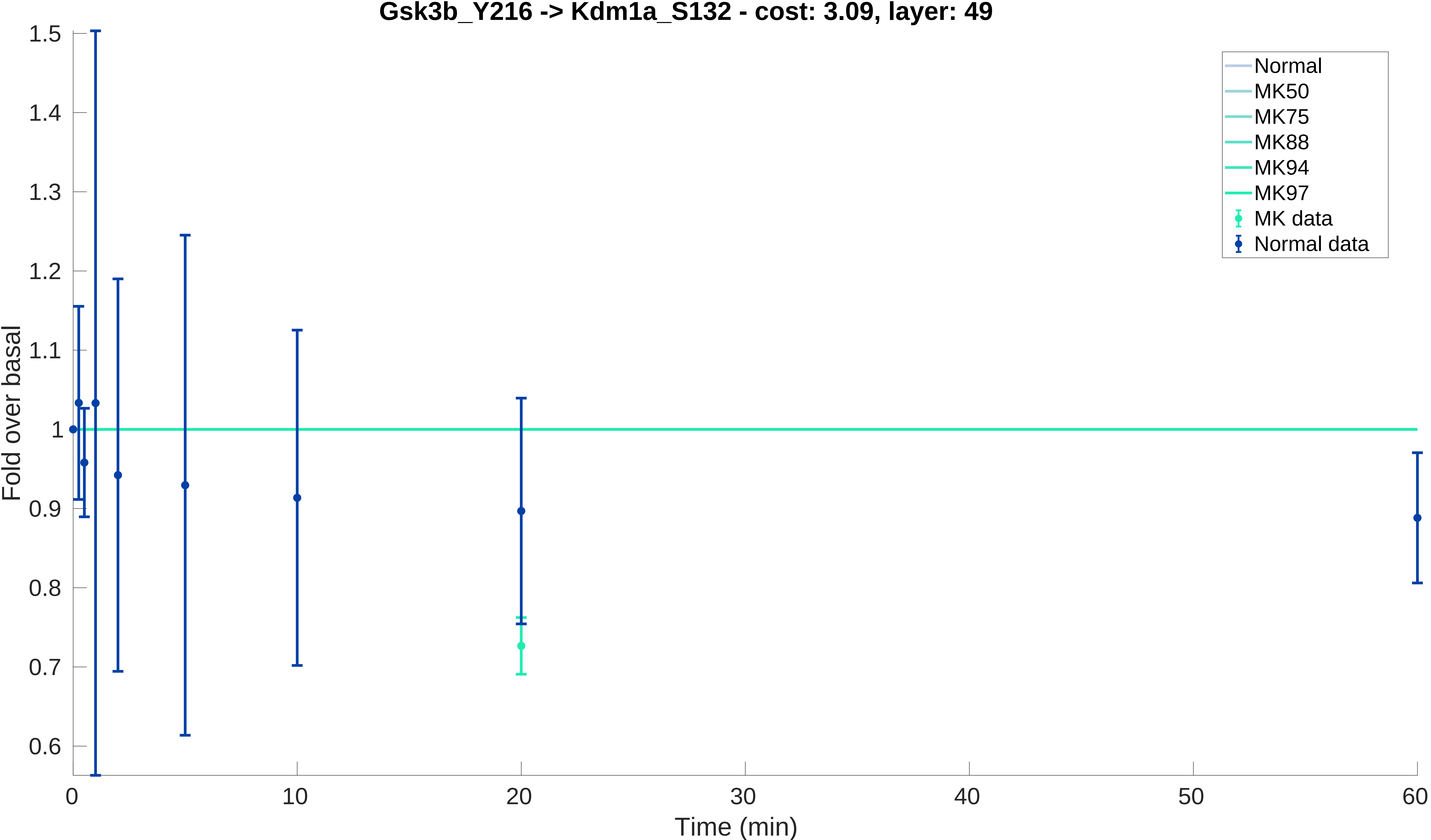
Gsk3b_Y216 -> Kdm1a_S132 - cost: 3.09, layer: 49
Fold over basal
1.5
1.4
1.3
1.2
1.1
1
0.9
0.8
0.7
0.6
0
10
20
30
40
50
60
Time (min)
Normal
MK50
MK75
MK88
MK94
MK97
MK data
Normal data

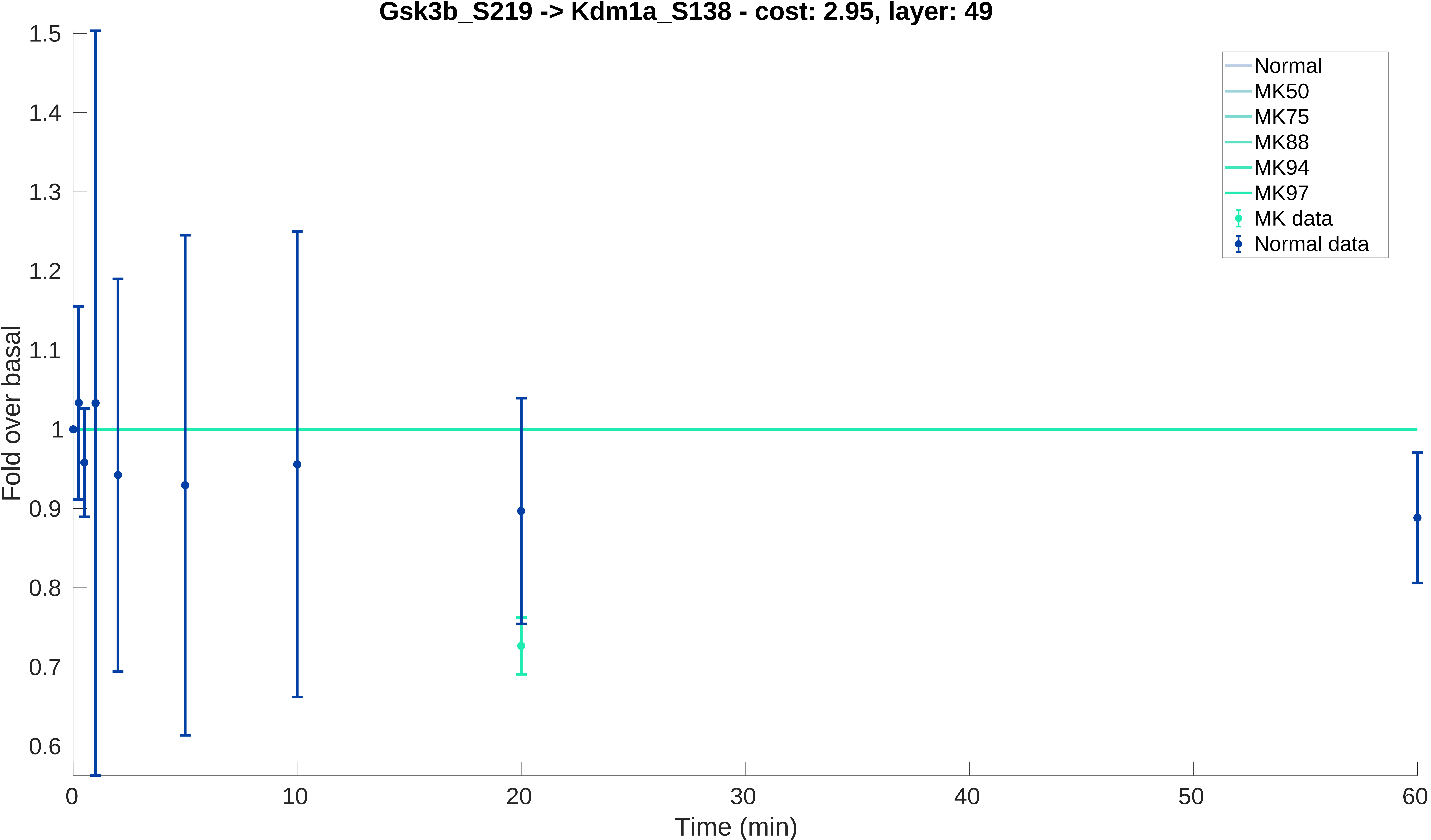
Gsk3b_S219 -> Kdm1a_S138 - cost: 2.95, layer: 49
Fold over basal
1.5
1.4
1.3
1.2
1.1
1
0.9
0.8
0.7
0.6
0
10
20
30
40
50
60
Time (min)
Normal
MK50
MK75
MK88
MK94
MK97
MK data
Normal data

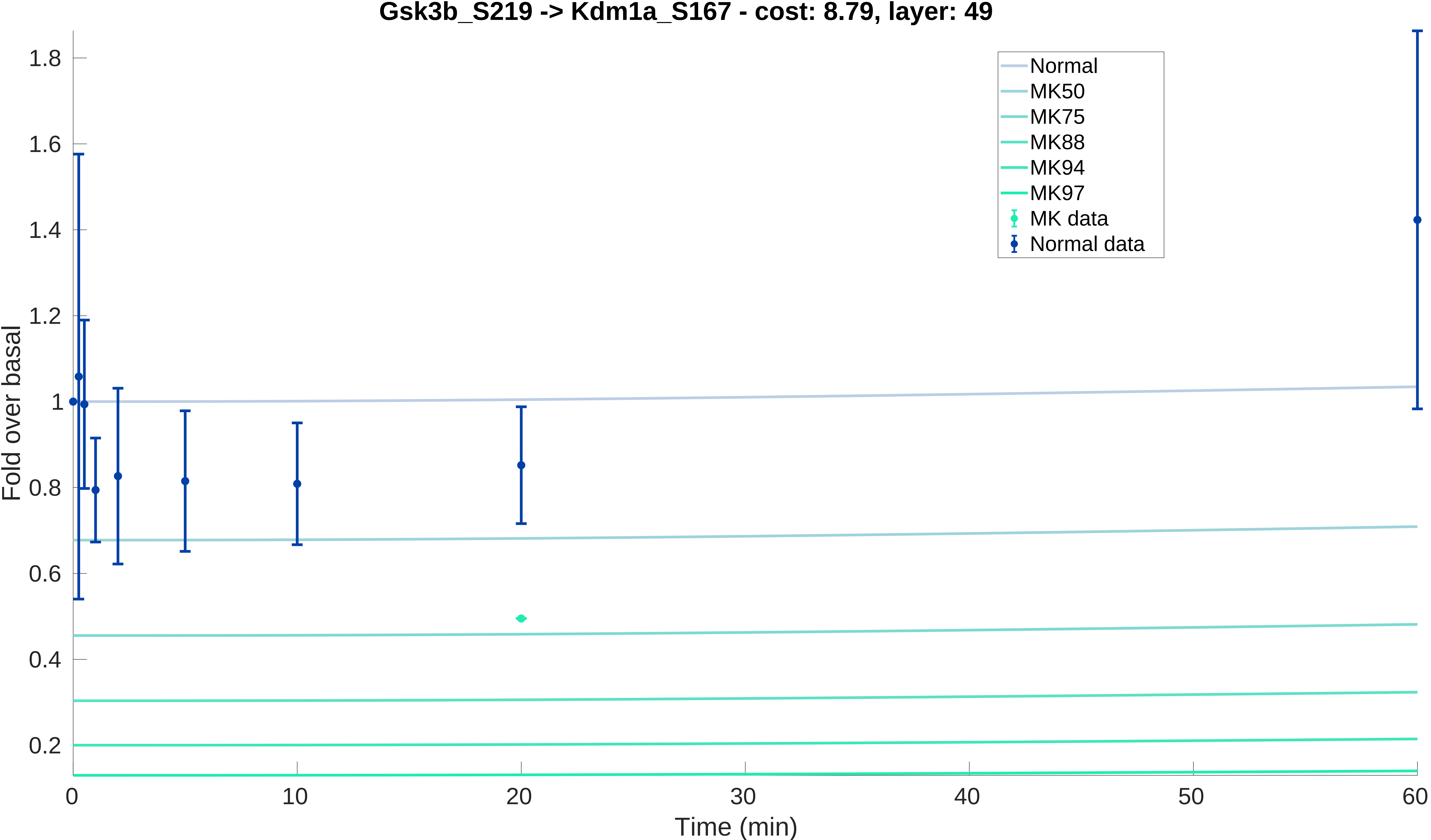

Gsk3b_S219 -> Kdm1a_S167 - cost: 8.79, layer: 49
Normal
MK50
MK75
MK88
MK94
MK97
MK data
Normal data
Fold over basal
1.8
1.6
1.4
1.2
1
0.8
0.6
0.4
0.2
0
10
20
30
40
50
60
Time (min)

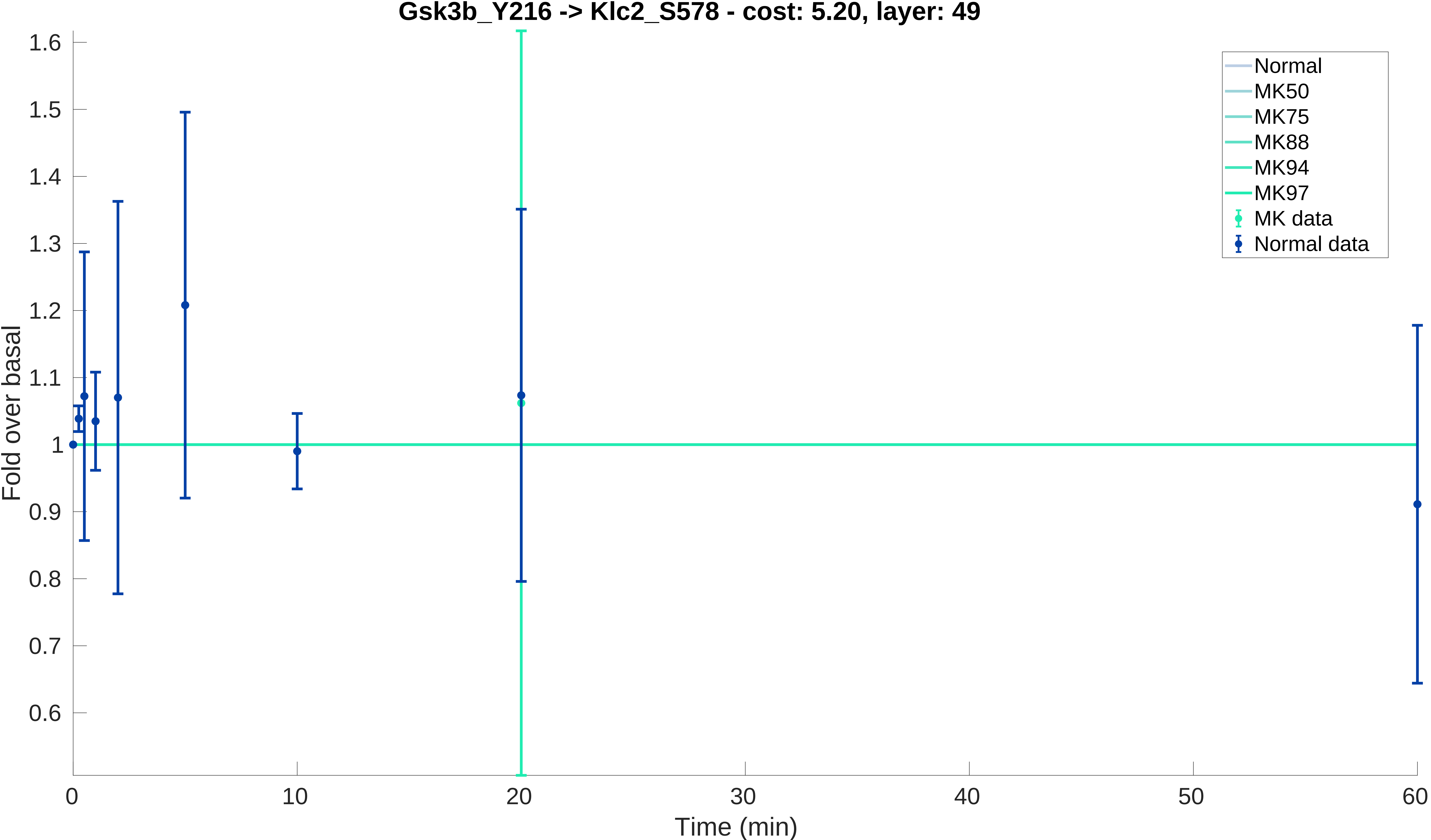

Gsk3b_Y216 -> Klc2_S578 - cost: 5.20, layer: 49
Fold over basal
Time (min)
1.6
1.5
1.4
1.3
1.2
1.1
1
0.9
0.8
0.7
0.6
0
10
20
30
40
50
60
Normal
MK50
MK75
MK88
MK94
MK97
MK data
Normal data

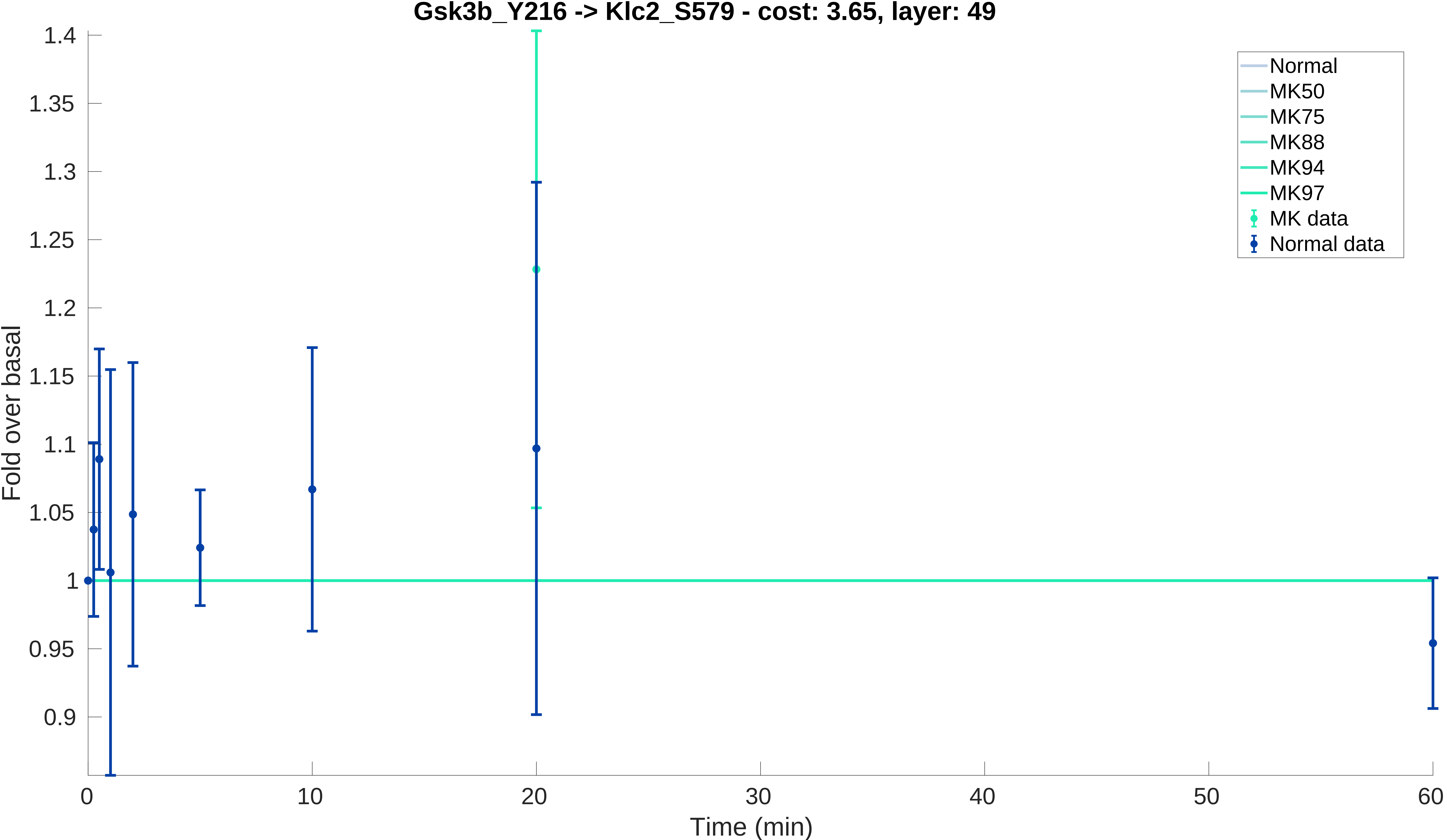
Gsk3b_Y216 -> Klc2_S579 - cost: 3.65, layer: 49
Fold over basal
Time (min)
1.4
1.35
1.3
1.25
1.2
1.15
1.1
1.05
1
0.95
0.9
0
10
20
30
40
50
60
Normal
MK50
MK75
MK88
MK94
MK97
MK data
Normal data

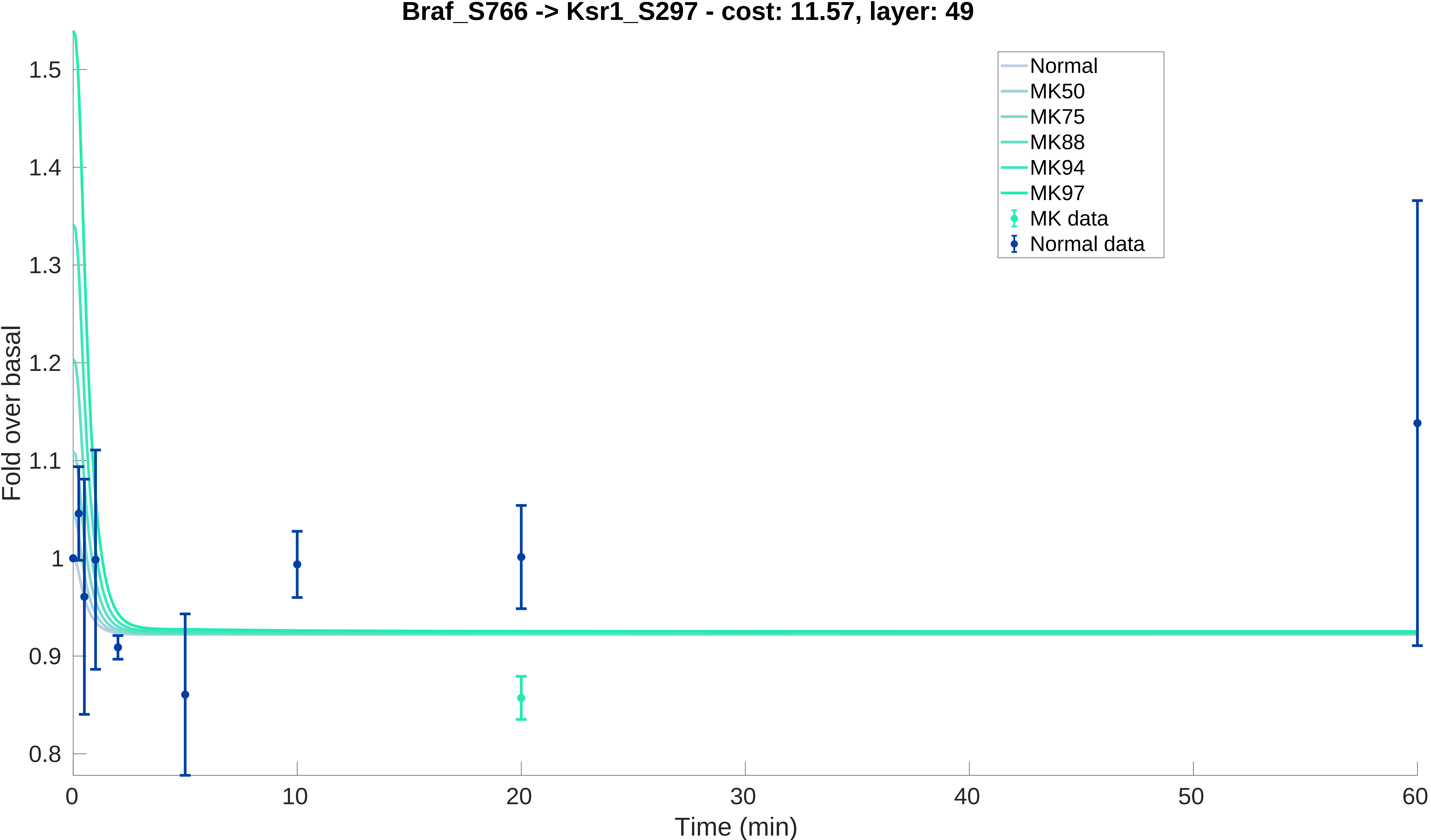

Braf_S766 -> Ksr1_S297 - cost: 11.57, layer: 49
Fold over basal
Time (min)
0.8
0.9
1
1.1
1.2
1.3
1.4
1.5
0
10
20
30
40
50
60
Normal
MK50
MK75
MK88
MK94
MK97
MK data
Normal data

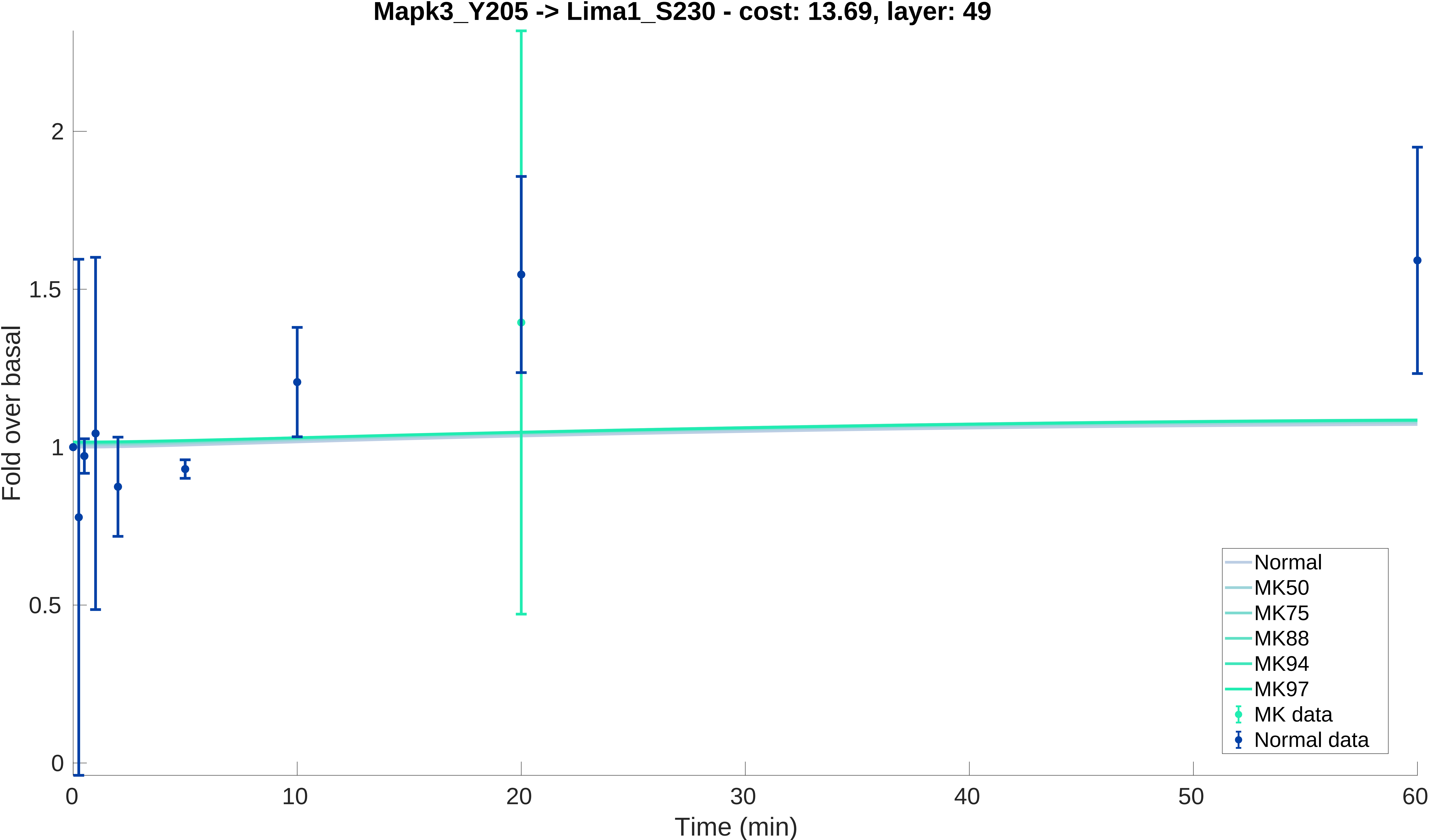

Mapk3_Y205 -> Lima1_S230 - cost: 13.69, layer: 49
Fold over basal
Time (min)
0
0.5
1
1.5
2
0
10
20
30
40
50
60
Normal
MK50
MK75
MK88
MK94
MK97
MK data
Normal data

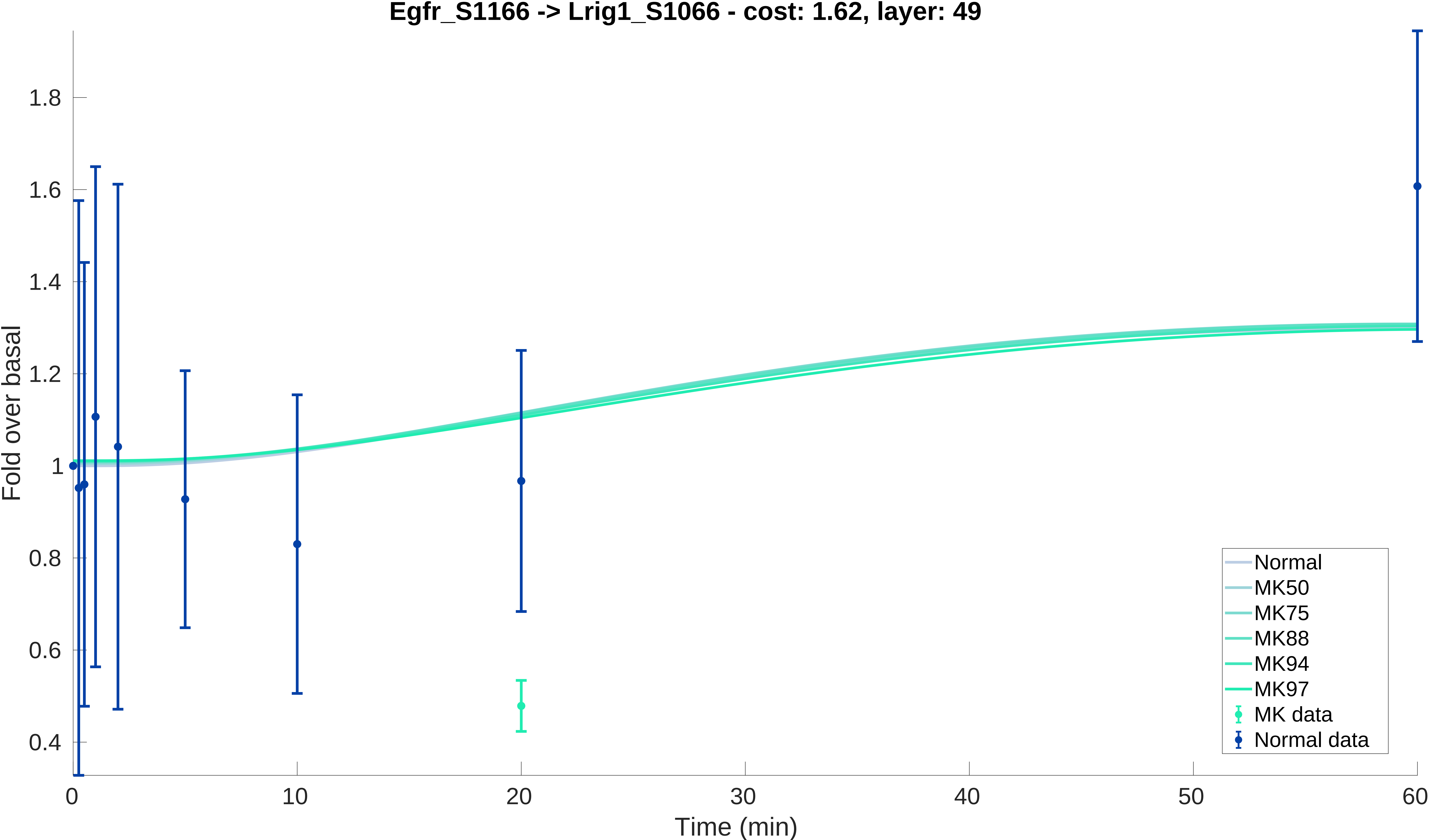
Egfr_S1166 -> Lrig1_S1066 - cost: 1.62, layer: 49
Fold over basal
Time (min)
1.8
1.6
1.4
1.2
1
0.8
0.6
0.4
0
10
20
30
40
50
60
Normal
MK50
MK75
MK88
MK94
MK97
MK data
Normal data

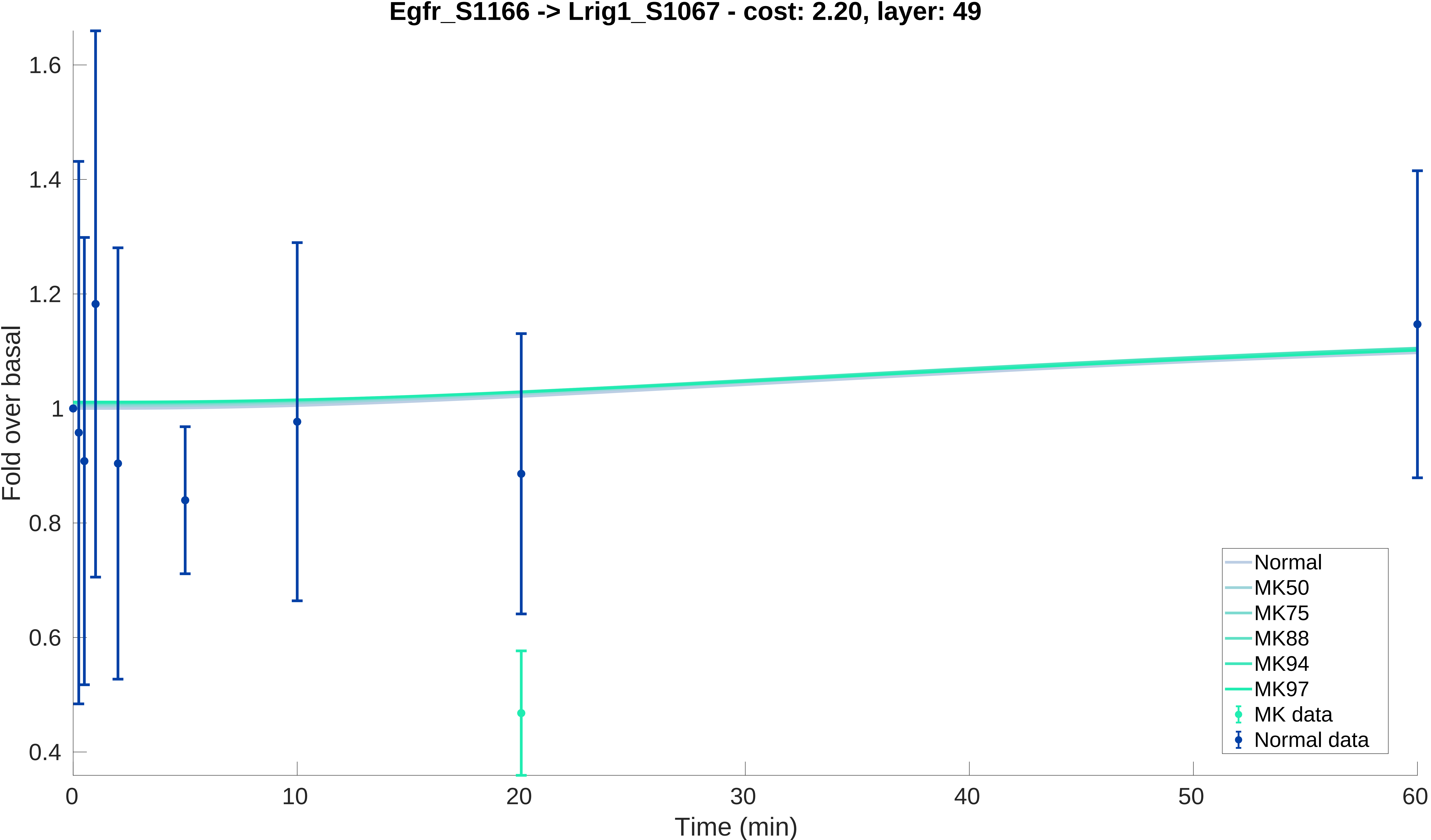

Egfr_S1166 -> Lrig1_S1067 - cost: 2.20, layer: 49
Fold over basal
1.6
1.4
1.2
1
0.8
0.6
0.4
0
10
20
30
40
50
60
Time (min)
Normal
MK50
MK75
MK88
MK94
MK97
MK data
Normal data

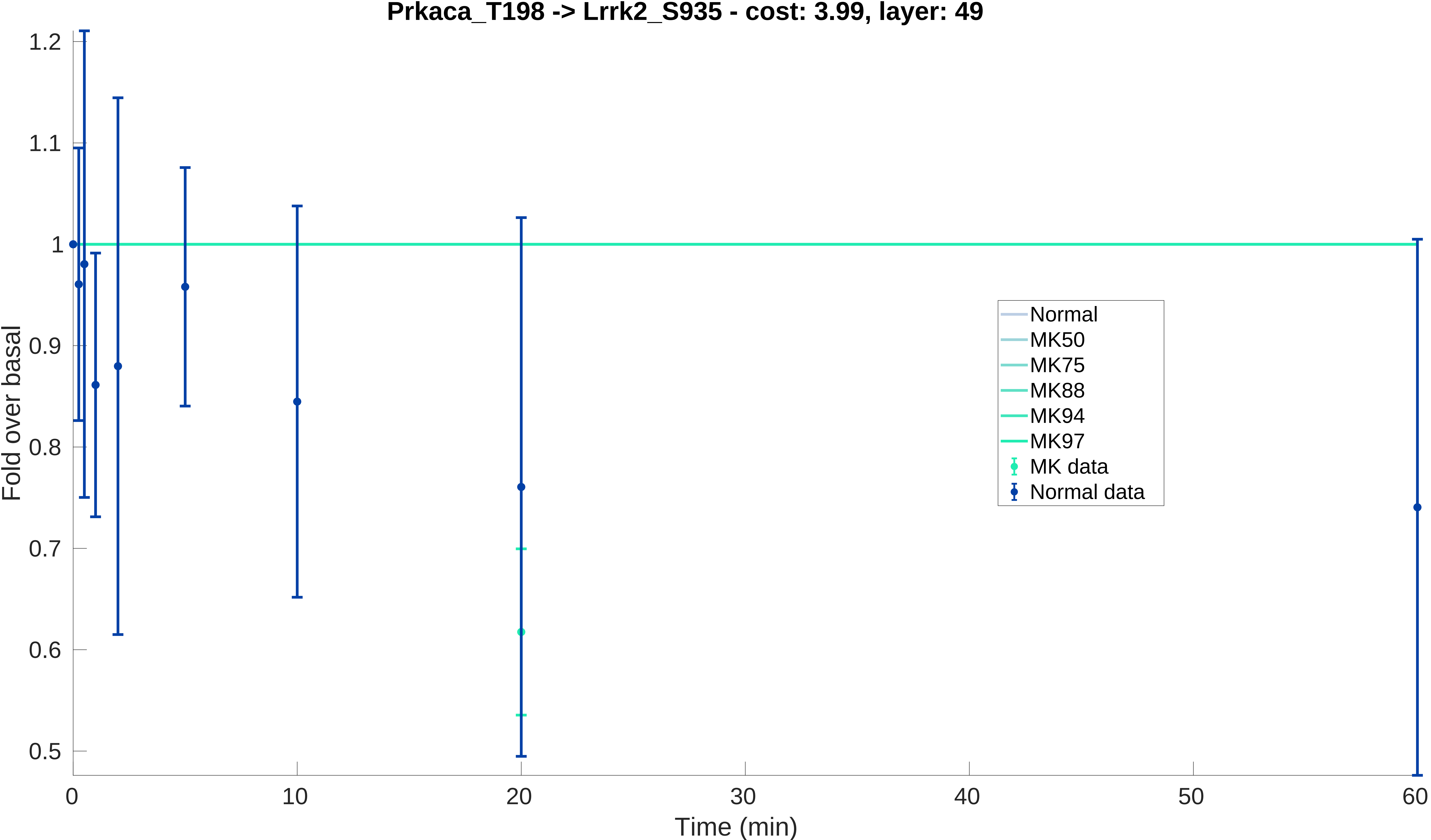
Prrkaca_T198 -> Lrrk2_S935 - cost: 3.99, layer: 49
Fold over basal
Time (min)
1.2
1.1
1
0.9
0.8
0.7
0.6
0.5
0
10
20
30
40
50
60
Normal
MK50
MK75
MK88
MK94
MK97
MK data
Normal data

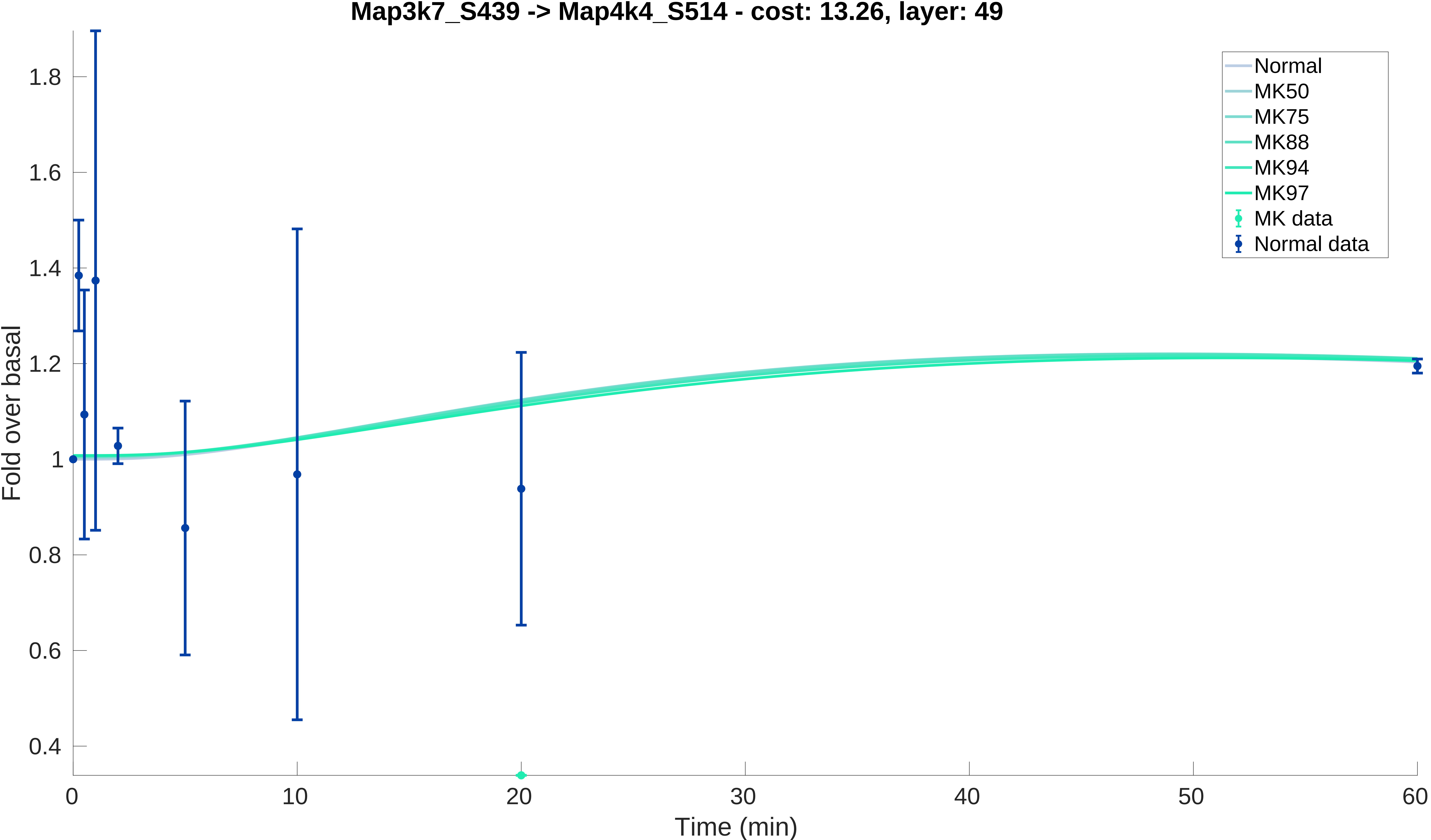

Map3k7_S439 -> Map4k4_S514 - cost: 13.26, layer: 49
Normal
MK50
MK75
MK88
MK94
MK97
MK data
Normal data
1.8
1.6
1.4
1.2
1
0.8
0.6
0.4
Fold over basal
0
10
20
30
40
50
60
Time (min)

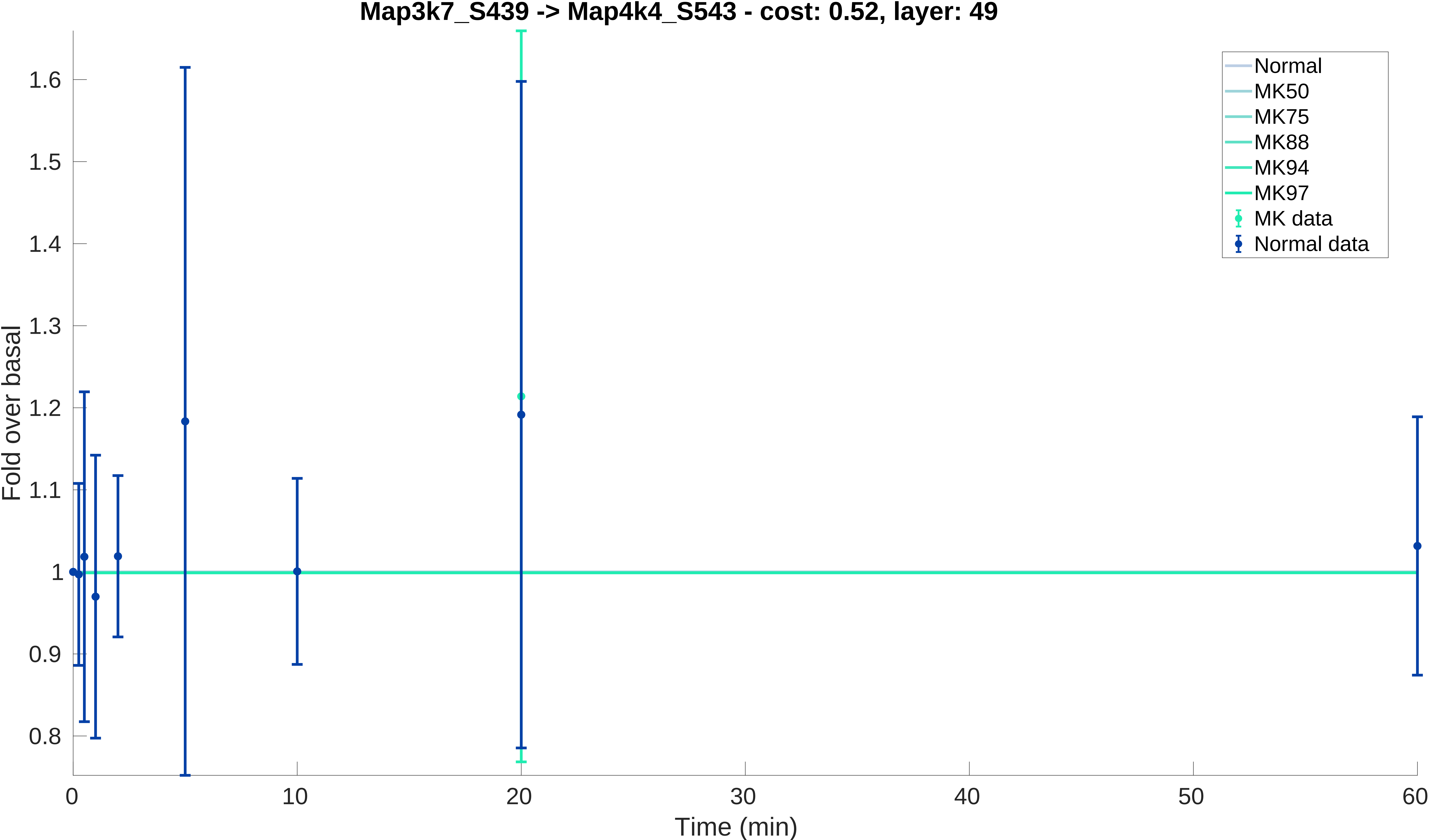

Map3k7_S439 -> Map4k4_S543 - cost: 0.52, layer: 49
Normal
MK50
MK75
MK88
MK94
MK97
MK data
Normal data
1.6
1.5
1.4
1.3
1.2
1.1
1
0.9
0.8
Fold over basal
0
10
20
30
40
50
60
Time (min)

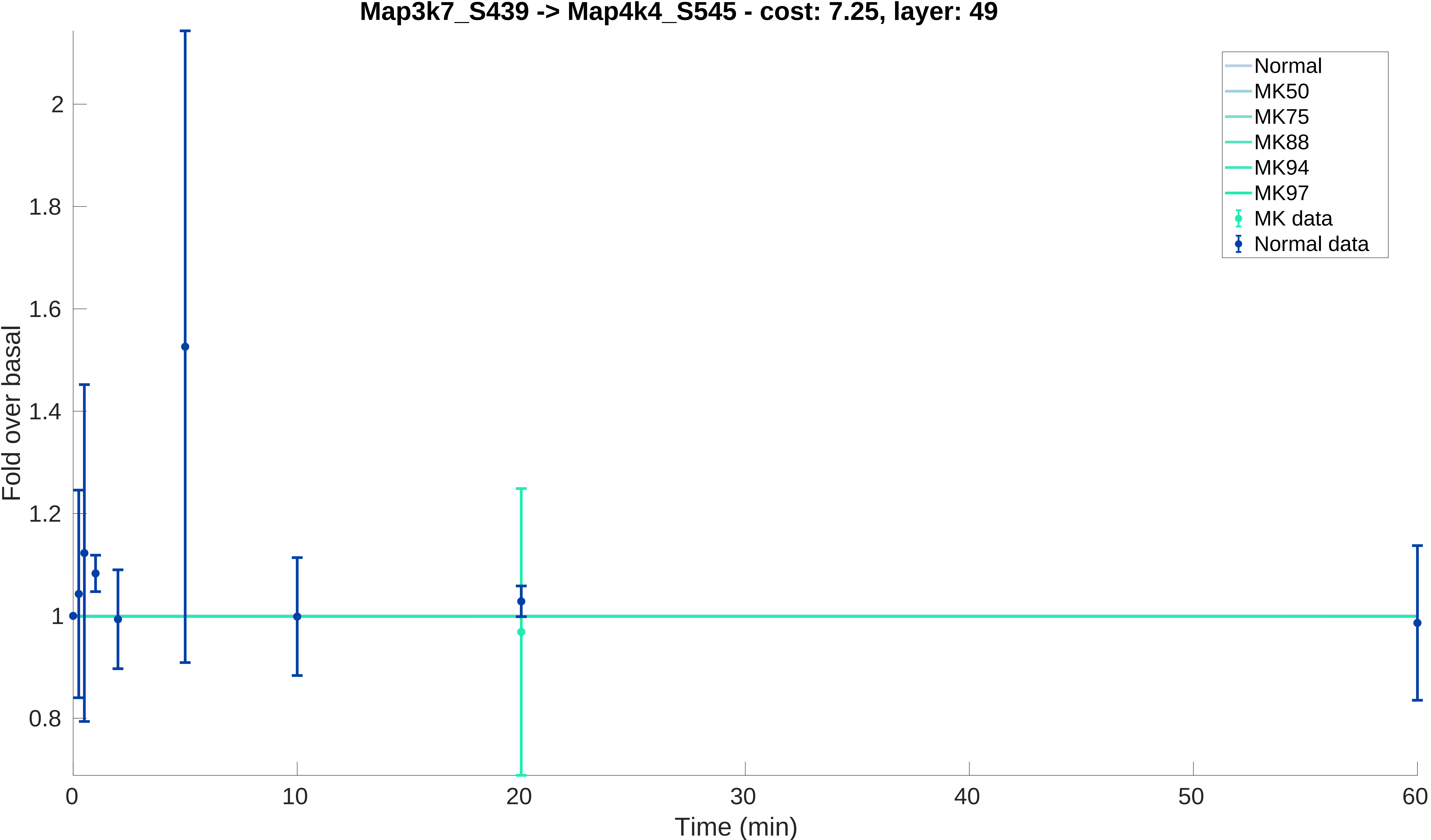

Map3k7_S439 -> Map4k4_S545 - cost: 7.25, layer: 49
Fold over basal
Time (min)
0.8
1
1.2
1.4
1.6
1.8
2
0
10
20
30
40
50
60
Normal
MK50
MK75
MK88
MK94
MK97
MK data
Normal data

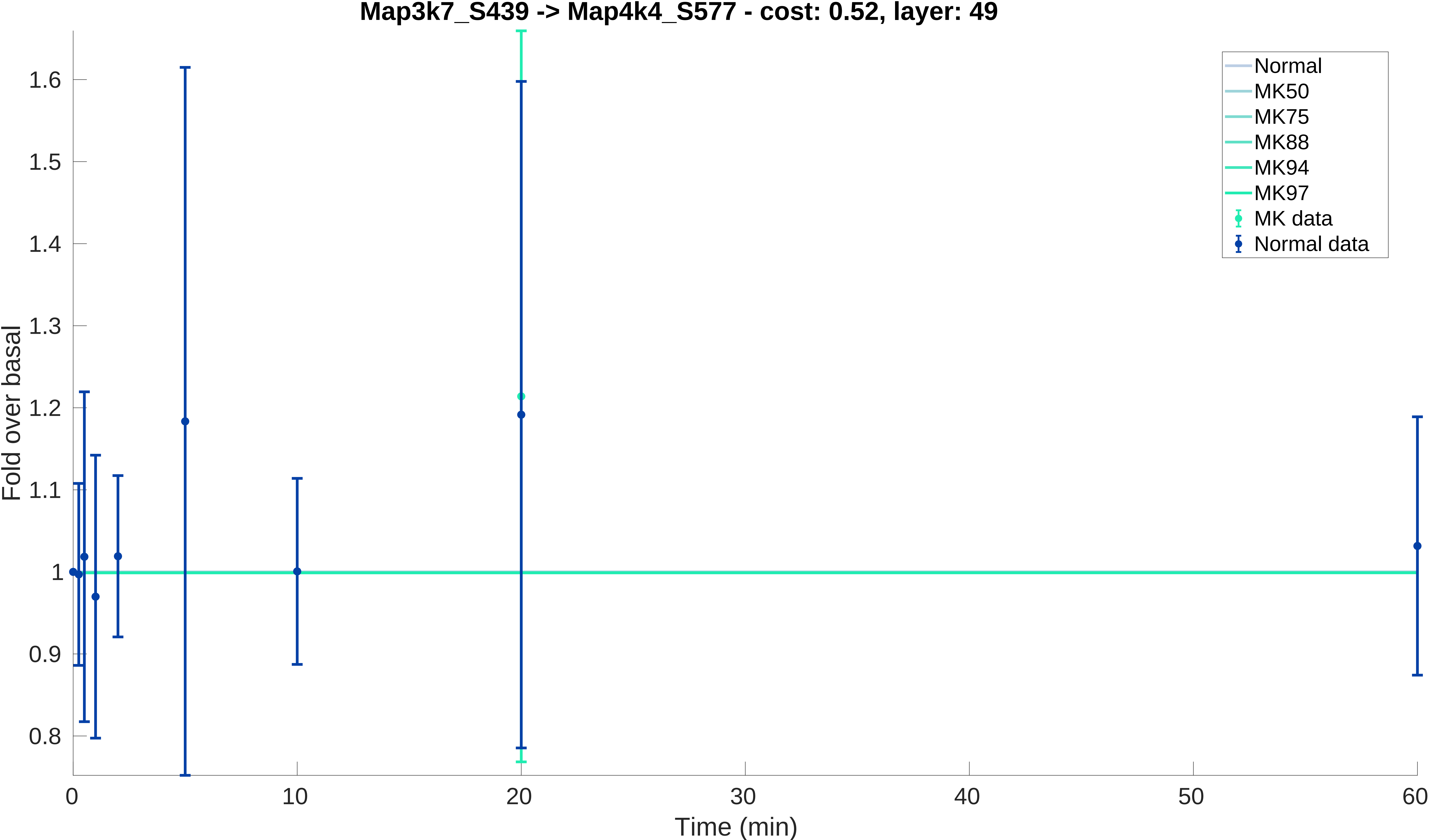

Map3k7_S439 -> Map4k4_S577 - cost: 0.52, layer: 49
Normal
MK50
MK75
MK88
MK94
MK97
MK data
Normal data
Fold over basal
1.6
1.5
1.4
1.3
1.2
1.1
1
0.9
0.8
0
10
20
30
40
50
60
Time (min)

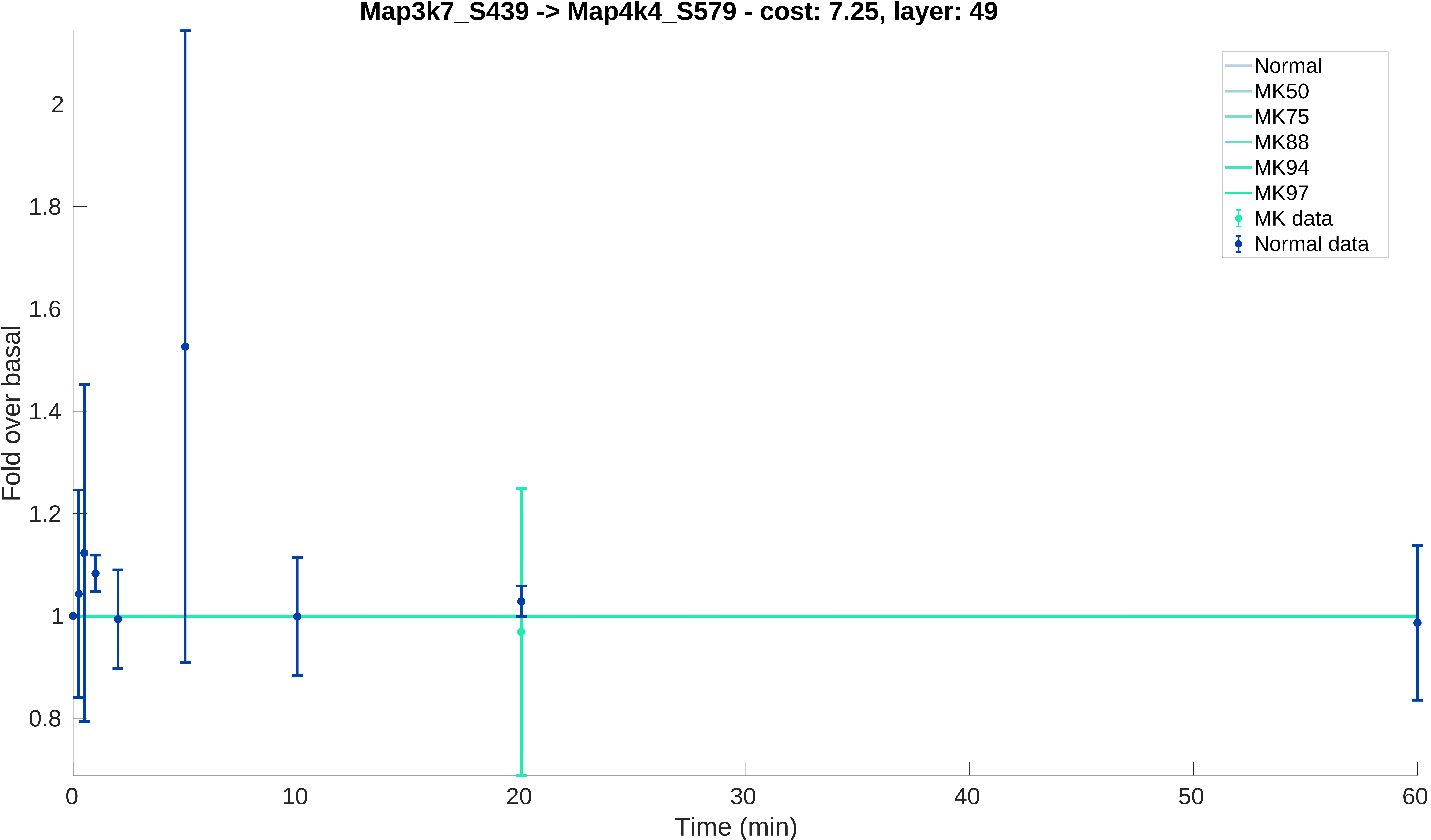

Map3k7_S439 -> Map4k4_S579 - cost: 7.25, layer: 49
Fold over basal
Time (min)
0.8
1
1.2
1.4
1.6
1.8
2
0
10
20
30
40
50
60
Normal
MK50
MK75
MK88
MK94
MK97
MK data
Normal data

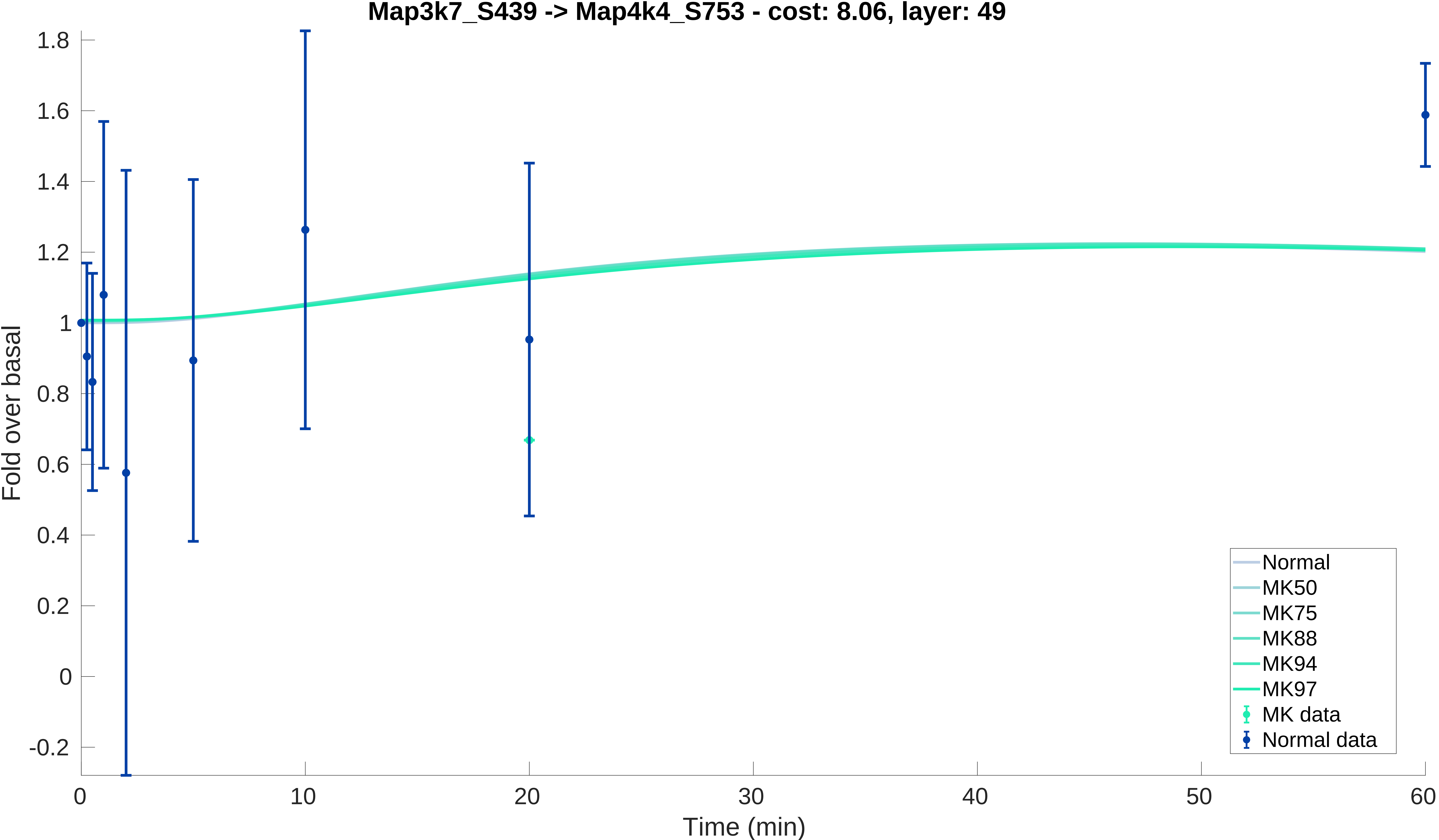

Map3k7_S439 -> Map4k4_S753 - cost: 8.06, layer: 49
Fold over basal
1.8
1.6
1.4
1.2
1
0.8
0.6
0.4
0.2
0
-0.2
Time (min)
0
10
20
30
40
50
60
Normal
MK50
MK75
MK88
MK94
MK97
MK data
Normal data

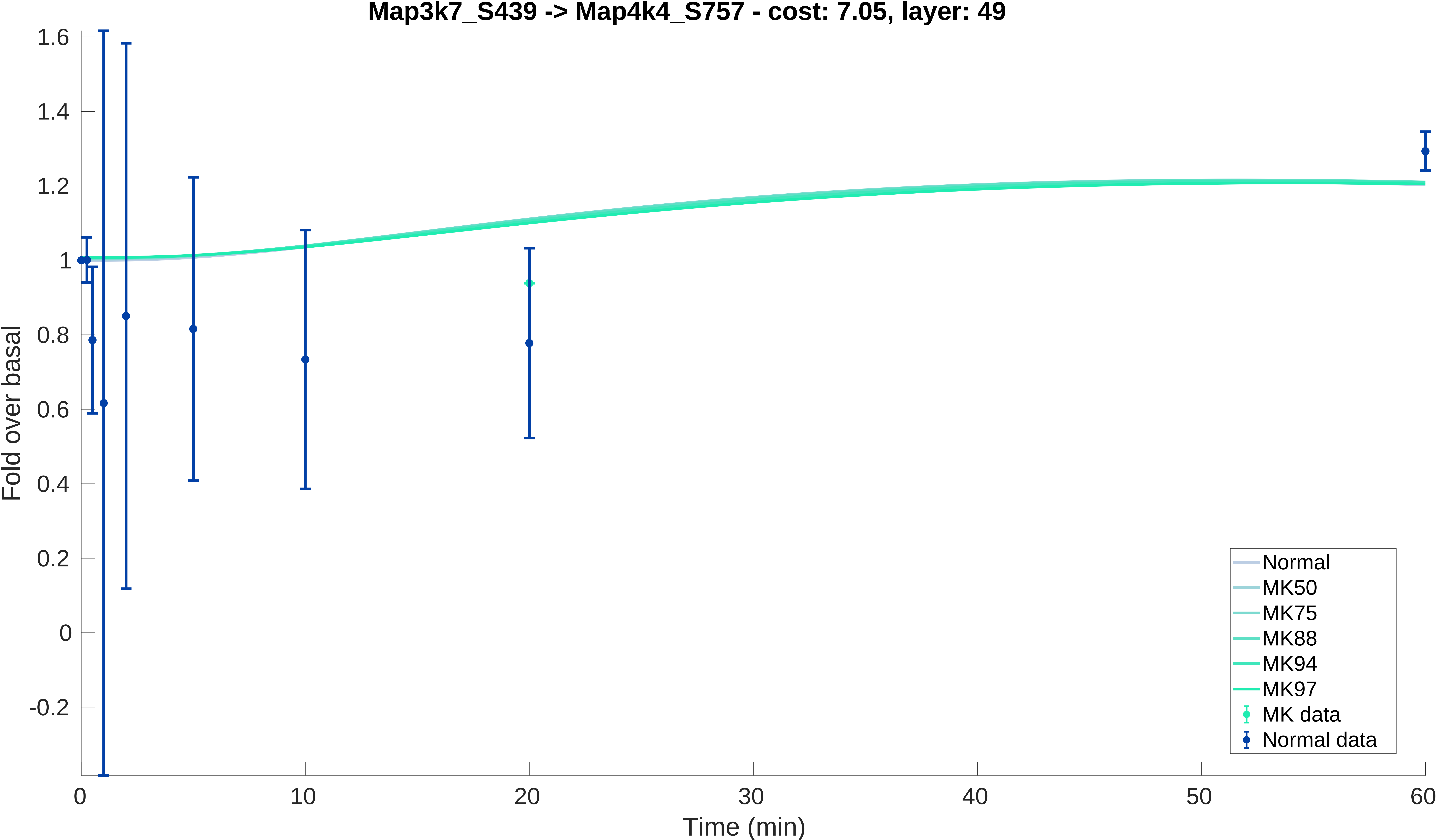

Map3k7_S439 -> Map4k4_S757 - cost: 7.05, layer: 49
Fold over basal
1.6
1.4
1.2
1
0.8
0.6
0.4
0.2
0
-0.2
0
10
20
30
40
50
60
Time (min)
Normal
MK50
MK75
MK88
MK94
MK97
MK data
Normal data

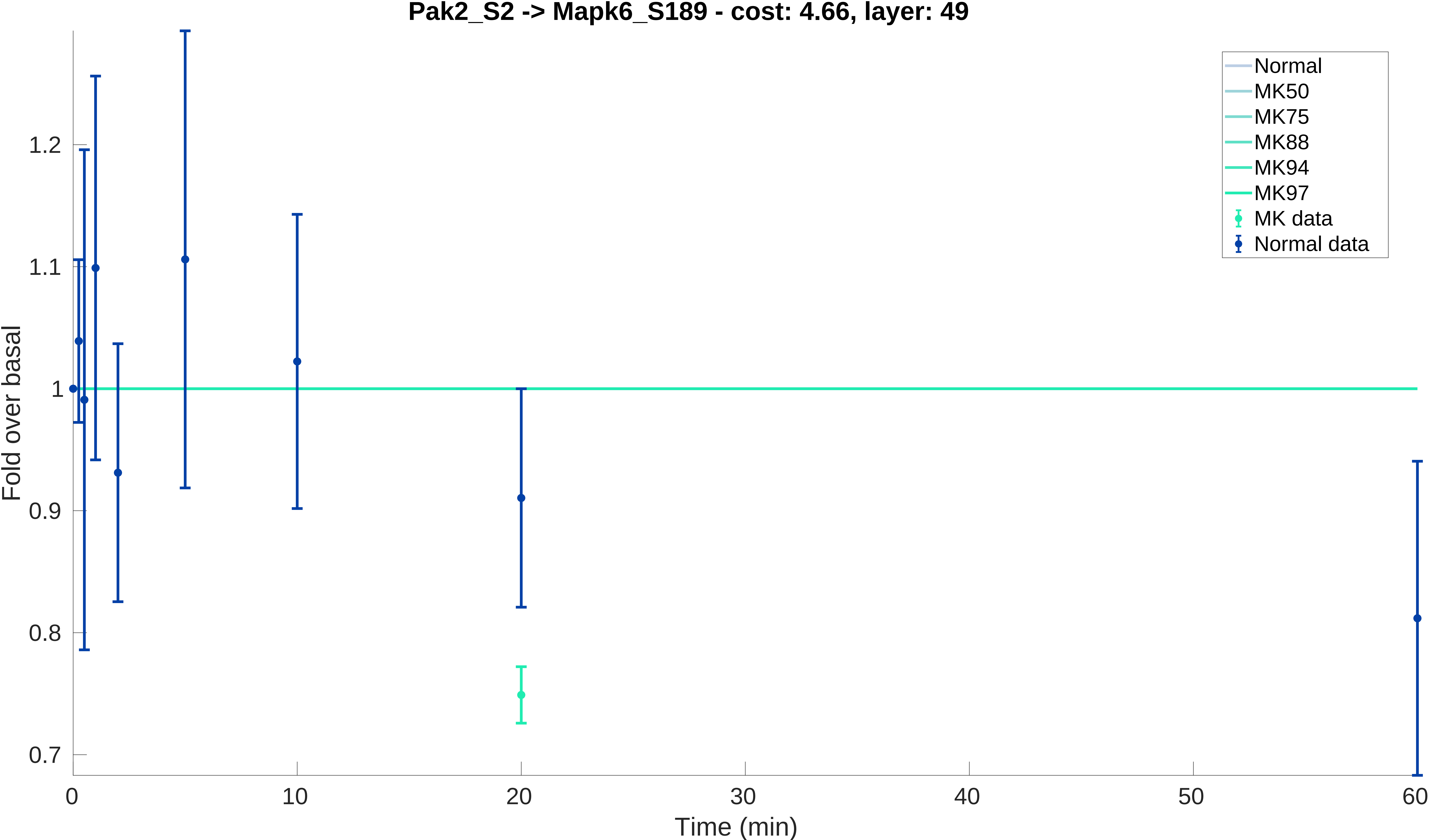

Pak2_S2 -> Mapk6_S189 - cost: 4.66, layer: 49
Normal
MK50
MK75
MK88
MK94
MK97
MK data
Normal data
Fold over basal
0.7
0.8
0.9
1
1.1
1.2
0
10
20
30
40
50
60
Time (min)

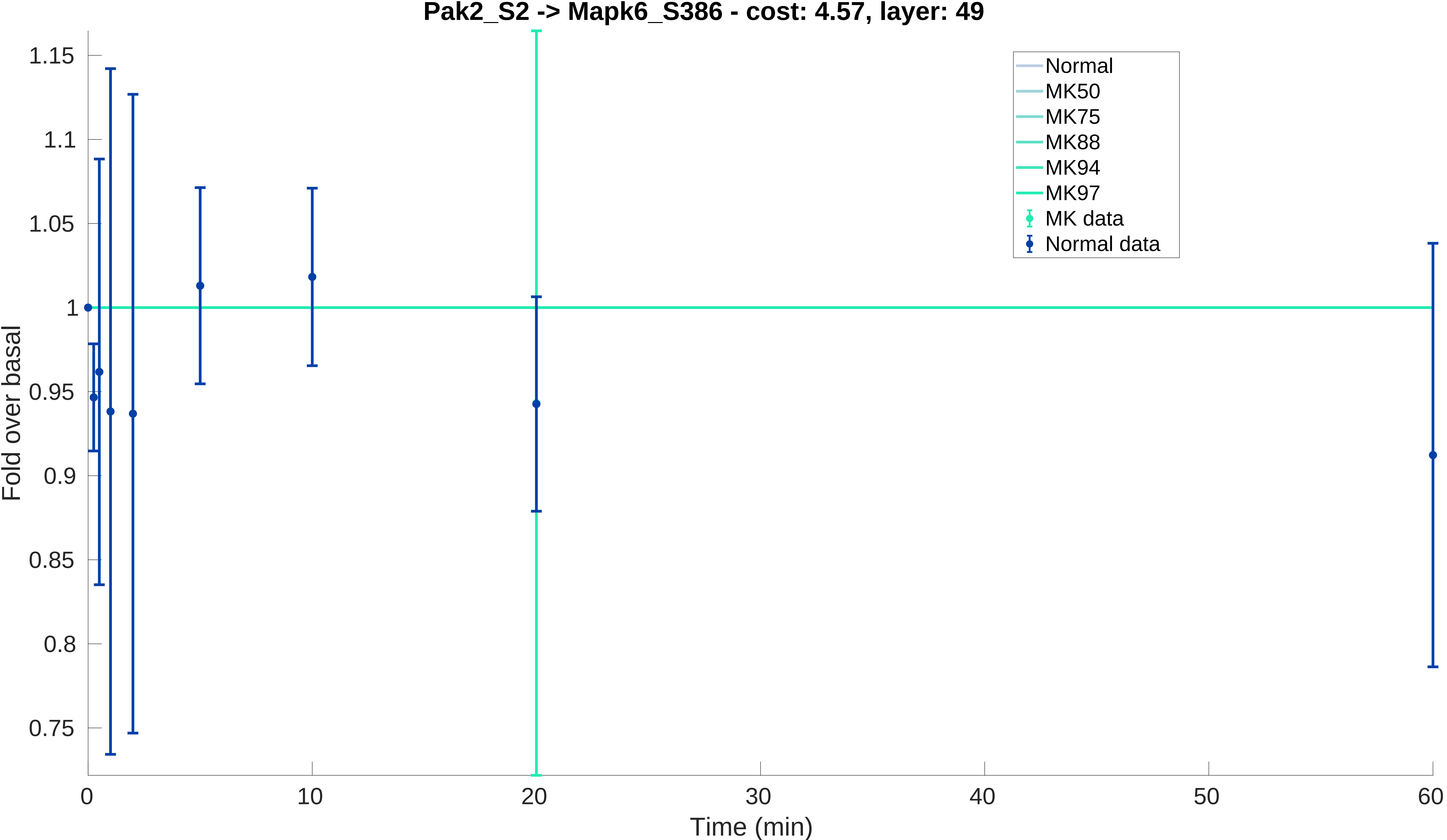

Pak2_S2 -> Mapk6_S386 - cost: 4.57, layer: 49
Normal
MK50
MK75
MK88
MK94
MK97
MK data
Normal data
Fold over basal
1.15
1.1
1.05
1
0.95
0.9
0.85
0.8
0.75
0
10
20
30
40
50
60
Time (min)

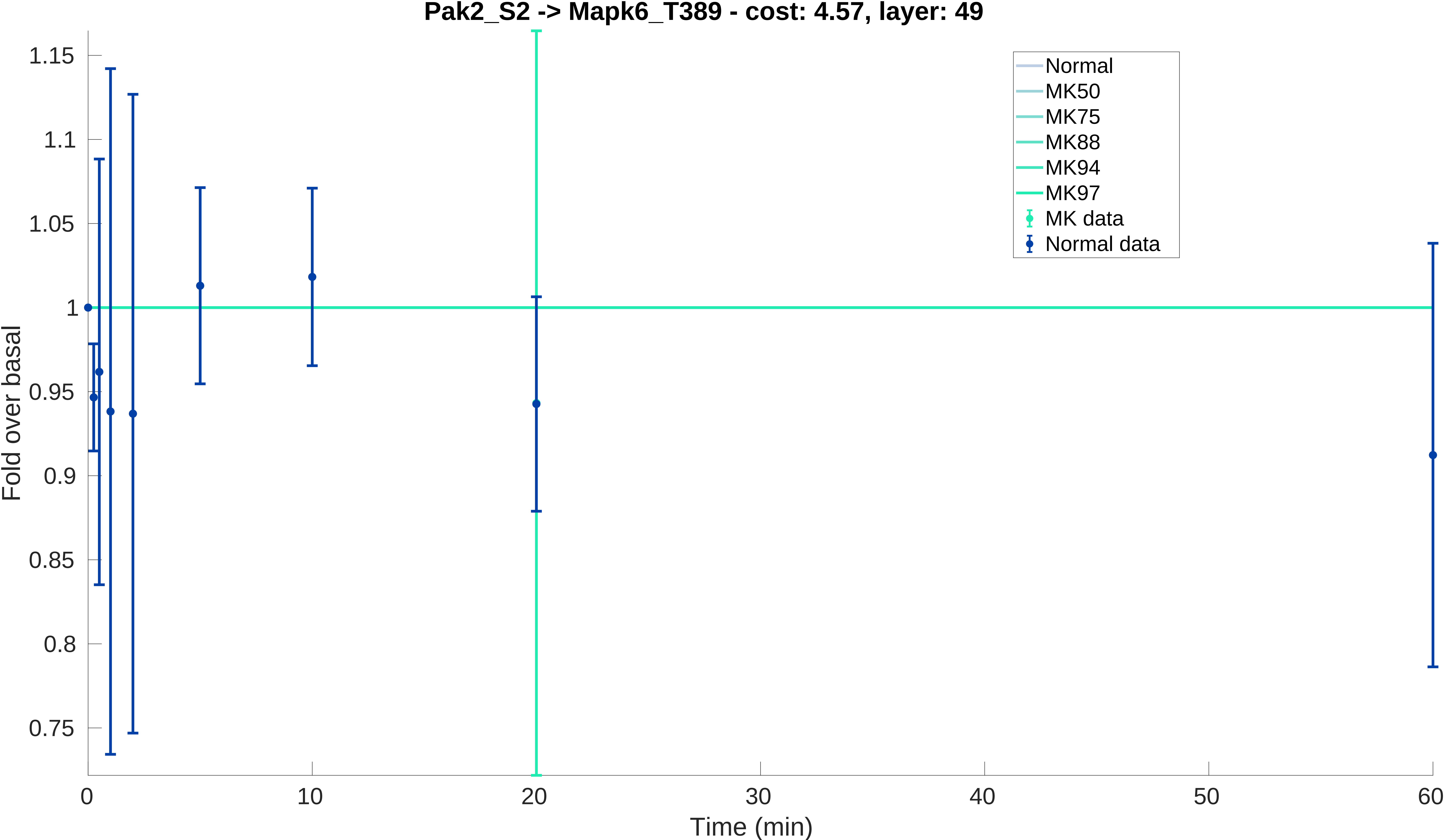

Pak2_S2 -> Mapk6_T389 - cost: 4.57, layer: 49
Normal
MK50
MK75
MK88
MK94
MK97
MK data
Normal data
Fold over basal
Time (min)
1.15
1.1
1.05
1
0.95
0.9
0.85
0.8
0.75
0
10
20
30
40
50
60

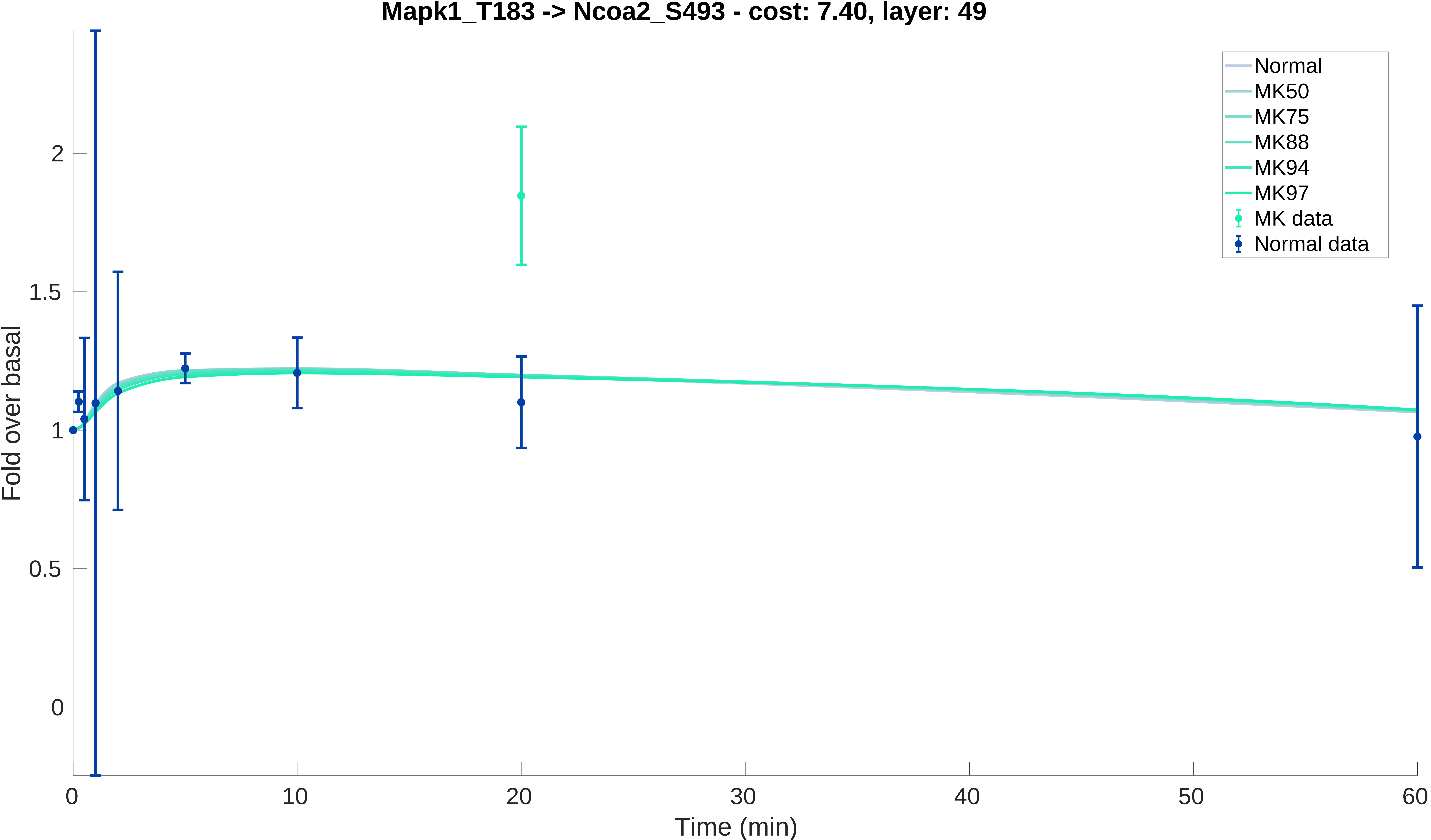

Mapk1_T183 -> Ncoa2_S493 - cost: 7.40, layer: 49
Fold over basal
Time (min)
0
0.5
1
1.5
2
0
10
20
30
40
50
60
Normal
MK50
MK75
MK88
MK94
MK97
MK data
Normal data

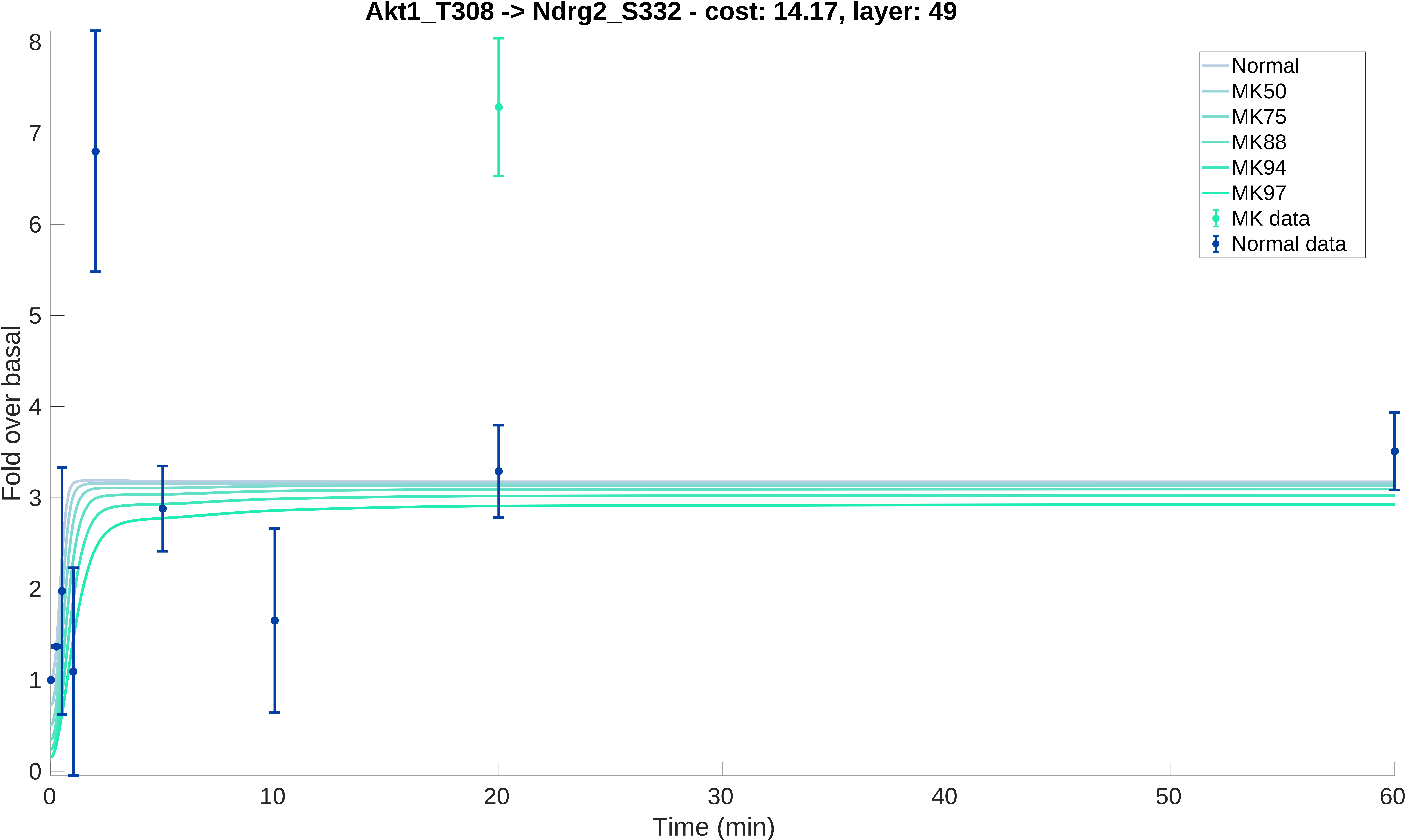
Akt1_T308 -> Ndrg2_S332 - cost: 14.17, layer: 49
Fold over basal
Time (min)
0
1
2
3
4
5
6
7
8
0
10
20
30
40
50
60
Normal
MK50
MK75
MK88
MK94
MK97
MK data
Normal data

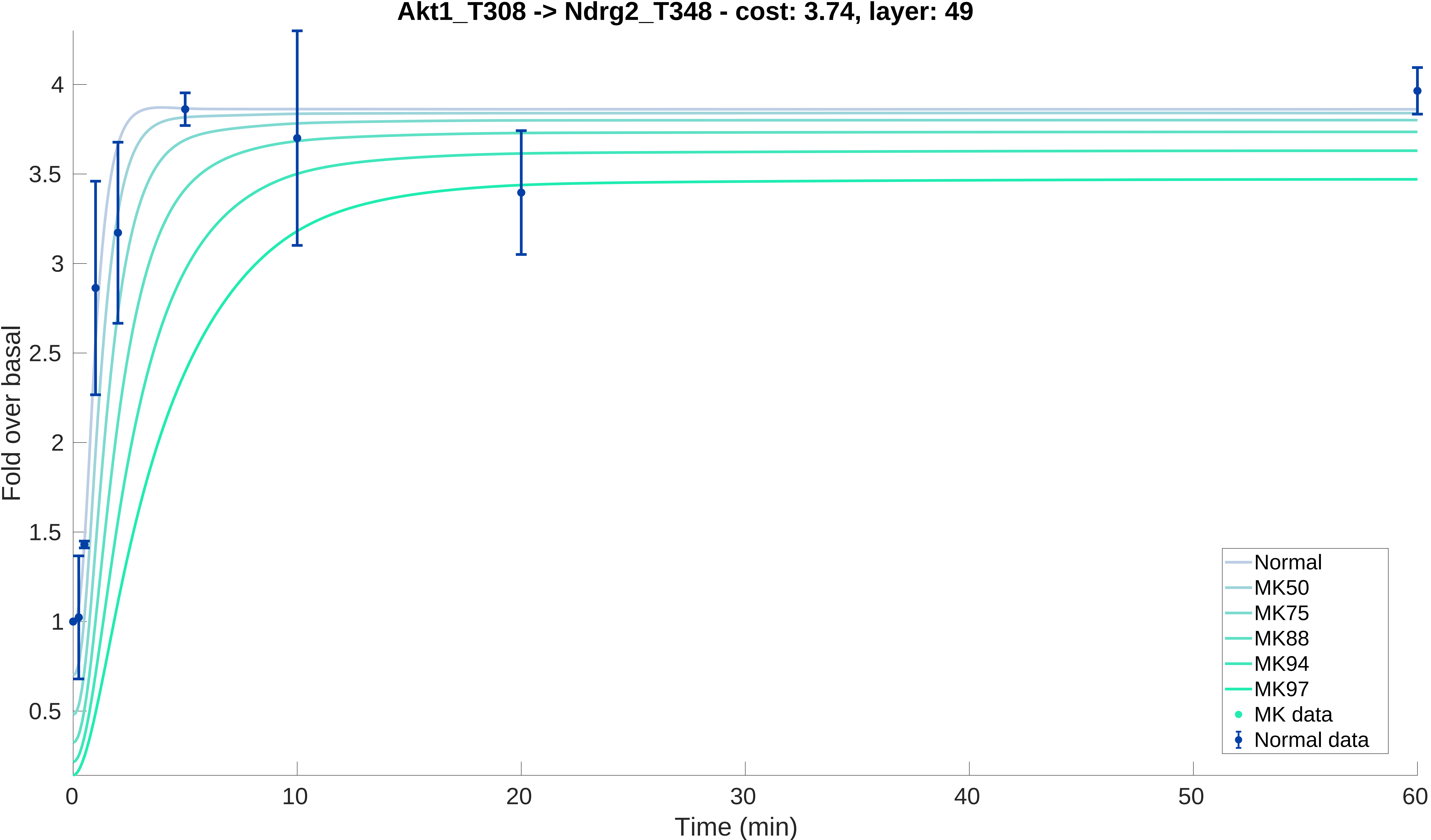

Akt1_T308 -> Ndrg2_T348 - cost: 3.74, layer: 49
Fold over basal
Time (min)
0.5
1
1.5
2
2.5
3
3.5
4
0
10
20
30
40
50
60
Normal
MK50
MK75
MK88
MK94
MK97
MK data
Normal data

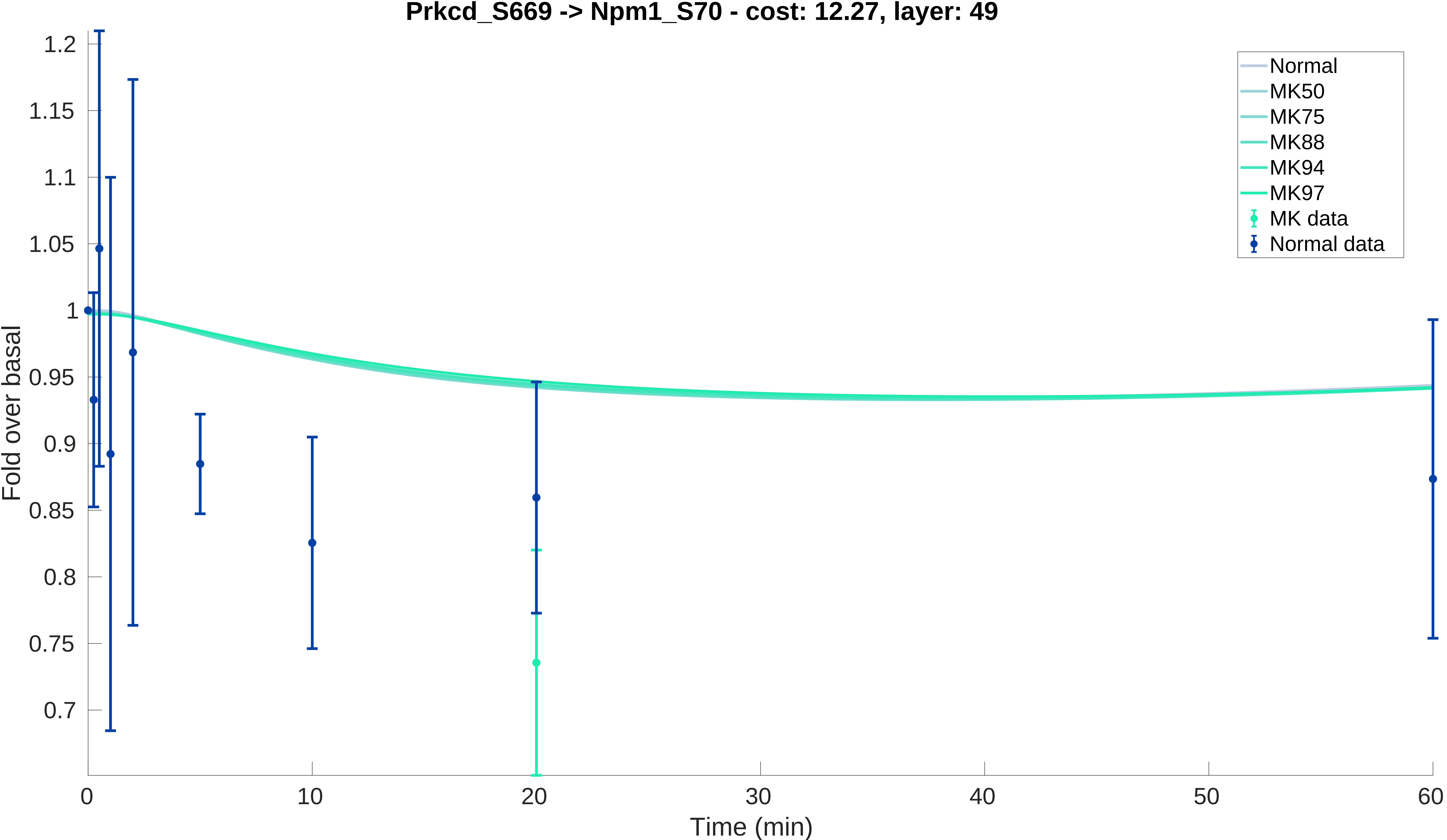
Prkcd_S669 -> Npm1_S70 - cost: 12.27, layer: 49
Fold over basal
Time (min)
1.2
1.15
1.1
1.05
1
0.95
0.9
0.85
0.8
0.75
0.7
0
10
20
30
40
50
60
Normal
MK50
MK75
MK88
MK94
MK97
MK data
Normal data

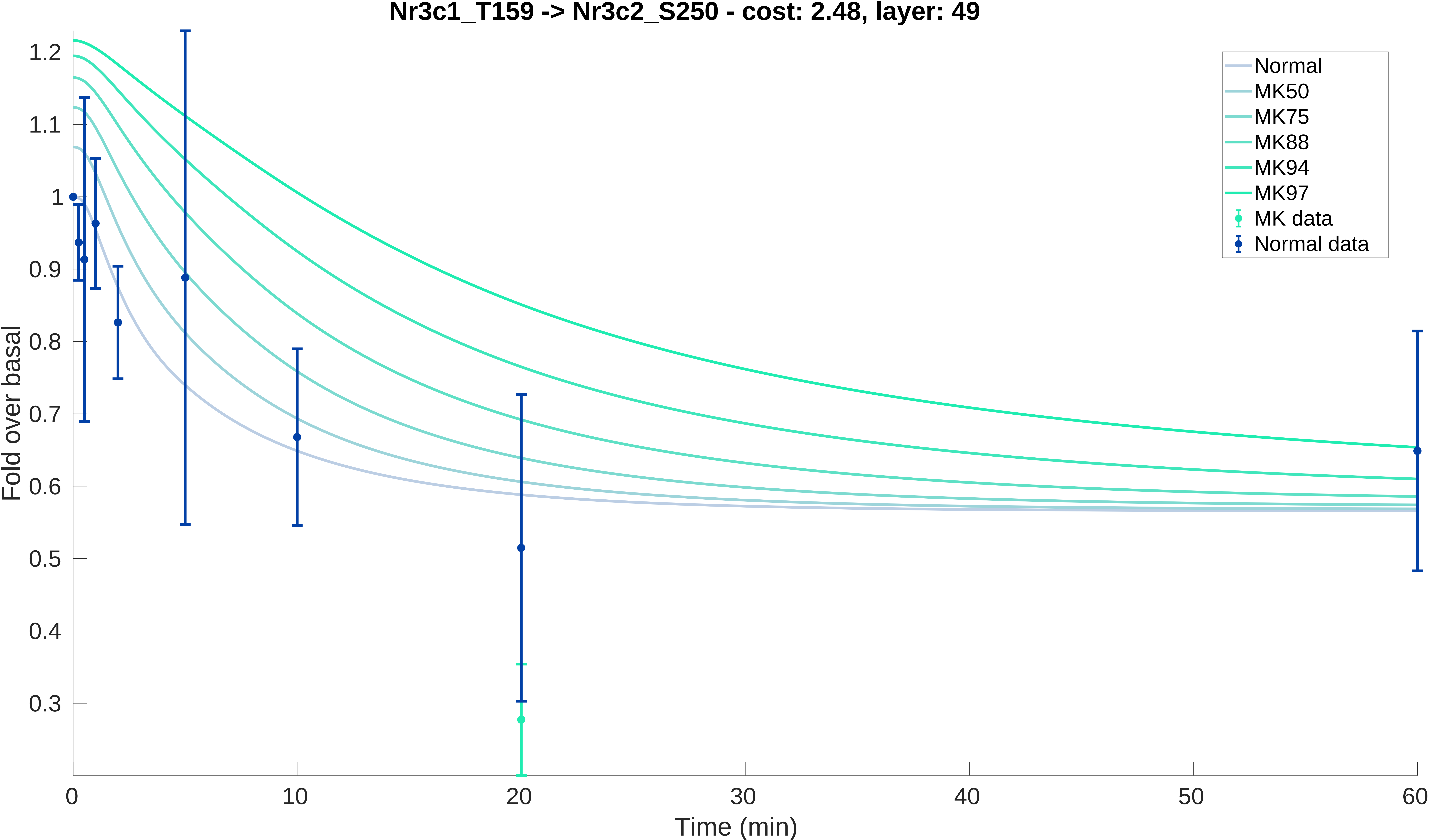
Nr3c1_T159 -> Nr3c2_S250 - cost: 2.48, layer: 49
Fold over basal
Time (min)
0
10
20
30
40
50
60
0.3
0.4
0.5
0.6
0.7
0.8
0.9
1
1.1
1.2
Normal
MK50
MK75
MK88
MK94
MK97
MK data
Normal data

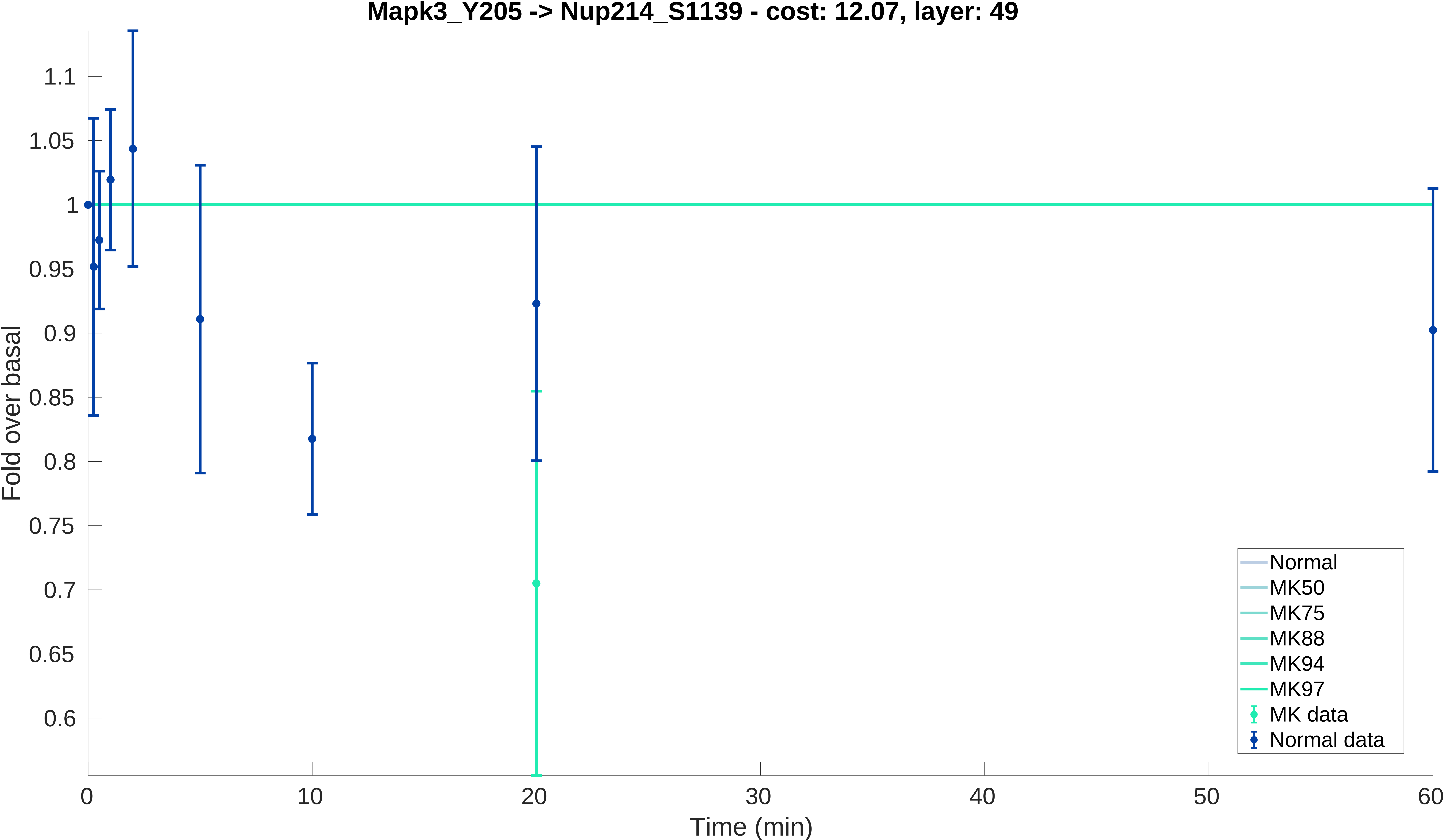

Mapk3_Y205 -> Nup214_S1139 - cost: 12.07, layer: 49
Fold over basal
1.1
1.05
1
0.95
0.9
0.85
0.8
0.75
0.7
0.65
0.6
0
10
20
30
40
50
60
Time (min)
Normal
MK50
MK75
MK88
MK94
MK97
MK data
Normal data

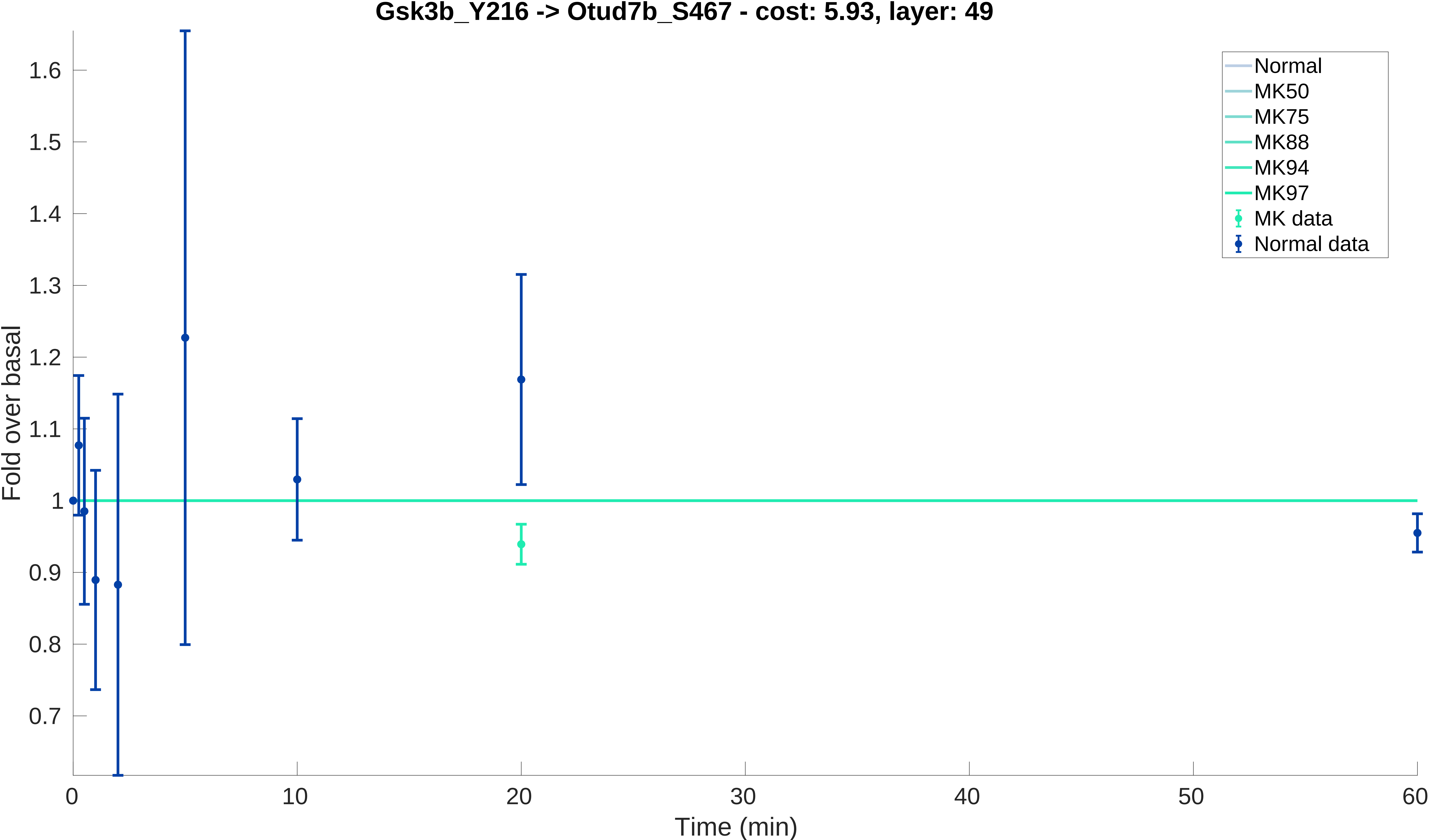

Gsk3b_Y216 -> Otud7b_S467 - cost: 5.93, layer: 49
Fold over basal
Time (min)
Normal
MK50
MK75
MK88
MK94
MK97
MK data
Normal data
1.6
1.5
1.4
1.3
1.2
1.1
1
0.9
0.8
0.7
0
10
20
30
40
50
60

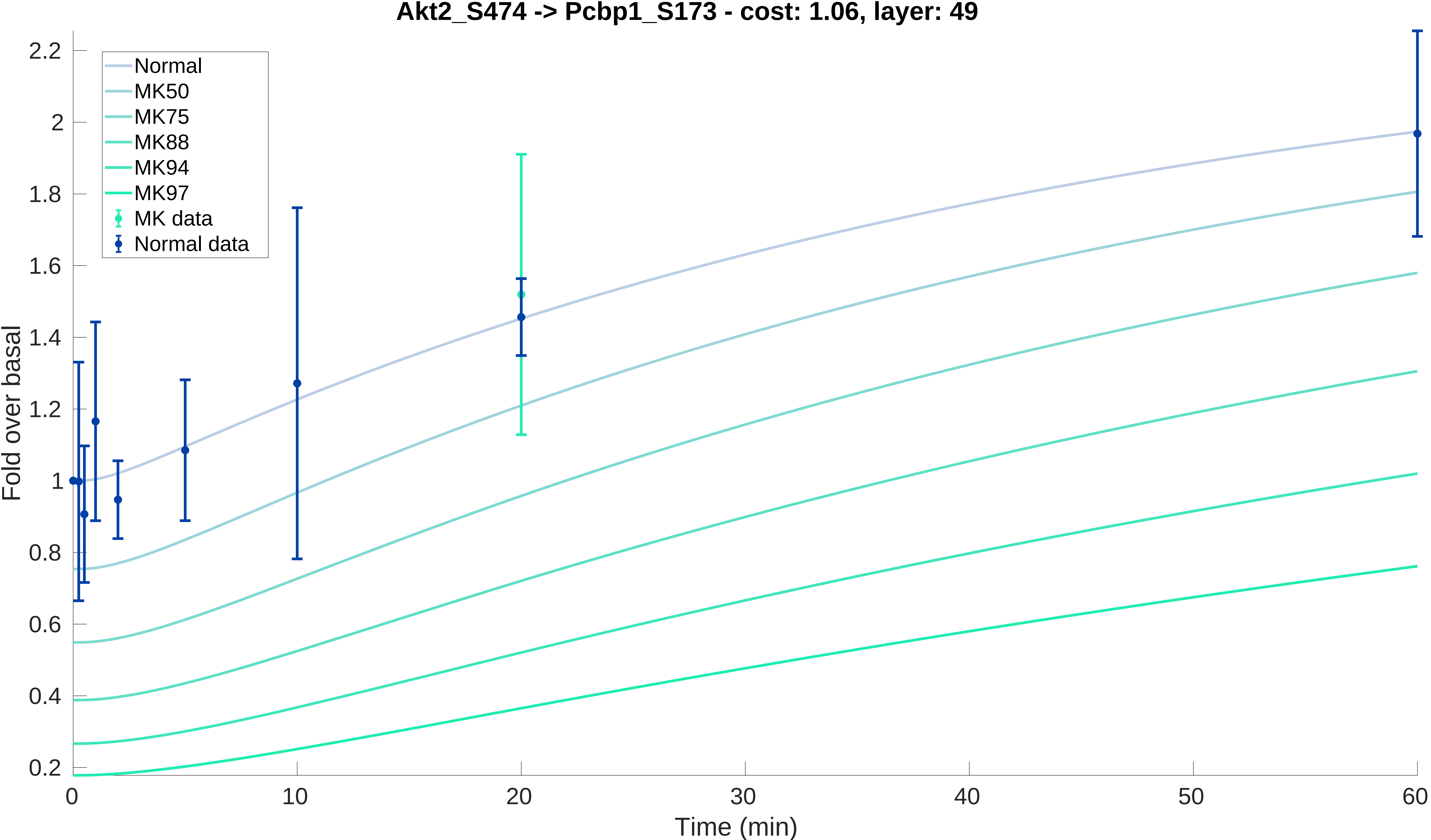

Akt2_S474 -> Pcbp1_S173 - cost: 1.06, layer: 49
Normal
MK50
MK75
MK88
MK94
MK97
MK data
Normal data
Fold over basal
Time (min)
0.2
0.4
0.6
0.8
1
1.2
1.4
1.6
1.8
2
2.2
0
10
20
30
40
50
60

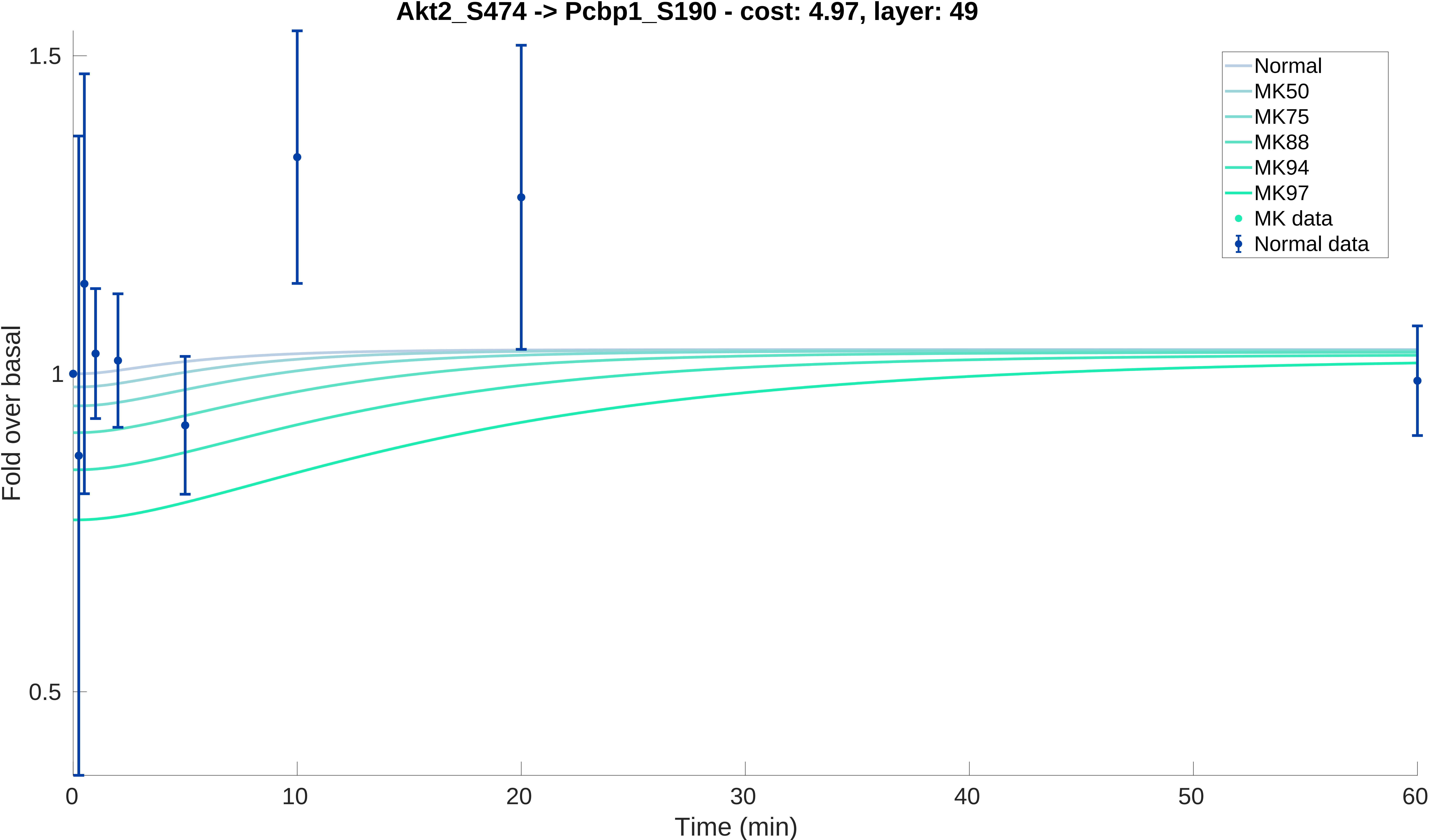
Akt2_S474 -> Pcbp1_S190 - cost: 4.97, layer: 49
Fold over basal
Time (min)
1.5
1
0.5
0
10
20
30
40
50
60
Normal
MK50
MK75
MK88
MK94
MK97
MK data
Normal data

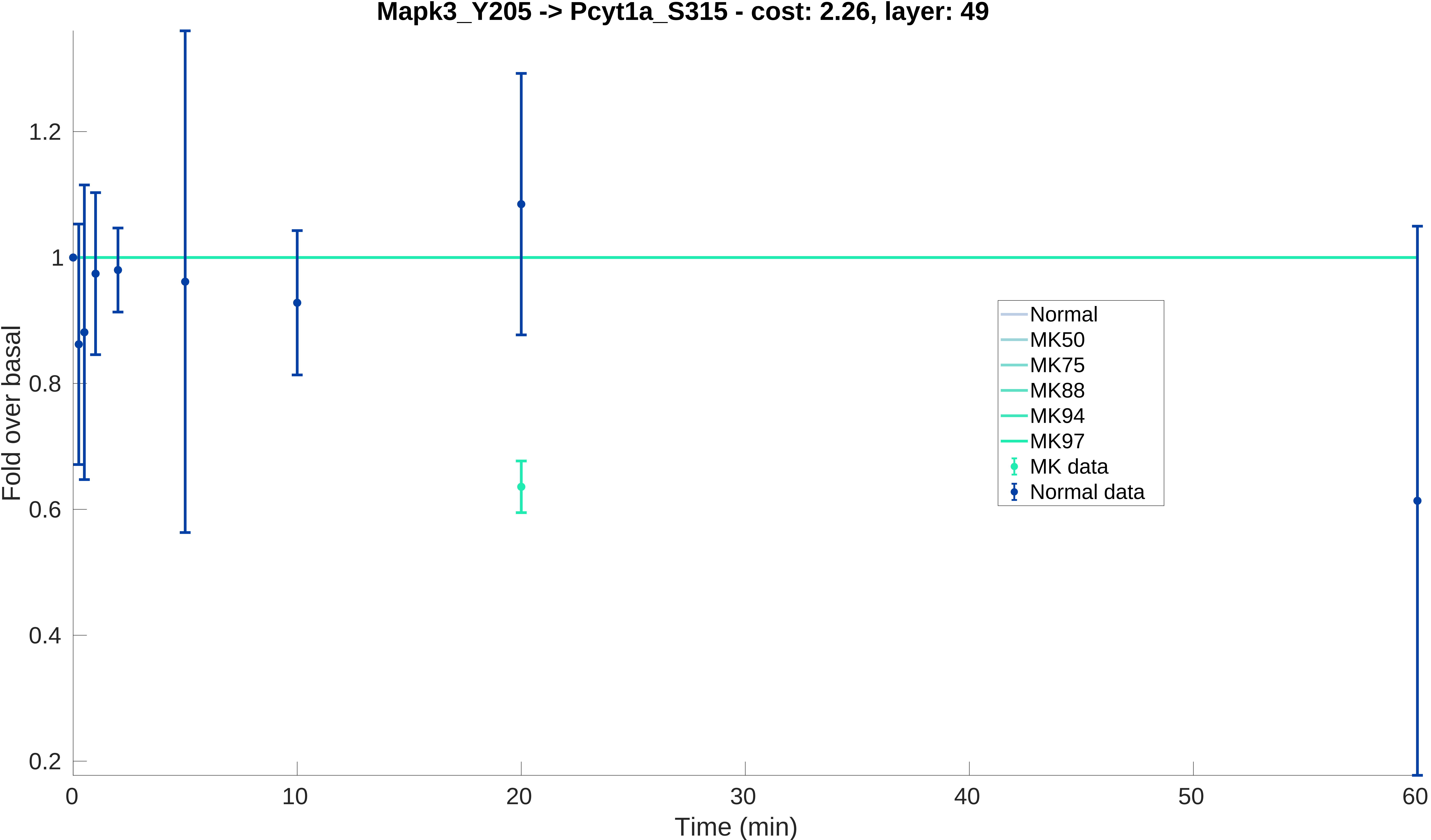
Mapk3_Y205 -> Pcyt1a_S315 - cost: 2.26, layer: 49
Fold over basal
1.2
1
0.8
0.6
0.4
0.2
0
10
20
30
40
50
60
Time (min)
Normal
MK50
MK75
MK88
MK94
MK97
MK data
Normal data

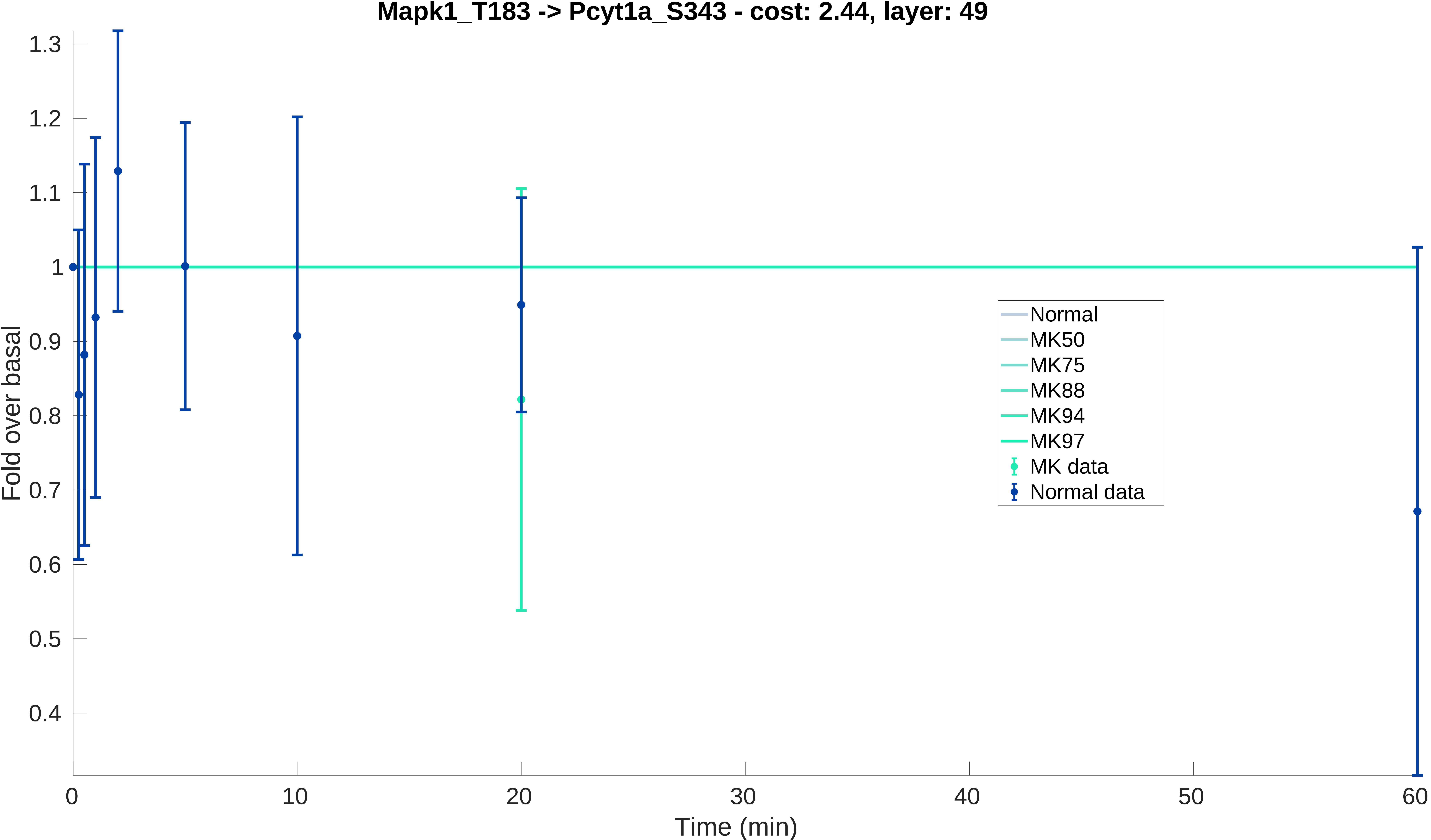

Mapk1_T183 -> Pcyt1a_S343 - cost: 2.44, layer: 49
Fold over basal
Time (min)
0
10
20
30
40
50
60
0.4
0.5
0.6
0.7
0.8
0.9
1
1.1
1.2
1.3
Normal
MK50
MK75
MK88
MK94
MK97
MK data
Normal data

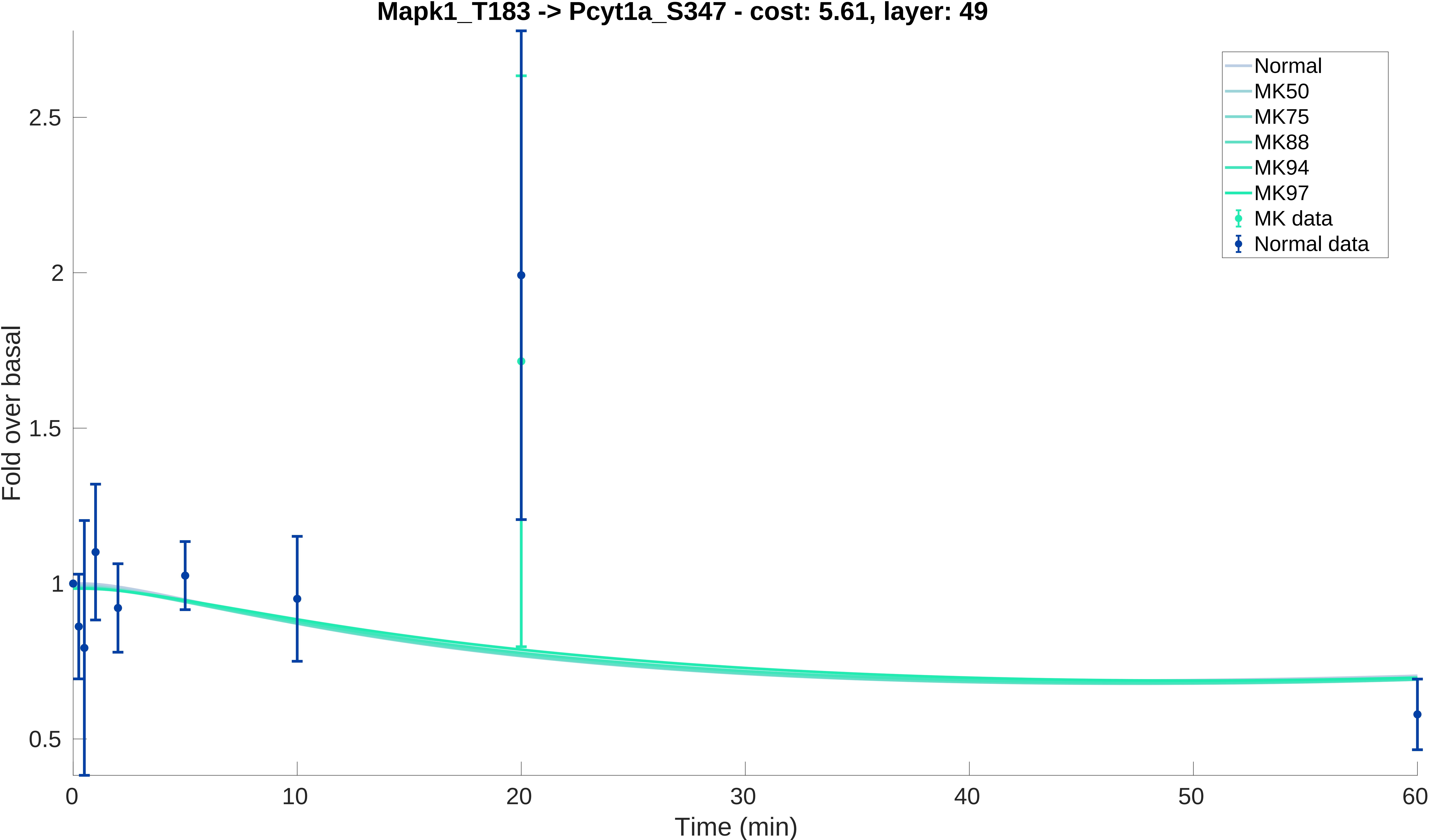
Mapk1_T183 -> Pcyt1a_S347 - cost: 5.61, layer: 49
Fold over basal
Time (min)
0.5
1
1.5
2
2.5
0
10
20
30
40
50
60
Normal
MK50
MK75
MK88
MK94
MK97
MK data
Normal data

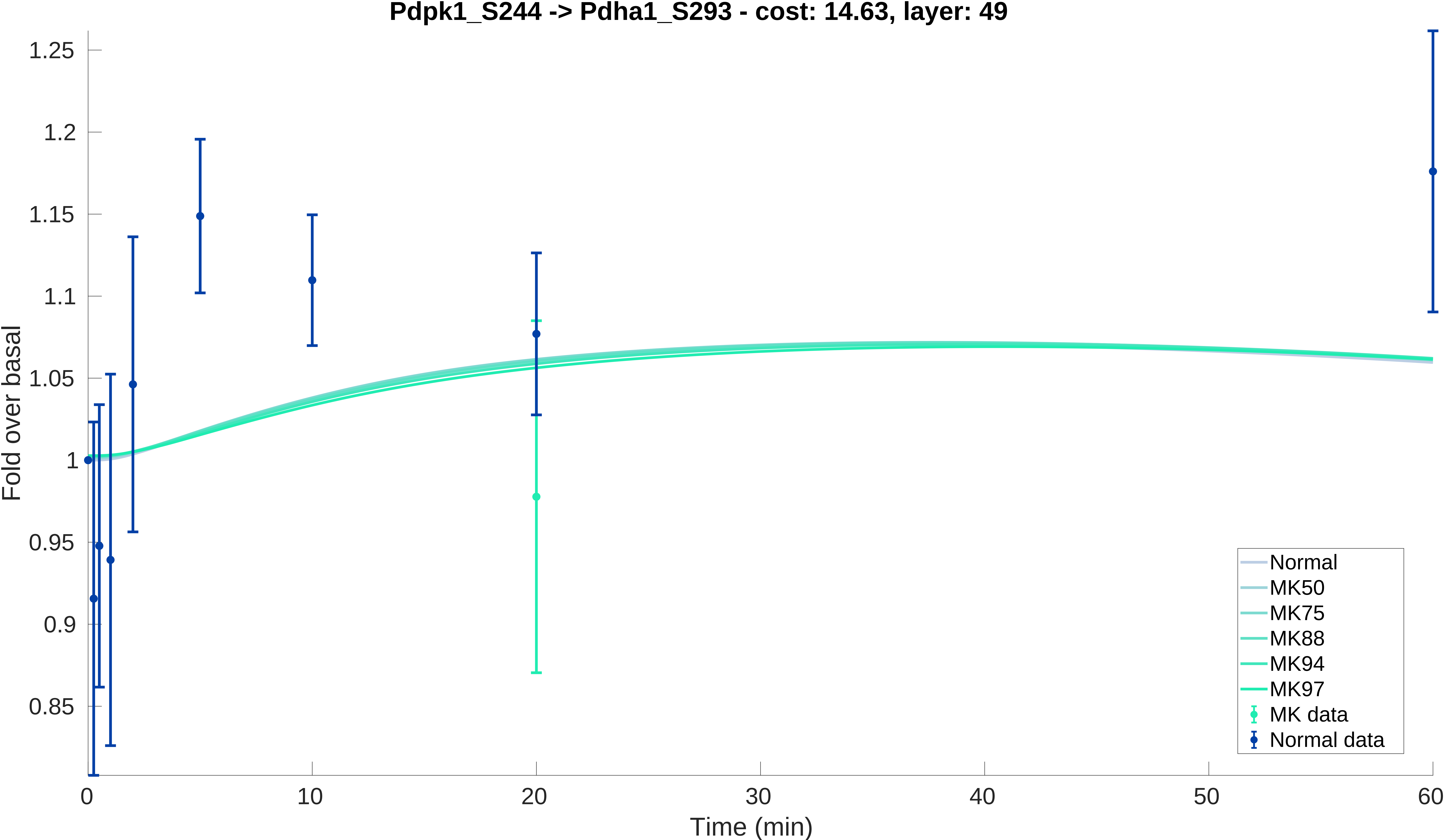

Pdpk1_S244 -> Pdha1_S293 - cost: 14.63, layer: 49
Fold over basal
Time (min)
1.25
1.2
1.15
1.1
1.05
1
0.95
0.9
0.85
0
10
20
30
40
50
60
Normal
MK50
MK75
MK88
MK94
MK97
MK data
Normal data

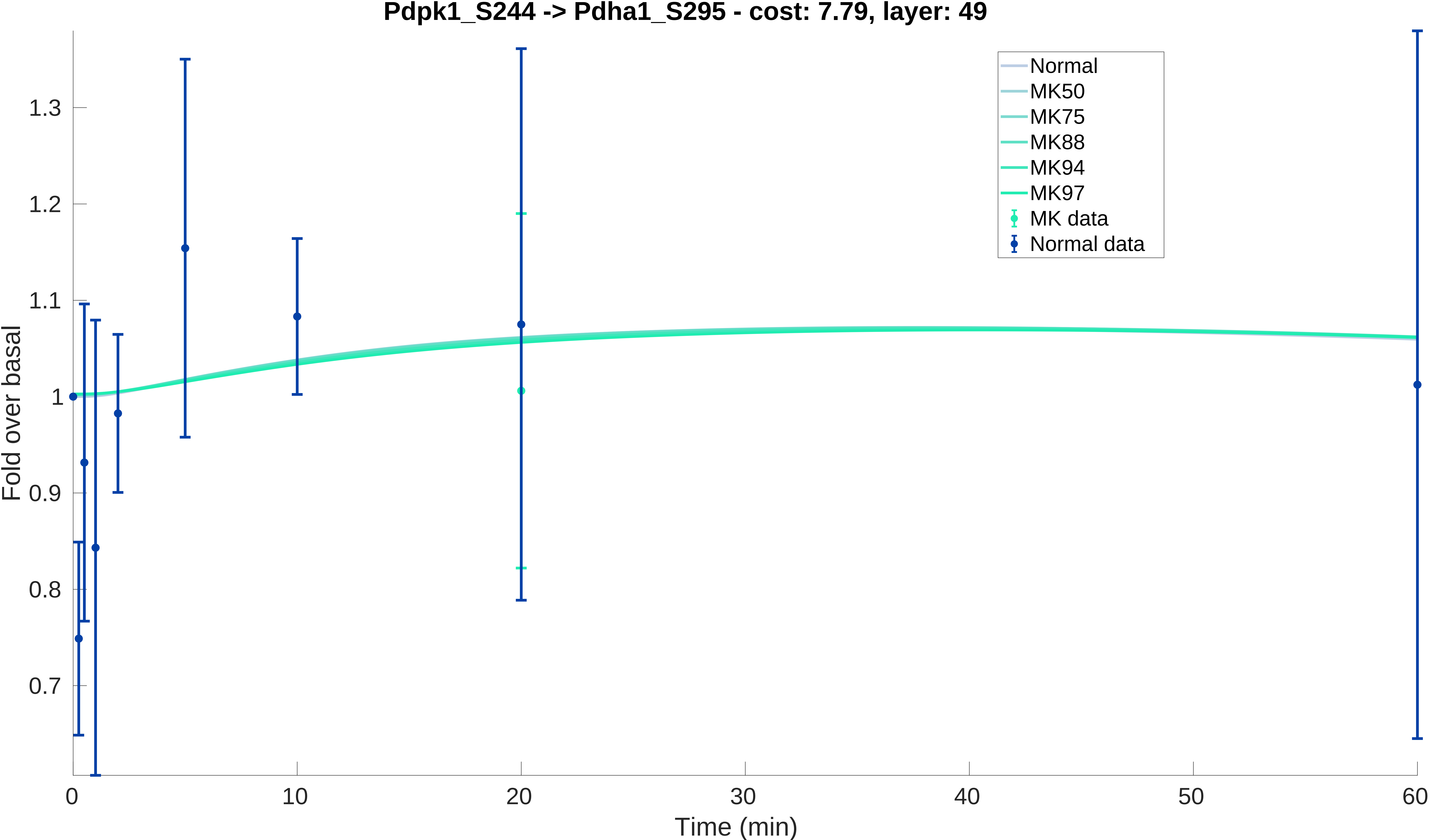

Pdpk1_S244 -> Pdha1_S295 - cost: 7.79, layer: 49
Fold over basal
Time (min)
1.3
1.2
1.1
1
0.9
0.8
0.7
0
10
20
30
40
50
60
Normal
MK50
MK75
MK88
MK94
MK97
MK data
Normal data

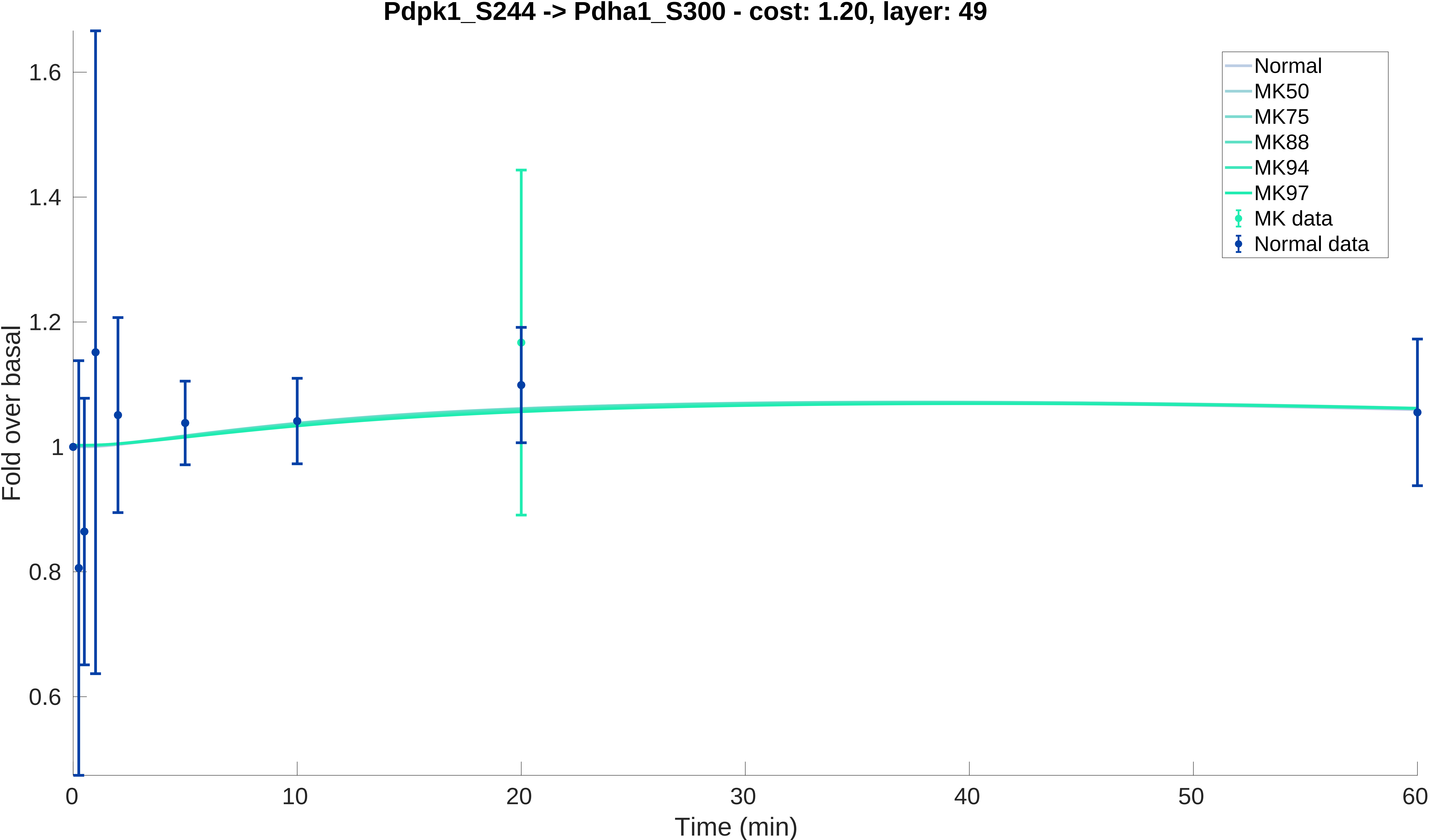

Pdpk1_S244 -> Pdha1_S300 - cost: 1.20, layer: 49
Fold over basal
Time (min)
1.6
1.4
1.2
1
0.8
0.6
0
10
20
30
40
50
60
Normal
MK50
MK75
MK88
MK94
MK97
MK data
Normal data

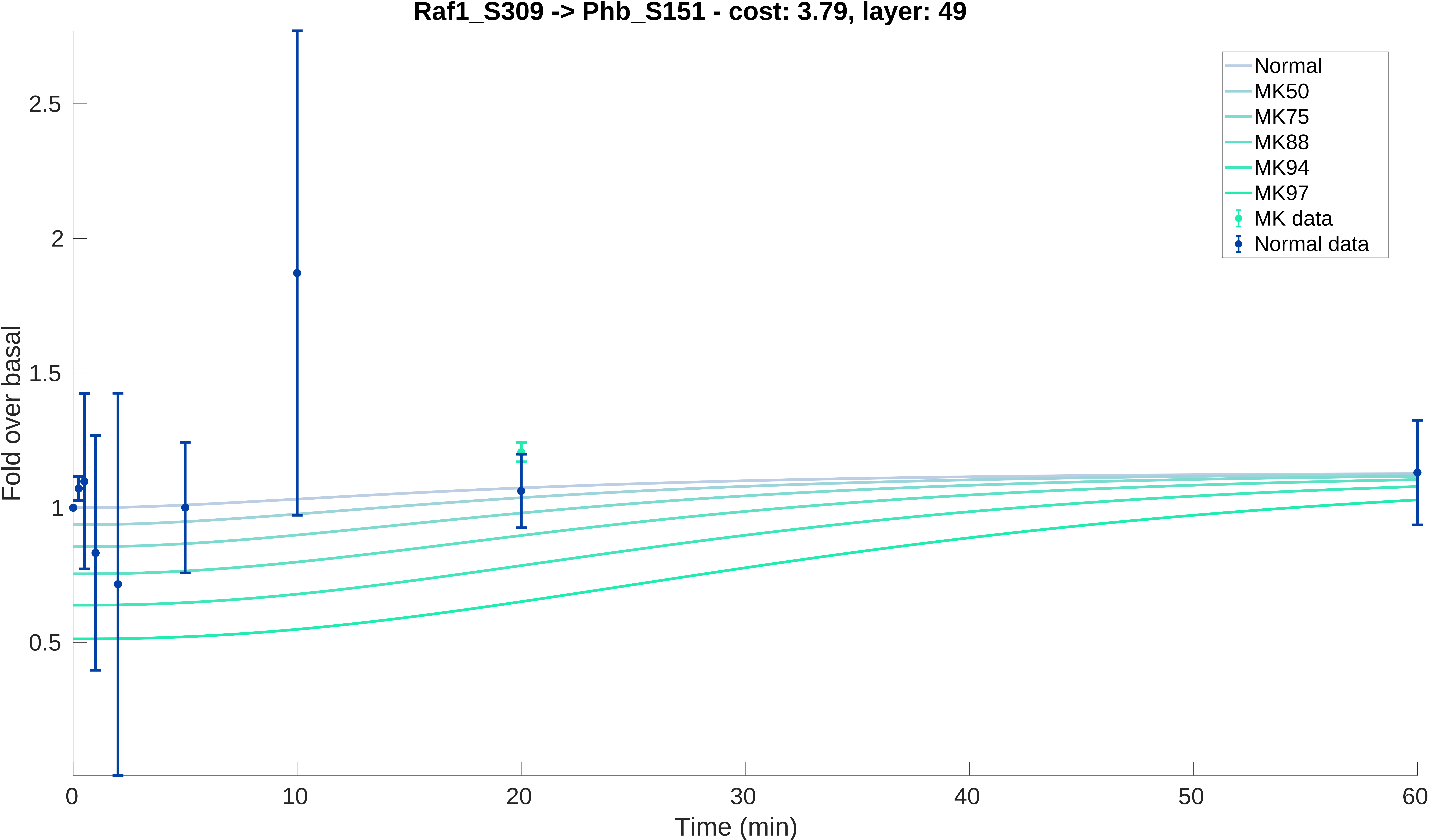

Raf1_S309 -> Phb_S151 - cost: 3.79, layer: 49
Normal
MK50
MK75
MK88
MK94
MK97
MK data
Normal data
Fold over basal
2.5
2
1.5
1
0.5
0
10
20
30
40
50
60
Time (min)

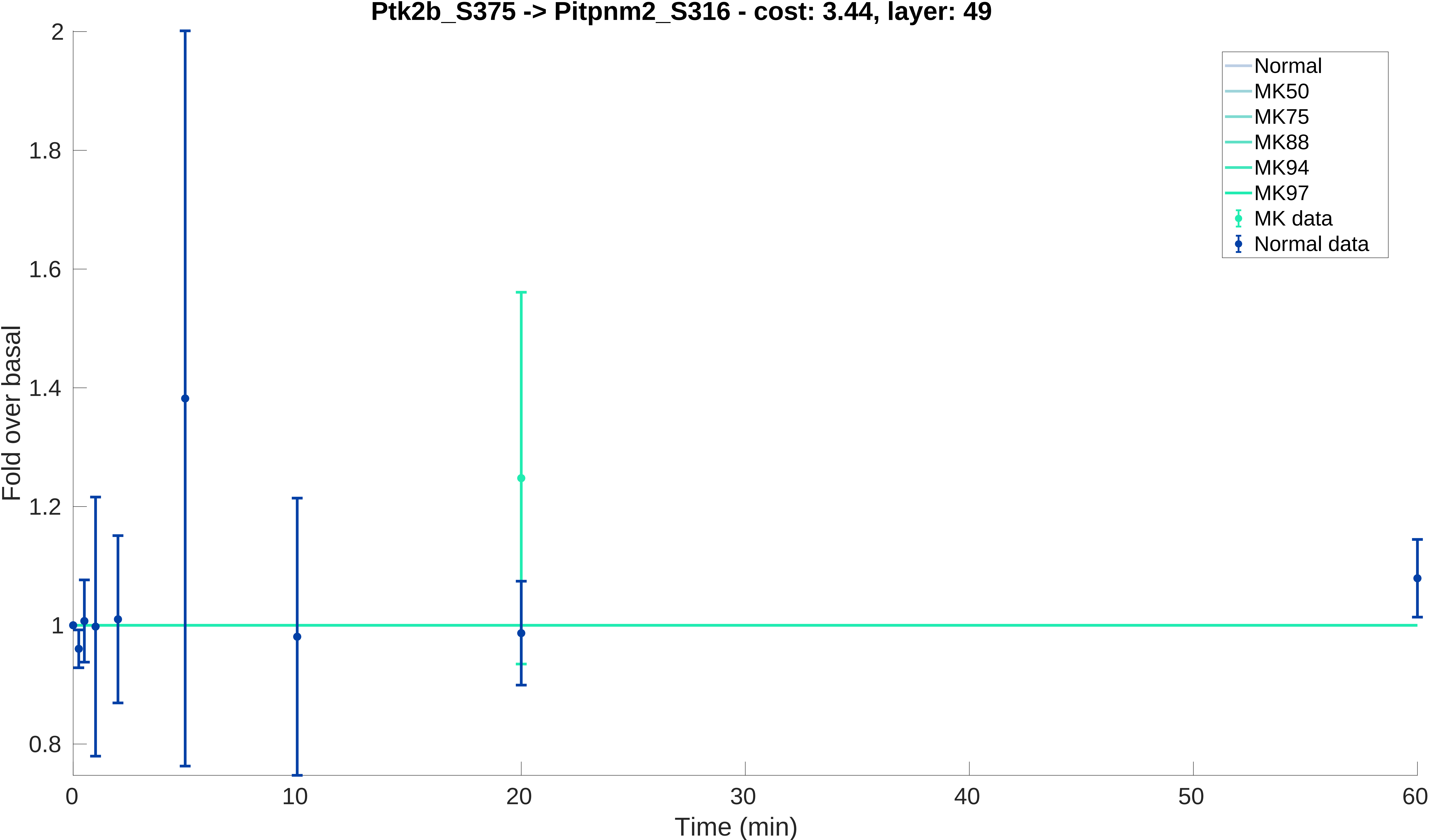

Ptk2b_S375 -> Pitpnm2_S316 - cost: 3.44, layer: 49
Fold over basal
2
1.8
1.6
1.4
1.2
1
0.8
0
10
20
30
40
50
60
Time (min)
Normal
MK50
MK75
MK88
MK94
MK97
MK data
Normal data

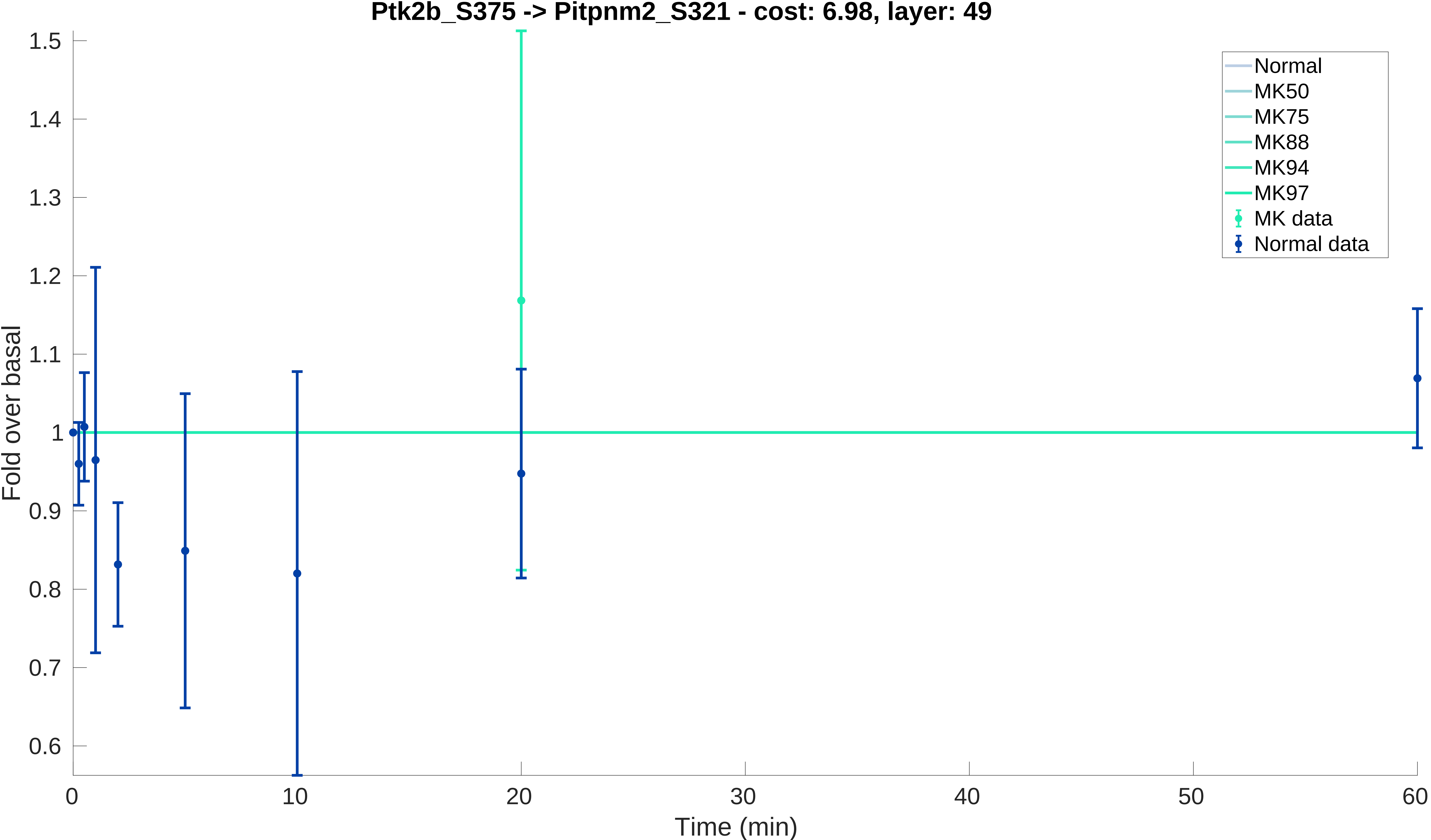

Ptk2b_S375 -> Pitpnm2_S321 - cost: 6.98, layer: 49
Fold over basal
Time (min)
1.5
1.4
1.3
1.2
1.1
1
0.9
0.8
0.7
0.6
0
10
20
30
40
50
60
Normal
MK50
MK75
MK88
MK94
MK97
MK data
Normal data

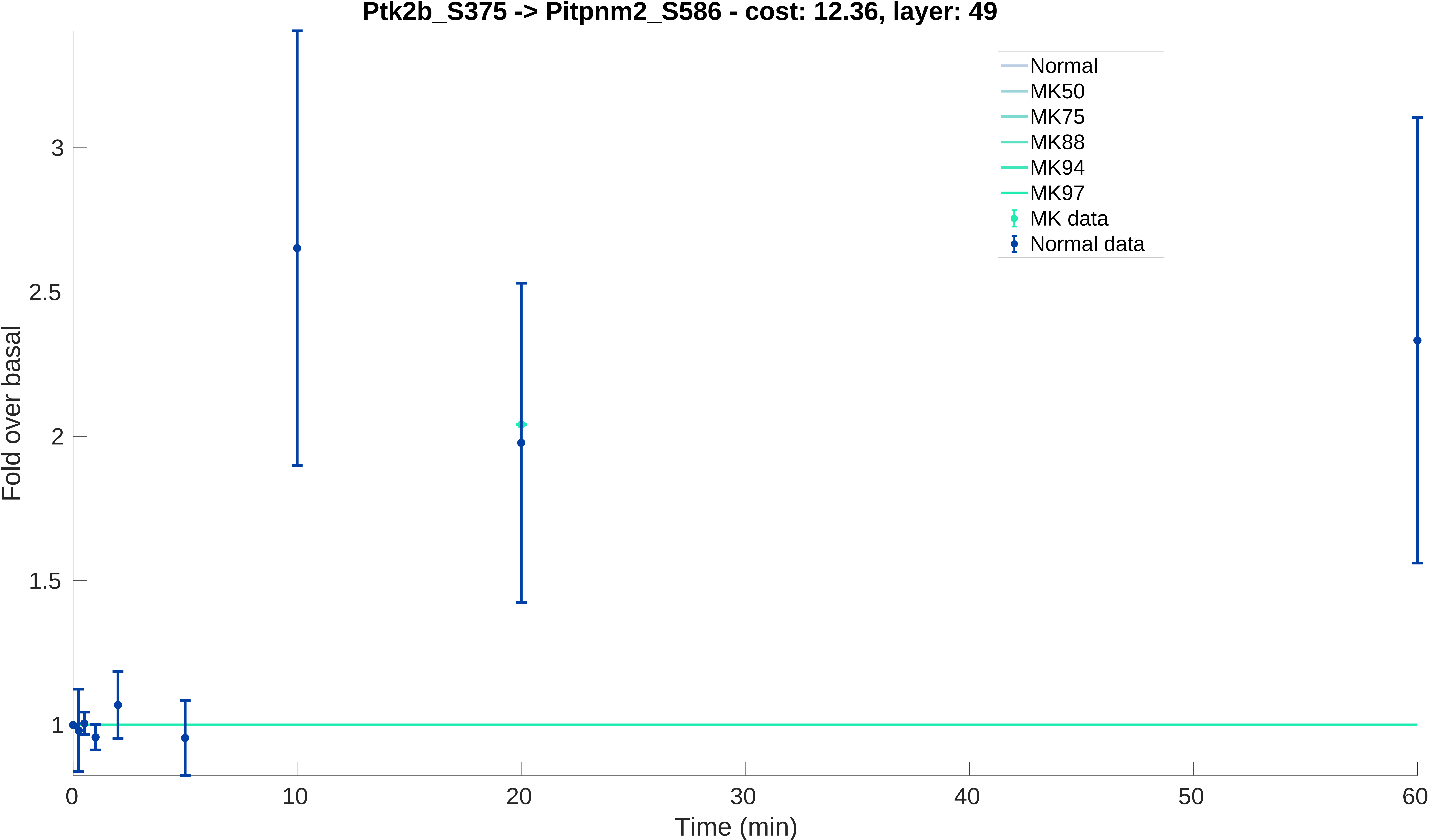

Ptk2b_S375 -> Pitpnm2_S586 - cost: 12.36, layer: 49
Normal
MK50
MK75
MK88
MK94
MK97
MK data
Normal data
Fold over basal
Time (min)
3
2.5
2
1.5
1
0
10
20
30
40
50
60

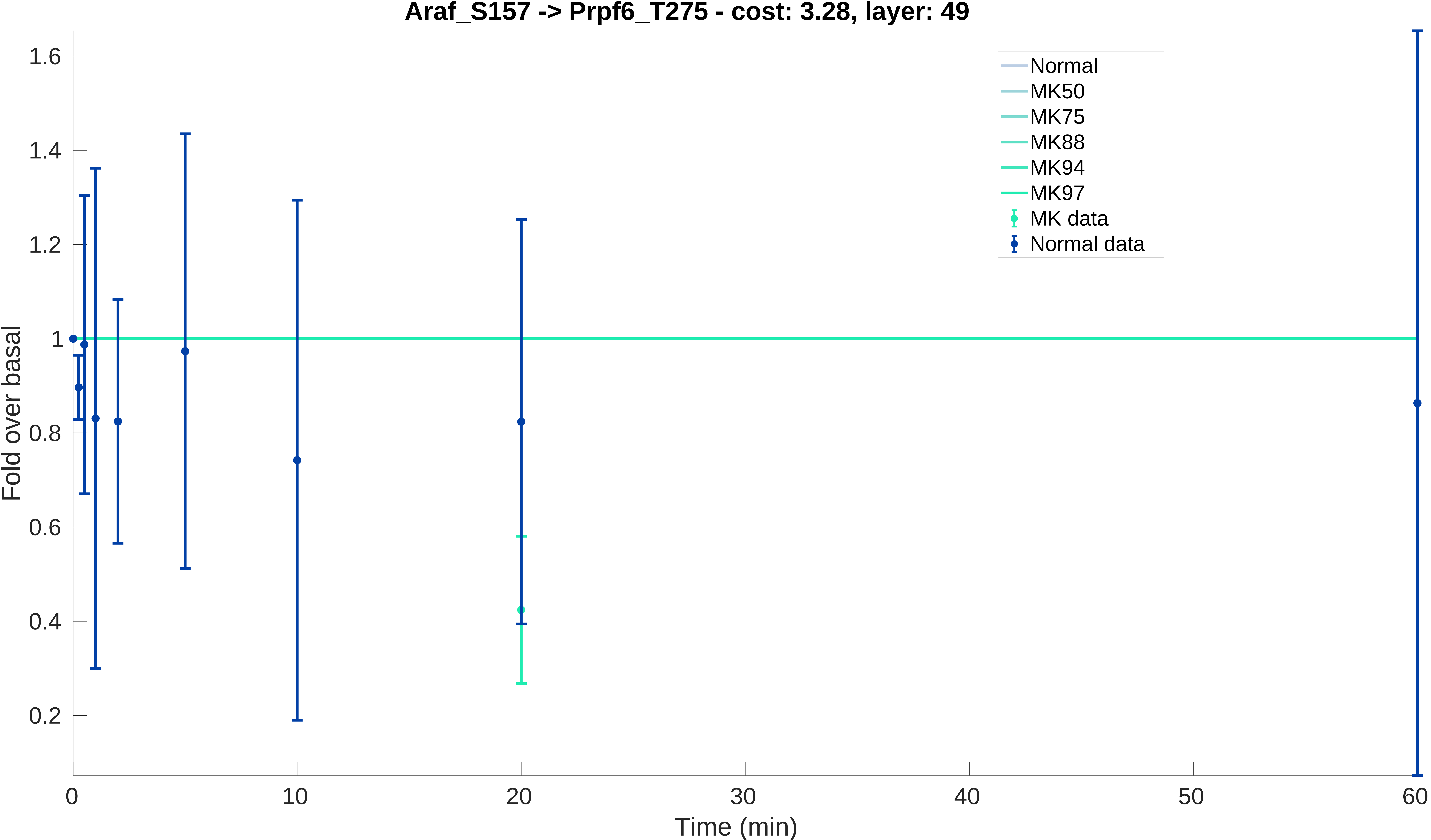
Araf_S157 -> Prpf6_T275 - cost: 3.28, layer: 49
Fold over basal
Time (min)
0
10
20
30
40
50
60
0.2
0.4
0.6
0.8
1
1.2
1.4
1.6
Normal
MK50
MK75
MK88
MK94
MK97
MK data
Normal data

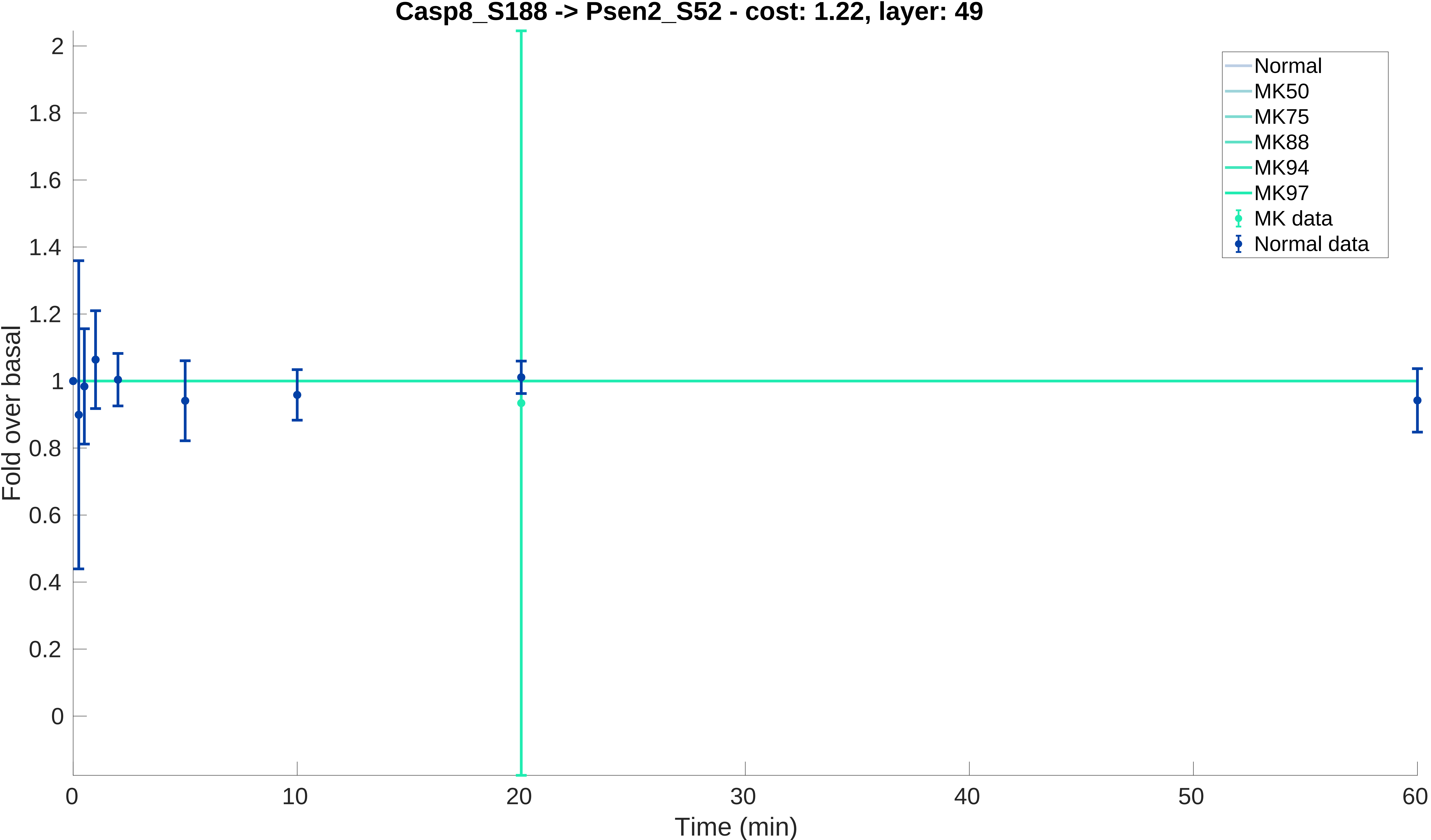

Casp8_S188 -> Psen2_S52 - cost: 1.22, layer: 49
Normal
MK50
MK75
MK88
MK94
MK97
MK data
Normal data
Fold over basal
Time (min)
0
0.2
0.4
0.6
0.8
1
1.2
1.4
1.6
1.8
2
0
10
20
30
40
50
60

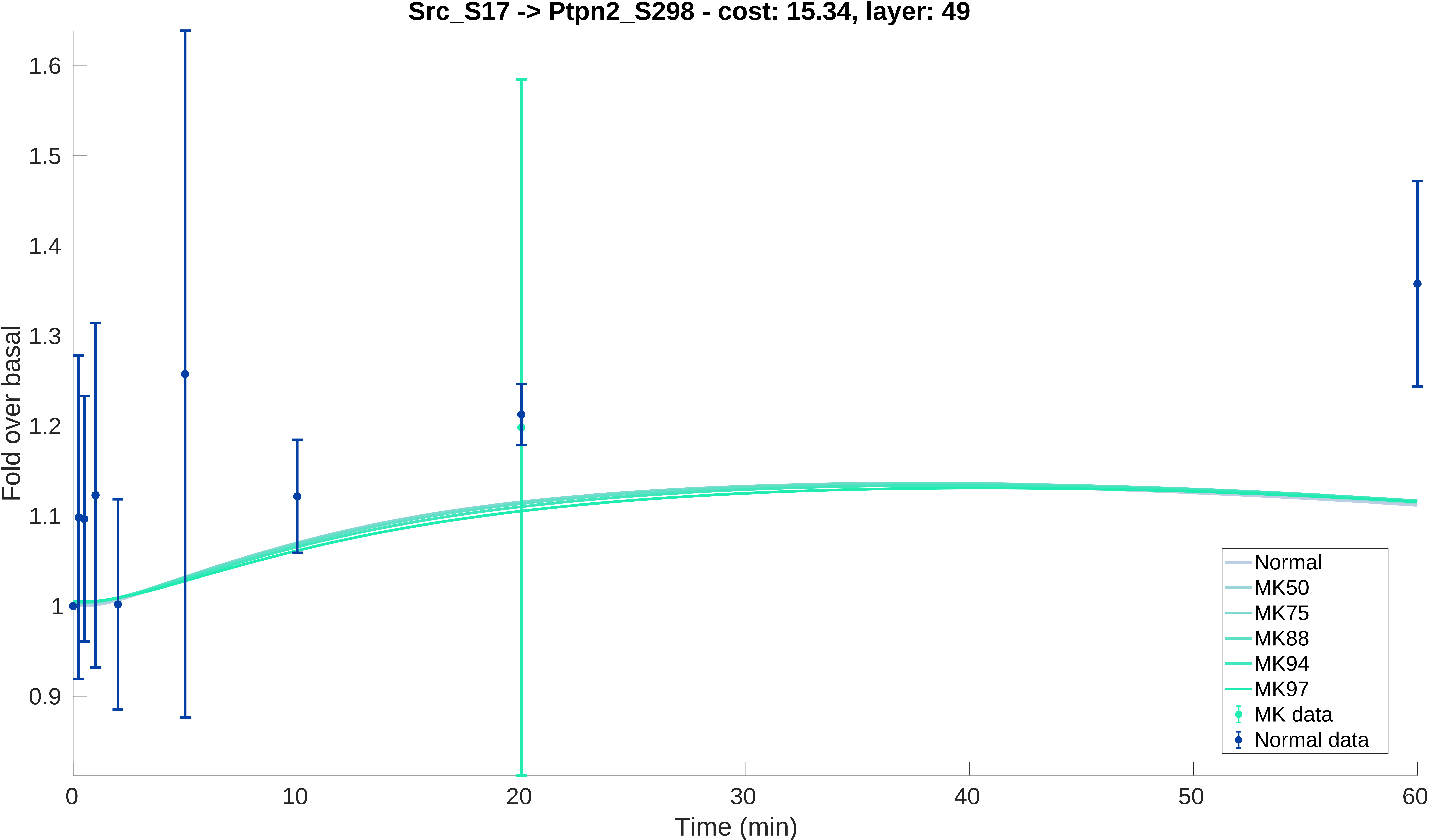

Src_S17 -> Ptpn2_S298 - cost: 15.34, layer: 49
Fold over basal
1.6
1.5
1.4
1.3
1.2
1.1
1
0.9
0
10
20
30
40
50
60
Time (min)
Normal
MK50
MK75
MK88
MK94
MK97
MK data
Normal data

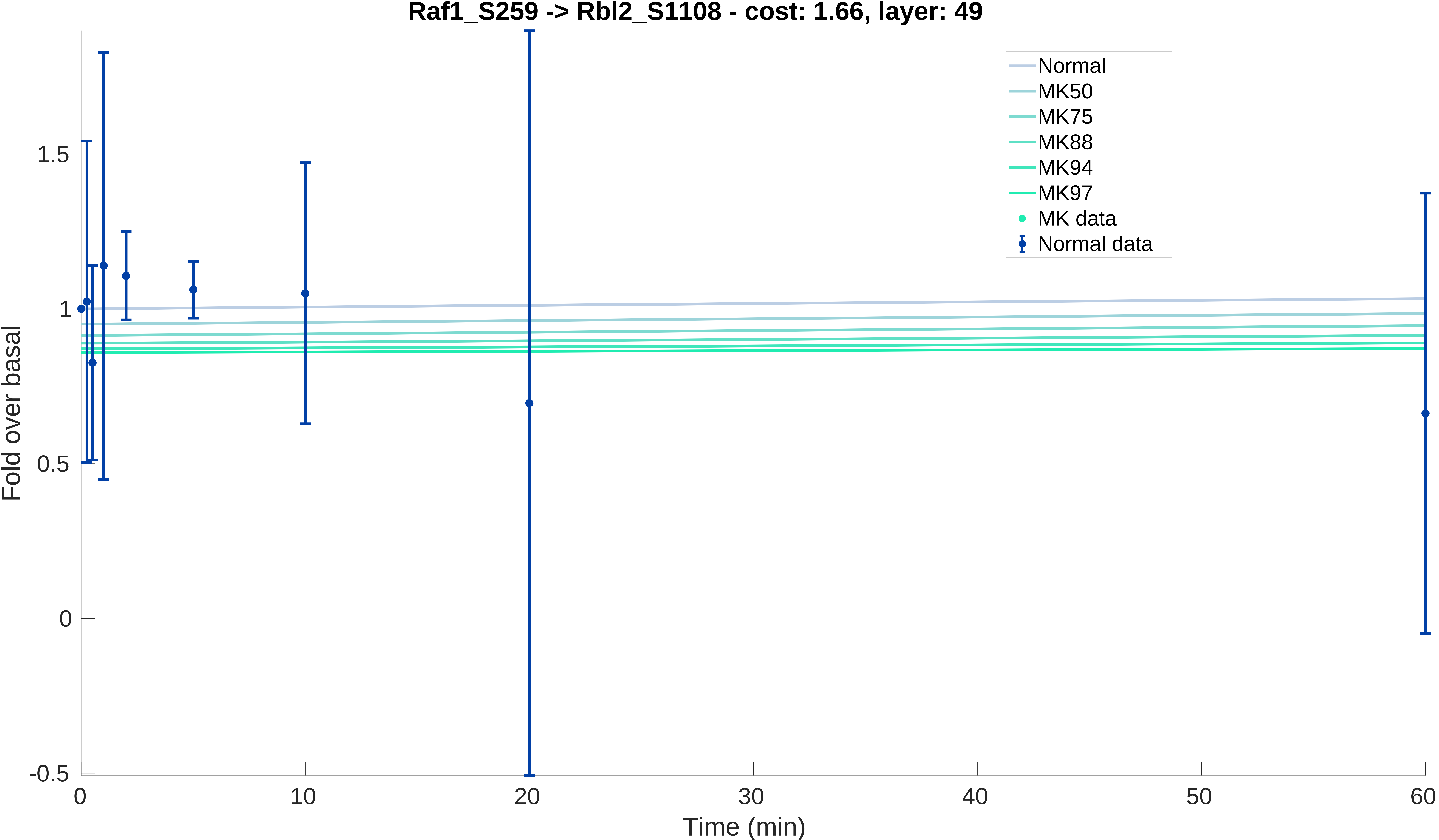

Raf1_S259 -> Rbl2_S1108 - cost: 1.66, layer: 49
Normal
MK50
MK75
MK88
MK94
MK97
MK data
Normal data
Fold over basal
1.5
1
0.5
0
-0.5
0
10
20
30
40
50
60
Time (min)

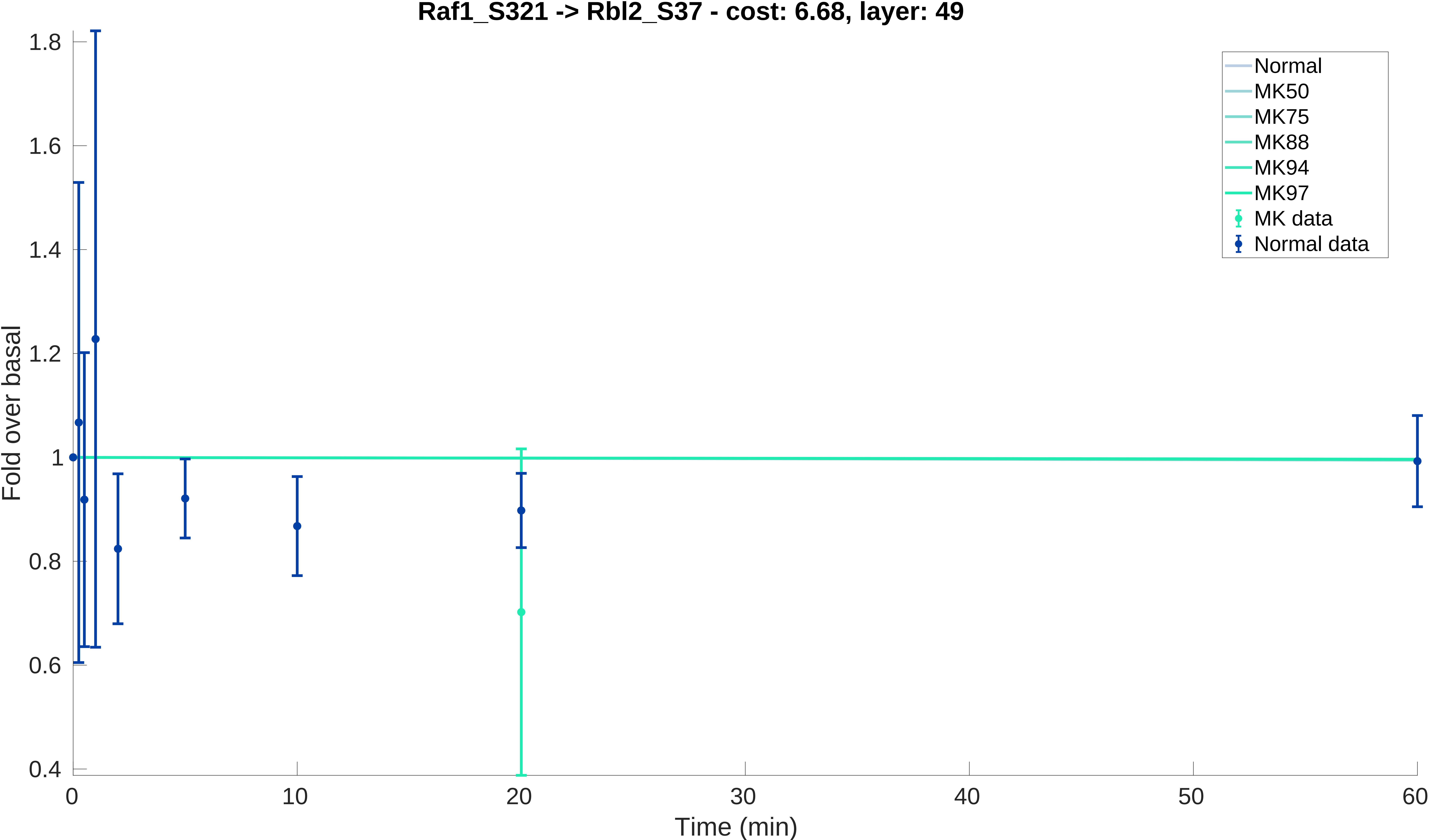

Raf1_S321 -> Rbl2_S37 - cost: 6.68, layer: 49
Fold over basal
1.8
1.6
1.4
1.2
1
0.8
0.6
0.4
0
10
20
30
40
50
60
Time (min)
Normal
MK50
MK75
MK88
MK94
MK97
MK data
Normal data

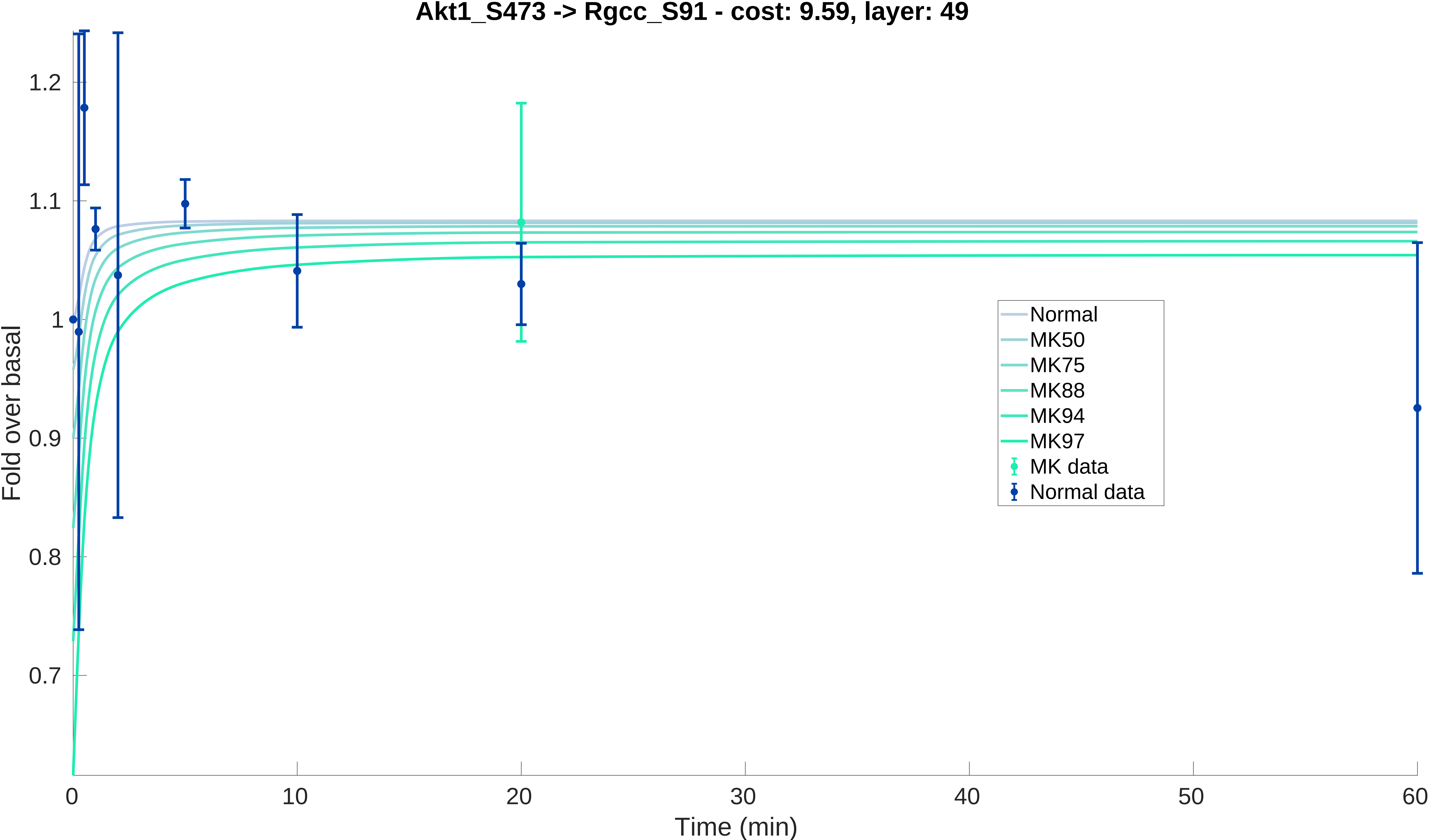
Akt1_S473 -> Rgcc_S91 - cost: 9.59, layer: 49
Fold over basal
1.2
1.1
1
0.9
0.8
0.7
0
10
20
30
40
50
60
Time (min)
Normal
MK50
MK75
MK88
MK94
MK97
MK data
Normal data

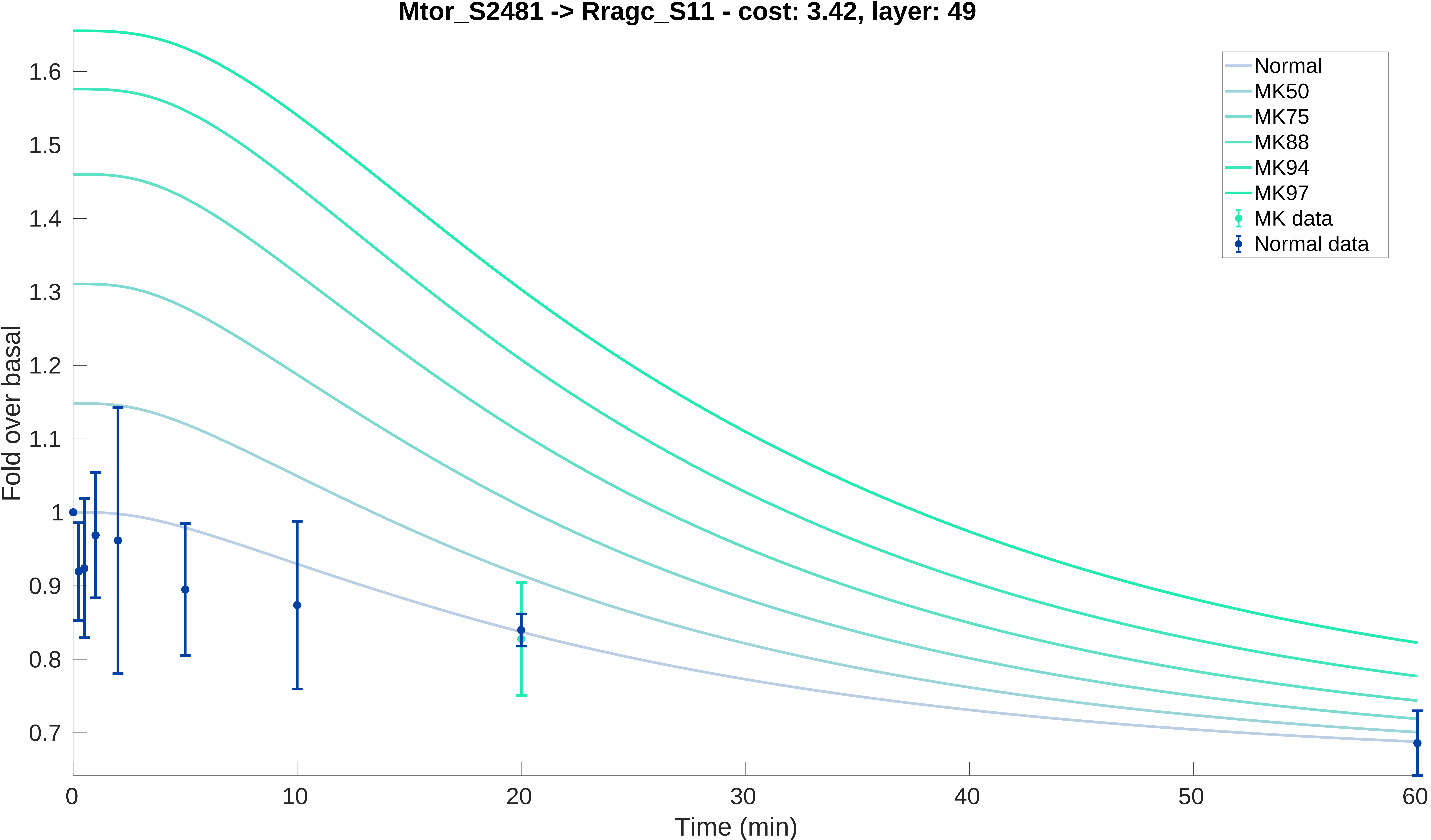

Mtor_S2481 -> Rragc_S11 - cost: 3.42, layer: 49
Fold over basal
Time (min)
0.7
0.8
0.9
1
1.1
1.2
1.3
1.4
1.5
1.6
0
10
20
30
40
50
60
Normal
MK50
MK75
MK88
MK94
MK97
MK data
Normal data

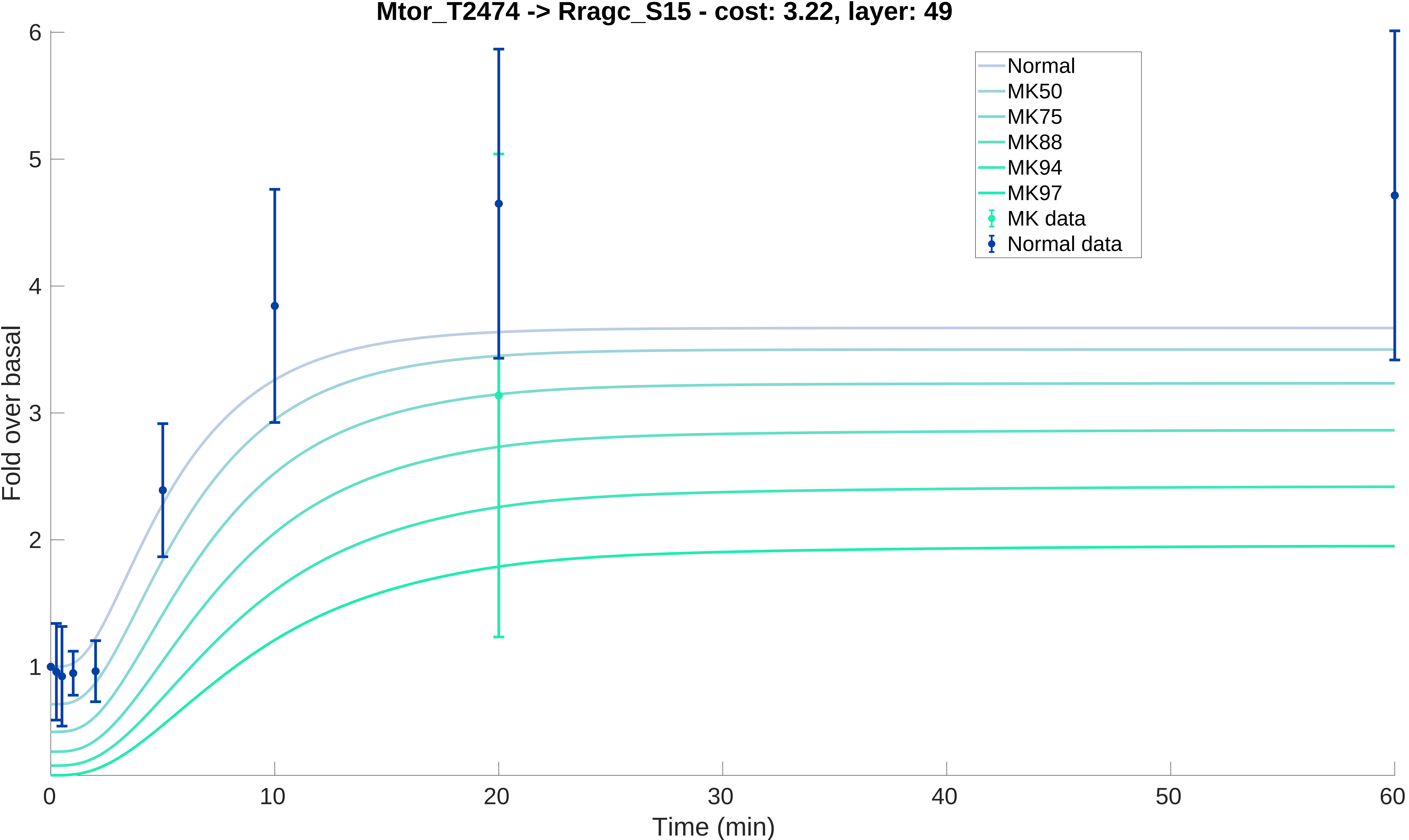

Mtor_T2474 -> Rragc_S15 - cost: 3.22, layer: 49
Fold over basal
Time (min)
0
10
20
30
40
50
60
1
2
3
4
5
6
Normal
MK50
MK75
MK88
MK94
MK97
MK data
Normal data

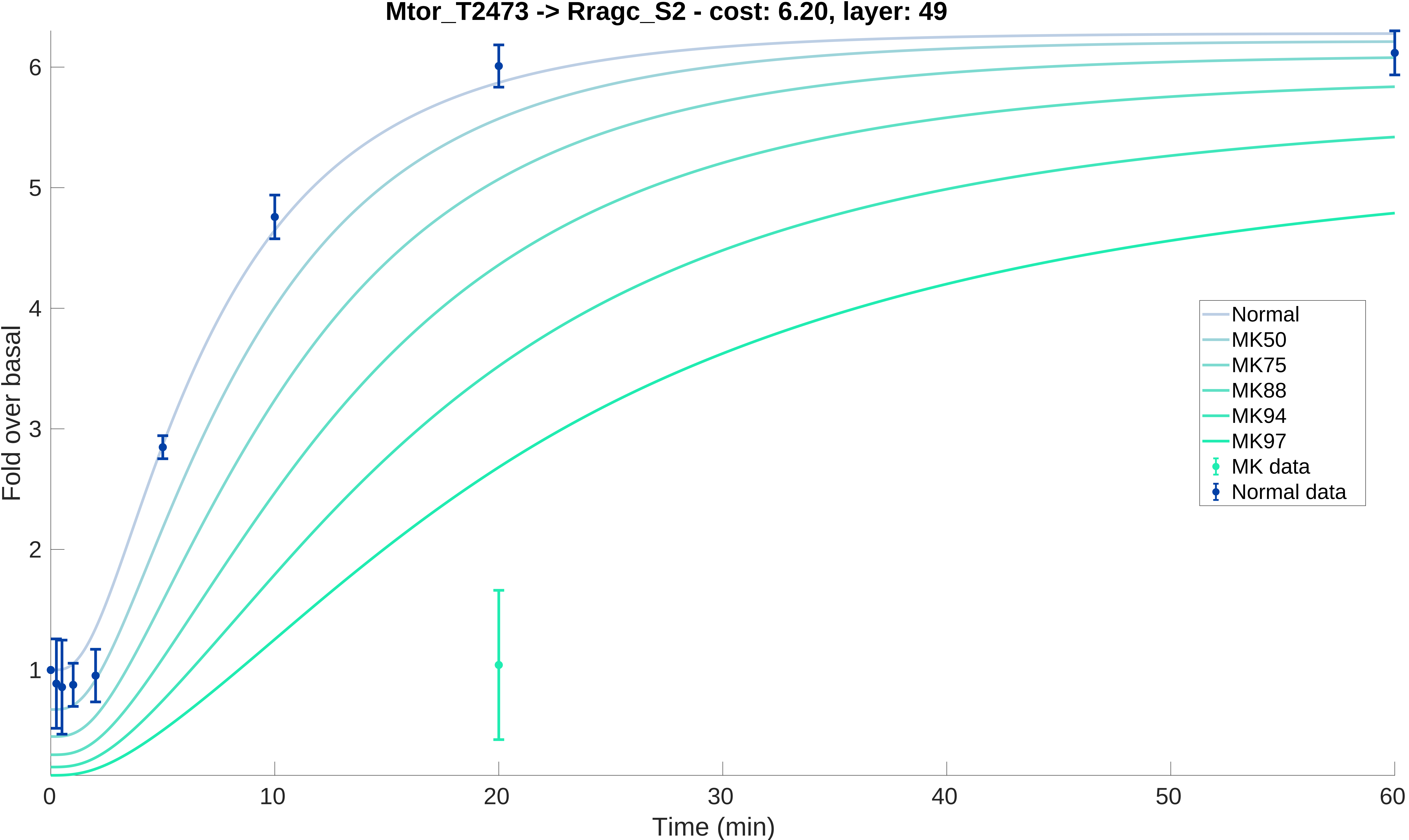
Mtor_T2473 -> Rragc_S2 - cost: 6.20, layer: 49
Fold over basal
Time (min)
0
10
20
30
40
50
60
1
2
3
4
5
6
Normal
MK50
MK75
MK88
MK94
MK97
MK data
Normal data

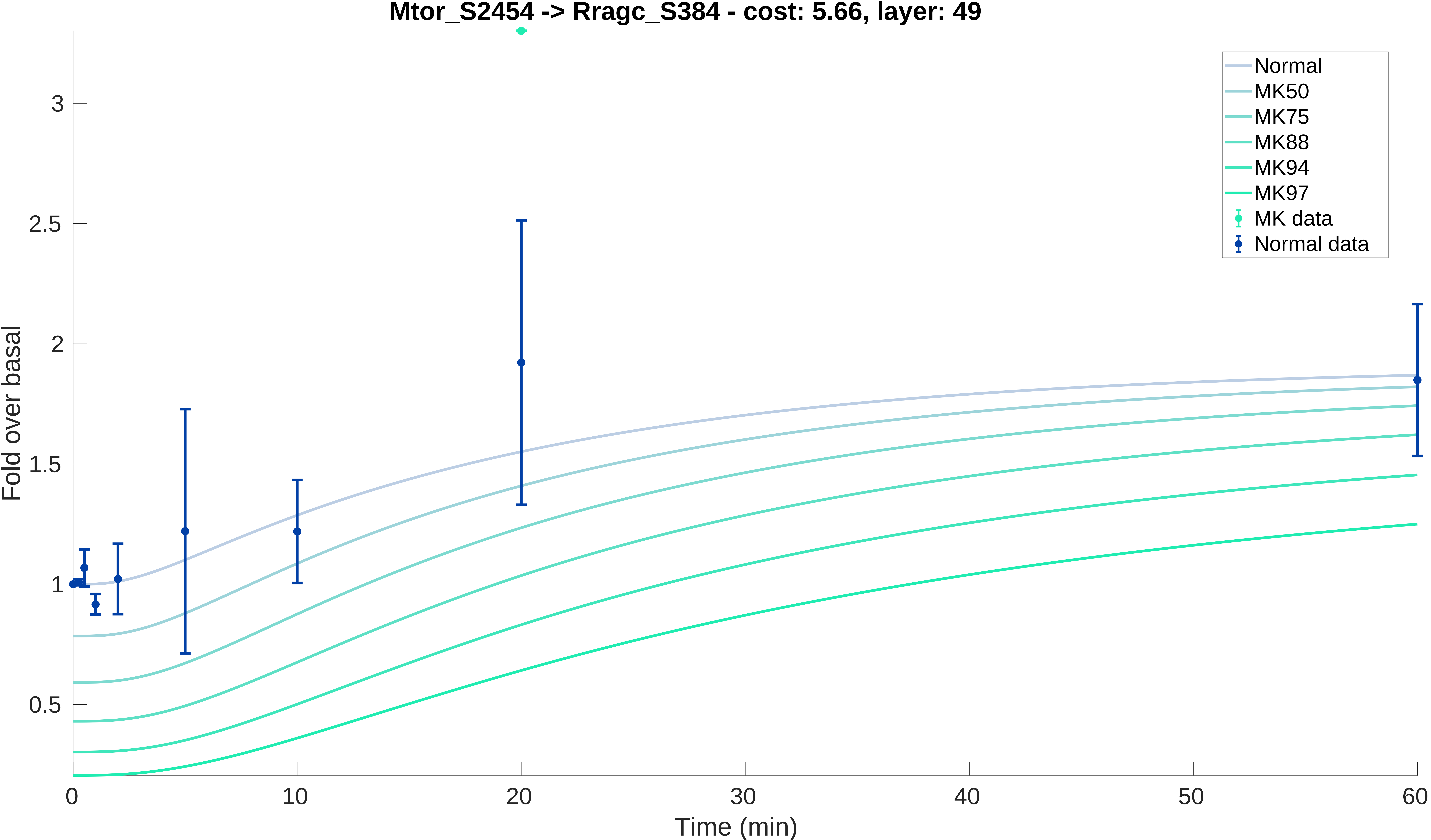

Mtor_S2454 -> Rragc_S384 - cost: 5.66, layer: 49
Normal
MK50
MK75
MK88
MK94
MK97
MK data
Normal data
Fold over basal
Time (min)
0.5
1
1.5
2
2.5
3
0
10
20
30
40
50
60

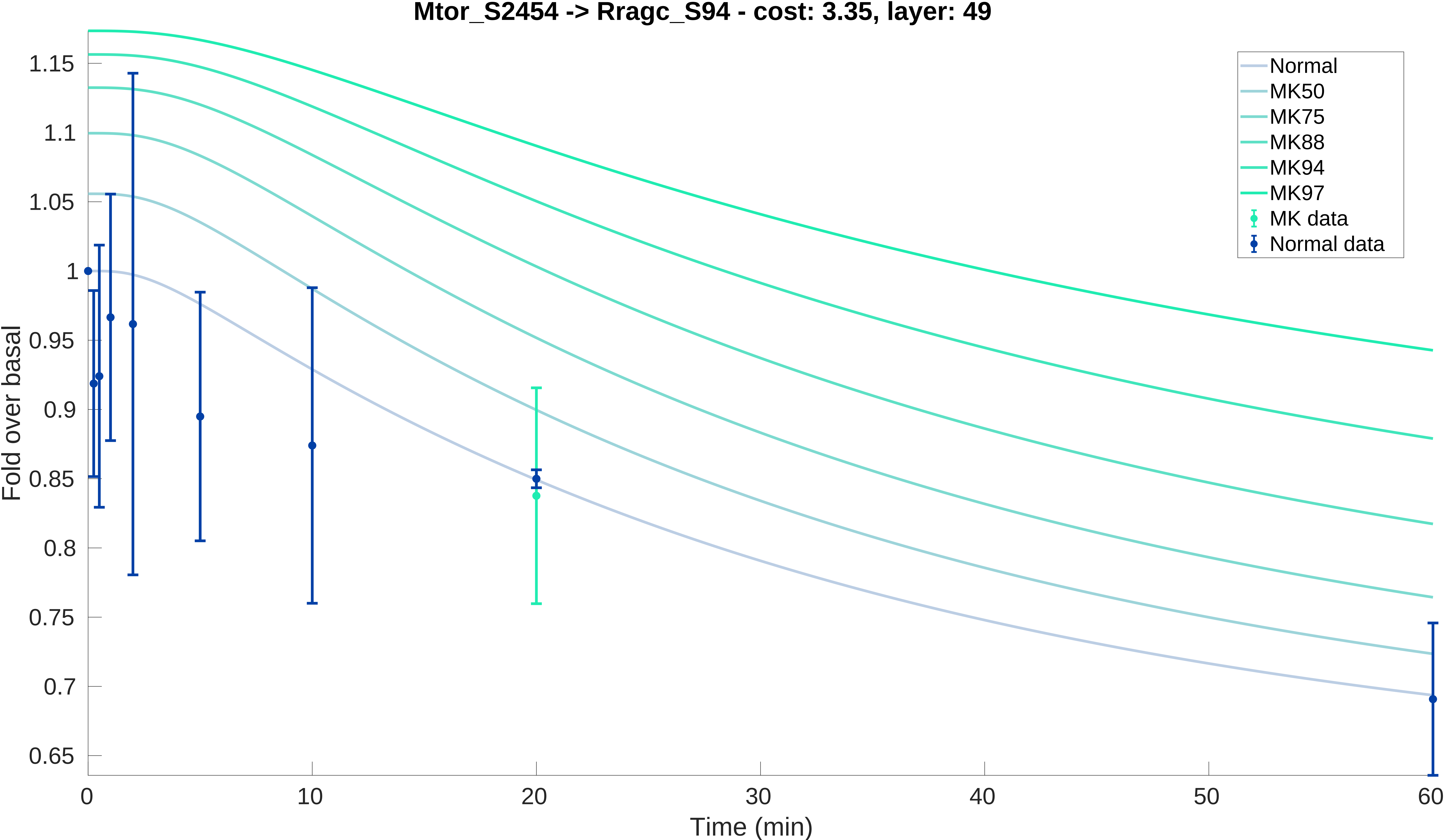
Mtor_S2454 -> Rragc_S94 - cost: 3.35, layer: 49
Fold over basal
Time (min)
0.65
0.7
0.75
0.8
0.85
0.9
0.95
1
1.05
1.1
1.15
0
10
20
30
40
50
60
Normal
MK50
MK75
MK88
MK94
MK97
MK data
Normal data

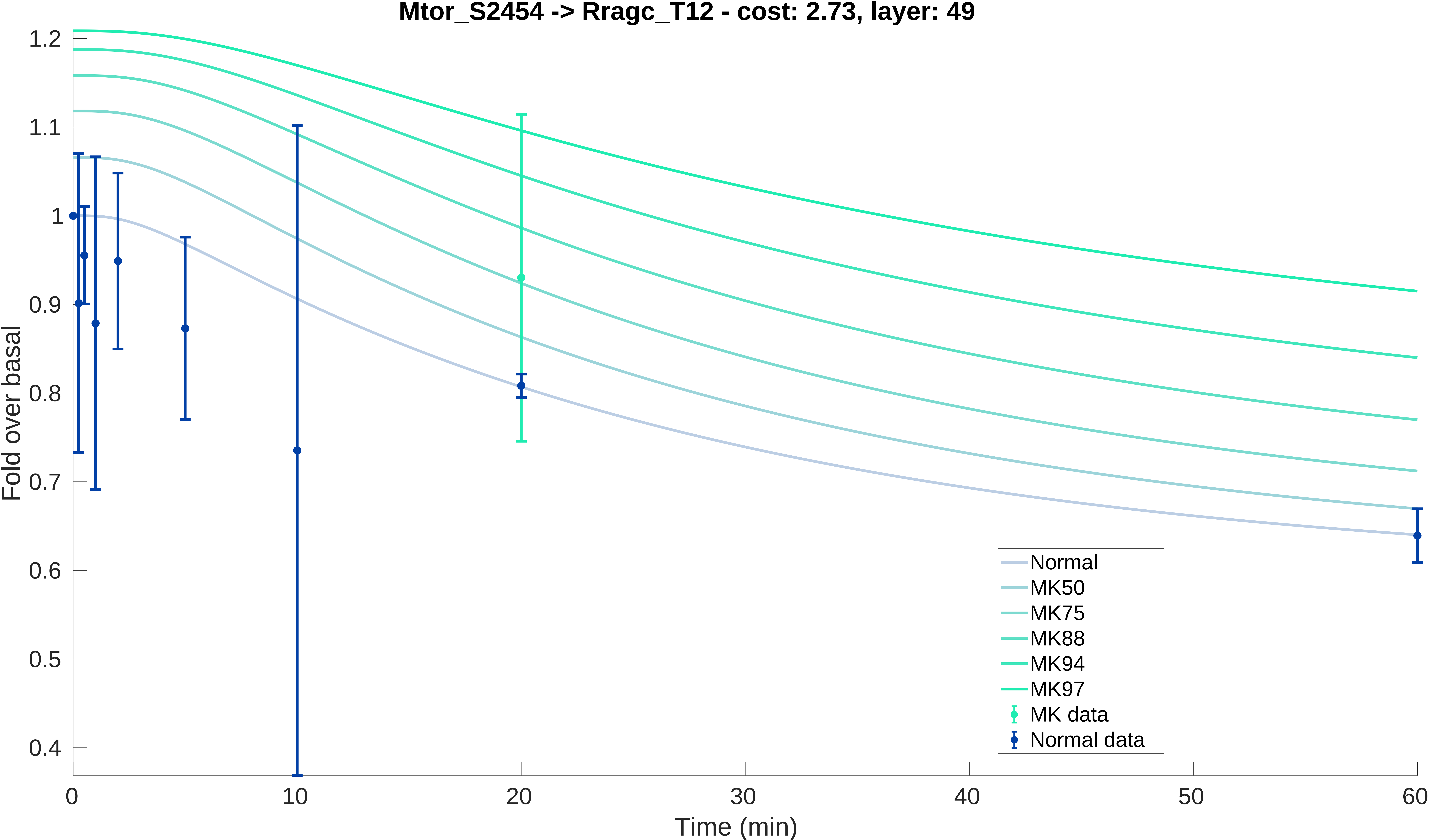

Mtor_S2454 -> Rragc_T12 - cost: 2.73, layer: 49
Fold over basal
Time (min)
0.4
0.5
0.6
0.7
0.8
0.9
1
1.1
1.2
0
10
20
30
40
50
60
Normal
MK50
MK75
MK88
MK94
MK97
MK data
Normal data

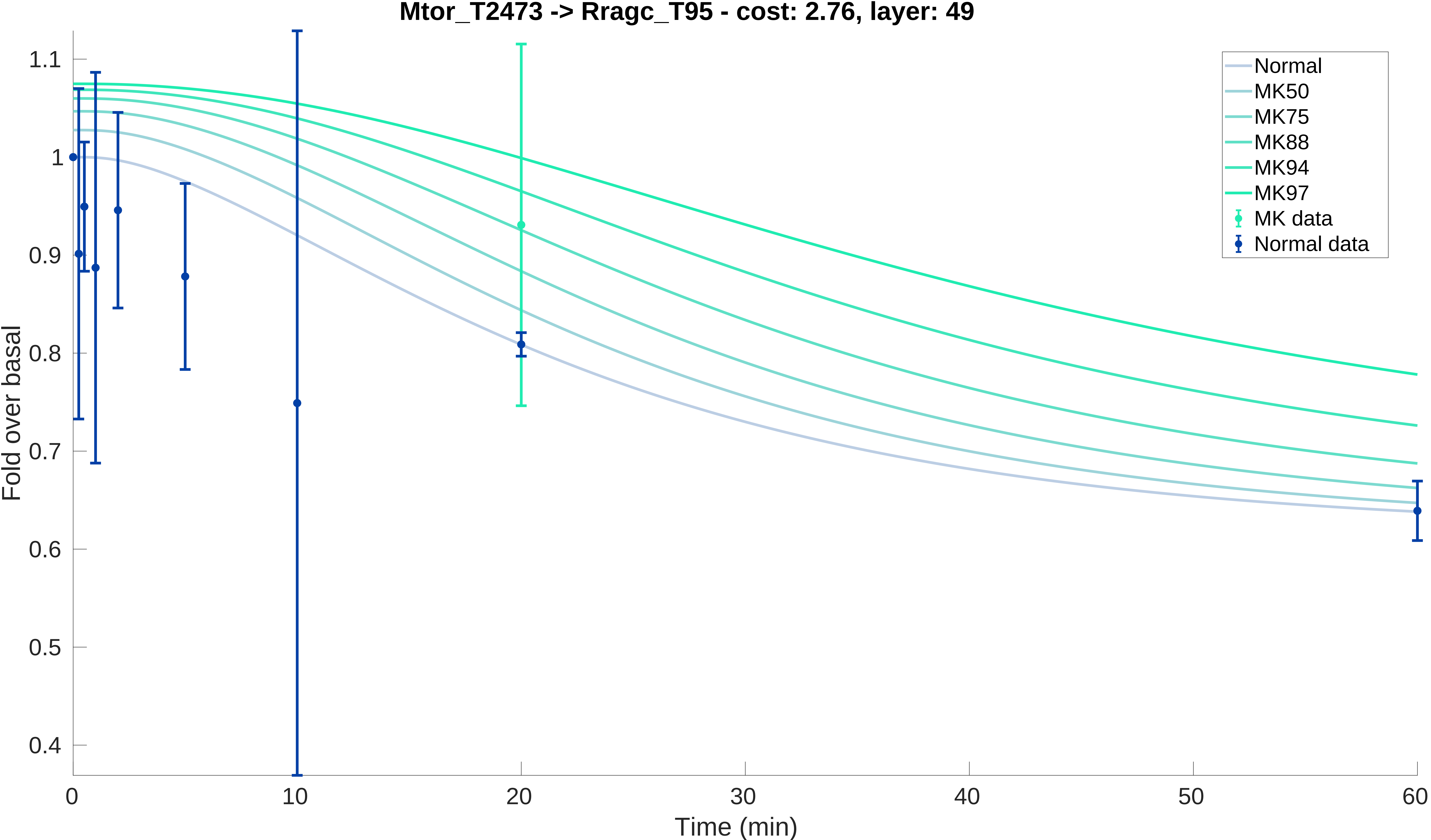

Mtor_T2473 -> Rragc_T95 - cost: 2.76, layer: 49
Fold over basal
Time (min)
1.1
1
0.9
0.8
0.7
0.6
0.5
0.4
0
10
20
30
40
50
60
Normal
MK50
MK75
MK88
MK94
MK97
MK data
Normal data

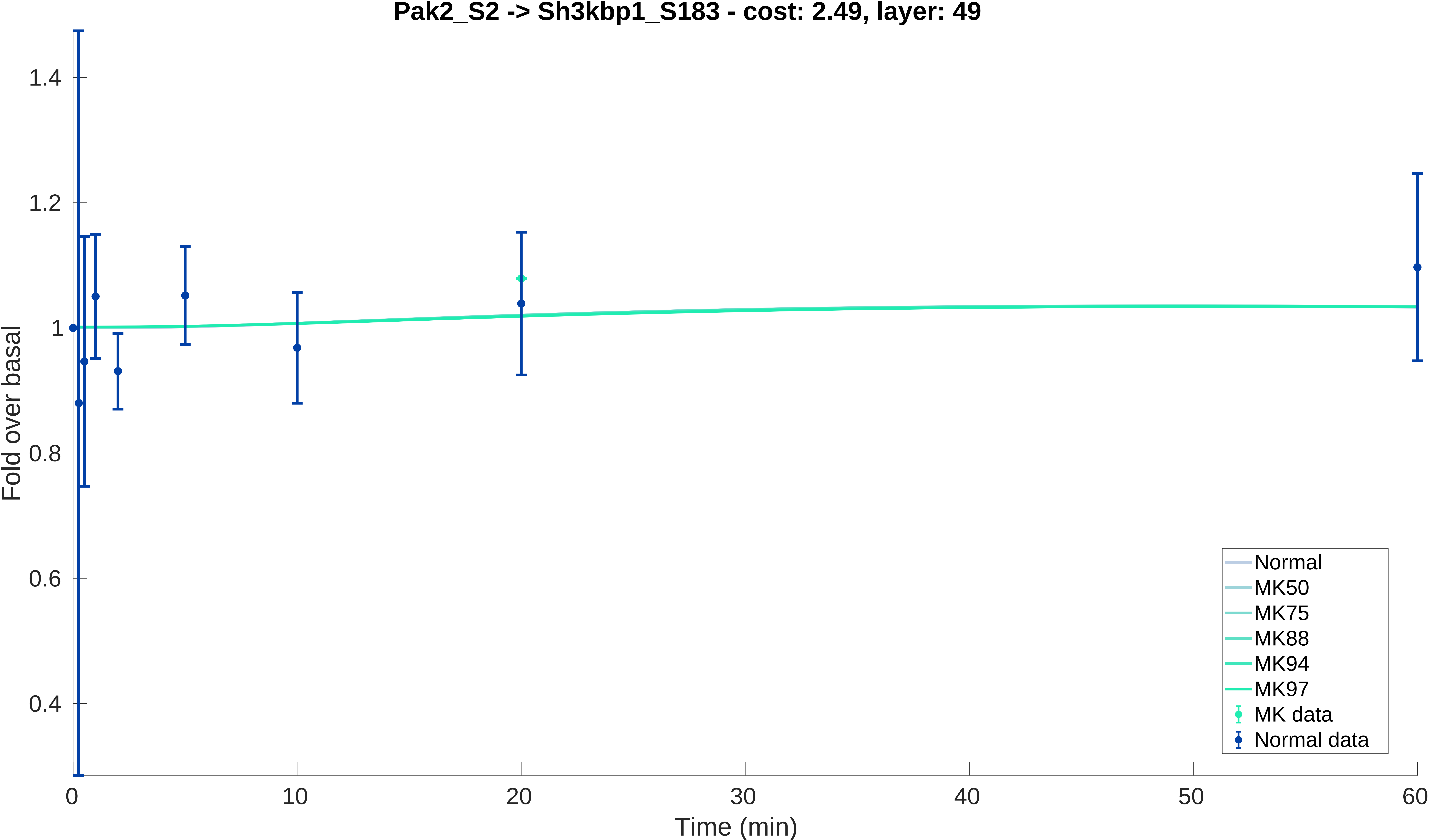

Pak2_S2 -> Sh3kbp1_S183 - cost: 2.49, layer: 49
Fold over basal
Time (min)
1.4
1.2
1
0.8
0.6
0.4
0
10
20
30
40
50
60
Normal
MK50
MK75
MK88
MK94
MK97
MK data
Normal data

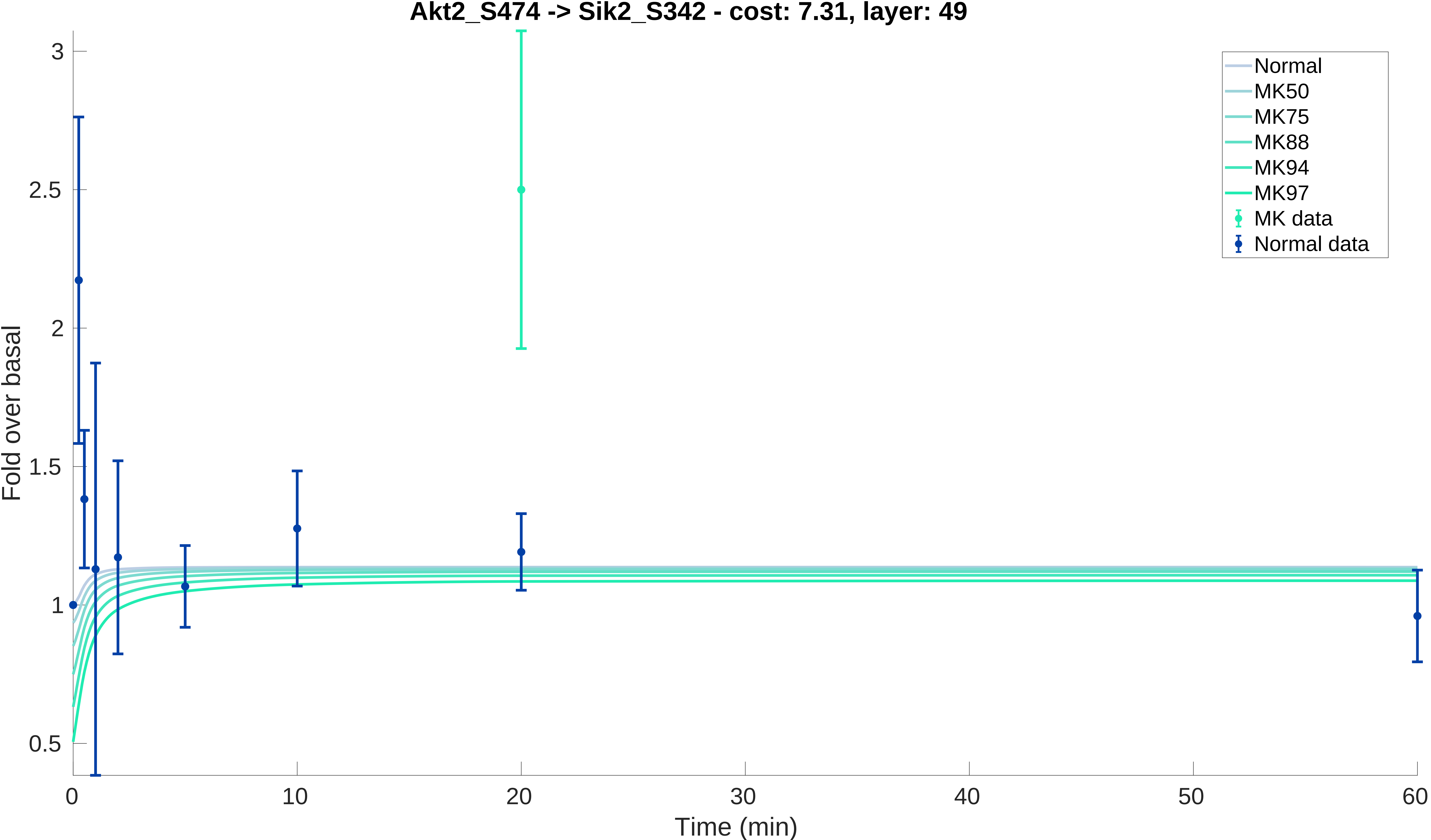
Akt2_S474 -> Sik2_S342 - cost: 7.31, layer: 49
Fold over basal
Time (min)
0
10
20
30
40
50
60
0.5
1
1.5
2
2.5
3
Normal
MK50
MK75
MK88
MK94
MK97
MK data
Normal data

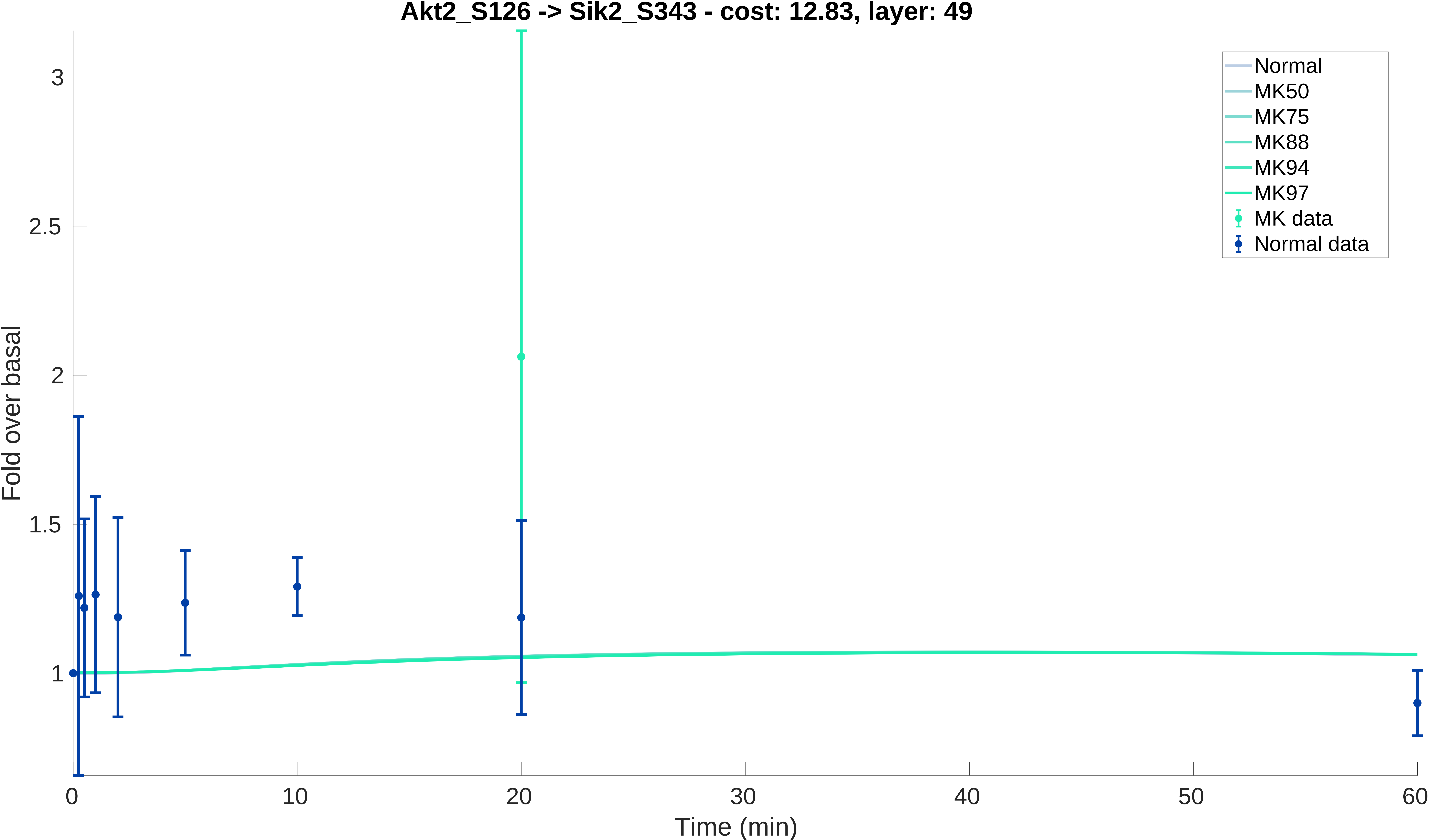

Akt2_S126 -> Sik2_S343 - cost: 12.83, layer: 49
Fold over basal
Time (min)
3
2.5
2
1.5
1
0
10
20
30
40
50
60
Normal
MK50
MK75
MK88
MK94
MK97
MK data
Normal data

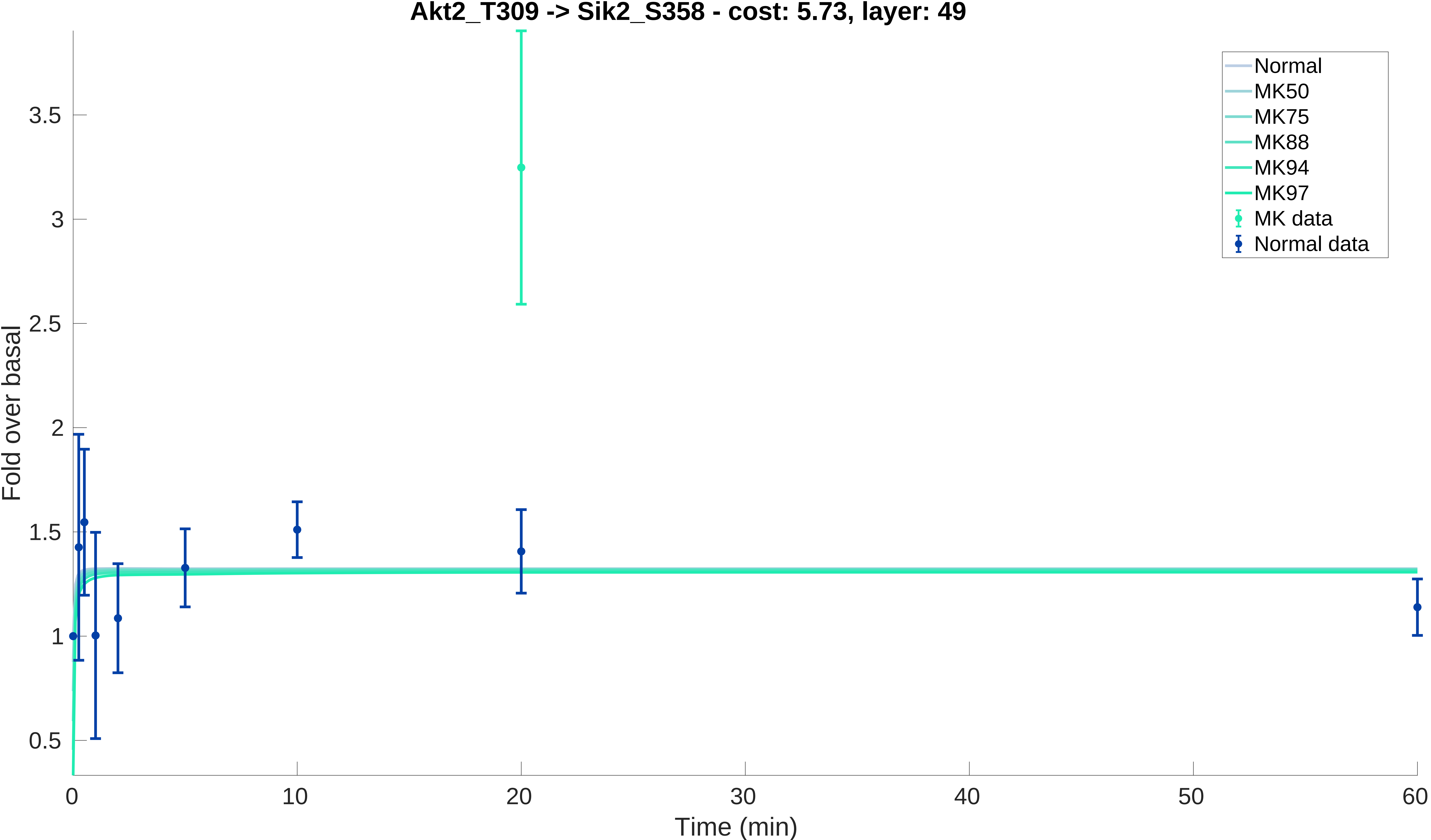

Akt2_T309 -> Sik2_S358 - cost: 5.73, layer: 49
Normal
MK50
MK75
MK88
MK94
MK97
MK data
Normal data
Fold over basal
3.5
3
2.5
2
1.5
1
0.5
0
10
20
30
40
50
60
Time (min)

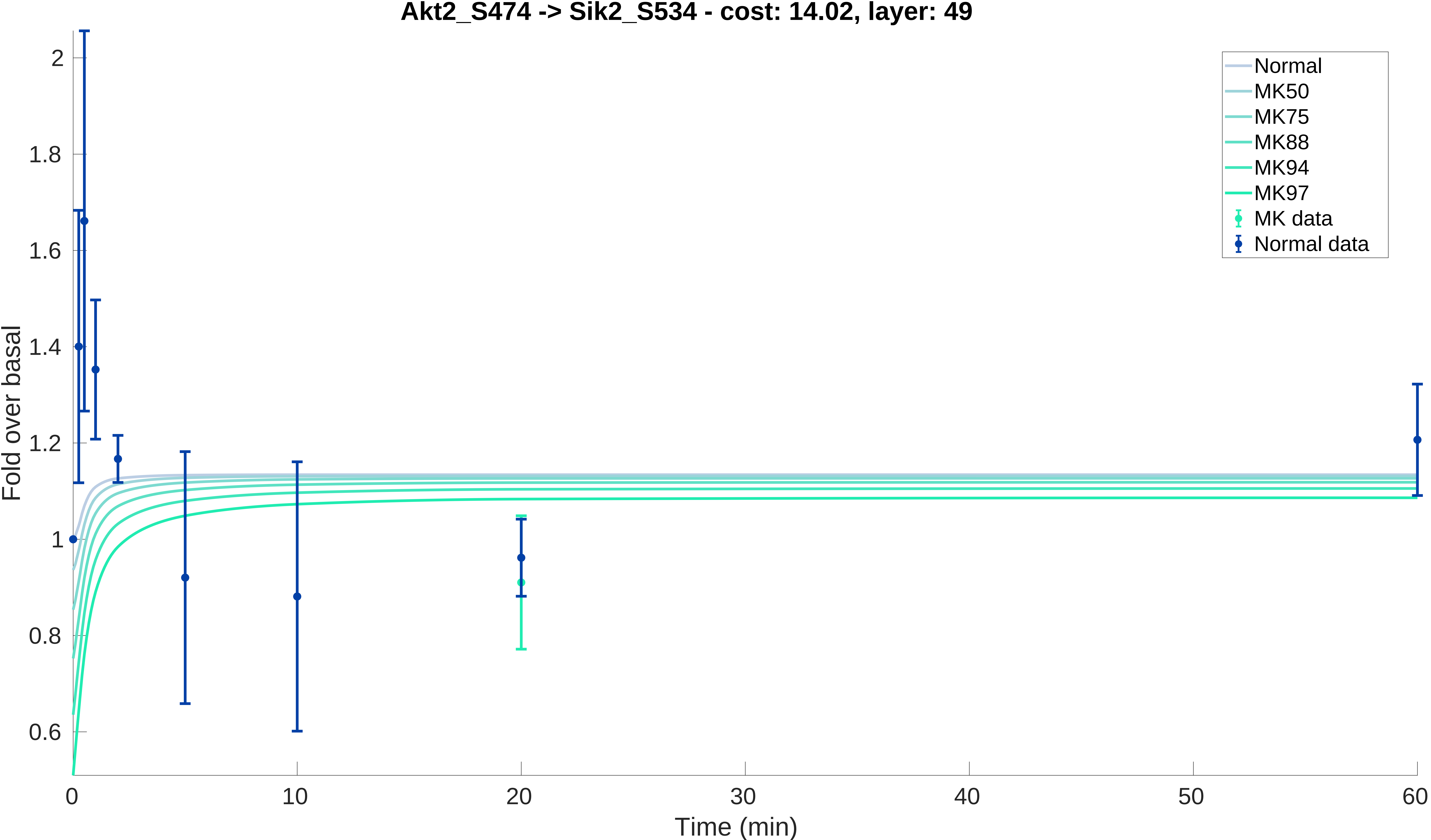

Akt2_S474 -> Sik2_S534 - cost: 14.02, layer: 49
Fold over basal
Time (min)
2
1.8
1.6
1.4
1.2
1
0.8
0.6
0
10
20
30
40
50
60
Normal
MK50
MK75
MK88
MK94
MK97
MK data
Normal data

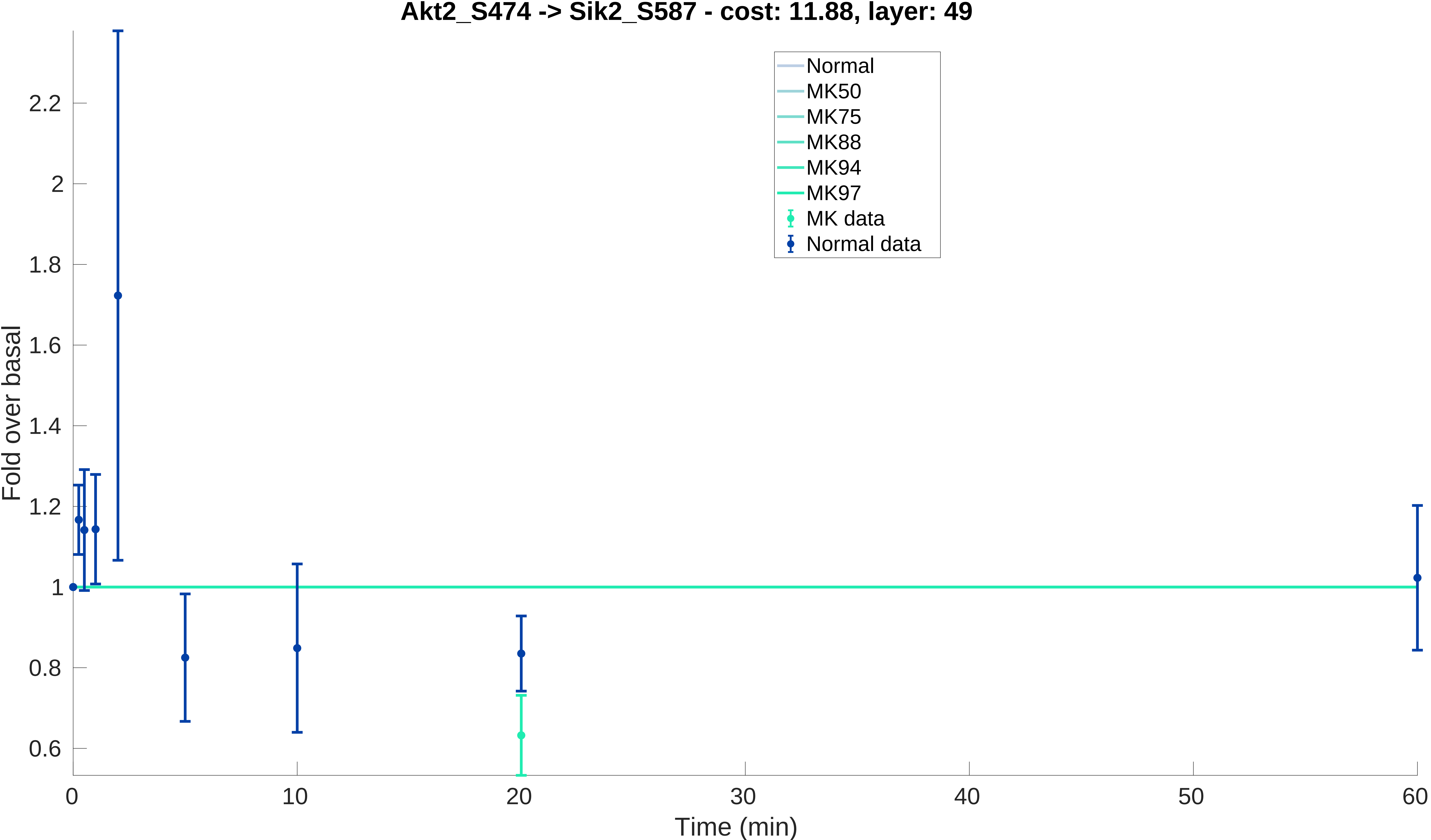
Akt2_S474 -> Sik2_S587 - cost: 11.88, layer: 49
Normal
MK50
MK75
MK88
MK94
MK97
MK data
Normal data
Fold over basal
2.2
2
1.8
1.6
1.4
1.2
1
0.8
0.6
0
10
20
30
40
50
60
Time (min)

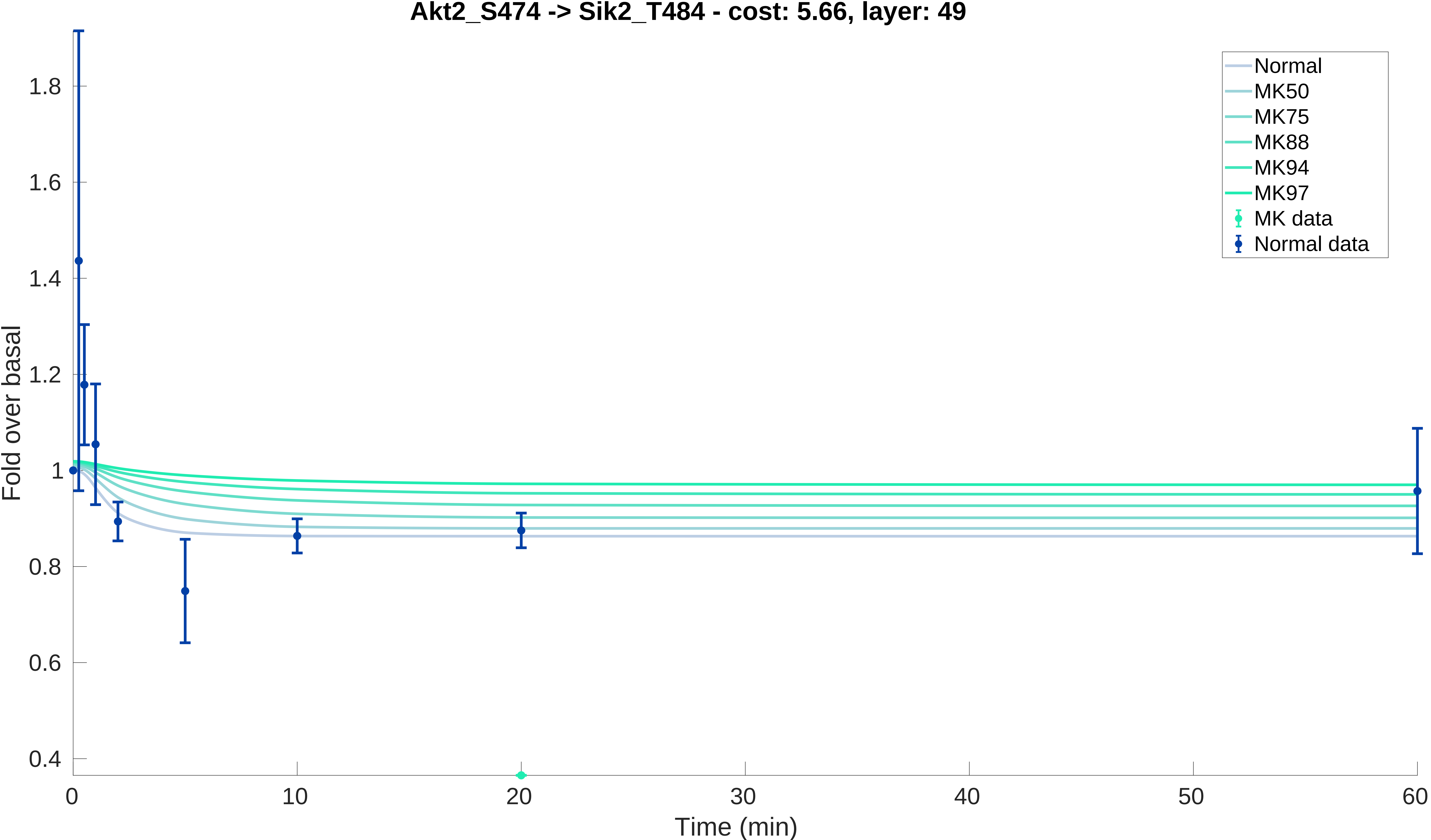

Akt2_S474 -> Sik2_T484 - cost: 5.66, layer: 49
Normal
MK50
MK75
MK88
MK94
MK97
MK data
Normal data
Fold over basal
0.4
0.6
0.8
1
1.2
1.4
1.6
1.8
0
10
20
30
40
50
60
Time (min)

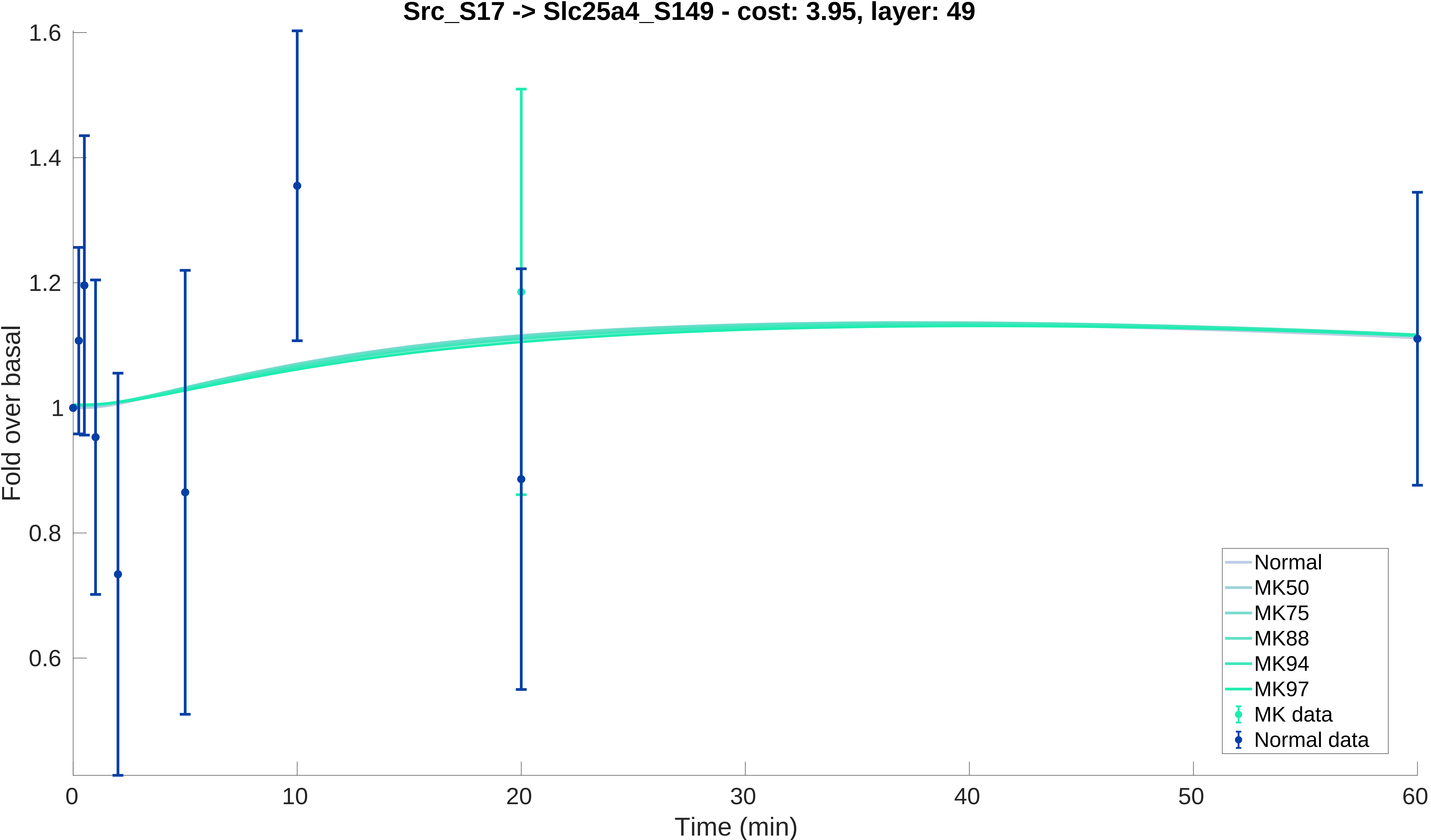

Src_S17 -> Slc25a4_S149 - cost: 3.95, layer: 49
Fold over basal
Time (min)
1.6
1.4
1.2
1
0.8
0.6
0
10
20
30
40
50
60
Normal
MK50
MK75
MK88
MK94
MK97
MK data
Normal data

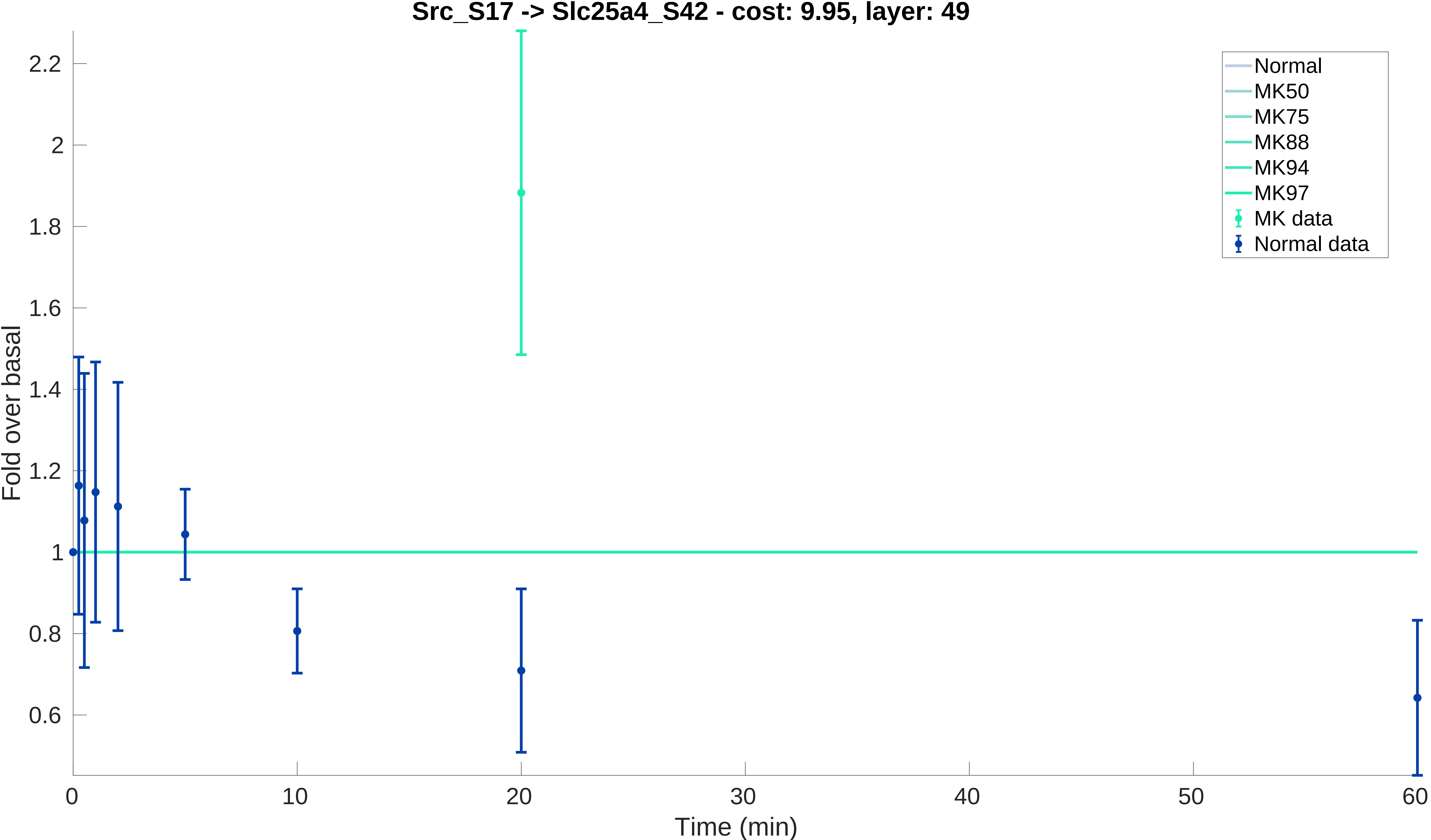

Src_S17 -> Slc25a4_S42 - cost: 9.95, layer: 49
Normal
MK50
MK75
MK88
MK94
MK97
MK data
Normal data
Fold over basal
2.2
2
1.8
1.6
1.4
1.2
1
0.8
0.6
0
10
20
30
40
50
60
Time (min)

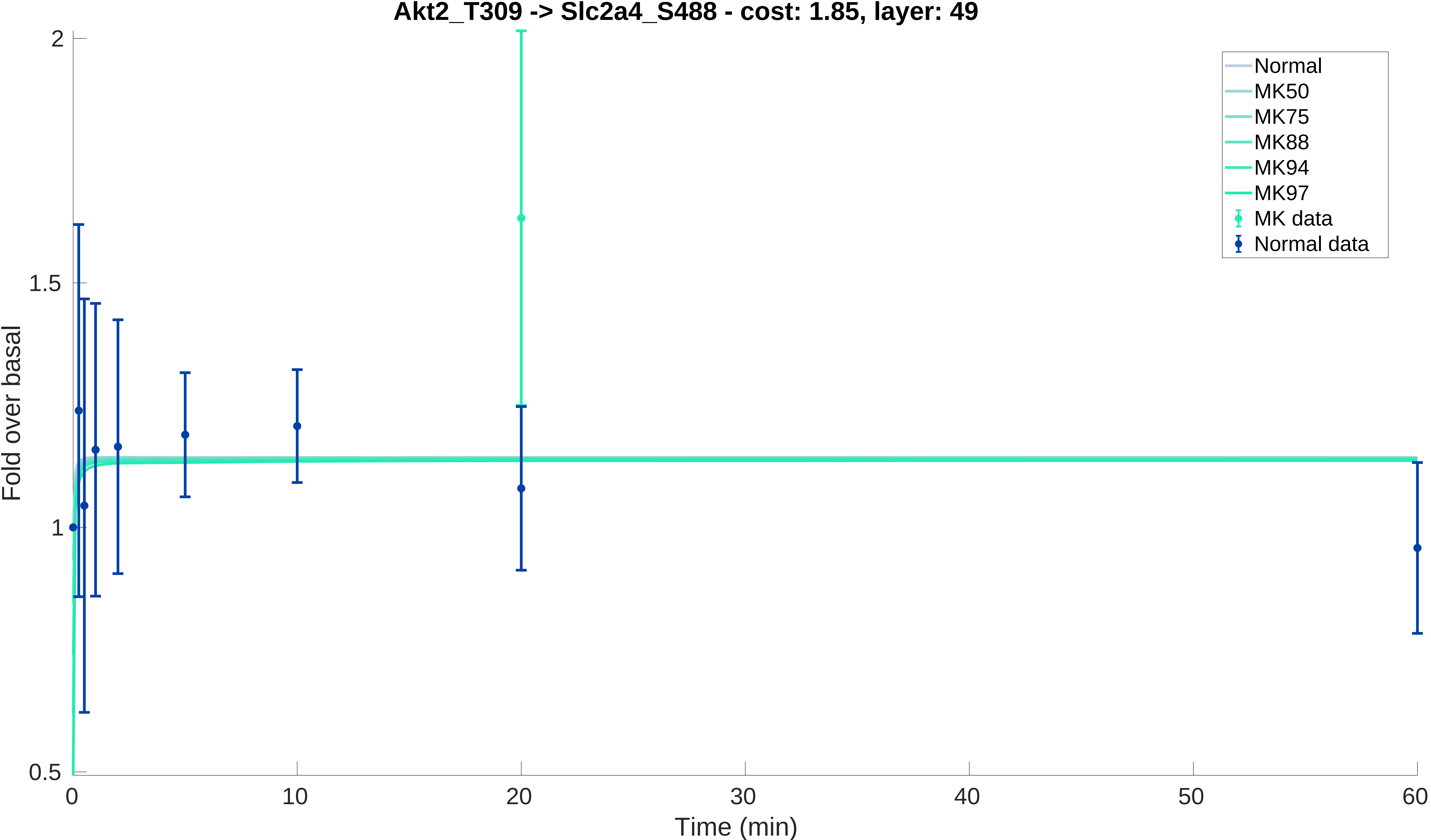

Akt2_T309 -> Slc2a4_S488 - cost: 1.85, layer: 49
Fold over basal
Time (min)
Normal
MK50
MK75
MK88
MK94
MK97
MK data
Normal data
0.5
1
1.5
2
0
10
20
30
40
50
60

#### Ncoa1_S702 -> Smarcc1_S327 - cost: 4.23, layer: 49

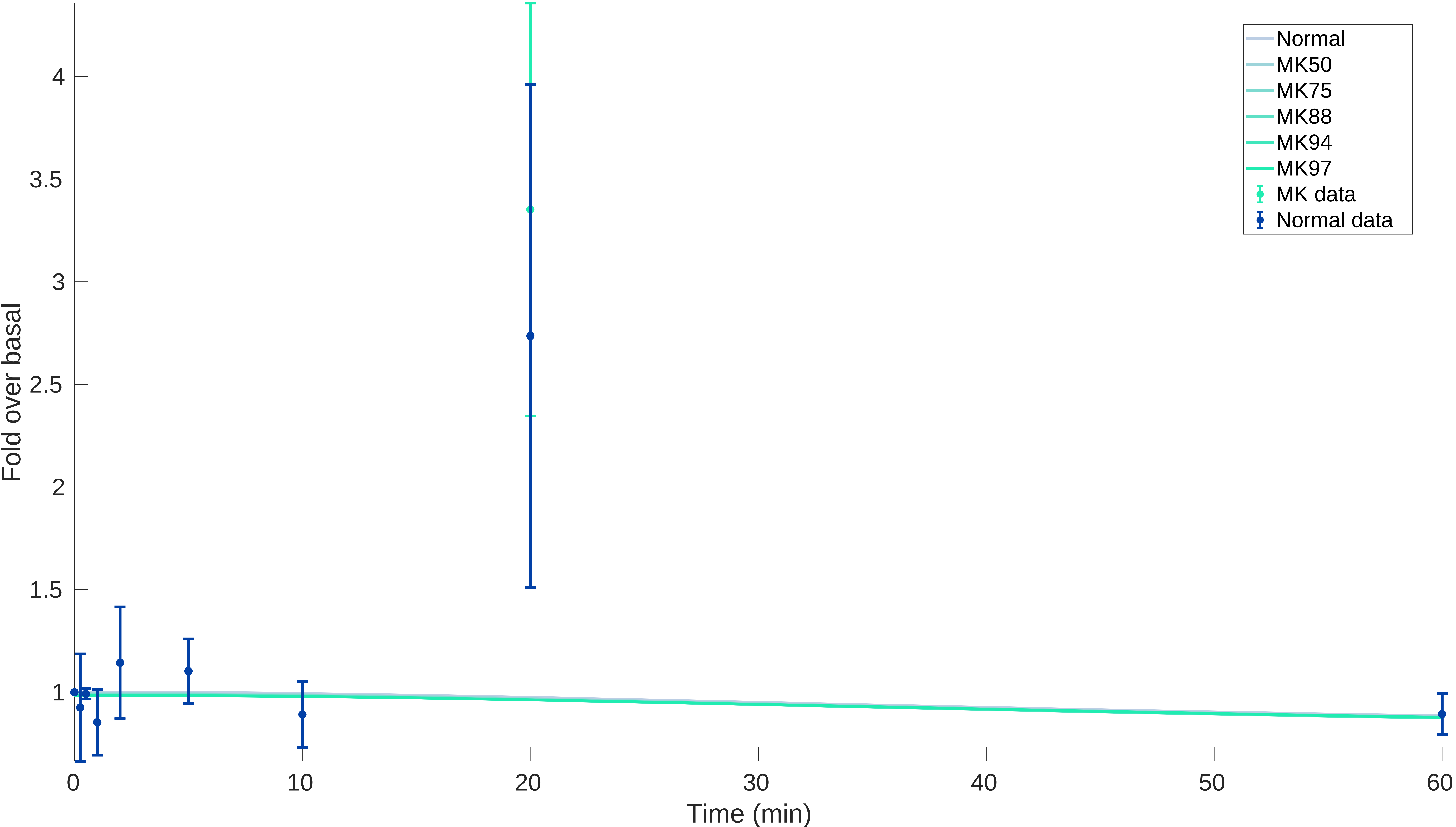

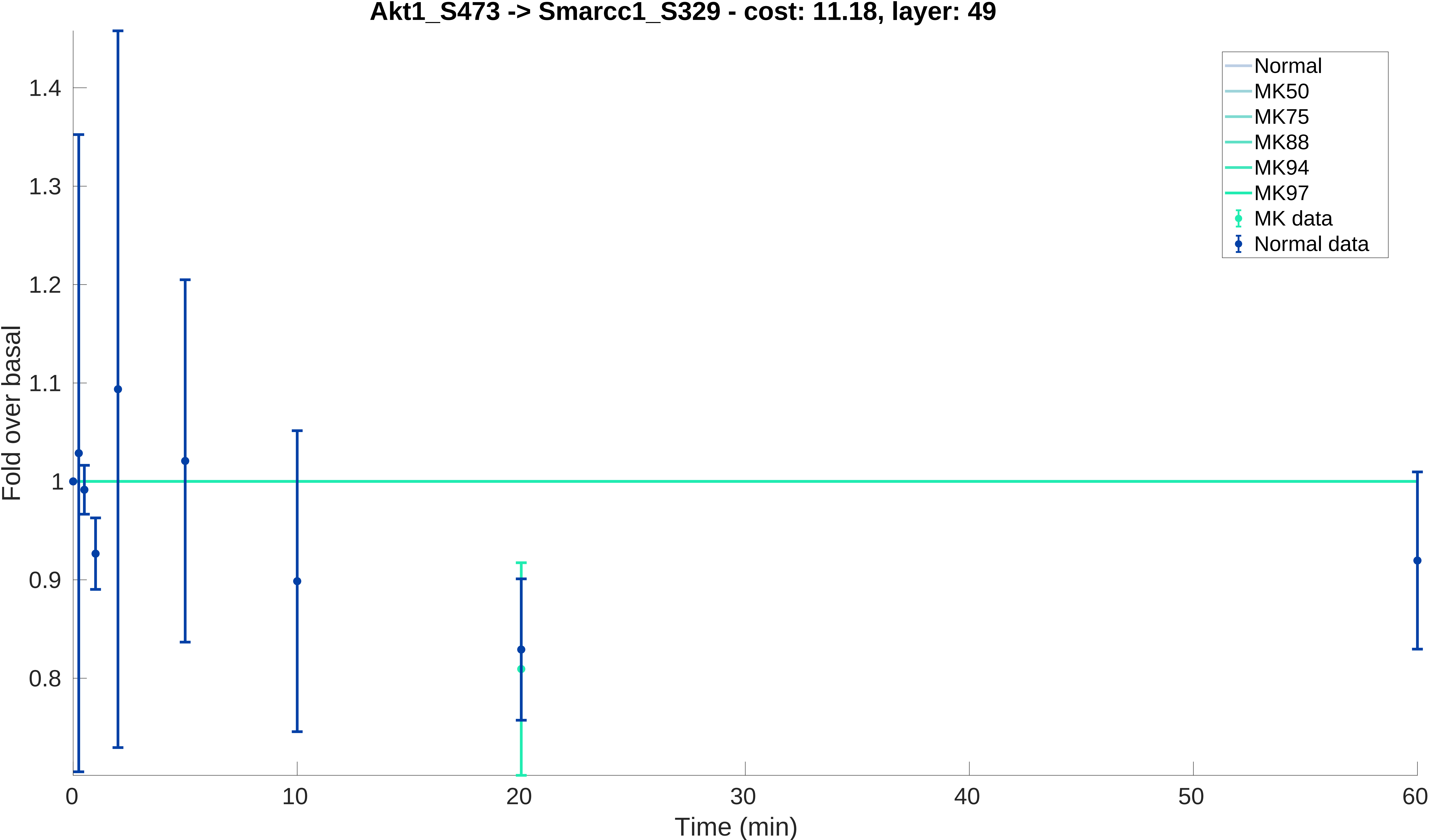
Akt1_S473 -> Smarcc1_S329 - cost: 11.18, layer: 49
Fold over basal
1.4
1.3
1.2
1.1
1
0.9
0.8
0
10
20
30
40
50
60
Time (min)
Normal
MK50
MK75
MK88
MK94
MK97
MK data
Normal data

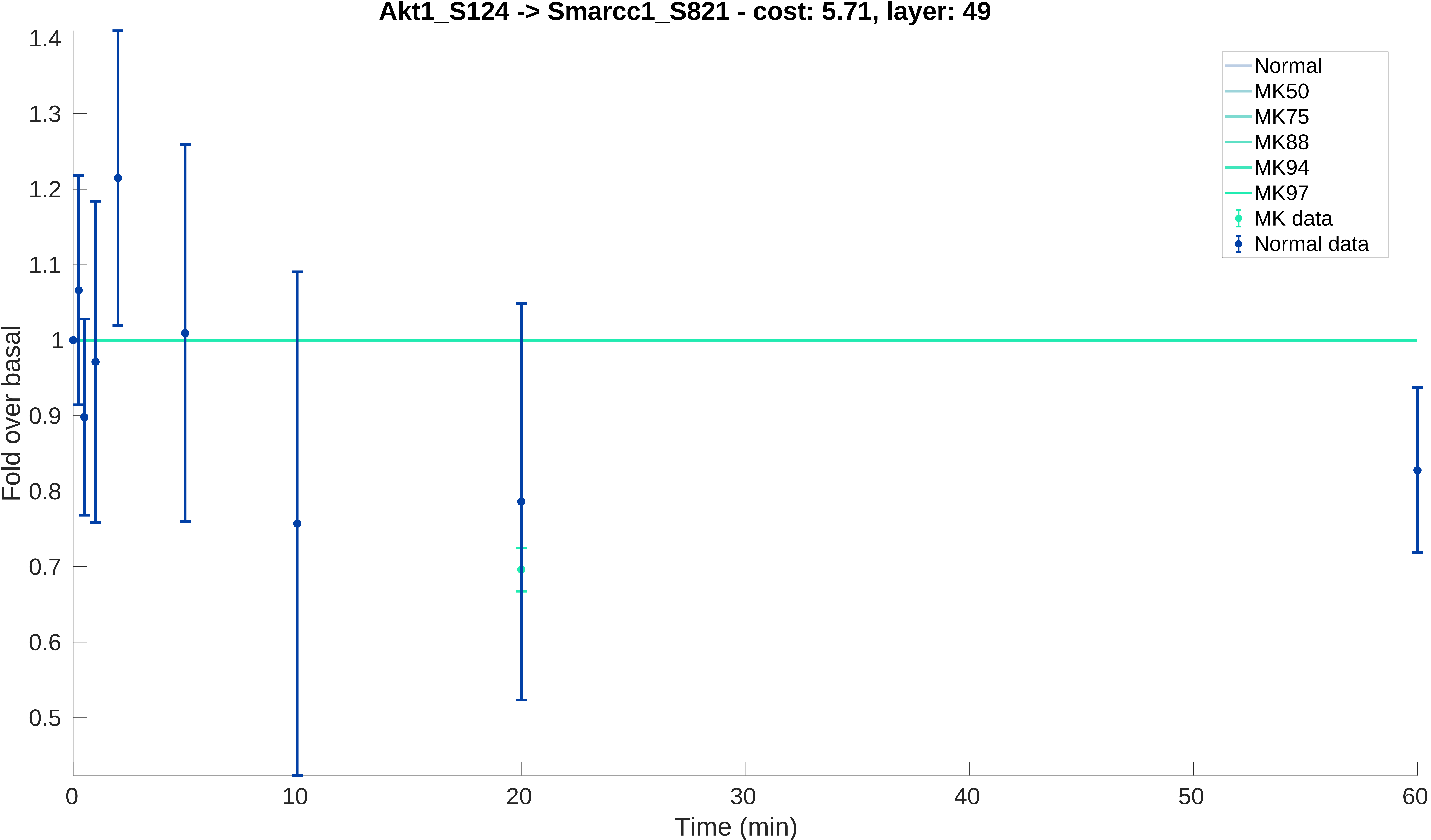

Akt1_S124 -> Smarcc1_S821 - cost: 5.71, layer: 49
Fold over basal
Time (min)
1.4
1.3
1.2
1.1
1
0.9
0.8
0.7
0.6
0.5
0
10
20
30
40
50
60
Normal
MK50
MK75
MK88
MK94
MK97
MK data
Normal data

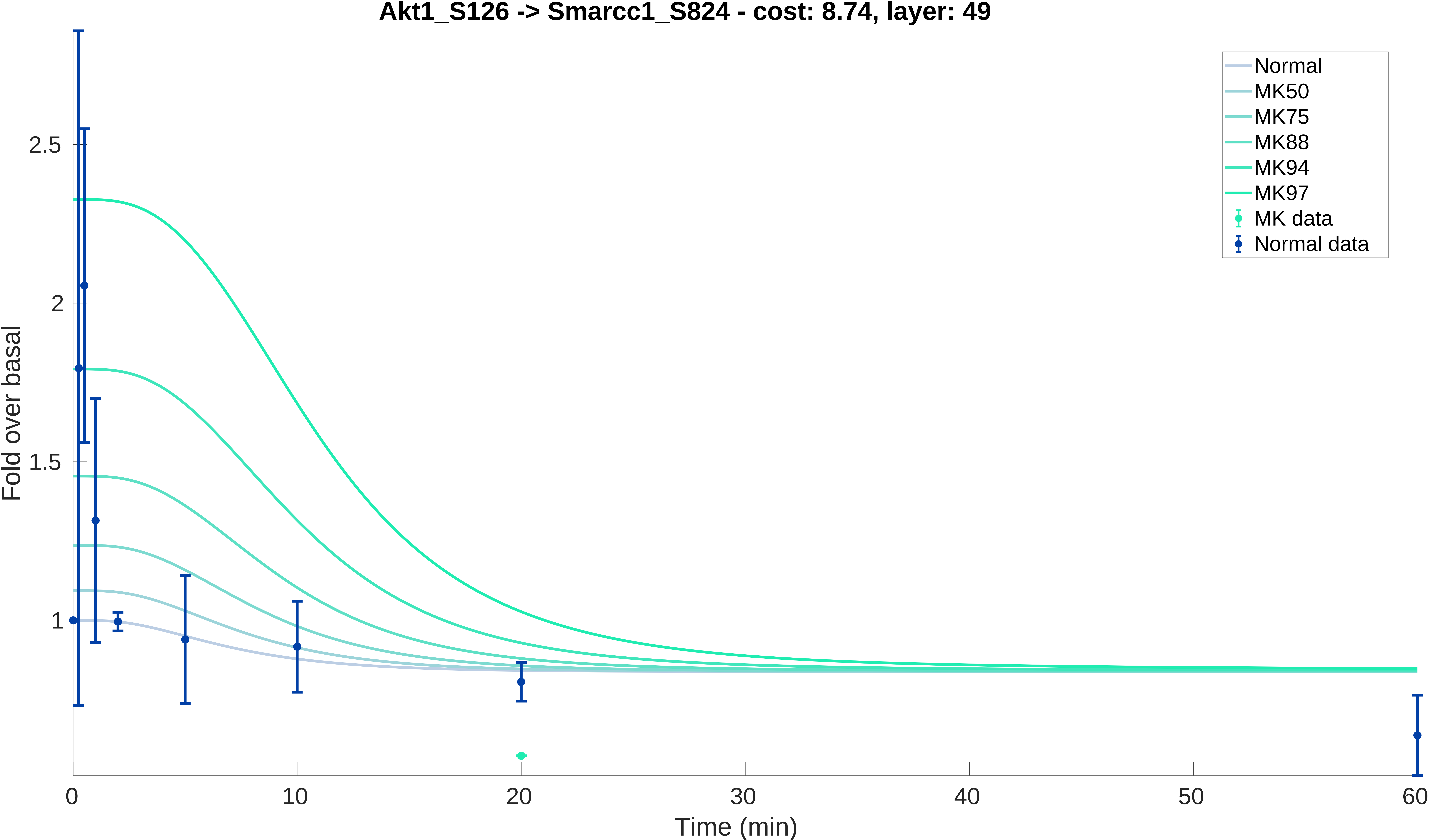

Akt1_S126 -> Smarcc1_S824 - cost: 8.74, layer: 49
Fold over basal
2.5
2
1.5
1
0
10
20
30
40
50
60
Time (min)
Normal
MK50
MK75
MK88
MK94
MK97
MK data
Normal data

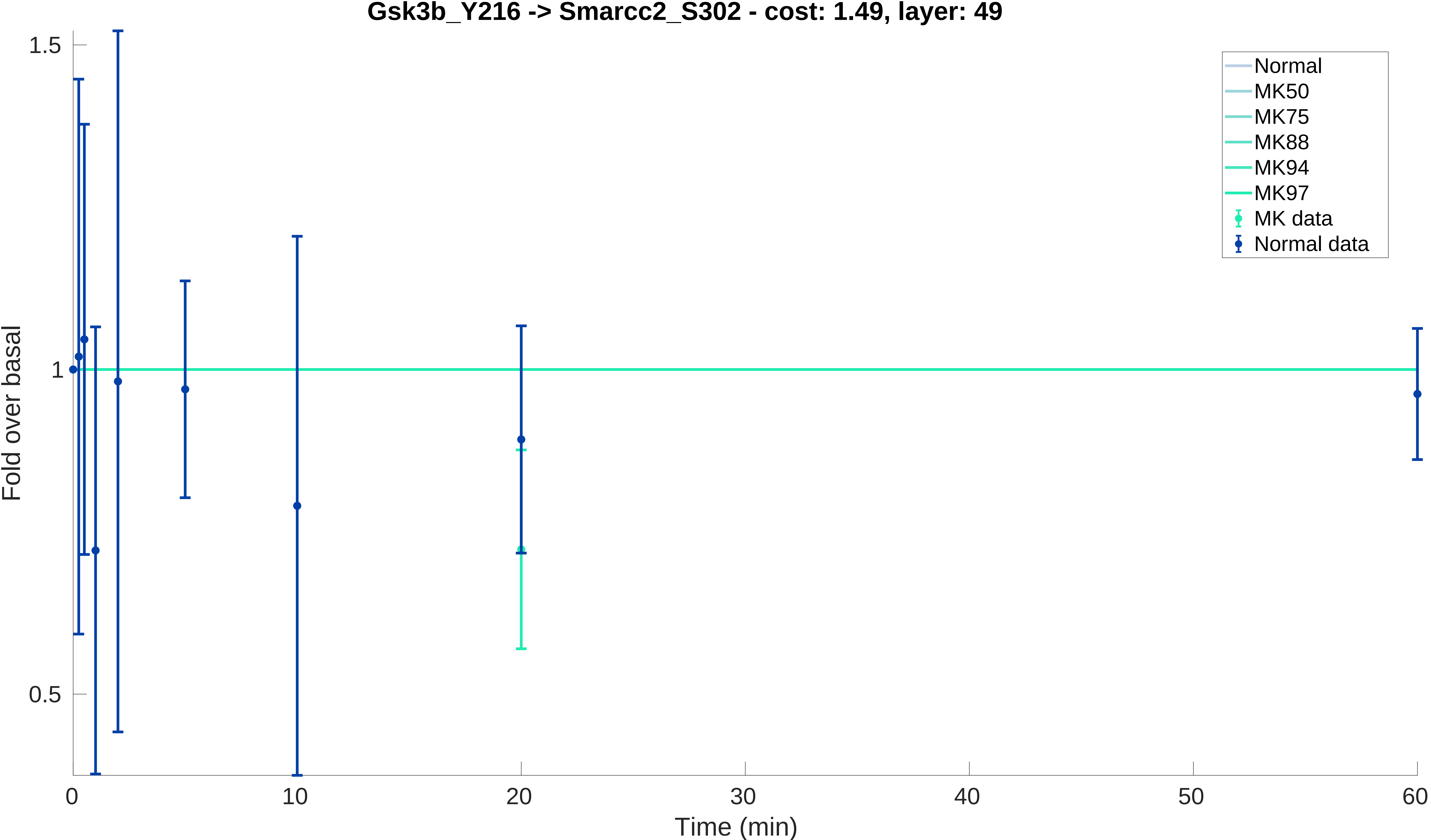

Gsk3b_Y216 -> Smarcc2_S302 - cost: 1.49, layer: 49
Fold over basal
1.5
1
0.5
0
10
20
30
40
50
60
Time (min)
Normal
MK50
MK75
MK88
MK94
MK97
MK data
Normal data

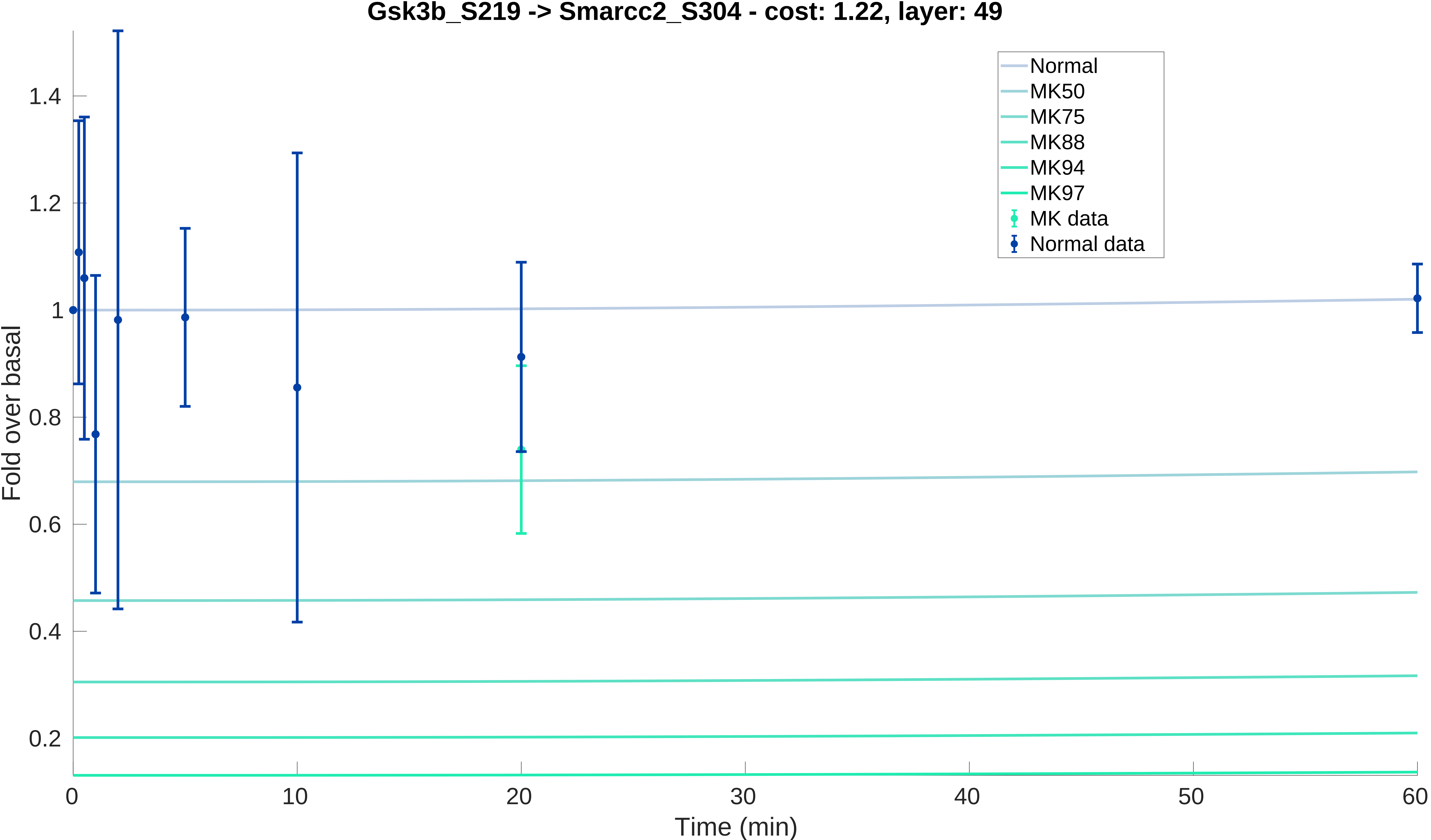
Gsk3b_S219 -> Smarcc2_S304 - cost: 1.22, layer: 49
Normal
MK50
MK75
MK88
MK94
MK97
MK data
Normal data
Fold over basal
Time (min)
0.2
0.4
0.6
0.8
1
1.2
1.4
0
10
20
30
40
50
60

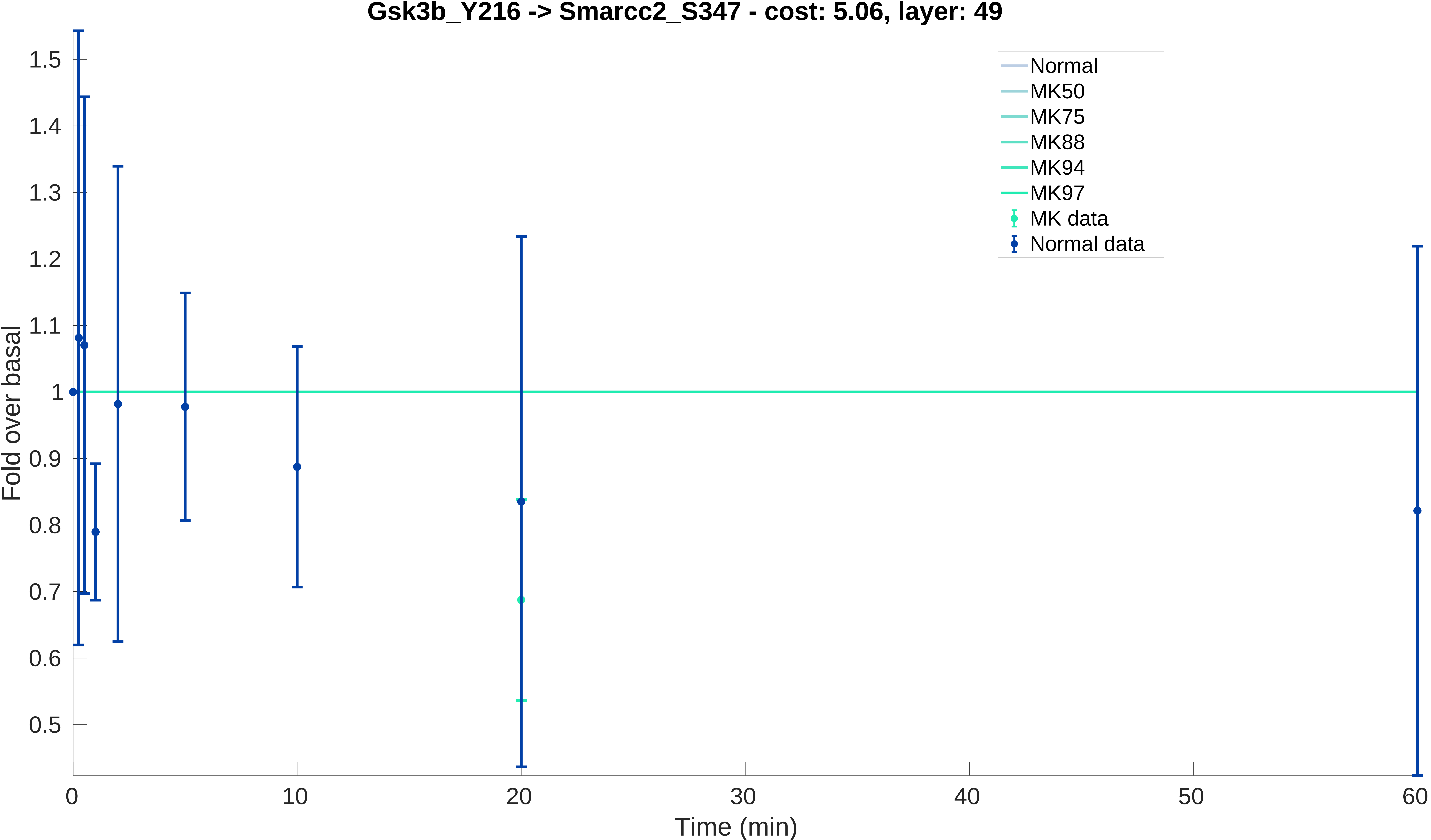

Gsk3b_Y216 -> Smarcc2_S347 - cost: 5.06, layer: 49
Normal
MK50
MK75
MK88
MK94
MK97
MK data
Normal data
Fold over basal
1.5
1.4
1.3
1.2
1.1
1
0.9
0.8
0.7
0.6
0.5
0
10
20
30
40
50
60
Time (min)

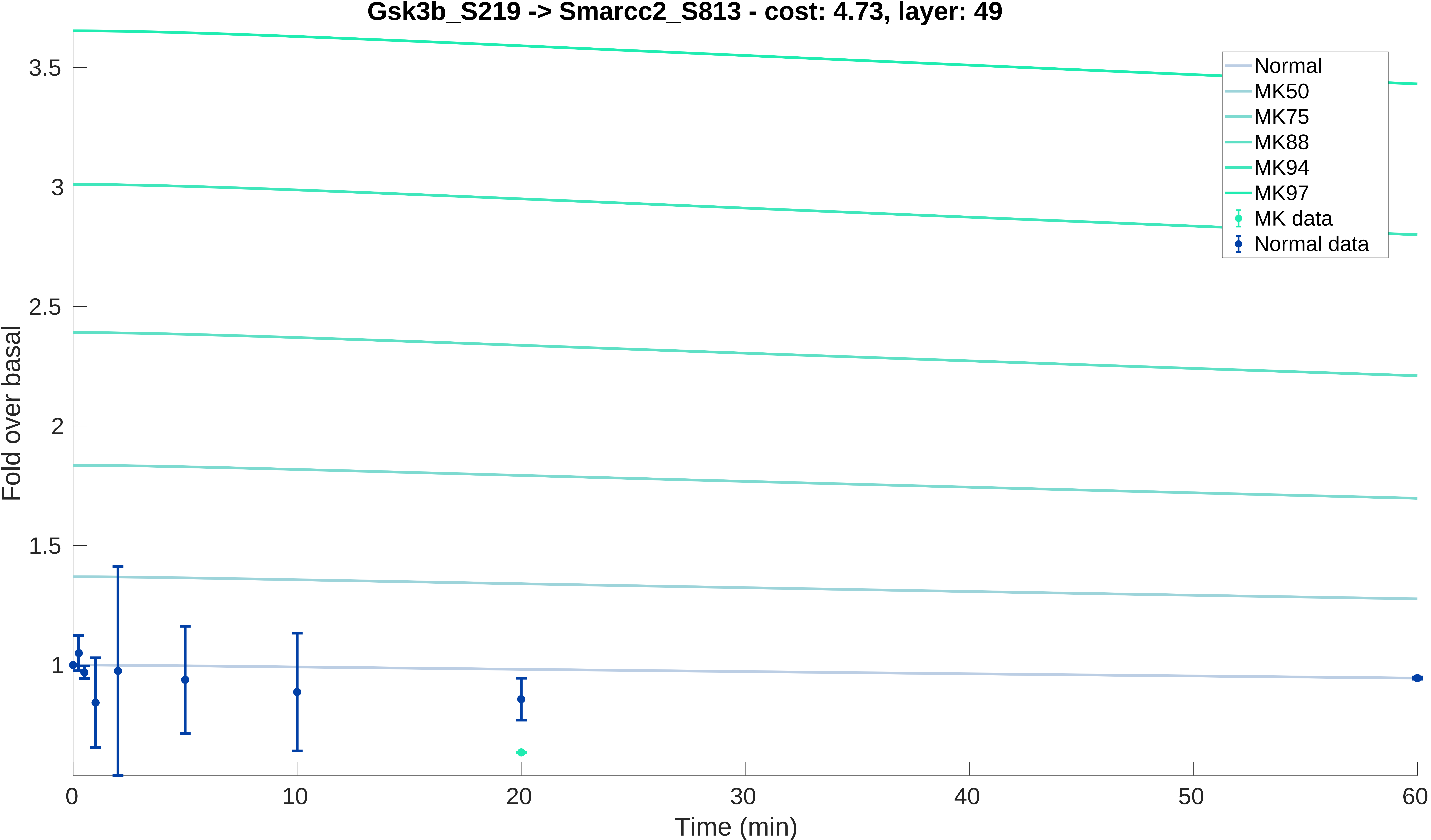

Gsk3b_S219 -> Smarcc2_S813 - cost: 4.73, layer: 49
Fold over basal
3.5
3
2.5
2
1.5
1
0
10
20
30
40
50
60
Time (min)
Normal
MK50
MK75
MK88
MK94
MK97
MK data
Normal data

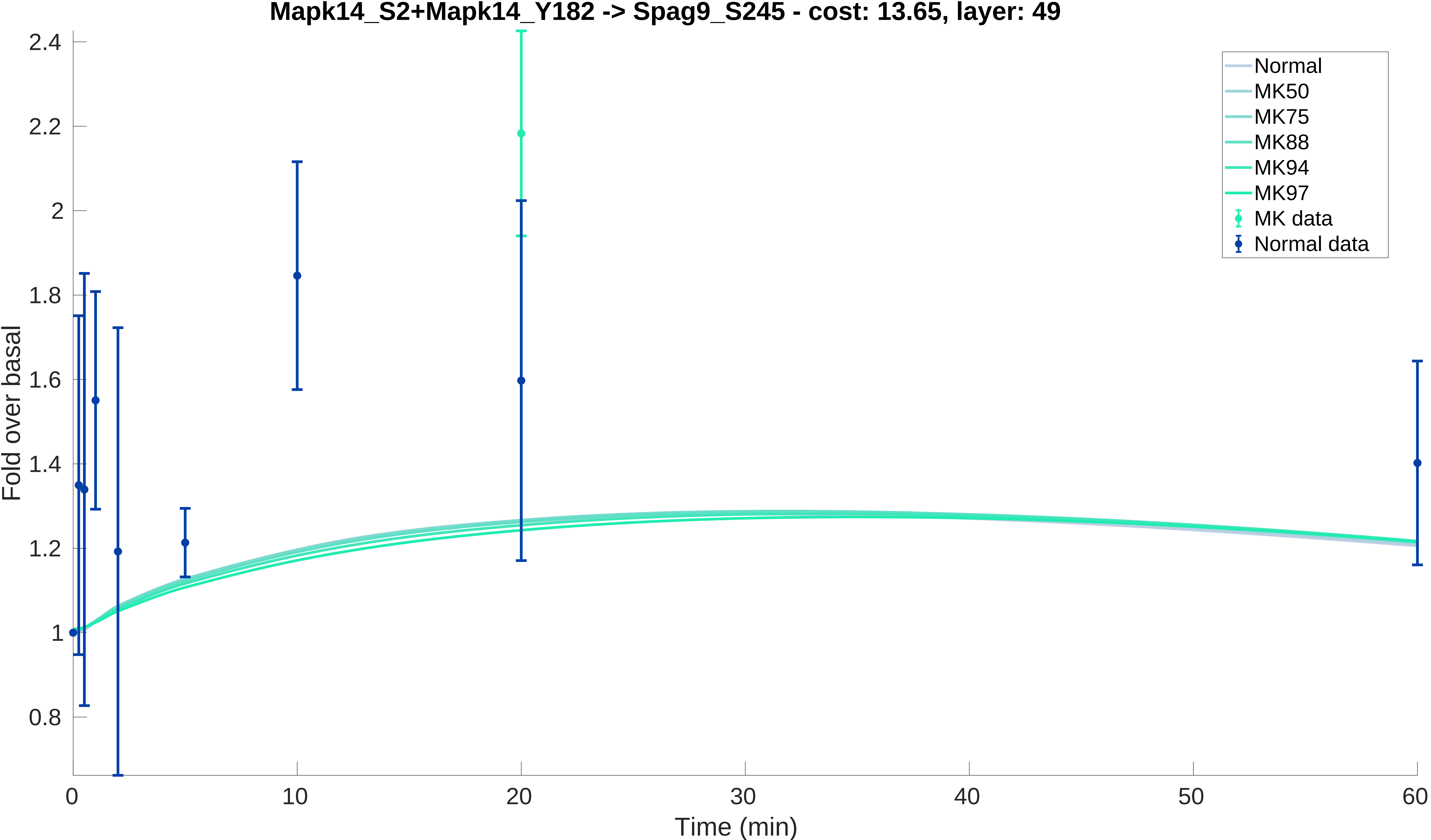

Mapk14_S2+Mapk14_Y182 -> Spag9_S245 - cost: 13.65, layer: 49
Fold over basal
Time (min)
2.4
2.2
2
1.8
1.6
1.4
1.2
1
0.8
0
10
20
30
40
50
60
Normal
MK50
MK75
MK88
MK94
MK97
MK data
Normal data

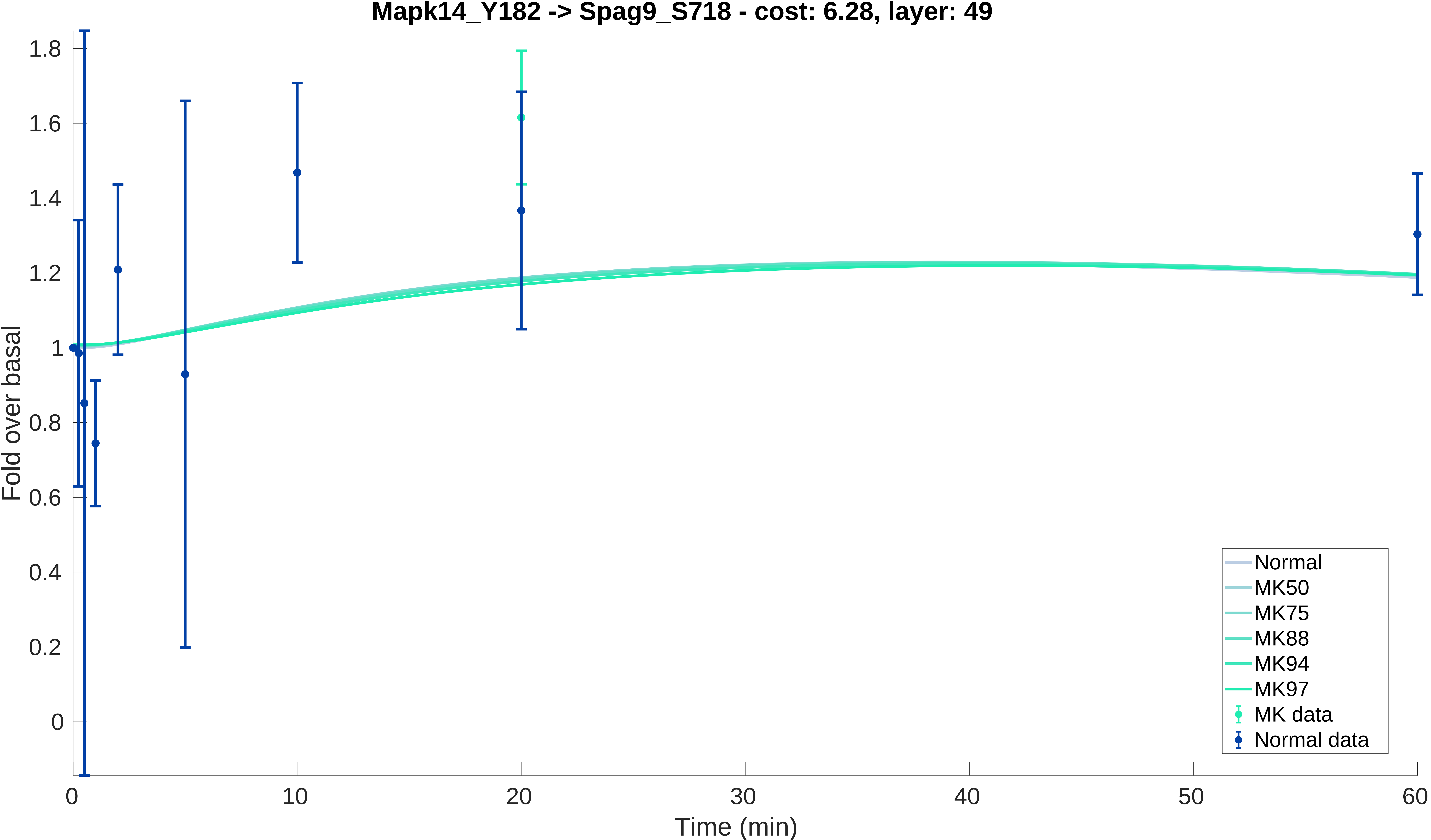

Mapk14_Y182 -> Spag9_S718 - cost: 6.28, layer: 49
Fold over basal
Time (min)
0
0.2
0.4
0.6
0.8
1
1.2
1.4
1.6
1.8
0
10
20
30
40
50
60
Normal
MK50
MK75
MK88
MK94
MK97
MK data
Normal data

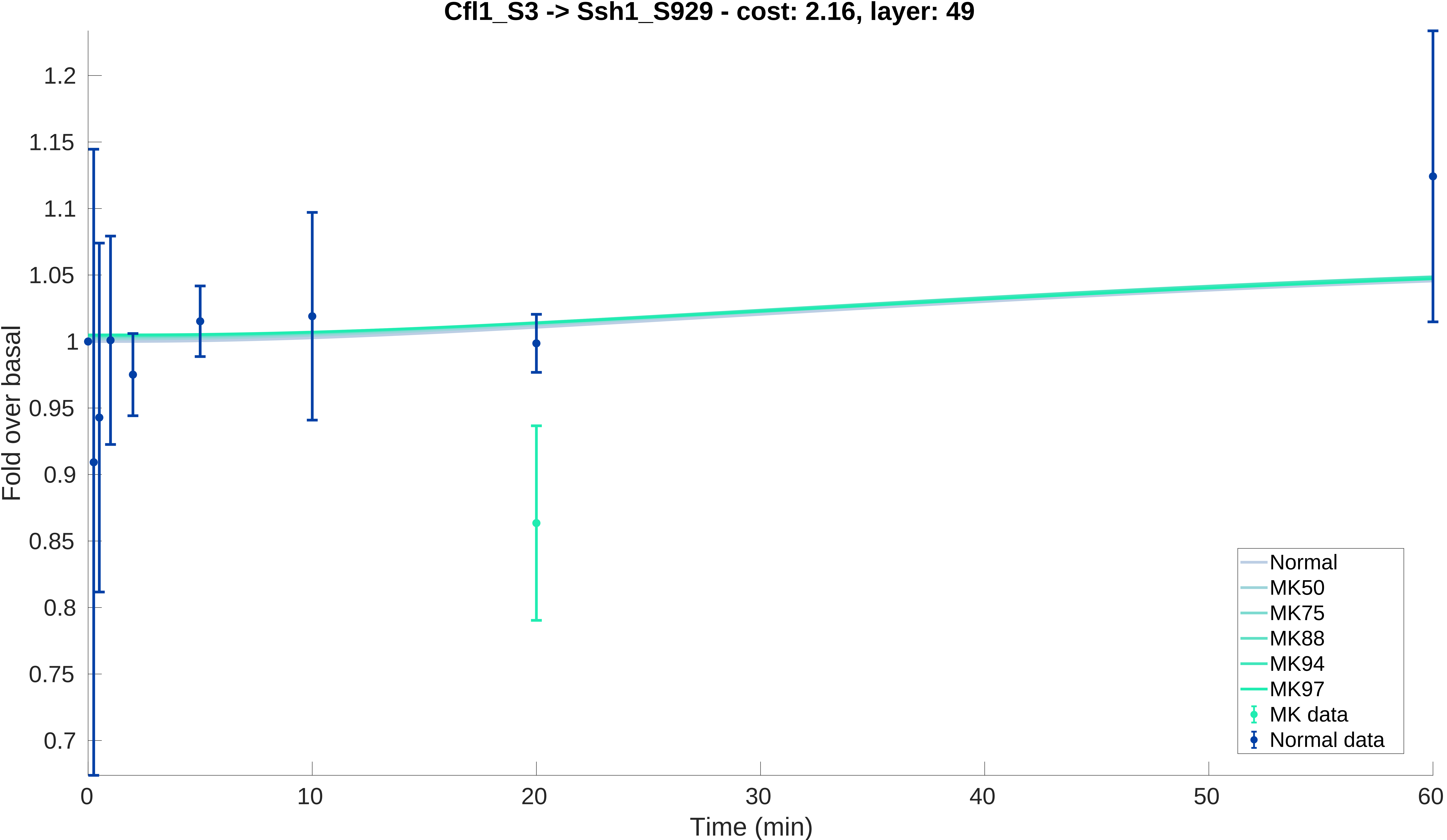

Cfl1_S3 -> Ssh1_S929 - cost: 2.16, layer: 49
Fold over basal
1.2
1.15
1.1
1.05
1
0.95
0.9
0.85
0.8
0.75
0.7
0
10
20
30
40
50
60
Time (min)
Normal
MK50
MK75
MK88
MK94
MK97
MK data
Normal data

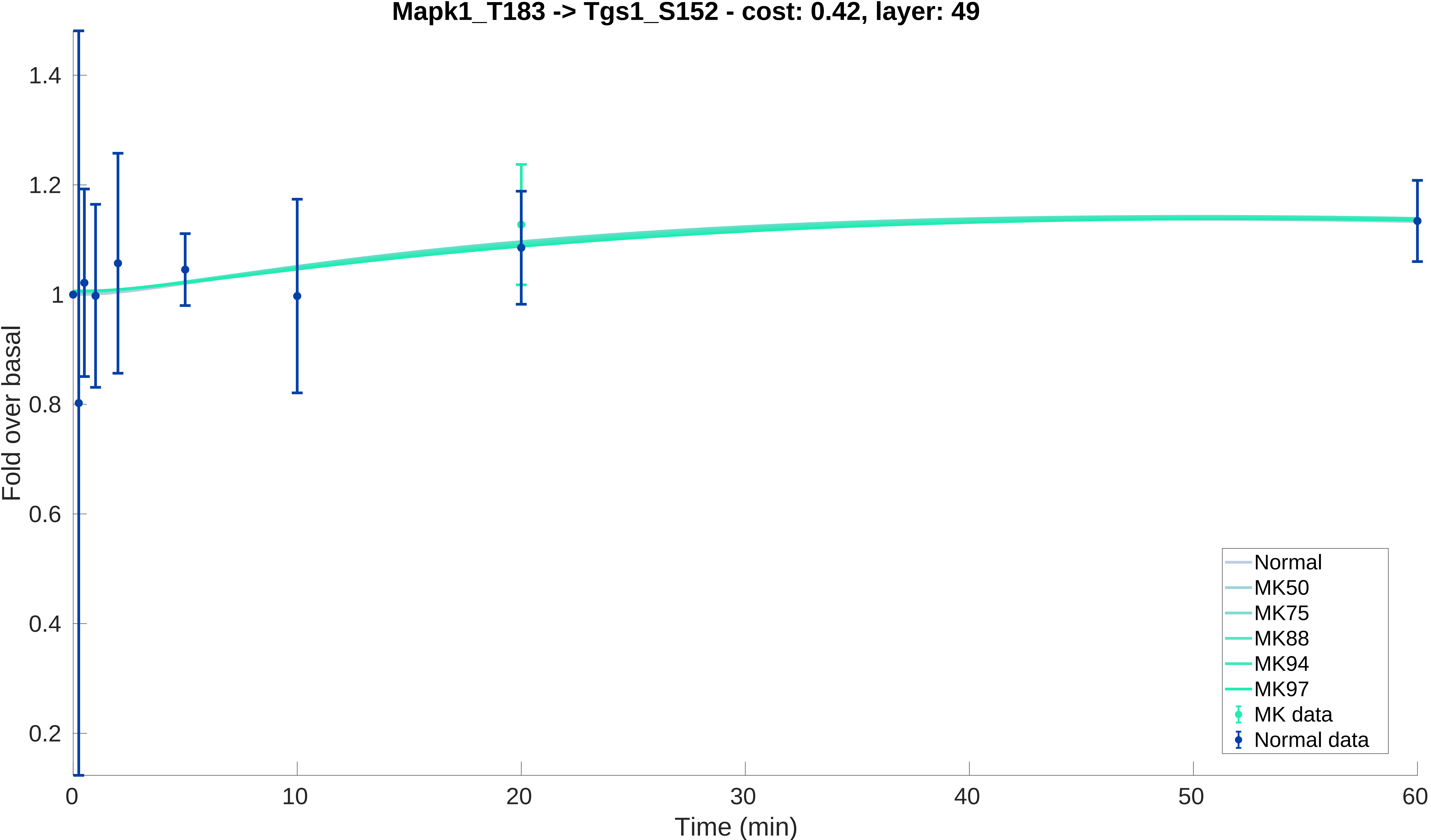

Mapk1_T183 -> Tgs1_S152 - cost: 0.42, layer: 49
Fold over basal
1.4
1.2
1
0.8
0.6
0.4
0.2
0
10
20
30
40
50
60
Time (min)
Normal
MK50
MK75
MK88
MK94
MK97
MK data
Normal data

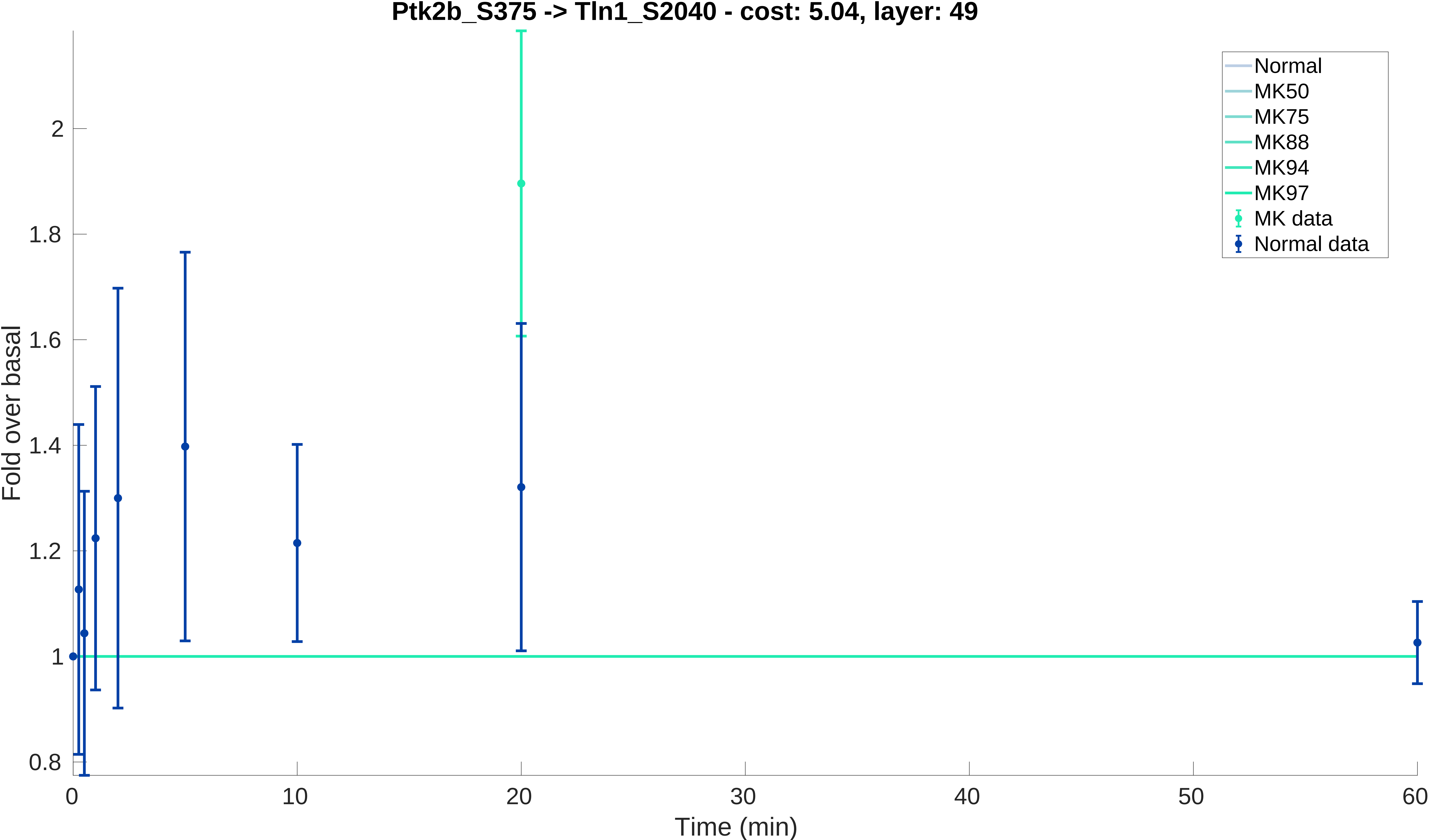

Ptk2b_S375 -> Tln1_S2040 - cost: 5.04, layer: 49
Normal
MK50
MK75
MK88
MK94
MK97
MK data
Normal data
Fold over basal
Time (min)
0.8
1
1.2
1.4
1.6
1.8
2
0
10
20
30
40
50
60

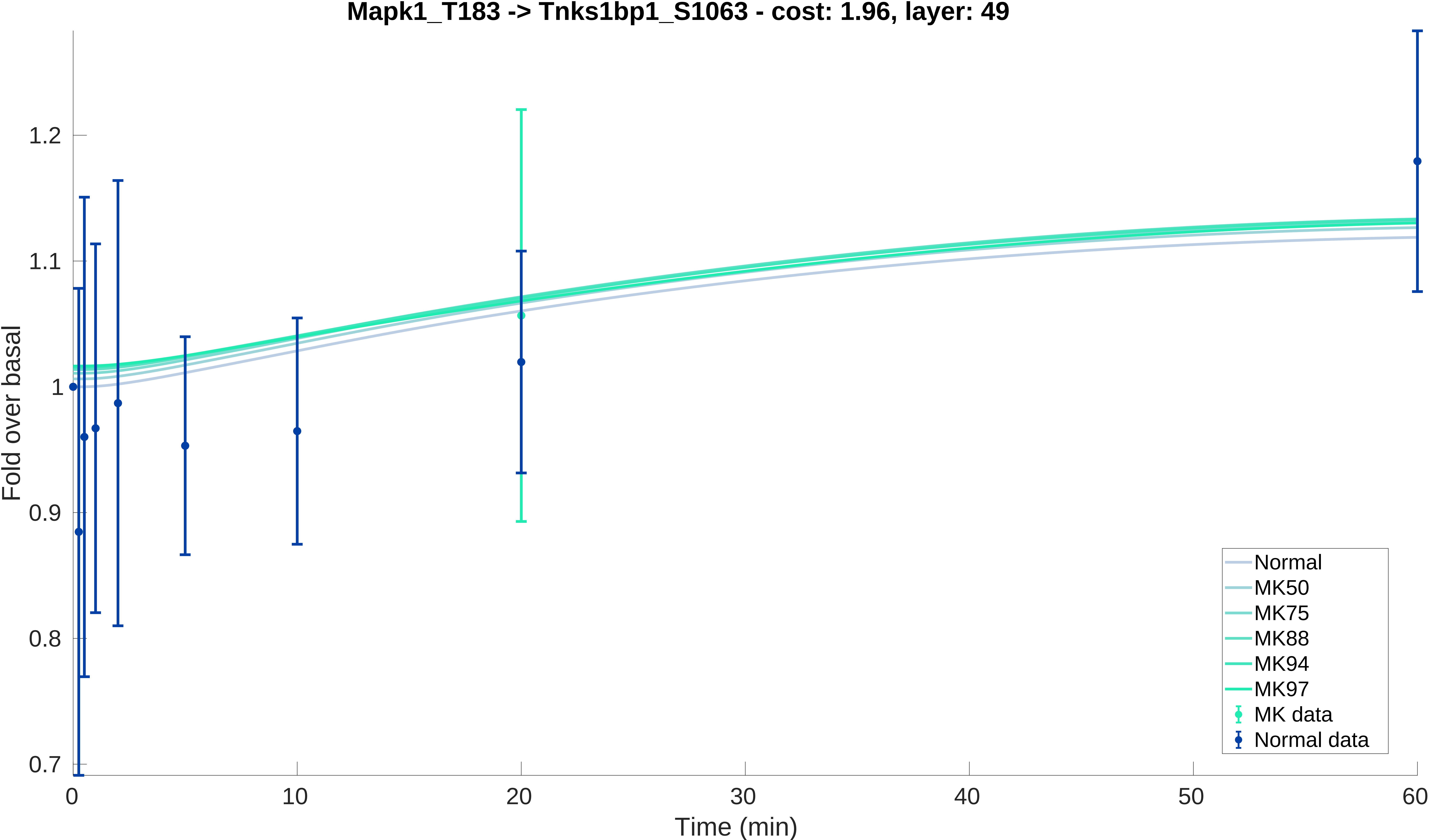
Mapk1_T183 -> Tnks1bp1_S1063 - cost: 1.96, layer: 49
Fold over basal
1.2
1.1
1
0.9
0.8
0.7
0
10
20
30
40
50
60
Time (min)
Normal
MK50
MK75
MK88
MK94
MK97
MK data
Normal data

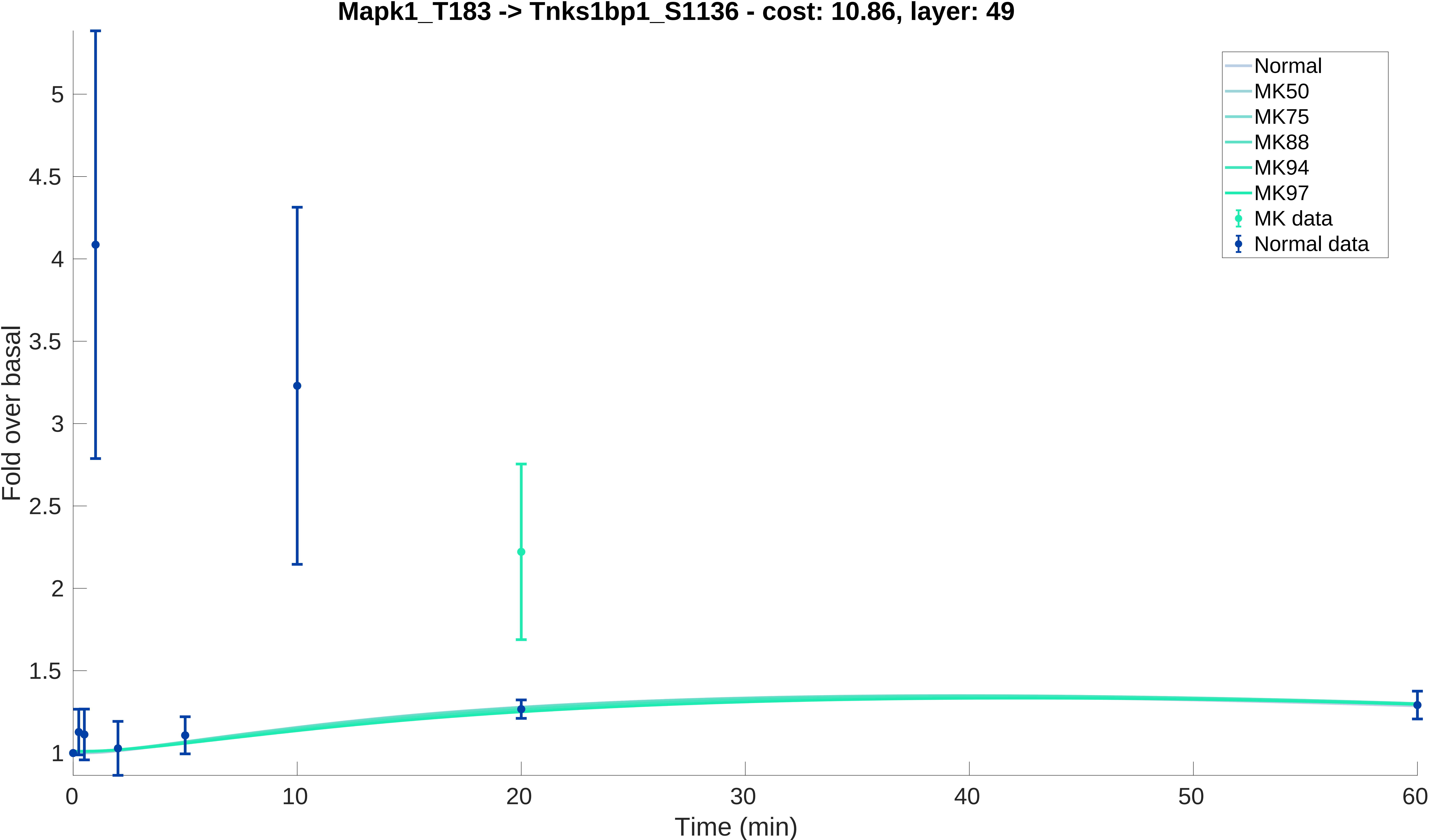

Mapk1_T183 -> Tnks1bp1_S1136 - cost: 10.86, layer: 49
Normal
MK50
MK75
MK88
MK94
MK97
MK data
Normal data
Fold over basal
5
4.5
4
3.5
3
2.5
2
1.5
1
0
10
20
30
40
50
60
Time (min)

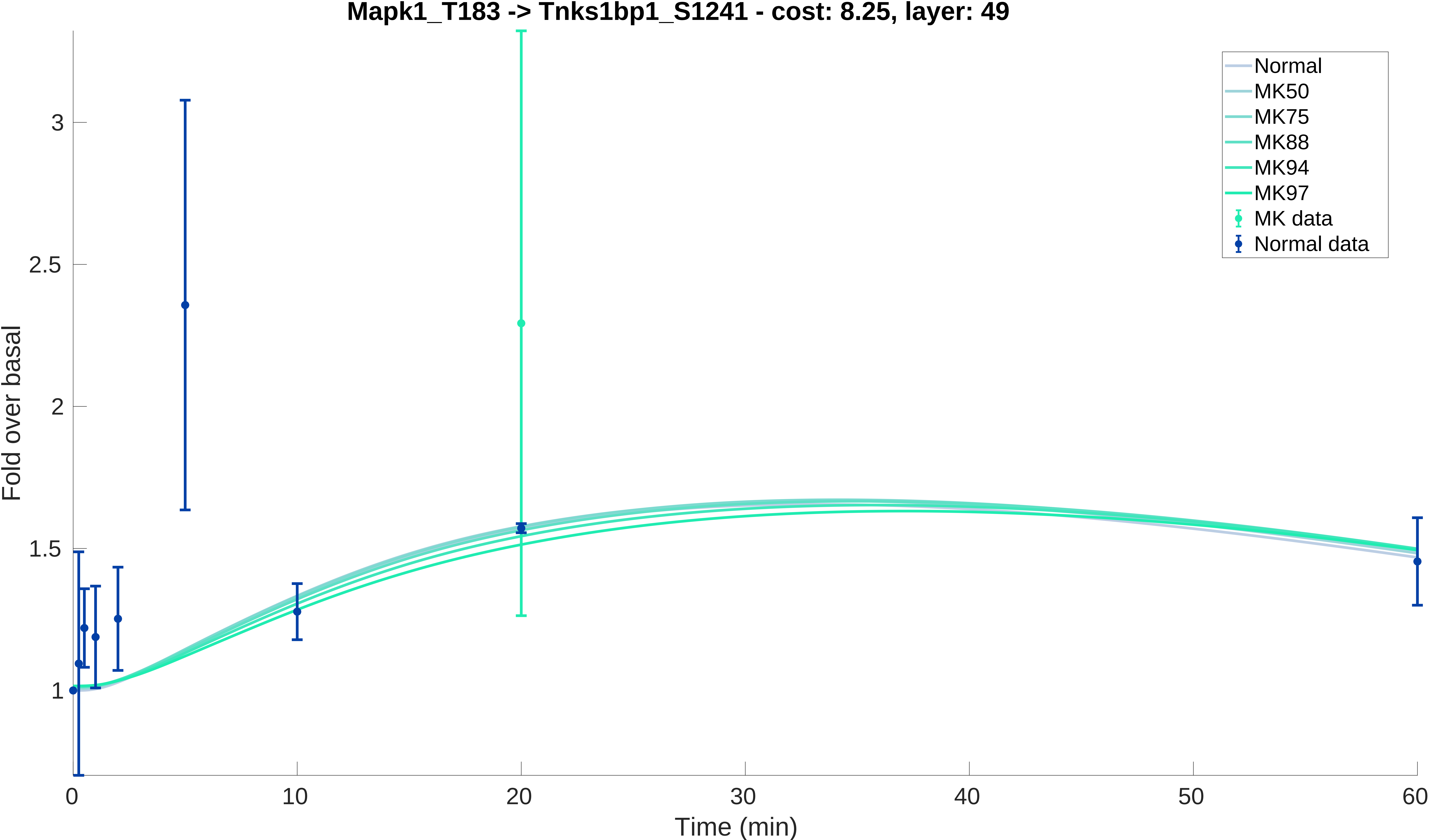

Mapk1_T183 -> Tnks1bp1_S1241 - cost: 8.25, layer: 49
Fold over basal
Time (min)
0
10
20
30
40
50
60
1
1.5
2
2.5
3
Normal
MK50
MK75
MK88
MK94
MK97
MK data
Normal data

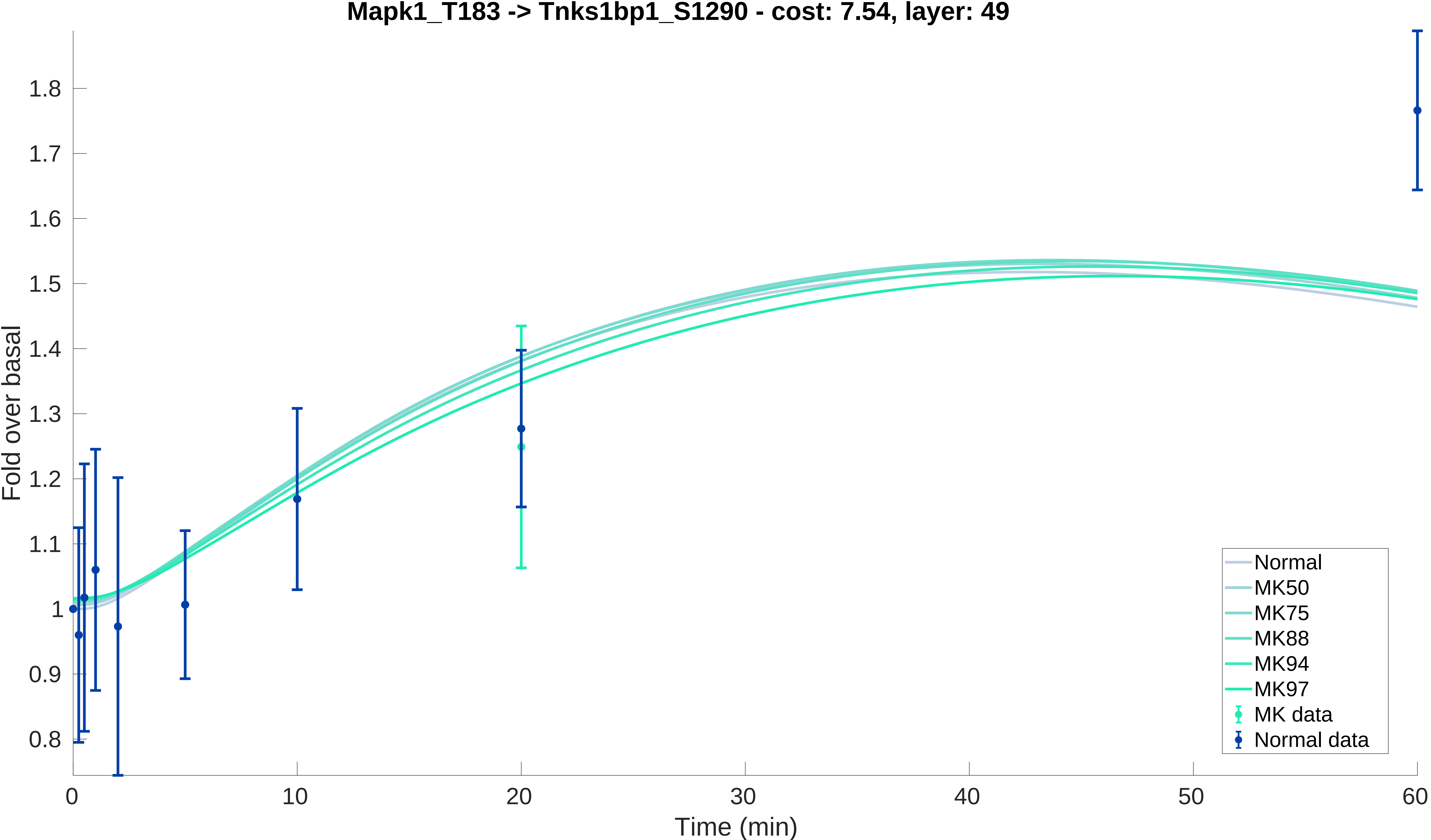
Mapk1_T183 -> Tnks1bp1_S1290 - cost: 7.54, layer: 49
Fold over basal
Time (min)
0.8
0.9
1
1.1
1.2
1.3
1.4
1.5
1.6
1.7
1.8
0
10
20
30
40
50
60
Normal
MK50
MK75
MK88
MK94
MK97
MK data
Normal data

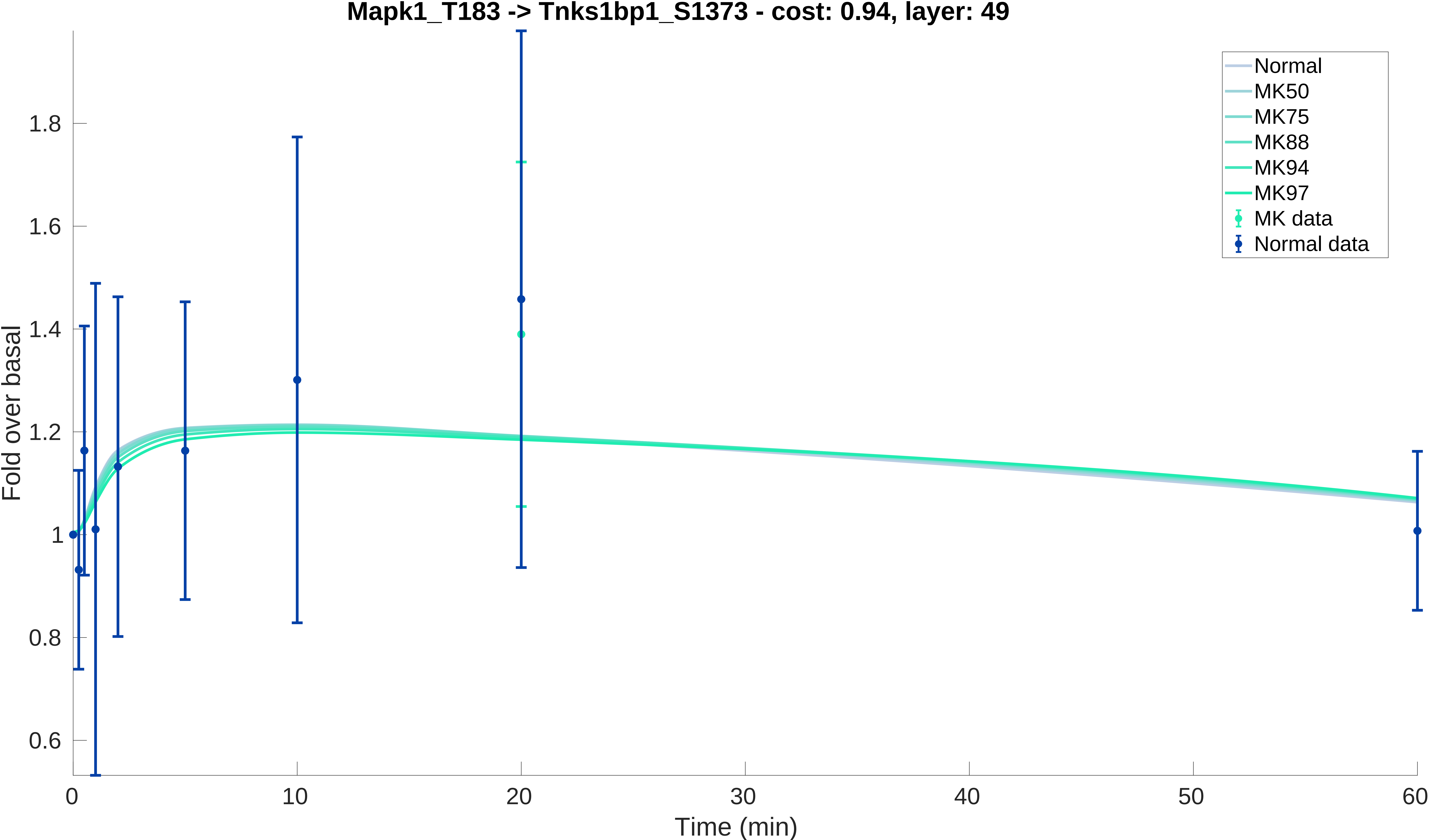

Mapk1_T183 -> Tnks1bp1_S1373 - cost: 0.94, layer: 49
Fold over basal
Time (min)
0.6
0.8
1
1.2
1.4
1.6
1.8
0
10
20
30
40
50
60
Normal
MK50
MK75
MK88
MK94
MK97
MK data
Normal data

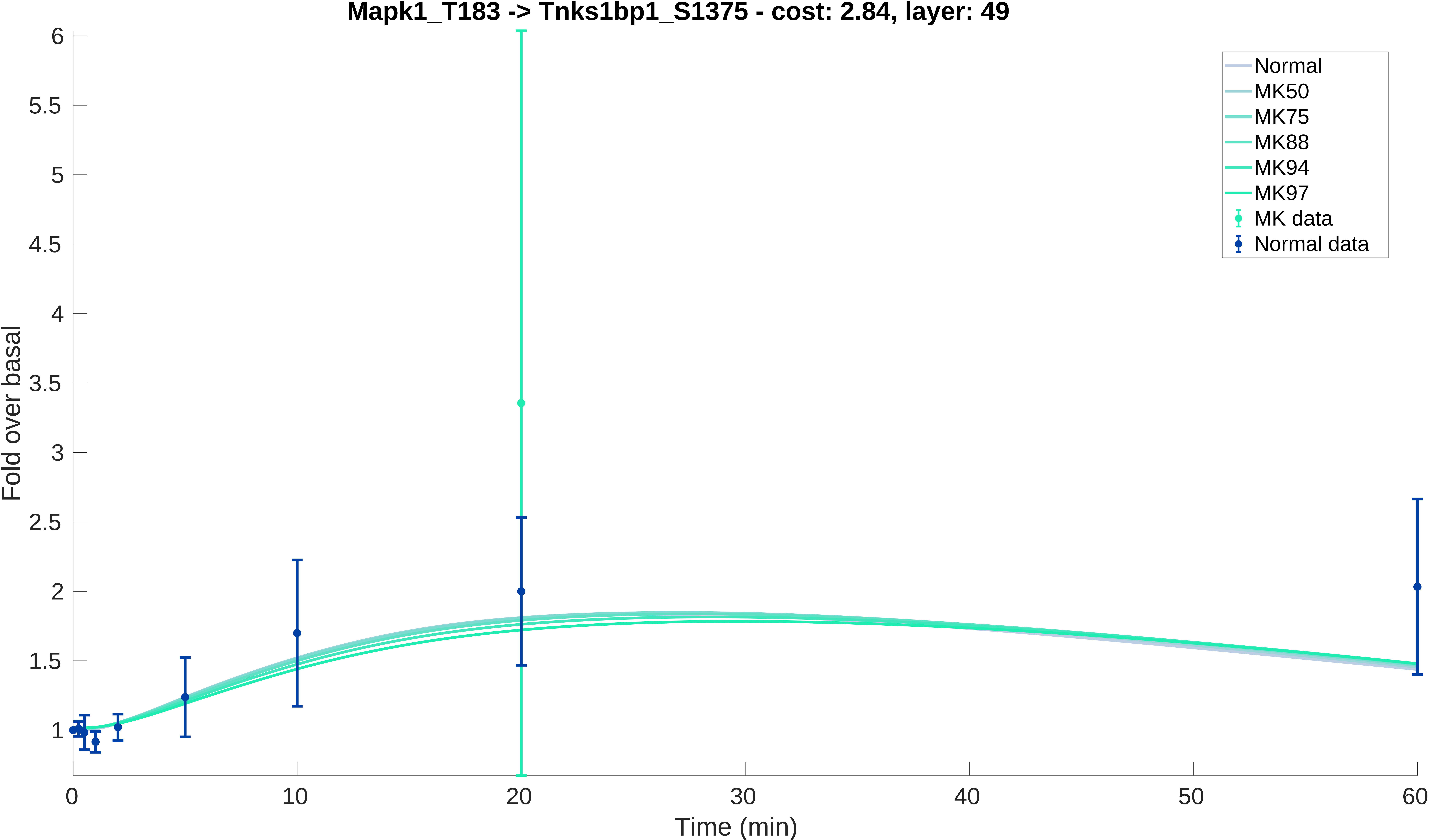
Mapk1_T183 -> Tnks1bp1_S1375 - cost: 2.84, layer: 49
Fold over basal
Time (min)
Normal
MK50
MK75
MK88
MK94
MK97
MK data
Normal data
0
10
20
30
40
50
60
1
1.5
2
2.5
3
3.5
4
4.5
5
5.5
6

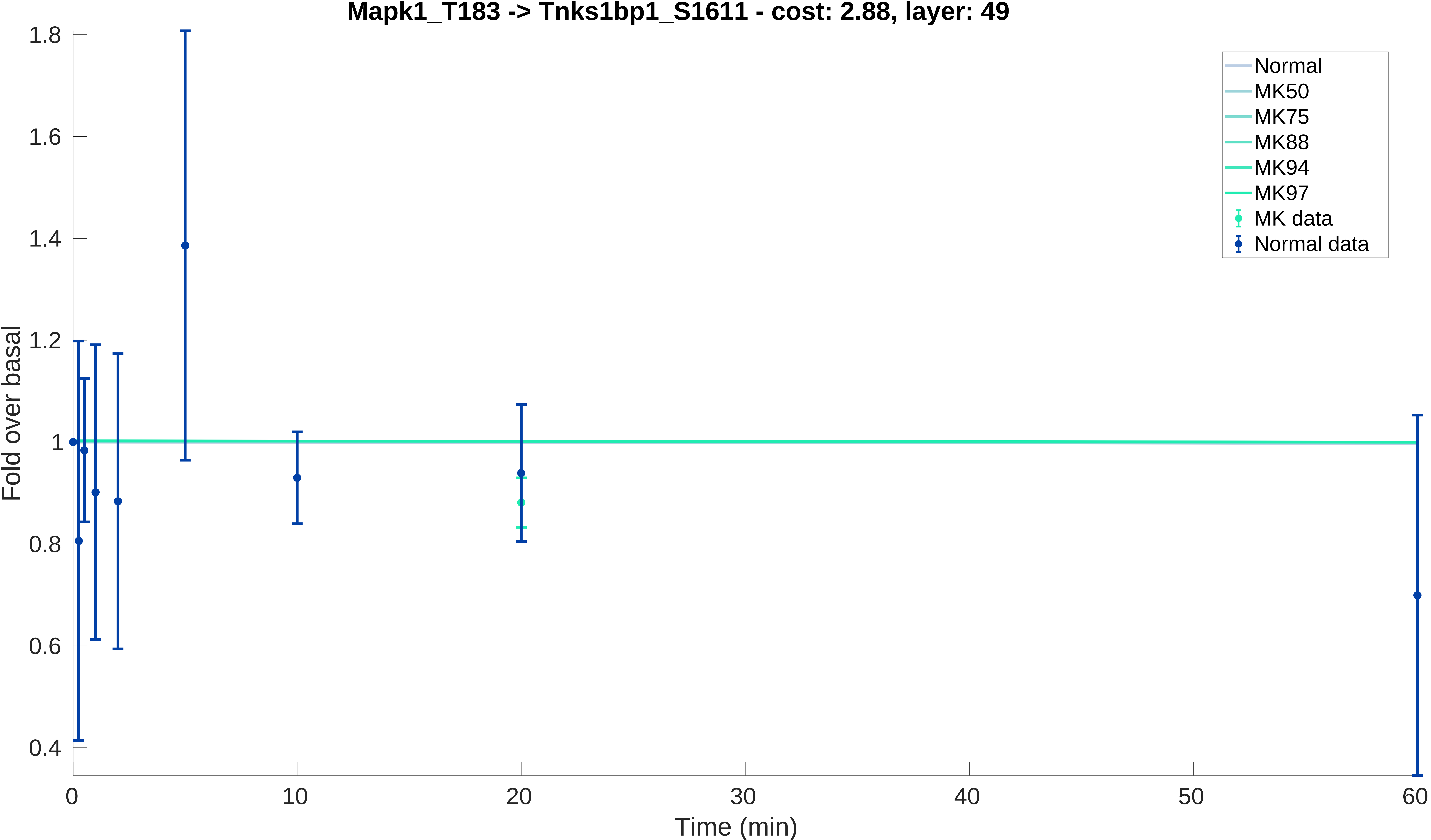

Mapk1_T183 -> Tnks1bp1_S1611 - cost: 2.88, layer: 49
Fold over basal
Time (min)
Normal
MK50
MK75
MK88
MK94
MK97
MK data
Normal data
1.8
1.6
1.4
1.2
1
0.8
0.6
0.4
0
10
20
30
40
50
60

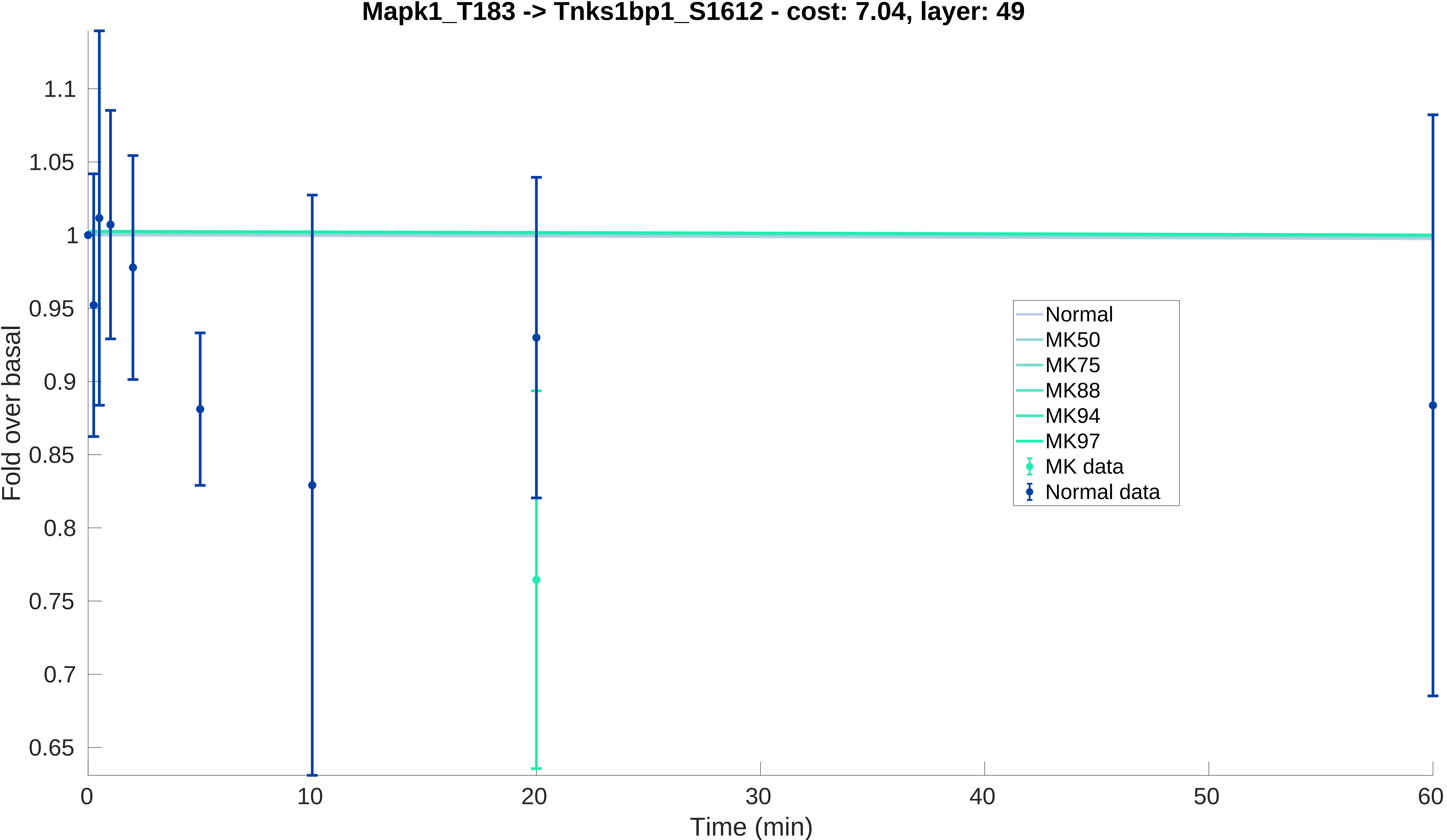

Mapk1_T183 -> Tnks1bp1_S1612 - cost: 7.04, layer: 49
Fold over basal
Time (min)
1.1
1.05
1
0.95
0.9
0.85
0.8
0.75
0.7
0.65
0
10
20
30
40
50
60
Normal
MK50
MK75
MK88
MK94
MK97
MK data
Normal data

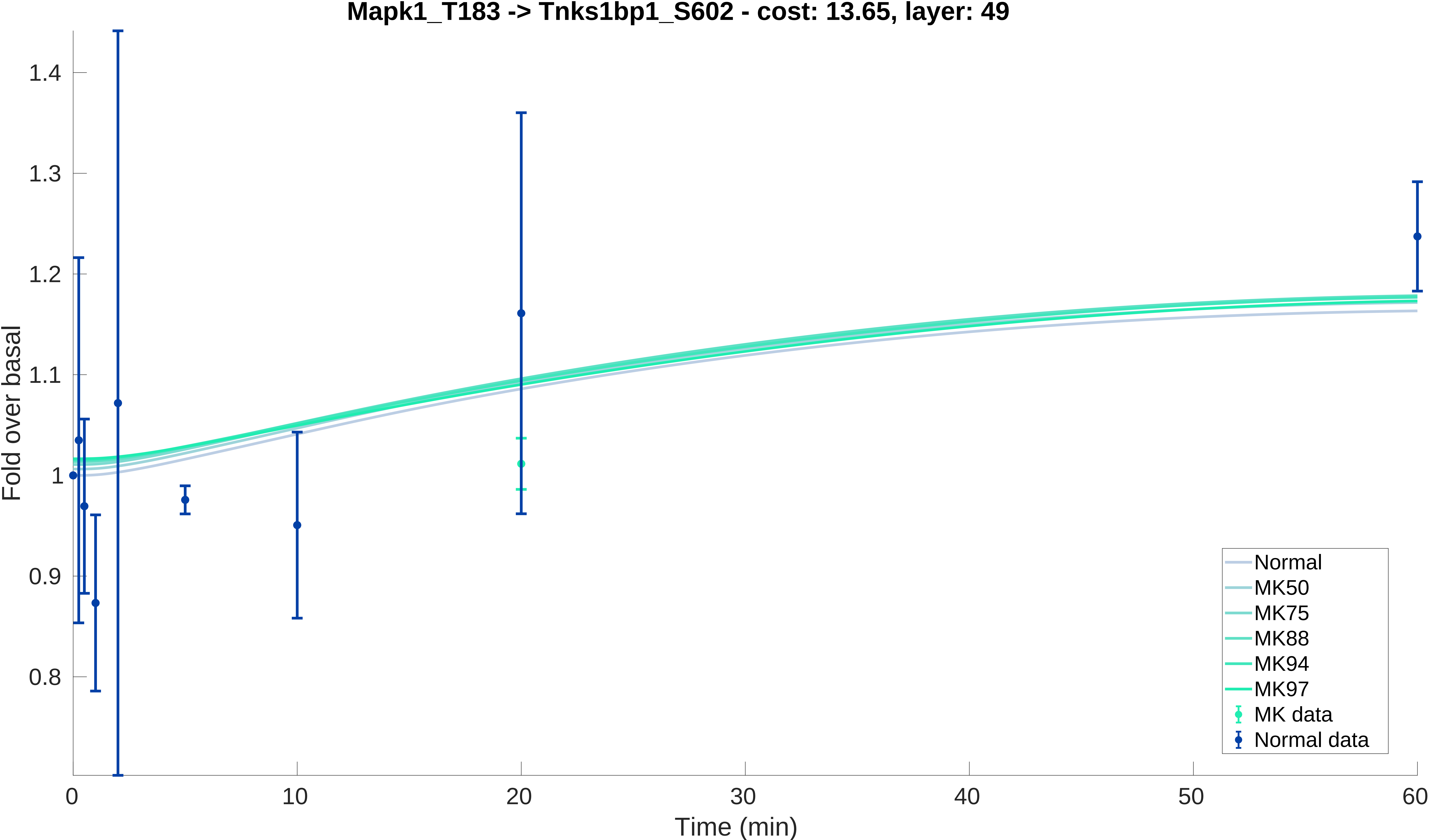
Mapk1_T183 -> Tnks1bp1_S602 - cost: 13.65, layer: 49
Fold over basal
Time (min)
1.4
1.3
1.2
1.1
1
0.9
0.8
0
10
20
30
40
50
60
Normal
MK50
MK75
MK88
MK94
MK97
MK data
Normal data

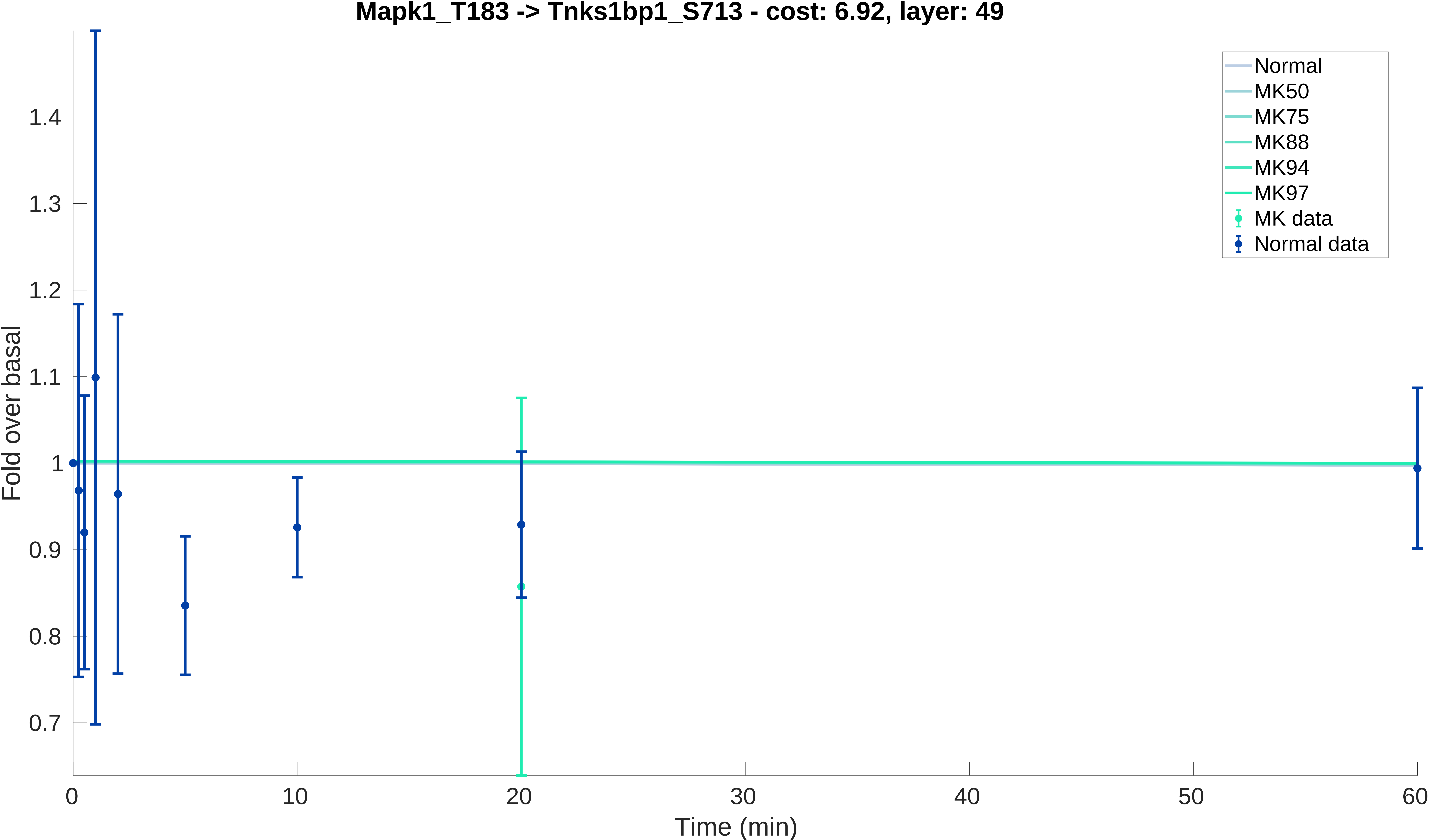

Mapk1_T183 -> Tnks1bp1_S713 - cost: 6.92, layer: 49
Fold over basal
Time (min)
1.4
1.3
1.2
1.1
1
0.9
0.8
0.7
0
10
20
30
40
50
60
Normal
MK50
MK75
MK88
MK94
MK97
MK data
Normal data

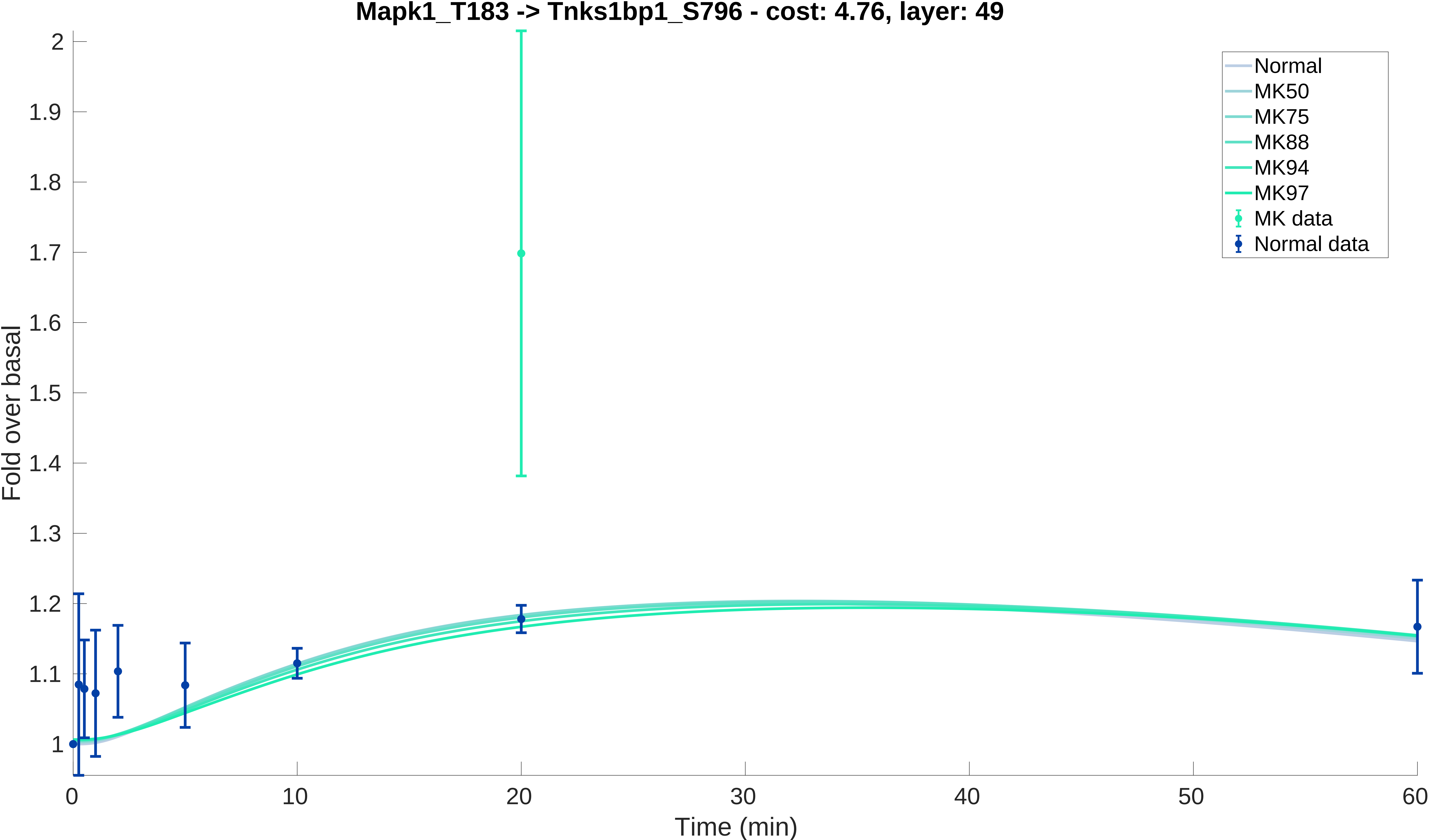

Mapk1_T183 -> Tnks1bp1_S796 - cost: 4.76, layer: 49
Fold over basal
Time (min)
2
1.9
1.8
1.7
1.6
1.5
1.4
1.3
1.2
1.1
1
0
10
20
30
40
50
60
Normal
MK50
MK75
MK88
MK94
MK97
MK data
Normal data

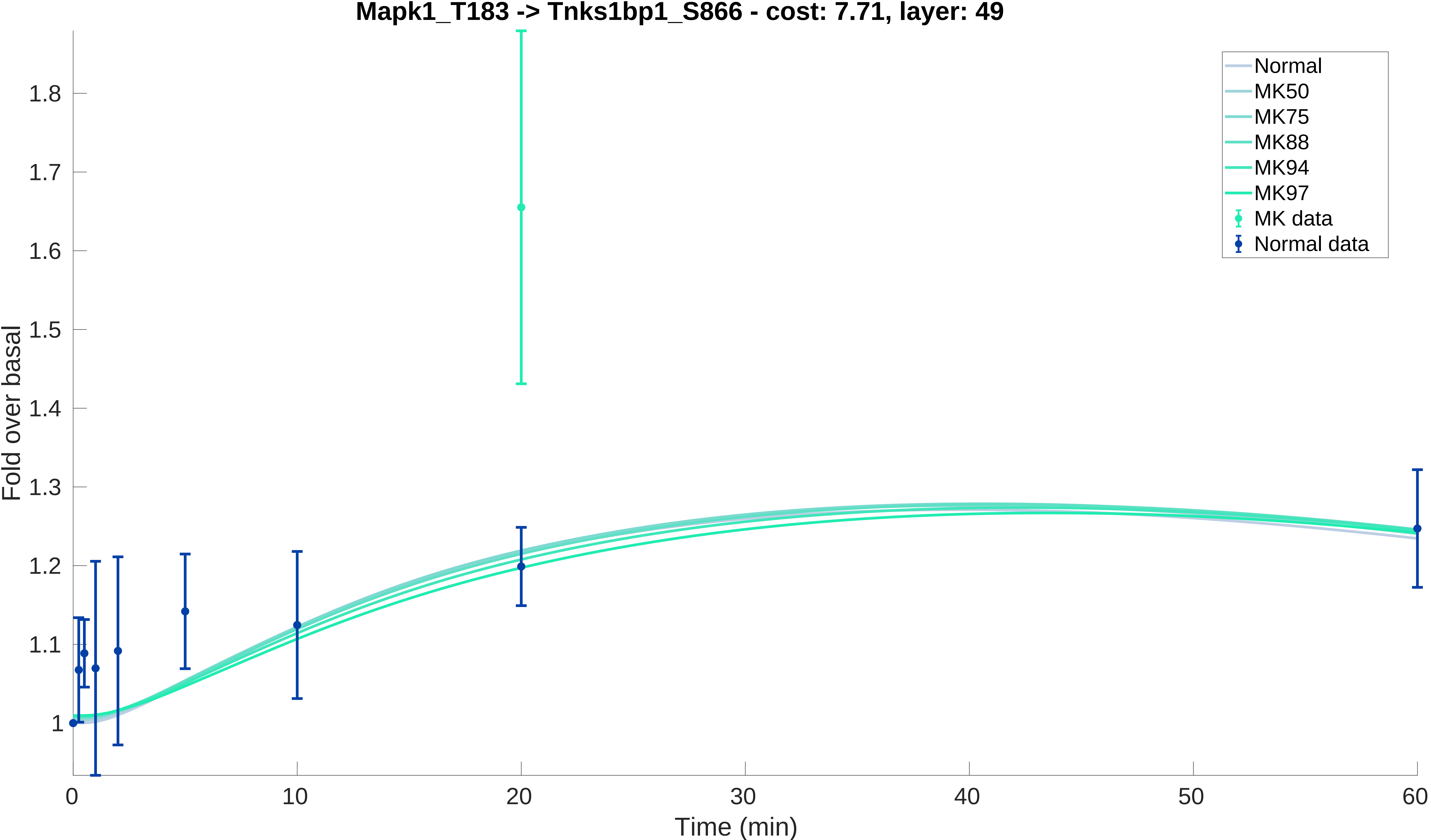
Mapk1_T183 -> Tnks1bp1_S866 - cost: 7.71, layer: 49
Fold over basal
Time (min)
Normal
MK50
MK75
MK88
MK94
MK97
MK data
Normal data
1
1.1
1.2
1.3
1.4
1.5
1.6
1.7
1.8
0
10
20
30
40
50
60

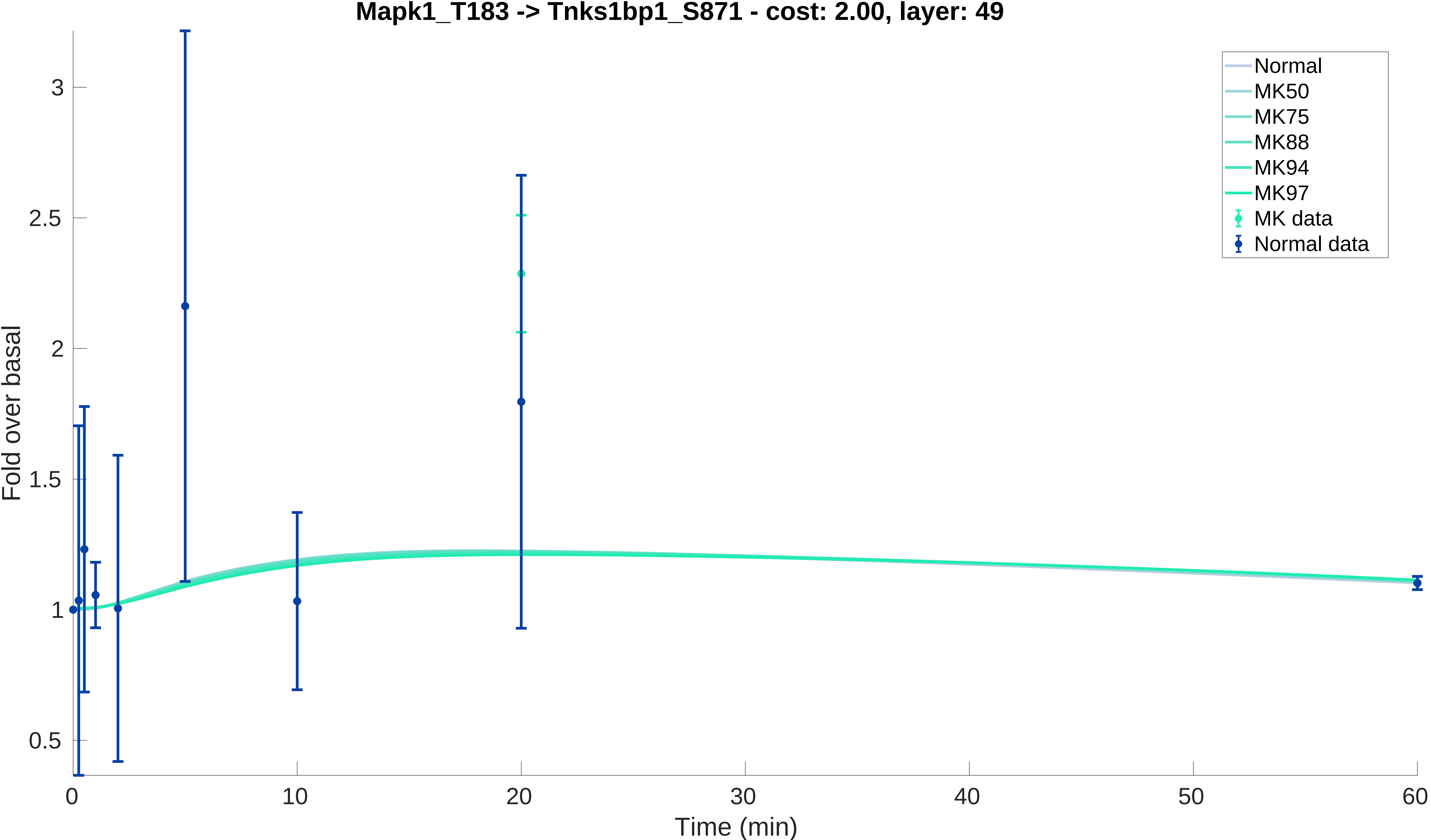

Mapk1_T183 -> Tnks1bp1_S871 - cost: 2.00, layer: 49
Fold over basal
Time (min)
0.5
1
1.5
2
2.5
3
0
10
20
30
40
50
60
Normal
MK50
MK75
MK88
MK94
MK97
MK data
Normal data

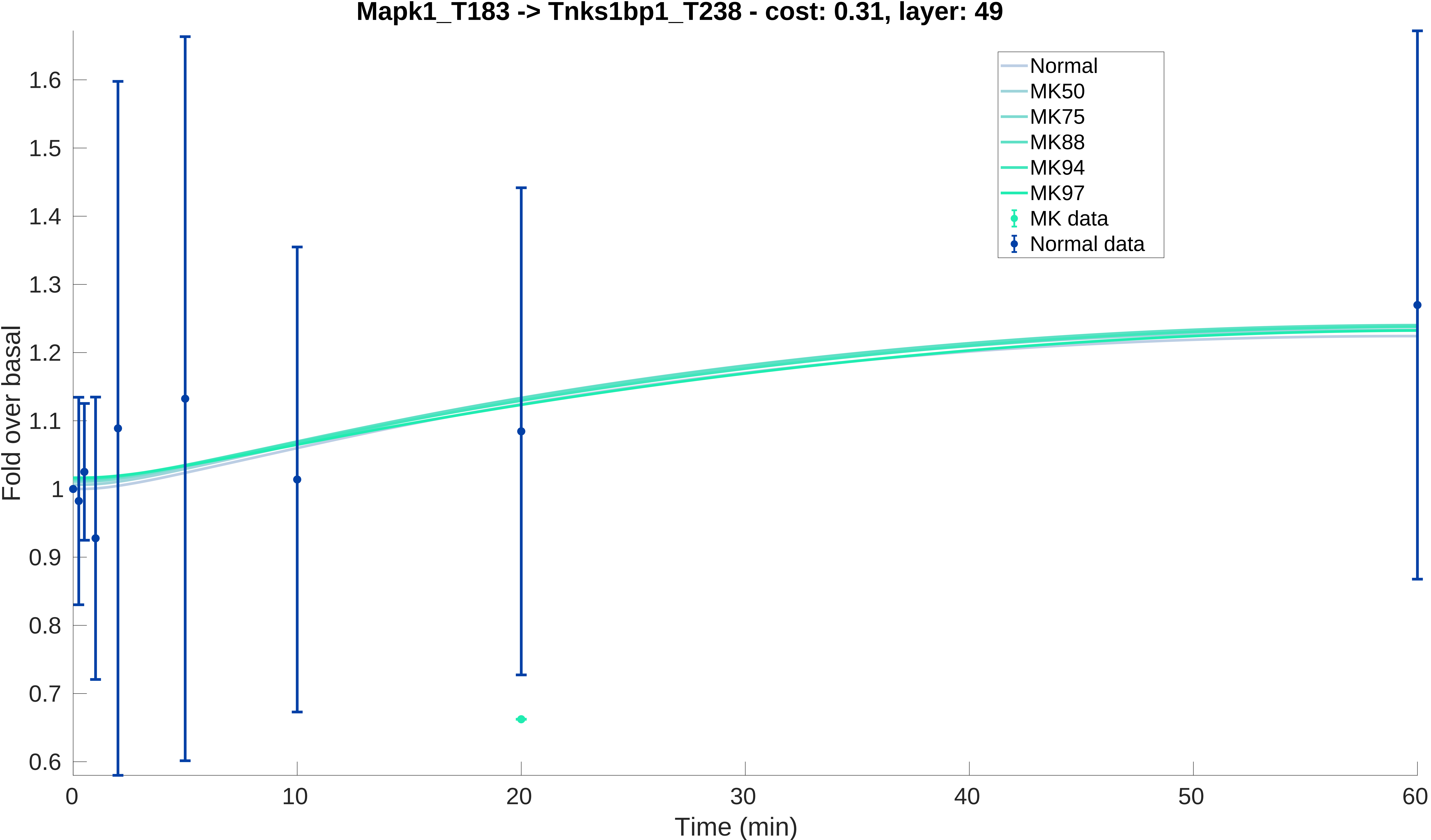

Mapk1_T183 -> Tnks1bp1_T238 - cost: 0.31, layer: 49
Normal
MK50
MK75
MK88
MK94
MK97
MK data
Normal data
Fold over basal
Time (min)
1.6
1.5
1.4
1.3
1.2
1.1
1
0.9
0.8
0.7
0.6
0
10
20
30
40
50
60

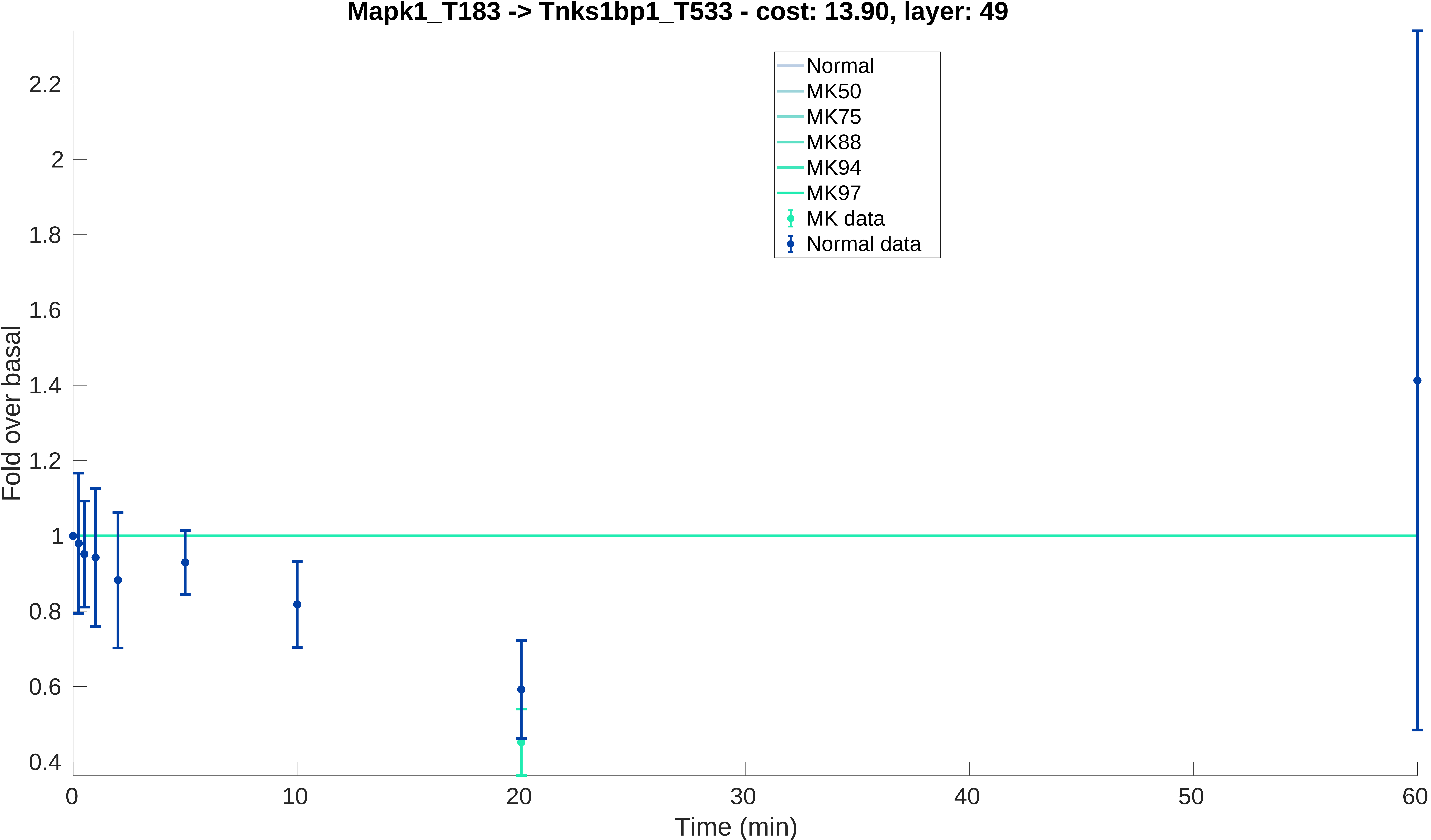

Mapk1_T183 -> Tnks1bp1_T533 - cost: 13.90, layer: 49
Normal
MK50
MK75
MK88
MK94
MK97
MK data
Normal data
Fold over basal
2.2
2
1.8
1.6
1.4
1.2
1
0.8
0.6
0.4
0
10
20
30
40
50
60
Time (min)

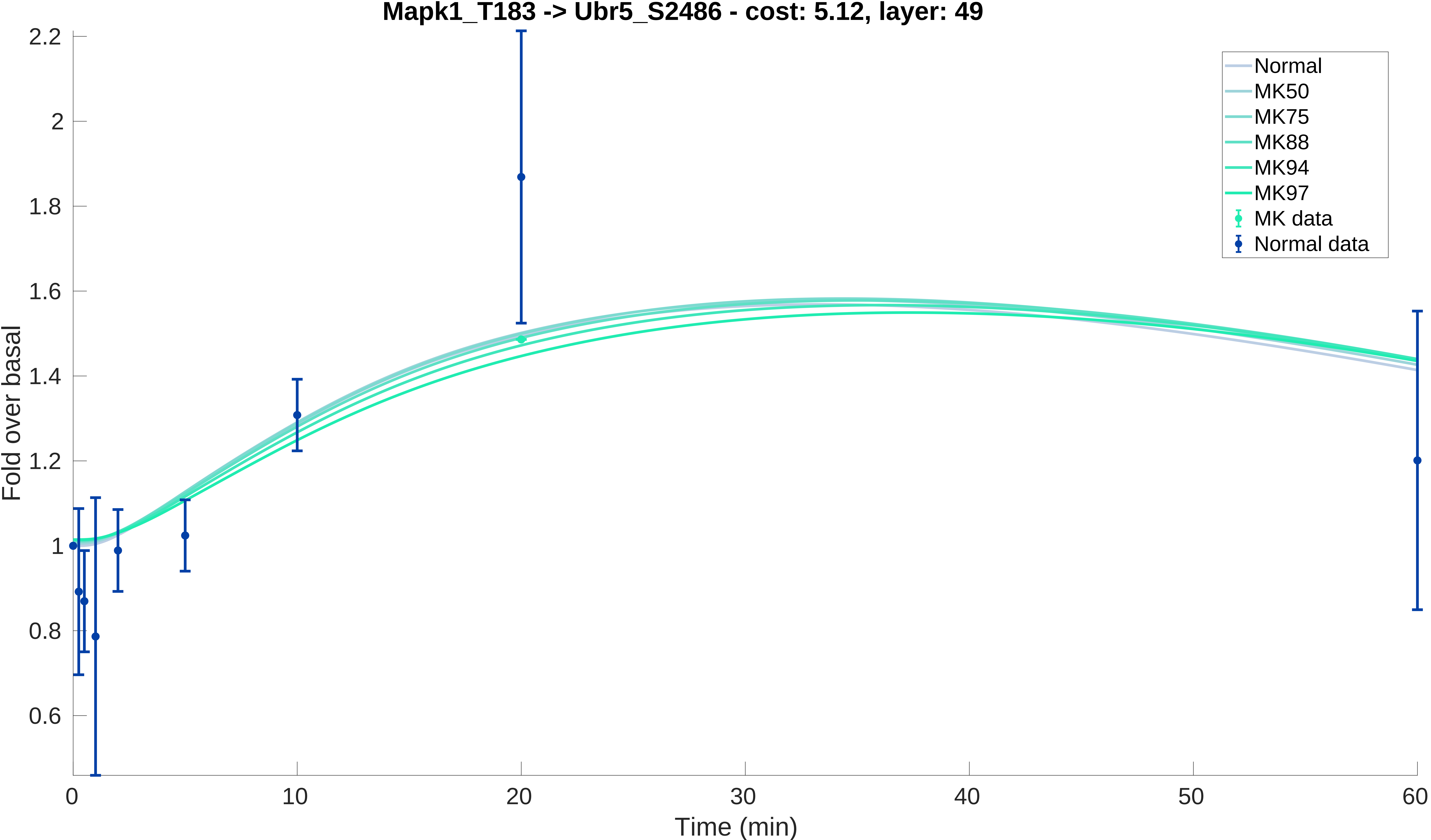
Mapk1_T183 -> Ubr5_S2486 - cost: 5.12, layer: 49
Fold over basal
Time (min)
0.6
0.8
1
1.2
1.4
1.6
1.8
2
2.2
0
10
20
30
40
50
60
Normal
MK50
MK75
MK88
MK94
MK97
MK data
Normal data

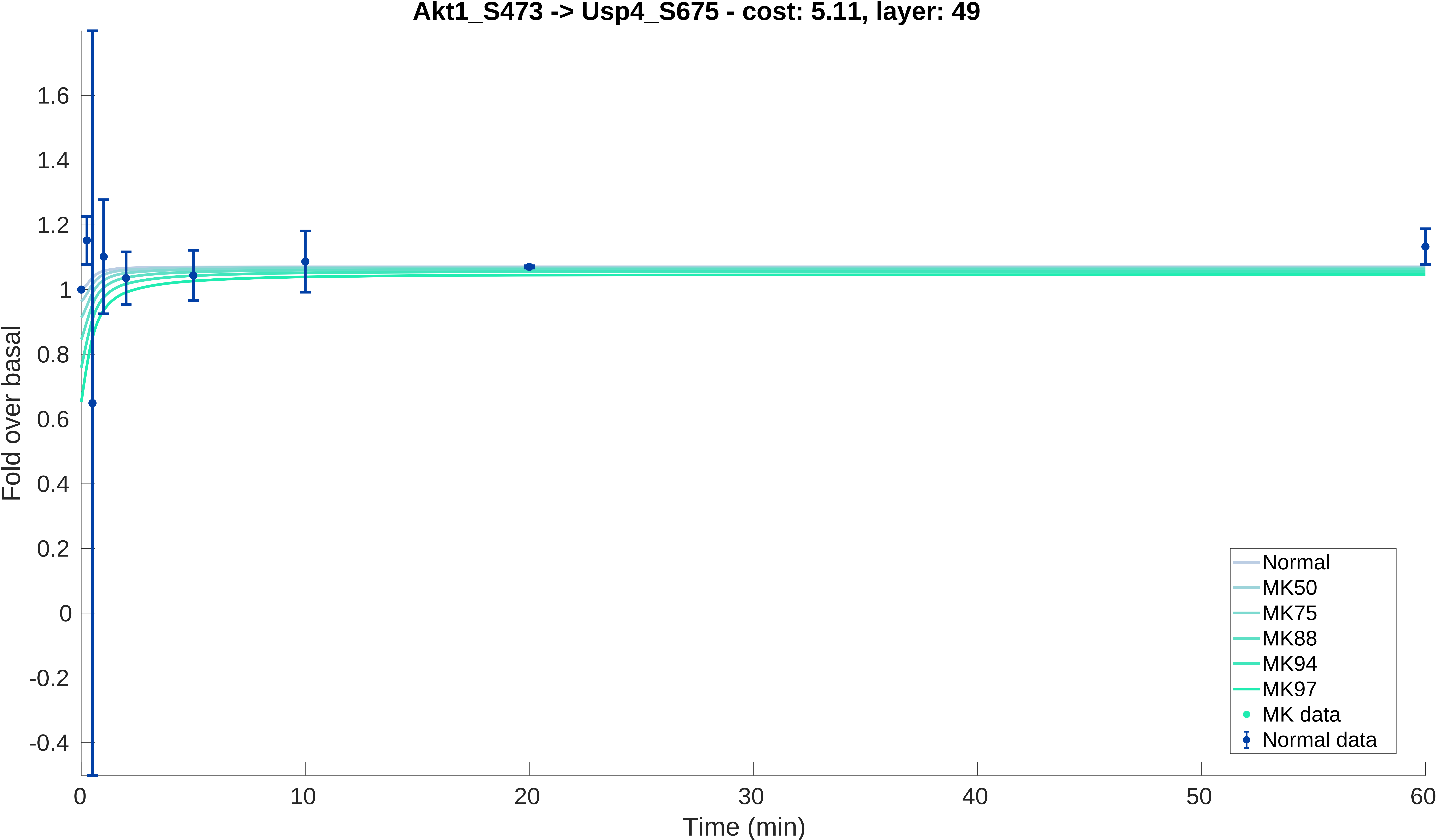

Akt1_S473 -> Usp4_S675 - cost: 5.11, layer: 49
Fold over basal
1.6
1.4
1.2
1
0.8
0.6
0.4
0.2
0
-0.2
-0.4
0
10
20
30
40
50
60
Time (min)
Normal
MK50
MK75
MK88
MK94
MK97
MK data
Normal data

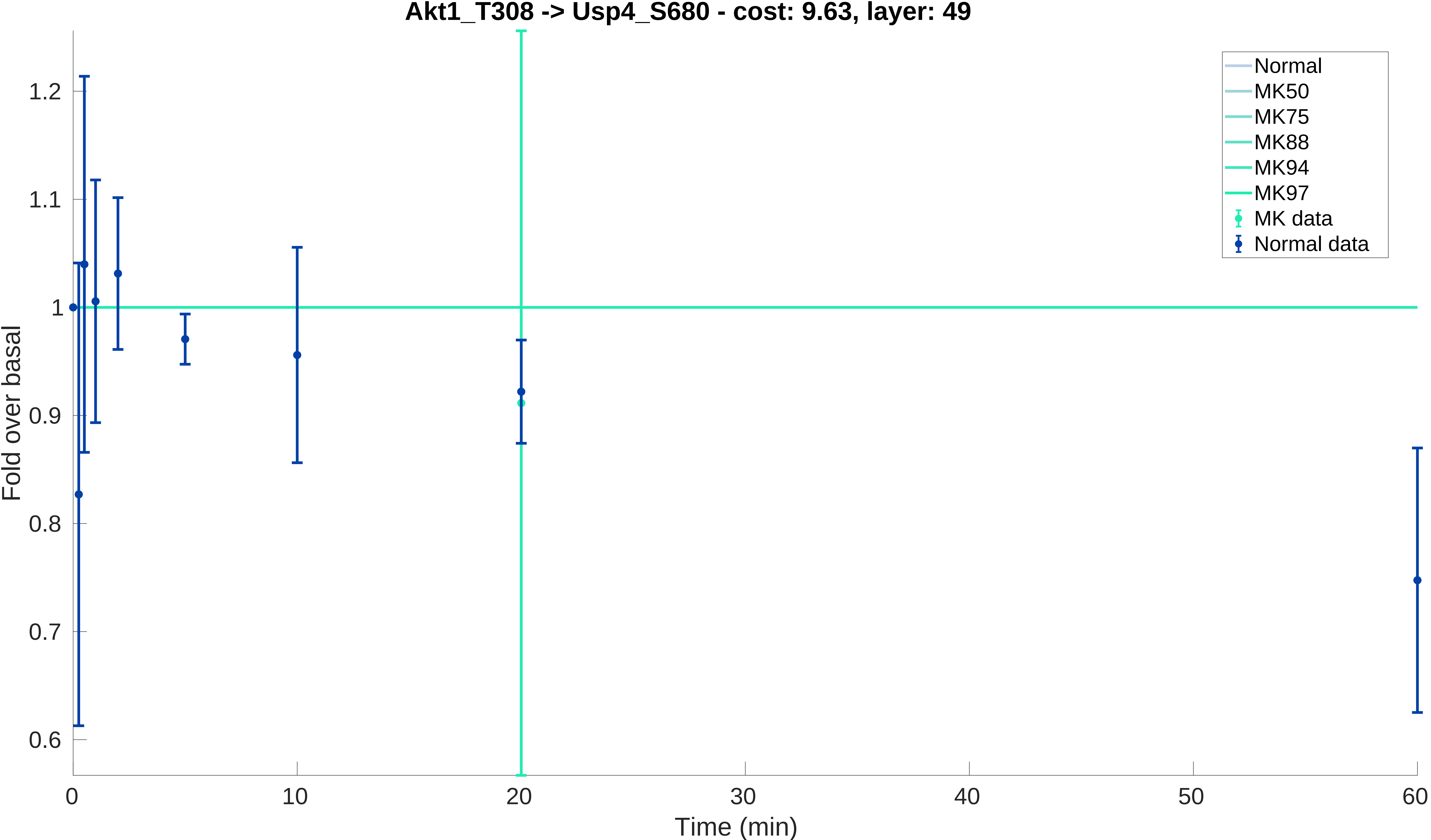

Akt1_T308 -> Usp4_S680 - cost: 9.63, layer: 49
Fold over basal
Time (min)
Normal
MK50
MK75
MK88
MK94
MK97
MK data
Normal data
0.6
0.7
0.8
0.9
1
1.1
1.2
0
10
20
30
40
50
60

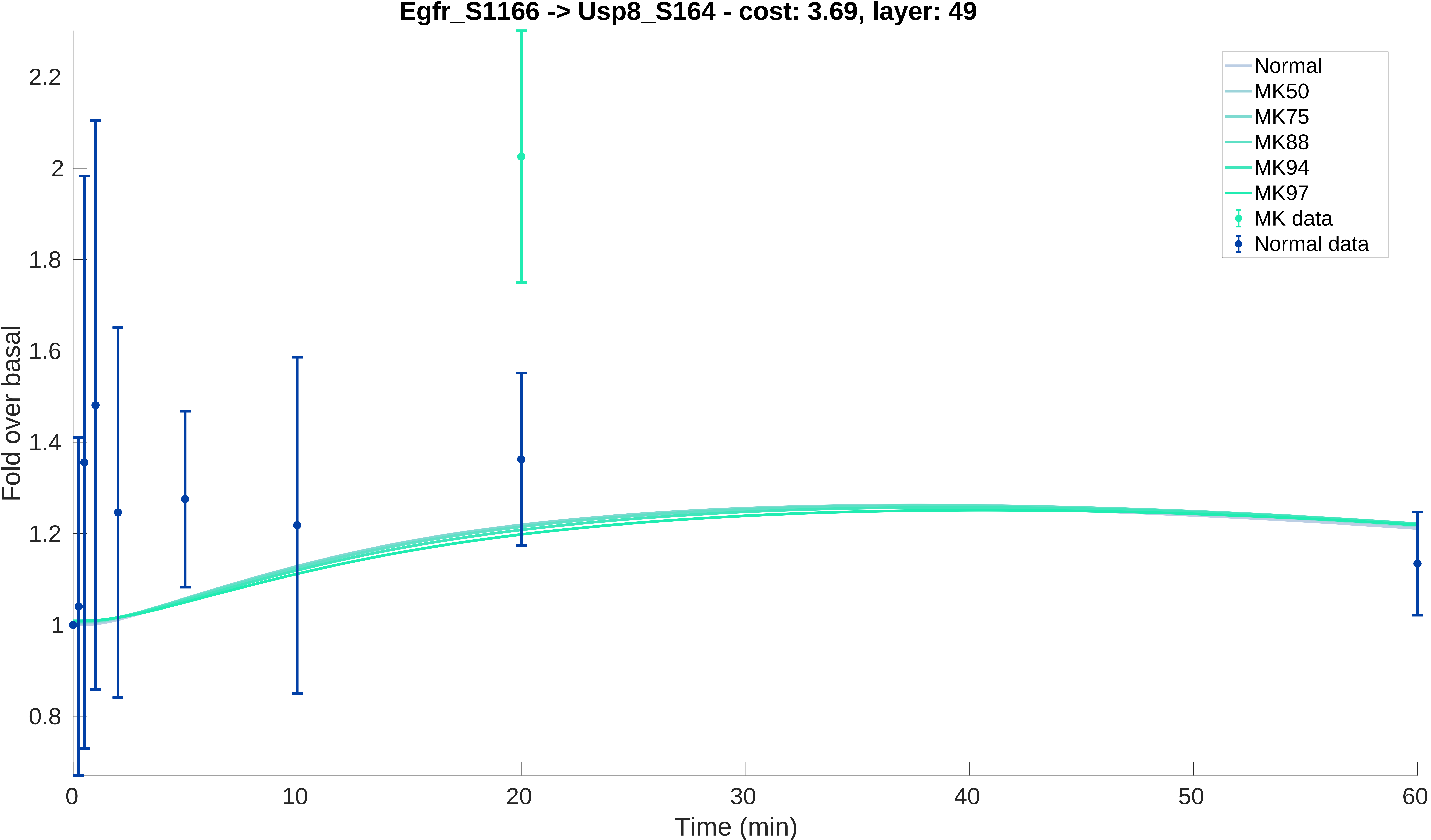

Egfr_S1166 -> Usp8_S164 - cost: 3.69, layer: 49
Fold over basal
Time (min)
0
10
20
30
40
50
60
0.8
1
1.2
1.4
1.6
1.8
2
2.2
Normal
MK50
MK75
MK88
MK94
MK97
MK data
Normal data

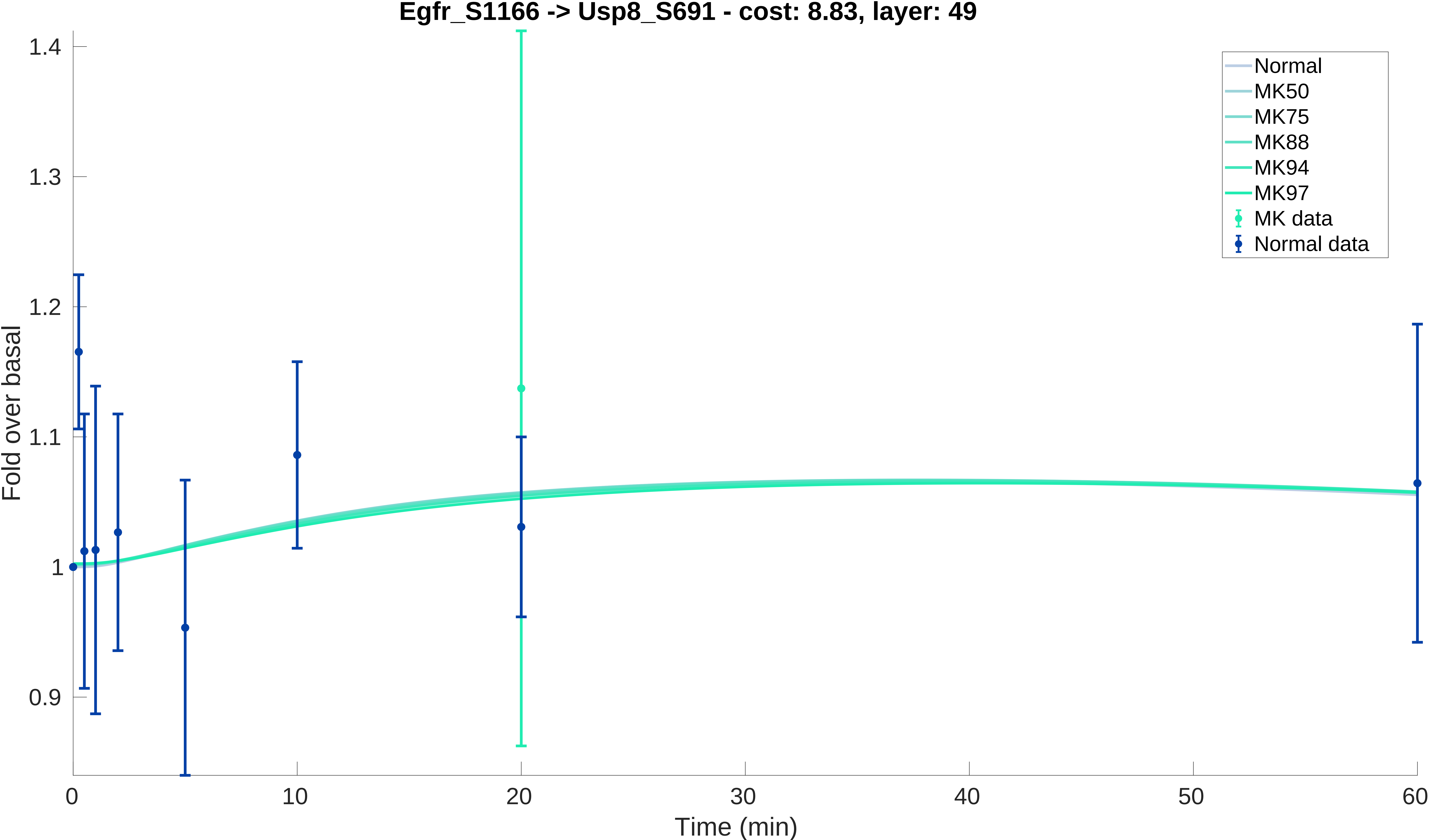

Egfr_S1166 -> Usp8_S691 - cost: 8.83, layer: 49
Fold over basal
Time (min)
1.4
1.3
1.2
1.1
1
0.9
0
10
20
30
40
50
60
Normal
MK50
MK75
MK88
MK94
MK97
MK data
Normal data

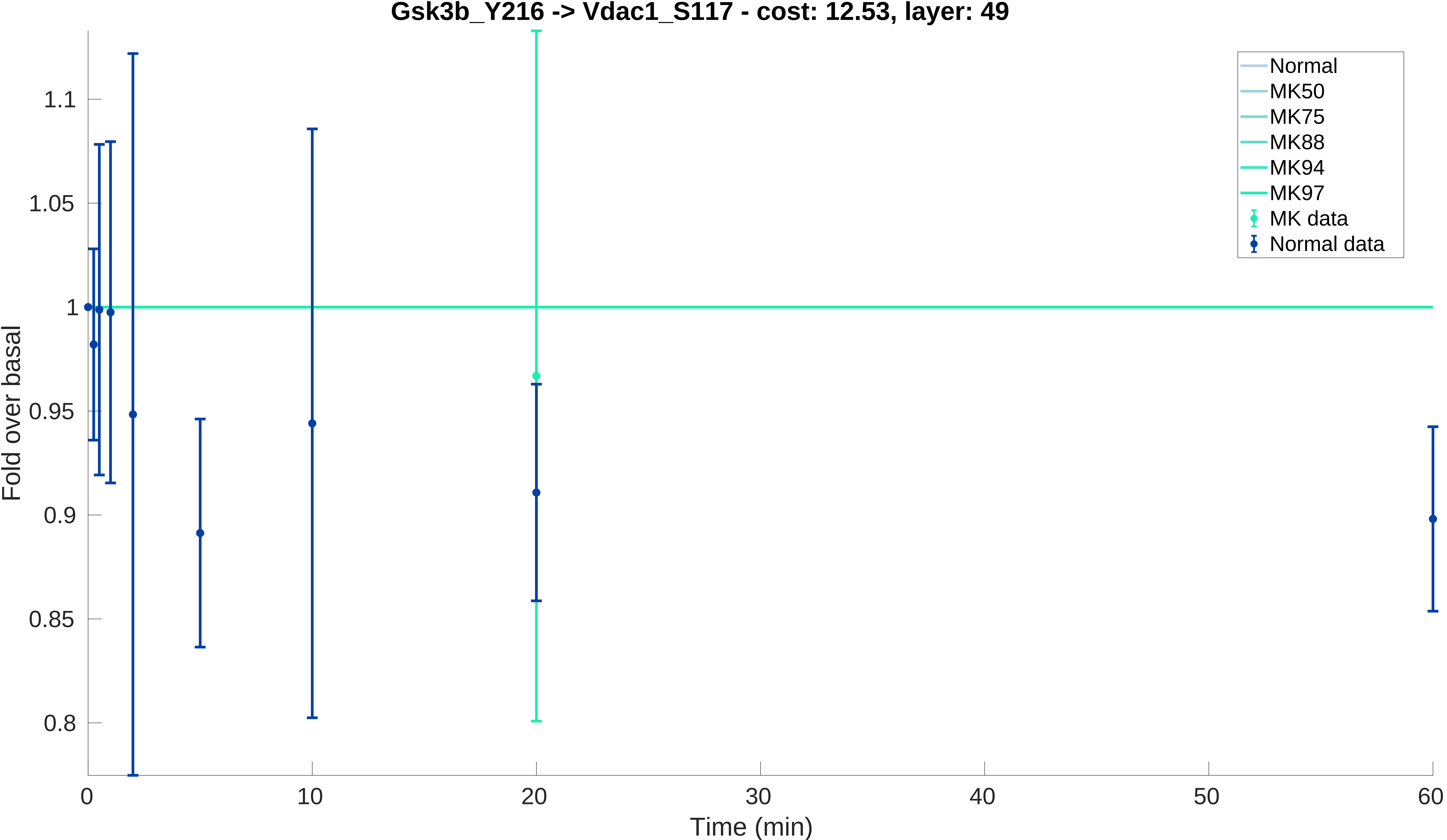
Gsk3b_Y216 -> Vdac1_S117 - cost: 12.53, layer: 49
Normal
MK50
MK75
MK88
MK94
MK97
MK data
Normal data
Fold over basal
1.1
1.05
1
0.95
0.9
0.85
0.8
0
10
20
30
40
50
60
Time (min)

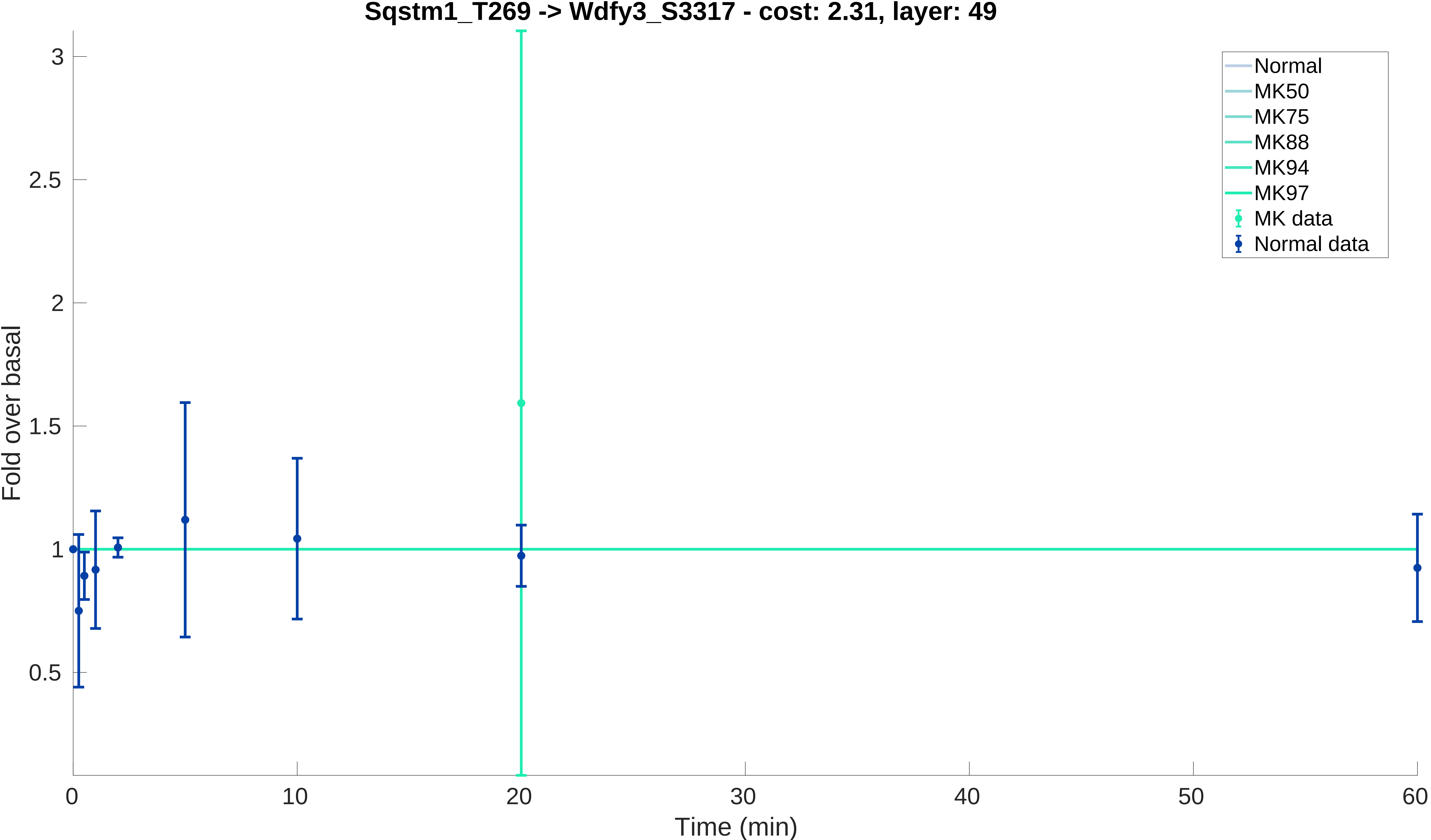
Sqstm1_T269 -> Wdfy3_S3317 - cost: 2.31, layer: 49
Fold over basal
Time (min)
0
10
20
30
40
50
60
0.5
1
1.5
2
2.5
3
Normal
MK50
MK75
MK88
MK94
MK97
MK data
Normal data

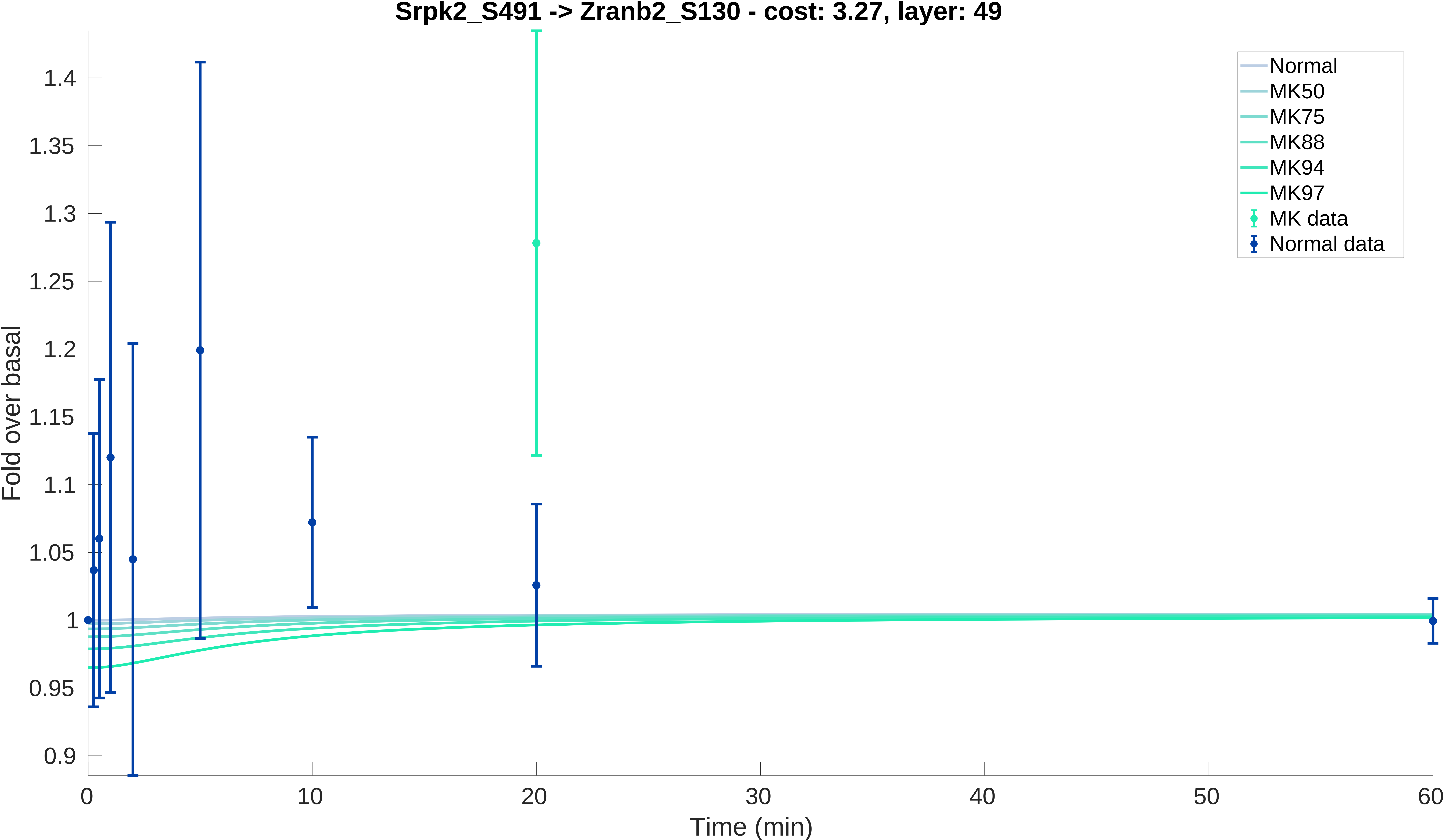

Srpk2_S491 -> Zranb2_S130 - cost: 3.27, layer: 49
Fold over basal
1.4
1.35
1.3
1.25
1.2
1.15
1.1
1.05
1
0.95
0.9
0
10
20
30
40
50
60
Time (min)
Normal
MK50
MK75
MK88
MK94
MK97
MK data
Normal data

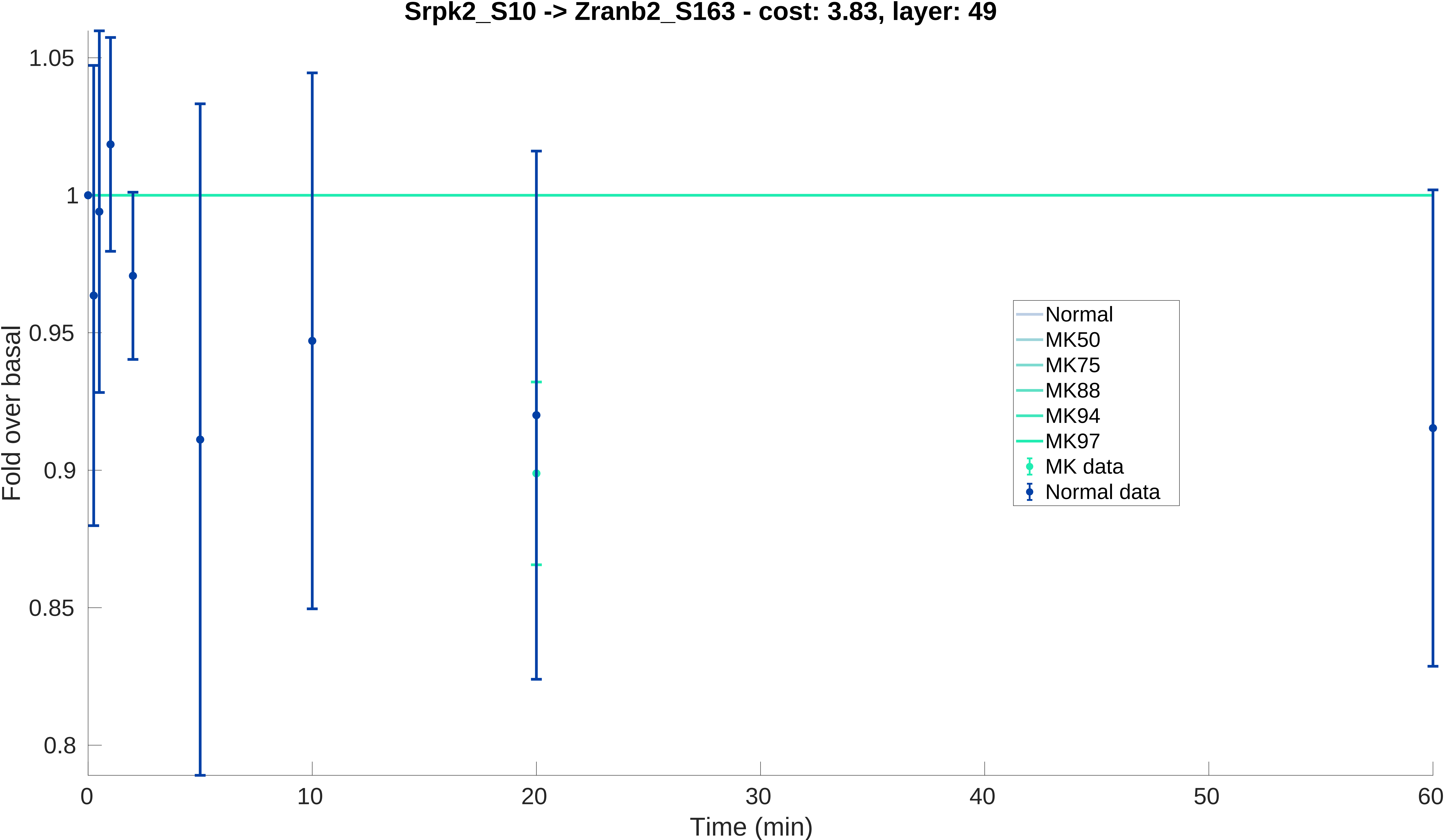
Srpk2_S10 -> Zranb2_S163 - cost: 3.83, layer: 49
Fold over basal
1.05
1
0.95
0.9
0.85
0.8
0
10
20
30
40
50
60
Time (min)
Normal
MK50
MK75
MK88
MK94
MK97
MK data
Normal data

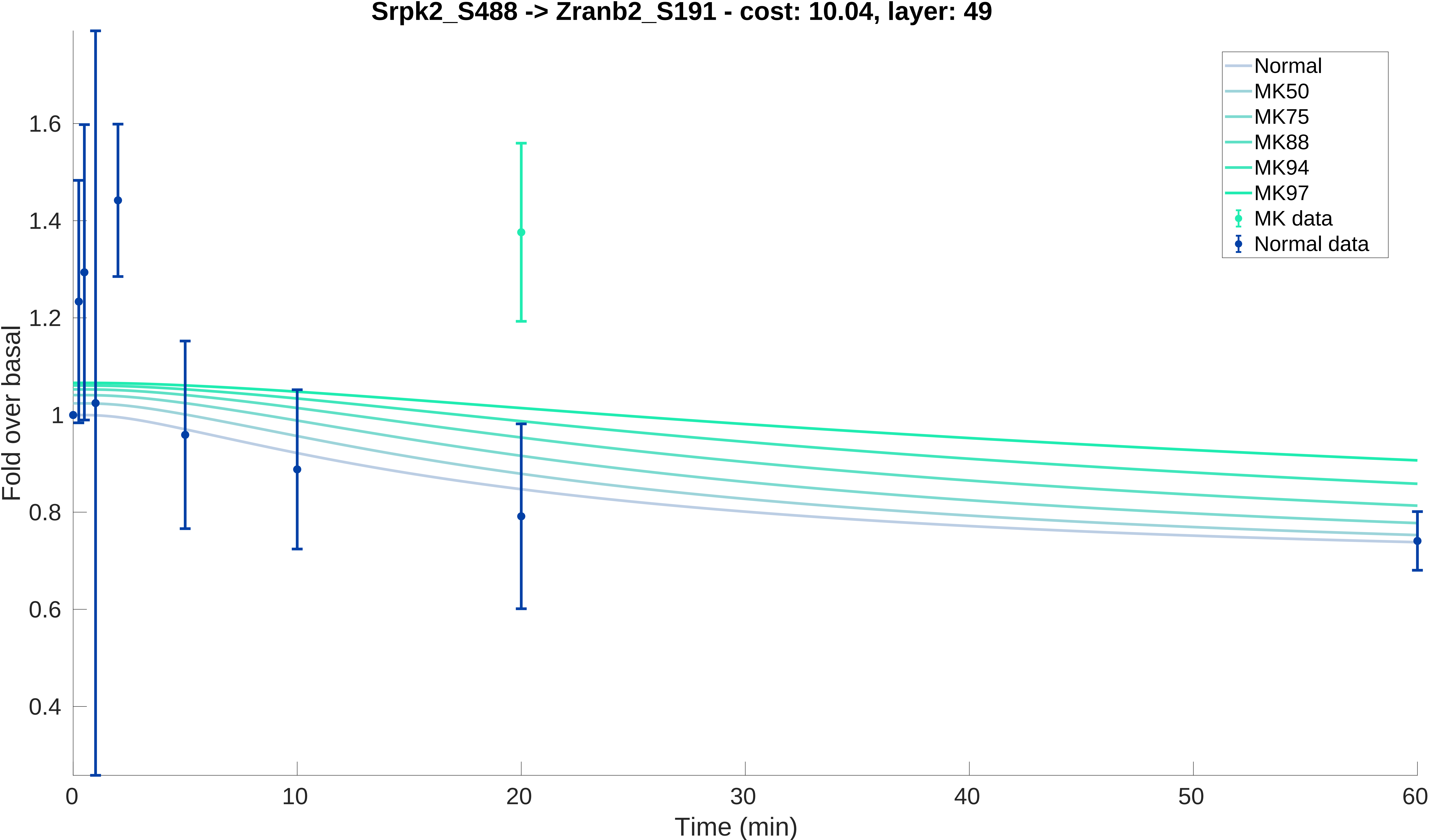
Srpk2_S488 -> Zranb2_S191 - cost: 10.04, layer: 49
Fold over basal
Time (min)
0
10
20
30
40
50
60
0.4
0.6
0.8
1
1.2
1.4
1.6
Normal
MK50
MK75
MK88
MK94
MK97
MK data
Normal data

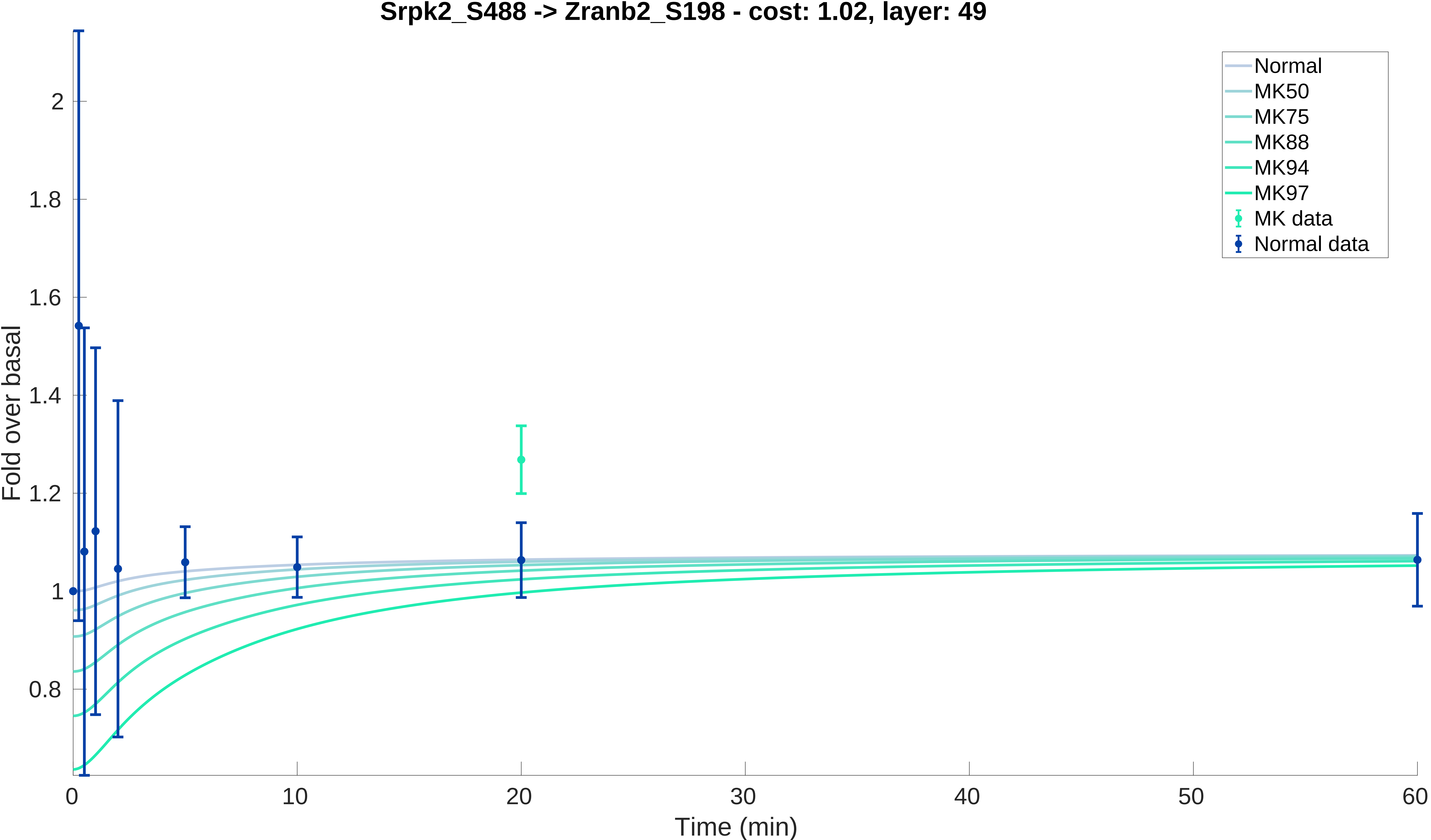

Srpk2_S488 -> Zranb2_S198 - cost: 1.02, layer: 49
Normal
MK50
MK75
MK88
MK94
MK97
MK data
Normal data
Fold over basal
0.8
1
1.2
1.4
1.6
1.8
2
0
10
20
30
40
50
60
Time (min)

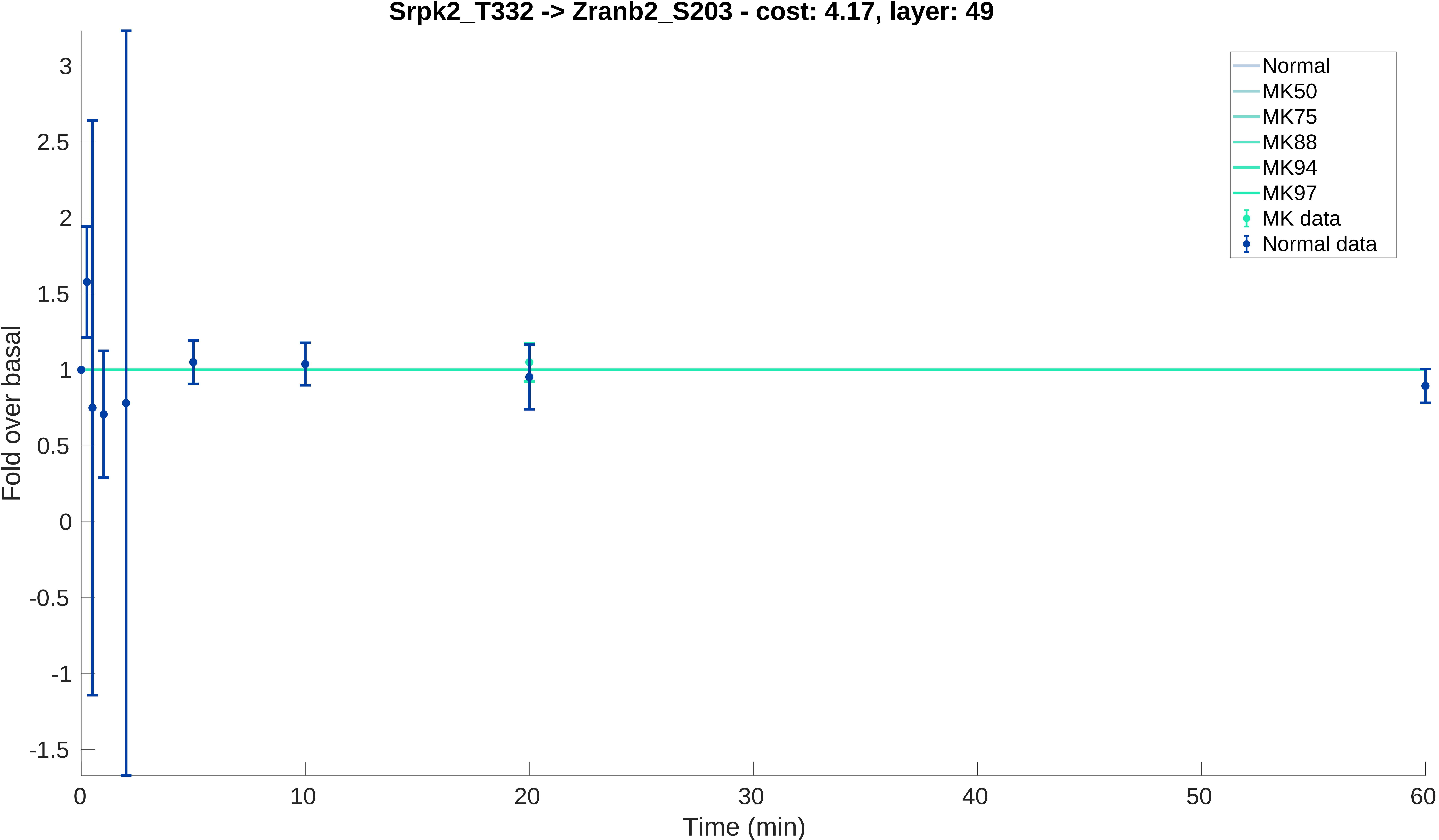
Srpk2_T332 -> Zranb2_S203 - cost: 4.17, layer: 49
Fold over basal
Time (min)
3
2.5
2
1.5
1
0.5
0
-0.5
-1
-1.5
0
10
20
30
40
50
60
Normal
MK50
MK75
MK88
MK94
MK97
MK data
Normal data

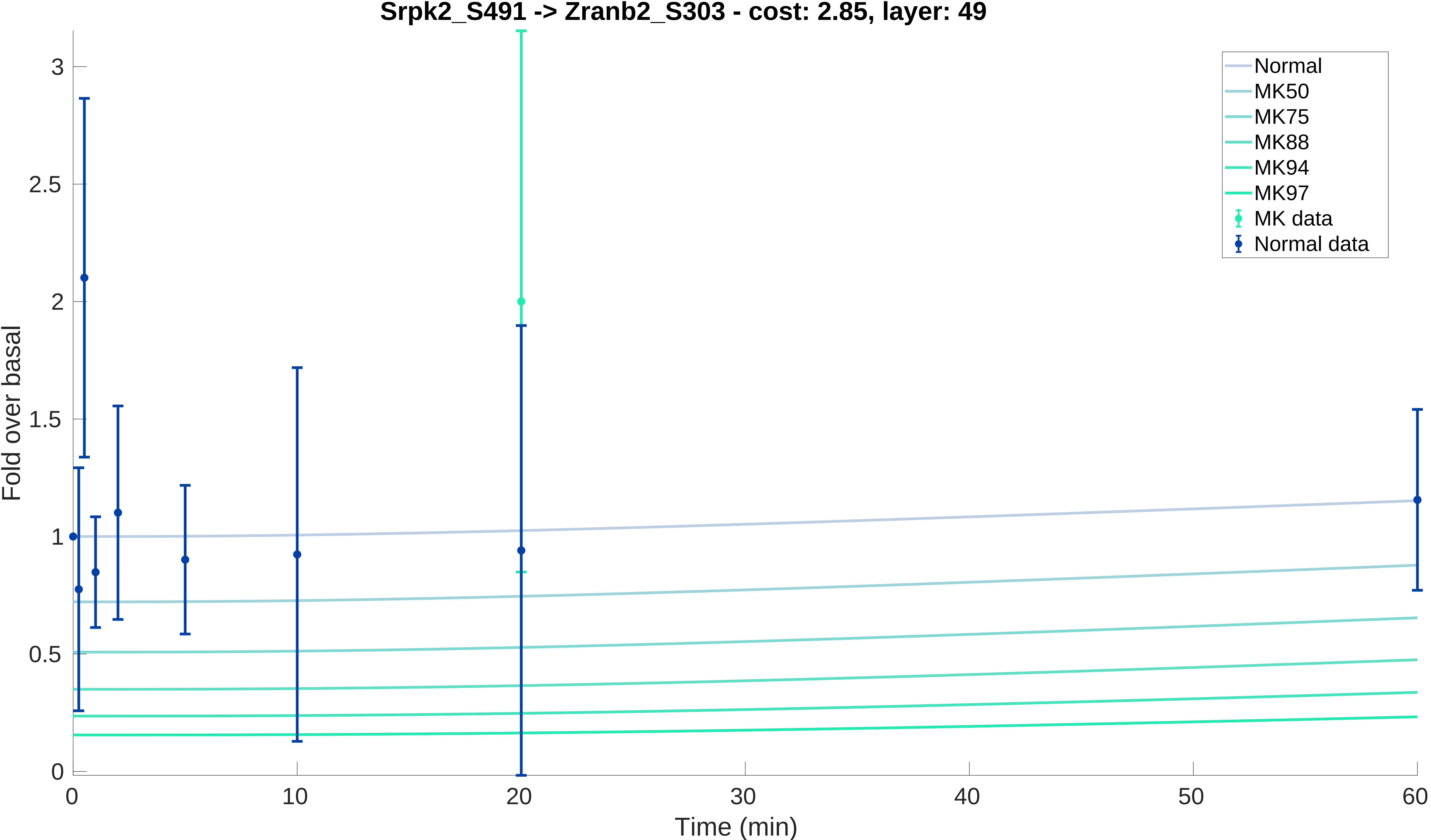
Srpk2_S491 -> Zranb2_S303 - cost: 2.85, layer: 49
Fold over basal
Time (min)
0
0.5
1
1.5
2
2.5
3
0
10
20
30
40
50
60
Normal
MK50
MK75
MK88
MK94
MK97
MK data
Normal data

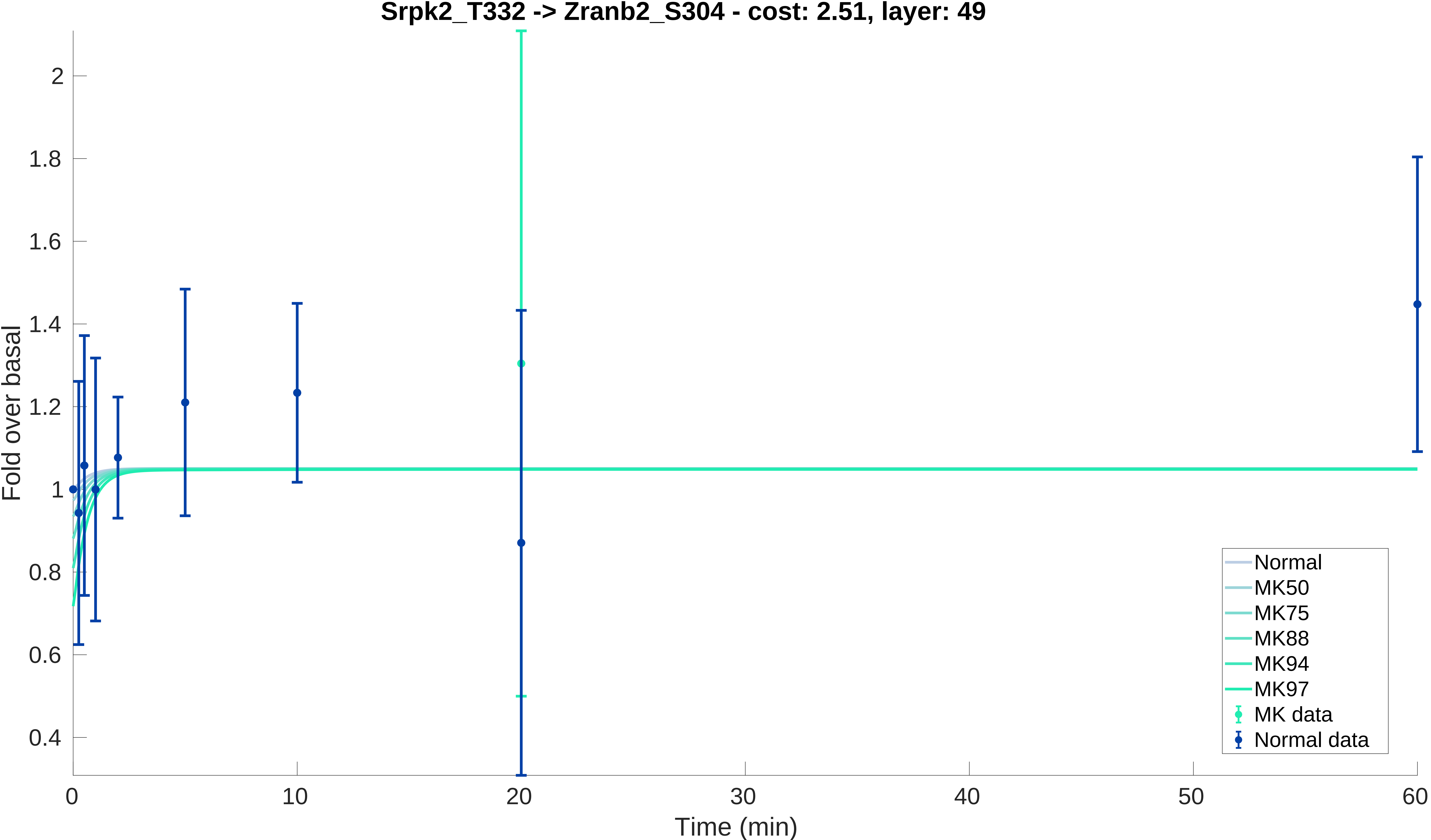

Srpk2_T332 -> Zranb2_S304 - cost: 2.51, layer: 49
Fold over basal
2
1.8
1.6
1.4
1.2
1
0.8
0.6
0.4
0
10
20
30
40
50
60
Time (min)
Normal
MK50
MK75
MK88
MK94
MK97
MK data
Normal data

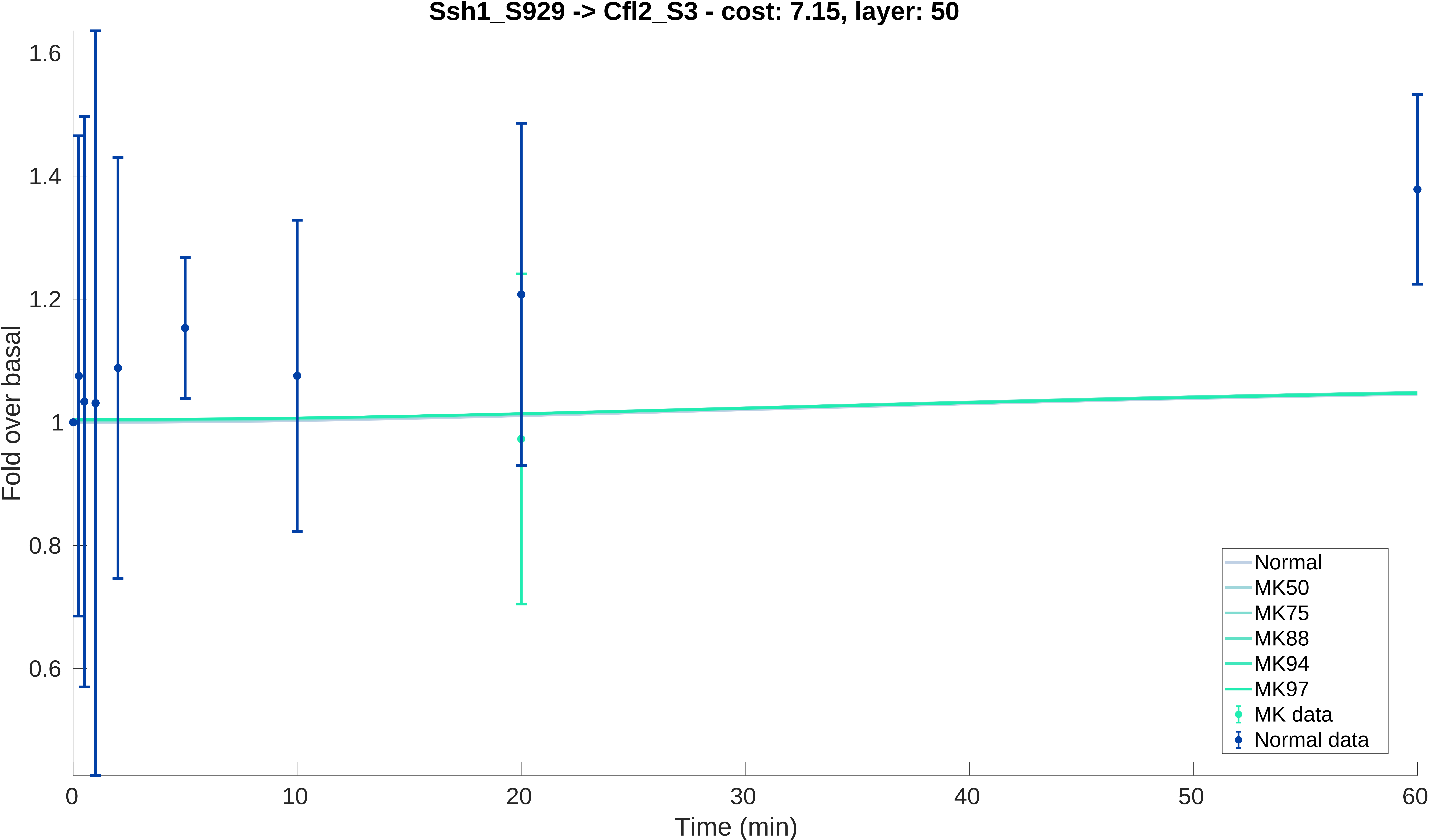

Ssh1_S929 -> Cfl2_S3 - cost: 7.15, layer: 50
Fold over basal
1.6
1.4
1.2
1
0.8
0.6
0
10
20
30
40
50
60
Time (min)
Normal
MK50
MK75
MK88
MK94
MK97
MK data
Normal data

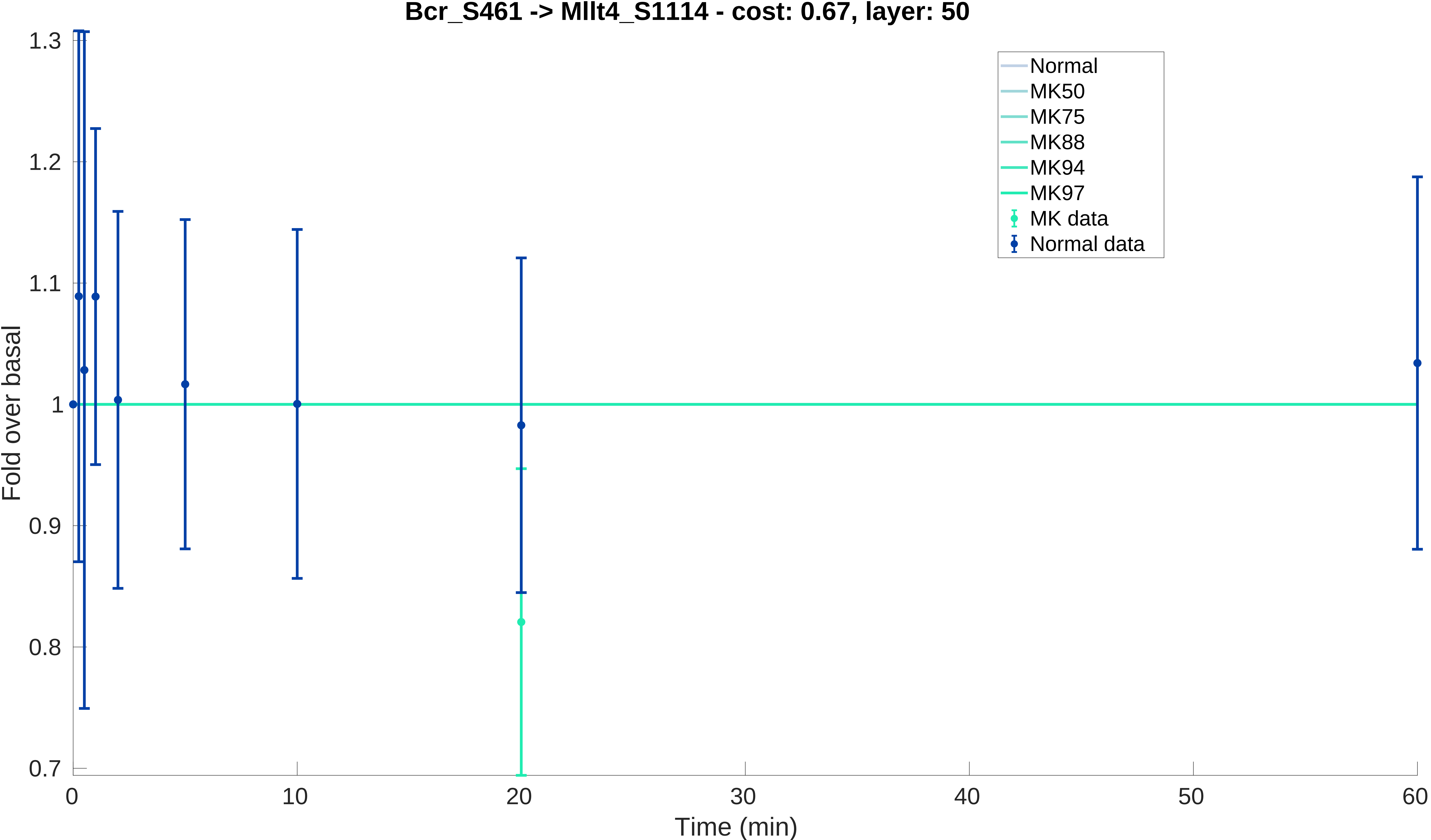

Bcr_S461 -> Mllt4_S1114 - cost: 0.67, layer: 50
Fold over basal
Time (min)
1.3
1.2
1.1
1
0.9
0.8
0.7
0
10
20
30
40
50
60
Normal
MK50
MK75
MK88
MK94
MK97
MK data
Normal data

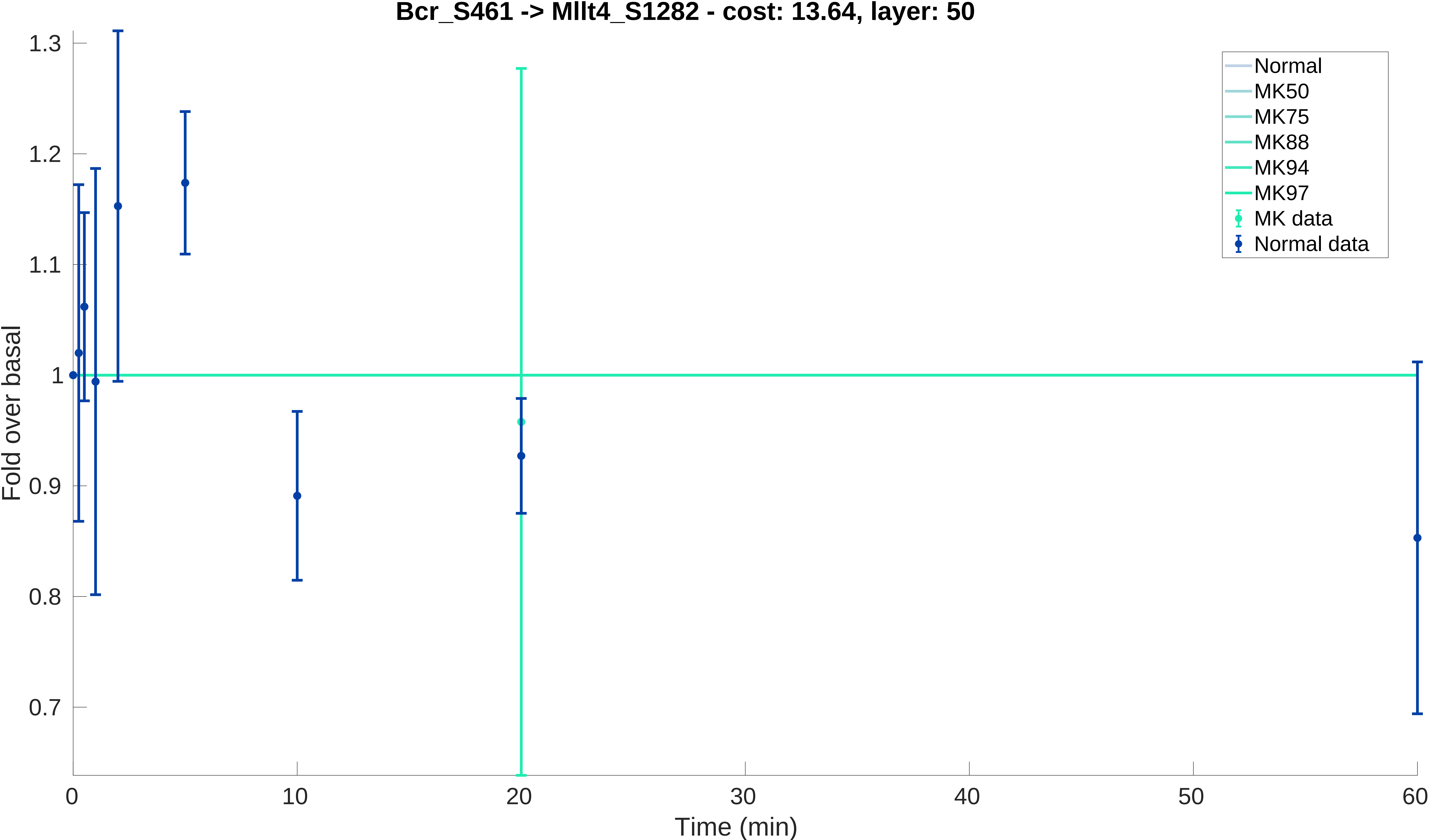

Bcr_S461 -> Mllt4_S1282 - cost: 13.64, layer: 50
Fold over basal
1.3
1.2
1.1
1
0.9
0.8
0.7
0
10
20
30
40
50
60
Time (min)
Normal
MK50
MK75
MK88
MK94
MK97
MK data
Normal data

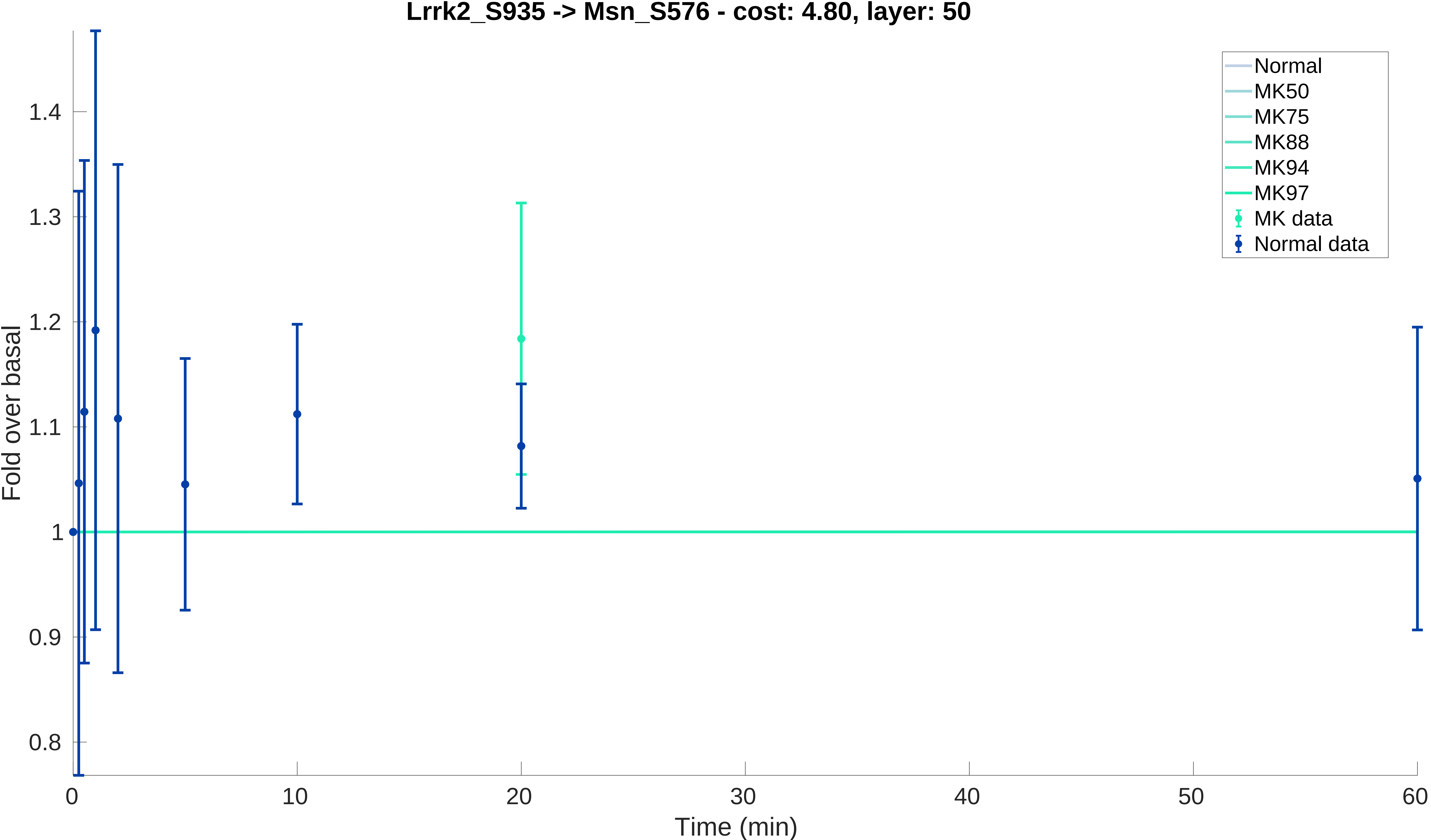

Lrrk2_S935 -> Msn_S576 - cost: 4.80, layer: 50
Normal
MK50
MK75
MK88
MK94
MK97
MK data
Normal data
Fold over basal
1.4
1.3
1.2
1.1
1
0.9
0.8
0
10
20
30
40
50
60
Time (min)

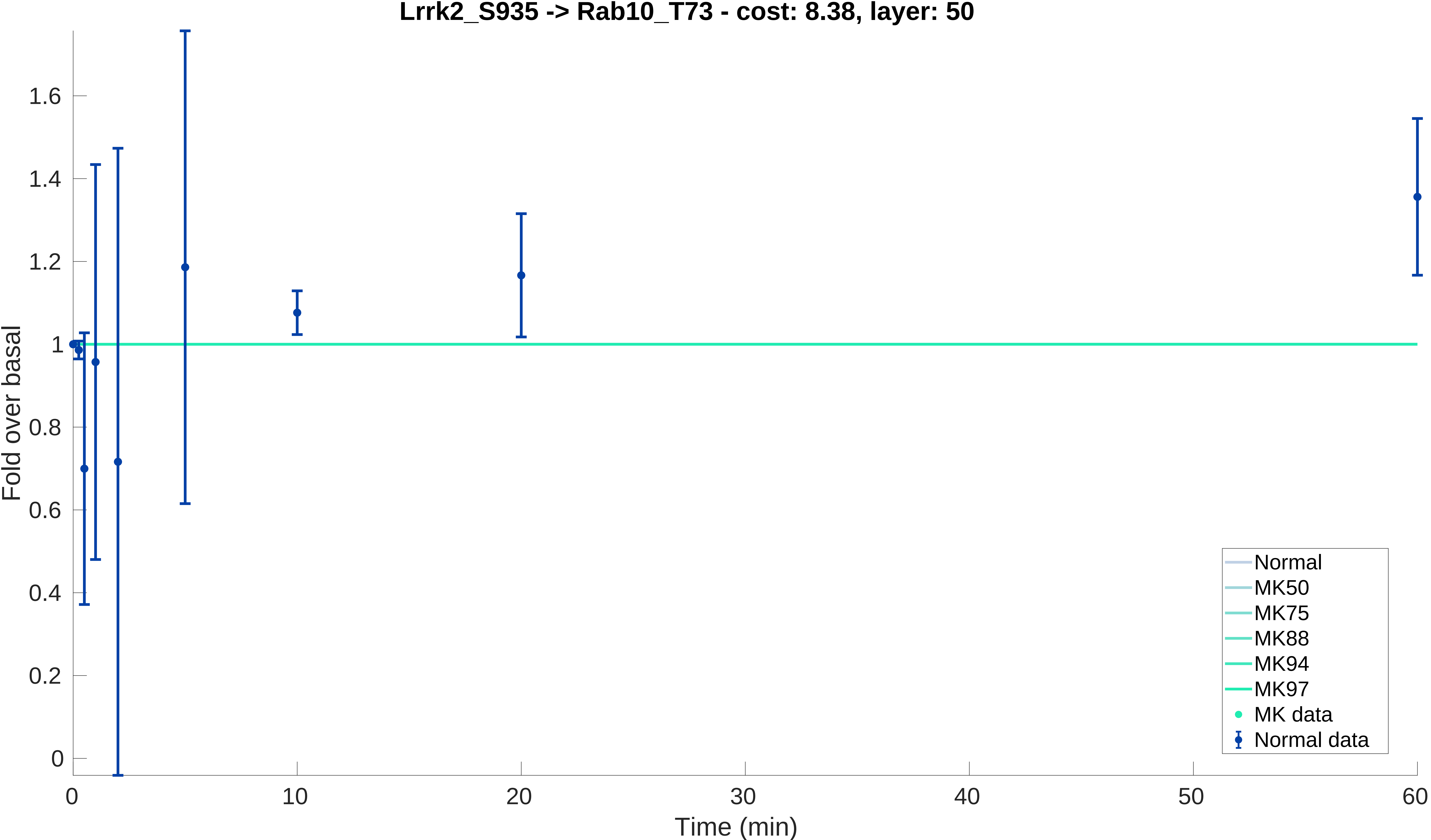
Lrrk2_S935 -> Rab10_T73 - cost: 8.38, layer: 50
Fold over basal
Time (min)
0
0.2
0.4
0.6
0.8
1
1.2
1.4
1.6
0
10
20
30
40
50
60
Normal
MK50
MK75
MK88
MK94
MK97
MK data
Normal data

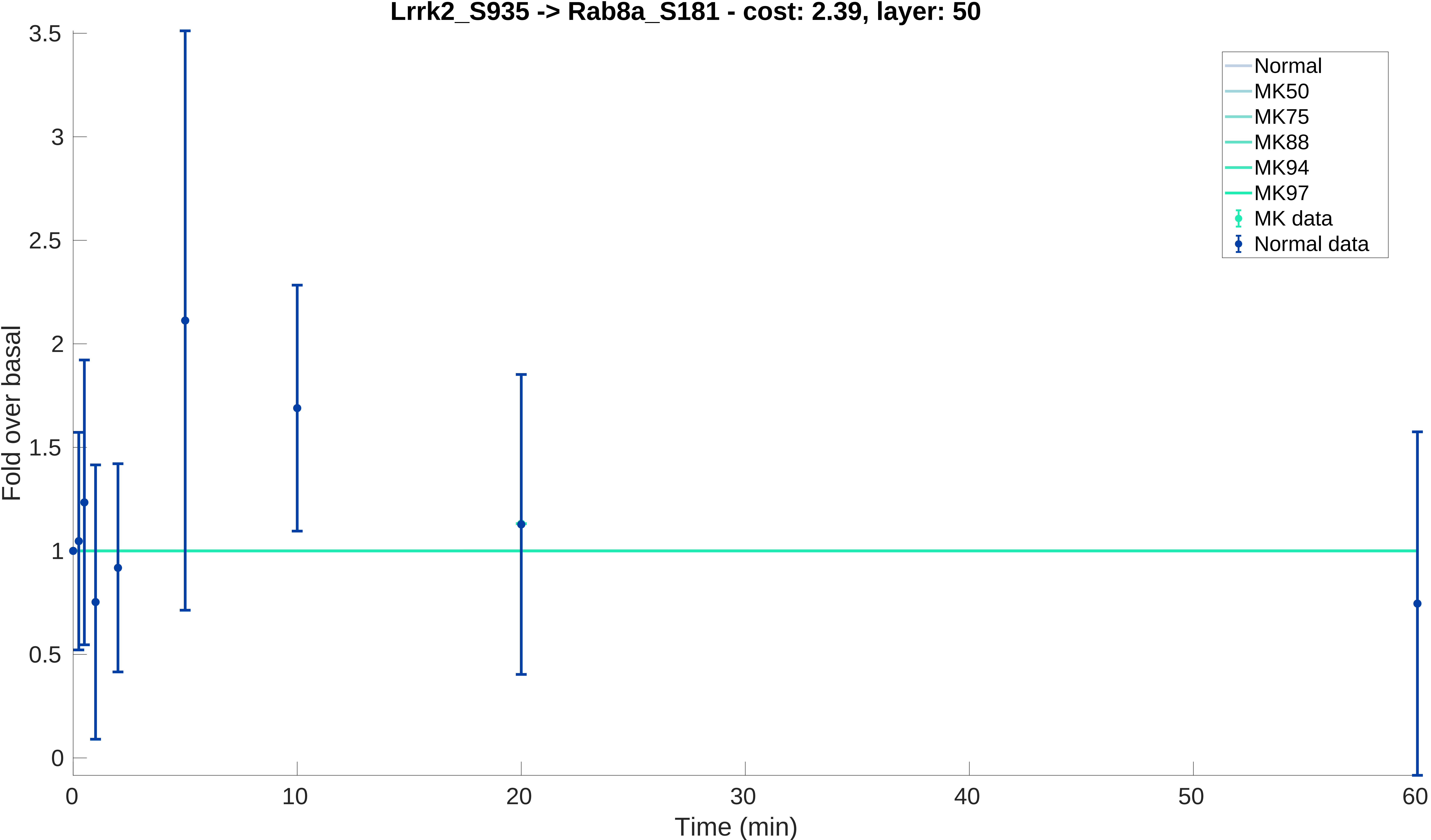

Lrrk2_S935 -> Rab8a_S181 - cost: 2.39, layer: 50
Fold over basal
Time (min)
0
0.5
1
1.5
2
2.5
3
3.5
0
10
20
30
40
50
60
Normal
MK50
MK75
MK88
MK94
MK97
MK data
Normal data

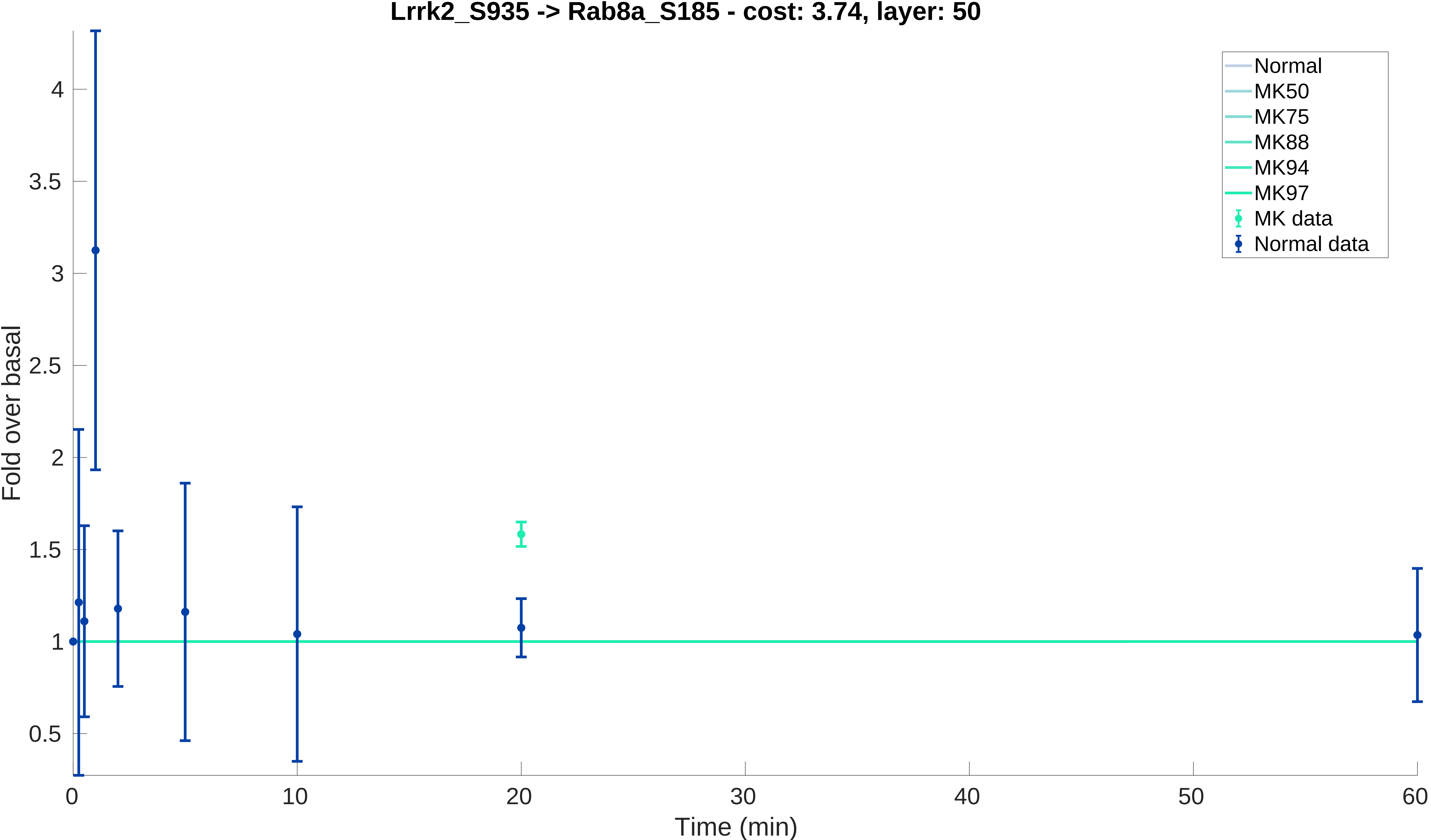

Lrrk2_S935 -> Rab8a_S185 - cost: 3.74, layer: 50
Fold over basal
Time (min)
0.5
1
1.5
2
2.5
3
3.5
4
0
10
20
30
40
50
60
Normal
MK50
MK75
MK88
MK94
MK97
MK data
Normal data

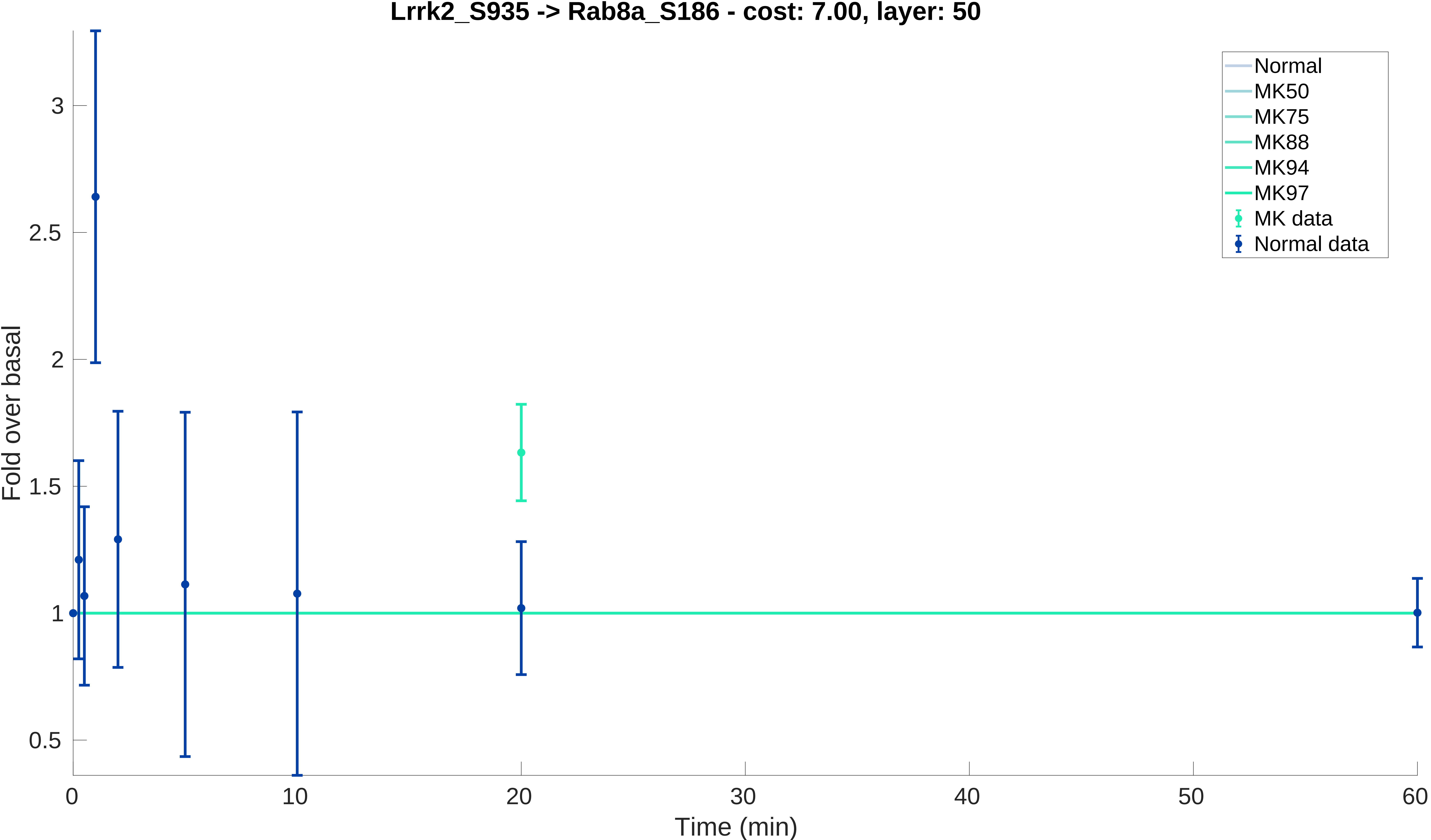

Lrrk2_S935 -> Rab8a_S186 - cost: 7.00, layer: 50
Normal
MK50
MK75
MK88
MK94
MK97
MK data
Normal data
Fold over basal
0.5
1
1.5
2
2.5
3
0
10
20
30
40
50
60
Time (min)

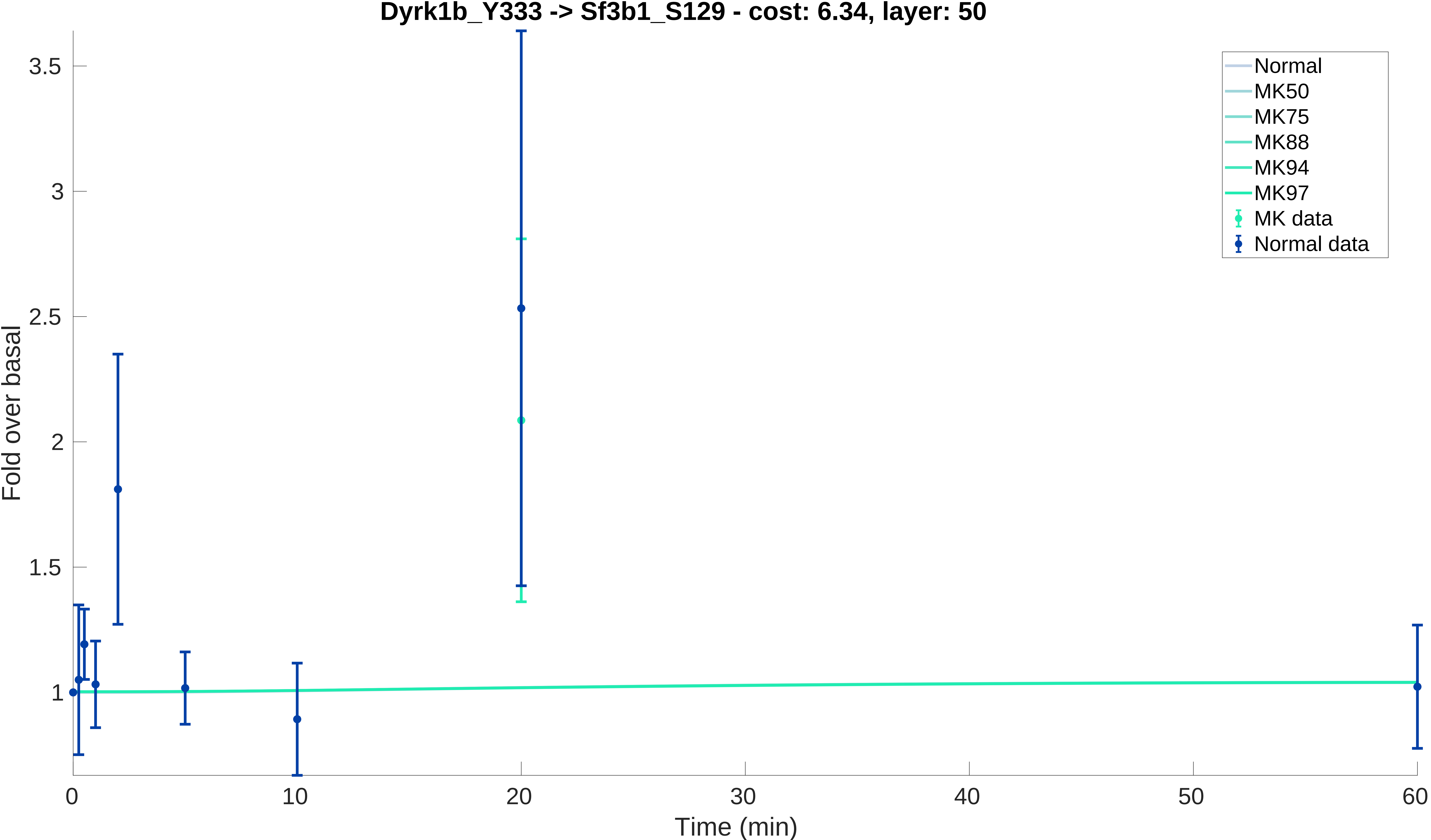

Dyrk1b_Y333 -> Sf3b1_S129 - cost: 6.34, layer: 50
Fold over basal
3.5
3
2.5
2
1.5
1
0
10
20
30
40
50
60
Time (min)
Normal
MK50
MK75
MK88
MK94
MK97
MK data
Normal data

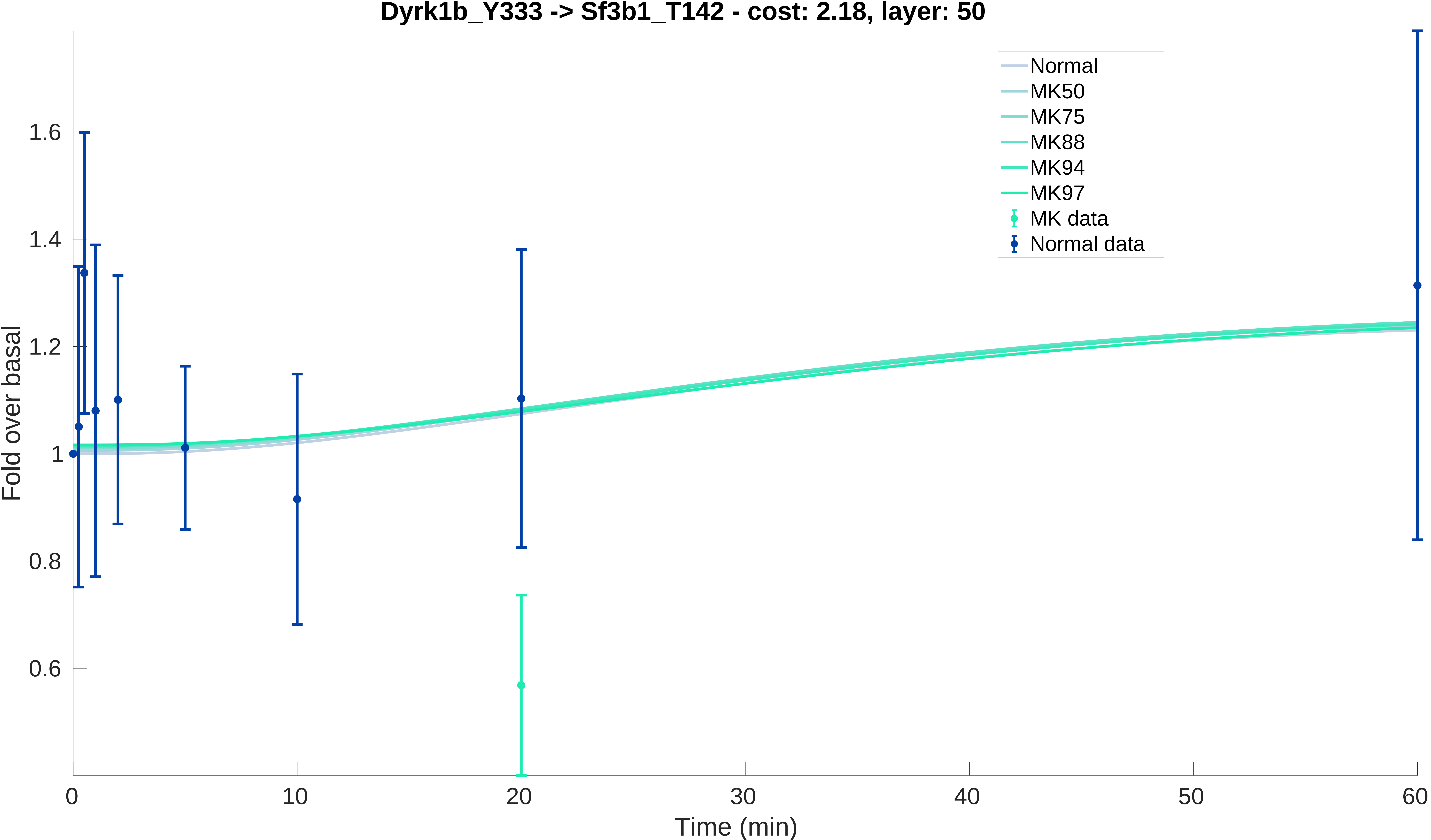
Dyrk1b_Y333 -> Sf3b1_T142 - cost: 2.18, layer: 50
Normal
MK50
MK75
MK88
MK94
MK97
MK data
Normal data
Fold over basal
1.6
1.4
1.2
1
0.8
0.6
0
10
20
30
40
50
60
Time (min)

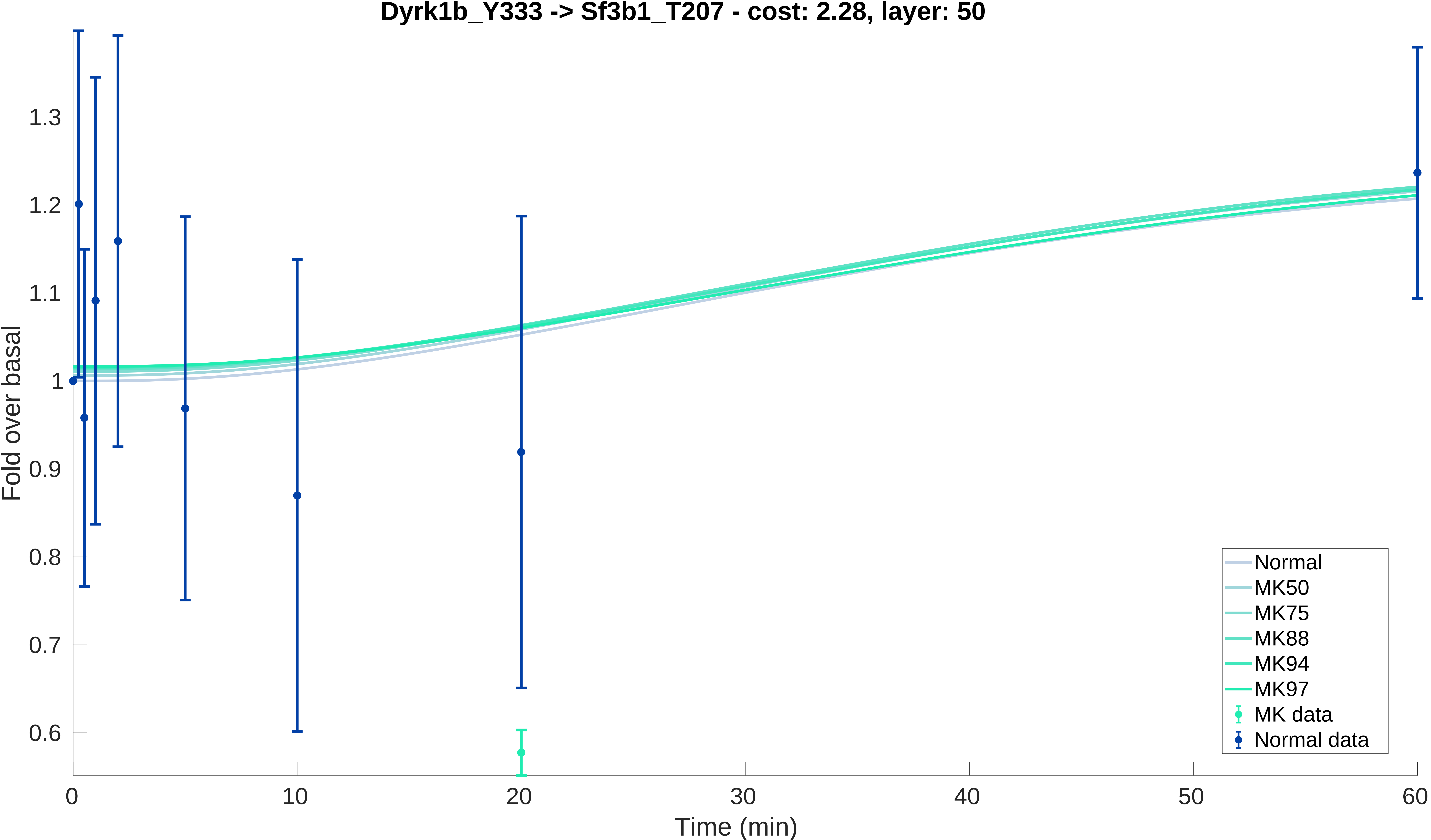

Dyrk1b_Y333 -> Sf3b1_T207 - cost: 2.28, layer: 50
Fold over basal
1.3
1.2
1.1
1
0.9
0.8
0.7
0.6
0
10
20
30
40
50
60
Time (min)
Normal
MK50
MK75
MK88
MK94
MK97
MK data
Normal data

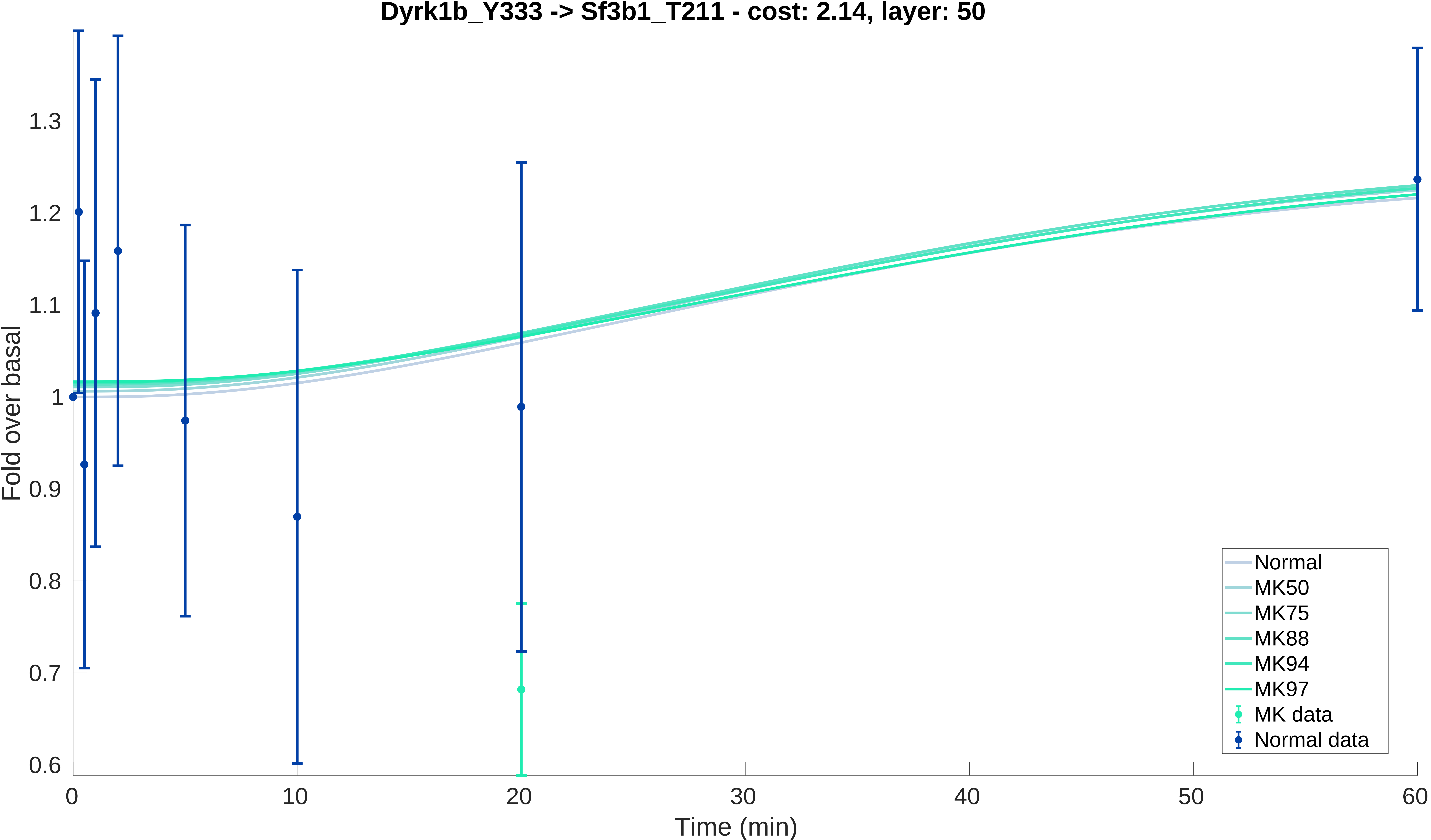

Dyrk1b_Y333 -> Sf3b1_T211 - cost: 2.14, layer: 50
Fold over basal
Time (min)
0.6
0.7
0.8
0.9
1
1.1
1.2
1.3
0
10
20
30
40
50
60
Normal
MK50
MK75
MK88
MK94
MK97
MK data
Normal data

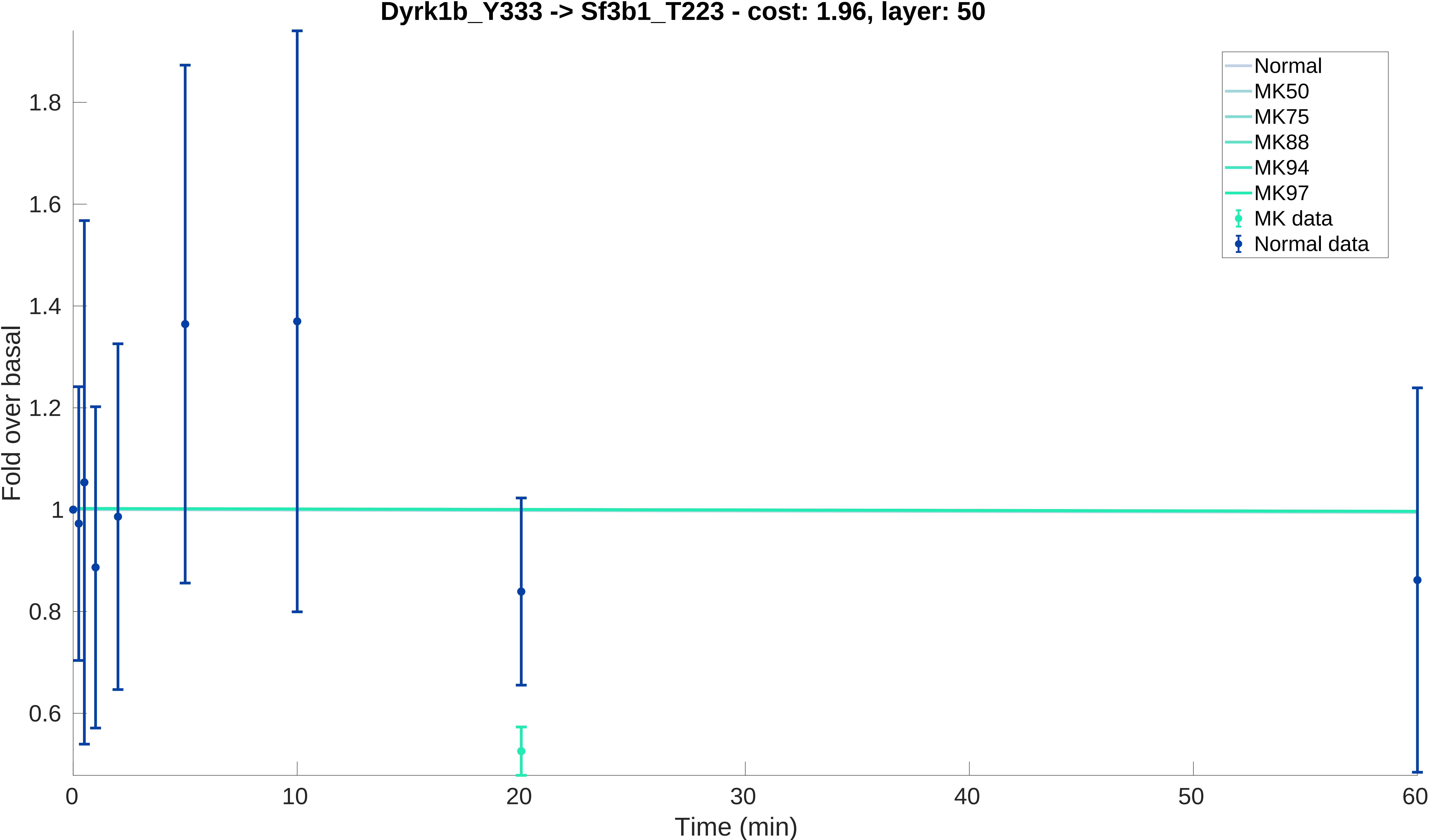

Dyrk1b_Y333 -> Sf3b1_T223 - cost: 1.96, layer: 50
Fold over basal
0.6
0.8
1
1.2
1.4
1.6
1.8
0
10
20
30
40
50
60
Time (min)
Normal
MK50
MK75
MK88
MK94
MK97
MK data
Normal data

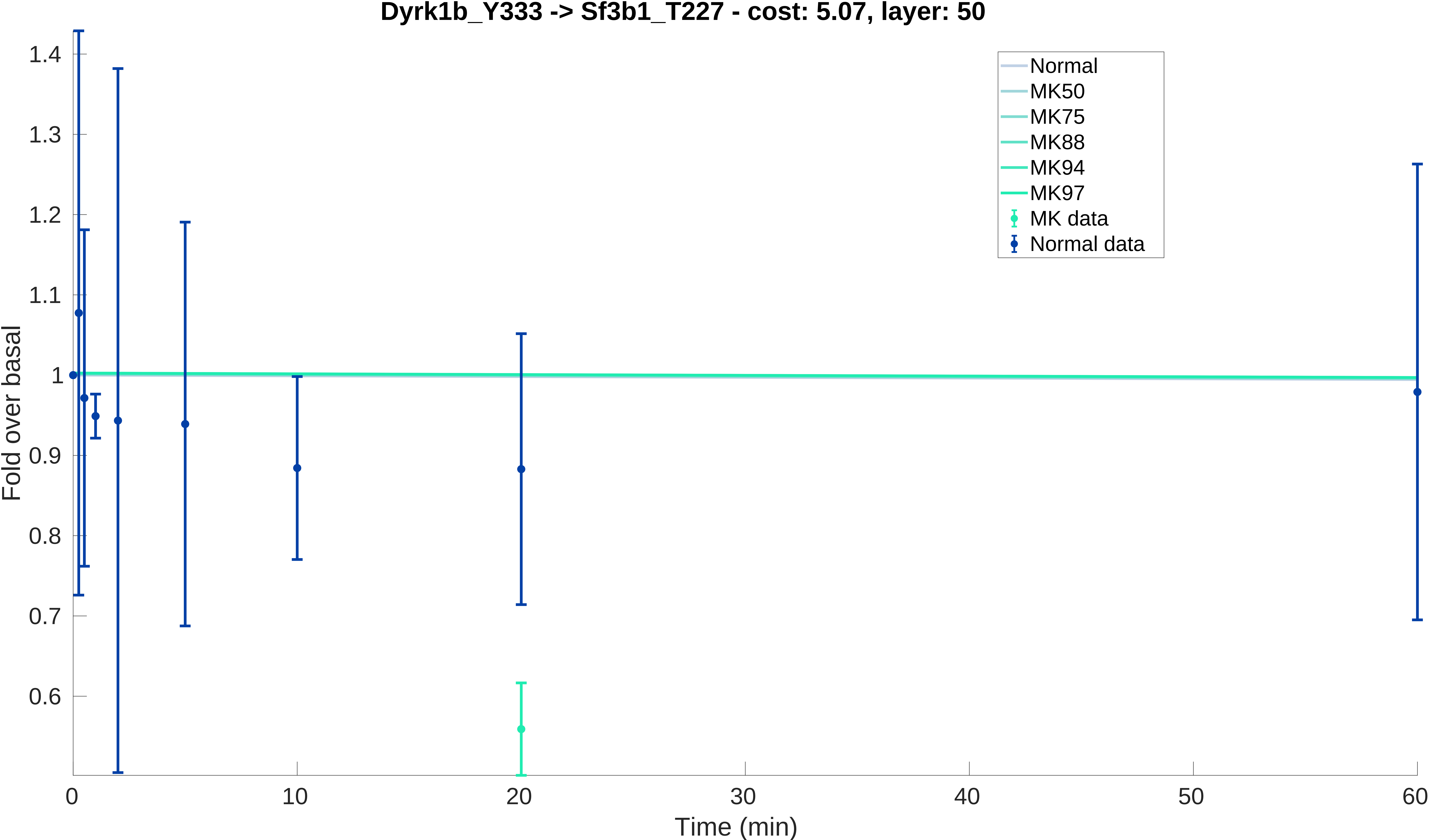

Dyrk1b_Y333 -> Sf3b1_T227 - cost: 5.07, layer: 50
Fold over basal
Time (min)
Normal
MK50
MK75
MK88
MK94
MK97
MK data
Normal data
1.4
1.3
1.2
1.1
1
0.9
0.8
0.7
0.6
0
10
20
30
40
50
60

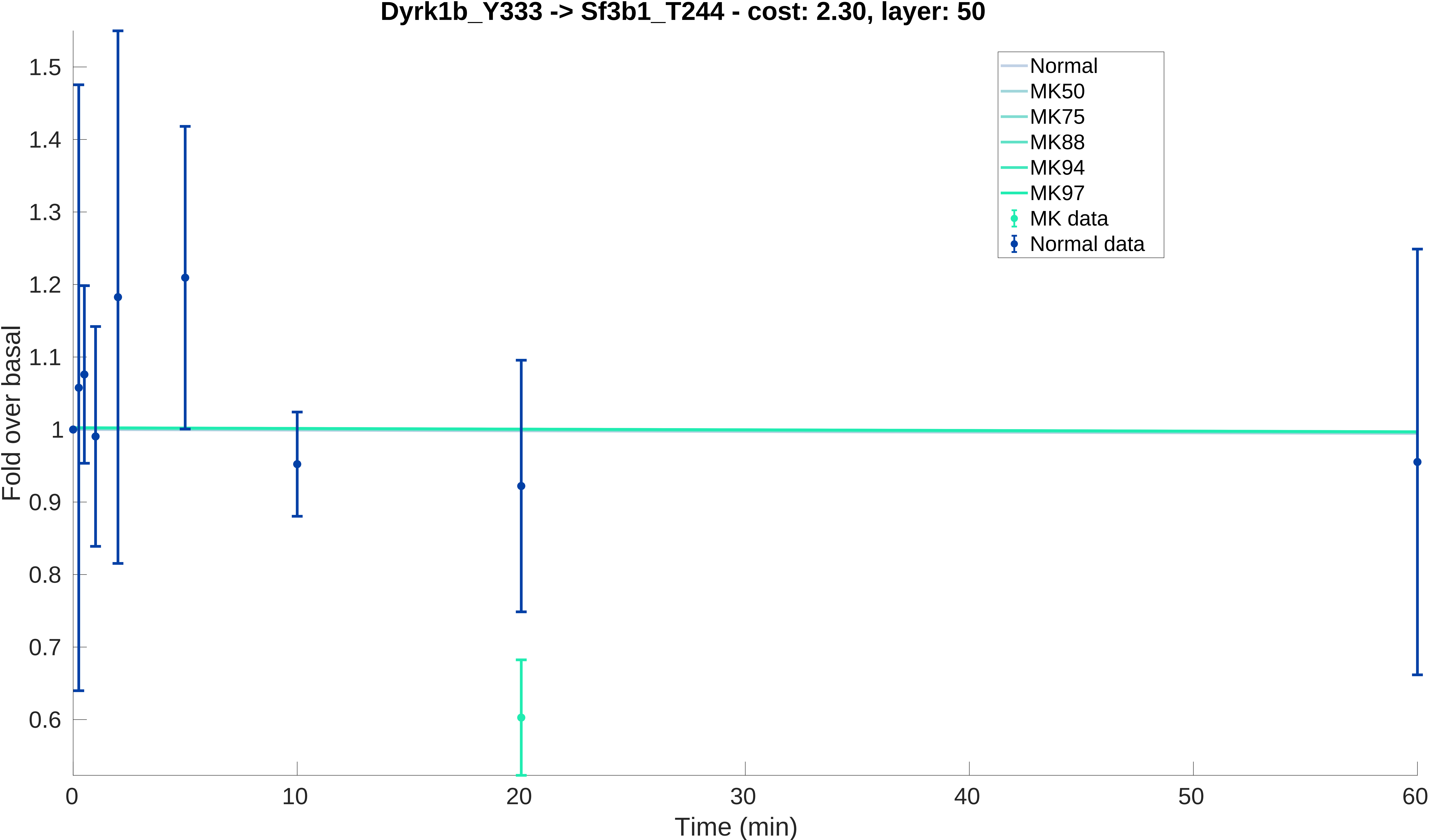

Dyrk1b_Y333 -> Sf3b1_T244 - cost: 2.30, layer: 50
Normal
MK50
MK75
MK88
MK94
MK97
MK data
Normal data
Fold over basal
1.5
1.4
1.3
1.2
1.1
1
0.9
0.8
0.7
0.6
0
10
20
30
40
50
60
Time (min)

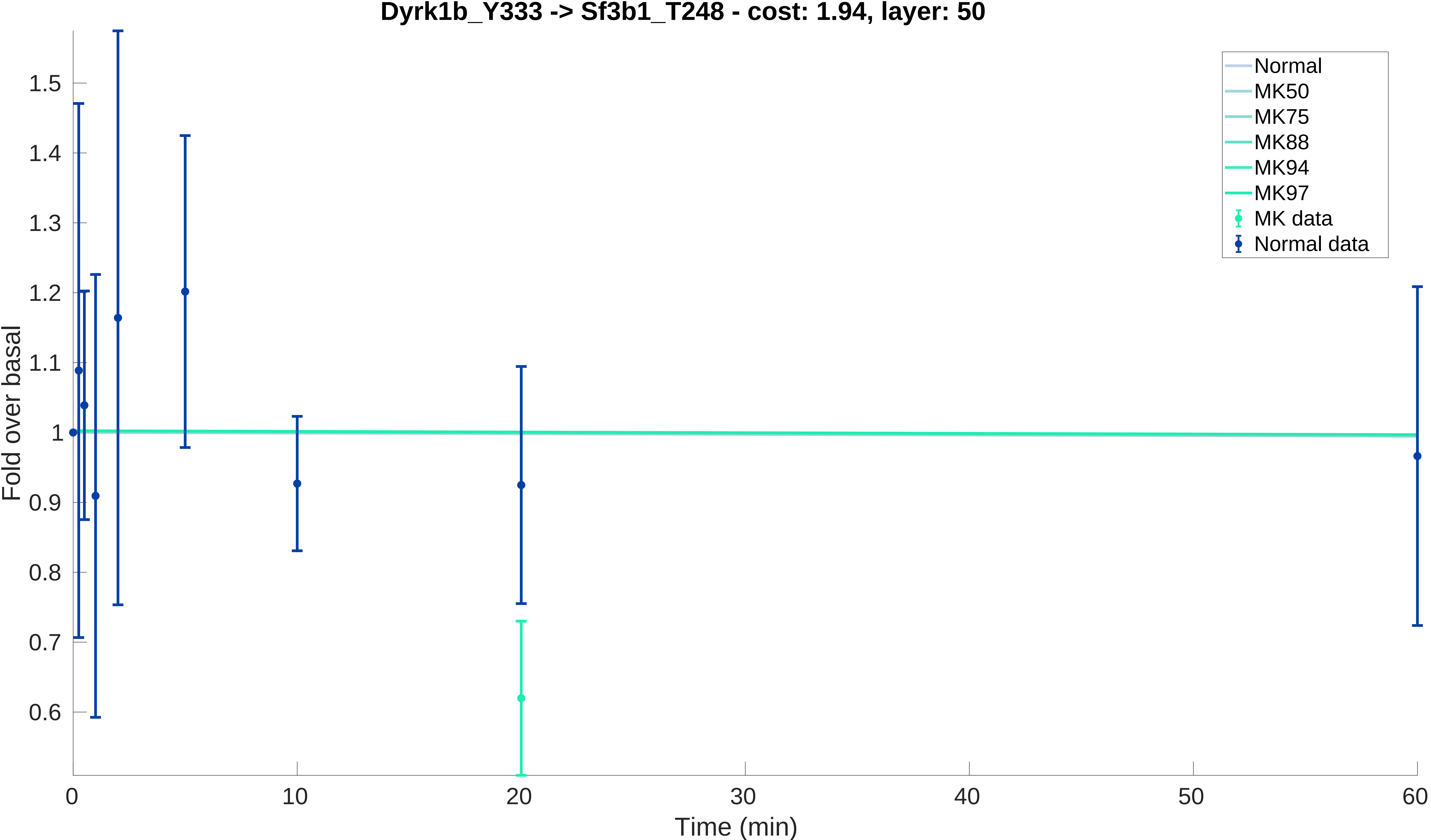

Dyrk1b_Y333 -> Sf3b1_T248 - cost: 1.94, layer: 50
Fold over basal
Time (min)
1.5
1.4
1.3
1.2
1.1
1
0.9
0.8
0.7
0.6
0
10
20
30
40
50
60
Normal
MK50
MK75
MK88
MK94
MK97
MK data
Normal data

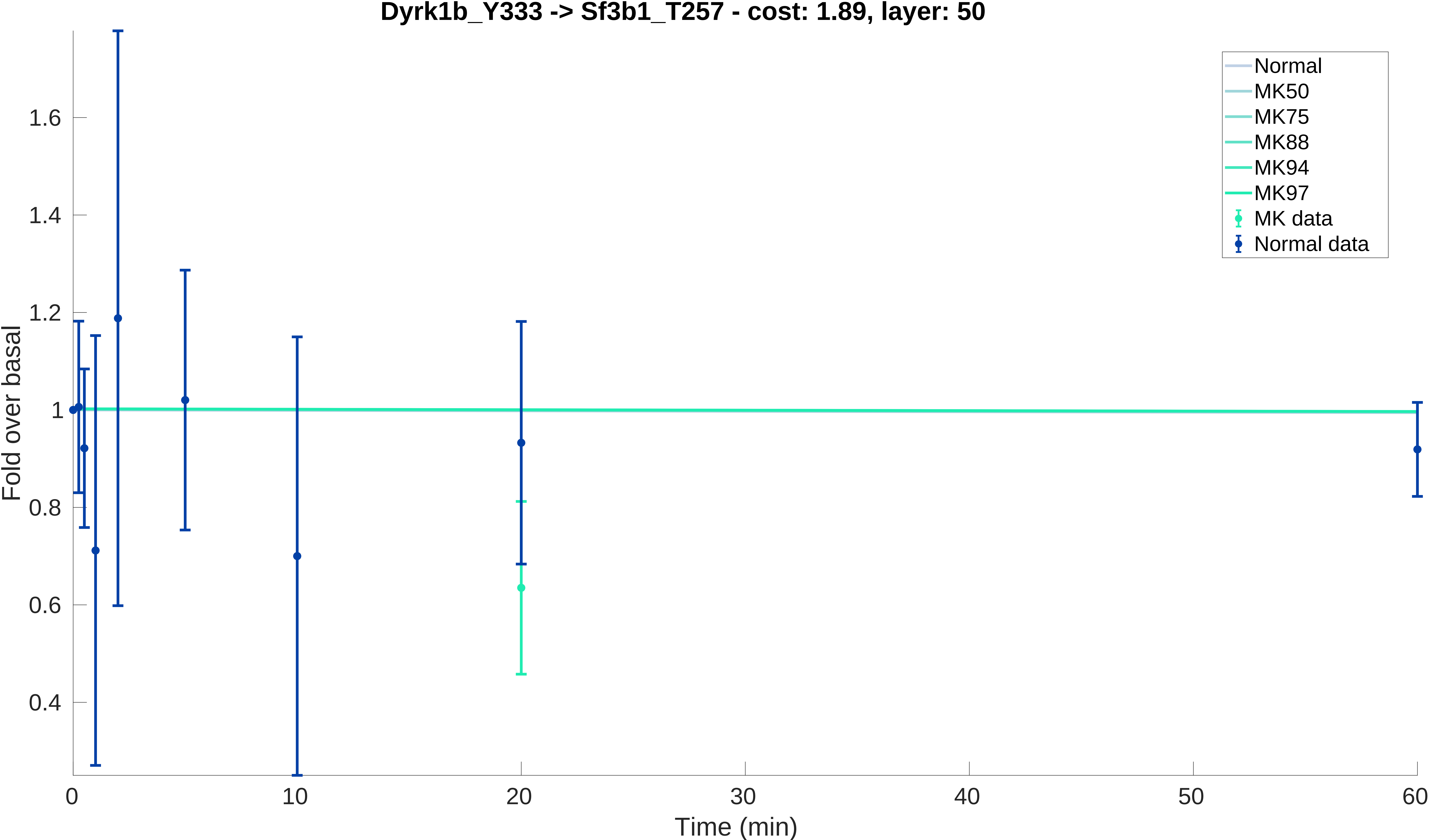
Dyrk1b_Y333 -> Sf3b1_T257 - cost: 1.89, layer: 50
Fold over basal
Time (min)
0.4
0.6
0.8
1
1.2
1.4
1.6
0
10
20
30
40
50
60
Normal
MK50
MK75
MK88
MK94
MK97
MK data
Normal data

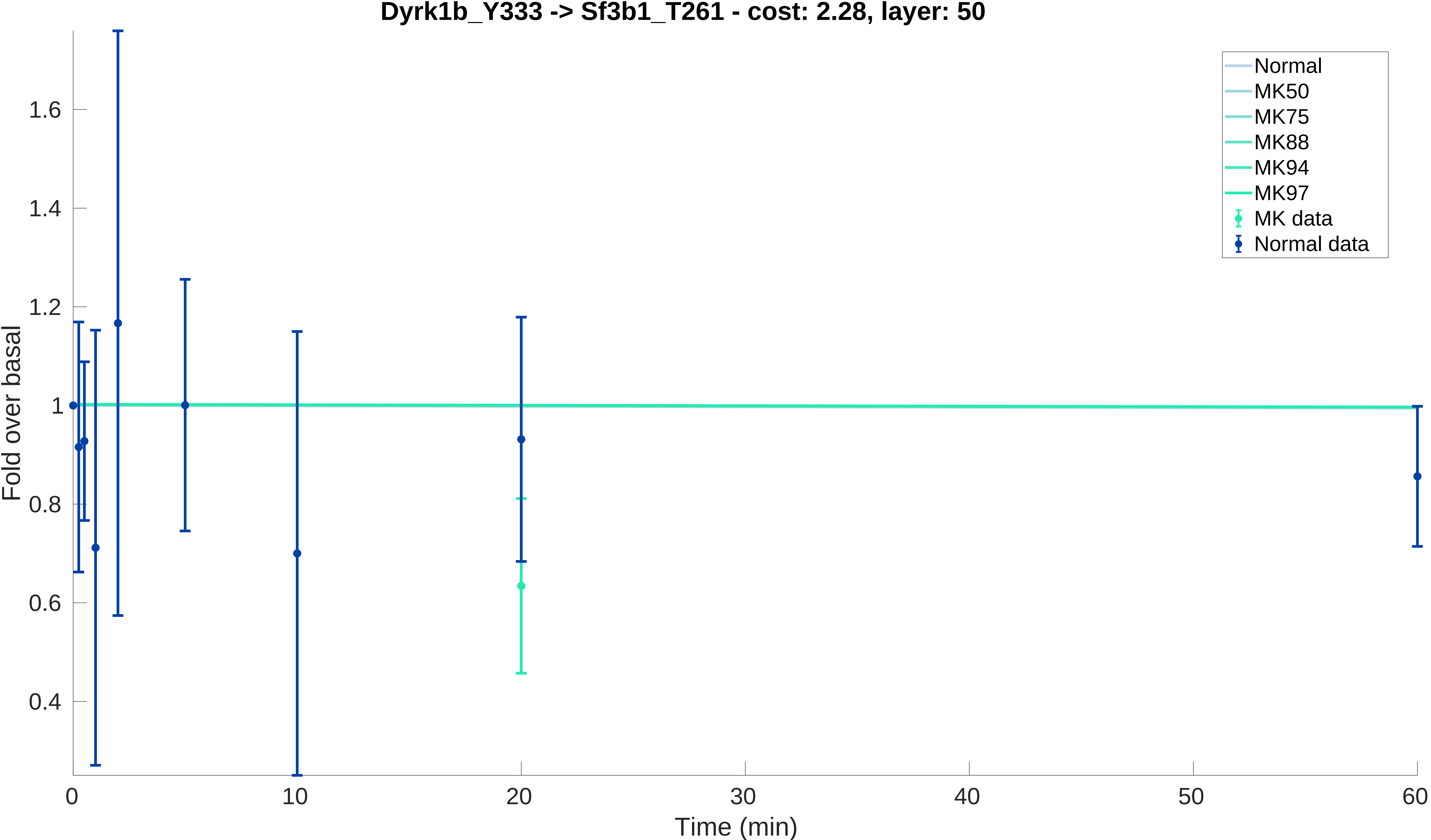
Dyrk1b_Y333 -> Sf3b1_T261 - cost: 2.28, layer: 50
Normal
MK50
MK75
MK88
MK94
MK97
MK data
Normal data
Fold over basal
1.6
1.4
1.2
1
0.8
0.6
0.4
0
10
20
30
40
50
60
Time (min)

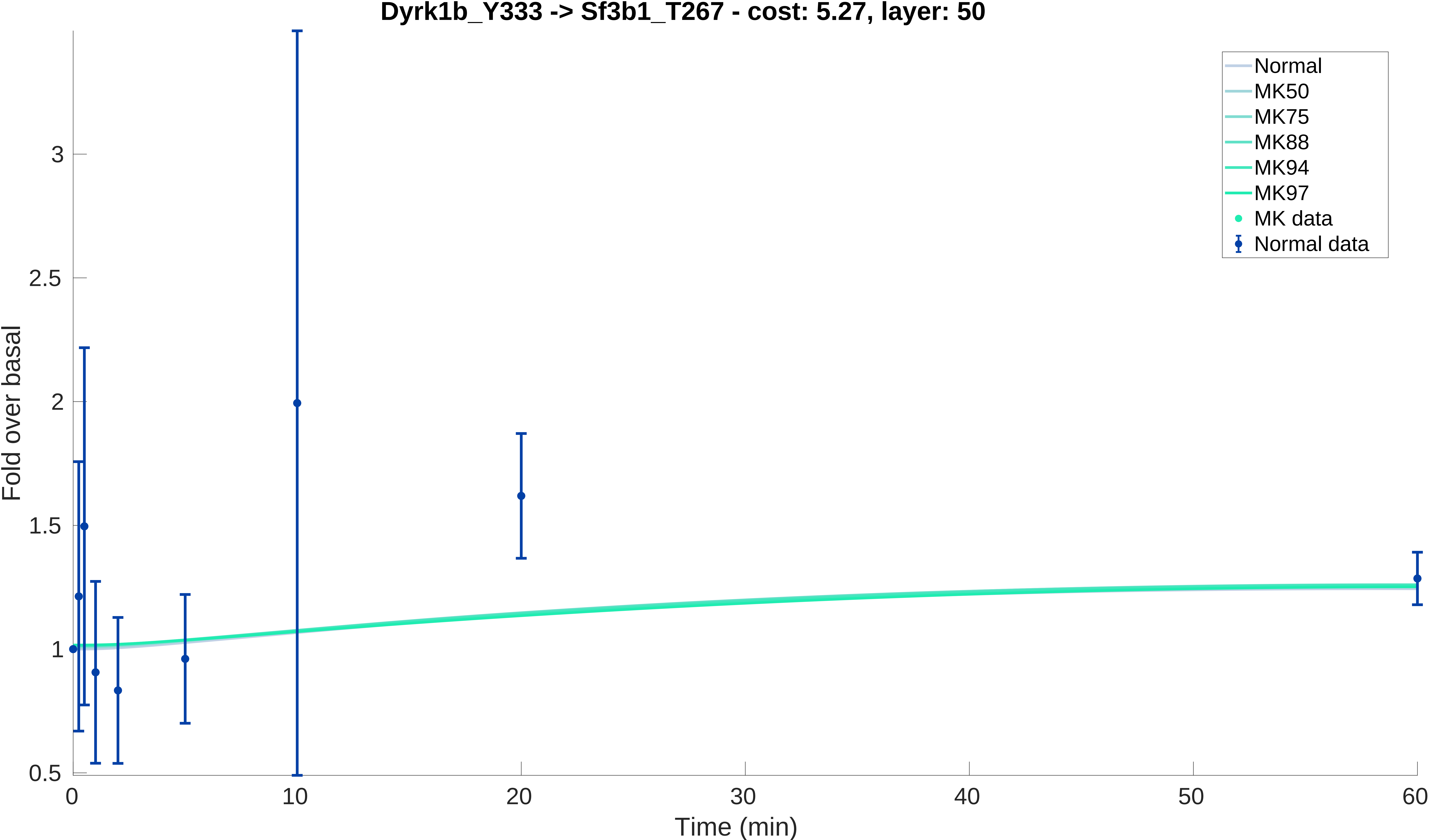
Dyrk1b_Y333 -> Sf3b1_T267 - cost: 5.27, layer: 50
Fold over basal
Time (min)
0.5
1
1.5
2
2.5
3
0
10
20
30
40
50
60
Normal
MK50
MK75
MK88
MK94
MK97
MK data
Normal data

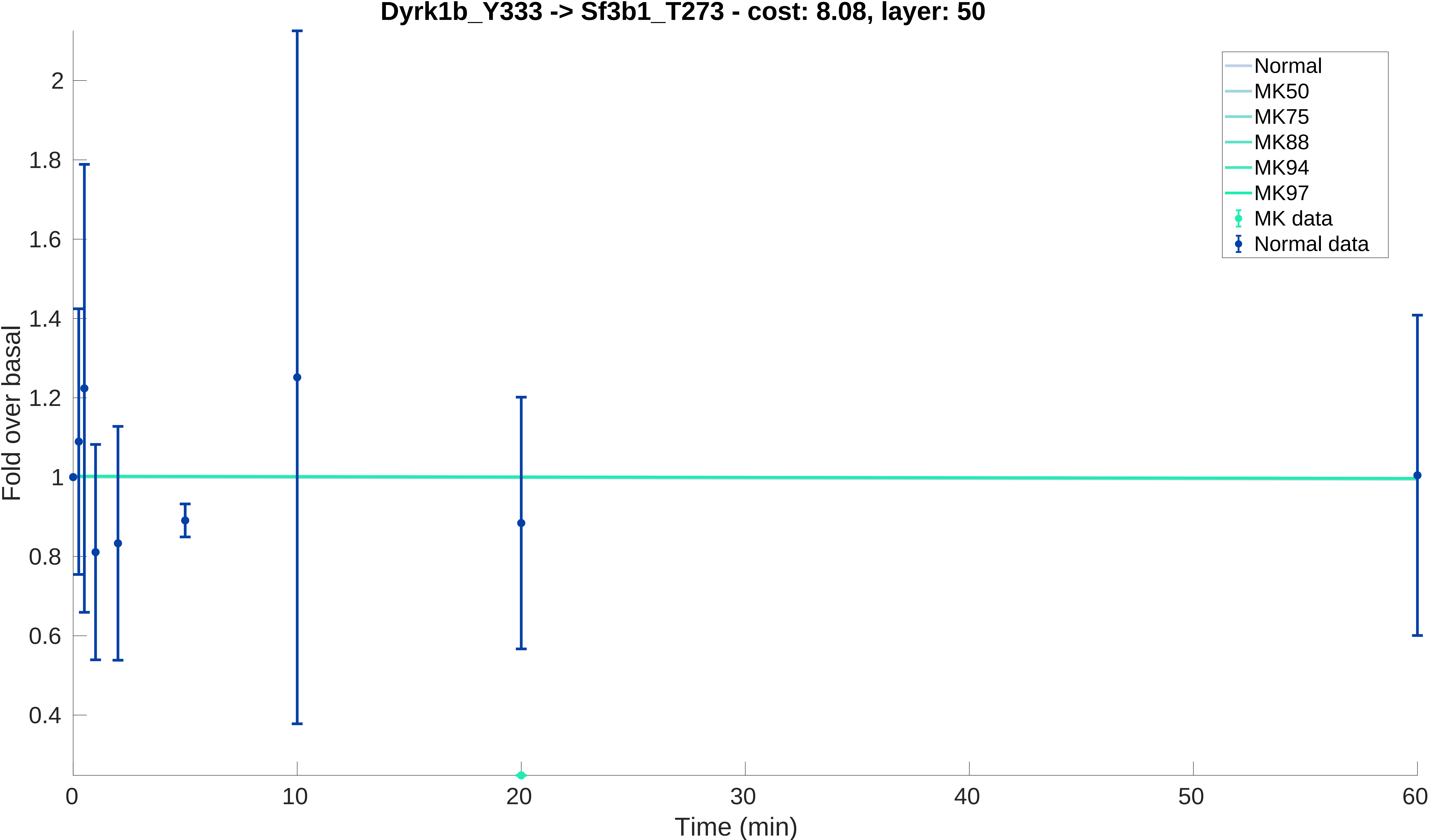

Dyrk1b_Y333 -> Sf3b1_T273 - cost: 8.08, layer: 50
Normal
MK50
MK75
MK88
MK94
MK97
MK data
Normal data
Fold over basal
2
1.8
1.6
1.4
1.2
1
0.8
0.6
0.4
0
10
20
30
40
50
60
Time (min)

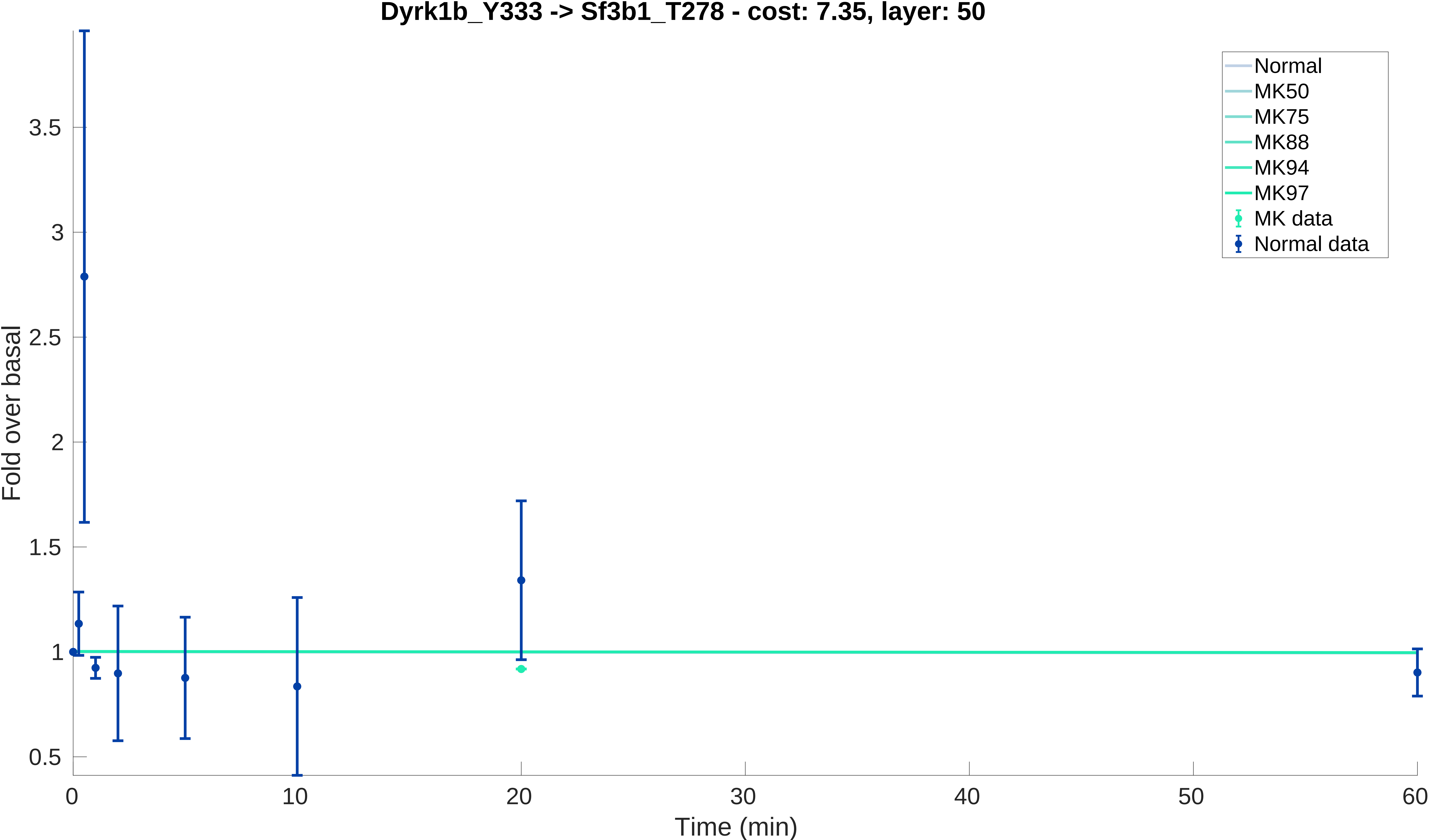

Dyrk1b_Y333 -> Sf3b1_T278 - cost: 7.35, layer: 50
Fold over basal
Time (min)
0.5
1
1.5
2
2.5
3
3.5
0
10
20
30
40
50
60
Normal
MK50
MK75
MK88
MK94
MK97
MK data
Normal data

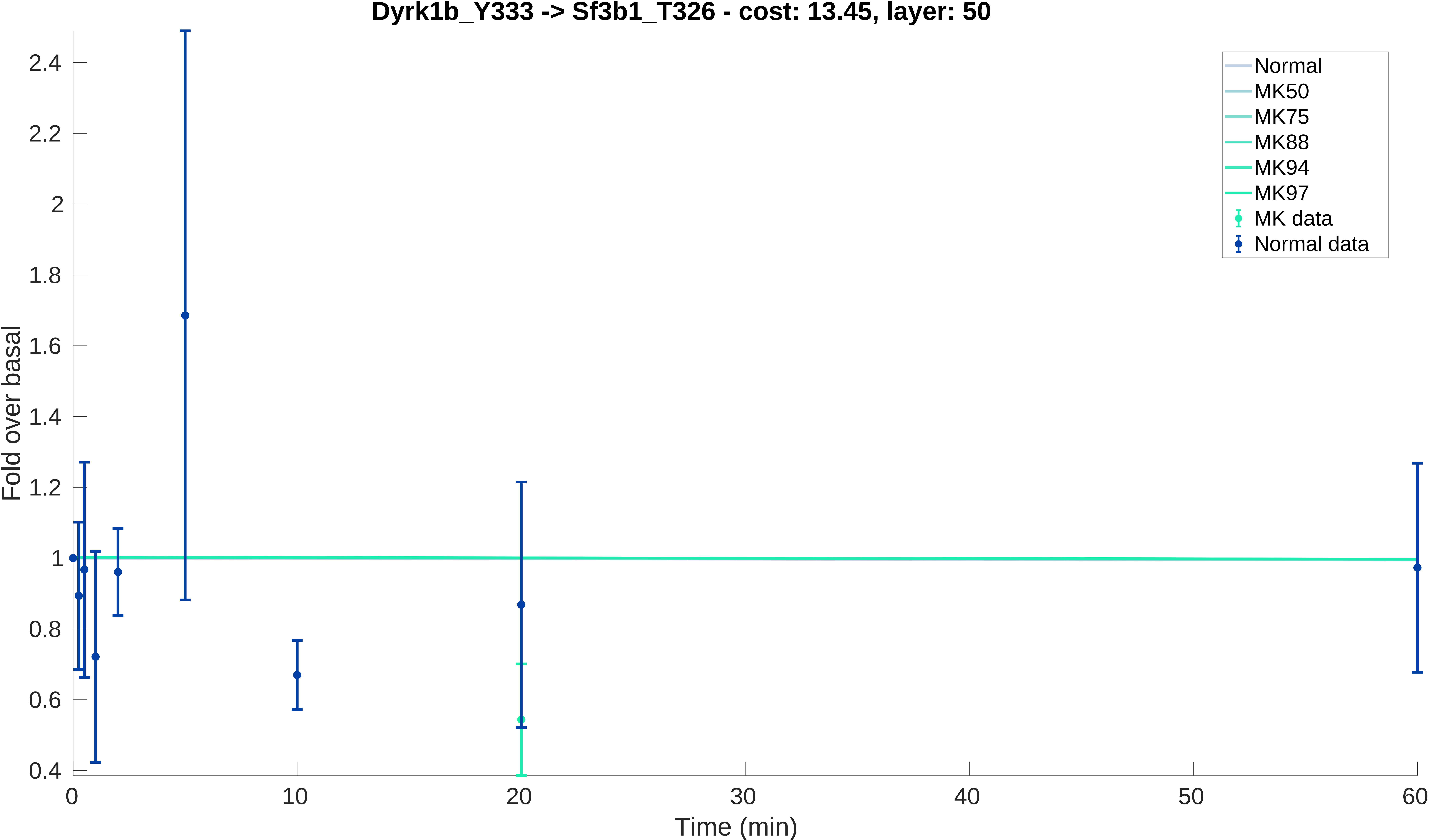

Dyrk1b_Y333 -> Sf3b1_T326 - cost: 13.45, layer: 50
Fold over basal
Time (min)
0.4
0.6
0.8
1
1.2
1.4
1.6
1.8
2
2.2
2.4
0
10
20
30
40
50
60
Normal
MK50
MK75
MK88
MK94
MK97
MK data
Normal data

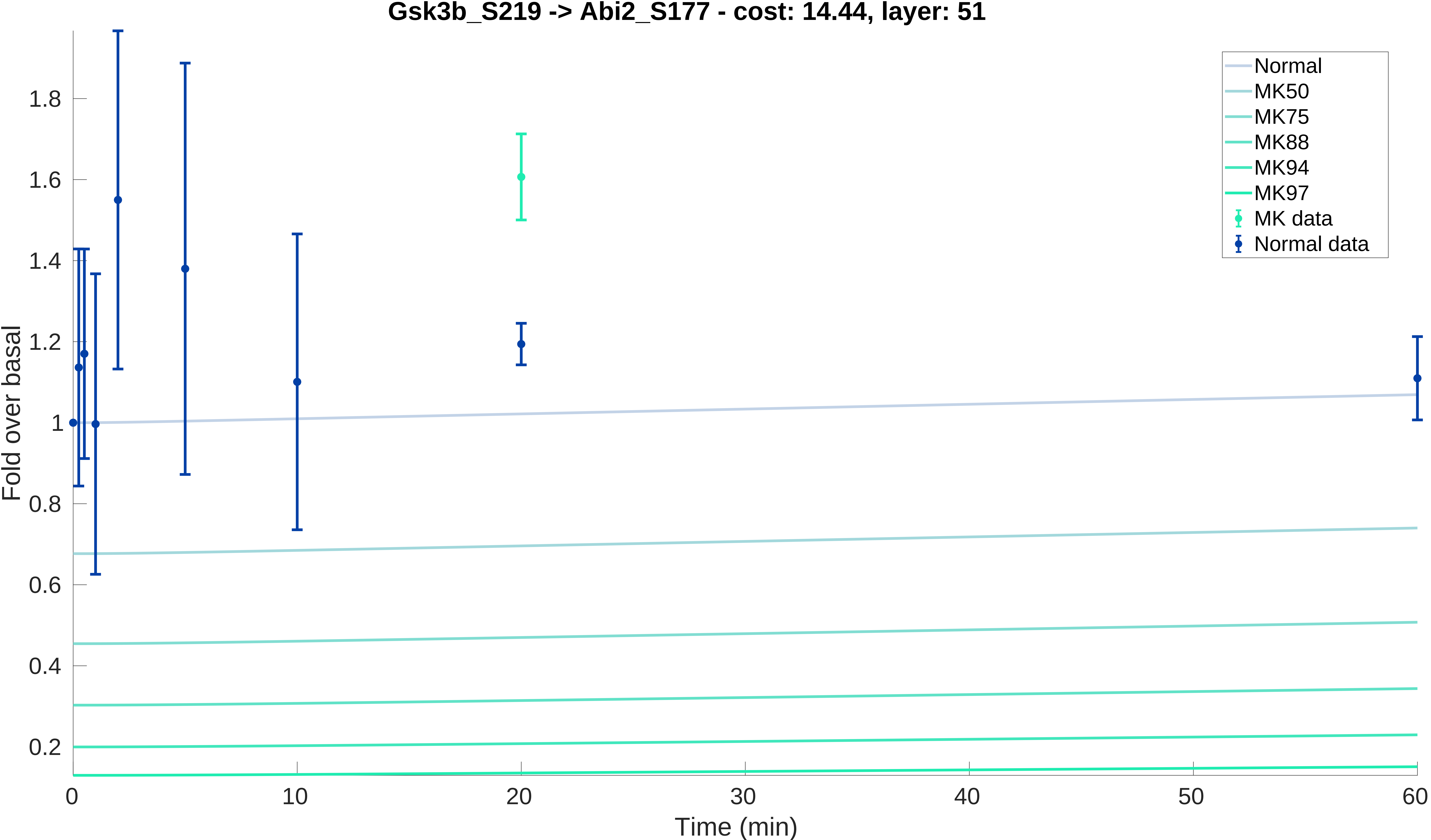

Gsk3b_S219 -> Abi2_S177 - cost: 14.44, layer: 51
Fold over basal
Time (min)
0.2
0.4
0.6
0.8
1
1.2
1.4
1.6
1.8
0
10
20
30
40
50
60
Normal
MK50
MK75
MK88
MK94
MK97
MK data
Normal data

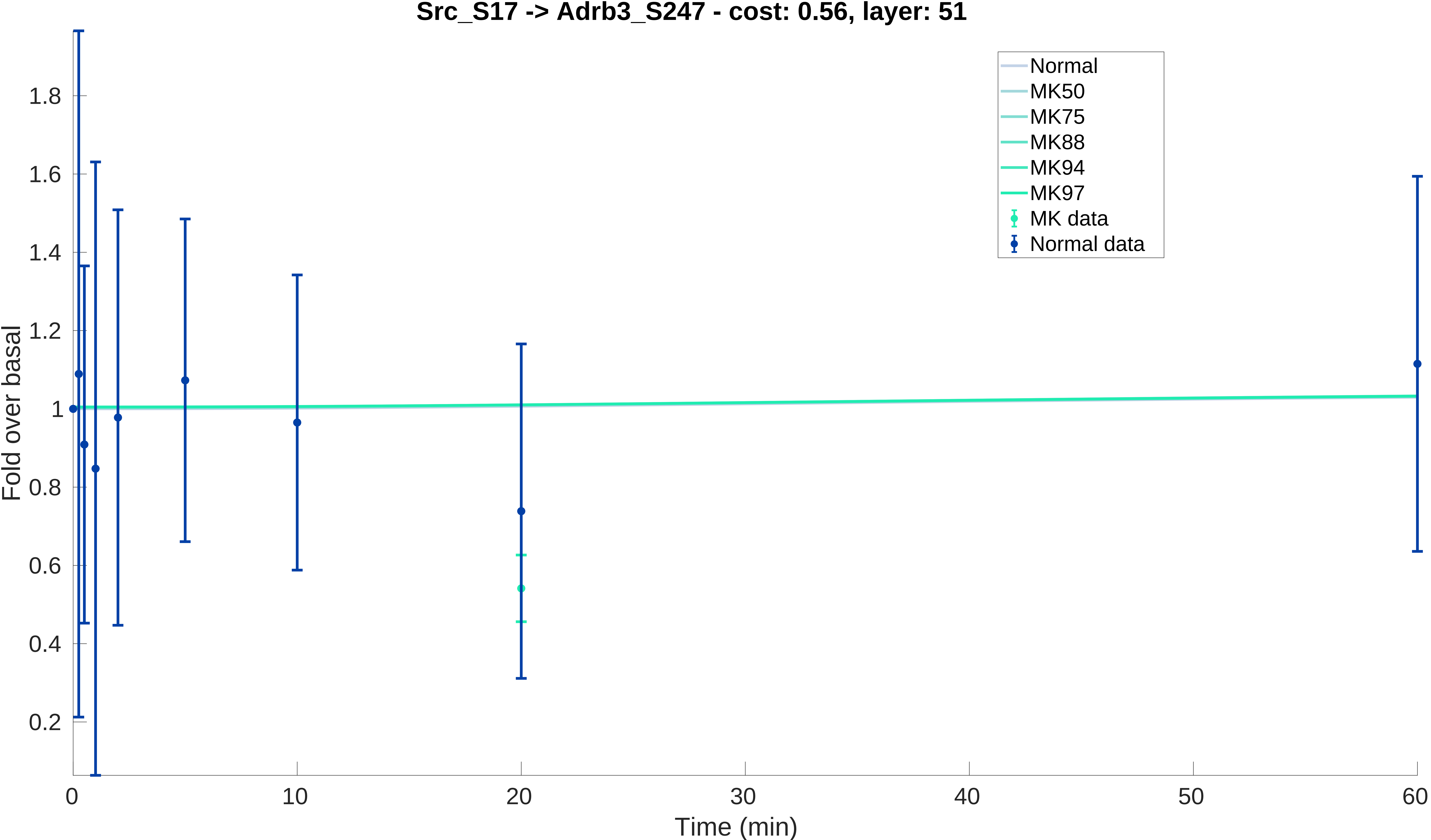

Src_S17 -> Adrb3_S247 - cost: 0.56, layer: 51
Normal
MK50
MK75
MK88
MK94
MK97
MK data
Normal data
Fold over basal
0.2
0.4
0.6
0.8
1
1.2
1.4
1.6
1.8
0
10
20
30
40
50
60
Time (min)

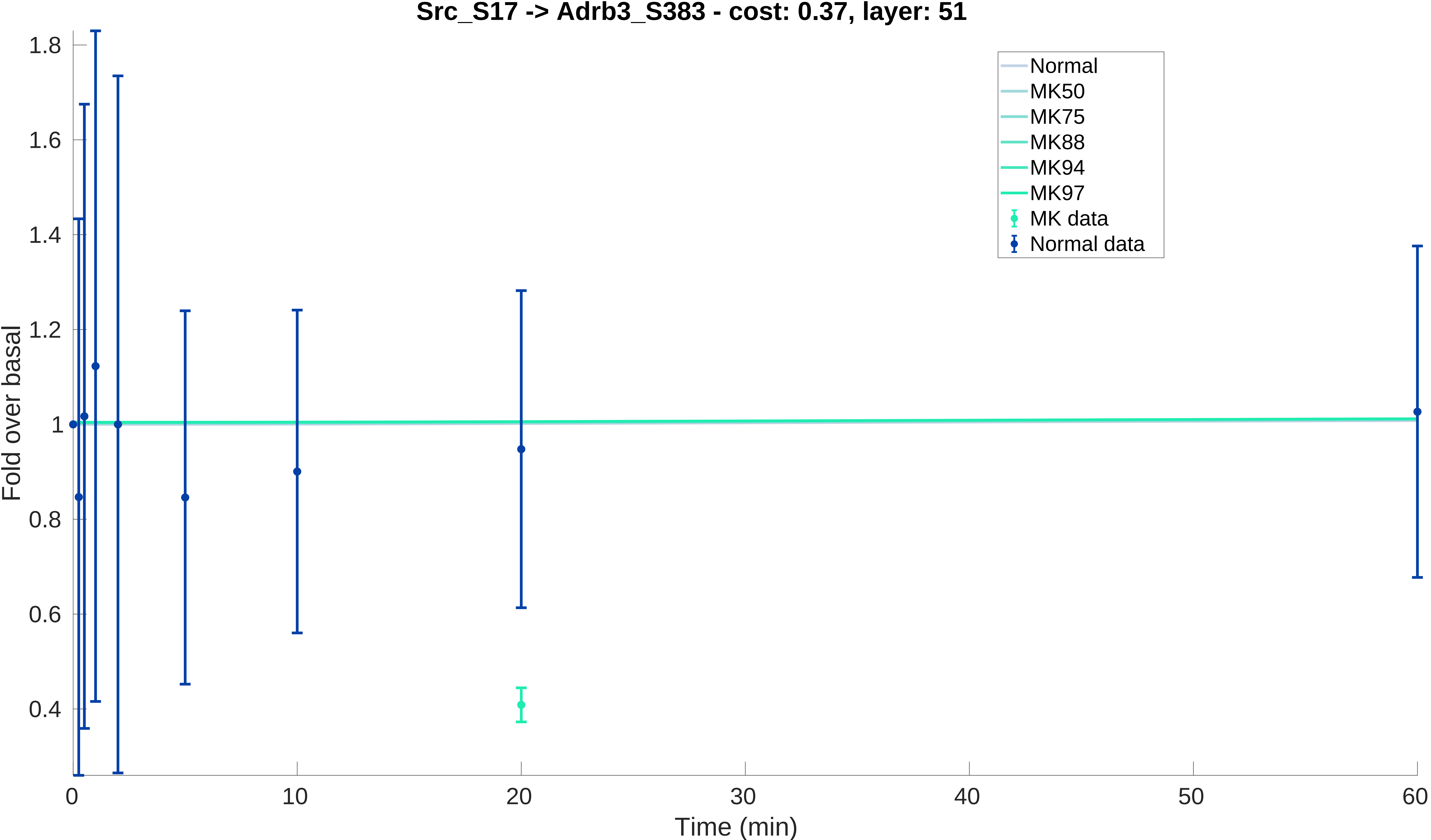

Src_S17 -> Adrb3_S383 - cost: 0.37, layer: 51
Fold over basal
Time (min)
Normal
MK50
MK75
MK88
MK94
MK97
MK data
Normal data
1.8
1.6
1.4
1.2
1
0.8
0.6
0.4
0
10
20
30
40
50
60

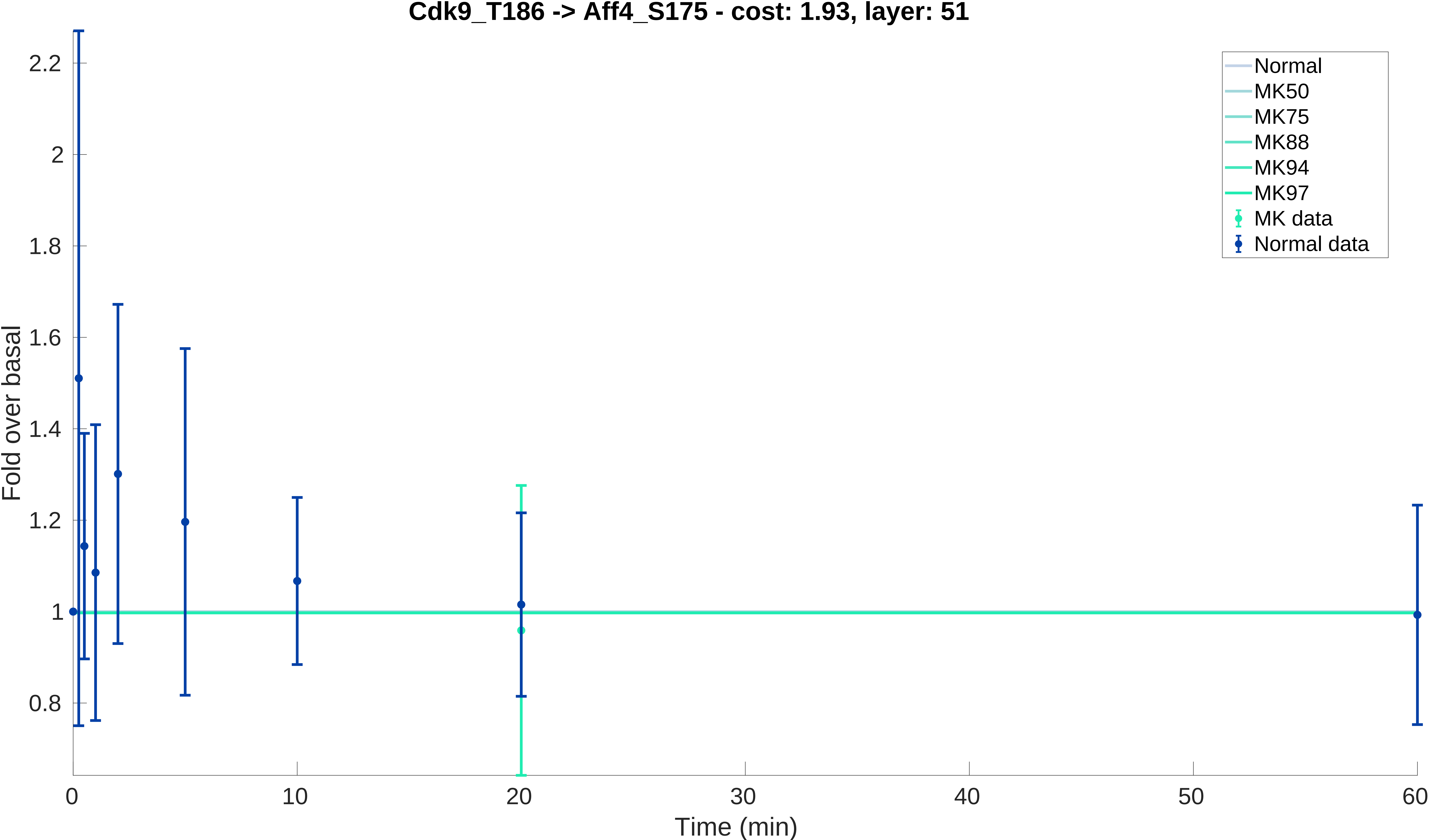
Cdk9_T186 -> Aff4_S175 - cost: 1.93, layer: 51
Normal
MK50
MK75
MK88
MK94
MK97
MK data
Normal data
Fold over basal
Time (min)
2.2
2
1.8
1.6
1.4
1.2
1
0.8
0
10
20
30
40
50
60

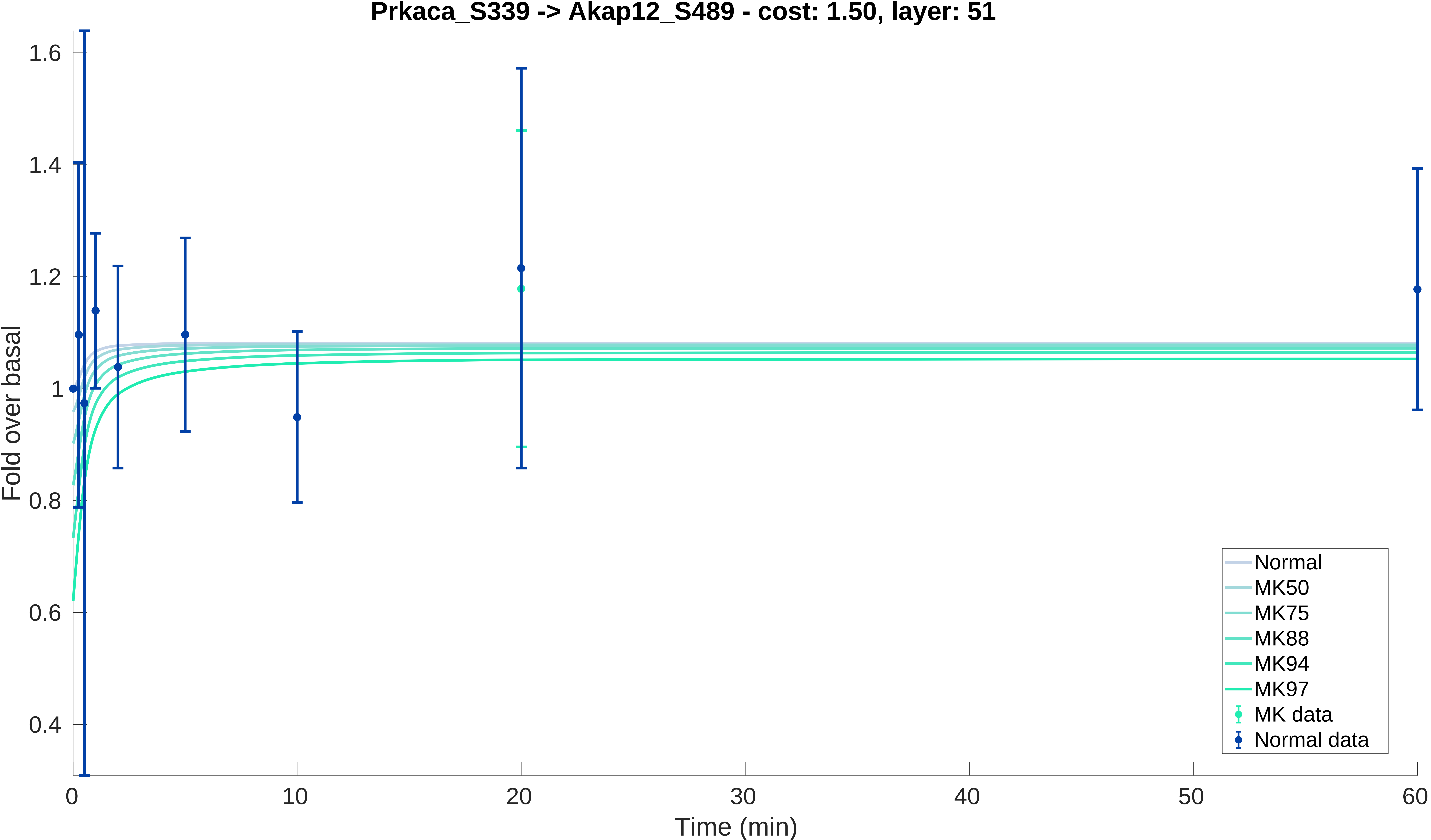

Prkaca_S339 -> Akap12_S489 - cost: 1.50, layer: 51
Fold over basal
Time (min)
1.6
1.4
1.2
1
0.8
0.6
0.4
0
10
20
30
40
50
60
Normal
MK50
MK75
MK88
MK94
MK97
MK data
Normal data

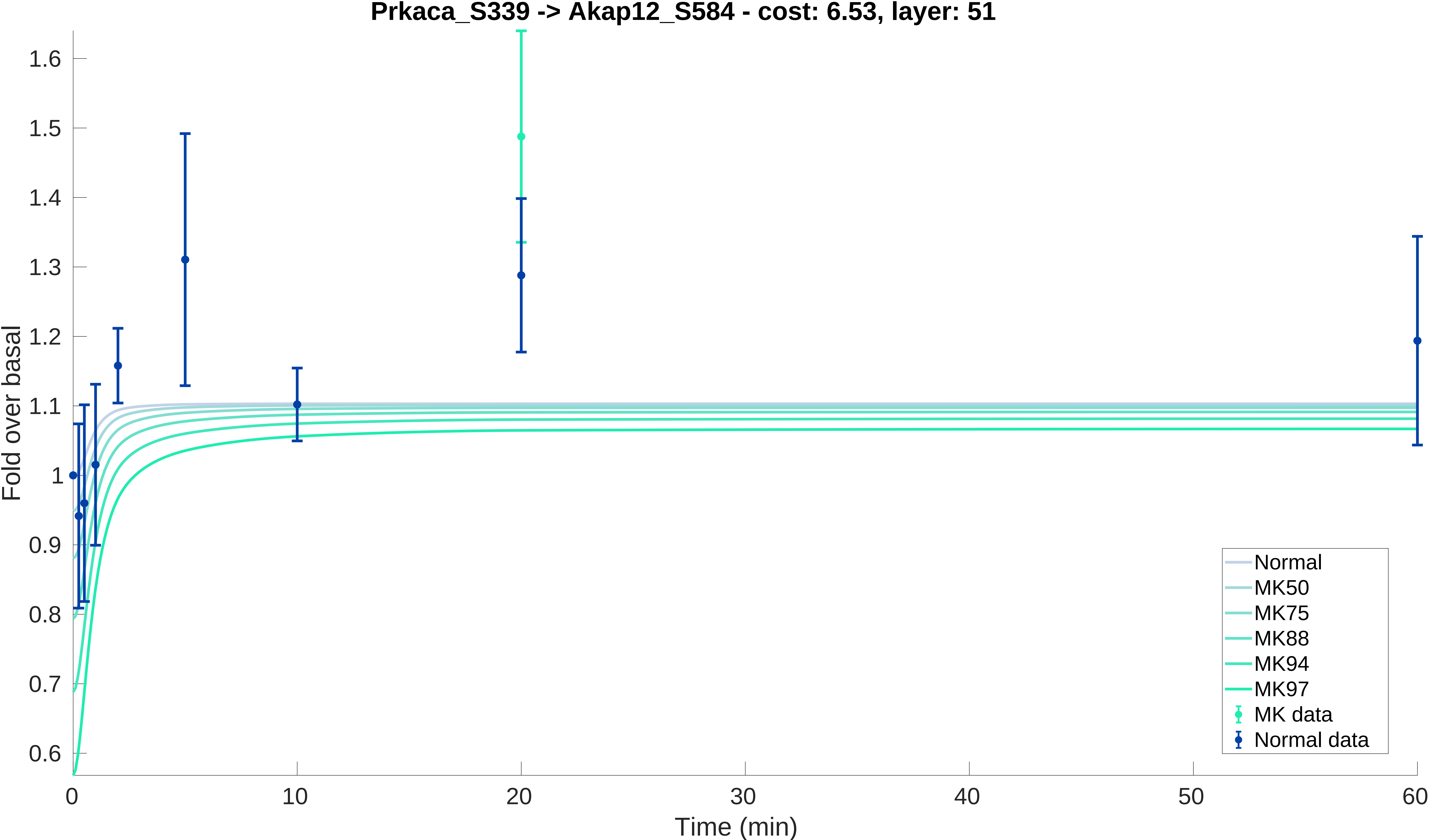

Prkaca_S339 -> Akap12_S584 - cost: 6.53, layer: 51
Fold over basal
1.6
1.5
1.4
1.3
1.2
1.1
1
0.9
0.8
0.7
0.6
0
10
20
30
40
50
60
Time (min)
Normal
MK50
MK75
MK88
MK94
MK97
MK data
Normal data

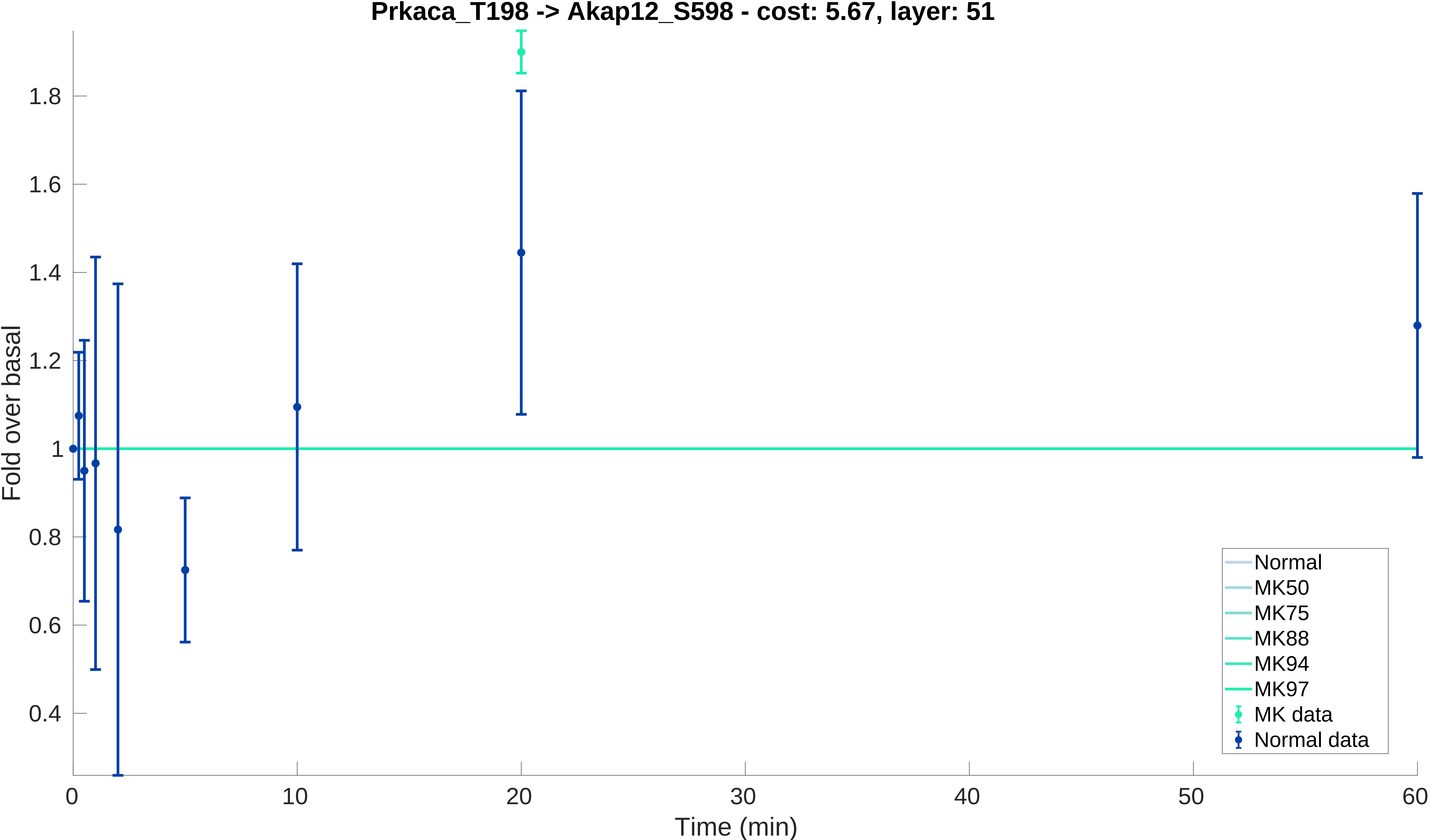

Prkaca_T198 -> Akap12_S598 - cost: 5.67, layer: 51
Fold over basal
1.8
1.6
1.4
1.2
1
0.8
0.6
0.4
0
10
20
30
40
50
60
Time (min)
Normal
MK50
MK75
MK88
MK94
MK97
MK data
Normal data

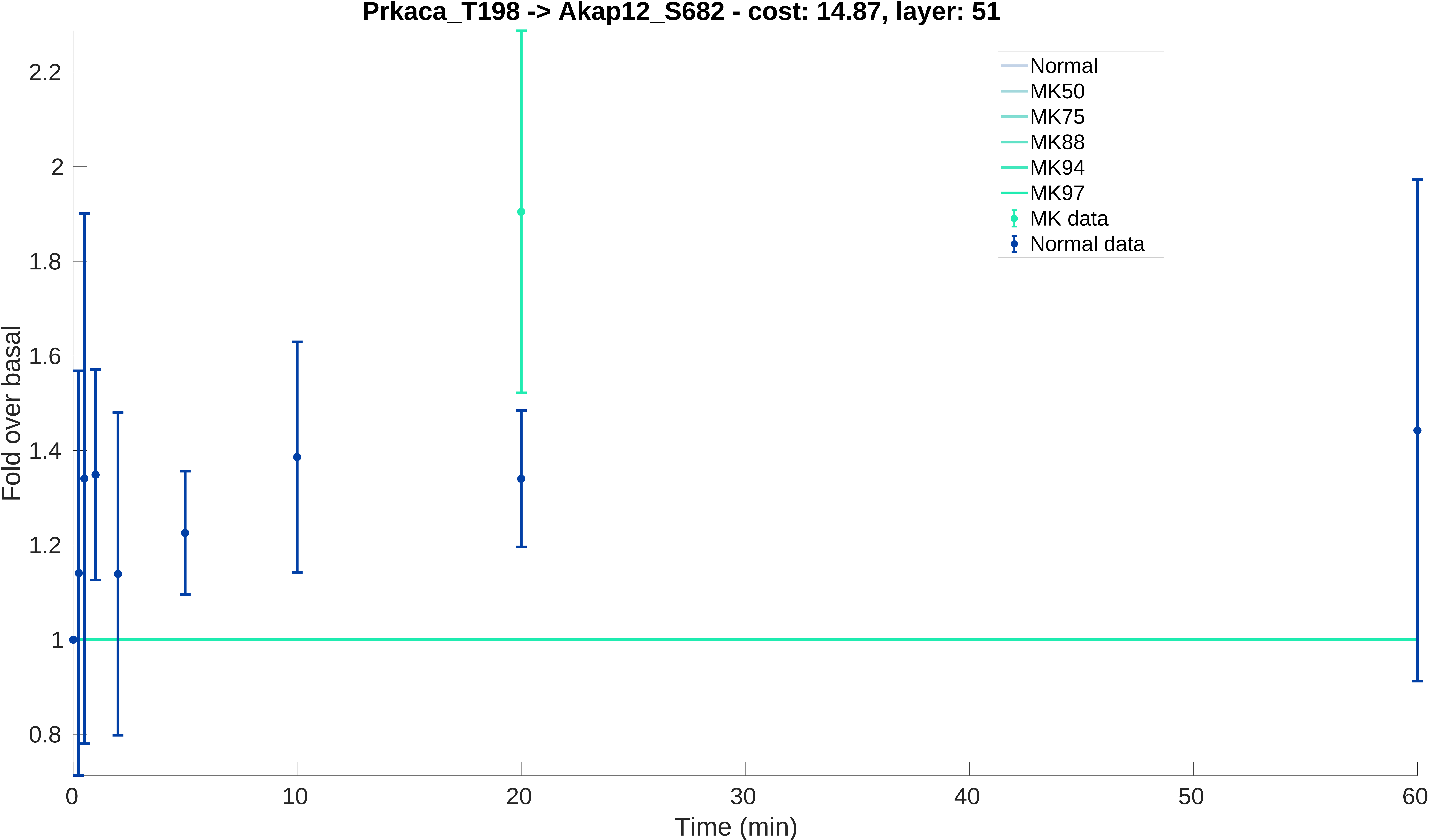

Prkaca_T198 -> Akap12_S682 - cost: 14.87, layer: 51
Fold over basal
Time (min)
0.8
1
1.2
1.4
1.6
1.8
2
2.2
0
10
20
30
40
50
60
Normal
MK50
MK75
MK88
MK94
MK97
MK data
Normal data

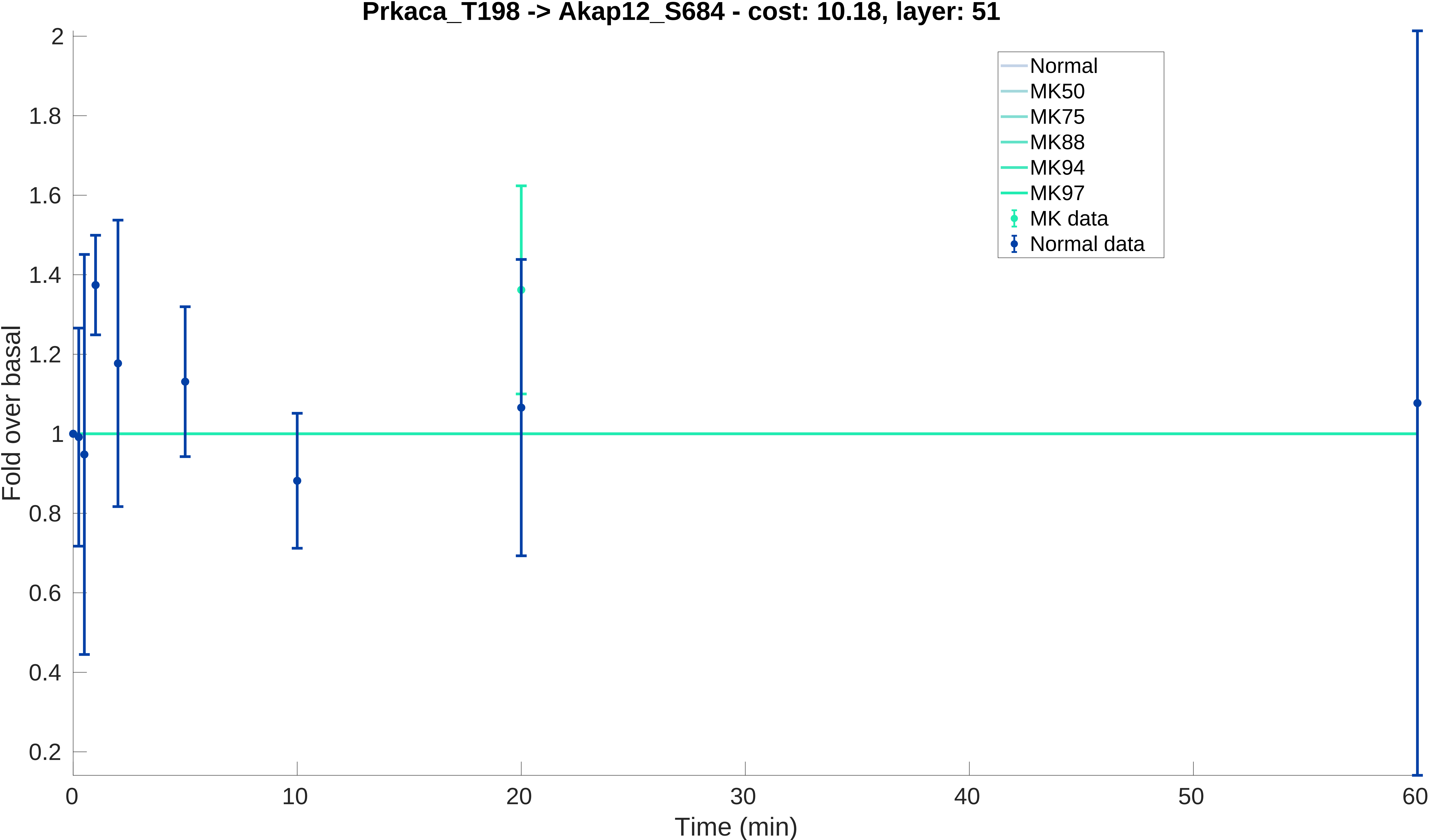

Prkaca_T198 -> Akap12_S684 - cost: 10.18, layer: 51
Fold over basal
2
1.8
1.6
1.4
1.2
1
0.8
0.6
0.4
0.2
0
10
20
30
40
50
60
Time (min)
Normal
MK50
MK75
MK88
MK94
MK97
MK data
Normal data

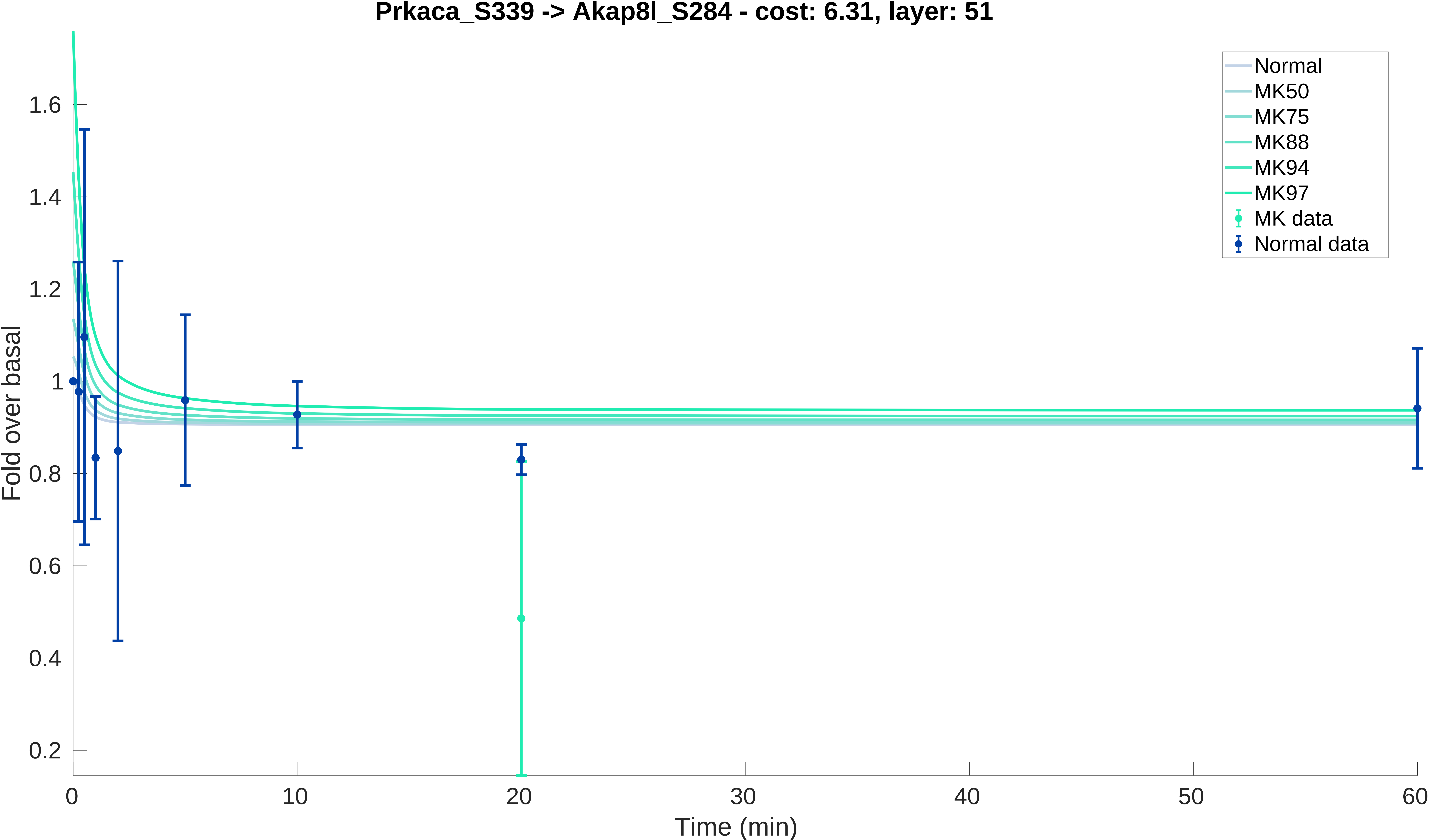

Prkaca_S339 -> Akap8l_S284 - cost: 6.31, layer: 51
Normal
MK50
MK75
MK88
MK94
MK97
MK data
Normal data
Fold over basal
1.6
1.4
1.2
1
0.8
0.6
0.4
0.2
0
10
20
30
40
50
60
Time (min)

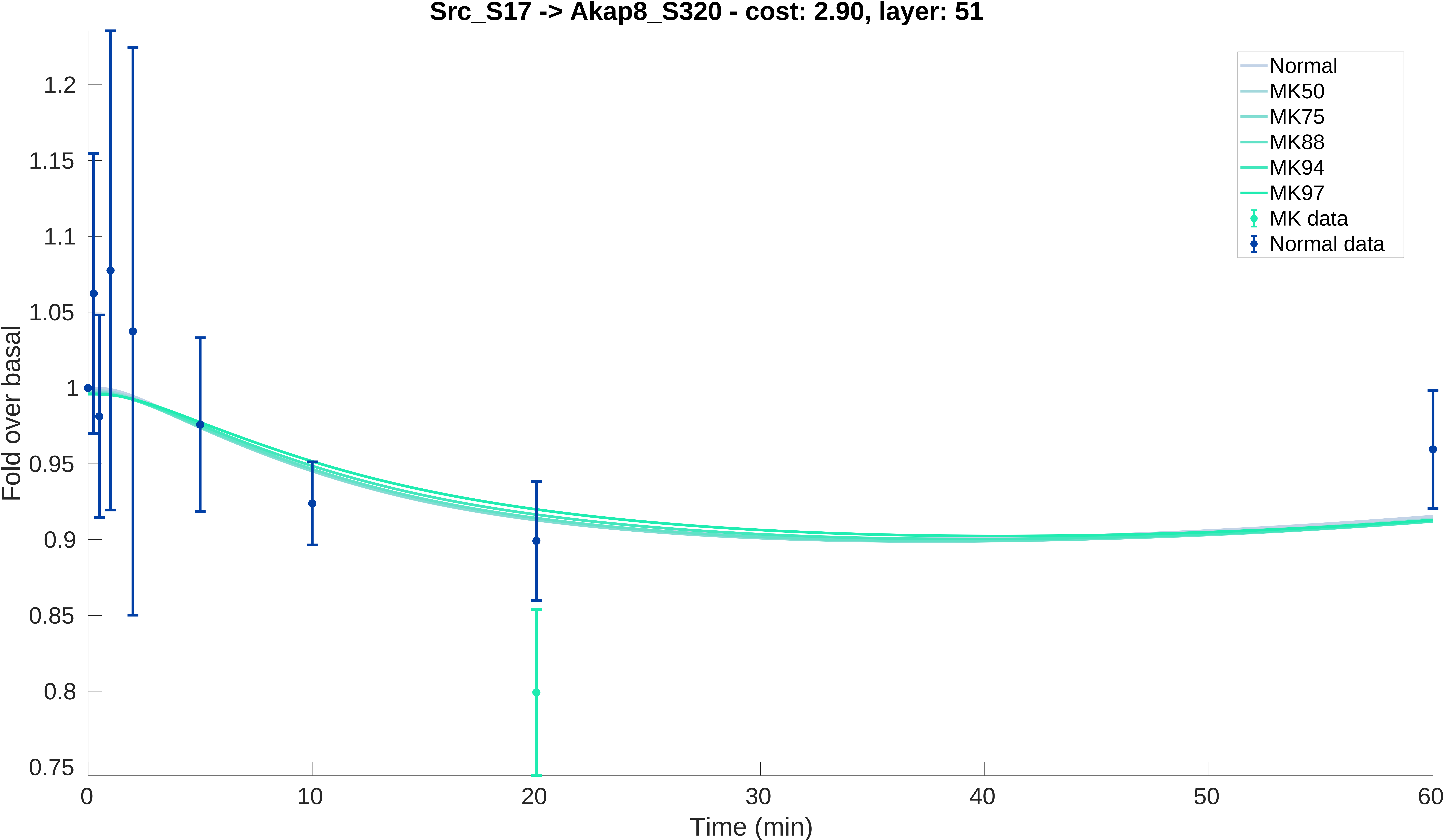
Src_S17 -> Akap8_S320 - cost: 2.90, layer: 51
Fold over basal
Time (min)
1.2
1.15
1.1
1.05
1
0.95
0.9
0.85
0.8
0.75
0
10
20
30
40
50
60
Normal
MK50
MK75
MK88
MK94
MK97
MK data
Normal data

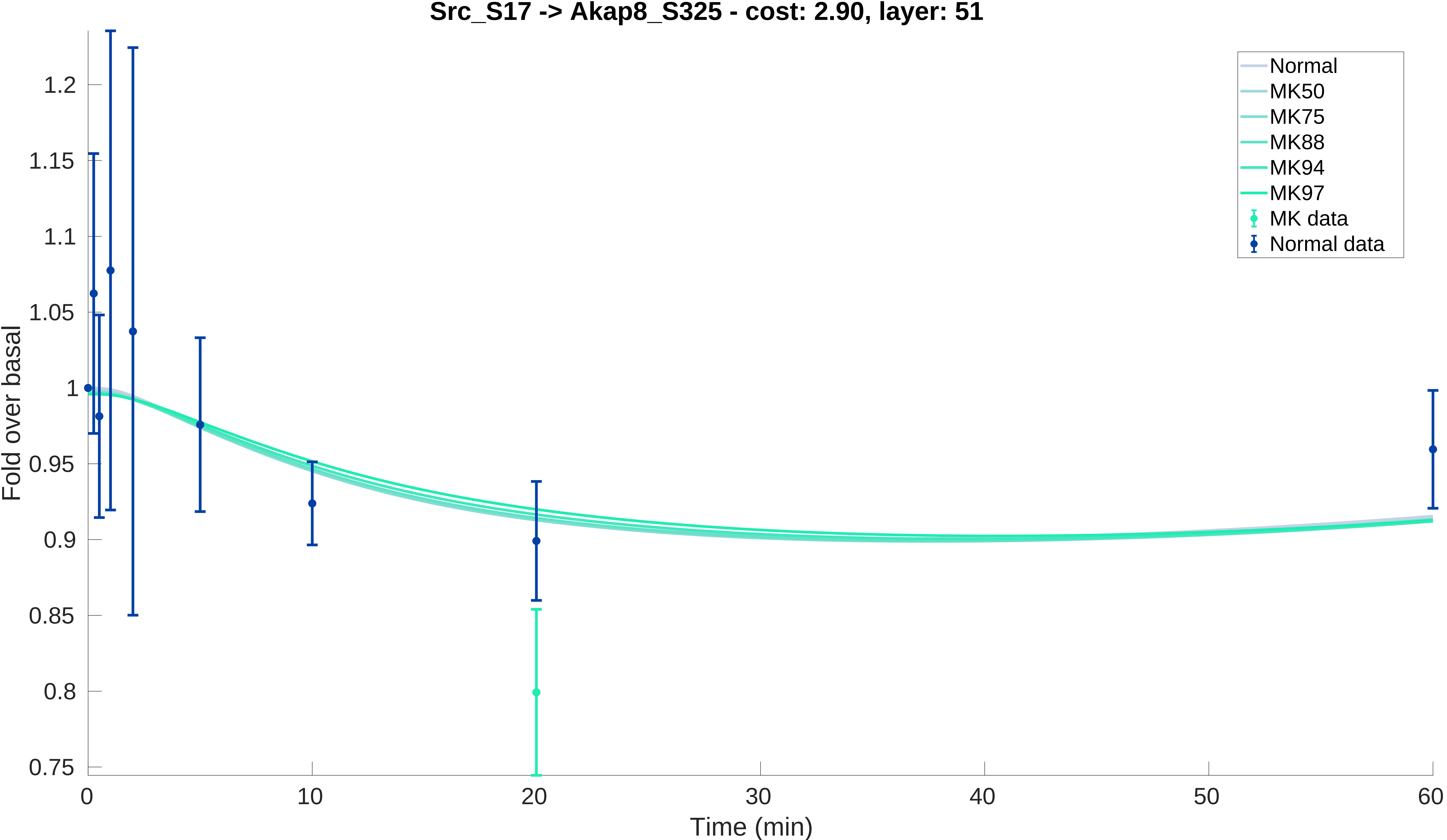

Src_S17 -> Akap8_S325 - cost: 2.90, layer: 51
Fold over basal
1.2
1.15
1.1
1.05
1
0.95
0.9
0.85
0.8
0.75
0
10
20
30
40
50
60
Time (min)
Normal
MK50
MK75
MK88
MK94
MK97
MK data
Normal data

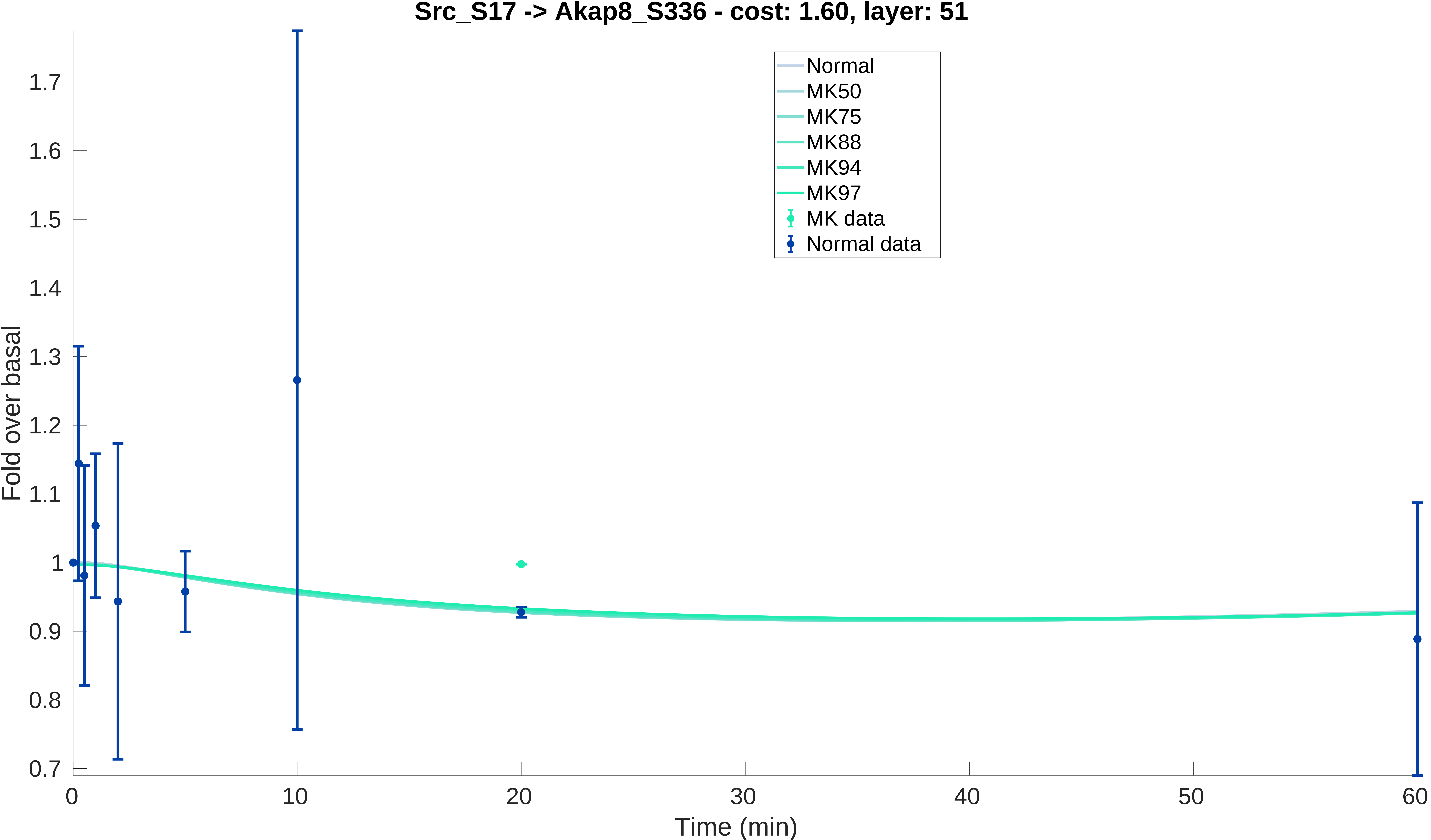
Src_S17 -> Akap8_S336 - cost: 1.60, layer: 51
Normal
MK50
MK75
MK88
MK94
MK97
MK data
Normal data
Fold over basal
1.7
1.6
1.5
1.4
1.3
1.2
1.1
1
0.9
0.8
0.7
0
10
20
30
40
50
60
Time (min)

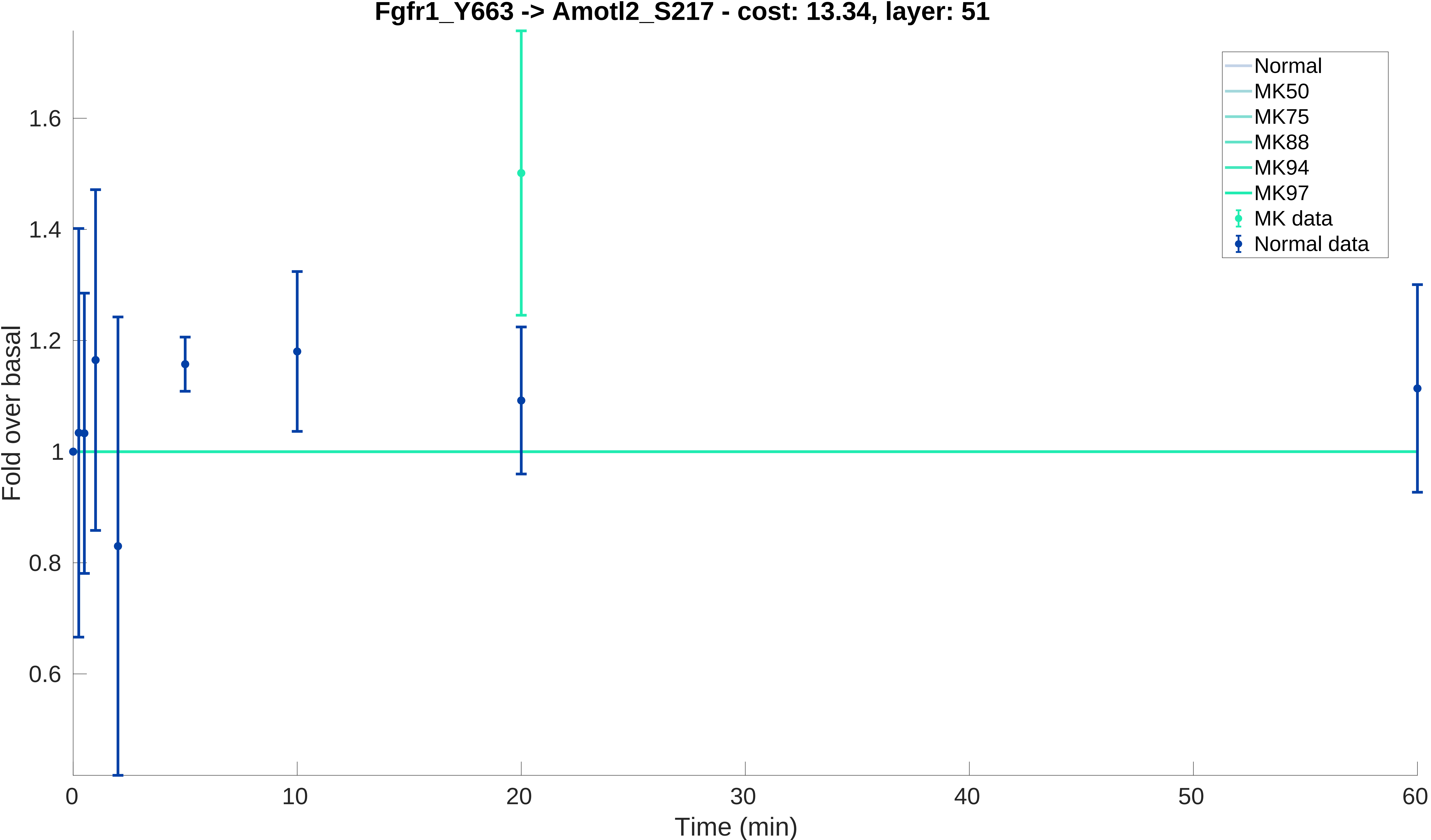

Fgfr1_Y663 -> Amotl2_S217 - cost: 13.34, layer: 51
Normal
MK50
MK75
MK88
MK94
MK97
MK data
Normal data
Fold over basal
1.6
1.4
1.2
1
0.8
0.6
0
10
20
30
40
50
60
Time (min)

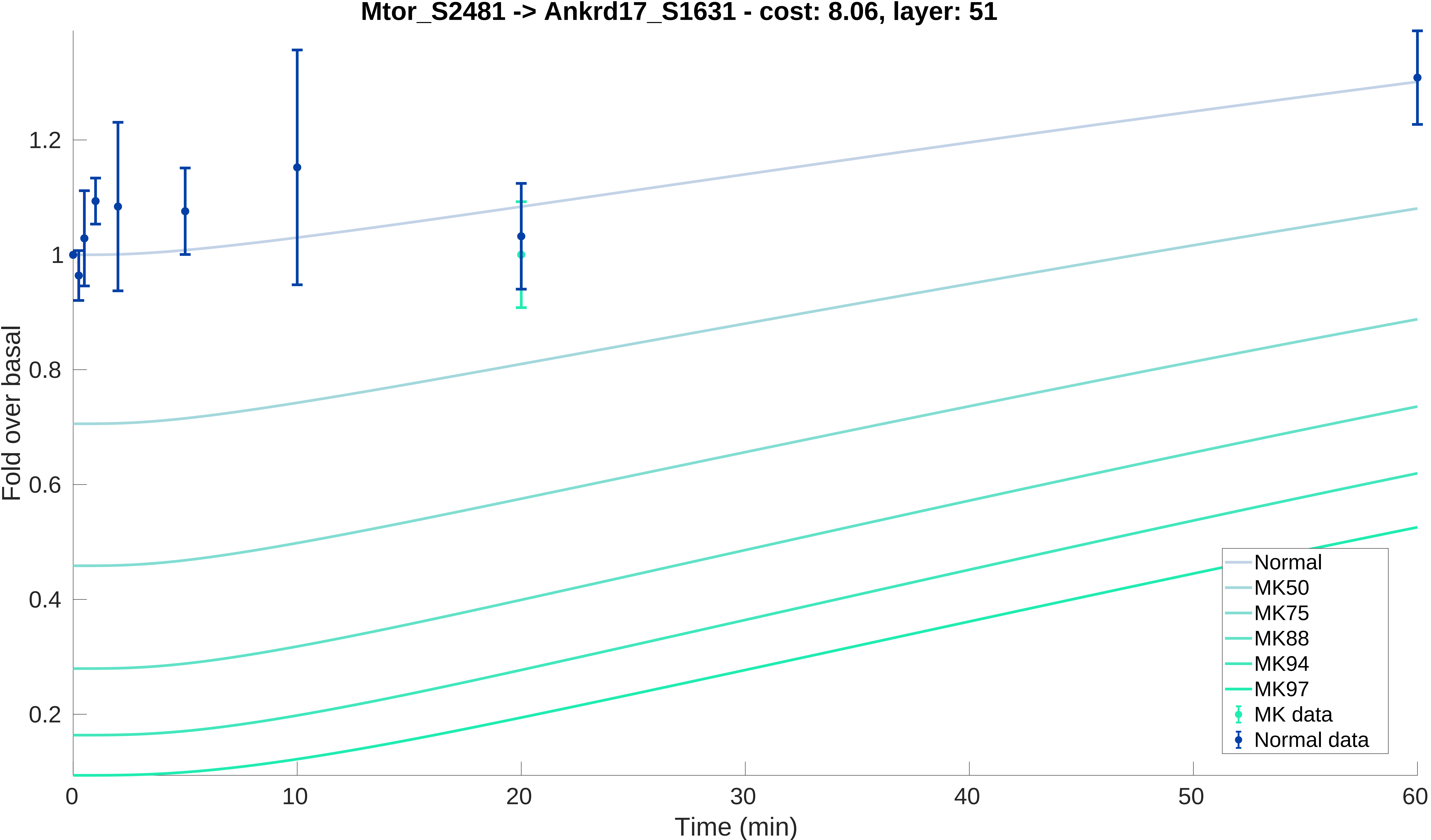

Mtor_S2481 -> Ankrd17_S1631 - cost: 8.06, layer: 51
Fold over basal
1.2
1
0.8
0.6
0.4
0.2
0
10
20
30
40
50
60
Time (min)
Normal
MK50
MK75
MK88
MK94
MK97
MK data
Normal data

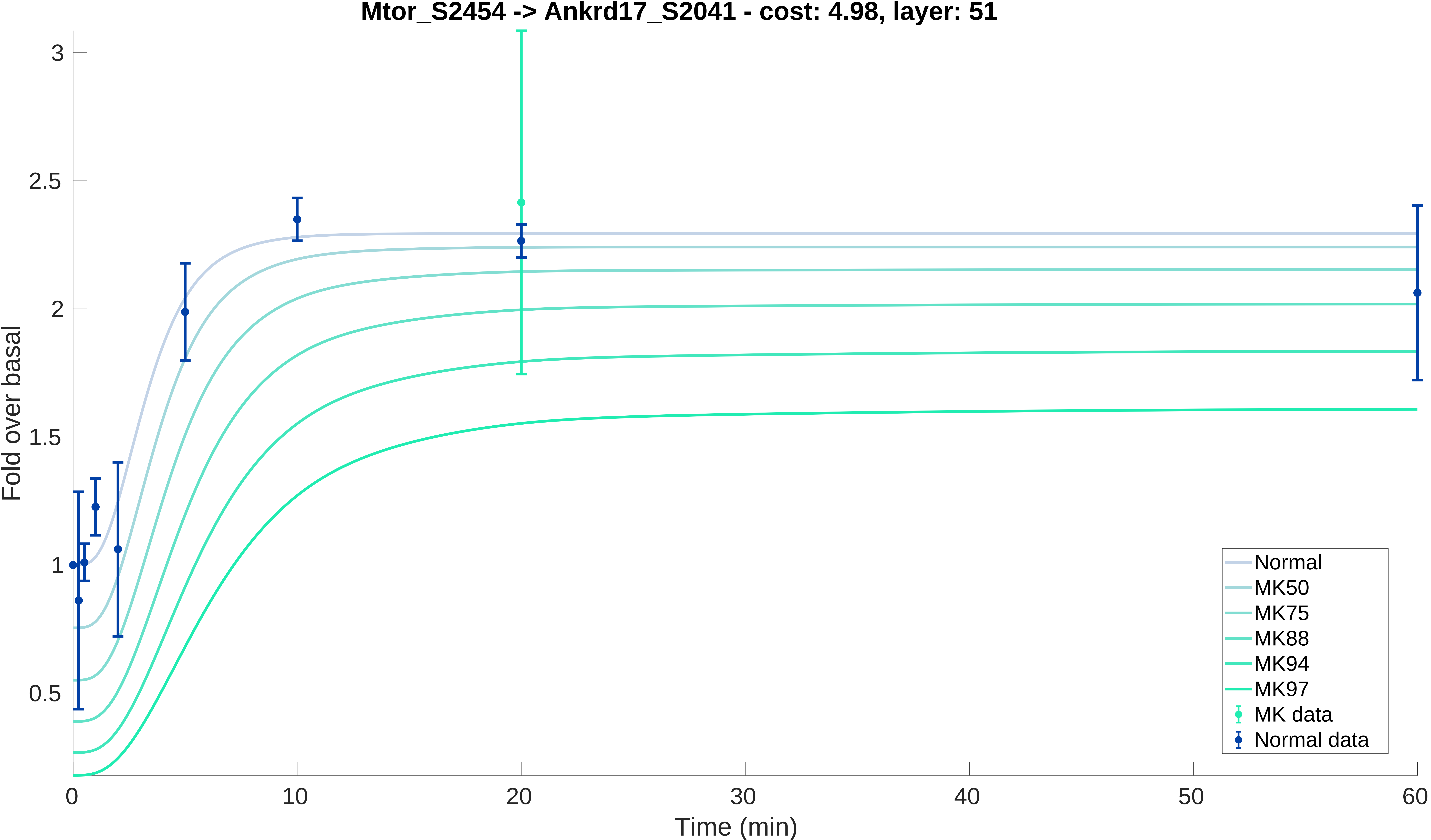
Mtor_S2454 -> Ankrd17_S2041 - cost: 4.98, layer: 51
Fold over basal
Time (min)
0
10
20
30
40
50
60
0.5
1
1.5
2
2.5
3
Normal
MK50
MK75
MK88
MK94
MK97
MK data
Normal data

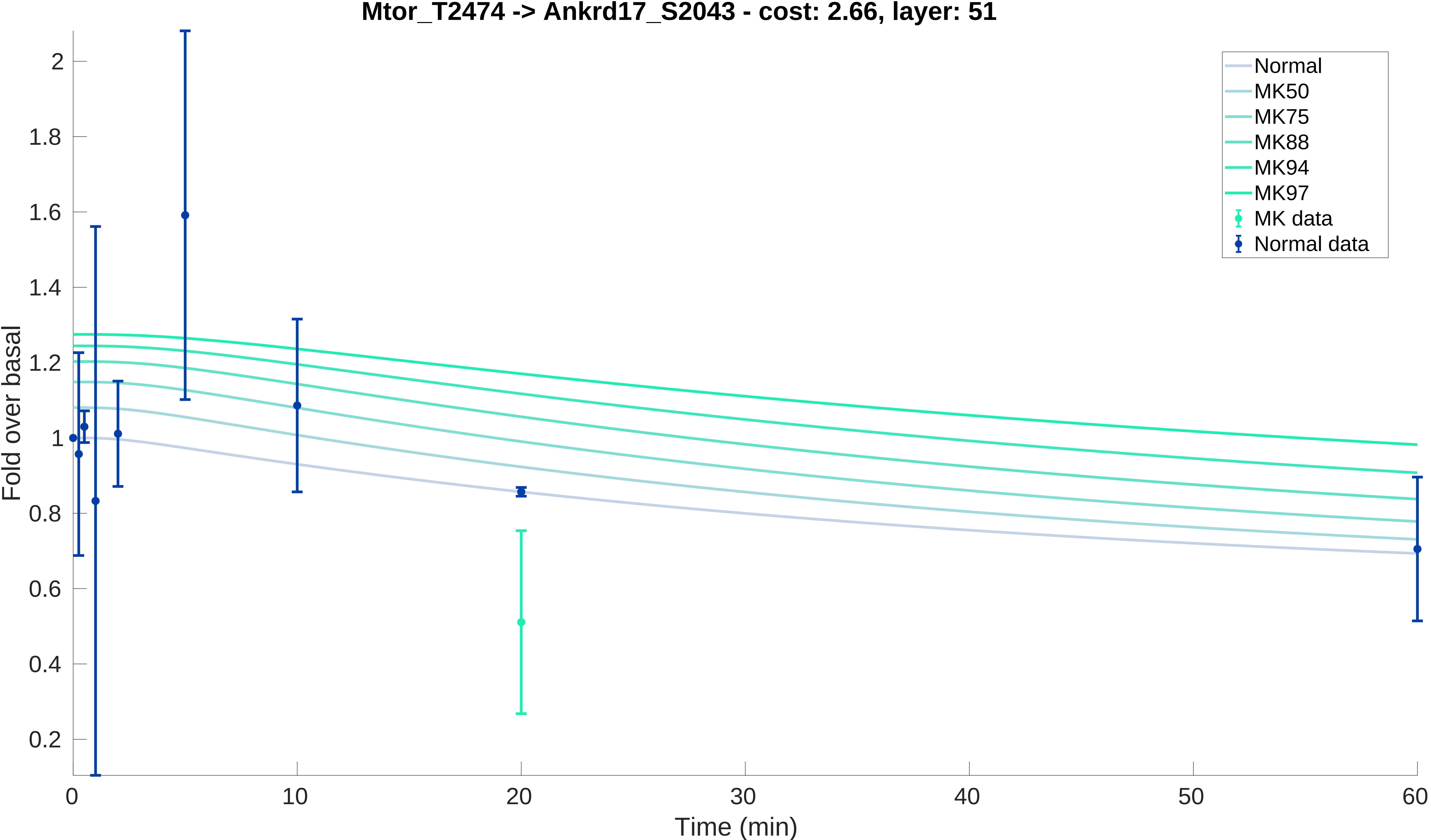

Mtor_T2474 -> Ankrd17_S2043 - cost: 2.66, layer: 51
Fold over basal
Time (min)
0
10
20
30
40
50
60
0.2
0.4
0.6
0.8
1
1.2
1.4
1.6
1.8
2
Normal
MK50
MK75
MK88
MK94
MK97
MK data
Normal data

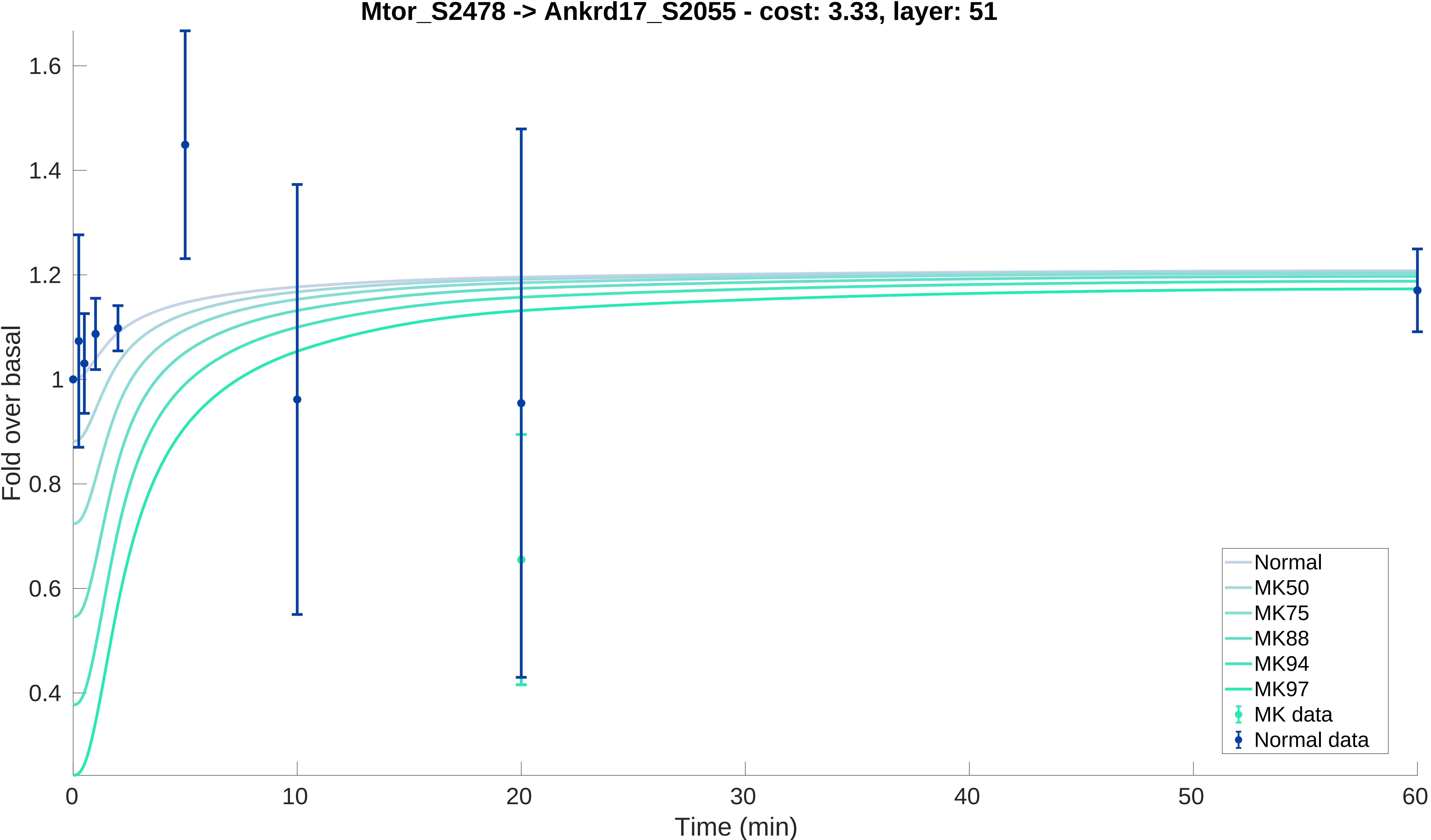

Mtor_S2478 -> Ankrd17_S2055 - cost: 3.33, layer: 51
Fold over basal
1.6
1.4
1.2
1
0.8
0.6
0.4
0
10
20
30
40
50
60
Time (min)
Normal
MK50
MK75
MK88
MK94
MK97
MK data
Normal data

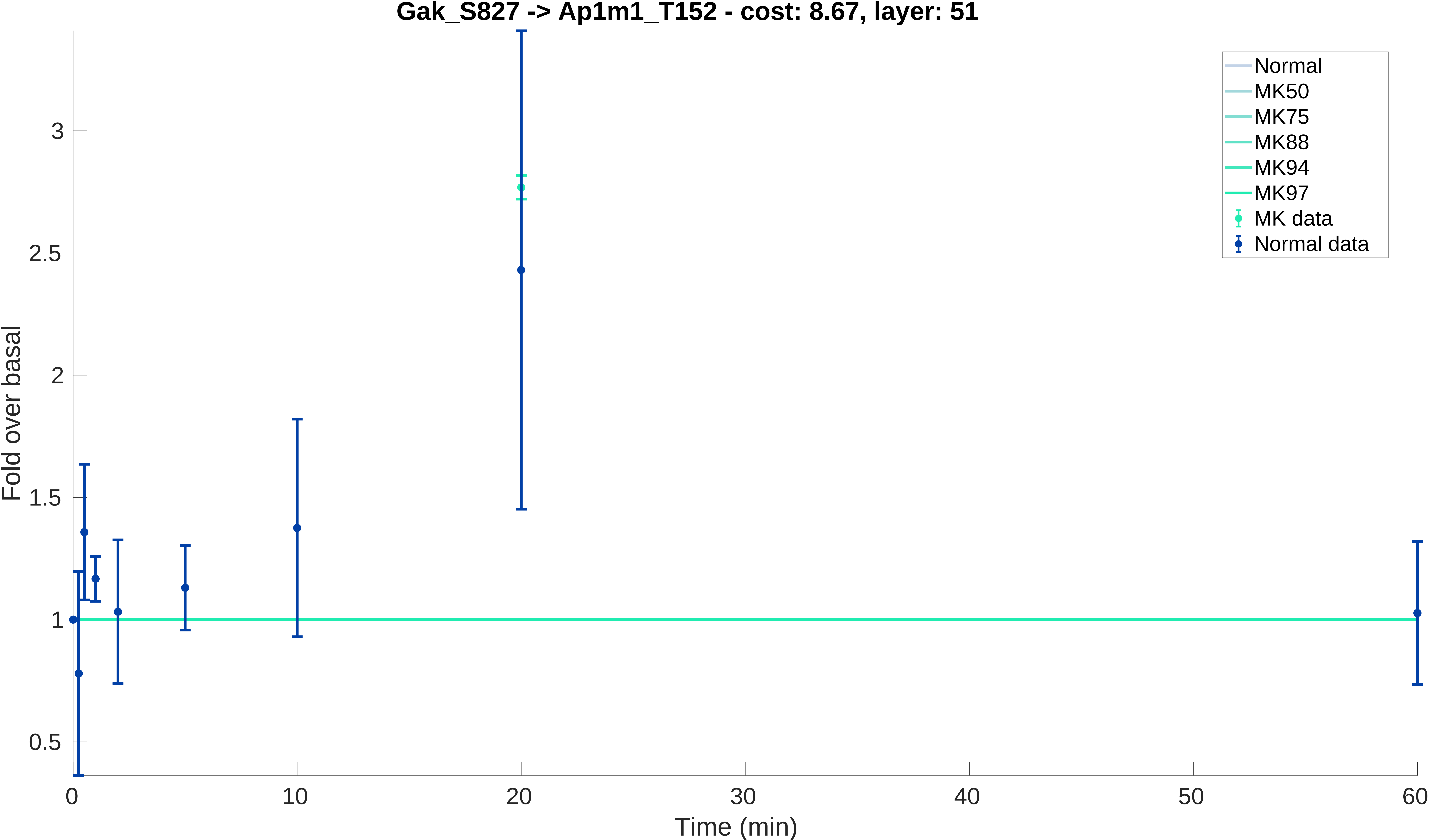

Gak_S827 -> Ap1m1_T152 - cost: 8.67, layer: 51
Normal
MK50
MK75
MK88
MK94
MK97
MK data
Normal data
Fold over basal
Time (min)
0.5
1
1.5
2
2.5
3
0
10
20
30
40
50
60

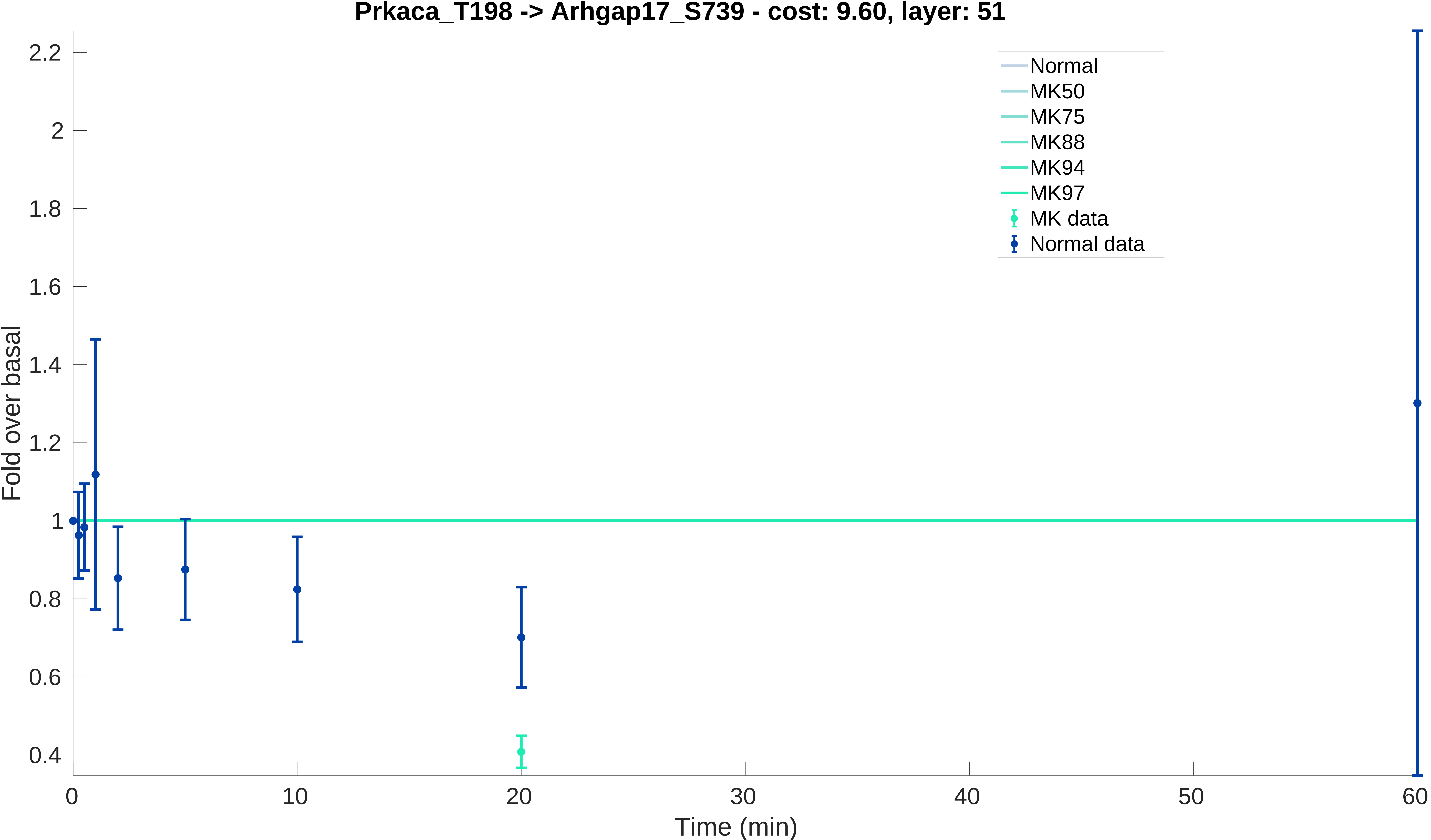

Prkaca_T198 -> Arhgap17_S739 - cost: 9.60, layer: 51
Fold over basal
Time (min)
2.2
2
1.8
1.6
1.4
1.2
1
0.8
0.6
0.4
0
10
20
30
40
50
60
Normal
MK50
MK75
MK88
MK94
MK97
MK data
Normal data

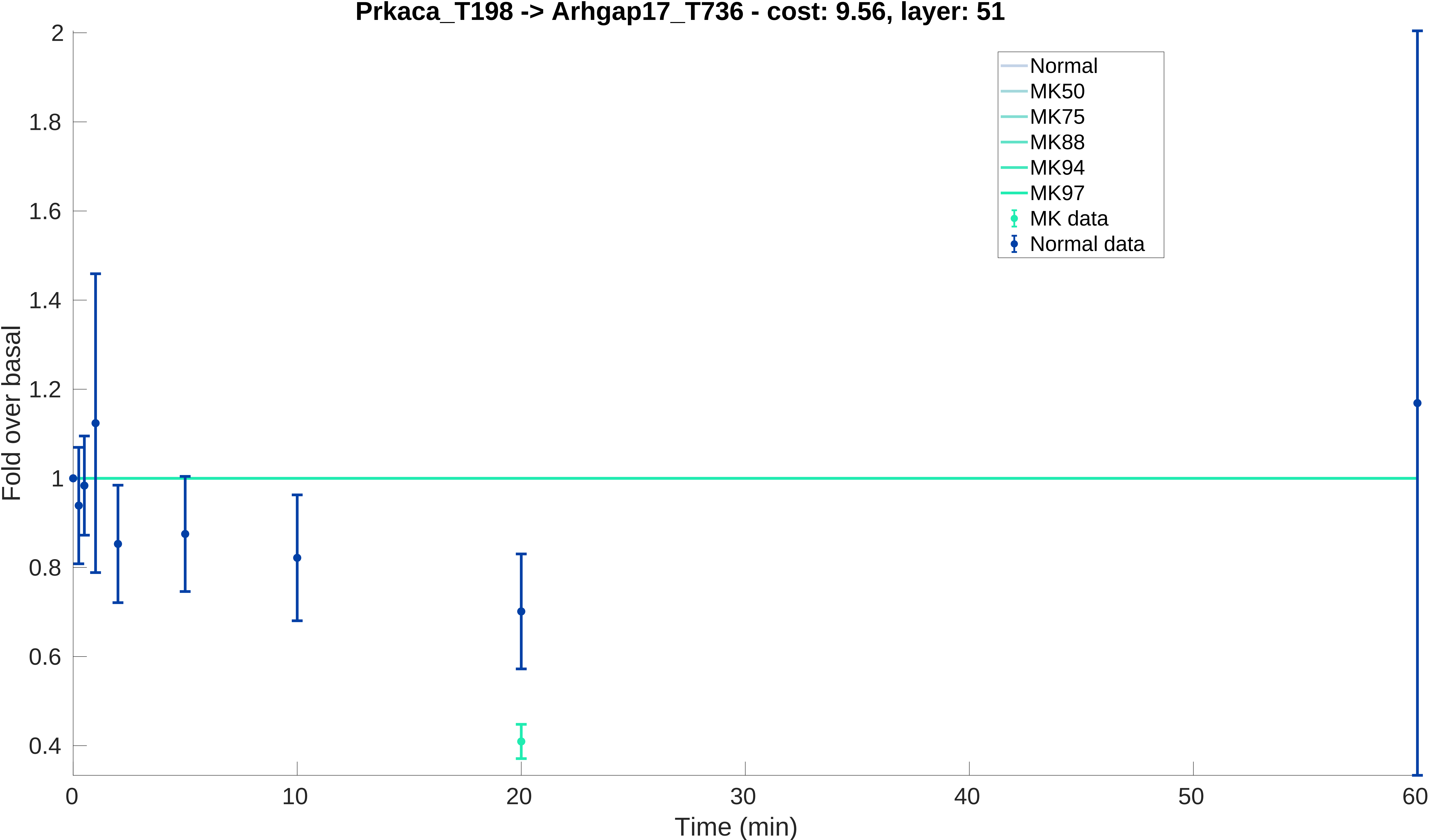

Prkaca_T198 -> Arhgap17_T736 - cost: 9.56, layer: 51
Fold over basal
Time (min)
0
10
20
30
40
50
60
0.4
0.6
0.8
1
1.2
1.4
1.6
1.8
2
Normal
MK50
MK75
MK88
MK94
MK97
MK data
Normal data

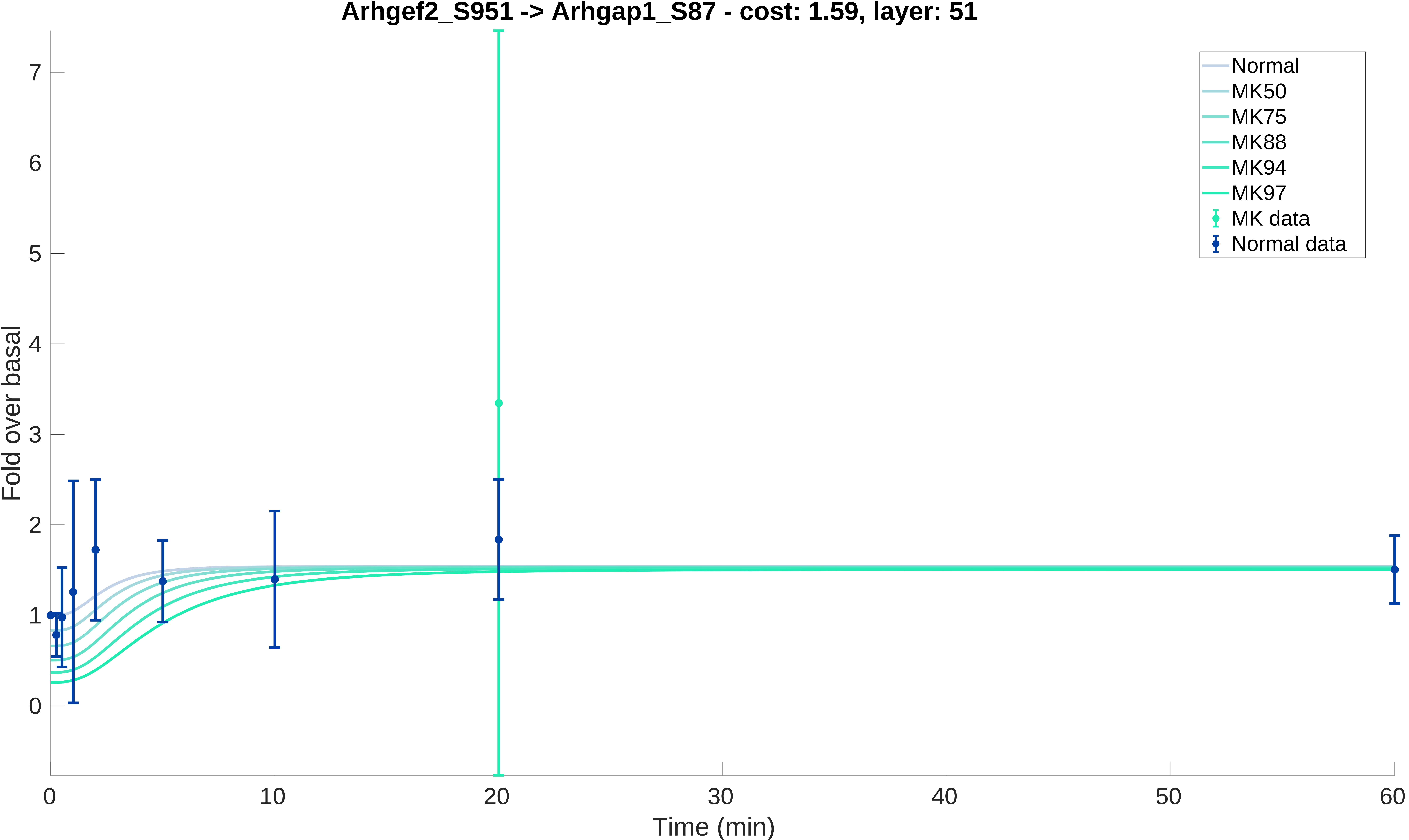
Arhgef2_S951 -> Arhgap1_S87 - cost: 1.59, layer: 51
Fold over basal
Time (min)
0
1
2
3
4
5
6
7
0
10
20
30
40
50
60
Normal
MK50
MK75
MK88
MK94
MK97
MK data
Normal data

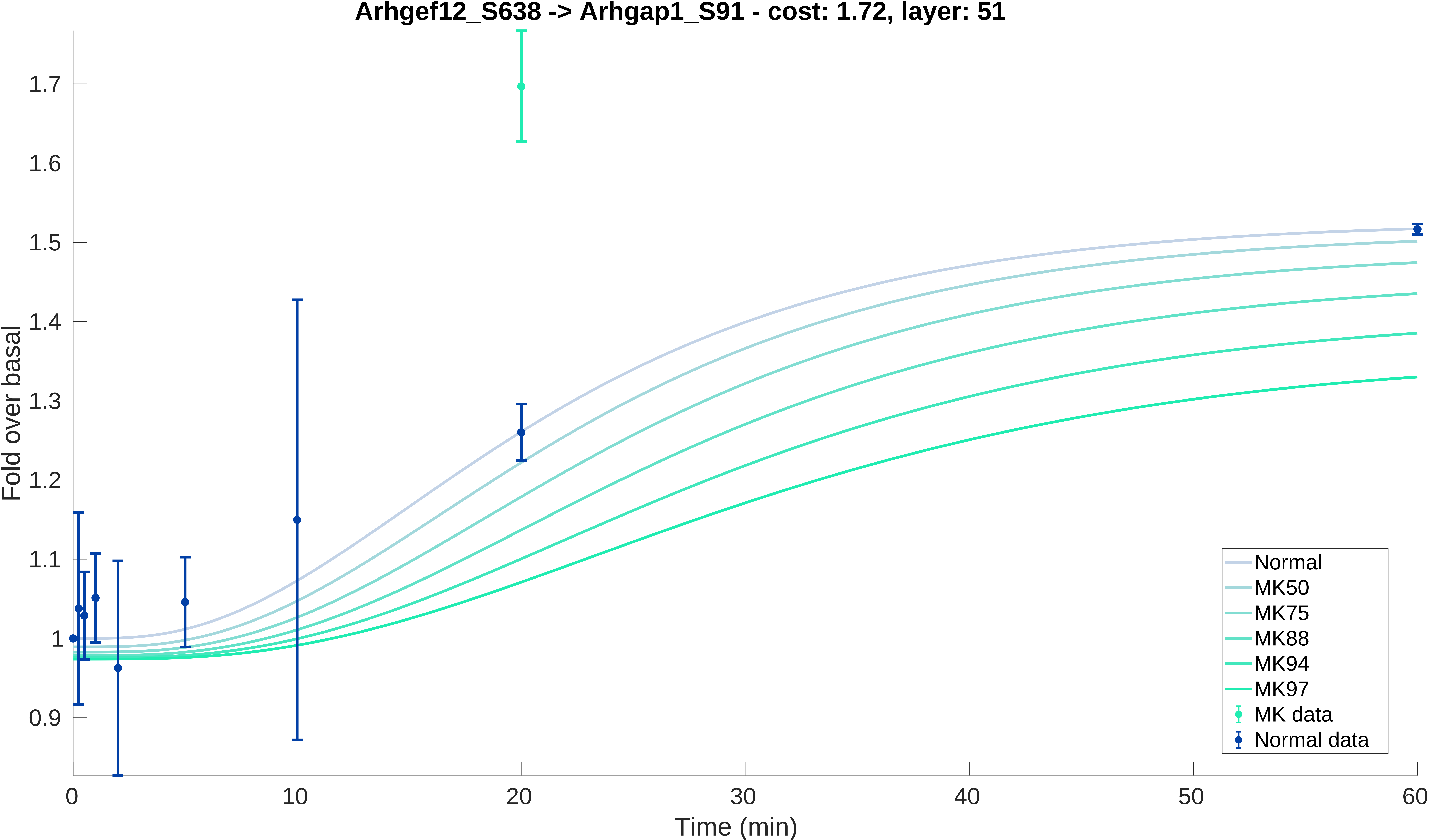

Arhgef12_S638 -> Arhgap1_S91 - cost: 1.72, layer: 51
Fold over basal
1.7
1.6
1.5
1.4
1.3
1.2
1.1
1
0.9
0
10
20
30
40
50
60
Time (min)
Normal
MK50
MK75
MK88
MK94
MK97
MK data
Normal data

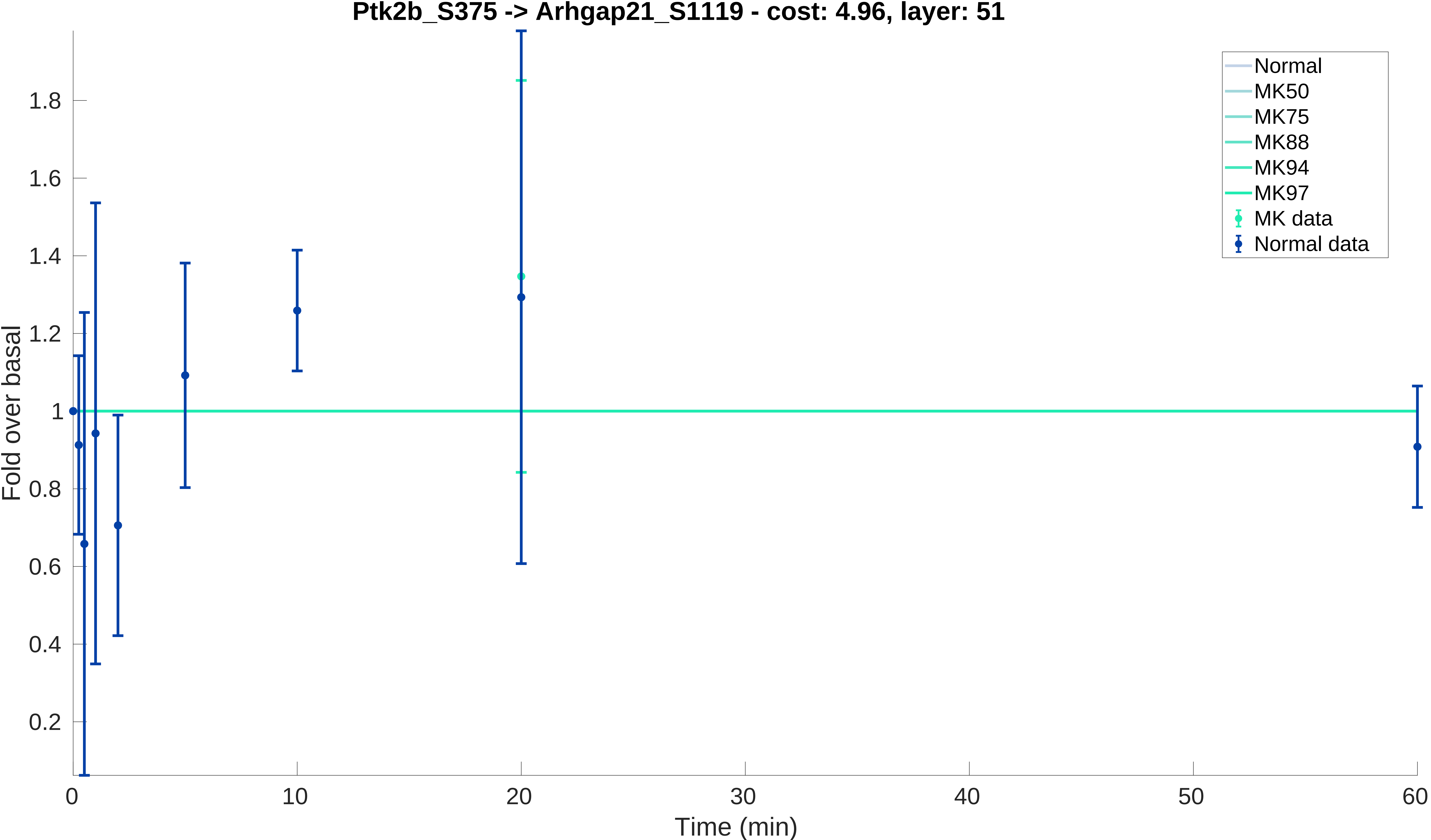

Ptk2b_S375 -> Arhgap21_S1119 - cost: 4.96, layer: 51
Fold over basal
Time (min)
Normal
MK50
MK75
MK88
MK94
MK97
MK data
Normal data
0.2
0.4
0.6
0.8
1
1.2
1.4
1.6
1.8
0
10
20
30
40
50
60

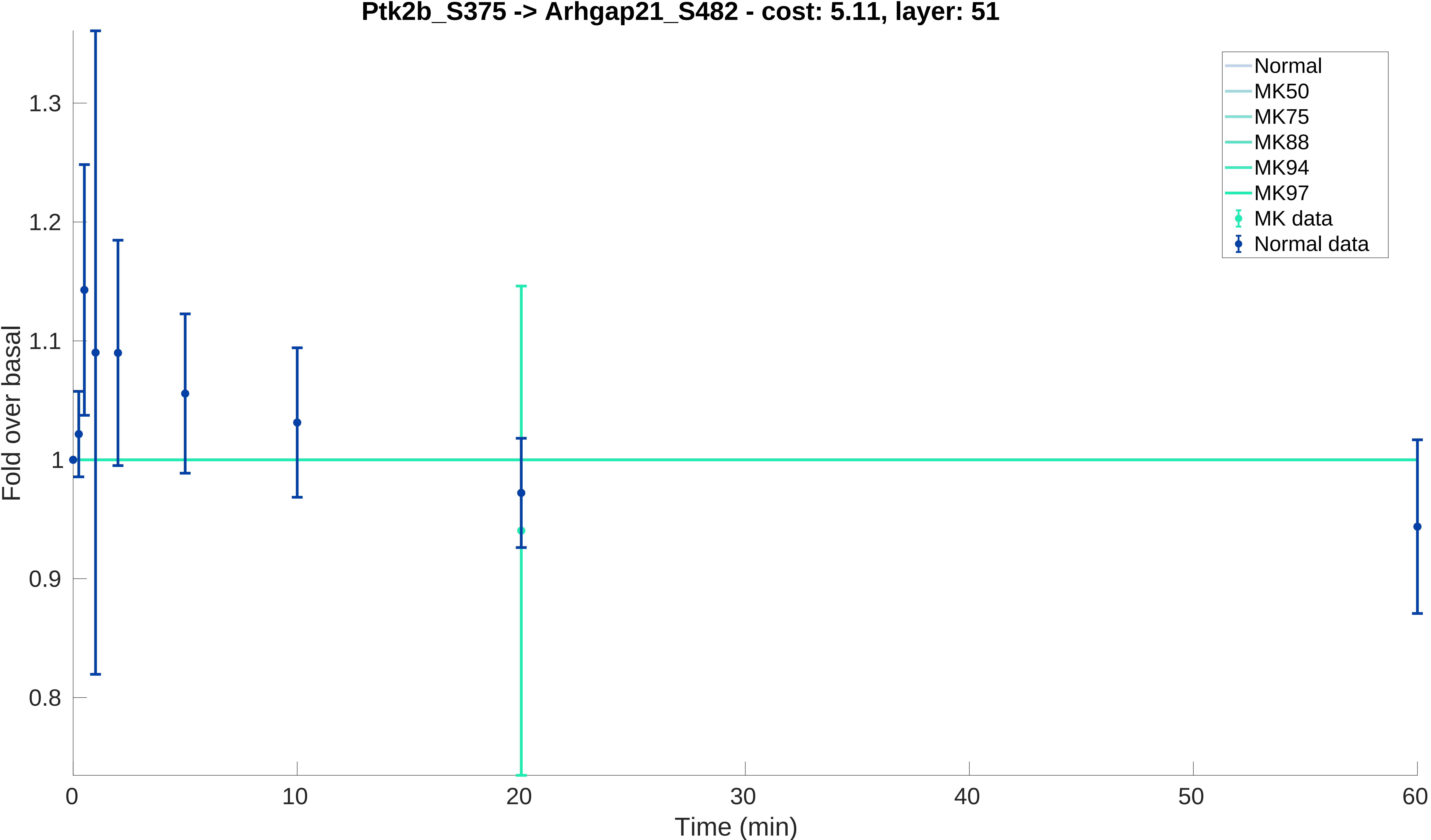

Ptk2b_S375 -> Arhgap21_S482 - cost: 5.11, layer: 51
Normal
MK50
MK75
MK88
MK94
MK97
MK data
Normal data
Fold over basal
1.3
1.2
1.1
1
0.9
0.8
0
10
20
30
40
50
60
Time (min)

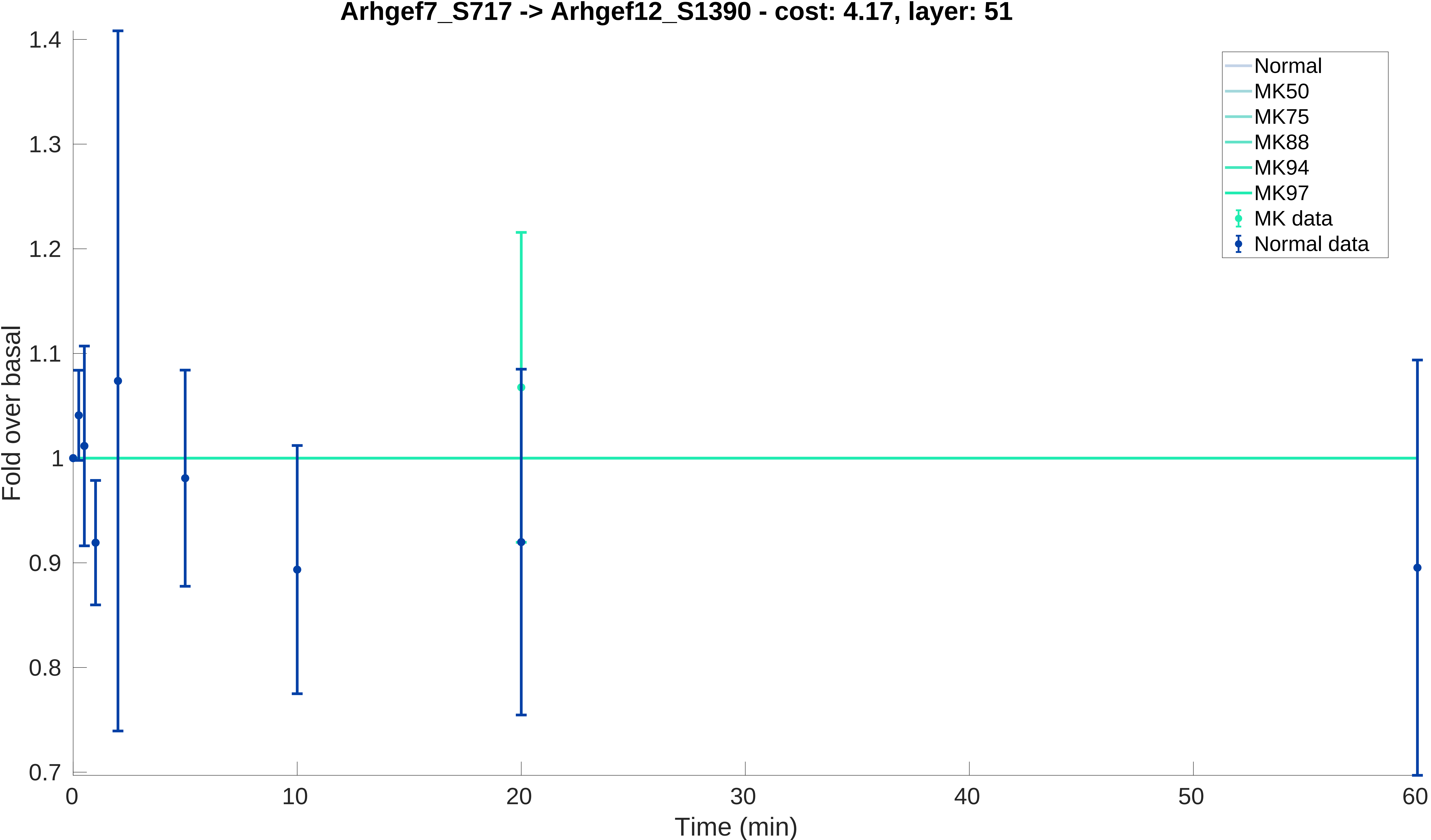

Arhgef7_S717 -> Arhgef12_S1390 - cost: 4.17, layer: 51
Fold over basal
1.4
1.3
1.2
1.1
1
0.9
0.8
0.7
0
10
20
30
40
50
60
Time (min)
Normal
MK50
MK75
MK88
MK94
MK97
MK data
Normal data

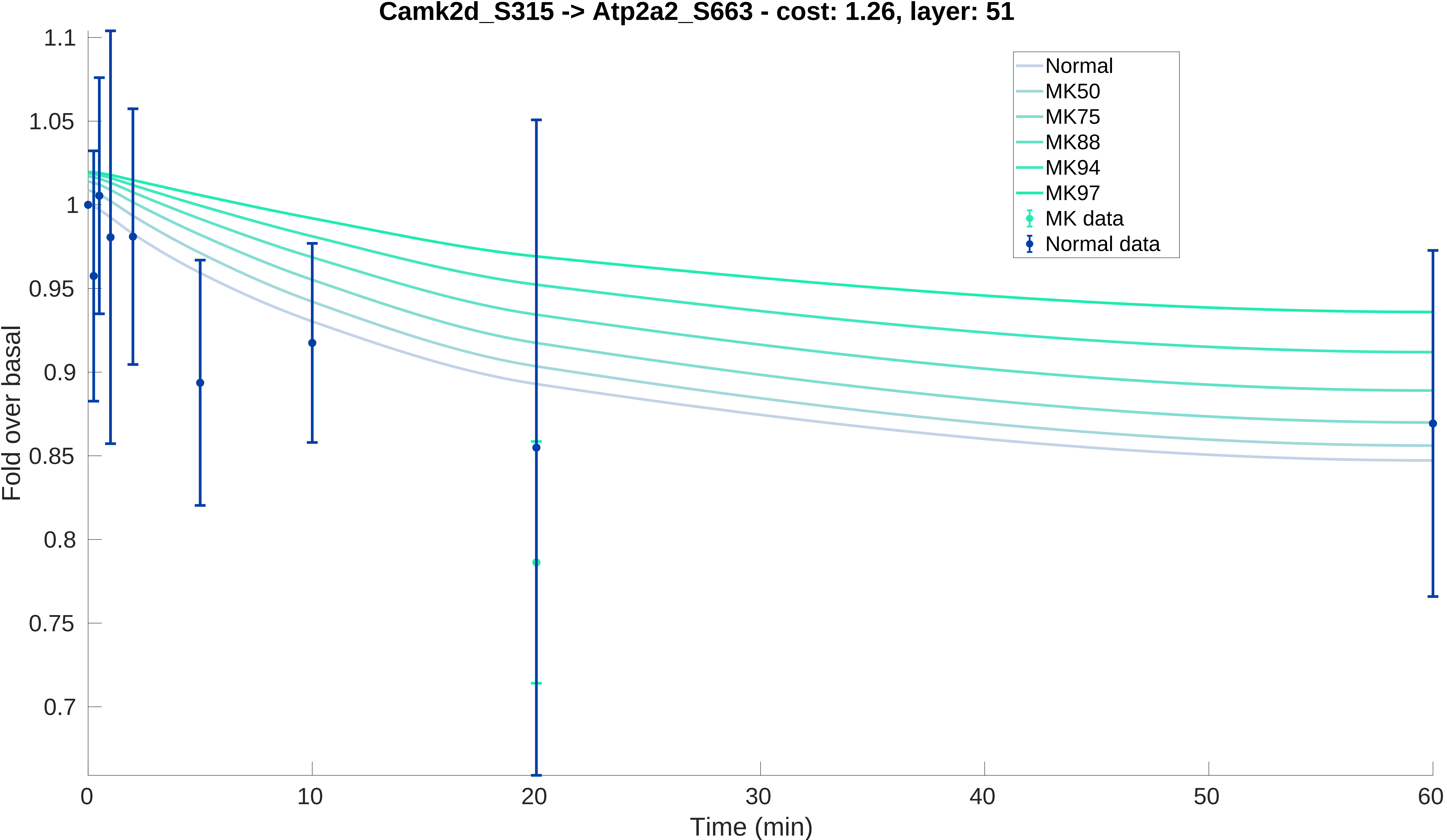

Camk2d_S315 -> Atp2a2_S663 - cost: 1.26, layer: 51
Fold over basal
Time (min)
1.1
1.05
1
0.95
0.9
0.85
0.8
0.75
0.7
0
10
20
30
40
50
60
Normal
MK50
MK75
MK88
MK94
MK97
MK data
Normal data

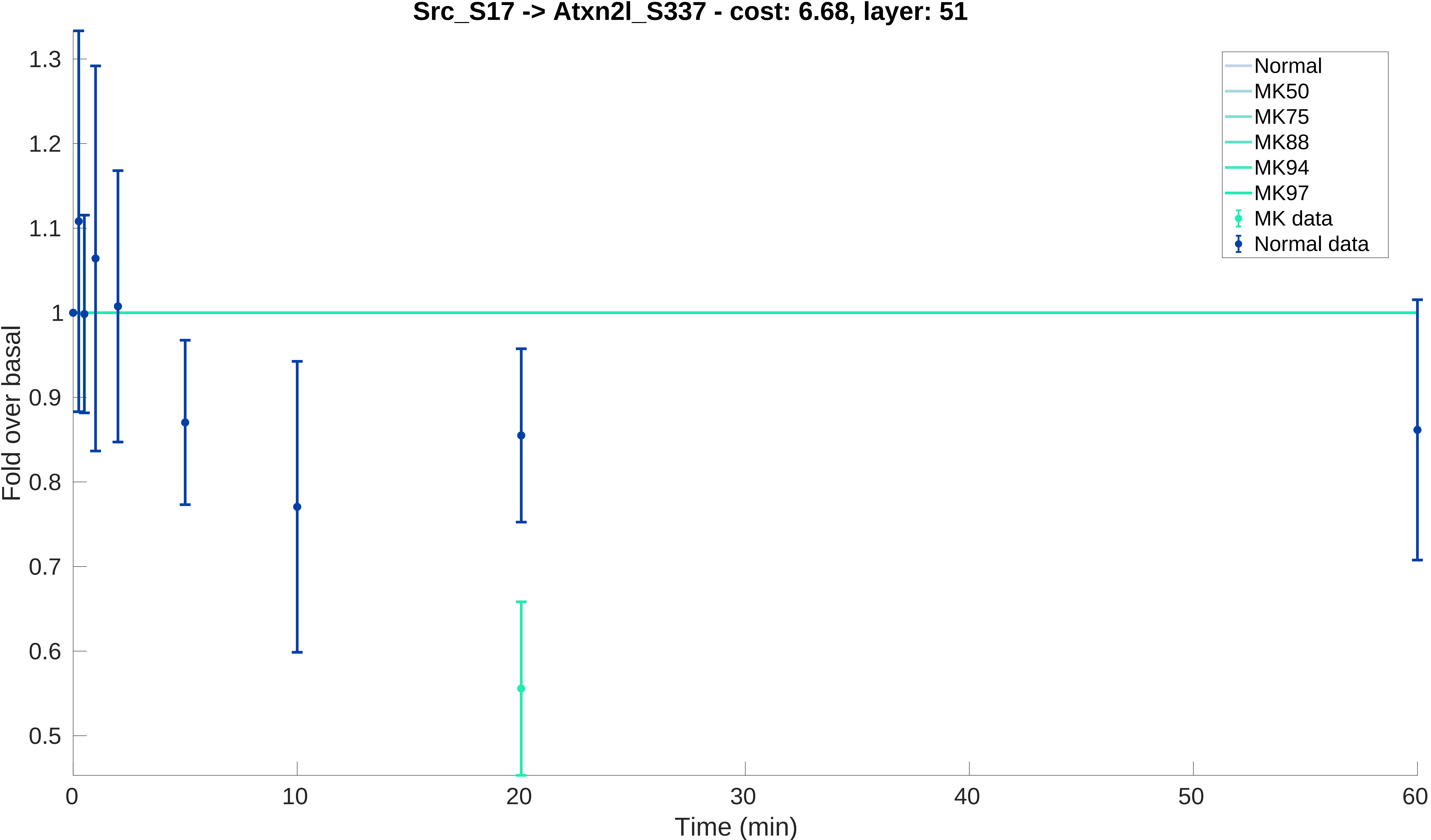

Src_S17 -> Atxn2l_S337 - cost: 6.68, layer: 51
Fold over basal
1.3
1.2
1.1
1
0.9
0.8
0.7
0.6
0.5
0
10
20
30
40
50
60
Time (min)
Normal
MK50
MK75
MK88
MK94
MK97
MK data
Normal data

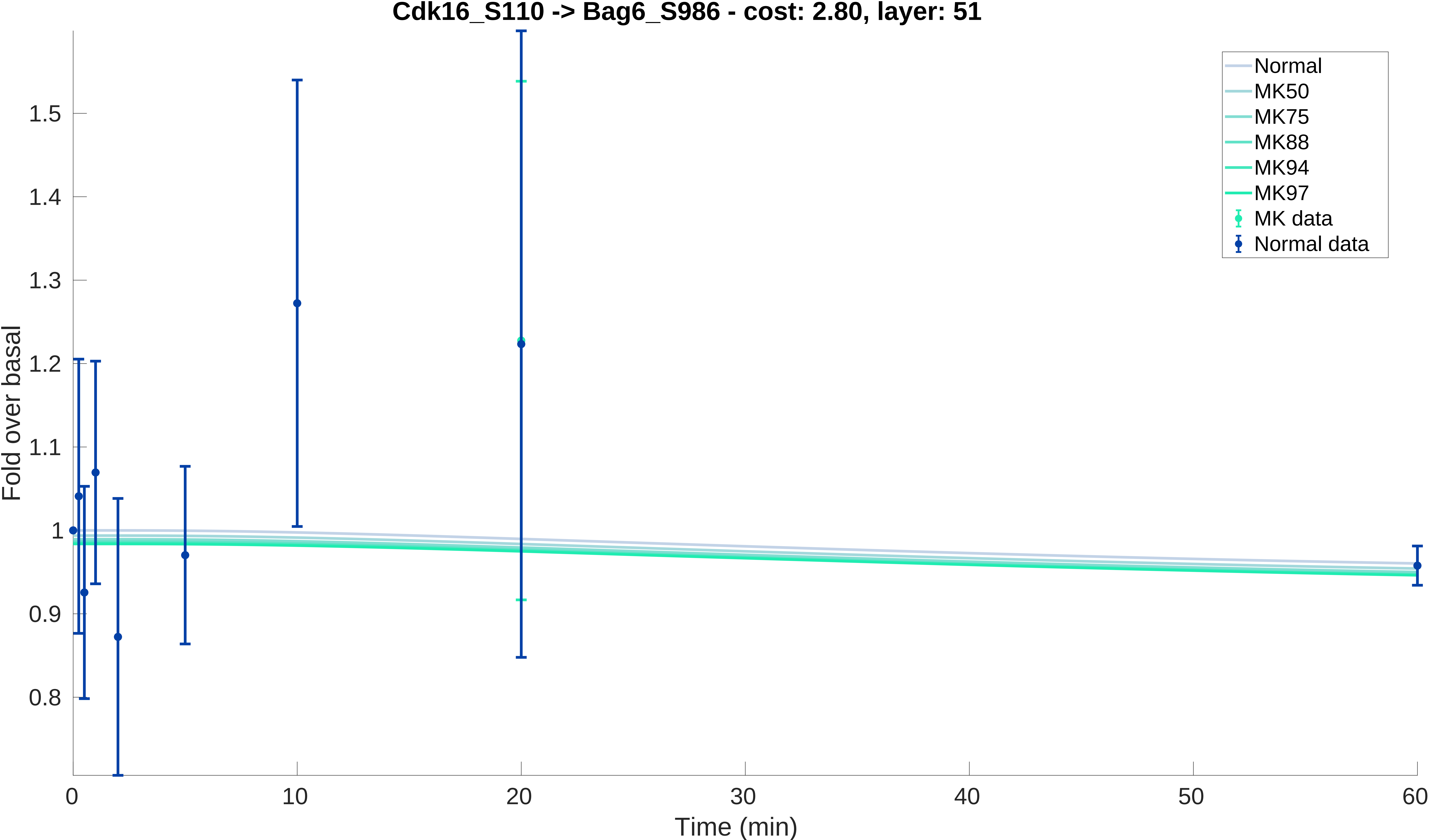

Cdk16_S110 -> Bag6_S986 - cost: 2.80, layer: 51
Normal
MK50
MK75
MK88
MK94
MK97
MK data
Normal data
Fold over basal
1.5
1.4
1.3
1.2
1.1
1
0.9
0.8
0
10
20
30
40
50
60
Time (min)

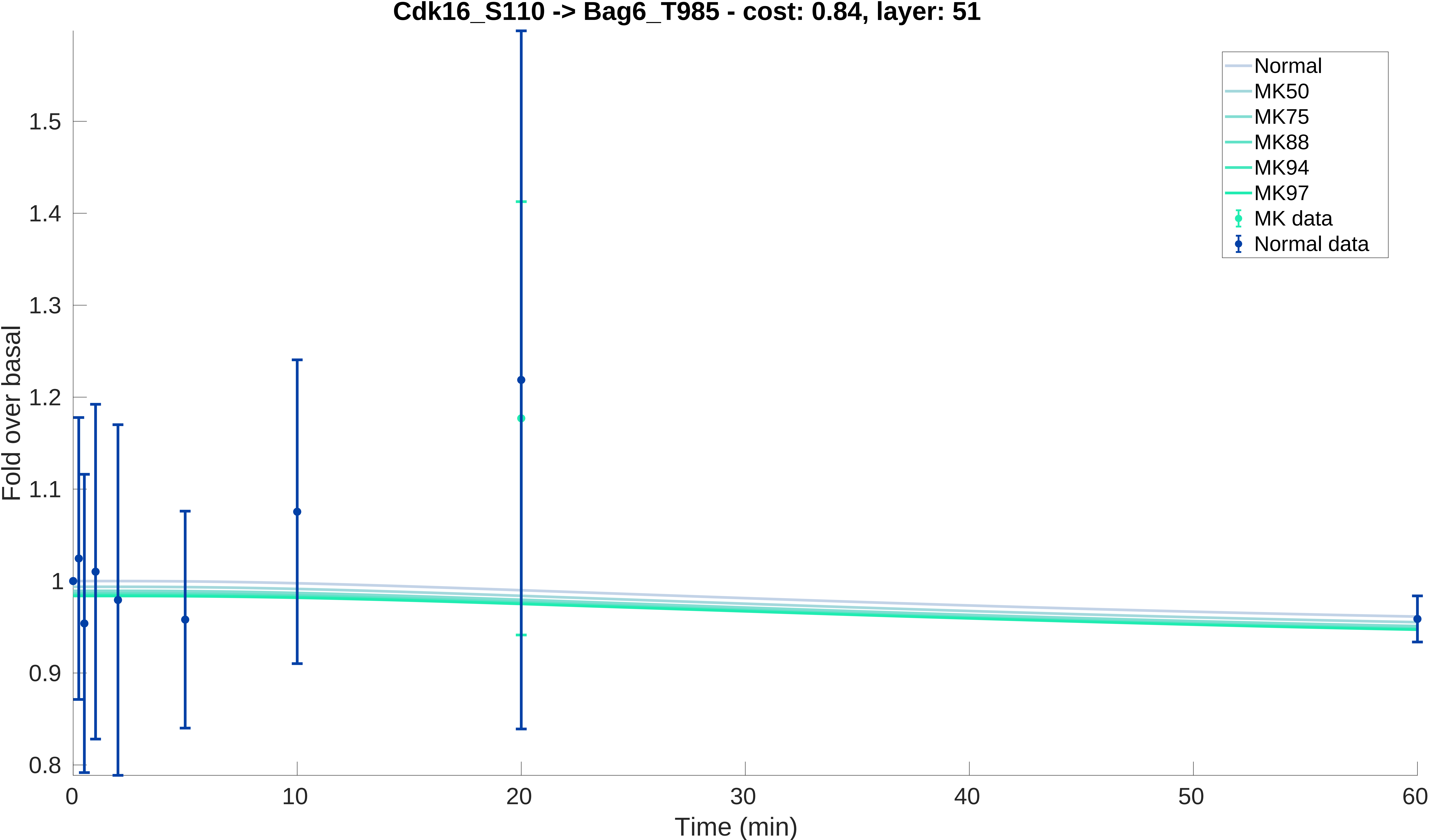

Cdk16_S110 -> Bag6_T985 - cost: 0.84, layer: 51
Fold over basal
Time (min)
0.8
0.9
1
1.1
1.2
1.3
1.4
1.5
0
10
20
30
40
50
60
Normal
MK50
MK75
MK88
MK94
MK97
MK data
Normal data

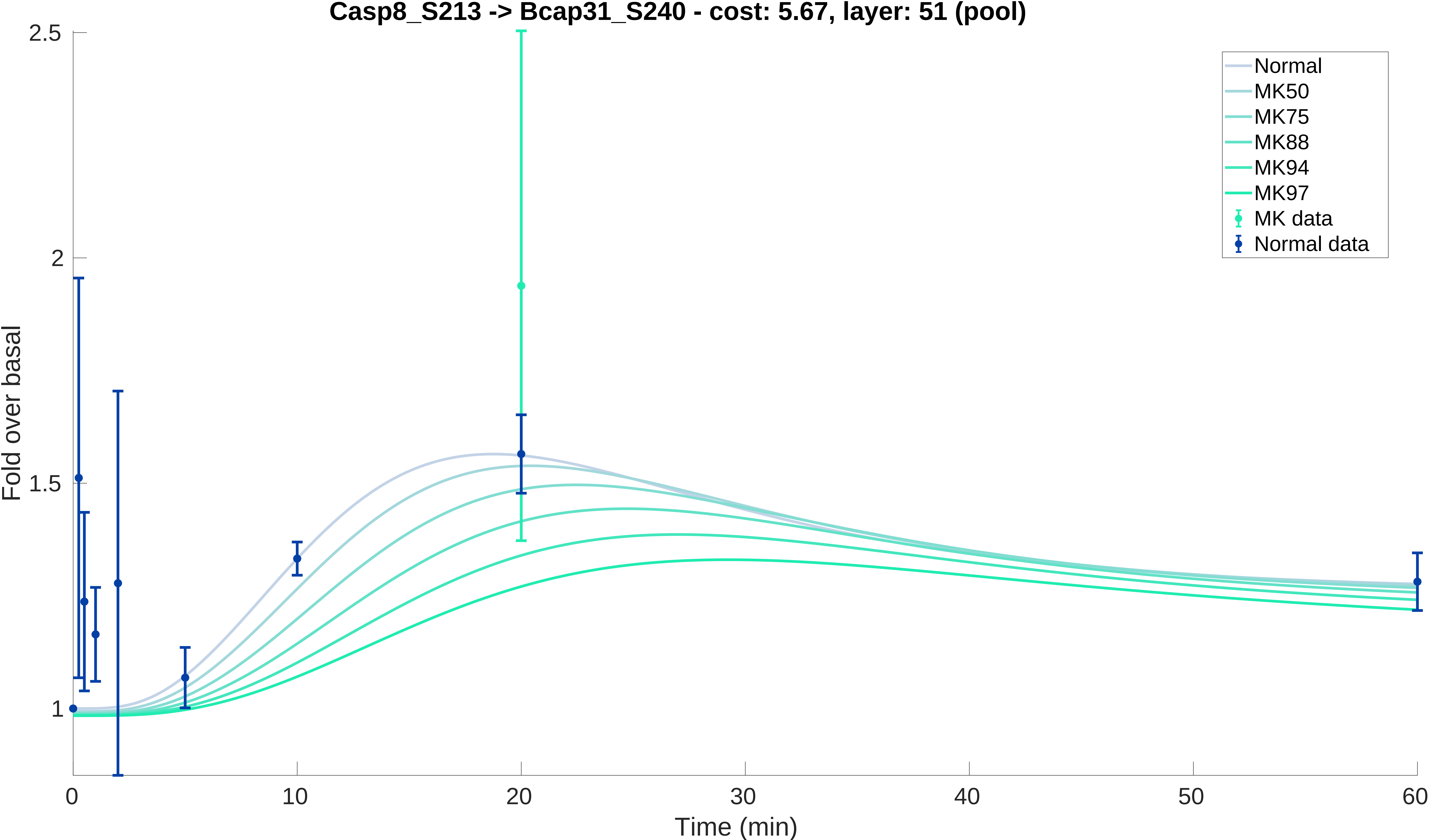
Casp8_S213 -> Bcap31_S240 - cost: 5.67, layer: 51 (pool)
Fold over basal
Time (min)
2.5
2
1.5
1
0
10
20
30
40
50
60
Normal
MK50
MK75
MK88
MK94
MK97
MK data
Normal data

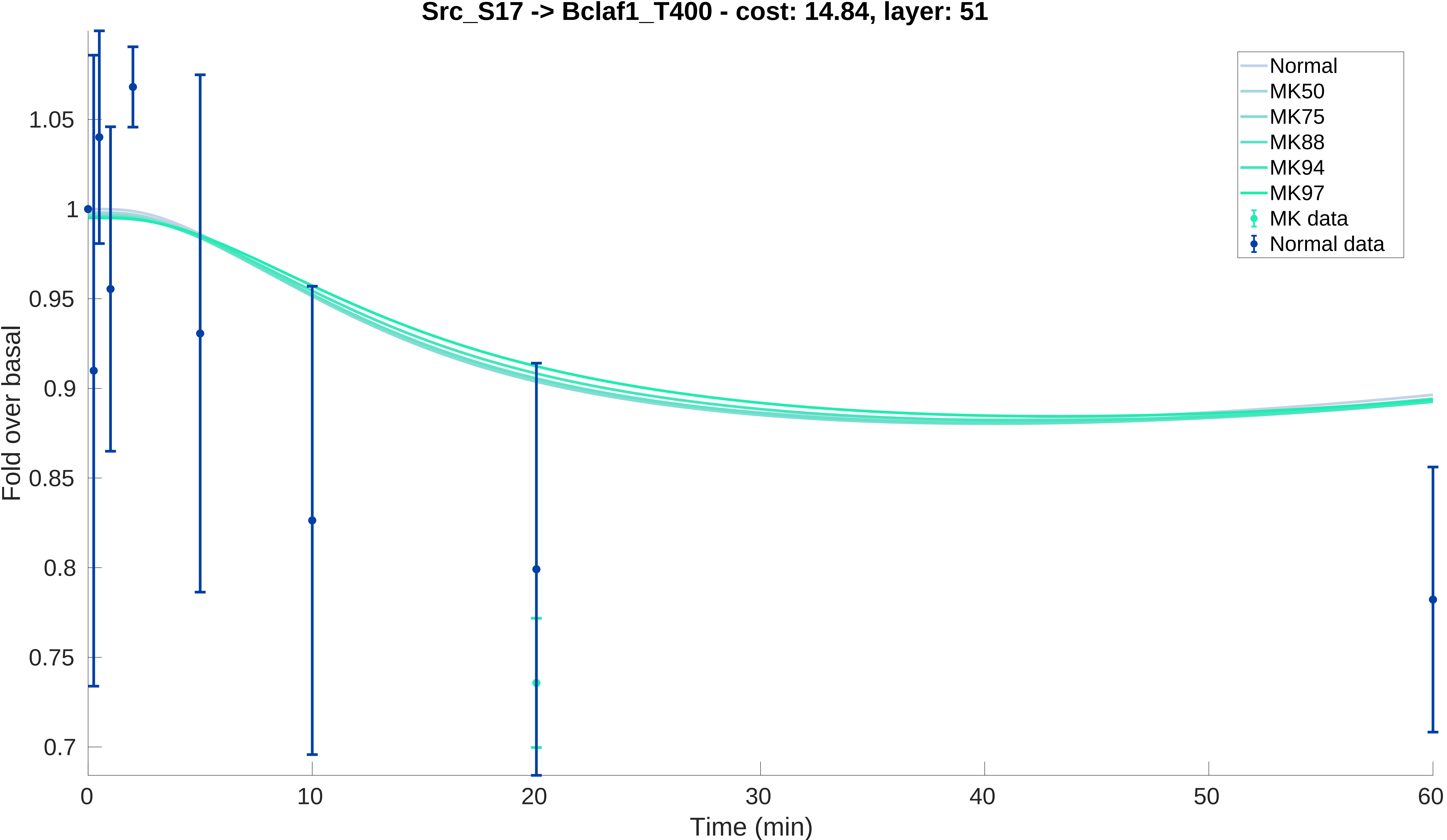

Src_S17 -> Bclaf1_T400 - cost: 14.84, layer: 51
Normal
MK50
MK75
MK88
MK94
MK97
MK data
Normal data
Fold over basal
1.05
1
0.95
0.9
0.85
0.8
0.75
0.7
0
10
20
30
40
50
60
Time (min)

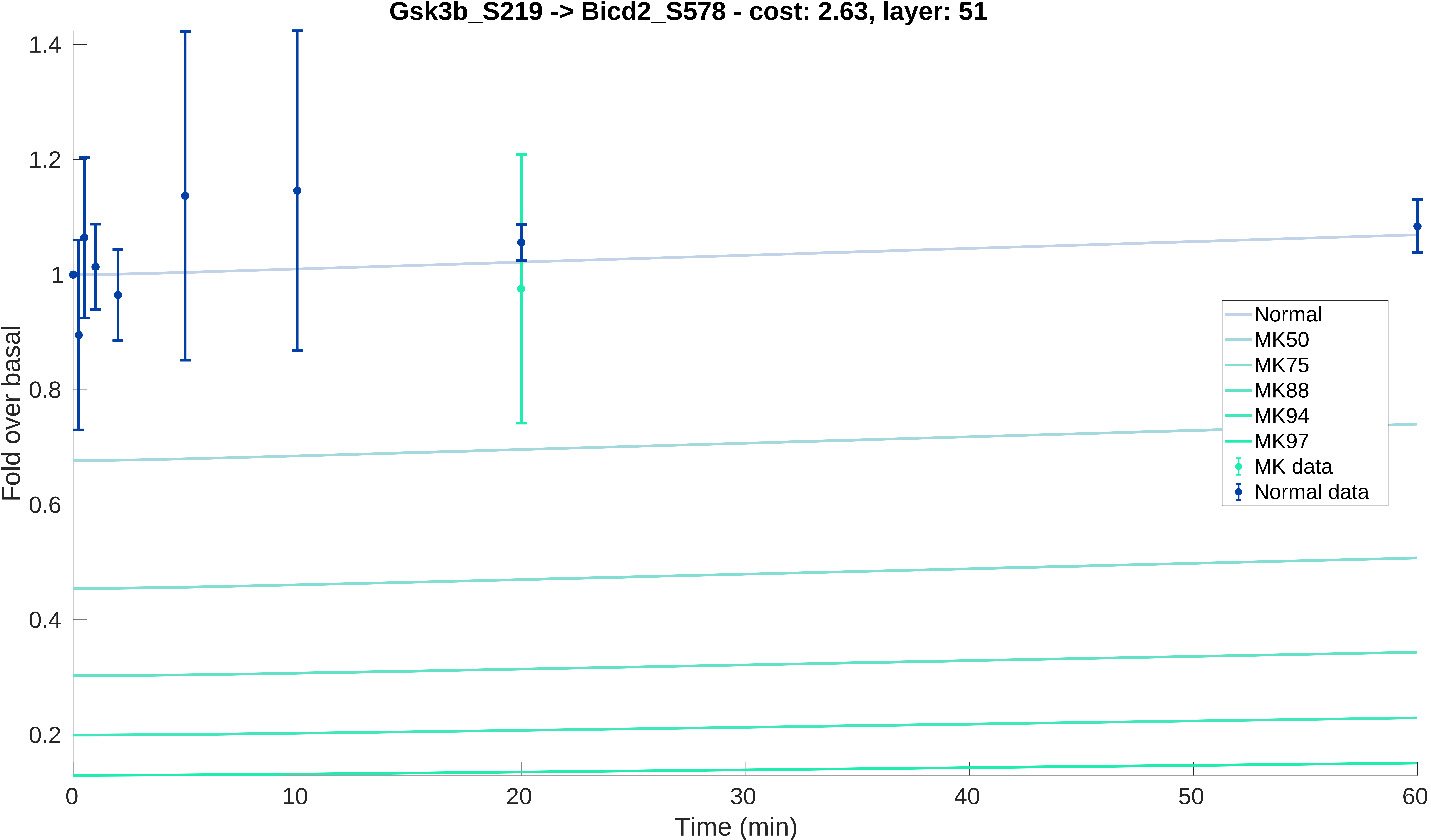

Gsk3b_S219 -> Bicd2_S578 - cost: 2.63, layer: 51
Fold over basal
Time (min)
0
10
20
30
40
50
60
0.2
0.4
0.6
0.8
1
1.2
1.4
Normal
MK50
MK75
MK88
MK94
MK97
MK data
Normal data

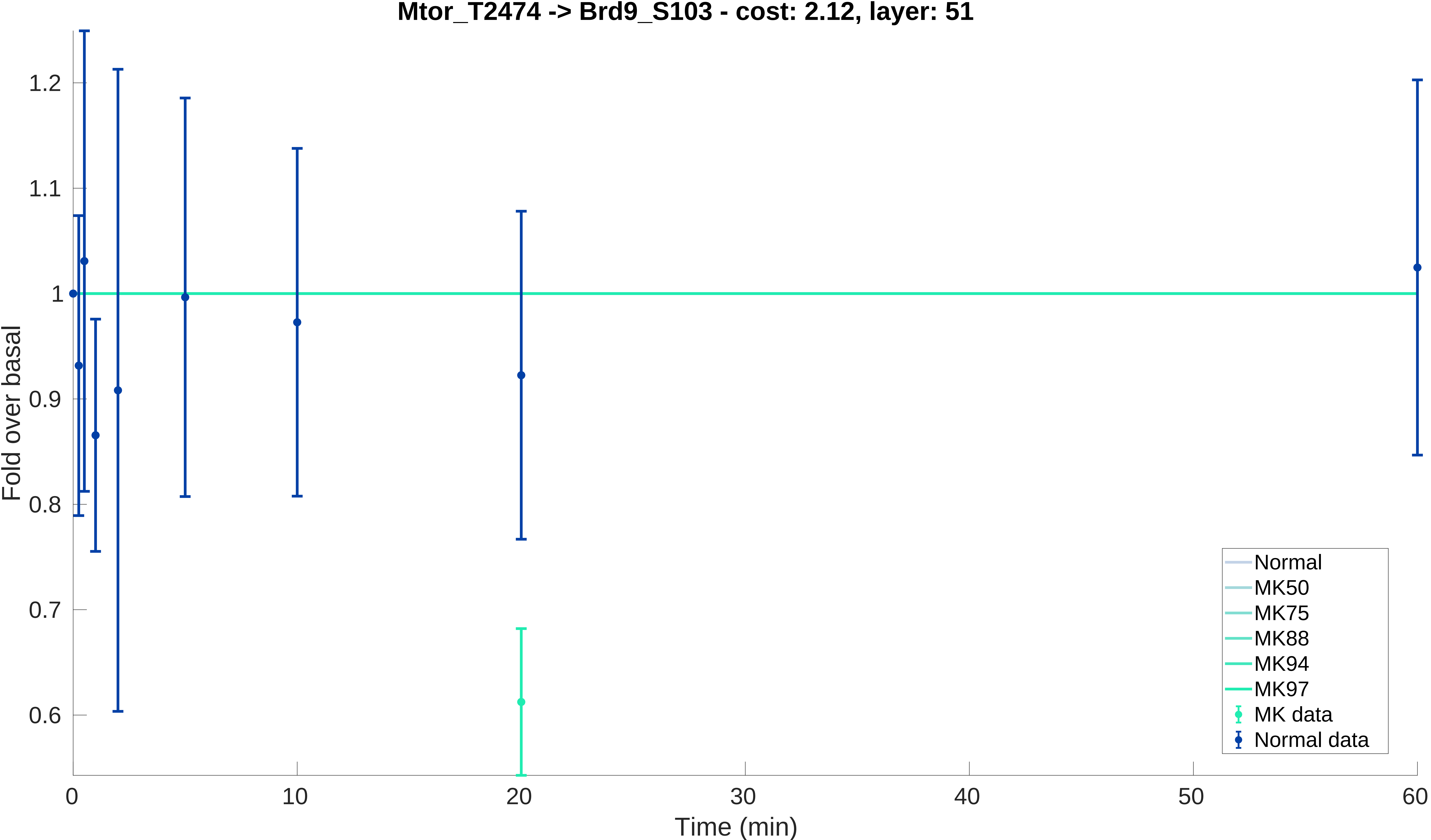

Mtor_T2474 -> Brd9_S103 - cost: 2.12, layer: 51
Fold over basal
1.2
1.1
1
0.9
0.8
0.7
0.6
0
10
20
30
40
50
60
Time (min)
Normal
MK50
MK75
MK88
MK94
MK97
MK data
Normal data

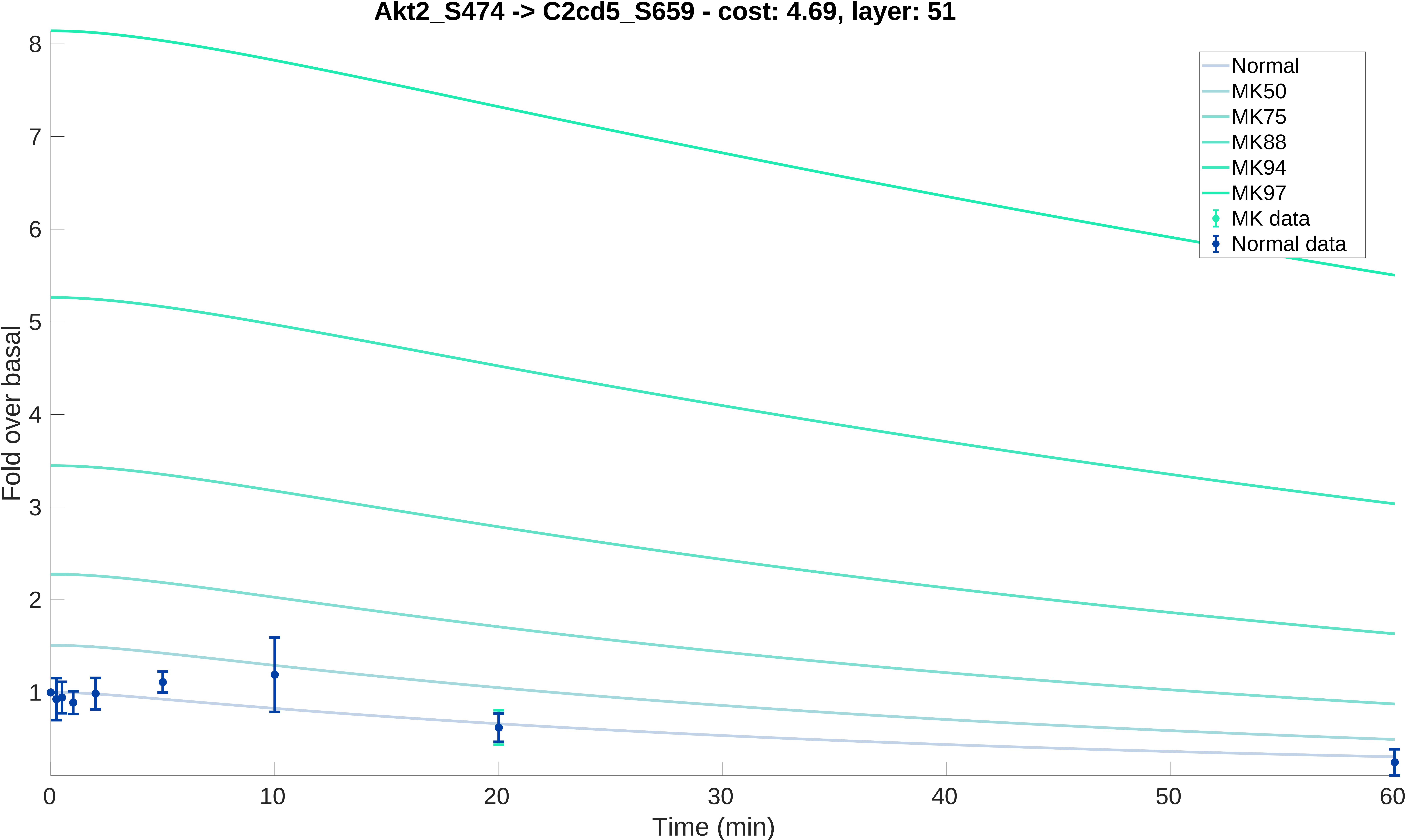

Akt2_S474 -> C2cd5_S659 - cost: 4.69, layer: 51
Fold over basal
Time (min)
0
10
20
30
40
50
60
1
2
3
4
5
6
7
8
Normal
MK50
MK75
MK88
MK94
MK97
MK data
Normal data

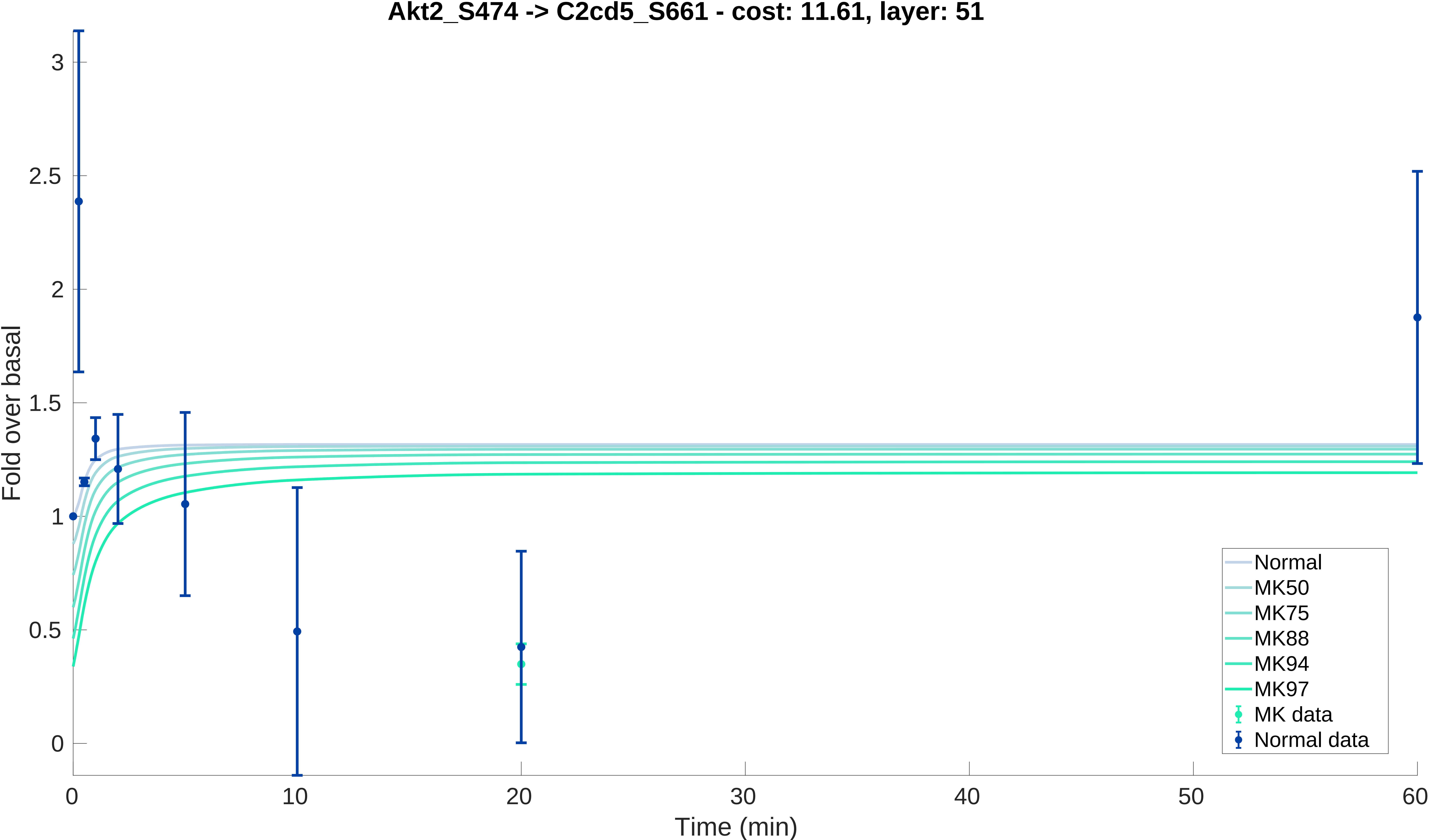

Akt2_S474 -> C2cd5_S661 - cost: 11.61, layer: 51
Fold over basal
Time (min)
0
0.5
1
1.5
2
2.5
3
0
10
20
30
40
50
60
Normal
MK50
MK75
MK88
MK94
MK97
MK data
Normal data

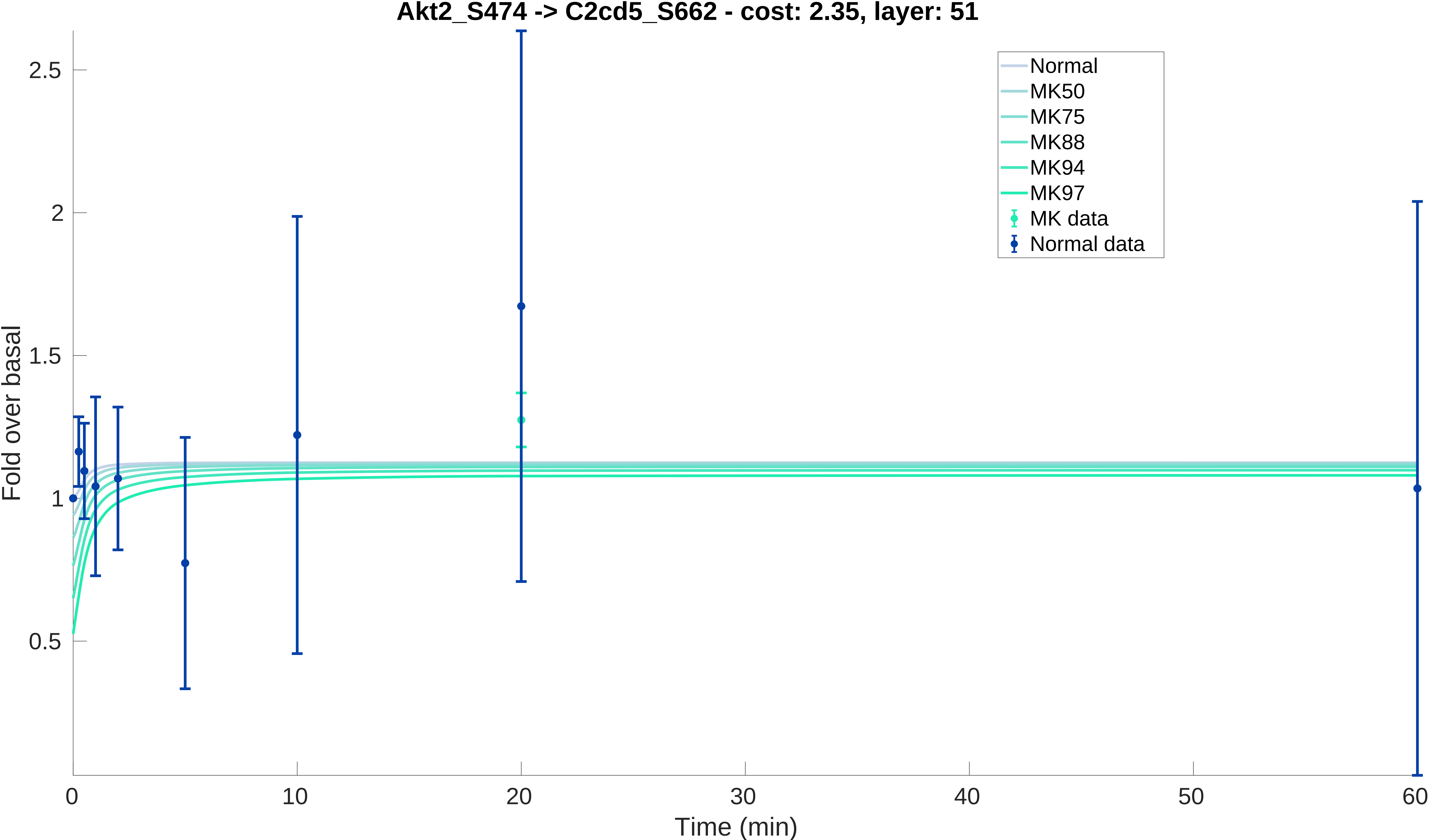

Akt2_S474 -> C2cd5_S662 - cost: 2.35, layer: 51
Normal
MK50
MK75
MK88
MK94
MK97
MK data
Normal data
Fold over basal
0.5
1
1.5
2
2.5
0
10
20
30
40
50
60
Time (min)

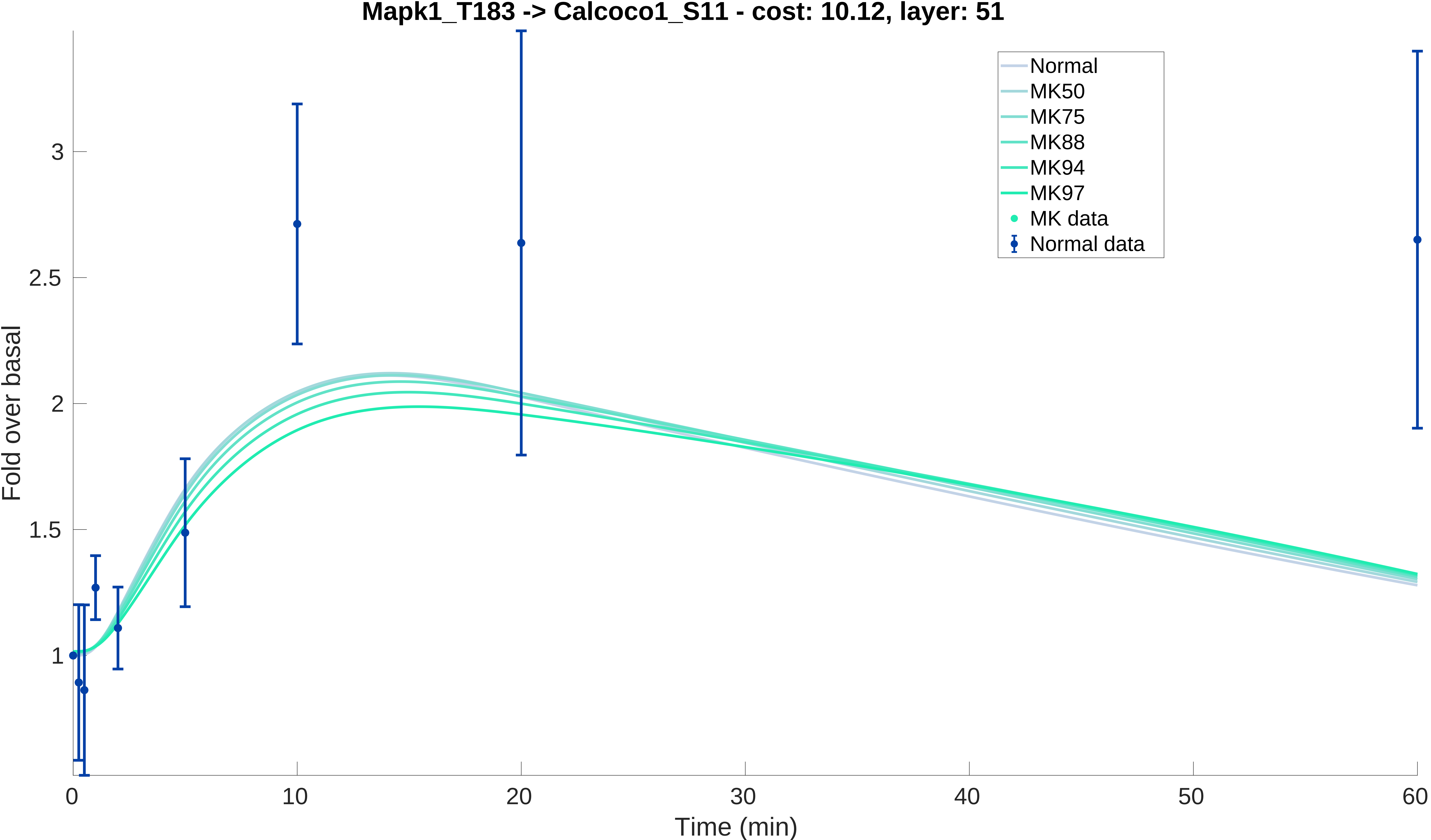

Mapk1_T183 -> Calcoco1_S11 - cost: 10.12, layer: 51
Normal
MK50
MK75
MK88
MK94
MK97
MK data
Normal data
Fold over basal
3
2.5
2
1.5
1
0
10
20
30
40
50
60
Time (min)

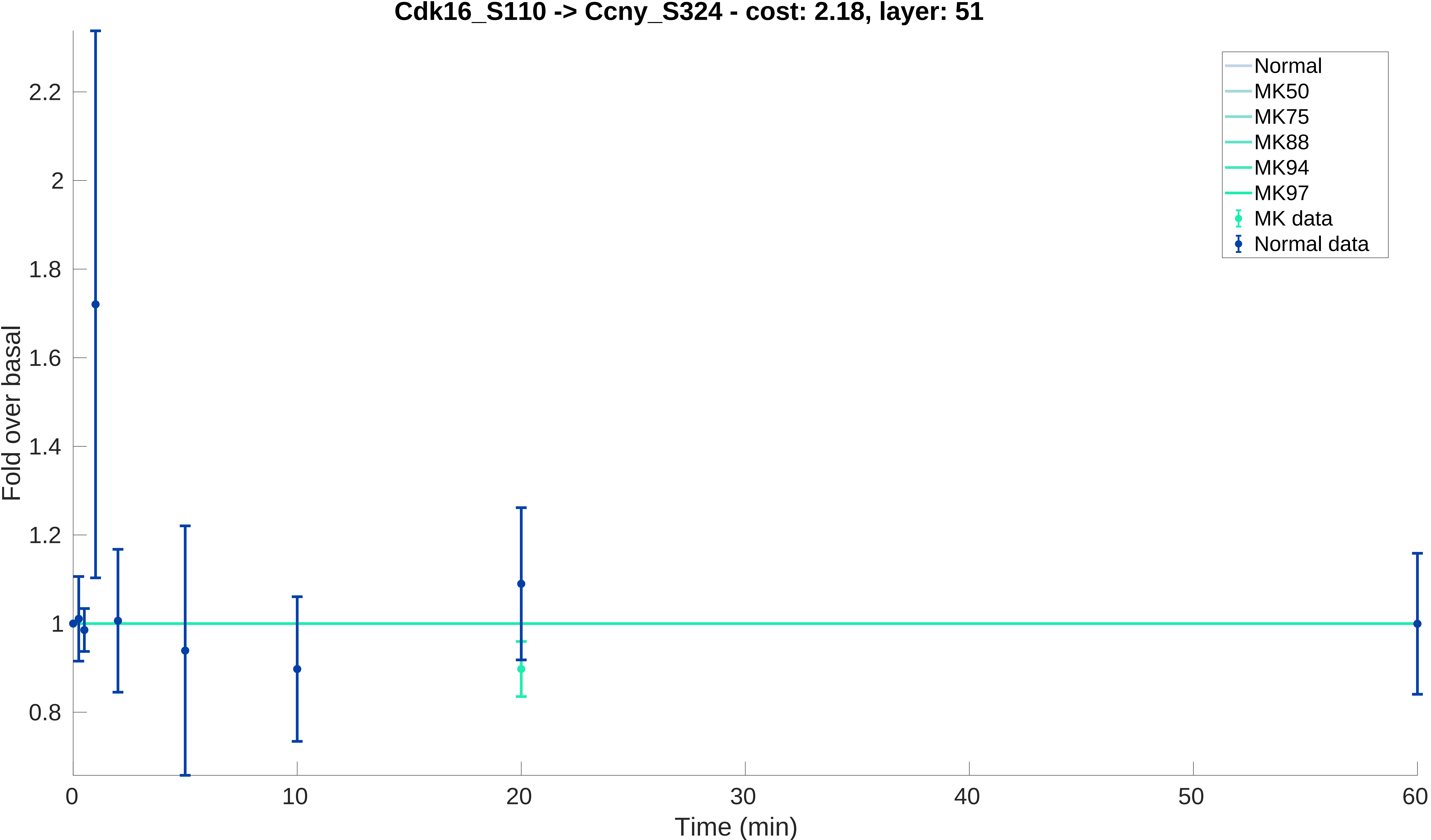

Cdk16_S110 -> Ccny_S324 - cost: 2.18, layer: 51
Fold over basal
2.2
2
1.8
1.6
1.4
1.2
1
0.8
0
10
20
30
40
50
60
Time (min)
Normal
MK50
MK75
MK88
MK94
MK97
MK data
Normal data

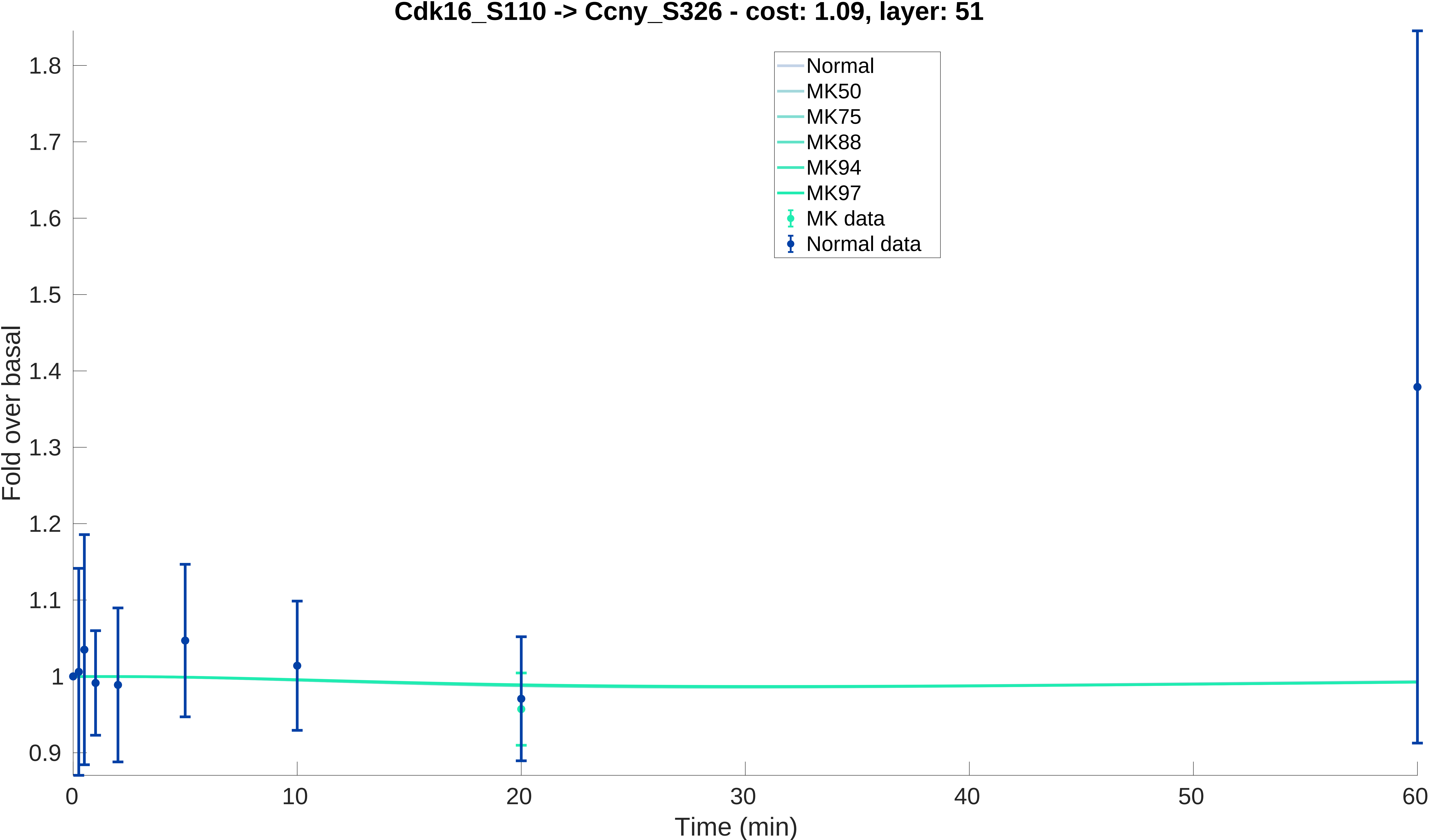
Cdk16_S110 -> Ccny_S326 - cost: 1.09, layer: 51
Normal
MK50
MK75
MK88
MK94
MK97
MK data
Normal data
Fold over basal
Time (min)
0.9
1
1.1
1.2
1.3
1.4
1.5
1.6
1.7
1.8
0
10
20
30
40
50
60

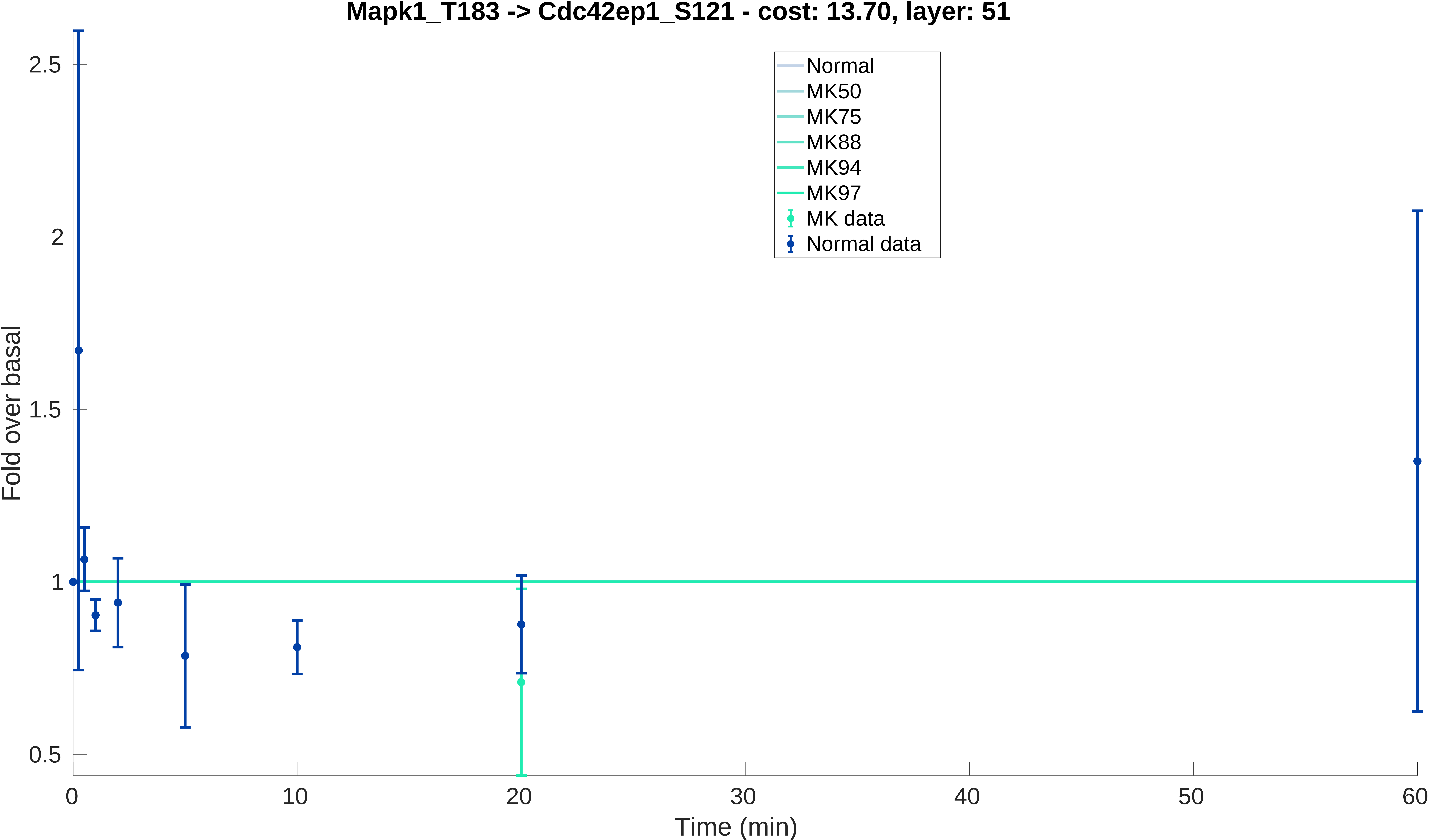

Mapk1_T183 -> Cdc42ep1_S121 - cost: 13.70, layer: 51
Normal
MK50
MK75
MK88
MK94
MK97
MK data
Normal data
Fold over basal
2.5
2
1.5
1
0.5
0
10
20
30
40
50
60
Time (min)

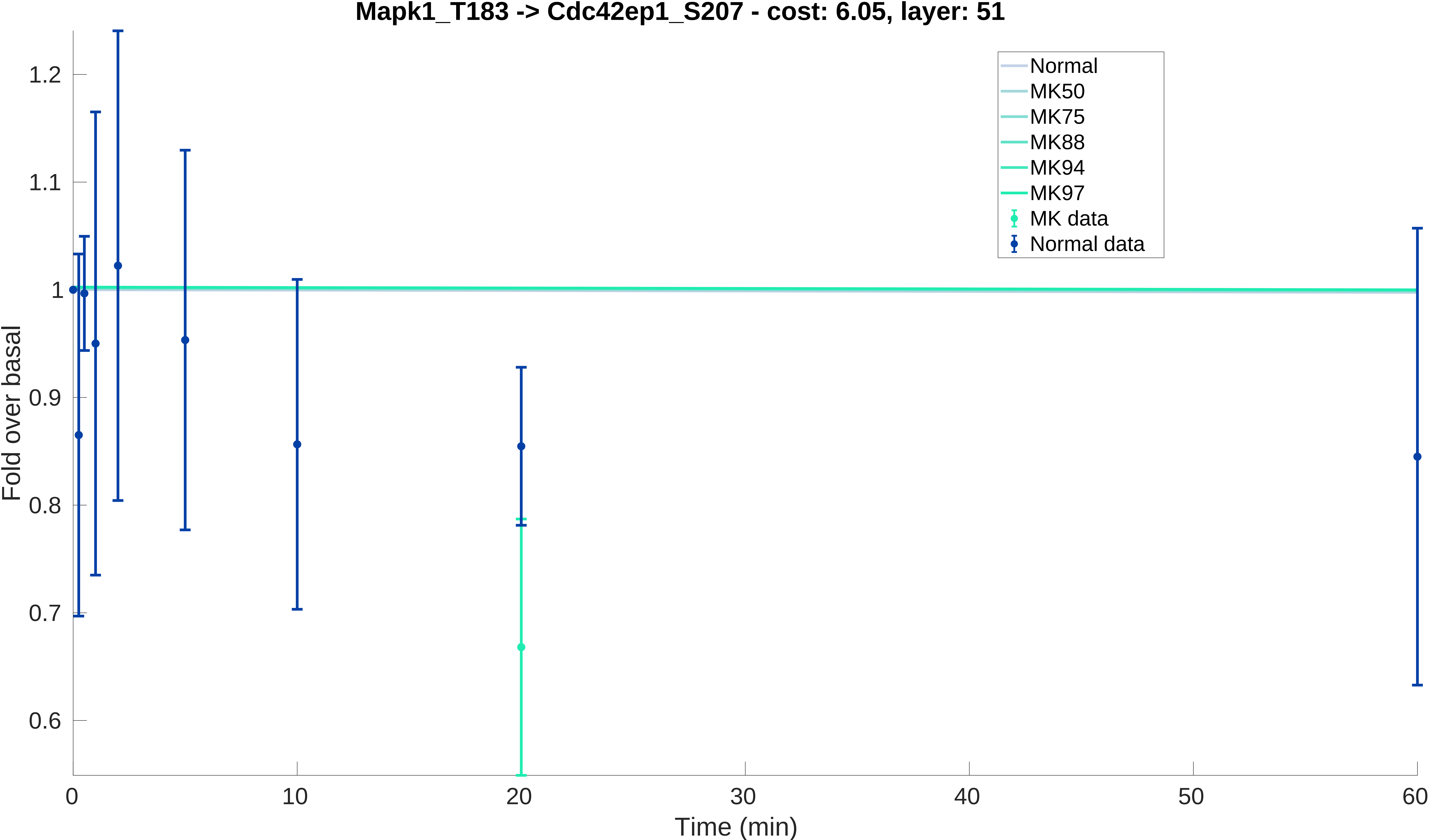

Mapk1_T183 -> Cdc42ep1_S207 - cost: 6.05, layer: 51
Normal
MK50
MK75
MK88
MK94
MK97
MK data
Normal data
Fold over basal
1.2
1.1
1
0.9
0.8
0.7
0.6
0
10
20
30
40
50
60
Time (min)

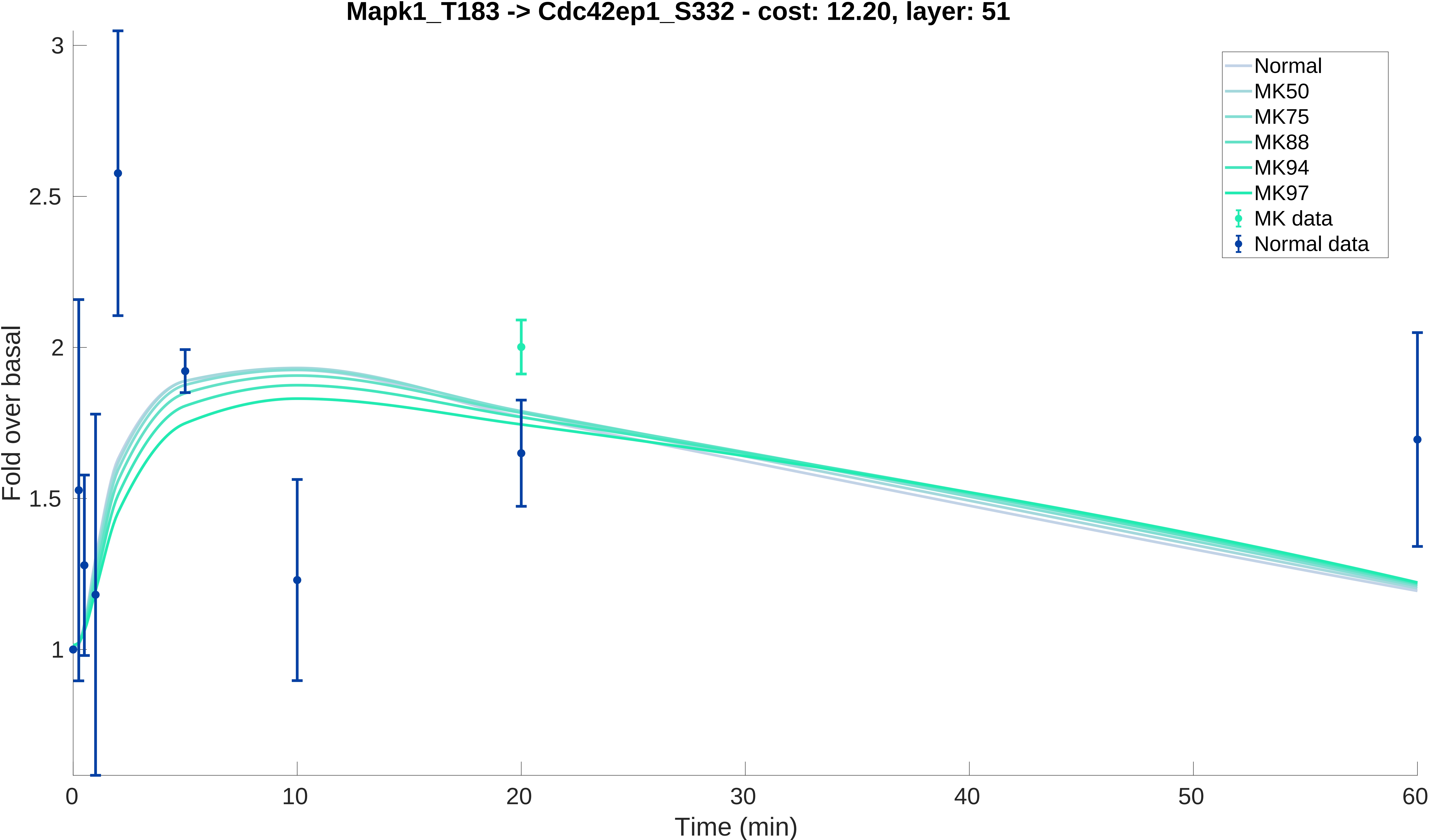

Mapk1_T183 -> Cdc42ep1_S332 - cost: 12.20, layer: 51
Fold over basal
Time (min)
Normal
MK50
MK75
MK88
MK94
MK97
MK data
Normal data
3
2.5
2
1.5
1
0
10
20
30
40
50
60

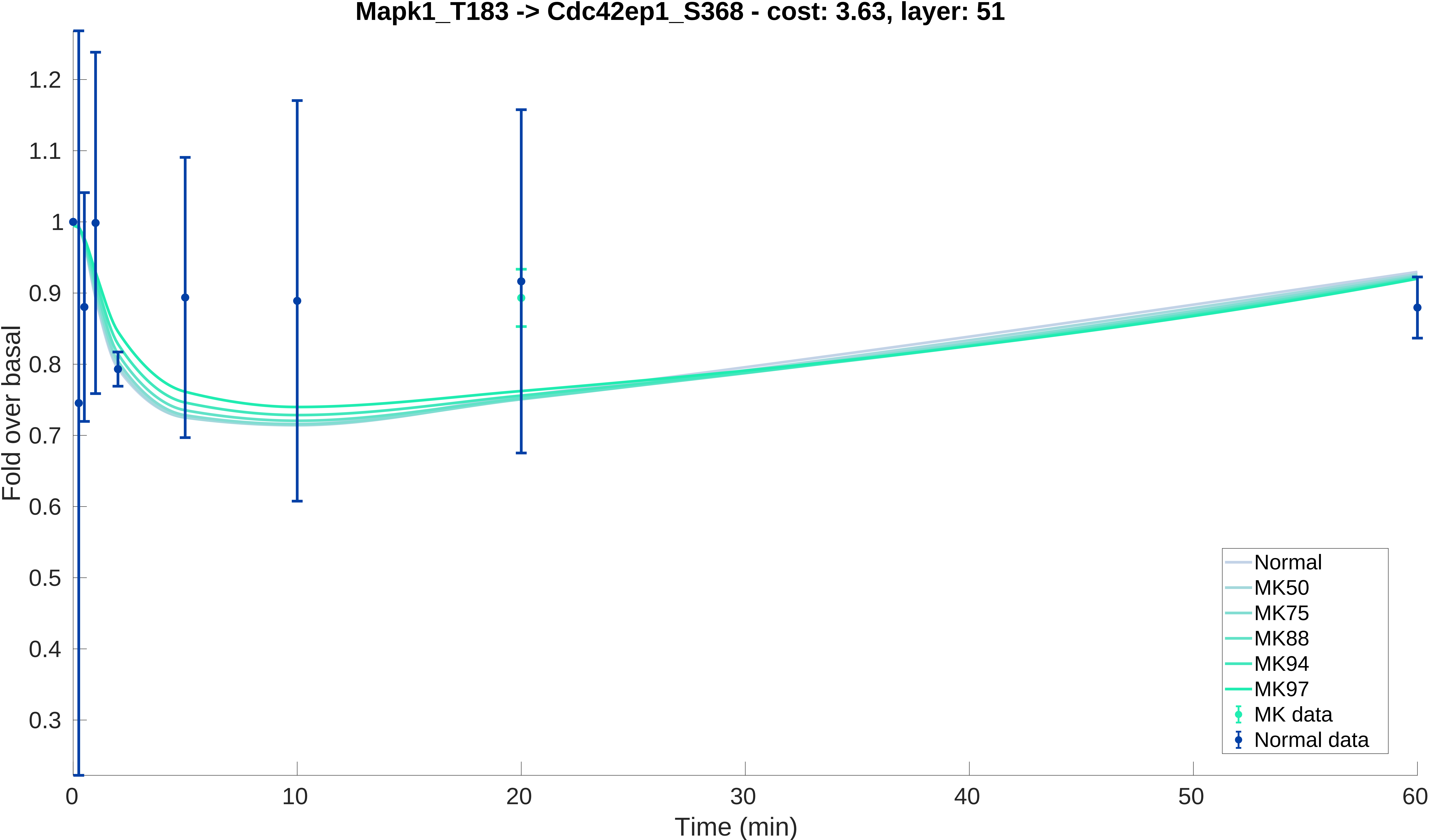

Mapk1_T183 -> Cdc42ep1_S368 - cost: 3.63, layer: 51
Fold over basal
1.2
1.1
1
0.9
0.8
0.7
0.6
0.5
0.4
0.3
0
10
20
30
40
50
60
Time (min)
Normal
MK50
MK75
MK88
MK94
MK97
MK data
Normal data

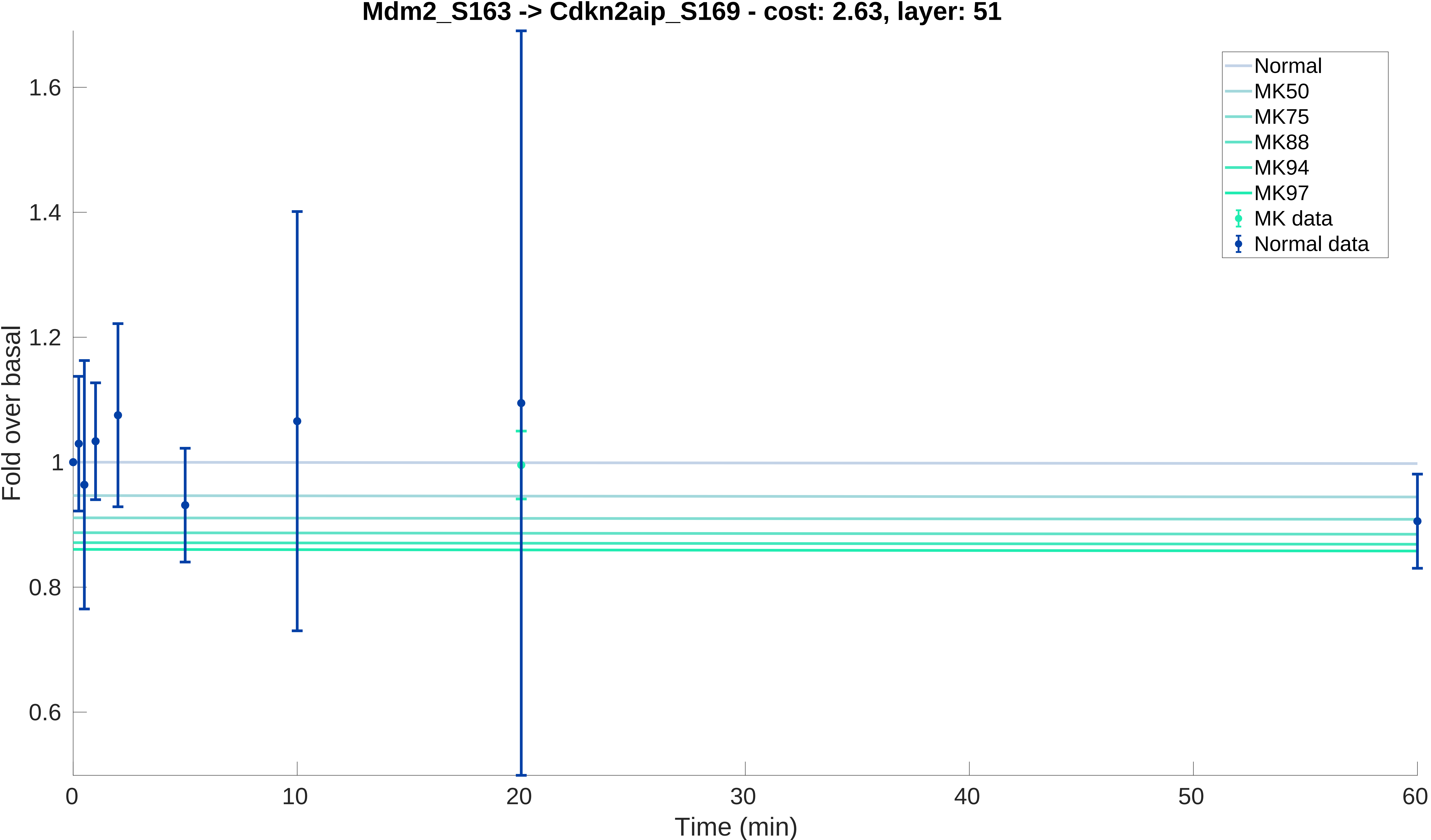
Mdm2_S163 -> Cdkn2aip_S169 - cost: 2.63, layer: 51
Fold over basal
Time (min)
Normal
MK50
MK75
MK88
MK94
MK97
MK data
Normal data
1.6
1.4
1.2
1
0.8
0.6
0
10
20
30
40
50
60

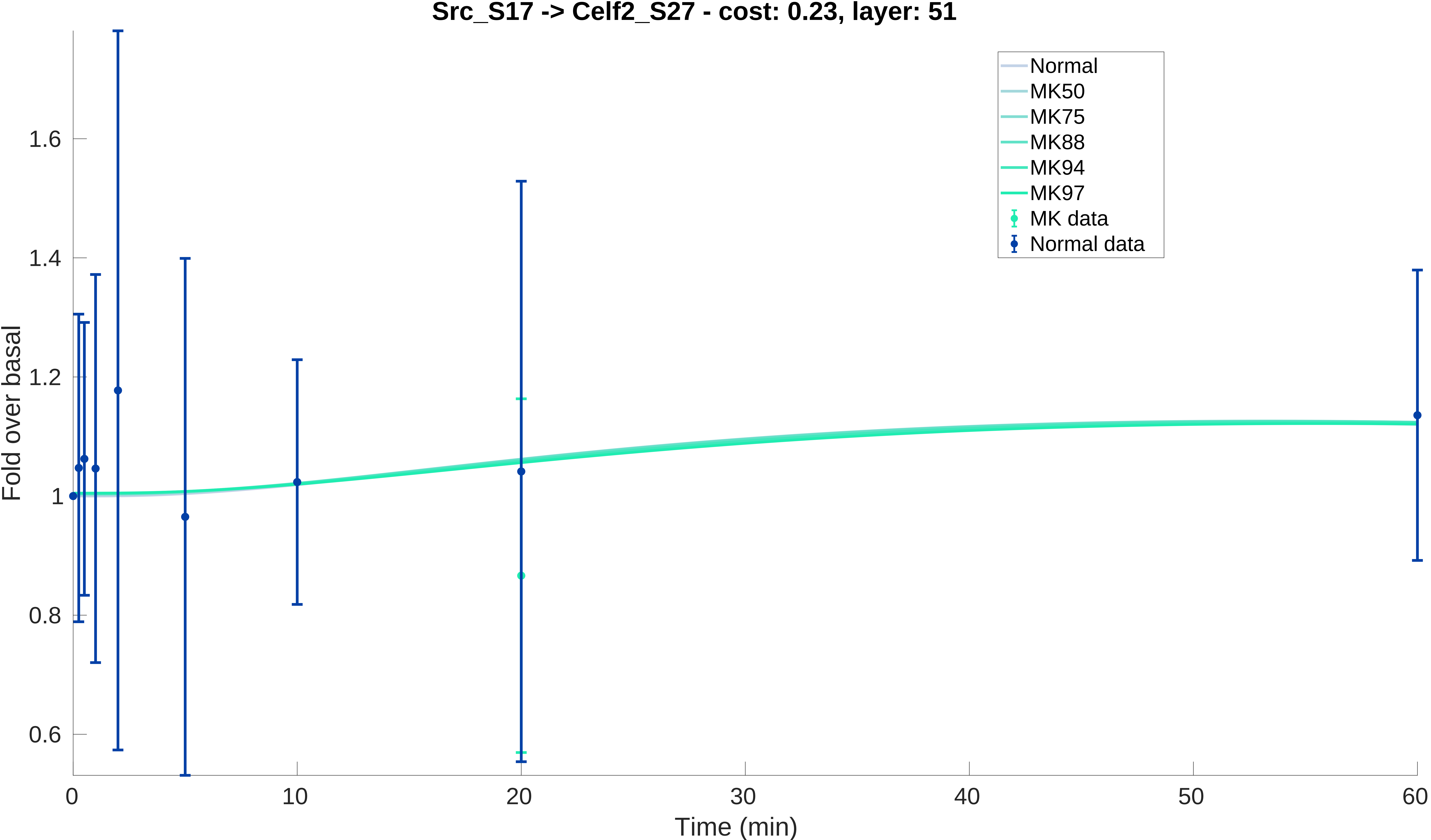

Src_S17 -> Celf2_S27 - cost: 0.23, layer: 51
Normal
MK50
MK75
MK88
MK94
MK97
MK data
Normal data
Fold over basal
1.6
1.4
1.2
1
0.8
0.6
0
10
20
30
40
50
60
Time (min)

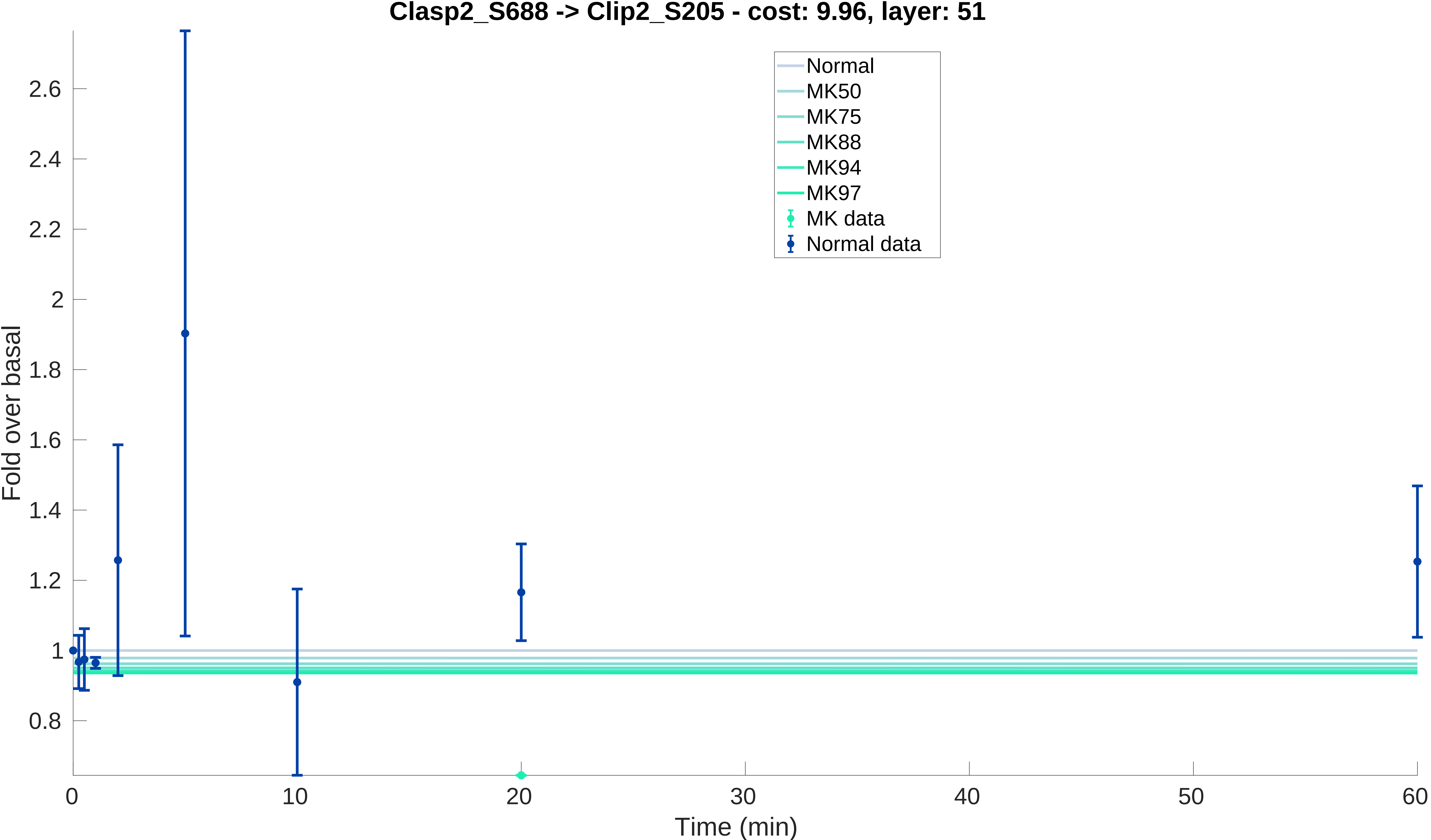

Clasp2_S688 -> Clip2_S205 - cost: 9.96, layer: 51
Normal
MK50
MK75
MK88
MK94
MK97
MK data
Normal data
Fold over basal
2.6
2.4
2.2
2
1.8
1.6
1.4
1.2
1
0.8
0
10
20
30
40
50
60
Time (min)

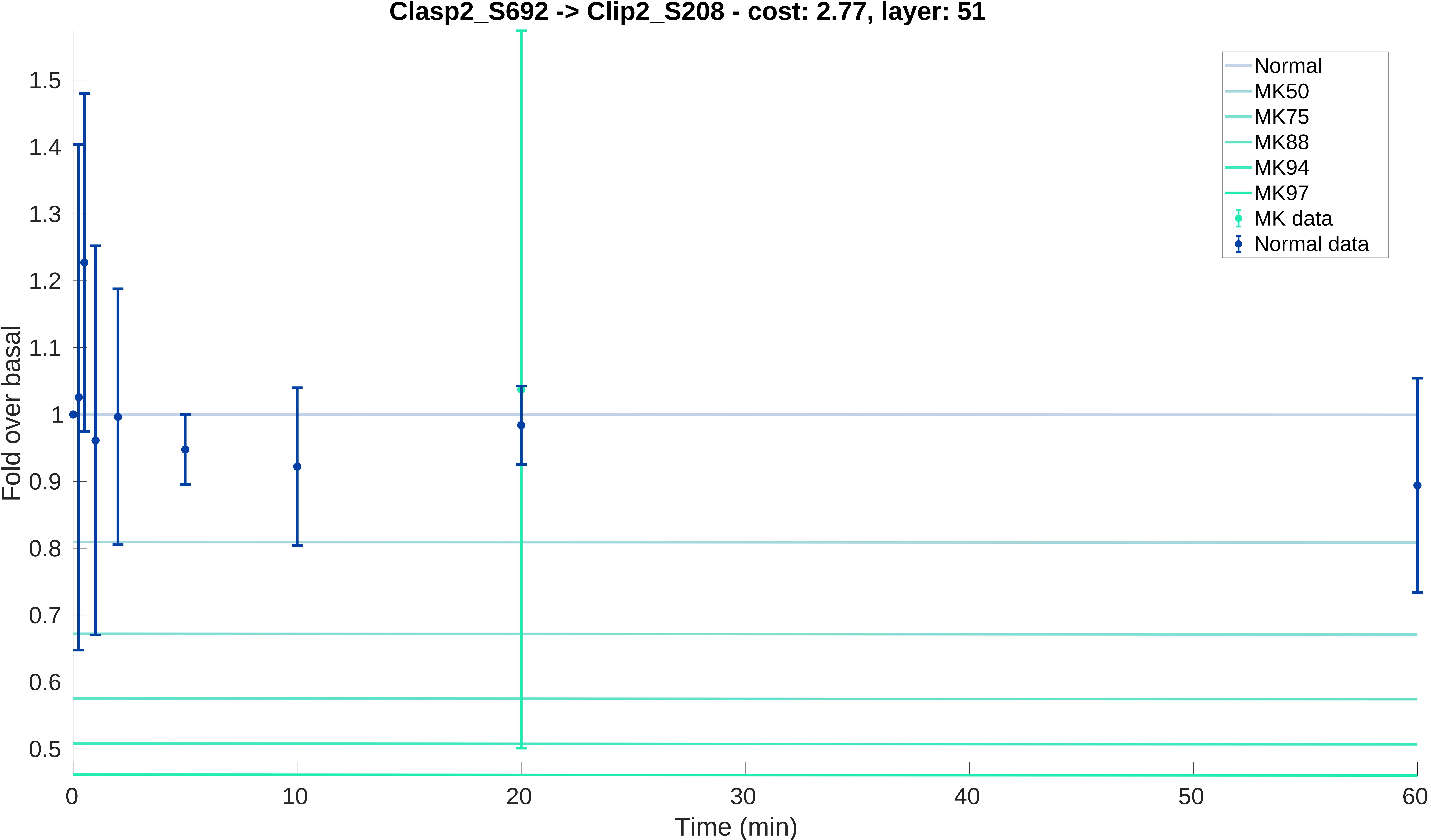

Clasp2_S692 -> Clip2_S208 - cost: 2.77, layer: 51
Fold over basal
Time (min)
1.5
1.4
1.3
1.2
1.1
1
0.9
0.8
0.7
0.6
0.5
0
10
20
30
40
50
60
Normal
MK50
MK75
MK88
MK94
MK97
MK data
Normal data

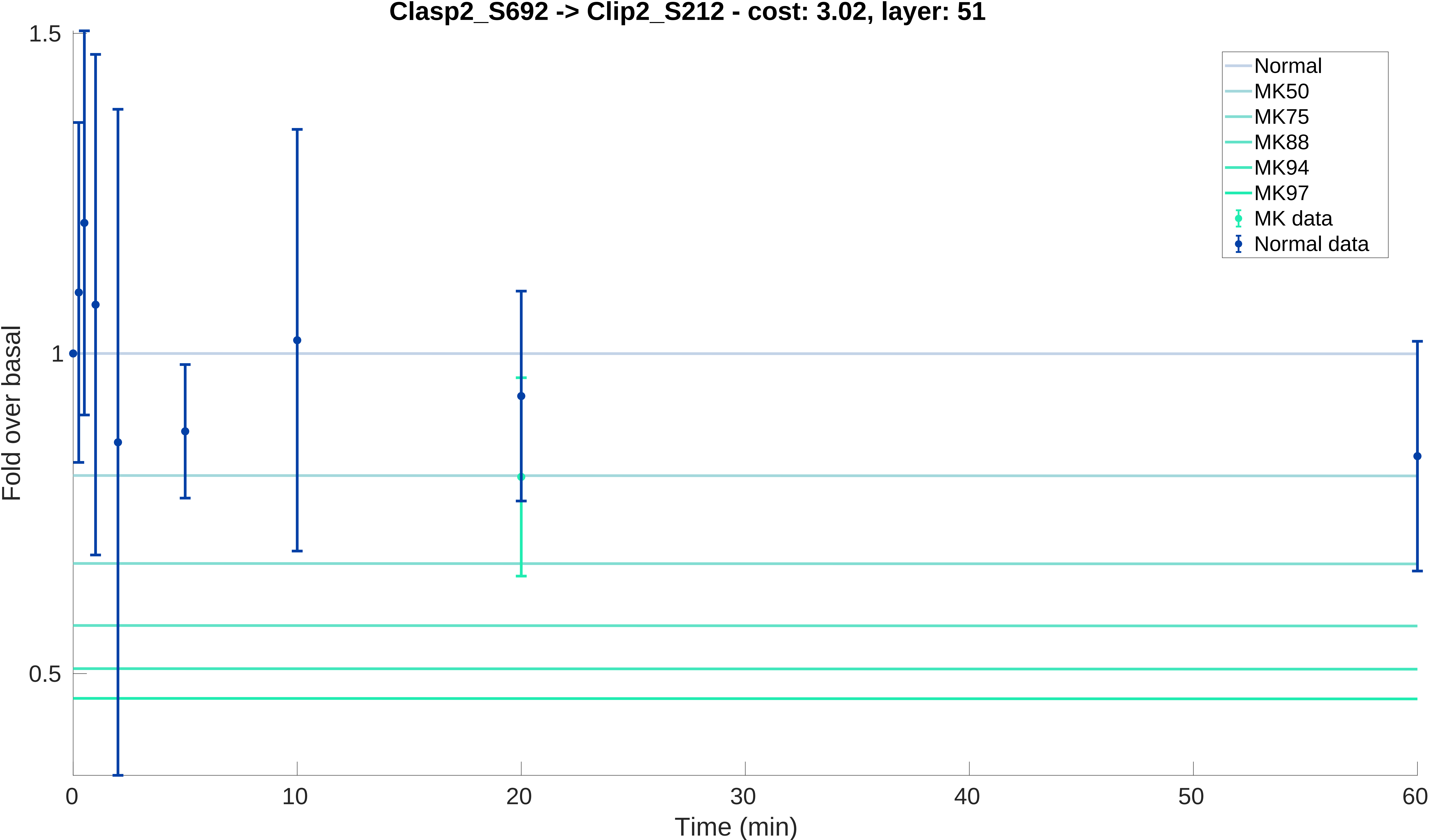

Clasp2_S692 -> Clip2_S212 - cost: 3.02, layer: 51
Fold over basal
1.5
1
0.5
Time (min)
0
10
20
30
40
50
60
Normal
MK50
MK75
MK88
MK94
MK97
MK data
Normal data

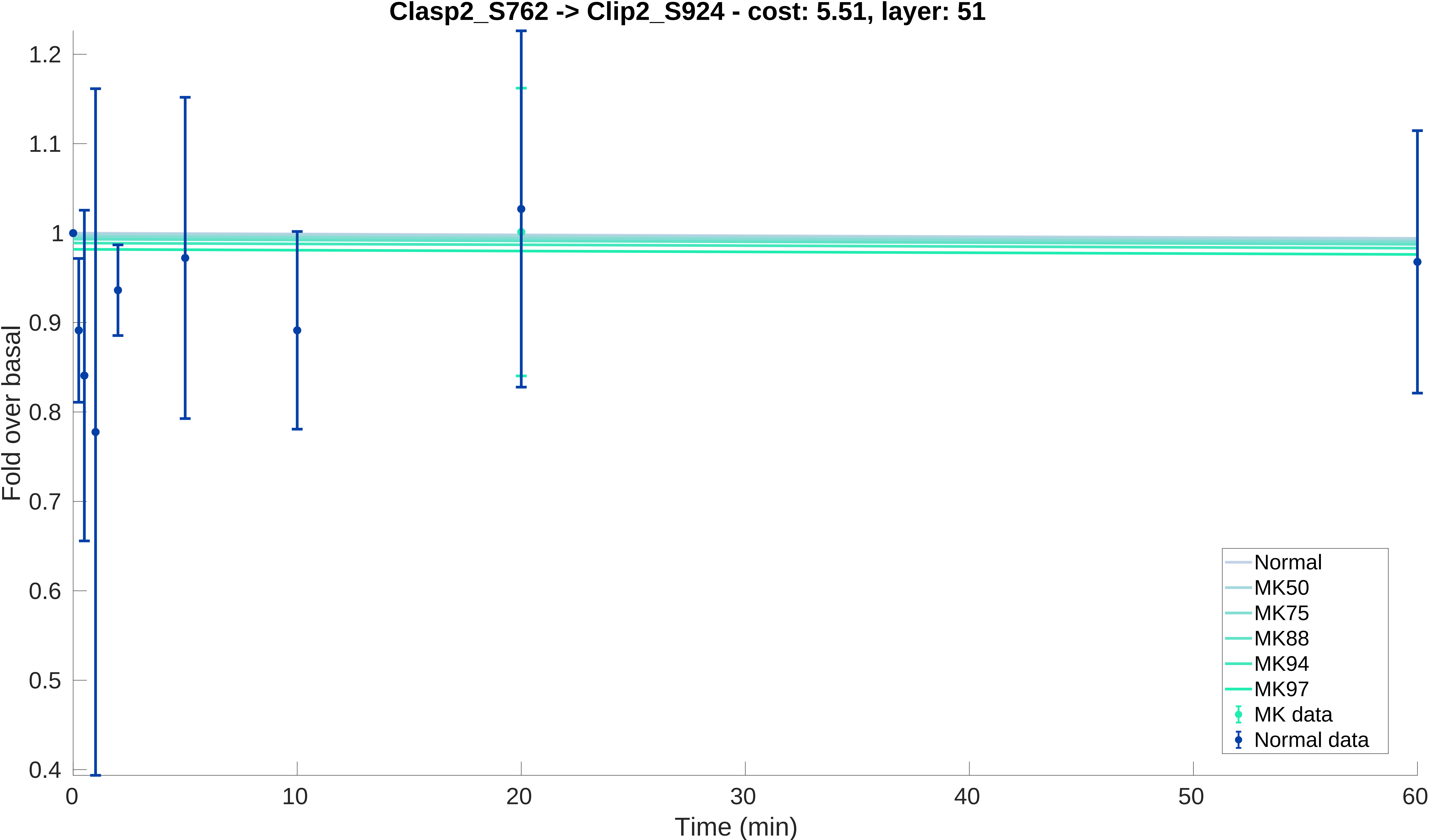
Clasp2_S762 -> Clip2_S924 - cost: 5.51, layer: 51
Fold over basal
1.2
1.1
1
0.9
0.8
0.7
0.6
0.5
0.4
0
10
20
30
40
50
60
Time (min)
Normal
MK50
MK75
MK88
MK94
MK97
MK data
Normal data

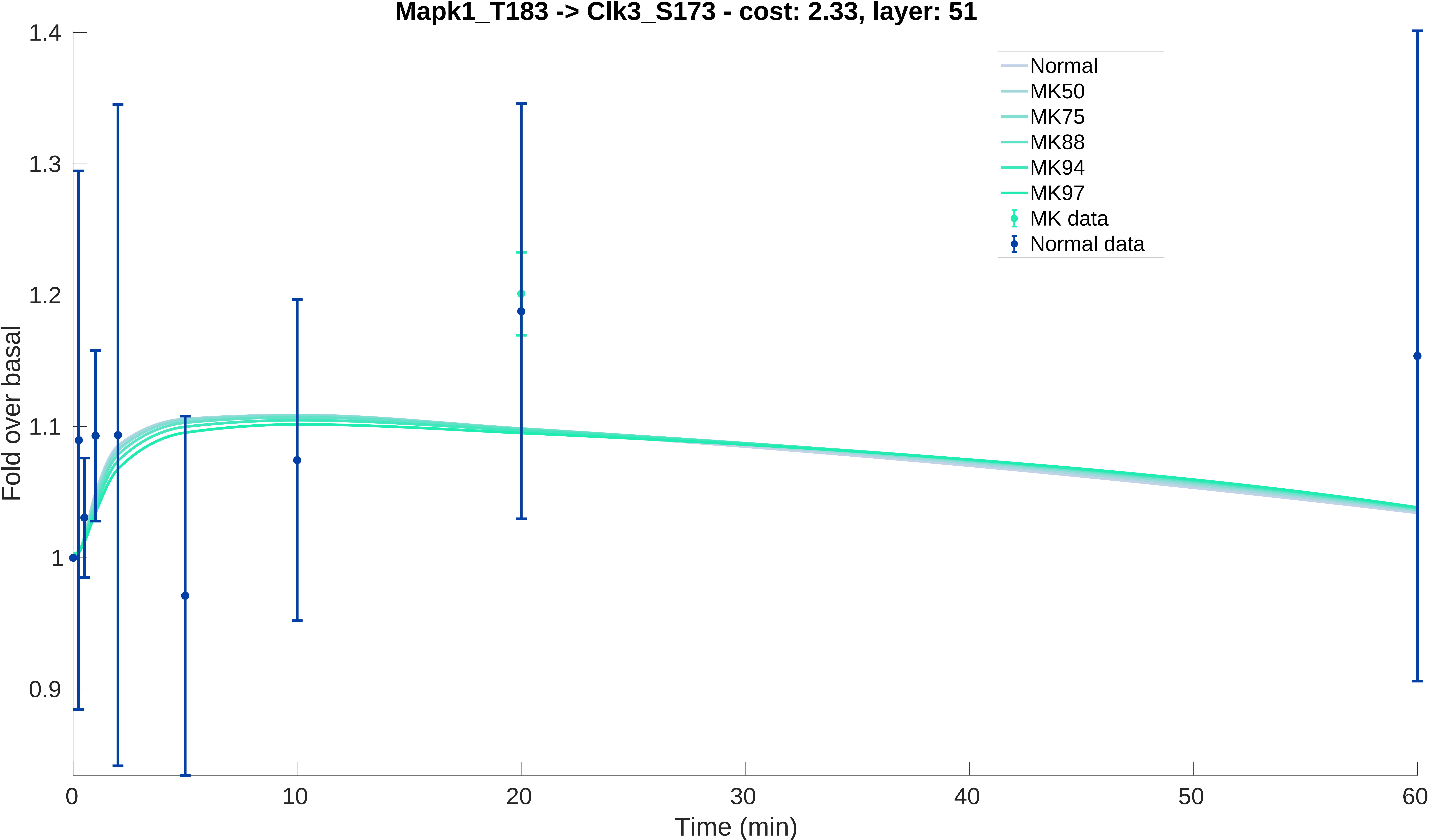
Mapk1_T183 -> Clk3_S173 - cost: 2.33, layer: 51
Fold over basal
1.4
1.3
1.2
1.1
1
0.9
0
10
20
30
40
50
60
Time (min)
Normal
MK50
MK75
MK88
MK94
MK97
MK data
Normal data

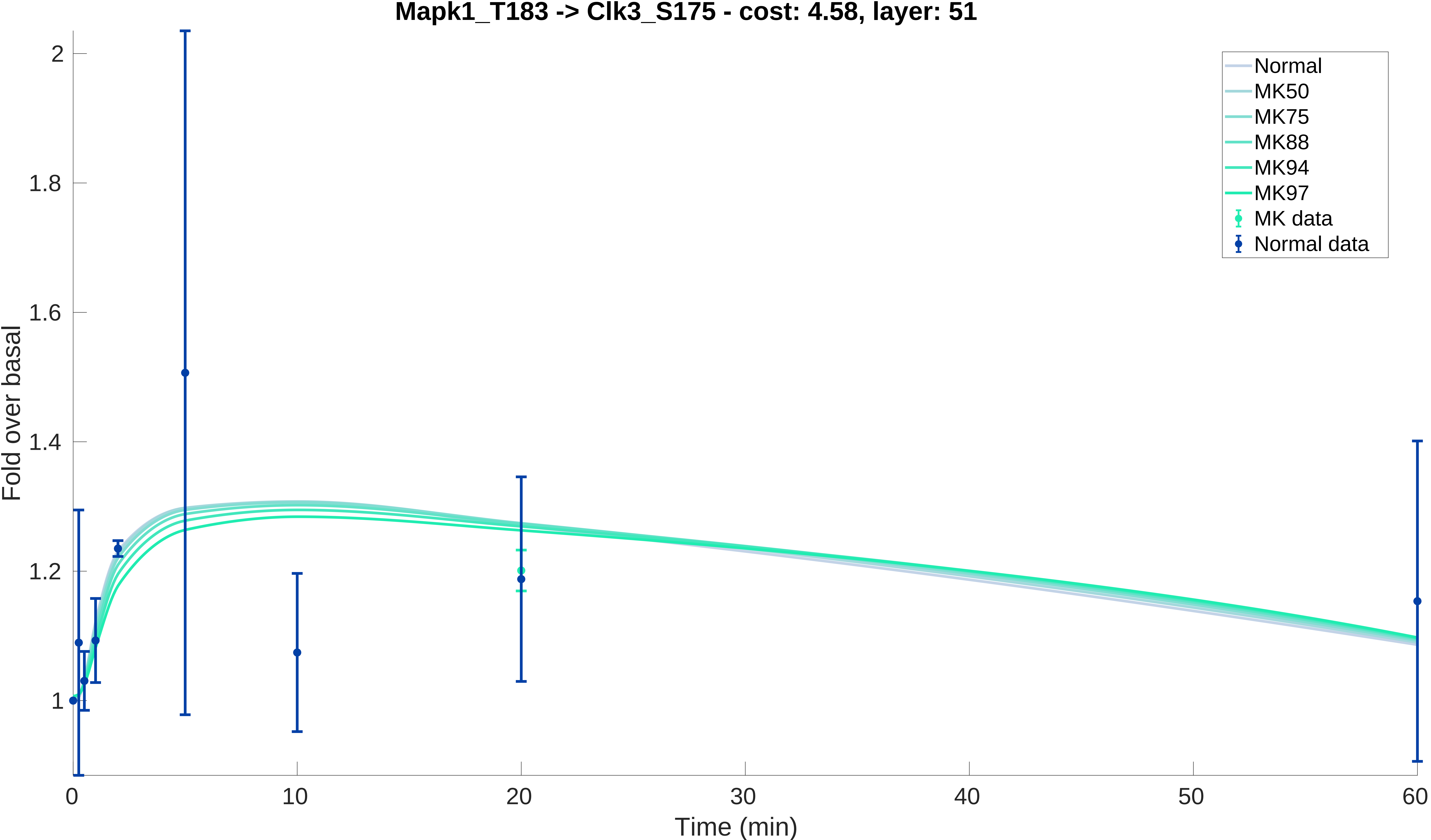
Mapk1_T183 -> Clk3_S175 - cost: 4.58, layer: 51
Fold over basal
Time (min)
2
1.8
1.6
1.4
1.2
1
0
10
20
30
40
50
60
Normal
MK50
MK75
MK88
MK94
MK97
MK data
Normal data

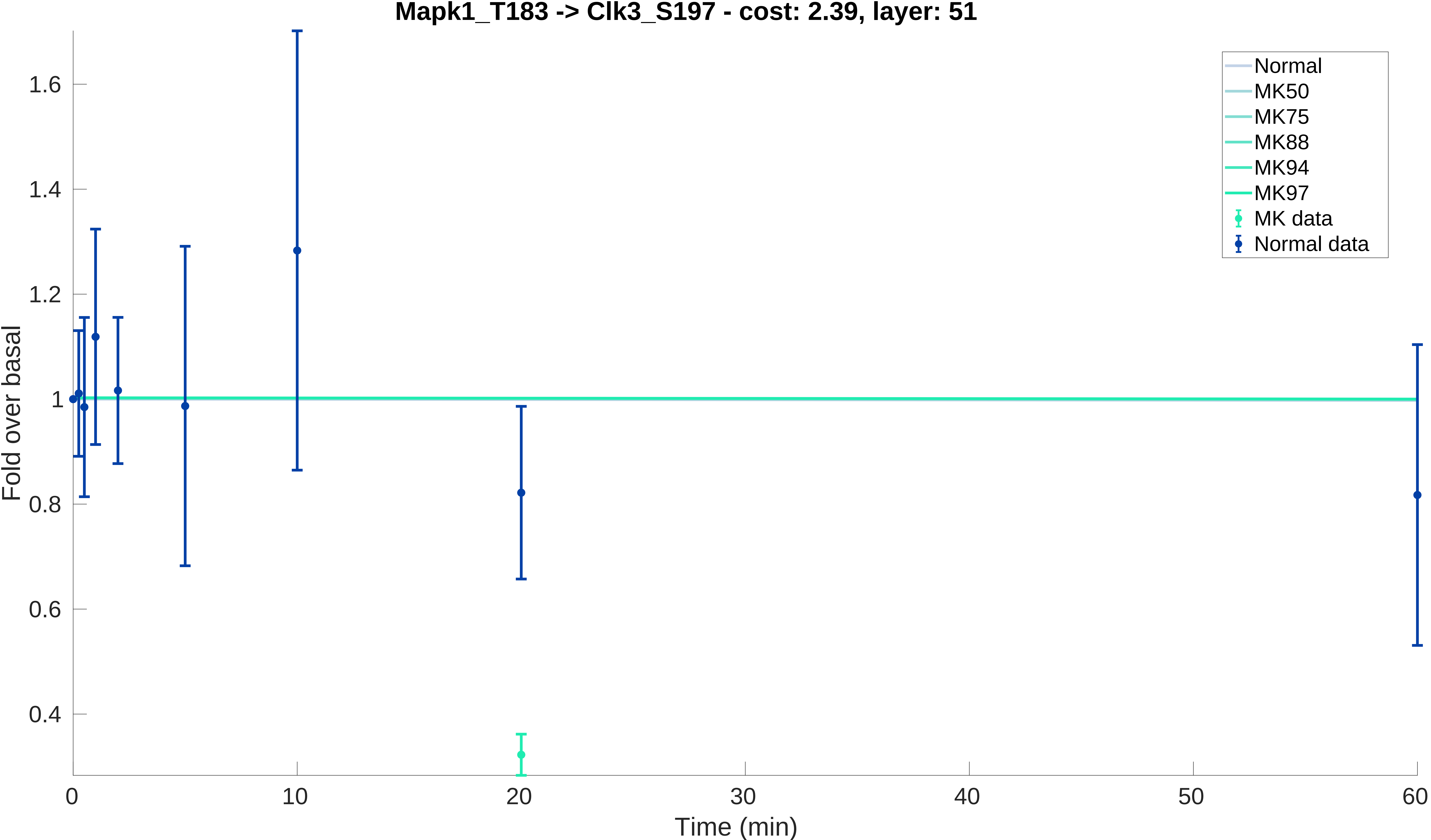

Mapk1_T183 -> Clk3_S197 - cost: 2.39, layer: 51
Fold over basal
Time (min)
0.4
0.6
0.8
1
1.2
1.4
1.6
0
10
20
30
40
50
60
Normal
MK50
MK75
MK88
MK94
MK97
MK data
Normal data

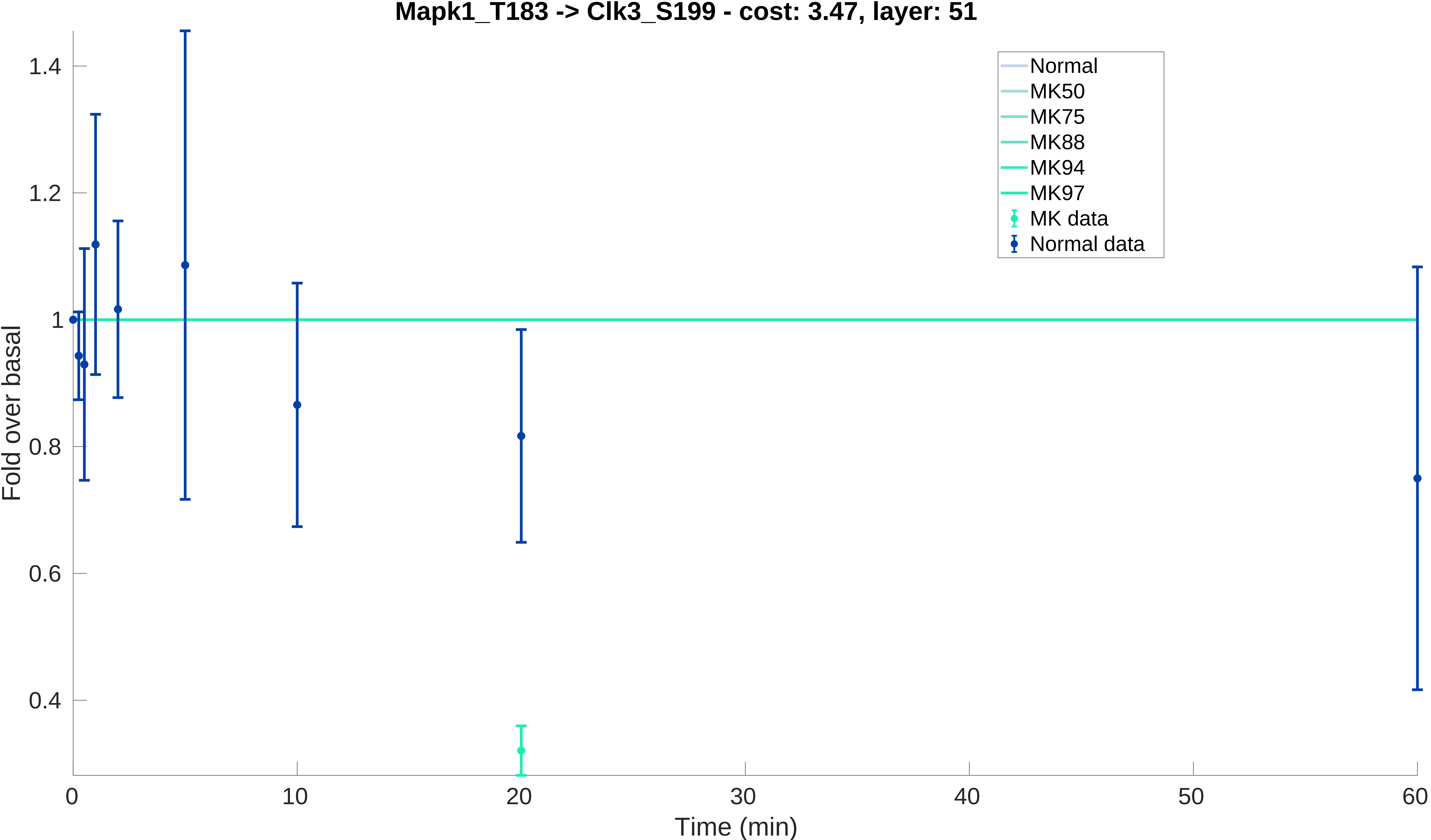

Mapk1_T183 -> Clk3_S199 - cost: 3.47, layer: 51
Fold over basal
Time (min)
0
10
20
30
40
50
60
0.4
0.6
0.8
1
1.2
1.4
Normal
MK50
MK75
MK88
MK94
MK97
MK data
Normal data

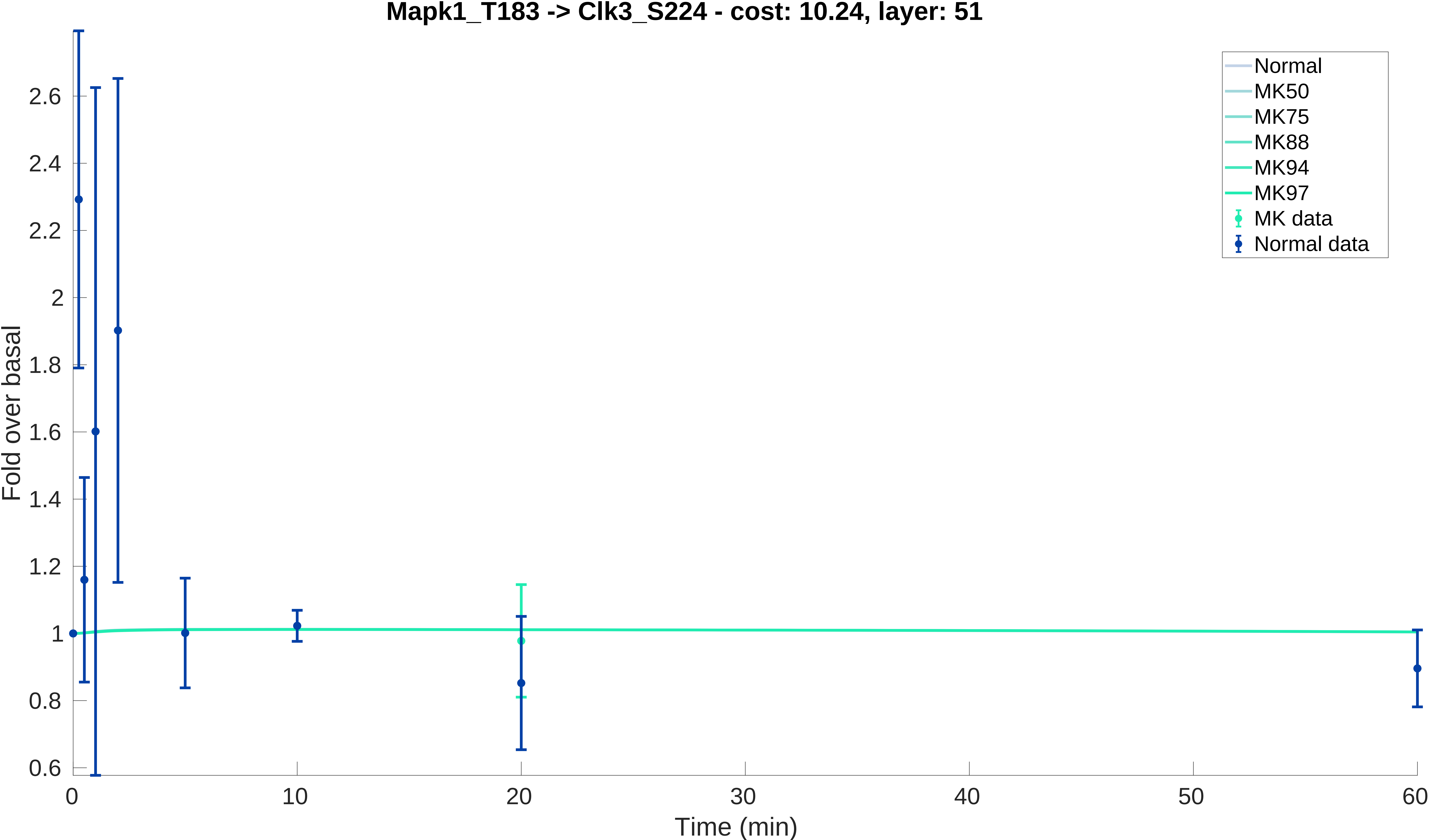

Mapk1_T183 -> Clk3_S224 - cost: 10.24, layer: 51
Fold over basal
Time (min)
Normal
MK50
MK75
MK88
MK94
MK97
MK data
Normal data
2.6
2.4
2.2
2
1.8
1.6
1.4
1.2
1
0.8
0.6
0
10
20
30
40
50
60

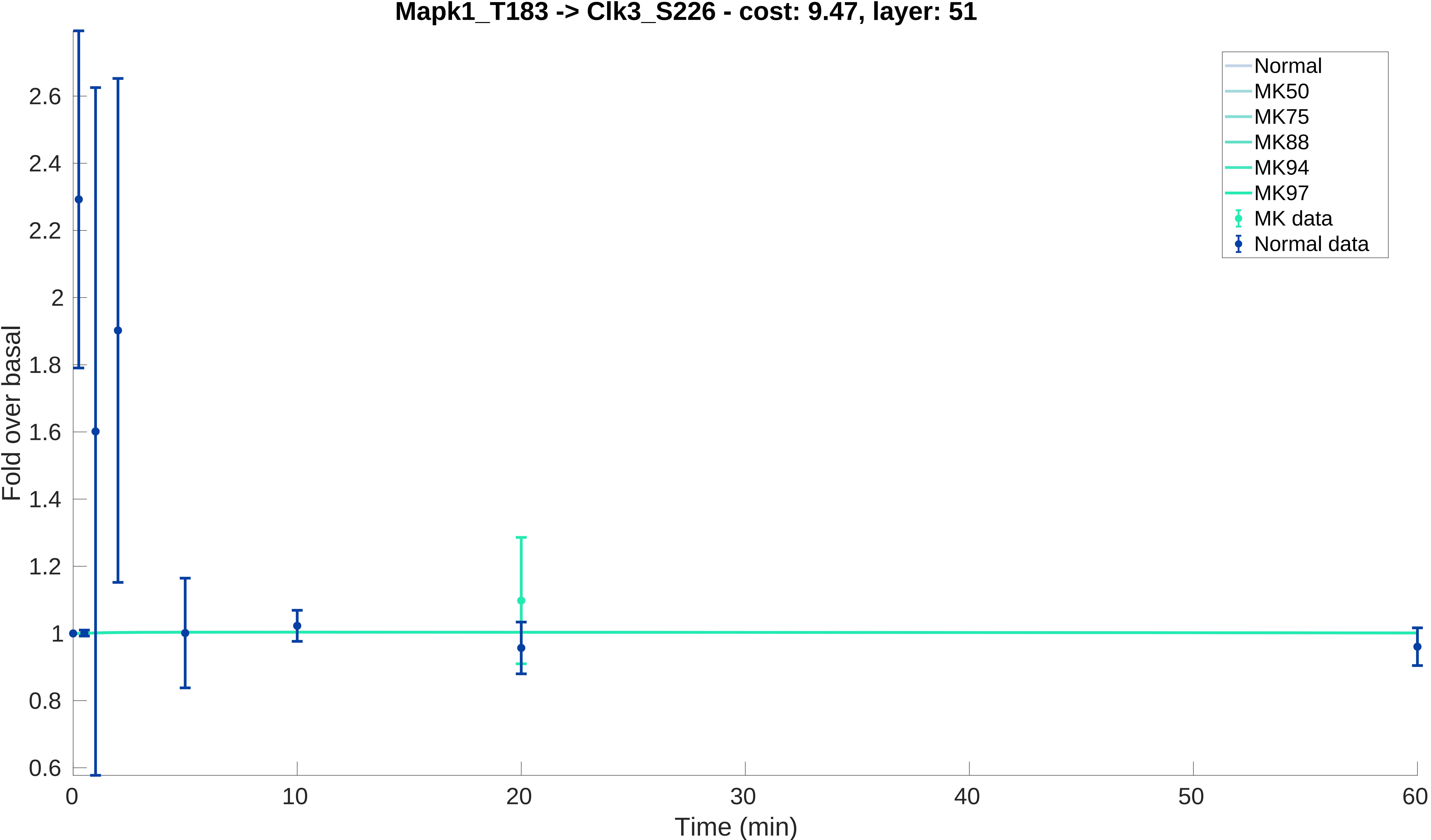

Mapk1_T183 -> Clk3_S226 - cost: 9.47, layer: 51
Normal
MK50
MK75
MK88
MK94
MK97
MK data
Normal data
Fold over basal
Time (min)
0.6
0.8
1
1.2
1.4
1.6
1.8
2
2.2
2.4
2.6
0
10
20
30
40
50
60

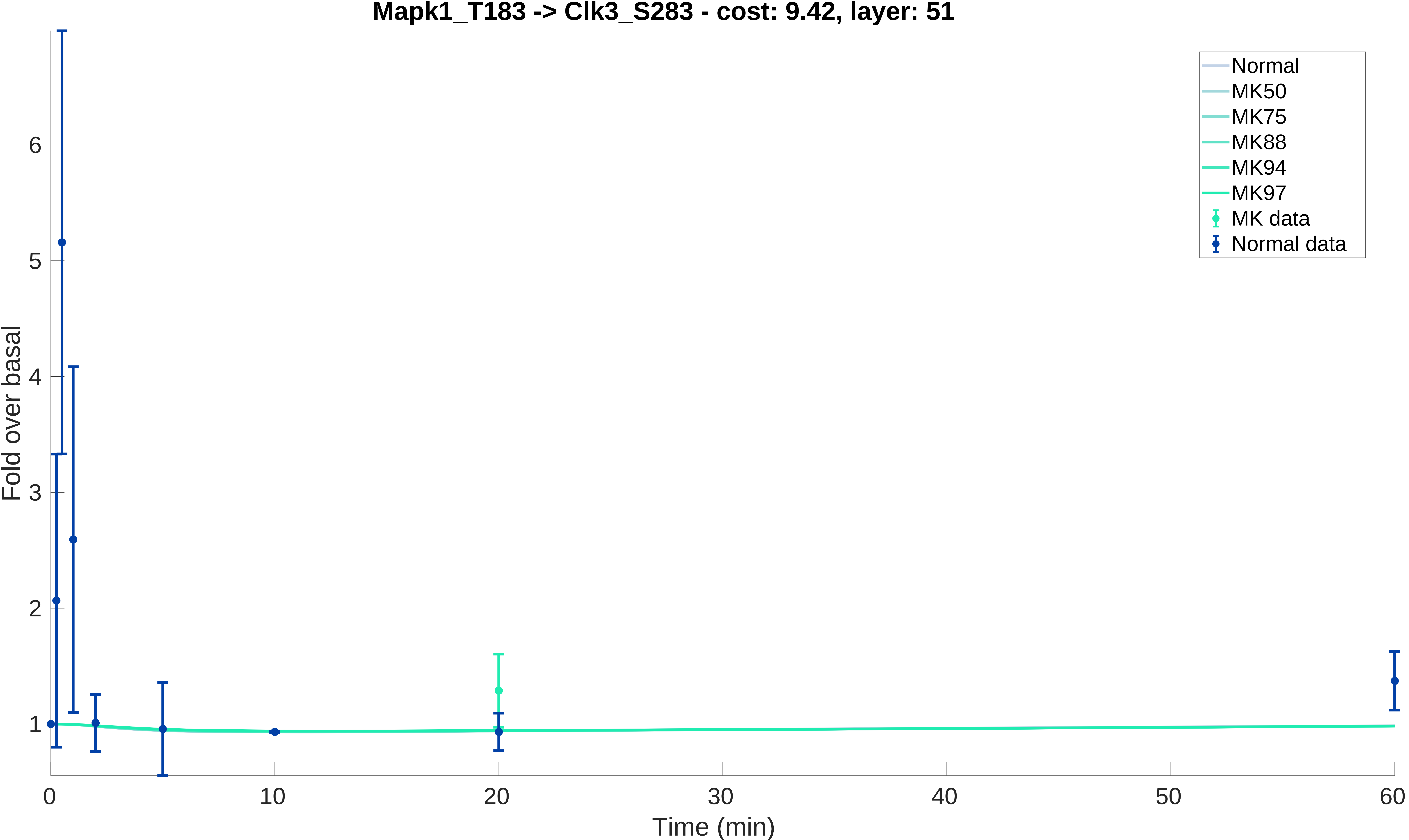

Mapk1_T183 -> Clk3_S283 - cost: 9.42, layer: 51
Fold over basal
Time (min)
1
2
3
4
5
6
0
10
20
30
40
50
60
Normal
MK50
MK75
MK88
MK94
MK97
MK data
Normal data

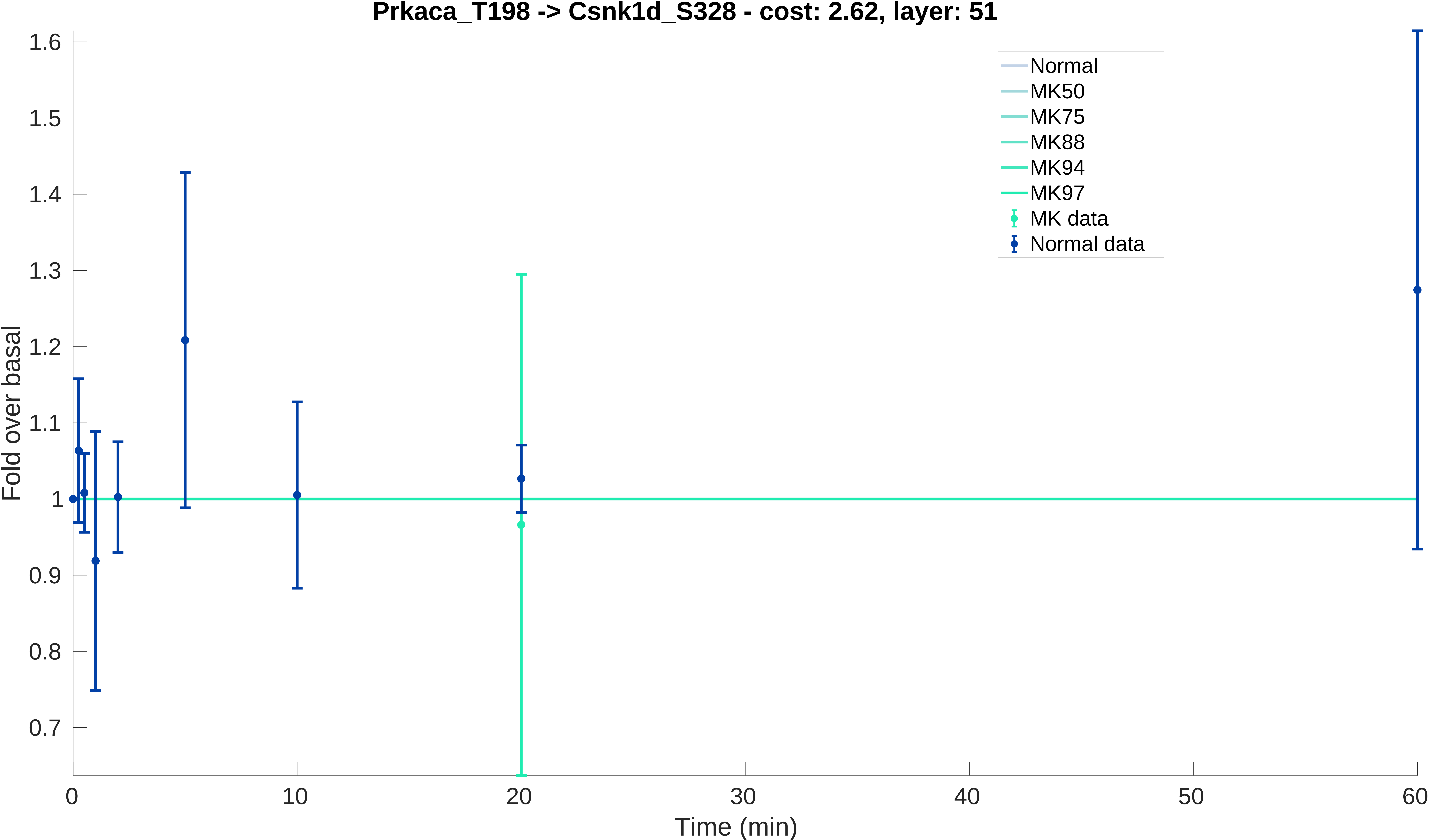

Prkaca_T198 -> Csnk1d_S328 - cost: 2.62, layer: 51
Normal
MK50
MK75
MK88
MK94
MK97
MK data
Normal data
Fold over basal
Time (min)
1.6
1.5
1.4
1.3
1.2
1.1
1
0.9
0.8
0.7
0
10
20
30
40
50
60

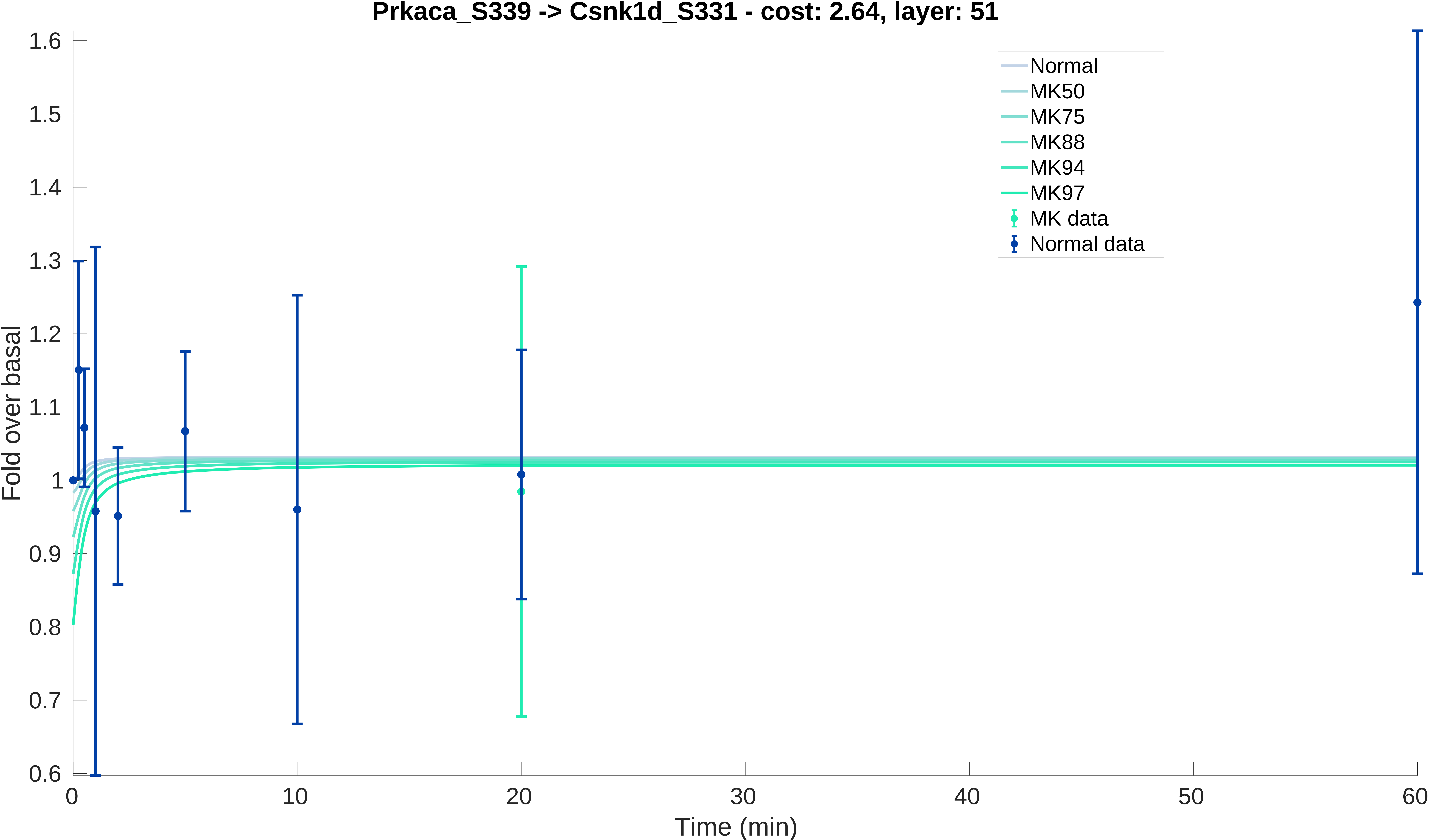

Prkaca_S339 -> Csnk1d_S331 - cost: 2.64, layer: 51
Fold over basal
Time (min)
0.6
0.7
0.8
0.9
1
1.1
1.2
1.3
1.4
1.5
1.6
0
10
20
30
40
50
60
Normal
MK50
MK75
MK88
MK94
MK97
MK data
Normal data

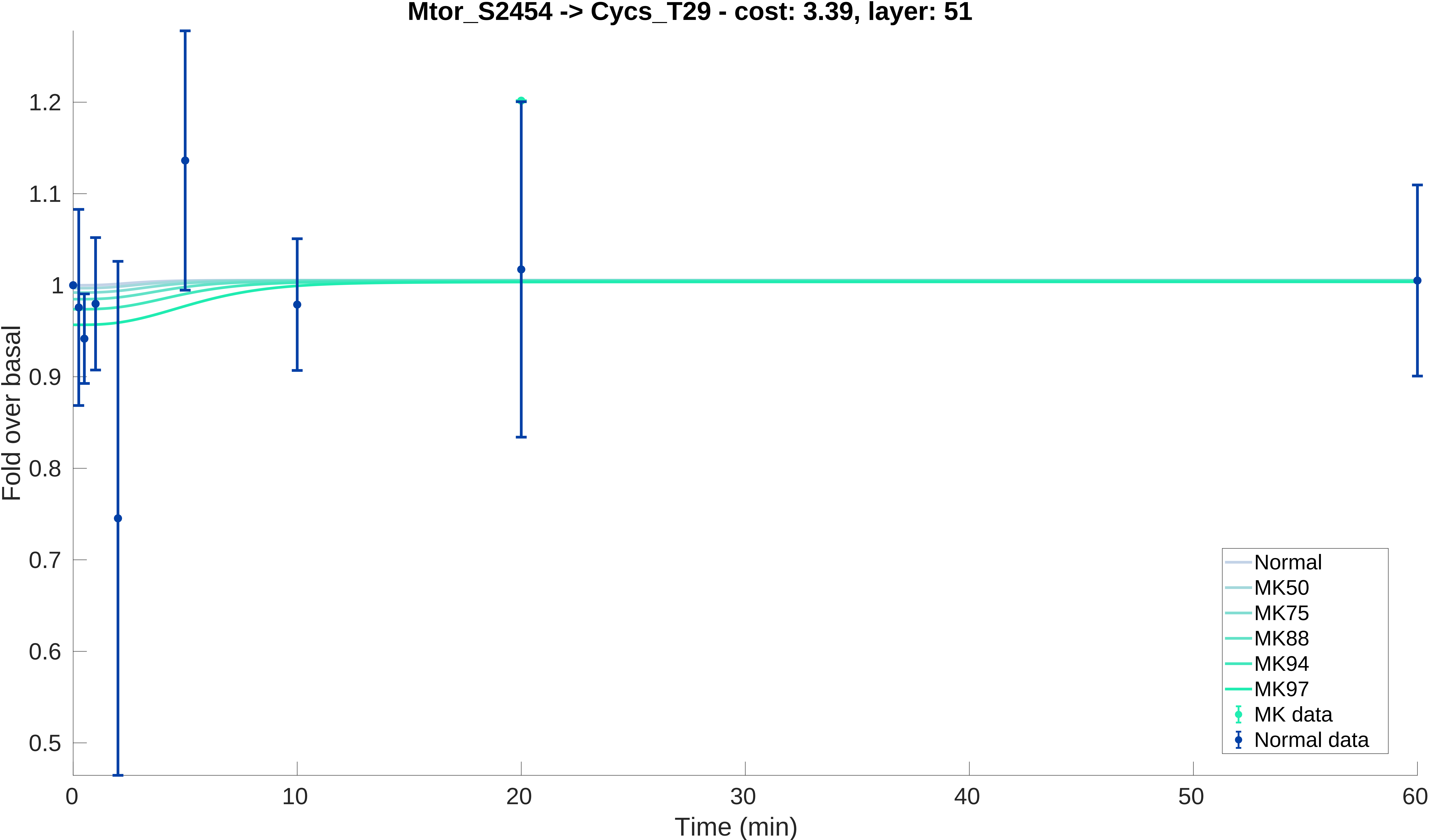
Mtor_S2454 -> Cycs_T29 - cost: 3.39, layer: 51
Fold over basal
Time (min)
0
10
20
30
40
50
60
0.5
0.6
0.7
0.8
0.9
1
1.1
1.2
Normal
MK50
MK75
MK88
MK94
MK97
MK data
Normal data

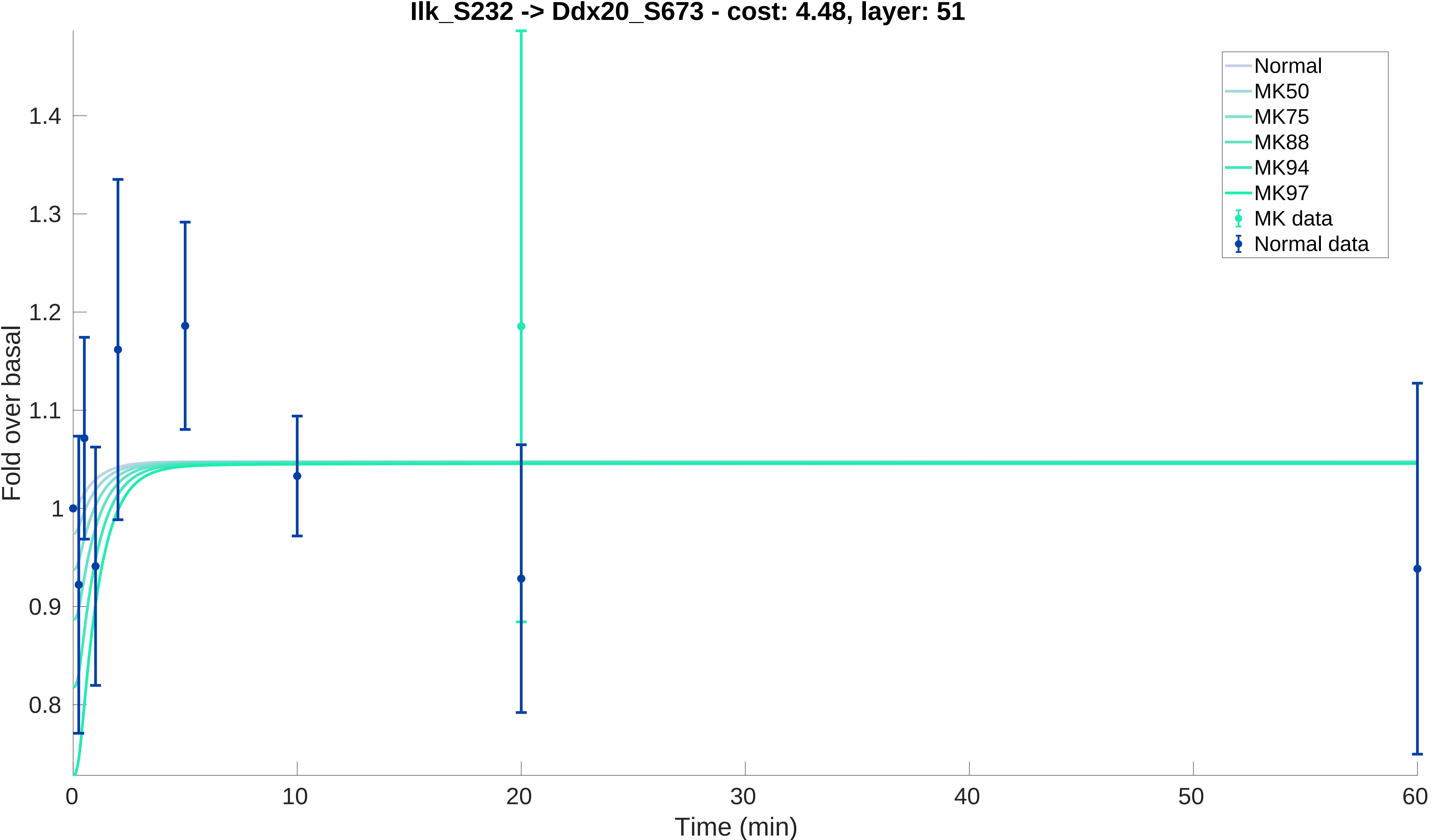

Ilk_S232 -> Ddx20_S673 - cost: 4.48, layer: 51
Normal
MK50
MK75
MK88
MK94
MK97
MK data
Normal data
Fold over basal
1.4
1.3
1.2
1.1
1
0.9
0.8
0
10
20
30
40
50
60
Time (min)

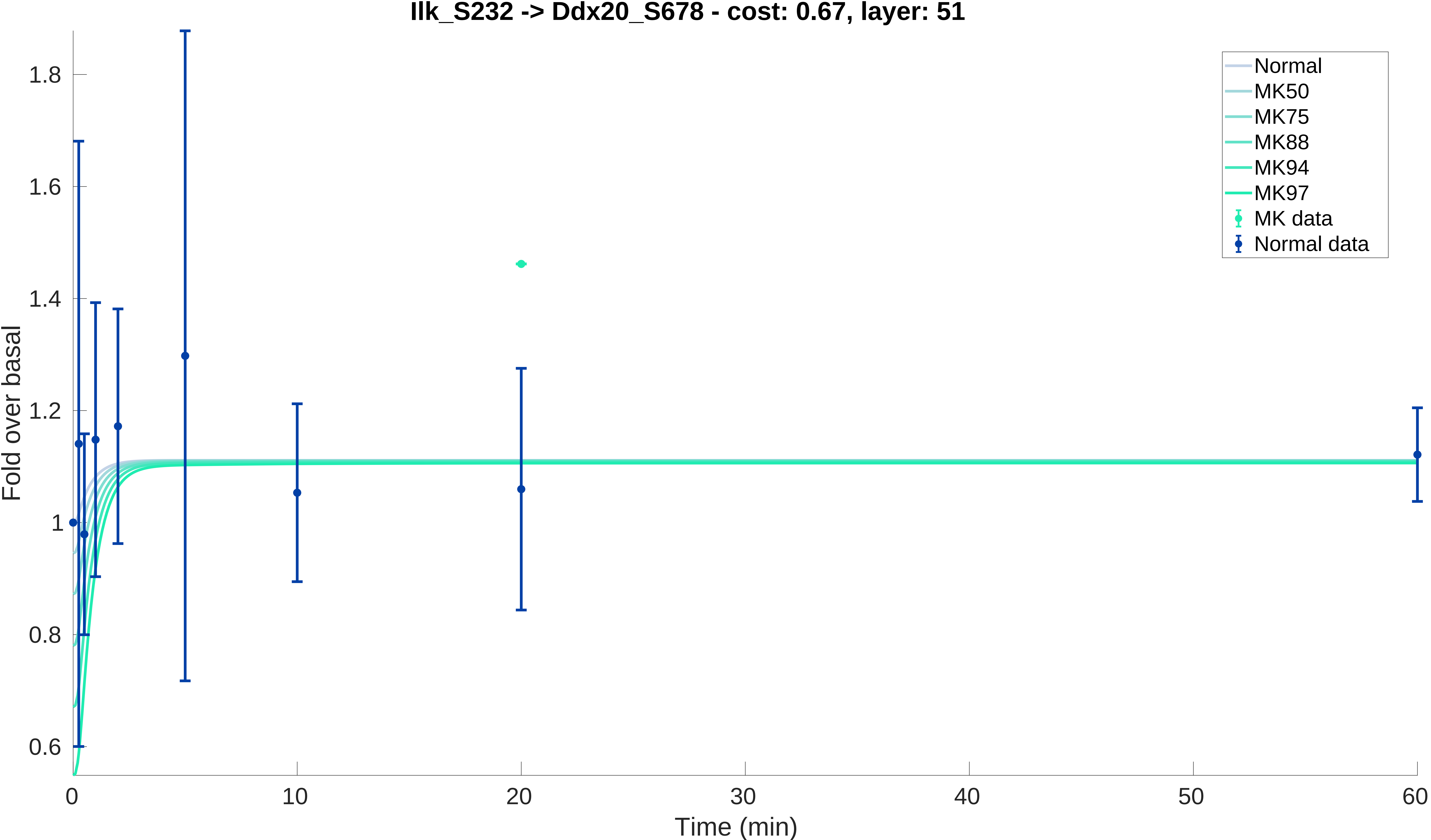
Ilk_S232 -> Ddx20_S678 - cost: 0.67, layer: 51
Fold over basal
1.8
1.6
1.4
1.2
1
0.8
0.6
0
10
20
30
40
50
60
Time (min)
Normal
MK50
MK75
MK88
MK94
MK97
MK data
Normal data

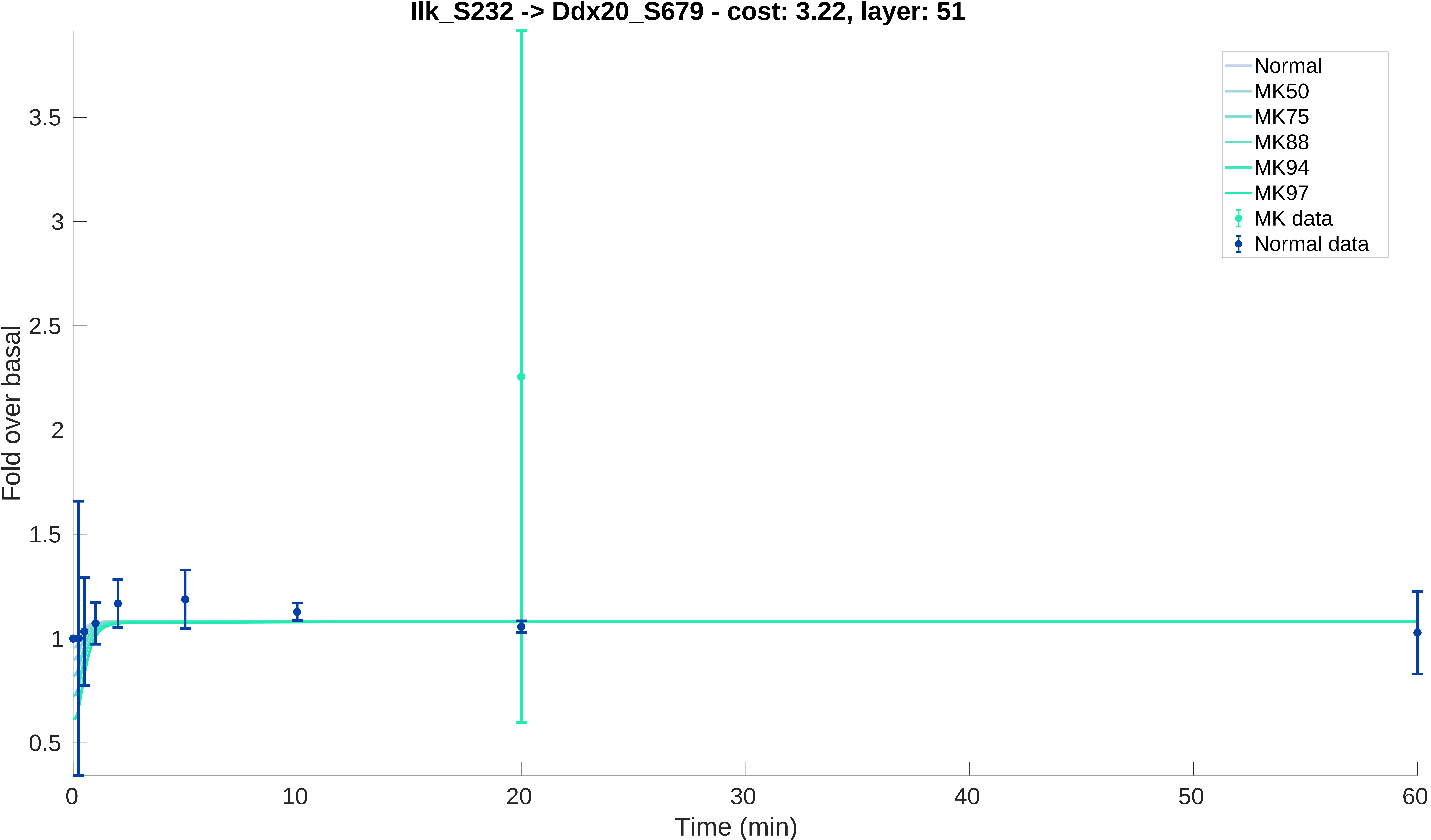

Ilk_S232 -> Ddx20_S679 - cost: 3.22, layer: 51
Fold over basal
Time (min)
0.5
1
1.5
2
2.5
3
3.5
0
10
20
30
40
50
60
Normal
MK50
MK75
MK88
MK94
MK97
MK data
Normal data

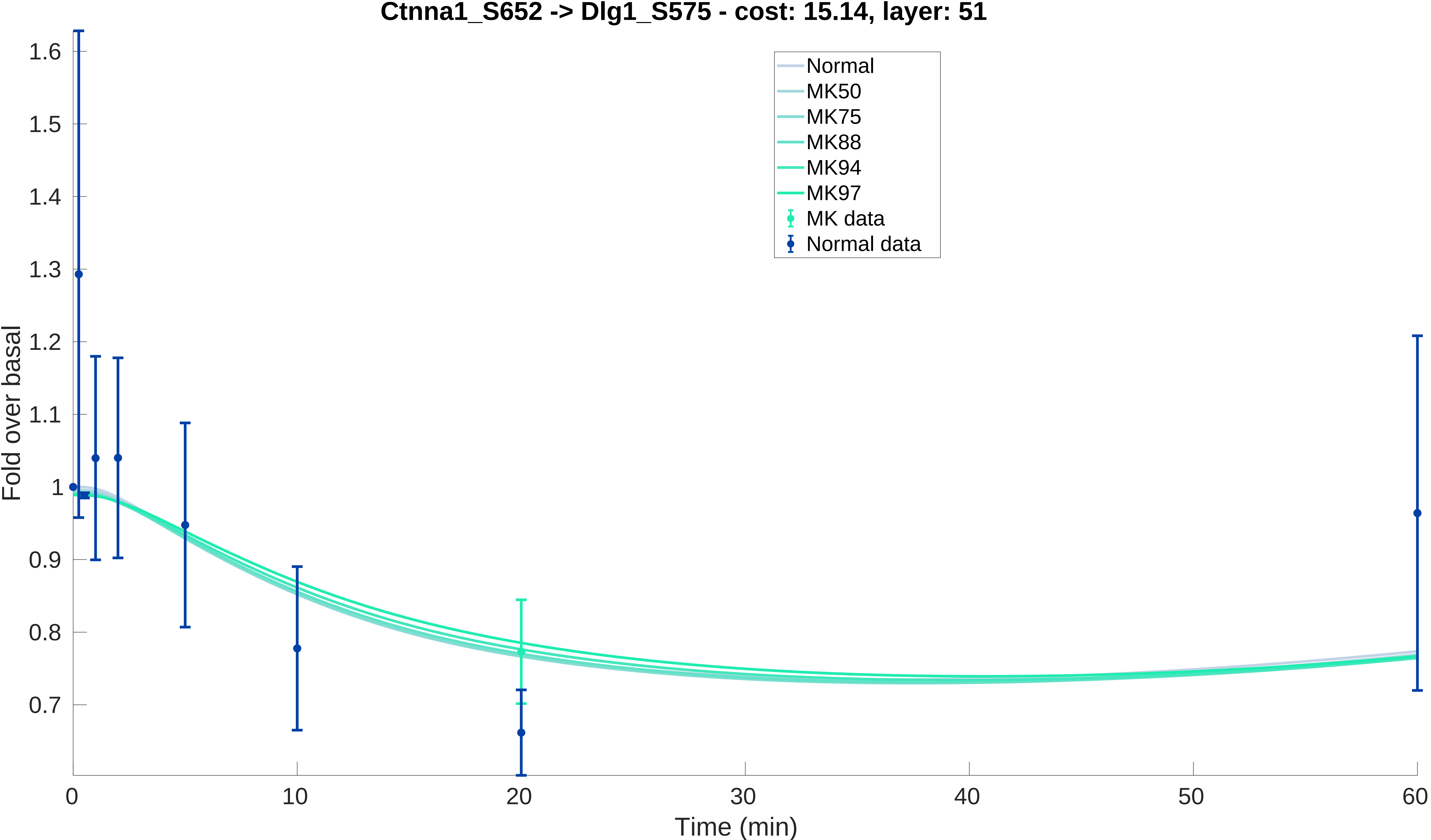

Ctnna1_S652 -> Dlg1_S575 - cost: 15.14, layer: 51
Normal
MK50
MK75
MK88
MK94
MK97
MK data
Normal data
Fold over basal
1.6
1.5
1.4
1.3
1.2
1.1
1
0.9
0.8
0.7
0
10
20
30
40
50
60
Time (min)

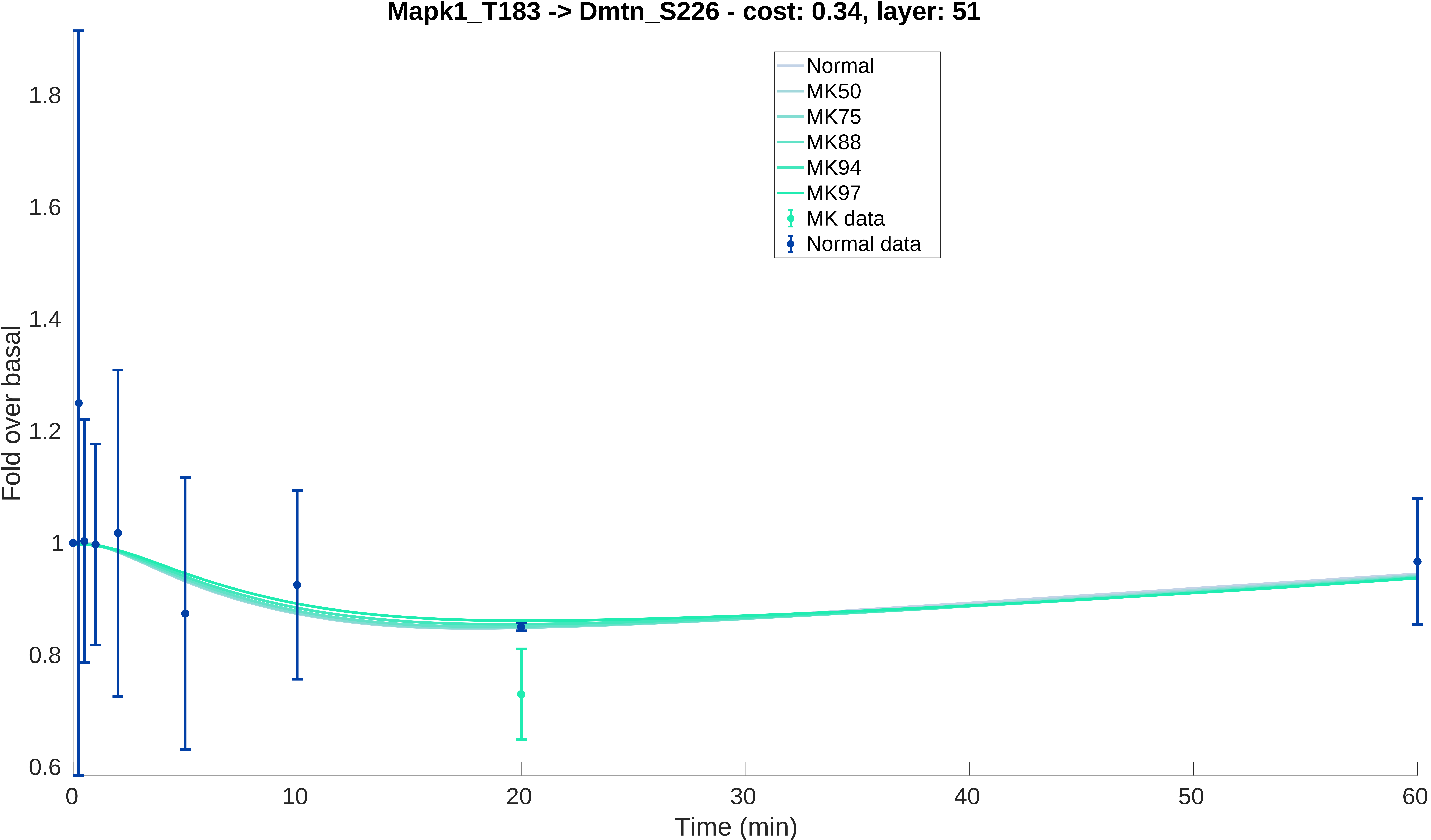

Mapk1_T183 -> Dmtn_S226 - cost: 0.34, layer: 51
Normal
MK50
MK75
MK88
MK94
MK97
MK data
Normal data
Fold over basal
1.8
1.6
1.4
1.2
1
0.8
0.6
0
10
20
30
40
50
60
Time (min)

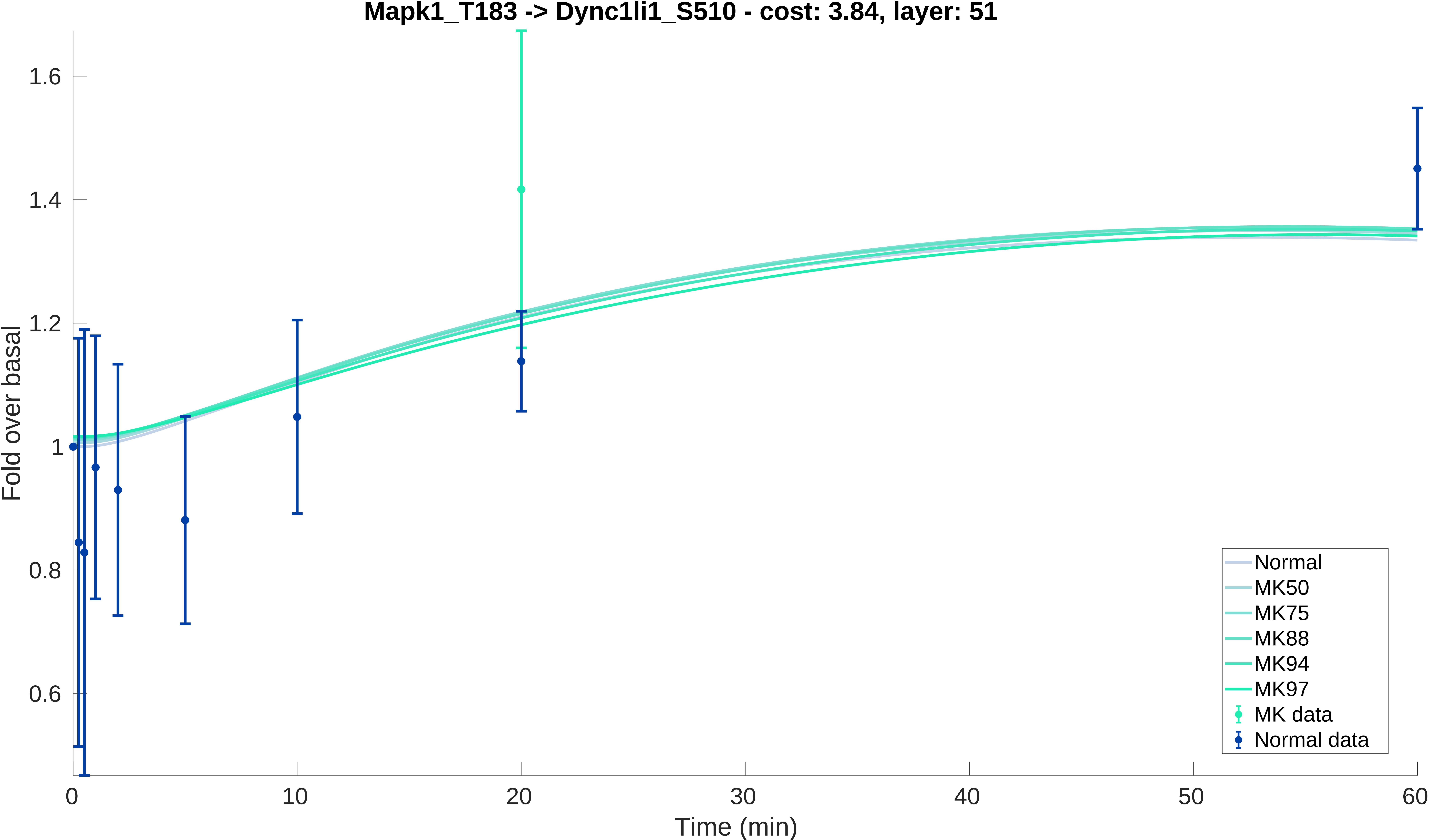

Mapk1_T183 -> Dync1li1_S510 - cost: 3.84, layer: 51
Fold over basal
1.6
1.4
1.2
1
0.8
0.6
0
10
20
30
40
50
60
Time (min)
Normal
MK50
MK75
MK88
MK94
MK97
MK data
Normal data

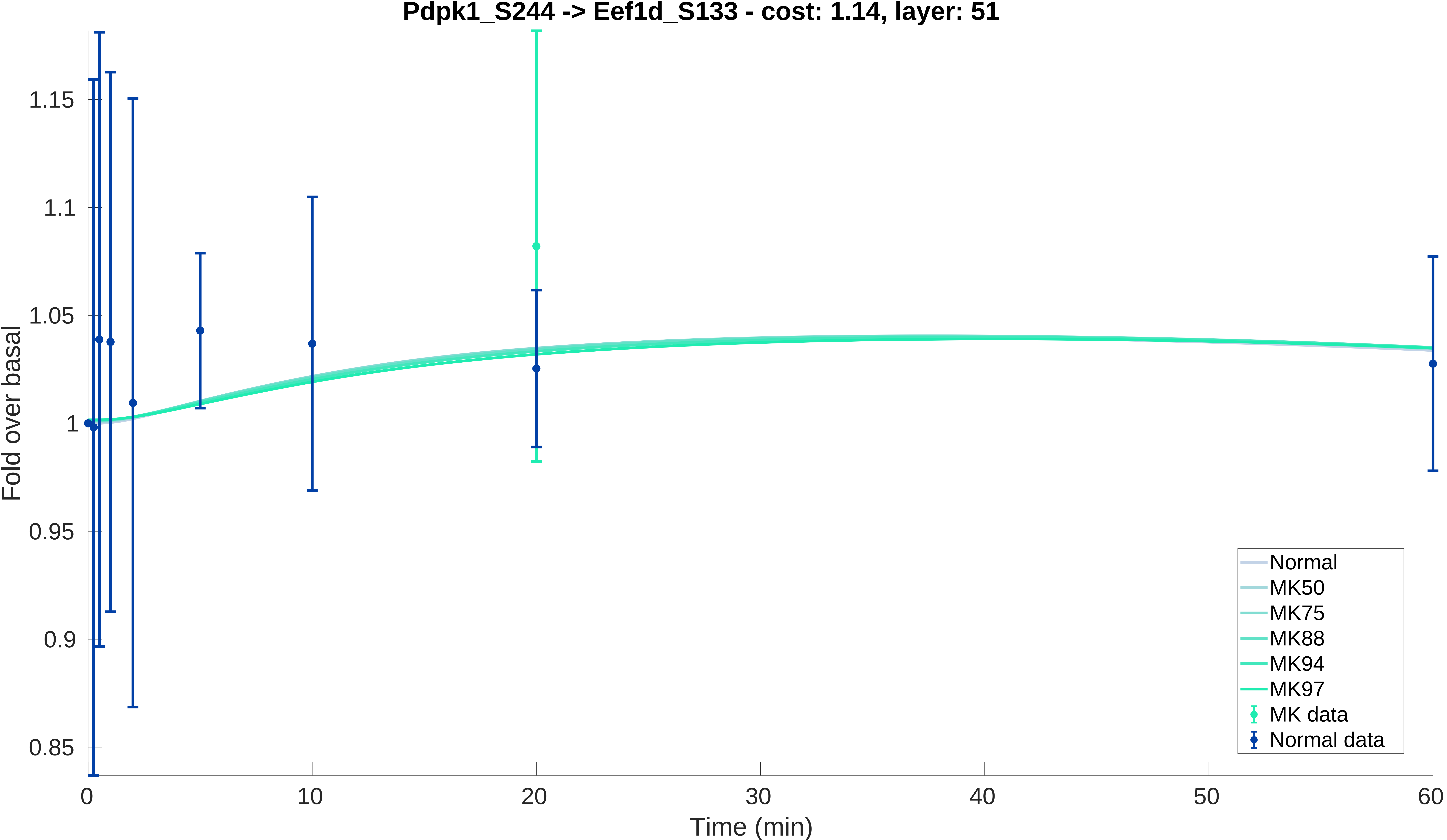

Pdpk1_S244 -> Eef1d_S133 - cost: 1.14, layer: 51
Fold over basal
1.15
1.1
1.05
1
0.95
0.9
0.85
Time (min)
0
10
20
30
40
50
60
Normal
MK50
MK75
MK88
MK94
MK97
MK data
Normal data

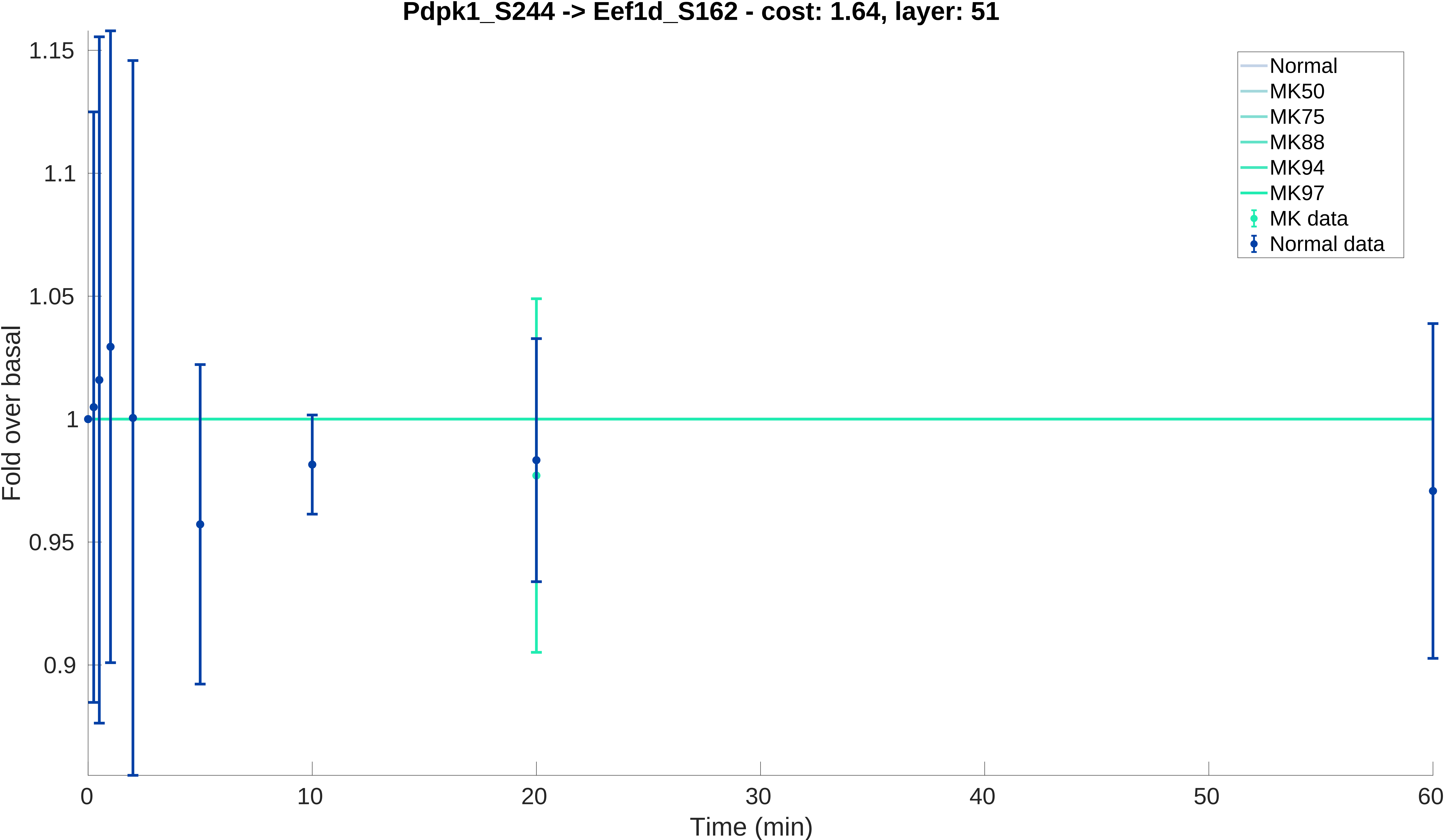

Pdpk1_S244 -> Eef1d_S162 - cost: 1.64, layer: 51
Fold over basal
1.15
1.1
1.05
1
0.95
0.9
0
10
20
30
40
50
60
Time (min)
Normal
MK50
MK75
MK88
MK94
MK97
MK data
Normal data

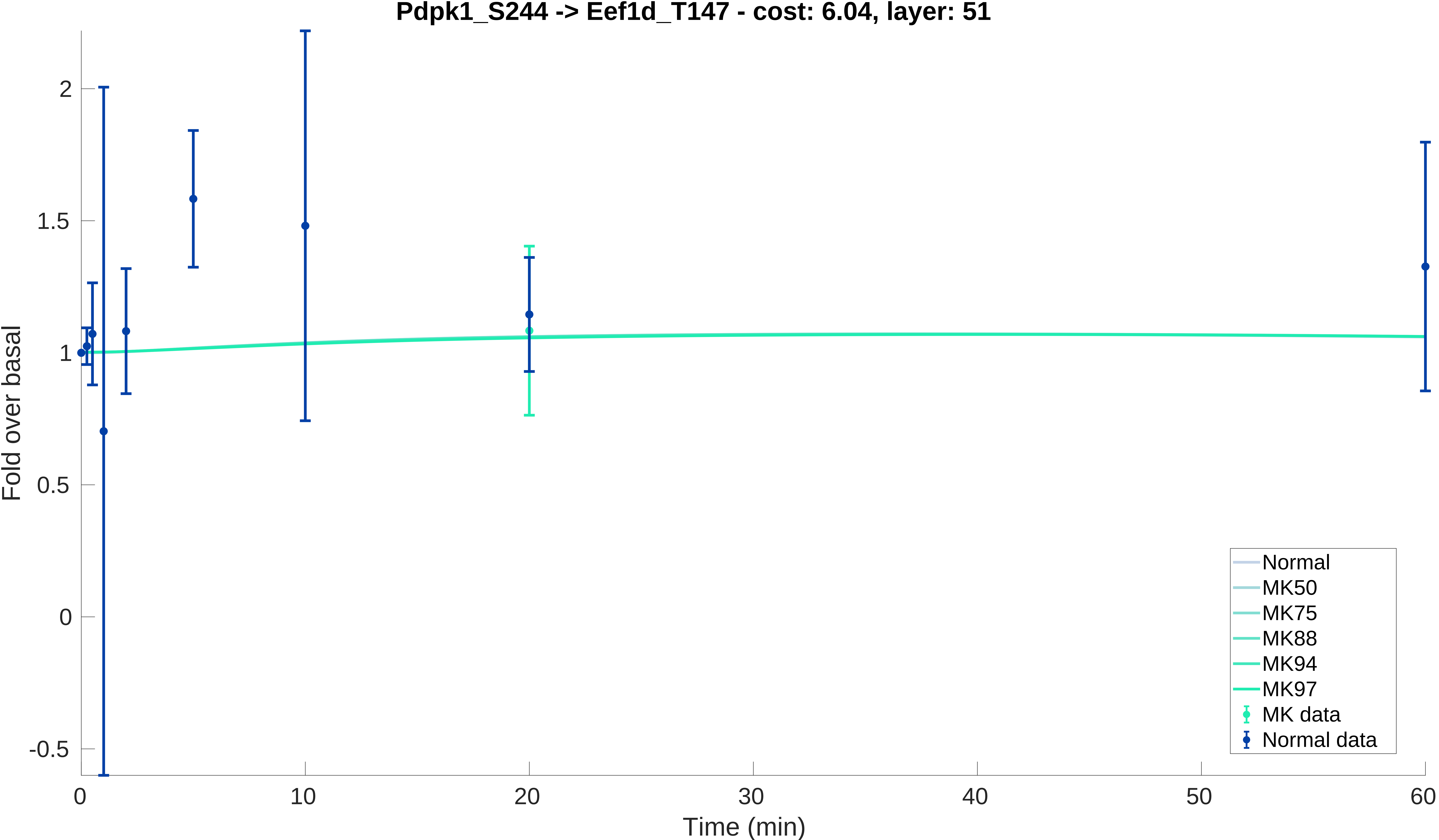

Pdpk1_S244 -> Eef1d_T147 - cost: 6.04, layer: 51
Fold over basal
Time (min)
2
1.5
1
0.5
0
-0.5
0
10
20
30
40
50
60
Normal
MK50
MK75
MK88
MK94
MK97
MK data
Normal data

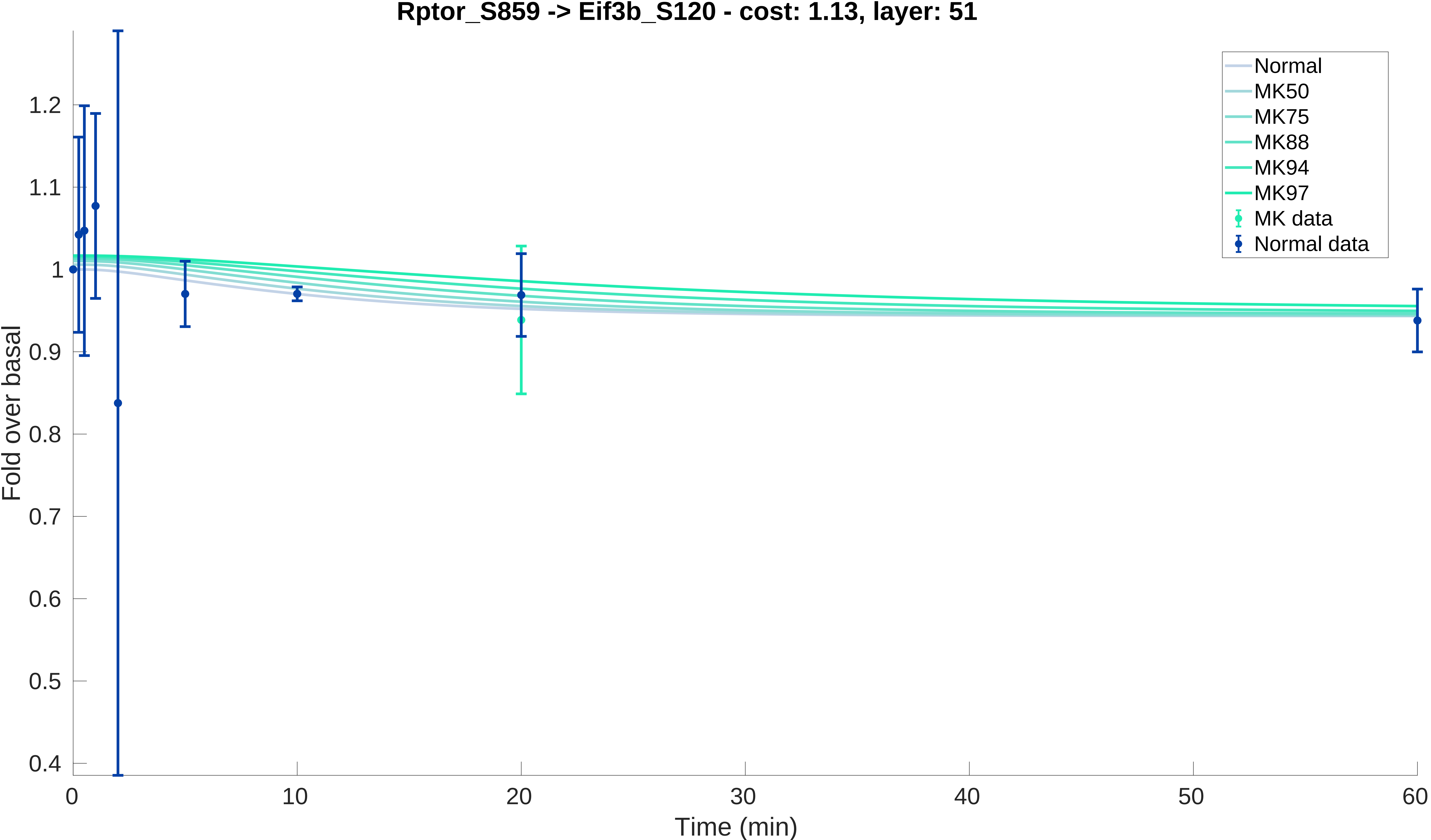
Rptor_S859 -> Eif3b_S120 - cost: 1.13, layer: 51
Normal
MK50
MK75
MK88
MK94
MK97
MK data
Normal data
Fold over basal
Time (min)
0.4
0.5
0.6
0.7
0.8
0.9
1
1.1
1.2
0
10
20
30
40
50
60

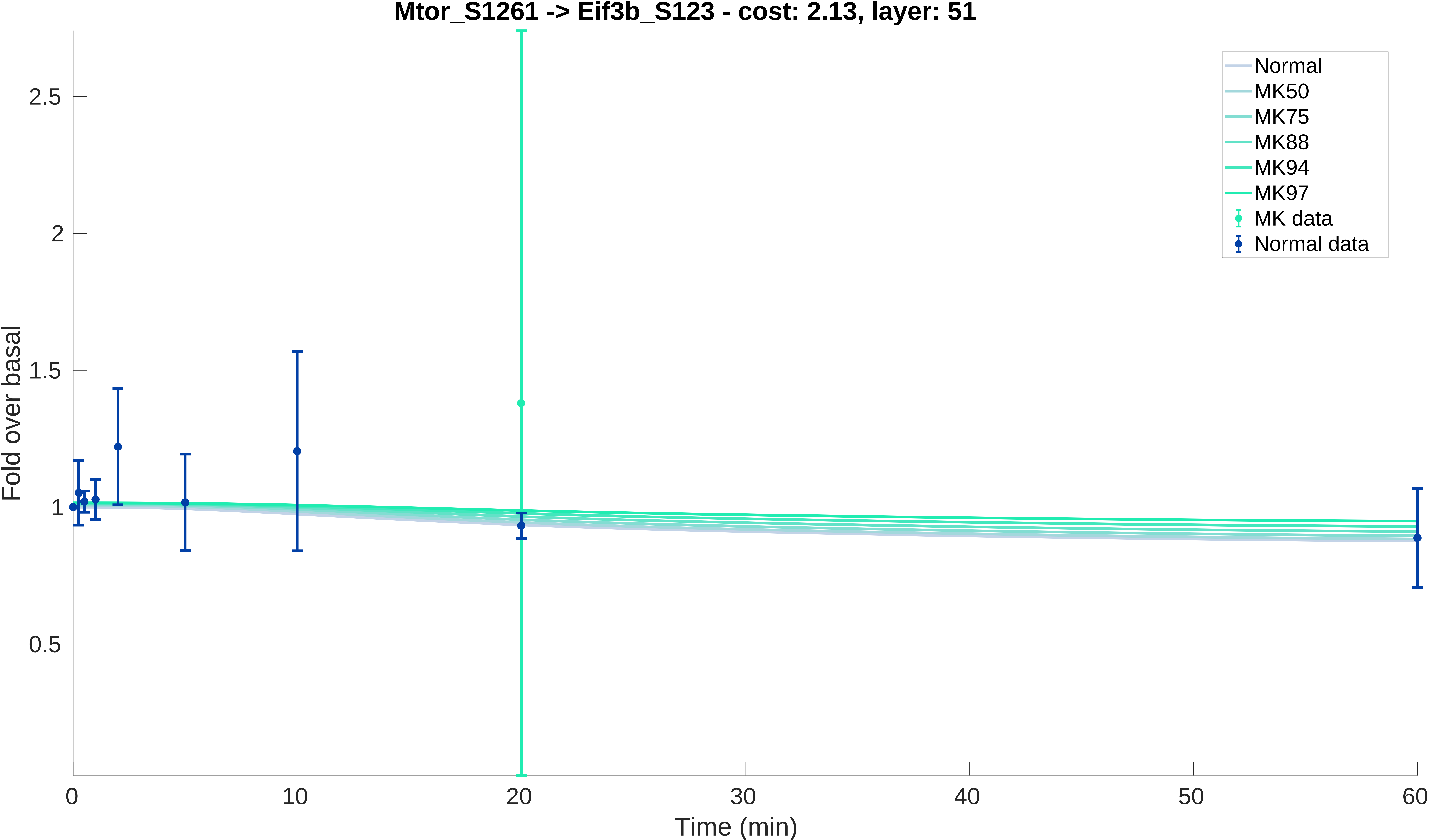

Mtor_S1261 -> Eif3b_S123 - cost: 2.13, layer: 51
Fold over basal
Time (min)
0
10
20
30
40
50
60
0.5
1
1.5
2
2.5
Normal
MK50
MK75
MK88
MK94
MK97
MK data
Normal data

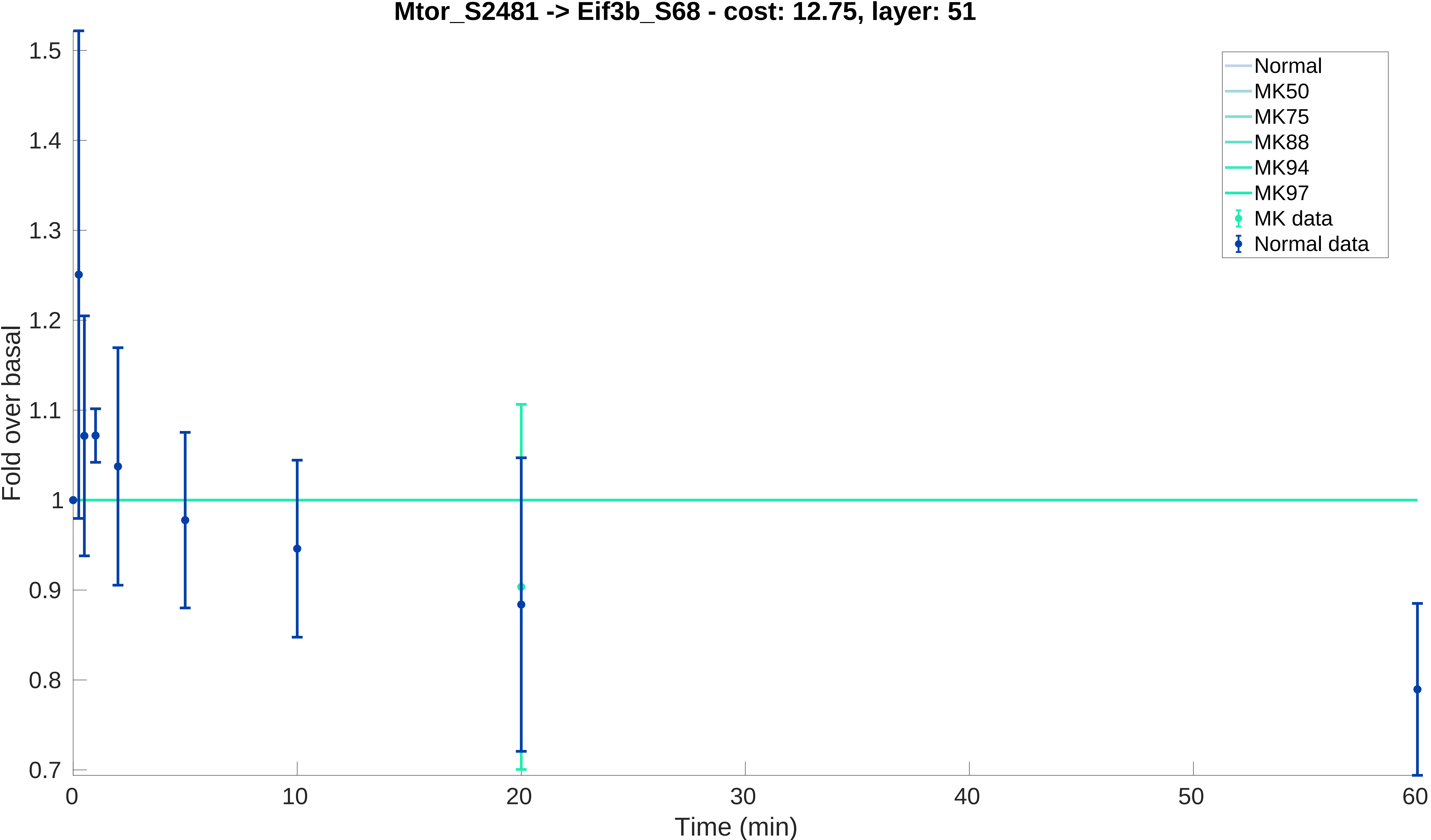
Mtor_S2481 -> Eif3b_S68 - cost: 12.75, layer: 51
Normal
MK50
MK75
MK88
MK94
MK97
MK data
Normal data
Fold over basal
1.5
1.4
1.3
1.2
1.1
1
0.9
0.8
0.7
0
10
20
30
40
50
60
Time (min)

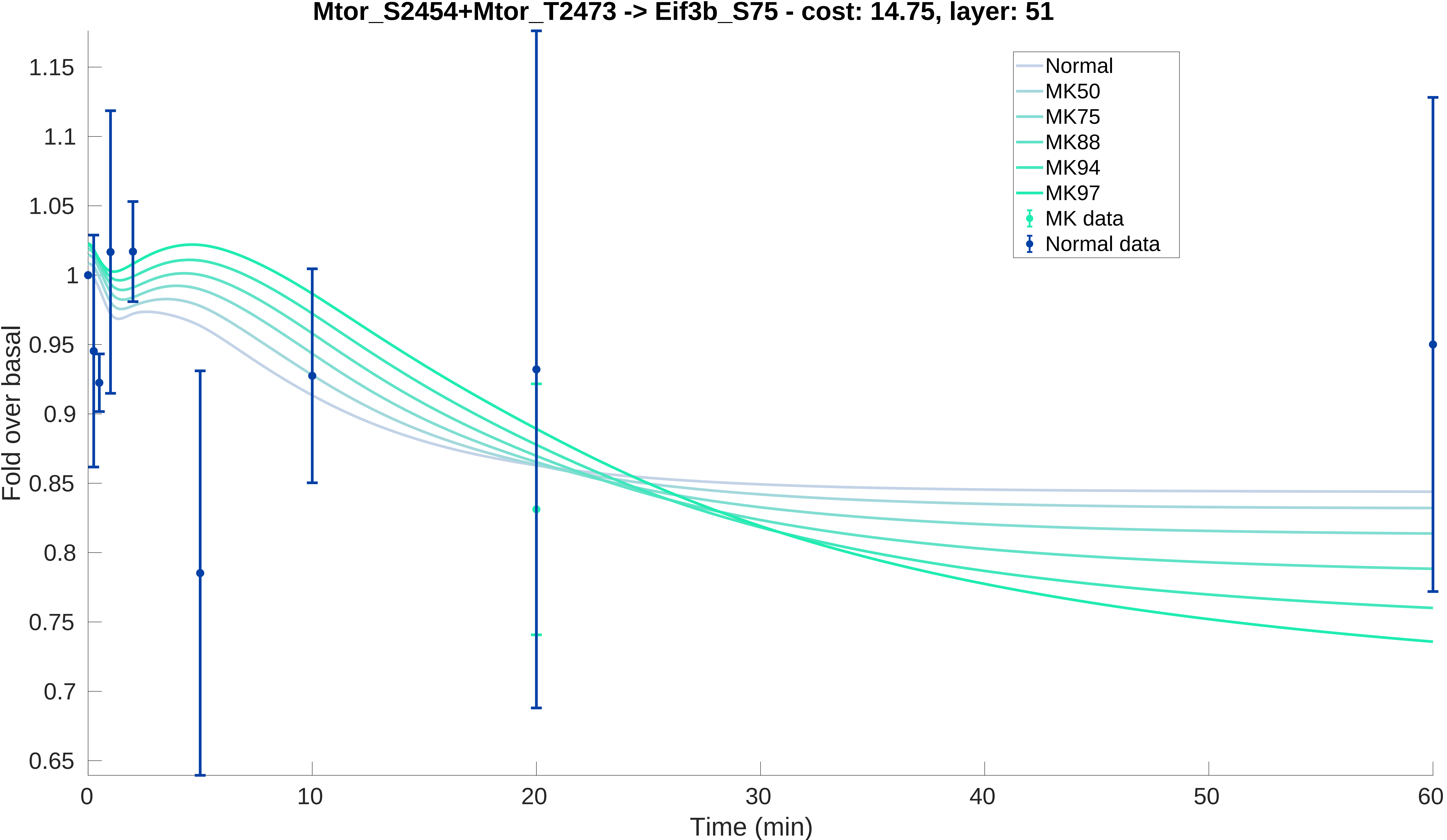

Mtor_S2454+Mtor_T2473 -> Eif3b_S75 - cost: 14.75, layer: 51
Fold over basal
Time (min)
1.15
1.1
1.05
1
0.95
0.9
0.85
0.8
0.75
0.7
0.65
0
10
20
30
40
50
60
Normal
MK50
MK75
MK88
MK94
MK97
MK data
Normal data

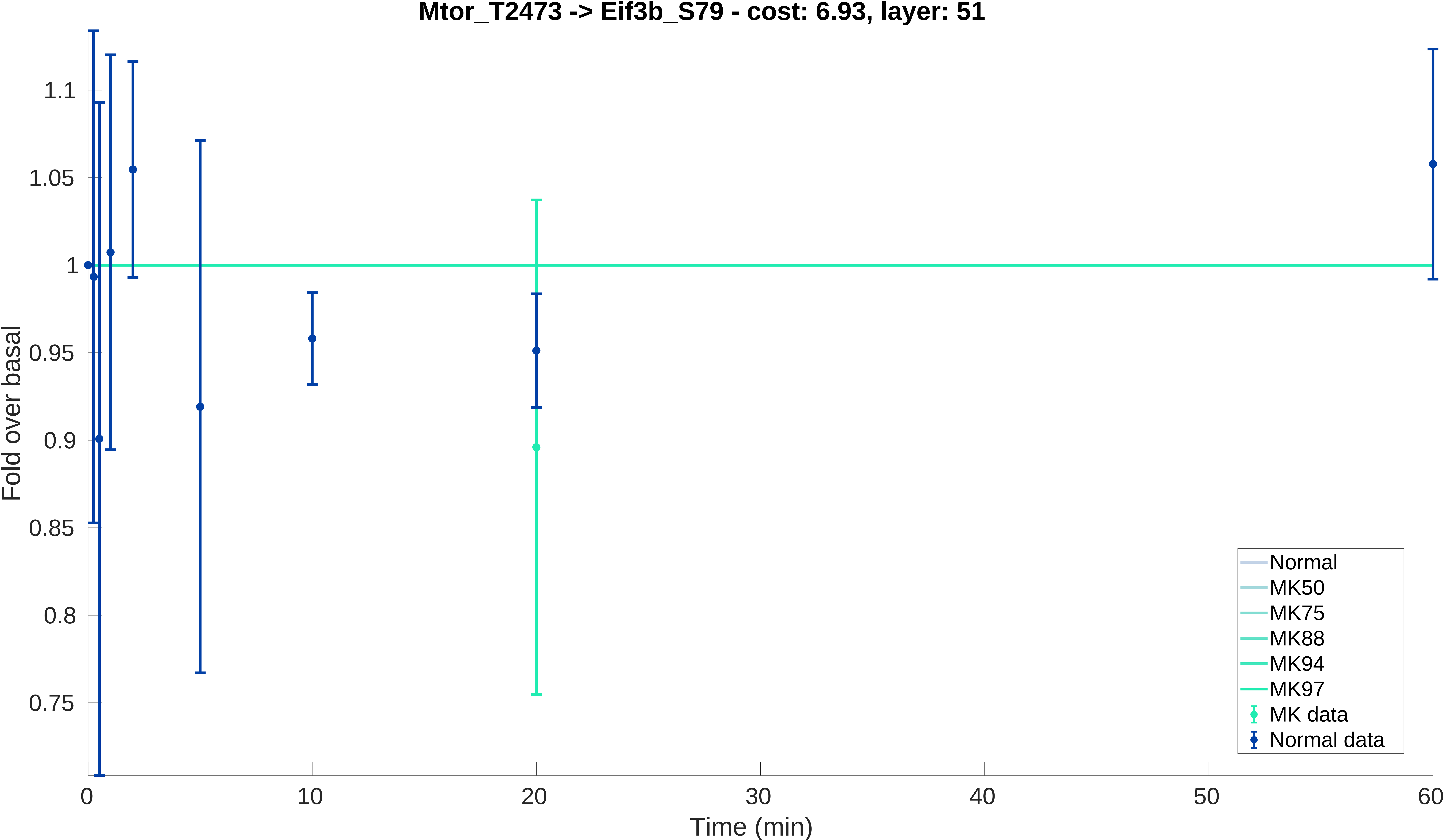

Mtor_T2473 -> Eif3b_S79 - cost: 6.93, layer: 51
Fold over basal
1.1
1.05
1
0.95
0.9
0.85
0.8
0.75
0
10
20
30
40
50
60
Time (min)
Normal
MK50
MK75
MK88
MK94
MK97
MK data
Normal data

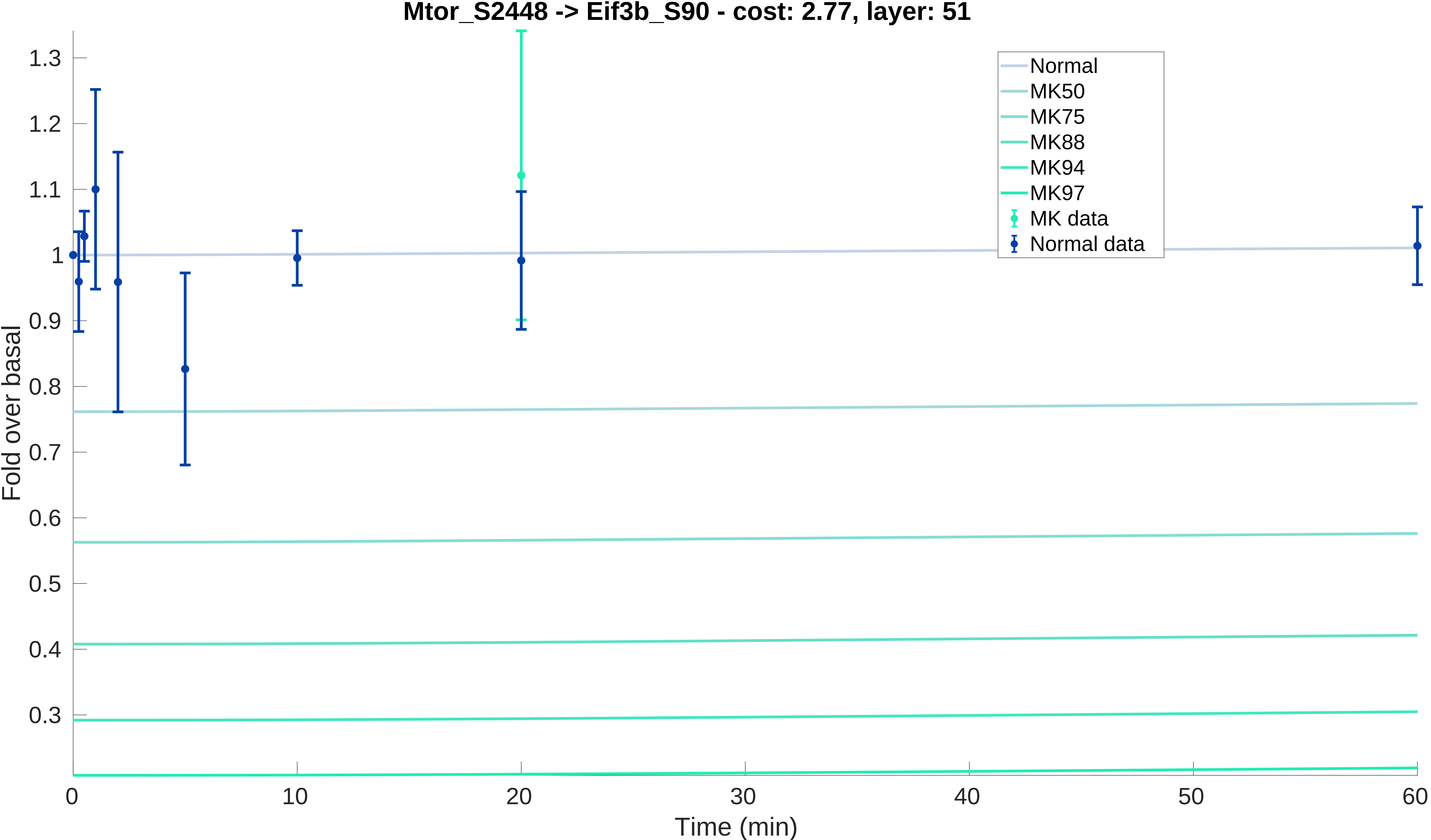

Mtor_S2448 -> Eif3b_S90 - cost: 2.77, layer: 51
Normal
MK50
MK75
MK88
MK94
MK97
MK data
Normal data
Fold over basal
Time (min)
1.3
1.2
1.1
1
0.9
0.8
0.7
0.6
0.5
0.4
0.3
0
10
20
30
40
50
60

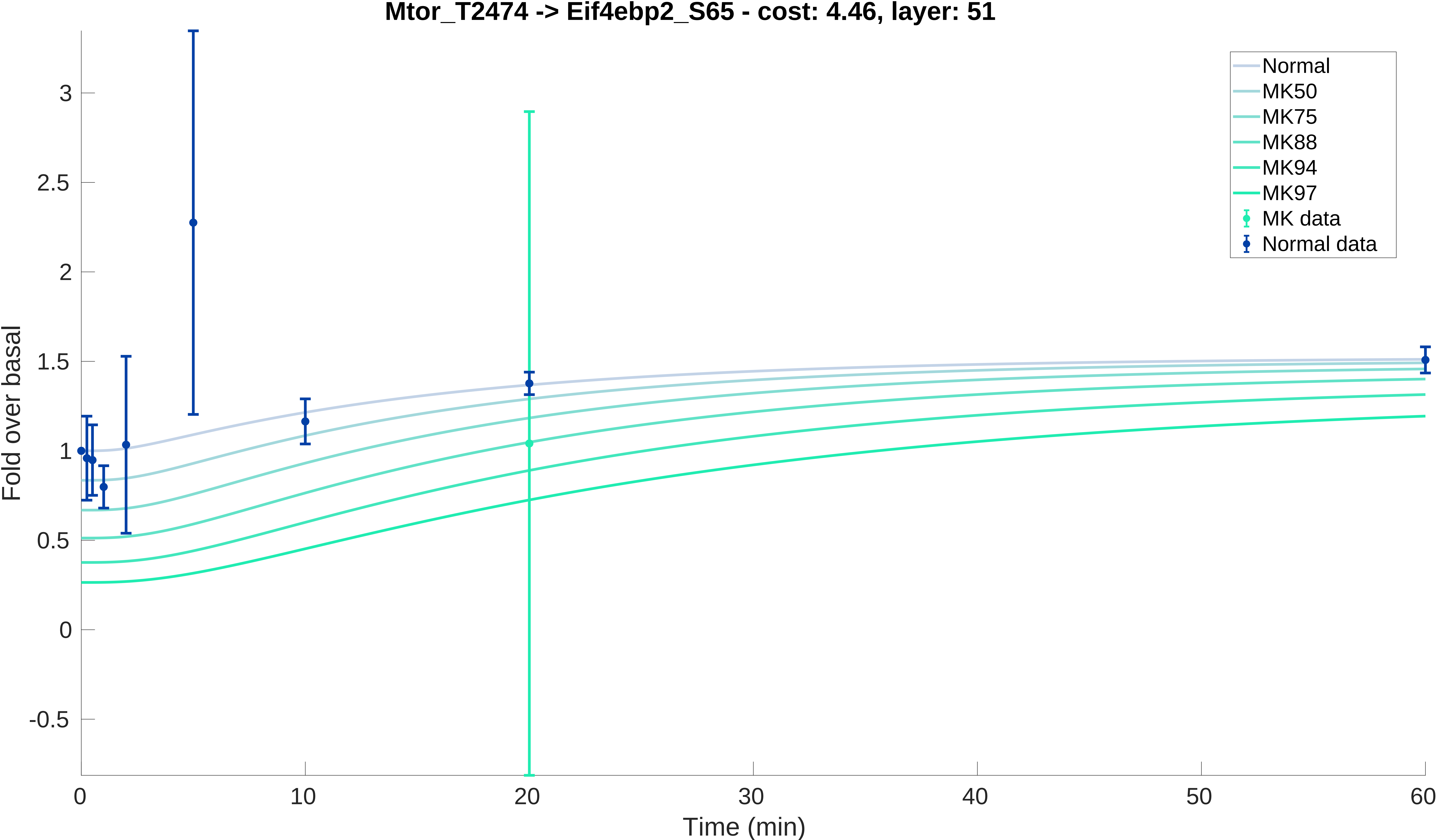

Mtor_T2474 -> Eif4ebp2_S65 - cost: 4.46, layer: 51
Fold over basal
Time (min)
3
2.5
2
1.5
1
0.5
0
-0.5
0
10
20
30
40
50
60
Normal
MK50
MK75
MK88
MK94
MK97
MK data
Normal data

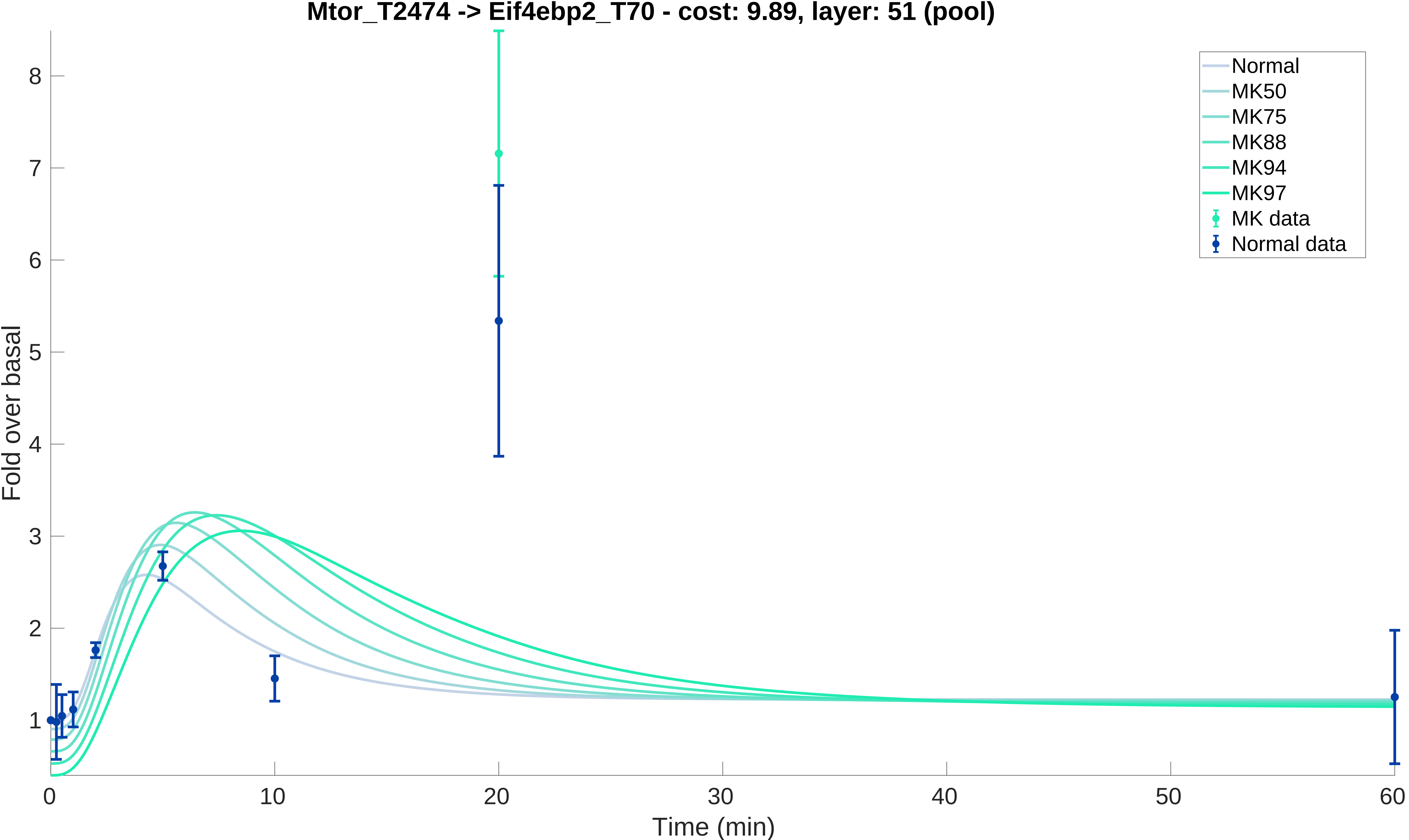

Mtor_T2474 -> Eif4ebp2_T70 - cost: 9.89, layer: 51 (pool)
Fold over basal
Time (min)
0
10
20
30
40
50
60
1
2
3
4
5
6
7
8
Normal
MK50
MK75
MK88
MK94
MK97
MK data
Normal data

#### Mapk1_T183 -> Eif4enif1_S563 - cost: 4.37, layer: 51

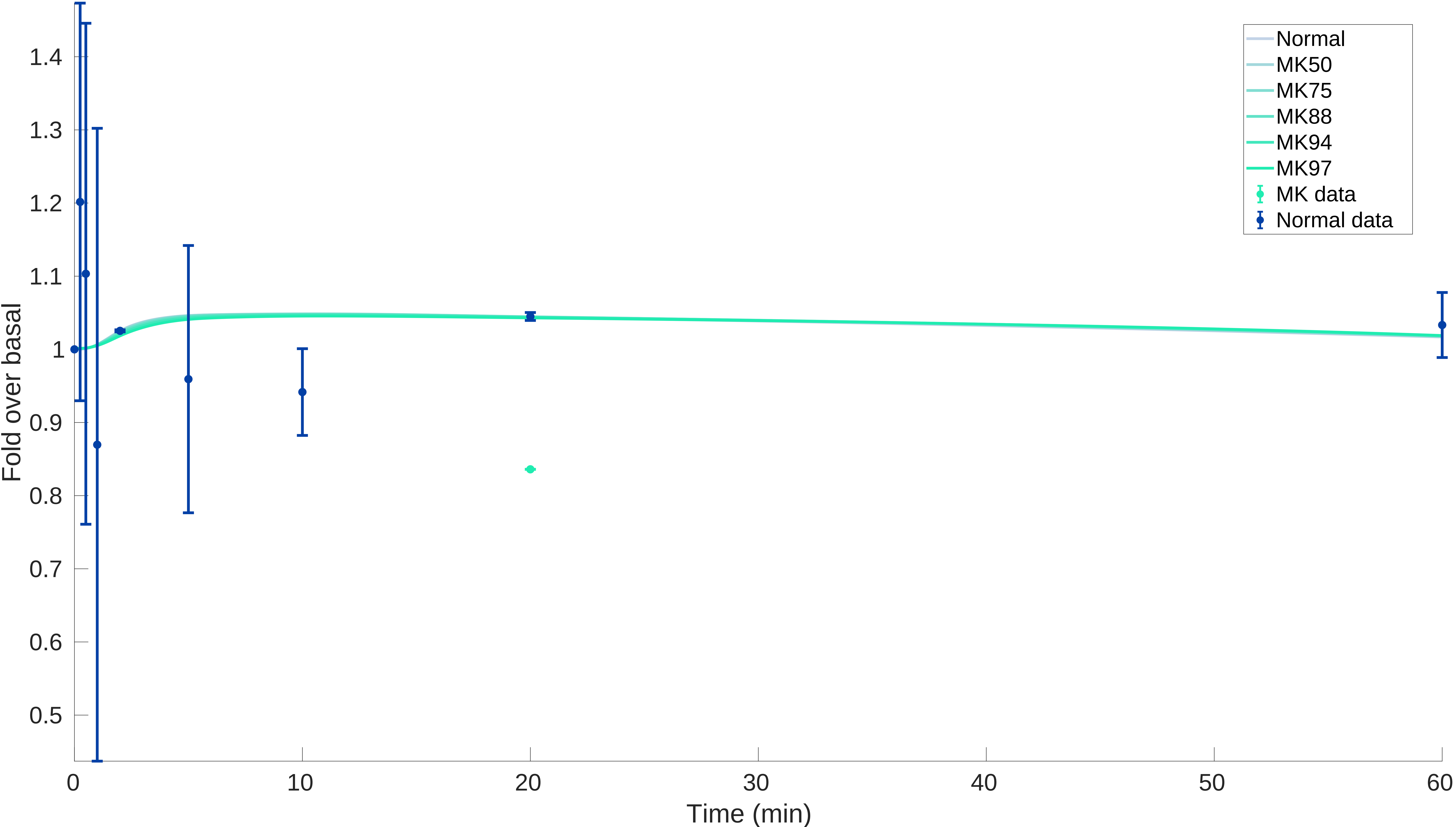

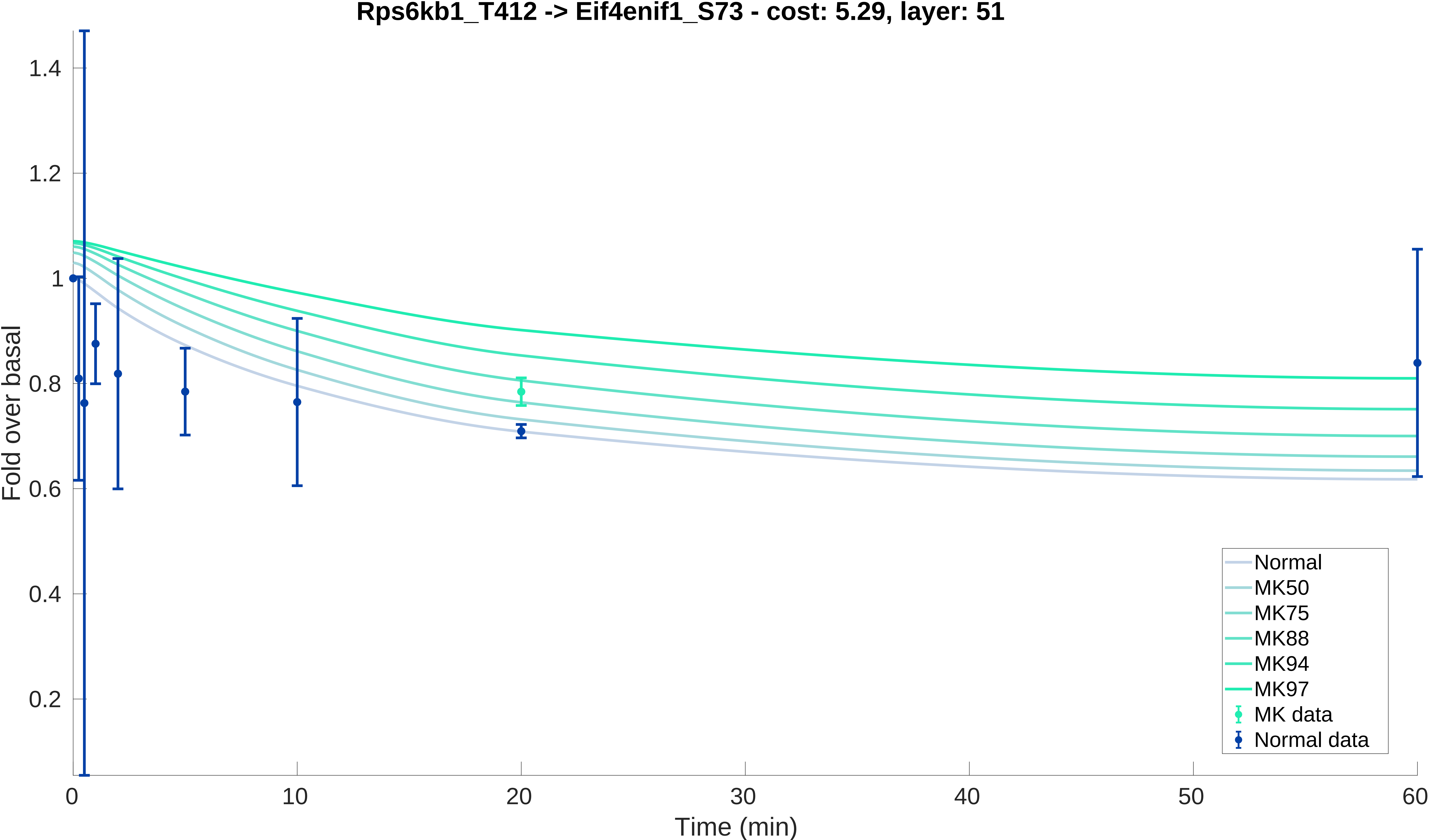

Rps6kb1_T412 -> Eif4enif1_S73 - cost: 5.29, layer: 51
Fold over basal
1.4
1.2
1
0.8
0.6
0.4
0.2
0
10
20
30
40
50
60
Time (min)
Normal
MK50
MK75
MK88
MK94
MK97
MK data
Normal data

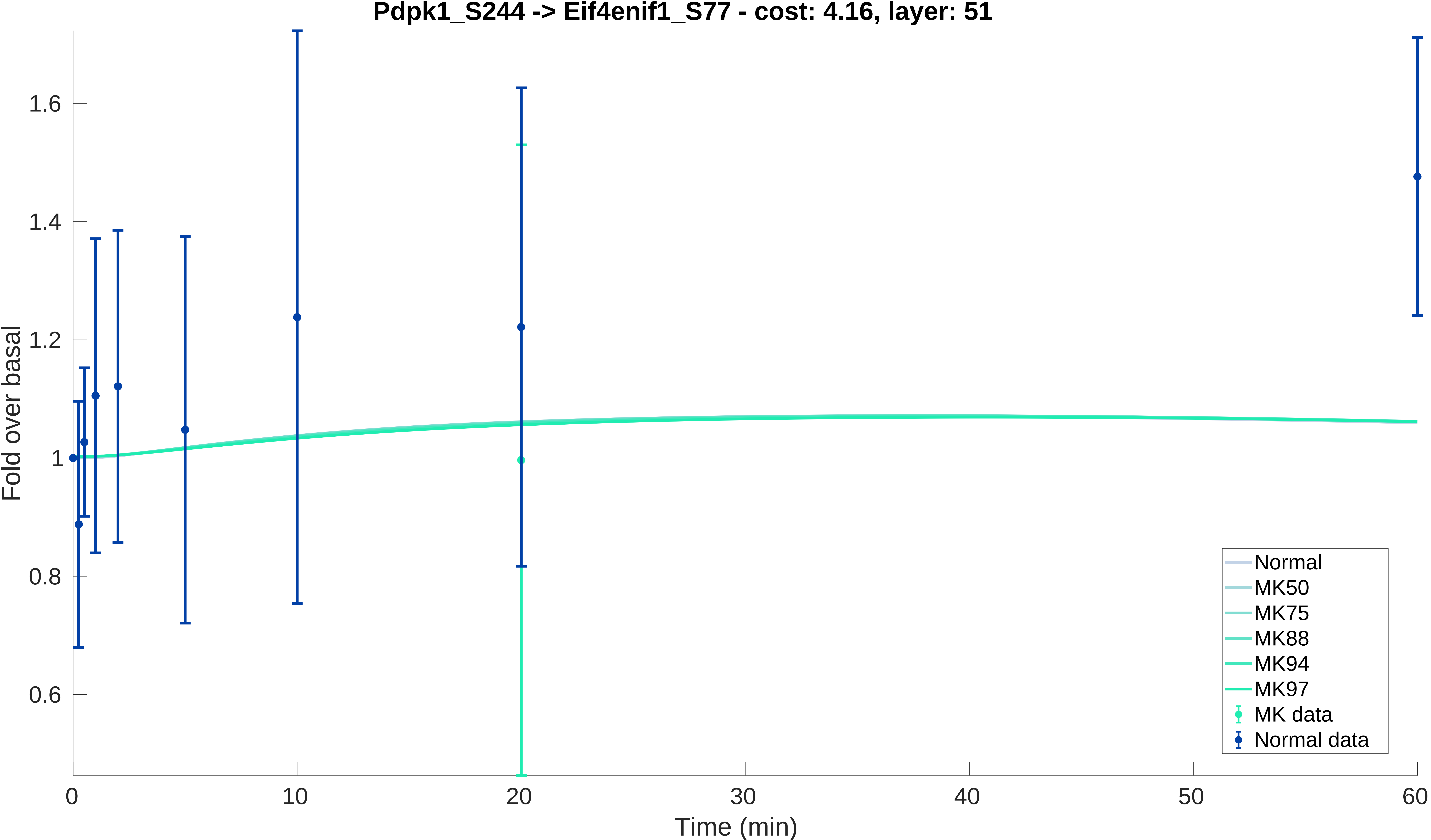

Pdpk1_S244 -> Eif4enif1_S77 - cost: 4.16, layer: 51
Fold over basal
Time (min)
0.6
0.8
1
1.2
1.4
1.6
0
10
20
30
40
50
60
Normal
MK50
MK75
MK88
MK94
MK97
MK data
Normal data

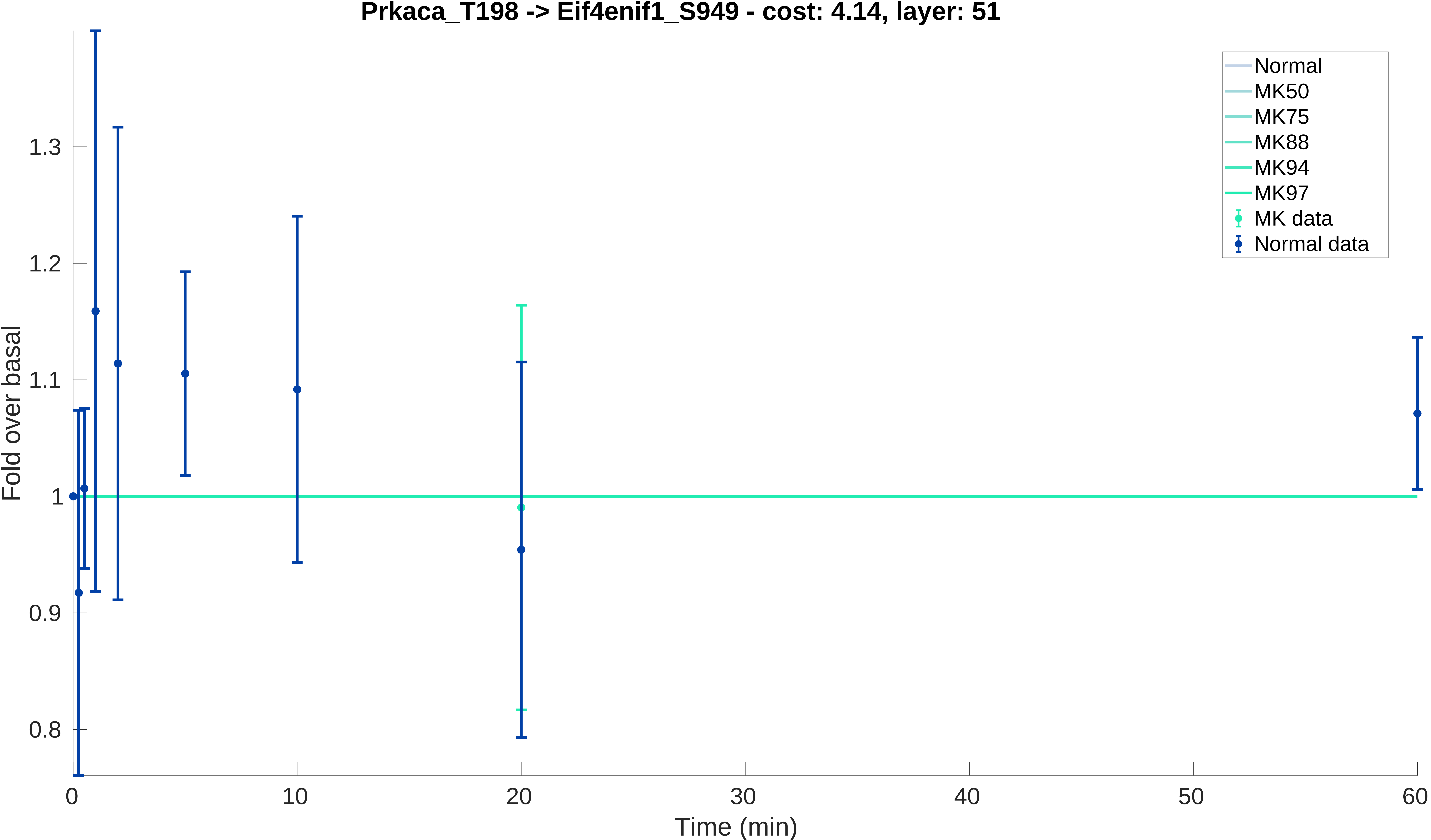

Prkaca_T198 -> Eif4enif1_S949 - cost: 4.14, layer: 51
Fold over basal
1.3
1.2
1.1
1
0.9
0.8
0
10
20
30
40
50
60
Time (min)
Normal
MK50
MK75
MK88
MK94
MK97
MK data
Normal data

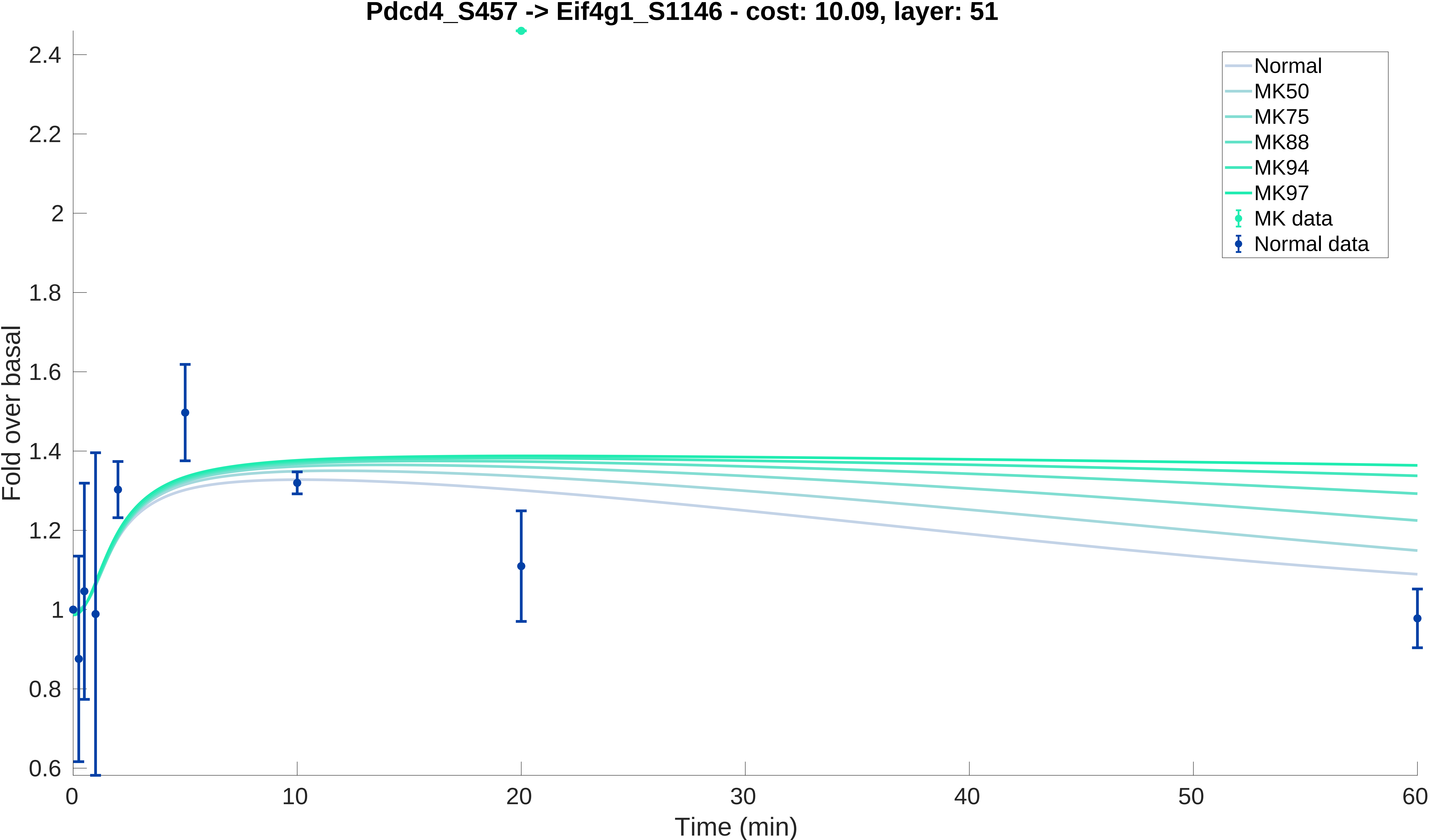

Pdcd4_S457 -> Eif4g1_S1146 - cost: 10.09, layer: 51
Fold over basal
Time (min)
2.4
2.2
2
1.8
1.6
1.4
1.2
1
0.8
0.6
0
10
20
30
40
50
60
Normal
MK50
MK75
MK88
MK94
MK97
MK data
Normal data

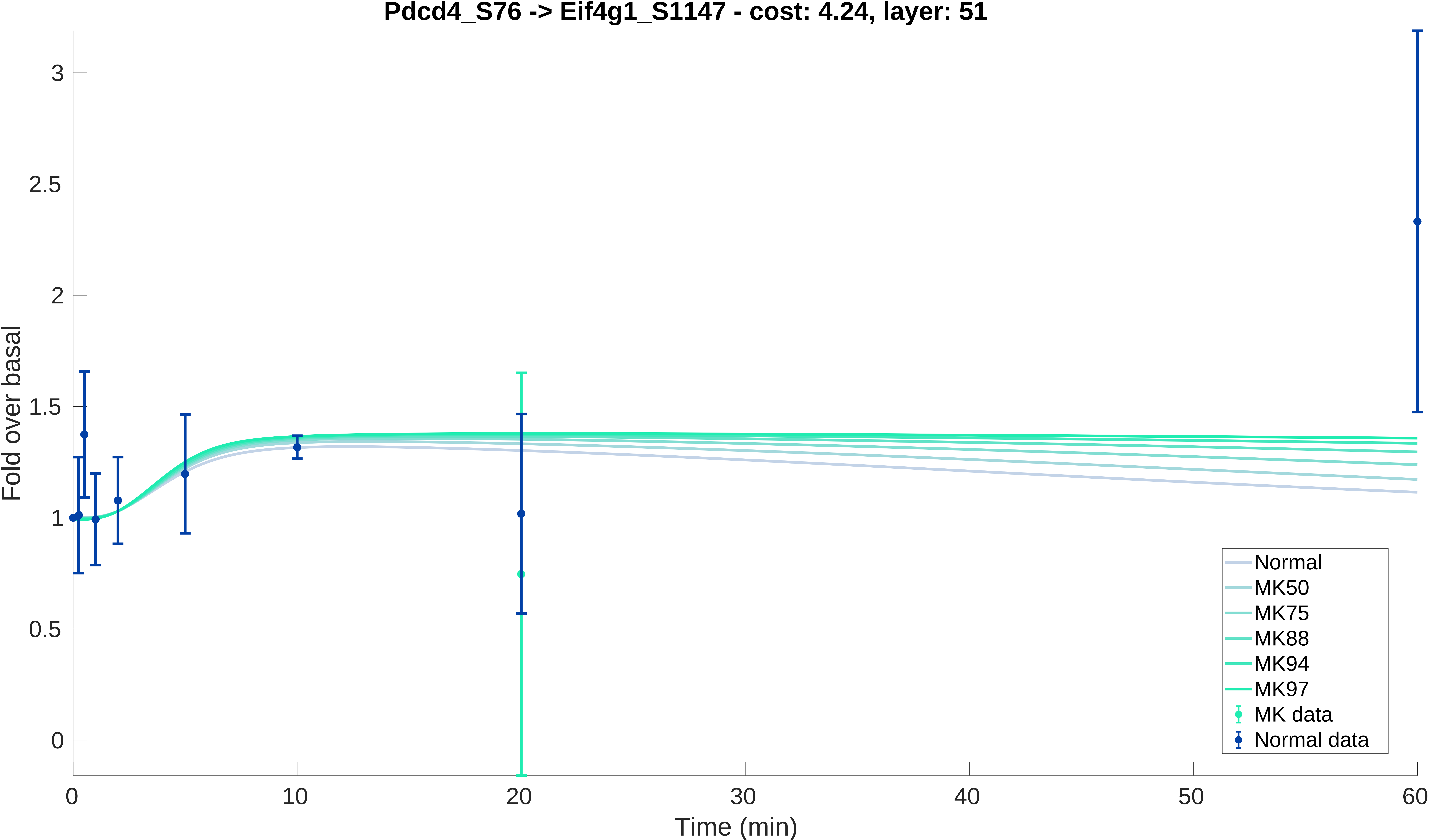
Pdcd4_S76 -> Eif4g1_S1147 - cost: 4.24, layer: 51
Fold over basal
Time (min)
0
0.5
1
1.5
2
2.5
3
0
10
20
30
40
50
60
Normal
MK50
MK75
MK88
MK94
MK97
MK data
Normal data

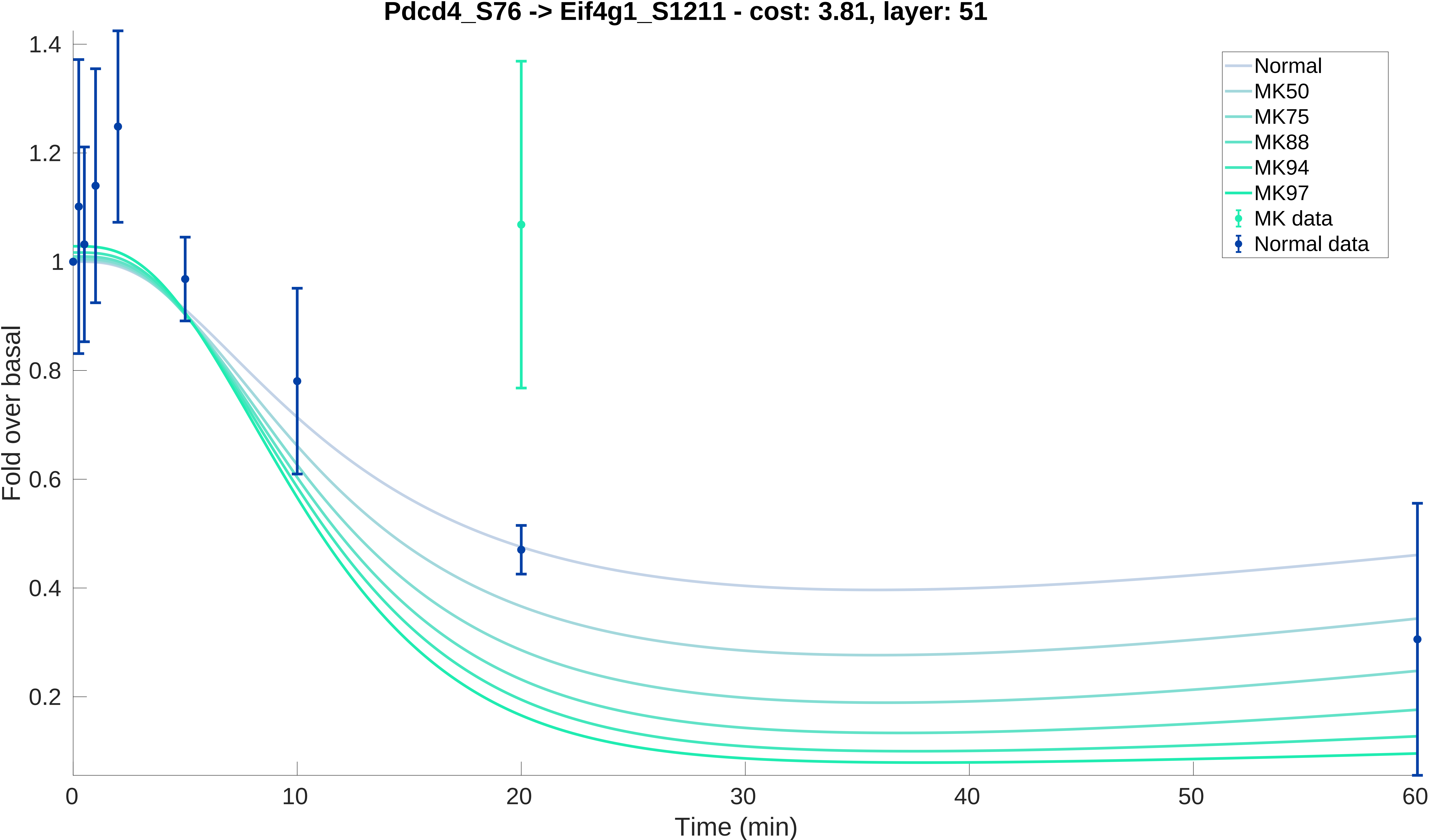

Pdcd4_S76 -> Eif4g1_S1211 - cost: 3.81, layer: 51
Fold over basal
Time (min)
0
10
20
30
40
50
60
0.2
0.4
0.6
0.8
1
1.2
1.4
Normal
MK50
MK75
MK88
MK94
MK97
MK data
Normal data

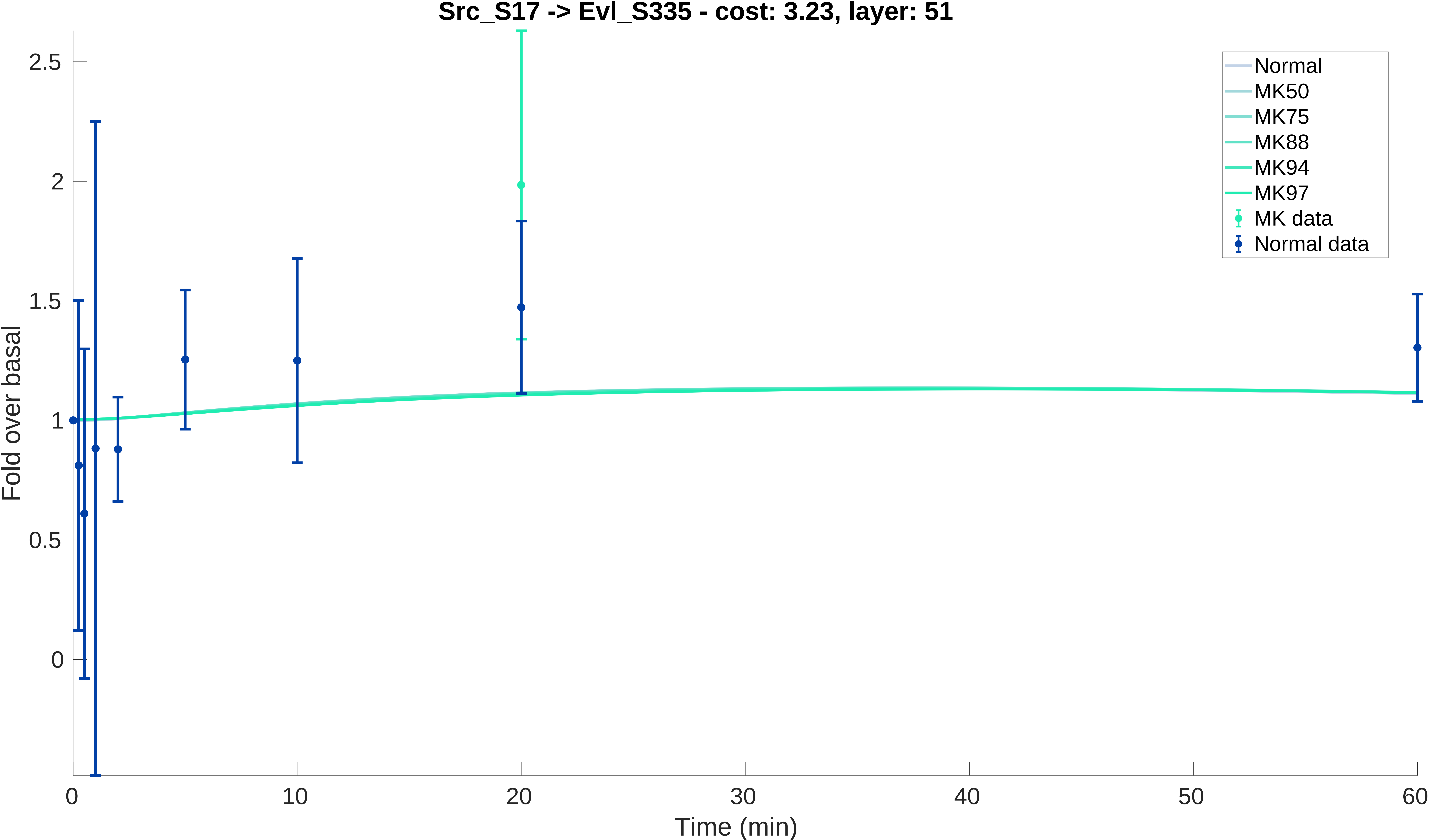
Src_S17 -> Evl_S335 - cost: 3.23, layer: 51
Fold over basal
Time (min)
Normal
MK50
MK75
MK88
MK94
MK97
MK data
Normal data
2.5
2
1.5
1
0.5
0
0
10
20
30
40
50
60

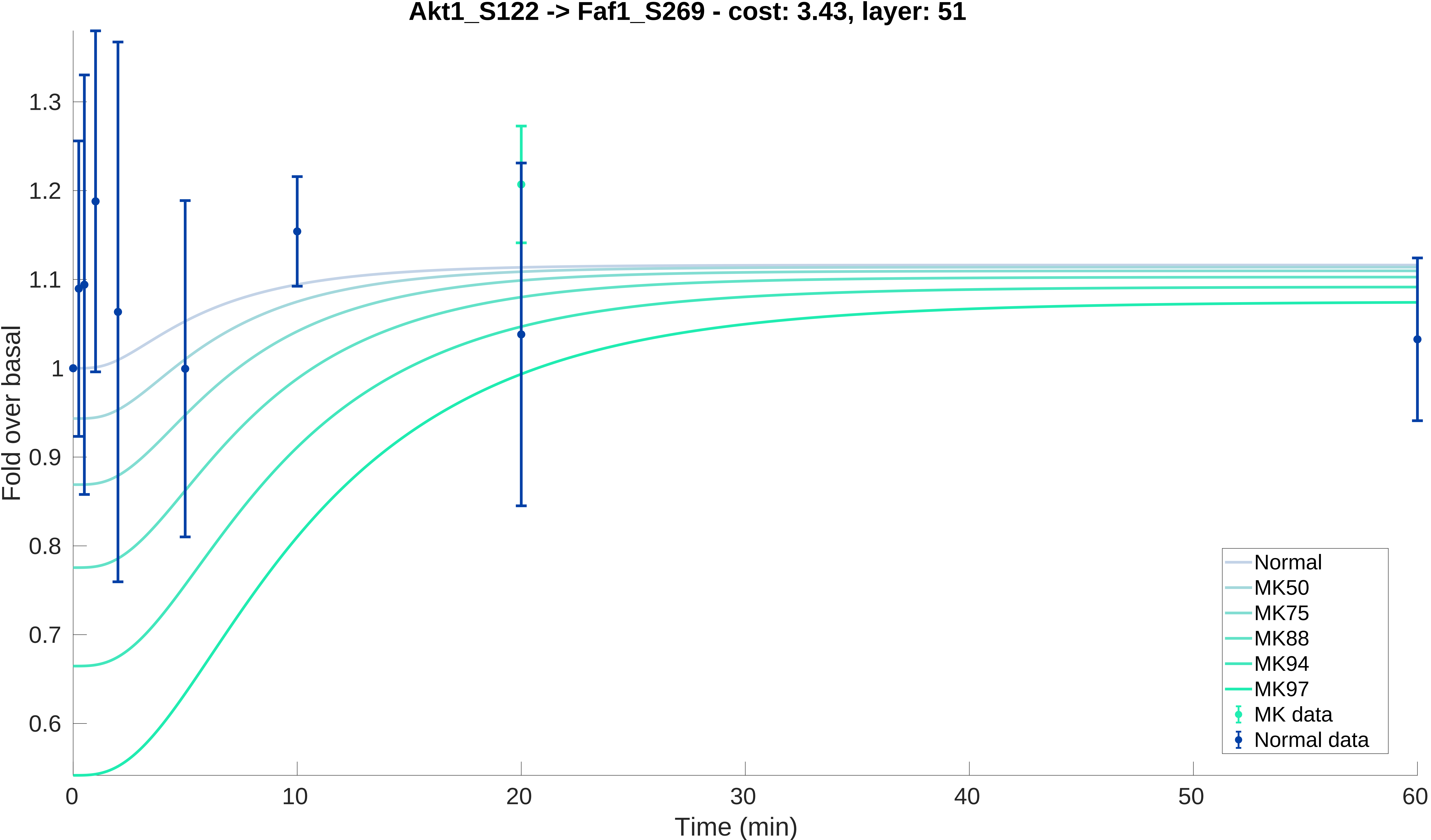

Akt1_S122 -> Faf1_S269 - cost: 3.43, layer: 51
Fold over basal
Time (min)
1.3
1.2
1.1
1
0.9
0.8
0.7
0.6
0
10
20
30
40
50
60
Normal
MK50
MK75
MK88
MK94
MK97
MK data
Normal data

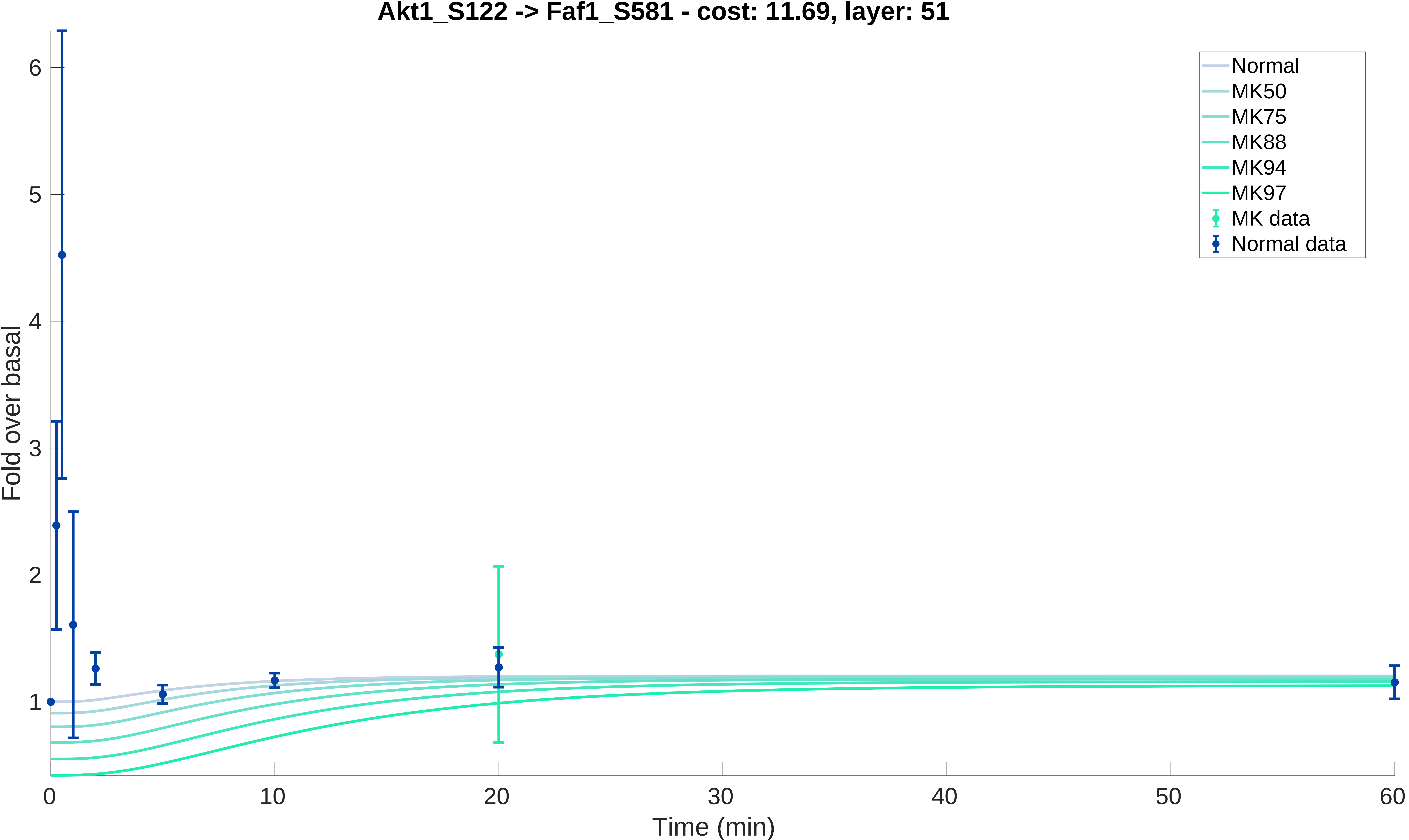

Akt1_S122 -> Faf1_S581 - cost: 11.69, layer: 51
Fold over basal
Time (min)
0
10
20
30
40
50
60
1
2
3
4
5
6
Normal
MK50
MK75
MK88
MK94
MK97
MK data
Normal data

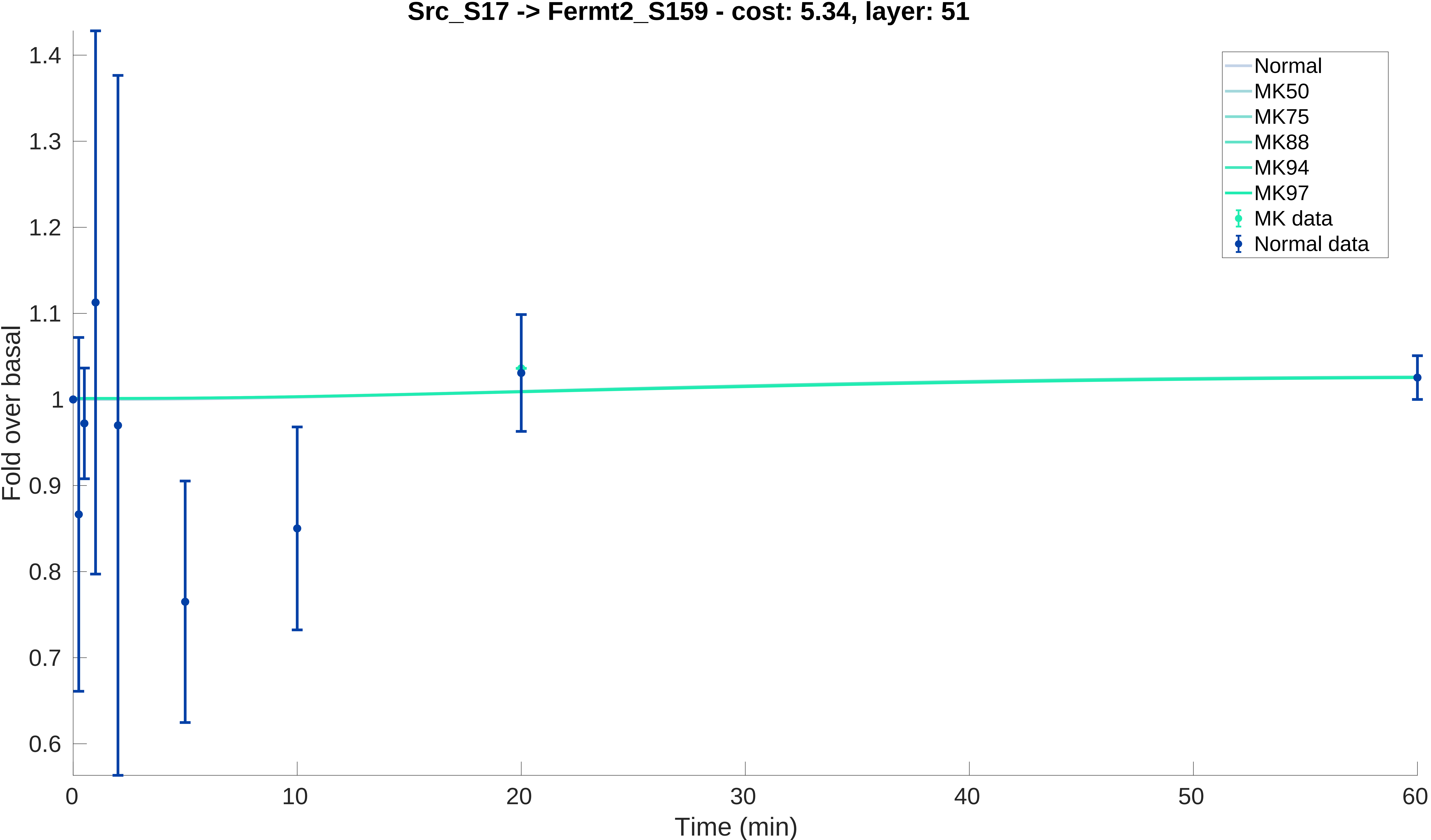

Src_S17 -> Fermt2_S159 - cost: 5.34, layer: 51
Fold over basal
1.4
1.3
1.2
1.1
1
0.9
0.8
0.7
0.6
Time (min)
0
10
20
30
40
50
60
Normal
MK50
MK75
MK88
MK94
MK97
MK data
Normal data

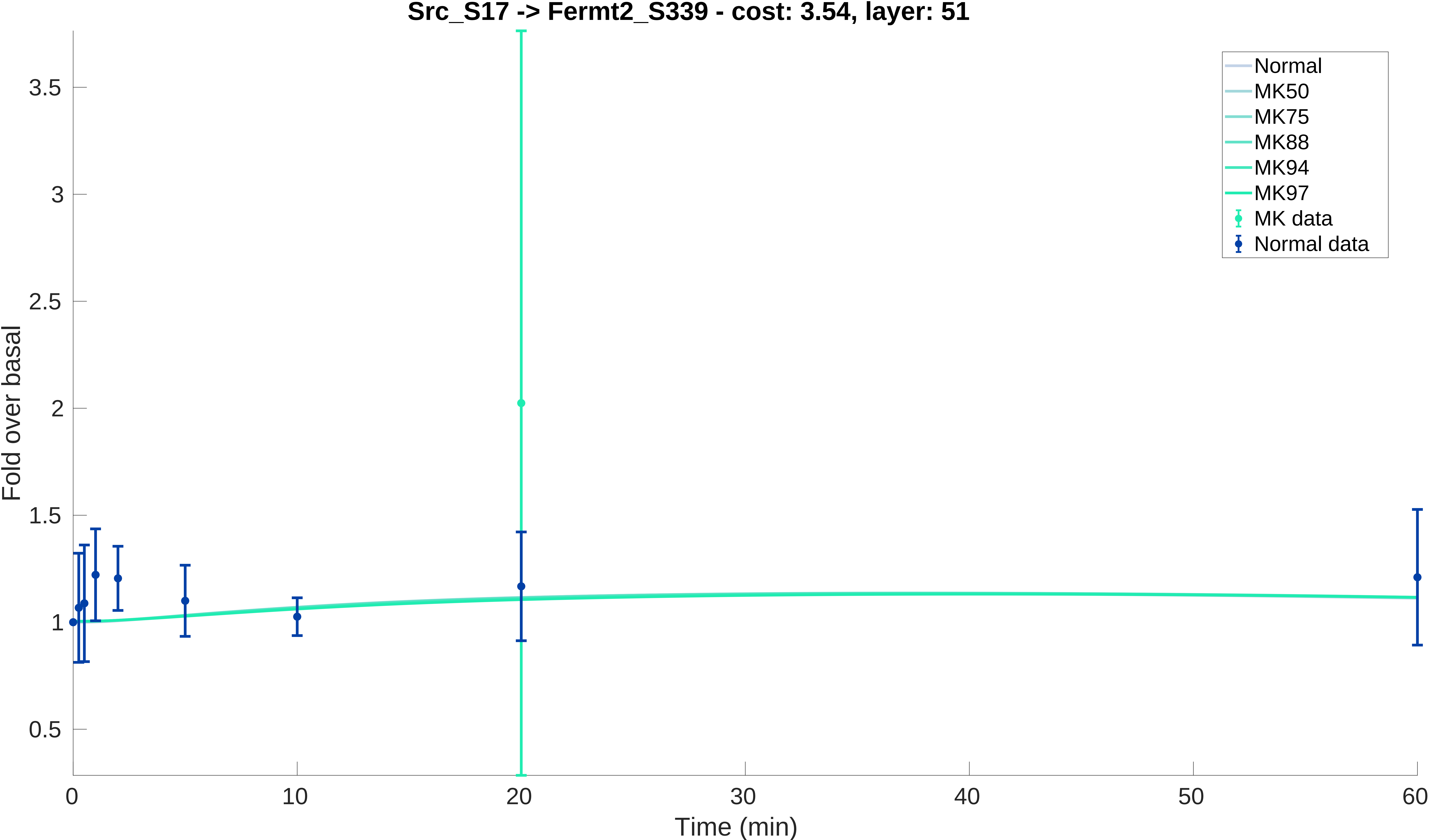

Src_S17 -> Fermt2_S339 - cost: 3.54, layer: 51
Fold over basal
Time (min)
Normal
MK50
MK75
MK88
MK94
MK97
MK data
Normal data
0.5
1
1.5
2
2.5
3
3.5
0
10
20
30
40
50
60

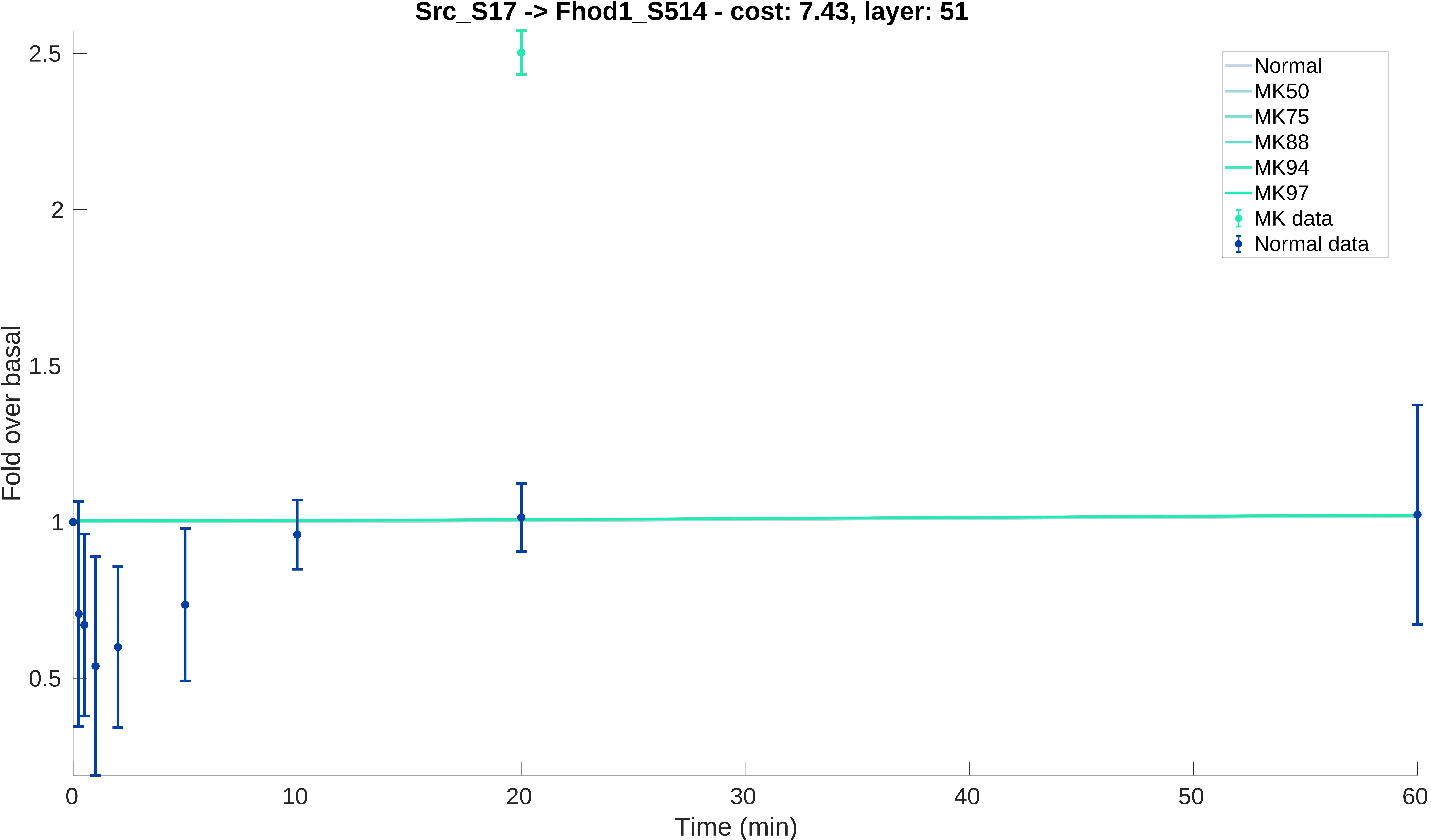
Src_S17 -> Fhod1_S514 - cost: 7.43, layer: 51
Fold over basal
2.5
2
1.5
1
0.5
0
10
20
30
40
50
60
Time (min)
Normal
MK50
MK75
MK88
MK94
MK97
MK data
Normal data

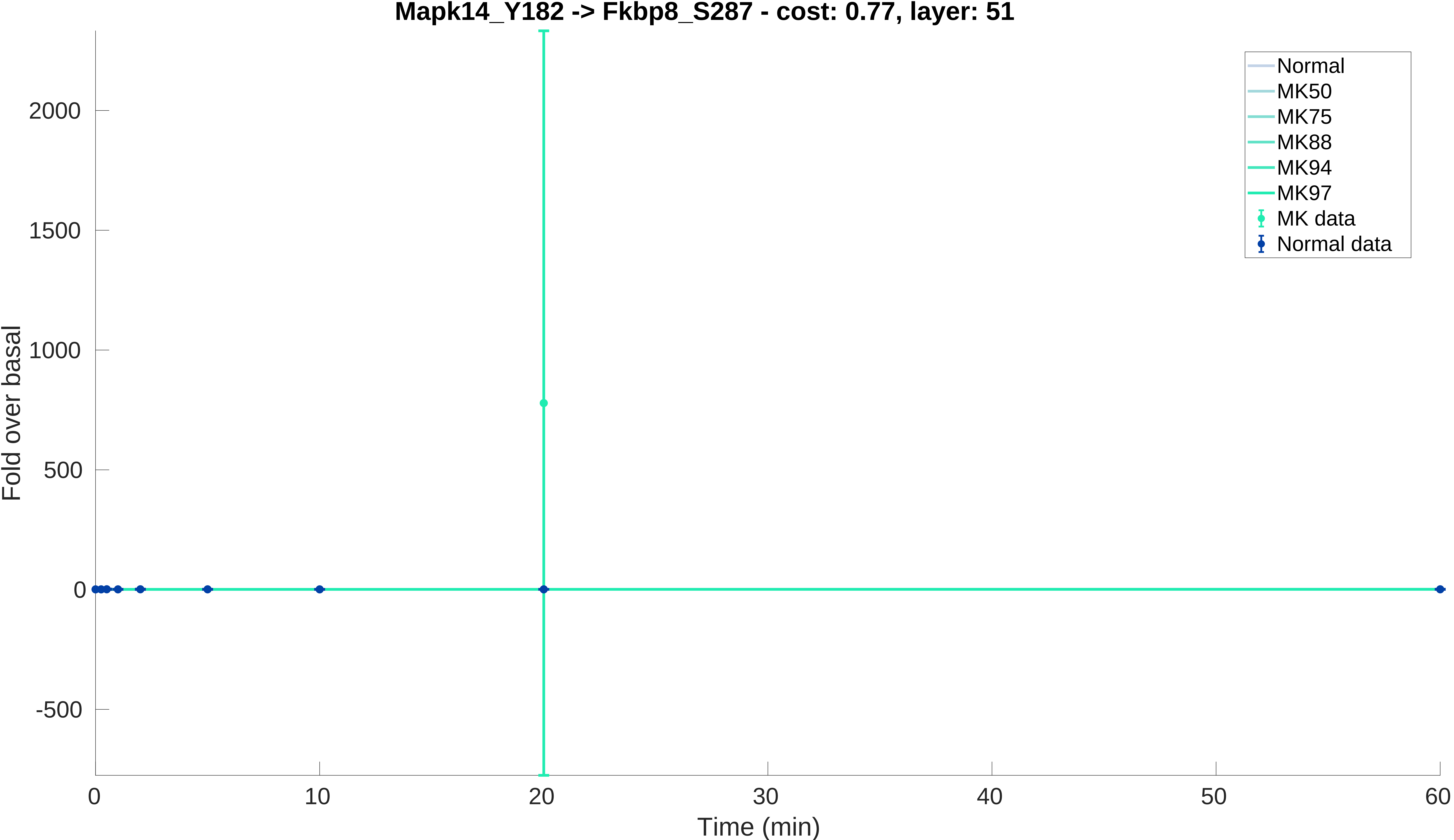

Mapk14_Y182 -> Fkbp8_S287 - cost: 0.77, layer: 51
Normal
MK50
MK75
MK88
MK94
MK97
MK data
Normal data
Fold over basal
2000
1500
1000
500
0
-500
0
10
20
30
40
50
60
Time (min)

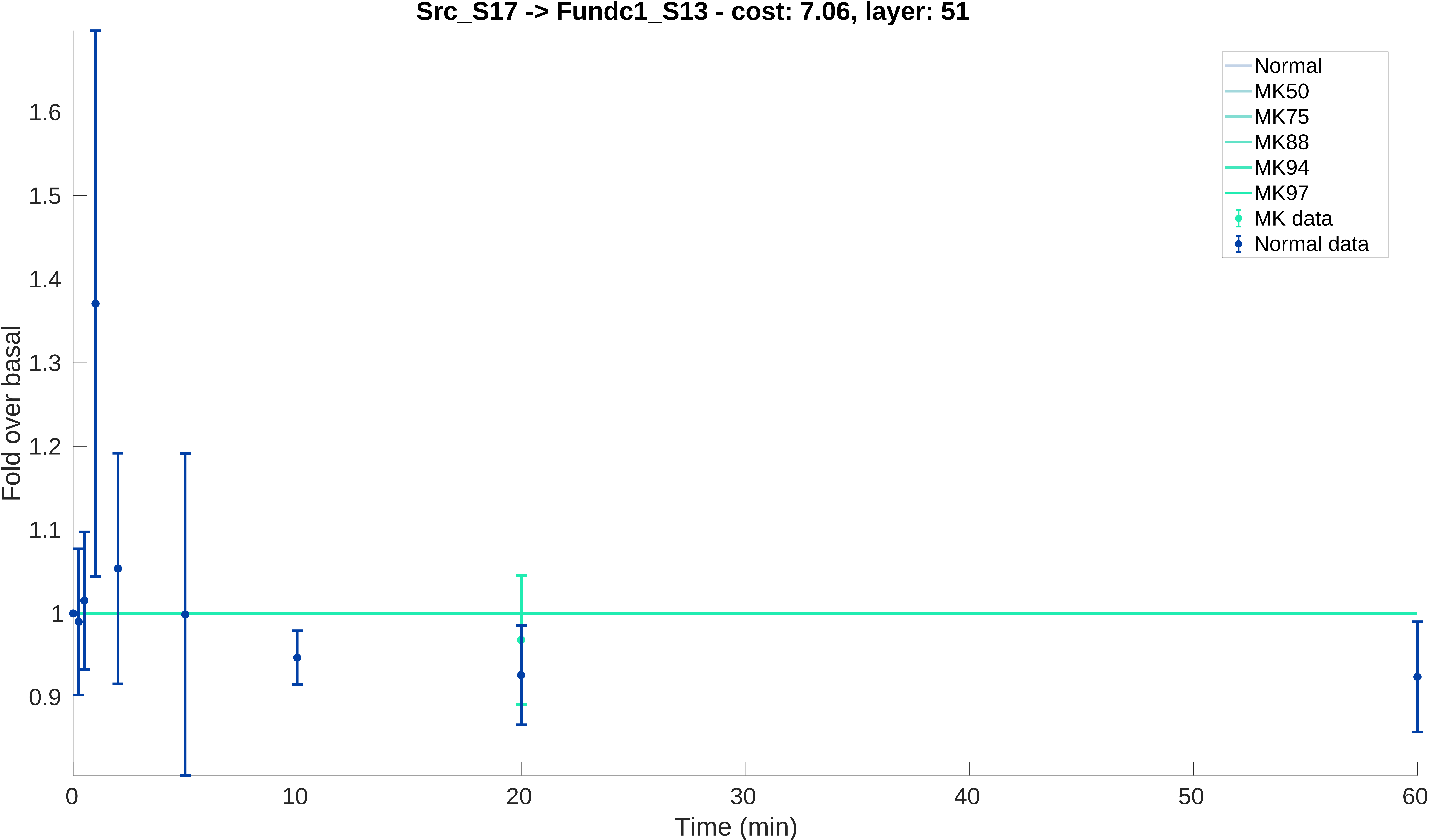
Src_S17 -> Fundc1_S13 - cost: 7.06, layer: 51
Fold over basal
Time (min)
Normal
MK50
MK75
MK88
MK94
MK97
MK data
Normal data
1.6
1.5
1.4
1.3
1.2
1.1
1
0.9
0
10
20
30
40
50
60

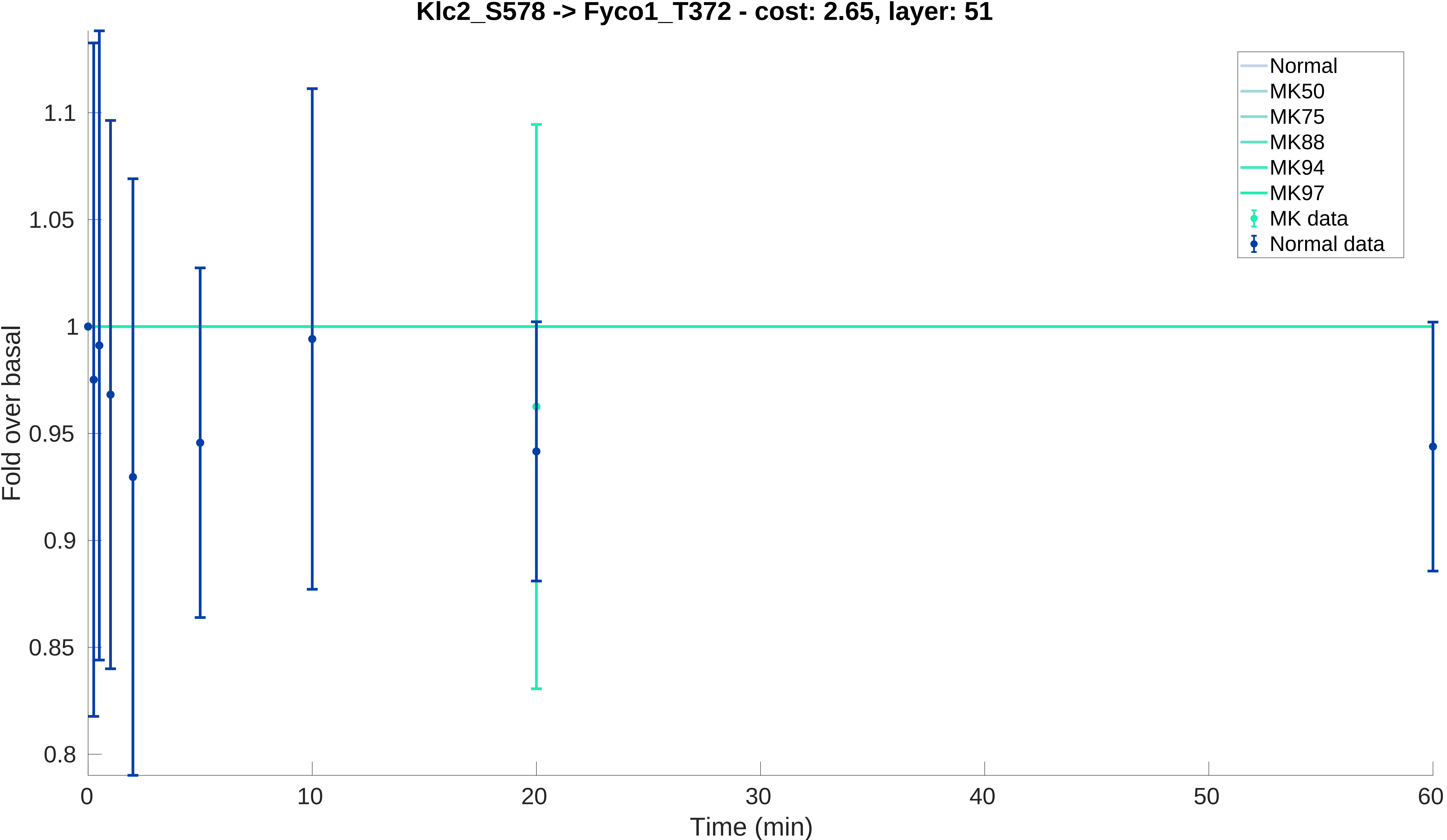

Klc2_S578 -> Fyco1_T372 - cost: 2.65, layer: 51
Fold over basal
Time (min)
0.8
0.85
0.9
0.95
1
1.05
1.1
0
10
20
30
40
50
60
Normal
MK50
MK75
MK88
MK94
MK97
MK data
Normal data

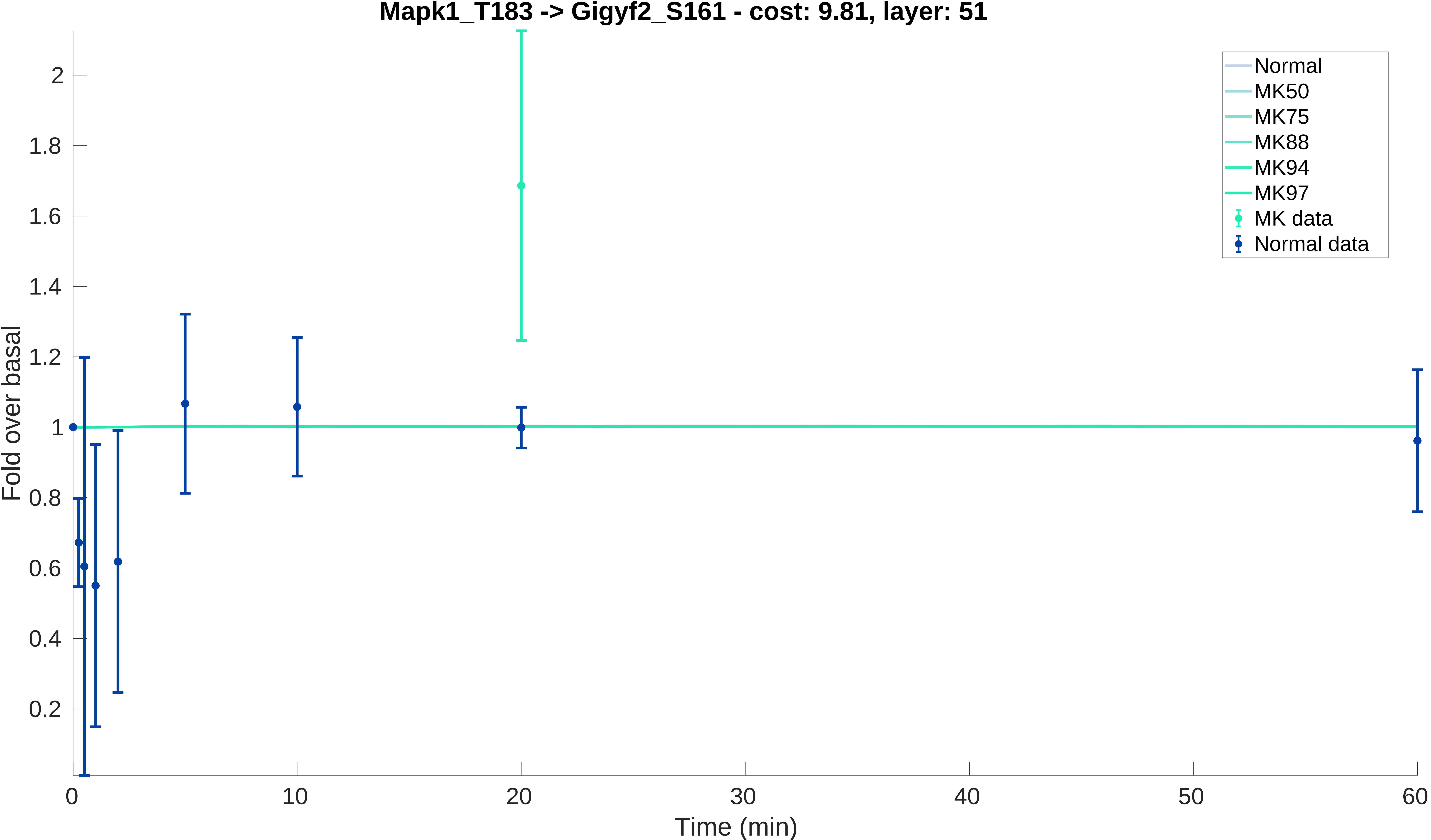

Mapk1_T183 -> Gigyf2_S161 - cost: 9.81, layer: 51
Fold over basal
Time (min)
0
10
20
30
40
50
60
0.2
0.4
0.6
0.8
1
1.2
1.4
1.6
1.8
2
Normal
MK50
MK75
MK88
MK94
MK97
MK data
Normal data

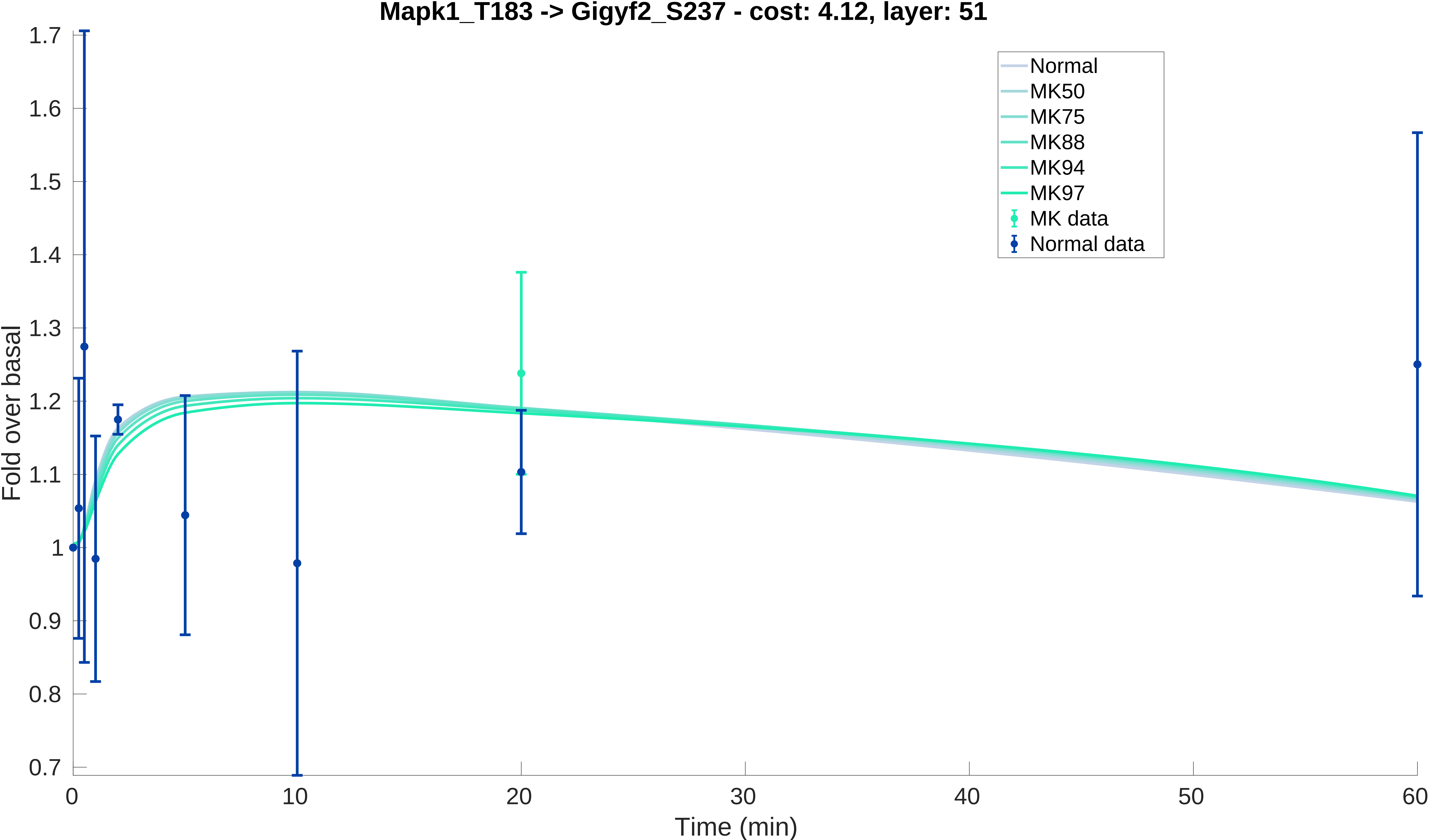

Mapk1_T183 -> Gigyf2_S237 - cost: 4.12, layer: 51
Fold over basal
Time (min)
1.7
1.6
1.5
1.4
1.3
1.2
1.1
1
0.9
0.8
0.7
0
10
20
30
40
50
60
Normal
MK50
MK75
MK88
MK94
MK97
MK data
Normal data

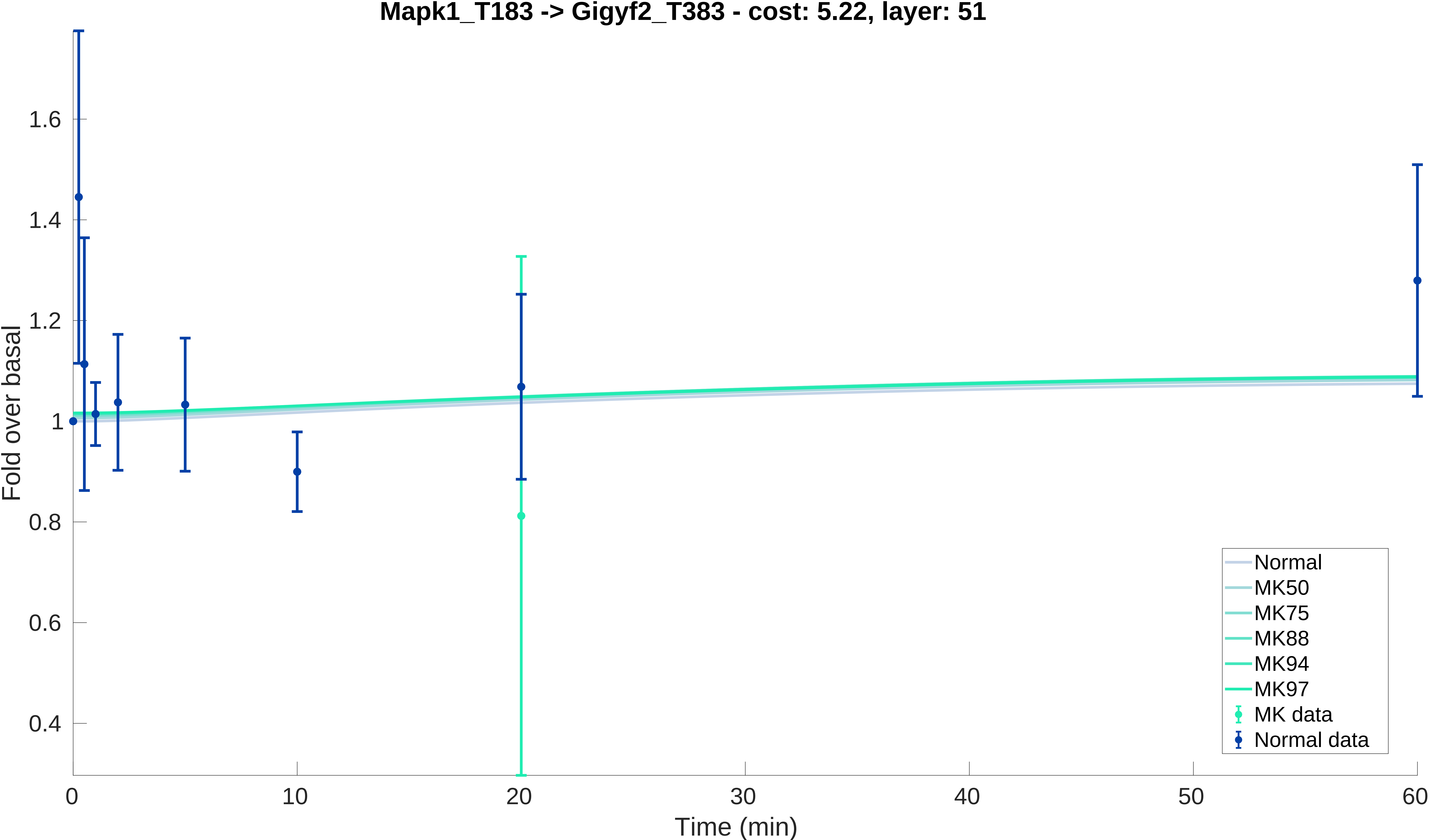
Mapk1_T183 -> Gigyf2_T383 - cost: 5.22, layer: 51
Fold over basal
1.6
1.4
1.2
1
0.8
0.6
0.4
0
10
20
30
40
50
60
Time (min)
Normal
MK50
MK75
MK88
MK94
MK97
MK data
Normal data

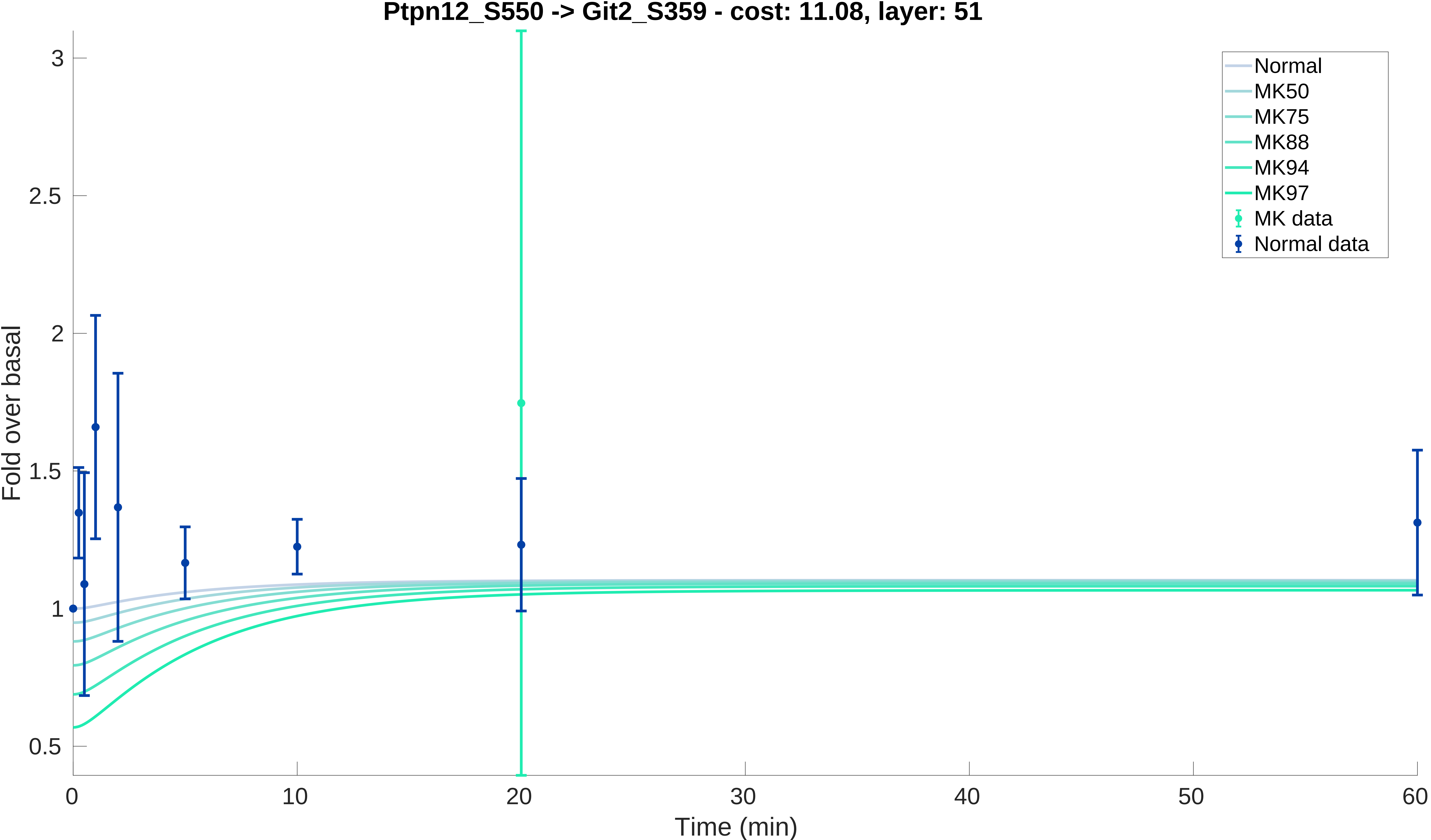

Ptpn12_S550 -> Git2_S359 - cost: 11.08, layer: 51
Fold over basal
Time (min)
Normal
MK50
MK75
MK88
MK94
MK97
MK data
Normal data
0.5
1
1.5
2
2.5
3
0
10
20
30
40
50
60

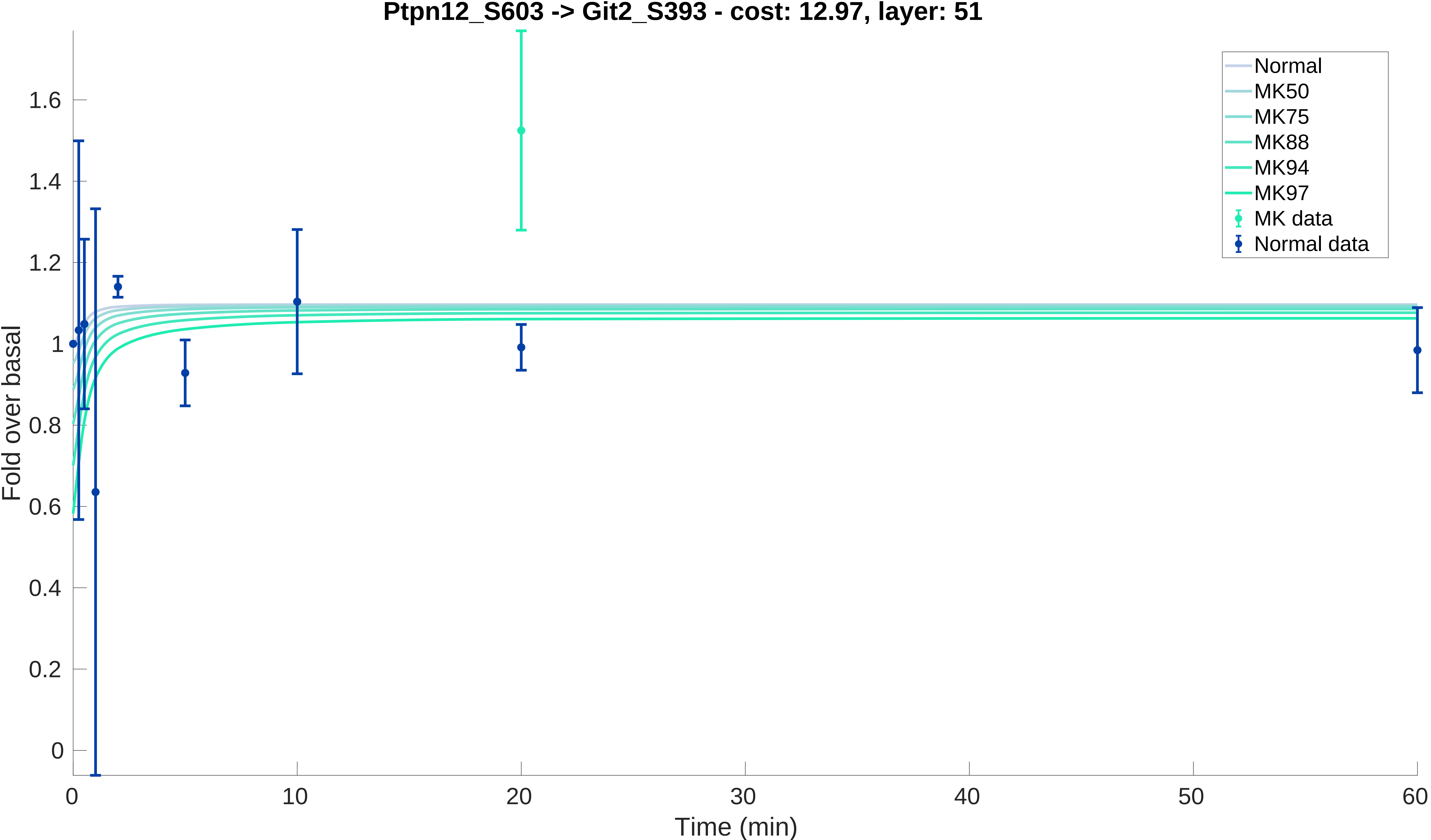

Ptpn12_S603 -> Git2_S393 - cost: 12.97, layer: 51
Fold over basal
Time (min)
1.6
1.4
1.2
1
0.8
0.6
0.4
0.2
0
0
10
20
30
40
50
60
Normal
MK50
MK75
MK88
MK94
MK97
MK data
Normal data

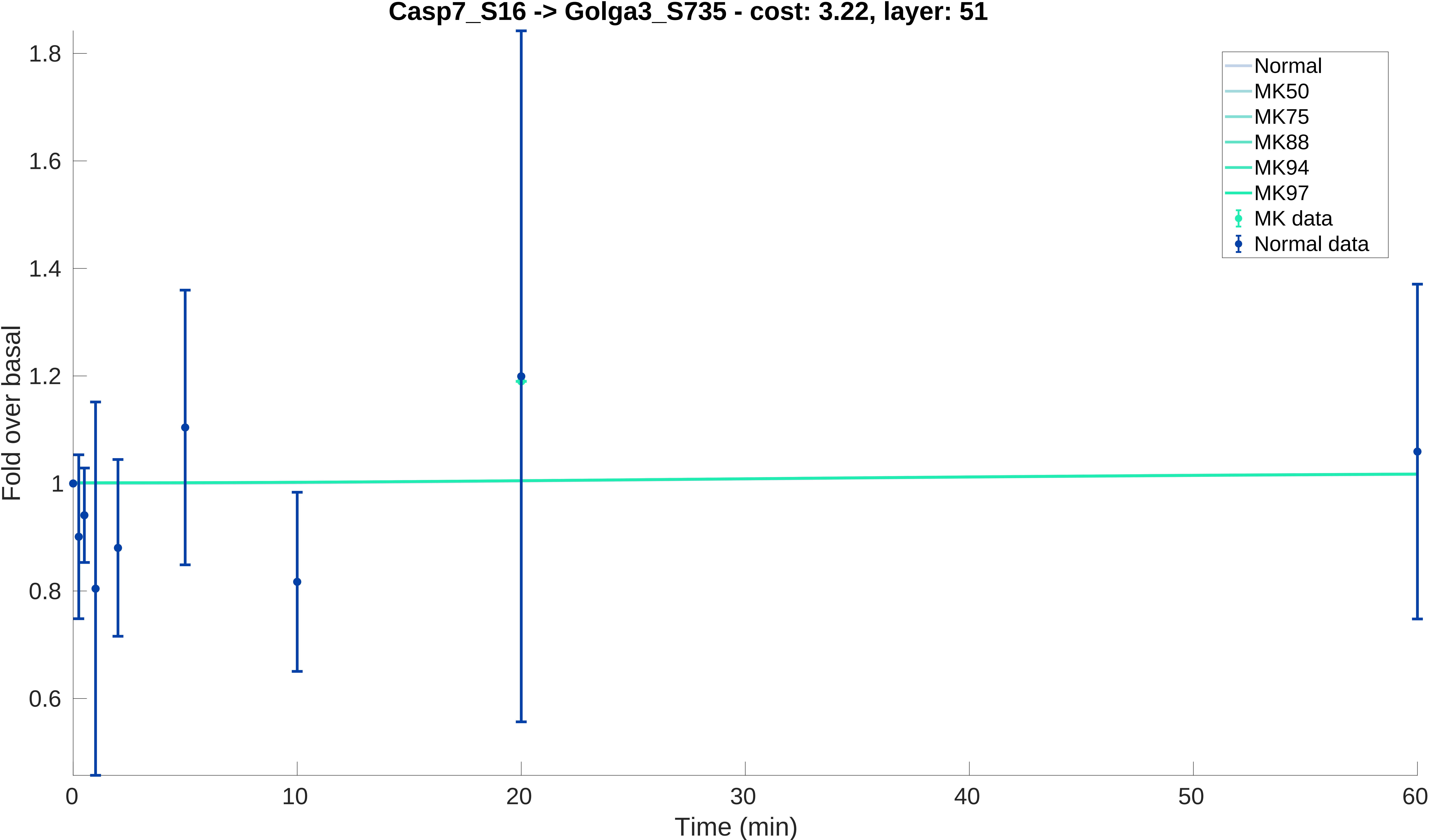

Casp7_S16 -> Golga3_S735 - cost: 3.22, layer: 51
Fold over basal
Time (min)
1.8
1.6
1.4
1.2
1
0.8
0.6
0
10
20
30
40
50
60
Normal
MK50
MK75
MK88
MK94
MK97
MK data
Normal data

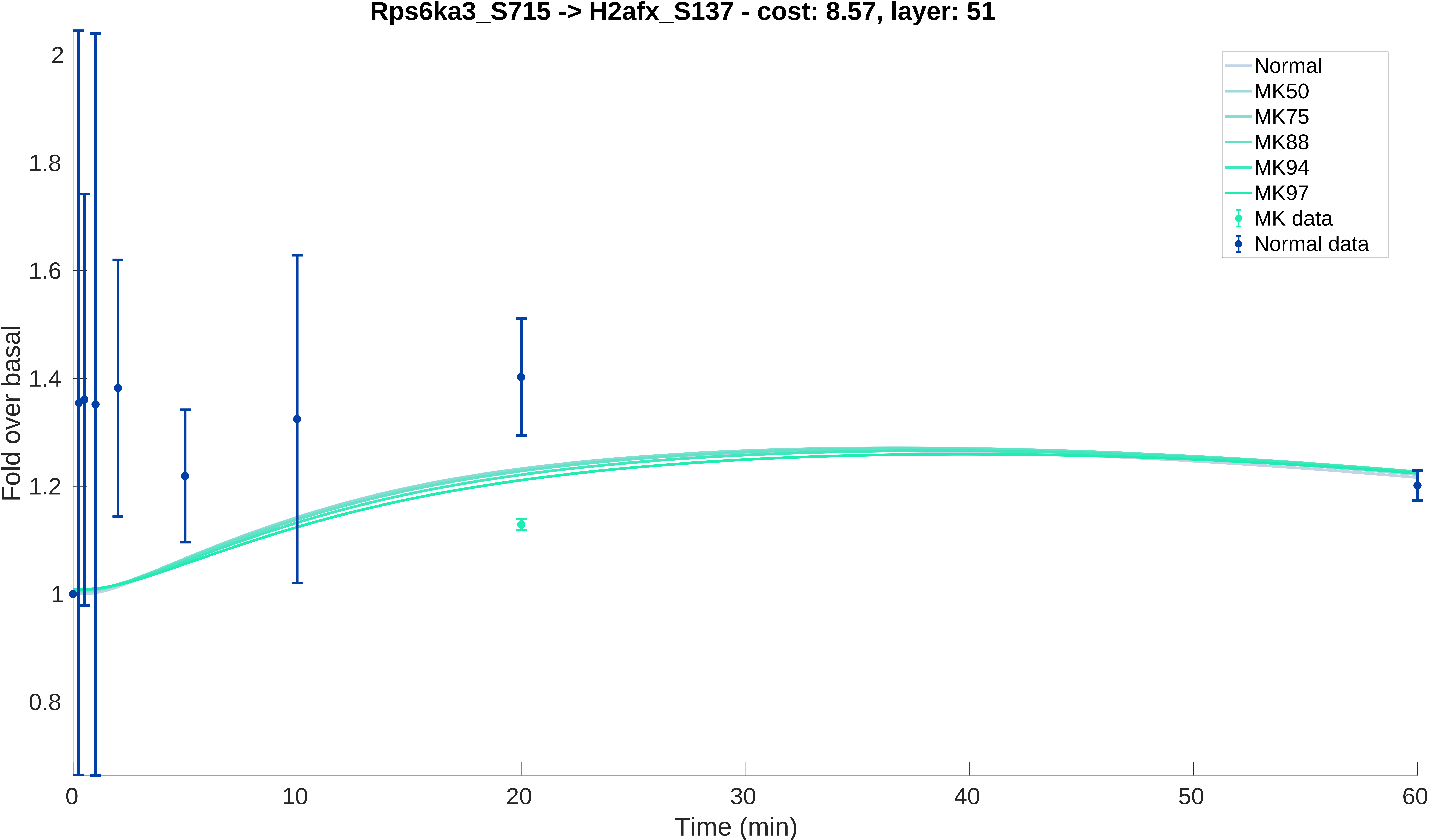

Rps6ka3_S715 -> H2afx_S137 - cost: 8.57, layer: 51
Normal
MK50
MK75
MK88
MK94
MK97
MK data
Normal data
Fold over basal
2
1.8
1.6
1.4
1.2
1
0.8
0
10
20
30
40
50
60
Time (min)

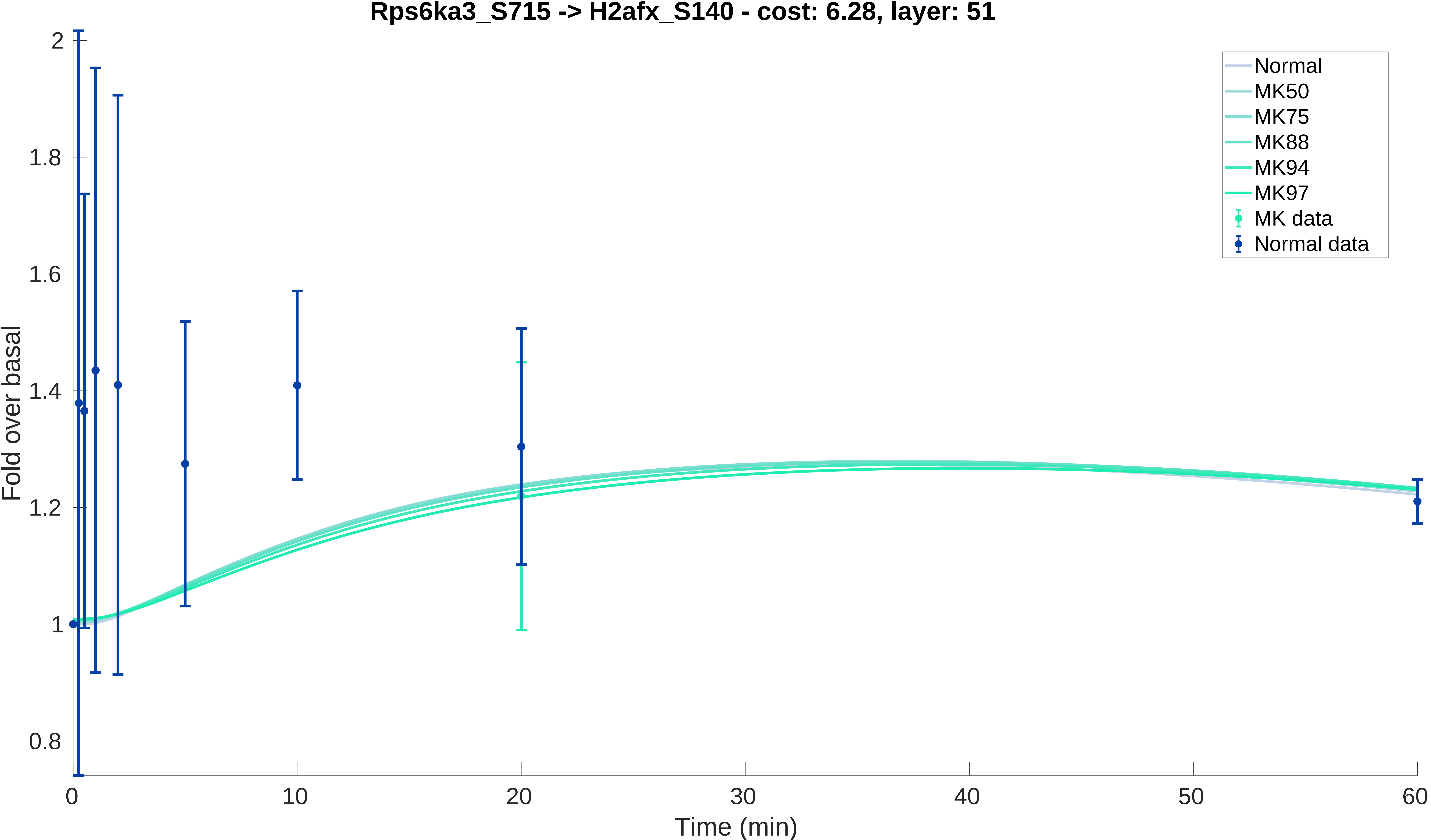
Rps6ka3_S715 -> H2afx_S140 - cost: 6.28, layer: 51
Fold over basal
Time (min)
0
10
20
30
40
50
60
0.8
1
1.2
1.4
1.6
1.8
2
Normal
MK50
MK75
MK88
MK94
MK97
MK data
Normal data

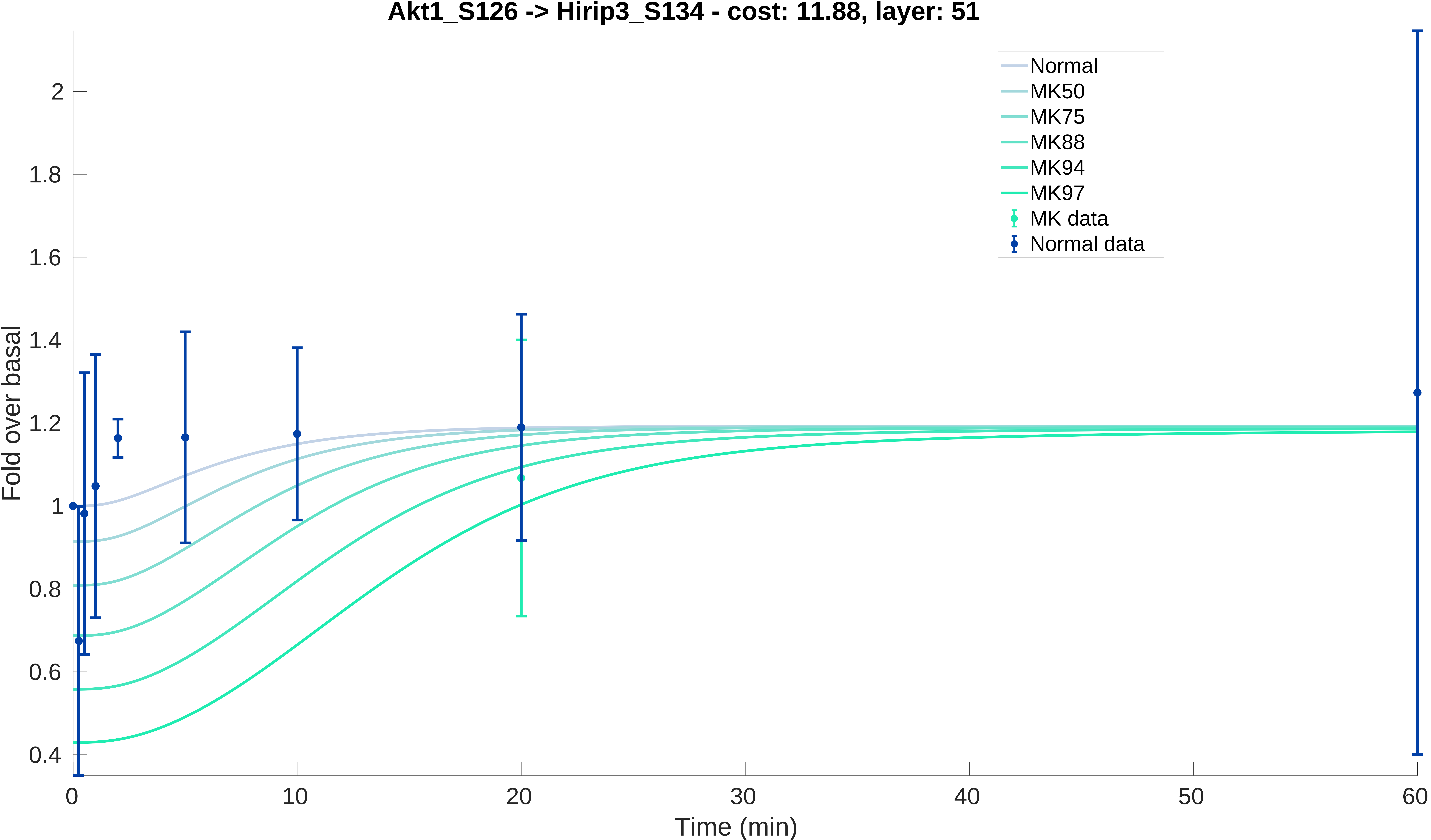
Akt1_S126 -> Hirip3_S134 - cost: 11.88, layer: 51
Fold over basal
Time (min)
Normal
MK50
MK75
MK88
MK94
MK97
MK data
Normal data
0.4
0.6
0.8
1
1.2
1.4
1.6
1.8
2
0
10
20
30
40
50
60

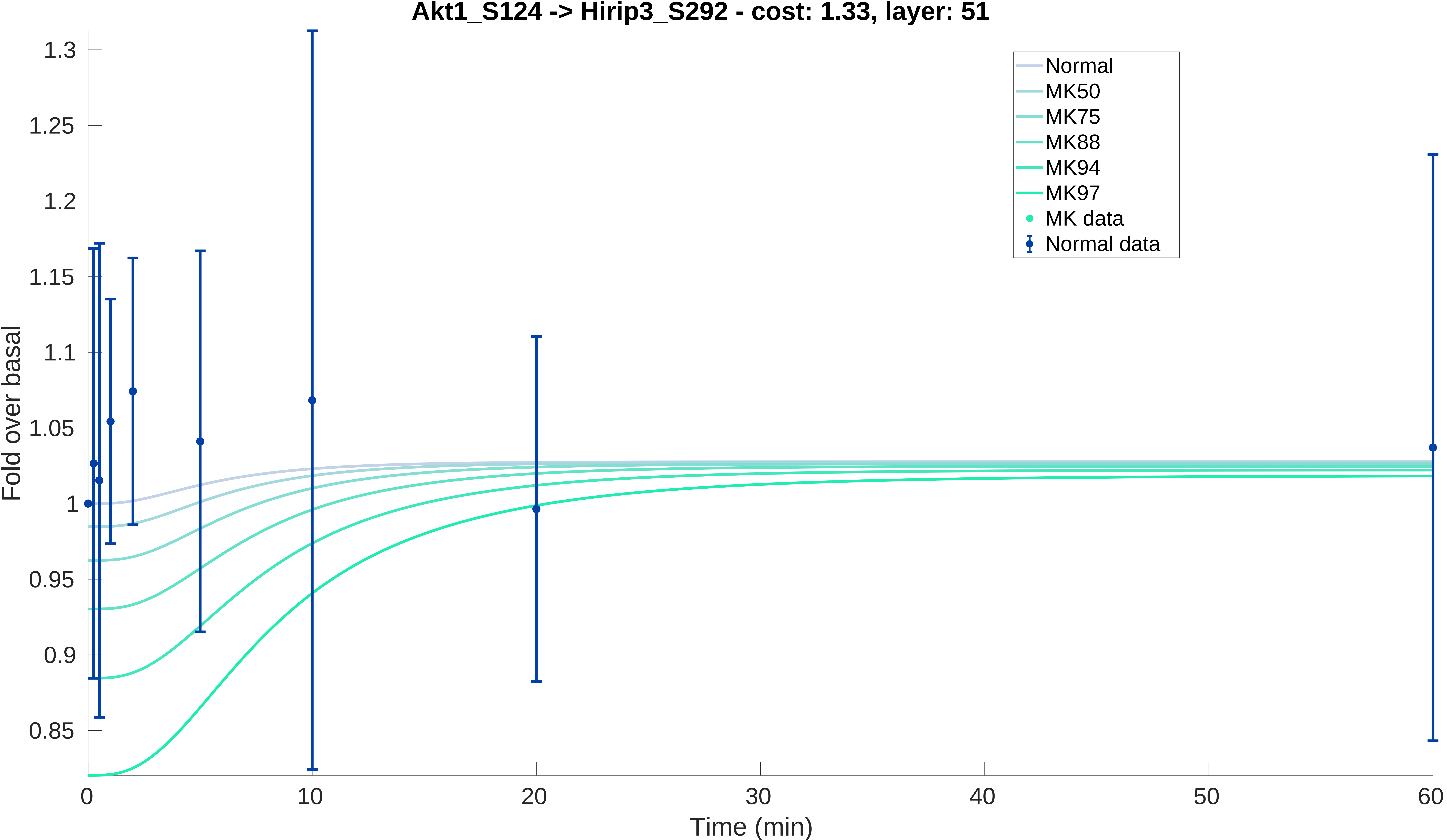

Akt1_S124 -> Hirip3_S292 - cost: 1.33, layer: 51
Fold over basal
1.3
1.25
1.2
1.15
1.1
1.05
1
0.95
0.9
0.85
0
10
20
30
40
50
60
Time (min)
Normal
MK50
MK75
MK88
MK94
MK97
MK data
Normal data

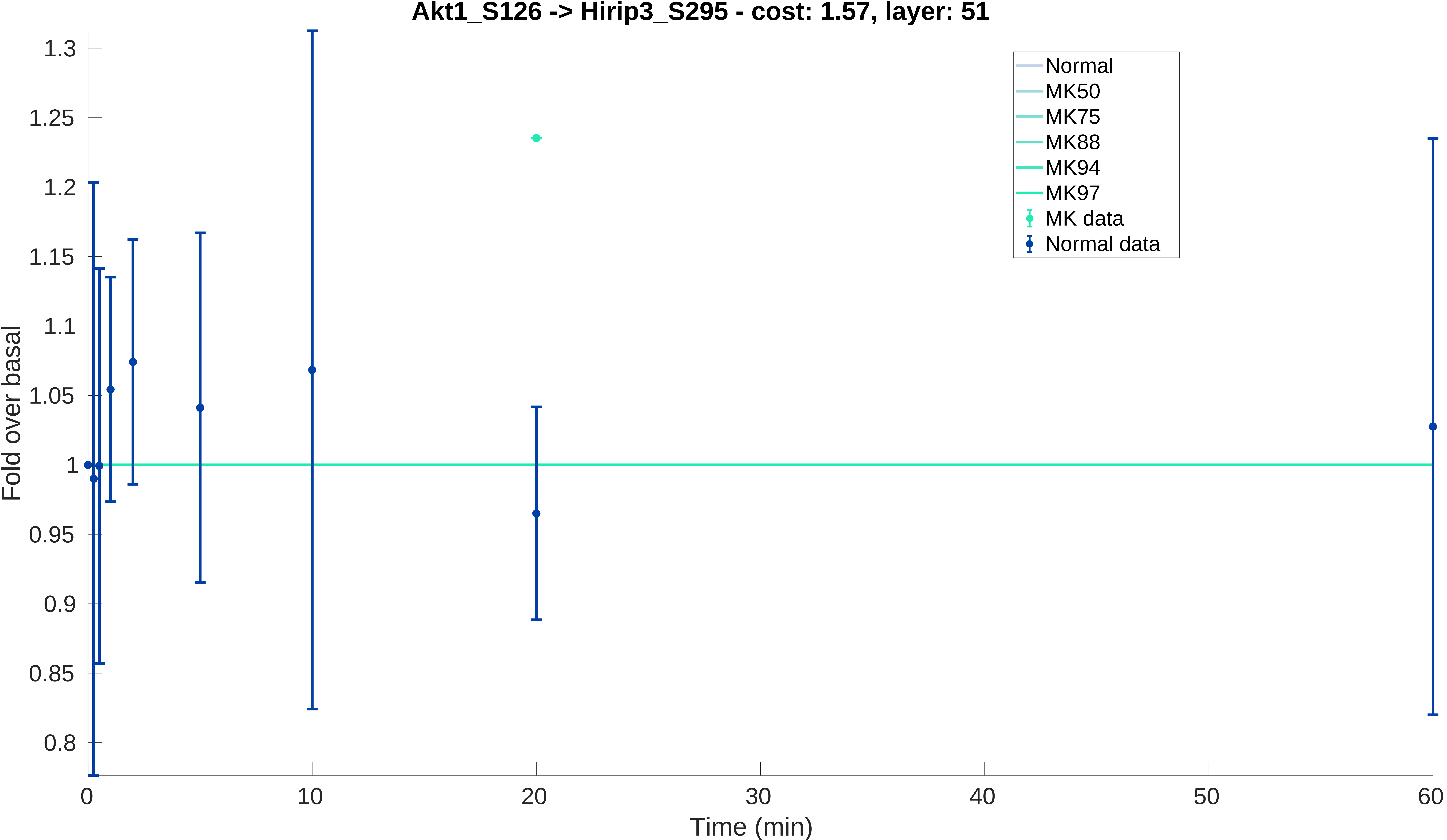

Akt1_S126 -> Hirip3_S295 - cost: 1.57, layer: 51
Fold over basal
Time (min)
1.3
1.25
1.2
1.15
1.1
1.05
1
0.95
0.9
0.85
0.8
0
10
20
30
40
50
60
Normal
MK50
MK75
MK88
MK94
MK97
MK data
Normal data

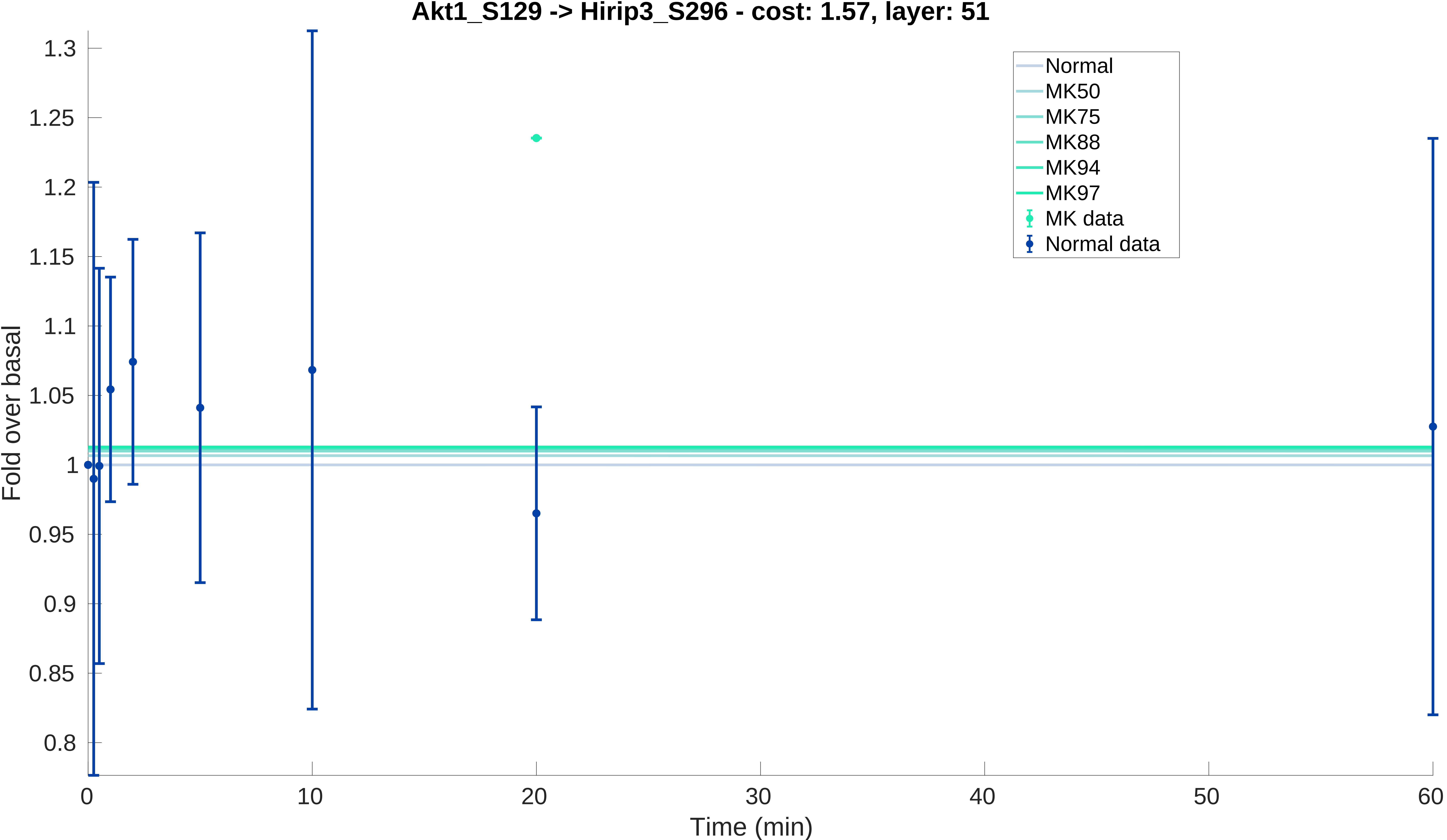
Akt1_S129 -> Hirip3_S296 - cost: 1.57, layer: 51
Fold over basal
1.3
1.25
1.2
1.15
1.1
1.05
1
0.95
0.9
0.85
0.8
0
10
20
30
40
50
60
Time (min)
Normal
MK50
MK75
MK88
MK94
MK97
MK data
Normal data

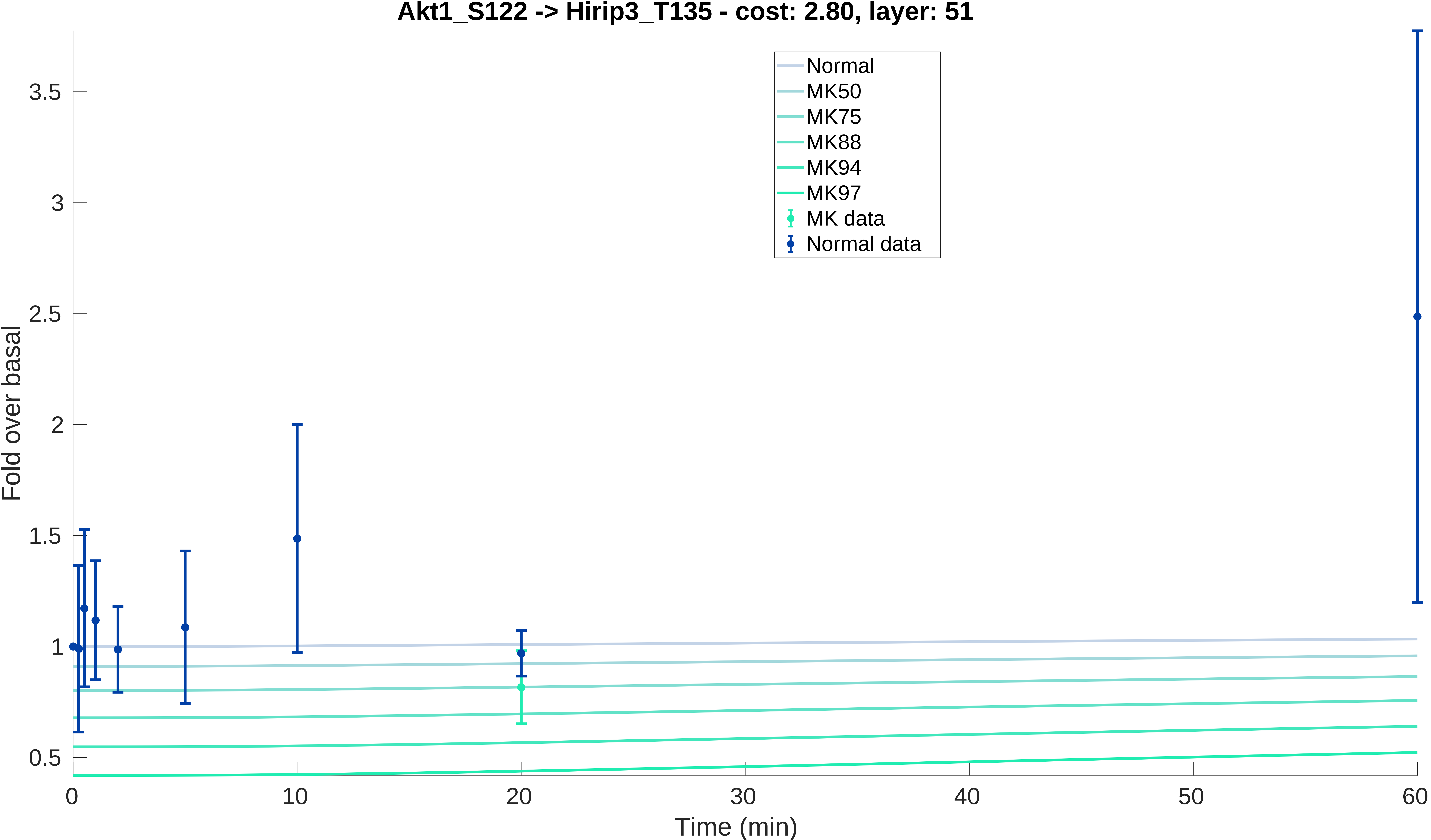

Akt1_S122 -> Hirip3_T135 - cost: 2.80, layer: 51
Normal
MK50
MK75
MK88
MK94
MK97
MK data
Normal data
Fold over basal
Time (min)
0.5
1
1.5
2
2.5
3
3.5
0
10
20
30
40
50
60

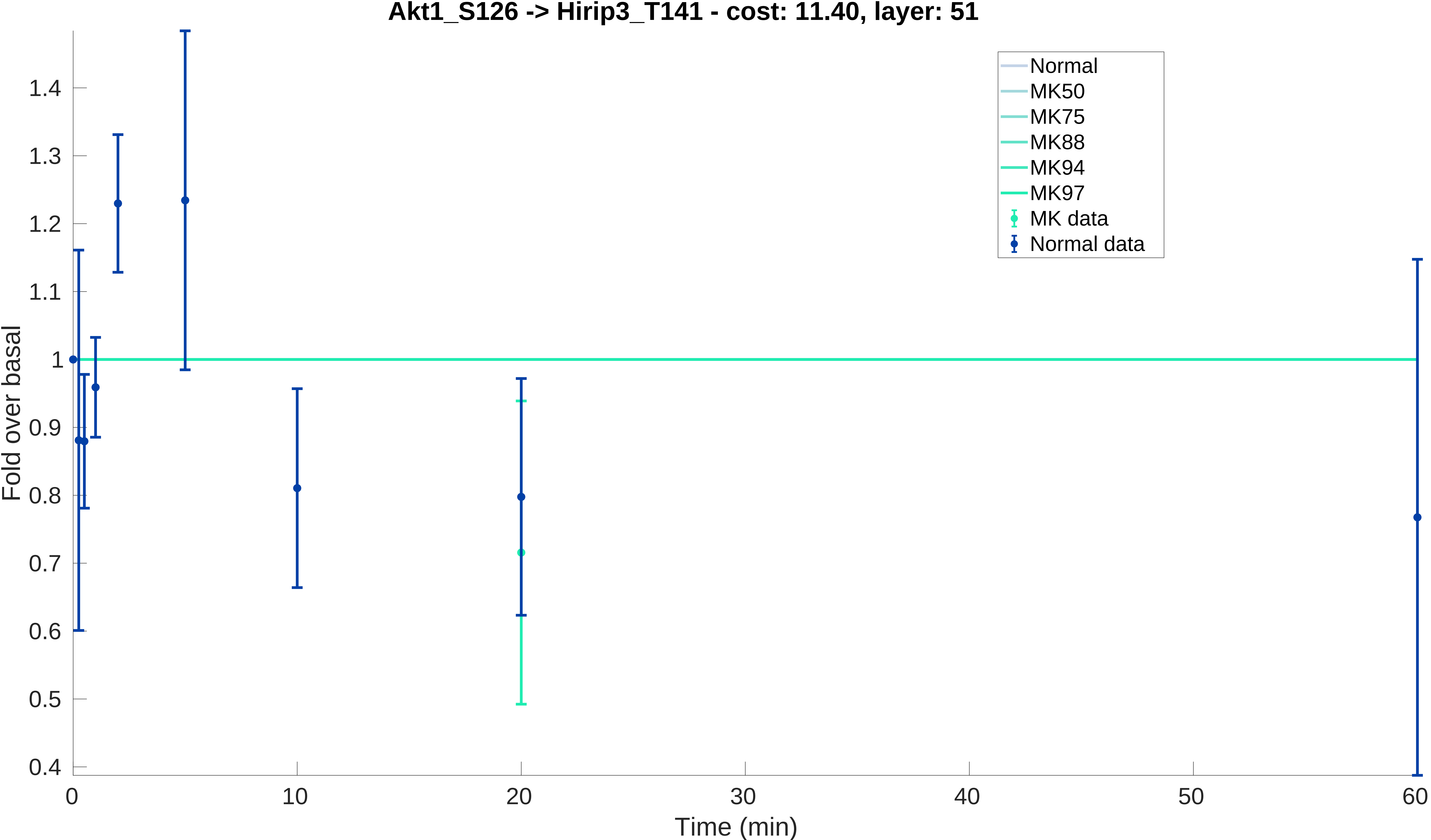

Akt1_S126 -> Hirip3_T141 - cost: 11.40, layer: 51
Normal
MK50
MK75
MK88
MK94
MK97
MK data
Normal data
Fold over basal
Time (min)
0.4
0.5
0.6
0.7
0.8
0.9
1
1.1
1.2
1.3
1.4
0
10
20
30
40
50
60

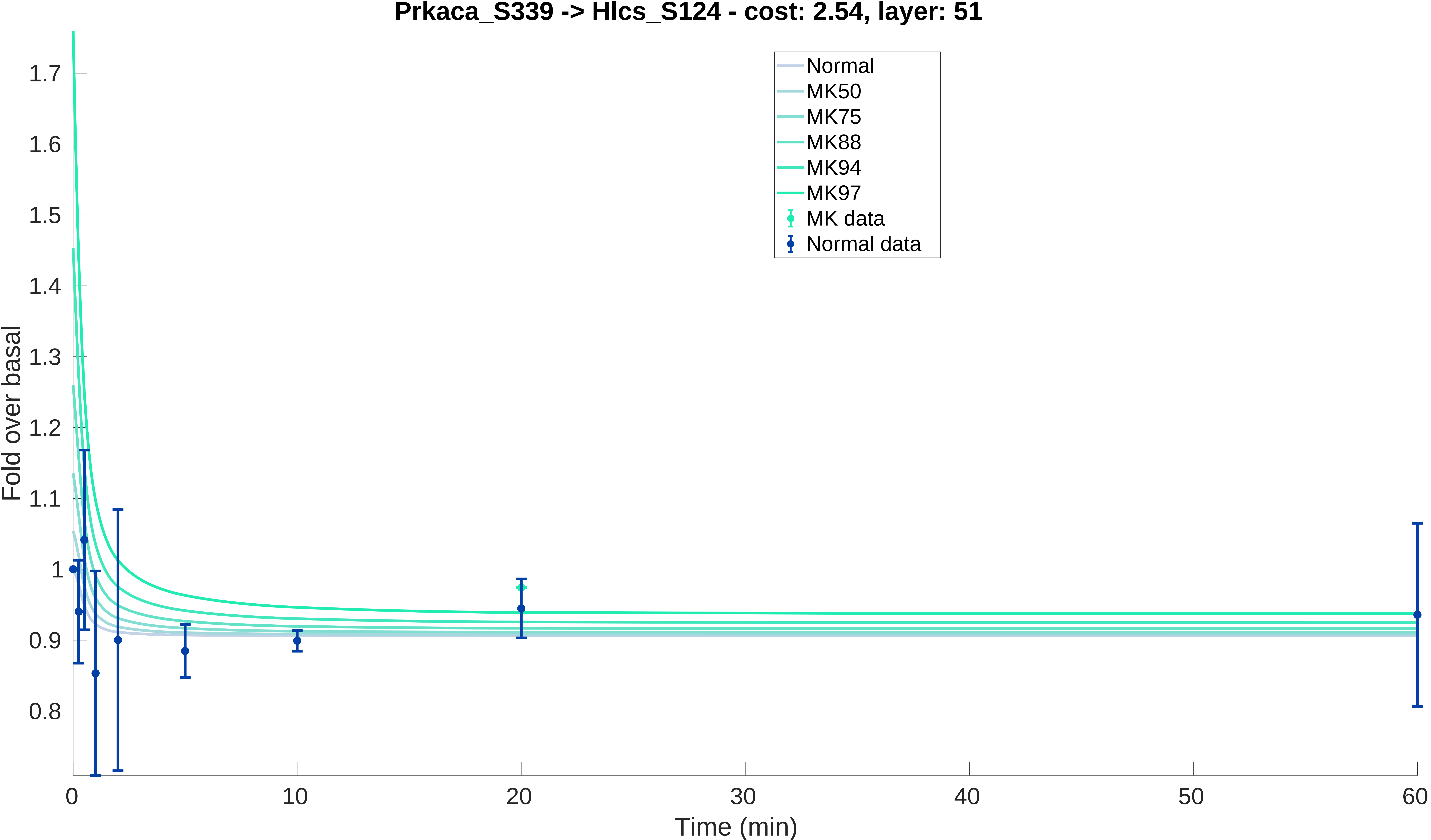

Prrkaca_S339 -> Hlcs_S124 - cost: 2.54, layer: 51
Normal
MK50
MK75
MK88
MK94
MK97
MK data
Normal data
Fold over basal
1.7
1.6
1.5
1.4
1.3
1.2
1.1
1
0.9
0.8
0
10
20
30
40
50
60
Time (min)

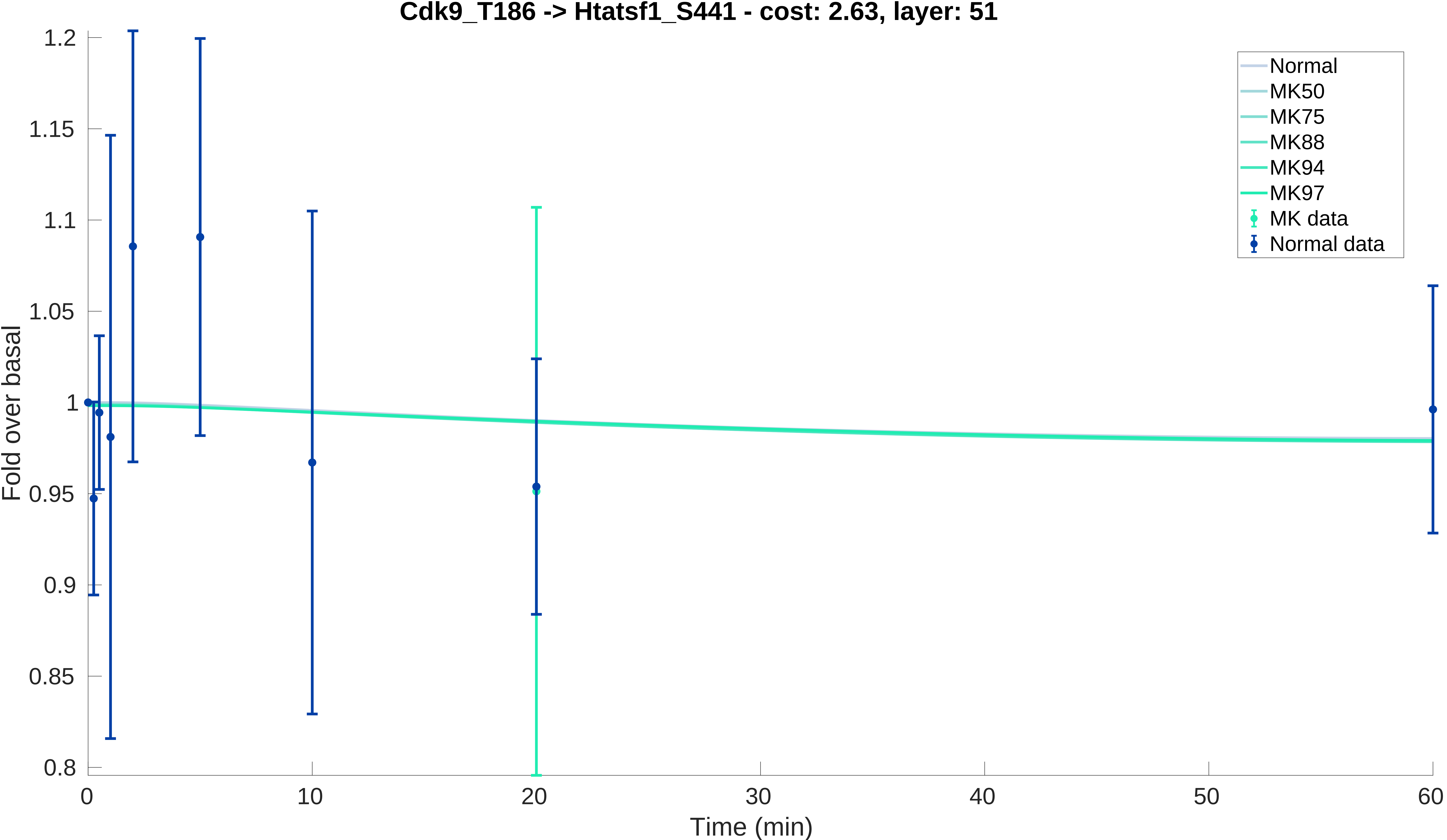

Cdk9_T186 -> Htatsf1_S441 - cost: 2.63, layer: 51
Fold over basal
Time (min)
1.2
1.15
1.1
1.05
1
0.95
0.9
0.85
0.8
0
10
20
30
40
50
60
Normal
MK50
MK75
MK88
MK94
MK97
MK data
Normal data

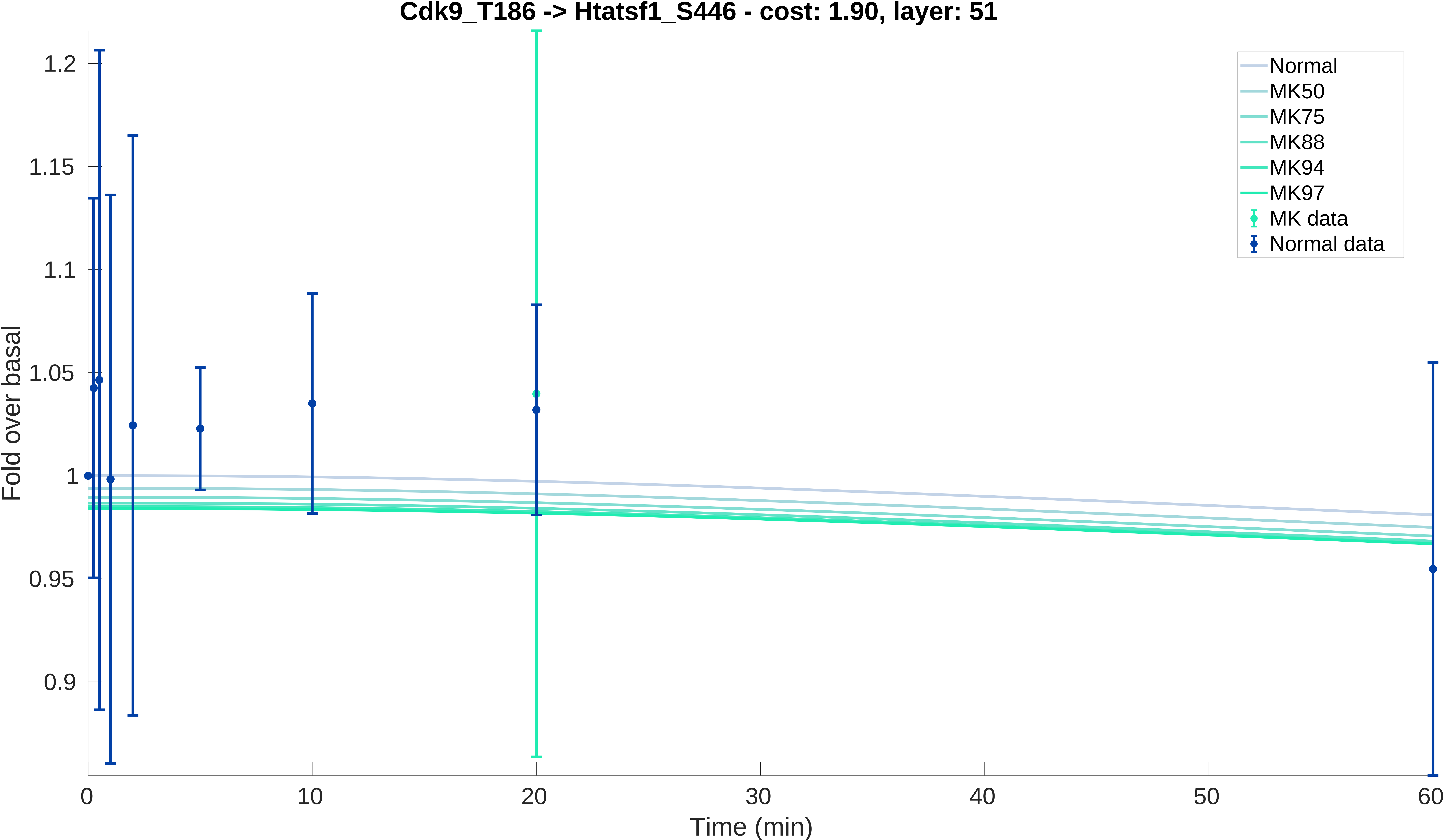
Cdk9_T186 -> Htatsf1_S446 - cost: 1.90, layer: 51
Fold over basal
1.2
1.15
1.1
1.05
1
0.95
0.9
0
10
20
30
40
50
60
Time (min)
Normal
MK50
MK75
MK88
MK94
MK97
MK data
Normal data

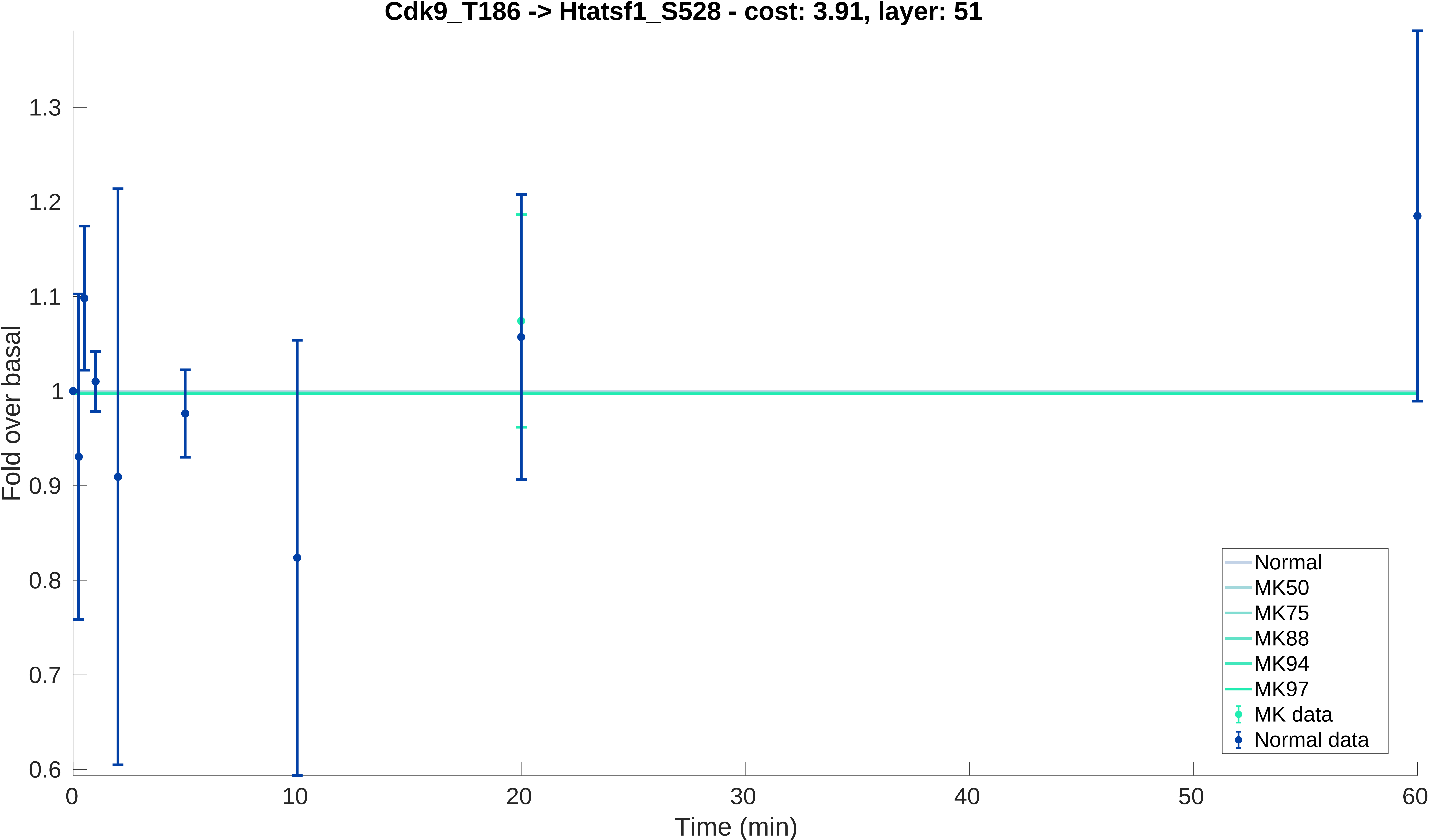
Cdk9_T186 -> Htatsf1_S528 - cost: 3.91, layer: 51
Fold over basal
Time (min)
1.3
1.2
1.1
1
0.9
0.8
0.7
0.6
0
10
20
30
40
50
60
Normal
MK50
MK75
MK88
MK94
MK97
MK data
Normal data

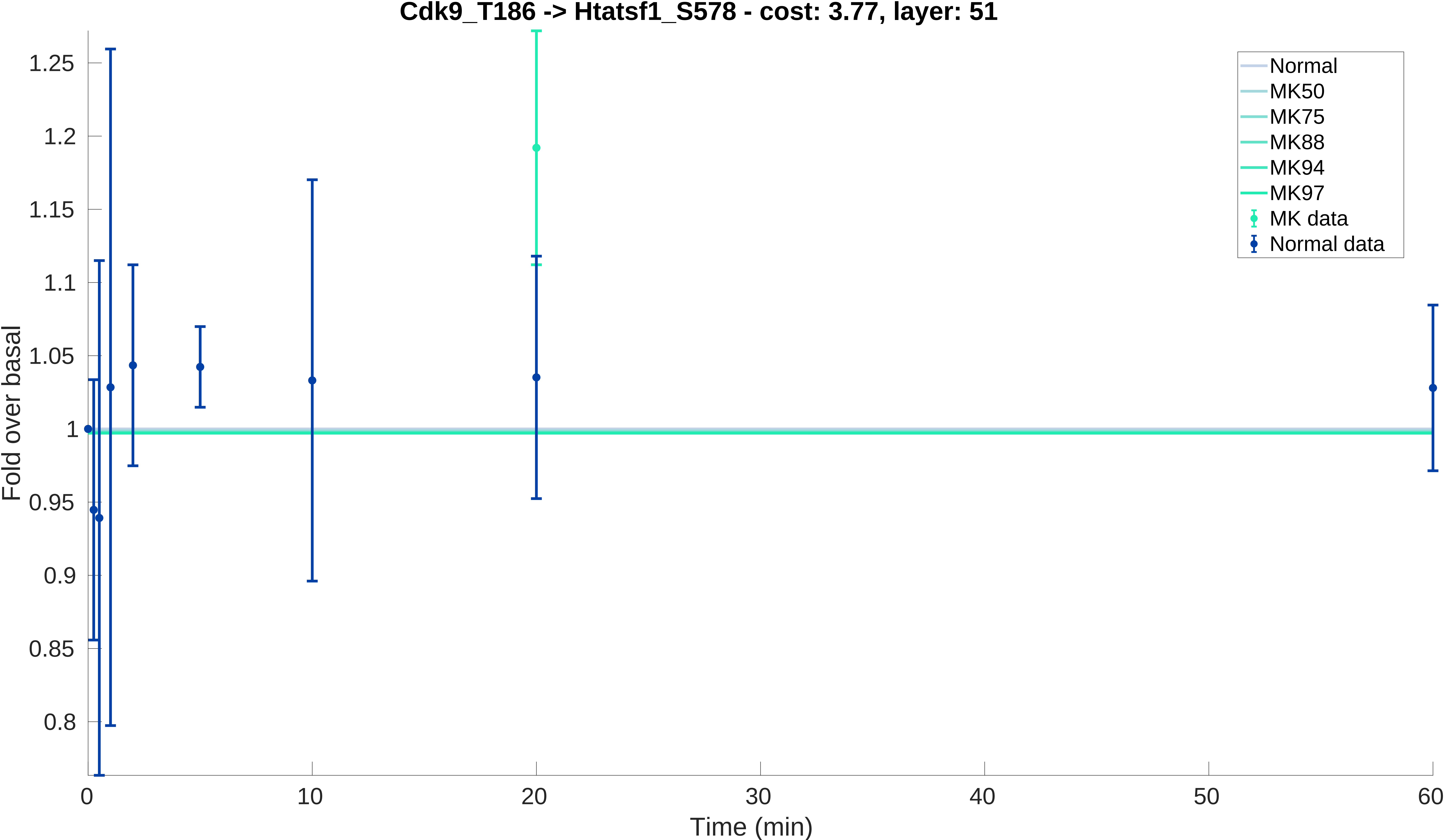

Cdk9_T186 -> Htatsf1_S578 - cost: 3.77, layer: 51
Normal
MK50
MK75
MK88
MK94
MK97
MK data
Normal data
Fold over basal
1.25
1.2
1.15
1.1
1.05
1
0.95
0.9
0.85
0.8
0
10
20
30
40
50
60
Time (min)

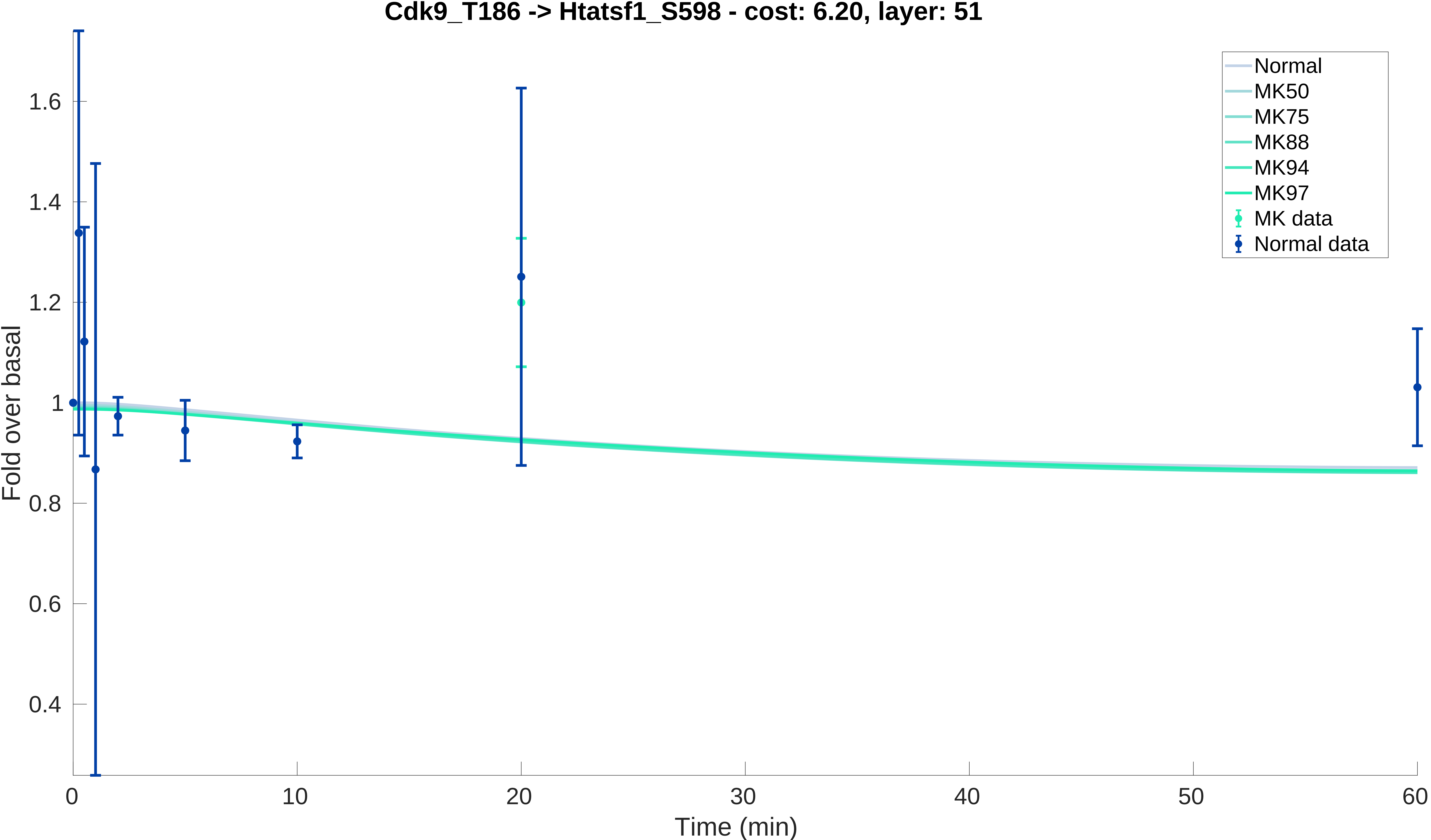

Cdk9_T186 -> Htatsf1_S598 - cost: 6.20, layer: 51
Fold over basal
Time (min)
0.4
0.6
0.8
1
1.2
1.4
1.6
0
10
20
30
40
50
60
Normal
MK50
MK75
MK88
MK94
MK97
MK data
Normal data

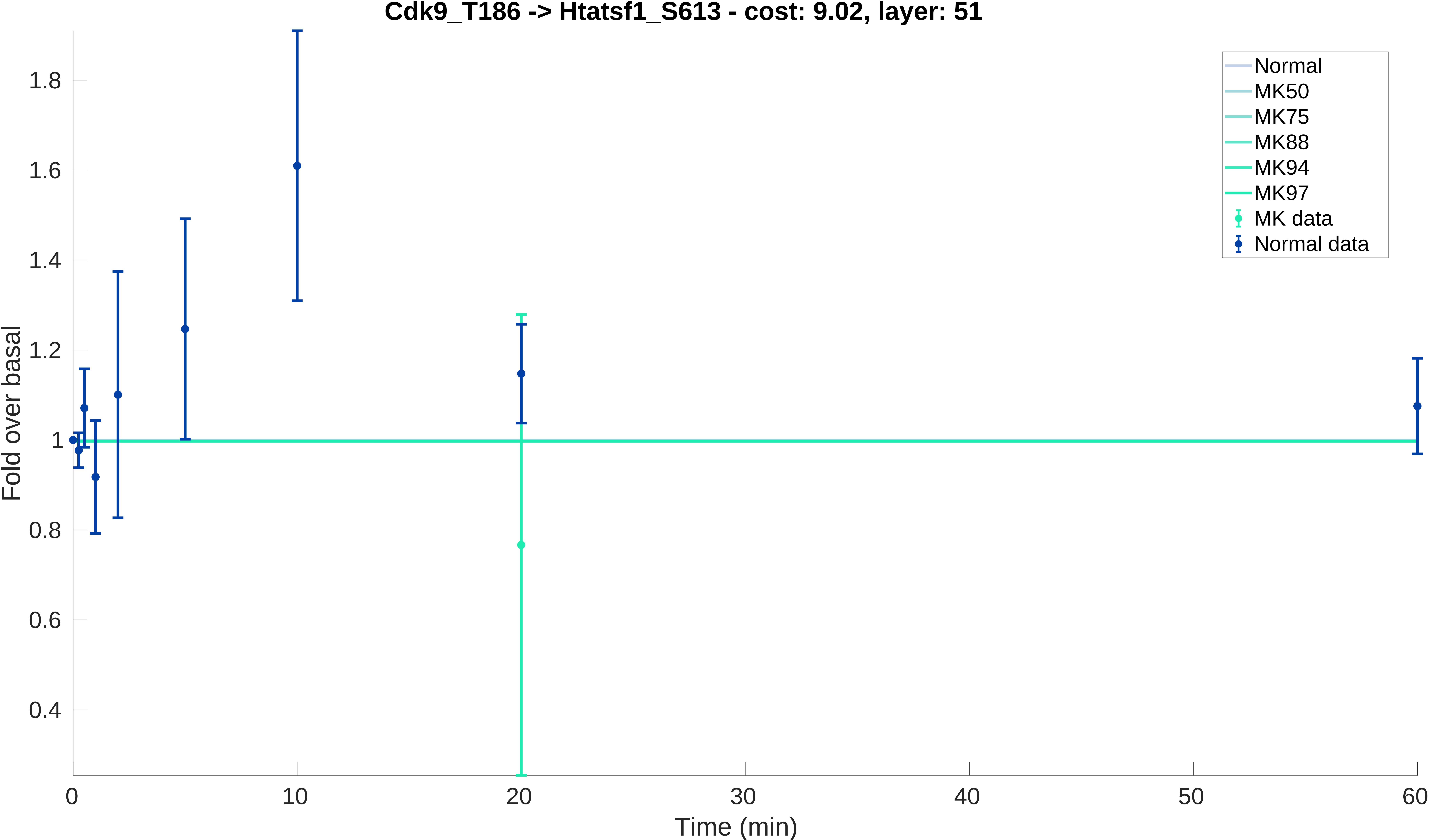

Cdk9_T186 -> Htatsf1_S613 - cost: 9.02, layer: 51
Fold over basal
Time (min)
Normal
MK50
MK75
MK88
MK94
MK97
MK data
Normal data
1.8
1.6
1.4
1.2
1
0.8
0.6
0.4
0
10
20
30
40
50
60

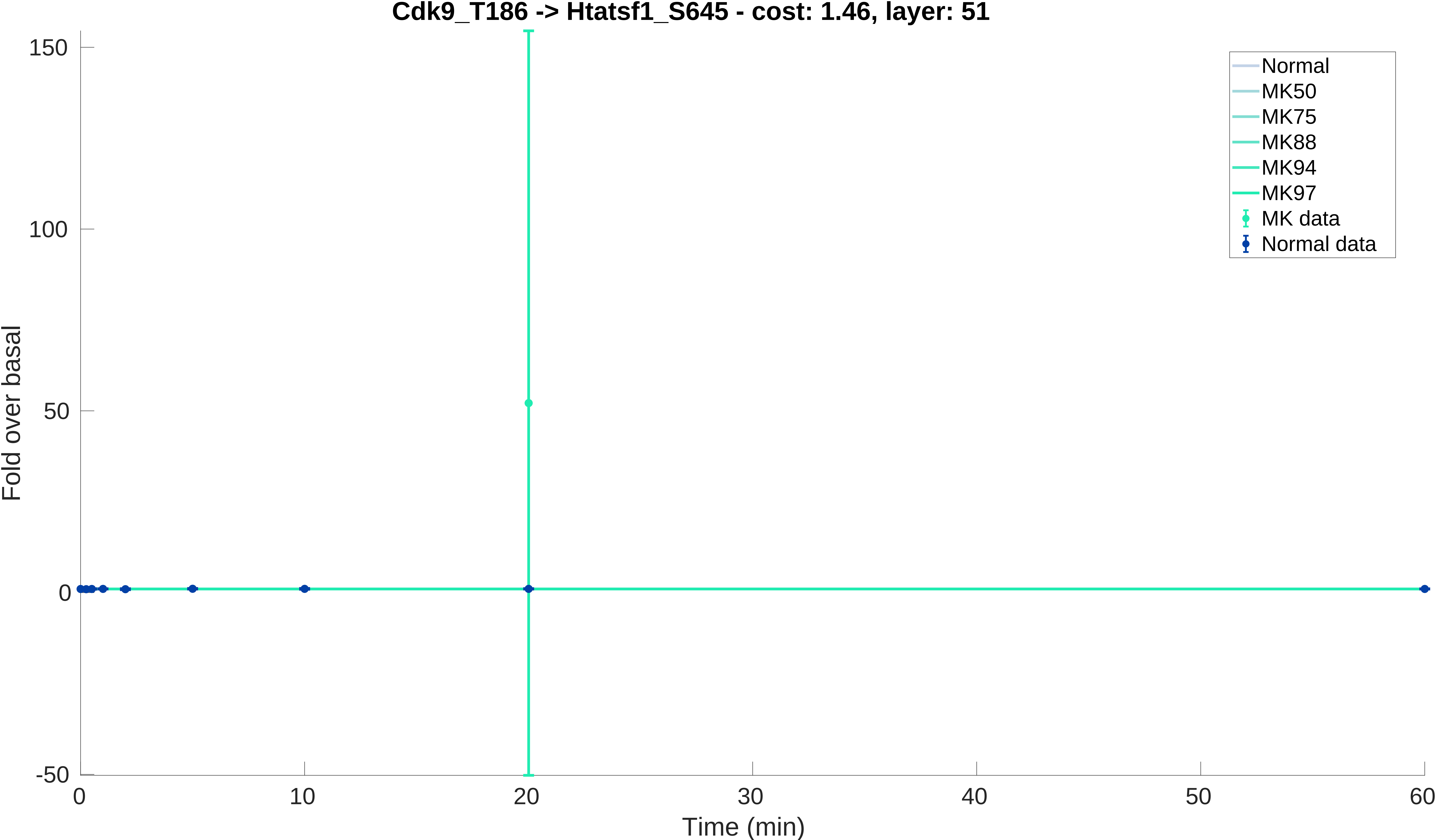

Cdk9_T186 -> Htatsf1_S645 - cost: 1.46, layer: 51
Fold over basal
Time (min)
150
100
50
0
-50
0
10
20
30
40
50
60
Normal
MK50
MK75
MK88
MK94
MK97
MK data
Normal data

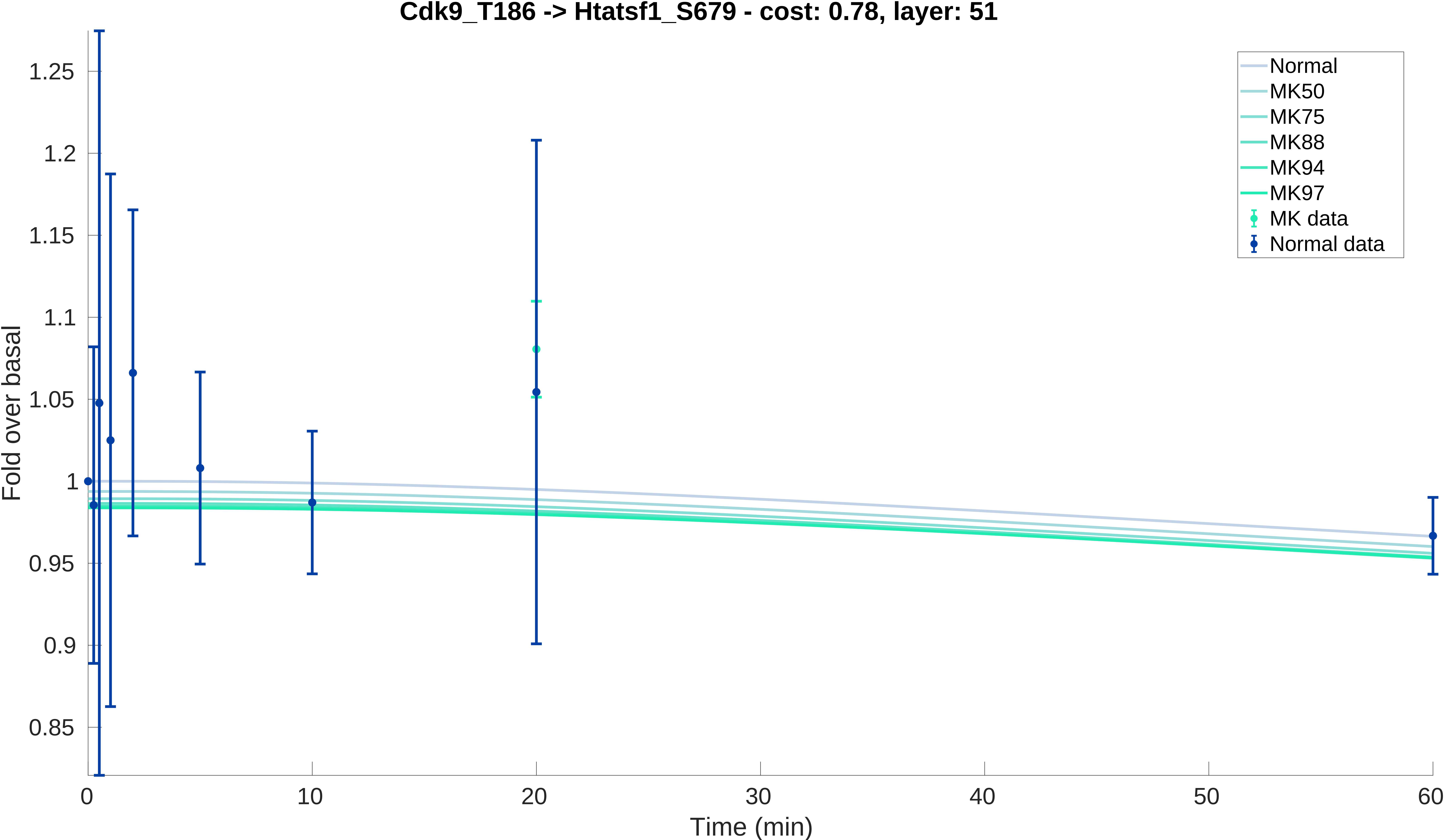

Cdk9_T186 -> Htatsf1_S679 - cost: 0.78, layer: 51
Normal
MK50
MK75
MK88
MK94
MK97
MK data
Normal data
Fold over basal
Time (min)
1.25
1.2
1.15
1.1
1.05
1
0.95
0.9
0.85
0
10
20
30
40
50
60

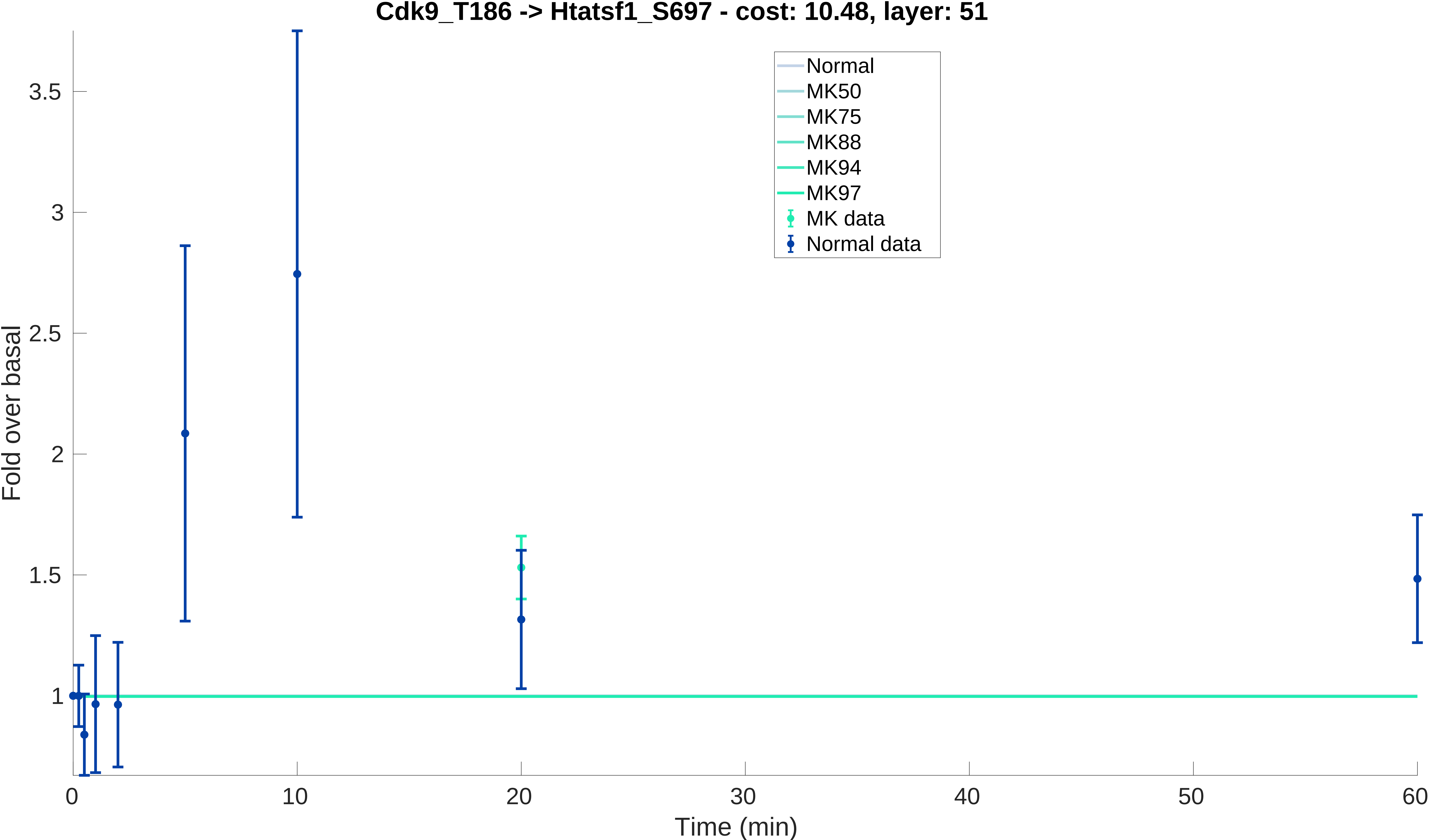

Cdk9_T186 -> Htatsf1_S697 - cost: 10.48, layer: 51
Normal
MK50
MK75
MK88
MK94
MK97
MK data
Normal data
Fold over basal
3.5
3
2.5
2
1.5
1
0
10
20
30
40
50
60
Time (min)

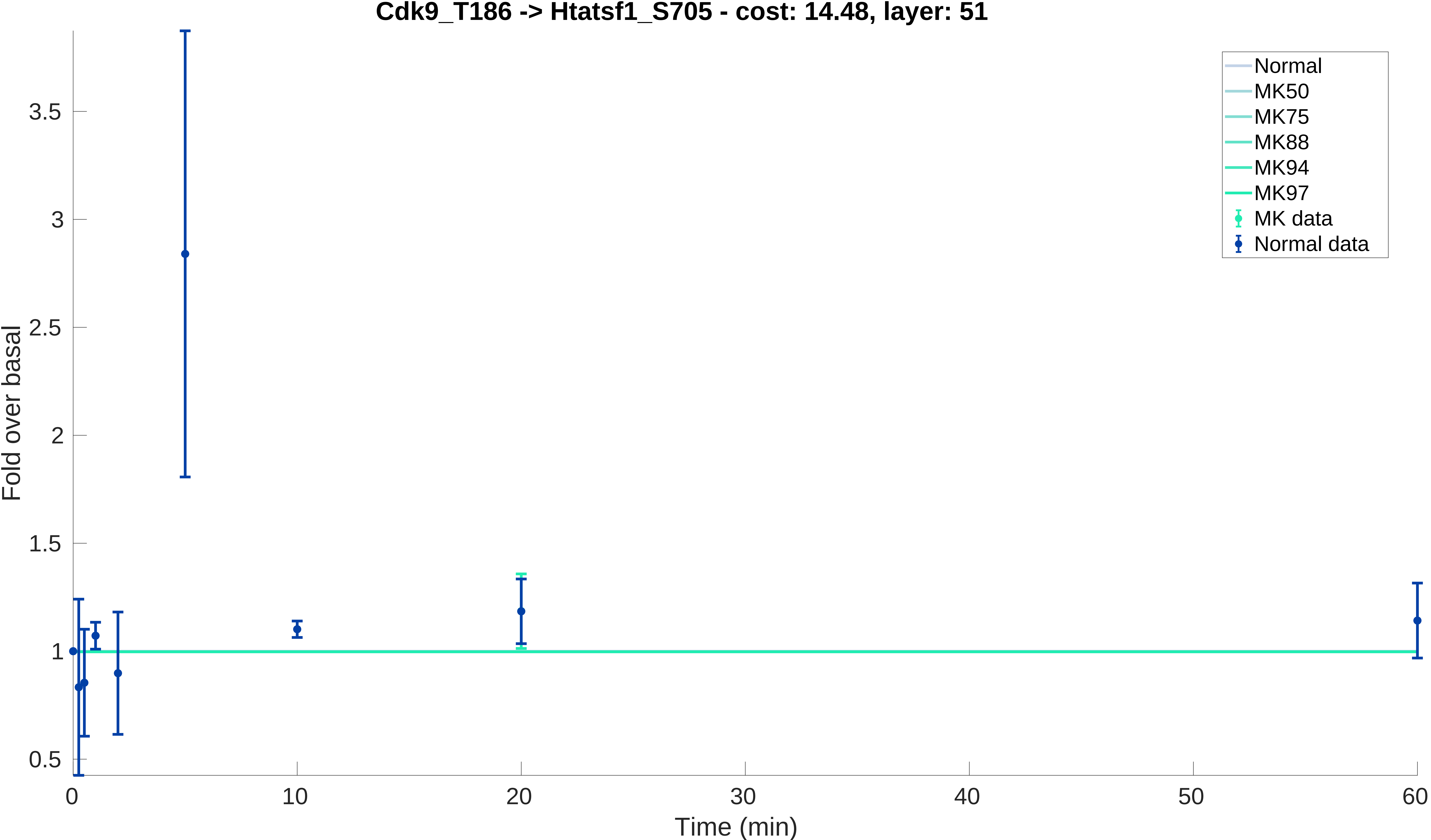

Cdk9_T186 -> Htatsf1_S705 - cost: 14.48, layer: 51
Fold over basal
Time (min)
0.5
1
1.5
2
2.5
3
3.5
0
10
20
30
40
50
60
Normal
MK50
MK75
MK88
MK94
MK97
MK data
Normal data

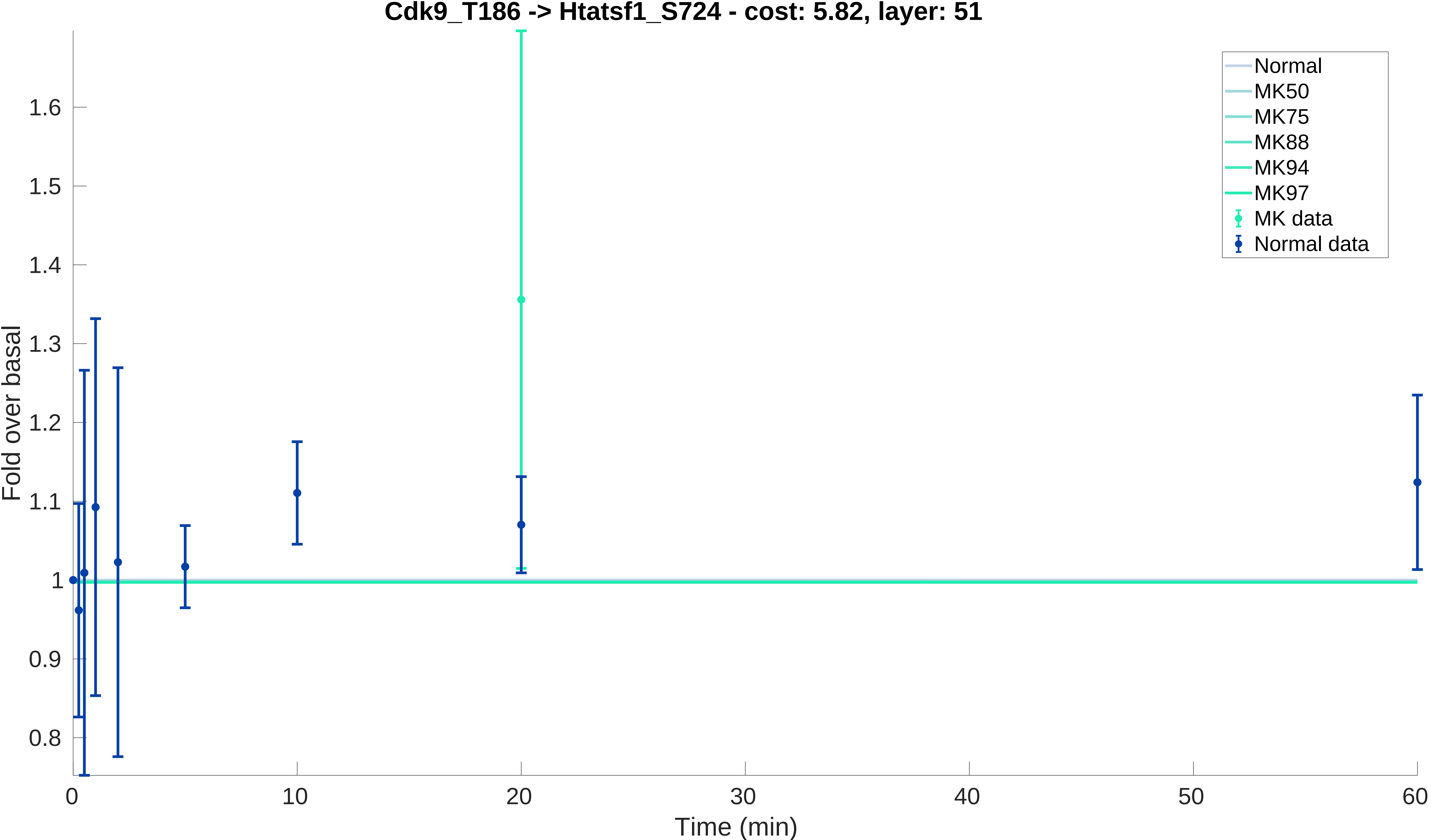

Cdk9_T186 -> Htatsf1_S724 - cost: 5.82, layer: 51
Fold over basal
Time (min)
Normal
MK50
MK75
MK88
MK94
MK97
MK data
Normal data
1.6
1.5
1.4
1.3
1.2
1.1
1
0.9
0.8
0
10
20
30
40
50
60

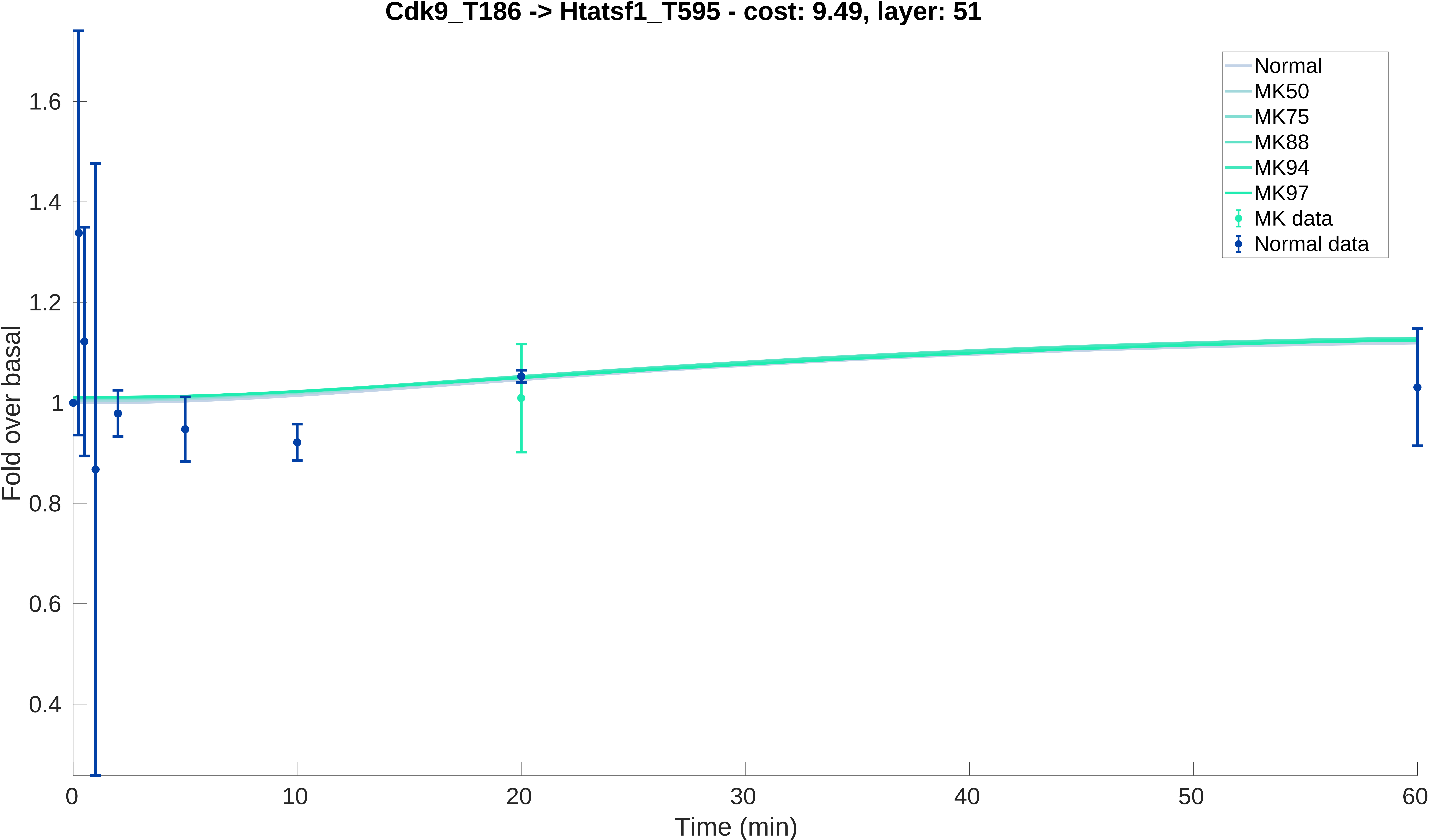

Cdk9_T186 -> Htatsf1_T595 - cost: 9.49, layer: 51
Normal
MK50
MK75
MK88
MK94
MK97
MK data
Normal data
Fold over basal
1.6
1.4
1.2
1
0.8
0.6
0.4
0
10
20
30
40
50
60
Time (min)

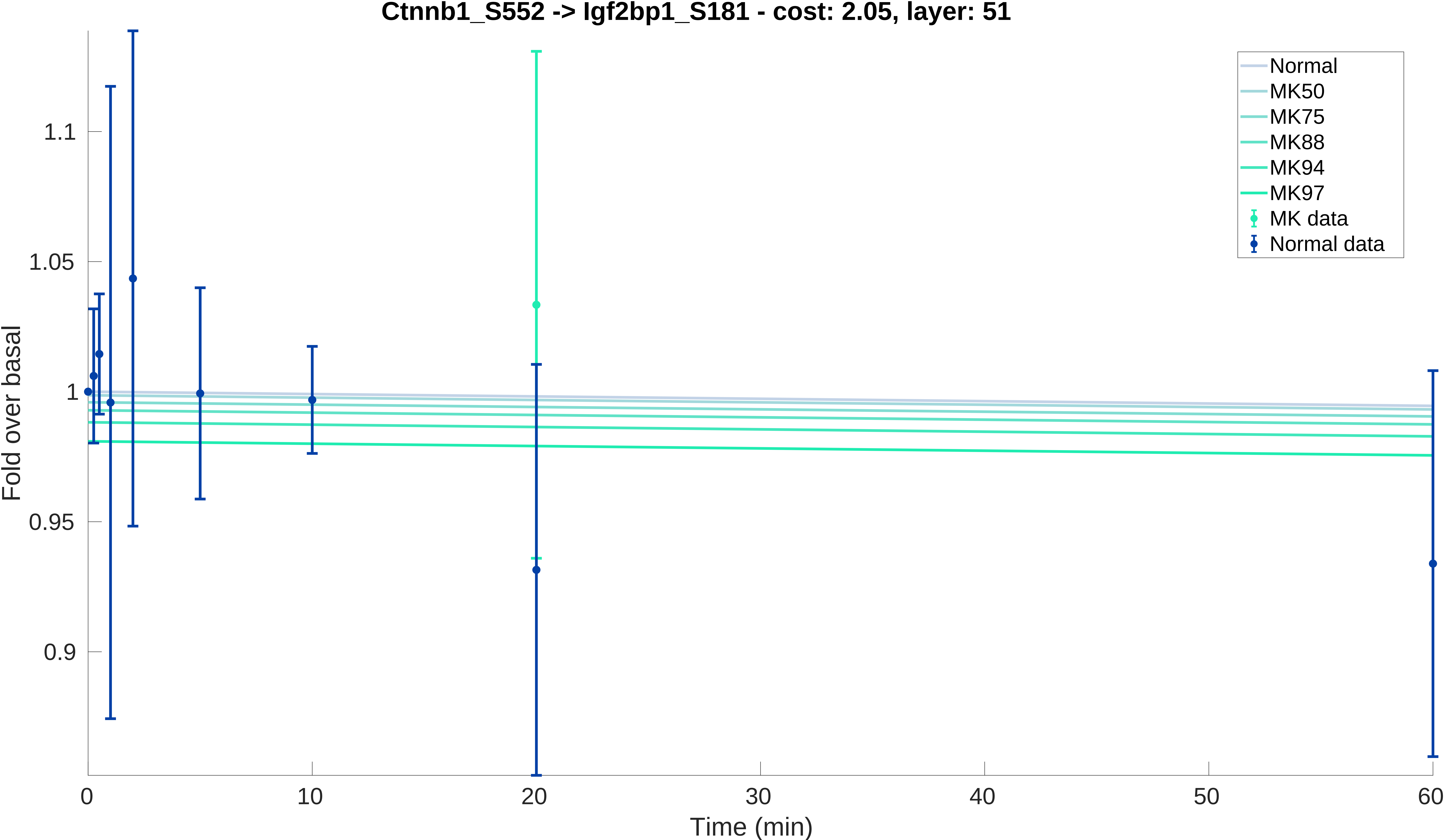
Ctnnb1_S552 -> Igf2bp1_S181 - cost: 2.05, layer: 51
Fold over basal
Time (min)
0
10
20
30
40
50
60
0.9
0.95
1
1.05
1.1
Normal
MK50
MK75
MK88
MK94
MK97
MK data
Normal data

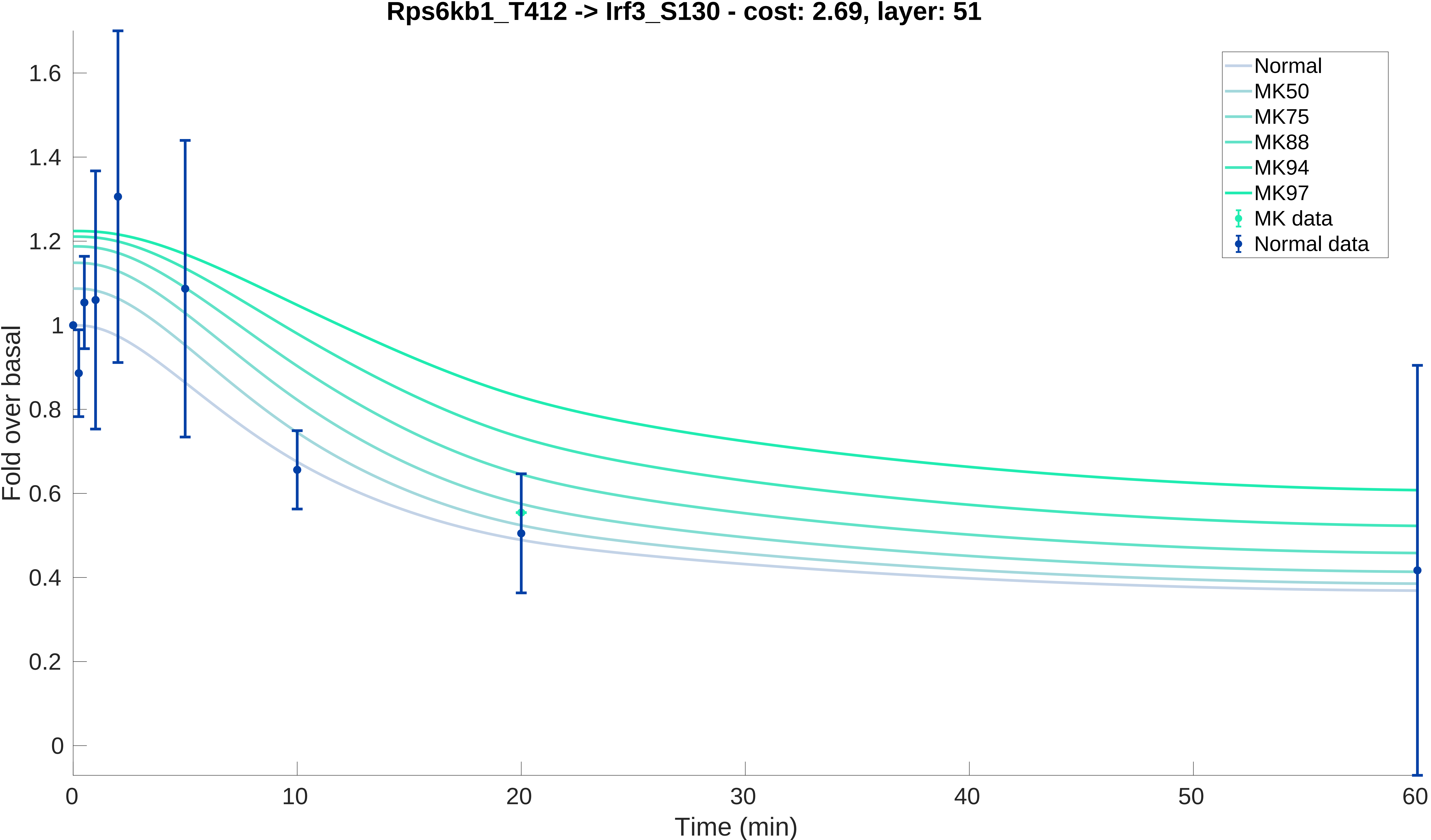
Rps6kb1_T412 -> Irf3_S130 - cost: 2.69, layer: 51
Fold over basal
Time (min)
0
0.2
0.4
0.6
0.8
1
1.2
1.4
1.6
0
10
20
30
40
50
60
Normal
MK50
MK75
MK88
MK94
MK97
MK data
Normal data

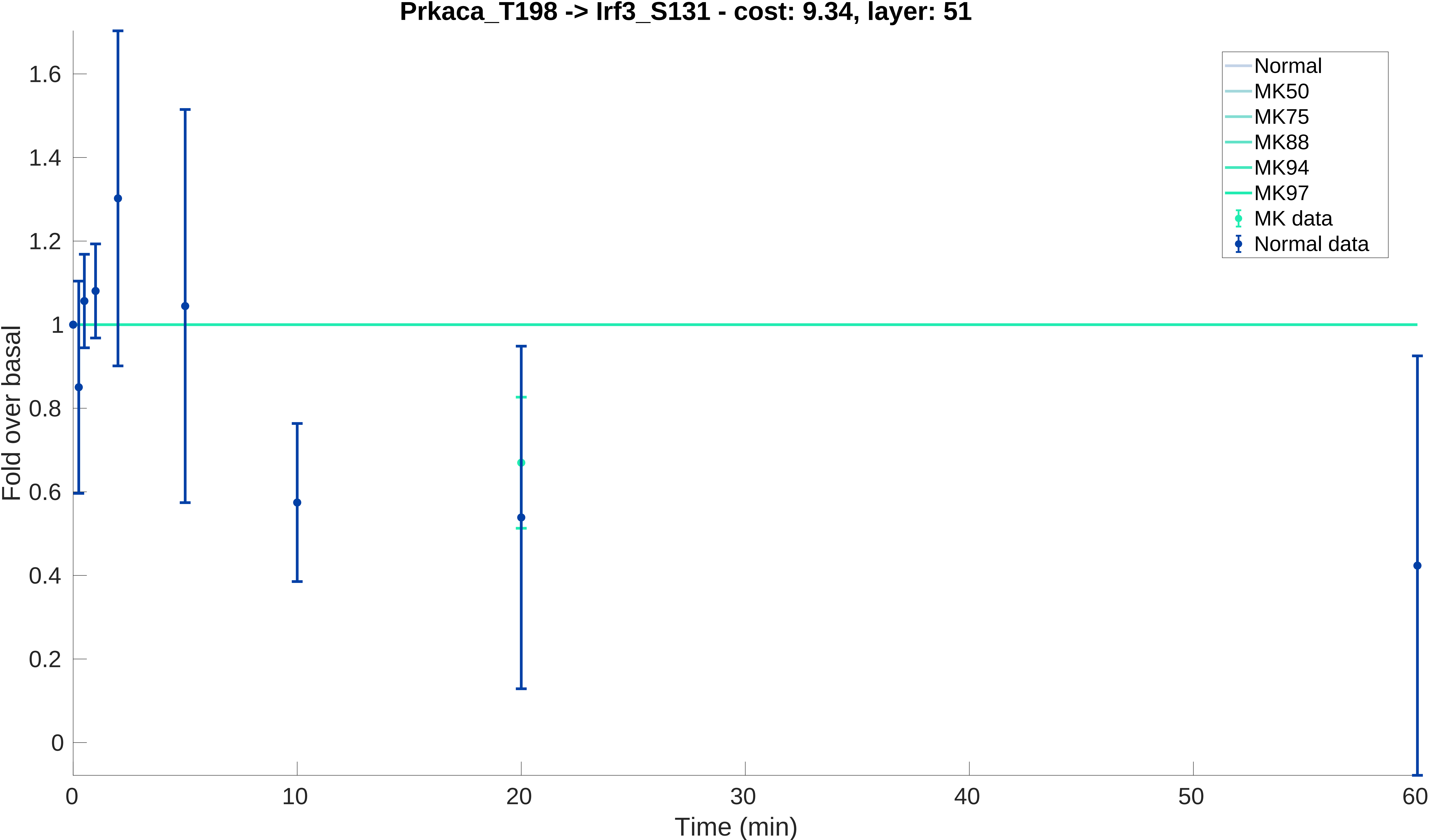
Prkaca_T198 -> Irf3_S131 - cost: 9.34, layer: 51
Normal
MK50
MK75
MK88
MK94
MK97
MK data
Normal data
Fold over basal
0
0.2
0.4
0.6
0.8
1
1.2
1.4
1.6
0
10
20
30
40
50
60
Time (min)

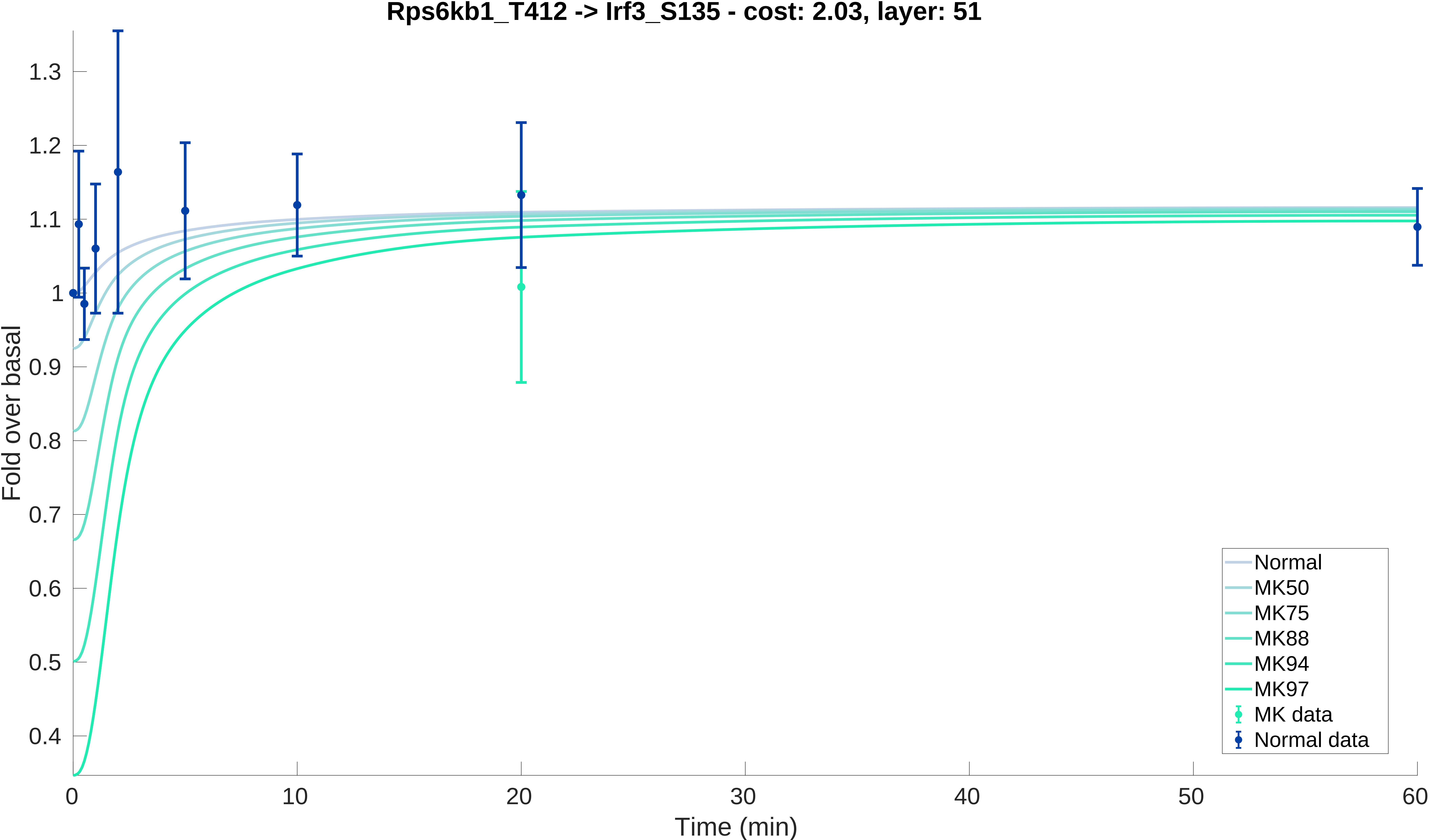
Rps6kb1_T412 -> Irf3_S135 - cost: 2.03, layer: 51
Fold over basal
Time (min)
0
10
20
30
40
50
60
0.4
0.5
0.6
0.7
0.8
0.9
1
1.1
1.2
1.3
Normal
MK50
MK75
MK88
MK94
MK97
MK data
Normal data

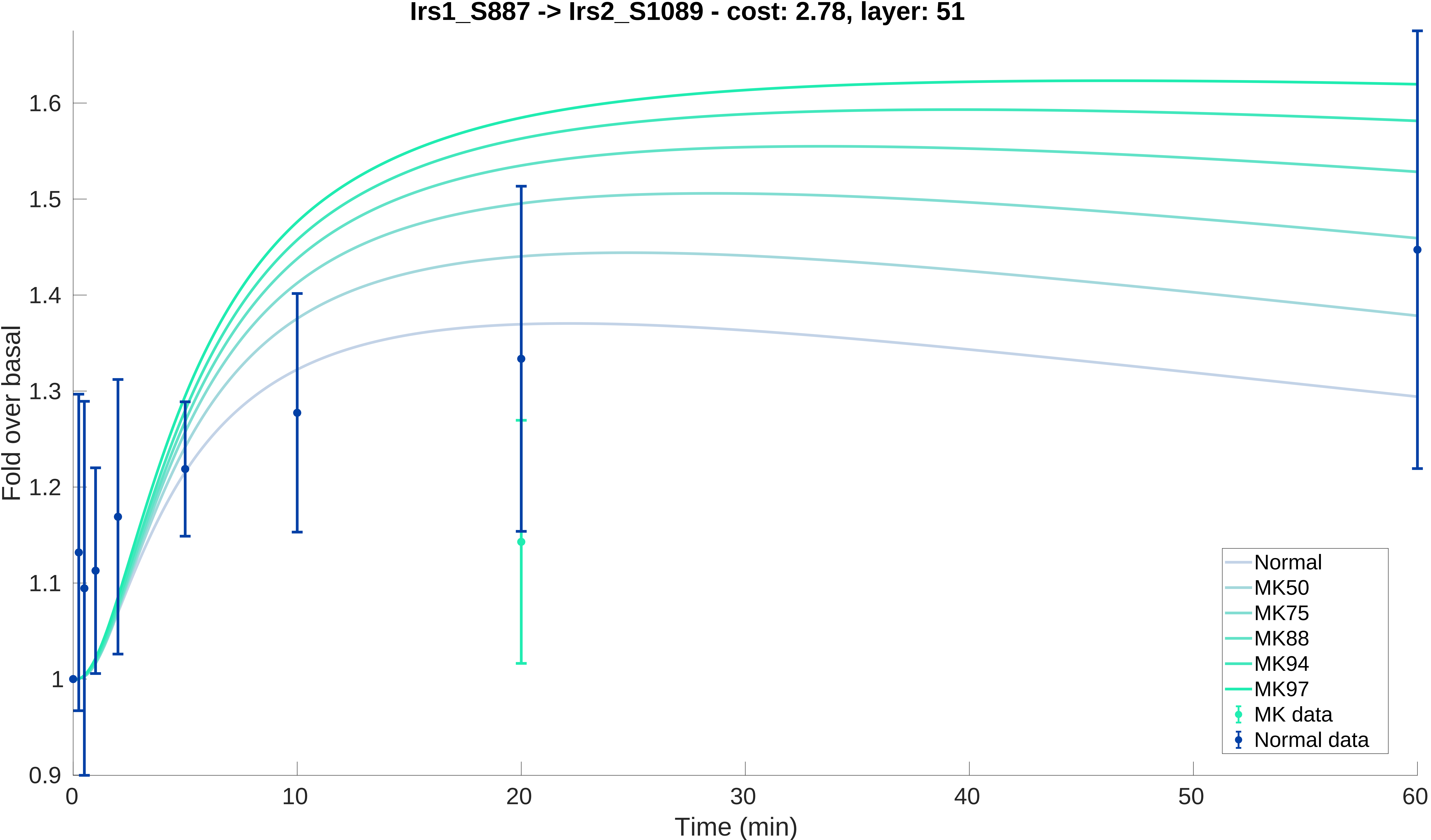

Irs1_S887 -> Irs2_S1089 - cost: 2.78, layer: 51
Fold over basal
1.6
1.5
1.4
1.3
1.2
1.1
1
0.9
0
10
20
30
40
50
60
Time (min)
Normal
MK50
MK75
MK88
MK94
MK97
MK data
Normal data

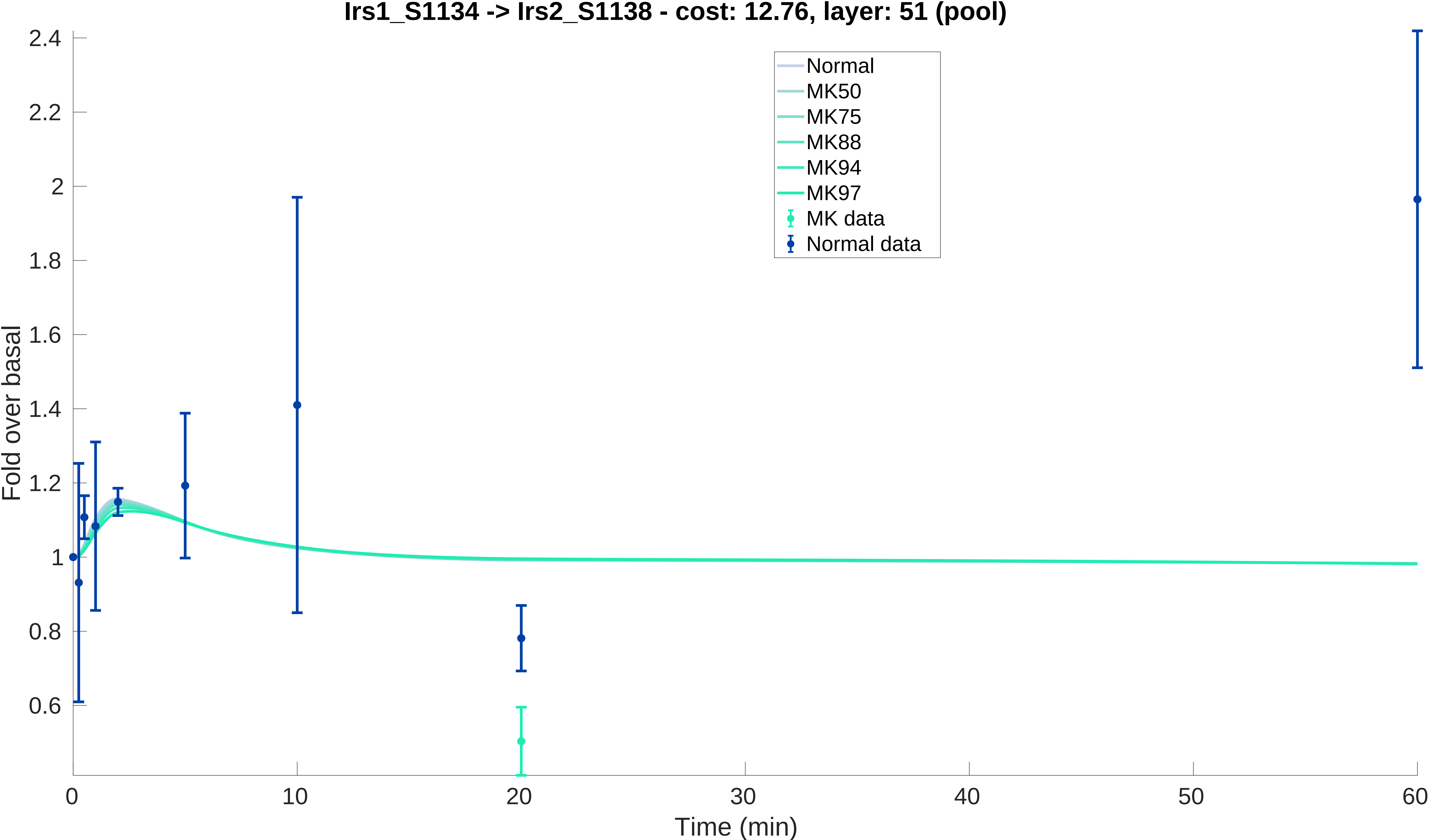

Irs1_S1134 -> Irs2_S1138 - cost: 12.76, layer: 51 (pool)
Fold over basal
Time (min)
2.4
2.2
2
1.8
1.6
1.4
1.2
1
0.8
0.6
0
10
20
30
40
50
60
Normal
MK50
MK75
MK88
MK94
MK97
MK data
Normal data

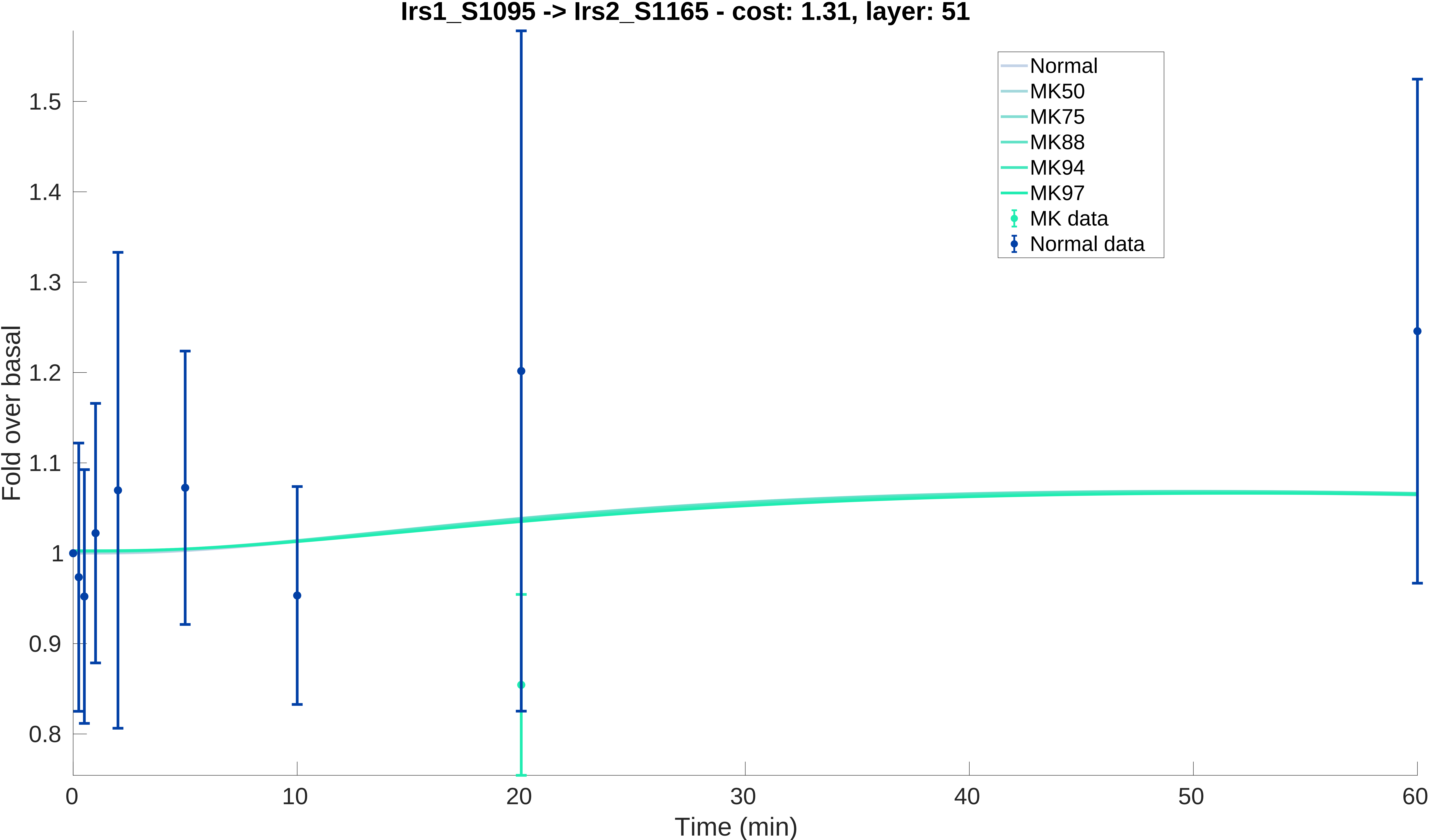

Irs1_S1095 -> Irs2_S1165 - cost: 1.31, layer: 51
Normal
MK50
MK75
MK88
MK94
MK97
MK data
Normal data
Fold over basal
Time (min)
0.8
0.9
1
1.1
1.2
1.3
1.4
1.5
0
10
20
30
40
50
60

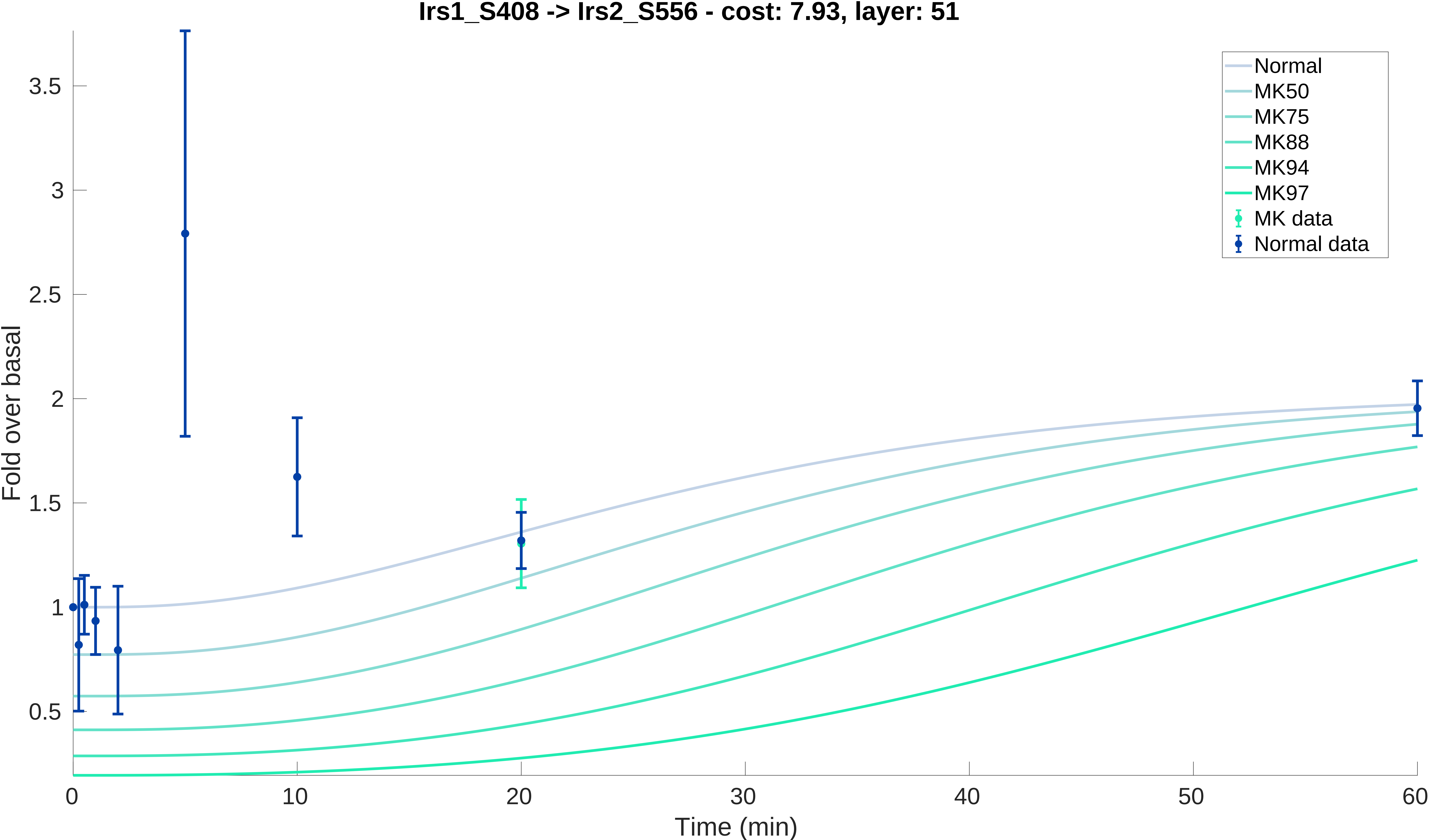
Irs1_S408 -> Irs2_S556 - cost: 7.93, layer: 51
Fold over basal
Time (min)
0
10
20
30
40
50
60
0.5
1
1.5
2
2.5
3
3.5
Normal
MK50
MK75
MK88
MK94
MK97
MK data
Normal data

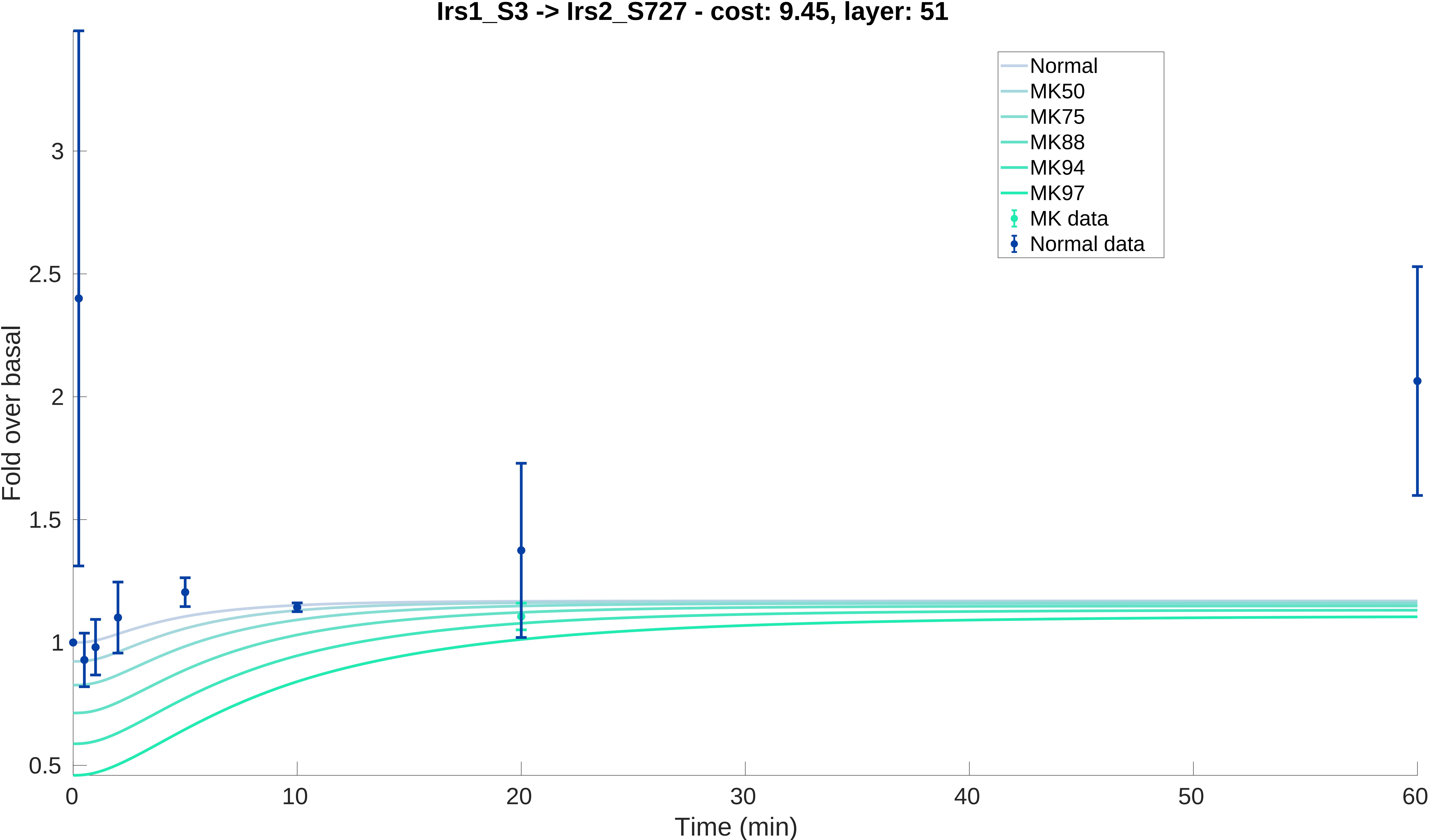

Irs1_S3 -> Irs2_S727 - cost: 9.45, layer: 51
Normal
MK50
MK75
MK88
MK94
MK97
MK data
Normal data
Fold over basal
3
2.5
2
1.5
1
0.5
0
10
20
30
40
50
60
Time (min)

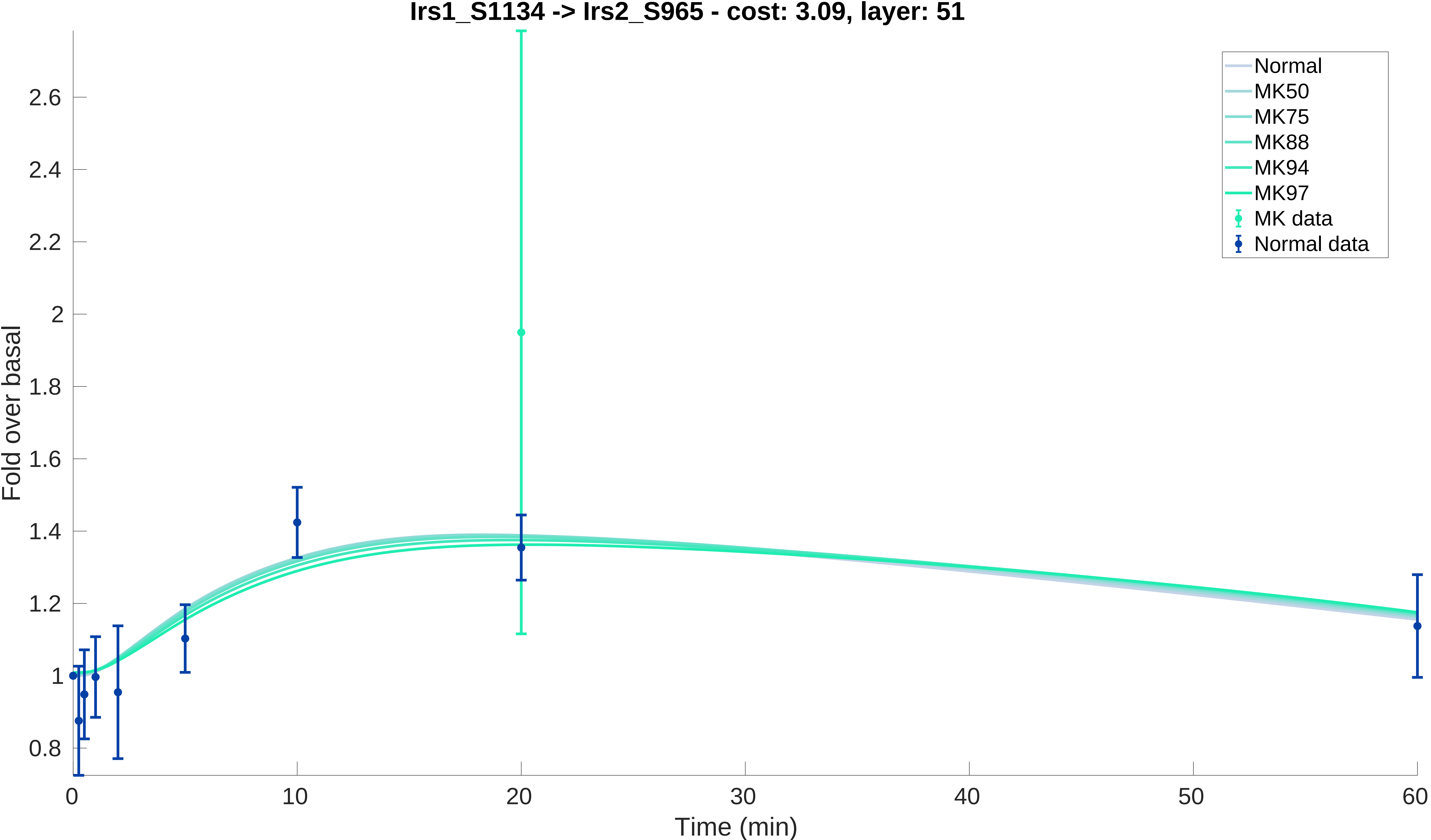

Irs1_S1134 -> Irs2_S965 - cost: 3.09, layer: 51
Normal
MK50
MK75
MK88
MK94
MK97
MK data
Normal data
Fold over basal
Time (min)
0
10
20
30
40
50
60
0.8
1
1.2
1.4
1.6
1.8
2
2.2
2.4
2.6

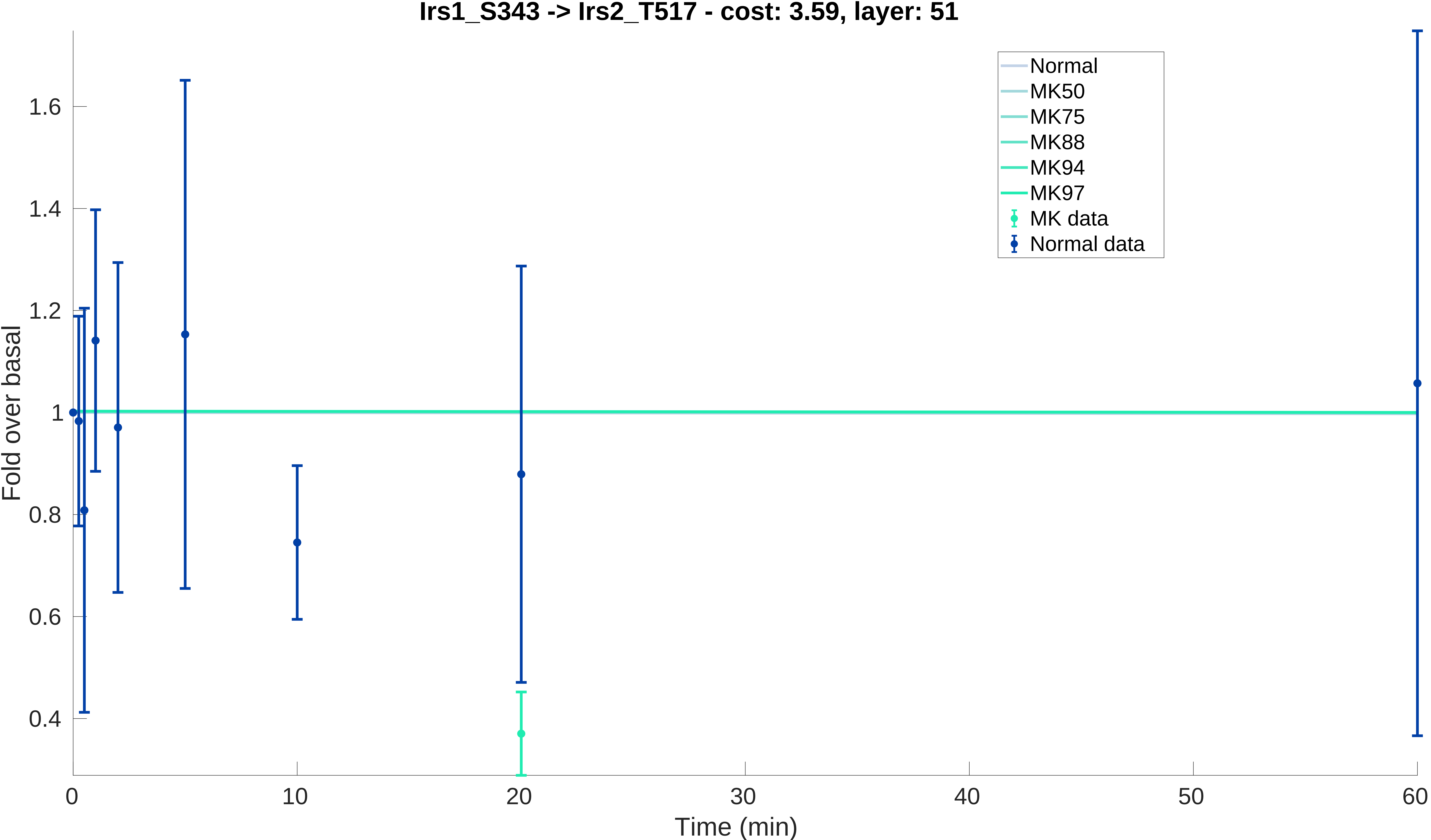
Irs1_S343 -> Irs2_T517 - cost: 3.59, layer: 51
Fold over basal
Time (min)
0.4
0.6
0.8
1
1.2
1.4
1.6
0
10
20
30
40
50
60
Normal
MK50
MK75
MK88
MK94
MK97
MK data
Normal data

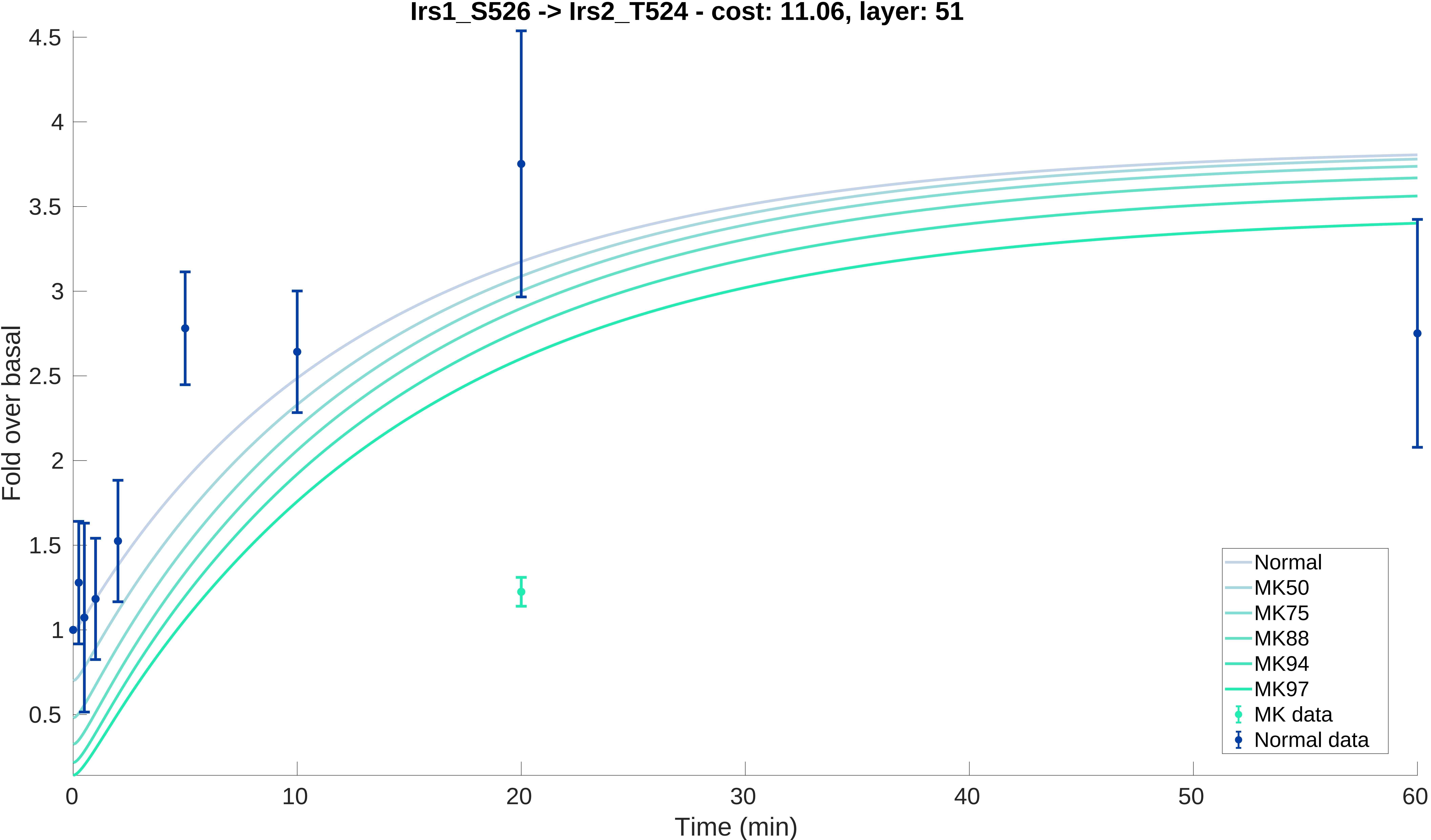
Irs1_S526 -> Irs2_T524 - cost: 11.06, layer: 51
Fold over basal
Time (min)
4.5
4
3.5
3
2.5
2
1.5
1
0.5
0
10
20
30
40
50
60
Normal
MK50
MK75
MK88
MK94
MK97
MK data
Normal data

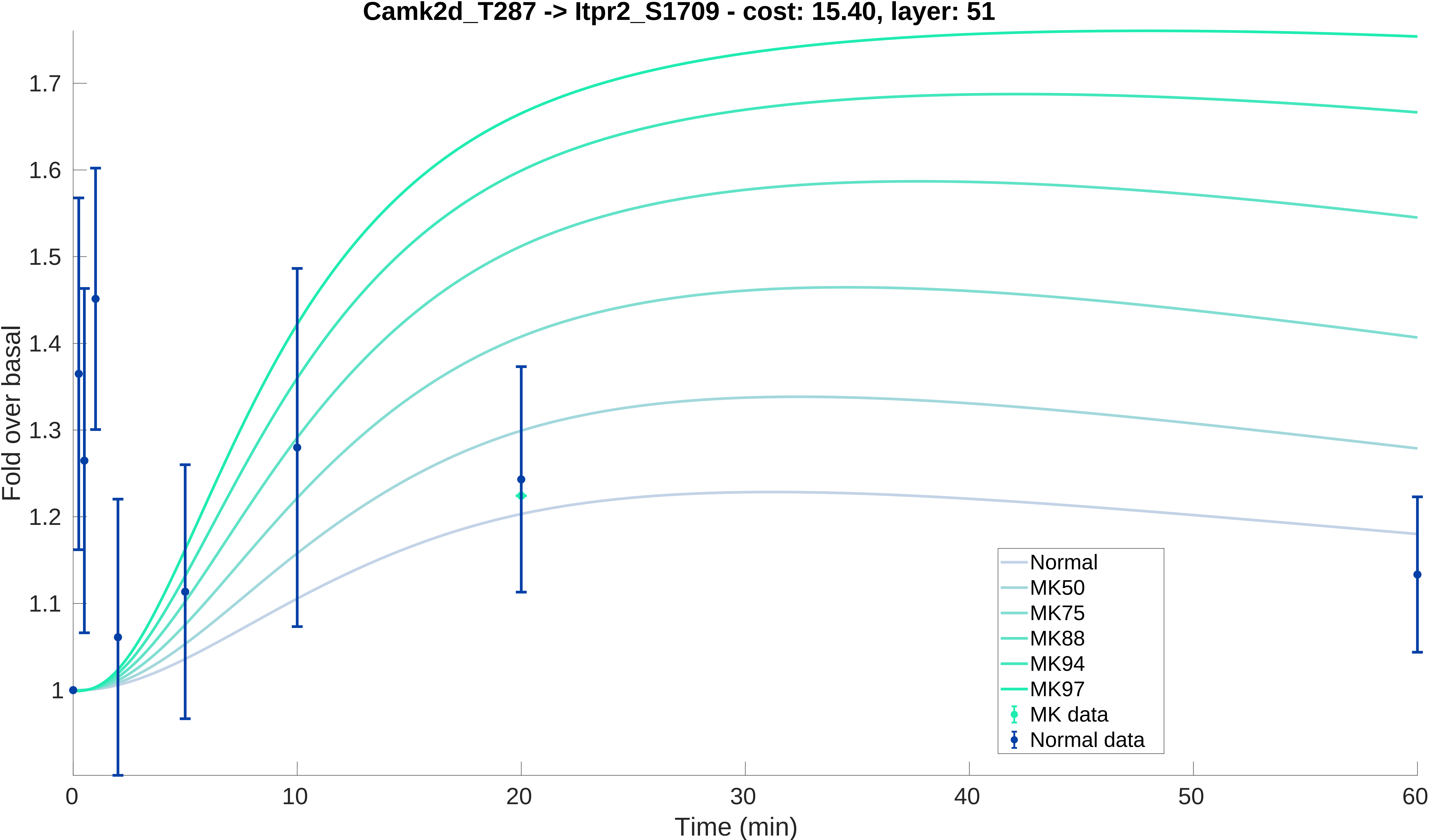

Camk2d_T287 -> Itpr2_S1709 - cost: 15.40, layer: 51
Fold over basal
1.7
1.6
1.5
1.4
1.3
1.2
1.1
1
0
10
20
30
40
50
60
Time (min)
Normal
MK50
MK75
MK88
MK94
MK97
MK data
Normal data

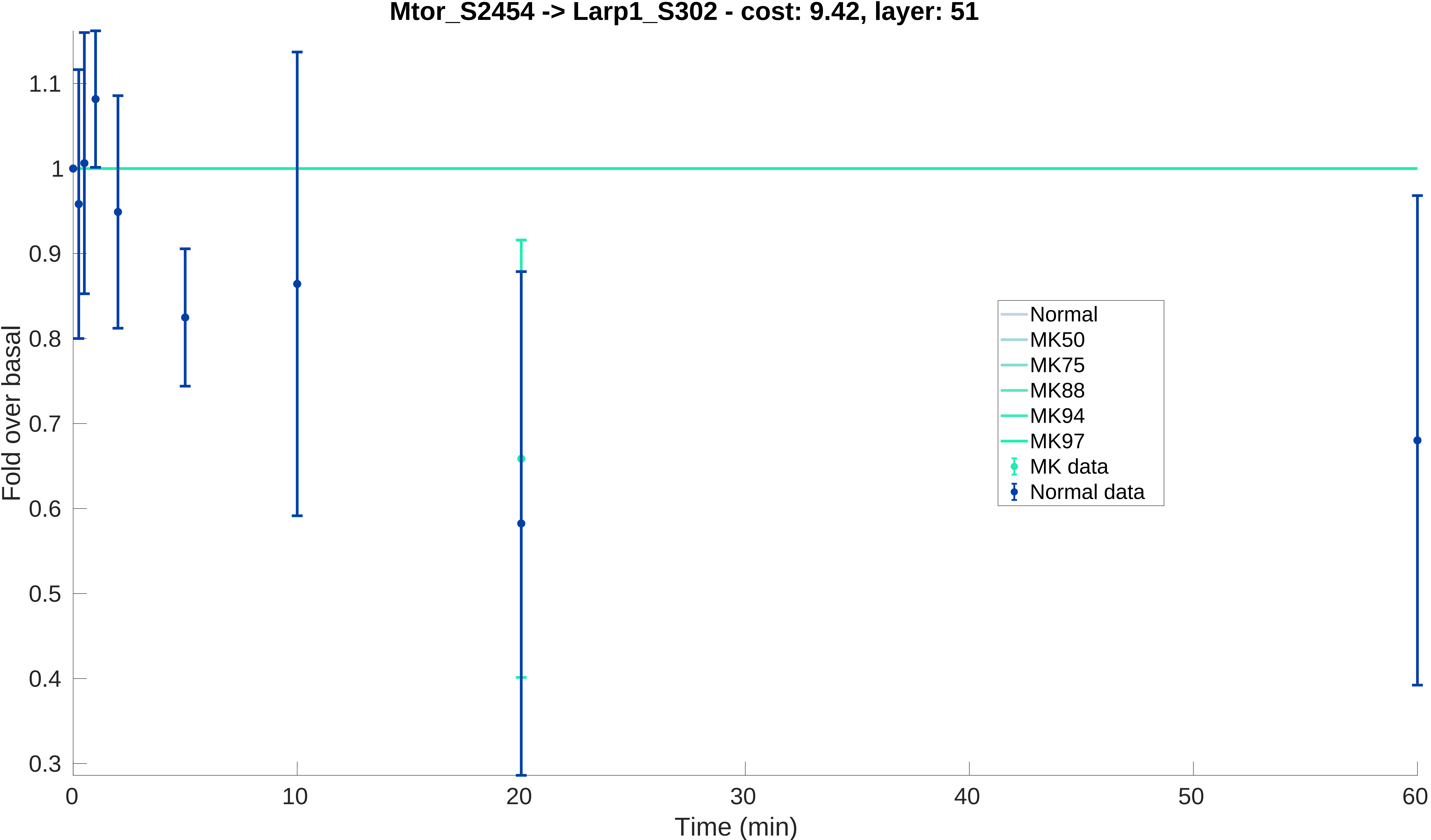

Mtor_S2454 -> Larp1_S302 - cost: 9.42, layer: 51
Fold over basal
1.1
1
0.9
0.8
0.7
0.6
0.5
0.4
0.3
0
10
20
30
40
50
60
Time (min)
Normal
MK50
MK75
MK88
MK94
MK97
MK data
Normal data

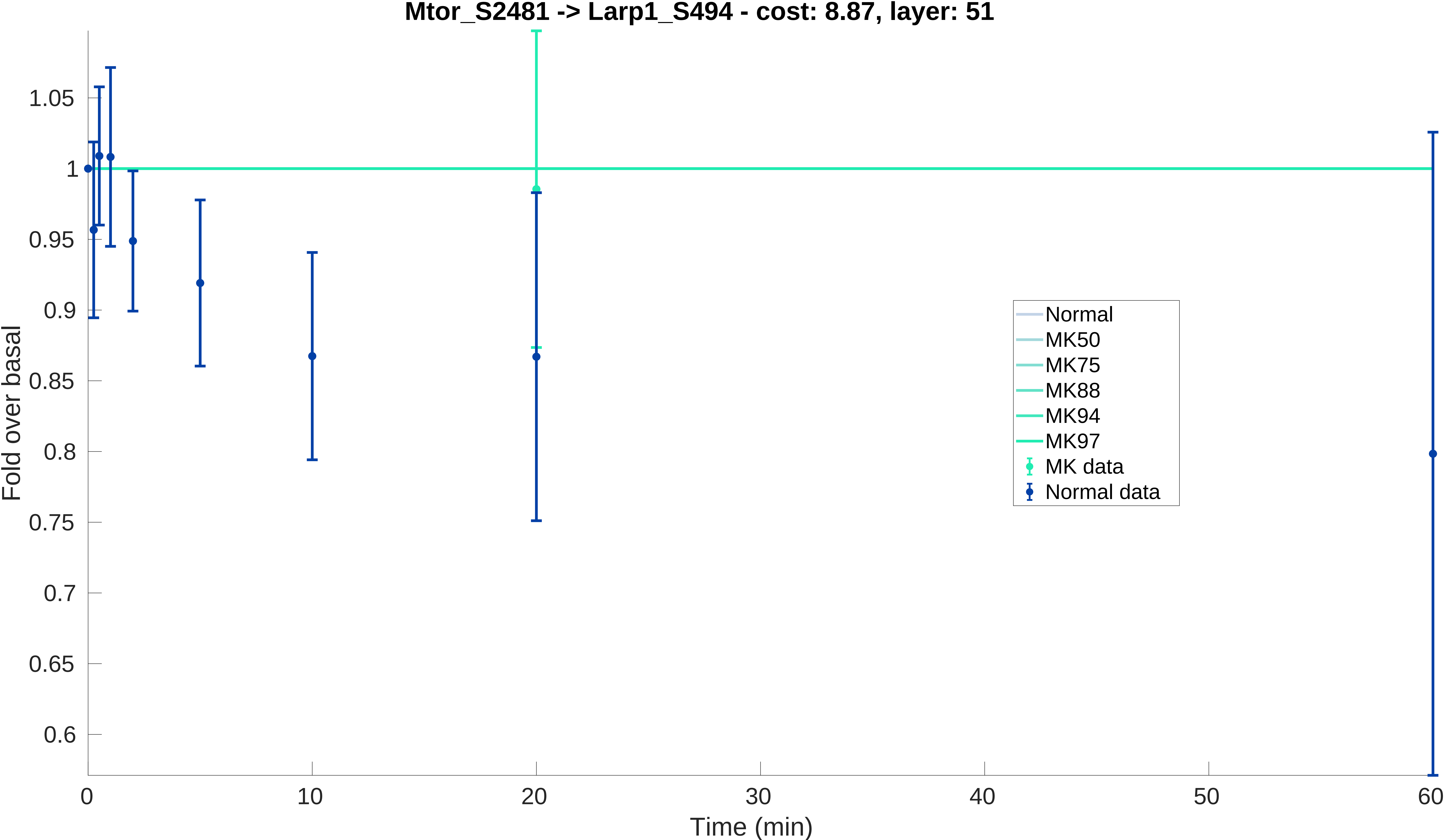
Mtor_S2481 -> Larp1_S494 - cost: 8.87, layer: 51
Fold over basal
Time (min)
1.05
1
0.95
0.9
0.85
0.8
0.75
0.7
0.65
0.6
0
10
20
30
40
50
60
Normal
MK50
MK75
MK88
MK94
MK97
MK data
Normal data

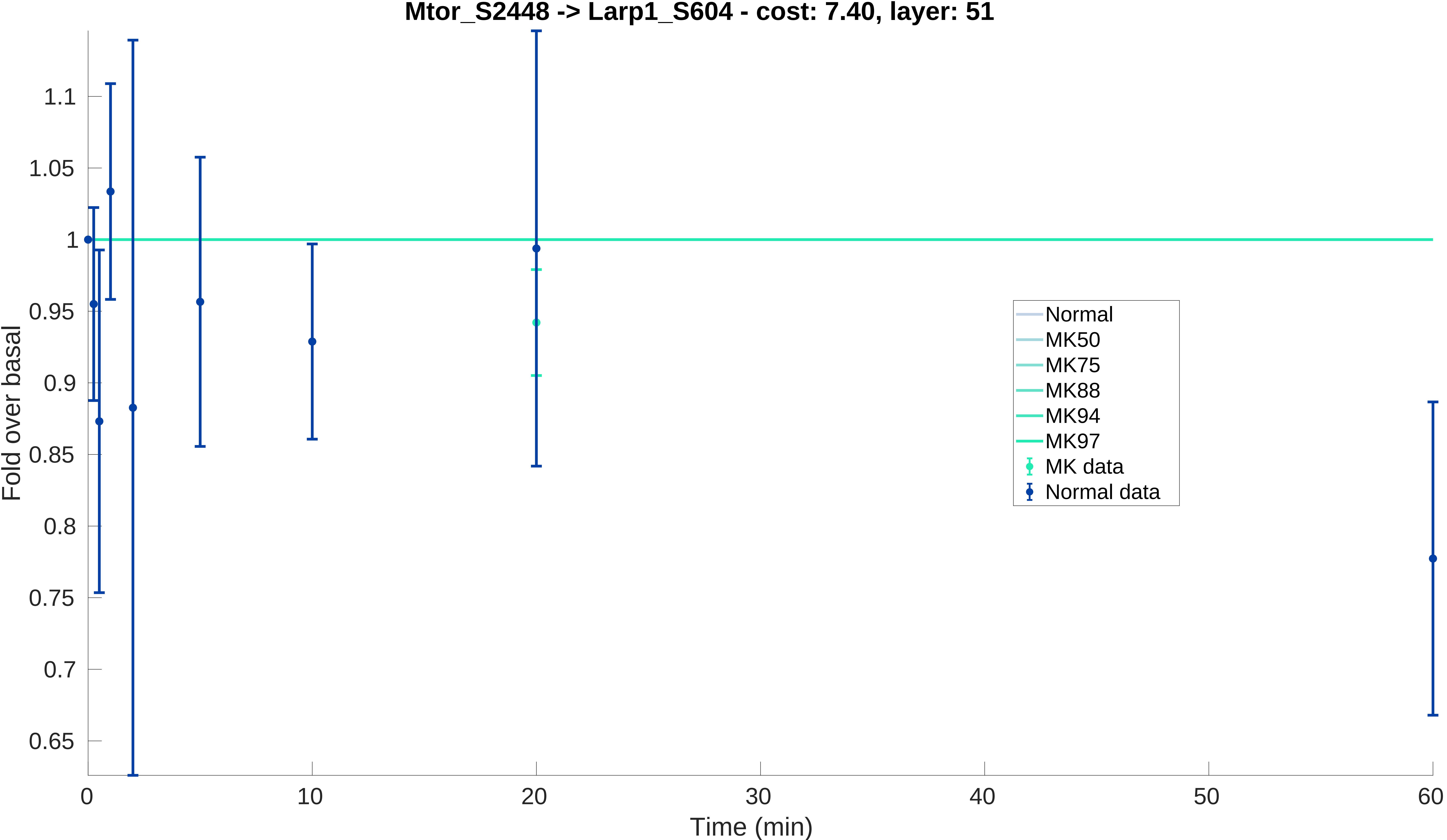
Mtor_S2448 -> Larp1_S604 - cost: 7.40, layer: 51
Fold over basal
1.1
1.05
1
0.95
0.9
0.85
0.8
0.75
0.7
0.65
0
10
20
30
40
50
60
Time (min)
Normal
MK50
MK75
MK88
MK94
MK97
MK data
Normal data

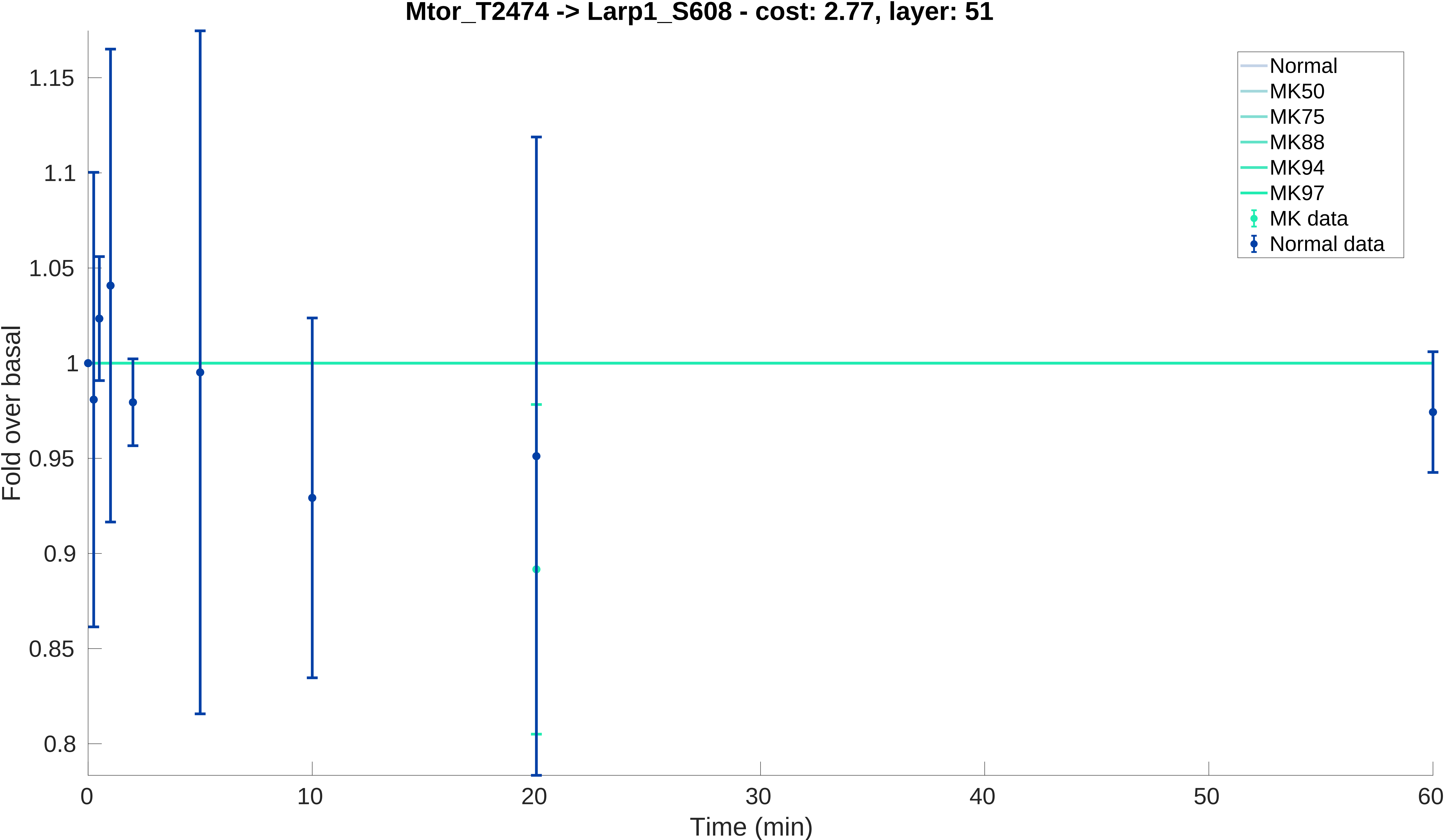
Mtor_T2474 -> Larp1_S608 - cost: 2.77, layer: 51
Fold over basal
Time (min)
0
10
20
30
40
50
60
0.8
0.85
0.9
0.95
1
1.05
1.1
1.15
Normal
MK50
MK75
MK88
MK94
MK97
MK data
Normal data

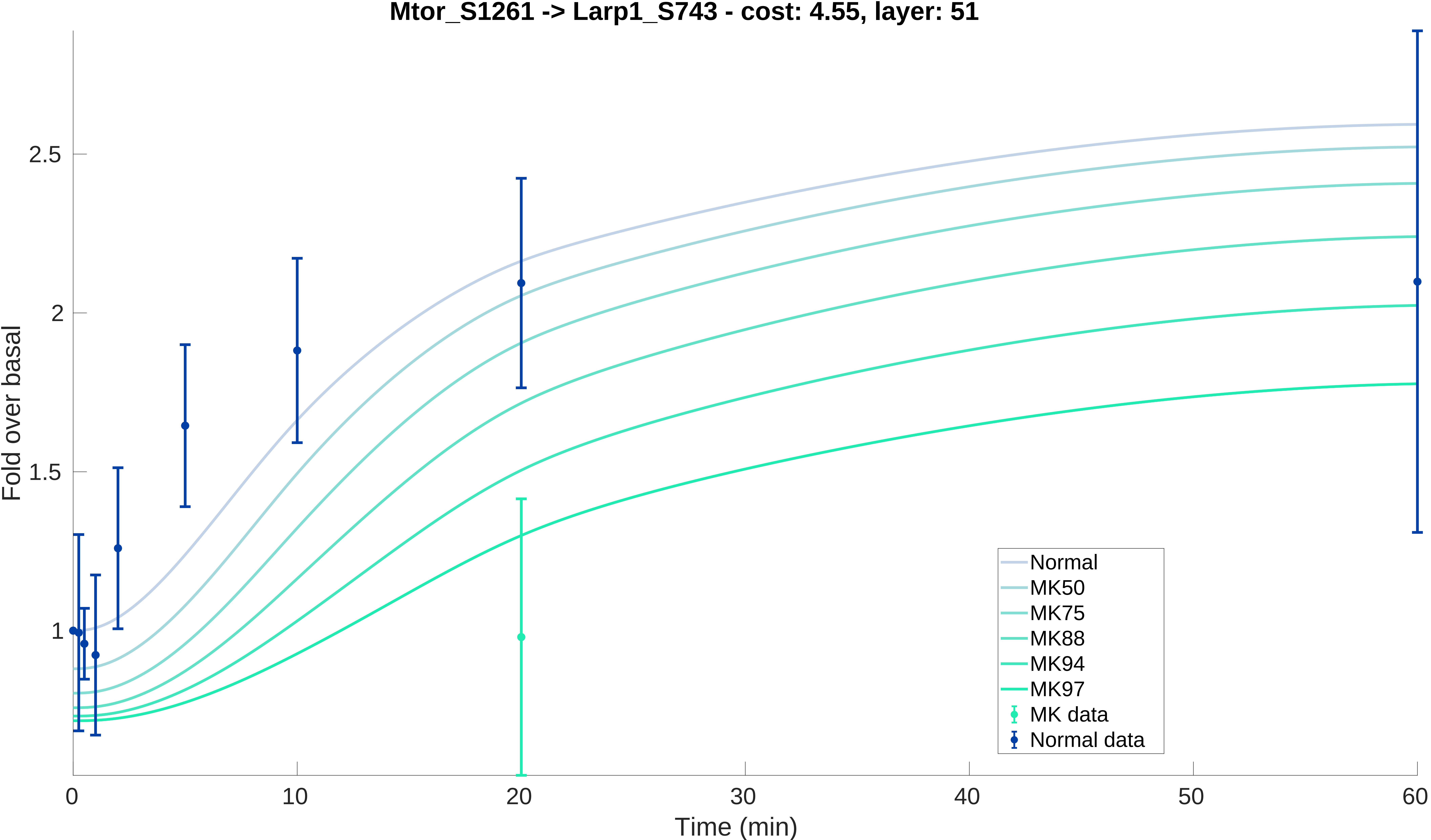

Mtor_S1261 -> Larp1_S743 - cost: 4.55, layer: 51
Fold over basal
2.5
2
1.5
1
0
10
20
30
40
50
60
Time (min)
Normal
MK50
MK75
MK88
MK94
MK97
MK data
Normal data

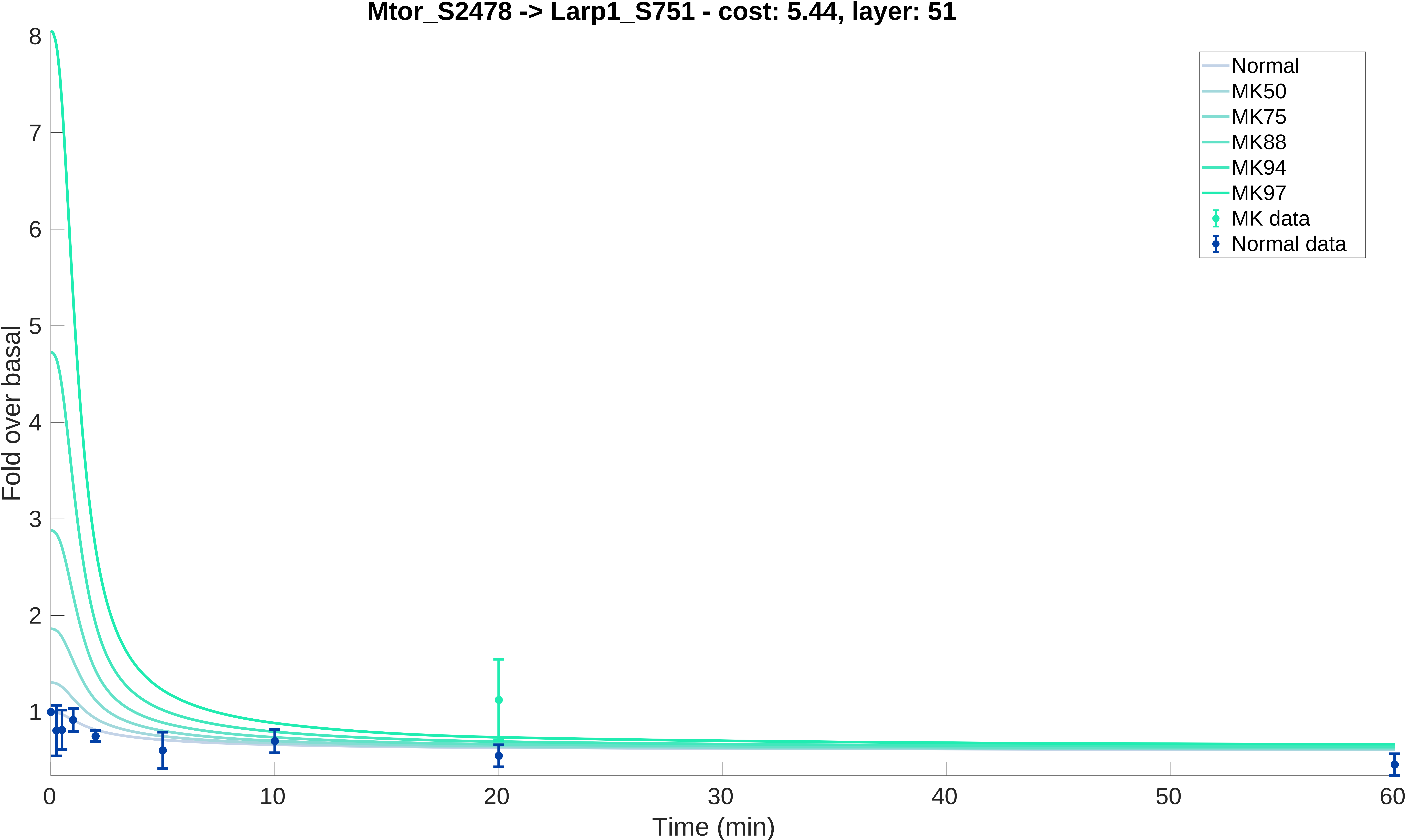

Mtor_S2478 -> Larp1_S751 - cost: 5.44, layer: 51
Fold over basal
Time (min)
Normal
MK50
MK75
MK88
MK94
MK97
MK data
Normal data
0
10
20
30
40
50
60
1
2
3
4
5
6
7
8

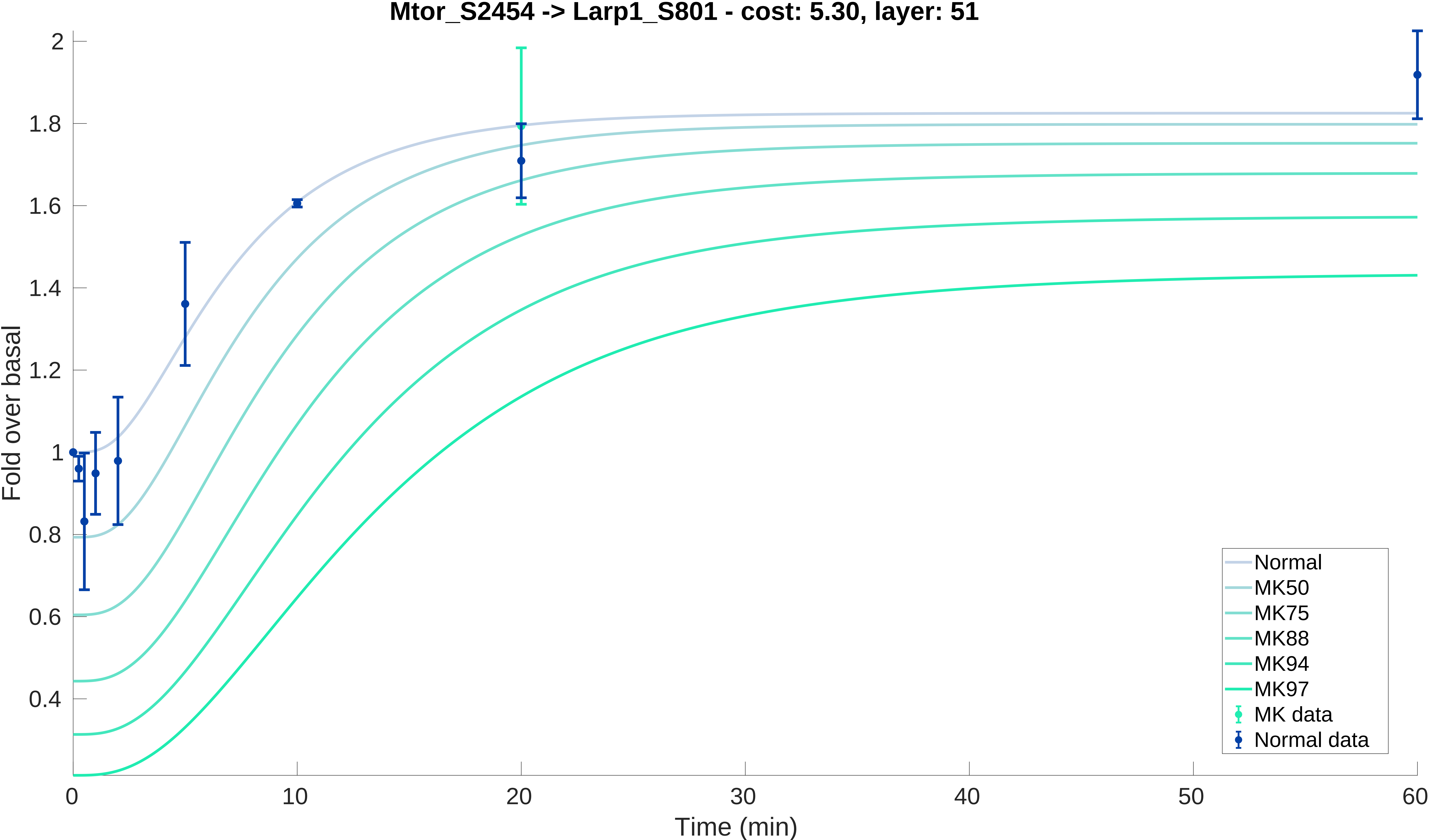
Mtor_S2454 -> Larp1_S801 - cost: 5.30, layer: 51
Fold over basal
2
1.8
1.6
1.4
1.2
1
0.8
0.6
0.4
0
10
20
30
40
50
60
Time (min)
Normal
MK50
MK75
MK88
MK94
MK97
MK data
Normal data

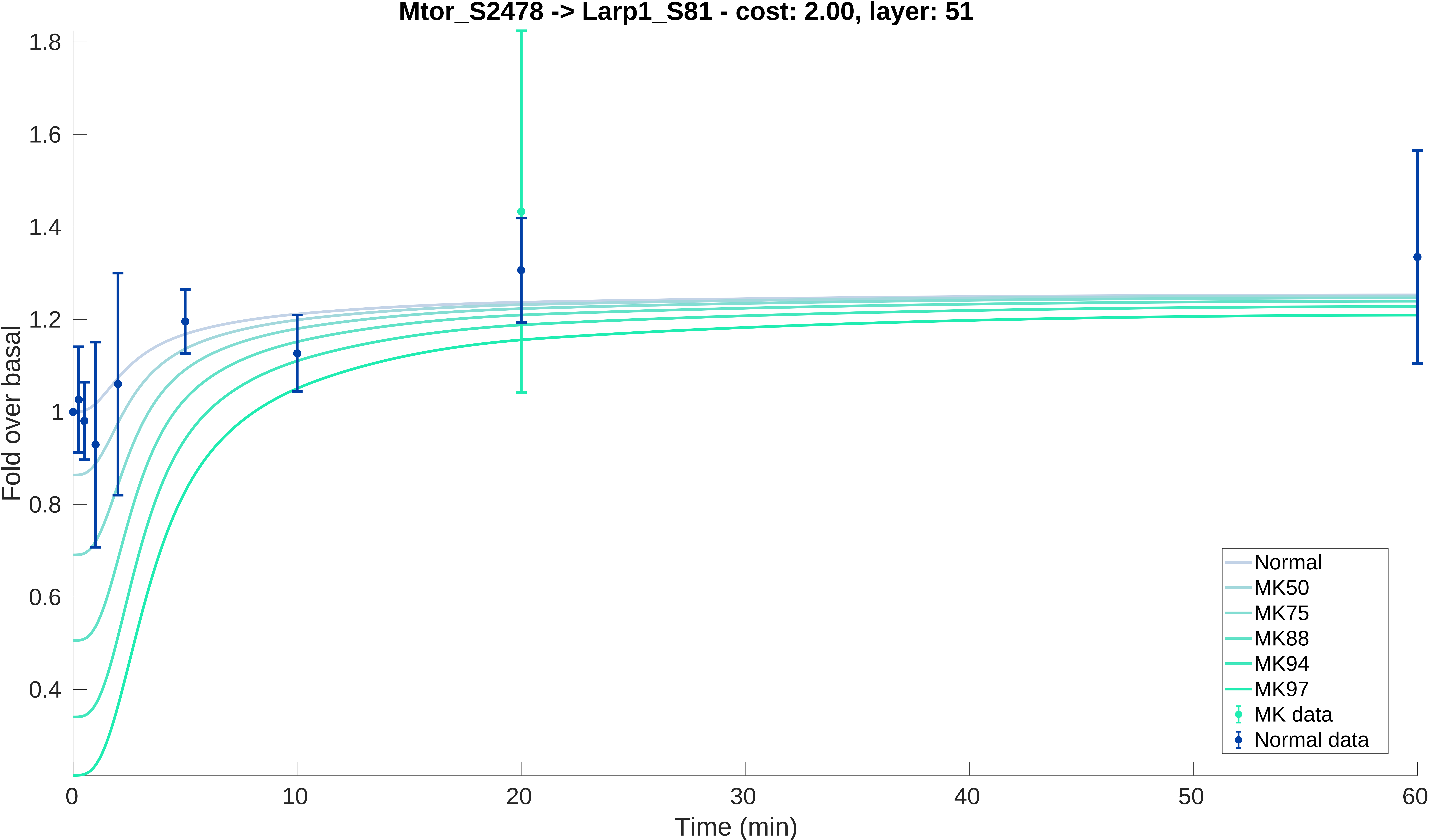

Mtor_S2478 -> Larp1_S81 - cost: 2.00, layer: 51
Fold over basal
1.8
1.6
1.4
1.2
1
0.8
0.6
0.4
0
10
20
30
40
50
60
Time (min)
Normal
MK50
MK75
MK88
MK94
MK97
MK data
Normal data

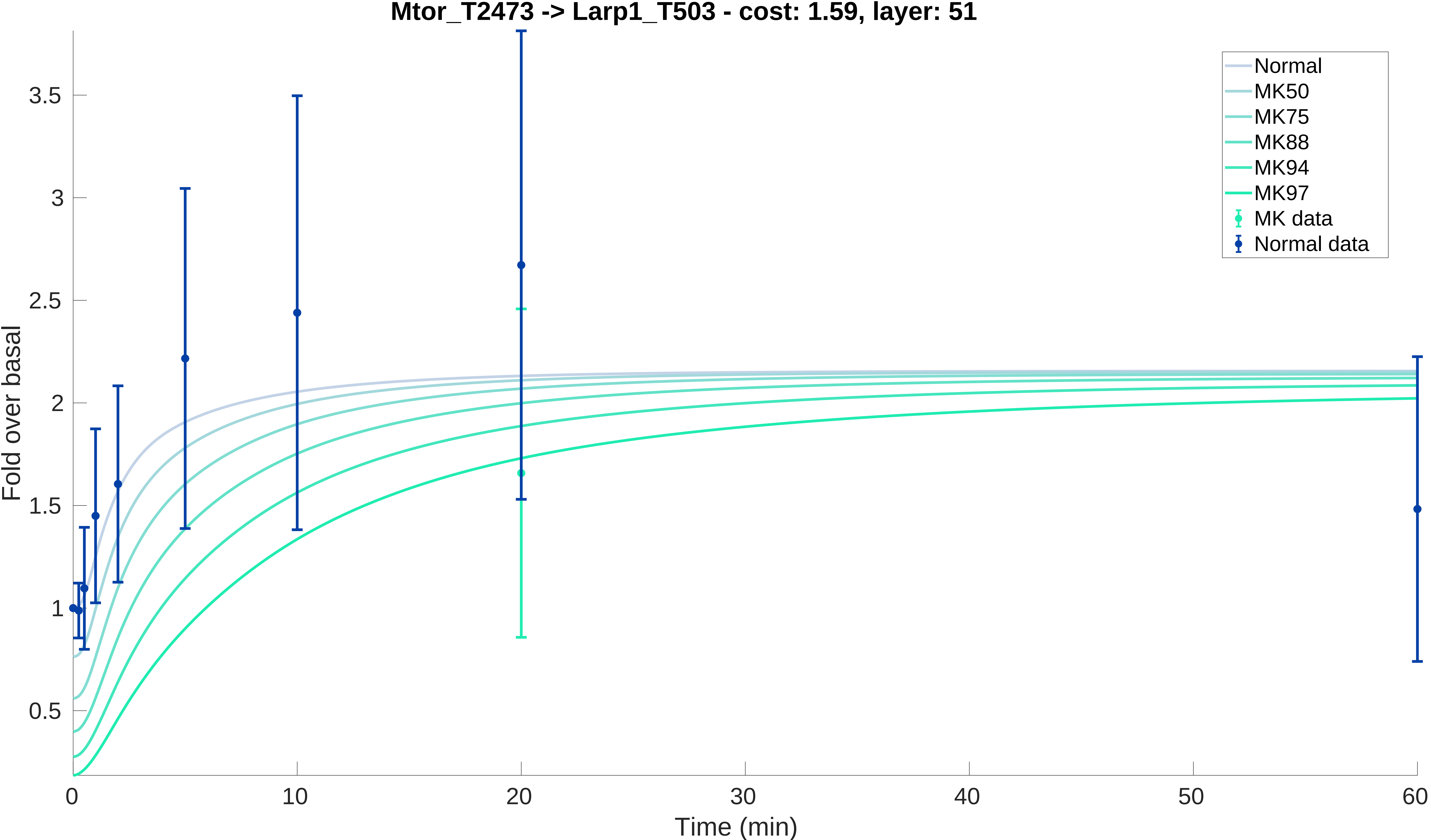

Mtor_T2473 -> Larp1_T503 - cost: 1.59, layer: 51
Fold over basal
Time (min)
0.5
1
1.5
2
2.5
3
3.5
0
10
20
30
40
50
60
Normal
MK50
MK75
MK88
MK94
MK97
MK data
Normal data

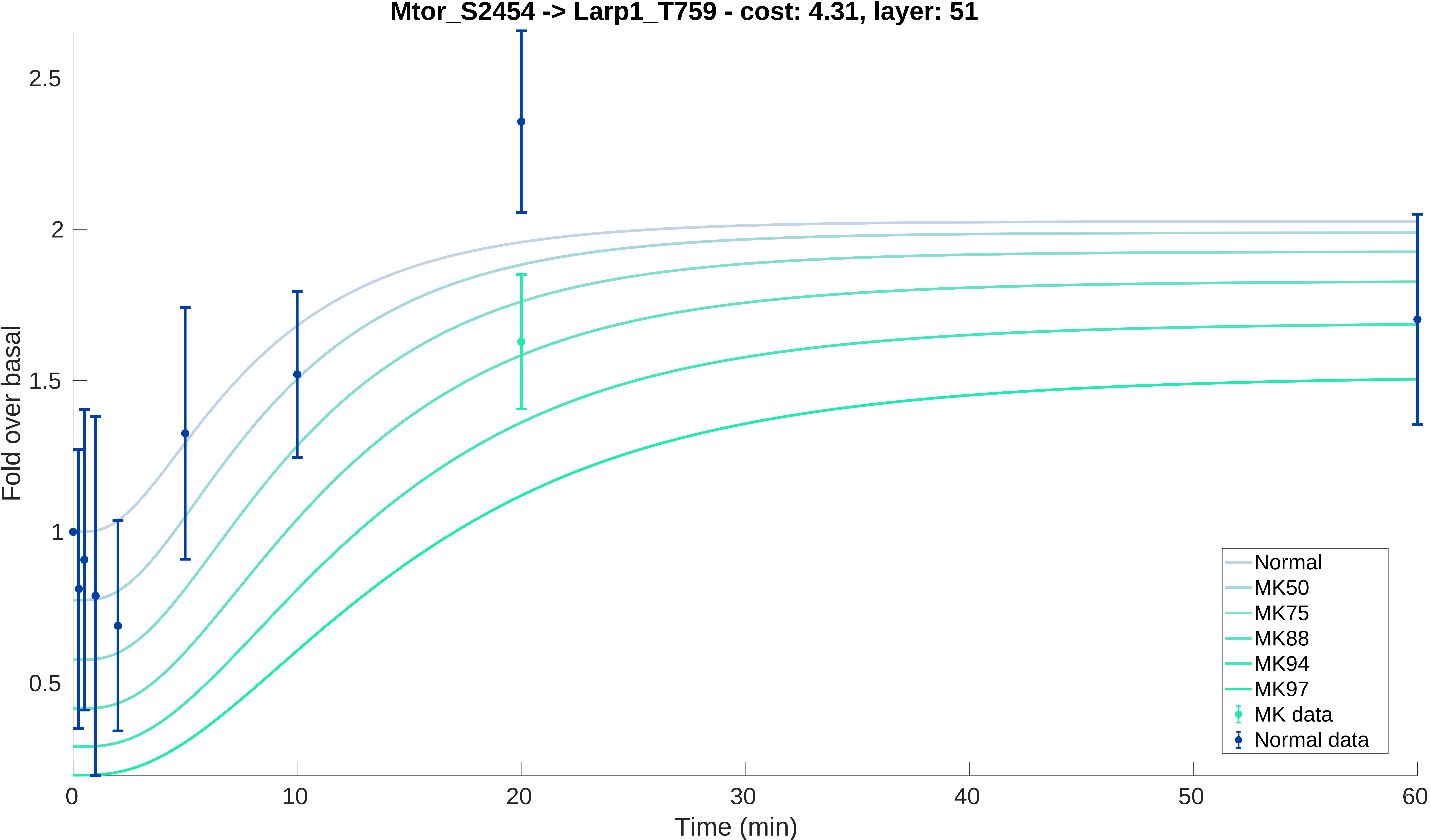

Mtor_S2454 -> Larp1_T759 - cost: 4.31, layer: 51
Fold over basal
Time (min)
0.5
1
1.5
2
2.5
0
10
20
30
40
50
60
Normal
MK50
MK75
MK88
MK94
MK97
MK data
Normal data

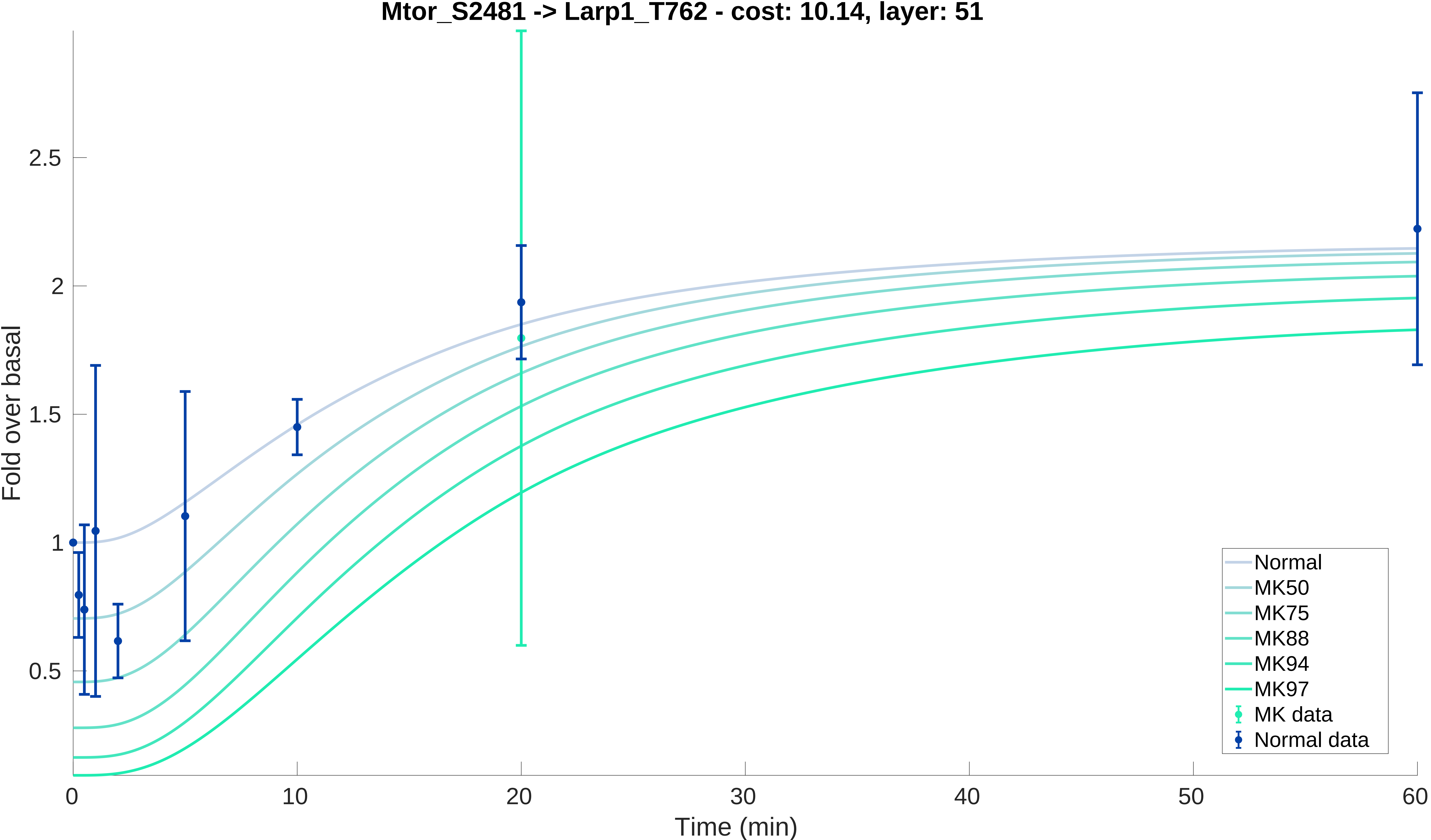

Mtor_S2481 -> Larp1_T762 - cost: 10.14, layer: 51
Fold over basal
Time (min)
0.5
1
1.5
2
2.5
0
10
20
30
40
50
60
Normal
MK50
MK75
MK88
MK94
MK97
MK data
Normal data

#### Prkcd_S671 -> Lmnb2_S427 - cost: 10.02, layer: 51

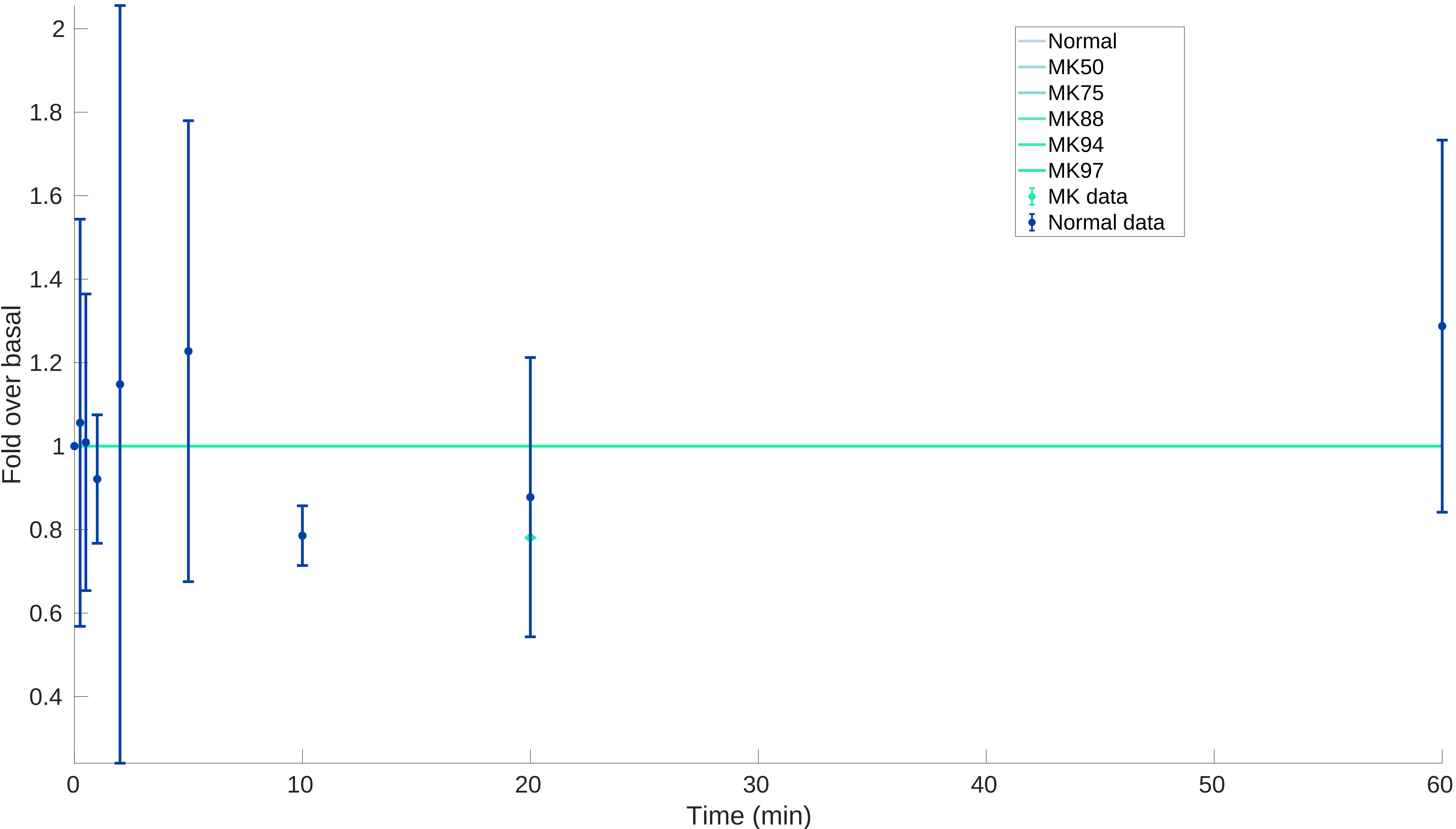

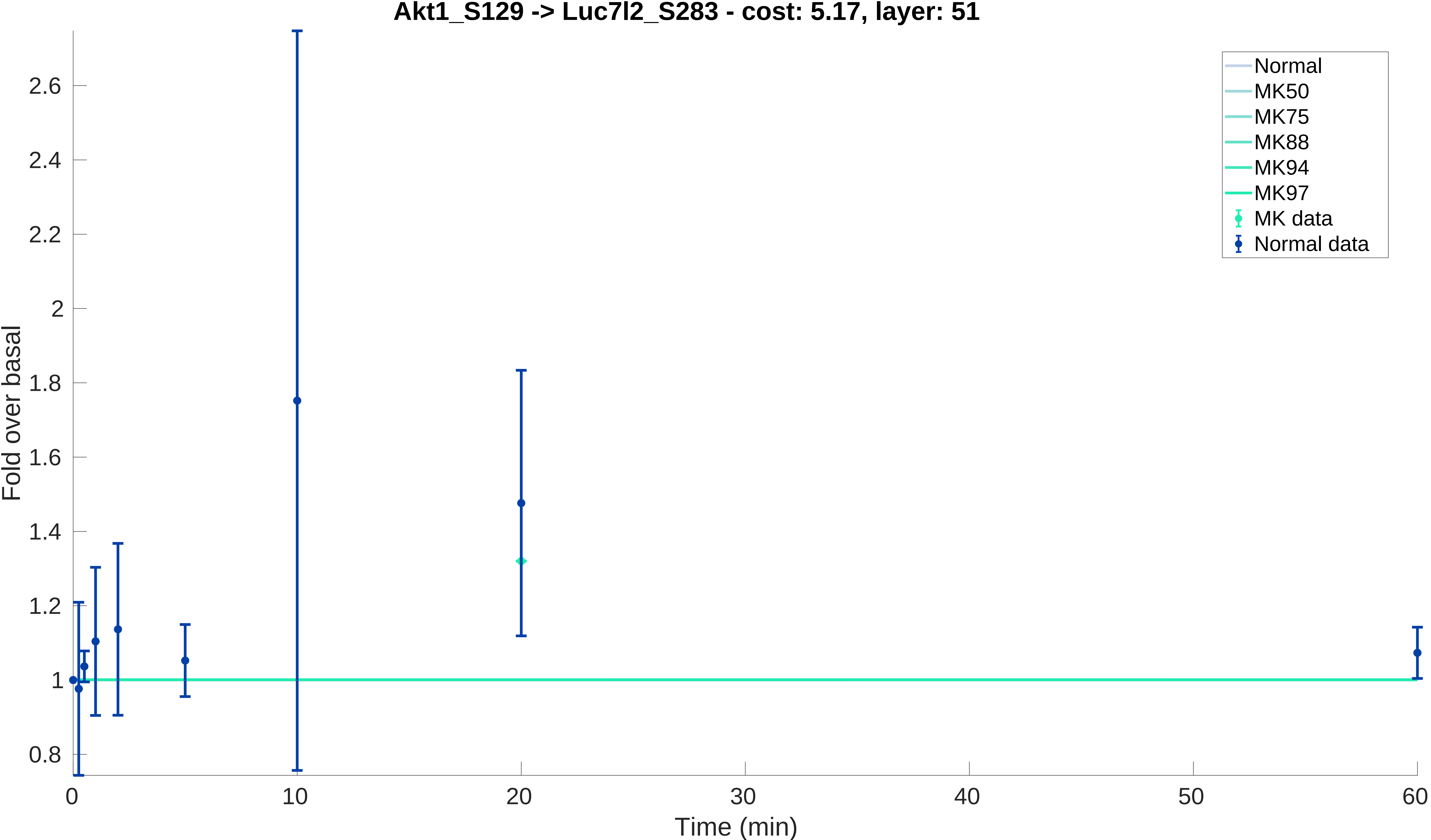

Akt1_S129 -> Luc7l2_S283 - cost: 5.17, layer: 51
Fold over basal
Time (min)
2.6
2.4
2.2
2
1.8
1.6
1.4
1.2
1
0.8
0
10
20
30
40
50
60
Normal
MK50
MK75
MK88
MK94
MK97
MK data
Normal data

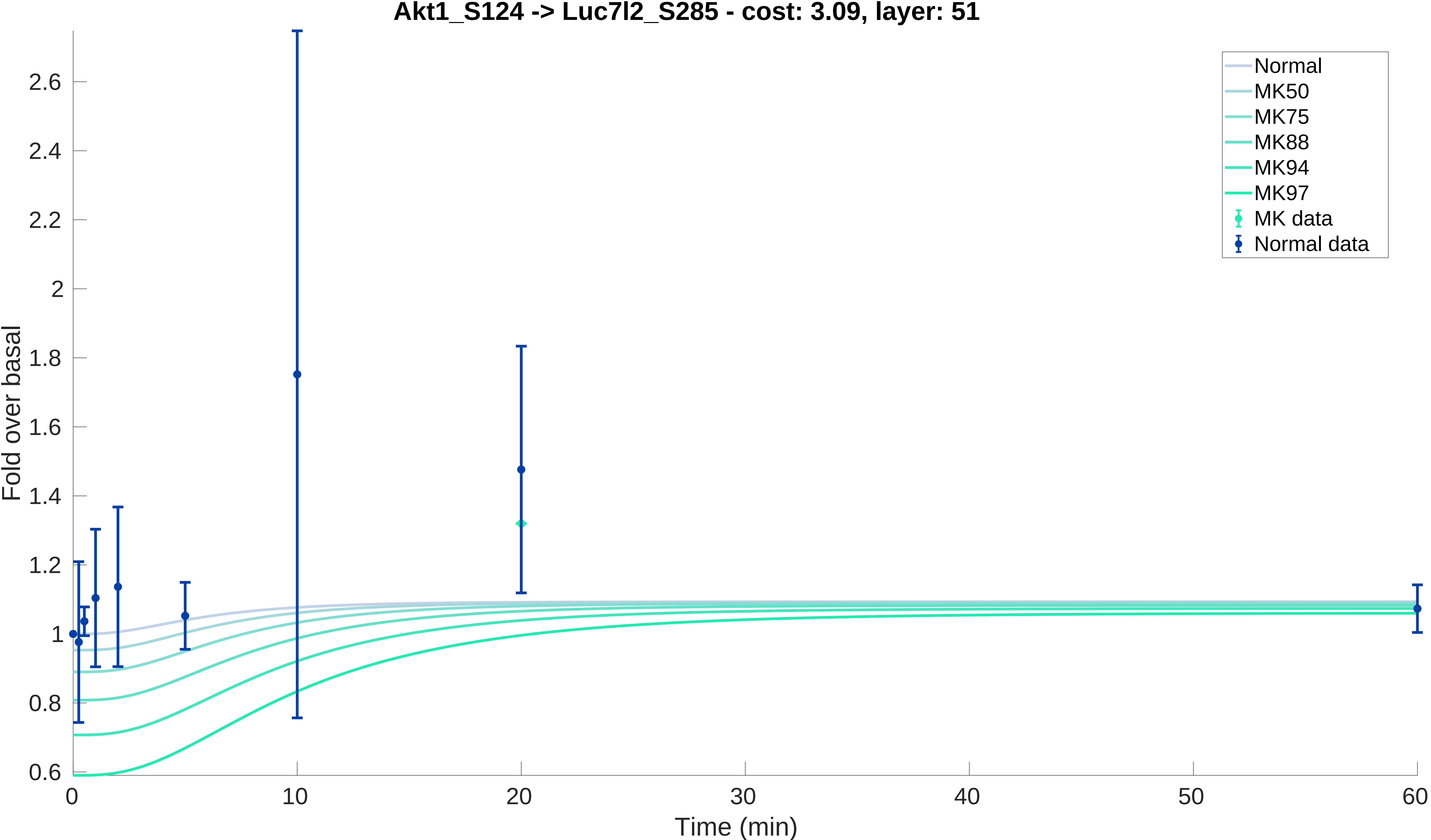

Akt1_S124 -> Luc7l2_S285 - cost: 3.09, layer: 51
Normal
MK50
MK75
MK88
MK94
MK97
MK data
Normal data
Fold over basal
Time (min)
0.6
0.8
1
1.2
1.4
1.6
1.8
2
2.2
2.4
2.6
0
10
20
30
40
50
60

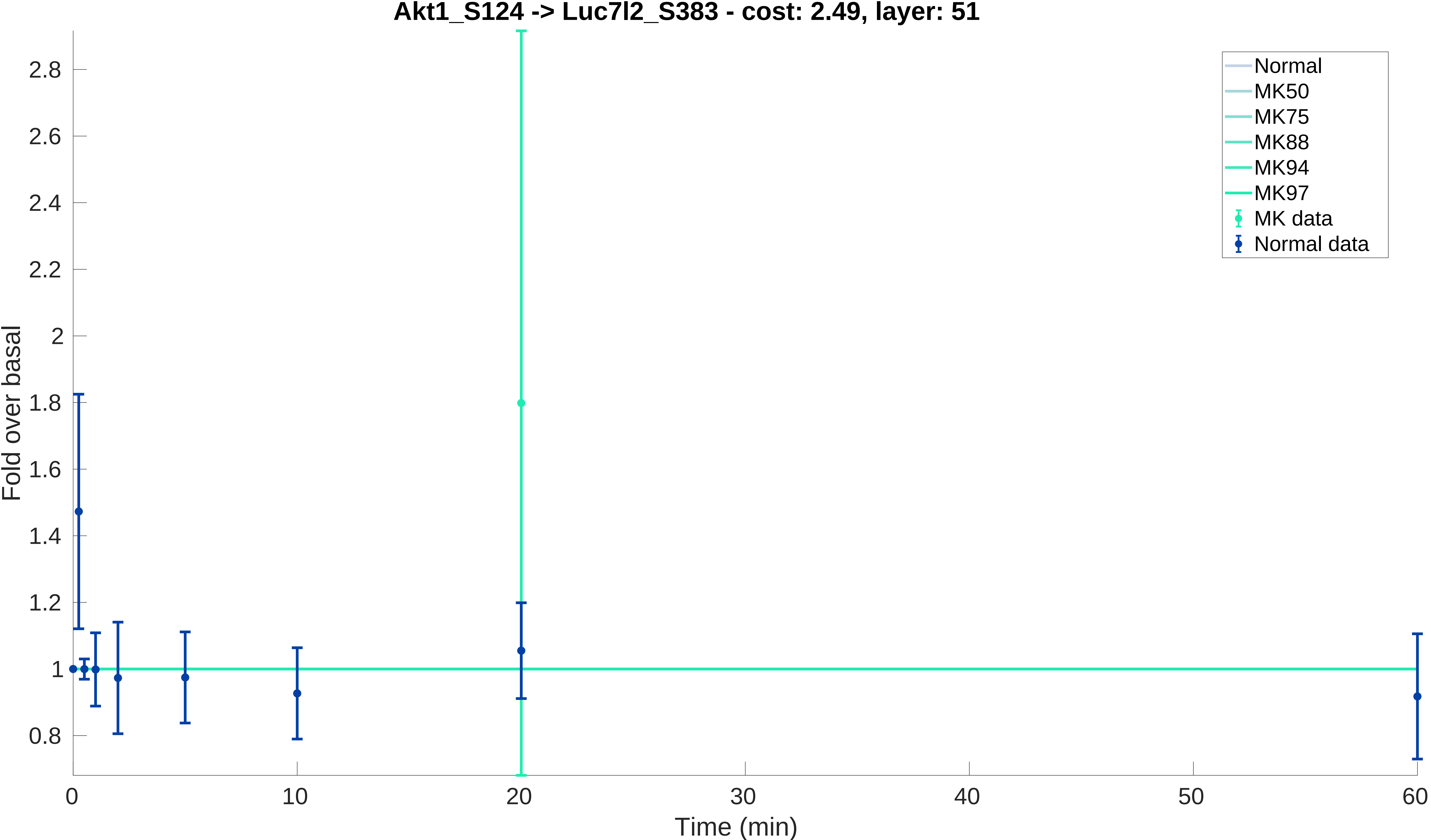

Akt1_S124 -> Luc7l2_S383 - cost: 2.49, layer: 51
Fold over basal
2.8
2.6
2.4
2.2
2
1.8
1.6
1.4
1.2
1
0.8
0
10
20
30
40
50
60
Time (min)
Normal
MK50
MK75
MK88
MK94
MK97
MK data
Normal data

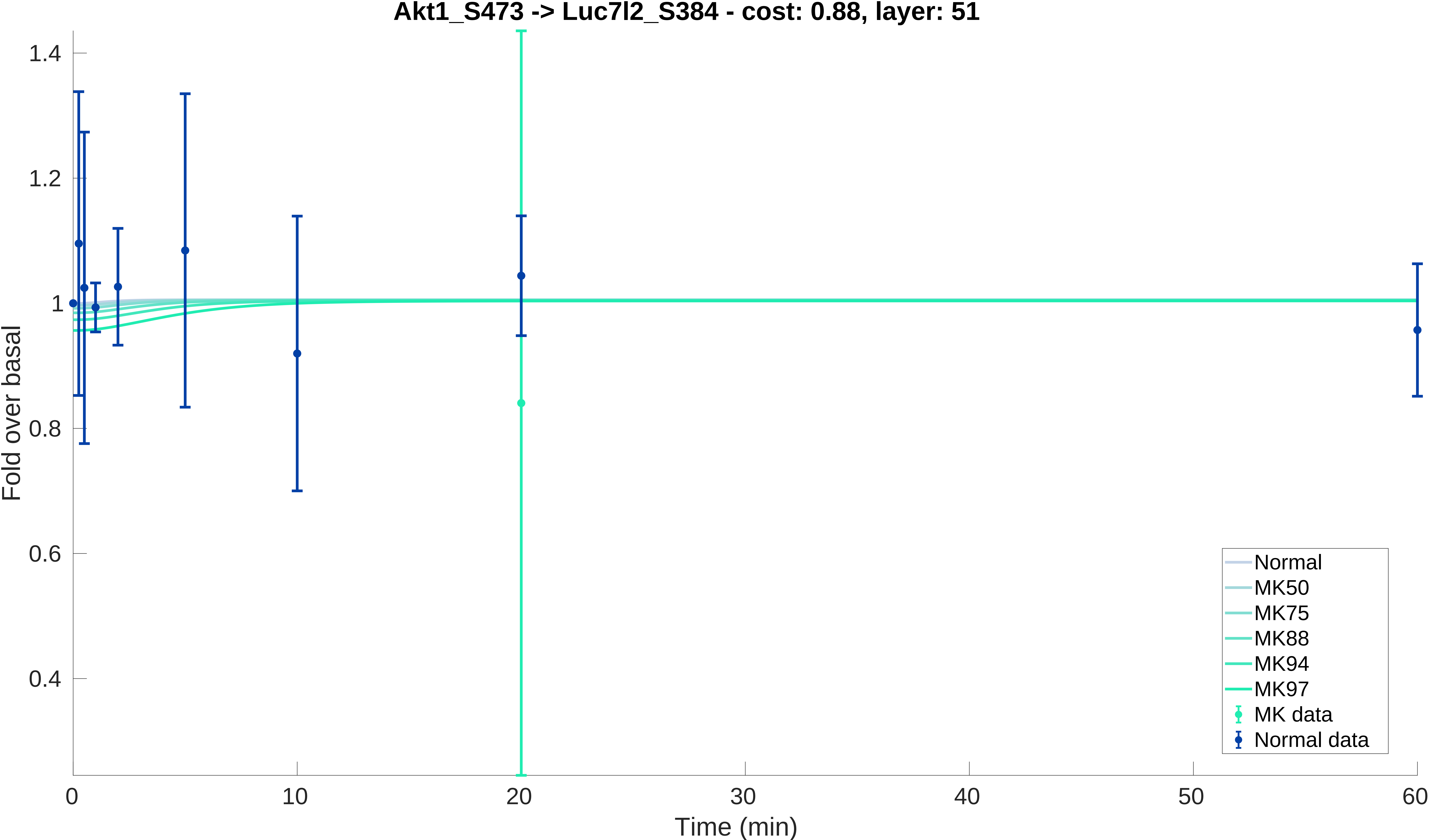

Akt1_S473 -> Luc7l2_S384 - cost: 0.88, layer: 51
Fold over basal
1.4
1.2
1
0.8
0.6
0.4
0
10
20
30
40
50
60
Time (min)
Normal
MK50
MK75
MK88
MK94
MK97
MK data
Normal data

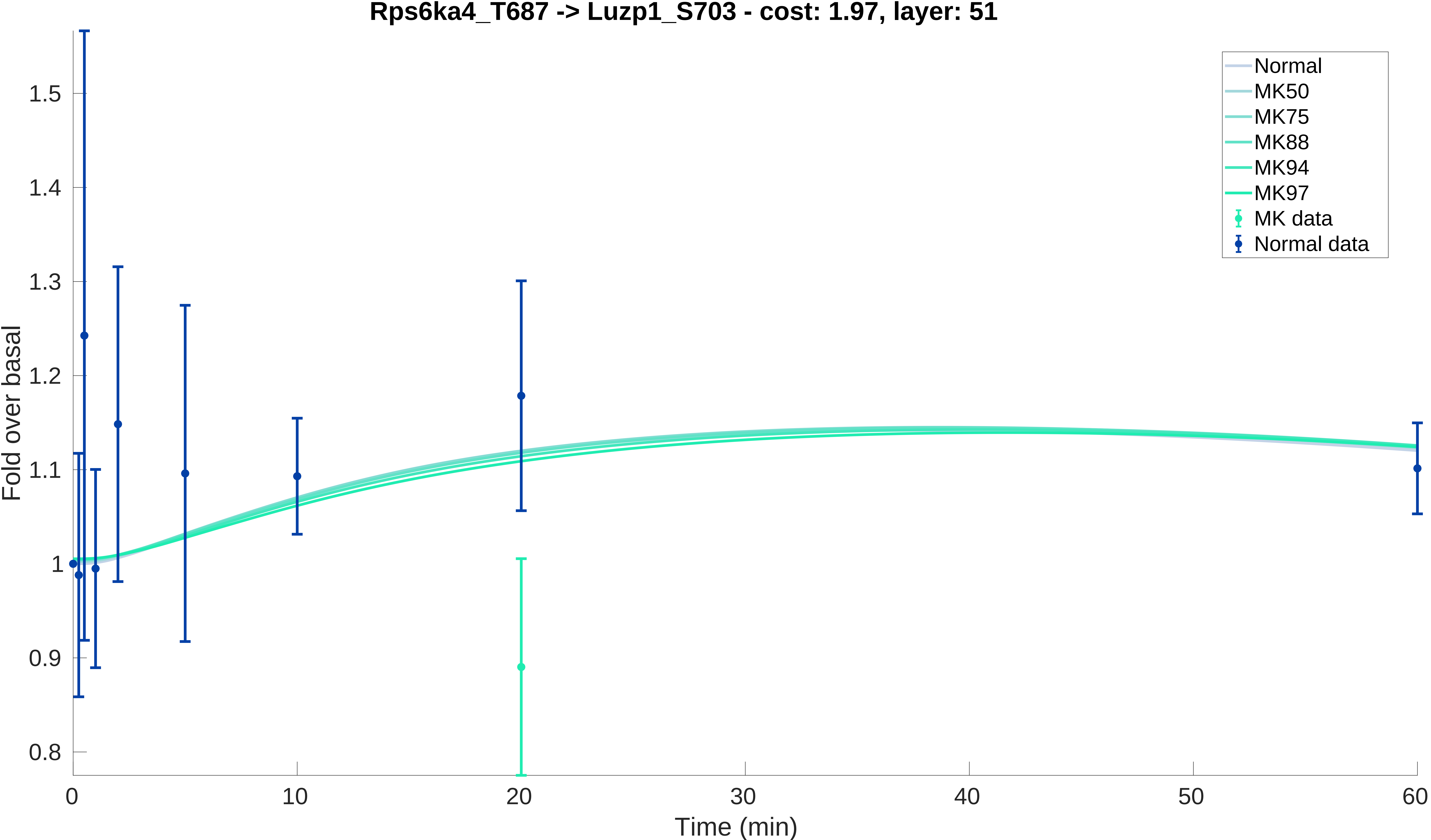
Rps6ka4_T687 -> Luzp1_S703 - cost: 1.97, layer: 51
Fold over basal
Time (min)
0.8
0.9
1
1.1
1.2
1.3
1.4
1.5
0
10
20
30
40
50
60
Normal
MK50
MK75
MK88
MK94
MK97
MK data
Normal data

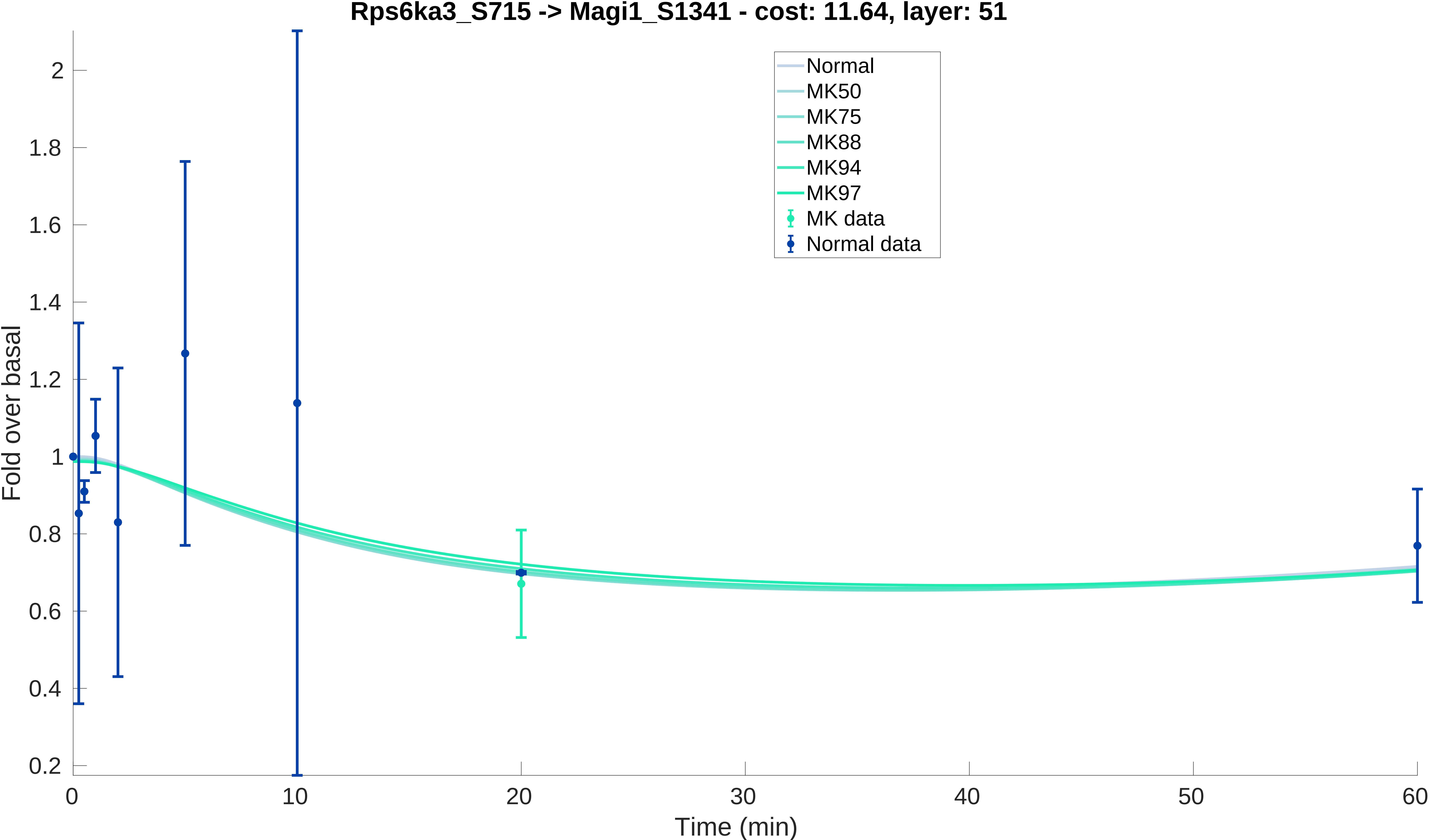

Rps6ka3_S715 -> Magi1_S1341 - cost: 11.64, layer: 51
Normal
MK50
MK75
MK88
MK94
MK97
MK data
Normal data
Fold over basal
Time (min)
0.2
0.4
0.6
0.8
1
1.2
1.4
1.6
1.8
2
0
10
20
30
40
50
60

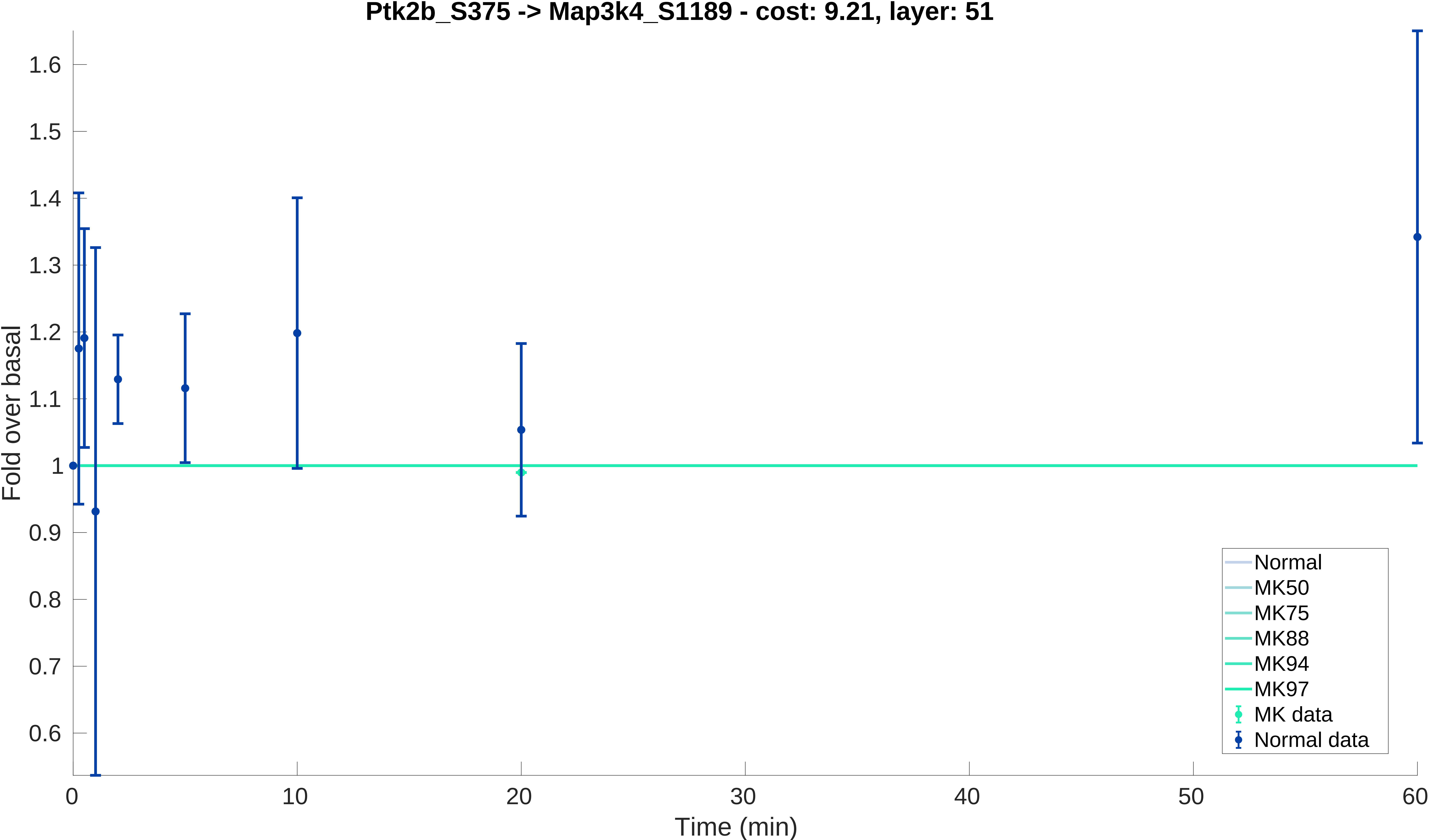
Ptk2b_S375 -> Map3k4_S1189 - cost: 9.21, layer: 51
Fold over basal
Time (min)
0.6
0.7
0.8
0.9
1
1.1
1.2
1.3
1.4
1.5
1.6
0
10
20
30
40
50
60
Normal
MK50
MK75
MK88
MK94
MK97
MK data
Normal data

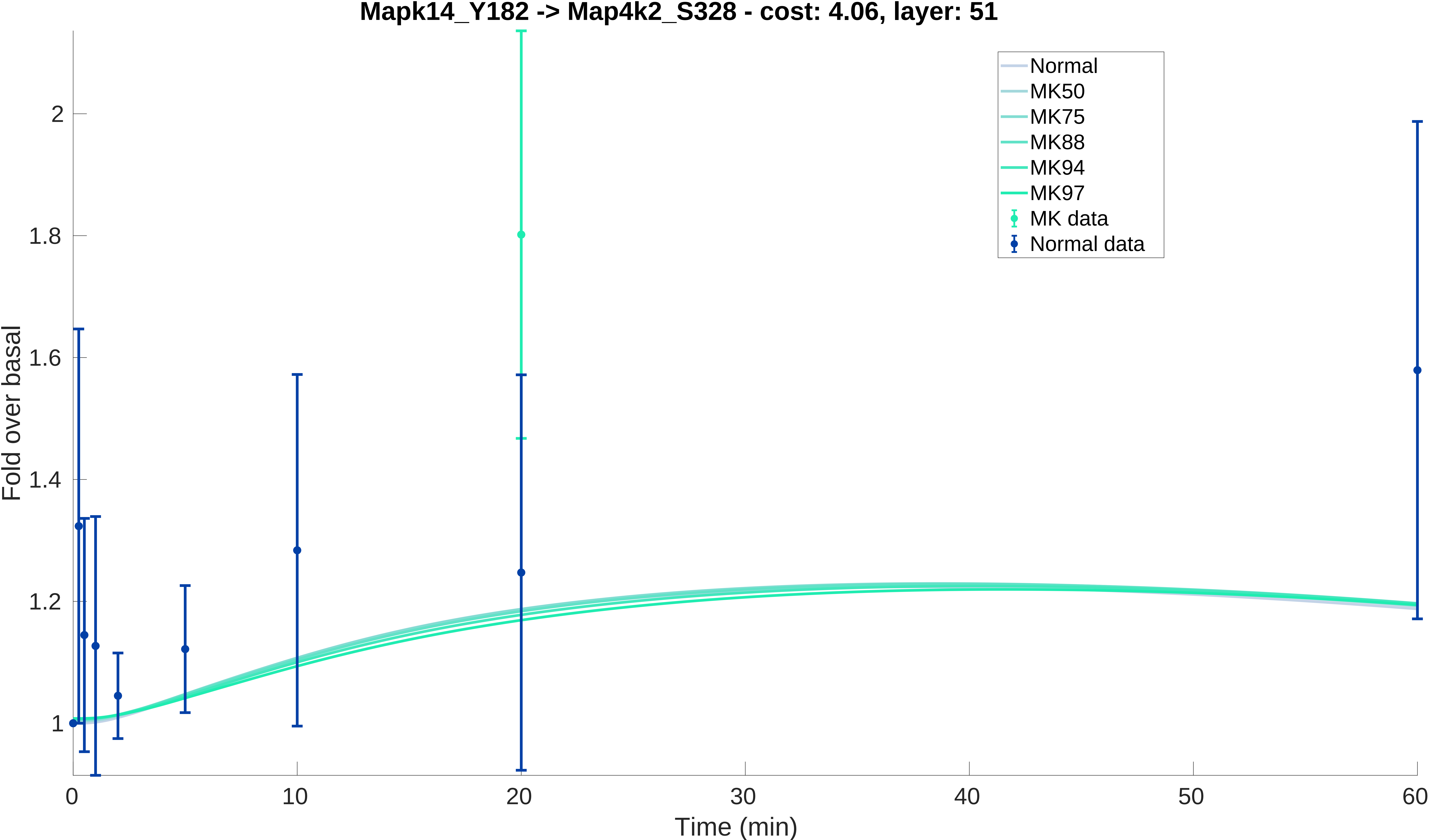

Mapk14_Y182 -> Map4k2_S328 - cost: 4.06, layer: 51
Normal
MK50
MK75
MK88
MK94
MK97
MK data
Normal data
Fold over basal
2
1.8
1.6
1.4
1.2
1
0
10
20
30
40
50
60
Time (min)

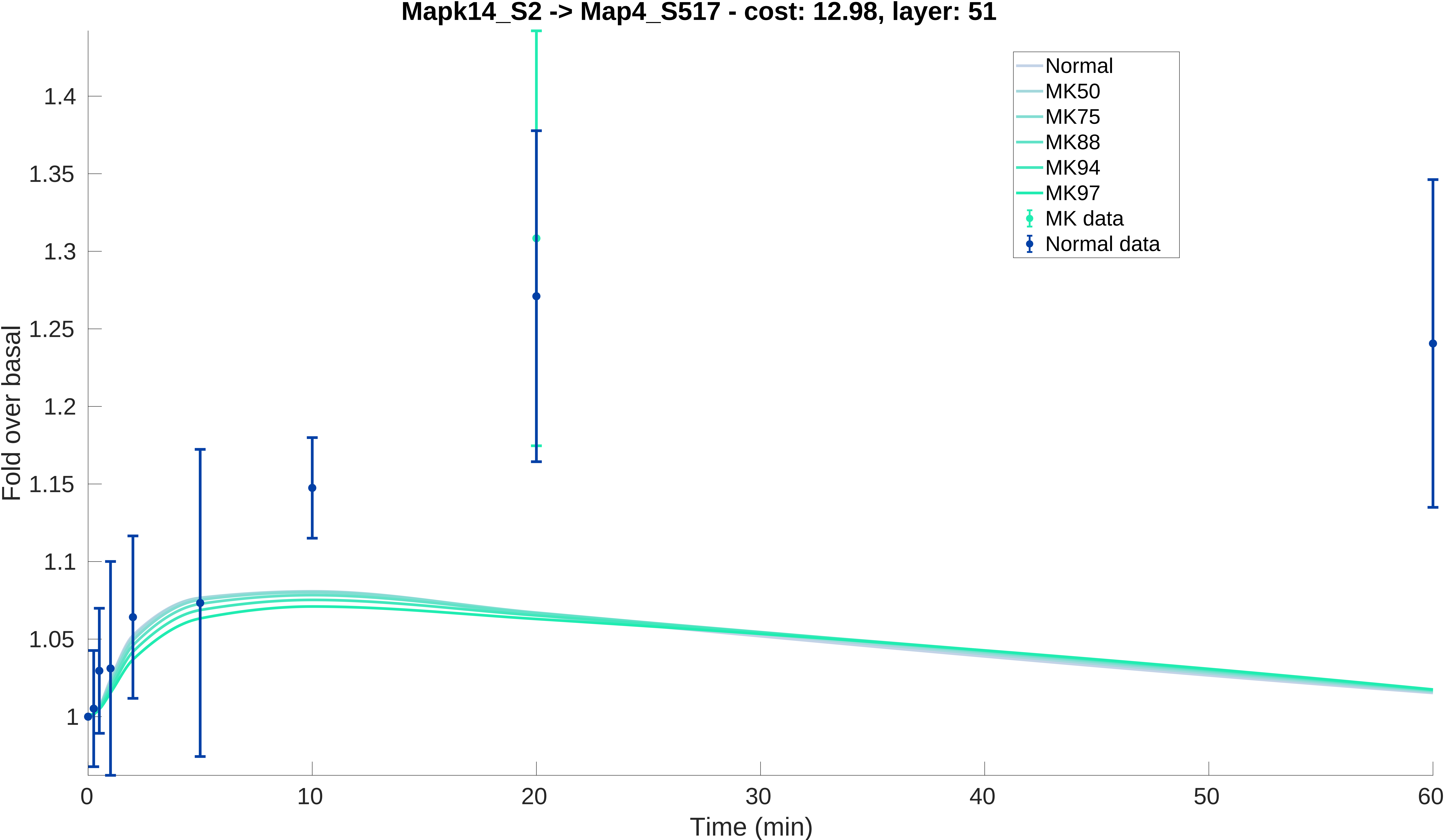

Mapk14_S2 -> Map4_S517 - cost: 12.98, layer: 51
Fold over basal
Time (min)
1.4
1.35
1.3
1.25
1.2
1.15
1.1
1.05
1
0
10
20
30
40
50
60
Normal
MK50
MK75
MK88
MK94
MK97
MK data
Normal data

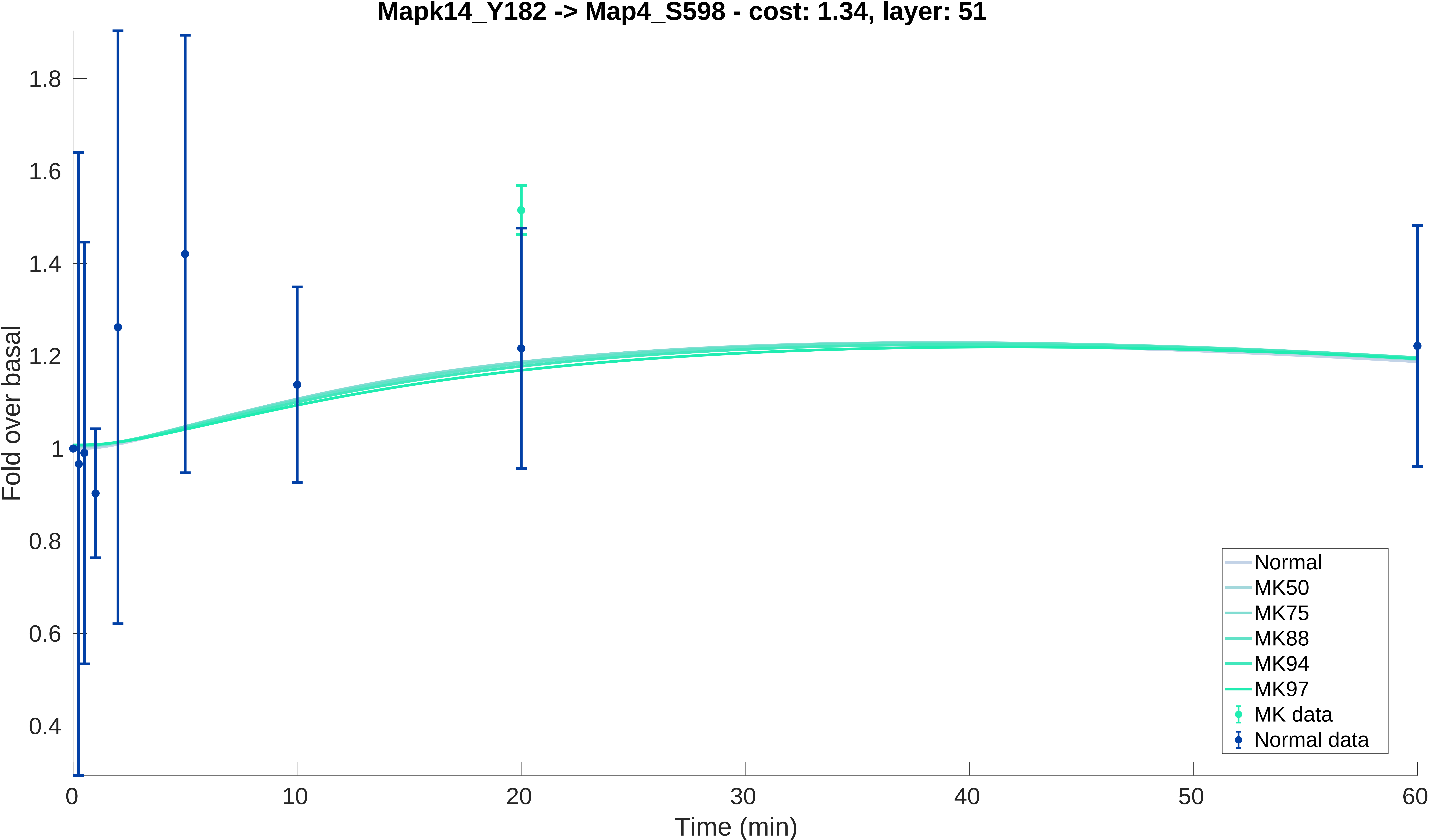

Mapk14_Y182 -> Map4_S598 - cost: 1.34, layer: 51
Fold over basal
1.8
1.6
1.4
1.2
1
0.8
0.6
0.4
0
10
20
30
40
50
60
Time (min)
Normal
MK50
MK75
MK88
MK94
MK97
MK data
Normal data

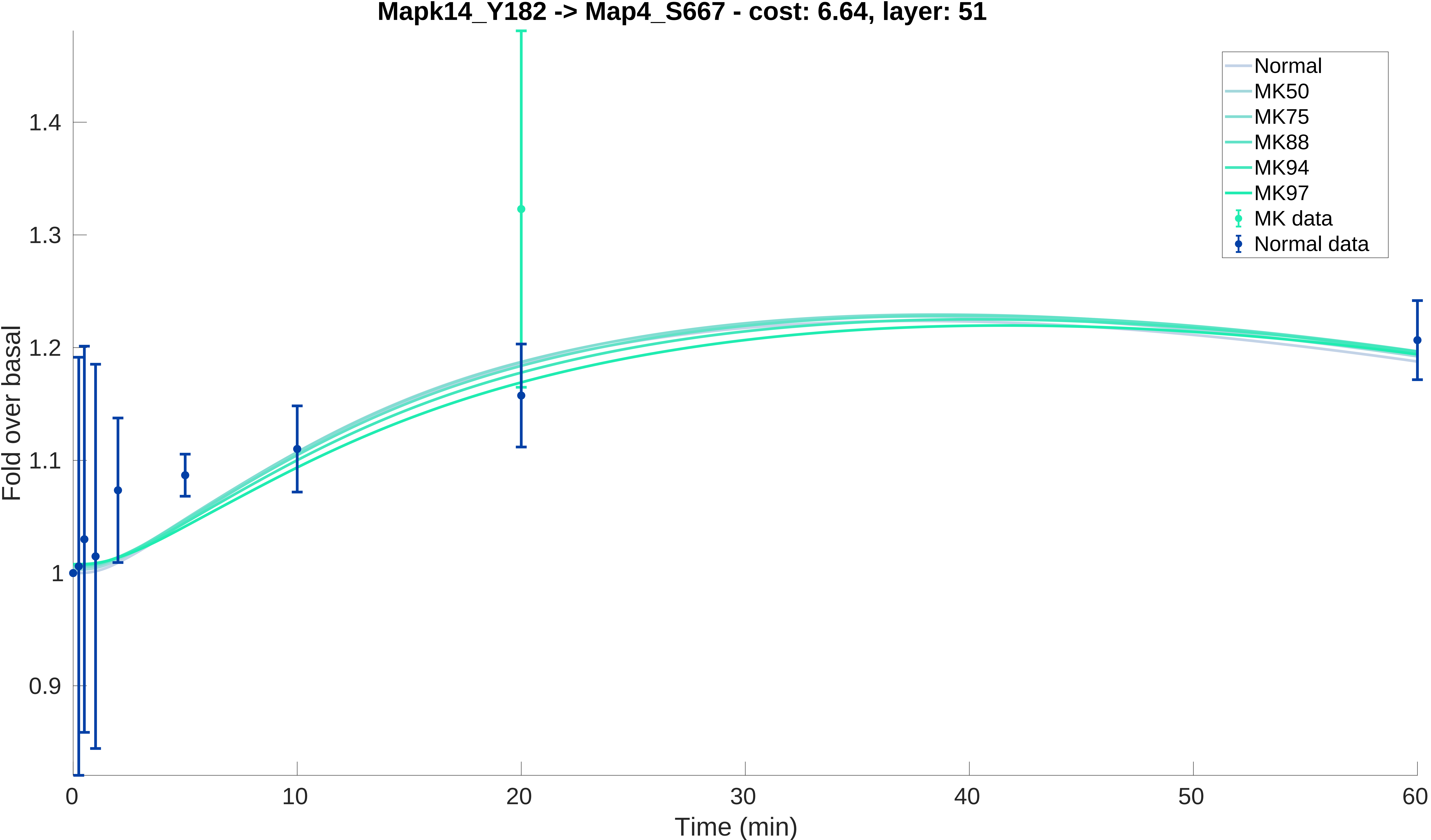

Mapk14_Y182 -> Map4_S667 - cost: 6.64, layer: 51
Fold over basal
Time (min)
0
10
20
30
40
50
60
0.9
1
1.1
1.2
1.3
1.4
Normal
MK50
MK75
MK88
MK94
MK97
MK data
Normal data

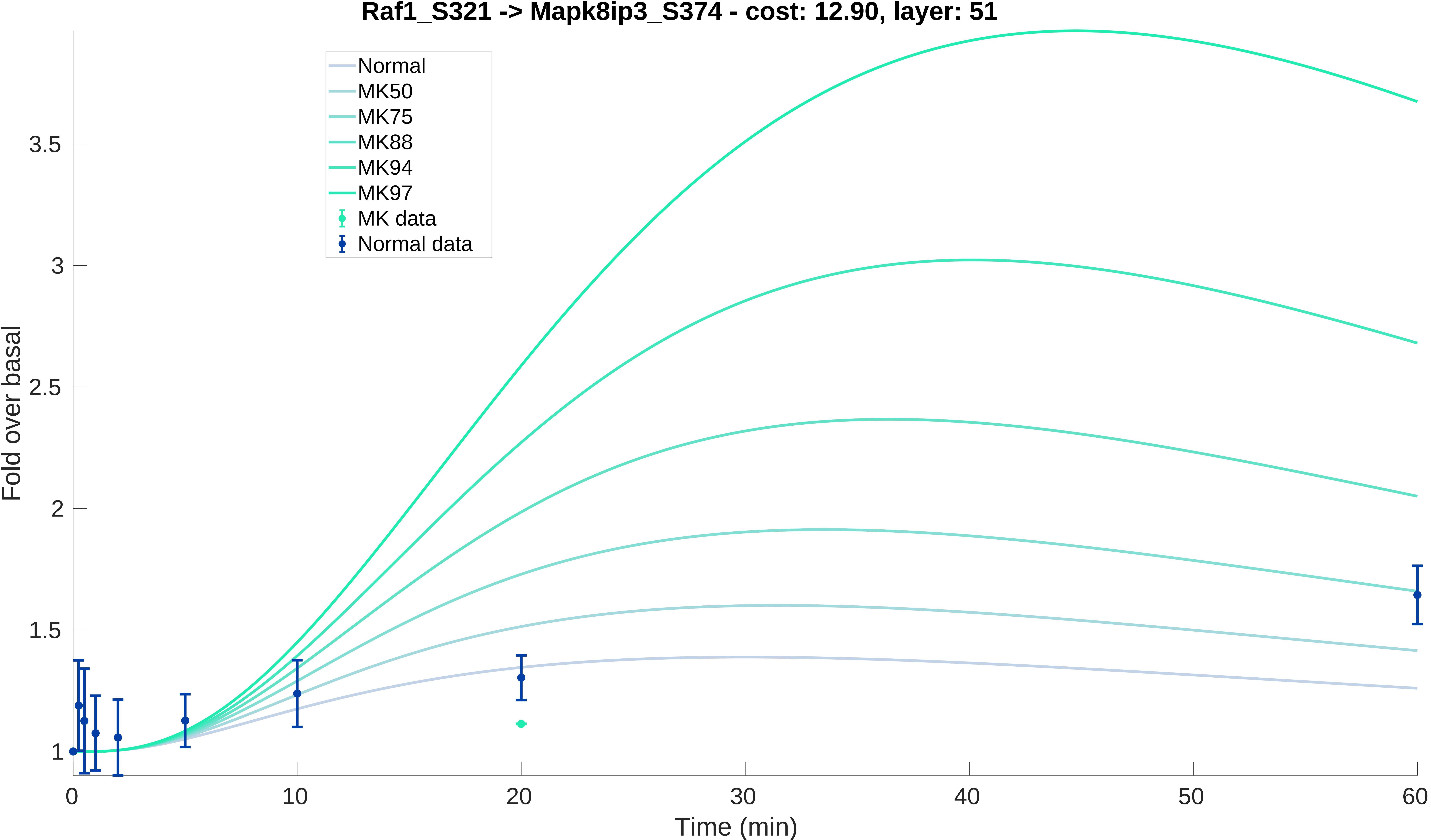

Raf1_S321 -> Mapk8ip3_S374 - cost: 12.90, layer: 51
Normal
MK50
MK75
MK88
MK94
MK97
MK data
Normal data
Fold over basal
3.5
3
2.5
2
1.5
1
0
10
20
30
40
50
60
Time (min)

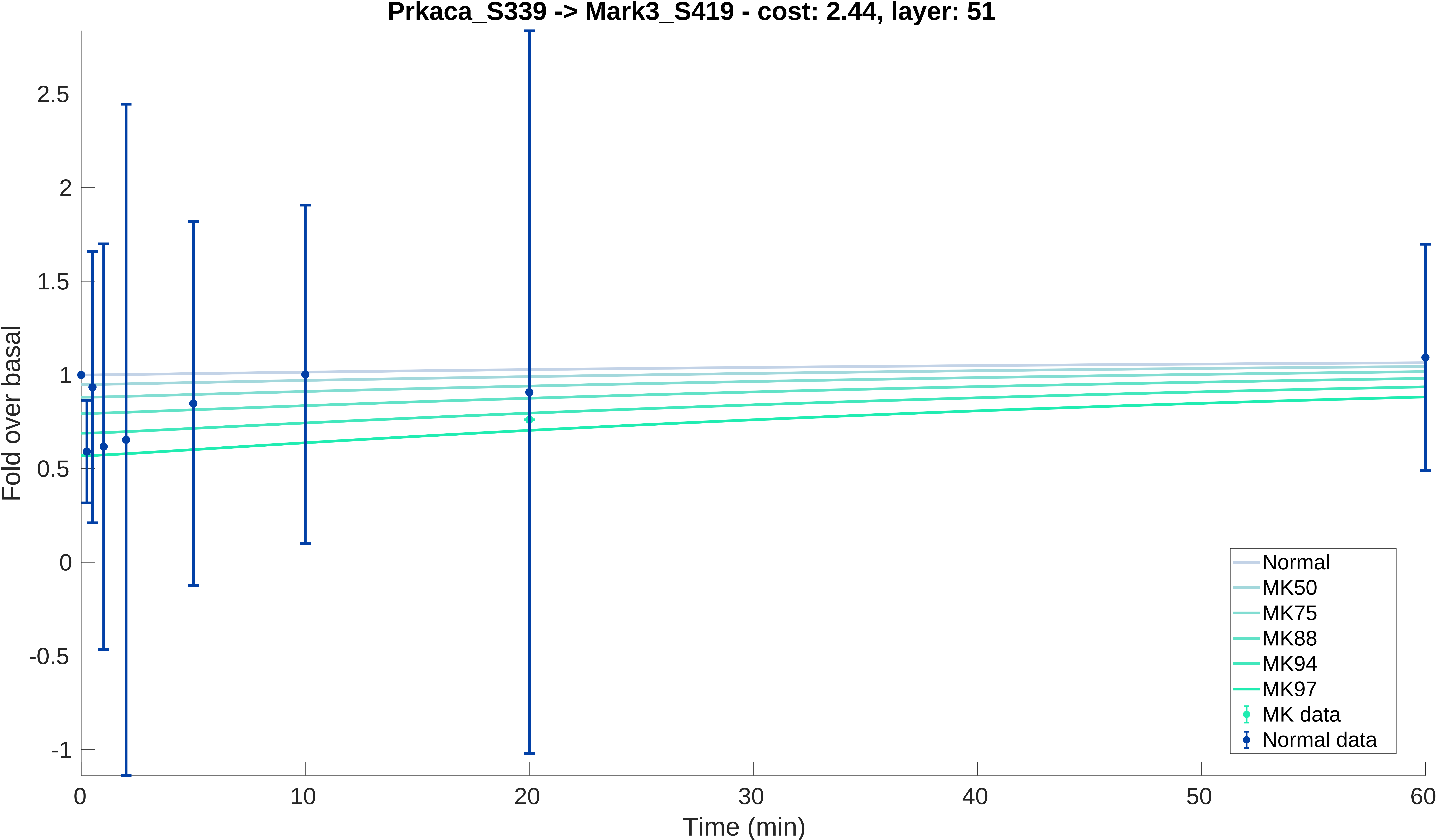

Prrkaca_S339 -> Mark3_S419 - cost: 2.44, layer: 51
Fold over basal
Time (min)
2.5
2
1.5
1
0.5
0
-0.5
-1
0
10
20
30
40
50
60
Normal
MK50
MK75
MK88
MK94
MK97
MK data
Normal data

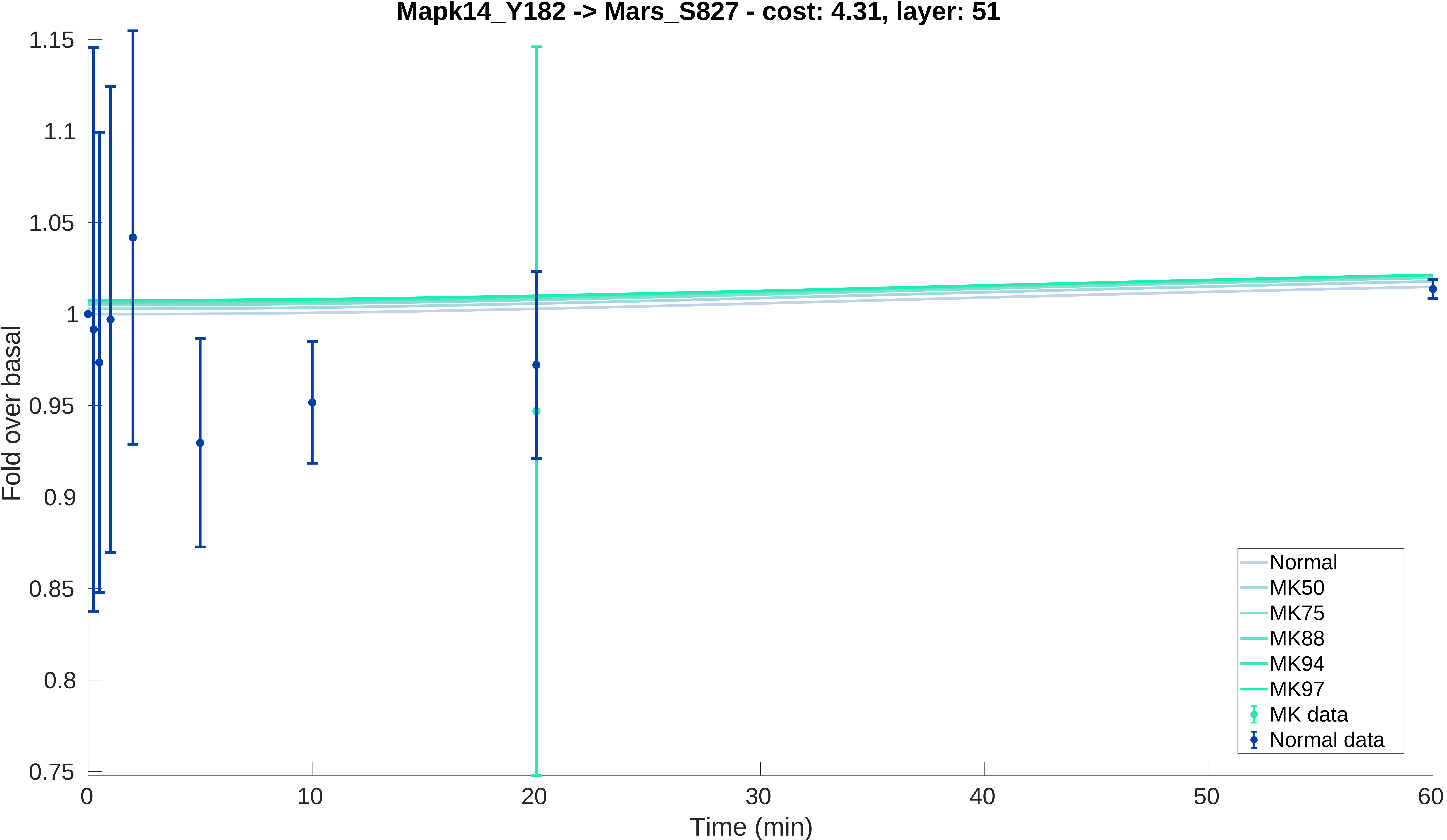

Mapk14_Y182 -> Mars_S827 - cost: 4.31, layer: 51
Fold over basal
1.15
1.1
1.05
1
0.95
0.9
0.85
0.8
0.75
0
10
20
30
40
50
60
Time (min)
Normal
MK50
MK75
MK88
MK94
MK97
MK data
Normal data

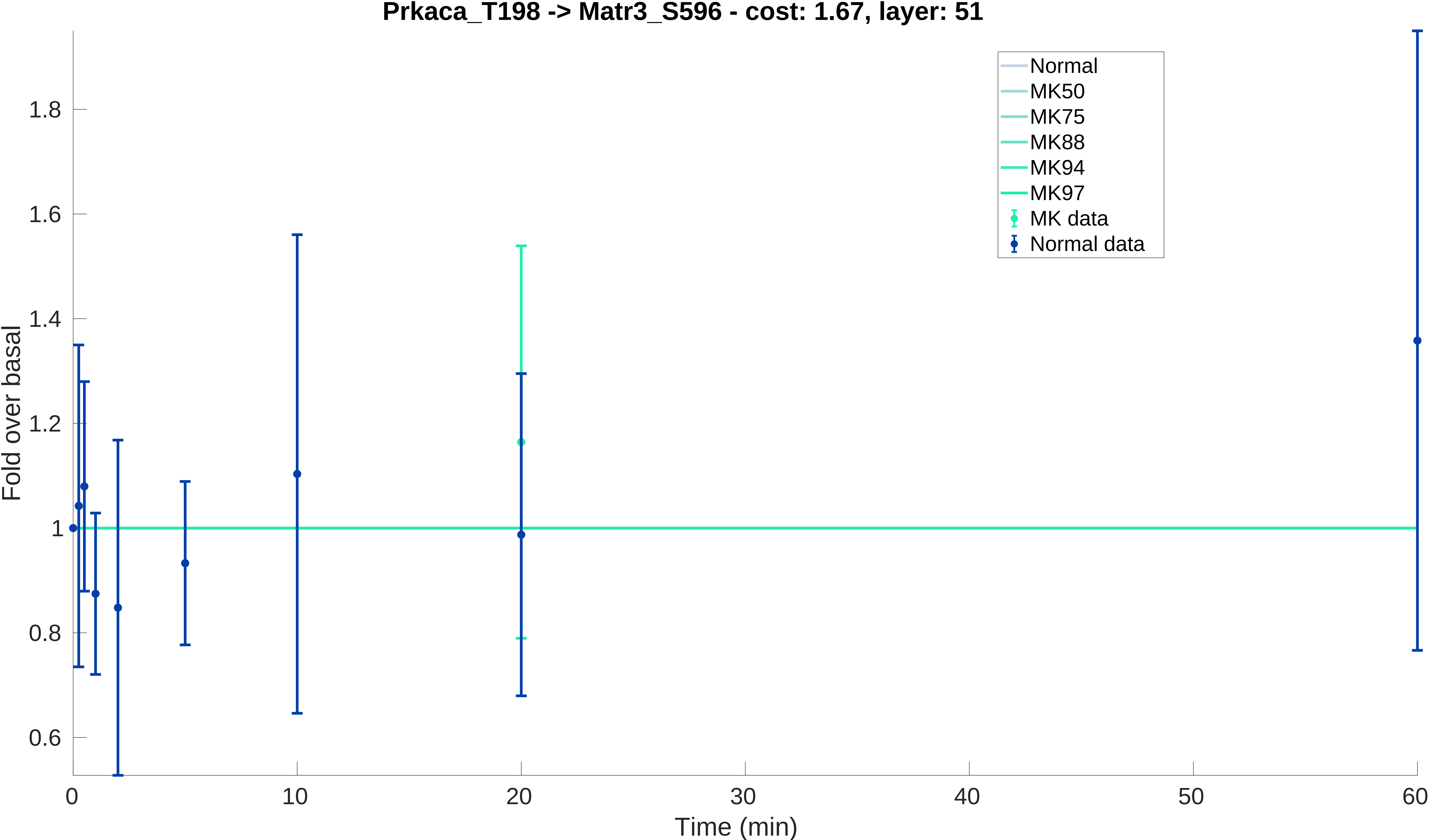

Prkaca_T198 -> Matr3_S596 - cost: 1.67, layer: 51
Normal
MK50
MK75
MK88
MK94
MK97
MK data
Normal data
Fold over basal
1.8
1.6
1.4
1.2
1
0.8
0.6
0
10
20
30
40
50
60
Time (min)

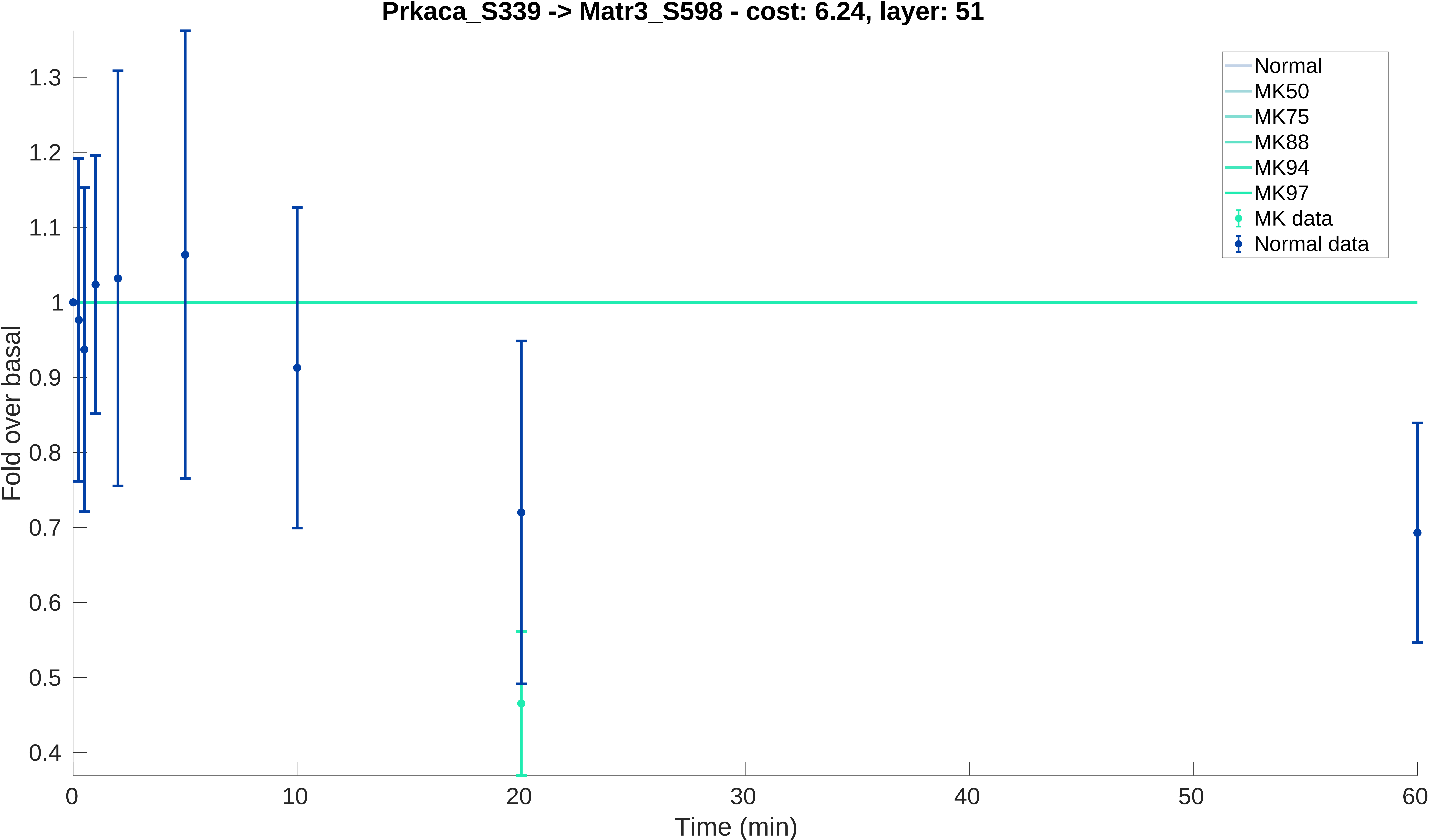

Prkaca_S339 -> Matr3_S598 - cost: 6.24, layer: 51
Fold over basal
Time (min)
0
10
20
30
40
50
60
0.4
0.5
0.6
0.7
0.8
0.9
1
1.1
1.2
1.3
Normal
MK50
MK75
MK88
MK94
MK97
MK data
Normal data

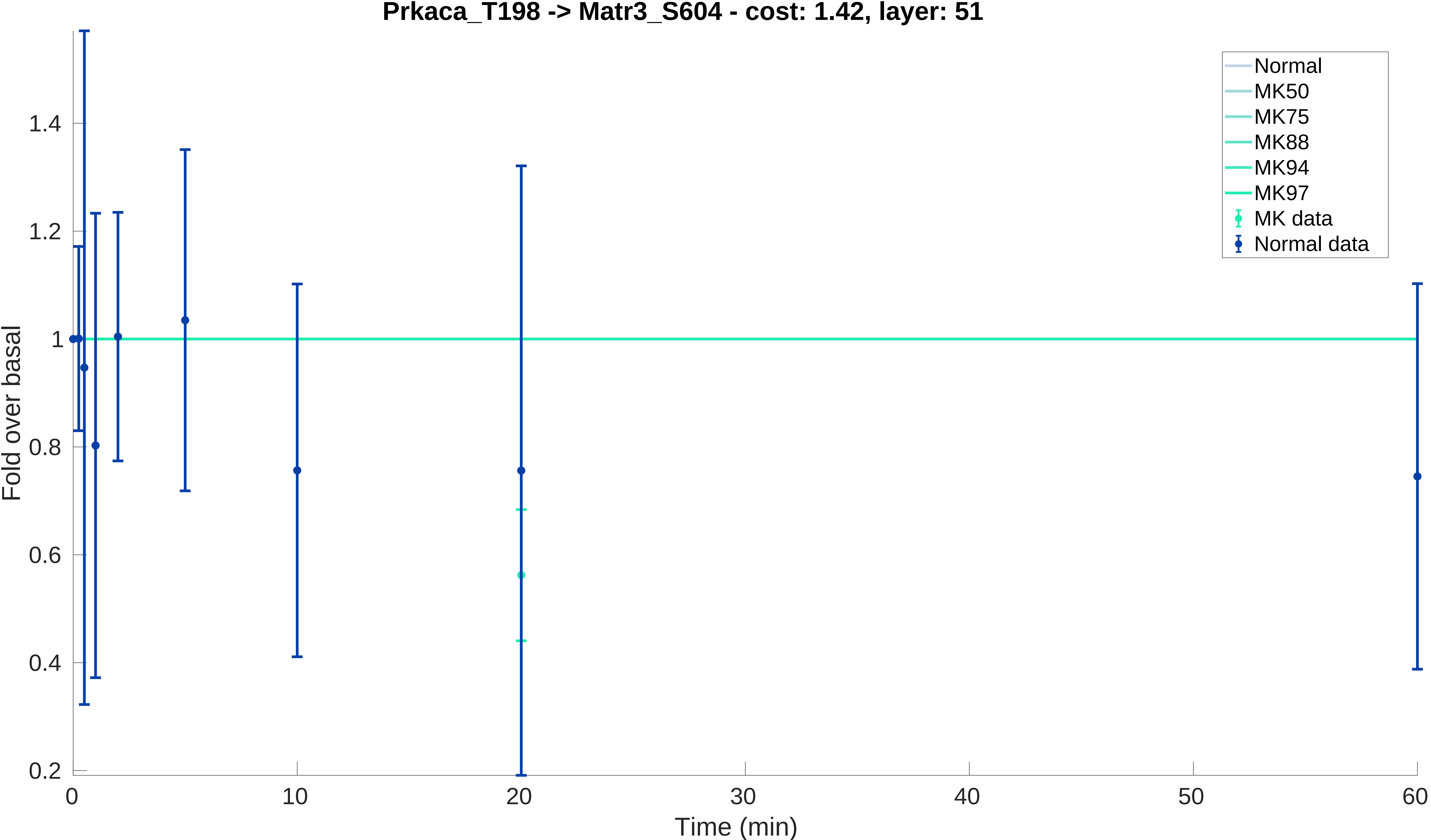

Prkaca_T198 -> Matr3_S604 - cost: 1.42, layer: 51
Normal
MK50
MK75
MK88
MK94
MK97
MK data
Normal data
Fold over basal
Time (min)
0.2
0.4
0.6
0.8
1
1.2
1.4
0
10
20
30
40
50
60

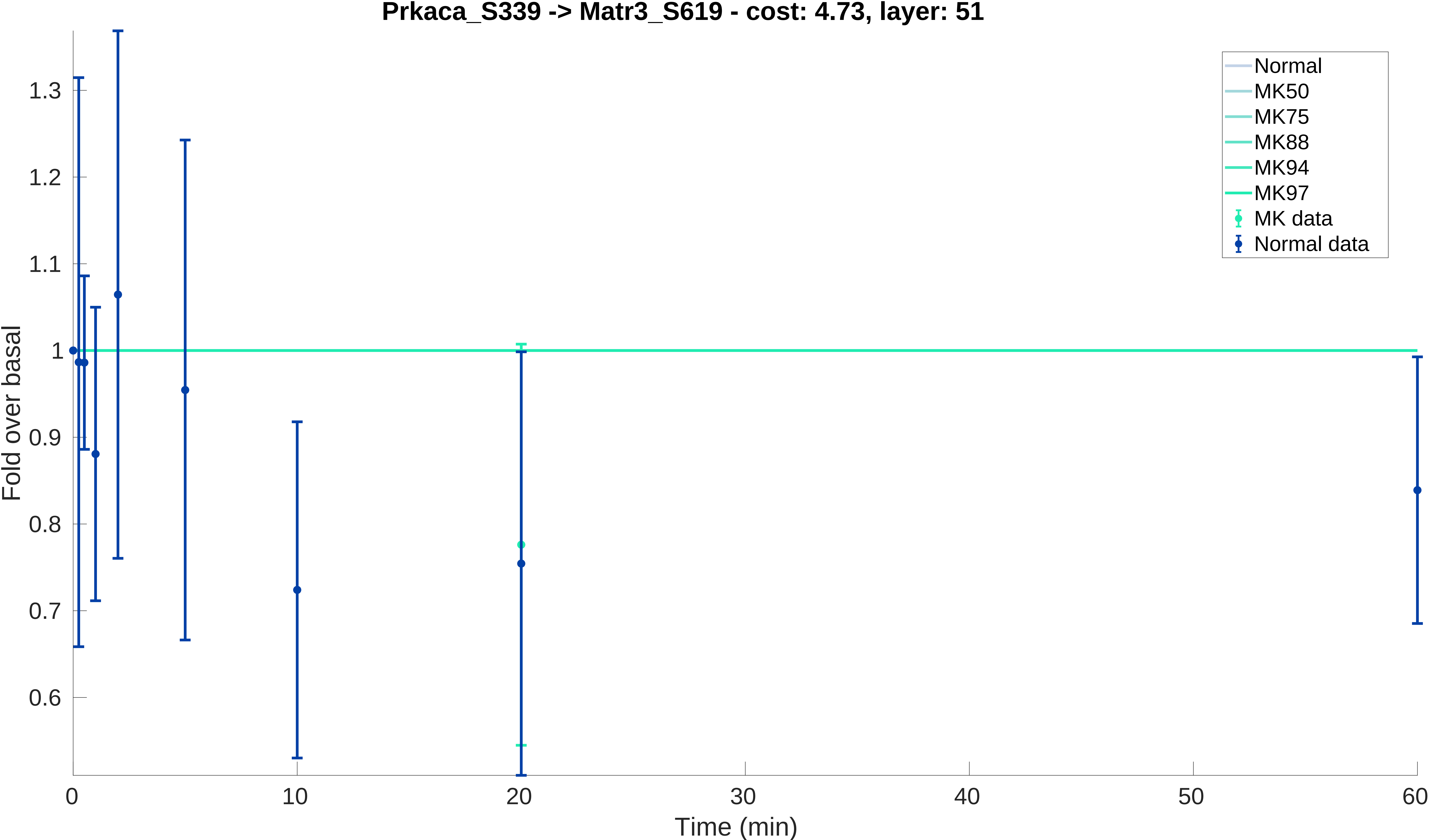

Prkaca_S339 -> Matr3_S619 - cost: 4.73, layer: 51
Fold over basal
Time (min)
1.3
1.2
1.1
1
0.9
0.8
0.7
0.6
0
10
20
30
40
50
60
Normal
MK50
MK75
MK88
MK94
MK97
MK data
Normal data

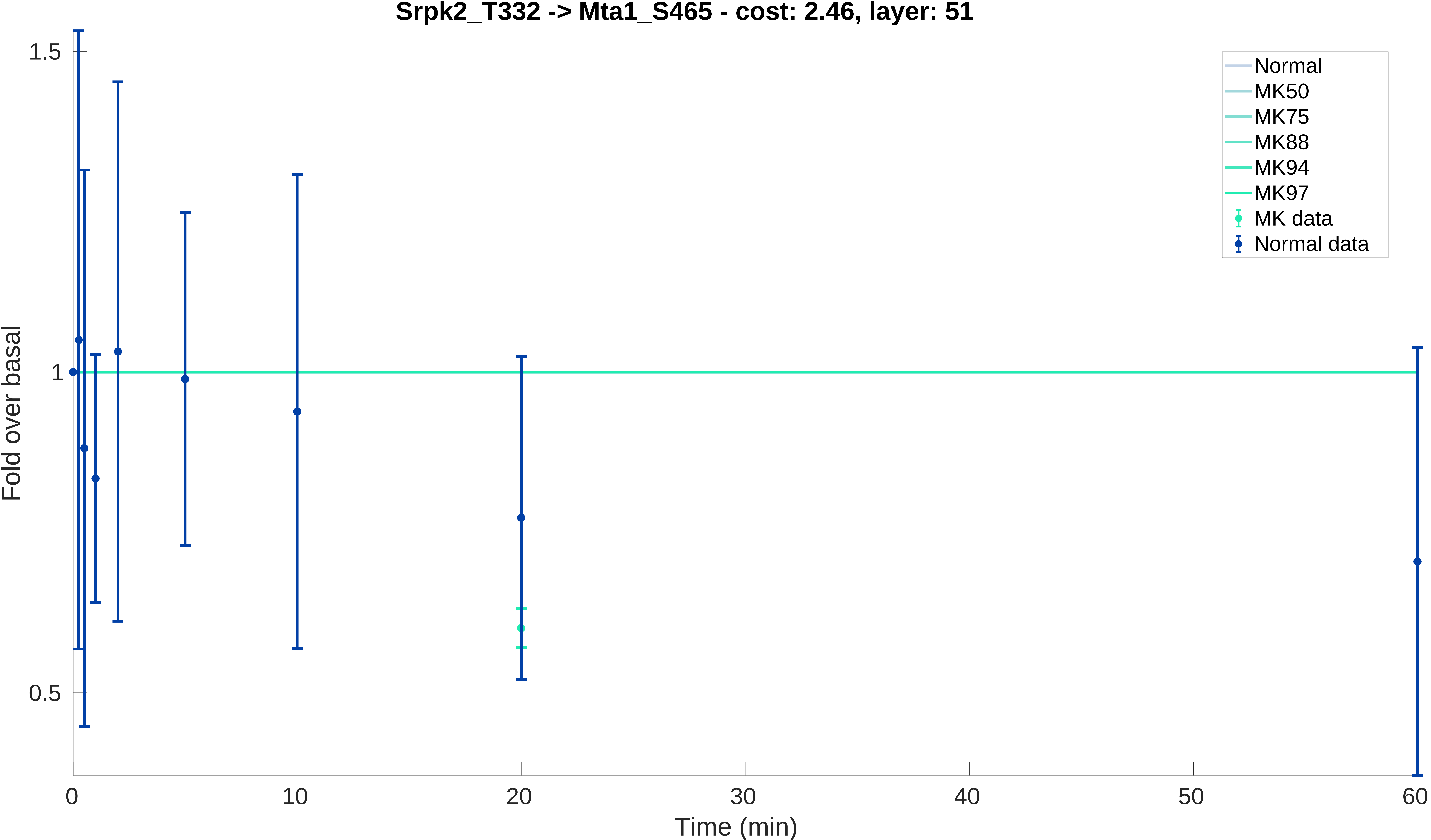
Srpk2_T332 -> Mta1_S465 - cost: 2.46, layer: 51
Fold over basal
1.5
1
0.5
0
10
20
30
40
50
60
Time (min)
Normal
MK50
MK75
MK88
MK94
MK97
MK data
Normal data

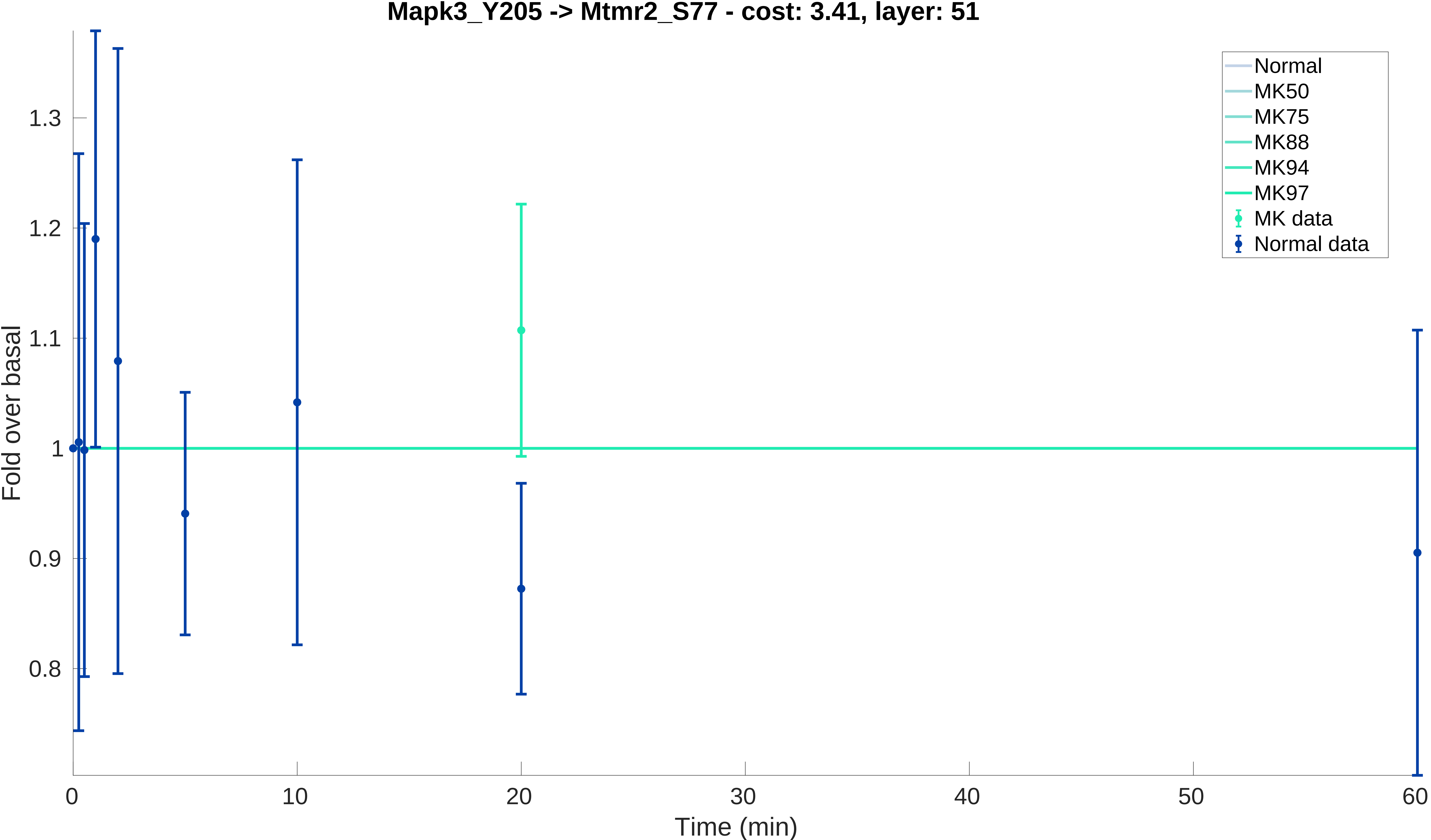

Mapk3_Y205 -> Mtmr2_S77 - cost: 3.41, layer: 51
Normal
MK50
MK75
MK88
MK94
MK97
MK data
Normal data
Fold over basal
1.3
1.2
1.1
1
0.9
0.8
0
10
20
30
40
50
60
Time (min)

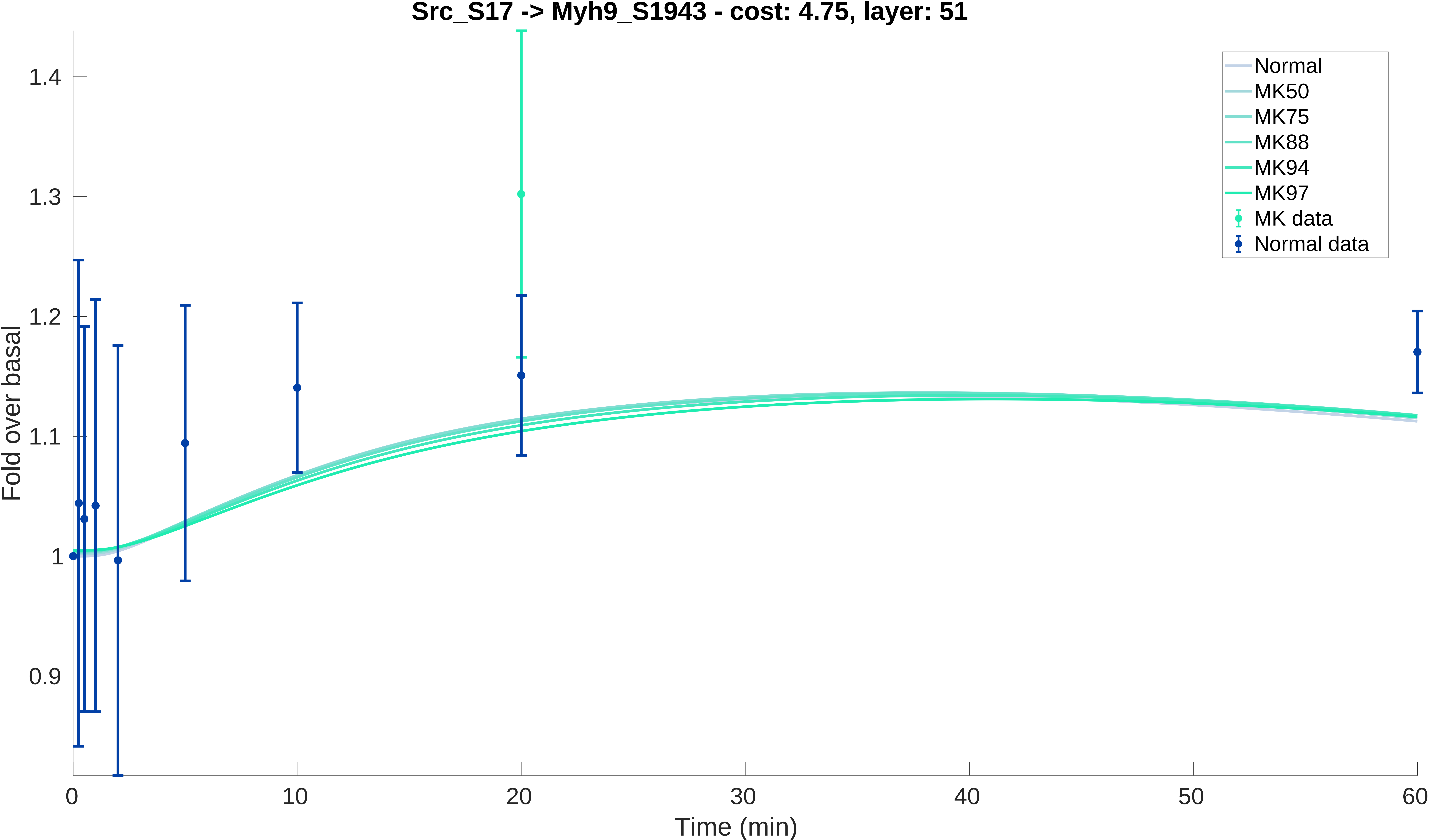

Src_S17 -> Myh9_S1943 - cost: 4.75, layer: 51
Normal
MK50
MK75
MK88
MK94
MK97
MK data
Normal data
Fold over basal
1.4
1.3
1.2
1.1
1
0.9
0
10
20
30
40
50
60
Time (min)

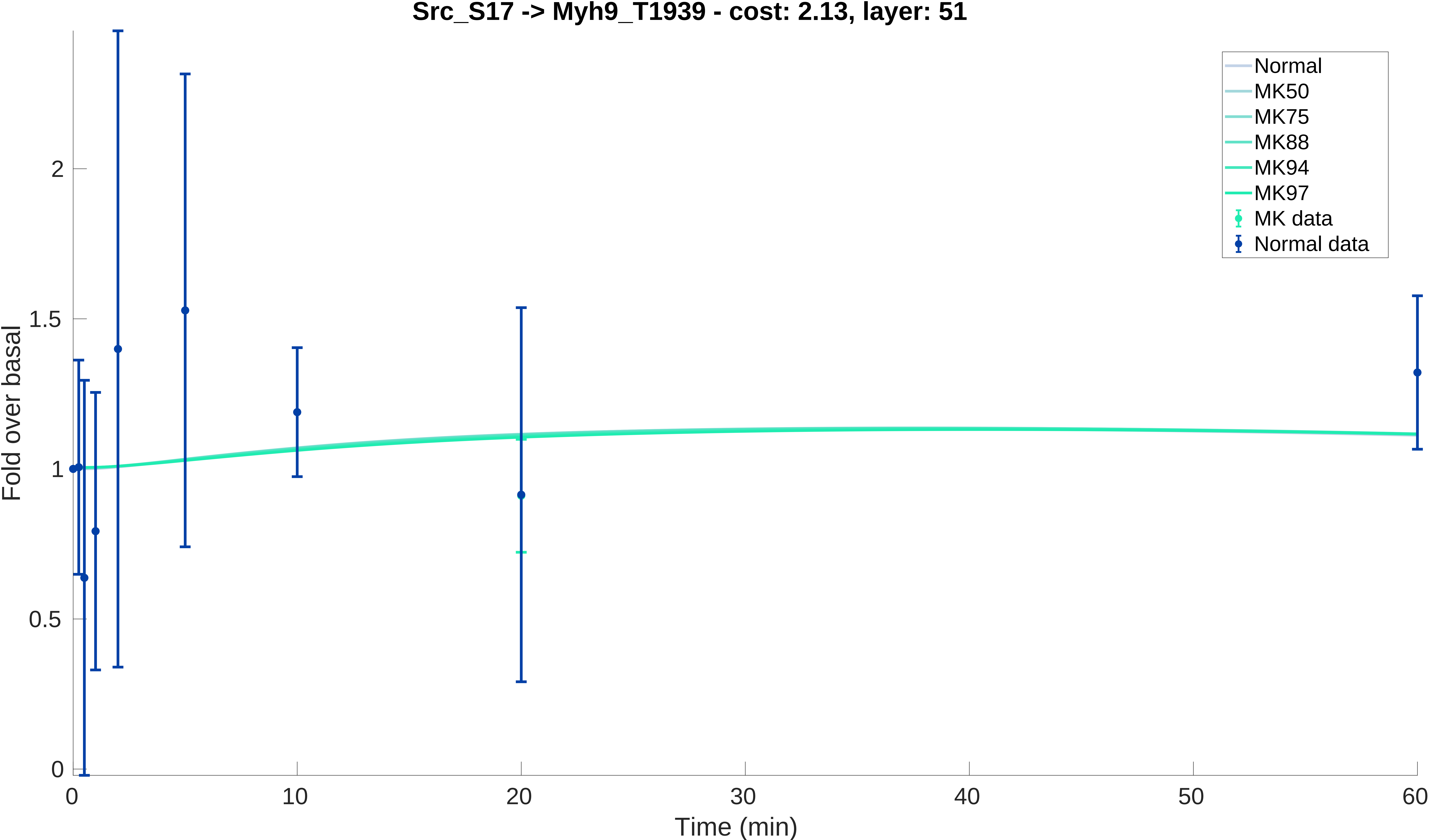

Src_S17 -> Myh9_T1939 - cost: 2.13, layer: 51
Normal
MK50
MK75
MK88
MK94
MK97
MK data
Normal data
Fold over basal
0
0.5
1
1.5
2
0
10
20
30
40
50
60
Time (min)

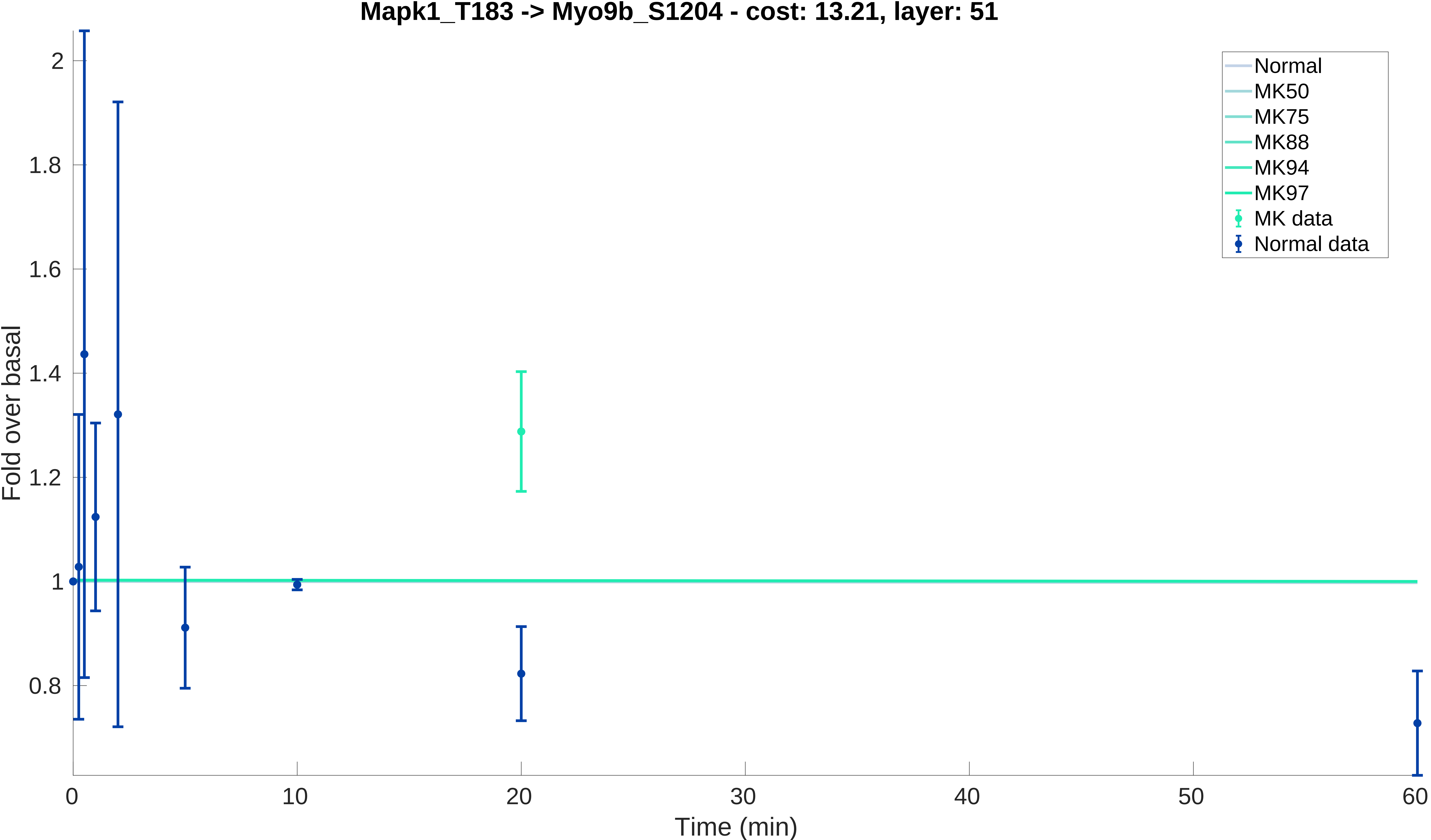

Mapk1_T183 -> Myo9b_S1204 - cost: 13.21, layer: 51
Fold over basal
Time (min)
2
1.8
1.6
1.4
1.2
1
0.8
0
10
20
30
40
50
60
Normal
MK50
MK75
MK88
MK94
MK97
MK data
Normal data

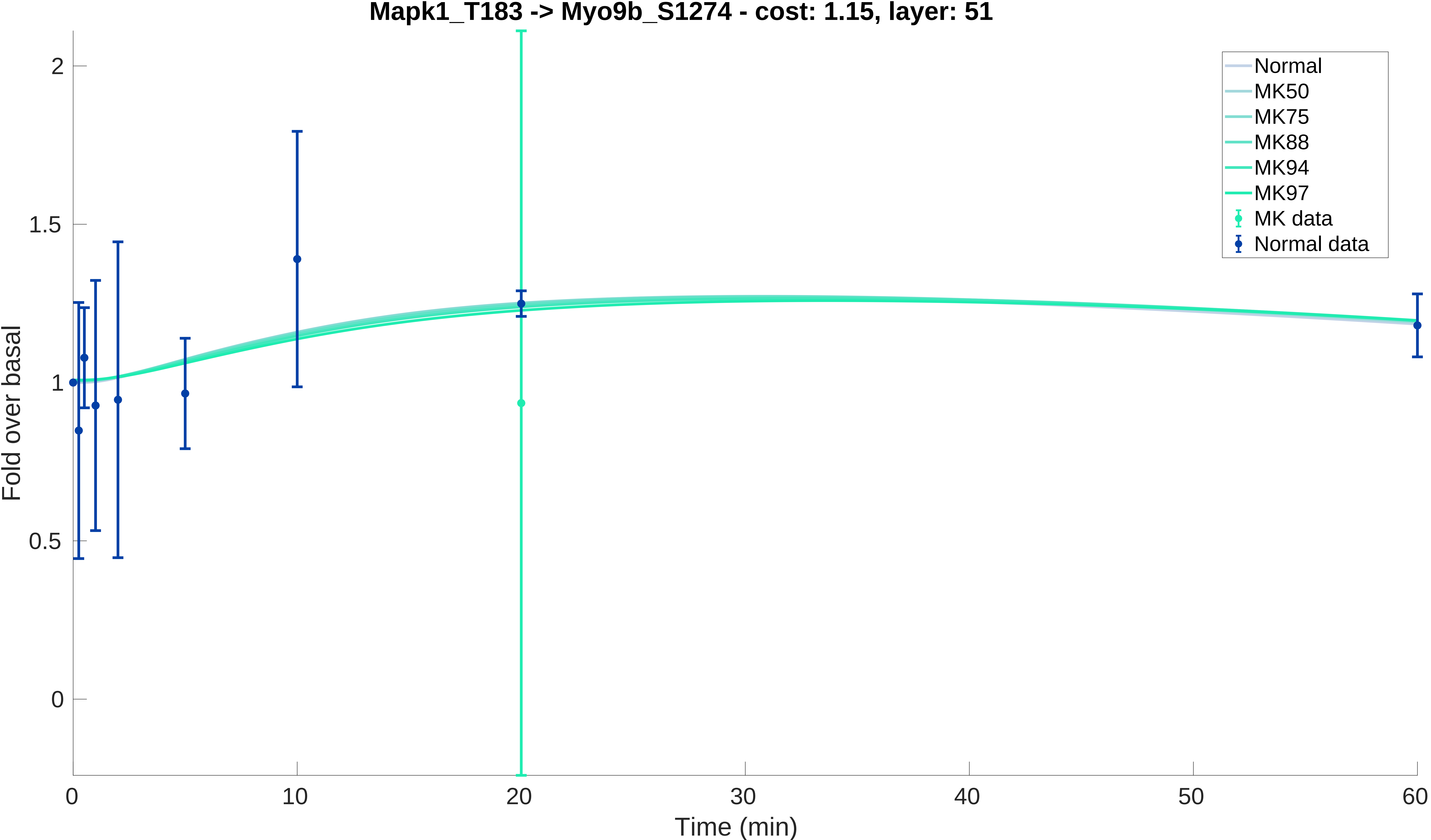

Mapk1_T183 -> Myo9b_S1274 - cost: 1.15, layer: 51
Fold over basal
Time (min)
0
0.5
1
1.5
2
0
10
20
30
40
50
60
Normal
MK50
MK75
MK88
MK94
MK97
MK data
Normal data

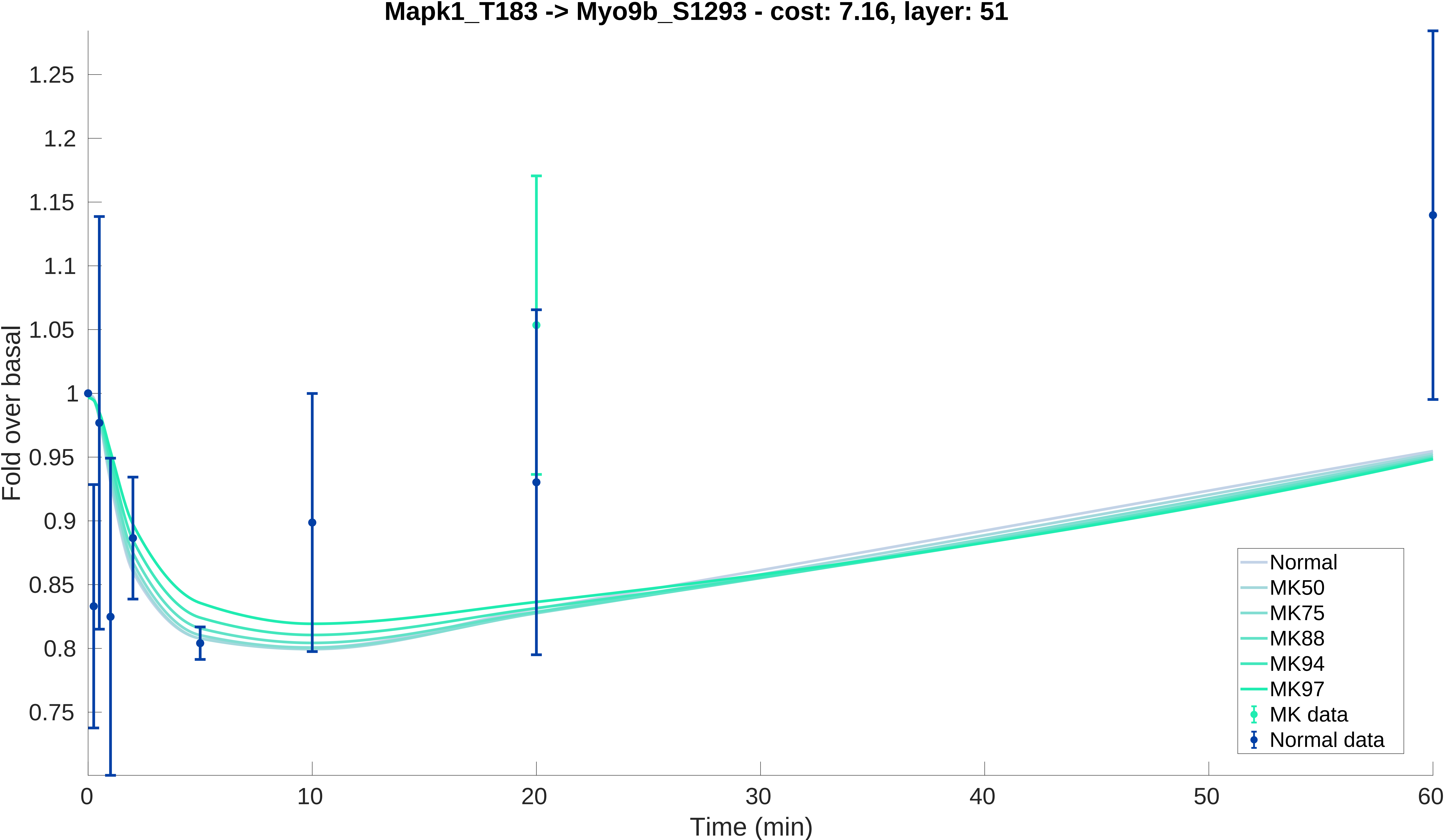
Mapk1_T183 -> Myo9b_S1293 - cost: 7.16, layer: 51
Fold over basal
1.25
1.2
1.15
1.1
1.05
1
0.95
0.9
0.85
0.8
0.75
0
10
20
30
40
50
60
Time (min)
Normal
MK50
MK75
MK88
MK94
MK97
MK data
Normal data

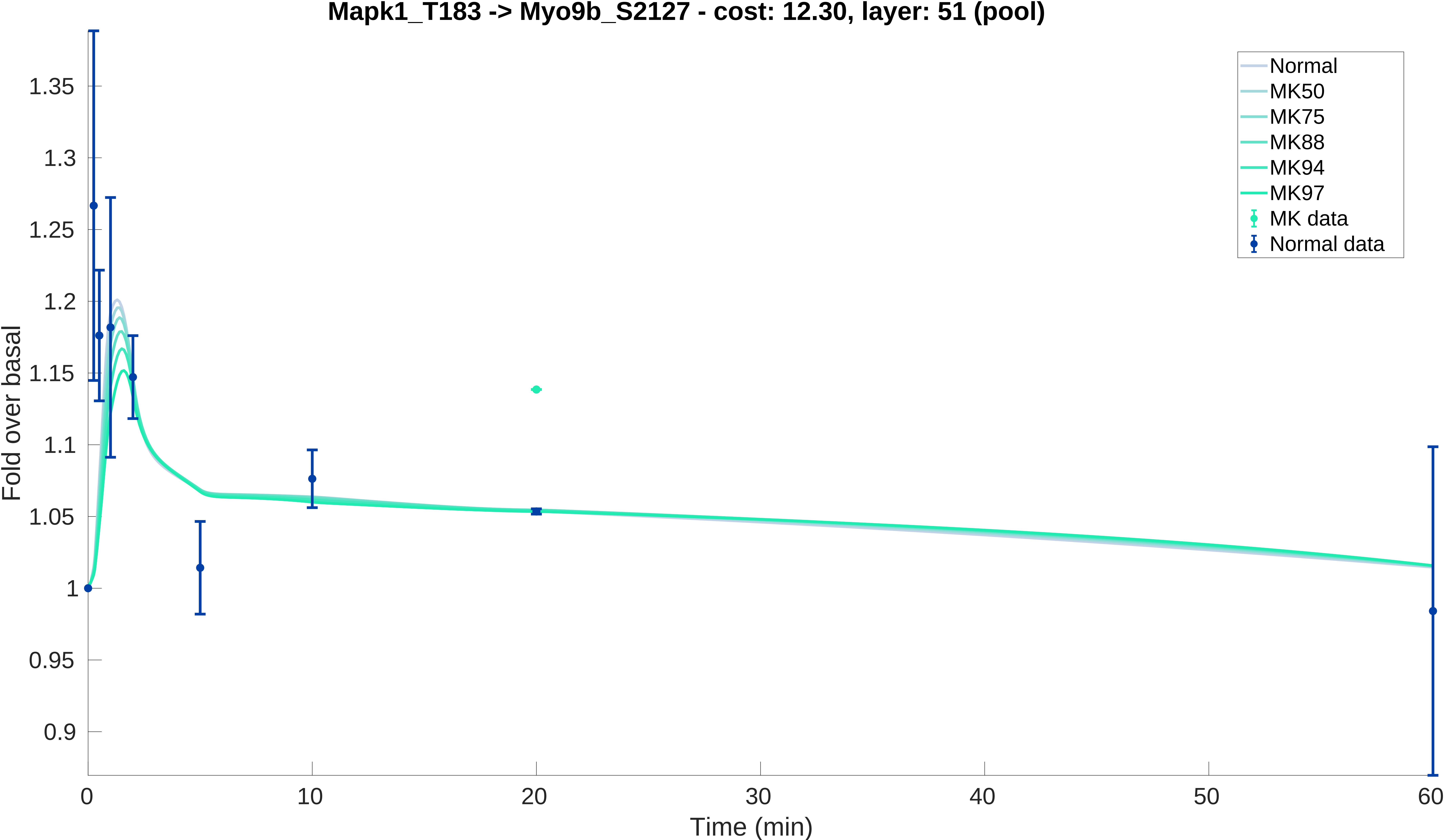
Mapk1_T183 -> Myo9b_S2127 - cost: 12.30, layer: 51 (pool)
Fold over basal
1.35
1.3
1.25
1.2
1.15
1.1
1.05
1
0.95
0.9
0
10
20
30
40
50
60
Time (min)
Normal
MK50
MK75
MK88
MK94
MK97
MK data
Normal data

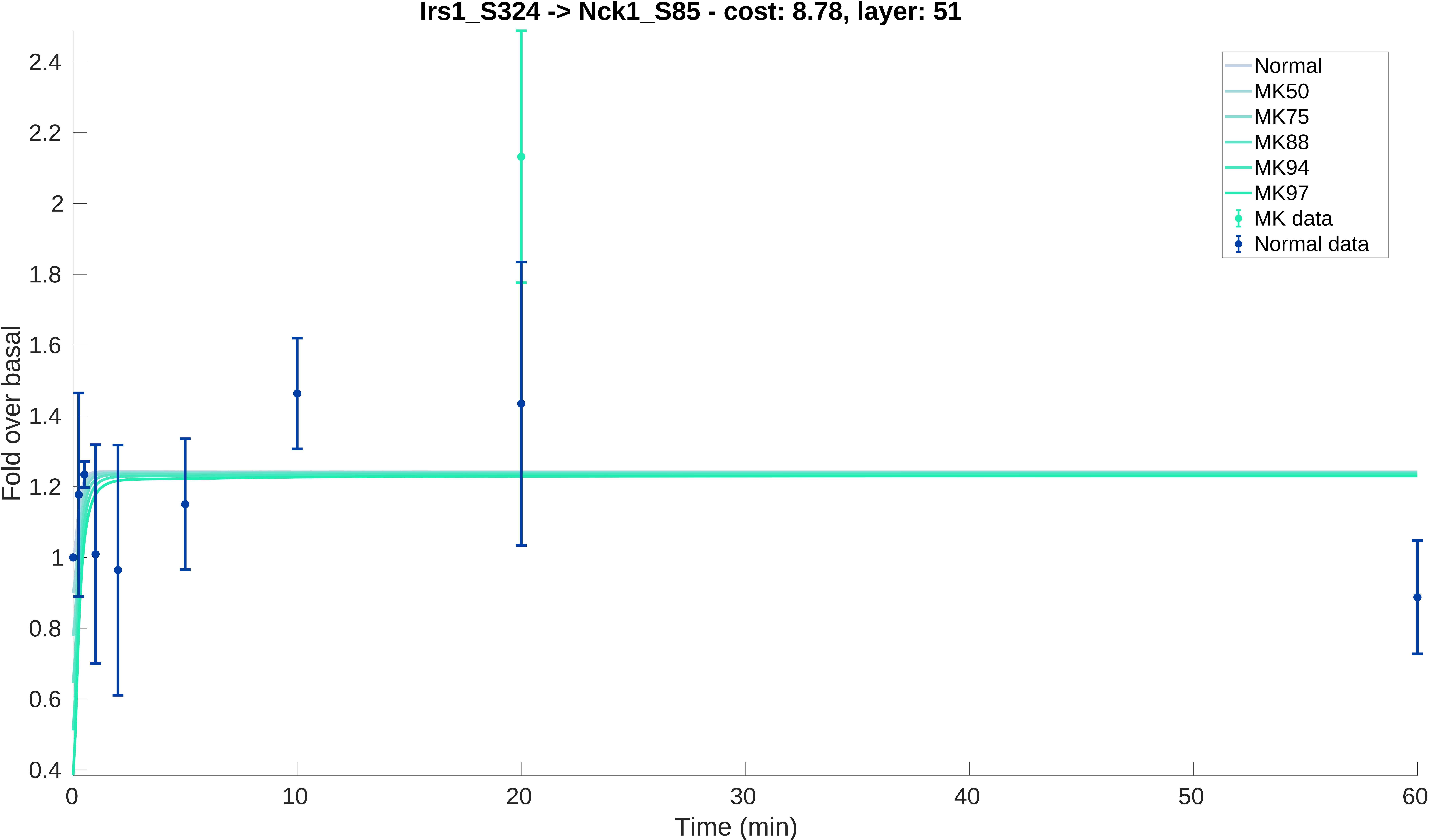
Irs1_S324 -> Nck1_S85 - cost: 8.78, layer: 51
Fold over basal
Time (min)
2.4
2.2
2
1.8
1.6
1.4
1.2
1
0.8
0.6
0.4
0
10
20
30
40
50
60
Normal
MK50
MK75
MK88
MK94
MK97
MK data
Normal data

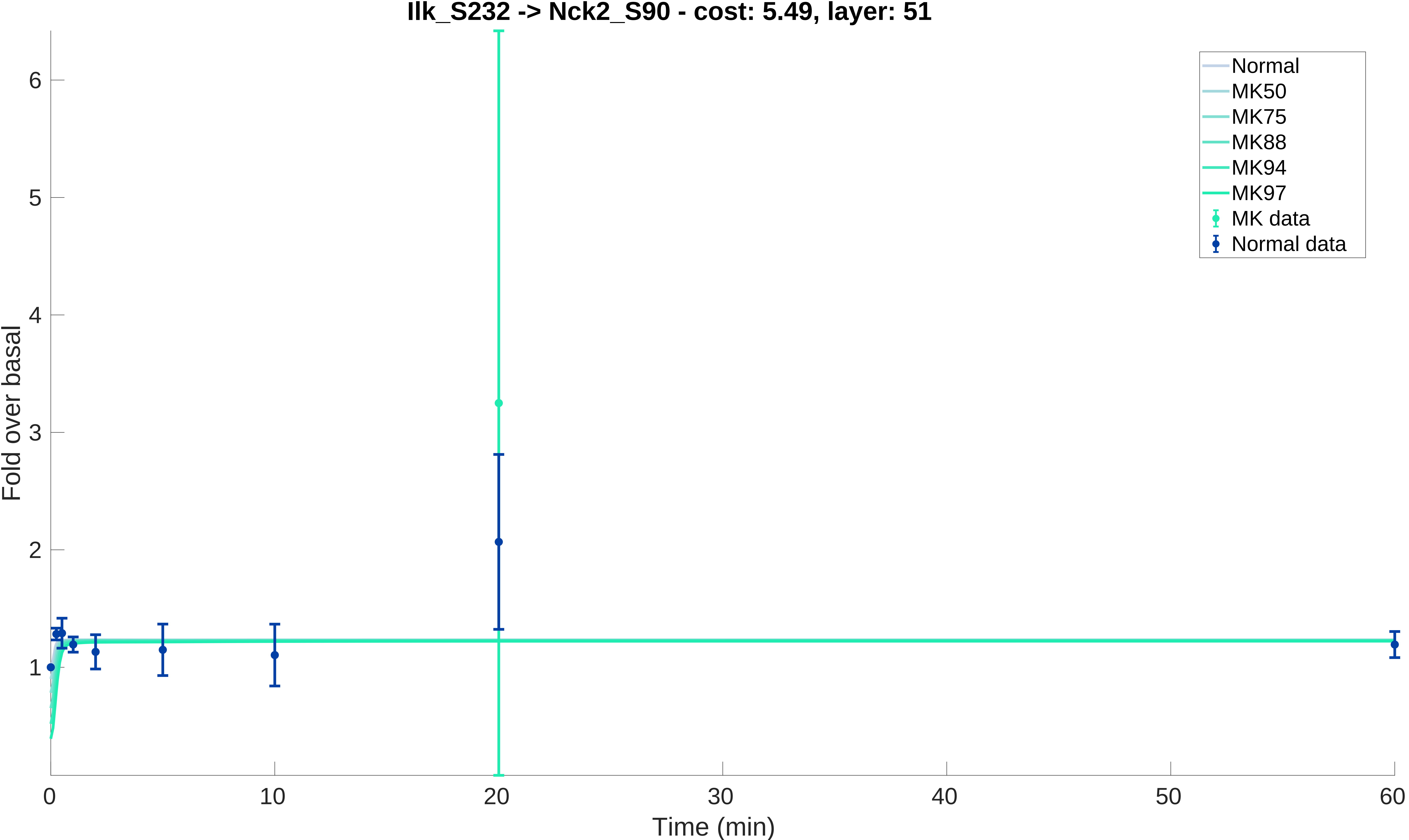
Ilk_S232 -> Nck2_S90 - cost: 5.49, layer: 51
Fold over basal
Time (min)
Normal
MK50
MK75
MK88
MK94
MK97
MK data
Normal data
0
10
20
30
40
50
60
1
2
3
4
5
6

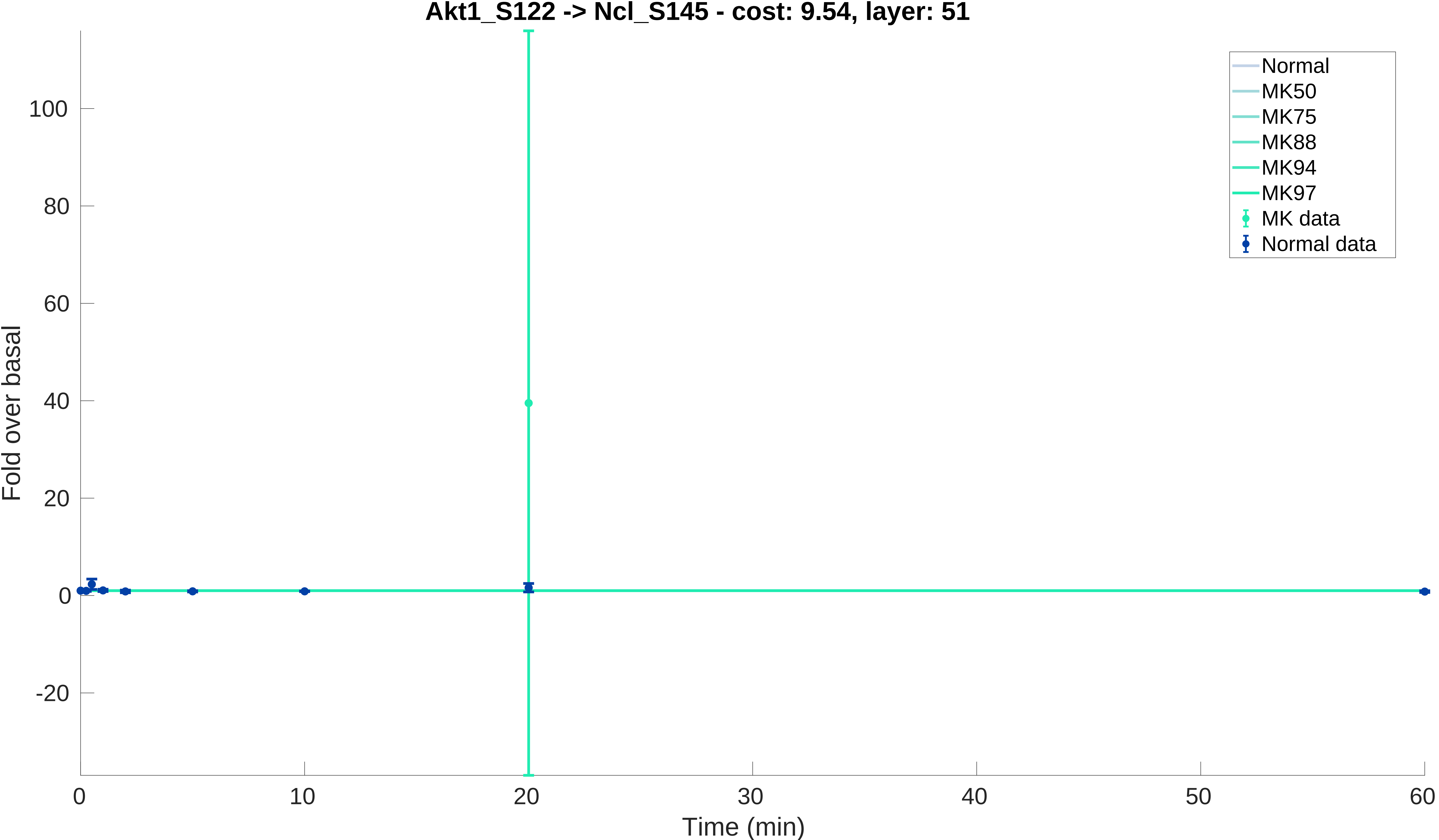

Akt1_S122 -> Ncl_S145 - cost: 9.54, layer: 51
Fold over basal
Time (min)
100
80
60
40
20
0
-20
0
10
20
30
40
50
60
Normal
MK50
MK75
MK88
MK94
MK97
MK data
Normal data

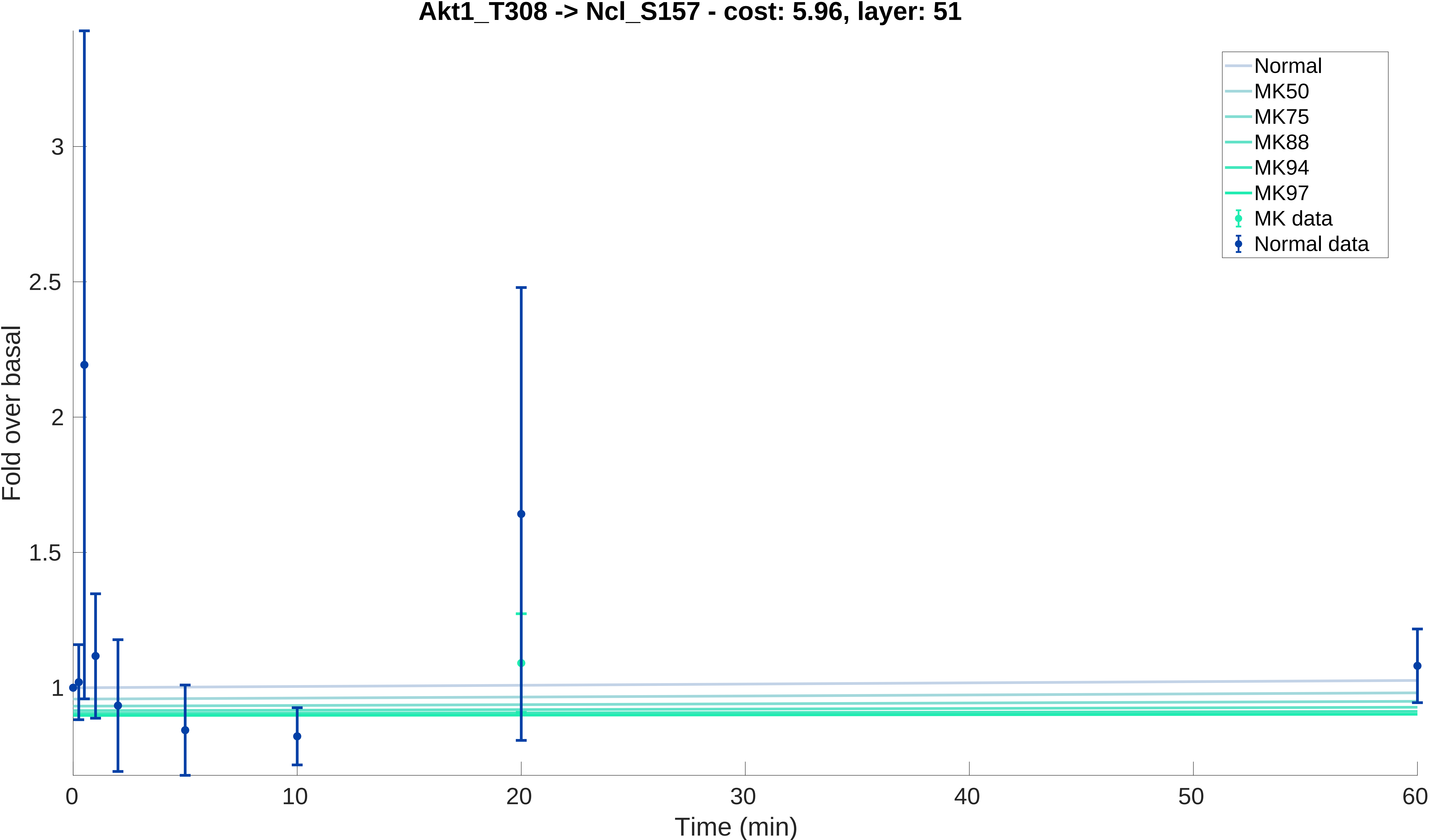
Akt1_T308 -> Ncl_S157 - cost: 5.96, layer: 51
Fold over basal
Time (min)
3
2.5
2
1.5
1
0
10
20
30
40
50
60
Normal
MK50
MK75
MK88
MK94
MK97
MK data
Normal data

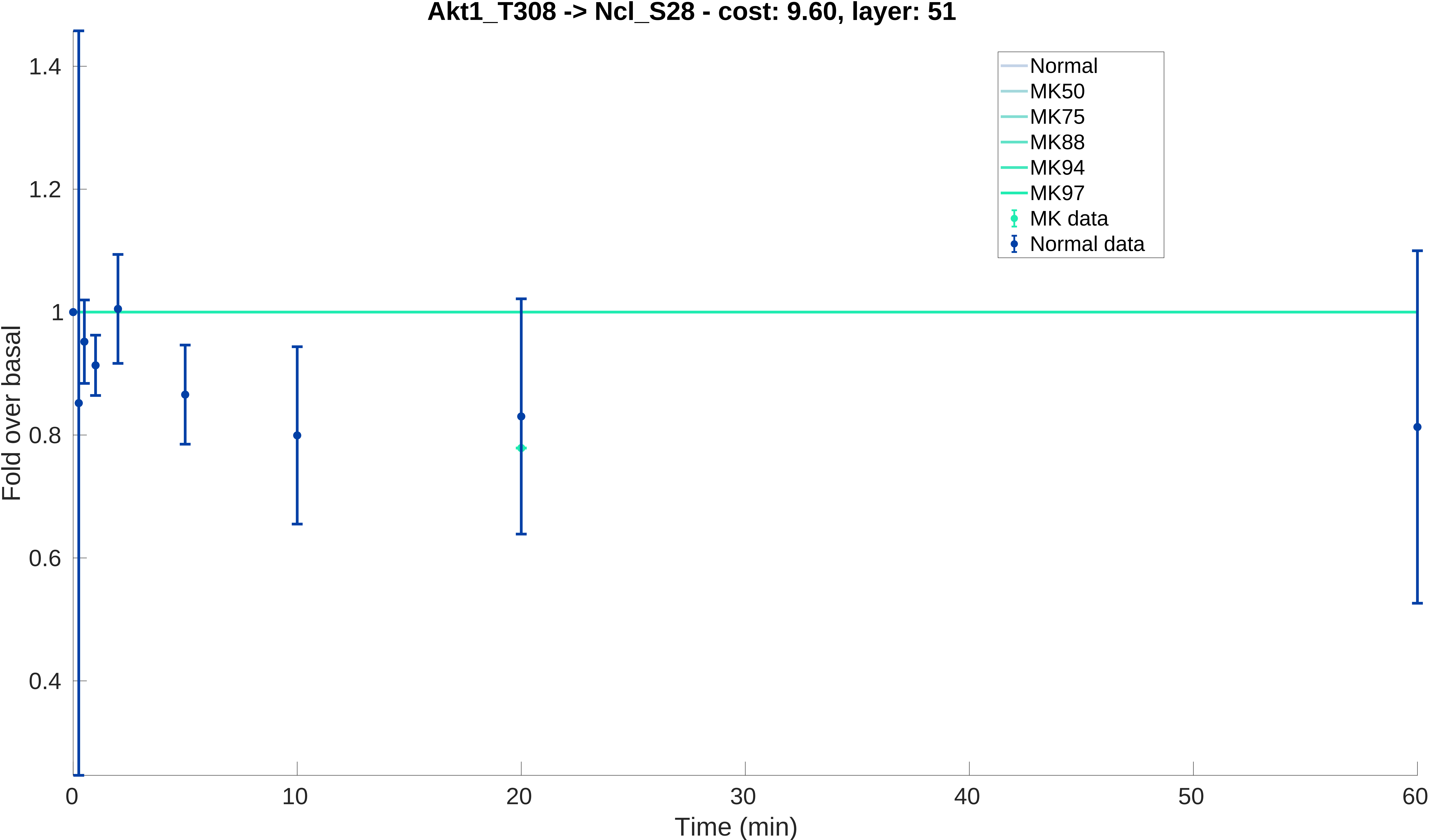

Akt1_T308 -> Ncl_S28 - cost: 9.60, layer: 51
Normal
MK50
MK75
MK88
MK94
MK97
MK data
Normal data
Fold over basal
Time (min)
1.4
1.2
1
0.8
0.6
0.4
0
10
20
30
40
50
60

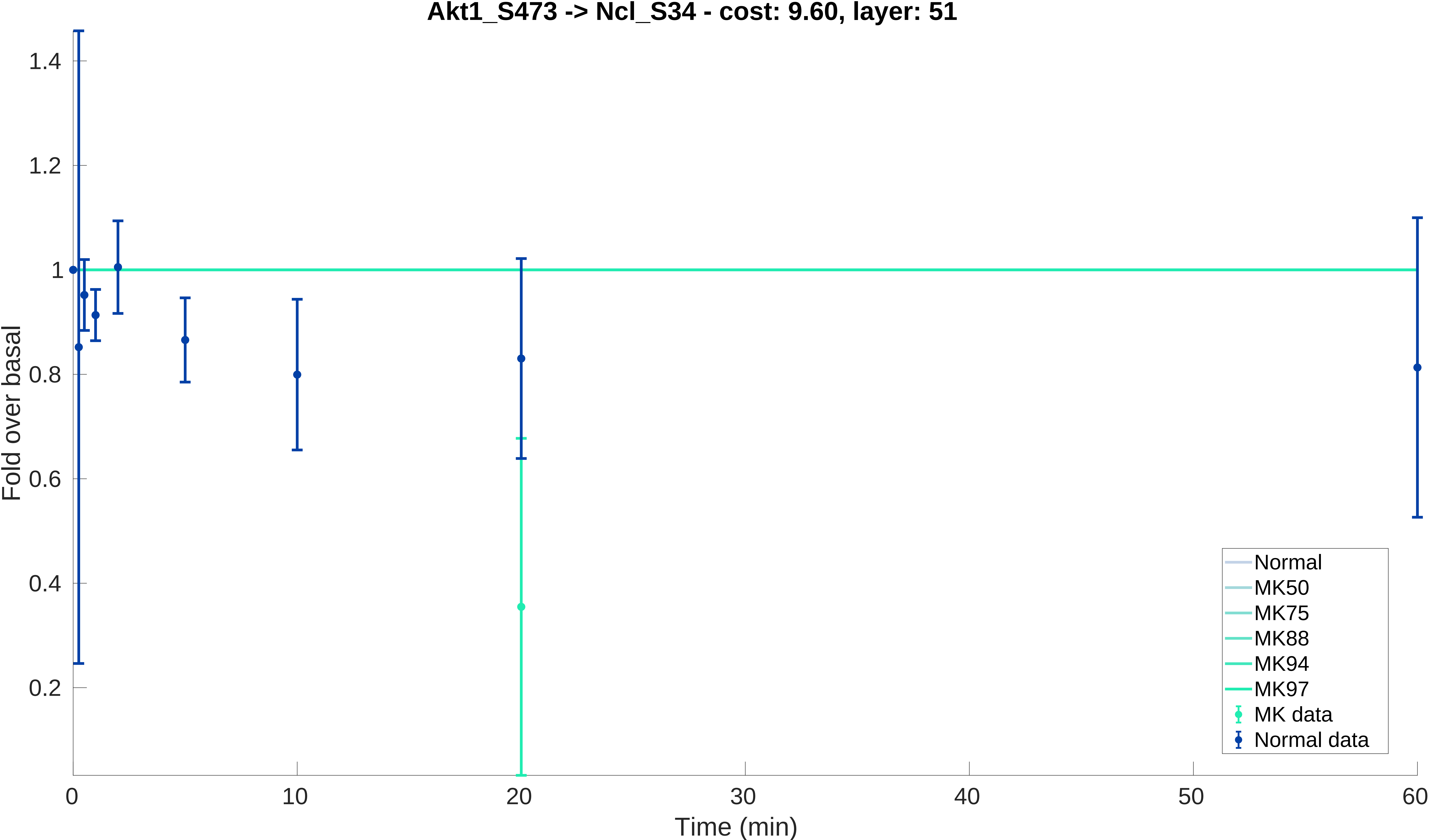

Akt1_S473 -> Ncl_S34 - cost: 9.60, layer: 51
Fold over basal
1.4
1.2
1
0.8
0.6
0.4
0.2
0
10
20
30
40
50
60
Time (min)
Normal
MK50
MK75
MK88
MK94
MK97
MK data
Normal data

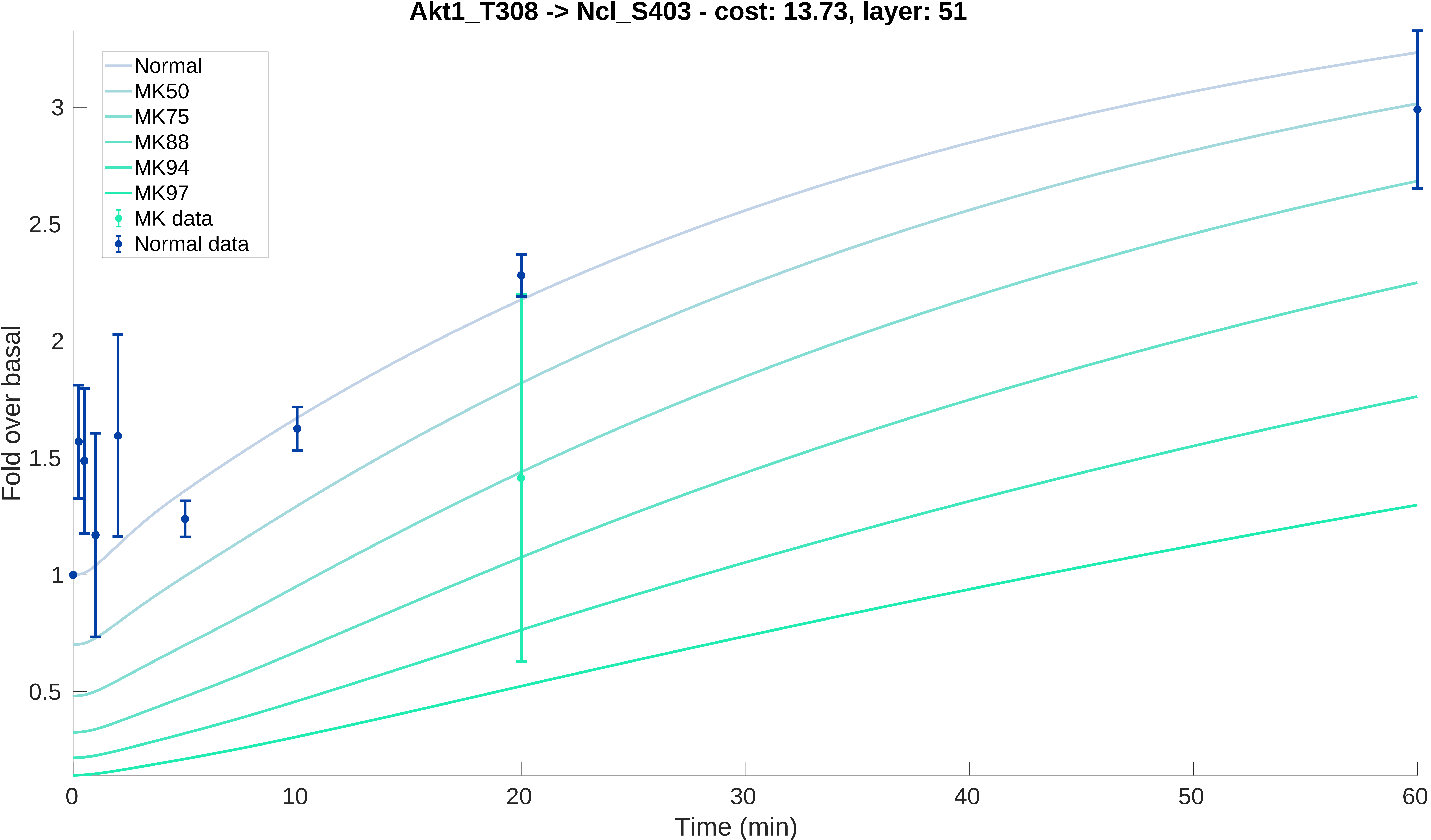

Akt1_T308 -> Ncl_S403 - cost: 13.73, layer: 51
Normal
MK50
MK75
MK88
MK94
MK97
MK data
Normal data
Fold over basal
Time (min)
0.5
1
1.5
2
2.5
3
0
10
20
30
40
50
60

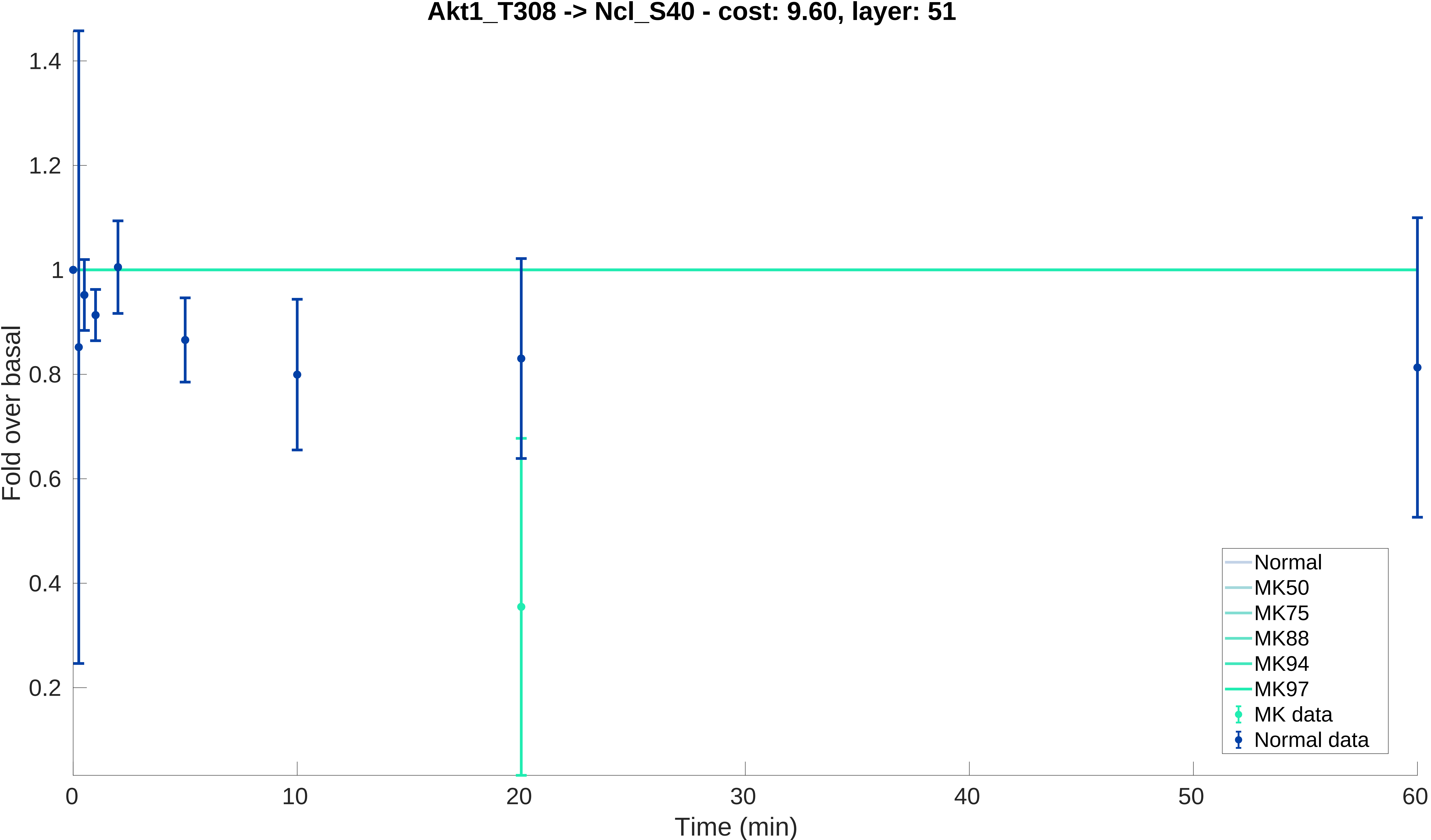

Akt1_T308 -> Ncl_S40 - cost: 9.60, layer: 51
1.4
1.2
1
0.8
0.6
0.4
0.2
Fold over basal
0
10
20
30
40
50
60
Time (min)
Normal
MK50
MK75
MK88
MK94
MK97
MK data
Normal data

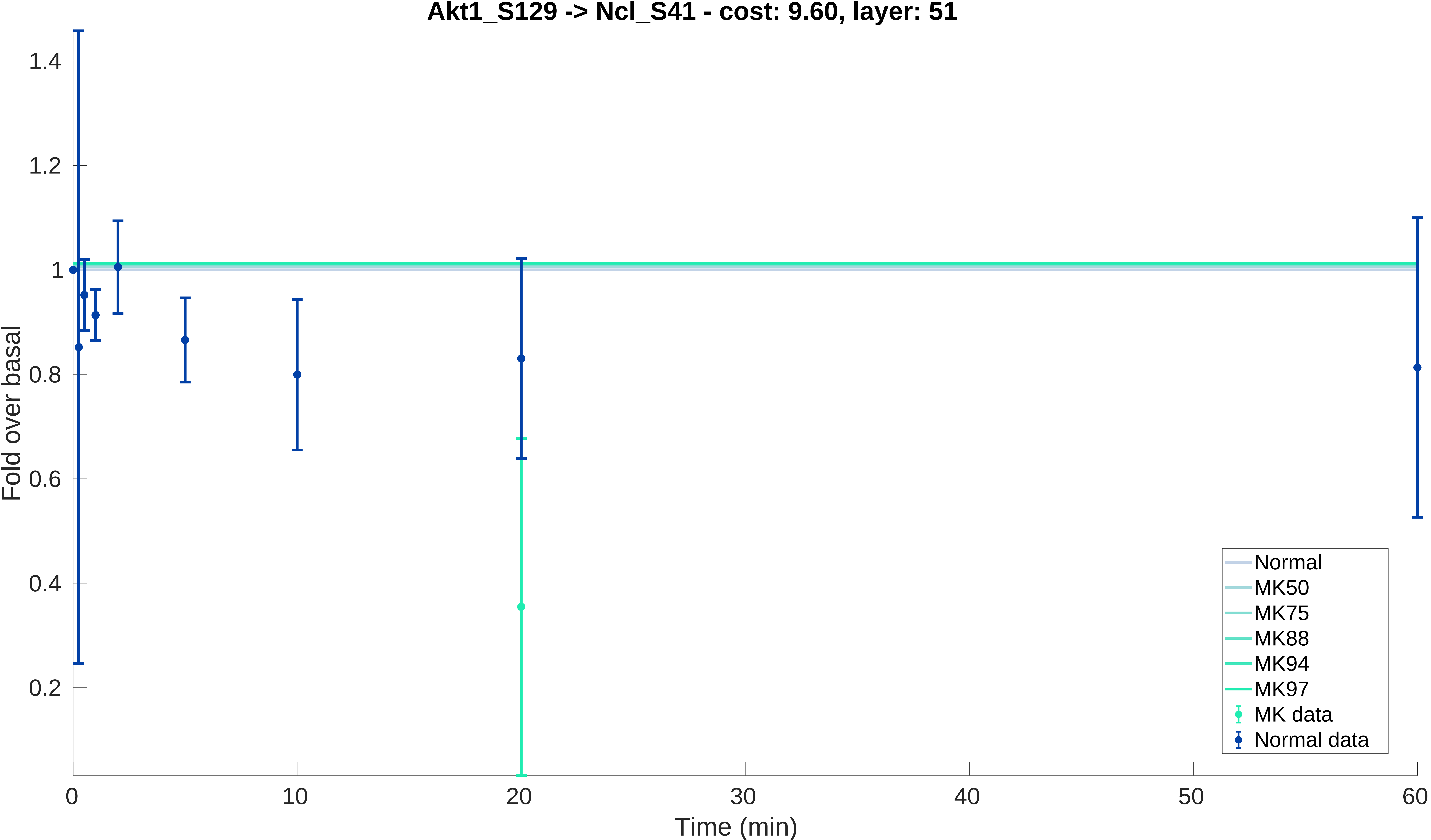

Akt1_S129 -> Ncl_S41 - cost: 9.60, layer: 51
Fold over basal
Time (min)
1.4
1.2
1
0.8
0.6
0.4
0.2
0
10
20
30
40
50
60
Normal
MK50
MK75
MK88
MK94
MK97
MK data
Normal data

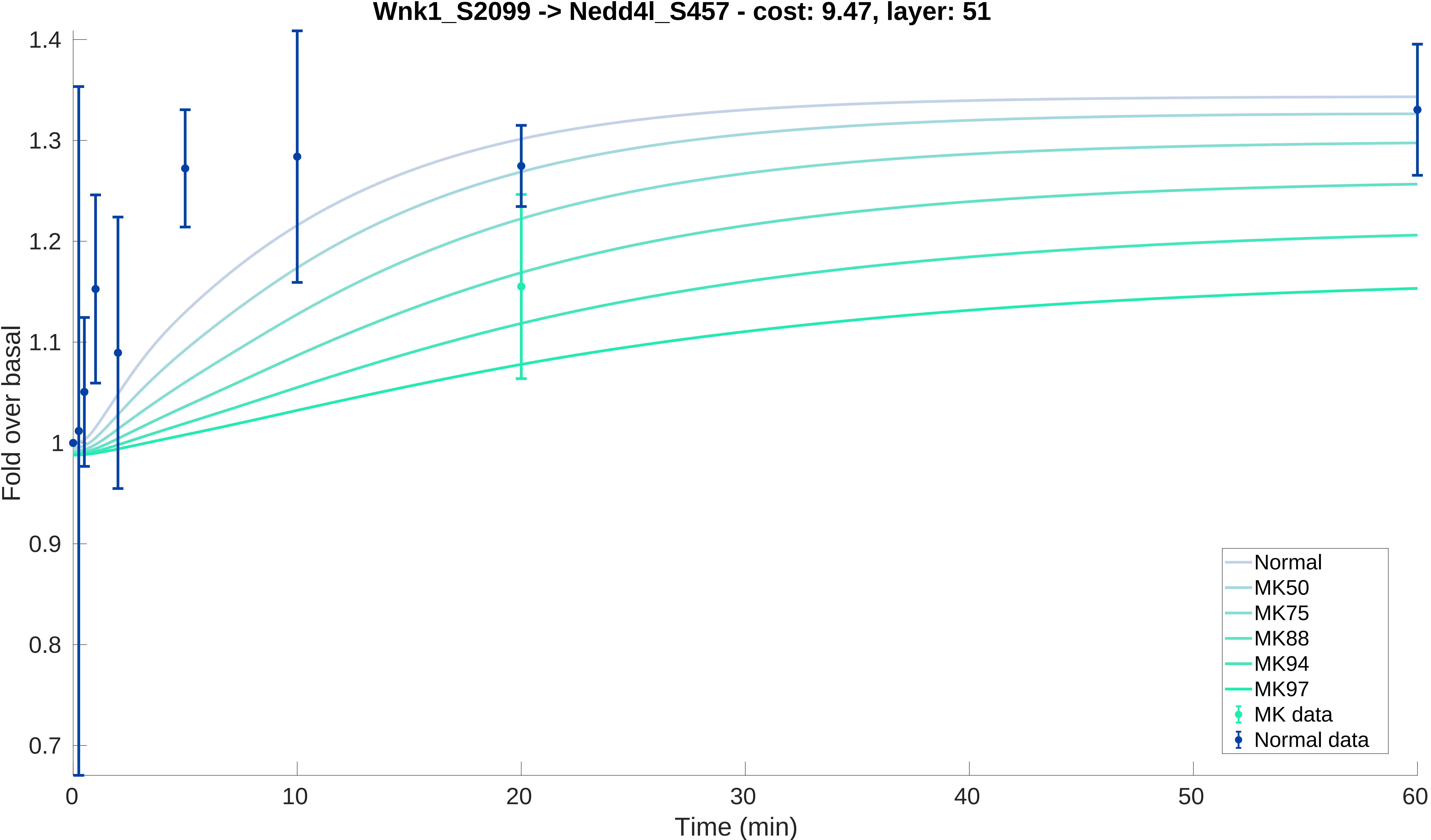

Wnk1_S2099 -> Nedd4l_S457 - cost: 9.47, layer: 51
Fold over basal
1.4
1.3
1.2
1.1
1
0.9
0.8
0.7
0
10
20
30
40
50
60
Time (min)
Normal
MK50
MK75
MK88
MK94
MK97
MK data
Normal data

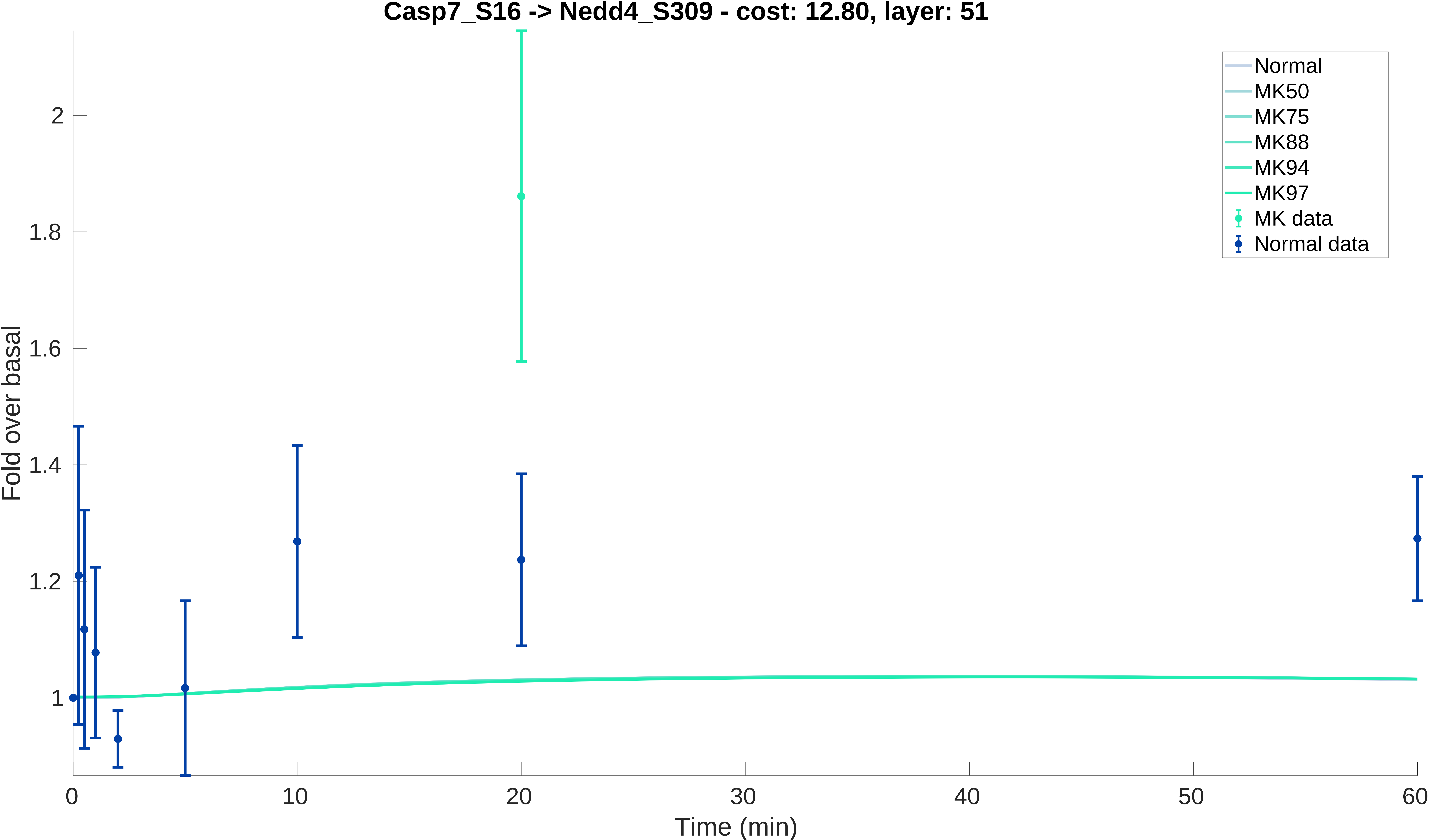

Casp7_S16 -> Nedd4_S309 - cost: 12.80, layer: 51
Normal
MK50
MK75
MK88
MK94
MK97
MK data
Normal data
Fold over basal
1
1.2
1.4
1.6
1.8
2
0
10
20
30
40
50
60
Time (min)

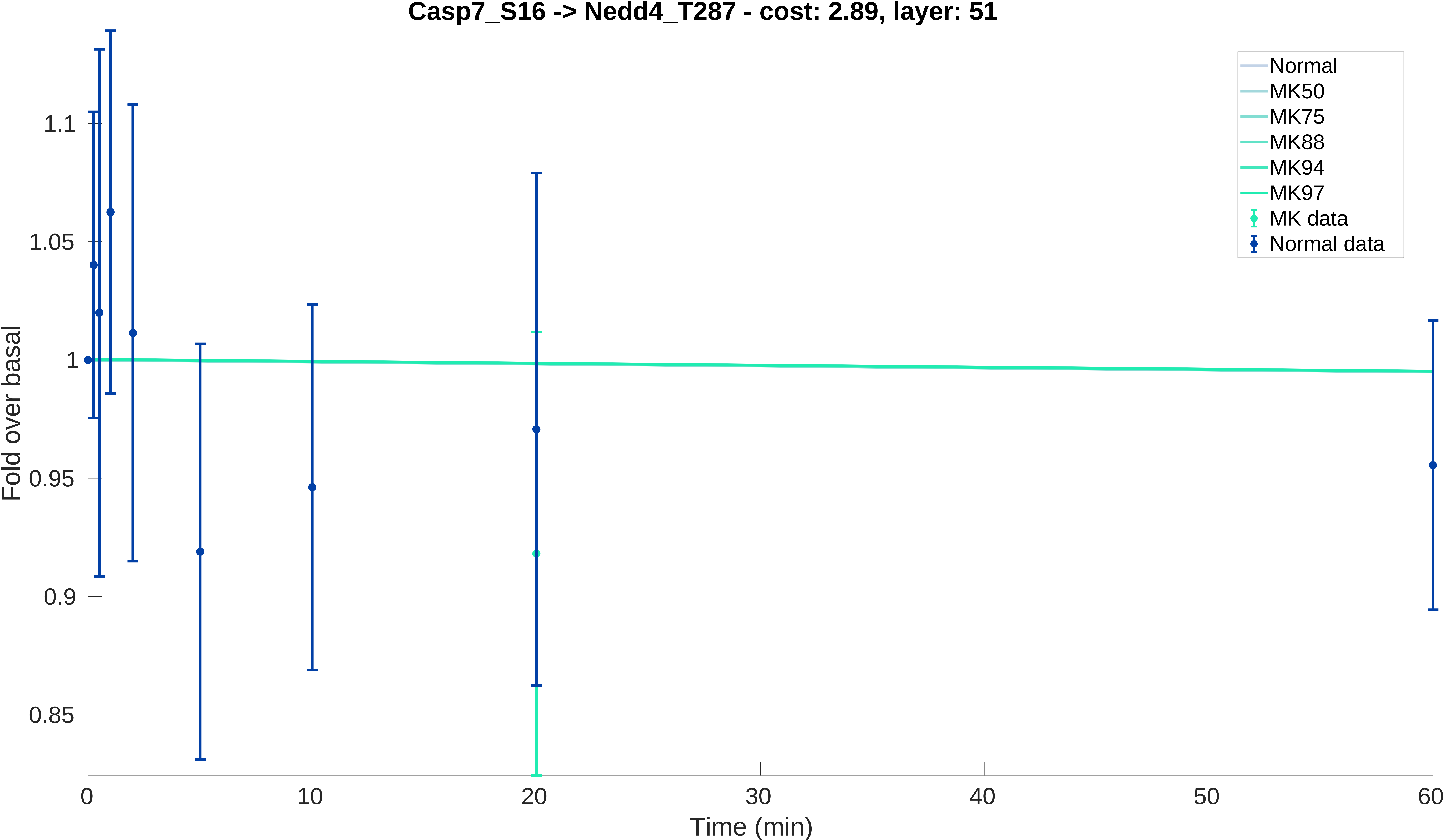
Casp7_S16 -> Nedd4_T287 - cost: 2.89, layer: 51
Normal
MK50
MK75
MK88
MK94
MK97
MK data
Normal data
Fold over basal
1.1
1.05
1
0.95
0.9
0.85
0
10
20
30
40
50
60
Time (min)

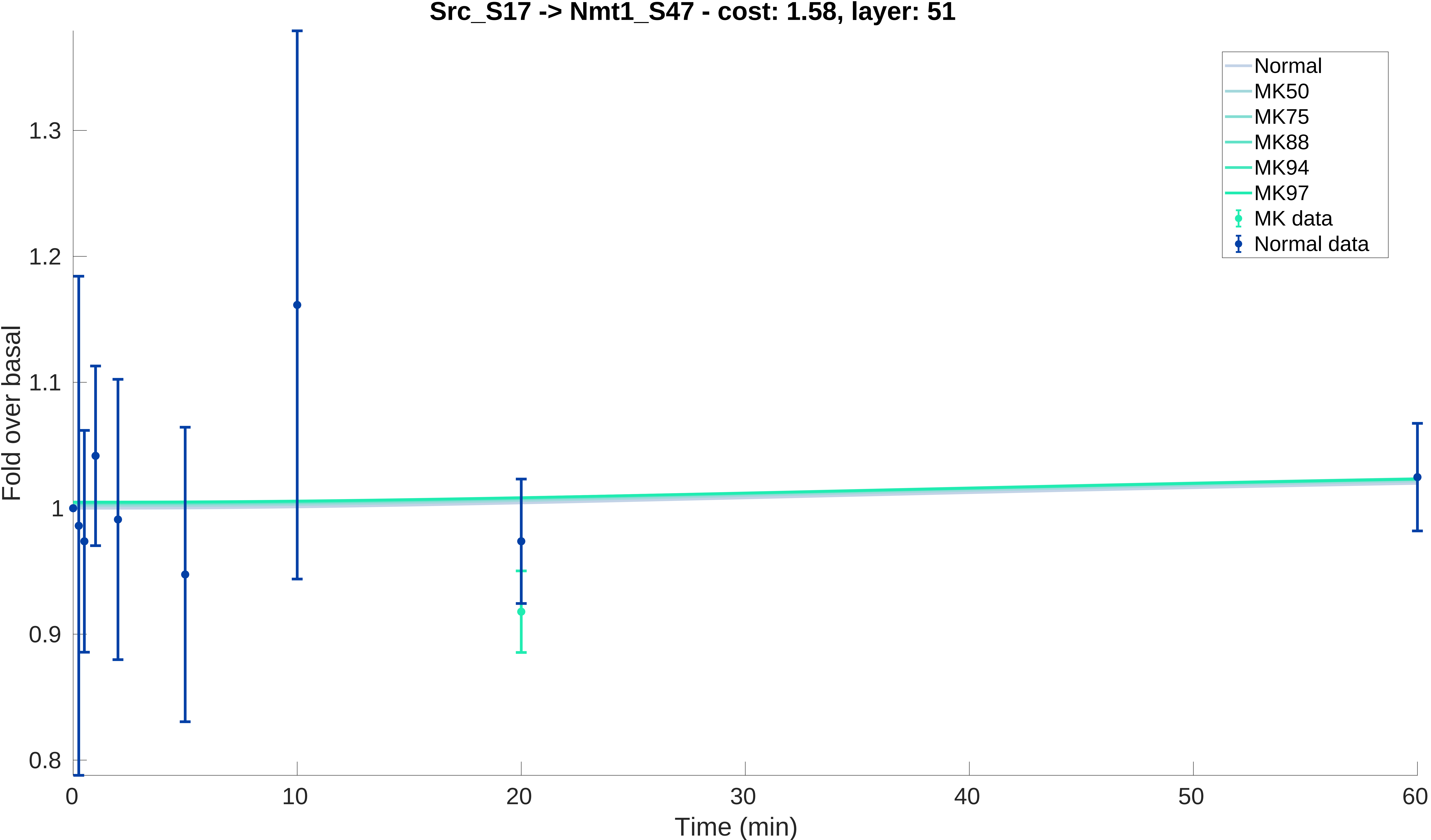

Src_S17 -> Nmt1_S47 - cost: 1.58, layer: 51
Normal
MK50
MK75
MK88
MK94
MK97
MK data
Normal data
Fold over basal
Time (min)
0.8
0.9
1
1.1
1.2
1.3
0
10
20
30
40
50
60

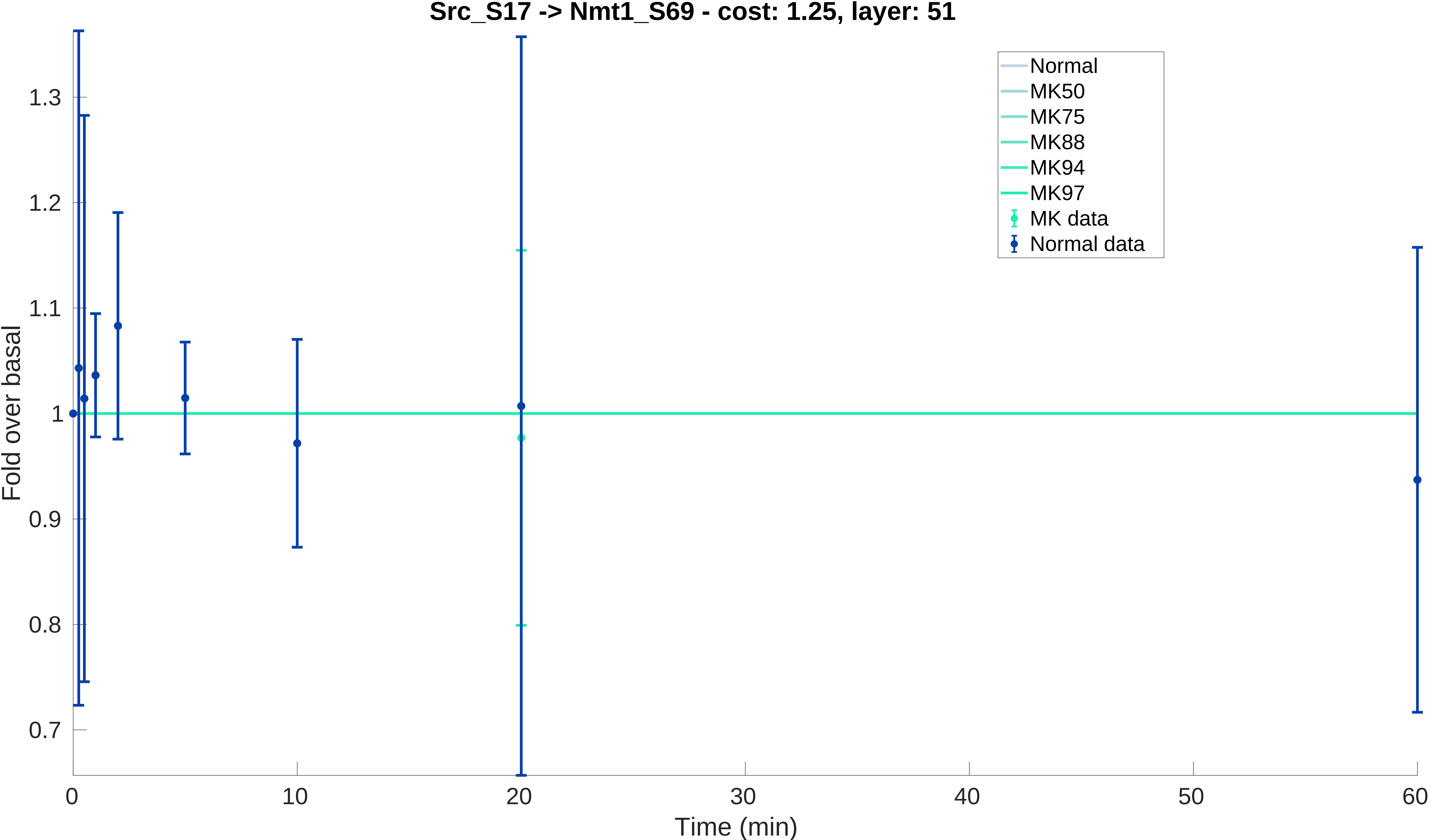
Src_S17 -> Nmt1_S69 - cost: 1.25, layer: 51
Normal
MK50
MK75
MK88
MK94
MK97
MK data
Normal data
Fold over basal
1.3
1.2
1.1
1
0.9
0.8
0.7
0
10
20
30
40
50
60
Time (min)

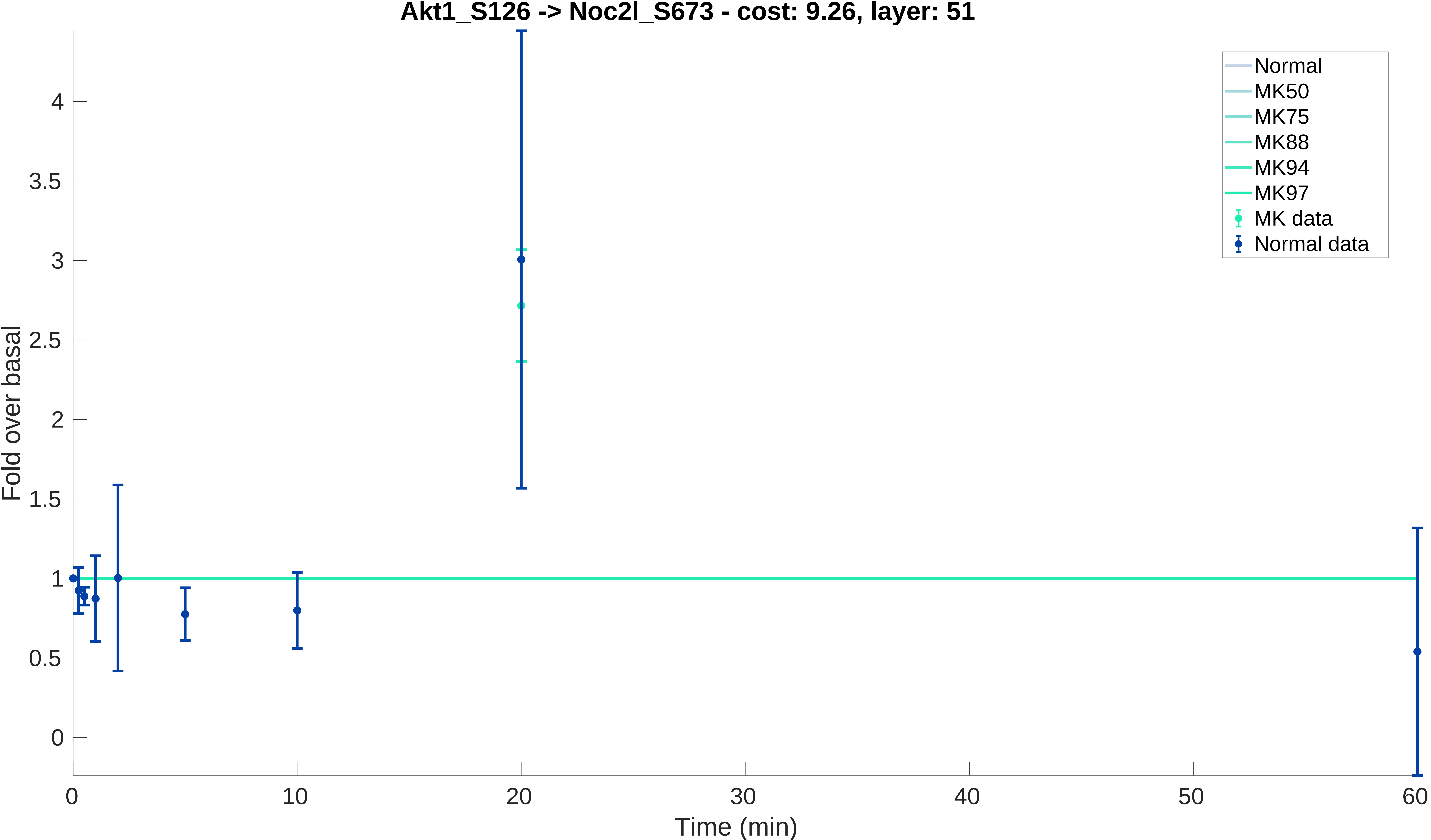

Akt1_S126 -> Noc2l_S673 - cost: 9.26, layer: 51
Normal
MK50
MK75
MK88
MK94
MK97
MK data
Normal data
Fold over basal
0
0.5
1
1.5
2
2.5
3
3.5
4
0
10
20
30
40
50
60
Time (min)

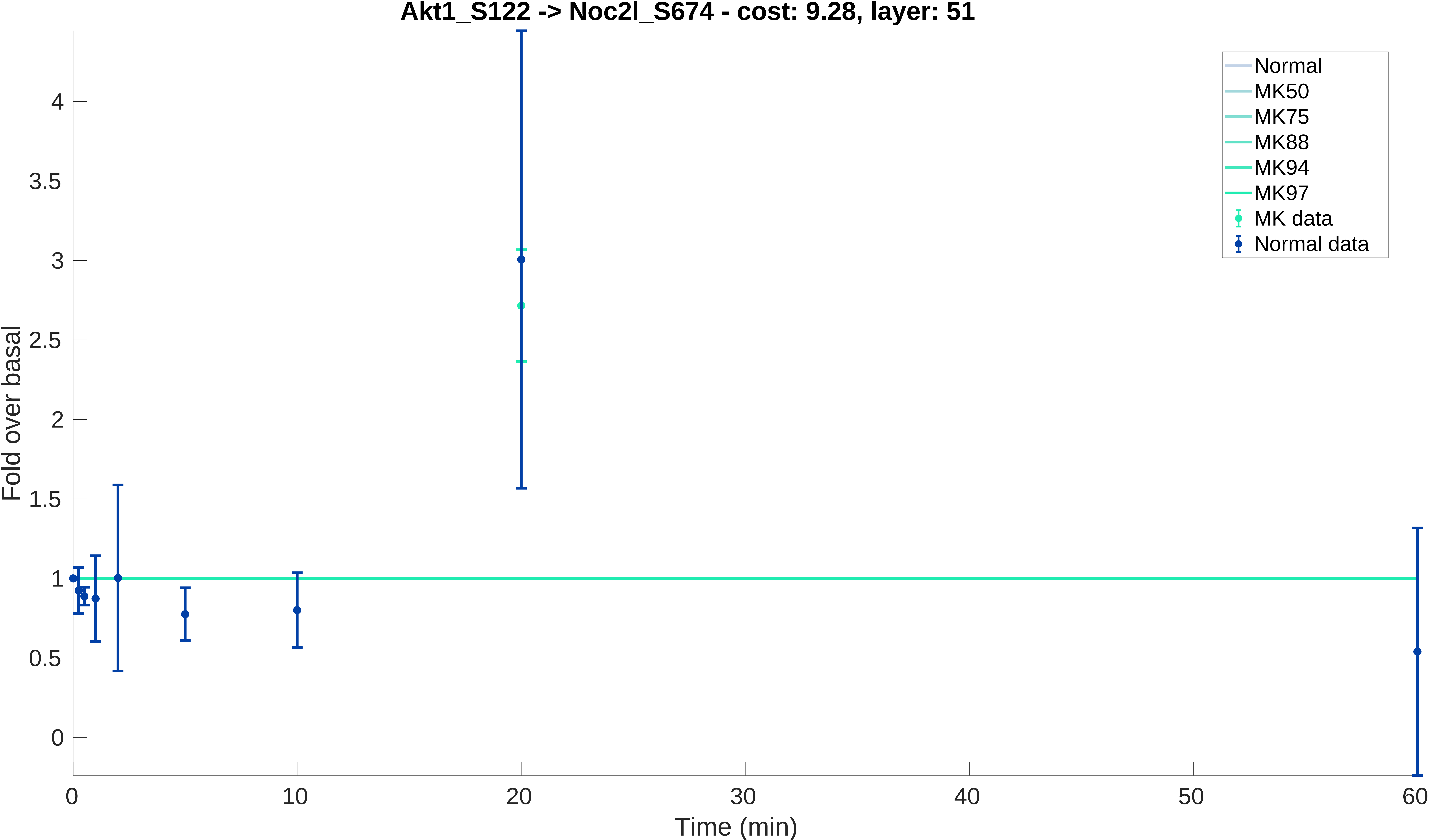

Akt1_S122 -> Noc2l_S674 - cost: 9.28, layer: 51
Fold over basal
Time (min)
0
0.5
1
1.5
2
2.5
3
3.5
4
0
10
20
30
40
50
60
Normal
MK50
MK75
MK88
MK94
MK97
MK data
Normal data

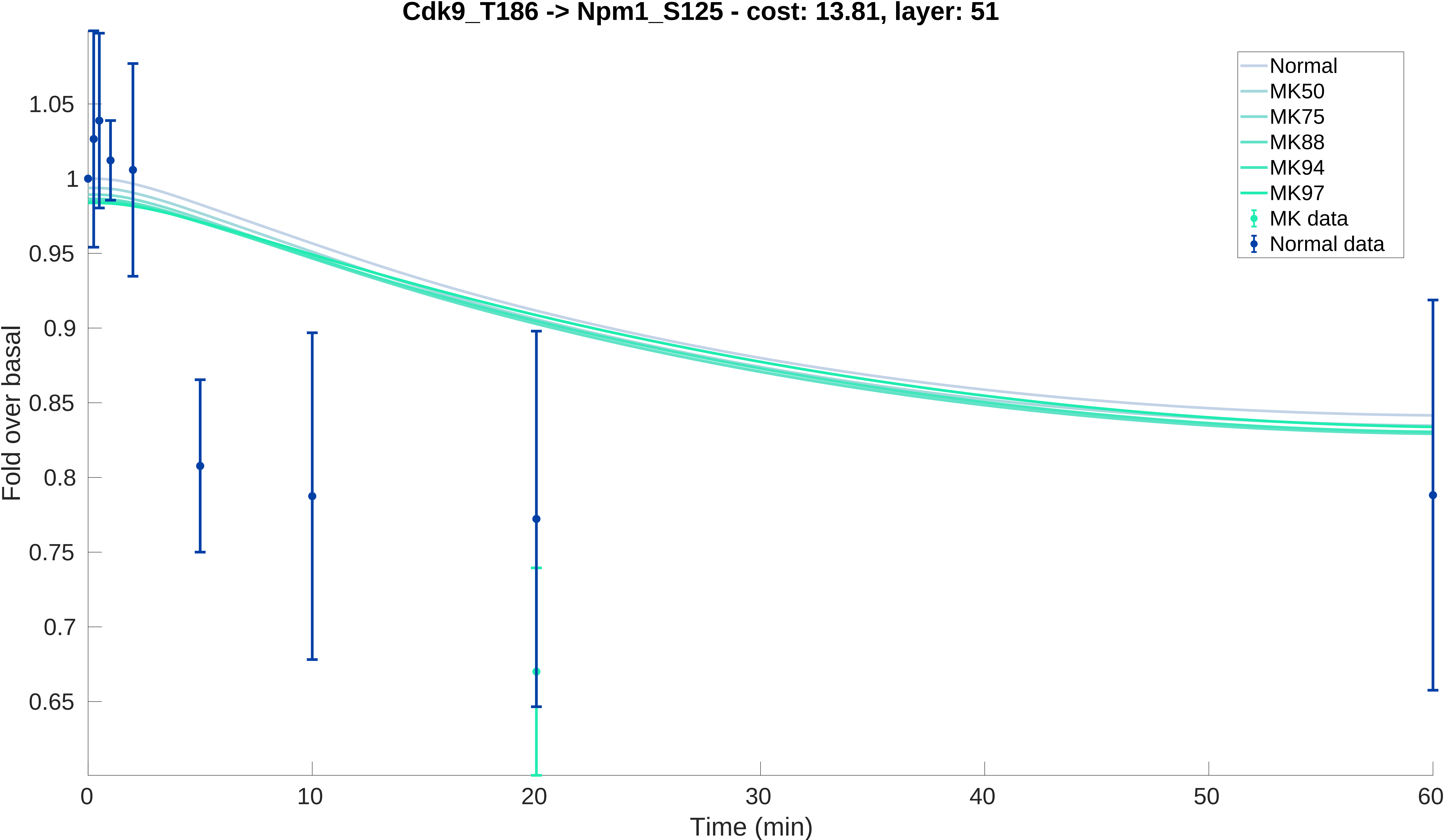
Cdk9_T186 -> Npm1_S125 - cost: 13.81, layer: 51
Normal
MK50
MK75
MK88
MK94
MK97
MK data
Normal data
Fold over basal
1.05
1
0.95
0.9
0.85
0.8
0.75
0.7
0.65
0
10
20
30
40
50
60
Time (min)

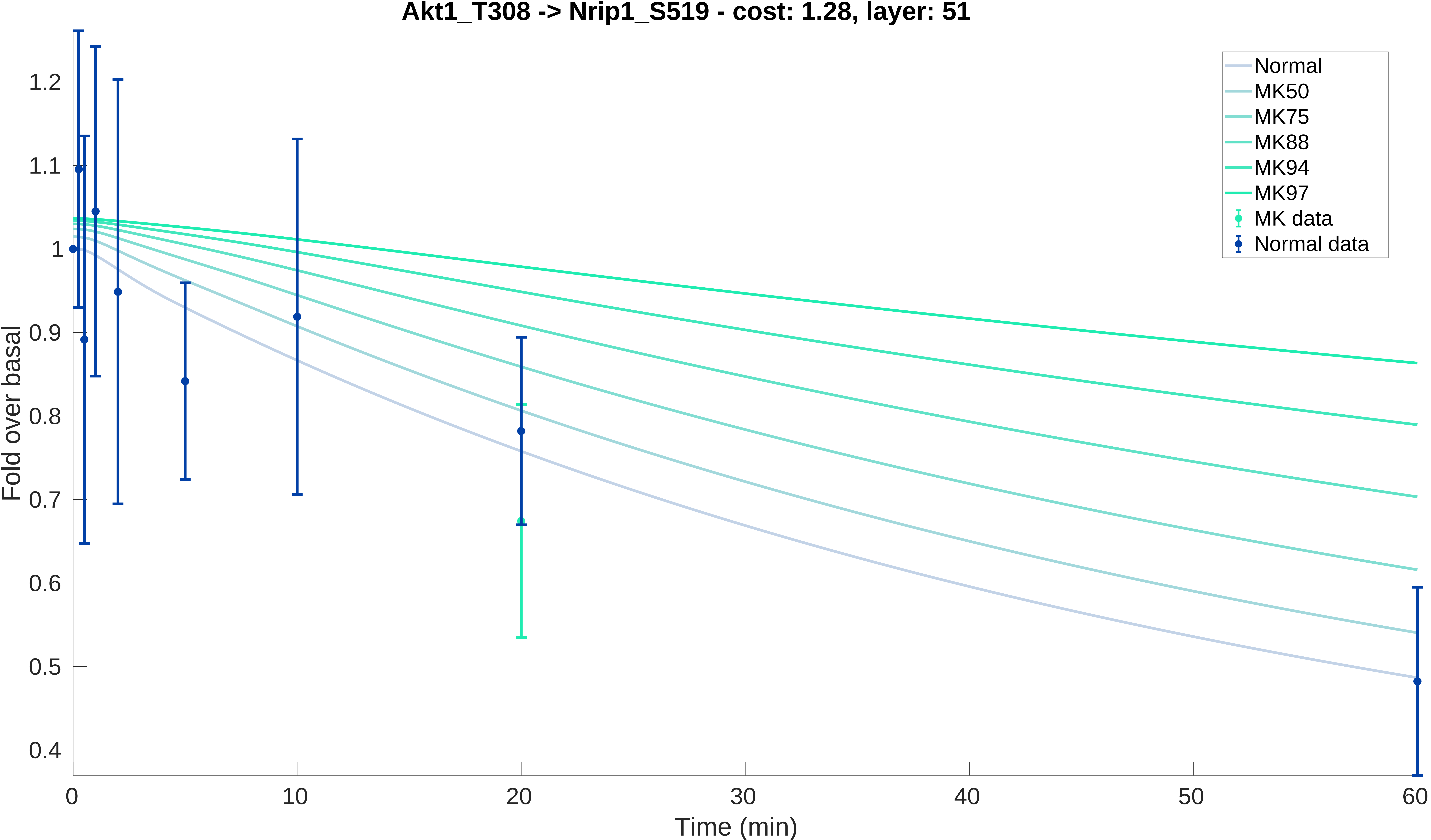

Akt1_T308 -> Nrip1_S519 - cost: 1.28, layer: 51
Normal
MK50
MK75
MK88
MK94
MK97
MK data
Normal data
Fold over basal
Time (min)
0.4
0.5
0.6
0.7
0.8
0.9
1
1.1
1.2
0
10
20
30
40
50
60

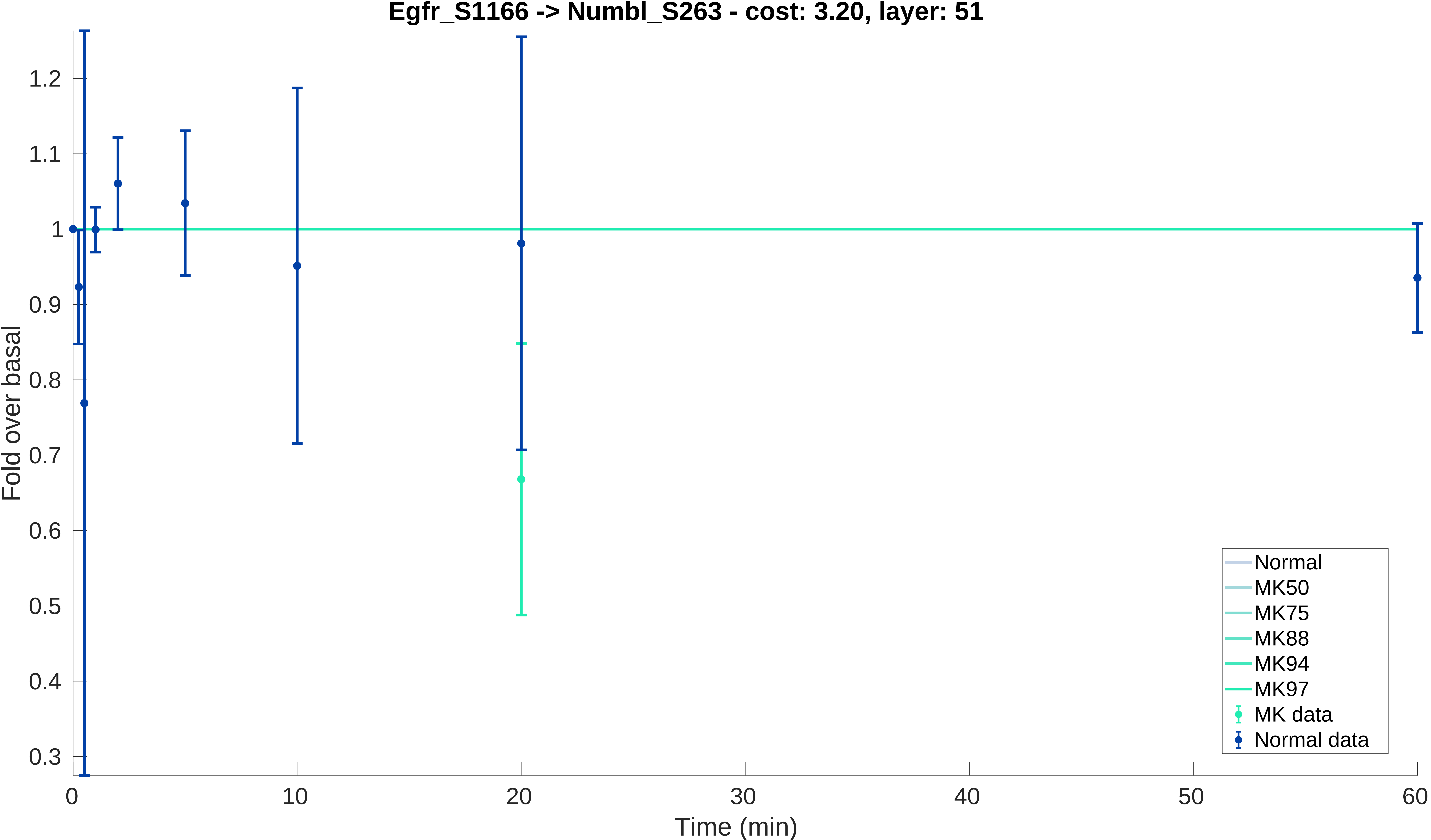
Egfr_S1166 -> Numbl_S263 - cost: 3.20, layer: 51
Fold over basal
1.2
1.1
1
0.9
0.8
0.7
0.6
0.5
0.4
0.3
0
10
20
30
40
50
60
Time (min)
Normal
MK50
MK75
MK88
MK94
MK97
MK data
Normal data

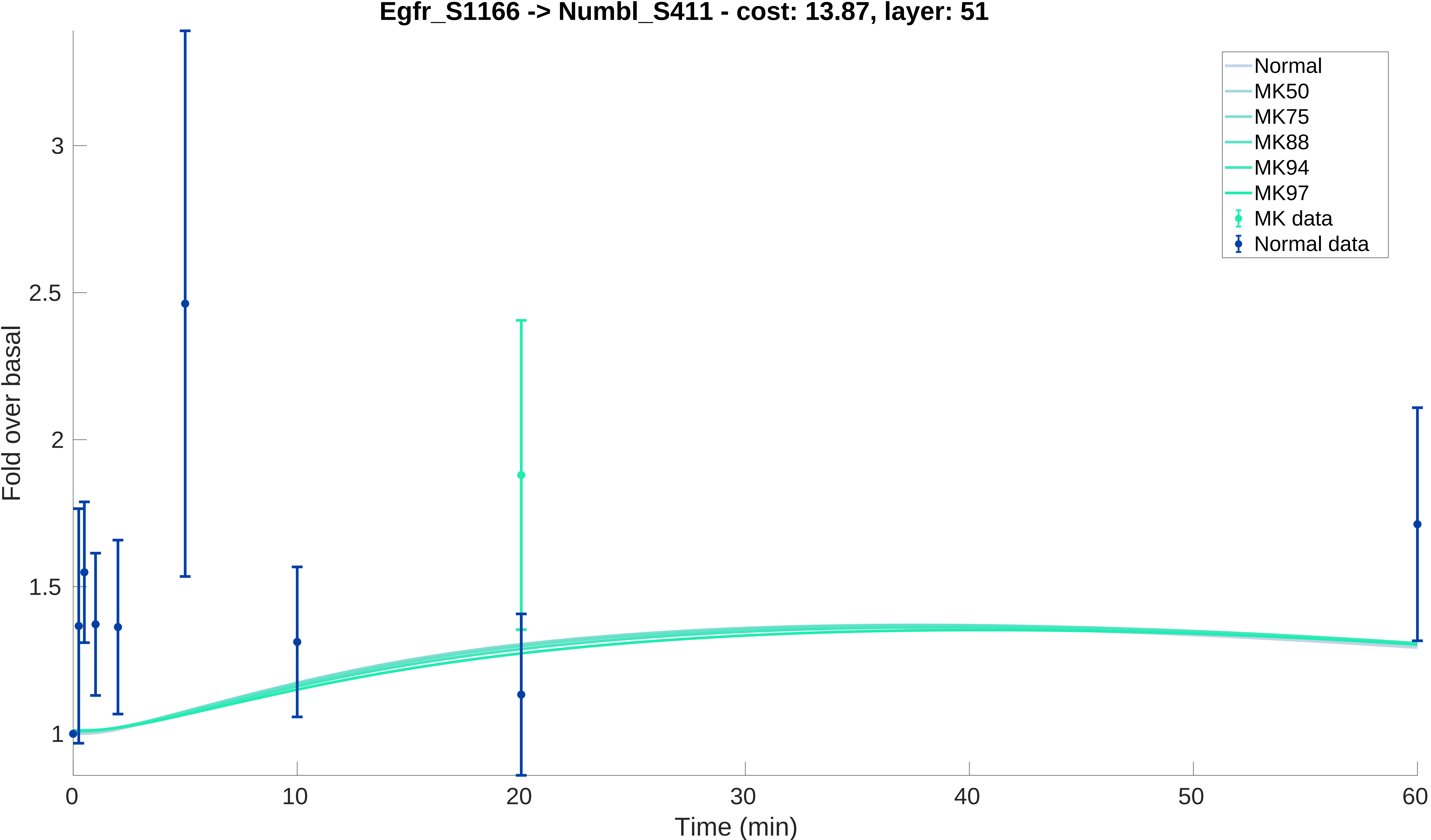

Egfr_S1166 -> Numbl_S411 - cost: 13.87, layer: 51
Normal
MK50
MK75
MK88
MK94
MK97
MK data
Normal data
Fold over basal
3
2.5
2
1.5
1
0
10
20
30
40
50
60
Time (min)

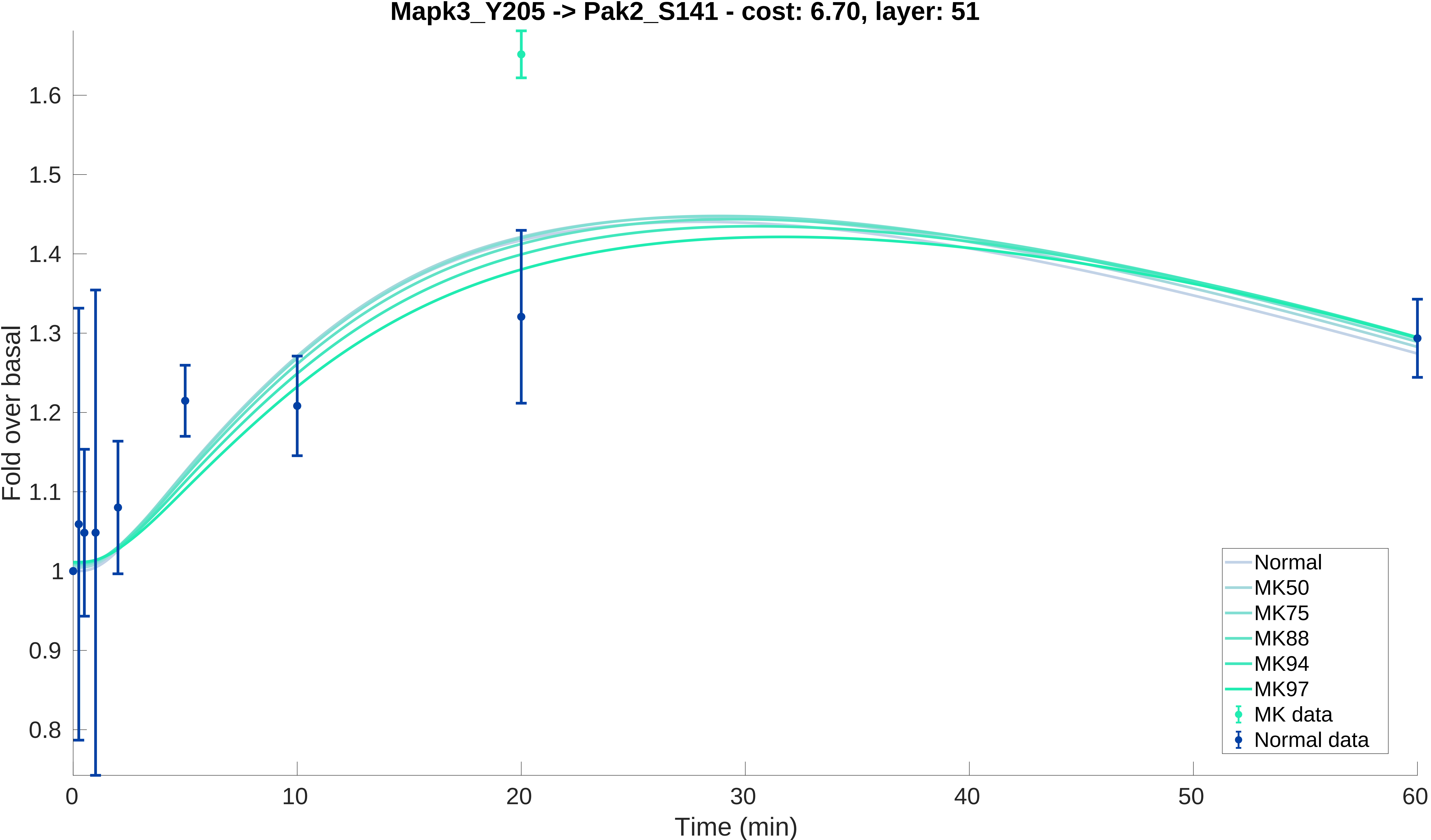

Mapk3_Y205 -> Pak2_S141 - cost: 6.70, layer: 51
Fold over basal
Time (min)
0.8
0.9
1
1.1
1.2
1.3
1.4
1.5
1.6
0
10
20
30
40
50
60
Normal
MK50
MK75
MK88
MK94
MK97
MK data
Normal data

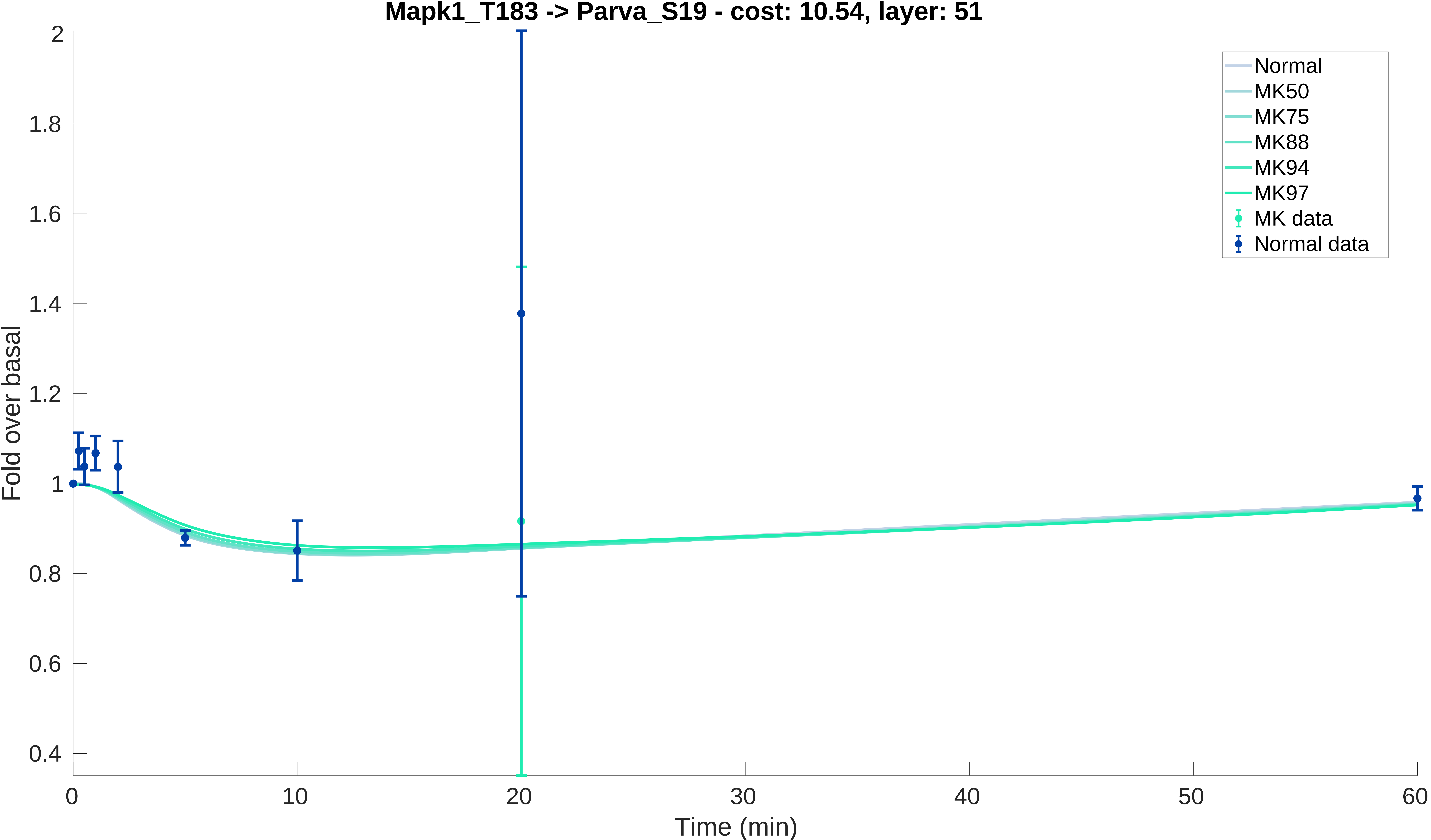

Mapk1_T183 -> Parva_S19 - cost: 10.54, layer: 51
Fold over basal
Time (min)
Normal
MK50
MK75
MK88
MK94
MK97
MK data
Normal data
2
1.8
1.6
1.4
1.2
1
0.8
0.6
0.4
0
10
20
30
40
50
60

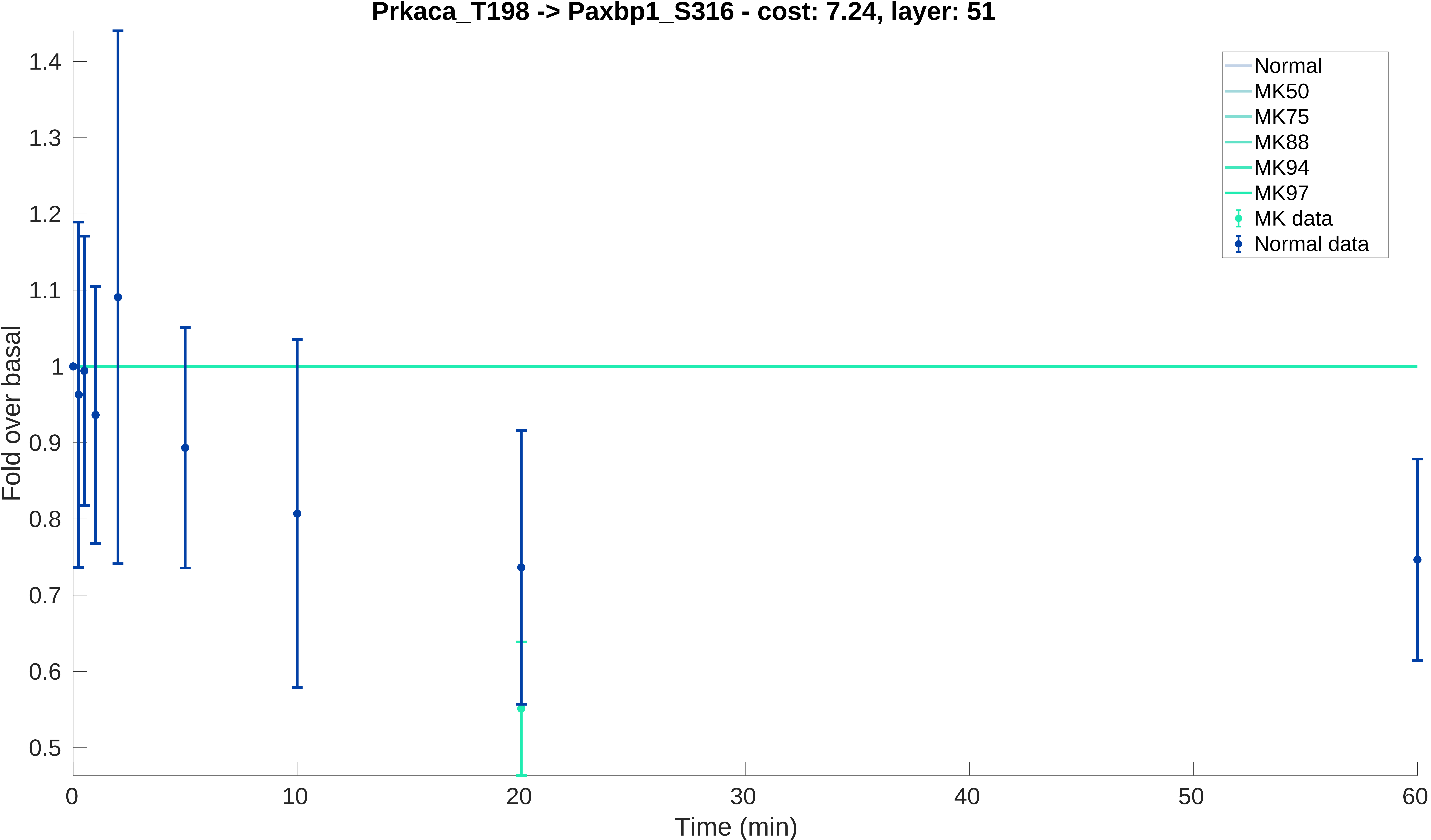

Prkaca_T198 -> Paxbp1_S316 - cost: 7.24, layer: 51
Fold over basal
1.4
1.3
1.2
1.1
1
0.9
0.8
0.7
0.6
0.5
0
10
20
30
40
50
60
Time (min)
Normal
MK50
MK75
MK88
MK94
MK97
MK data
Normal data

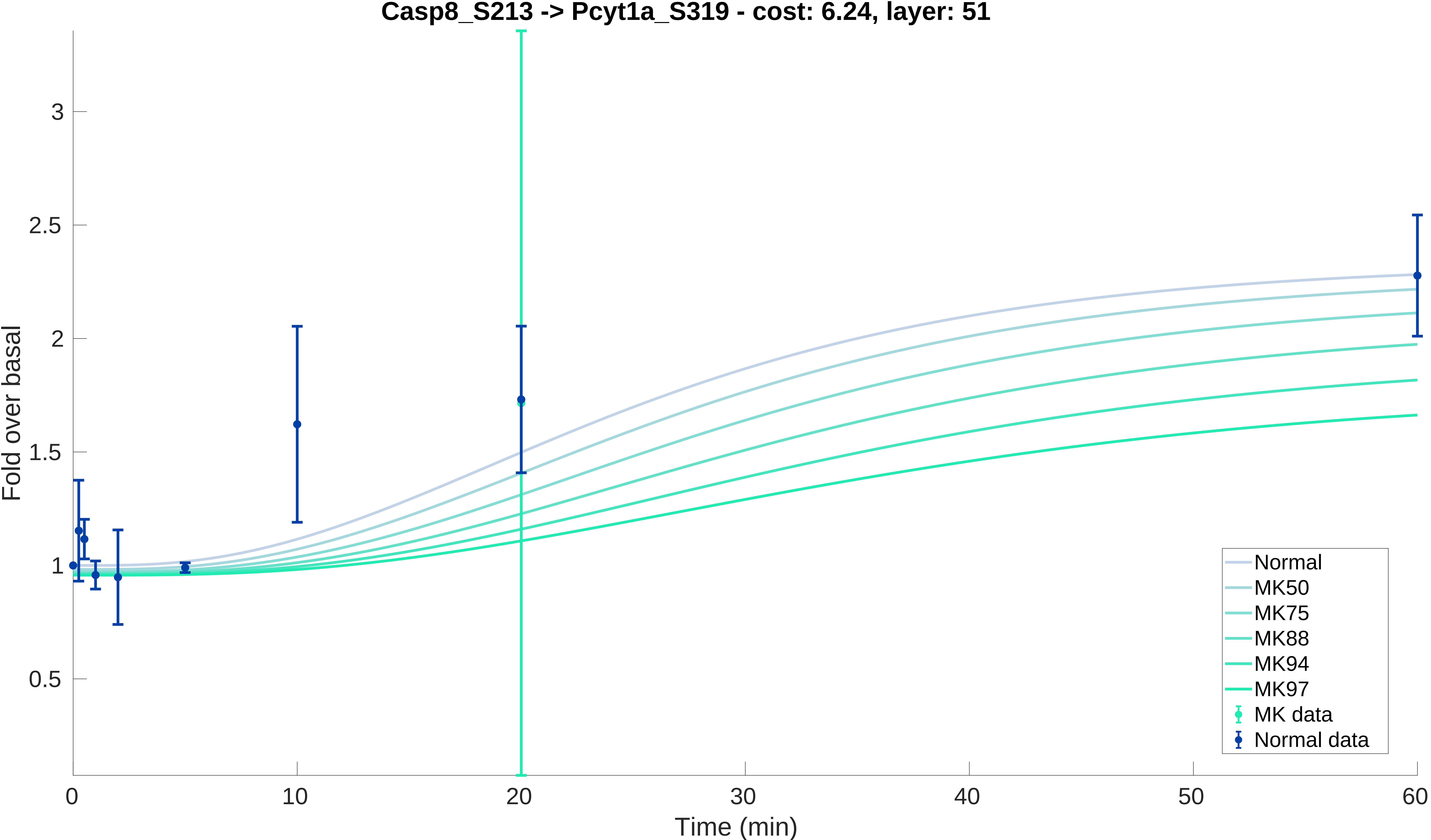

Casp8_S213 -> Pcyt1a_S319 - cost: 6.24, layer: 51
Fold over basal
Time (min)
0.5
1
1.5
2
2.5
3
0
10
20
30
40
50
60
Normal
MK50
MK75
MK88
MK94
MK97
MK data
Normal data

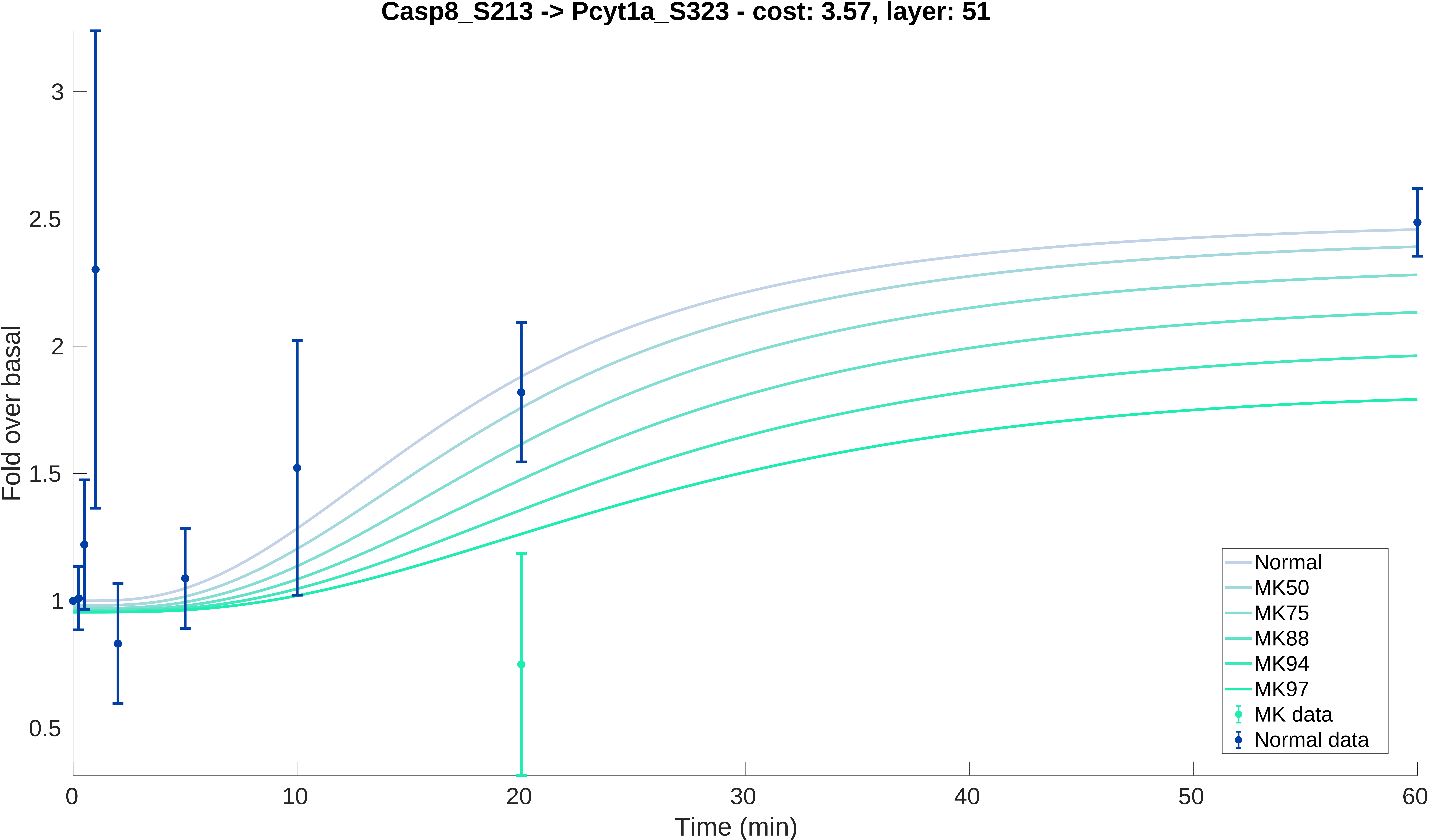

Casp8_S213 -> Pcyt1a_S323 - cost: 3.57, layer: 51
Fold over basal
Time (min)
0.5
1
1.5
2
2.5
3
0
10
20
30
40
50
60
Normal
MK50
MK75
MK88
MK94
MK97
MK data
Normal data

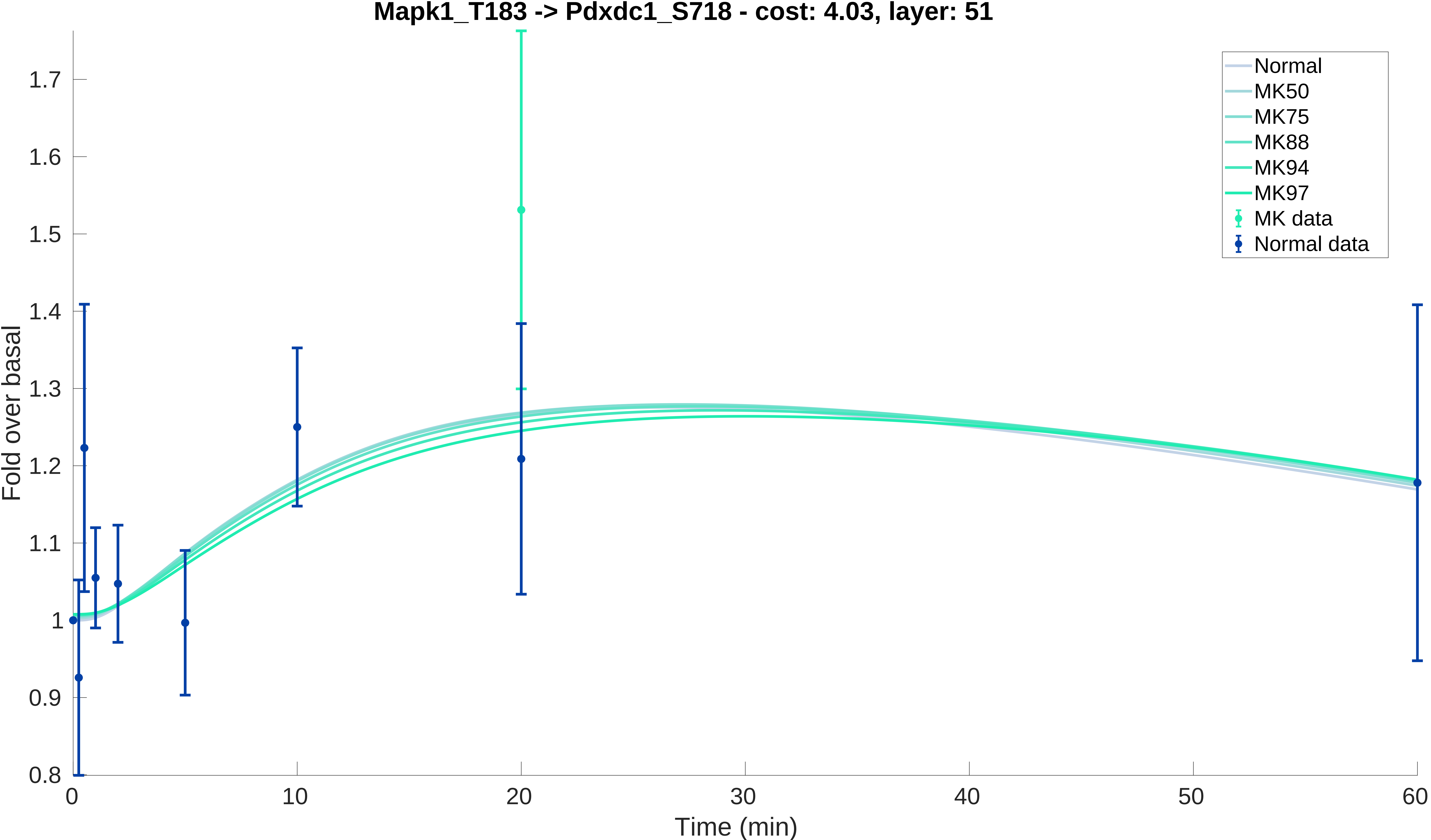

Mapk1_T183 -> Pdxdc1_S718 - cost: 4.03, layer: 51
Normal
MK50
MK75
MK88
MK94
MK97
MK data
Normal data
Fold over basal
1.7
1.6
1.5
1.4
1.3
1.2
1.1
1
0.9
0.8
0
10
20
30
40
50
60
Time (min)

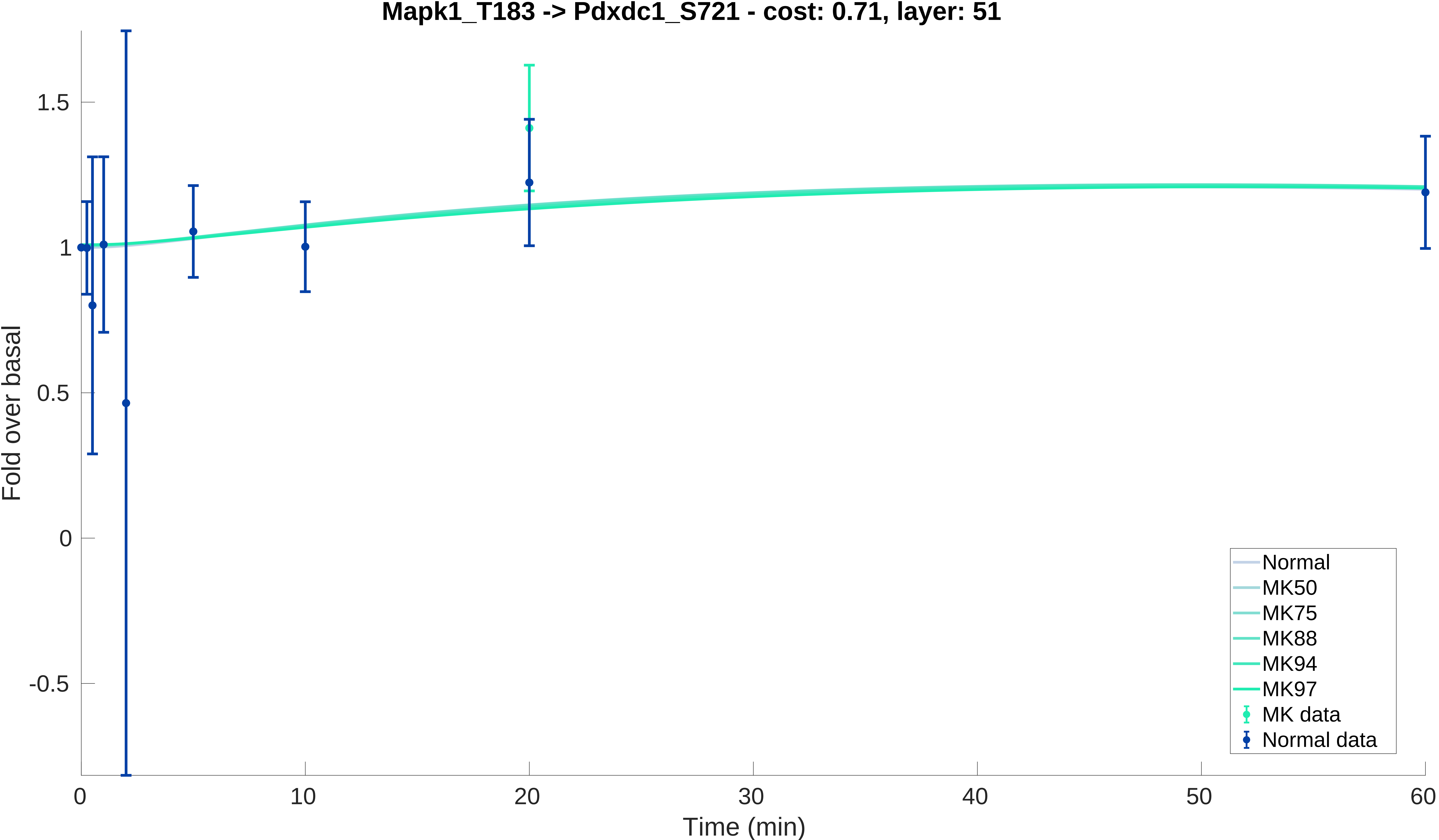

Mapk1_T183 -> Pdxdc1_S721 - cost: 0.71, layer: 51
Fold over basal
1.5
1
0.5
0
-0.5
0
10
20
30
40
50
60
Time (min)
Normal
MK50
MK75
MK88
MK94
MK97
MK data
Normal data

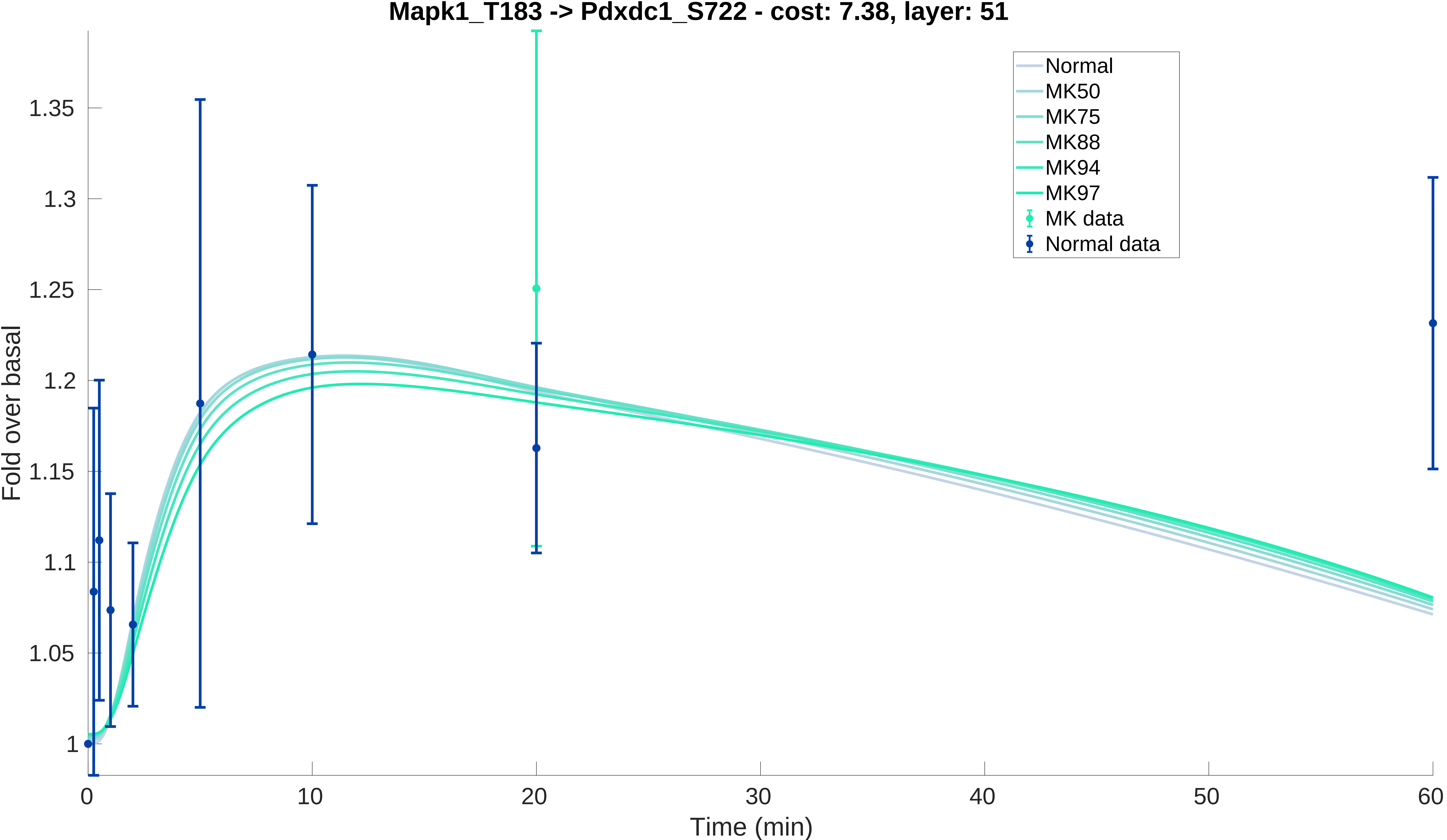

Mapk1_T183 -> Pdxdc1_S722 - cost: 7.38, layer: 51
Fold over basal
Time (min)
1
1.05
1.1
1.15
1.2
1.25
1.3
1.35
0
10
20
30
40
50
60
Normal
MK50
MK75
MK88
MK94
MK97
MK data
Normal data

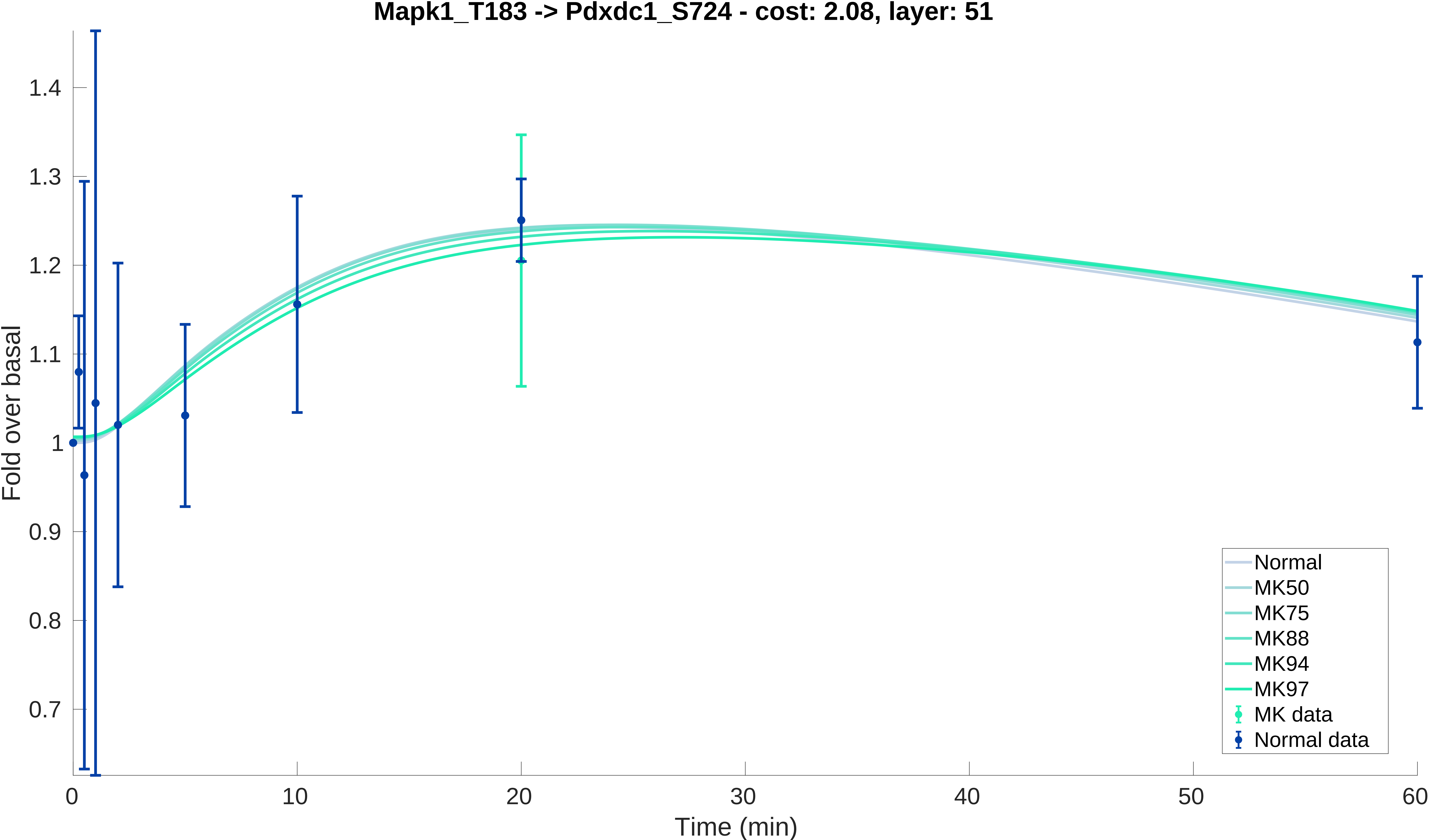

Mapk1_T183 -> Pdxdc1_S724 - cost: 2.08, layer: 51
Fold over basal
1.4
1.3
1.2
1.1
1
0.9
0.8
0.7
0
10
20
30
40
50
60
Time (min)
Normal
MK50
MK75
MK88
MK94
MK97
MK data
Normal data

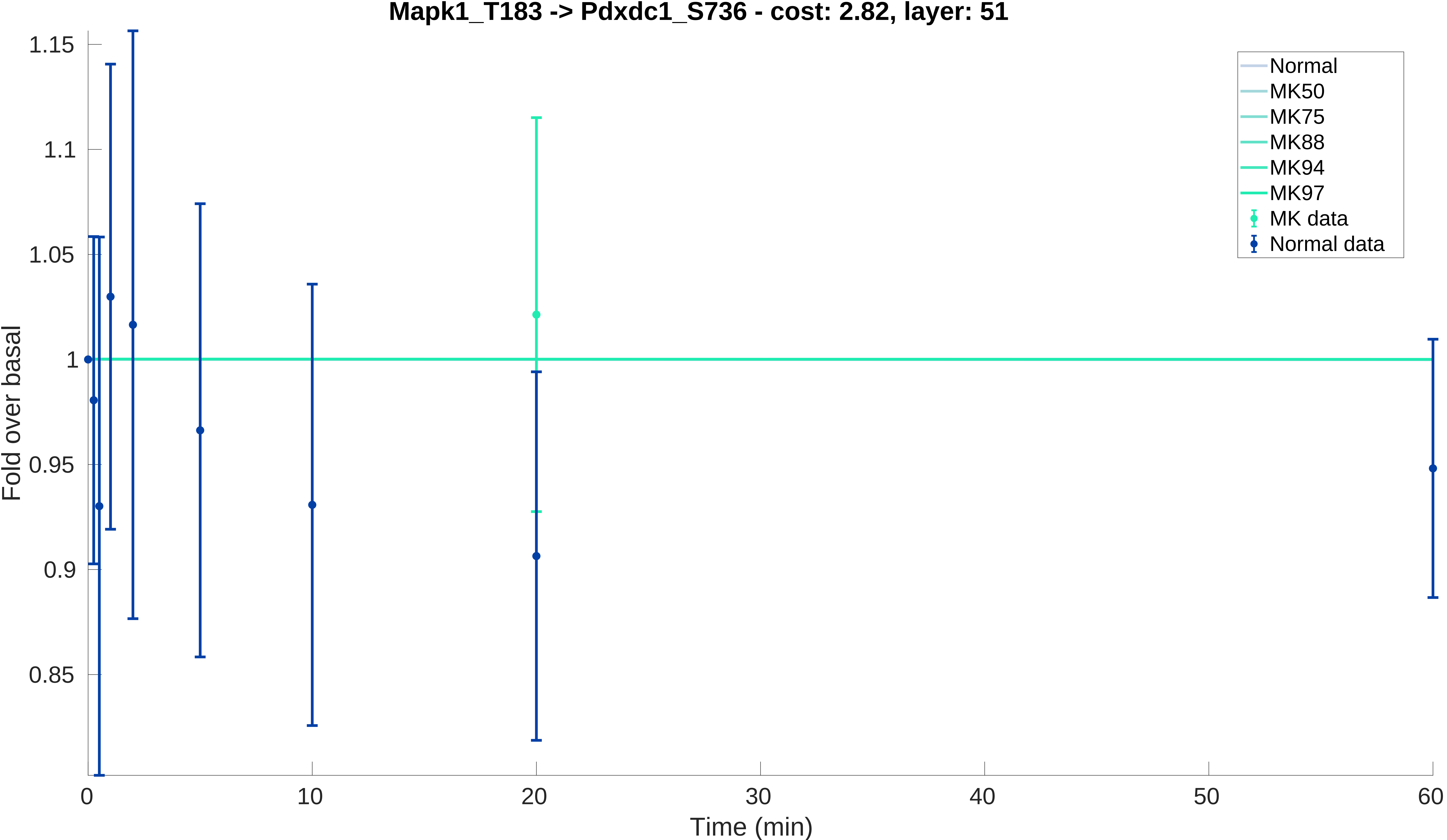
Mapk1_T183 -> Pdxdc1_S736 - cost: 2.82, layer: 51
1.15
1.1
1.05
1
0.95
0.9
0.85
Fold over basal
0
10
20
30
40
50
60
Time (min)
Normal
MK50
MK75
MK88
MK94
MK97
MK data
Normal data

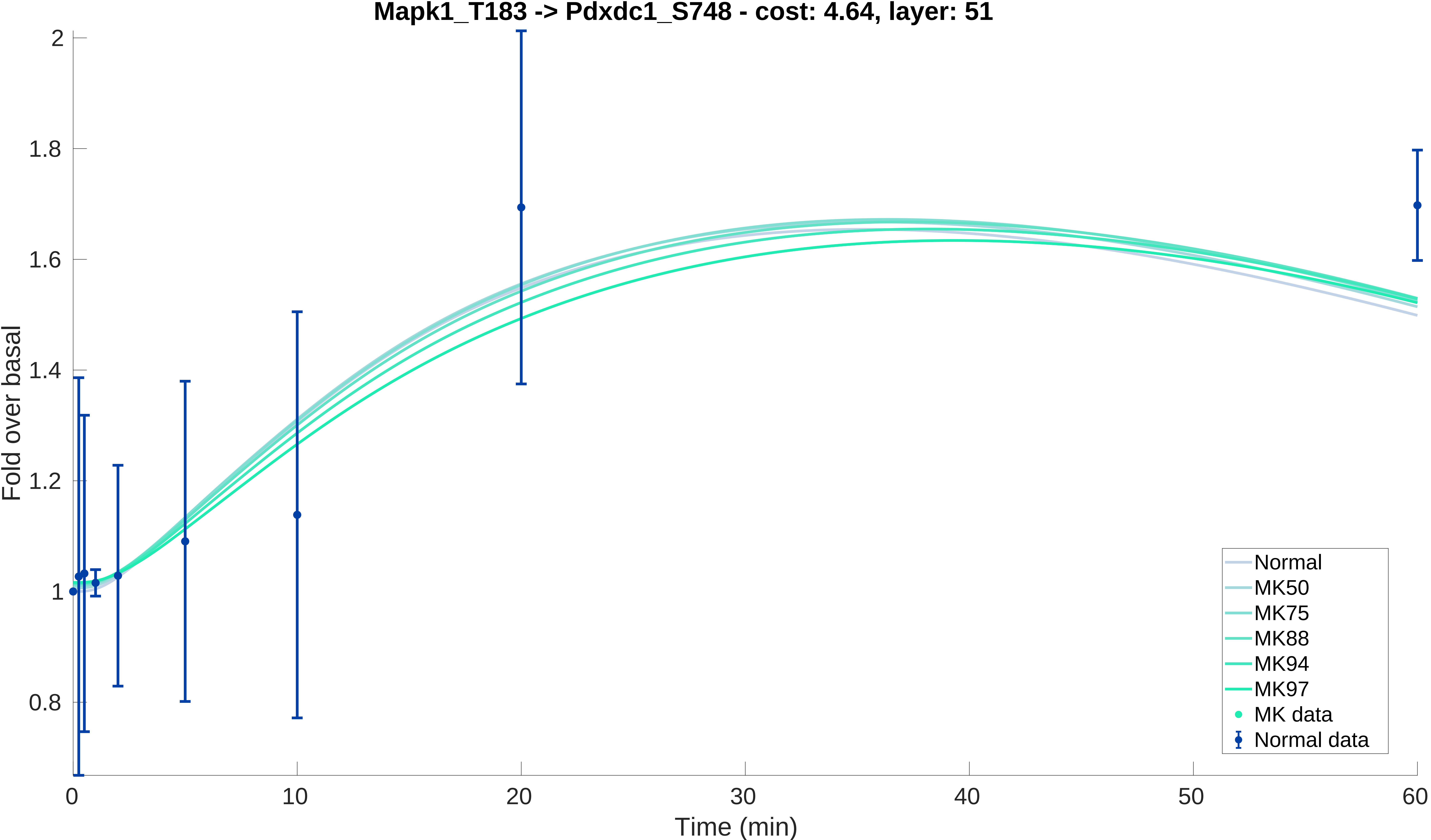

Mapk1_T183 -> Pdxdc1_S748 - cost: 4.64, layer: 51
Fold over basal
Time (min)
0.8
1
1.2
1.4
1.6
1.8
2
0
10
20
30
40
50
60
Normal
MK50
MK75
MK88
MK94
MK97
MK data
Normal data

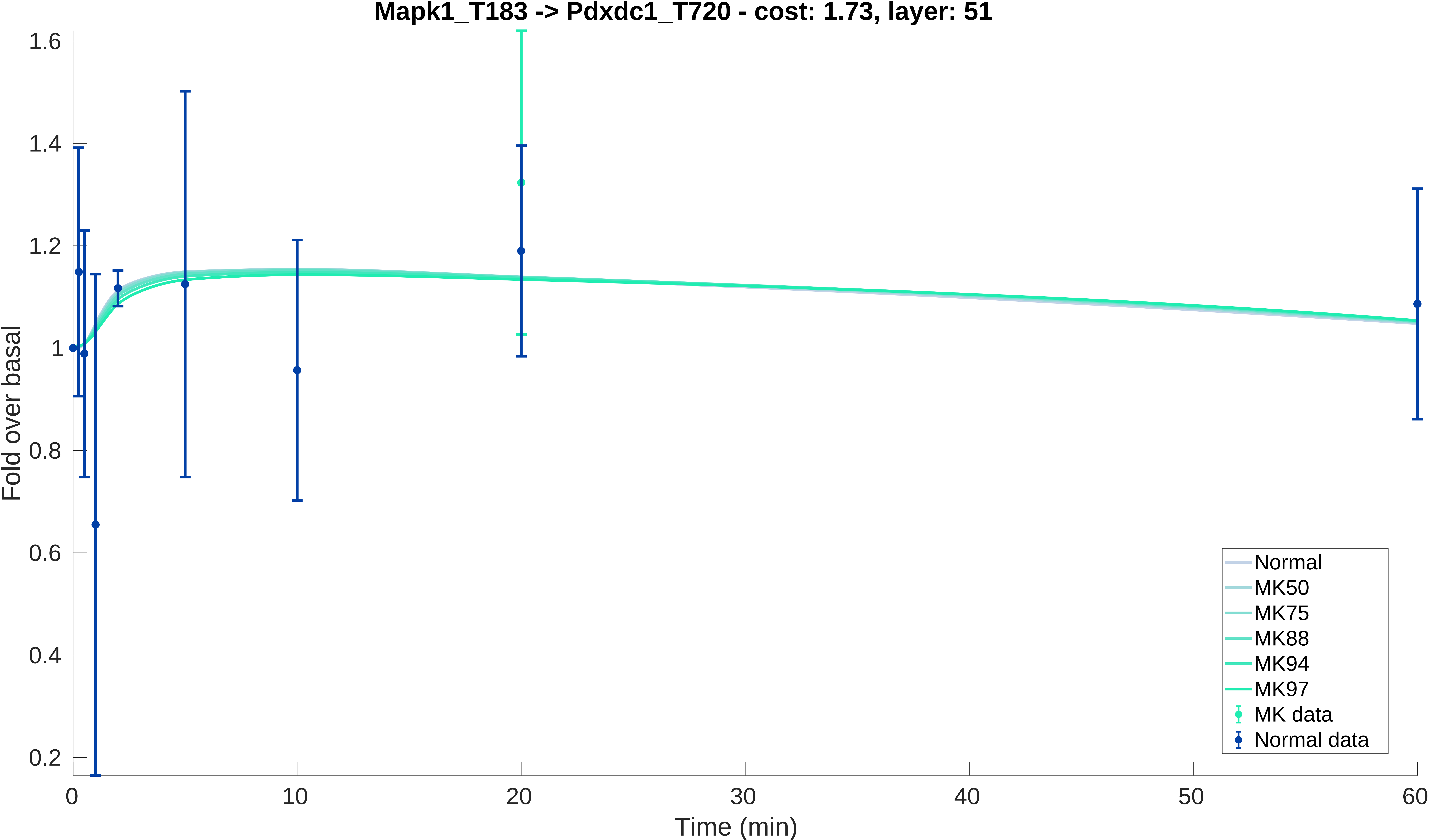

Mapk1_T183 -> Pdxdc1_T720 - cost: 1.73, layer: 51
Fold over basal
Time (min)
0.2
0.4
0.6
0.8
1
1.2
1.4
1.6
0
10
20
30
40
50
60
Normal
MK50
MK75
MK88
MK94
MK97
MK data
Normal data

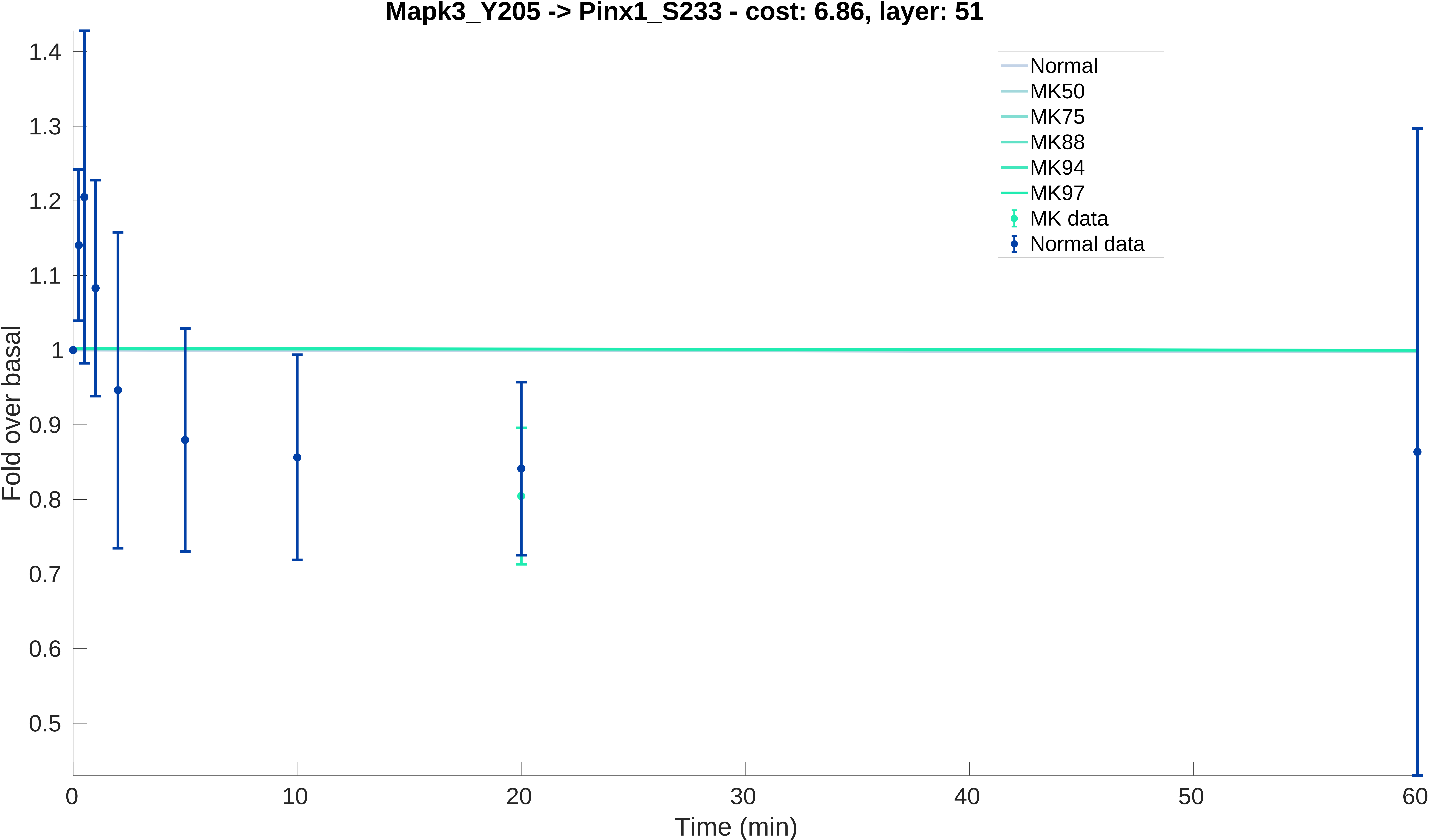

Mapk3_Y205 -> Pinx1_S233 - cost: 6.86, layer: 51
Fold over basal
Time (min)
0
10
20
30
40
50
60
0.5
0.6
0.7
0.8
0.9
1
1.1
1.2
1.3
1.4
Normal
MK50
MK75
MK88
MK94
MK97
MK data
Normal data

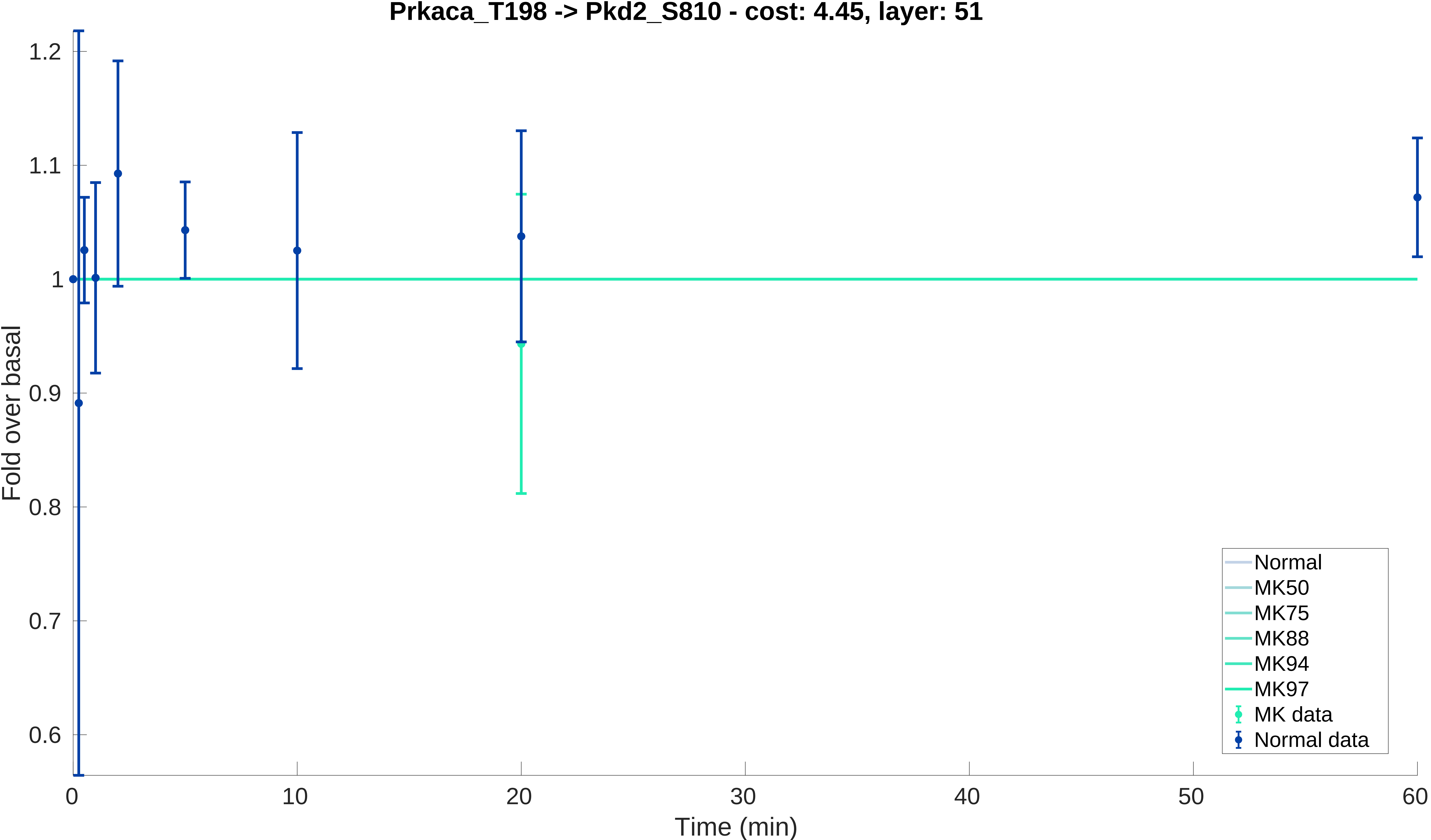
Prkaca_T198 -> Pkd2_S810 - cost: 4.45, layer: 51
Fold over basal
1.2
1.1
1
0.9
0.8
0.7
0.6
0
10
20
30
40
50
60
Time (min)
Normal
MK50
MK75
MK88
MK94
MK97
MK data
Normal data

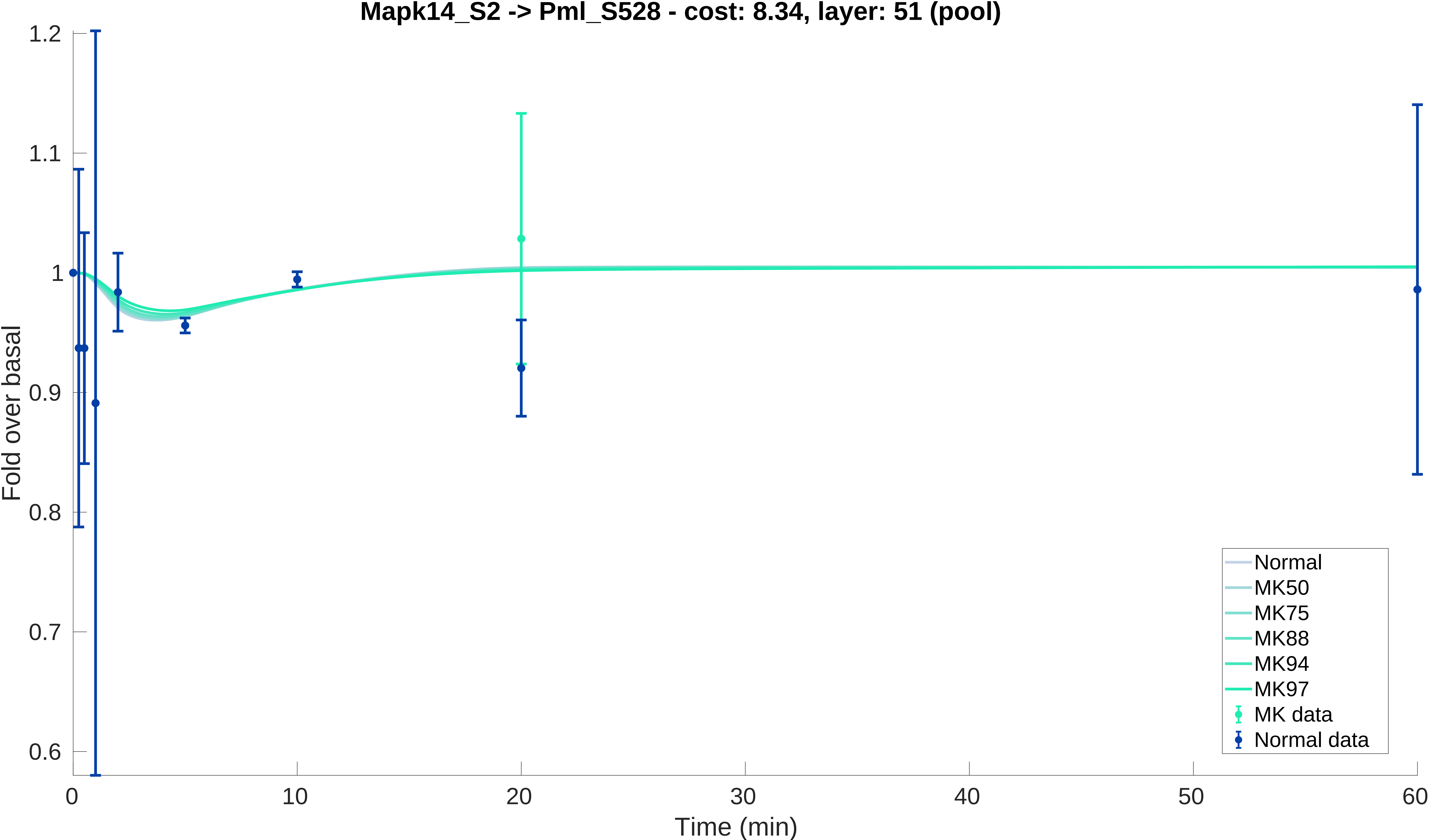

Mapk14_S2 -> Pml_S528 - cost: 8.34, layer: 51 (pool)
Fold over basal
1.2
1.1
1
0.9
0.8
0.7
0.6
0
10
20
30
40
50
60
Time (min)
Normal
MK50
MK75
MK88
MK94
MK97
MK data
Normal data

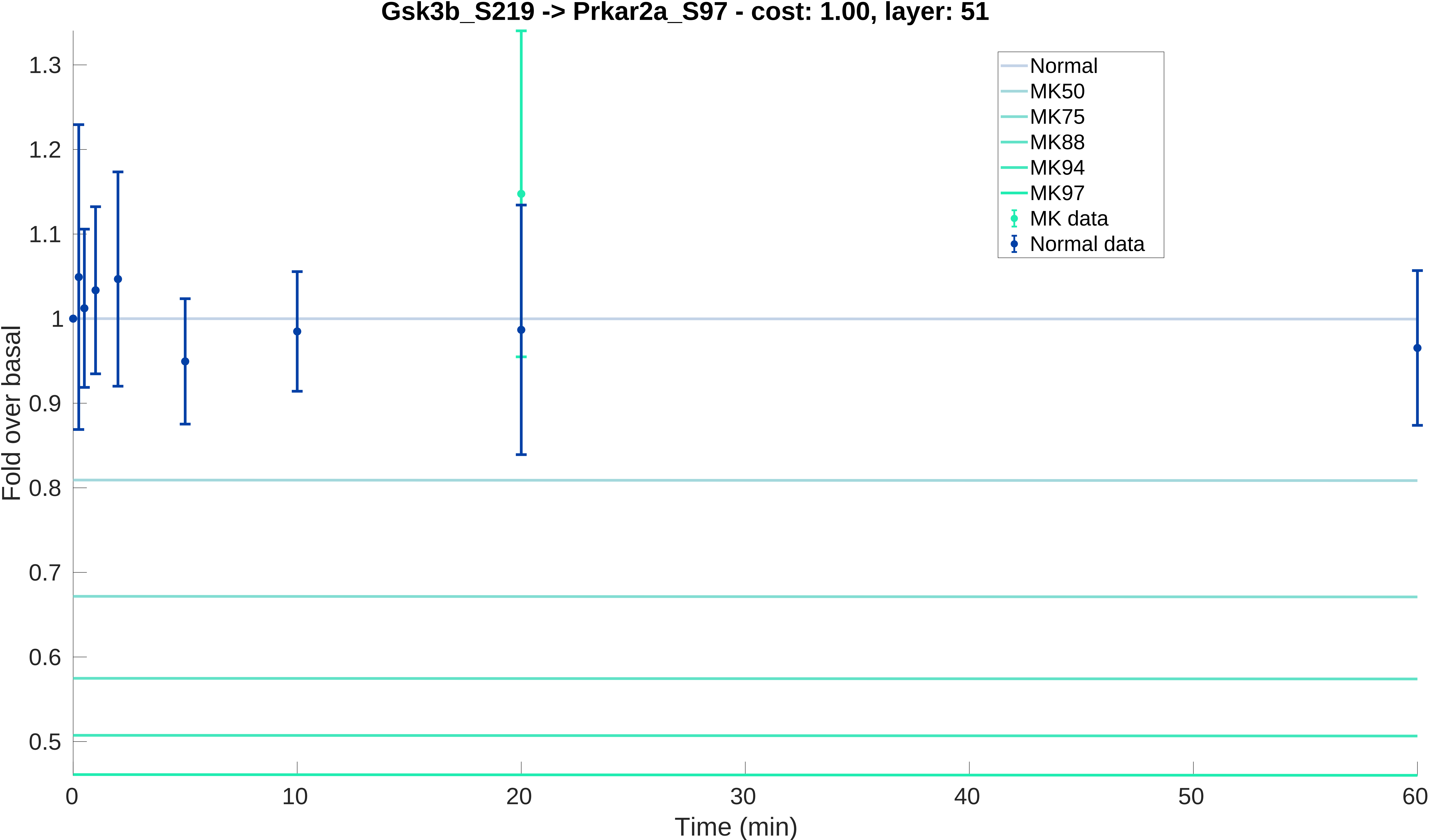

Gsk3b_S219 -> Prkar2a_S97 - cost: 1.00, layer: 51
Normal
MK50
MK75
MK88
MK94
MK97
MK data
Normal data
Fold over basal
Time (min)
1.3
1.2
1.1
1
0.9
0.8
0.7
0.6
0.5
0
10
20
30
40
50
60

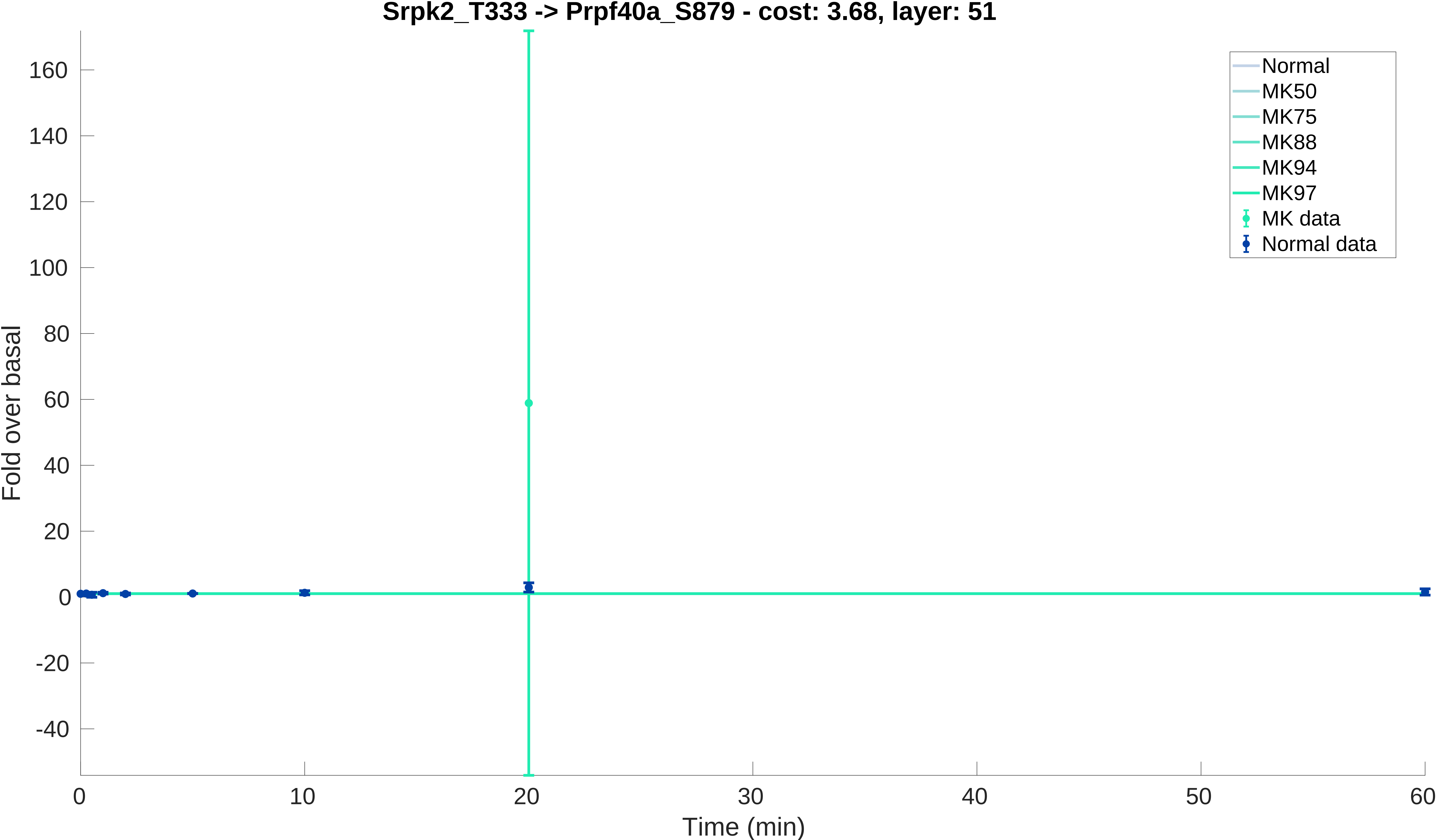

Srpk2_T333 -> Prpf40a_S879 - cost: 3.68, layer: 51
Normal
MK50
MK75
MK88
MK94
MK97
MK data
Normal data
Fold over basal
160
140
120
100
80
60
40
20
0
-20
-40
0
10
20
30
40
50
60
Time (min)

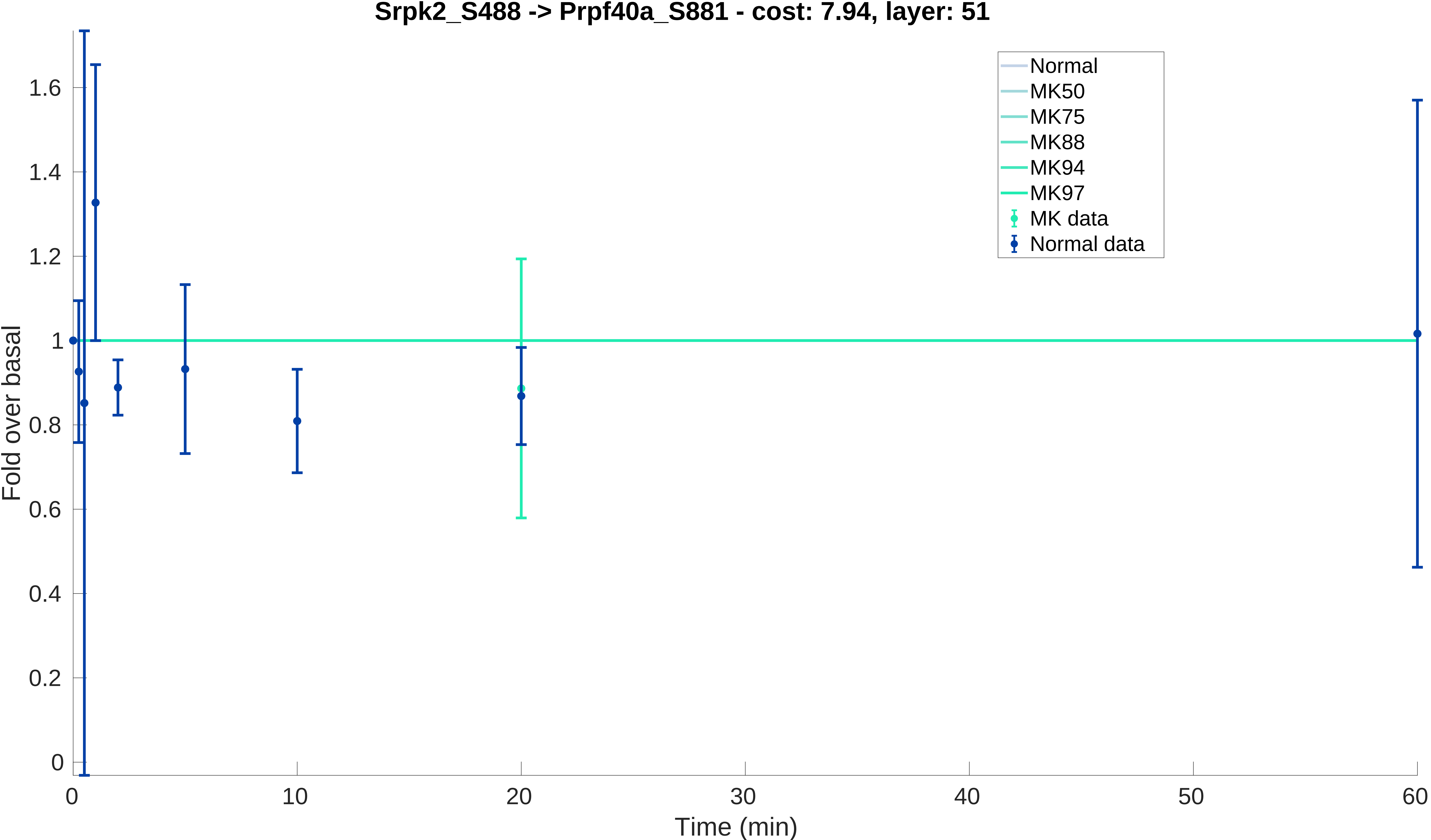

Srpk2_S488 -> Prpf40a_S881 - cost: 7.94, layer: 51
Normal
MK50
MK75
MK88
MK94
MK97
MK data
Normal data
Fold over basal
1.6
1.4
1.2
1
0.8
0.6
0.4
0.2
0
0
10
20
30
40
50
60
Time (min)

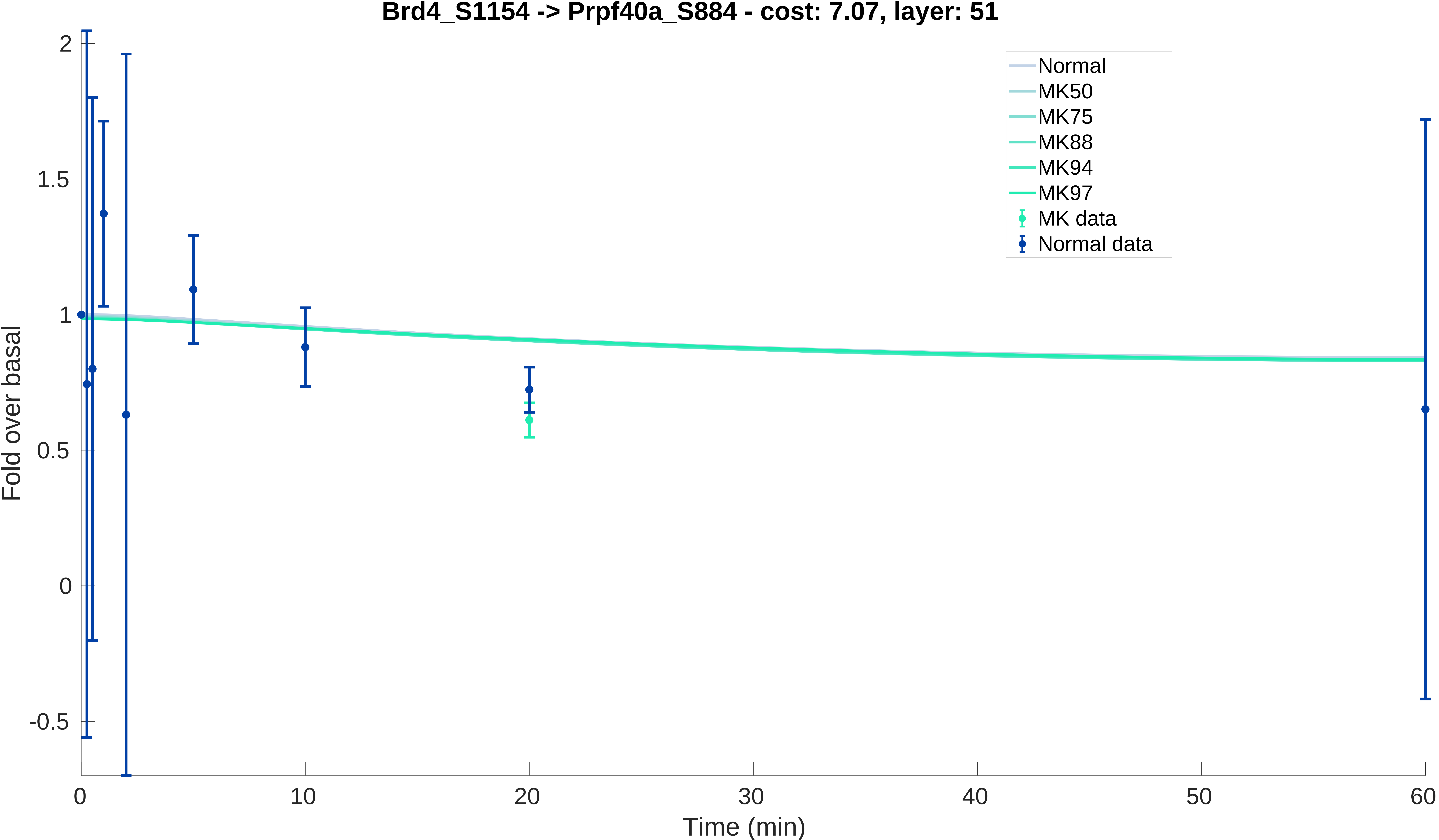

Brd4_S1154 -> Prpf40a_S884 - cost: 7.07, layer: 51
Normal
MK50
MK75
MK88
MK94
MK97
MK data
Normal data
Fold over basal
Time (min)
2
1.5
1
0.5
0
-0.5
0
10
20
30
40
50
60

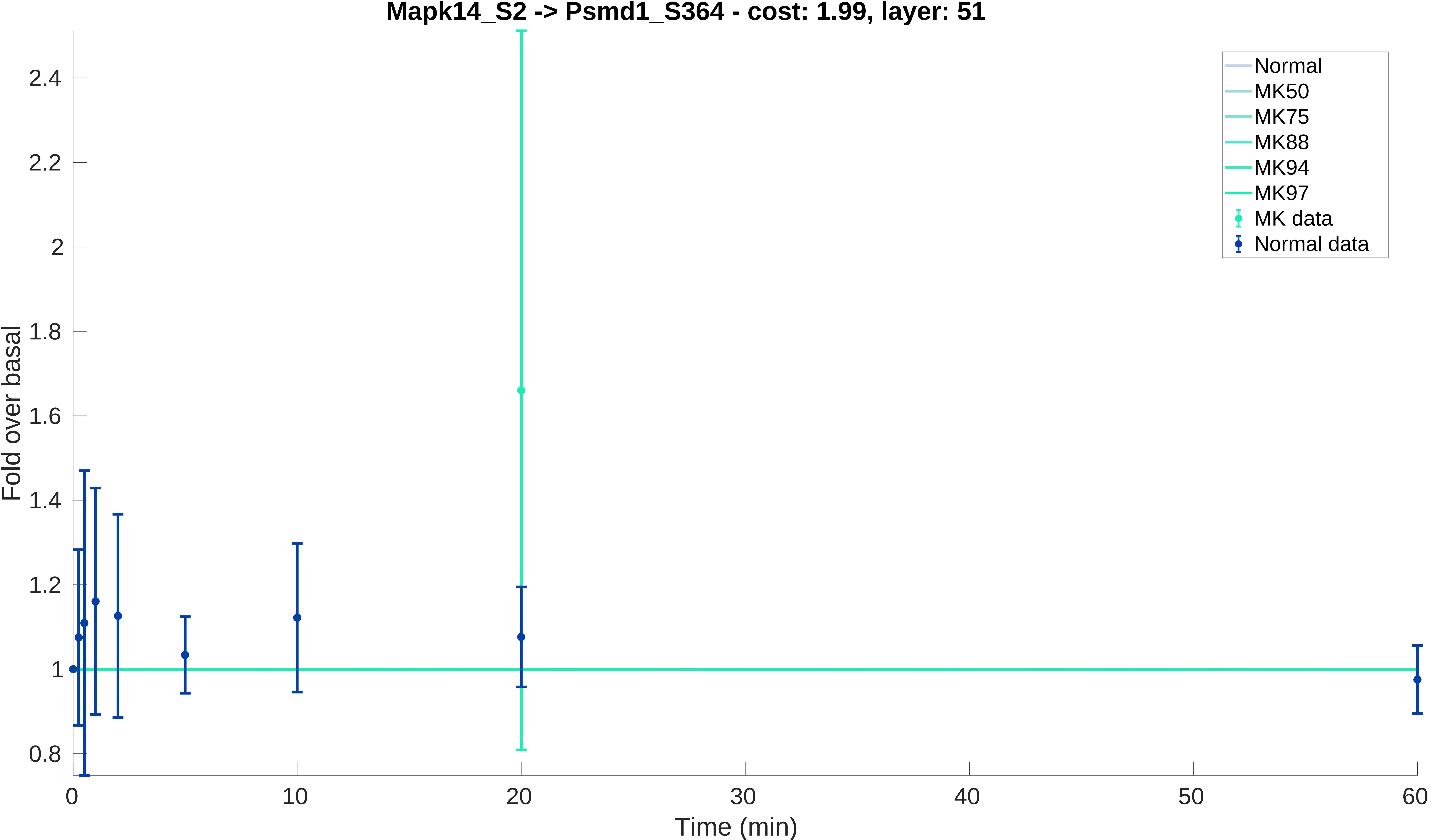
Mapk14_S2 -> Psmd1_S364 - cost: 1.99, layer: 51
Fold over basal
Time (min)
0.8
1
1.2
1.4
1.6
1.8
2
2.2
2.4
0
10
20
30
40
50
60
Normal
MK50
MK75
MK88
MK94
MK97
MK data
Normal data

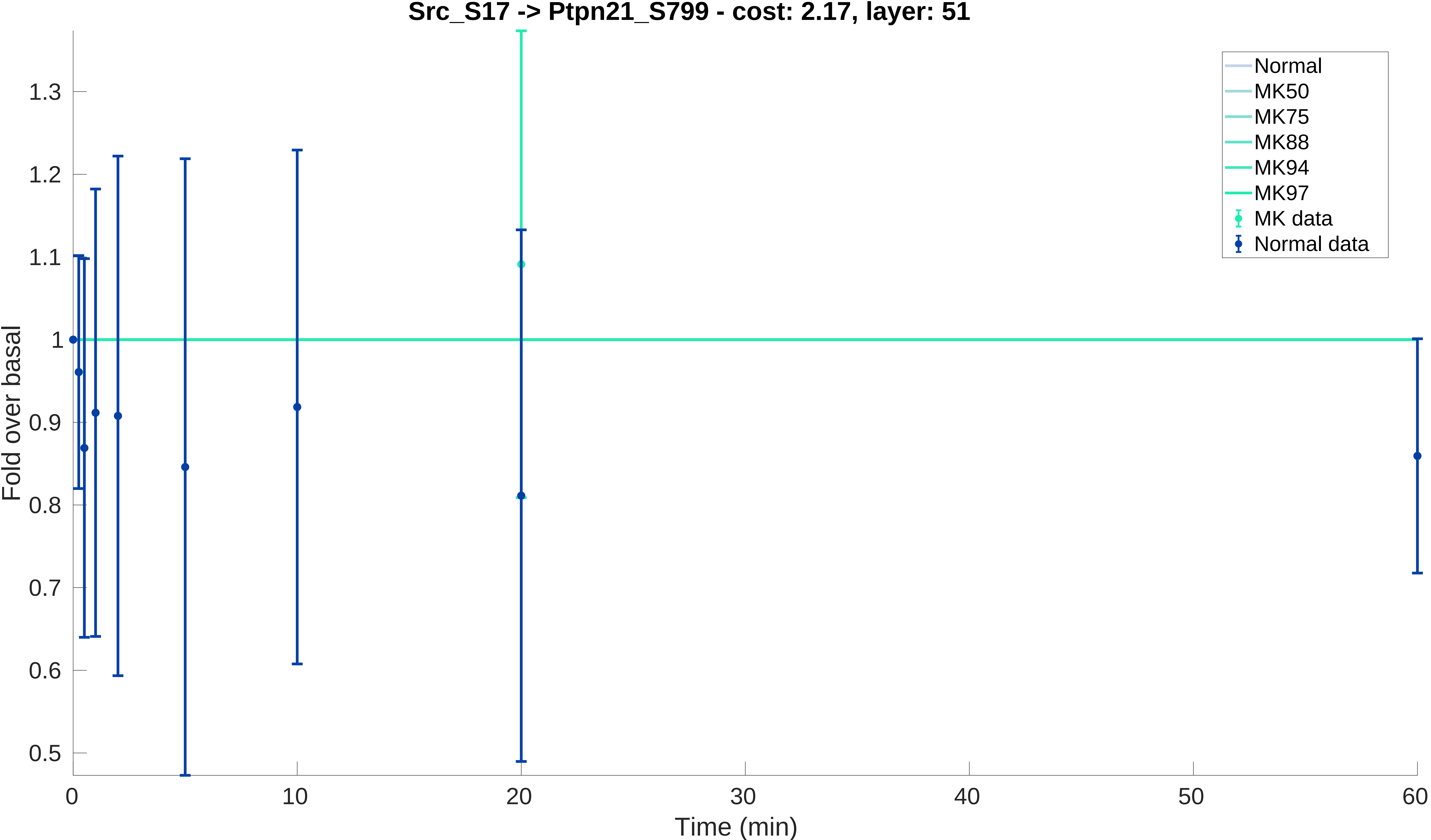

Src_S17 -> Ptpn21_S799 - cost: 2.17, layer: 51
Fold over basal
1.3
1.2
1.1
1
0.9
0.8
0.7
0.6
0.5
0
10
20
30
40
50
60
Time (min)
Normal
MK50
MK75
MK88
MK94
MK97
MK data
Normal data

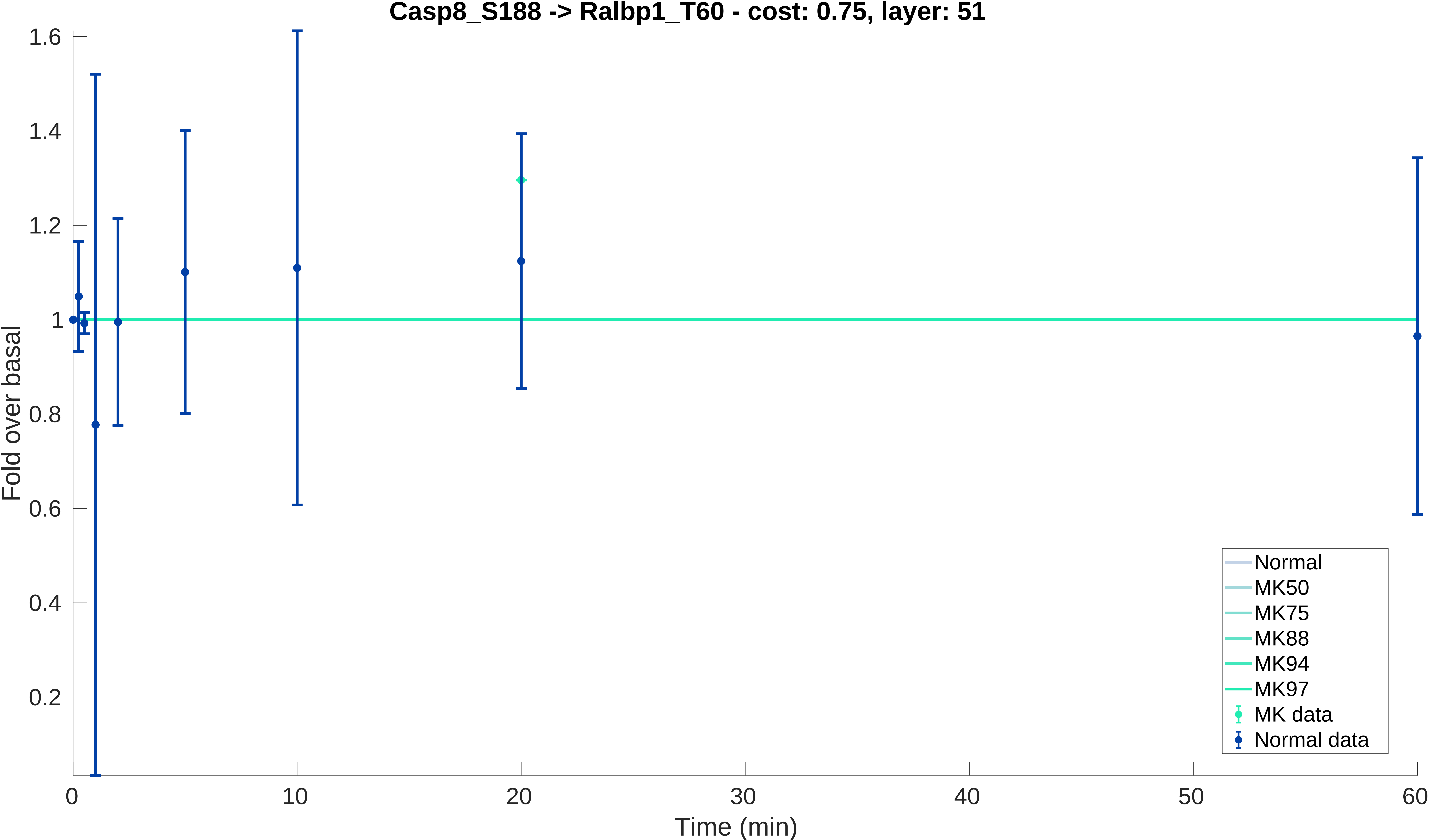

Casp8_S188 -> Ralbp1_T60 - cost: 0.75, layer: 51
Fold over basal
Time (min)
1.6
1.4
1.2
1
0.8
0.6
0.4
0.2
0
10
20
30
40
50
60
Normal
MK50
MK75
MK88
MK94
MK97
MK data
Normal data

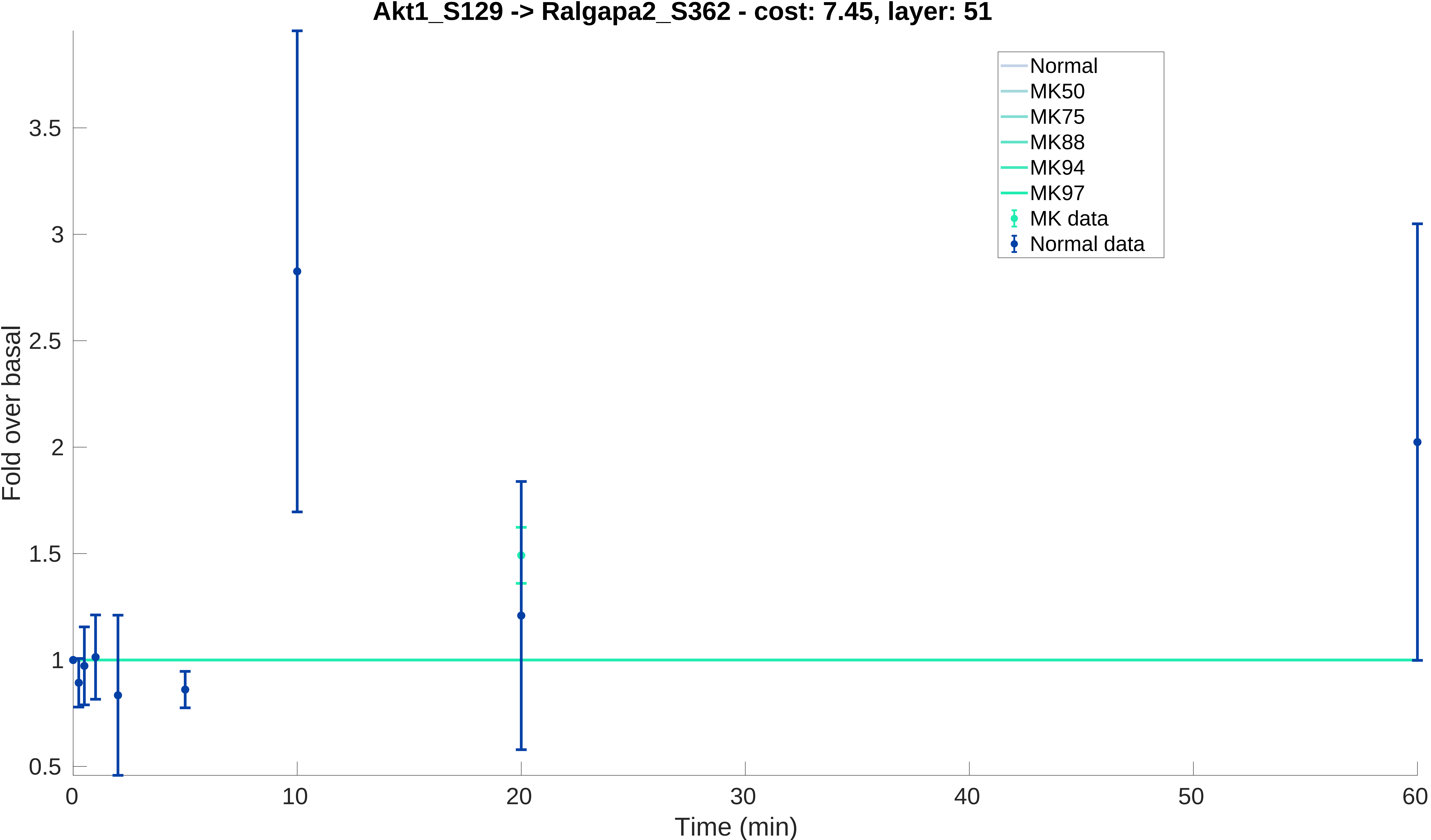

Akt1_S129 -> Ralgapa2_S362 - cost: 7.45, layer: 51
Fold over basal
0.5
1
1.5
2
2.5
3
3.5
0
10
20
30
40
50
60
Time (min)
Normal
MK50
MK75
MK88
MK94
MK97
MK data
Normal data

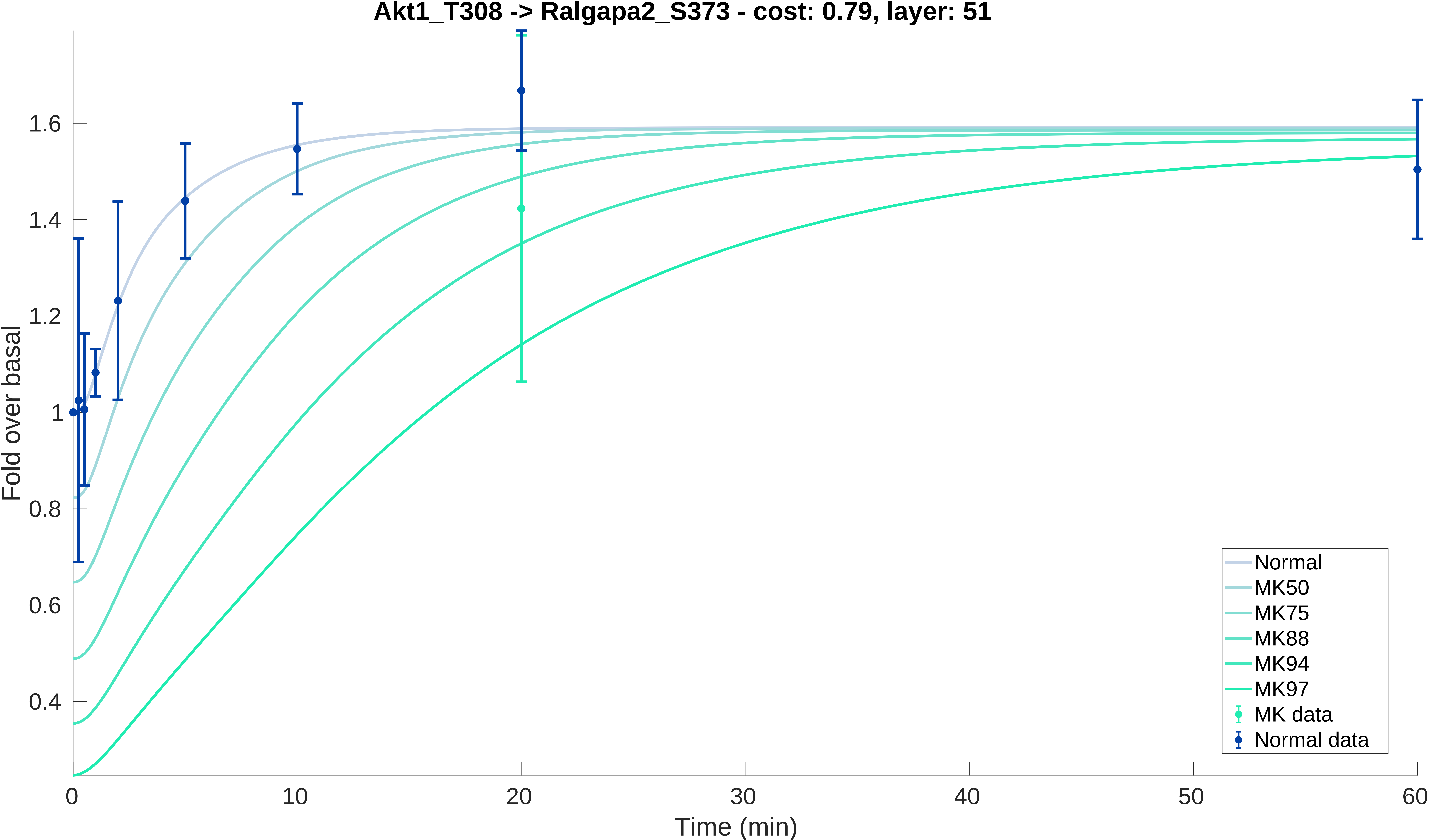
Akt1_T308 -> Ralgapa2_S373 - cost: 0.79, layer: 51
Fold over basal
1.6
1.4
1.2
1
0.8
0.6
0.4
0
10
20
30
40
50
60
Time (min)
Normal
MK50
MK75
MK88
MK94
MK97
MK data
Normal data

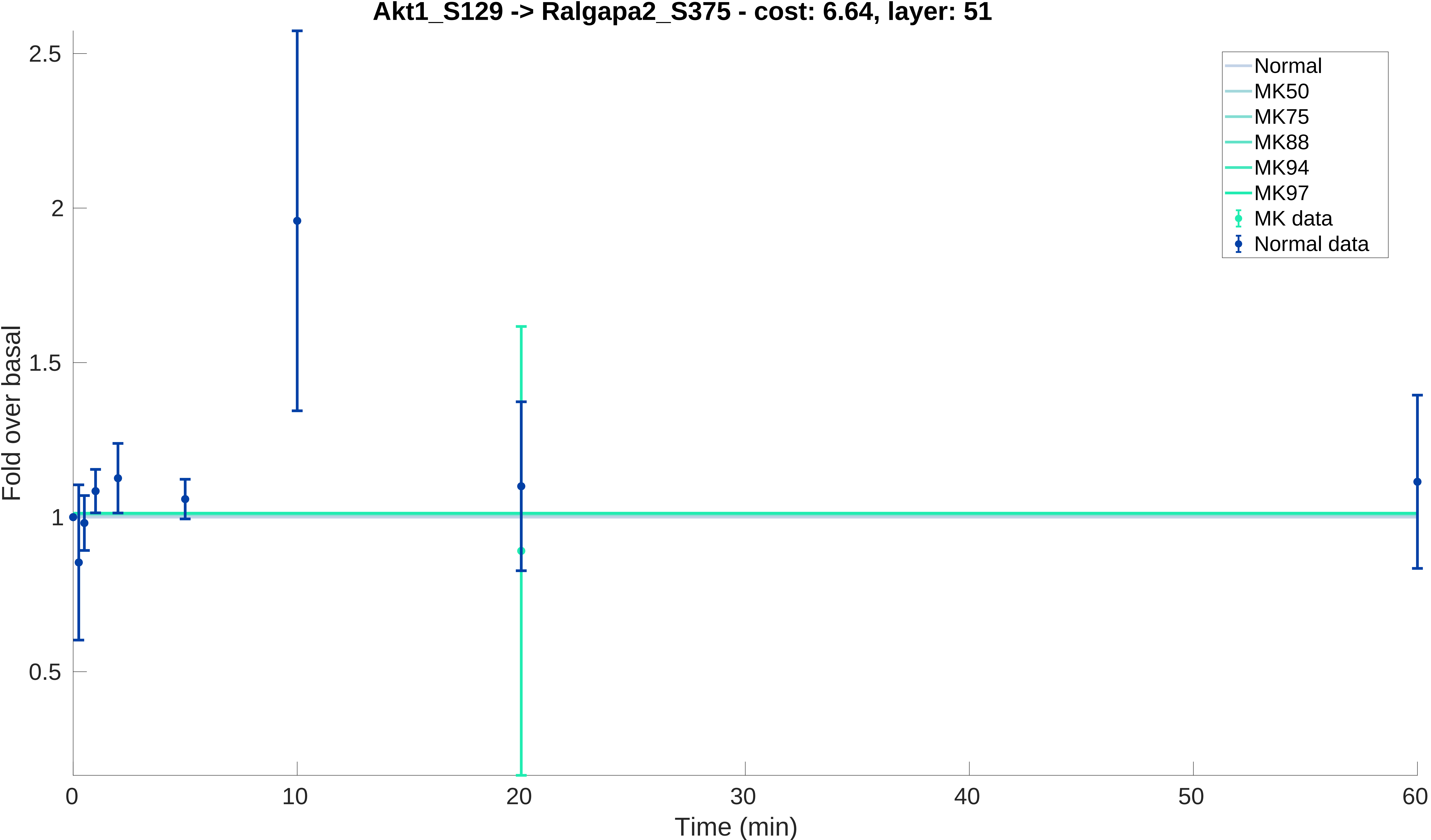

Akt1_S129 -> Ralgapa2_S375 - cost: 6.64, layer: 51
2.5
2
1.5
1
0.5
0
10
20
30
40
50
60
Time (min)
Fold over basal
Normal
MK50
MK75
MK88
MK94
MK97
MK data
Normal data

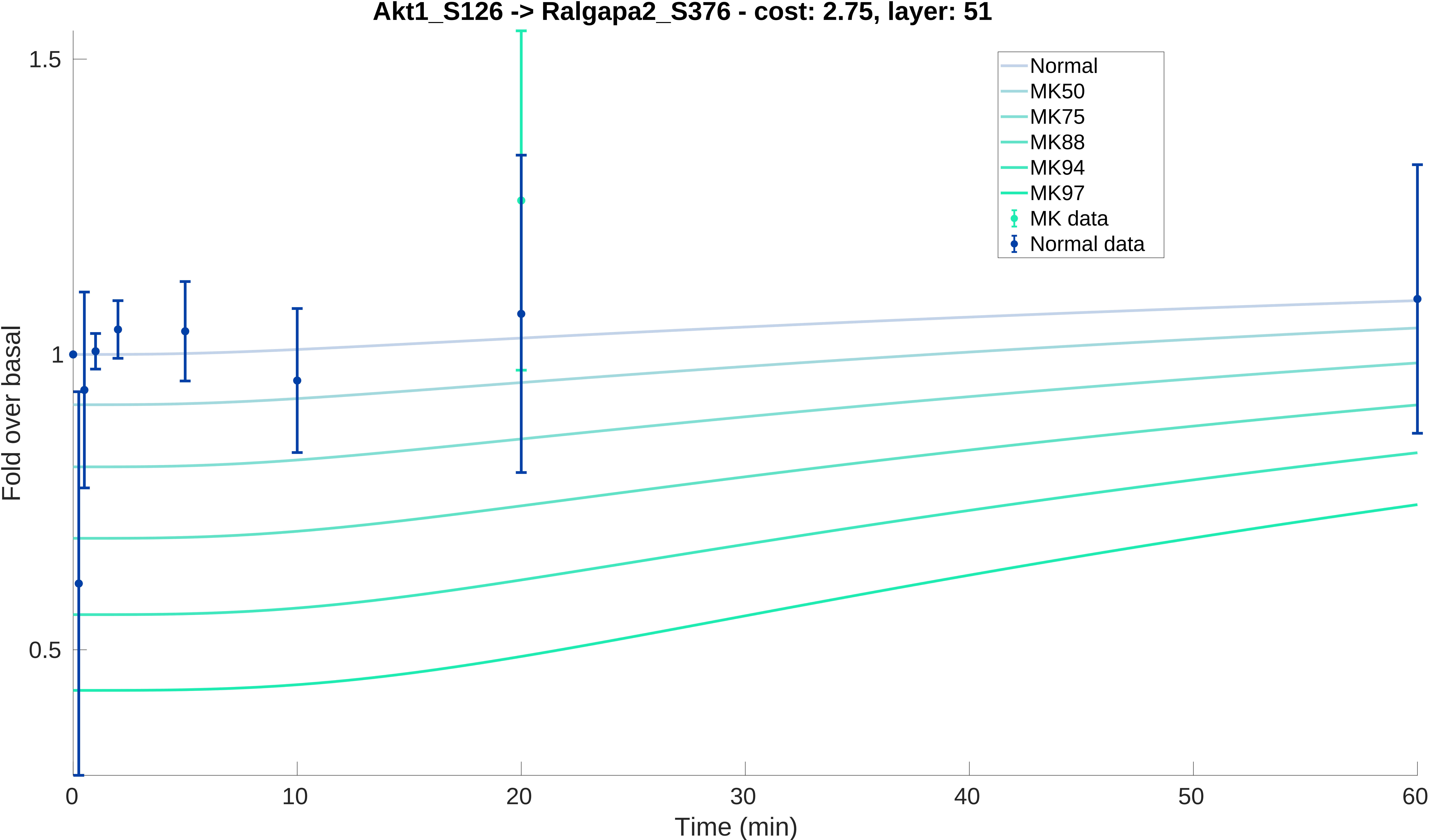

Akt1_S126 -> Ralgapa2_S376 - cost: 2.75, layer: 51
Normal
MK50
MK75
MK88
MK94
MK97
MK data
Normal data
Fold over basal
1.5
1
0.5
0
10
20
30
40
50
60
Time (min)

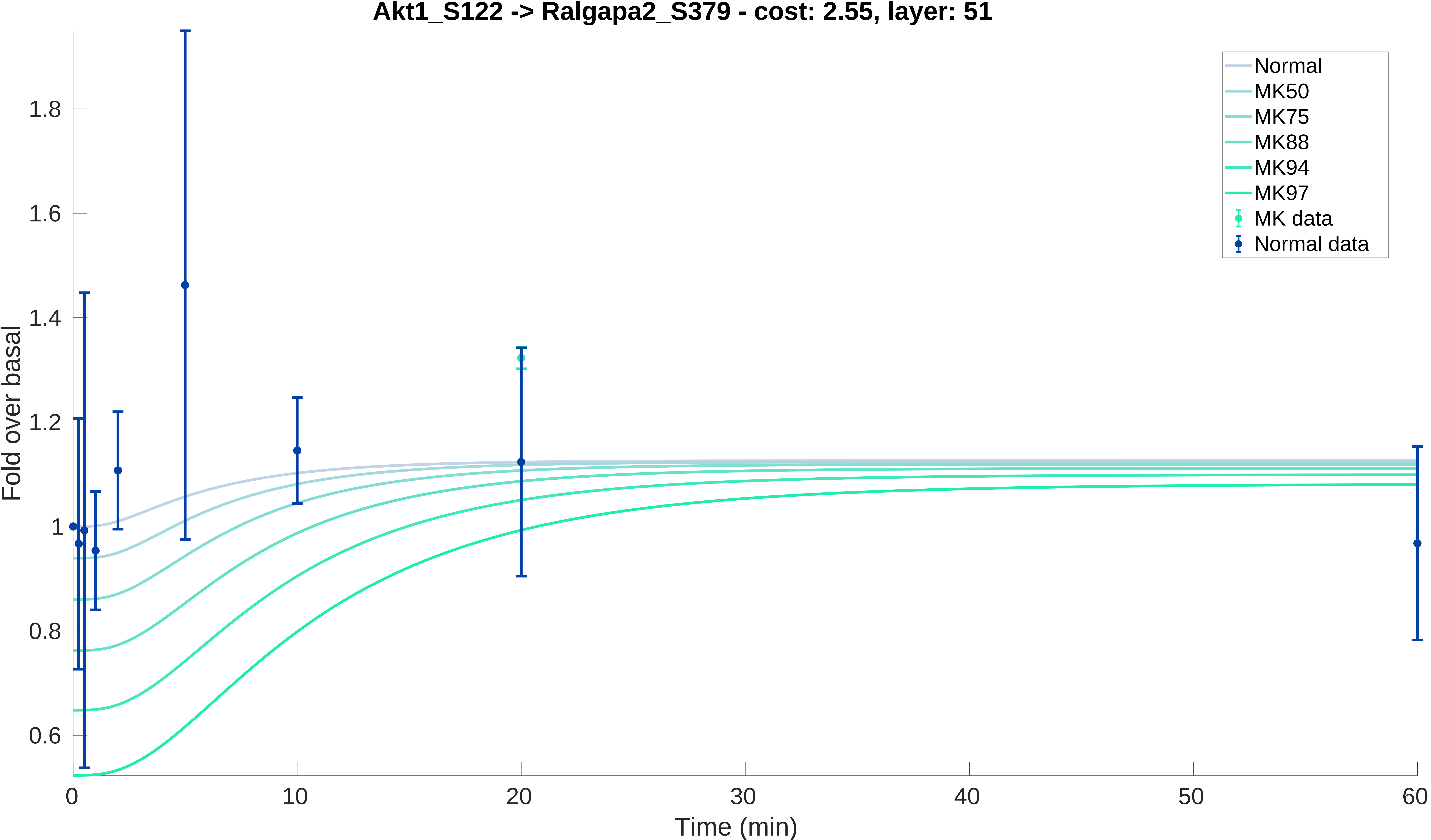

Akt1_S122 -> Ralgapa2_S379 - cost: 2.55, layer: 51
Normal
MK50
MK75
MK88
MK94
MK97
MK data
Normal data
Fold over basal
1.8
1.6
1.4
1.2
1
0.8
0.6
0
10
20
30
40
50
60
Time (min)

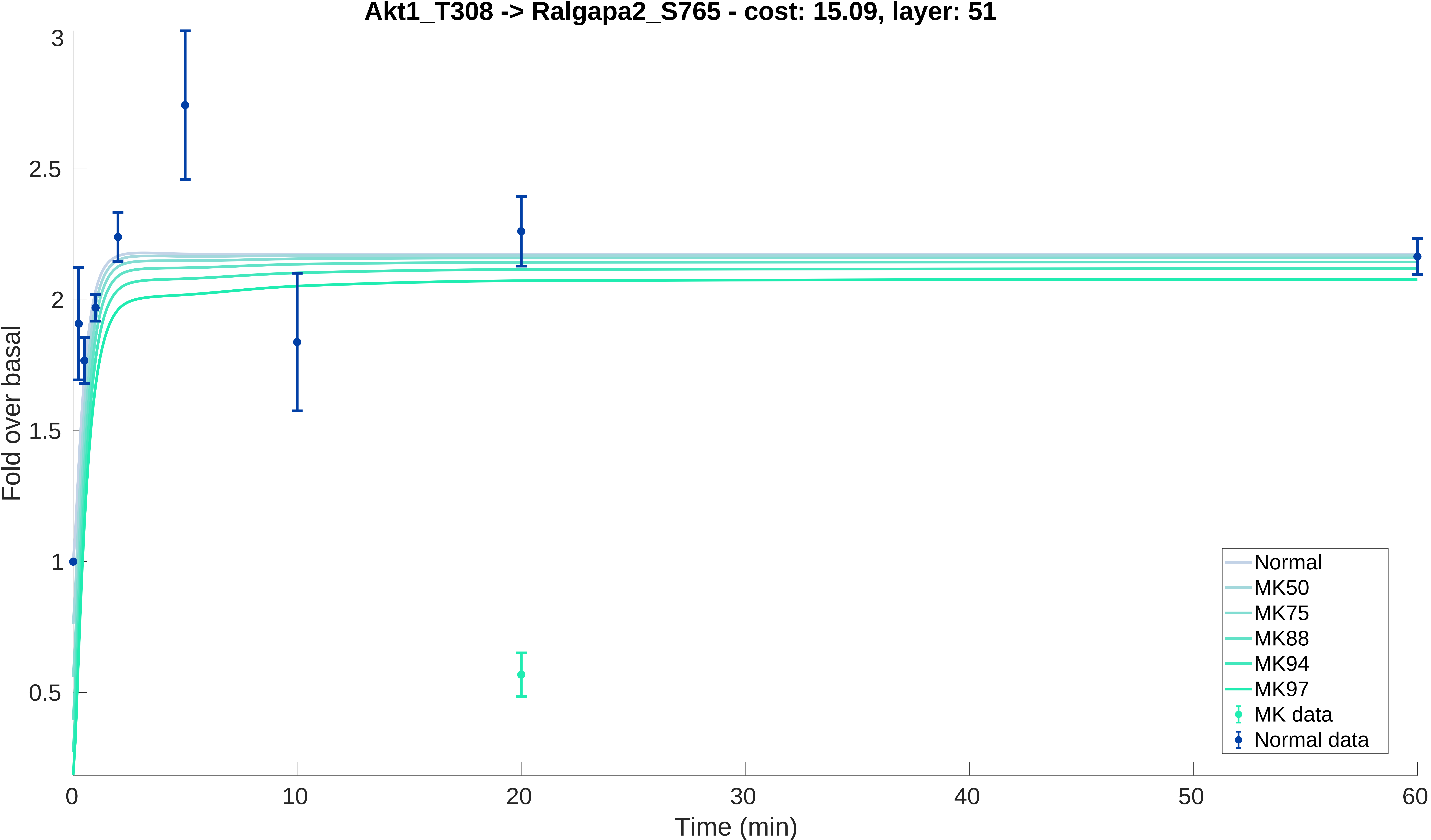

Akt1_T308 -> Ralgapa2_S765 - cost: 15.09, layer: 51
Fold over basal
Time (min)
0
10
20
30
40
50
60
0.5
1
1.5
2
2.5
3
Normal
MK50
MK75
MK88
MK94
MK97
MK data
Normal data

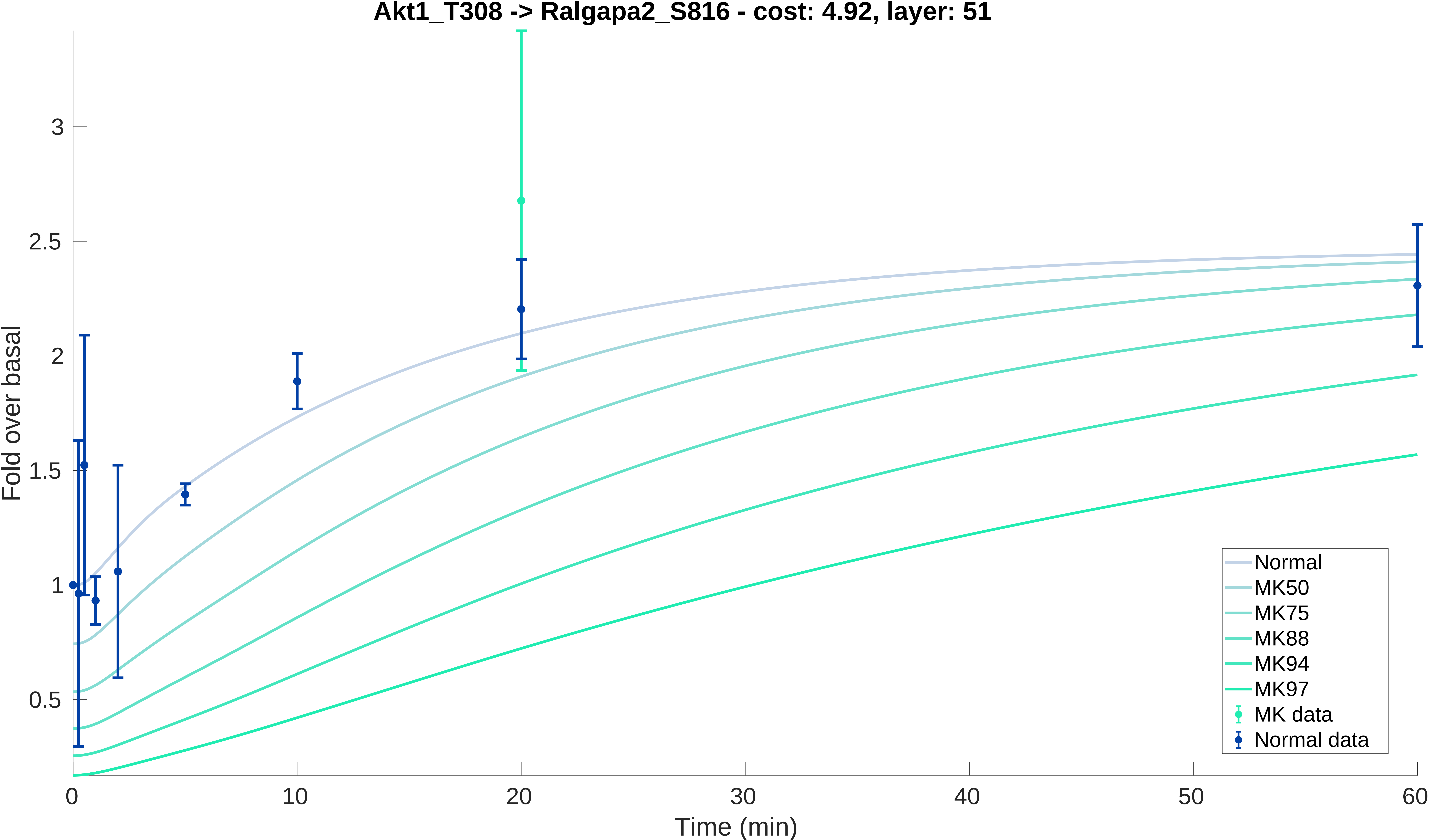

Akt1_T308 -> Ralgapa2_S816 - cost: 4.92, layer: 51
Fold over basal
Time (min)
3
2.5
2
1.5
1
0.5
0
10
20
30
40
50
60
Normal
MK50
MK75
MK88
MK94
MK97
MK data
Normal data

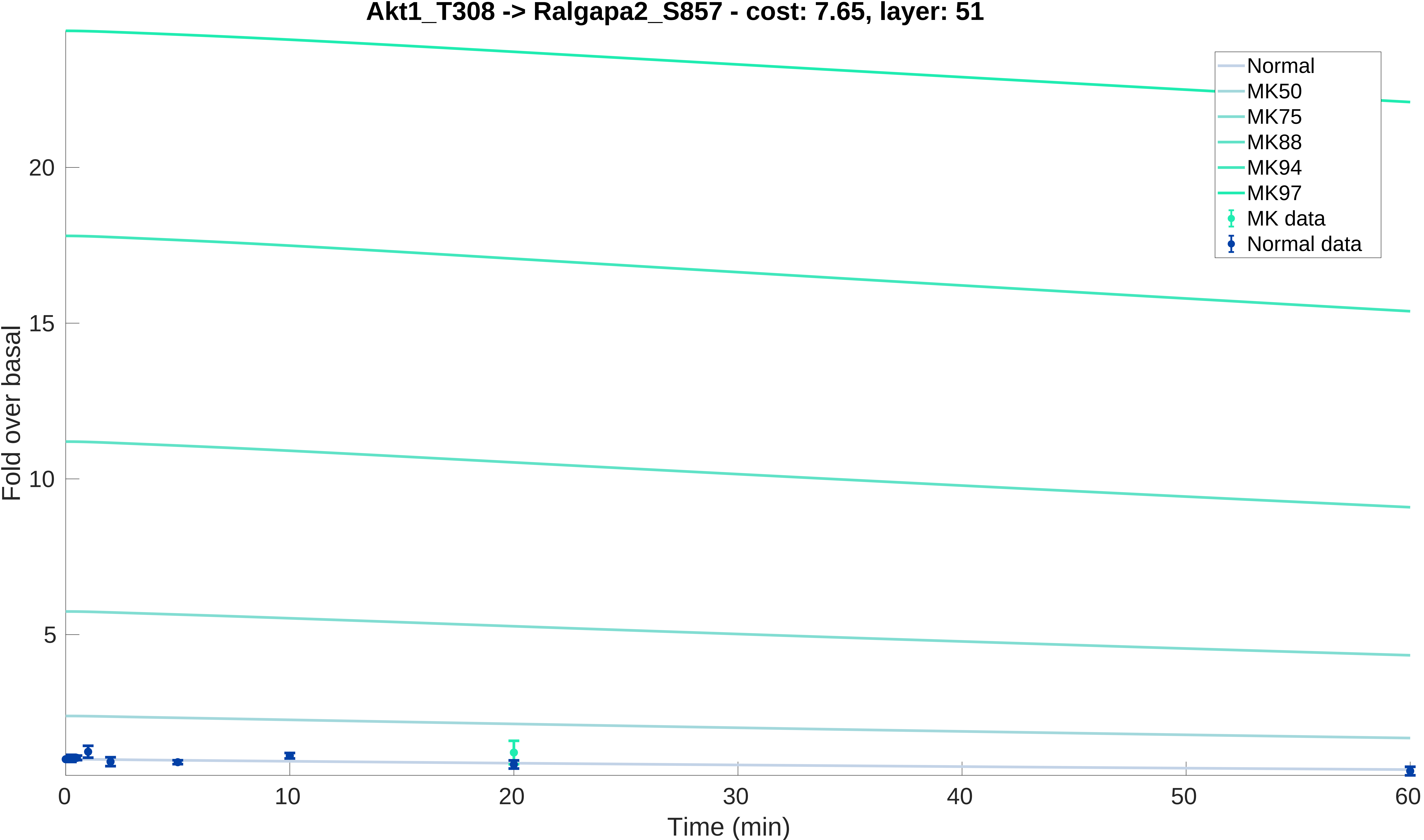

Akt1_T308 -> Ralgapa2_S857 - cost: 7.65, layer: 51
Normal
MK50
MK75
MK88
MK94
MK97
MK data
Normal data
Fold over basal
20
15
10
5
0
10
20
30
40
50
60
Time (min)

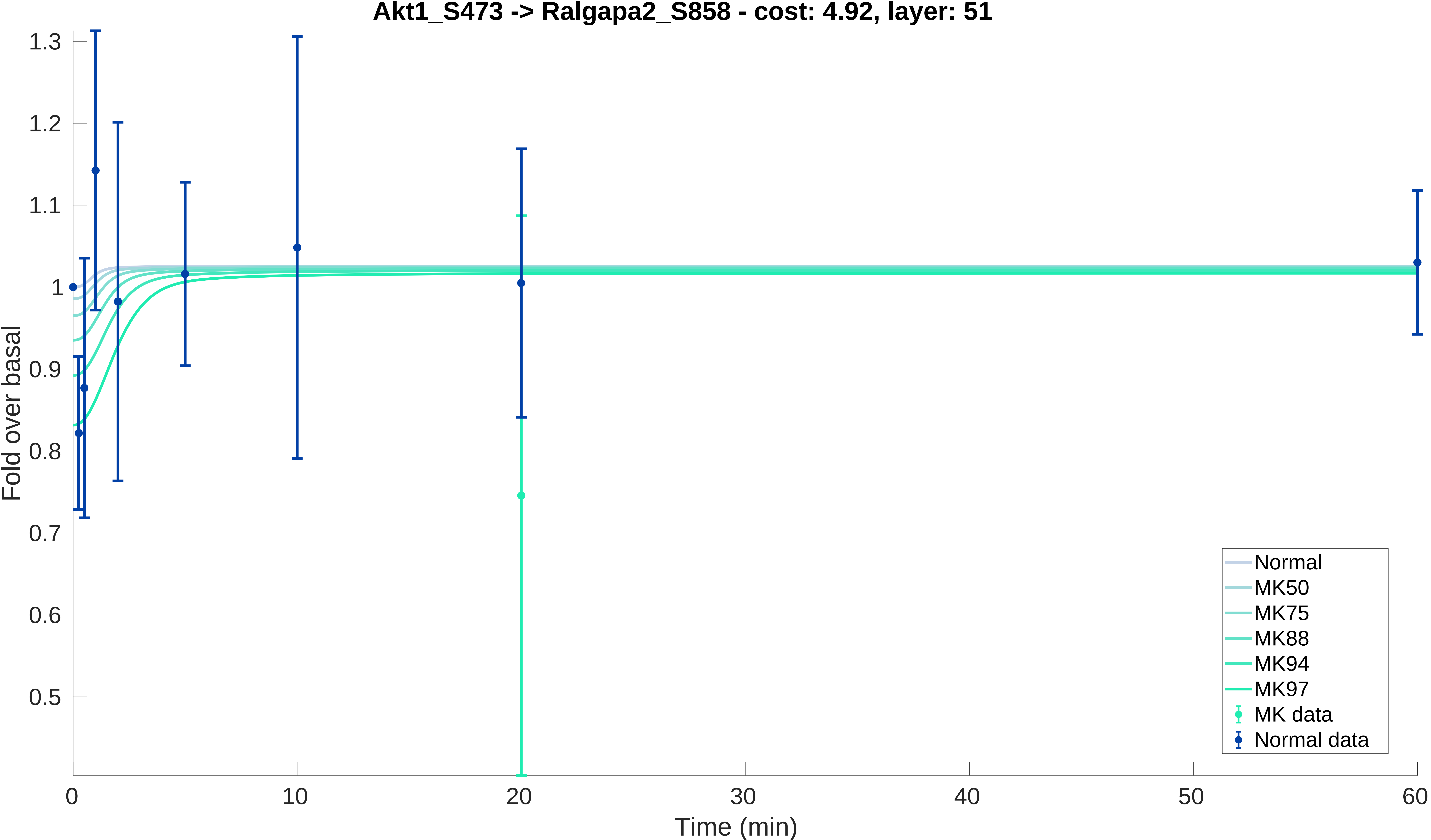

Akt1_S473 -> Ralgapa2_S858 - cost: 4.92, layer: 51
Fold over basal
Time (min)
1.3
1.2
1.1
1
0.9
0.8
0.7
0.6
0.5
0
10
20
30
40
50
60
Normal
MK50
MK75
MK88
MK94
MK97
MK data
Normal data

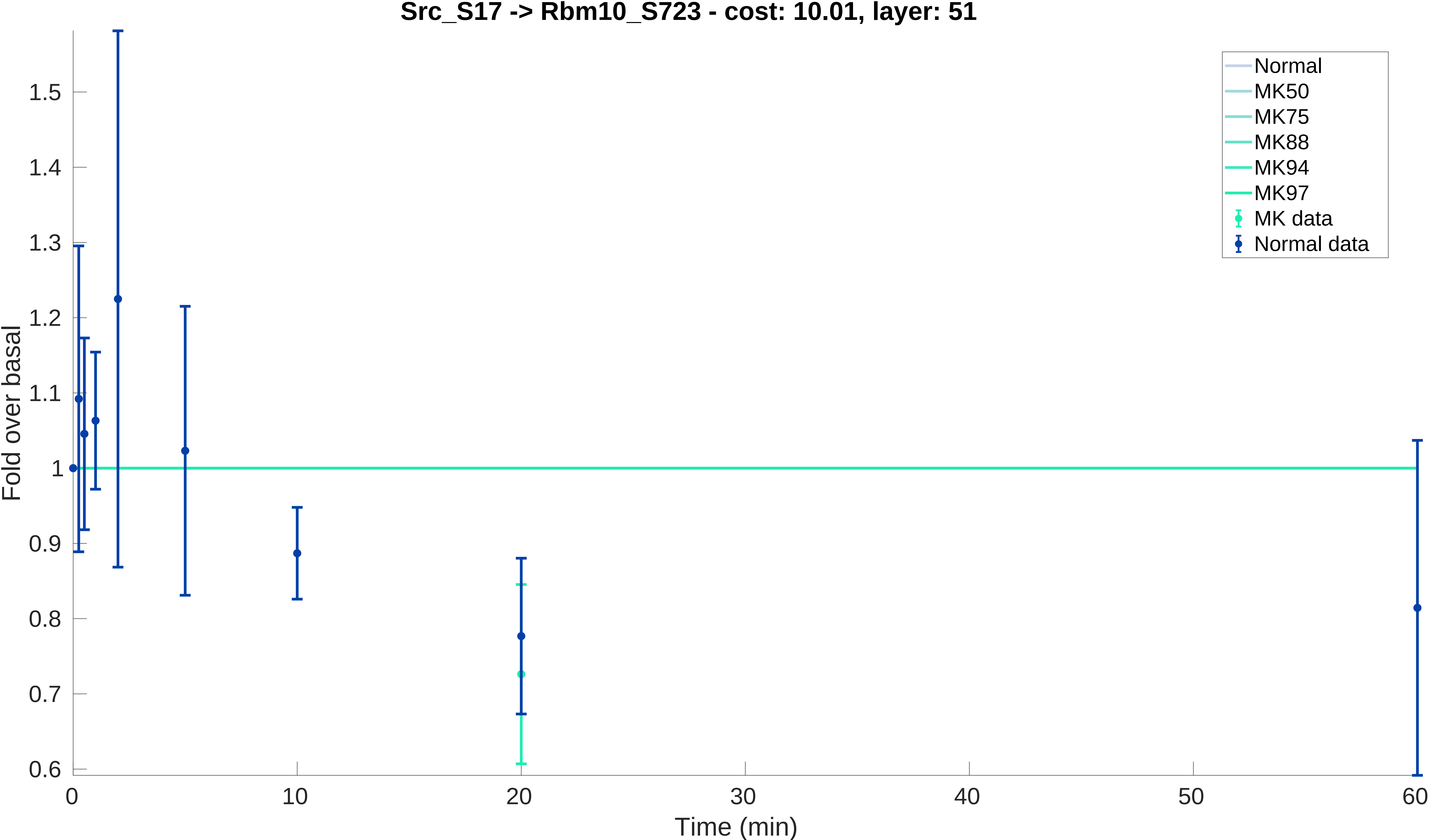
Src_S17 -> Rbm10_S723 - cost: 10.01, layer: 51
Fold over basal
Time (min)
1.5
1.4
1.3
1.2
1.1
1
0.9
0.8
0.7
0.6
0
10
20
30
40
50
60
Normal
MK50
MK75
MK88
MK94
MK97
MK data
Normal data

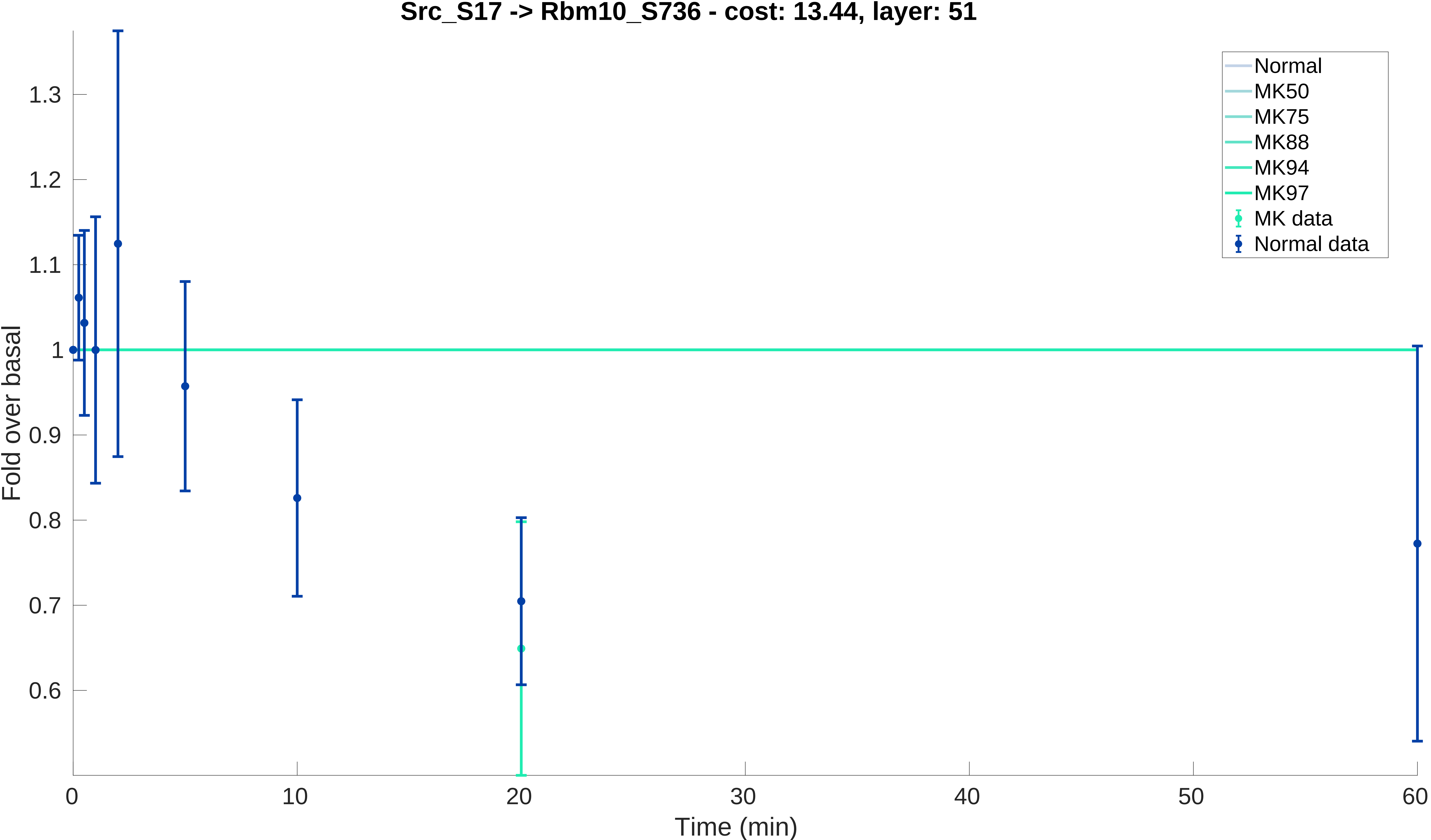

Src_S17 -> Rbm10_S736 - cost: 13.44, layer: 51
Normal
MK50
MK75
MK88
MK94
MK97
MK data
Normal data
Fold over basal
Time (min)
1.3
1.2
1.1
1
0.9
0.8
0.7
0.6
0
10
20
30
40
50
60

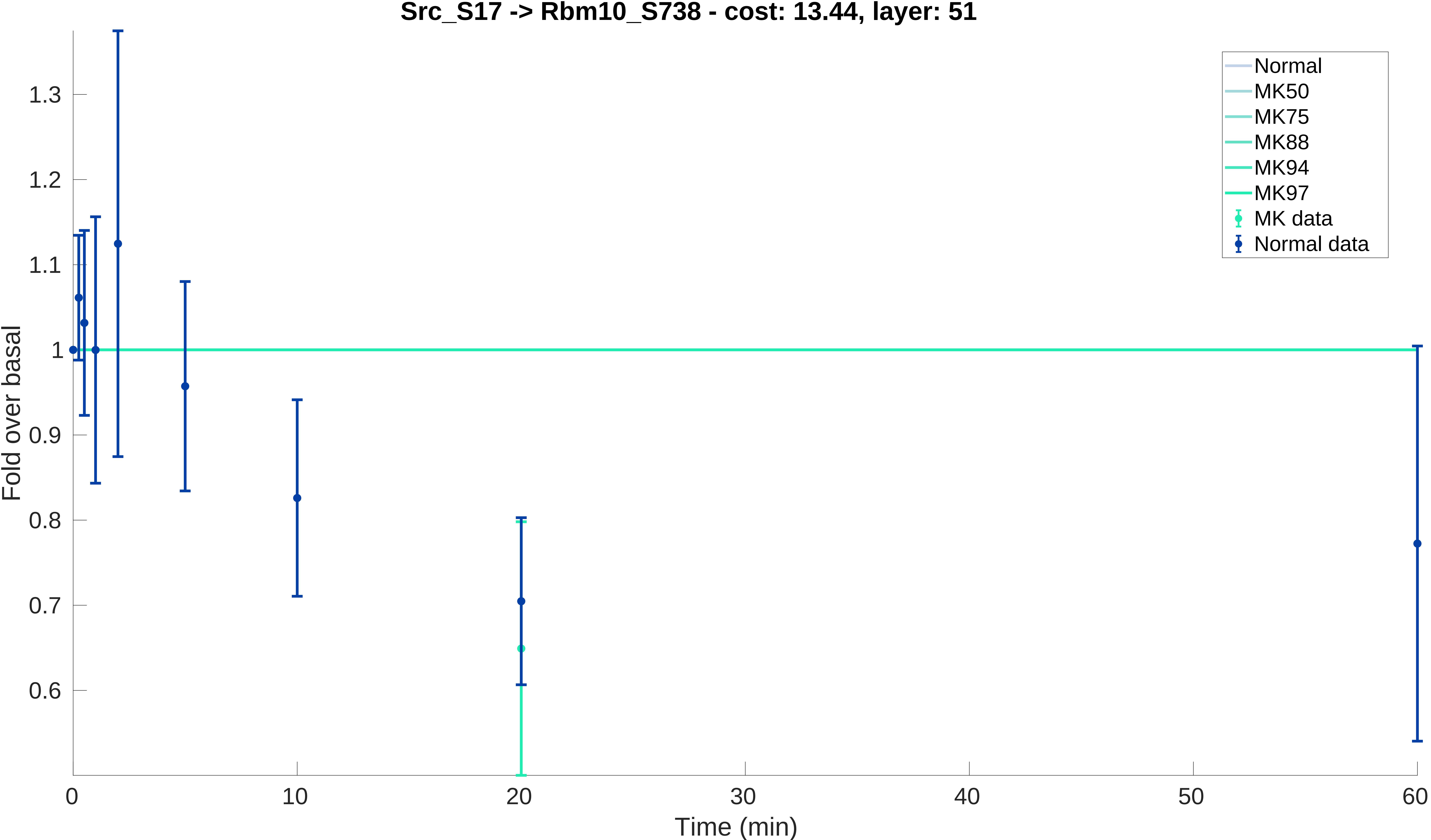

Src_S17 -> Rbm10_S738 - cost: 13.44, layer: 51
Fold over basal
Time (min)
1.3
1.2
1.1
1
0.9
0.8
0.7
0.6
0
10
20
30
40
50
60
Normal
MK50
MK75
MK88
MK94
MK97
MK data
Normal data

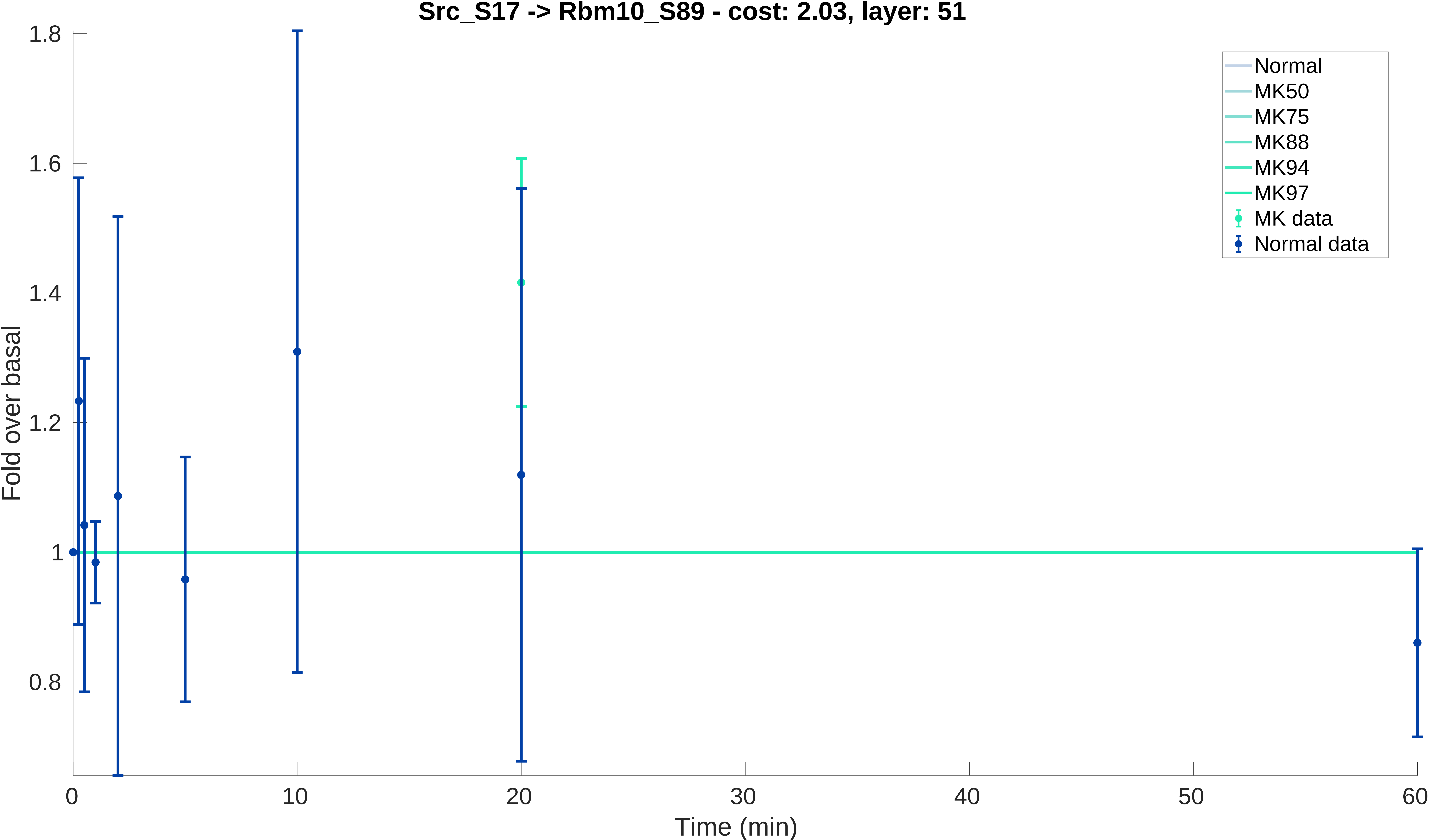

Src_S17 -> Rbm10_S89 - cost: 2.03, layer: 51
Fold over basal
Time (min)
1.8
1.6
1.4
1.2
1
0.8
0
10
20
30
40
50
60
Normal
MK50
MK75
MK88
MK94
MK97
MK data
Normal data

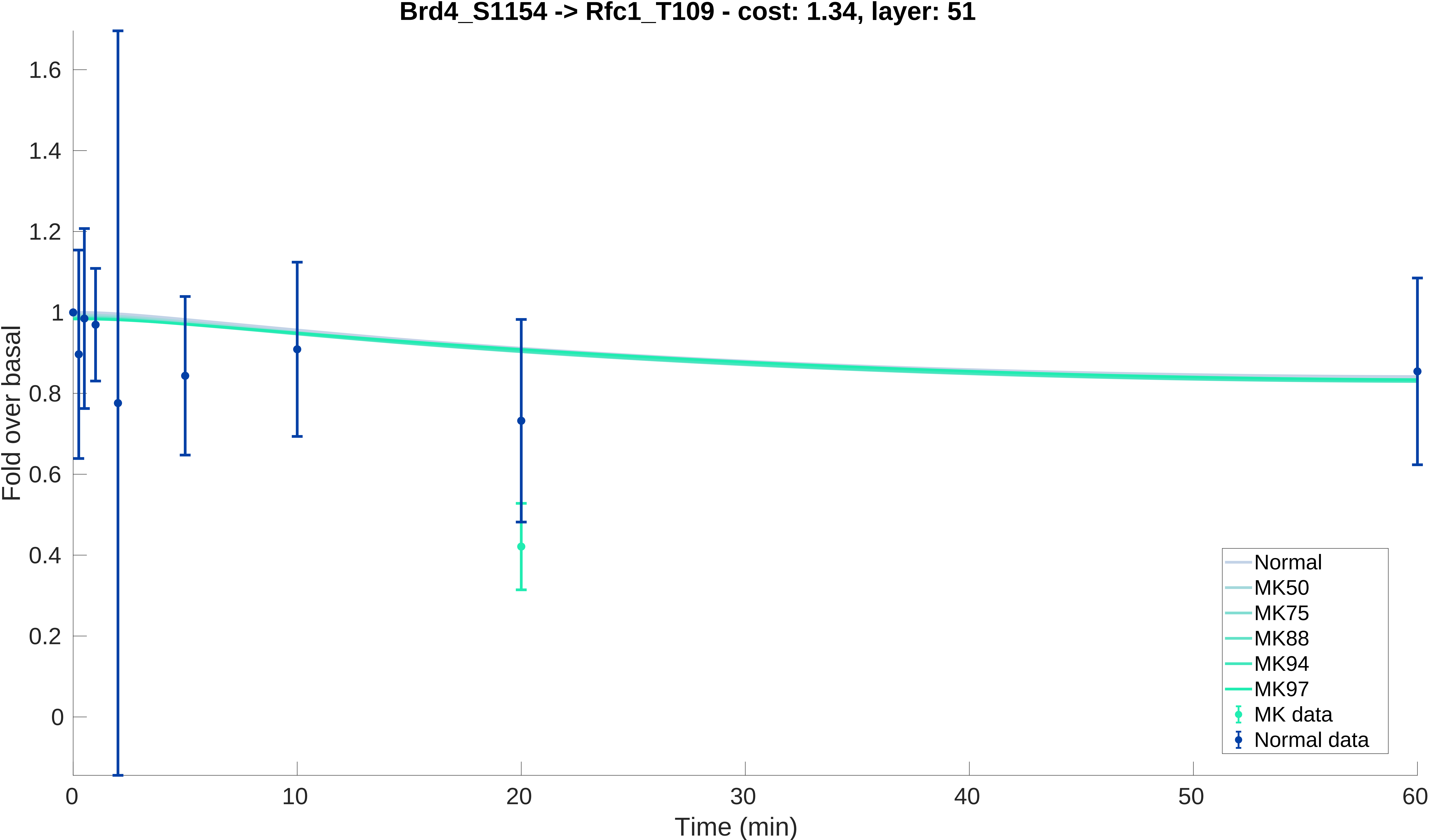

Brd4_S1154 -> Rfc1_T109 - cost: 1.34, layer: 51
Fold over basal
Time (min)
0
0.2
0.4
0.6
0.8
1
1.2
1.4
1.6
0
10
20
30
40
50
60
Normal
MK50
MK75
MK88
MK94
MK97
MK data
Normal data

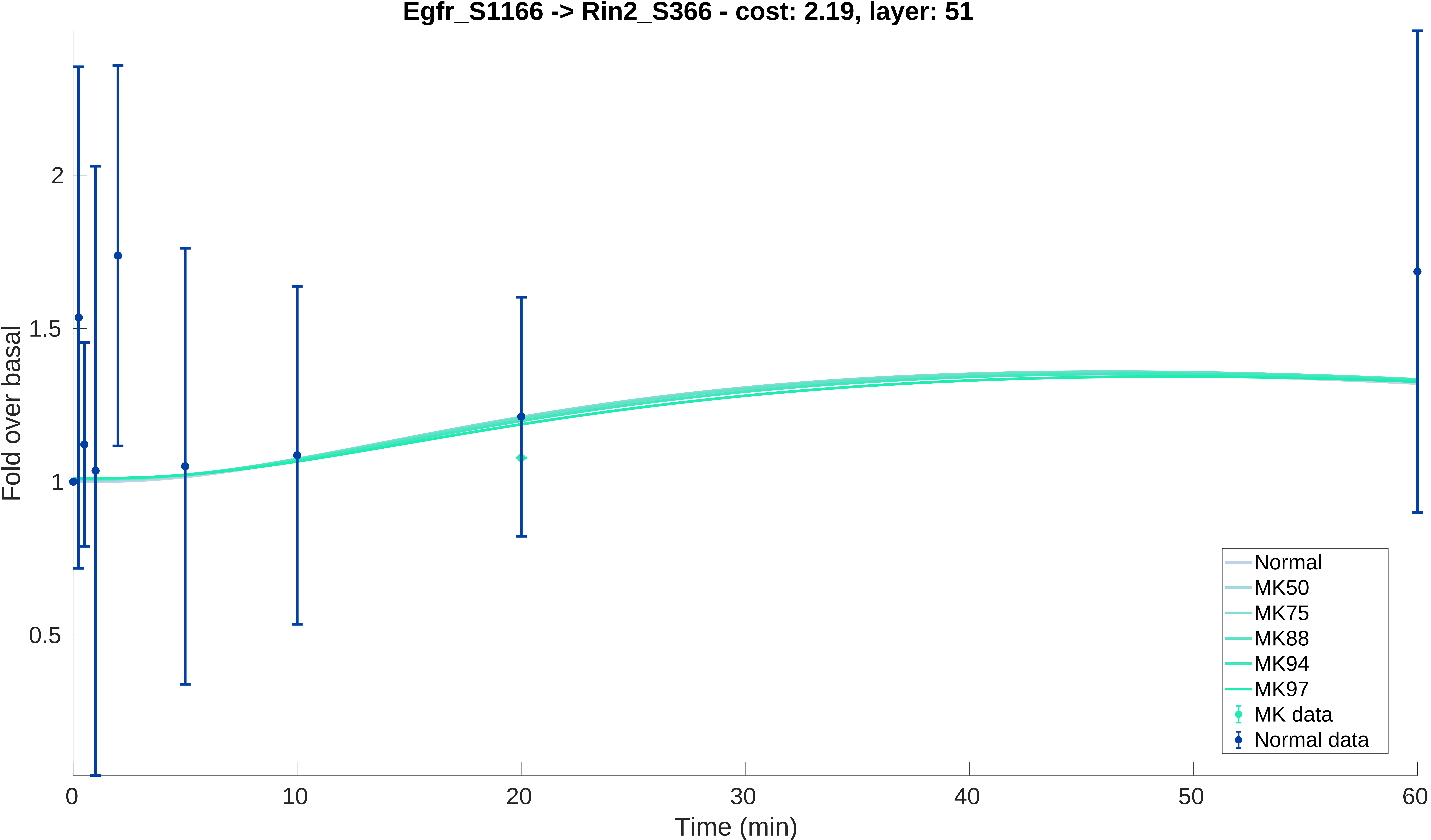

Egfr_S1166 -> Rin2_S366 - cost: 2.19, layer: 51
Fold over basal
0.5
1
1.5
2
0
10
20
30
40
50
60
Time (min)
Normal
MK50
MK75
MK88
MK94
MK97
MK data
Normal data

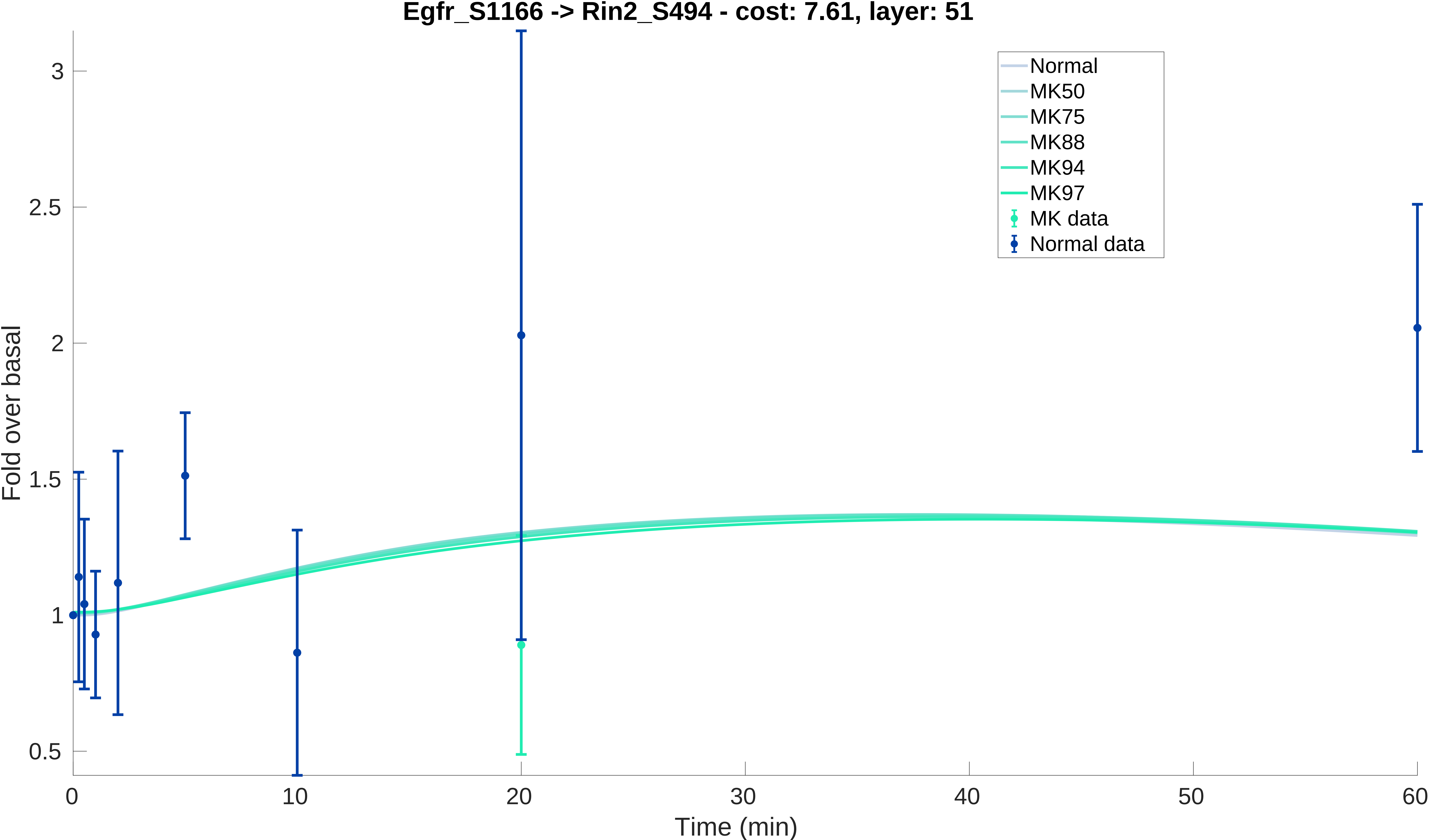
Egfr_S1166 -> Rin2_S494 - cost: 7.61, layer: 51
Fold over basal
Time (min)
0.5
1
1.5
2
2.5
3
0
10
20
30
40
50
60
Normal
MK50
MK75
MK88
MK94
MK97
MK data
Normal data

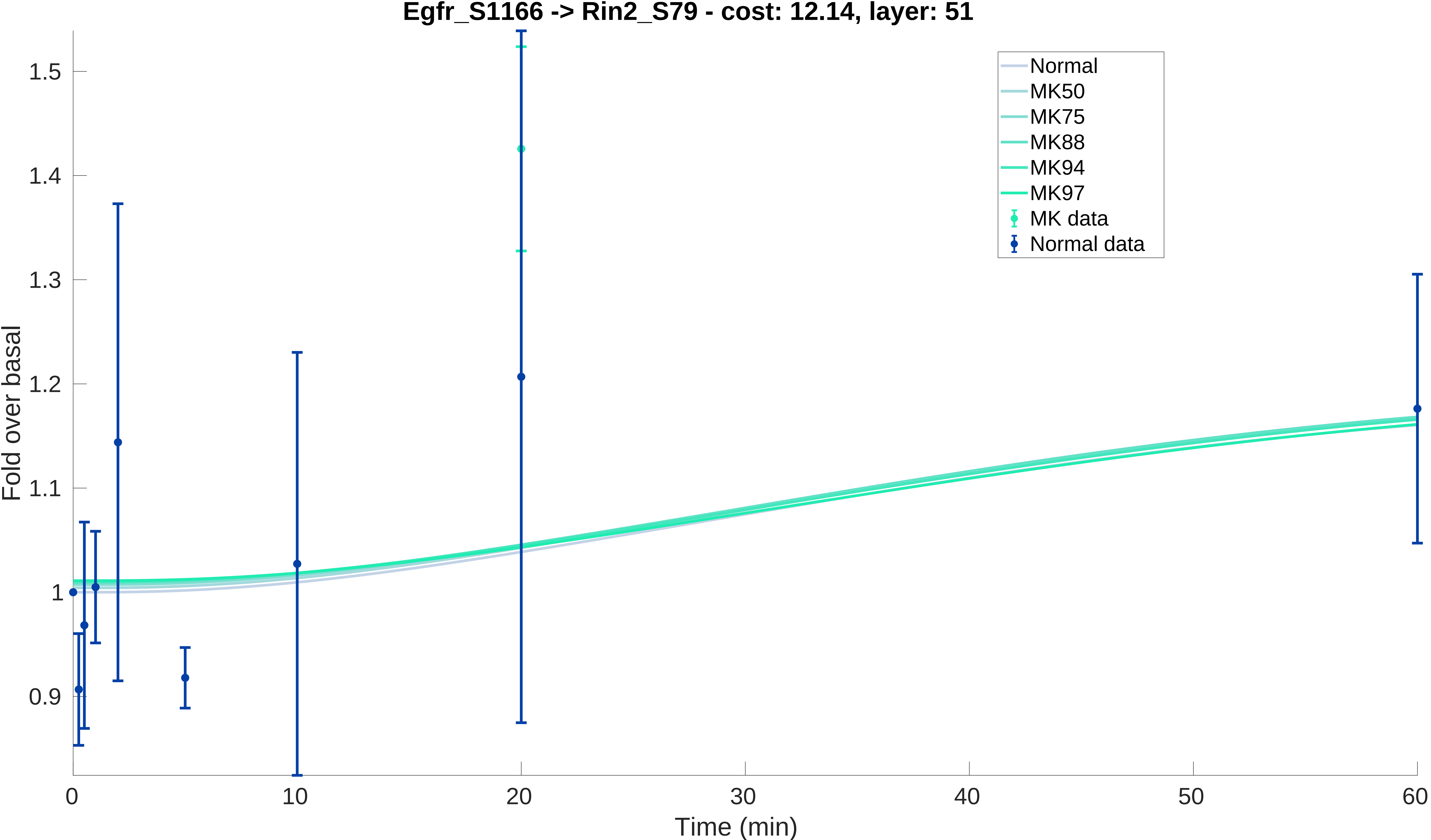

Egfr_S1166 -> Rin2_S79 - cost: 12.14, layer: 51
Fold over basal
Time (min)
1.5
1.4
1.3
1.2
1.1
1
0.9
0
10
20
30
40
50
60
Normal
MK50
MK75
MK88
MK94
MK97
MK data
Normal data

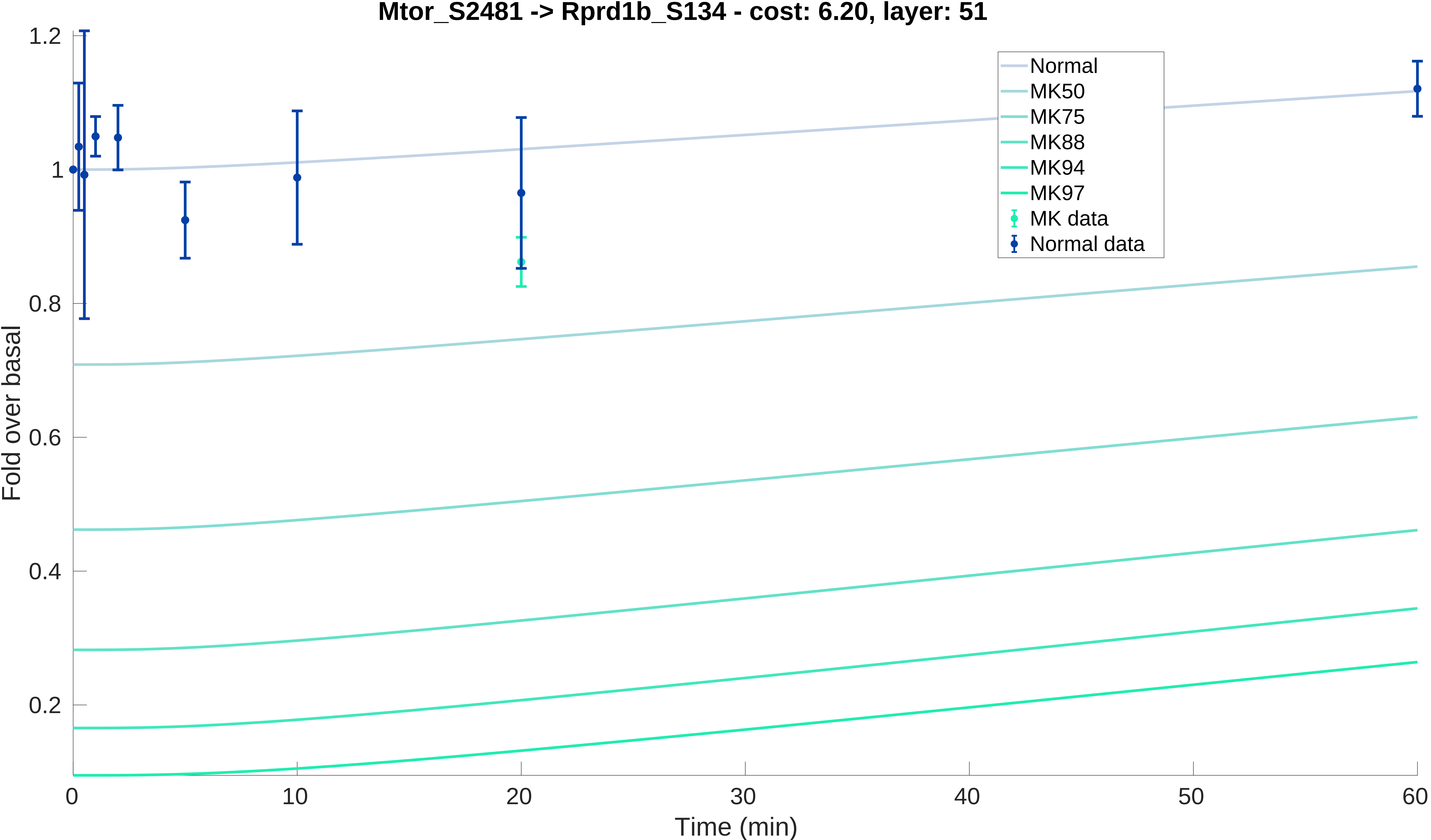

Mtor_S2481 -> Rprd1b_S134 - cost: 6.20, layer: 51
Fold over basal
Time (min)
Normal
MK50
MK75
MK88
MK94
MK97
MK data
Normal data
1.2
1
0.8
0.6
0.4
0.2
0
10
20
30
40
50
60

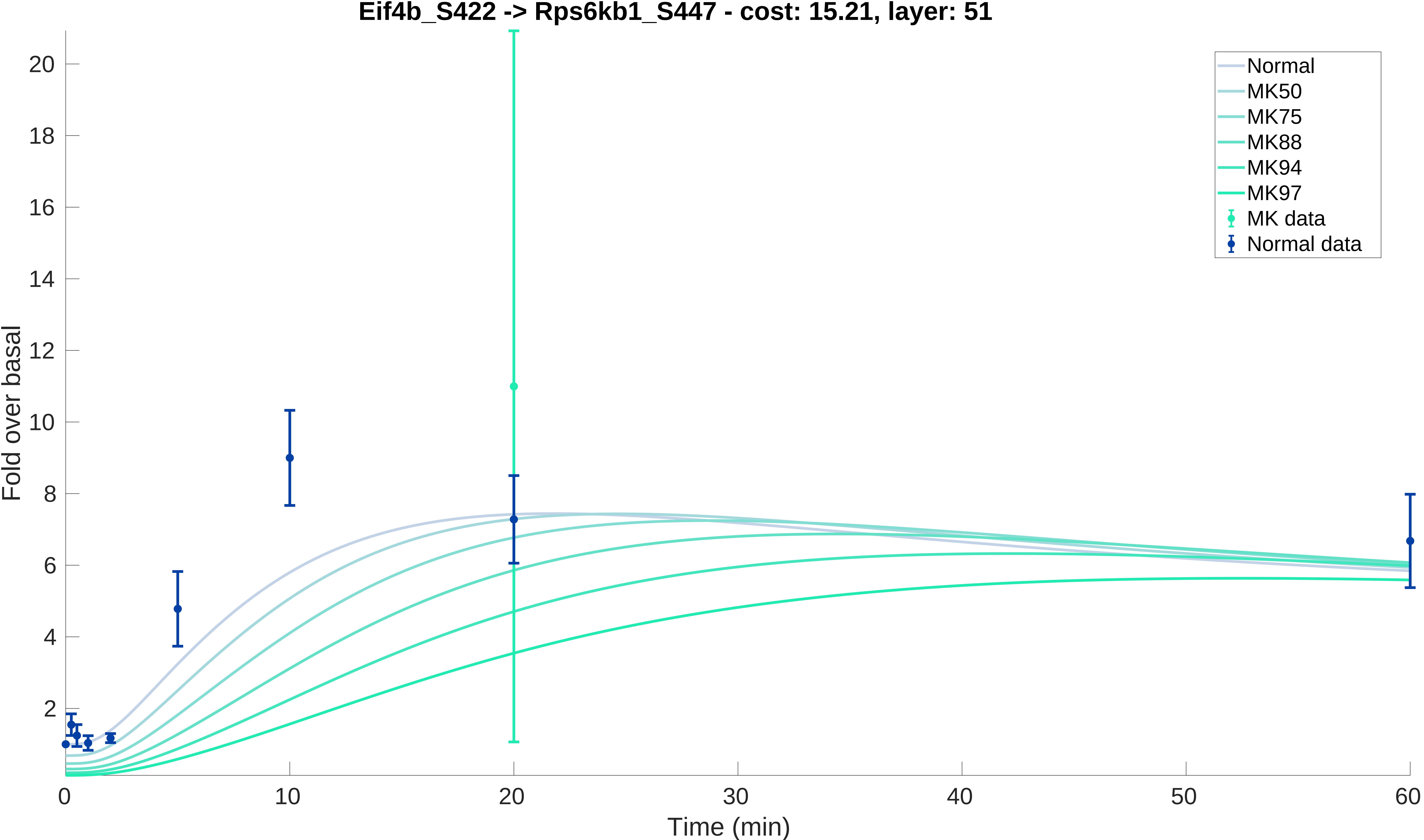

Eif4b_S422 -> Rps6kb1_S447 - cost: 15.21, layer: 51
Fold over basal
Time (min)
0
10
20
30
40
50
60
2
4
6
8
10
12
14
16
18
20
Normal
MK50
MK75
MK88
MK94
MK97
MK data
Normal data

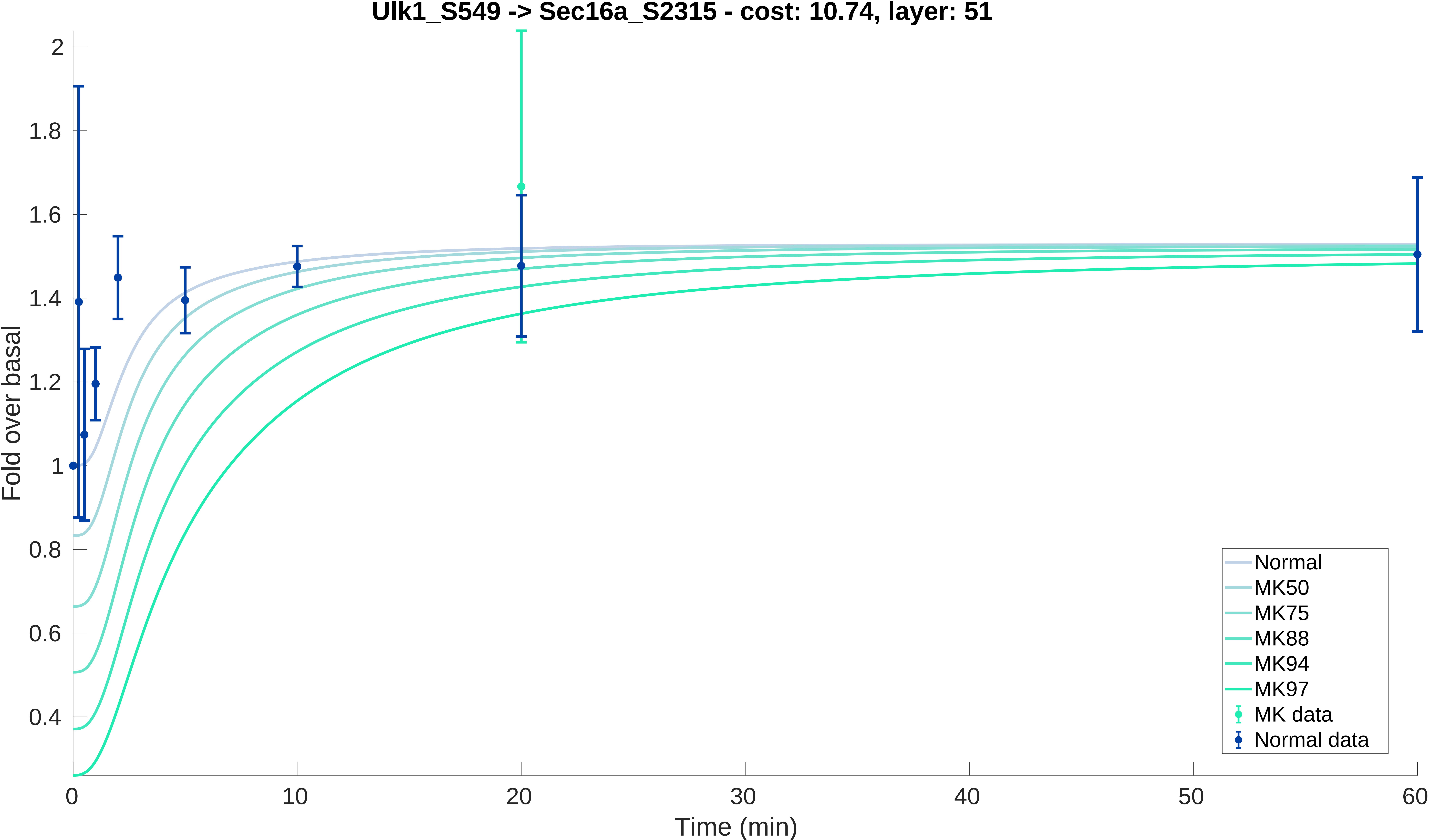

Ulk1_S549 -> Sec16a_S2315 - cost: 10.74, layer: 51
Fold over basal
Time (min)
2
1.8
1.6
1.4
1.2
1
0.8
0.6
0.4
0
10
20
30
40
50
60
Normal
MK50
MK75
MK88
MK94
MK97
MK data
Normal data

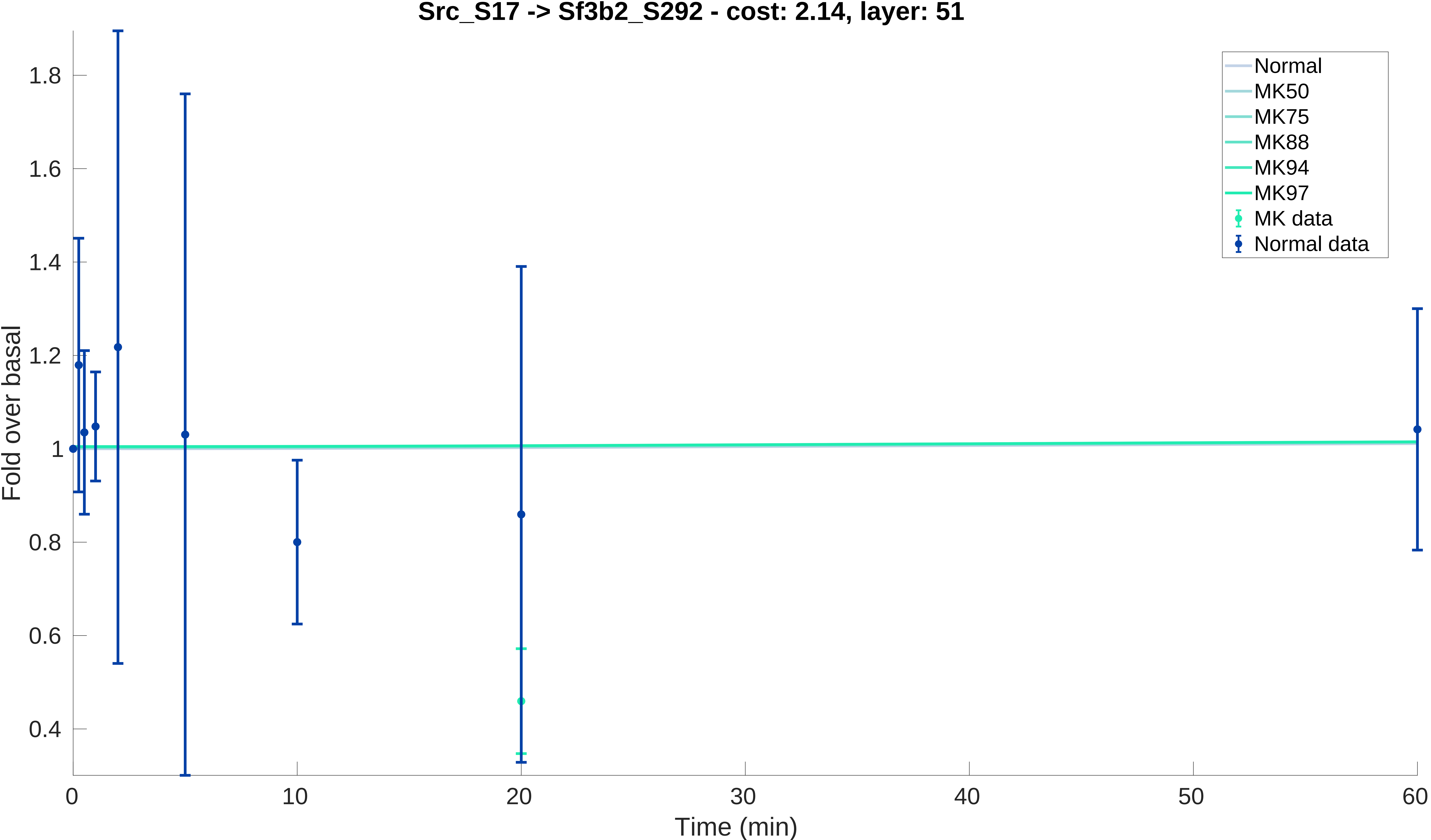

Src_S17 -> Sf3b2_S292 - cost: 2.14, layer: 51
Normal
MK50
MK75
MK88
MK94
MK97
MK data
Normal data
Fold over basal
1.8
1.6
1.4
1.2
1
0.8
0.6
0.4
0
10
20
30
40
50
60
Time (min)

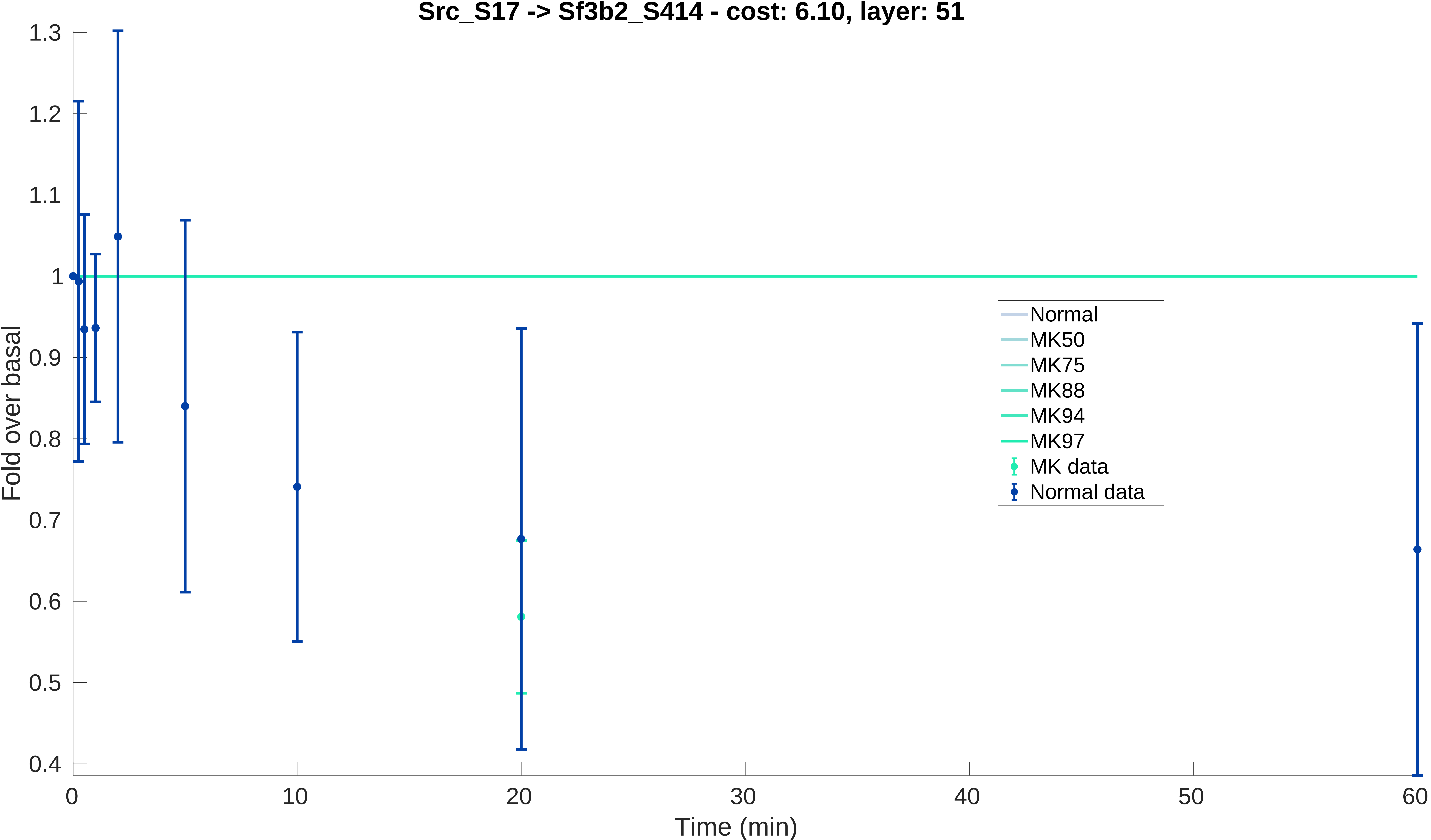

Src_S17 -> Sf3b2_S414 - cost: 6.10, layer: 51
Fold over basal
Time (min)
Normal
MK50
MK75
MK88
MK94
MK97
MK data
Normal data
1.3
1.2
1.1
1
0.9
0.8
0.7
0.6
0.5
0.4
0
10
20
30
40
50
60

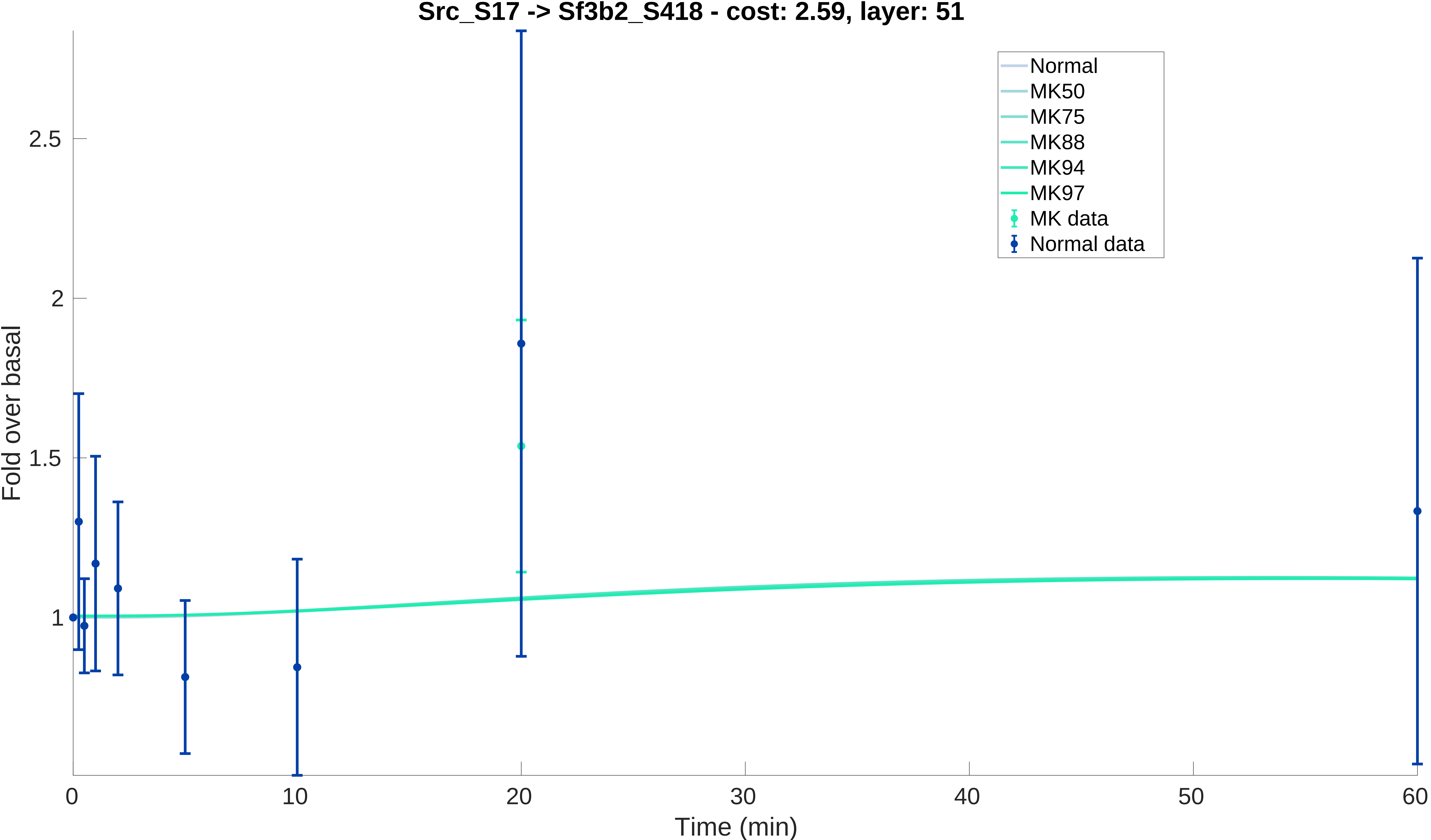

Src_S17 -> Sf3b2_S418 - cost: 2.59, layer: 51
Fold over basal
2.5
2
1.5
1
Time (min)
0
10
20
30
40
50
60
Normal
MK50
MK75
MK88
MK94
MK97
MK data
Normal data

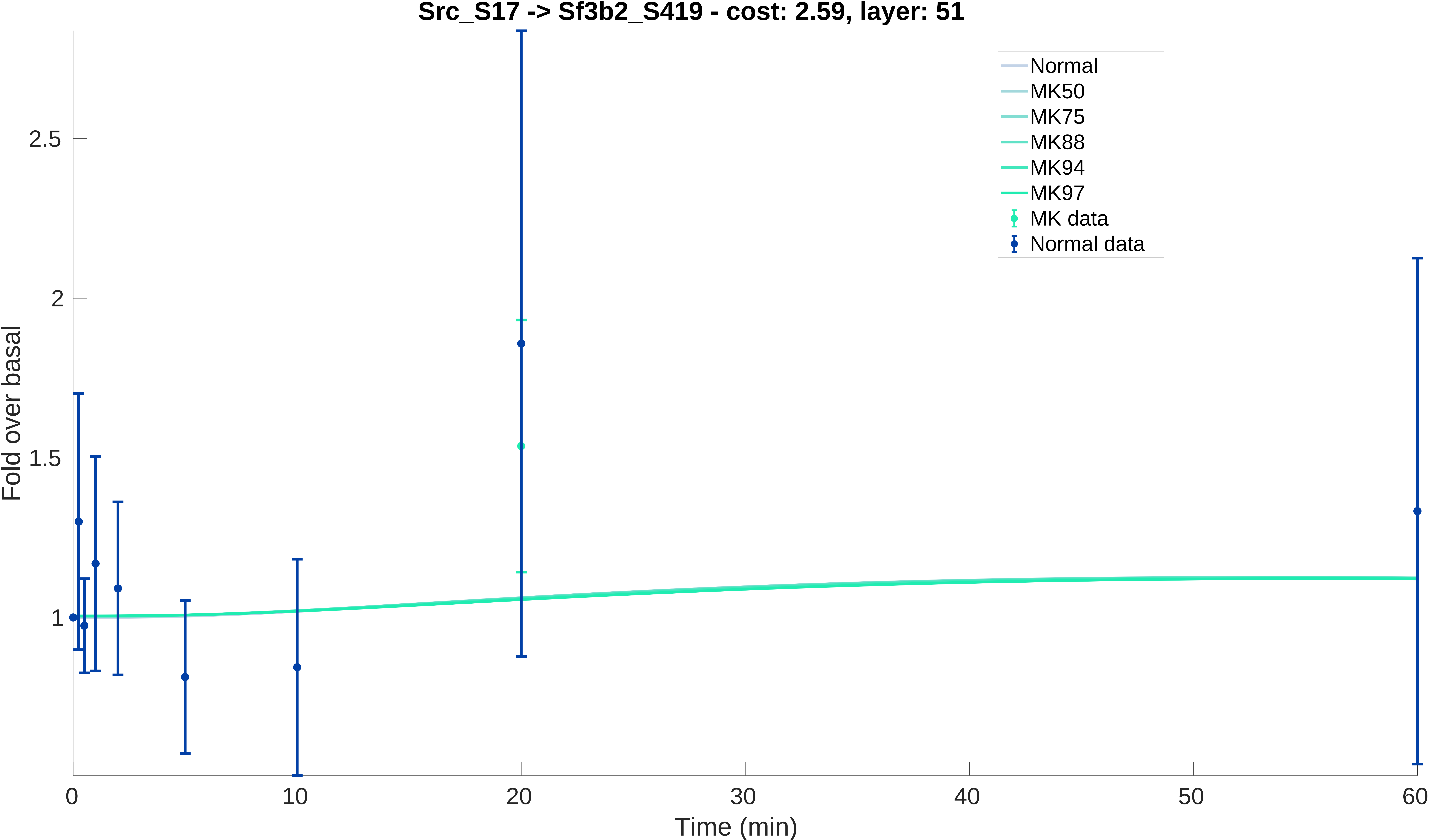
Src_S17 -> Sf3b2_S419 - cost: 2.59, layer: 51
Normal
MK50
MK75
MK88
MK94
MK97
MK data
Normal data
Fold over basal
2.5
2
1.5
1
0
10
20
30
40
50
60
Time (min)

#### Src_S17 -> Sf3b2_S844 - cost: 15.21, layer: 51

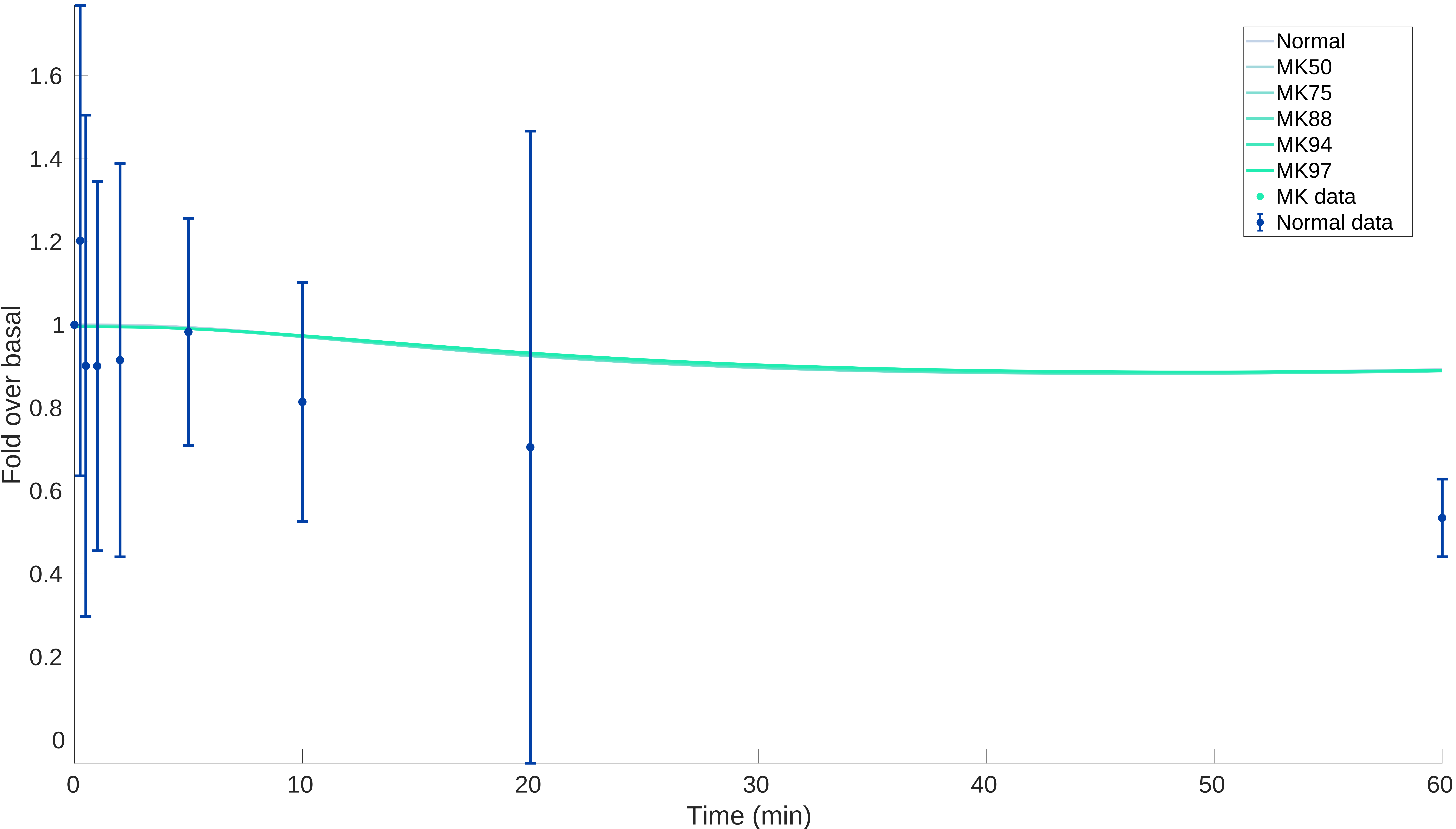

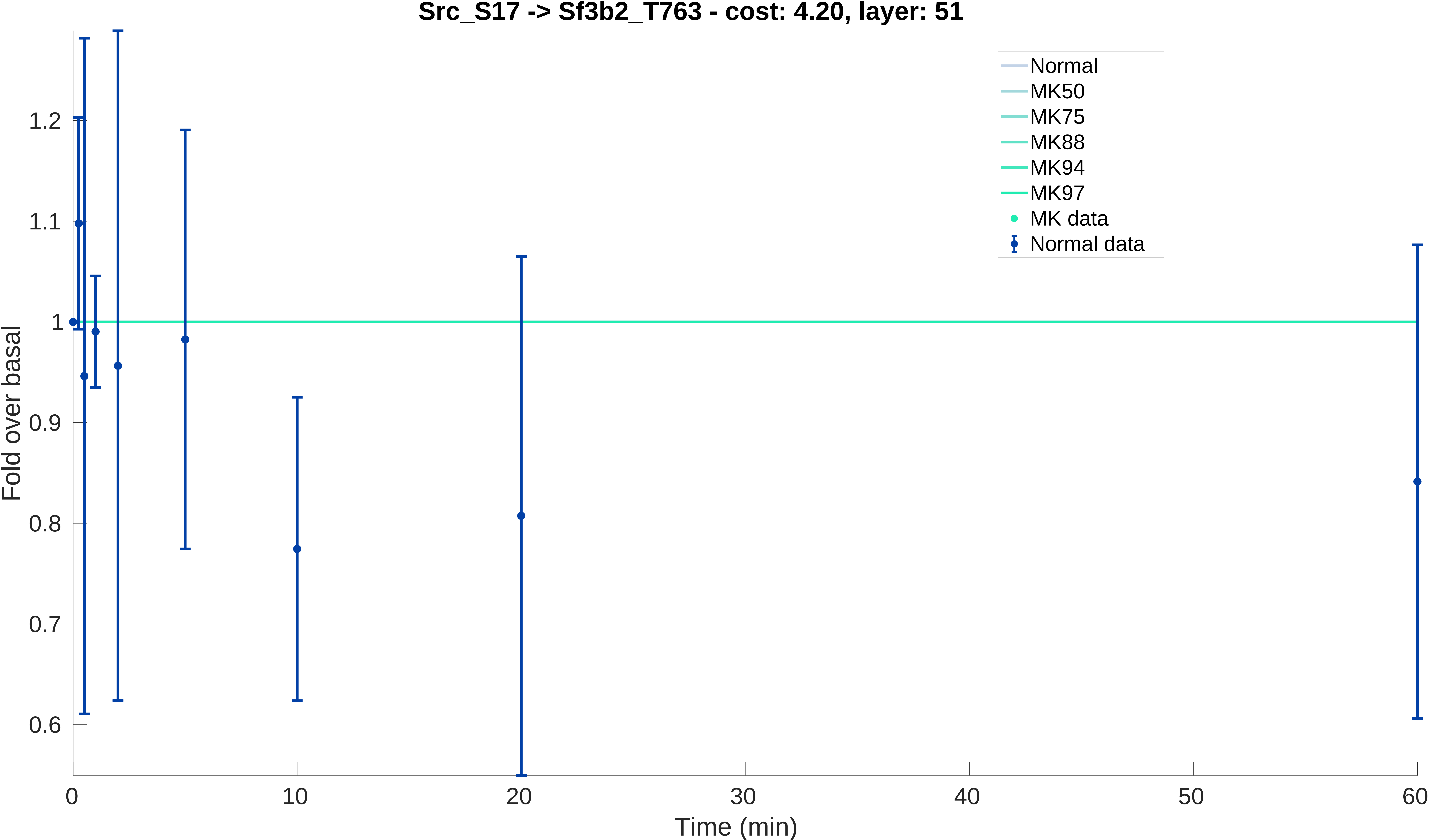

Src_S17 -> Sf3b2_T763 - cost: 4.20, layer: 51
Normal
MK50
MK75
MK88
MK94
MK97
MK data
Normal data
Fold over basal
1.2
1.1
1
0.9
0.8
0.7
0.6
0
10
20
30
40
50
60
Time (min)

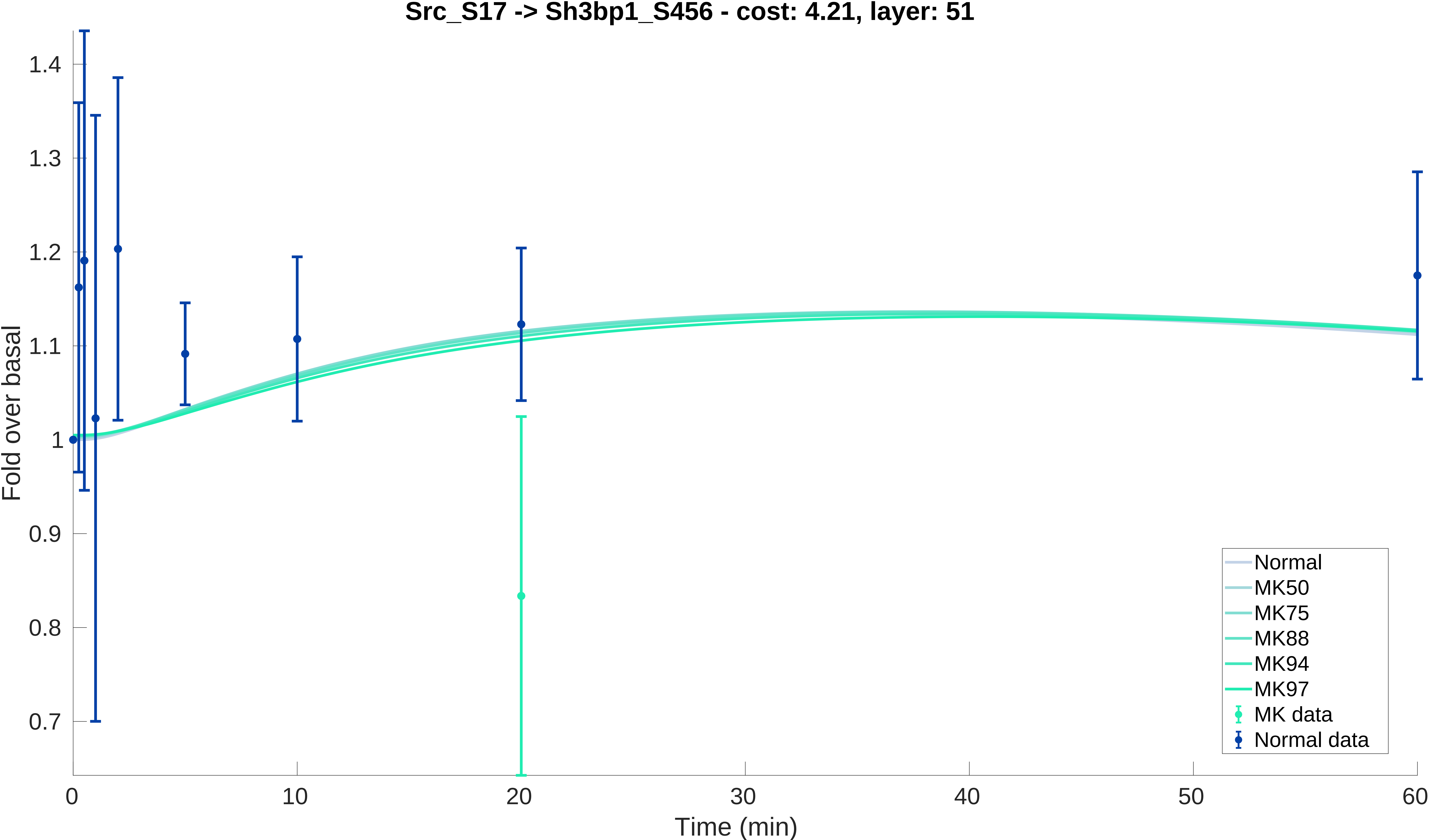

Src_S17 -> Sh3bp1_S456 - cost: 4.21, layer: 51
Fold over basal
1.4
1.3
1.2
1.1
1
0.9
0.8
0.7
0
10
20
30
40
50
60
Time (min)
Normal
MK50
MK75
MK88
MK94
MK97
MK data
Normal data

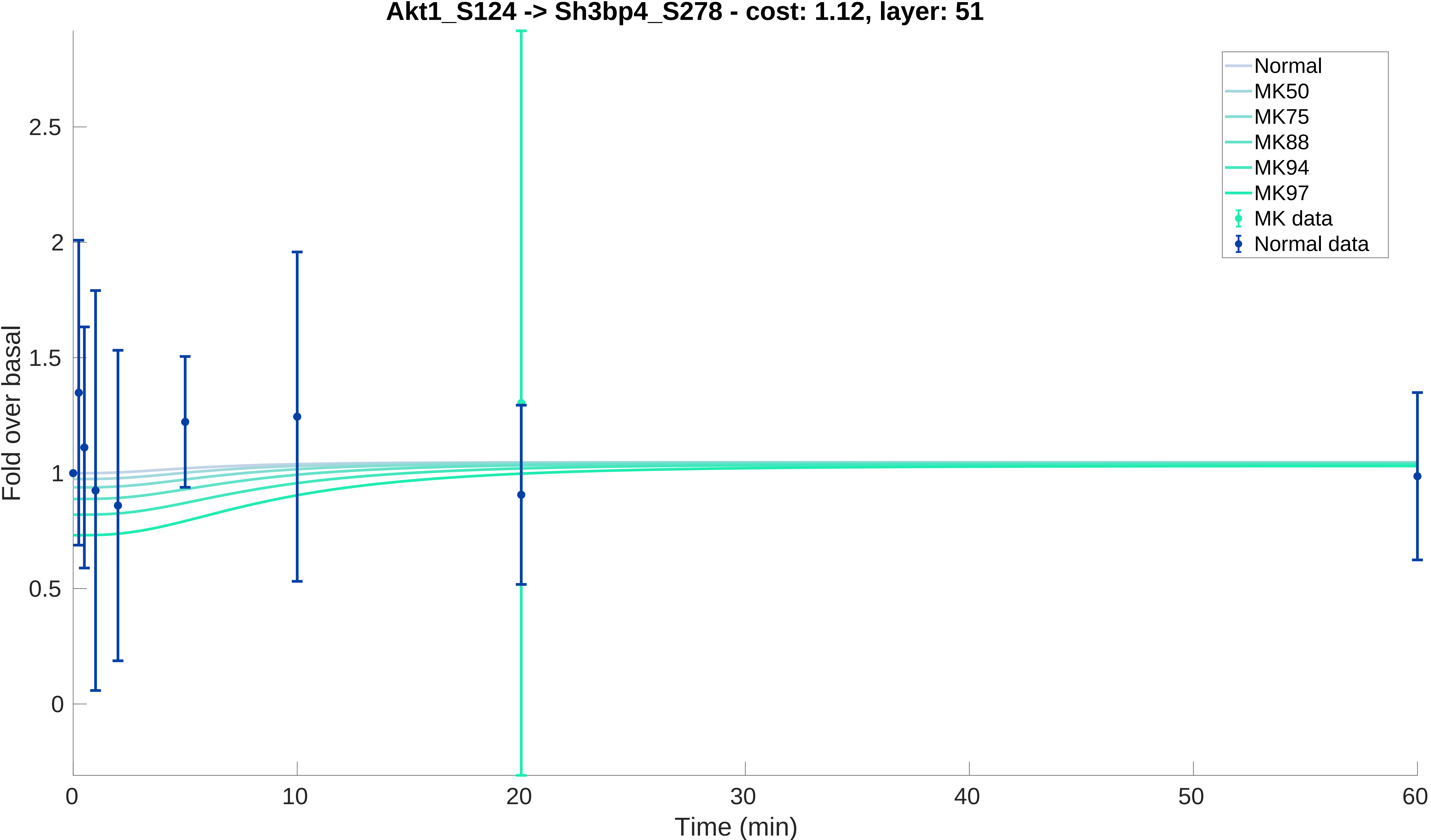

Akt1_S124 -> Sh3bp4_S278 - cost: 1.12, layer: 51
Normal
MK50
MK75
MK88
MK94
MK97
MK data
Normal data
Fold over basal
2.5
2
1.5
1
0.5
0
0
10
20
30
40
50
60
Time (min)

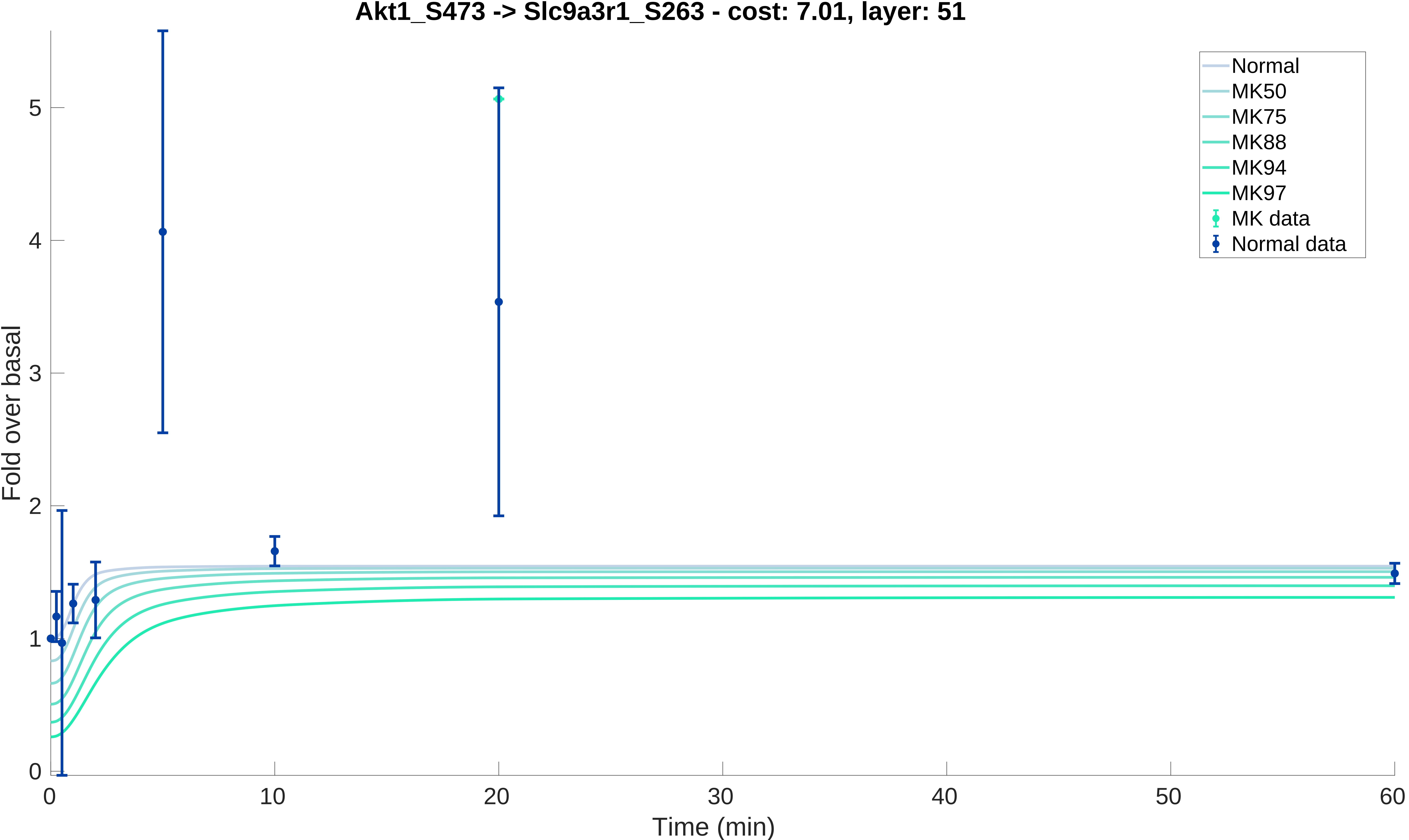

Akt1_S473 -> Slc9a3r1_S263 - cost: 7.01, layer: 51
Fold over basal
Time (min)
0
1
2
3
4
5
0
10
20
30
40
50
60
Normal
MK50
MK75
MK88
MK94
MK97
MK data
Normal data

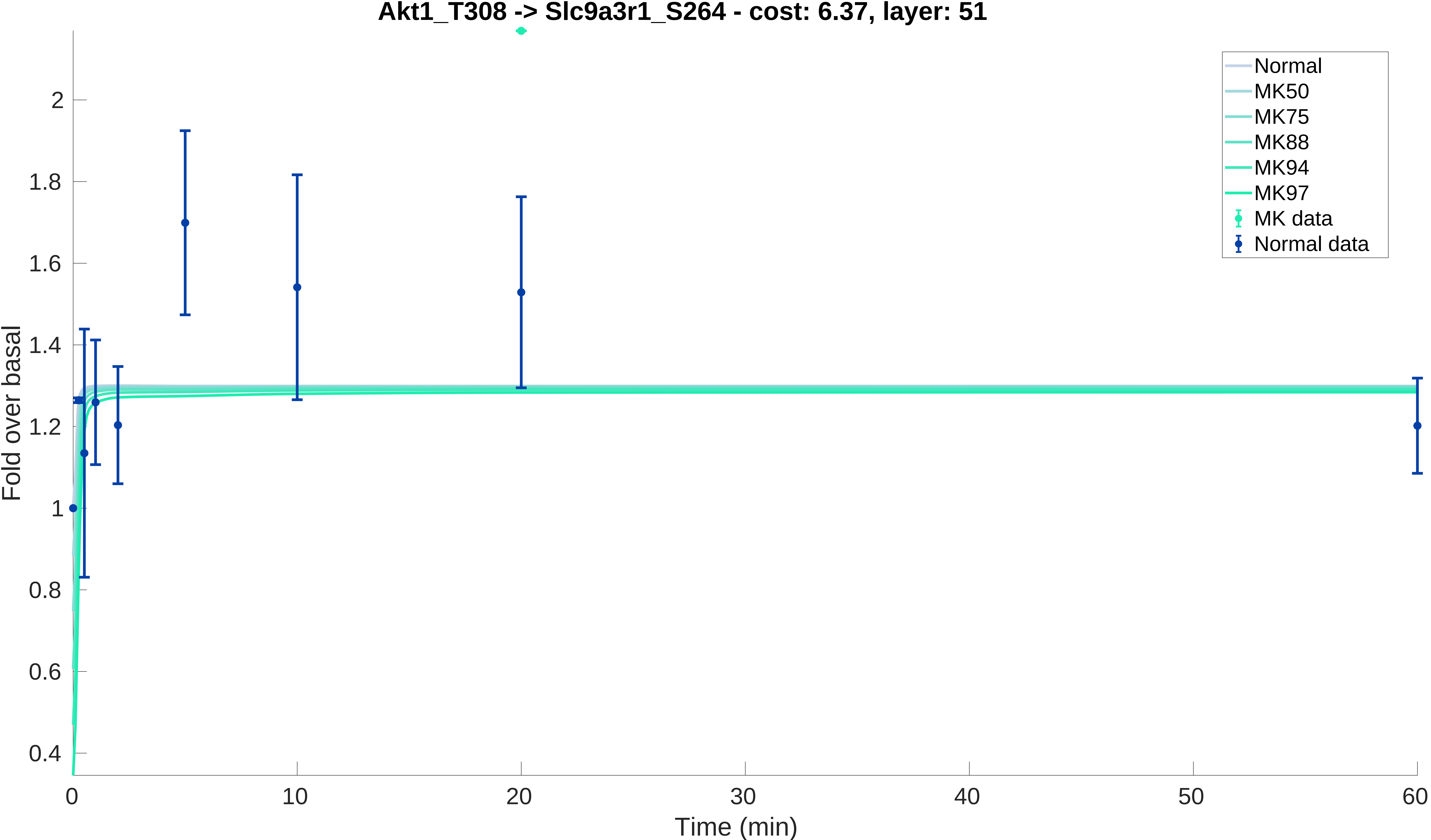

Akt1_T308 -> Slc9a3r1_S264 - cost: 6.37, layer: 51
Fold over basal
Time (min)
Normal
MK50
MK75
MK88
MK94
MK97
MK data
Normal data
0.4
0.6
0.8
1
1.2
1.4
1.6
1.8
2
0
10
20
30
40
50
60

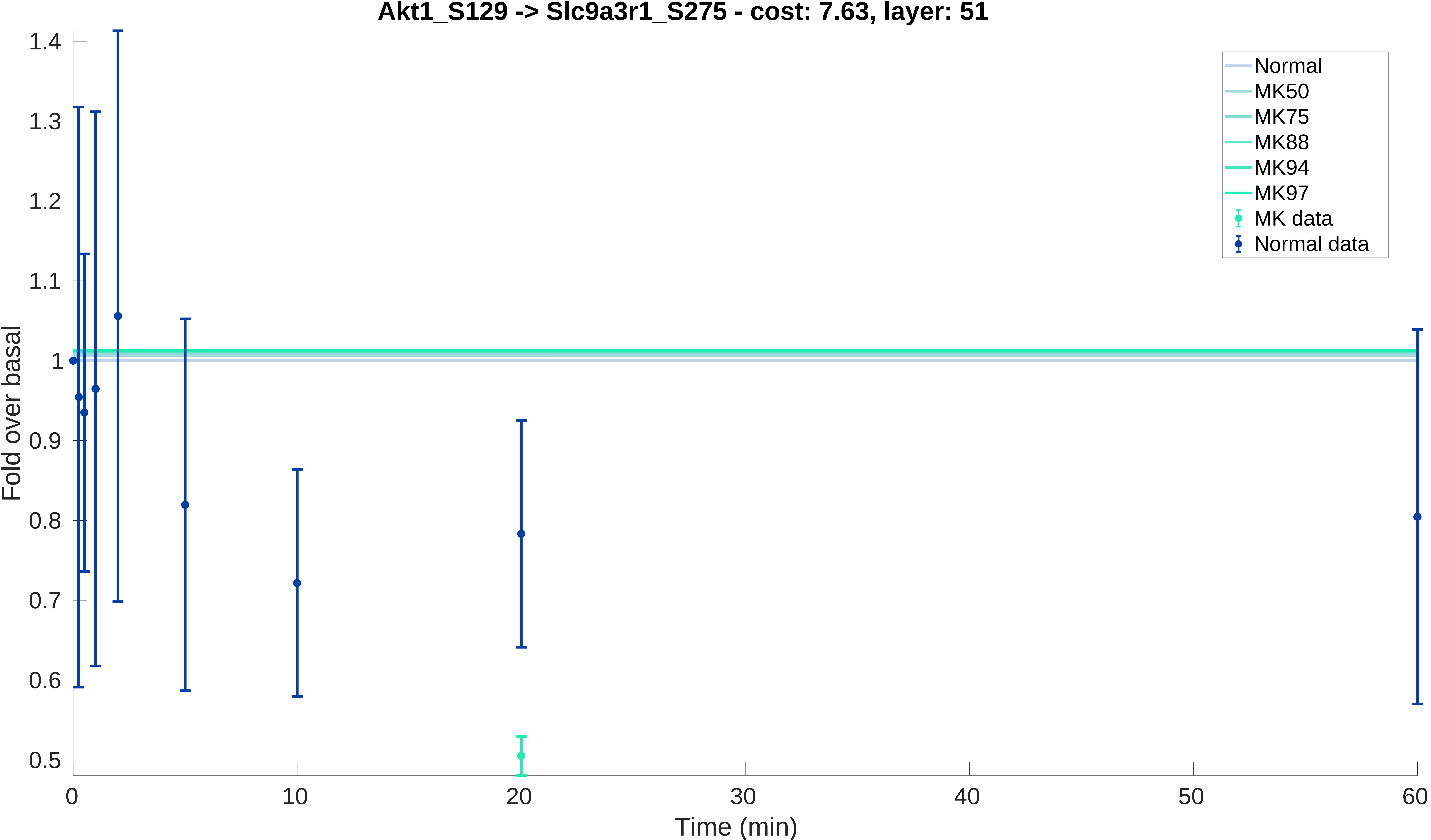

Akt1_S129 -> Slc9a3r1_S275 - cost: 7.63, layer: 51
Fold over basal
Time (min)
Normal
MK50
MK75
MK88
MK94
MK97
MK data
Normal data
0.5
0.6
0.7
0.8
0.9
1
1.1
1.2
1.3
1.4
0
10
20
30
40
50
60

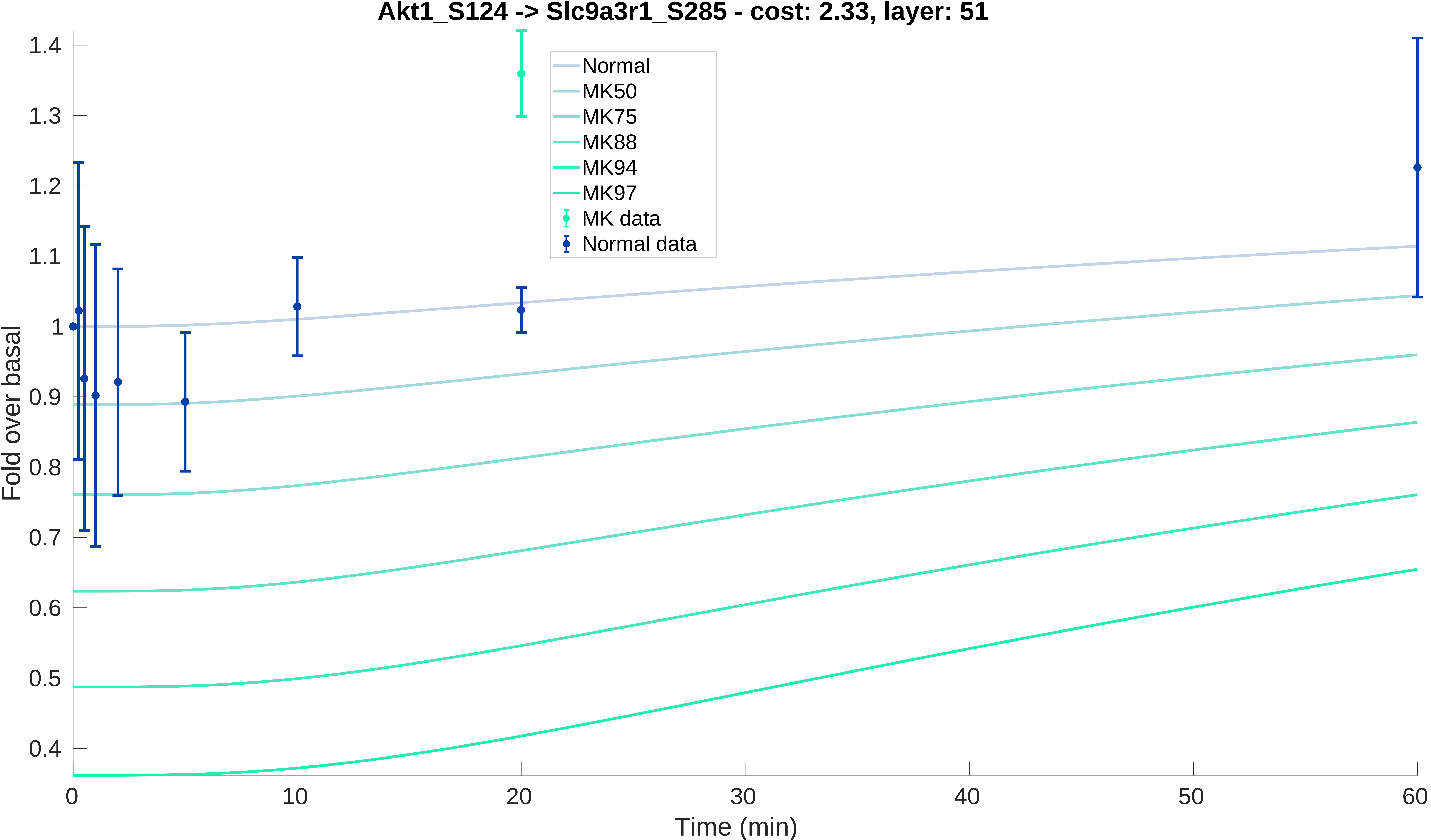

Akt1_S124 -> Slc9a3r1_S285 - cost: 2.33, layer: 51
Normal
MK50
MK75
MK88
MK94
MK97
MK data
Normal data
Fold over basal
Time (min)
1.4
1.3
1.2
1.1
1
0.9
0.8
0.7
0.6
0.5
0.4
0
10
20
30
40
50
60

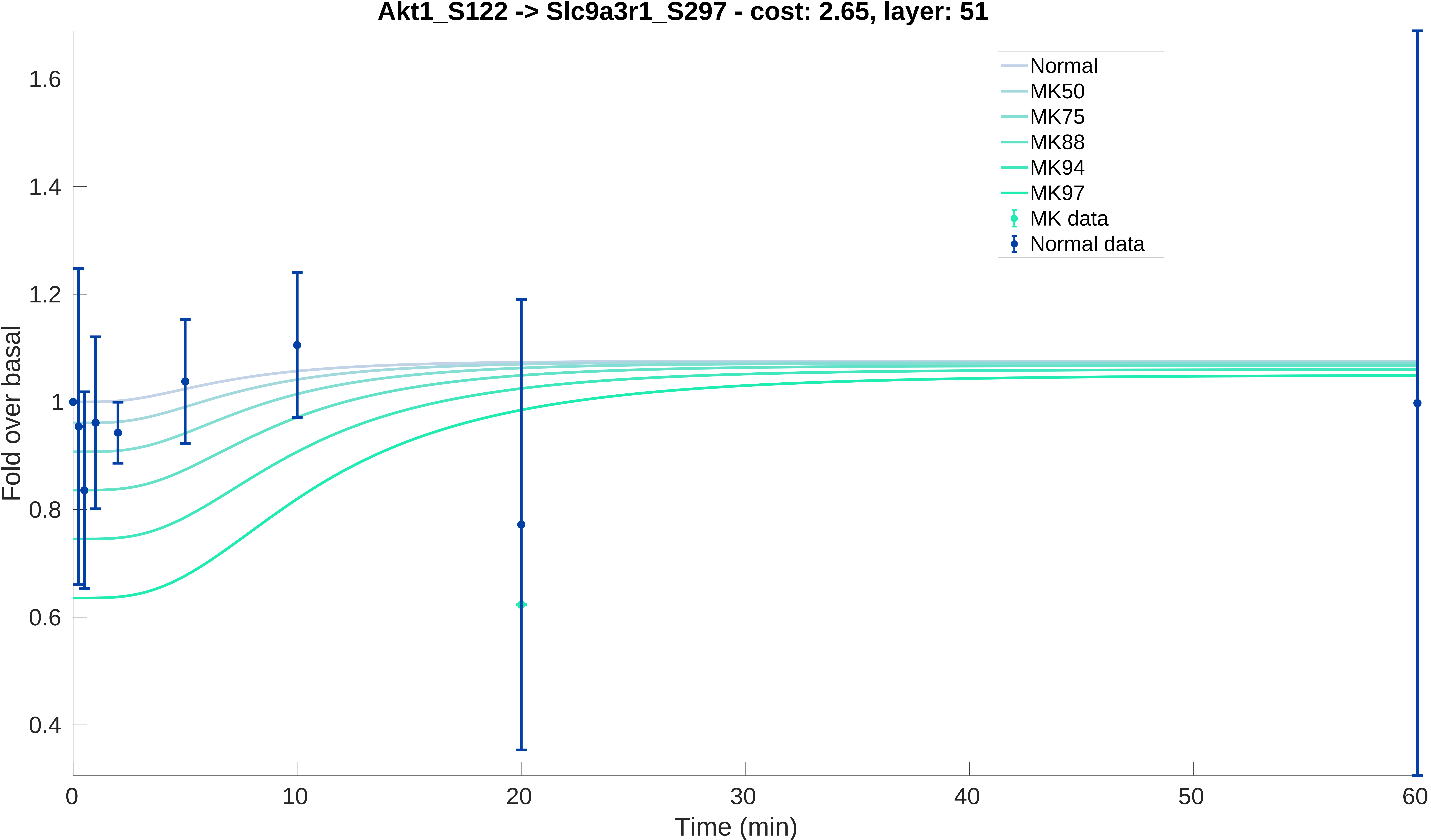

Akt1_S122 -> Slc9a3r1_S297 - cost: 2.65, layer: 51
Normal
MK50
MK75
MK88
MK94
MK97
MK data
Normal data
Fold over basal
1.6
1.4
1.2
1
0.8
0.6
0.4
0
10
20
30
40
50
60
Time (min)

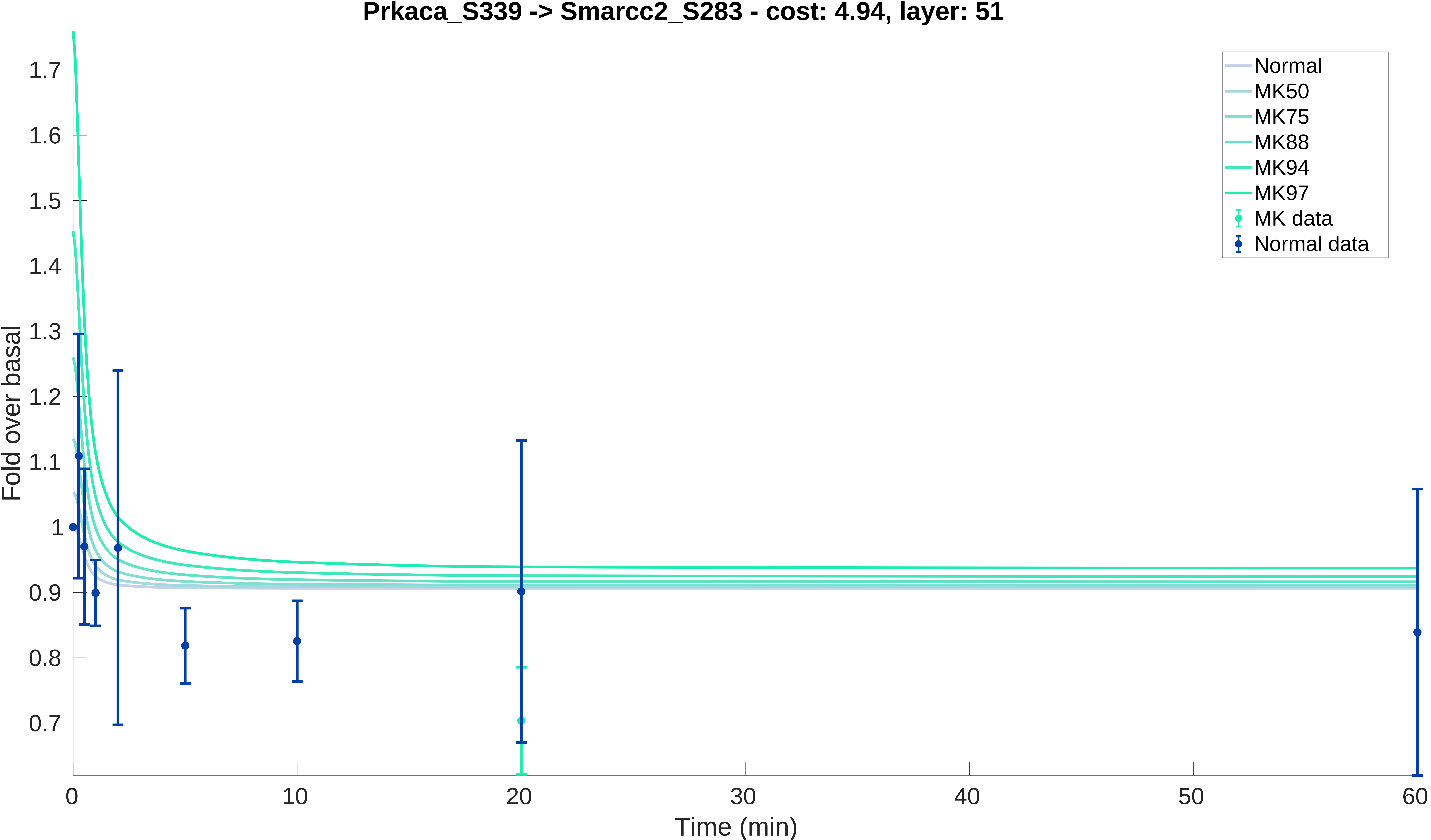

Prkaca_S339 -> Smarcc2_S283 - cost: 4.94, layer: 51
Fold over basal
1.7
1.6
1.5
1.4
1.3
1.2
1.1
1
0.9
0.8
0.7
0
10
20
30
40
50
60
Time (min)
Normal
MK50
MK75
MK88
MK94
MK97
MK data
Normal data

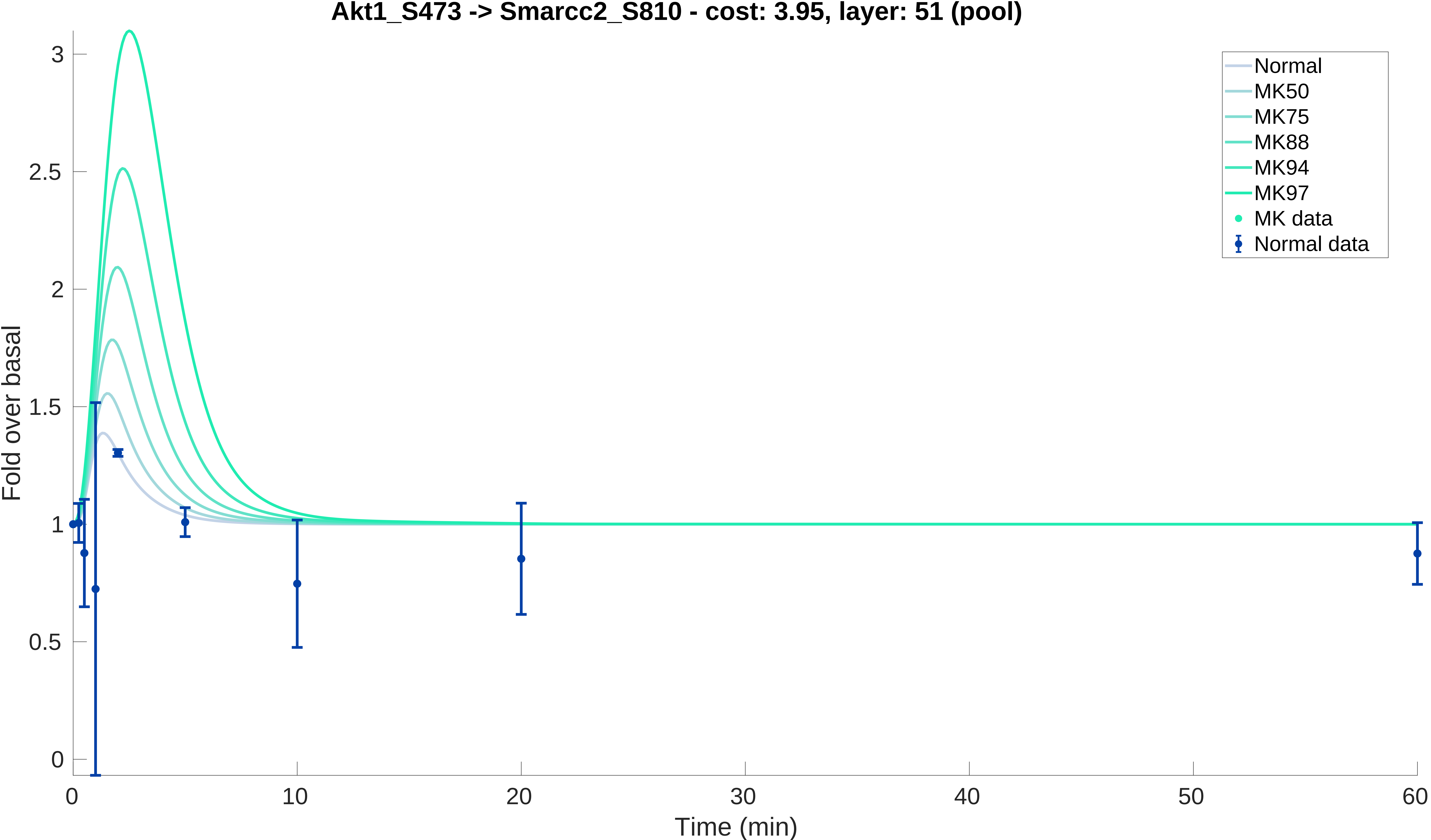
Akt1_S473 -> Smarcc2_S810 - cost: 3.95, layer: 51 (pool)
Fold over basal
Time (min)
0
0.5
1
1.5
2
2.5
3
0
10
20
30
40
50
60
Normal
MK50
MK75
MK88
MK94
MK97
MK data
Normal data

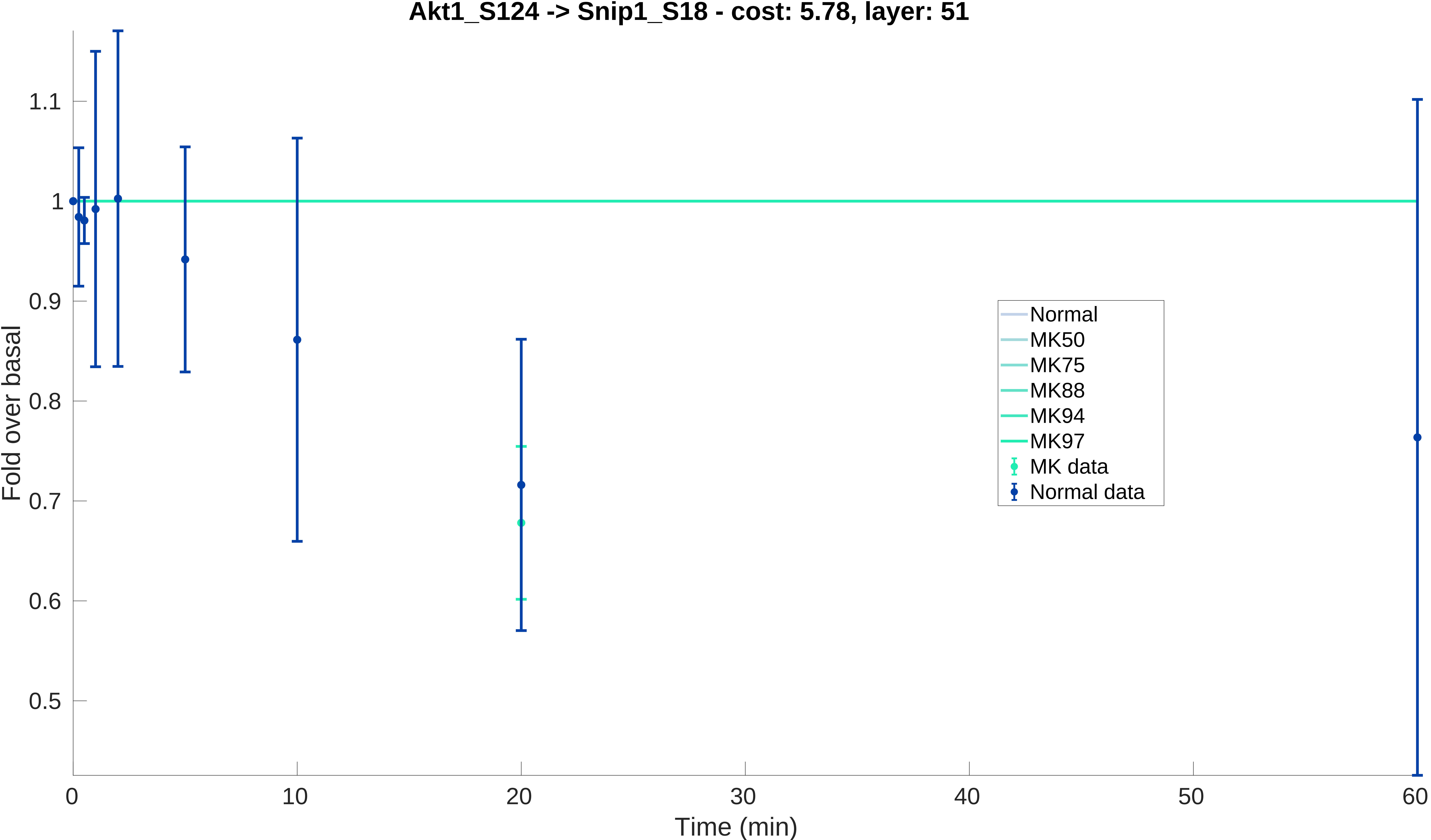

Akt1_S124 -> Snip1_S18 - cost: 5.78, layer: 51
Fold over basal
1.1
1
0.9
0.8
0.7
0.6
0.5
0
10
20
30
40
50
60
Time (min)
Normal
MK50
MK75
MK88
MK94
MK97
MK data
Normal data

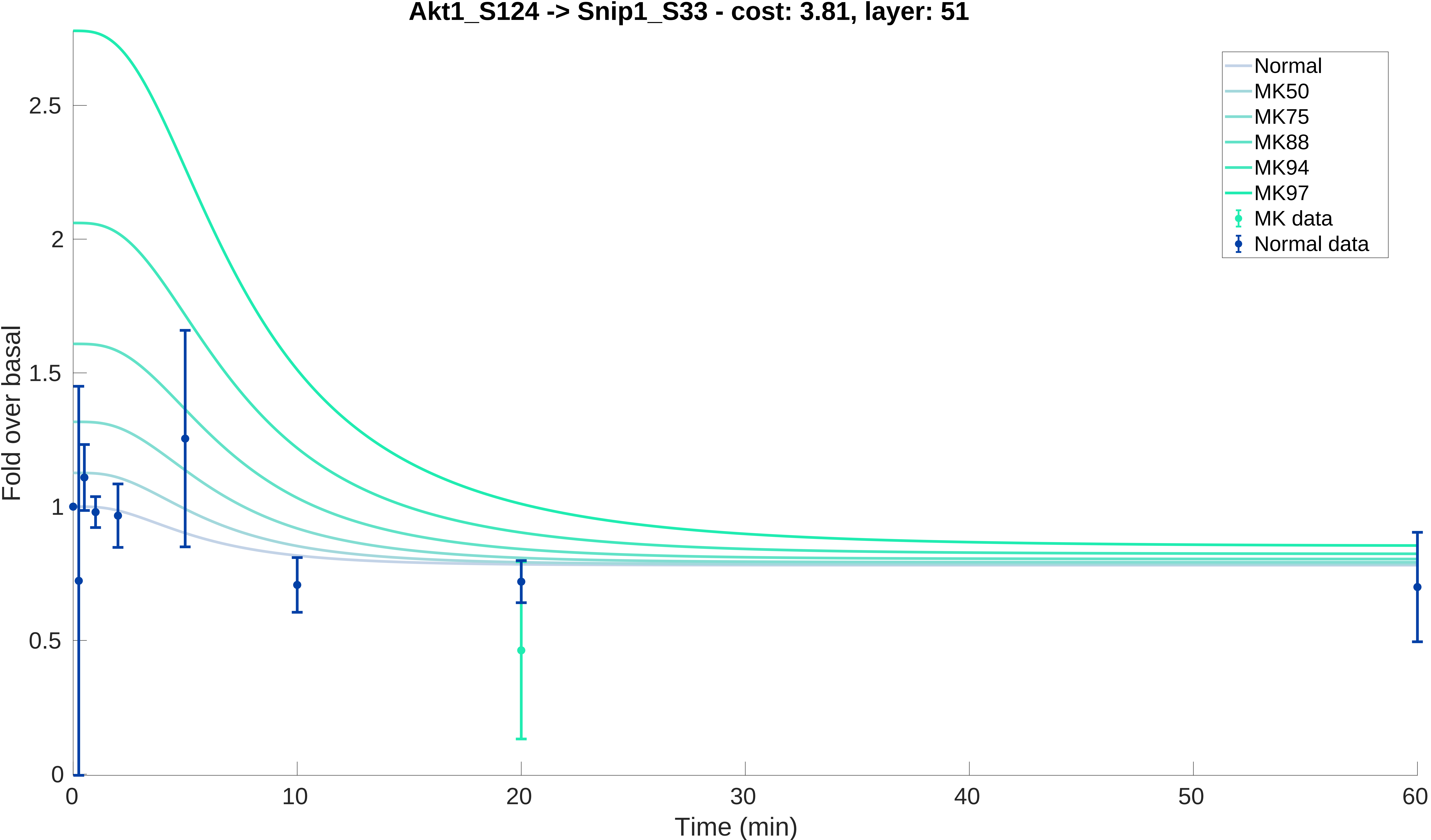

Akt1_S124 -> Snip1_S33 - cost: 3.81, layer: 51
Normal
MK50
MK75
MK88
MK94
MK97
MK data
Normal data
Fold over basal
Time (min)
0
0.5
1
1.5
2
2.5
0
10
20
30
40
50
60

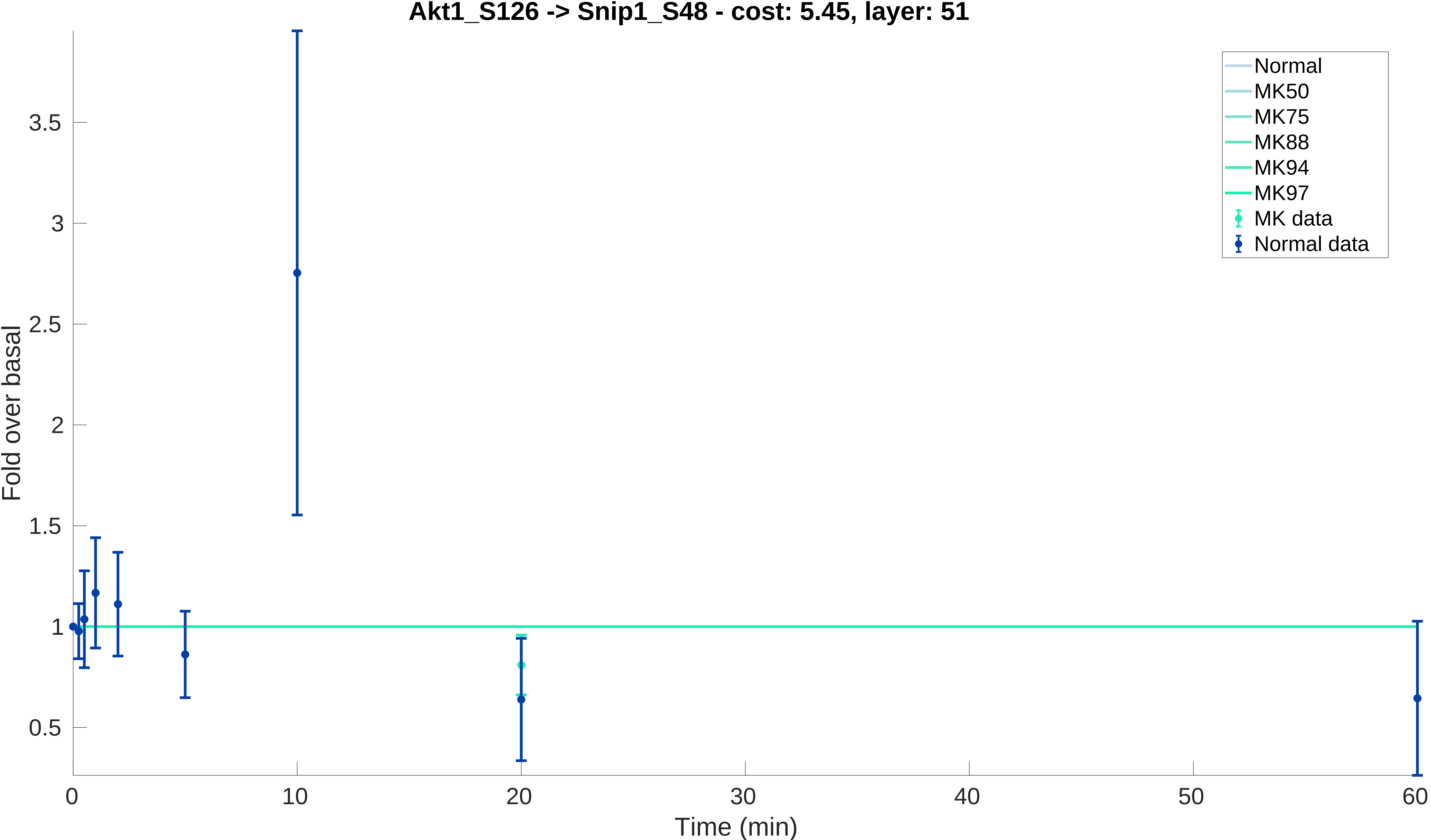
Akt1_S126 -> Snip1_S48 - cost: 5.45, layer: 51
Fold over basal
3.5
3
2.5
2
1.5
1
0.5
0
10
20
30
40
50
60
Time (min)
Normal
MK50
MK75
MK88
MK94
MK97
MK data
Normal data

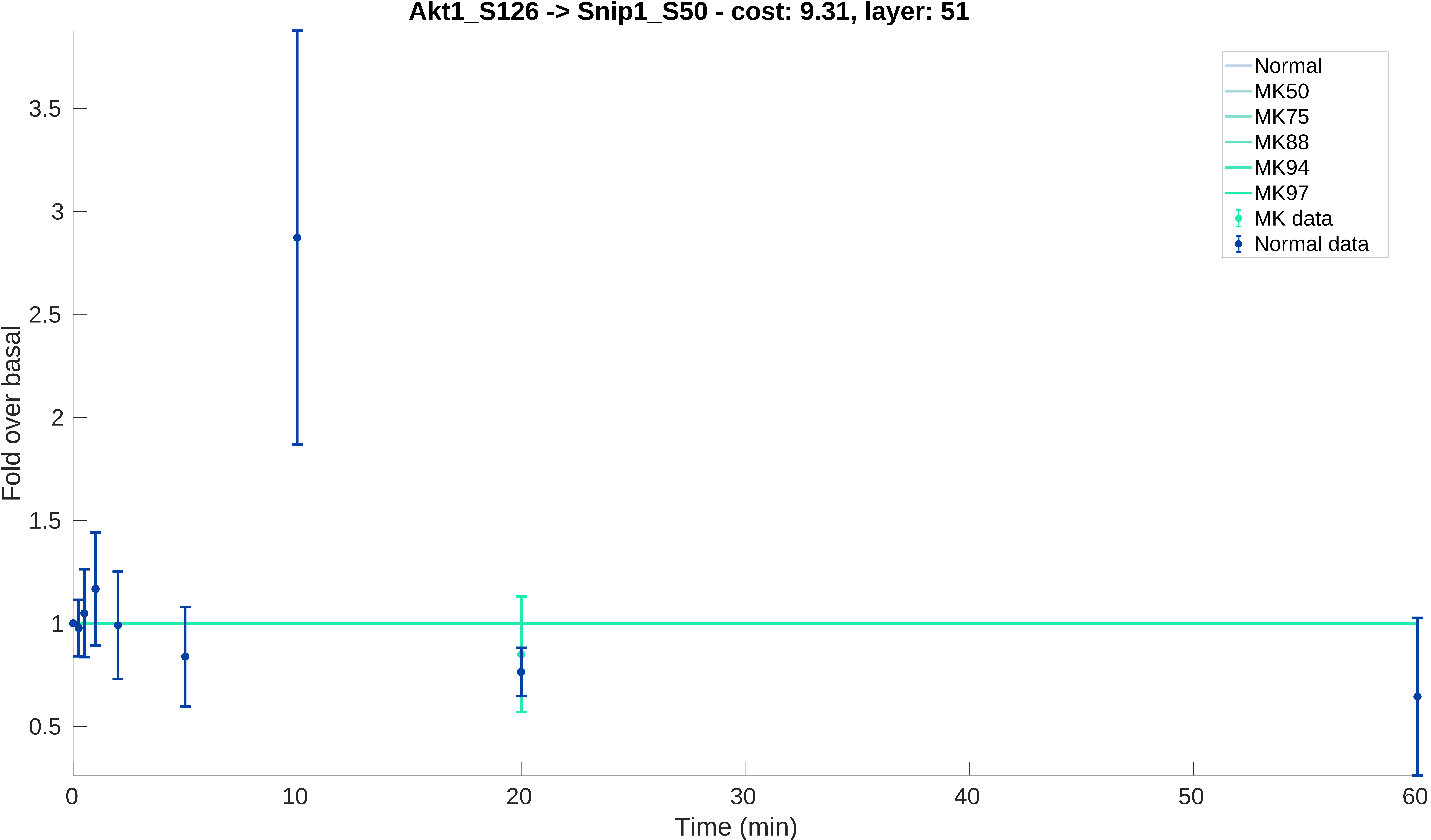

Akt1_S126 -> Snip1_S50 - cost: 9.31, layer: 51
Normal
MK50
MK75
MK88
MK94
MK97
MK data
Normal data
Fold over basal
3.5
3
2.5
2
1.5
1
0.5
0
10
20
30
40
50
60
Time (min)

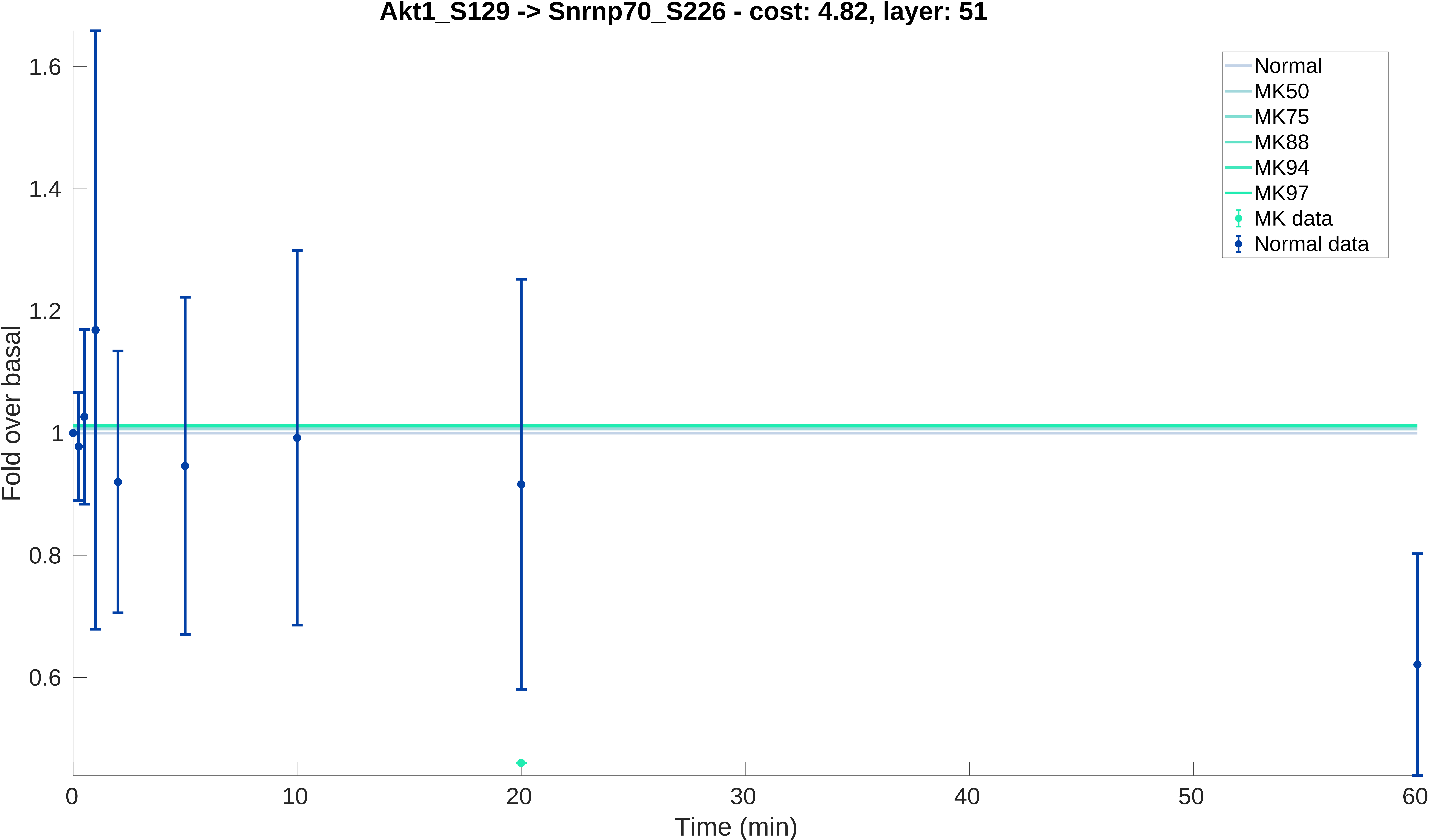
Akt1_S129 -> Snrnp70_S226 - cost: 4.82, layer: 51
Fold over basal
1.6
1.4
1.2
1
0.8
0.6
0
10
20
30
40
50
60
Time (min)
Normal
MK50
MK75
MK88
MK94
MK97
MK data
Normal data

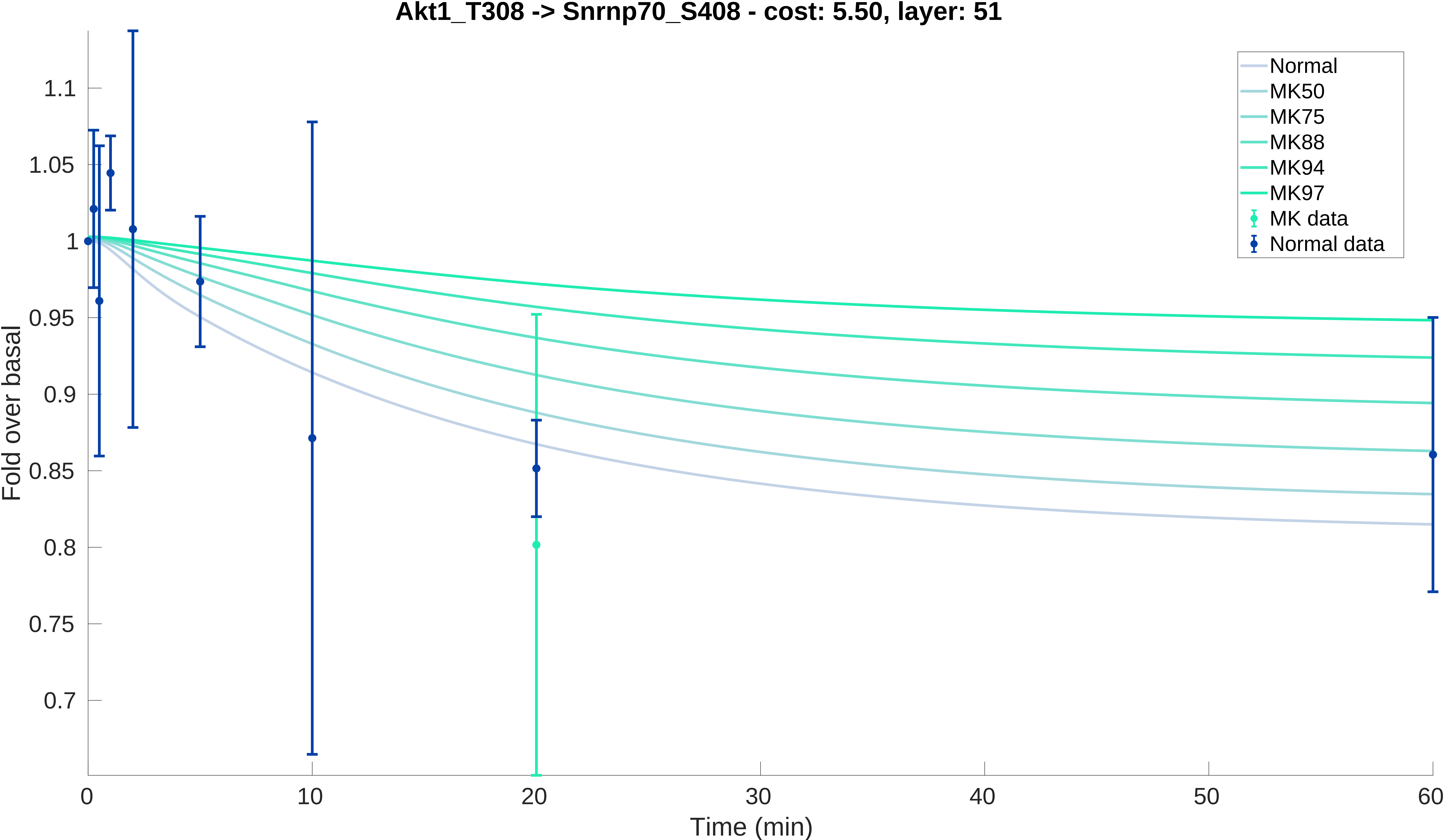
Akt1_T308 -> Snrnp70_S408 - cost: 5.50, layer: 51
Fold over basal
Time (min)
1.1
1.05
1
0.95
0.9
0.85
0.8
0.75
0.7
0
10
20
30
40
50
60
Normal
MK50
MK75
MK88
MK94
MK97
MK data
Normal data

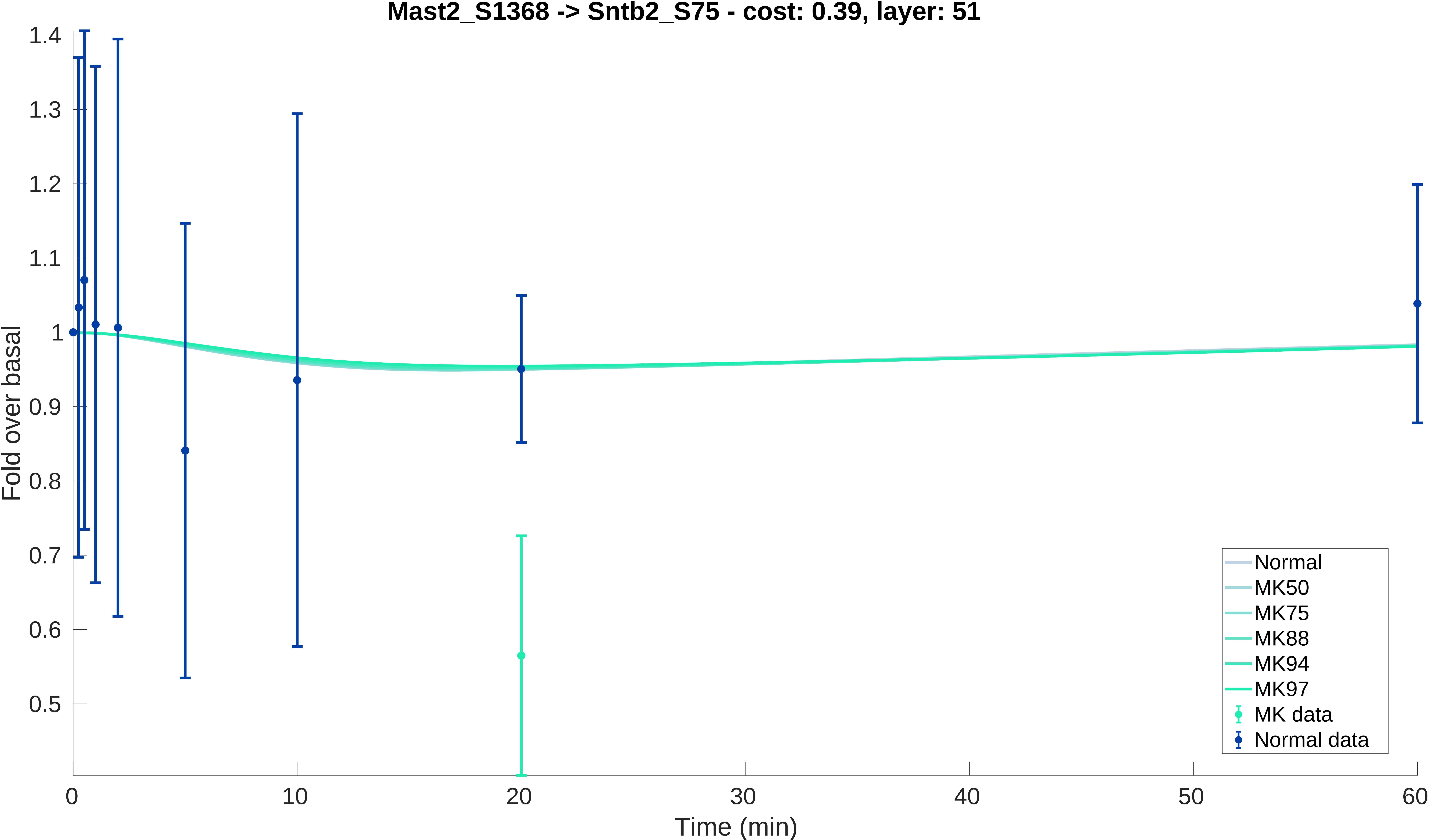

Mast2_S1368 -> Sntb2_S75 - cost: 0.39, layer: 51
Fold over basal
Time (min)
1.4
1.3
1.2
1.1
1
0.9
0.8
0.7
0.6
0.5
0
10
20
30
40
50
60
Normal
MK50
MK75
MK88
MK94
MK97
MK data
Normal data

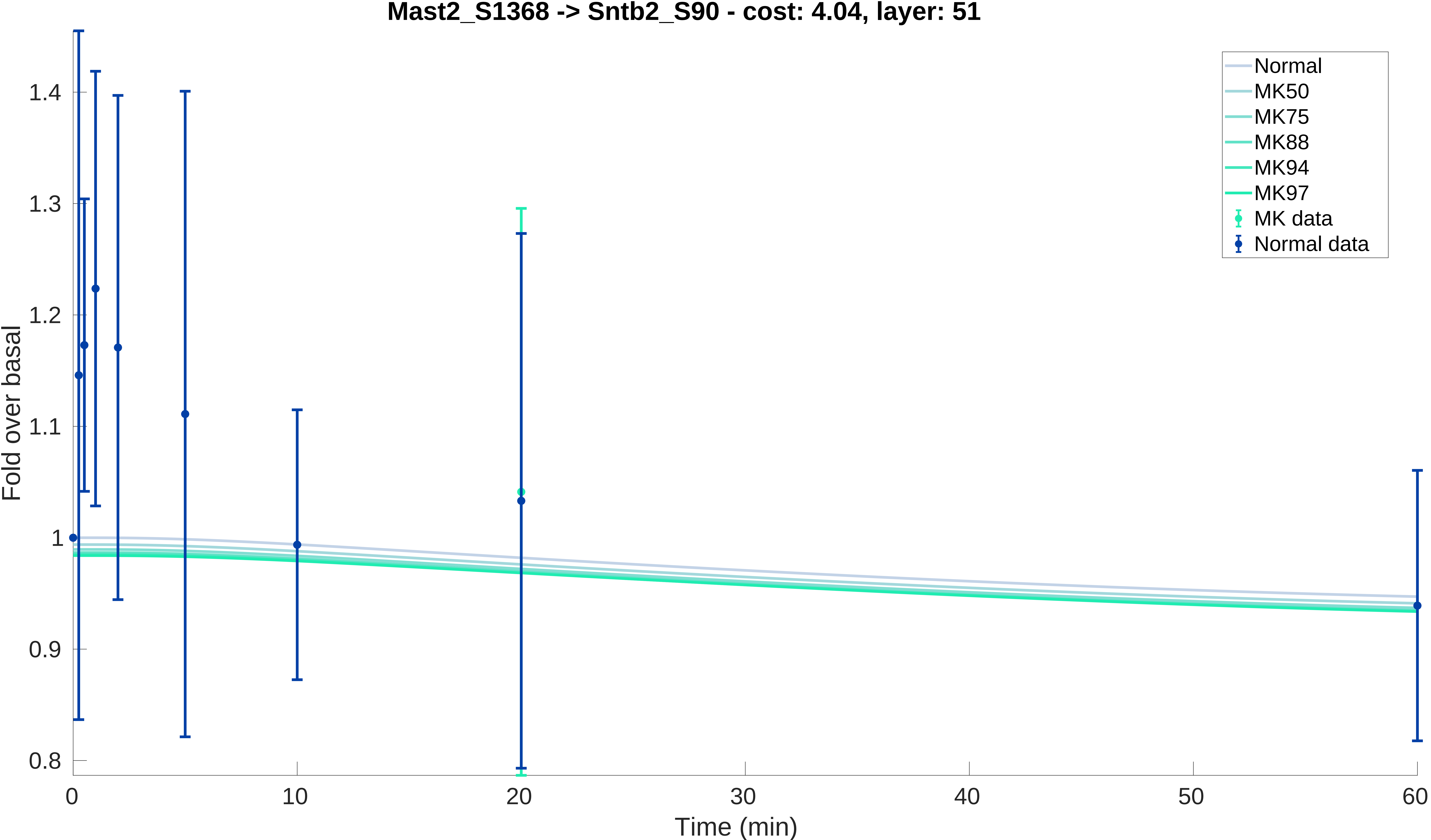
Mast2_S1368 -> Sntb2_S90 - cost: 4.04, layer: 51
Fold over basal
Time (min)
0.8
0.9
1
1.1
1.2
1.3
1.4
0
10
20
30
40
50
60
Normal
MK50
MK75
MK88
MK94
MK97
MK data
Normal data

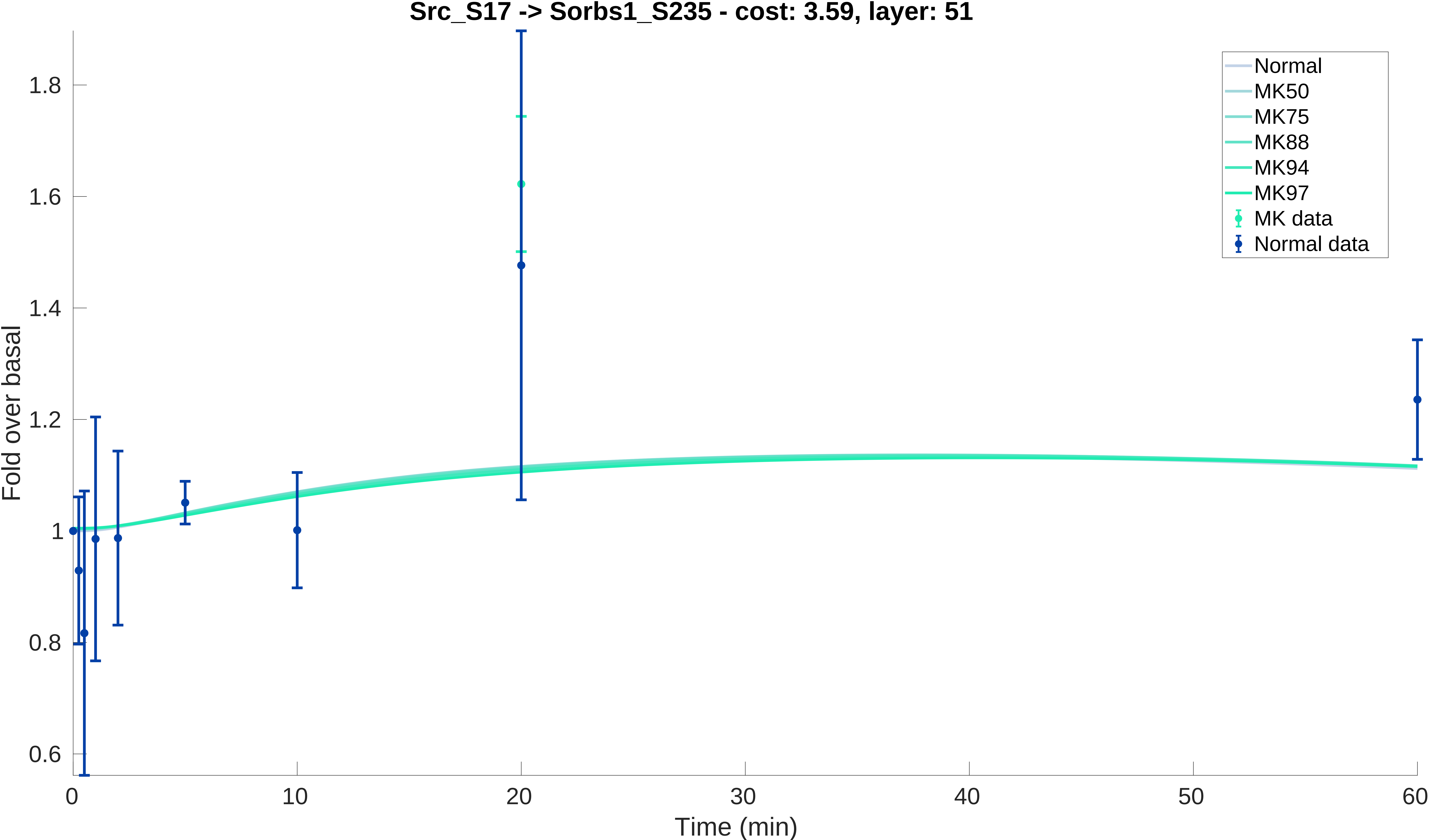

Src_S17 -> Sorbs1_S235 - cost: 3.59, layer: 51
Fold over basal
Time (min)
0.6
0.8
1
1.2
1.4
1.6
1.8
0
10
20
30
40
50
60
Normal
MK50
MK75
MK88
MK94
MK97
MK data
Normal data

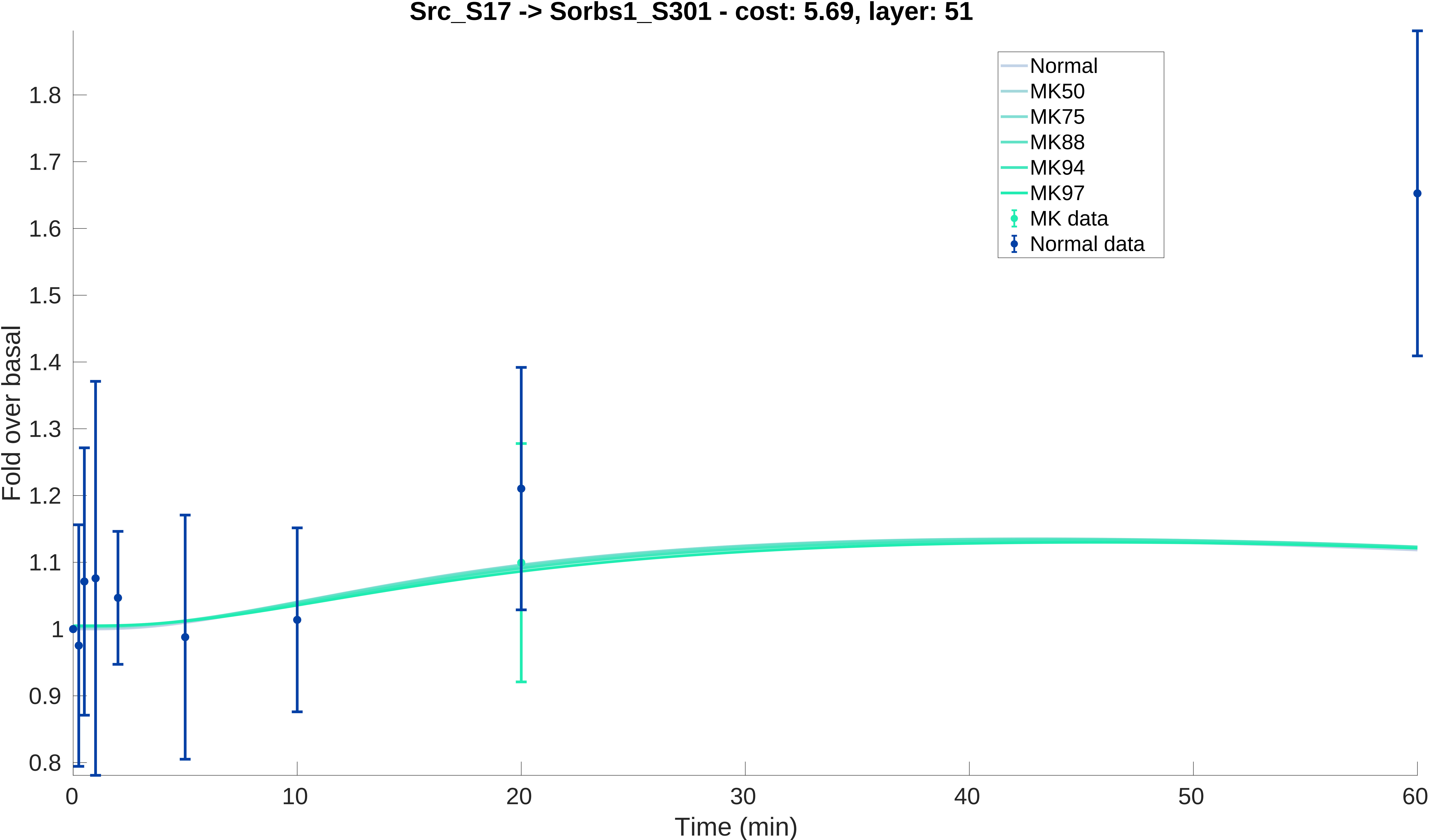

Src_S17 -> Sorbs1_S301 - cost: 5.69, layer: 51
Normal
MK50
MK75
MK88
MK94
MK97
MK data
Normal data
Fold over basal
1.8
1.7
1.6
1.5
1.4
1.3
1.2
1.1
1
0.9
0.8
0
10
20
30
40
50
60
Time (min)

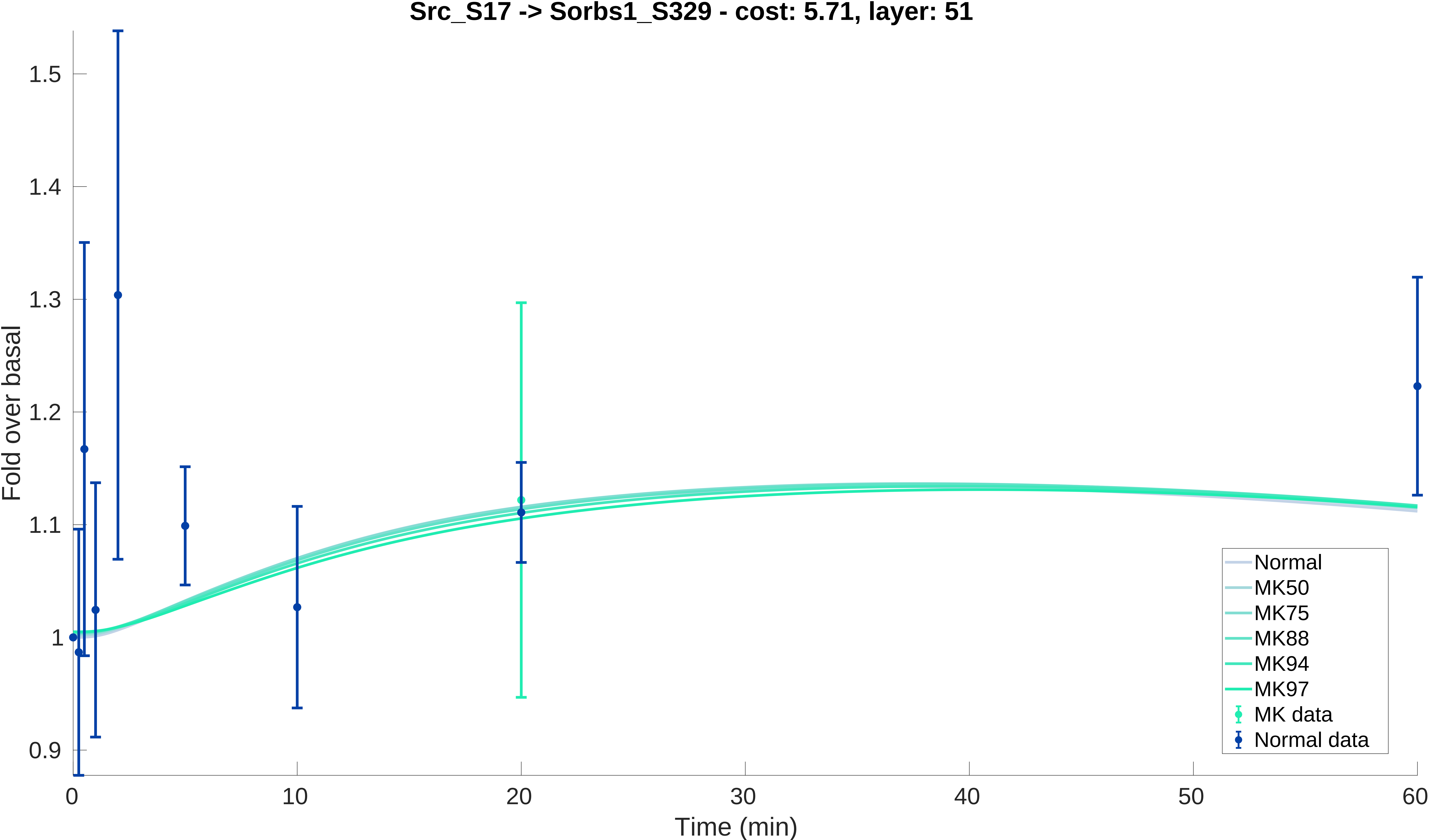
Src_S17 -> Sorbs1_S329 - cost: 5.71, layer: 51
Fold over basal
0.9
1
1.1
1.2
1.3
1.4
1.5
0
10
20
30
40
50
60
Time (min)
Normal
MK50
MK75
MK88
MK94
MK97
MK data
Normal data

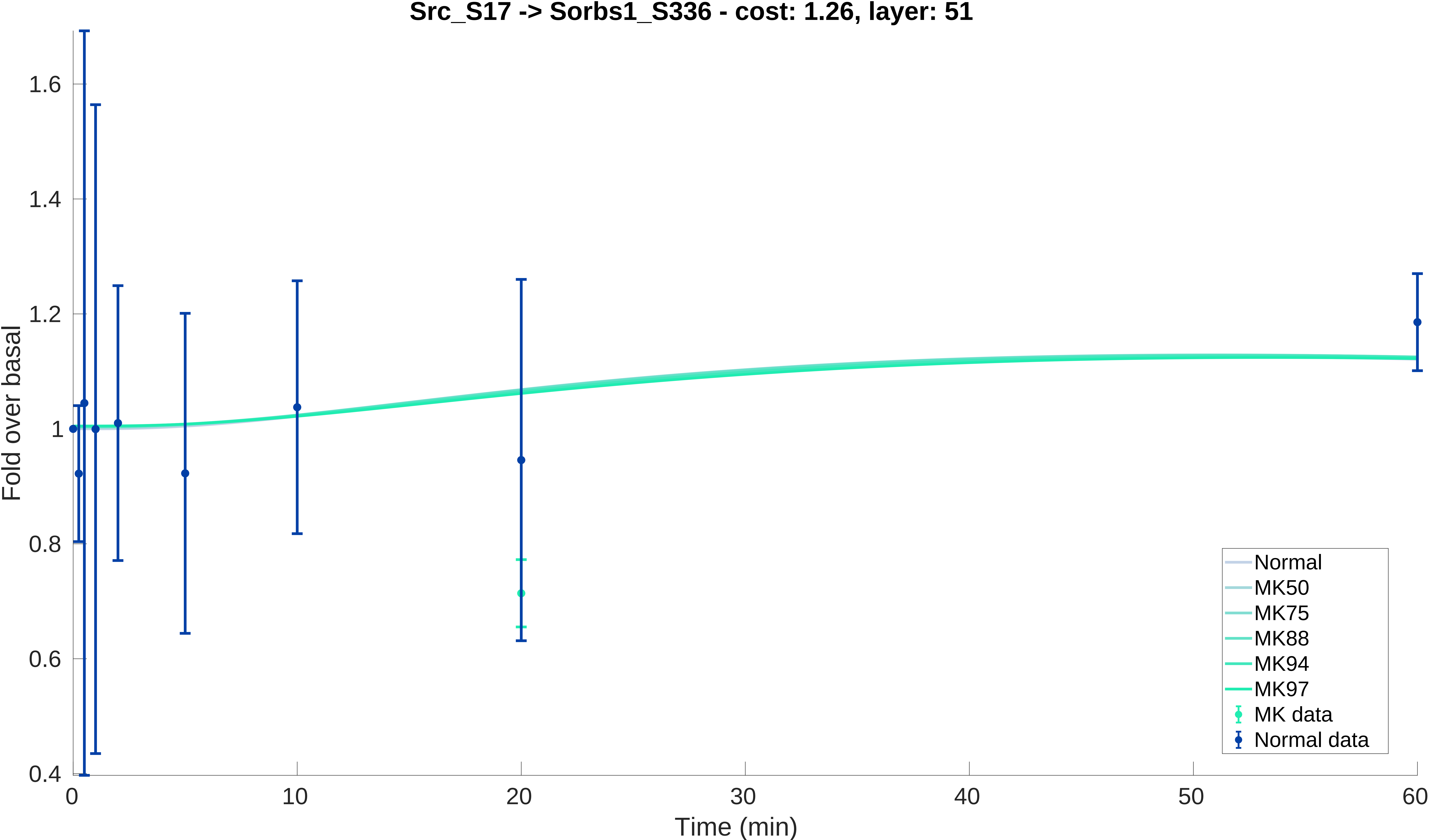

Src_S17 -> Sorbs1_S336 - cost: 1.26, layer: 51
Fold over basal
1.6
1.4
1.2
1
0.8
0.6
0.4
0
10
20
30
40
50
60
Time (min)
Normal
MK50
MK75
MK88
MK94
MK97
MK data
Normal data

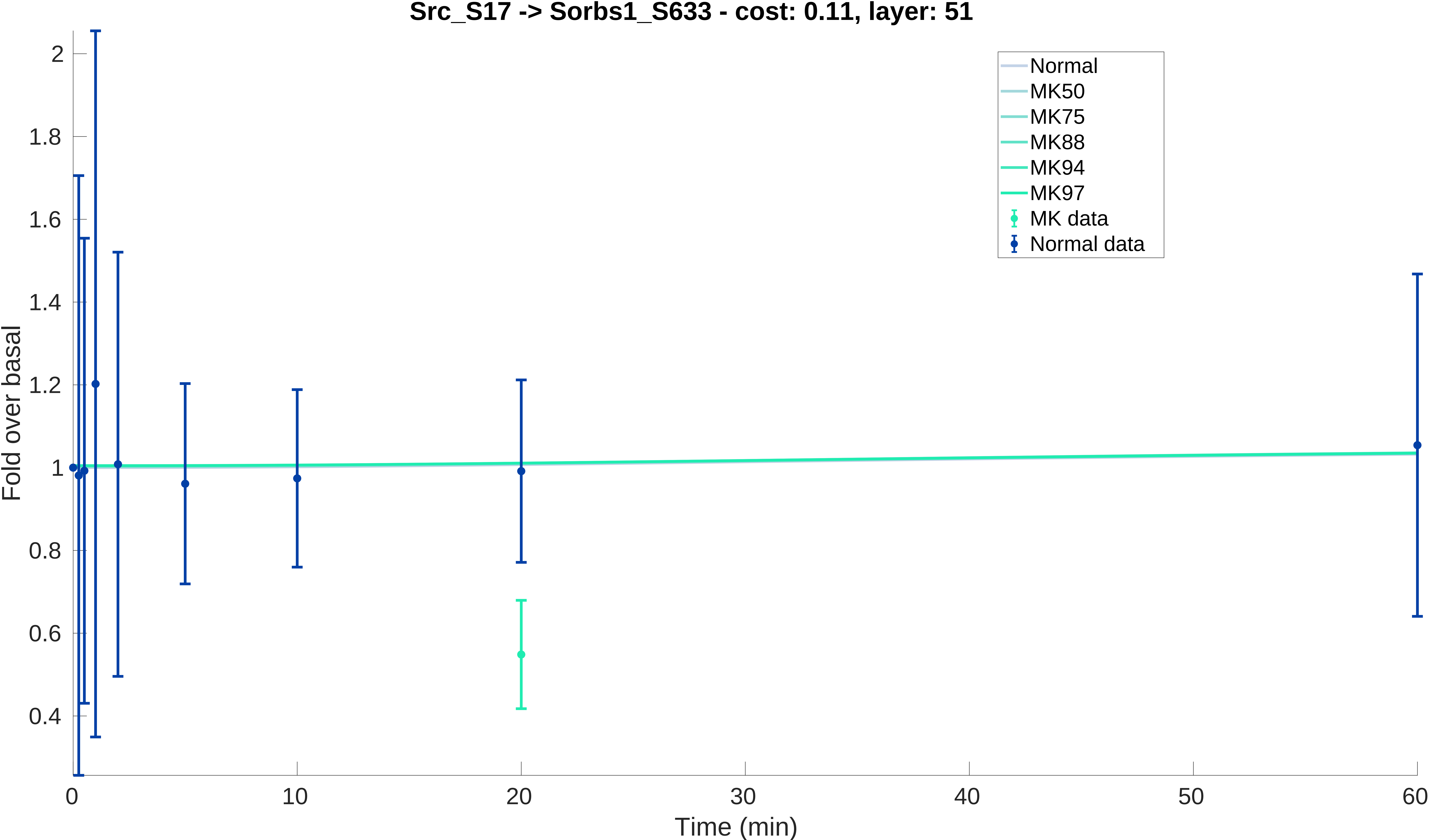

Src_S17 -> Sorbs1_S633 - cost: 0.11, layer: 51
Normal
MK50
MK75
MK88
MK94
MK97
MK data
Normal data
Fold over basal
Time (min)
0.4
0.6
0.8
1
1.2
1.4
1.6
1.8
2
0
10
20
30
40
50
60

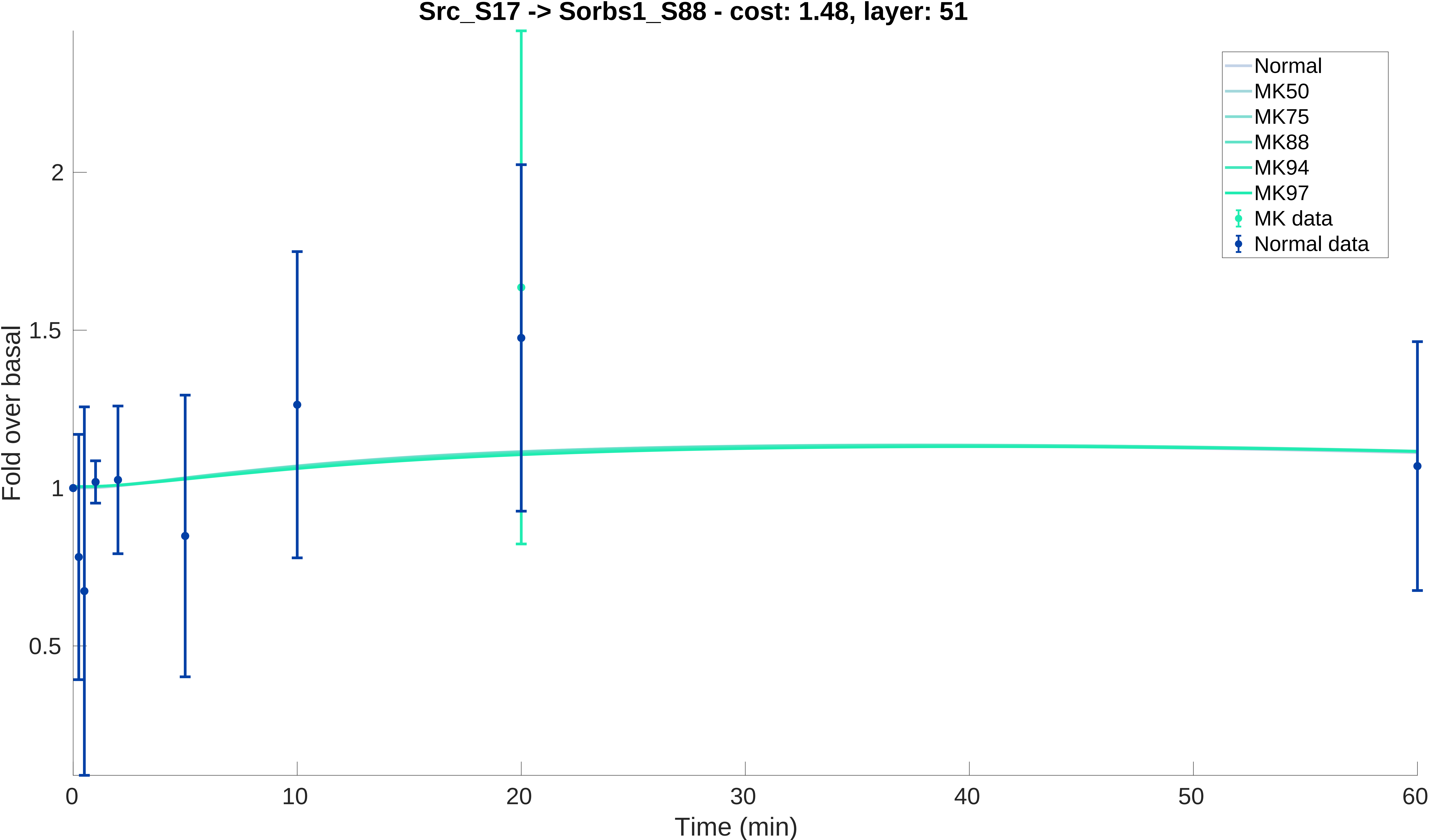

Src_S17 -> Sorbs1_S88 - cost: 1.48, layer: 51
Normal
MK50
MK75
MK88
MK94
MK97
MK data
Normal data
Fold over basal
0.5
1
1.5
2
0
10
20
30
40
50
60
Time (min)

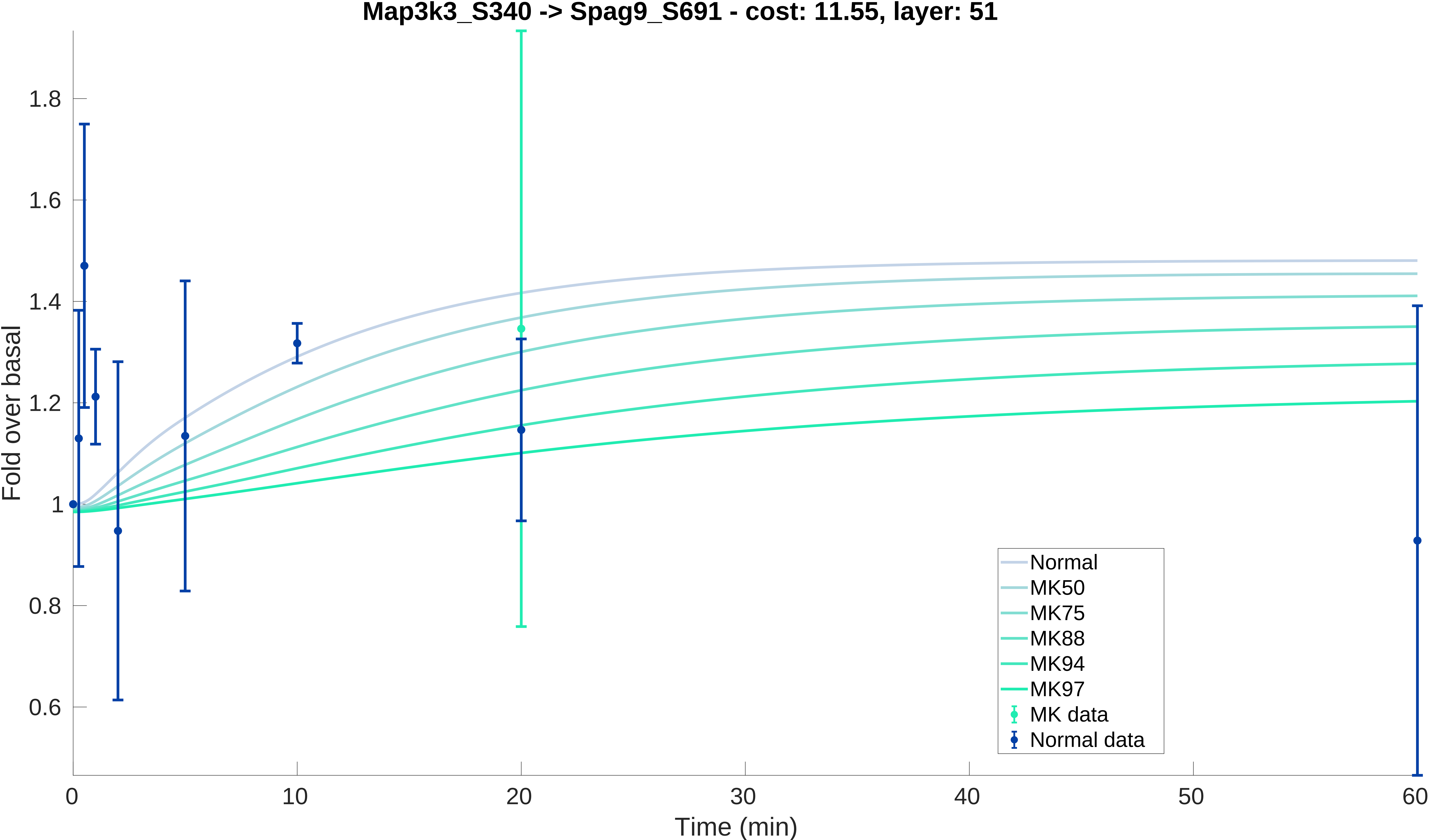

Map3k3_S340 -> Spag9_S691 - cost: 11.55, layer: 51
Fold over basal
1.8
1.6
1.4
1.2
1
0.8
0.6
0
10
20
30
40
50
60
Time (min)
Normal
MK50
MK75
MK88
MK94
MK97
MK data
Normal data

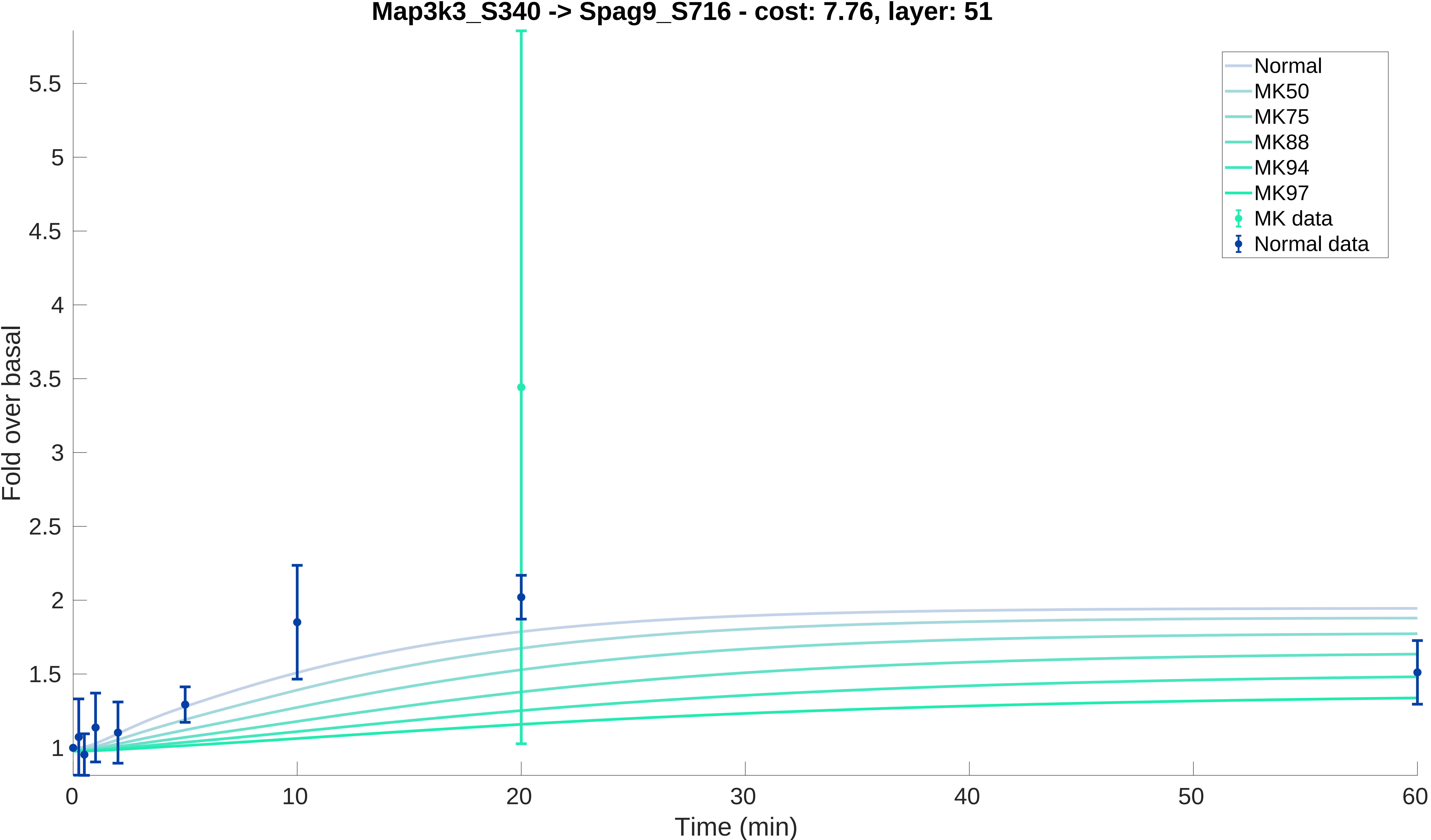
Map3k3_S340 -> Spag9_S716 - cost: 7.76, layer: 51
Normal
MK50
MK75
MK88
MK94
MK97
MK data
Normal data
Fold over basal
Time (min)
5.5
5
4.5
4
3.5
3
2.5
2
1.5
1
0
10
20
30
40
50
60

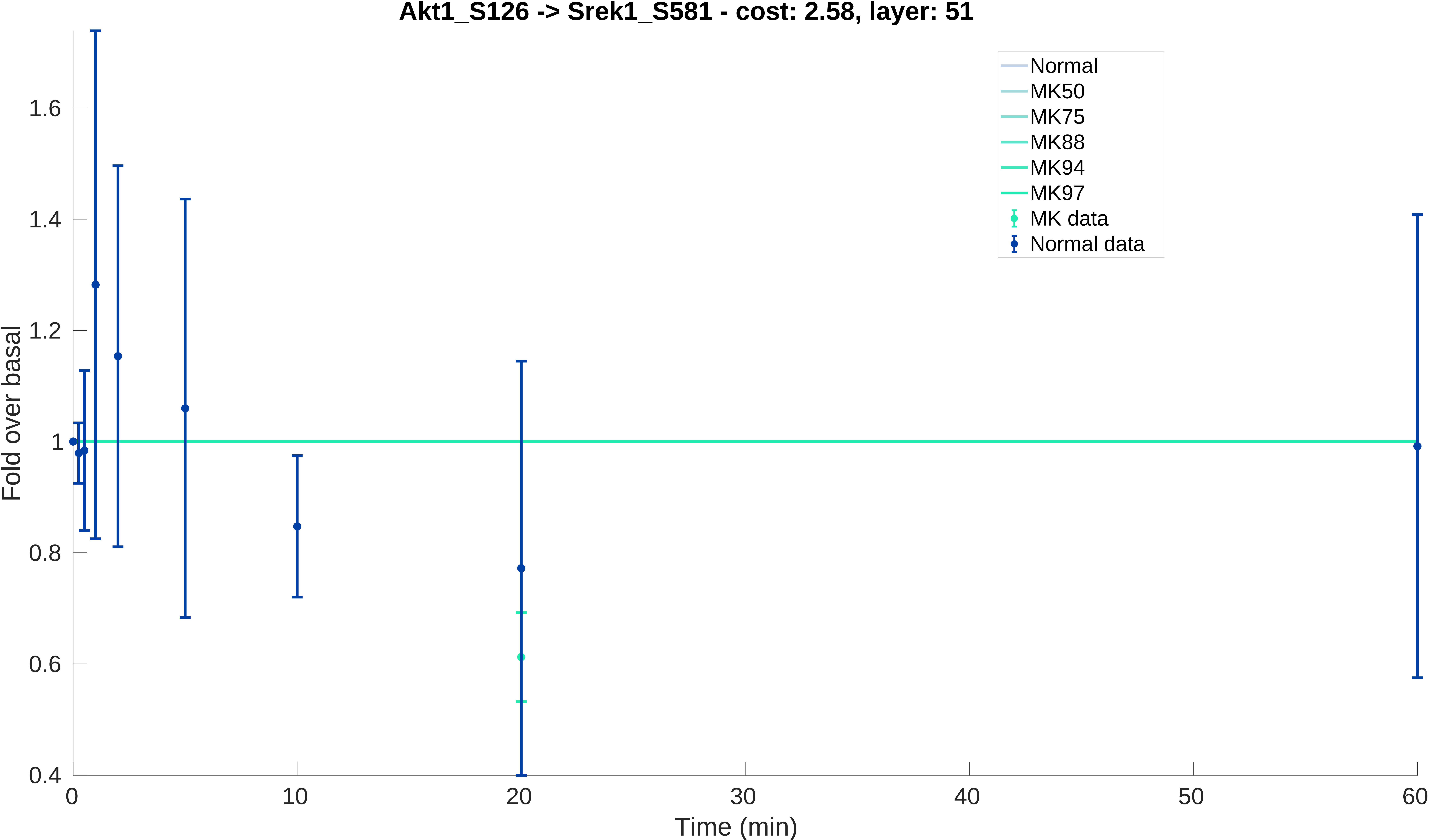

Akt1_S126 -> Srek1_S581 - cost: 2.58, layer: 51
Normal
MK50
MK75
MK88
MK94
MK97
MK data
Normal data
Fold over basal
Time (min)
1.6
1.4
1.2
1
0.8
0.6
0.4
0
10
20
30
40
50
60

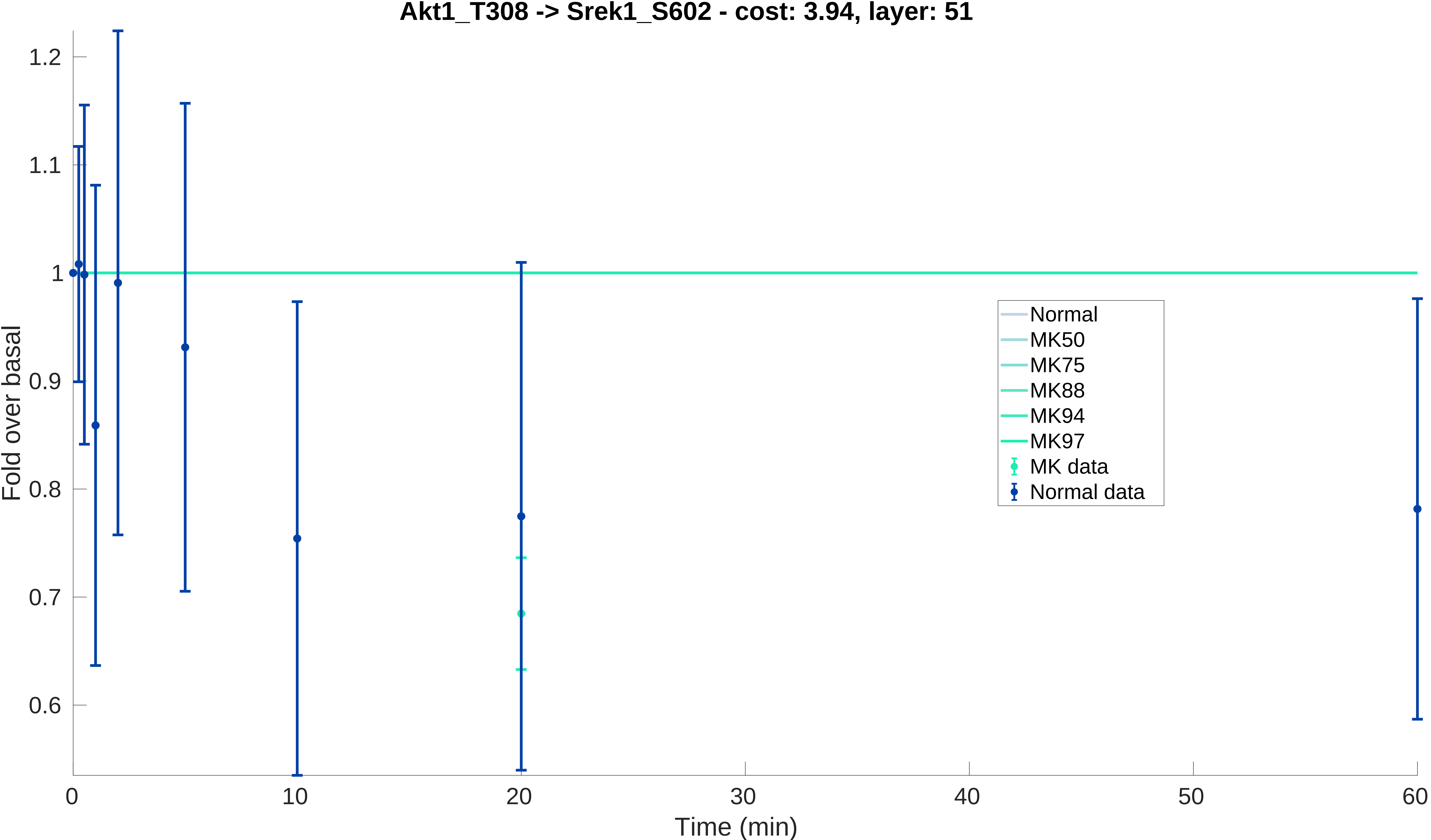

Akt1_T308 -> Srek1_S602 - cost: 3.94, layer: 51
Fold over basal
Time (min)
1.2
1.1
1
0.9
0.8
0.7
0.6
0
10
20
30
40
50
60
Normal
MK50
MK75
MK88
MK94
MK97
MK data
Normal data

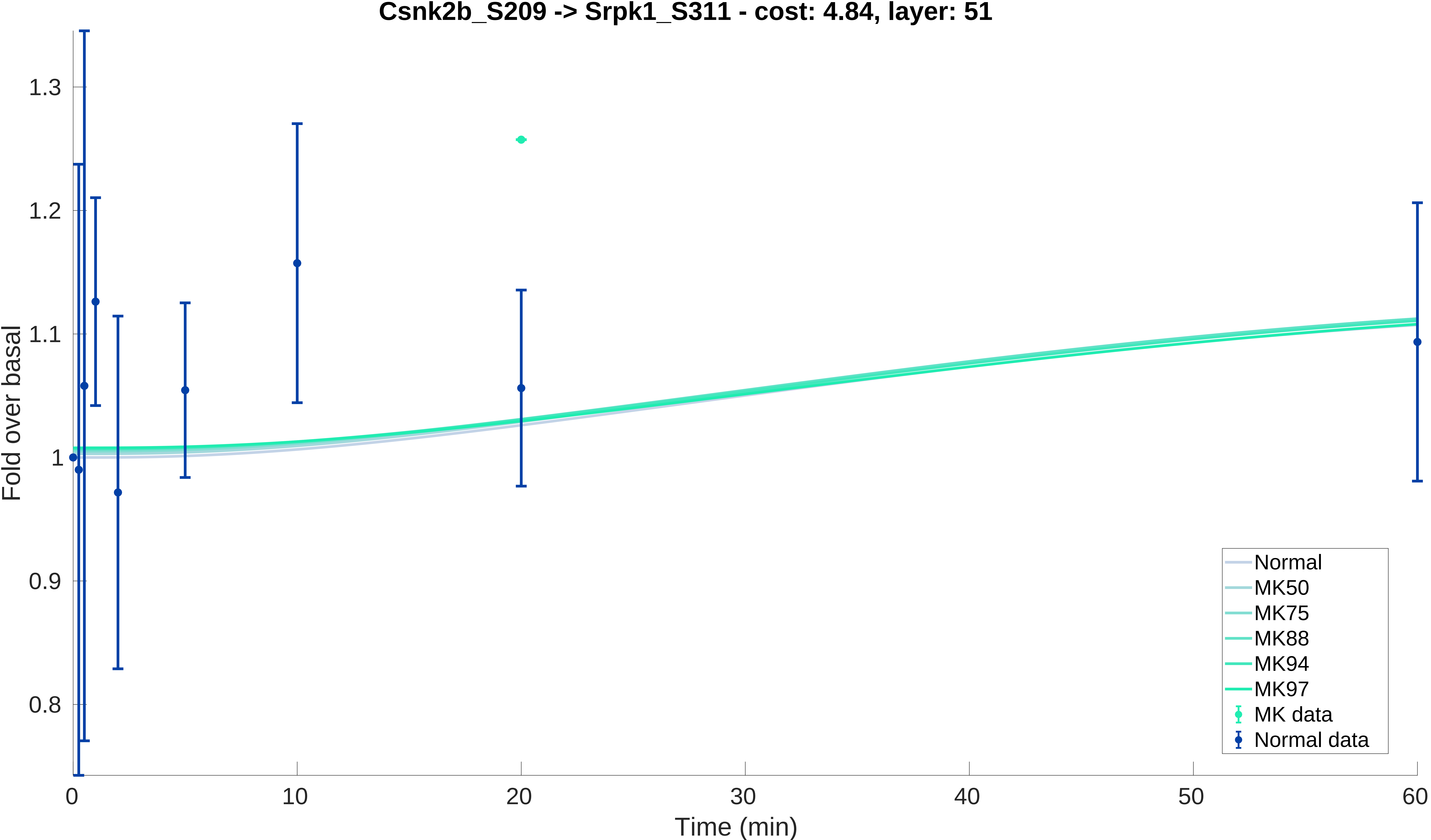

Csnk2b_S209 -> Srpk1_S311 - cost: 4.84, layer: 51
Fold over basal
1.3
1.2
1.1
1
0.9
0.8
0
10
20
30
40
50
60
Time (min)
Normal
MK50
MK75
MK88
MK94
MK97
MK data
Normal data

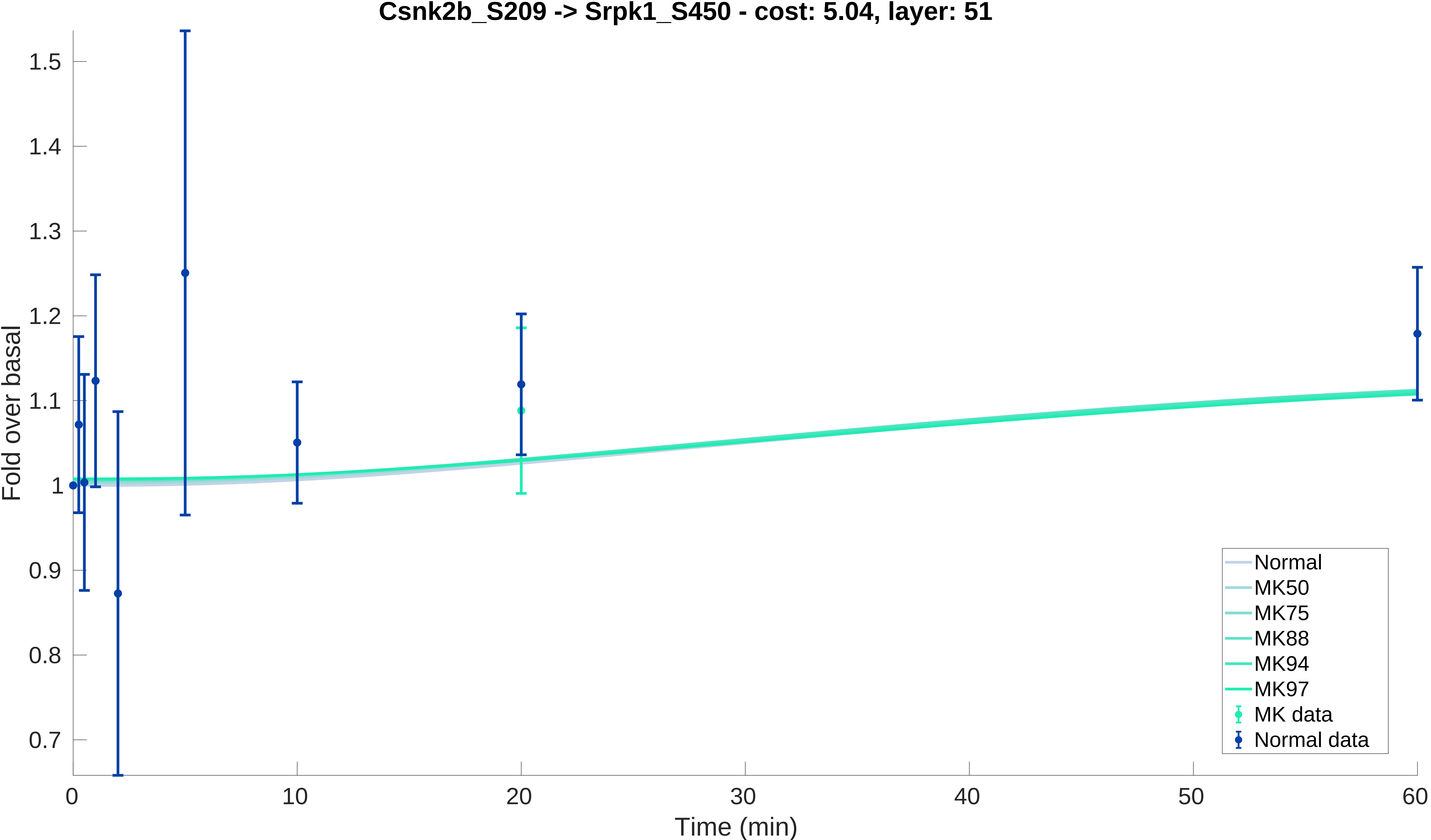
Csnk2b_S209 -> Srpk1_S450 - cost: 5.04, layer: 51
Fold over basal
1.5
1.4
1.3
1.2
1.1
1
0.9
0.8
0.7
0
10
20
30
40
50
60
Time (min)
Normal
MK50
MK75
MK88
MK94
MK97
MK data
Normal data

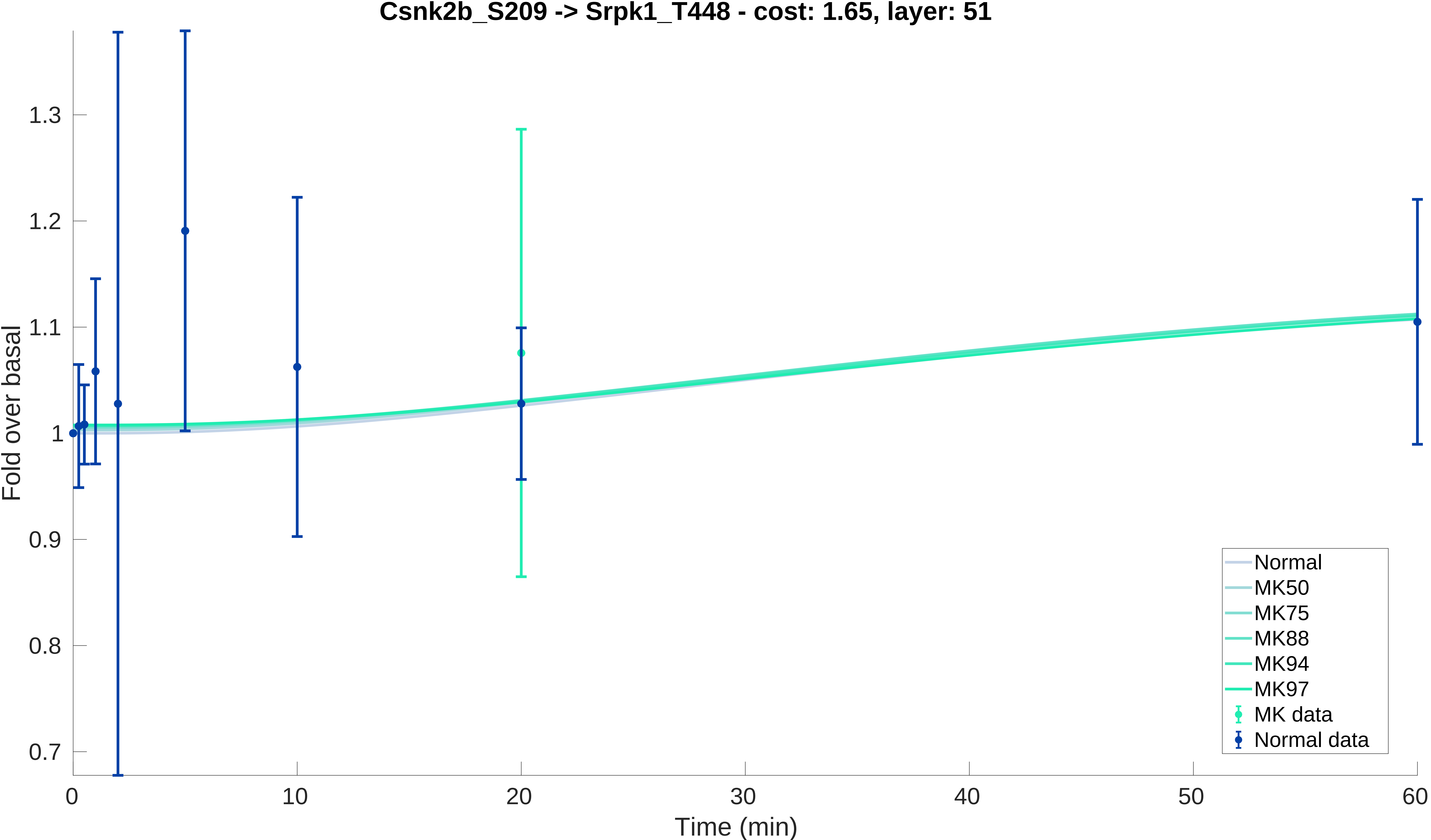

Csnk2b_S209 -> Srpk1_T448 - cost: 1.65, layer: 51
Fold over basal
1.3
1.2
1.1
1
0.9
0.8
0.7
0
10
20
30
40
50
60
Time (min)
Normal
MK50
MK75
MK88
MK94
MK97
MK data
Normal data

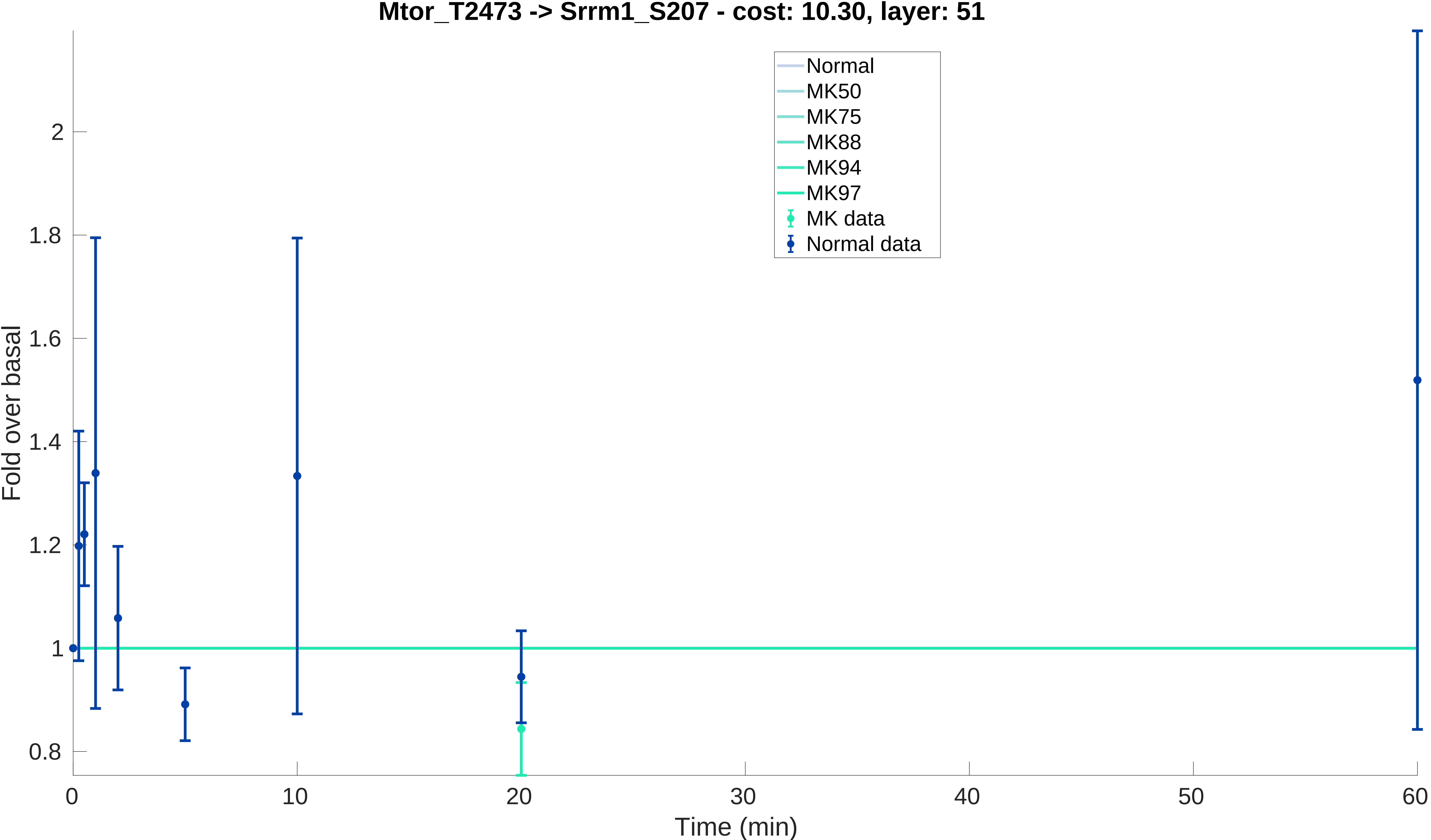

Mtor_T2473 -> Srrm1_S207 - cost: 10.30, layer: 51
Normal
MK50
MK75
MK88
MK94
MK97
MK data
Normal data
Fold over basal
2
1.8
1.6
1.4
1.2
1
0.8
0
10
20
30
40
50
60
Time (min)

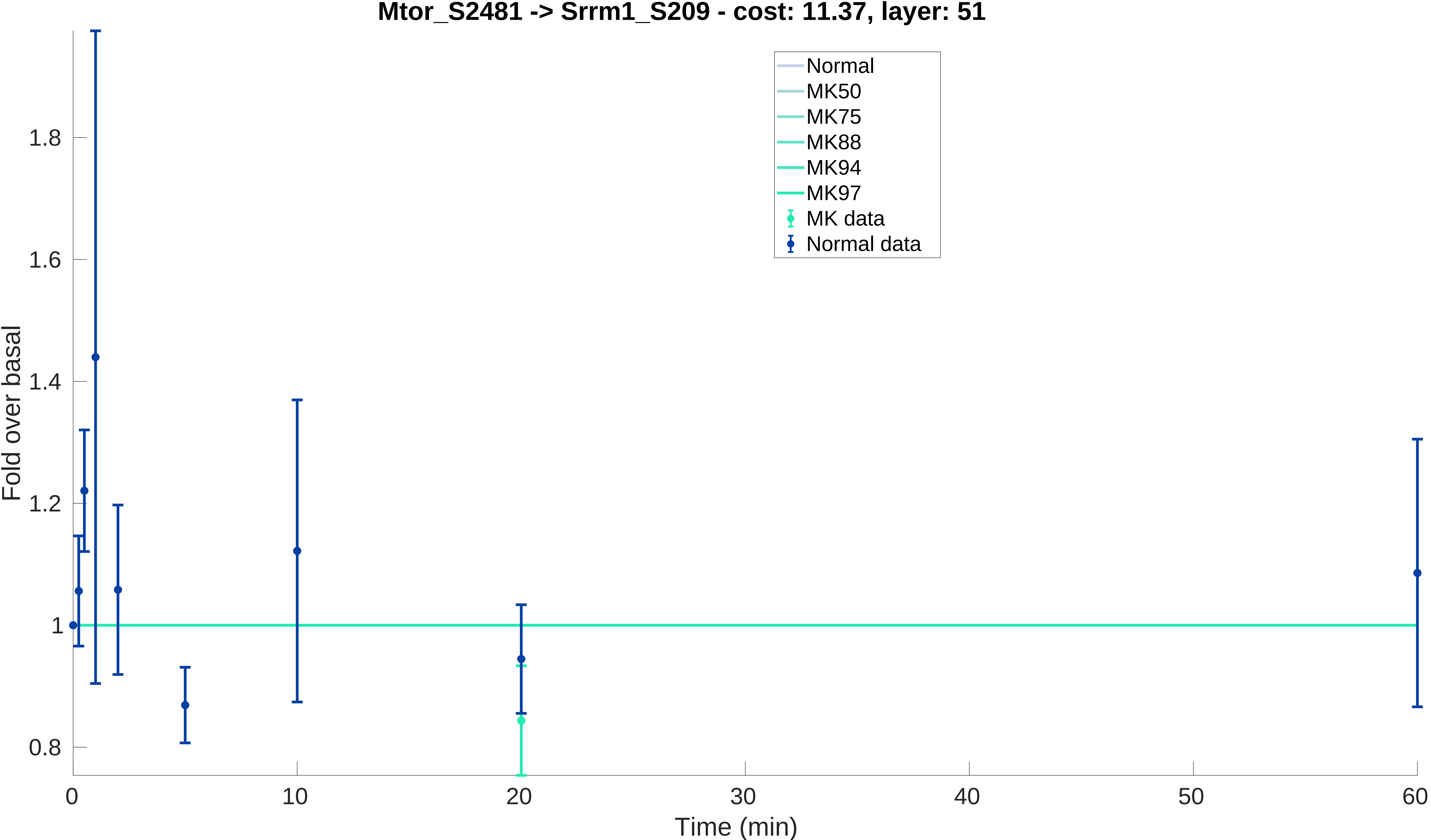

Mtor_S2481 -> Srrm1_S209 - cost: 11.37, layer: 51
Normal
MK50
MK75
MK88
MK94
MK97
MK data
Normal data
Fold over basal
1.8
1.6
1.4
1.2
1
0.8
0
10
20
30
40
50
60
Time (min)

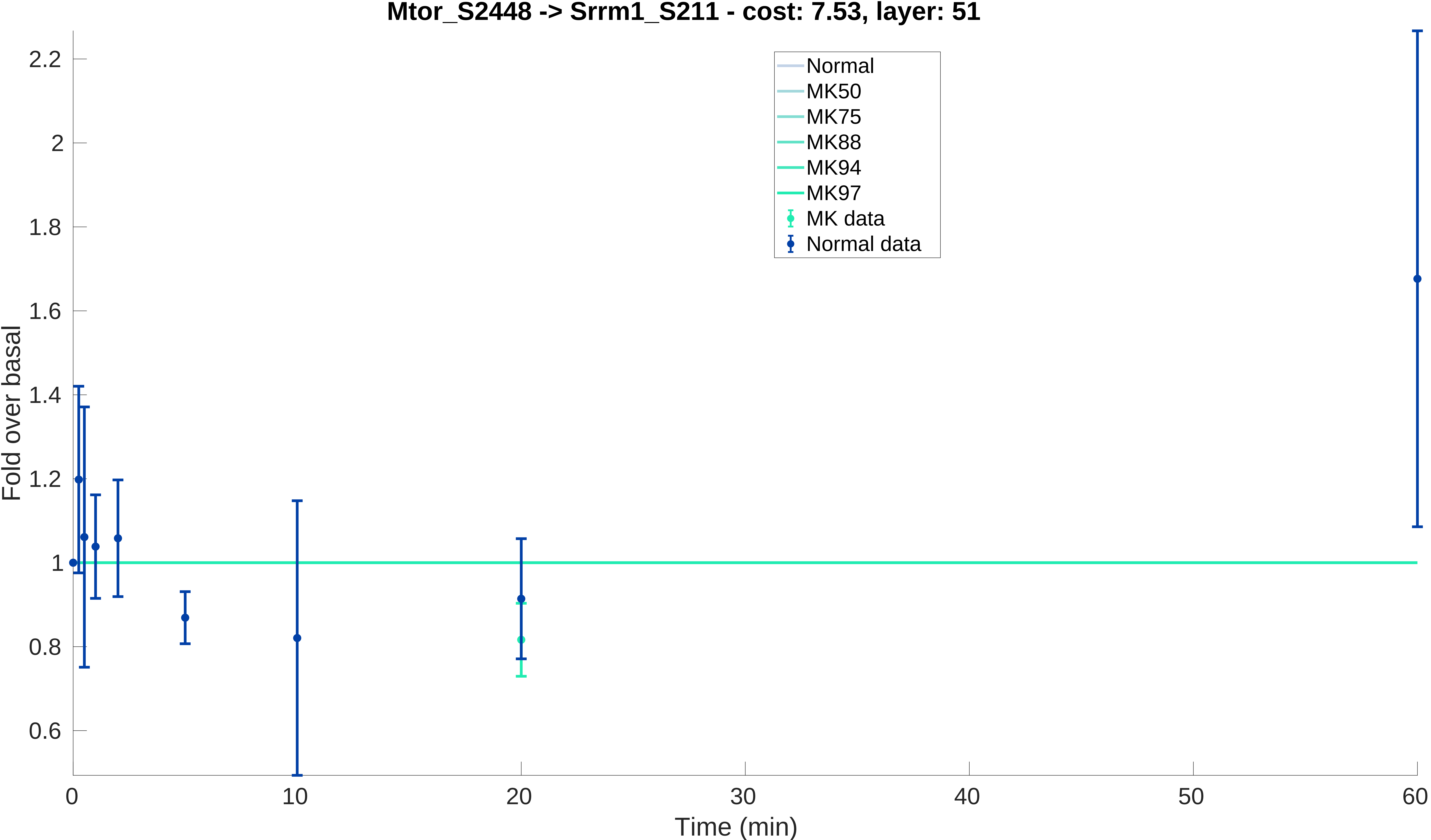
Mtor_S2448 -> Srrm1_S211 - cost: 7.53, layer: 51
Fold over basal
Time (min)
0.6
0.8
1
1.2
1.4
1.6
1.8
2
2.2
0
10
20
30
40
50
60
Normal
MK50
MK75
MK88
MK94
MK97
MK data
Normal data

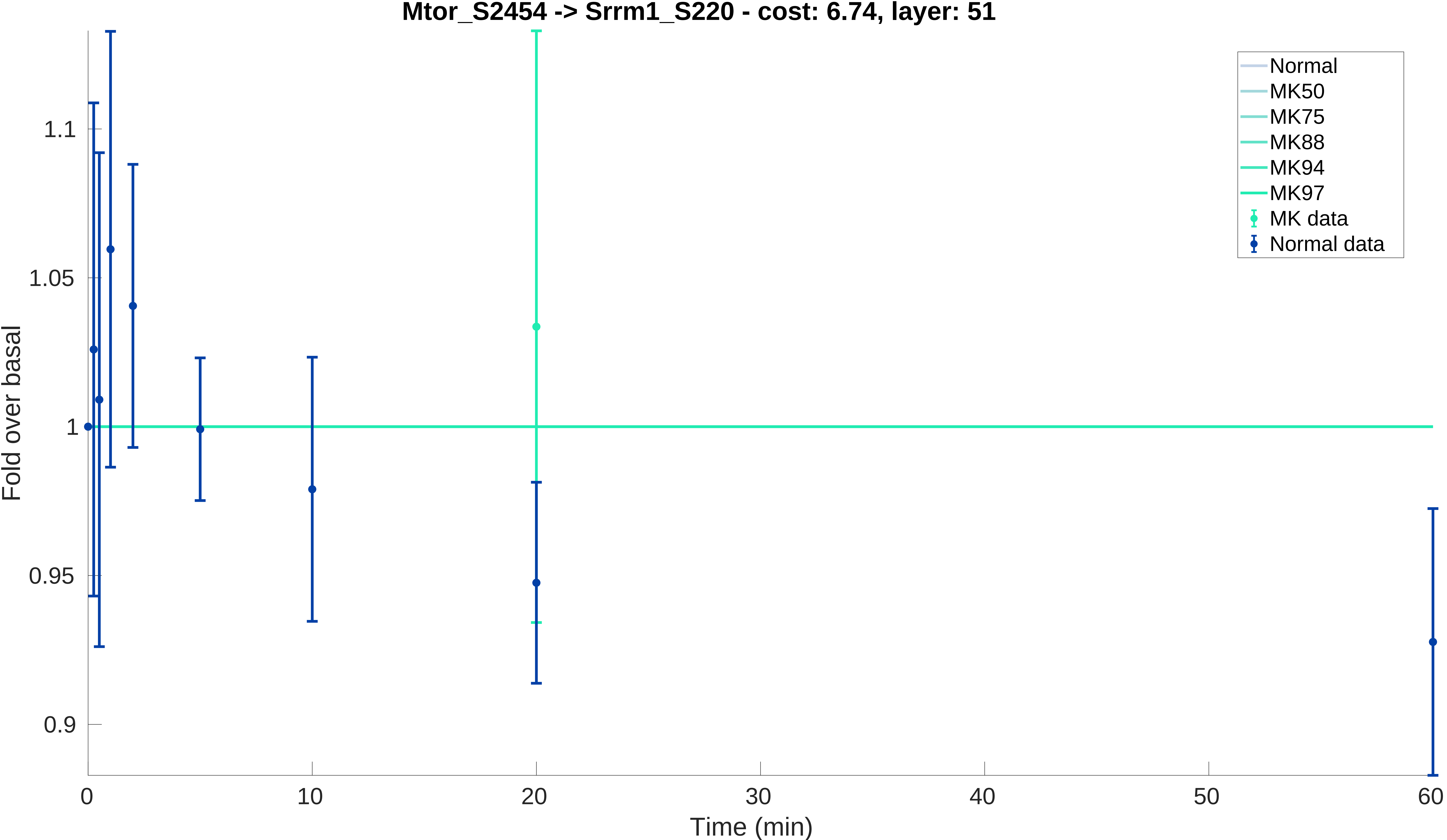

Mtor_S2454 -> Srrm1_S220 - cost: 6.74, layer: 51
Fold over basal
1.1
1.05
1
0.95
0.9
0
10
20
30
40
50
60
Time (min)
Normal
MK50
MK75
MK88
MK94
MK97
MK data
Normal data

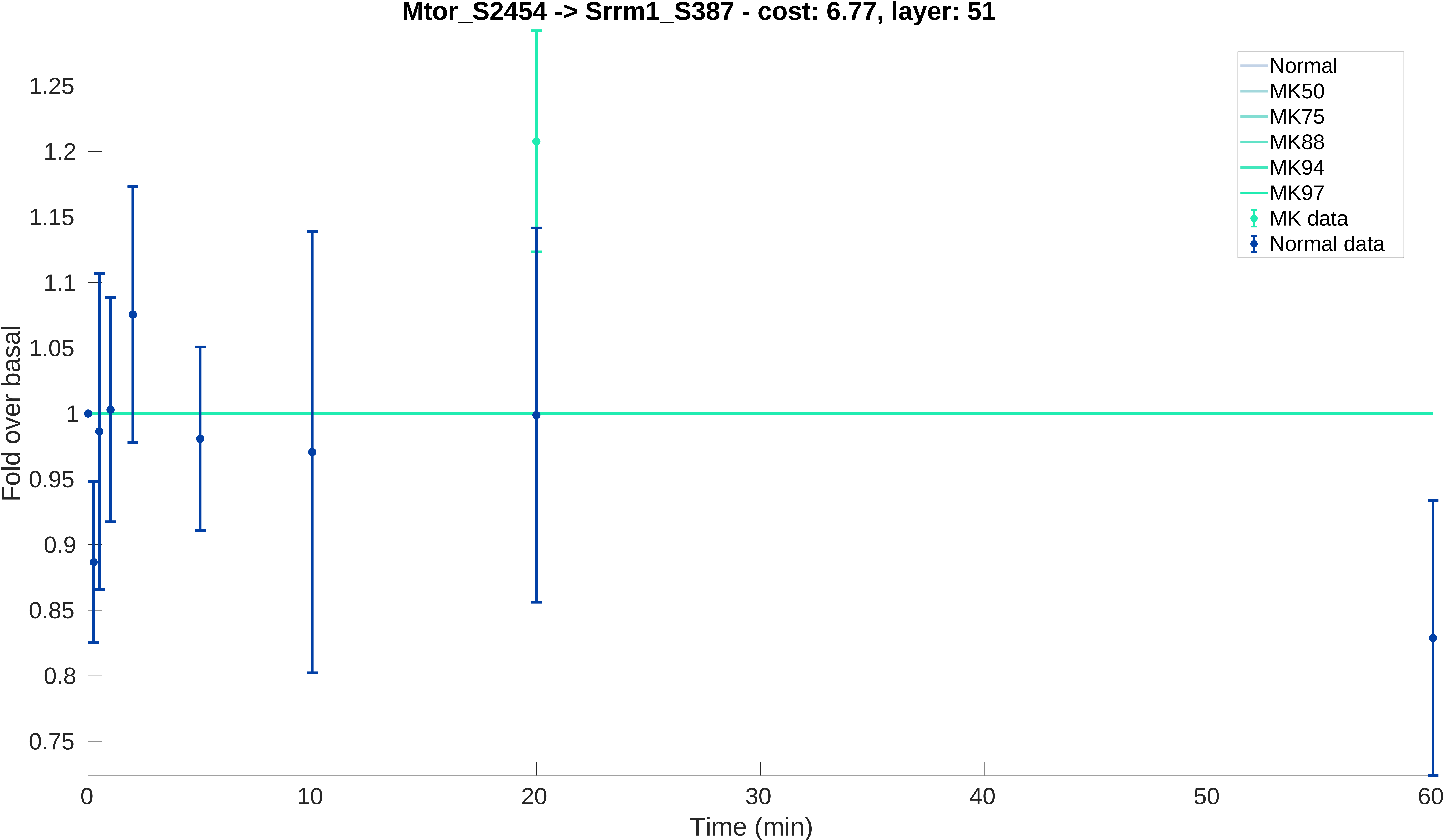

Mtor_S2454 -> Srrm1_S387 - cost: 6.77, layer: 51
Fold over basal
Time (min)
Normal
MK50
MK75
MK88
MK94
MK97
MK data
Normal data
1.25
1.2
1.15
1.1
1.05
1
0.95
0.9
0.85
0.8
0.75
0
10
20
30
40
50
60

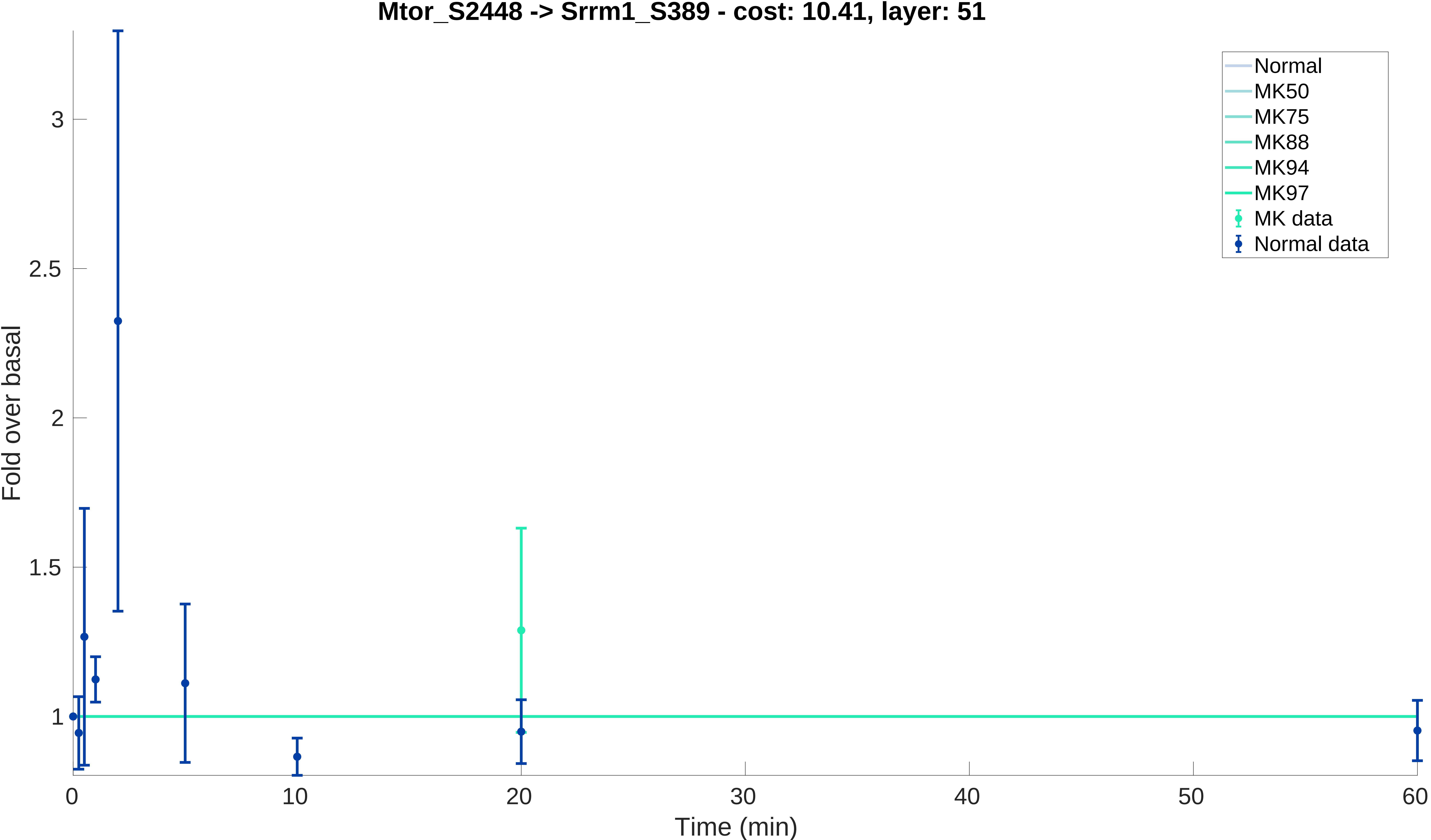
Mtor_S2448 -> Srrm1_S389 - cost: 10.41, layer: 51
Fold over basal
Time (min)
3
2.5
2
1.5
1
0
10
20
30
40
50
60
Normal
MK50
MK75
MK88
MK94
MK97
MK data
Normal data

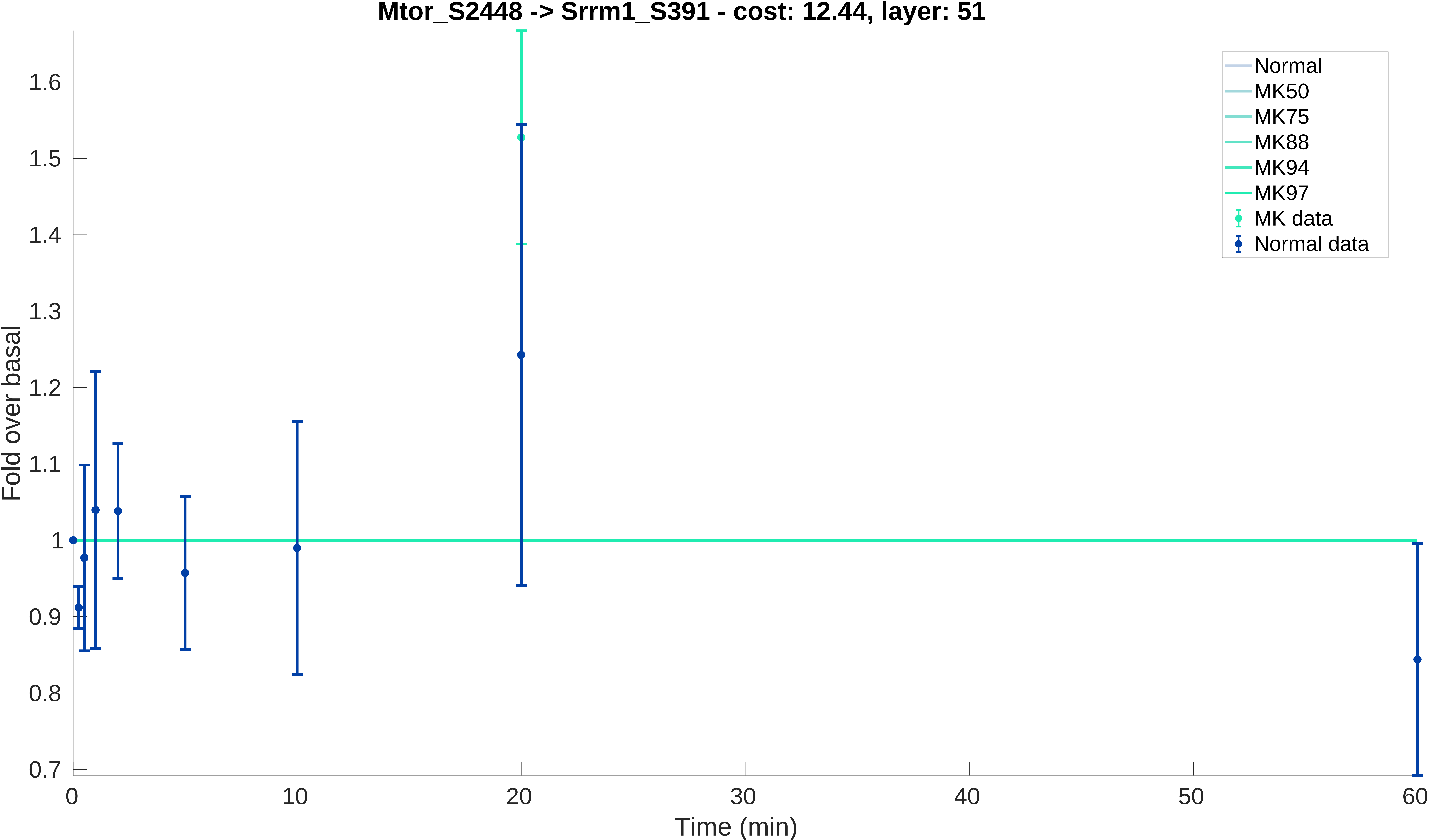

Mtor_S2448 -> Srrm1_S391 - cost: 12.44, layer: 51
Fold over basal
Time (min)
1.6
1.5
1.4
1.3
1.2
1.1
1
0.9
0.8
0.7
0
10
20
30
40
50
60
Normal
MK50
MK75
MK88
MK94
MK97
MK data
Normal data

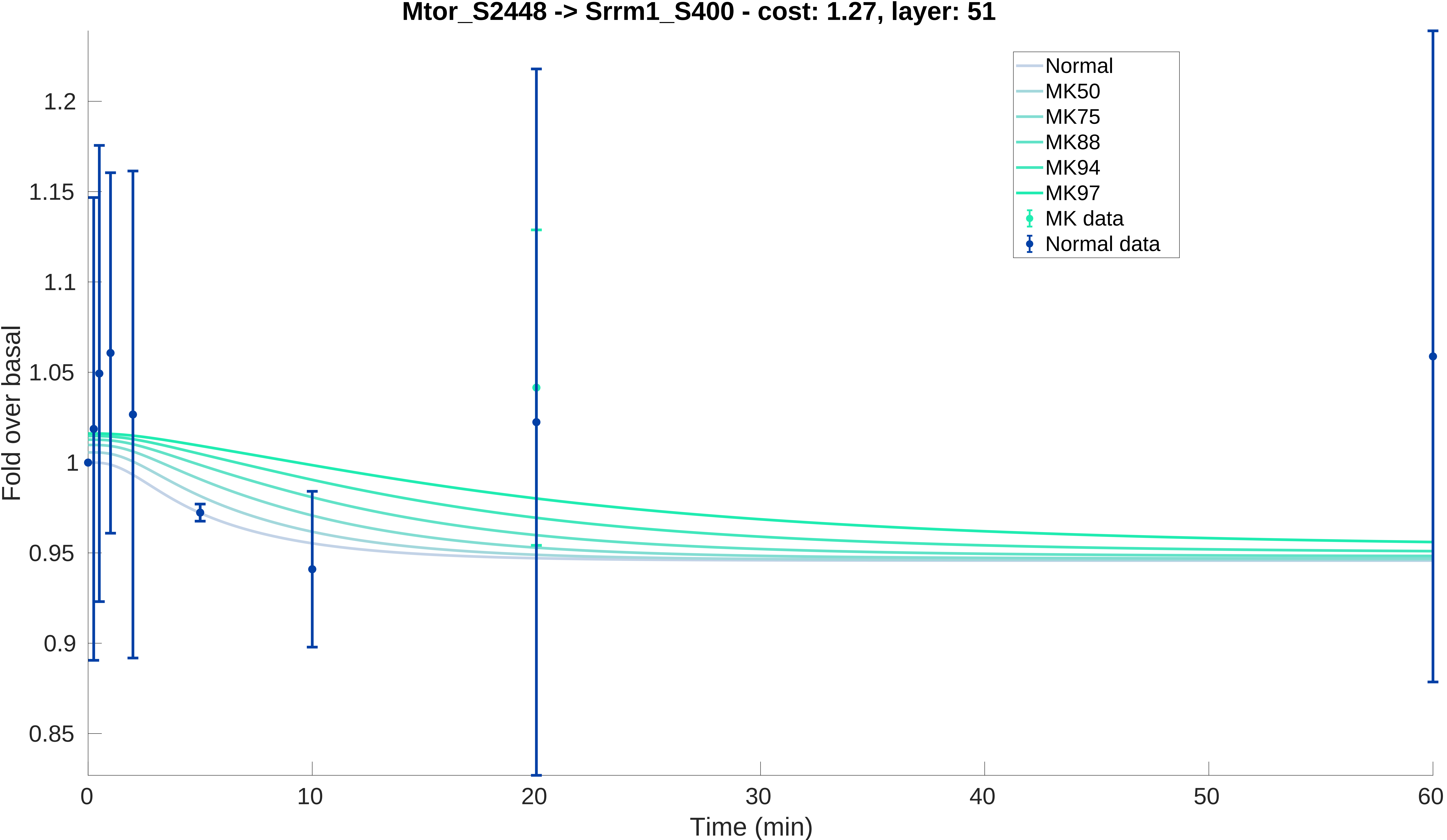

Mtor_S2448 -> Srrm1_S400 - cost: 1.27, layer: 51
Normal
MK50
MK75
MK88
MK94
MK97
MK data
Normal data
Fold over basal
1.2
1.15
1.1
1.05
1
0.95
0.9
0.85
0
10
20
30
40
50
60
Time (min)

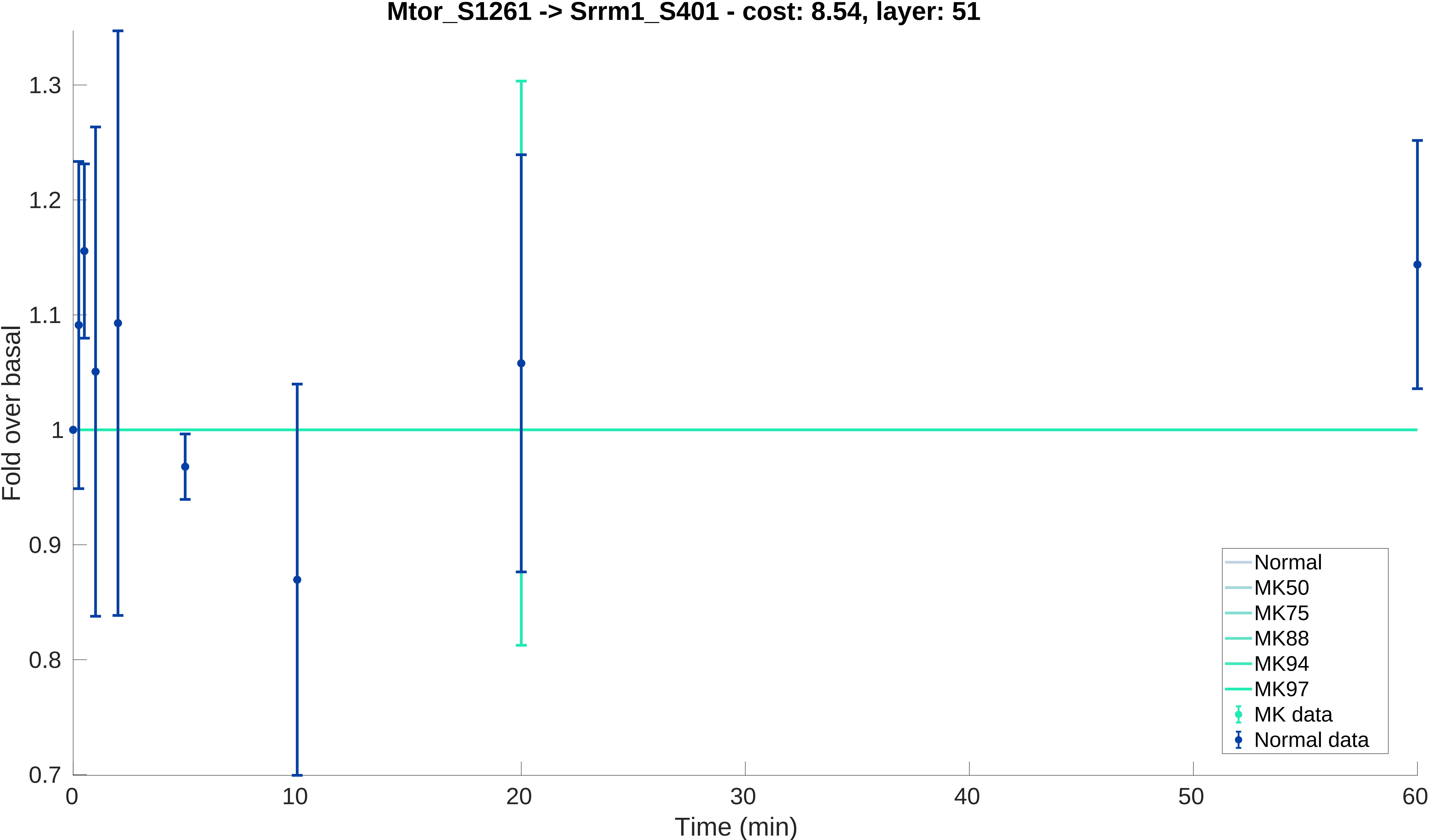

Mtor_S1261 -> Srrm1_S401 - cost: 8.54, layer: 51
Fold over basal
Time (min)
1.3
1.2
1.1
1
0.9
0.8
0.7
0
10
20
30
40
50
60
Normal
MK50
MK75
MK88
MK94
MK97
MK data
Normal data

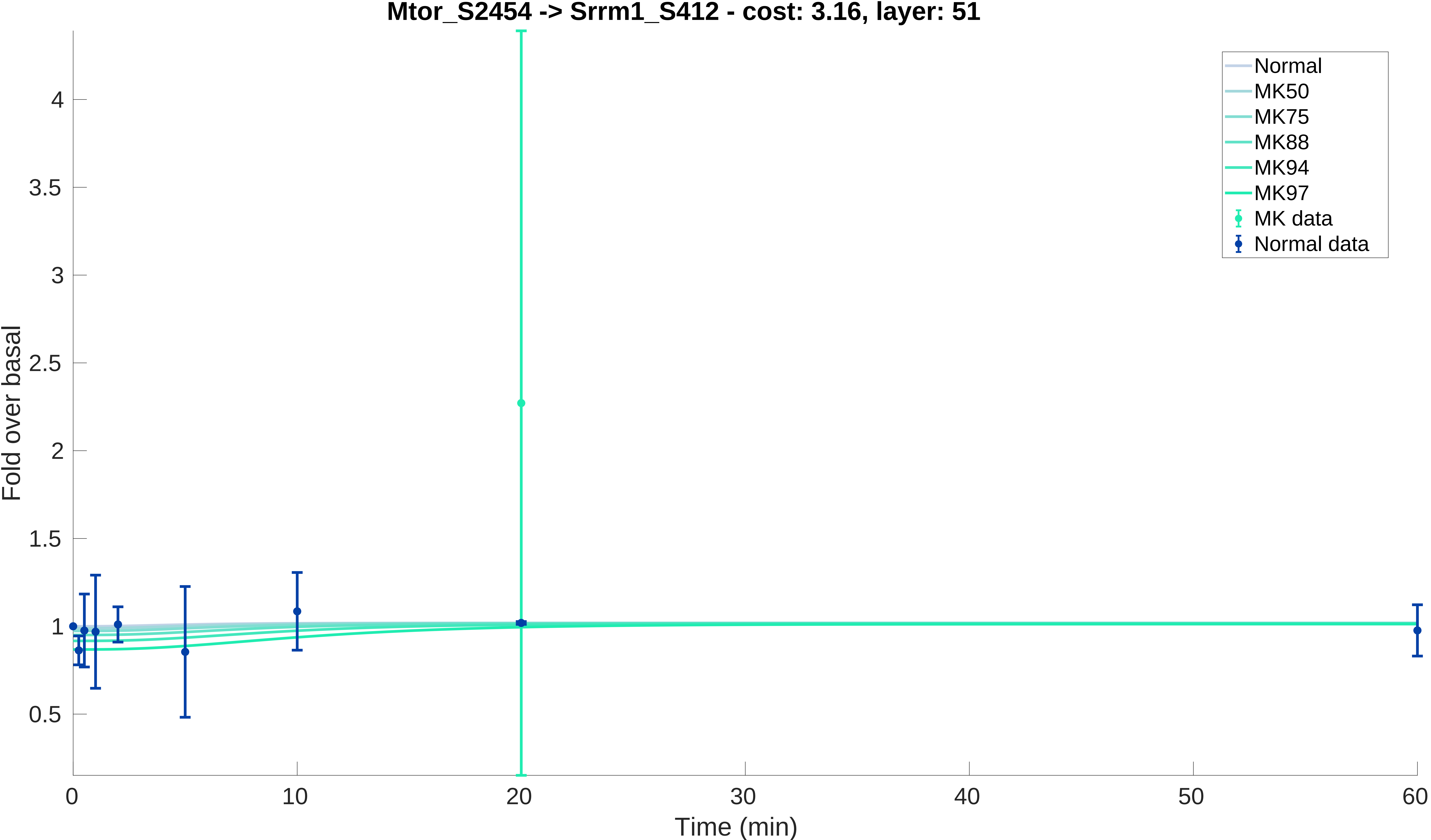
Mtor_S2454 -> Srrm1_S412 - cost: 3.16, layer: 51
Fold over basal
Time (min)
4
3.5
3
2.5
2
1.5
1
0.5
0
10
20
30
40
50
60
Normal
MK50
MK75
MK88
MK94
MK97
MK data
Normal data

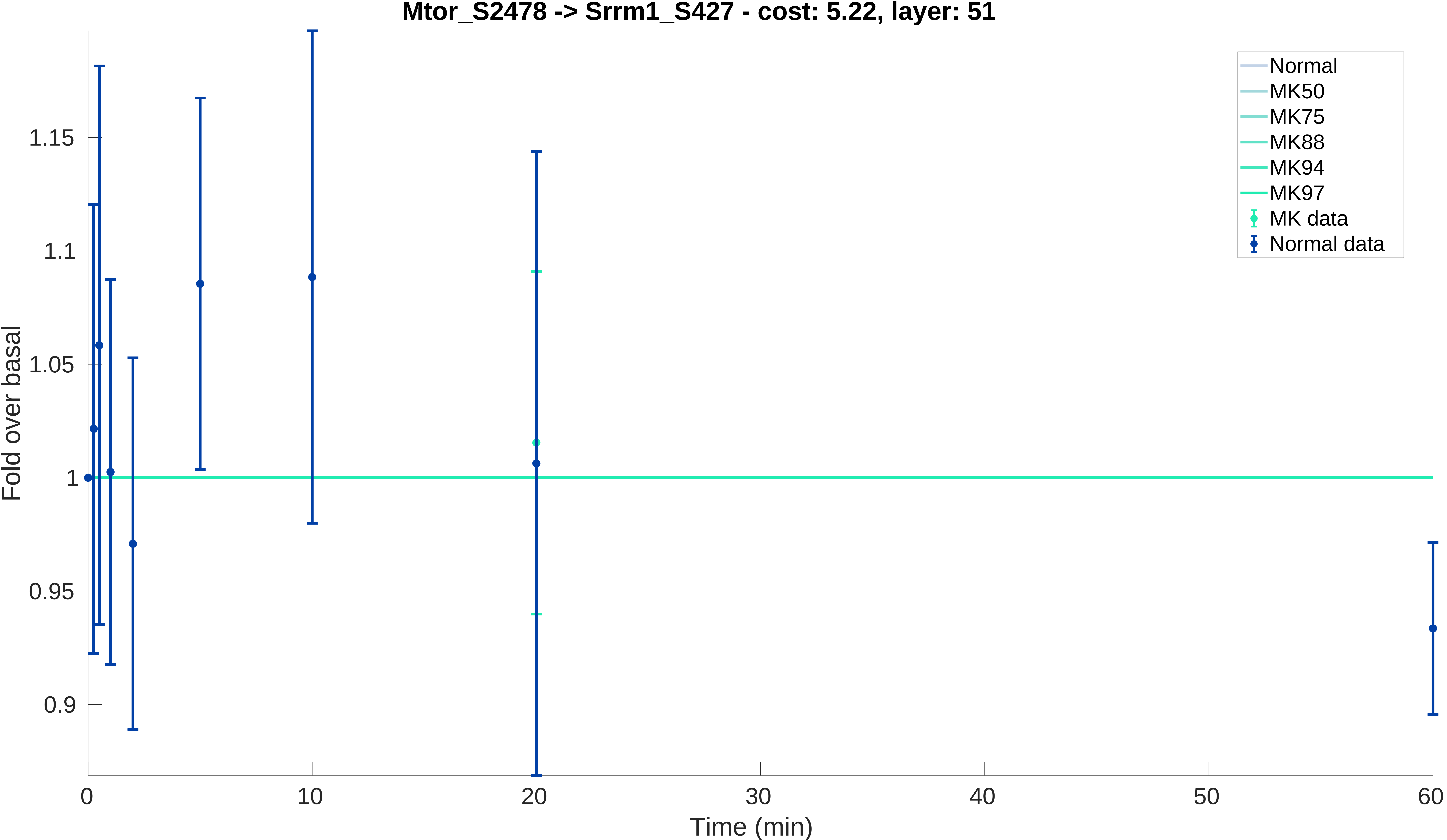
Mtor_S2478 -> Srrm1_S427 - cost: 5.22, layer: 51
Fold over basal
Time (min)
0
10
20
30
40
50
60
0.9
0.95
1
1.05
1.1
1.15
Normal
MK50
MK75
MK88
MK94
MK97
MK data
Normal data

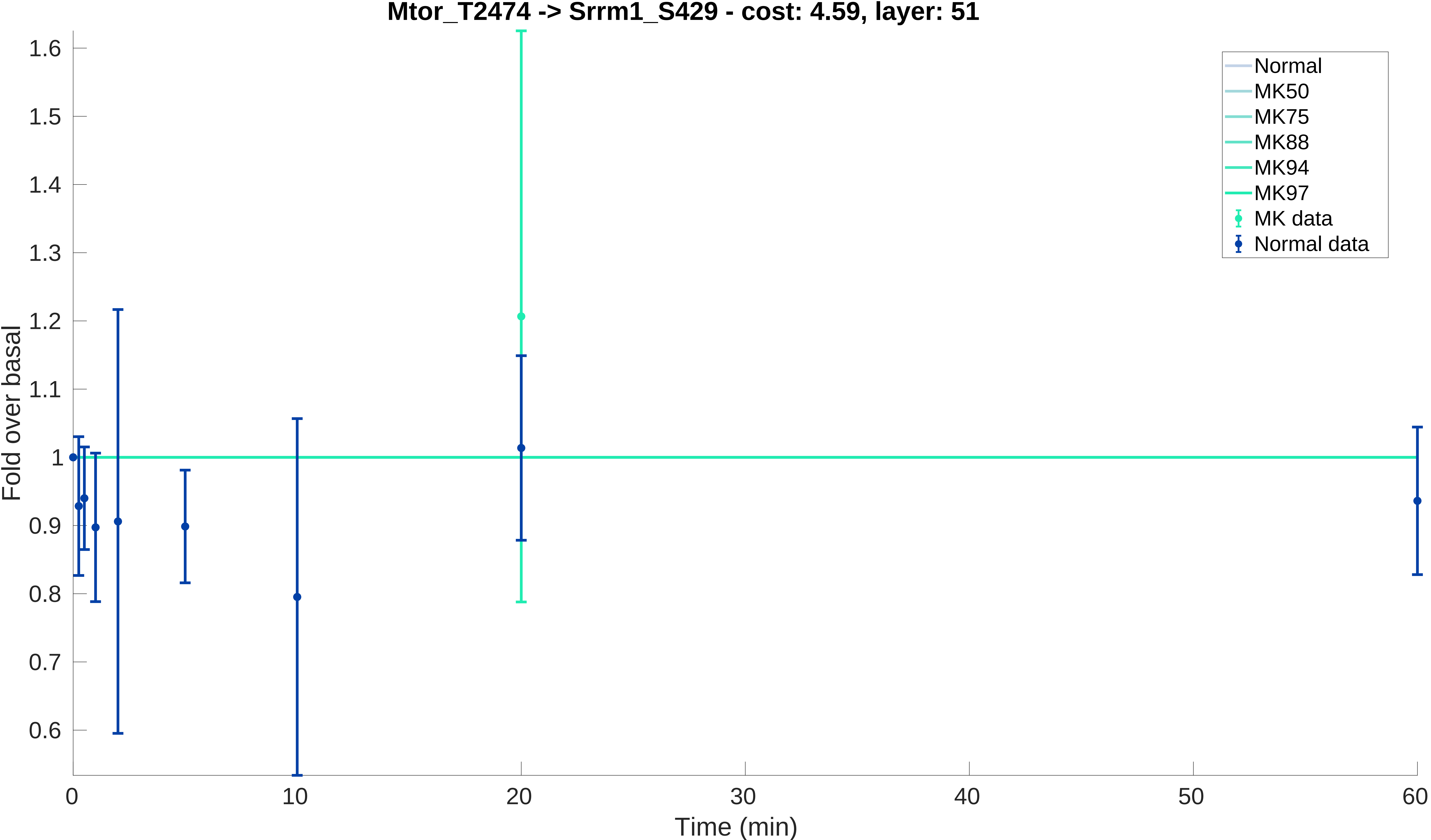

Mtor_T2474 -> Srrm1_S429 - cost: 4.59, layer: 51
Fold over basal
Time (min)
Normal
MK50
MK75
MK88
MK94
MK97
MK data
Normal data
1.6
1.5
1.4
1.3
1.2
1.1
1
0.9
0.8
0.7
0.6
0
10
20
30
40
50
60

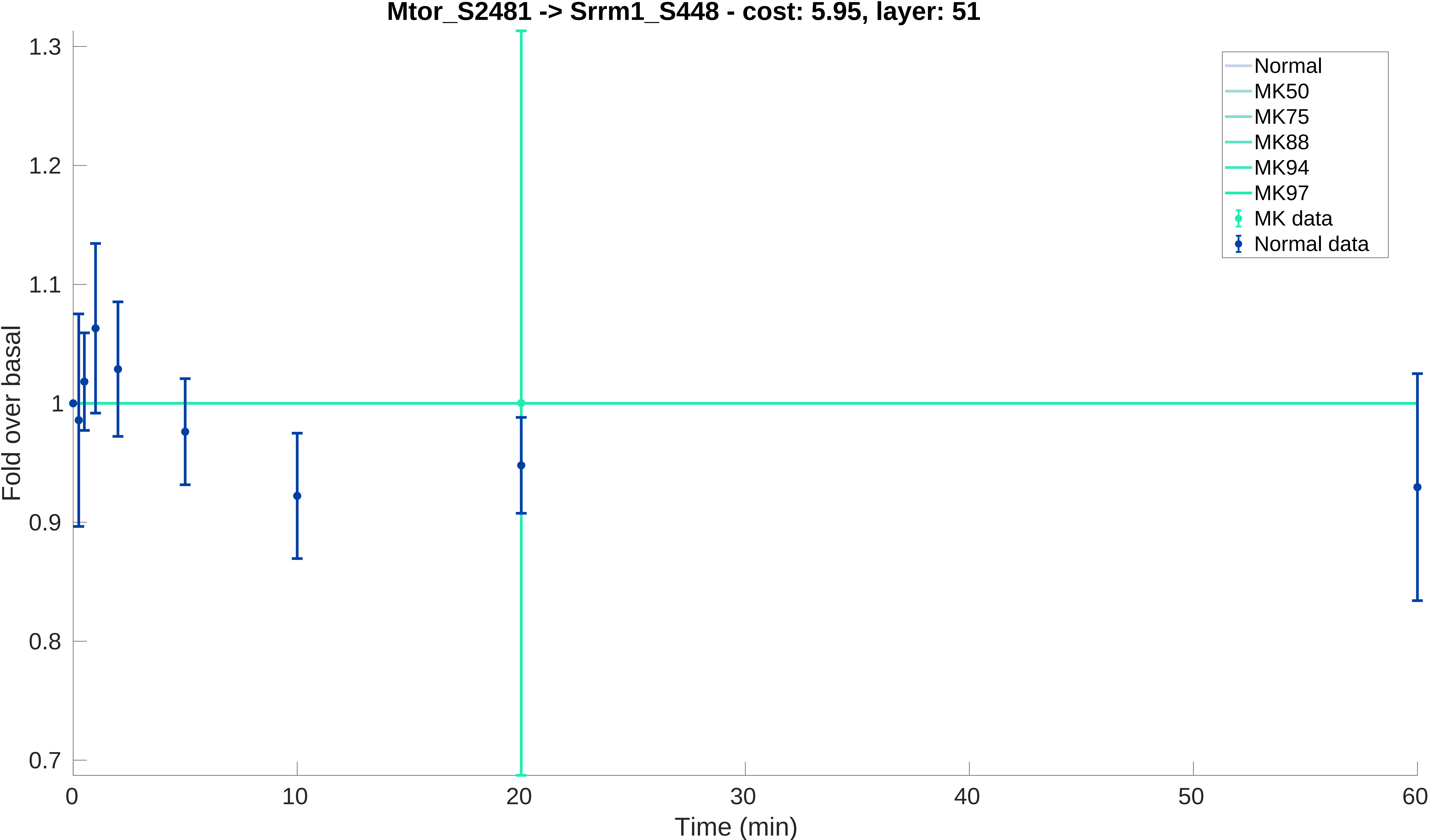

Mtor_S2481 -> Srrm1_S448 - cost: 5.95, layer: 51
Fold over basal
Time (min)
1.3
1.2
1.1
1
0.9
0.8
0.7
0
10
20
30
40
50
60
Normal
MK50
MK75
MK88
MK94
MK97
MK data
Normal data

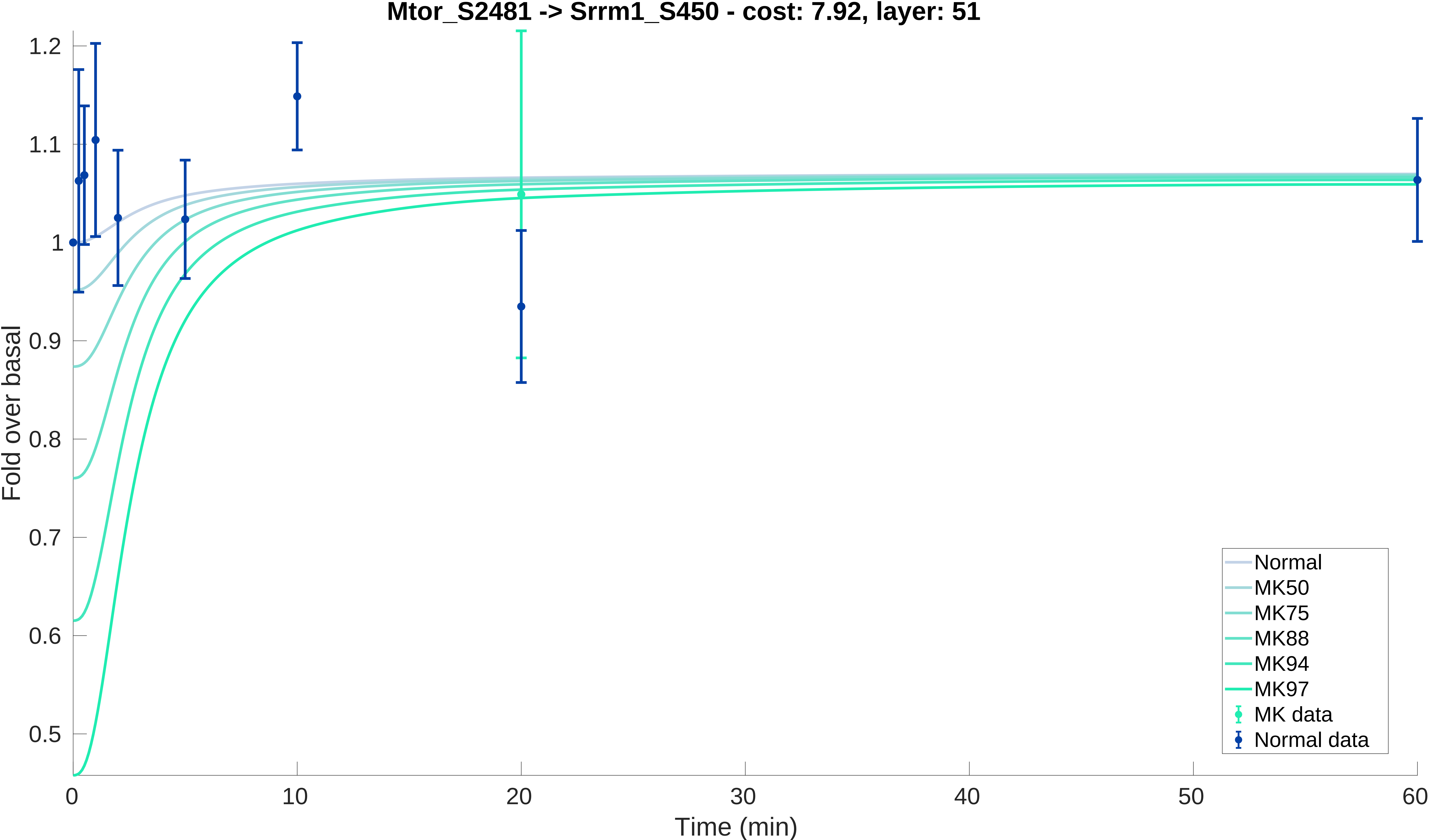

Mtor_S2481 -> Srrm1_S450 - cost: 7.92, layer: 51
1.2
1.1
1
0.9
0.8
0.7
0.6
0.5
Fold over basal
0
10
20
30
40
50
60
Time (min)
Normal
MK50
MK75
MK88
MK94
MK97
MK data
Normal data

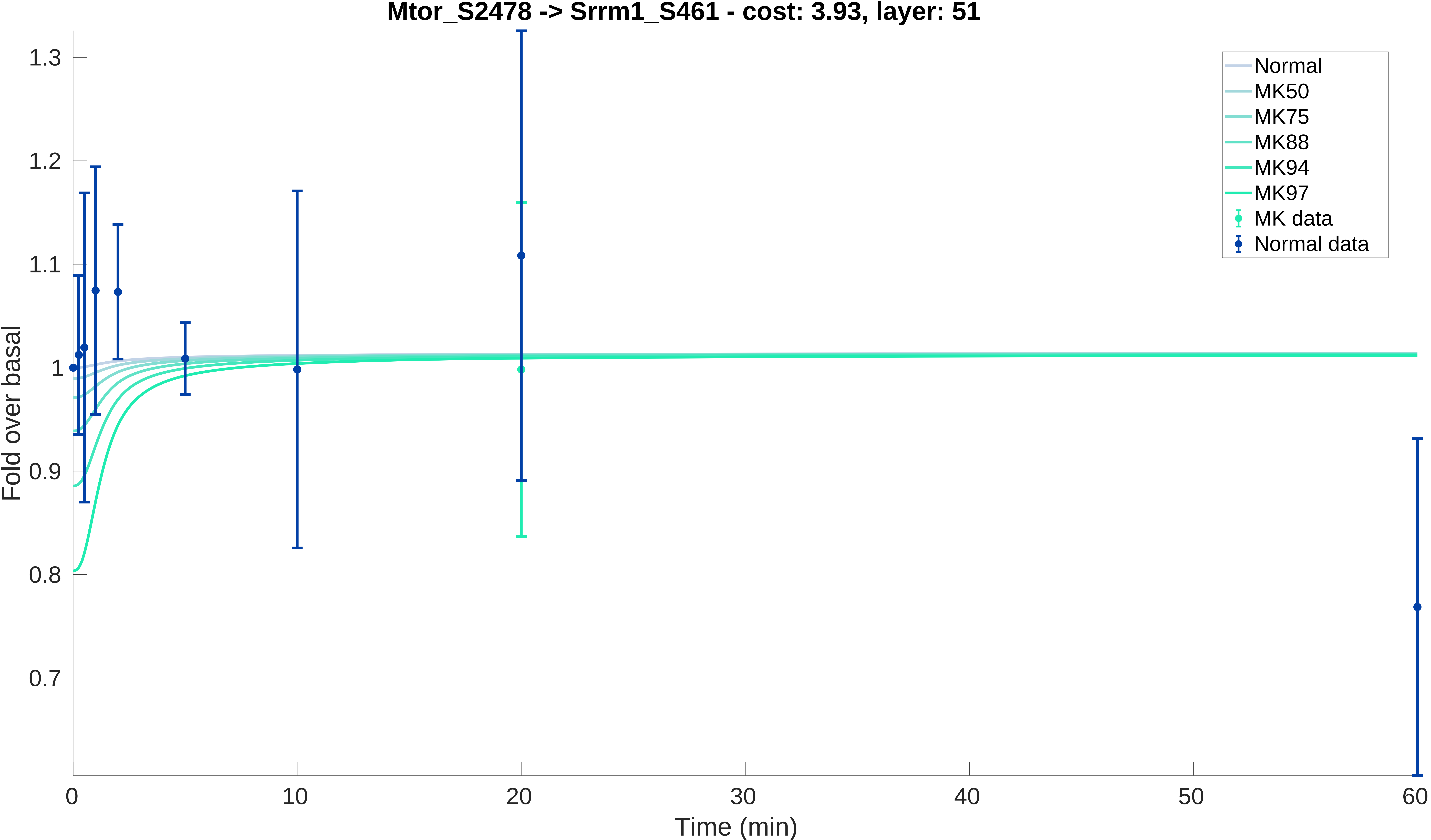

Mtor_S2478 -> Srrm1_S461 - cost: 3.93, layer: 51
Fold over basal
Time (min)
1.3
1.2
1.1
1
0.9
0.8
0.7
0
10
20
30
40
50
60
Normal
MK50
MK75
MK88
MK94
MK97
MK data
Normal data

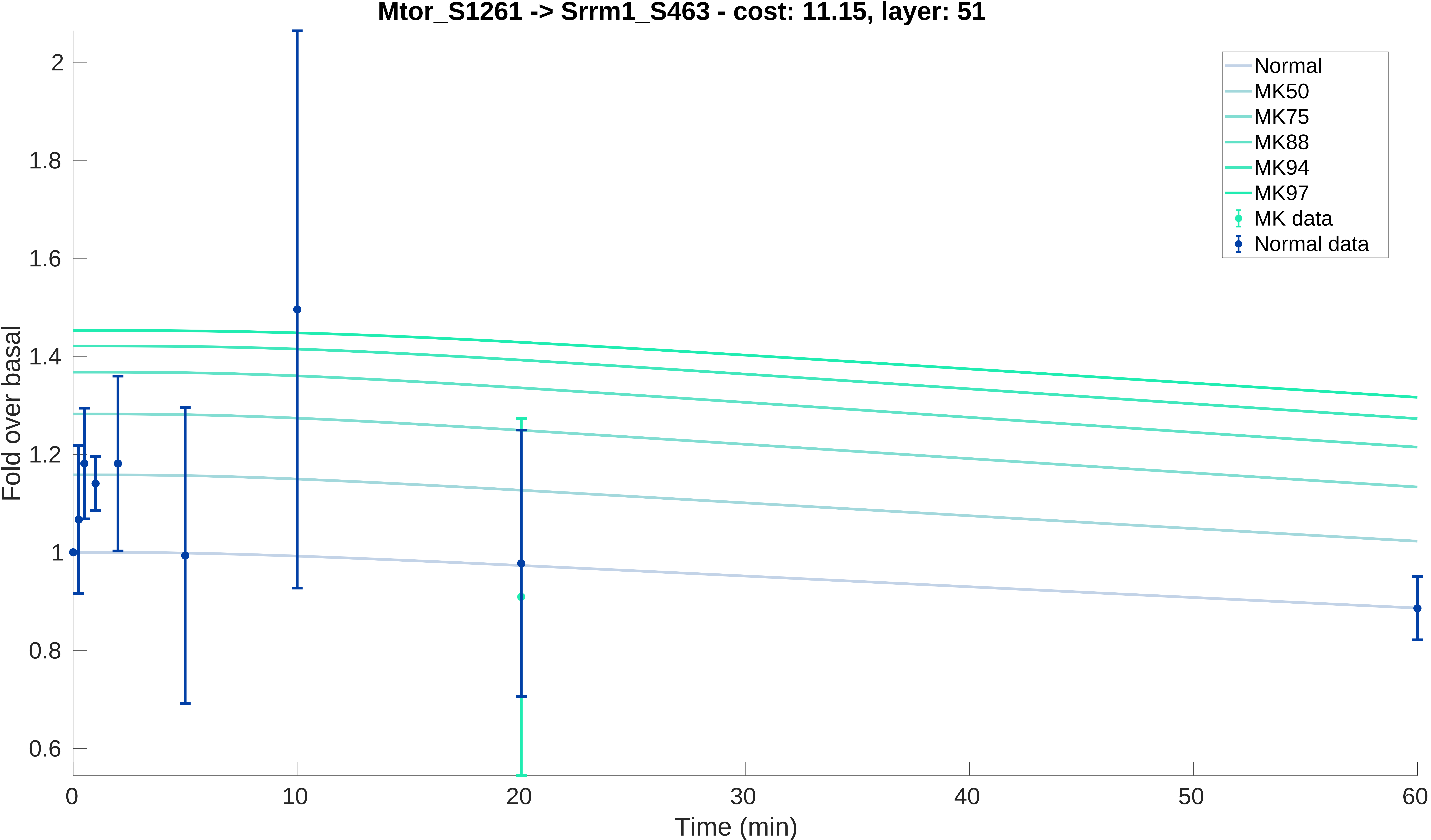
Mtor_S1261 -> Srrm1_S463 - cost: 11.15, layer: 51
Fold over basal
Time (min)
Normal
MK50
MK75
MK88
MK94
MK97
MK data
Normal data
0.6
0.8
1
1.2
1.4
1.6
1.8
2
0
10
20
30
40
50
60

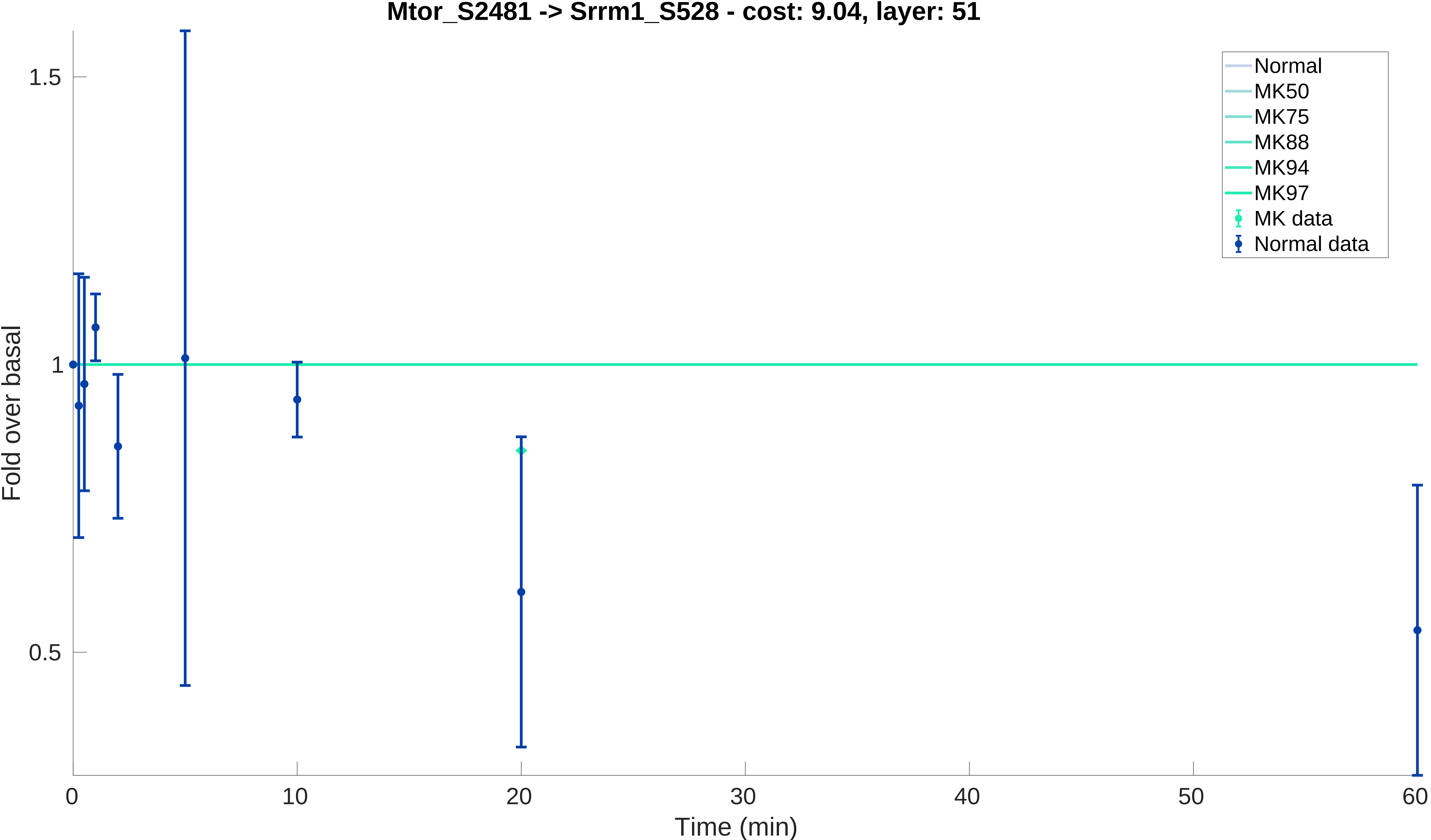

Mtor_S2481 -> Srrm1_S528 - cost: 9.04, layer: 51
Fold over basal
1.5
1
0.5
Time (min)
0
10
20
30
40
50
60
Normal
MK50
MK75
MK88
MK94
MK97
MK data
Normal data

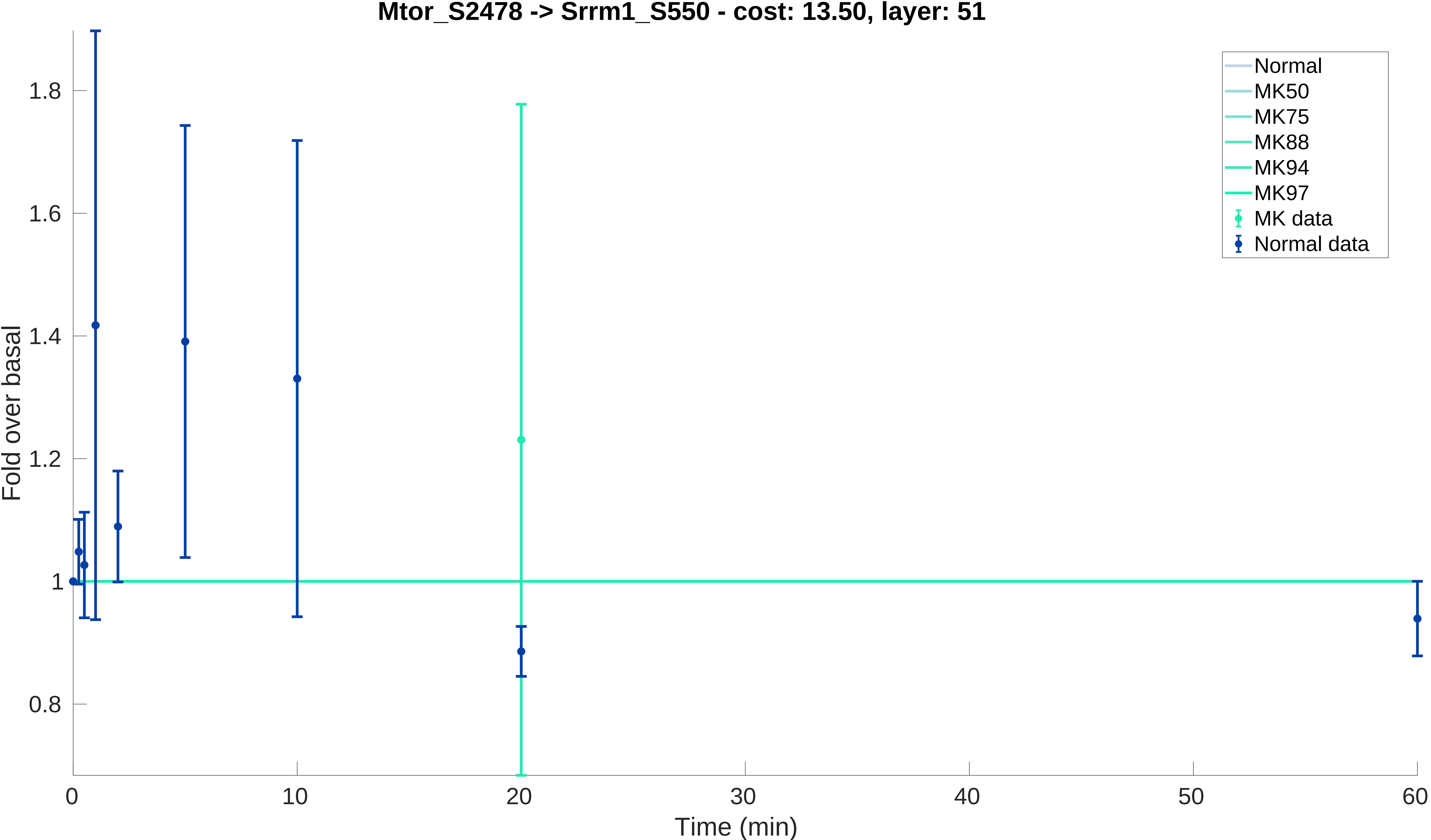

Mtor_S2478 -> Srrm1_S550 - cost: 13.50, layer: 51
Fold over basal
Time (min)
0.8
1
1.2
1.4
1.6
1.8
0
10
20
30
40
50
60
Normal
MK50
MK75
MK88
MK94
MK97
MK data
Normal data

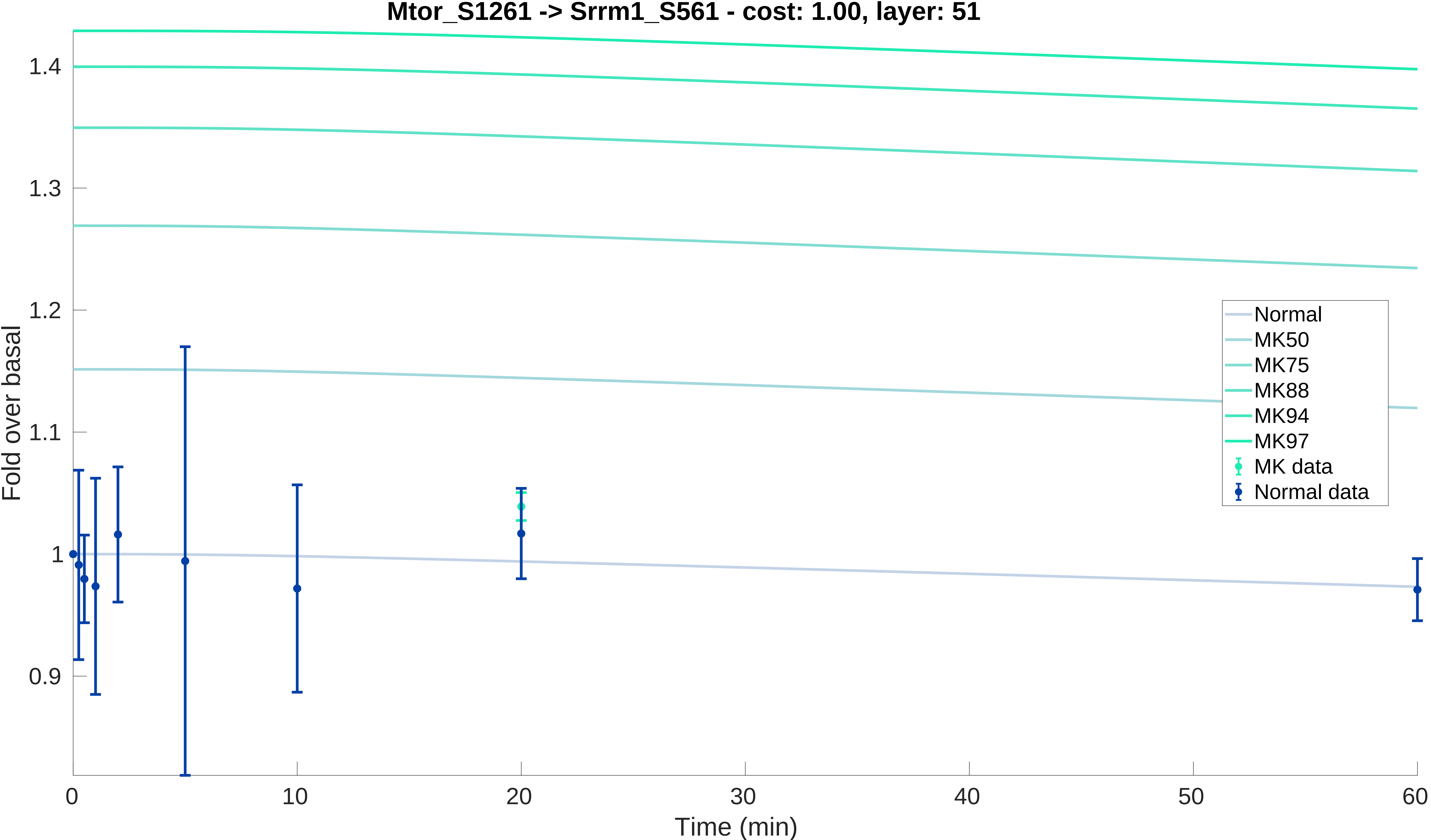
Mtor_S1261 -> Srrm1_S561 - cost: 1.00, layer: 51
Fold over basal
1.4
1.3
1.2
1.1
1
0.9
Time (min)
0
10
20
30
40
50
60
Normal
MK50
MK75
MK88
MK94
MK97
MK data
Normal data

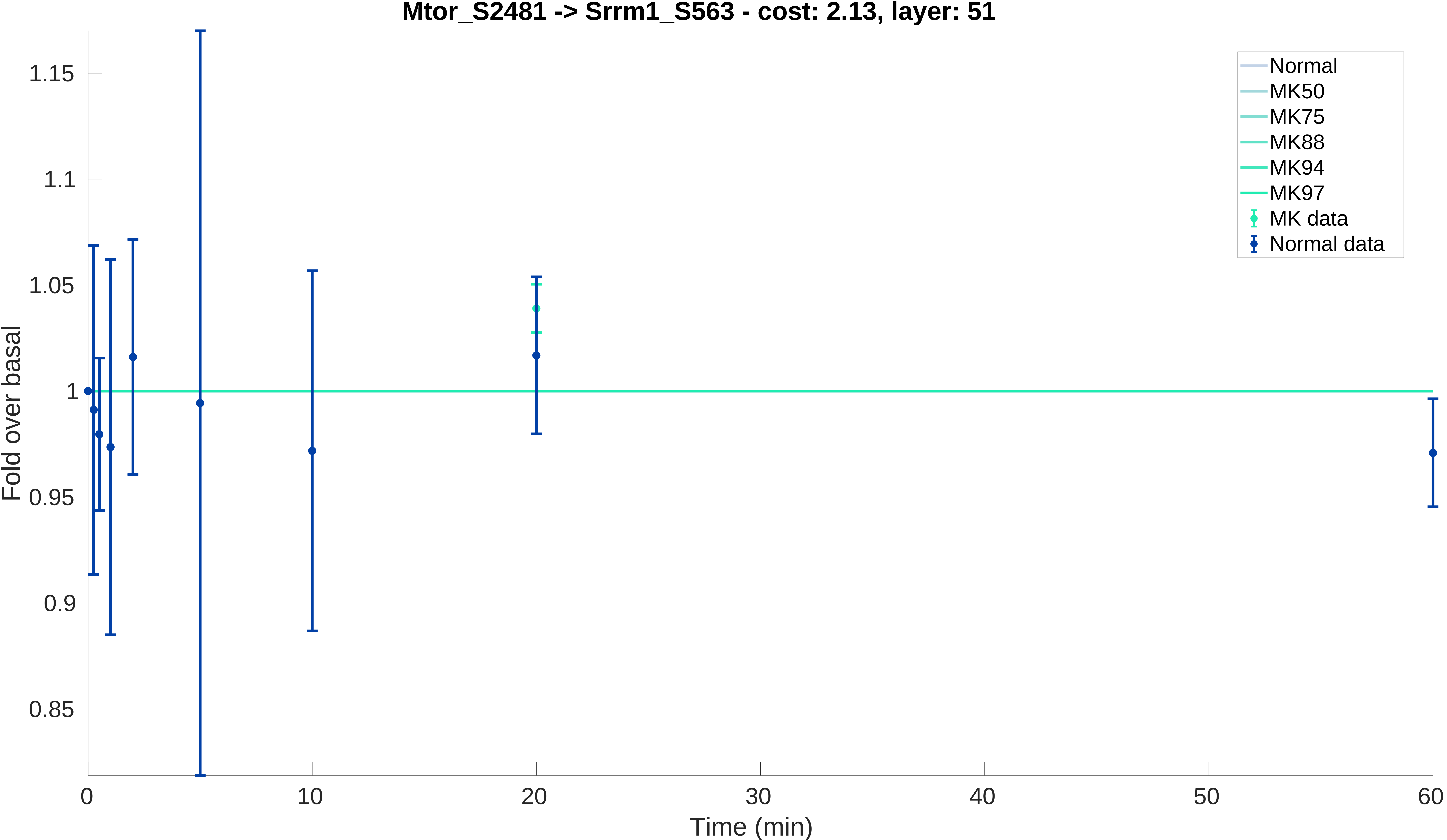

Mtor_S2481 -> Srrm1_S563 - cost: 2.13, layer: 51
Fold over basal
Time (min)
1.15
1.1
1.05
1
0.95
0.9
0.85
0
10
20
30
40
50
60
Normal
MK50
MK75
MK88
MK94
MK97
MK data
Normal data

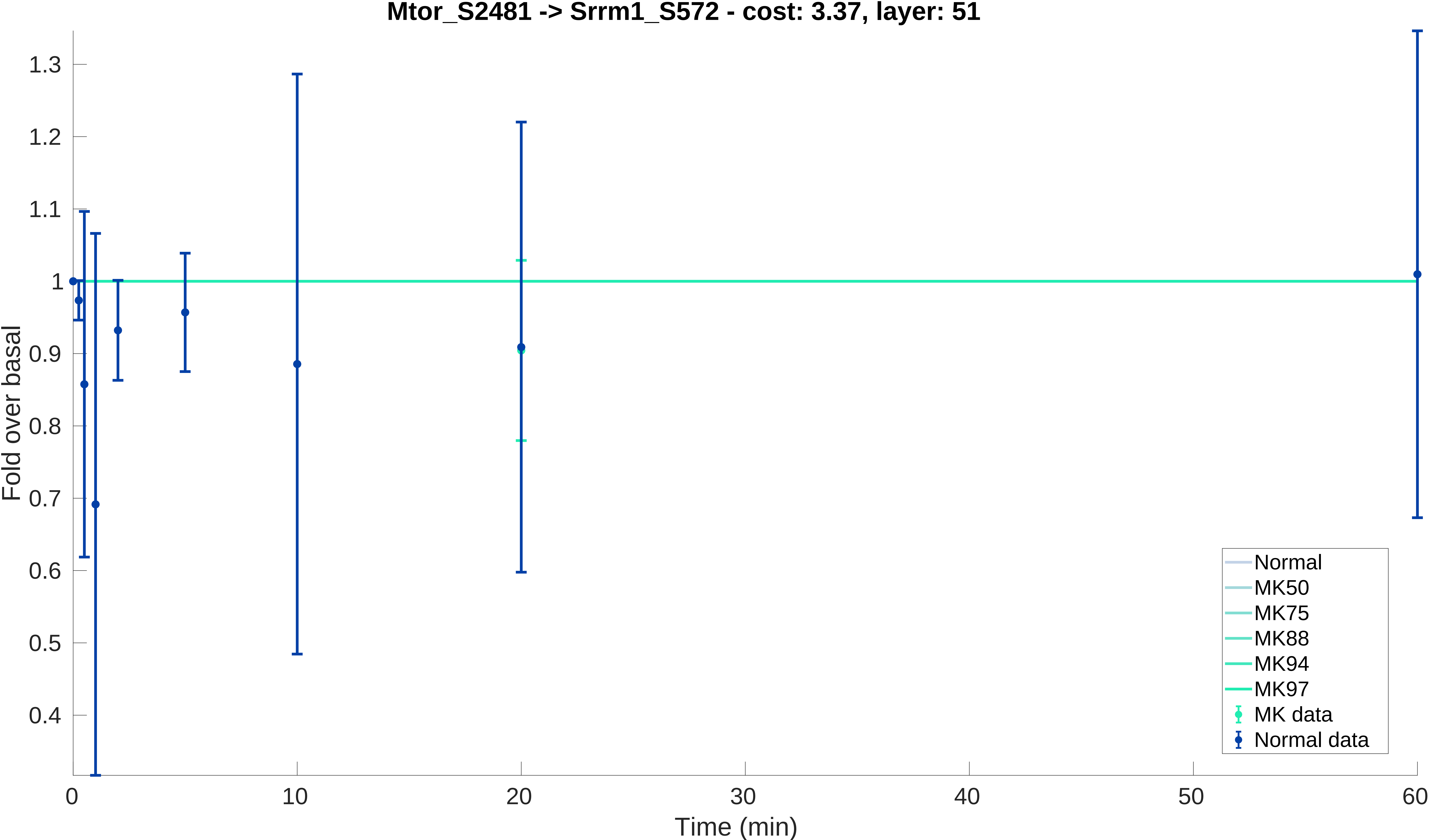

Mtor_S2481 -> Srrm1_S572 - cost: 3.37, layer: 51
Fold over basal
Time (min)
1.3
1.2
1.1
1
0.9
0.8
0.7
0.6
0.5
0.4
0
10
20
30
40
50
60
Normal
MK50
MK75
MK88
MK94
MK97
MK data
Normal data

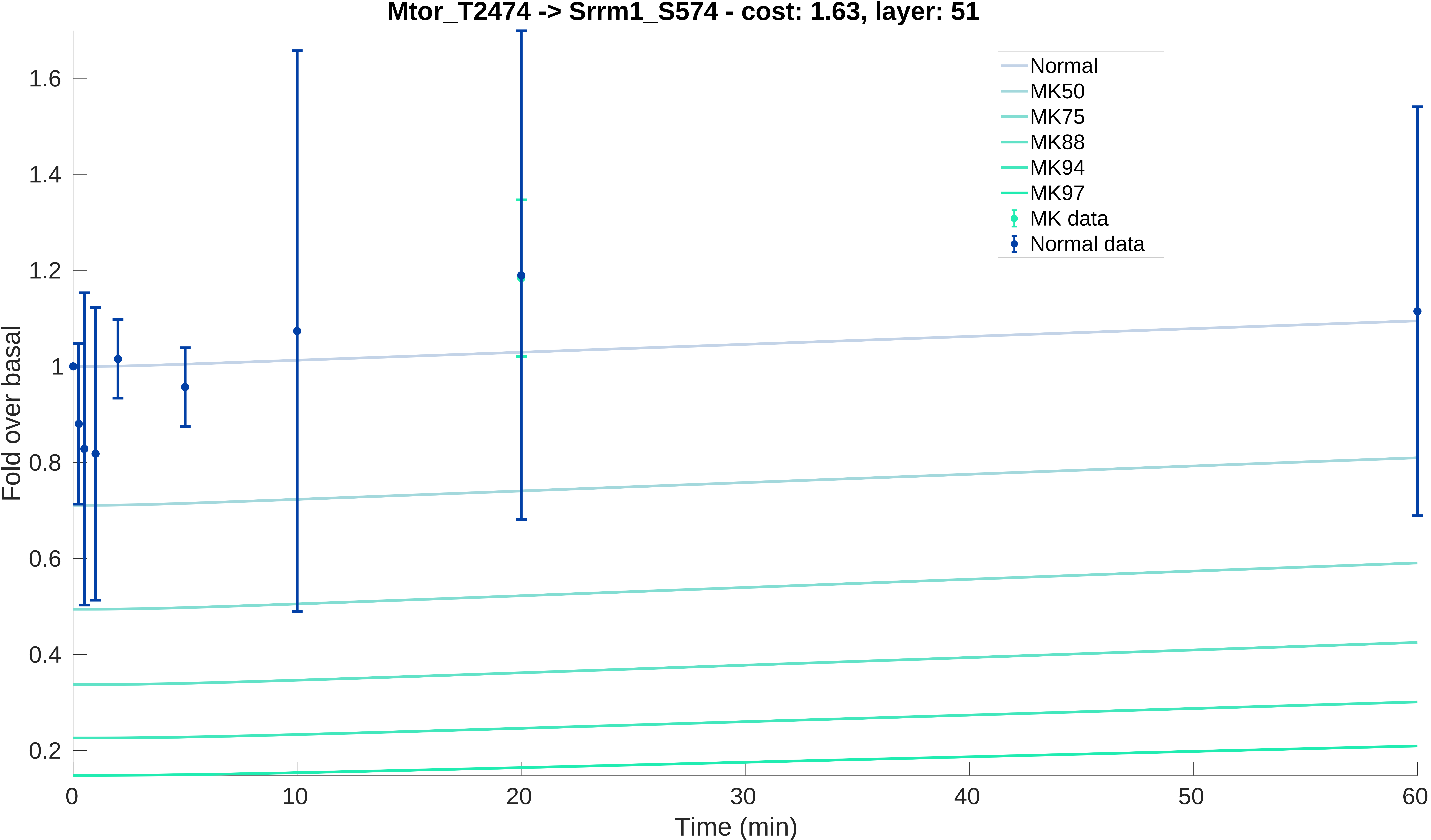

Mtor_T2474 -> Srrm1_S574 - cost: 1.63, layer: 51
Normal
MK50
MK75
MK88
MK94
MK97
MK data
Normal data
Fold over basal
1.6
1.4
1.2
1
0.8
0.6
0.4
0.2
0
10
20
30
40
50
60
Time (min)

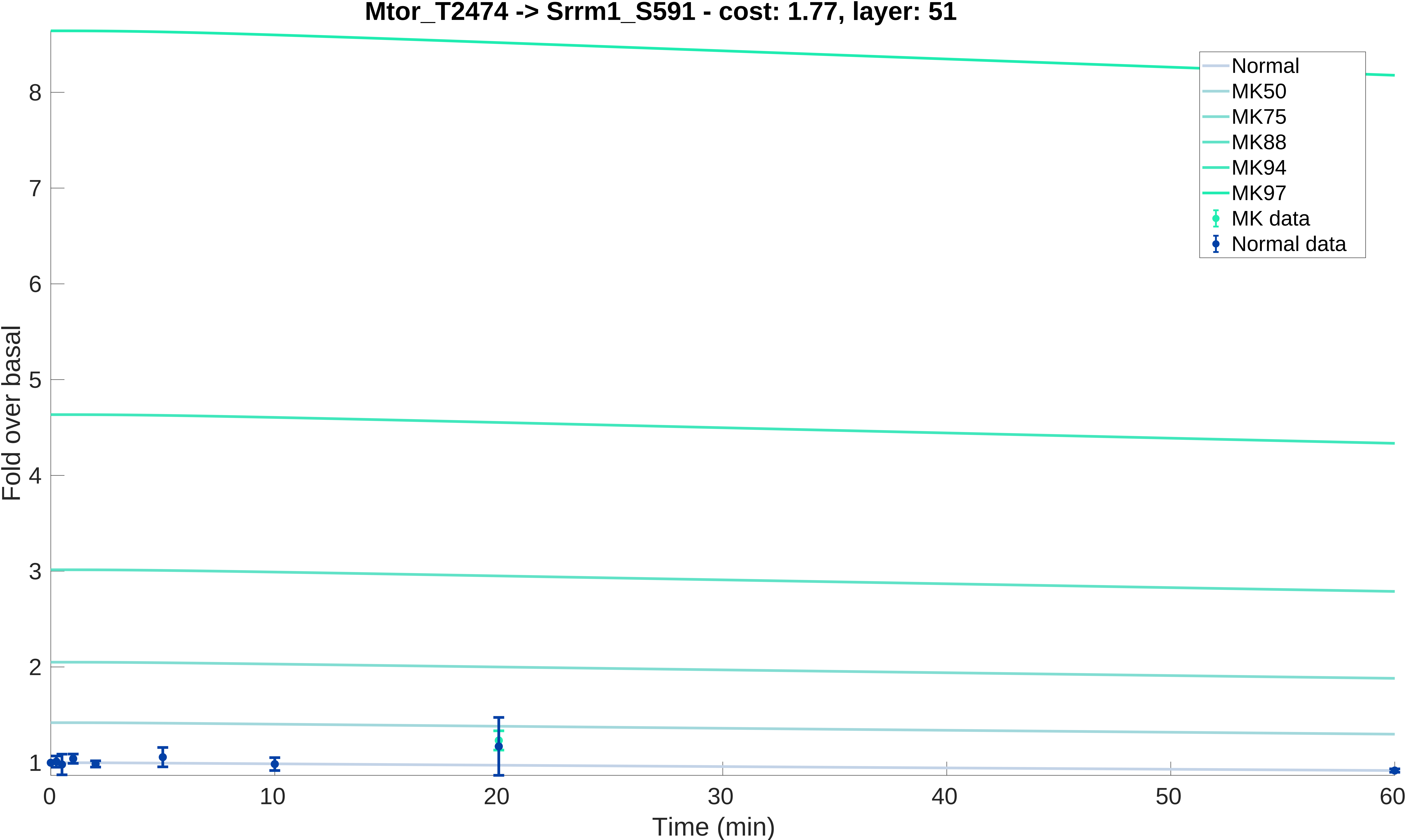

Mtor_T2474 -> Srrm1_S591 - cost: 1.77, layer: 51
Fold over basal
Time (min)
Normal
MK50
MK75
MK88
MK94
MK97
MK data
Normal data
1
2
3
4
5
6
7
8
0
10
20
30
40
50
60

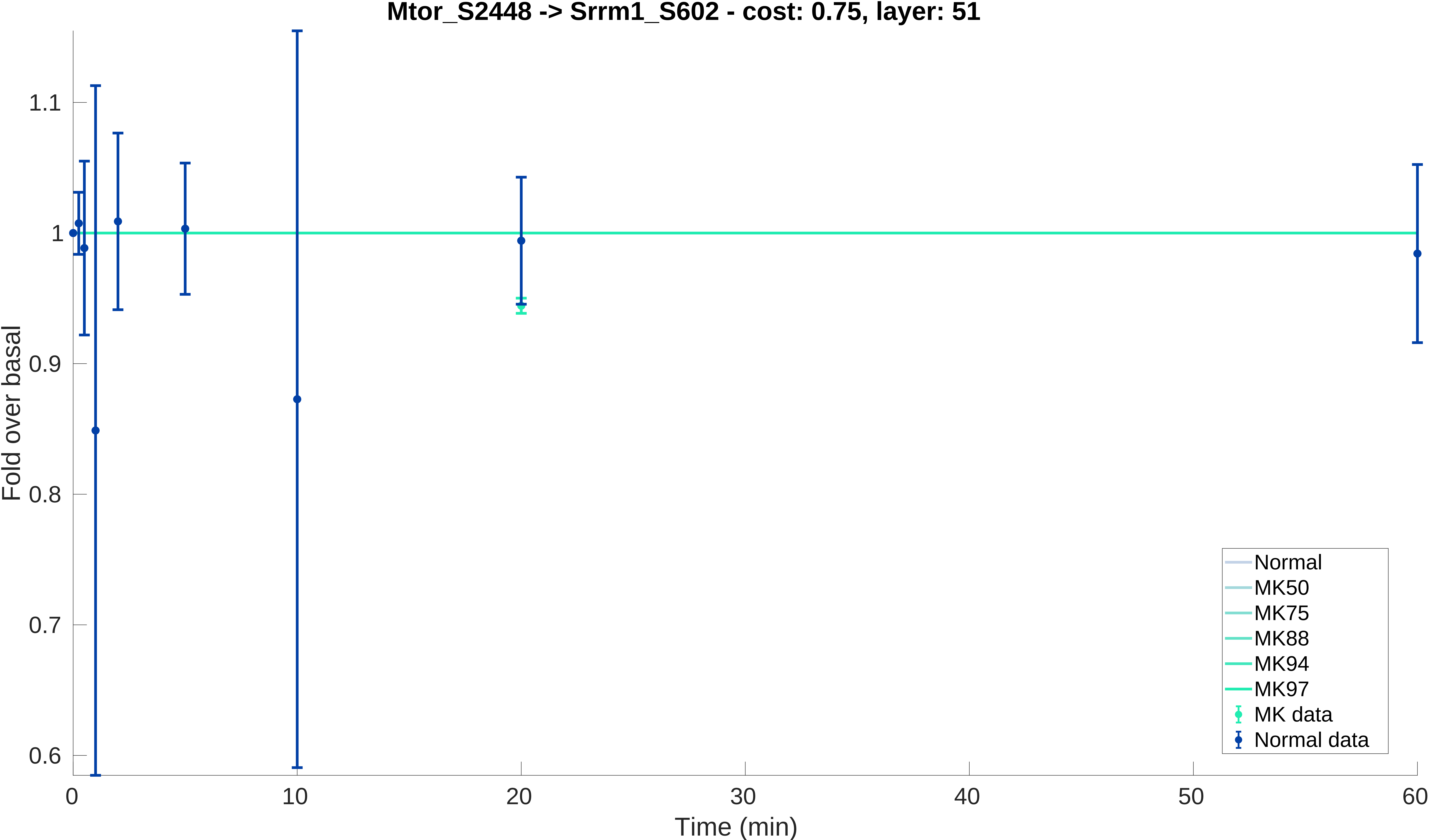

Mtor_S2448 -> Srrm1_S602 - cost: 0.75, layer: 51
Fold over basal
1.1
1
0.9
0.8
0.7
0.6
0
10
20
30
40
50
60
Time (min)
Normal
MK50
MK75
MK88
MK94
MK97
MK data
Normal data

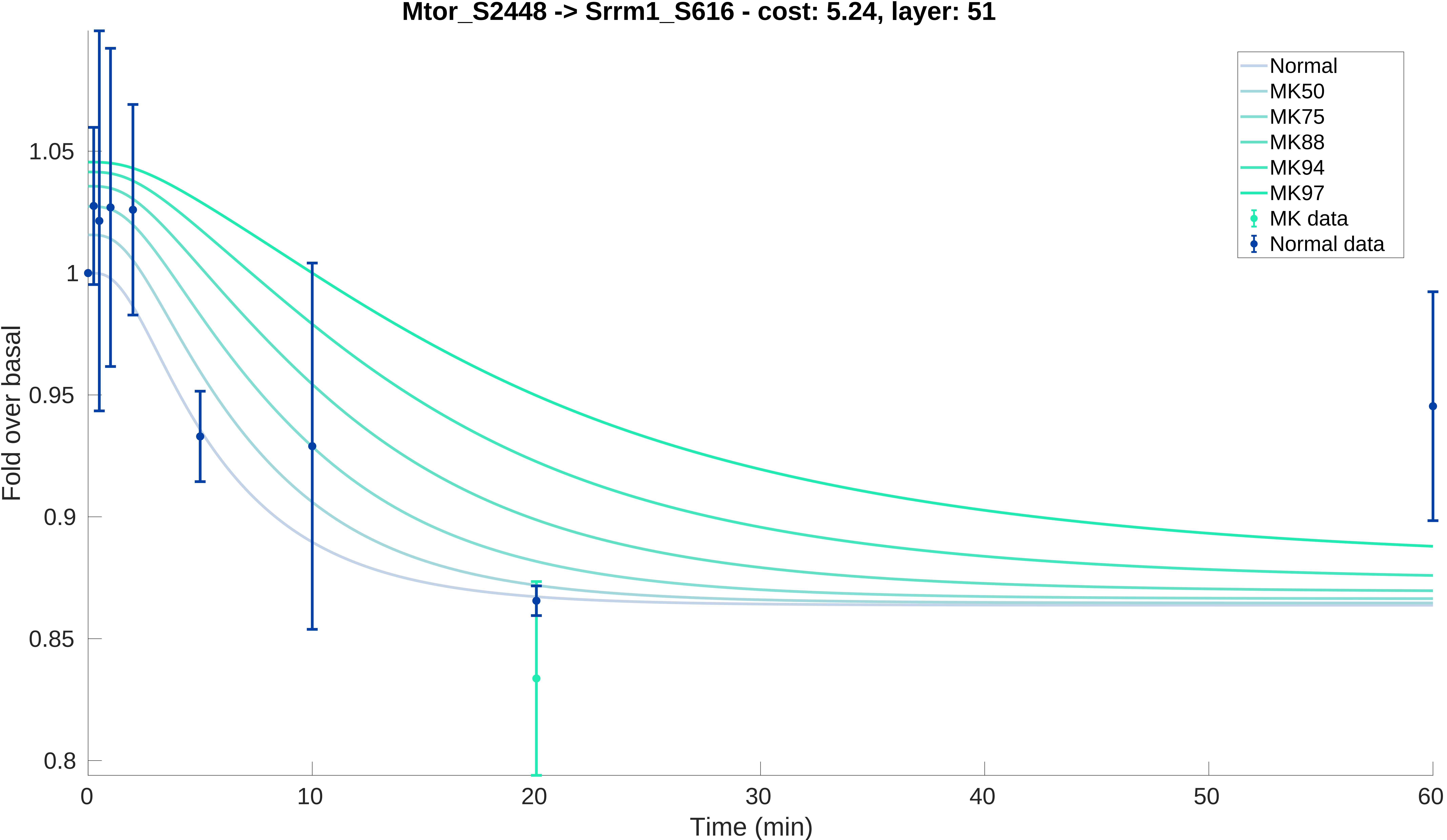

Mtor_S2448 -> Srrm1_S616 - cost: 5.24, layer: 51
Fold over basal
Time (min)
1.05
1
0.95
0.9
0.85
0.8
0
10
20
30
40
50
60
Normal
MK50
MK75
MK88
MK94
MK97
MK data
Normal data

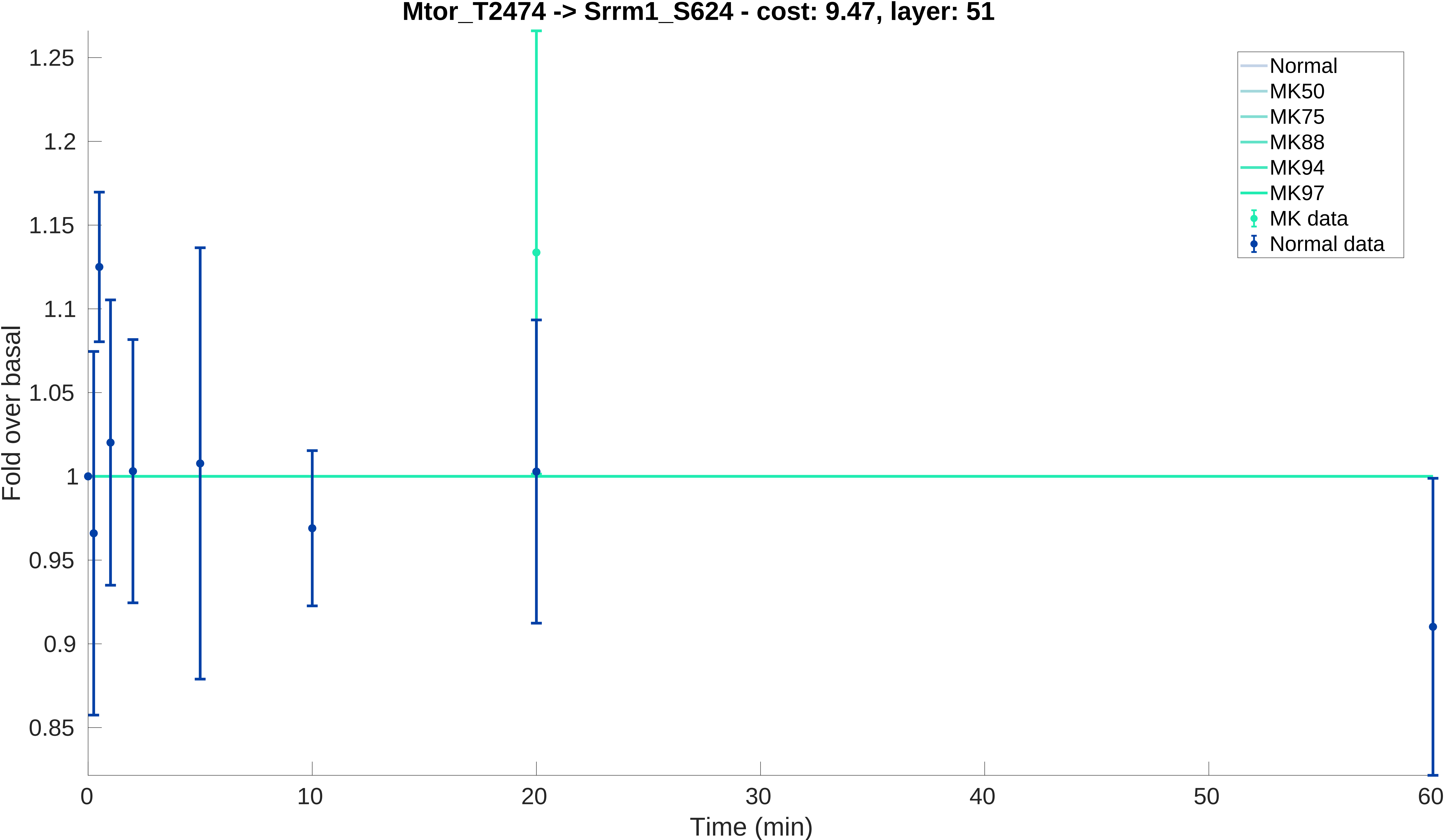

Mtor_T2474 -> Srrm1_S624 - cost: 9.47, layer: 51
Fold over basal
Time (min)
1.25
1.2
1.15
1.1
1.05
1
0.95
0.9
0.85
0
10
20
30
40
50
60
Normal
MK50
MK75
MK88
MK94
MK97
MK data
Normal data

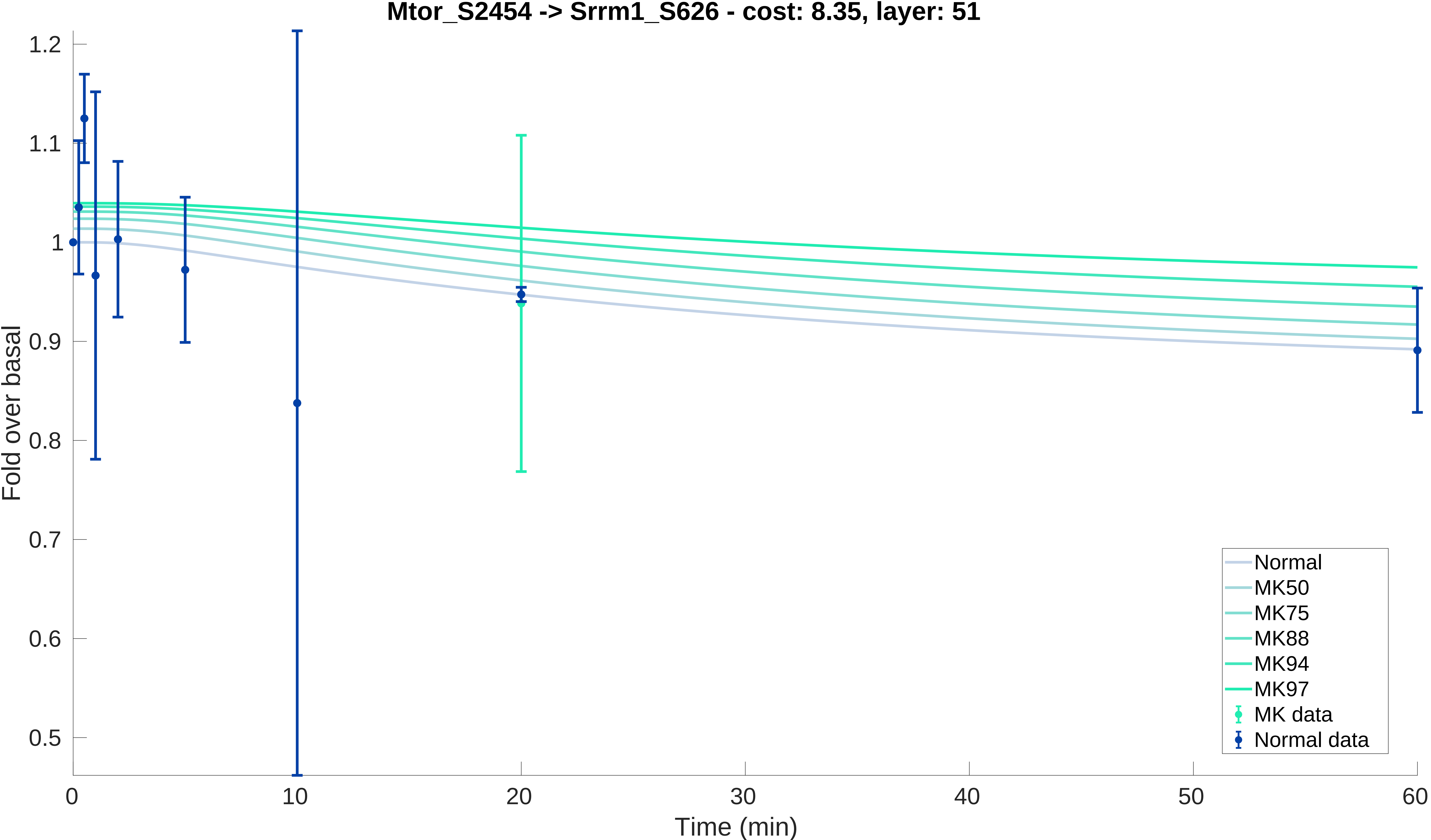

Mtor_S2454 -> Srrm1_S626 - cost: 8.35, layer: 51
Fold over basal
Time (min)
1.2
1.1
1
0.9
0.8
0.7
0.6
0.5
0
10
20
30
40
50
60
Normal
MK50
MK75
MK88
MK94
MK97
MK data
Normal data

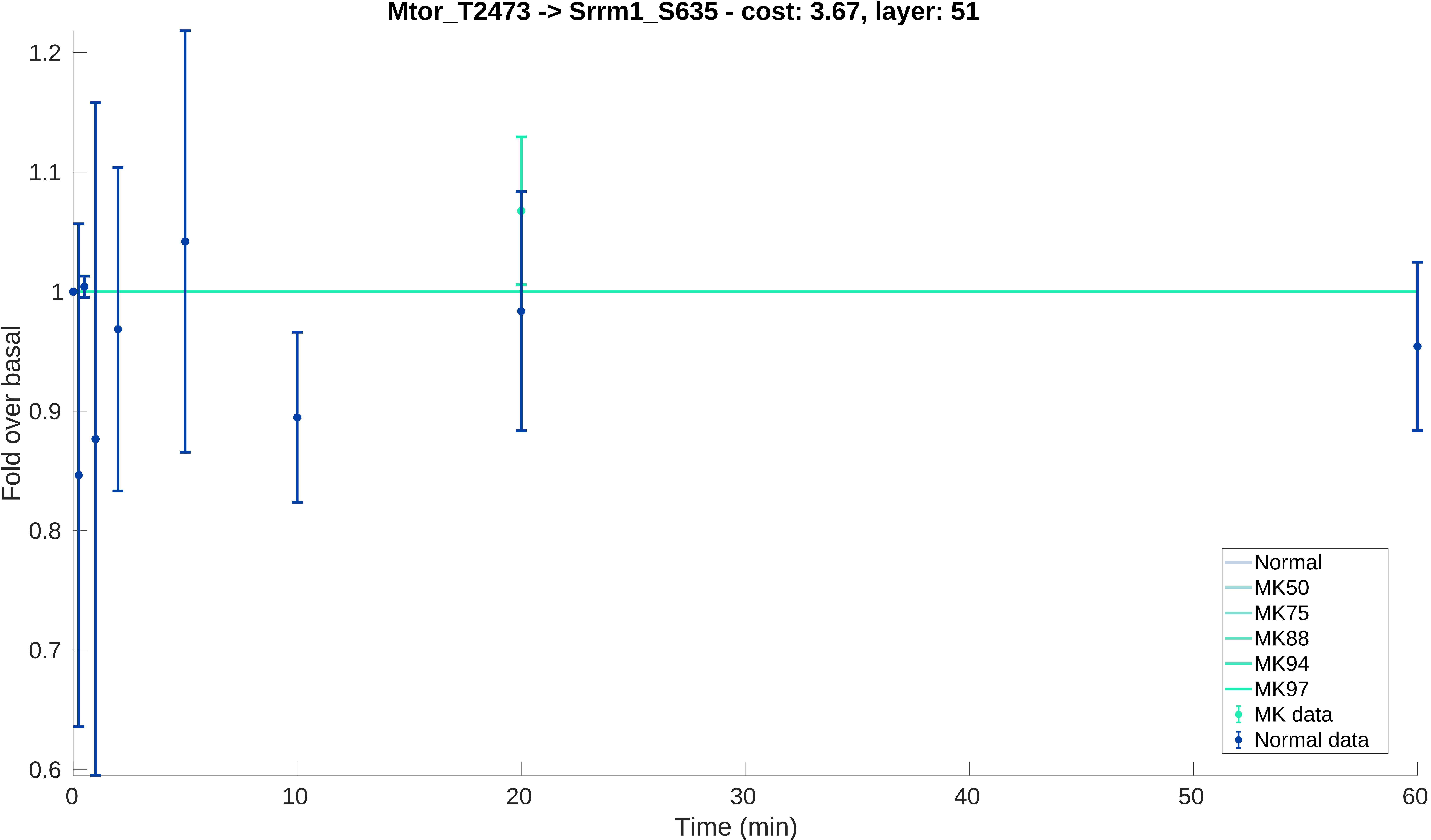

Mtor_T2473 -> Srrm1_S635 - cost: 3.67, layer: 51
Fold over basal
1.2
1.1
1
0.9
0.8
0.7
0.6
0
10
20
30
40
50
60
Time (min)
Normal
MK50
MK75
MK88
MK94
MK97
MK data
Normal data

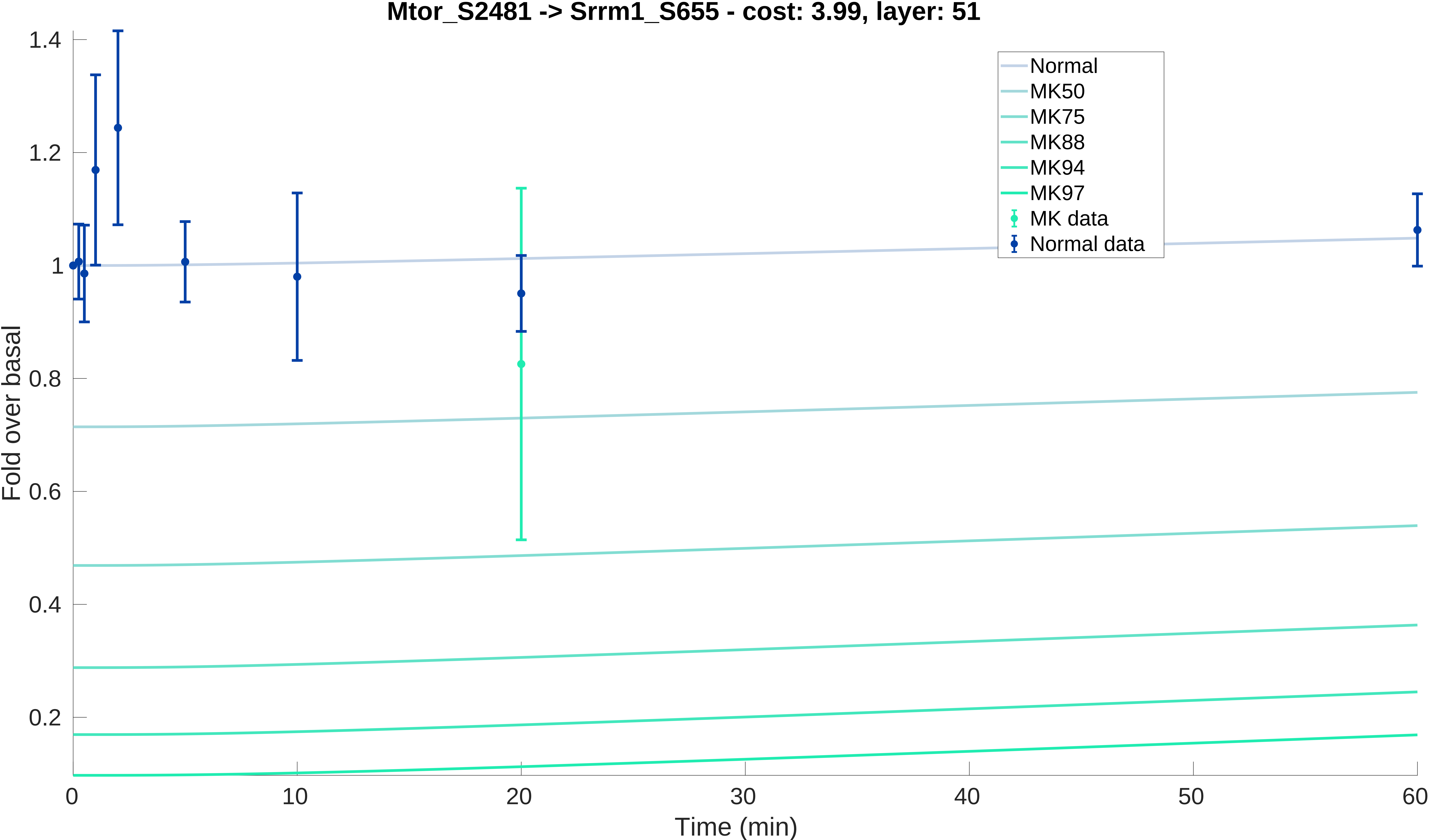

Mtor_S2481 -> Srrm1_S655 - cost: 3.99, layer: 51
Fold over basal
1.4
1.2
1
0.8
0.6
0.4
0.2
0
10
20
30
40
50
60
Time (min)
Normal
MK50
MK75
MK88
MK94
MK97
MK data
Normal data

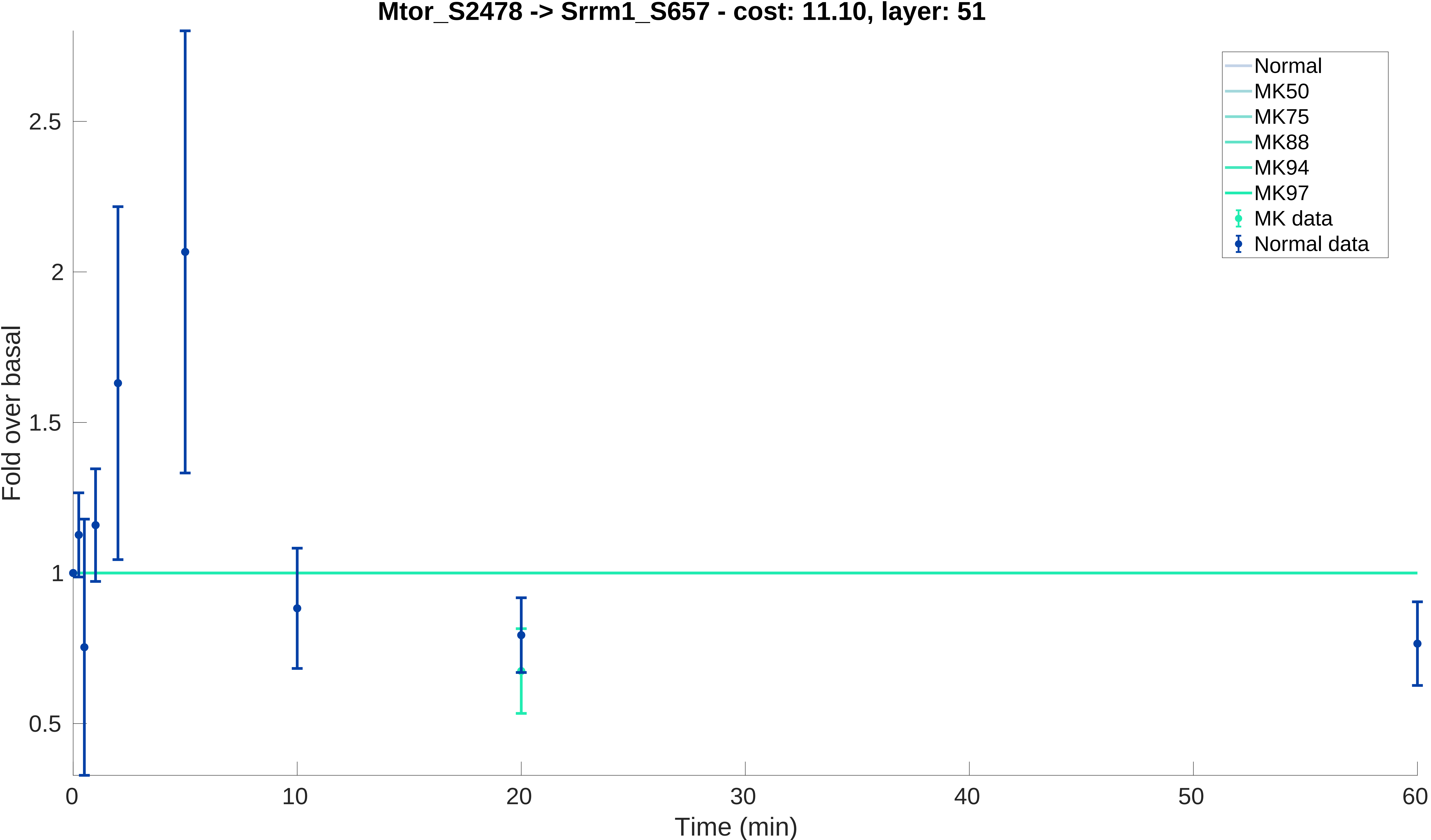

Mtor_S2478 -> Srrm1_S657 - cost: 11.10, layer: 51
Fold over basal
Time (min)
0.5
1
1.5
2
2.5
0
10
20
30
40
50
60
Normal
MK50
MK75
MK88
MK94
MK97
MK data
Normal data

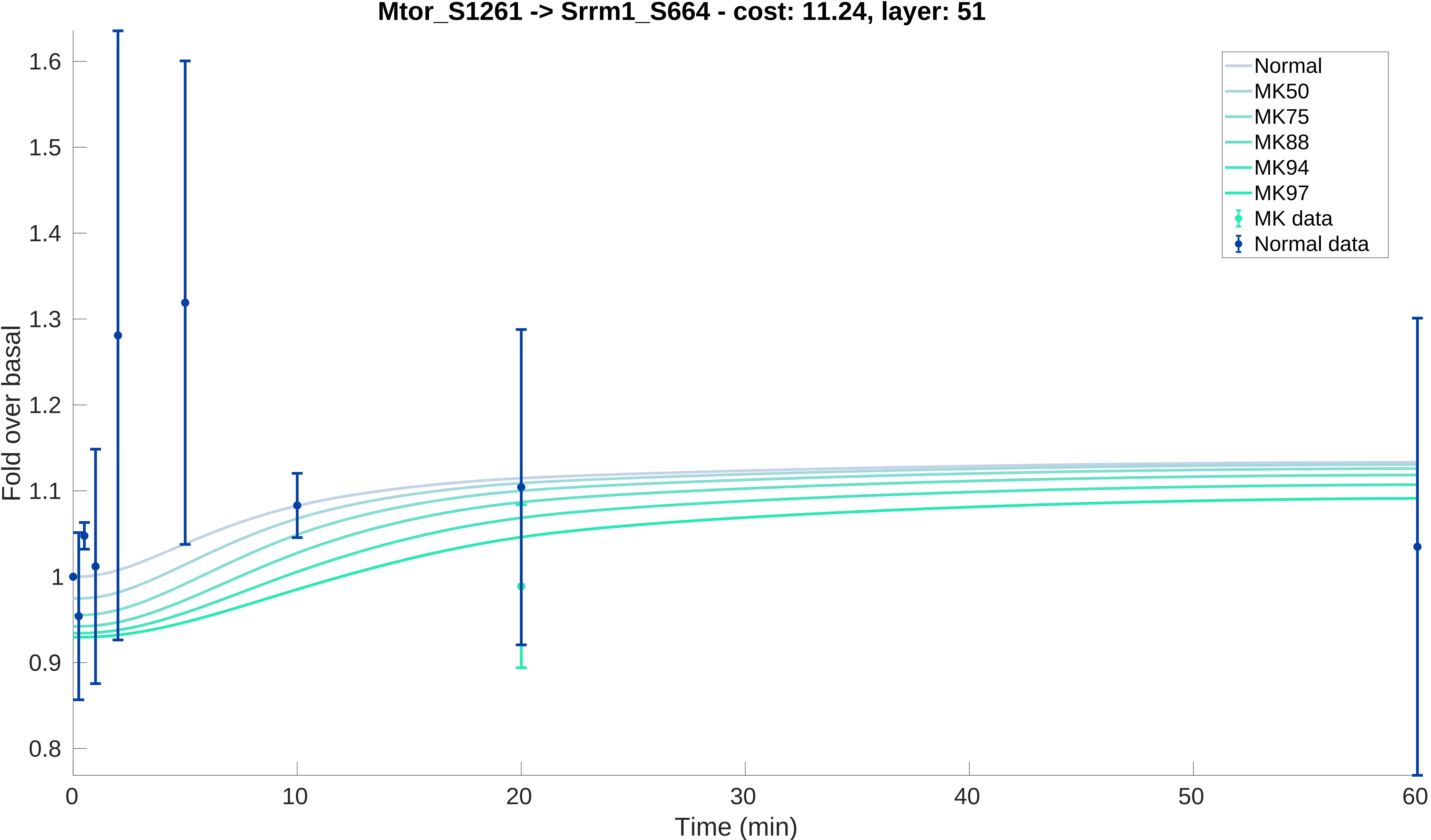

Mtor_S1261 -> Srrm1_S664 - cost: 11.24, layer: 51
Fold over basal
Time (min)
0.8
0.9
1
1.1
1.2
1.3
1.4
1.5
1.6
0
10
20
30
40
50
60
Normal
MK50
MK75
MK88
MK94
MK97
MK data
Normal data

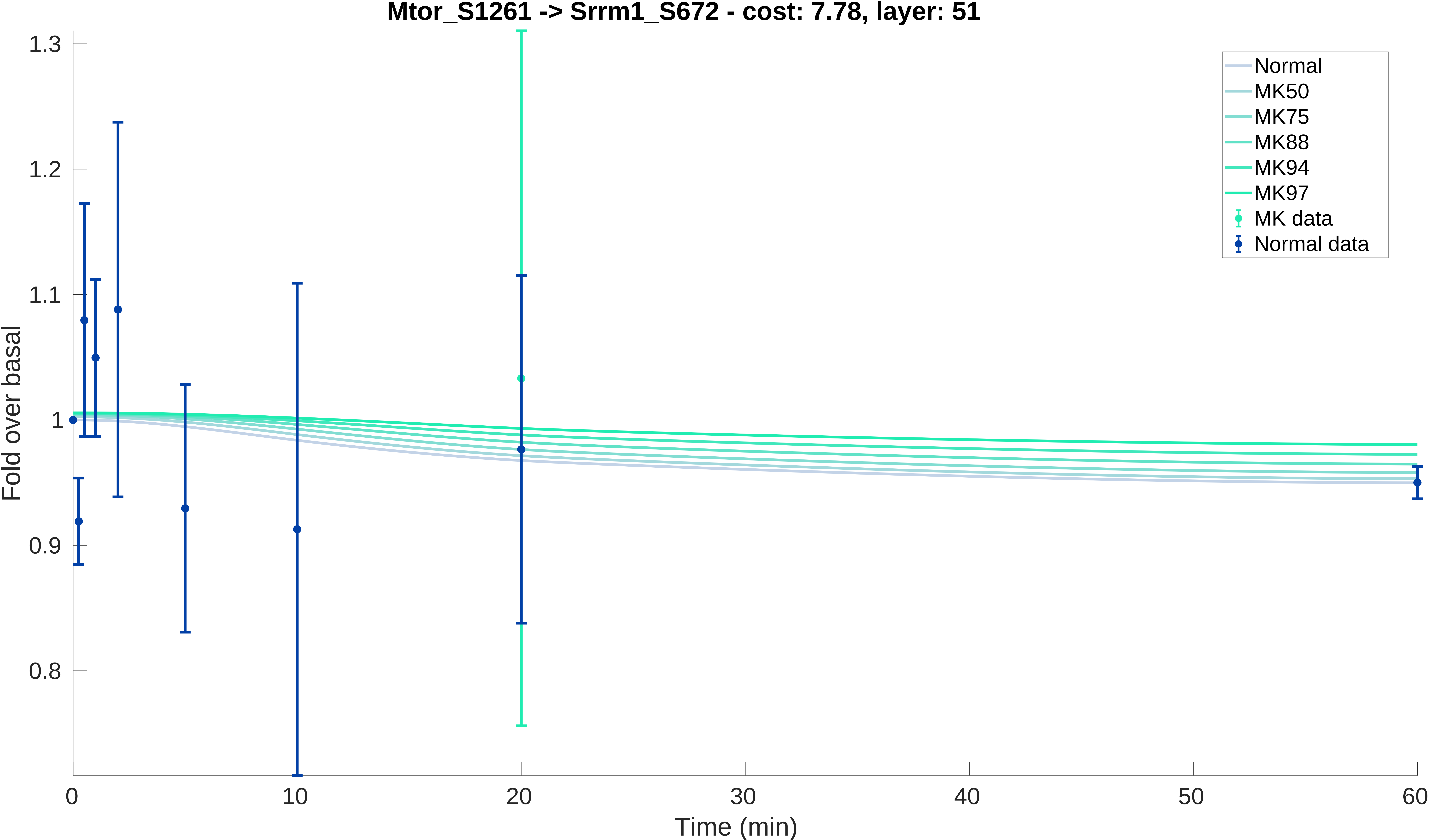

Mtor_S1261 -> Srrm1_S672 - cost: 7.78, layer: 51
Fold over basal
Time (min)
1.3
1.2
1.1
1
0.9
0.8
0
10
20
30
40
50
60
Normal
MK50
MK75
MK88
MK94
MK97
MK data
Normal data

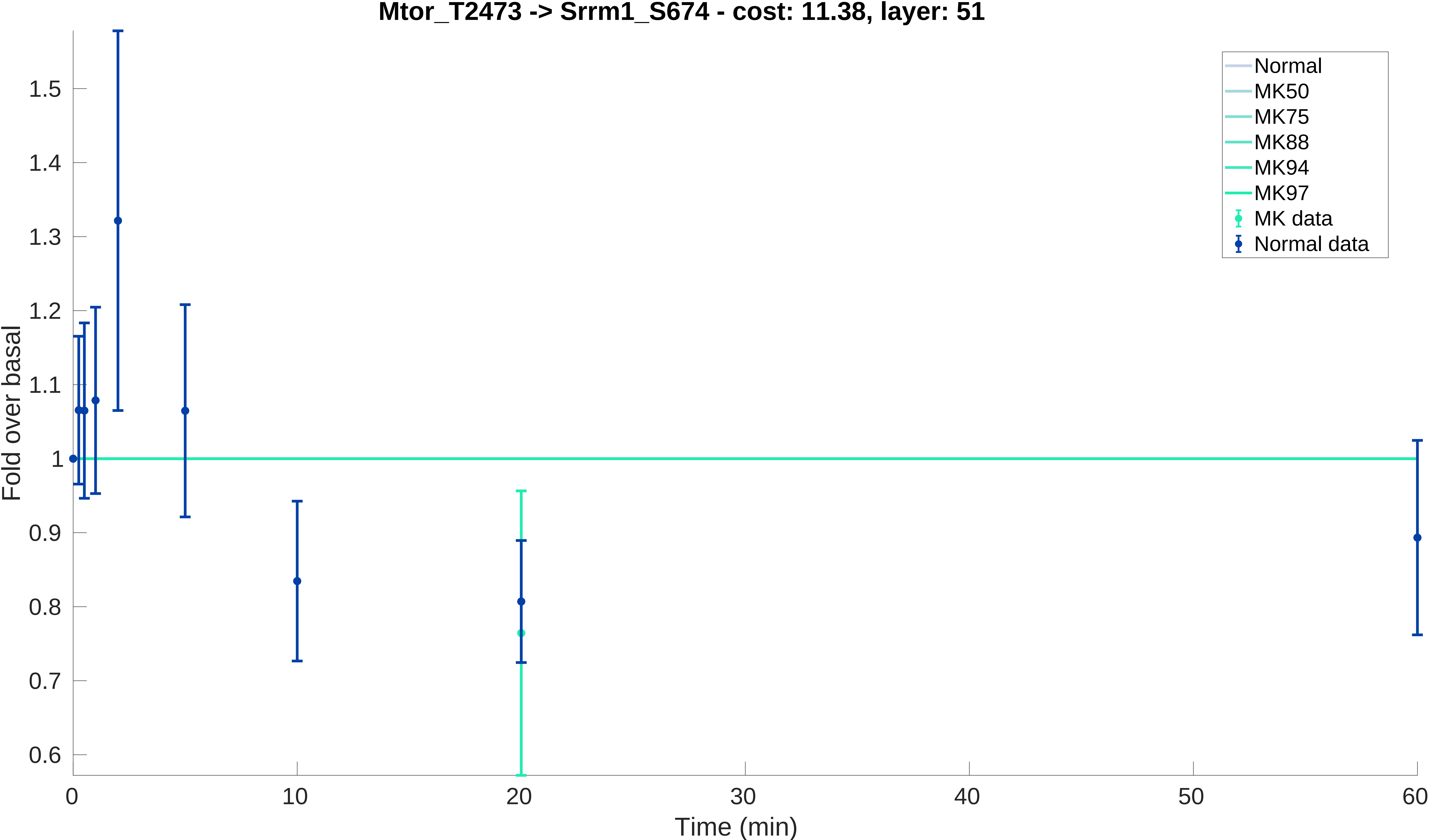

Mtor_T2473 -> Srrm1_S674 - cost: 11.38, layer: 51
Fold over basal
Time (min)
0.6
0.7
0.8
0.9
1
1.1
1.2
1.3
1.4
1.5
0
10
20
30
40
50
60
Normal
MK50
MK75
MK88
MK94
MK97
MK data
Normal data

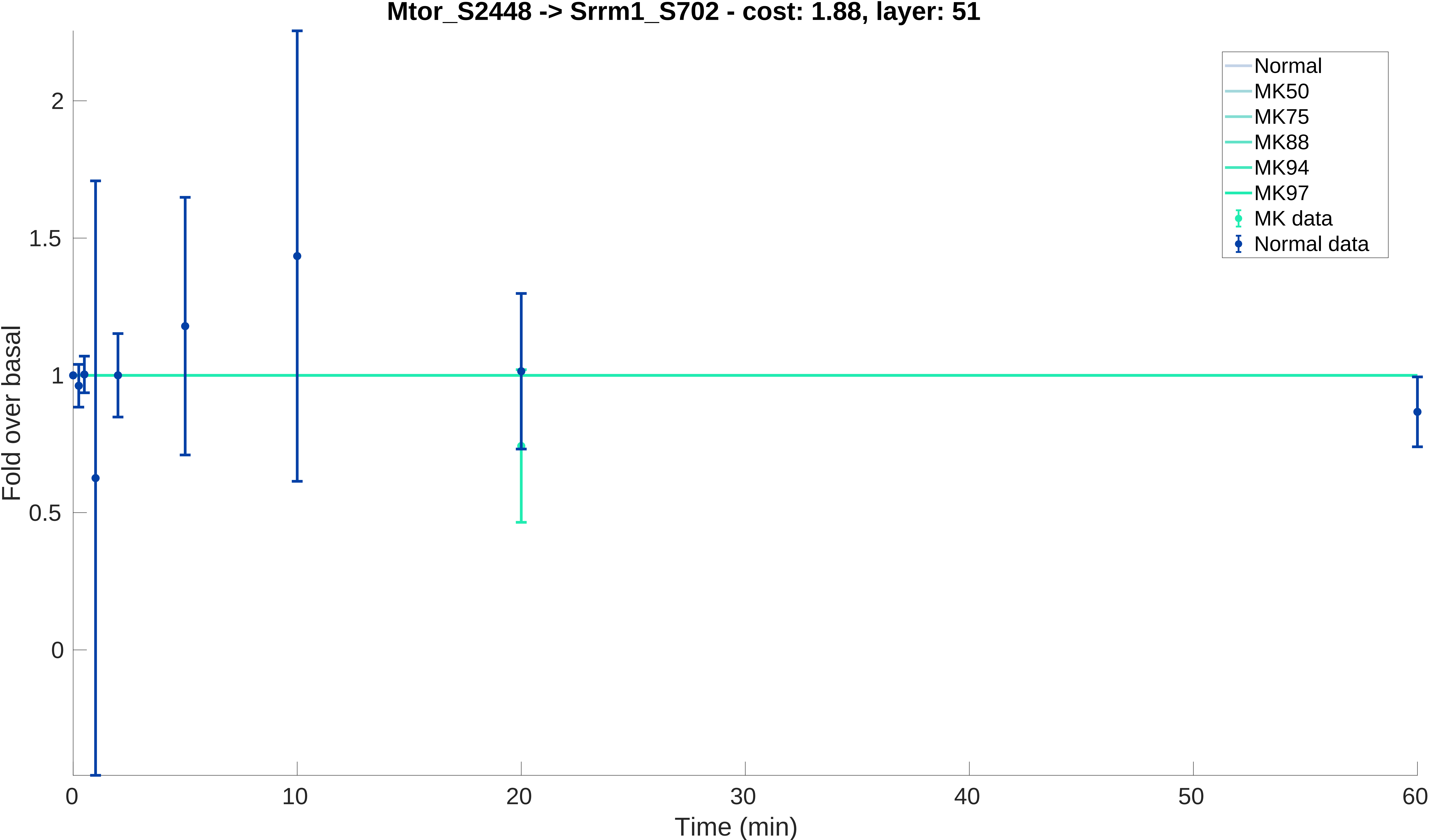

Mtor_S2448 -> Srrm1_S702 - cost: 1.88, layer: 51
Normal
MK50
MK75
MK88
MK94
MK97
MK data
Normal data
Fold over basal
Time (min)
0
10
20
30
40
50
60
0
0.5
1
1.5
2

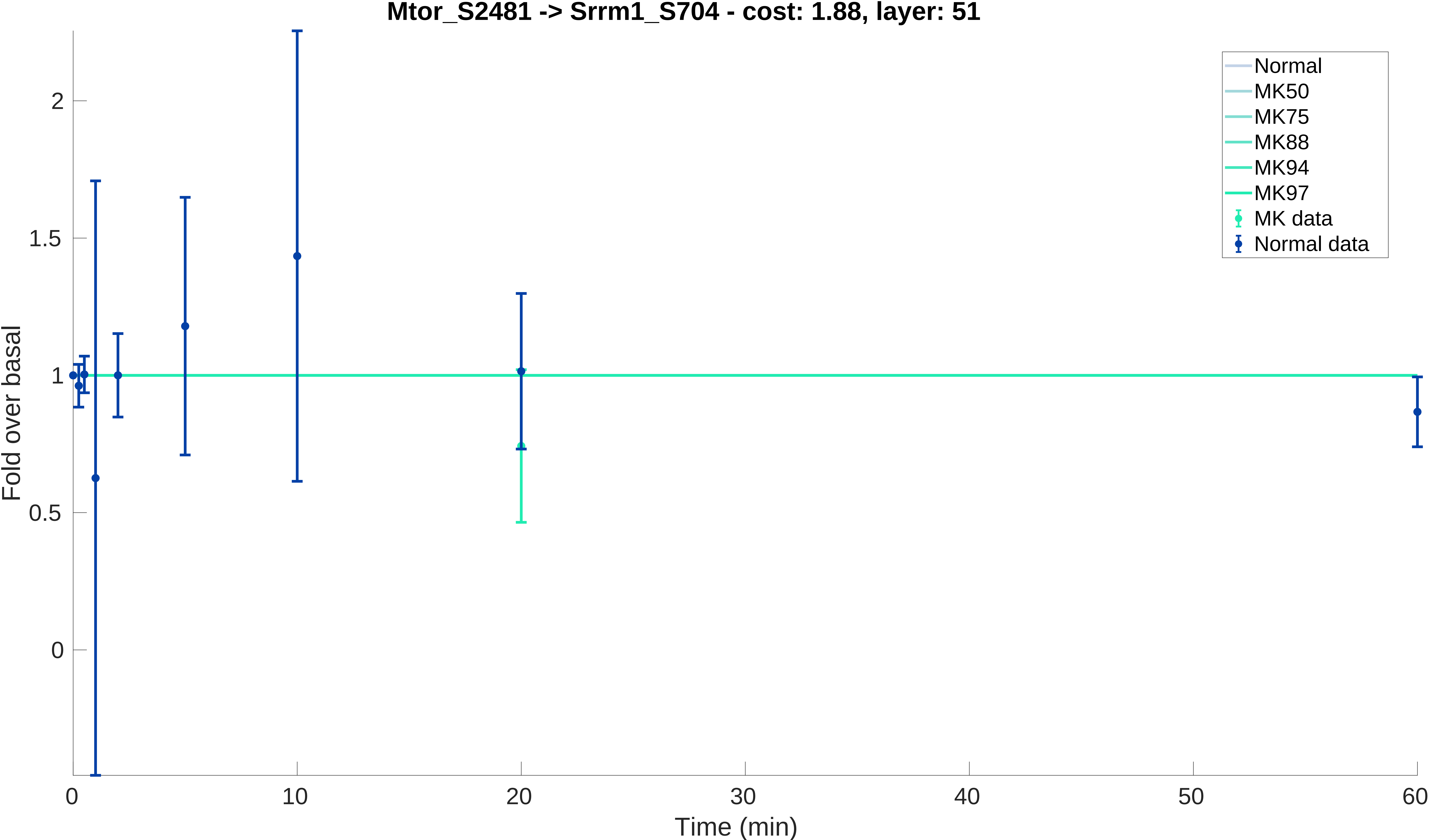
Mtor_S2481 -> Srrm1_S704 - cost: 1.88, layer: 51
Normal
MK50
MK75
MK88
MK94
MK97
MK data
Normal data
Fold over basal
Time (min)
2
1.5
1
0.5
0
0
10
20
30
40
50
60

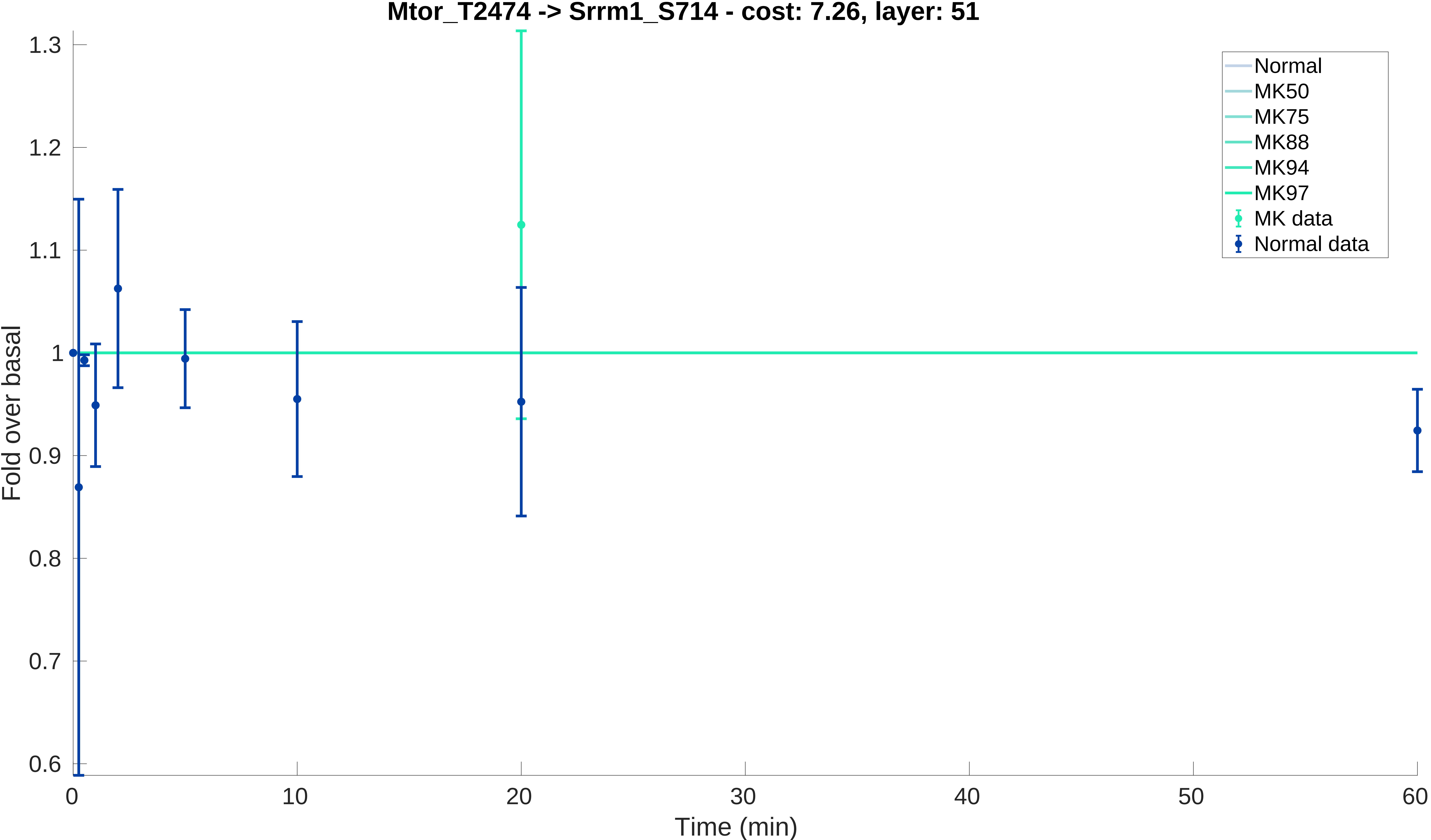

Mtor_T2474 -> Srrm1_S714 - cost: 7.26, layer: 51
Fold over basal
Time (min)
0.6
0.7
0.8
0.9
1
1.1
1.2
1.3
0
10
20
30
40
50
60
Normal
MK50
MK75
MK88
MK94
MK97
MK data
Normal data

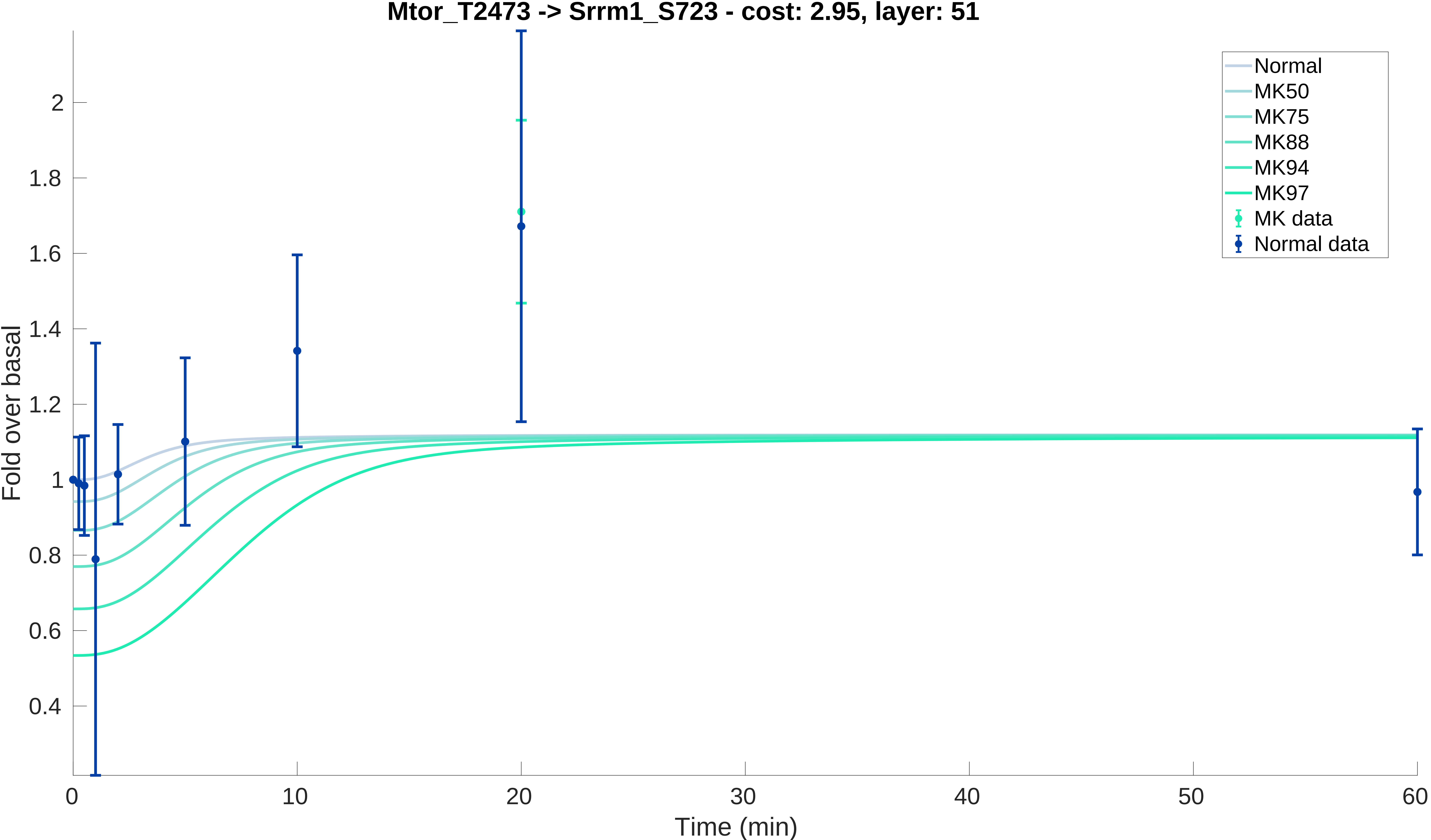

Mtor_T2473 -> Srrm1_S723 - cost: 2.95, layer: 51
Fold over basal
Time (min)
0.4
0.6
0.8
1
1.2
1.4
1.6
1.8
2
0
10
20
30
40
50
60
Normal
MK50
MK75
MK88
MK94
MK97
MK data
Normal data

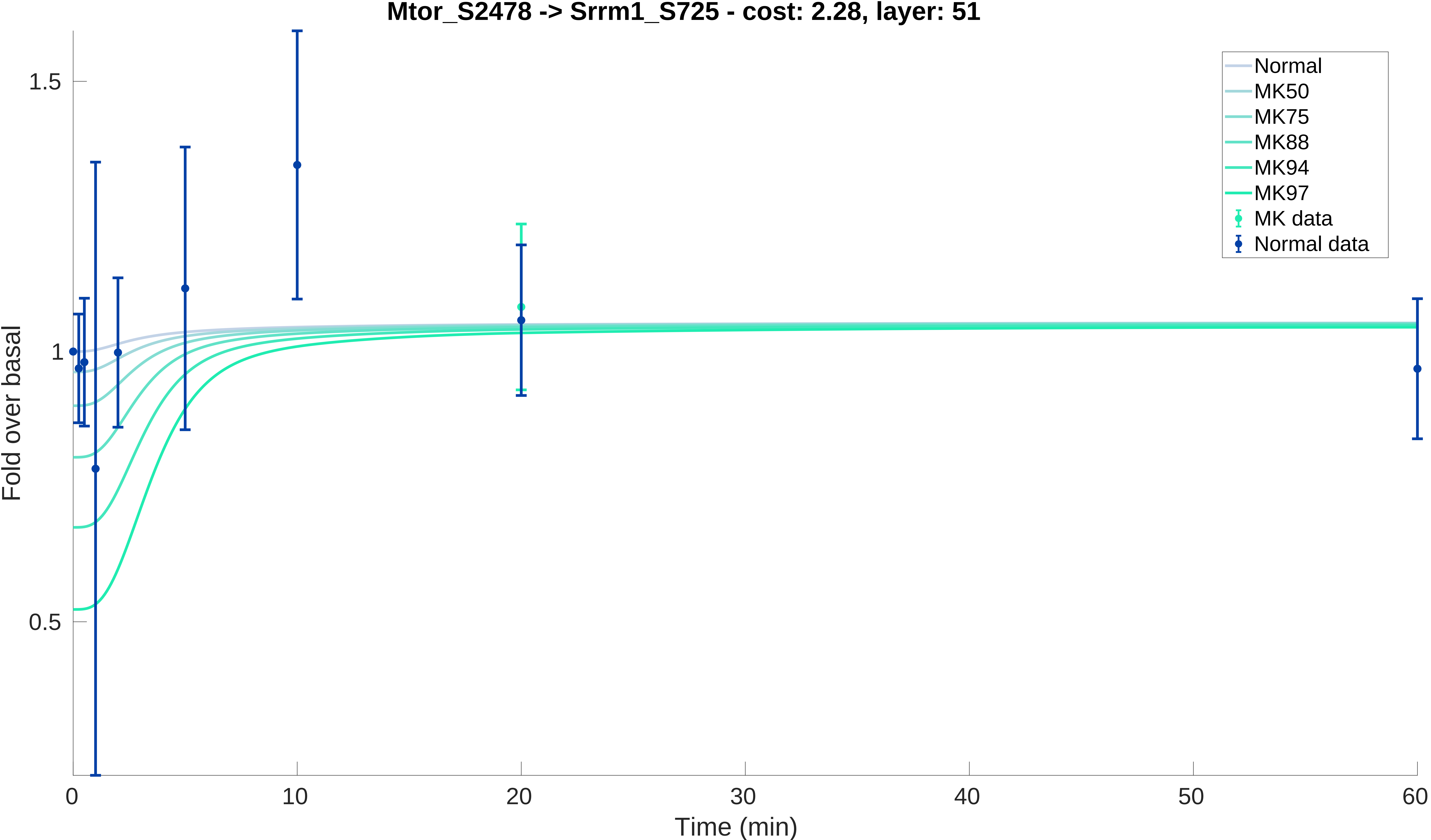

Mtor_S2478 -> Srrm1_S725 - cost: 2.28, layer: 51
Fold over basal
Time (min)
1.5
1
0.5
0
10
20
30
40
50
60
Normal
MK50
MK75
MK88
MK94
MK97
MK data
Normal data

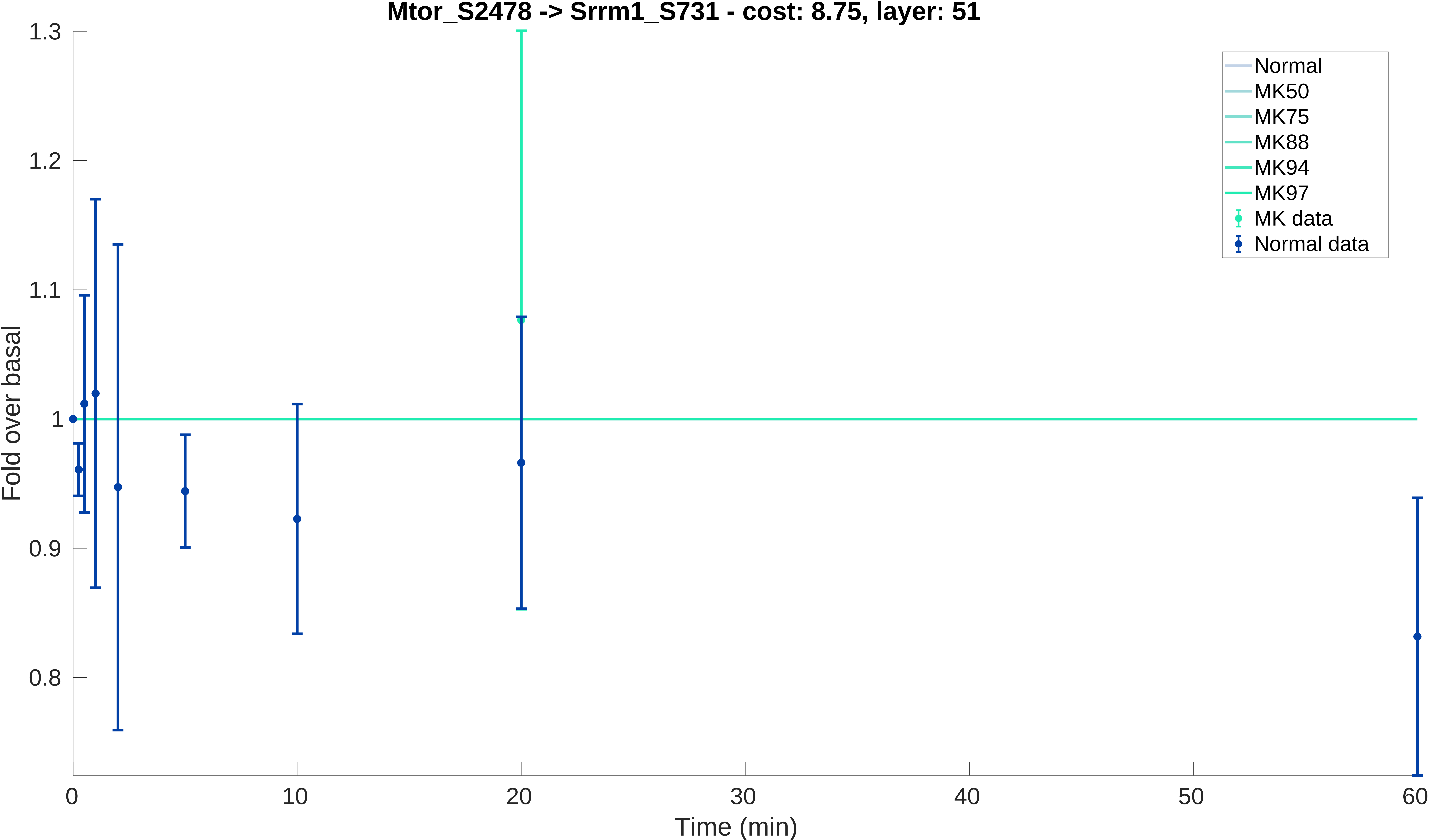

Mtor_S2478 -> Srrm1_S731 - cost: 8.75, layer: 51
1.3
1.2
1.1
1
0.9
0.8
Fold over basal
0
10
20
30
40
50
60
Time (min)
Normal
MK50
MK75
MK88
MK94
MK97
MK data
Normal data

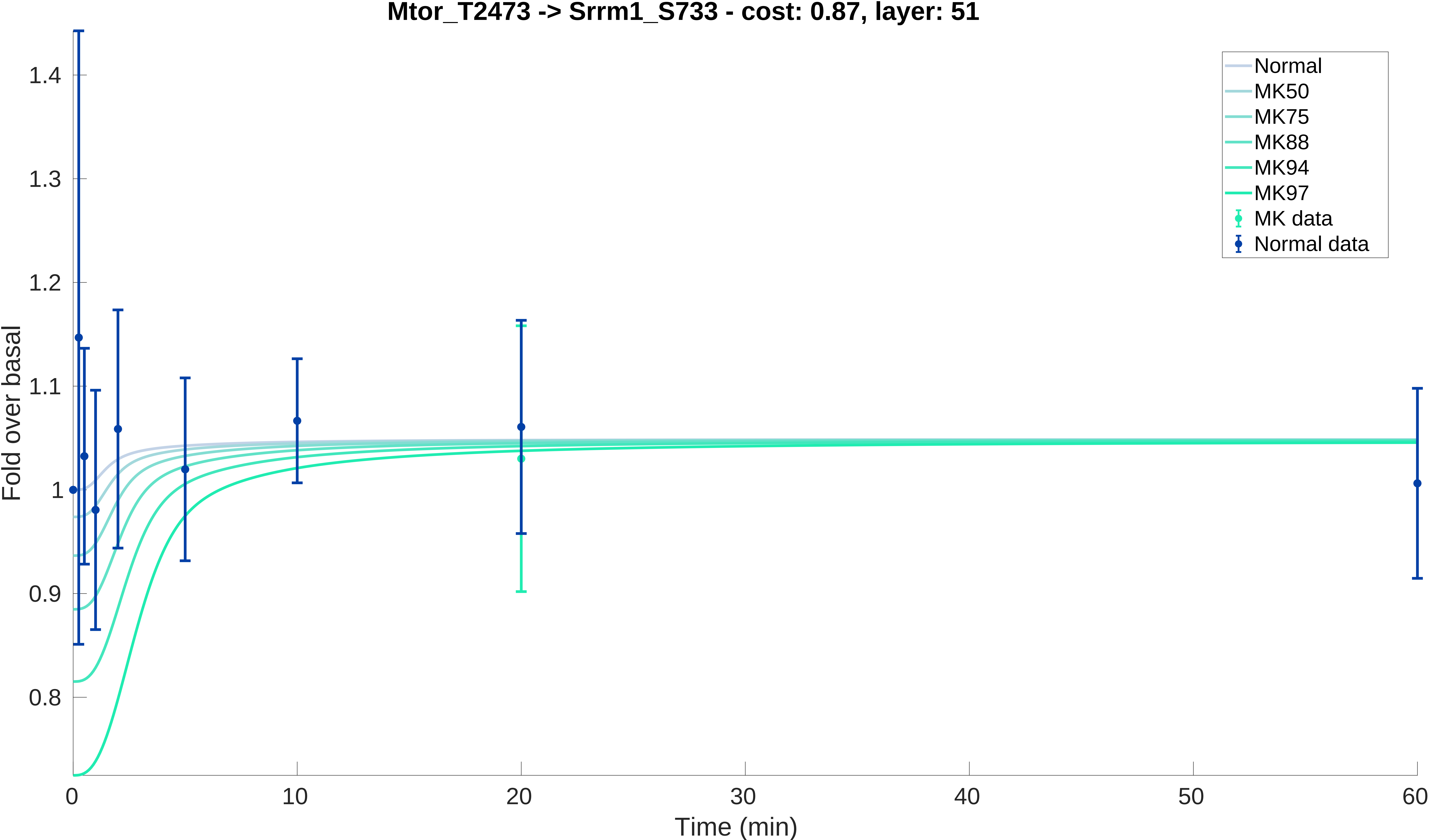
Mtor_T2473 -> Srrm1_S733 - cost: 0.87, layer: 51
Normal
MK50
MK75
MK88
MK94
MK97
MK data
Normal data
Fold over basal
1.4
1.3
1.2
1.1
1
0.9
0.8
0
10
20
30
40
50
60
Time (min)

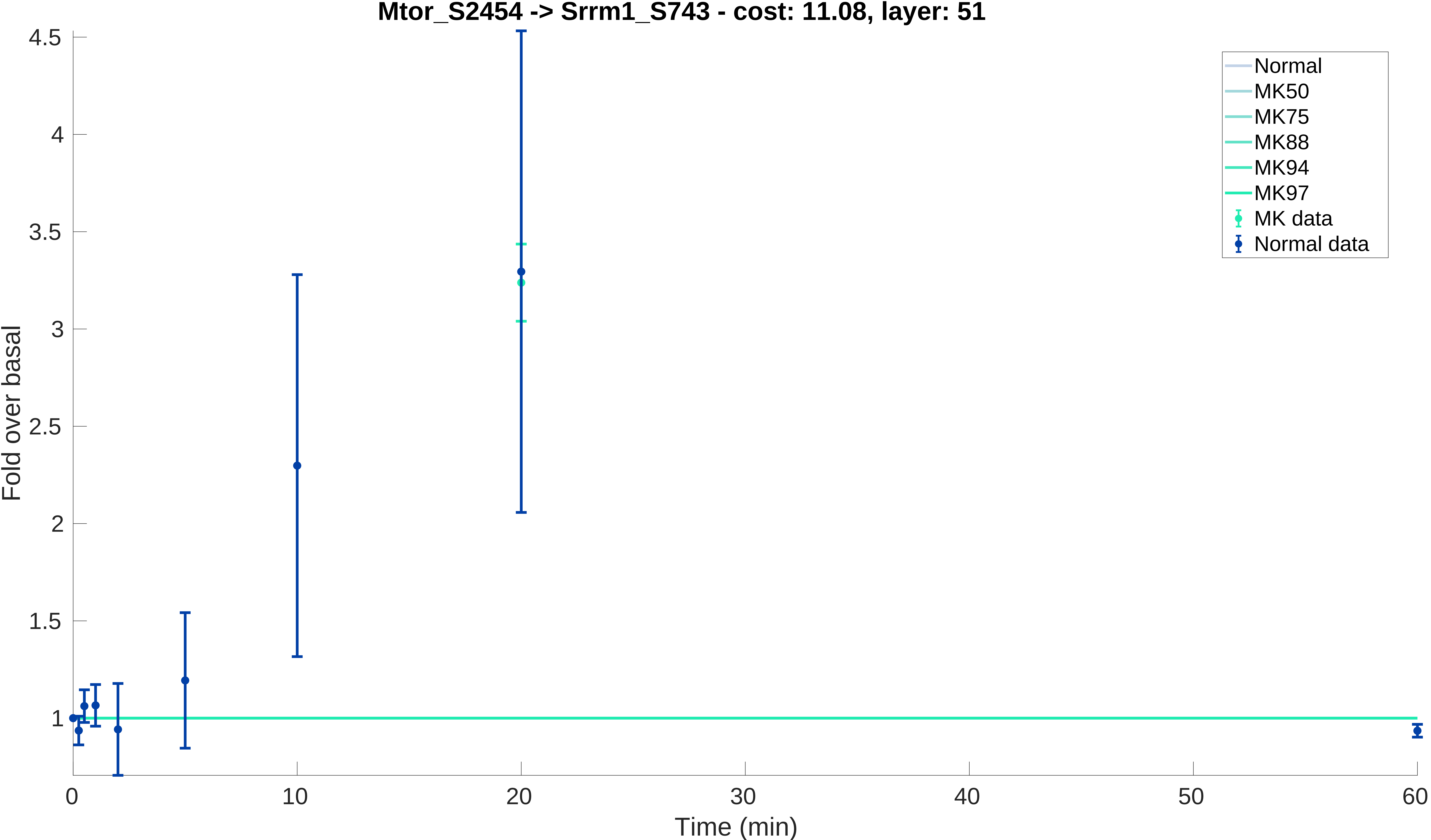
Mtor_S2454 -> Srrm1_S743 - cost: 11.08, layer: 51
Fold over basal
4.5
4
3.5
3
2.5
2
1.5
1
Time (min)
0
10
20
30
40
50
60
Normal
MK50
MK75
MK88
MK94
MK97
MK data
Normal data

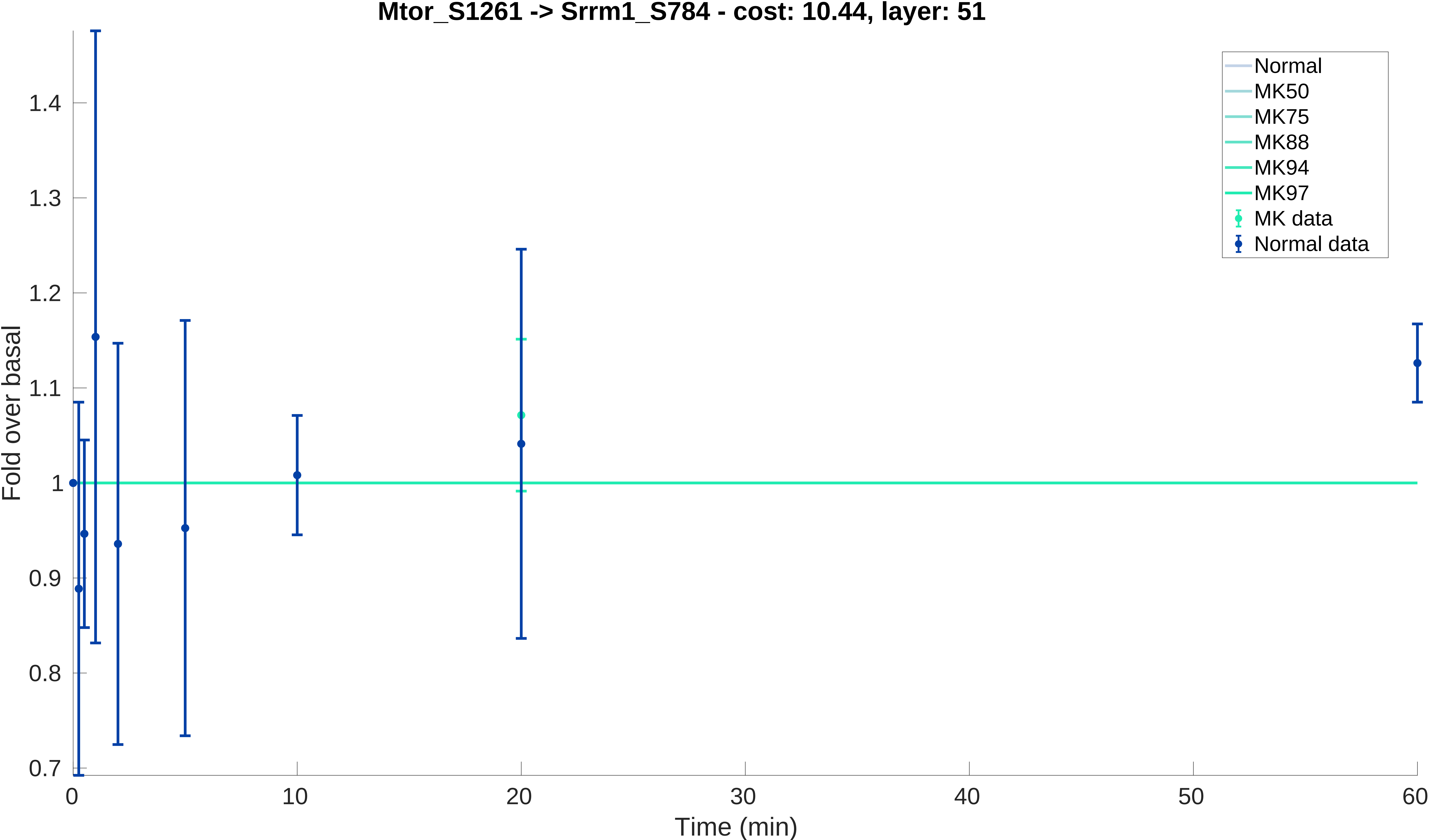

Mtor_S1261 -> Srrm1_S784 - cost: 10.44, layer: 51
Fold over basal
Time (min)
1.4
1.3
1.2
1.1
1
0.9
0.8
0.7
0
10
20
30
40
50
60
Normal
MK50
MK75
MK88
MK94
MK97
MK data
Normal data

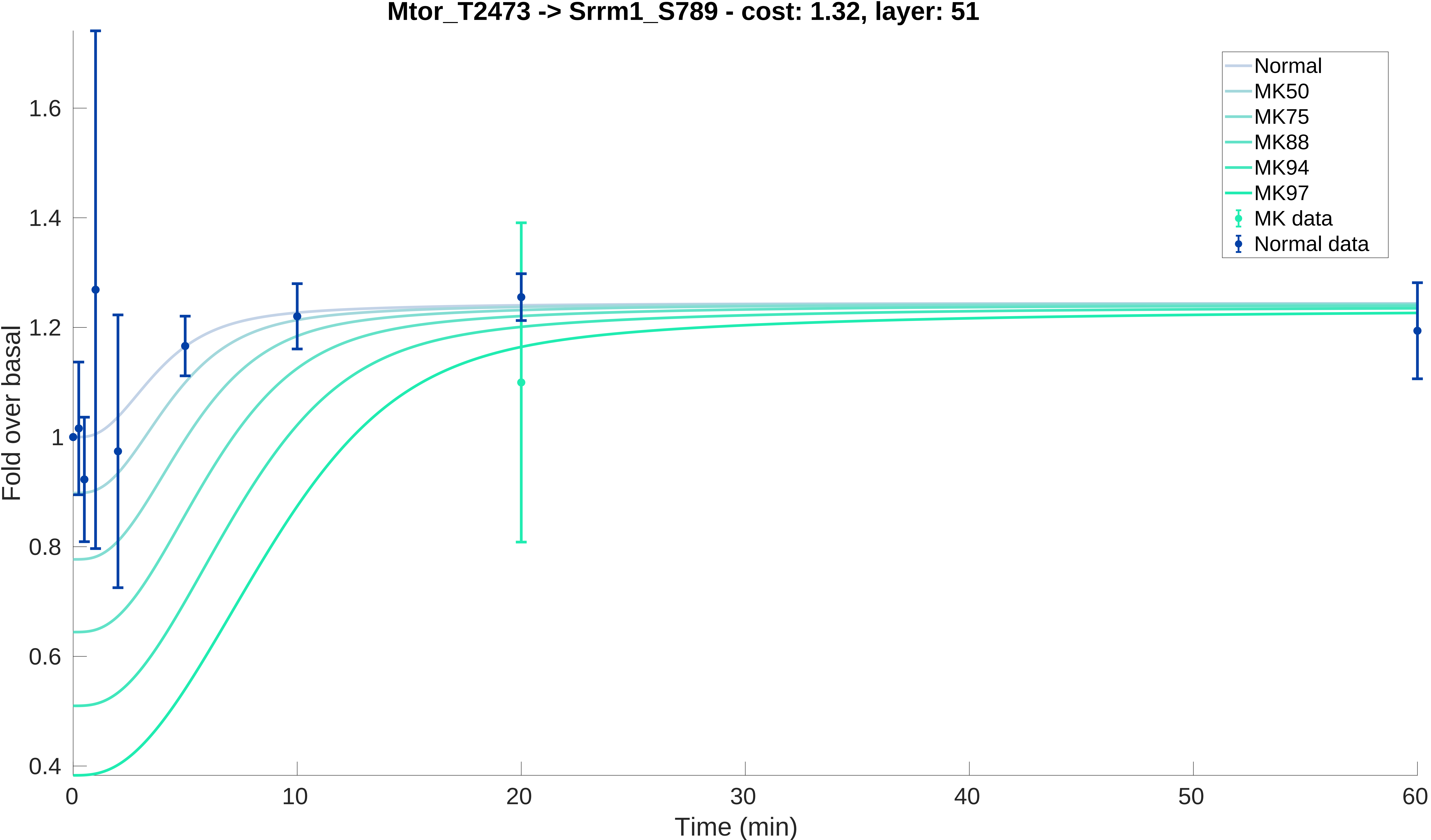
Mtor_T2473 -> Srrm1_S789 - cost: 1.32, layer: 51
Normal
MK50
MK75
MK88
MK94
MK97
MK data
Normal data
Fold over basal
1.6
1.4
1.2
1
0.8
0.6
0.4
0
10
20
30
40
50
60
Time (min)

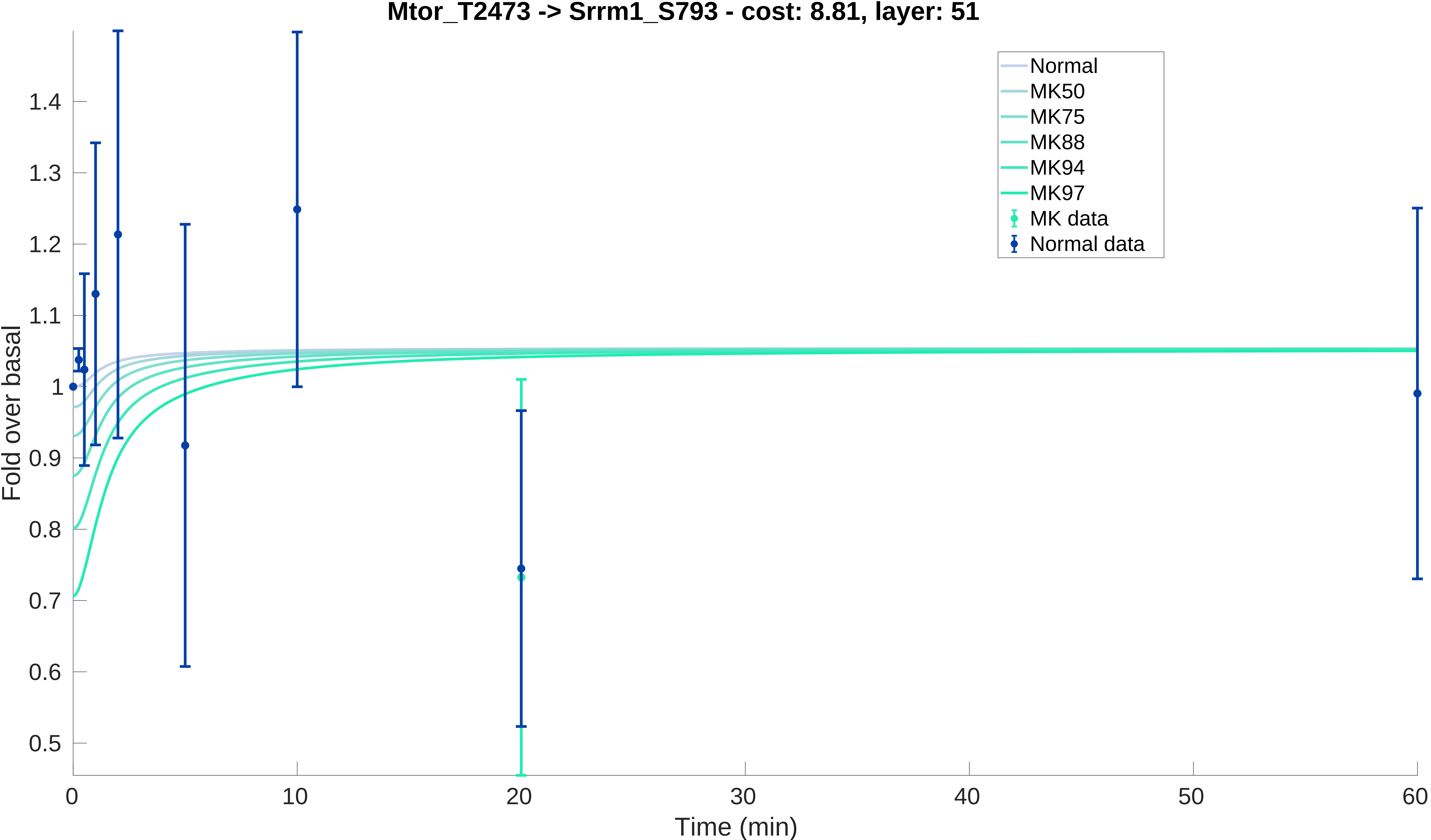
Mtor_T2473 -> Srrm1_S793 - cost: 8.81, layer: 51
Fold over basal
Time (min)
Normal
MK50
MK75
MK88
MK94
MK97
MK data
Normal data
1.4
1.3
1.2
1.1
1
0.9
0.8
0.7
0.6
0.5
0
10
20
30
40
50
60

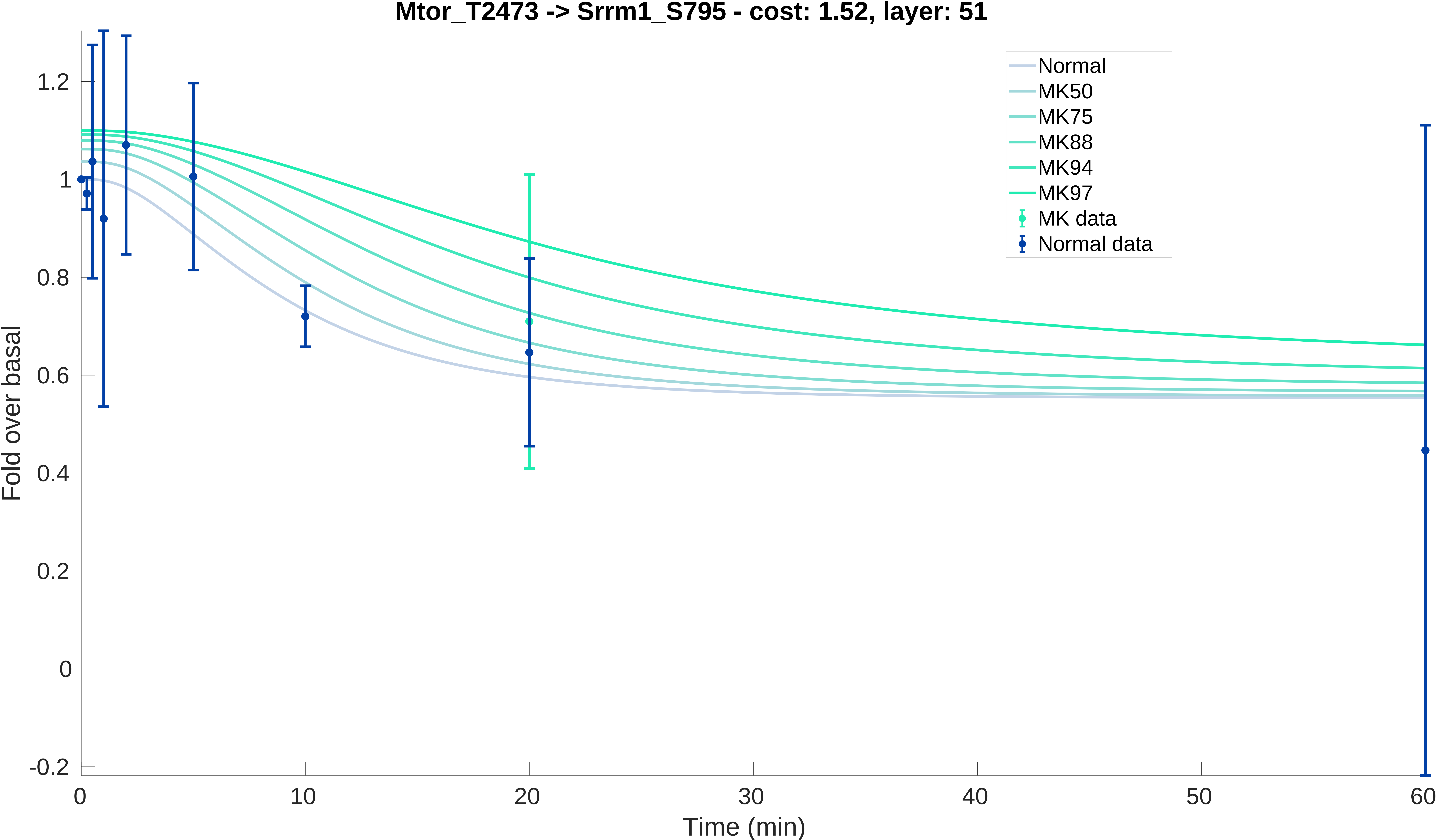

Mtor_T2473 -> Srrm1_S795 - cost: 1.52, layer: 51
Normal
MK50
MK75
MK88
MK94
MK97
MK data
Normal data
Fold over basal
1.2
1
0.8
0.6
0.4
0.2
0
-0.2
0
10
20
30
40
50
60
Time (min)

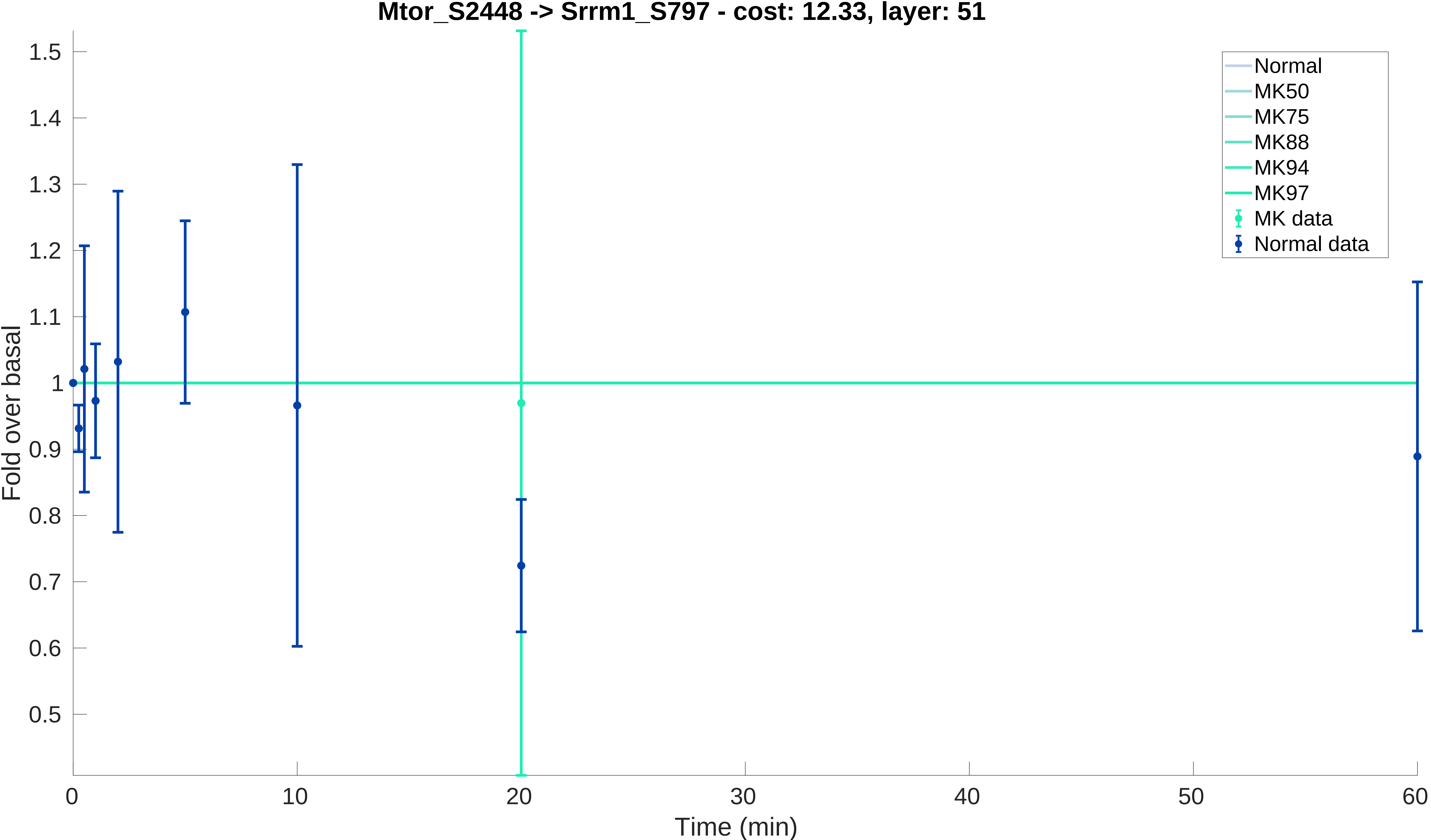

Mtor_S2448 -> Srrm1_S797 - cost: 12.33, layer: 51
Fold over basal
Time (min)
1.5
1.4
1.3
1.2
1.1
1
0.9
0.8
0.7
0.6
0.5
0
10
20
30
40
50
60
Normal
MK50
MK75
MK88
MK94
MK97
MK data
Normal data

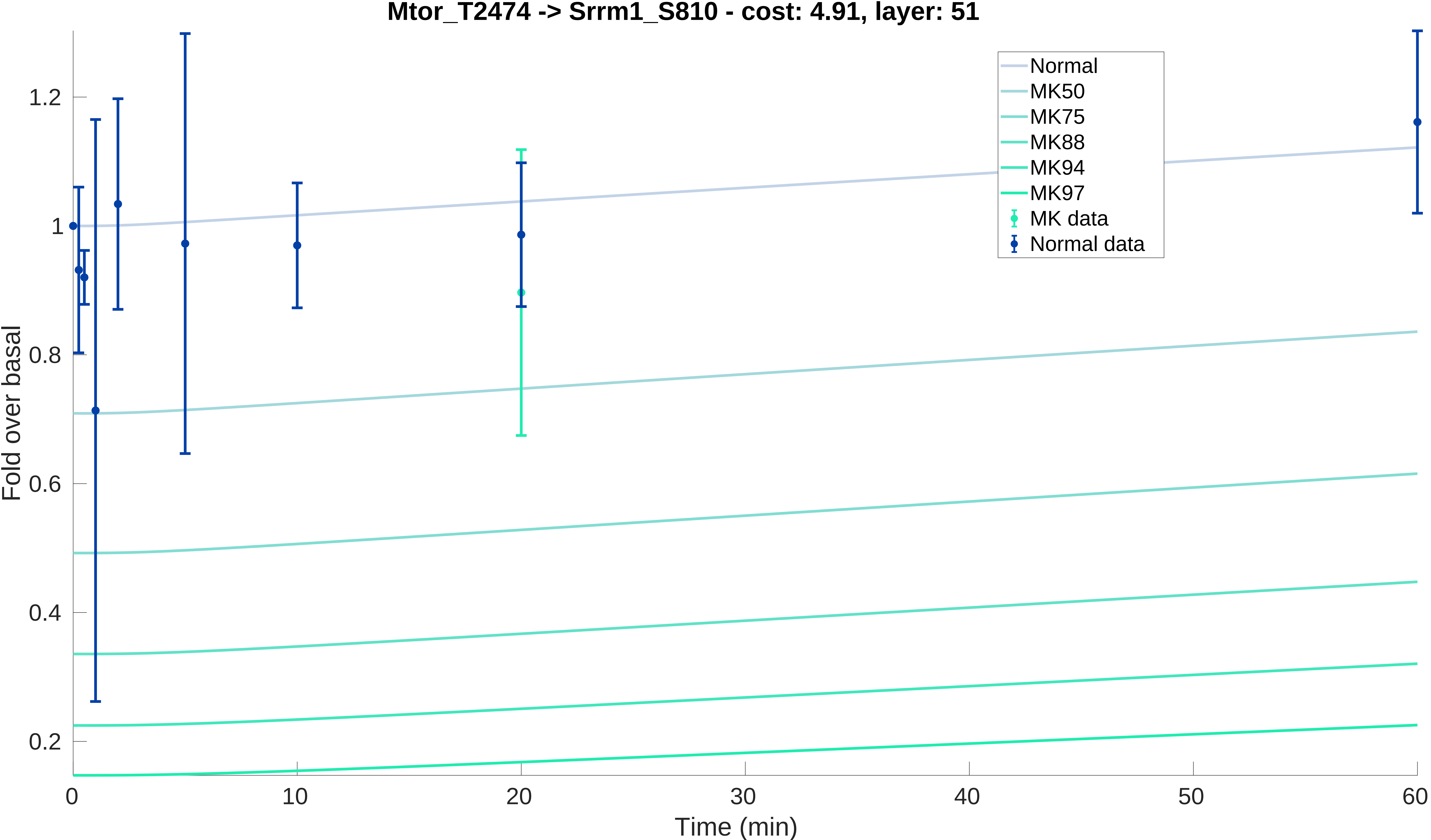
Mtor_T2474 -> Srrm1_S810 - cost: 4.91, layer: 51
Normal
MK50
MK75
MK88
MK94
MK97
MK data
Normal data
Fold over basal
1.2
1
0.8
0.6
0.4
0.2
0
10
20
30
40
50
60
Time (min)

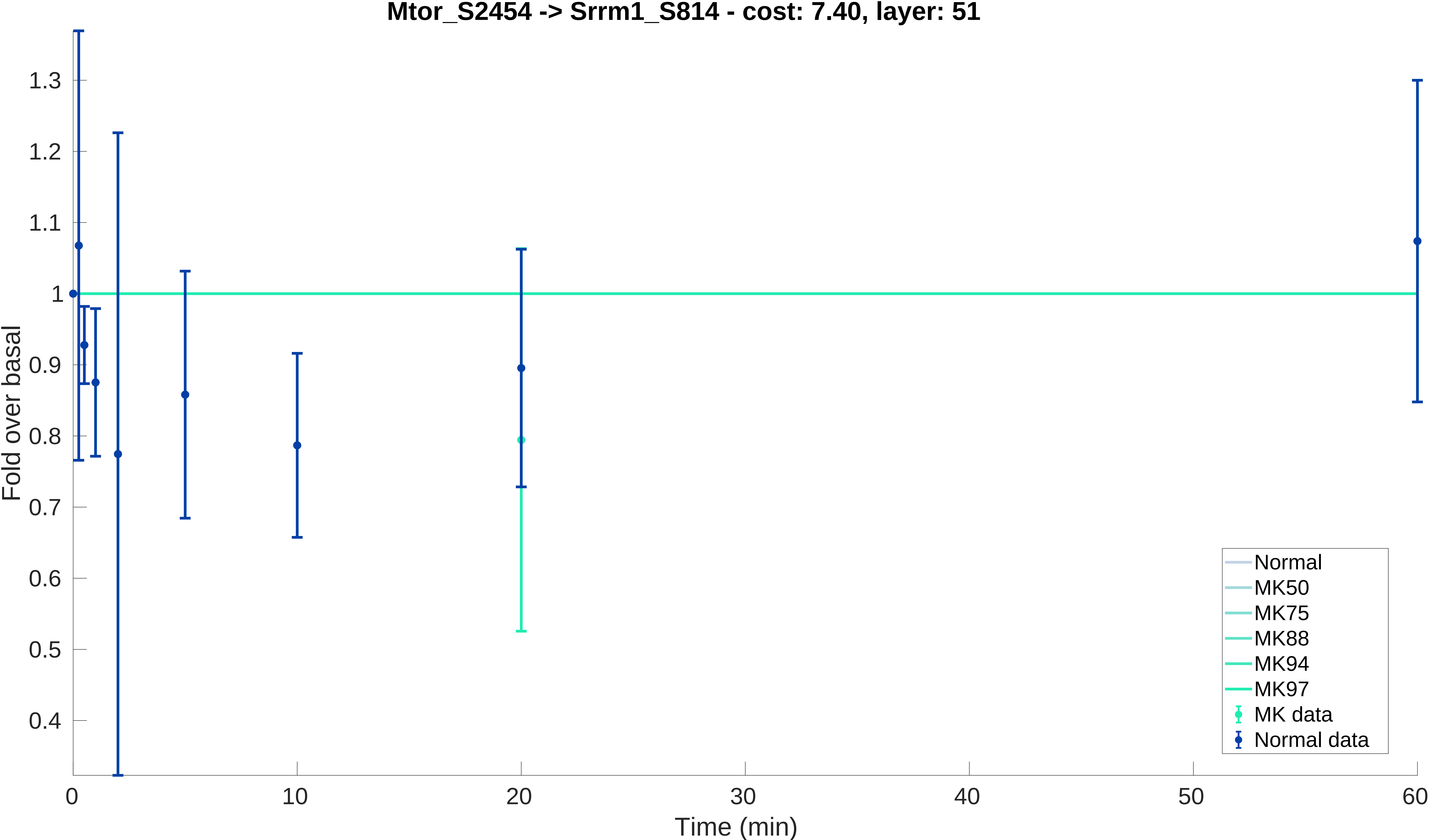

Mtor_S2454 -> Srrm1_S814 - cost: 7.40, layer: 51
Fold over basal
1.3
1.2
1.1
1
0.9
0.8
0.7
0.6
0.5
0.4
0
10
20
30
40
50
60
Time (min)
Normal
MK50
MK75
MK88
MK94
MK97
MK data
Normal data

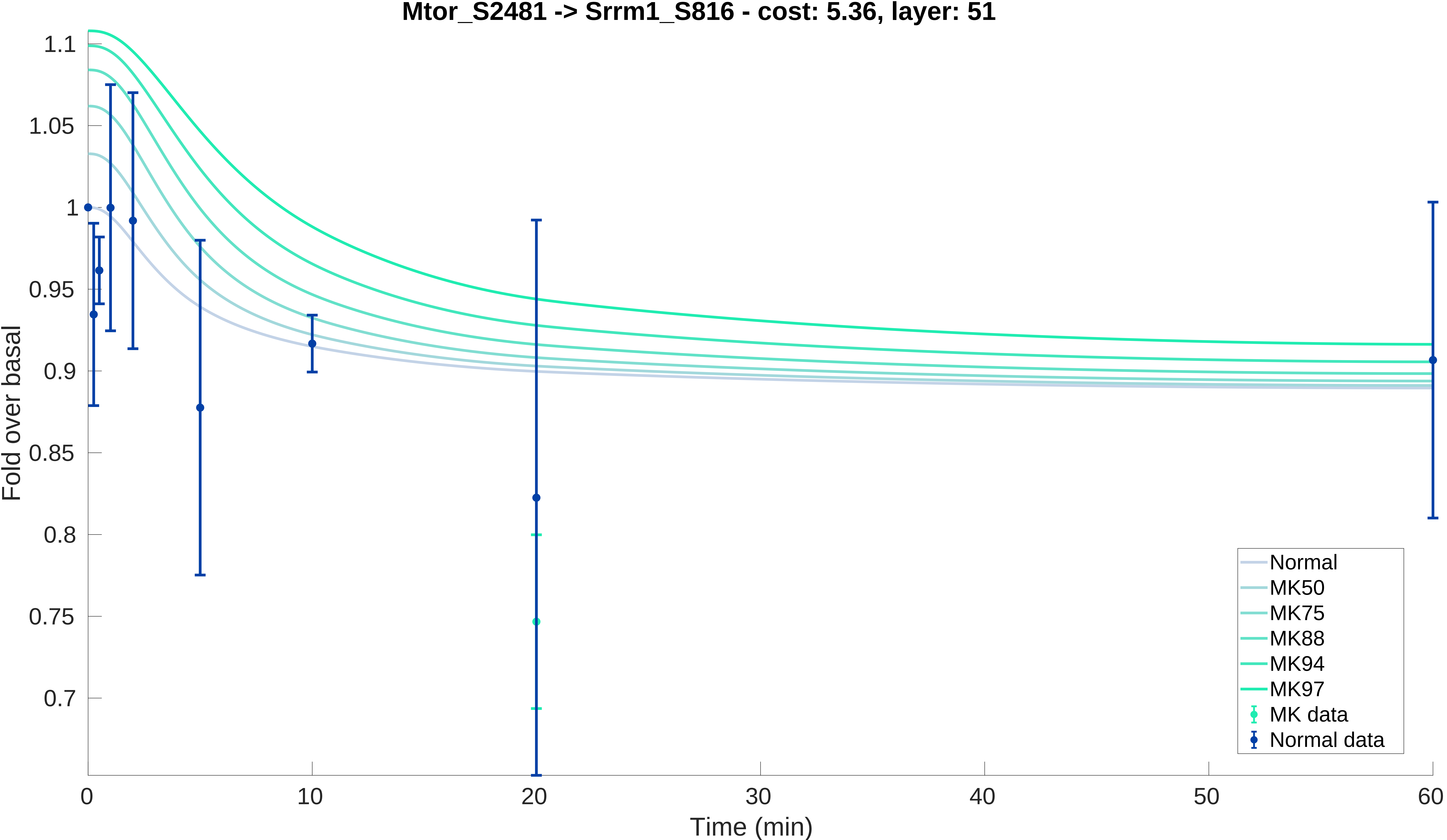

Mtor_S2481 -> Srrm1_S816 - cost: 5.36, layer: 51
Fold over basal
1.1
1.05
1
0.95
0.9
0.85
0.8
0.75
0.7
0
10
20
30
40
50
60
Time (min)
Normal
MK50
MK75
MK88
MK94
MK97
MK data
Normal data

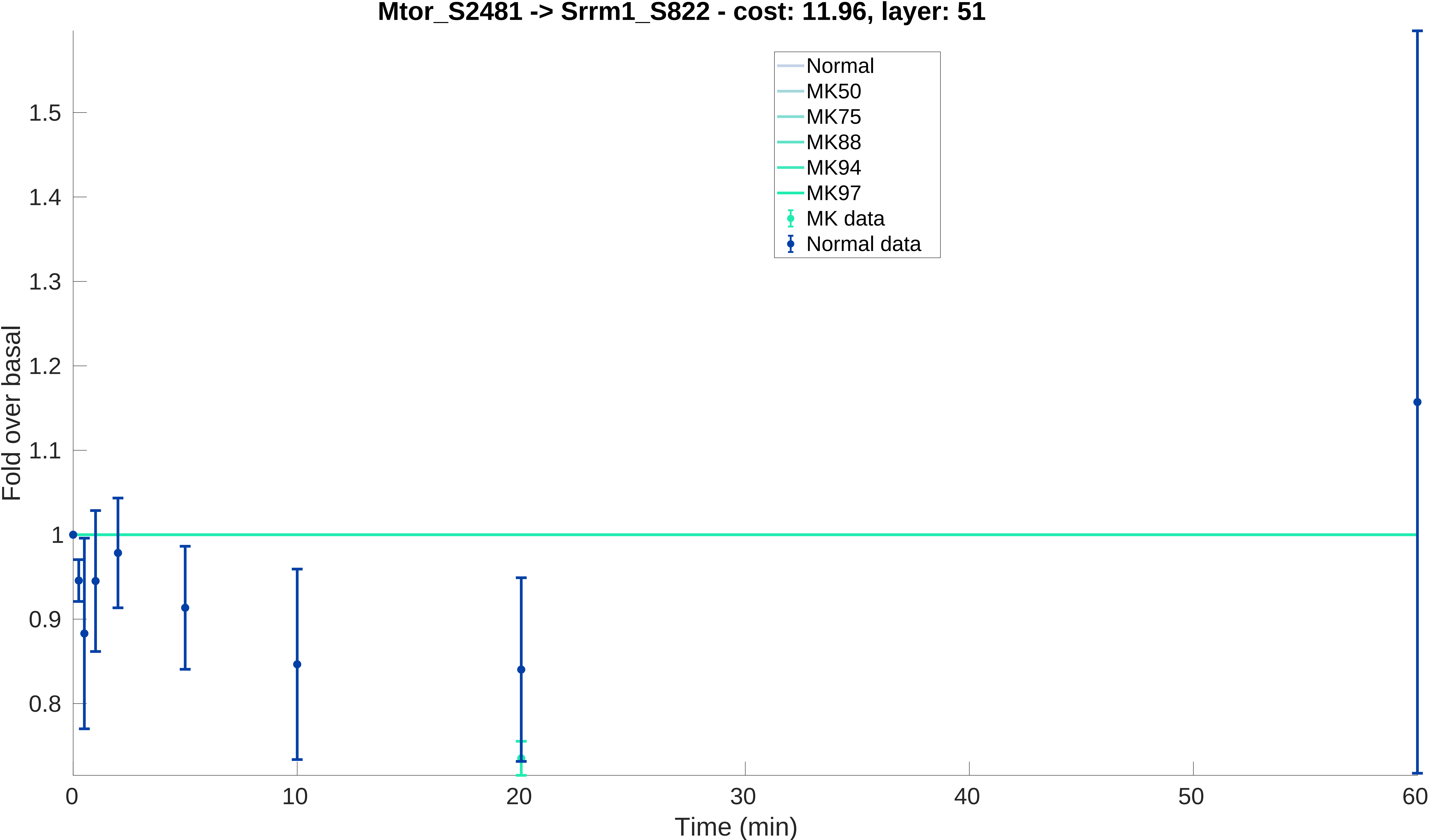

Mtor_S2481 -> Srrm1_S822 - cost: 11.96, layer: 51
Normal
MK50
MK75
MK88
MK94
MK97
MK data
Normal data
Fold over basal
1.5
1.4
1.3
1.2
1.1
1
0.9
0.8
0
10
20
30
40
50
60
Time (min)

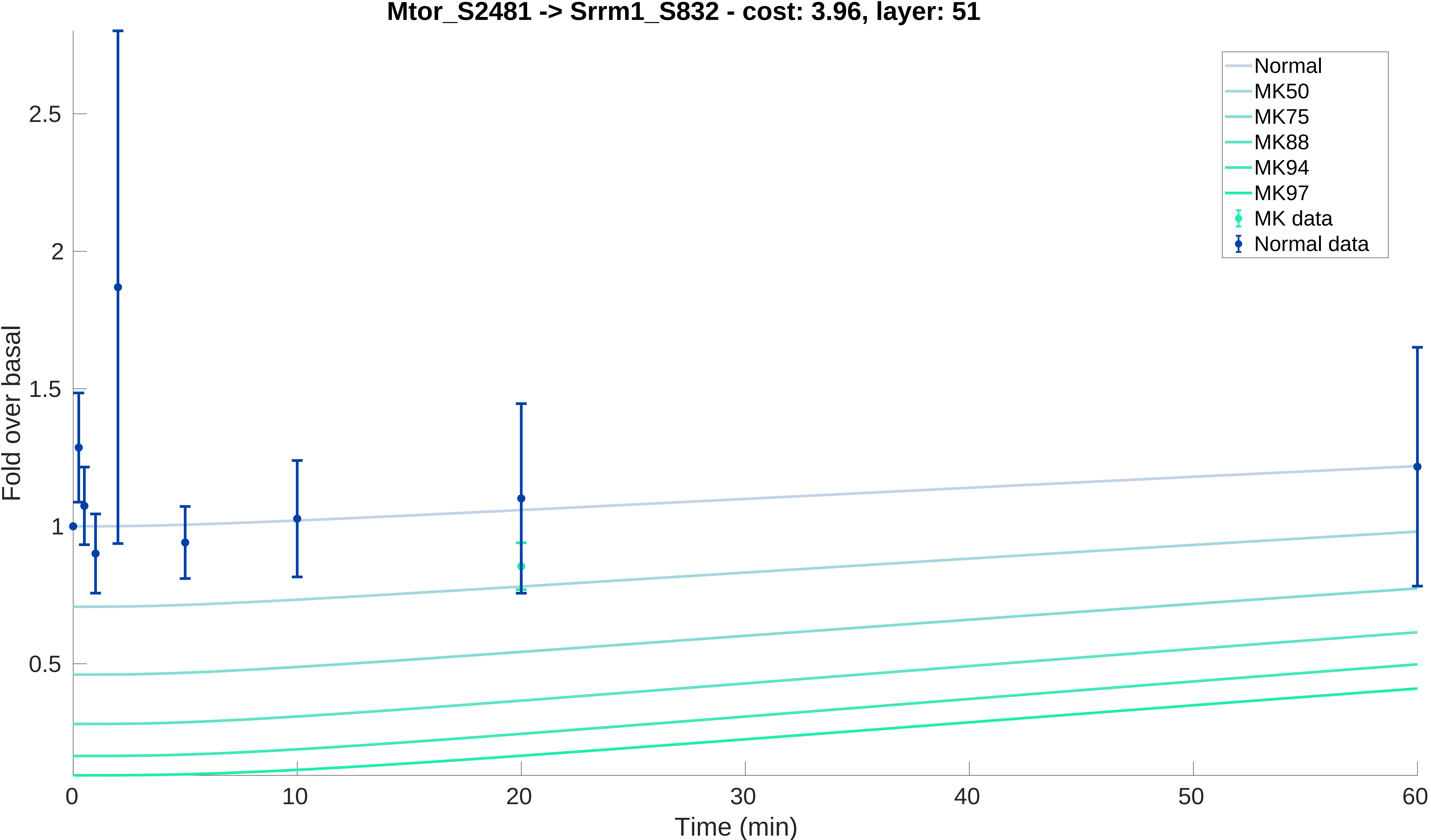

Mtor_S2481 -> Srrm1_S832 - cost: 3.96, layer: 51
Fold over basal
Time (min)
0
10
20
30
40
50
60
0.5
1
1.5
2
2.5
Normal
MK50
MK75
MK88
MK94
MK97
MK data
Normal data

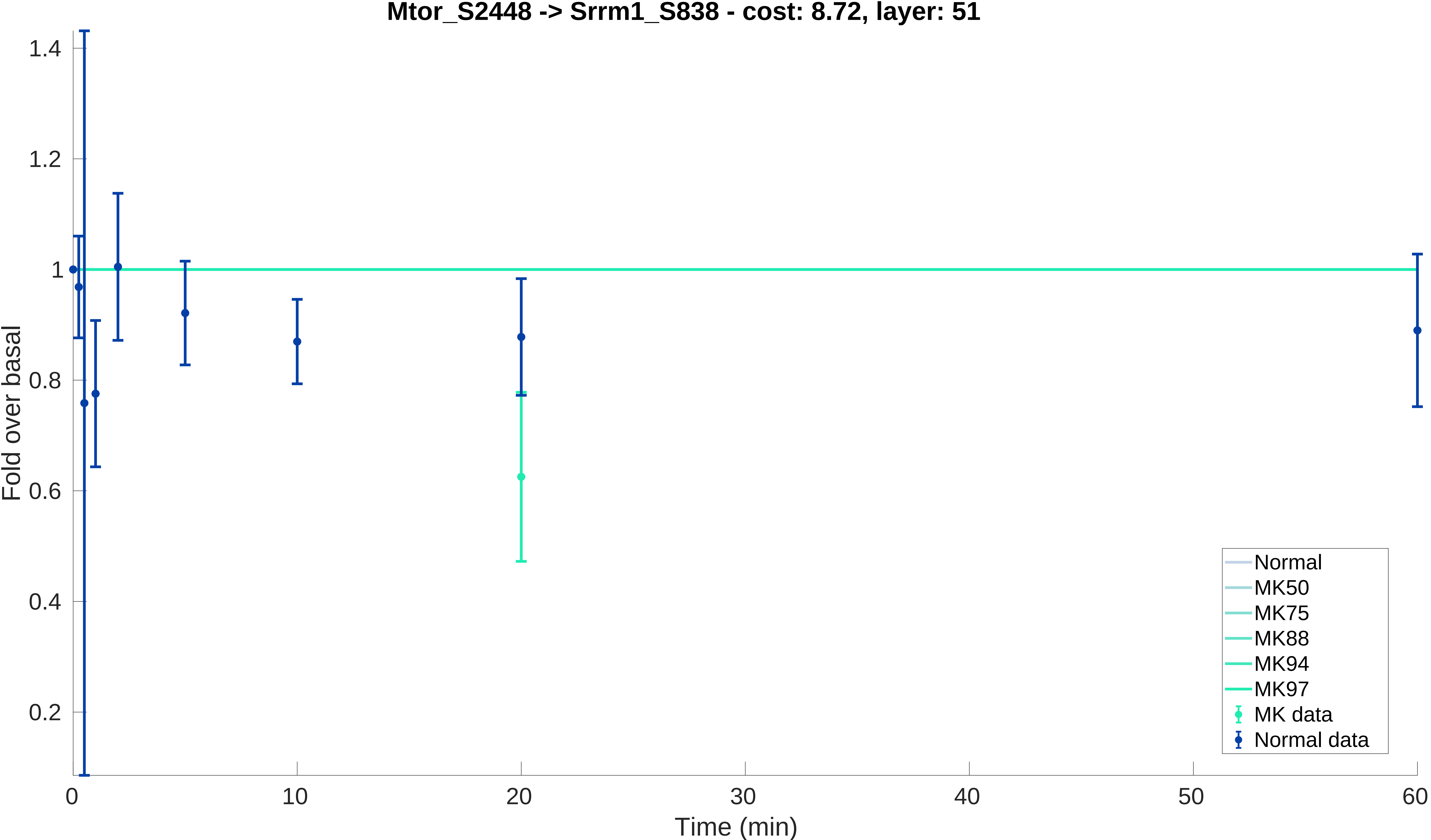
Mtor_S2448 -> Srrm1_S838 - cost: 8.72, layer: 51
Fold over basal
1.4
1.2
1
0.8
0.6
0.4
0.2
0
10
20
30
40
50
60
Time (min)
Normal
MK50
MK75
MK88
MK94
MK97
MK data
Normal data

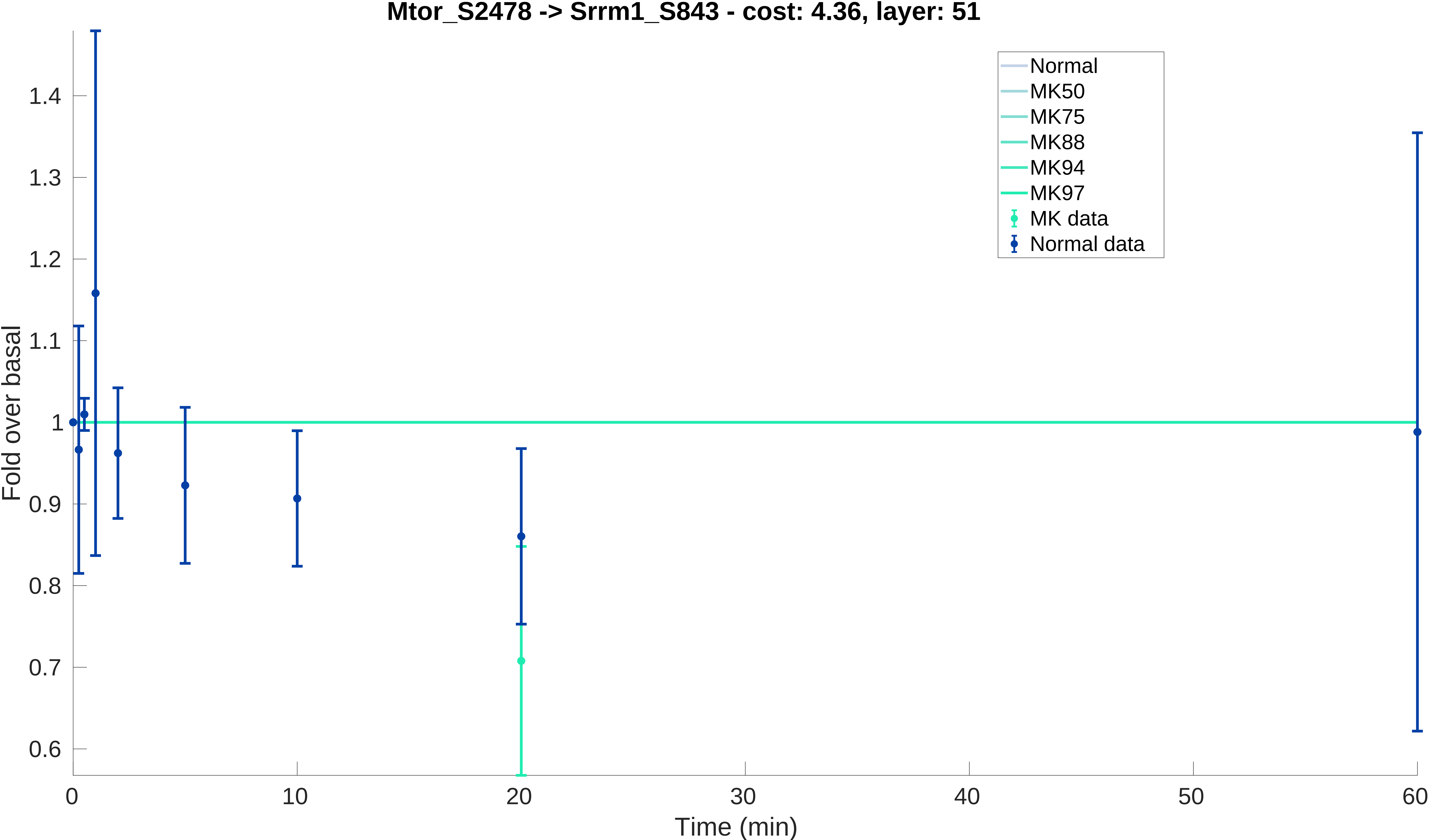
Mtor_S2478 -> Srrm1_S843 - cost: 4.36, layer: 51
Fold over basal
Time (min)
0.6
0.7
0.8
0.9
1
1.1
1.2
1.3
1.4
0
10
20
30
40
50
60
Normal
MK50
MK75
MK88
MK94
MK97
MK data
Normal data

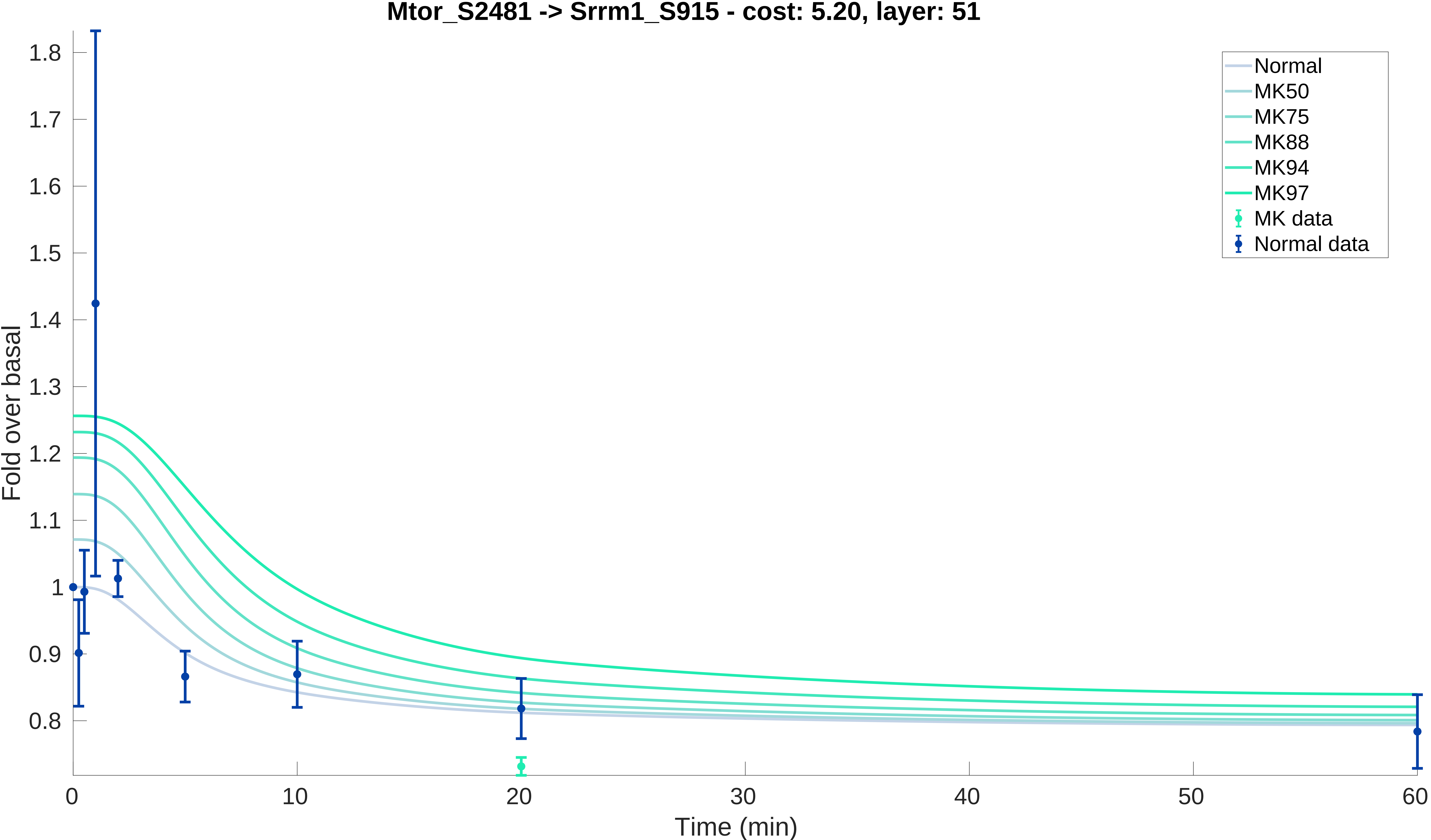
Mtor_S2481 -> Srrm1_S915 - cost: 5.20, layer: 51
Fold over basal
Time (min)
0
10
20
30
40
50
60
0.8
0.9
1
1.1
1.2
1.3
1.4
1.5
1.6
1.7
1.8
Normal
MK50
MK75
MK88
MK94
MK97
MK data
Normal data

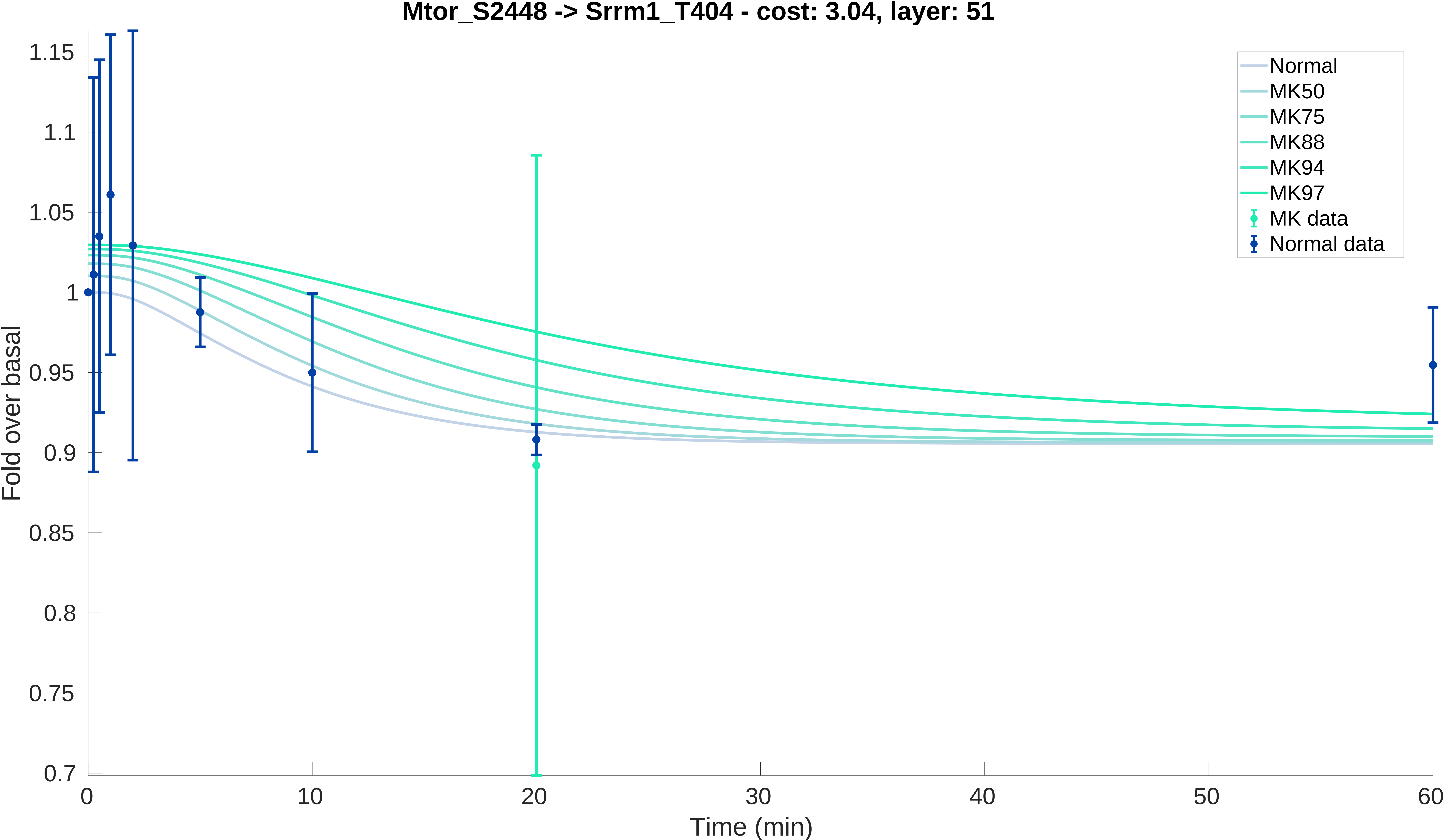

Mtor_S2448 -> Srrm1_T404 - cost: 3.04, layer: 51
Fold over basal
Time (min)
1.15
1.1
1.05
1
0.95
0.9
0.85
0.8
0.75
0.7
0
10
20
30
40
50
60
Normal
MK50
MK75
MK88
MK94
MK97
MK data
Normal data

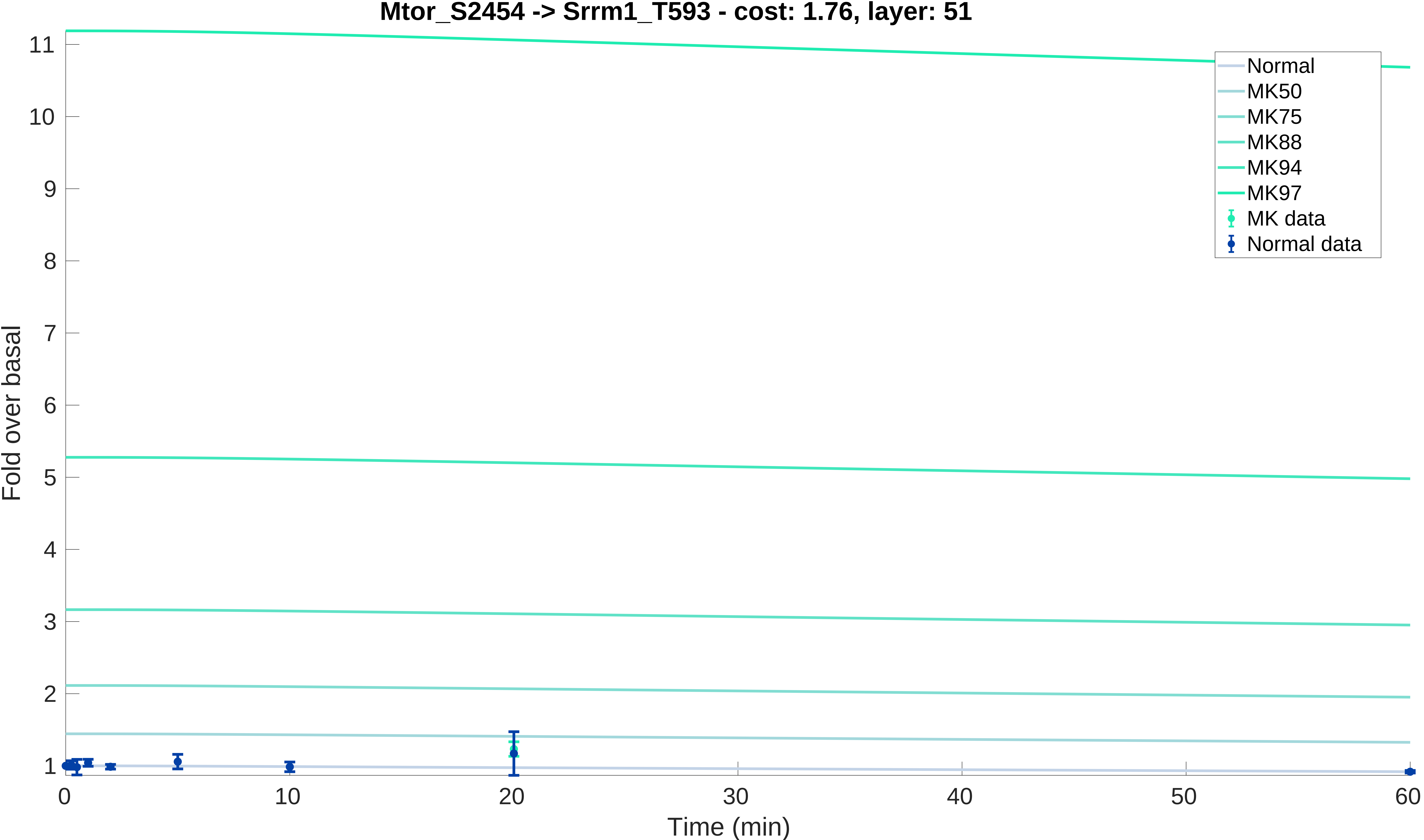
Mtor_S2454 -> Srrm1_T593 - cost: 1.76, layer: 51
Fold over basal
Time (min)
Normal
MK50
MK75
MK88
MK94
MK97
MK data
Normal data
1
2
3
4
5
6
7
8
9
10
11
0
10
20
30
40
50
60

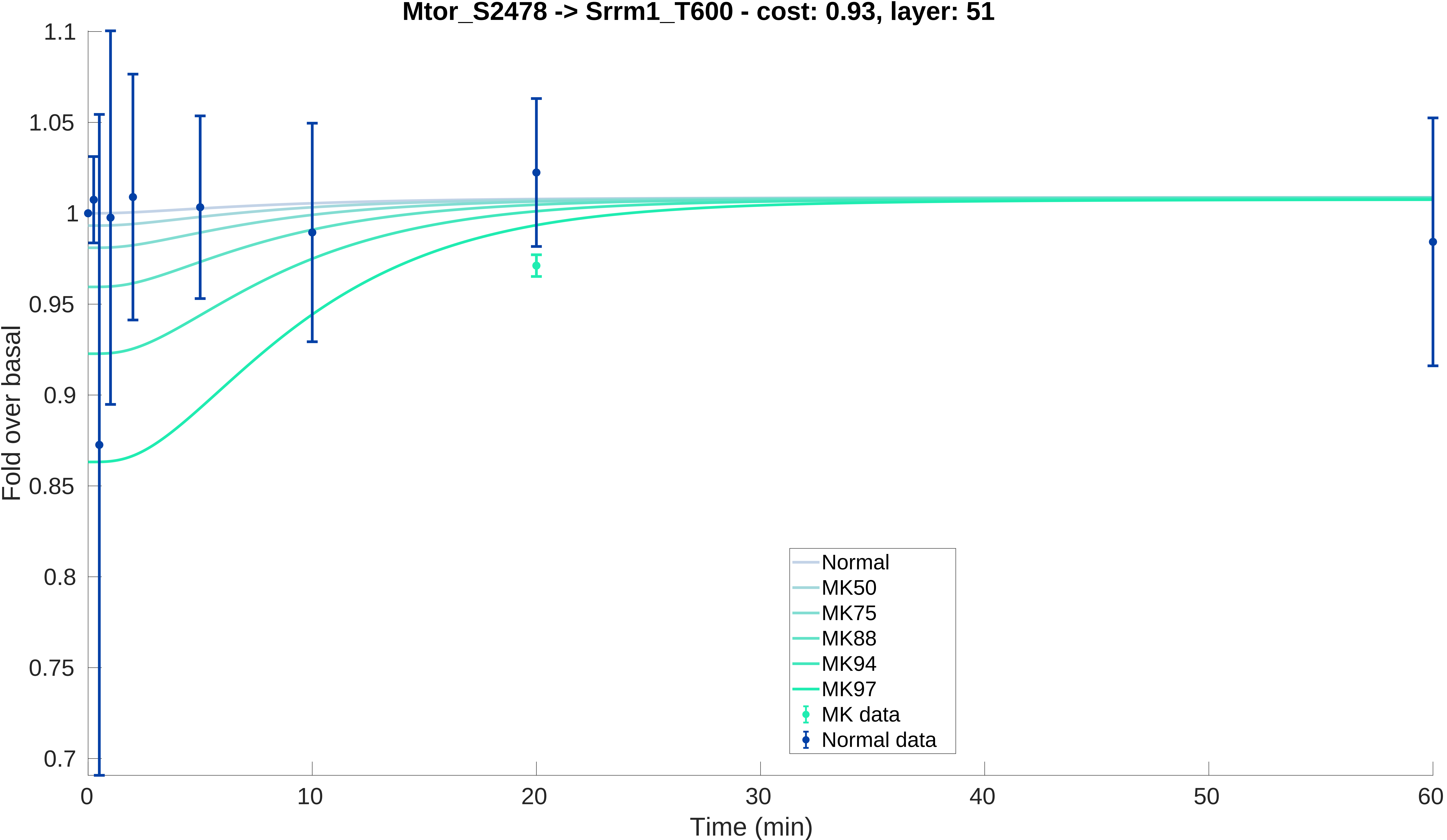

Mtor_S2478 -> Srrm1_T600 - cost: 0.93, layer: 51
Fold over basal
Time (min)
1.1
1.05
1
0.95
0.9
0.85
0.8
0.75
0.7
0
10
20
30
40
50
60
Normal
MK50
MK75
MK88
MK94
MK97
MK data
Normal data

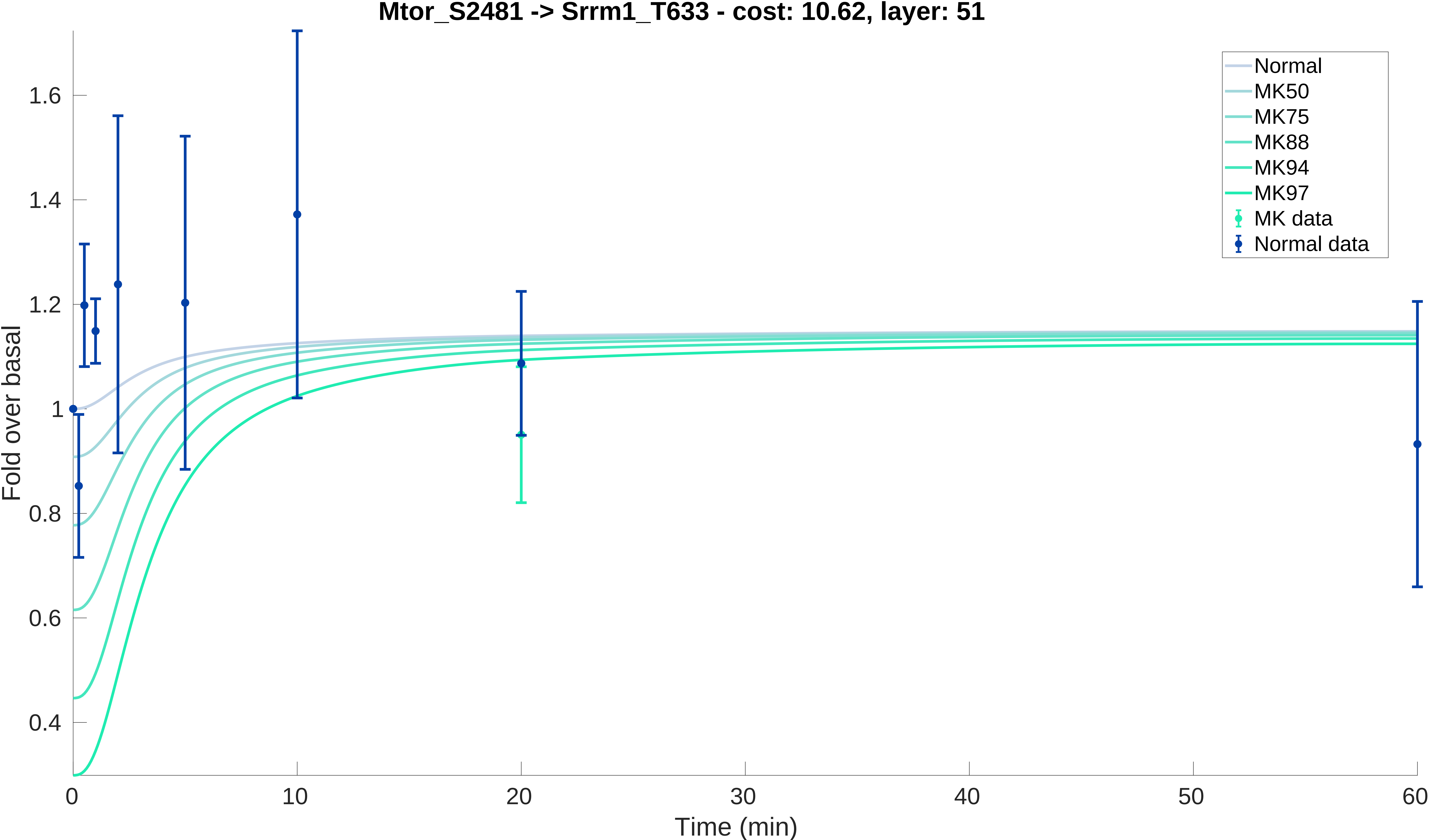

Mtor_S2481 -> Srrm1_T633 - cost: 10.62, layer: 51
Normal
MK50
MK75
MK88
MK94
MK97
MK data
Normal data
Fold over basal
1.6
1.4
1.2
1
0.8
0.6
0.4
0
10
20
30
40
50
60
Time (min)

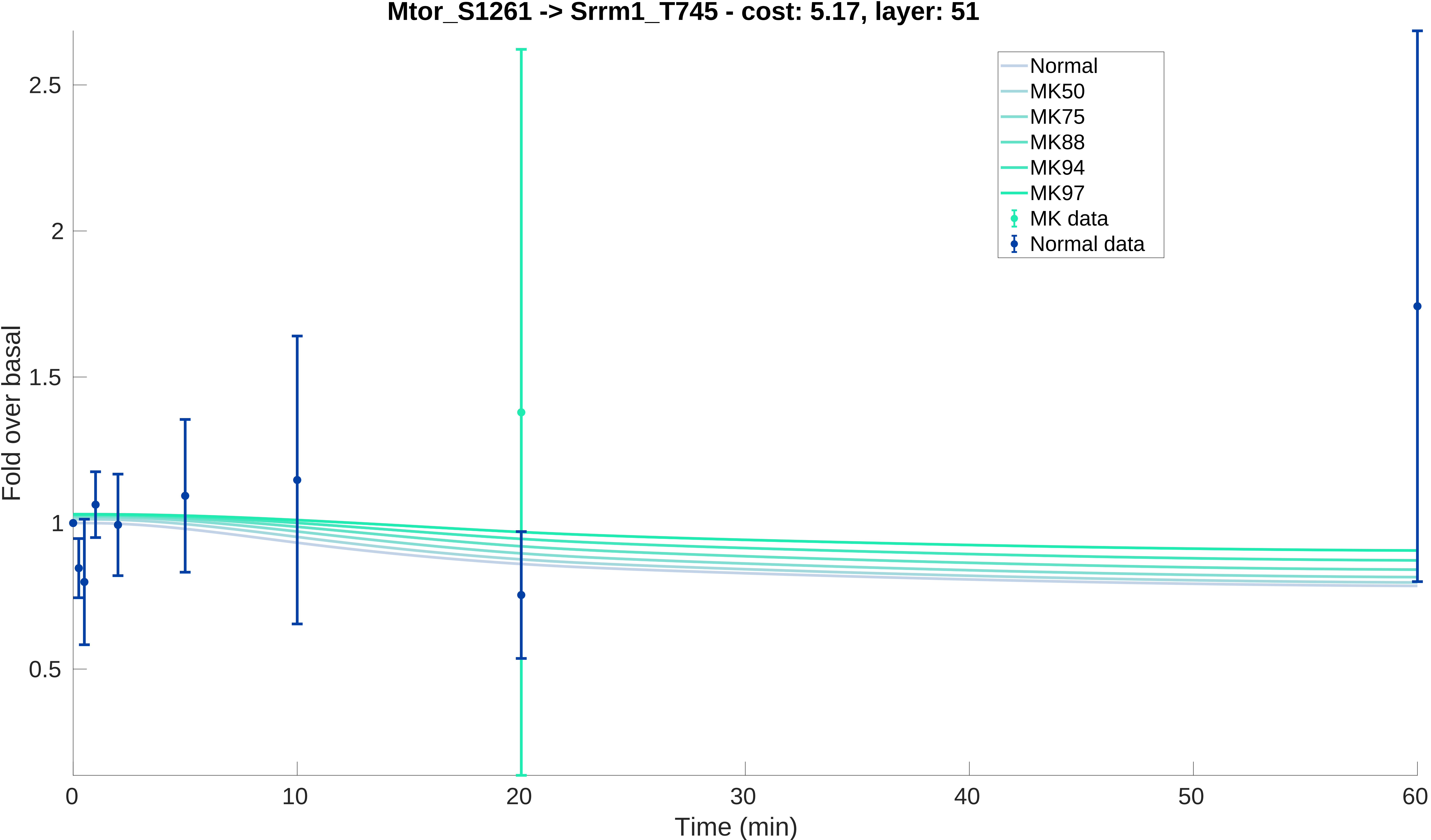

Mtor_S1261 -> Srrm1_T745 - cost: 5.17, layer: 51
Normal
MK50
MK75
MK88
MK94
MK97
MK data
Normal data
Fold over basal
Time (min)
0.5
1
1.5
2
2.5
0
10
20
30
40
50
60

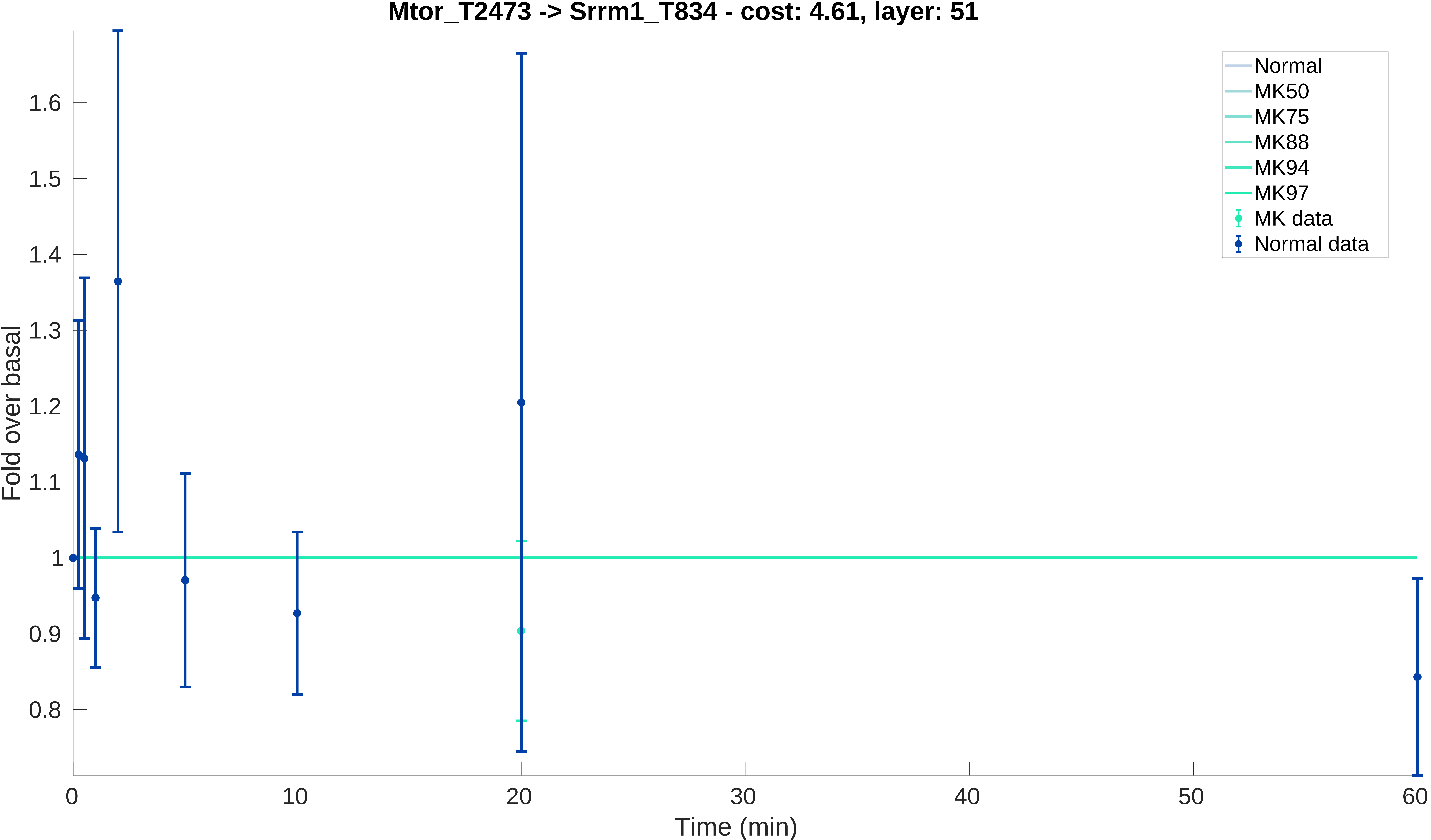

Mtor_T2473 -> Srrm1_T834 - cost: 4.61, layer: 51
Fold over basal
Time (min)
1.6
1.5
1.4
1.3
1.2
1.1
1
0.9
0.8
0
10
20
30
40
50
60
Normal
MK50
MK75
MK88
MK94
MK97
MK data
Normal data

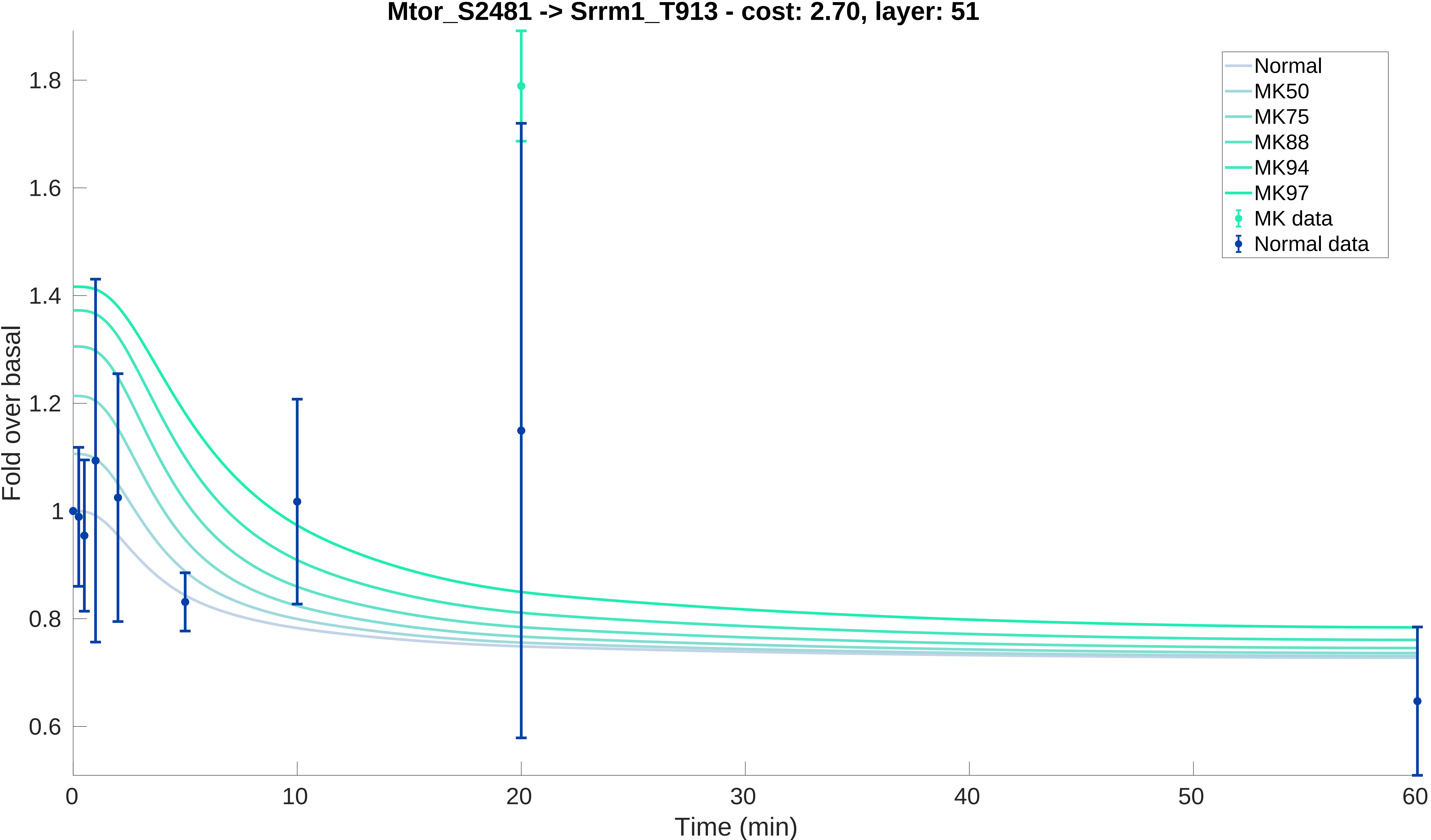

Mtor_S2481 -> Srrm1_T913 - cost: 2.70, layer: 51
Fold over basal
Time (min)
0.6
0.8
1
1.2
1.4
1.6
1.8
0
10
20
30
40
50
60
Normal
MK50
MK75
MK88
MK94
MK97
MK data
Normal data

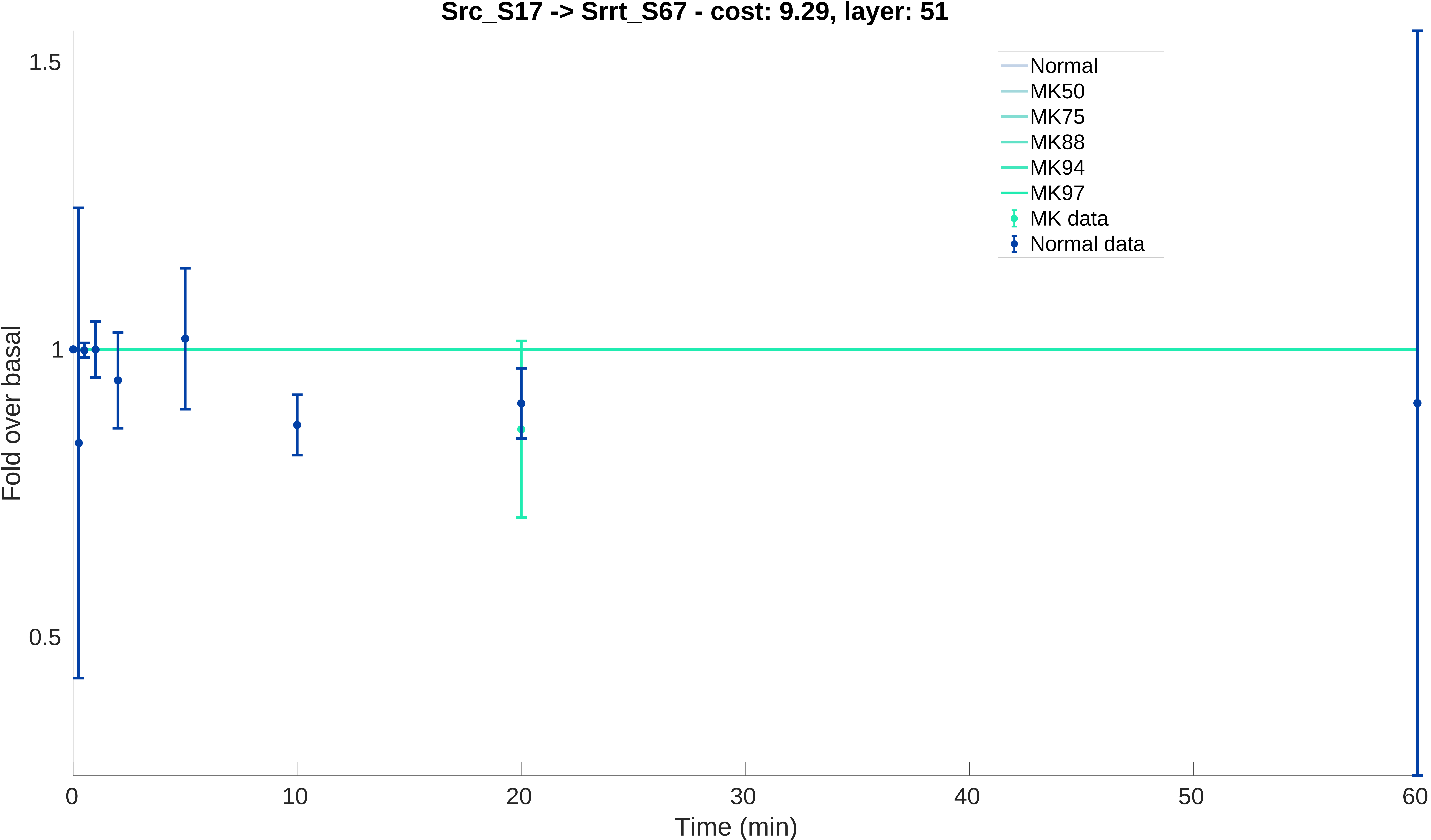
Src_S17 -> Srrt_S67 - cost: 9.29, layer: 51
Normal
MK50
MK75
MK88
MK94
MK97
MK data
Normal data
Fold over basal
1.5
1
0.5
0
10
20
30
40
50
60
Time (min)

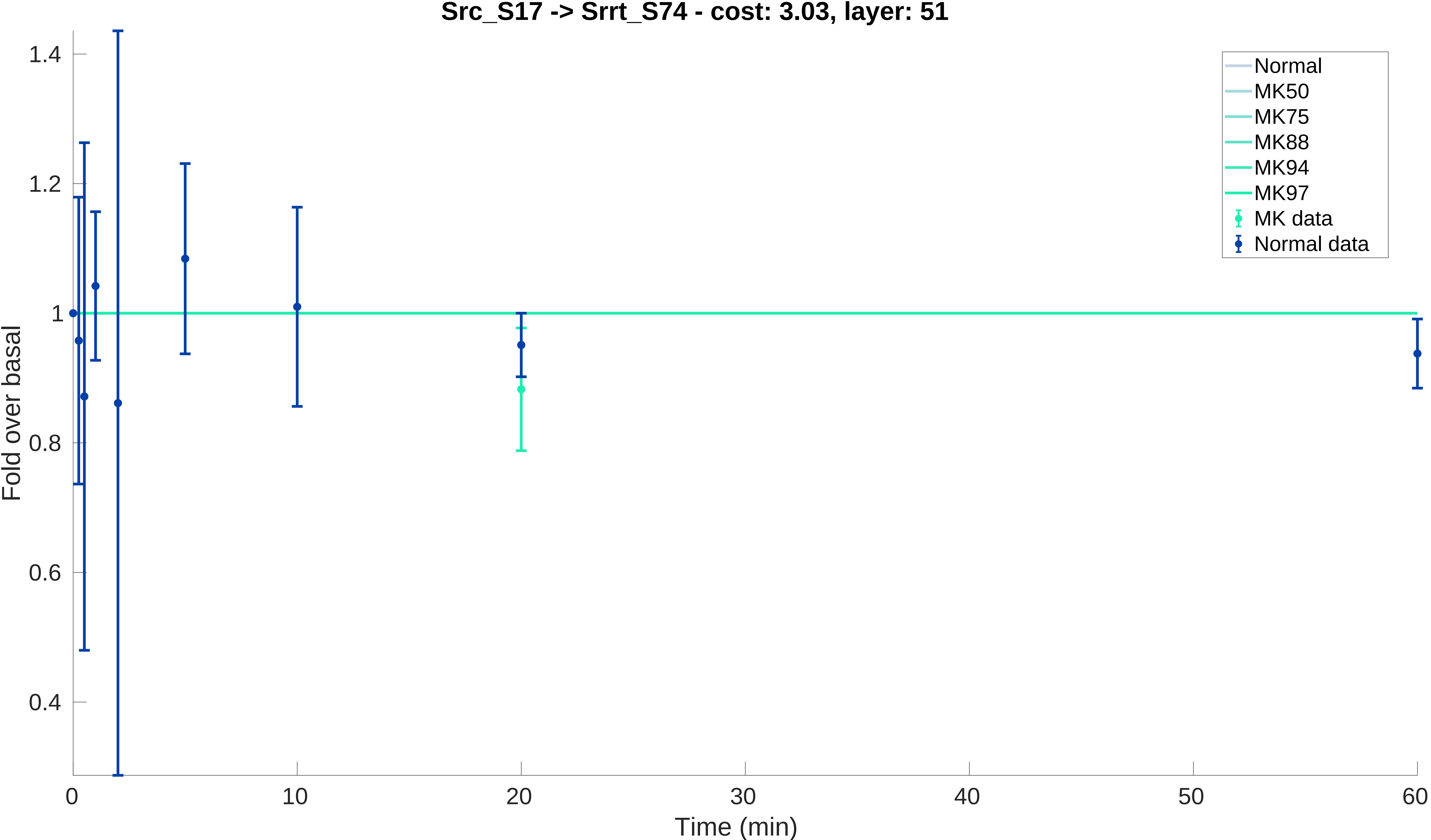

Src_S17 -> Srrt_S74 - cost: 3.03, layer: 51
Fold over basal
Time (min)
0.4
0.6
0.8
1
1.2
1.4
0
10
20
30
40
50
60
Normal
MK50
MK75
MK88
MK94
MK97
MK data
Normal data

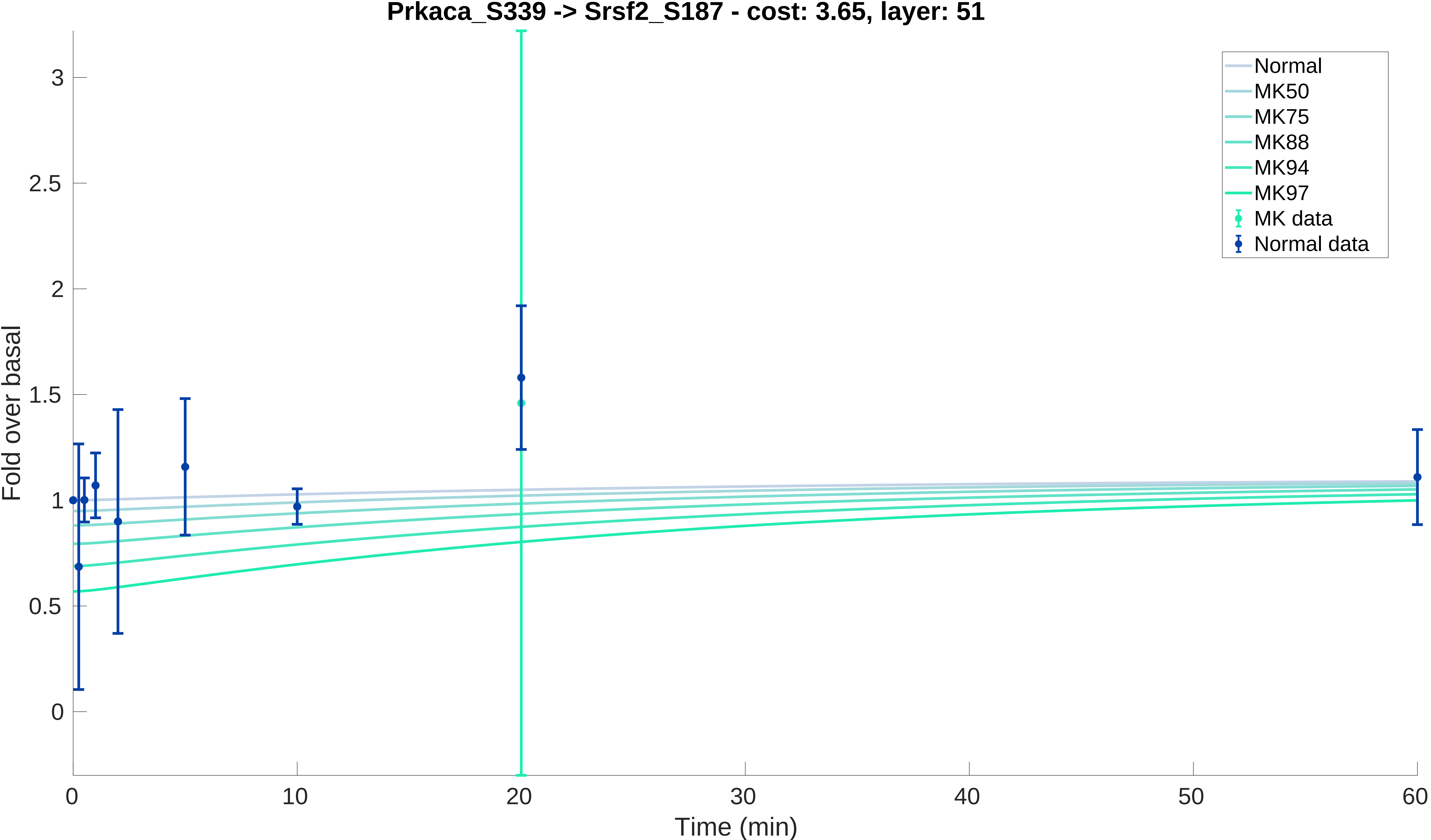

Prkaca_S339 -> Srsf2_S187 - cost: 3.65, layer: 51
Normal
MK50
MK75
MK88
MK94
MK97
MK data
Normal data
Fold over basal
Time (min)
0
0.5
1
1.5
2
2.5
3
0
10
20
30
40
50
60

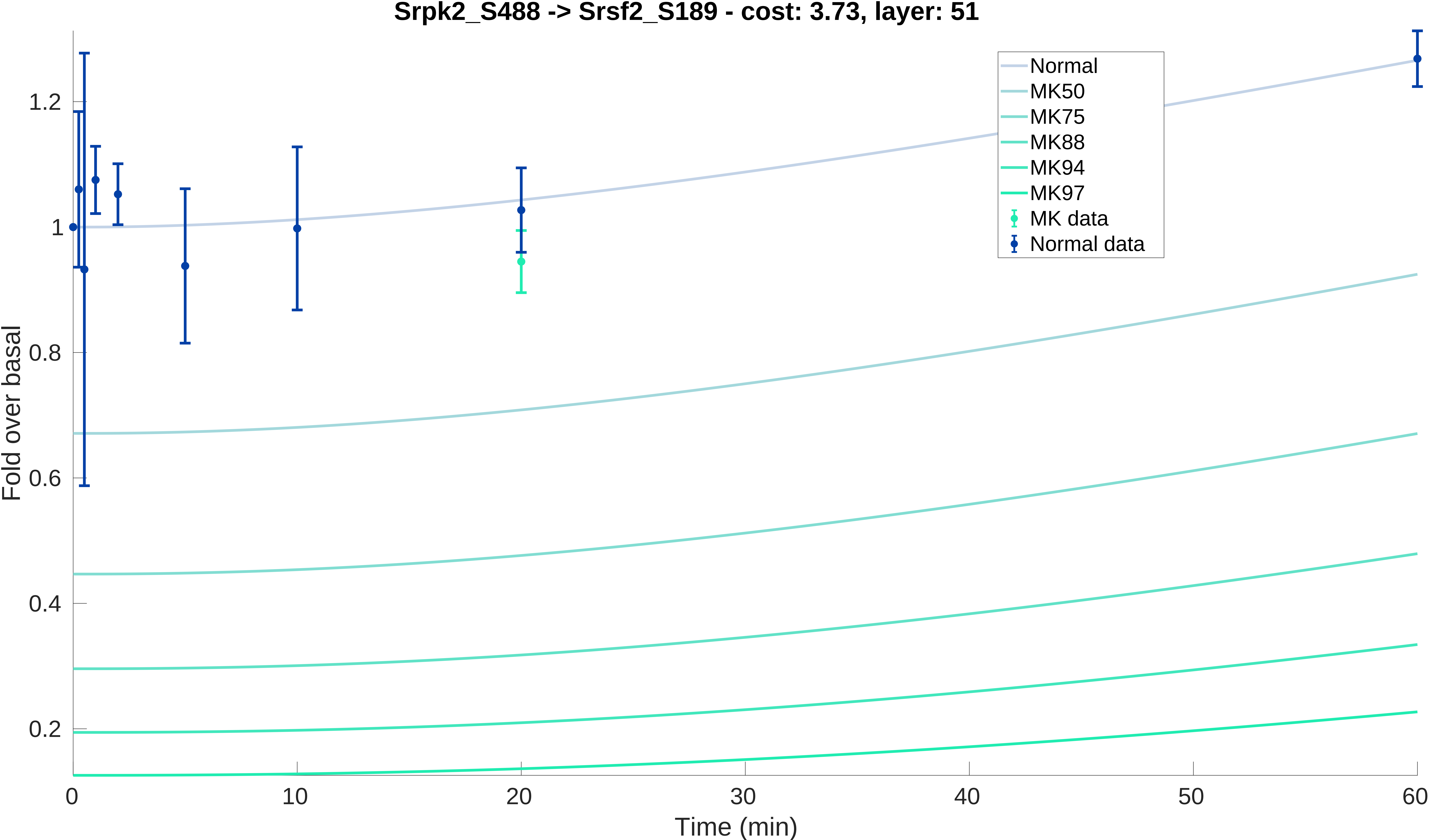

Srpk2_S488 -> Srsf2_S189 - cost: 3.73, layer: 51
Normal
MK50
MK75
MK88
MK94
MK97
MK data
Normal data
Fold over basal
0.2
0.4
0.6
0.8
1
1.2
0
10
20
30
40
50
60
Time (min)

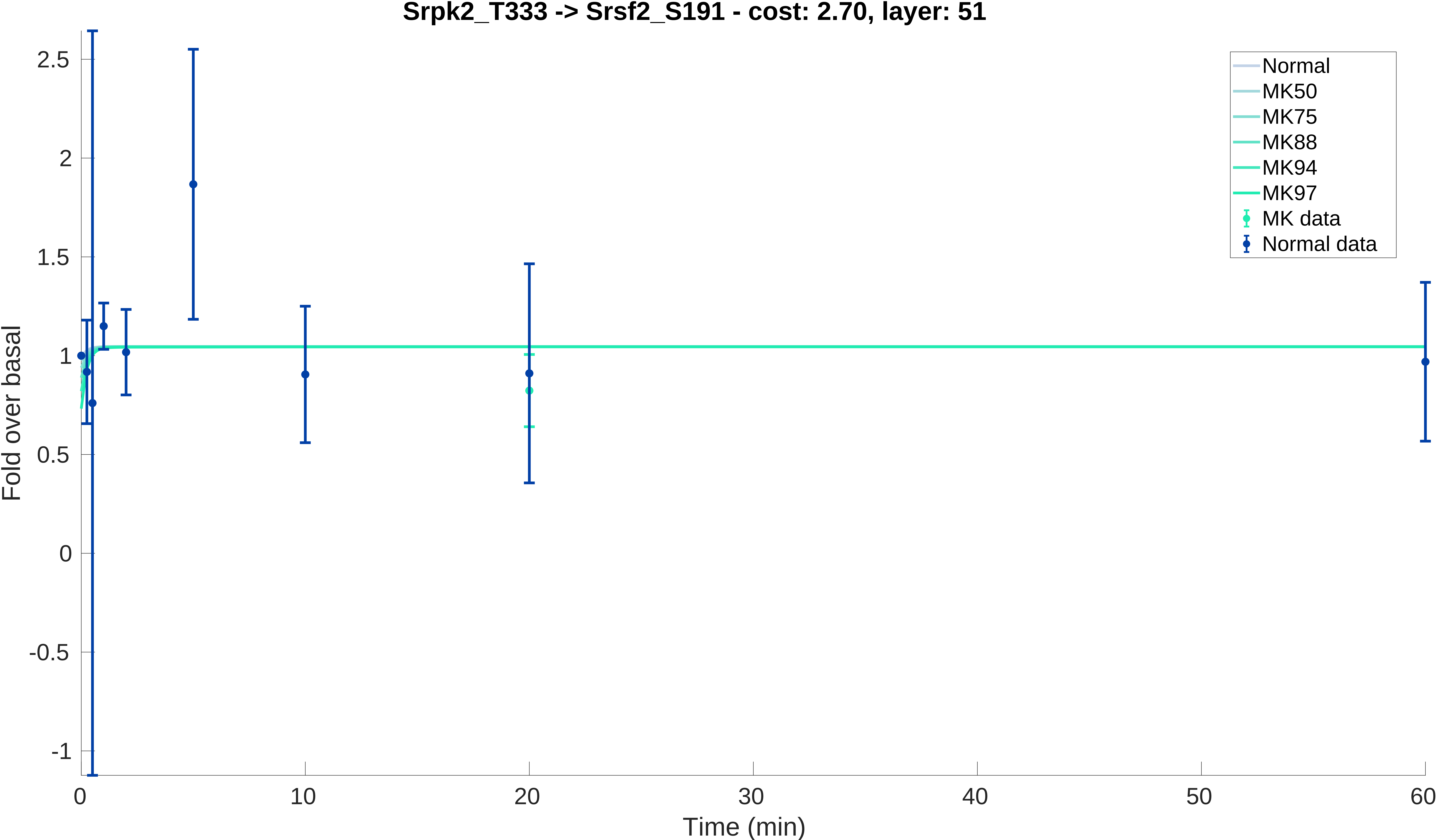
Srpk2_T333 -> Srsf2_S191 - cost: 2.70, layer: 51
Normal
MK50
MK75
MK88
MK94
MK97
MK data
Normal data
Fold over basal
Time (min)
2.5
2
1.5
1
0.5
0
-0.5
-1
0
10
20
30
40
50
60

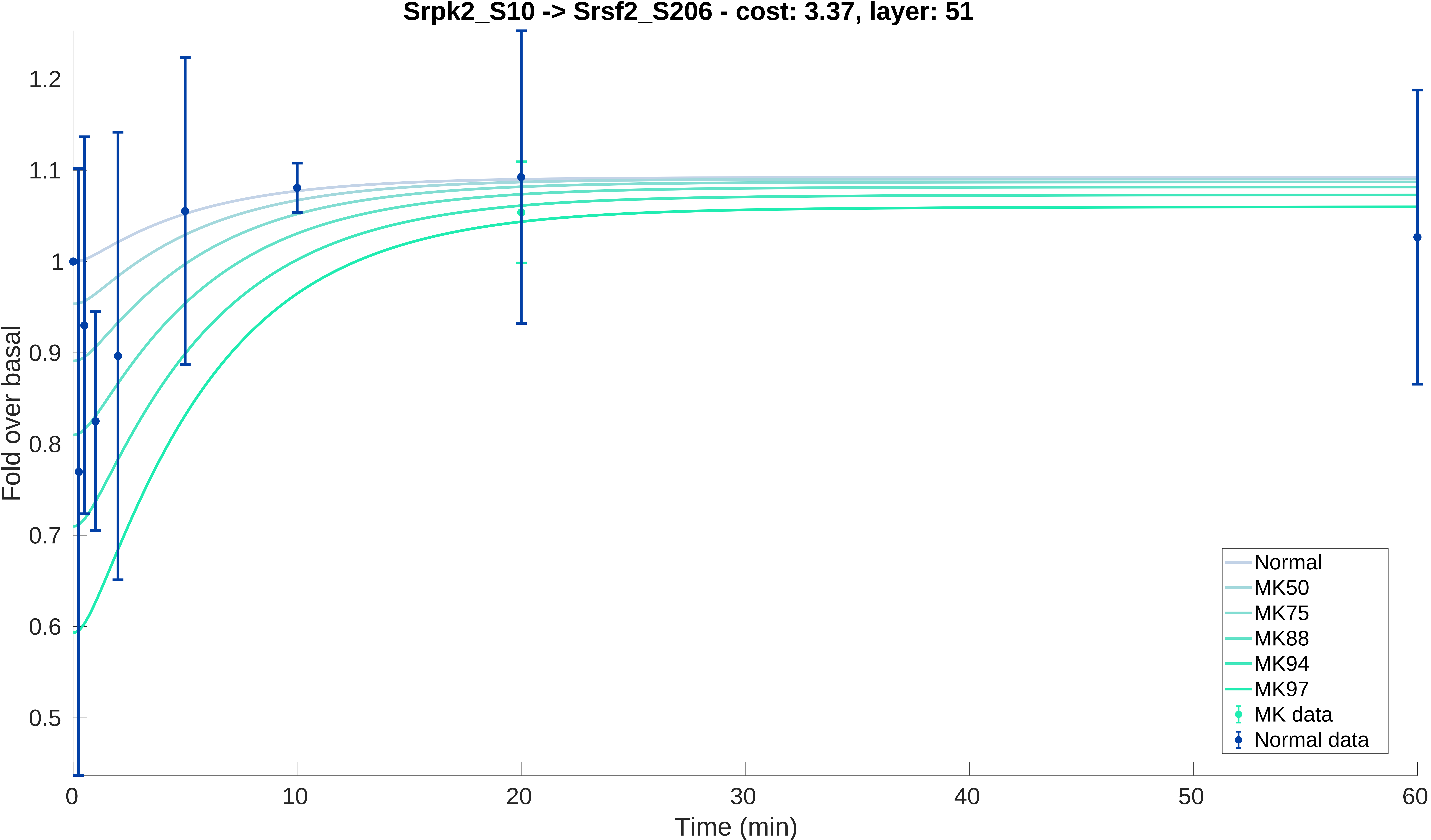
Srpk2_S10 -> Srsf2_S206 - cost: 3.37, layer: 51
Fold over basal
Time (min)
0.5
0.6
0.7
0.8
0.9
1
1.1
1.2
0
10
20
30
40
50
60
Normal
MK50
MK75
MK88
MK94
MK97
MK data
Normal data

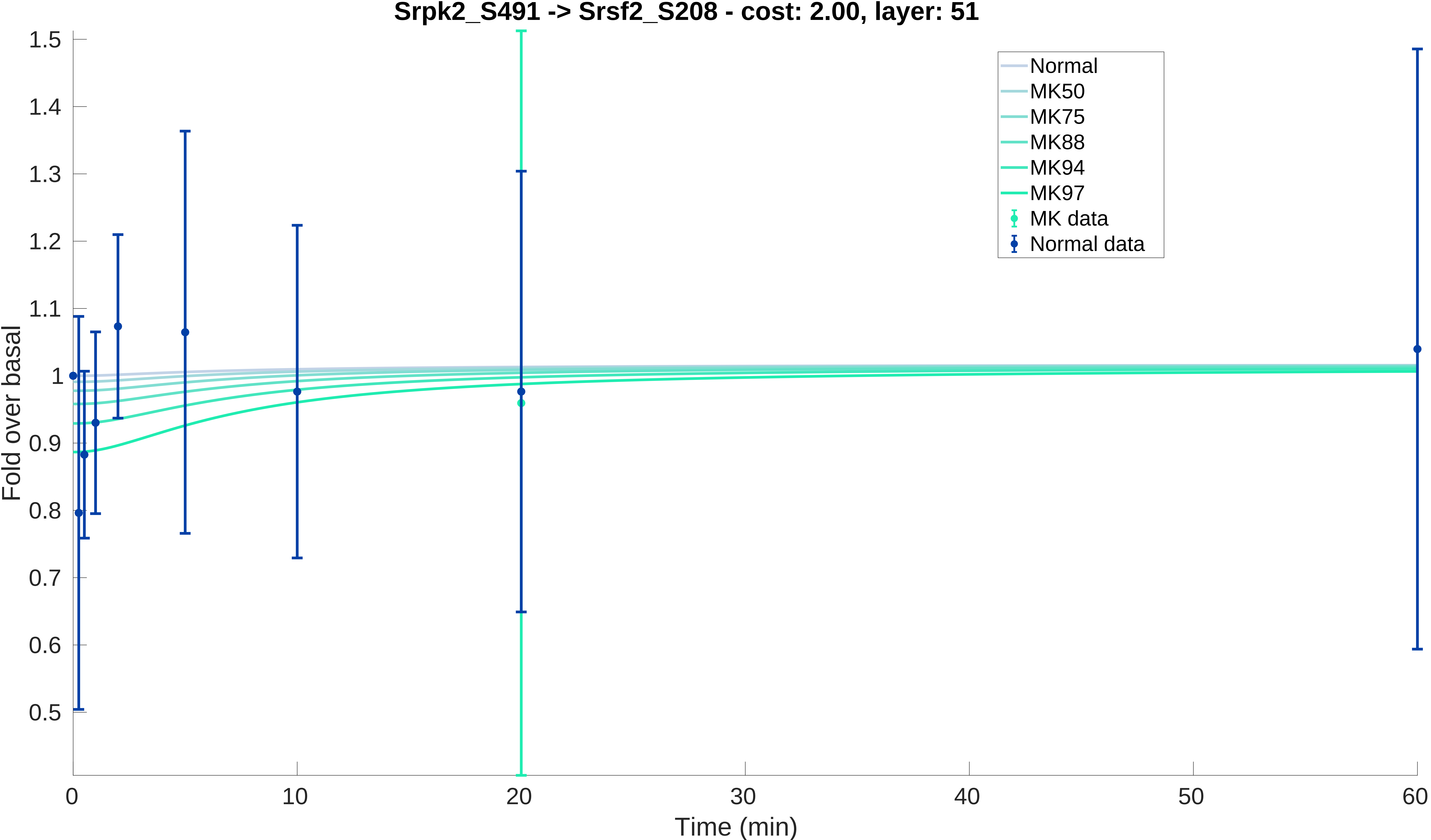

Srpk2_S491 -> Srsf2_S208 - cost: 2.00, layer: 51
Fold over basal
Time (min)
0.5
0.6
0.7
0.8
0.9
1
1.1
1.2
1.3
1.4
1.5
0
10
20
30
40
50
60
Normal
MK50
MK75
MK88
MK94
MK97
MK data
Normal data

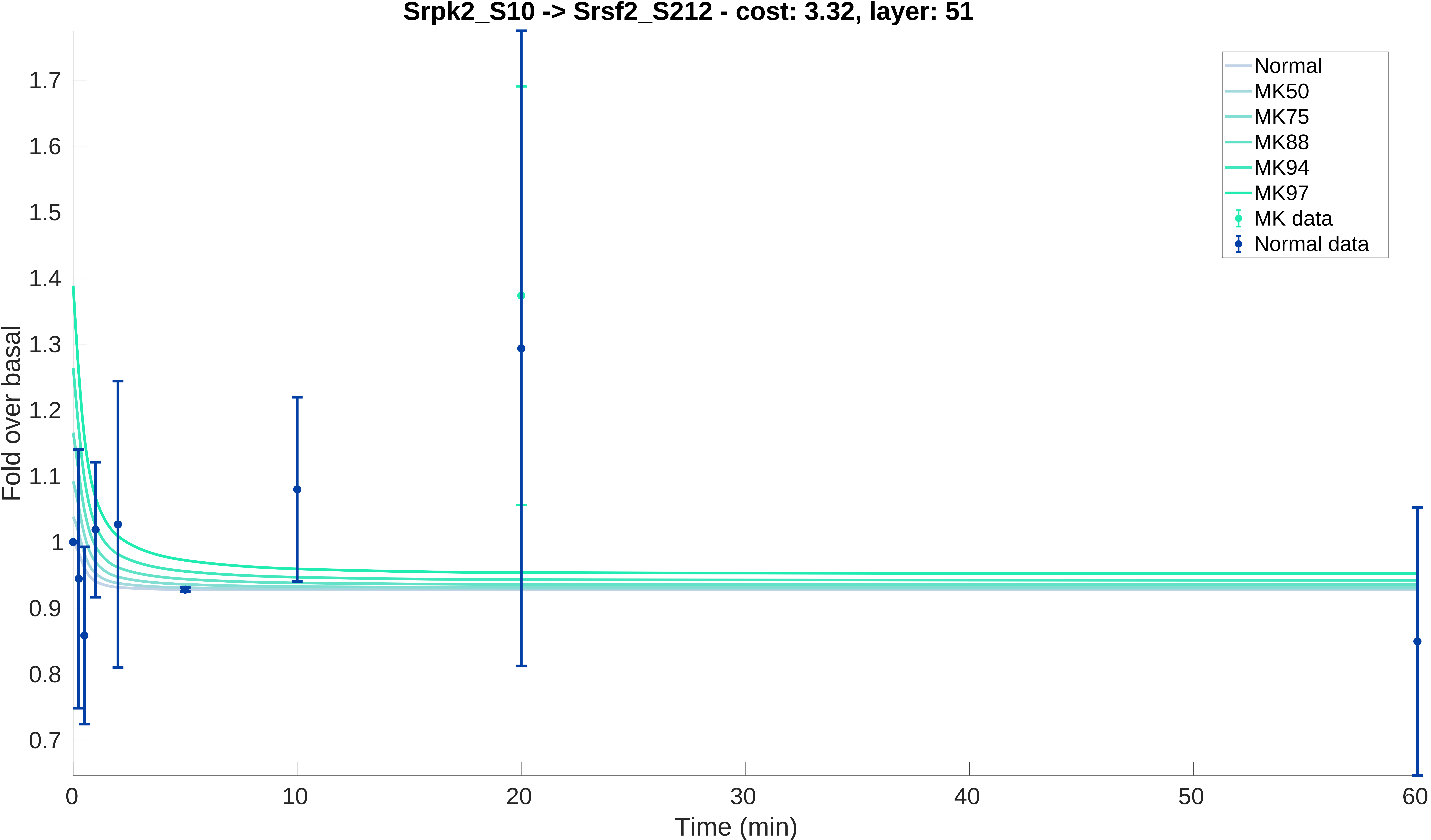

Srpk2_S10 -> Srsf2_S212 - cost: 3.32, layer: 51
Fold over basal
Time (min)
0.7
0.8
0.9
1
1.1
1.2
1.3
1.4
1.5
1.6
1.7
0
10
20
30
40
50
60
Normal
MK50
MK75
MK88
MK94
MK97
MK data
Normal data

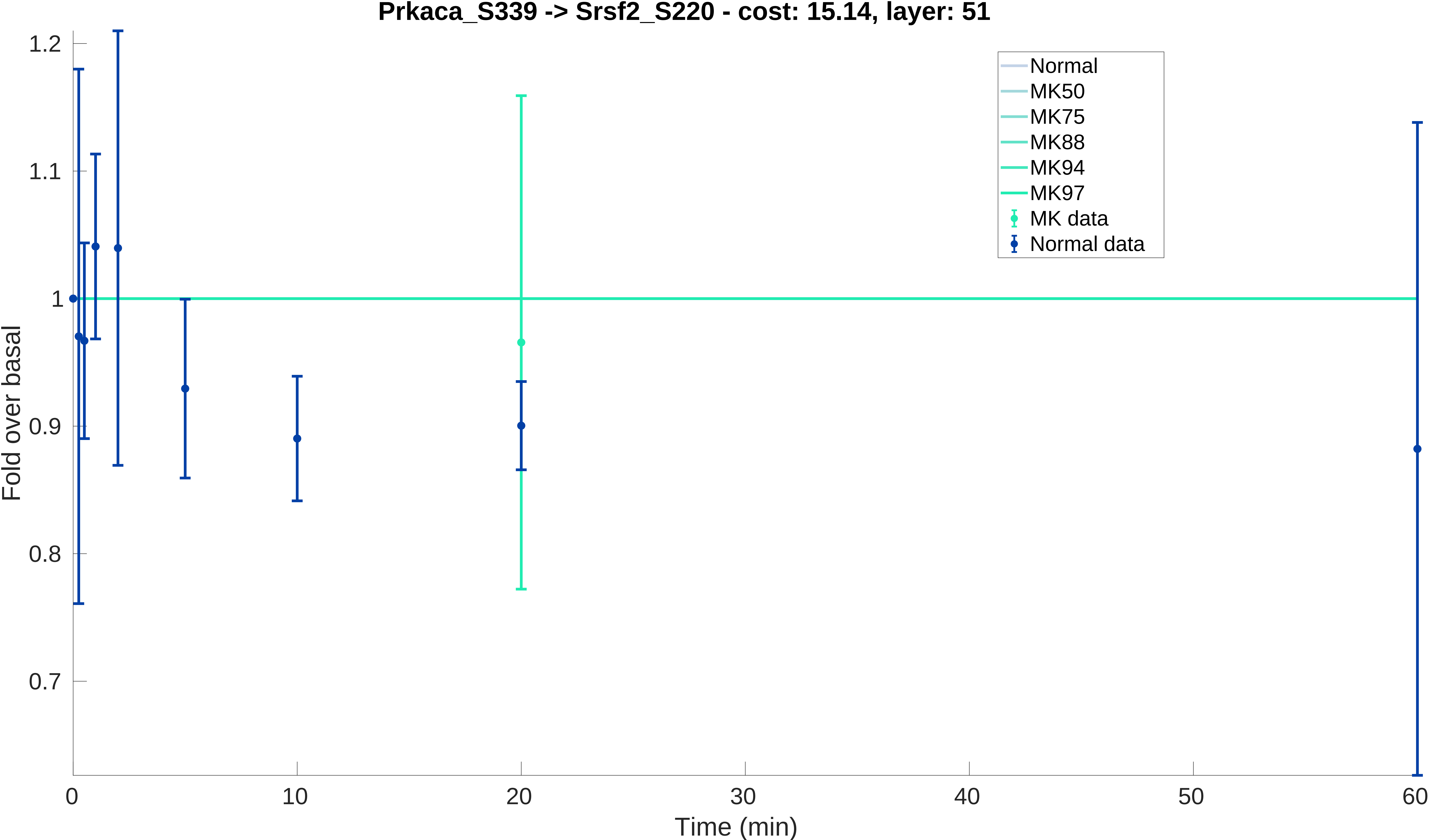
Prkaca_S339 -> Srsf2_S220 - cost: 15.14, layer: 51
Normal
MK50
MK75
MK88
MK94
MK97
MK data
Normal data
Fold over basal
1.2
1.1
1
0.9
0.8
0.7
0
10
20
30
40
50
60
Time (min)

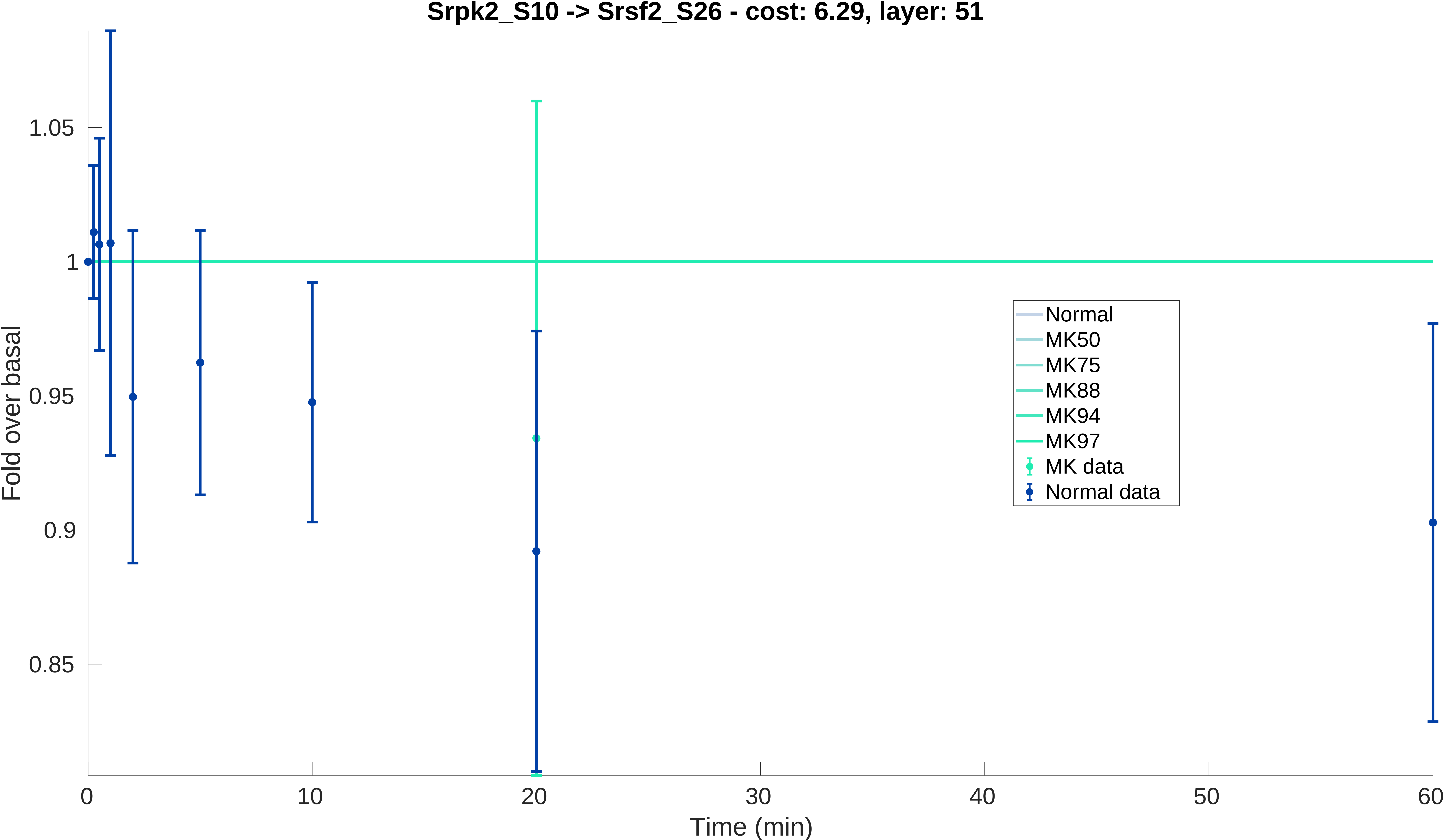

Srpk2_S10 -> Srsf2_S26 - cost: 6.29, layer: 51
Fold over basal
1.05
1
0.95
0.9
0.85
0
10
20
30
40
50
60
Time (min)
Normal
MK50
MK75
MK88
MK94
MK97
MK data
Normal data

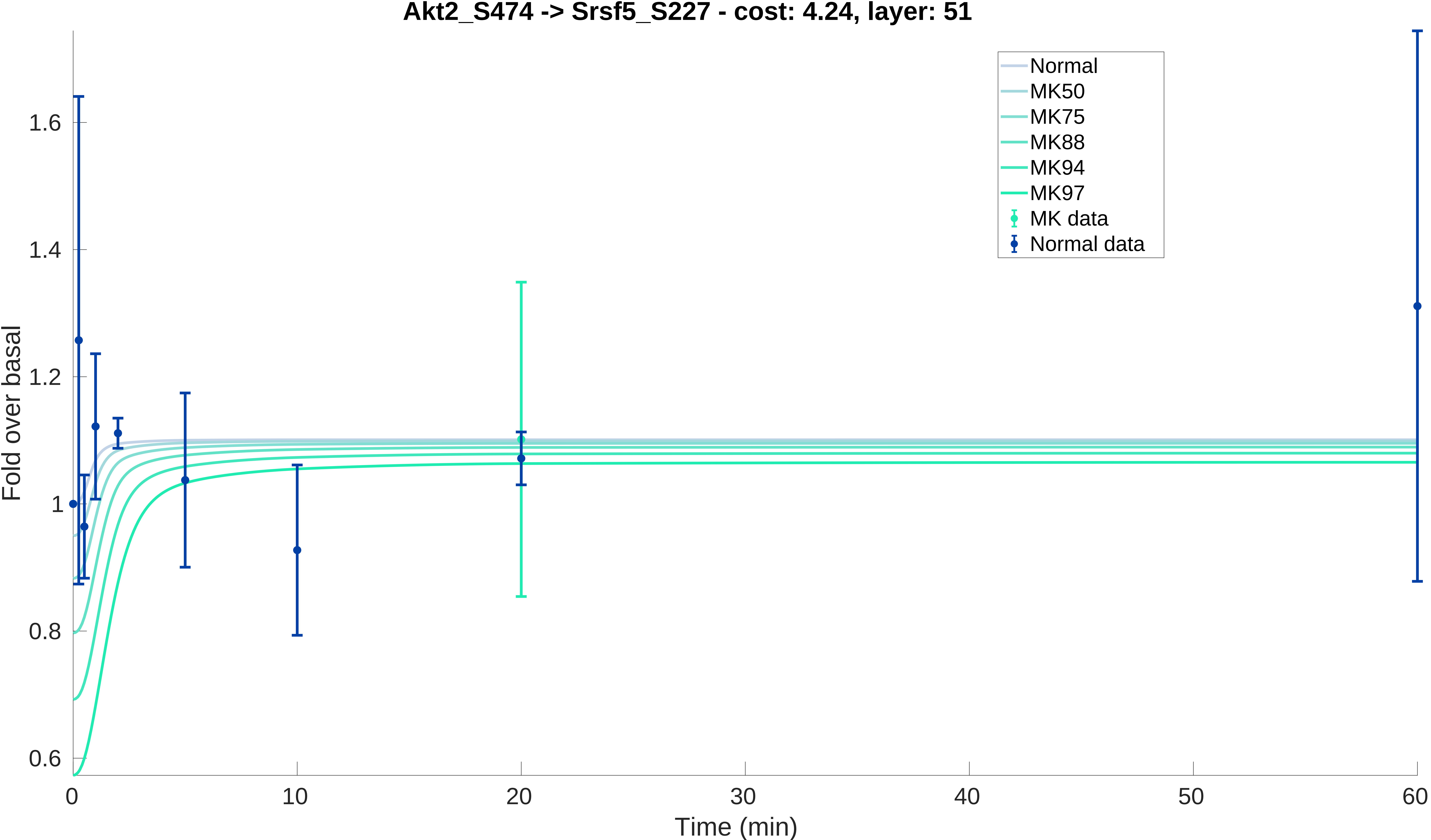

Akt2_S474 -> Srsf5_S227 - cost: 4.24, layer: 51
Normal
MK50
MK75
MK88
MK94
MK97
MK data
Normal data
Fold over basal
1.6
1.4
1.2
1
0.8
0.6
0
10
20
30
40
50
60
Time (min)

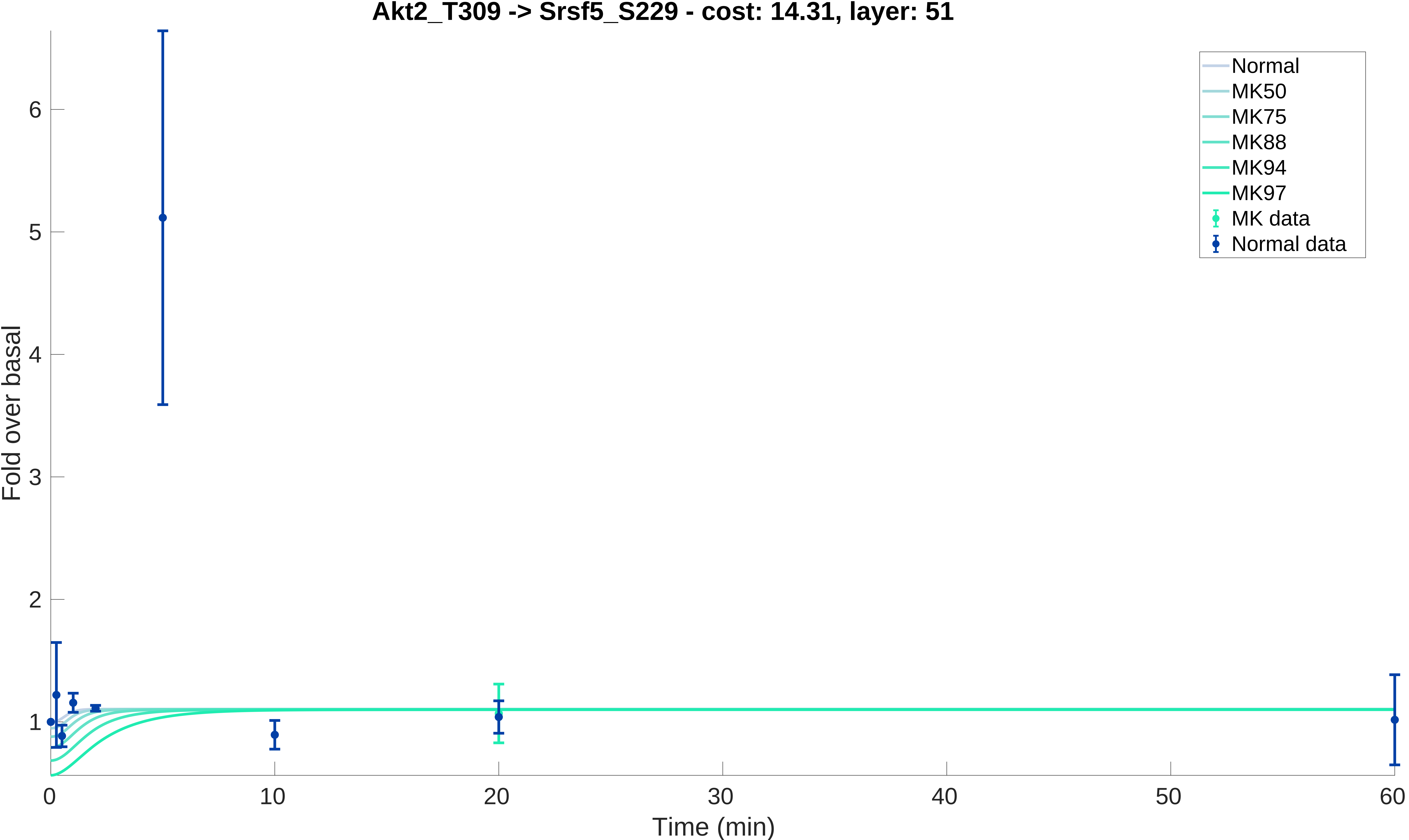

Akt2_T309 -> Srsf5_S229 - cost: 14.31, layer: 51
Fold over basal
Time (min)
0
10
20
30
40
50
60
1
2
3
4
5
6
Normal
MK50
MK75
MK88
MK94
MK97
MK data
Normal data

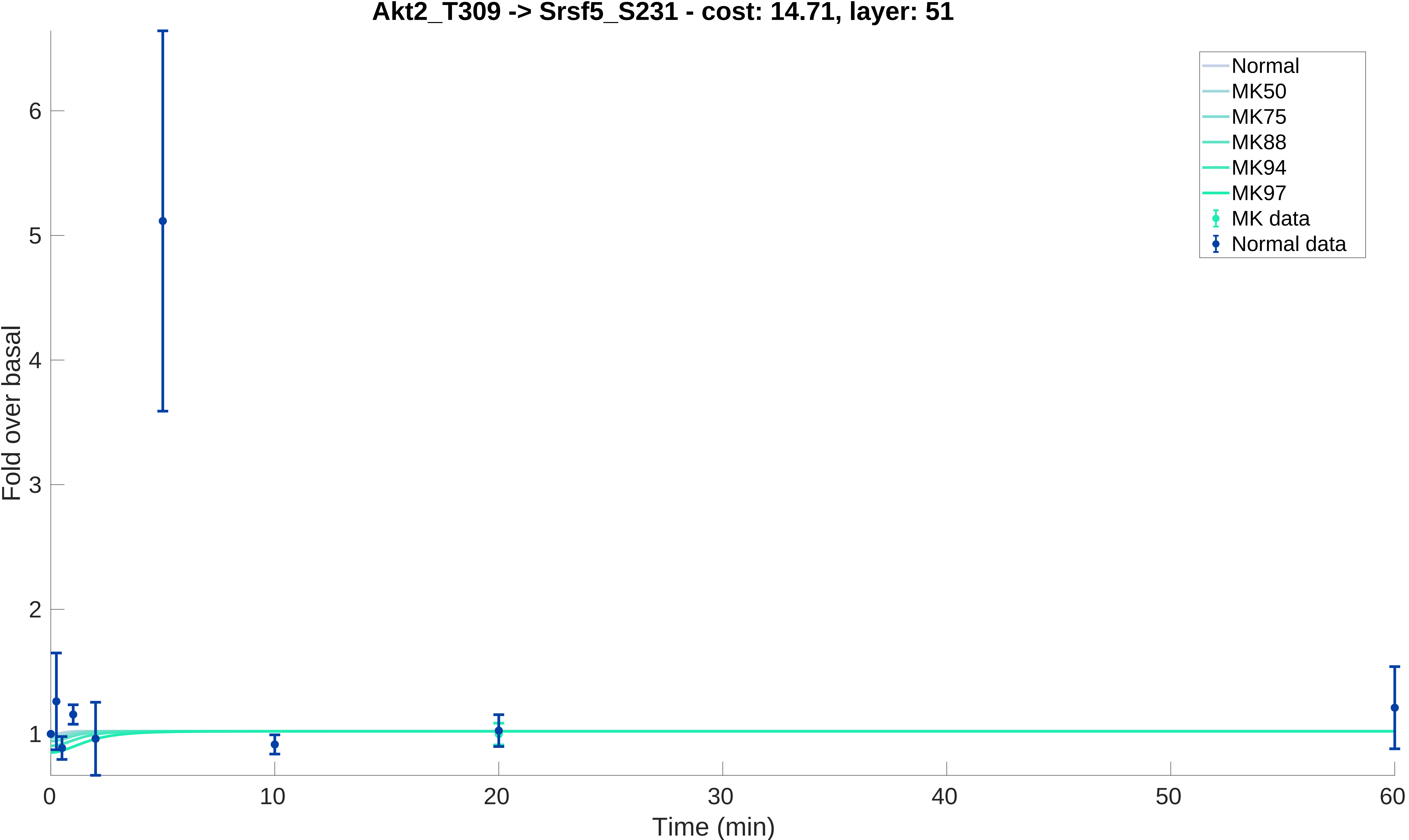

Akt2_T309 -> Srsf5_S231 - cost: 14.71, layer: 51
Fold over basal
Time (min)
0
10
20
30
40
50
60
1
2
3
4
5
6
Normal
MK50
MK75
MK88
MK94
MK97
MK data
Normal data

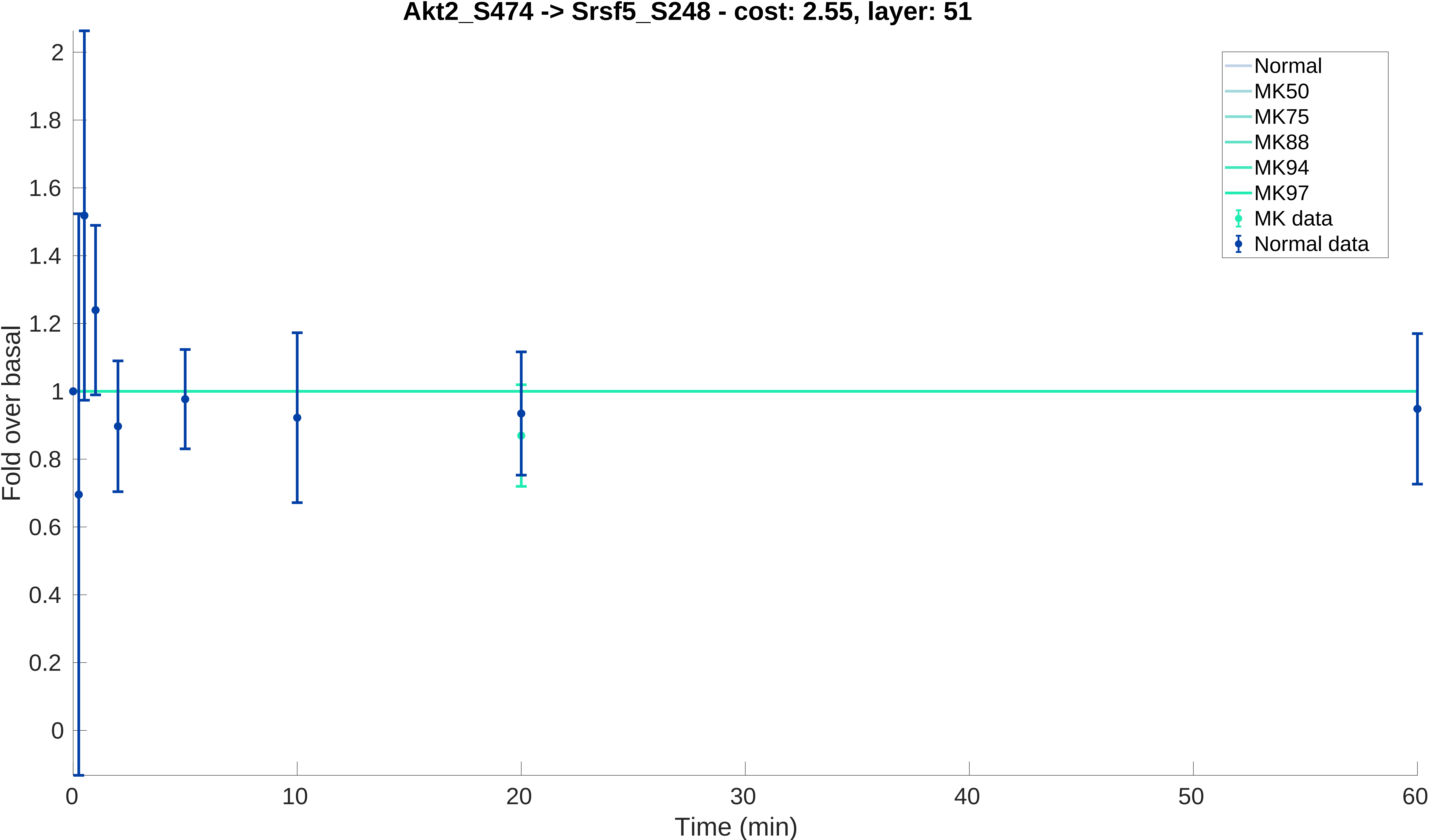

Akt2_S474 -> Srsf5_S248 - cost: 2.55, layer: 51
Normal
MK50
MK75
MK88
MK94
MK97
MK data
Normal data
Fold over basal
Time (min)
2
1.8
1.6
1.4
1.2
1
0.8
0.6
0.4
0.2
0
0
10
20
30
40
50
60

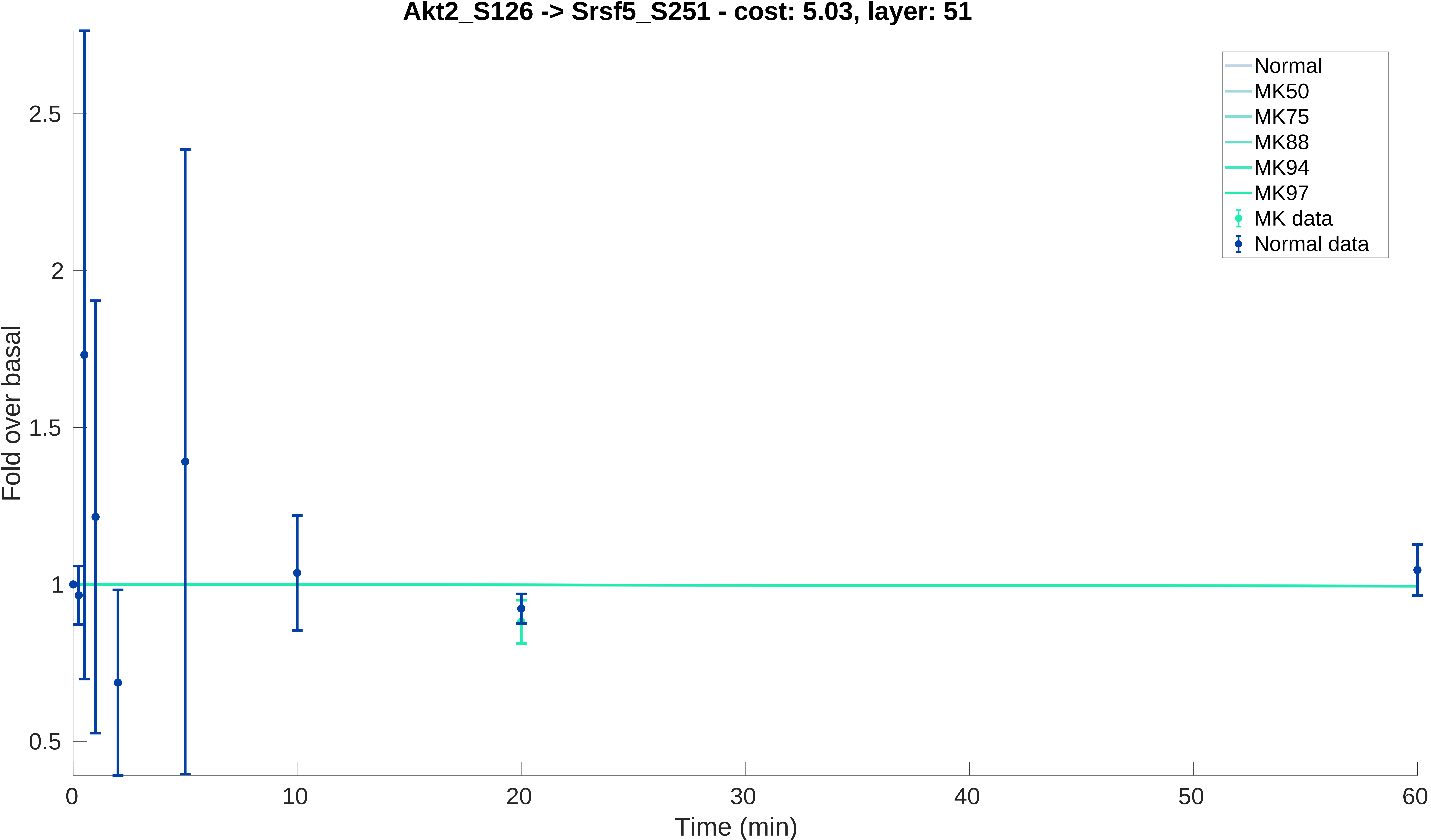

Akt2_S126 -> Srsf5_S251 - cost: 5.03, layer: 51
Fold over basal
Time (min)
0.5
1
1.5
2
2.5
0
10
20
30
40
50
60
Normal
MK50
MK75
MK88
MK94
MK97
MK data
Normal data

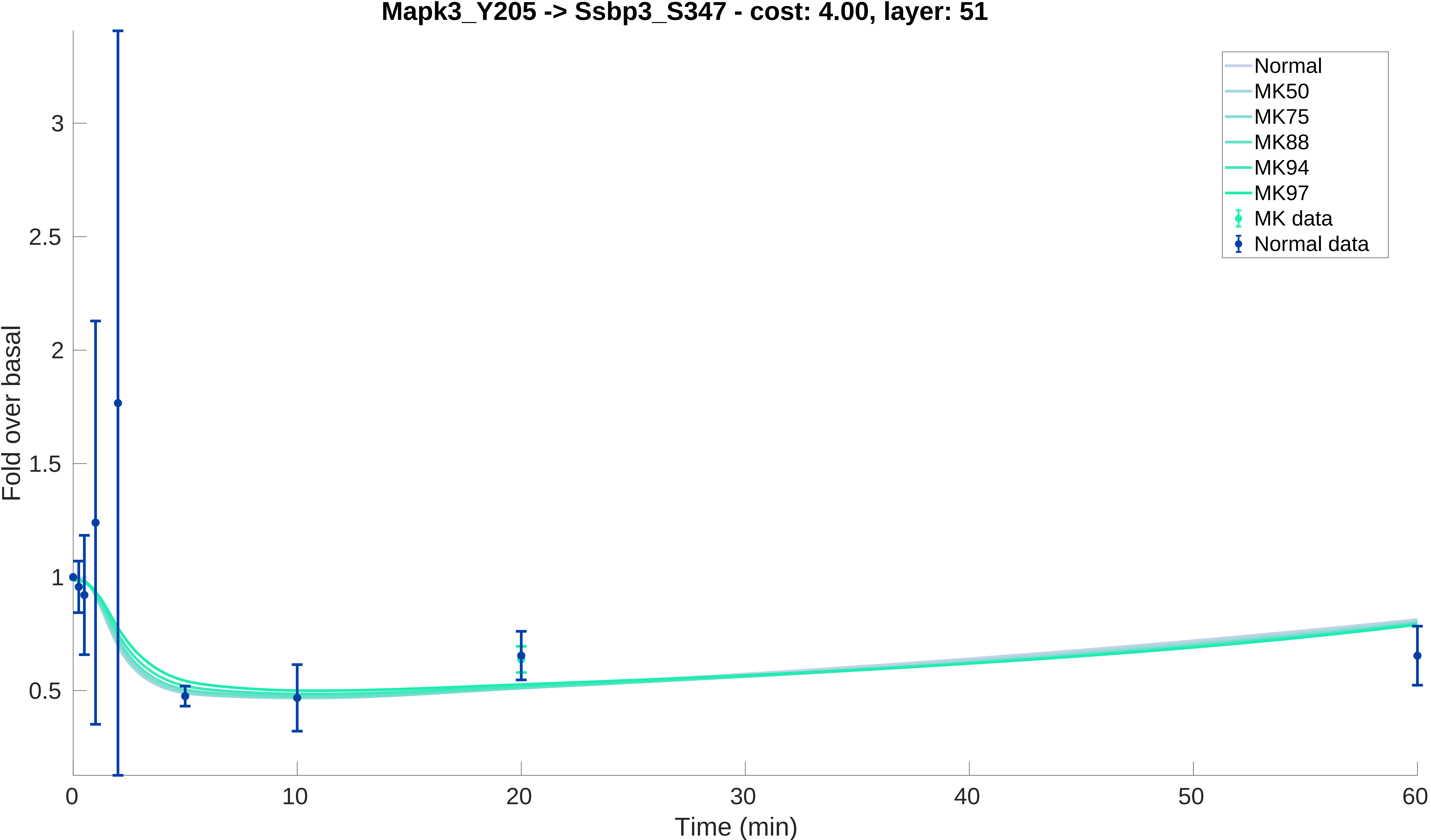

Mapk3_Y205 -> Ssbp3_S347 - cost: 4.00, layer: 51
Normal
MK50
MK75
MK88
MK94
MK97
MK data
Normal data
Fold over basal
3
2.5
2
1.5
1
0.5
0
10
20
30
40
50
60
Time (min)

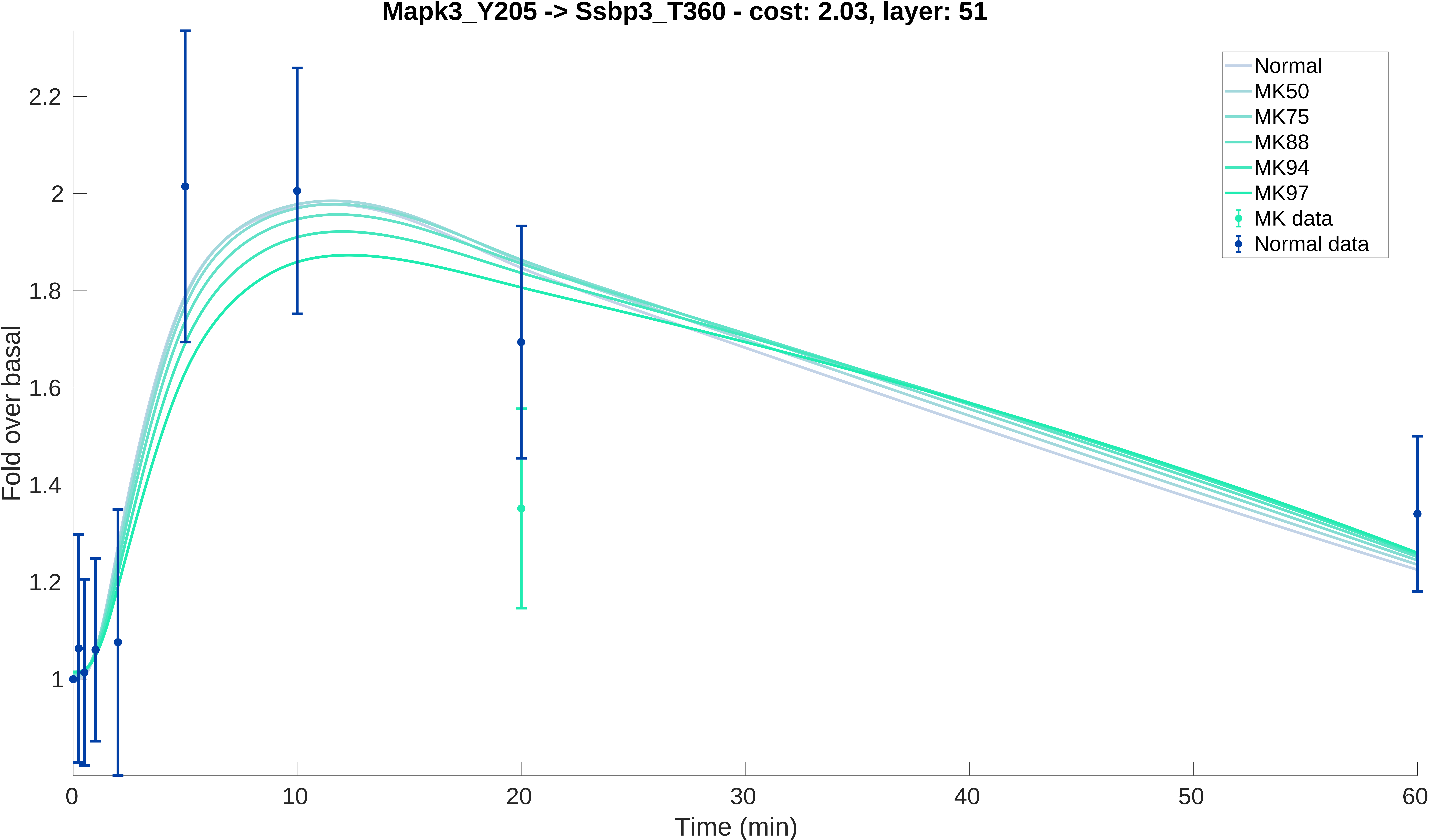

Mapk3_Y205 -> Ssbp3_T360 - cost: 2.03, layer: 51
Normal
MK50
MK75
MK88
MK94
MK97
MK data
Normal data
Fold over basal
2.2
2
1.8
1.6
1.4
1.2
1
0
10
20
30
40
50
60
Time (min)

#### Akt1_T308 -> Ssb_S92 - cost: 4.99, layer: 51

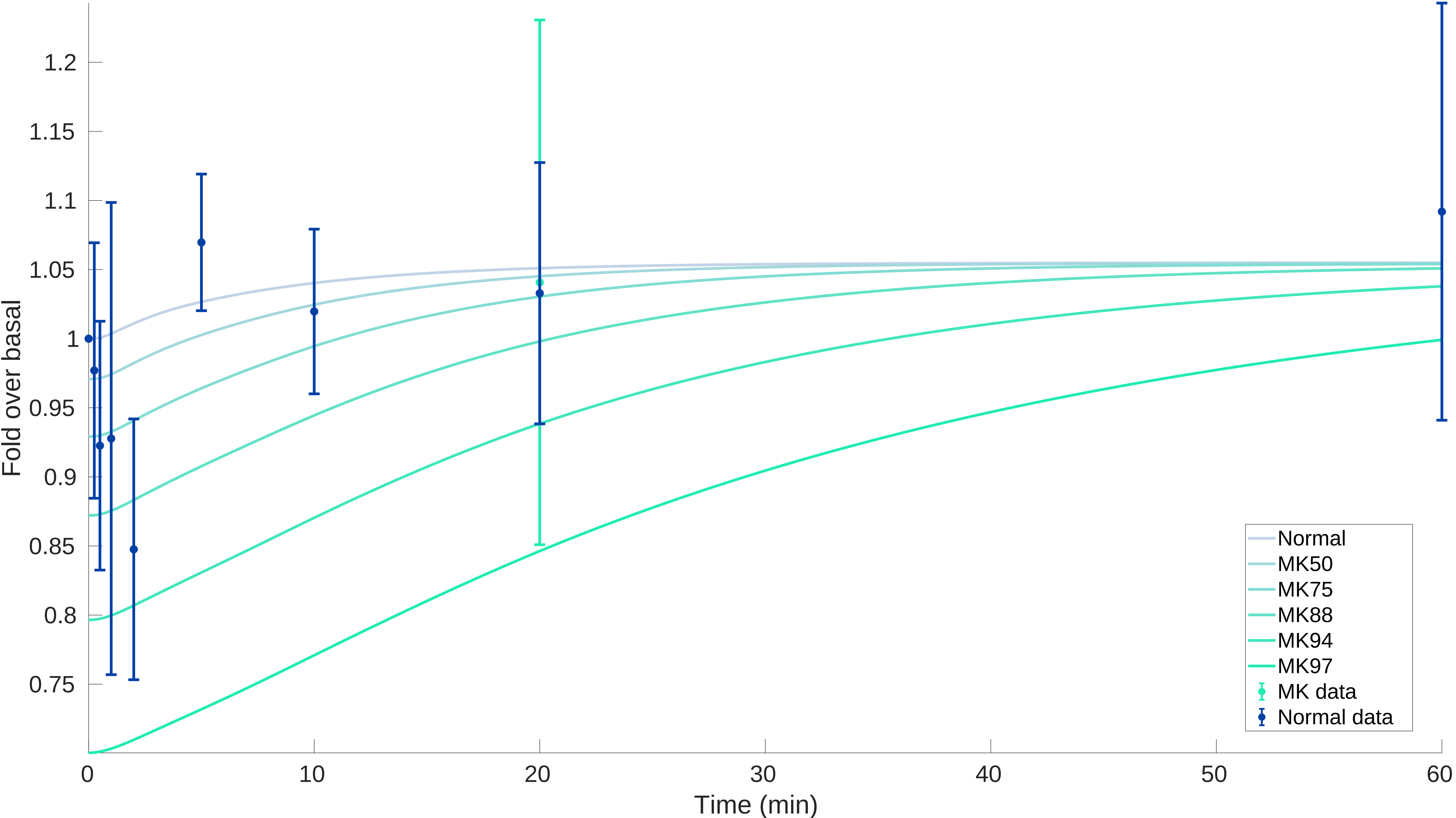

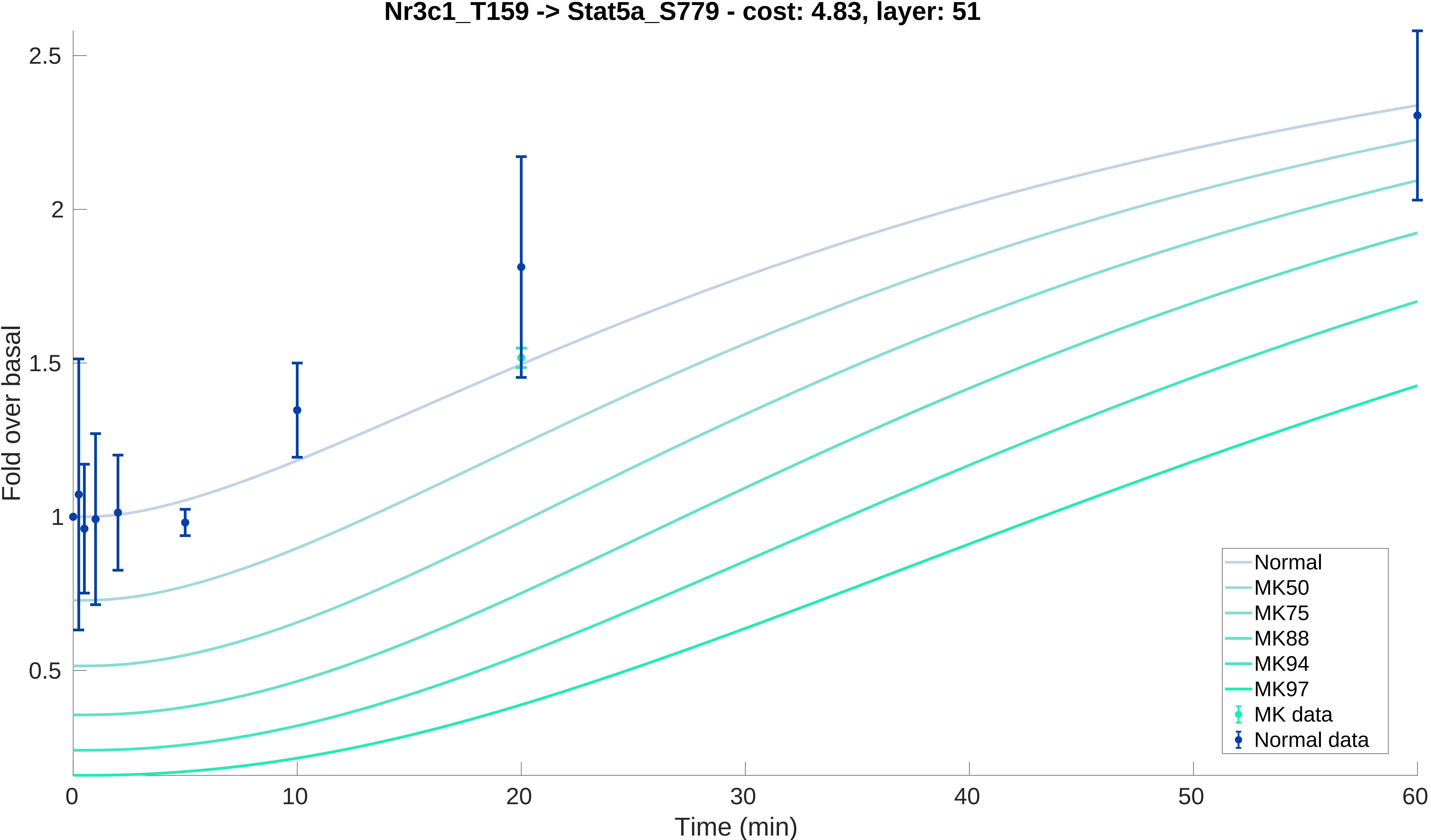

Nr3c1_T159 -> Stat5a_S779 - cost: 4.83, layer: 51
Fold over basal
2.5
2
1.5
1
0.5
0
10
20
30
40
50
60
Time (min)
Normal
MK50
MK75
MK88
MK94
MK97
MK data
Normal data

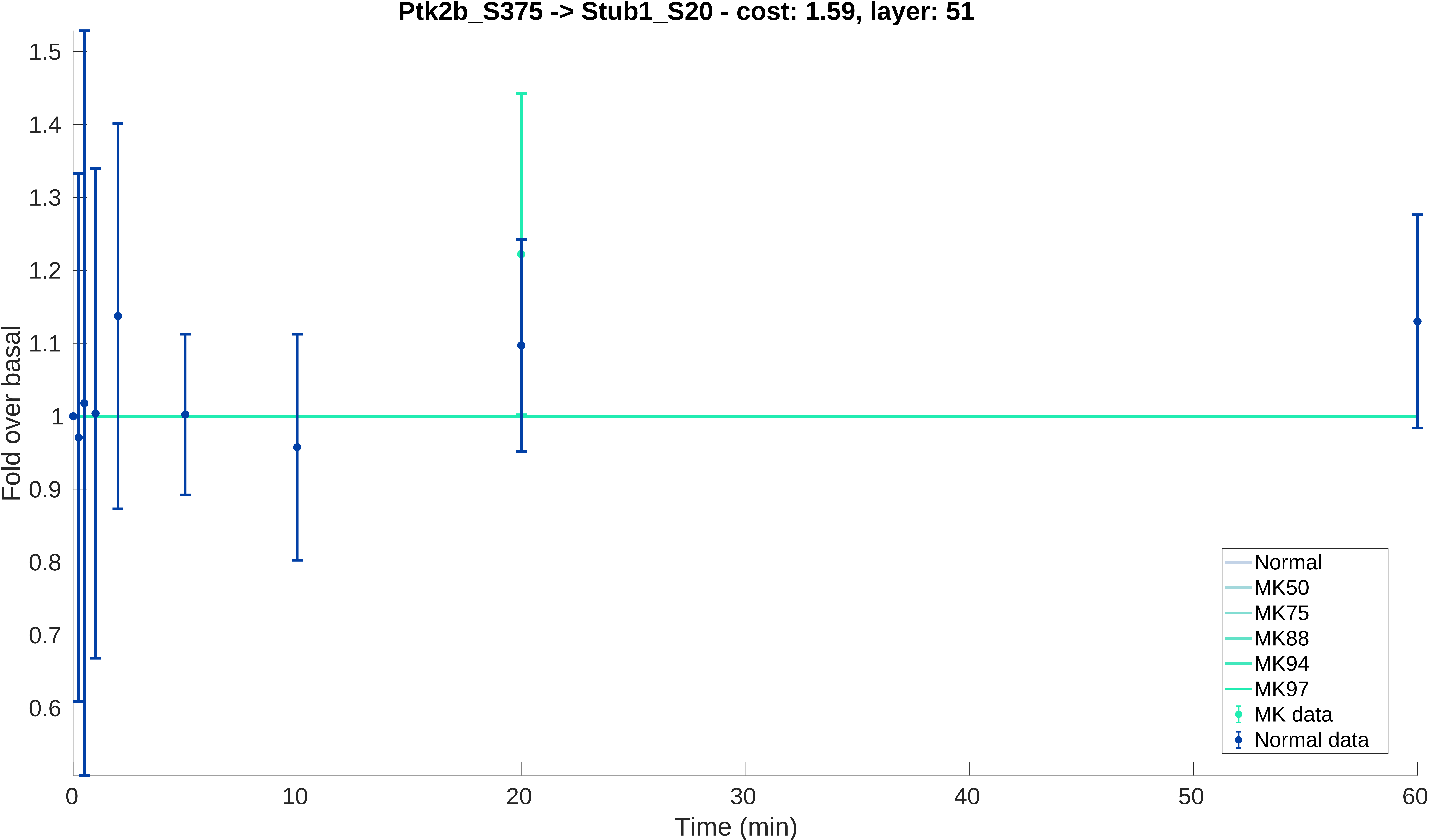

Ptk2b_S375 -> Stub1_S20 - cost: 1.59, layer: 51
Fold over basal
1.5
1.4
1.3
1.2
1.1
1
0.9
0.8
0.7
0.6
0
10
20
30
40
50
60
Time (min)
Normal
MK50
MK75
MK88
MK94
MK97
MK data
Normal data

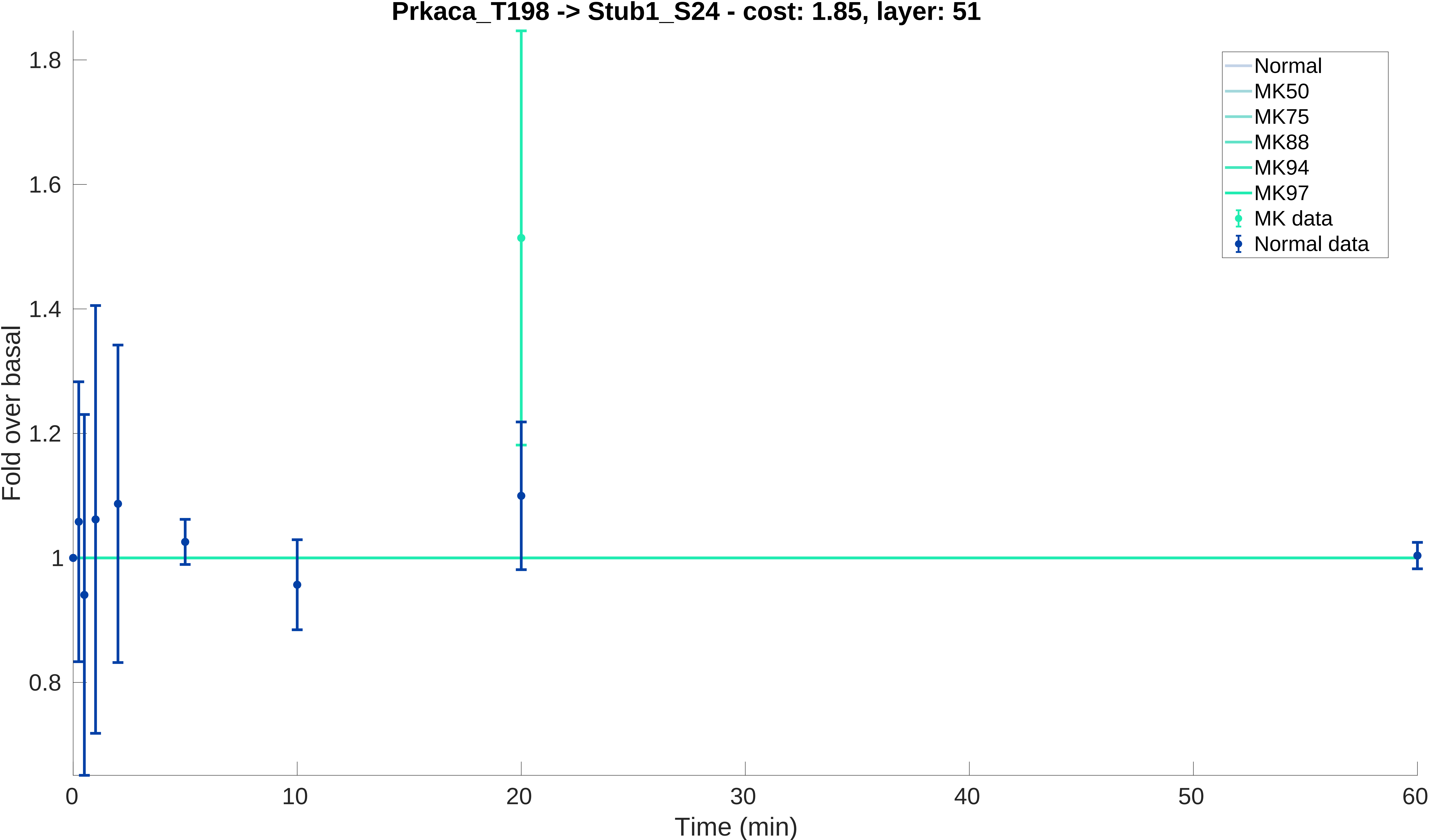
Prrkaca_T198 -> Stub1_S24 - cost: 1.85, layer: 51
Fold over basal
Time (min)
1.8
1.6
1.4
1.2
1
0.8
0
10
20
30
40
50
60
Normal
MK50
MK75
MK88
MK94
MK97
MK data
Normal data

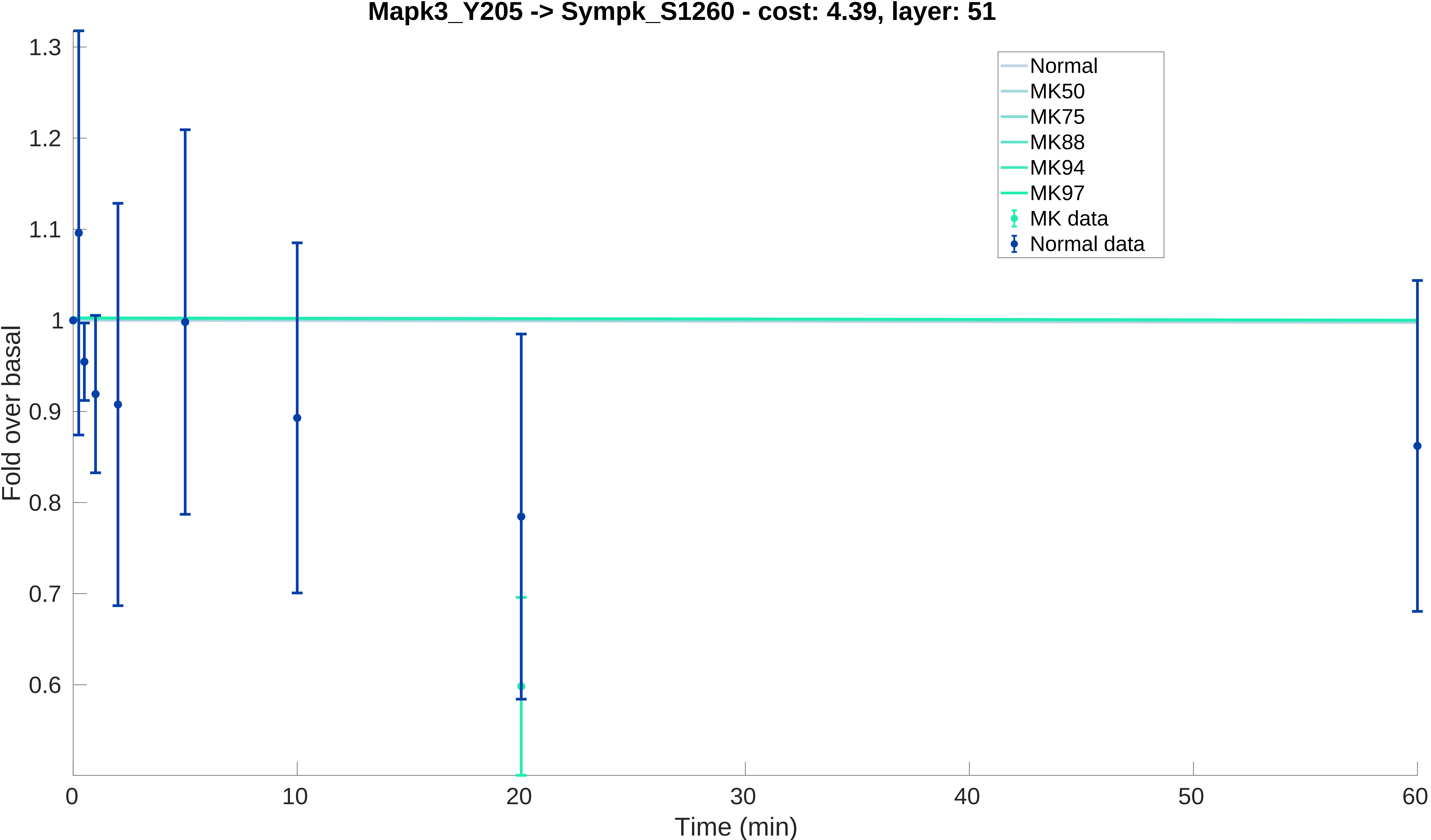

Mapk3_Y205 -> Sympk_S1260 - cost: 4.39, layer: 51
Fold over basal
Time (min)
Normal
MK50
MK75
MK88
MK94
MK97
MK data
Normal data
1.3
1.2
1.1
1
0.9
0.8
0.7
0.6
0
10
20
30
40
50
60

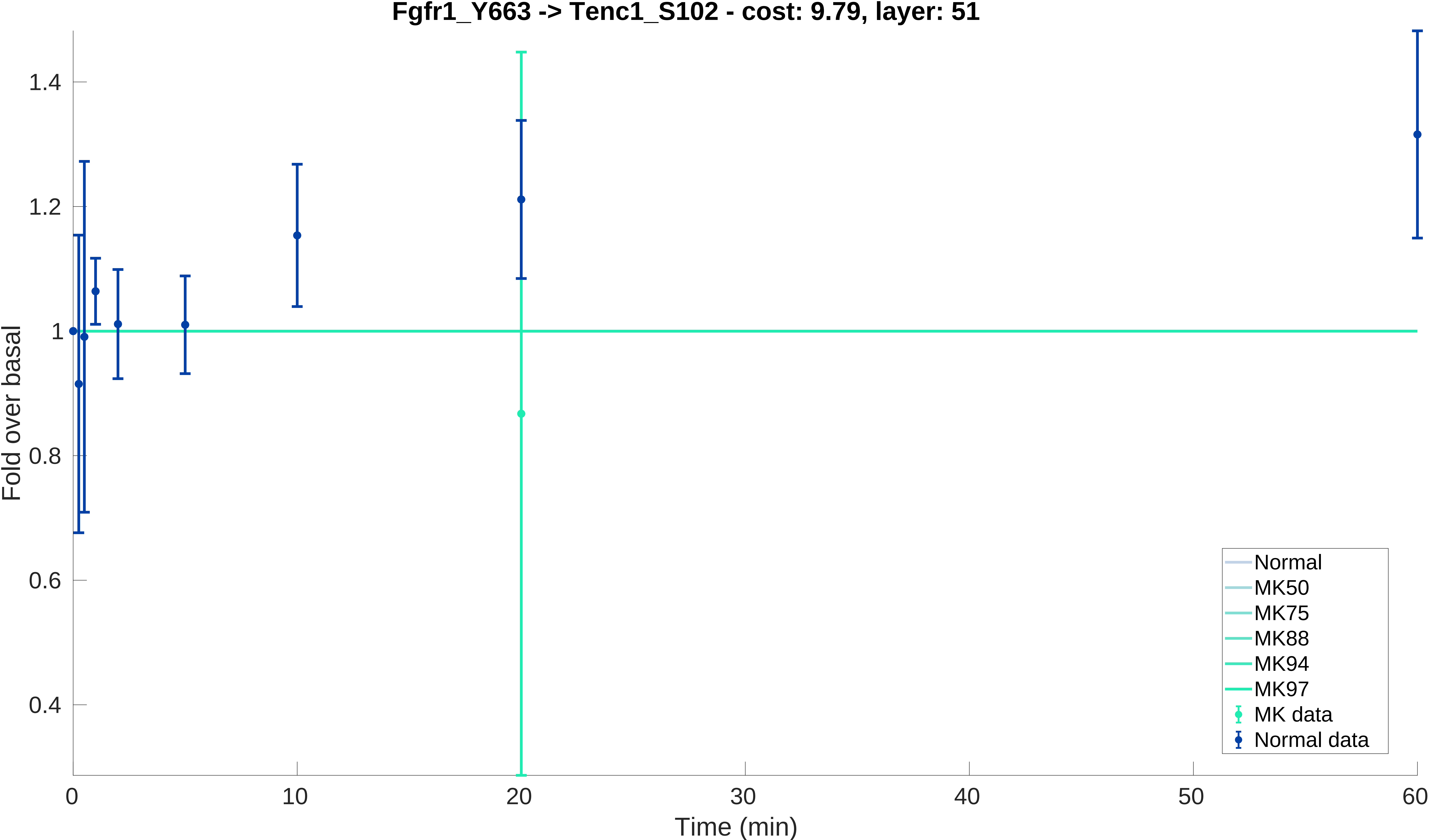

Fgfr1_Y663 -> Tenc1_S102 - cost: 9.79, layer: 51
Fold over basal
1.4
1.2
1
0.8
0.6
0.4
0
10
20
30
40
50
60
Time (min)
Normal
MK50
MK75
MK88
MK94
MK97
MK data
Normal data

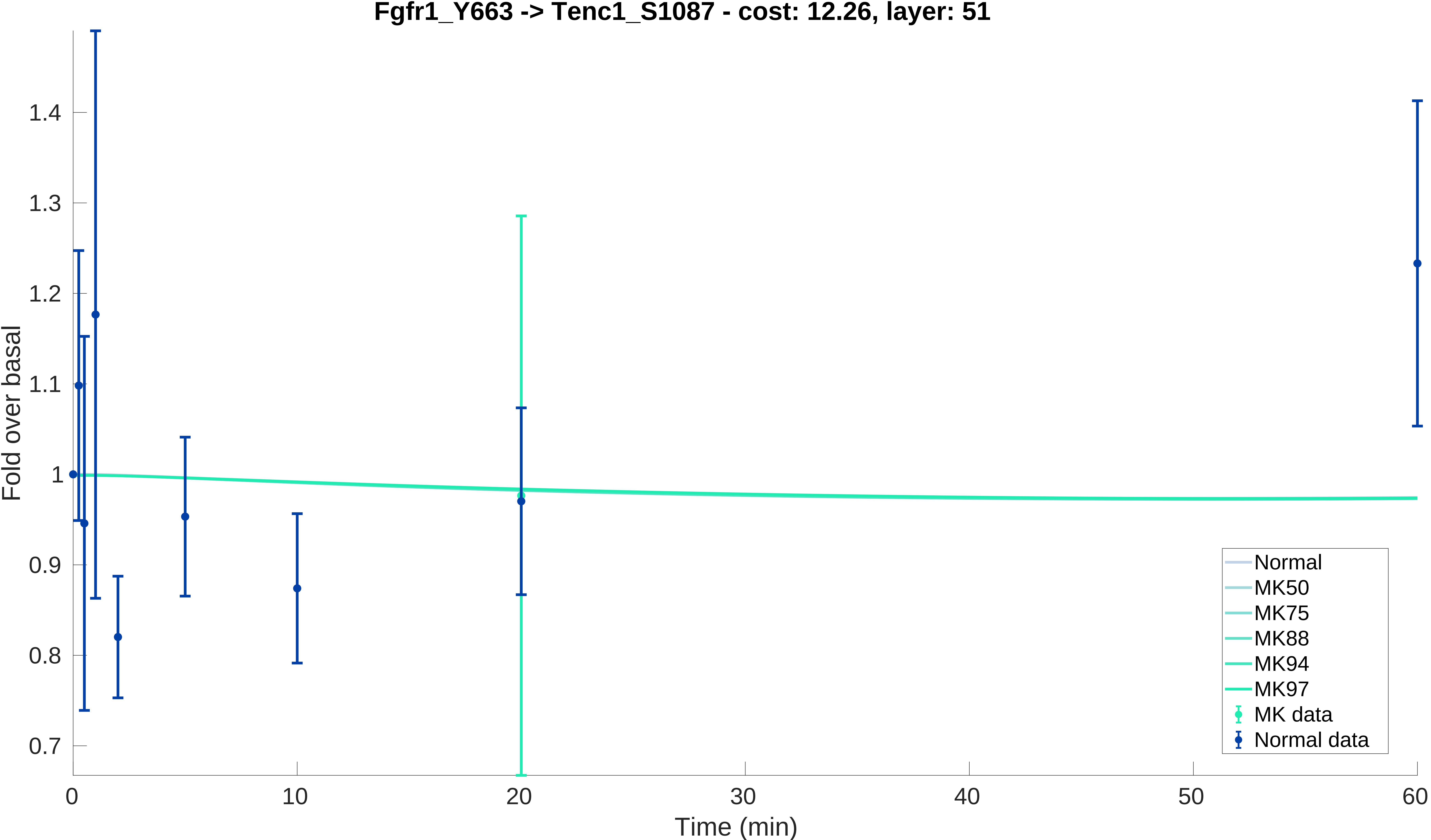

Fgfr1_Y663 -> Tenc1_S1087 - cost: 12.26, layer: 51
Fold over basal
1.4
1.3
1.2
1.1
1
0.9
0.8
0.7
0
10
20
30
40
50
60
Time (min)
Normal
MK50
MK75
MK88
MK94
MK97
MK data
Normal data

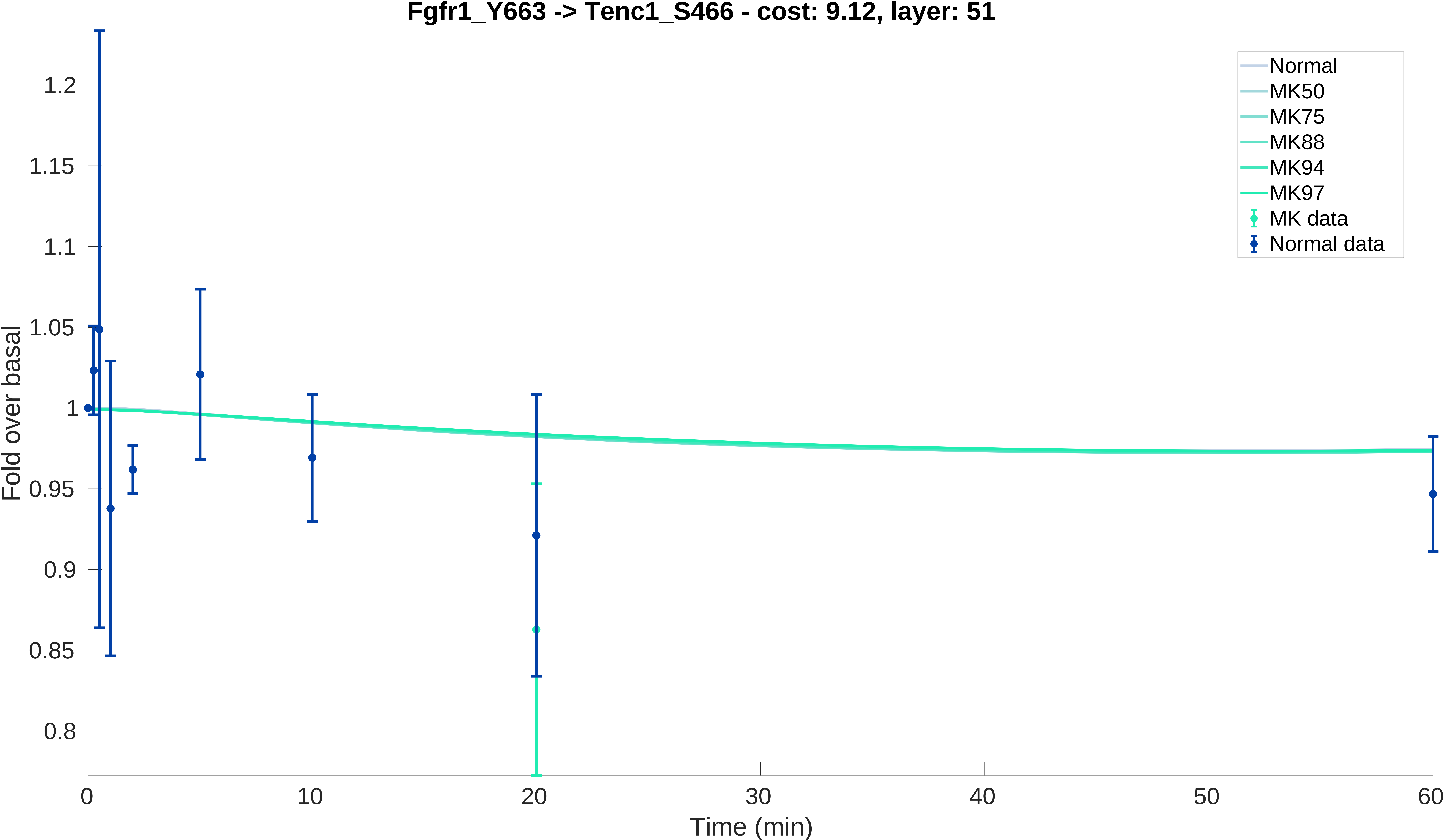
Fgfr1_Y663 -> Tenc1_S466 - cost: 9.12, layer: 51
Normal
MK50
MK75
MK88
MK94
MK97
MK data
Normal data
Fold over basal
1.2
1.15
1.1
1.05
1
0.95
0.9
0.85
0.8
0
10
20
30
40
50
60
Time (min)

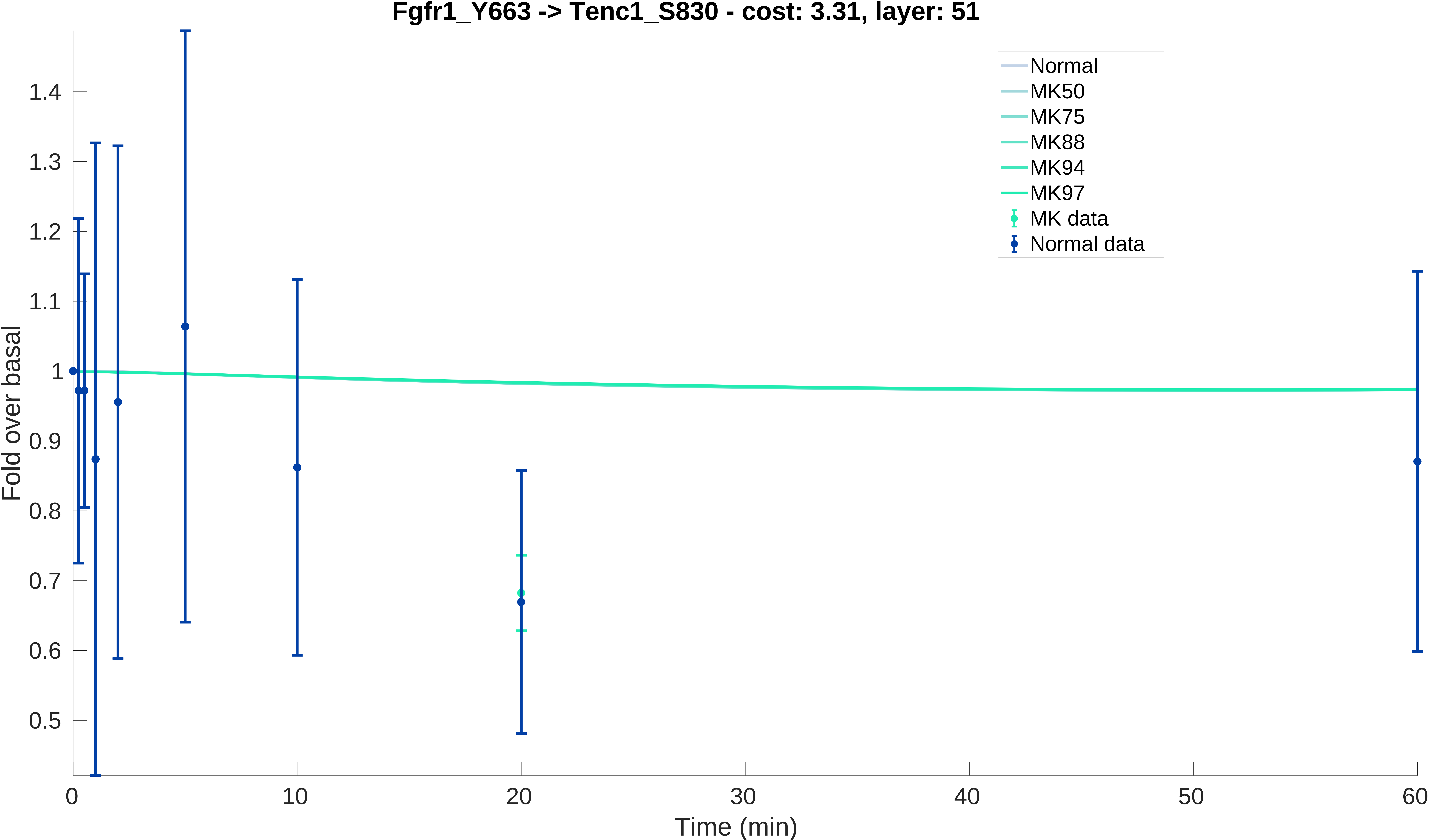

Fgfr1_Y663 -> Tenc1_S830 - cost: 3.31, layer: 51
Fold over basal
Time (min)
1.4
1.3
1.2
1.1
1
0.9
0.8
0.7
0.6
0.5
0
10
20
30
40
50
60
Normal
MK50
MK75
MK88
MK94
MK97
MK data
Normal data

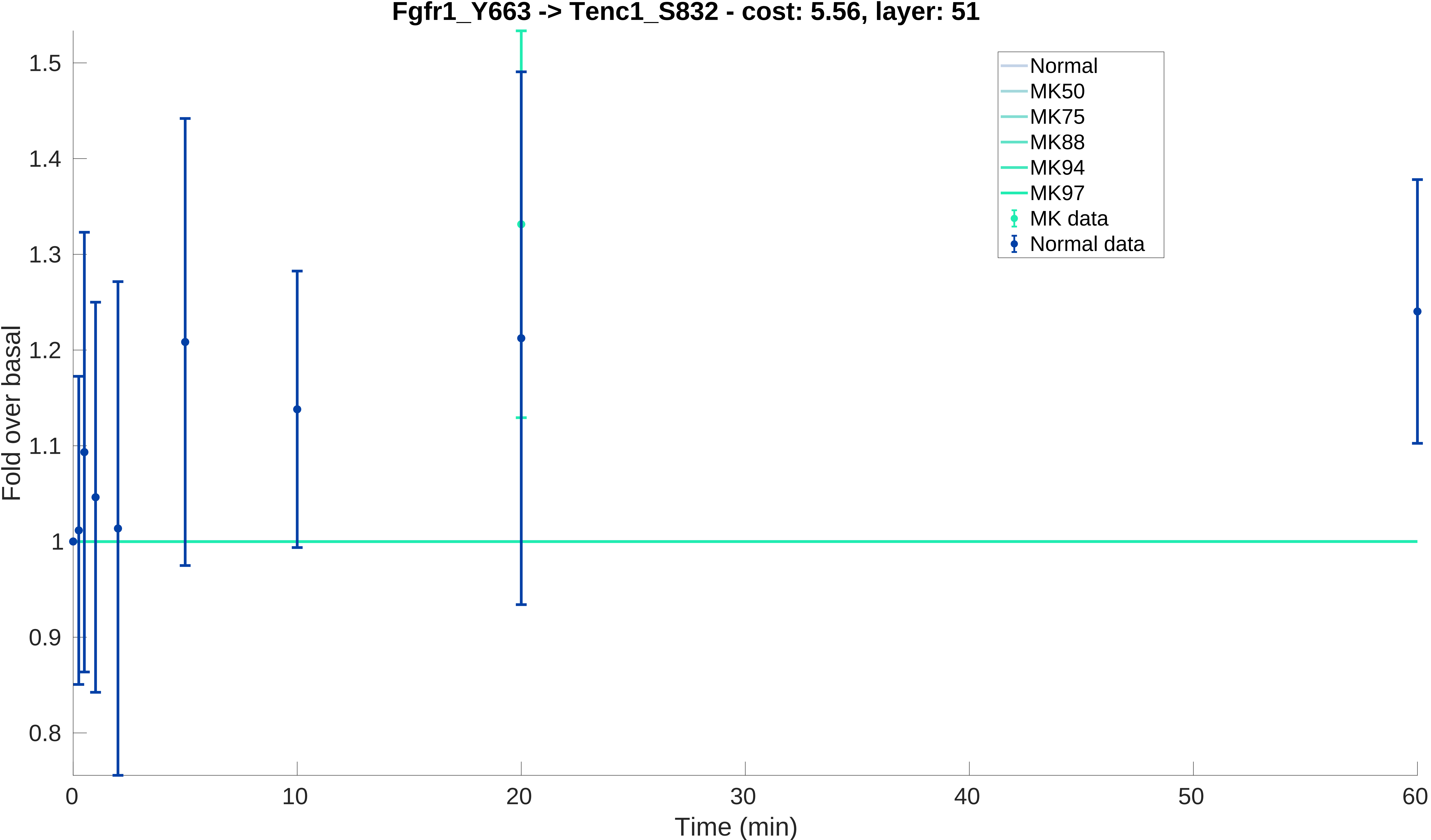

Fgfr1_Y663 -> Tenc1_S832 - cost: 5.56, layer: 51
Normal
MK50
MK75
MK88
MK94
MK97
MK data
Normal data
Fold over basal
1.5
1.4
1.3
1.2
1.1
1
0.9
0.8
0
10
20
30
40
50
60
Time (min)

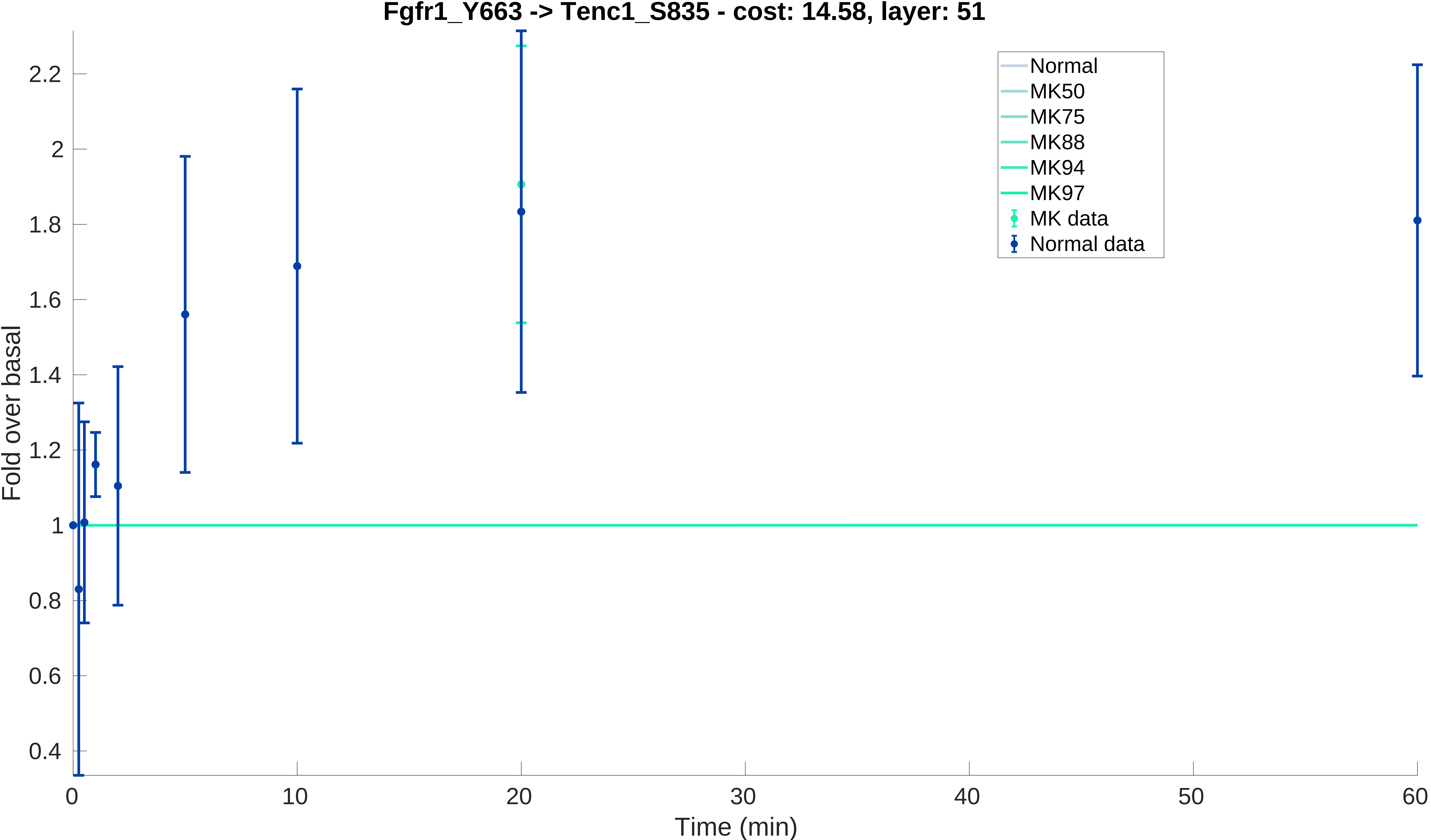

Fgfr1_Y663 -> Tenc1_S835 - cost: 14.58, layer: 51
Normal
MK50
MK75
MK88
MK94
MK97
MK data
Normal data
Fold over basal
0.4
0.6
0.8
1
1.2
1.4
1.6
1.8
2
2.2
0
10
20
30
40
50
60
Time (min)

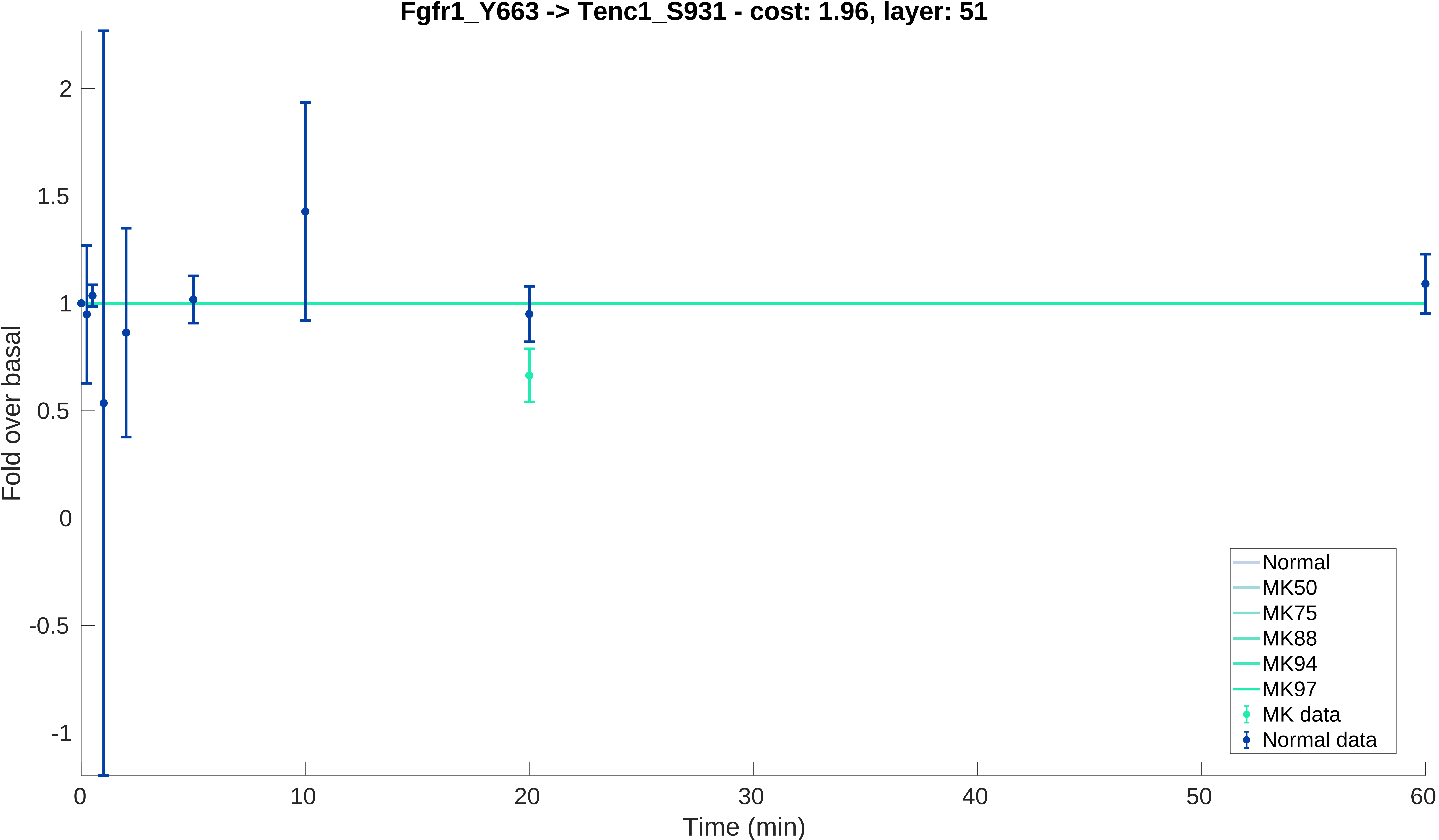

Fgfr1_Y663 -> Tenc1_S931 - cost: 1.96, layer: 51
Fold over basal
Time (min)
2
1.5
1
0.5
0
-0.5
-1
0
10
20
30
40
50
60
Normal
MK50
MK75
MK88
MK94
MK97
MK data
Normal data

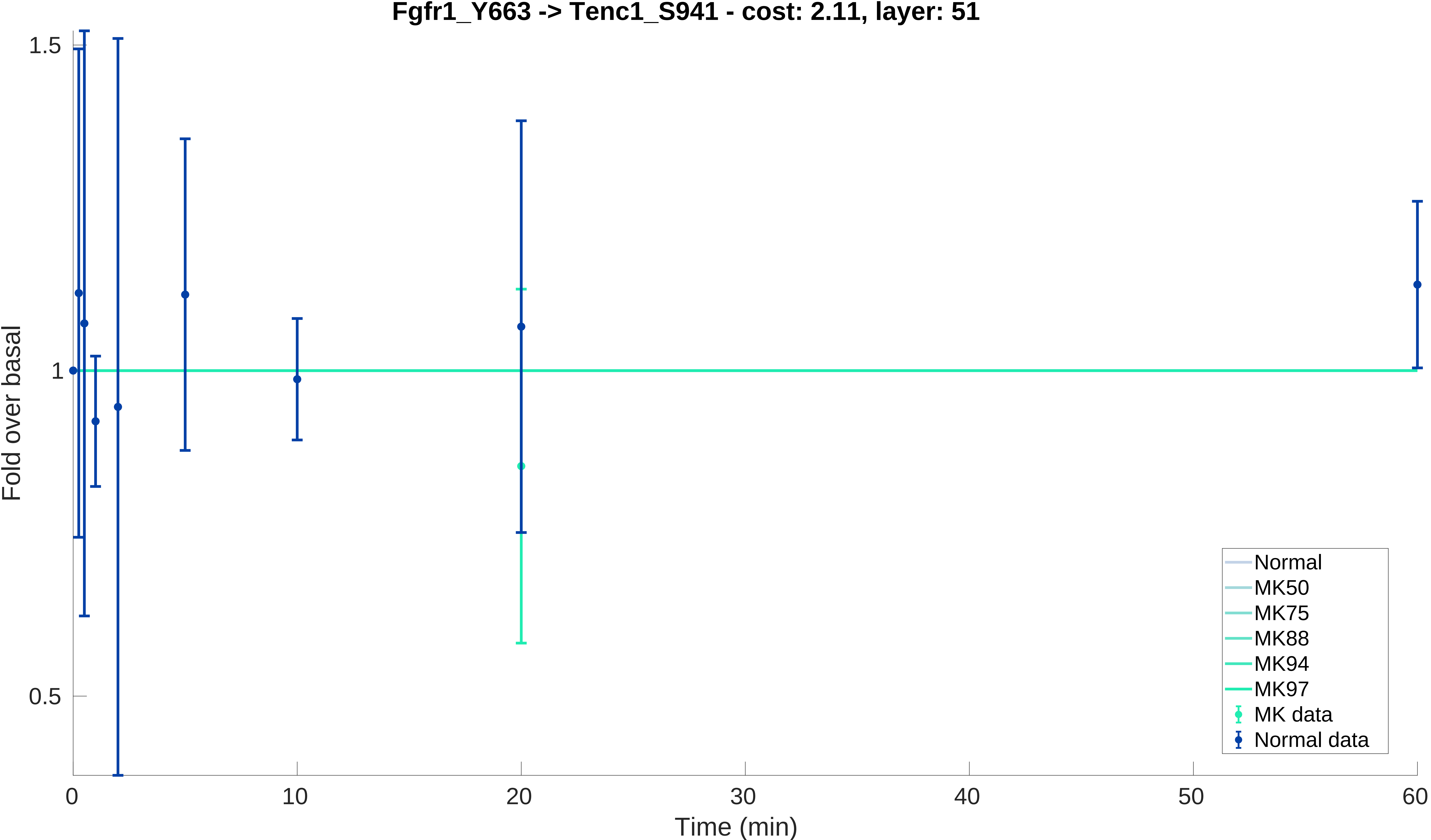

Fgfr1_Y663 -> Tenc1_S941 - cost: 2.11, layer: 51
Fold over basal
1.5
1
0.5
0
10
20
30
40
50
60
Time (min)
Normal
MK50
MK75
MK88
MK94
MK97
MK data
Normal data

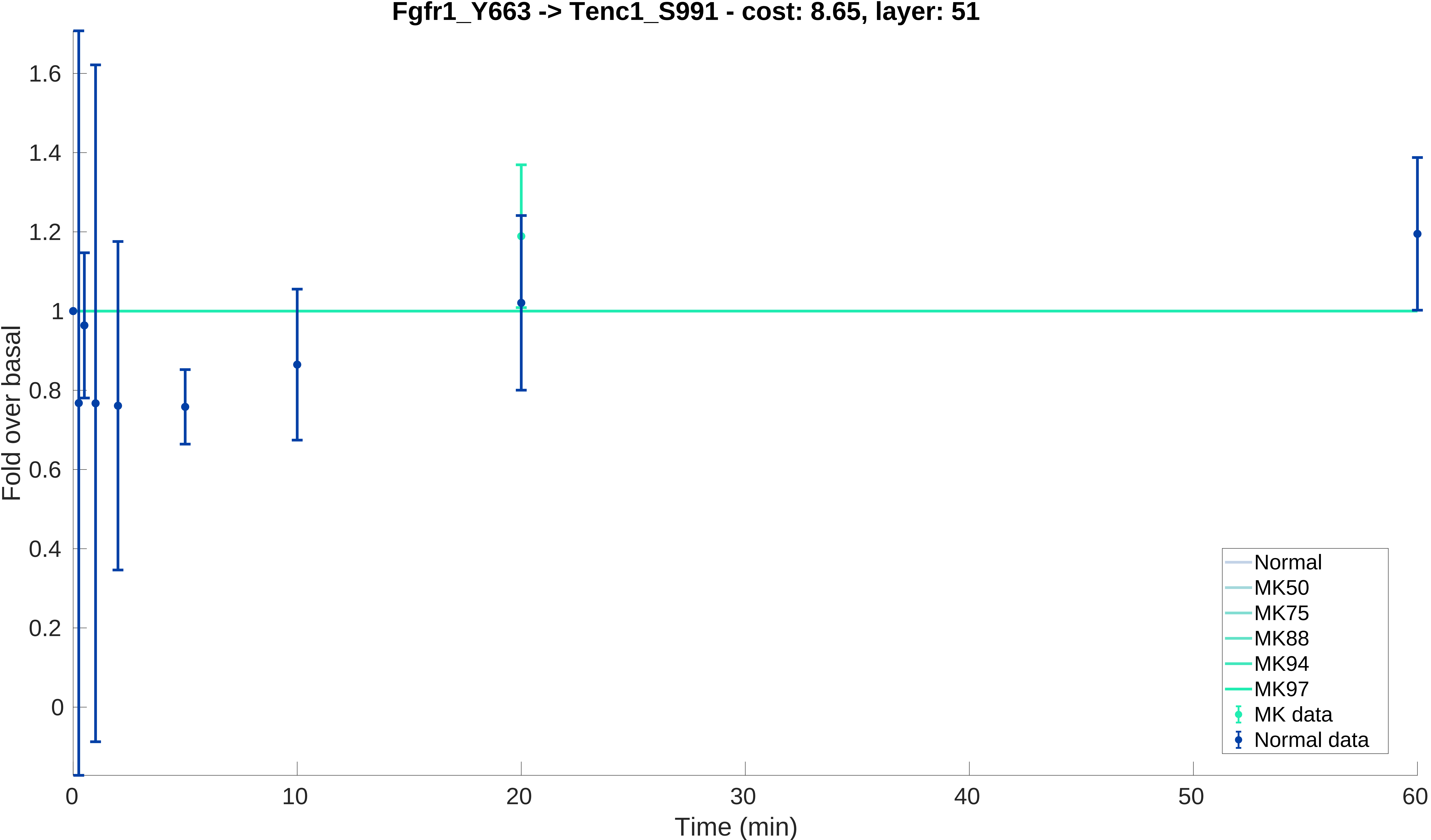

Fgfr1_Y663 -> Tenc1_S991 - cost: 8.65, layer: 51
Fold over basal
1.6
1.4
1.2
1
0.8
0.6
0.4
0.2
0
0
10
20
30
40
50
60
Time (min)
Normal
MK50
MK75
MK88
MK94
MK97
MK data
Normal data

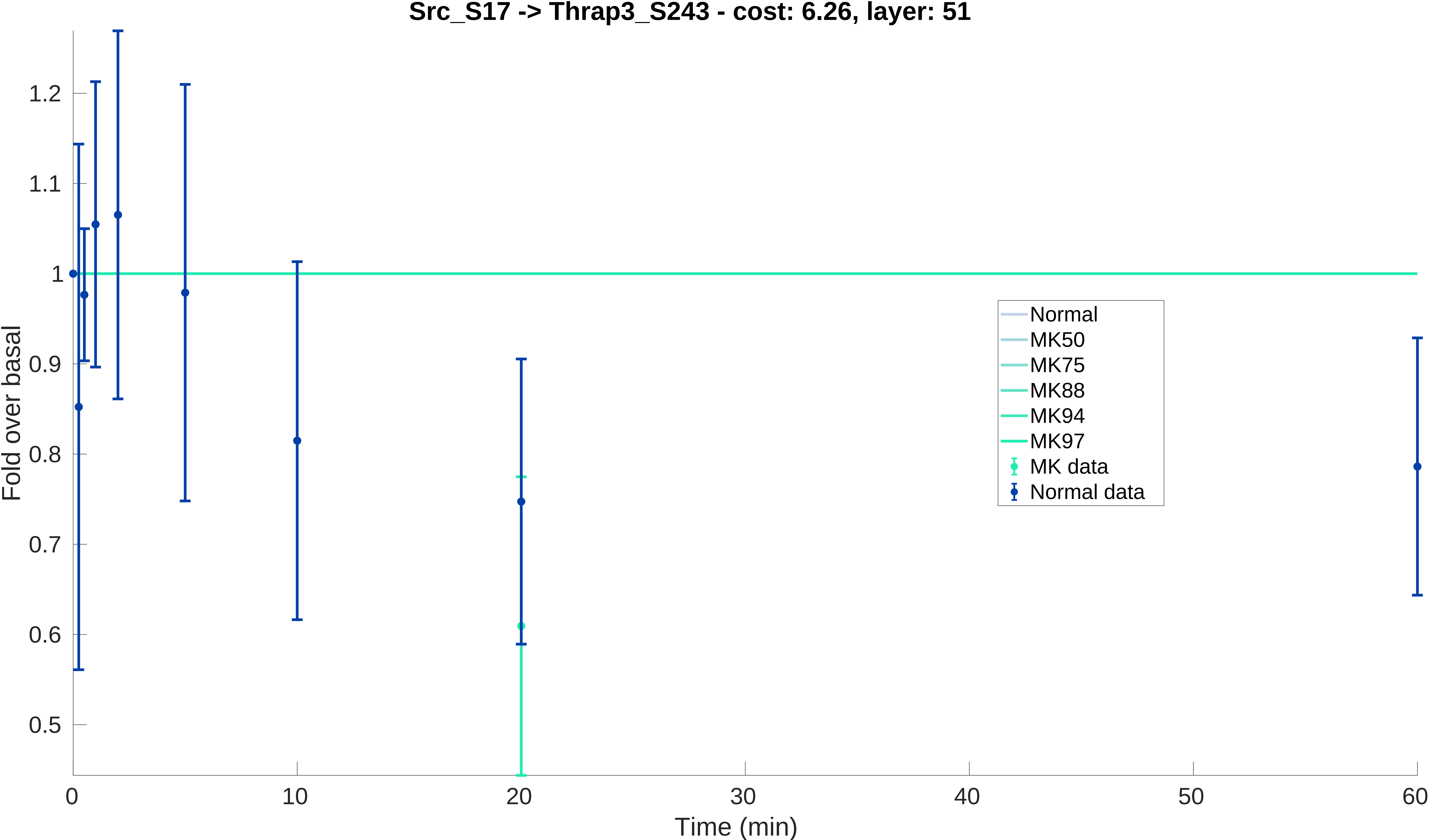

Src_S17 -> Thrap3_S243 - cost: 6.26, layer: 51
Fold over basal
1.2
1.1
1
0.9
0.8
0.7
0.6
0.5
0
10
20
30
40
50
60
Time (min)
Normal
MK50
MK75
MK88
MK94
MK97
MK data
Normal data

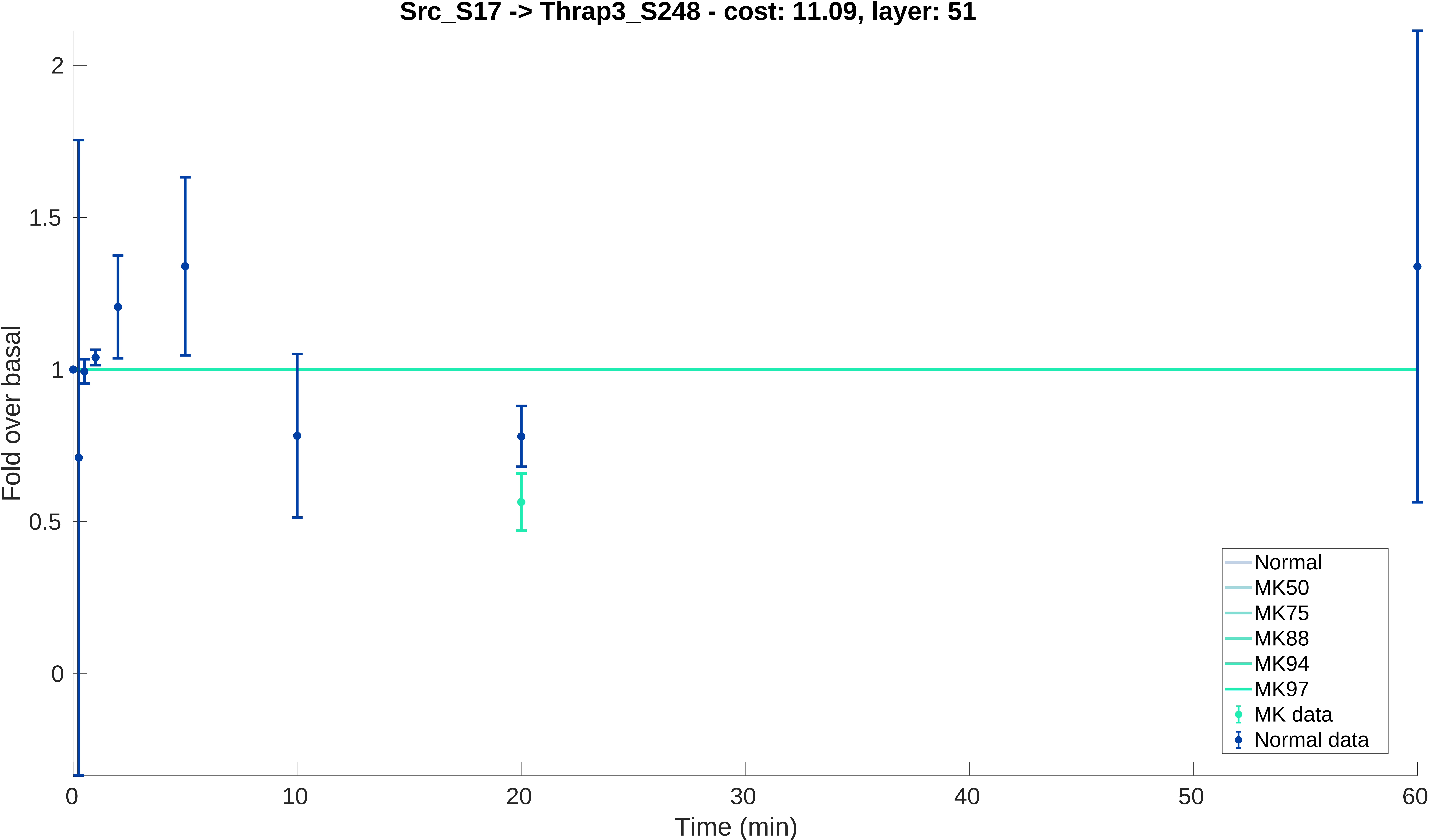
Src_S17 -> Thrap3_S248 - cost: 11.09, layer: 51
Fold over basal
Time (min)
0
0.5
1
1.5
2
0
10
20
30
40
50
60
Normal
MK50
MK75
MK88
MK94
MK97
MK data
Normal data

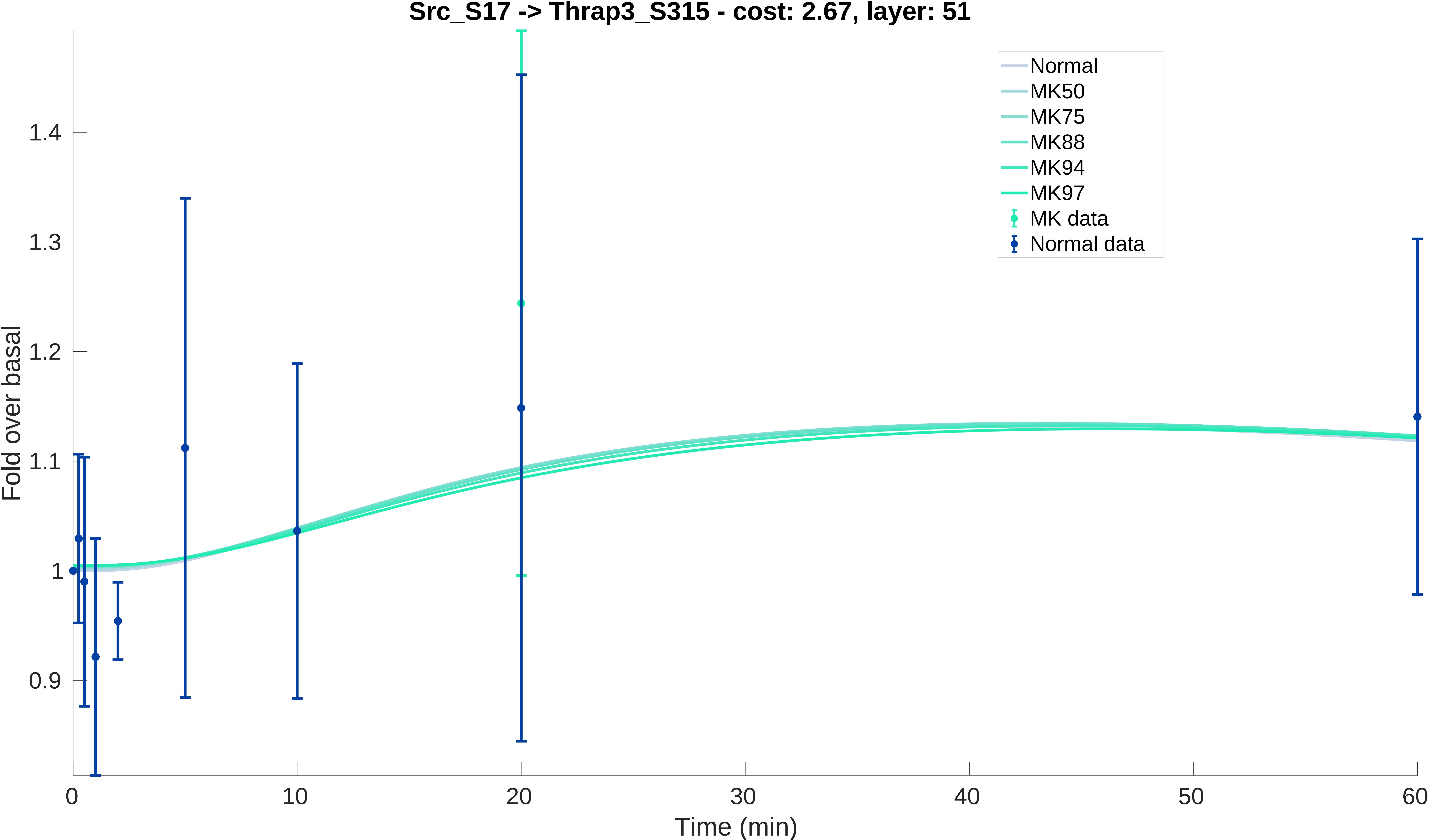
Src_S17 -> Thrap3_S315 - cost: 2.67, layer: 51
Fold over basal
Time (min)
0.9
1
1.1
1.2
1.3
1.4
0
10
20
30
40
50
60
Normal
MK50
MK75
MK88
MK94
MK97
MK data
Normal data

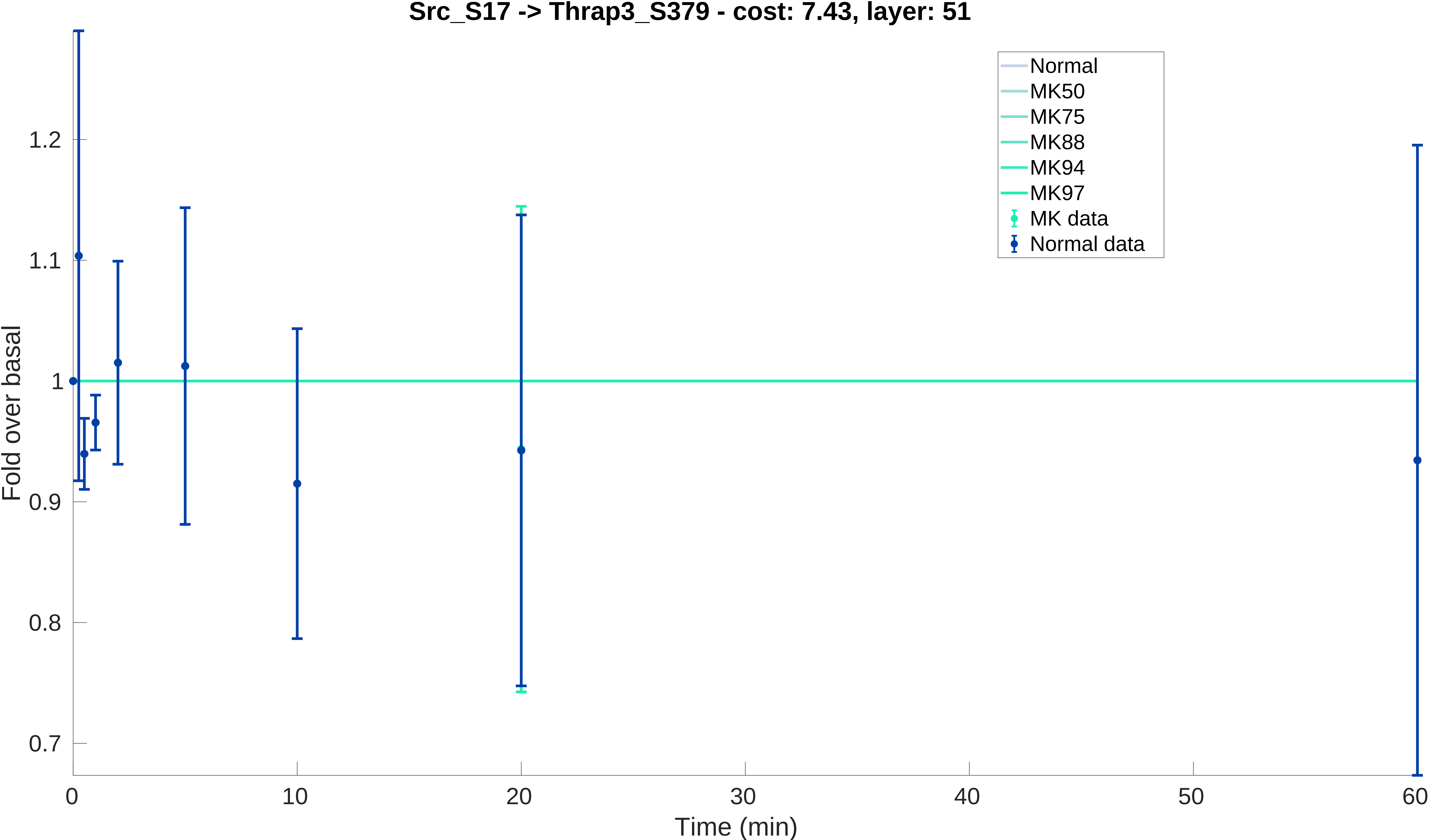

Src_S17 -> Thrap3_S379 - cost: 7.43, layer: 51
Normal
MK50
MK75
MK88
MK94
MK97
MK data
Normal data
Fold over basal
1.2
1.1
1
0.9
0.8
0.7
0
10
20
30
40
50
60
Time (min)

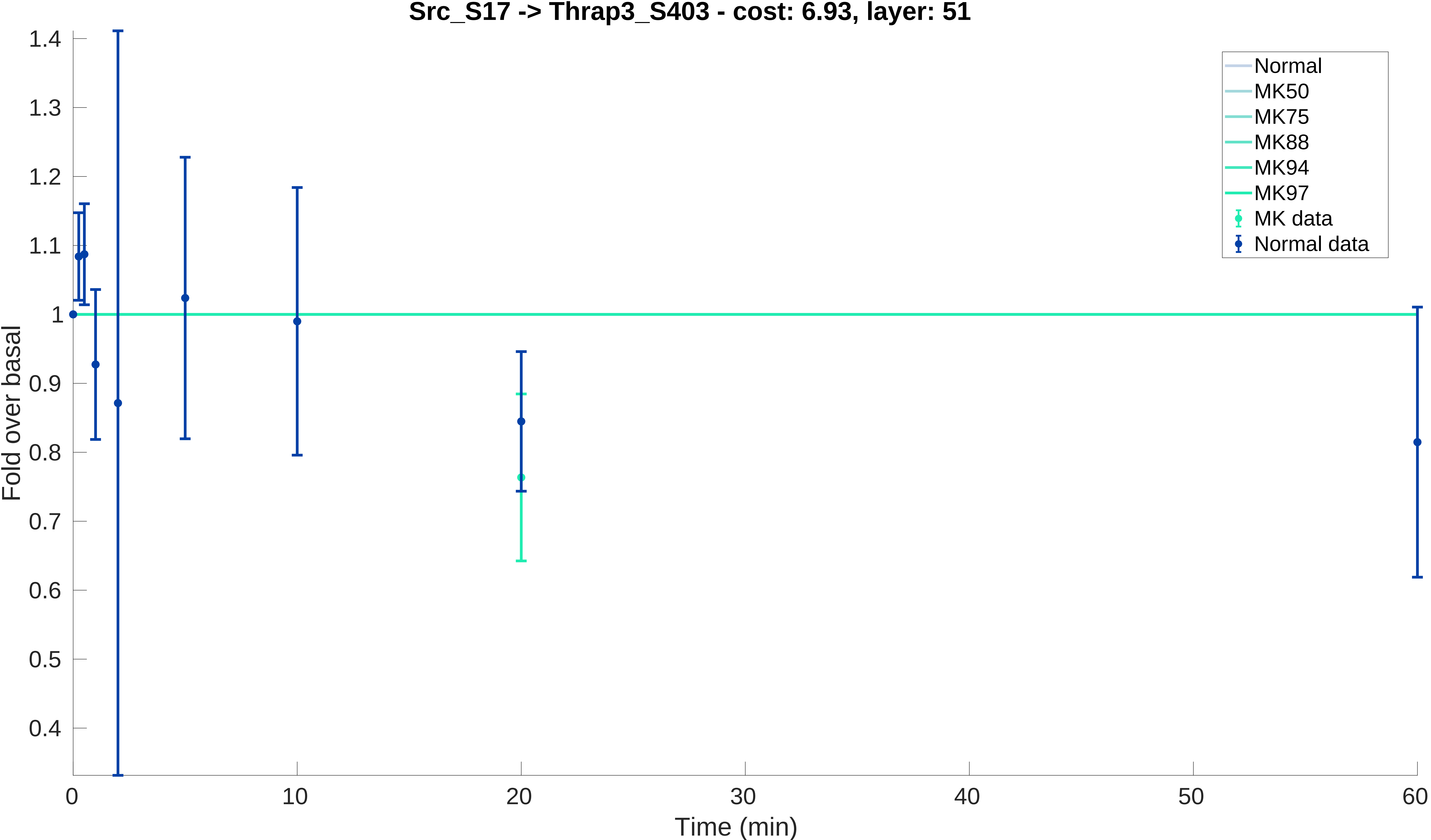

Src_S17 -> Thrap3_S403 - cost: 6.93, layer: 51
Fold over basal
Time (min)
1.4
1.3
1.2
1.1
1
0.9
0.8
0.7
0.6
0.5
0.4
0
10
20
30
40
50
60
Normal
MK50
MK75
MK88
MK94
MK97
MK data
Normal data

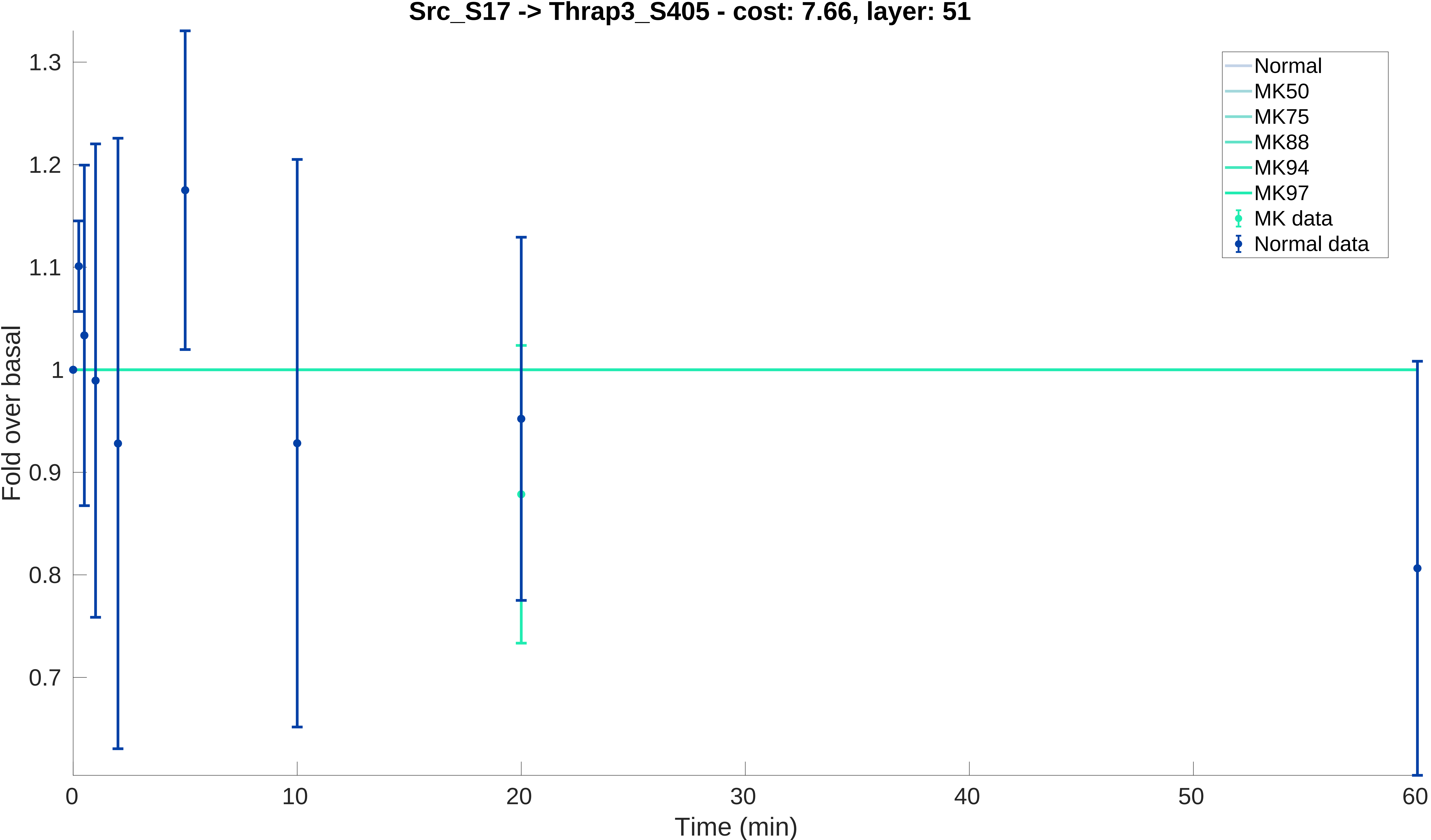

Src_S17 -> Thrap3_S405 - cost: 7.66, layer: 51
Fold over basal
Time (min)
1.3
1.2
1.1
1
0.9
0.8
0.7
0
10
20
30
40
50
60
Normal
MK50
MK75
MK88
MK94
MK97
MK data
Normal data

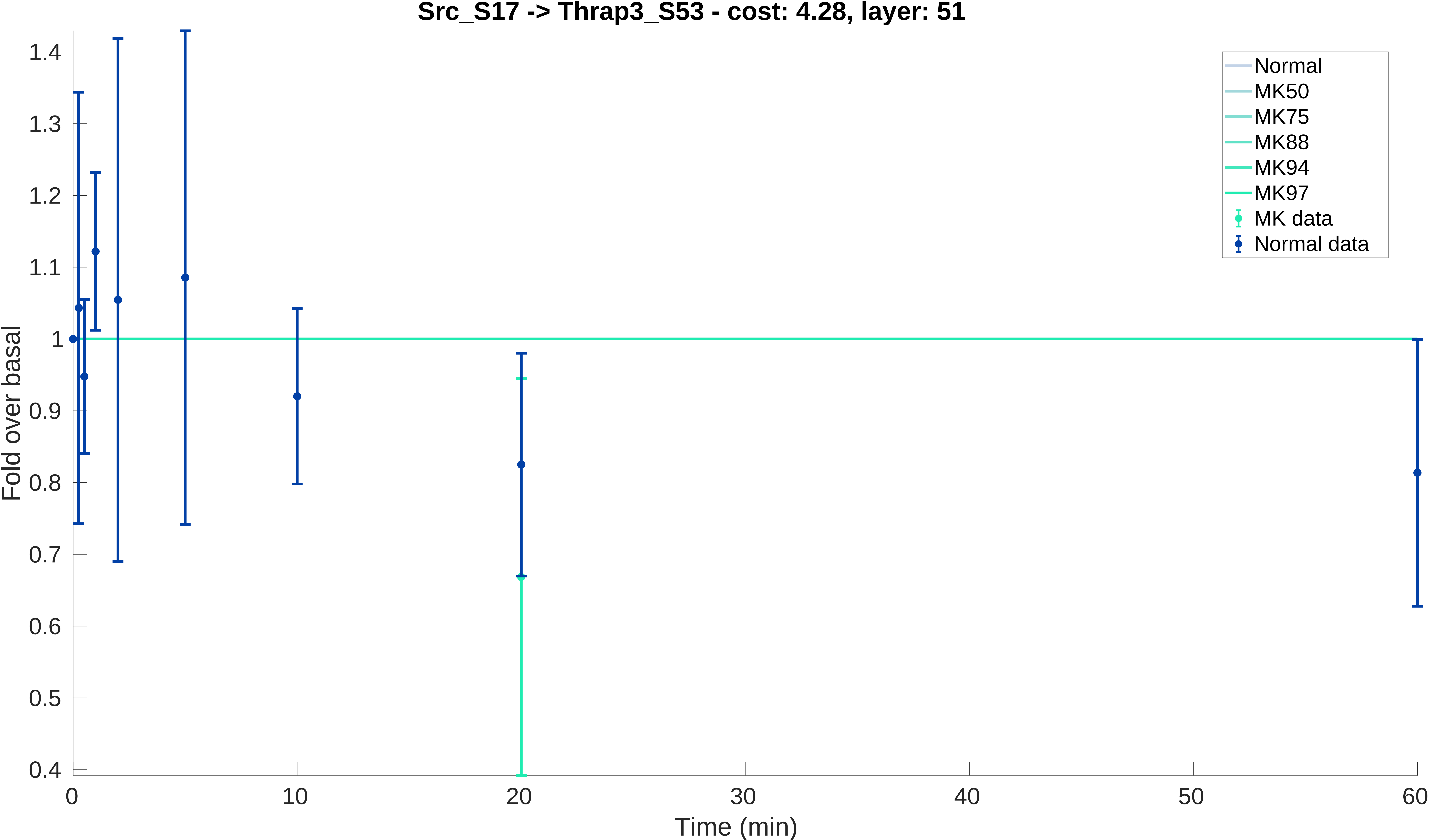

Src_S17 -> Thrap3_S53 - cost: 4.28, layer: 51
Fold over basal
1.4
1.3
1.2
1.1
1
0.9
0.8
0.7
0.6
0.5
0.4
0
10
20
30
40
50
60
Time (min)
Normal
MK50
MK75
MK88
MK94
MK97
MK data
Normal data

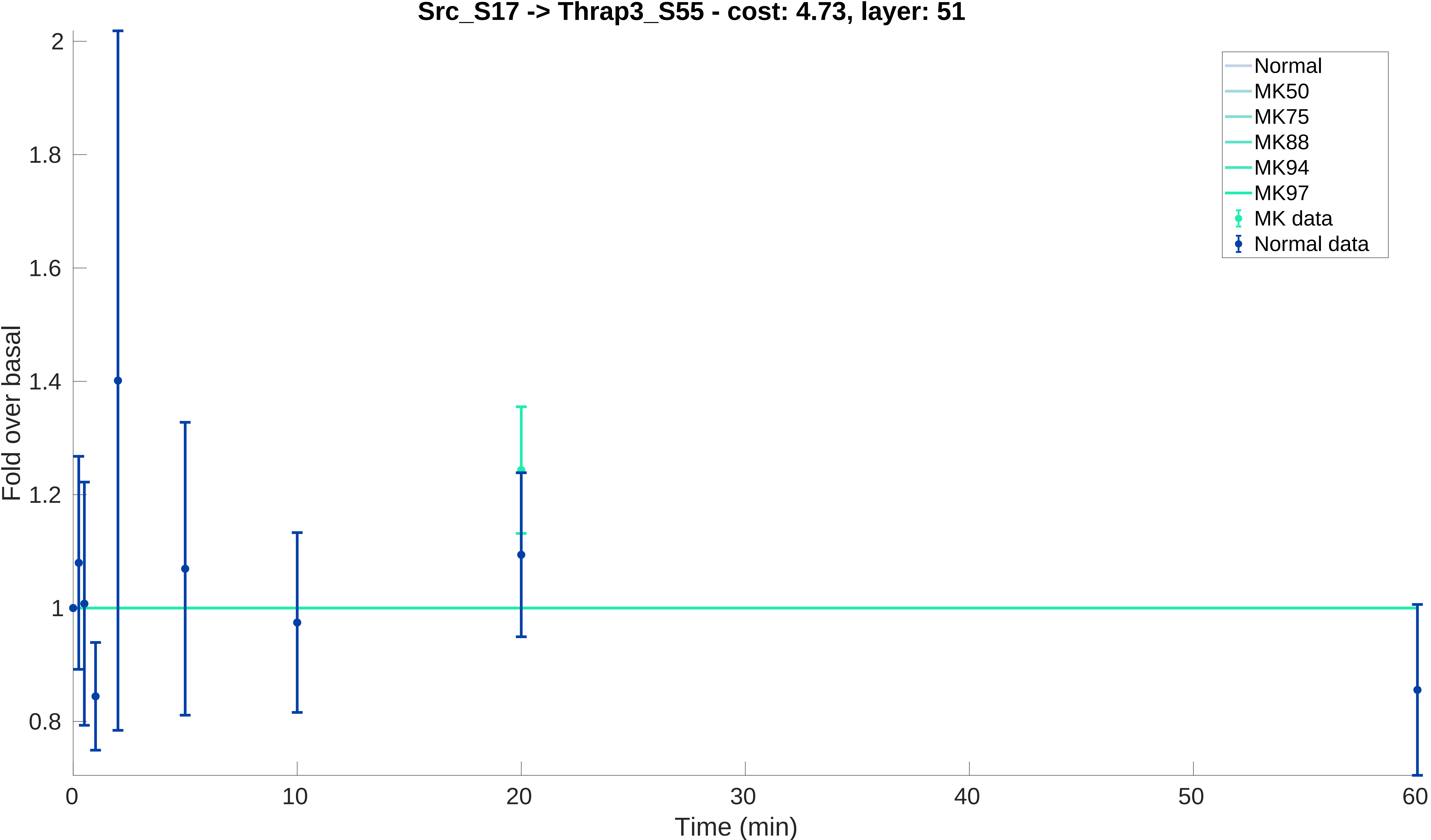

Src_S17 -> Thrap3_S55 - cost: 4.73, layer: 51
Fold over basal
Time (min)
2
1.8
1.6
1.4
1.2
1
0.8
0
10
20
30
40
50
60
Normal
MK50
MK75
MK88
MK94
MK97
MK data
Normal data

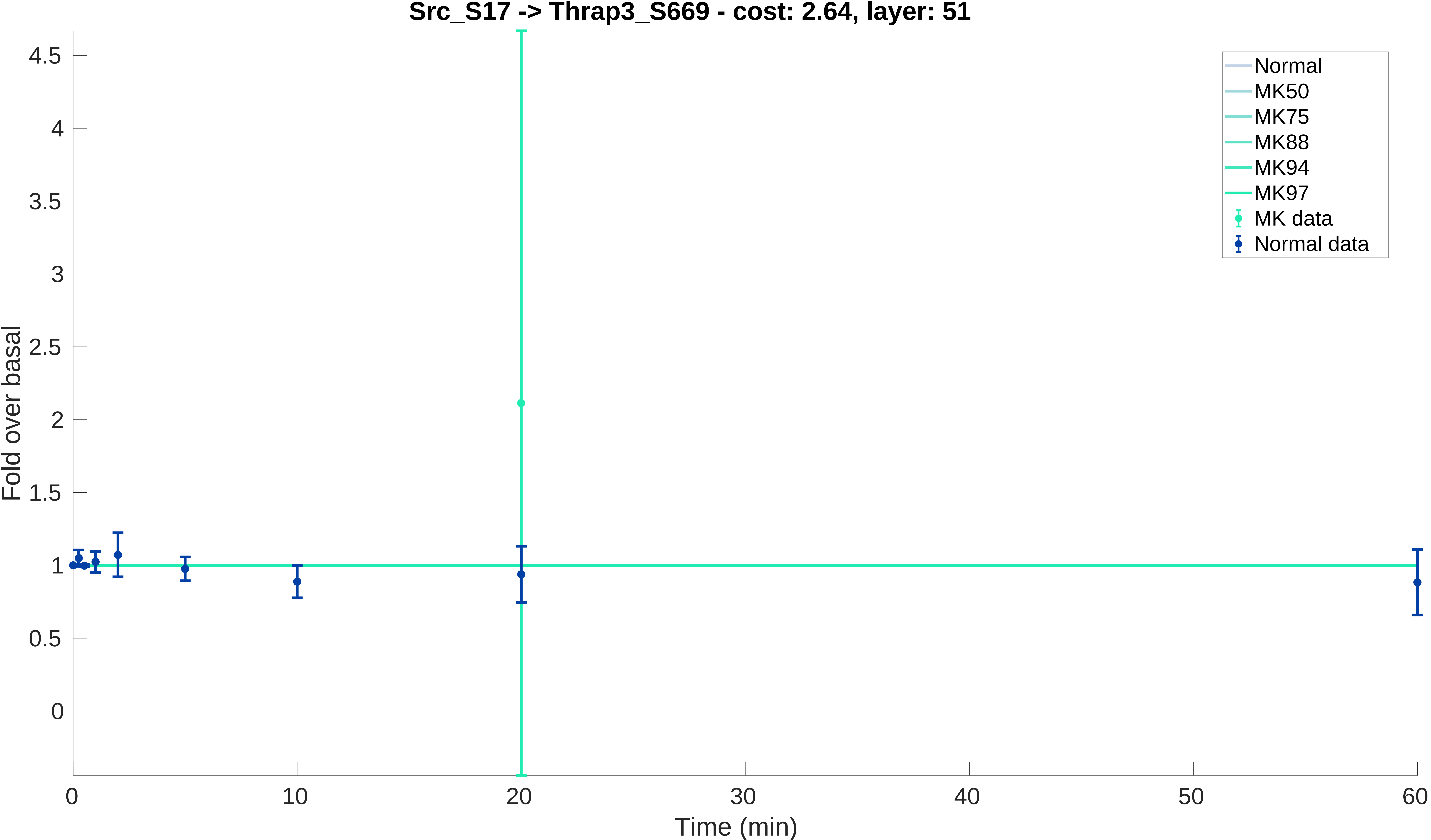

Src_S17 -> Thrap3_S669 - cost: 2.64, layer: 51
Normal
MK50
MK75
MK88
MK94
MK97
MK data
Normal data
Fold over basal
4.5
4
3.5
3
2.5
2
1.5
1
0.5
0
0
10
20
30
40
50
60
Time (min)

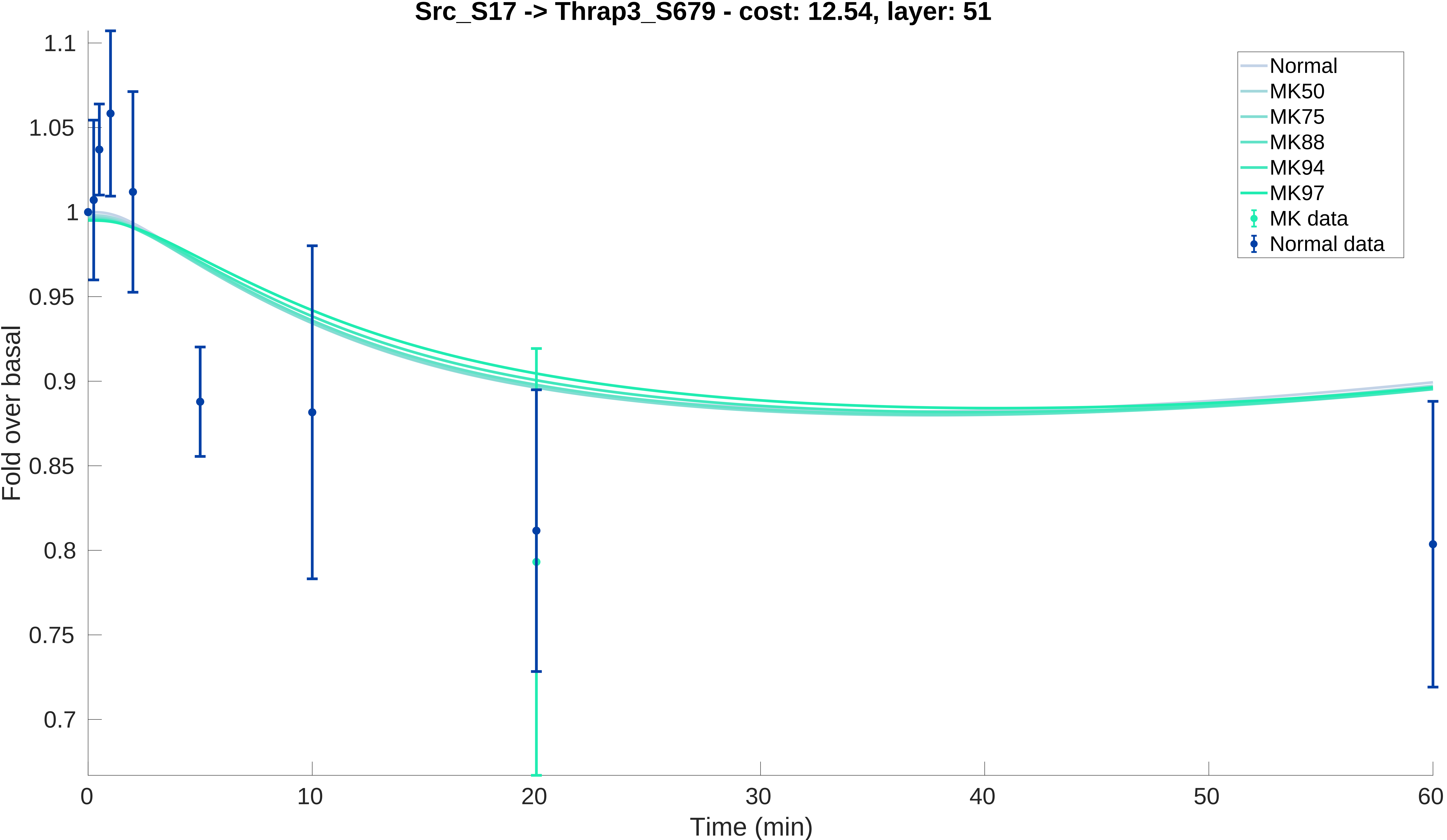

Src_S17 -> Thrap3_S679 - cost: 12.54, layer: 51
Fold over basal
Time (min)
1.1
1.05
1
0.95
0.9
0.85
0.8
0.75
0.7
0
10
20
30
40
50
60
Normal
MK50
MK75
MK88
MK94
MK97
MK data
Normal data

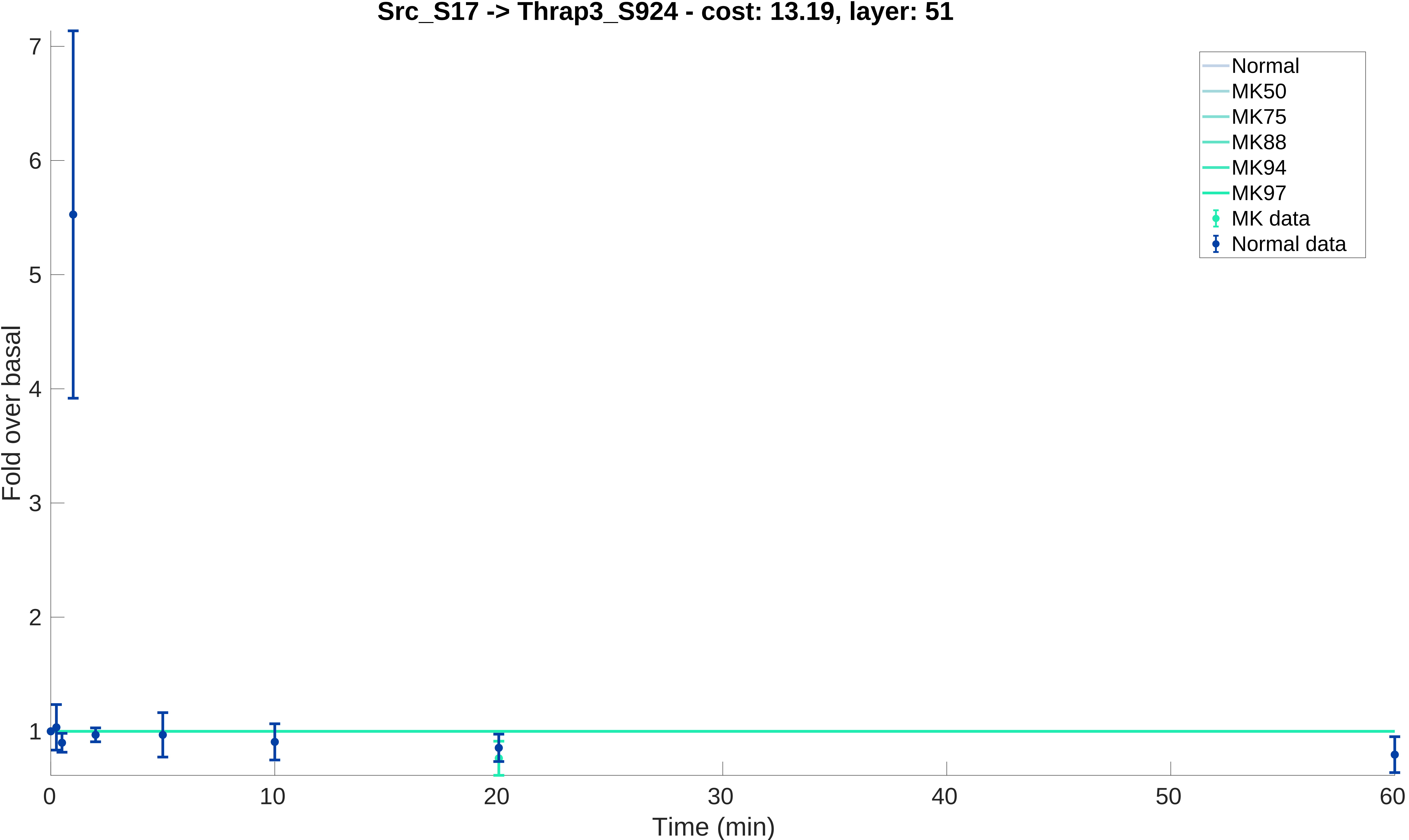

Src_S17 -> Thrap3_S924 - cost: 13.19, layer: 51
Fold over basal
Time (min)
1
2
3
4
5
6
7
0
10
20
30
40
50
60
Normal
MK50
MK75
MK88
MK94
MK97
MK data
Normal data

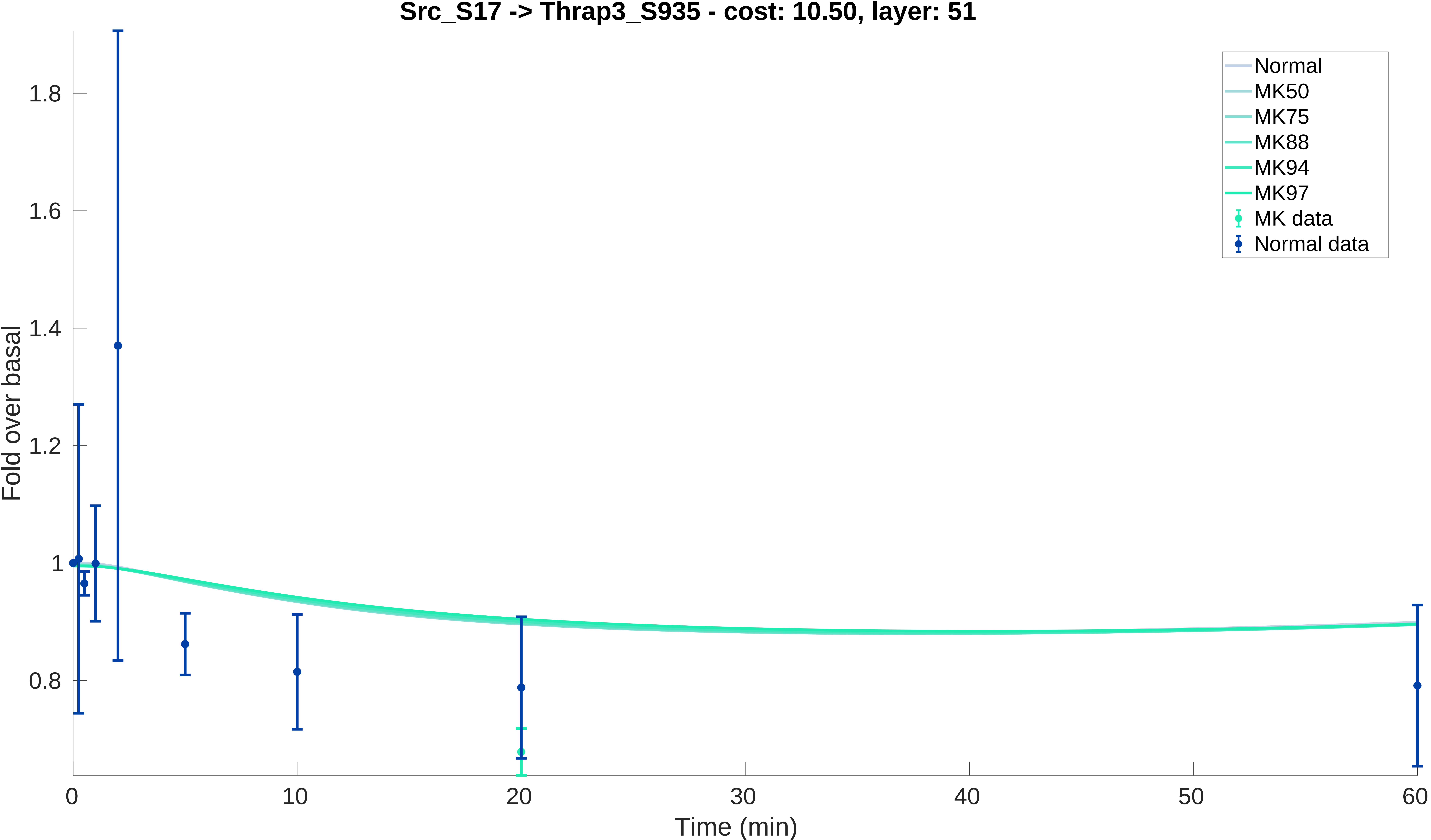
Src_S17 -> Thrap3_S935 - cost: 10.50, layer: 51
Normal
MK50
MK75
MK88
MK94
MK97
MK data
Normal data
Fold over basal
1.8
1.6
1.4
1.2
1
0.8
0
10
20
30
40
50
60
Time (min)

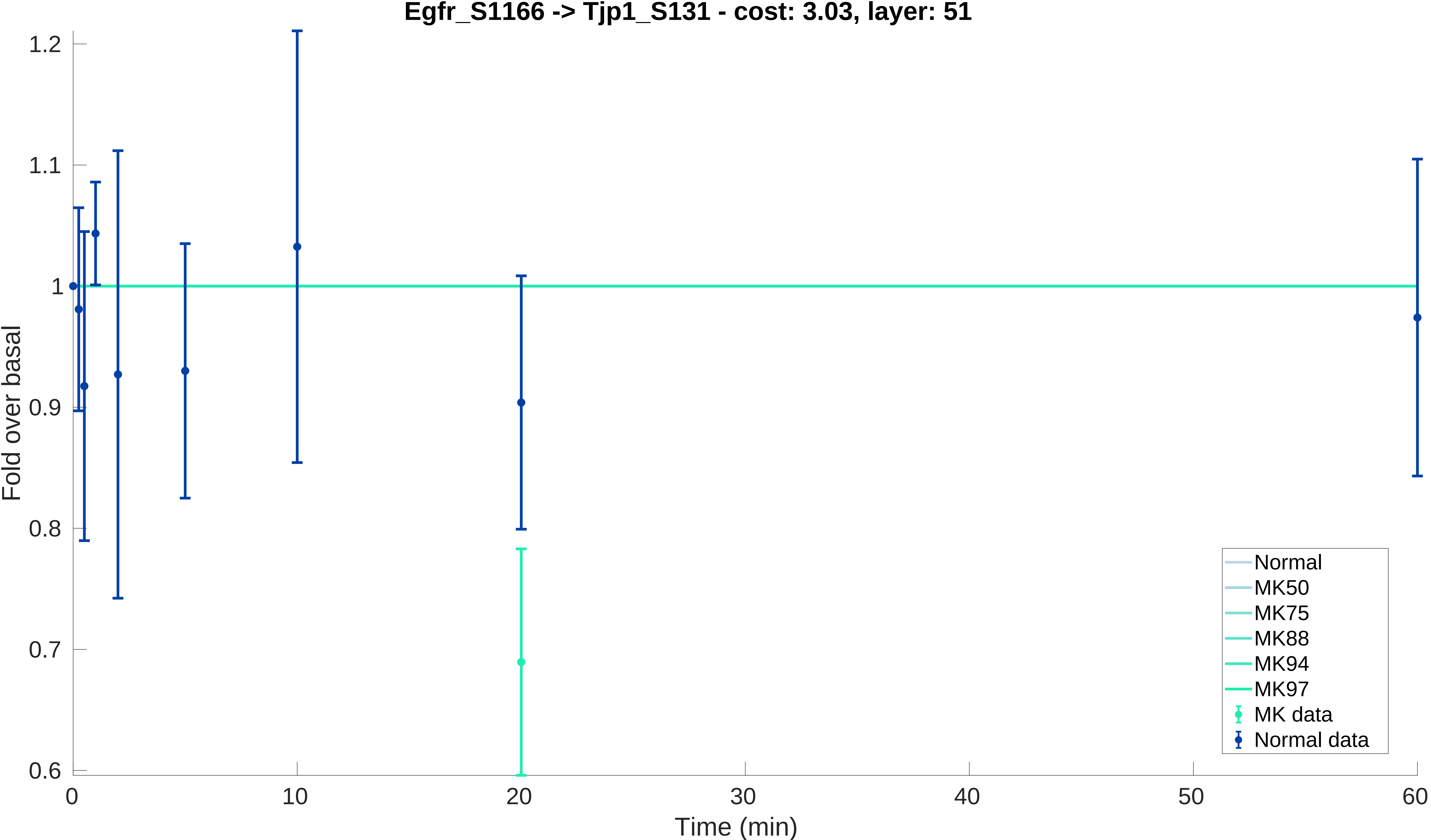

Egfr_S1166 -> Tjp1_S131 - cost: 3.03, layer: 51
1.2
1.1
1
0.9
0.8
0.7
0.6
Fold over basal
0
10
20
30
40
50
60
Time (min)
Normal
MK50
MK75
MK88
MK94
MK97
MK data
Normal data

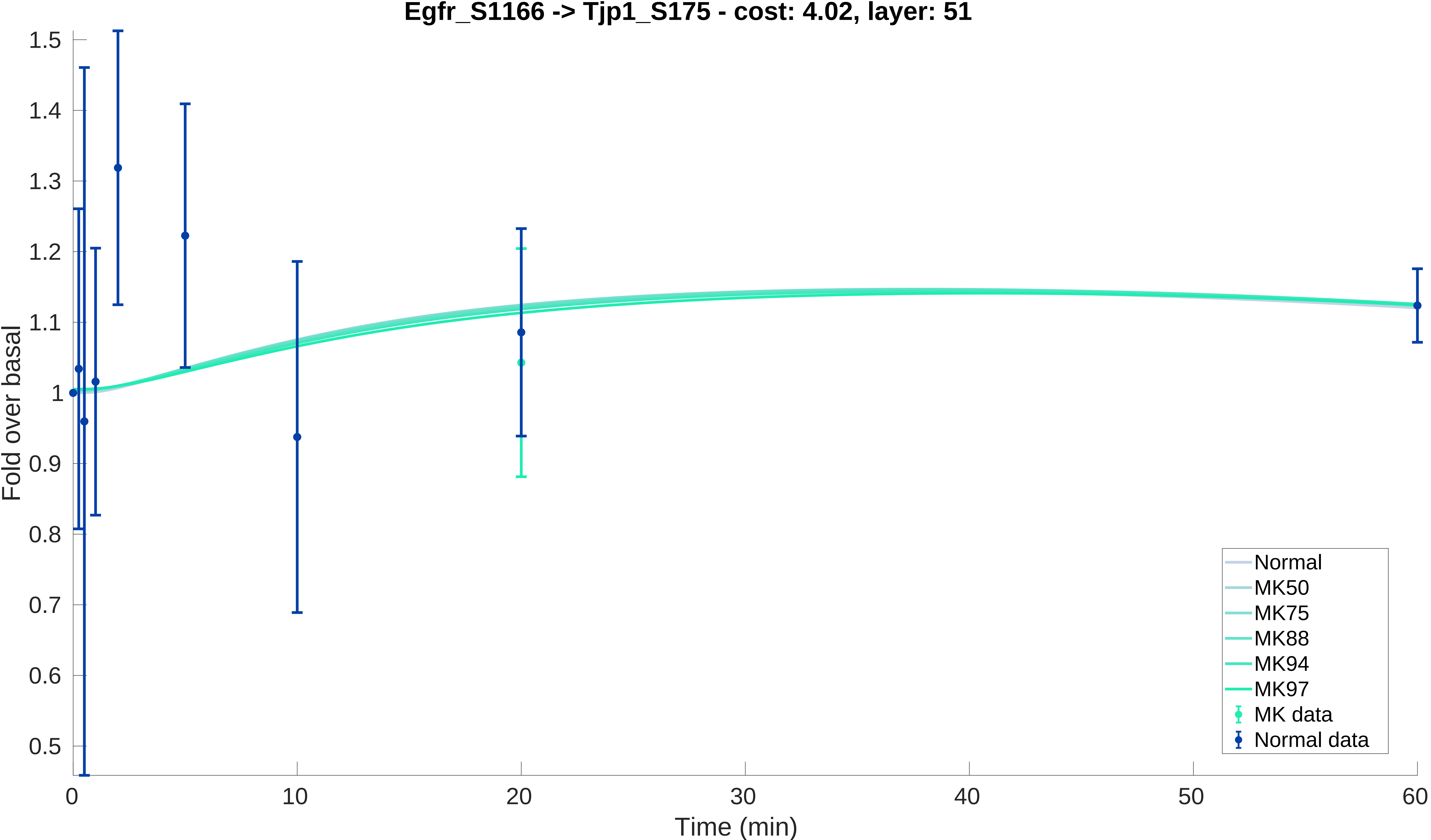
Egfr_S1166 -> Tjp1_S175 - cost: 4.02, layer: 51
Fold over basal
Time (min)
1.5
1.4
1.3
1.2
1.1
1
0.9
0.8
0.7
0.6
0.5
0
10
20
30
40
50
60
Normal
MK50
MK75
MK88
MK94
MK97
MK data
Normal data

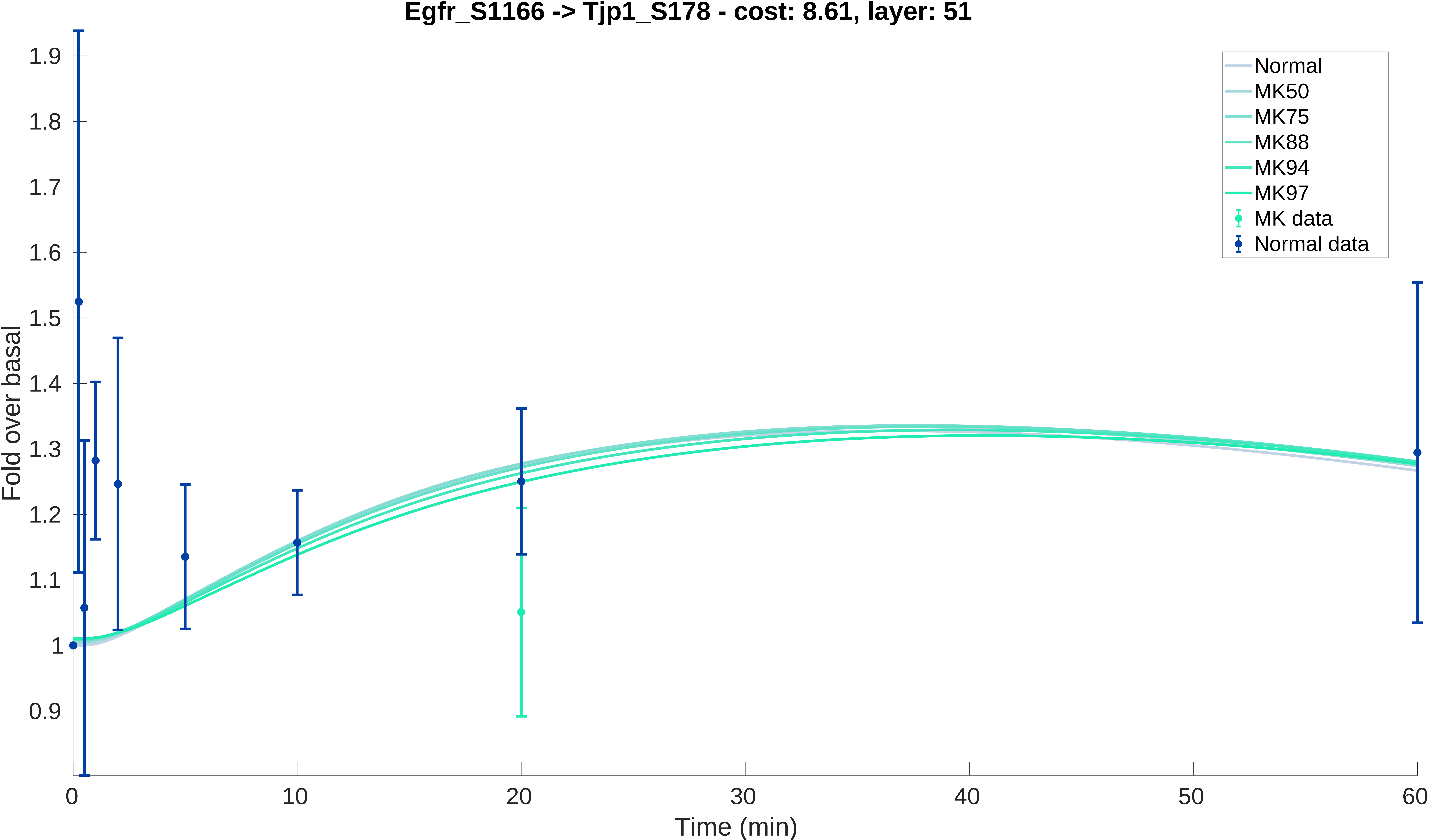

Egfr_S1166 -> Tjp1_S178 - cost: 8.61, layer: 51
Normal
MK50
MK75
MK88
MK94
MK97
MK data
Normal data
Fold over basal
0.9
1
1.1
1.2
1.3
1.4
1.5
1.6
1.7
1.8
1.9
0
10
20
30
40
50
60
Time (min)

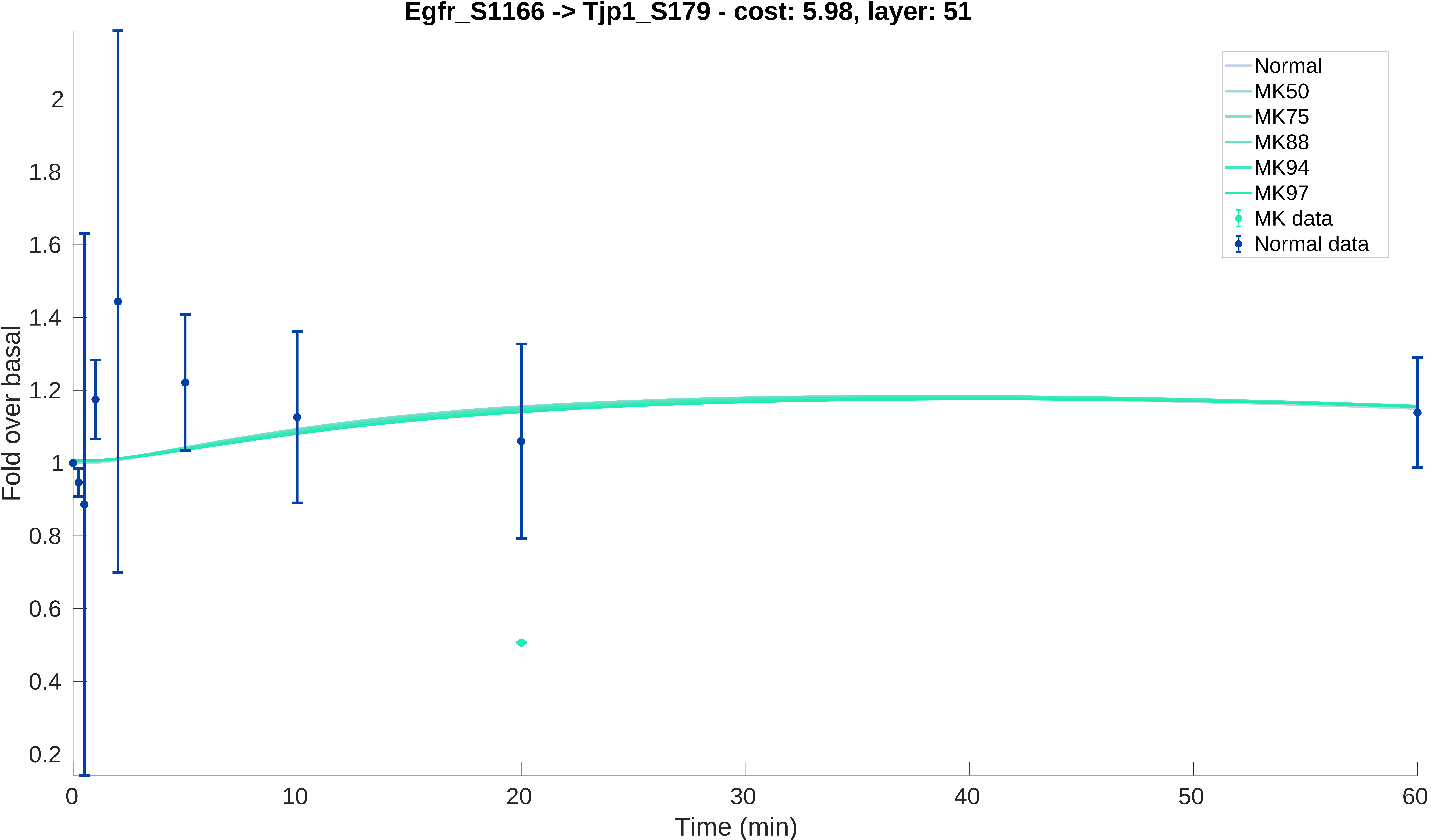

Egfr_S1166 -> Tjp1_S179 - cost: 5.98, layer: 51
Fold over basal
Time (min)
0
10
20
30
40
50
60
0.2
0.4
0.6
0.8
1
1.2
1.4
1.6
1.8
2
Normal
MK50
MK75
MK88
MK94
MK97
MK data
Normal data

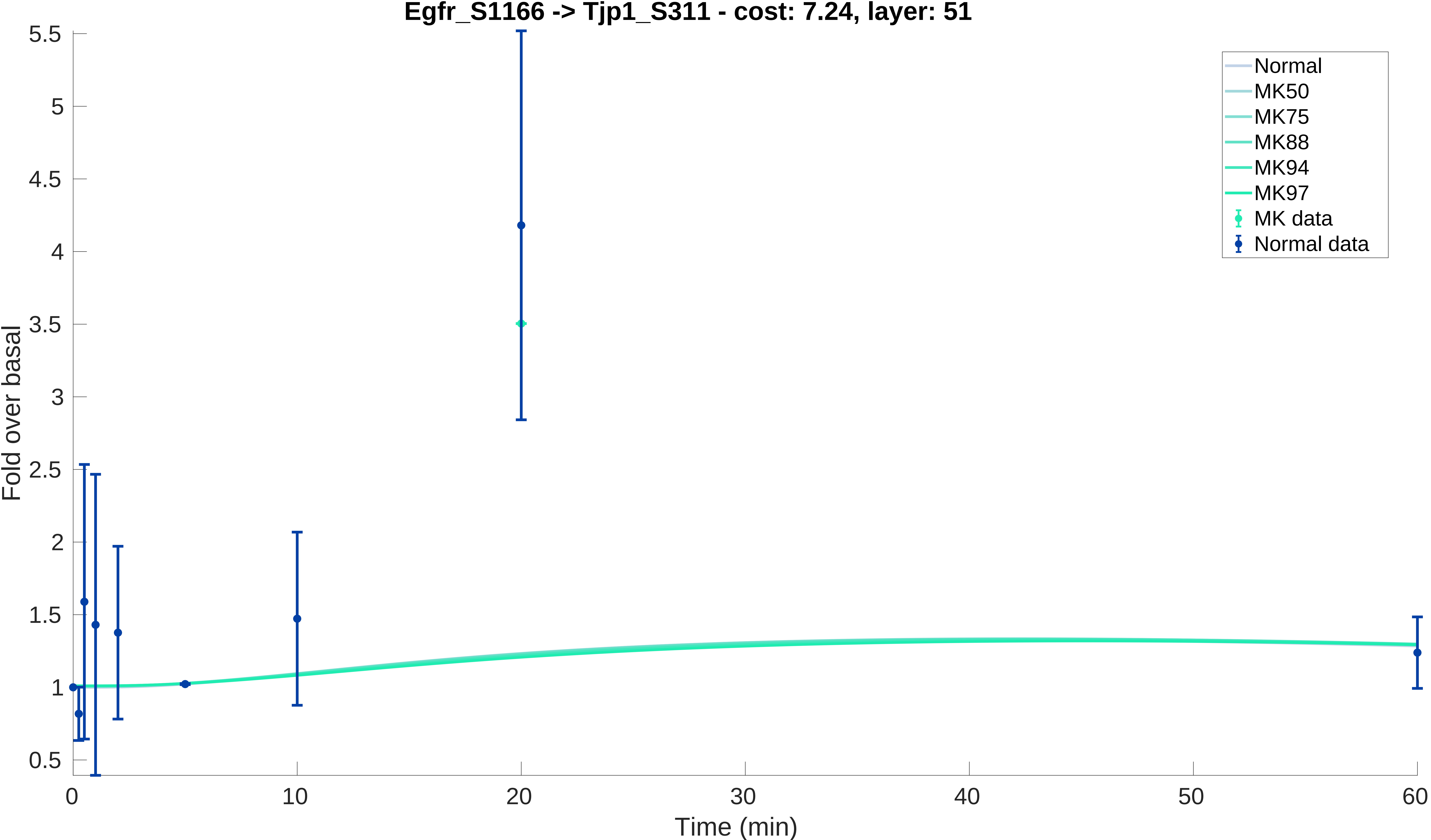

Egfr_S1166 -> Tjp1_S311 - cost: 7.24, layer: 51
Fold over basal
Time (min)
0.5
1
1.5
2
2.5
3
3.5
4
4.5
5
5.5
0
10
20
30
40
50
60
Normal
MK50
MK75
MK88
MK94
MK97
MK data
Normal data

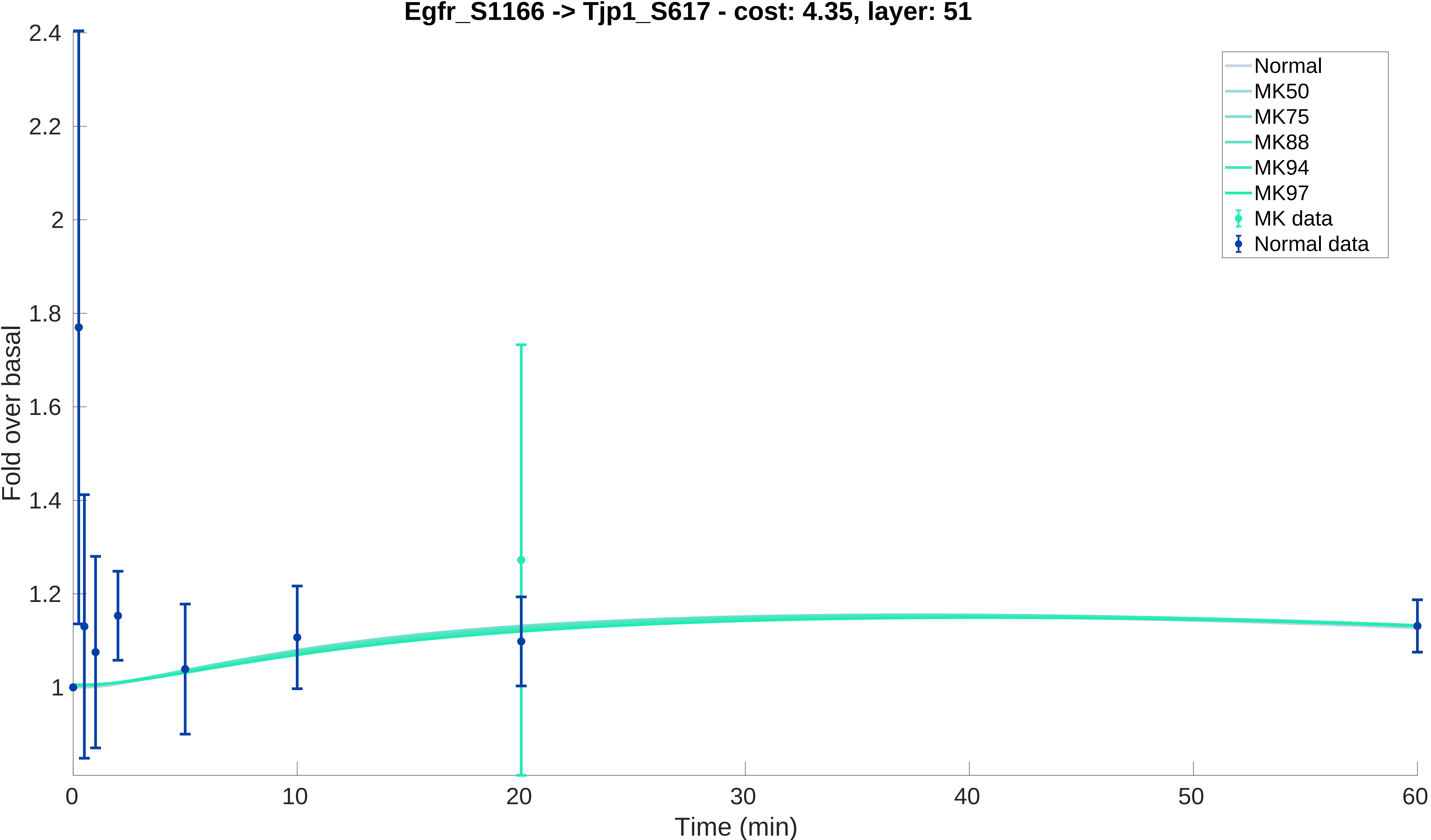
Egfr_S1166 -> Tjp1_S617 - cost: 4.35, layer: 51
Fold over basal
Time (min)
2.4
2.2
2
1.8
1.6
1.4
1.2
1
0
10
20
30
40
50
60
Normal
MK50
MK75
MK88
MK94
MK97
MK data
Normal data

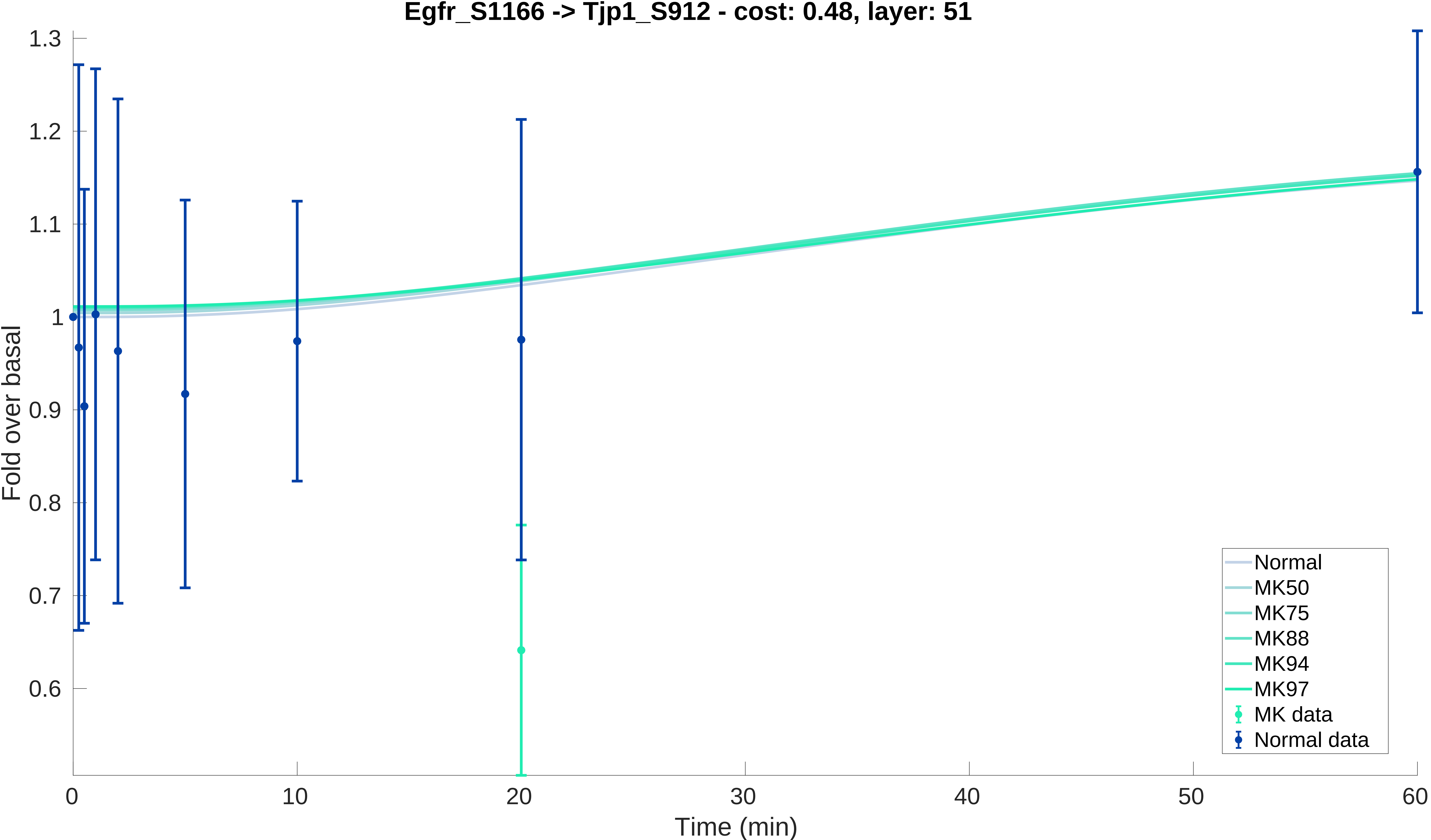

Egfr_S1166 -> Tjp1_S912 - cost: 0.48, layer: 51
Fold over basal
Time (min)
1.3
1.2
1.1
1
0.9
0.8
0.7
0.6
0
10
20
30
40
50
60
Normal
MK50
MK75
MK88
MK94
MK97
MK data
Normal data

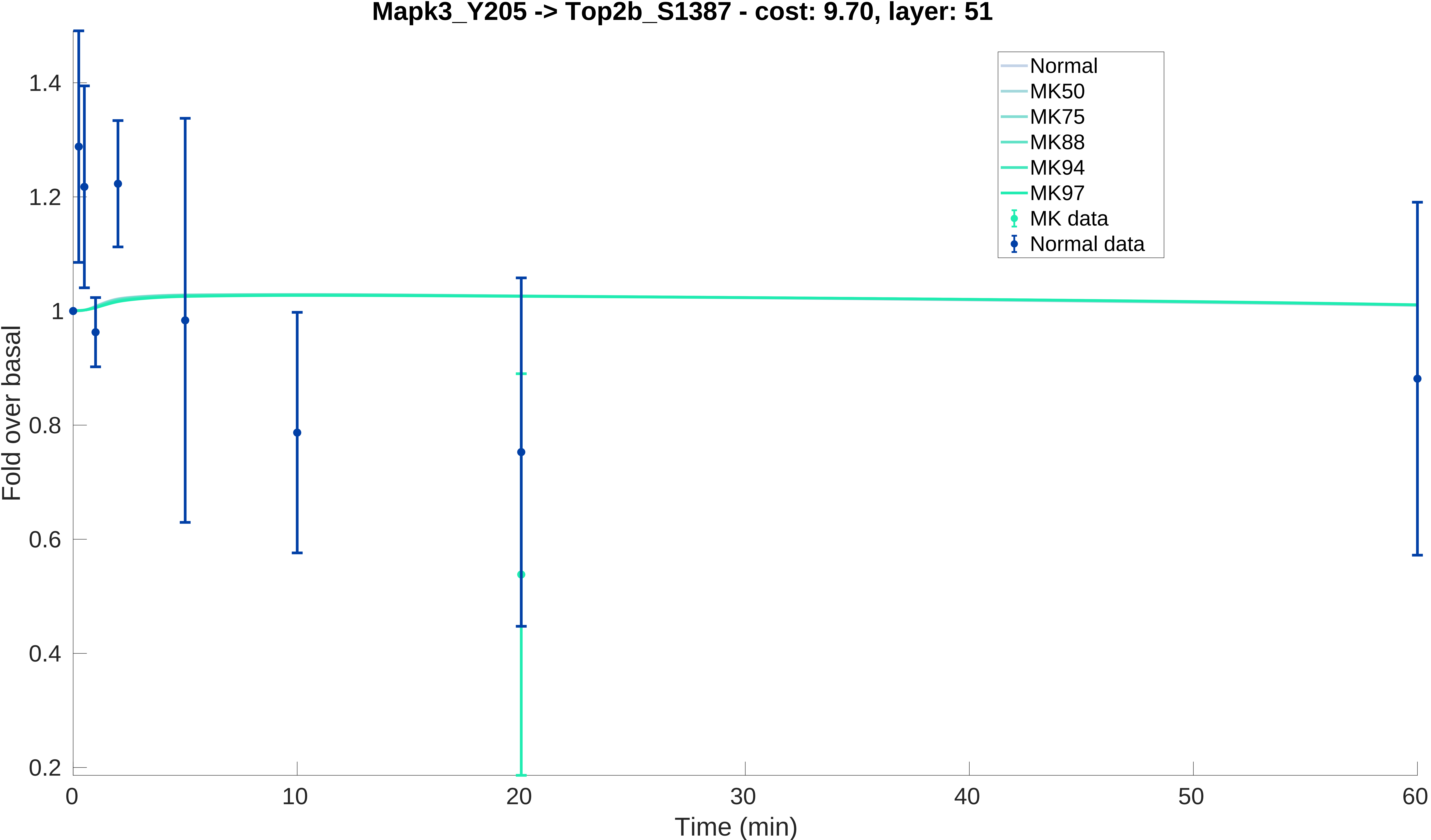

Mapk3_Y205 -> Top2b_S1387 - cost: 9.70, layer: 51
Normal
MK50
MK75
MK88
MK94
MK97
MK data
Normal data
Fold over basal
0.2
0.4
0.6
0.8
1
1.2
1.4
0
10
20
30
40
50
60
Time (min)

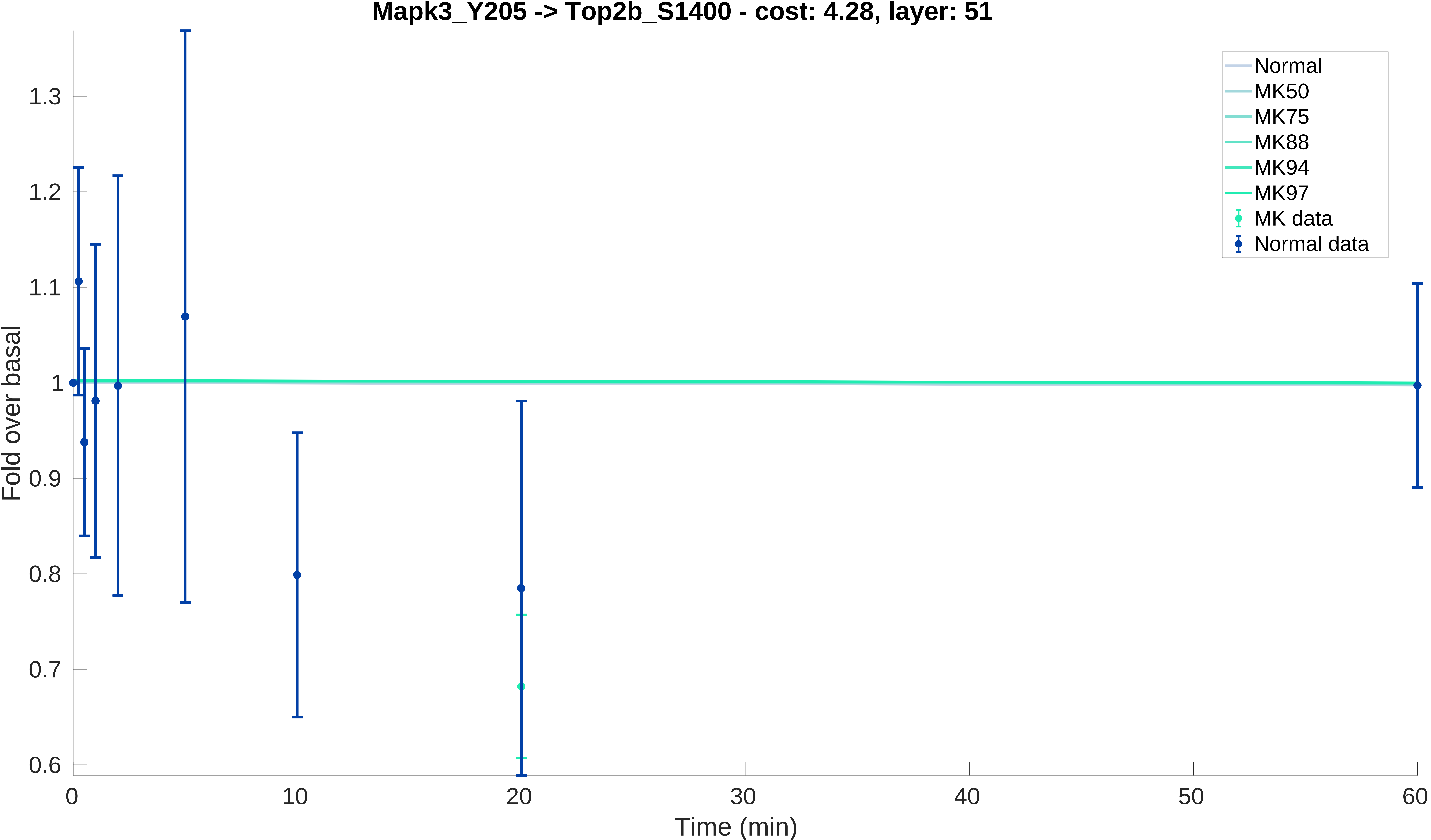

Mapk3_Y205 -> Top2b_S1400 - cost: 4.28, layer: 51
Fold over basal
Time (min)
1.3
1.2
1.1
1
0.9
0.8
0.7
0.6
0
10
20
30
40
50
60
Normal
MK50
MK75
MK88
MK94
MK97
MK data
Normal data

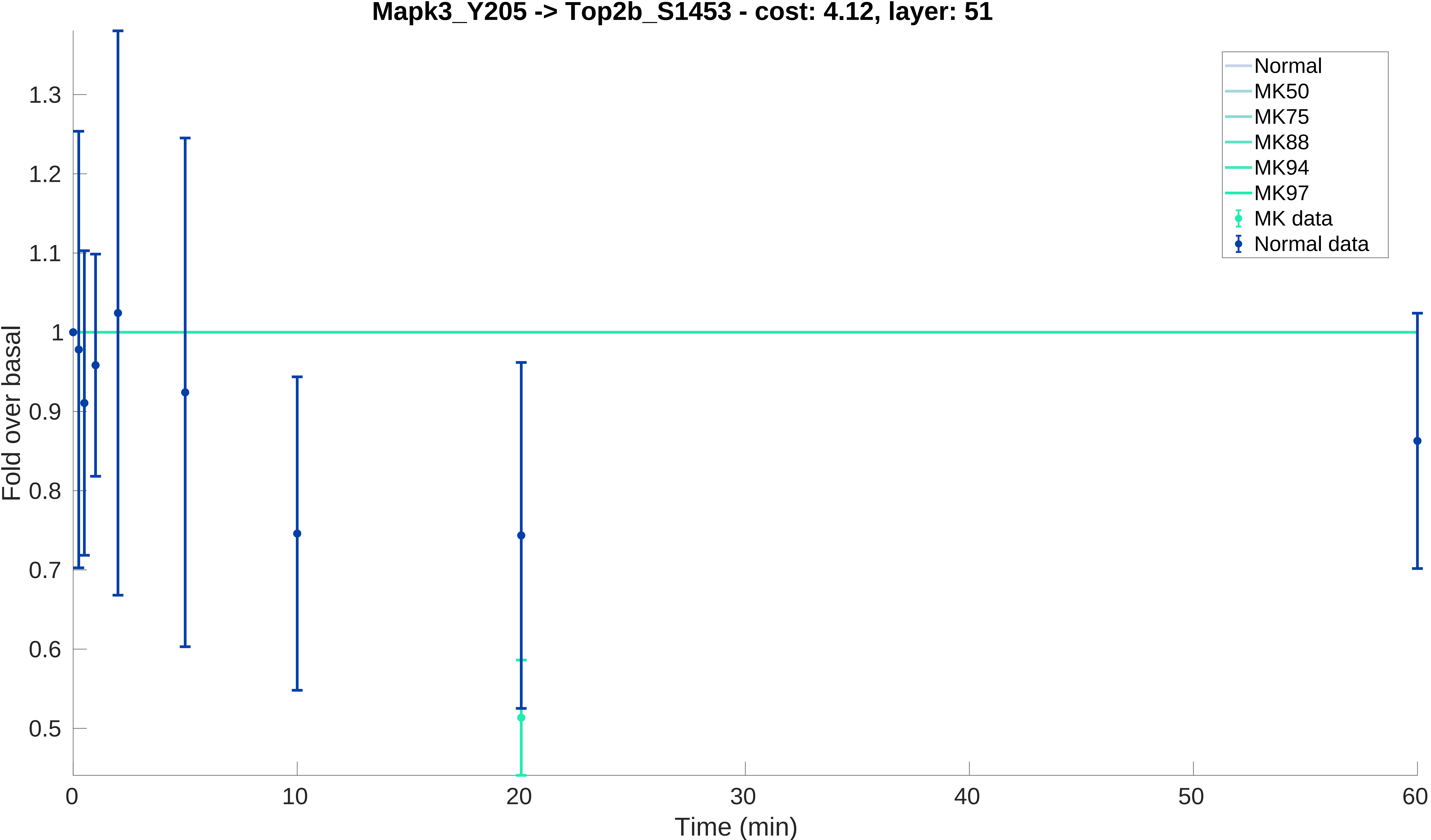

Mapk3_Y205 -> Top2b_S1453 - cost: 4.12, layer: 51
Fold over basal
Time (min)
0.5
0.6
0.7
0.8
0.9
1
1.1
1.2
1.3
0
10
20
30
40
50
60
Normal
MK50
MK75
MK88
MK94
MK97
MK data
Normal data

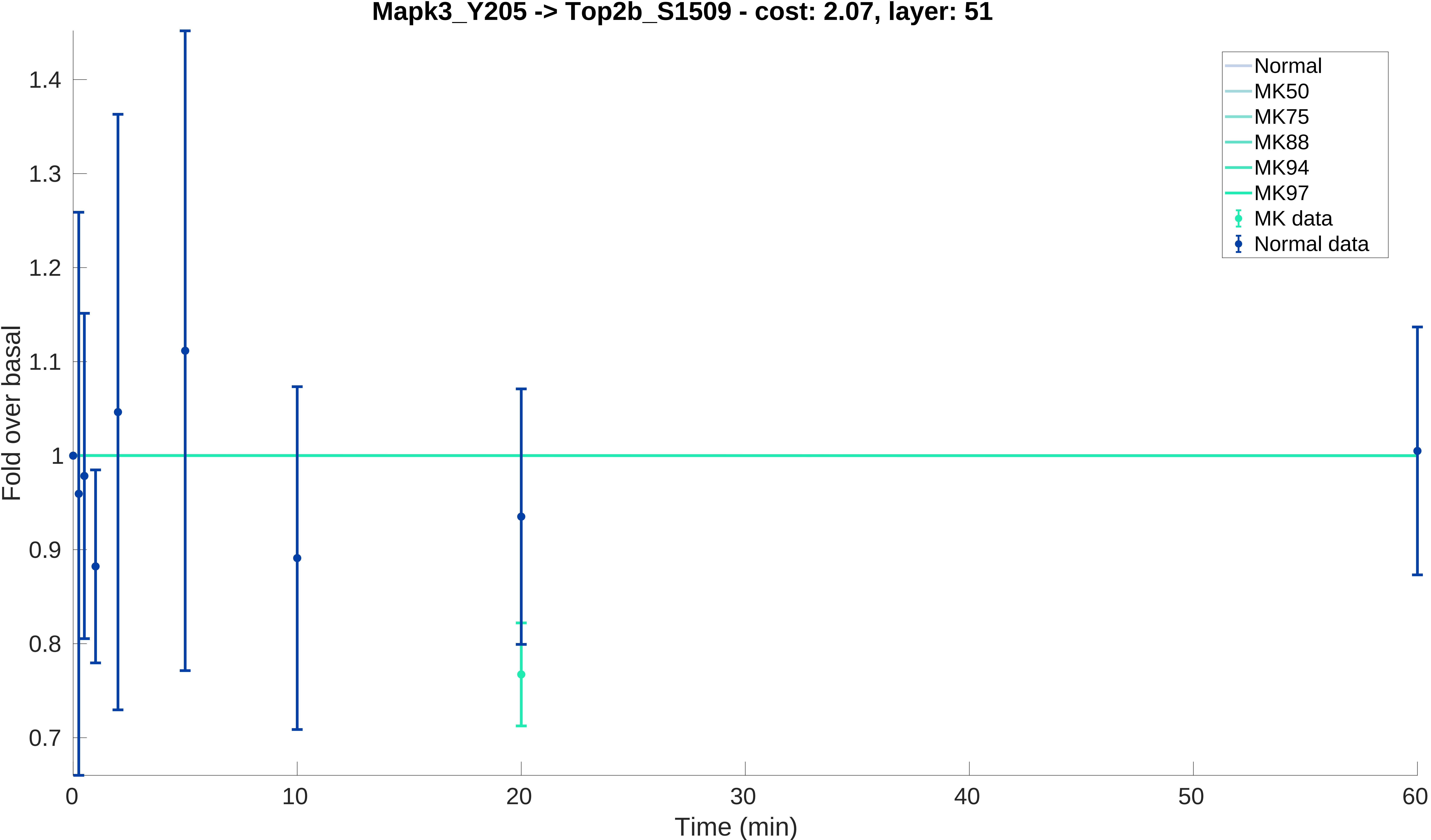

Mapk3_Y205 -> Top2b_S1509 - cost: 2.07, layer: 51
Fold over basal
Time (min)
1.4
1.3
1.2
1.1
1
0.9
0.8
0.7
0
10
20
30
40
50
60
Normal
MK50
MK75
MK88
MK94
MK97
MK data
Normal data

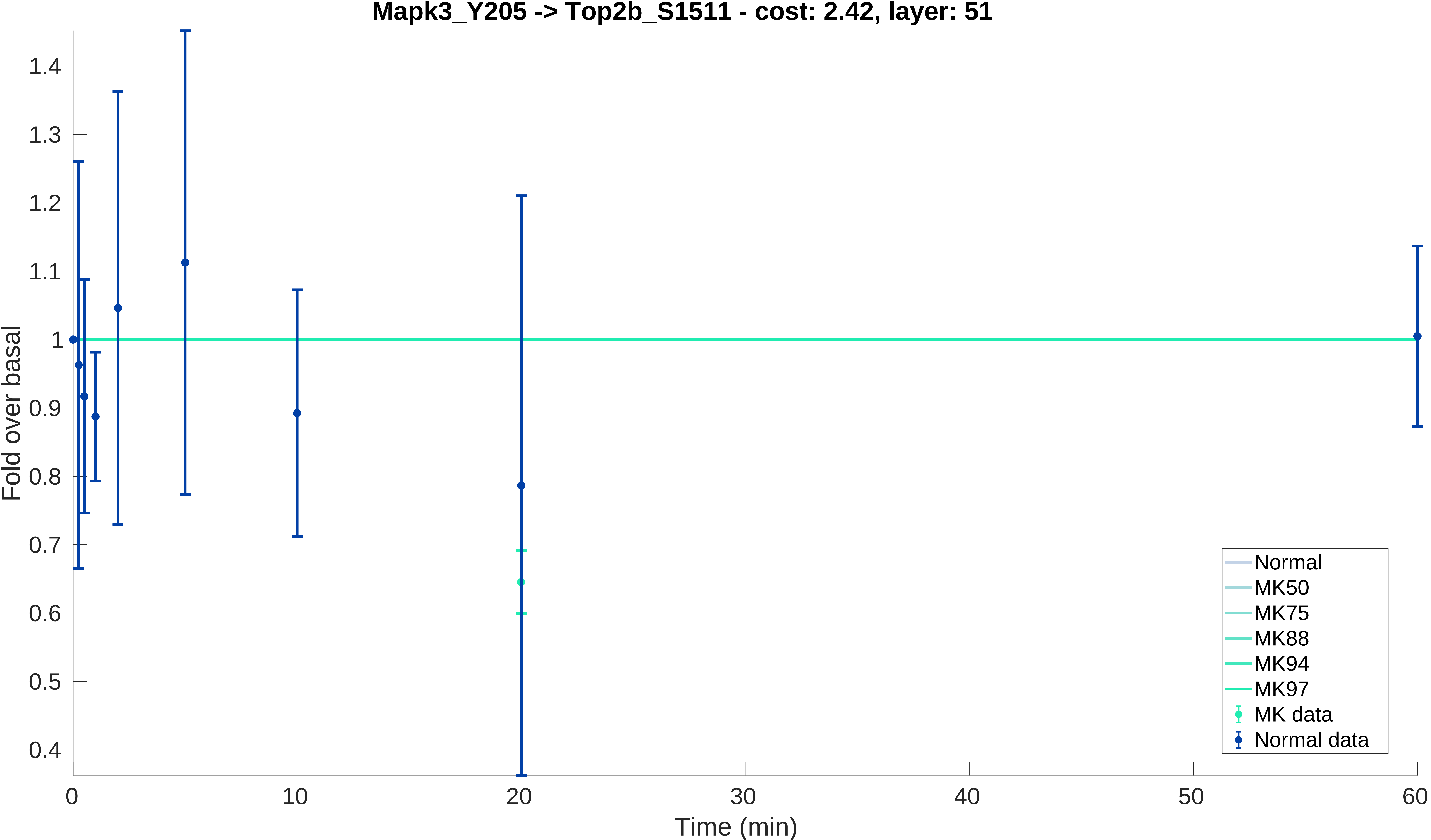

Mapk3_Y205 -> Top2b_S1511 - cost: 2.42, layer: 51
Fold over basal
Time (min)
1.4
1.3
1.2
1.1
1
0.9
0.8
0.7
0.6
0.5
0.4
0
10
20
30
40
50
60
Normal
MK50
MK75
MK88
MK94
MK97
MK data
Normal data

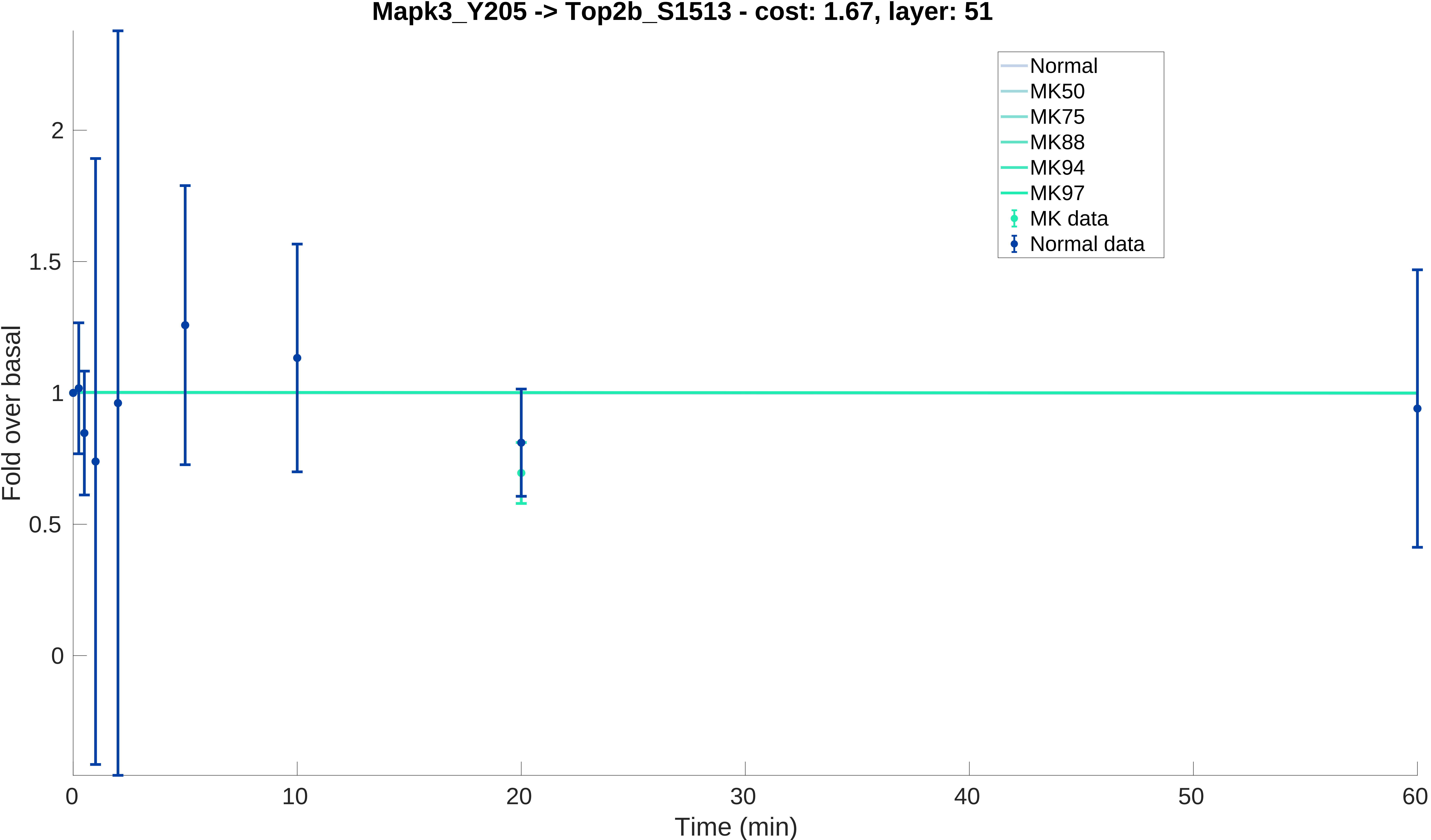
Mapk3_Y205 -> Top2b_S1513 - cost: 1.67, layer: 51
Normal
MK50
MK75
MK88
MK94
MK97
MK data
Normal data
Fold over basal
Time (min)
2
1.5
1
0.5
0
0
10
20
30
40
50
60

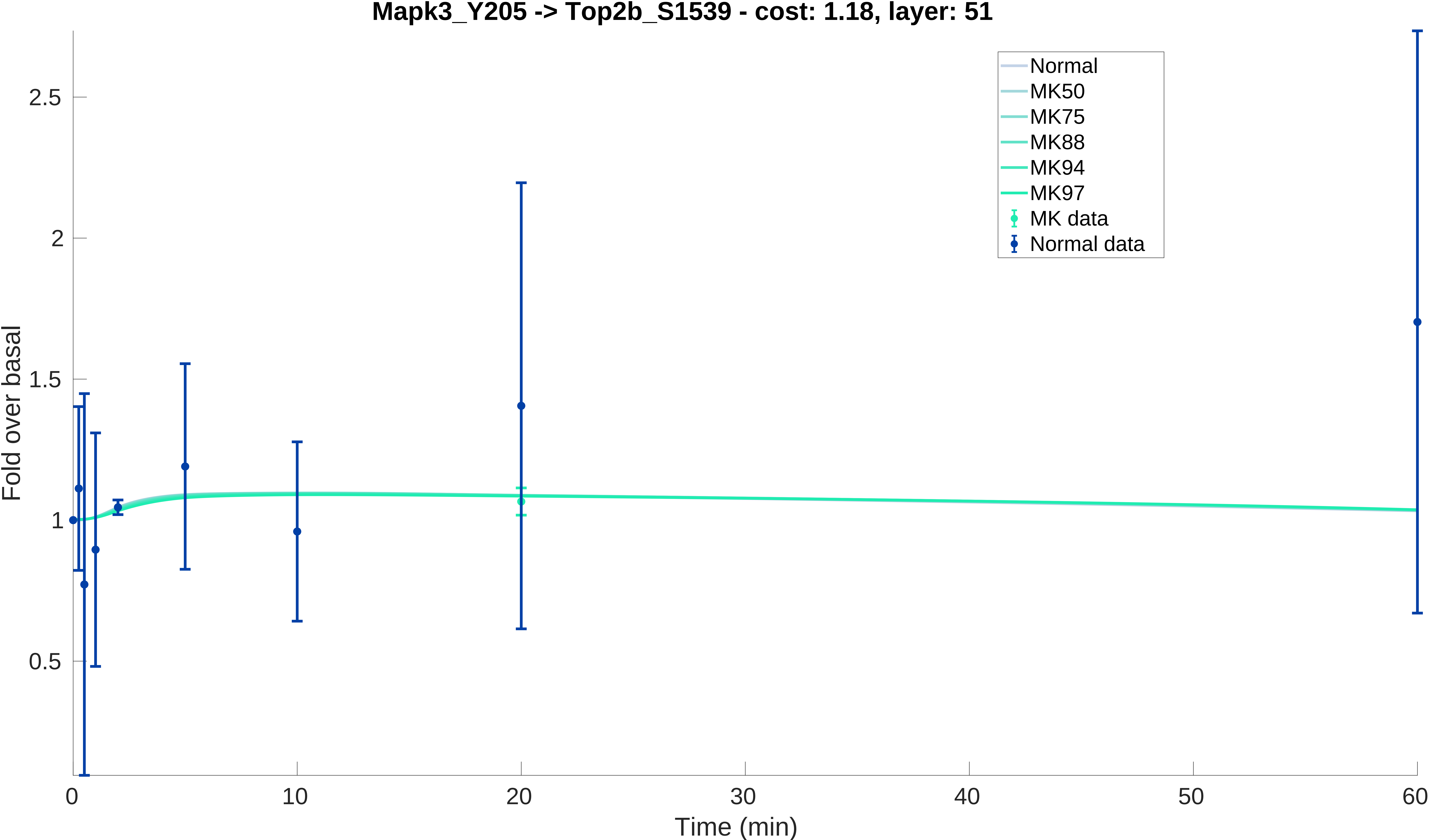
Mapk3_Y205 -> Top2b_S1539 - cost: 1.18, layer: 51
Fold over basal
Time (min)
0.5
1
1.5
2
2.5
0
10
20
30
40
50
60
Normal
MK50
MK75
MK88
MK94
MK97
MK data
Normal data

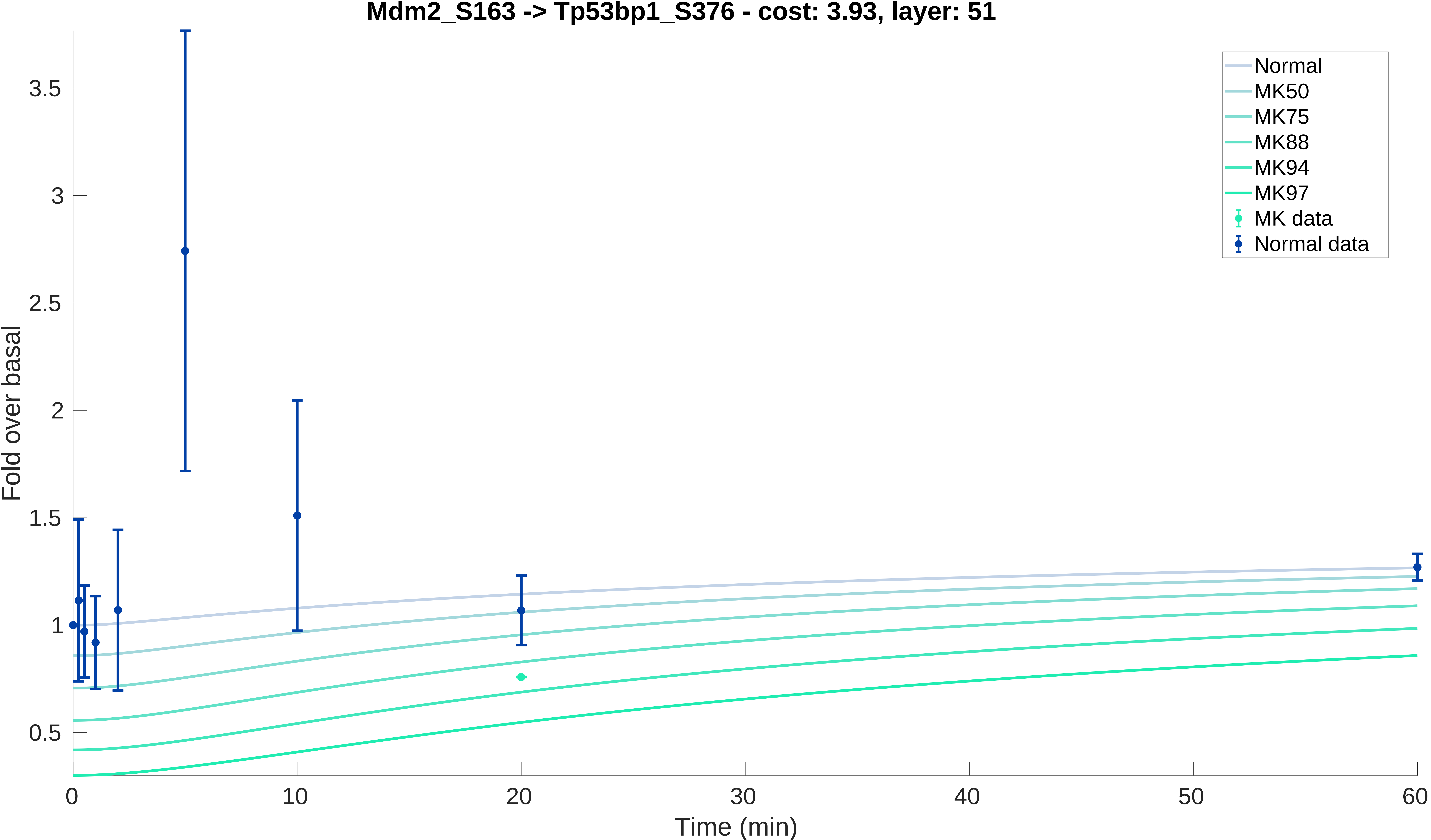

Mdm2_S163 -> Tp53bp1_S376 - cost: 3.93, layer: 51
Fold over basal
Time (min)
0.5
1
1.5
2
2.5
3
3.5
0
10
20
30
40
50
60
Normal
MK50
MK75
MK88
MK94
MK97
MK data
Normal data

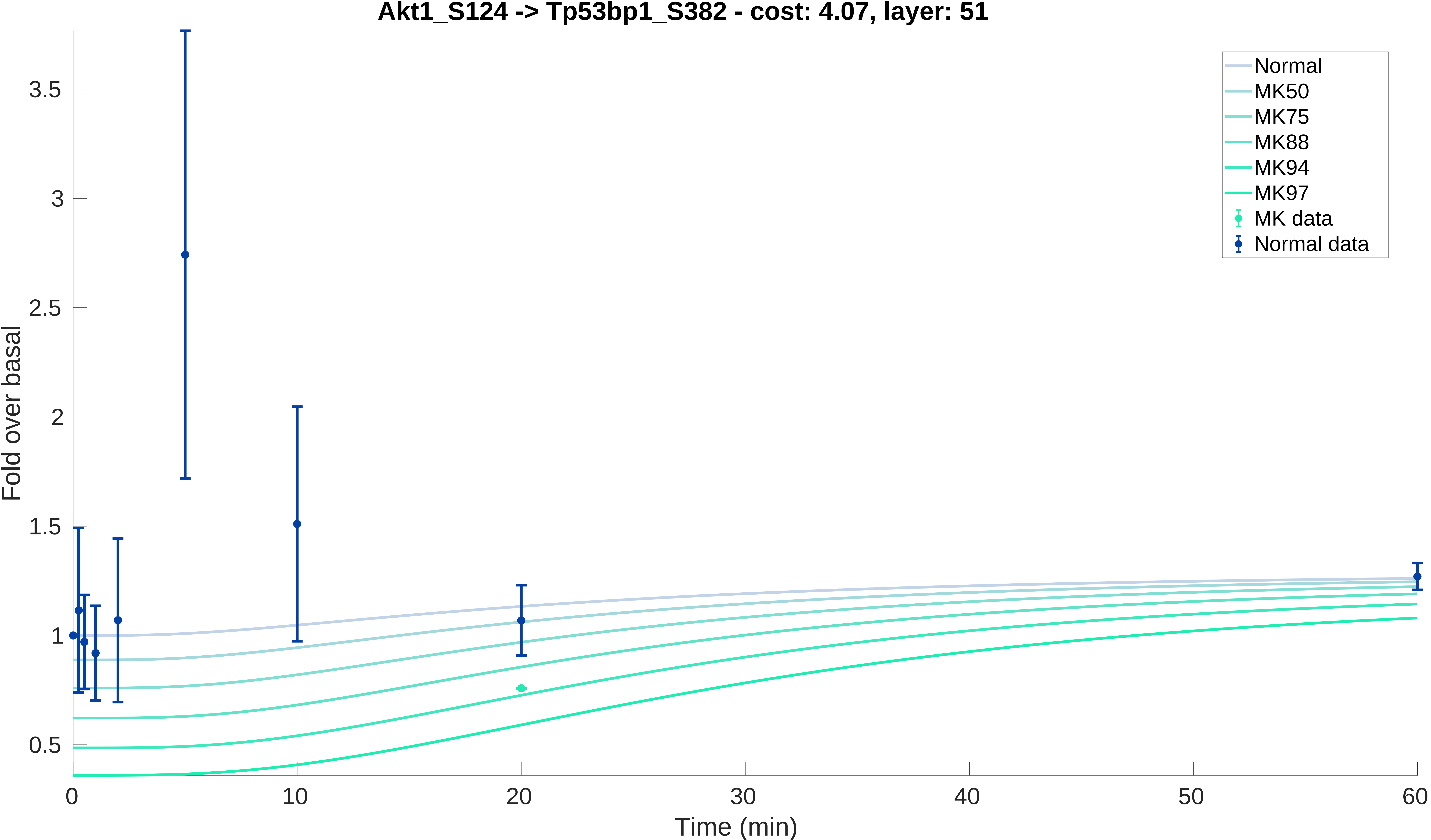

Akt1_S124 -> Tp53bp1_S382 - cost: 4.07, layer: 51
Fold over basal
Time (min)
0
10
20
30
40
50
60
0.5
1
1.5
2
2.5
3
3.5
Normal
MK50
MK75
MK88
MK94
MK97
MK data
Normal data

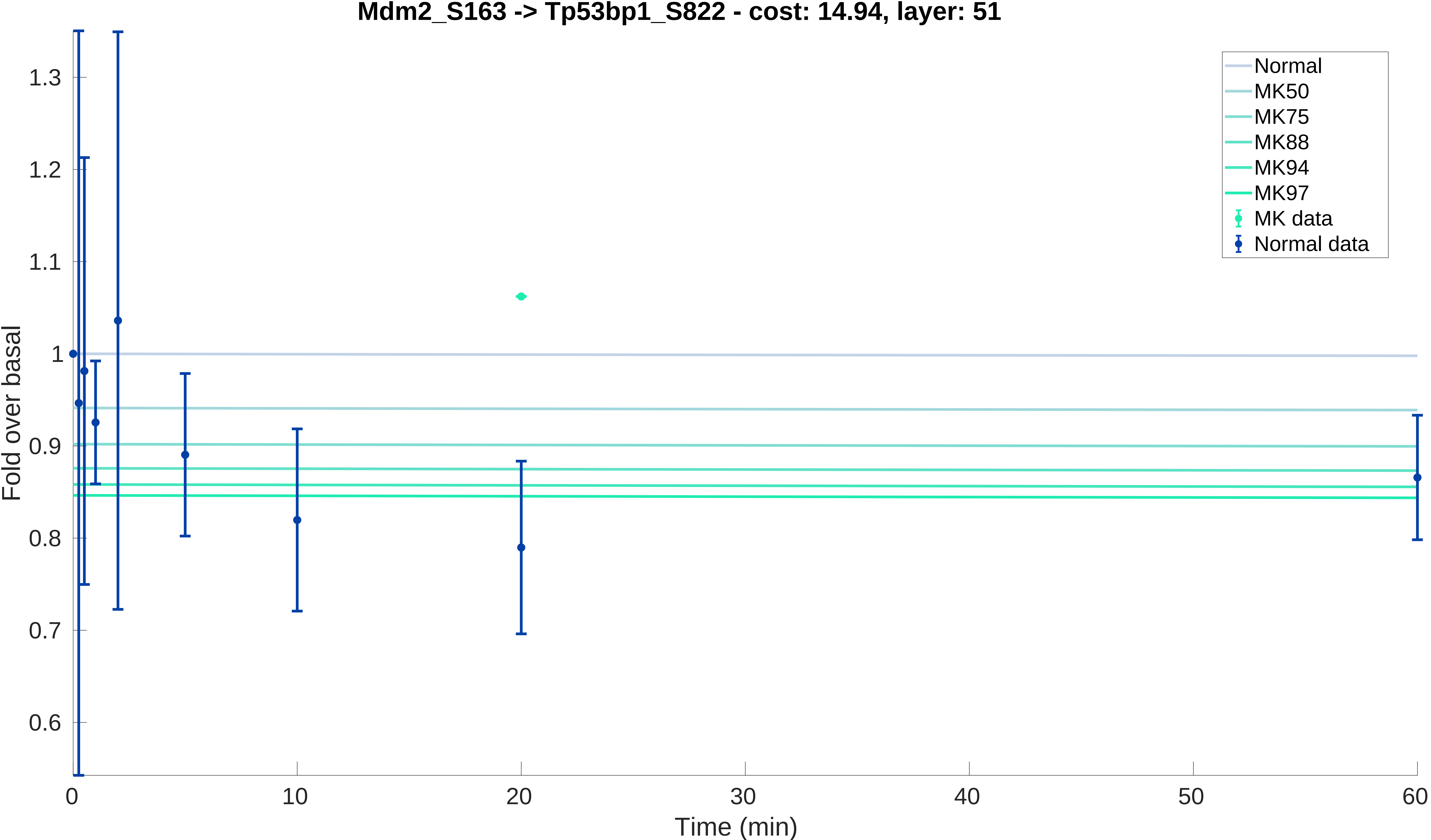

Mdm2_S163 -> Tp53bp1_S822 - cost: 14.94, layer: 51
Fold over basal
Time (min)
0.6
0.7
0.8
0.9
1
1.1
1.2
1.3
0
10
20
30
40
50
60
Normal
MK50
MK75
MK88
MK94
MK97
MK data
Normal data

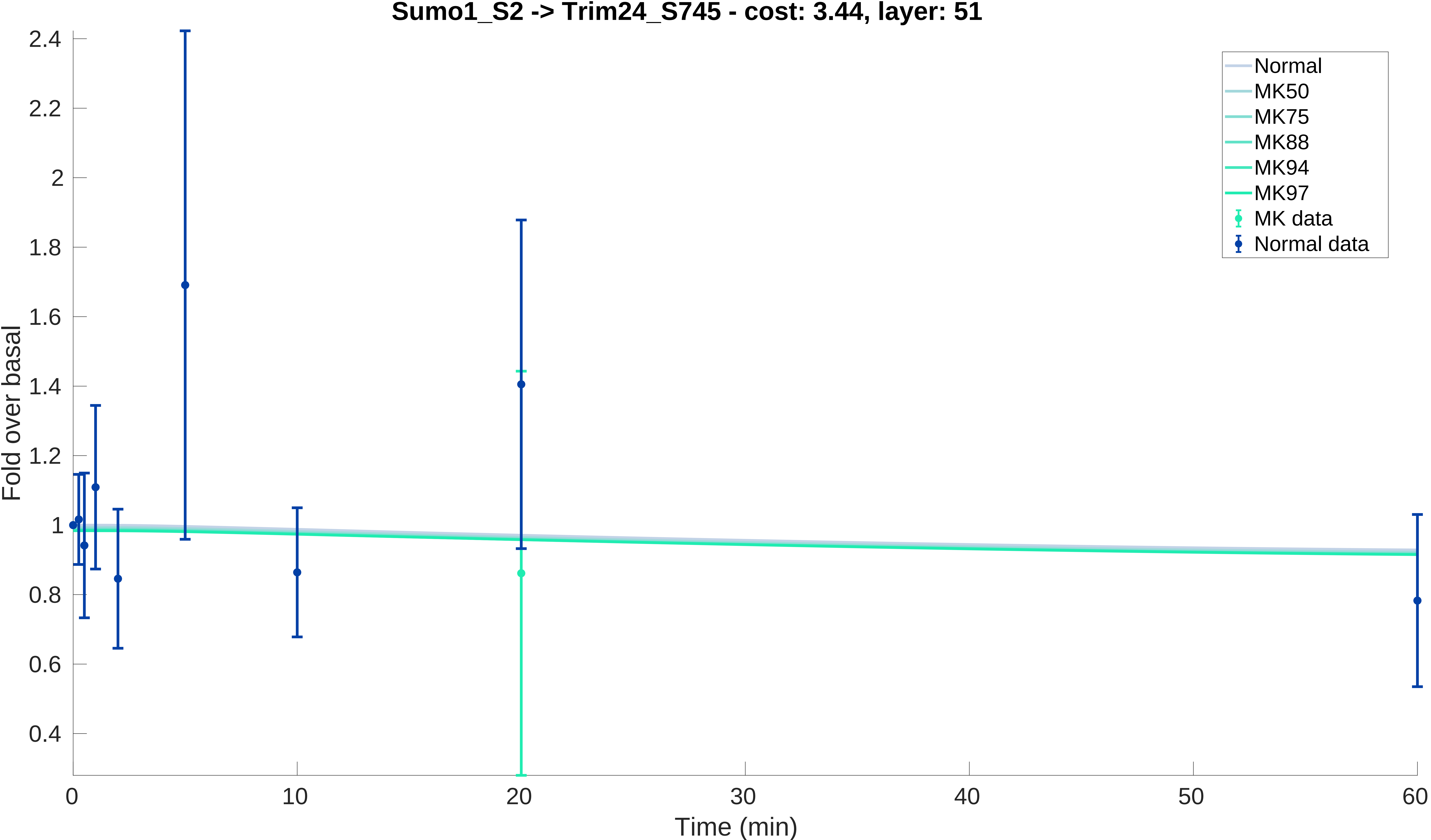

Sumo1_S2 -> Trim24_S745 - cost: 3.44, layer: 51
Normal
MK50
MK75
MK88
MK94
MK97
MK data
Normal data
Fold over basal
Time (min)
0
10
20
30
40
50
60
0.4
0.6
0.8
1
1.2
1.4
1.6
1.8
2
2.2
2.4

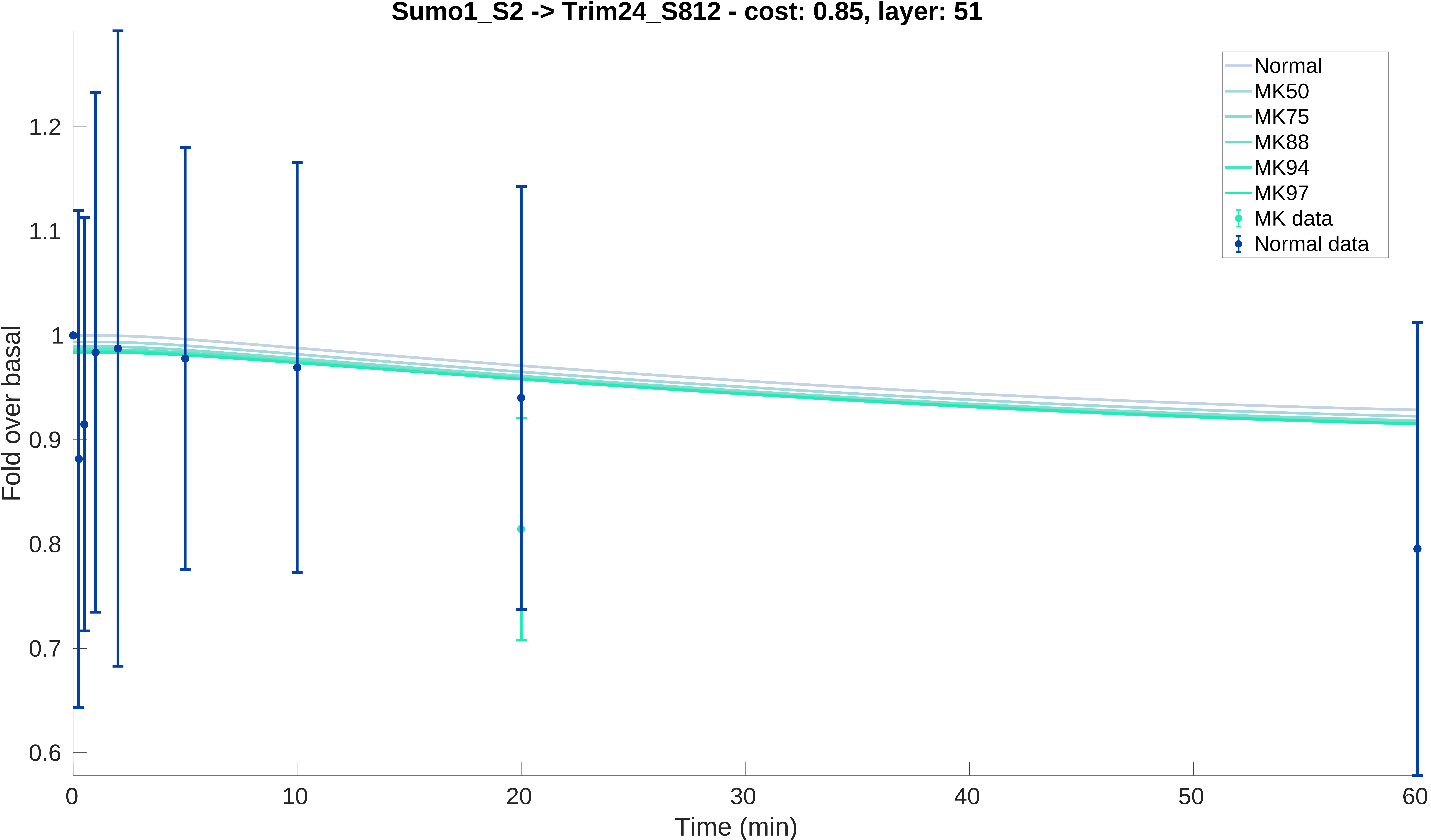

Sumo1_S2 -> Trim24_S812 - cost: 0.85, layer: 51
Normal
MK50
MK75
MK88
MK94
MK97
MK data
Normal data
Fold over basal
Time (min)
0.6
0.7
0.8
0.9
1
1.1
1.2
0
10
20
30
40
50
60

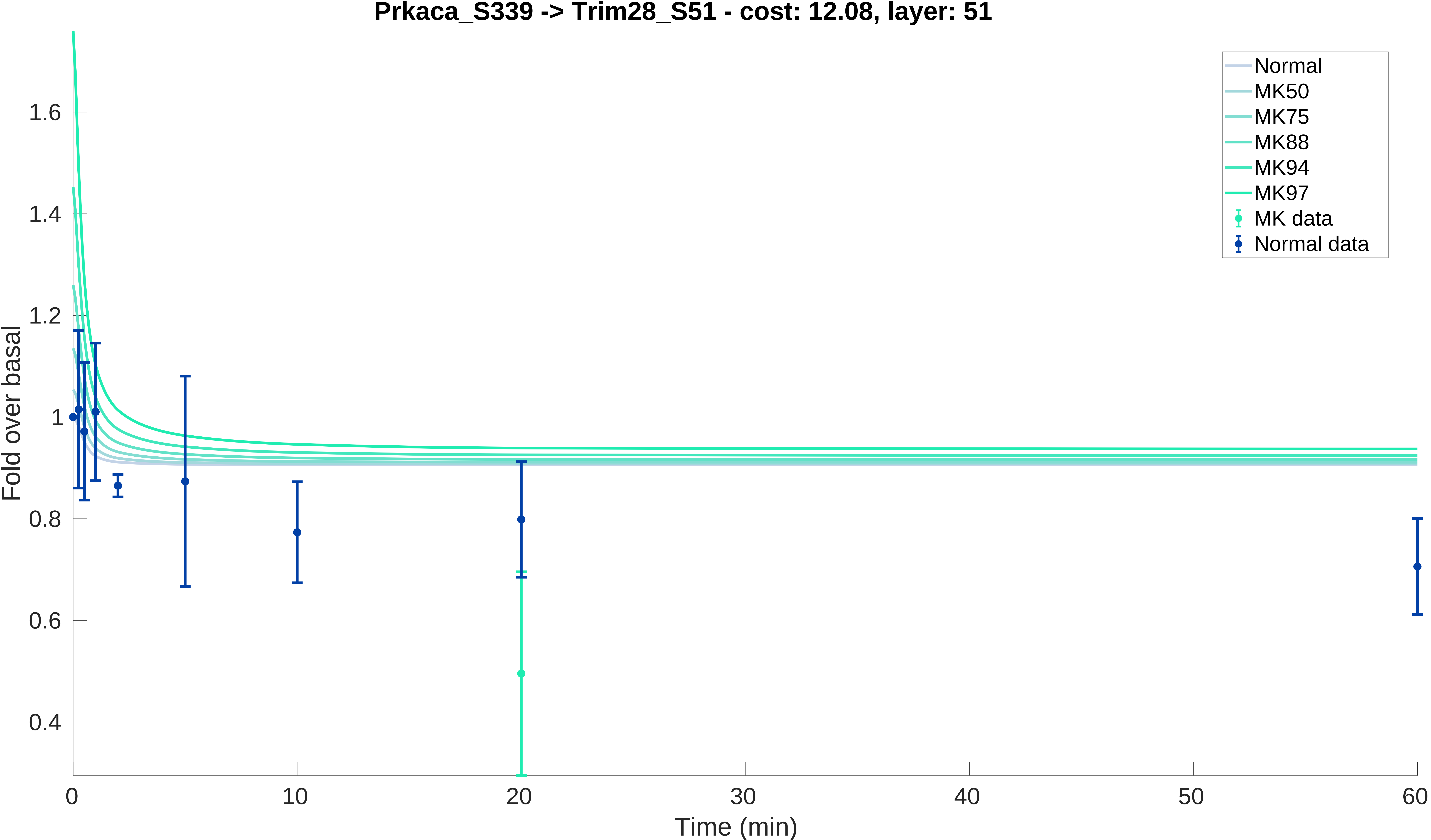

Prkaca_S339 -> Trim28_S51 - cost: 12.08, layer: 51
Fold over basal
1.6
1.4
1.2
1
0.8
0.6
0.4
0
10
20
30
40
50
60
Time (min)
Normal
MK50
MK75
MK88
MK94
MK97
MK data
Normal data

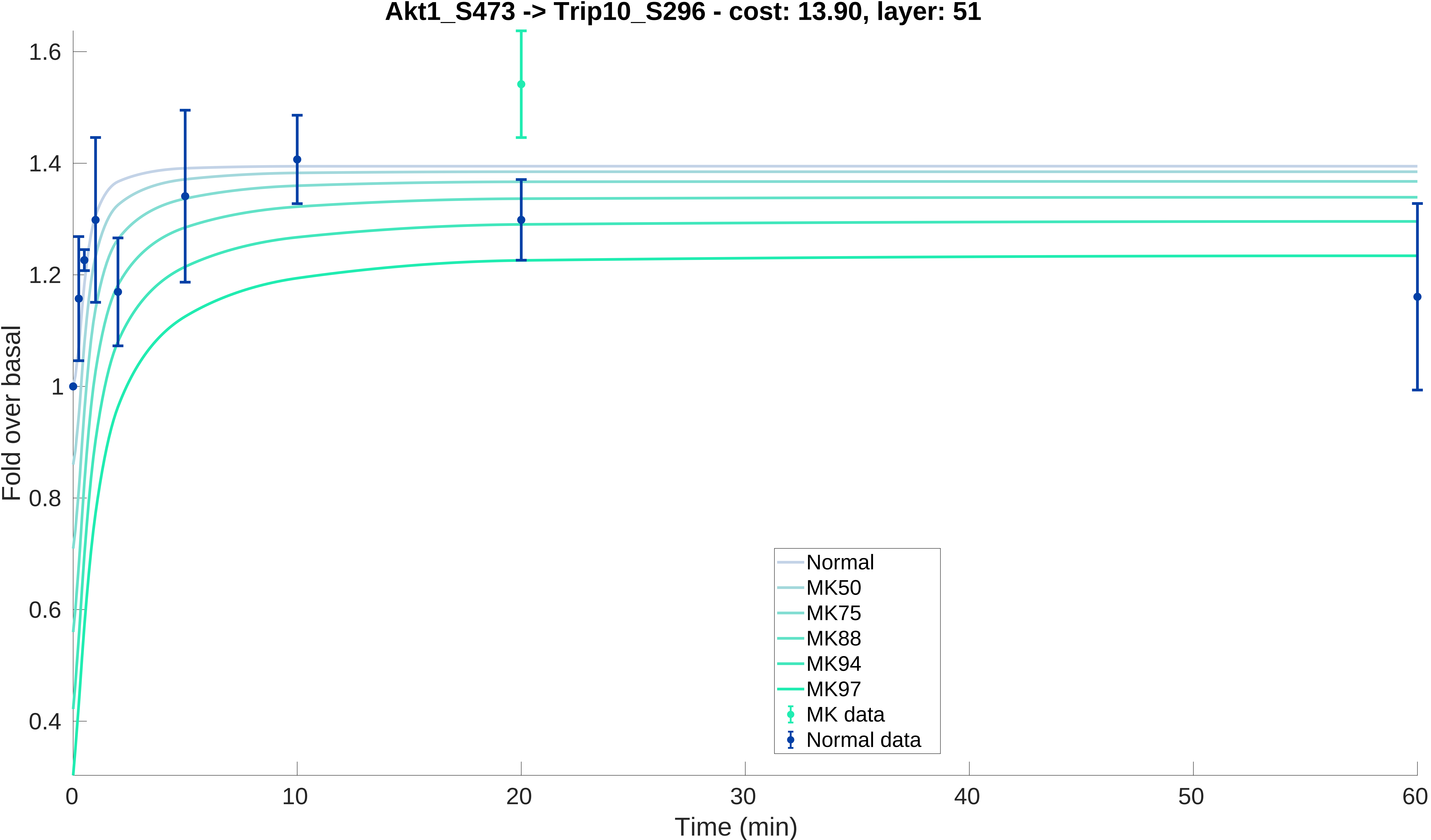

Akt1_S473 -> Trip10_S296 - cost: 13.90, layer: 51
Fold over basal
1.6
1.4
1.2
1
0.8
0.6
0.4
0
10
20
30
40
50
60
Time (min)
Normal
MK50
MK75
MK88
MK94
MK97
MK data
Normal data

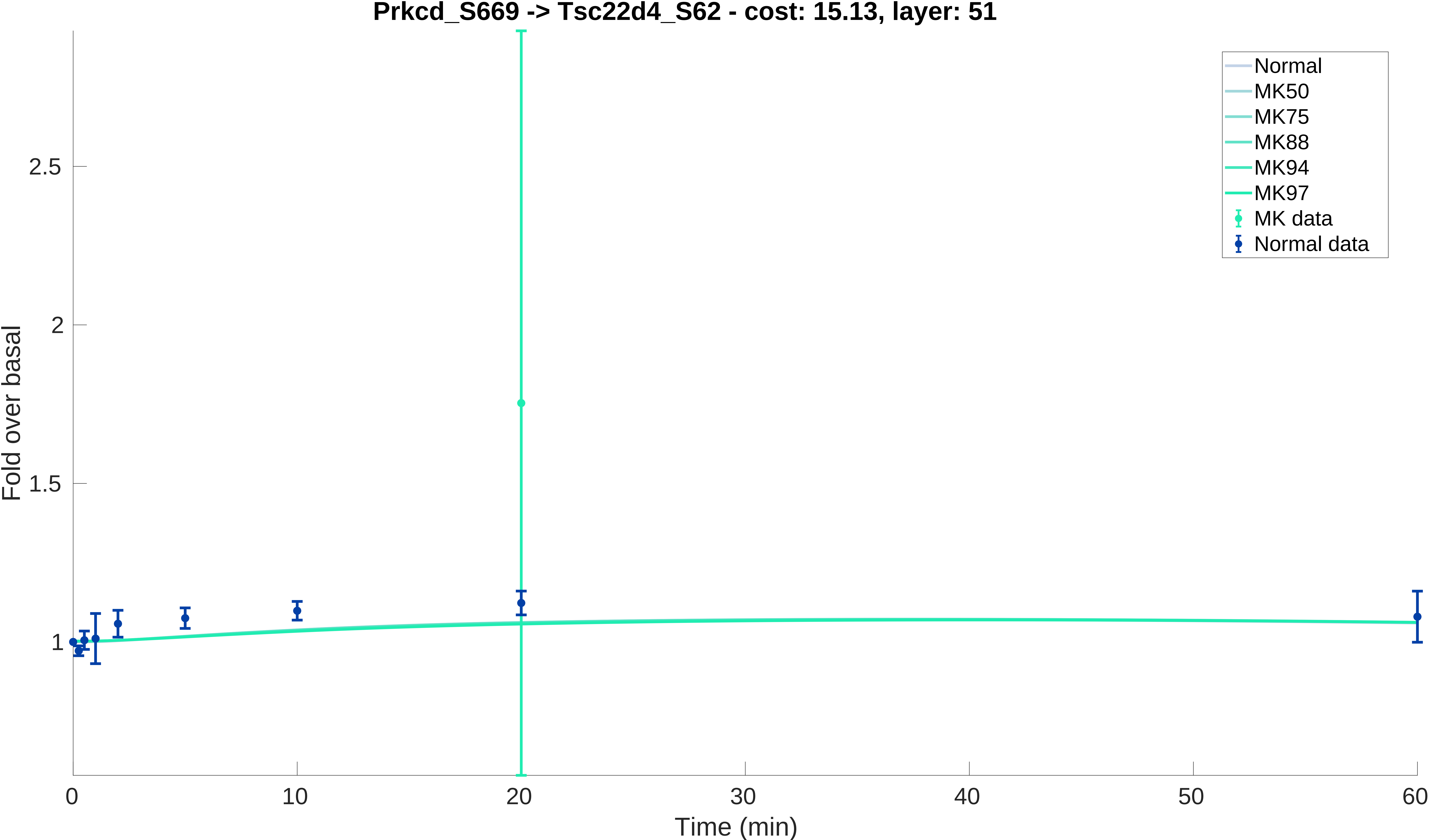

Prkcd_S669 -> Tsc22d4_S62 - cost: 15.13, layer: 51
Normal
MK50
MK75
MK88
MK94
MK97
MK data
Normal data
Fold over basal
2.5
2
1.5
1
0
10
20
30
40
50
60
Time (min)

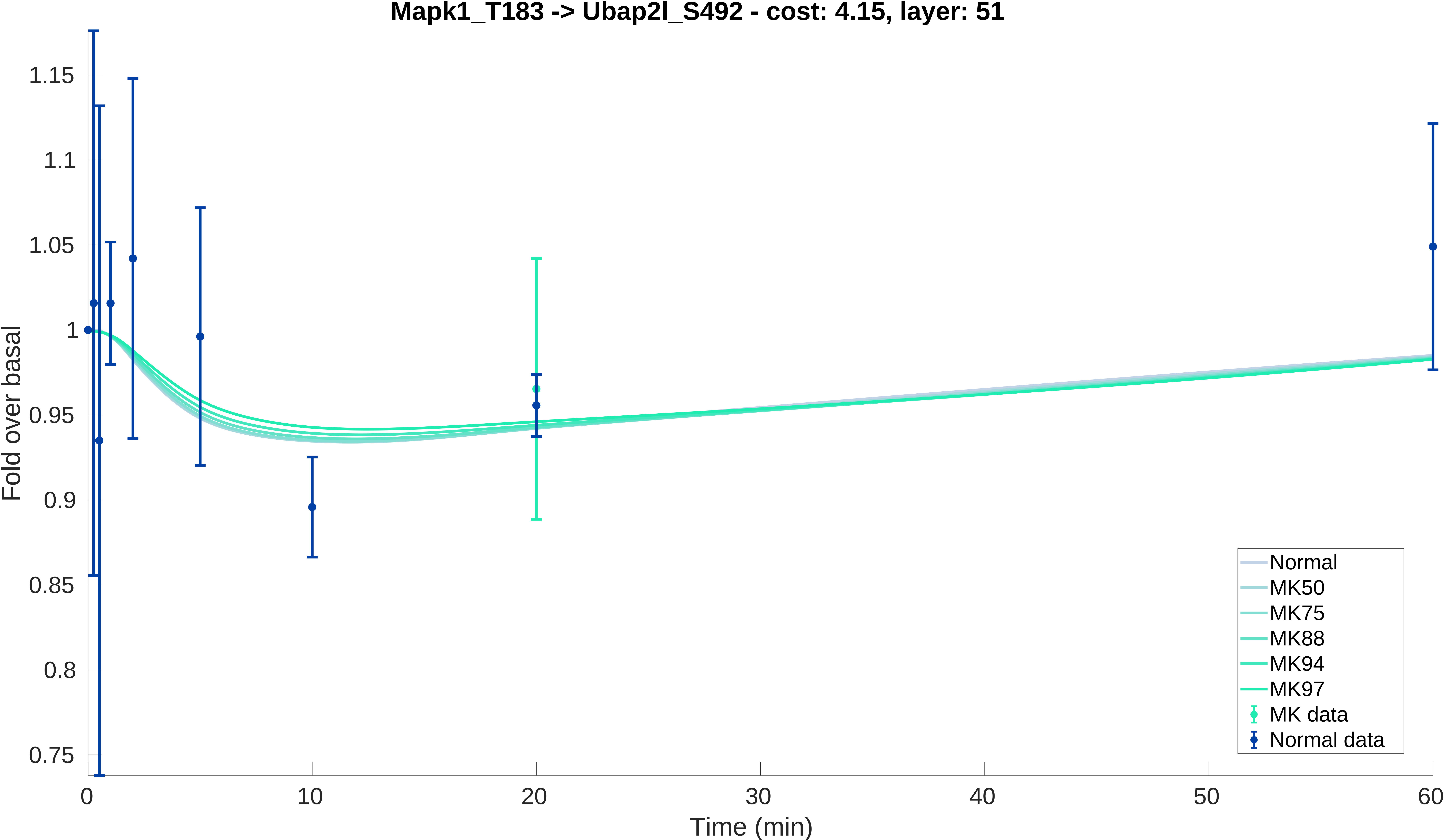

Mapk1_T183 -> Ubap2l_S492 - cost: 4.15, layer: 51
Fold over basal
1.15
1.1
1.05
1
0.95
0.9
0.85
0.8
0.75
0
10
20
30
40
50
60
Time (min)
Normal
MK50
MK75
MK88
MK94
MK97
MK data
Normal data

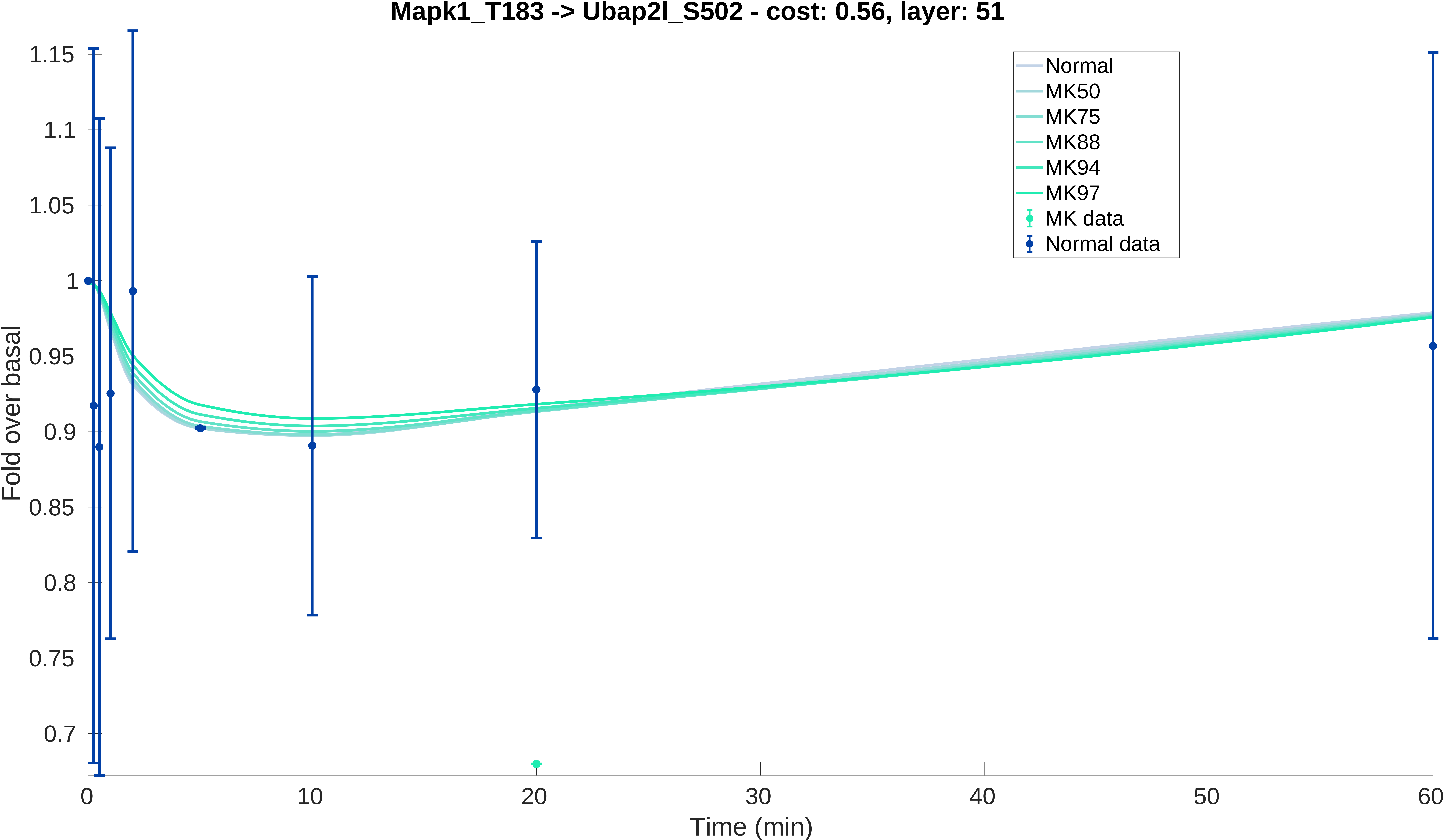
Mapk1_T183 -> Ubap2l_S502 - cost: 0.56, layer: 51
Fold over basal
Time (min)
1.15
1.1
1.05
1
0.95
0.9
0.85
0.8
0.75
0.7
0
10
20
30
40
50
60
Normal
MK50
MK75
MK88
MK94
MK97
MK data
Normal data

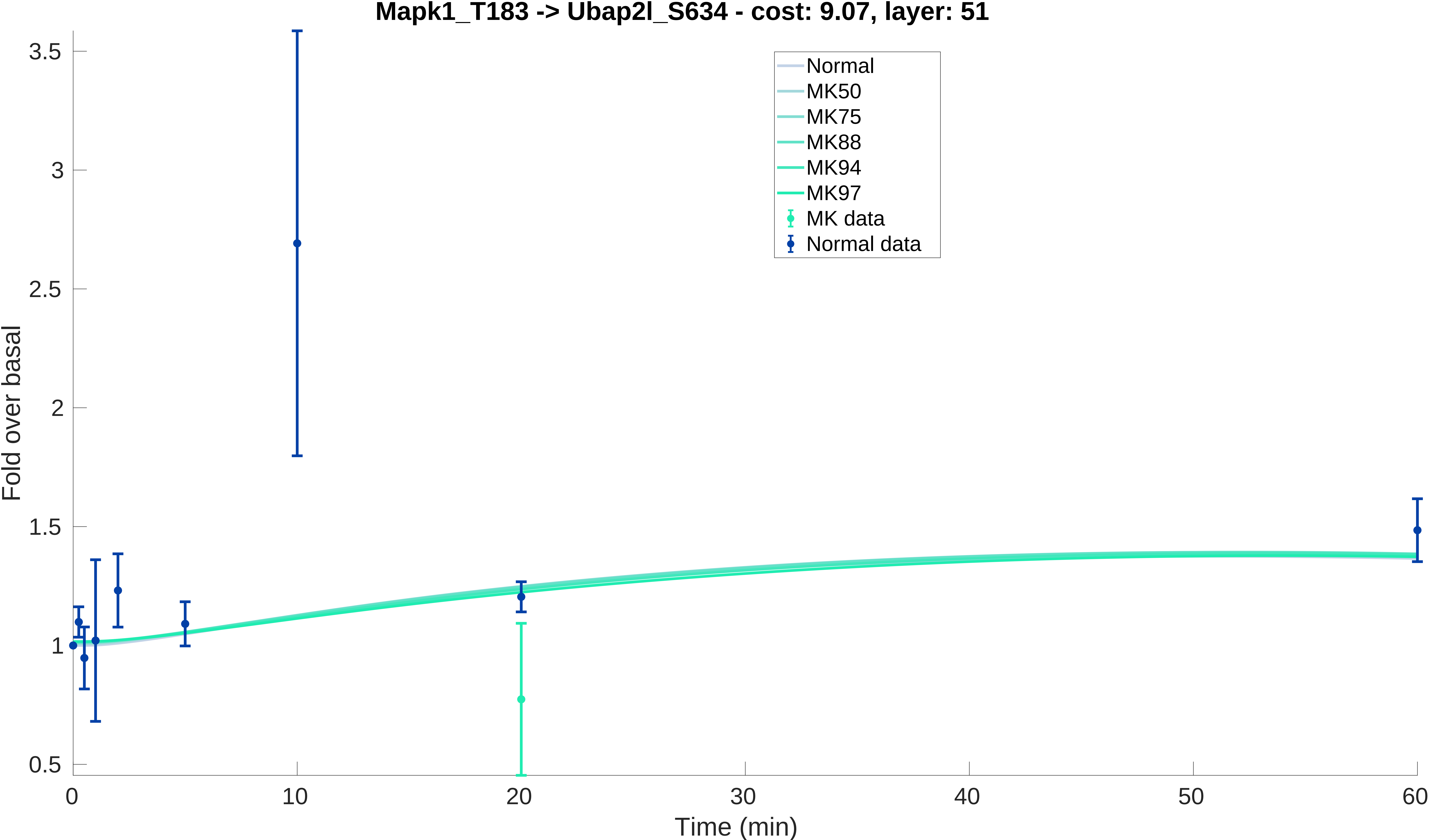

Mapk1_T183 -> Ubap2l_S634 - cost: 9.07, layer: 51
Normal
MK50
MK75
MK88
MK94
MK97
MK data
Normal data
Fold over basal
3.5
3
2.5
2
1.5
1
0.5
0
10
20
30
40
50
60
Time (min)

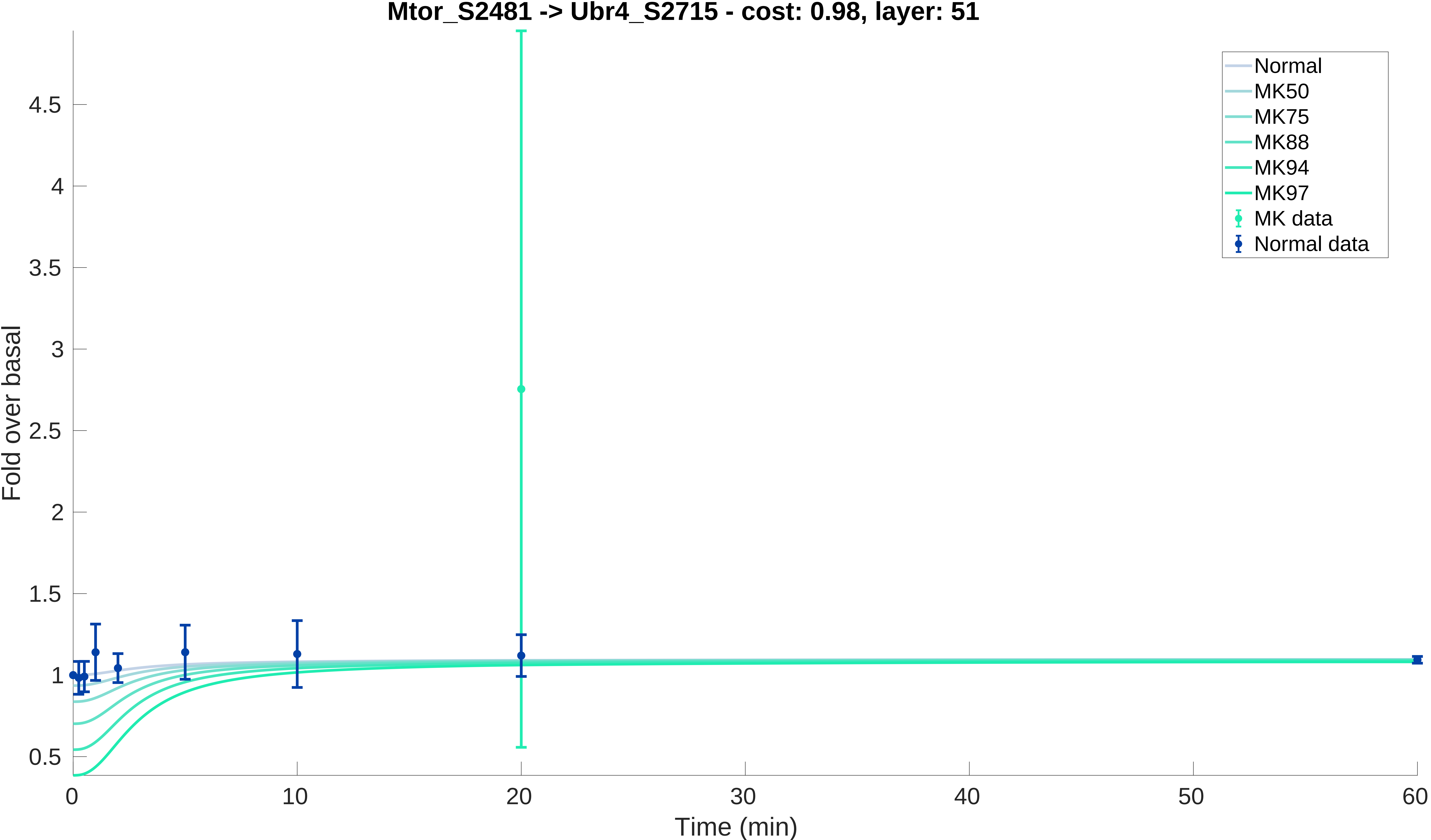
Mtor_S2481 -> Ubr4_S2715 - cost: 0.98, layer: 51
Normal
MK50
MK75
MK88
MK94
MK97
MK data
Normal data
Fold over basal
0.5
1
1.5
2
2.5
3
3.5
4
4.5
0
10
20
30
40
50
60
Time (min)

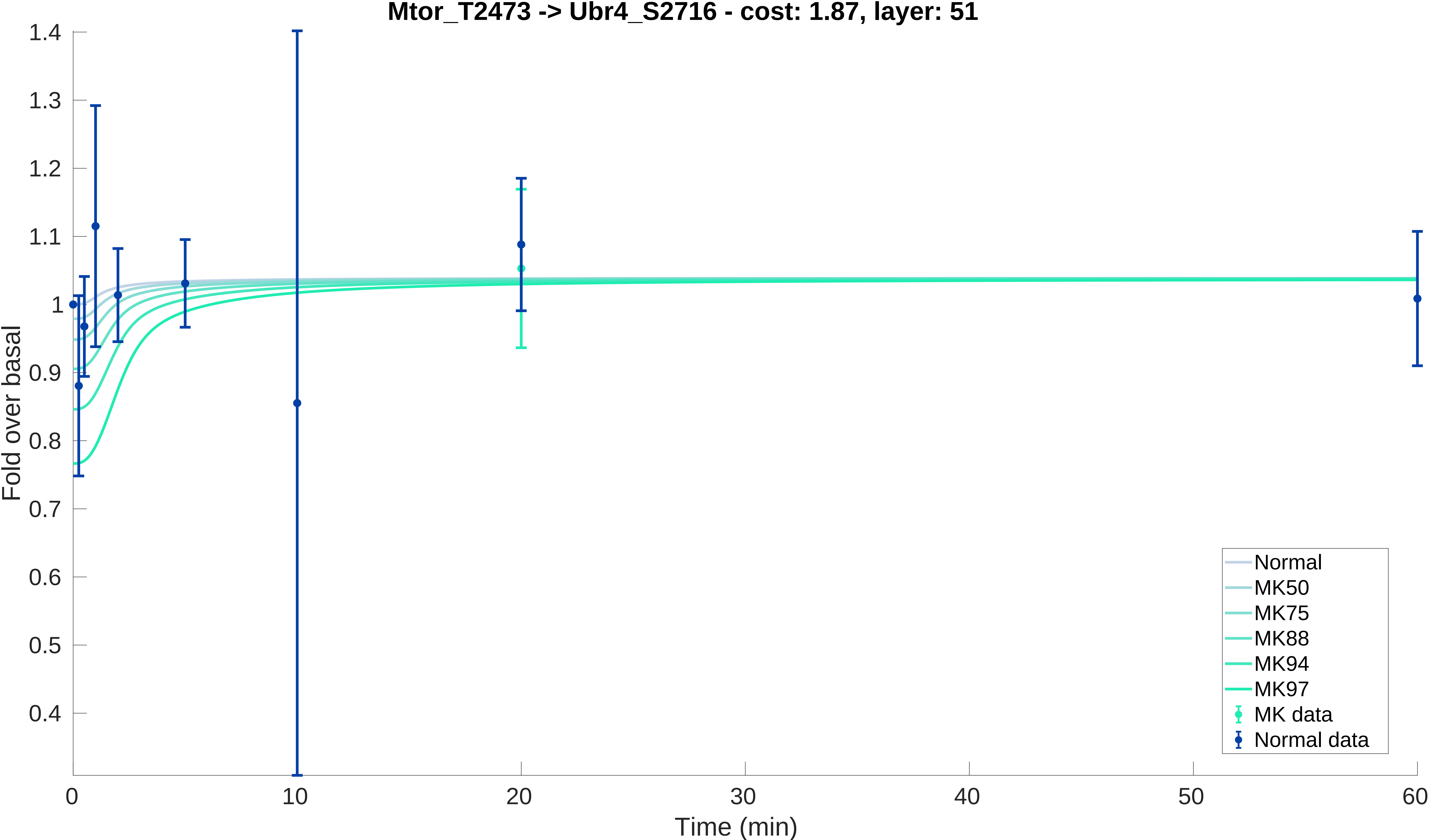

Mtor_T2473 -> Ubr4_S2716 - cost: 1.87, layer: 51
Fold over basal
1.4
1.3
1.2
1.1
1
0.9
0.8
0.7
0.6
0.5
0.4
0
10
20
30
40
50
60
Time (min)
Normal
MK50
MK75
MK88
MK94
MK97
MK data
Normal data

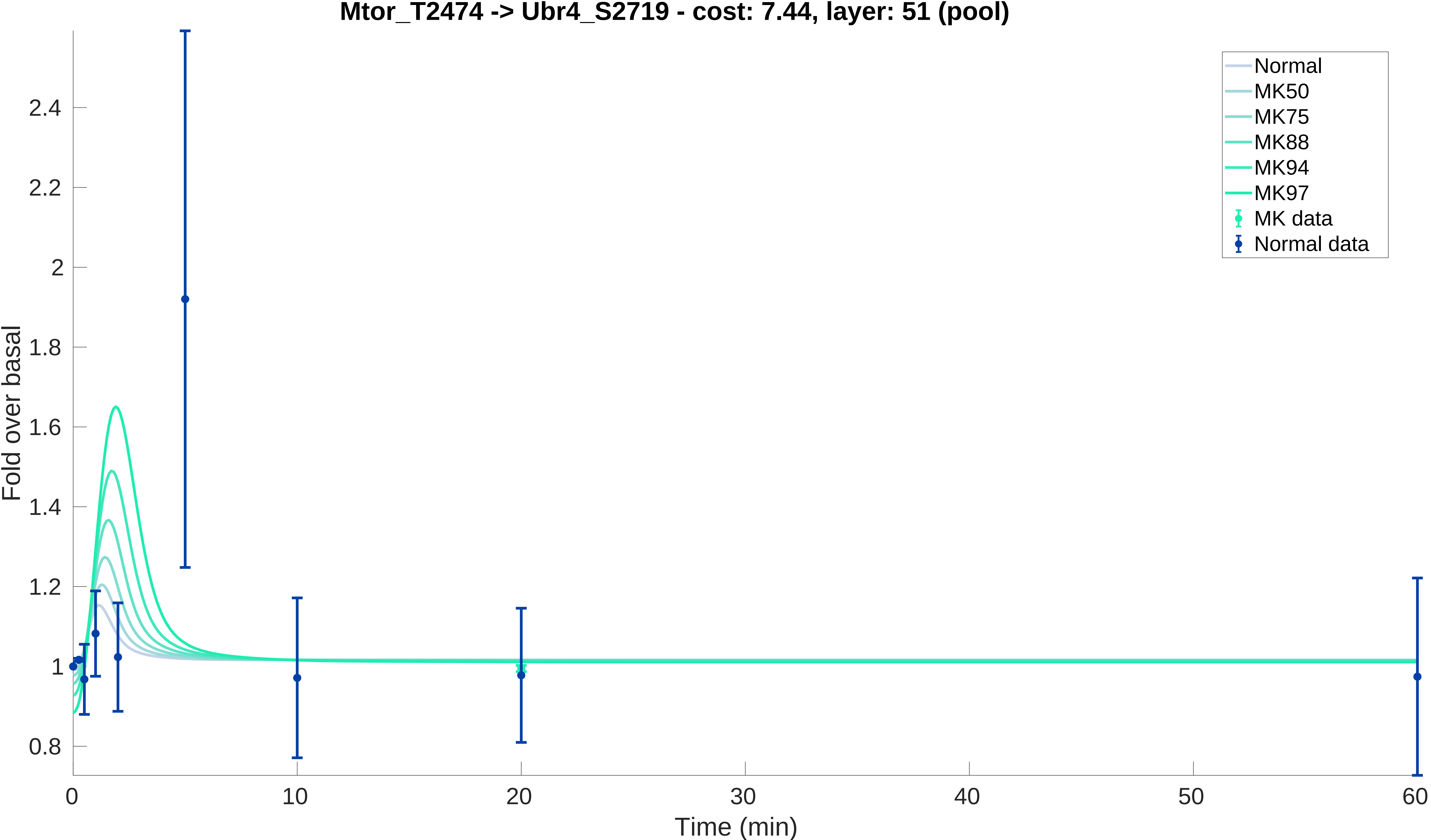
Mtor_T2474 -> Ubr4_S2719 - cost: 7.44, layer: 51 (pool)
Fold over basal
Time (min)
0.8
1
1.2
1.4
1.6
1.8
2
2.2
2.4
0
10
20
30
40
50
60
Normal
MK50
MK75
MK88
MK94
MK97
MK data
Normal data

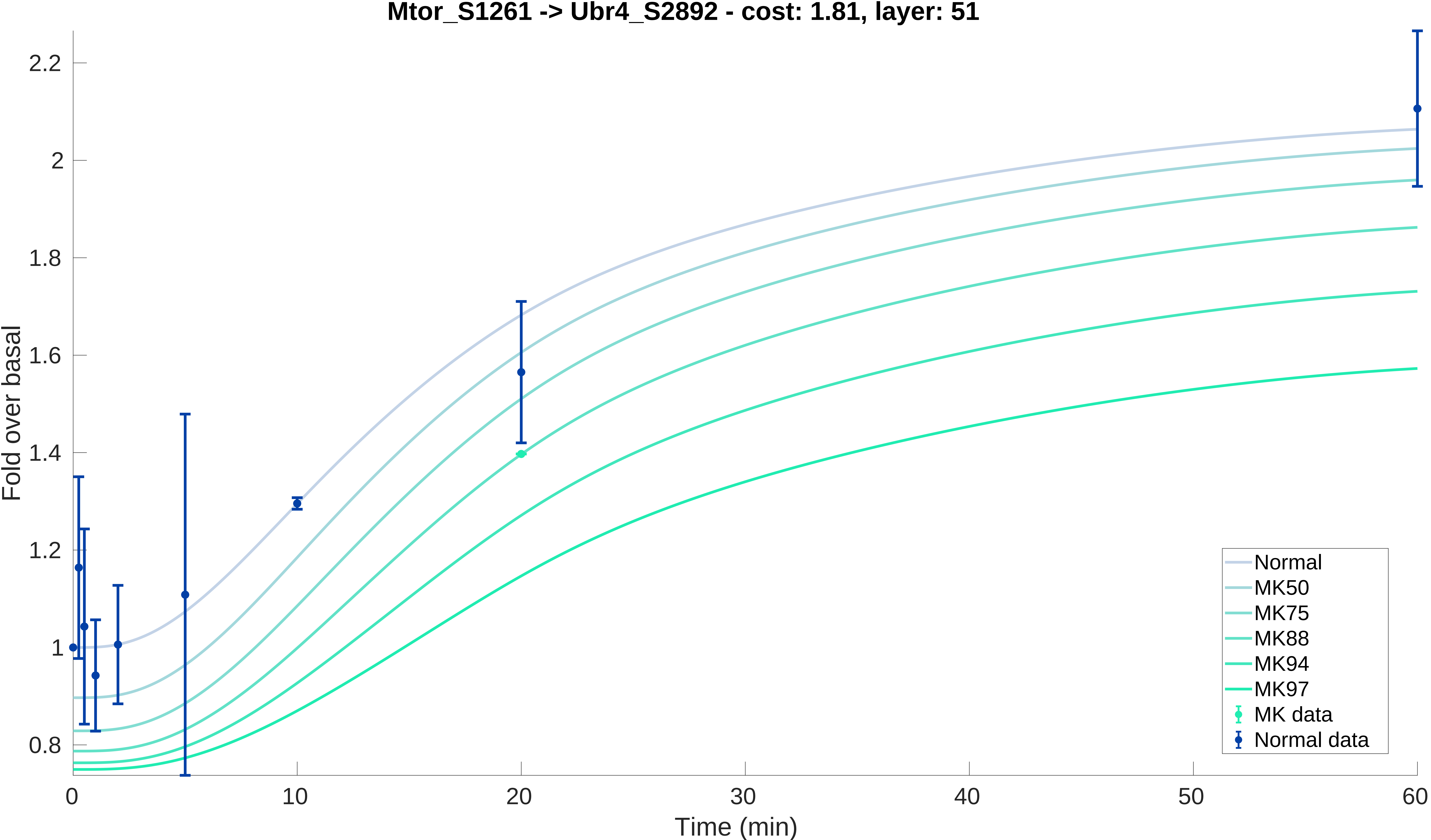

Mtor_S1261 -> Ubr4_S2892 - cost: 1.81, layer: 51
Fold over basal
Time (min)
2.2
2
1.8
1.6
1.4
1.2
1
0.8
0
10
20
30
40
50
60
Normal
MK50
MK75
MK88
MK94
MK97
MK data
Normal data

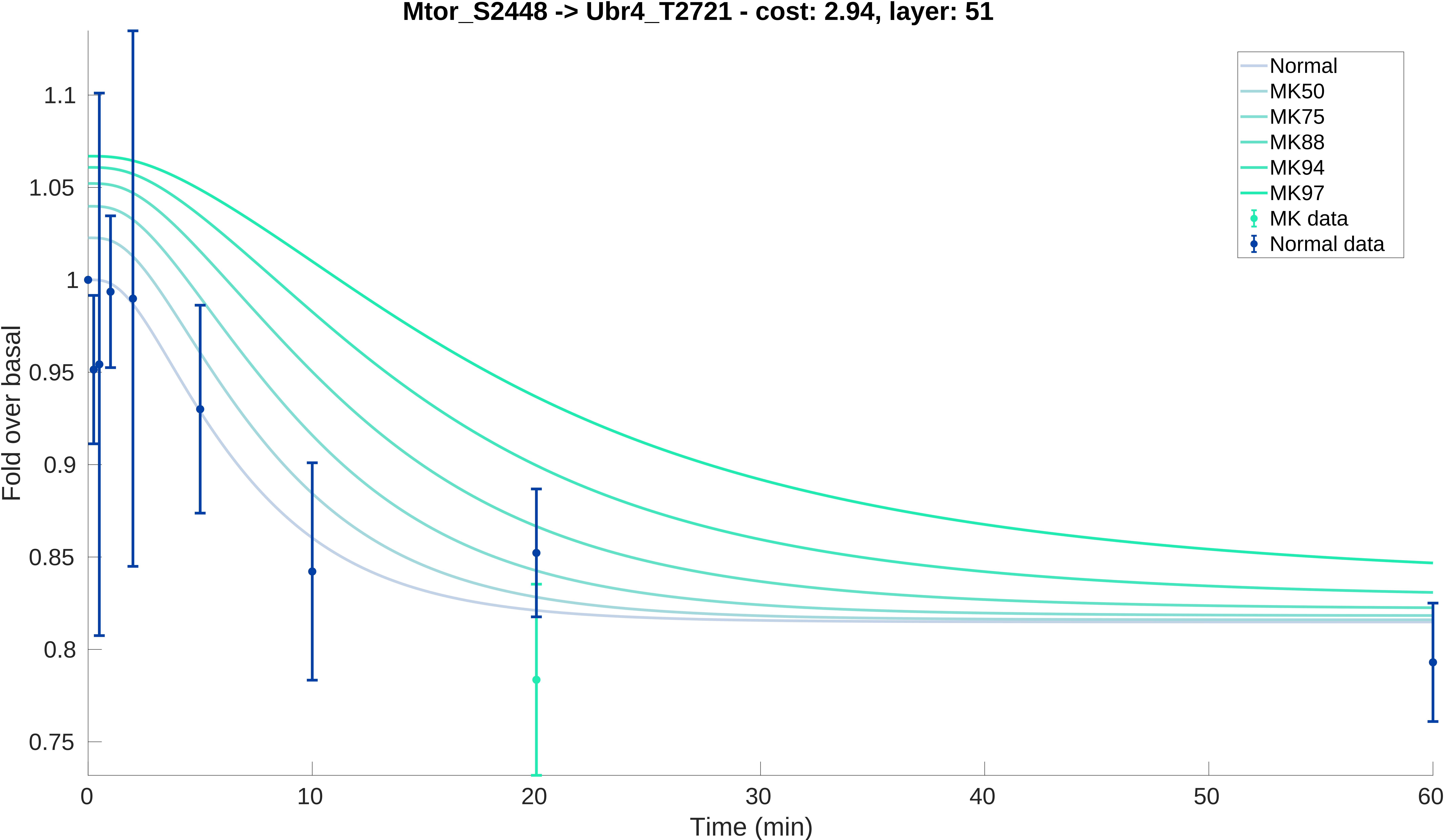

Mtor_S2448 -> Ubr4_T2721 - cost: 2.94, layer: 51
Fold over basal
1.1
1.05
1
0.95
0.9
0.85
0.8
0.75
0
10
20
30
40
50
60
Time (min)
Normal
MK50
MK75
MK88
MK94
MK97
MK data
Normal data

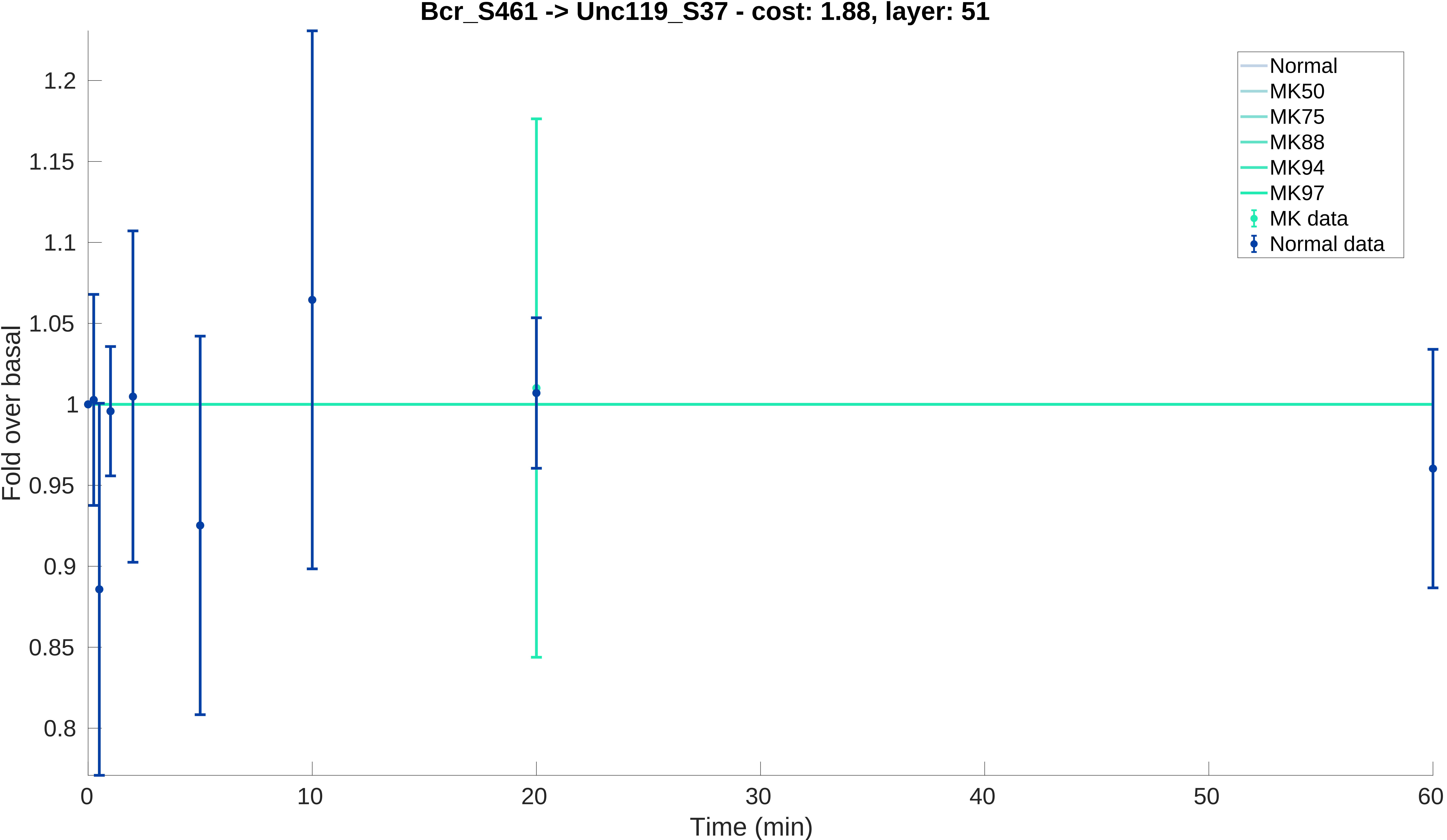

Bcr_S461 -> Unc119_S37 - cost: 1.88, layer: 51
Fold over basal
Time (min)
1.2
1.15
1.1
1.05
1
0.95
0.9
0.85
0.8
0
10
20
30
40
50
60
Normal
MK50
MK75
MK88
MK94
MK97
MK data
Normal data

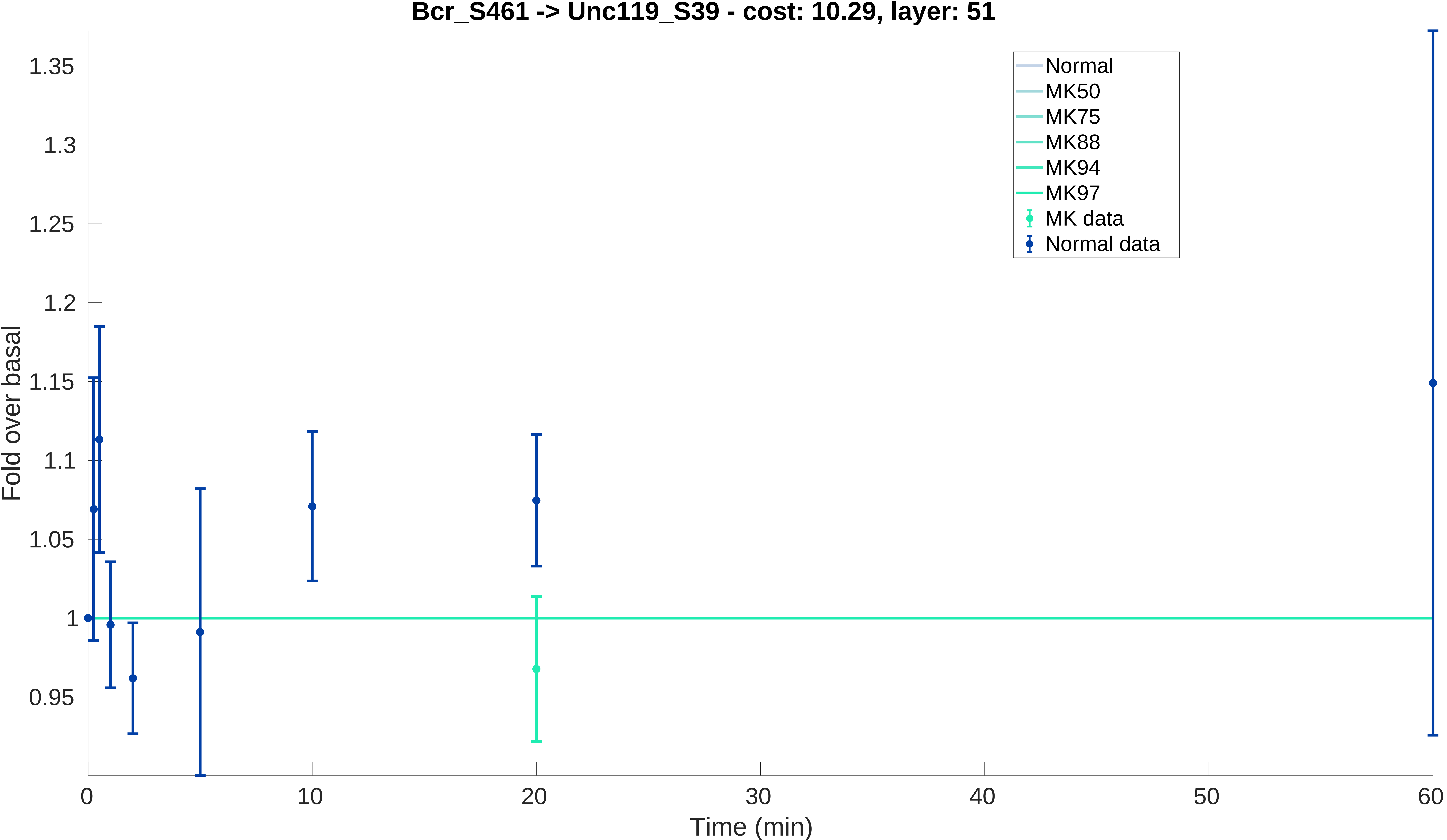

Bcr_S461 -> Unc119_S39 - cost: 10.29, layer: 51
Normal
MK50
MK75
MK88
MK94
MK97
MK data
Normal data
Fold over basal
Time (min)
1.35
1.3
1.25
1.2
1.15
1.1
1.05
1
0.95
0
10
20
30
40
50
60

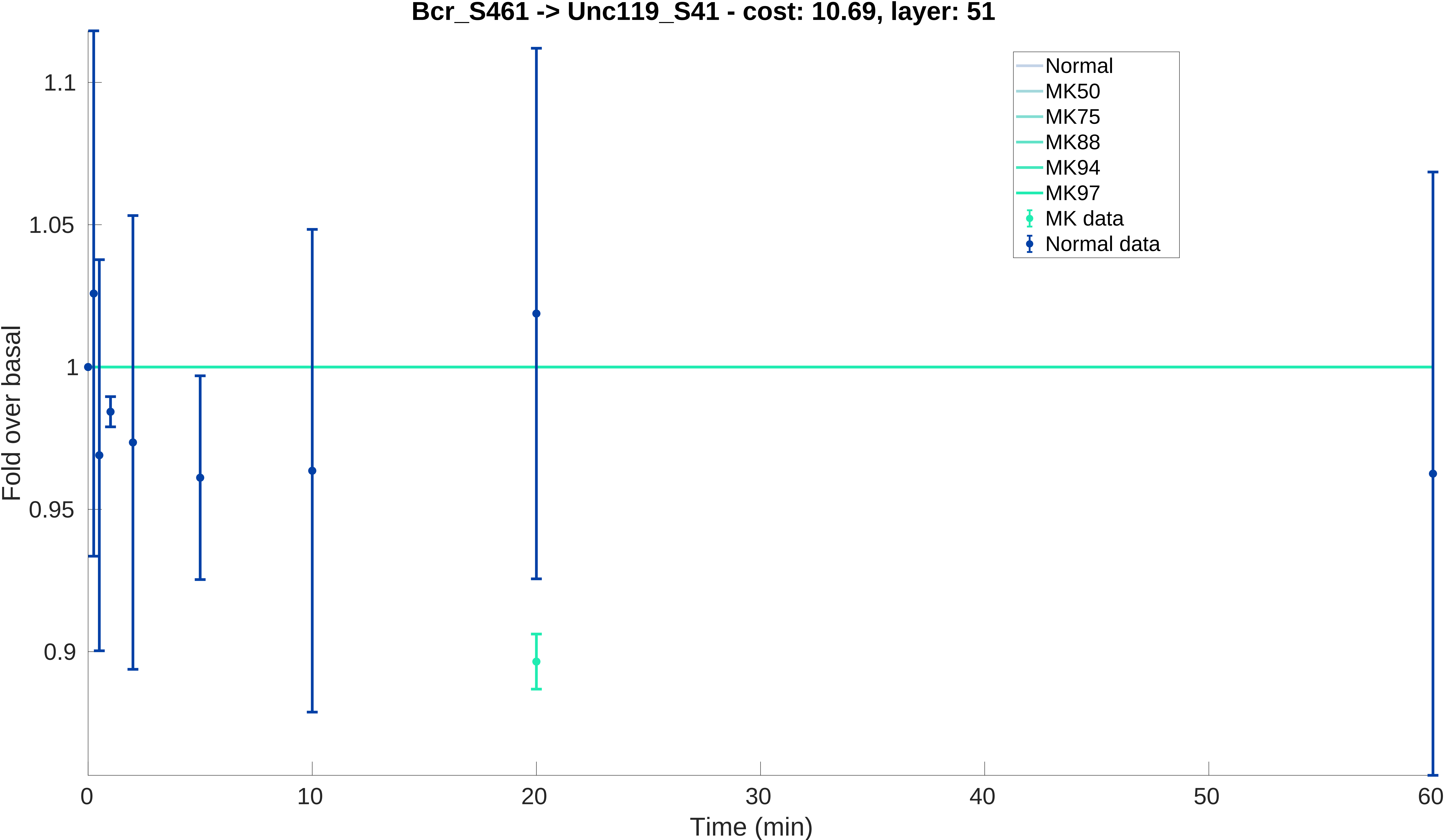

Bcr_S461 -> Unc119_S41 - cost: 10.69, layer: 51
Normal
MK50
MK75
MK88
MK94
MK97
MK data
Normal data
Fold over basal
1.1
1.05
1
0.95
0.9
0
10
20
30
40
50
60
Time (min)

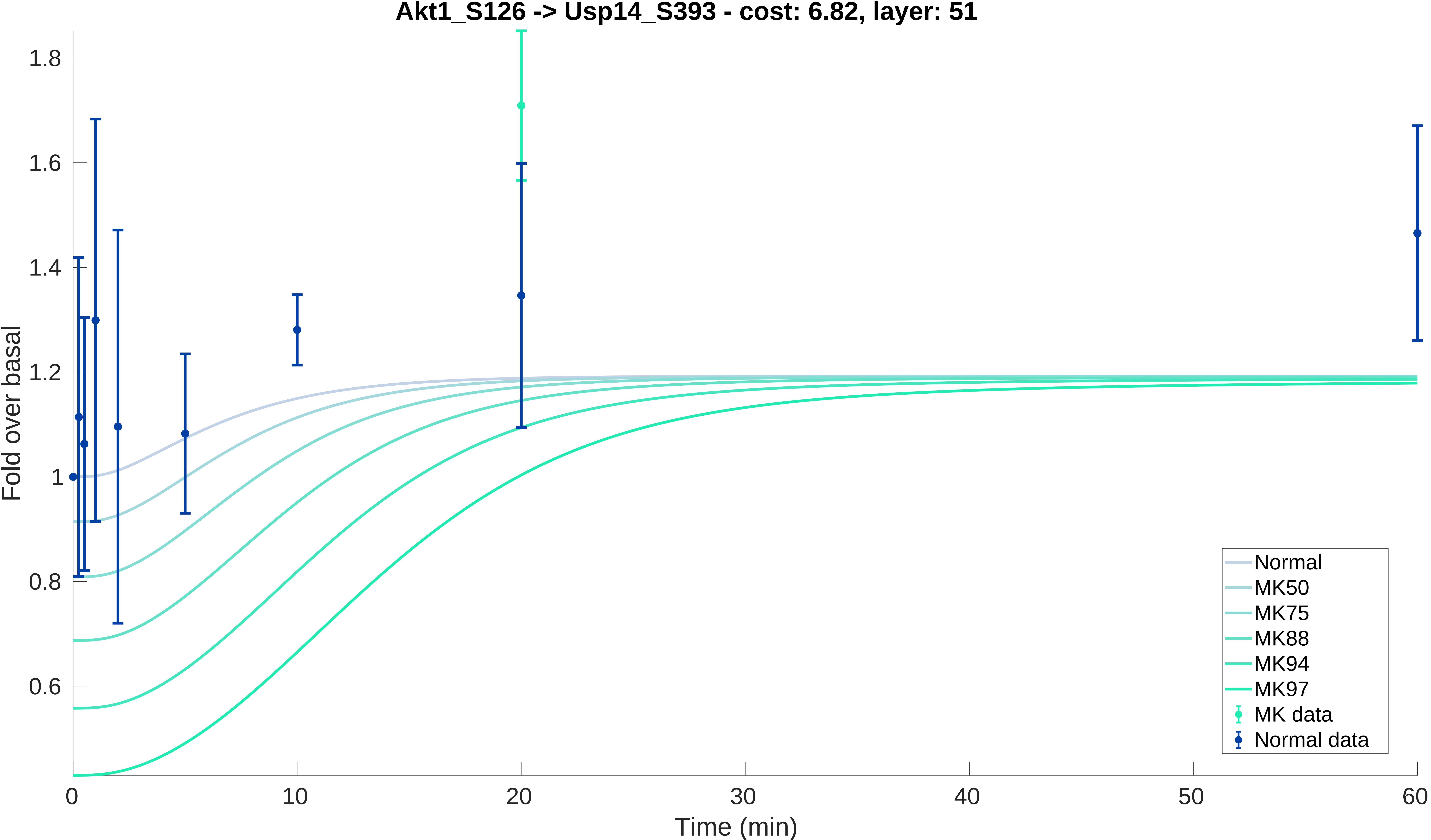
Akt1_S126 -> Usp14_S393 - cost: 6.82, layer: 51
Fold over basal
Time (min)
0.6
0.8
1
1.2
1.4
1.6
1.8
0
10
20
30
40
50
60
Normal
MK50
MK75
MK88
MK94
MK97
MK data
Normal data

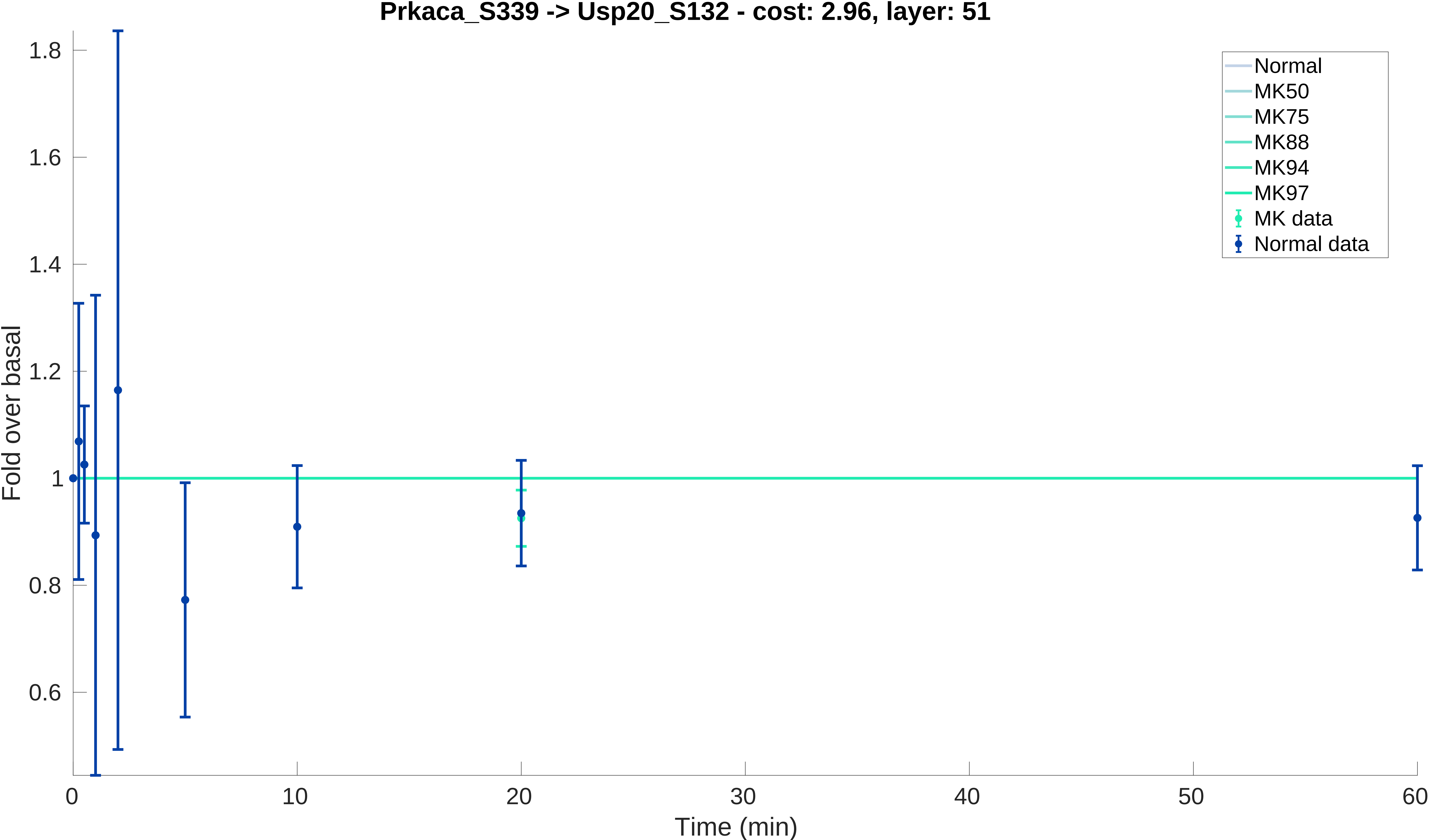

Prkaca_S339 -> Usp20_S132 - cost: 2.96, layer: 51
Fold over basal
1.8
1.6
1.4
1.2
1
0.8
0.6
0
10
20
30
40
50
60
Time (min)
Normal
MK50
MK75
MK88
MK94
MK97
MK data
Normal data

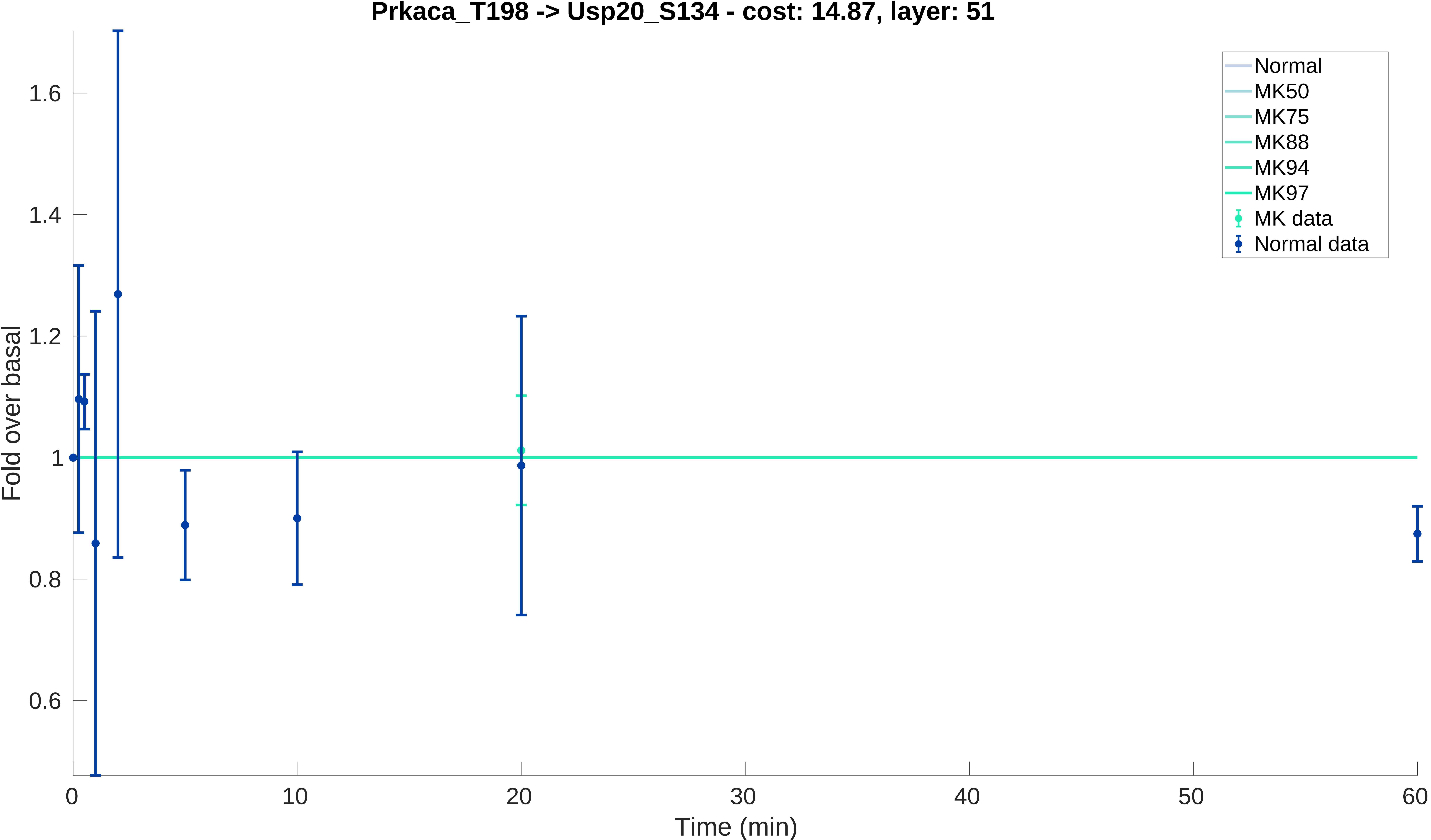

Prkaca_T198 -> Usp20_S134 - cost: 14.87, layer: 51
Fold over basal
1.6
1.4
1.2
1
0.8
0.6
0
10
20
30
40
50
60
Time (min)
Normal
MK50
MK75
MK88
MK94
MK97
MK data
Normal data

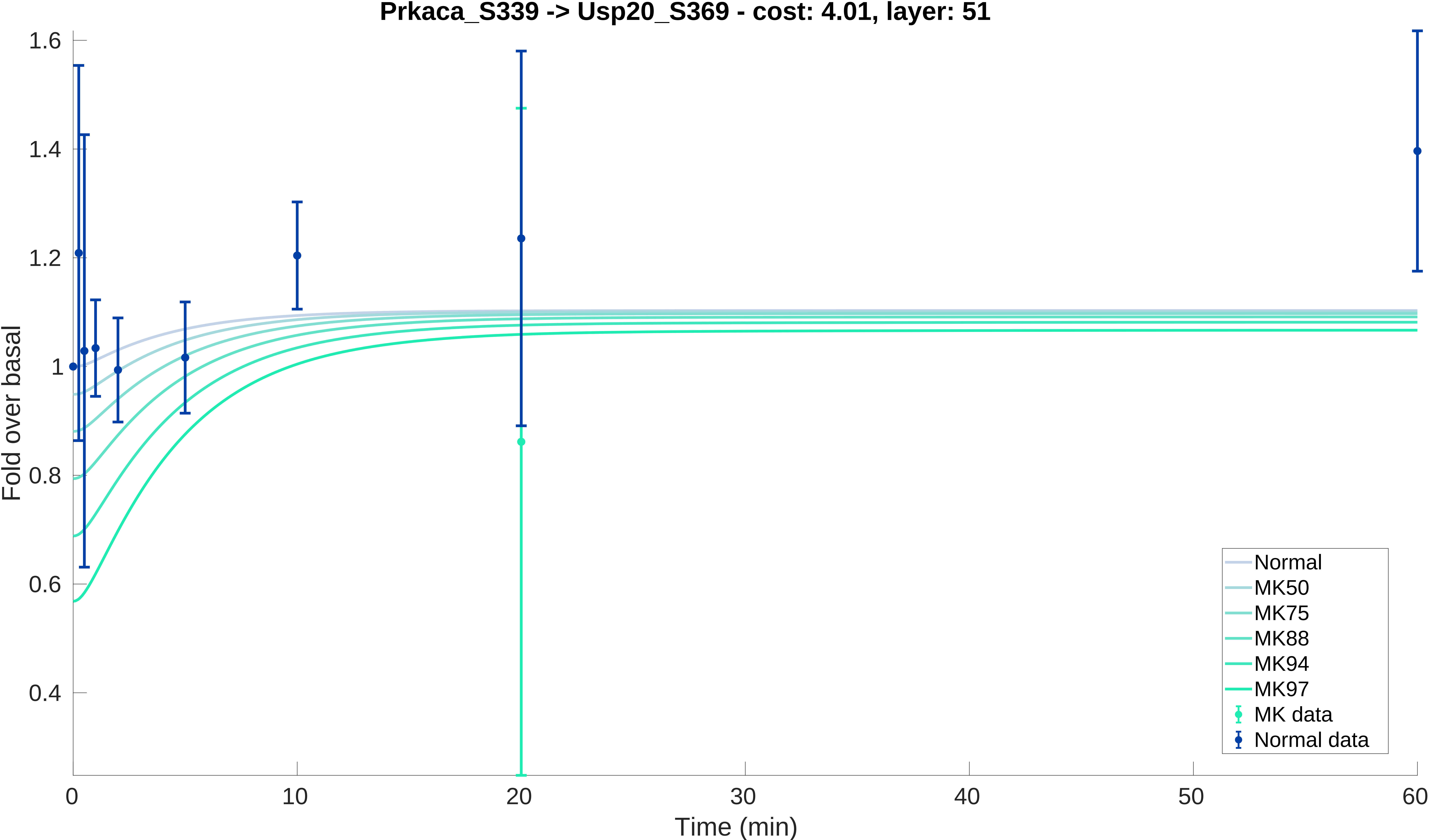

Prkaca_S339 -> Usp20_S369 - cost: 4.01, layer: 51
Fold over basal
Time (min)
1.6
1.4
1.2
1
0.8
0.6
0.4
0
10
20
30
40
50
60
Normal
MK50
MK75
MK88
MK94
MK97
MK data
Normal data

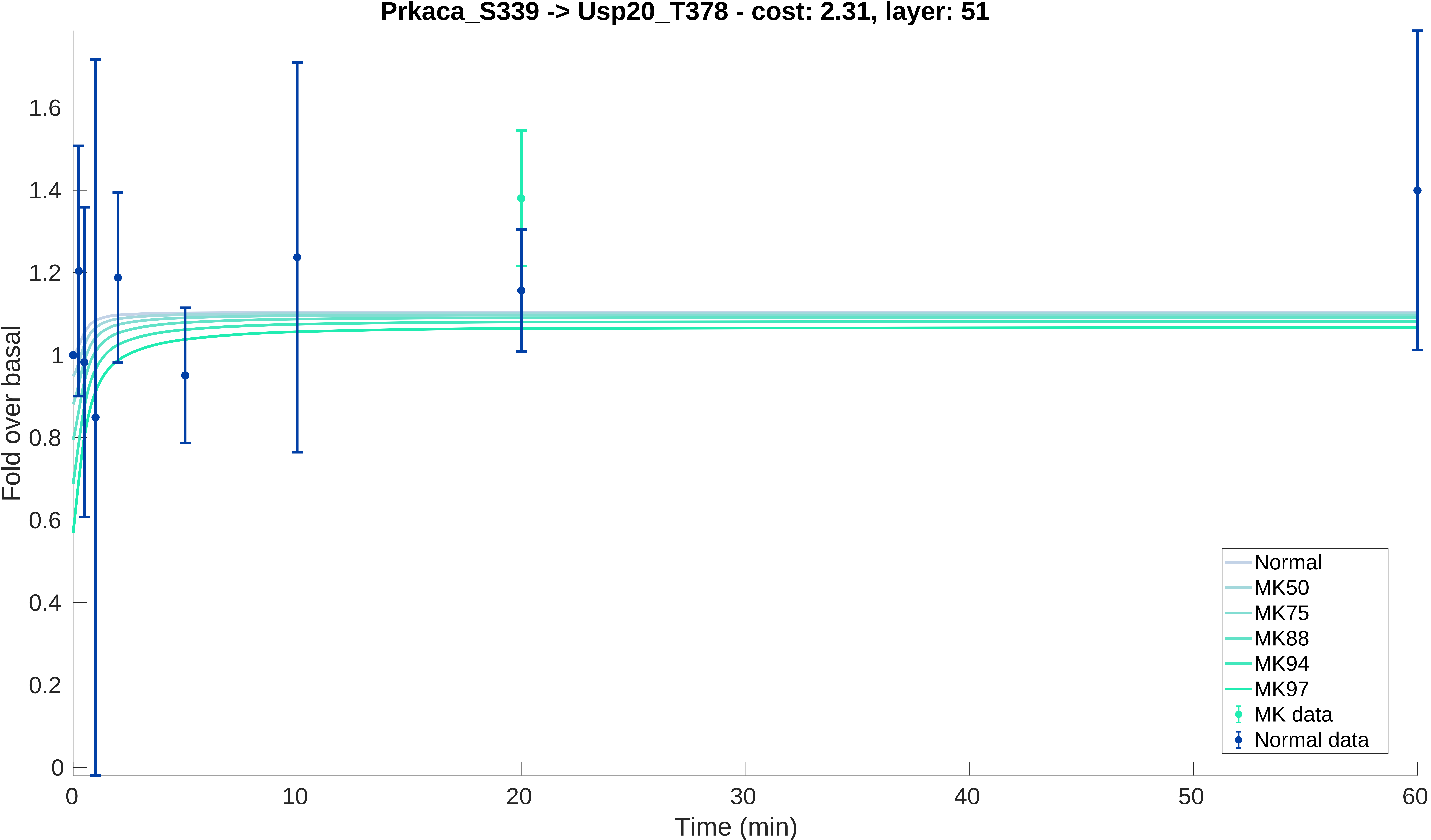
Prkaca_S339 -> Usp20_T378 - cost: 2.31, layer: 51
Fold over basal
Time (min)
0
0.2
0.4
0.6
0.8
1
1.2
1.4
1.6
0
10
20
30
40
50
60
Normal
MK50
MK75
MK88
MK94
MK97
MK data
Normal data

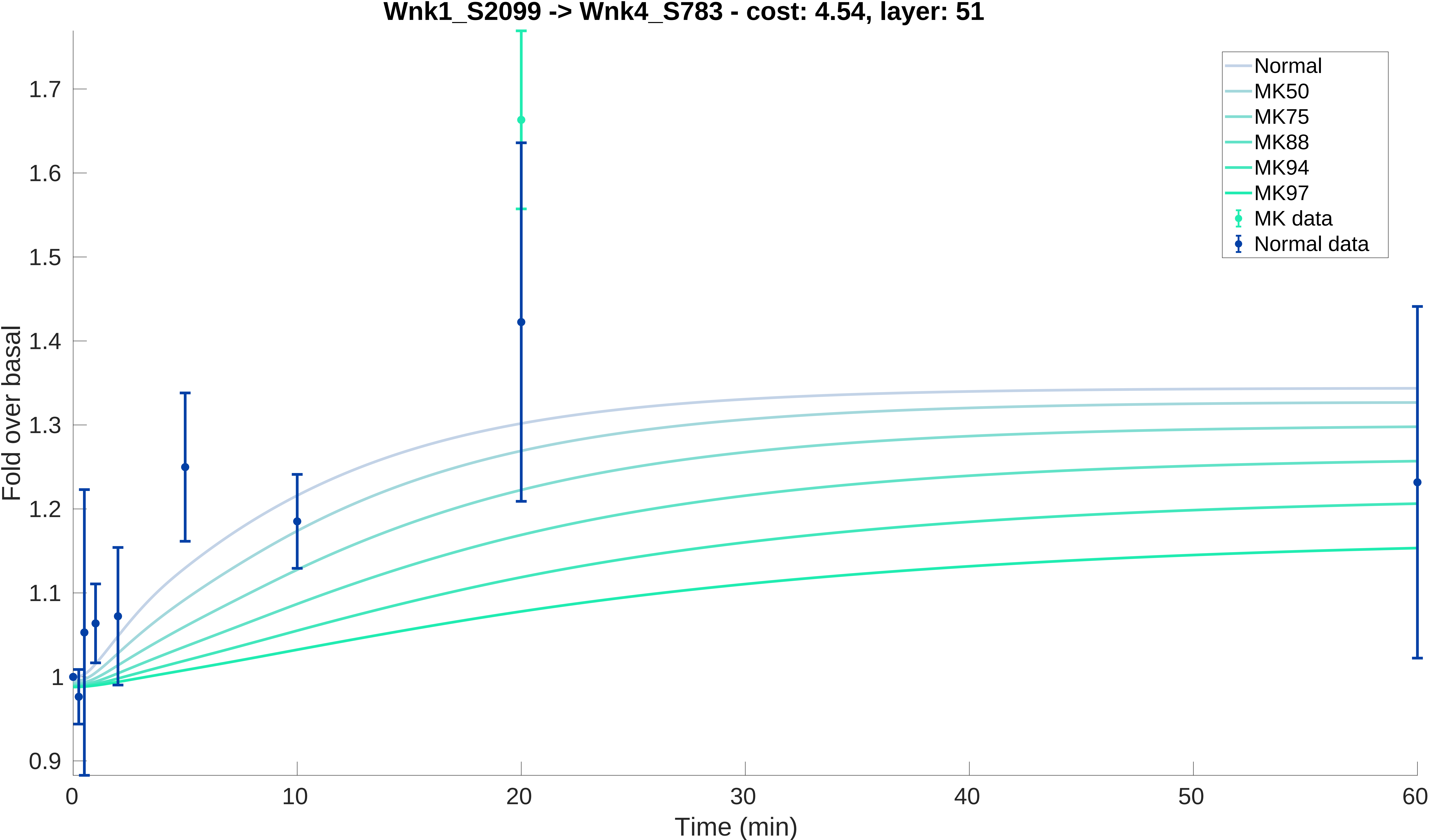

Wnk1_S2099 -> Wnk4_S783 - cost: 4.54, layer: 51
Fold over basal
1.7
1.6
1.5
1.4
1.3
1.2
1.1
1
0.9
0
10
20
30
40
50
60
Time (min)
Normal
MK50
MK75
MK88
MK94
MK97
MK data
Normal data

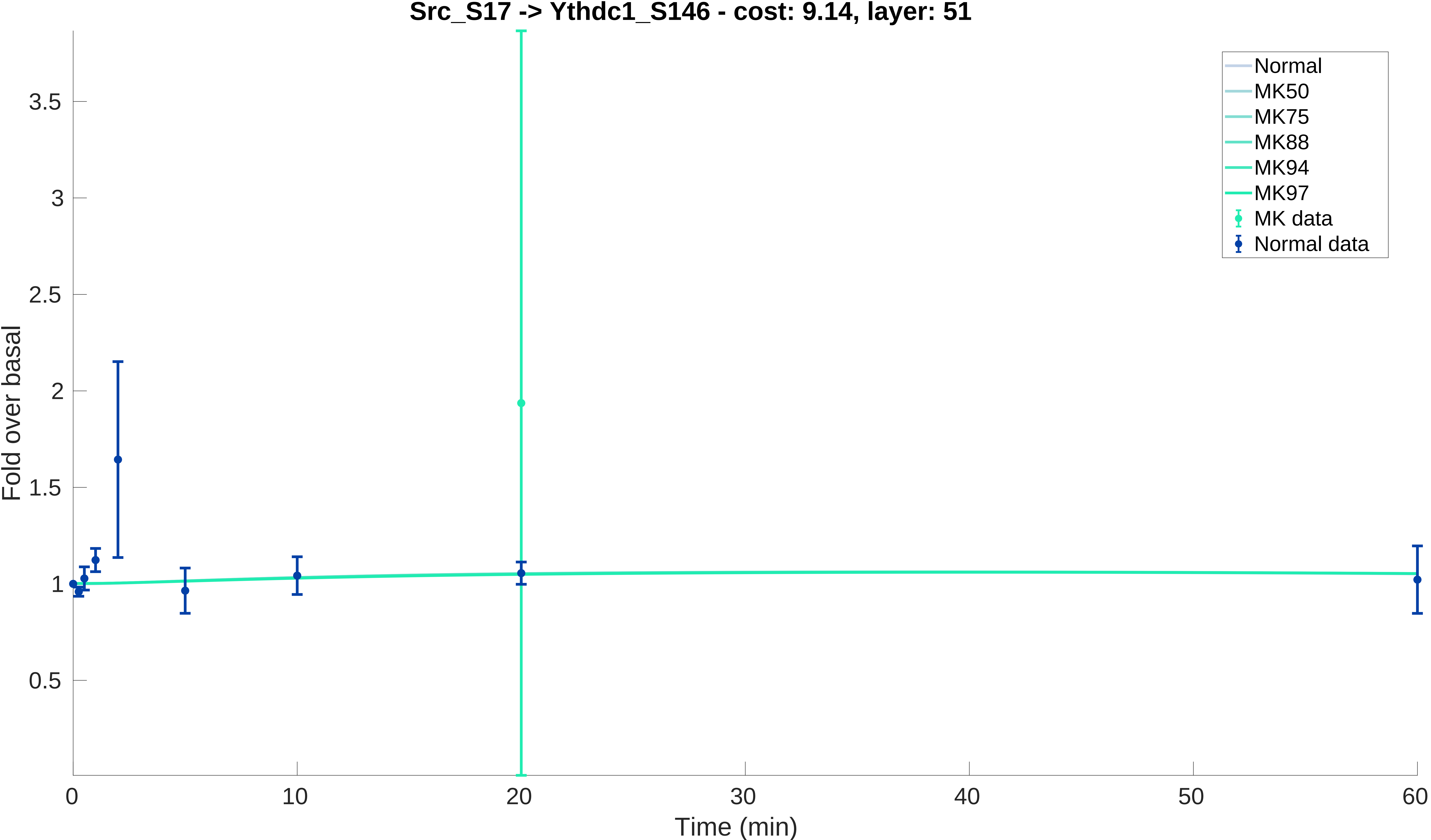

Src_S17 -> Ythdc1_S146 - cost: 9.14, layer: 51
Fold over basal
Time (min)
3.5
3
2.5
2
1.5
1
0.5
0
10
20
30
40
50
60
Normal
MK50
MK75
MK88
MK94
MK97
MK data
Normal data

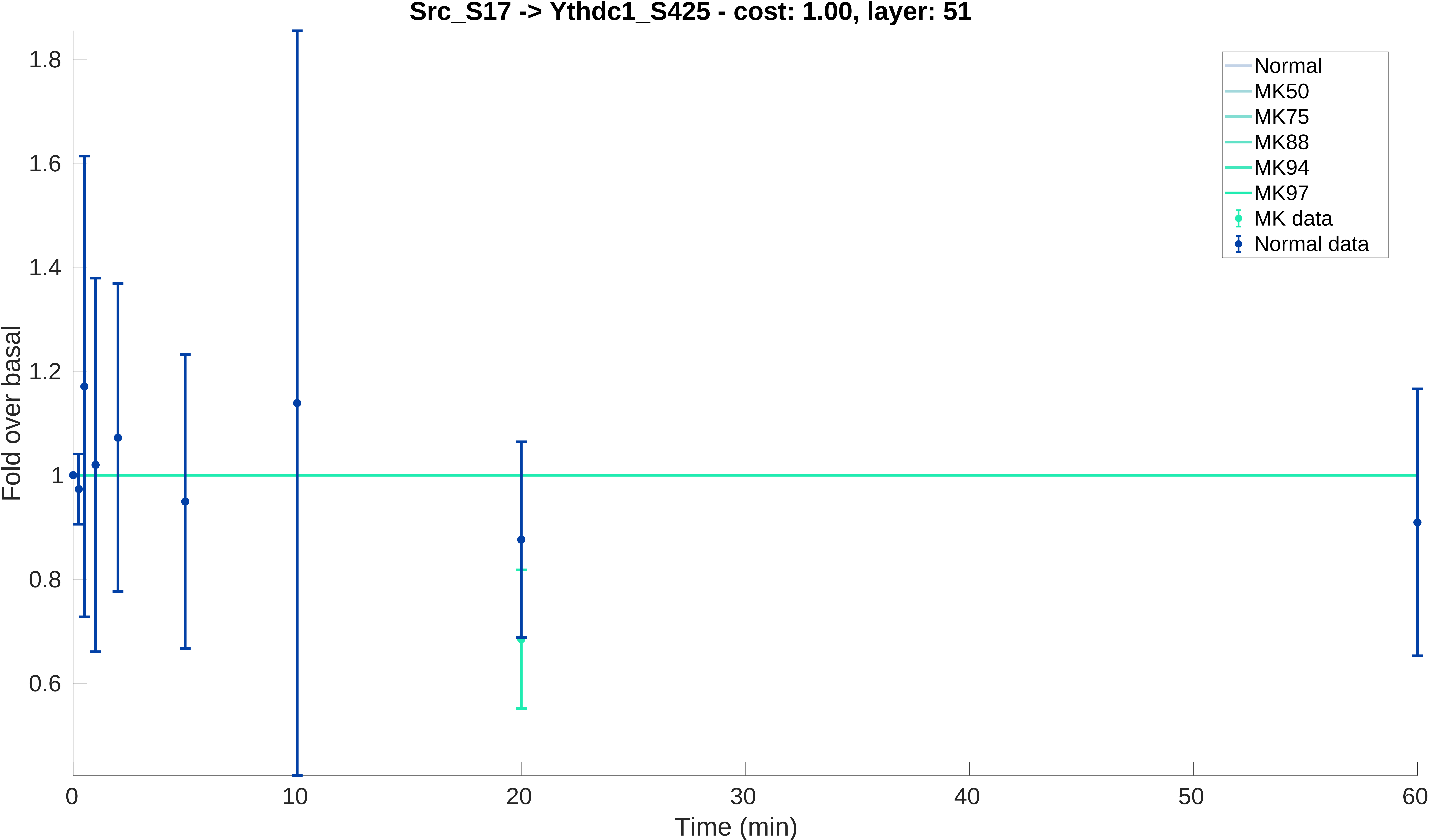

Src_S17 -> Ythdc1_S425 - cost: 1.00, layer: 51
Fold over basal
Time (min)
0
10
20
30
40
50
60
0.6
0.8
1
1.2
1.4
1.6
1.8
Normal
MK50
MK75
MK88
MK94
MK97
MK data
Normal data

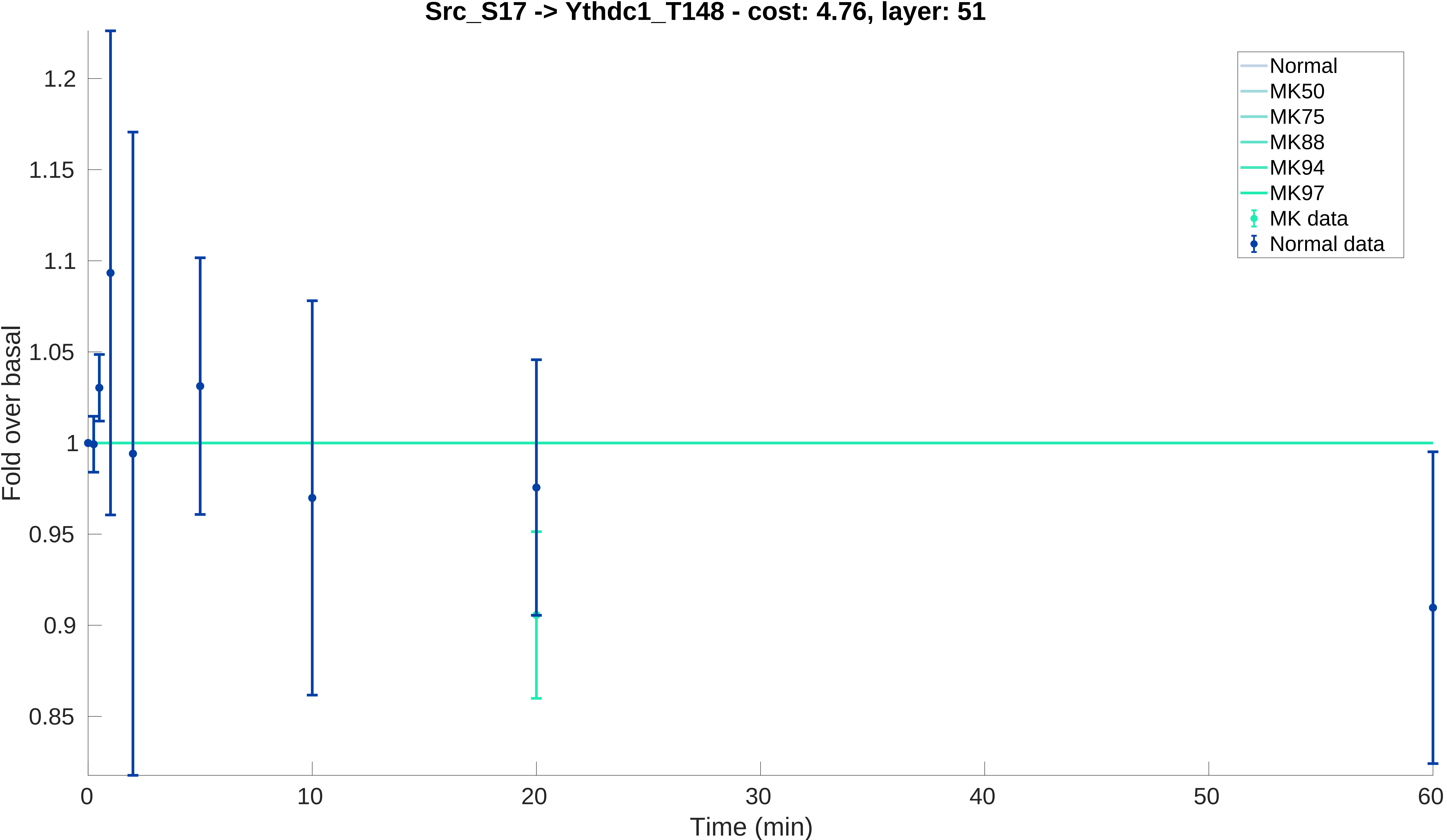

Src_S17 -> Ythdc1_T148 - cost: 4.76, layer: 51
Fold over basal
1.2
1.15
1.1
1.05
1
0.95
0.9
0.85
0
10
20
30
40
50
60
Time (min)
Normal
MK50
MK75
MK88
MK94
MK97
MK data
Normal data

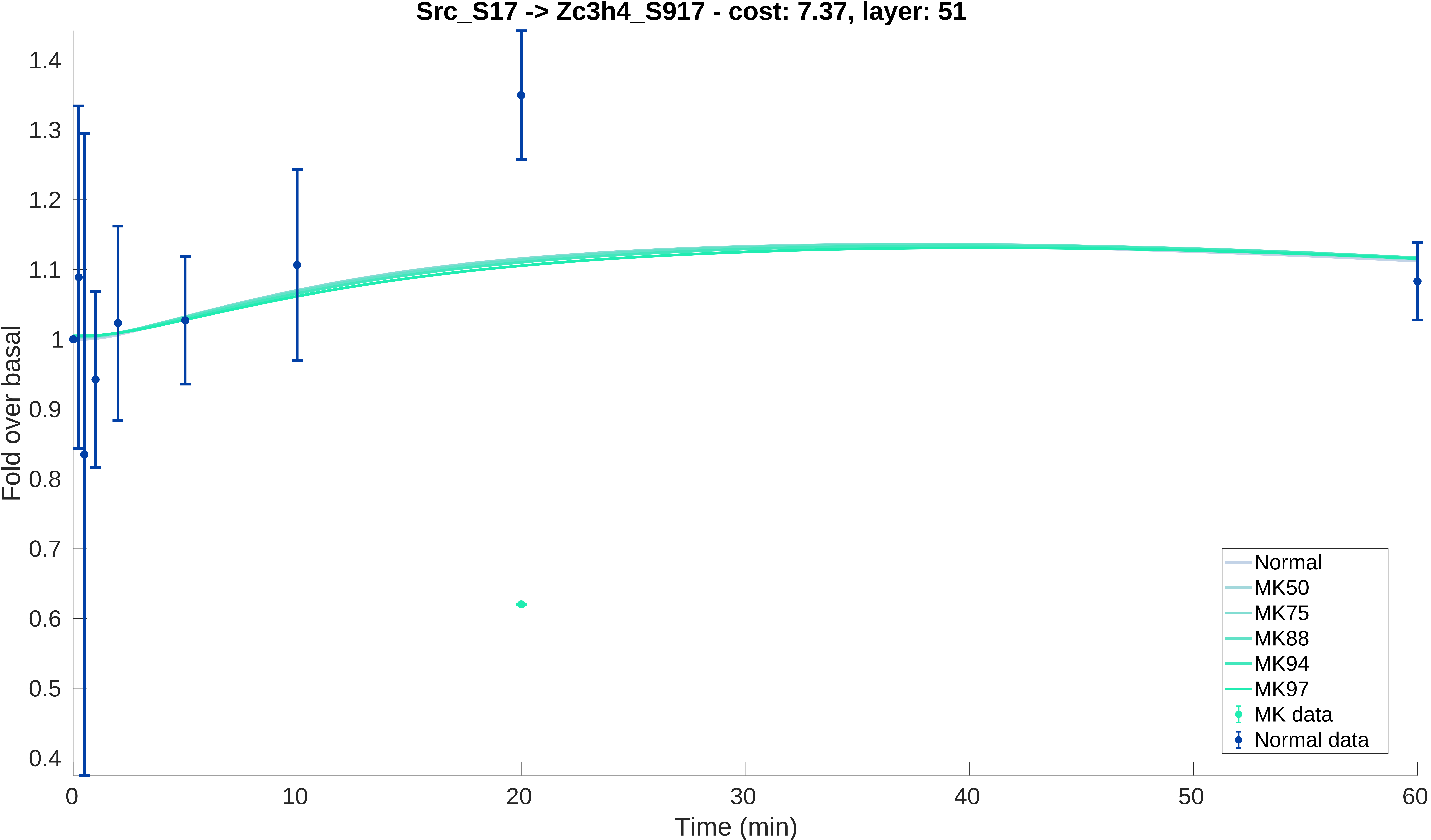

Src_S17 -> Zc3h4_S917 - cost: 7.37, layer: 51
Fold over basal
1.4
1.3
1.2
1.1
1
0.9
0.8
0.7
0.6
0.5
0.4
0
10
20
30
40
50
60
Time (min)
Normal
MK50
MK75
MK88
MK94
MK97
MK data
Normal data

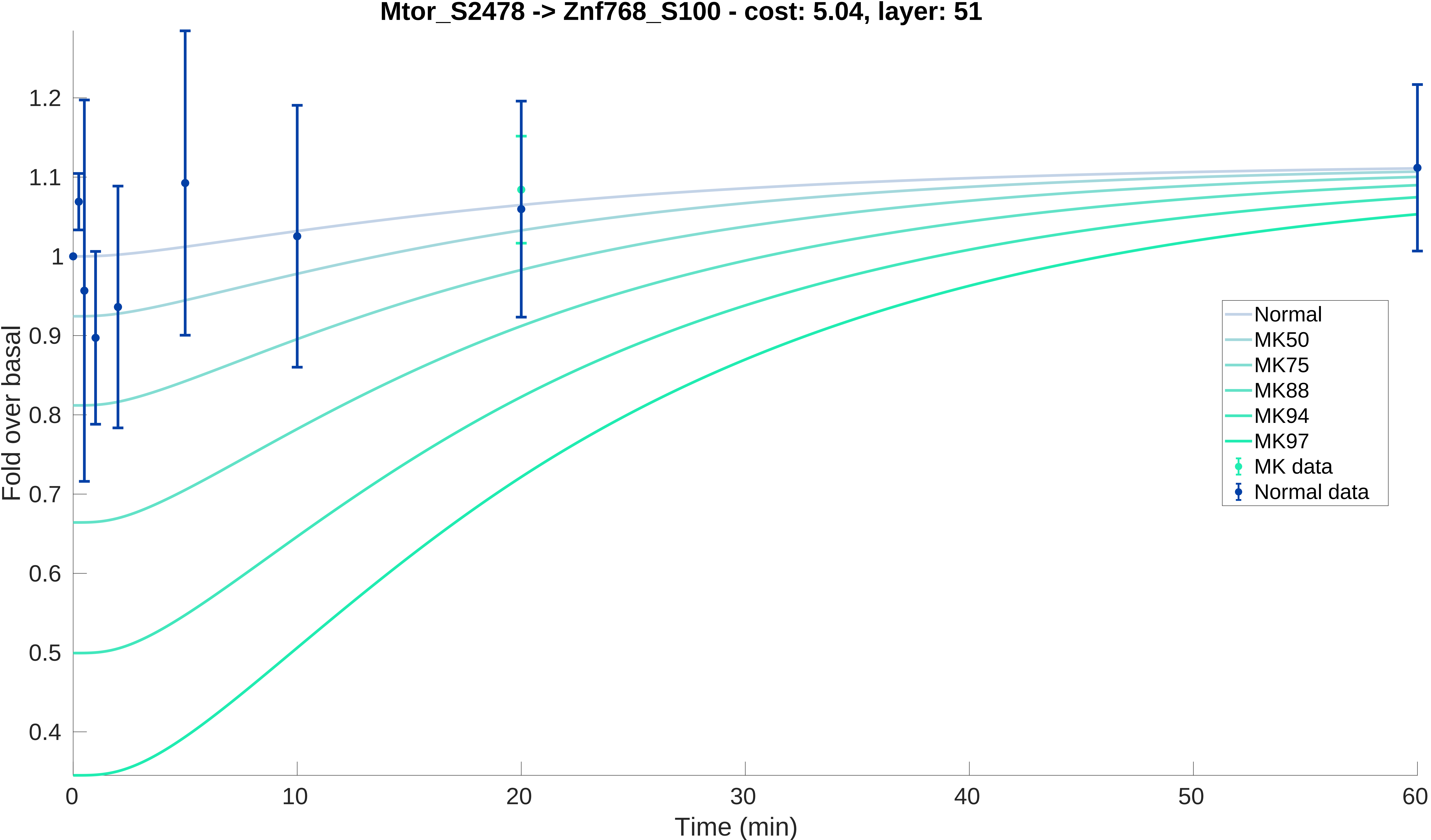
Mtor_S2478 -> Znf768_S100 - cost: 5.04, layer: 51
Fold over basal
Time (min)
0.4
0.5
0.6
0.7
0.8
0.9
1
1.1
1.2
0
10
20
30
40
50
60
Normal
MK50
MK75
MK88
MK94
MK97
MK data
Normal data

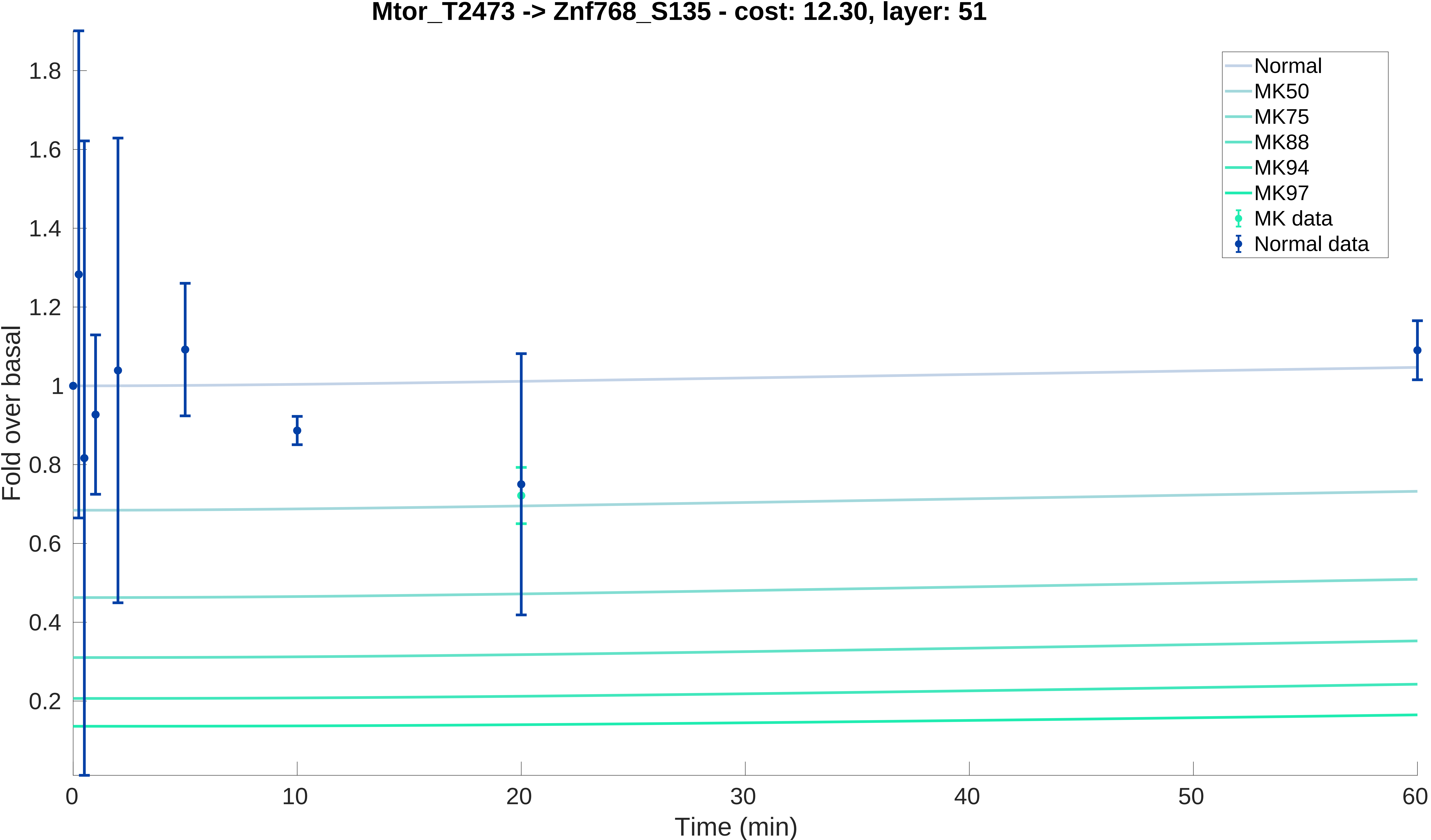

Mtor_T2473 -> Znf768_S135 - cost: 12.30, layer: 51
Fold over basal
Time (min)
0
10
20
30
40
50
60
0.2
0.4
0.6
0.8
1
1.2
1.4
1.6
1.8
Normal
MK50
MK75
MK88
MK94
MK97
MK data
Normal data

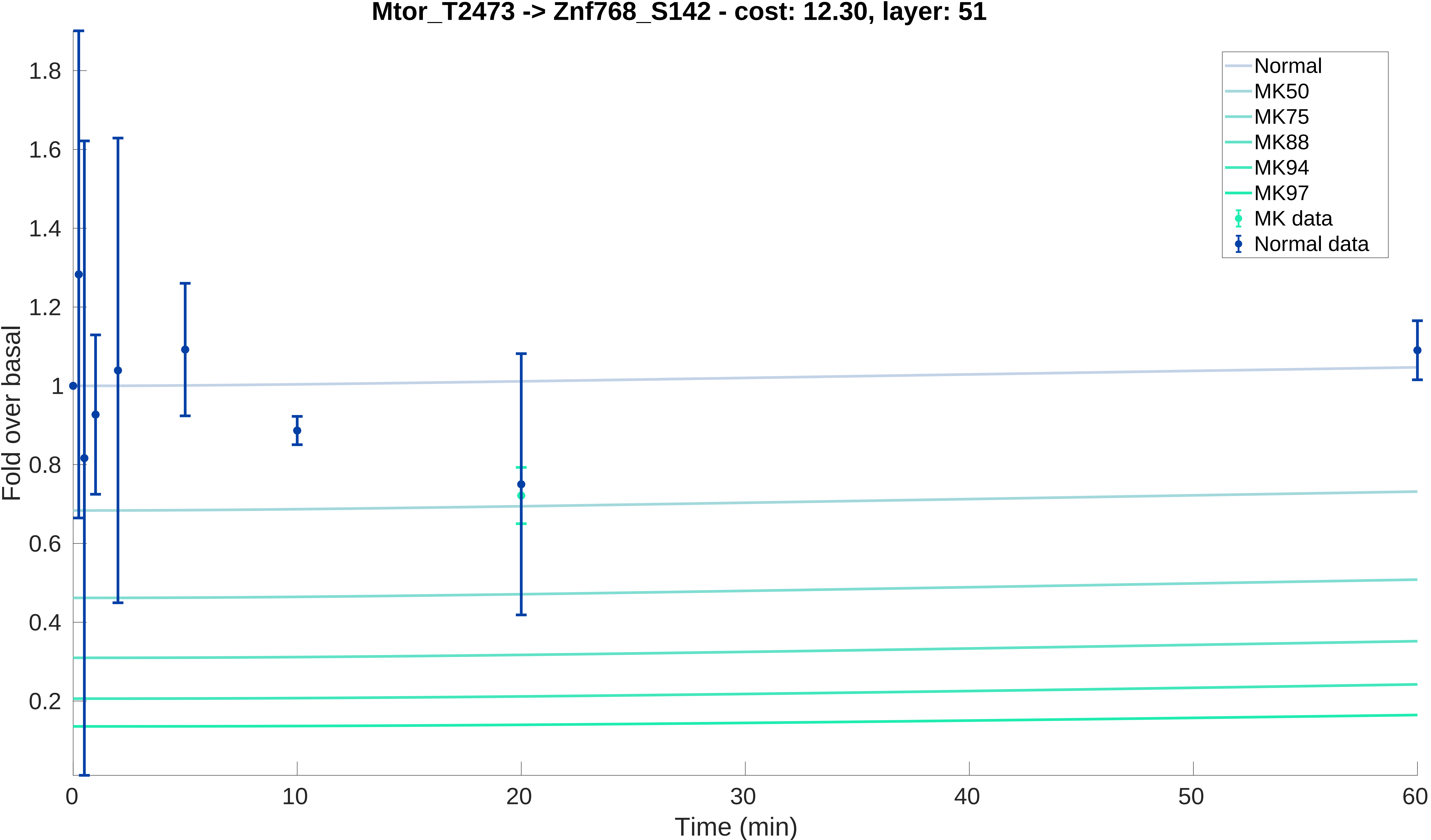
Mtor_T2473 -> Znf768_S142 - cost: 12.30, layer: 51
Fold over basal
Time (min)
0
10
20
30
40
50
60
0.2
0.4
0.6
0.8
1
1.2
1.4
1.6
1.8
Normal
MK50
MK75
MK88
MK94
MK97
MK data
Normal data

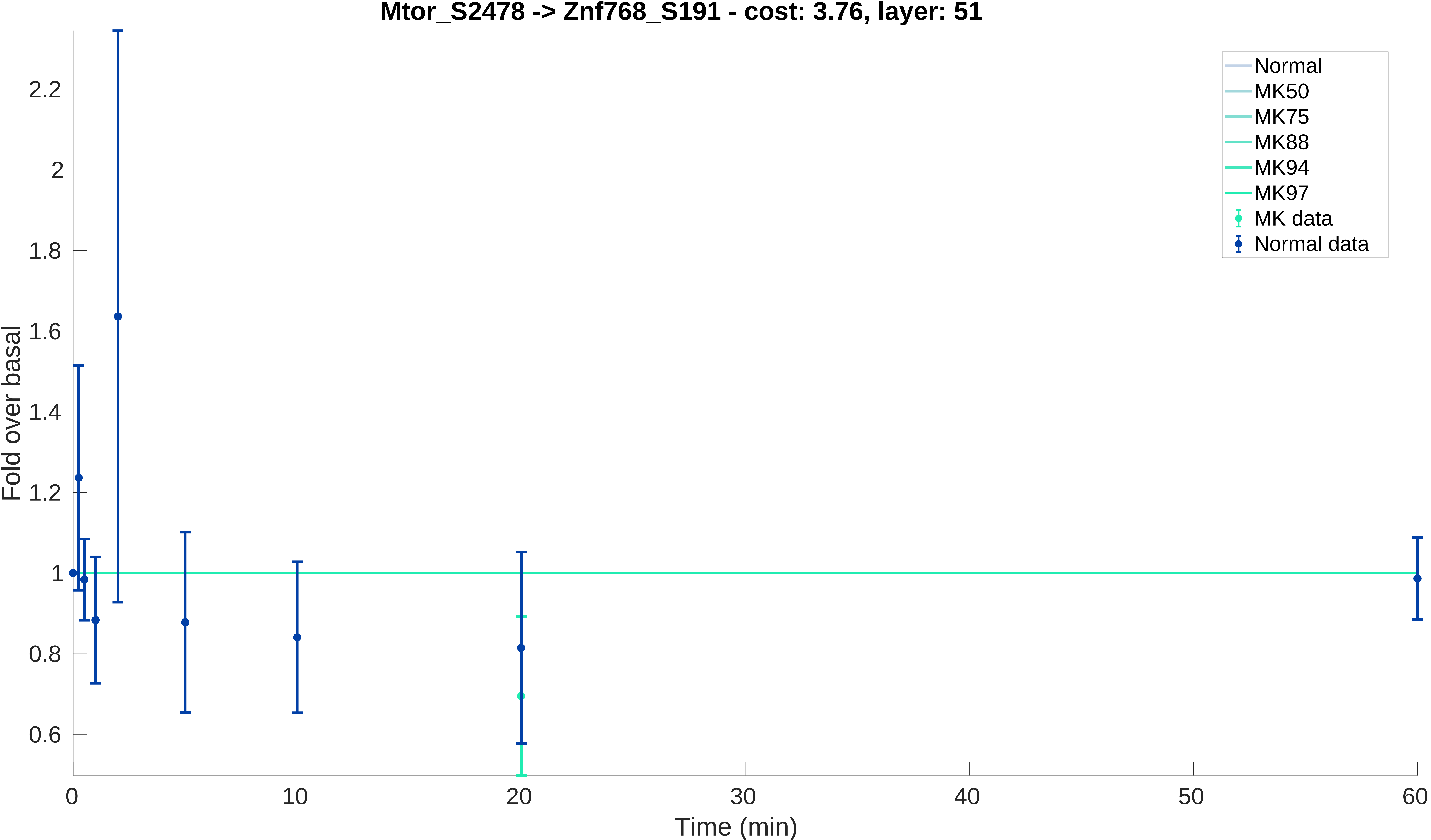

Mtor_S2478 -> Znf768_S191 - cost: 3.76, layer: 51
Fold over basal
Time (min)
0.6
0.8
1
1.2
1.4
1.6
1.8
2
2.2
0
10
20
30
40
50
60
Normal
MK50
MK75
MK88
MK94
MK97
MK data
Normal data

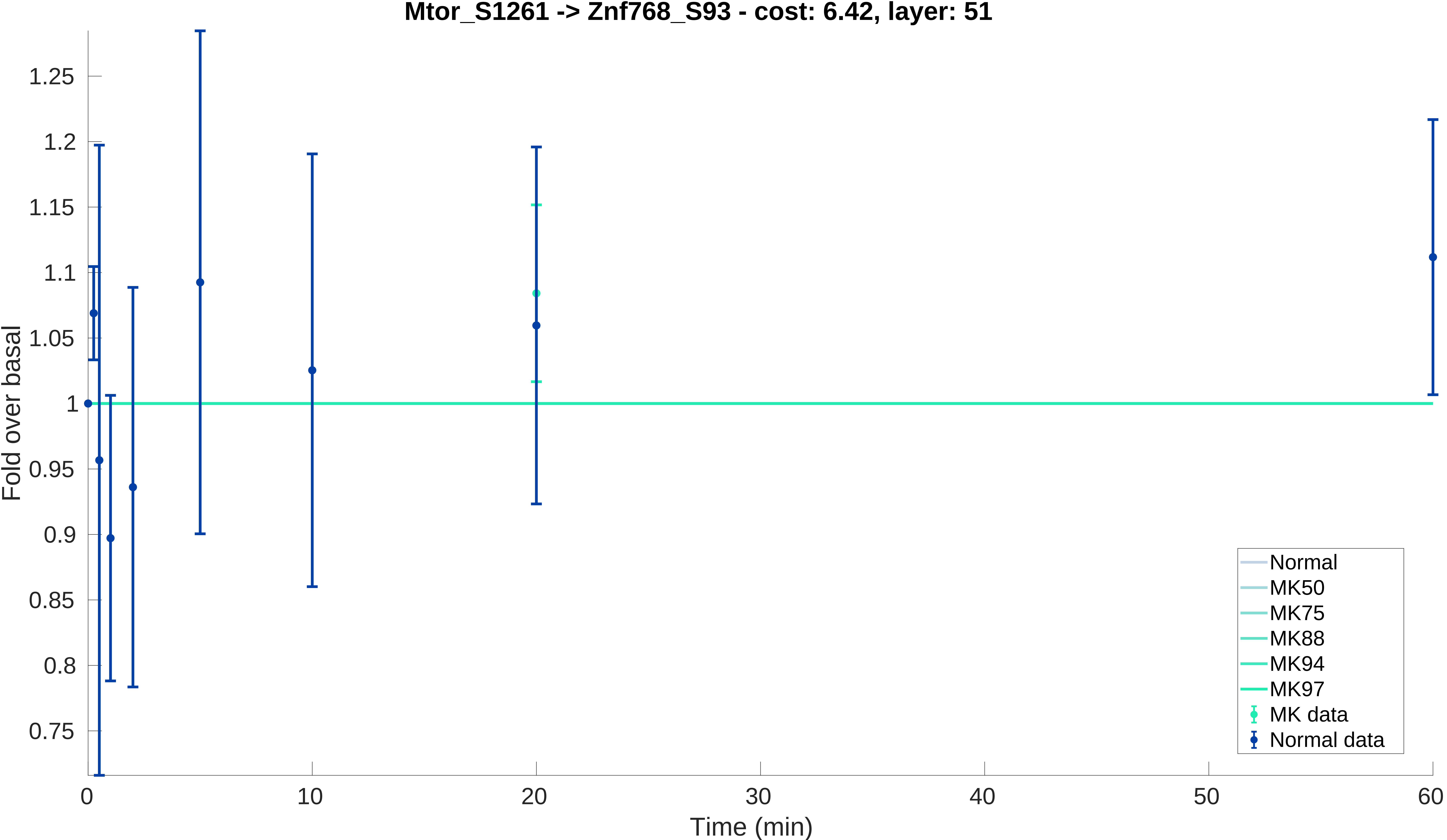

Mtor_S1261 -> Znf768_S93 - cost: 6.42, layer: 51
Fold over basal
1.25
1.2
1.15
1.1
1.05
1
0.95
0.9
0.85
0.8
0.75
0
10
20
30
40
50
60
Time (min)
Normal
MK50
MK75
MK88
MK94
MK97
MK data
Normal data

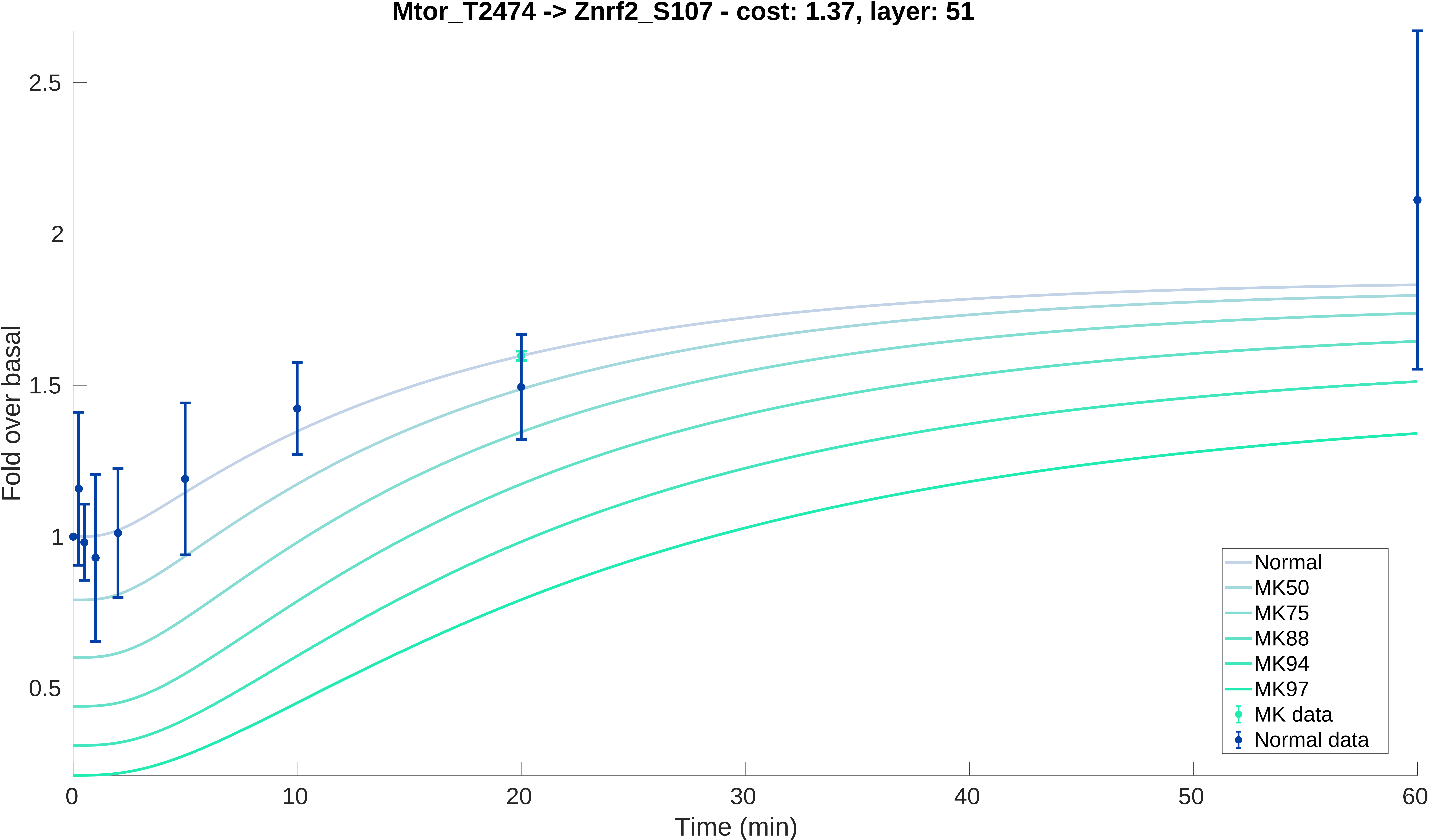

Mtor_T2474 -> Znrf2_S107 - cost: 1.37, layer: 51
Fold over basal
Time (min)
0
10
20
30
40
50
60
0.5
1
1.5
2
2.5
Normal
MK50
MK75
MK88
MK94
MK97
MK data
Normal data

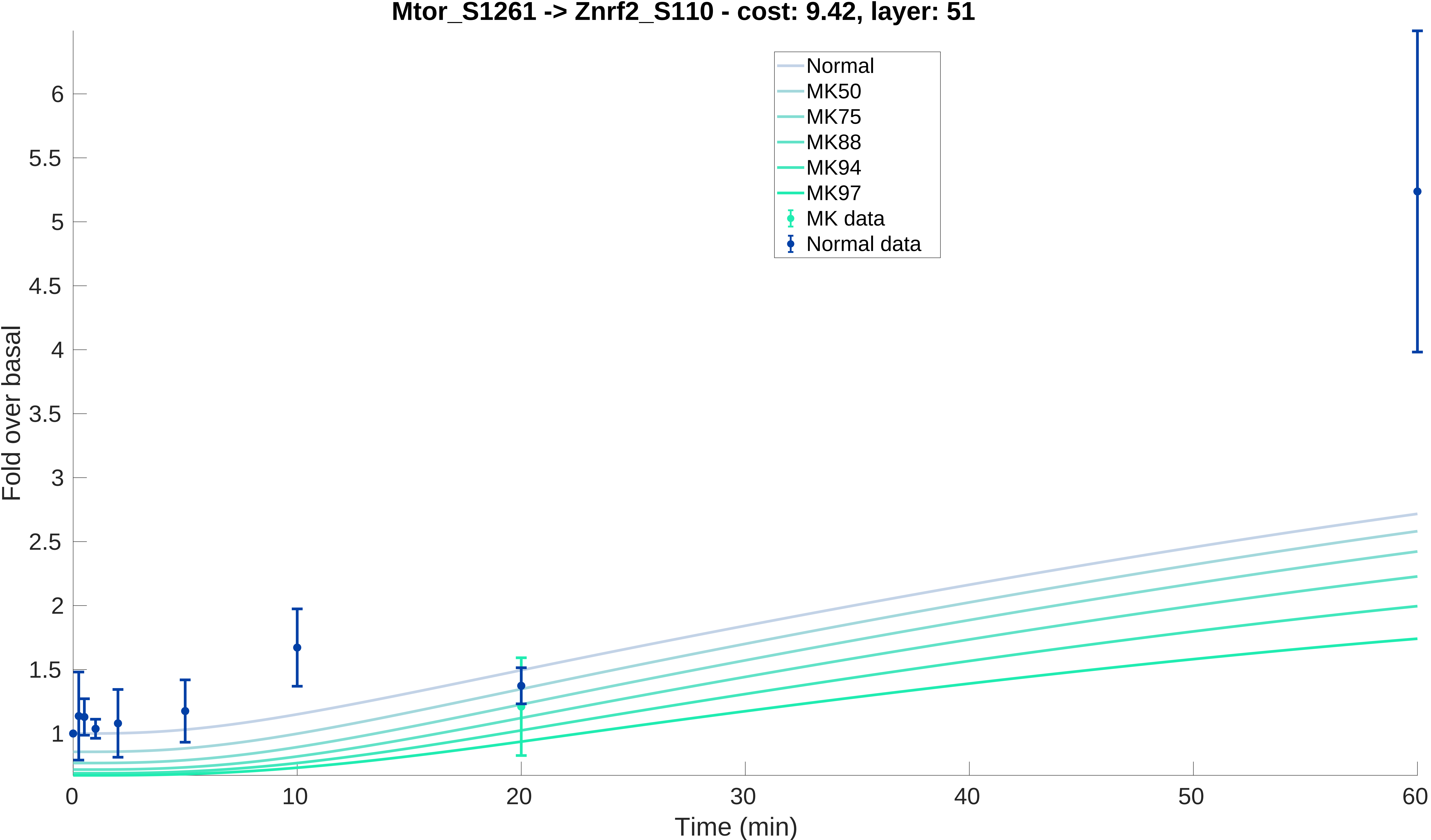

Mtor_S1261 -> Znrf2_S110 - cost: 9.42, layer: 51
Normal
MK50
MK75
MK88
MK94
MK97
MK data
Normal data
Fold over basal
6
5.5
5
4.5
4
3.5
3
2.5
2
1.5
1
0
10
20
30
40
50
60
Time (min)

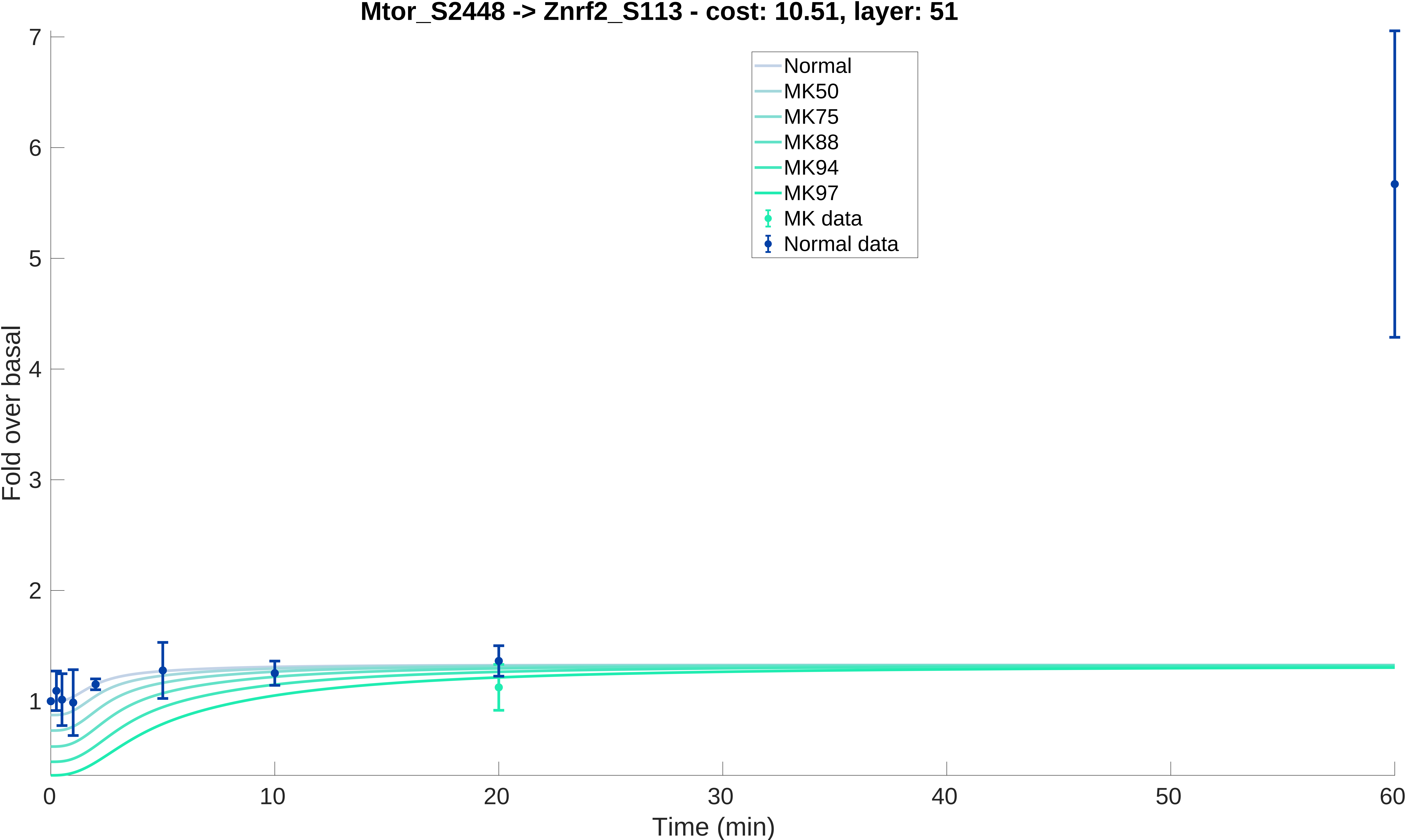

Mtor_S2448 -> Znrf2_S113 - cost: 10.51, layer: 51
Normal
MK50
MK75
MK88
MK94
MK97
MK data
Normal data
Fold over basal
Time (min)
0
10
20
30
40
50
60
1
2
3
4
5
6
7

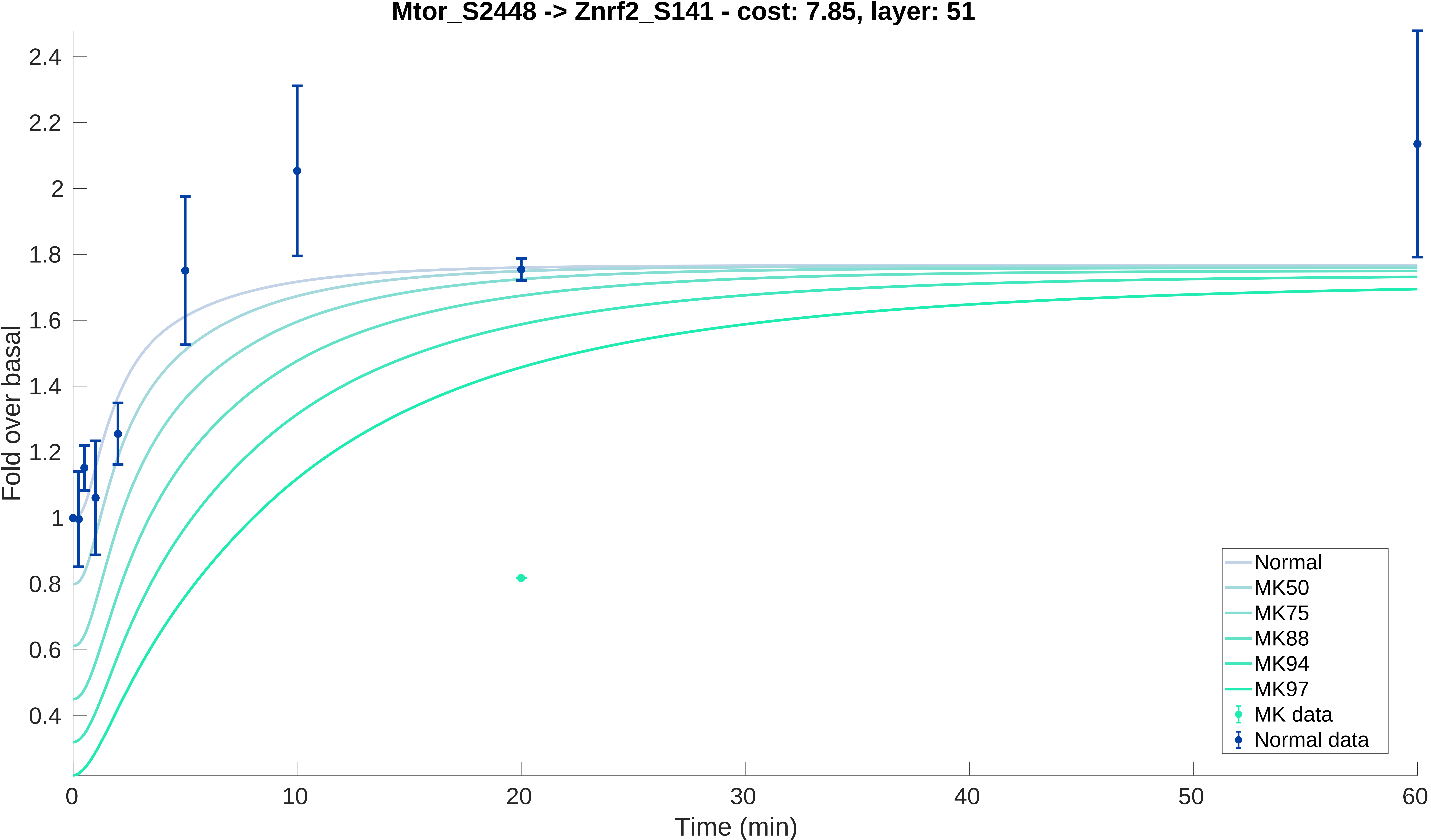
Mtor_S2448 -> Znrf2_S141 - cost: 7.85, layer: 51
Fold over basal
Time (min)
2.4
2.2
2
1.8
1.6
1.4
1.2
1
0.8
0.6
0.4
0
10
20
30
40
50
60
Normal
MK50
MK75
MK88
MK94
MK97
MK data
Normal data

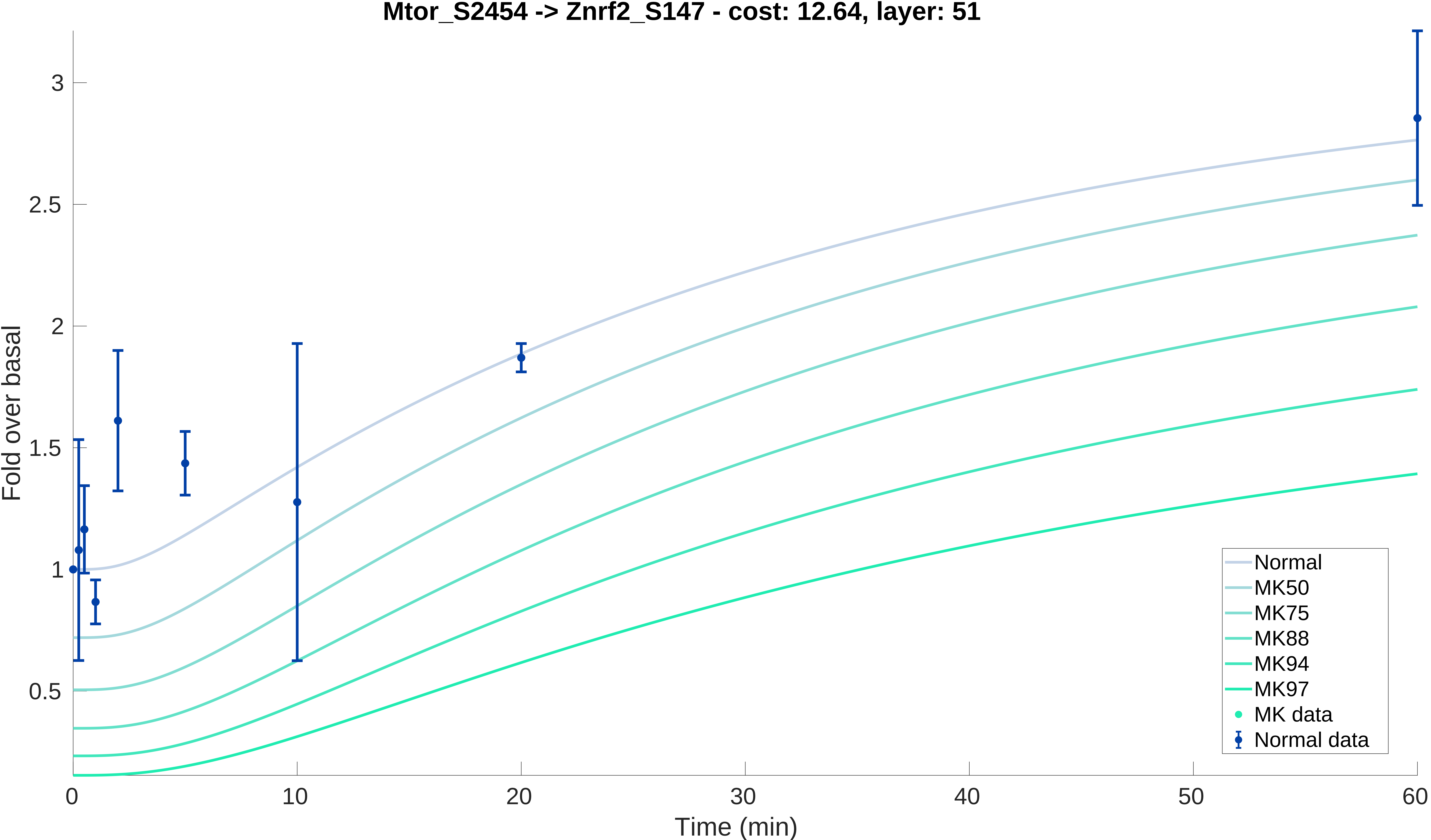

Mtor_S2454 -> Znrf2_S147 - cost: 12.64, layer: 51
Fold over basal
Time (min)
0.5
1
1.5
2
2.5
3
0
10
20
30
40
50
60
Normal
MK50
MK75
MK88
MK94
MK97
MK data
Normal data

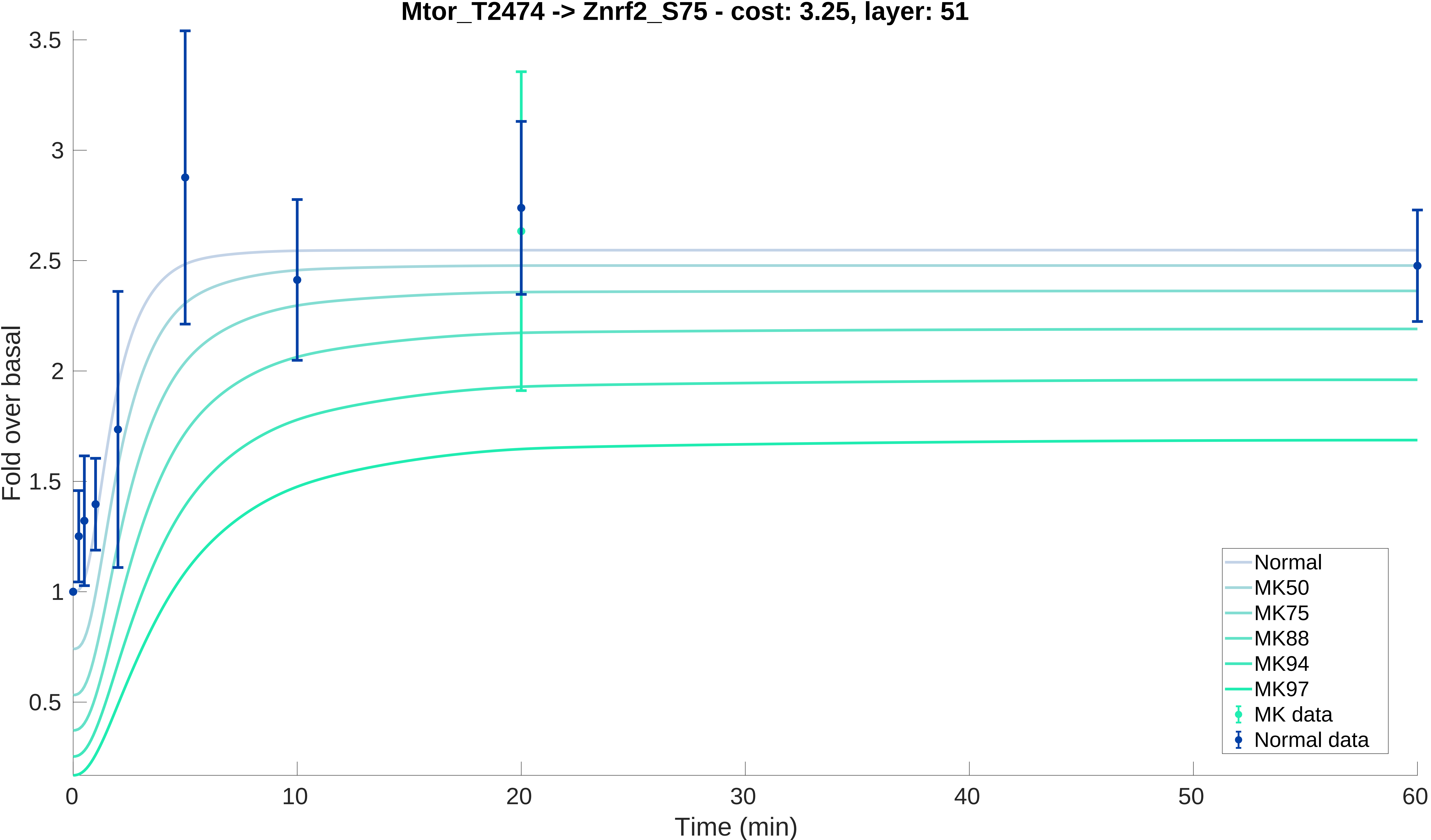

Mtor_T2474 -> Znrf2_S75 - cost: 3.25, layer: 51
Fold over basal
Time (min)
3.5
3
2.5
2
1.5
1
0.5
0
10
20
30
40
50
60
Normal
MK50
MK75
MK88
MK94
MK97
MK data
Normal data

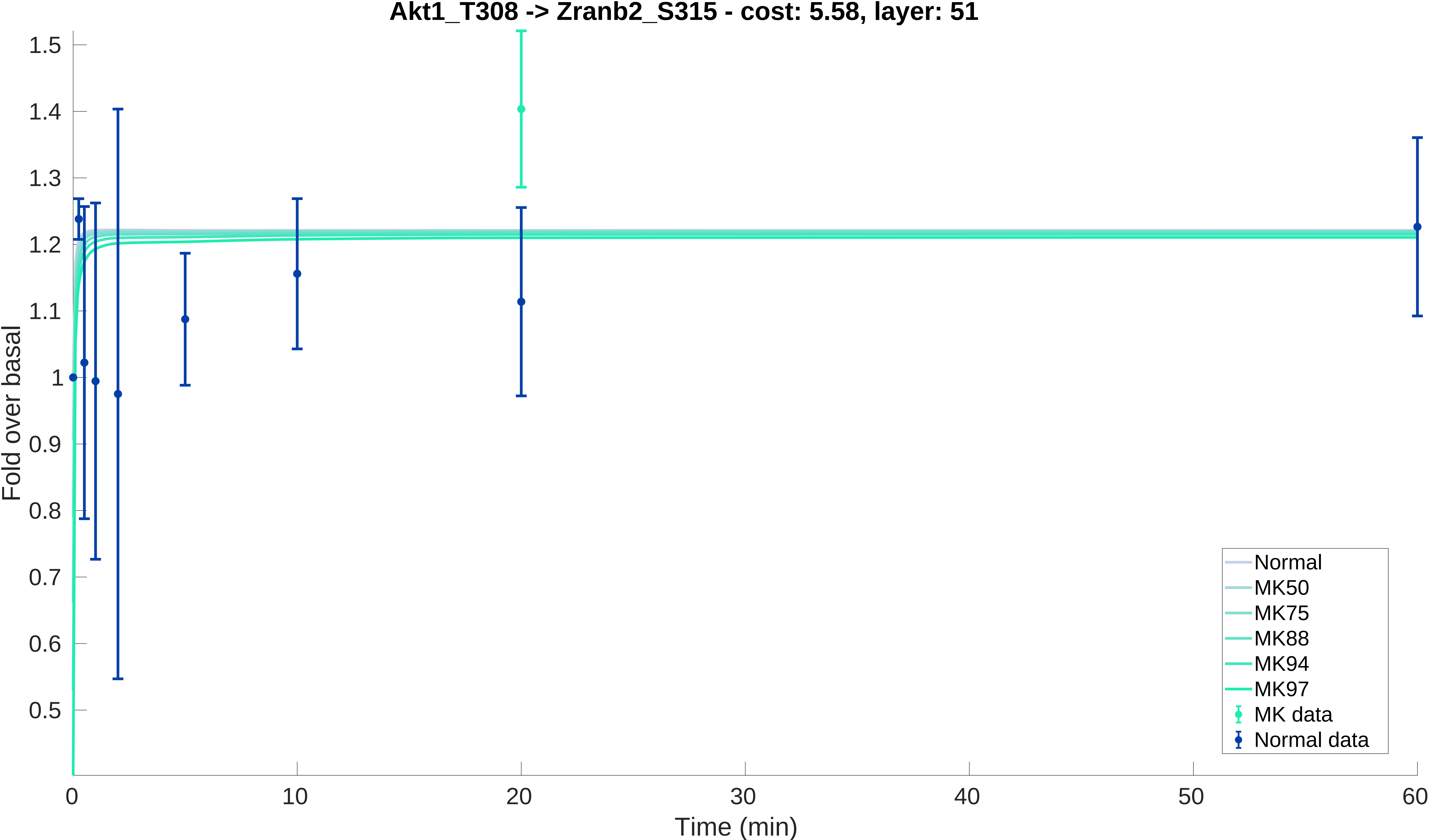

Akt1_T308 -> Zranb2_S315 - cost: 5.58, layer: 51
Fold over basal
Time (min)
1.5
1.4
1.3
1.2
1.1
1
0.9
0.8
0.7
0.6
0.5
0
10
20
30
40
50
60
Normal
MK50
MK75
MK88
MK94
MK97
MK data
Normal data

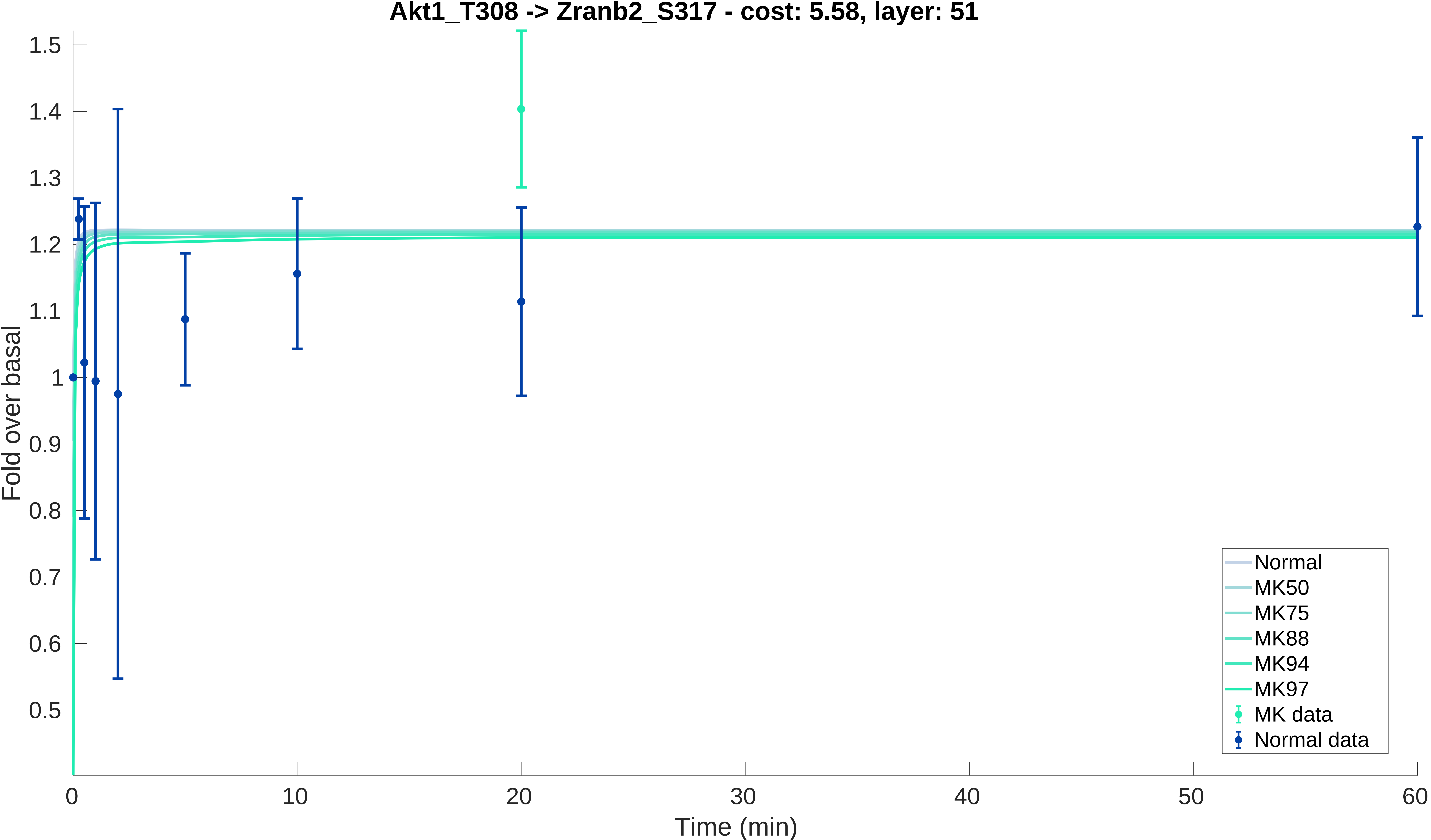

Akt1_T308 -> Zranb2_S317 - cost: 5.58, layer: 51
Fold over basal
Time (min)
1.5
1.4
1.3
1.2
1.1
1
0.9
0.8
0.7
0.6
0.5
0
10
20
30
40
50
60
Normal
MK50
MK75
MK88
MK94
MK97
MK data
Normal data

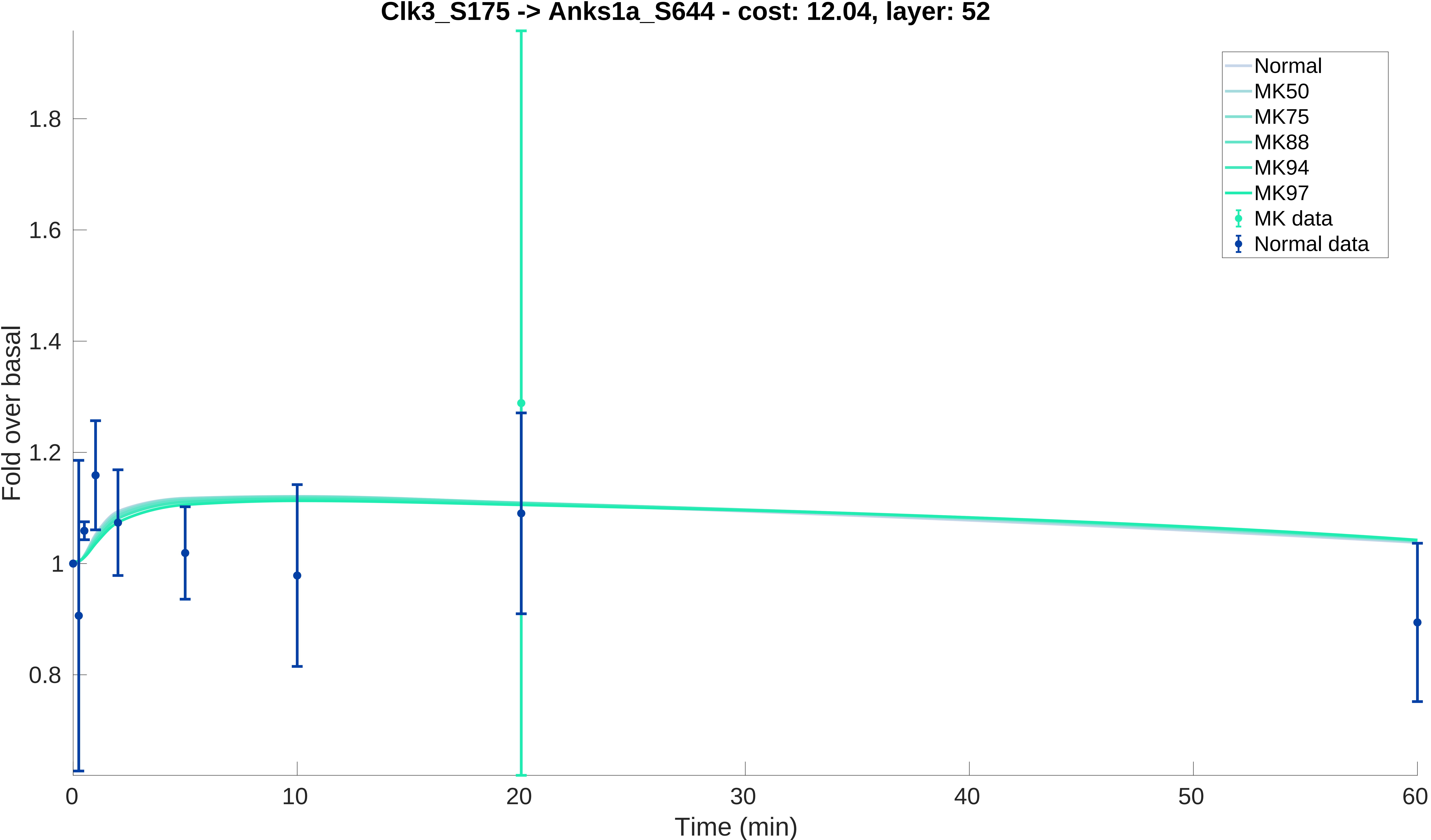
Clk3_S175 -> Anks1a_S644 - cost: 12.04, layer: 52
Normal
MK50
MK75
MK88
MK94
MK97
MK data
Normal data
Fold over basal
1.8
1.6
1.4
1.2
1
0.8
0
10
20
30
40
50
60
Time (min)

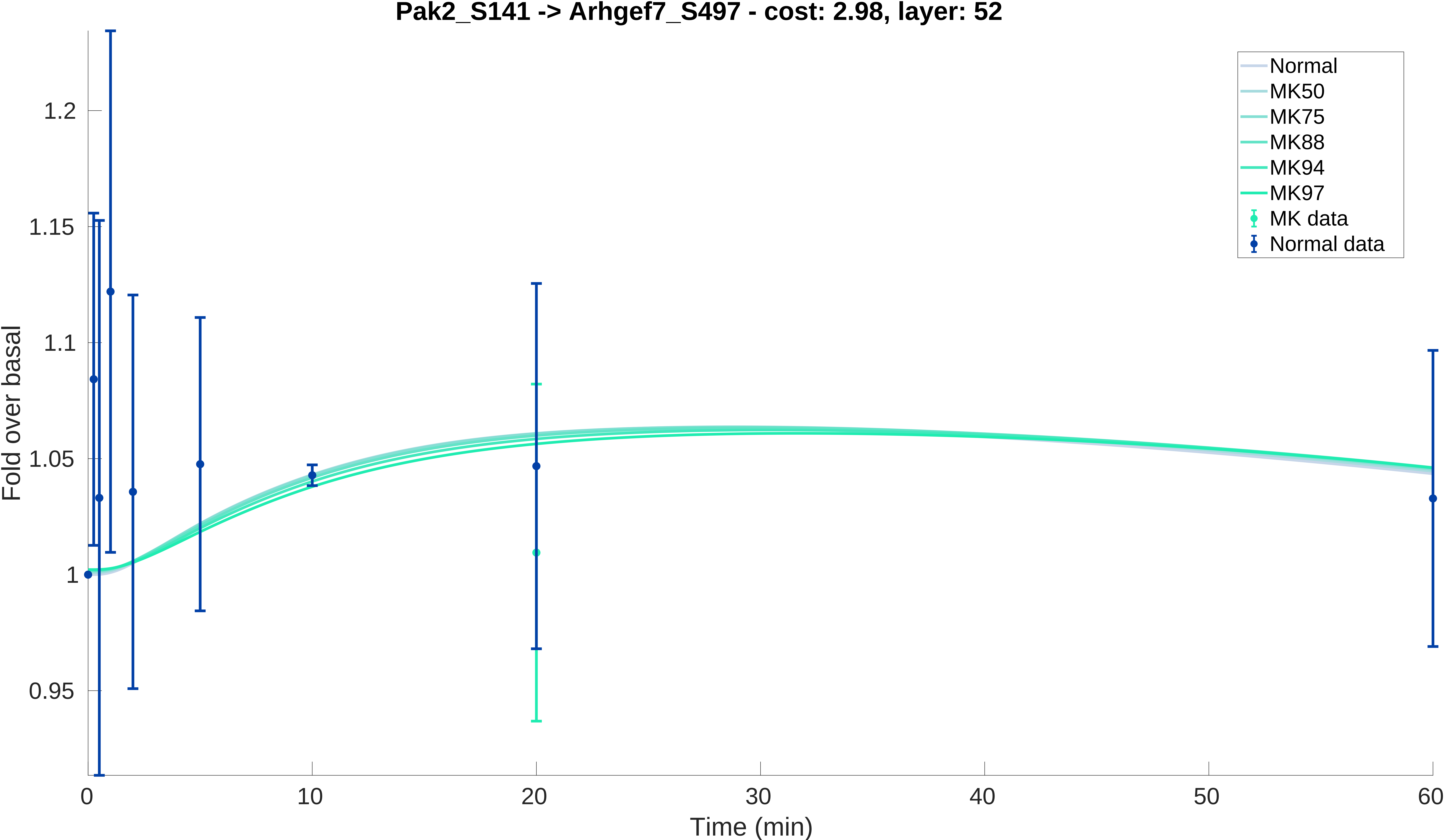

Pak2_S141 -> Arhgef7_S497 - cost: 2.98, layer: 52
Fold over basal
Time (min)
1.2
1.15
1.1
1.05
1
0.95
0
10
20
30
40
50
60
Normal
MK50
MK75
MK88
MK94
MK97
MK data
Normal data

#### Ccny_S326 -> Cdk14_S120 - cost: 7.66, layer: 52

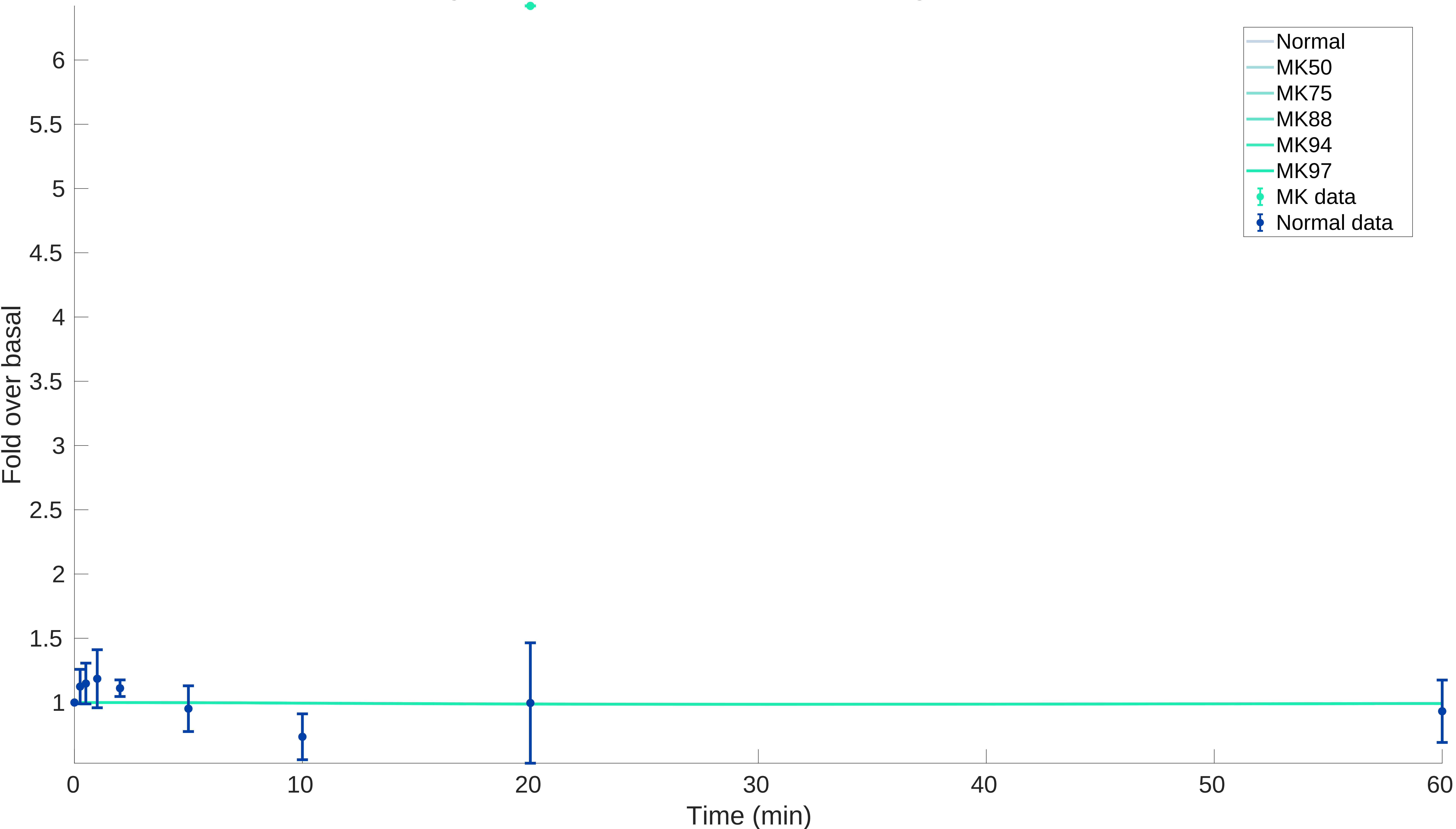

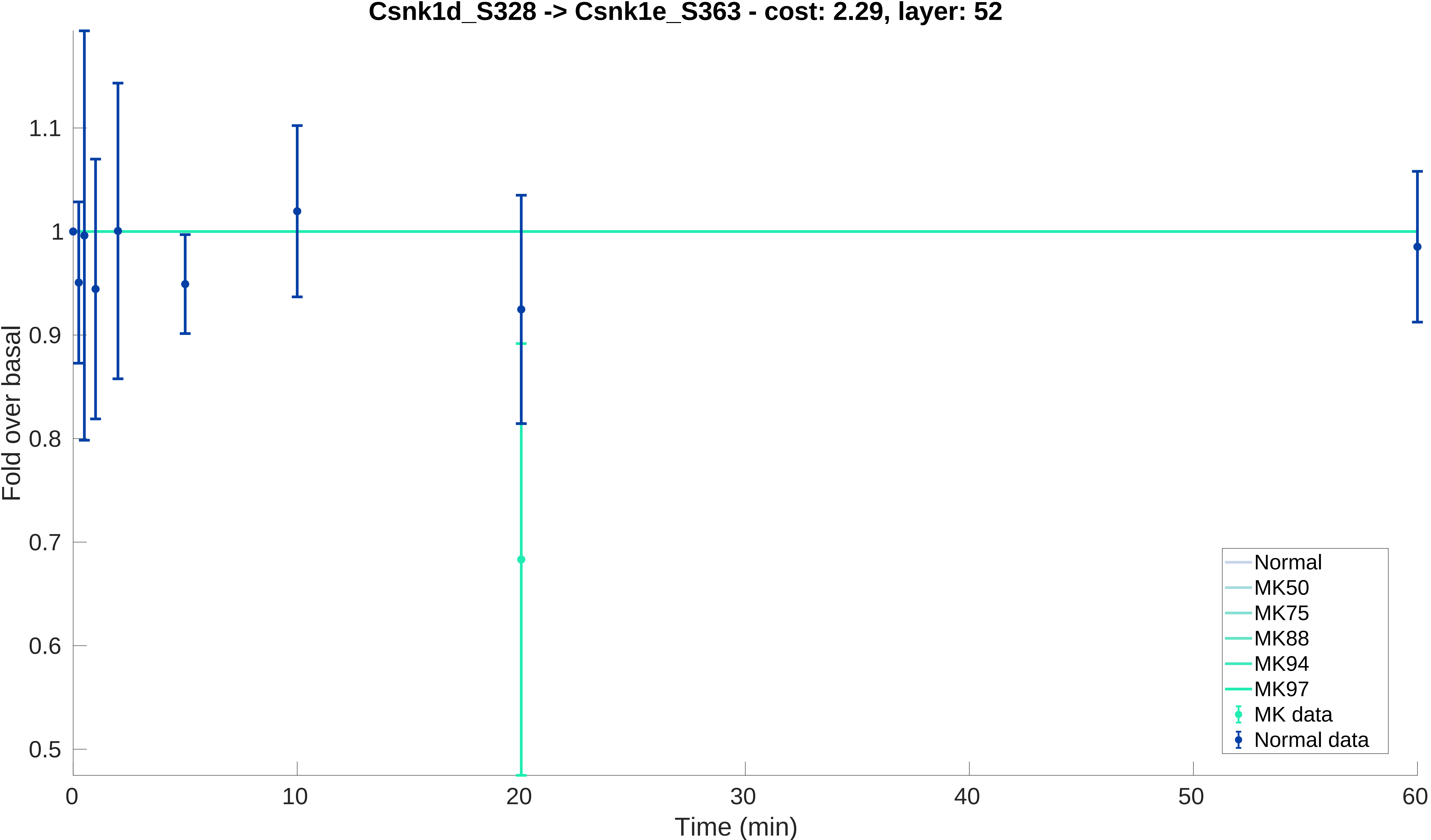

Csnk1d_S328 -> Csnk1e_S363 - cost: 2.29, layer: 52
Fold over basal
Time (min)
0.5
0.6
0.7
0.8
0.9
1
1.1
0
10
20
30
40
50
60
Normal
MK50
MK75
MK88
MK94
MK97
MK data
Normal data

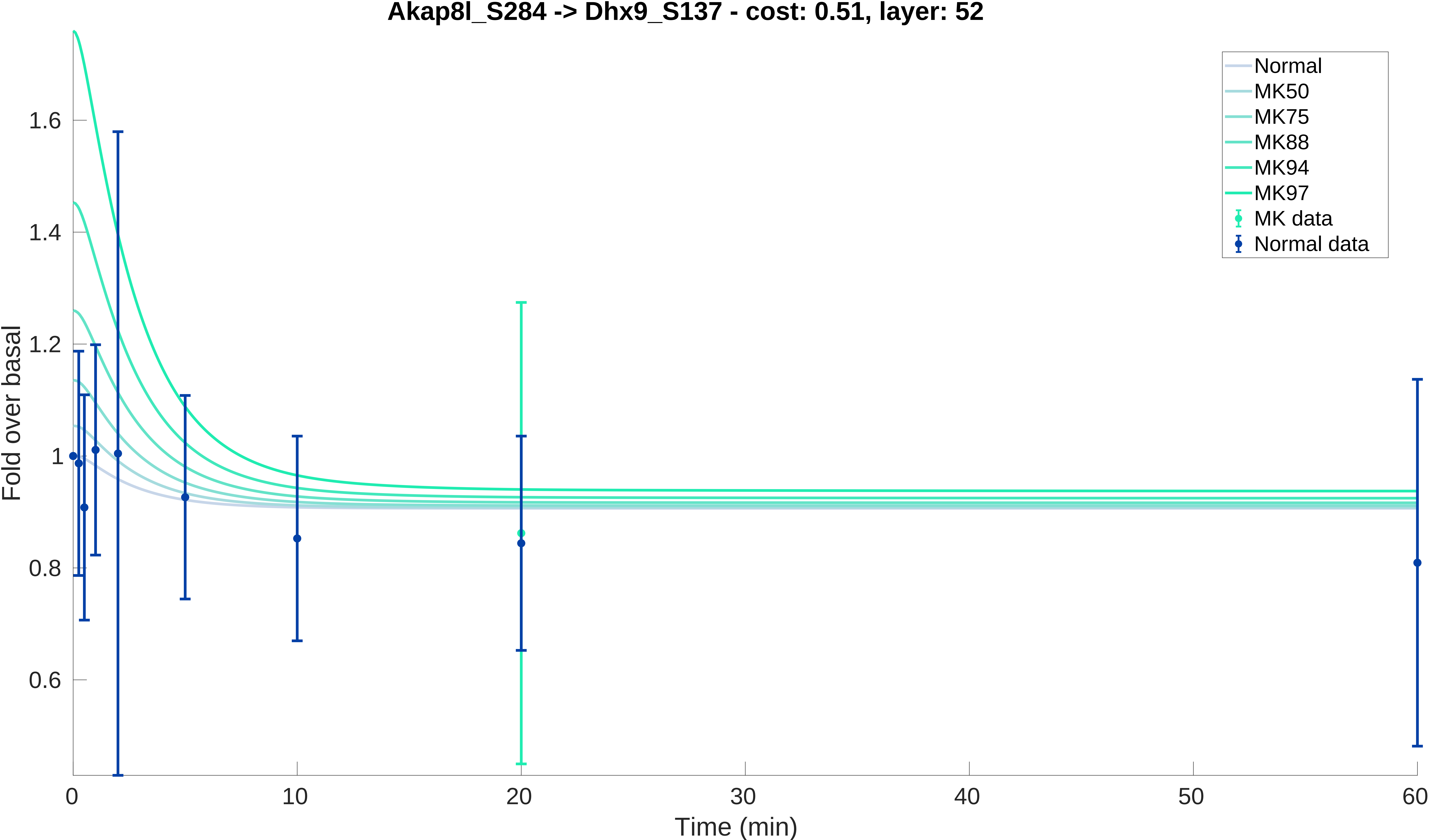
Akap8l_S284 -> Dhx9_S137 - cost: 0.51, layer: 52
Fold over basal
Time (min)
1.6
1.4
1.2
1
0.8
0.6
0
10
20
30
40
50
60
Normal
MK50
MK75
MK88
MK94
MK97
MK data
Normal data

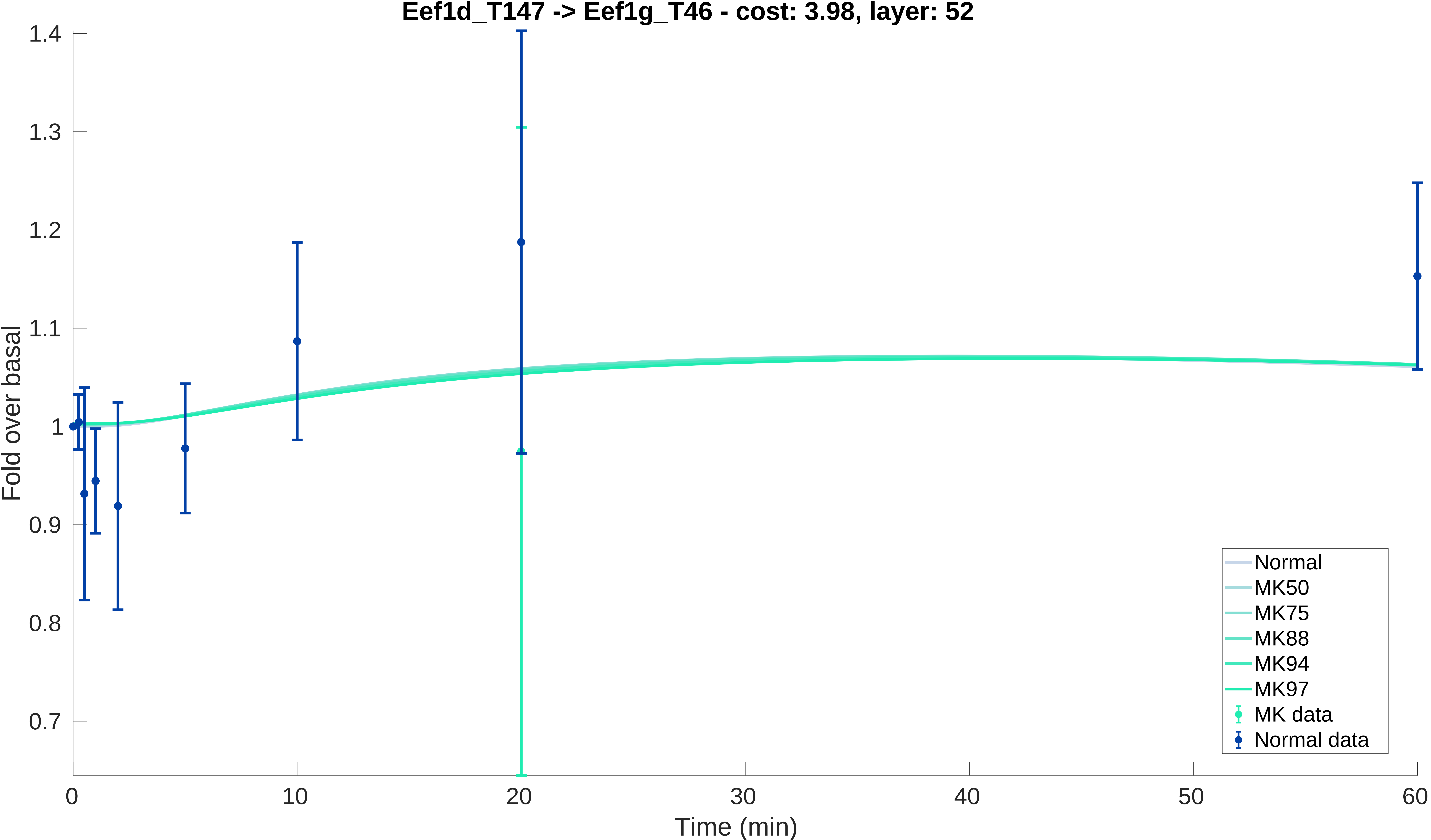
Eef1d_T147 -> Eef1g_T46 - cost: 3.98, layer: 52
Fold over basal
Time (min)
1.4
1.3
1.2
1.1
1
0.9
0.8
0.7
0
10
20
30
40
50
60
Normal
MK50
MK75
MK88
MK94
MK97
MK data
Normal data

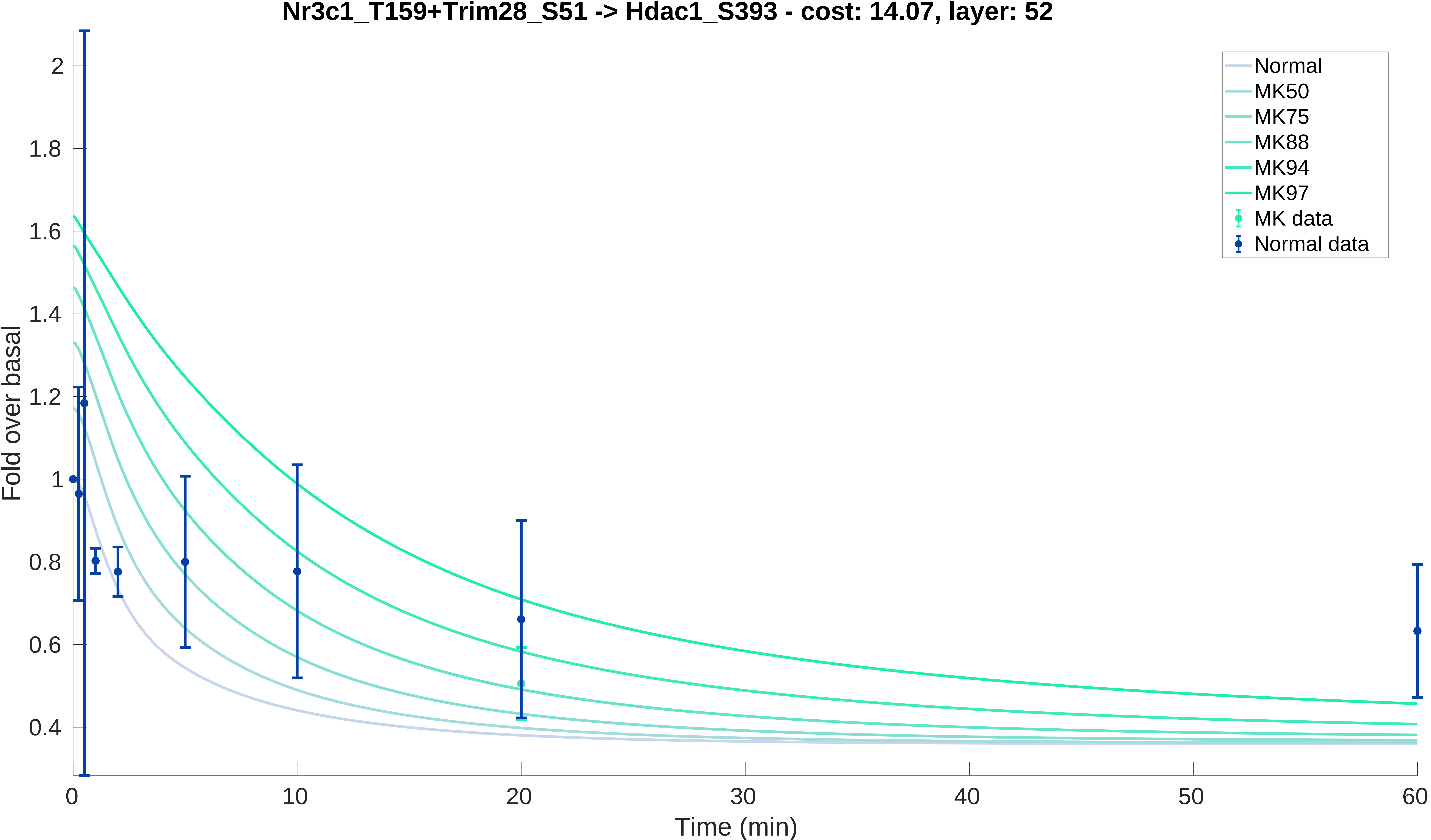

Nr3c1_T159+Trim28_S51 -> Hdac1_S393 - cost: 14.07, layer: 52
Normal
MK50
MK75
MK88
MK94
MK97
MK data
Normal data
Fold over basal
2
1.8
1.6
1.4
1.2
1
0.8
0.6
0.4
0
10
20
30
40
50
60
Time (min)

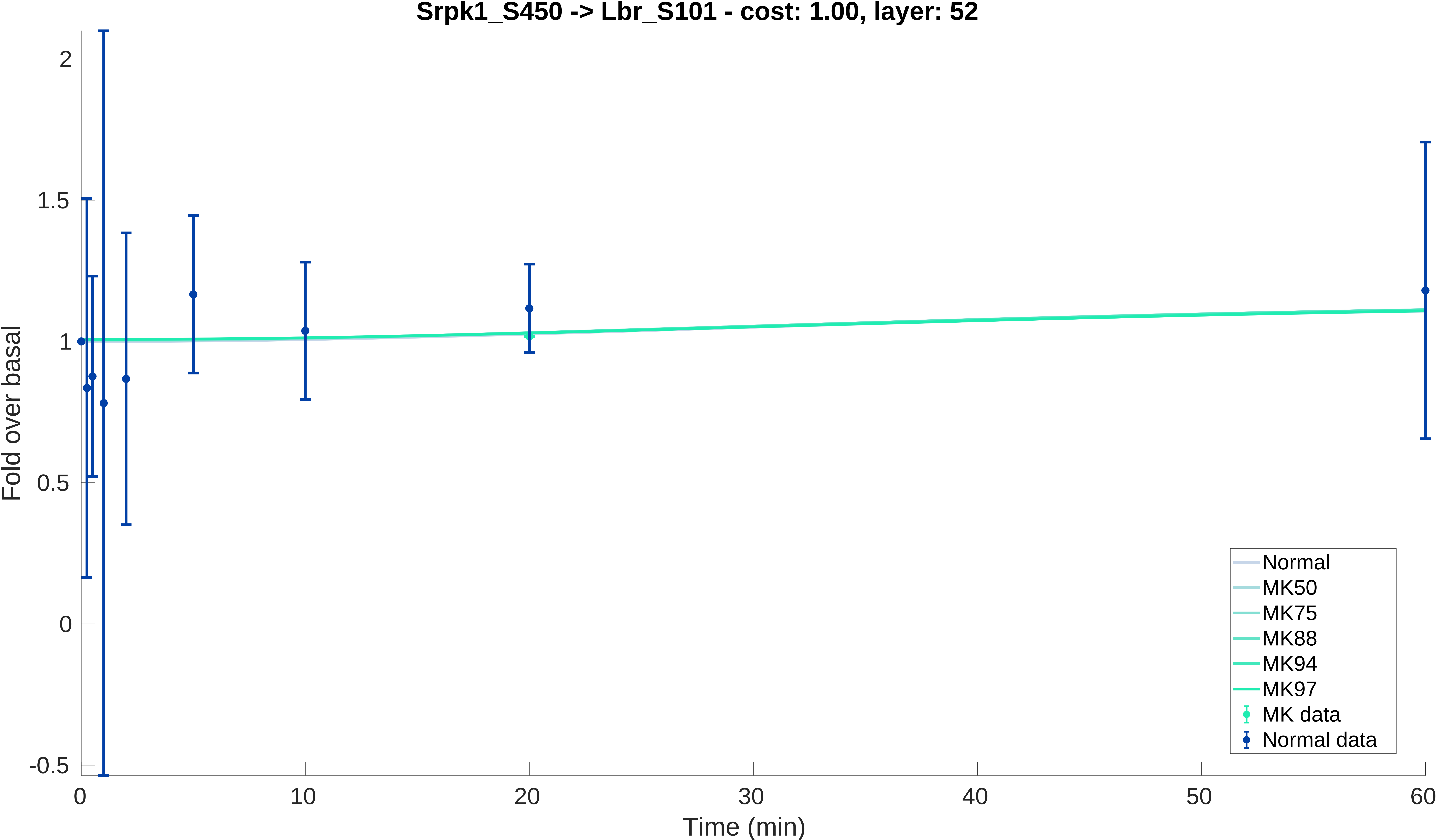
Srpk1_S450 -> Lbr_S101 - cost: 1.00, layer: 52
Fold over basal
Time (min)
2
1.5
1
0.5
0
-0.5
0
10
20
30
40
50
60
Normal
MK50
MK75
MK88
MK94
MK97
MK data
Normal data

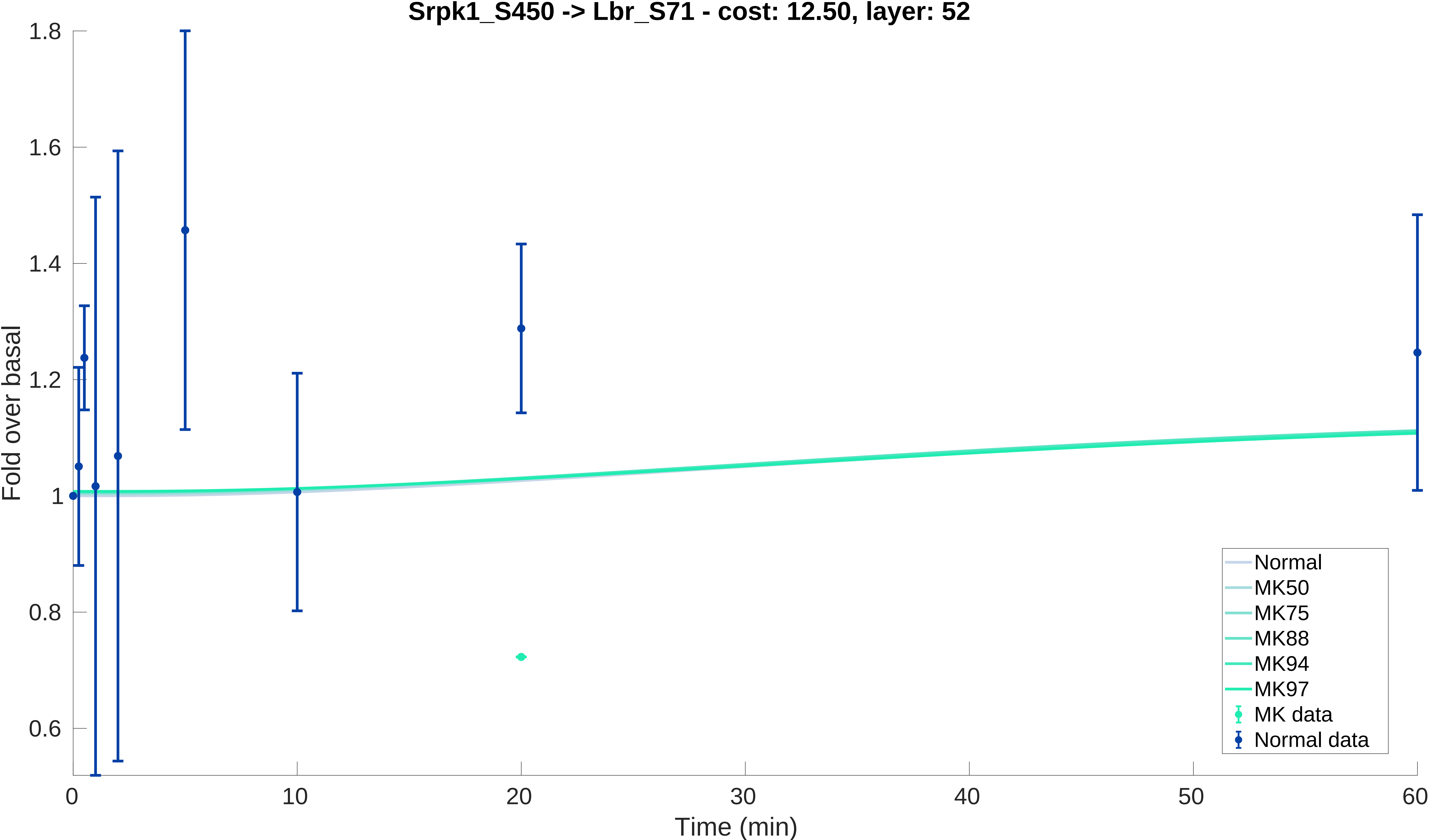
Srpk1_S450 -> Lbr_S71 - cost: 12.50, layer: 52
1.8
1.6
1.4
1.2
1
0.8
0.6
Fold over basal
0
10
20
30
40
50
60
Time (min)
Normal
MK50
MK75
MK88
MK94
MK97
MK data
Normal data

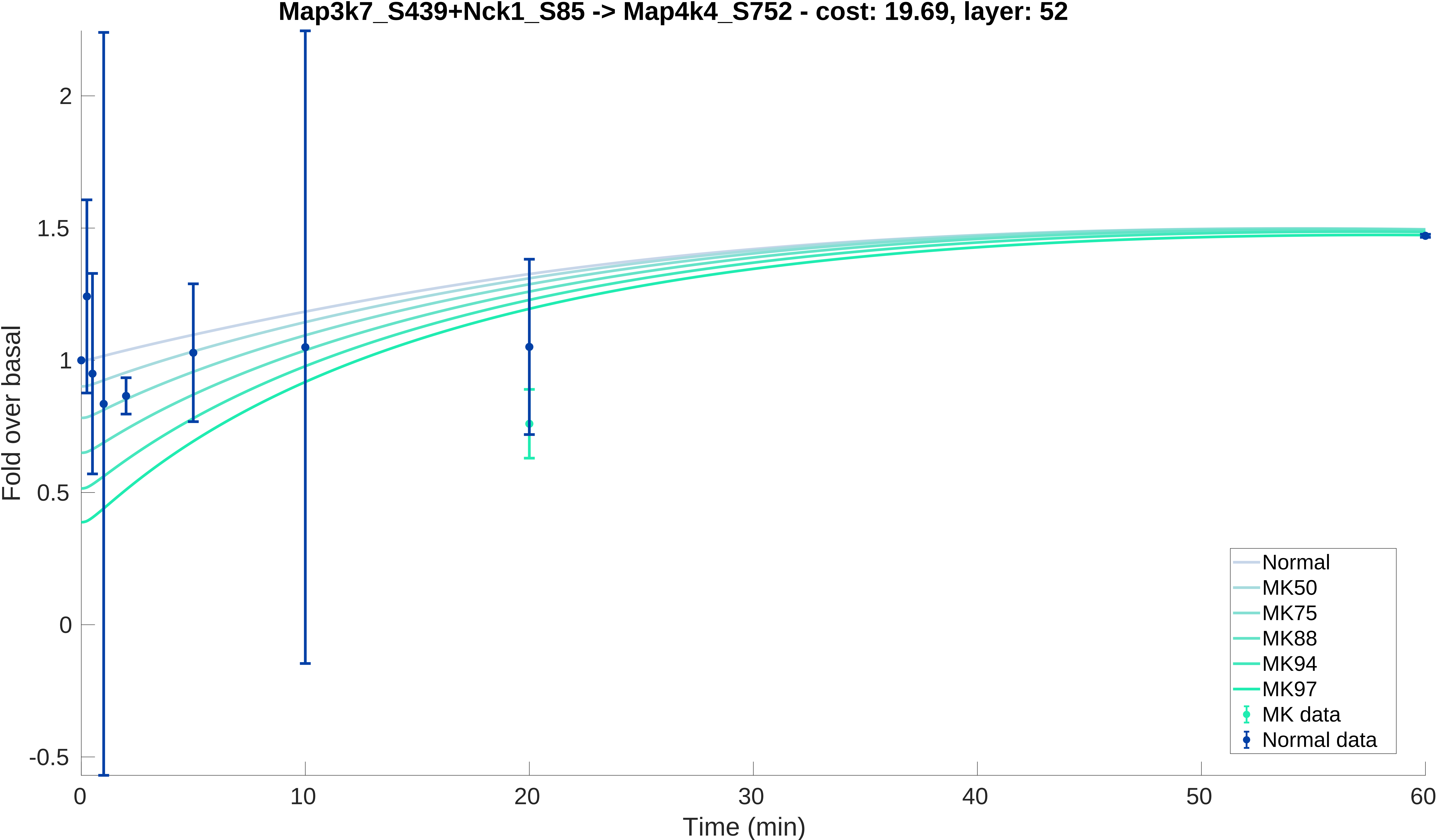
Map3k7_S439+Nck1_S85 -> Map4k4_S752 - cost: 19.69, layer: 52
Fold over basal
Time (min)
2
1.5
1
0.5
0
-0.5
0
10
20
30
40
50
60
Normal
MK50
MK75
MK88
MK94
MK97
MK data
Normal data

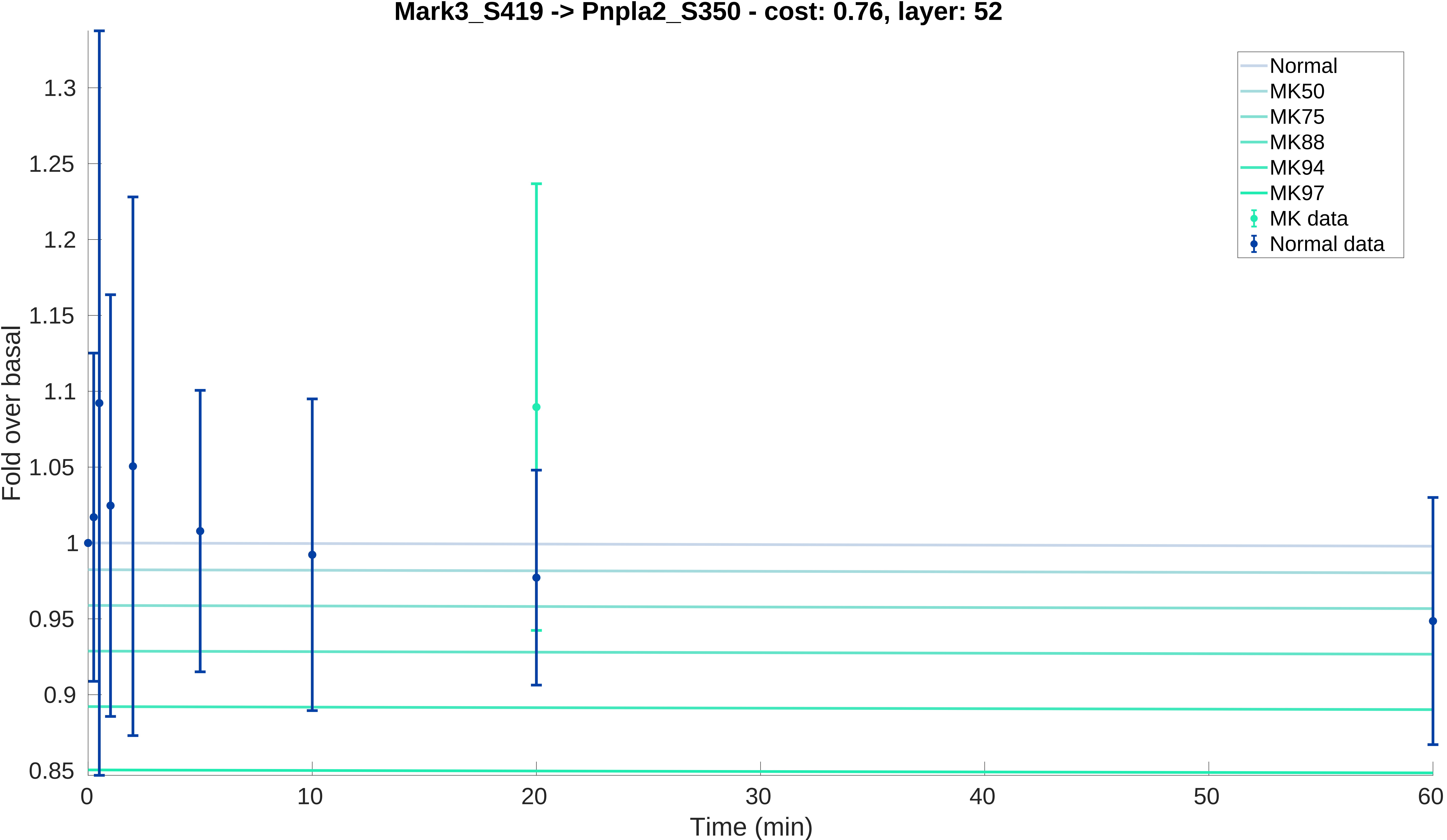

Mark3_S419 -> Pnpla2_S350 - cost: 0.76, layer: 52
Normal
MK50
MK75
MK88
MK94
MK97
MK data
Normal data
Fold over basal
1.3
1.25
1.2
1.15
1.1
1.05
1
0.95
0.9
0.85
0
10
20
30
40
50
60
Time (min)

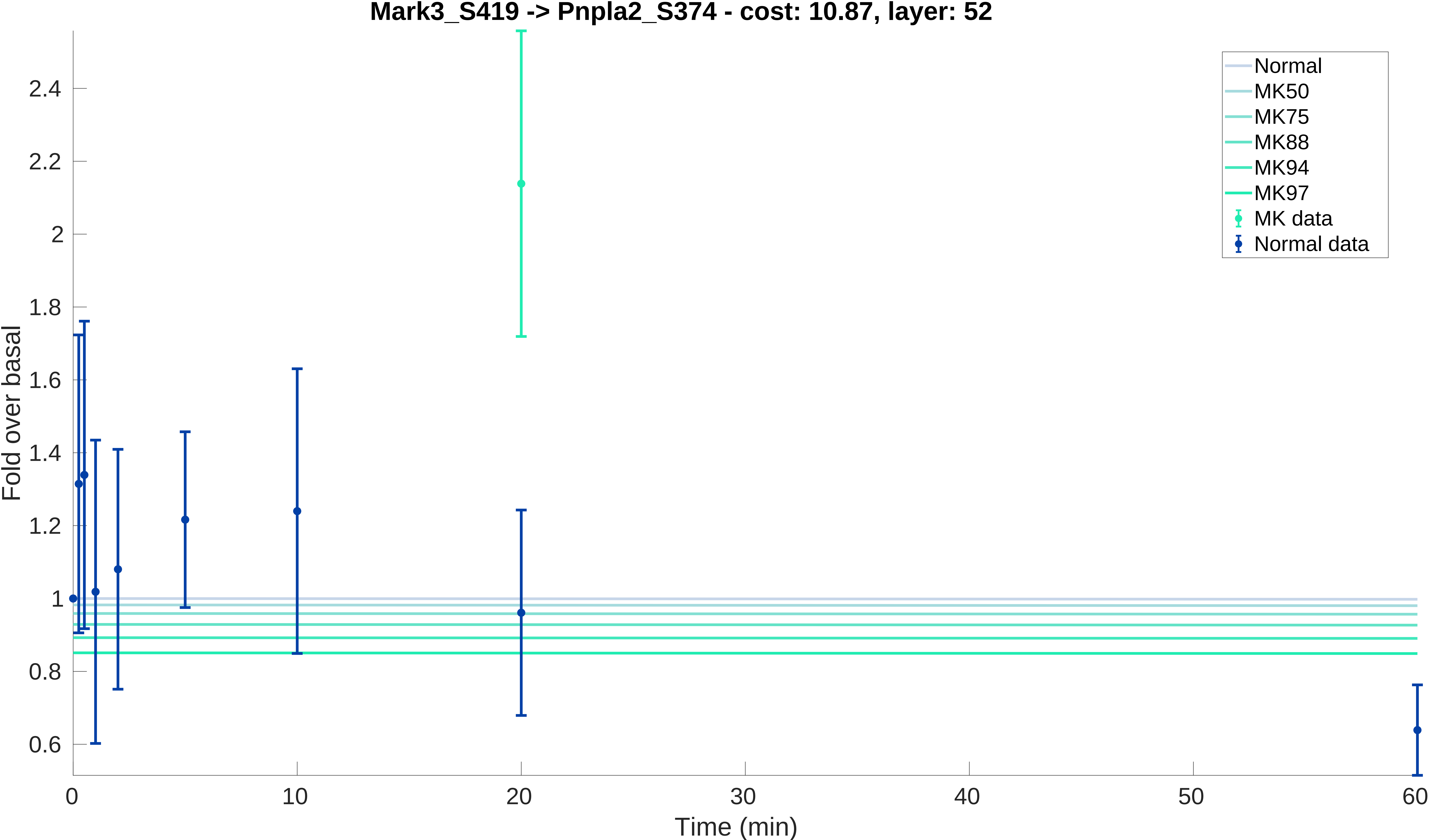

Mark3_S419 -> Pnpla2_S374 - cost: 10.87, layer: 52
Fold over basal
Time (min)
0
10
20
30
40
50
60
0.6
0.8
1
1.2
1.4
1.6
1.8
2
2.2
2.4
Normal
MK50
MK75
MK88
MK94
MK97
MK data
Normal data

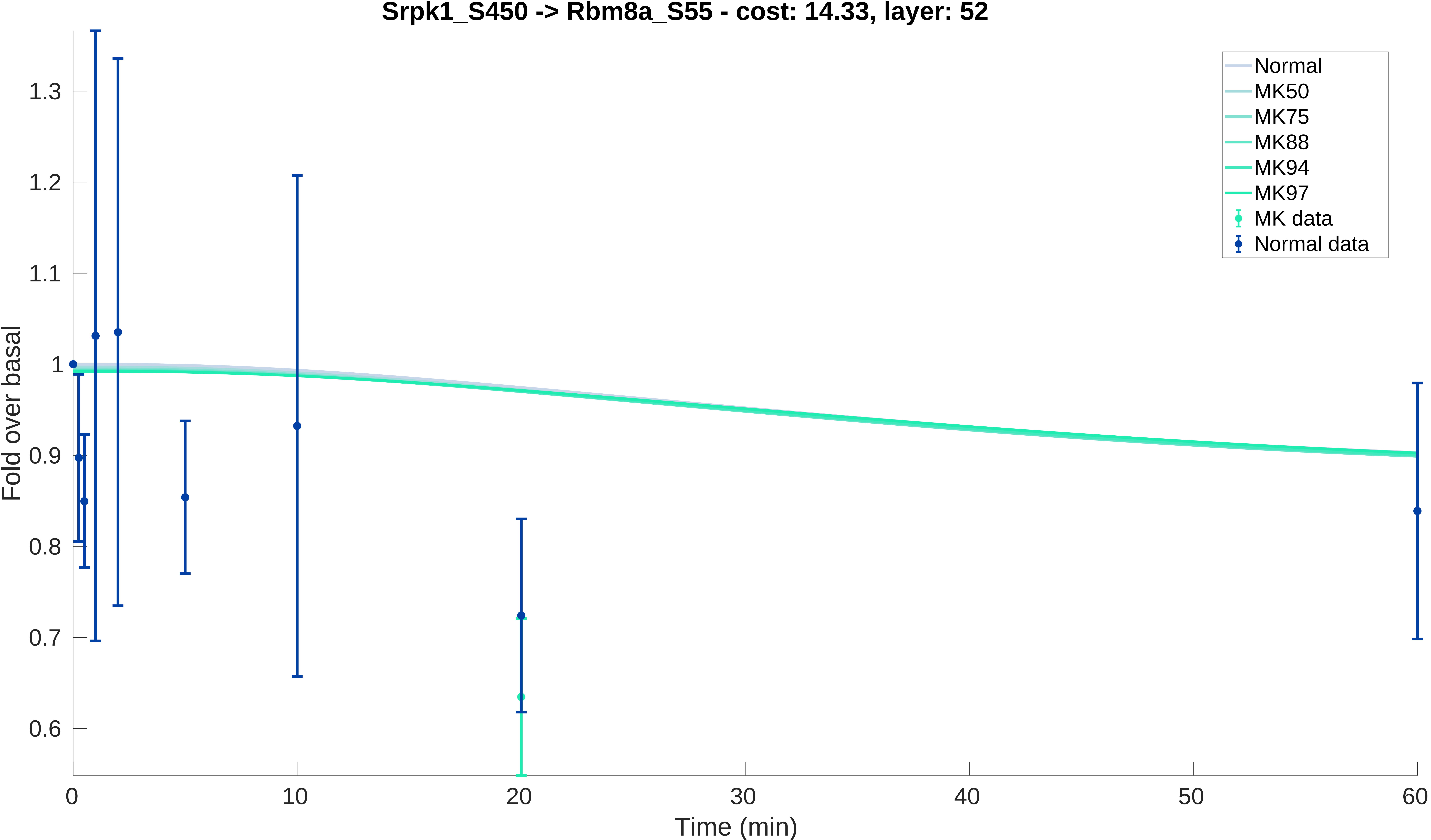

Srpk1_S450 -> Rbm8a_S55 - cost: 14.33, layer: 52
Fold over basal
Time (min)
0.6
0.7
0.8
0.9
1
1.1
1.2
1.3
0
10
20
30
40
50
60
Normal
MK50
MK75
MK88
MK94
MK97
MK data
Normal data

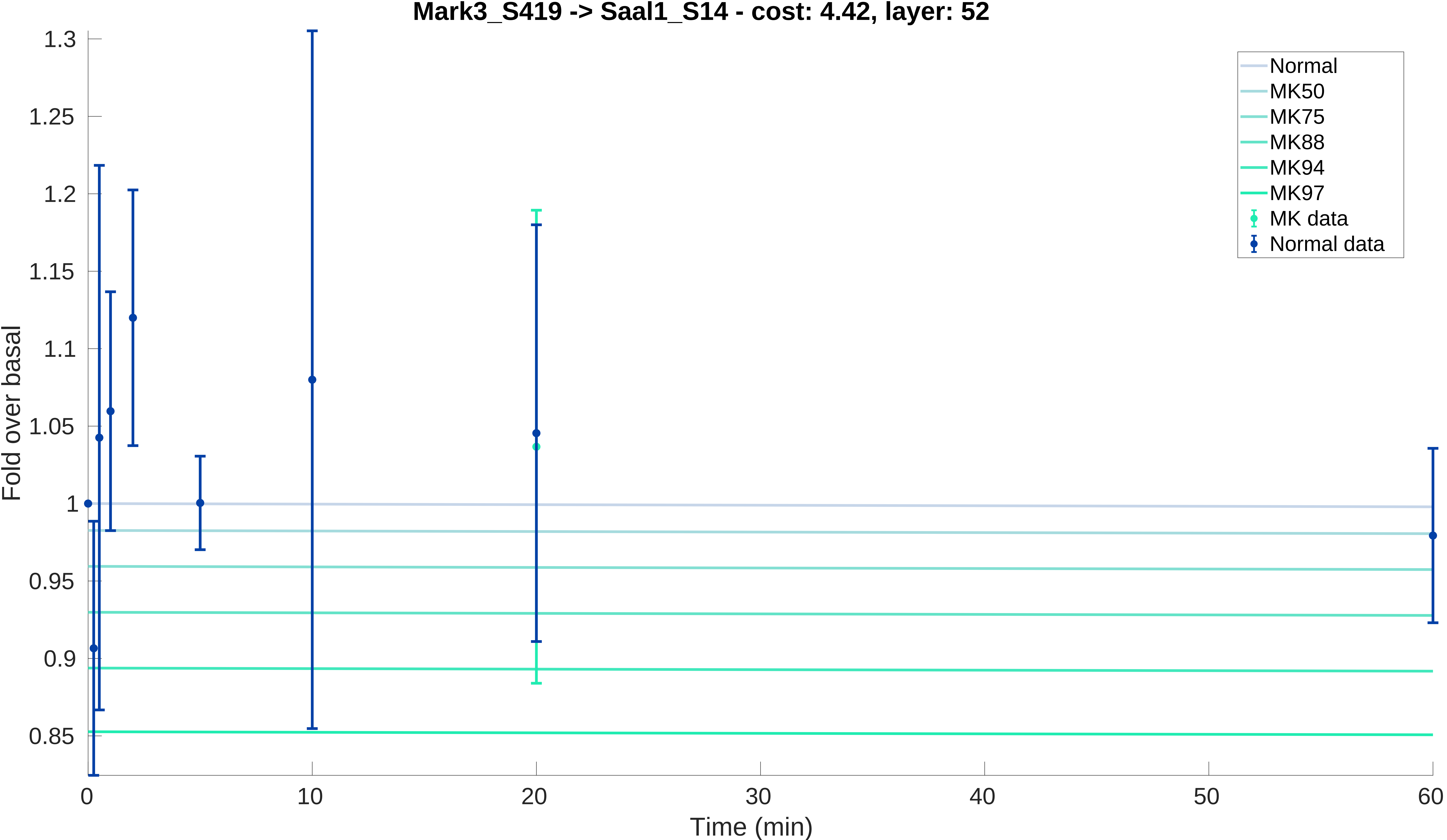
Mark3_S419 -> Saal1_S14 - cost: 4.42, layer: 52
Fold over basal
1.3
1.25
1.2
1.15
1.1
1.05
1
0.95
0.9
0.85
0
10
20
30
40
50
60
Time (min)
Normal
MK50
MK75
MK88
MK94
MK97
MK data
Normal data

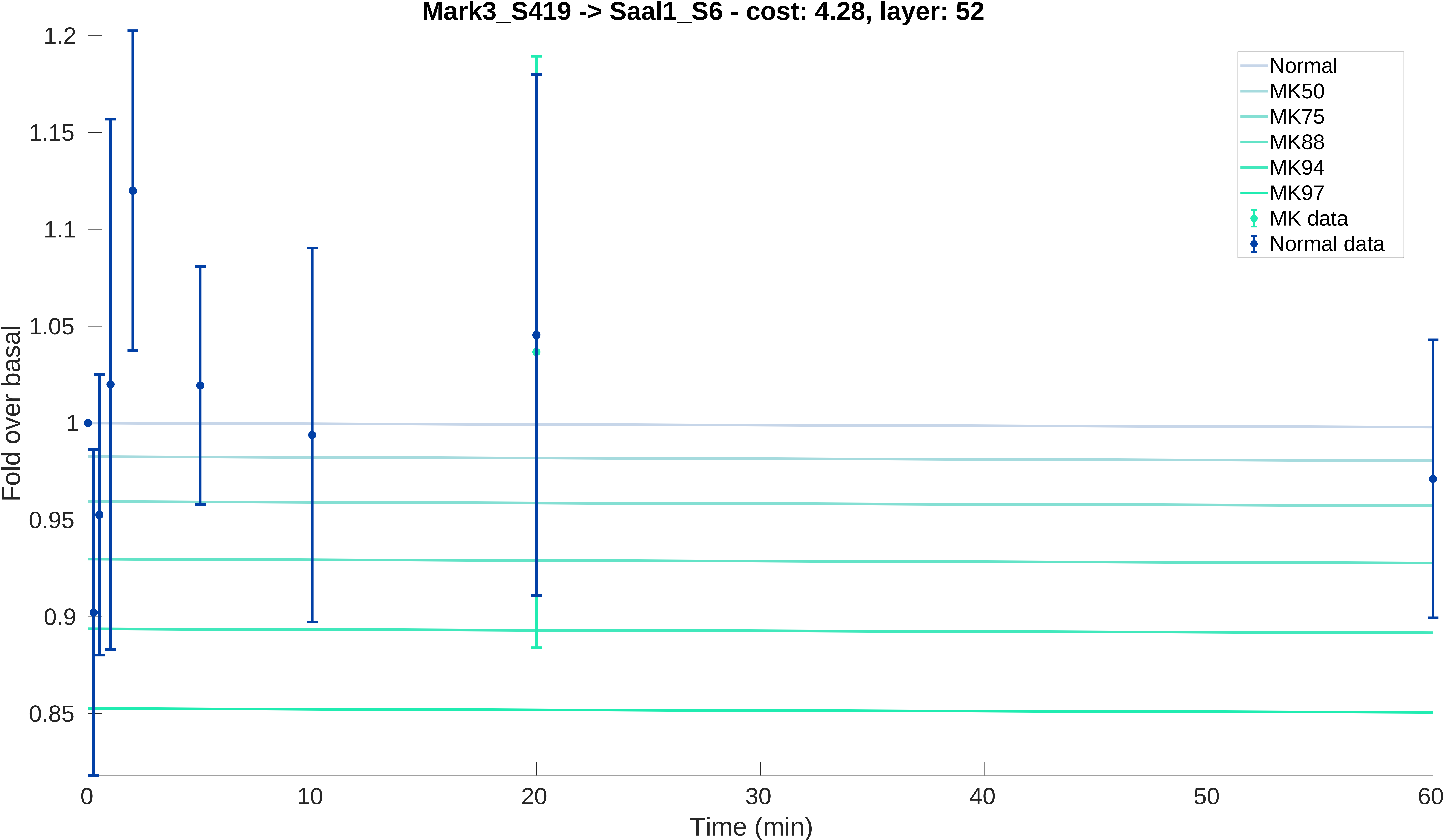
Mark3_S419 -> Saal1_S6 - cost: 4.28, layer: 52
Fold over basal
Time (min)
0.85
0.9
0.95
1
1.05
1.1
1.15
1.2
0
10
20
30
40
50
60
Normal
MK50
MK75
MK88
MK94
MK97
MK data
Normal data

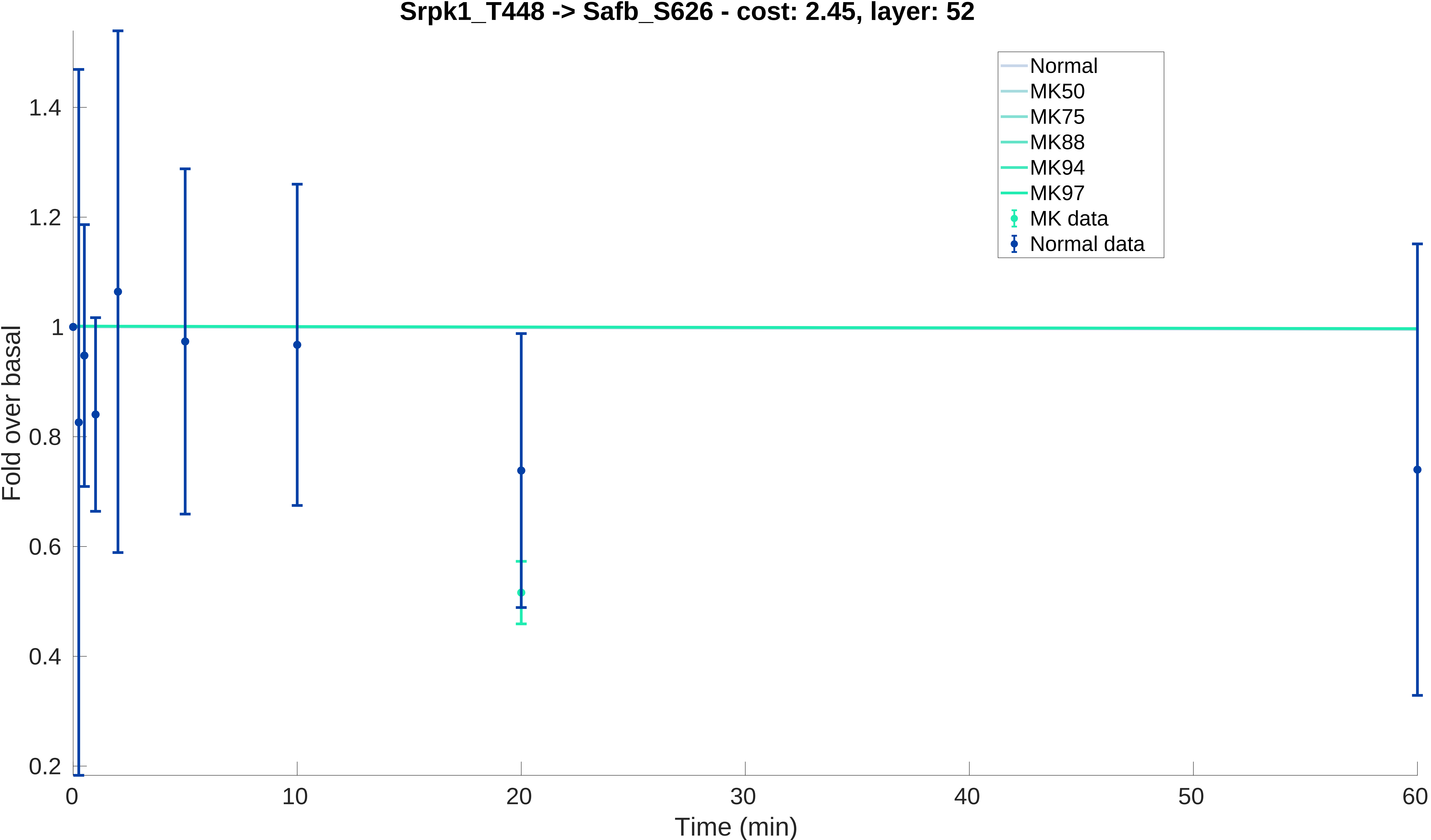

Srpk1_T448 -> Safb_S626 - cost: 2.45, layer: 52
Normal
MK50
MK75
MK88
MK94
MK97
MK data
Normal data
Fold over basal
1.4
1.2
1
0.8
0.6
0.4
0.2
0
10
20
30
40
50
60
Time (min)

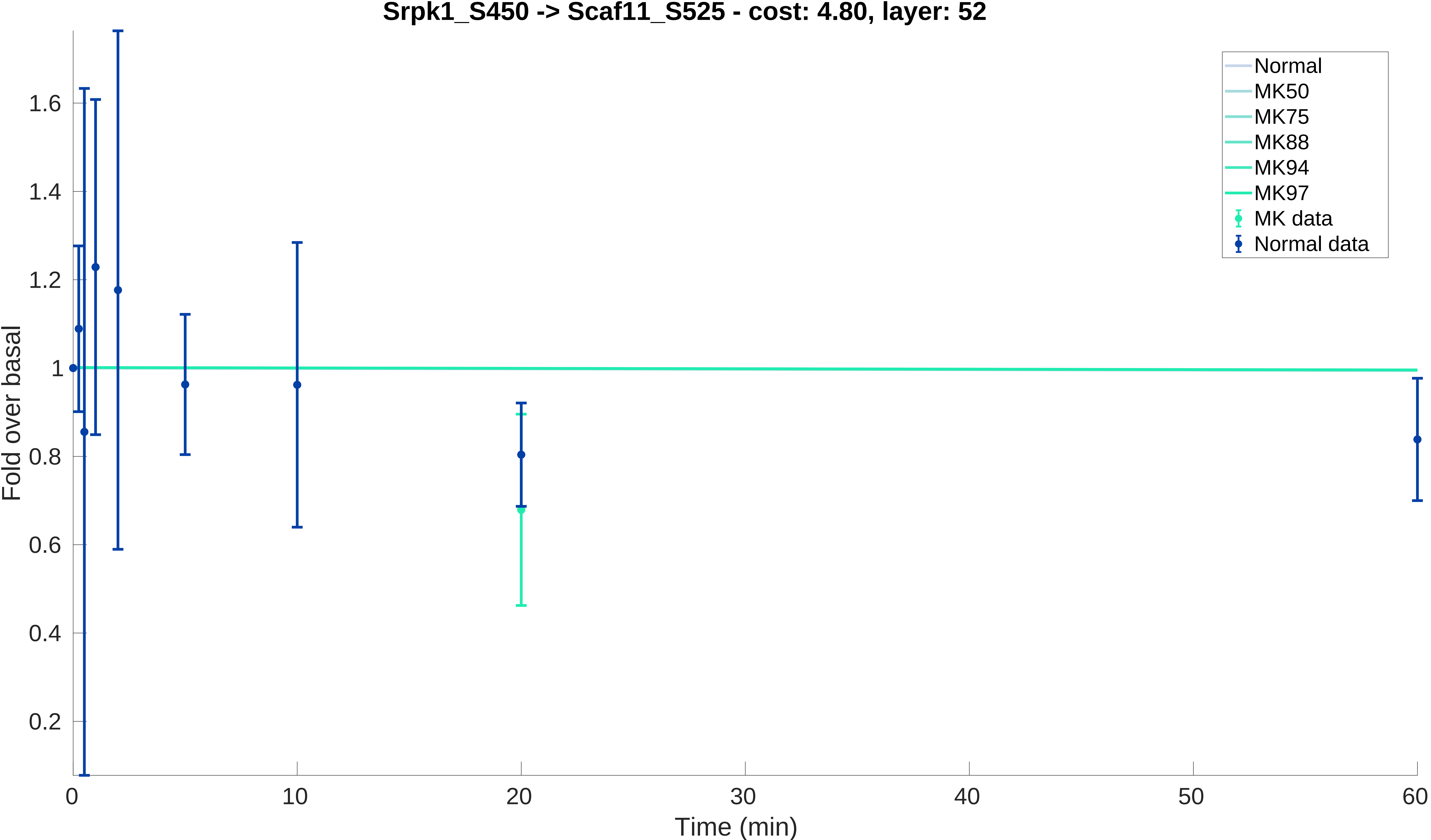
Srpk1_S450 -> Scaf11_S525 - cost: 4.80, layer: 52
Fold over basal
1.6
1.4
1.2
1
0.8
0.6
0.4
0.2
0
10
20
30
40
50
60
Time (min)
Normal
MK50
MK75
MK88
MK94
MK97
MK data
Normal data

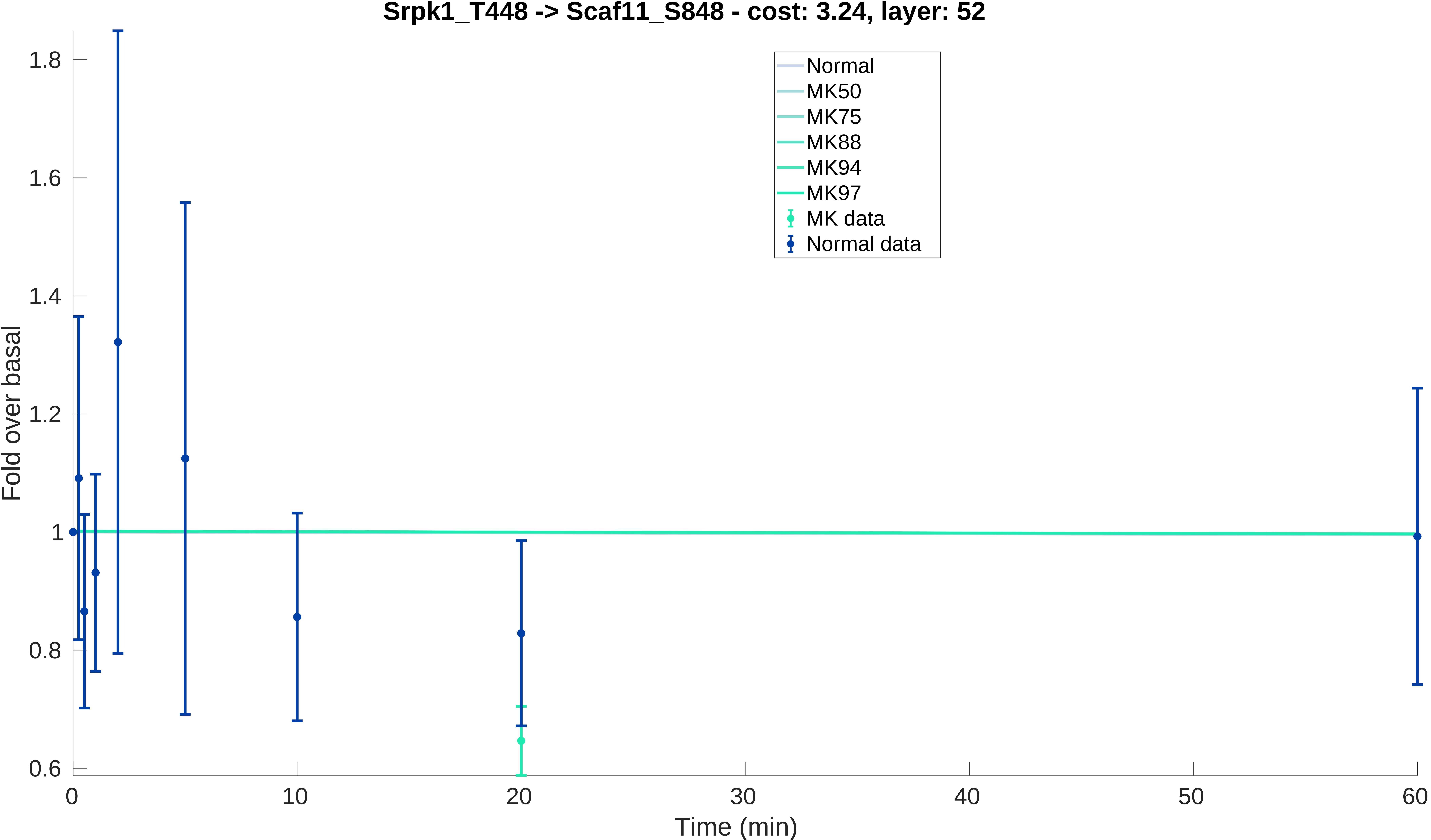

Srpk1_T448 -> Scaf11_S848 - cost: 3.24, layer: 52
Normal
MK50
MK75
MK88
MK94
MK97
MK data
Normal data
Fold over basal
Time (min)
0.6
0.8
1
1.2
1.4
1.6
1.8
0
10
20
30
40
50
60

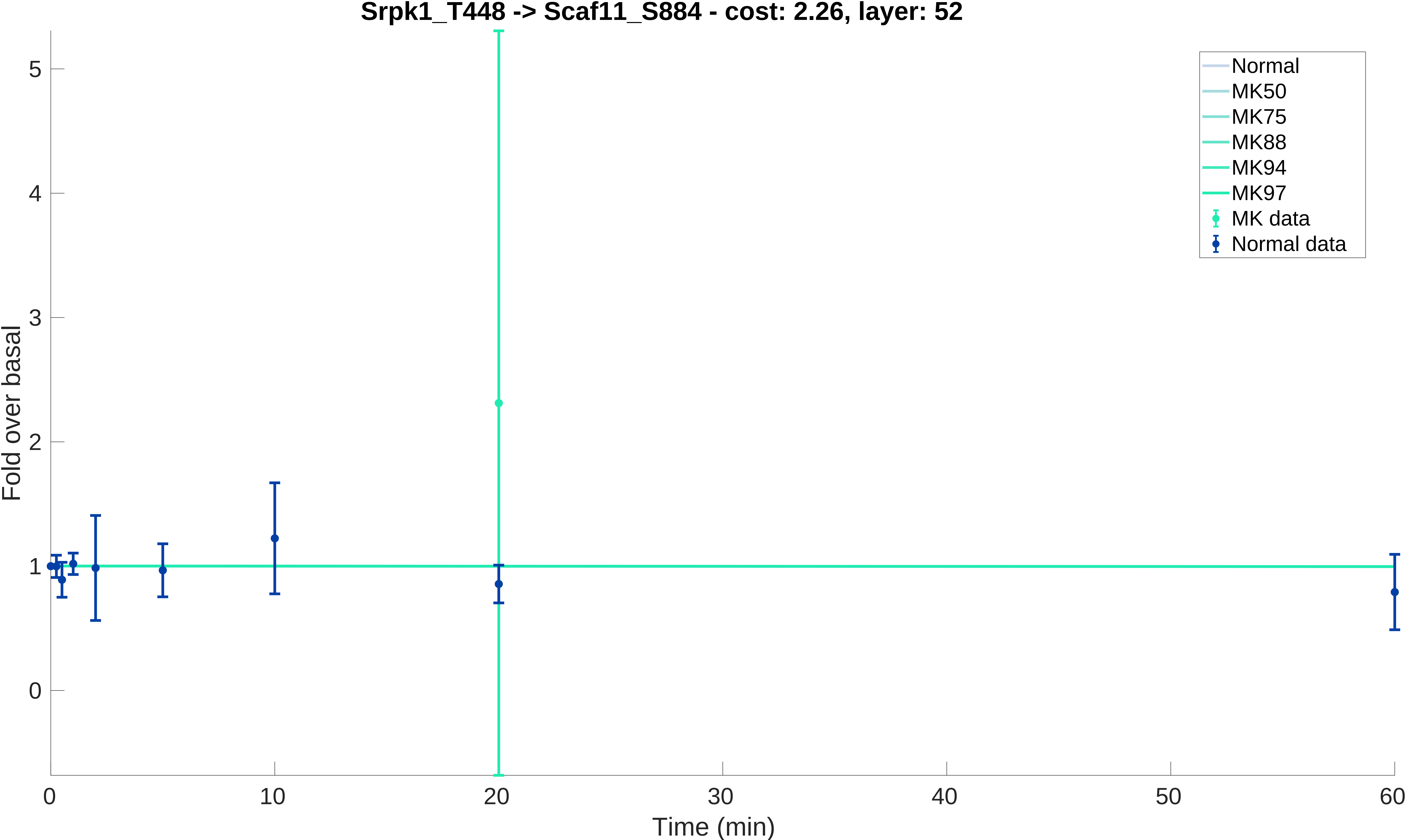

Srpk1_T448 -> Scaf11_S884 - cost: 2.26, layer: 52
Fold over basal
Time (min)
0
1
2
3
4
5
0
10
20
30
40
50
60
Normal
MK50
MK75
MK88
MK94
MK97
MK data
Normal data

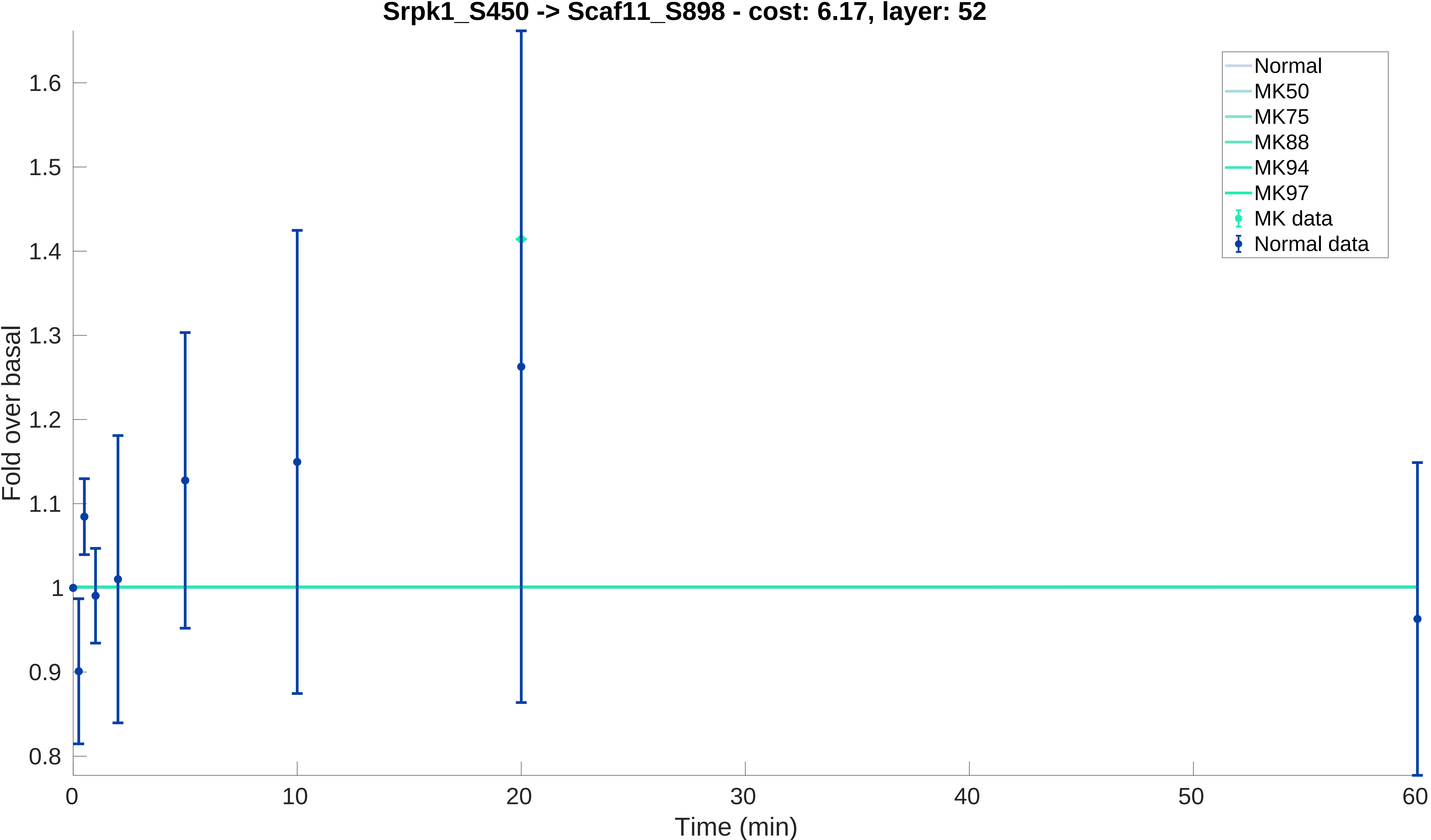
Srpk1_S450 -> Scaf11_S898 - cost: 6.17, layer: 52
Fold over basal
Time (min)
0.8
0.9
1
1.1
1.2
1.3
1.4
1.5
1.6
0
10
20
30
40
50
60
Normal
MK50
MK75
MK88
MK94
MK97
MK data
Normal data

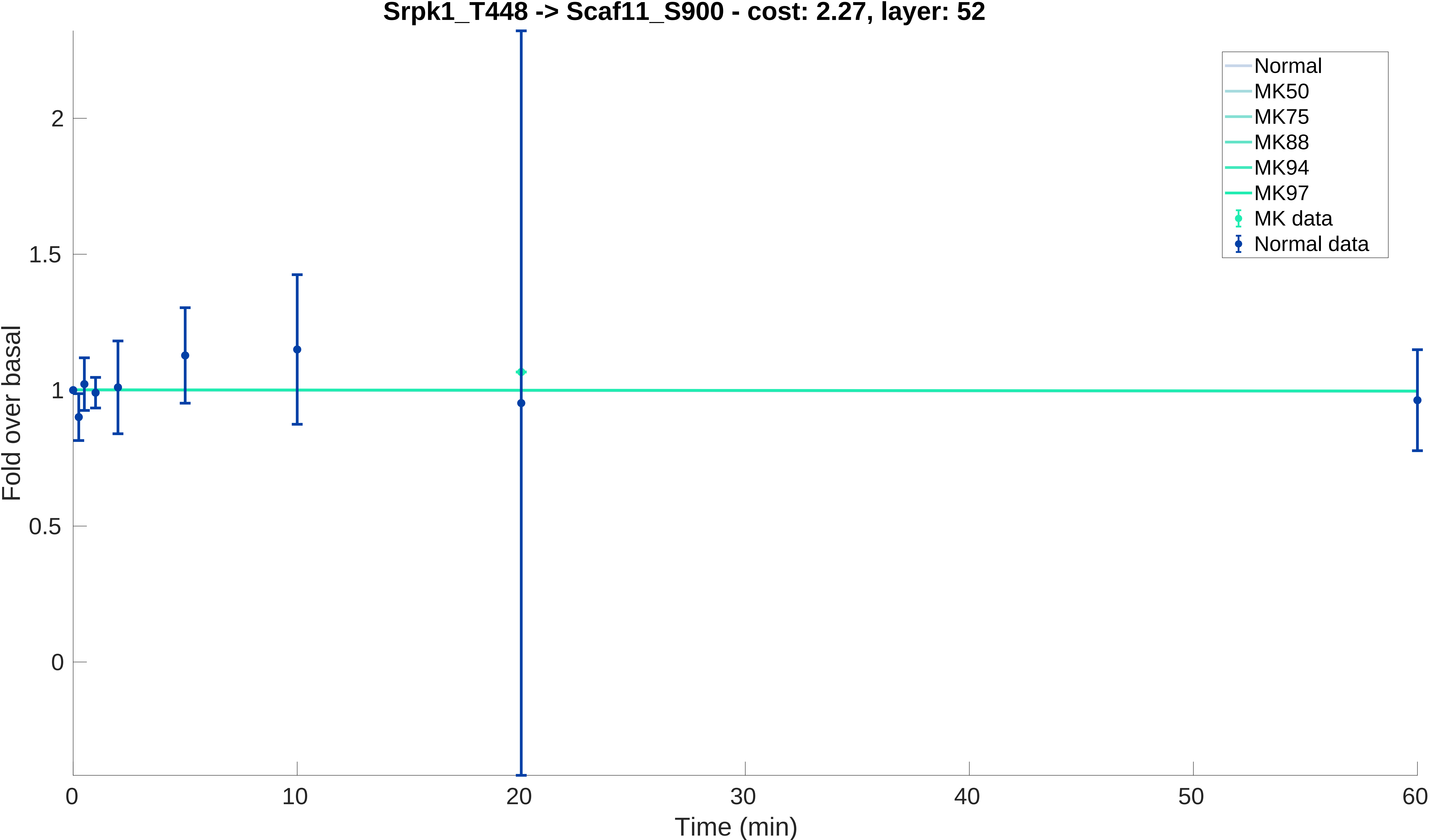

Srpk1_T448 -> Scaf11_S900 - cost: 2.27, layer: 52
Normal
MK50
MK75
MK88
MK94
MK97
MK data
Normal data
2
1.5
1
0.5
0
Fold over basal
0
10
20
30
40
50
60
Time (min)

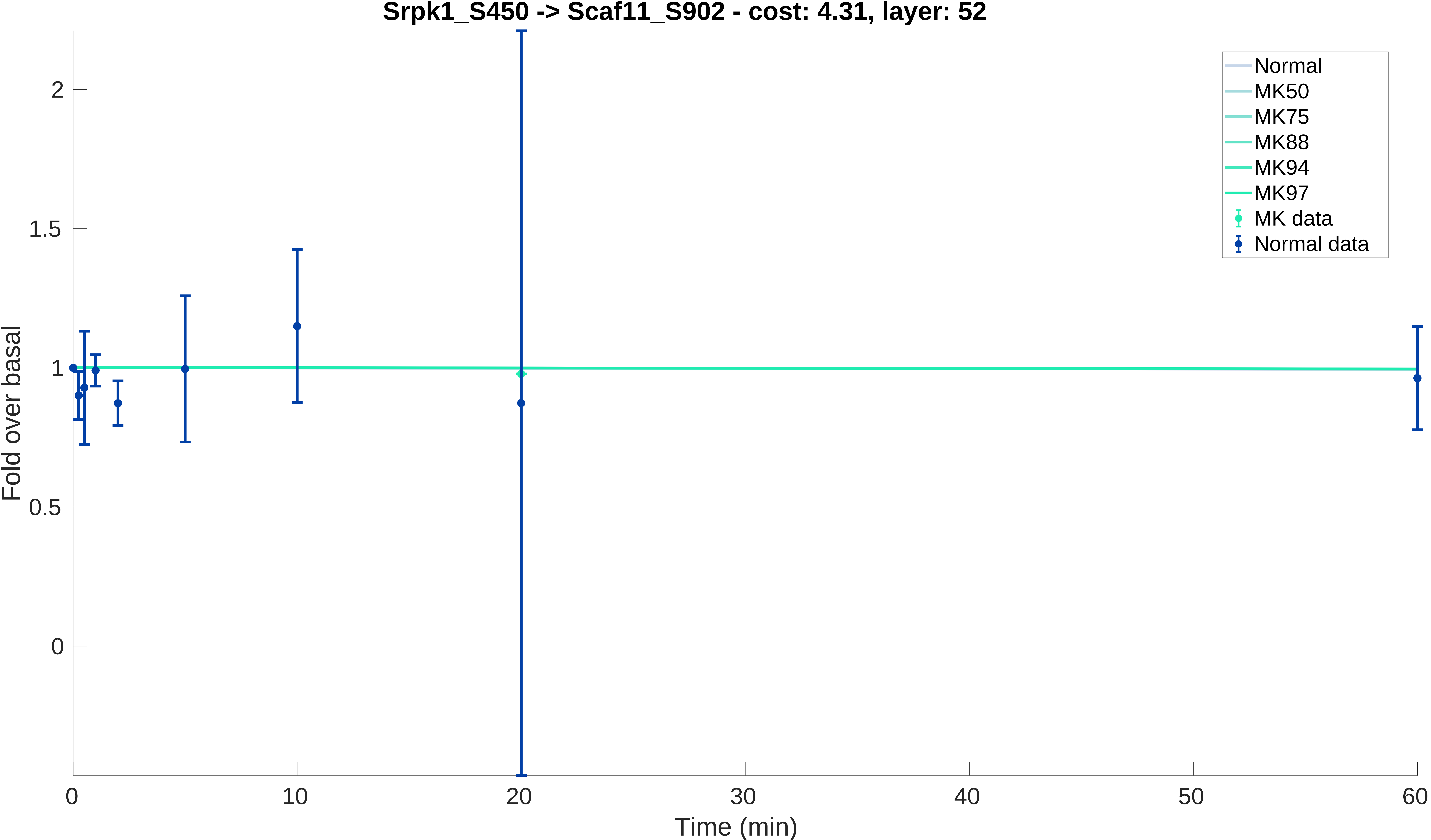

Srpk1_S450 -> Scaf11_S902 - cost: 4.31, layer: 52
Normal
MK50
MK75
MK88
MK94
MK97
MK data
Normal data
Fold over basal
Time (min)
0
0.5
1
1.5
2
0
10
20
30
40
50
60

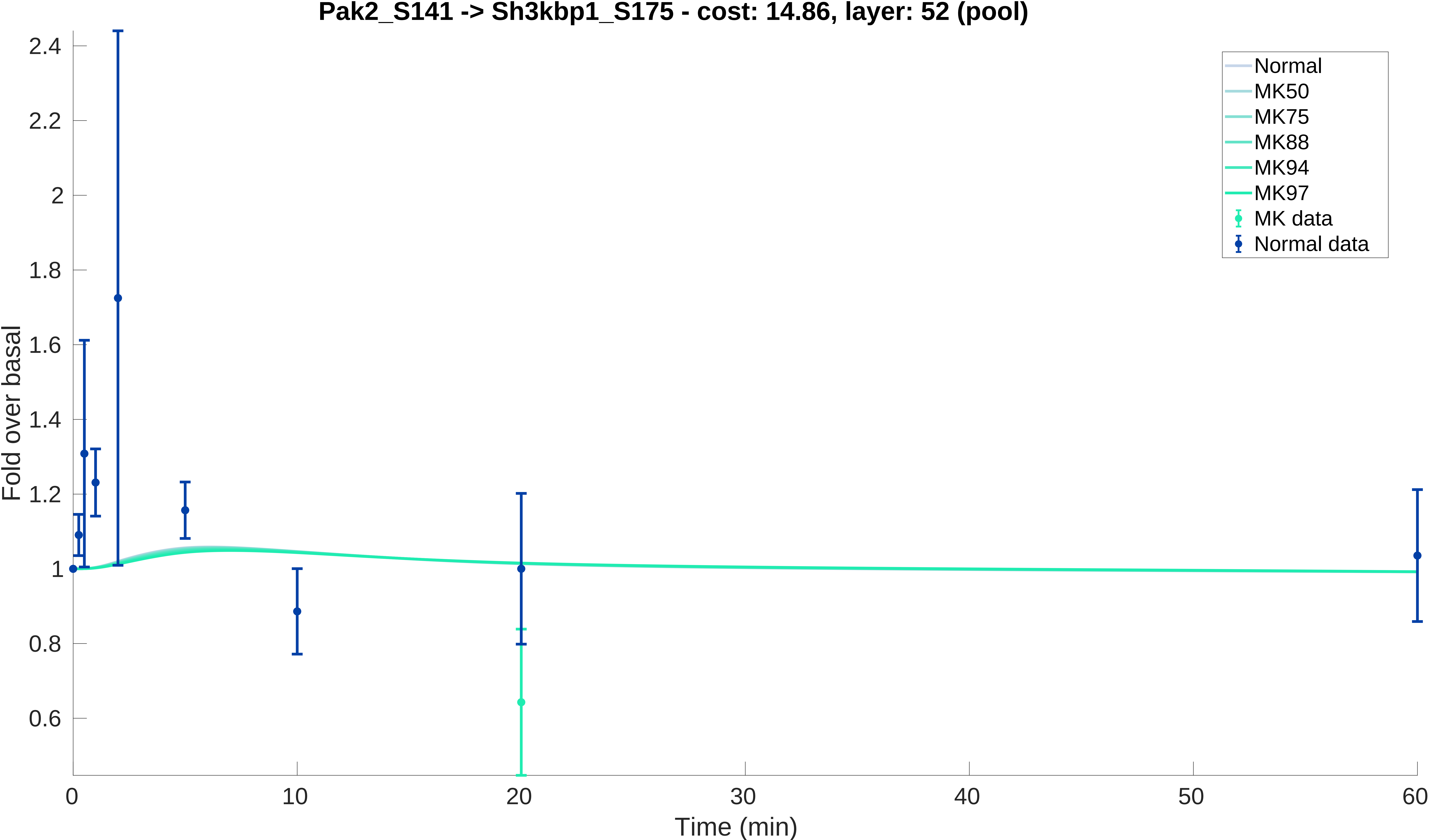

Pak2_S141 -> Sh3kbp1_S175 - cost: 14.86, layer: 52 (pool)
Fold over basal
2.4
2.2
2
1.8
1.6
1.4
1.2
1
0.8
0.6
0
10
20
30
40
50
60
Time (min)
Normal
MK50
MK75
MK88
MK94
MK97
MK data
Normal data

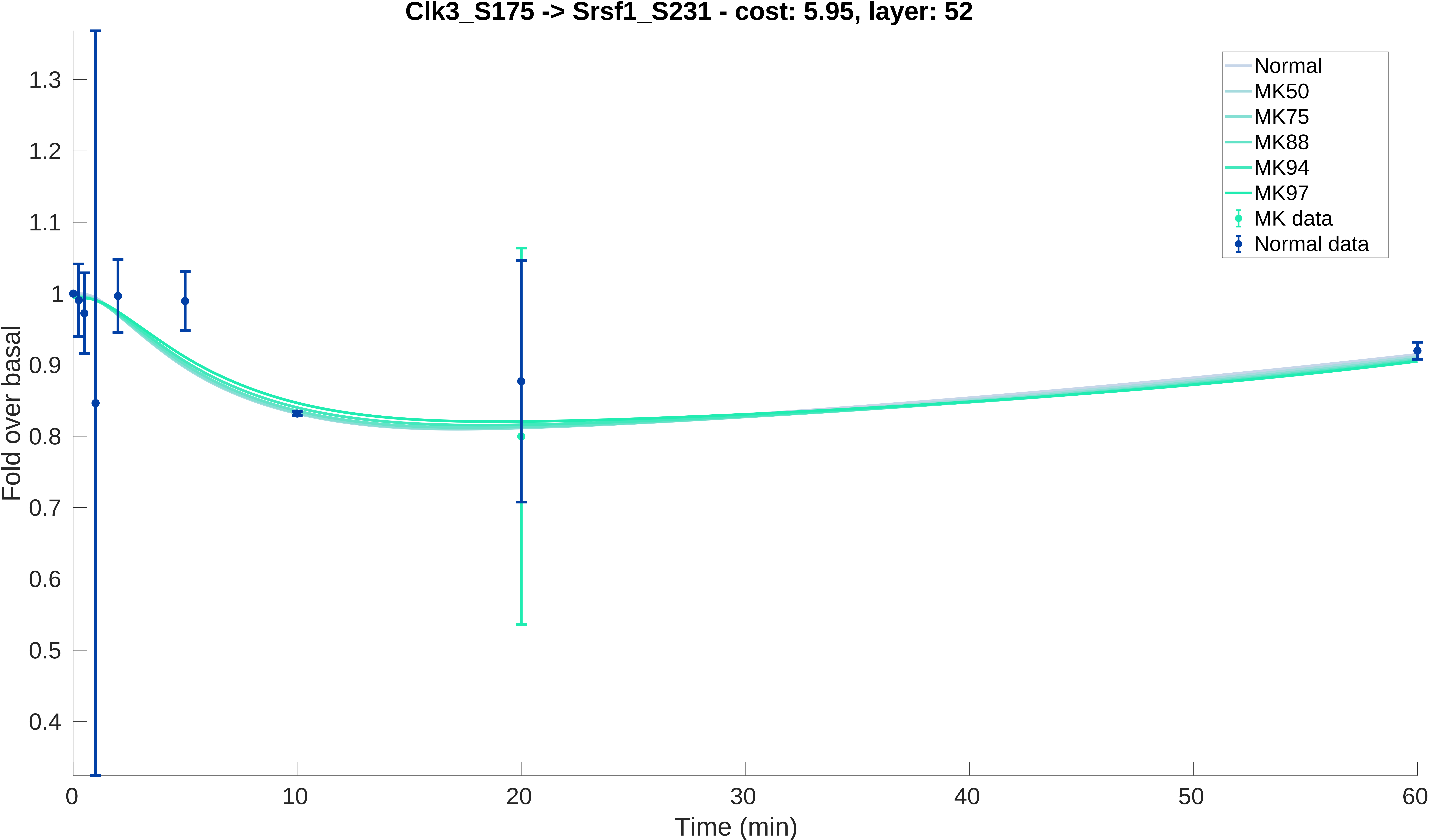

Clk3_S175 -> Srsf1_S231 - cost: 5.95, layer: 52
Normal
MK50
MK75
MK88
MK94
MK97
MK data
Normal data
Fold over basal
1.3
1.2
1.1
1
0.9
0.8
0.7
0.6
0.5
0.4
0
10
20
30
40
50
60
Time (min)

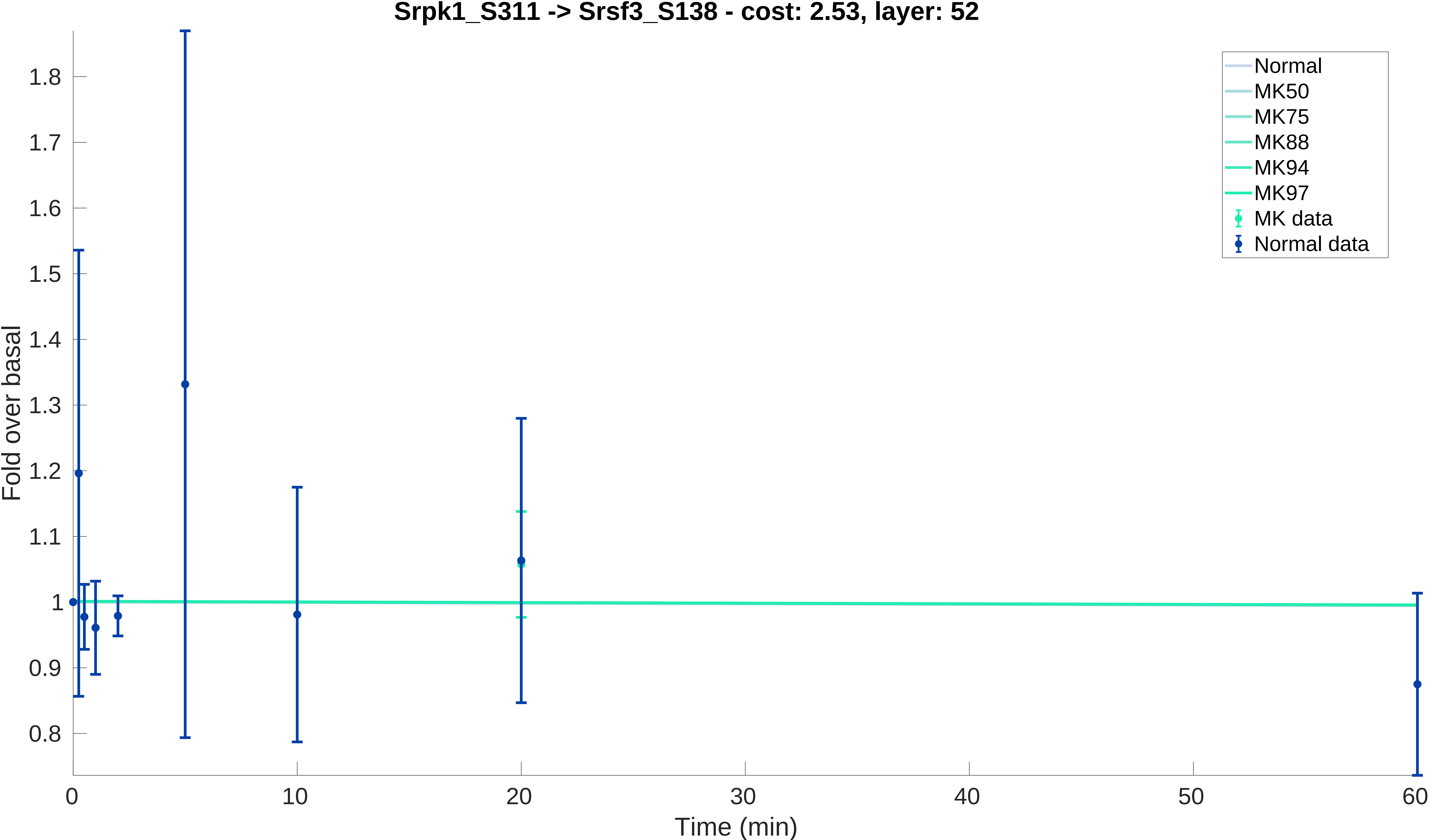

Srpk1_S311 -> Srsf3_S138 - cost: 2.53, layer: 52
Fold over basal
Time (min)
Normal
MK50
MK75
MK88
MK94
MK97
MK data
Normal data
0.8
0.9
1
1.1
1.2
1.3
1.4
1.5
1.6
1.7
1.8
0
10
20
30
40
50
60

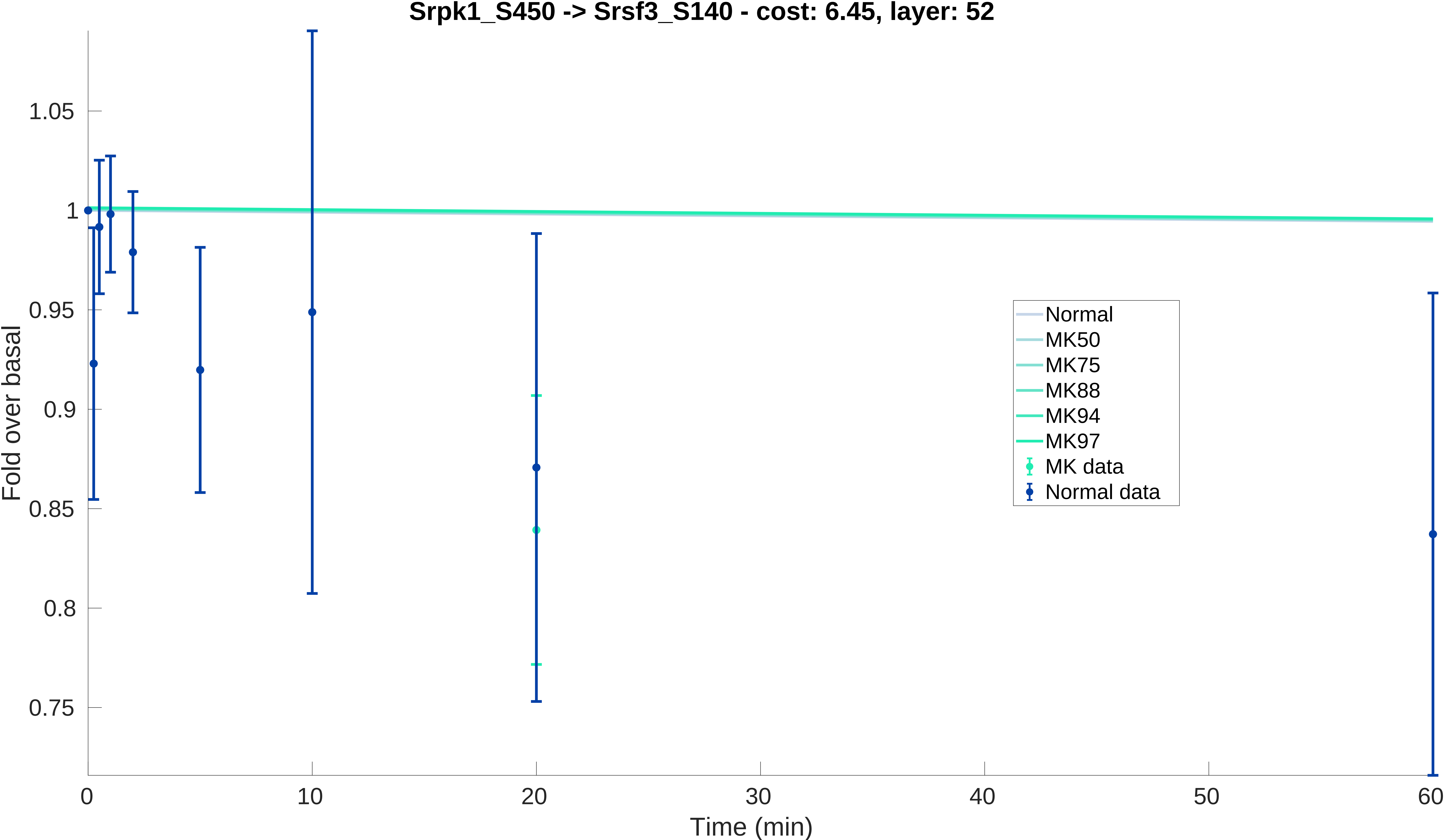

Srpk1_S450 -> Srsf3_S140 - cost: 6.45, layer: 52
Fold over basal
Time (min)
1.05
1
0.95
0.9
0.85
0.8
0.75
0
10
20
30
40
50
60
Normal
MK50
MK75
MK88
MK94
MK97
MK data
Normal data

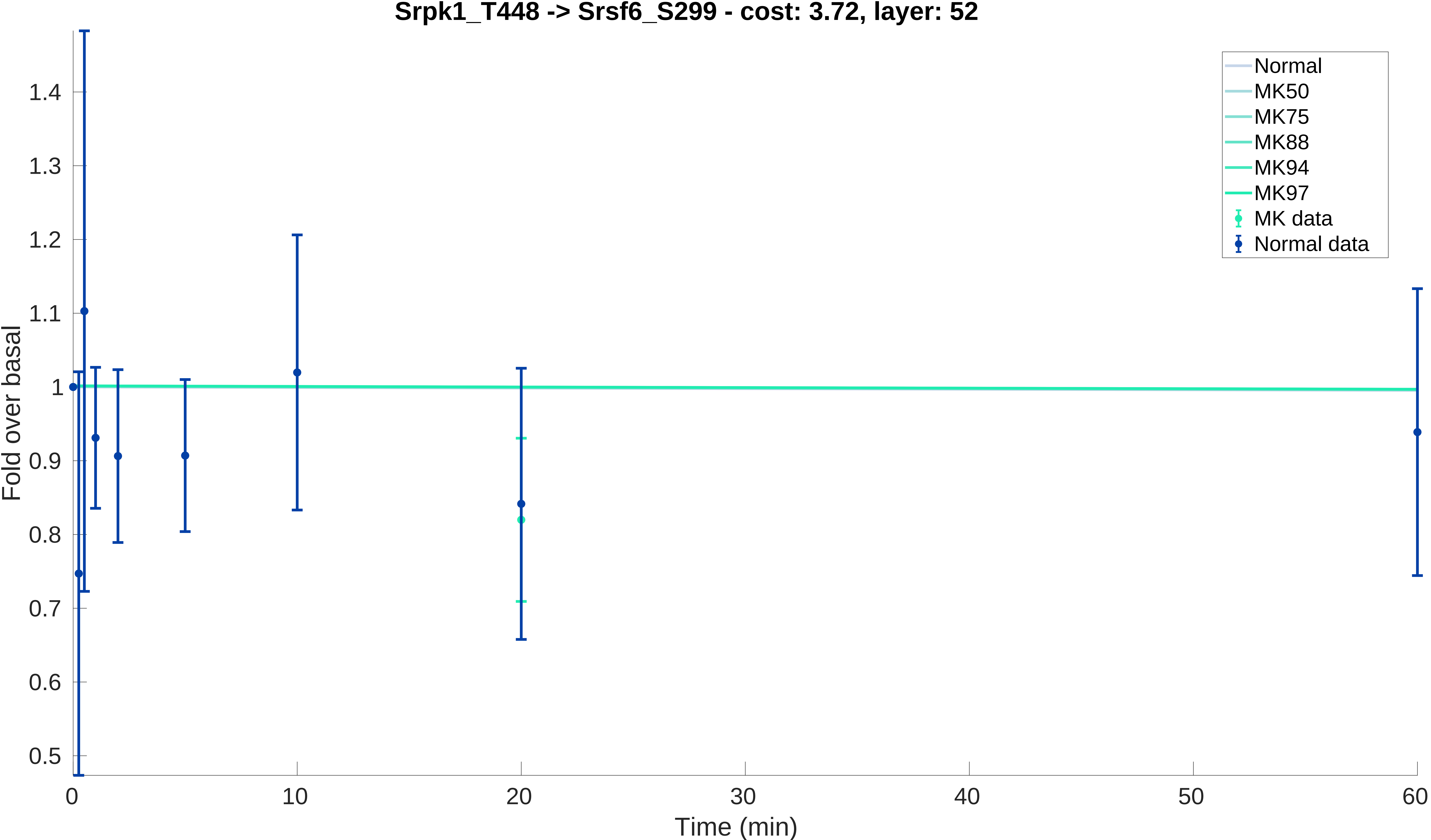

Srpk1_T448 -> Srsf6_S299 - cost: 3.72, layer: 52
Fold over basal
Time (min)
1.4
1.3
1.2
1.1
1
0.9
0.8
0.7
0.6
0.5
0
10
20
30
40
50
60
Normal
MK50
MK75
MK88
MK94
MK97
MK data
Normal data

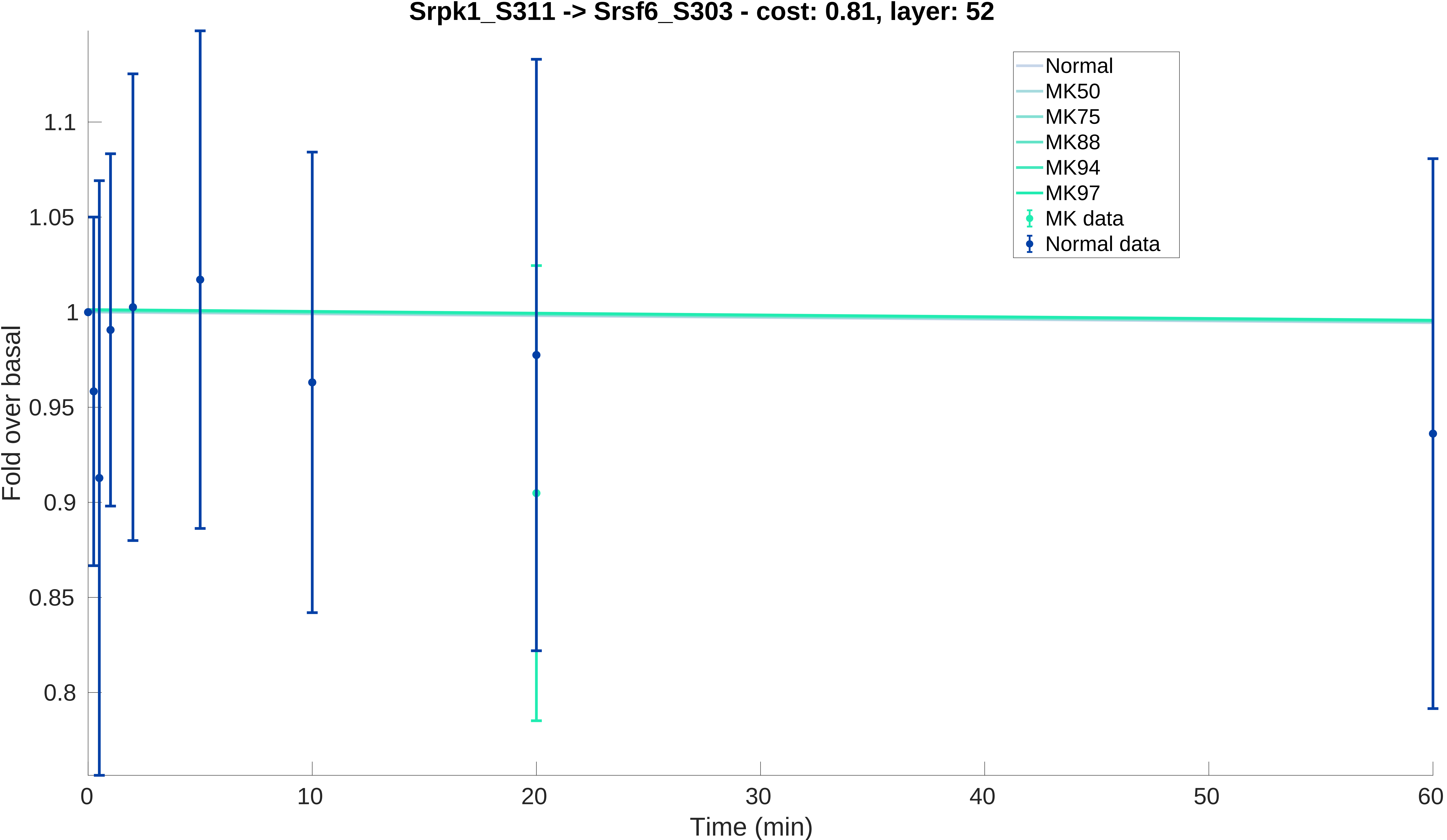

Srpk1_S311 -> Srsf6_S303 - cost: 0.81, layer: 52
Normal
MK50
MK75
MK88
MK94
MK97
MK data
Normal data
Fold over basal
1.1
1.05
1
0.95
0.9
0.85
0.8
0
10
20
30
40
50
60
Time (min)

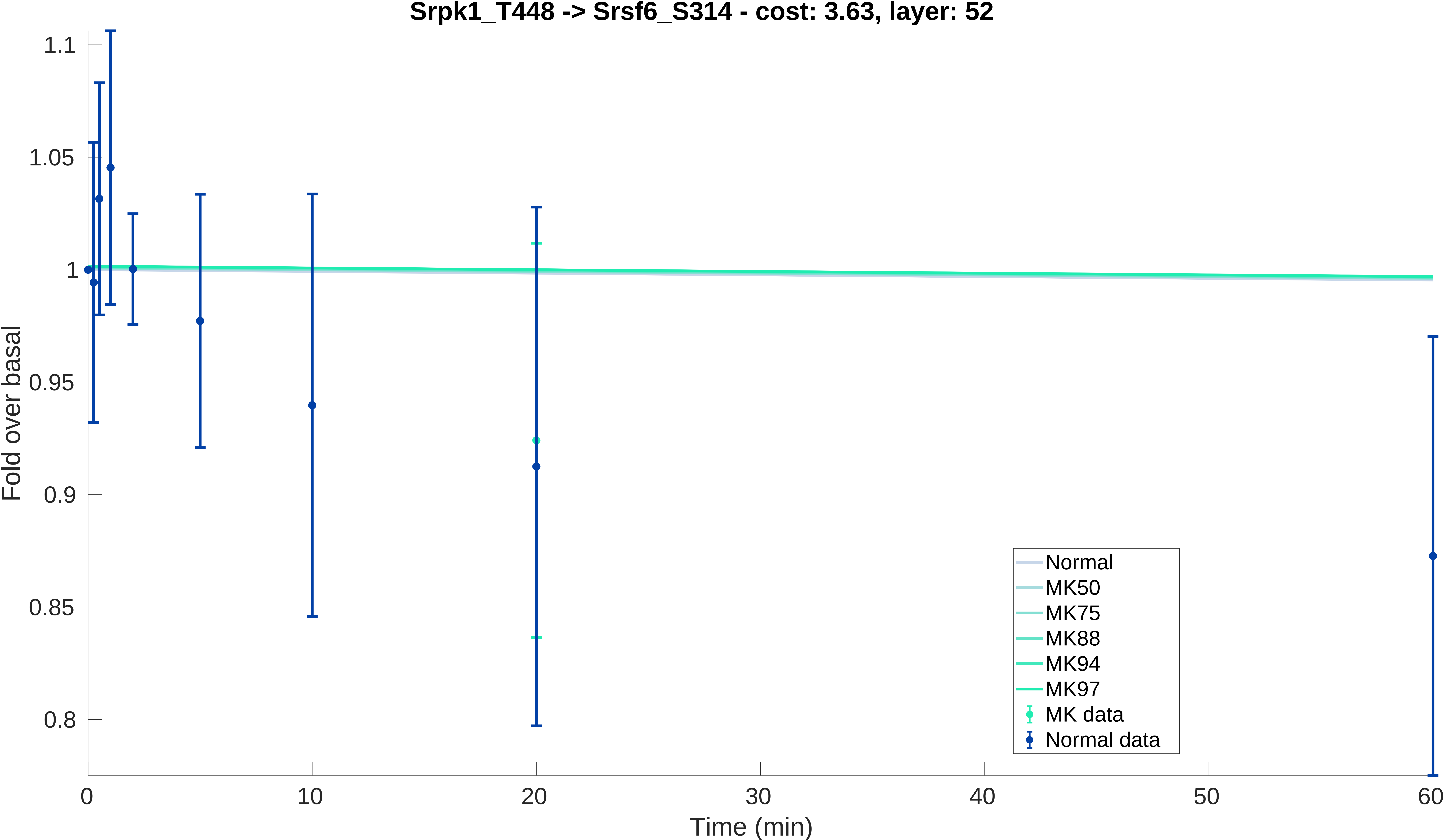

Srpk1_T448 -> Srsf6_S314 - cost: 3.63, layer: 52
Fold over basal
1.1
1.05
1
0.95
0.9
0.85
0.8
0
10
20
30
40
50
60
Time (min)
Normal
MK50
MK75
MK88
MK94
MK97
MK data
Normal data

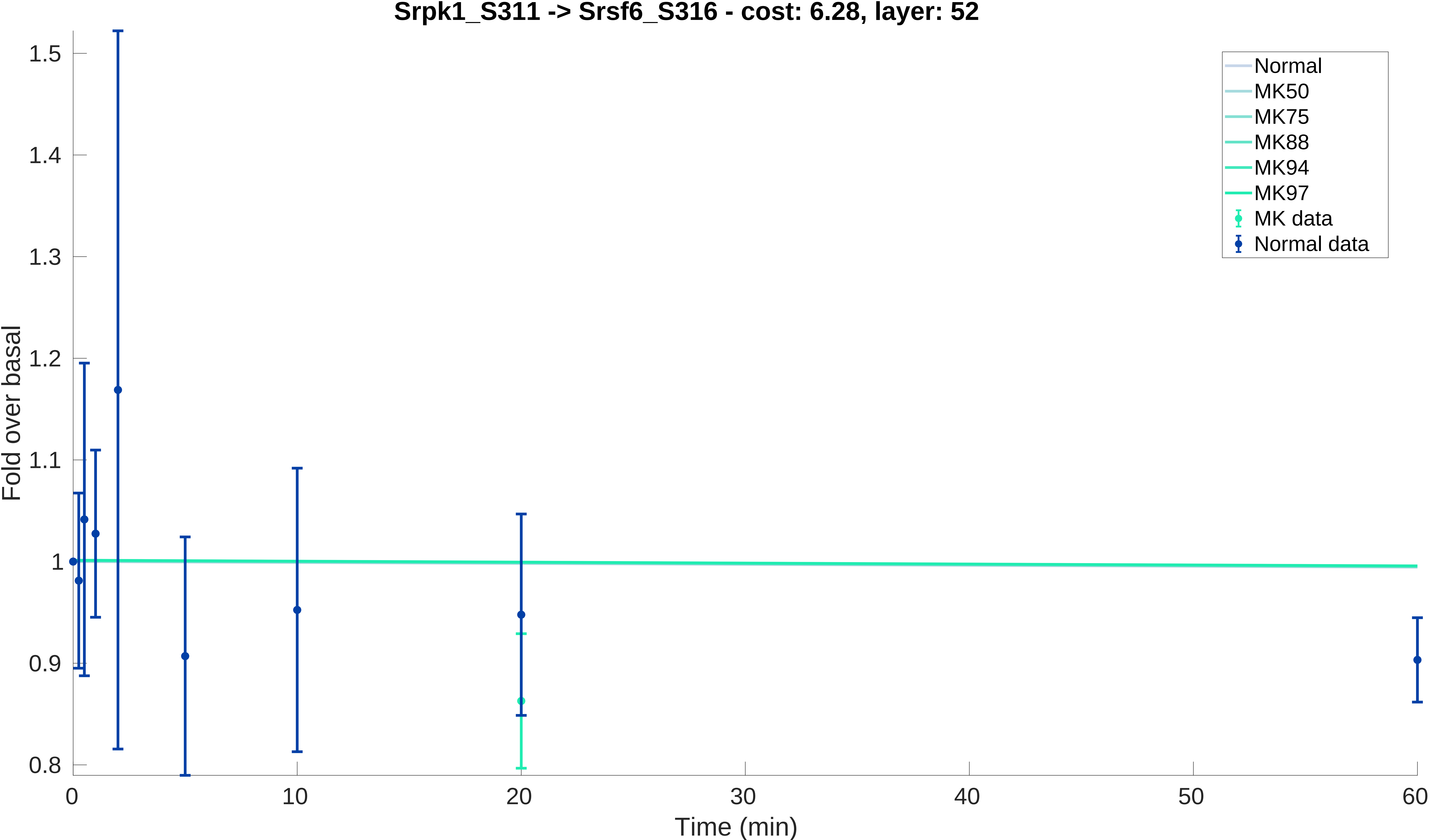

Srpk1_S311 -> Srsf6_S316 - cost: 6.28, layer: 52
Fold over basal
1.5
1.4
1.3
1.2
1.1
1
0.9
0.8
0
10
20
30
40
50
60
Time (min)
Normal
MK50
MK75
MK88
MK94
MK97
MK data
Normal data

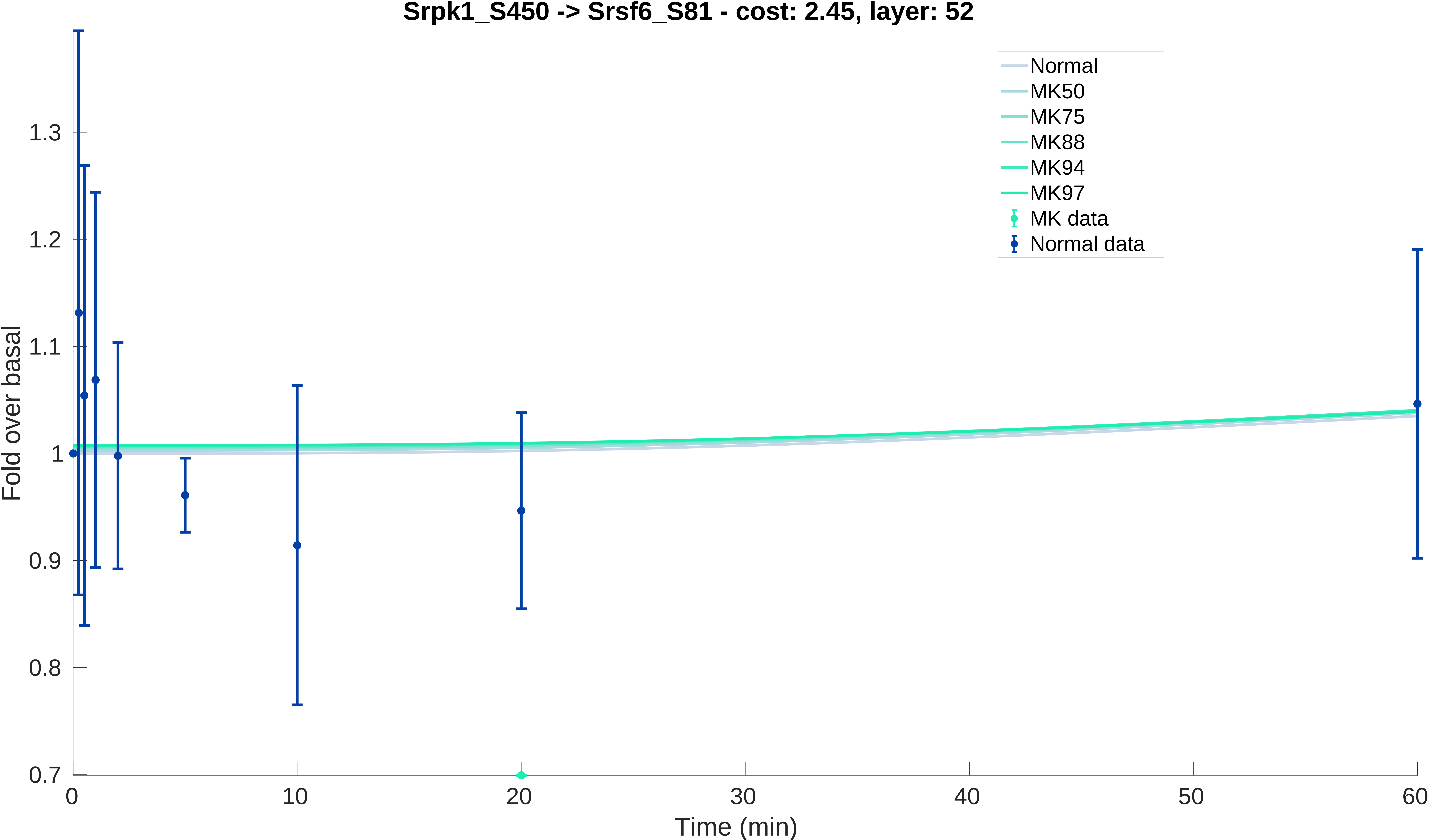

Srpk1_S450 -> Srsf6_S81 - cost: 2.45, layer: 52
Normal
MK50
MK75
MK88
MK94
MK97
MK data
Normal data
Fold over basal
1.3
1.2
1.1
1
0.9
0.8
0.7
0
10
20
30
40
50
60
Time (min)

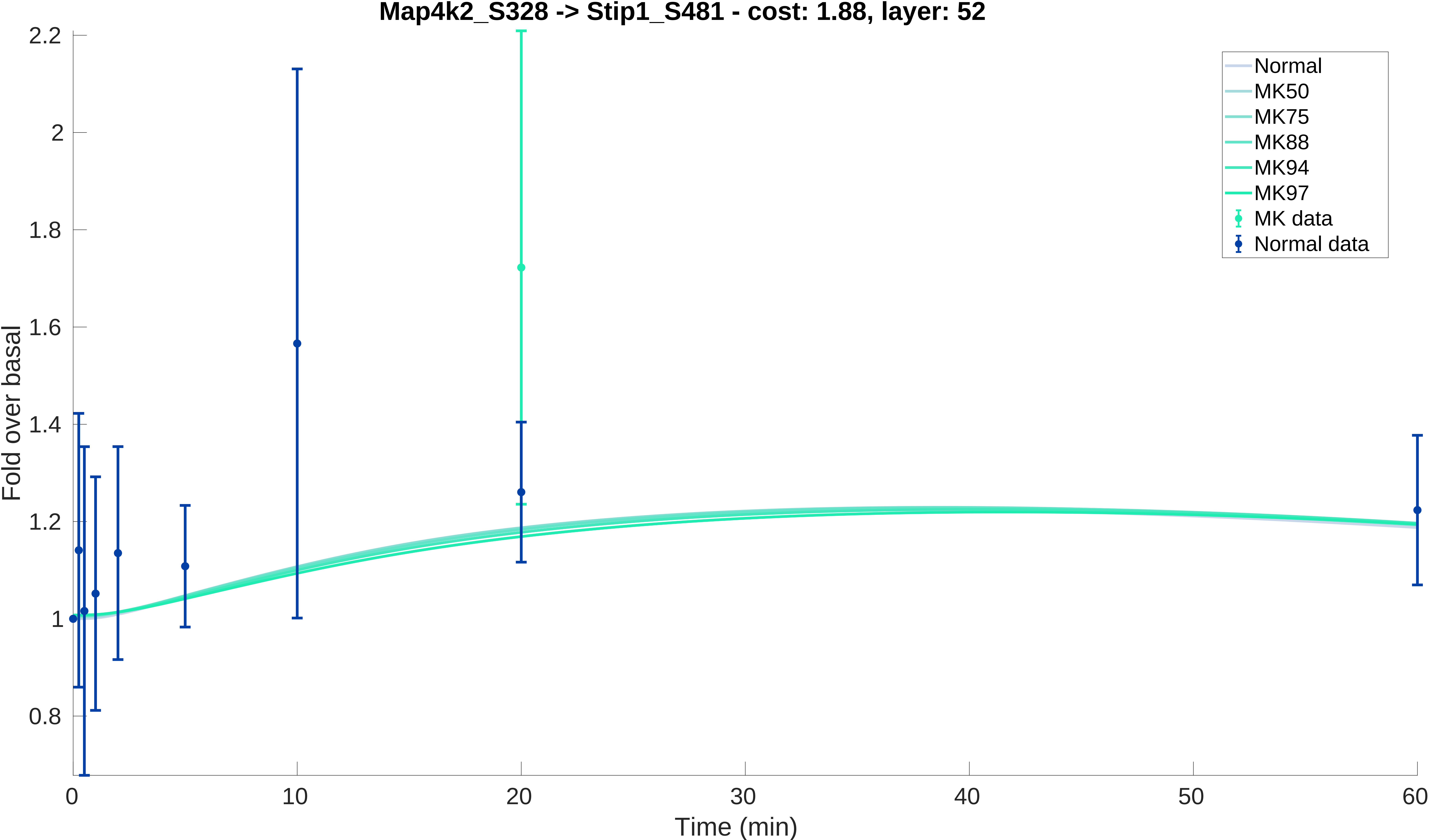

Map4k2_S328 -> Stip1_S481 - cost: 1.88, layer: 52
Fold over basal
Time (min)
Normal
MK50
MK75
MK88
MK94
MK97
MK data
Normal data
2.2
2
1.8
1.6
1.4
1.2
1
0.8
0
10
20
30
40
50
60

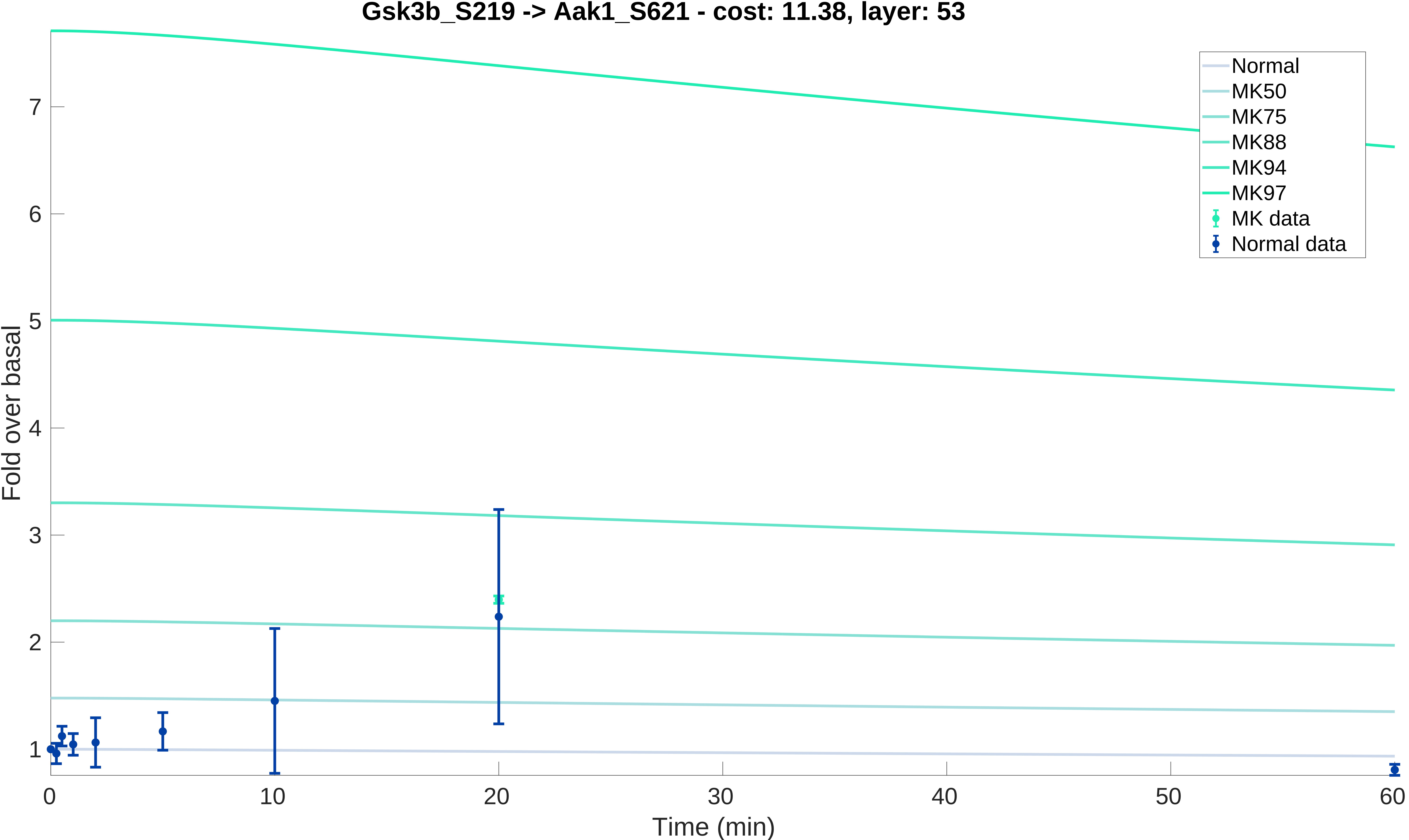
Gsk3b_S219 -> Aak1_S621 - cost: 11.38, layer: 53
Fold over basal
Time (min)
0
10
20
30
40
50
60
1
2
3
4
5
6
7
Normal
MK50
MK75
MK88
MK94
MK97
MK data
Normal data

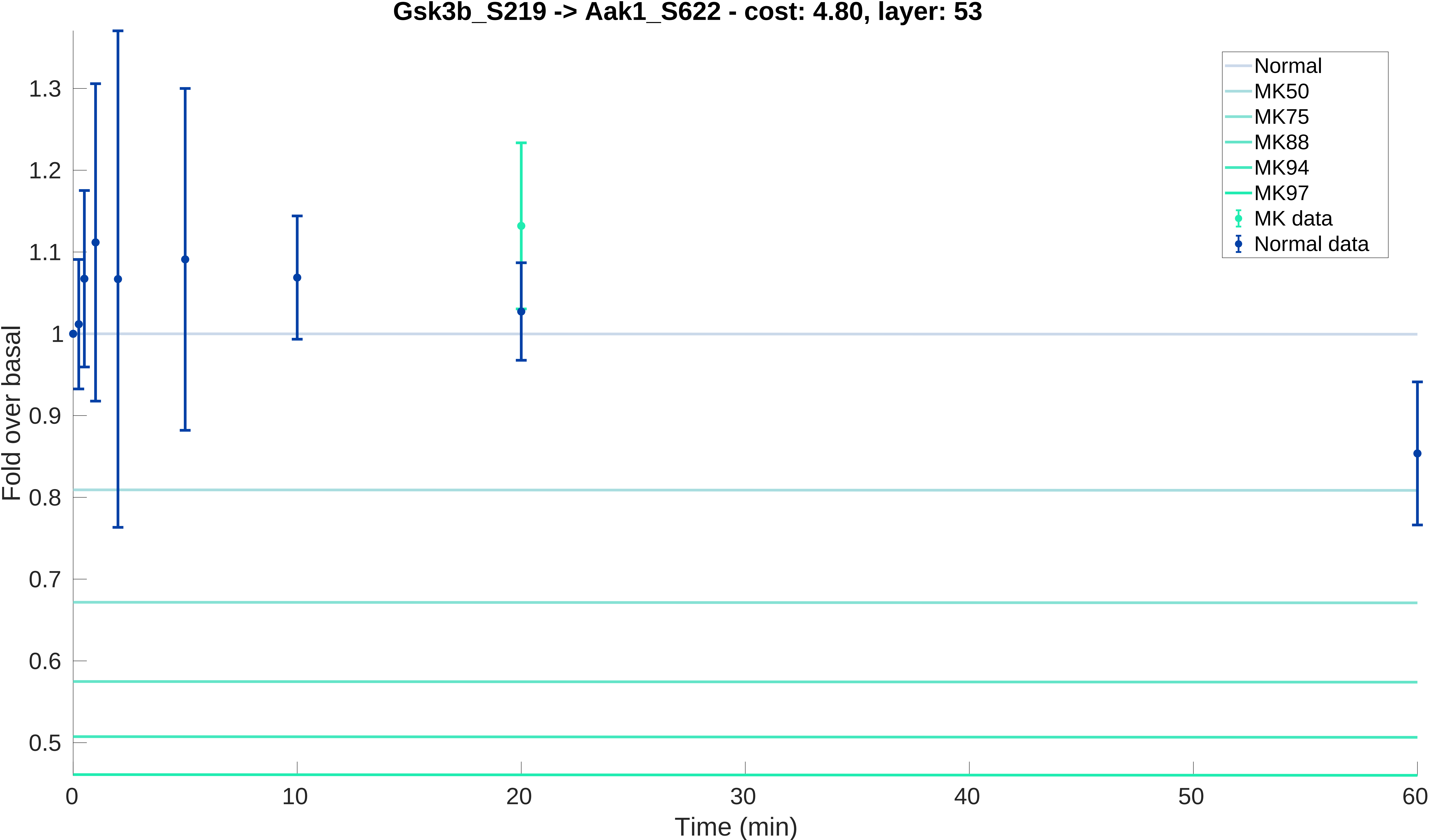

Gsk3b_S219 -> Aak1_S622 - cost: 4.80, layer: 53
Fold over basal
Time (min)
Normal
MK50
MK75
MK88
MK94
MK97
MK data
Normal data
1.3
1.2
1.1
1
0.9
0.8
0.7
0.6
0.5
0
10
20
30
40
50
60

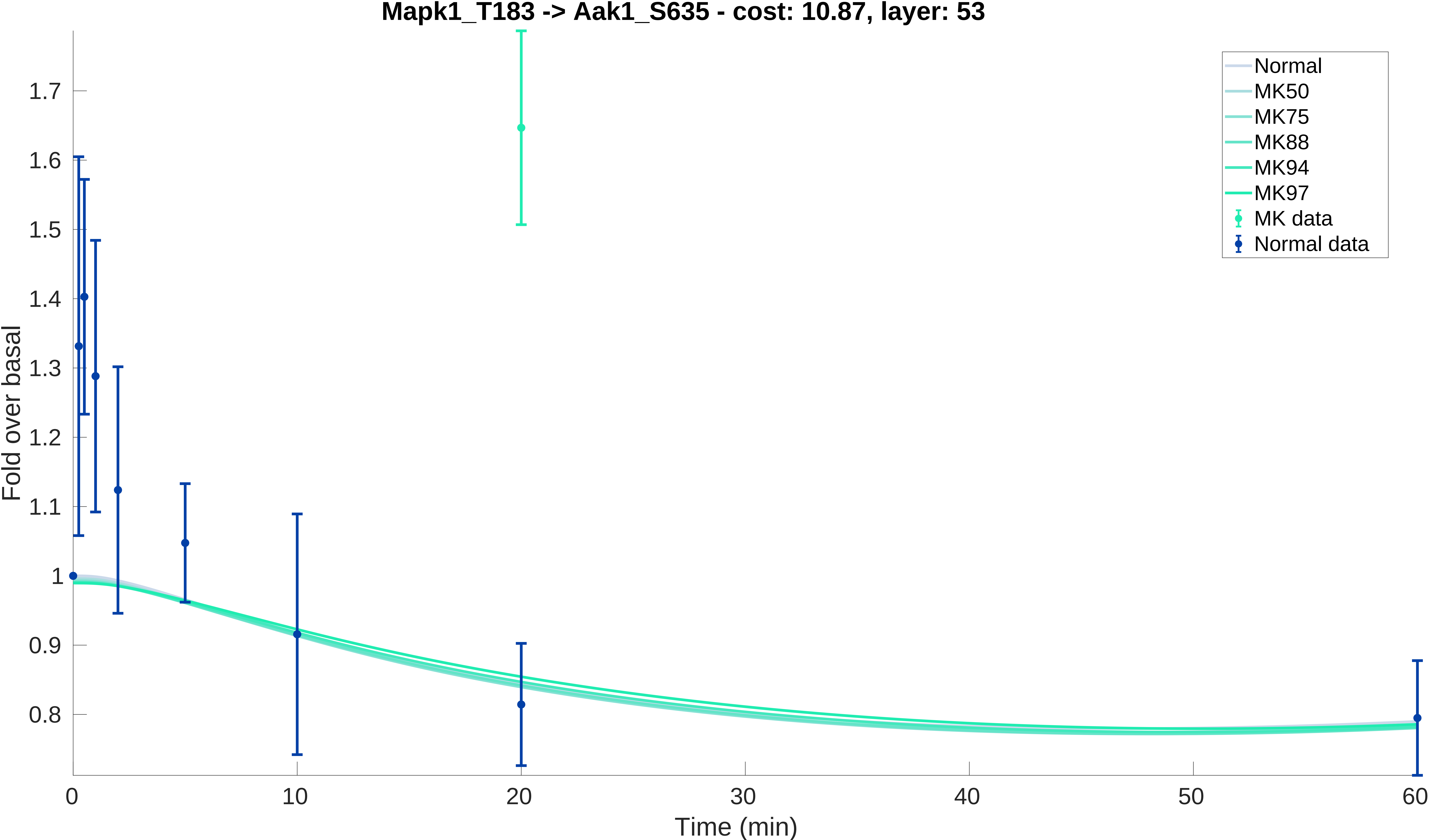

Mapk1_T183 -> Aak1_S635 - cost: 10.87, layer: 53
Fold over basal
Time (min)
0.8
0.9
1
1.1
1.2
1.3
1.4
1.5
1.6
1.7
0
10
20
30
40
50
60
Normal
MK50
MK75
MK88
MK94
MK97
MK data
Normal data

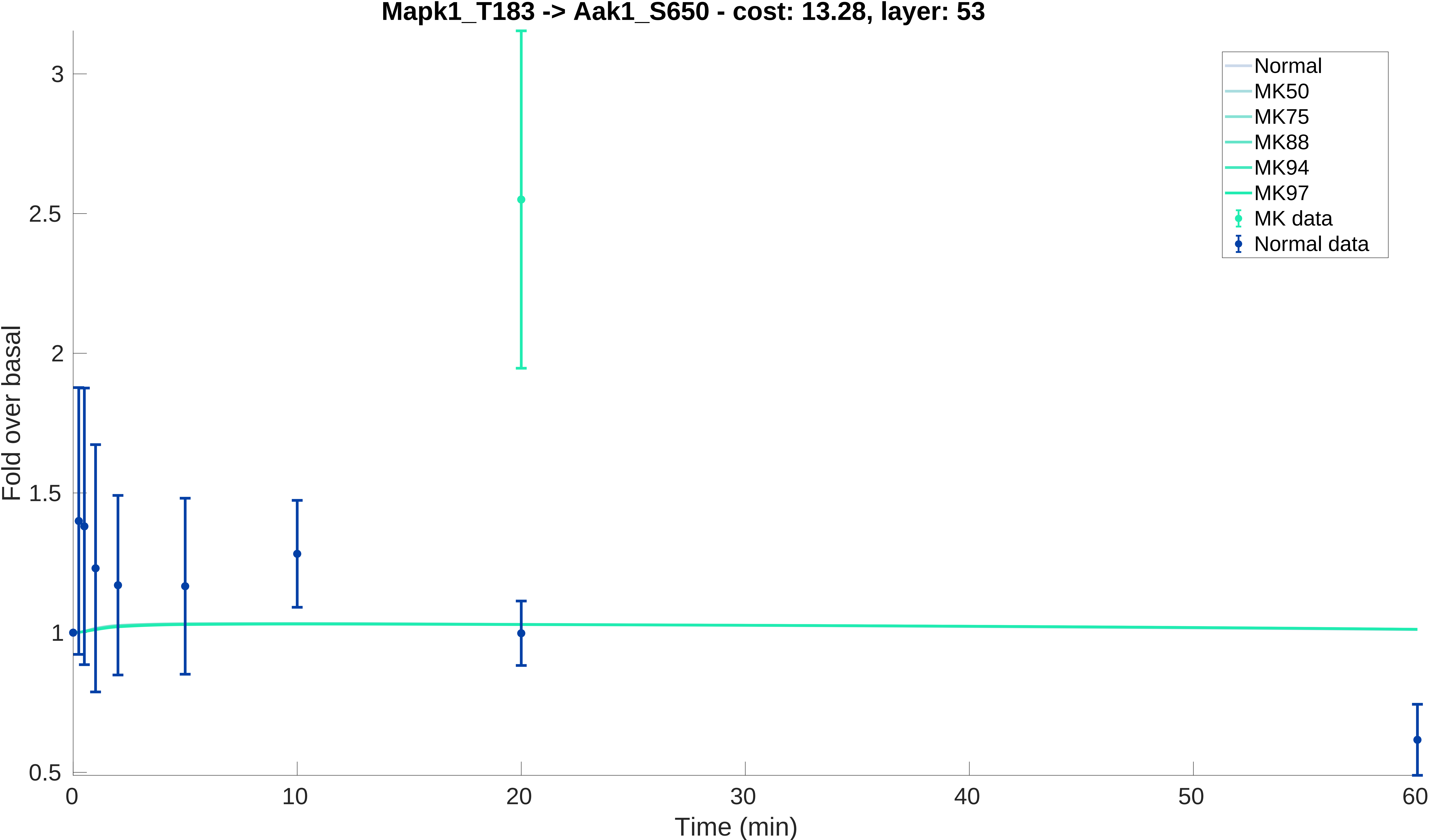

Mapk1_T183 -> Aak1_S650 - cost: 13.28, layer: 53
Normal
MK50
MK75
MK88
MK94
MK97
MK data
Normal data
Fold over basal
Time (min)
0.5
1
1.5
2
2.5
3
0
10
20
30
40
50
60

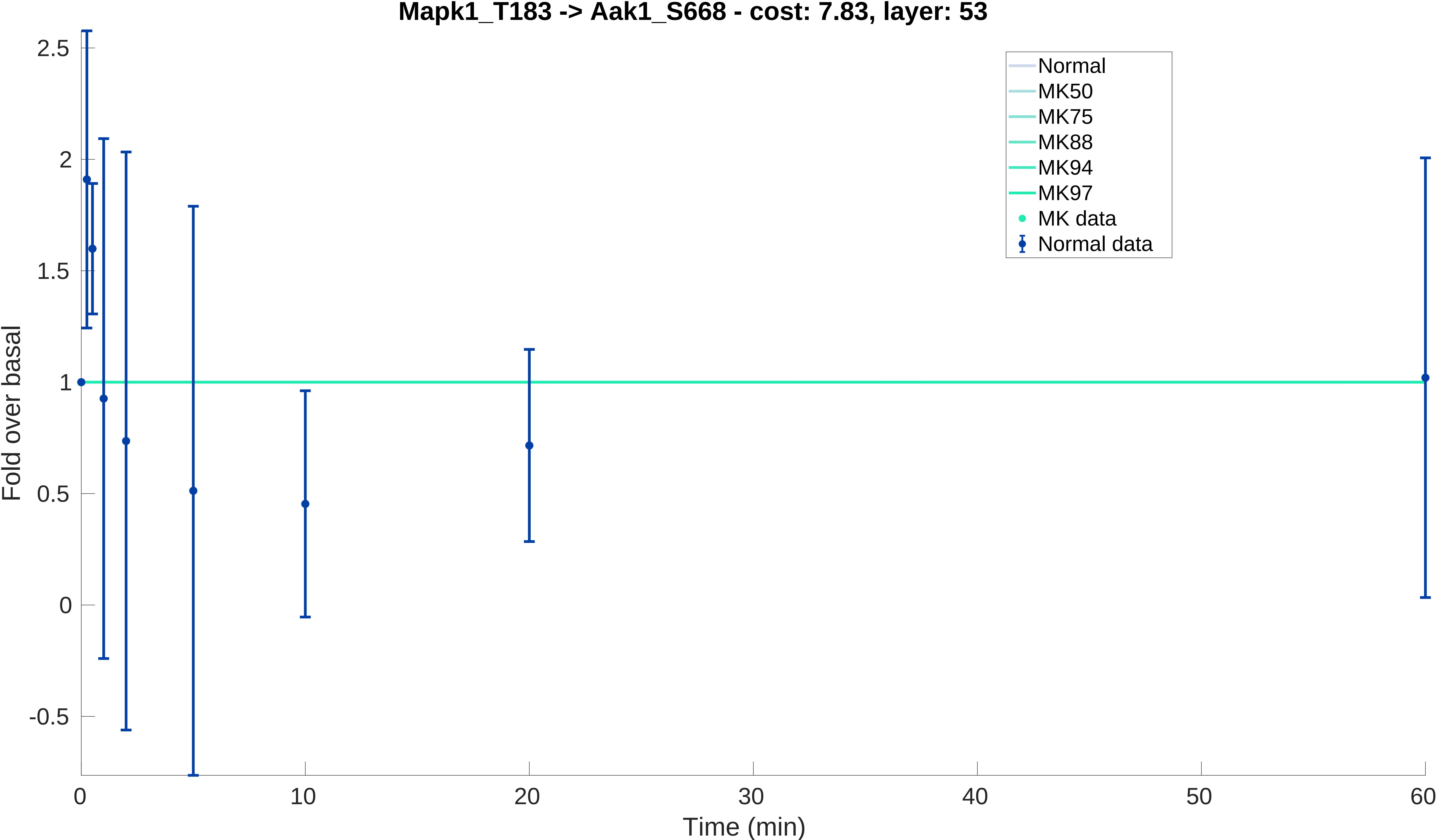

Mapk1_T183 -> Aak1_S668 - cost: 7.83, layer: 53
Fold over basal
Time (min)
2.5
2
1.5
1
0.5
0
-0.5
0
10
20
30
40
50
60
Normal
MK50
MK75
MK88
MK94
MK97
MK data
Normal data

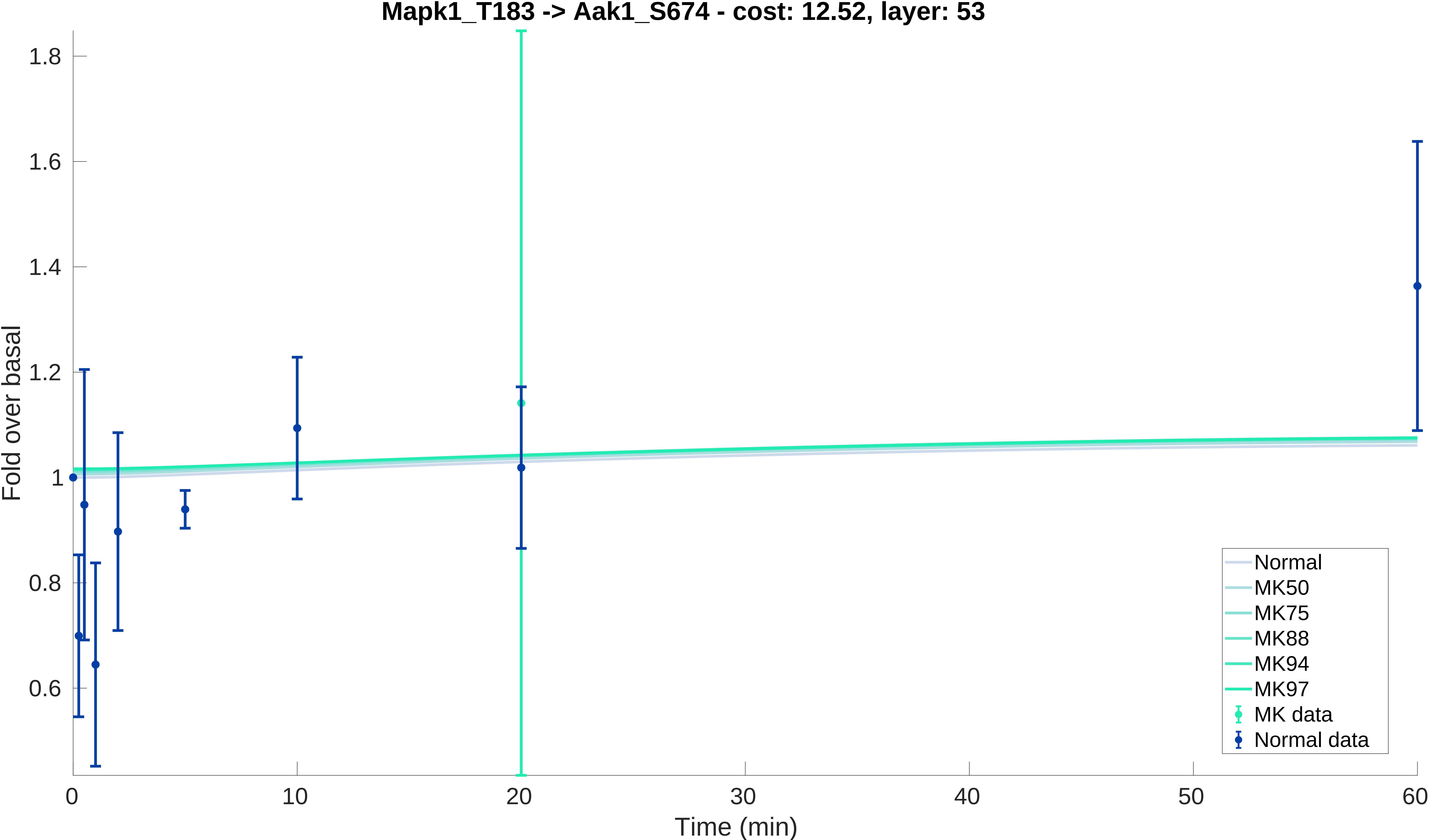

Mapk1_T183 -> Aak1_S674 - cost: 12.52, layer: 53
Fold over basal
1.8
1.6
1.4
1.2
1
0.8
0.6
0
10
20
30
40
50
60
Time (min)
Normal
MK50
MK75
MK88
MK94
MK97
MK data
Normal data

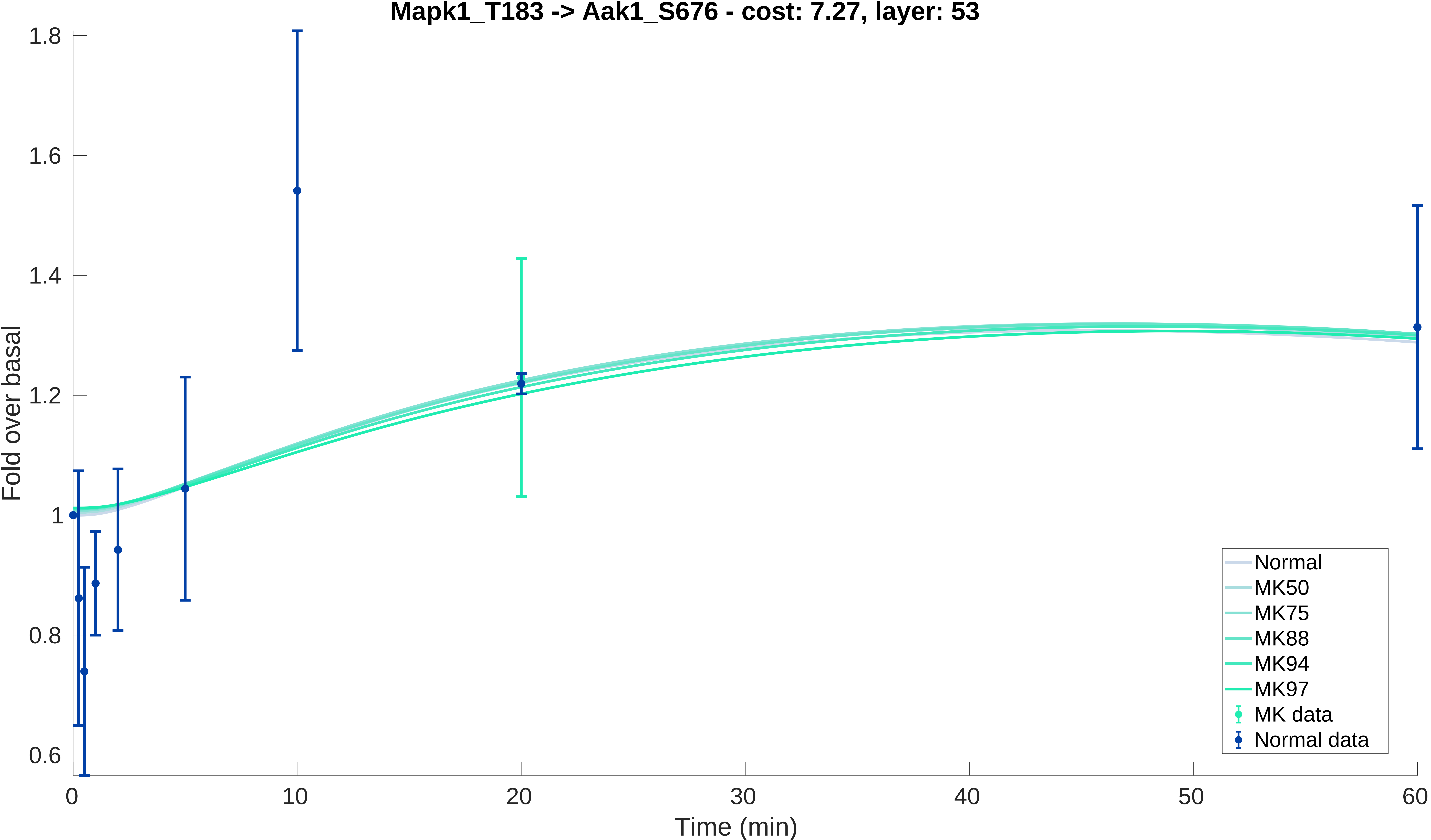

Mapk1_T183 -> Aak1_S676 - cost: 7.27, layer: 53
Fold over basal
Time (min)
1.8
1.6
1.4
1.2
1
0.8
0.6
0
10
20
30
40
50
60
Normal
MK50
MK75
MK88
MK94
MK97
MK data
Normal data

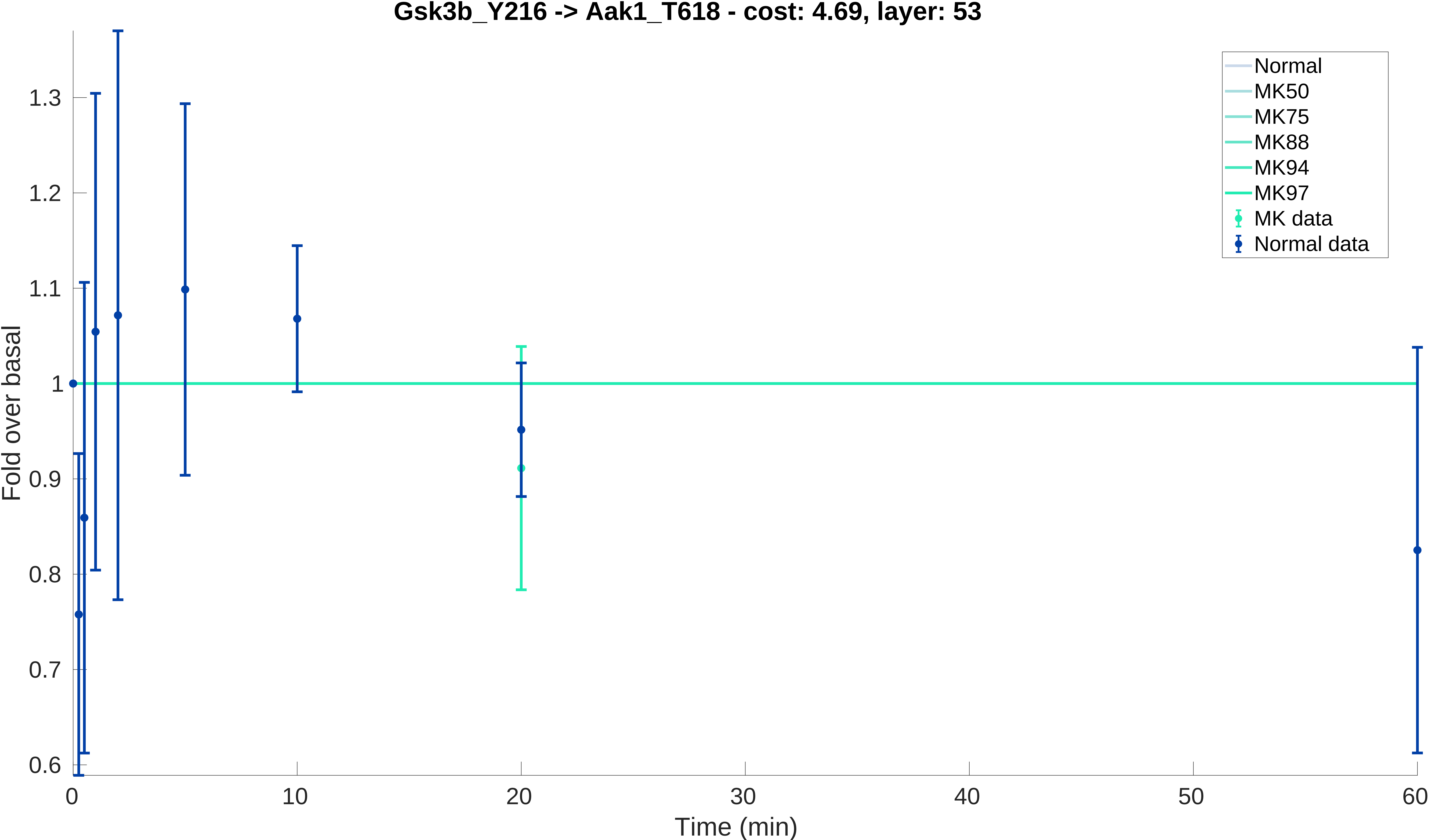

Gsk3b_Y216 -> Aak1_T618 - cost: 4.69, layer: 53
Fold over basal
Time (min)
0.6
0.7
0.8
0.9
1
1.1
1.2
1.3
0
10
20
30
40
50
60
Normal
MK50
MK75
MK88
MK94
MK97
MK data
Normal data

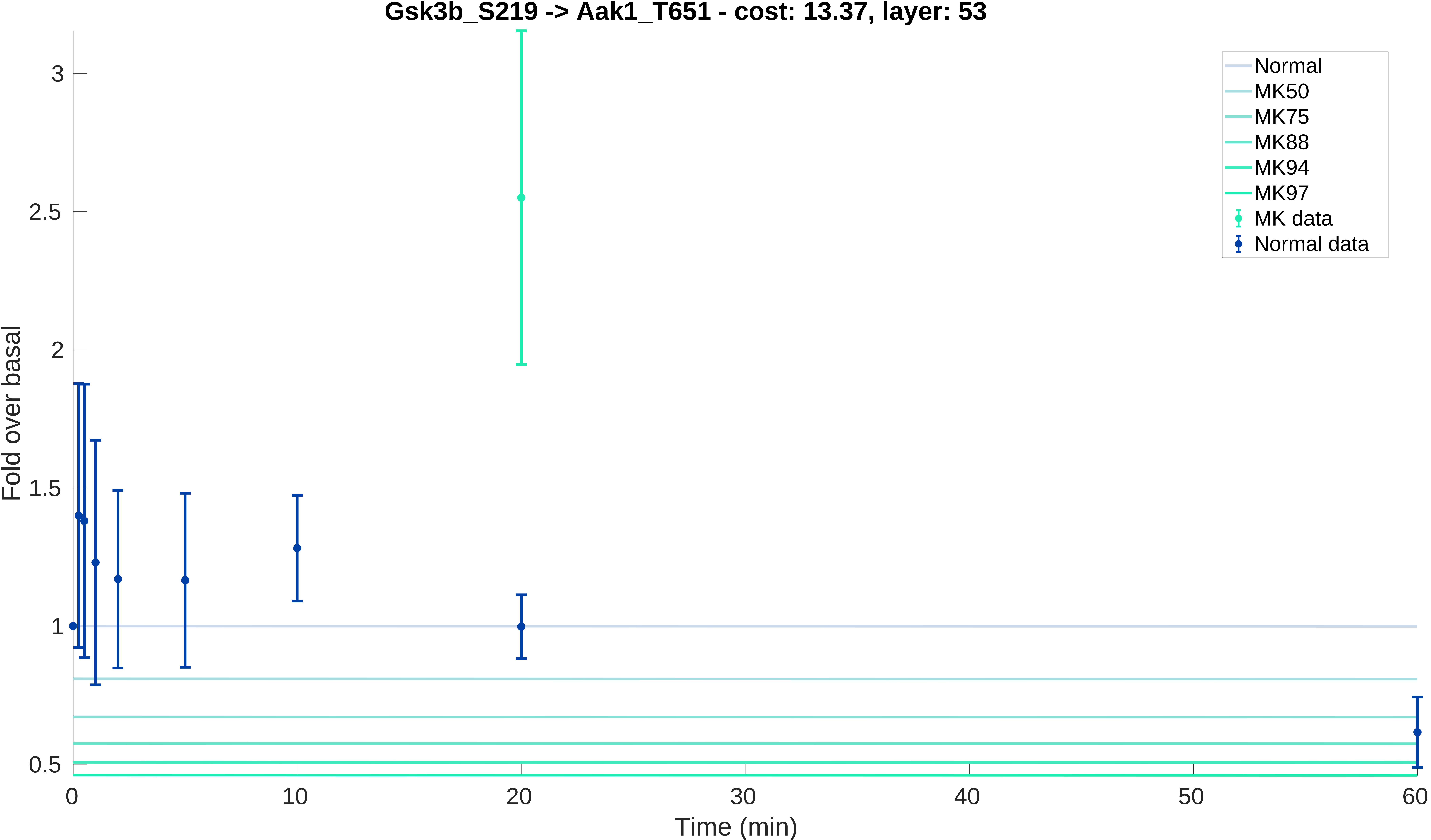

Gsk3b_S219 -> Aak1_T651 - cost: 13.37, layer: 53
Normal
MK50
MK75
MK88
MK94
MK97
MK data
Normal data
Fold over basal
0.5
1
1.5
2
2.5
3
0
10
20
30
40
50
60
Time (min)

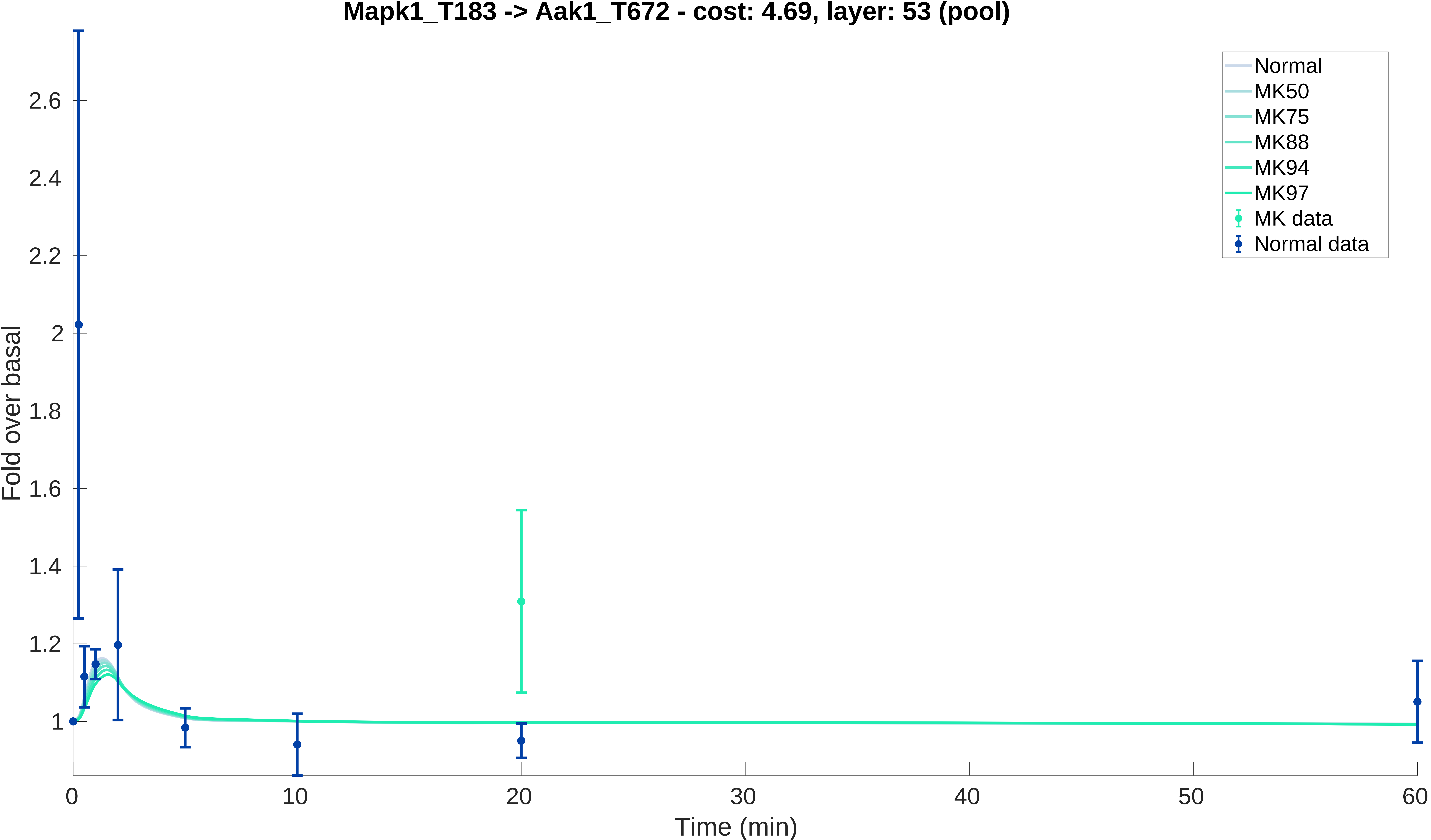

Mapk1_T183 -> Aak1_T672 - cost: 4.69, layer: 53 (pool)
Fold over basal
2.6
2.4
2.2
2
1.8
1.6
1.4
1.2
1
0
10
20
30
40
50
60
Time (min)
Normal
MK50
MK75
MK88
MK94
MK97
MK data
Normal data

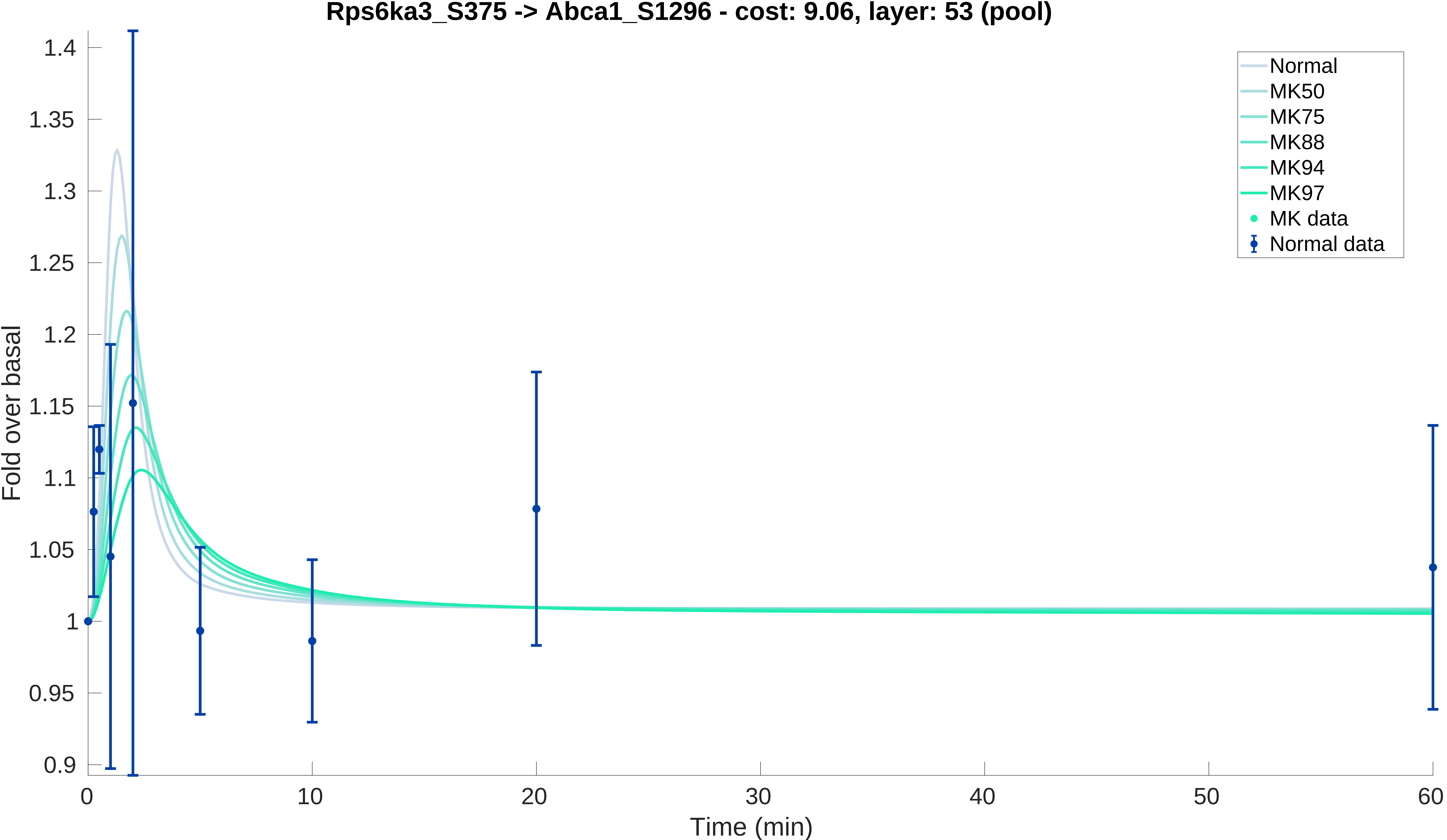

Rps6ka3_S375 -> Abca1_S1296 - cost: 9.06, layer: 53 (pool)
Fold over basal
Time (min)
1.4
1.35
1.3
1.25
1.2
1.15
1.1
1.05
1
0.95
0.9
0
10
20
30
40
50
60
Normal
MK50
MK75
MK88
MK94
MK97
MK data
Normal data

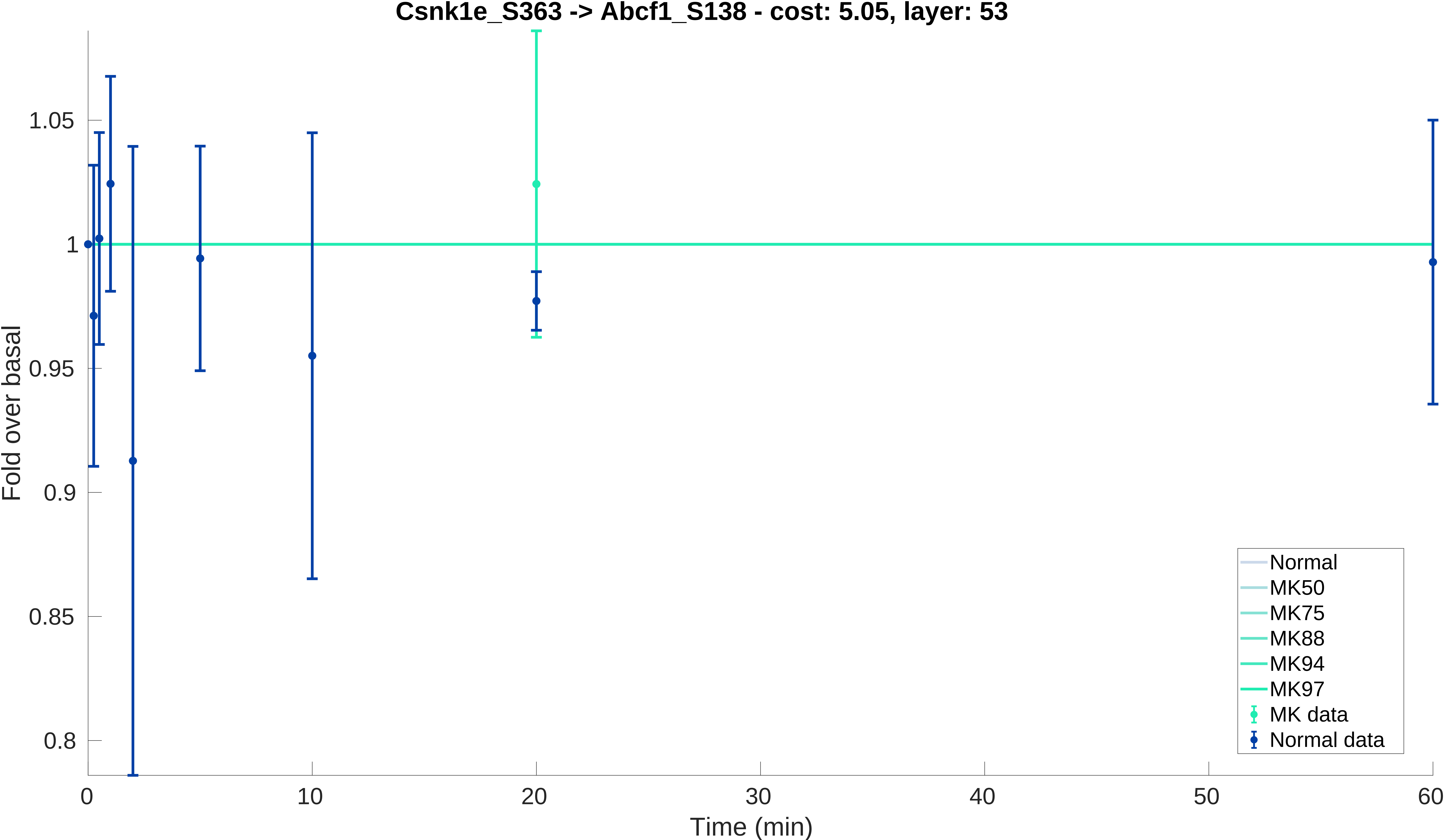

Csnk1e_S363 -> Abcf1_S138 - cost: 5.05, layer: 53
Fold over basal
1.05
1
0.95
0.9
0.85
0.8
0
10
20
30
40
50
60
Time (min)
Normal
MK50
MK75
MK88
MK94
MK97
MK data
Normal data

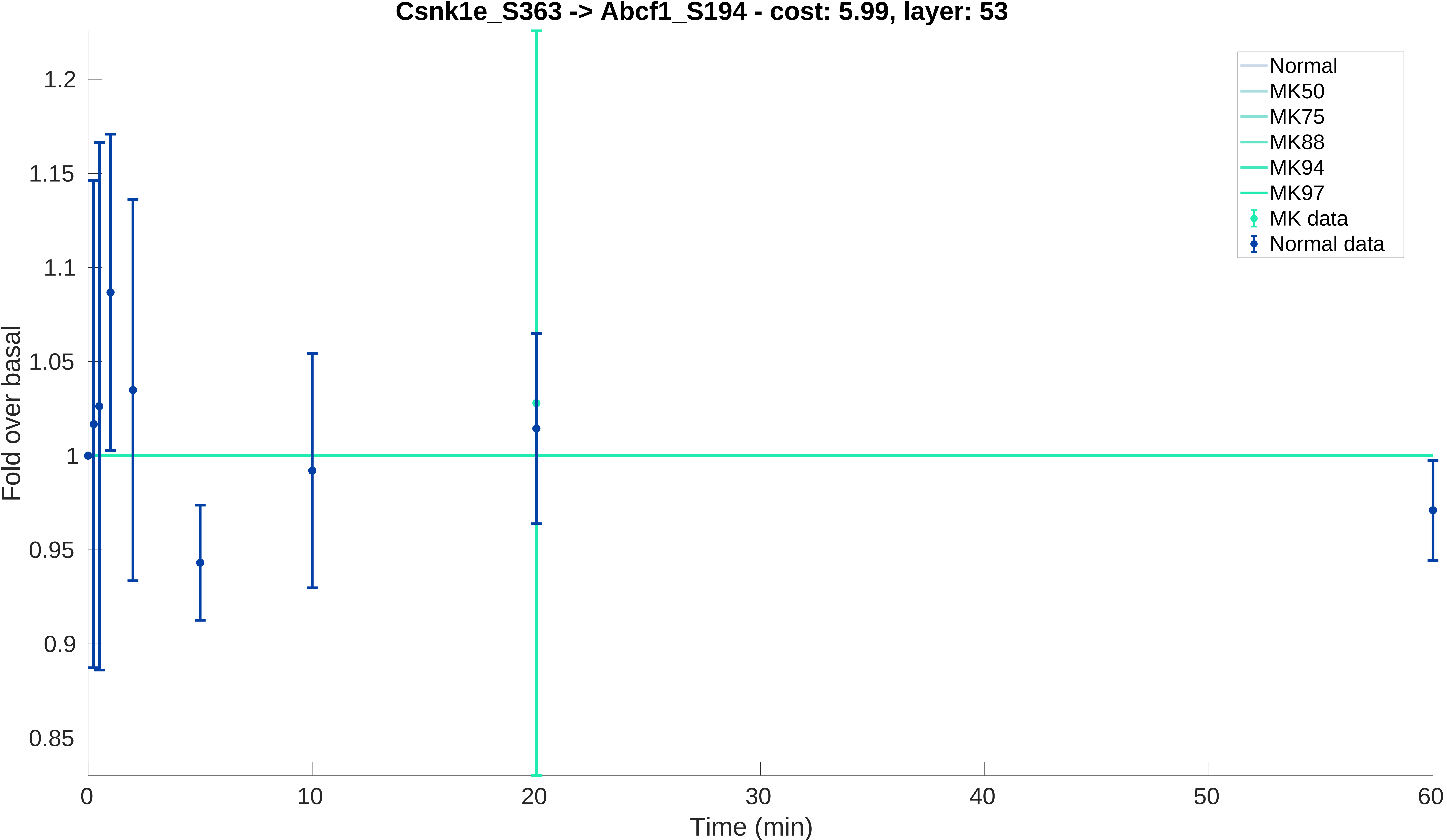
Csnk1e_S363 -> Abcf1_S194 - cost: 5.99, layer: 53
Fold over basal
Time (min)
1.2
1.15
1.1
1.05
1
0.95
0.9
0.85
0
10
20
30
40
50
60
Normal
MK50
MK75
MK88
MK94
MK97
MK data
Normal data

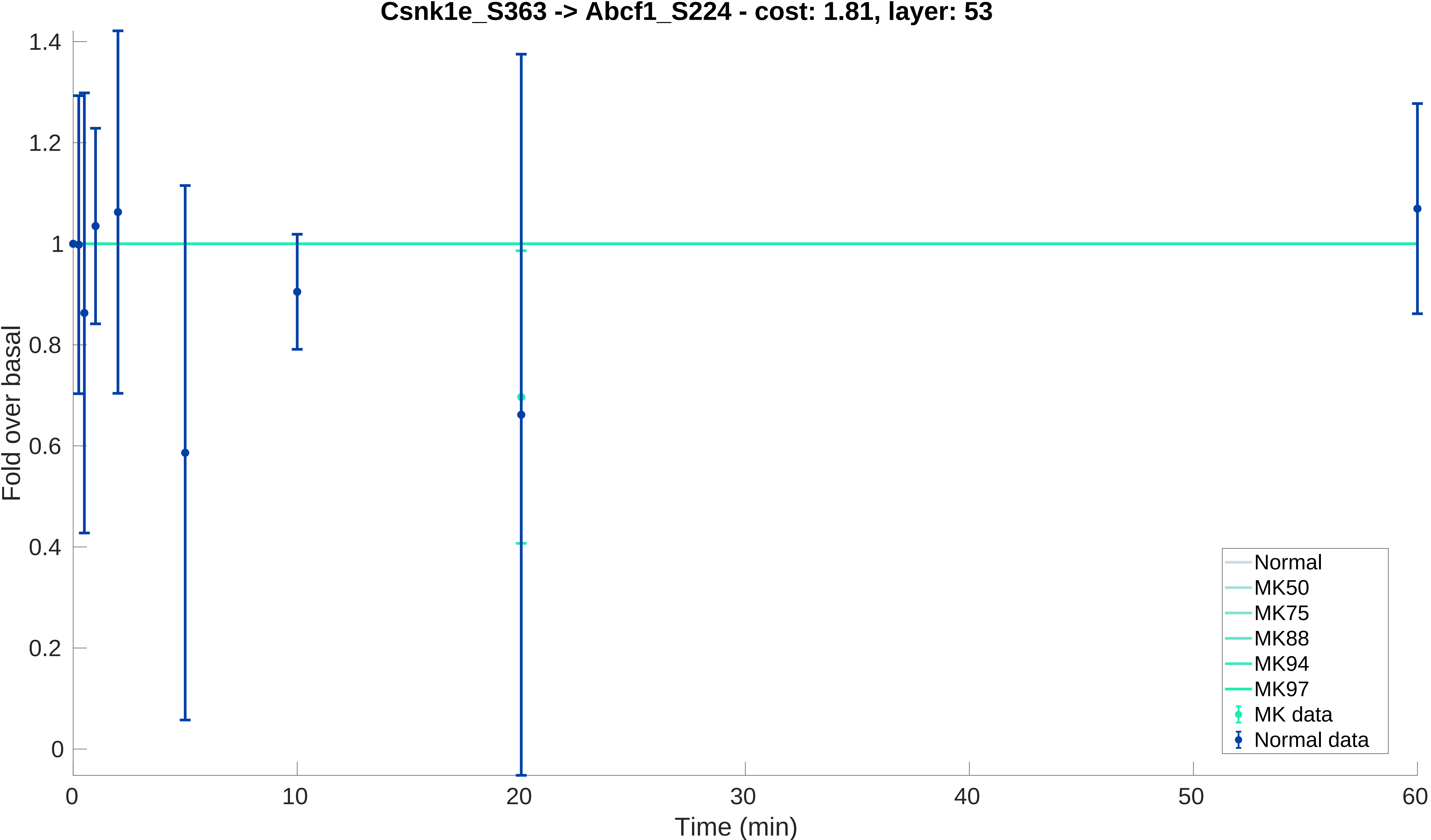

Csnk1e_S363 -> Abcf1_S224 - cost: 1.81, layer: 53
Fold over basal
Time (min)
0
0.2
0.4
0.6
0.8
1
1.2
1.4
0
10
20
30
40
50
60
Normal
MK50
MK75
MK88
MK94
MK97
MK data
Normal data

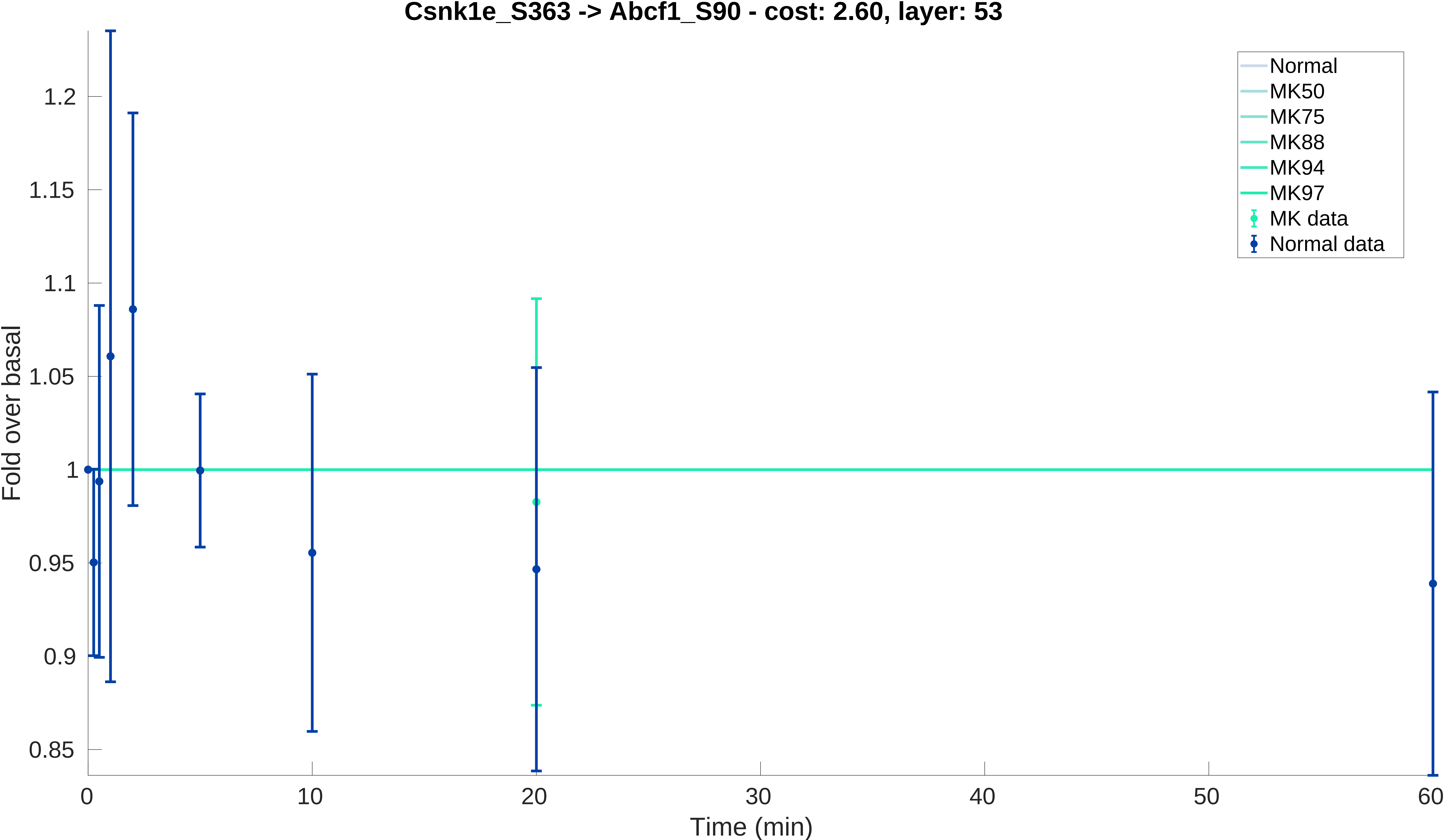

Csnk1e_S363 -> Abcf1_S90 - cost: 2.60, layer: 53
Normal
MK50
MK75
MK88
MK94
MK97
MK data
Normal data
Fold over basal
1.2
1.15
1.1
1.05
1
0.95
0.9
0.85
0
10
20
30
40
50
60
Time (min)

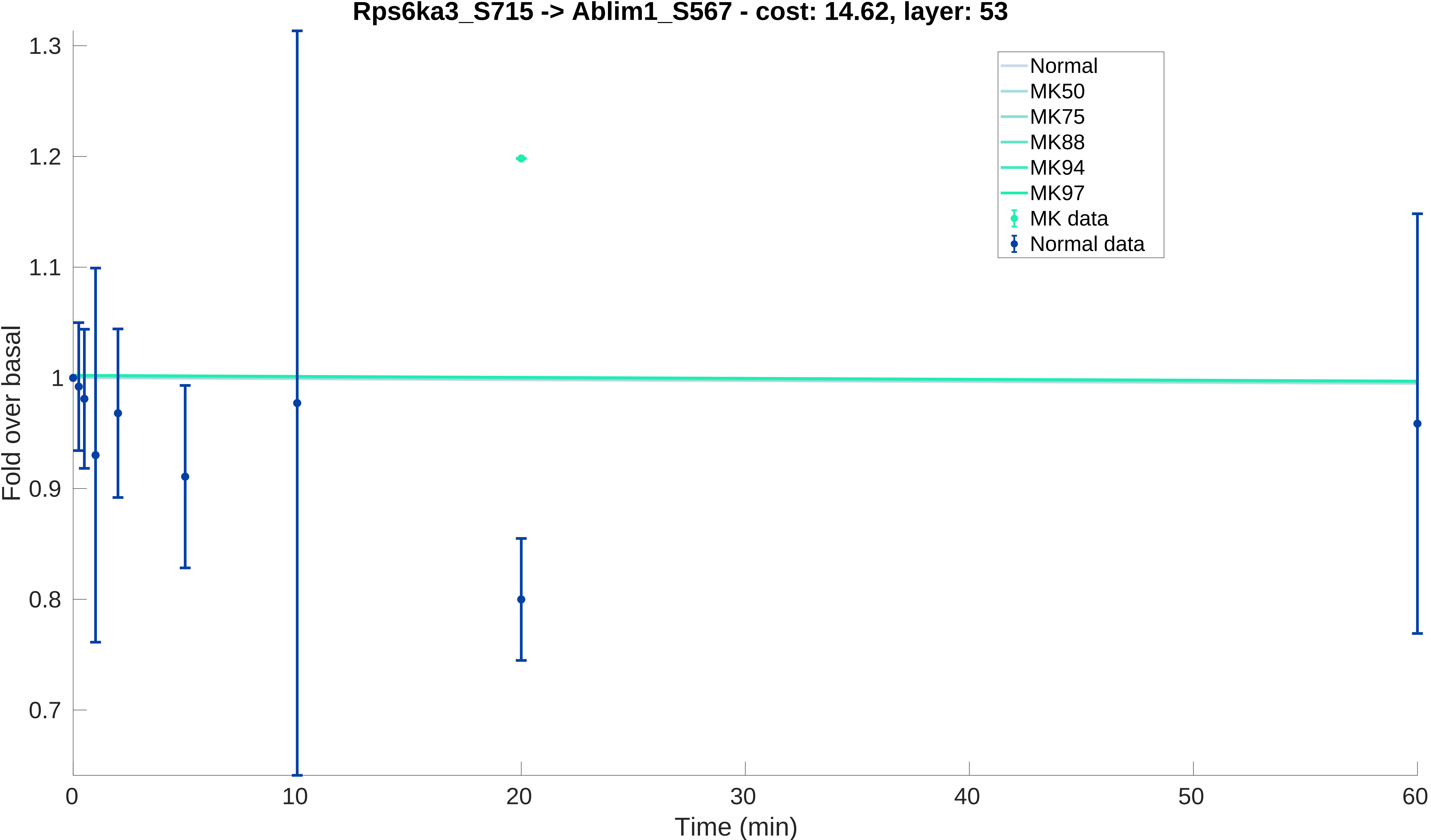

Rps6ka3_S715 -> Ablim1_S567 - cost: 14.62, layer: 53
Fold over basal
Time (min)
1.3
1.2
1.1
1
0.9
0.8
0.7
0
10
20
30
40
50
60
Normal
MK50
MK75
MK88
MK94
MK97
MK data
Normal data

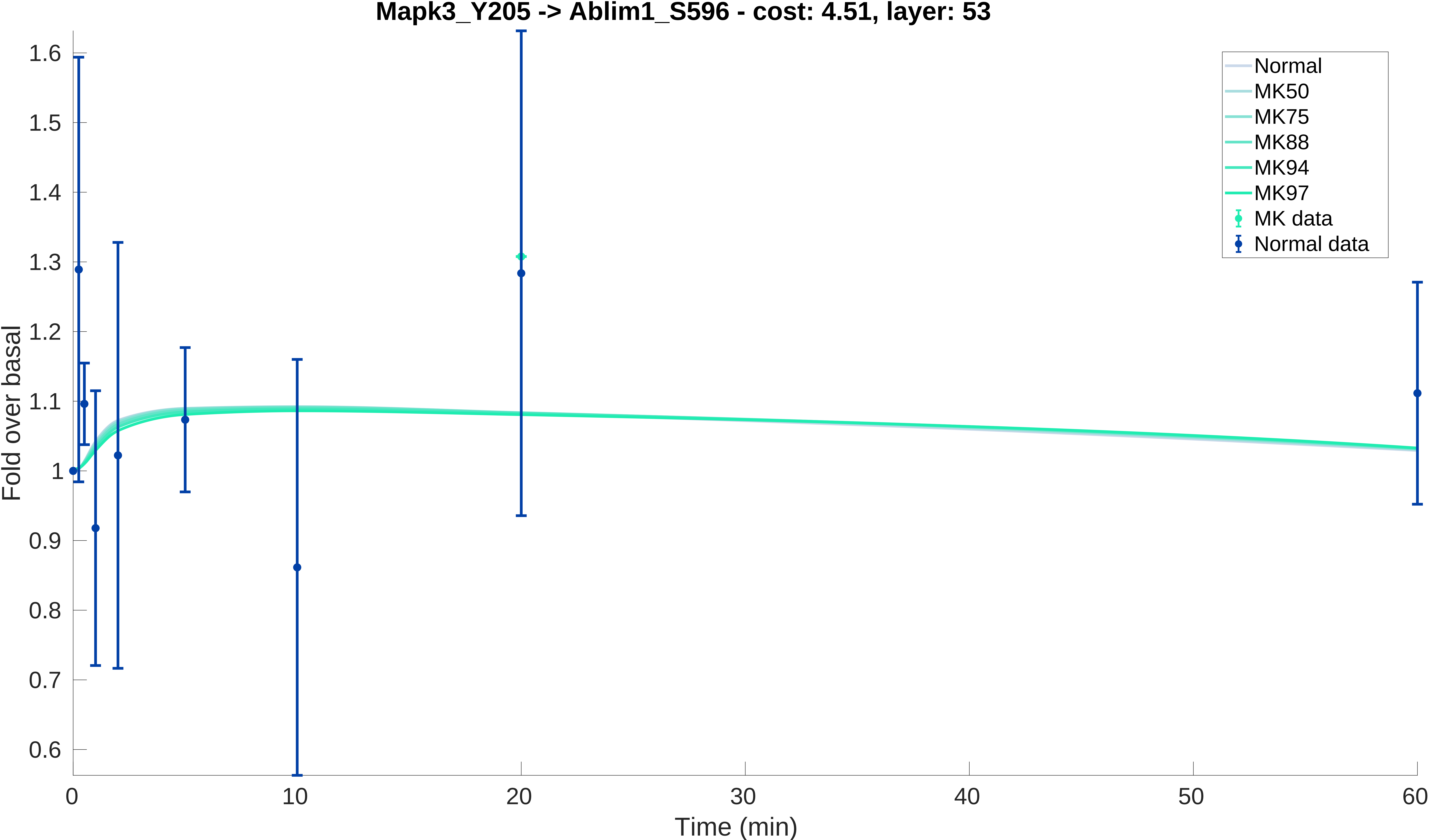

Mapk3_Y205 -> Ablim1_S596 - cost: 4.51, layer: 53
Fold over basal
Time (min)
Normal
MK50
MK75
MK88
MK94
MK97
MK data
Normal data
1.6
1.5
1.4
1.3
1.2
1.1
1
0.9
0.8
0.7
0.6
0
10
20
30
40
50
60

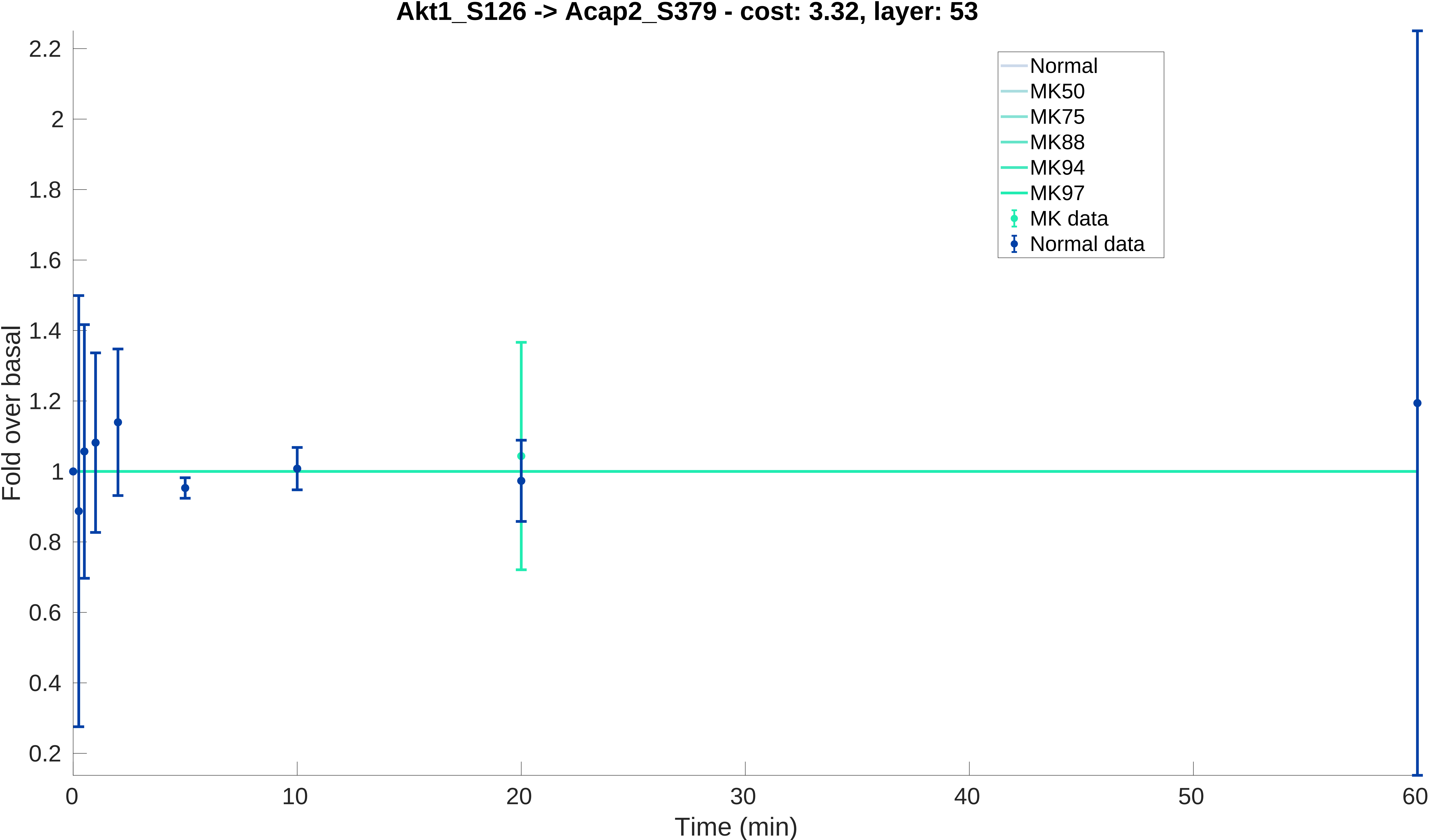

Akt1_S126 -> Acap2_S379 - cost: 3.32, layer: 53
Fold over basal
2.2
2
1.8
1.6
1.4
1.2
1
0.8
0.6
0.4
0.2
0
10
20
30
40
50
60
Time (min)
Normal
MK50
MK75
MK88
MK94
MK97
MK data
Normal data

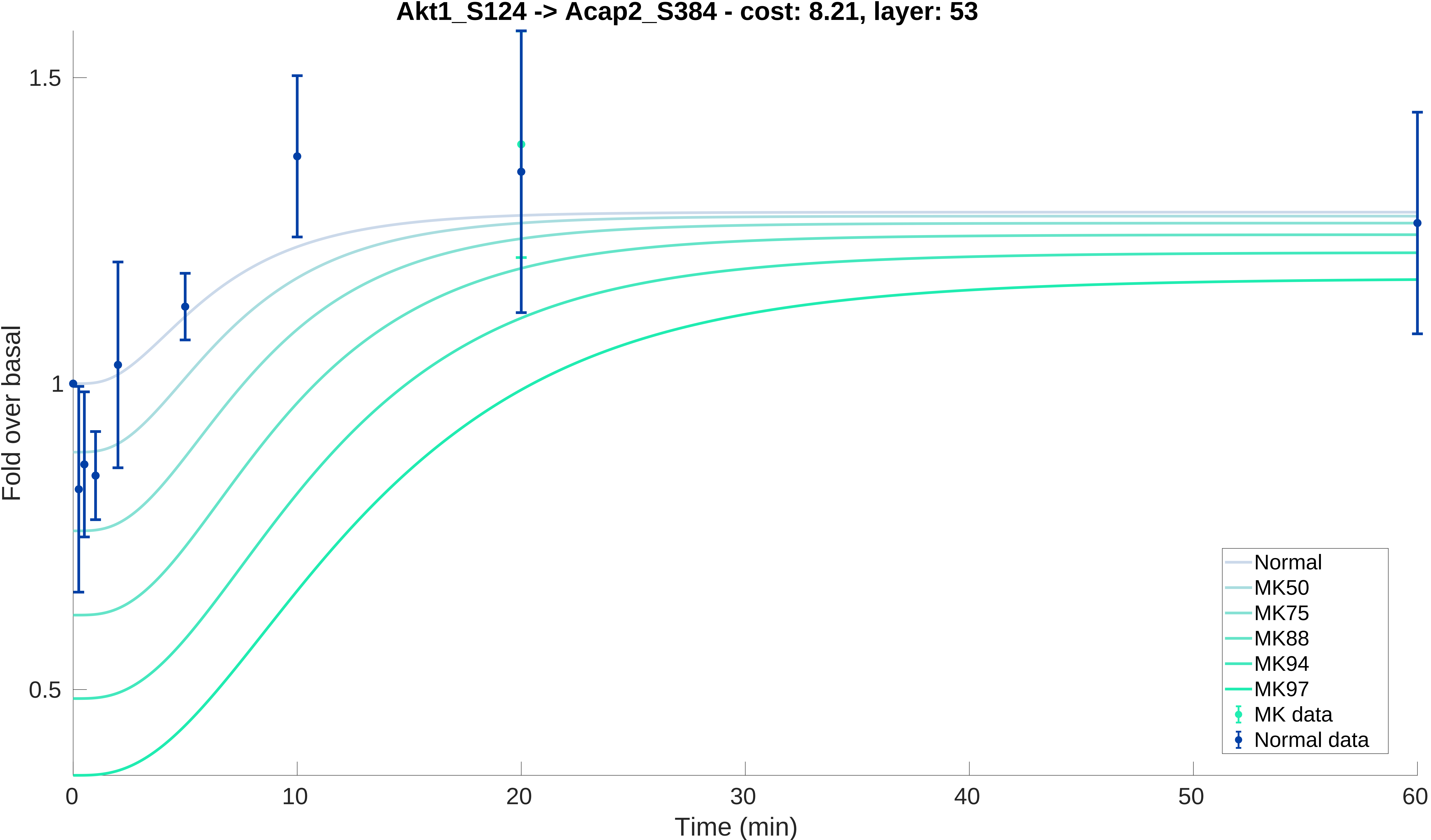

Akt1_S124 -> Acap2_S384 - cost: 8.21, layer: 53
Fold over basal
1.5
1
0.5
0
10
20
30
40
50
60
Time (min)
Normal
MK50
MK75
MK88
MK94
MK97
MK data
Normal data

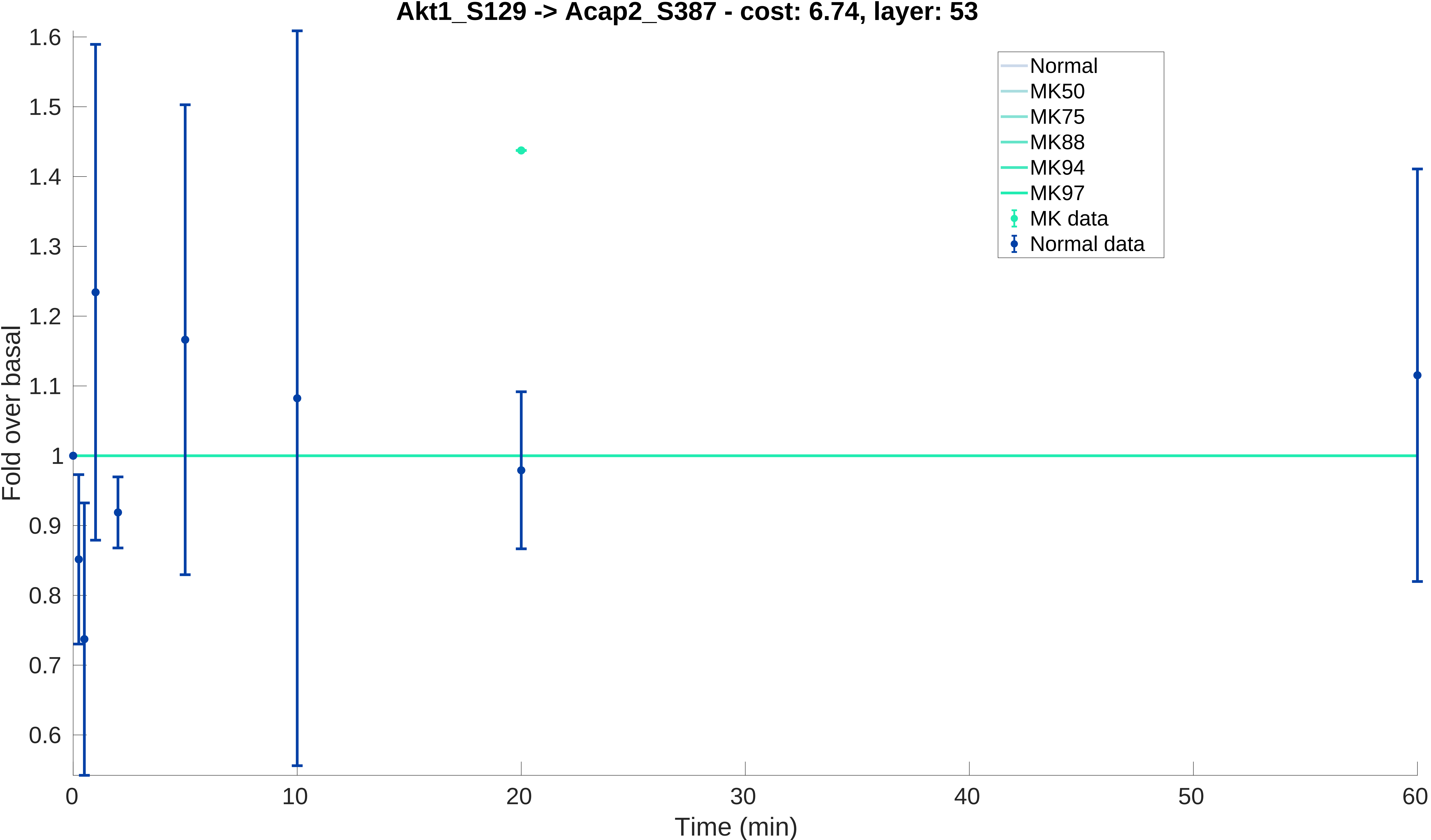

Akt1_S129 -> Acap2_S387 - cost: 6.74, layer: 53
Fold over basal
Time (min)
1.6
1.5
1.4
1.3
1.2
1.1
1
0.9
0.8
0.7
0.6
0
10
20
30
40
50
60
Normal
MK50
MK75
MK88
MK94
MK97
MK data
Normal data

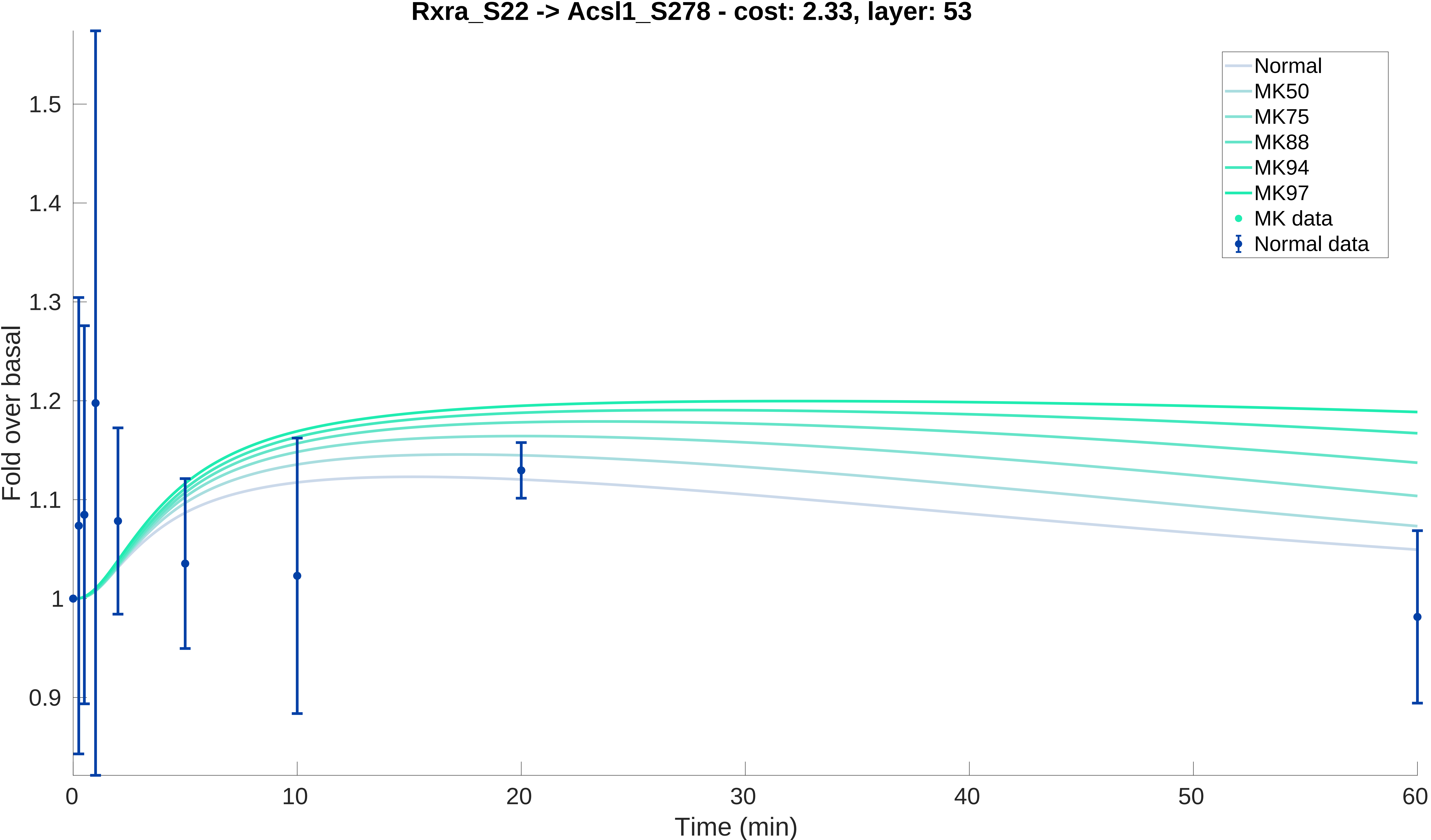

Rxra_S22 -> Acsl1_S278 - cost: 2.33, layer: 53
Normal
MK50
MK75
MK88
MK94
MK97
MK data
Normal data
Fold over basal
Time (min)
0
10
20
30
40
50
60
0.9
1
1.1
1.2
1.3
1.4
1.5

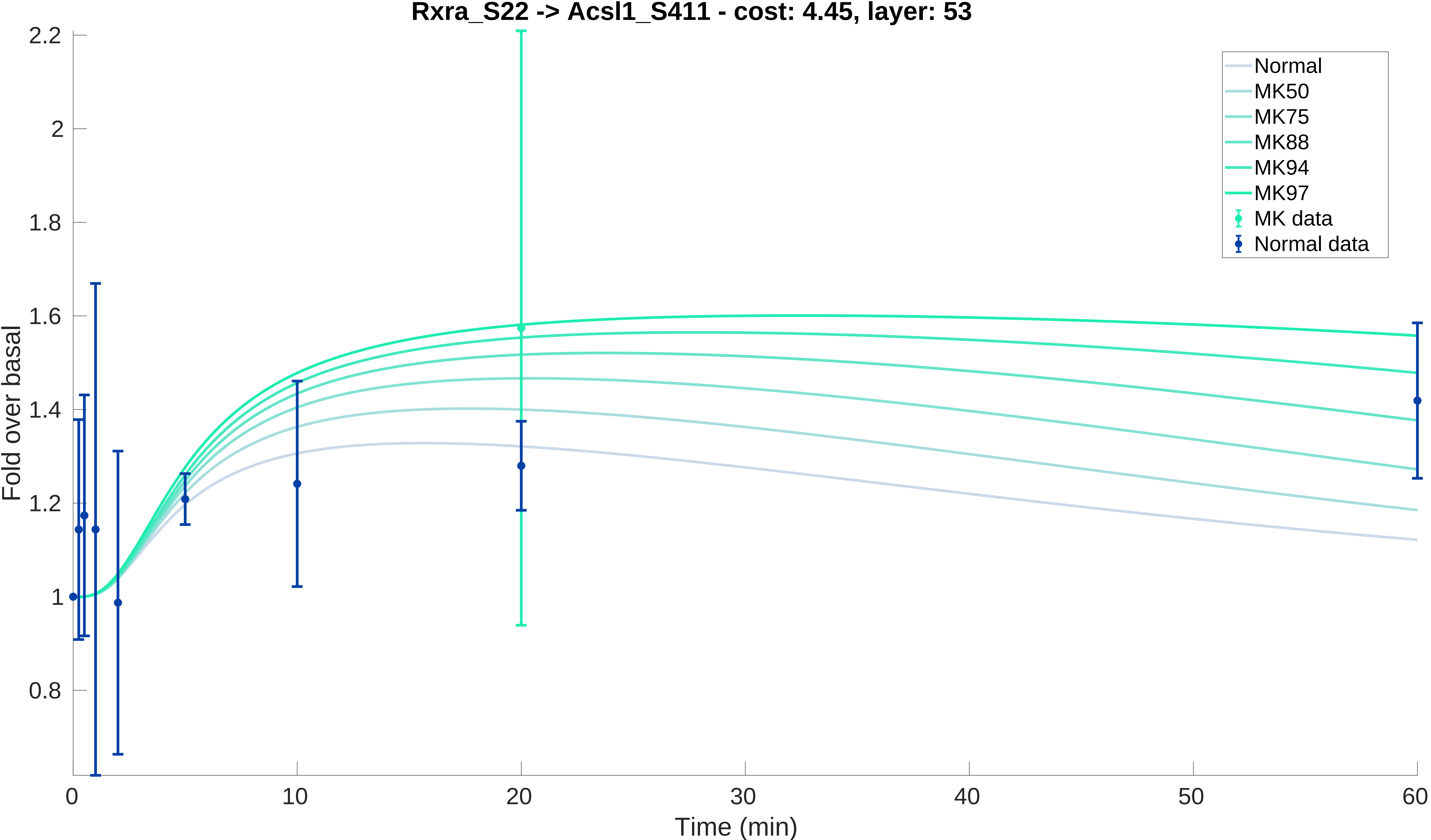

Rxra_S22 -> Acsl1_S411 - cost: 4.45, layer: 53
Fold over basal
Time (min)
0.8
1
1.2
1.4
1.6
1.8
2
2.2
0
10
20
30
40
50
60
Normal
MK50
MK75
MK88
MK94
MK97
MK data
Normal data

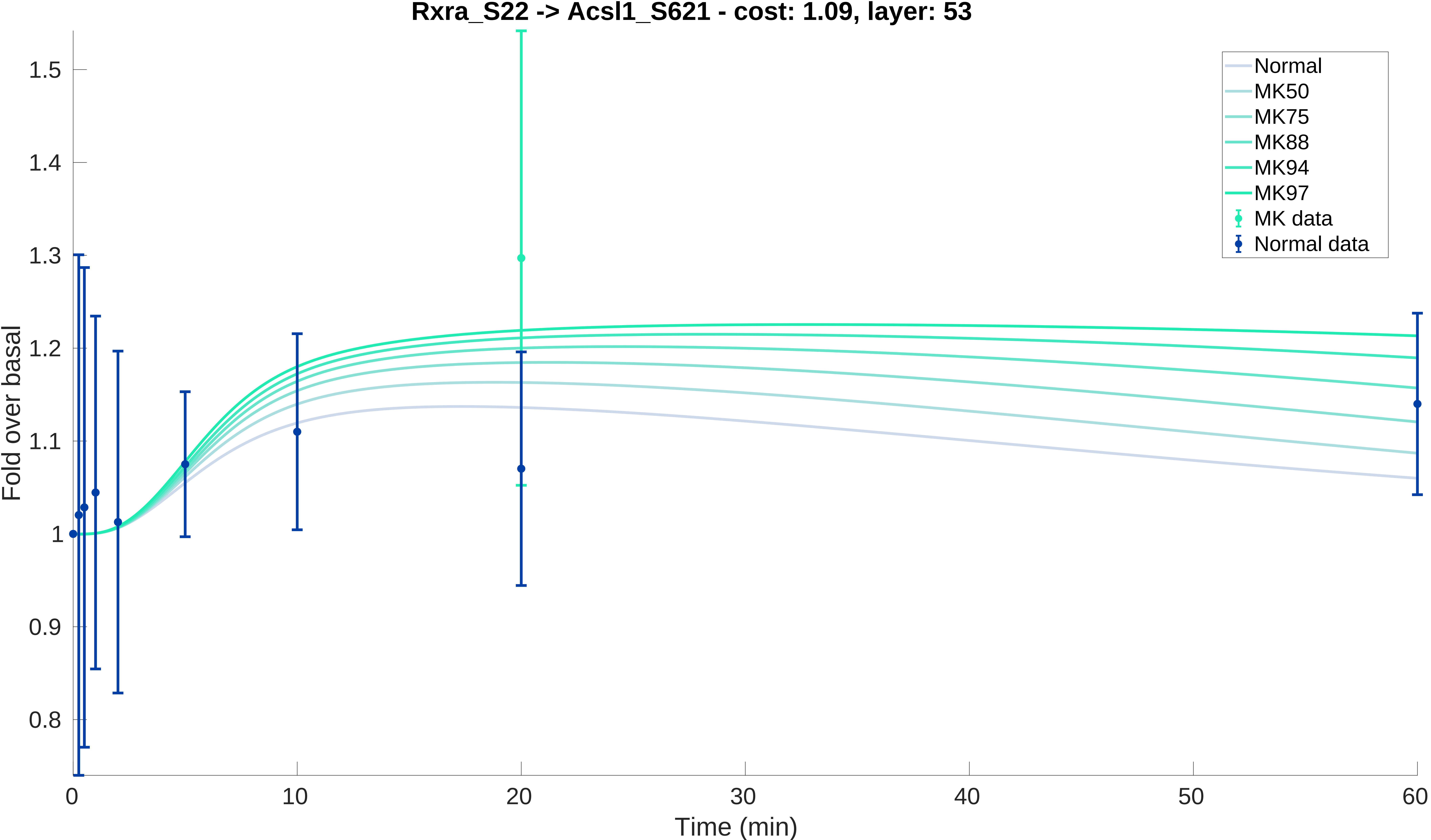
Rxra_S22 -> Acsl1_S621 - cost: 1.09, layer: 53
Fold over basal
Time (min)
0
10
20
30
40
50
60
0.8
0.9
1
1.1
1.2
1.3
1.4
1.5
Normal
MK50
MK75
MK88
MK94
MK97
MK data
Normal data

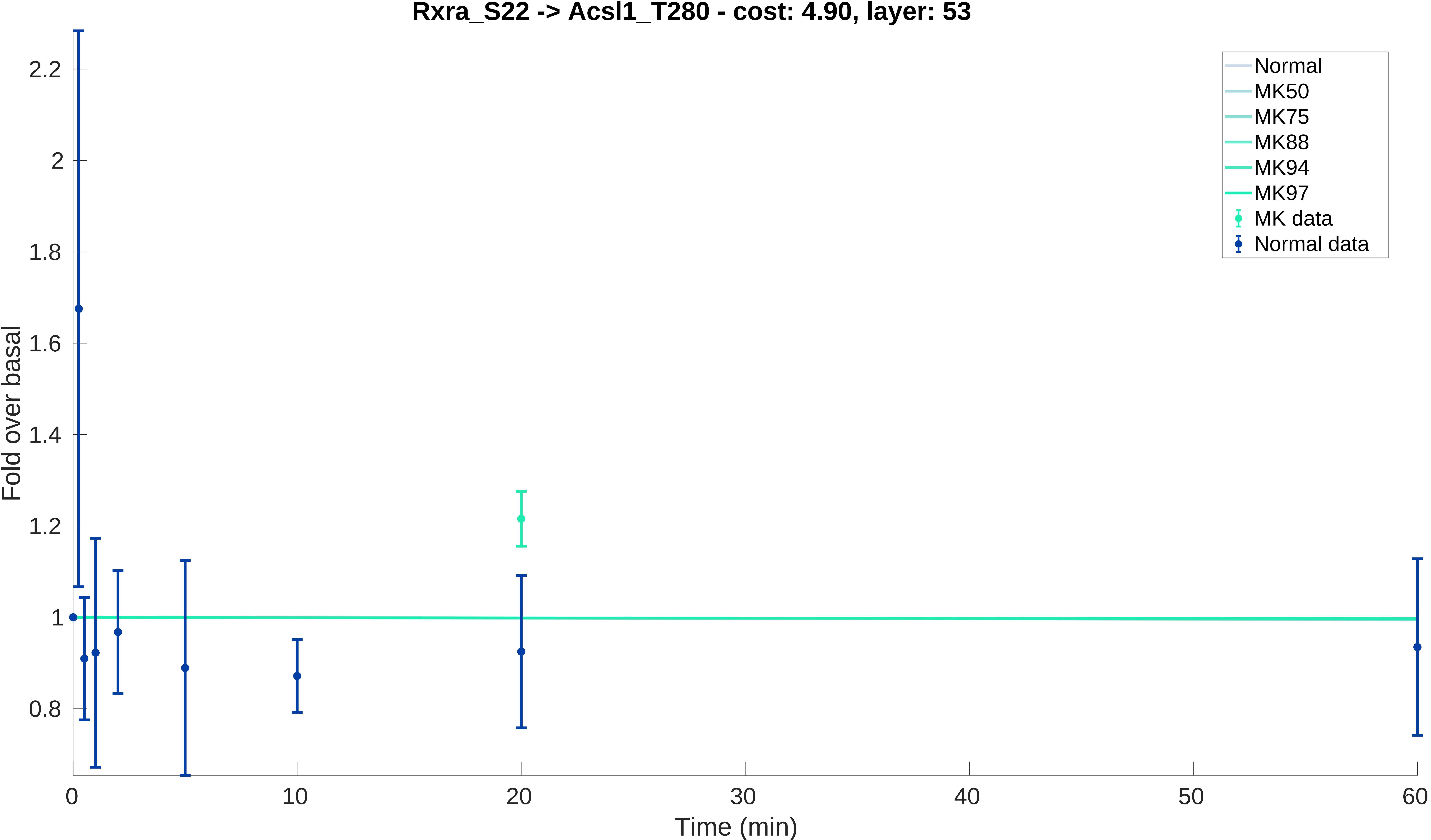
Rxra_S22 -> Acsl1_T280 - cost: 4.90, layer: 53
Fold over basal
Time (min)
0.8
1
1.2
1.4
1.6
1.8
2
2.2
0
10
20
30
40
50
60
Normal
MK50
MK75
MK88
MK94
MK97
MK data
Normal data

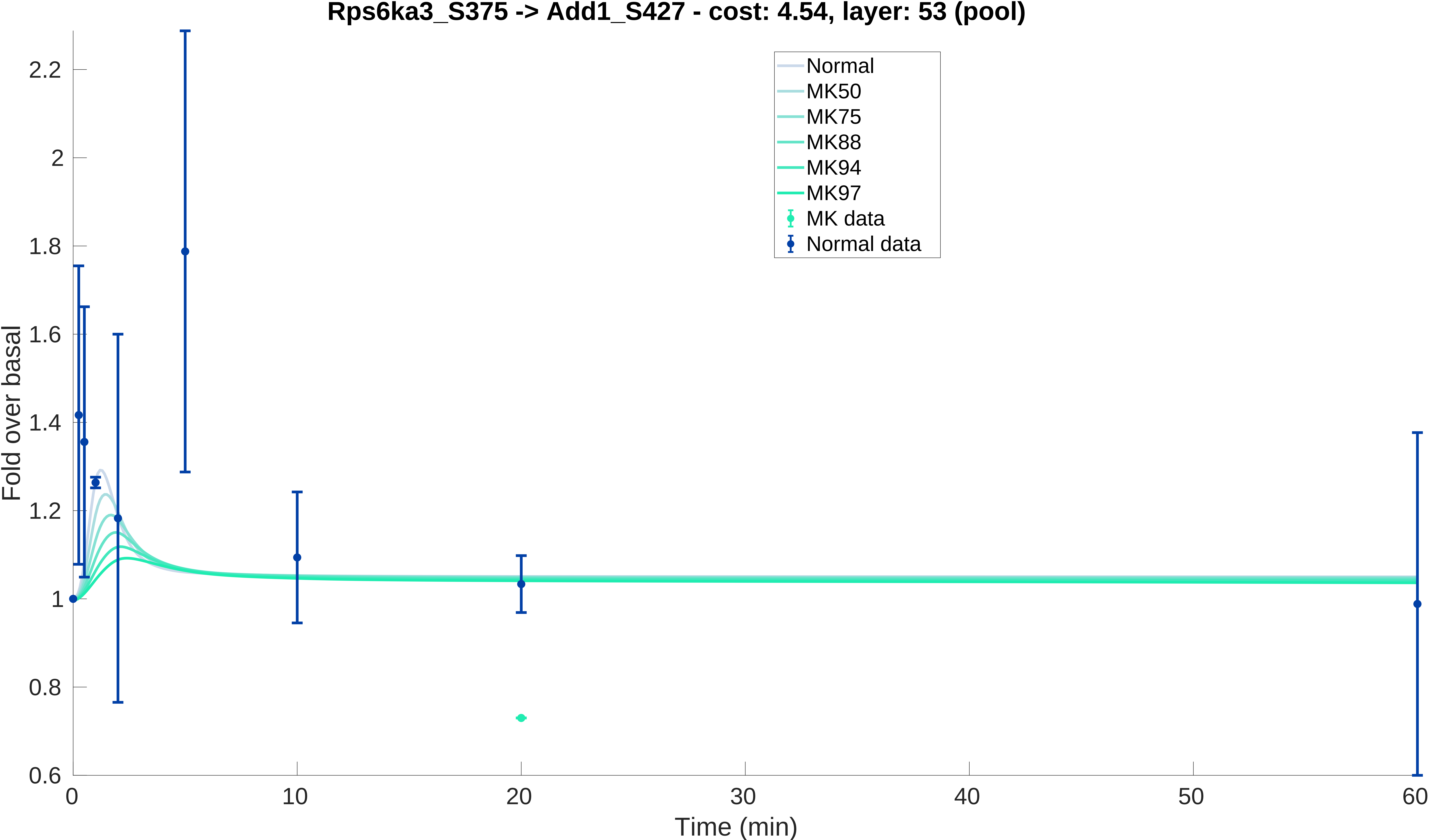

Rps6ka3_S375 -> Add1_S427 - cost: 4.54, layer: 53 (pool)
Normal
MK50
MK75
MK88
MK94
MK97
MK data
Normal data
Fold over basal
2.2
2
1.8
1.6
1.4
1.2
1
0.8
0.6
0
10
20
30
40
50
60
Time (min)

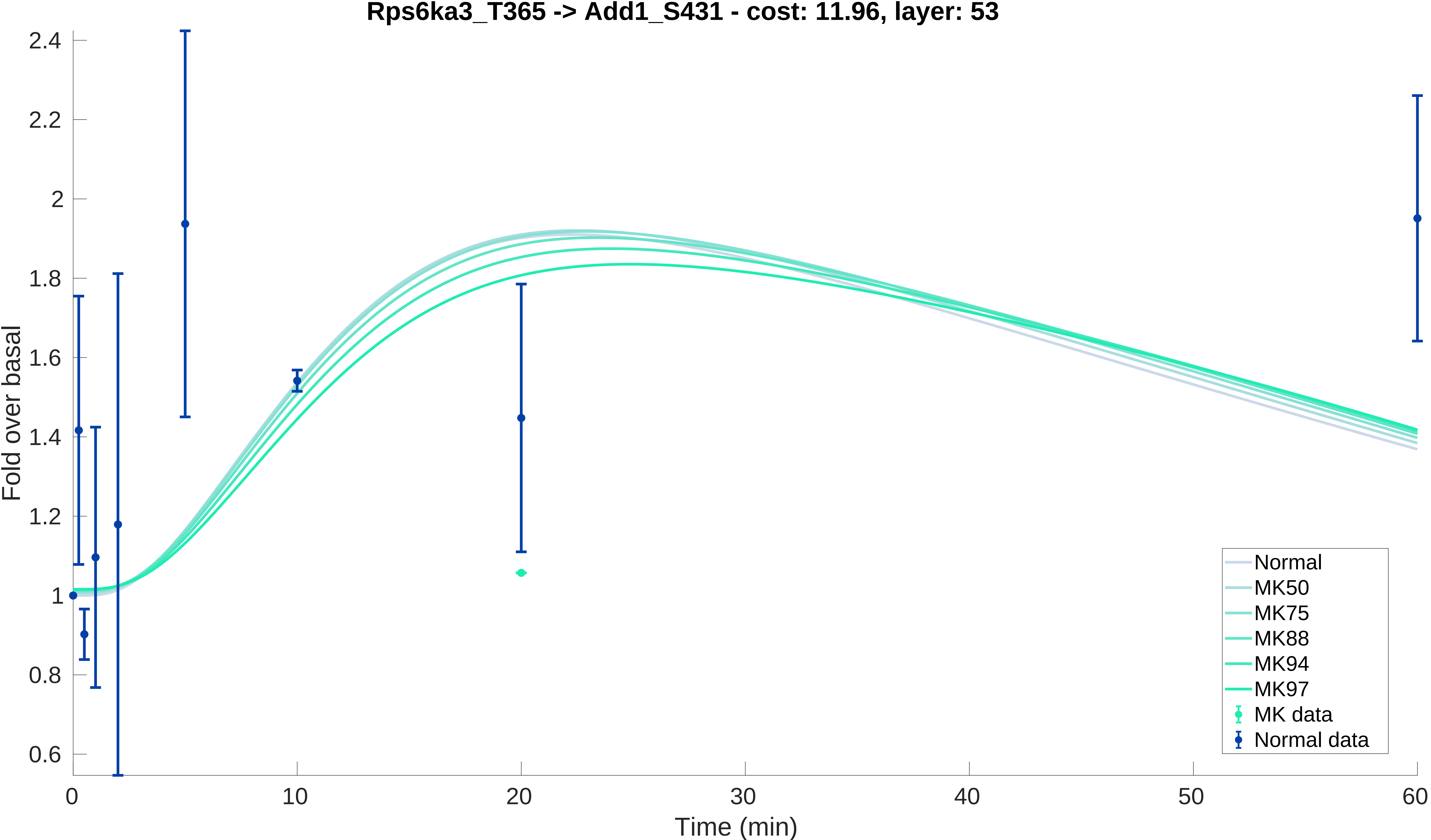

Rps6ka3_T365 -> Add1_S431 - cost: 11.96, layer: 53
Fold over basal
Time (min)
0.6
0.8
1
1.2
1.4
1.6
1.8
2
2.2
2.4
0
10
20
30
40
50
60
Normal
MK50
MK75
MK88
MK94
MK97
MK data
Normal data

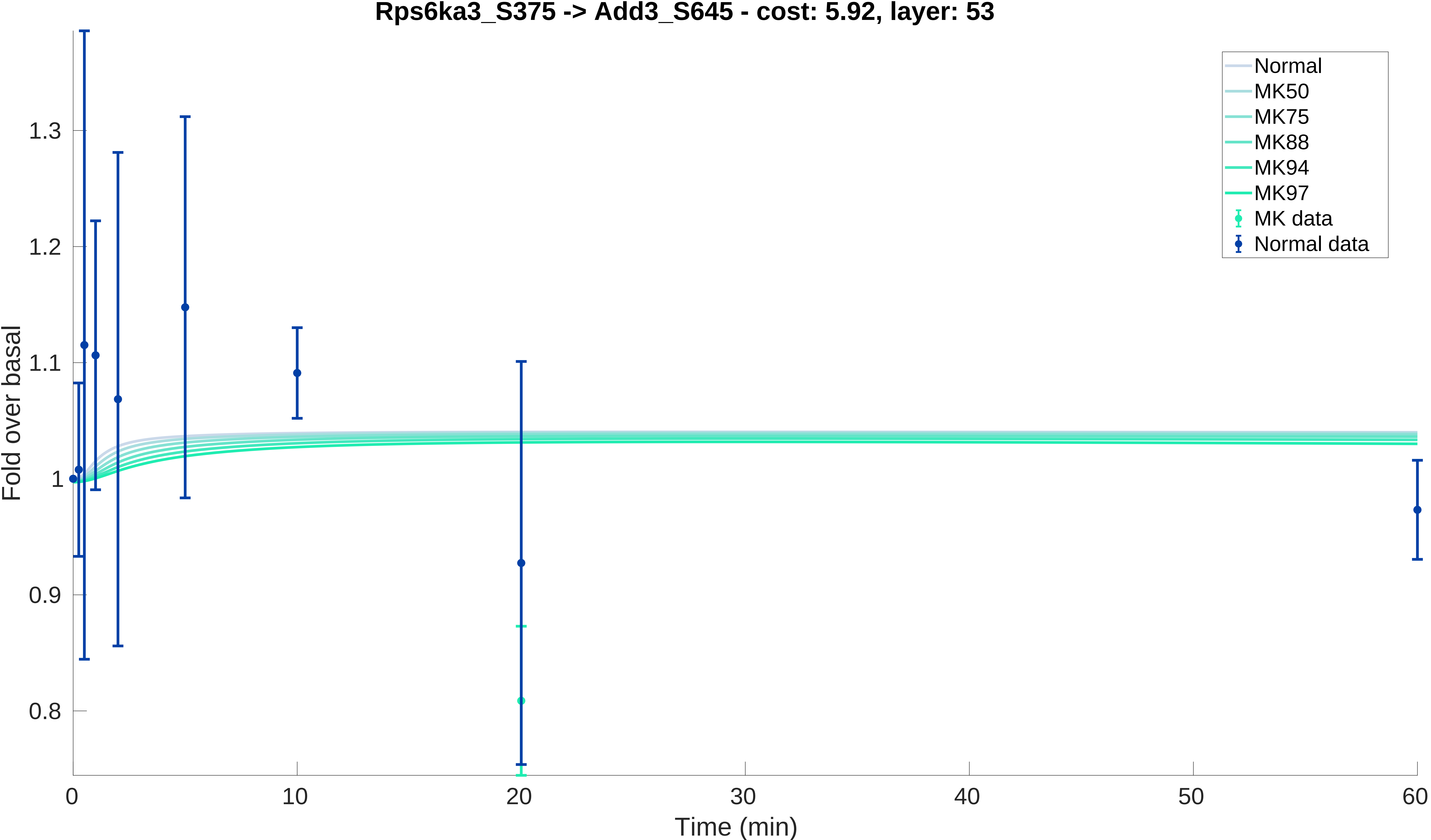

Rps6ka3_S375 -> Add3_S645 - cost: 5.92, layer: 53
Normal
MK50
MK75
MK88
MK94
MK97
MK data
Normal data
Fold over basal
1.3
1.2
1.1
1
0.9
0.8
0
10
20
30
40
50
60
Time (min)

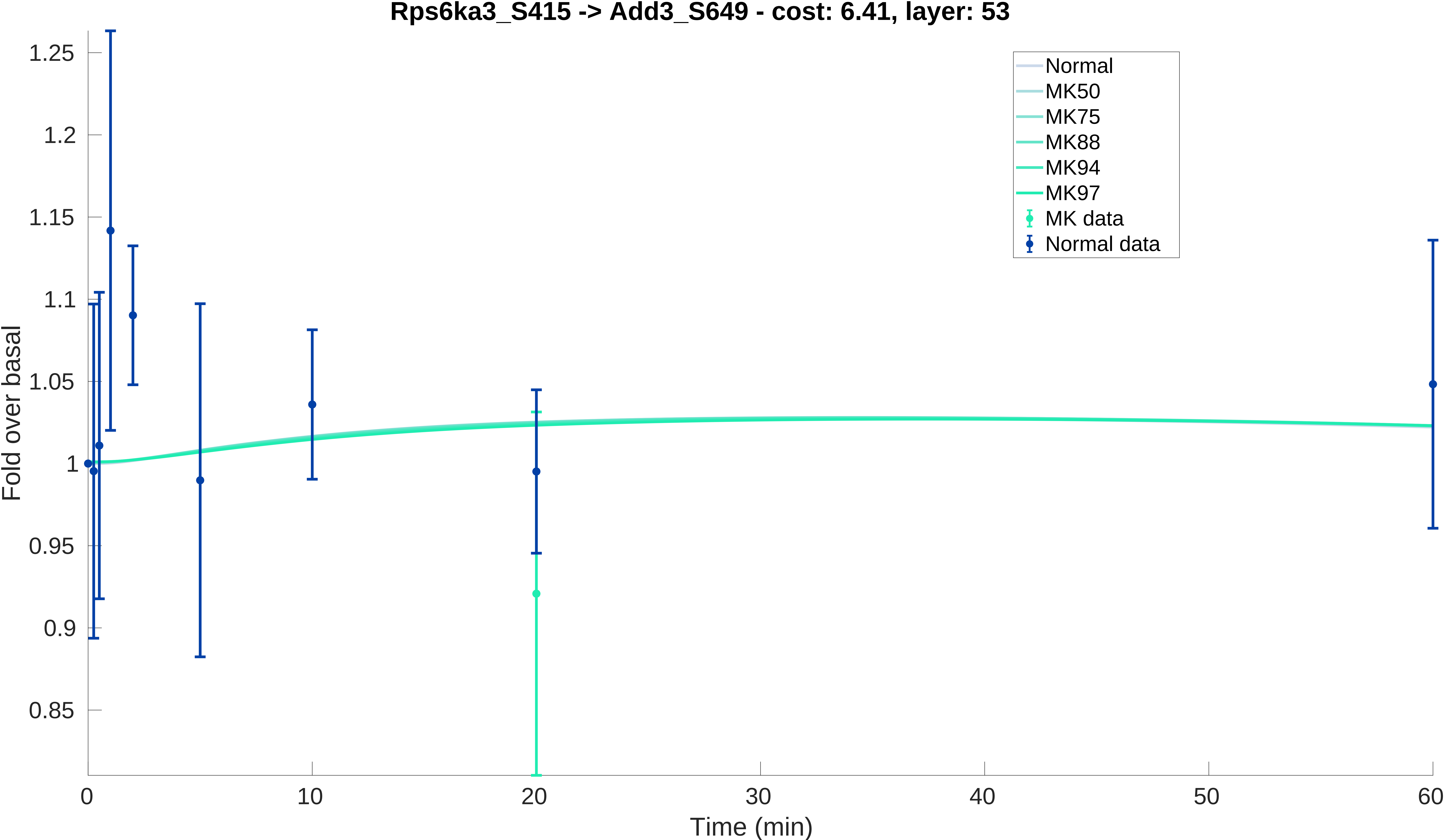

Rps6ka3_S415 -> Add3_S649 - cost: 6.41, layer: 53
Normal
MK50
MK75
MK88
MK94
MK97
MK data
Normal data
Fold over basal
Time (min)
1.25
1.2
1.15
1.1
1.05
1
0.95
0.9
0.85
0
10
20
30
40
50
60

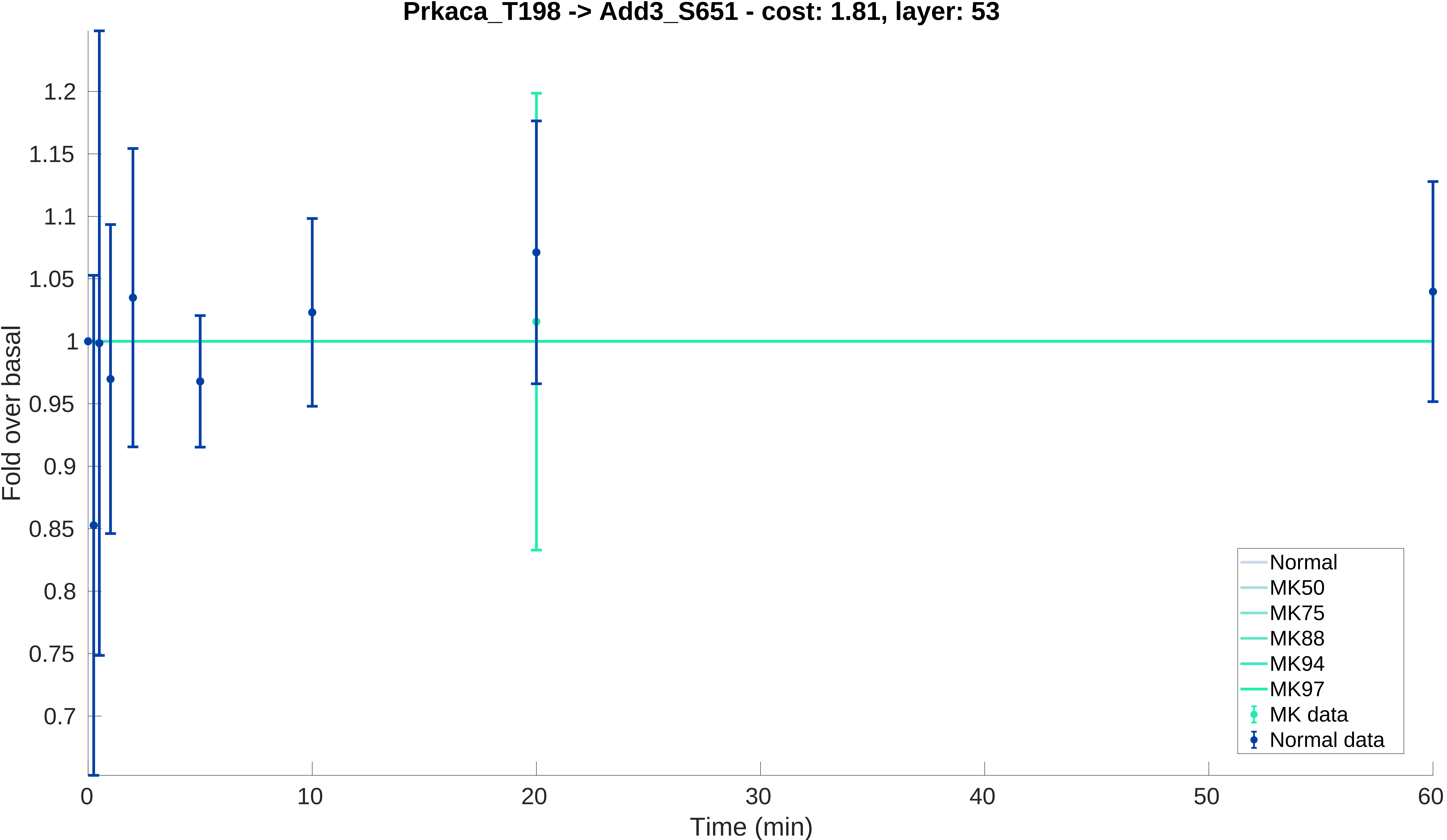

Prkaca_T198 -> Add3_S651 - cost: 1.81, layer: 53
Fold over basal
Time (min)
0
10
20
30
40
50
60
1.2
1.15
1.1
1.05
1
0.95
0.9
0.85
0.8
0.75
0.7
Normal
MK50
MK75
MK88
MK94
MK97
MK data
Normal data

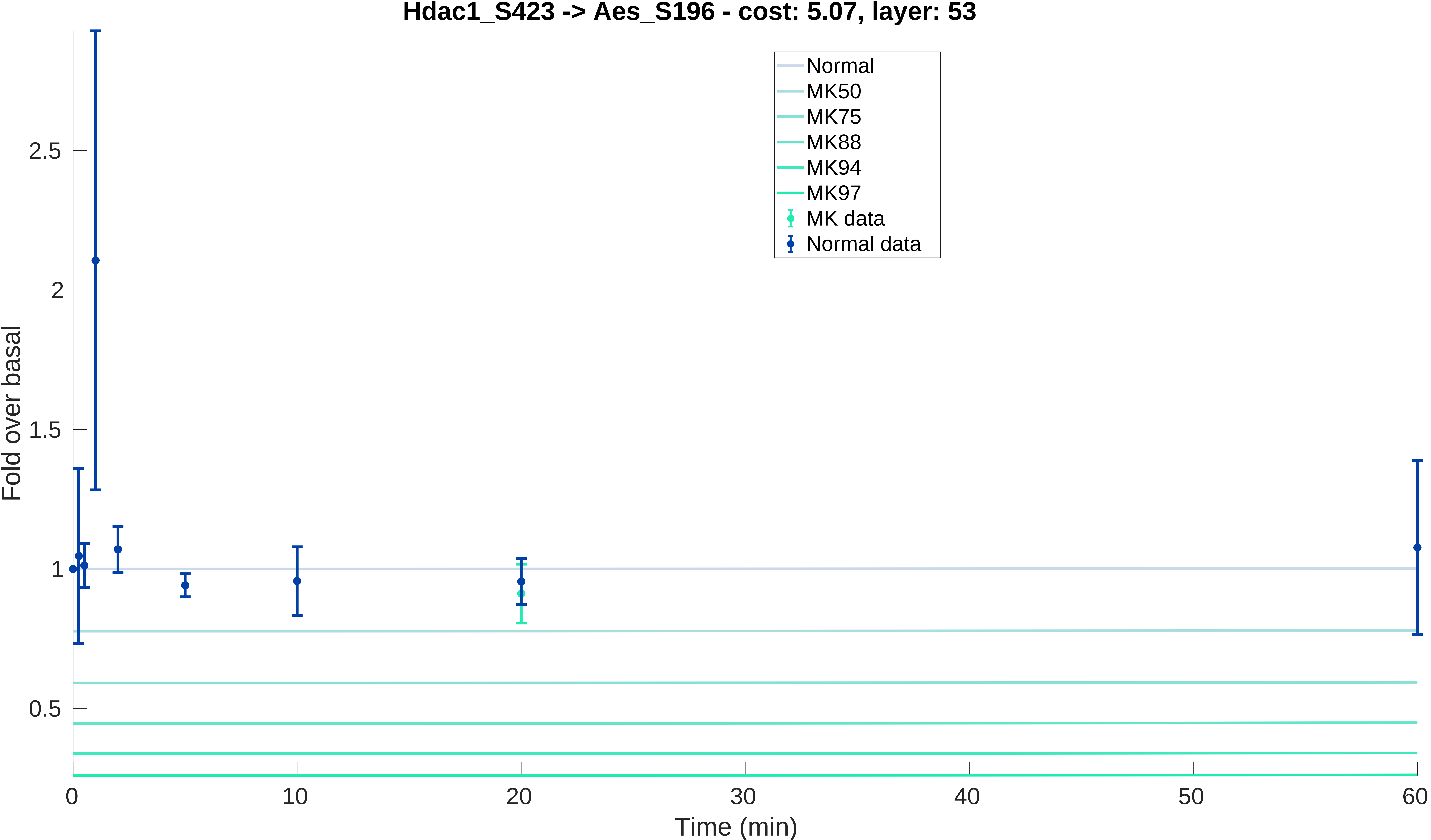
Hdac1_S423 -> Aes_S196 - cost: 5.07, layer: 53
Fold over basal
Time (min)
0
10
20
30
40
50
60
0.5
1
1.5
2
2.5
Normal
MK50
MK75
MK88
MK94
MK97
MK data
Normal data

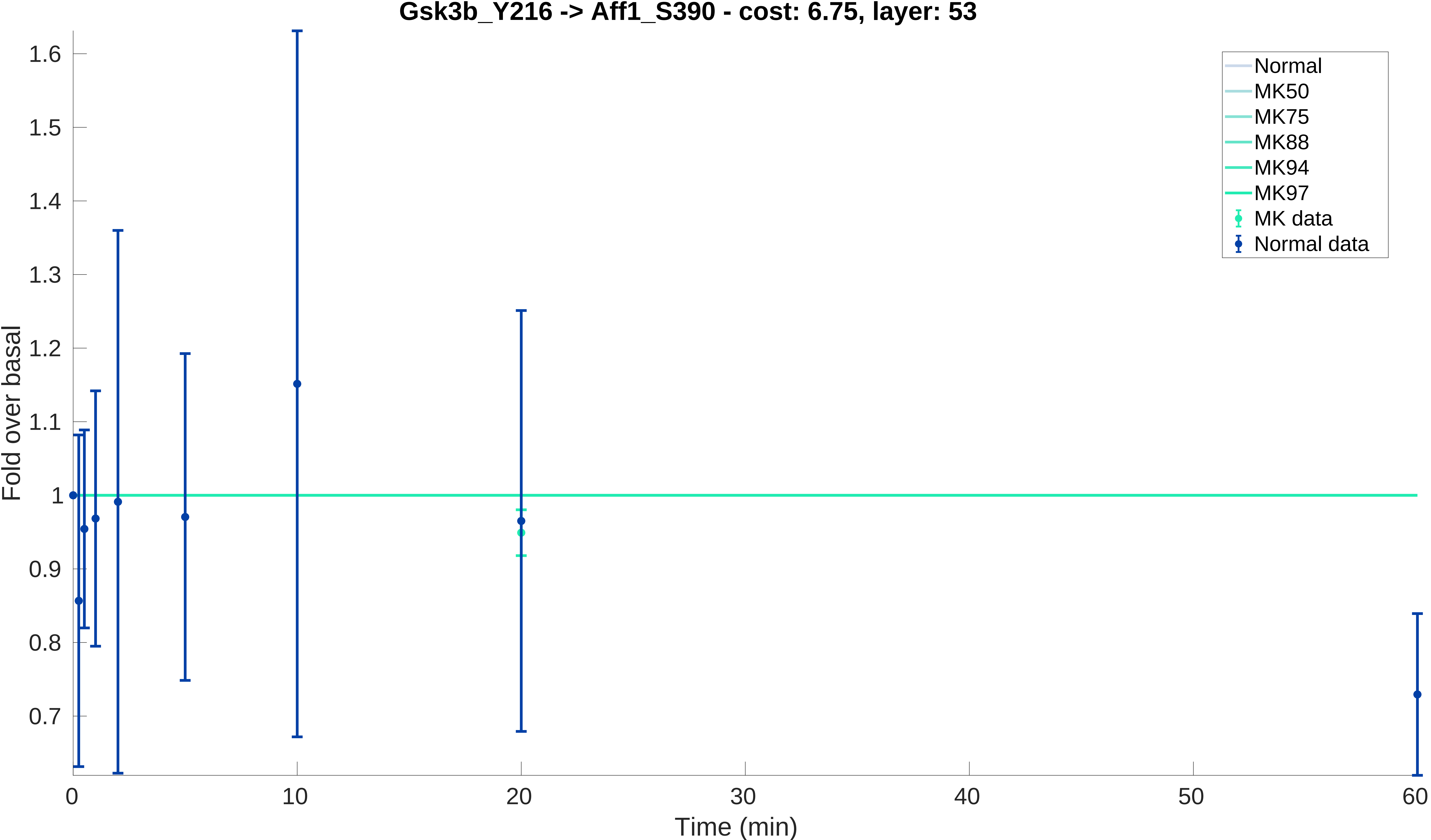

Gsk3b_Y216 -> Aff1_S390 - cost: 6.75, layer: 53
Fold over basal
Time (min)
1.6
1.5
1.4
1.3
1.2
1.1
1
0.9
0.8
0.7
0
10
20
30
40
50
60
Normal
MK50
MK75
MK88
MK94
MK97
MK data
Normal data

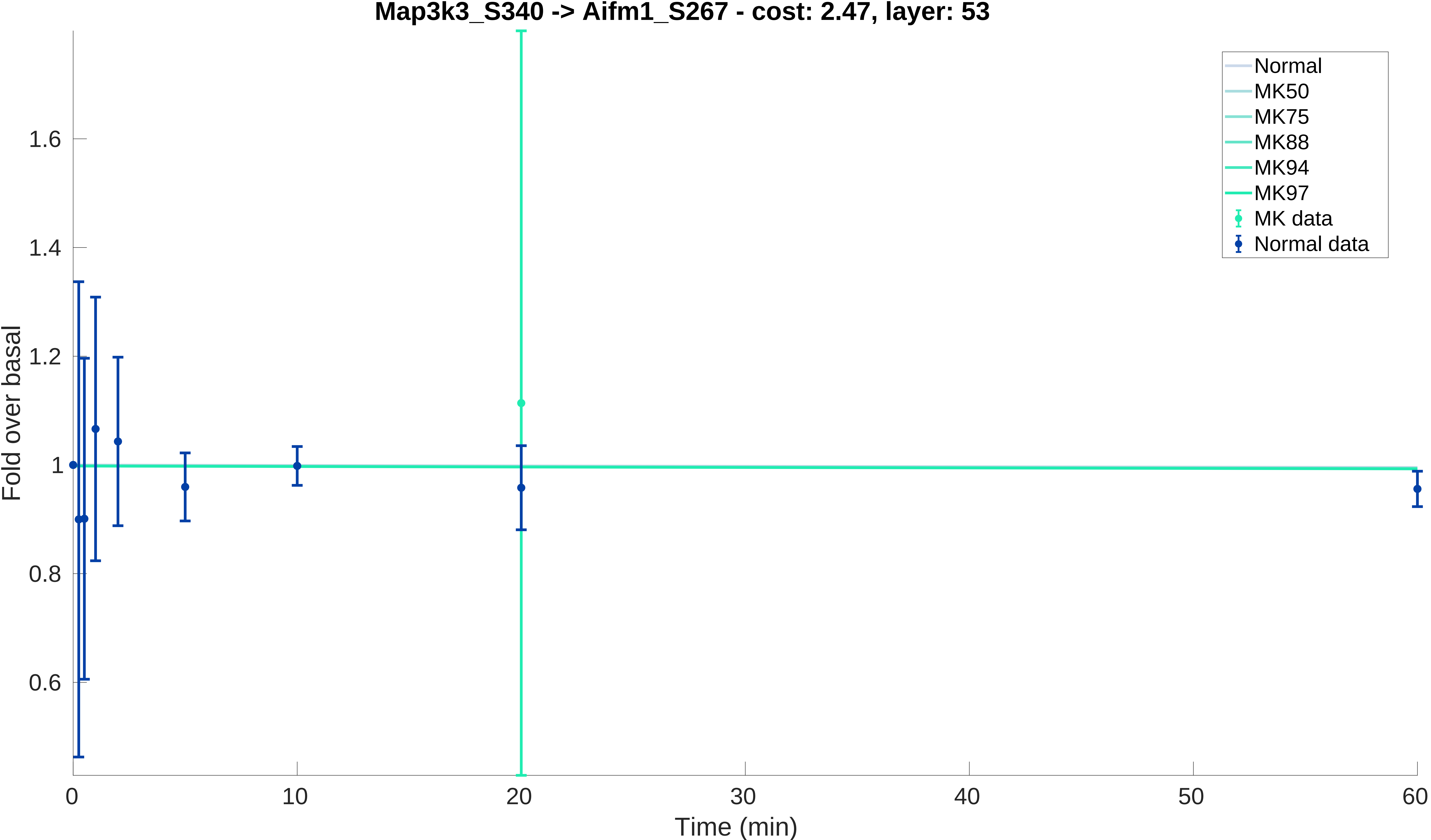

Map3k3_S340 -> Aifm1_S267 - cost: 2.47, layer: 53
Fold over basal
1.6
1.4
1.2
1
0.8
0.6
0
10
20
30
40
50
60
Time (min)
Normal
MK50
MK75
MK88
MK94
MK97
MK data
Normal data

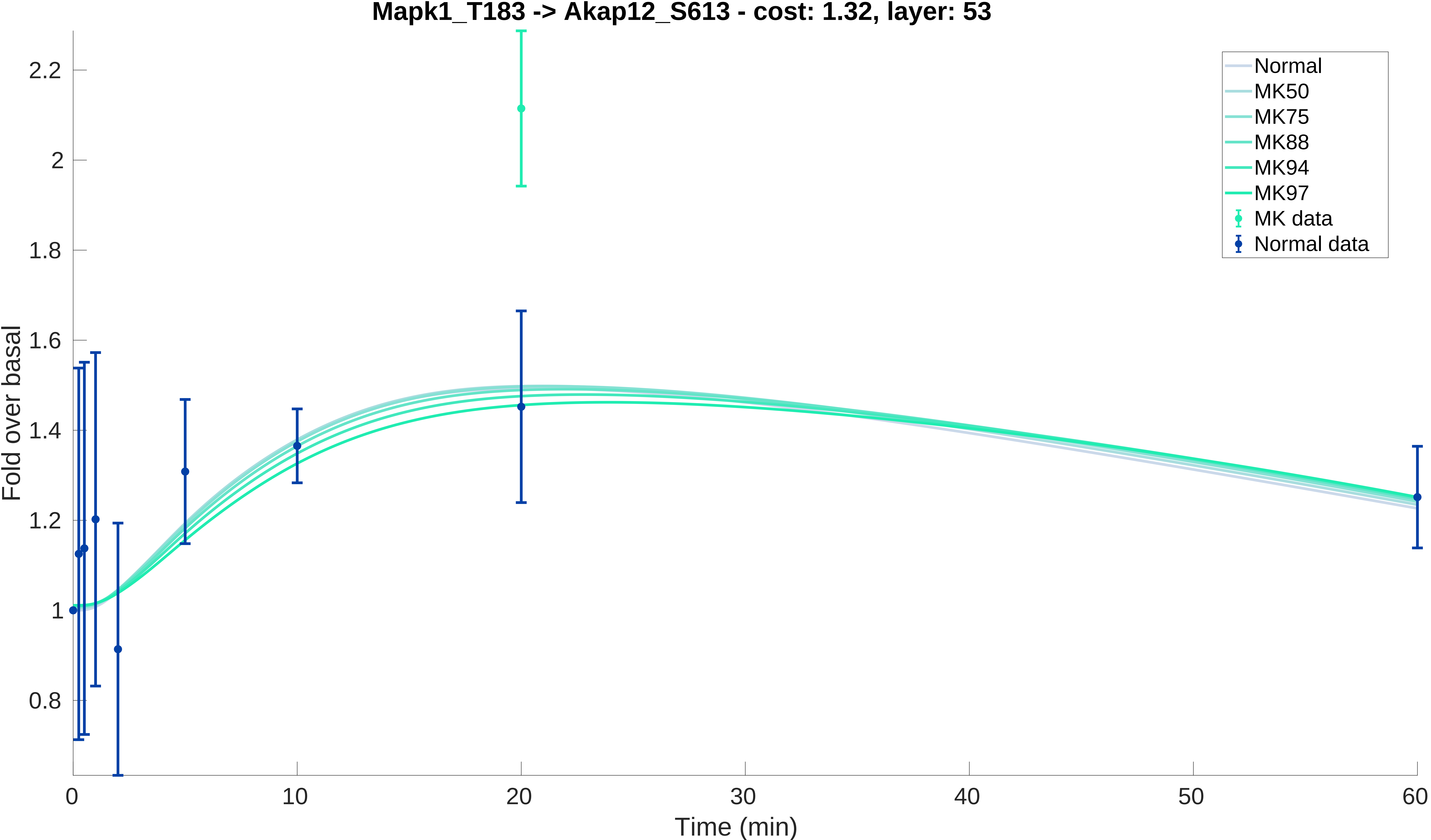

Mapk1_T183 -> Akap12_S613 - cost: 1.32, layer: 53
Normal
MK50
MK75
MK88
MK94
MK97
MK data
Normal data
Fold over basal
2.2
2
1.8
1.6
1.4
1.2
1
0.8
0
10
20
30
40
50
60
Time (min)

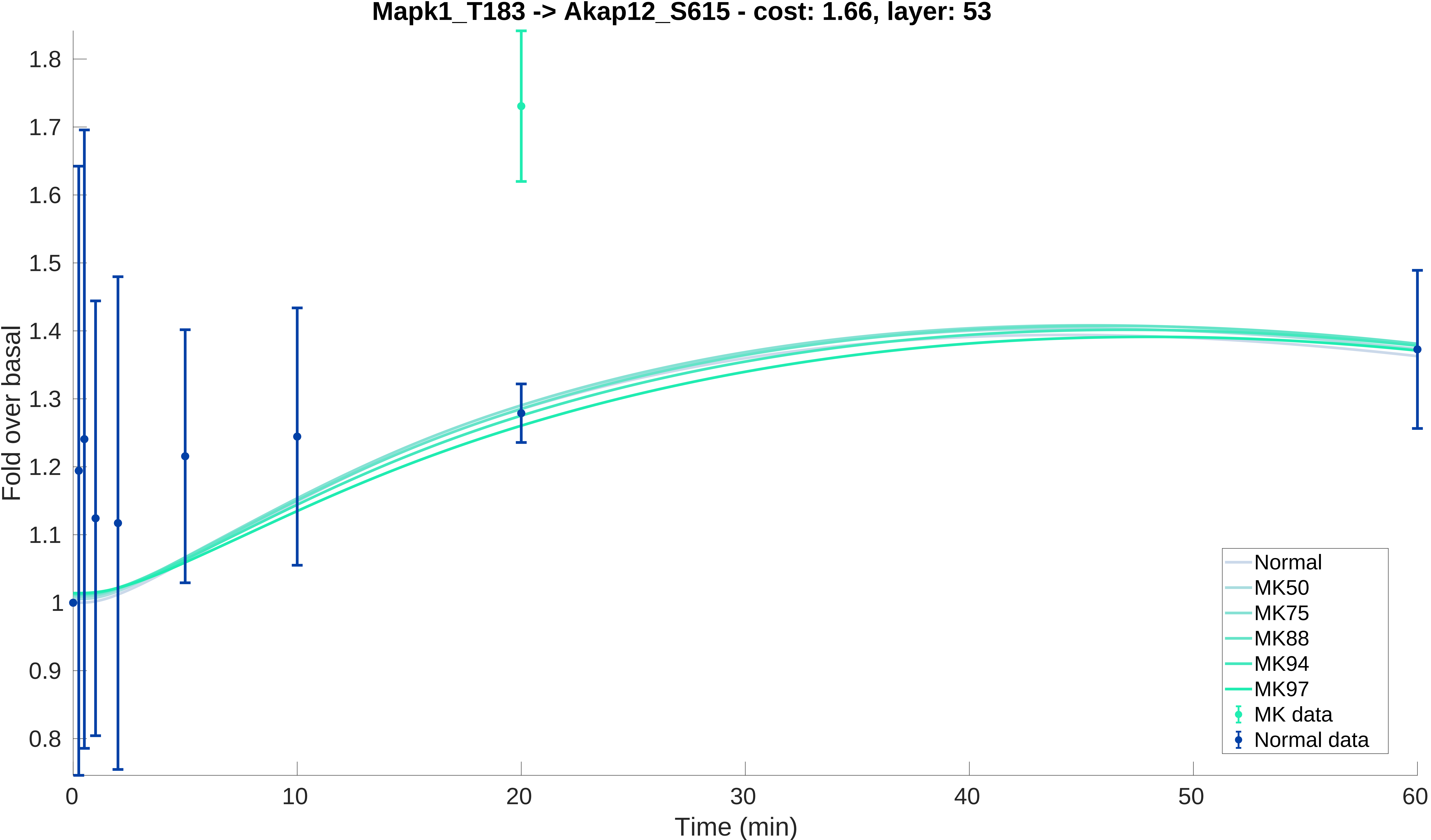
Mapk1_T183 -> Akap12_S615 - cost: 1.66, layer: 53
Fold over basal
Time (min)
1.8
1.7
1.6
1.5
1.4
1.3
1.2
1.1
1
0.9
0.8
0
10
20
30
40
50
60
Normal
MK50
MK75
MK88
MK94
MK97
MK data
Normal data

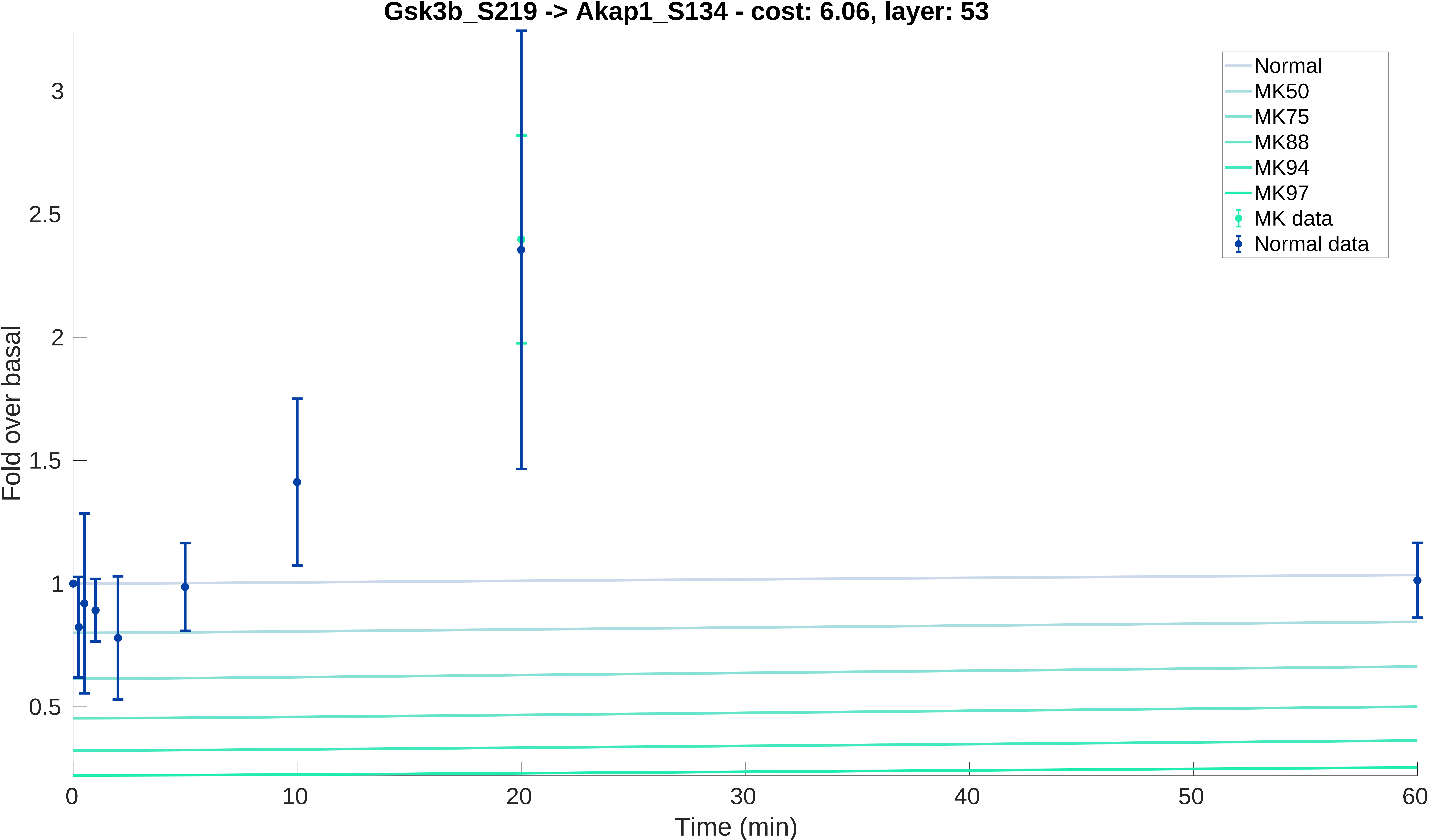

Gsk3b_S219 -> Akap1_S134 - cost: 6.06, layer: 53
Fold over basal
Time (min)
0
10
20
30
40
50
60
0.5
1
1.5
2
2.5
3
Normal
MK50
MK75
MK88
MK94
MK97
MK data
Normal data

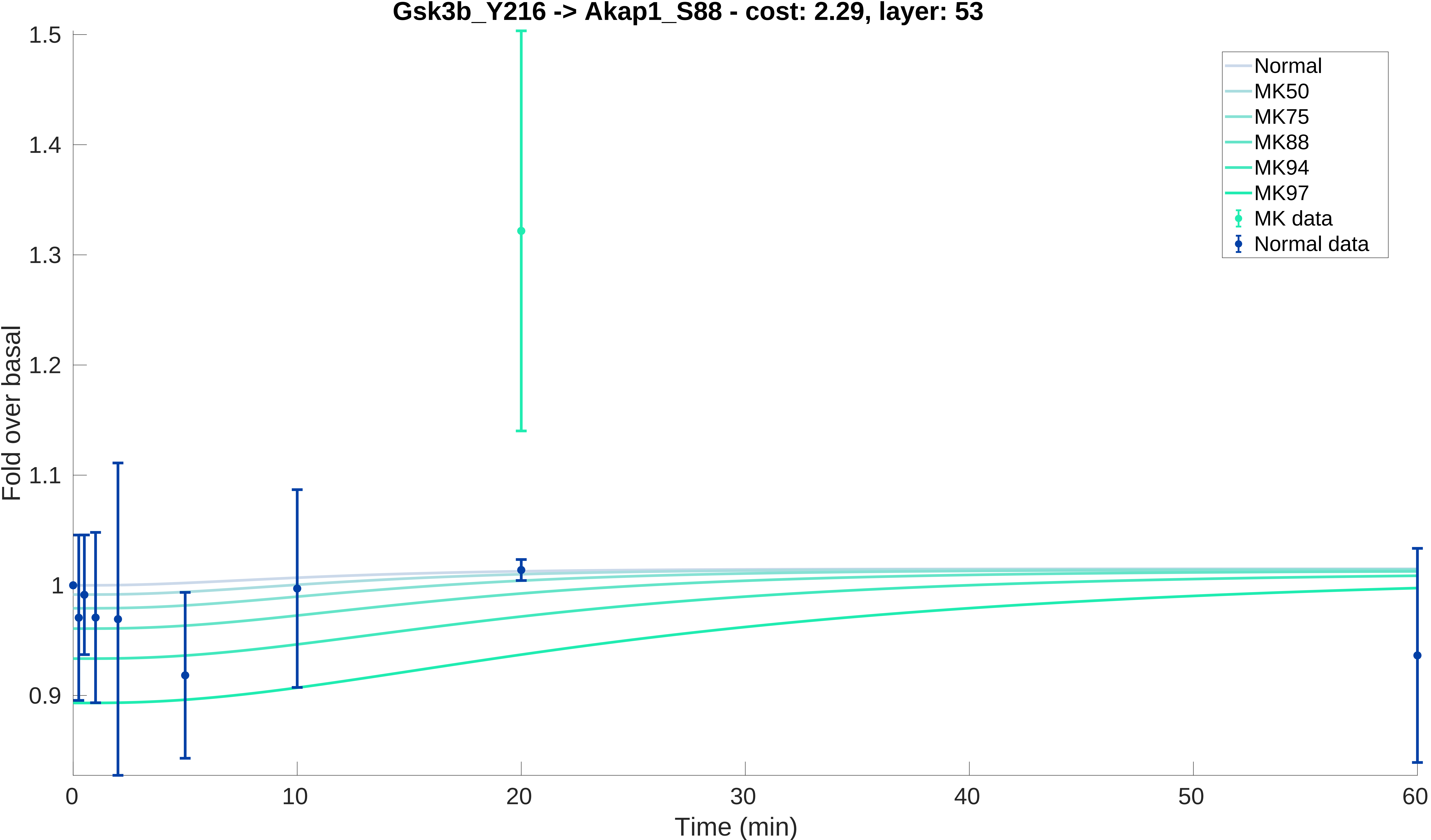

Gsk3b_Y216 -> Akap1_S88 - cost: 2.29, layer: 53
Fold over basal
Time (min)
1.5
1.4
1.3
1.2
1.1
1
0.9
0
10
20
30
40
50
60
Normal
MK50
MK75
MK88
MK94
MK97
MK data
Normal data

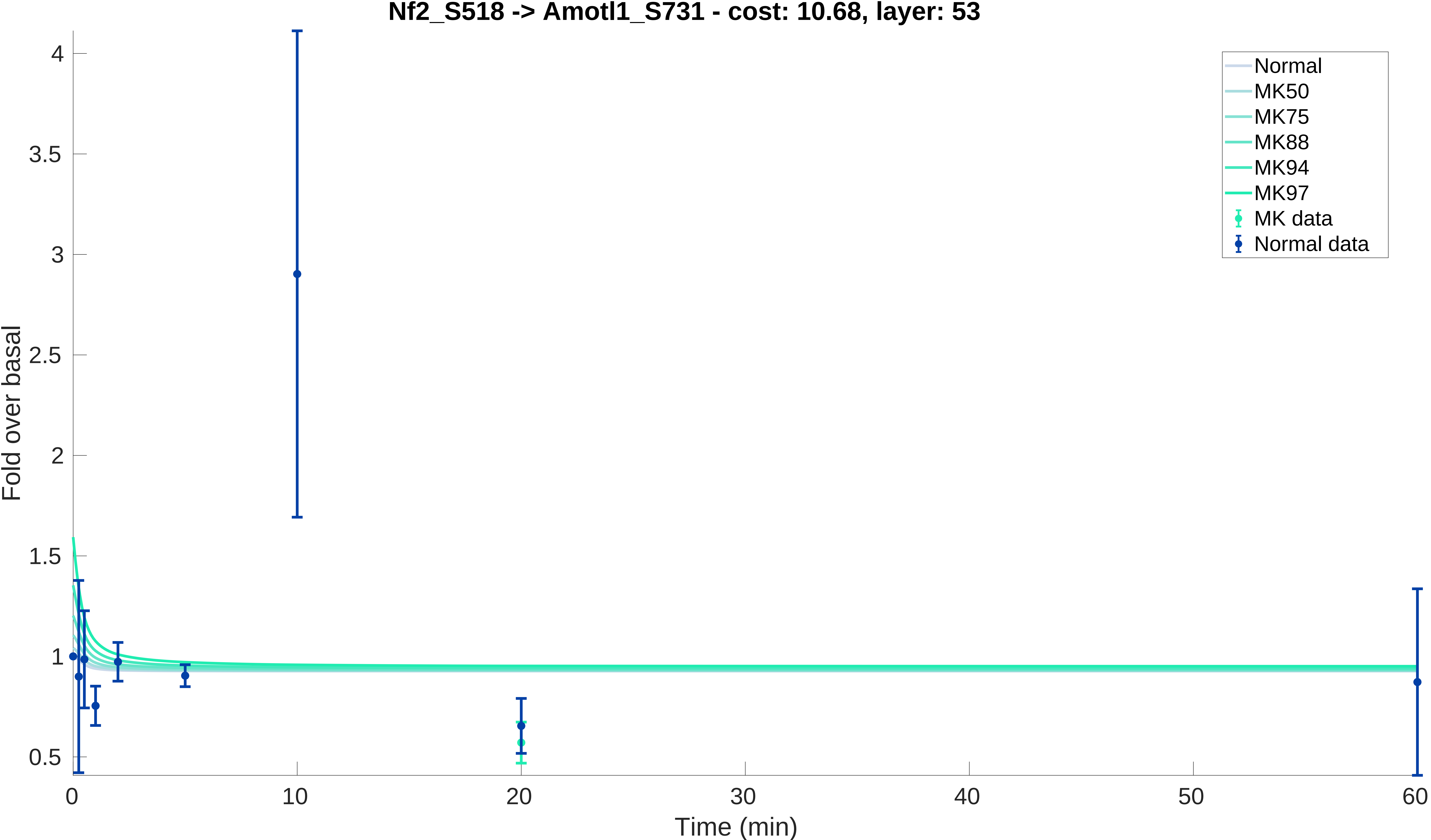

Nf2_S518 -> Amotl1_S731 - cost: 10.68, layer: 53
Fold over basal
4
3.5
3
2.5
2
1.5
1
0.5
0
10
20
30
40
50
60
Time (min)
Normal
MK50
MK75
MK88
MK94
MK97
MK data
Normal data

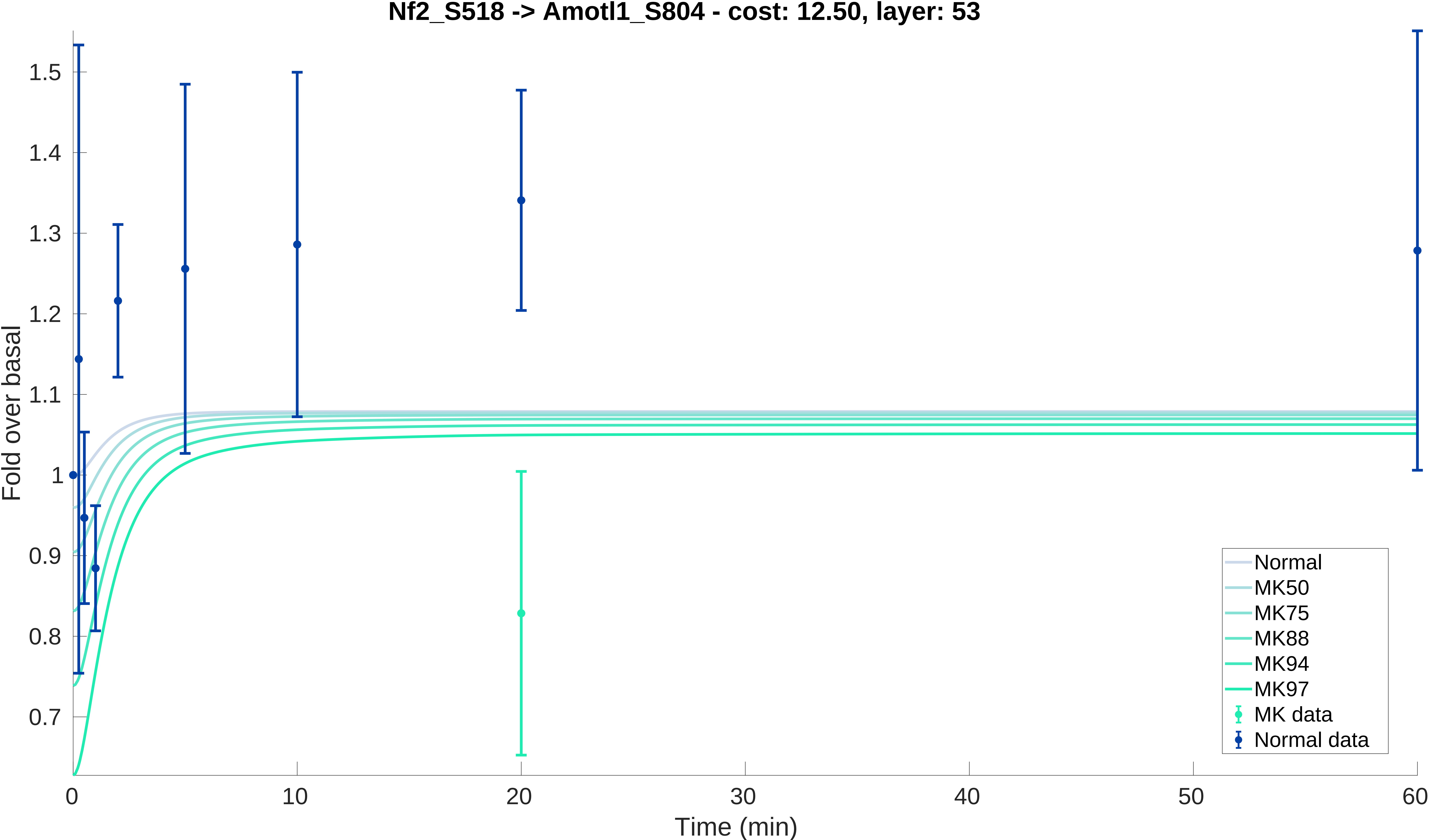

Nf2_S518 -> Amotl1_S804 - cost: 12.50, layer: 53
Fold over basal
Time (min)
0
10
20
30
40
50
60
0.7
0.8
0.9
1
1.1
1.2
1.3
1.4
1.5
Normal
MK50
MK75
MK88
MK94
MK97
MK data
Normal data

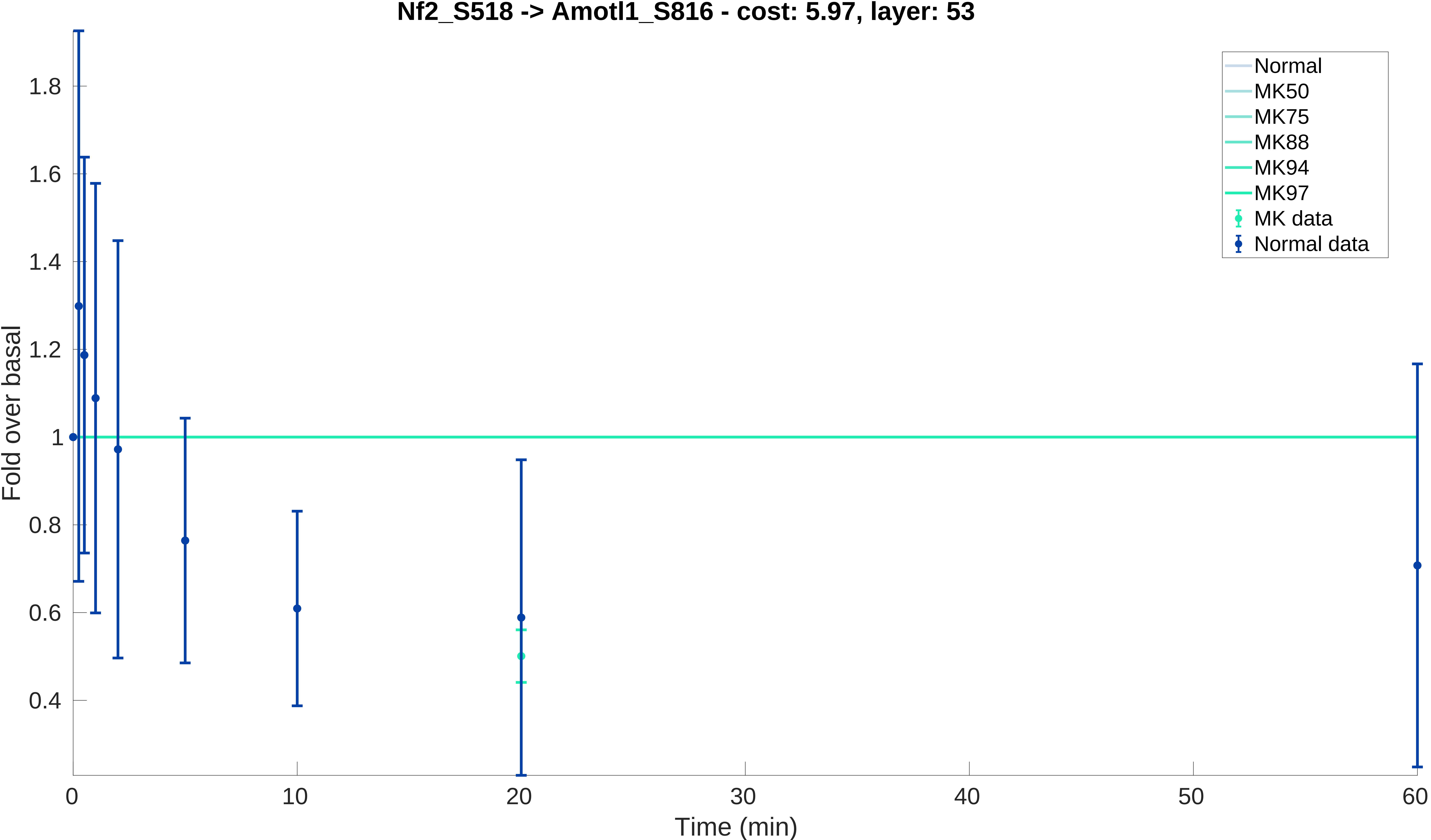
Nf2_S518 -> Amotl1_S816 - cost: 5.97, layer: 53
Fold over basal
1.8
1.6
1.4
1.2
1
0.8
0.6
0.4
0
10
20
30
40
50
60
Time (min)
Normal
MK50
MK75
MK88
MK94
MK97
MK data
Normal data

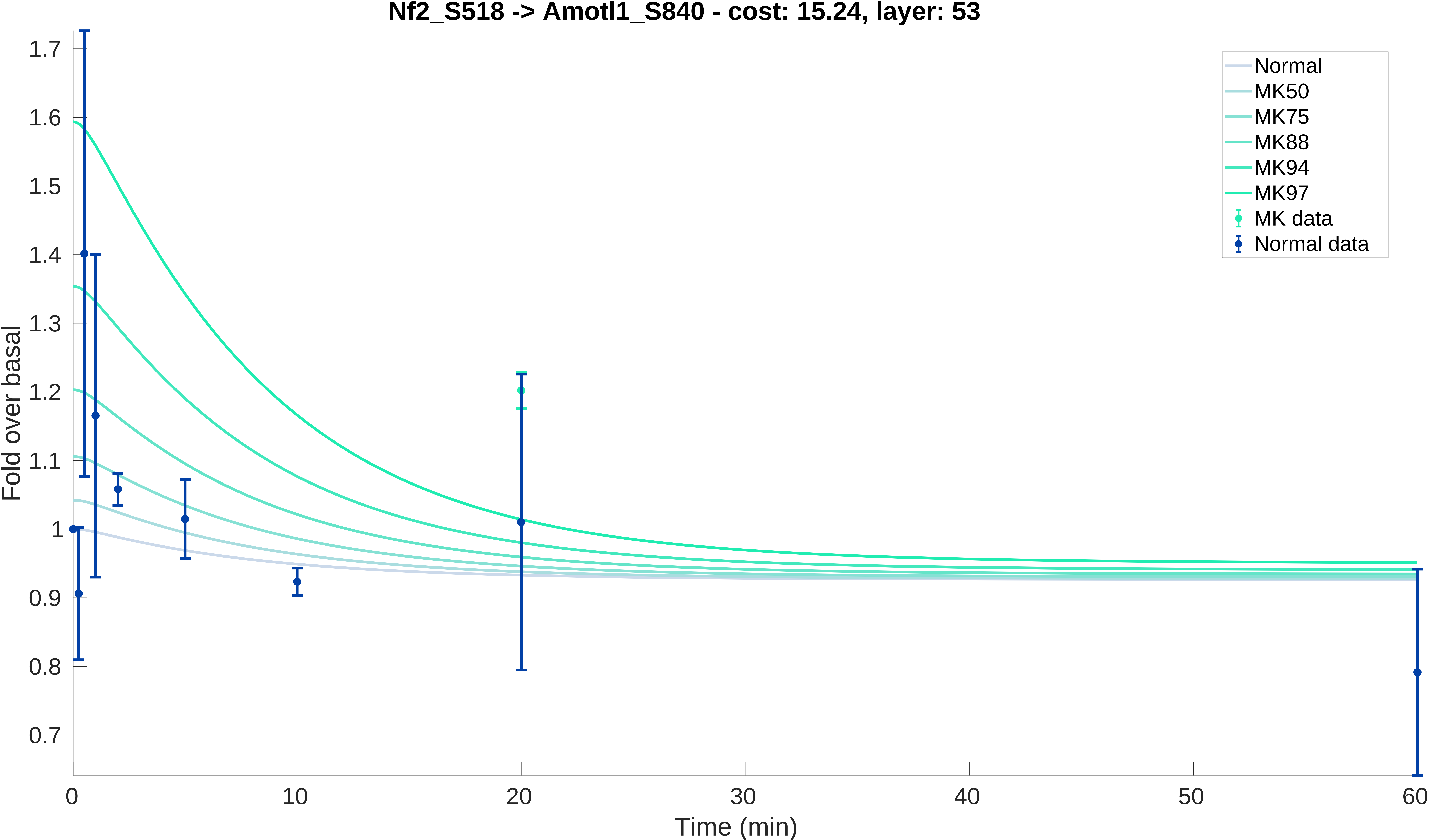

Nf2_S518 -> Amotl1_S840 - cost: 15.24, layer: 53
Fold over basal
Time (min)
Normal
MK50
MK75
MK88
MK94
MK97
MK data
Normal data
1.7
1.6
1.5
1.4
1.3
1.2
1.1
1
0.9
0.8
0.7
0
10
20
30
40
50
60

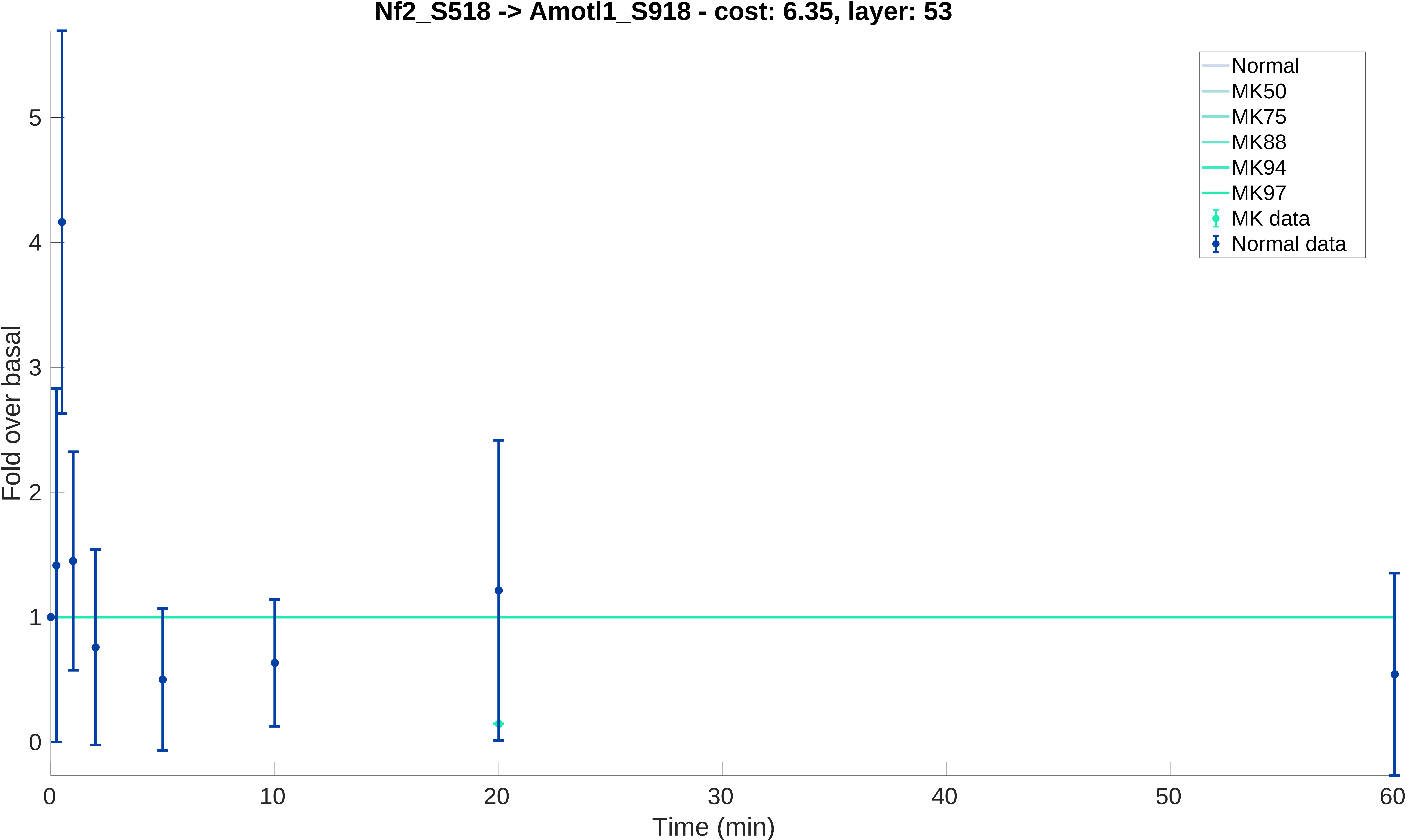

Nf2_S518 -> Amotl1_S918 - cost: 6.35, layer: 53
Normal
MK50
MK75
MK88
MK94
MK97
MK data
Normal data
Fold over basal
Time (min)
0
1
2
3
4
5
0
10
20
30
40
50
60

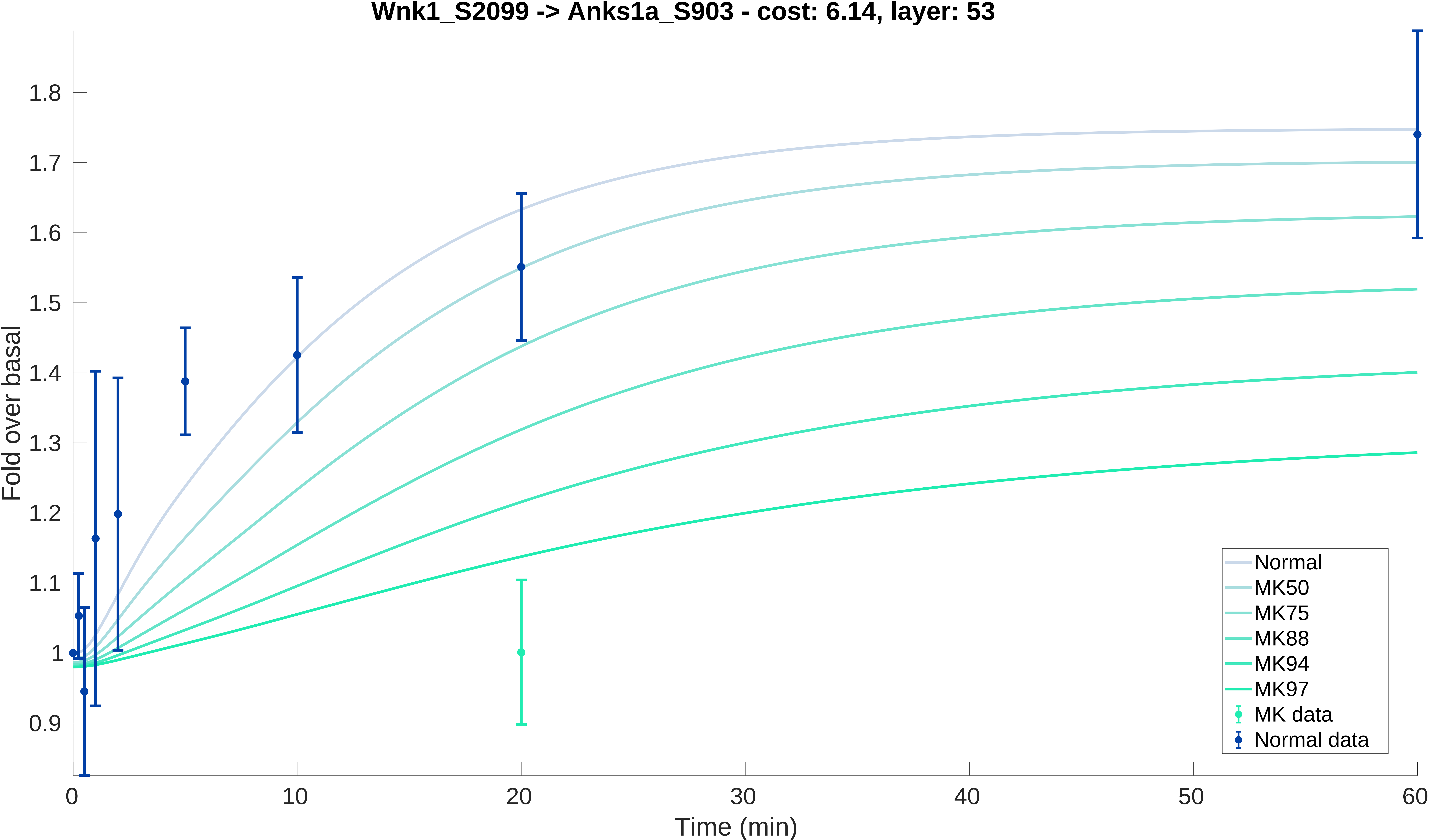

Wnk1_S2099 -> Anks1a_S903 - cost: 6.14, layer: 53
Fold over basal
Time (min)
0.9
1
1.1
1.2
1.3
1.4
1.5
1.6
1.7
1.8
0
10
20
30
40
50
60
Normal
MK50
MK75
MK88
MK94
MK97
MK data
Normal data

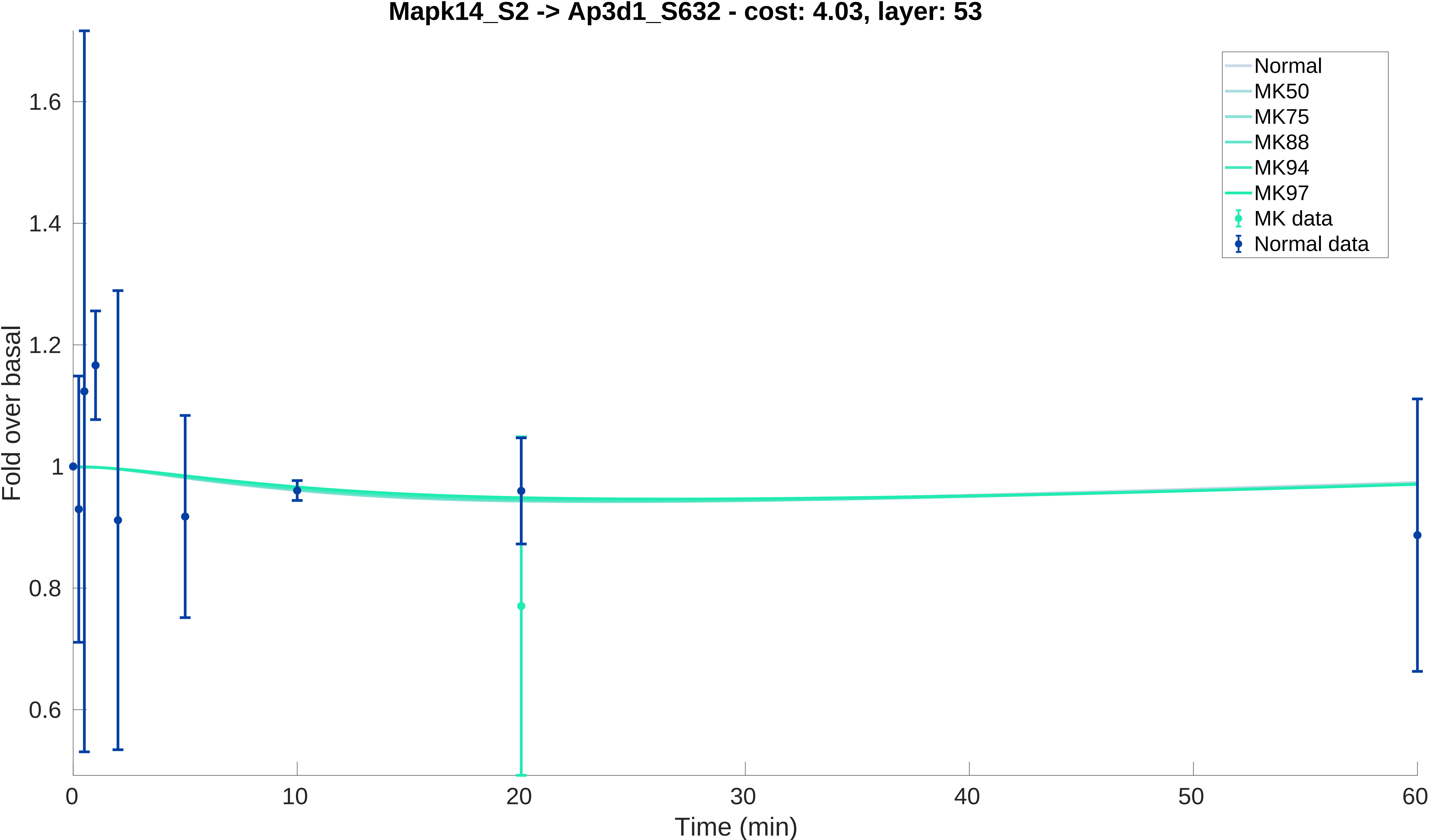

Mapk14_S2 -> Ap3d1_S632 - cost: 4.03, layer: 53
Normal
MK50
MK75
MK88
MK94
MK97
MK data
Normal data
Fold over basal
1.6
1.4
1.2
1
0.8
0.6
0
10
20
30
40
50
60
Time (min)

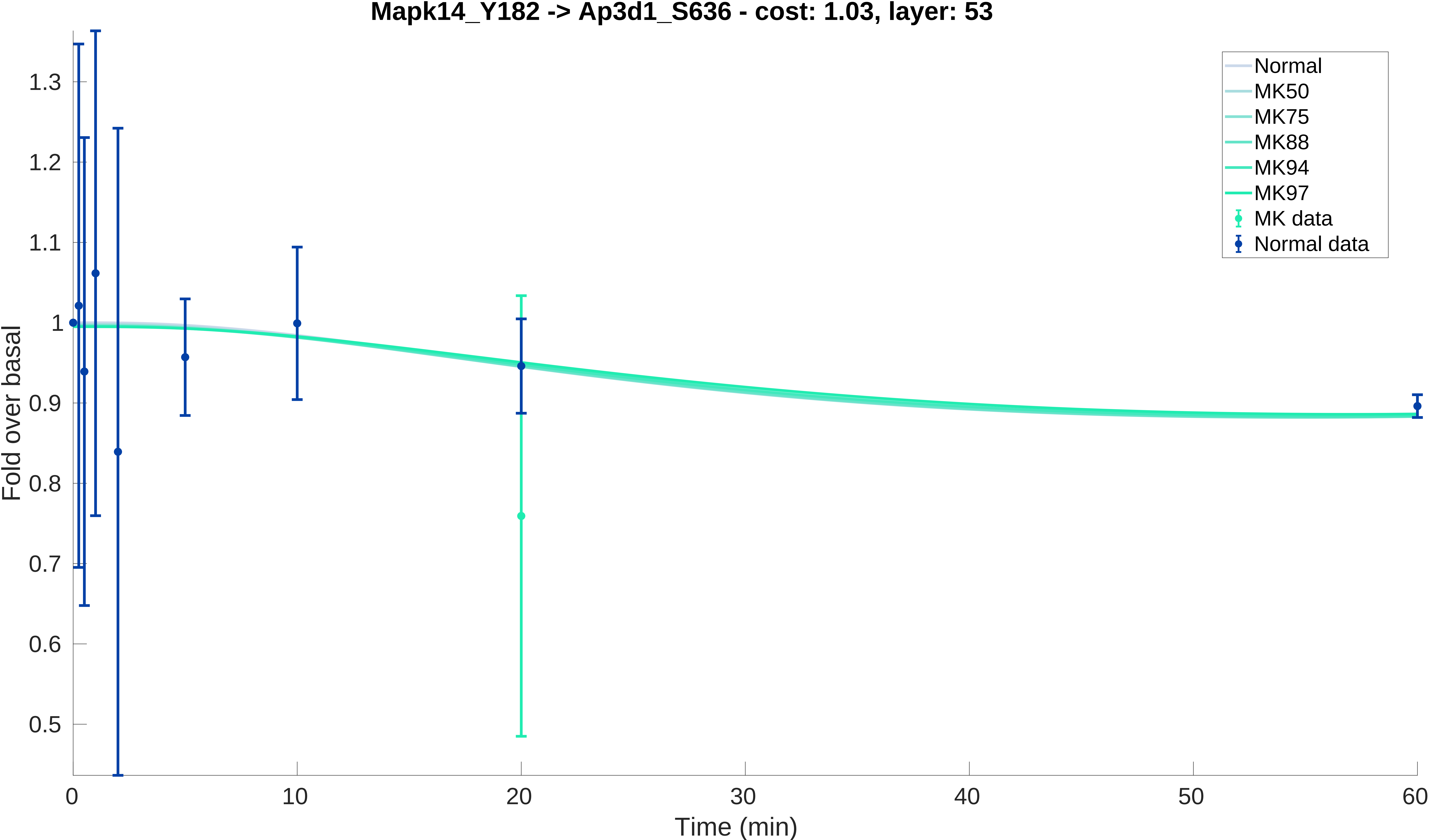

Mapk14_Y182 -> Ap3d1_S636 - cost: 1.03, layer: 53
Normal
MK50
MK75
MK88
MK94
MK97
MK data
Normal data
Fold over basal
Time (min)
1.3
1.2
1.1
1
0.9
0.8
0.7
0.6
0.5
0
10
20
30
40
50
60

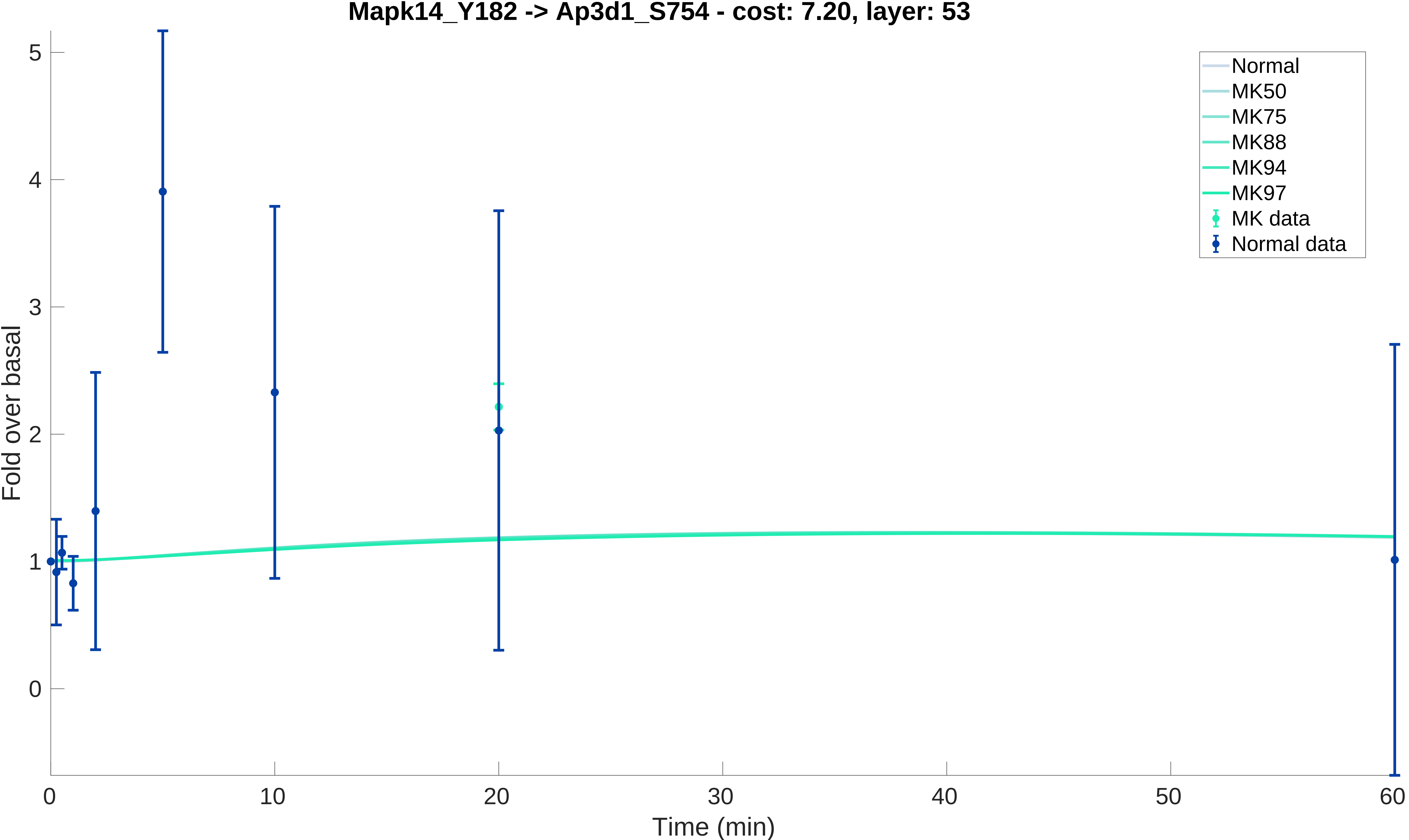

Mapk14_Y182 -> Ap3d1_S754 - cost: 7.20, layer: 53
Normal
MK50
MK75
MK88
MK94
MK97
MK data
Normal data
Fold over basal
Time (min)
0
1
2
3
4
5
0
10
20
30
40
50
60

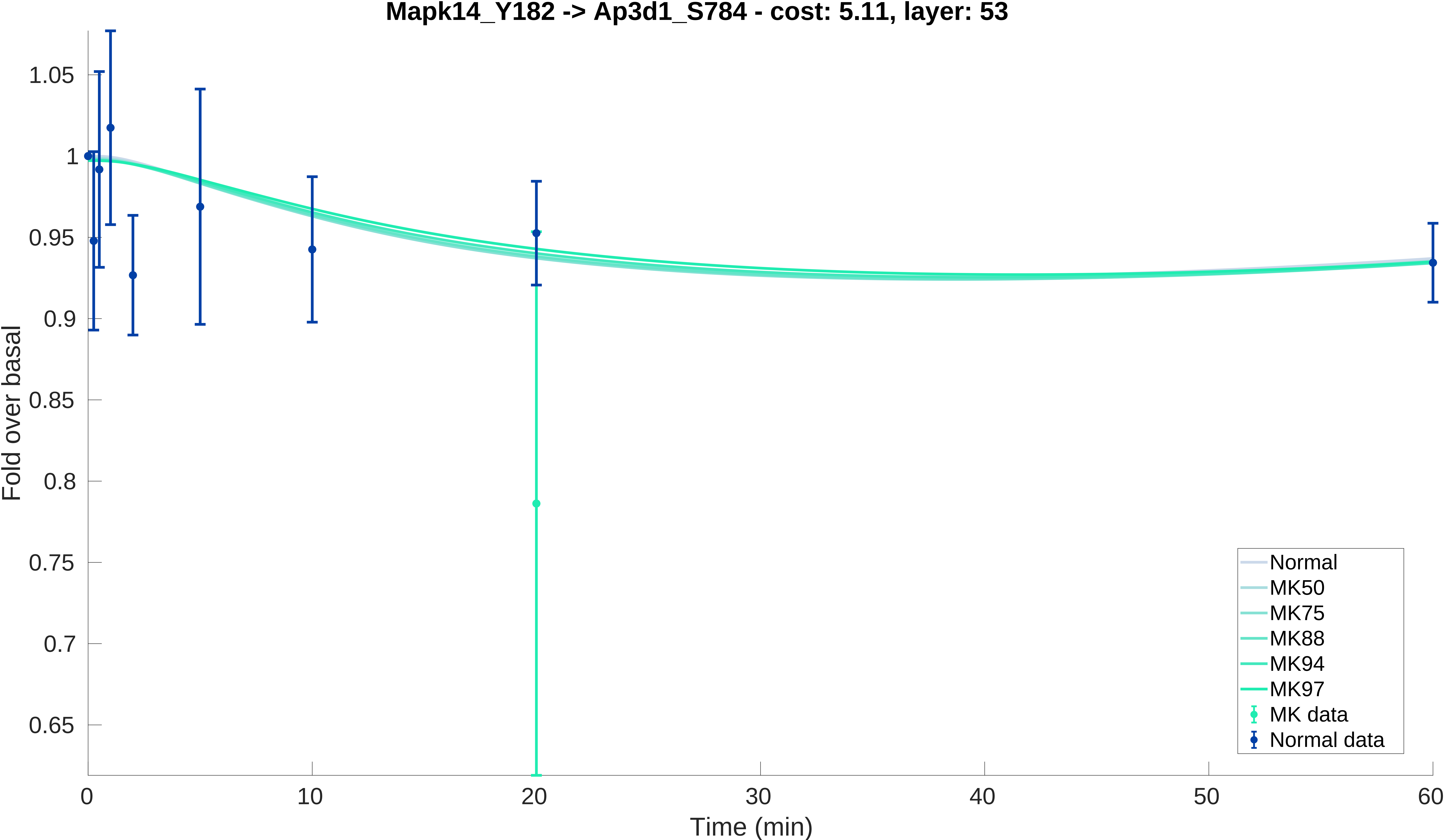

Mapk14_Y182 -> Ap3d1_S784 - cost: 5.11, layer: 53
Fold over basal
1.05
1
0.95
0.9
0.85
0.8
0.75
0.7
0.65
0
10
20
30
40
50
60
Time (min)
Normal
MK50
MK75
MK88
MK94
MK97
MK data
Normal data

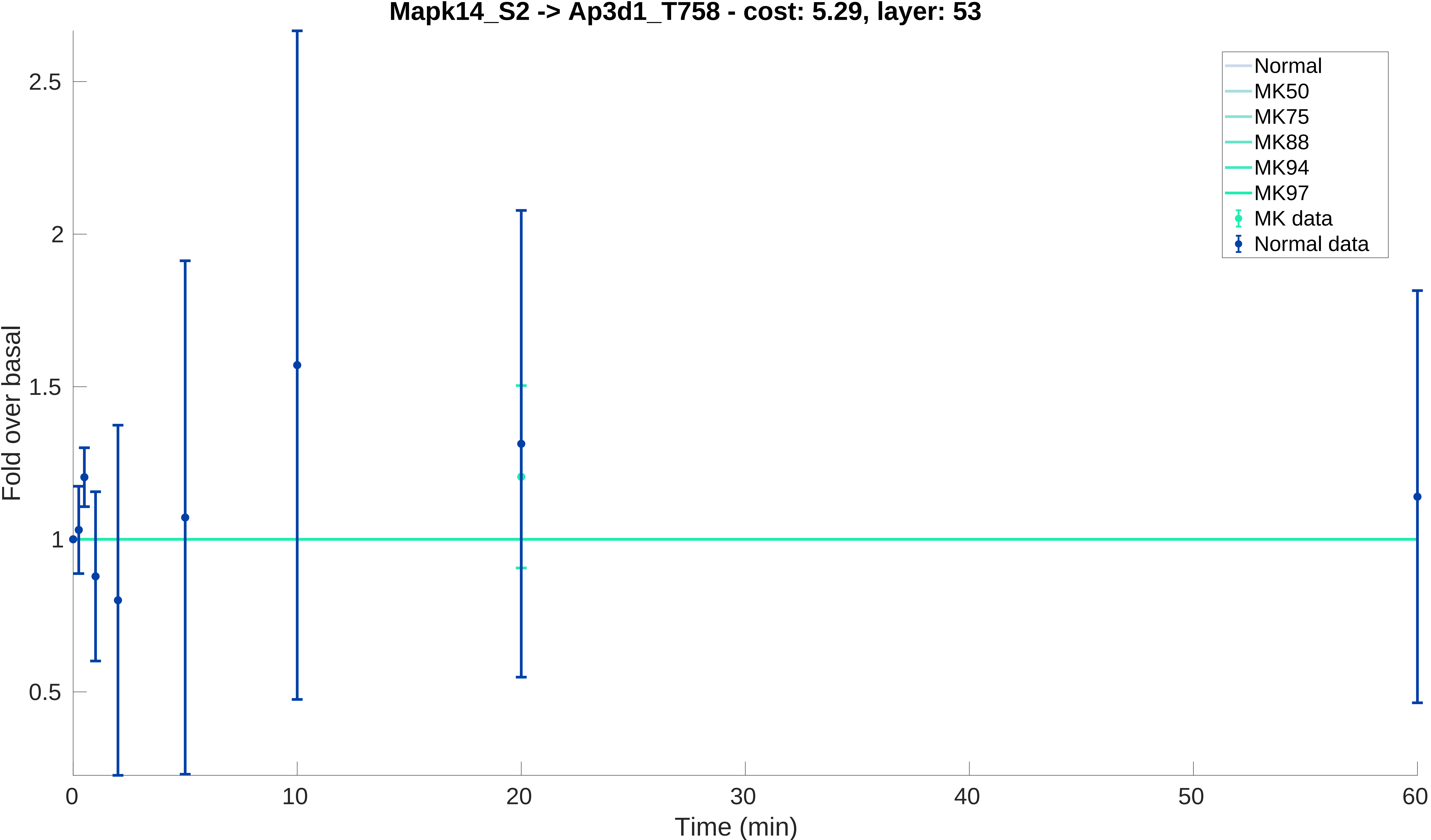

Mapk14_S2 -> Ap3d1_T758 - cost: 5.29, layer: 53
Fold over basal
Time (min)
2.5
2
1.5
1
0.5
0
10
20
30
40
50
60
Normal
MK50
MK75
MK88
MK94
MK97
MK data
Normal data

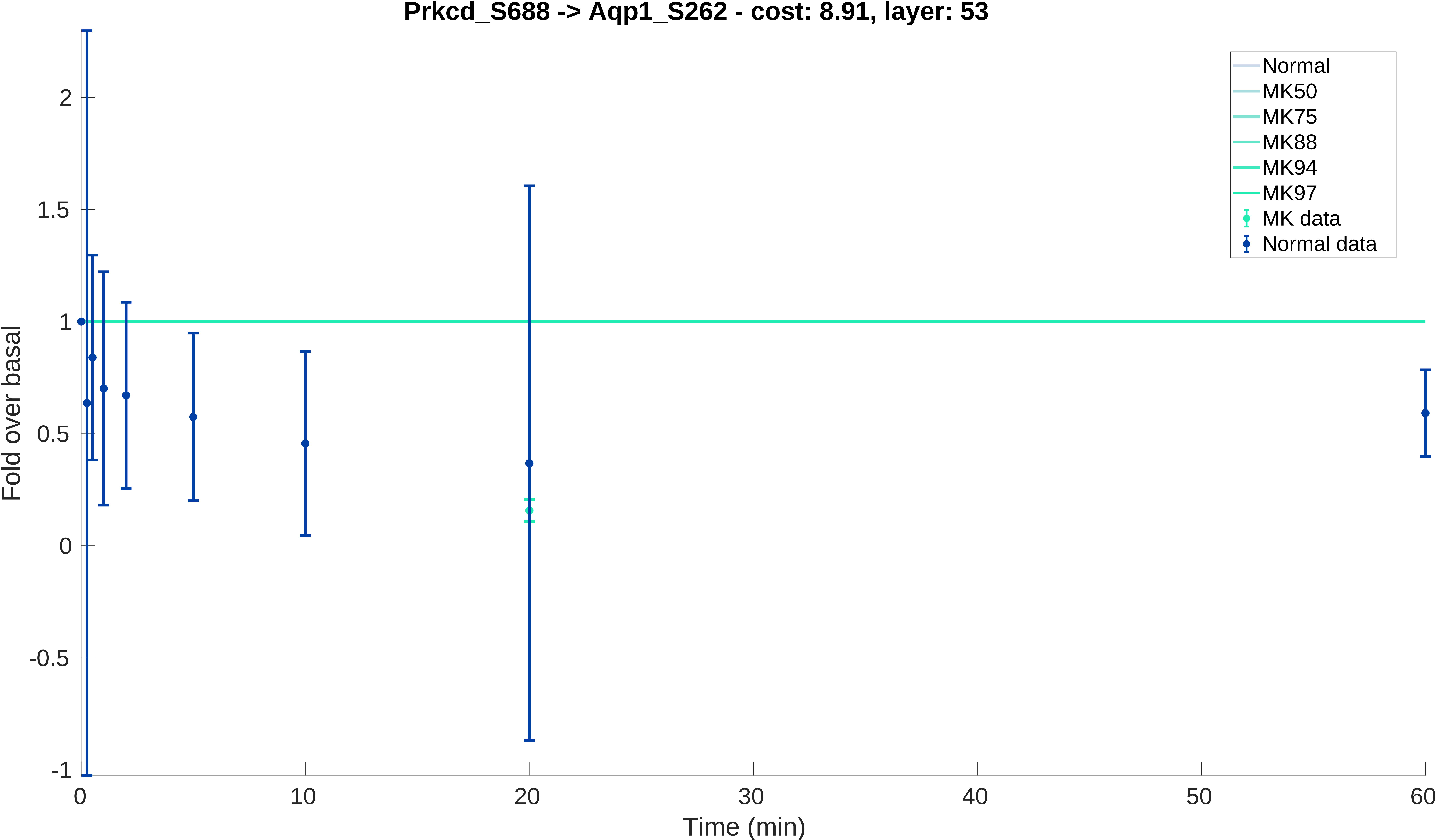

Prkcd_S688 -> Aqp1_S262 - cost: 8.91, layer: 53
Normal
MK50
MK75
MK88
MK94
MK97
MK data
Normal data
Fold over basal
Time (min)
2
1.5
1
0.5
0
-0.5
-1
0
10
20
30
40
50
60

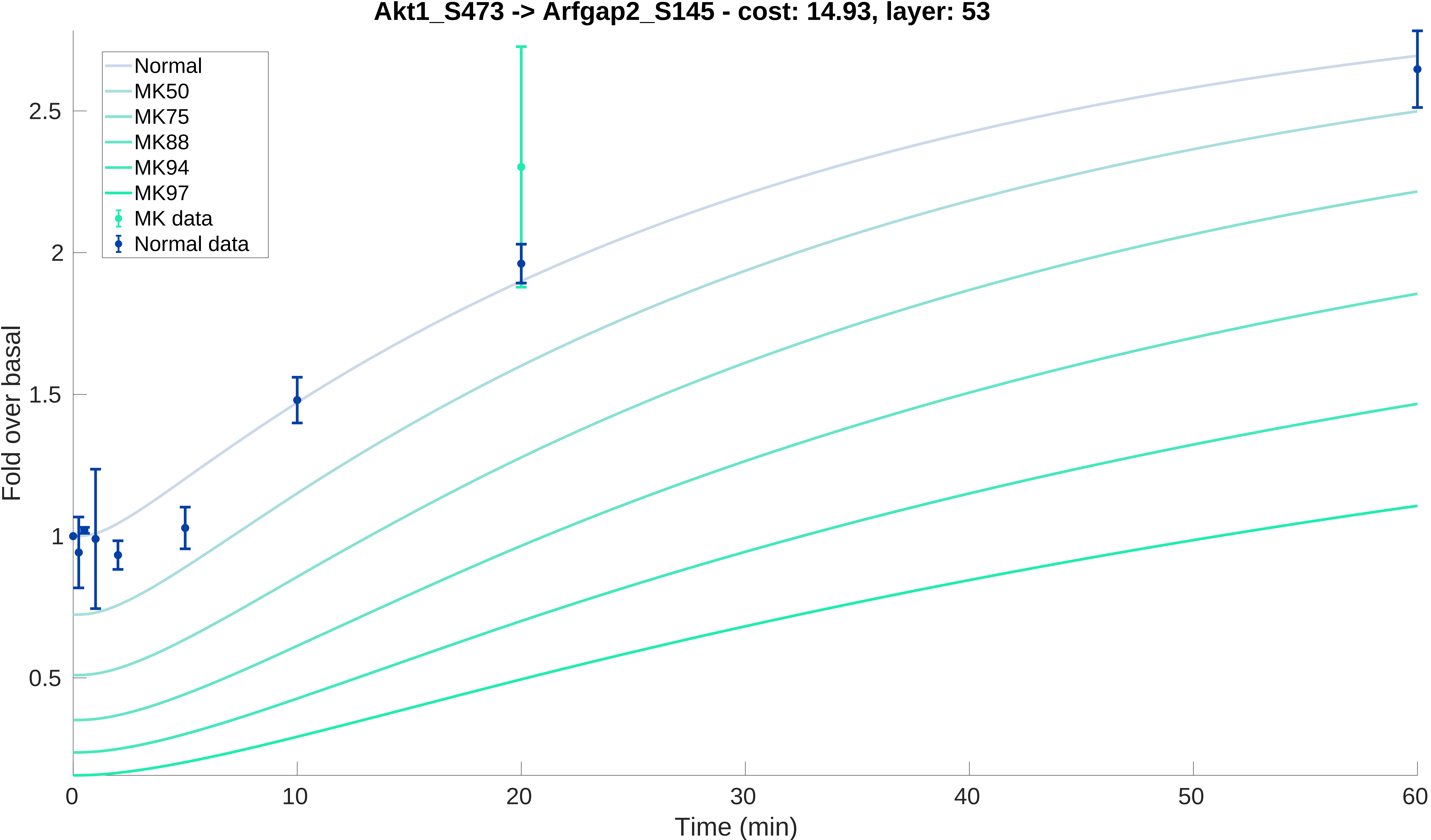

Akt1_S473 -> Arfgap2_S145 - cost: 14.93, layer: 53
Normal
MK50
MK75
MK88
MK94
MK97
MK data
Normal data
Fold over basal
2.5
2
1.5
1
0.5
0
10
20
30
40
50
60
Time (min)

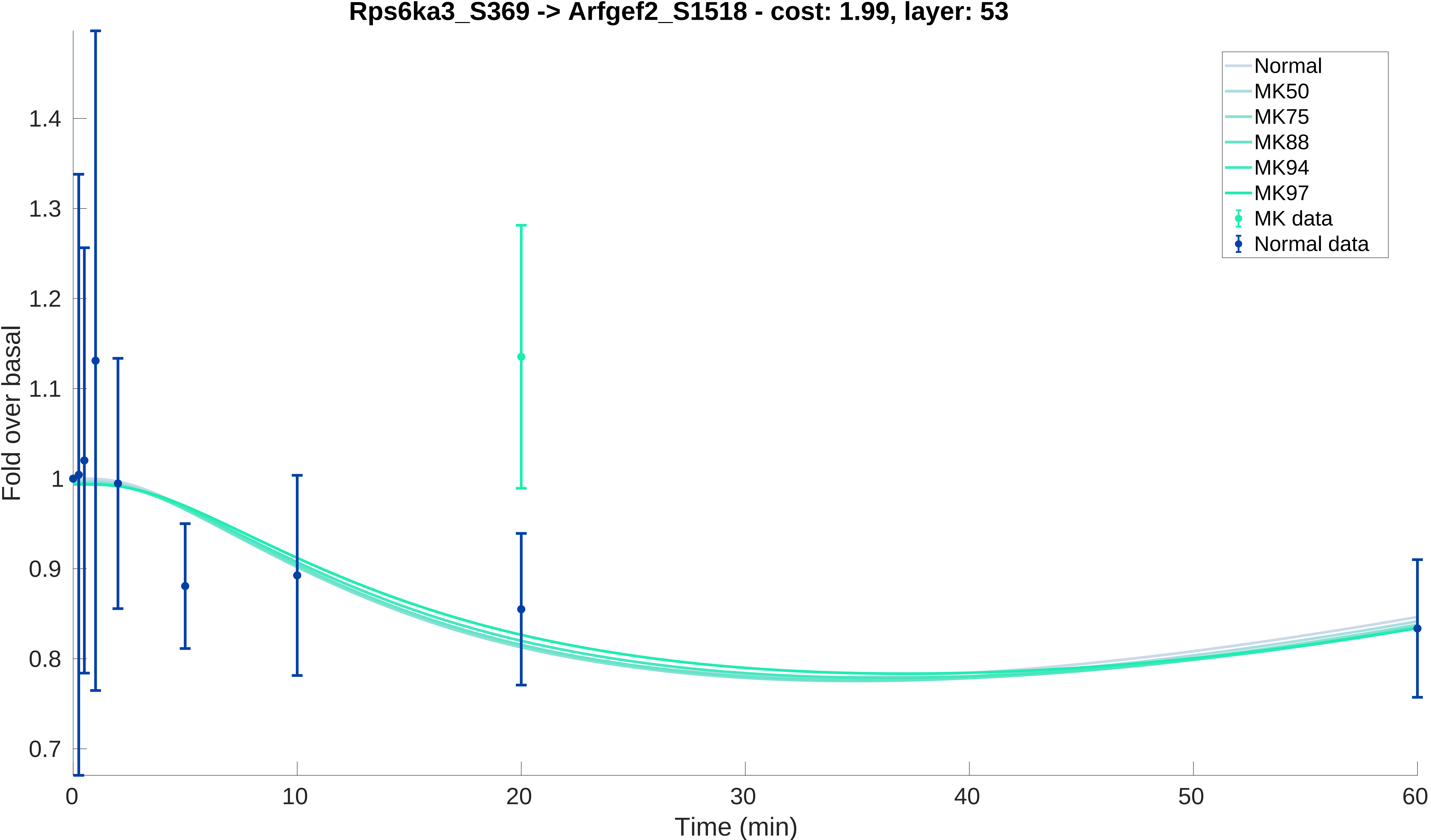
Rps6ka3_S369 -> Arfgef2_S1518 - cost: 1.99, layer: 53
Fold over basal
Time (min)
Normal
MK50
MK75
MK88
MK94
MK97
MK data
Normal data
0.7
0.8
0.9
1
1.1
1.2
1.3
1.4
0
10
20
30
40
50
60

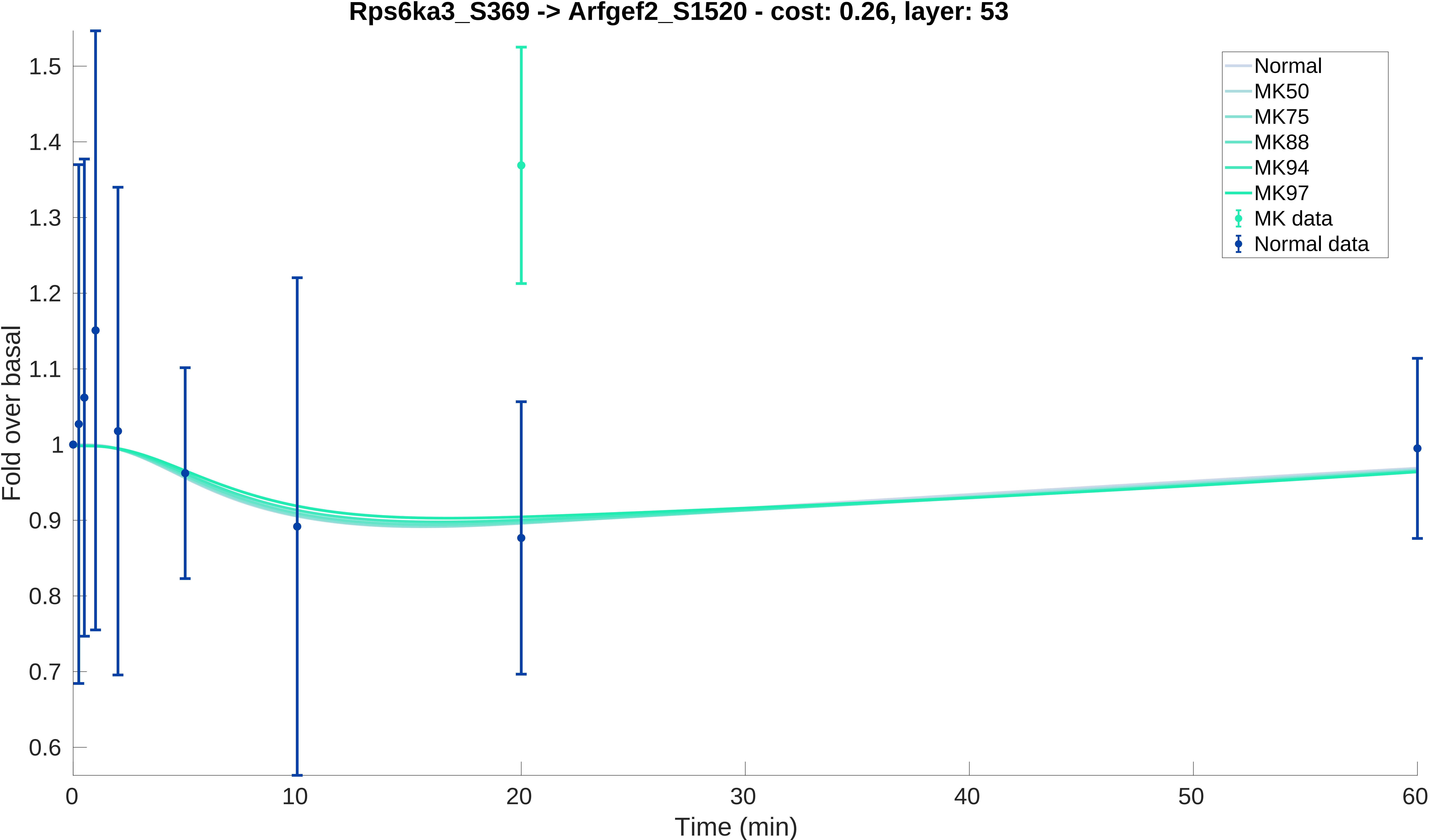

Rps6ka3_S369 -> Arfgef2_S1520 - cost: 0.26, layer: 53
Fold over basal
Time (min)
0.6
0.7
0.8
0.9
1
1.1
1.2
1.3
1.4
1.5
0
10
20
30
40
50
60
Normal
MK50
MK75
MK88
MK94
MK97
MK data
Normal data

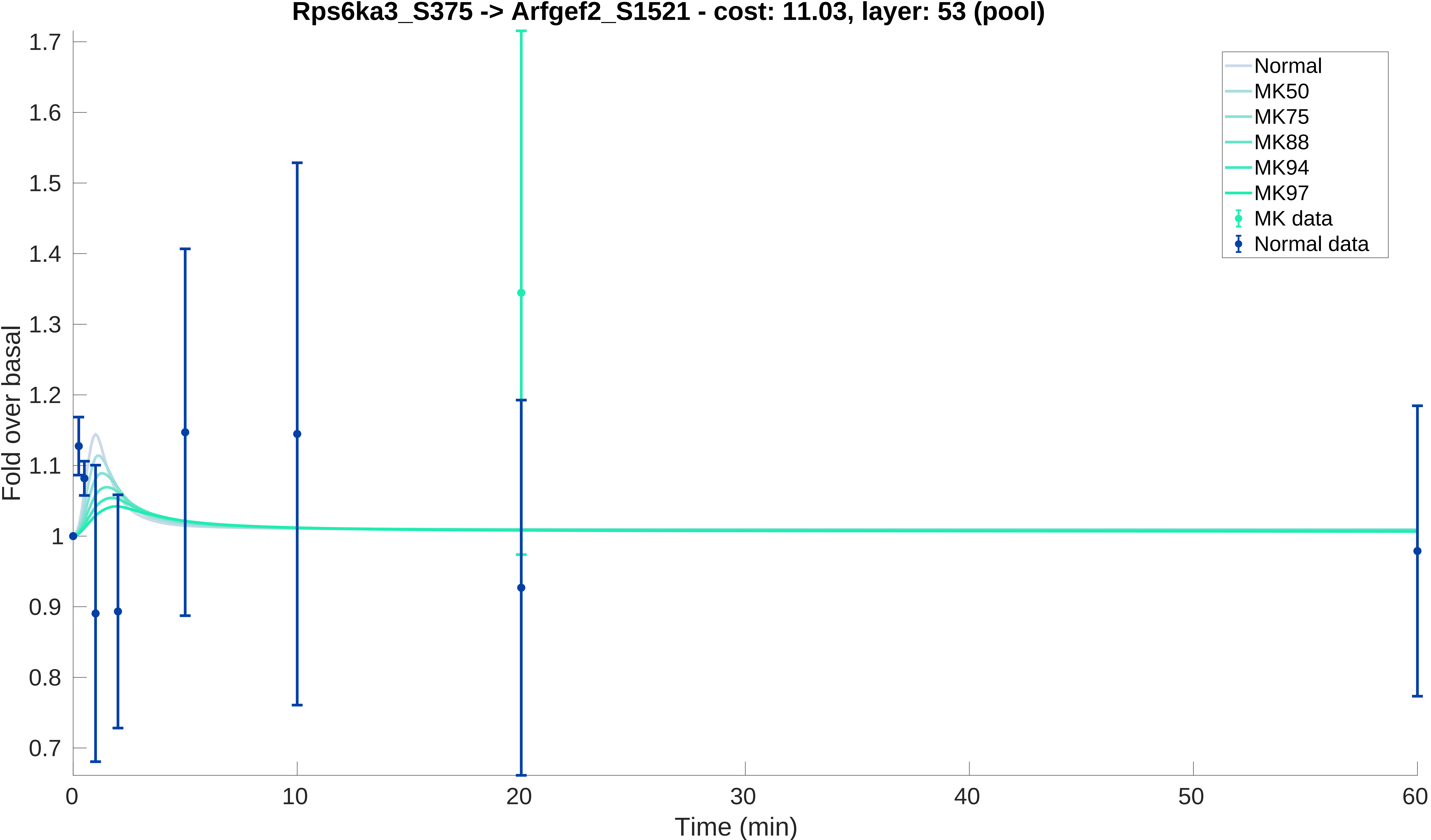

Rps6ka3_S375 -> Arfgef2_S1521 - cost: 11.03, layer: 53 (pool)
Fold over basal
Time (min)
0.7
0.8
0.9
1
1.1
1.2
1.3
1.4
1.5
1.6
1.7
0
10
20
30
40
50
60
Normal
MK50
MK75
MK88
MK94
MK97
MK data
Normal data

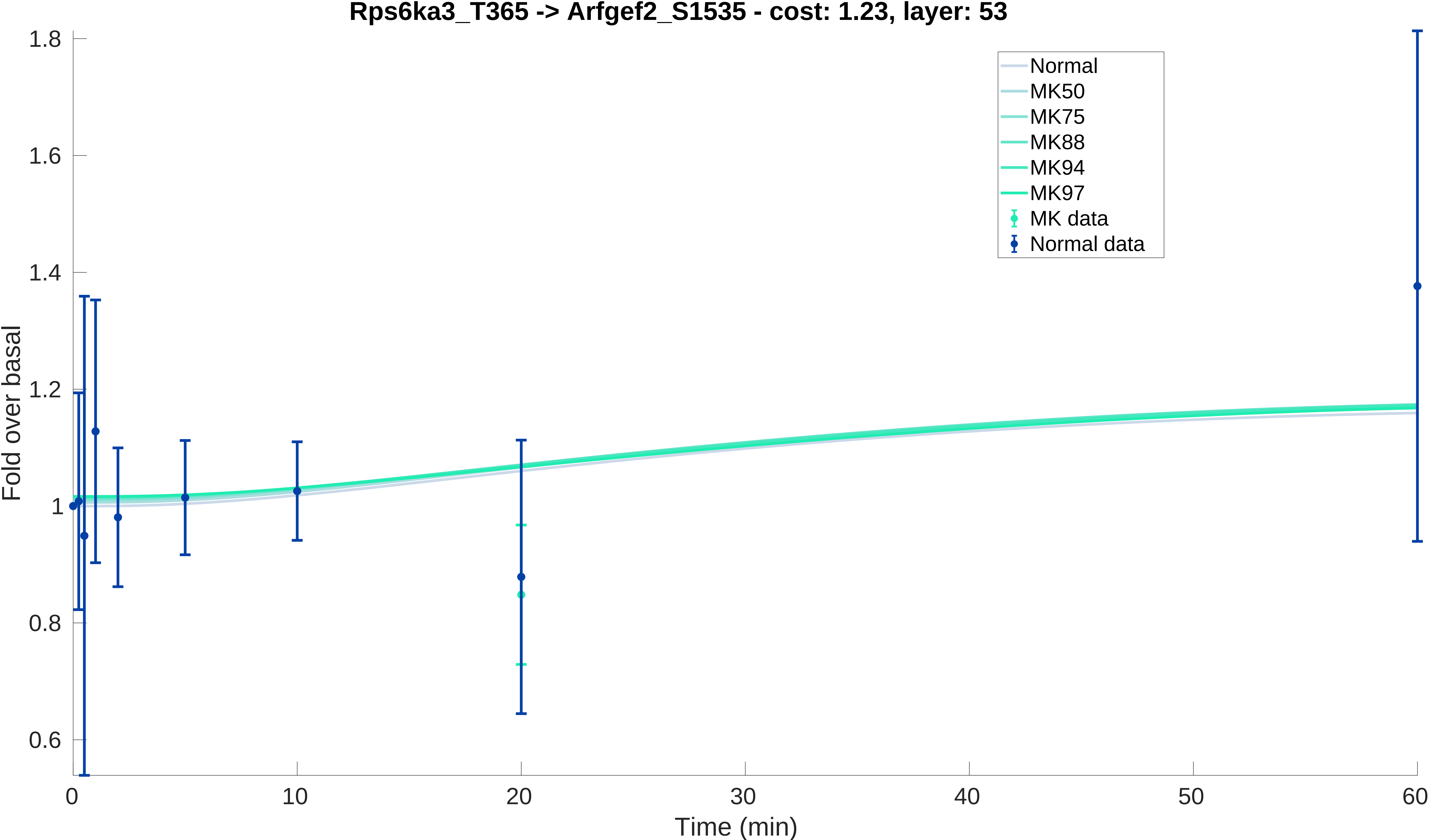

Rps6ka3_T365 -> Arfgef2_S1535 - cost: 1.23, layer: 53
Fold over basal
Time (min)
0.6
0.8
1
1.2
1.4
1.6
1.8
0
10
20
30
40
50
60
Normal
MK50
MK75
MK88
MK94
MK97
MK data
Normal data

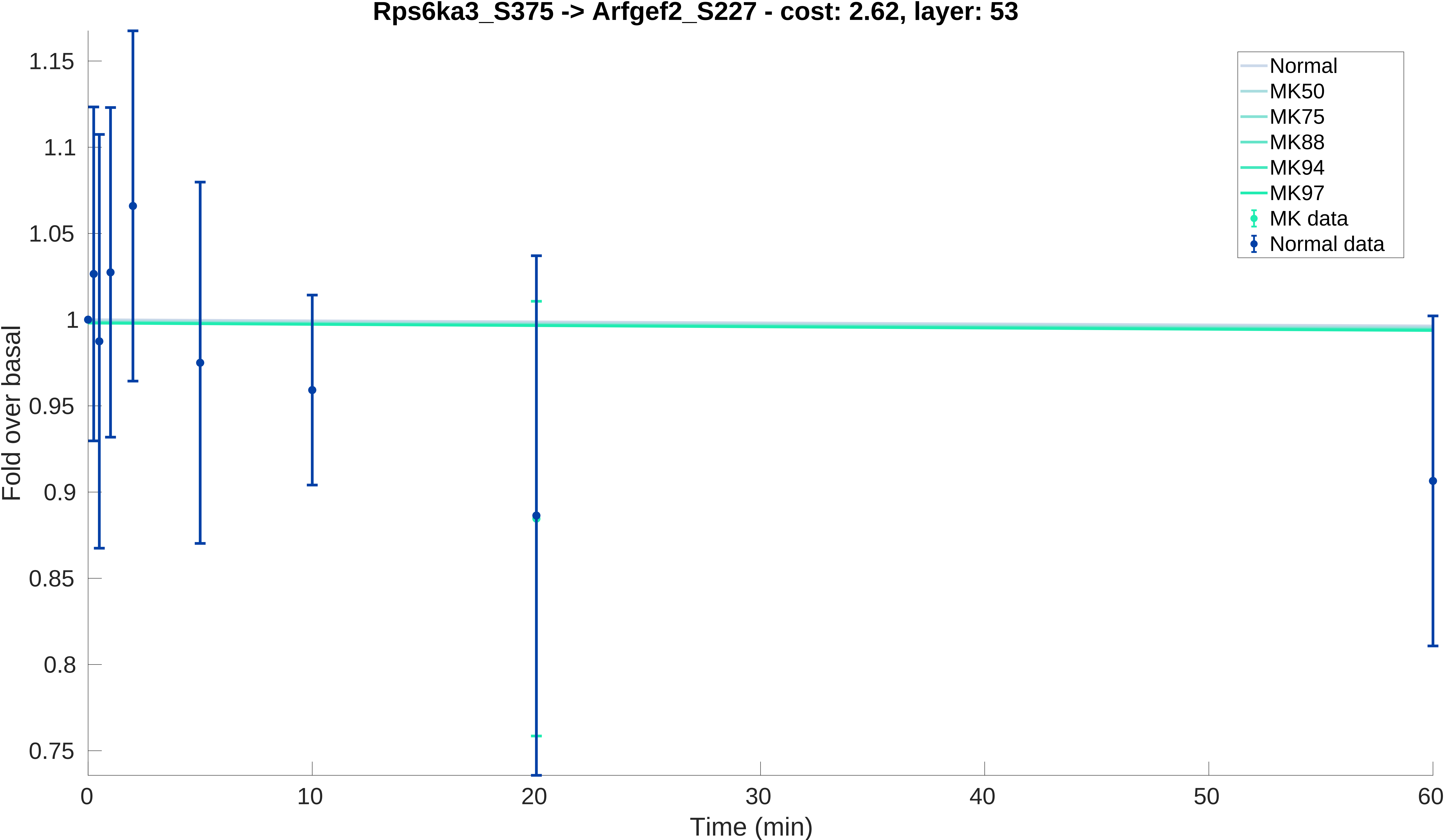

Rps6ka3_S375 -> Arfgef2_S227 - cost: 2.62, layer: 53
Fold over basal
Time (min)
Normal
MK50
MK75
MK88
MK94
MK97
MK data
Normal data
1.15
1.1
1.05
1
0.95
0.9
0.85
0.8
0.75
0
10
20
30
40
50
60

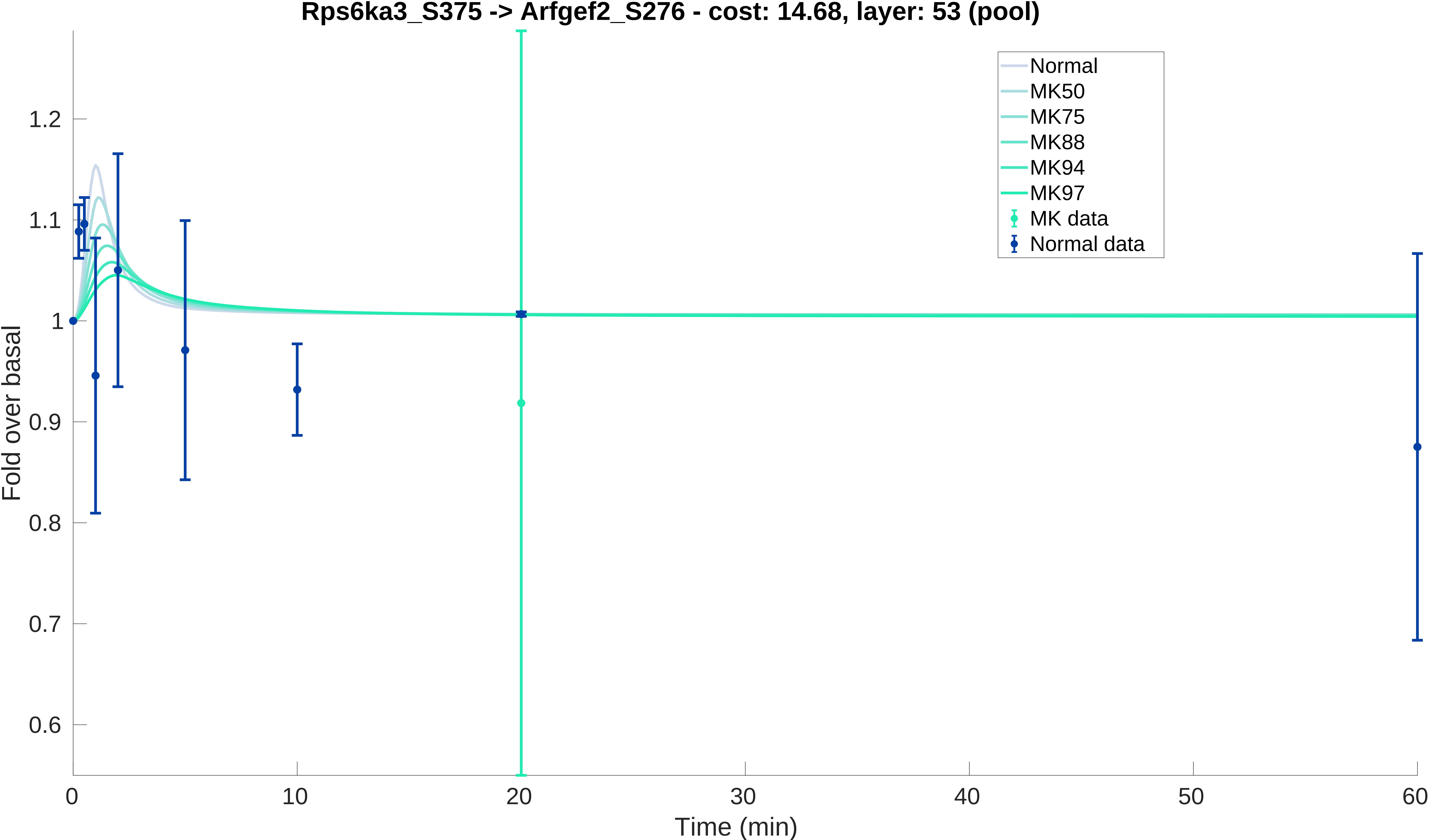

Rps6ka3_S375 -> Arfgef2_S276 - cost: 14.68, layer: 53 (pool)
Normal
MK50
MK75
MK88
MK94
MK97
MK data
Normal data
Fold over basal
1.2
1.1
1
0.9
0.8
0.7
0.6
0
10
20
30
40
50
60
Time (min)

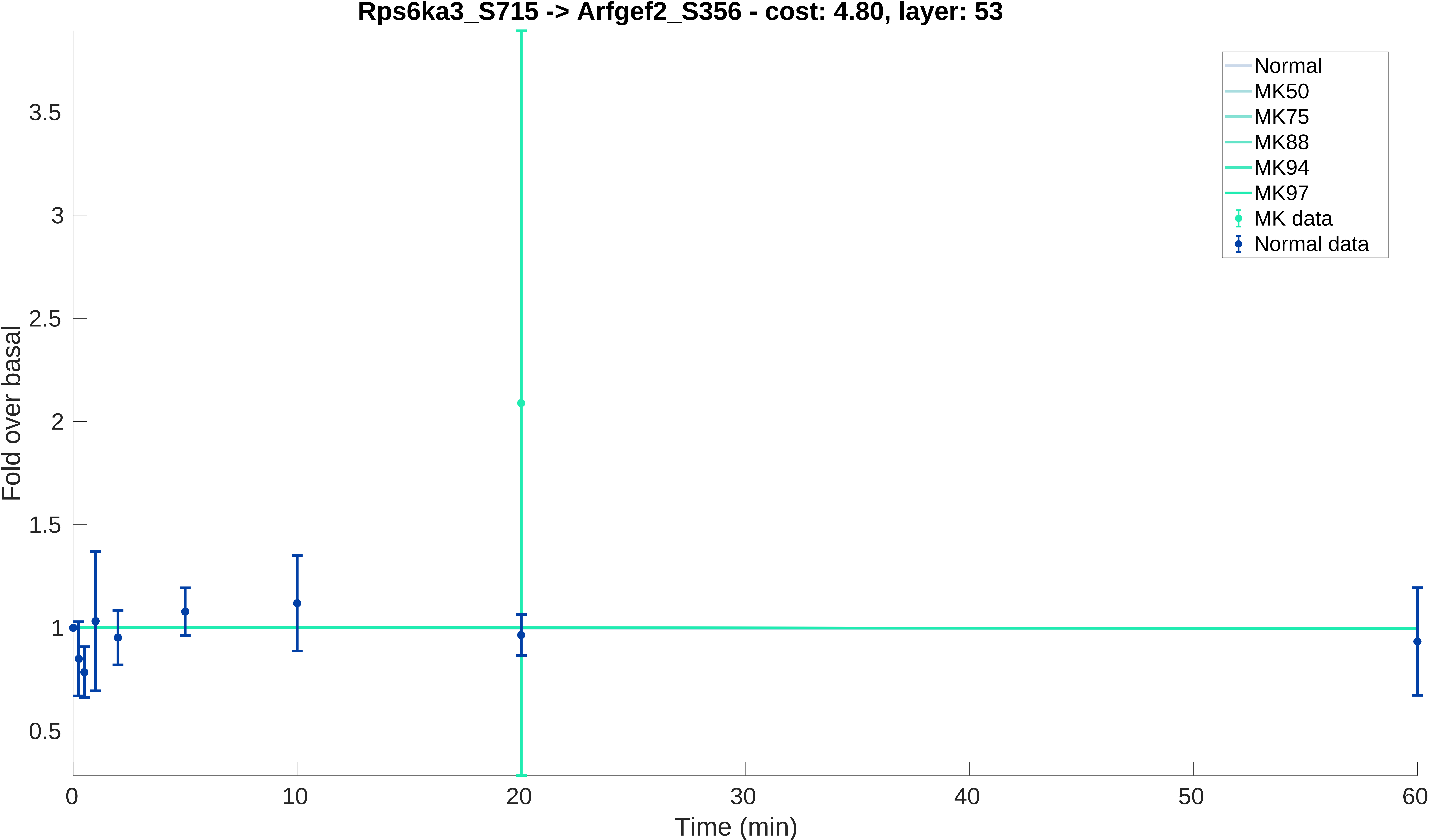
Rps6ka3_S715 -> Arfgef2_S356 - cost: 4.80, layer: 53
Fold over basal
Time (min)
0.5
1
1.5
2
2.5
3
3.5
0
10
20
30
40
50
60
Normal
MK50
MK75
MK88
MK94
MK97
MK data
Normal data

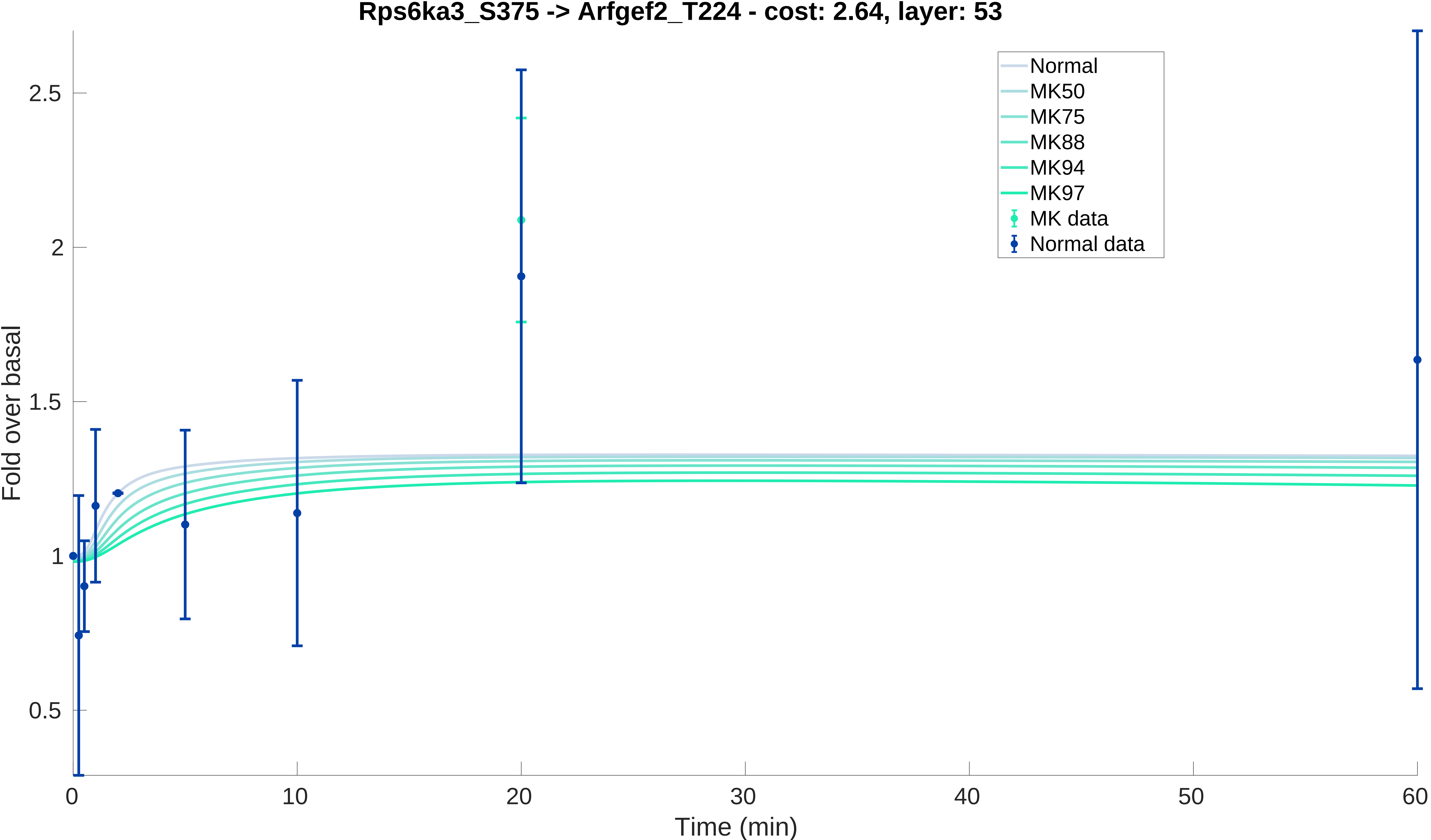
Rps6ka3_S375 -> Arfgef2_T224 - cost: 2.64, layer: 53
Fold over basal
Time (min)
0.5
1
1.5
2
2.5
0
10
20
30
40
50
60
Normal
MK50
MK75
MK88
MK94
MK97
MK data
Normal data

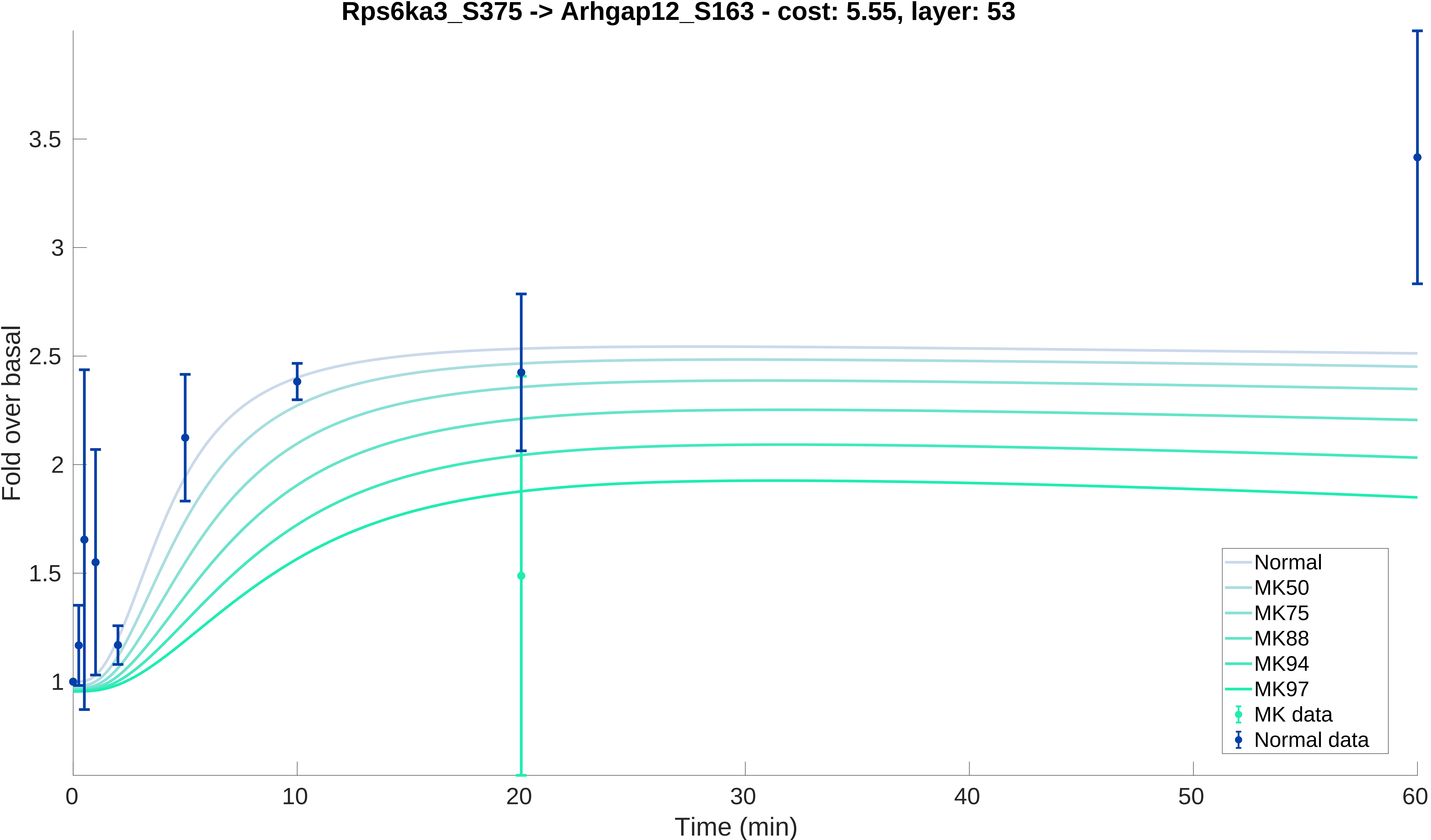
Rps6ka3_S375 -> Arhgap12_S163 - cost: 5.55, layer: 53
Fold over basal
Time (min)
1
1.5
2
2.5
3
3.5
0
10
20
30
40
50
60
Normal
MK50
MK75
MK88
MK94
MK97
MK data
Normal data

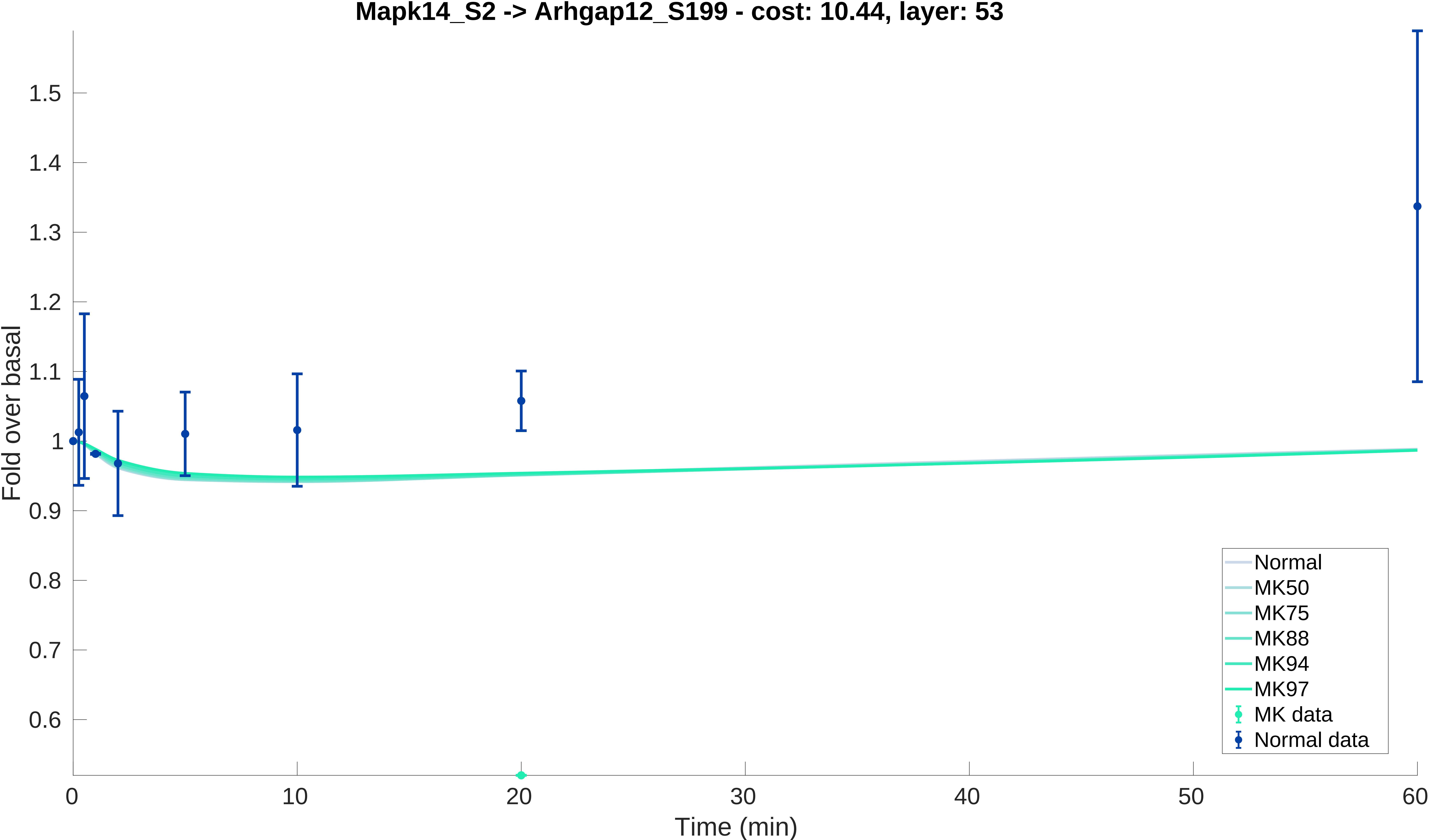

Mapk14_S2 -> Arhgap12_S199 - cost: 10.44, layer: 53
Fold over basal
1.5
1.4
1.3
1.2
1.1
1
0.9
0.8
0.7
0.6
0
10
20
30
40
50
60
Time (min)
Normal
MK50
MK75
MK88
MK94
MK97
MK data
Normal data

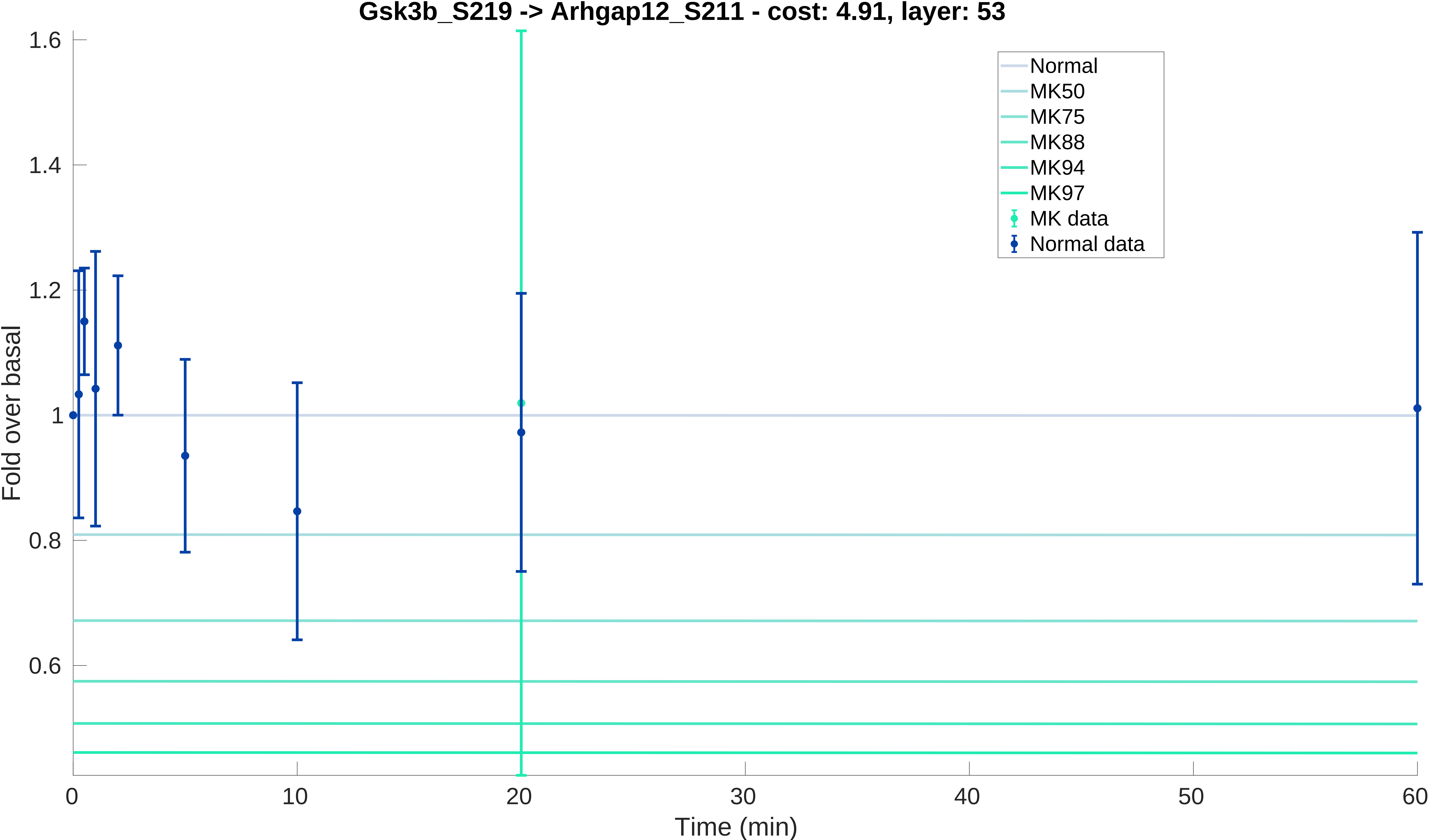

Gsk3b_S219 -> Arhgap12_S211 - cost: 4.91, layer: 53
Fold over basal
Time (min)
1.6
1.4
1.2
1
0.8
0.6
0
10
20
30
40
50
60
Normal
MK50
MK75
MK88
MK94
MK97
MK data
Normal data

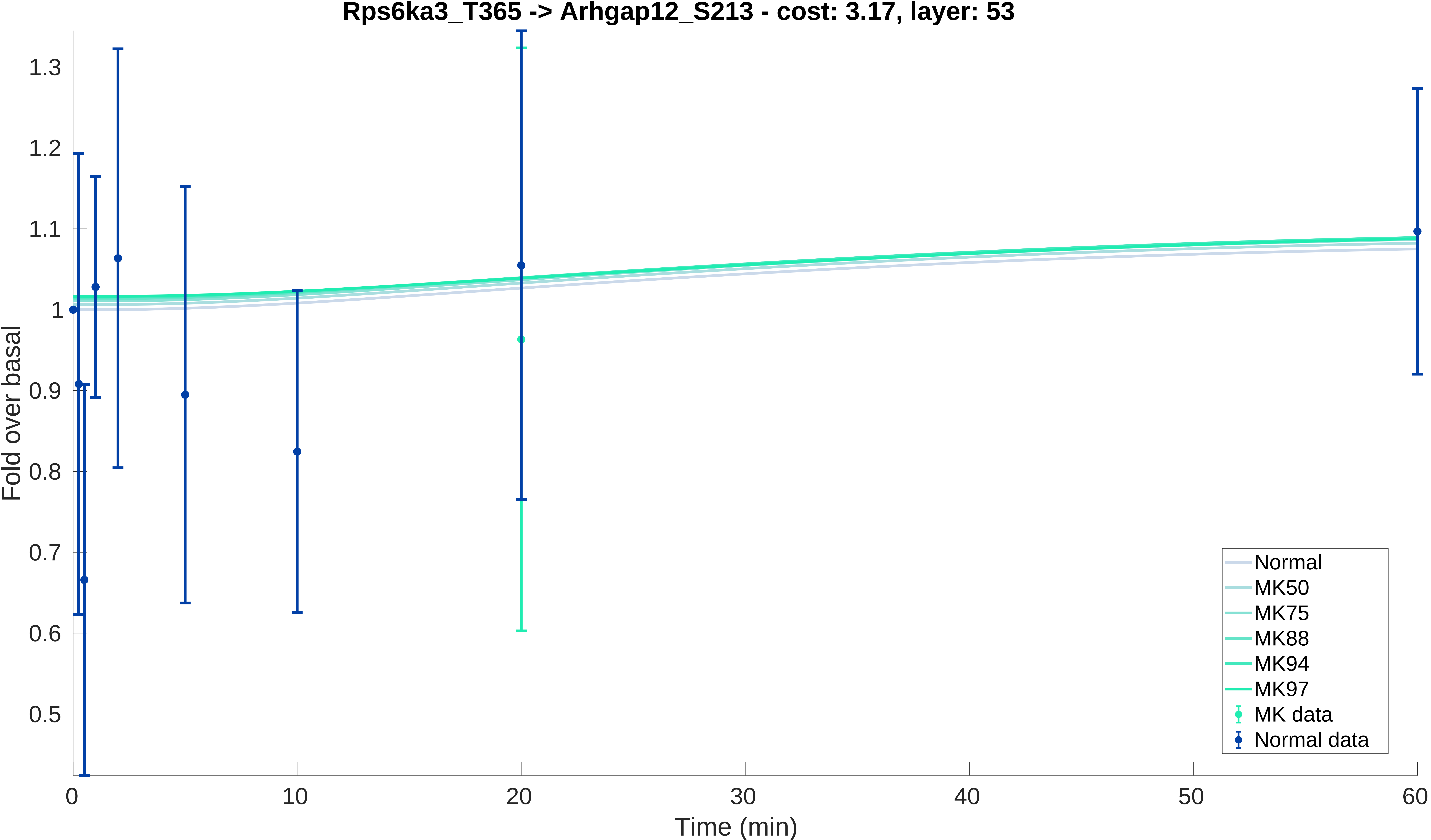

Rps6ka3_T365 -> Arhgap12_S213 - cost: 3.17, layer: 53
Fold over basal
Time (min)
0.5
0.6
0.7
0.8
0.9
1
1.1
1.2
1.3
0
10
20
30
40
50
60
Normal
MK50
MK75
MK88
MK94
MK97
MK data
Normal data

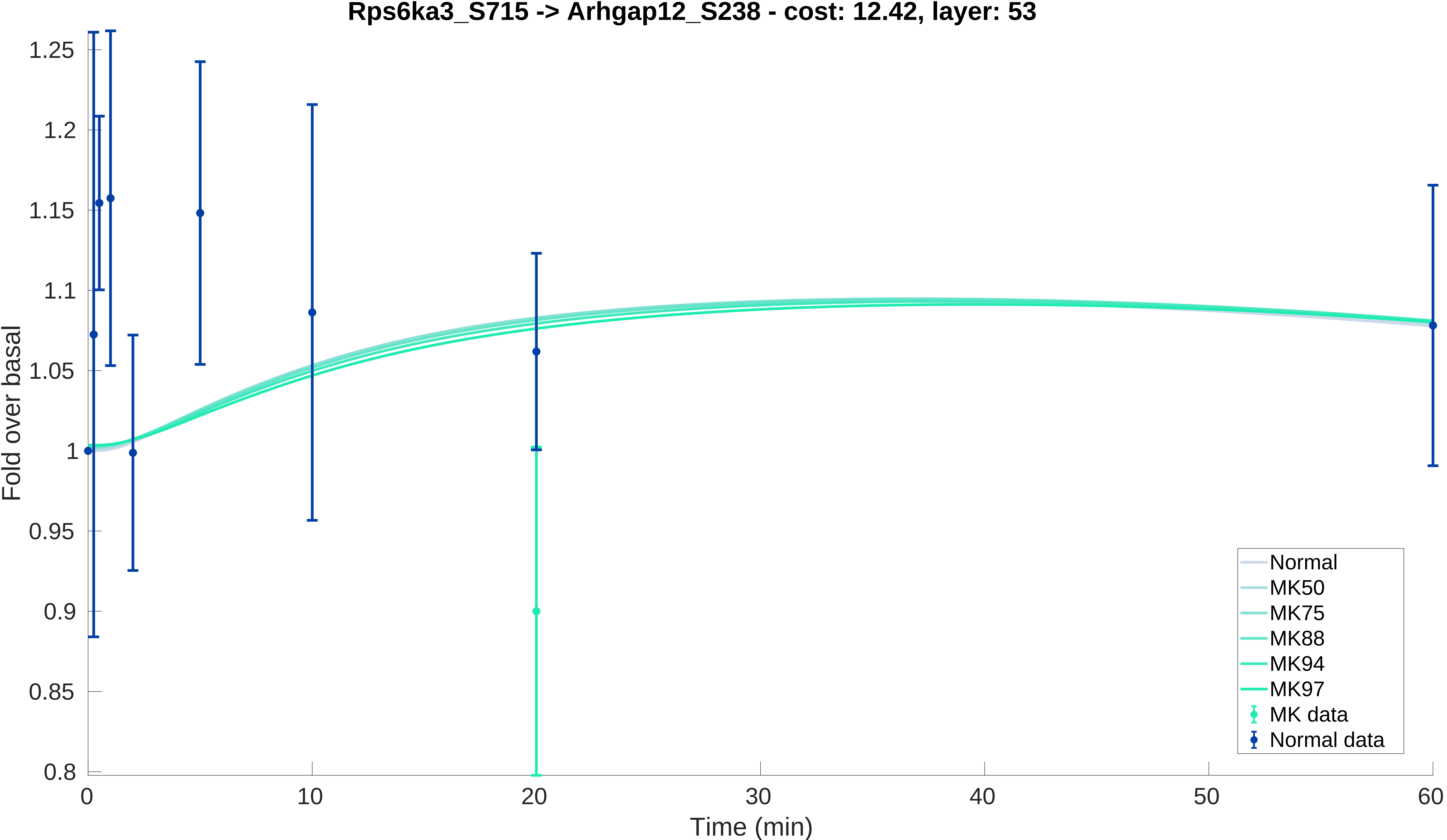

Rps6ka3_S715 -> Arhgap12_S238 - cost: 12.42, layer: 53
Fold over basal
1.25
1.2
1.15
1.1
1.05
1
0.95
0.9
0.85
0.8
0
10
20
30
40
50
60
Time (min)
Normal
MK50
MK75
MK88
MK94
MK97
MK data
Normal data

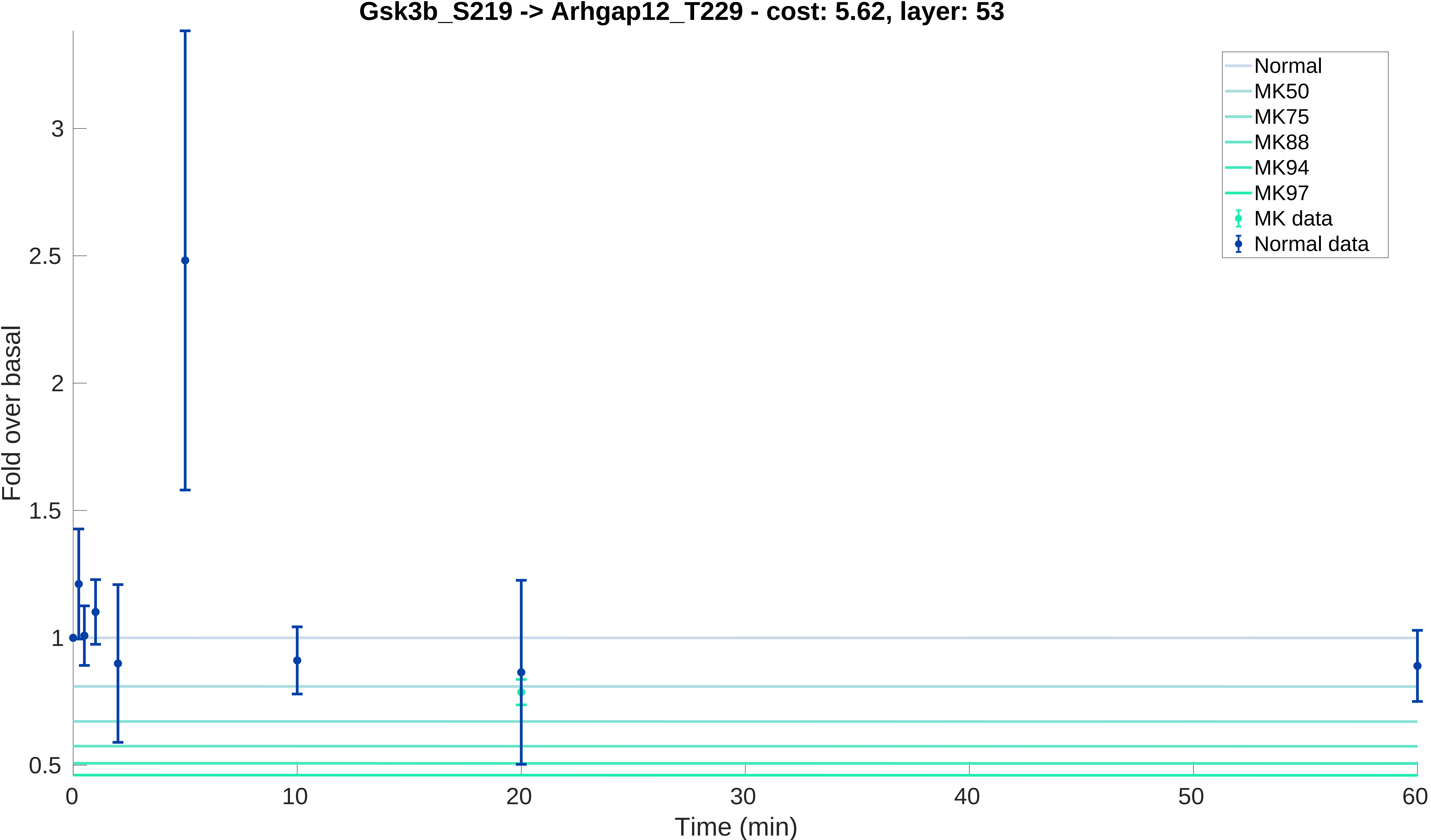

Gsk3b_S219 -> Arhgap12_T229 - cost: 5.62, layer: 53
Fold over basal
Time (min)
0.5
1
1.5
2
2.5
3
0
10
20
30
40
50
60
Normal
MK50
MK75
MK88
MK94
MK97
MK data
Normal data

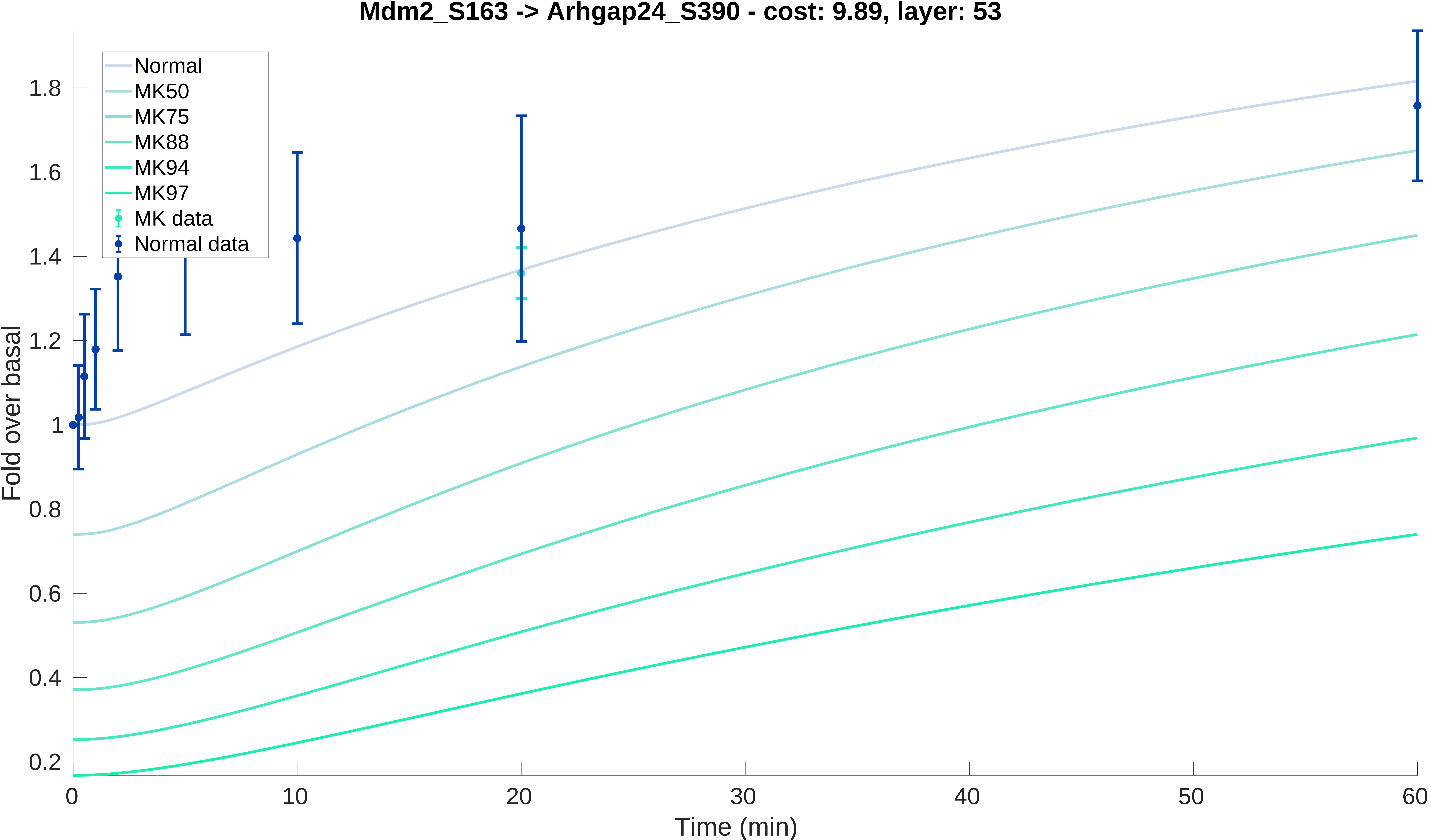

Mdm2_S163 -> Arhgap24_S390 - cost: 9.89, layer: 53
Normal
MK50
MK75
MK88
MK94
MK97
MK data
Normal data
Fold over basal
Time (min)
0.2
0.4
0.6
0.8
1
1.2
1.4
1.6
1.8
0
10
20
30
40
50
60

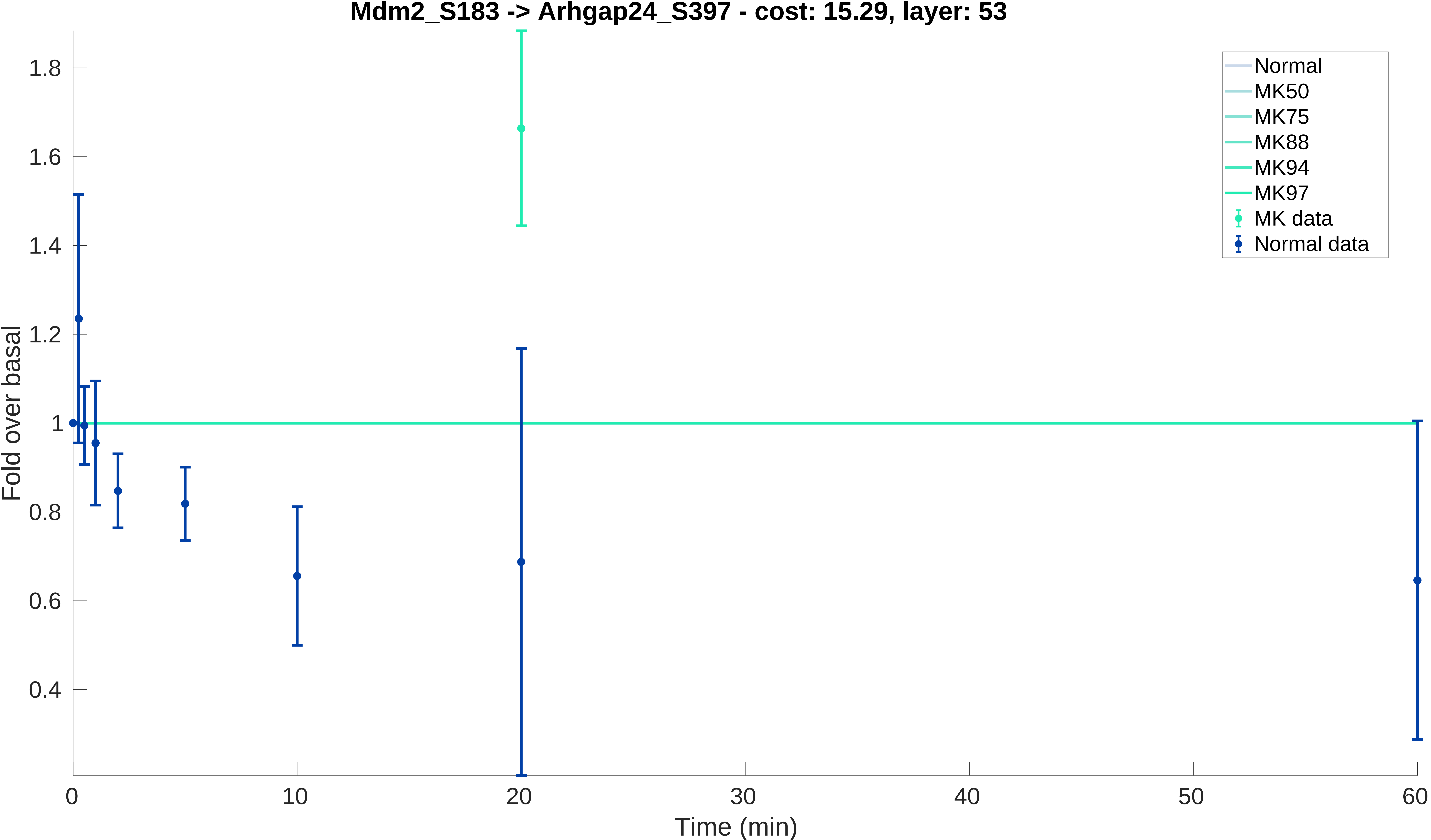

Mdm2_S183 -> Arhgap24_S397 - cost: 15.29, layer: 53
Normal
MK50
MK75
MK88
MK94
MK97
MK data
Normal data
Fold over basal
1.8
1.6
1.4
1.2
1
0.8
0.6
0.4
0
10
20
30
40
50
60
Time (min)

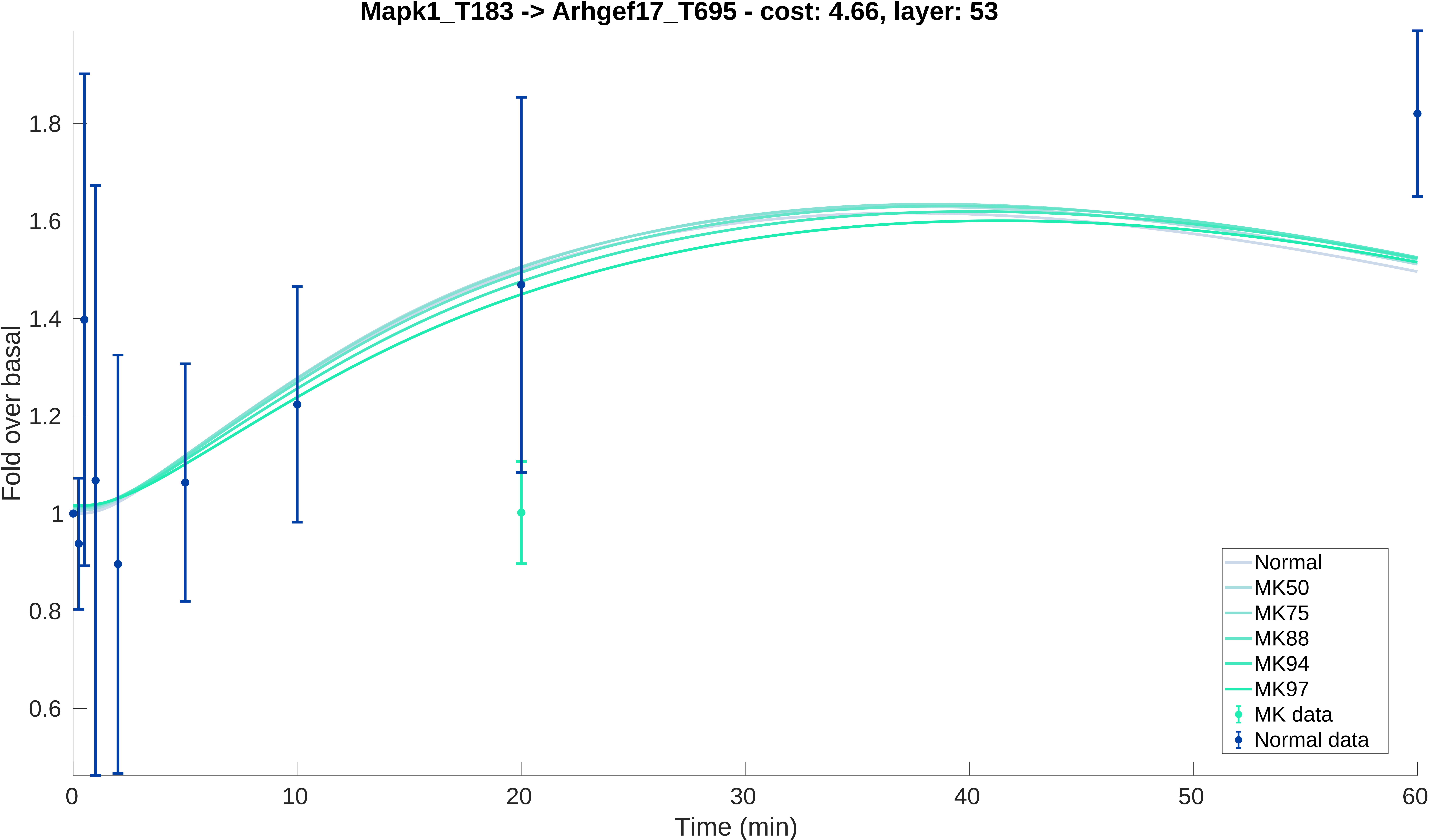

Mapk1_T183 -> Arhgef17_T695 - cost: 4.66, layer: 53
Fold over basal
Time (min)
0.6
0.8
1
1.2
1.4
1.6
1.8
0
10
20
30
40
50
60
Normal
MK50
MK75
MK88
MK94
MK97
MK data
Normal data

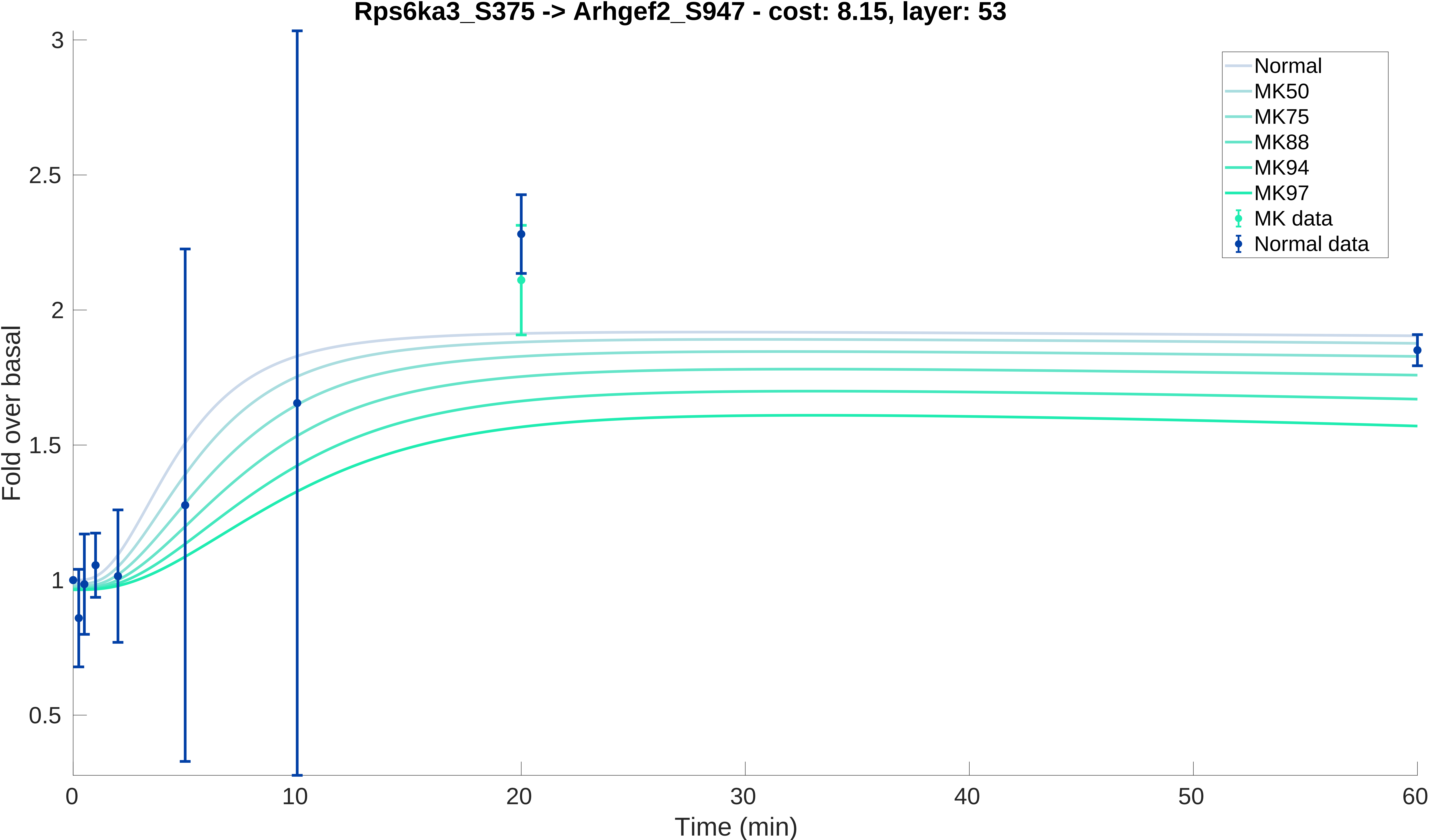

Rps6ka3_S375 -> Arhgef2_S947 - cost: 8.15, layer: 53
Fold over basal
Time (min)
0.5
1
1.5
2
2.5
3
0
10
20
30
40
50
60
Normal
MK50
MK75
MK88
MK94
MK97
MK data
Normal data

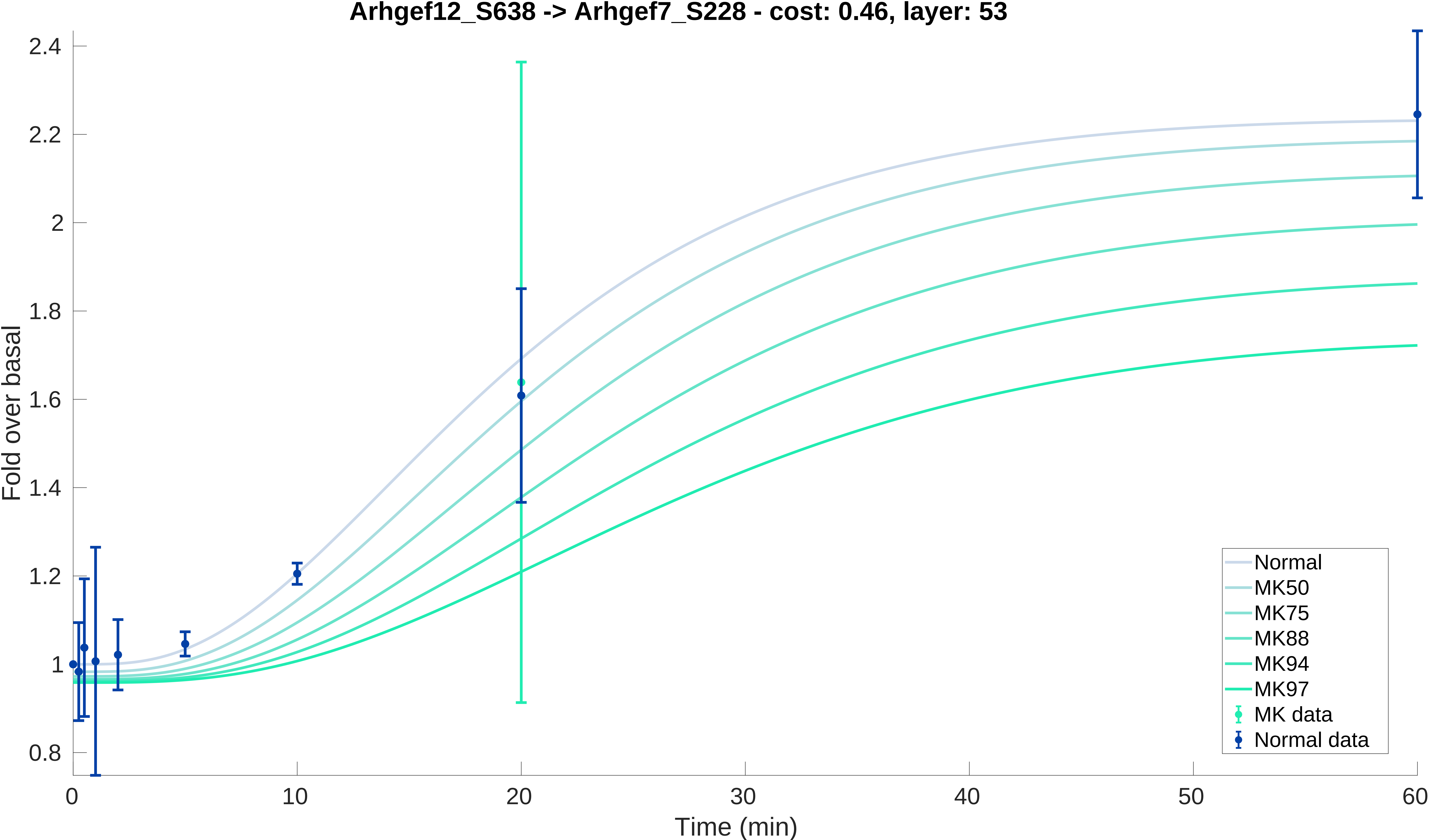

Arhgef12_S638 -> Arhgef7_S228 - cost: 0.46, layer: 53
Fold over basal
Time (min)
2.4
2.2
2
1.8
1.6
1.4
1.2
1
0.8
0
10
20
30
40
50
60
Normal
MK50
MK75
MK88
MK94
MK97
MK data
Normal data

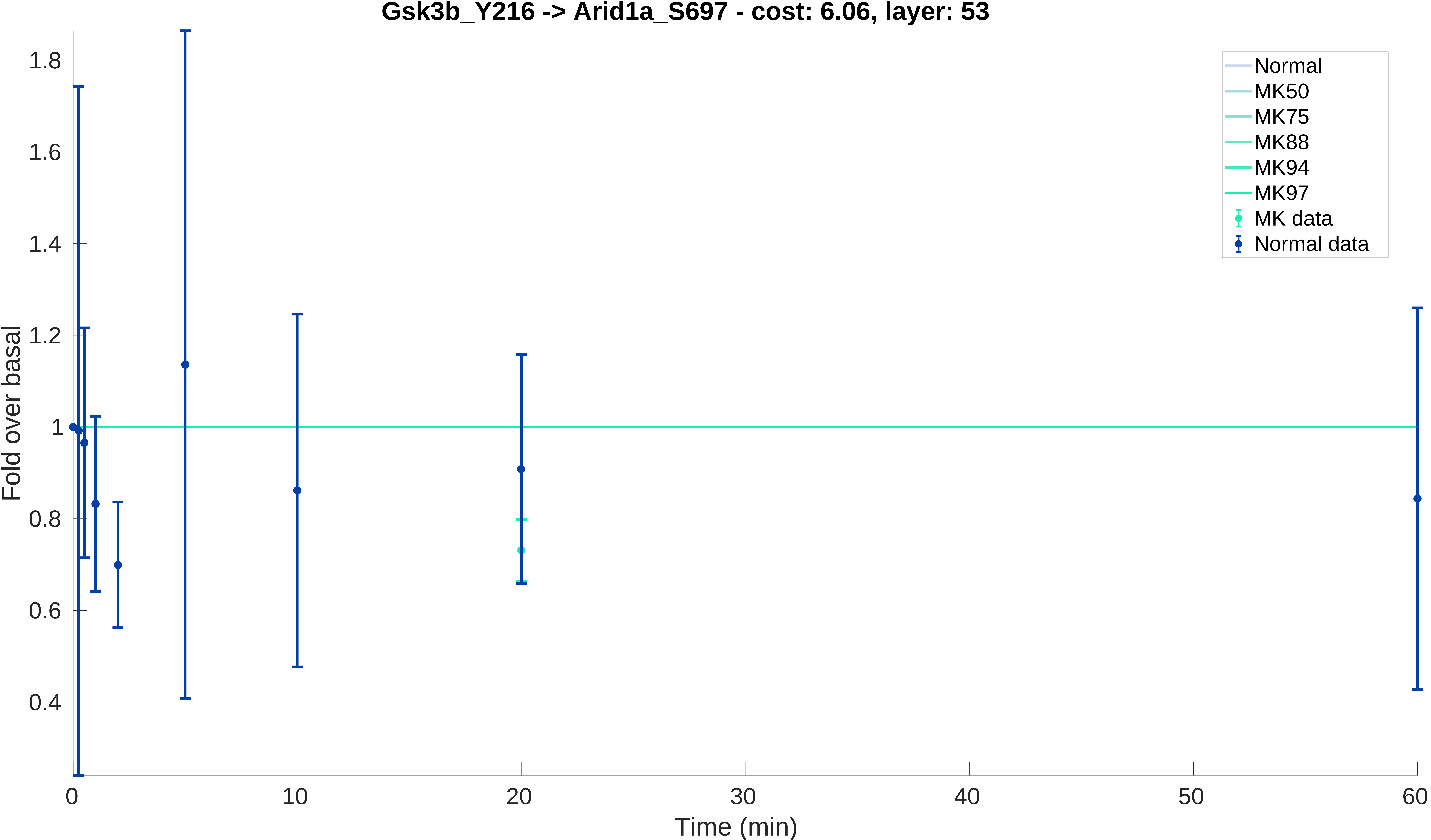

Gsk3b_Y216 -> Arid1a_S697 - cost: 6.06, layer: 53
Fold over basal
Time (min)
1.8
1.6
1.4
1.2
1
0.8
0.6
0.4
0
10
20
30
40
50
60
Normal
MK50
MK75
MK88
MK94
MK97
MK data
Normal data

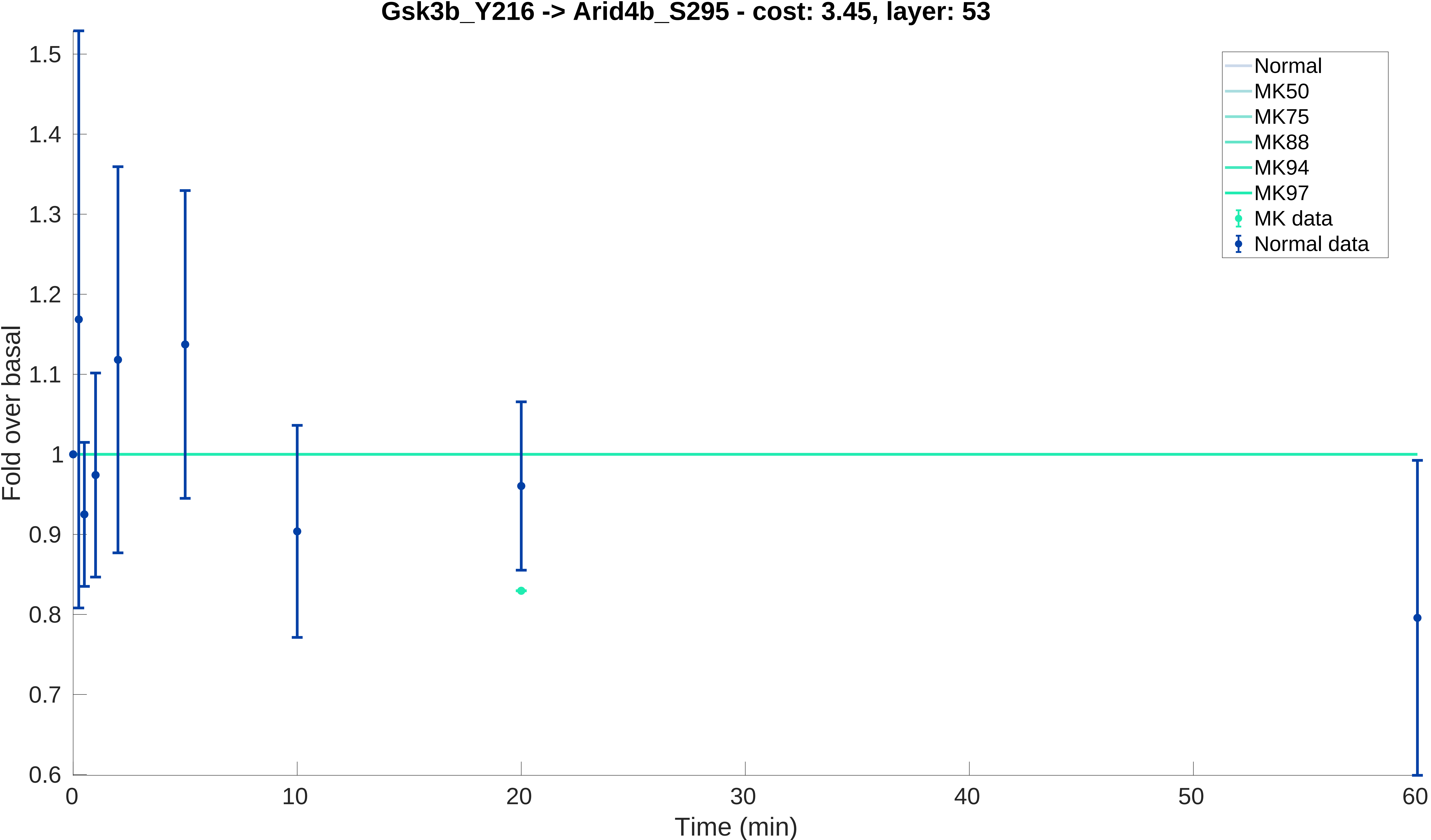
Gsk3b_Y216 -> Arid4b_S295 - cost: 3.45, layer: 53
Fold over basal
Time (min)
Normal
MK50
MK75
MK88
MK94
MK97
MK data
Normal data
1.5
1.4
1.3
1.2
1.1
1
0.9
0.8
0.7
0.6
0
10
20
30
40
50
60

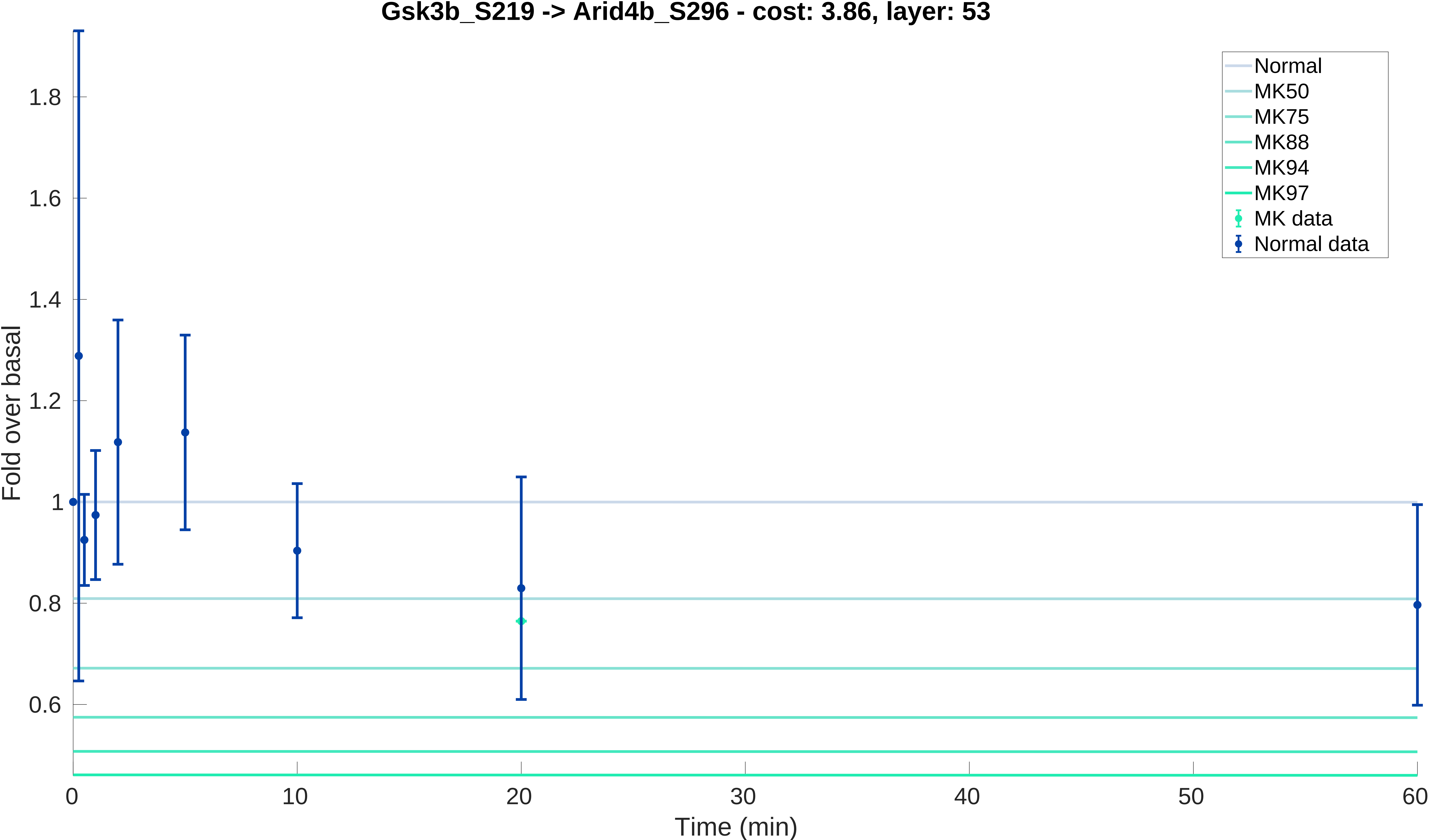

Gsk3b_S219 -> Arid4b_S296 - cost: 3.86, layer: 53
Normal
MK50
MK75
MK88
MK94
MK97
MK data
Normal data
Fold over basal
1.8
1.6
1.4
1.2
1
0.8
0.6
0
10
20
30
40
50
60
Time (min)

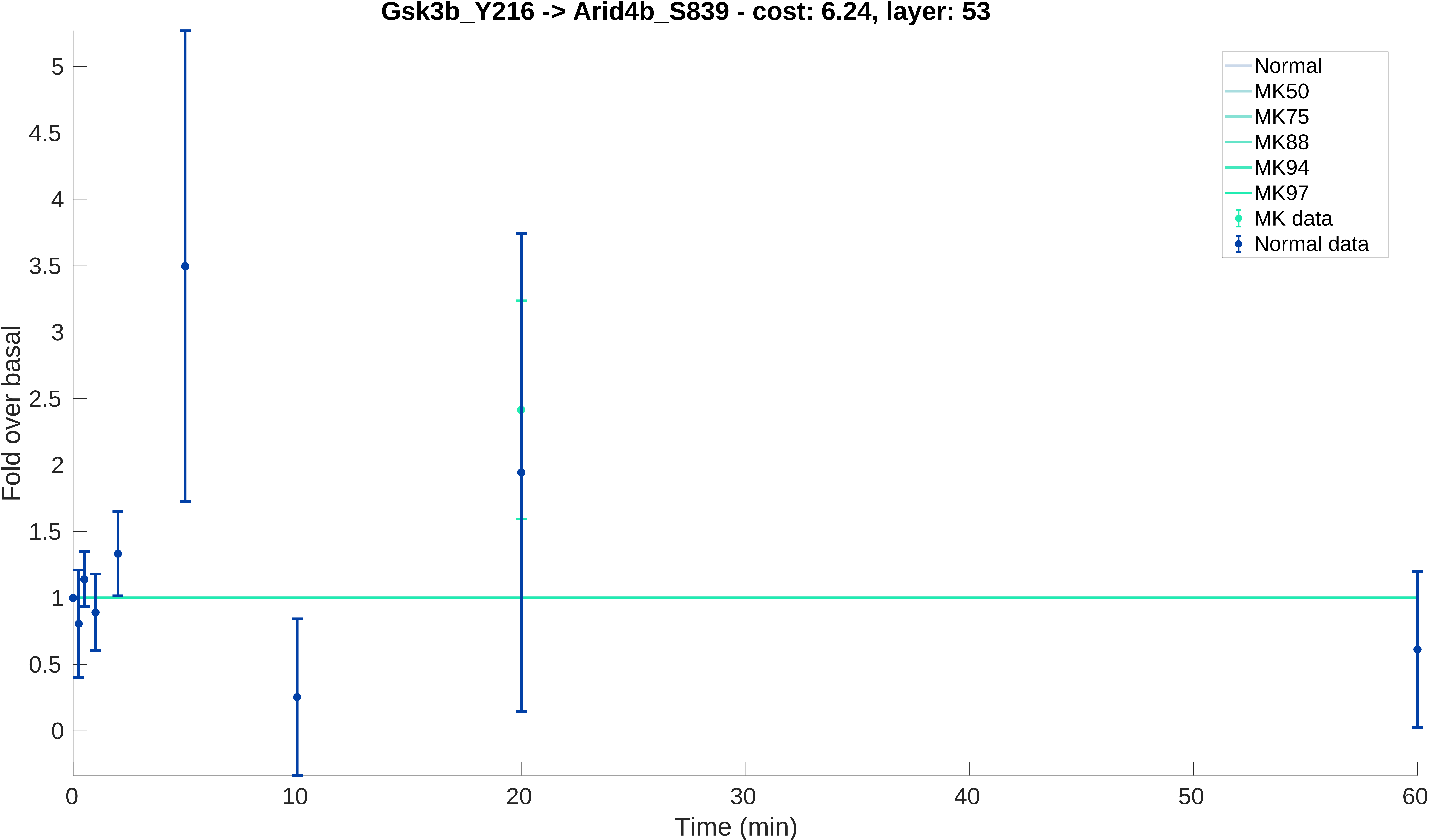

Gsk3b_Y216 -> Arid4b_S839 - cost: 6.24, layer: 53
Fold over basal
Time (min)
Normal
MK50
MK75
MK88
MK94
MK97
MK data
Normal data
0
0.5
1
1.5
2
2.5
3
3.5
4
4.5
5
0
10
20
30
40
50
60

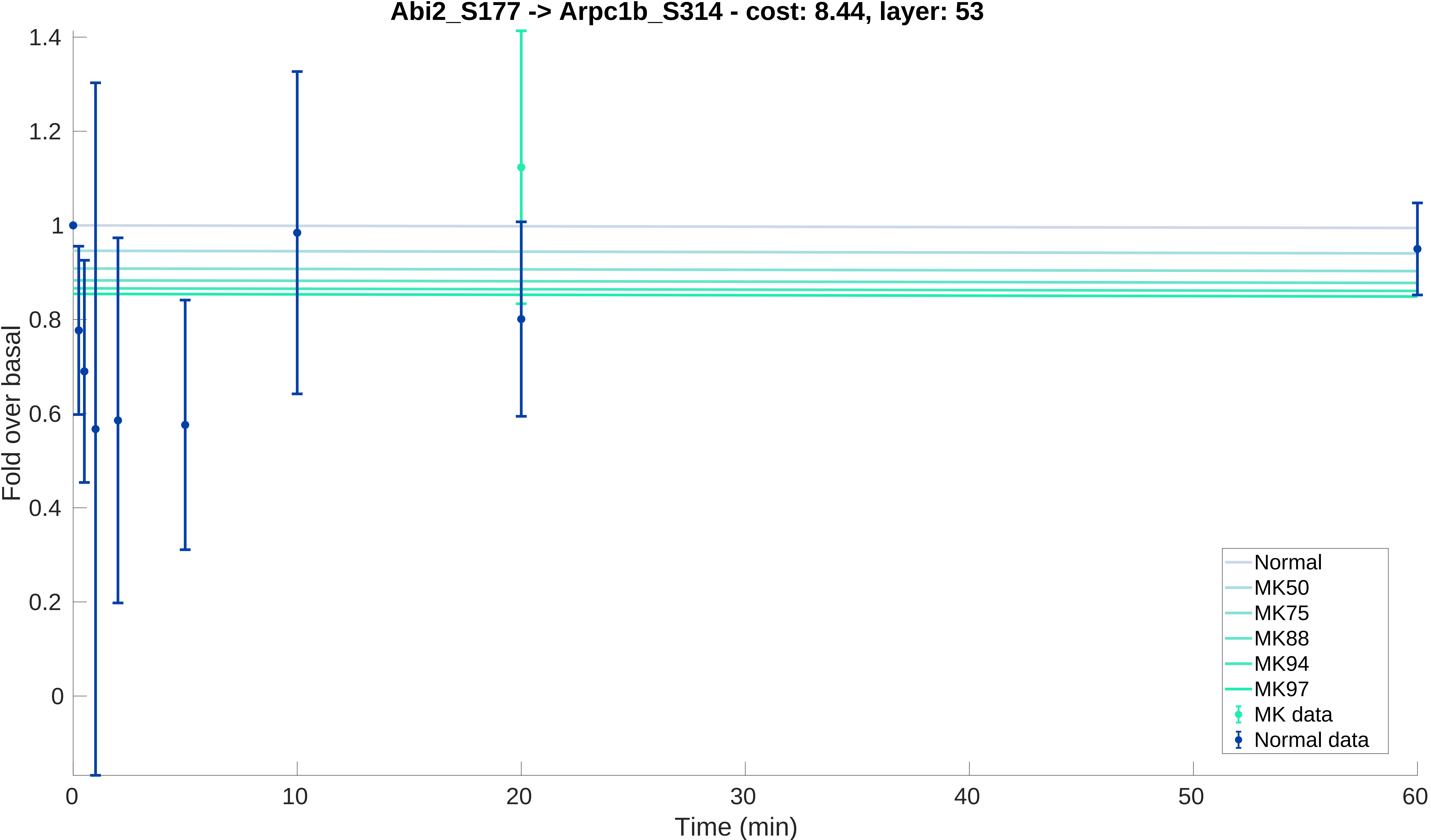

Abi2_S177 -> Arpc1b_S314 - cost: 8.44, layer: 53
Fold over basal
1.4
1.2
1
0.8
0.6
0.4
0.2
0
Time (min)
0
10
20
30
40
50
60
Normal
MK50
MK75
MK88
MK94
MK97
MK data
Normal data

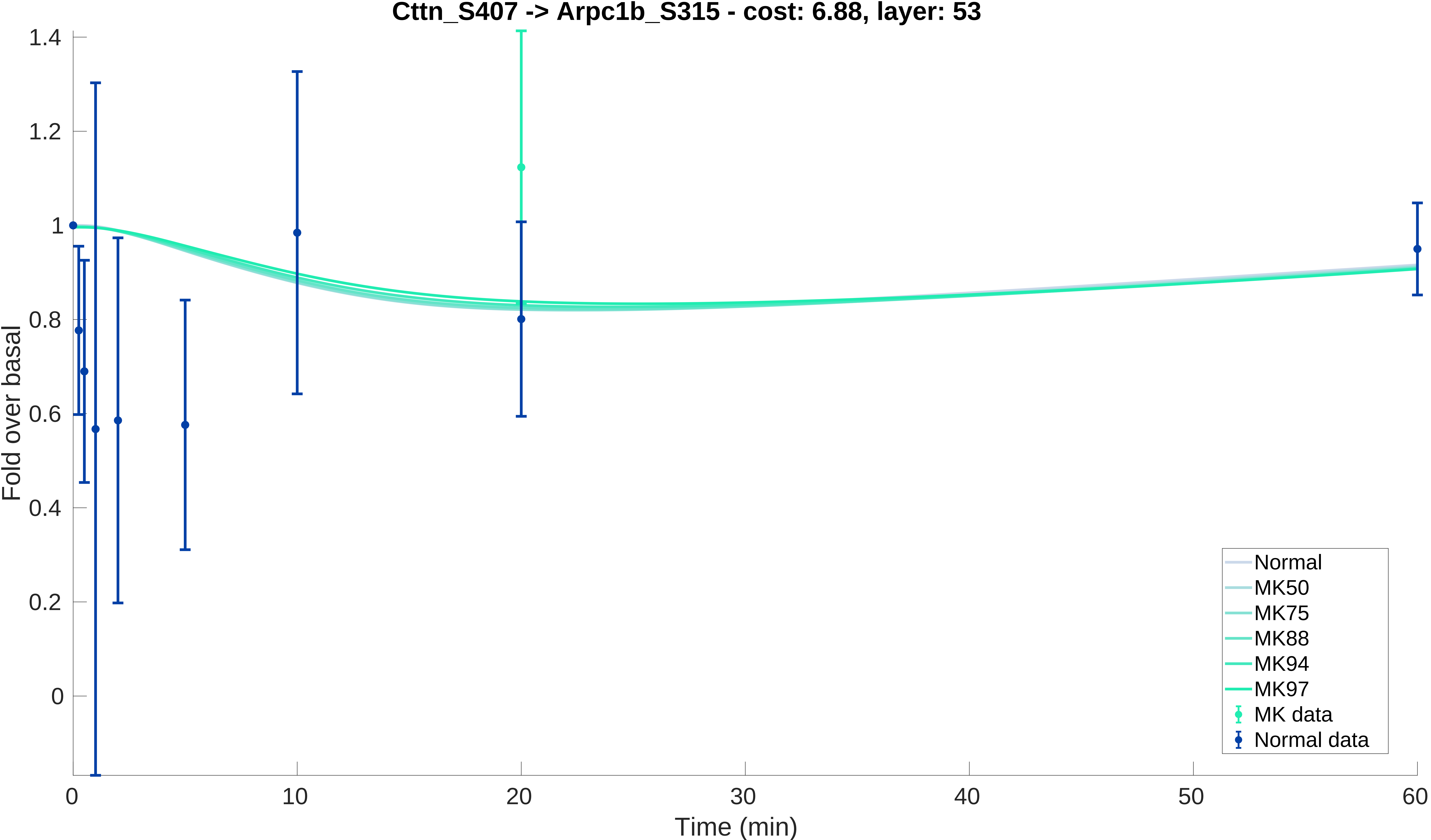
Cttn_S407 -> Arpc1b_S315 - cost: 6.88, layer: 53
Fold over basal
Time (min)
1.4
1.2
1
0.8
0.6
0.4
0.2
0
0
10
20
30
40
50
60
Normal
MK50
MK75
MK88
MK94
MK97
MK data
Normal data

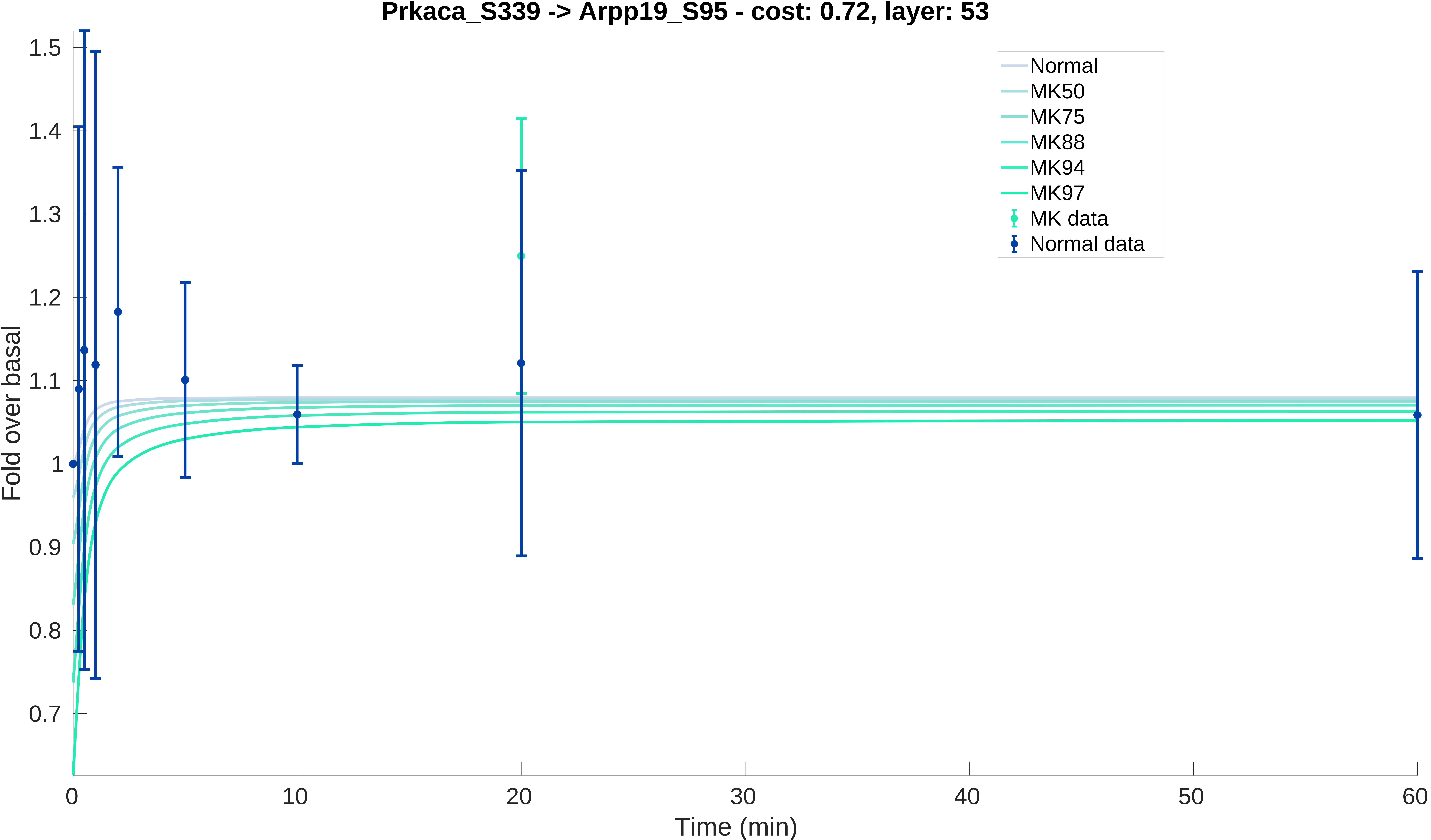
Prrkaca_S339 -> Arpp19_S95 - cost: 0.72, layer: 53
Fold over basal
Time (min)
Normal
MK50
MK75
MK88
MK94
MK97
MK data
Normal data
0.7
0.8
0.9
1
1.1
1.2
1.3
1.4
1.5
0
10
20
30
40
50
60

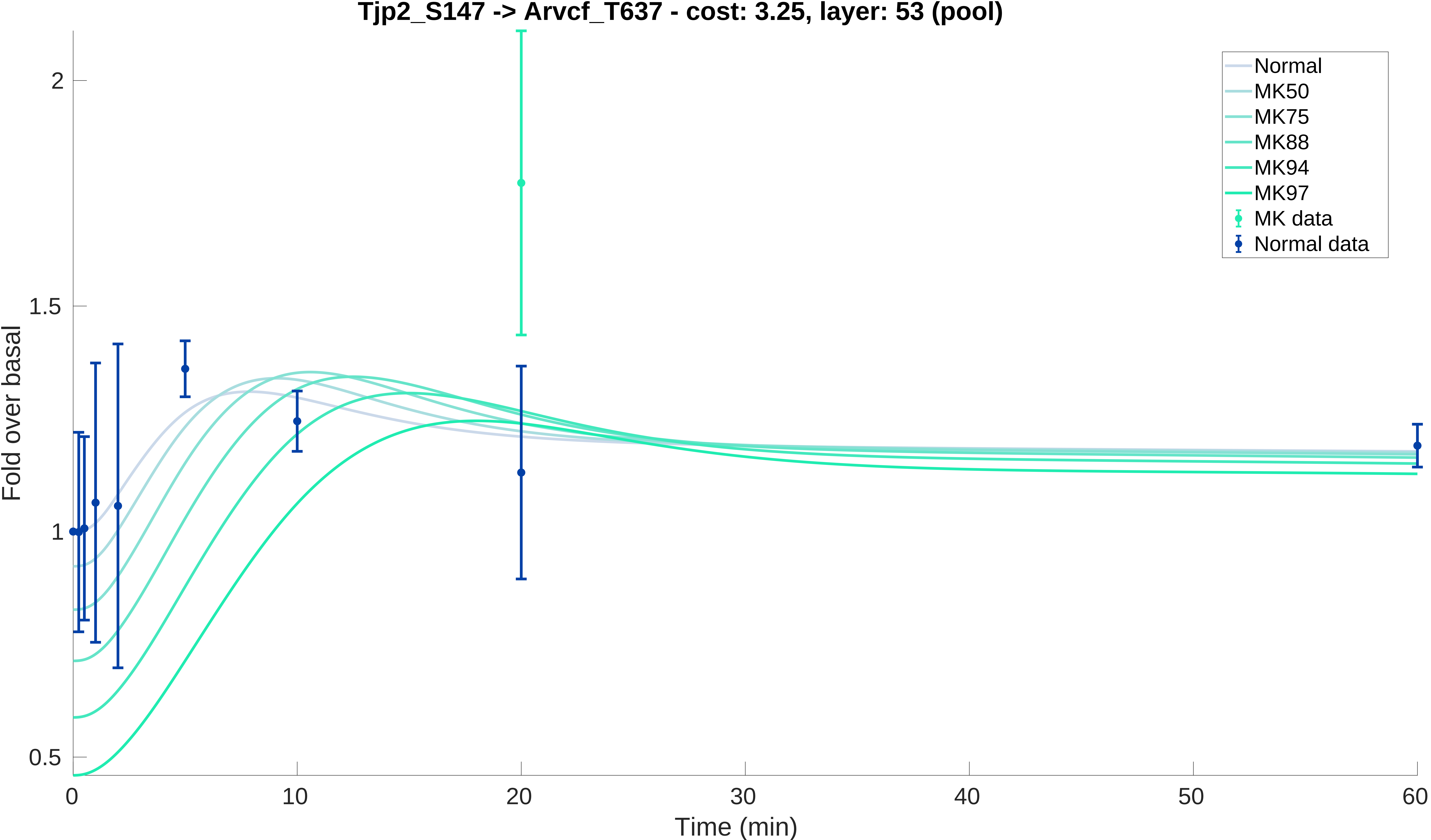

Tjp2_S147 -> Arvcf_T637 - cost: 3.25, layer: 53 (pool)
Normal
MK50
MK75
MK88
MK94
MK97
MK data
Normal data
Fold over basal
2
1.5
1
0.5
0
10
20
30
40
50
60
Time (min)

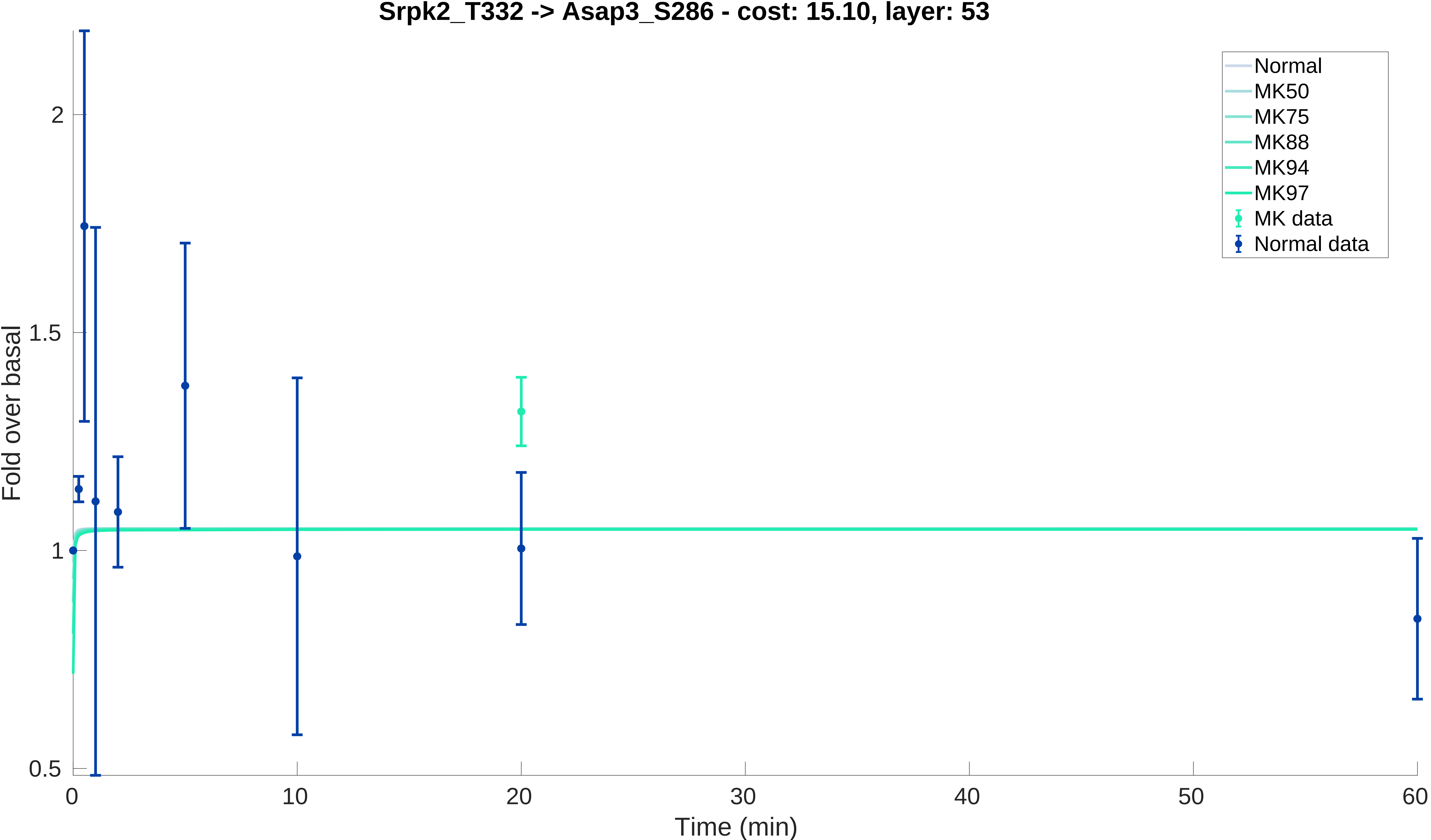

Srpk2_T332 -> Asap3_S286 - cost: 15.10, layer: 53
Normal
MK50
MK75
MK88
MK94
MK97
MK data
Normal data
Fold over basal
2
1.5
1
0.5
0
10
20
30
40
50
60
Time (min)

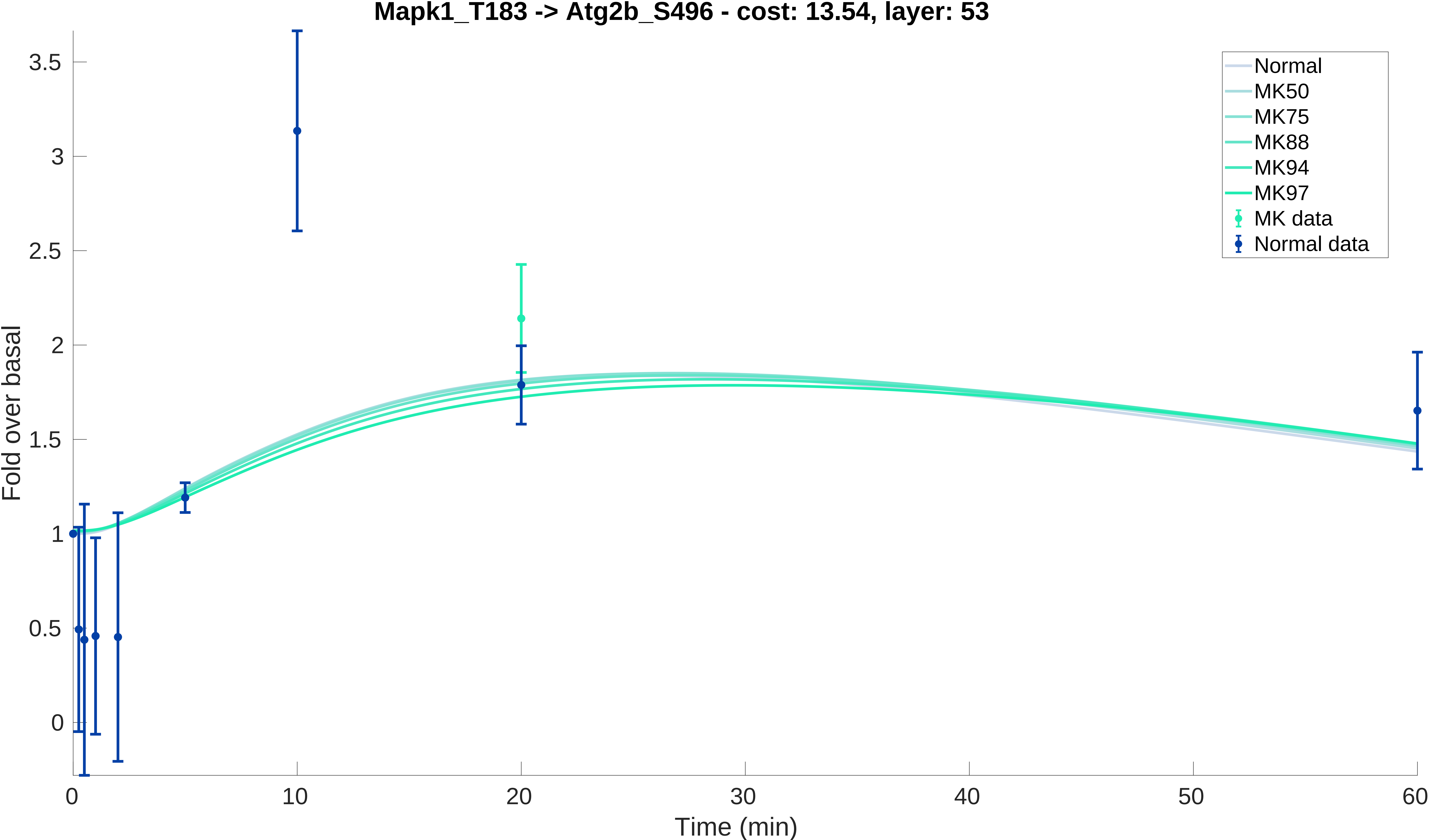

Mapk1_T183 -> Atg2b_S496 - cost: 13.54, layer: 53
Fold over basal
Time (min)
0
0.5
1
1.5
2
2.5
3
3.5
0
10
20
30
40
50
60
Normal
MK50
MK75
MK88
MK94
MK97
MK data
Normal data

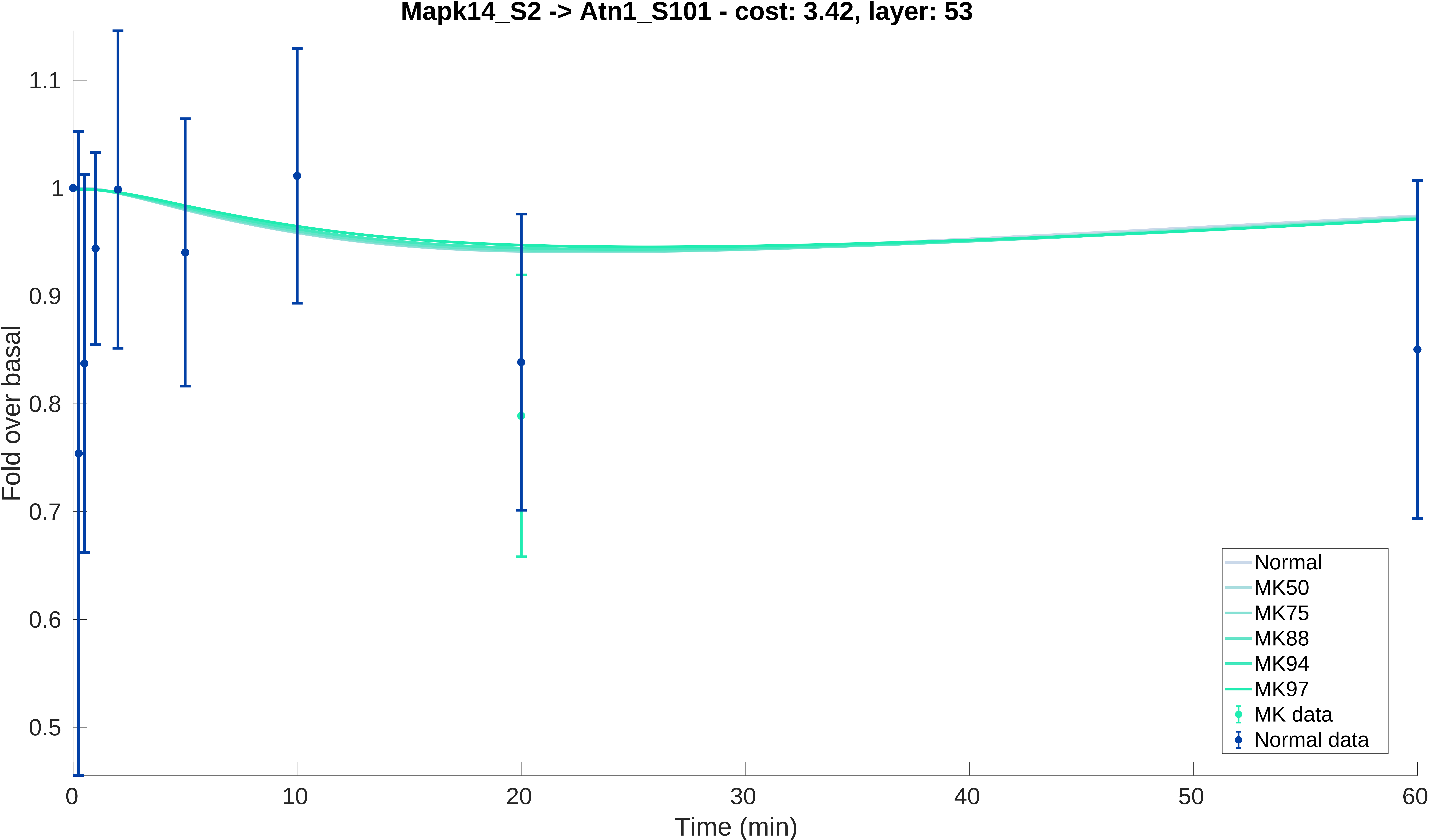

Mapk14_S2 -> Atn1_S101 - cost: 3.42, layer: 53
Fold over basal
Time (min)
1.1
1
0.9
0.8
0.7
0.6
0.5
0
10
20
30
40
50
60
Normal
MK50
MK75
MK88
MK94
MK97
MK data
Normal data

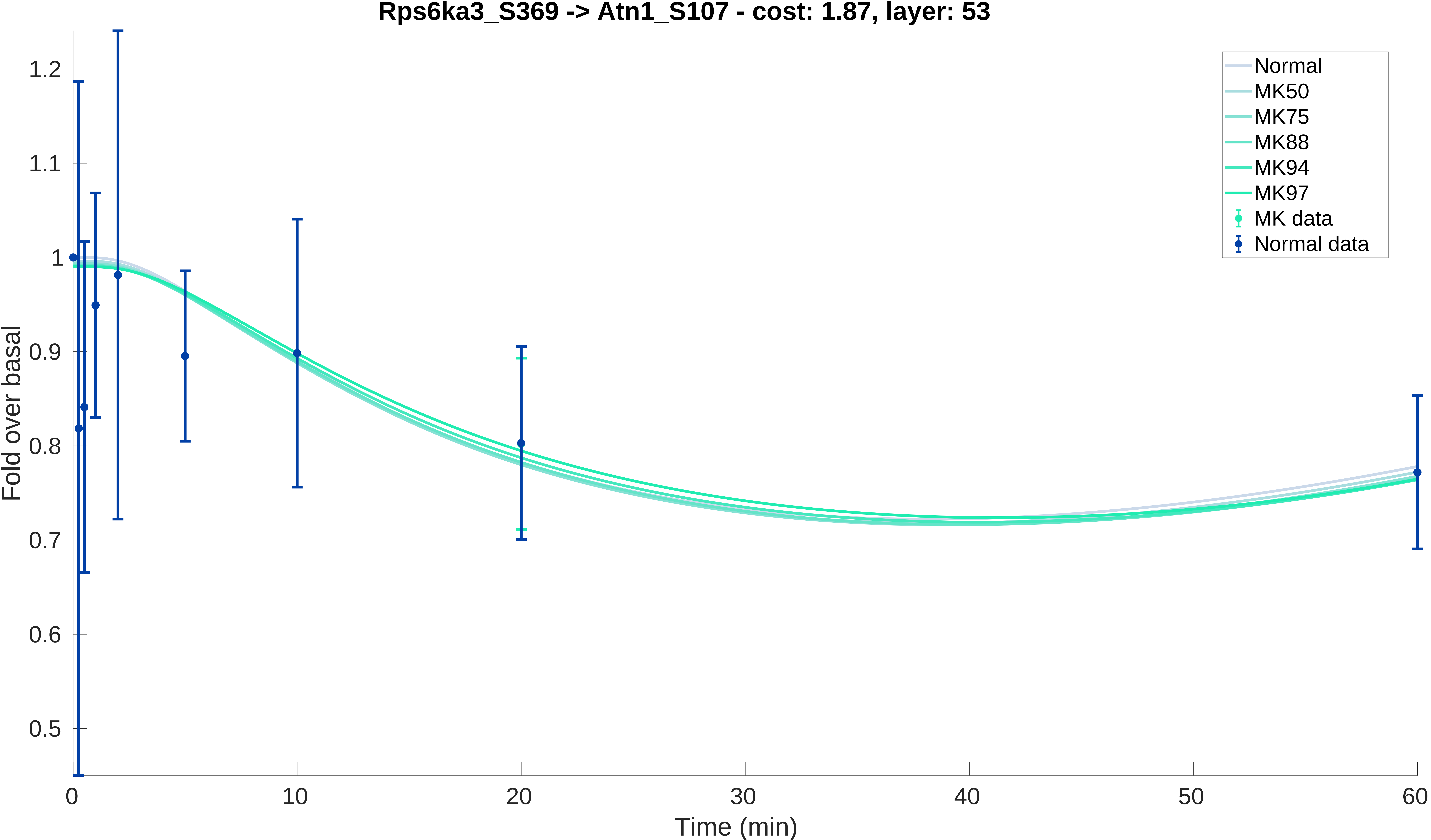

Rps6ka3_S369 -> Atn1_S107 - cost: 1.87, layer: 53
Fold over basal
Time (min)
0.5
0.6
0.7
0.8
0.9
1
1.1
1.2
0
10
20
30
40
50
60
Normal
MK50
MK75
MK88
MK94
MK97
MK data
Normal data

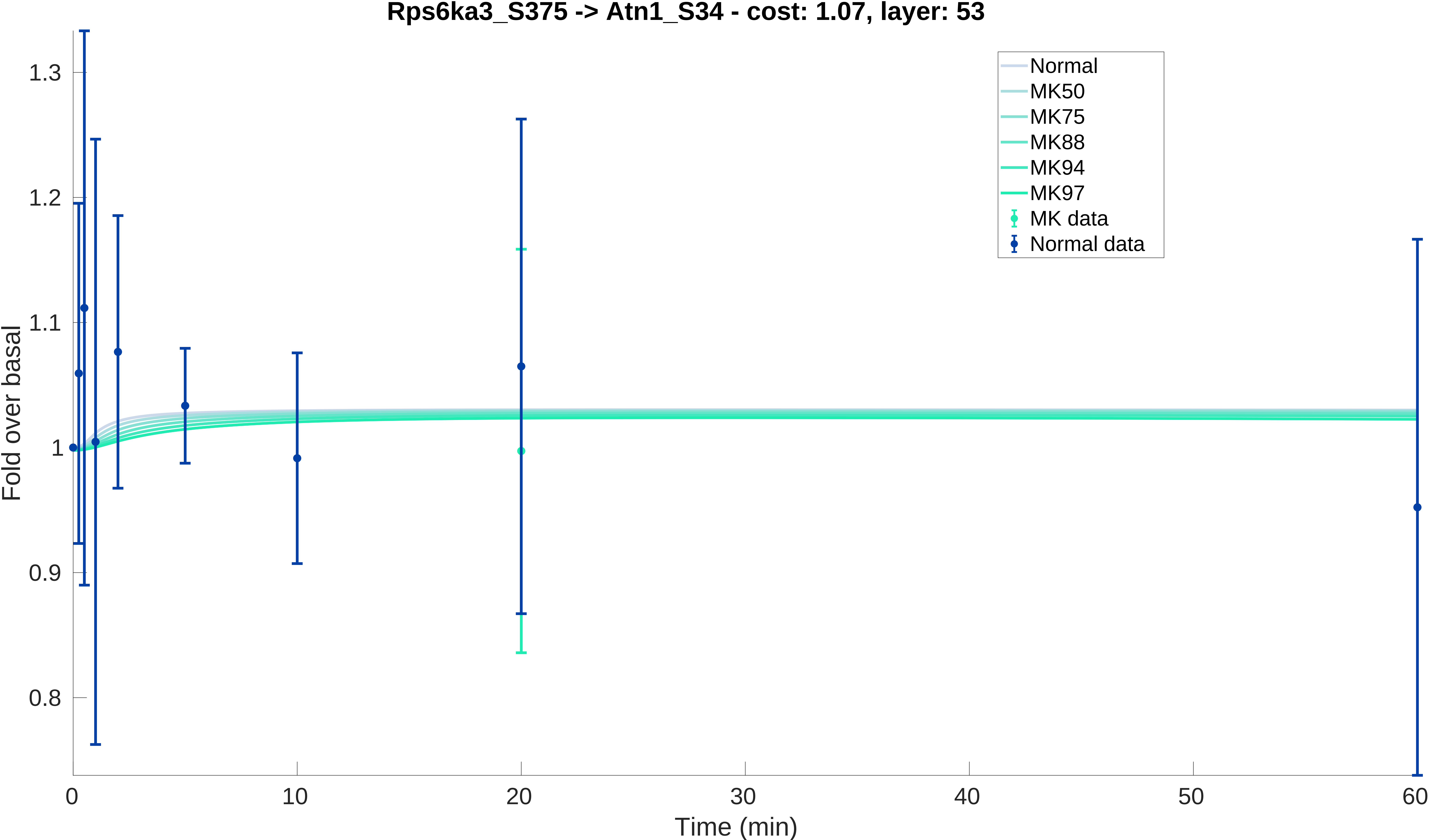

Rps6ka3_S375 -> Atn1_S34 - cost: 1.07, layer: 53
Normal
MK50
MK75
MK88
MK94
MK97
MK data
Normal data
Fold over basal
1.3
1.2
1.1
1
0.9
0.8
0
10
20
30
40
50
60
Time (min)

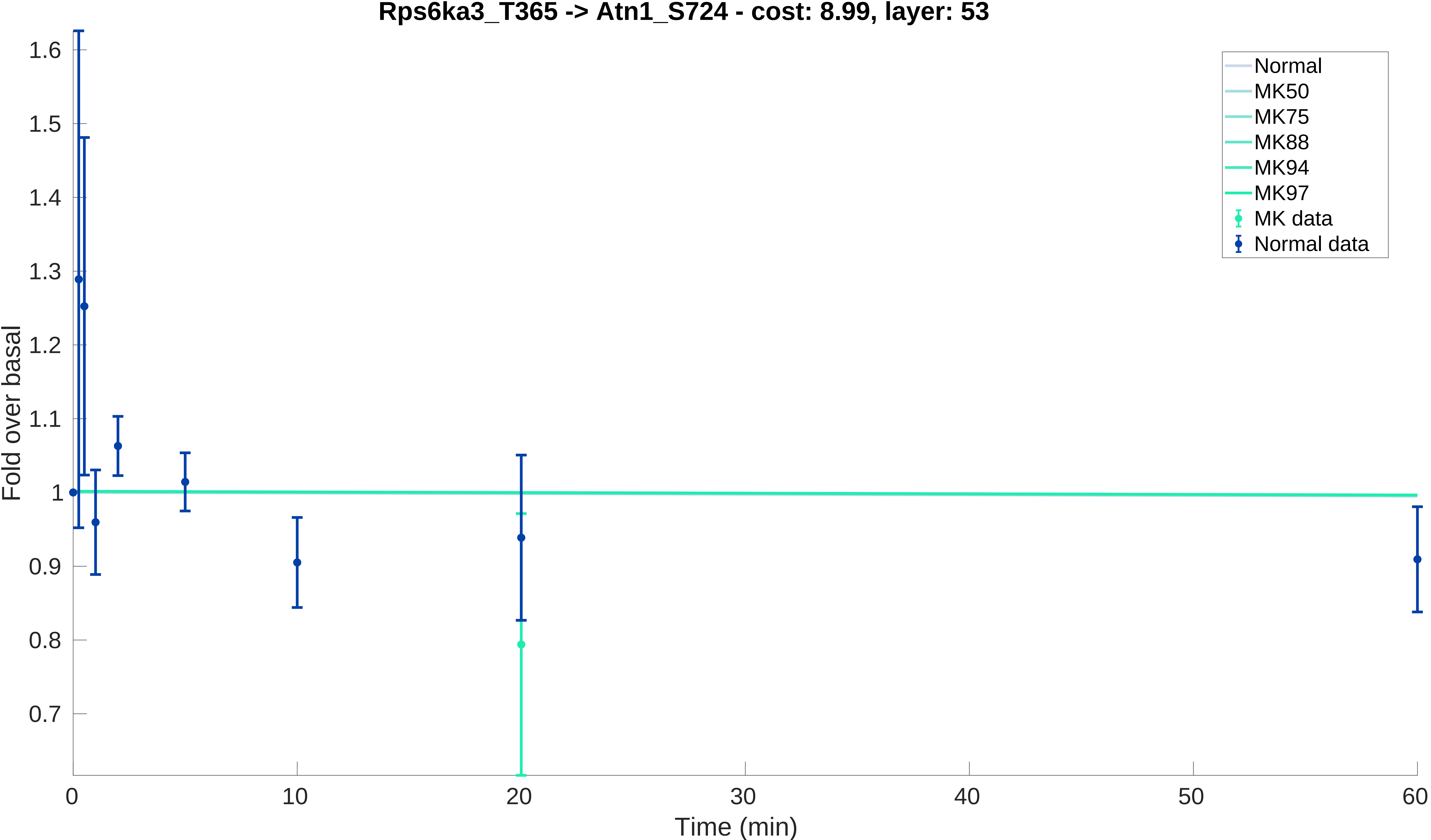

Rps6ka3_T365 -> Atn1_S724 - cost: 8.99, layer: 53
Fold over basal
Time (min)
1.6
1.5
1.4
1.3
1.2
1.1
1
0.9
0.8
0.7
0
10
20
30
40
50
60
Normal
MK50
MK75
MK88
MK94
MK97
MK data
Normal data

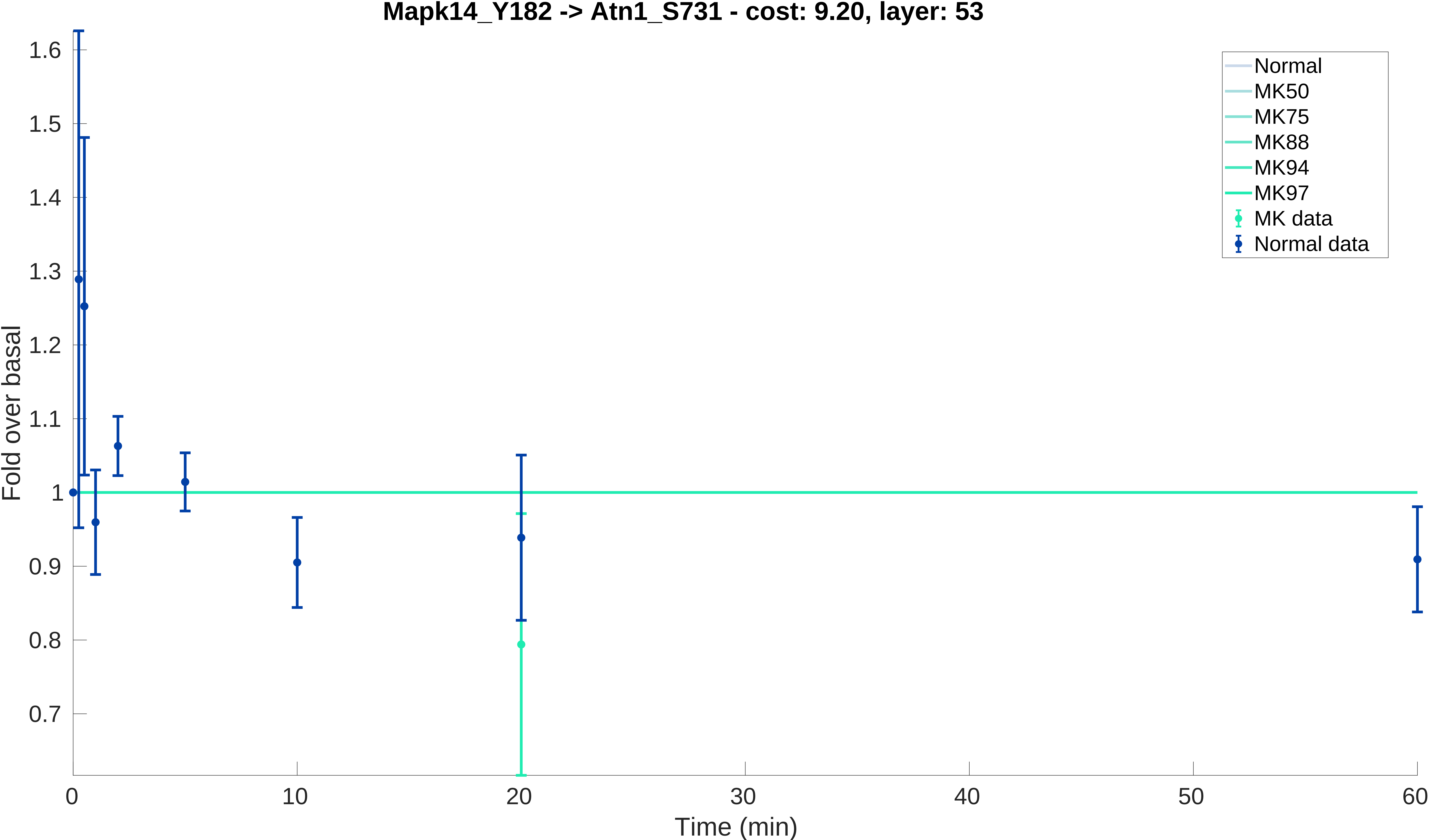

Mapk14_Y182 -> Atn1_S731 - cost: 9.20, layer: 53
Fold over basal
1.6
1.5
1.4
1.3
1.2
1.1
1
0.9
0.8
0.7
0
10
20
30
40
50
60
Time (min)
Normal
MK50
MK75
MK88
MK94
MK97
MK data
Normal data

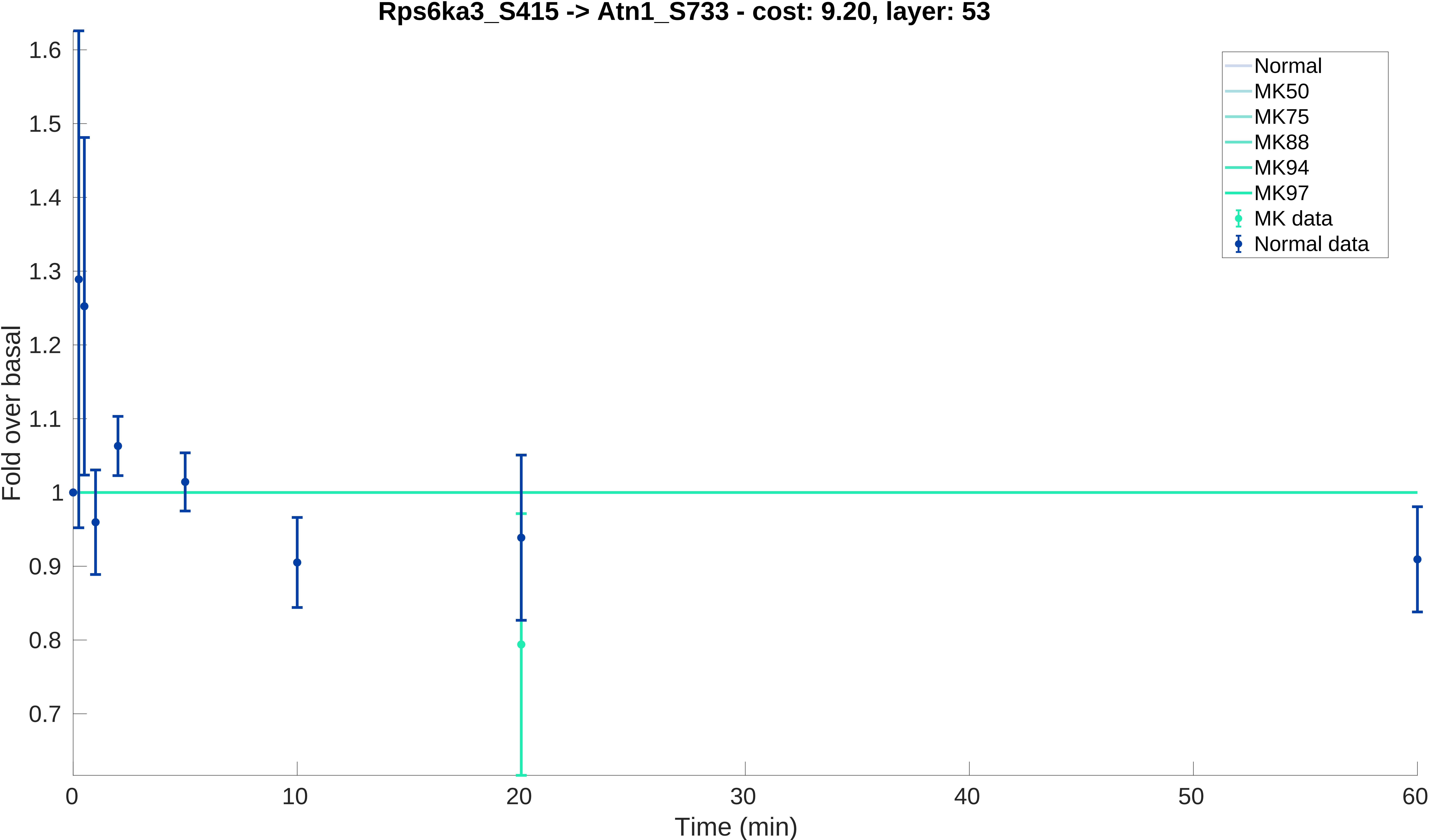

Rps6ka3_S415 -> Atn1_S733 - cost: 9.20, layer: 53
Fold over basal
Time (min)
1.6
1.5
1.4
1.3
1.2
1.1
1
0.9
0.8
0.7
0
10
20
30
40
50
60
Normal
MK50
MK75
MK88
MK94
MK97
MK data
Normal data

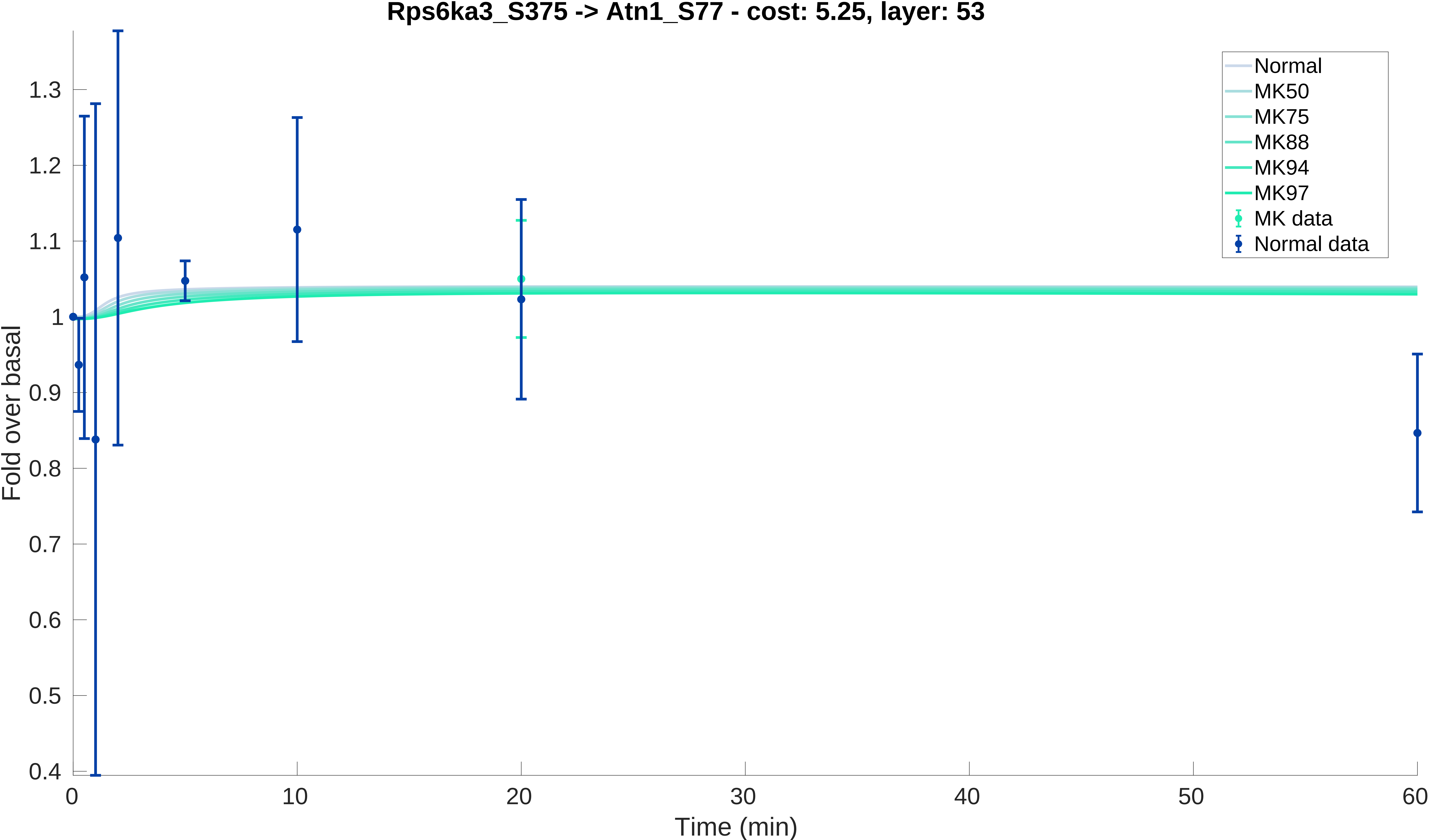
Rps6ka3_S375 -> Atn1_S77 - cost: 5.25, layer: 53
Fold over basal
Time (min)
1.3
1.2
1.1
1
0.9
0.8
0.7
0.6
0.5
0.4
0
10
20
30
40
50
60
Normal
MK50
MK75
MK88
MK94
MK97
MK data
Normal data

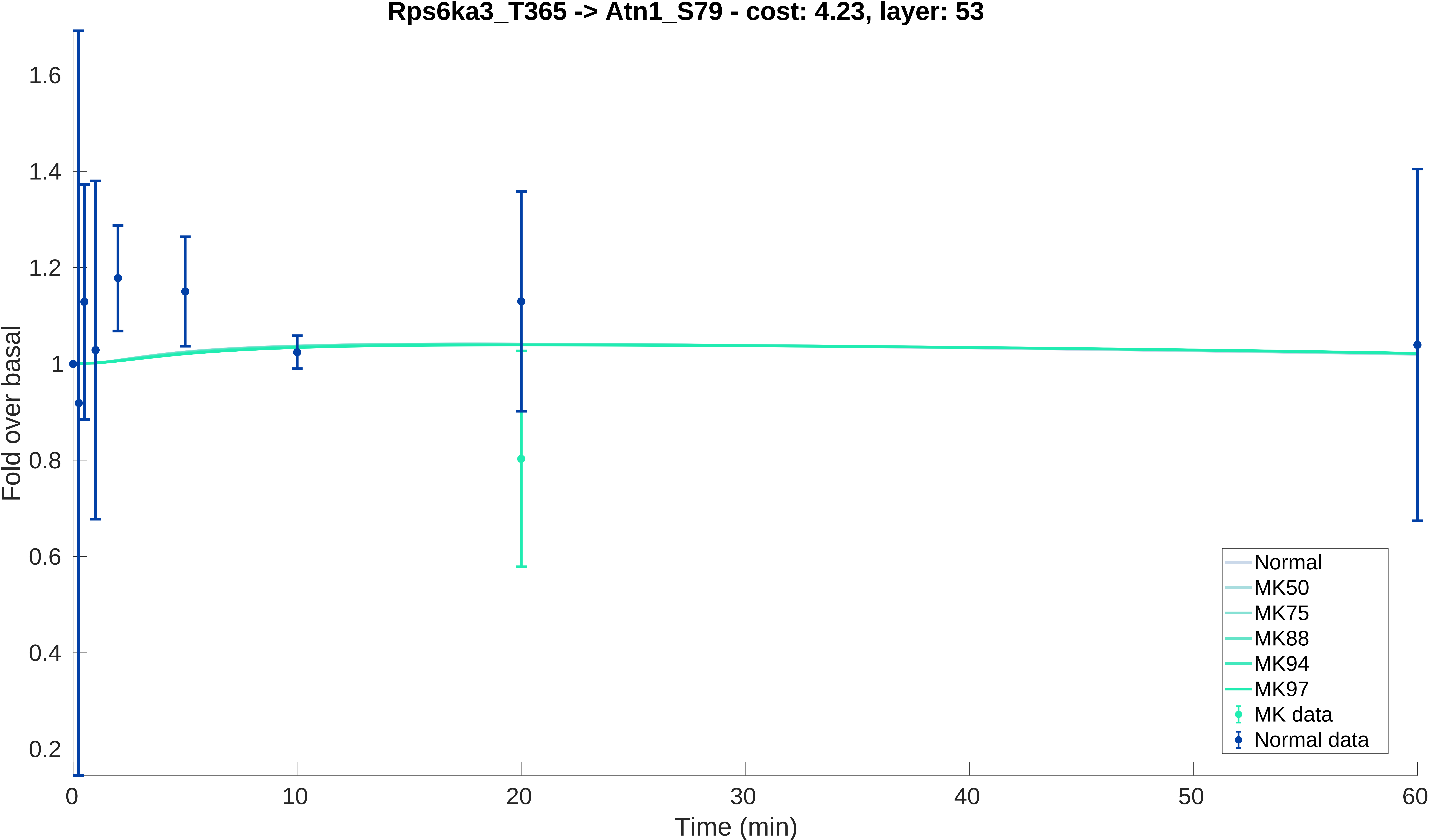
Rps6ka3_T365 -> Atn1_S79 - cost: 4.23, layer: 53
Fold over basal
1.6
1.4
1.2
1
0.8
0.6
0.4
0.2
0
10
20
30
40
50
60
Time (min)
Normal
MK50
MK75
MK88
MK94
MK97
MK data
Normal data

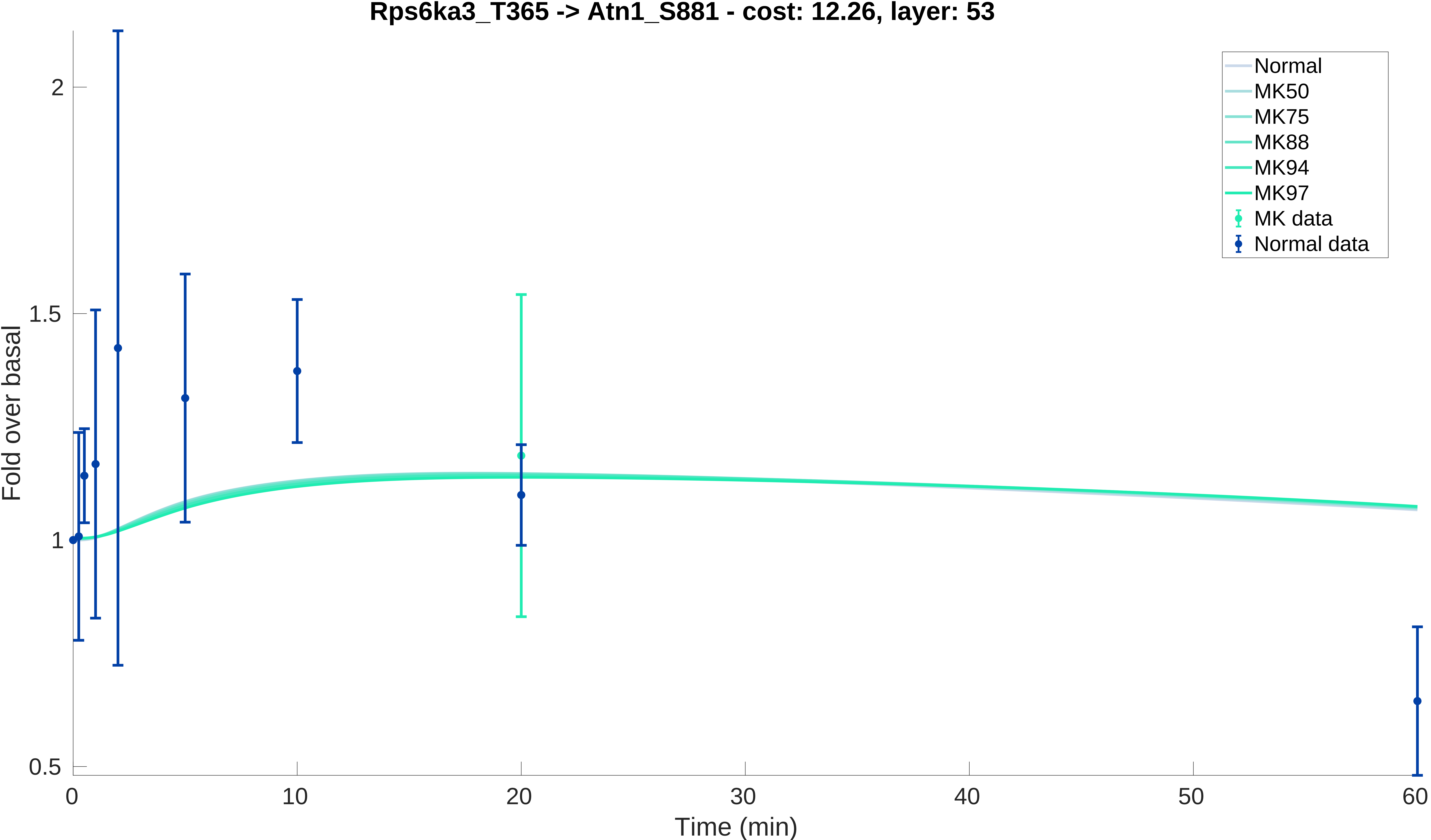

Rps6ka3_T365 -> Atn1_S881 - cost: 12.26, layer: 53
Normal
MK50
MK75
MK88
MK94
MK97
MK data
Normal data
Fold over basal
0.5
1
1.5
2
0
10
20
30
40
50
60
Time (min)

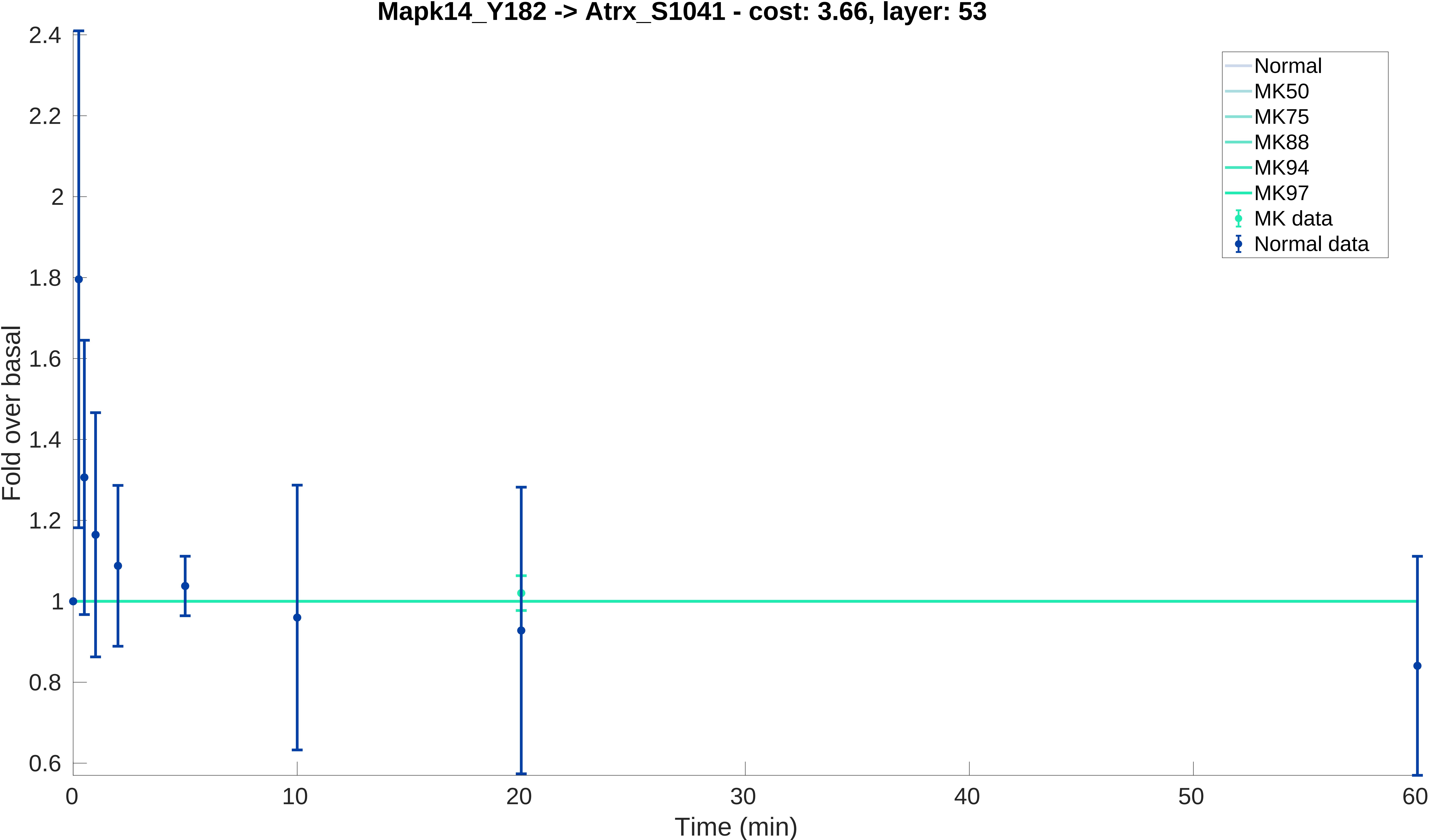

Mapk14_Y182 -> Atrx_S1041 - cost: 3.66, layer: 53
Fold over basal
Time (min)
2.4
2.2
2
1.8
1.6
1.4
1.2
1
0.8
0.6
0
10
20
30
40
50
60
Normal
MK50
MK75
MK88
MK94
MK97
MK data
Normal data

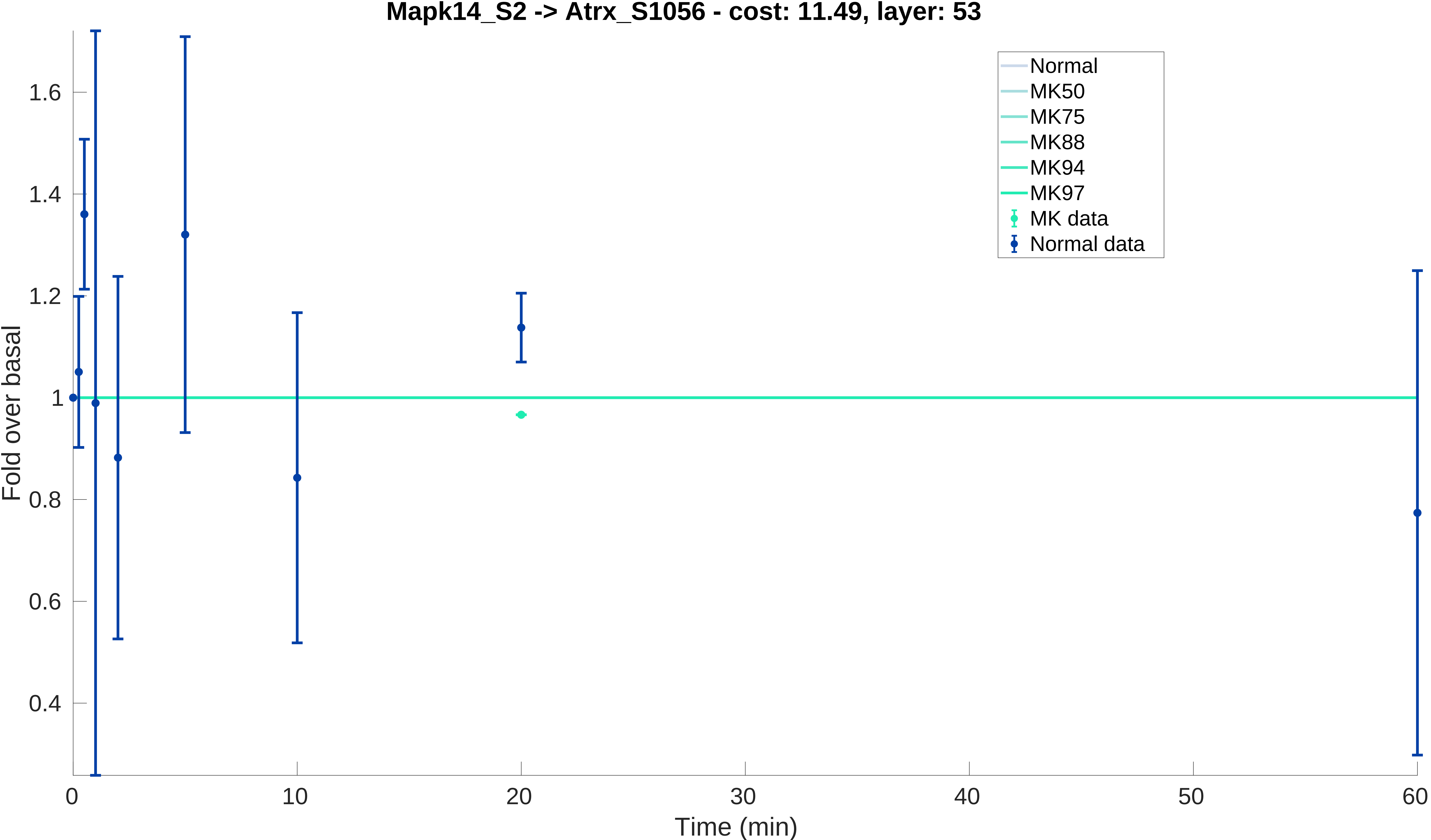

Mapk14_S2 -> Atrx_S1056 - cost: 11.49, layer: 53
Normal
MK50
MK75
MK88
MK94
MK97
MK data
Normal data
Fold over basal
Time (min)
0.4
0.6
0.8
1
1.2
1.4
1.6
0
10
20
30
40
50
60

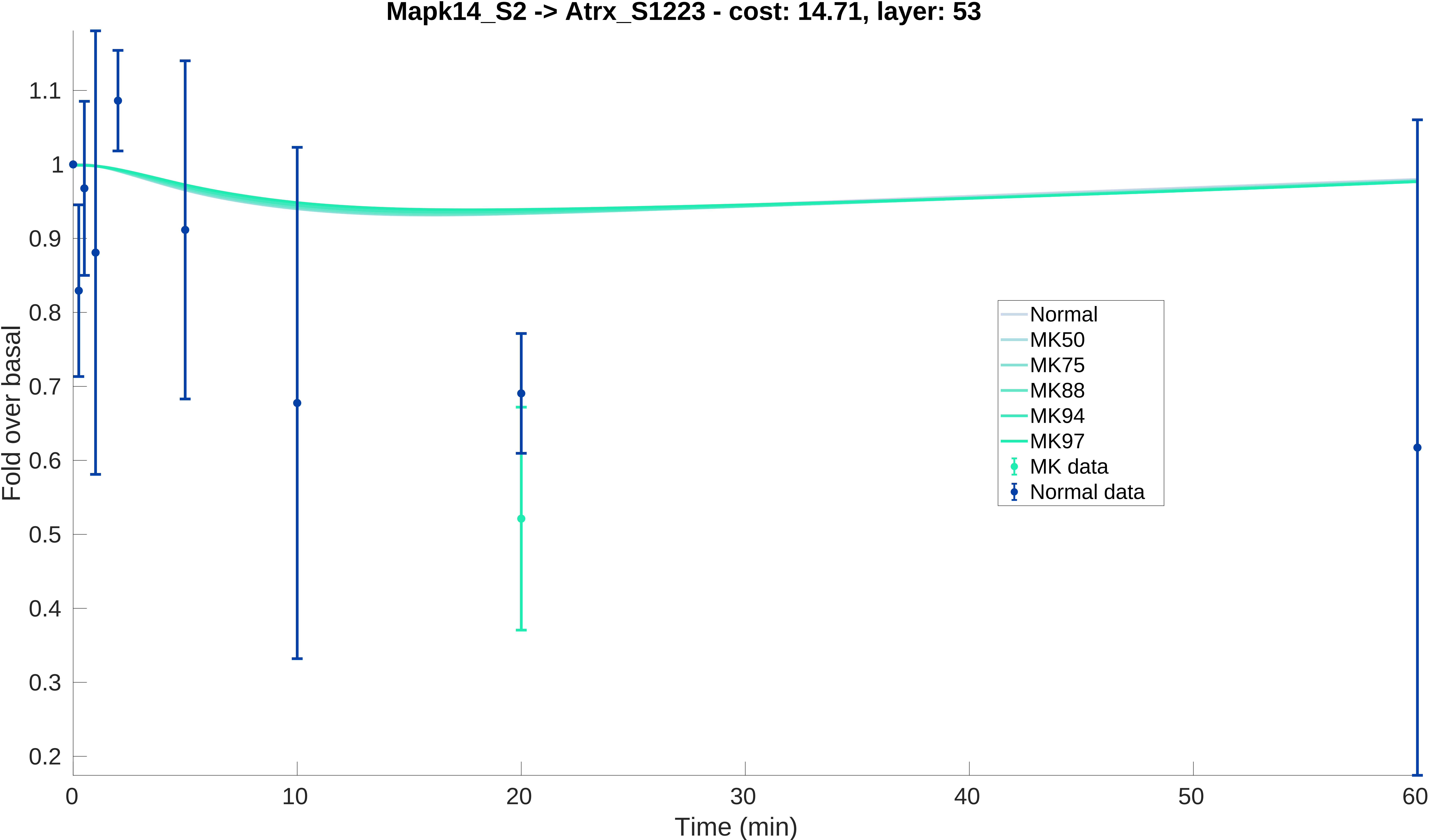
Mapk14_S2 -> Atrx_S1223 - cost: 14.71, layer: 53
Fold over basal
1.1
1
0.9
0.8
0.7
0.6
0.5
0.4
0.3
0.2
0
10
20
30
40
50
60
Time (min)
Normal
MK50
MK75
MK88
MK94
MK97
MK data
Normal data

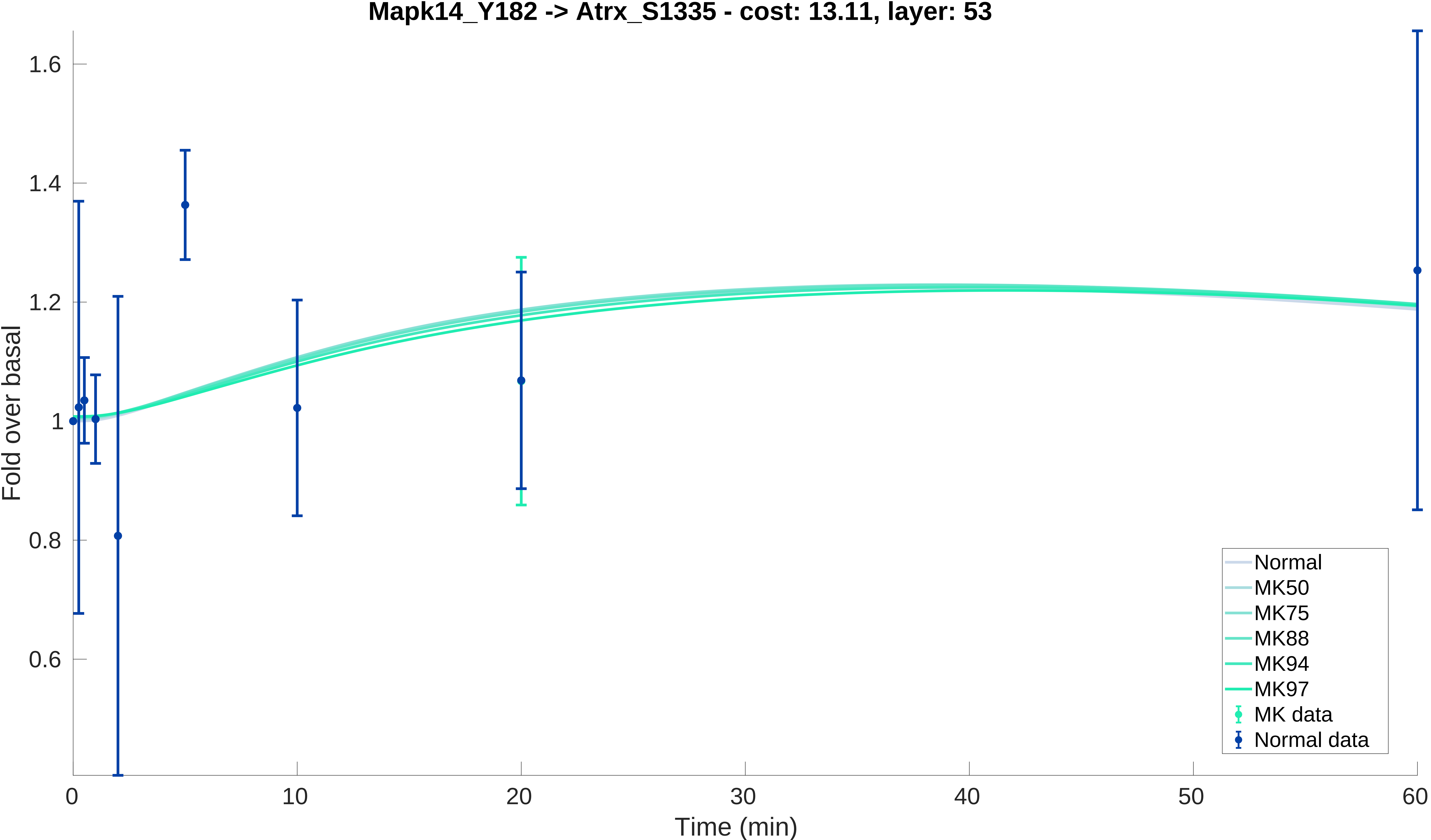

Mapk14_Y182 -> Atrx_S1335 - cost: 13.11, layer: 53
Fold over basal
Time (min)
1.6
1.4
1.2
1
0.8
0.6
0
10
20
30
40
50
60
Normal
MK50
MK75
MK88
MK94
MK97
MK data
Normal data

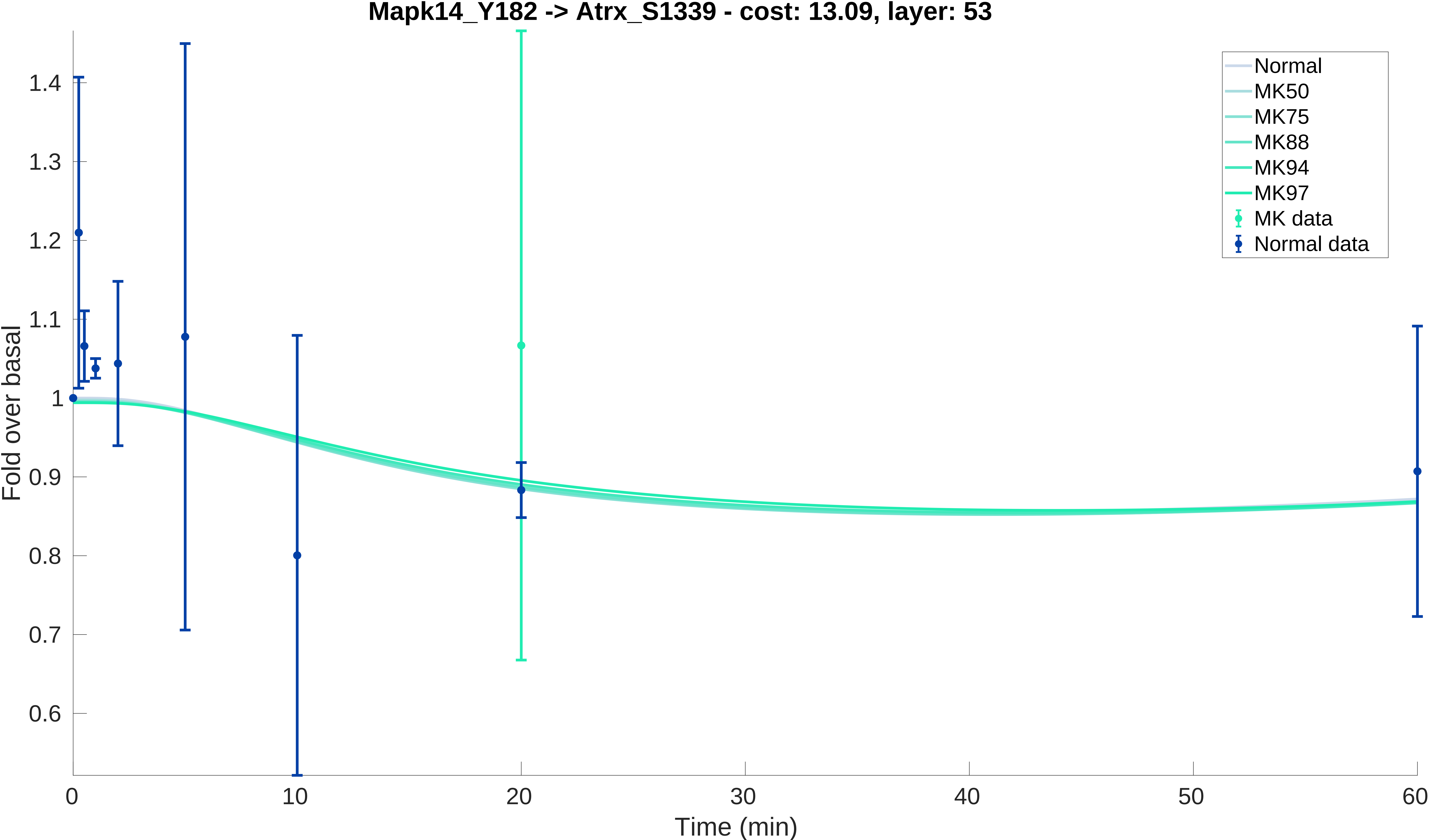

Mapk14_Y182 -> Atrx_S1339 - cost: 13.09, layer: 53
Fold over basal
1.4
1.3
1.2
1.1
1
0.9
0.8
0.7
0.6
0
10
20
30
40
50
60
Time (min)
Normal
MK50
MK75
MK88
MK94
MK97
MK data
Normal data

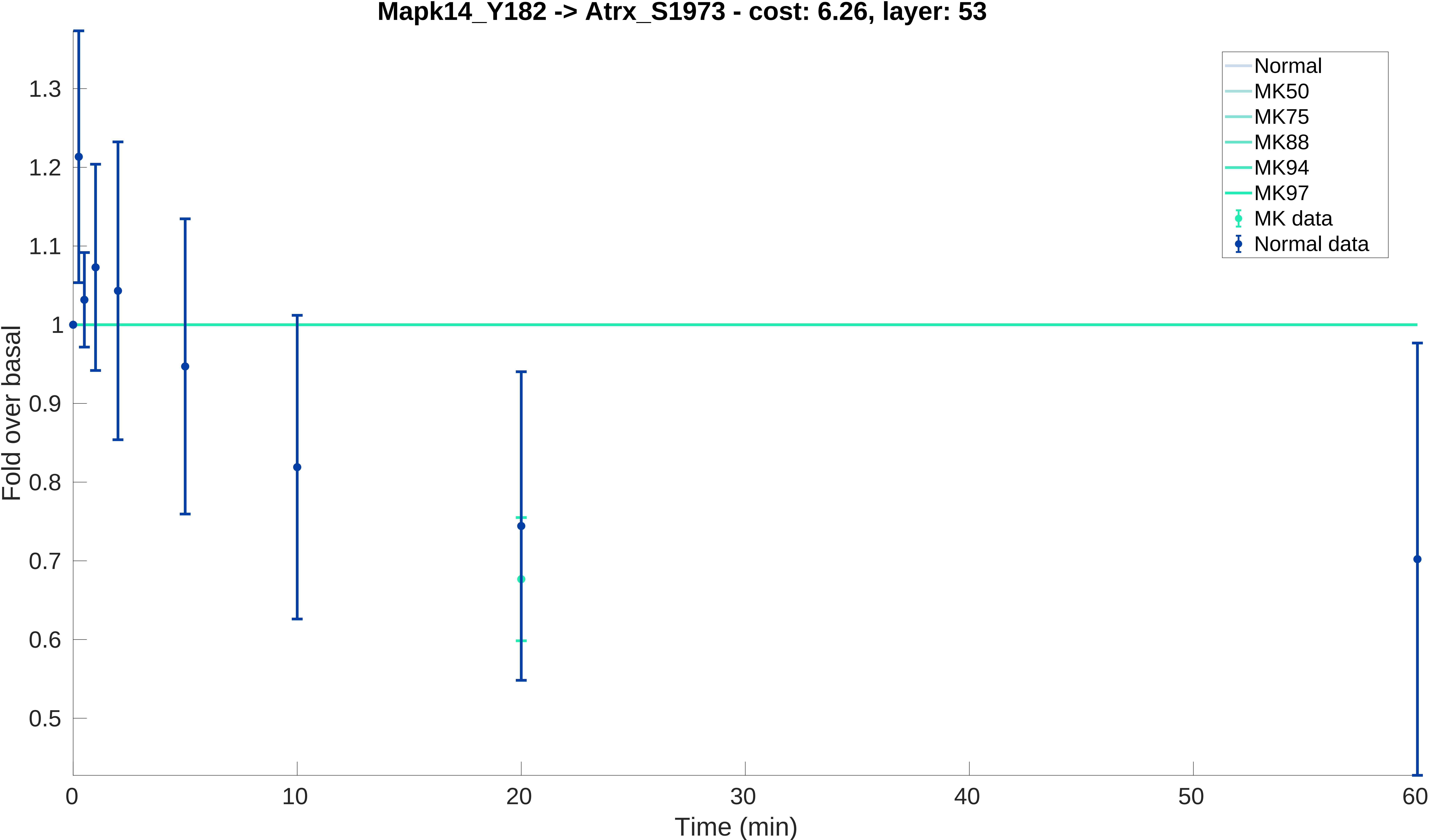

Mapk14_Y182 -> Atrx_S1973 - cost: 6.26, layer: 53
Fold over basal
1.3
1.2
1.1
1
0.9
0.8
0.7
0.6
0.5
0
10
20
30
40
50
60
Time (min)
Normal
MK50
MK75
MK88
MK94
MK97
MK data
Normal data

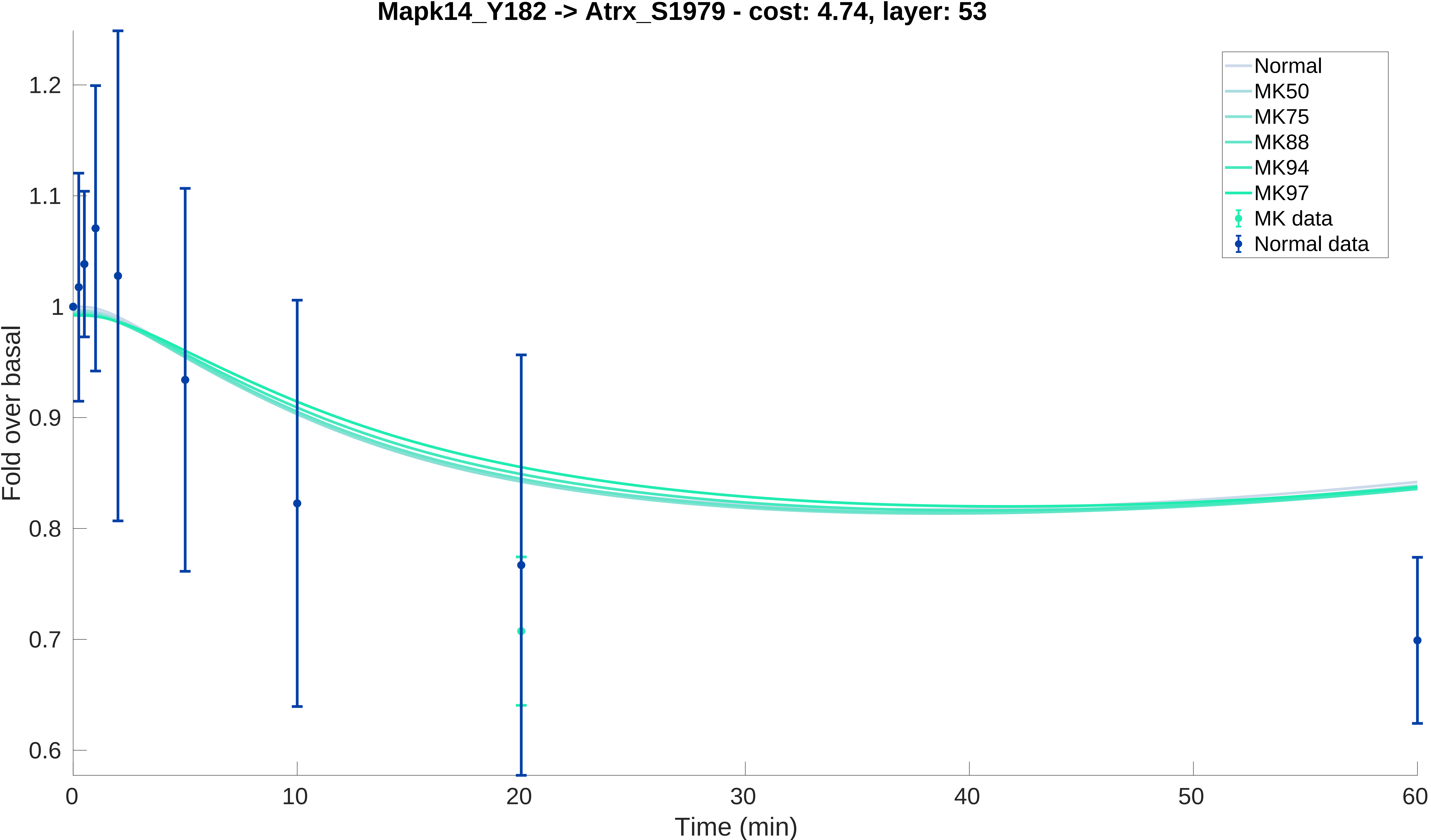

Mapk14_Y182 -> Atrx_S1979 - cost: 4.74, layer: 53
Fold over basal
Time (min)
1.2
1.1
1
0.9
0.8
0.7
0.6
0
10
20
30
40
50
60
Normal
MK50
MK75
MK88
MK94
MK97
MK data
Normal data

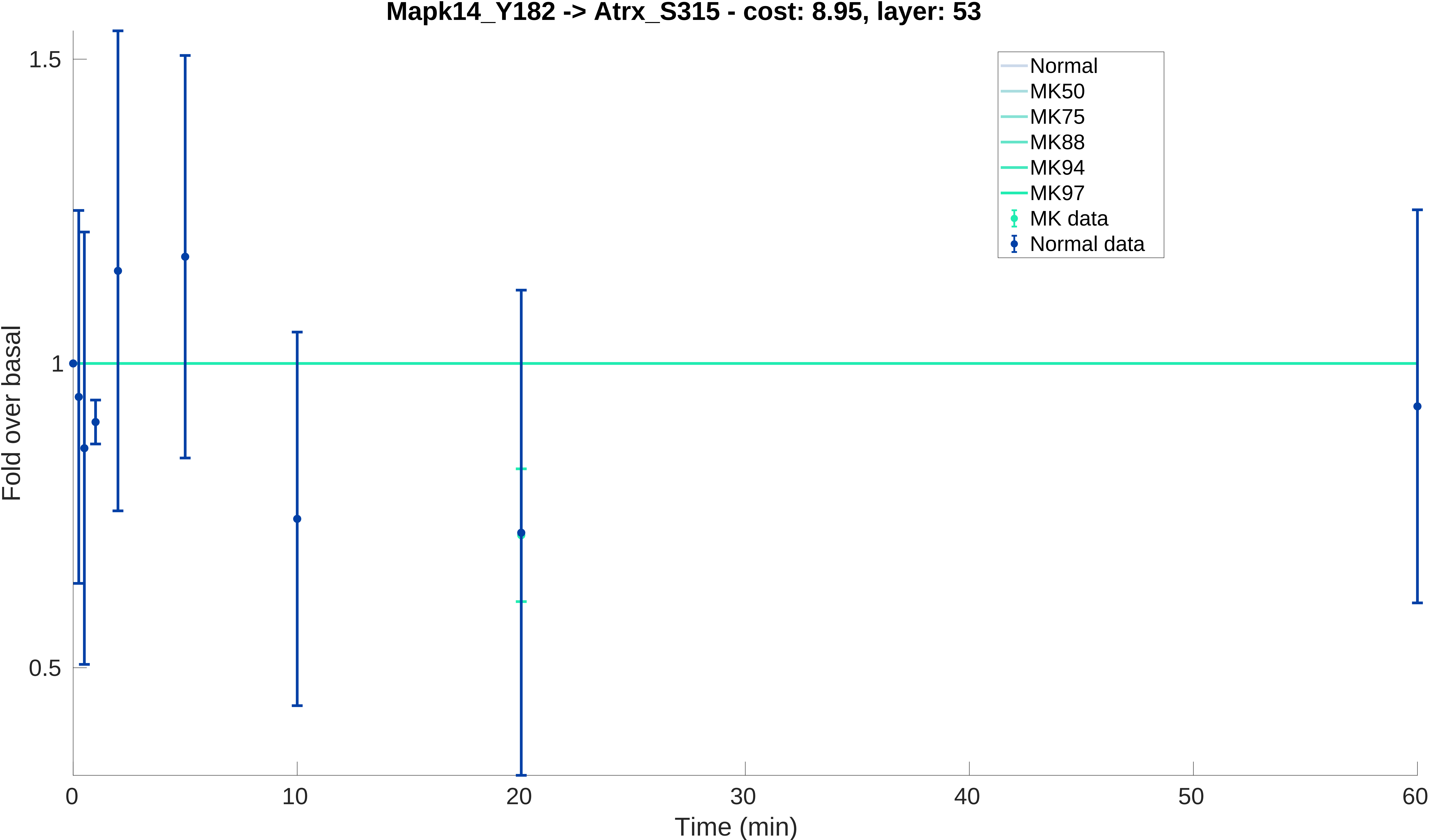
Mapk14_Y182 -> Atrx_S315 - cost: 8.95, layer: 53
Normal
MK50
MK75
MK88
MK94
MK97
MK data
Normal data
Fold over basal
1.5
1
0.5
Time (min)
0
10
20
30
40
50
60

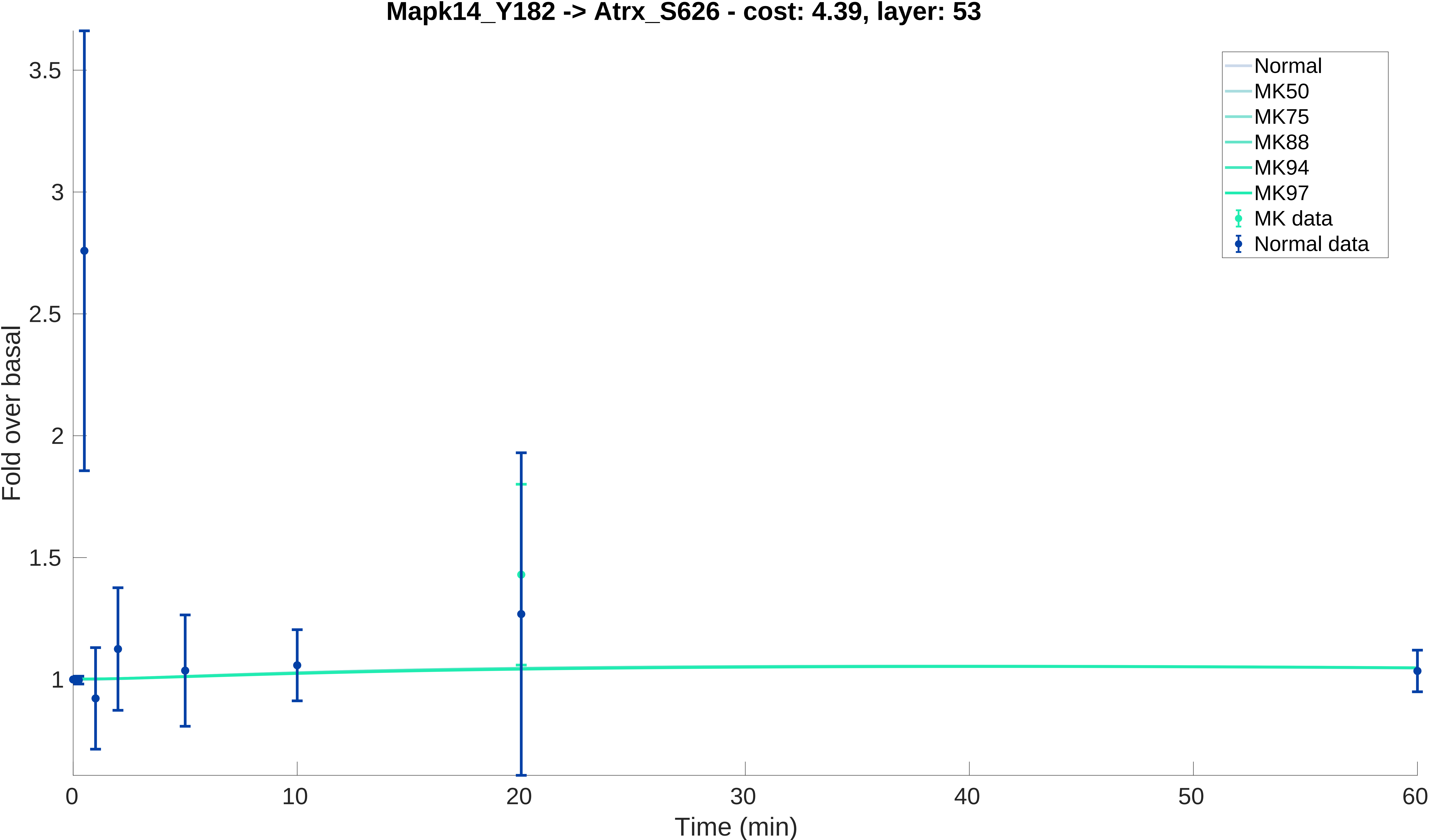

Mapk14_Y182 -> Atrx_S626 - cost: 4.39, layer: 53
Fold over basal
3.5
3
2.5
2
1.5
1
0
10
20
30
40
50
60
Time (min)
Normal
MK50
MK75
MK88
MK94
MK97
MK data
Normal data

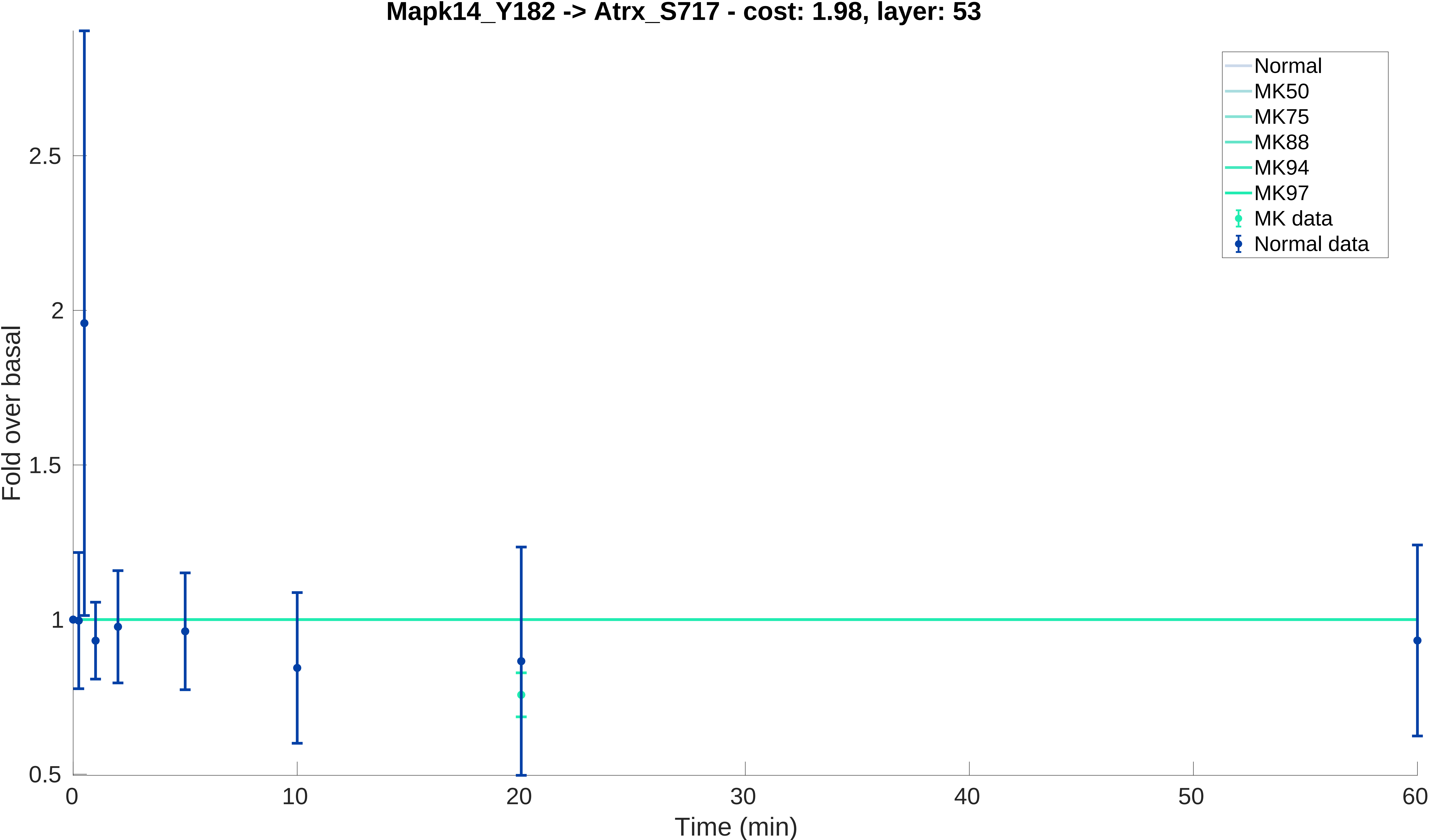

Mapk14_Y182 -> Atrx_S717 - cost: 1.98, layer: 53
Fold over basal
2.5
2
1.5
1
0.5
0
10
20
30
40
50
60
Time (min)
Normal
MK50
MK75
MK88
MK94
MK97
MK data
Normal data

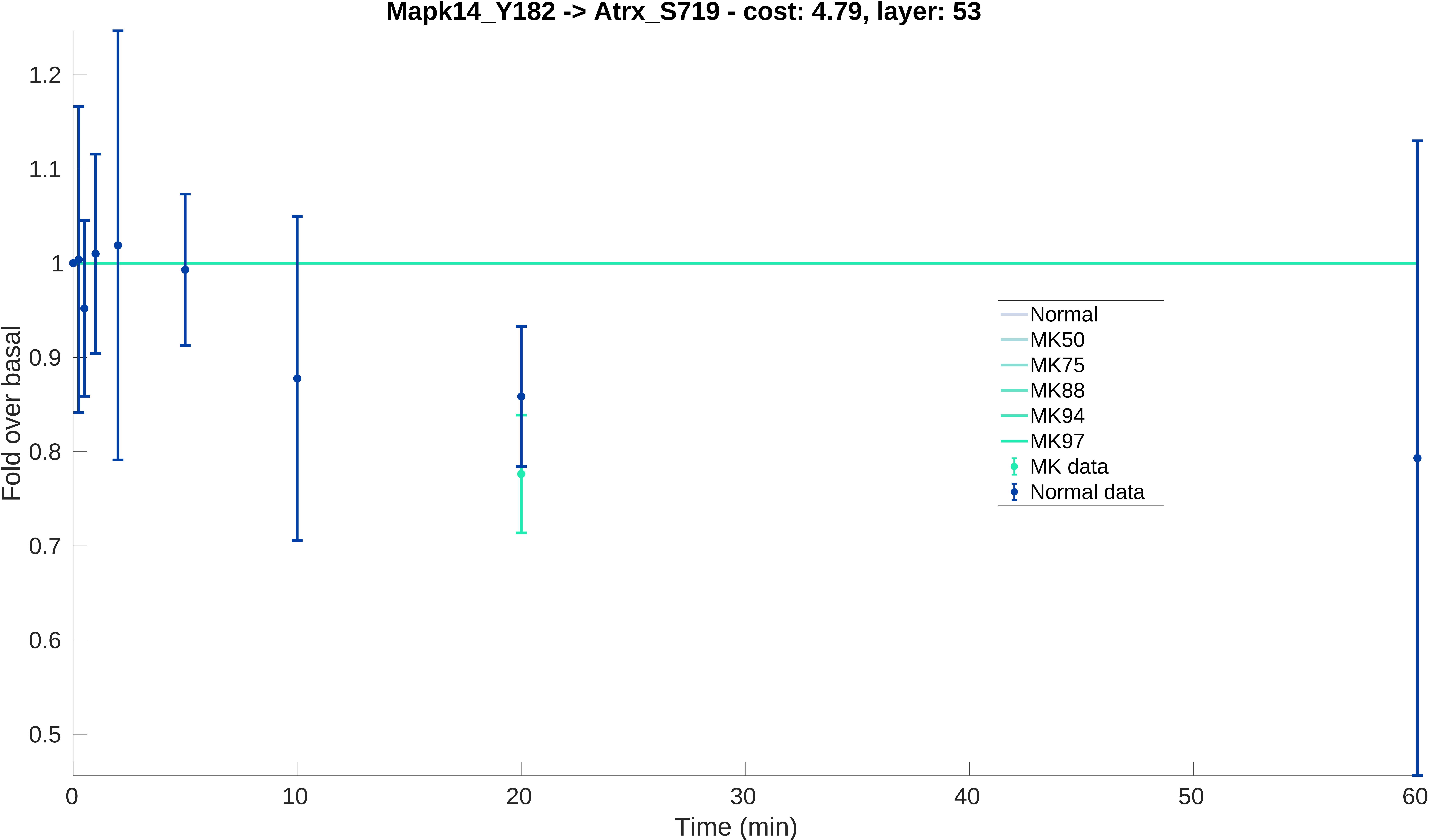

Mapk14_Y182 -> Atrx_S719 - cost: 4.79, layer: 53
Fold over basal
1.2
1.1
1
0.9
0.8
0.7
0.6
0.5
0
10
20
30
40
50
60
Time (min)
Normal
MK50
MK75
MK88
MK94
MK97
MK data
Normal data

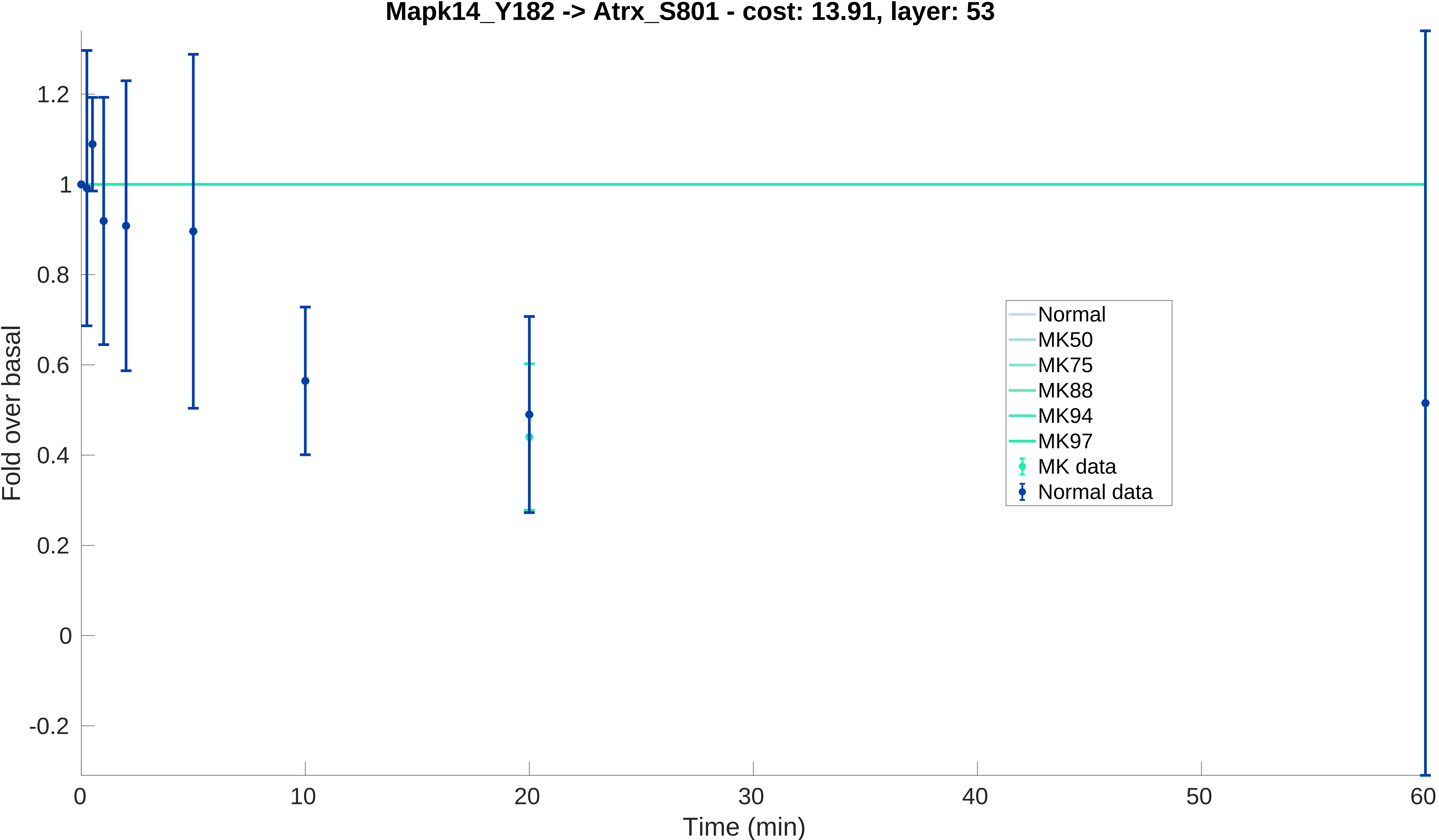

Mapk14_Y182 -> Atrx_S801 - cost: 13.91, layer: 53
Fold over basal
Time (min)
1.2
1
0.8
0.6
0.4
0.2
0
-0.2
0
10
20
30
40
50
60
Normal
MK50
MK75
MK88
MK94
MK97
MK data
Normal data

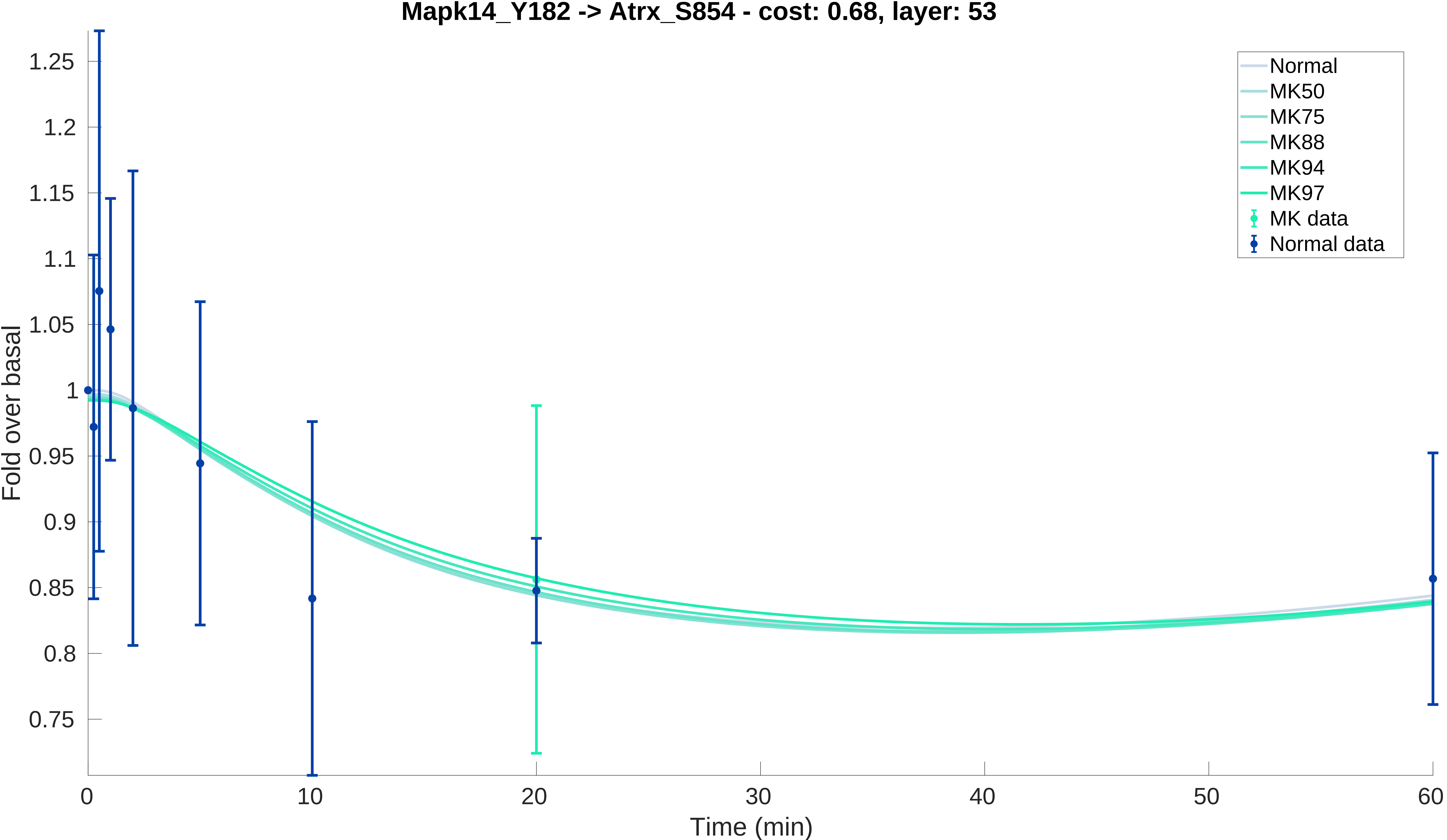

Mapk14_Y182 -> Atrx_S854 - cost: 0.68, layer: 53
Fold over basal
1.25
1.2
1.15
1.1
1.05
1
0.95
0.9
0.85
0.8
0.75
0
10
20
30
40
50
60
Time (min)
Normal
MK50
MK75
MK88
MK94
MK97
MK data
Normal data

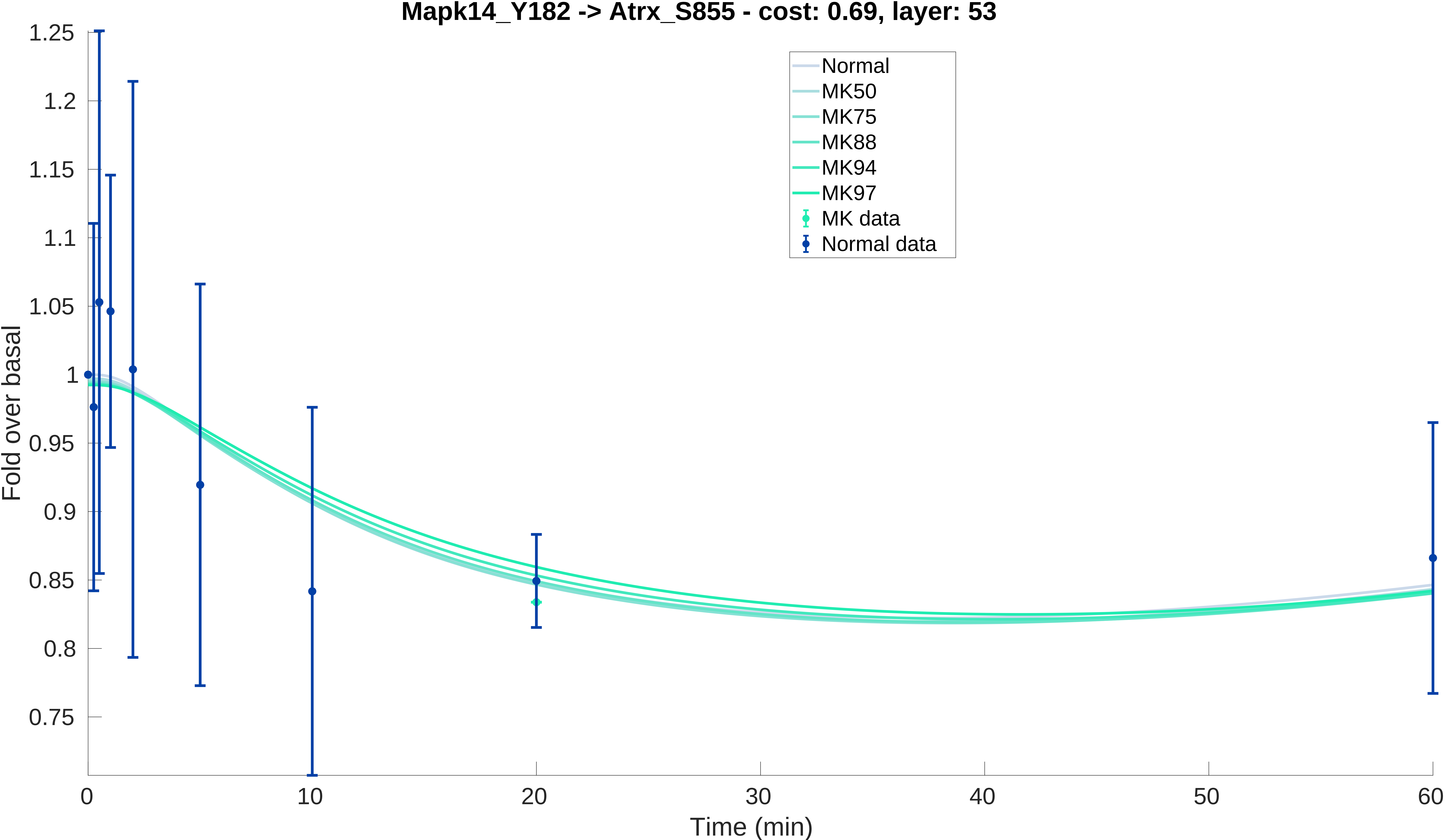

Mapk14_Y182 -> Atrx_S855 - cost: 0.69, layer: 53
Normal
MK50
MK75
MK88
MK94
MK97
MK data
Normal data
Fold over basal
1.25
1.2
1.15
1.1
1.05
1
0.95
0.9
0.85
0.8
0.75
0
10
20
30
40
50
60
Time (min)

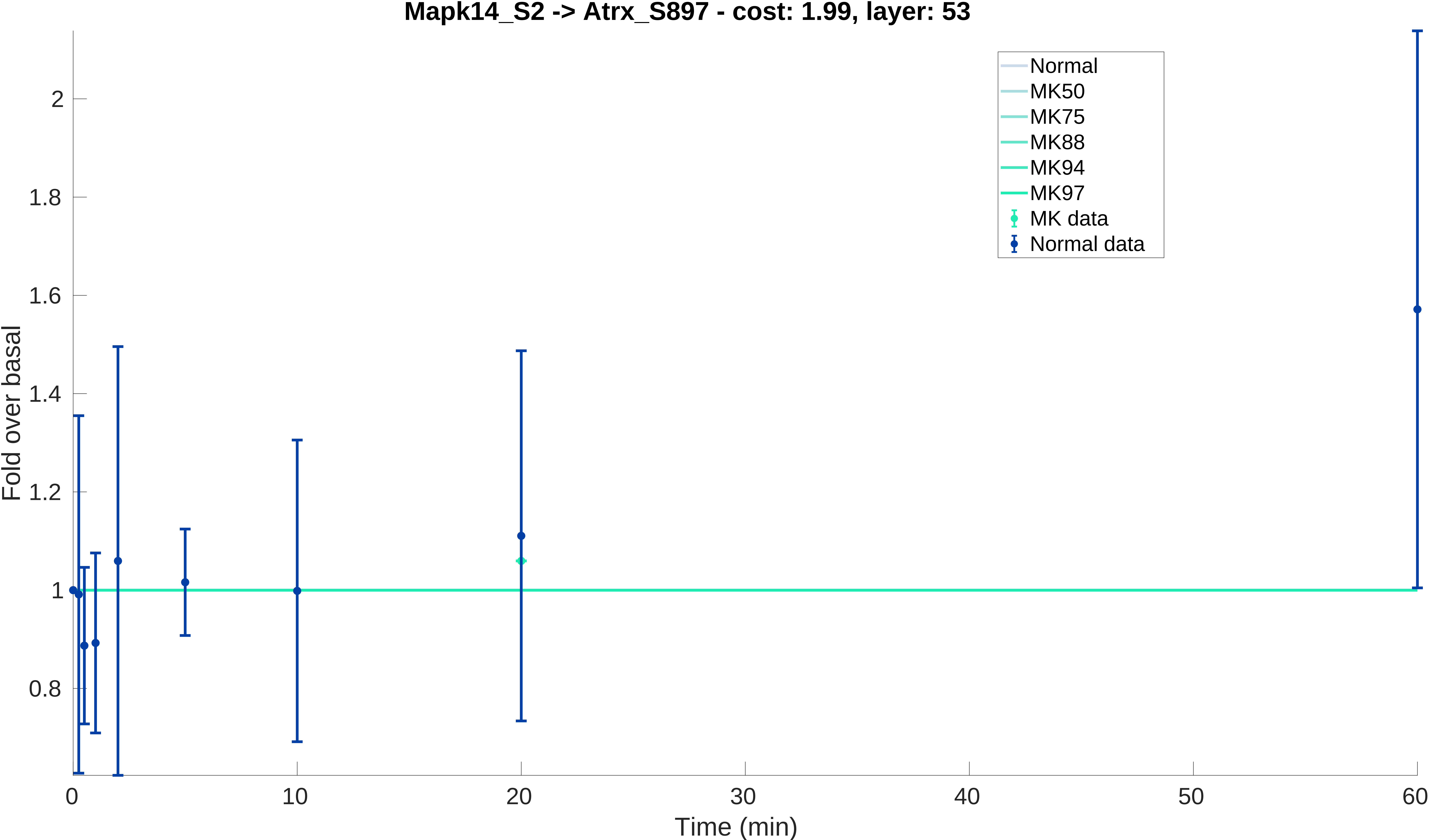

Mapk14_S2 -> Atrx_S897 - cost: 1.99, layer: 53
Fold over basal
2
1.8
1.6
1.4
1.2
1
0.8
Time (min)
0
10
20
30
40
50
60
Normal
MK50
MK75
MK88
MK94
MK97
MK data
Normal data

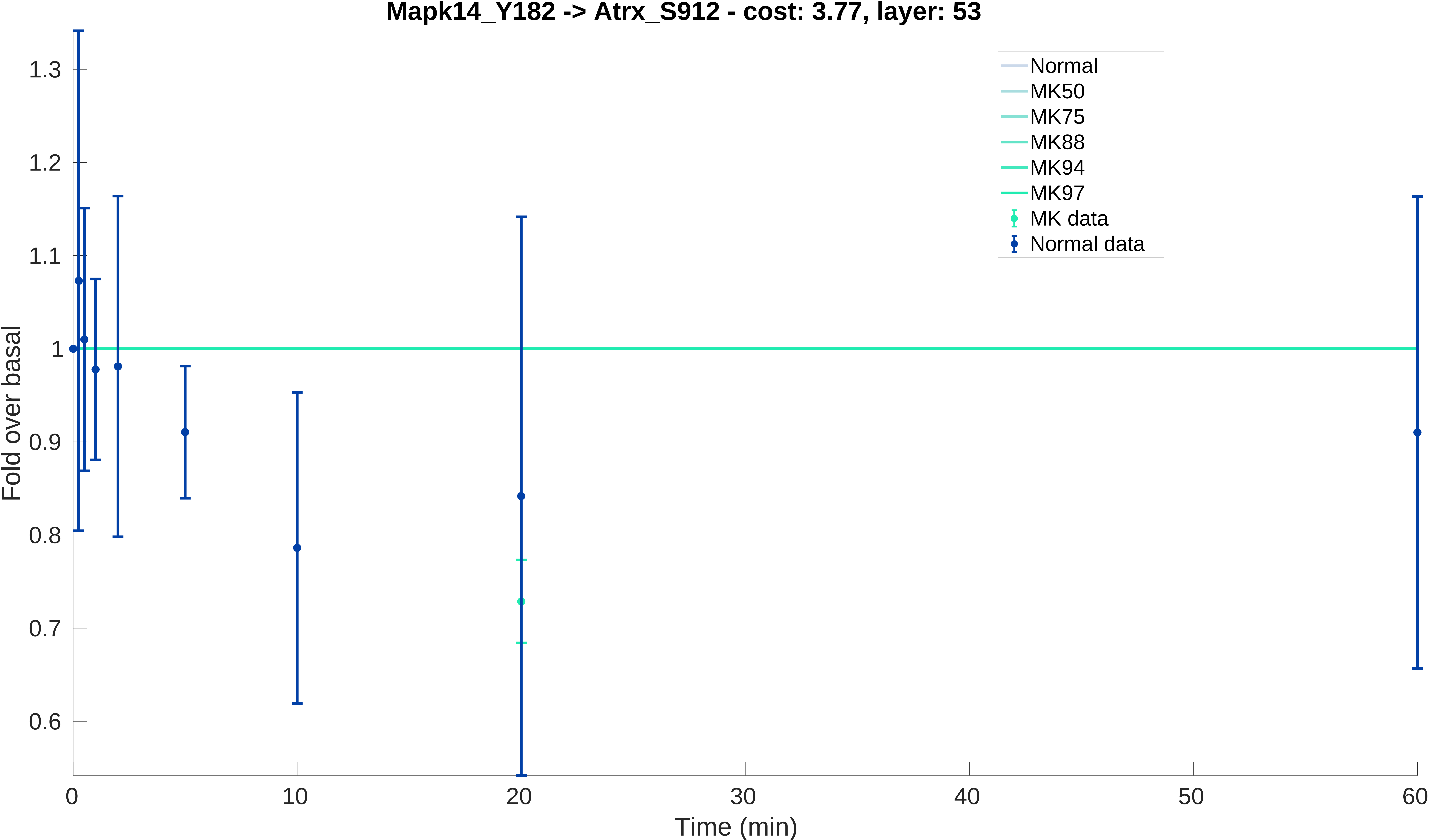
Mapk14_Y182 -> Atrx_S912 - cost: 3.77, layer: 53
Fold over basal
Time (min)
Normal
MK50
MK75
MK88
MK94
MK97
MK data
Normal data
0
10
20
30
40
50
60
0.6
0.7
0.8
0.9
1
1.1
1.2
1.3

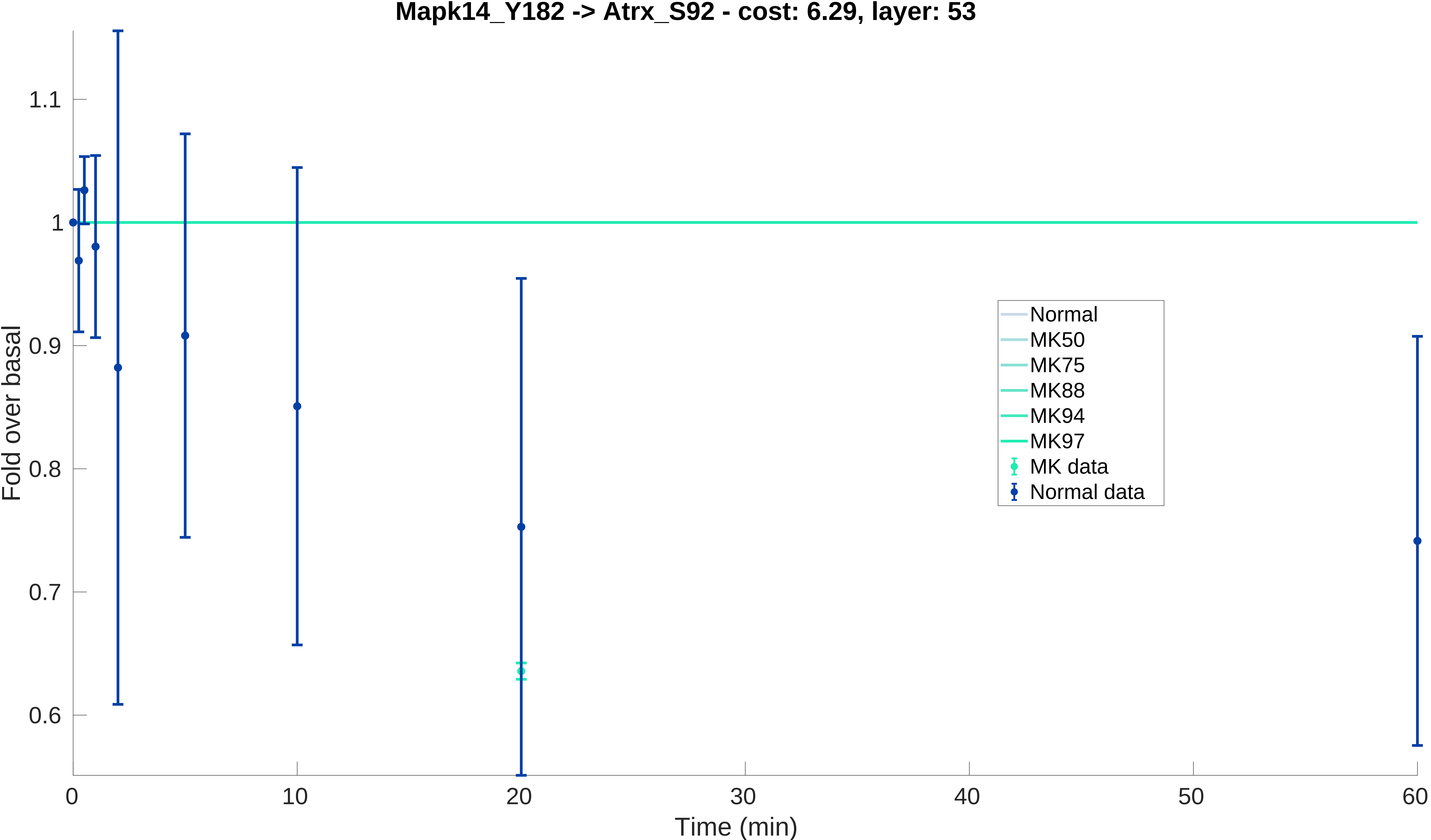

Mapk14_Y182 -> Atrx_S92 - cost: 6.29, layer: 53
Fold over basal
1.1
1
0.9
0.8
0.7
0.6
0
10
20
30
40
50
60
Time (min)
Normal
MK50
MK75
MK88
MK94
MK97
MK data
Normal data

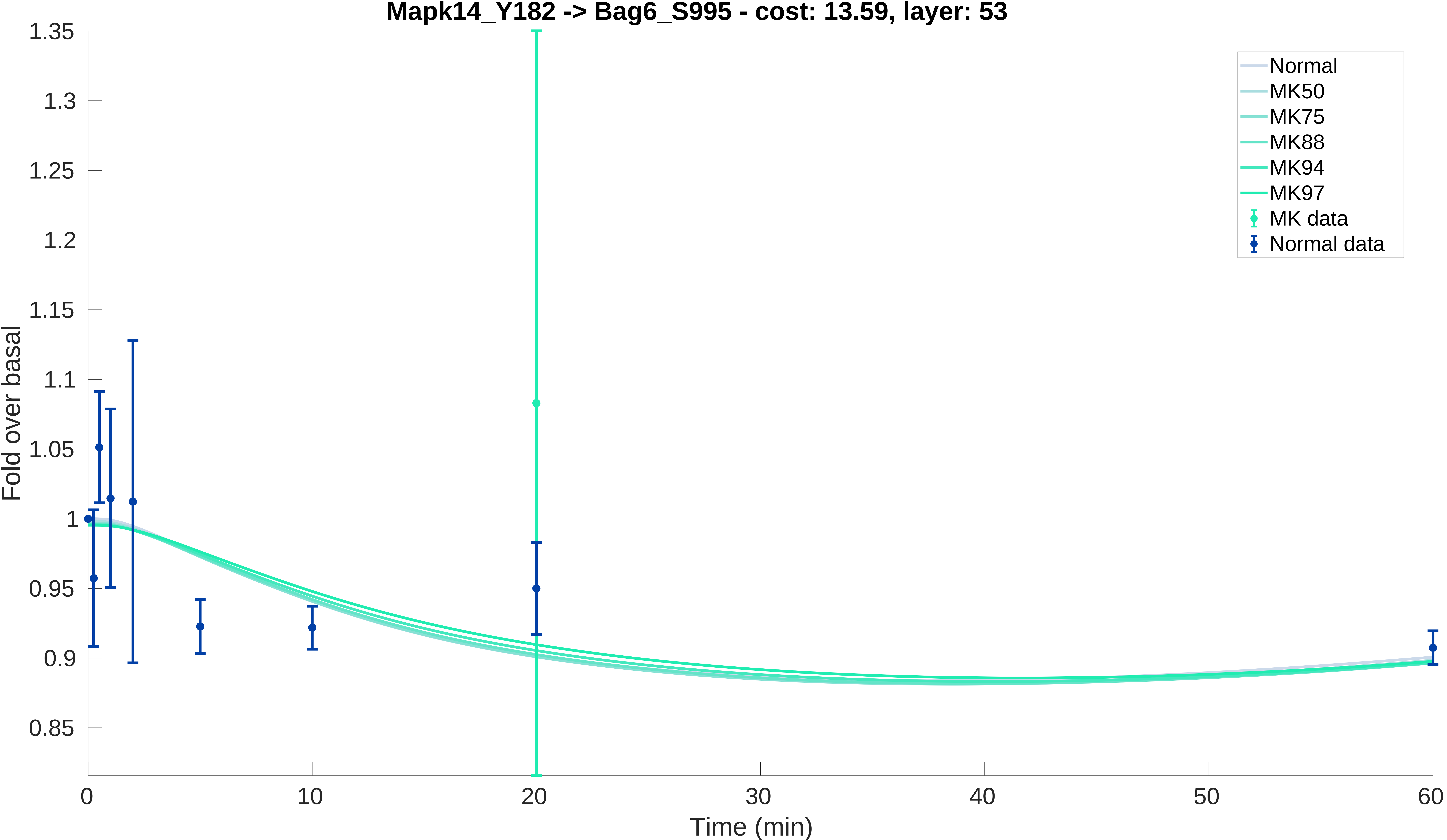

Mapk14_Y182 -> Bag6_S995 - cost: 13.59, layer: 53
1.35
1.3
1.25
1.2
1.15
1.1
1.05
1
0.95
0.9
0.85
Fold over basal
0
10
20
30
40
50
60
Time (min)
Normal
MK50
MK75
MK88
MK94
MK97
MK data
Normal data

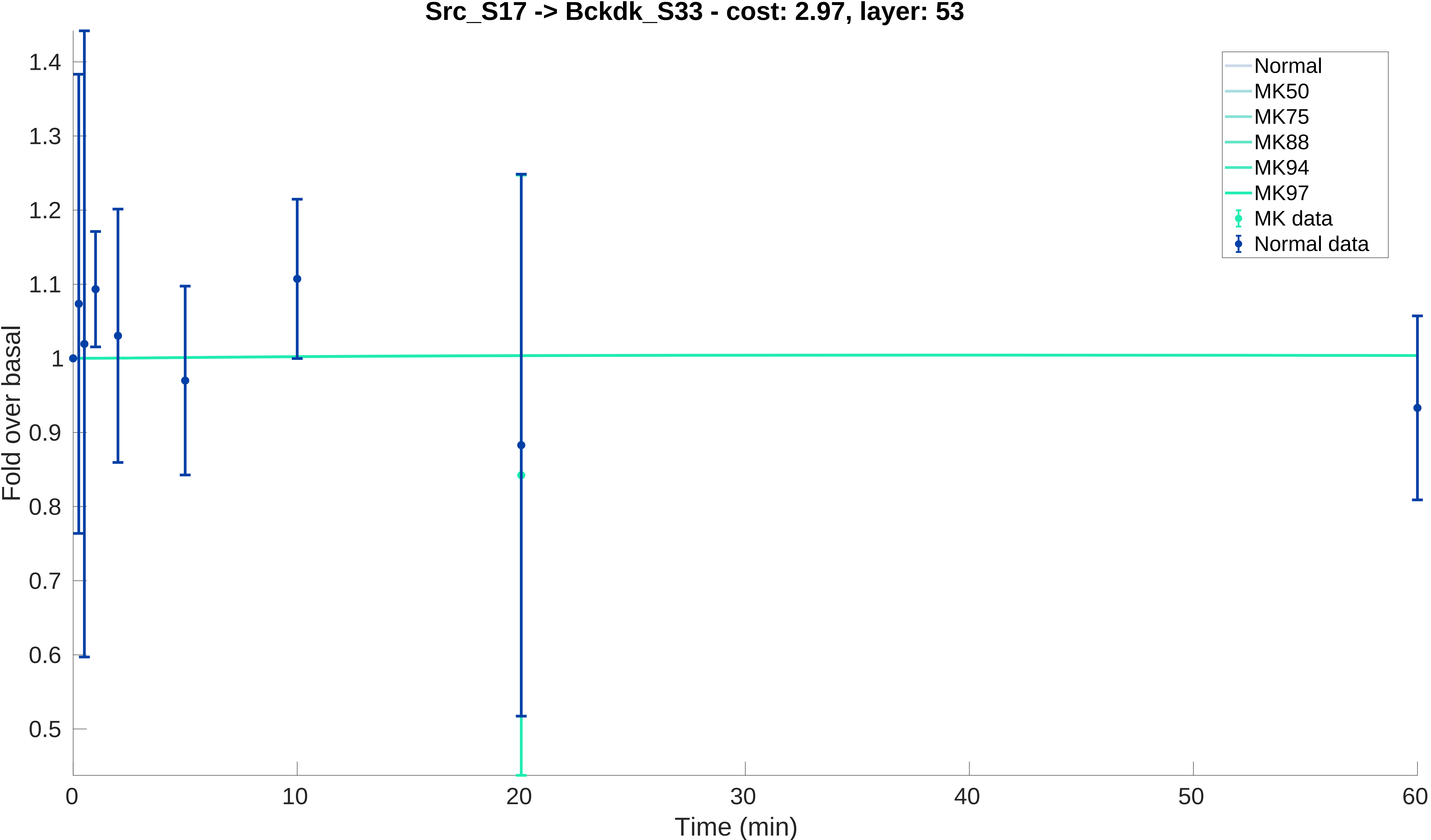

Src_S17 -> Bckdk_S33 - cost: 2.97, layer: 53
Normal
MK50
MK75
MK88
MK94
MK97
MK data
Normal data
Fold over basal
Time (min)
0
10
20
30
40
50
60
0.5
0.6
0.7
0.8
0.9
1
1.1
1.2
1.3
1.4

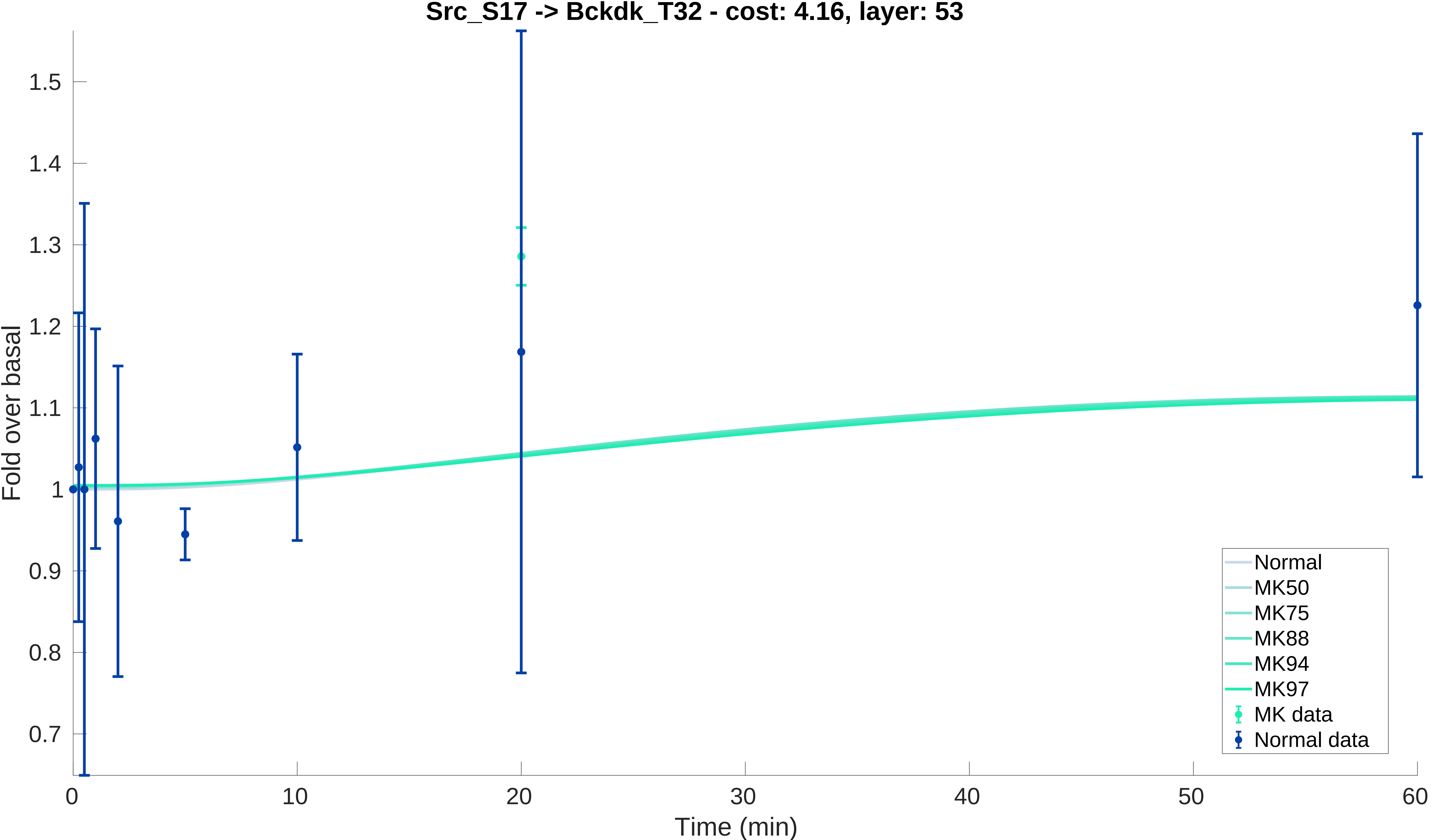
Src_S17 -> Bckdk_T32 - cost: 4.16, layer: 53
Fold over basal
Time (min)
1.5
1.4
1.3
1.2
1.1
1
0.9
0.8
0.7
0
10
20
30
40
50
60
Normal
MK50
MK75
MK88
MK94
MK97
MK data
Normal data

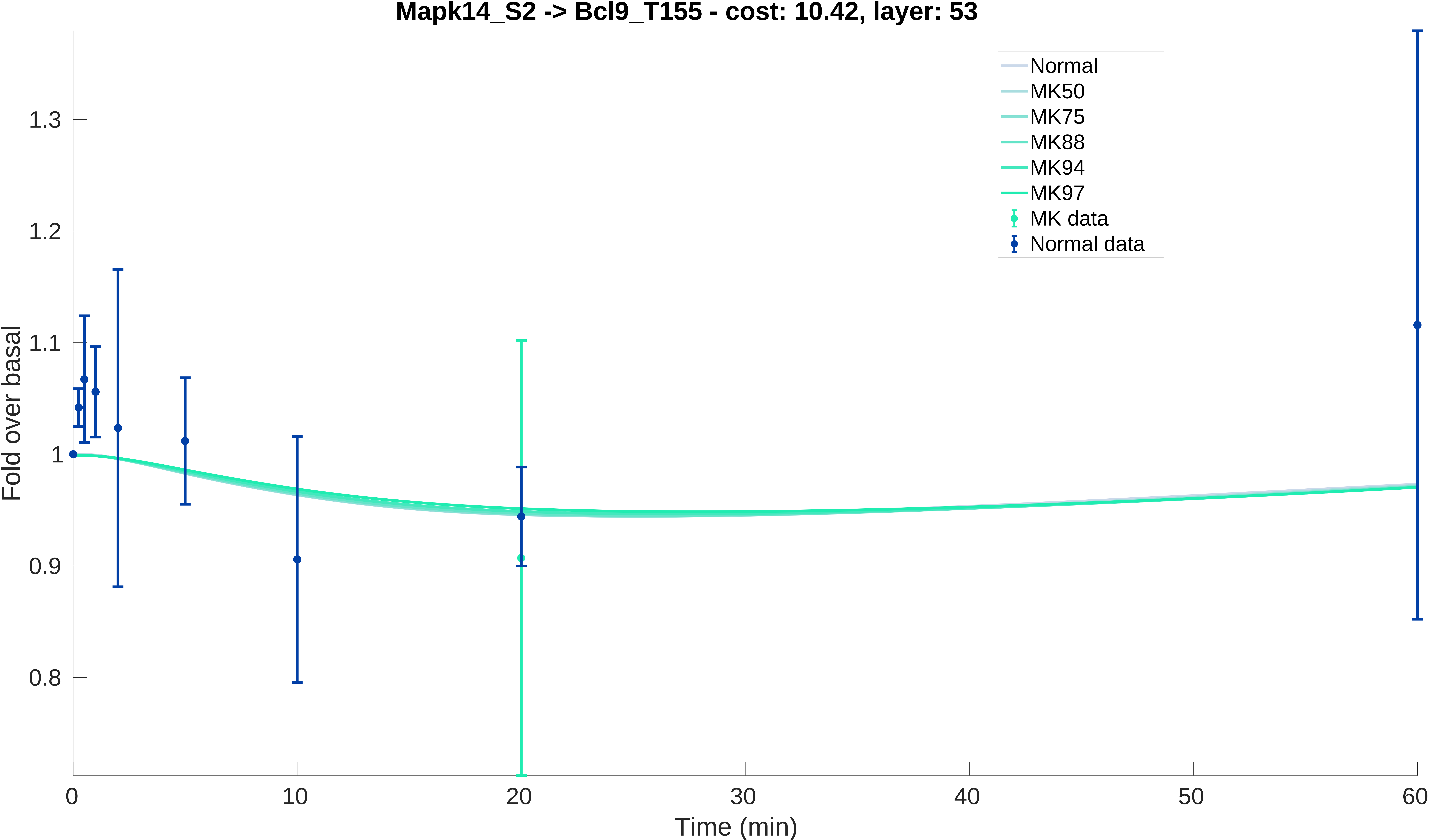
Mapk14_S2 -> Bcl9_T155 - cost: 10.42, layer: 53
Fold over basal
1.3
1.2
1.1
1
0.9
0.8
0
10
20
30
40
50
60
Time (min)
Normal
MK50
MK75
MK88
MK94
MK97
MK data
Normal data

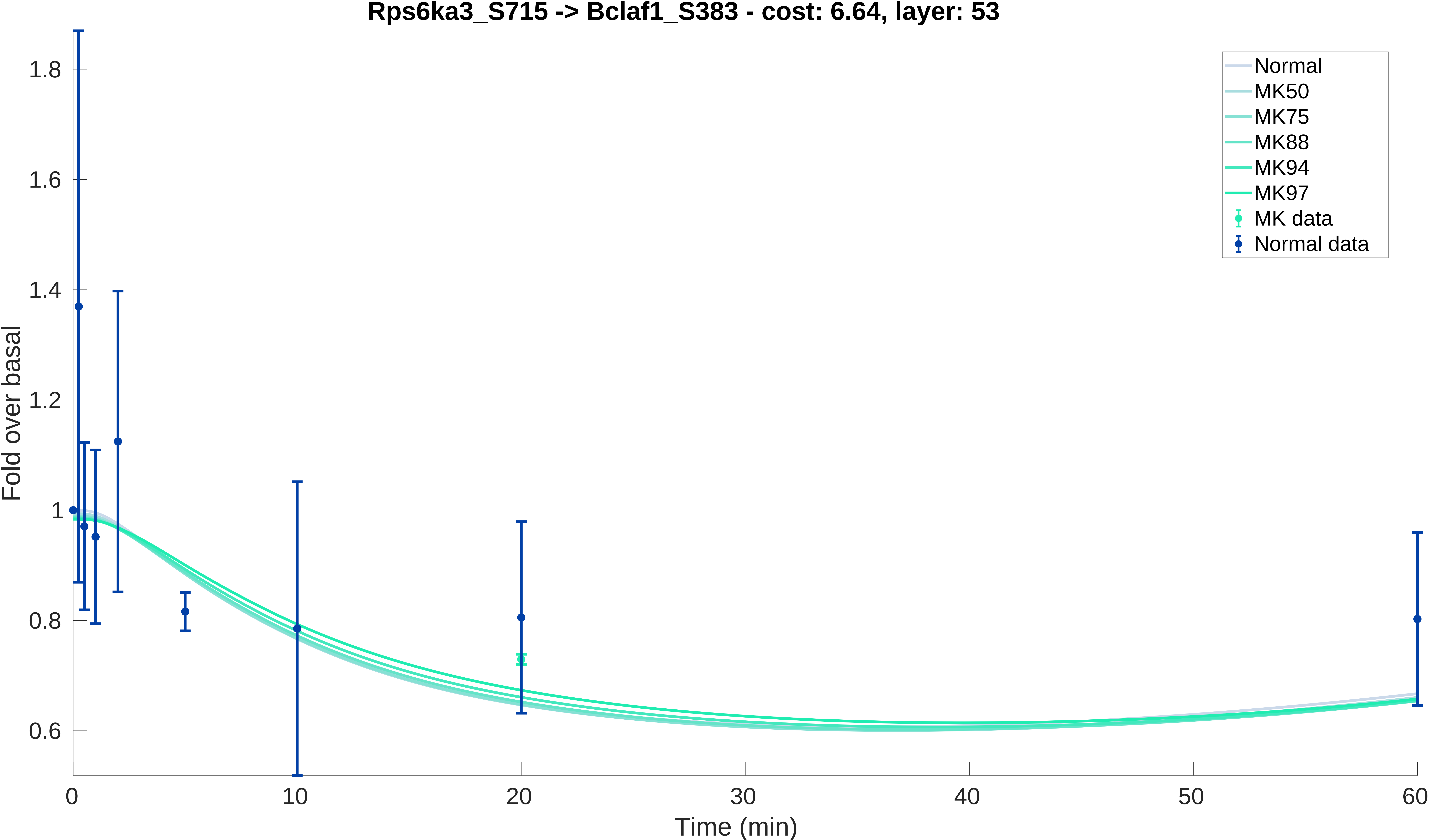

Rps6ka3_S715 -> Bclaf1_S383 - cost: 6.64, layer: 53
Fold over basal
Time (min)
0.6
0.8
1
1.2
1.4
1.6
1.8
0
10
20
30
40
50
60
Normal
MK50
MK75
MK88
MK94
MK97
MK data
Normal data

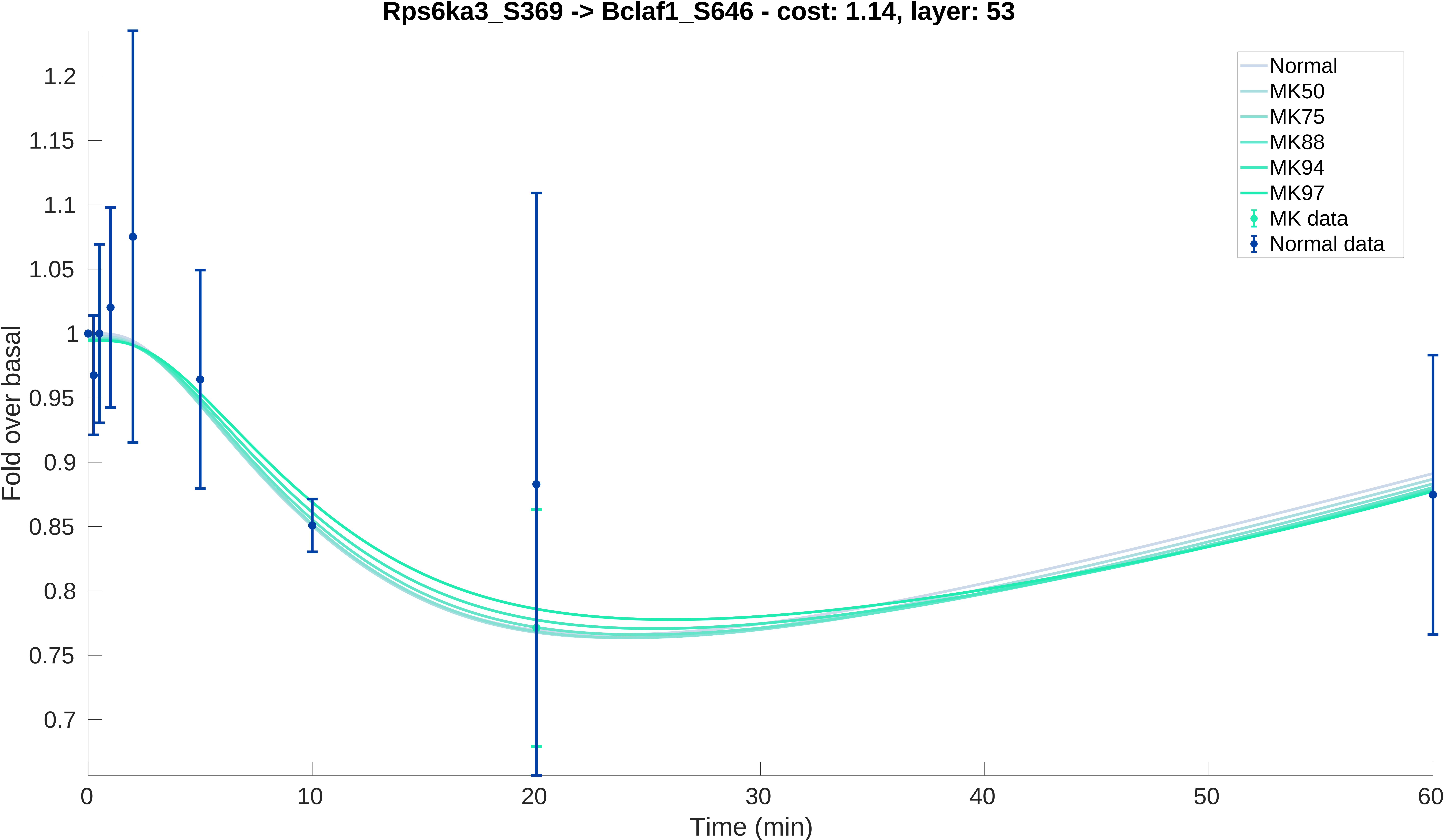

Rps6ka3_S369 -> Bclaf1_S646 - cost: 1.14, layer: 53
Fold over basal
Time (min)
Normal
MK50
MK75
MK88
MK94
MK97
MK data
Normal data
1.2
1.15
1.1
1.05
1
0.95
0.9
0.85
0.8
0.75
0.7
0
10
20
30
40
50
60

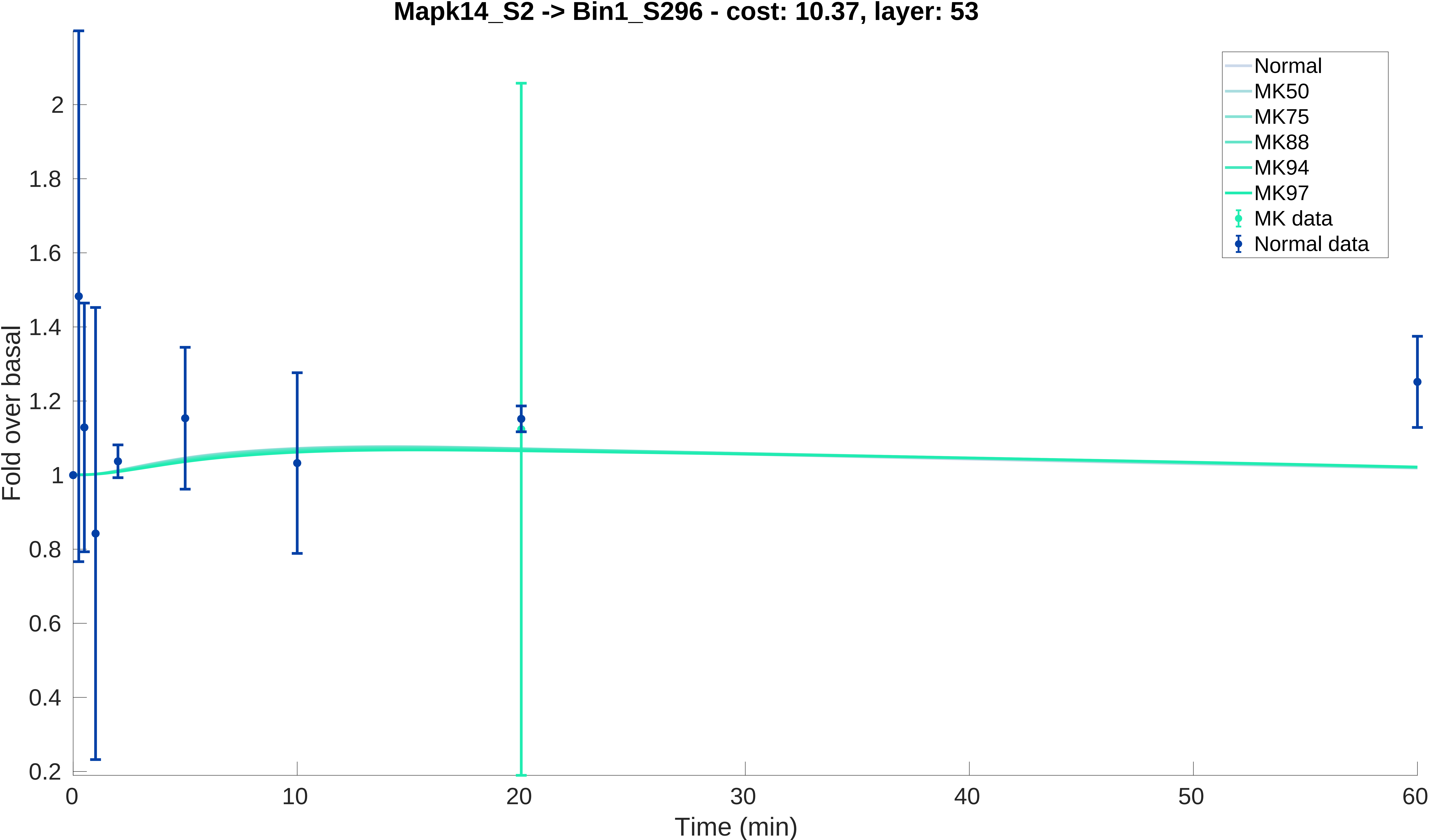

Mapk14_S2 -> Bin1_S296 - cost: 10.37, layer: 53
Normal
MK50
MK75
MK88
MK94
MK97
MK data
Normal data
Fold over basal
Time (min)
0.2
0.4
0.6
0.8
1
1.2
1.4
1.6
1.8
2
0
10
20
30
40
50
60

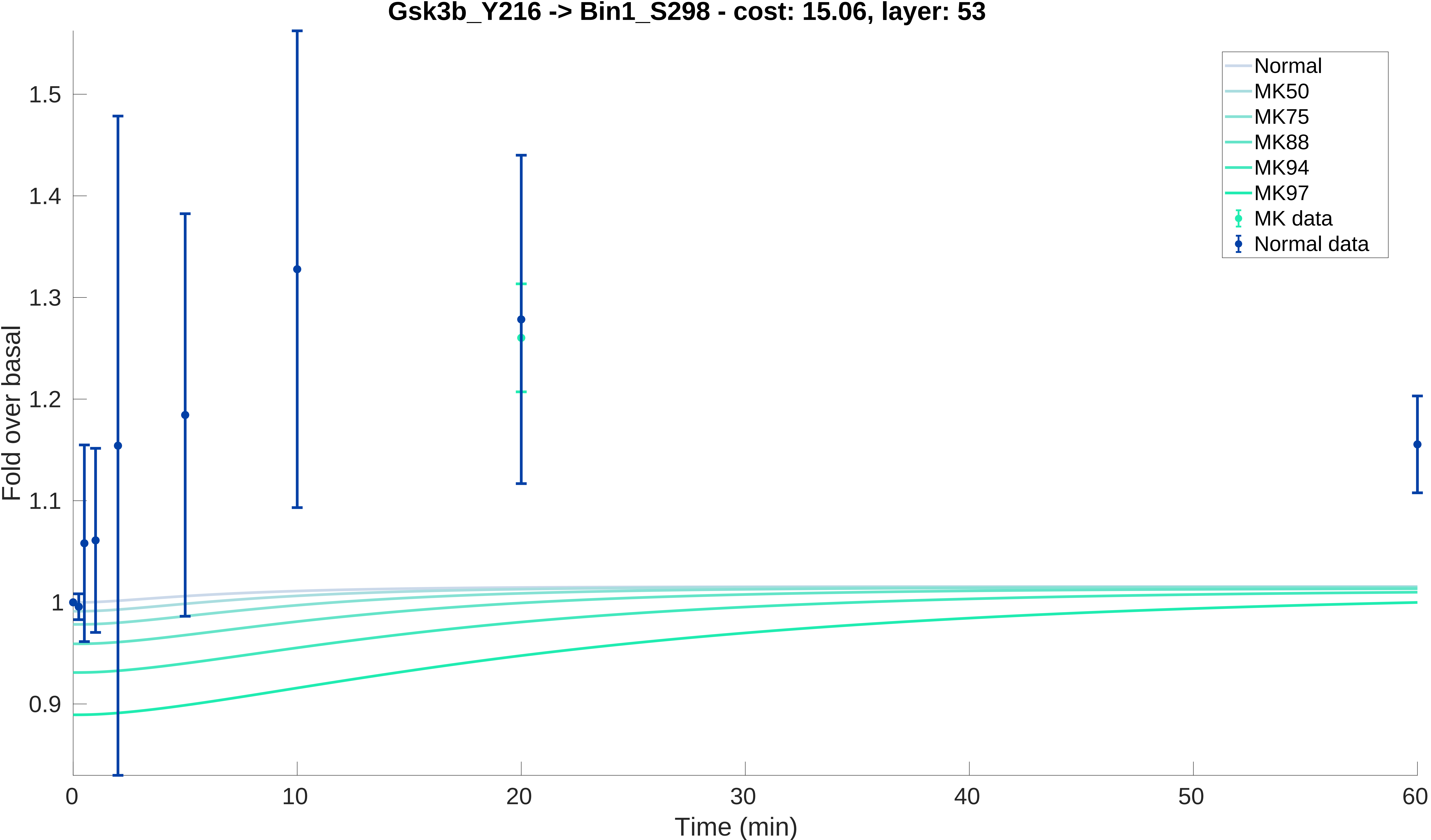

Gsk3b_Y216 -> Bin1_S298 - cost: 15.06, layer: 53
Normal
MK50
MK75
MK88
MK94
MK97
MK data
Normal data
Fold over basal
Time (min)
0.9
1
1.1
1.2
1.3
1.4
1.5
0
10
20
30
40
50
60

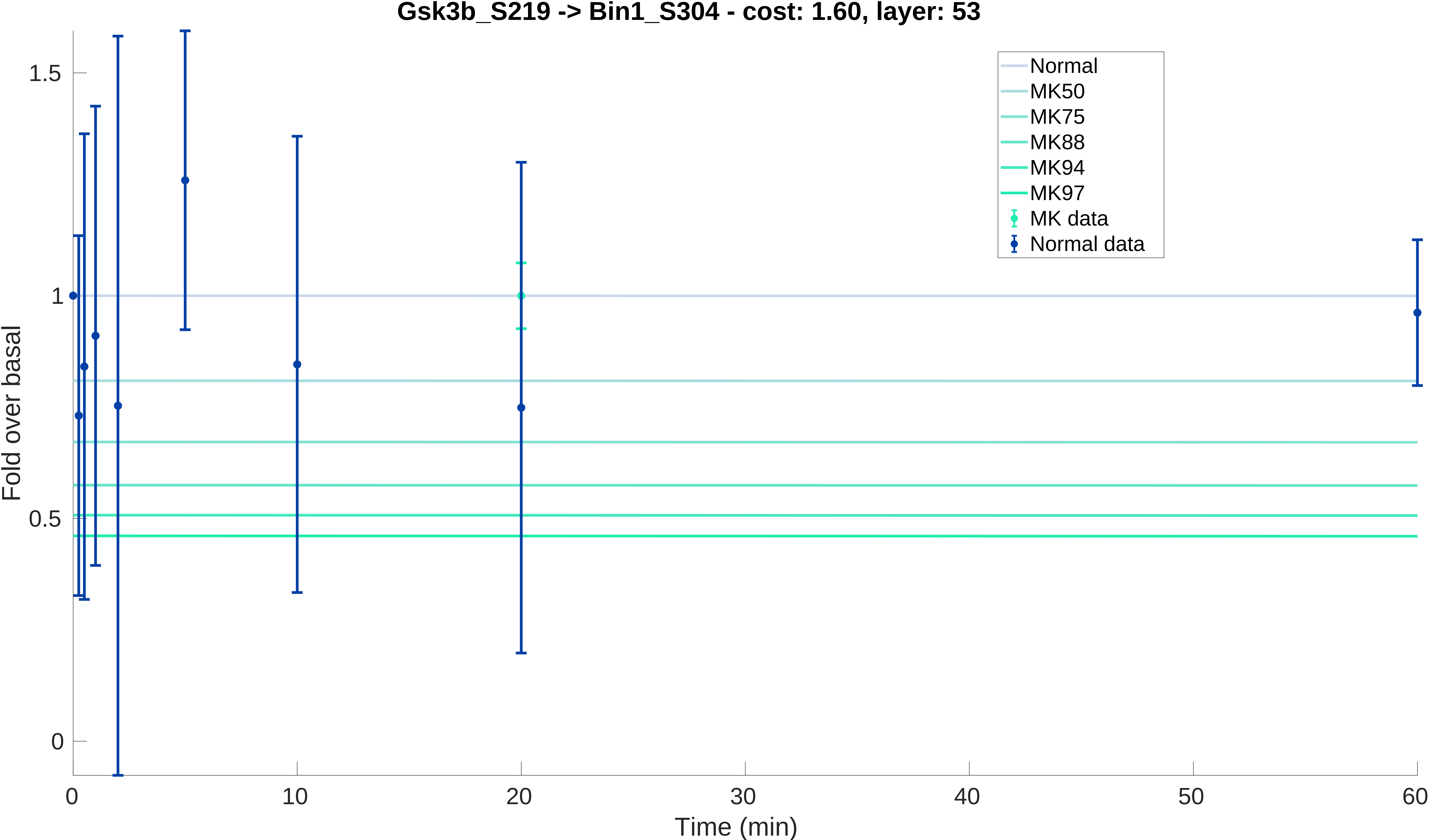

Gsk3b_S219 -> Bin1_S304 - cost: 1.60, layer: 53
Fold over basal
Time (min)
0
0.5
1
1.5
0
10
20
30
40
50
60
Normal
MK50
MK75
MK88
MK94
MK97
MK data
Normal data

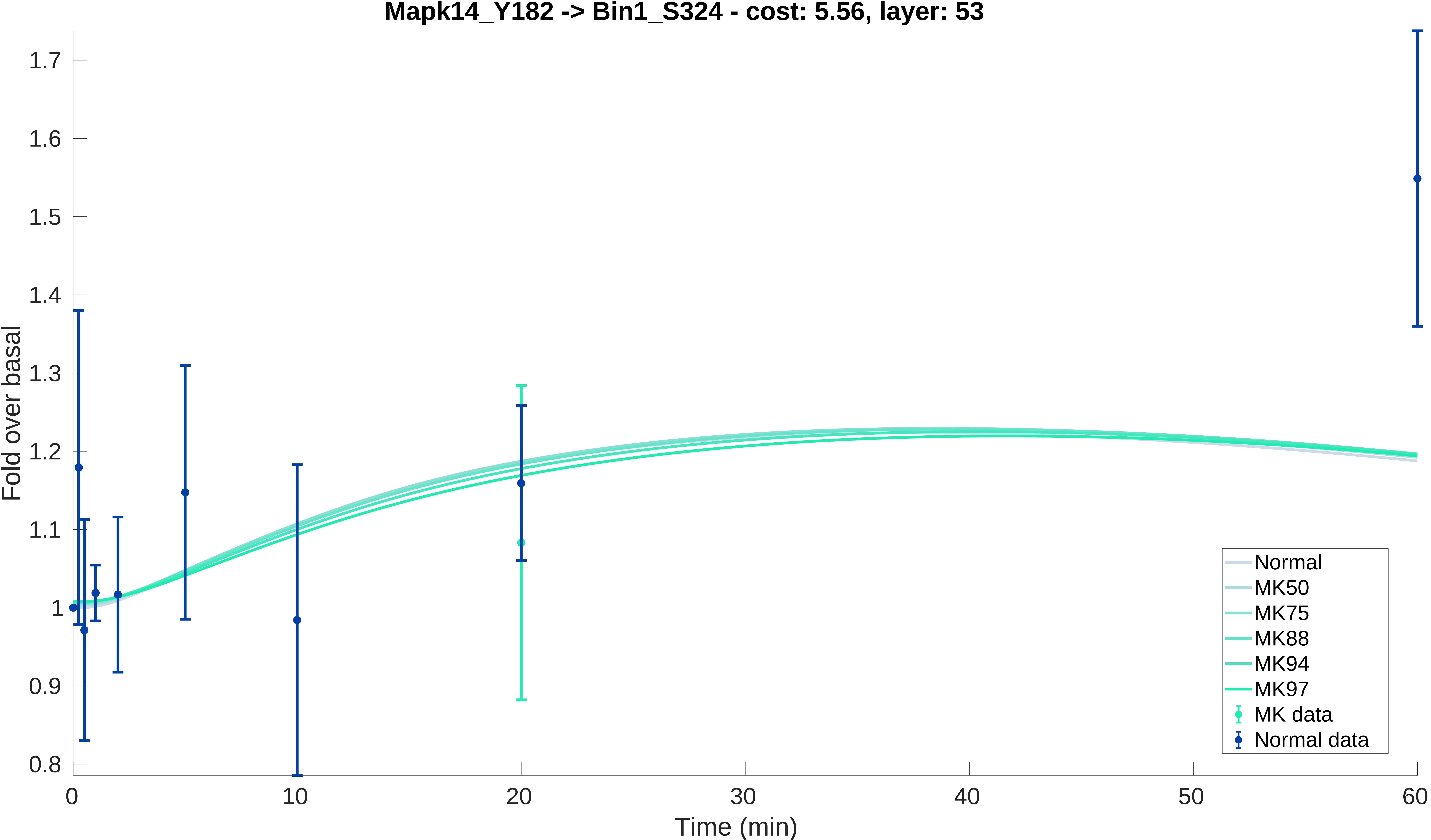

Mapk14_Y182 -> Bin1_S324 - cost: 5.56, layer: 53
Fold over basal
1.7
1.6
1.5
1.4
1.3
1.2
1.1
1
0.9
0.8
0
10
20
30
40
50
60
Time (min)
Normal
MK50
MK75
MK88
MK94
MK97
MK data
Normal data

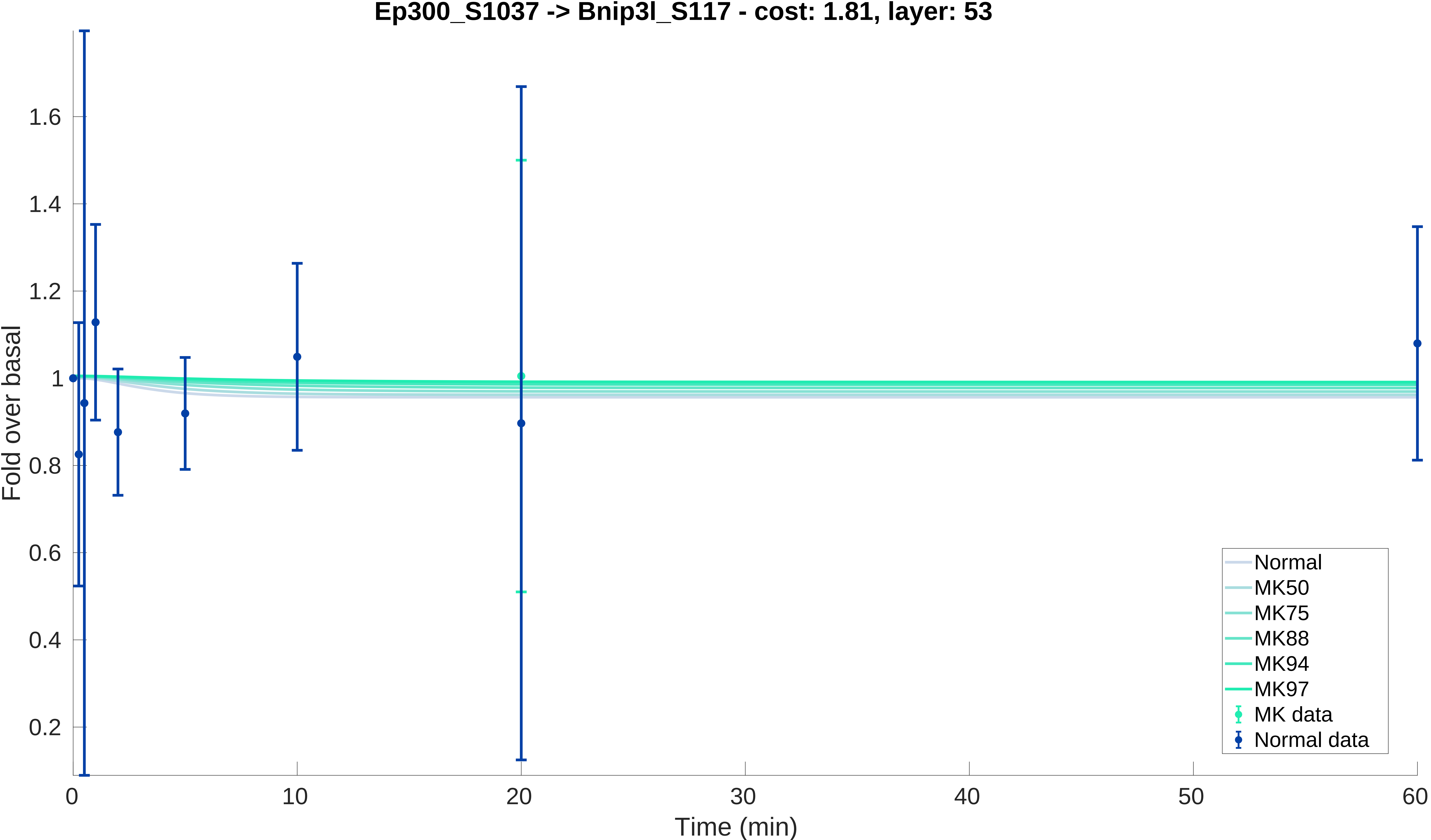

Ep300_S1037 -> Bnip3l_S117 - cost: 1.81, layer: 53
Fold over basal
Time (min)
1.6
1.4
1.2
1
0.8
0.6
0.4
0.2
0
10
20
30
40
50
60
Normal
MK50
MK75
MK88
MK94
MK97
MK data
Normal data

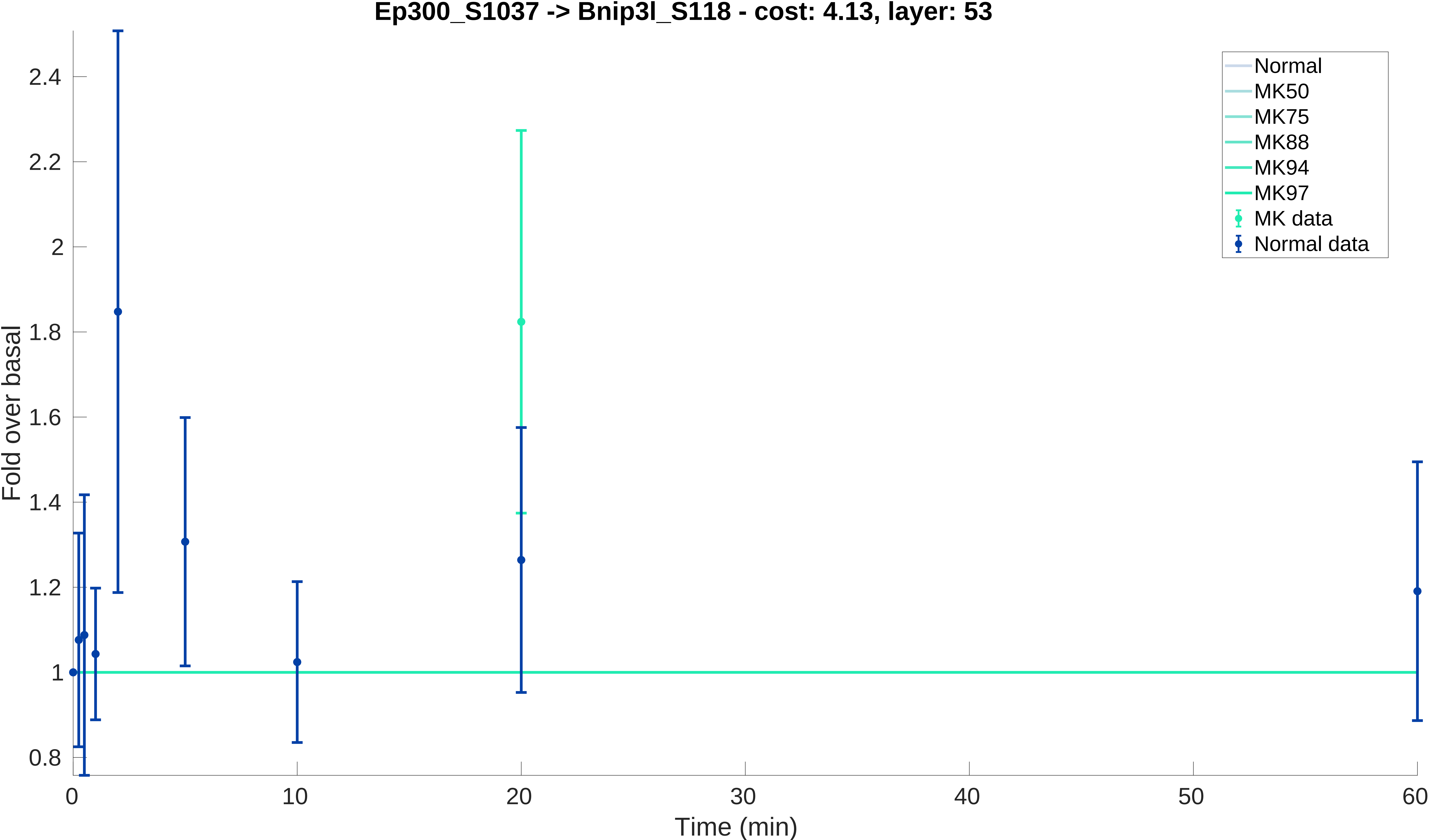

Ep300_S1037 -> Bnip3l_S118 - cost: 4.13, layer: 53
Fold over basal
2.4
2.2
2
1.8
1.6
1.4
1.2
1
0.8
0
10
20
30
40
50
60
Time (min)
Normal
MK50
MK75
MK88
MK94
MK97
MK data
Normal data

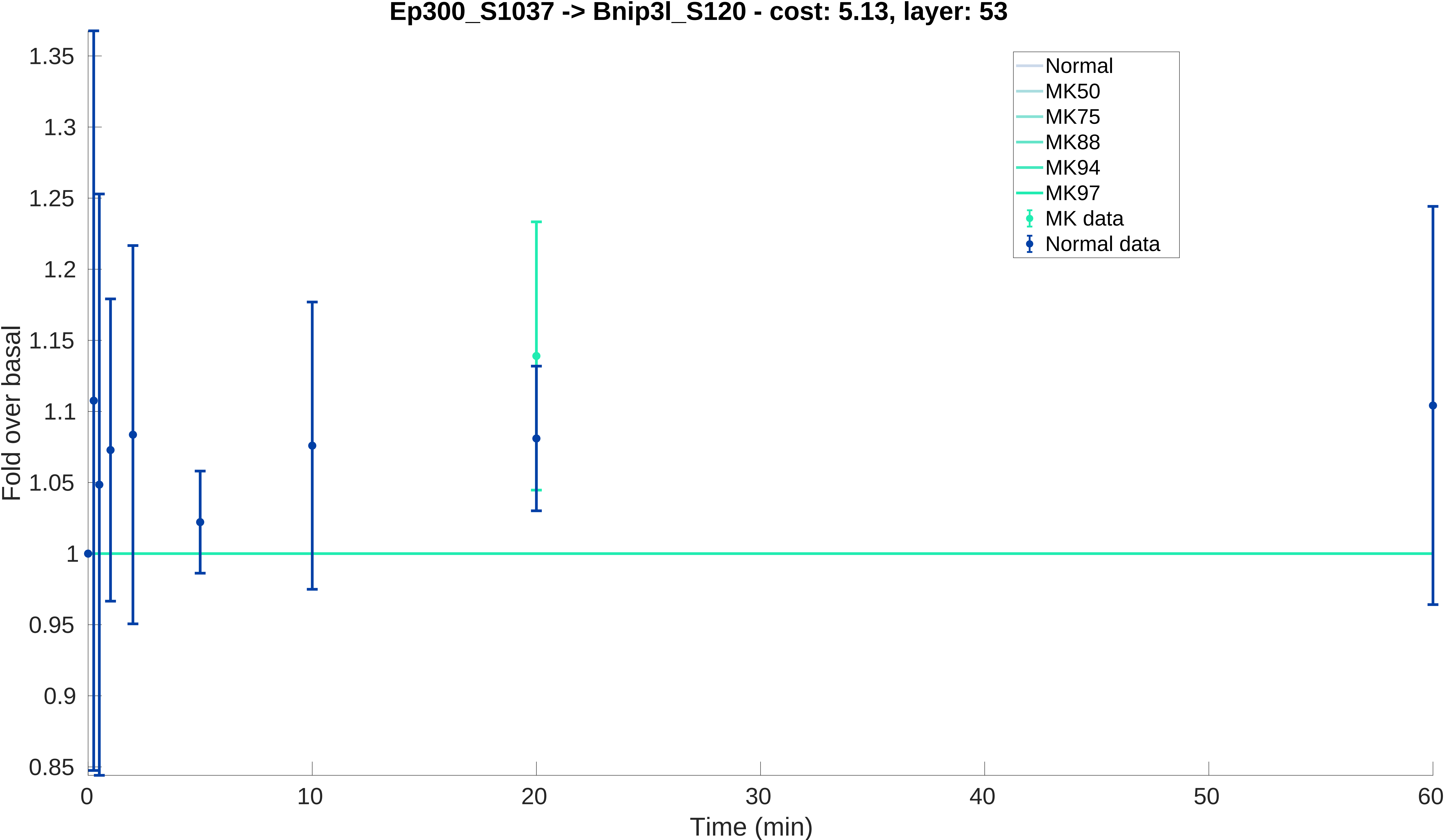

Ep300_S1037 -> Bnip3l_S120 - cost: 5.13, layer: 53
Fold over basal
Time (min)
1.35
1.3
1.25
1.2
1.15
1.1
1.05
1
0.95
0.9
0.85
0
10
20
30
40
50
60
Normal
MK50
MK75
MK88
MK94
MK97
MK data
Normal data

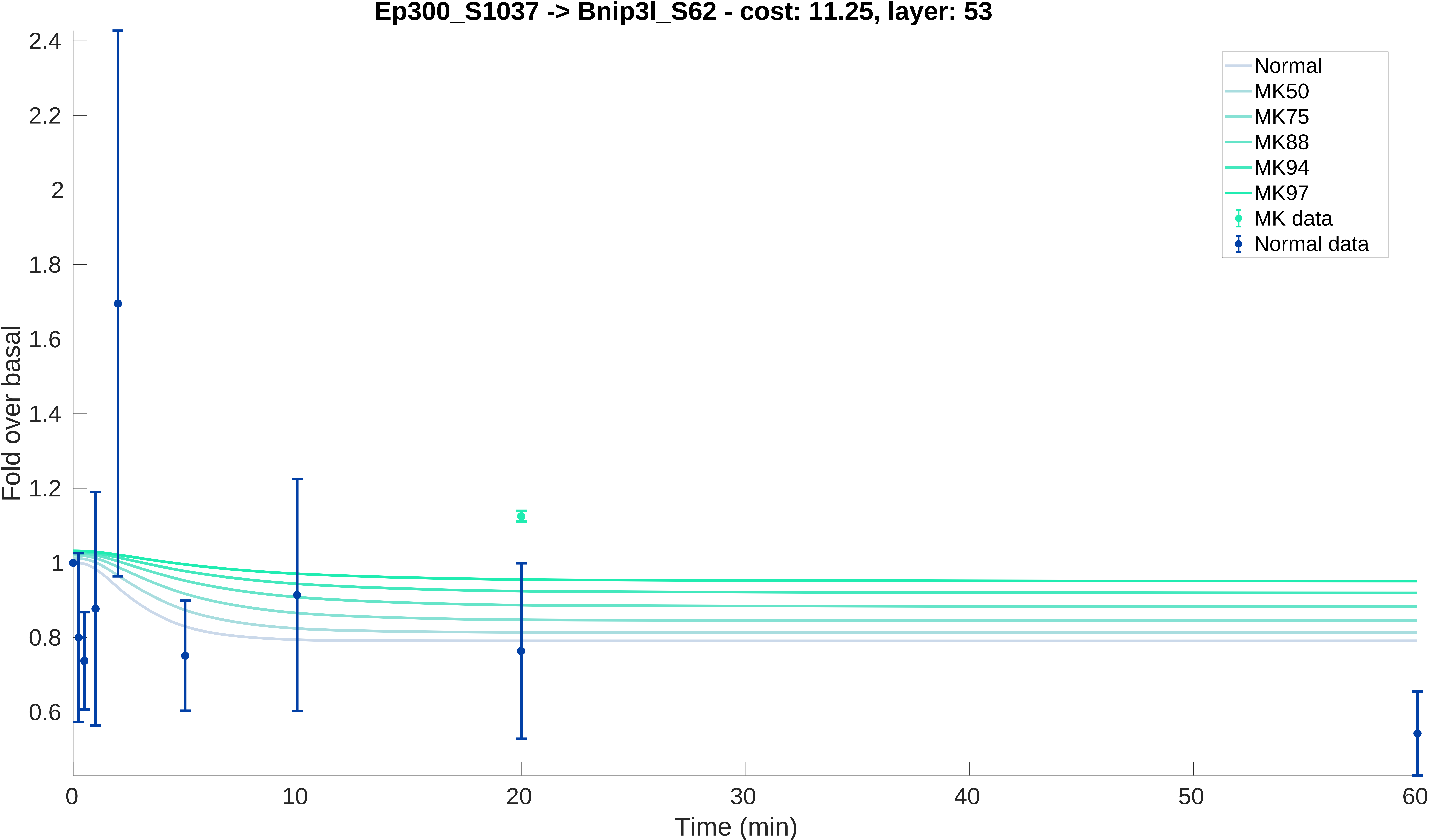
Ep300_S1037 -> Bnip3l_S62 - cost: 11.25, layer: 53
Fold over basal
Time (min)
2.4
2.2
2
1.8
1.6
1.4
1.2
1
0.8
0.6
0
10
20
30
40
50
60
Normal
MK50
MK75
MK88
MK94
MK97
MK data
Normal data

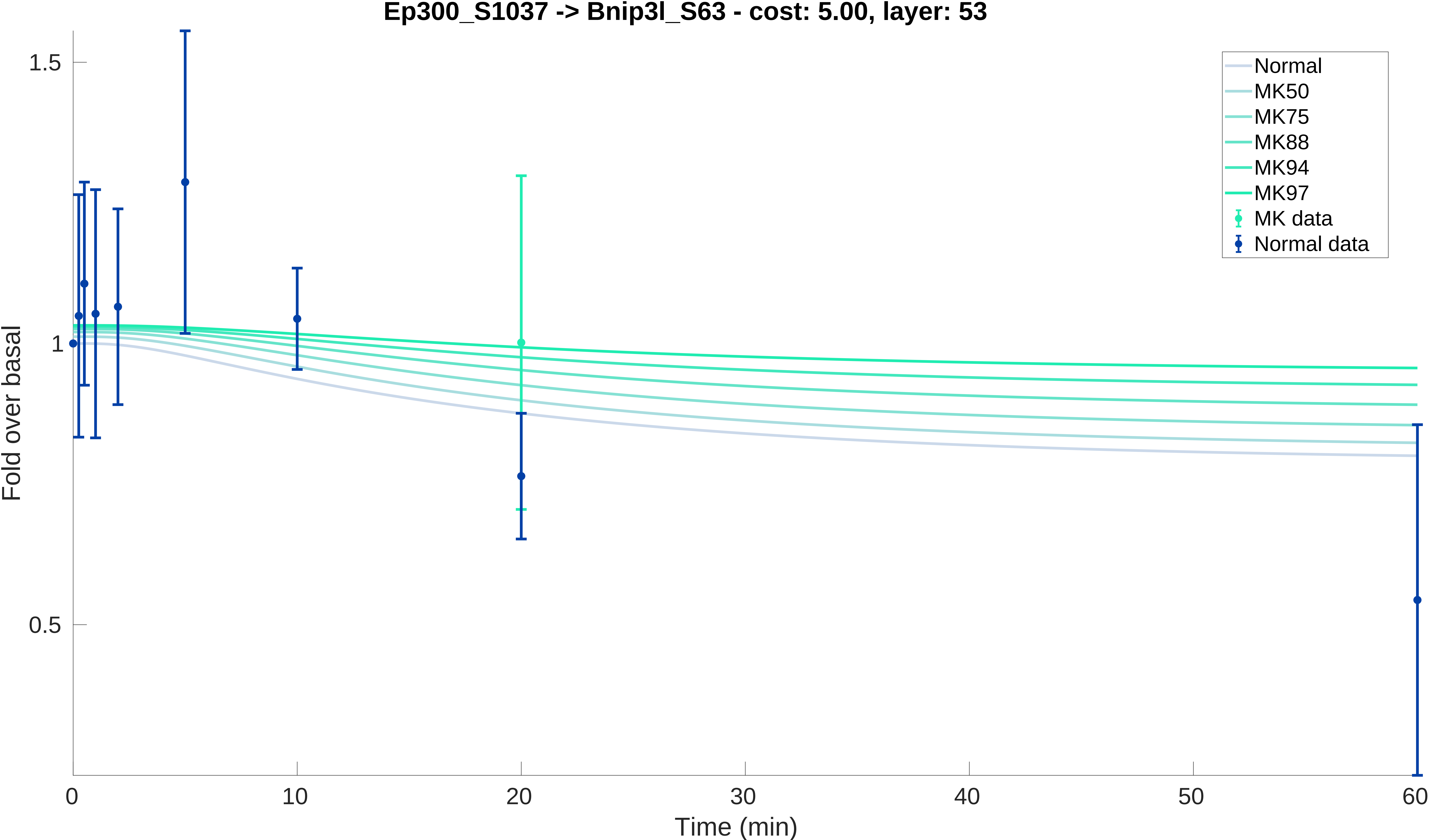

Ep300_S1037 -> Bnip3l_S63 - cost: 5.00, layer: 53
Normal
MK50
MK75
MK88
MK94
MK97
MK data
Normal data
Fold over basal
Time (min)
1.5
1
0.5
0
10
20
30
40
50
60

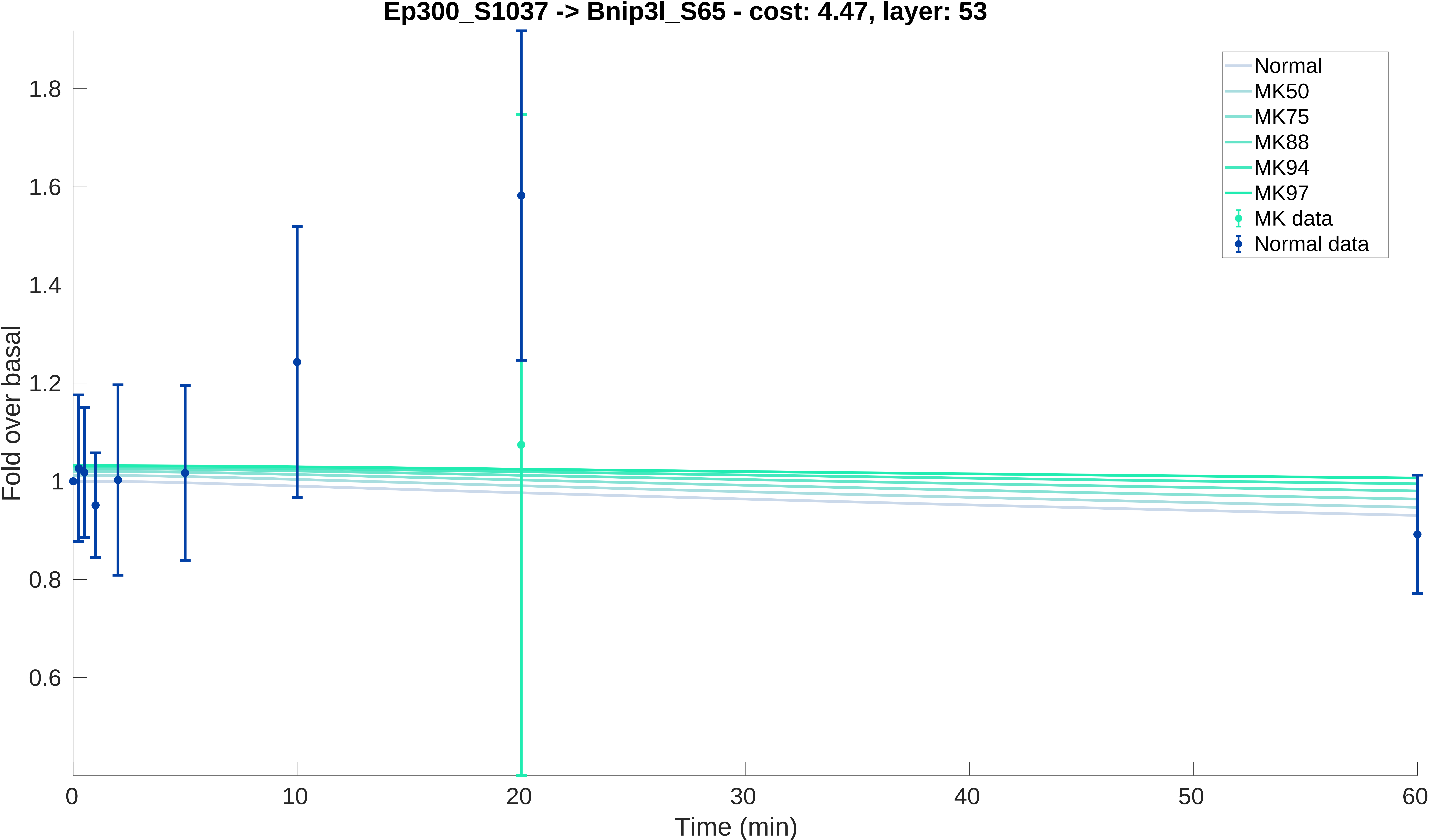

Ep300_S1037 -> Bnip3l_S65 - cost: 4.47, layer: 53
Fold over basal
1.8
1.6
1.4
1.2
1
0.8
0.6
0
10
20
30
40
50
60
Time (min)
Normal
MK50
MK75
MK88
MK94
MK97
MK data
Normal data

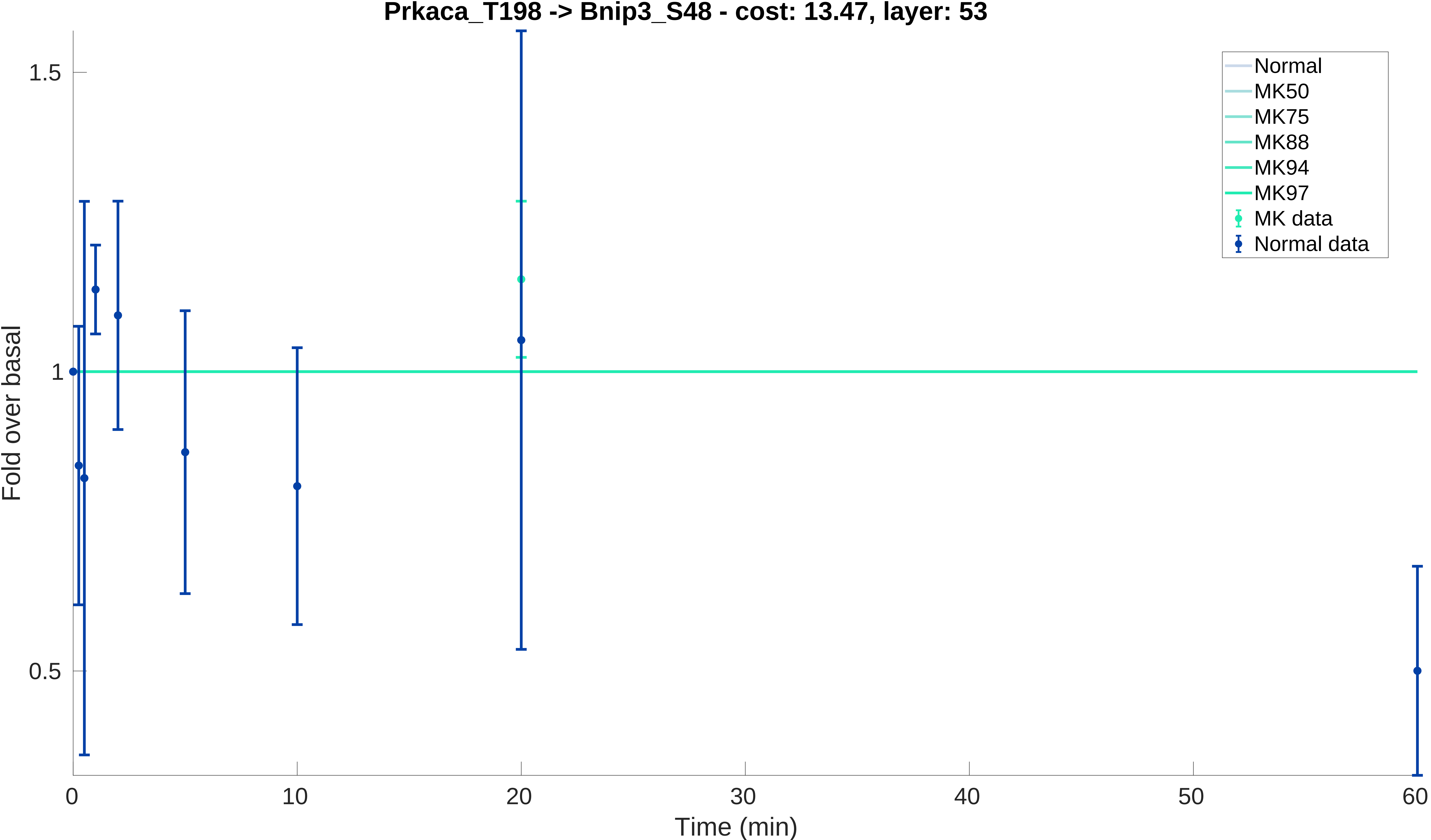

Prkaca_T198 -> Bnip3_S48 - cost: 13.47, layer: 53
Fold over basal
1.5
1
0.5
Time (min)
0
10
20
30
40
50
60
Normal
MK50
MK75
MK88
MK94
MK97
MK data
Normal data

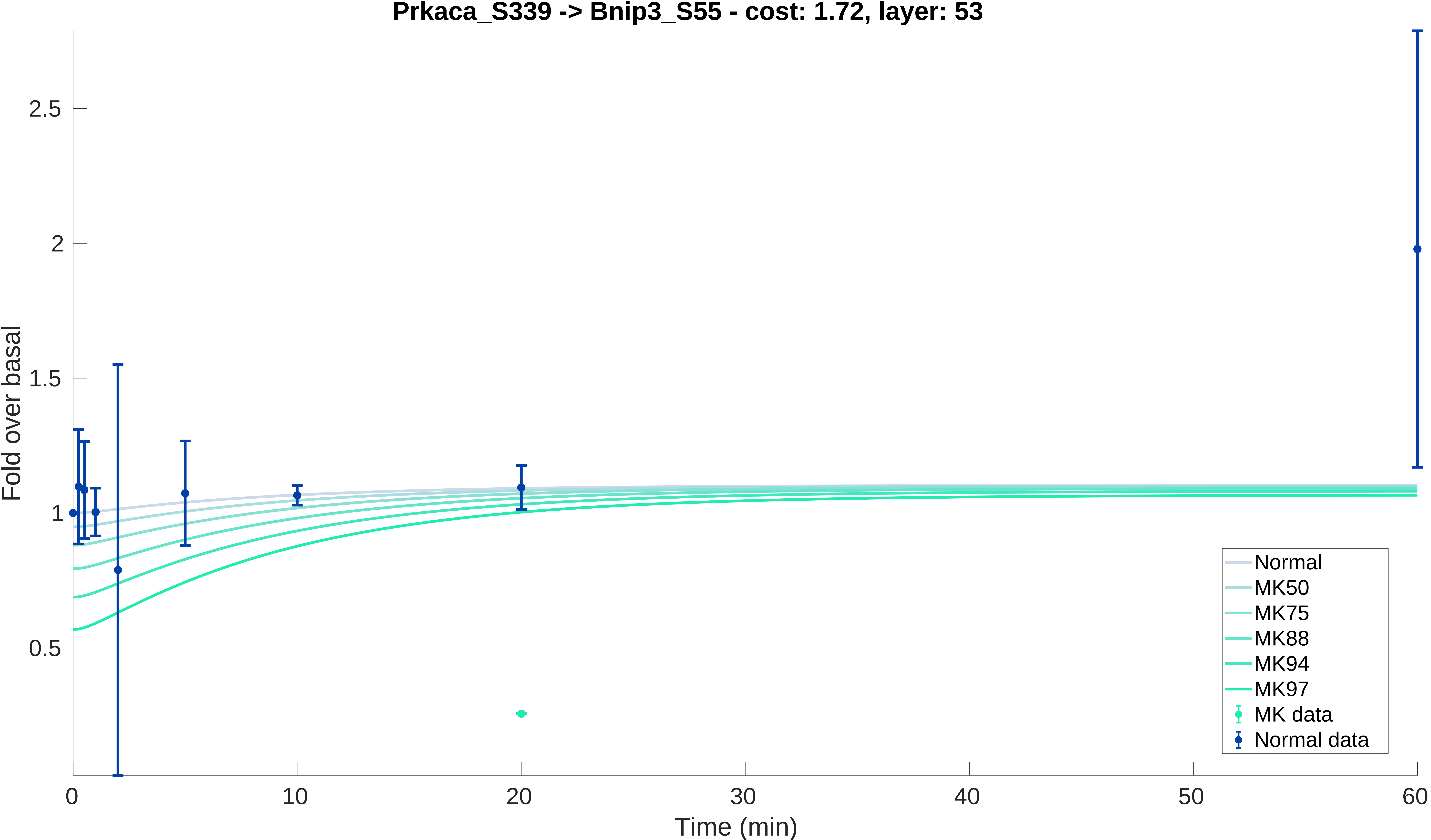
Prrkaca_S339 -> Bnip3_S55 - cost: 1.72, layer: 53
Fold over basal
2.5
2
1.5
1
0.5
0
10
20
30
40
50
60
Time (min)
Normal
MK50
MK75
MK88
MK94
MK97
MK data
Normal data

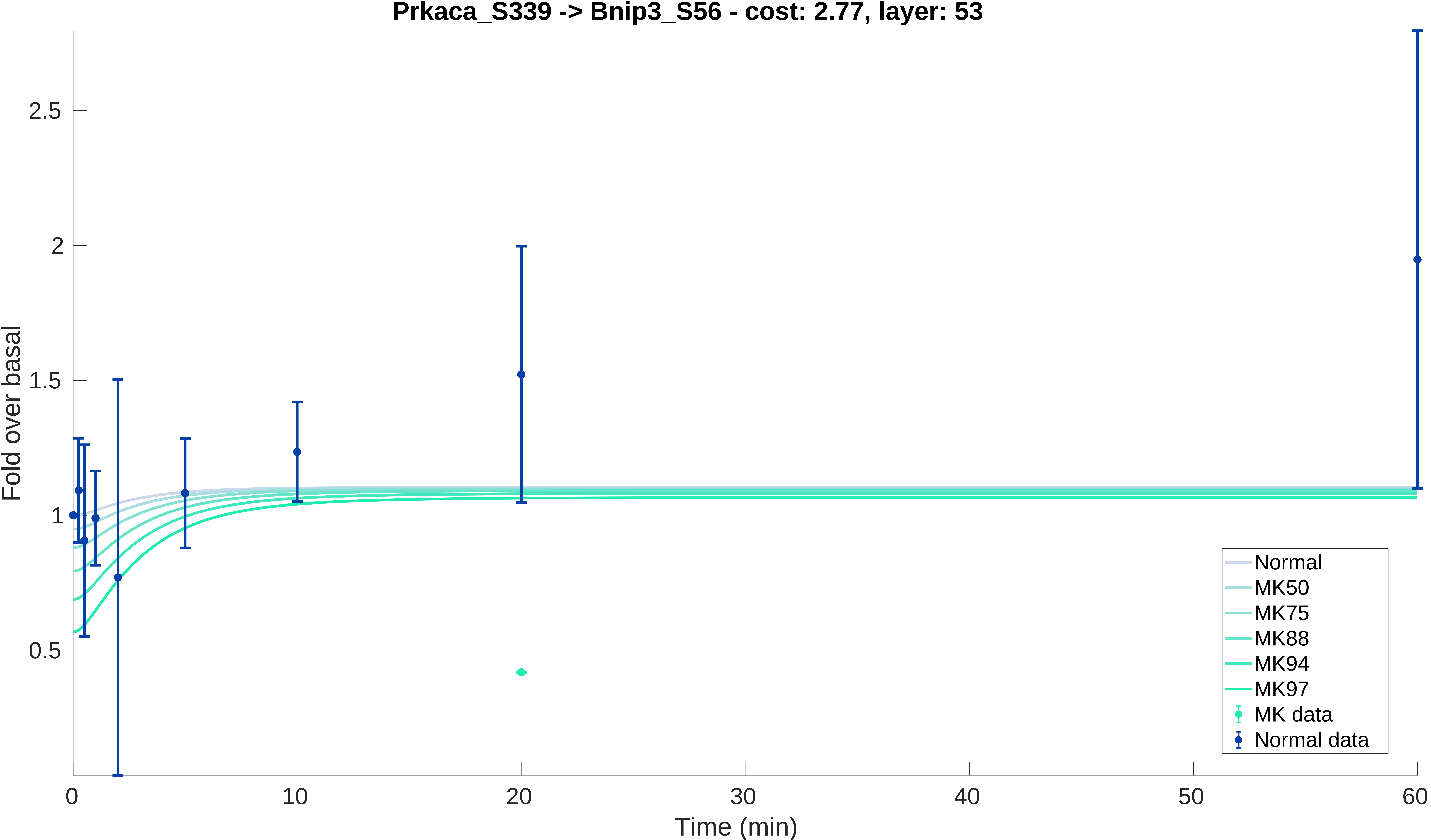

Prkaca_S339 -> Bnip3_S56 - cost: 2.77, layer: 53
Fold over basal
Time (min)
0.5
1
1.5
2
2.5
0
10
20
30
40
50
60
Normal
MK50
MK75
MK88
MK94
MK97
MK data
Normal data

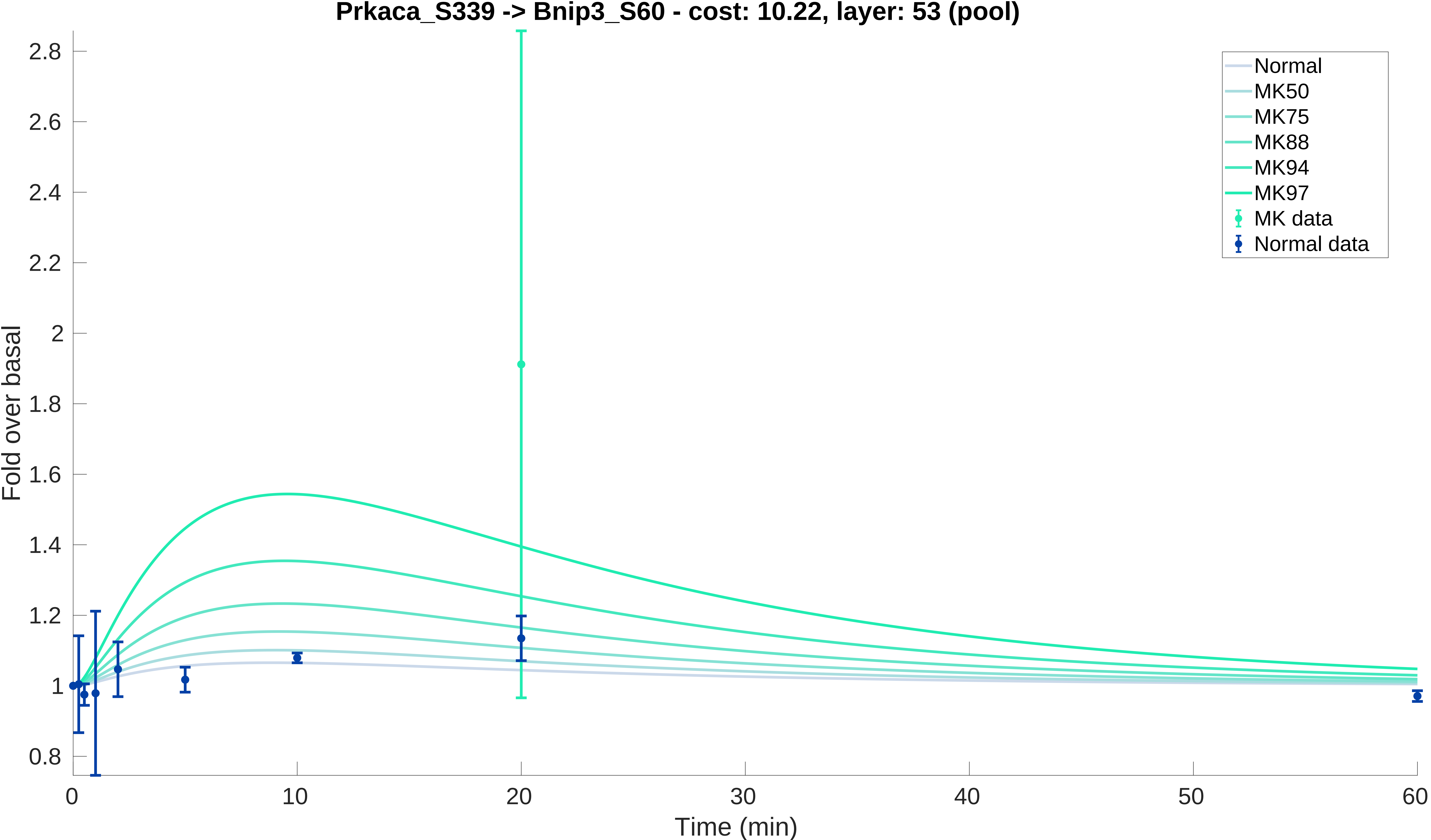
Prkaca_S339 -> Bnip3_S60 - cost: 10.22, layer: 53 (pool)
Fold over basal
Time (min)
2.8
2.6
2.4
2.2
2
1.8
1.6
1.4
1.2
1
0.8
0
10
20
30
40
50
60
Normal
MK50
MK75
MK88
MK94
MK97
MK data
Normal data

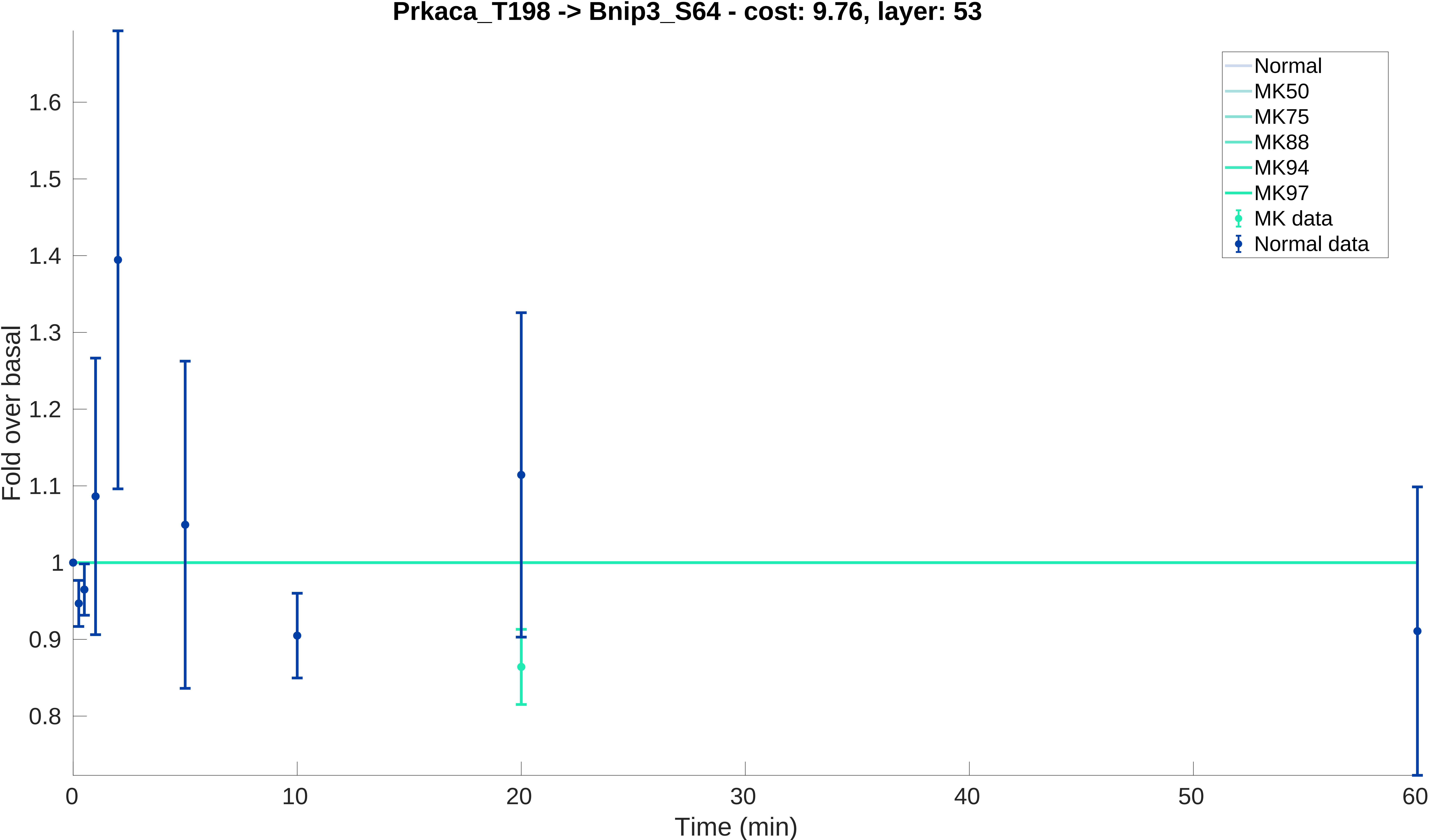

Prkaca_T198 -> Bnip3_S64 - cost: 9.76, layer: 53
Fold over basal
Time (min)
0.8
0.9
1
1.1
1.2
1.3
1.4
1.5
1.6
0
10
20
30
40
50
60
Normal
MK50
MK75
MK88
MK94
MK97
MK data
Normal data

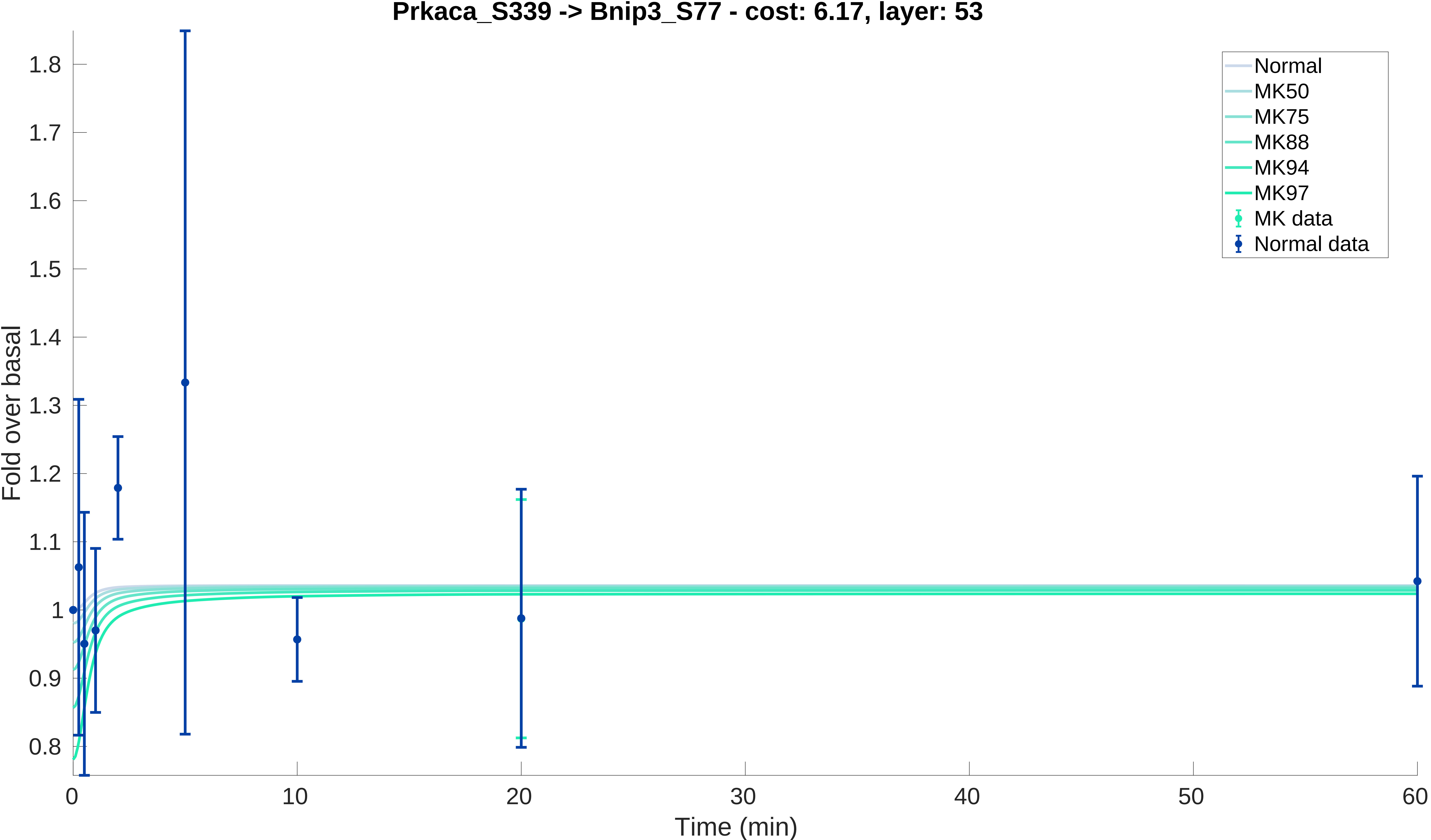

Prrkaca_S339 -> Bnip3_S77 - cost: 6.17, layer: 53
Fold over basal
Time (min)
1.8
1.7
1.6
1.5
1.4
1.3
1.2
1.1
1
0.9
0.8
0
10
20
30
40
50
60
Normal
MK50
MK75
MK88
MK94
MK97
MK data
Normal data

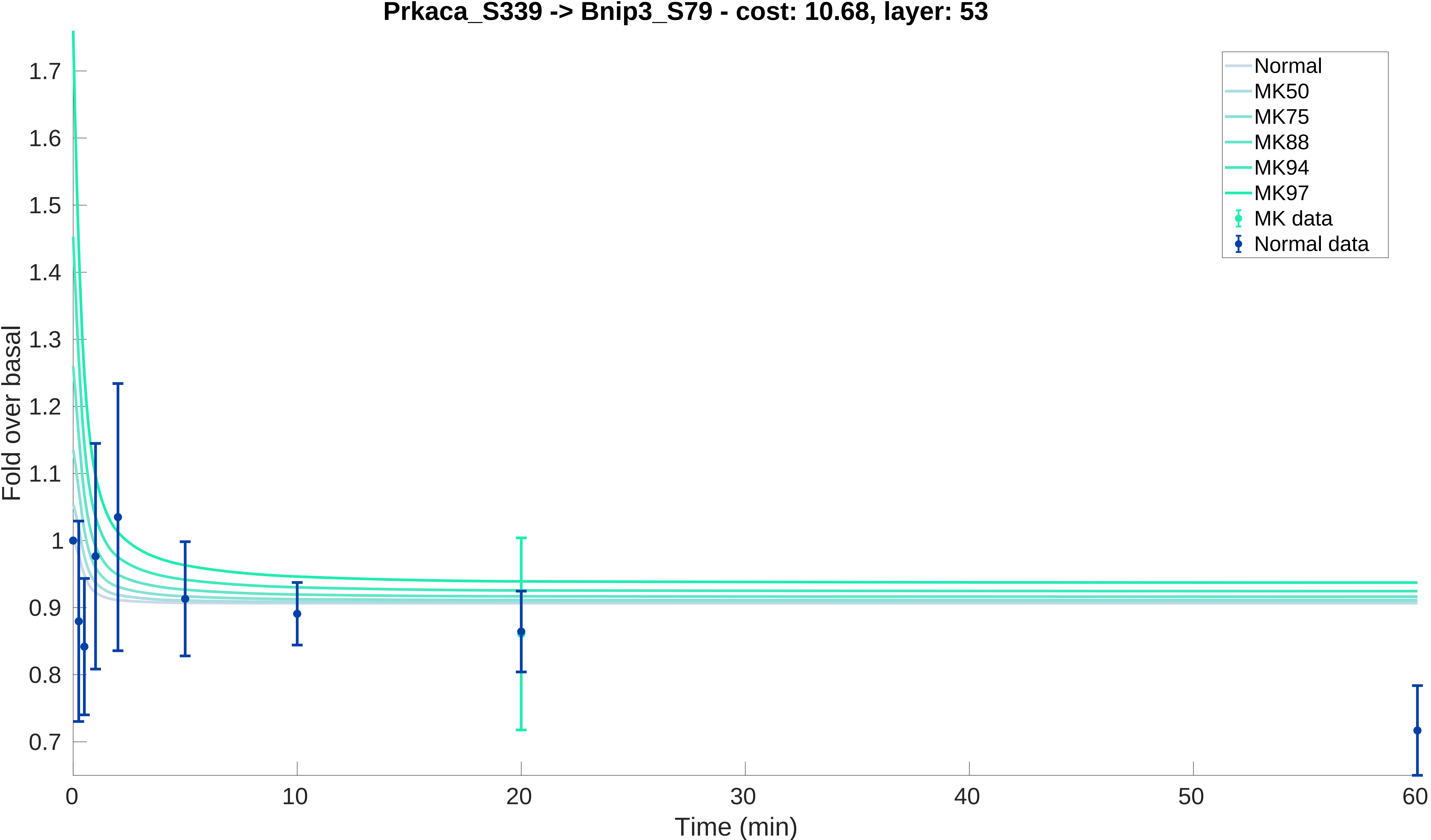
Prkaca_S339 -> Bnip3_S79 - cost: 10.68, layer: 53
Fold over basal
Time (min)
0.7
0.8
0.9
1
1.1
1.2
1.3
1.4
1.5
1.6
1.7
0
10
20
30
40
50
60
Normal
MK50
MK75
MK88
MK94
MK97
MK data
Normal data

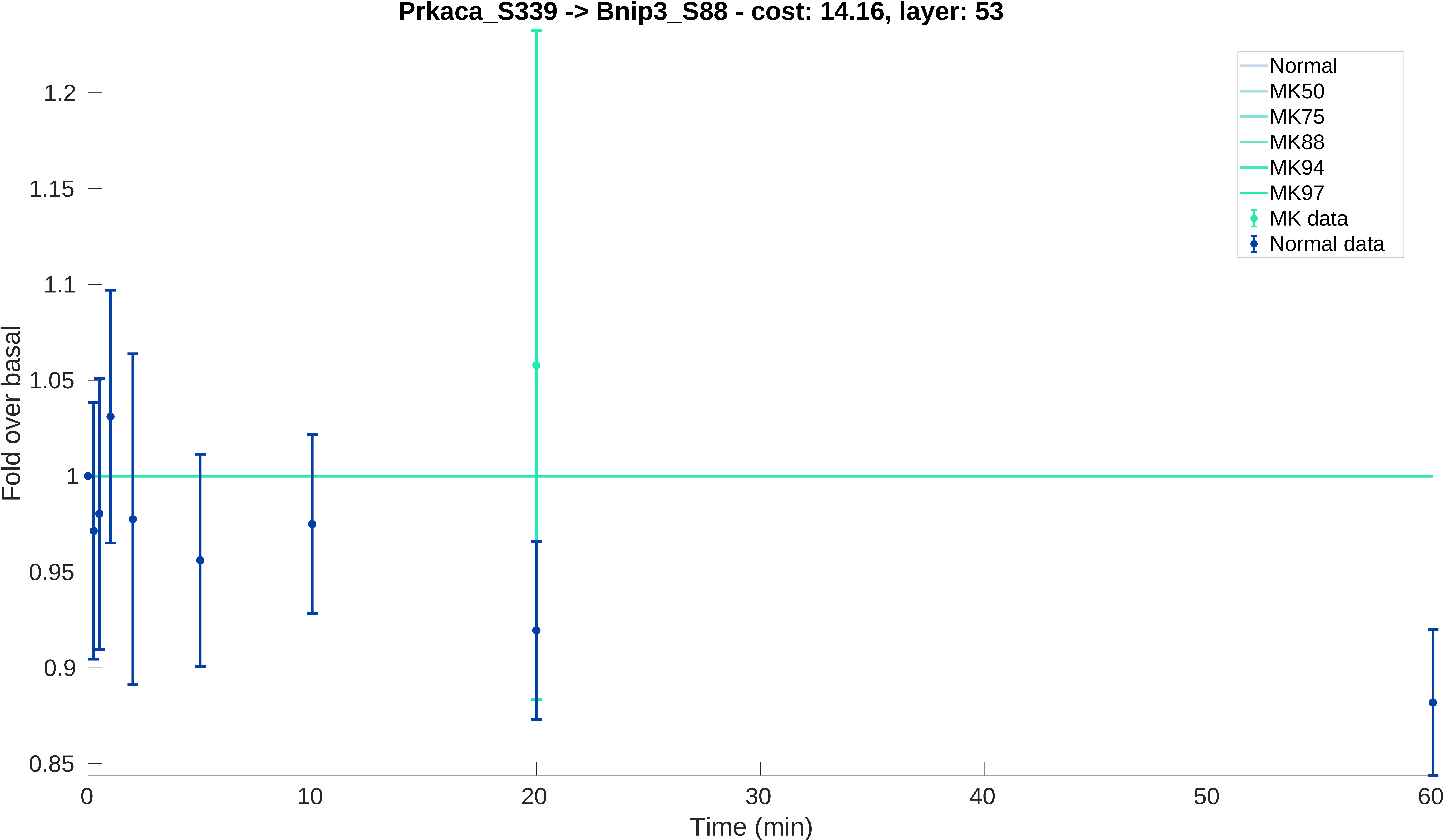

Prkaca_S339 -> Bnip3_S88 - cost: 14.16, layer: 53
Normal
MK50
MK75
MK88
MK94
MK97
MK data
Normal data
Fold over basal
1.2
1.15
1.1
1.05
1
0.95
0.9
0.85
0
10
20
30
40
50
60
Time (min)

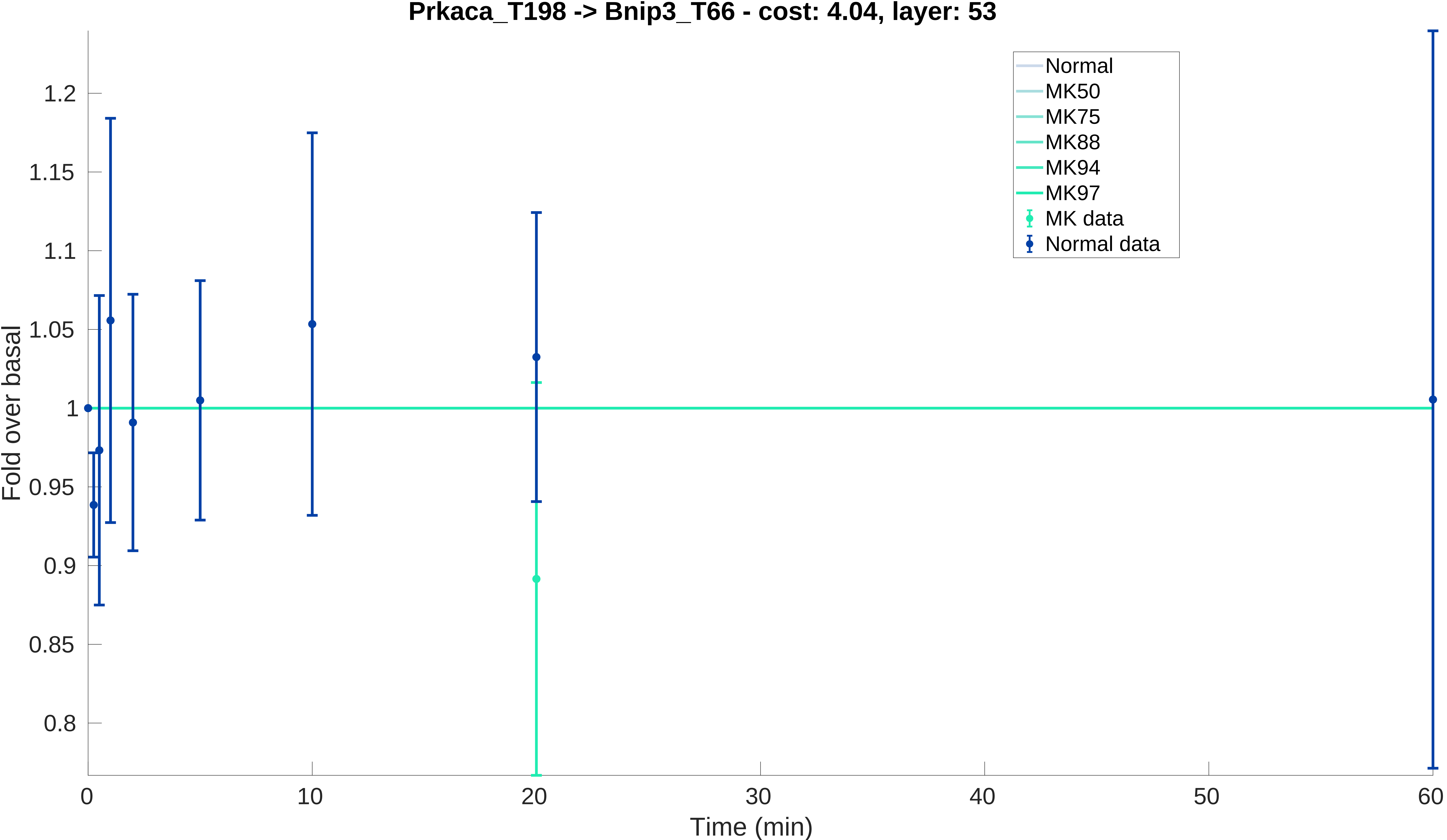

Prkaca_T198 -> Bnip3_T66 - cost: 4.04, layer: 53
Fold over basal
Time (min)
0
10
20
30
40
50
60
0.8
0.85
0.9
0.95
1
1.05
1.1
1.15
1.2
Normal
MK50
MK75
MK88
MK94
MK97
MK data
Normal data

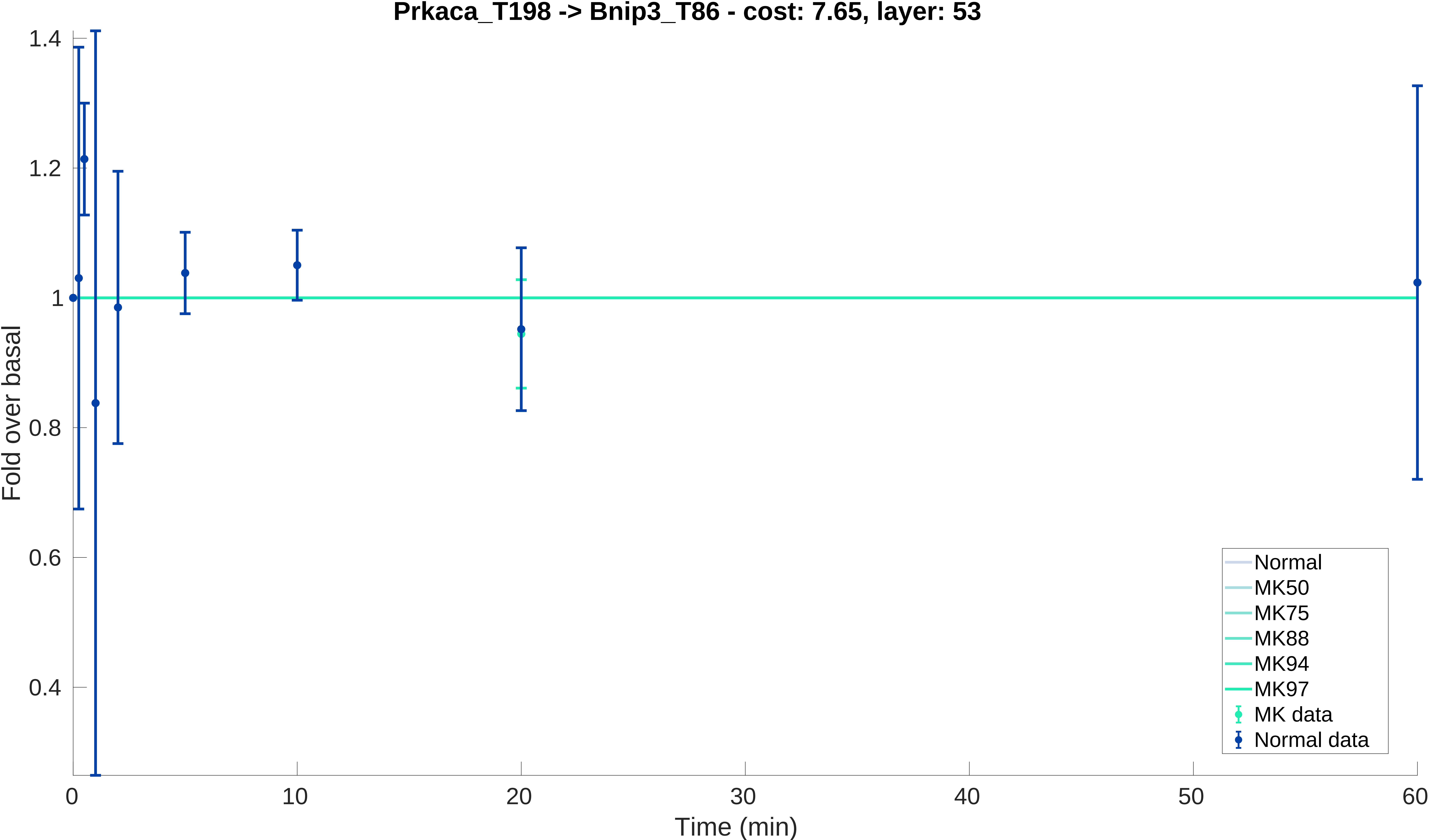
Prkaca_T198 -> Bnip3_T86 - cost: 7.65, layer: 53
Fold over basal
Time (min)
1.4
1.2
1
0.8
0.6
0.4
0
10
20
30
40
50
60
Normal
MK50
MK75
MK88
MK94
MK97
MK data
Normal data

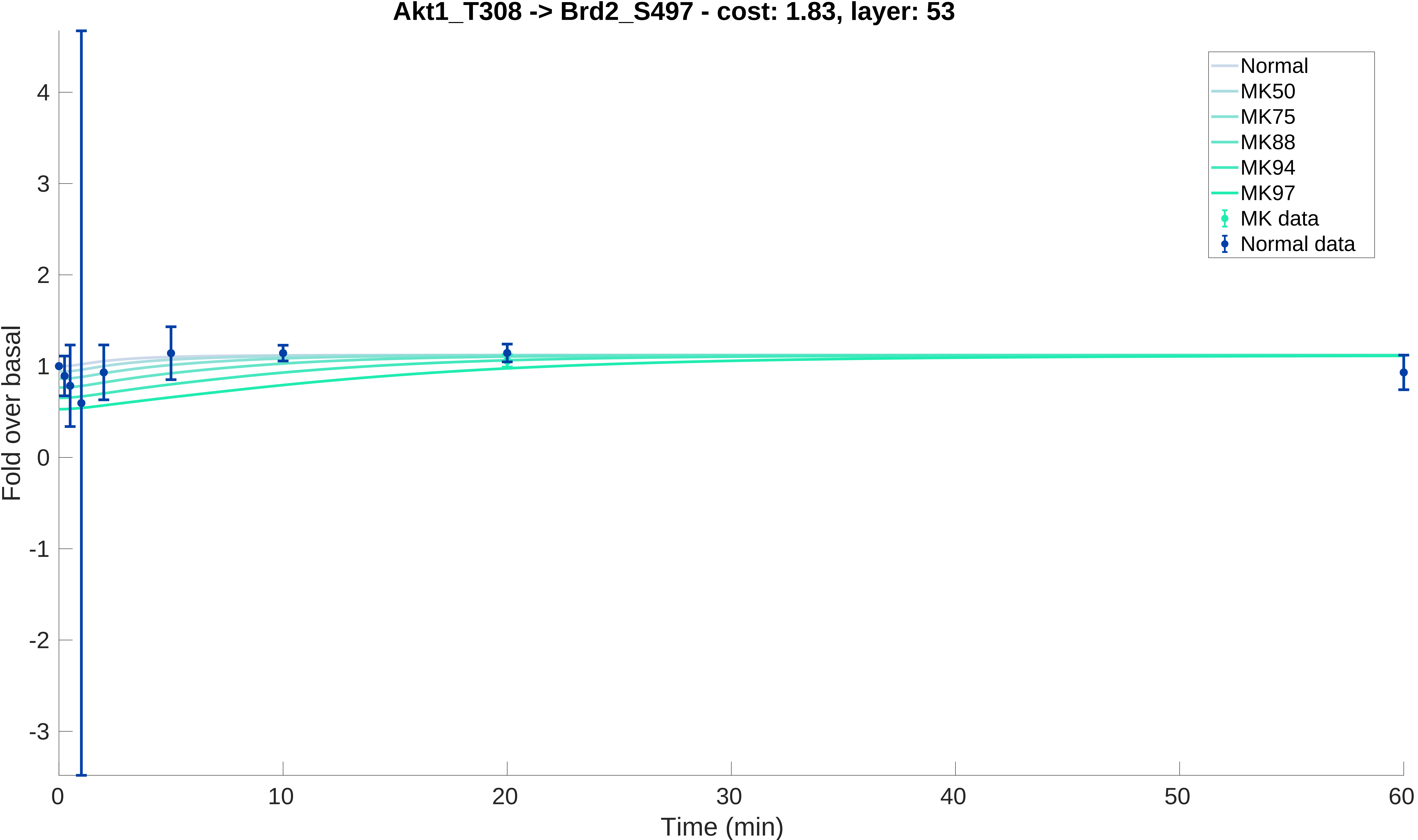
Akt1_T308 -> Brd2_S497 - cost: 1.83, layer: 53
Fold over basal
Time (min)
Normal
MK50
MK75
MK88
MK94
MK97
MK data
Normal data
4
3
2
1
0
-1
-2
-3
0
10
20
30
40
50
60

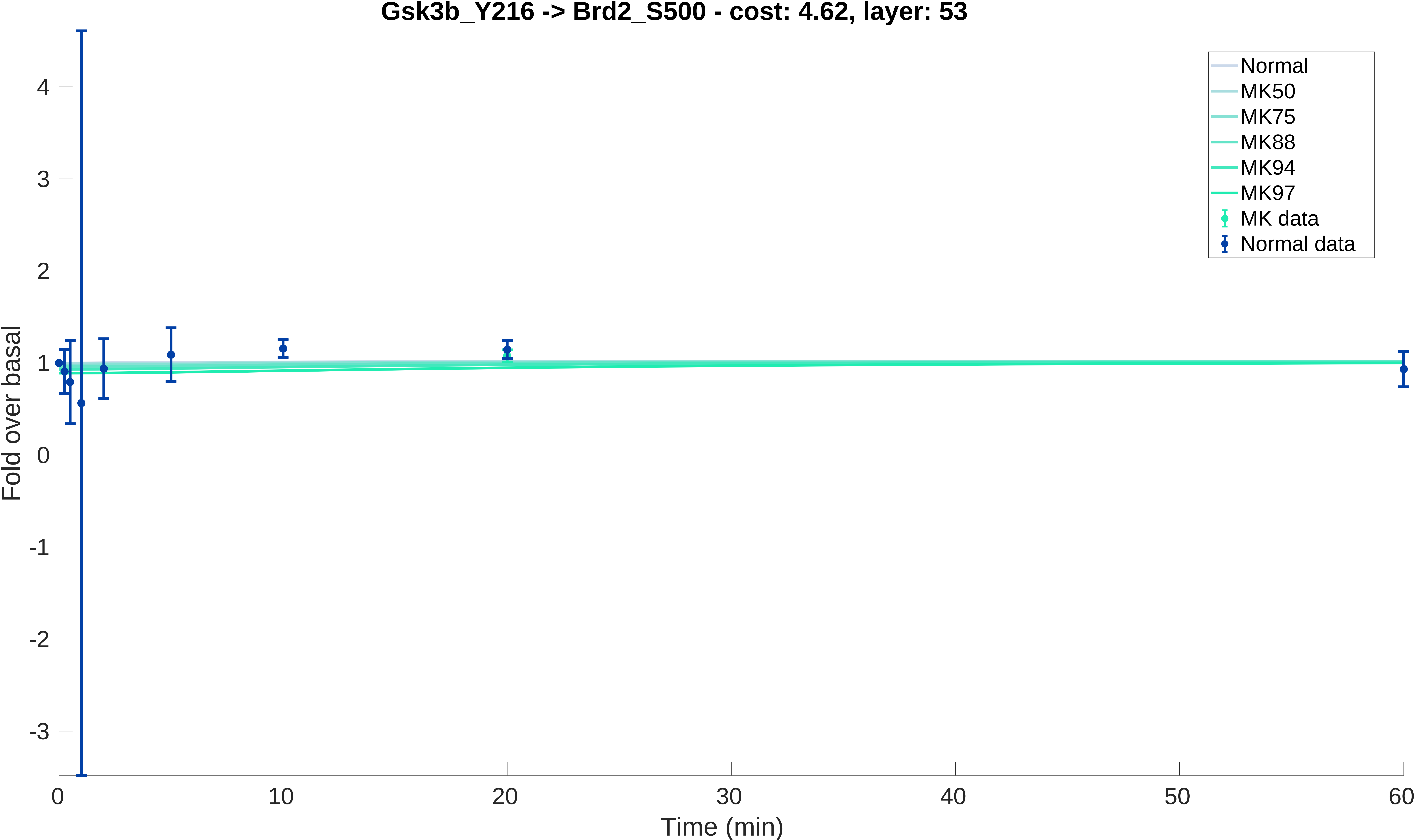
Gsk3b_Y216 -> Brd2_S500 - cost: 4.62, layer: 53
Normal
MK50
MK75
MK88
MK94
MK97
MK data
Normal data
4
3
2
1
0
-1
-2
-3
Fold over basal
0
10
20
30
40
50
60
Time (min)

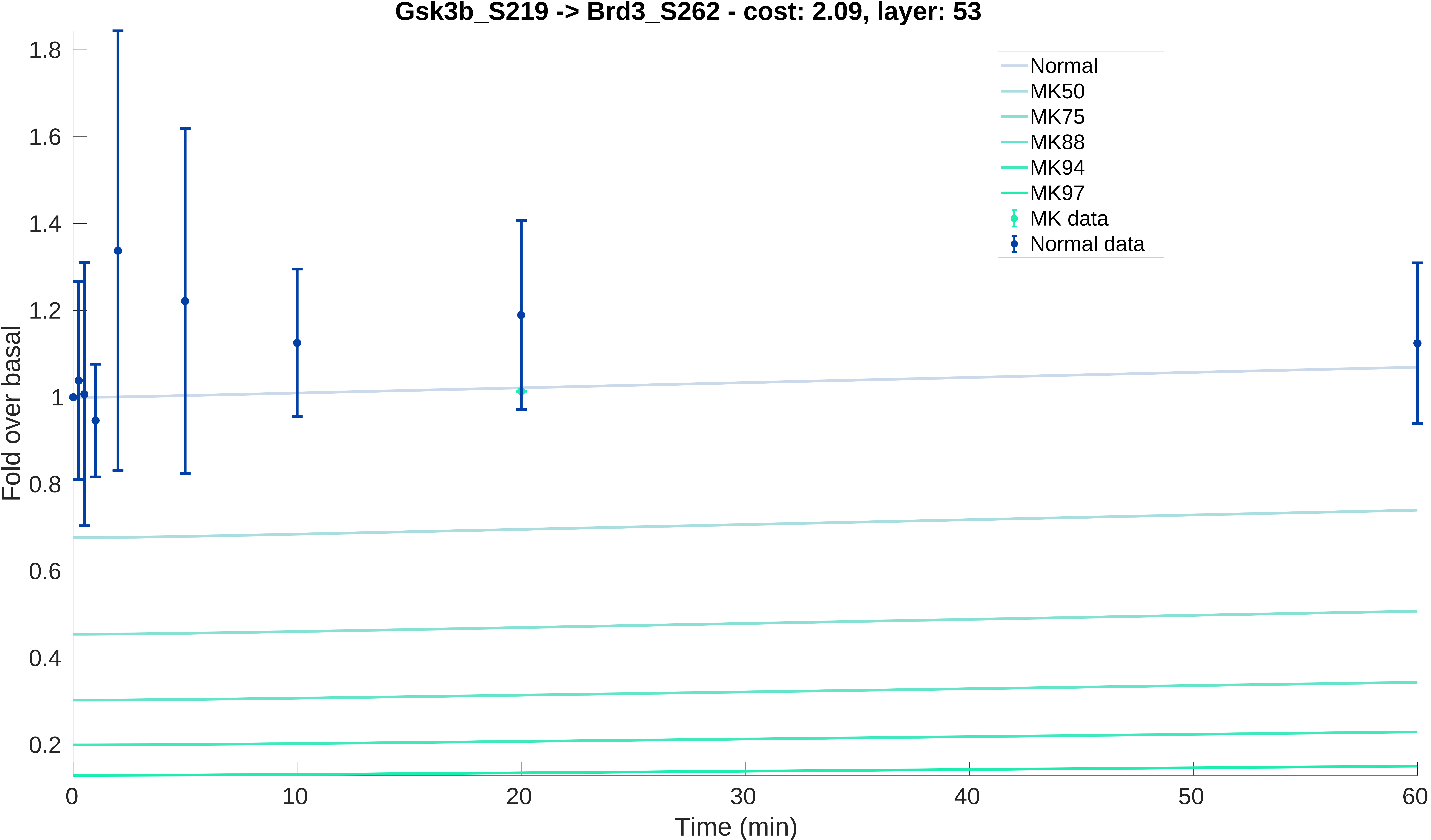
Gsk3b_S219 -> Brd3_S262 - cost: 2.09, layer: 53
Fold over basal
1.8
1.6
1.4
1.2
1
0.8
0.6
0.4
0.2
0
10
20
30
40
50
60
Time (min)
Normal
MK50
MK75
MK88
MK94
MK97
MK data
Normal data

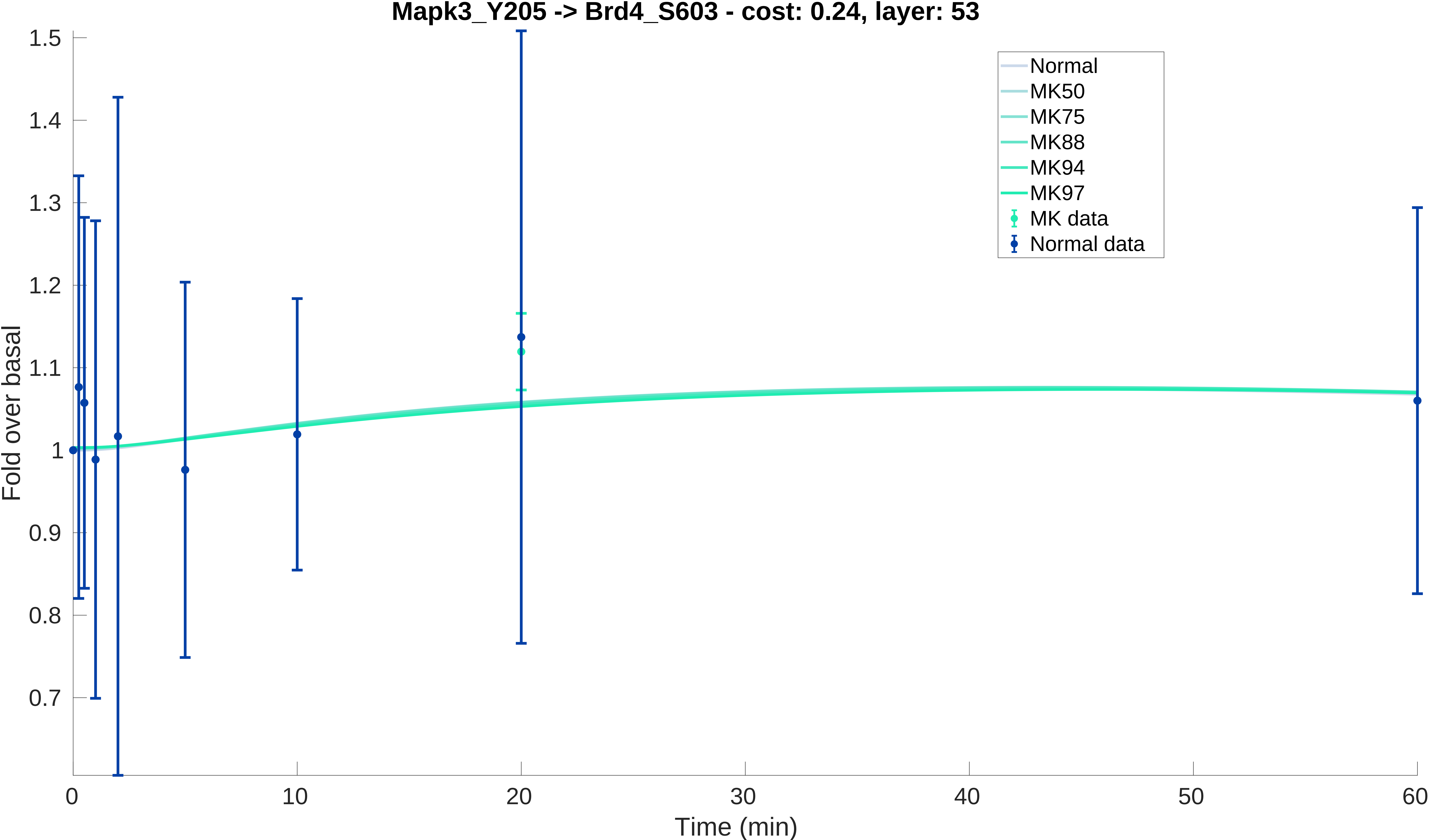

Mapk3_Y205 -> Brd4_S603 - cost: 0.24, layer: 53
Fold over basal
1.5
1.4
1.3
1.2
1.1
1
0.9
0.8
0.7
0
10
20
30
40
50
60
Time (min)
Normal
MK50
MK75
MK88
MK94
MK97
MK data
Normal data

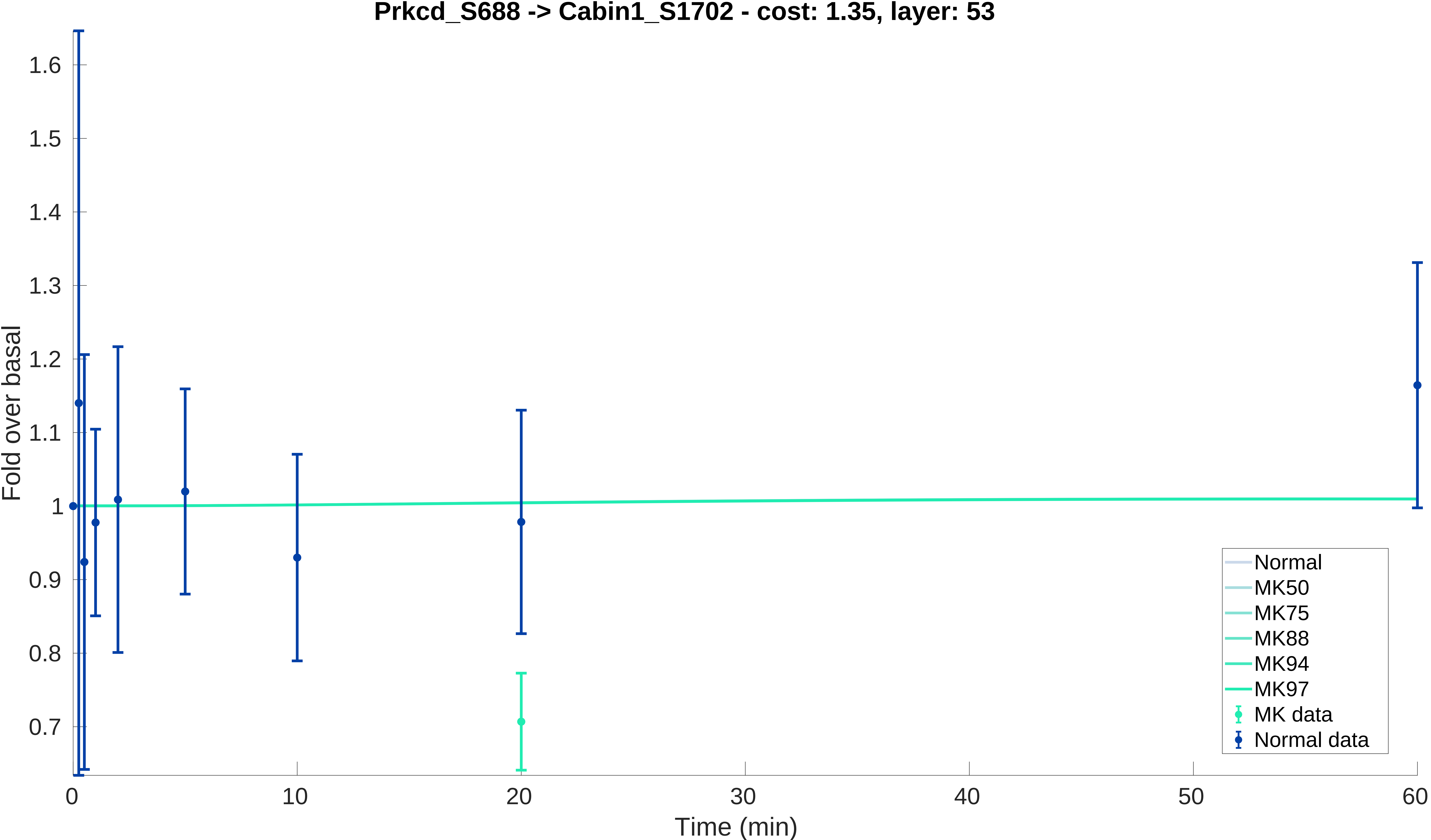
Prkcd_S688 -> Cabin1_S1702 - cost: 1.35, layer: 53
Fold over basal
1.6
1.5
1.4
1.3
1.2
1.1
1
0.9
0.8
0.7
0
10
20
30
40
50
60
Time (min)
Normal
MK50
MK75
MK88
MK94
MK97
MK data
Normal data

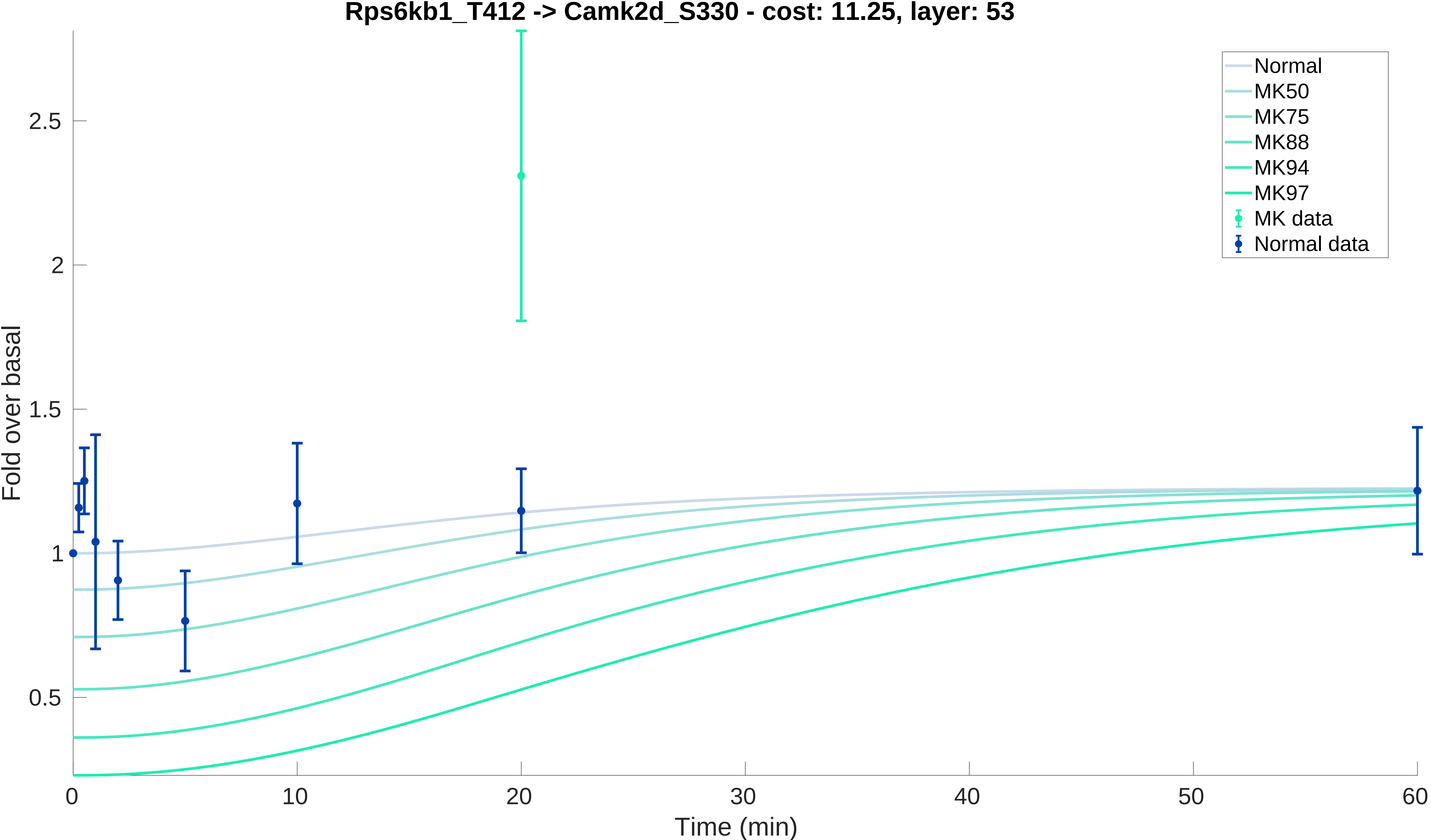

Rps6kb1_T412 -> Camk2d_S330 - cost: 11.25, layer: 53
Fold over basal
Time (min)
0.5
1
1.5
2
2.5
0
10
20
30
40
50
60
Normal
MK50
MK75
MK88
MK94
MK97
MK data
Normal data

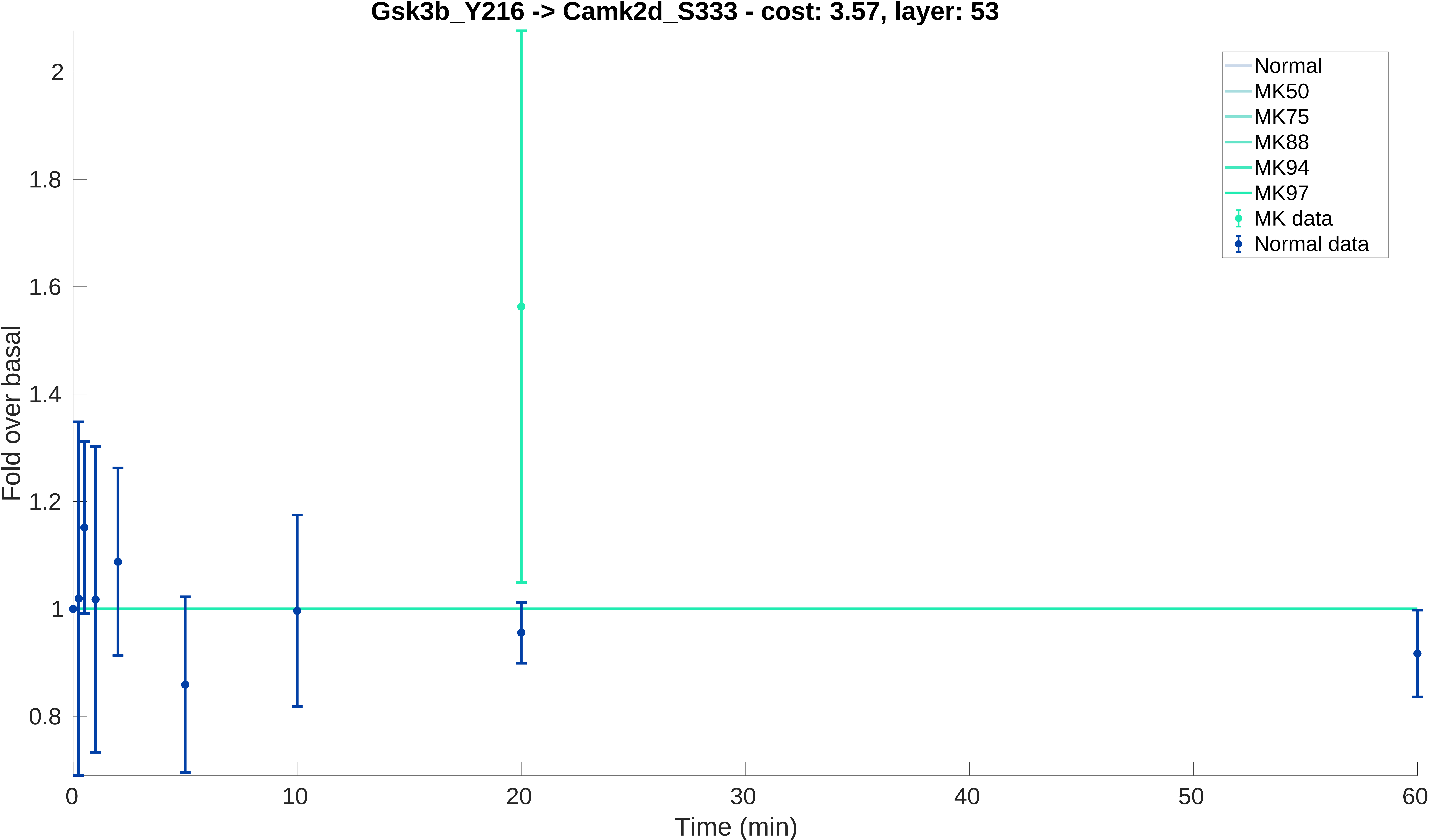

Gsk3b_Y216 -> Camk2d_S333 - cost: 3.57, layer: 53
Fold over basal
2
1.8
1.6
1.4
1.2
1
0.8
0
10
20
30
40
50
60
Time (min)
Normal
MK50
MK75
MK88
MK94
MK97
MK data
Normal data

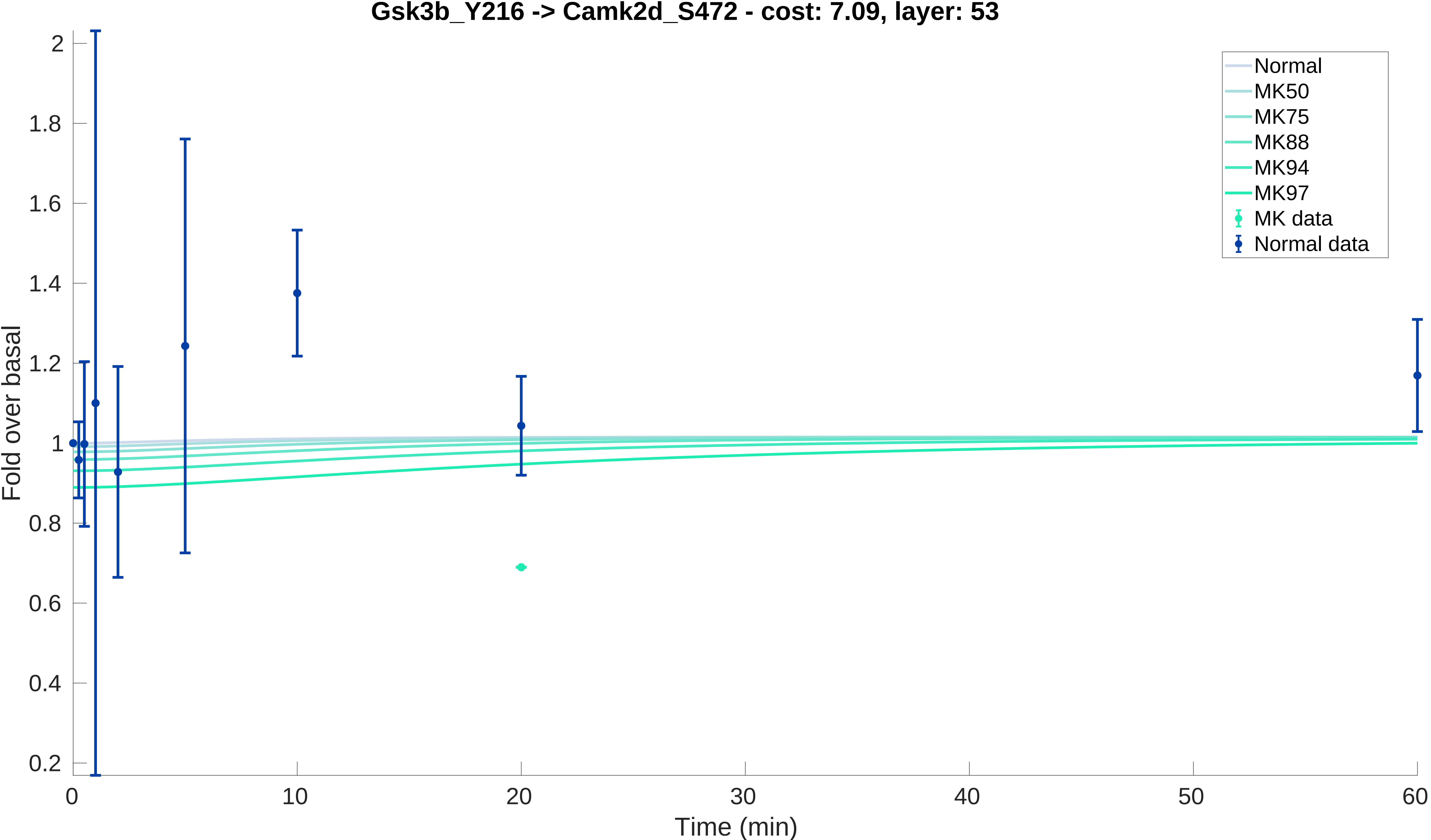

Gsk3b_Y216 -> Camk2d_S472 - cost: 7.09, layer: 53
Fold over basal
Time (min)
0.2
0.4
0.6
0.8
1
1.2
1.4
1.6
1.8
2
0
10
20
30
40
50
60
Normal
MK50
MK75
MK88
MK94
MK97
MK data
Normal data

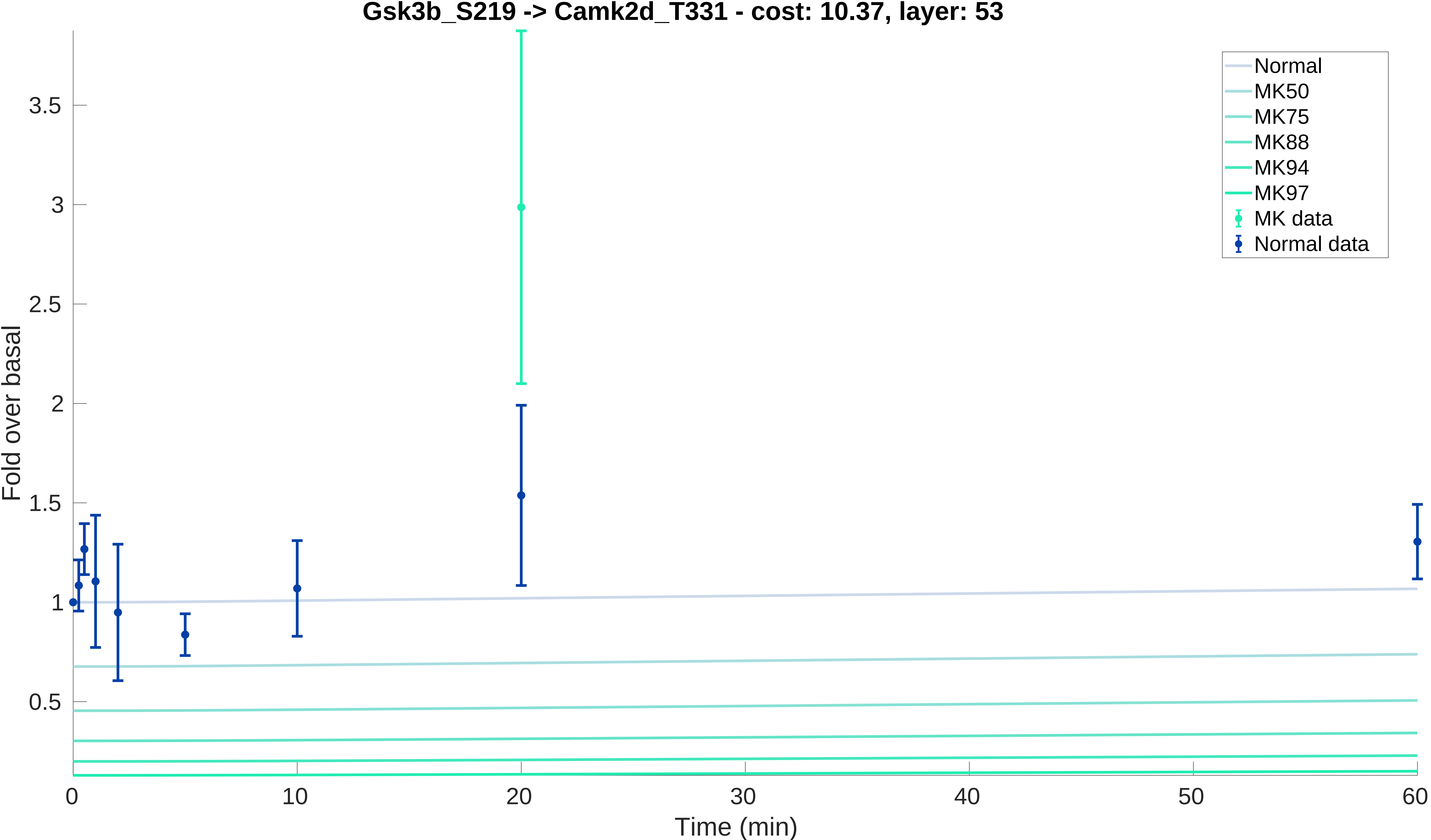

Gsk3b_S219 -> Camk2d_T331 - cost: 10.37, layer: 53
Normal
MK50
MK75
MK88
MK94
MK97
MK data
Normal data
Fold over basal
3.5
3
2.5
2
1.5
1
0.5
0
10
20
30
40
50
60
Time (min)

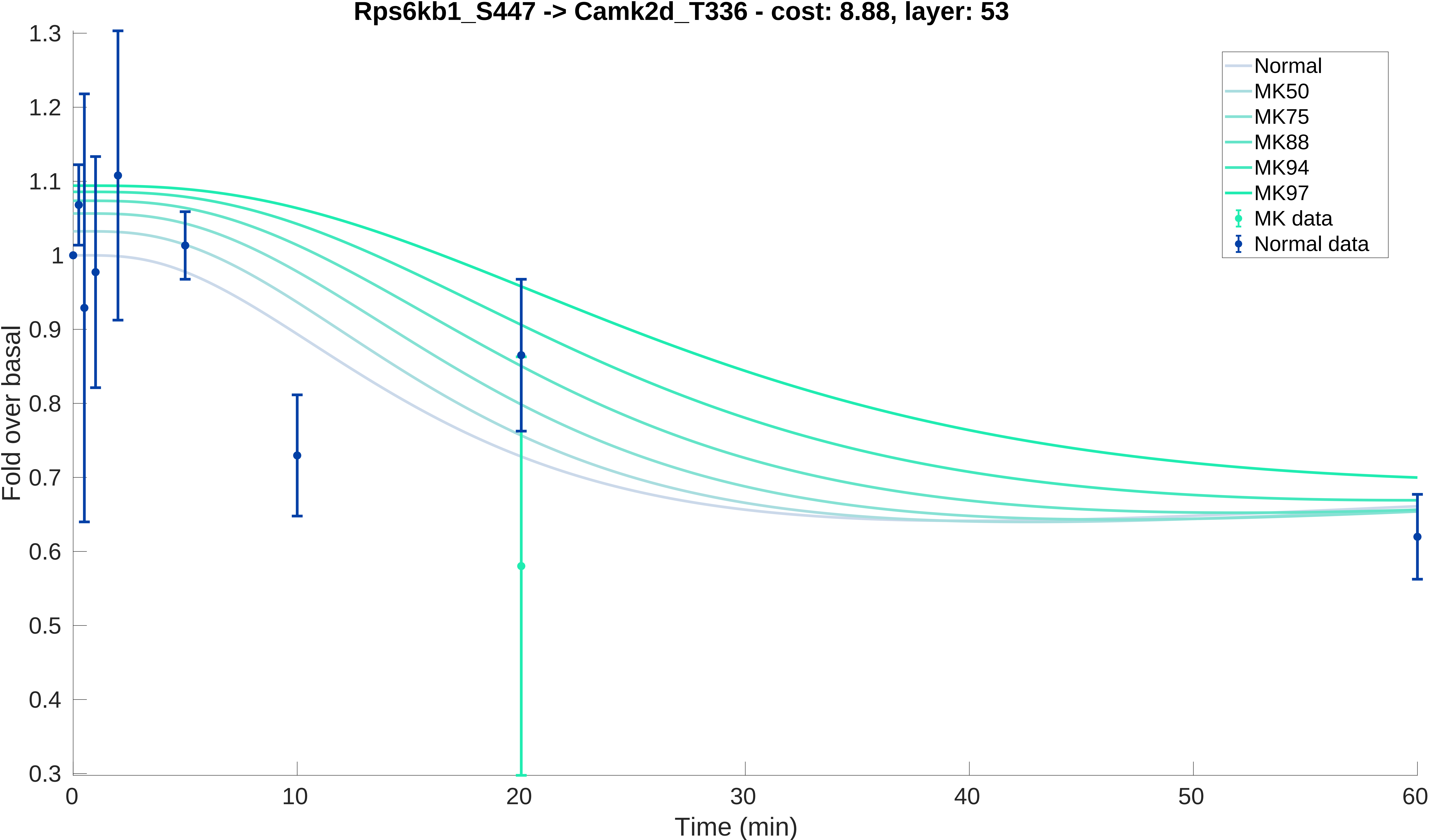

Rps6kb1_S447 -> Camk2d_T336 - cost: 8.88, layer: 53
Fold over basal
Time (min)
1.3
1.2
1.1
1
0.9
0.8
0.7
0.6
0.5
0.4
0.3
0
10
20
30
40
50
60
Normal
MK50
MK75
MK88
MK94
MK97
MK data
Normal data

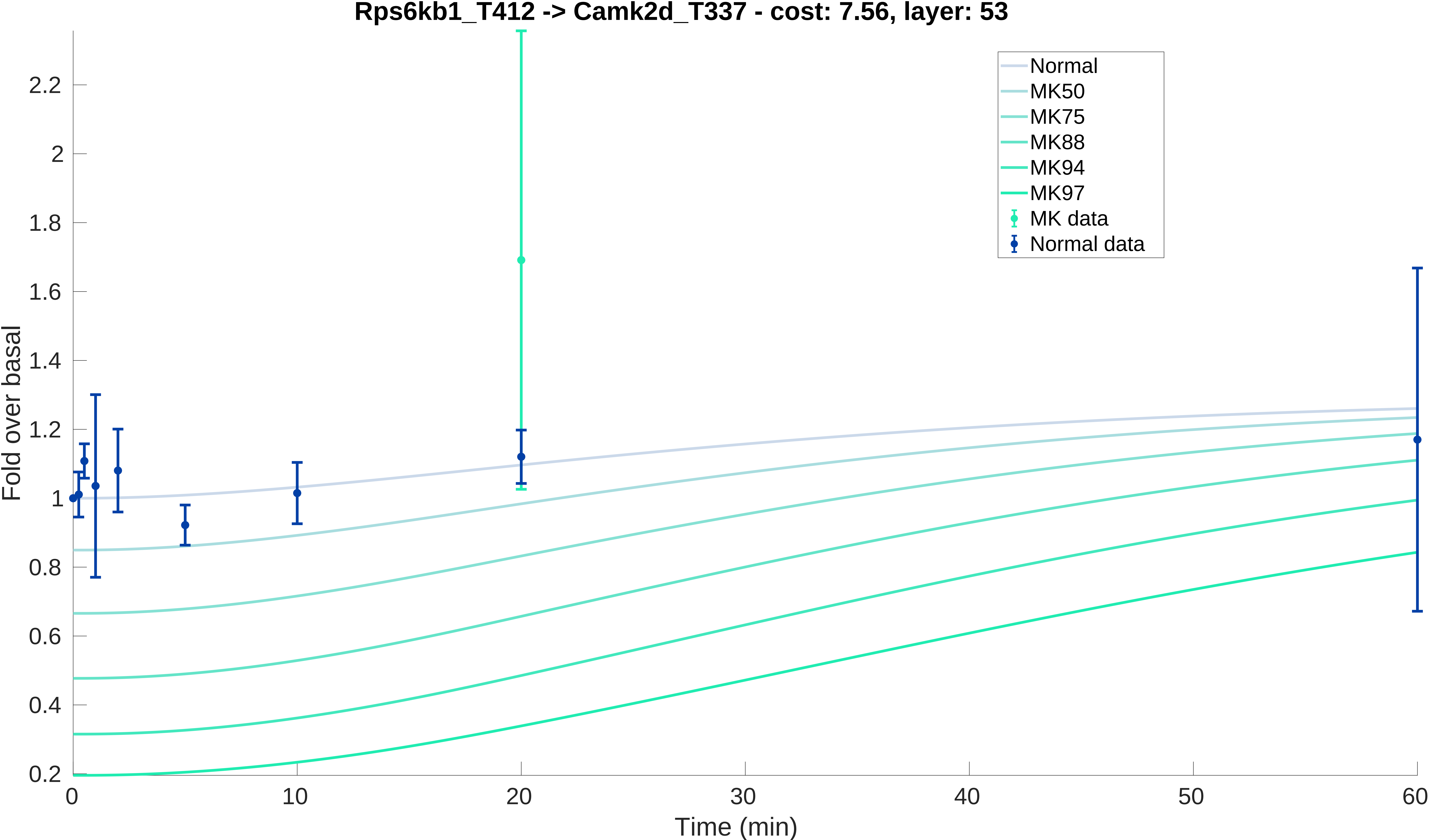
Rps6kb1_T412 -> Camk2d_T337 - cost: 7.56, layer: 53
Fold over basal
Time (min)
0
10
20
30
40
50
60
0.2
0.4
0.6
0.8
1
1.2
1.4
1.6
1.8
2
2.2
Normal
MK50
MK75
MK88
MK94
MK97
MK data
Normal data

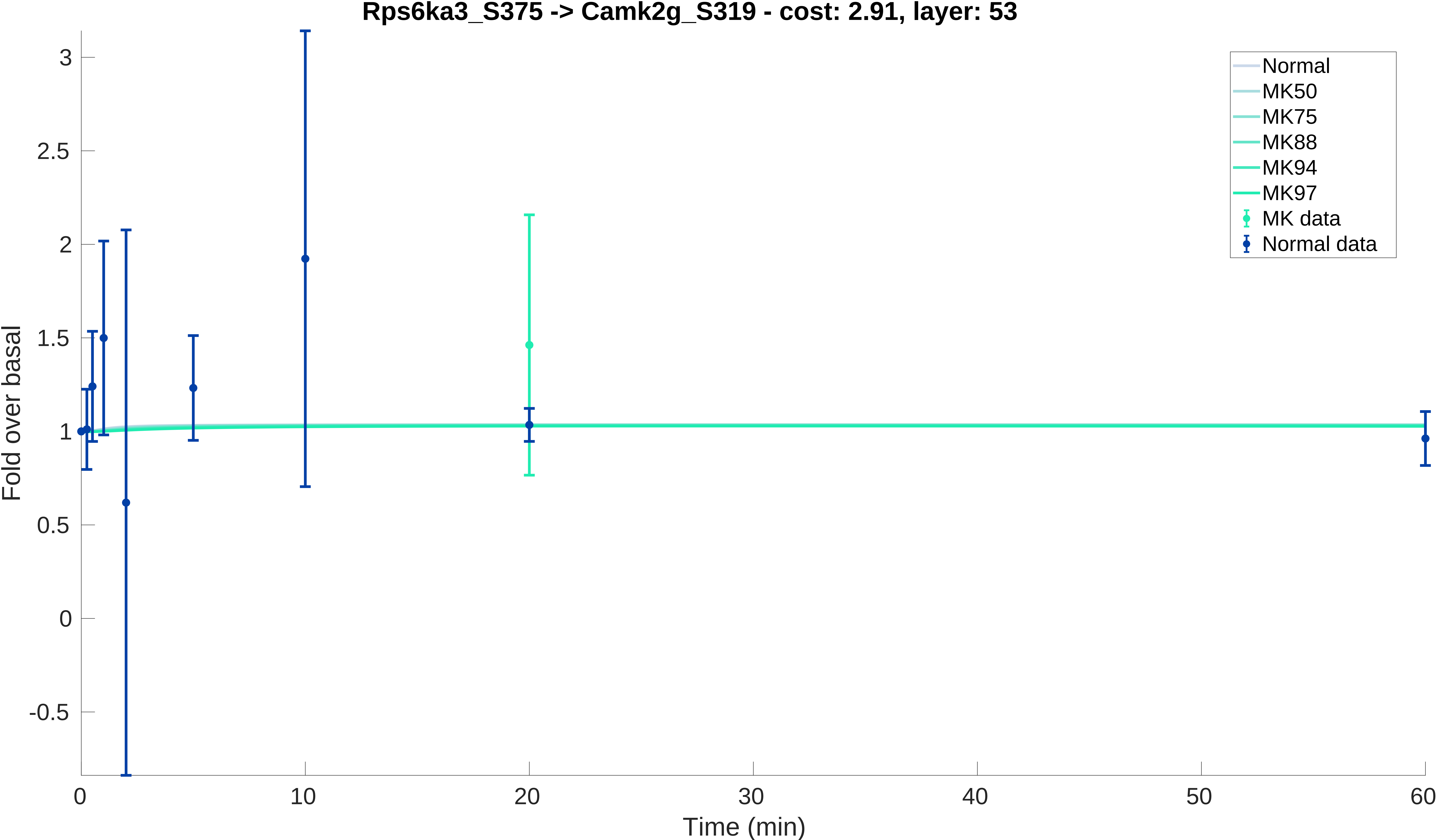
Rps6ka3_S375 -> Camk2g_S319 - cost: 2.91, layer: 53
Fold over basal
Time (min)
3
2.5
2
1.5
1
0.5
0
-0.5
0
10
20
30
40
50
60
Normal
MK50
MK75
MK88
MK94
MK97
MK data
Normal data

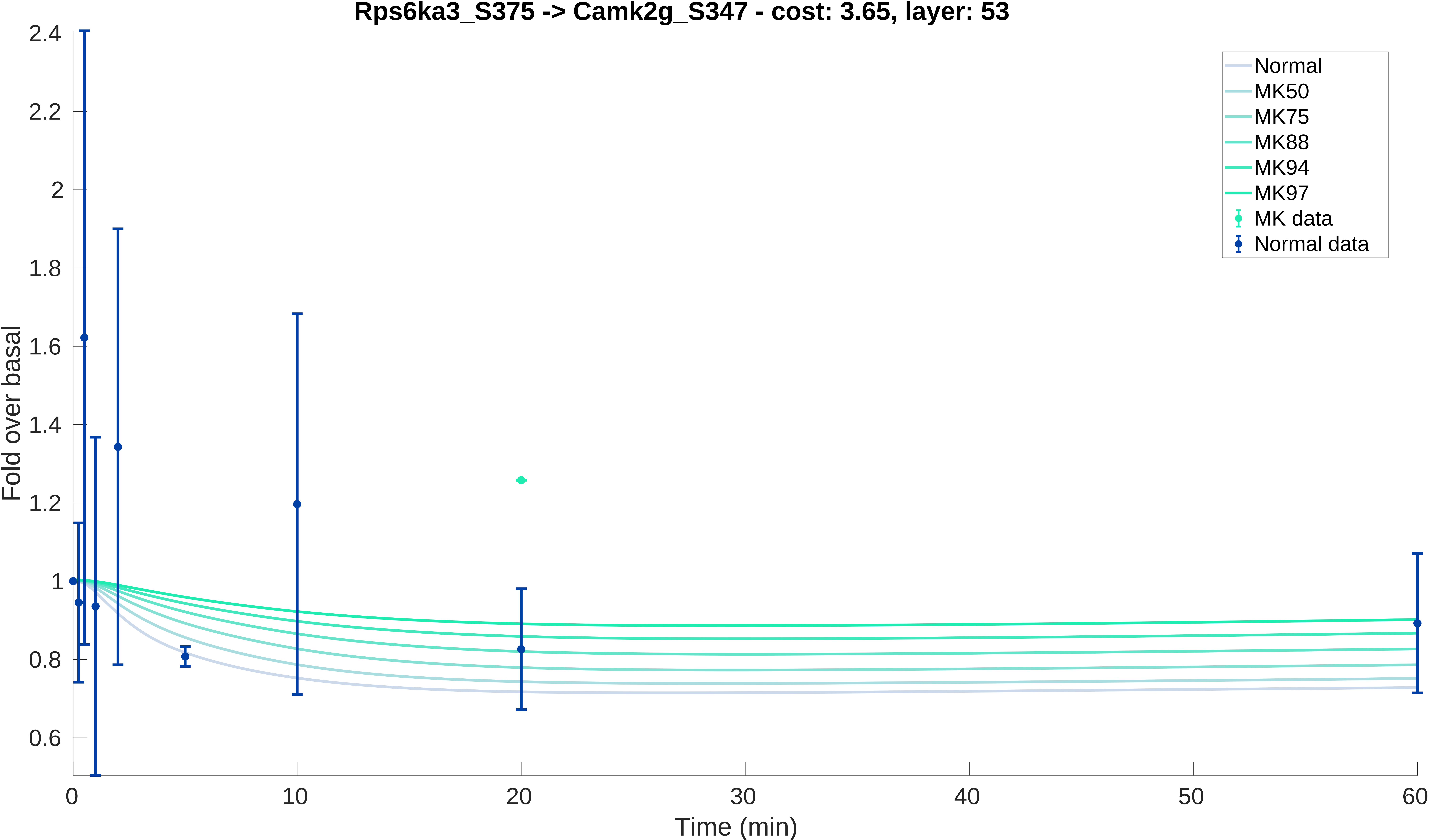

Rps6ka3_S375 -> Camk2g_S347 - cost: 3.65, layer: 53
Fold over basal
Time (min)
0.6
0.8
1
1.2
1.4
1.6
1.8
2
2.2
2.4
0
10
20
30
40
50
60
Normal
MK50
MK75
MK88
MK94
MK97
MK data
Normal data

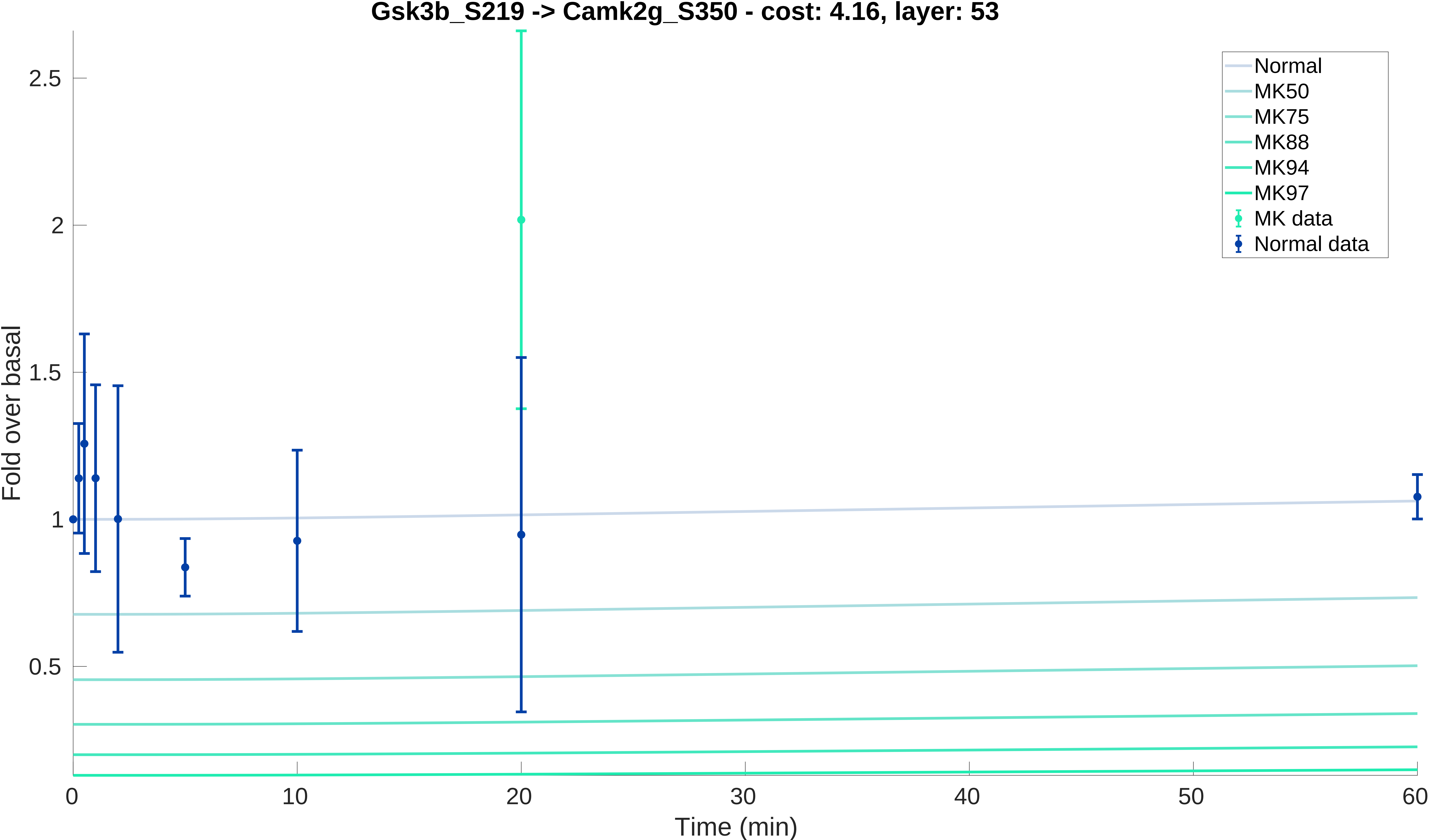
Gsk3b_S219 -> Camk2g_S350 - cost: 4.16, layer: 53
Fold over basal
Time (min)
2.5
2
1.5
1
0.5
0
10
20
30
40
50
60
Normal
MK50
MK75
MK88
MK94
MK97
MK data
Normal data

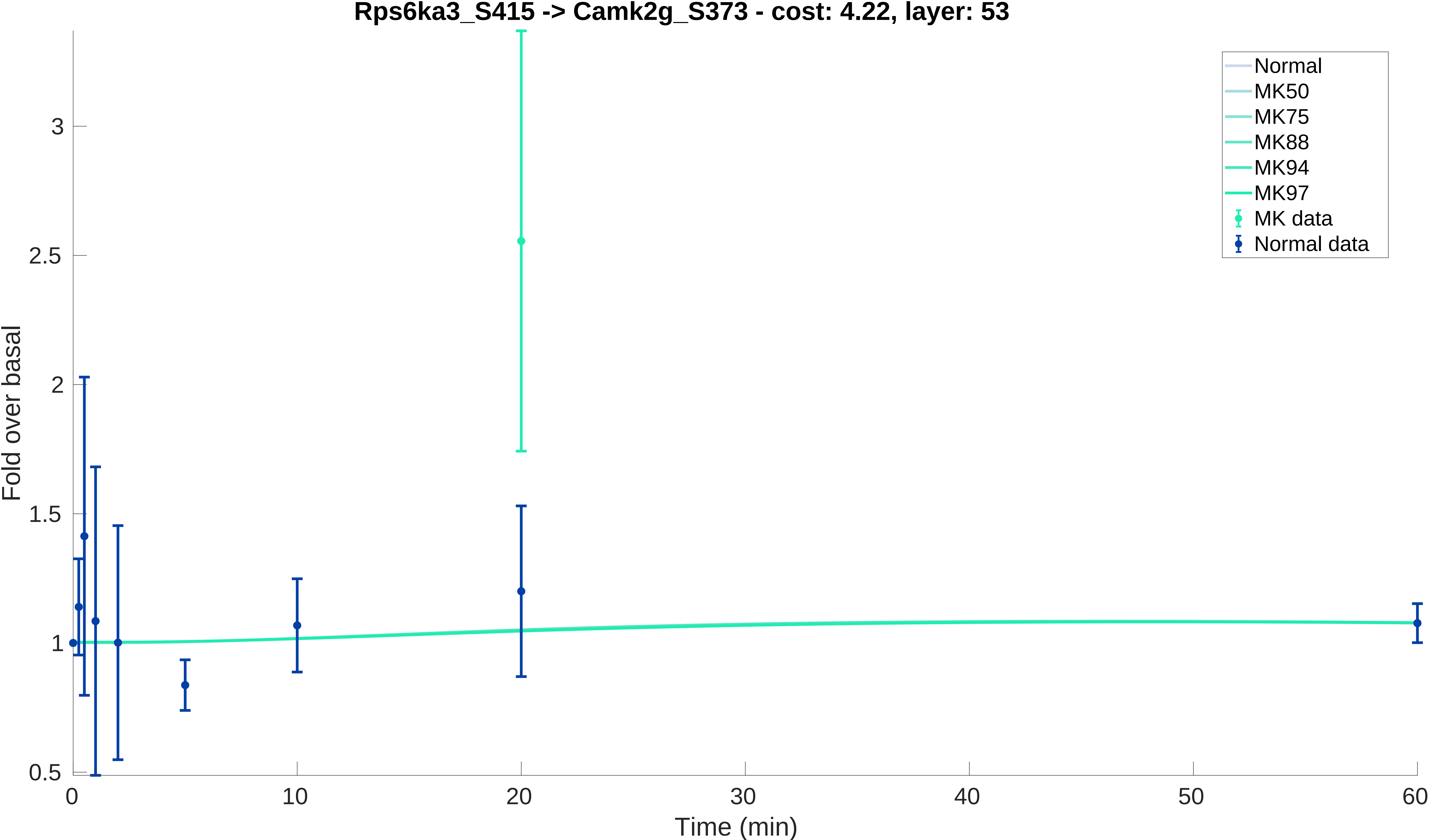

Rps6ka3_S415 -> Camk2g_S373 - cost: 4.22, layer: 53
Normal
MK50
MK75
MK88
MK94
MK97
MK data
Normal data
Fold over basal
Time (min)
0.5
1
1.5
2
2.5
3
0
10
20
30
40
50
60

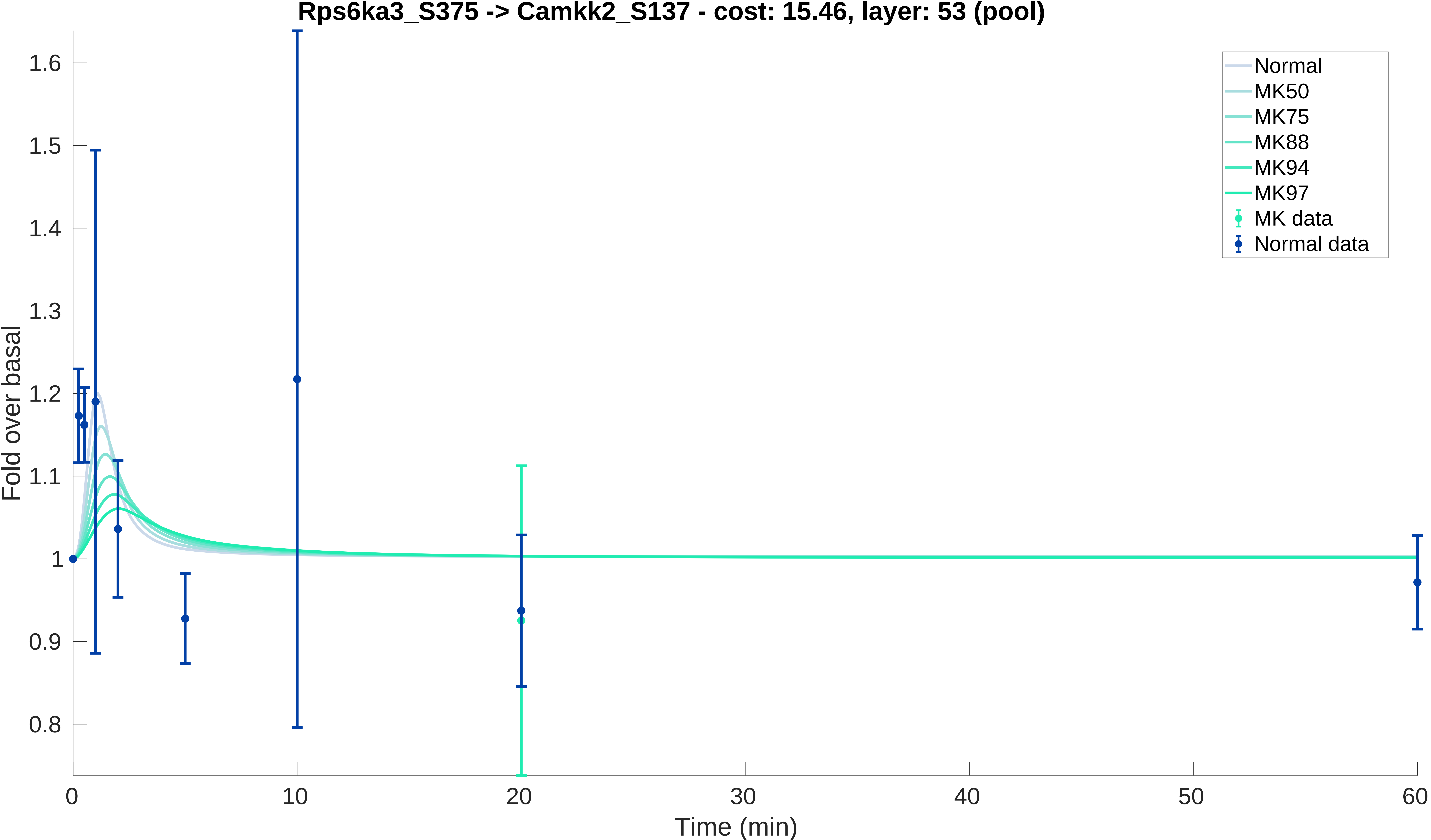

Rps6ka3_S375 -> Camkk2_S137 - cost: 15.46, layer: 53 (pool)
Fold over basal
Time (min)
1.6
1.5
1.4
1.3
1.2
1.1
1
0.9
0.8
0
10
20
30
40
50
60
Normal
MK50
MK75
MK88
MK94
MK97
MK data
Normal data

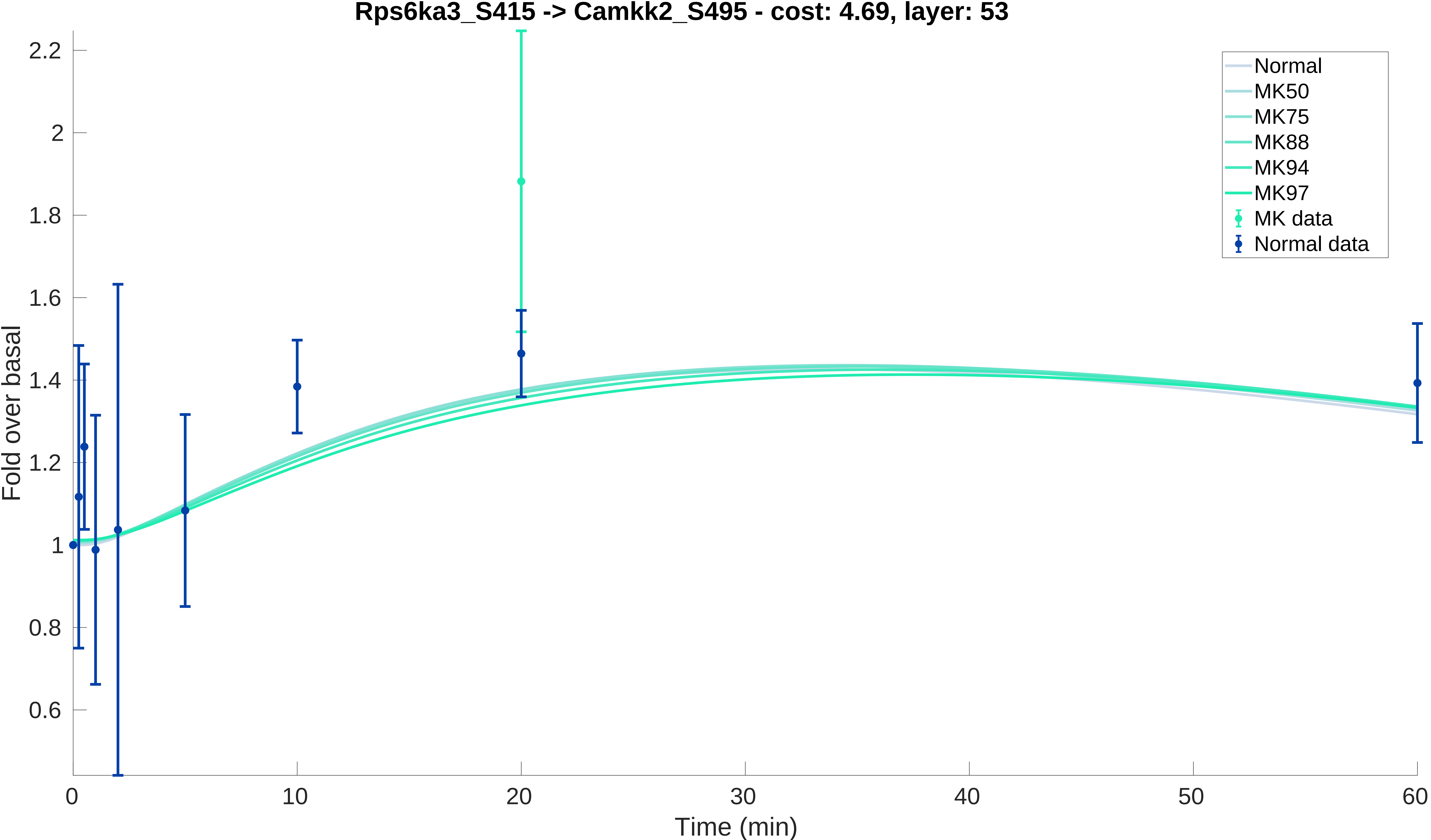

Rps6ka3_S415 -> Camkk2_S495 - cost: 4.69, layer: 53
Fold over basal
Time (min)
0.6
0.8
1
1.2
1.4
1.6
1.8
2
2.2
0
10
20
30
40
50
60
Normal
MK50
MK75
MK88
MK94
MK97
MK data
Normal data

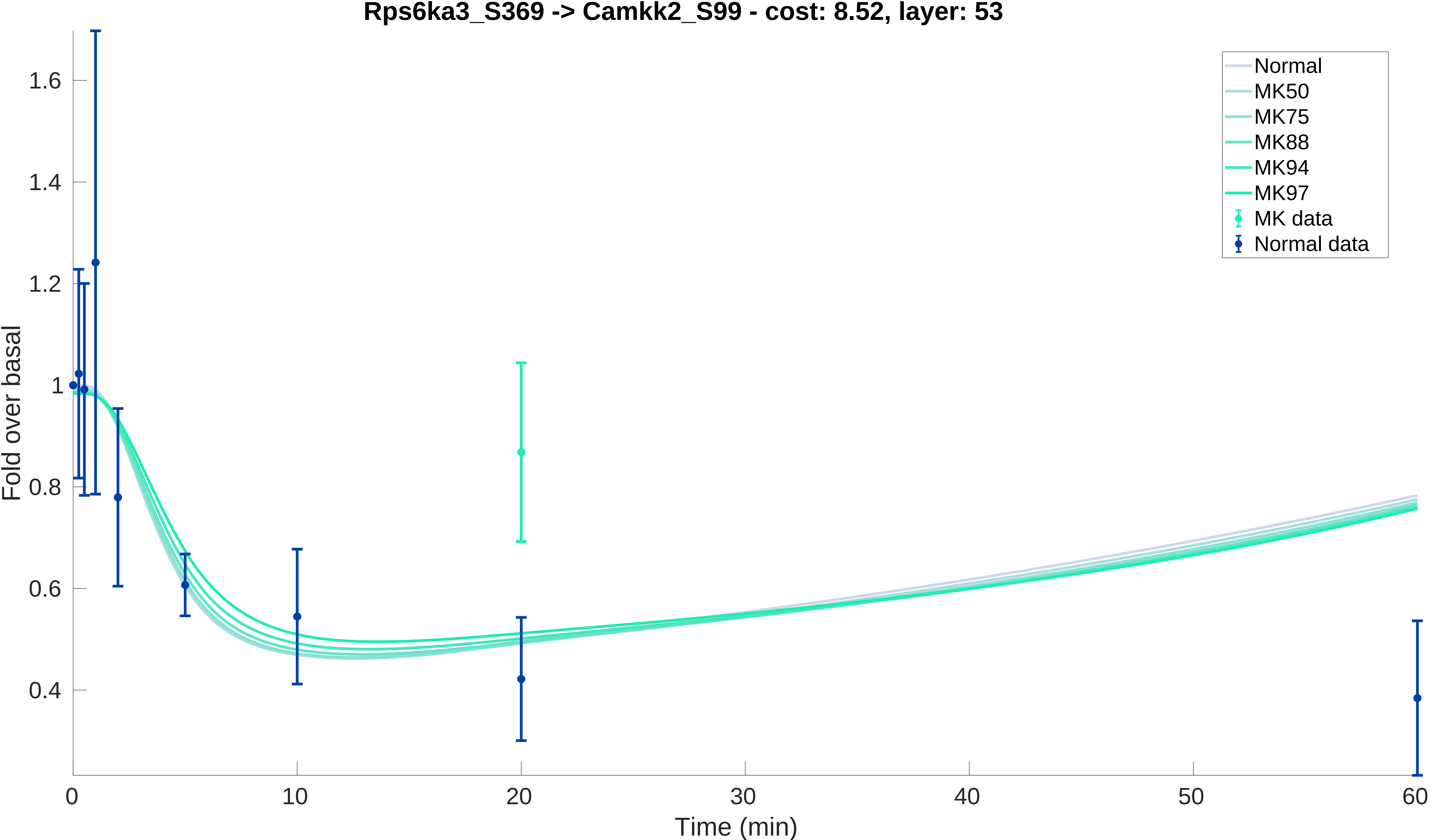

Rps6ka3_S369 -> Camkk2_S99 - cost: 8.52, layer: 53
Fold over basal
Time (min)
0.4
0.6
0.8
1
1.2
1.4
1.6
0
10
20
30
40
50
60
Normal
MK50
MK75
MK88
MK94
MK97
MK data
Normal data

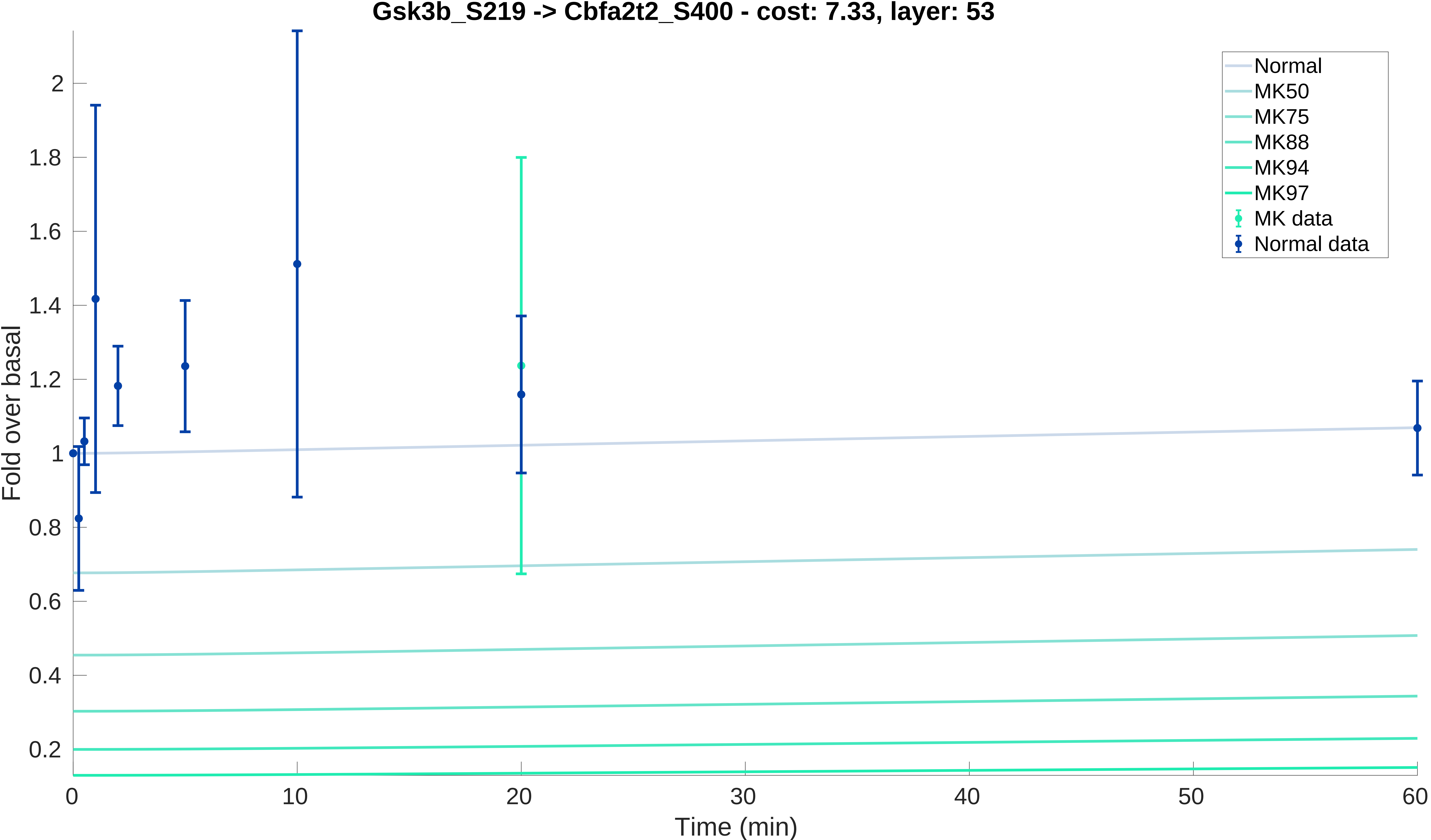

Gsk3b_S219 -> Cbfa2t2_S400 - cost: 7.33, layer: 53
Fold over basal
Time (min)
Normal
MK50
MK75
MK88
MK94
MK97
MK data
Normal data
0
10
20
30
40
50
60
0.2
0.4
0.6
0.8
1
1.2
1.4
1.6
1.8
2

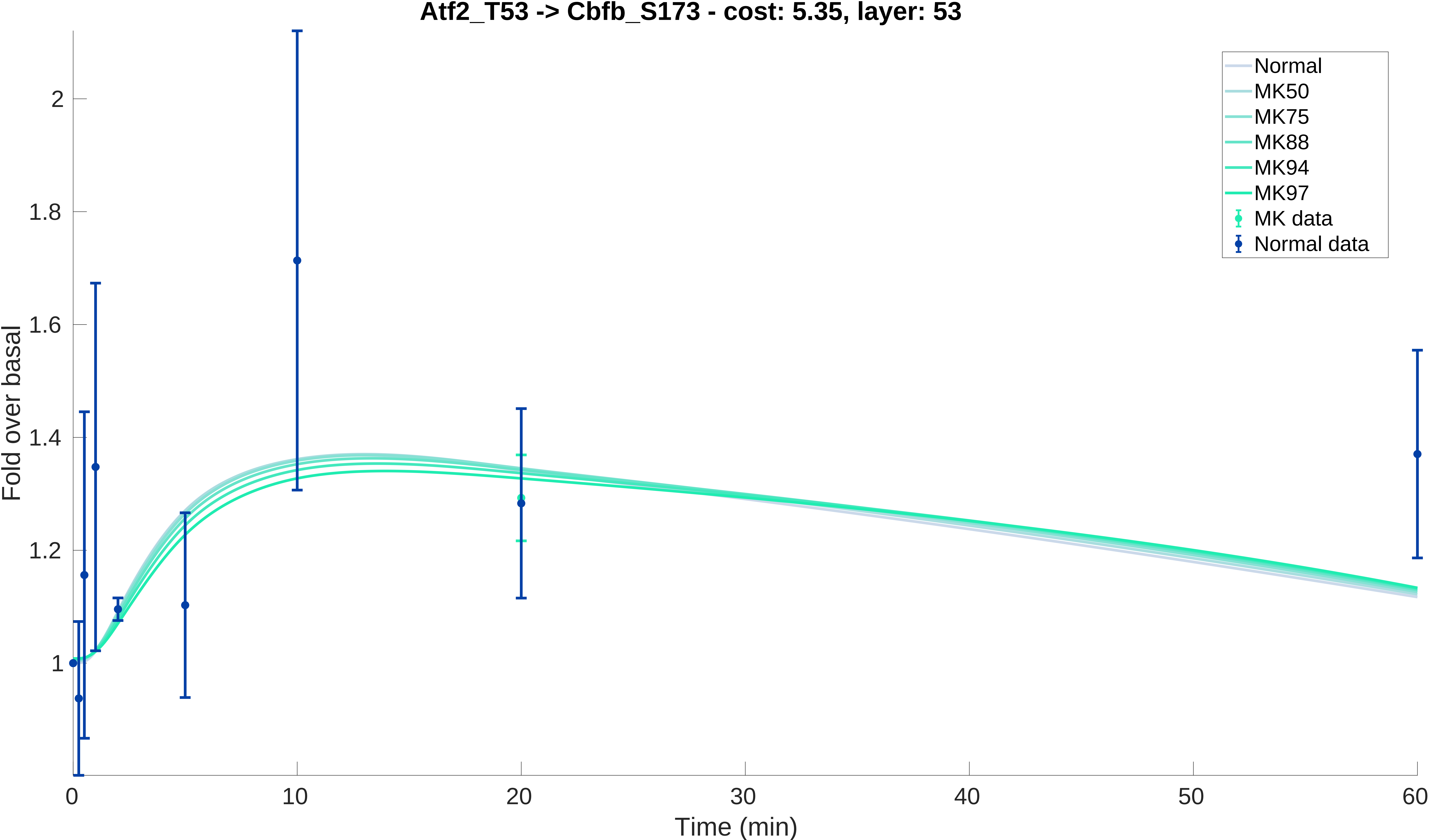
Atf2_T53 -> Cbfb_S173 - cost: 5.35, layer: 53
Fold over basal
Time (min)
0
10
20
30
40
50
60
1
1.2
1.4
1.6
1.8
2
Normal
MK50
MK75
MK88
MK94
MK97
MK data
Normal data

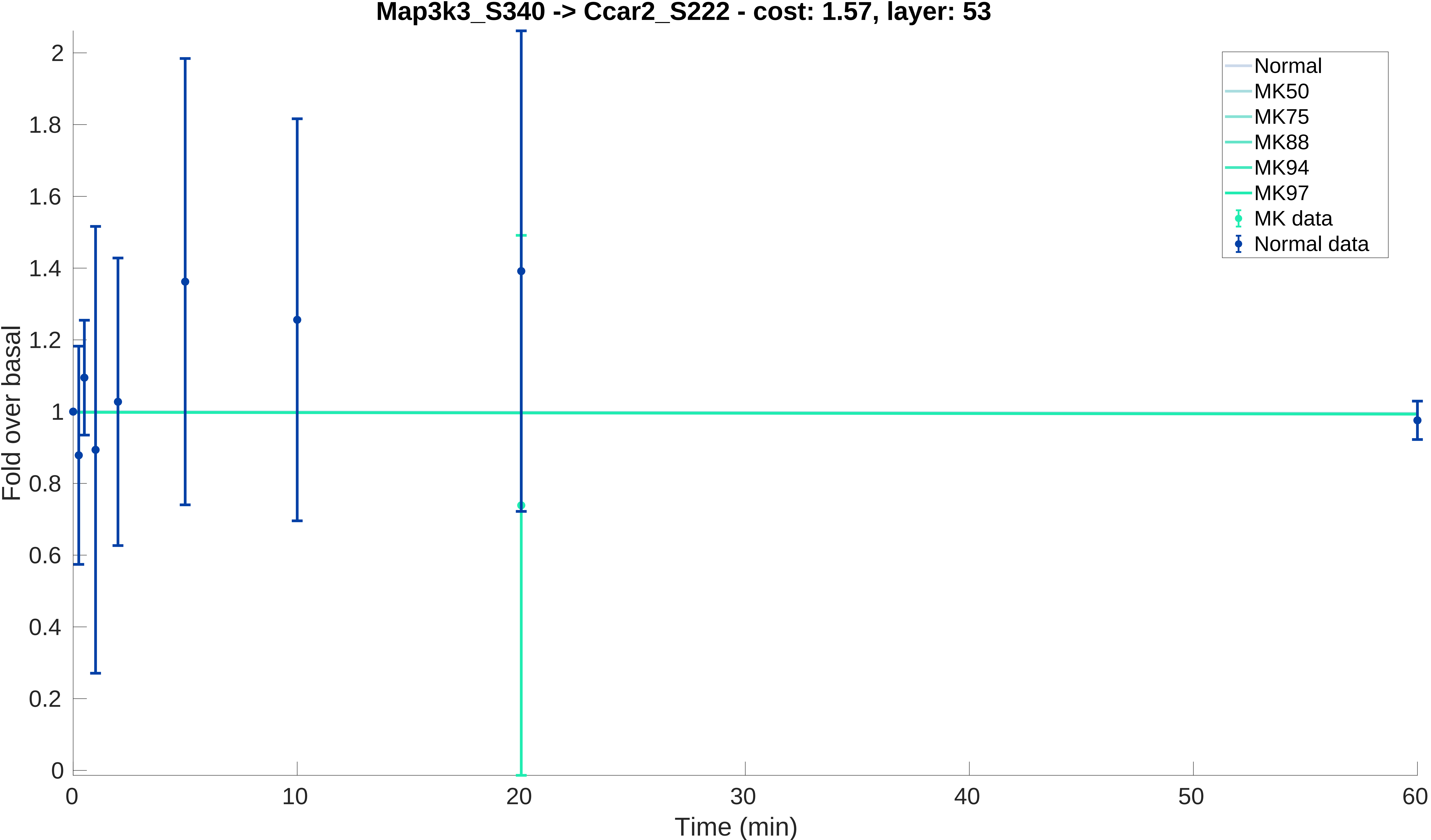

Map3k3_S340 -> Ccar2_S222 - cost: 1.57, layer: 53
Fold over basal
Time (min)
0
0.2
0.4
0.6
0.8
1
1.2
1.4
1.6
1.8
2
0
10
20
30
40
50
60
Normal
MK50
MK75
MK88
MK94
MK97
MK data
Normal data

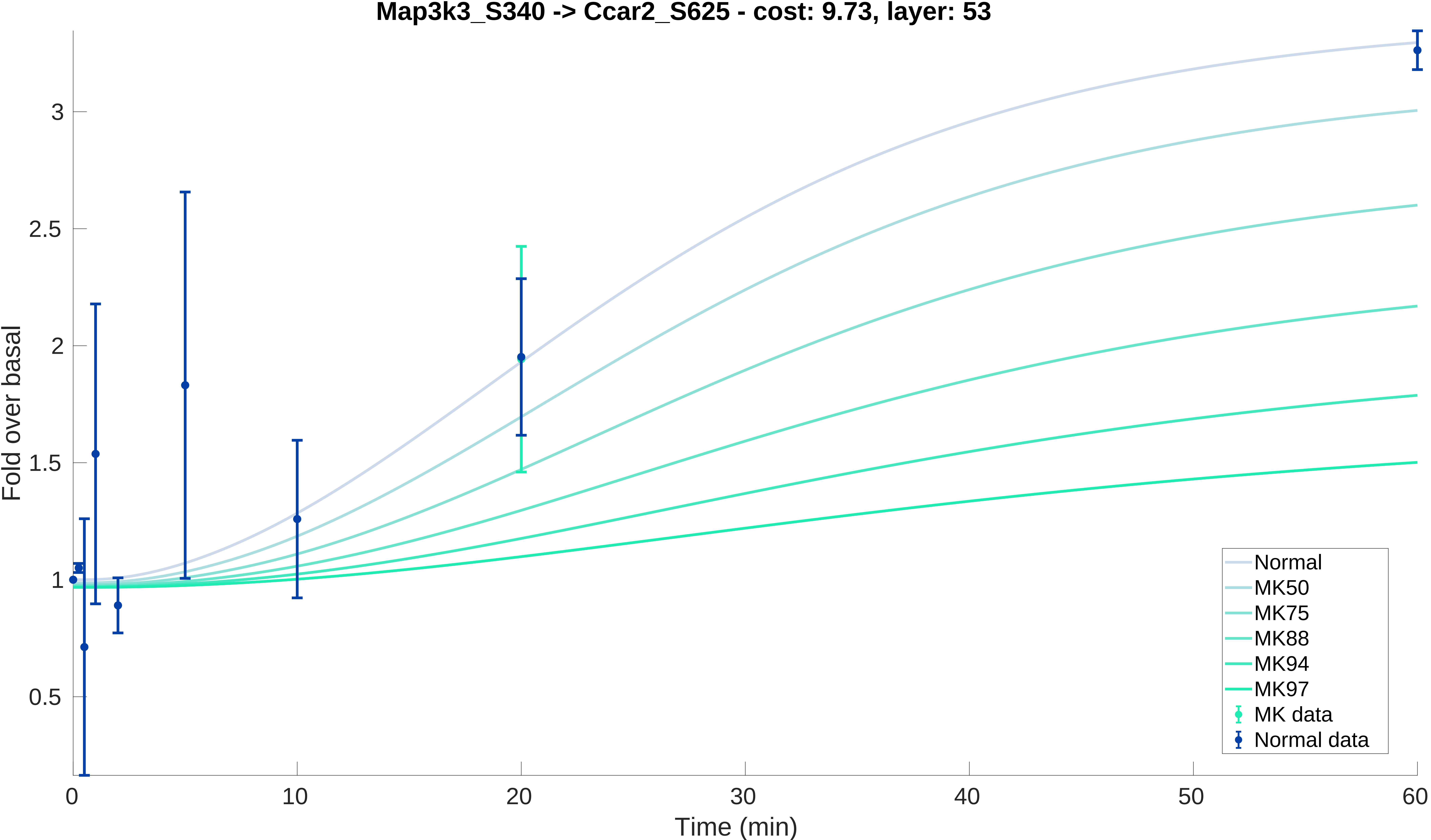
Map3k3_S340 -> Ccar2_S625 - cost: 9.73, layer: 53
Fold over basal
3
2.5
2
1.5
1
0.5
0
10
20
30
40
50
60
Time (min)
Normal
MK50
MK75
MK88
MK94
MK97
MK data
Normal data

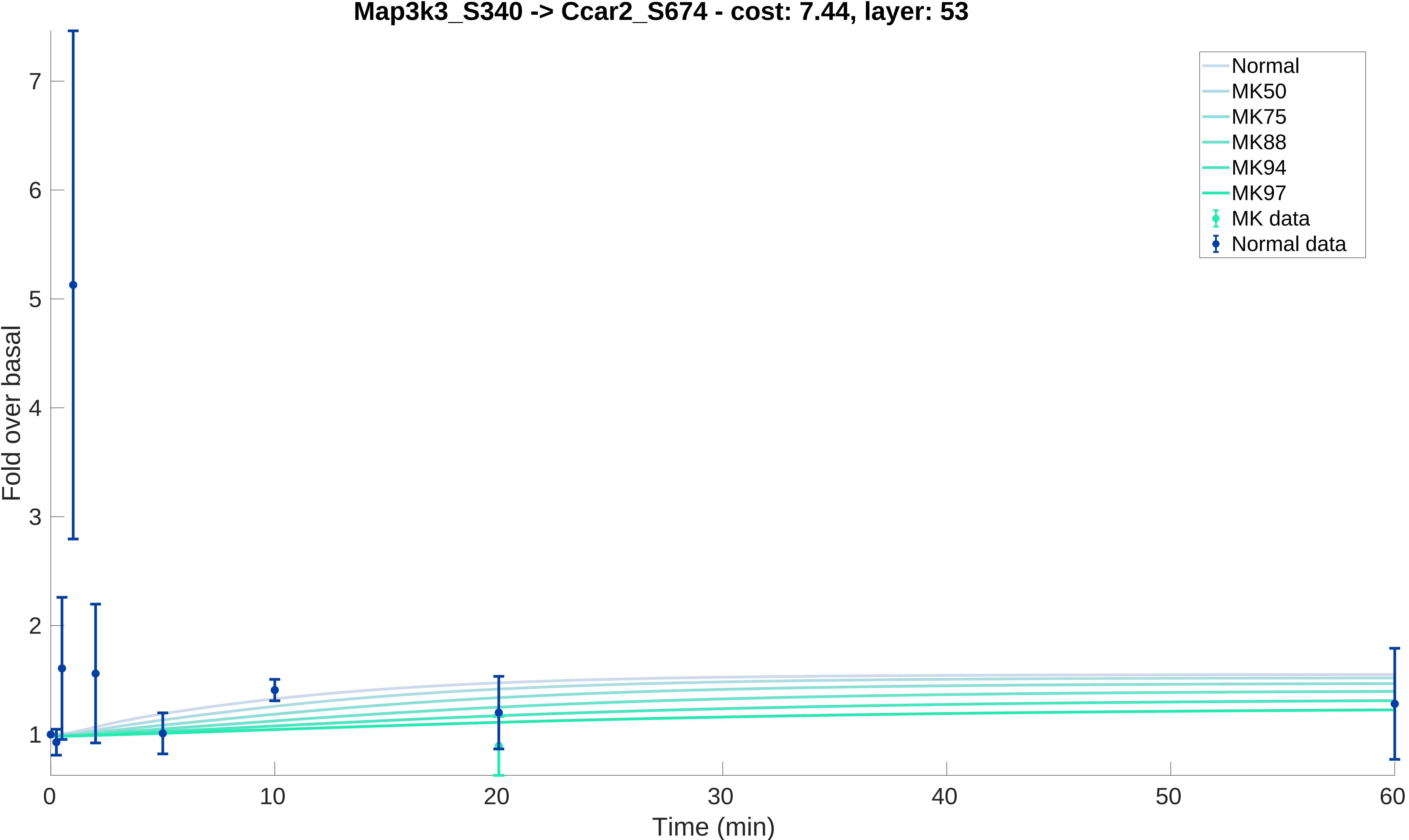

Map3k3_S340 -> Ccar2_S674 - cost: 7.44, layer: 53
Fold over basal
Time (min)
0
10
20
30
40
50
60
1
2
3
4
5
6
7
Normal
MK50
MK75
MK88
MK94
MK97
MK data
Normal data

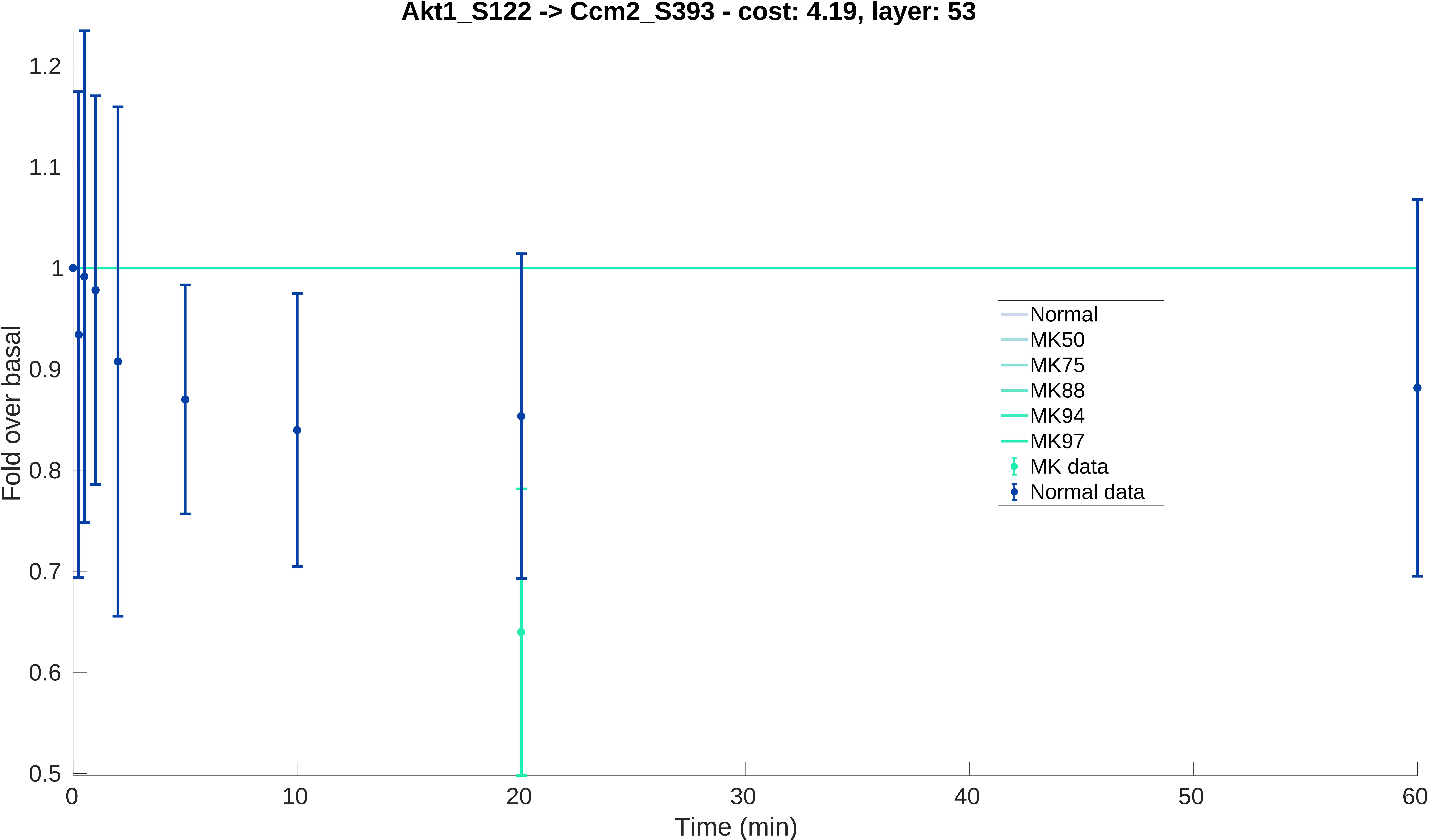

Akt1_S122 -> Ccm2_S393 - cost: 4.19, layer: 53
Fold over basal
Time (min)
1.2
1.1
1
0.9
0.8
0.7
0.6
0.5
0
10
20
30
40
50
60
Normal
MK50
MK75
MK88
MK94
MK97
MK data
Normal data

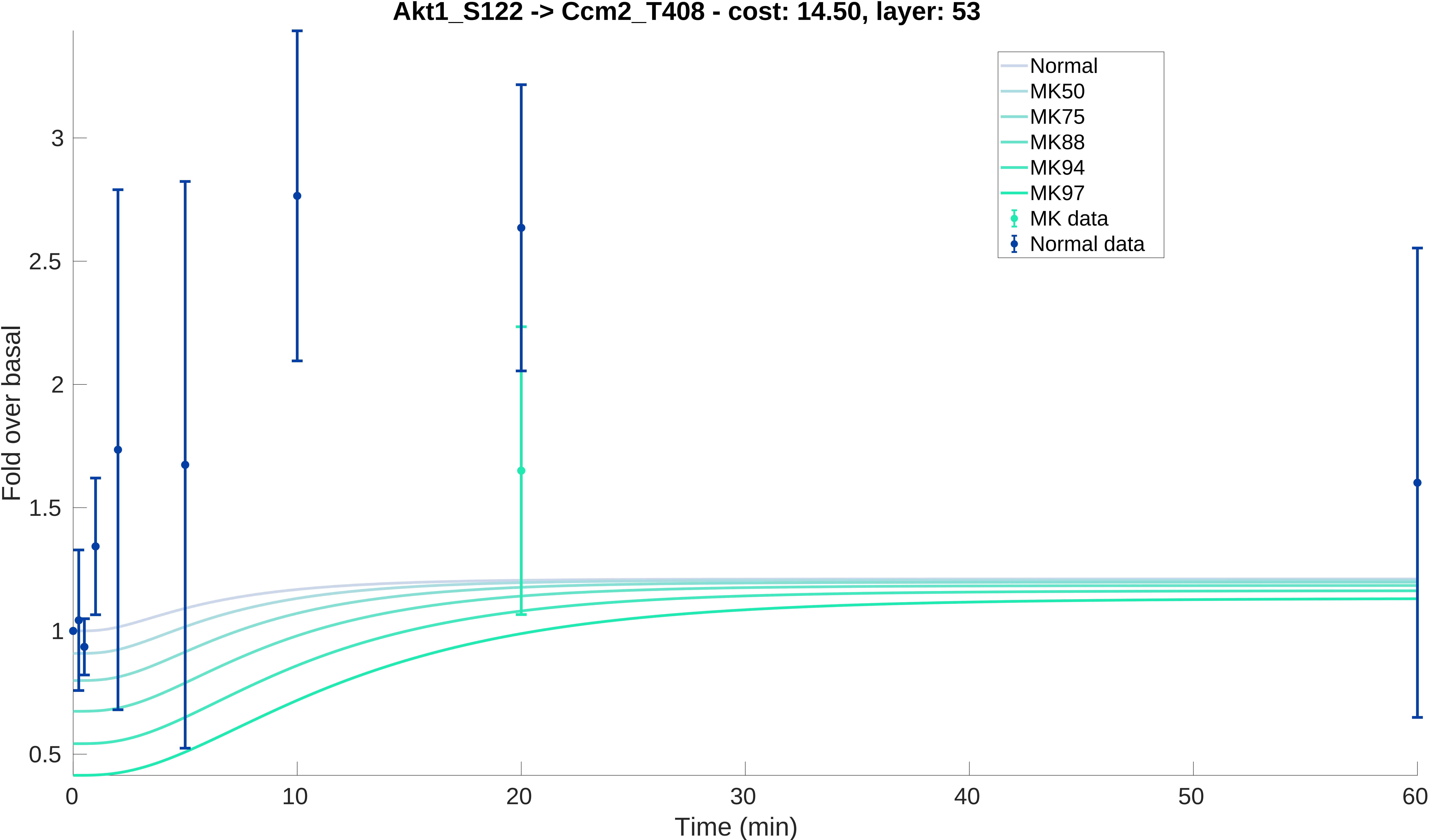

Akt1_S122 -> Ccm2_T408 - cost: 14.50, layer: 53
Fold over basal
Time (min)
0
10
20
30
40
50
60
0.5
1
1.5
2
2.5
3
Normal
MK50
MK75
MK88
MK94
MK97
MK data
Normal data

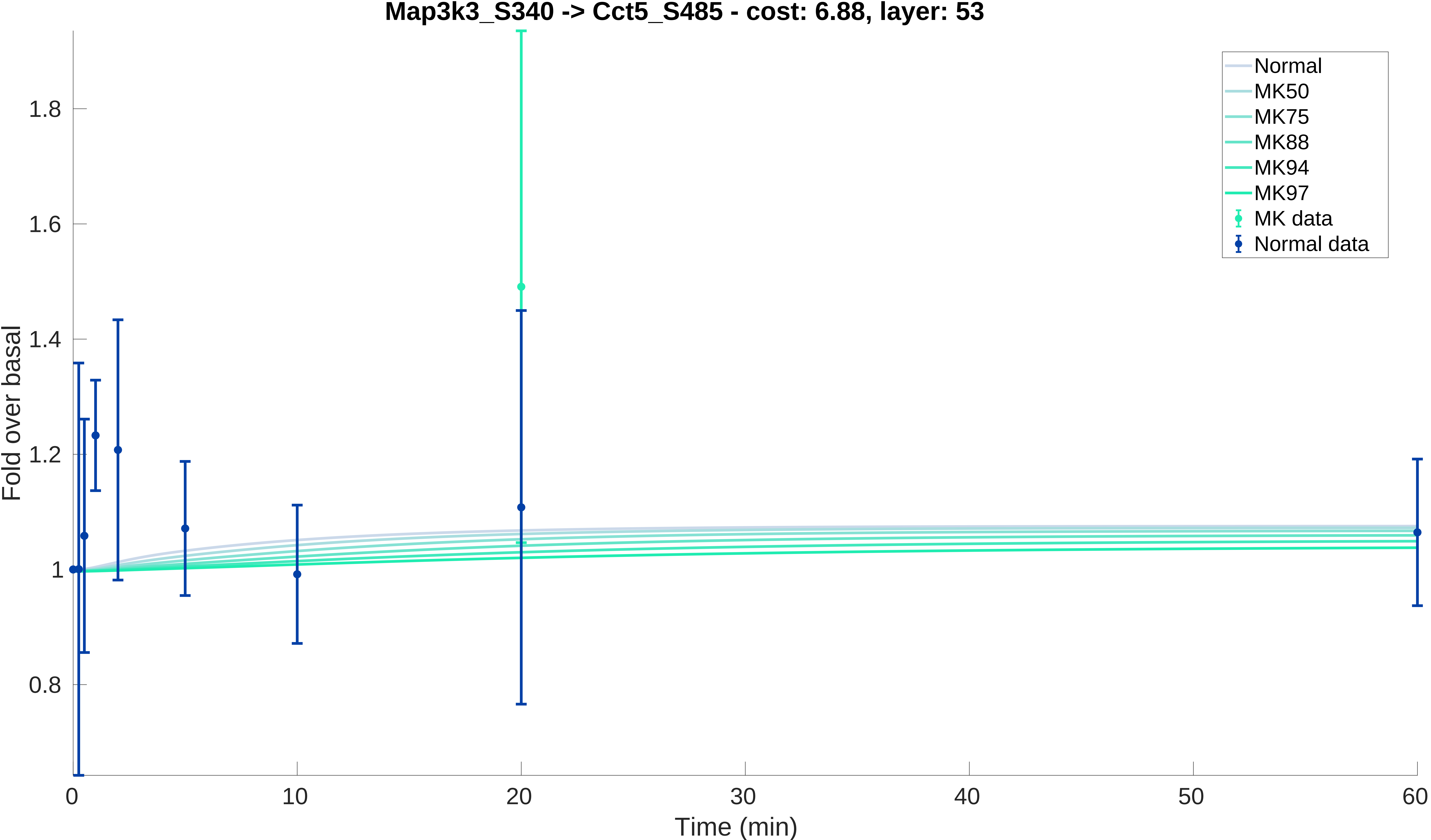
Map3k3_S340 -> Cct5_S485 - cost: 6.88, layer: 53
Fold over basal
Time (min)
0.8
1
1.2
1.4
1.6
1.8
0
10
20
30
40
50
60
Normal
MK50
MK75
MK88
MK94
MK97
MK data
Normal data

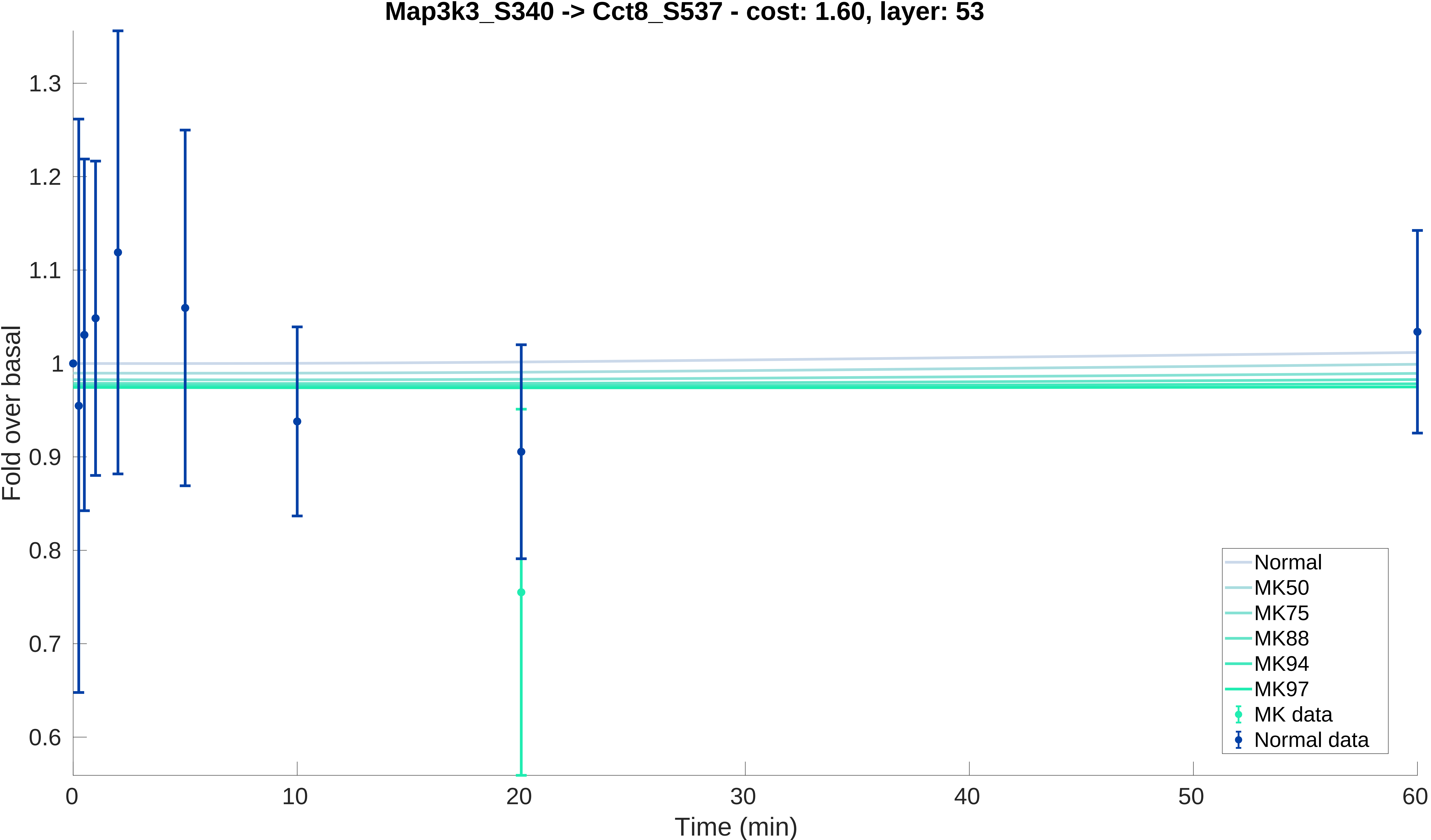
Map3k3_S340 -> Cct8_S537 - cost: 1.60, layer: 53
Fold over basal
Time (min)
0.6
0.7
0.8
0.9
1
1.1
1.2
1.3
0
10
20
30
40
50
60
Normal
MK50
MK75
MK88
MK94
MK97
MK data
Normal data

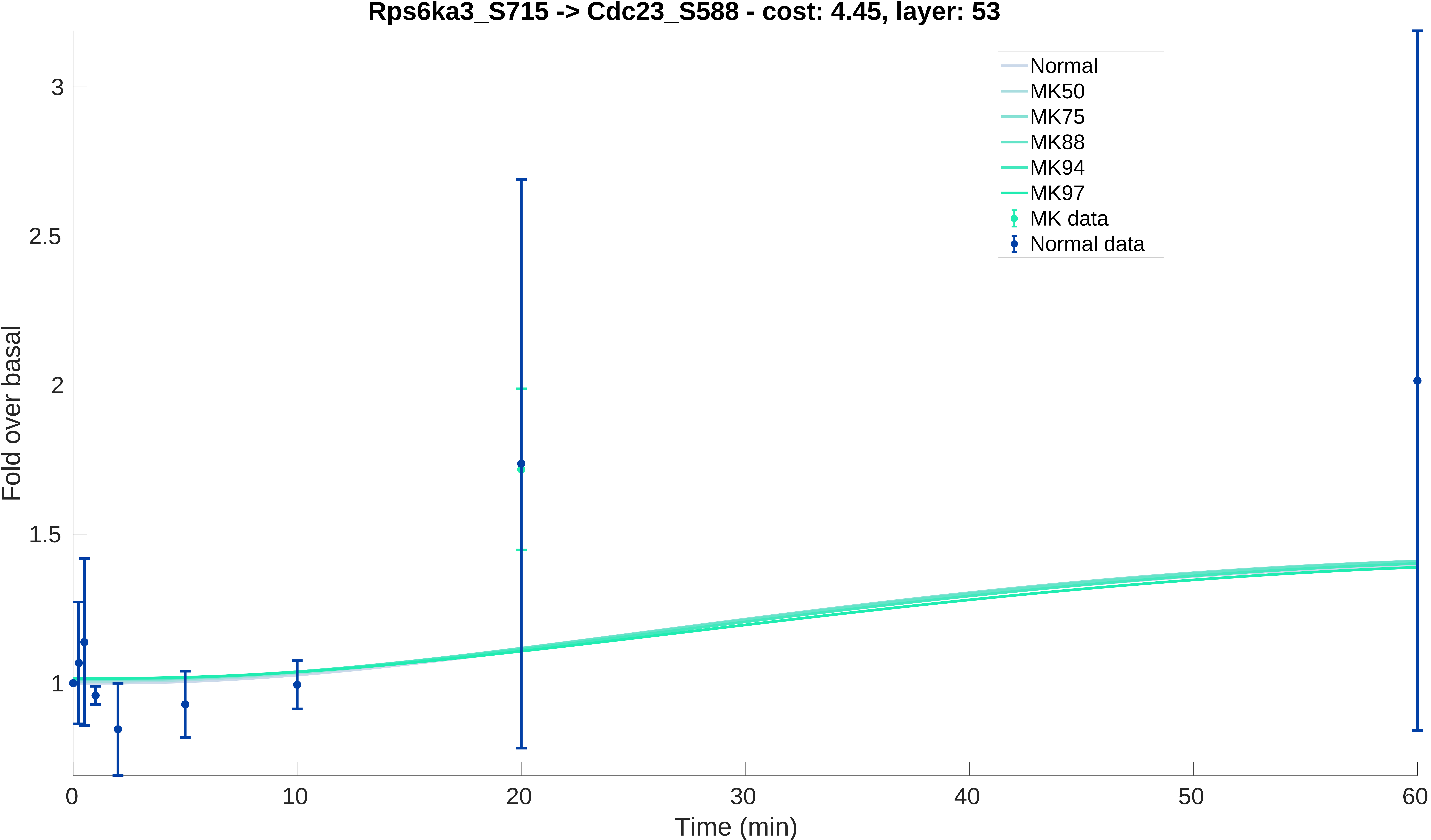

Rps6ka3_S715 -> Cdc23_S588 - cost: 4.45, layer: 53
Fold over basal
Time (min)
0
10
20
30
40
50
60
1
1.5
2
2.5
3
Normal
MK50
MK75
MK88
MK94
MK97
MK data
Normal data

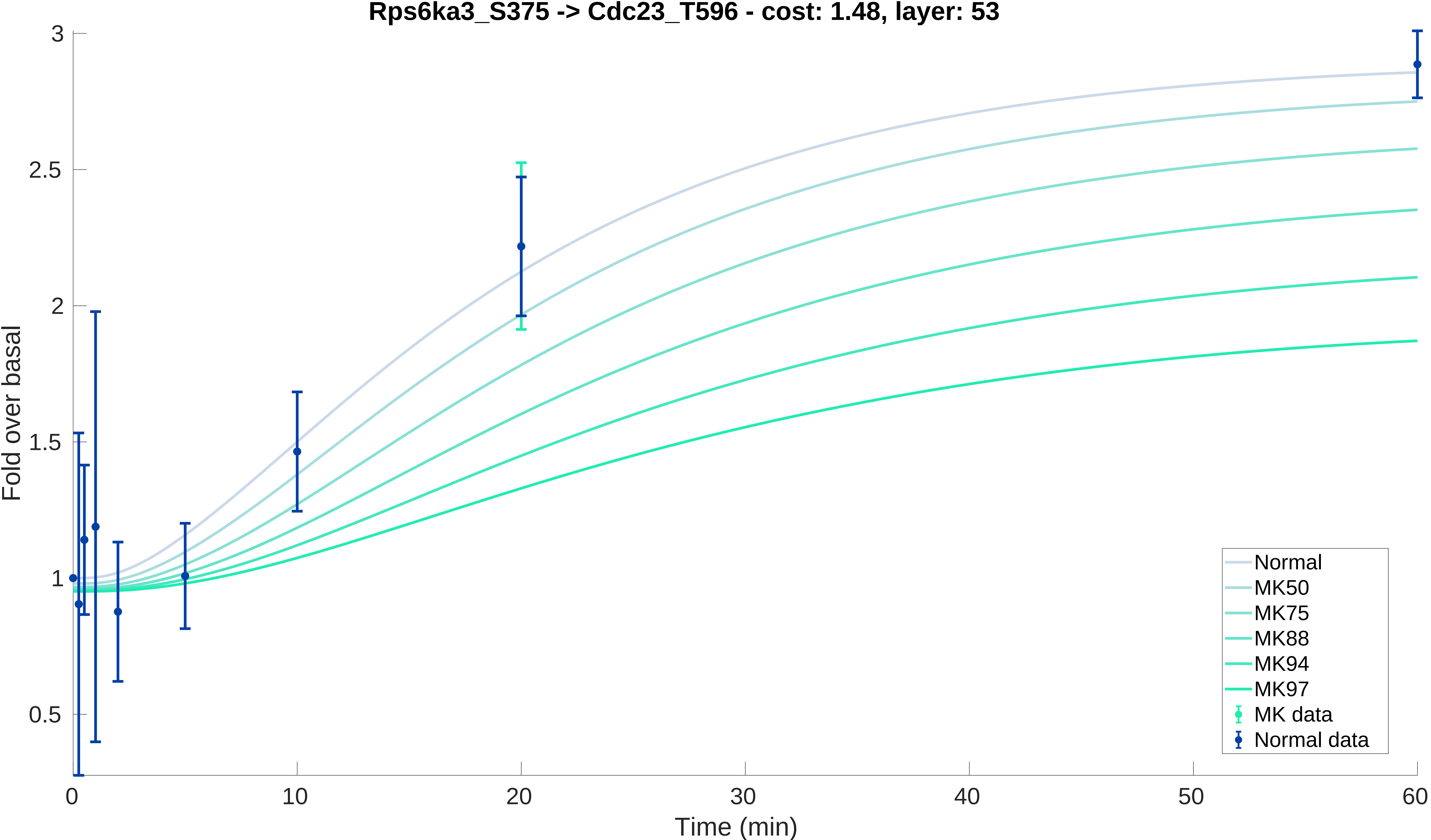

Rps6ka3_S375 -> Cdc23_T596 - cost: 1.48, layer: 53
Fold over basal
Time (min)
3
2.5
2
1.5
1
0.5
0
10
20
30
40
50
60
Normal
MK50
MK75
MK88
MK94
MK97
MK data
Normal data

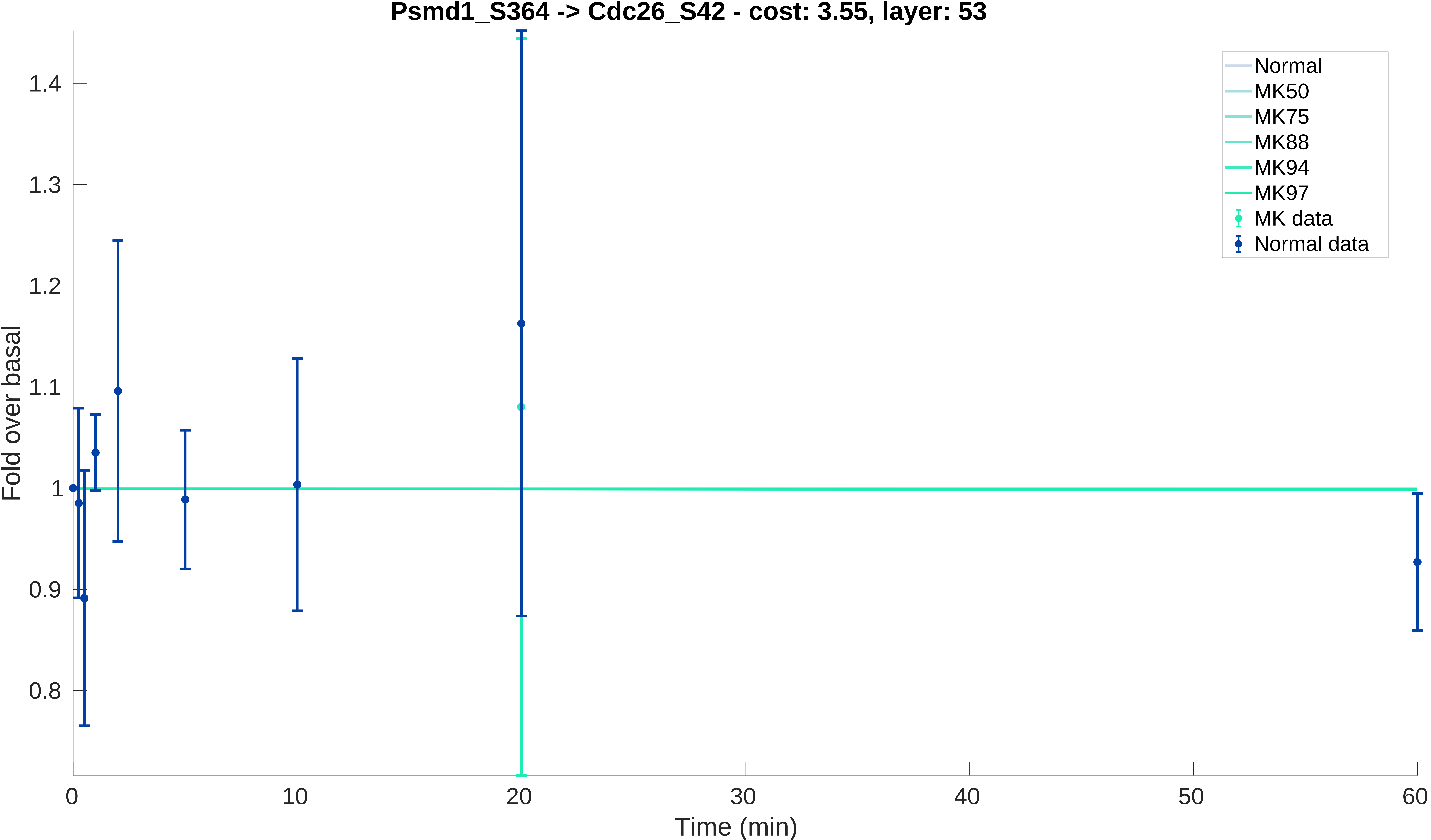
Psmd1_S364 -> Cdc26_S42 - cost: 3.55, layer: 53
Fold over basal
Time (min)
0.8
0.9
1
1.1
1.2
1.3
1.4
0
10
20
30
40
50
60
Normal
MK50
MK75
MK88
MK94
MK97
MK data
Normal data

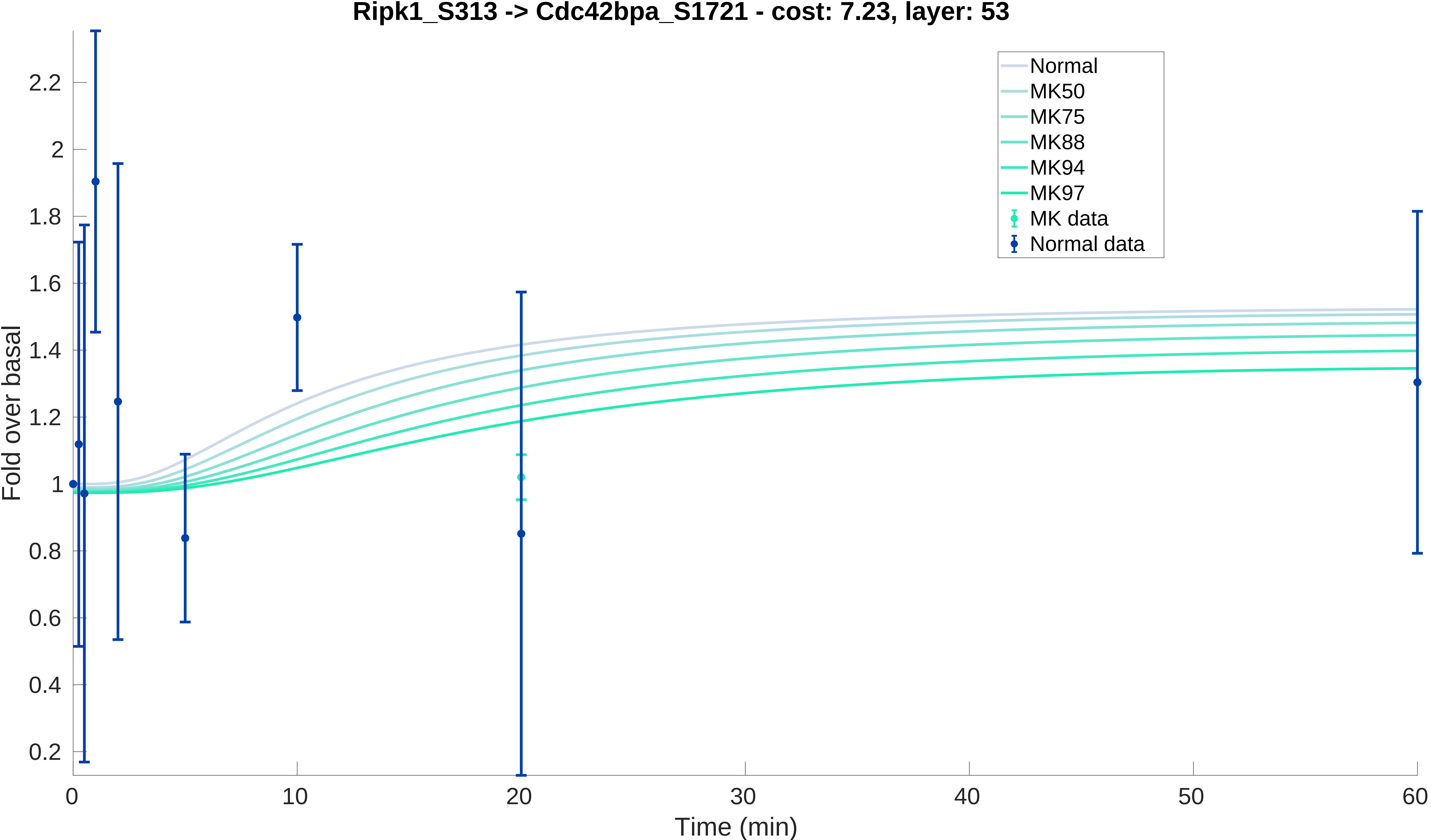

Ripk1_S313 -> Cdc42bpa_S1721 - cost: 7.23, layer: 53
Normal
MK50
MK75
MK88
MK94
MK97
MK data
Normal data
Fold over basal
Time (min)
0.2
0.4
0.6
0.8
1
1.2
1.4
1.6
1.8
2
2.2
0
10
20
30
40
50
60

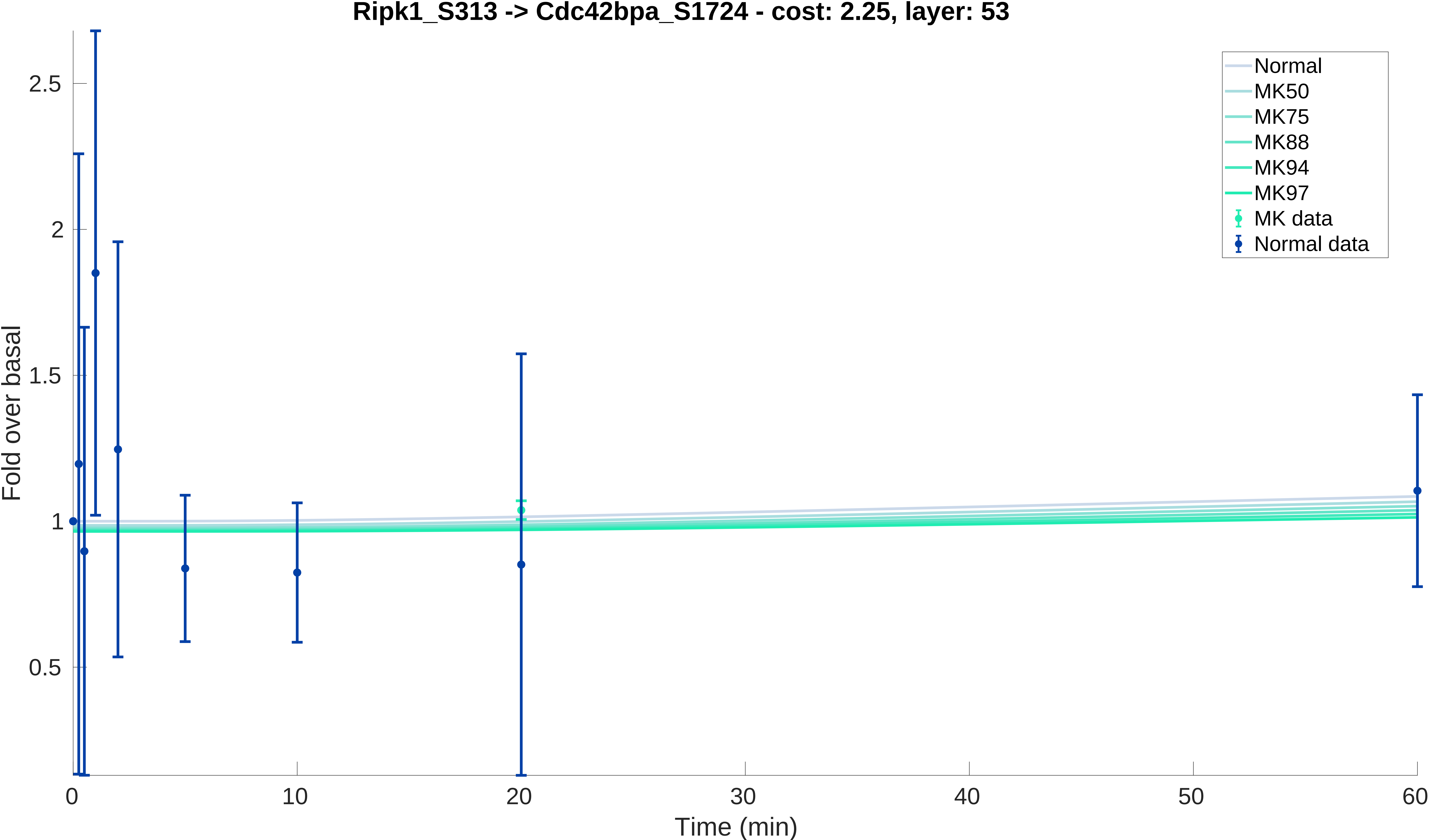

Ripk1_S313 -> Cdc42bpa_S1724 - cost: 2.25, layer: 53
Fold over basal
Time (min)
Normal
MK50
MK75
MK88
MK94
MK97
MK data
Normal data
0.5
1
1.5
2
2.5
0
10
20
30
40
50
60

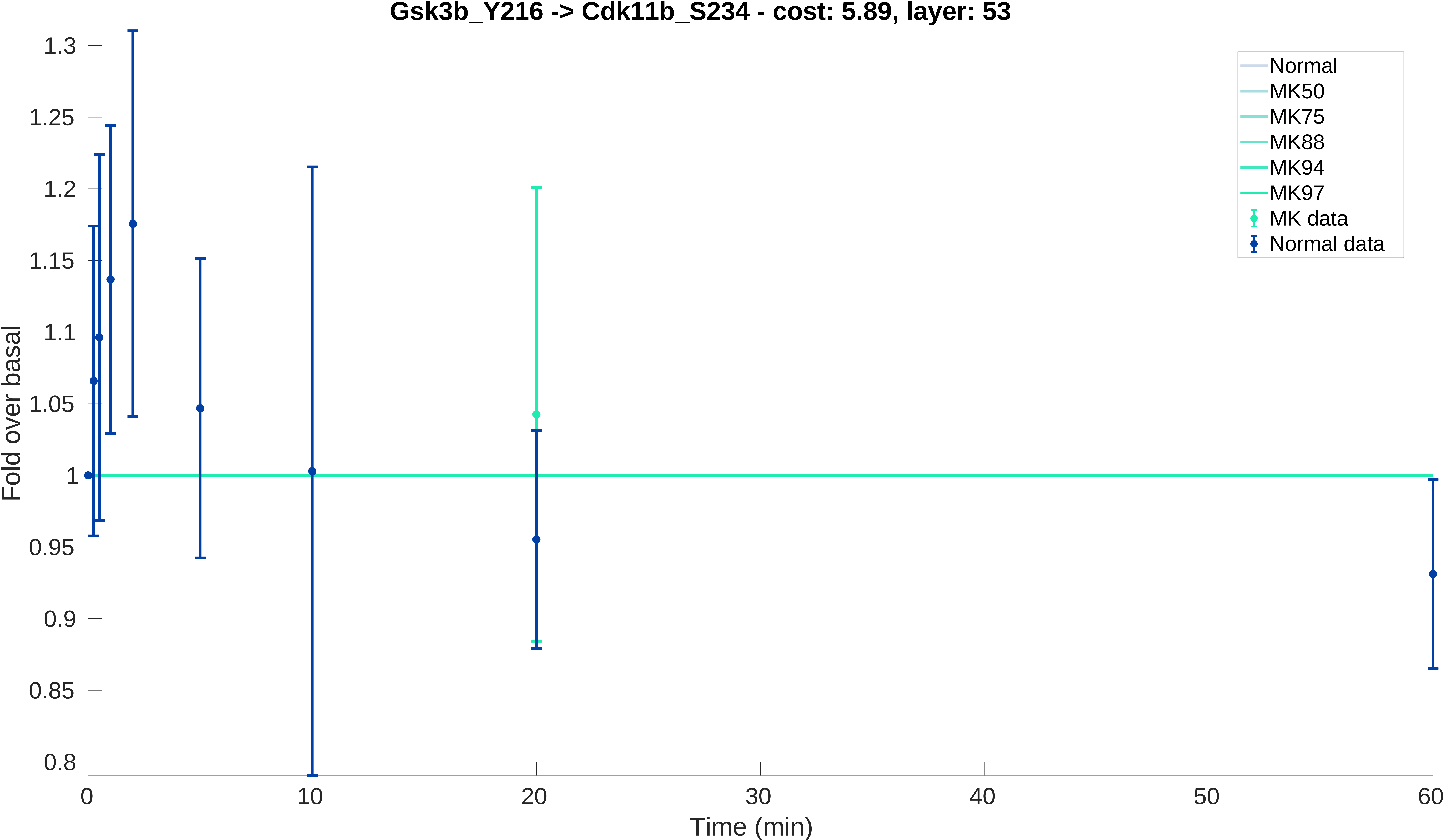

Gsk3b_Y216 -> Cdk11b_S234 - cost: 5.89, layer: 53
Fold over basal
Time (min)
1.3
1.25
1.2
1.15
1.1
1.05
1
0.95
0.9
0.85
0.8
0
10
20
30
40
50
60
Normal
MK50
MK75
MK88
MK94
MK97
MK data
Normal data

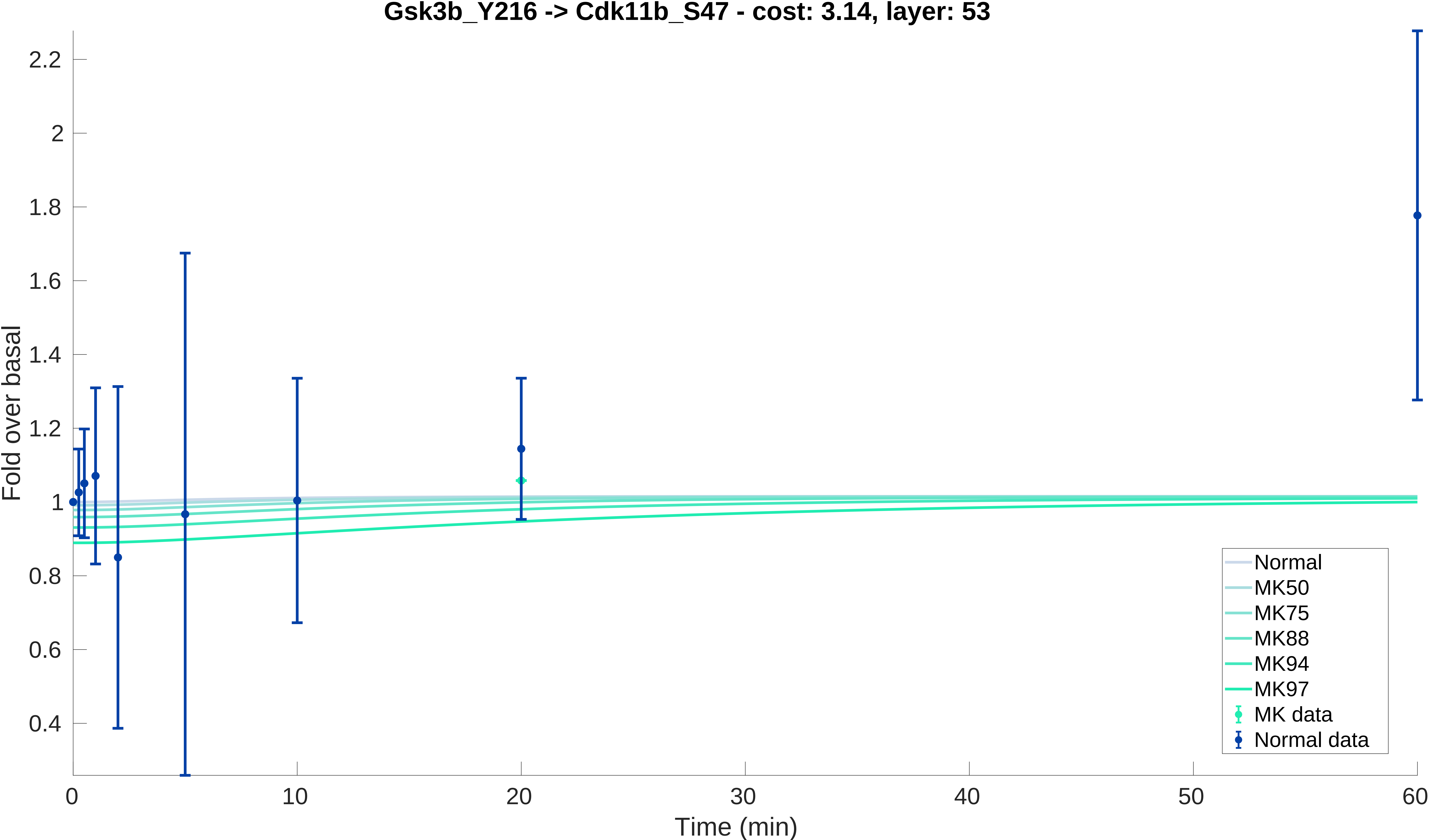

Gsk3b_Y216 -> Cdk11b_S47 - cost: 3.14, layer: 53
Fold over basal
2.2
2
1.8
1.6
1.4
1.2
1
0.8
0.6
0.4
0
10
20
30
40
50
60
Time (min)
Normal
MK50
MK75
MK88
MK94
MK97
MK data
Normal data

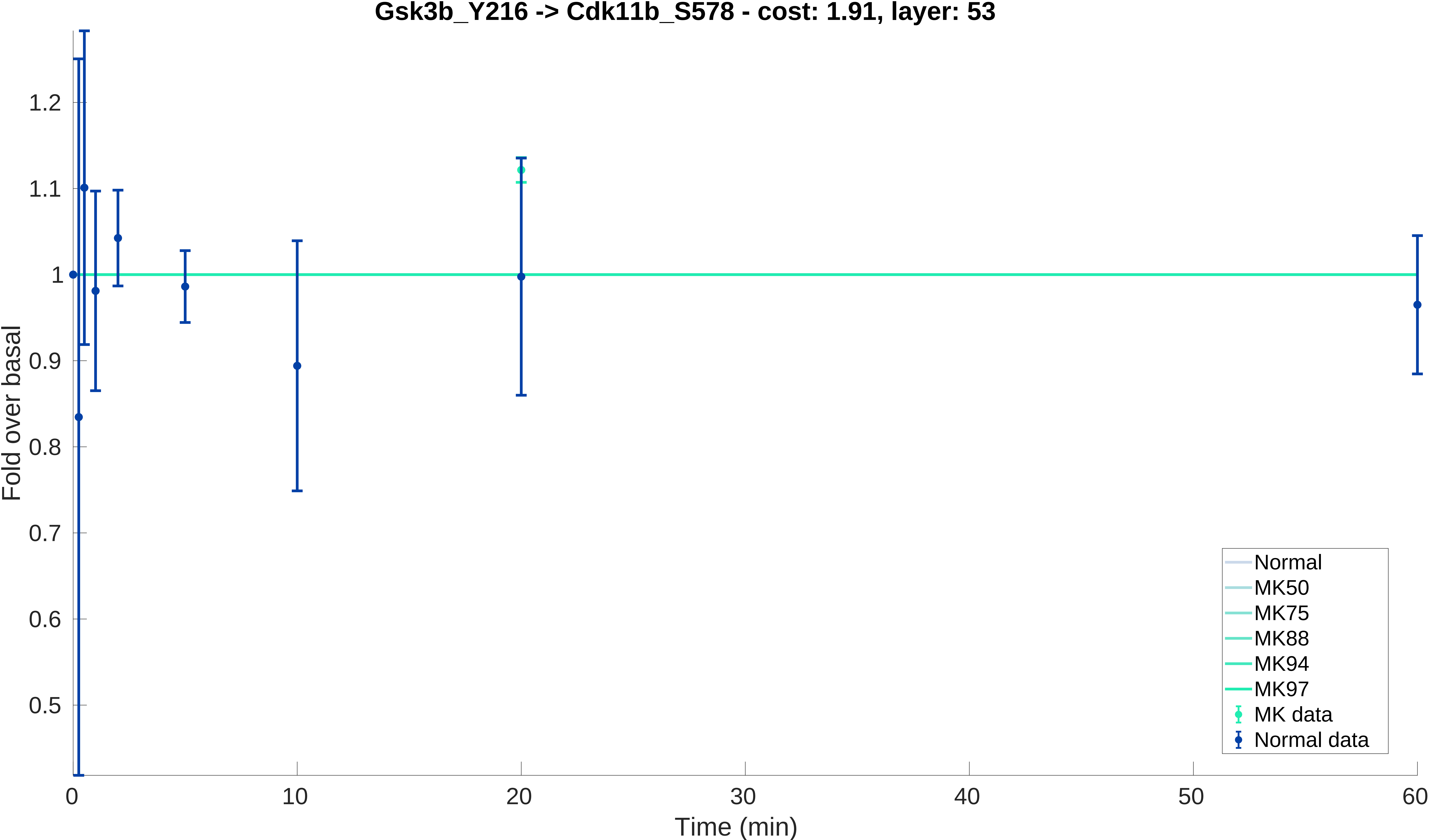

Gsk3b_Y216 -> Cdk11b_S578 - cost: 1.91, layer: 53
Fold over basal
1.2
1.1
1
0.9
0.8
0.7
0.6
0.5
0
10
20
30
40
50
60
Time (min)
Normal
MK50
MK75
MK88
MK94
MK97
MK data
Normal data

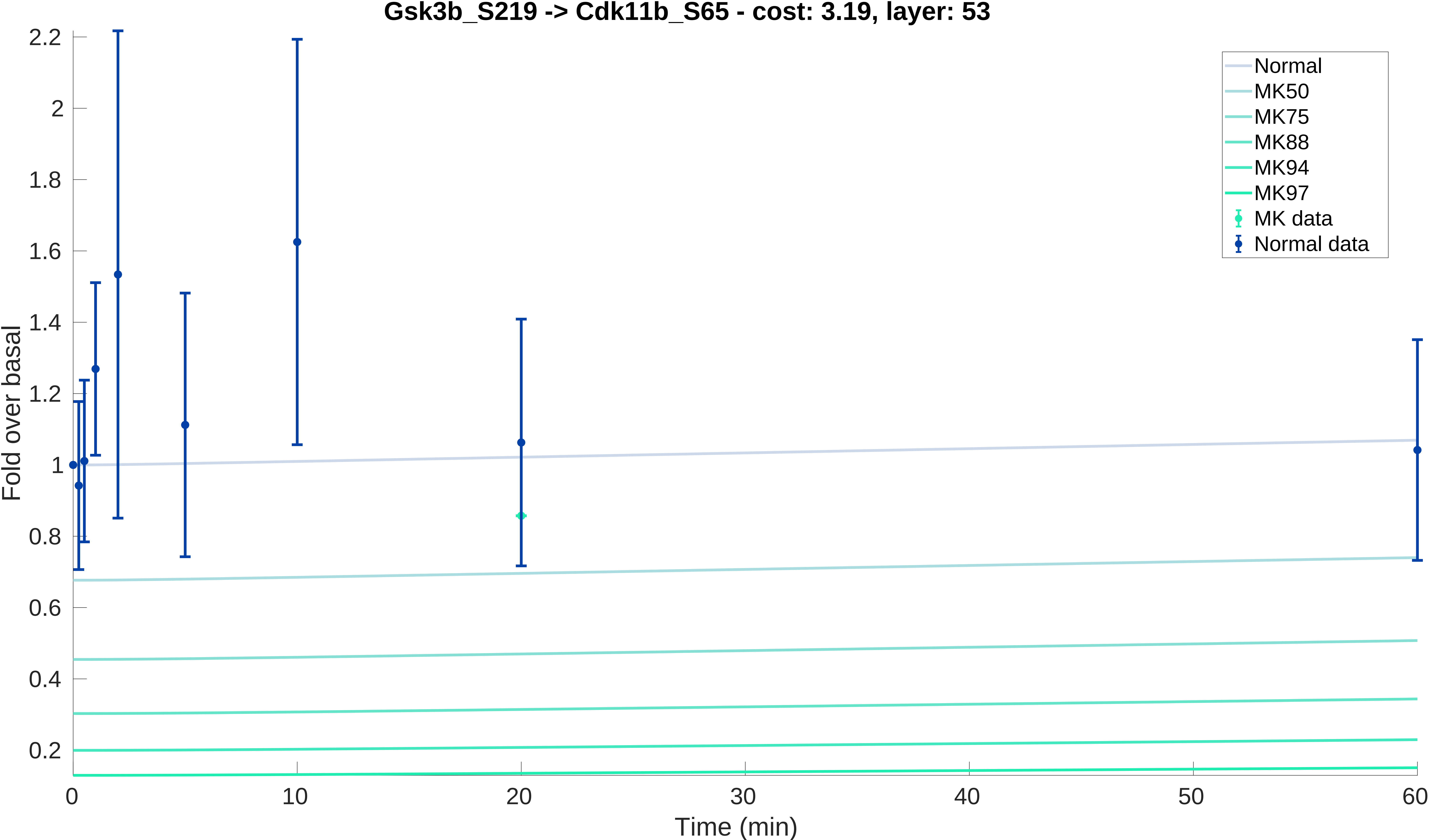

Gsk3b_S219 -> Cdk11b_S65 - cost: 3.19, layer: 53
Fold over basal
Time (min)
2.2
2
1.8
1.6
1.4
1.2
1
0.8
0.6
0.4
0.2
0
10
20
30
40
50
60
Normal
MK50
MK75
MK88
MK94
MK97
MK data
Normal data

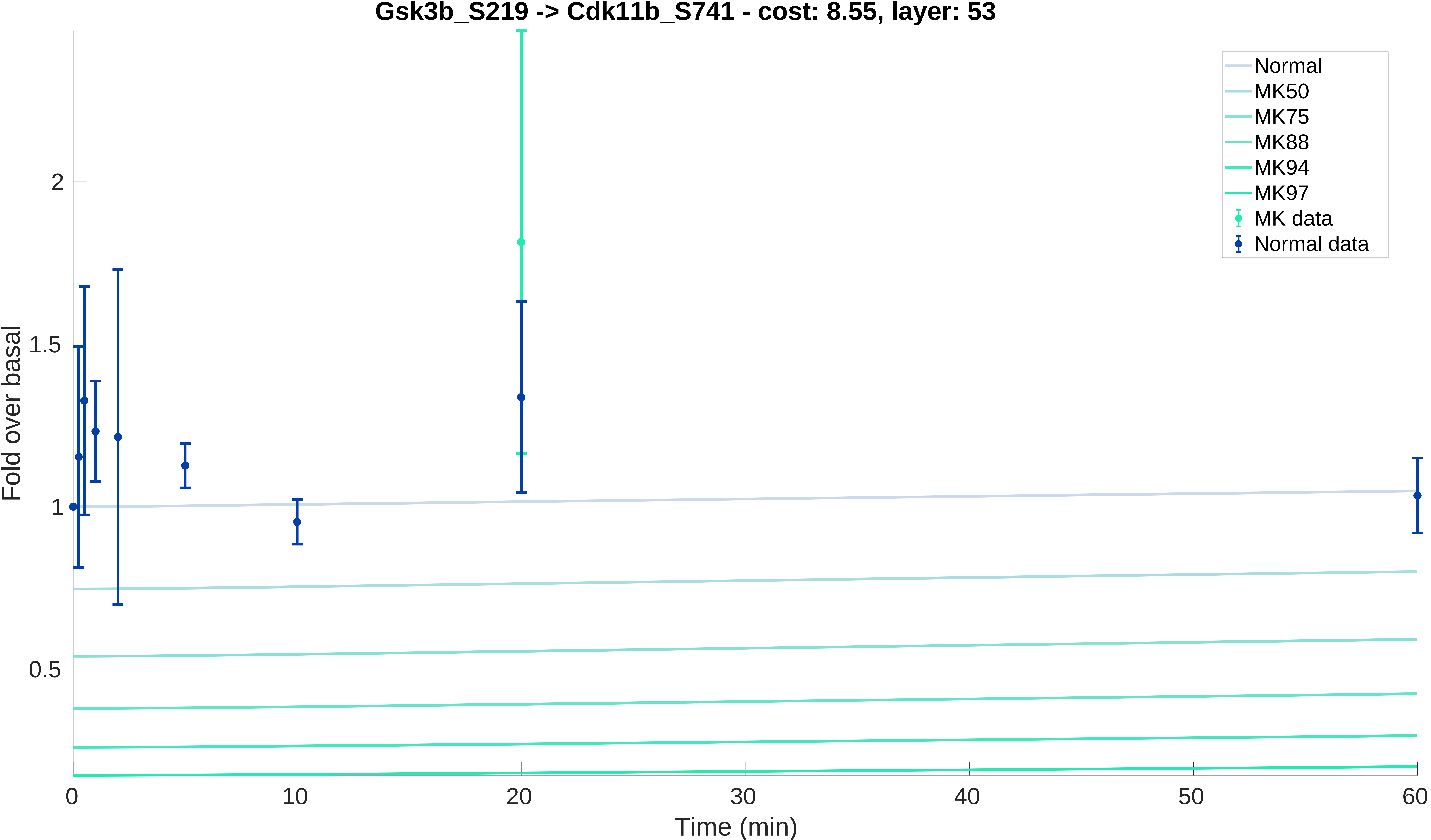

Gsk3b_S219 -> Cdk11b_S741 - cost: 8.55, layer: 53
Fold over basal
Time (min)
0
10
20
30
40
50
60
0.5
1
1.5
2
Normal
MK50
MK75
MK88
MK94
MK97
MK data
Normal data

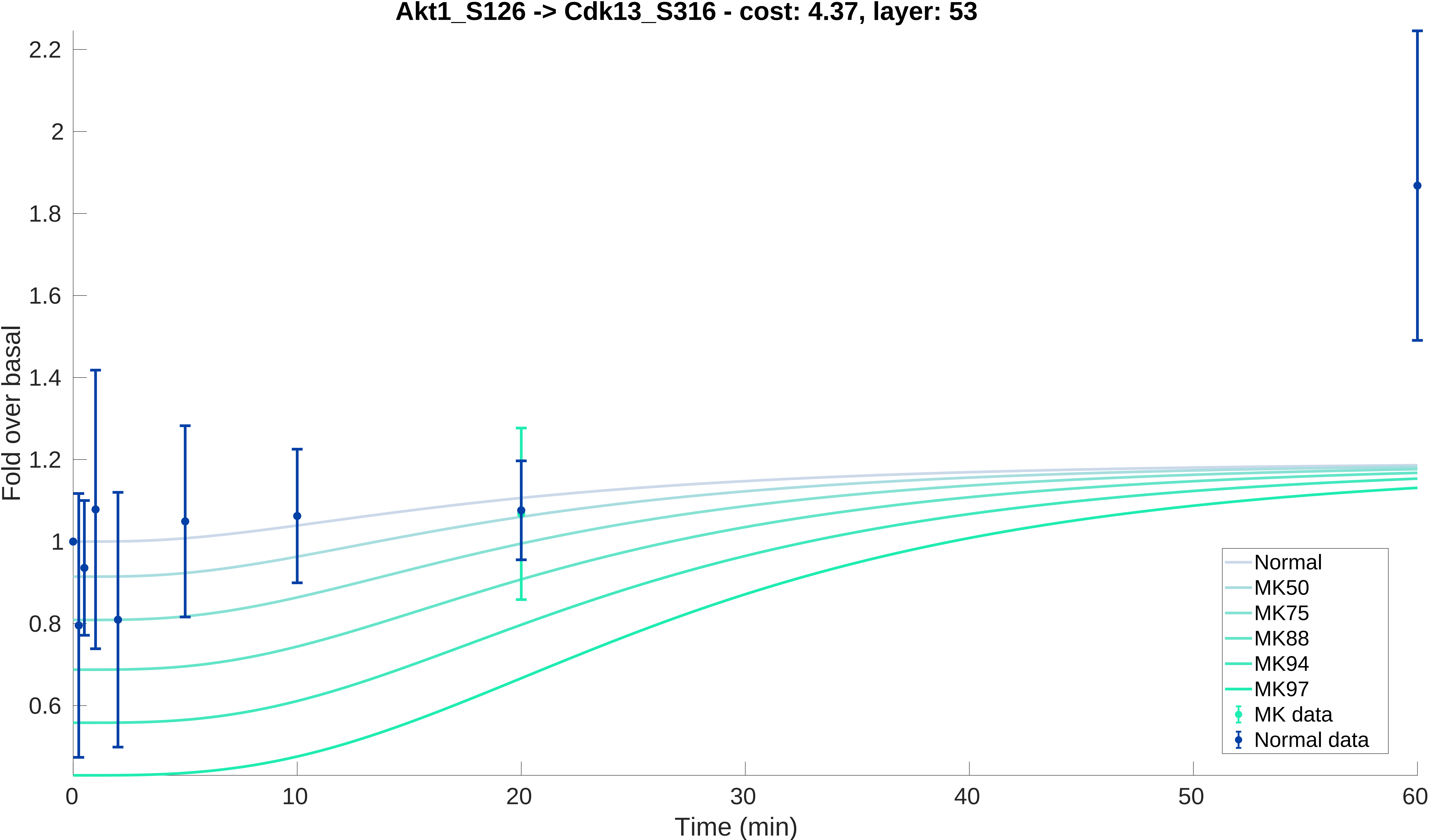

Akt1_S126 -> Cdk13_S316 - cost: 4.37, layer: 53
Fold over basal
2.2
2
1.8
1.6
1.4
1.2
1
0.8
0.6
Time (min)
0
10
20
30
40
50
60
Normal
MK50
MK75
MK88
MK94
MK97
MK data
Normal data

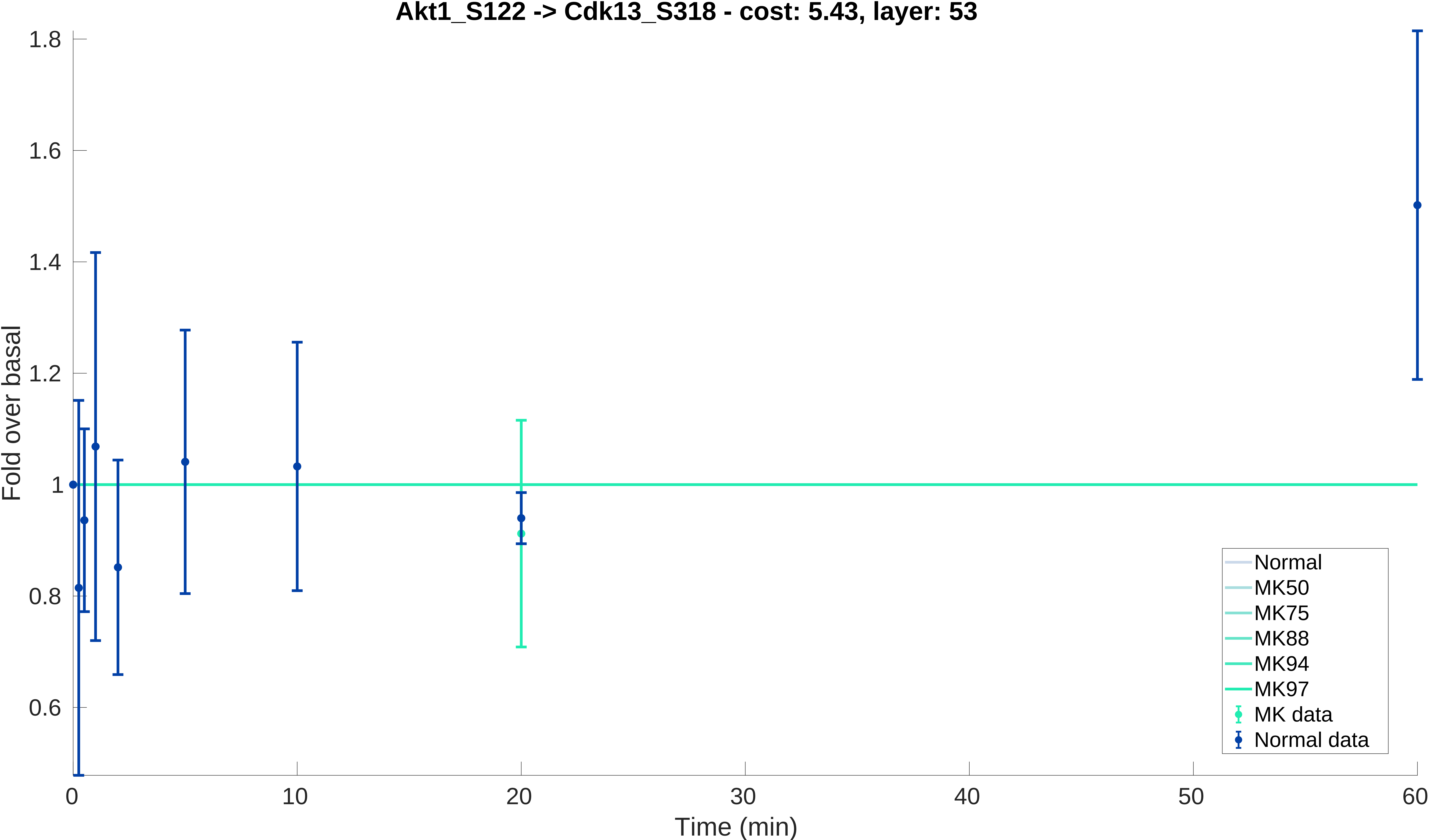

Akt1_S122 -> Cdk13_S318 - cost: 5.43, layer: 53
Fold over basal
1.8
1.6
1.4
1.2
1
0.8
0.6
0
10
20
30
40
50
60
Time (min)
Normal
MK50
MK75
MK88
MK94
MK97
MK data
Normal data

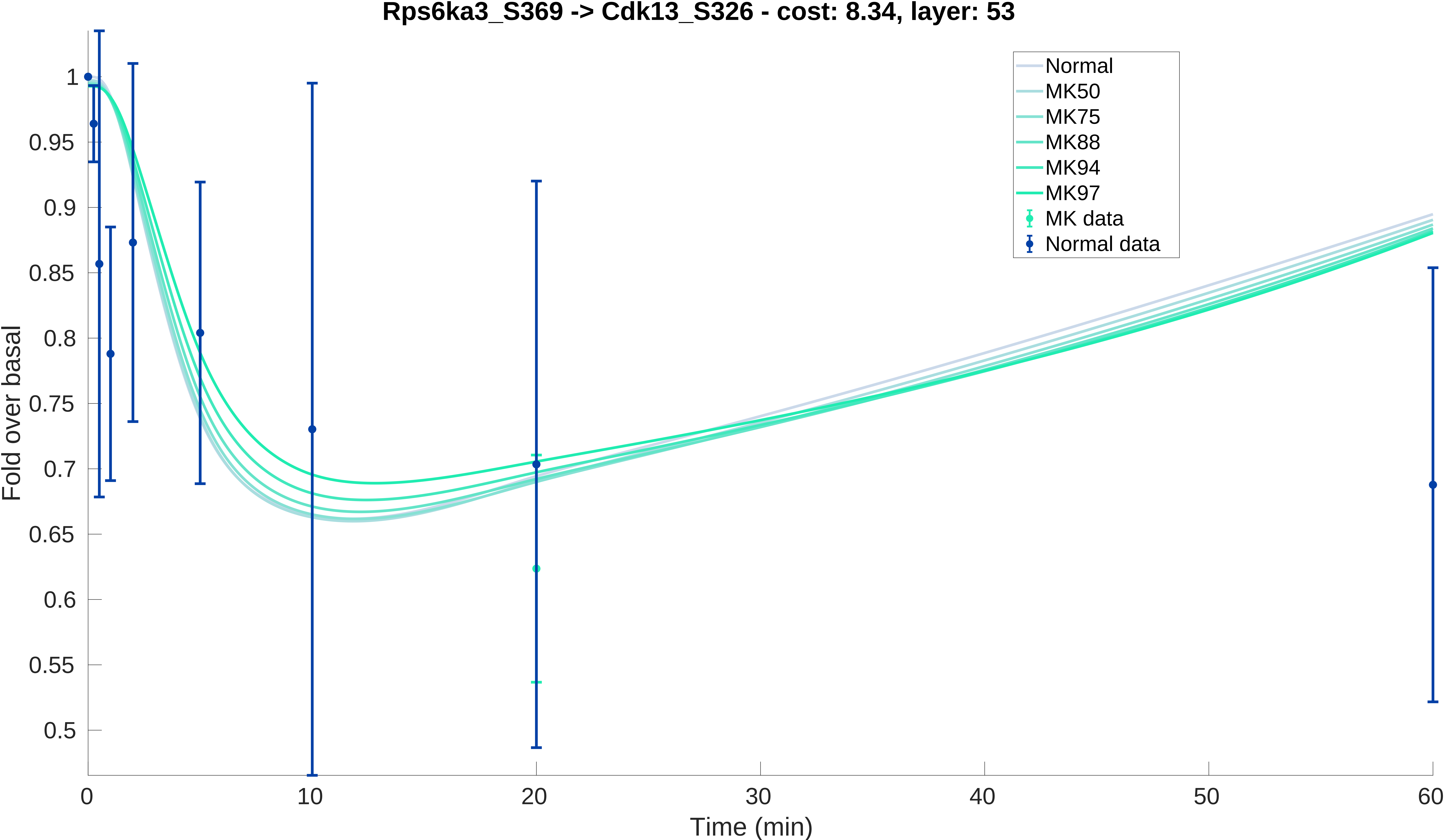

Rps6ka3_S369 -> Cdk13_S326 - cost: 8.34, layer: 53
Normal
MK50
MK75
MK88
MK94
MK97
MK data
Normal data
Fold over basal
Time (min)
1
0.95
0.9
0.85
0.8
0.75
0.7
0.65
0.6
0.55
0.5
0
10
20
30
40
50
60

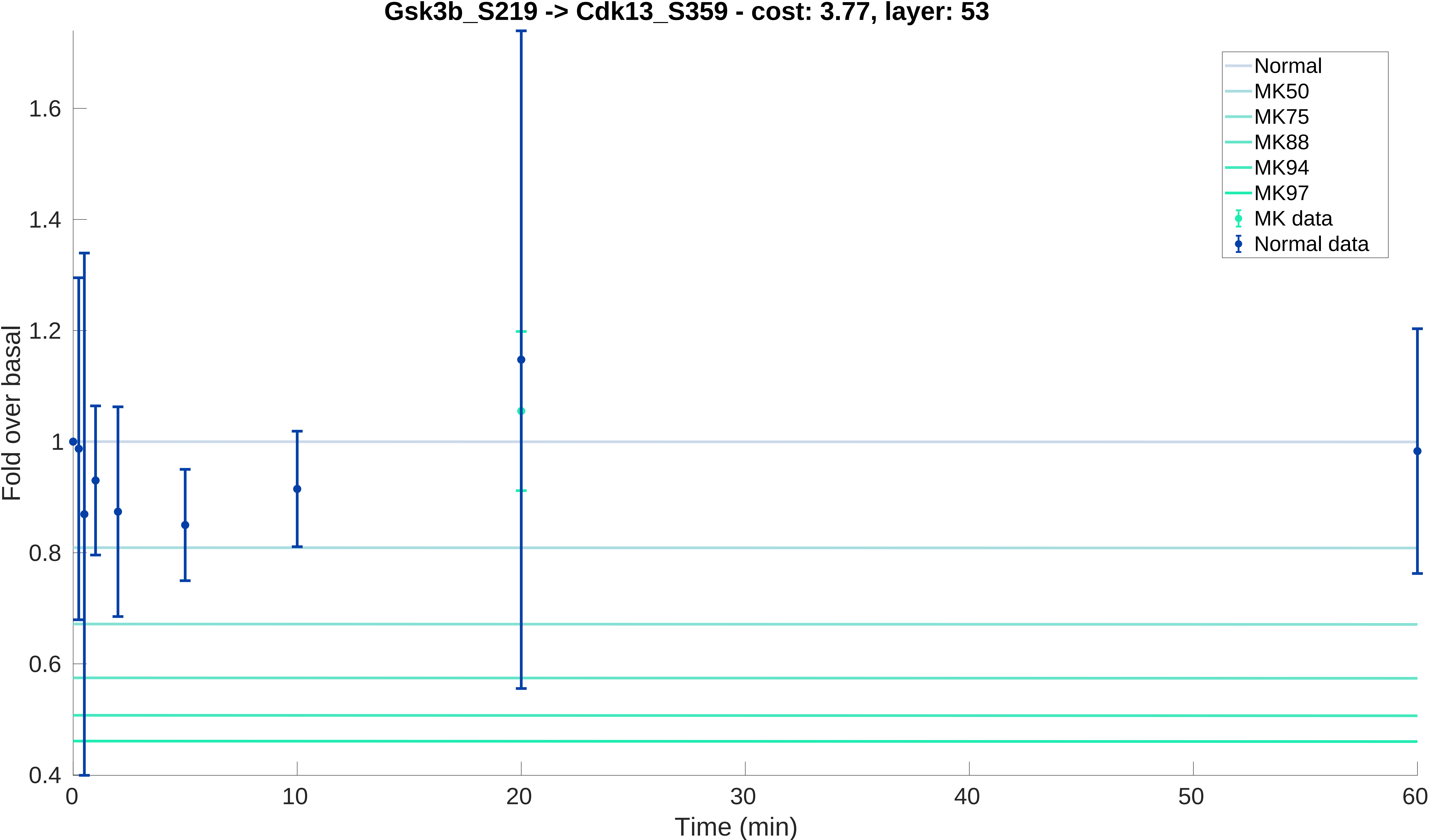

Gsk3b_S219 -> Cdk13_S359 - cost: 3.77, layer: 53
Fold over basal
Time (min)
0.4
0.6
0.8
1
1.2
1.4
1.6
0
10
20
30
40
50
60
Normal
MK50
MK75
MK88
MK94
MK97
MK data
Normal data

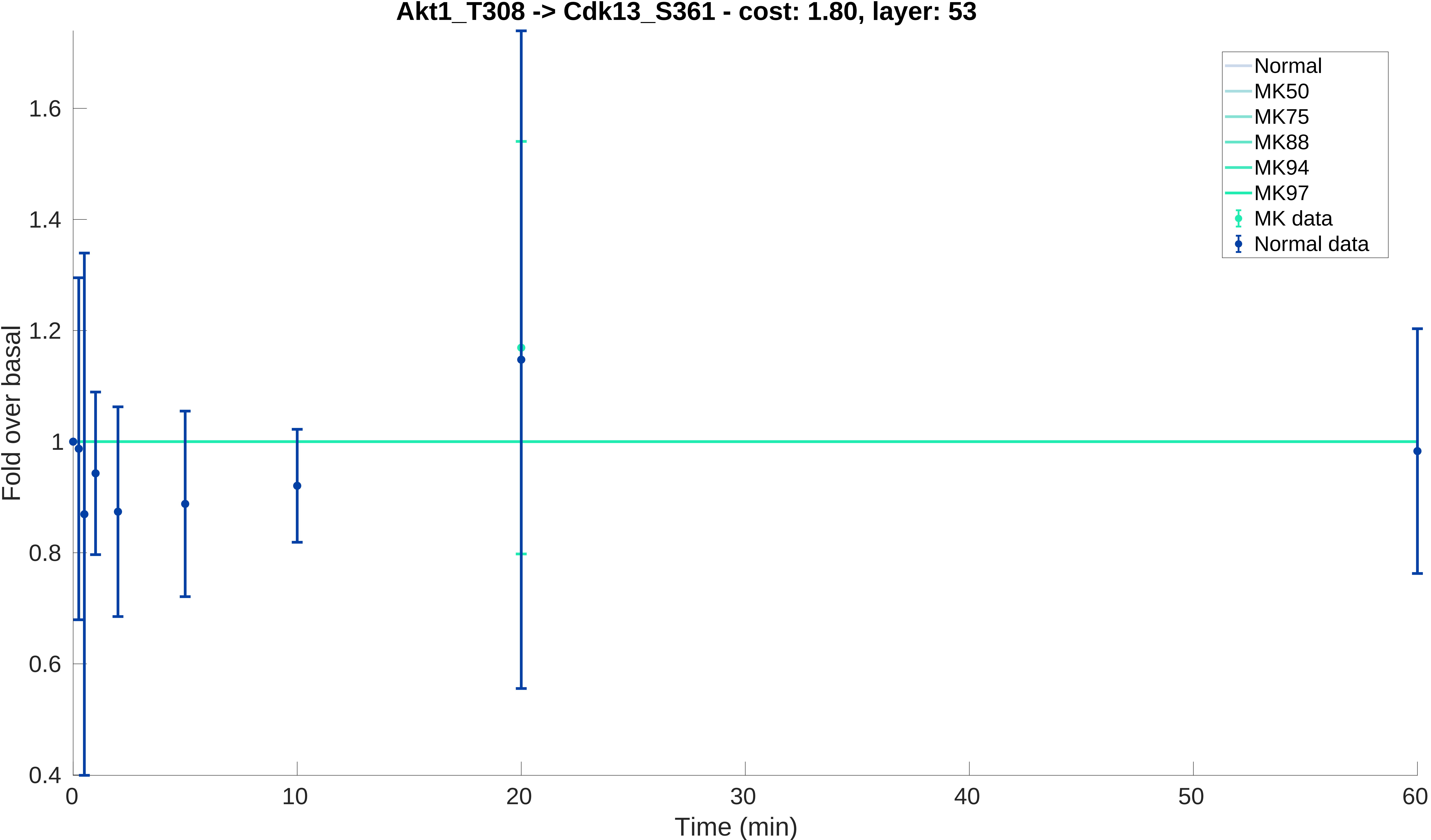
Akt1_T308 -> Cdk13_S361 - cost: 1.80, layer: 53
Fold over basal
Time (min)
0.4
0.6
0.8
1
1.2
1.4
1.6
0
10
20
30
40
50
60
Normal
MK50
MK75
MK88
MK94
MK97
MK data
Normal data

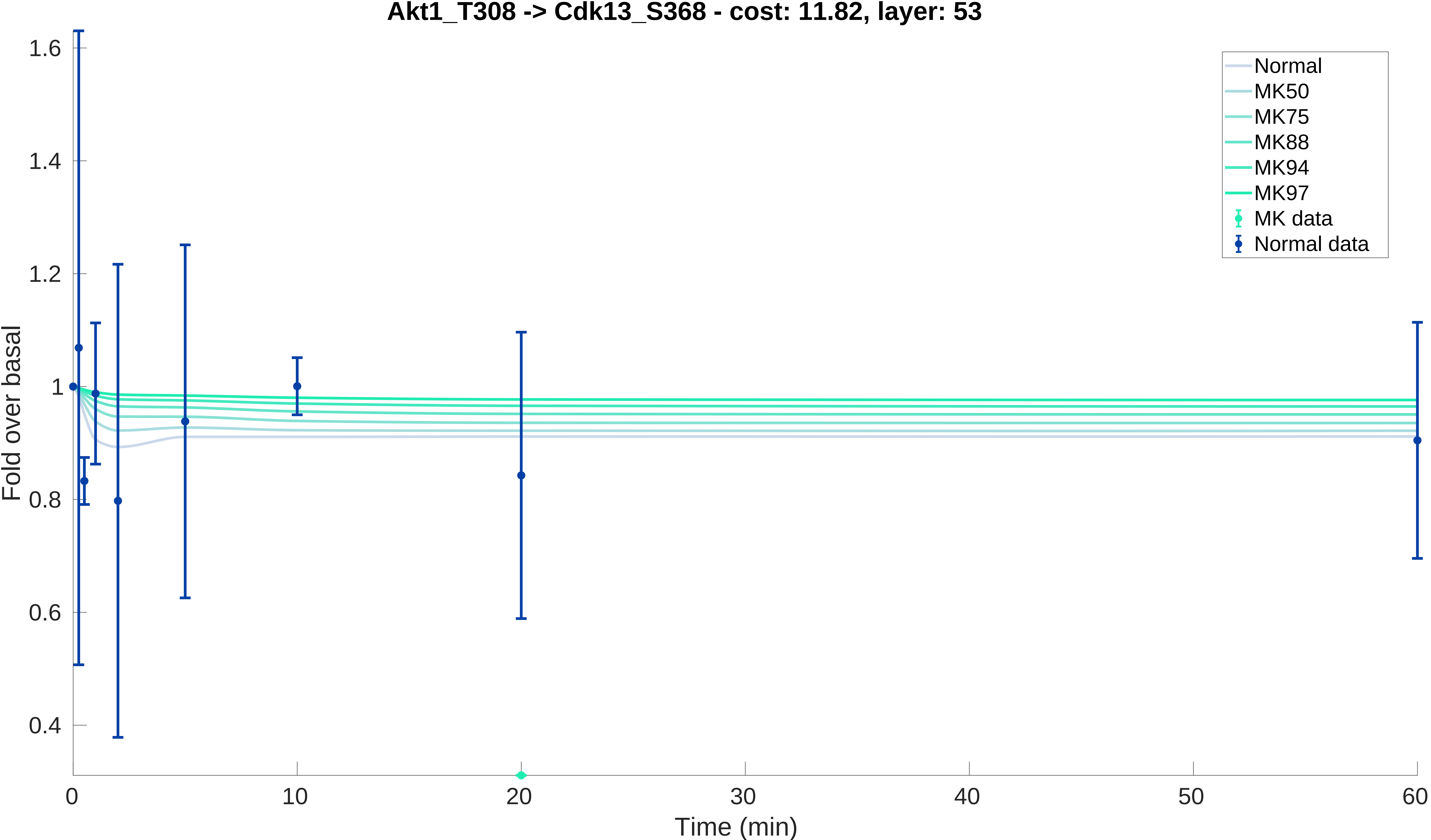

Akt1_T308 -> Cdk13_S368 - cost: 11.82, layer: 53
Fold over basal
Time (min)
1.6
1.4
1.2
1
0.8
0.6
0.4
0
10
20
30
40
50
60
Normal
MK50
MK75
MK88
MK94
MK97
MK data
Normal data

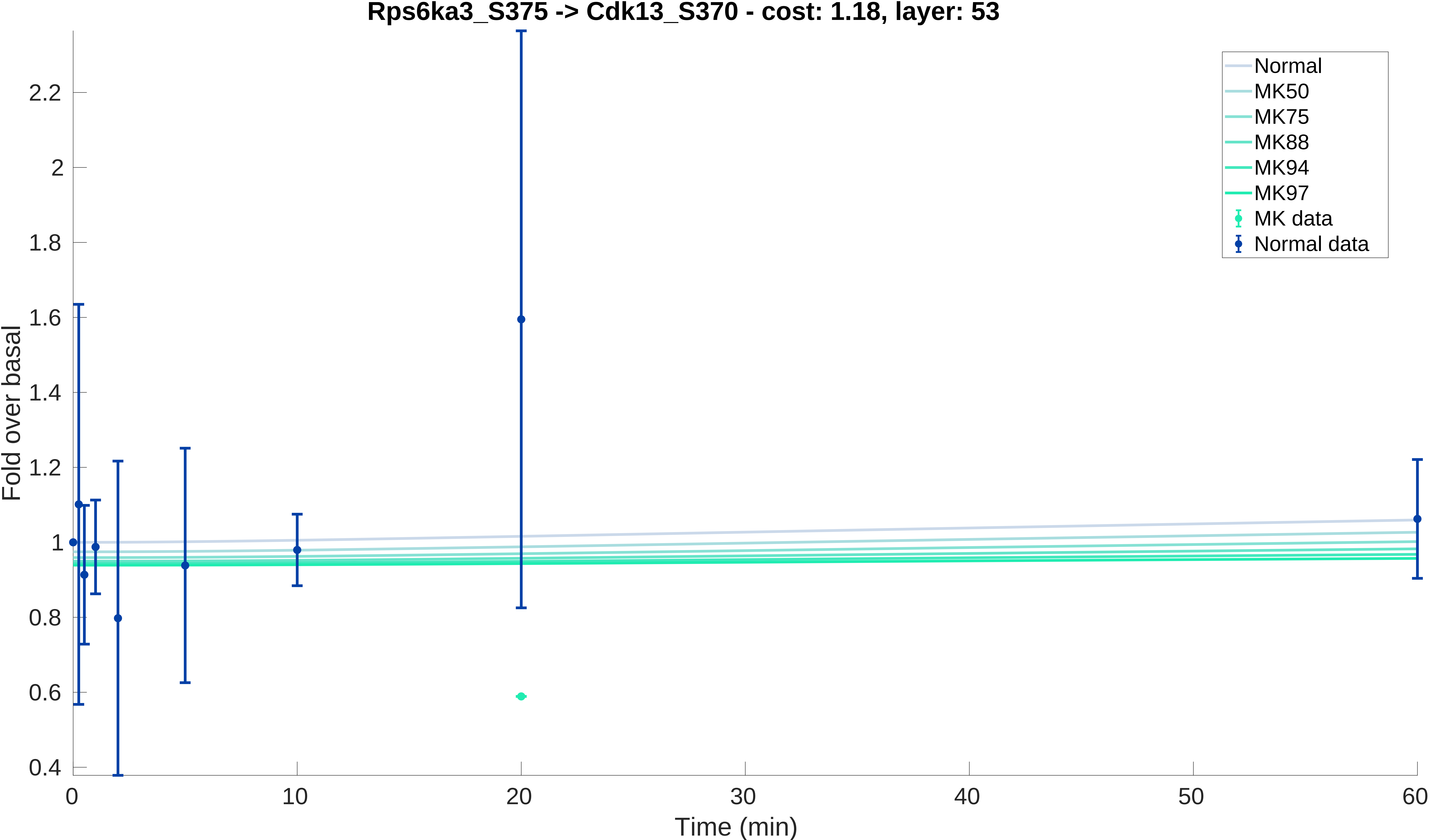
Rps6ka3_S375 -> Cdk13_S370 - cost: 1.18, layer: 53
Fold over basal
Time (min)
Normal
MK50
MK75
MK88
MK94
MK97
MK data
Normal data
0.4
0.6
0.8
1
1.2
1.4
1.6
1.8
2
2.2
0
10
20
30
40
50
60

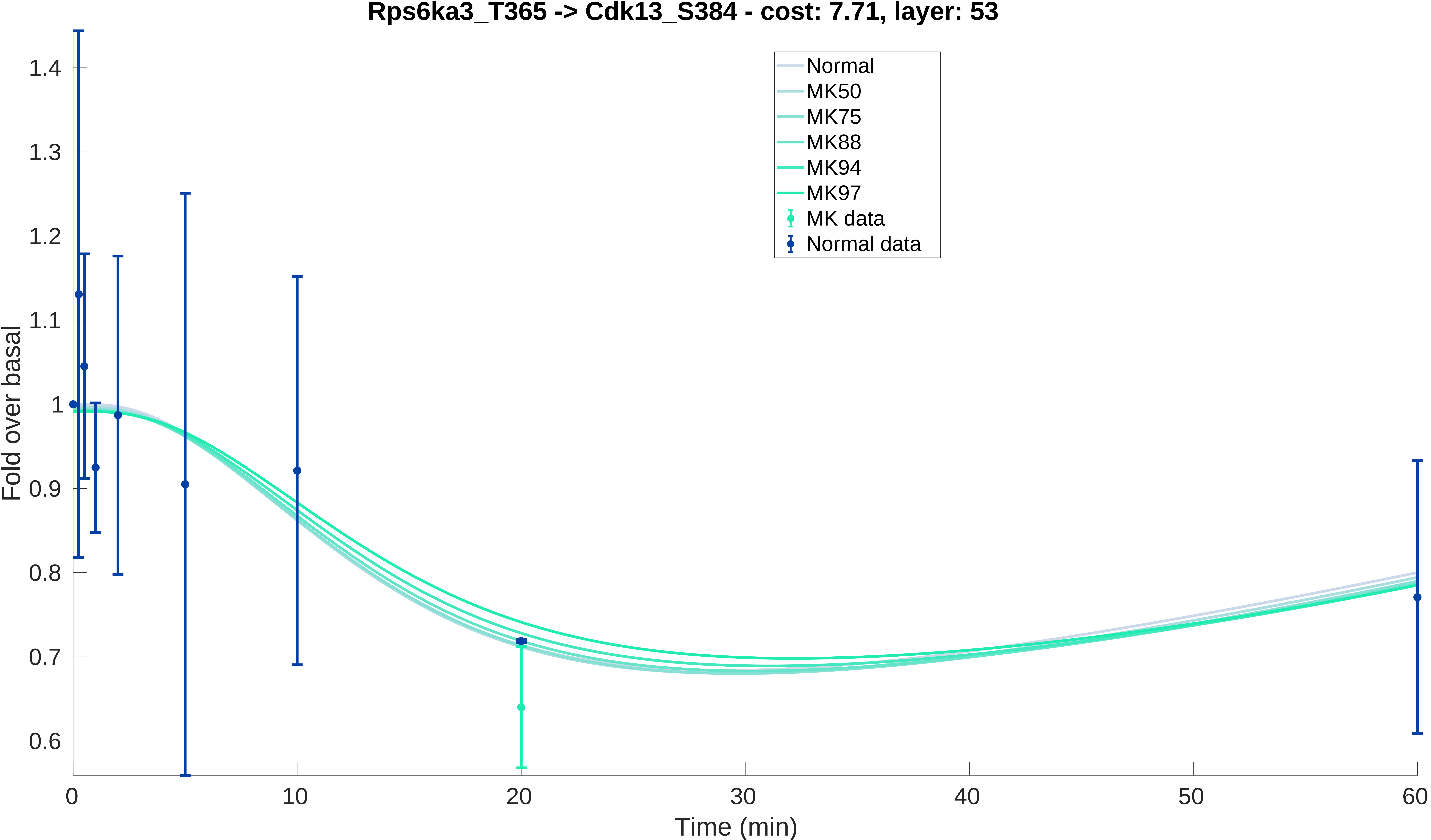

Rps6ka3_T365 -> Cdk13_S384 - cost: 7.71, layer: 53
Normal
MK50
MK75
MK88
MK94
MK97
MK data
Normal data
Fold over basal
1.4
1.3
1.2
1.1
1
0.9
0.8
0.7
0.6
0
10
20
30
40
50
60
Time (min)

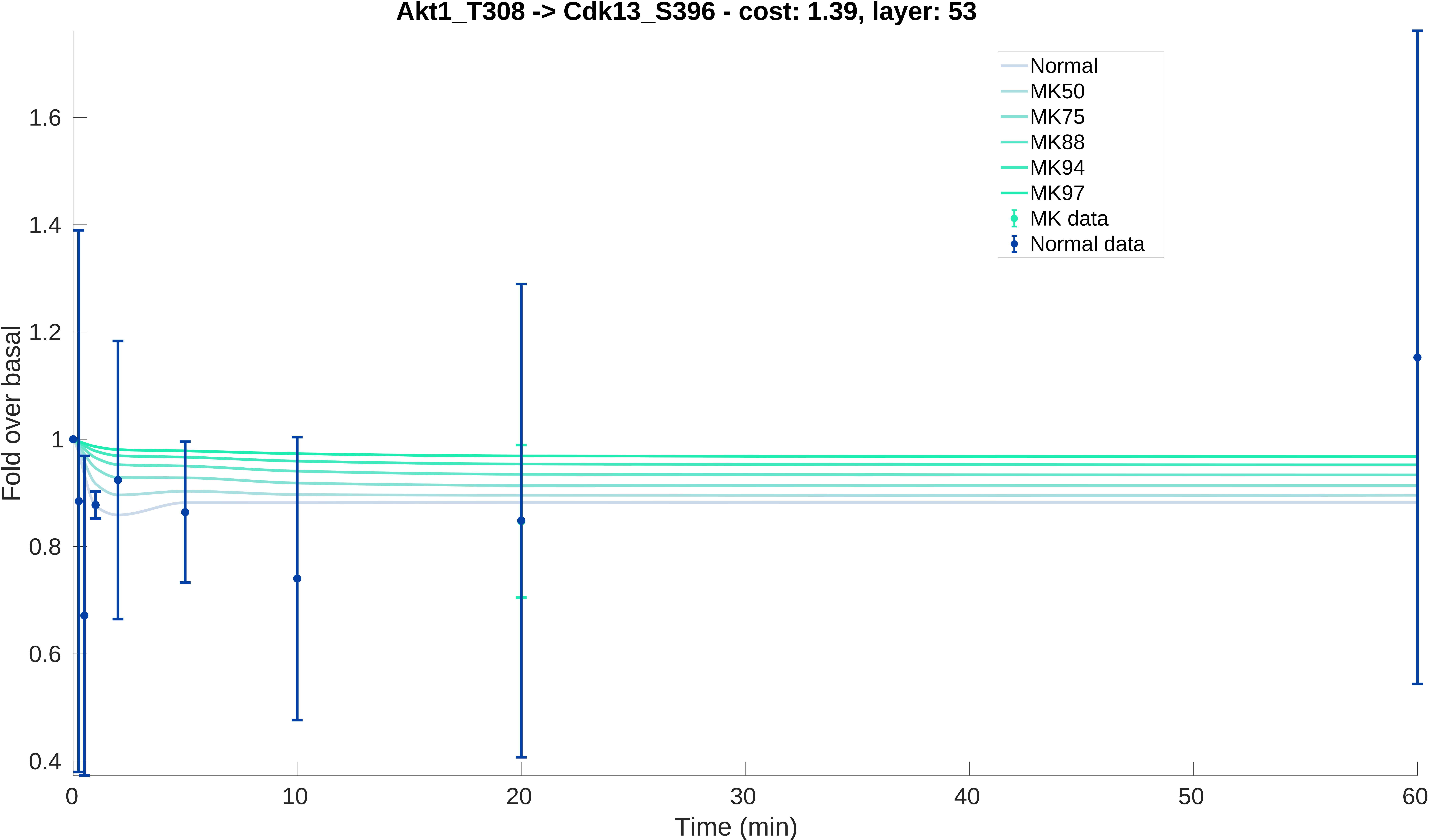
Akt1_T308 -> Cdk13_S396 - cost: 1.39, layer: 53
Fold over basal
Time (min)
Normal
MK50
MK75
MK88
MK94
MK97
MK data
Normal data
0.4
0.6
0.8
1
1.2
1.4
1.6
0
10
20
30
40
50
60

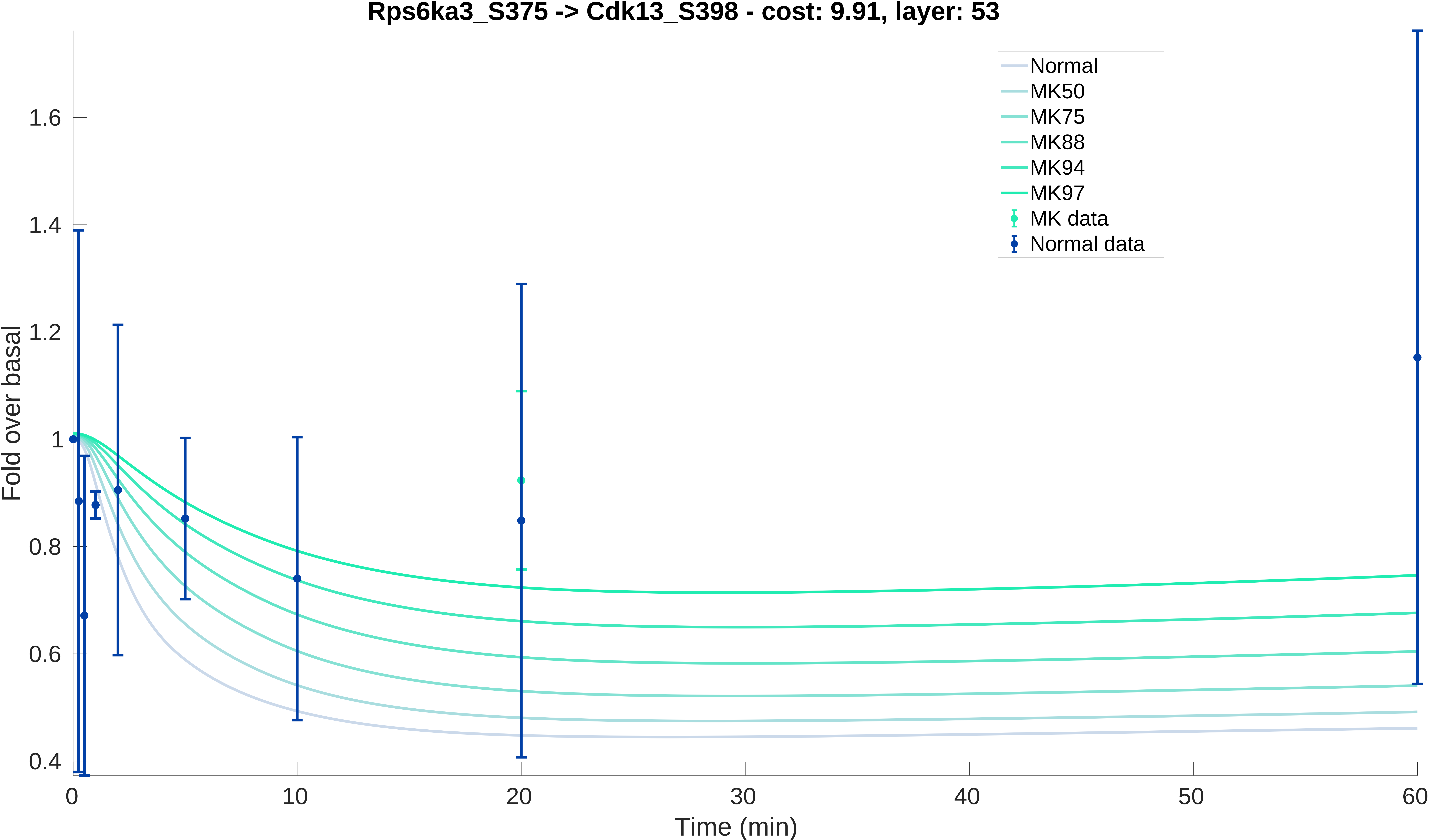

Rps6ka3_S375 -> Cdk13_S398 - cost: 9.91, layer: 53
Normal
MK50
MK75
MK88
MK94
MK97
MK data
Normal data
Fold over basal
Time (min)
0.4
0.6
0.8
1
1.2
1.4
1.6
0
10
20
30
40
50
60

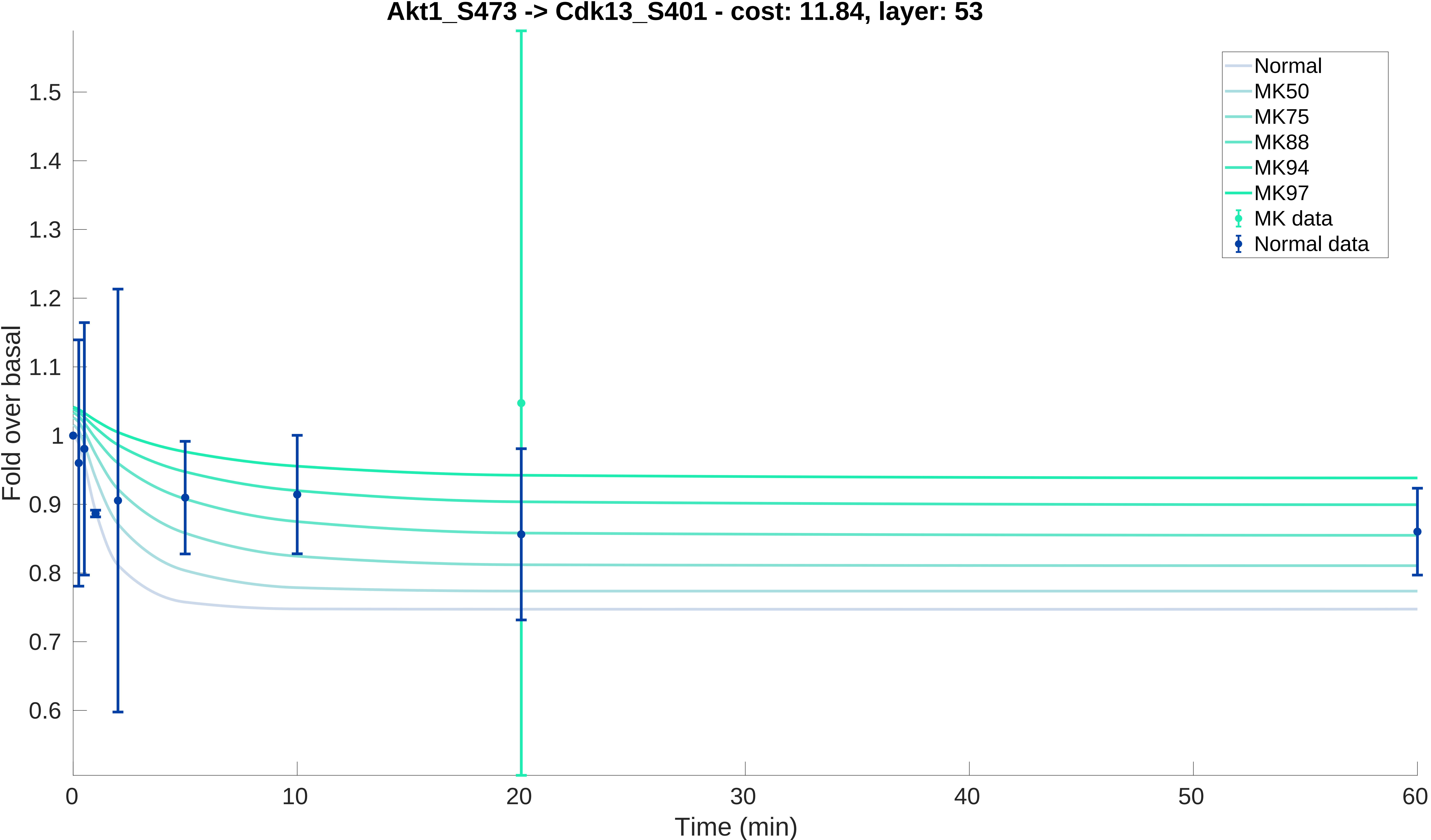

Akt1_S473 -> Cdk13_S401 - cost: 11.84, layer: 53
Fold over basal
Time (min)
0
10
20
30
40
50
60
0.6
0.7
0.8
0.9
1
1.1
1.2
1.3
1.4
1.5
Normal
MK50
MK75
MK88
MK94
MK97
MK data
Normal data

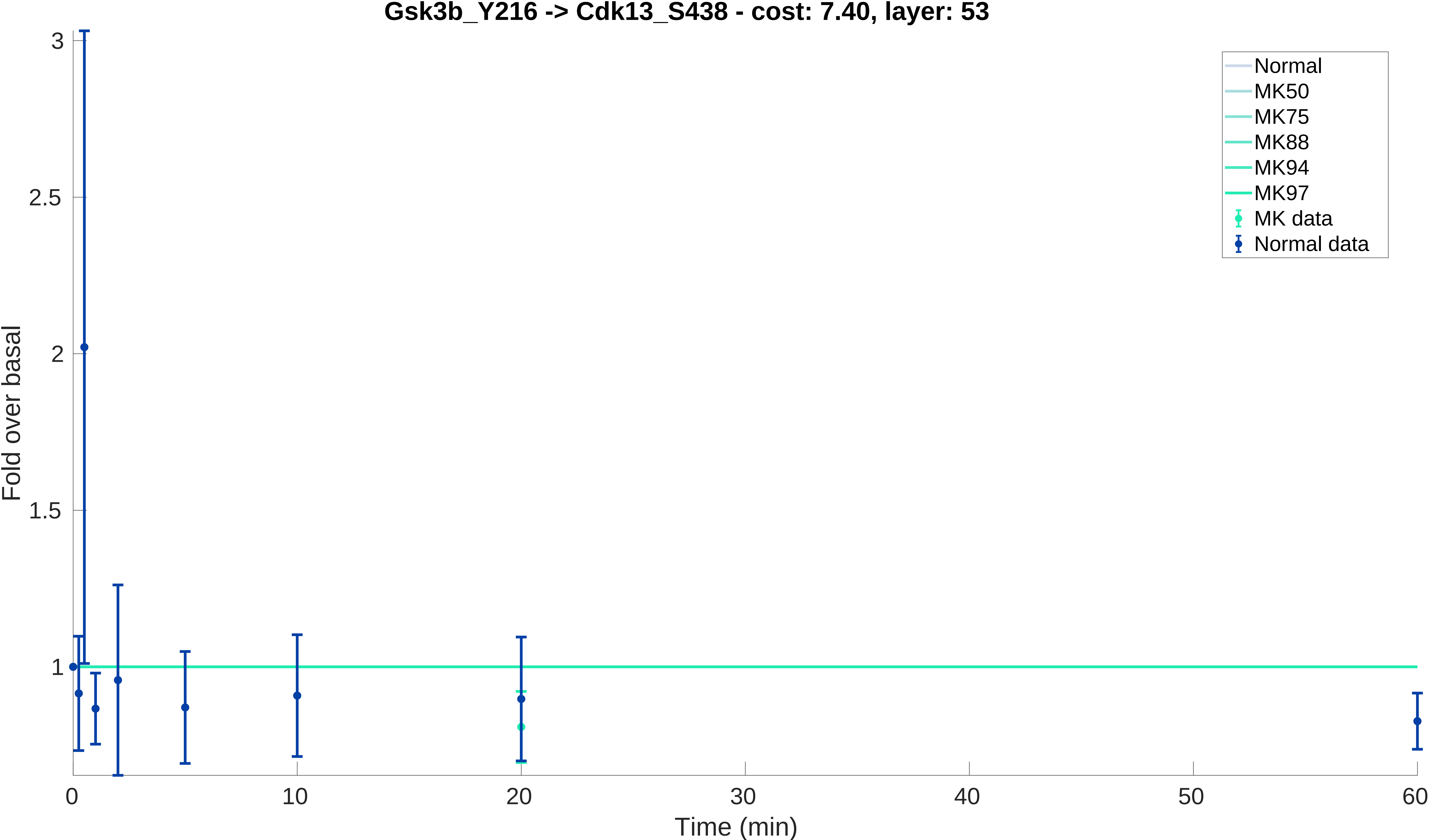
Gsk3b_Y216 -> Cdk13_S438 - cost: 7.40, layer: 53
Fold over basal
Time (min)
0
10
20
30
40
50
60
1
1.5
2
2.5
3
Normal
MK50
MK75
MK88
MK94
MK97
MK data
Normal data

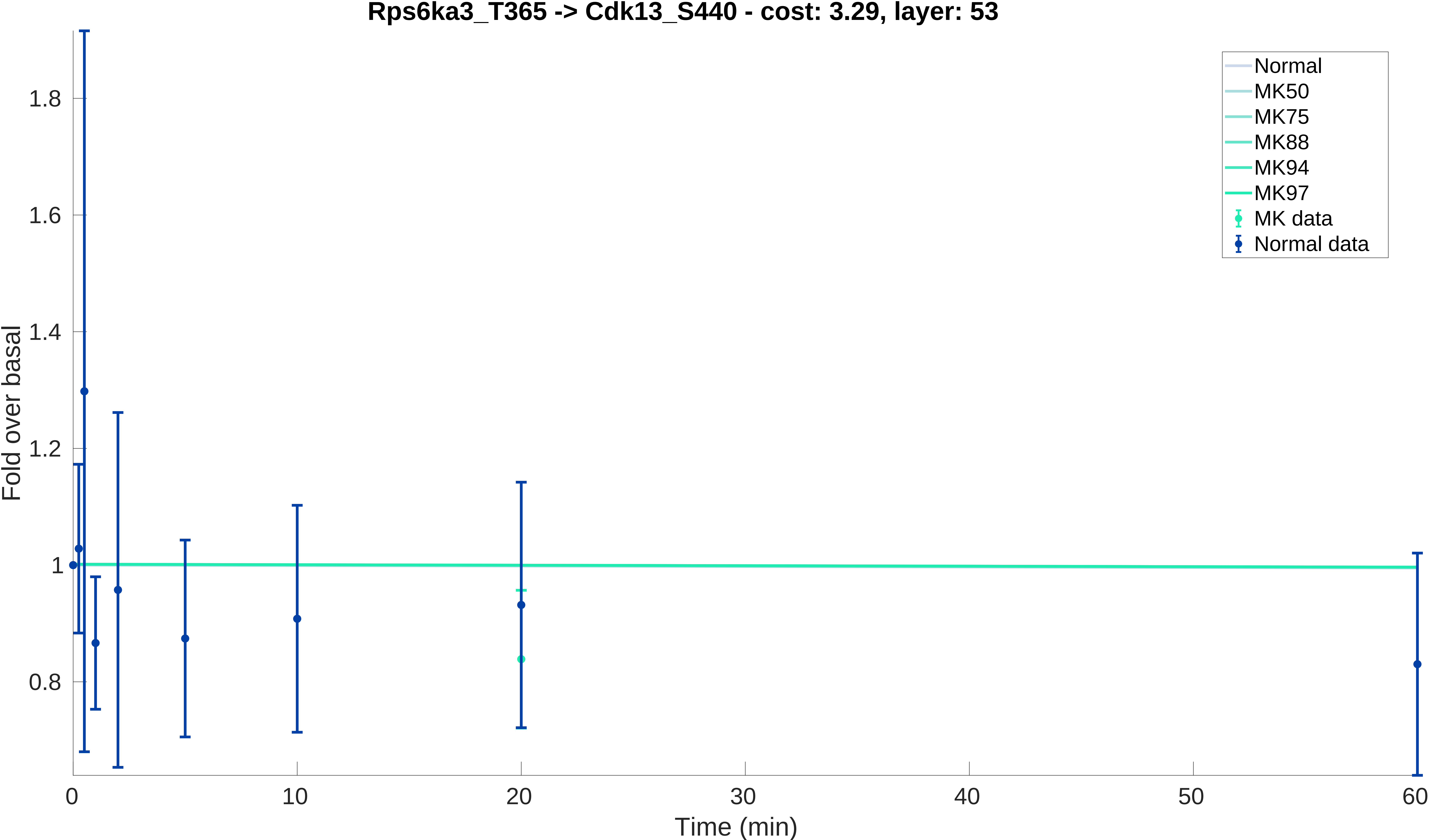

Rps6ka3_T365 -> Cdk13_S440 - cost: 3.29, layer: 53
Fold over basal
1.8
1.6
1.4
1.2
1
0.8
0
10
20
30
40
50
60
Time (min)
Normal
MK50
MK75
MK88
MK94
MK97
MK data
Normal data

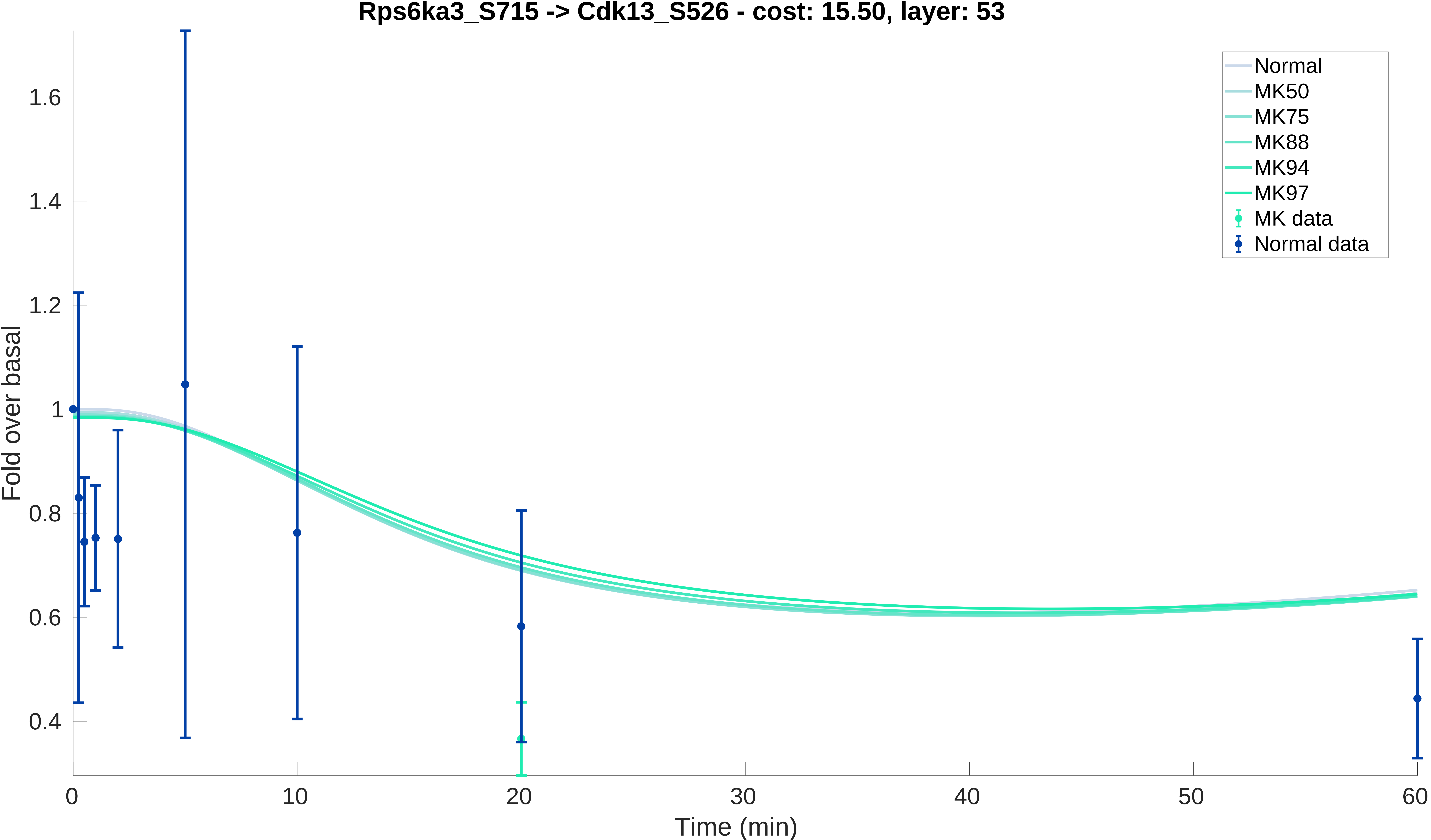

Rps6ka3_S715 -> Cdk13_S526 - cost: 15.50, layer: 53
Fold over basal
Time (min)
0.4
0.6
0.8
1
1.2
1.4
1.6
0
10
20
30
40
50
60
Normal
MK50
MK75
MK88
MK94
MK97
MK data
Normal data

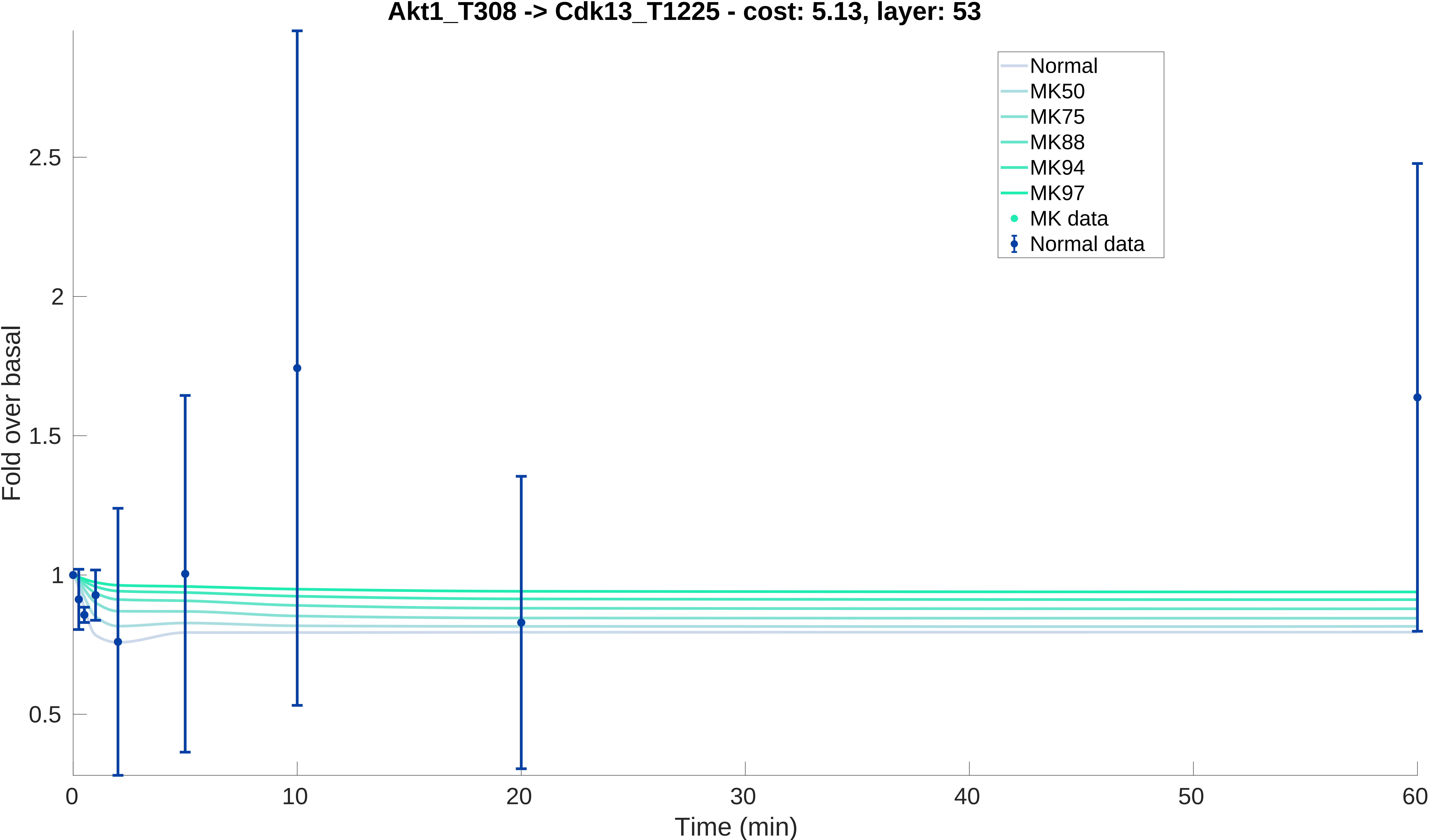

Akt1_T308 -> Cdk13_T1225 - cost: 5.13, layer: 53
Fold over basal
2.5
2
1.5
1
0.5
0
10
20
30
40
50
60
Time (min)
Normal
MK50
MK75
MK88
MK94
MK97
MK data
Normal data

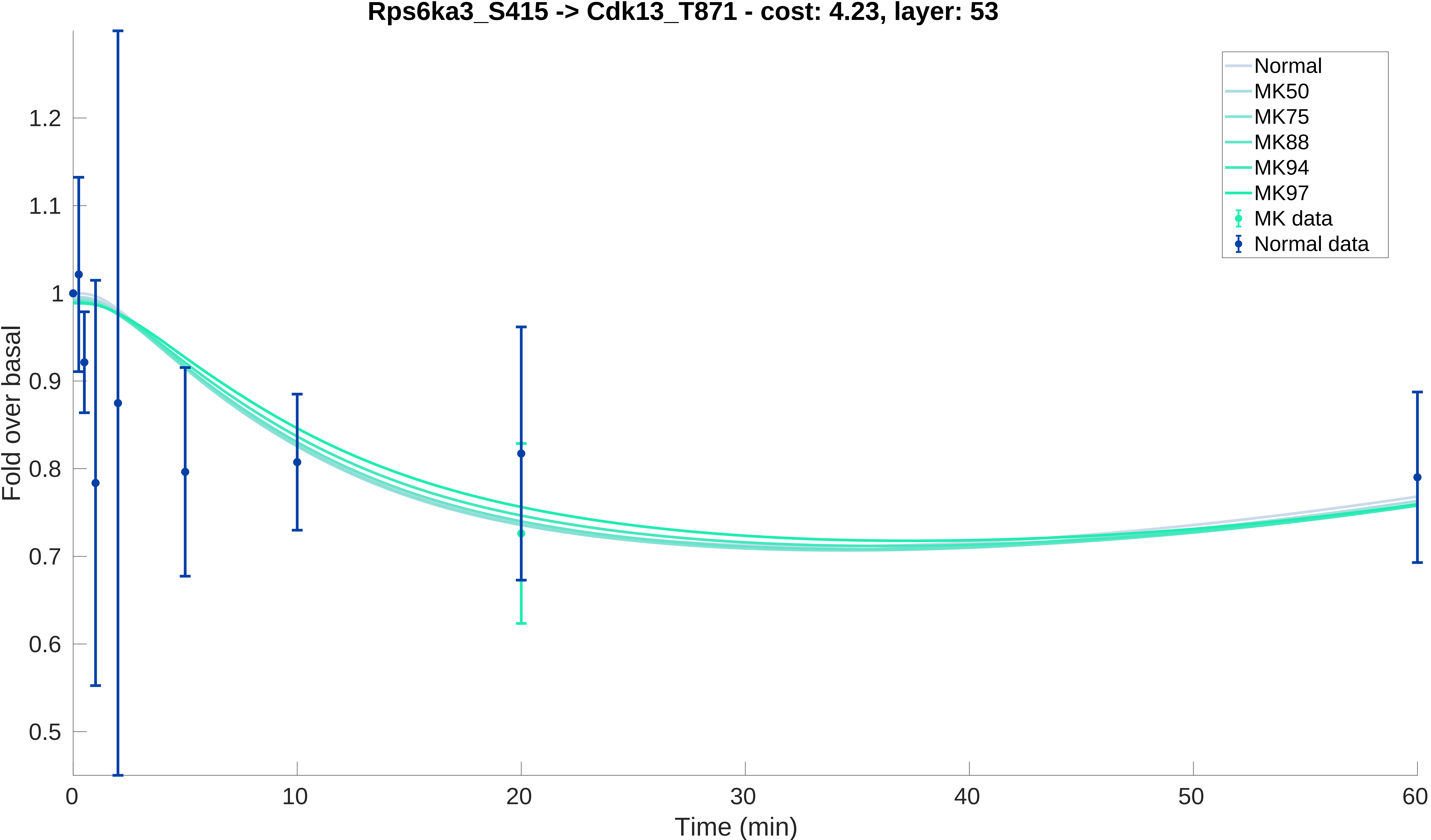

Rps6ka3_S415 -> Cdk13_T871 - cost: 4.23, layer: 53
Fold over basal
Time (min)
1.2
1.1
1
0.9
0.8
0.7
0.6
0.5
0
10
20
30
40
50
60
Normal
MK50
MK75
MK88
MK94
MK97
MK data
Normal data

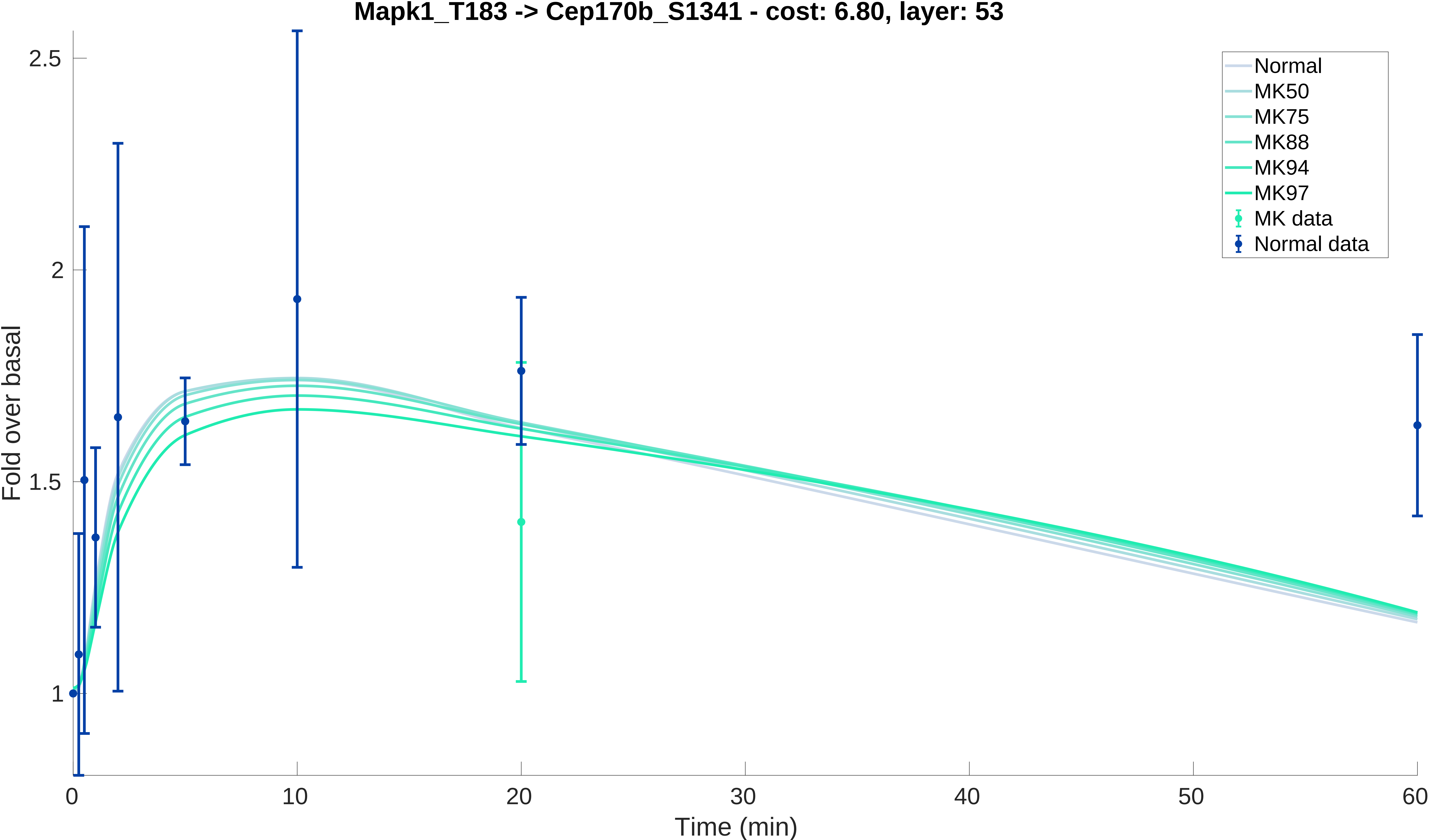

Mapk1_T183 -> Cep170b_S1341 - cost: 6.80, layer: 53
Fold over basal
2.5
2
1.5
1
Time (min)
0
10
20
30
40
50
60
Normal
MK50
MK75
MK88
MK94
MK97
MK data
Normal data

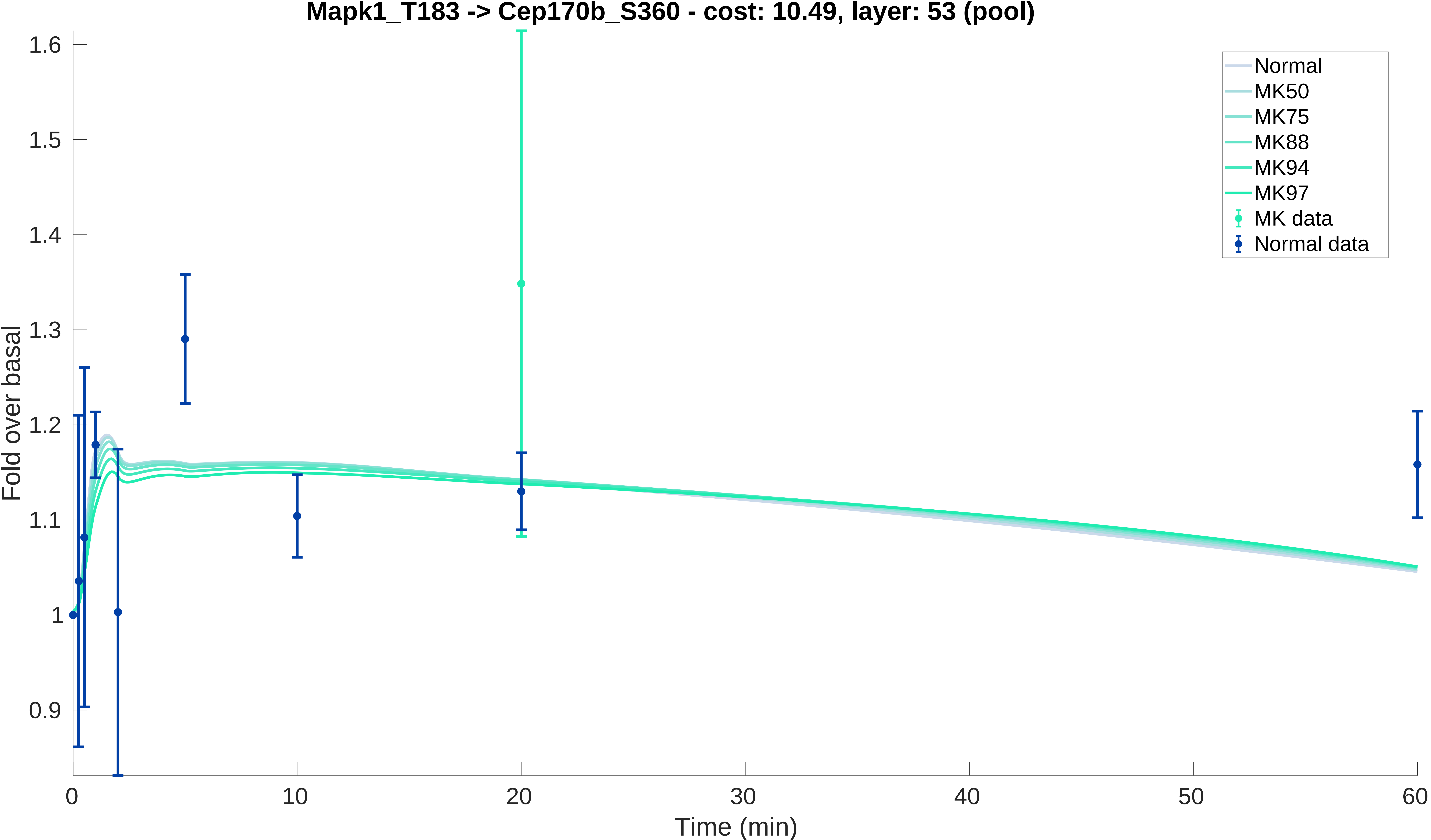
Mapk1_T183 -> Cep170b_S360 - cost: 10.49, layer: 53 (pool)
Fold over basal
Time (min)
0.9
1
1.1
1.2
1.3
1.4
1.5
1.6
0
10
20
30
40
50
60
Normal
MK50
MK75
MK88
MK94
MK97
MK data
Normal data

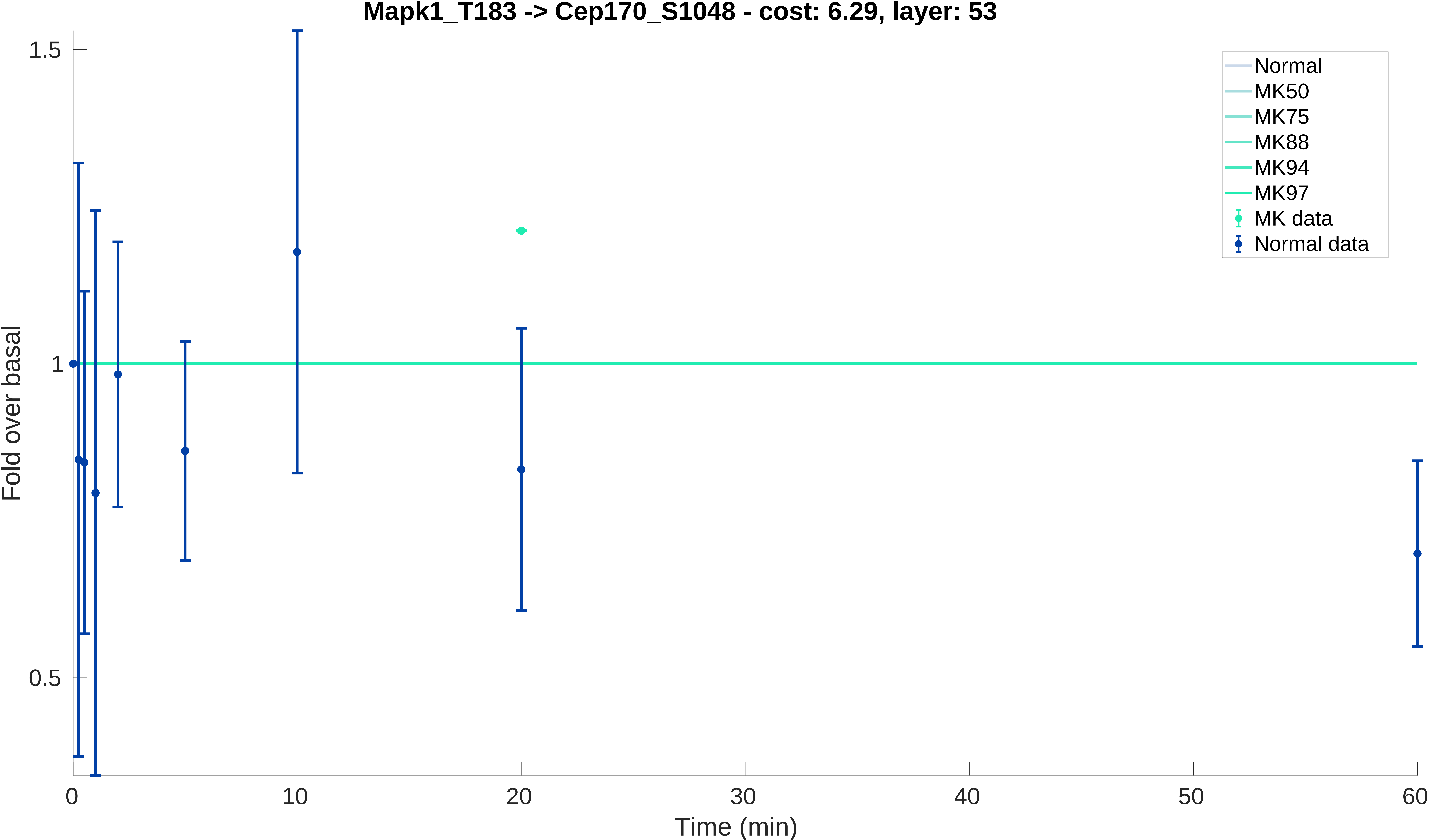
Mapk1_T183 -> Cep170_S1048 - cost: 6.29, layer: 53
Fold over basal
1.5
1
0.5
0
10
20
30
40
50
60
Time (min)
Normal
MK50
MK75
MK88
MK94
MK97
MK data
Normal data

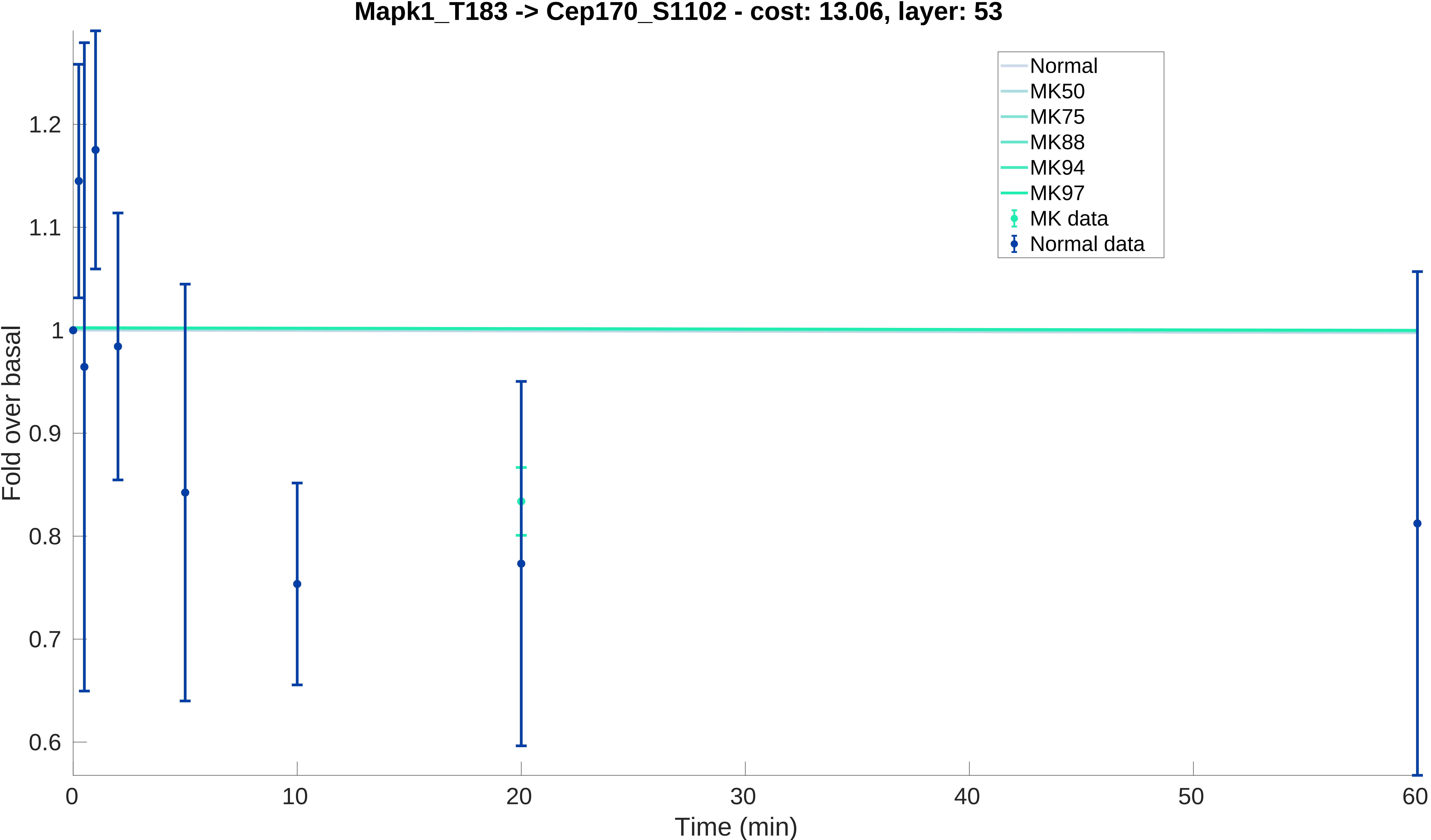

Mapk1_T183 -> Cep170_S1102 - cost: 13.06, layer: 53
Normal
MK50
MK75
MK88
MK94
MK97
MK data
Normal data
Fold over basal
1.2
1.1
1
0.9
0.8
0.7
0.6
0
10
20
30
40
50
60
Time (min)

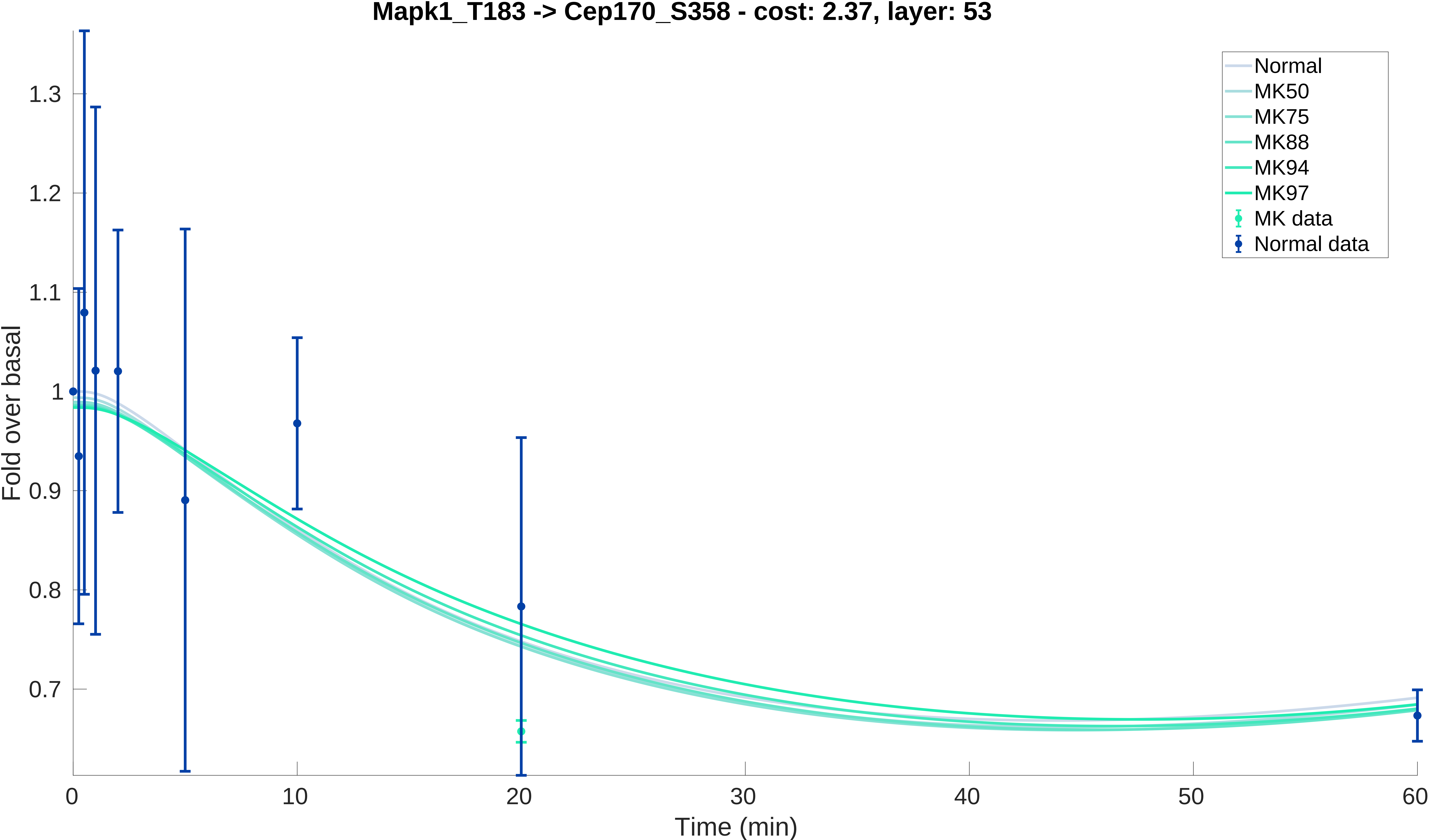

Mapk1_T183 -> Cep170_S358 - cost: 2.37, layer: 53
Normal
MK50
MK75
MK88
MK94
MK97
MK data
Normal data
Fold over basal
1.3
1.2
1.1
1
0.9
0.8
0.7
0
10
20
30
40
50
60
Time (min)

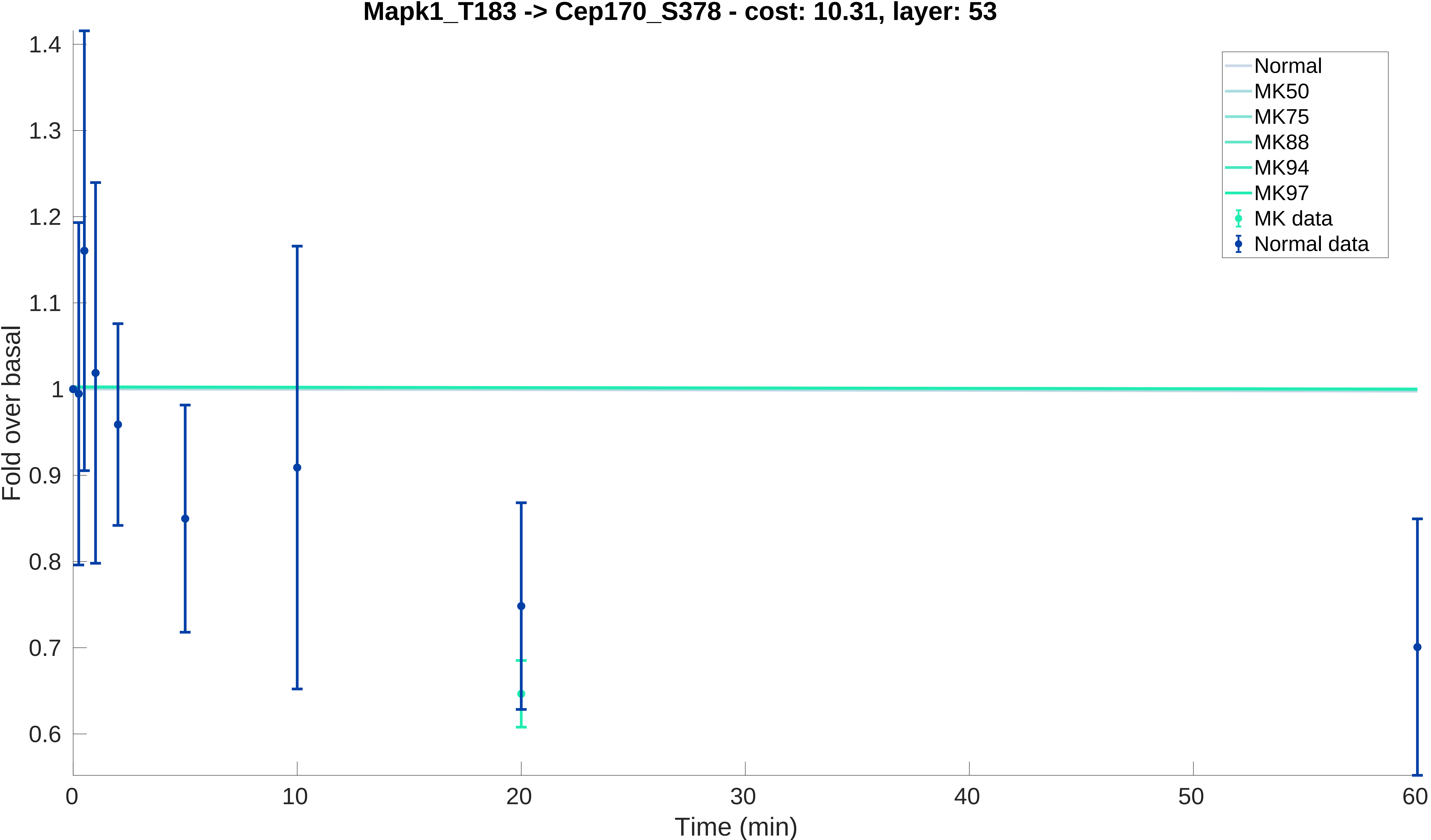
Mapk1_T183 -> Cep170_S378 - cost: 10.31, layer: 53
Fold over basal
Time (min)
1.4
1.3
1.2
1.1
1
0.9
0.8
0.7
0.6
0
10
20
30
40
50
60
Normal
MK50
MK75
MK88
MK94
MK97
MK data
Normal data

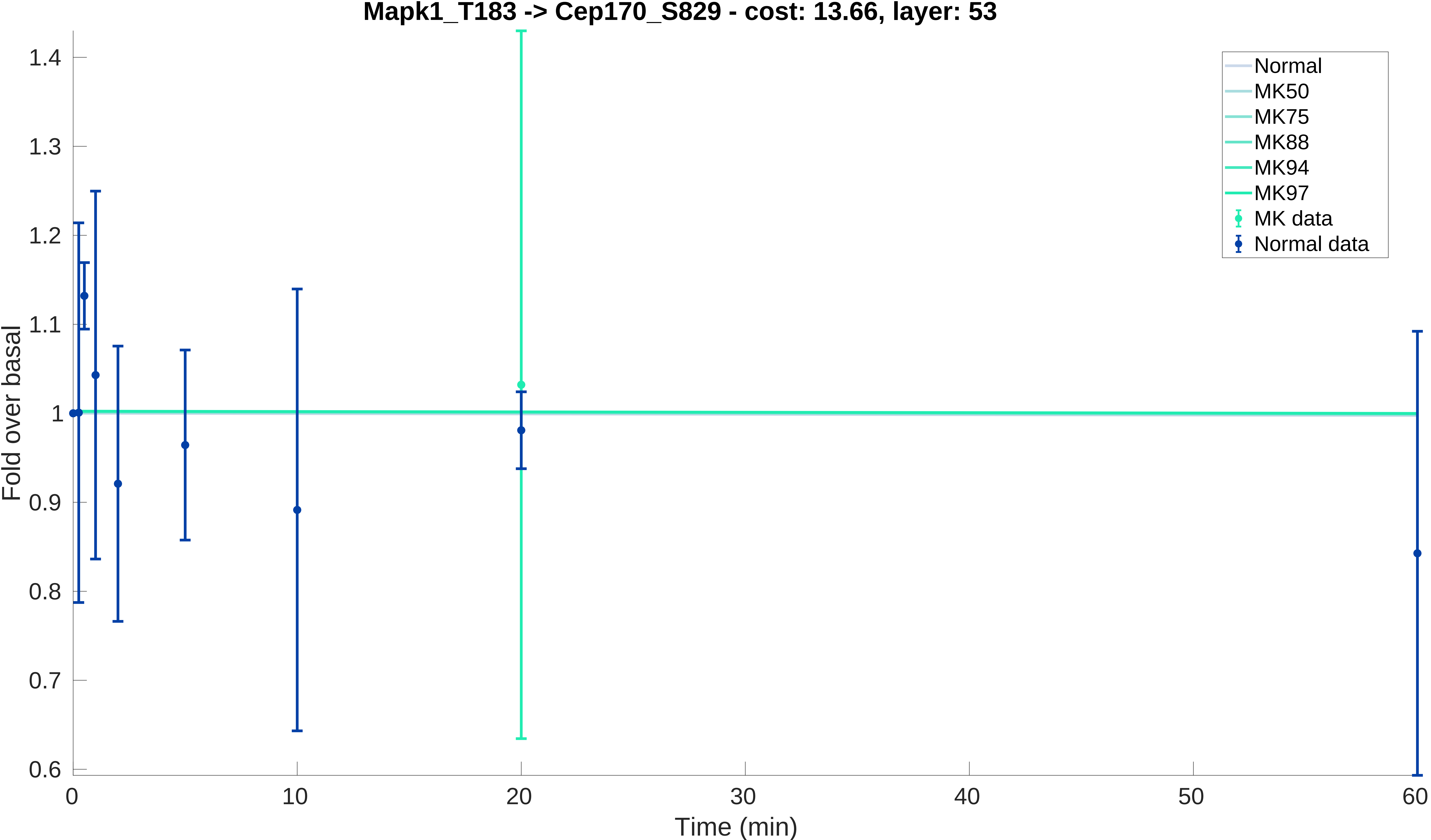

Mapk1_T183 -> Cep170_S829 - cost: 13.66, layer: 53
Fold over basal
Time (min)
0.6
0.7
0.8
0.9
1
1.1
1.2
1.3
1.4
0
10
20
30
40
50
60
Normal
MK50
MK75
MK88
MK94
MK97
MK data
Normal data

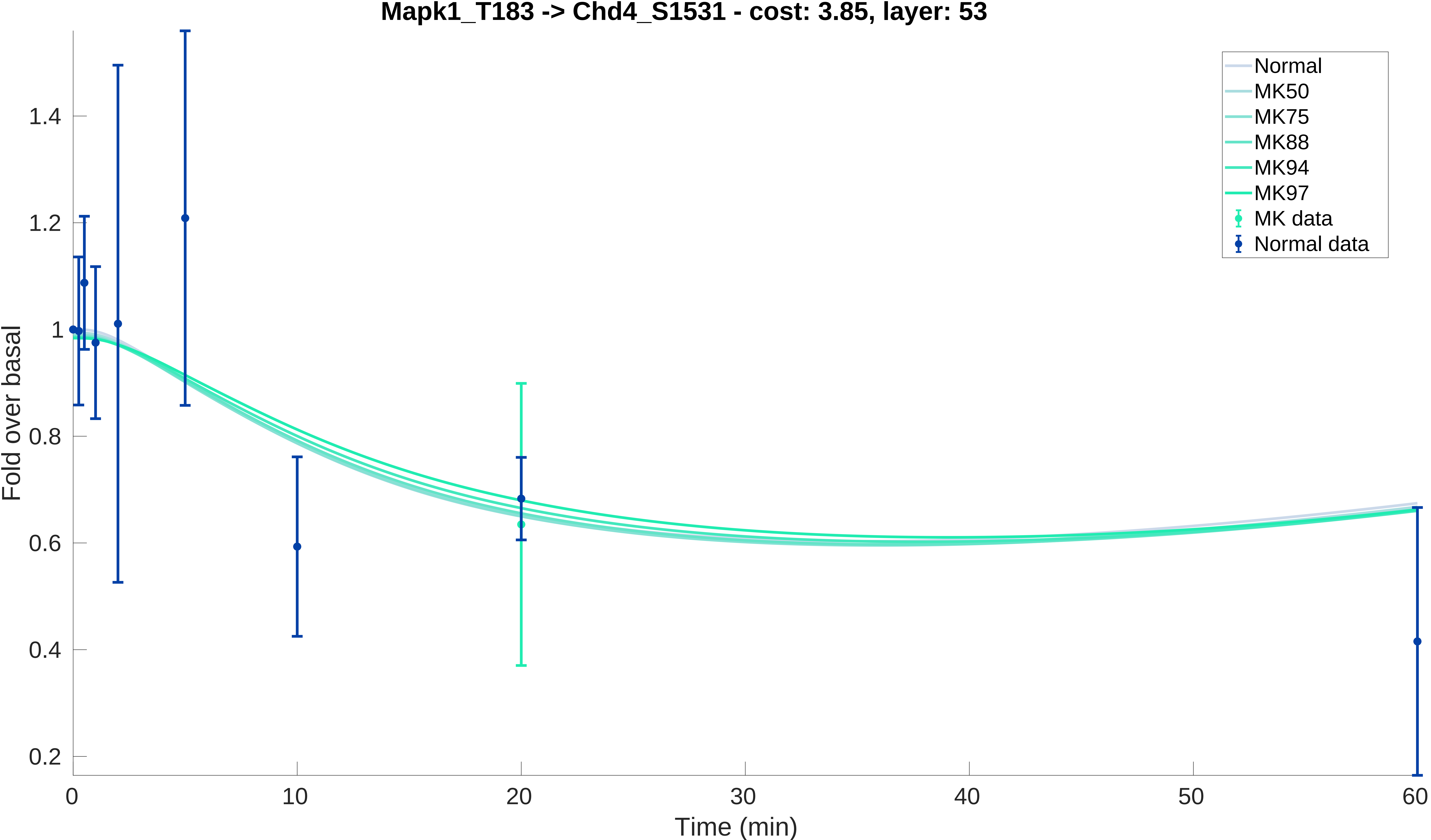

Mapk1_T183 -> Chd4_S1531 - cost: 3.85, layer: 53
Normal
MK50
MK75
MK88
MK94
MK97
MK data
Normal data
Fold over basal
Time (min)
1.4
1.2
1
0.8
0.6
0.4
0.2
0
10
20
30
40
50
60

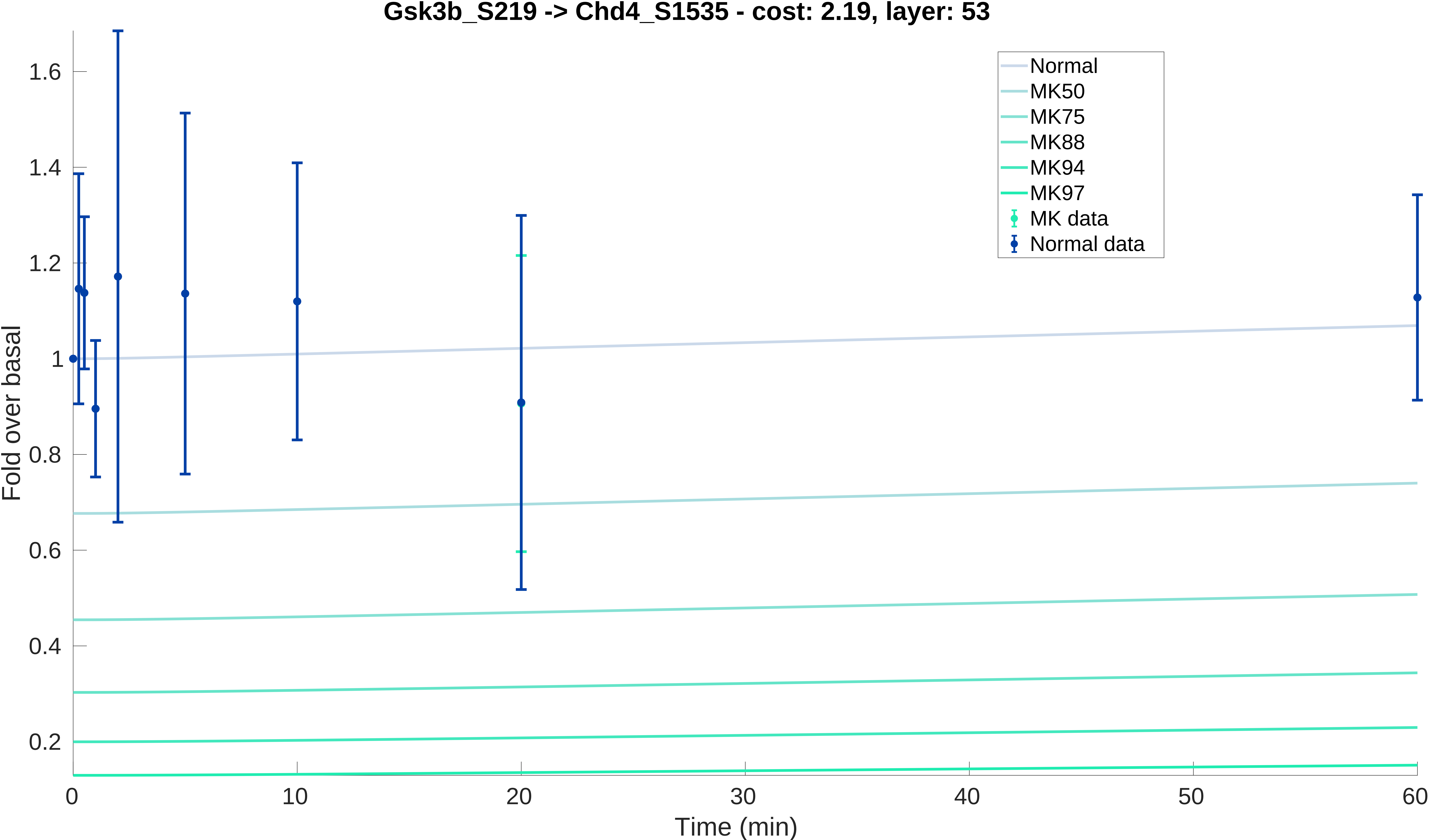

Gsk3b_S219 -> Chd4_S1535 - cost: 2.19, layer: 53
Normal
MK50
MK75
MK88
MK94
MK97
MK data
Normal data
Fold over basal
1.6
1.4
1.2
1
0.8
0.6
0.4
0.2
0
10
20
30
40
50
60
Time (min)

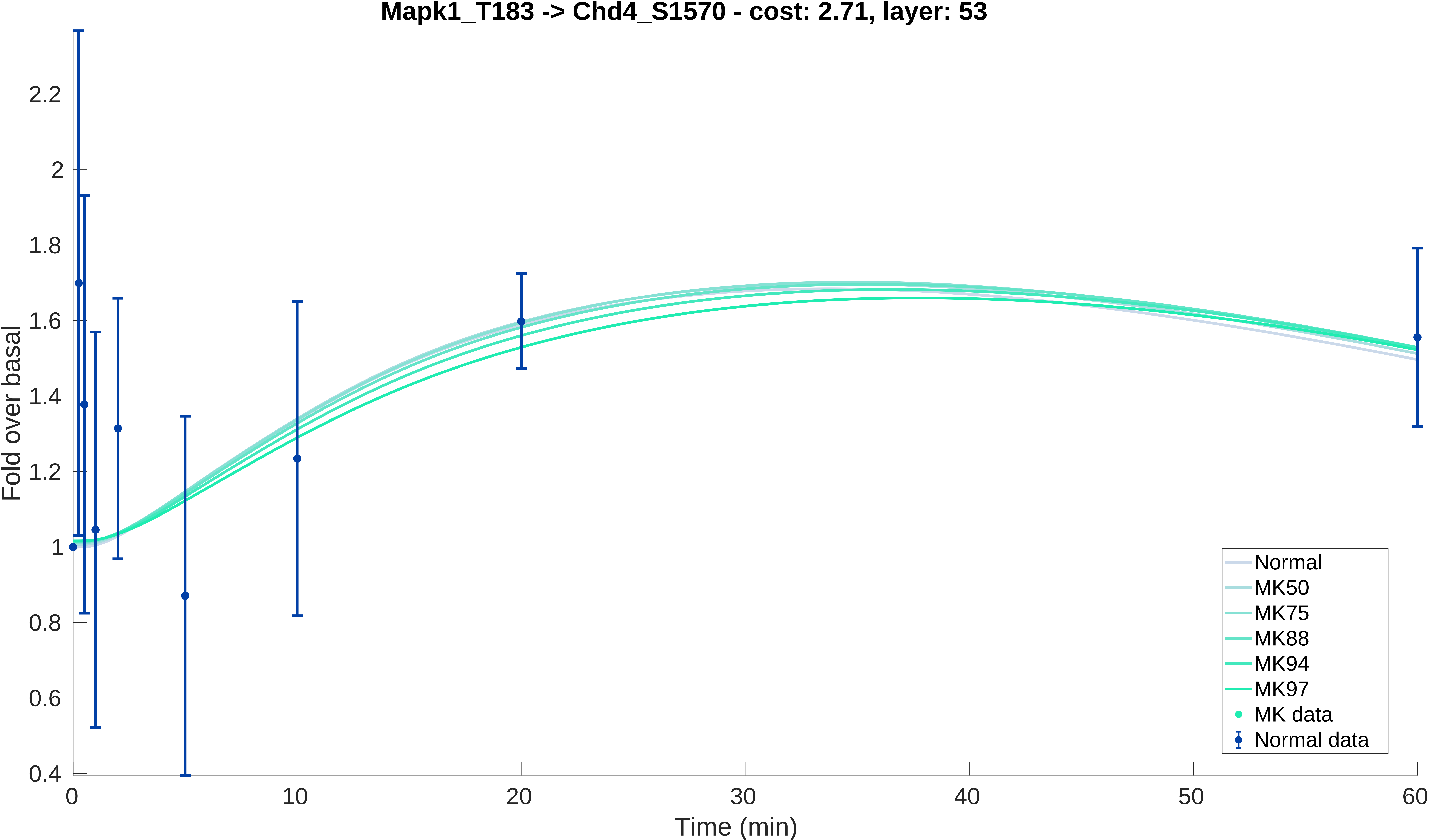

Mapk1_T183 -> Chd4_S1570 - cost: 2.71, layer: 53
Fold over basal
Time (min)
2.2
2
1.8
1.6
1.4
1.2
1
0.8
0.6
0.4
0
10
20
30
40
50
60
Normal
MK50
MK75
MK88
MK94
MK97
MK data
Normal data

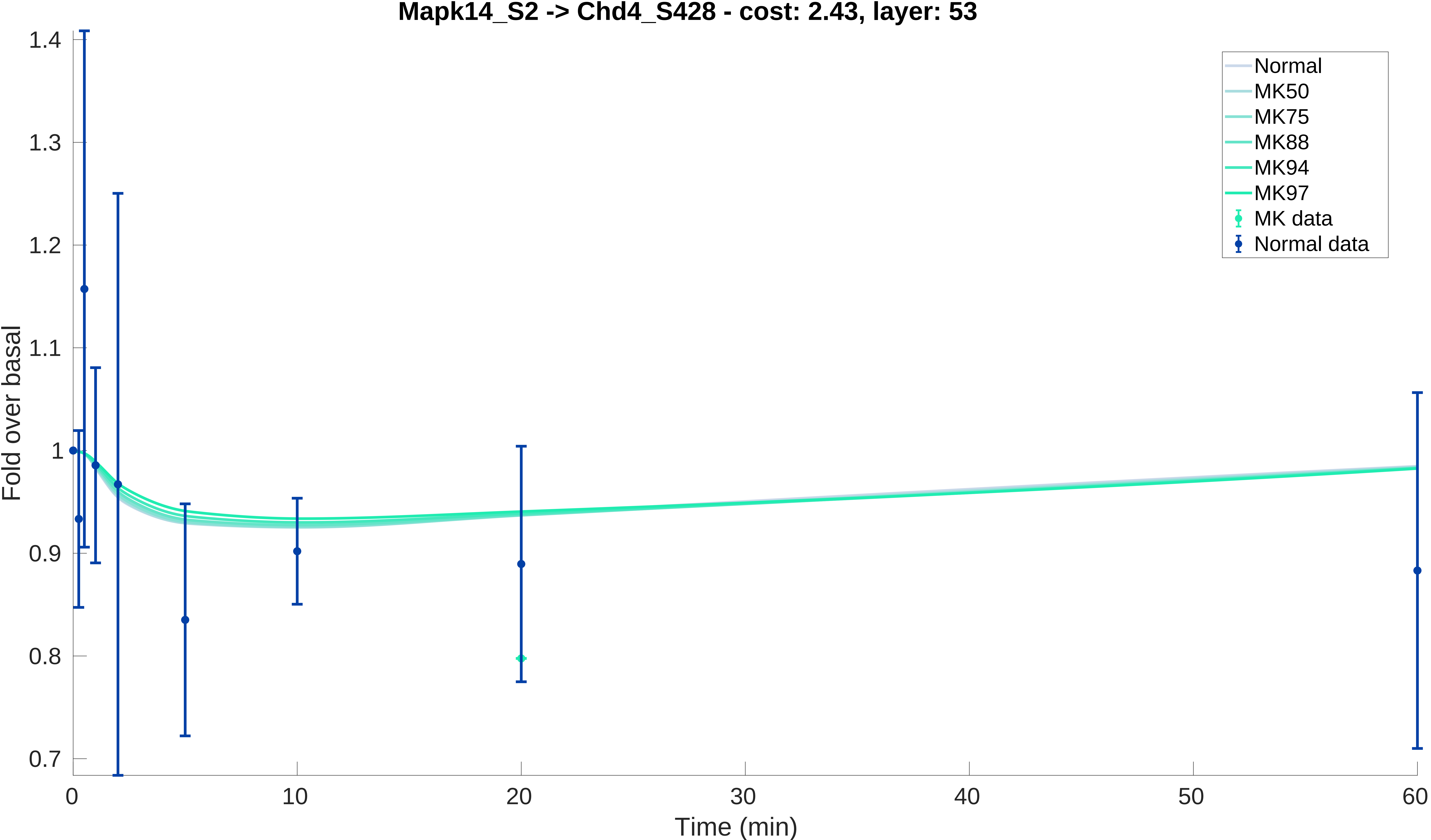

Mapk14_S2 -> Chd4_S428 - cost: 2.43, layer: 53
Normal
MK50
MK75
MK88
MK94
MK97
MK data
Normal data
Fold over basal
1.4
1.3
1.2
1.1
1
0.9
0.8
0.7
0
10
20
30
40
50
60
Time (min)

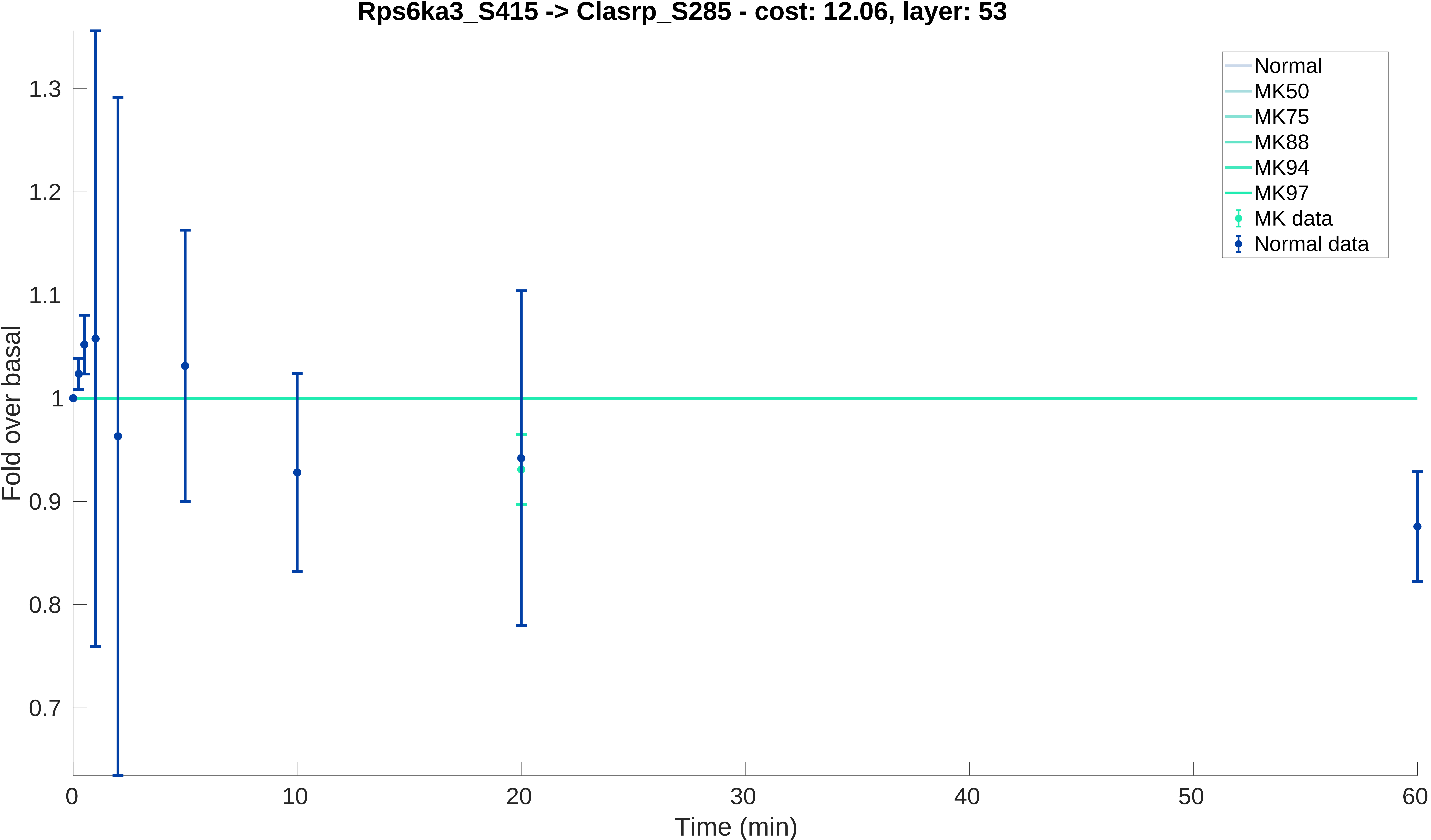
Rps6ka3_S415 -> Clasrp_S285 - cost: 12.06, layer: 53
Fold over basal
Time (min)
1.3
1.2
1.1
1
0.9
0.8
0.7
0
10
20
30
40
50
60
Normal
MK50
MK75
MK88
MK94
MK97
MK data
Normal data

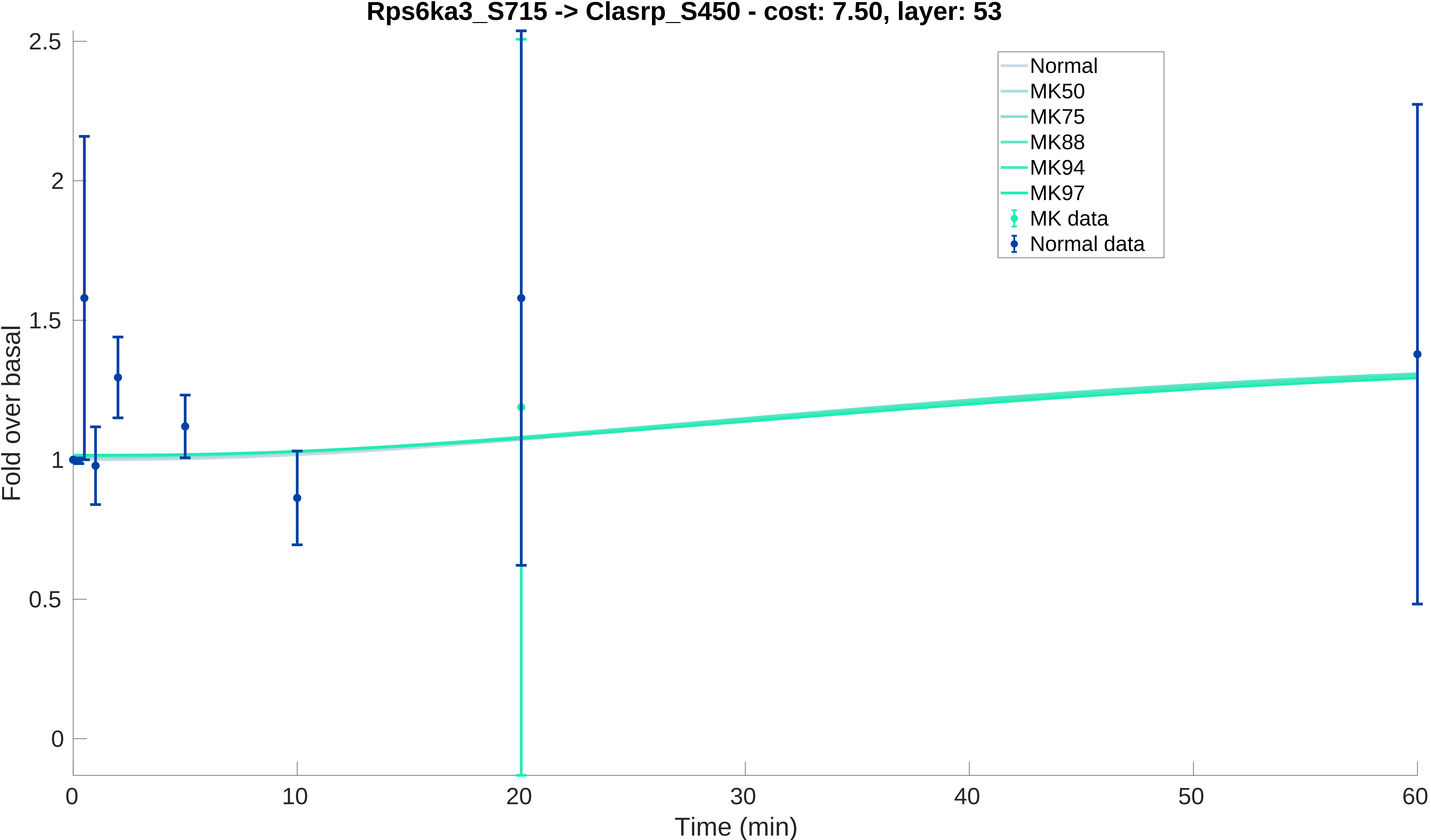

Rps6ka3_S715 -> Clasrp_S450 - cost: 7.50, layer: 53
Fold over basal
Time (min)
0
0.5
1
1.5
2
2.5
0
10
20
30
40
50
60
Normal
MK50
MK75
MK88
MK94
MK97
MK data
Normal data

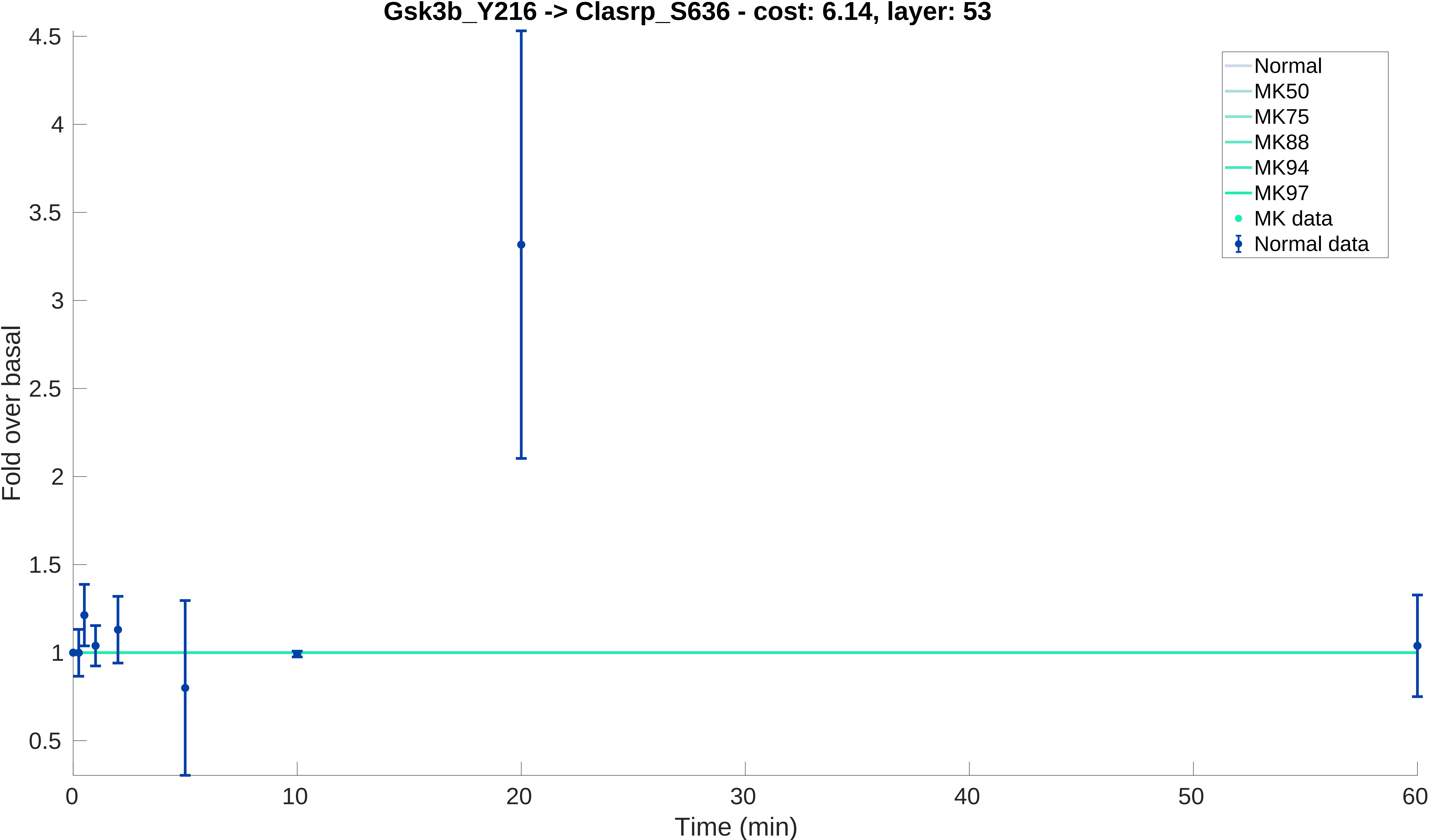

Gsk3b_Y216 -> Clasrp_S636 - cost: 6.14, layer: 53
Fold over basal
Time (min)
0.5
1
1.5
2
2.5
3
3.5
4
4.5
0
10
20
30
40
50
60
Normal
MK50
MK75
MK88
MK94
MK97
MK data
Normal data

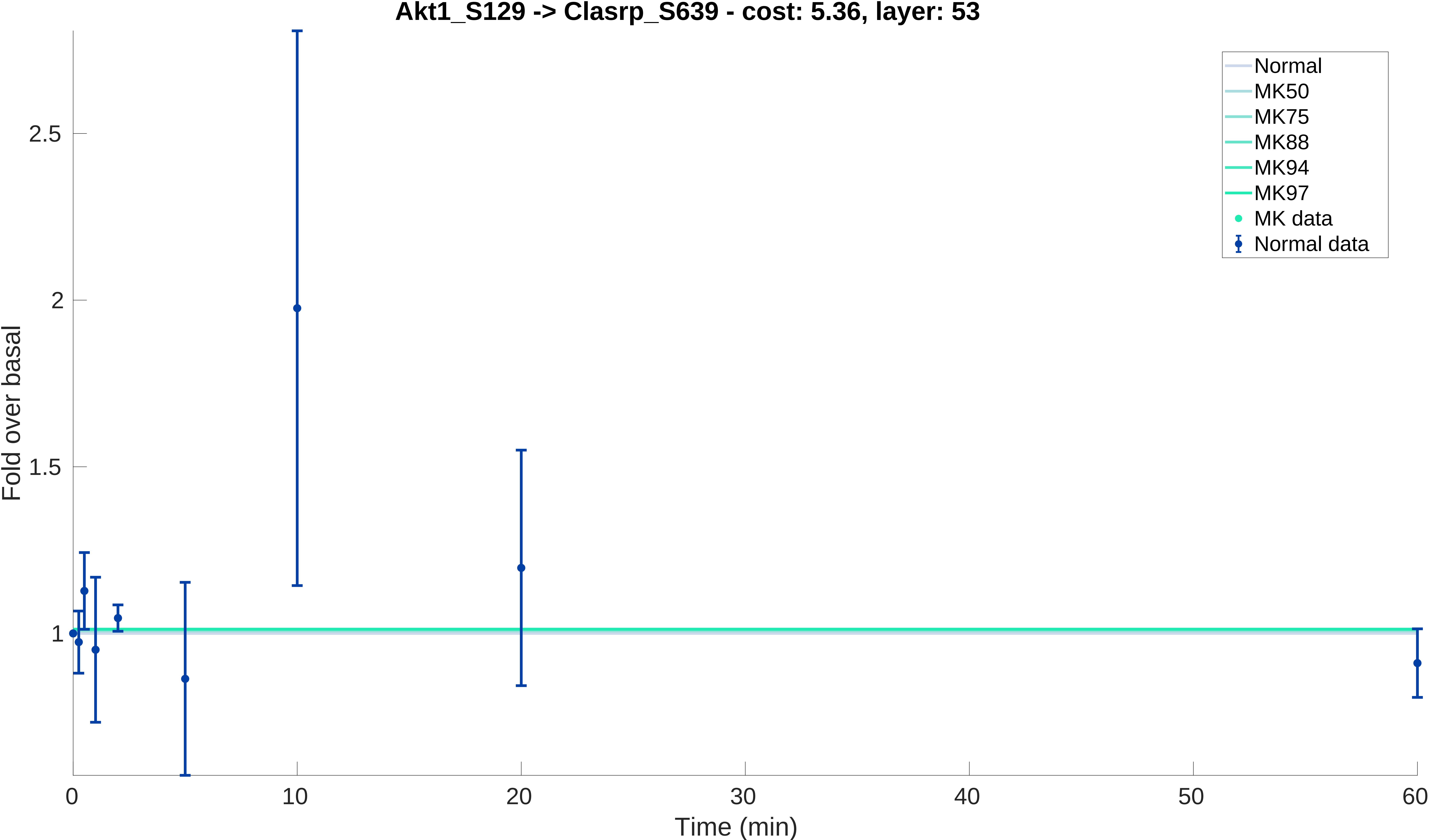

Akt1_S129 -> Clasrp_S639 - cost: 5.36, layer: 53
Normal
MK50
MK75
MK88
MK94
MK97
MK data
Normal data
2.5
2
1.5
1
Fold over basal
0
10
20
30
40
50
60
Time (min)

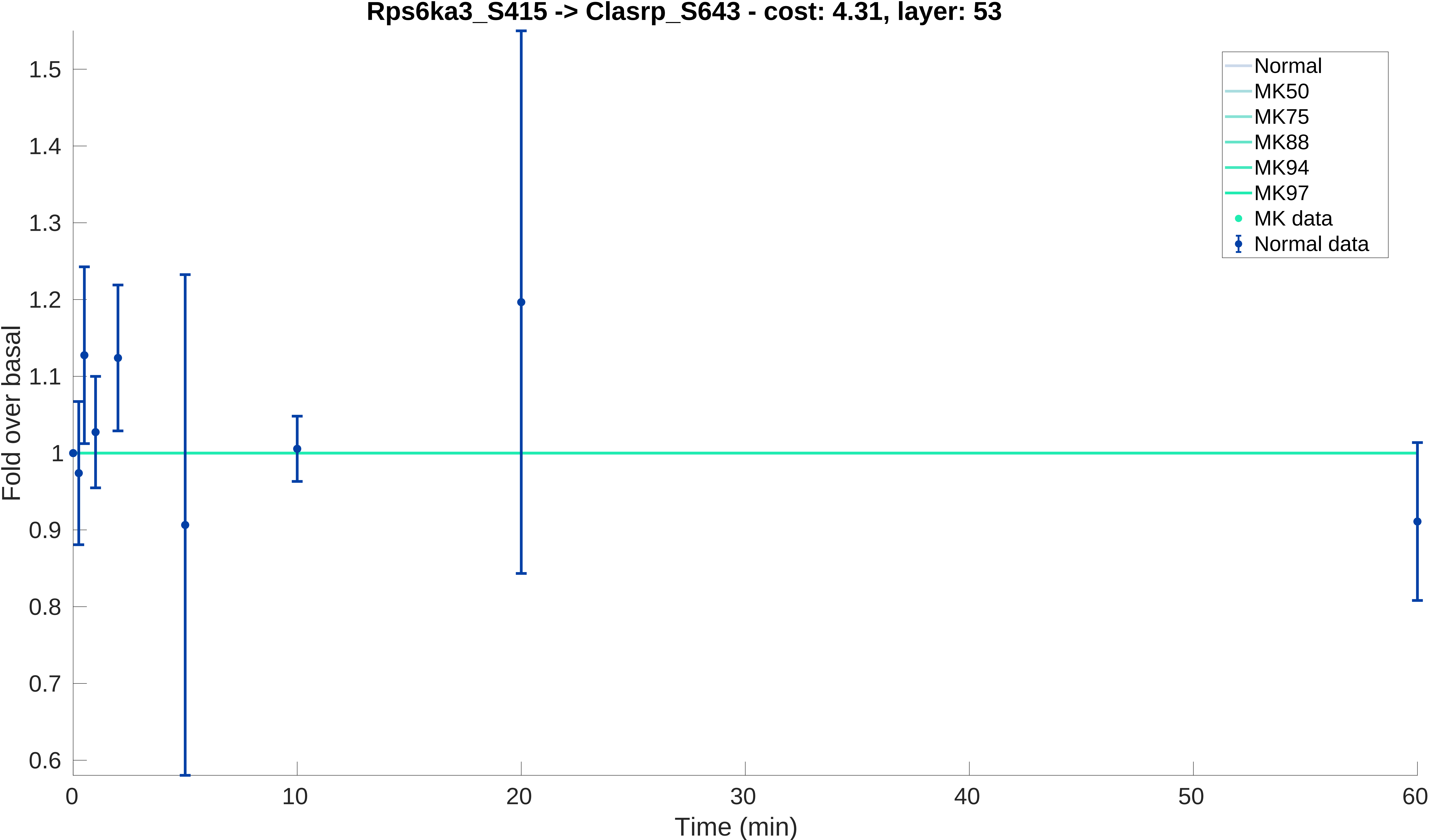

Rps6ka3_S415 -> Clasrp_S643 - cost: 4.31, layer: 53
Normal
MK50
MK75
MK88
MK94
MK97
MK data
Normal data
Fold over basal
1.5
1.4
1.3
1.2
1.1
1
0.9
0.8
0.7
0.6
0
10
20
30
40
50
60
Time (min)

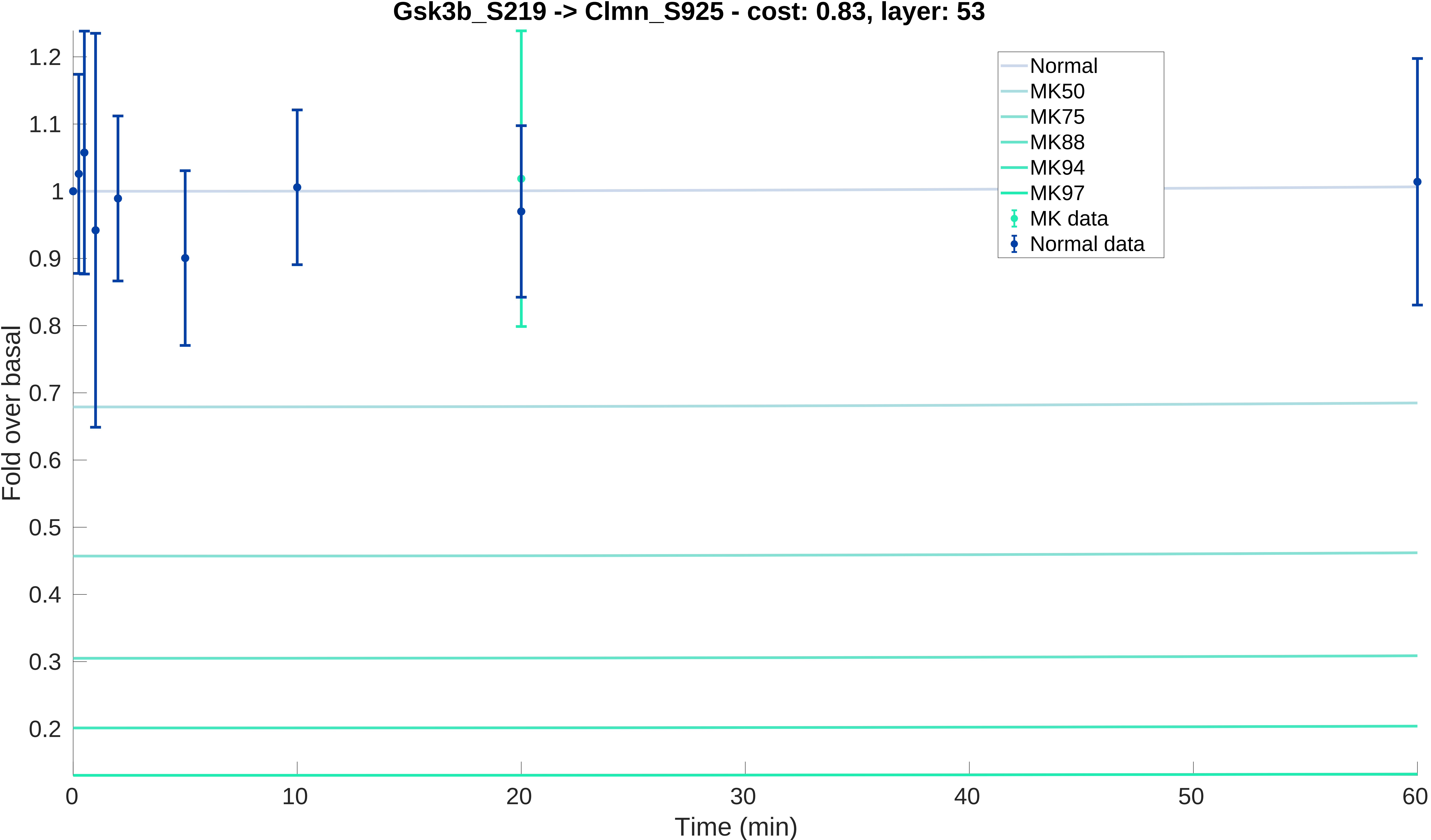
Gsk3b_S219 -> Clmn_S925 - cost: 0.83, layer: 53
Fold over basal
Time (min)
0
10
20
30
40
50
60
0.2
0.3
0.4
0.5
0.6
0.7
0.8
0.9
1
1.1
1.2
Normal
MK50
MK75
MK88
MK94
MK97
MK data
Normal data

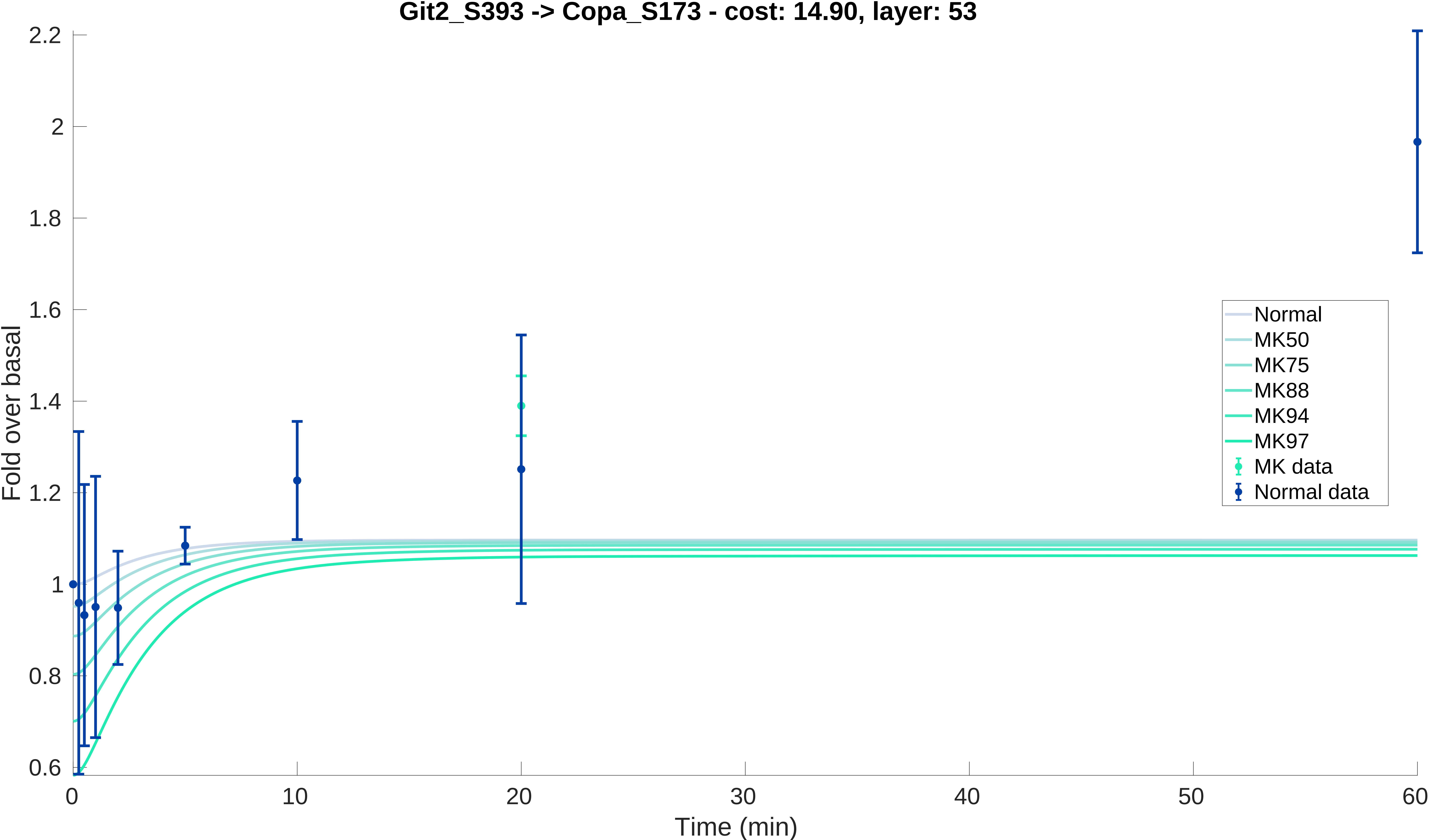
Git2_S393 -> Copa_S173 - cost: 14.90, layer: 53
Fold over basal
Time (min)
0.6
0.8
1
1.2
1.4
1.6
1.8
2
2.2
0
10
20
30
40
50
60
Normal
MK50
MK75
MK88
MK94
MK97
MK data
Normal data

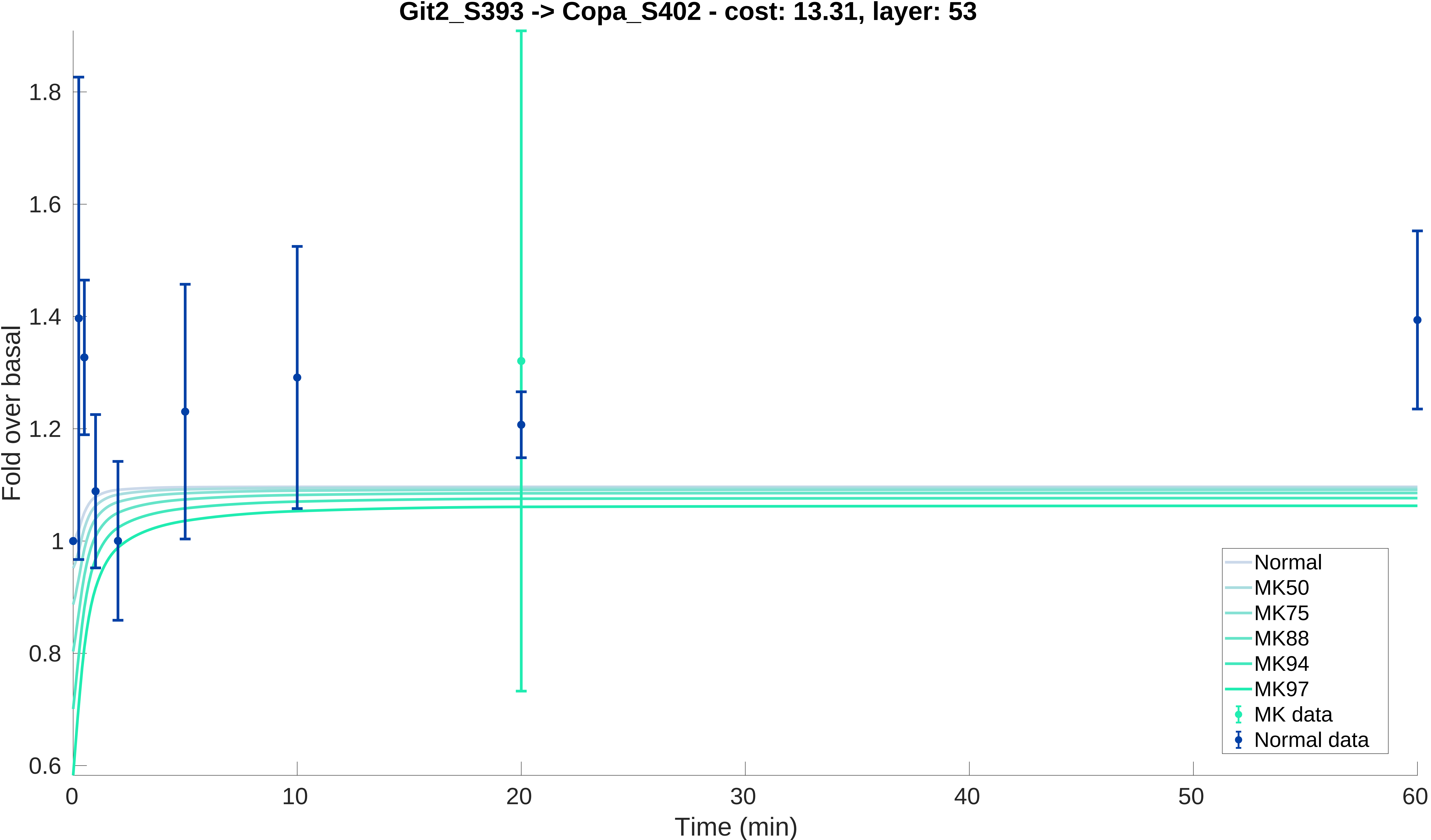

Git2_S393 -> Copa_S402 - cost: 13.31, layer: 53
Fold over basal
1.8
1.6
1.4
1.2
1
0.8
0.6
0
10
20
30
40
50
60
Time (min)
Normal
MK50
MK75
MK88
MK94
MK97
MK data
Normal data

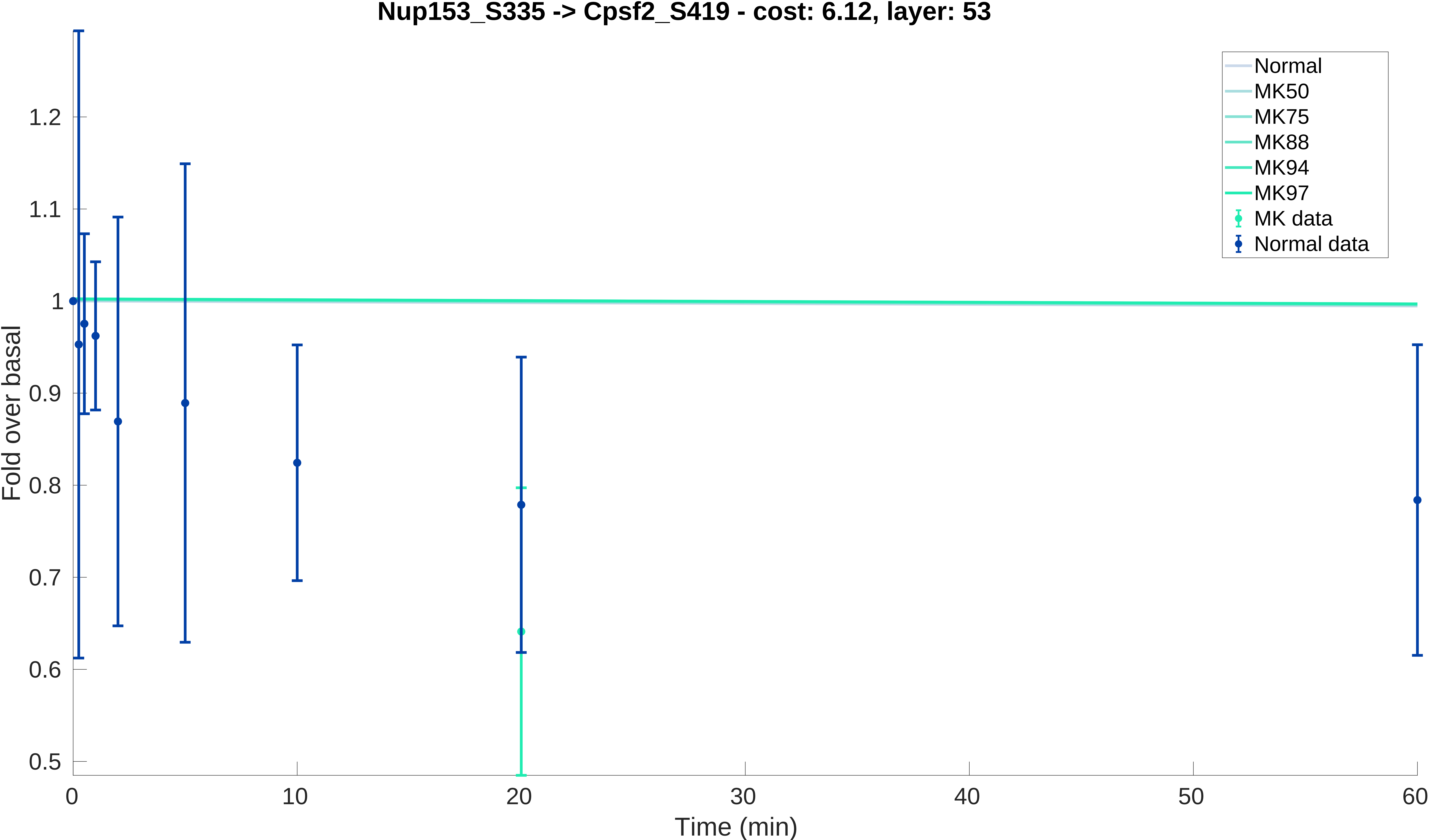
Nup153_S335 -> Cpsf2_S419 - cost: 6.12, layer: 53
Fold over basal
Time (min)
0.5
0.6
0.7
0.8
0.9
1
1.1
1.2
0
10
20
30
40
50
60
Normal
MK50
MK75
MK88
MK94
MK97
MK data
Normal data

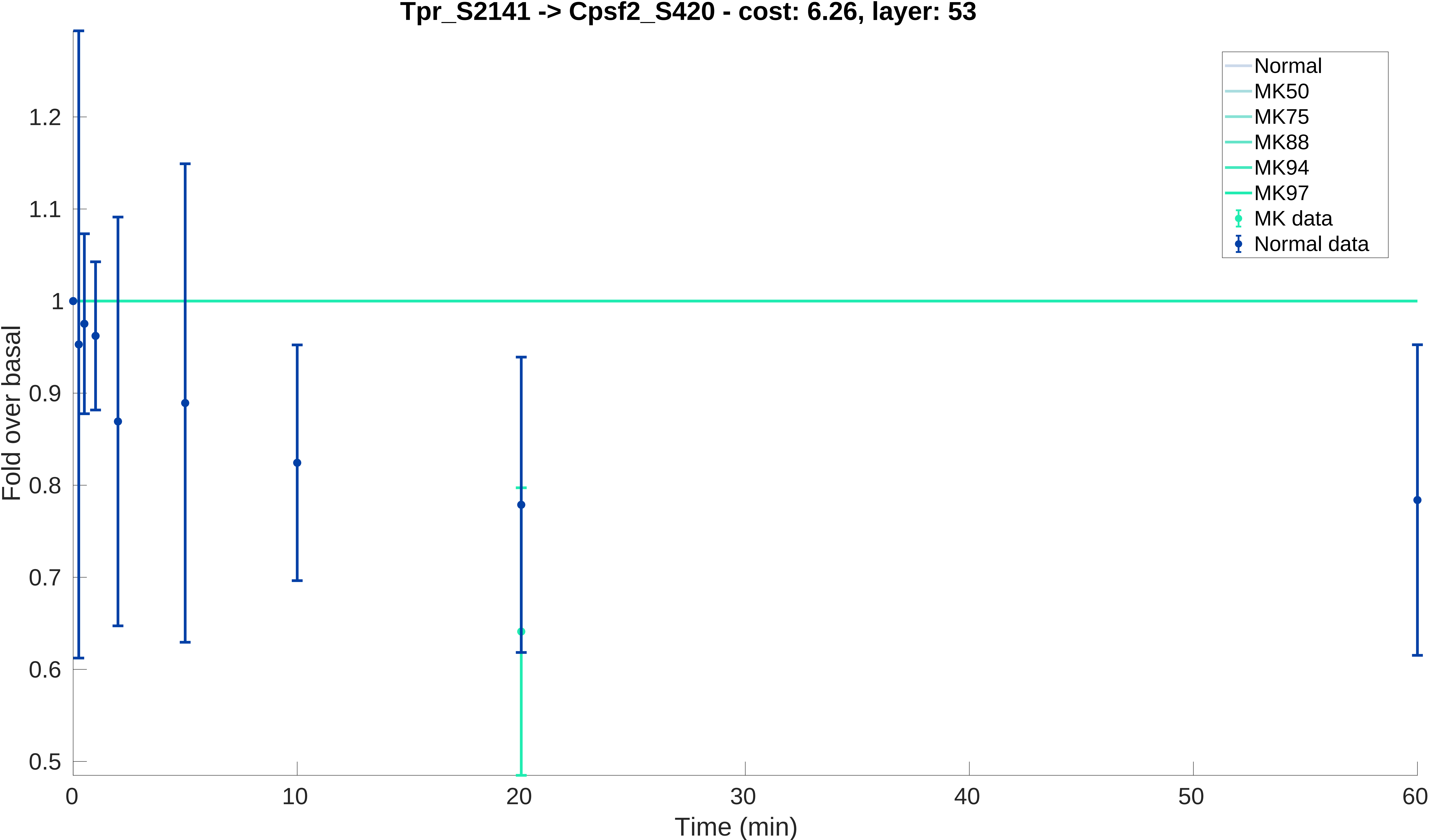

Tpr_S2141 -> Cpsf2_S420 - cost: 6.26, layer: 53
Normal
MK50
MK75
MK88
MK94
MK97
MK data
Normal data
Fold over basal
1.2
1.1
1
0.9
0.8
0.7
0.6
0.5
0
10
20
30
40
50
60
Time (min)

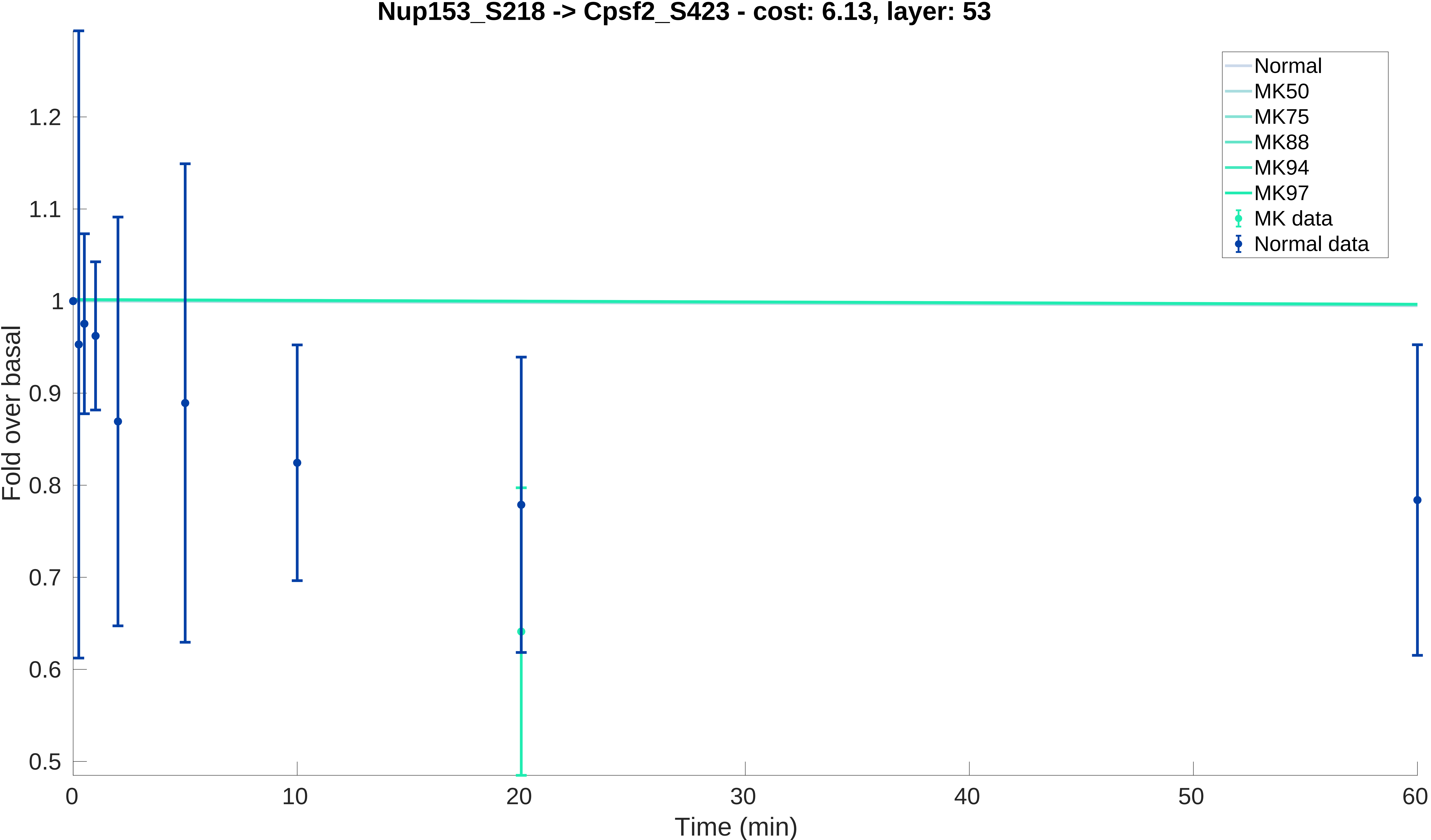

Nup153_S218 -> Cpsf2_S423 - cost: 6.13, layer: 53
Normal
MK50
MK75
MK88
MK94
MK97
MK data
Normal data
Fold over basal
1.2
1.1
1
0.9
0.8
0.7
0.6
0.5
0
10
20
30
40
50
60
Time (min)

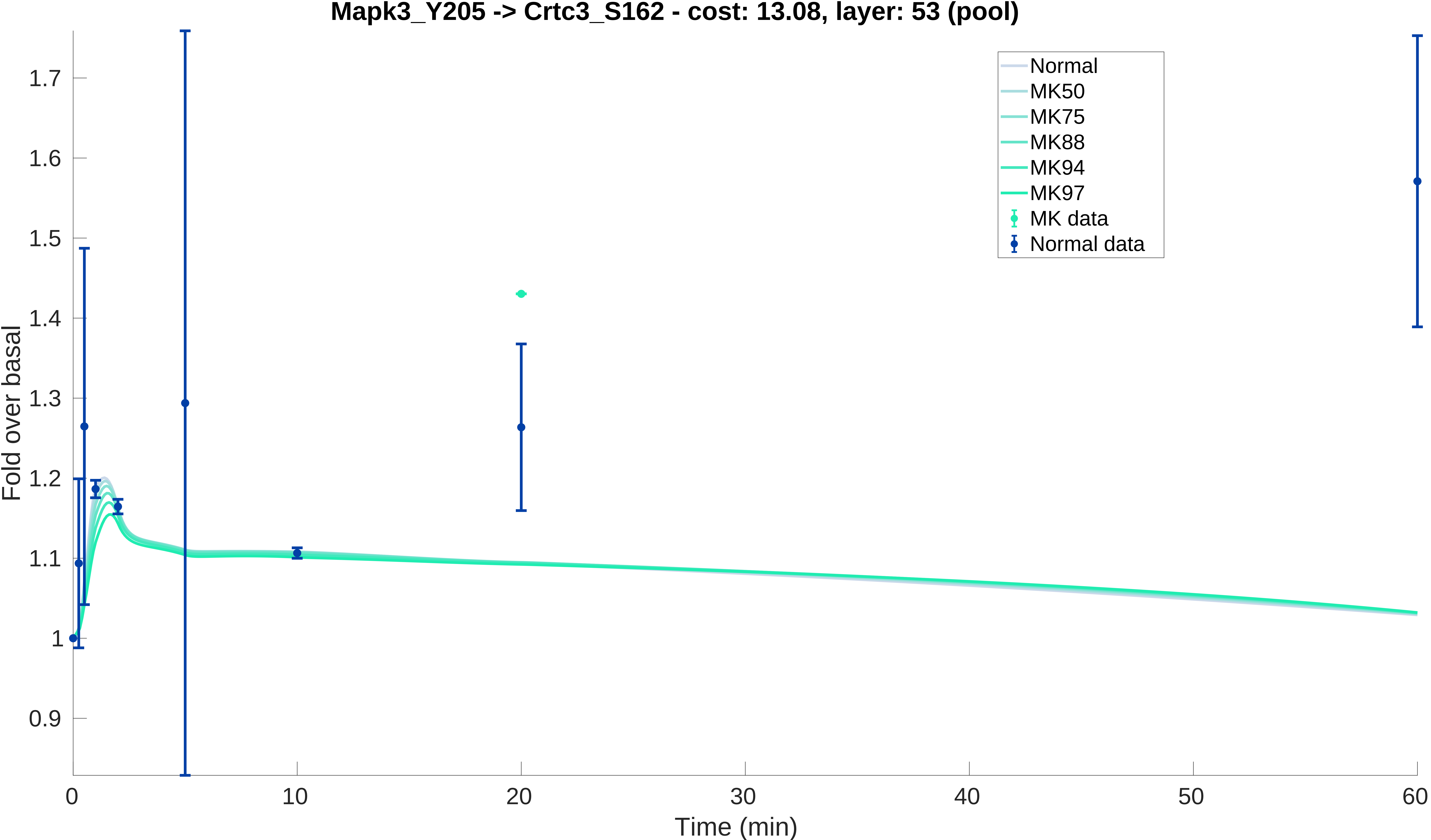

Mapk3_Y205 -> Crtc3_S162 - cost: 13.08, layer: 53 (pool)
Fold over basal
Time (min)
Normal
MK50
MK75
MK88
MK94
MK97
MK data
Normal data
1.7
1.6
1.5
1.4
1.3
1.2
1.1
1
0.9
0
10
20
30
40
50
60

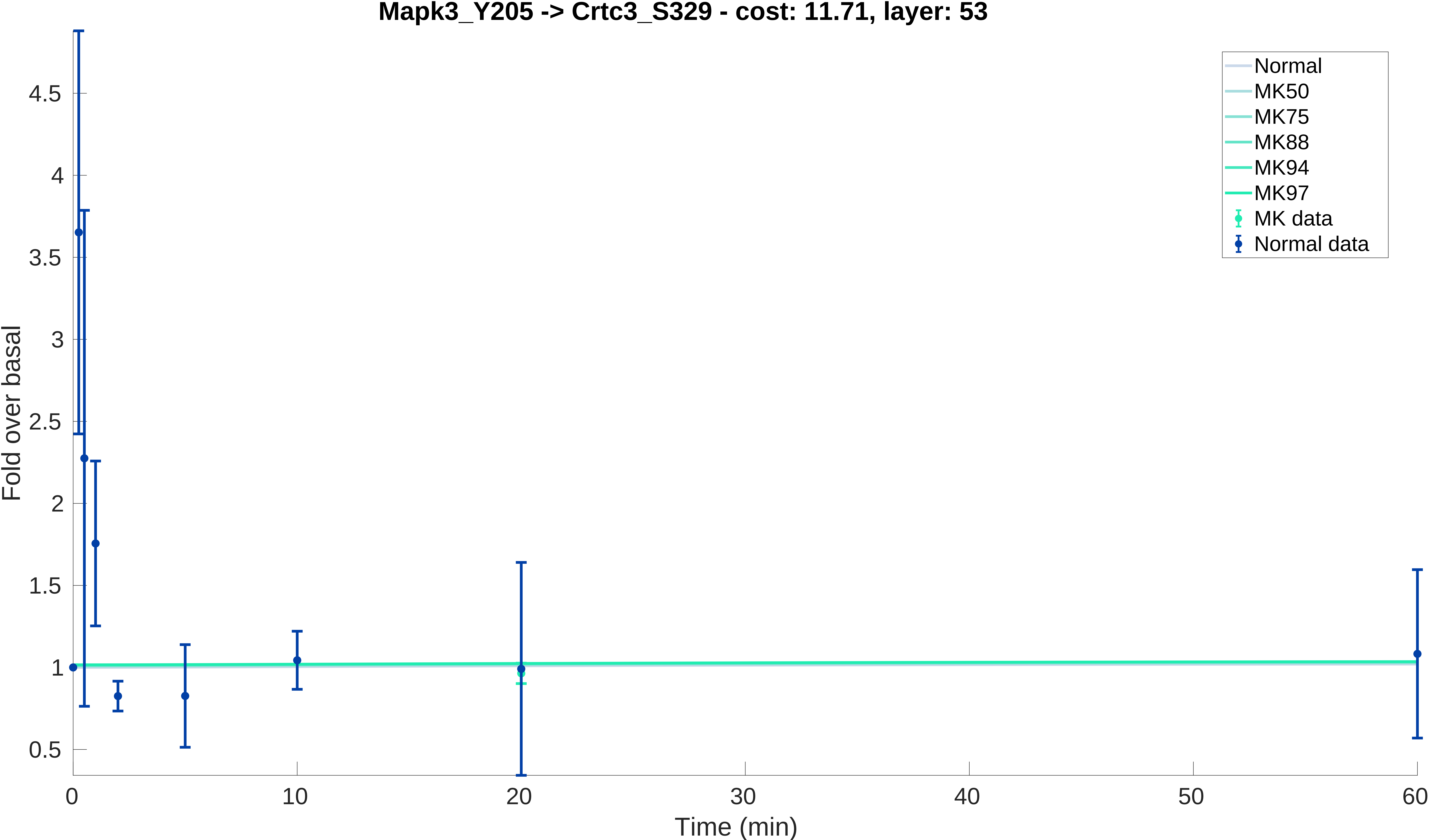
Mapk3_Y205 -> Crtc3_S329 - cost: 11.71, layer: 53
Fold over basal
Time (min)
0.5
1
1.5
2
2.5
3
3.5
4
4.5
0
10
20
30
40
50
60
Normal
MK50
MK75
MK88
MK94
MK97
MK data
Normal data

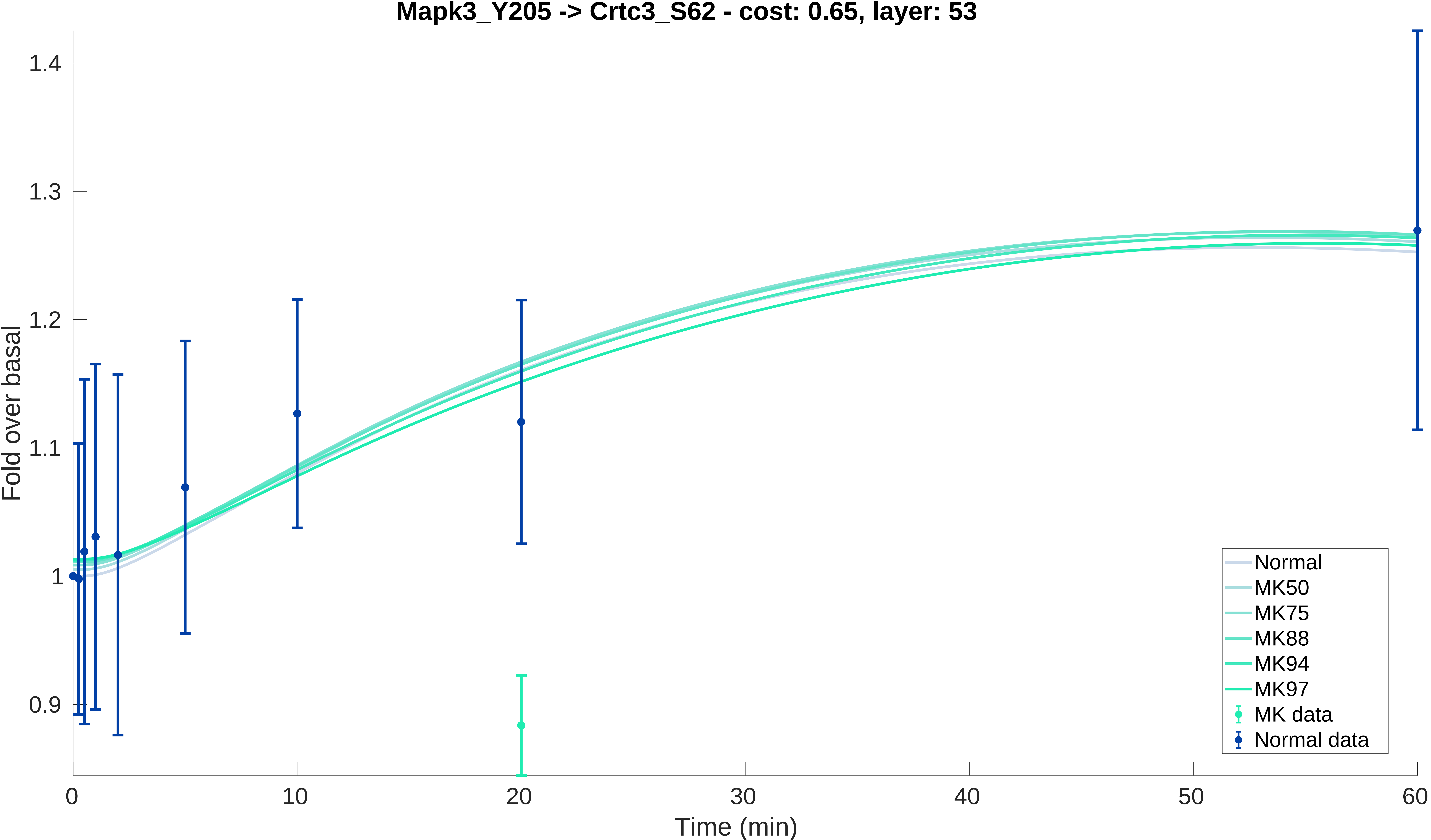

Mapk3_Y205 -> Crtc3_S62 - cost: 0.65, layer: 53
Fold over basal
Time (min)
0.9
1
1.1
1.2
1.3
1.4
0
10
20
30
40
50
60
Normal
MK50
MK75
MK88
MK94
MK97
MK data
Normal data

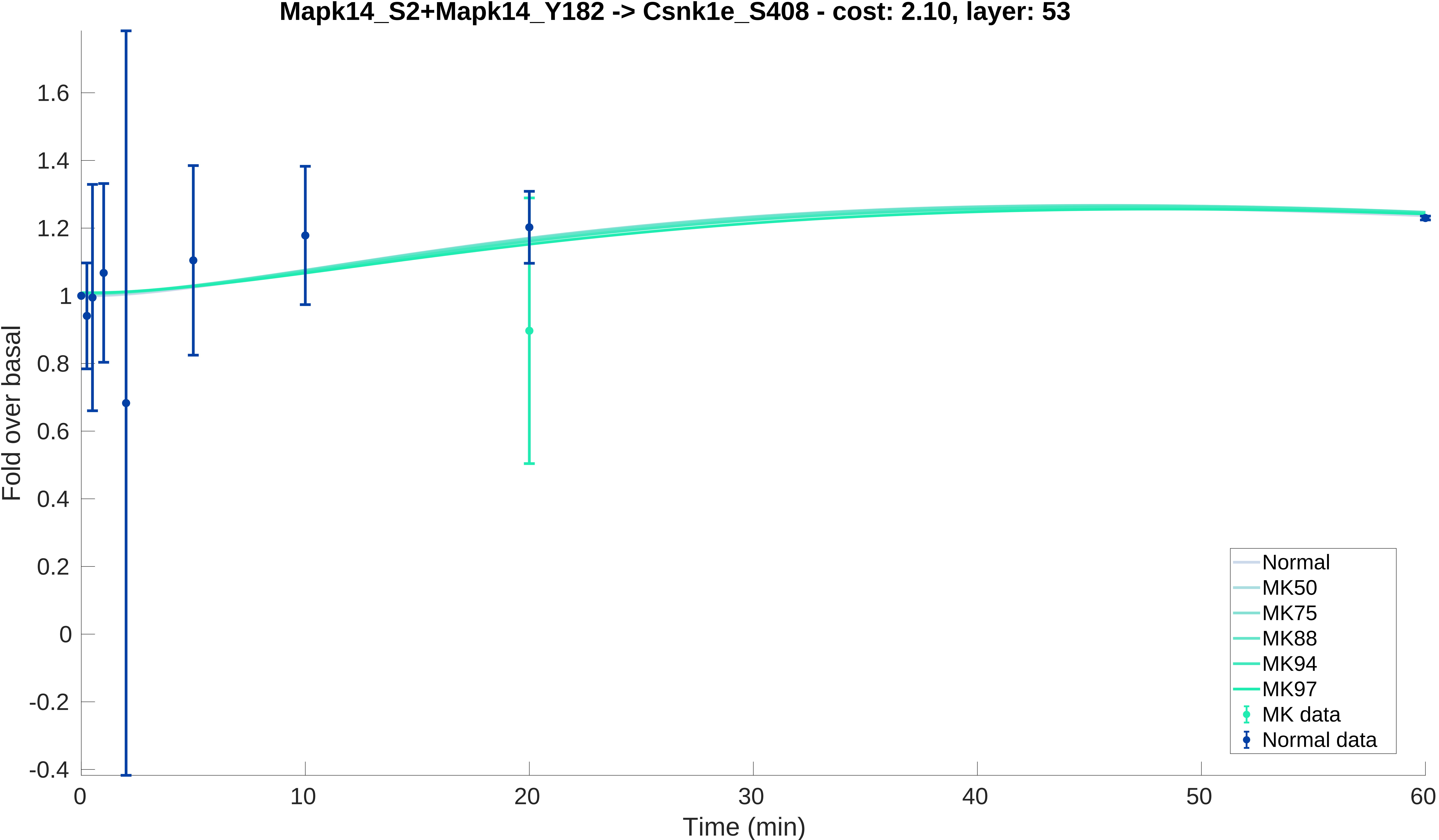

Mapk14_S2+Mapk14_Y182 -> Csnk1e_S408 - cost: 2.10, layer: 53
Fold over basal
Time (min)
1.6
1.4
1.2
1
0.8
0.6
0.4
0.2
0
-0.2
-0.4
0
10
20
30
40
50
60
Normal
MK50
MK75
MK88
MK94
MK97
MK data
Normal data

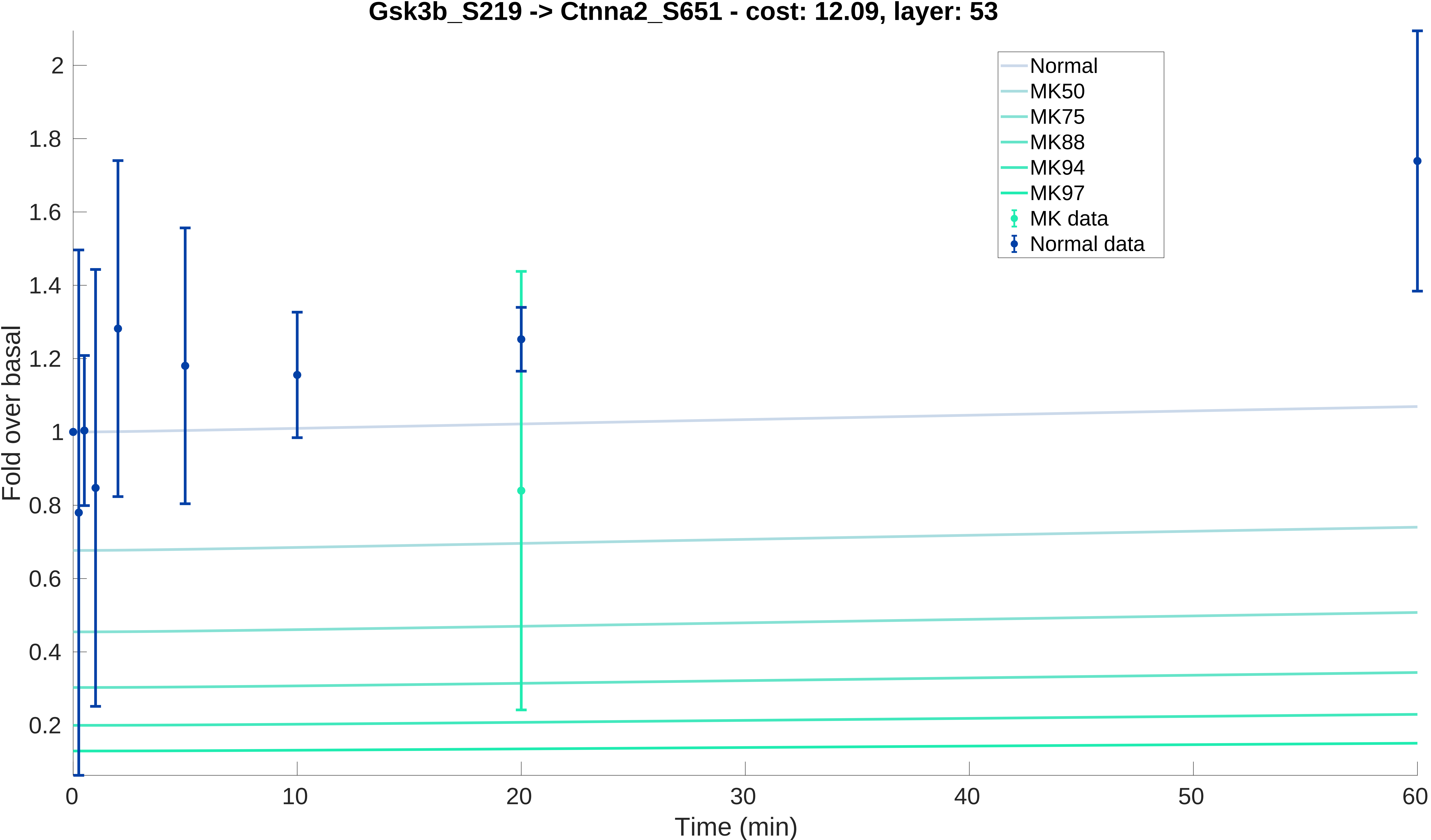

Gsk3b_S219 -> Ctnna2_S651 - cost: 12.09, layer: 53
Fold over basal
Time (min)
0
10
20
30
40
50
60
0.2
0.4
0.6
0.8
1
1.2
1.4
1.6
1.8
2
Normal
MK50
MK75
MK88
MK94
MK97
MK data
Normal data

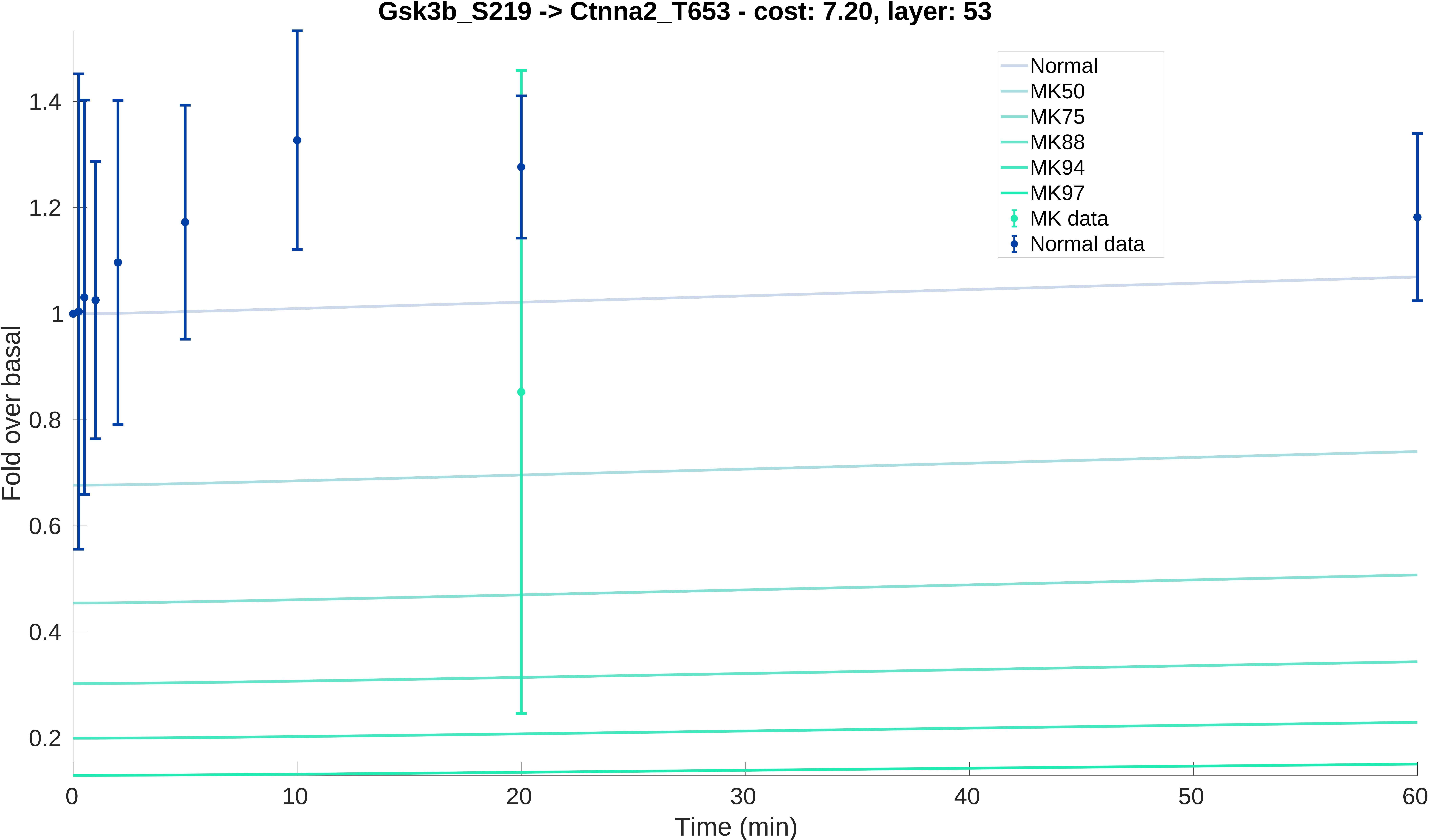

Gsk3b_S219 -> Ctnna2_T653 - cost: 7.20, layer: 53
Normal
MK50
MK75
MK88
MK94
MK97
MK data
Normal data
Fold over basal
Time (min)
0.2
0.4
0.6
0.8
1
1.2
1.4
0
10
20
30
40
50
60

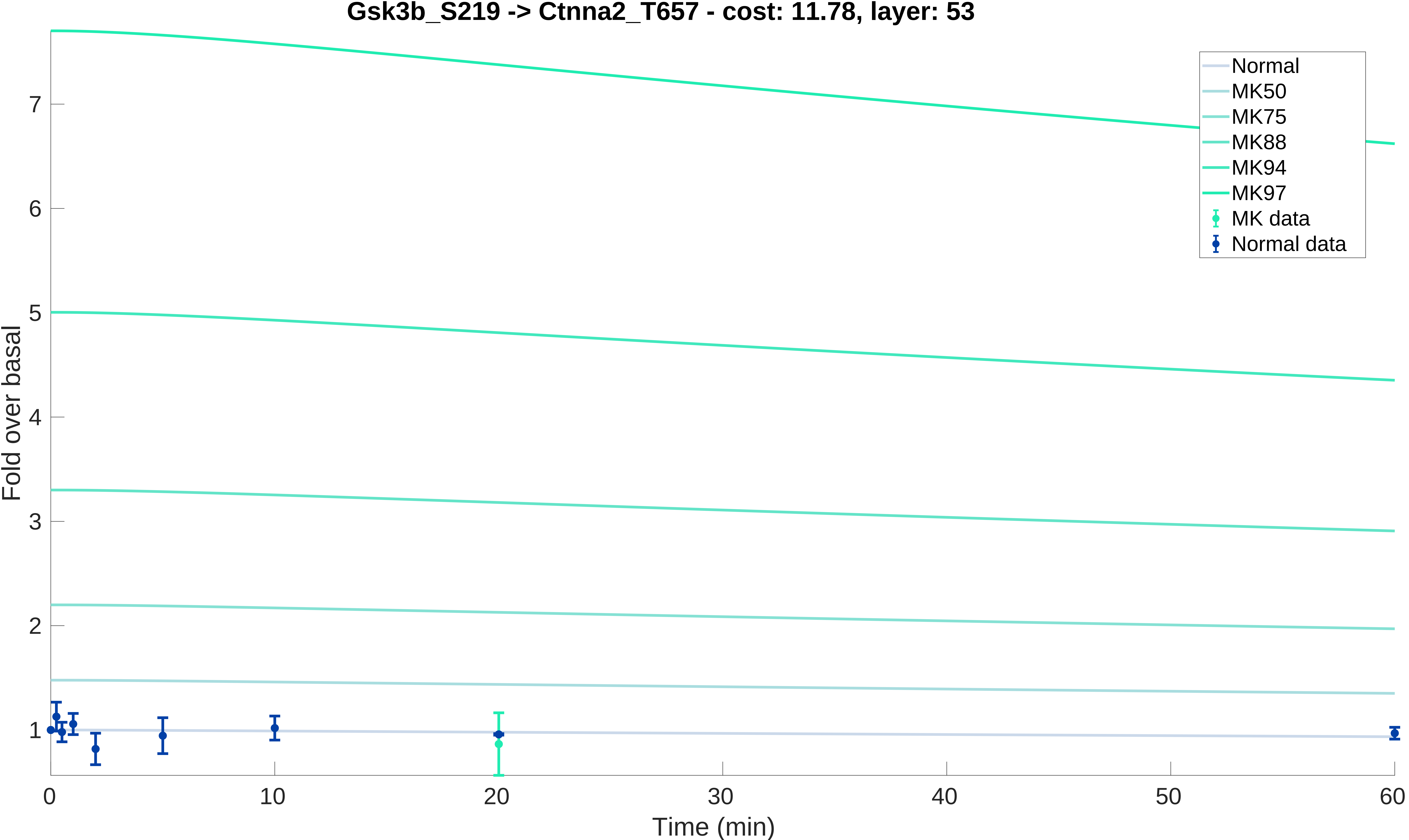

Gsk3b_S219 -> Ctnna2_T657 - cost: 11.78, layer: 53
Fold over basal
Time (min)
Normal
MK50
MK75
MK88
MK94
MK97
MK data
Normal data
1
2
3
4
5
6
7
0
10
20
30
40
50
60

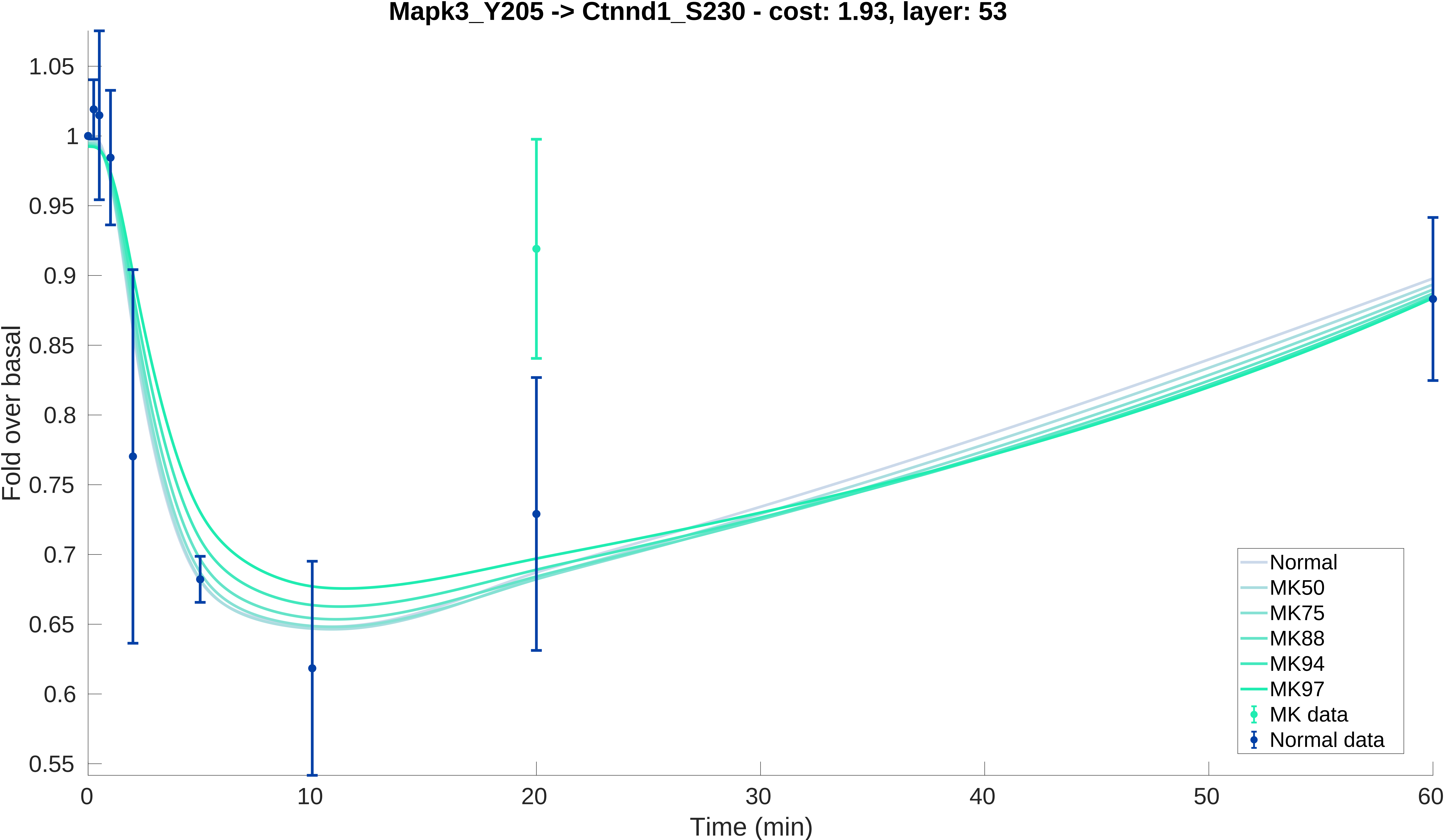
Mapk3_Y205 -> Ctnnd1_S230 - cost: 1.93, layer: 53
Fold over basal
1.05
1
0.95
0.9
0.85
0.8
0.75
0.7
0.65
0.6
0.55
0
10
20
30
40
50
60
Time (min)
Normal
MK50
MK75
MK88
MK94
MK97
MK data
Normal data

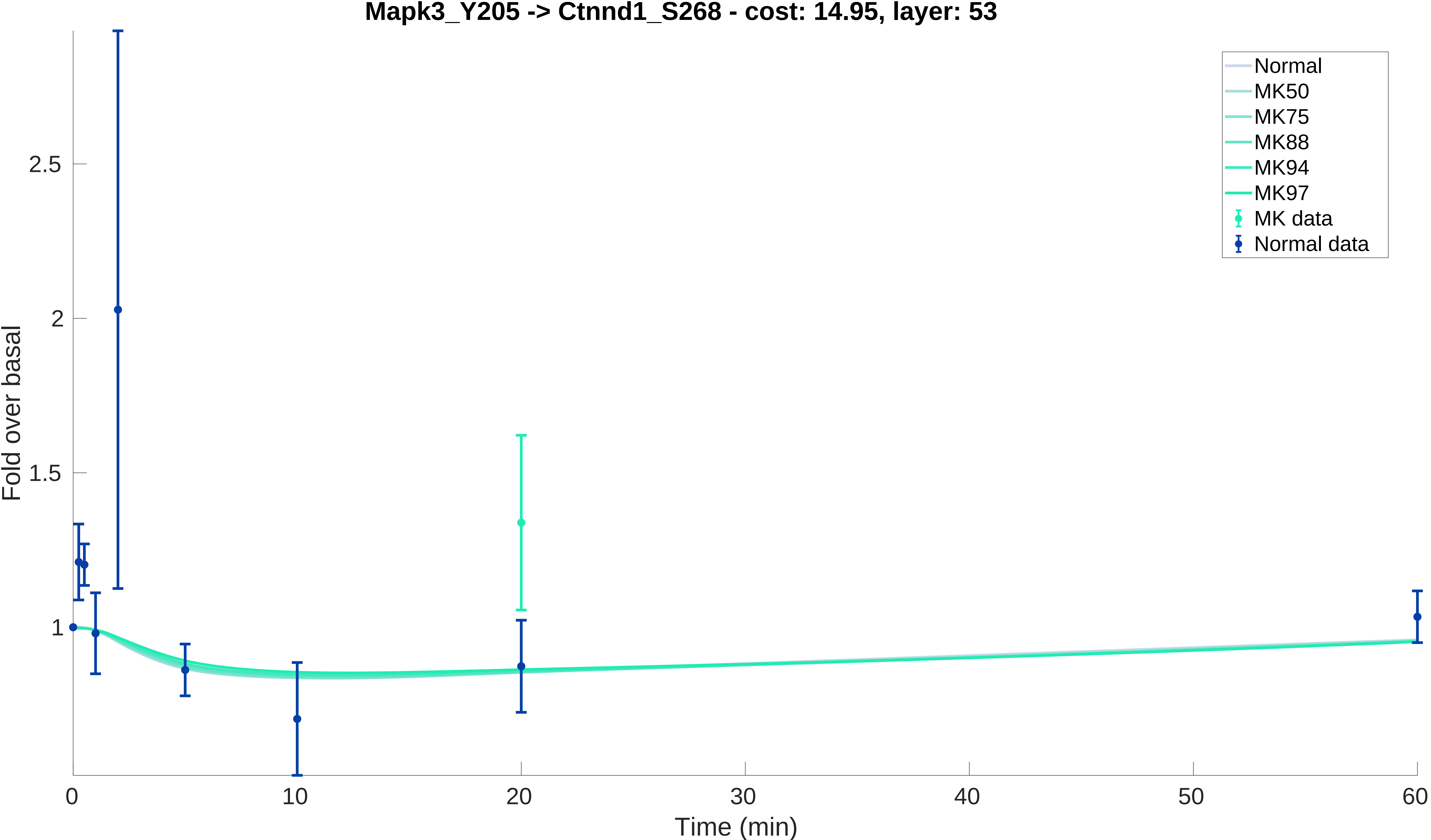
Mapk3_Y205 -> Ctnnd1_S268 - cost: 14.95, layer: 53
Fold over basal
Time (min)
0
10
20
30
40
50
60
1
1.5
2
2.5
Normal
MK50
MK75
MK88
MK94
MK97
MK data
Normal data

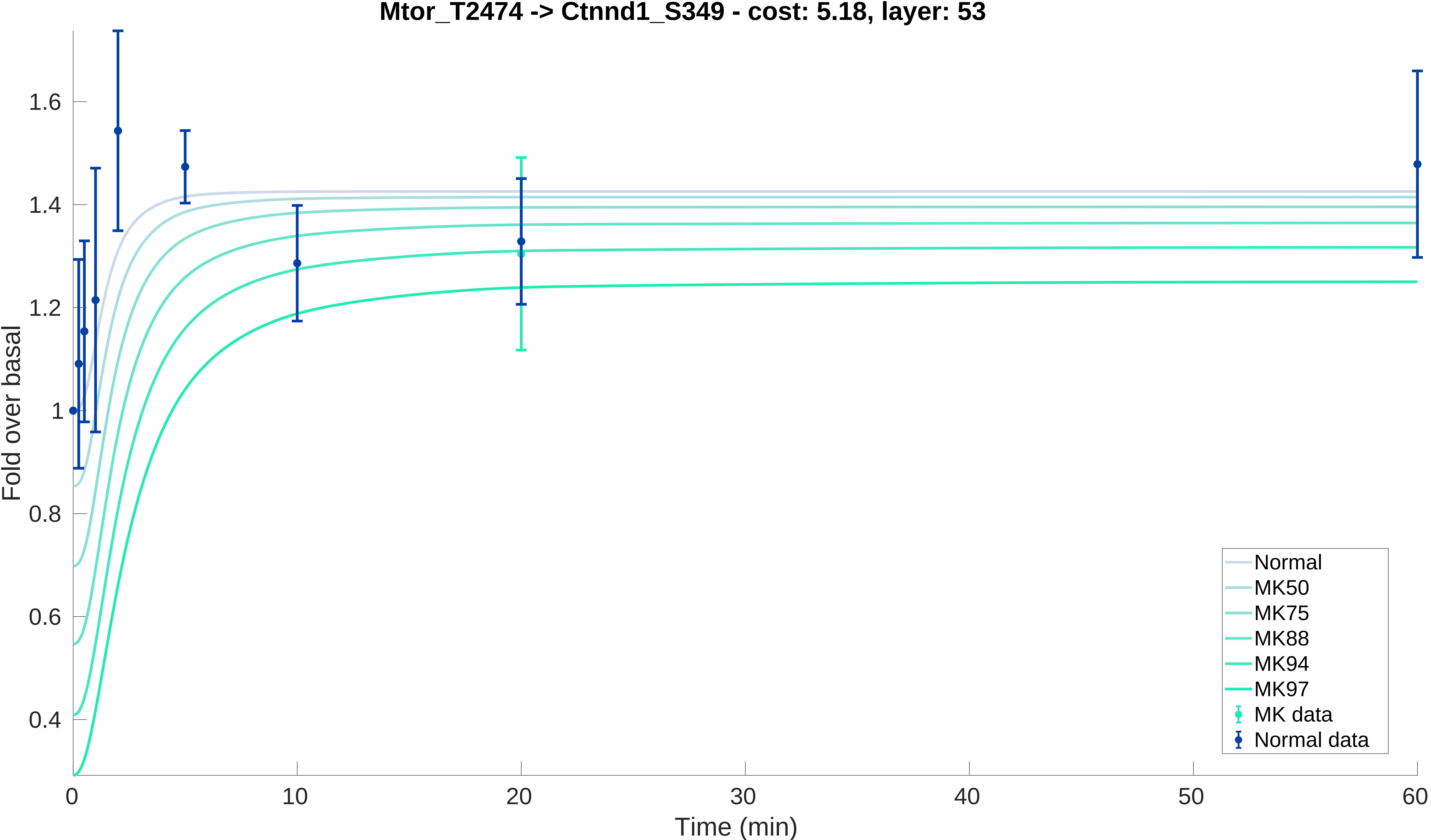
Mtor_T2474 -> Ctnnd1_S349 - cost: 5.18, layer: 53
Fold over basal
Time (min)
0.4
0.6
0.8
1
1.2
1.4
1.6
0
10
20
30
40
50
60
Normal
MK50
MK75
MK88
MK94
MK97
MK data
Normal data

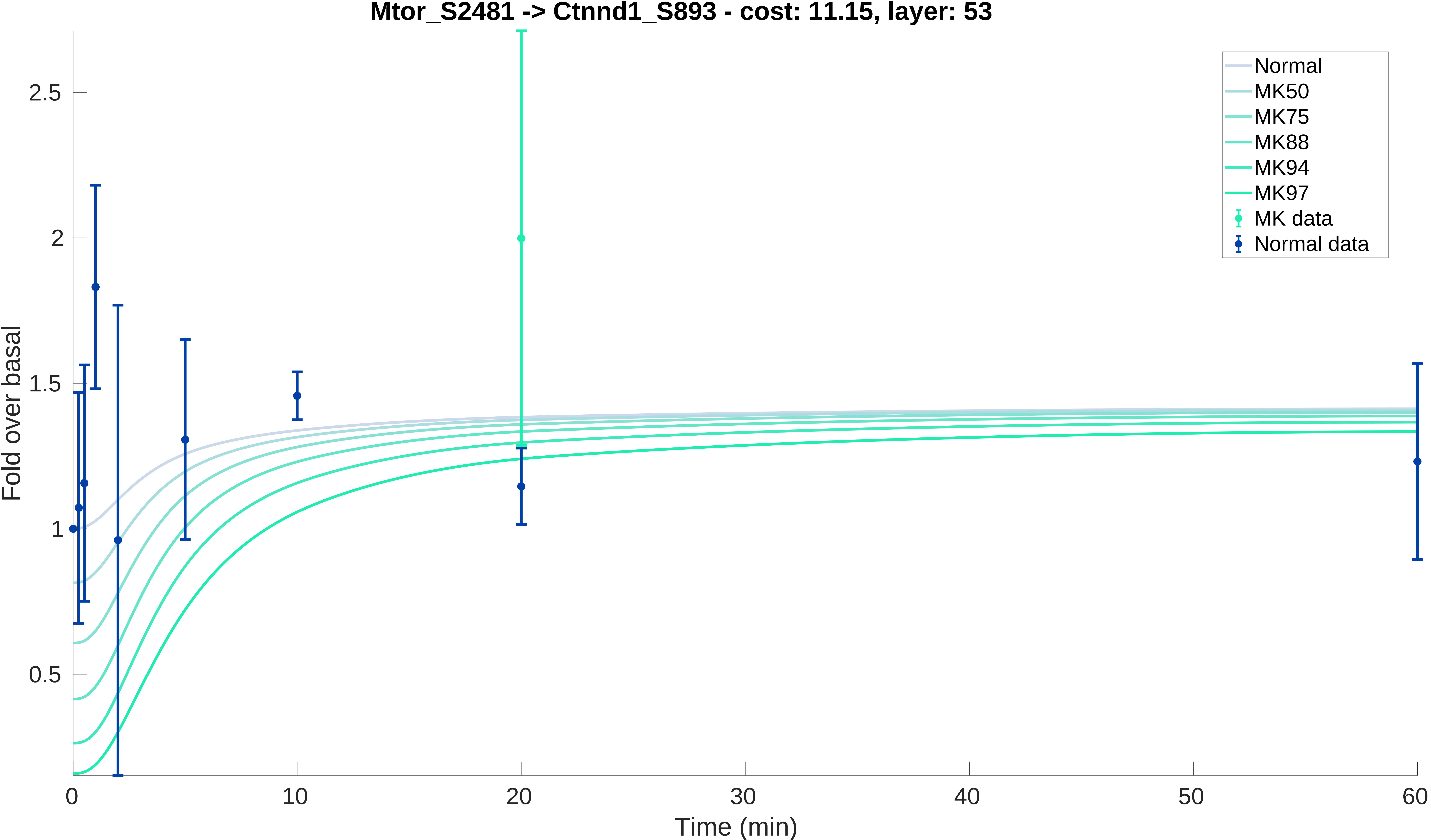

Mtor_S2481 -> Ctnnd1_S893 - cost: 11.15, layer: 53
Fold over basal
Time (min)
2.5
2
1.5
1
0.5
0
10
20
30
40
50
60
Normal
MK50
MK75
MK88
MK94
MK97
MK data
Normal data

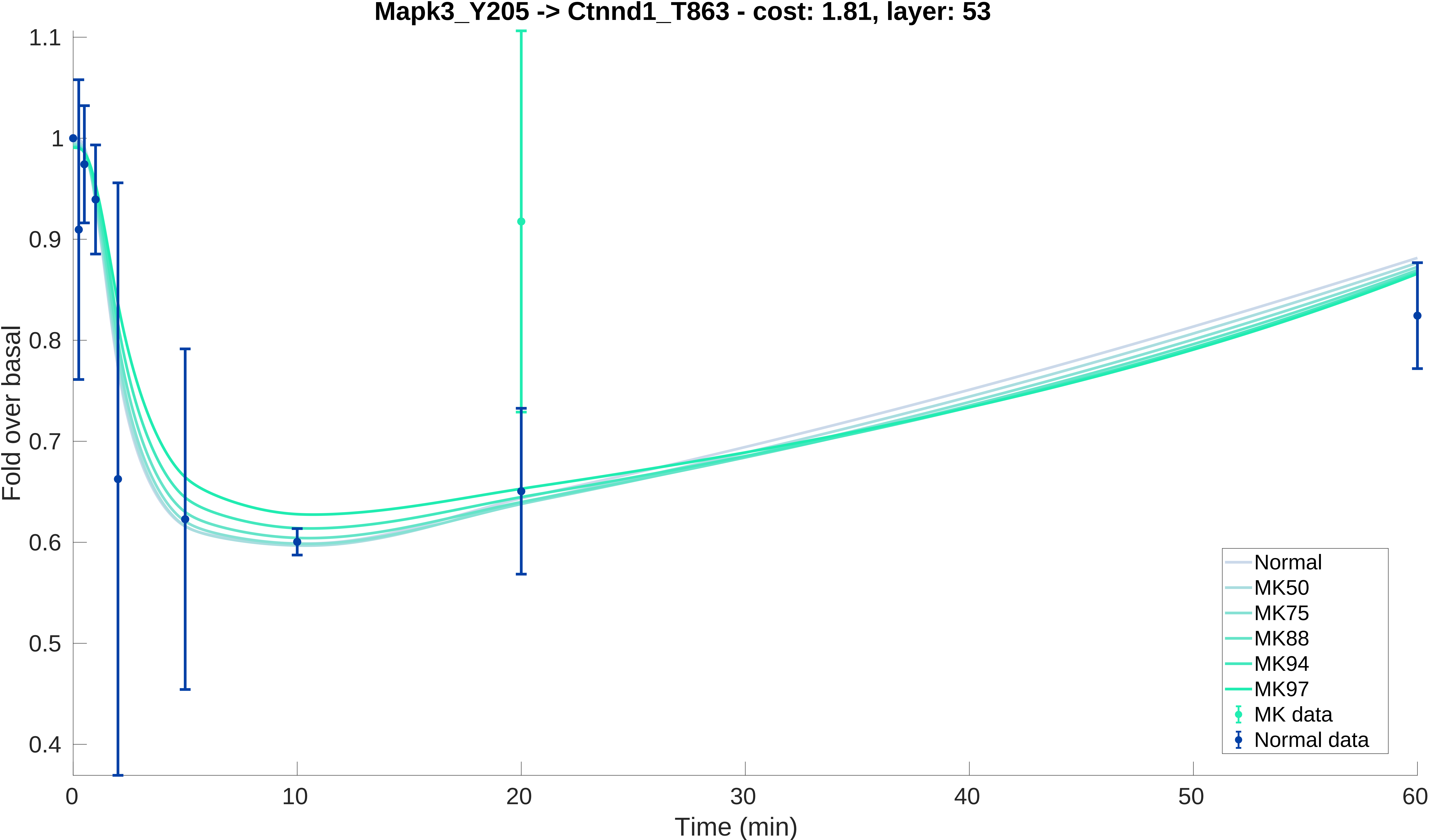

Mapk3_Y205 -> Ctnnd1_T863 - cost: 1.81, layer: 53
Fold over basal
1.1
1
0.9
0.8
0.7
0.6
0.5
0.4
0
10
20
30
40
50
60
Time (min)
Normal
MK50
MK75
MK88
MK94
MK97
MK data
Normal data

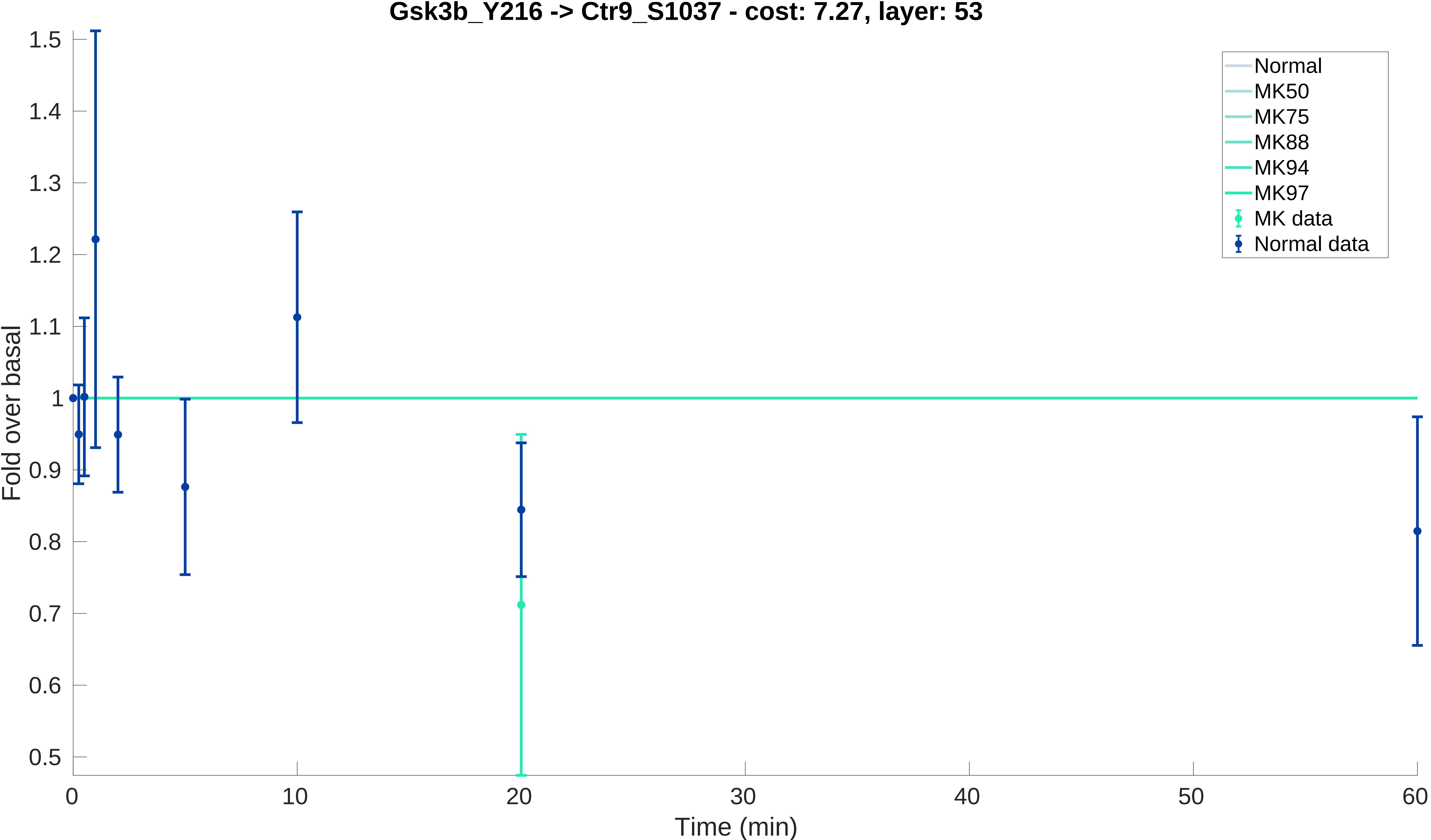

Gsk3b_Y216 -> Ctr9_S1037 - cost: 7.27, layer: 53
Fold over basal
Time (min)
1.5
1.4
1.3
1.2
1.1
1
0.9
0.8
0.7
0.6
0.5
0
10
20
30
40
50
60
Normal
MK50
MK75
MK88
MK94
MK97
MK data
Normal data

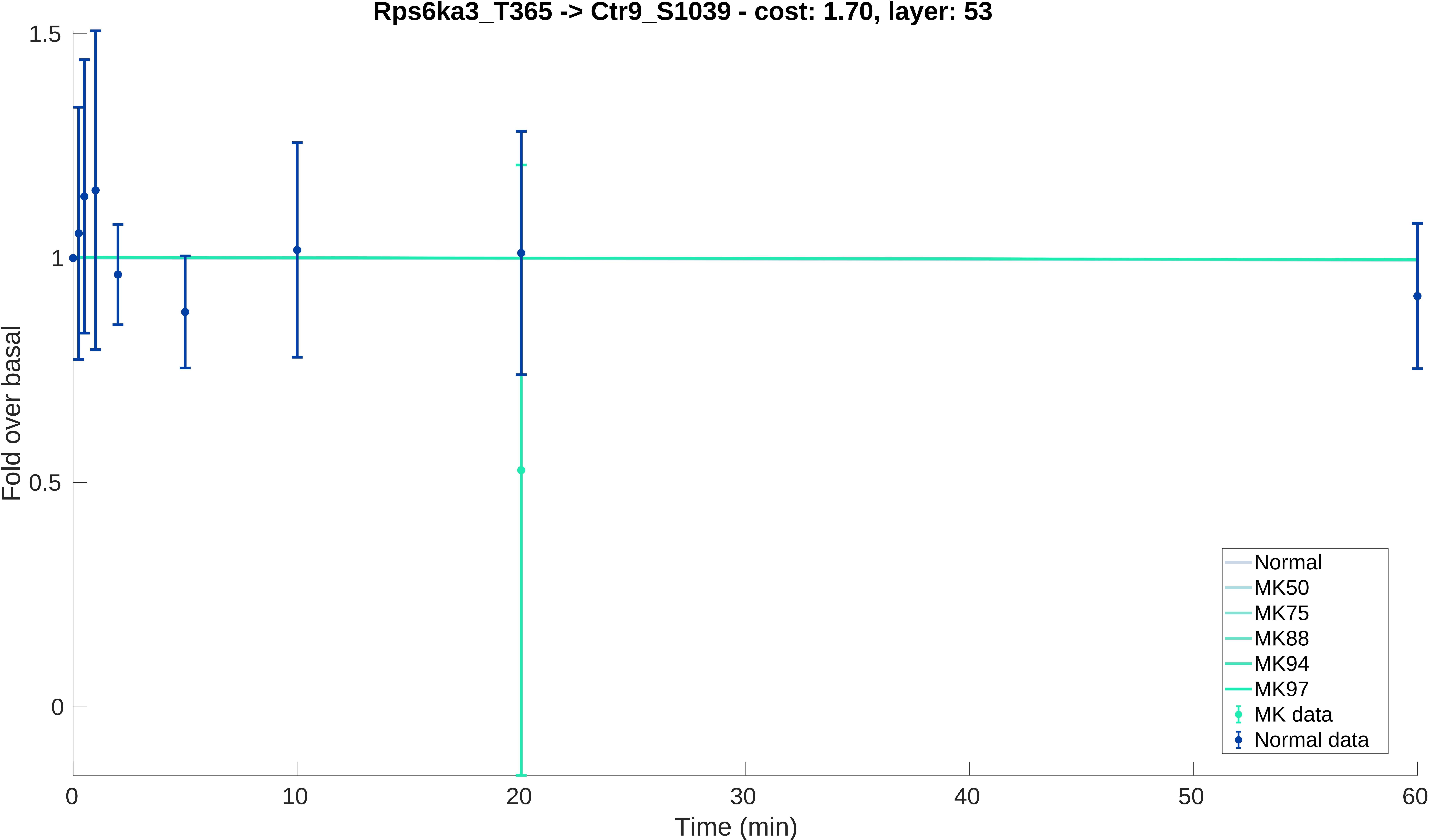

Rps6ka3_T365 -> Ctr9_S1039 - cost: 1.70, layer: 53
1.5
1
0.5
0
Fold over basal
0
10
20
30
40
50
60
Time (min)
Normal
MK50
MK75
MK88
MK94
MK97
MK data
Normal data

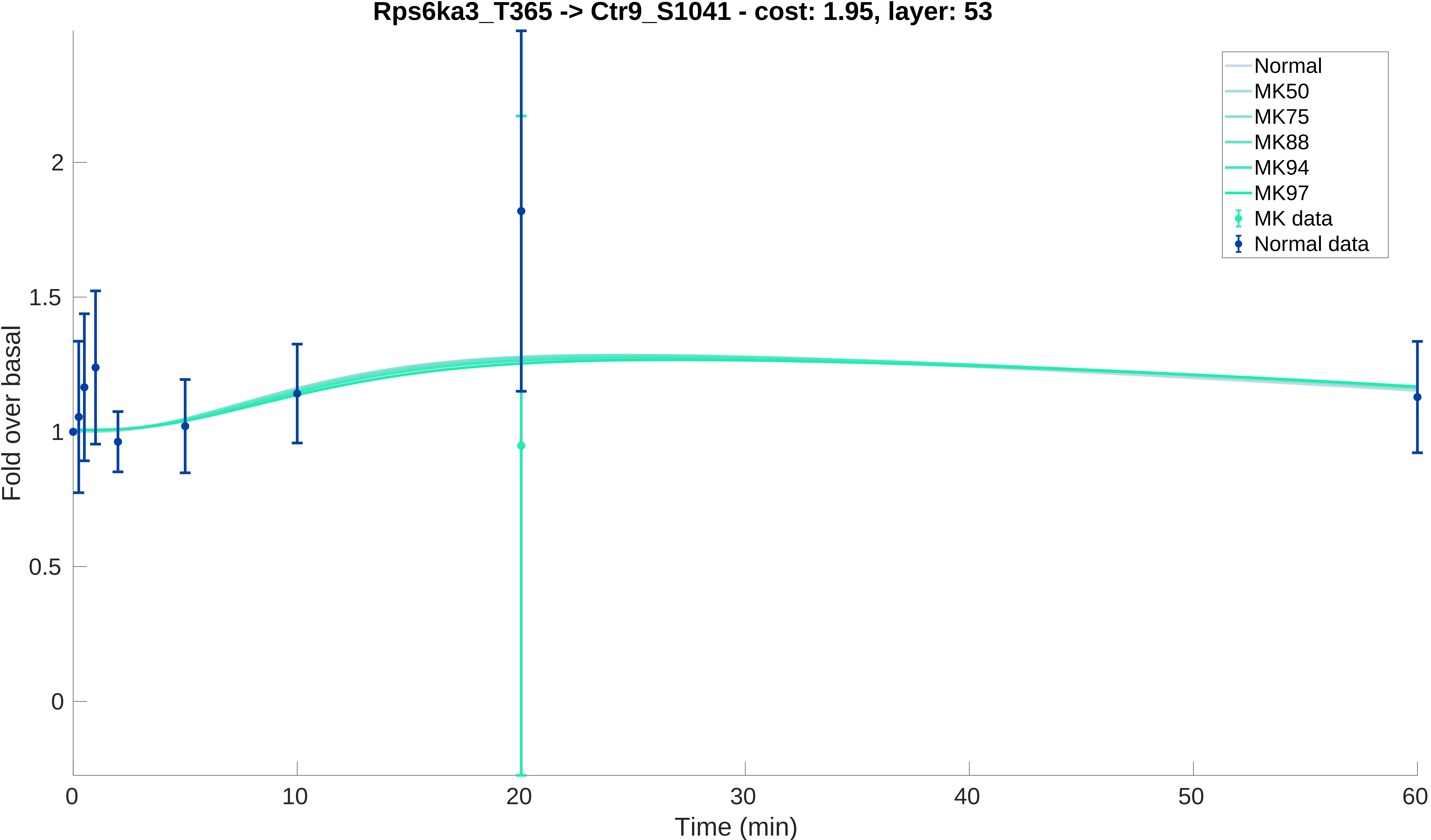

Rps6ka3_T365 -> Ctr9_S1041 - cost: 1.95, layer: 53
Normal
MK50
MK75
MK88
MK94
MK97
MK data
Normal data
Fold over basal
0
0.5
1
1.5
2
0
10
20
30
40
50
60
Time (min)

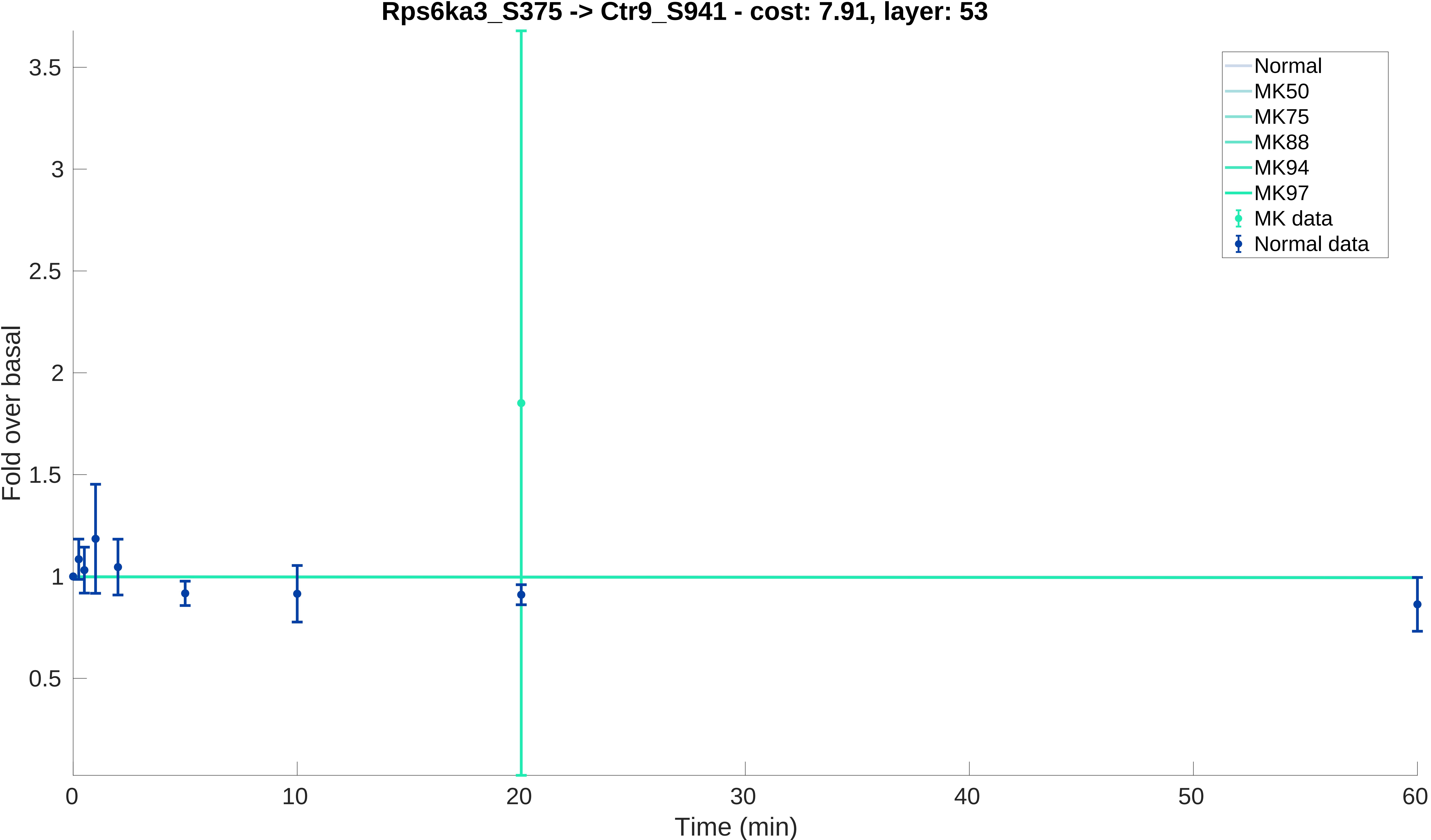
Rps6ka3_S375 -> Ctr9_S941 - cost: 7.91, layer: 53
Fold over basal
Time (min)
0.5
1
1.5
2
2.5
3
3.5
0
10
20
30
40
50
60
Normal
MK50
MK75
MK88
MK94
MK97
MK data
Normal data

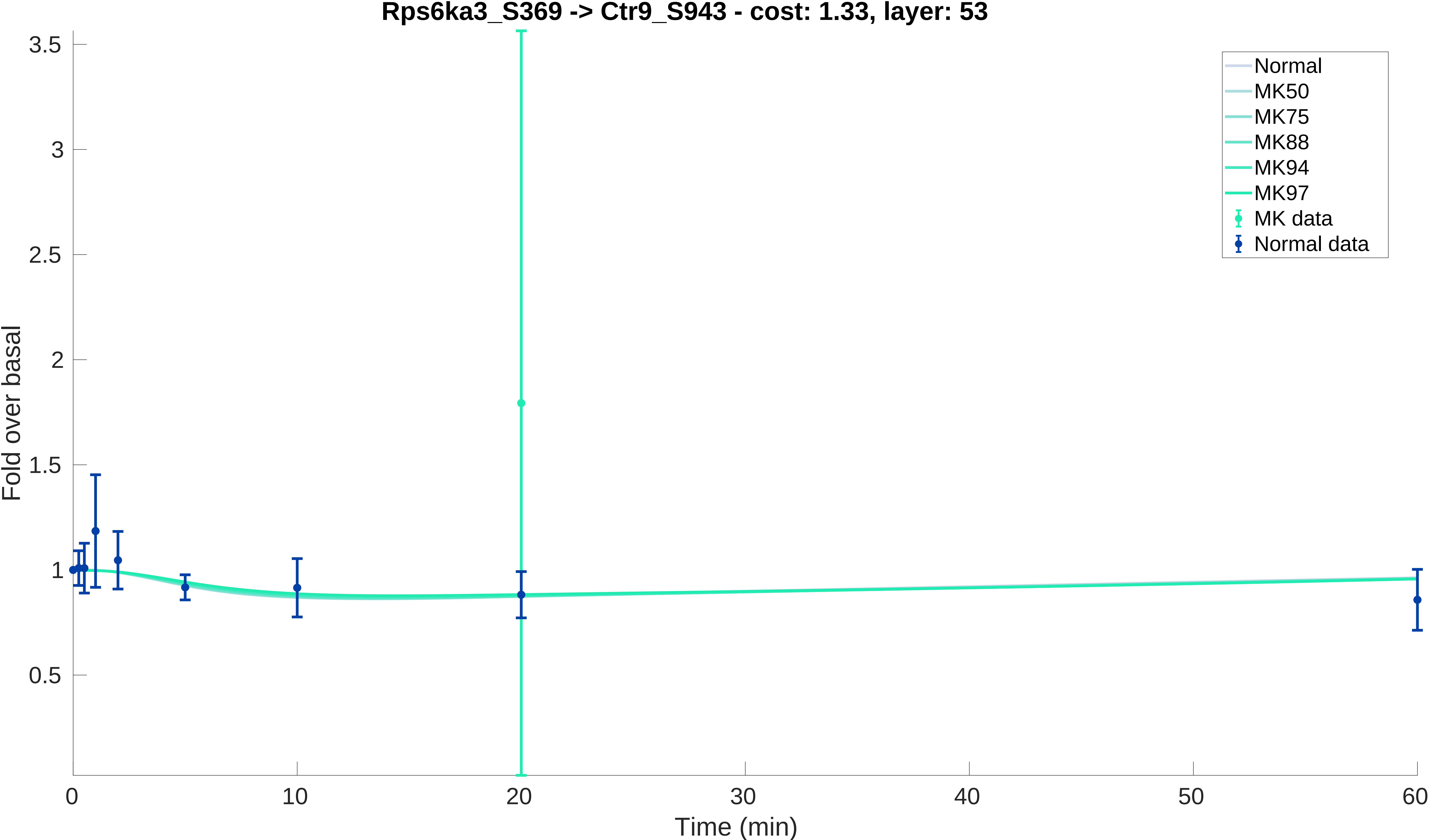
Rps6ka3_S369 -> Ctr9_S943 - cost: 1.33, layer: 53
Fold over basal
Time (min)
0.5
1
1.5
2
2.5
3
3.5
0
10
20
30
40
50
60
Normal
MK50
MK75
MK88
MK94
MK97
MK data
Normal data

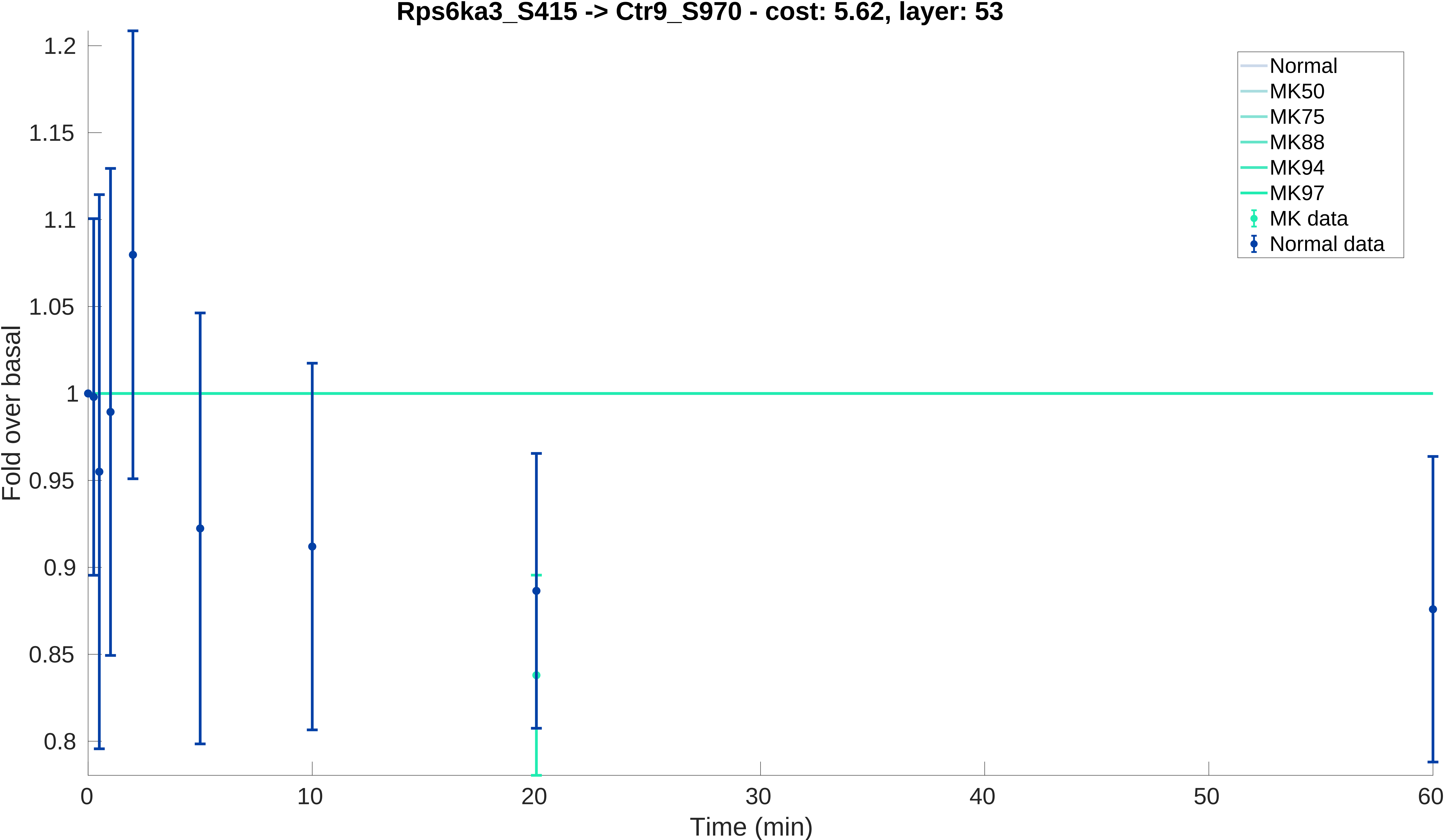

Rps6ka3_S415 -> Ctr9_S970 - cost: 5.62, layer: 53
Fold over basal
Time (min)
1.2
1.15
1.1
1.05
1
0.95
0.9
0.85
0.8
0
10
20
30
40
50
60
Normal
MK50
MK75
MK88
MK94
MK97
MK data
Normal data

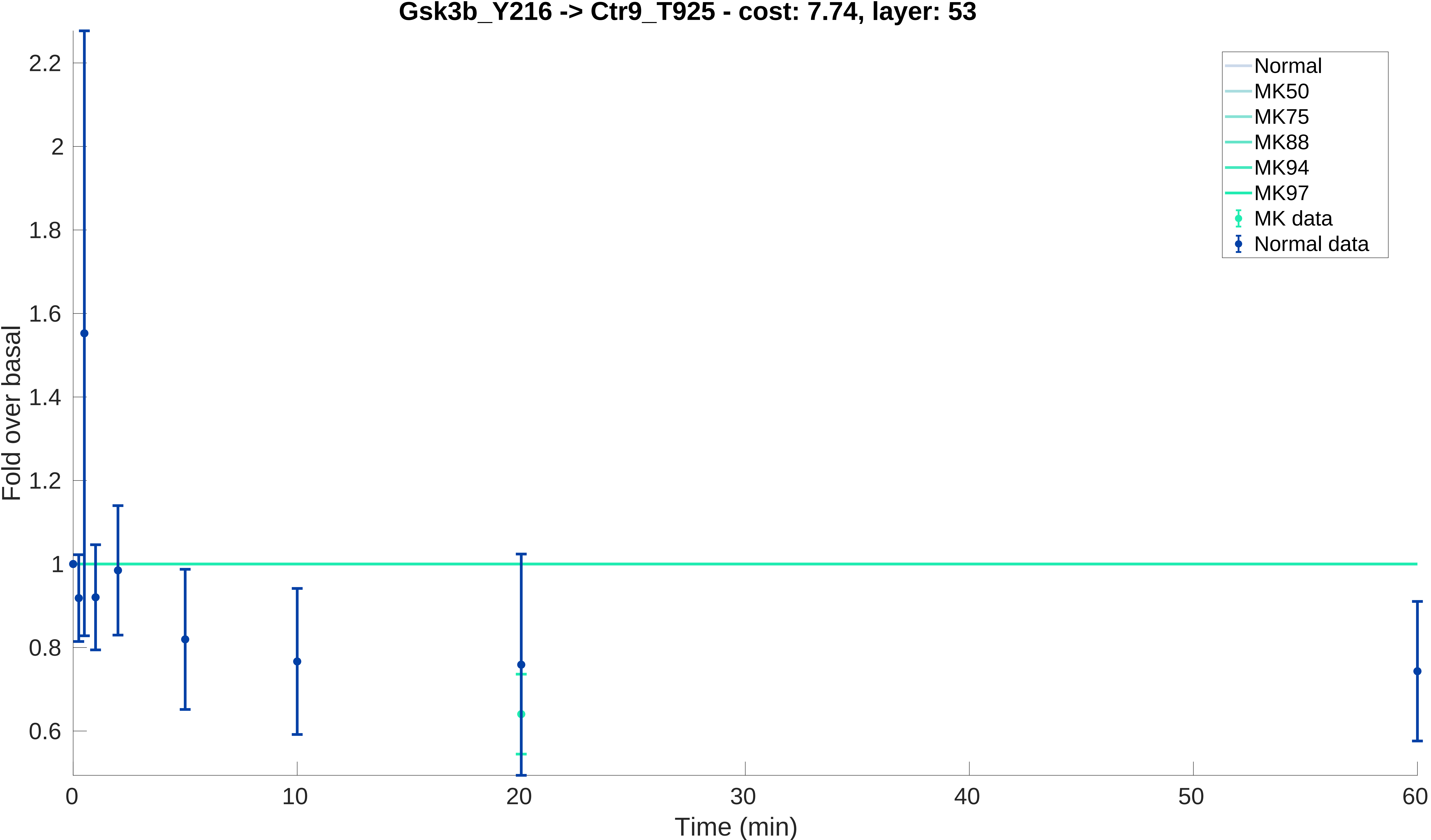

Gsk3b_Y216 -> Ctr9_T925 - cost: 7.74, layer: 53
Fold over basal
Time (min)
Normal
MK50
MK75
MK88
MK94
MK97
MK data
Normal data
0.6
0.8
1
1.2
1.4
1.6
1.8
2
2.2
0
10
20
30
40
50
60

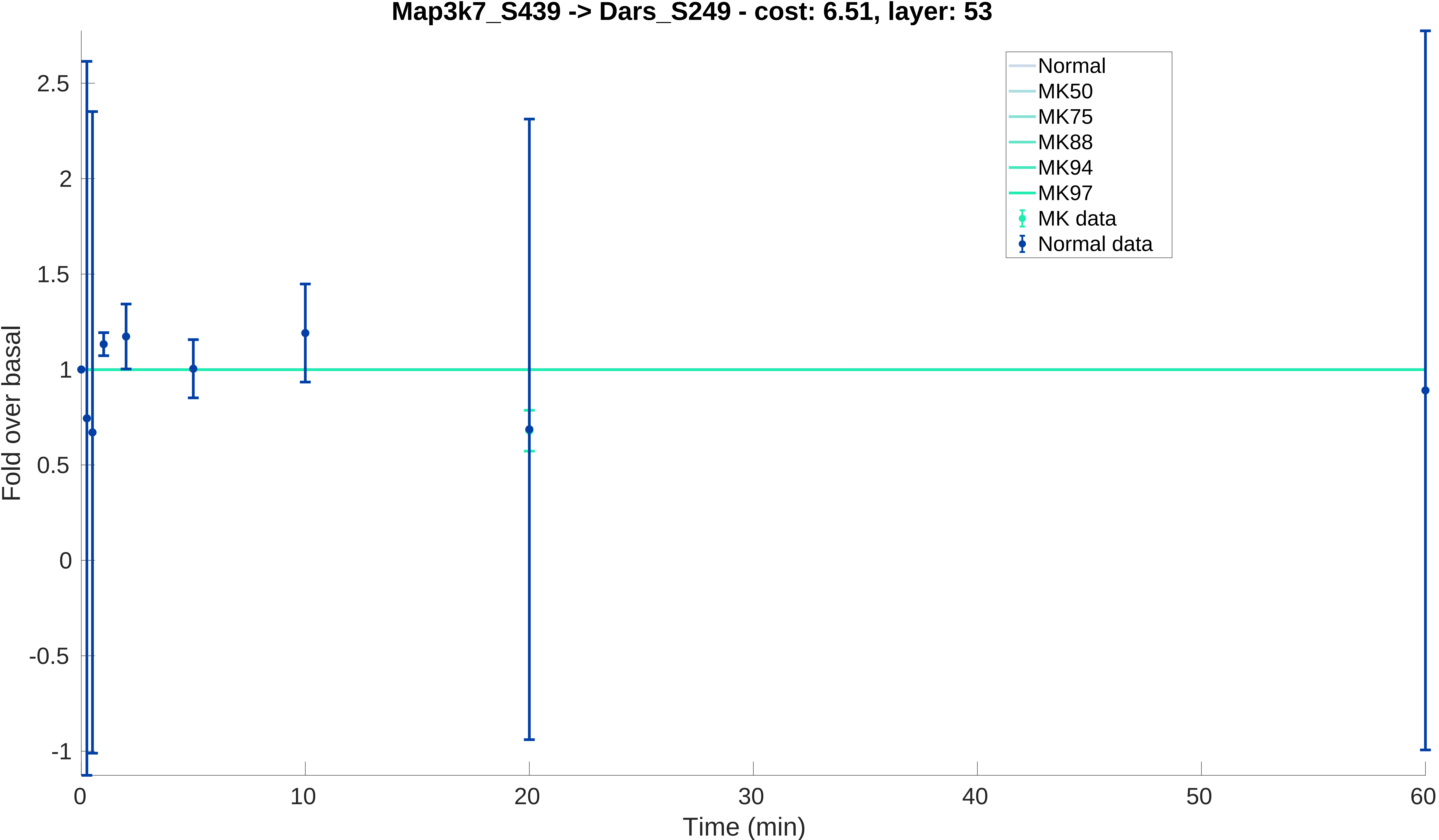

Map3k7_S439 -> Dars_S249 - cost: 6.51, layer: 53
Normal
MK50
MK75
MK88
MK94
MK97
MK data
Normal data
Fold over basal
Time (min)
2.5
2
1.5
1
0.5
0
-0.5
-1
0
10
20
30
40
50
60

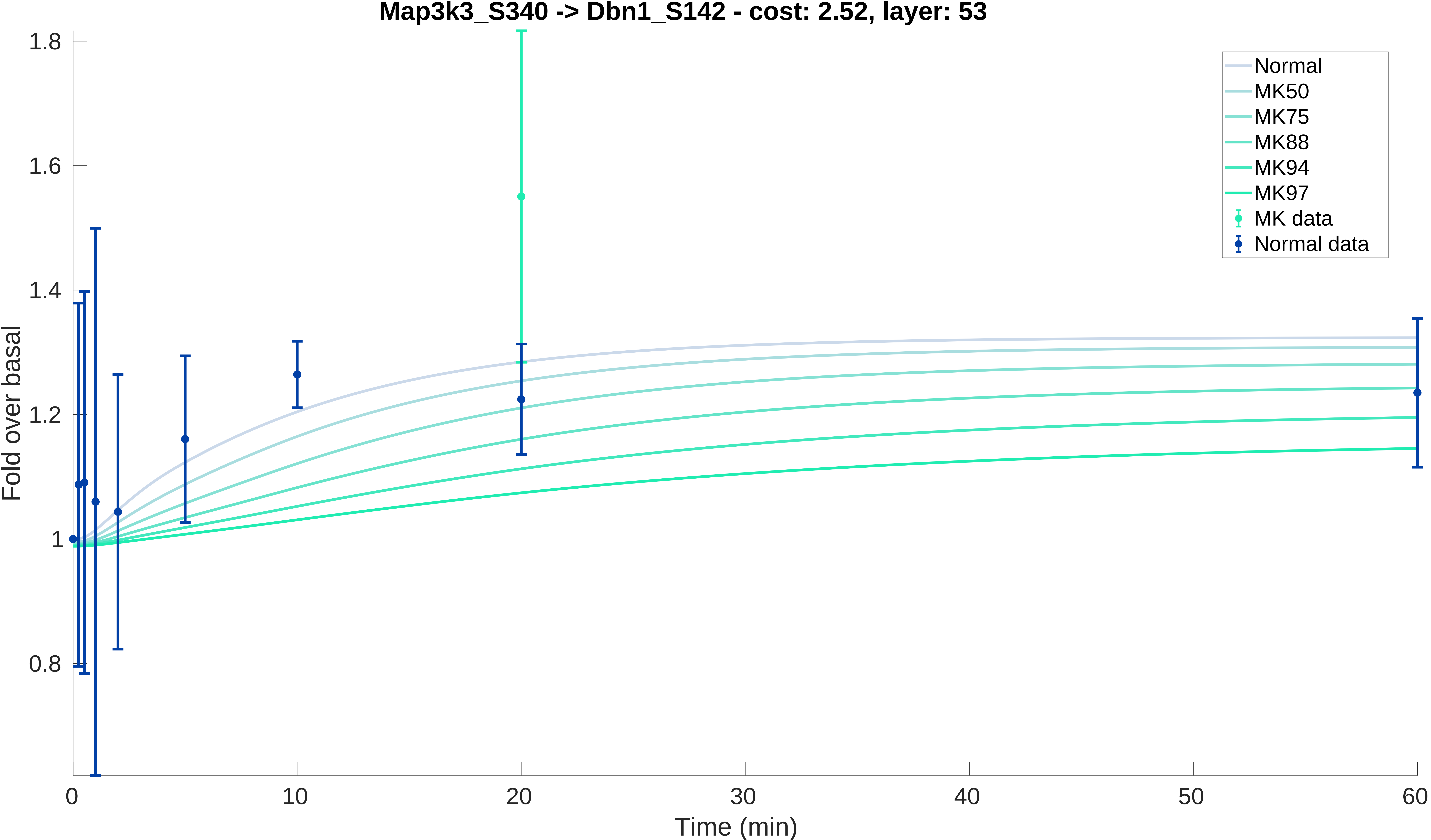

Map3k3_S340 -> Dbn1_S142 - cost: 2.52, layer: 53
Fold over basal
1.8
1.6
1.4
1.2
1
0.8
0
10
20
30
40
50
60
Time (min)
Normal
MK50
MK75
MK88
MK94
MK97
MK data
Normal data

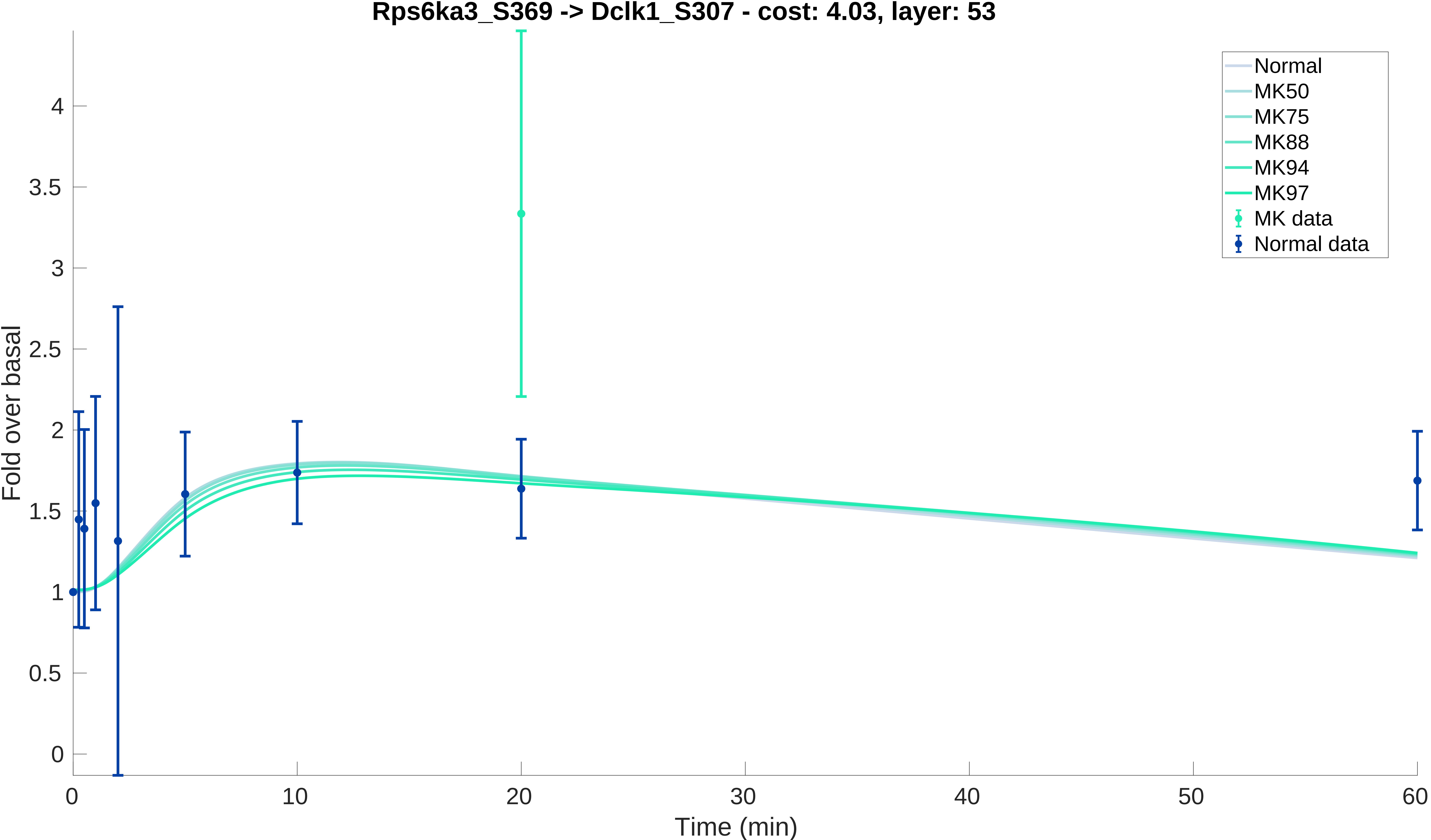

Rps6ka3_S369 -> Dclk1_S307 - cost: 4.03, layer: 53
Fold over basal
Time (min)
0
0.5
1
1.5
2
2.5
3
3.5
4
0
10
20
30
40
50
60
Normal
MK50
MK75
MK88
MK94
MK97
MK data
Normal data

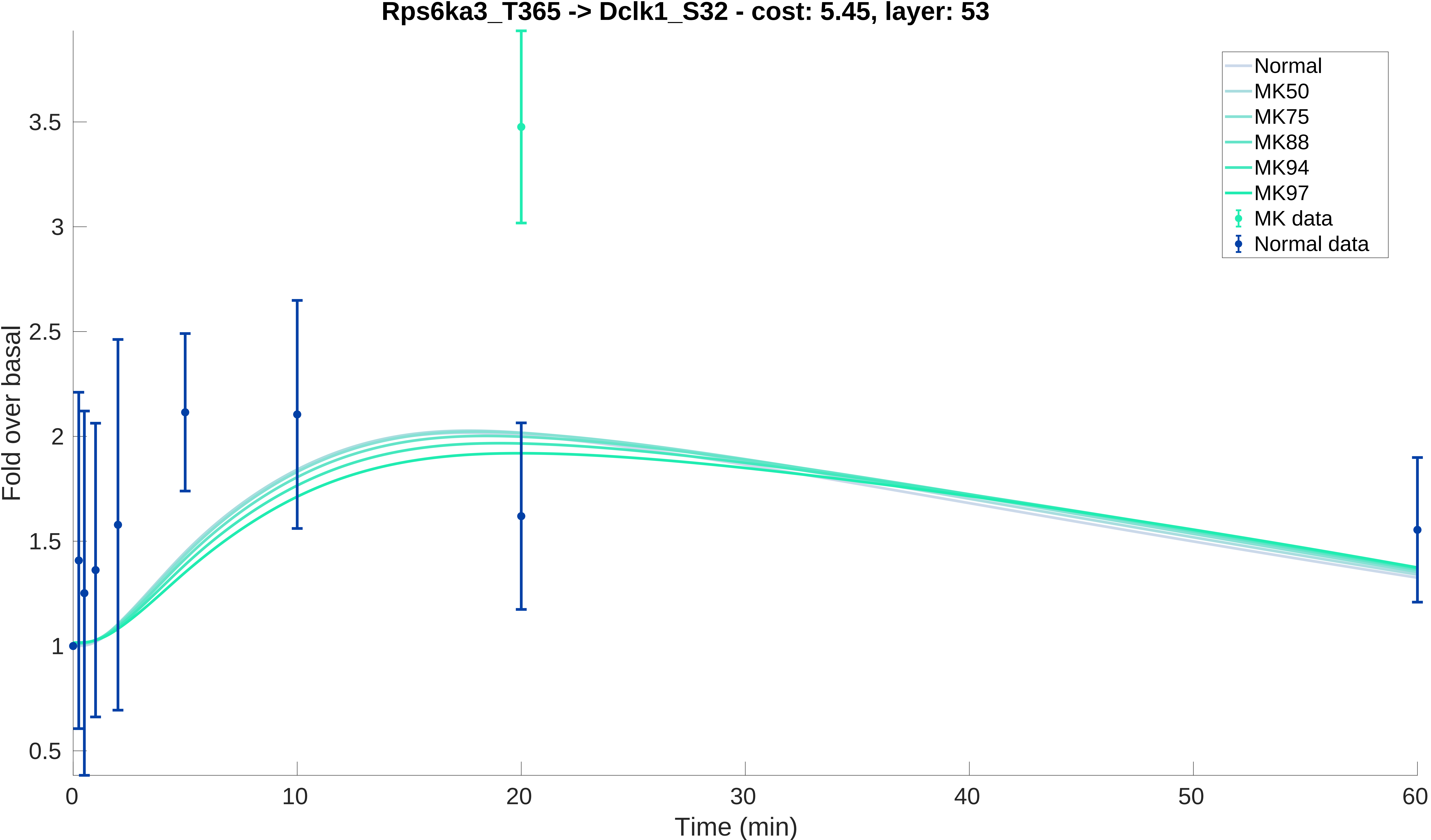

Rps6ka3_T365 -> Dclk1_S32 - cost: 5.45, layer: 53
Normal
MK50
MK75
MK88
MK94
MK97
MK data
Normal data
Fold over basal
0.5
1
1.5
2
2.5
3
3.5
0
10
20
30
40
50
60
Time (min)

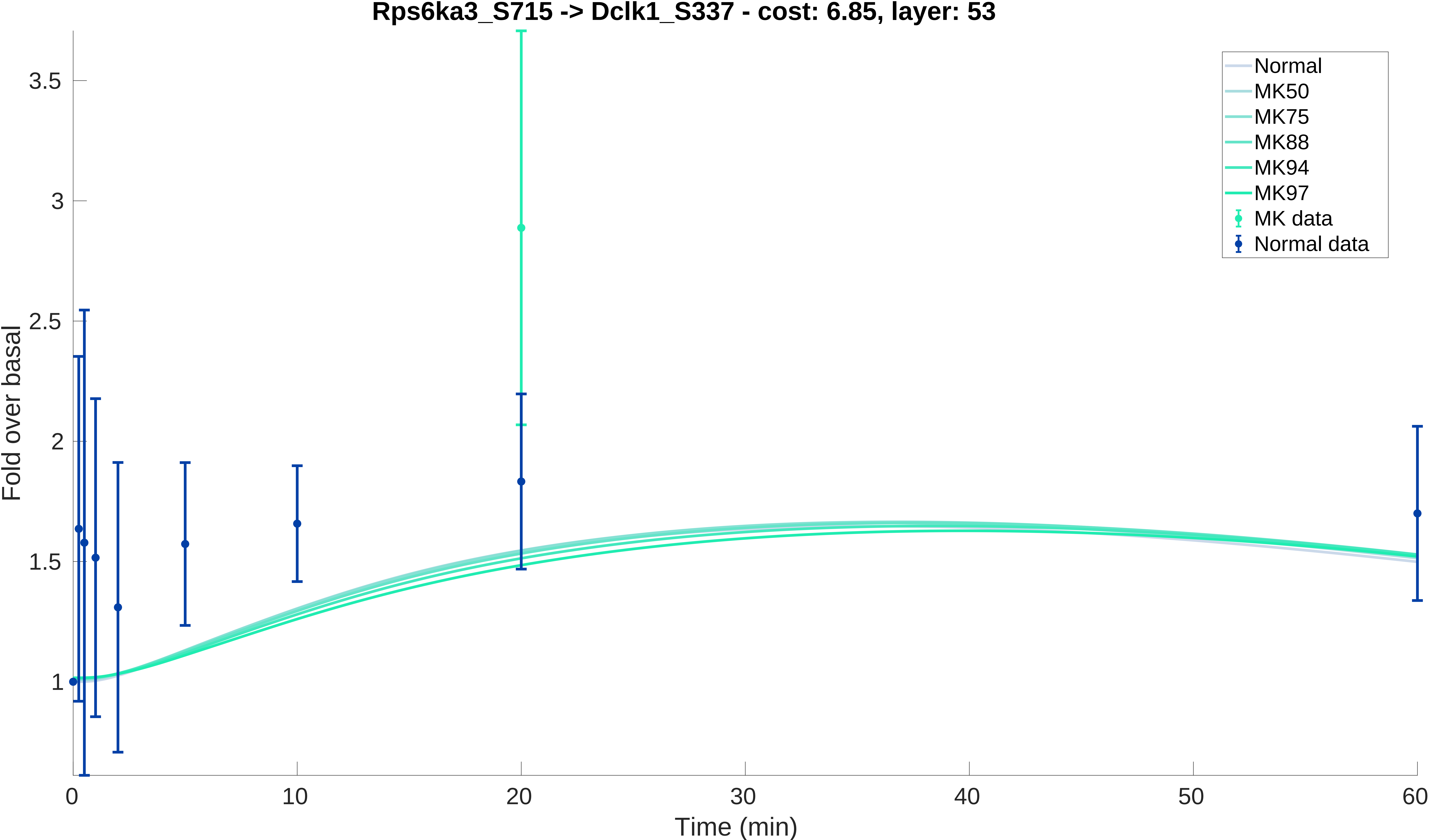

Rps6ka3_S715 -> Dclk1_S337 - cost: 6.85, layer: 53
Fold over basal
3.5
3
2.5
2
1.5
1
0
10
20
30
40
50
60
Time (min)
Normal
MK50
MK75
MK88
MK94
MK97
MK data
Normal data

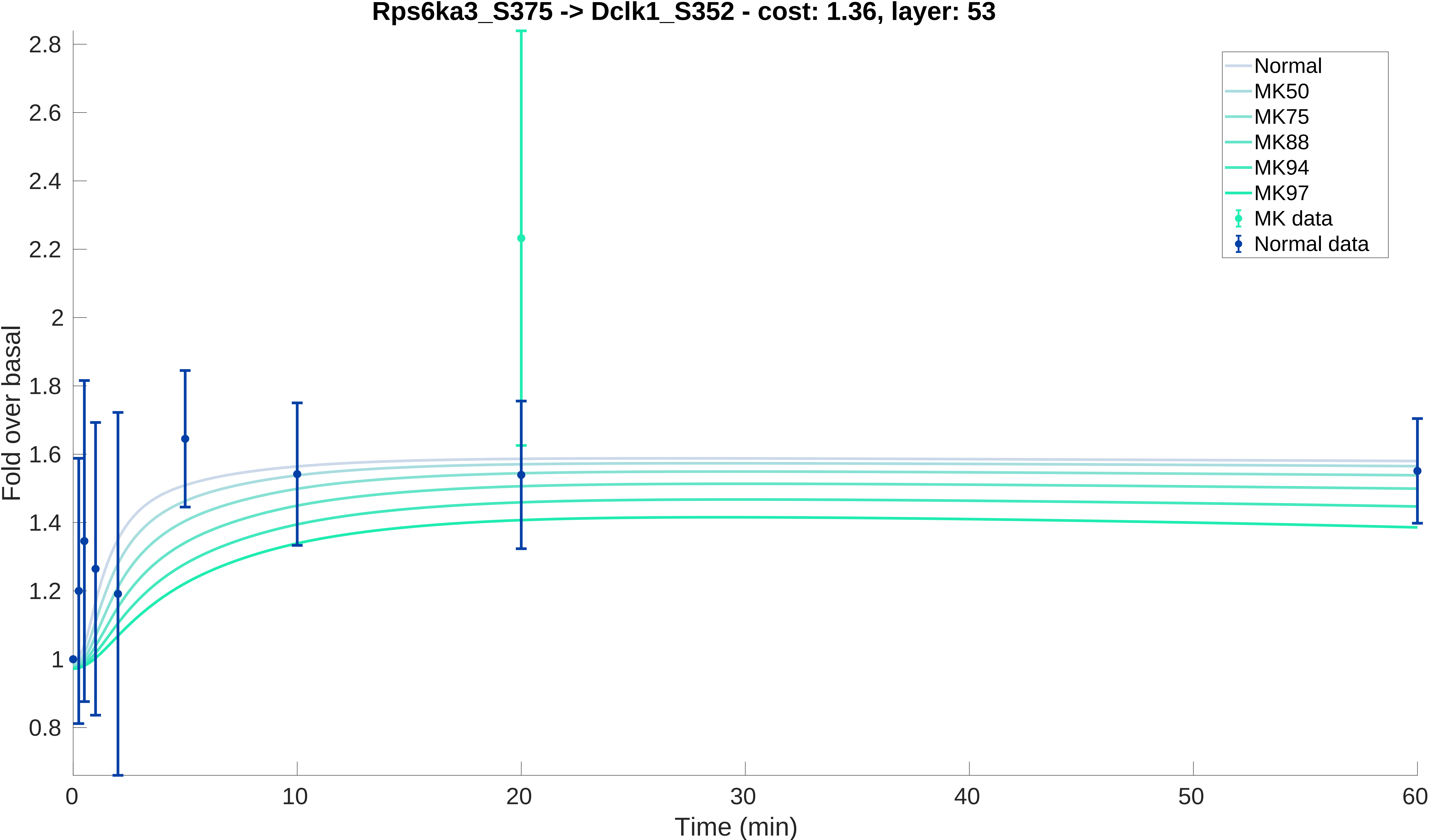

Rps6ka3_S375 -> Dclk1_S352 - cost: 1.36, layer: 53
Fold over basal
Time (min)
2.8
2.6
2.4
2.2
2
1.8
1.6
1.4
1.2
1
0.8
0
10
20
30
40
50
60
Normal
MK50
MK75
MK88
MK94
MK97
MK data
Normal data

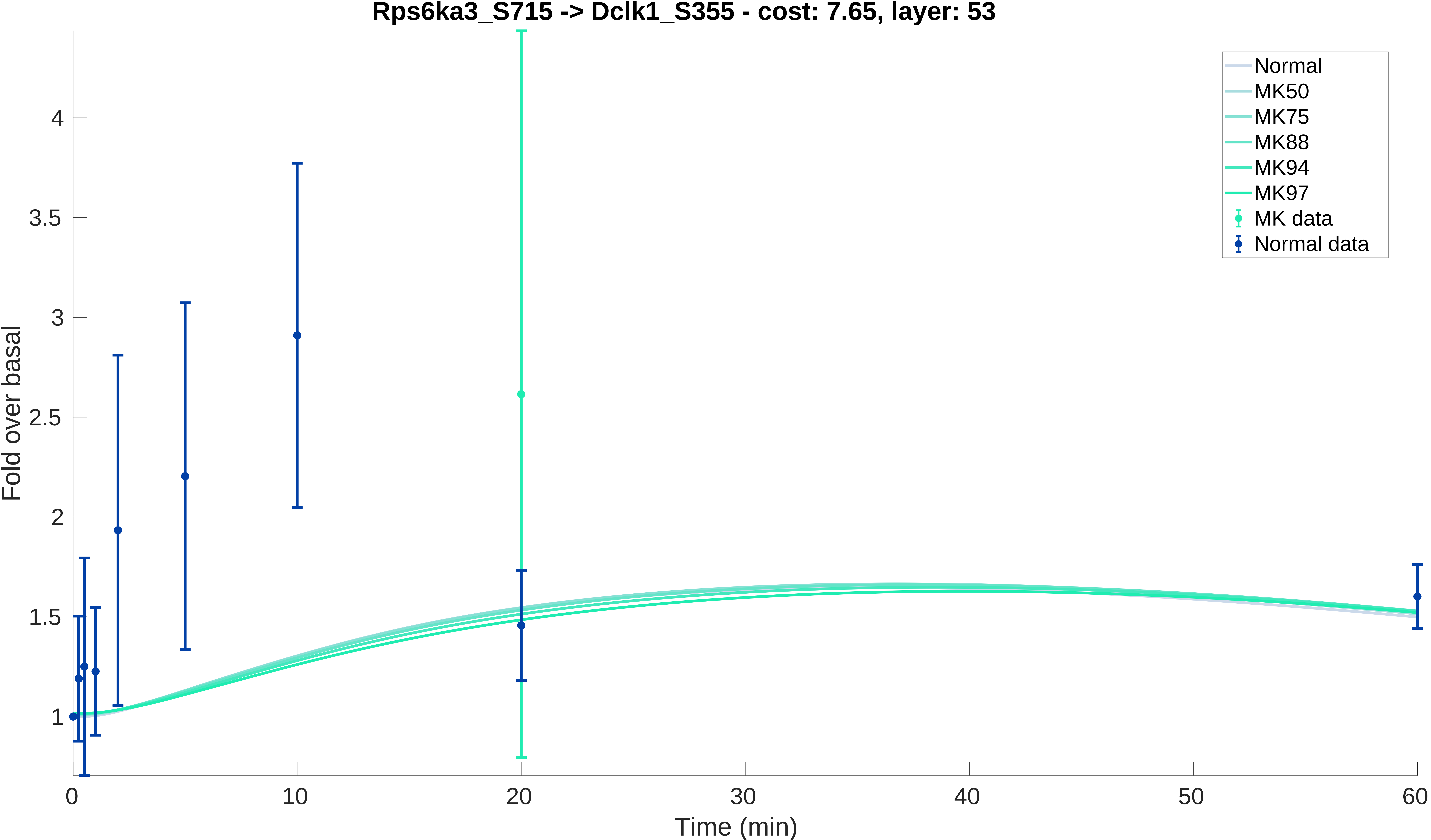
Rps6ka3_S715 -> Dclk1_S355 - cost: 7.65, layer: 53
Fold over basal
Time (min)
Normal
MK50
MK75
MK88
MK94
MK97
MK data
Normal data
4
3.5
3
2.5
2
1.5
1
0
10
20
30
40
50
60

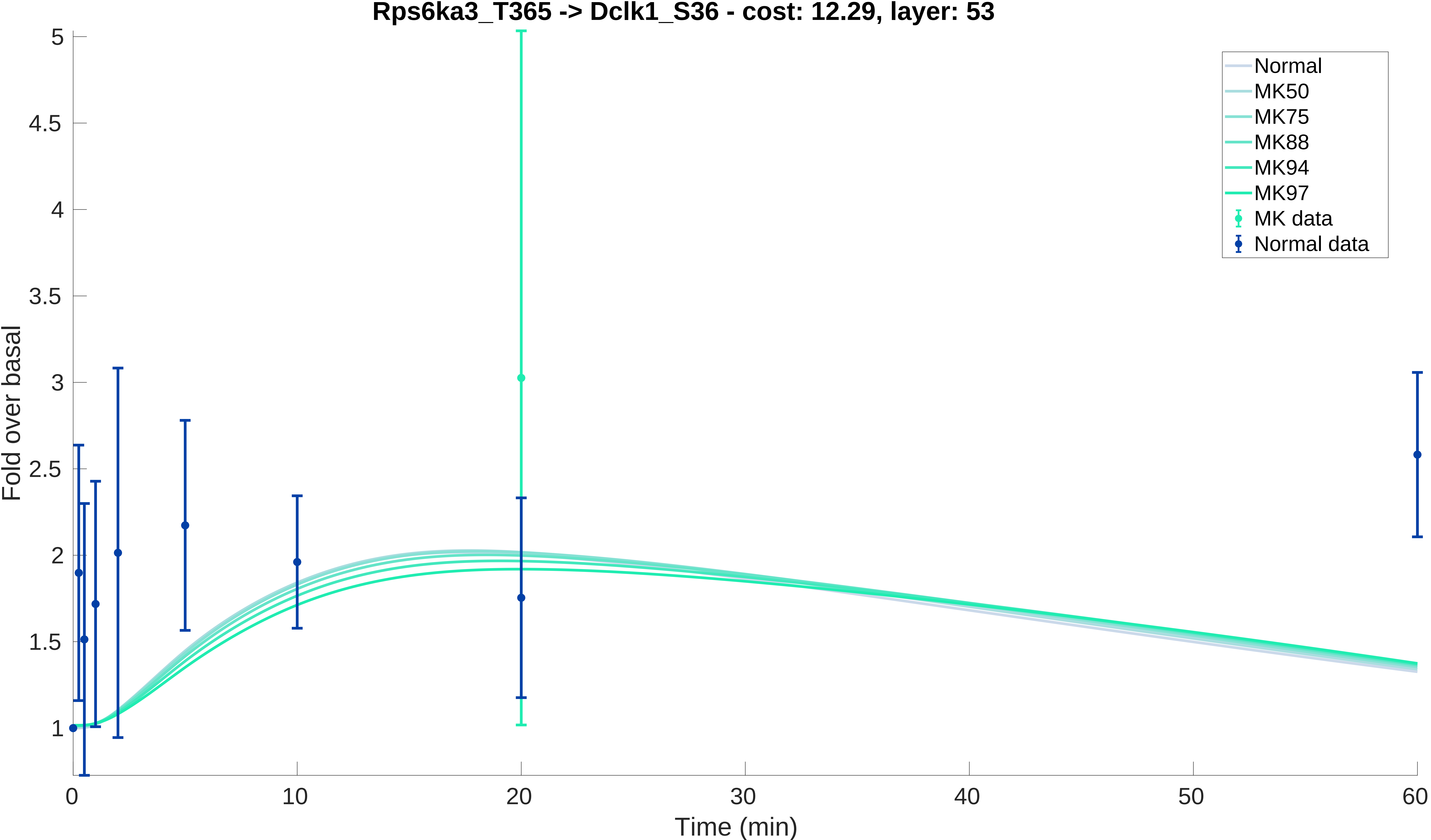

Rps6ka3_T365 -> Dclk1_S36 - cost: 12.29, layer: 53
Fold over basal
Time (min)
5
4.5
4
3.5
3
2.5
2
1.5
1
0
10
20
30
40
50
60
Normal
MK50
MK75
MK88
MK94
MK97
MK data
Normal data

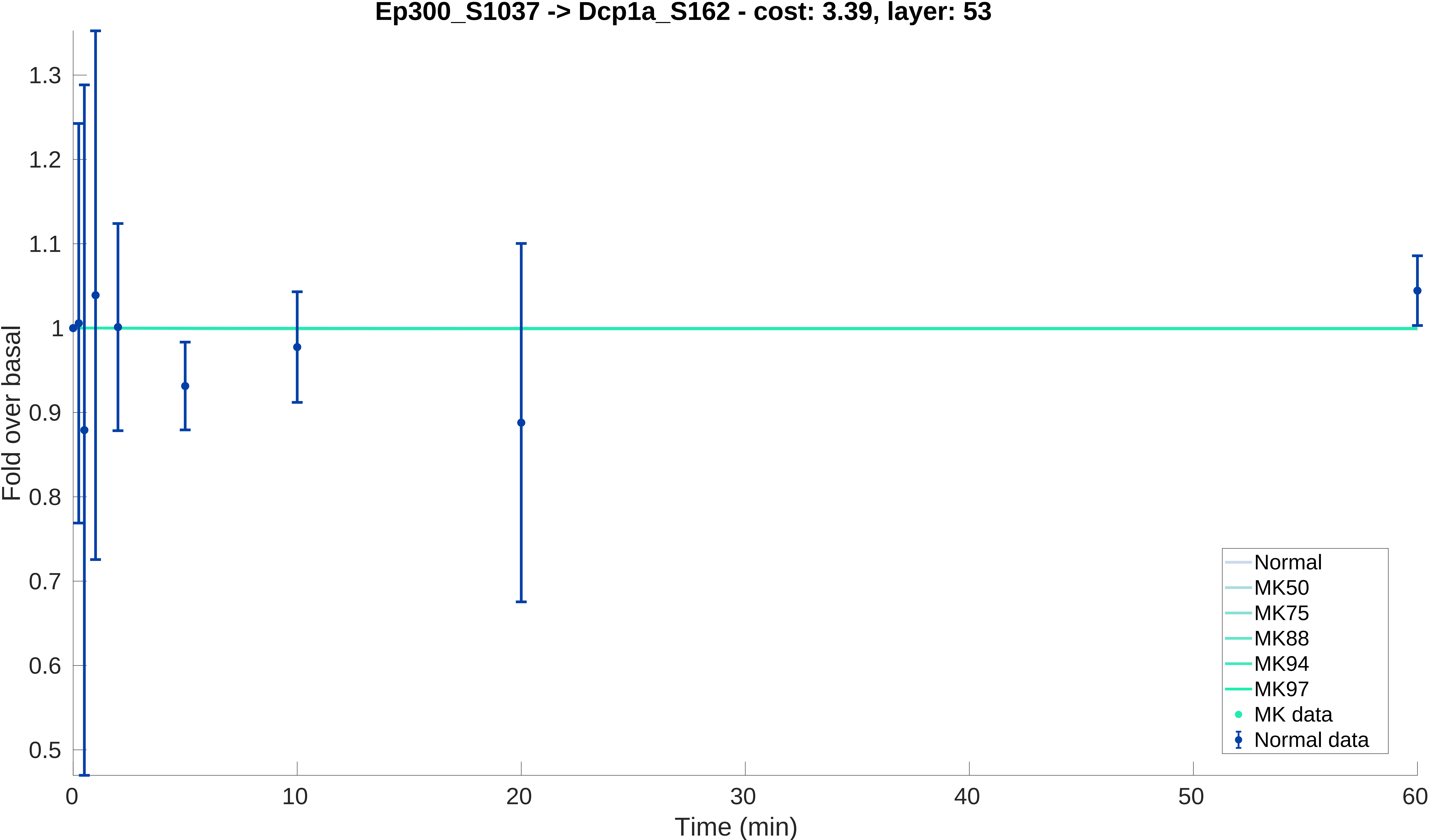

Ep300_S1037 -> Dcp1a_S162 - cost: 3.39, layer: 53
Fold over basal
Time (min)
0
10
20
30
40
50
60
0.5
0.6
0.7
0.8
0.9
1
1.1
1.2
1.3
Normal
MK50
MK75
MK88
MK94
MK97
MK data
Normal data

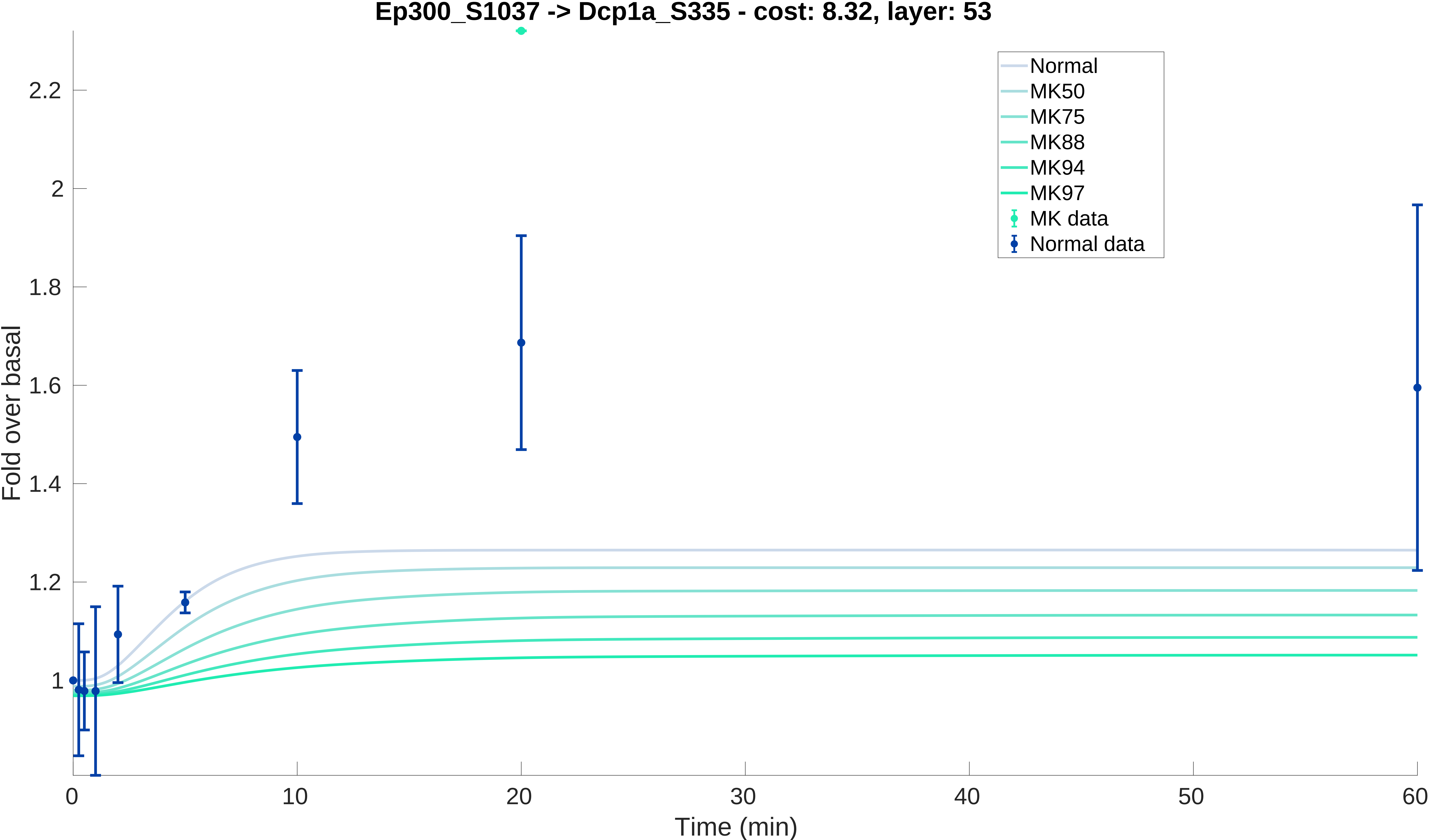

Ep300_S1037 -> Dcp1a_S335 - cost: 8.32, layer: 53
Normal
MK50
MK75
MK88
MK94
MK97
MK data
Normal data
Fold over basal
Time (min)
2.2
2
1.8
1.6
1.4
1.2
1
0
10
20
30
40
50
60

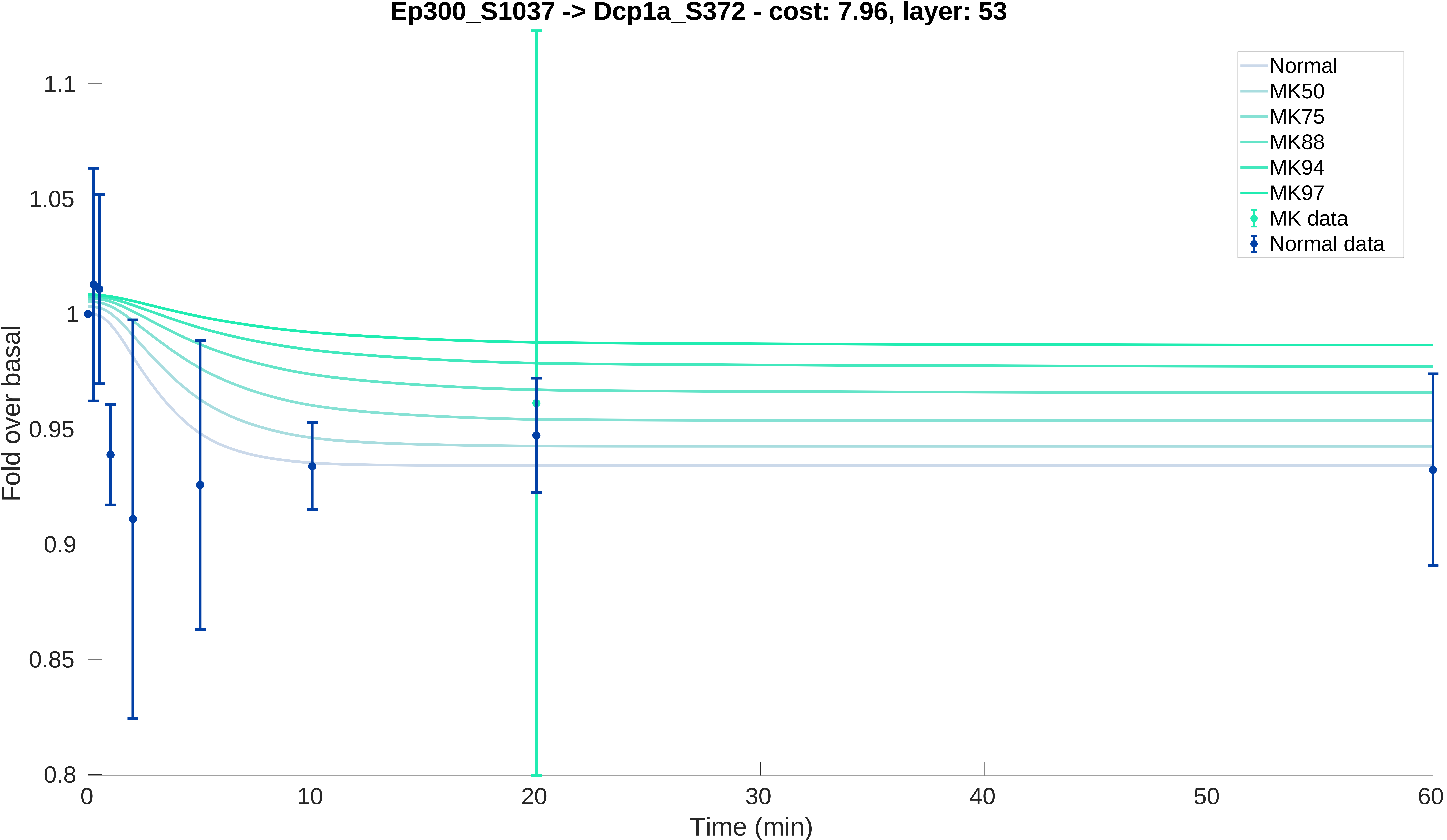
Ep300_S1037 -> Dcp1a_S372 - cost: 7.96, layer: 53
Fold over basal
Time (min)
0.8
0.85
0.9
0.95
1
1.05
1.1
0
10
20
30
40
50
60
Normal
MK50
MK75
MK88
MK94
MK97
MK data
Normal data

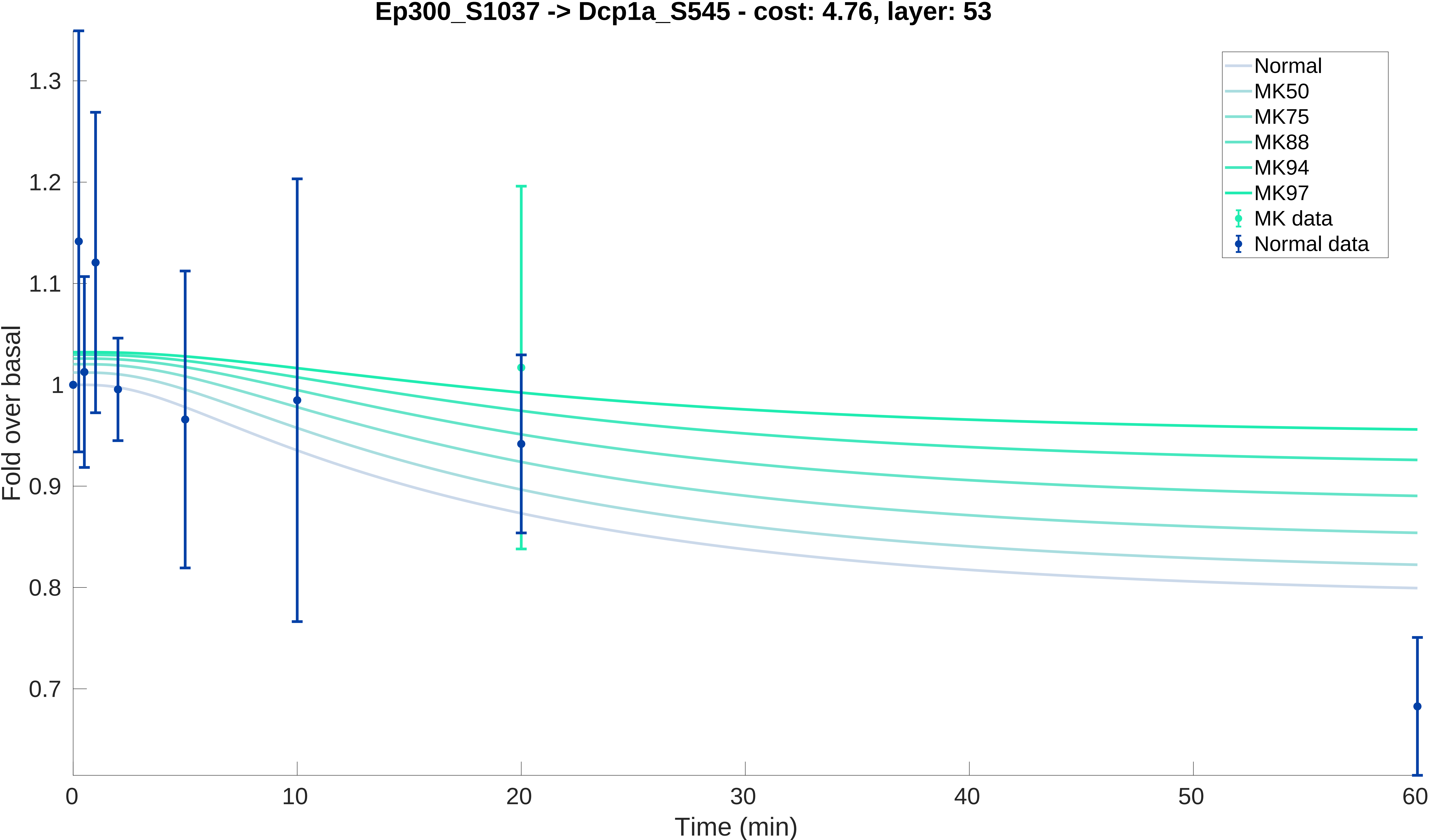

Ep300_S1037 -> Dcp1a_S545 - cost: 4.76, layer: 53
Fold over basal
1.3
1.2
1.1
1
0.9
0.8
0.7
0
10
20
30
40
50
60
Time (min)
Normal
MK50
MK75
MK88
MK94
MK97
MK data
Normal data

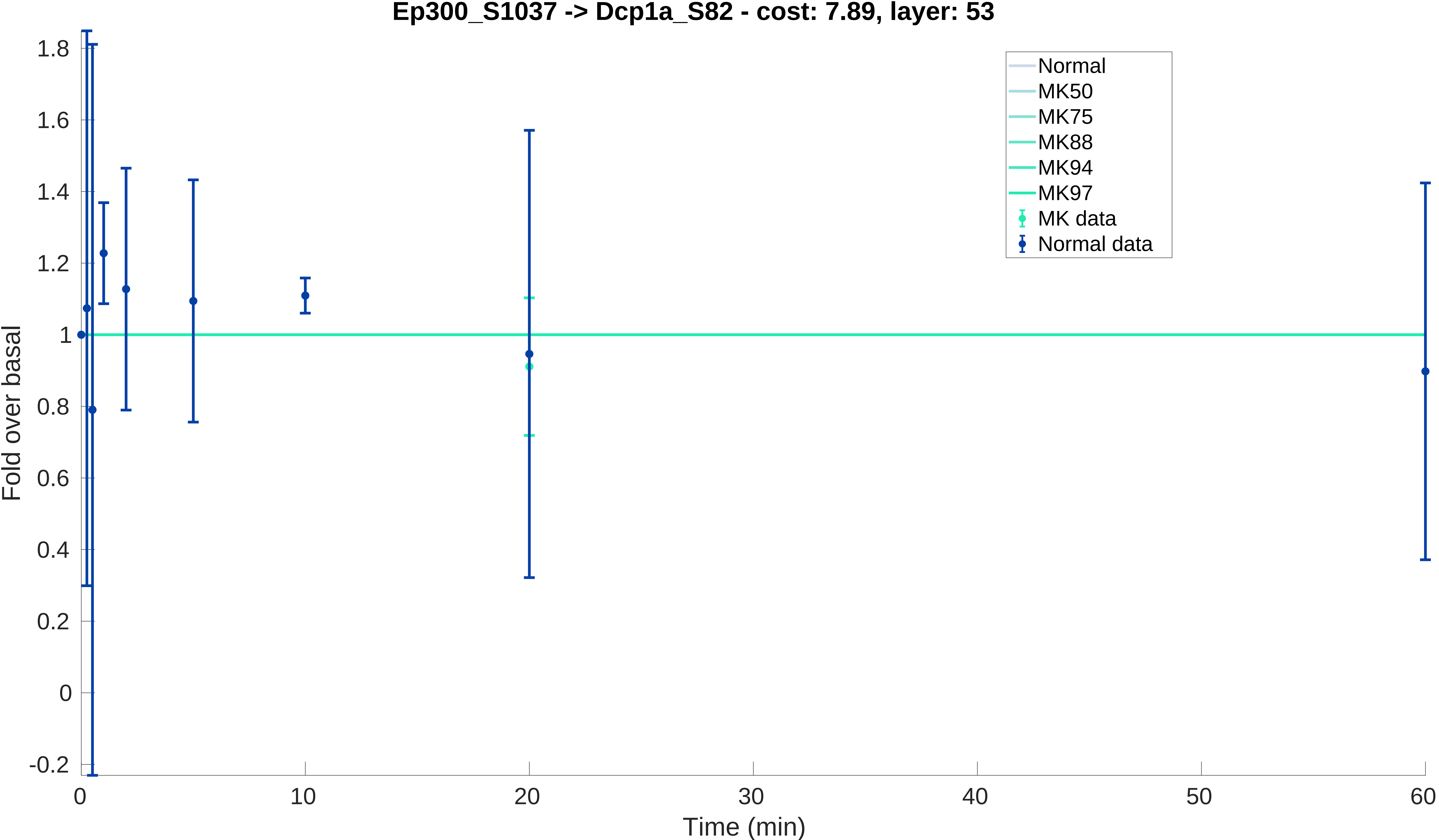

Ep300_S1037 -> Dcp1a_S82 - cost: 7.89, layer: 53
Normal
MK50
MK75
MK88
MK94
MK97
MK data
Normal data
Fold over basal
Time (min)
1.8
1.6
1.4
1.2
1
0.8
0.6
0.4
0.2
0
-0.2
0
10
20
30
40
50
60

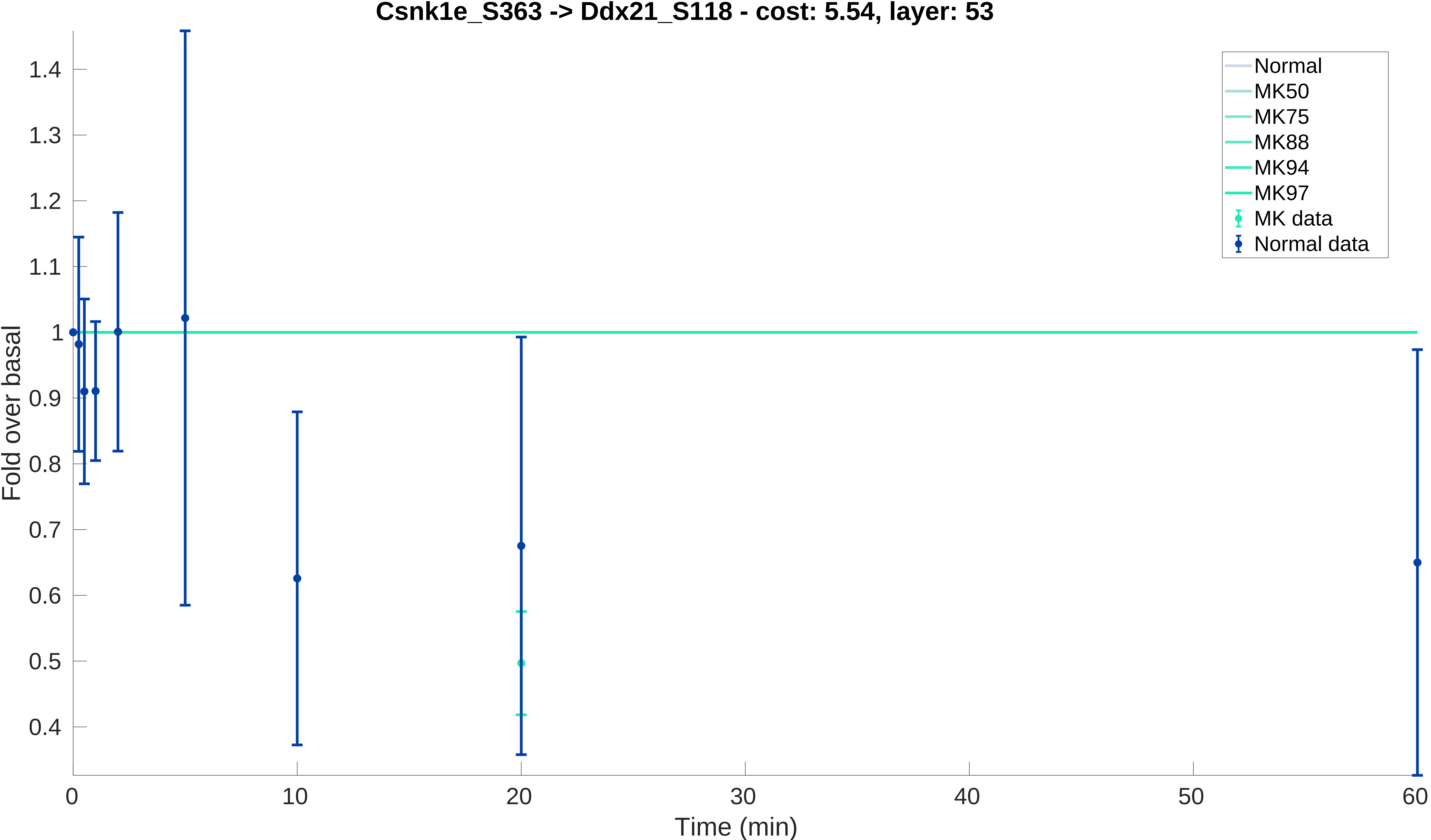
Csnk1e_S363 -> Ddx21_S118 - cost: 5.54, layer: 53
Fold over basal
Time (min)
1.4
1.3
1.2
1.1
1
0.9
0.8
0.7
0.6
0.5
0.4
0
10
20
30
40
50
60
Normal
MK50
MK75
MK88
MK94
MK97
MK data
Normal data

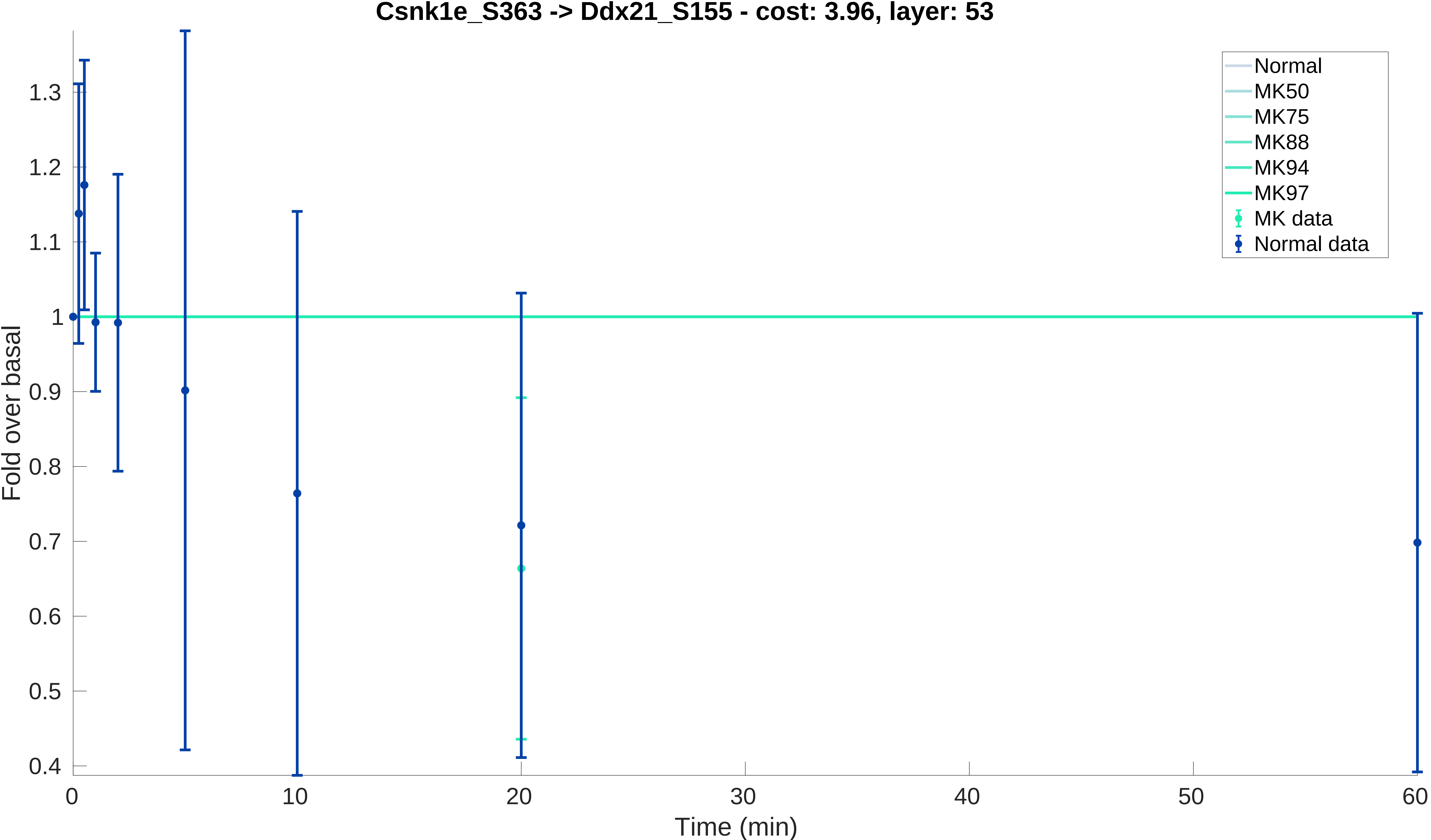

Csnk1e_S363 -> Ddx21_S155 - cost: 3.96, layer: 53
Normal
MK50
MK75
MK88
MK94
MK97
MK data
Normal data
Fold over basal
Time (min)
1.3
1.2
1.1
1
0.9
0.8
0.7
0.6
0.5
0.4
0
10
20
30
40
50
60

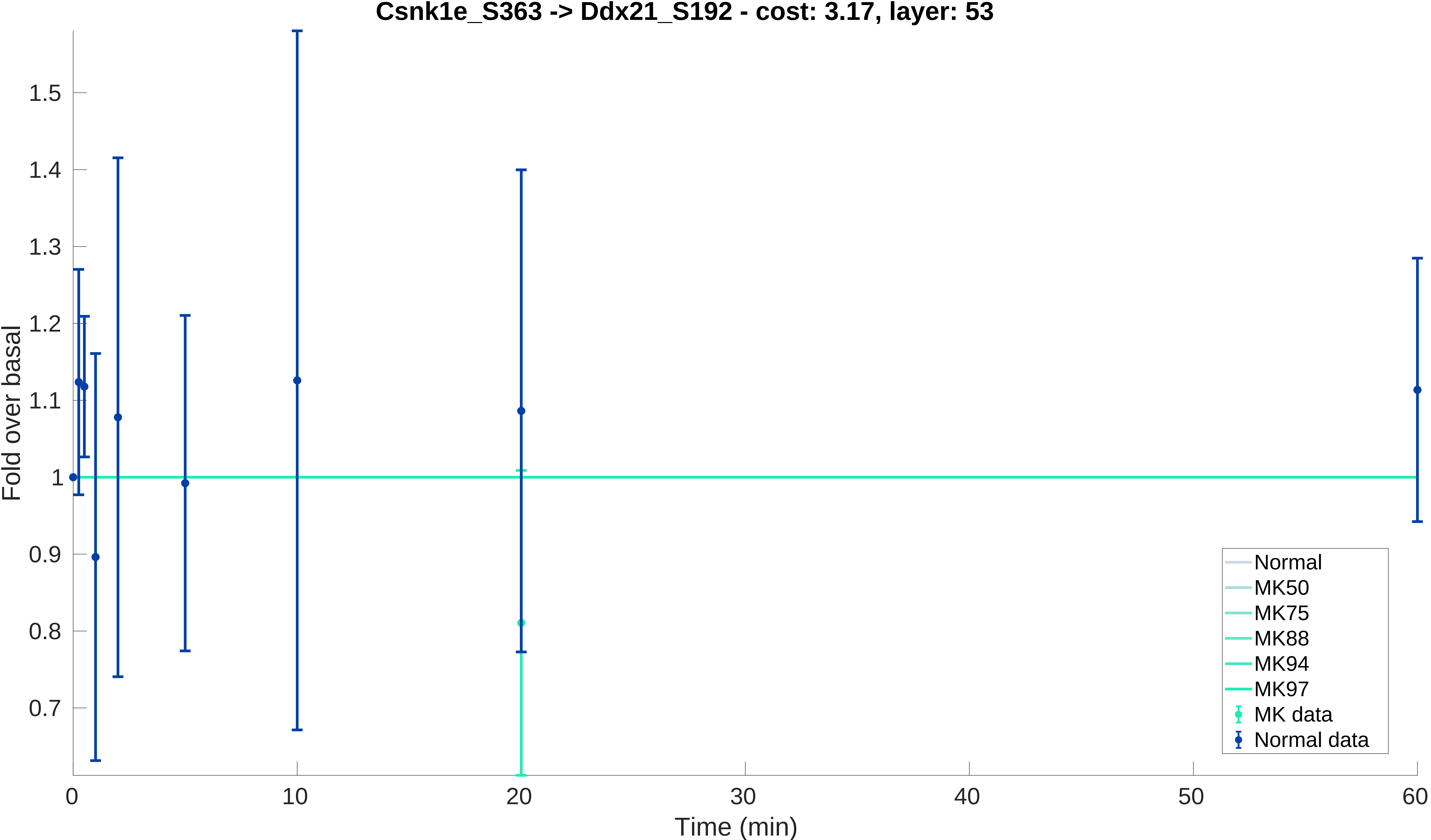
Csnk1e_S363 -> Ddx21_S192 - cost: 3.17, layer: 53
Fold over basal
Time (min)
1.5
1.4
1.3
1.2
1.1
1
0.9
0.8
0.7
0
10
20
30
40
50
60
Normal
MK50
MK75
MK88
MK94
MK97
MK data
Normal data

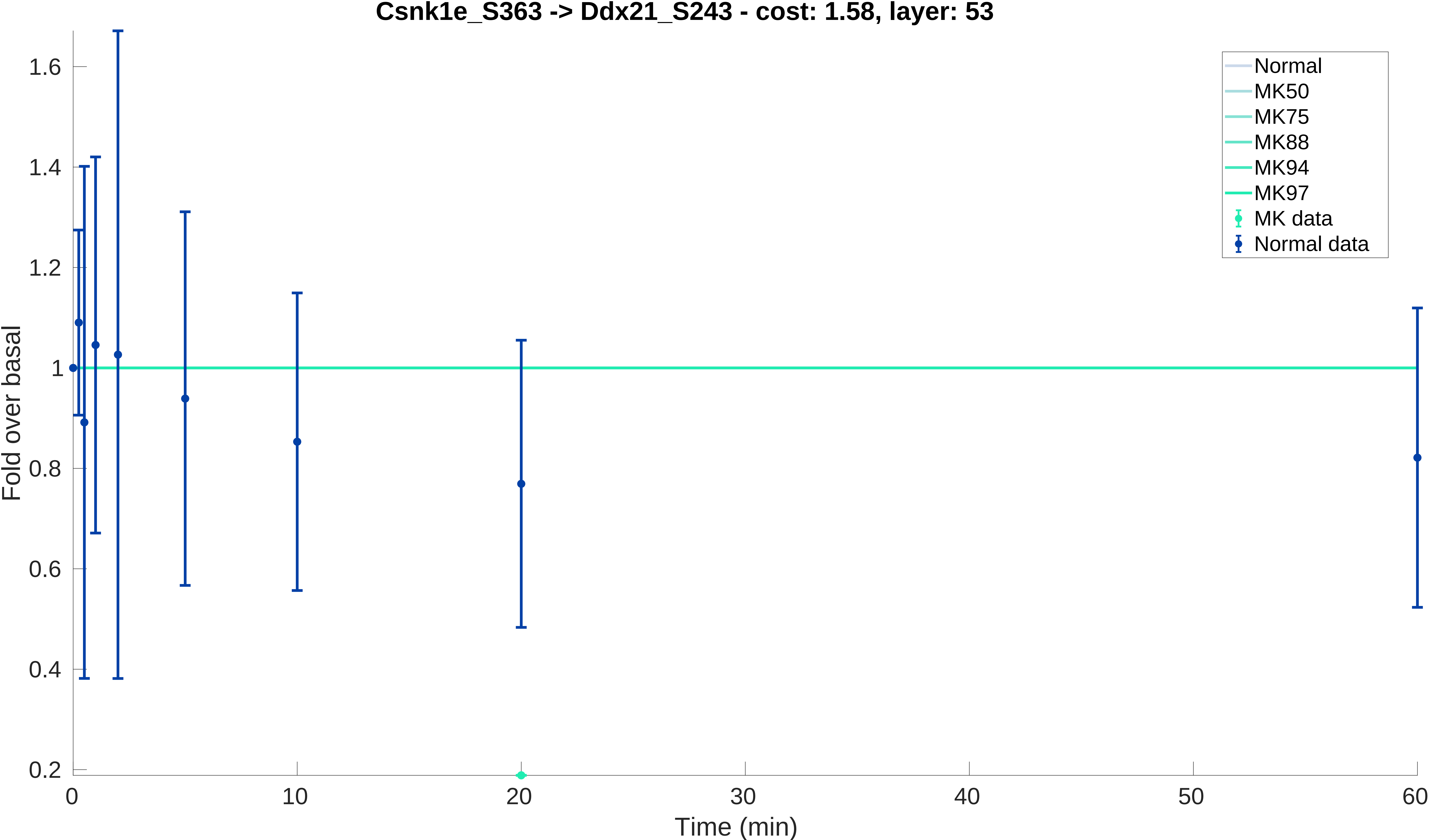
Csnk1e_S363 -> Ddx21_S243 - cost: 1.58, layer: 53
Fold over basal
Time (min)
1.6
1.4
1.2
1
0.8
0.6
0.4
0.2
0
10
20
30
40
50
60
Normal
MK50
MK75
MK88
MK94
MK97
MK data
Normal data

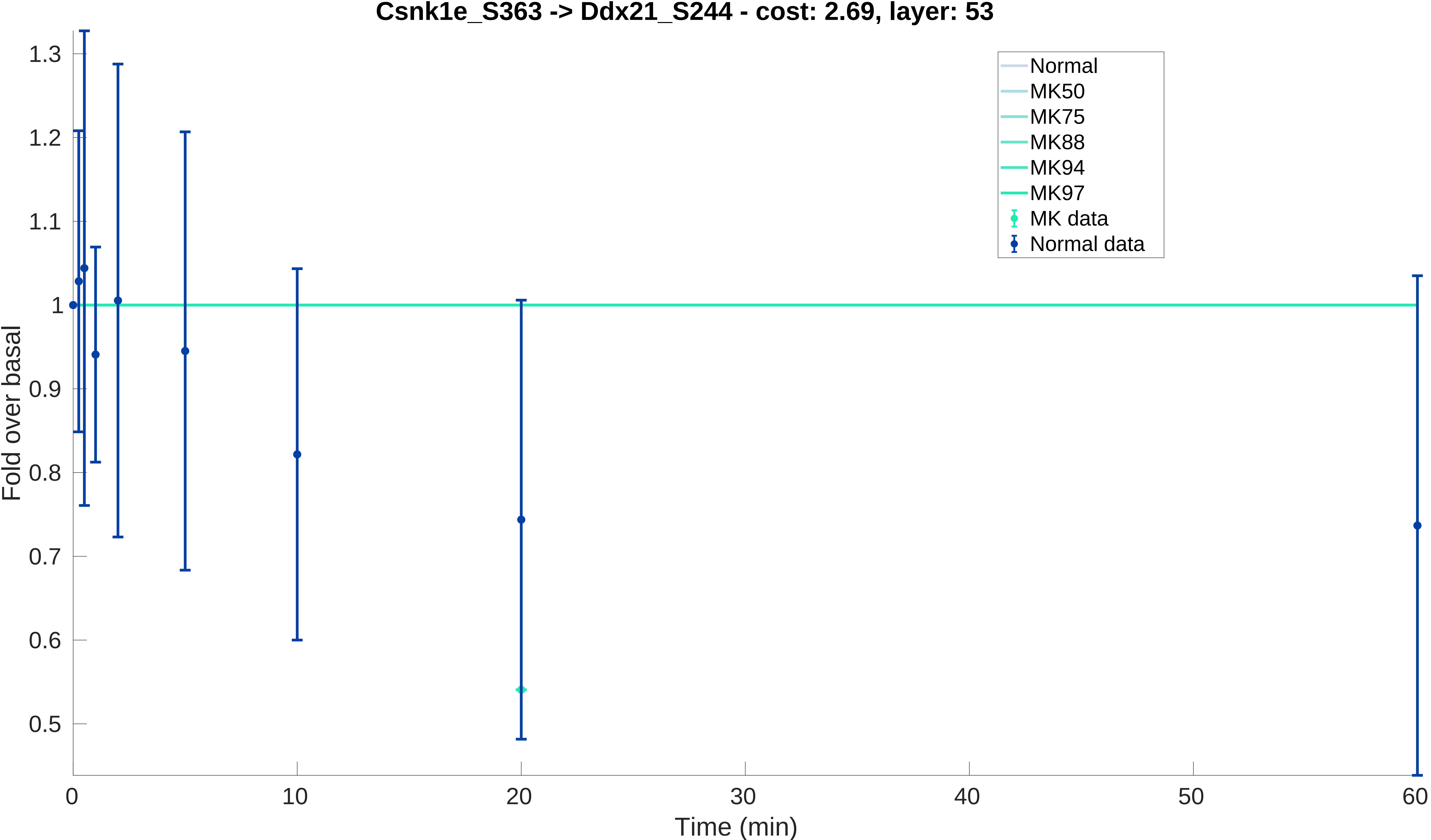

Csnk1e_S363 -> Ddx21_S244 - cost: 2.69, layer: 53
Fold over basal
Time (min)
1.3
1.2
1.1
1
0.9
0.8
0.7
0.6
0.5
0
10
20
30
40
50
60
Normal
MK50
MK75
MK88
MK94
MK97
MK data
Normal data

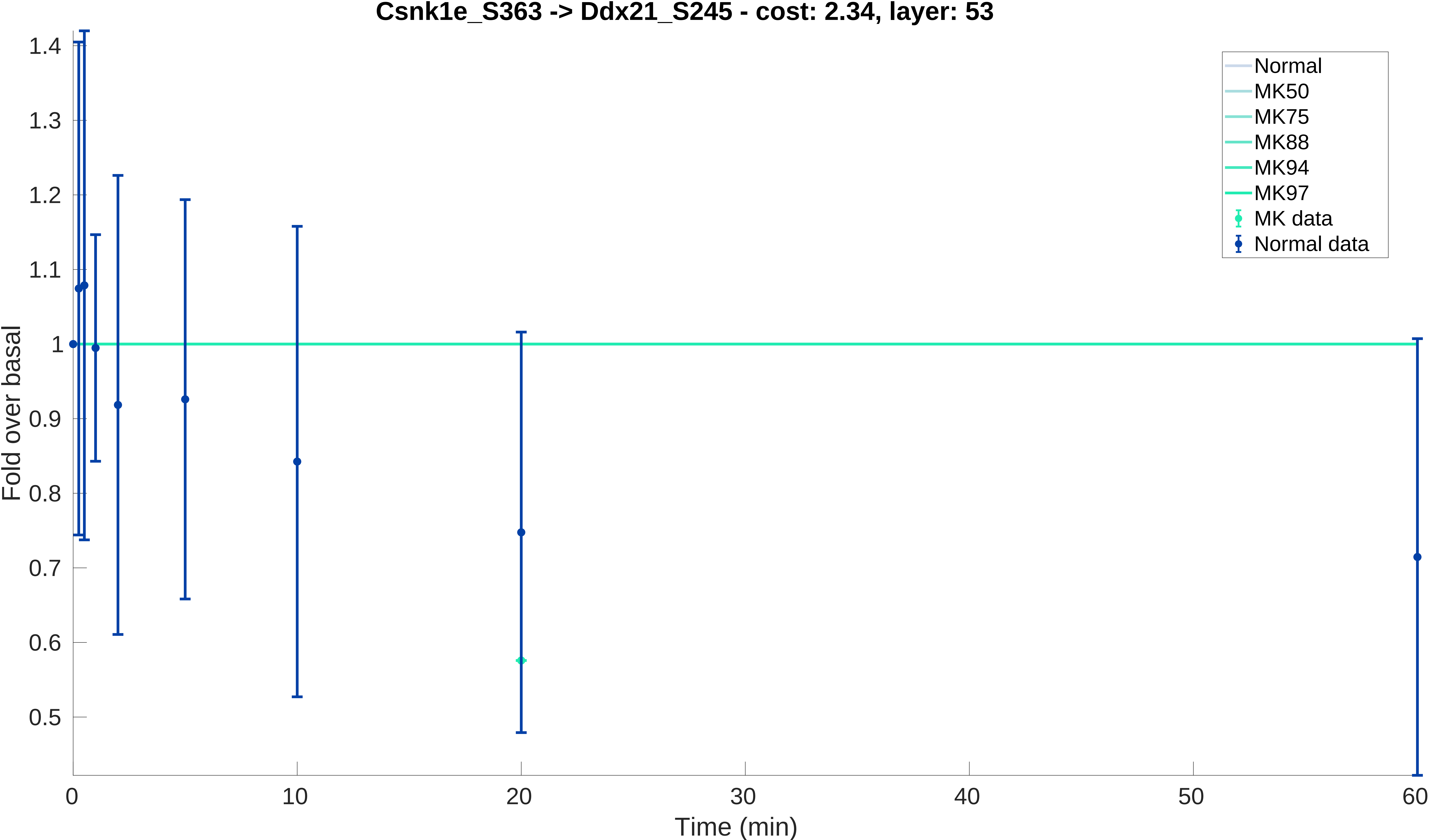

Csnk1e_S363 -> Ddx21_S245 - cost: 2.34, layer: 53
Fold over basal
1.4
1.3
1.2
1.1
1
0.9
0.8
0.7
0.6
0.5
0
10
20
30
40
50
60
Time (min)
Normal
MK50
MK75
MK88
MK94
MK97
MK data
Normal data

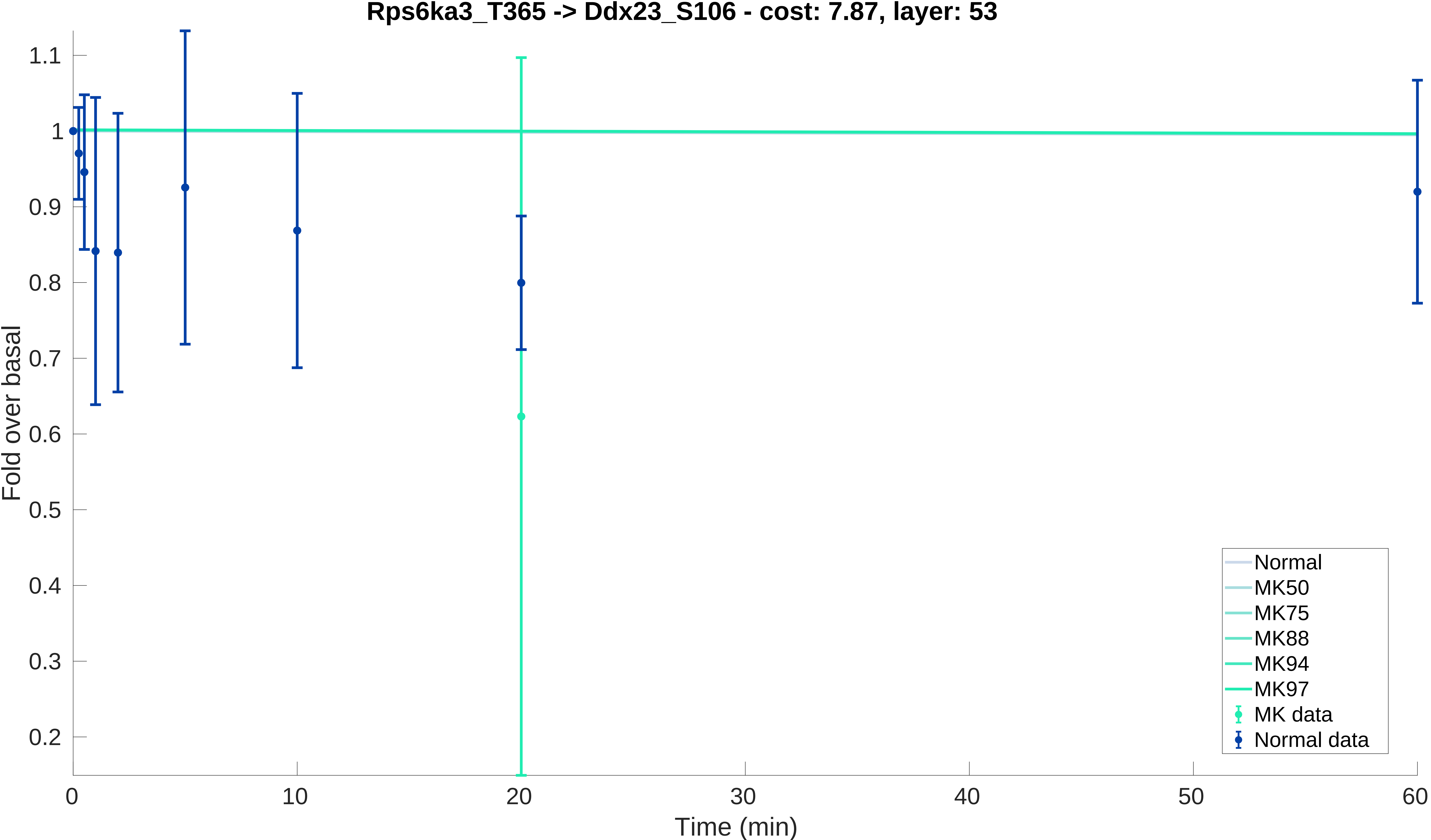

Rps6ka3_T365 -> Ddx23_S106 - cost: 7.87, layer: 53
Fold over basal
1.1
1
0.9
0.8
0.7
0.6
0.5
0.4
0.3
0.2
0
10
20
30
40
50
60
Time (min)
Normal
MK50
MK75
MK88
MK94
MK97
MK data
Normal data

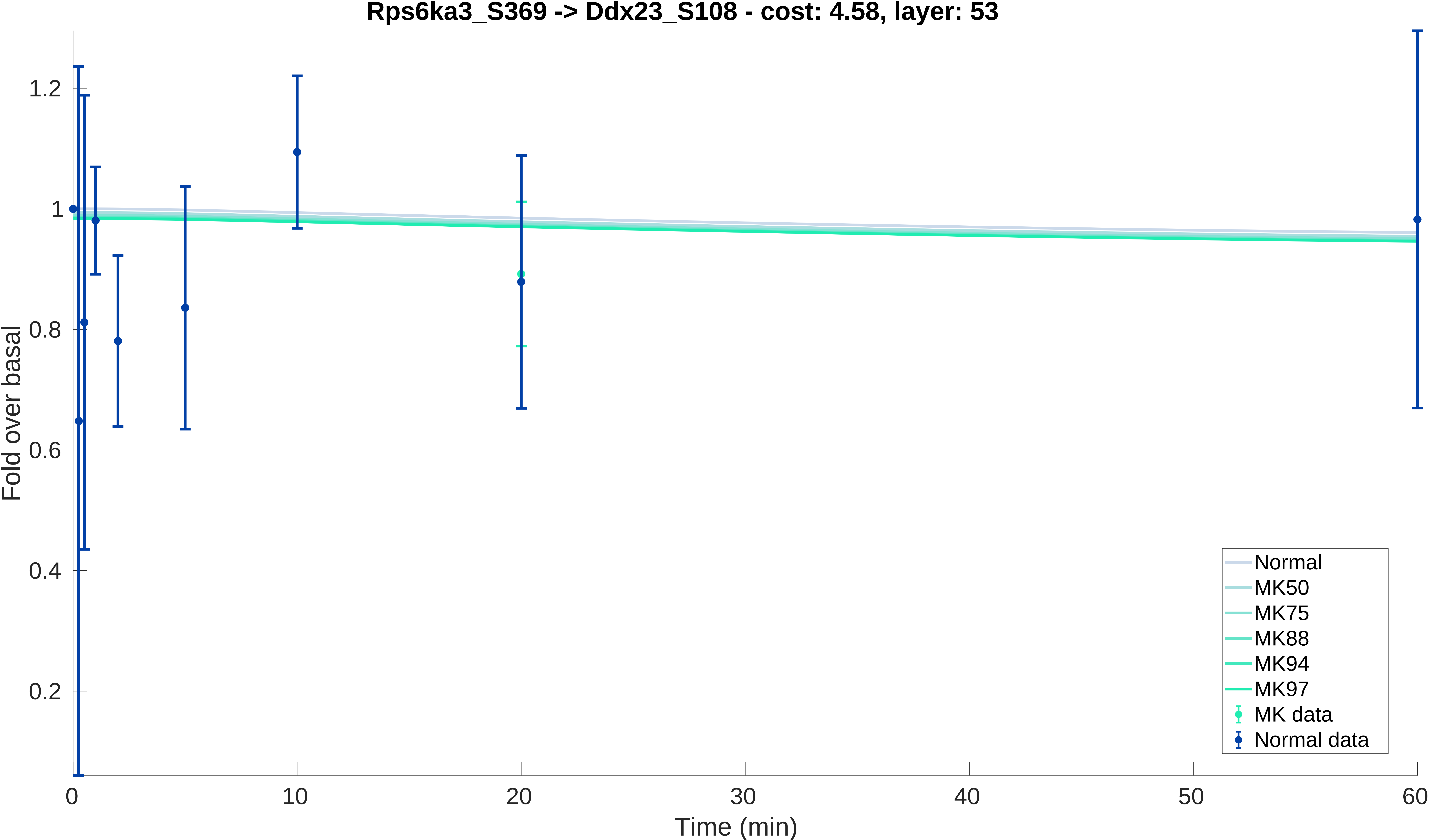

Rps6ka3_S369 -> Ddx23_S108 - cost: 4.58, layer: 53
Fold over basal
Time (min)
0
10
20
30
40
50
60
0.2
0.4
0.6
0.8
1
1.2
Normal
MK50
MK75
MK88
MK94
MK97
MK data
Normal data

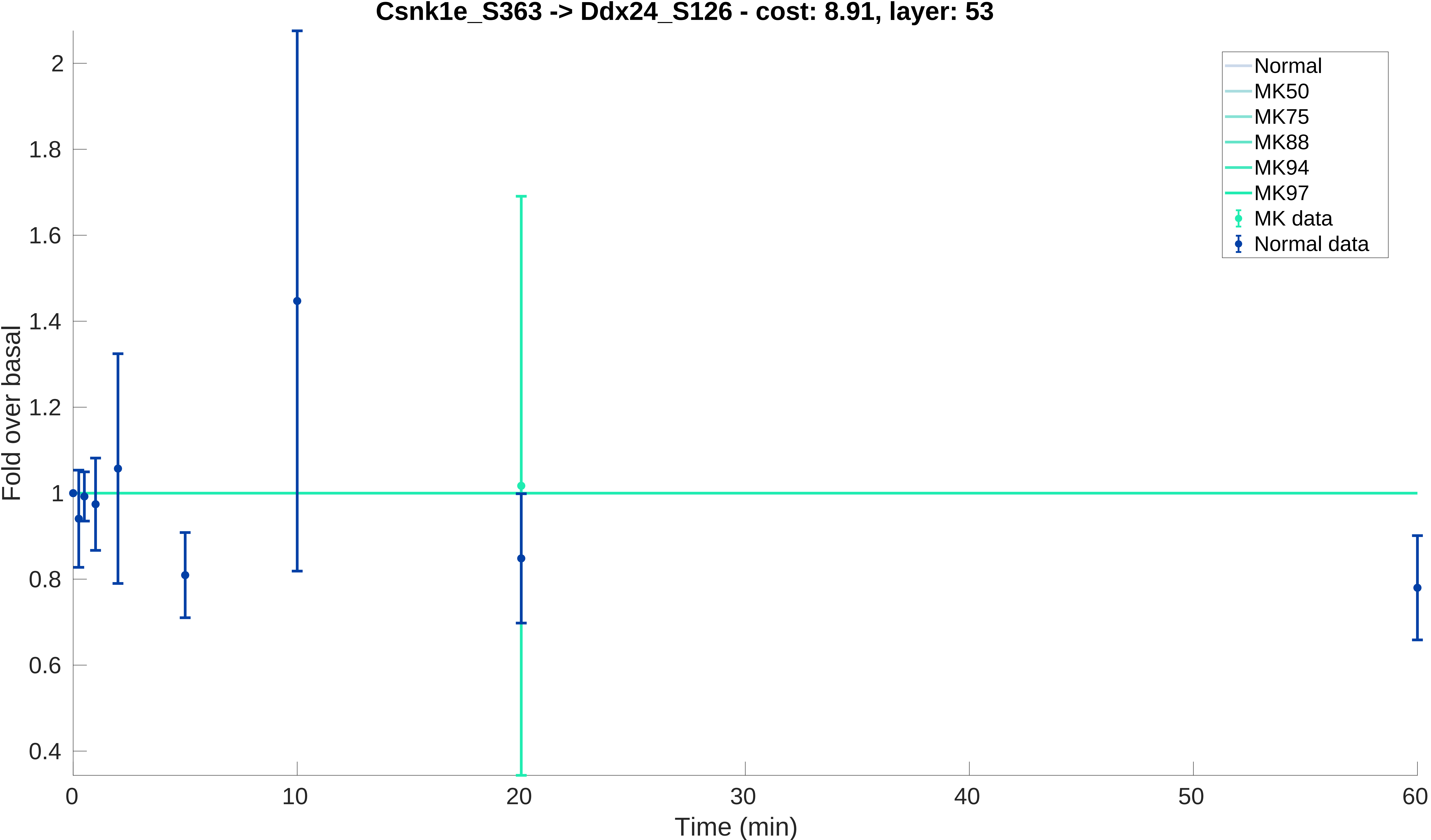

Csnk1e_S363 -> Ddx24_S126 - cost: 8.91, layer: 53
Fold over basal
2
1.8
1.6
1.4
1.2
1
0.8
0.6
0.4
0
10
20
30
40
50
60
Time (min)
Normal
MK50
MK75
MK88
MK94
MK97
MK data
Normal data

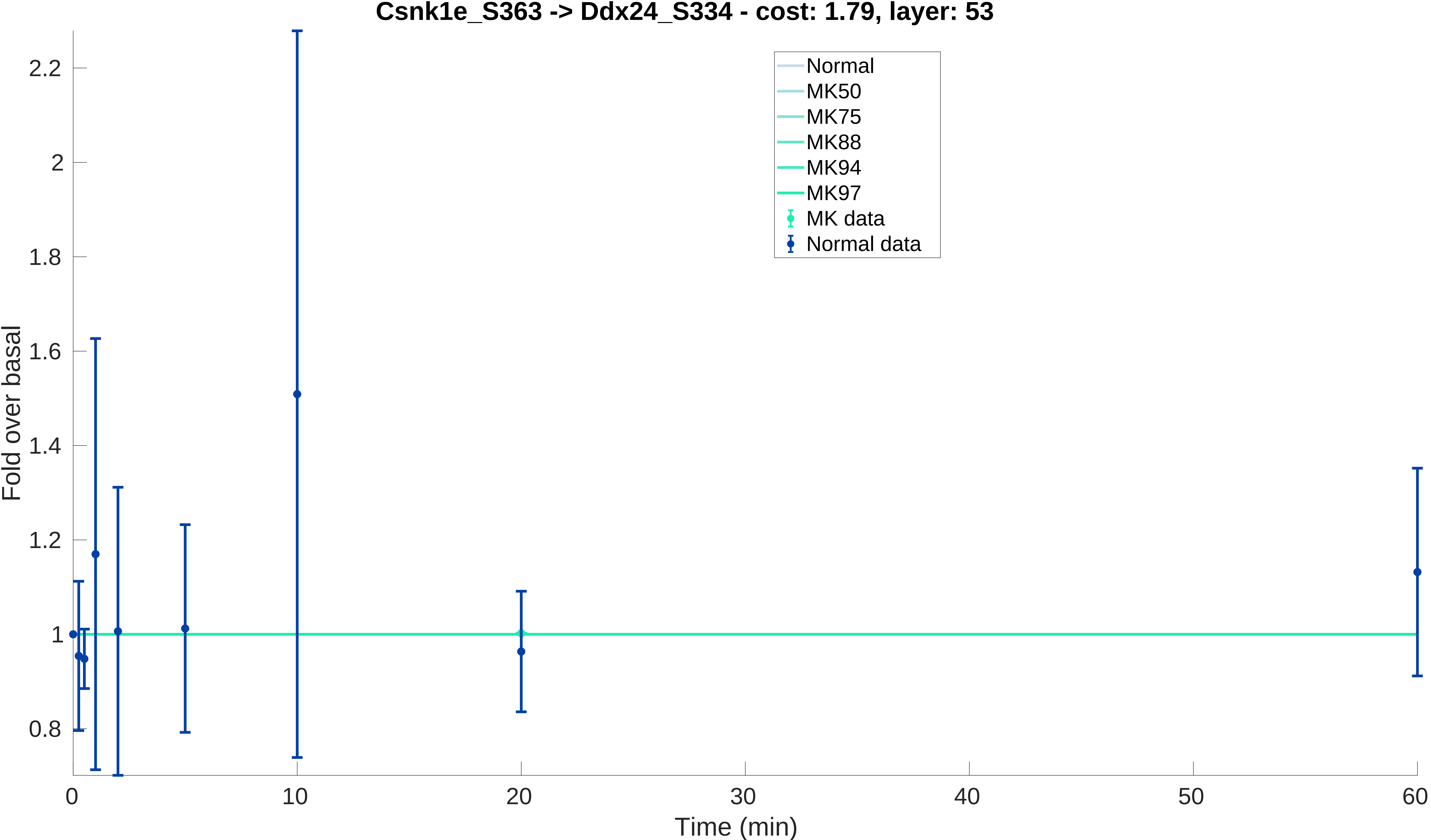

Csnk1e_S363 -> Ddx24_S334 - cost: 1.79, layer: 53
Normal
MK50
MK75
MK88
MK94
MK97
MK data
Normal data
Fold over basal
Time (min)
0.8
1
1.2
1.4
1.6
1.8
2
2.2
0
10
20
30
40
50
60

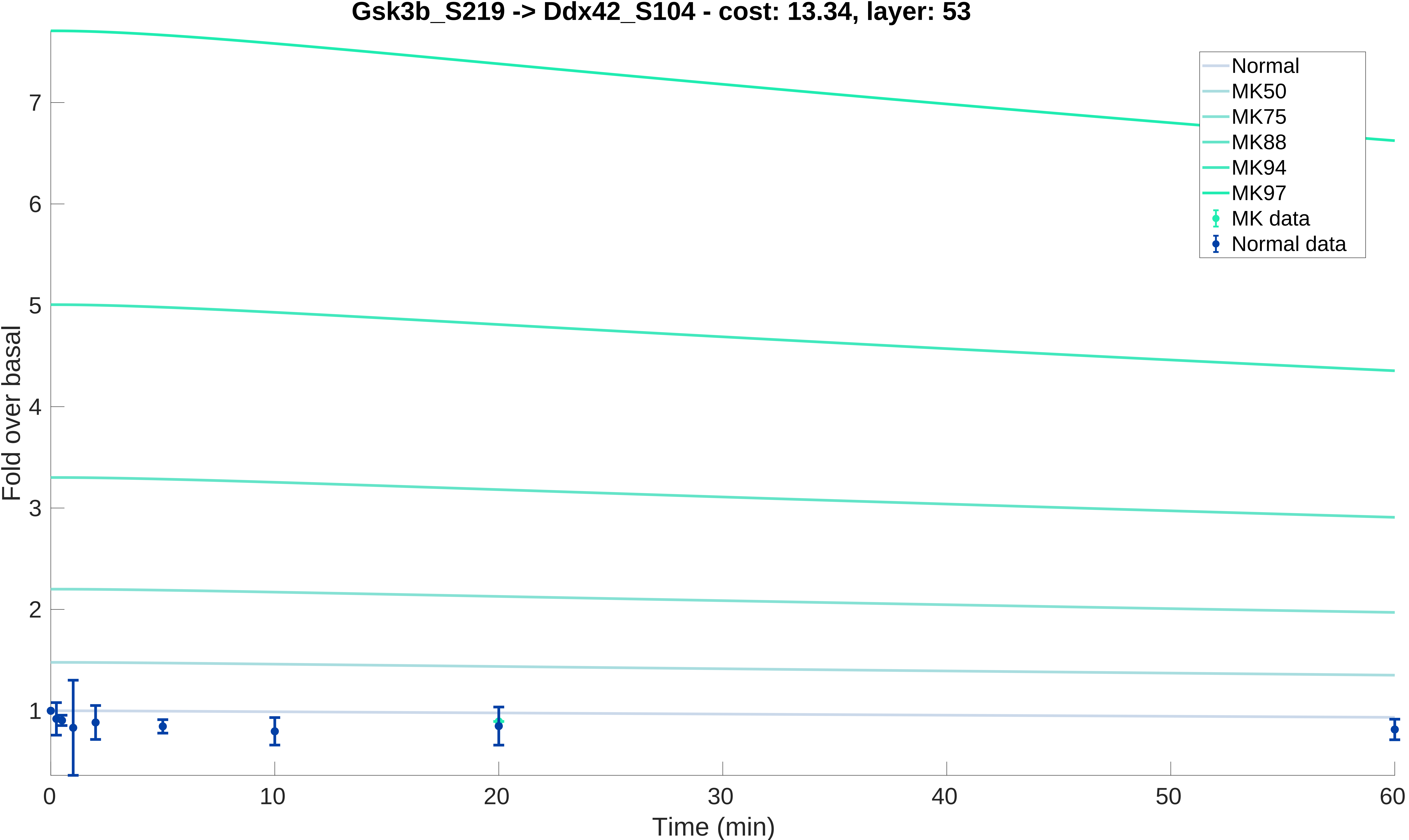

Gsk3b_S219 -> Ddx42_S104 - cost: 13.34, layer: 53
Fold over basal
Time (min)
Normal
MK50
MK75
MK88
MK94
MK97
MK data
Normal data
0
10
20
30
40
50
60
1
2
3
4
5
6
7

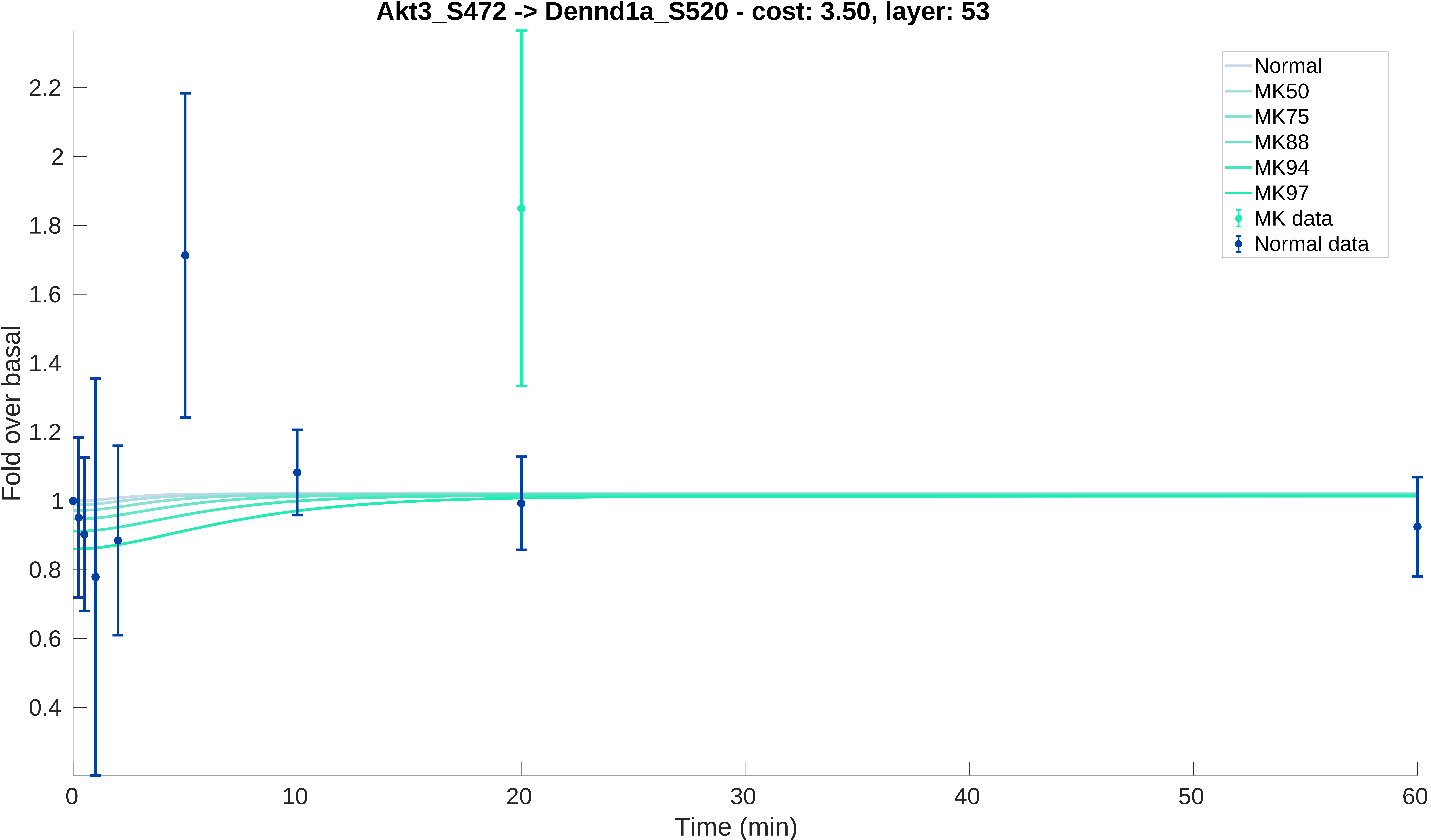
Akt3_S472 -> Dennd1a_S520 - cost: 3.50, layer: 53
Fold over basal
Time (min)
0
10
20
30
40
50
60
0.4
0.6
0.8
1
1.2
1.4
1.6
1.8
2
2.2
Normal
MK50
MK75
MK88
MK94
MK97
MK data
Normal data

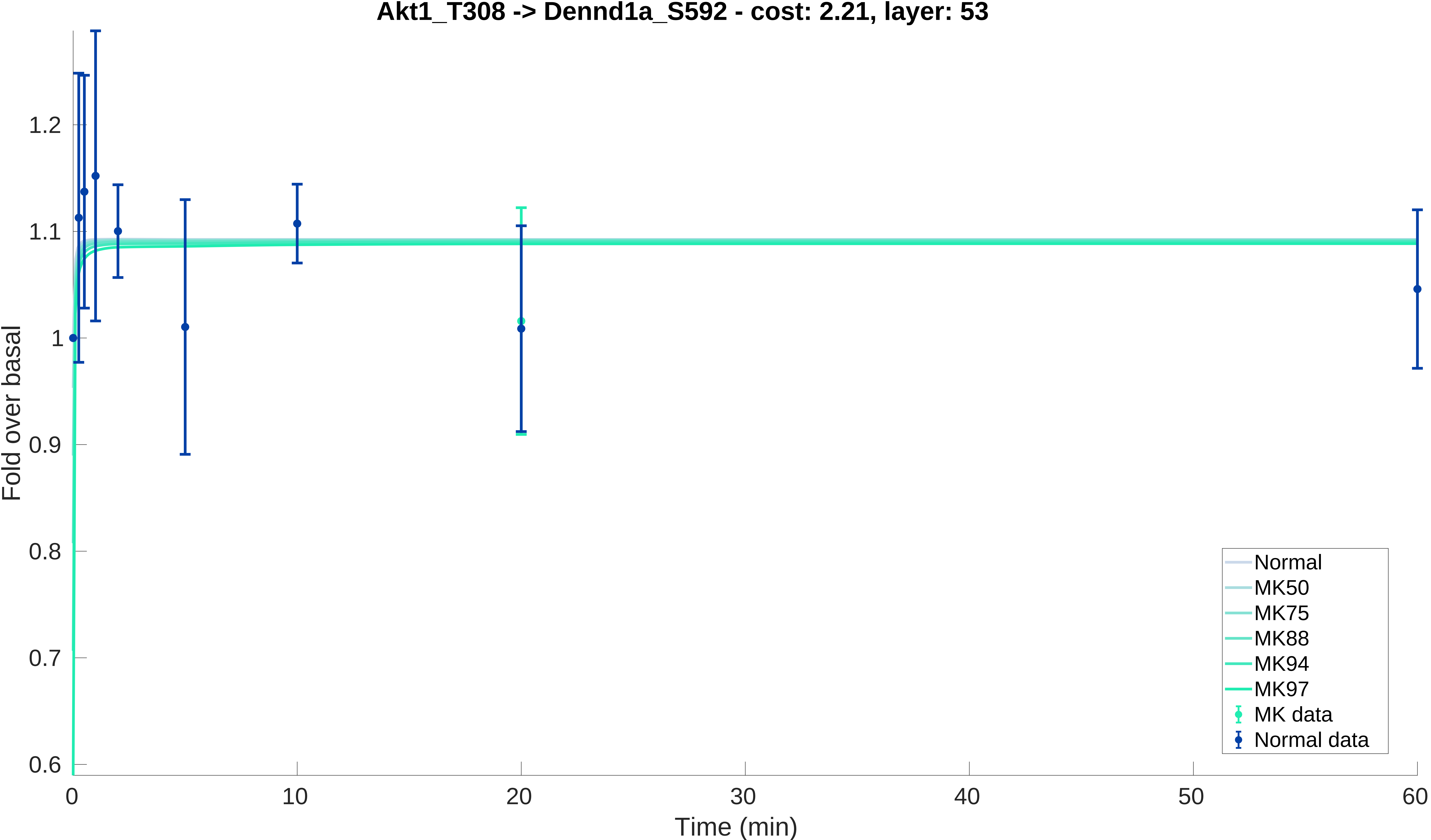

Akt1_T308 -> Dennd1a_S592 - cost: 2.21, layer: 53
Fold over basal
Time (min)
1.2
1.1
1
0.9
0.8
0.7
0.6
0
10
20
30
40
50
60
Normal
MK50
MK75
MK88
MK94
MK97
MK data
Normal data

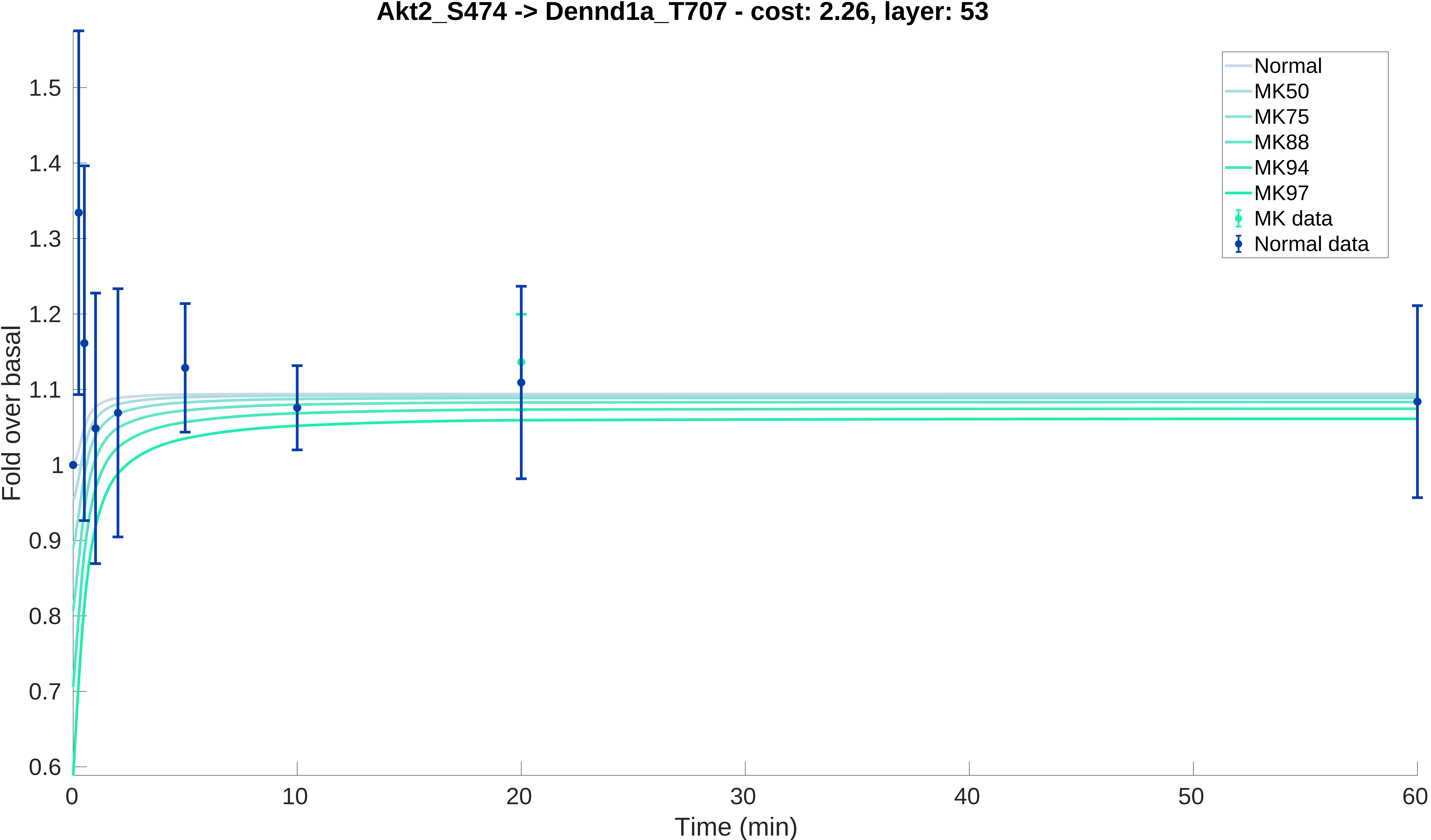

Akt2_S474 -> Dennd1a_T707 - cost: 2.26, layer: 53
Normal
MK50
MK75
MK88
MK94
MK97
MK data
Normal data
Fold over basal
1.5
1.4
1.3
1.2
1.1
1
0.9
0.8
0.7
0.6
0
10
20
30
40
50
60
Time (min)

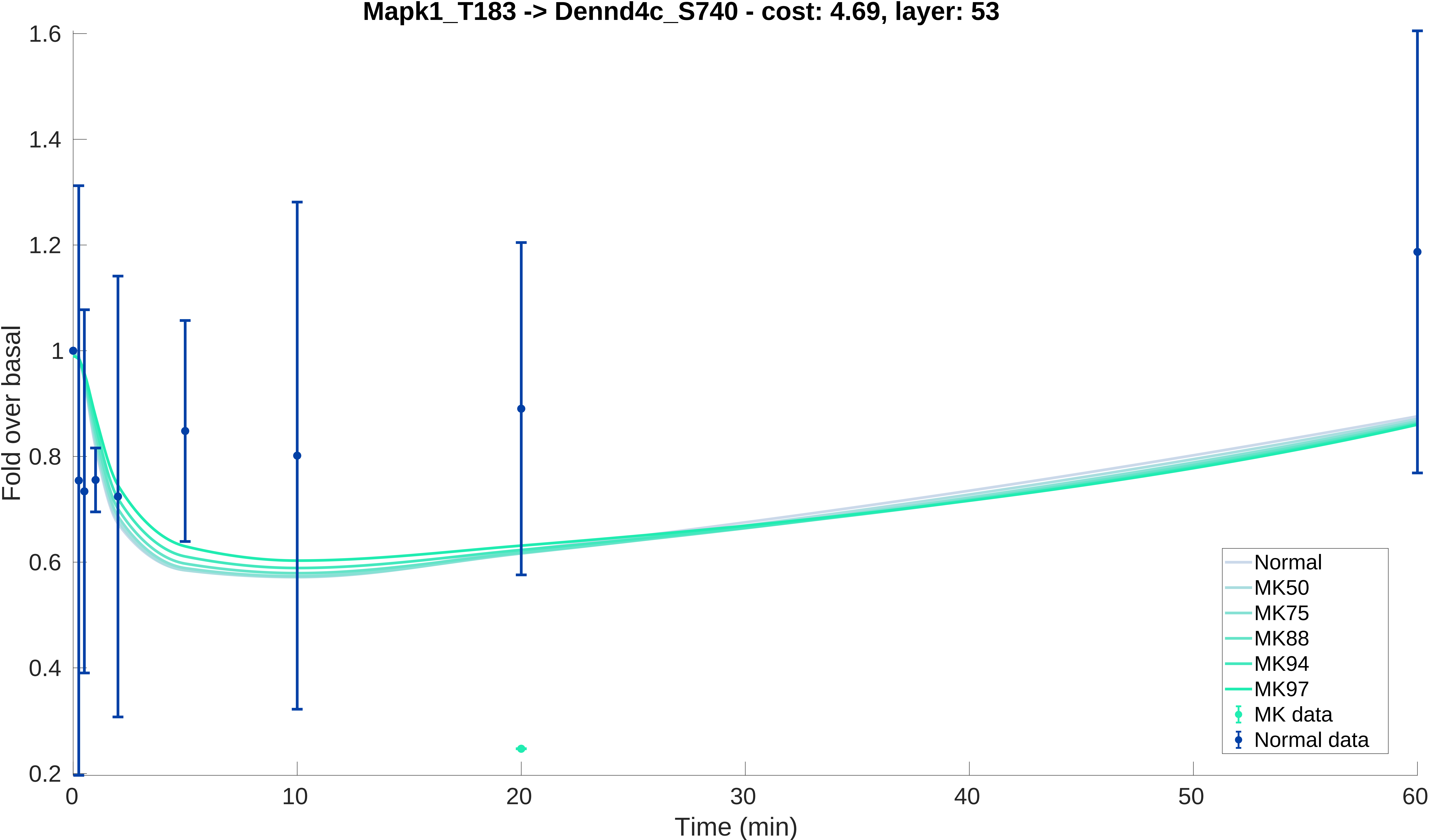

Mapk1_T183 -> Dennd4c_S740 - cost: 4.69, layer: 53
Fold over basal
Time (min)
1.6
1.4
1.2
1
0.8
0.6
0.4
0.2
0
10
20
30
40
50
60
Normal
MK50
MK75
MK88
MK94
MK97
MK data
Normal data

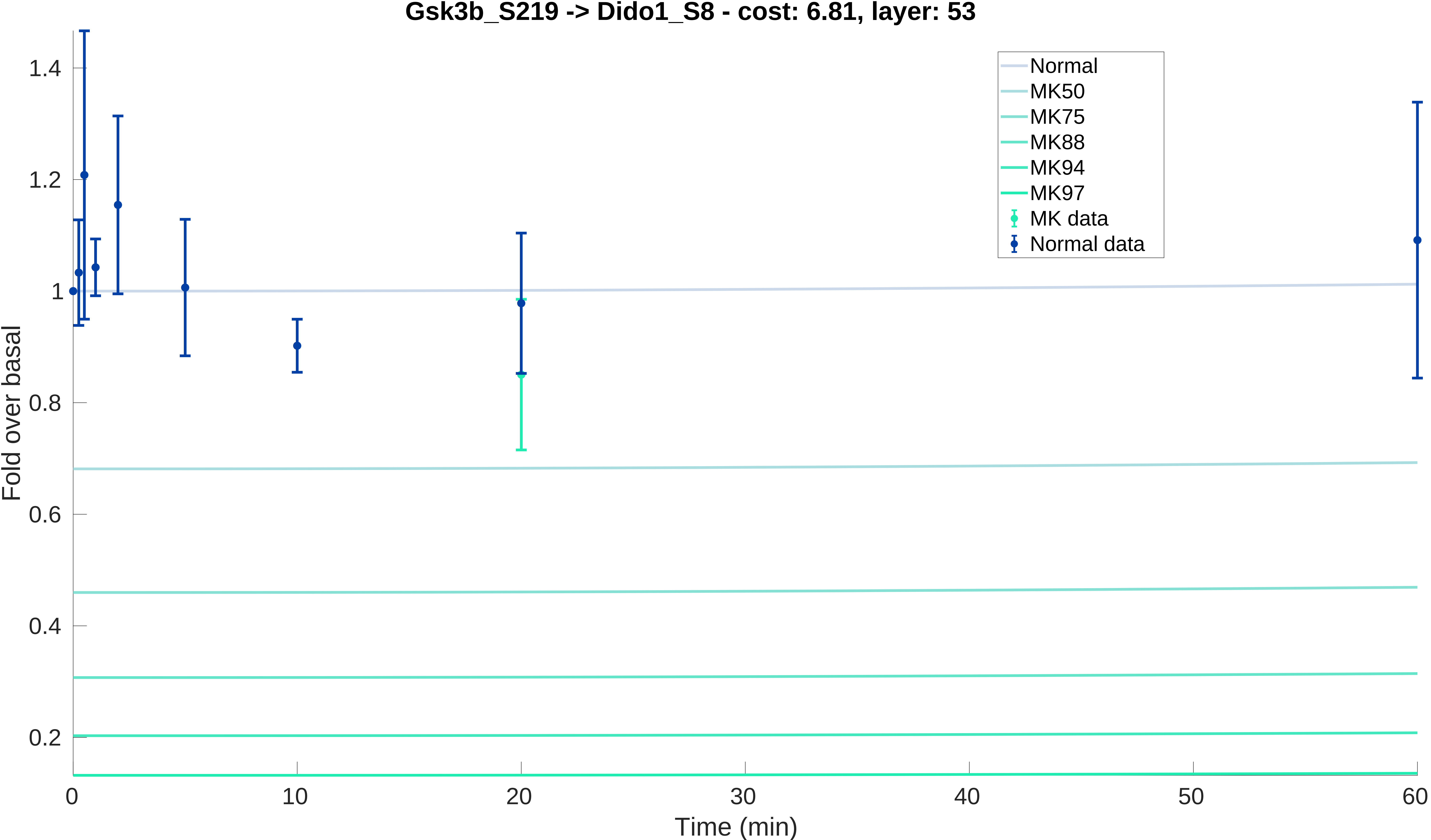

Gsk3b_S219 -> Dido1_S8 - cost: 6.81, layer: 53
Normal
MK50
MK75
MK88
MK94
MK97
MK data
Normal data
Fold over basal
Time (min)
1.4
1.2
1
0.8
0.6
0.4
0.2
0
10
20
30
40
50
60

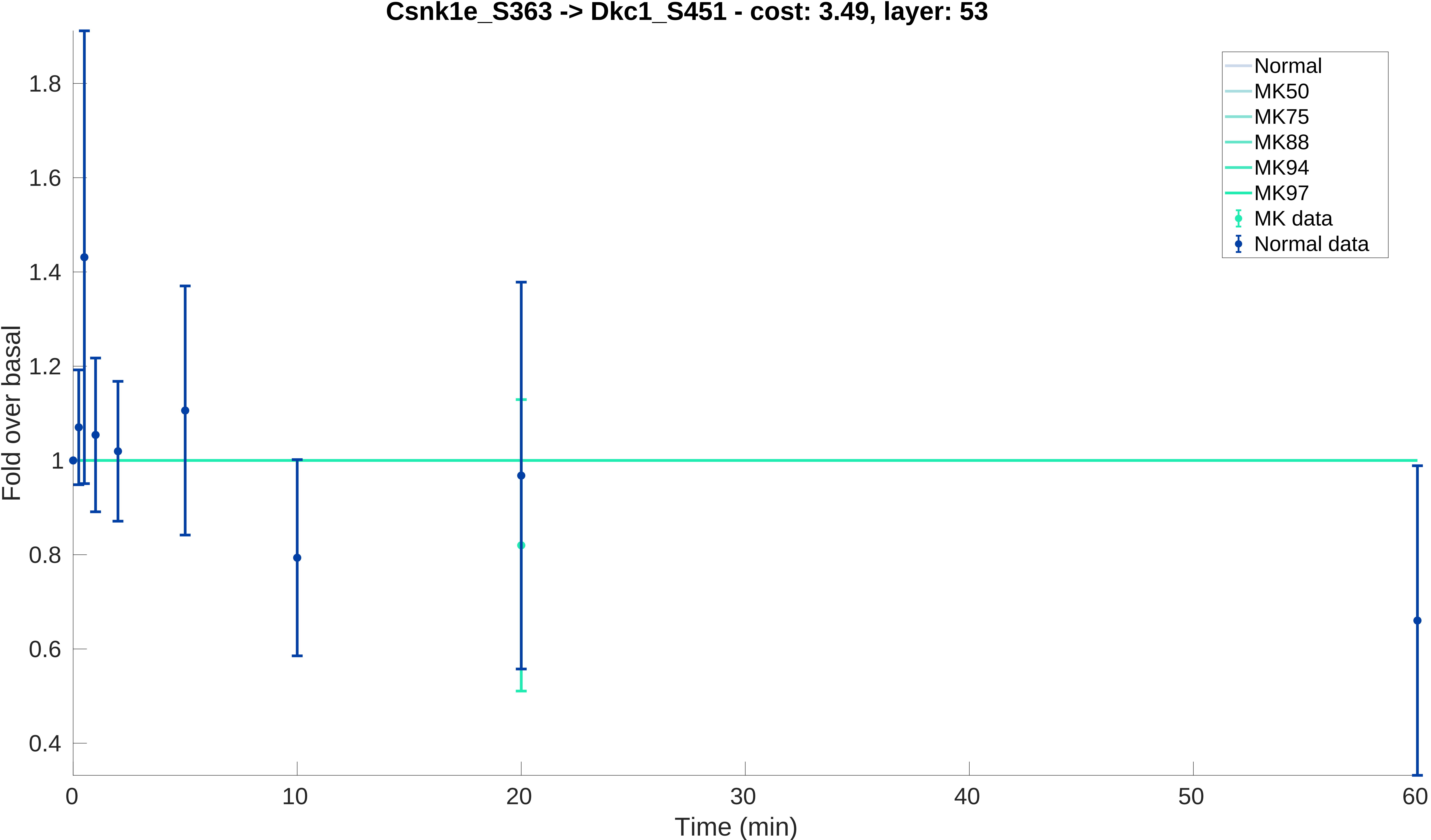

Csnk1e_S363 -> Dkc1_S451 - cost: 3.49, layer: 53
Fold over basal
Time (min)
0.4
0.6
0.8
1
1.2
1.4
1.6
1.8
0
10
20
30
40
50
60
Normal
MK50
MK75
MK88
MK94
MK97
MK data
Normal data

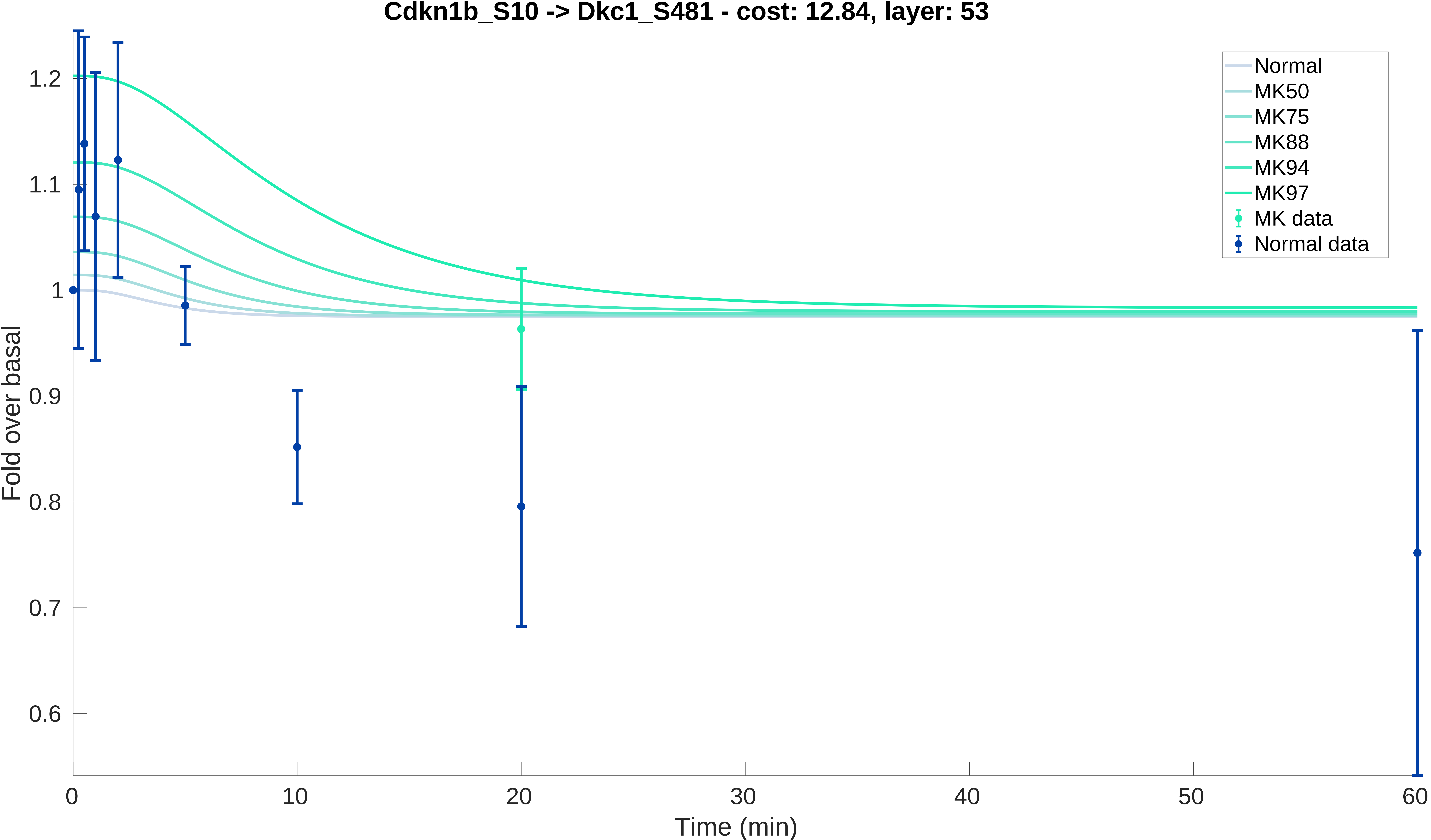
Cdkn1b_S10 -> Dkc1_S481 - cost: 12.84, layer: 53
Fold over basal
Time (min)
0.6
0.7
0.8
0.9
1
1.1
1.2
0
10
20
30
40
50
60
Normal
MK50
MK75
MK88
MK94
MK97
MK data
Normal data

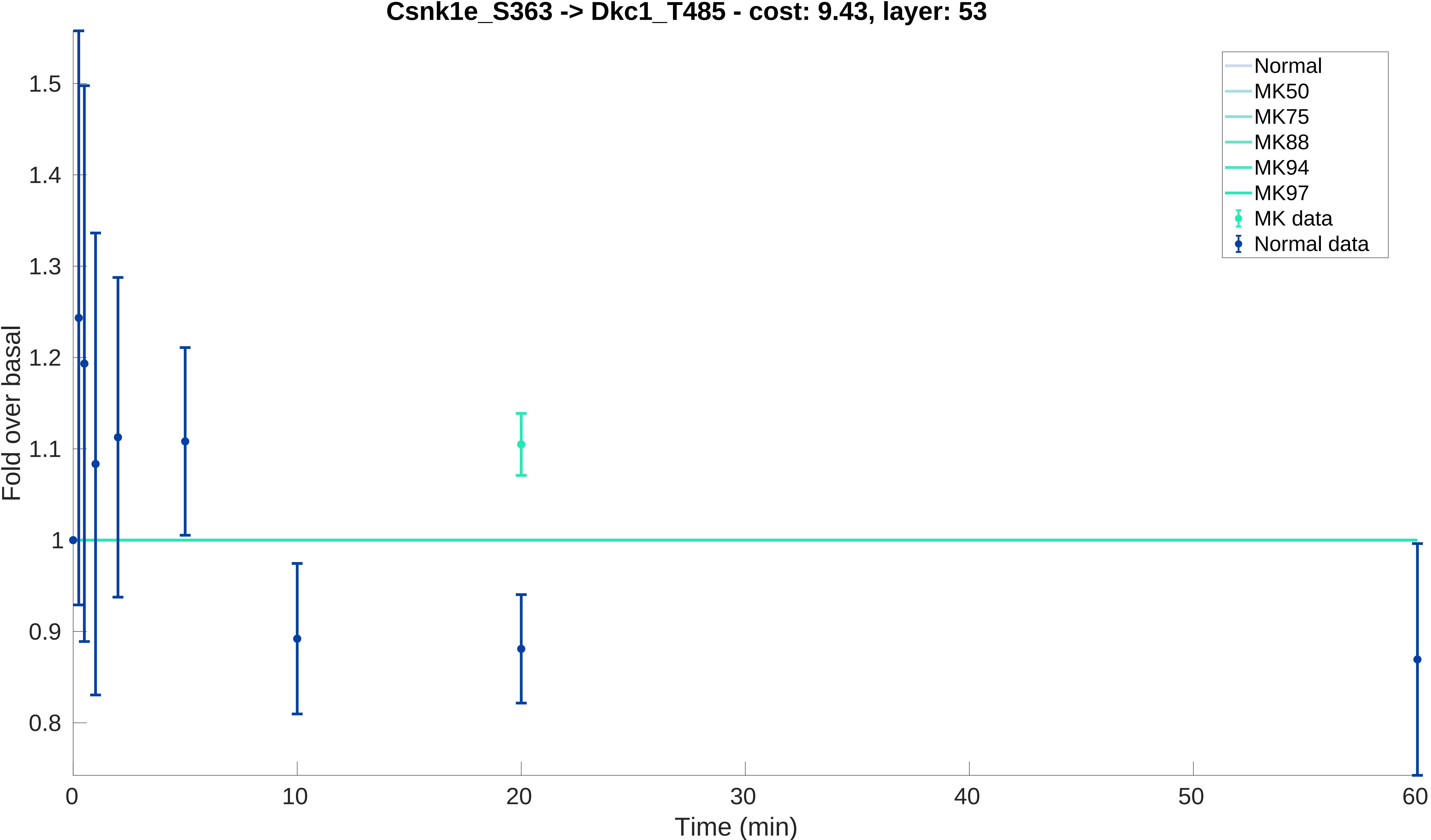
Csnk1e_S363 -> Dkc1_T485 - cost: 9.43, layer: 53
Fold over basal
Time (min)
1.5
1.4
1.3
1.2
1.1
1
0.9
0.8
0
10
20
30
40
50
60
Normal
MK50
MK75
MK88
MK94
MK97
MK data
Normal data

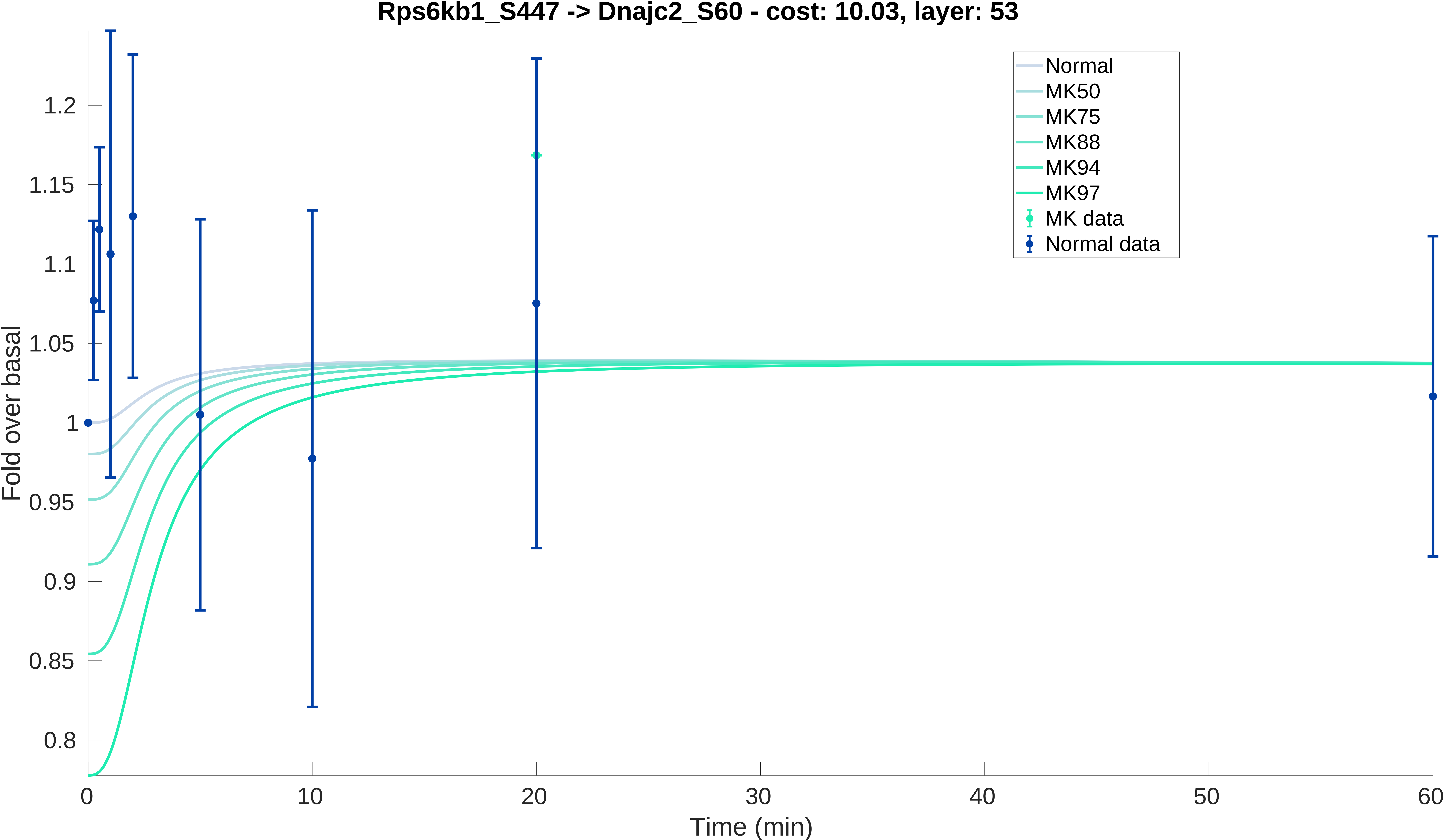

Rps6kb1_S447 -> Dnajc2_S60 - cost: 10.03, layer: 53
Normal
MK50
MK75
MK88
MK94
MK97
MK data
Normal data
Fold over basal
1.2
1.15
1.1
1.05
1
0.95
0.9
0.85
0.8
0
10
20
30
40
50
60
Time (min)

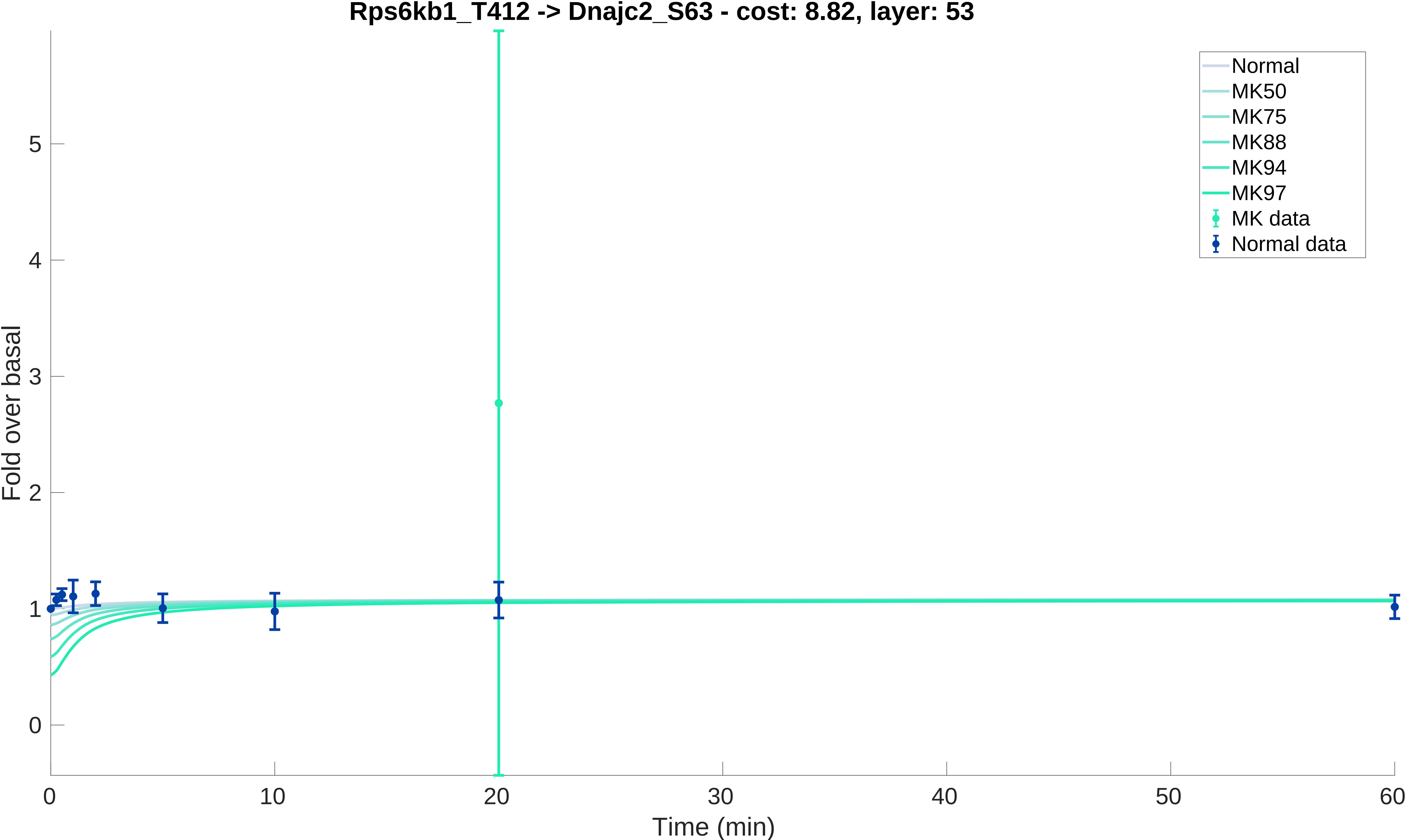

Rps6kb1_T412 -> Dnajc2_S63 - cost: 8.82, layer: 53
Fold over basal
Time (min)
0
1
2
3
4
5
0
10
20
30
40
50
60
Normal
MK50
MK75
MK88
MK94
MK97
MK data
Normal data

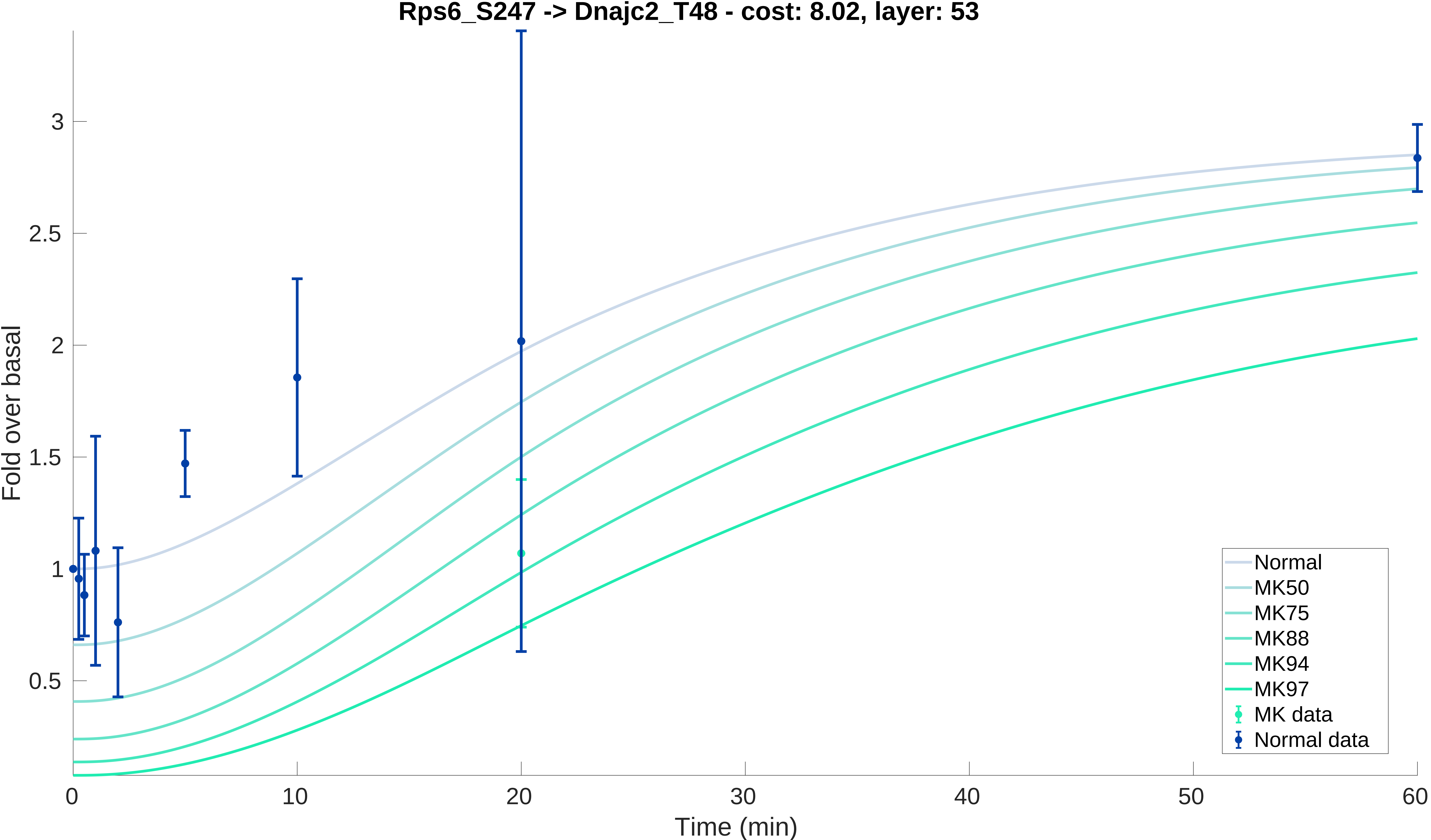

Rps6_S247 -> Dnajc2_T48 - cost: 8.02, layer: 53
Fold over basal
Time (min)
0.5
1
1.5
2
2.5
3
0
10
20
30
40
50
60
Normal
MK50
MK75
MK88
MK94
MK97
MK data
Normal data

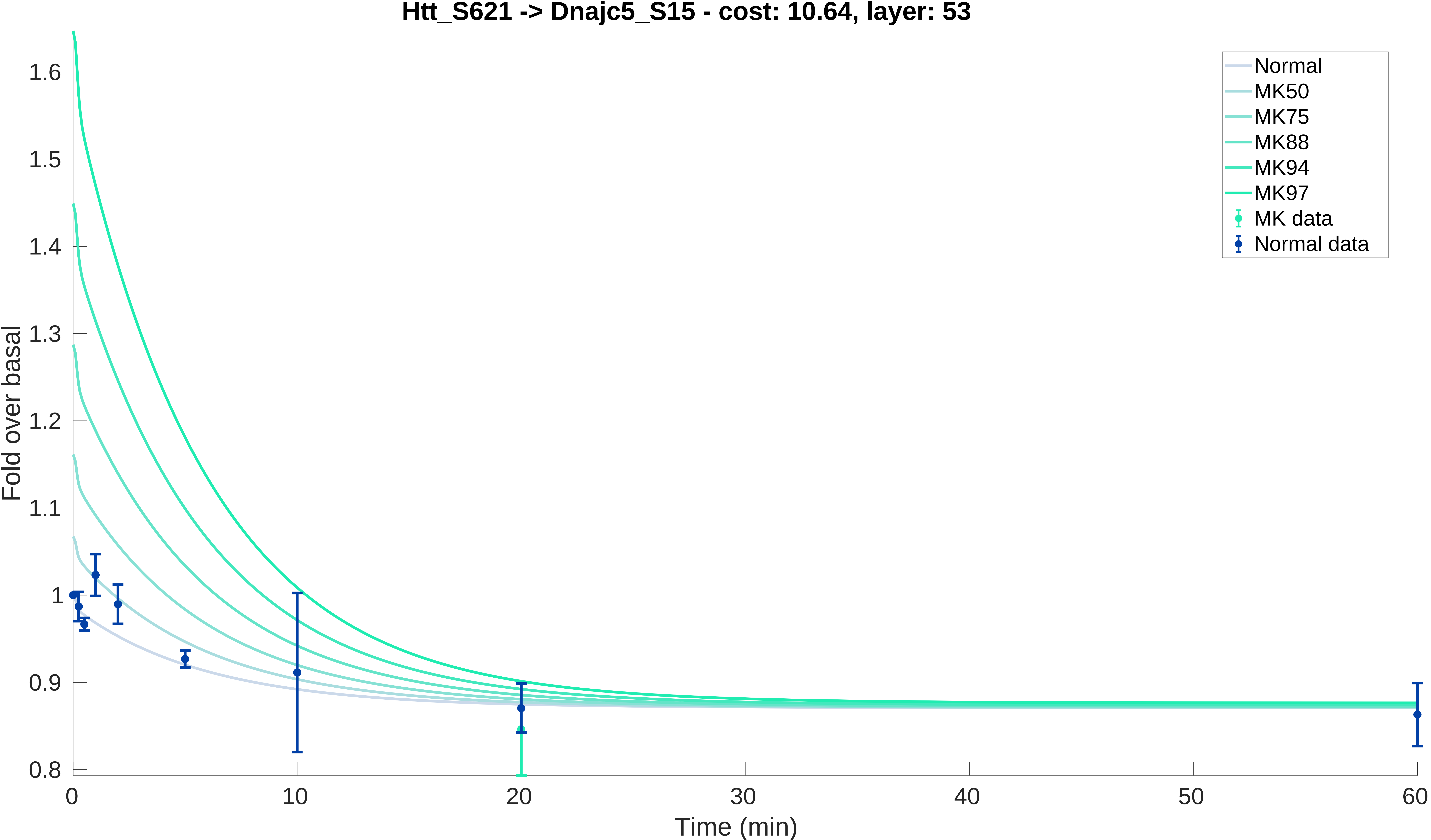

Htt_S621 -> Dnajc5_S15 - cost: 10.64, layer: 53
Fold over basal
1.6
1.5
1.4
1.3
1.2
1.1
1
0.9
0.8
0
10
20
30
40
50
60
Time (min)
Normal
MK50
MK75
MK88
MK94
MK97
MK data
Normal data

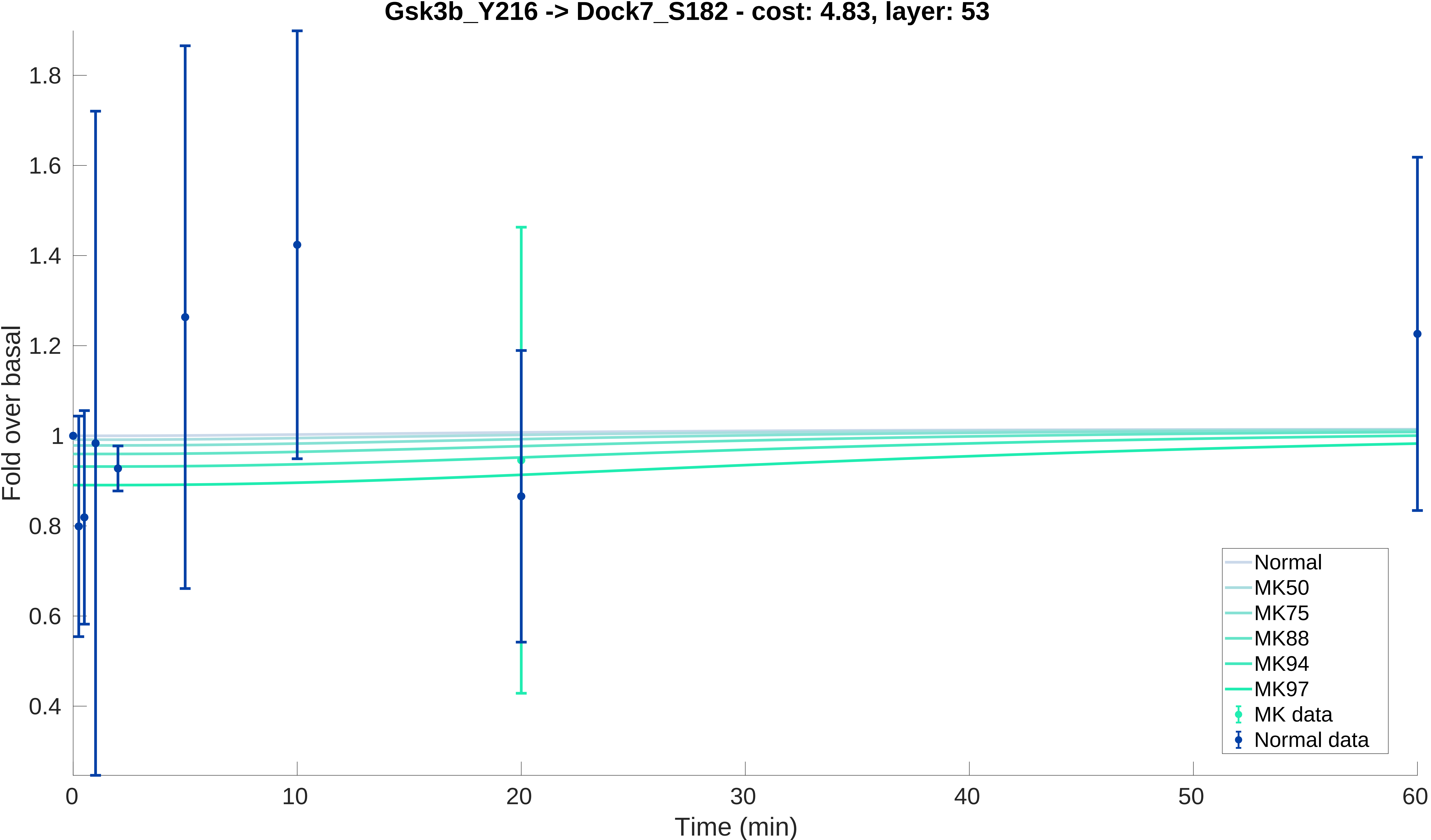

Gsk3b_Y216 -> Dock7_S182 - cost: 4.83, layer: 53
Fold over basal
1.8
1.6
1.4
1.2
1
0.8
0.6
0.4
0
10
20
30
40
50
60
Time (min)
Normal
MK50
MK75
MK88
MK94
MK97
MK data
Normal data

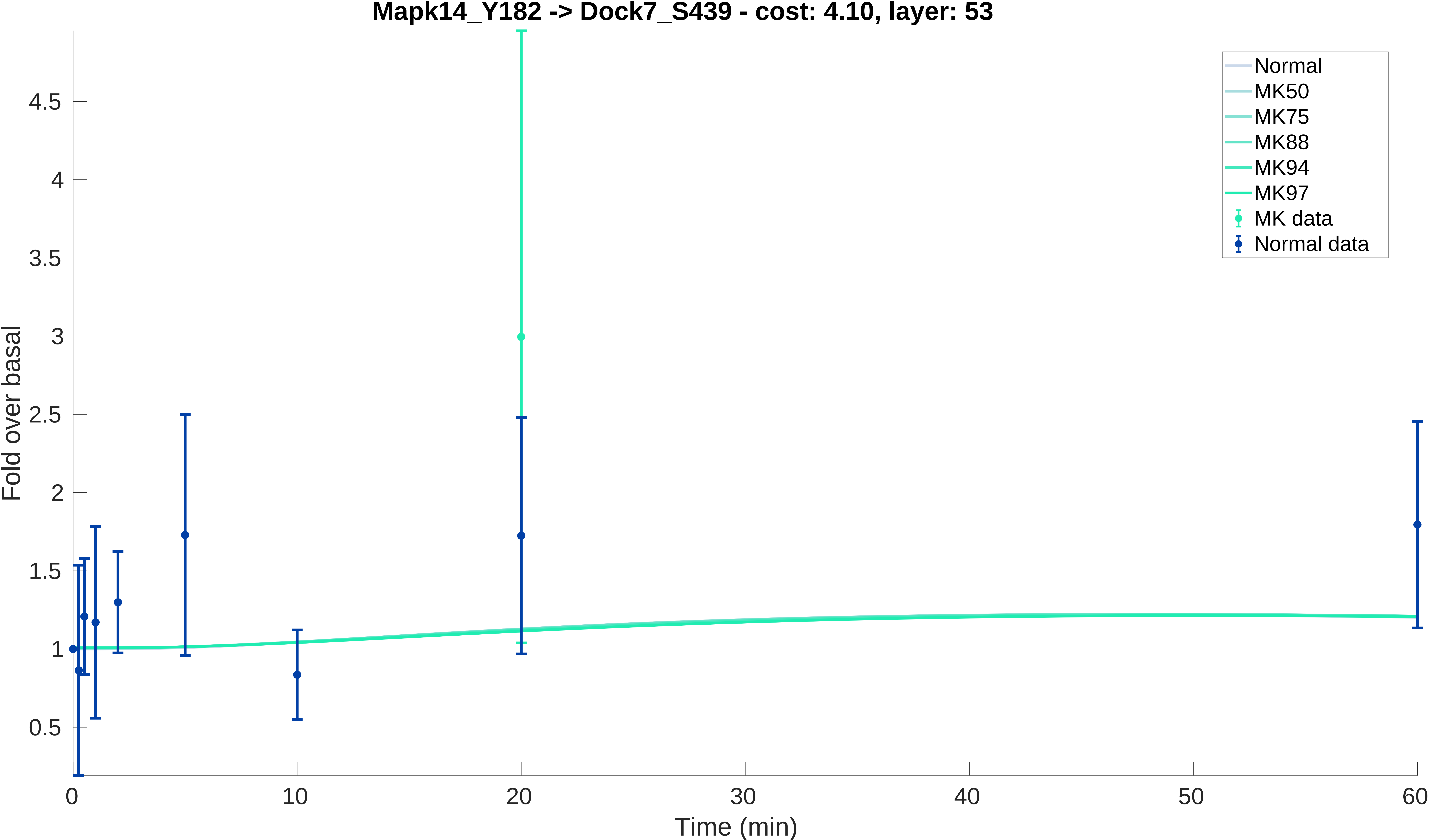

Mapk14_Y182 -> Dock7_S439 - cost: 4.10, layer: 53
Fold over basal
Time (min)
Normal
MK50
MK75
MK88
MK94
MK97
MK data
Normal data
4.5
4
3.5
3
2.5
2
1.5
1
0.5
0
10
20
30
40
50
60

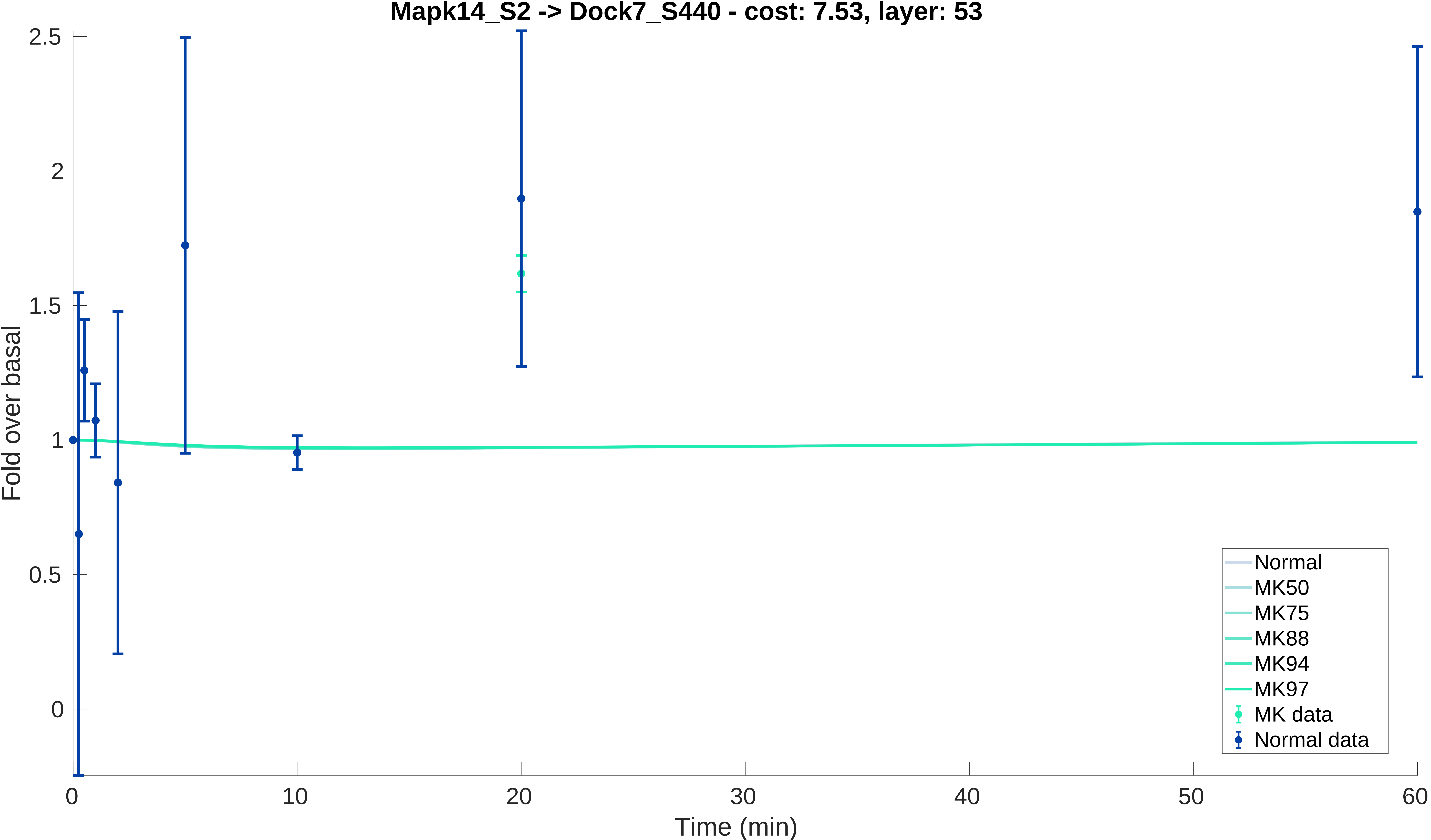
Mapk14_S2 -> Dock7_S440 - cost: 7.53, layer: 53
Fold over basal
2.5
2
1.5
1
0.5
0
Time (min)
0
10
20
30
40
50
60
Normal
MK50
MK75
MK88
MK94
MK97
MK data
Normal data

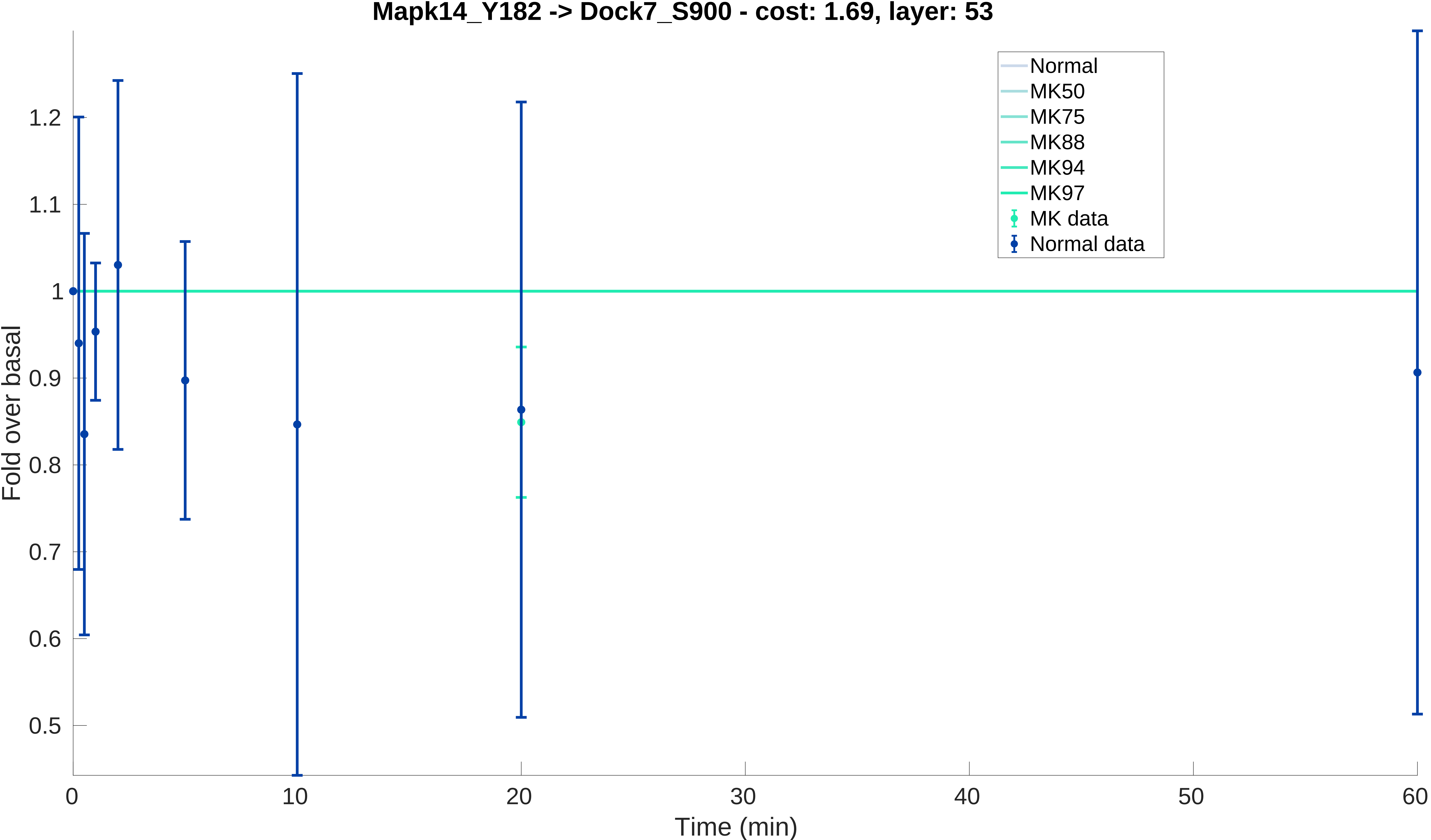

Mapk14_Y182 -> Dock7_S900 - cost: 1.69, layer: 53
Normal
MK50
MK75
MK88
MK94
MK97
MK data
Normal data
Fold over basal
Time (min)
1.2
1.1
1
0.9
0.8
0.7
0.6
0.5
0
10
20
30
40
50
60

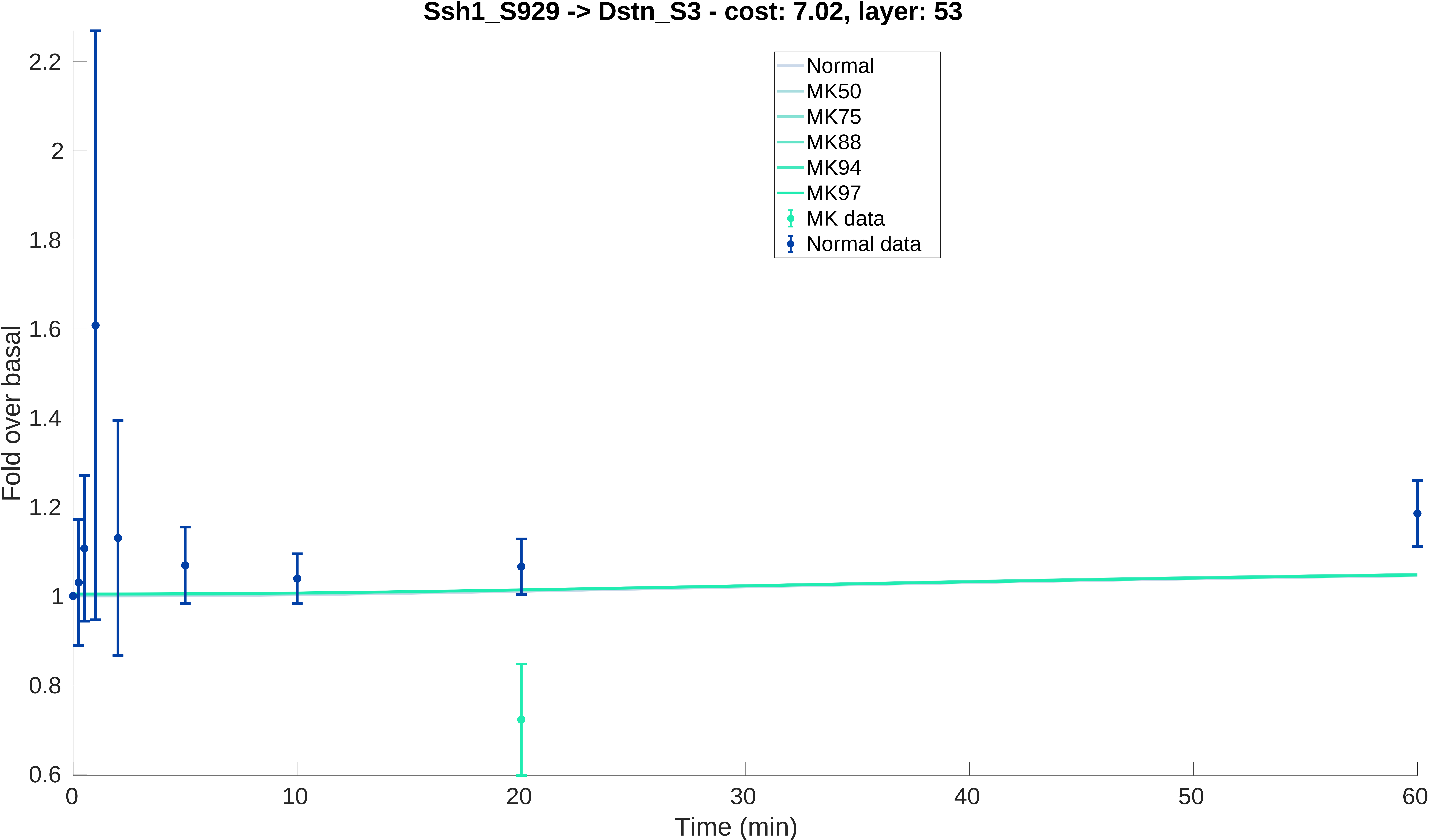
Ssh1_S929 -> Dstn_S3 - cost: 7.02, layer: 53
Normal
MK50
MK75
MK88
MK94
MK97
MK data
Normal data
Fold over basal
2.2
2
1.8
1.6
1.4
1.2
1
0.8
0.6
0
10
20
30
40
50
60
Time (min)

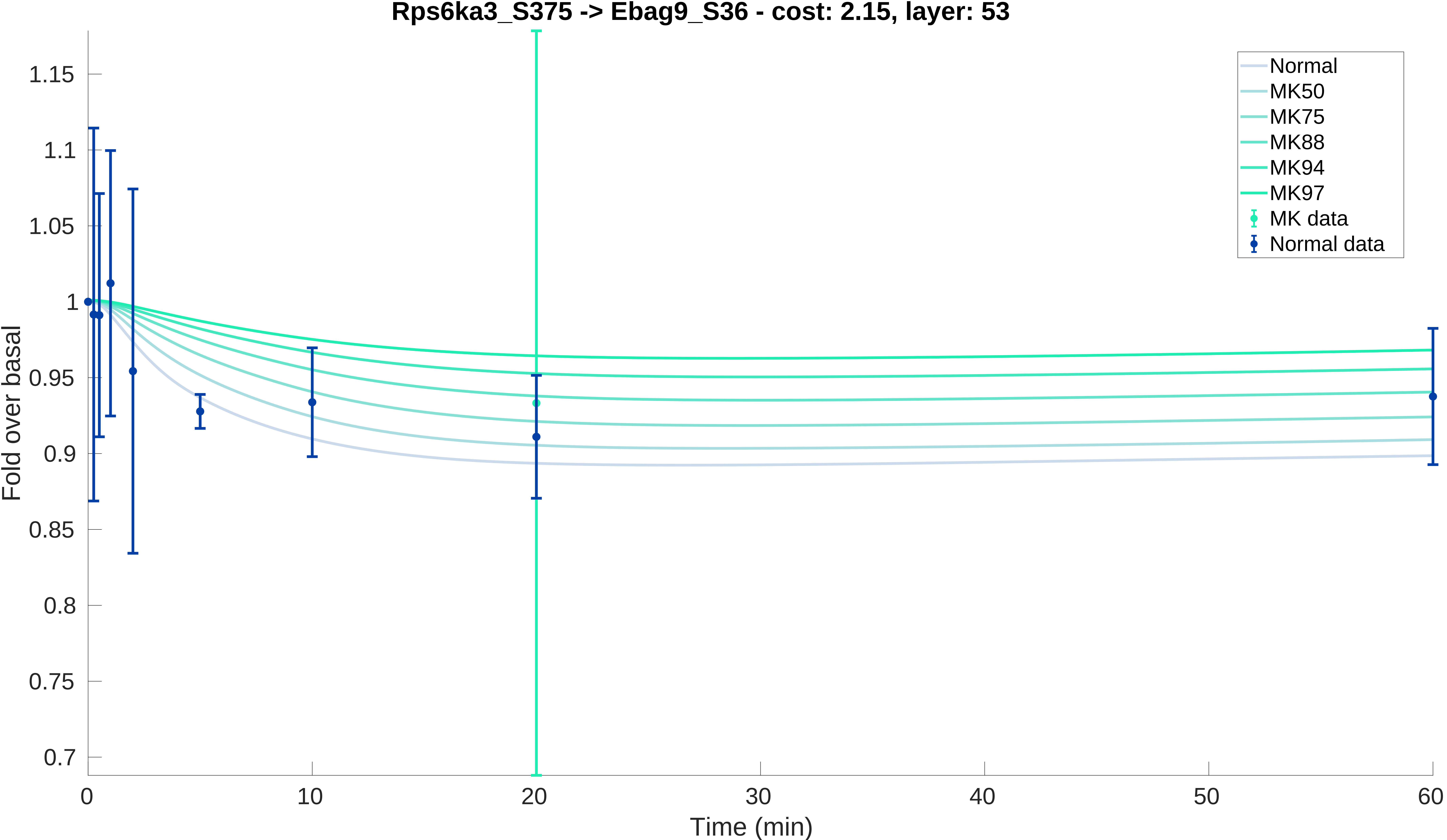
Rps6ka3_S375 -> Ebag9_S36 - cost: 2.15, layer: 53
Fold over basal
1.15
1.1
1.05
1
0.95
0.9
0.85
0.8
0.75
0.7
0
10
20
30
40
50
60
Time (min)
Normal
MK50
MK75
MK88
MK94
MK97
MK data
Normal data

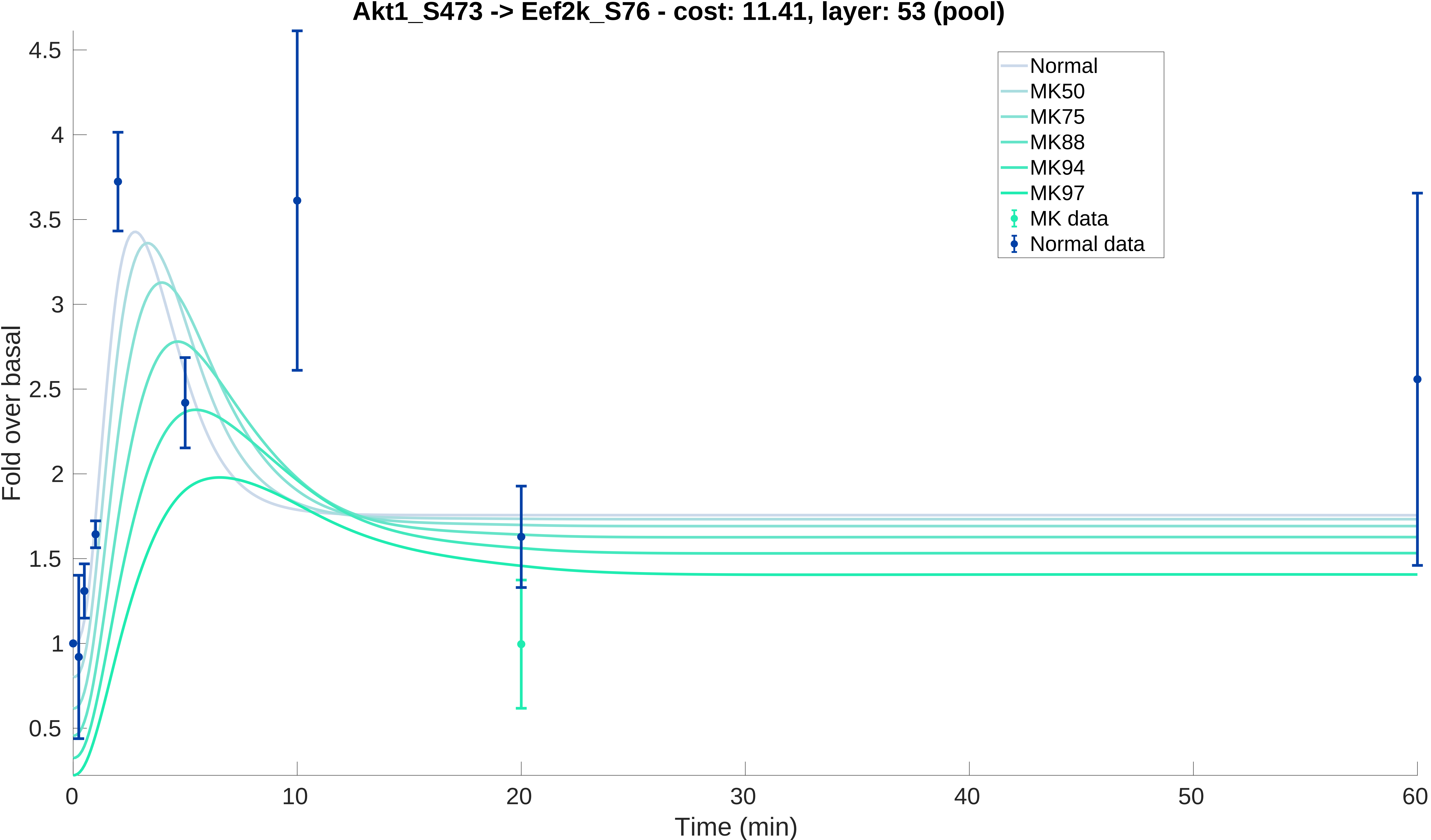

Akt1_S473 -> Eef2k_S76 - cost: 11.41, layer: 53 (pool)
Normal
MK50
MK75
MK88
MK94
MK97
MK data
Normal data
Fold over basal
Time (min)
4.5
4
3.5
3
2.5
2
1.5
1
0.5
0
10
20
30
40
50
60

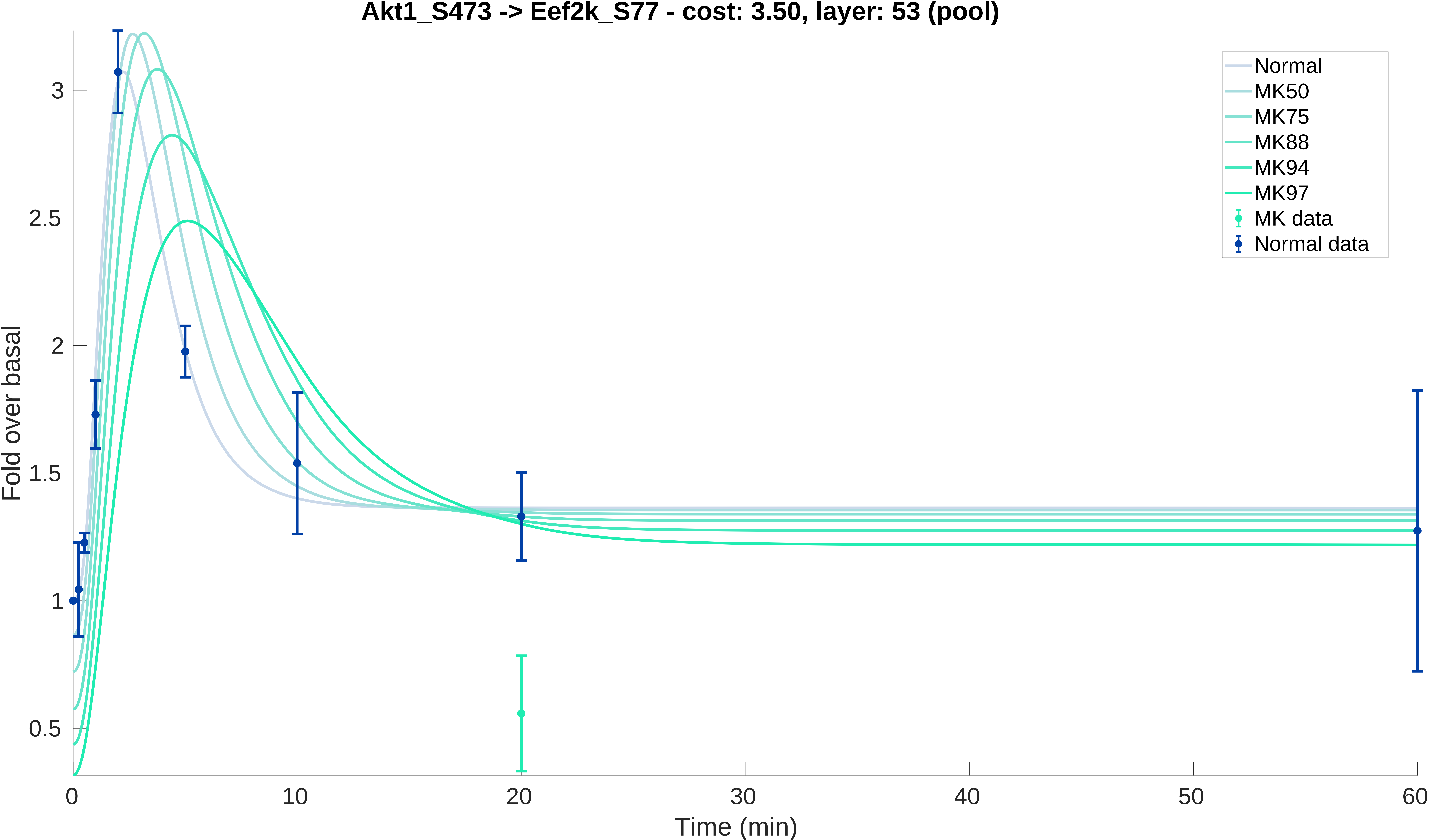

Akt1_S473 -> Eef2k_S77 - cost: 3.50, layer: 53 (pool)
Fold over basal
Time (min)
0.5
1
1.5
2
2.5
3
0
10
20
30
40
50
60
Normal
MK50
MK75
MK88
MK94
MK97
MK data
Normal data

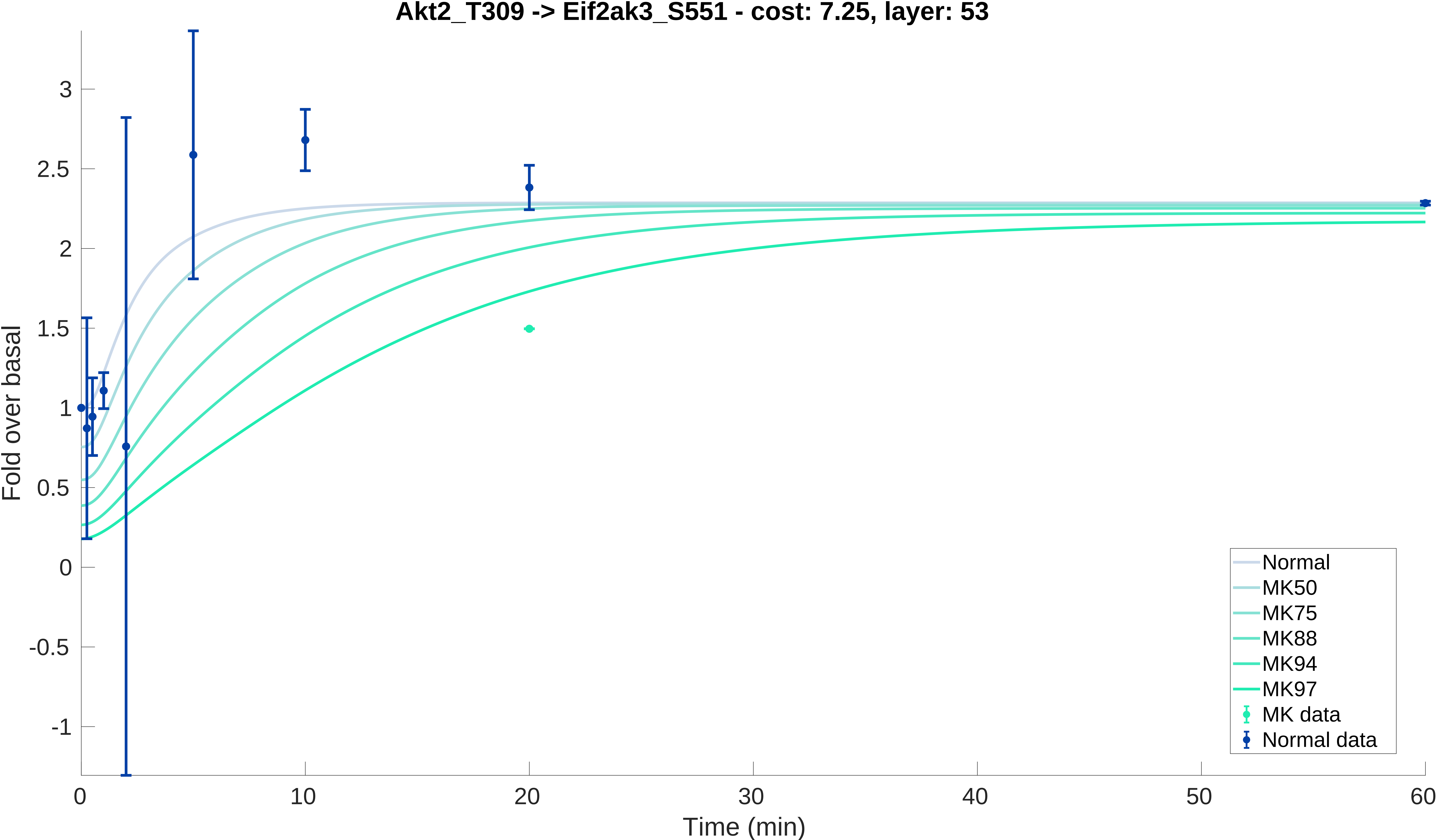

Akt2_T309 -> Eif2ak3_S551 - cost: 7.25, layer: 53
Fold over basal
Time (min)
3
2.5
2
1.5
1
0.5
0
-0.5
-1
0
10
20
30
40
50
60
Normal
MK50
MK75
MK88
MK94
MK97
MK data
Normal data

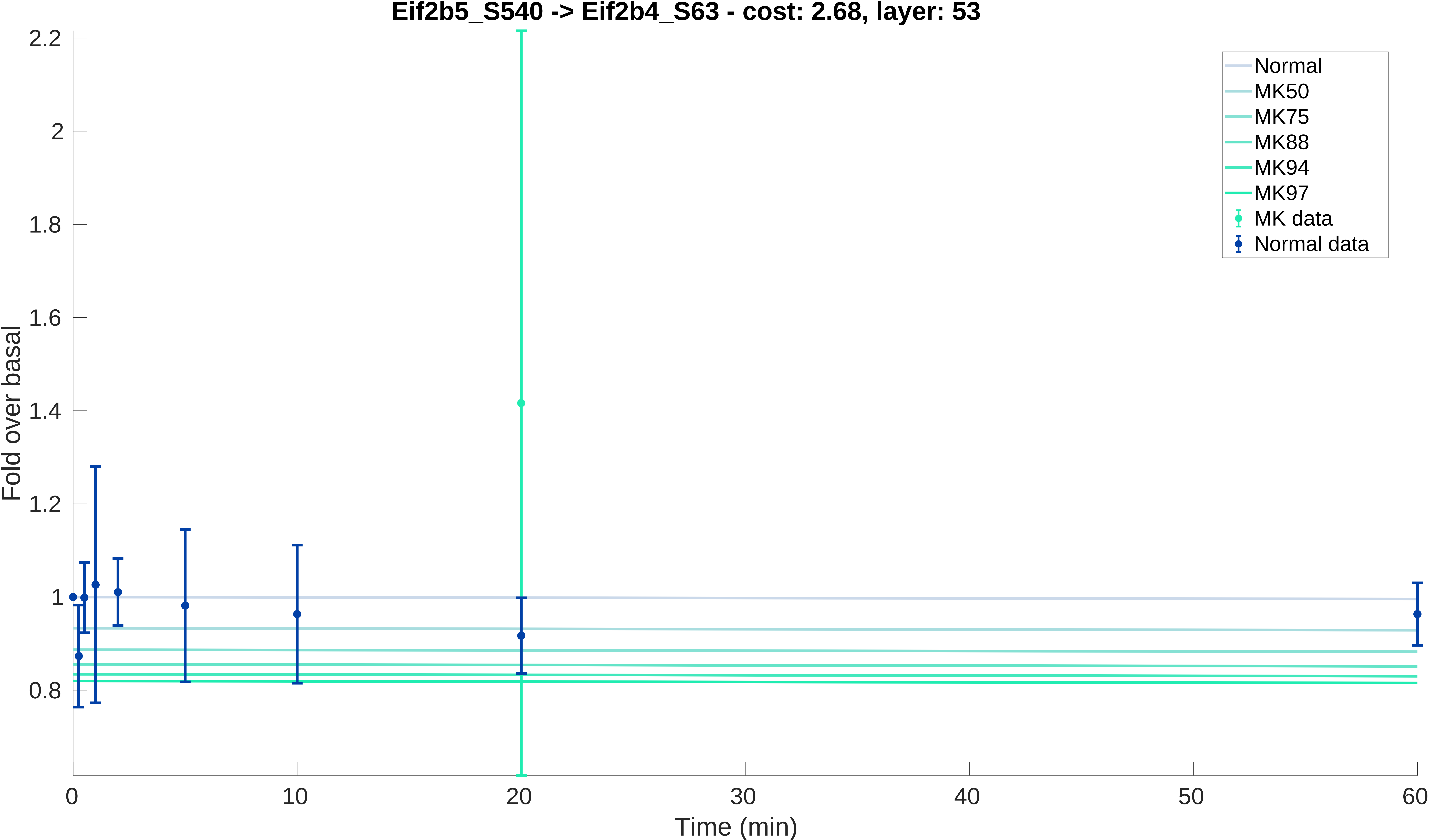

Eif2b5_S540 -> Eif2b4_S63 - cost: 2.68, layer: 53
Fold over basal
2.2
2
1.8
1.6
1.4
1.2
1
0.8
0
10
20
30
40
50
60
Time (min)
Normal
MK50
MK75
MK88
MK94
MK97
MK data
Normal data

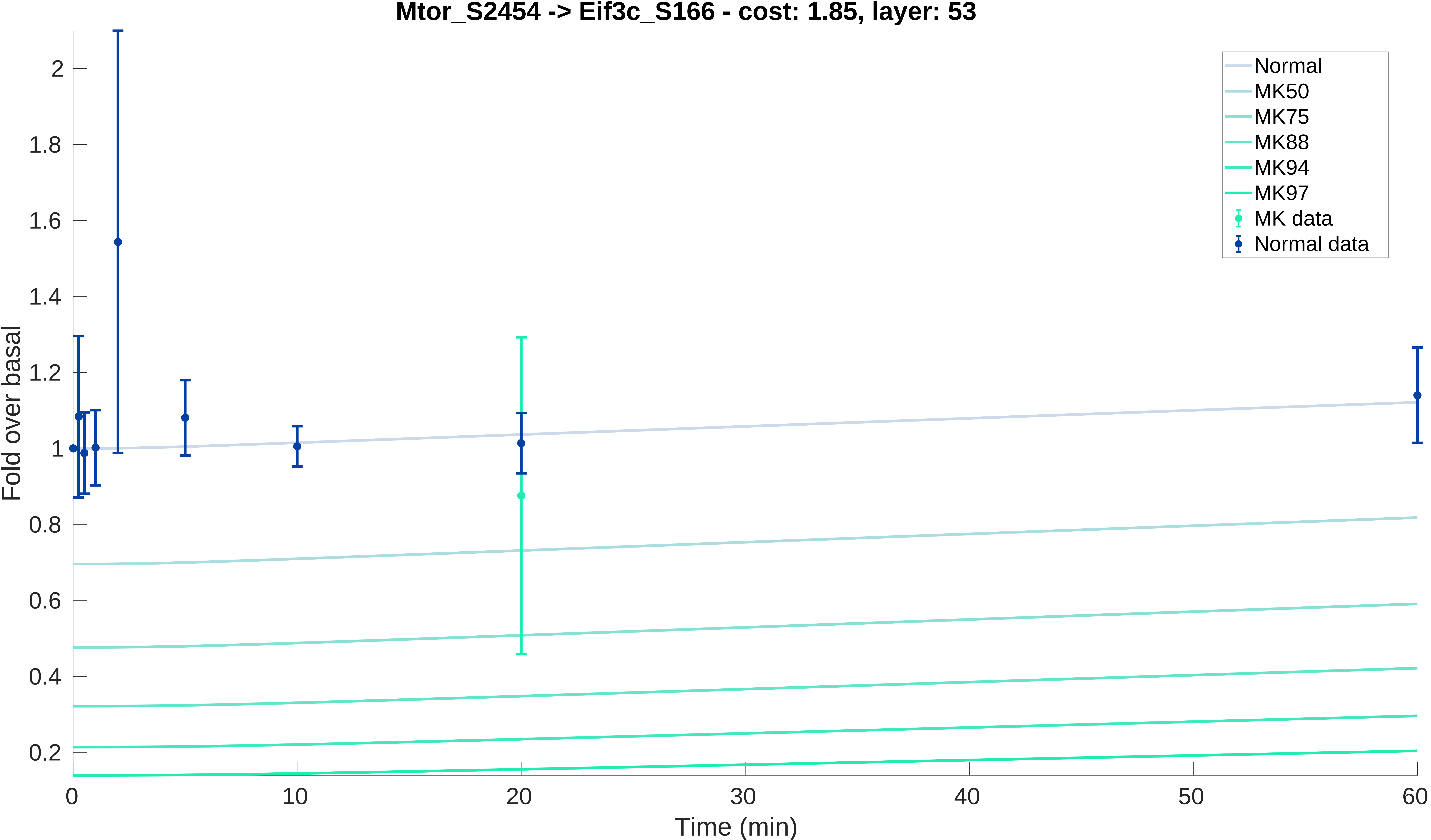

Mtor_S2454 -> Eif3c_S166 - cost: 1.85, layer: 53
Fold over basal
Time (min)
Normal
MK50
MK75
MK88
MK94
MK97
MK data
Normal data
2
1.8
1.6
1.4
1.2
1
0.8
0.6
0.4
0.2
0
10
20
30
40
50
60

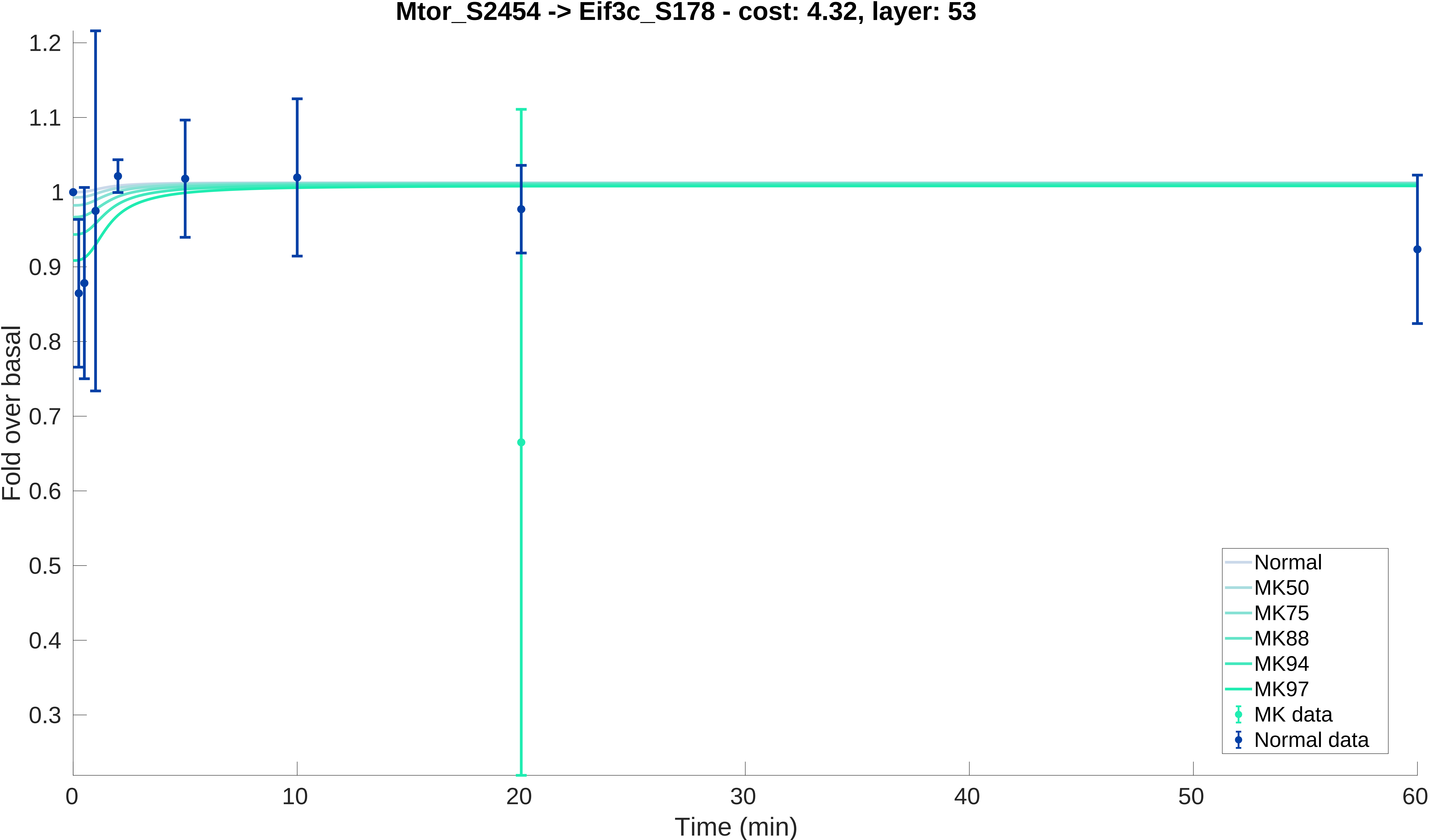

Mtor_S2454 -> Eif3c_S178 - cost: 4.32, layer: 53
Fold over basal
1.2
1.1
1
0.9
0.8
0.7
0.6
0.5
0.4
0.3
0
10
20
30
40
50
60
Time (min)
Normal
MK50
MK75
MK88
MK94
MK97
MK data
Normal data

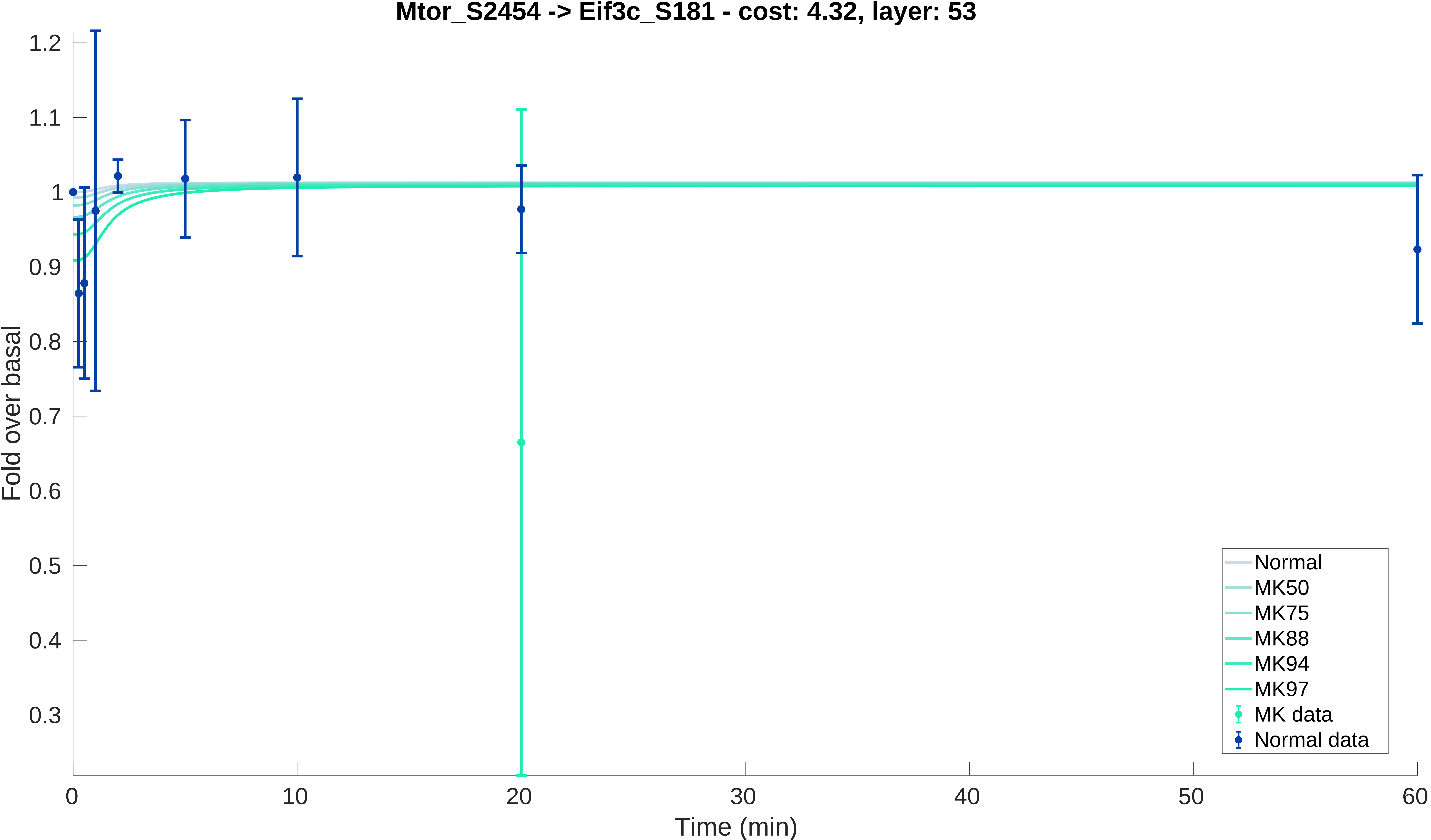

Mtor_S2454 -> Eif3c_S181 - cost: 4.32, layer: 53
Fold over basal
Time (min)
1.2
1.1
1
0.9
0.8
0.7
0.6
0.5
0.4
0.3
0
10
20
30
40
50
60
Normal
MK50
MK75
MK88
MK94
MK97
MK data
Normal data

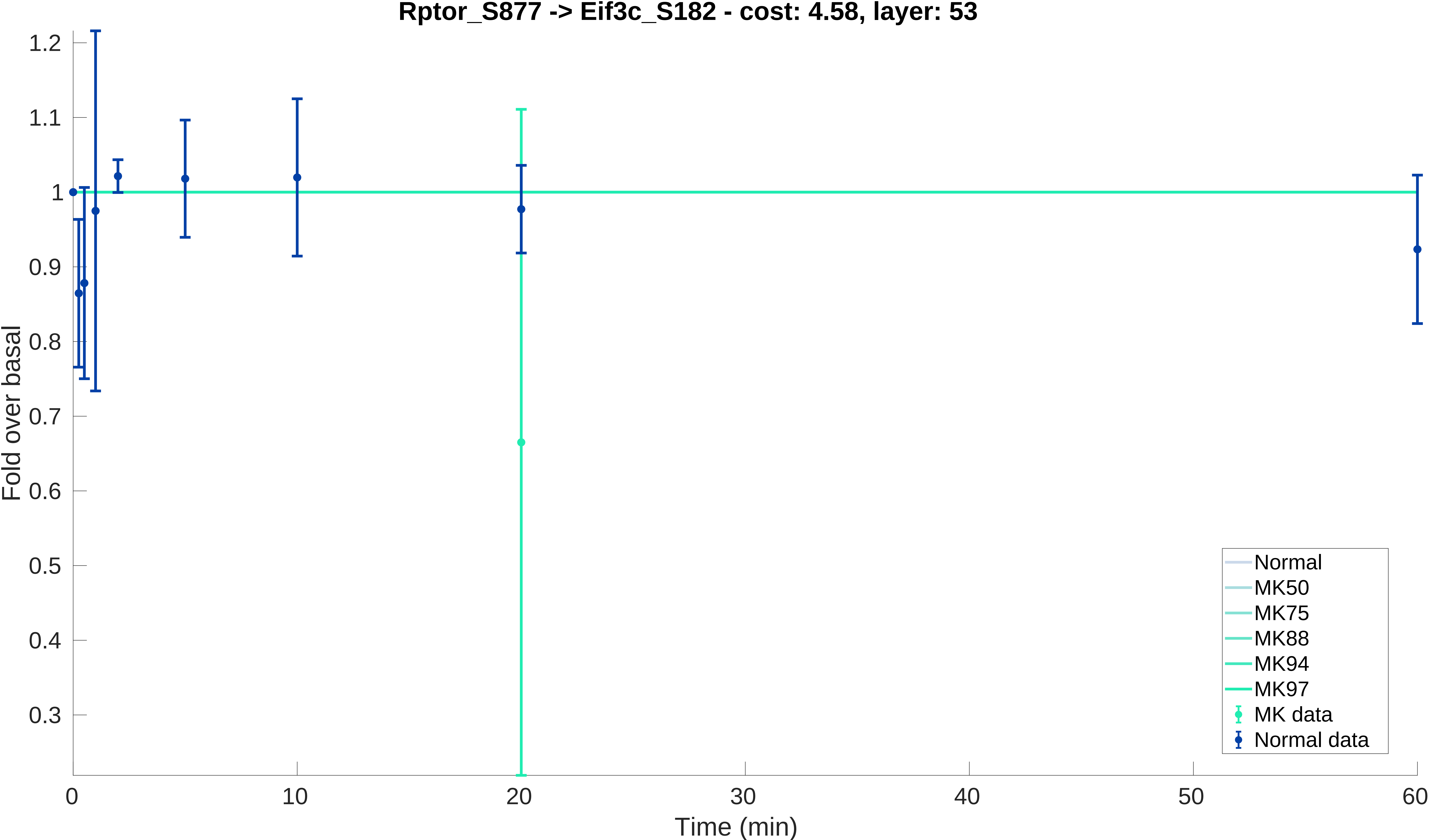
Rptor_S877 -> Eif3c_S182 - cost: 4.58, layer: 53
Fold over basal
1.2
1.1
1
0.9
0.8
0.7
0.6
0.5
0.4
0.3
0
10
20
30
40
50
60
Time (min)
Normal
MK50
MK75
MK88
MK94
MK97
MK data
Normal data

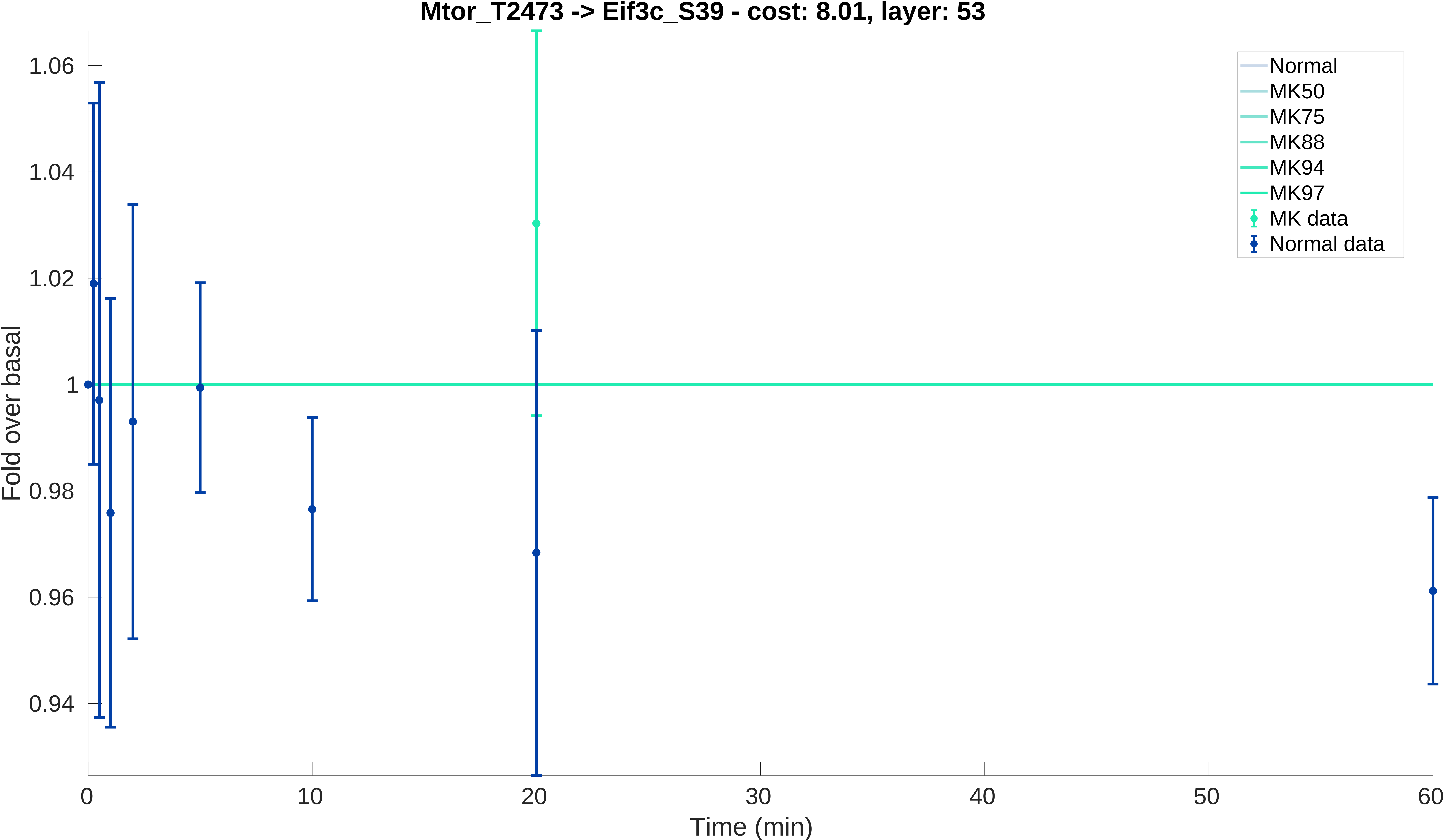
Mtor_T2473 -> Eif3c_S39 - cost: 8.01, layer: 53
Fold over basal
Time (min)
1.06
1.04
1.02
1
0.98
0.96
0.94
0
10
20
30
40
50
60
Normal
MK50
MK75
MK88
MK94
MK97
MK data
Normal data

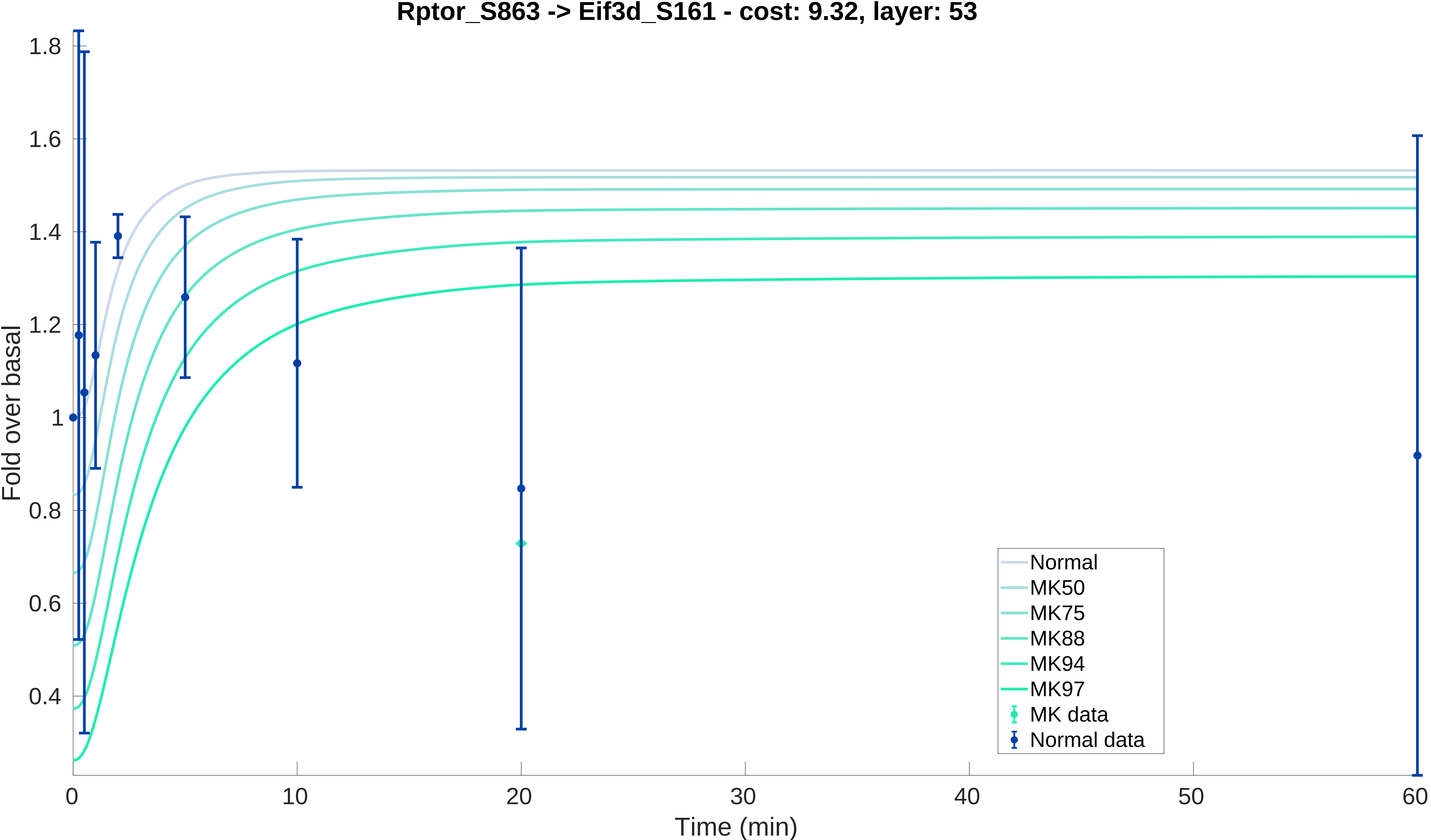

Rptor_S863 -> Eif3d_S161 - cost: 9.32, layer: 53
Fold over basal
1.8
1.6
1.4
1.2
1
0.8
0.6
0.4
0
10
20
30
40
50
60
Time (min)
Normal
MK50
MK75
MK88
MK94
MK97
MK data
Normal data

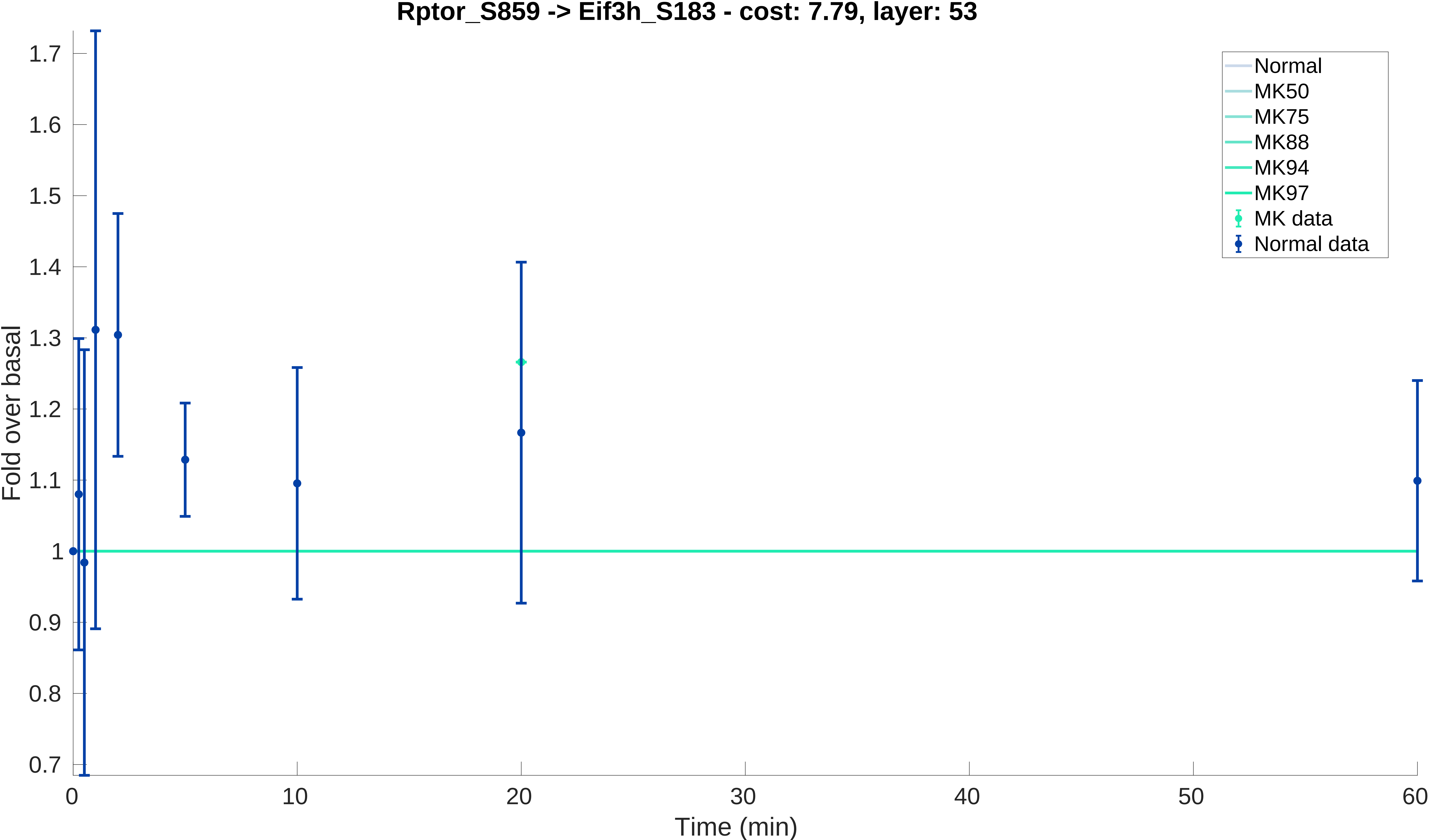
Rptor_S859 -> Eif3h_S183 - cost: 7.79, layer: 53
Fold over basal
Time (min)
0.7
0.8
0.9
1
1.1
1.2
1.3
1.4
1.5
1.6
1.7
0
10
20
30
40
50
60
Normal
MK50
MK75
MK88
MK94
MK97
MK data
Normal data

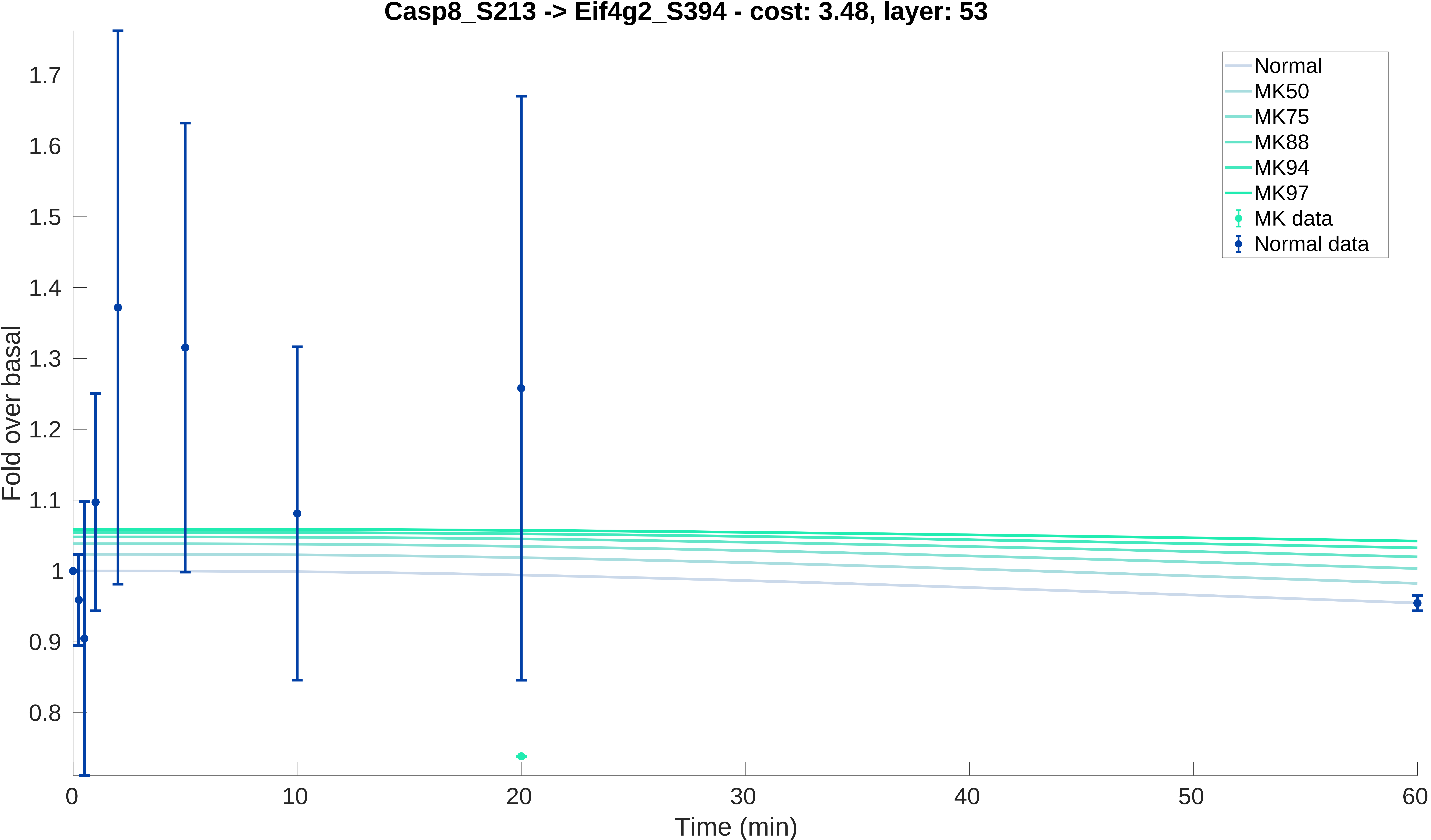

Casp8_S213 -> Eif4g2_S394 - cost: 3.48, layer: 53
Fold over basal
1.7
1.6
1.5
1.4
1.3
1.2
1.1
1
0.9
0.8
0
10
20
30
40
50
60
Time (min)
Normal
MK50
MK75
MK88
MK94
MK97
MK data
Normal data

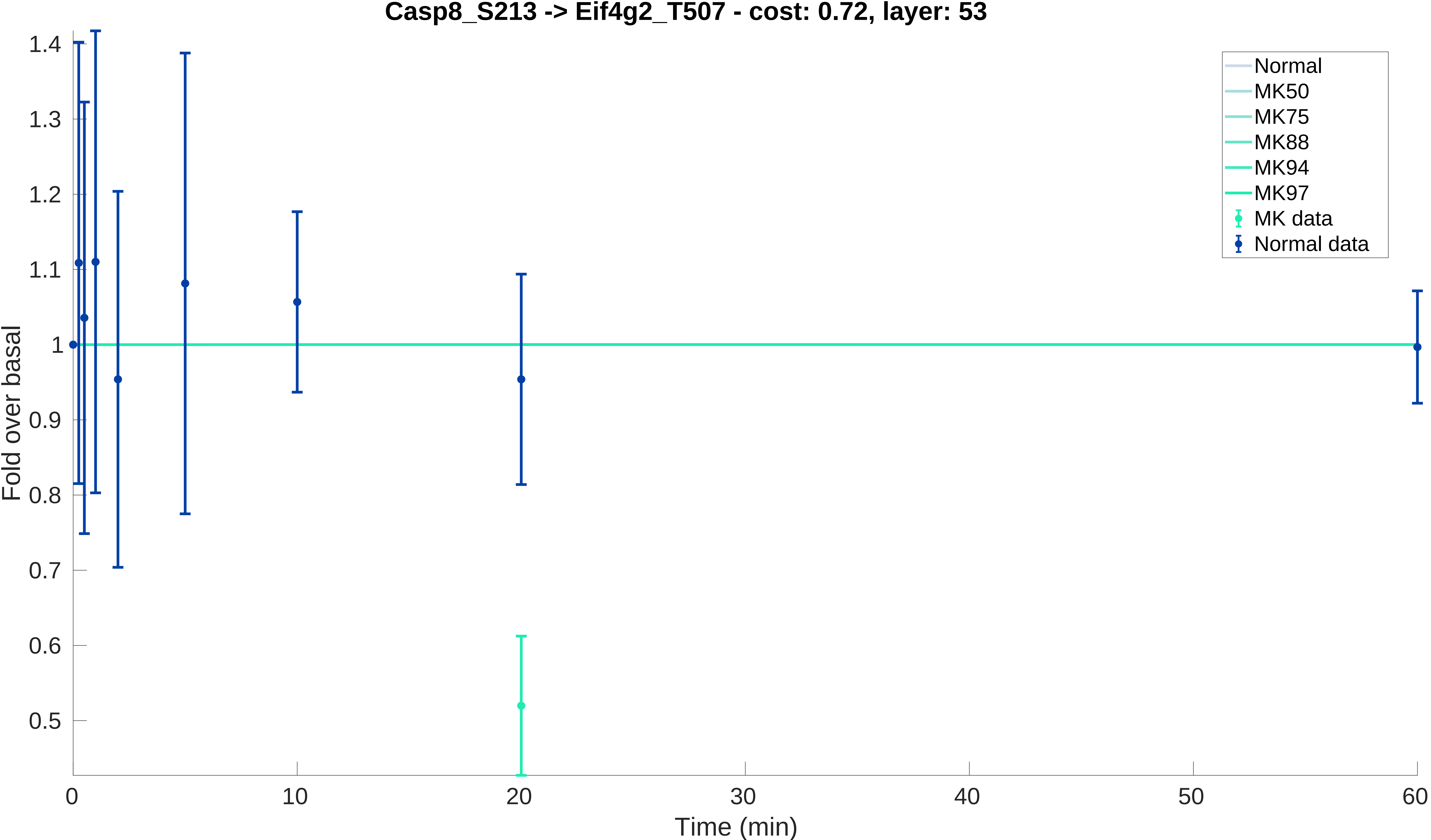

Casp8_S213 -> Eif4g2_T507 - cost: 0.72, layer: 53
Normal
MK50
MK75
MK88
MK94
MK97
MK data
Normal data
Fold over basal
Time (min)
0
10
20
30
40
50
60
0.5
0.6
0.7
0.8
0.9
1
1.1
1.2
1.3
1.4

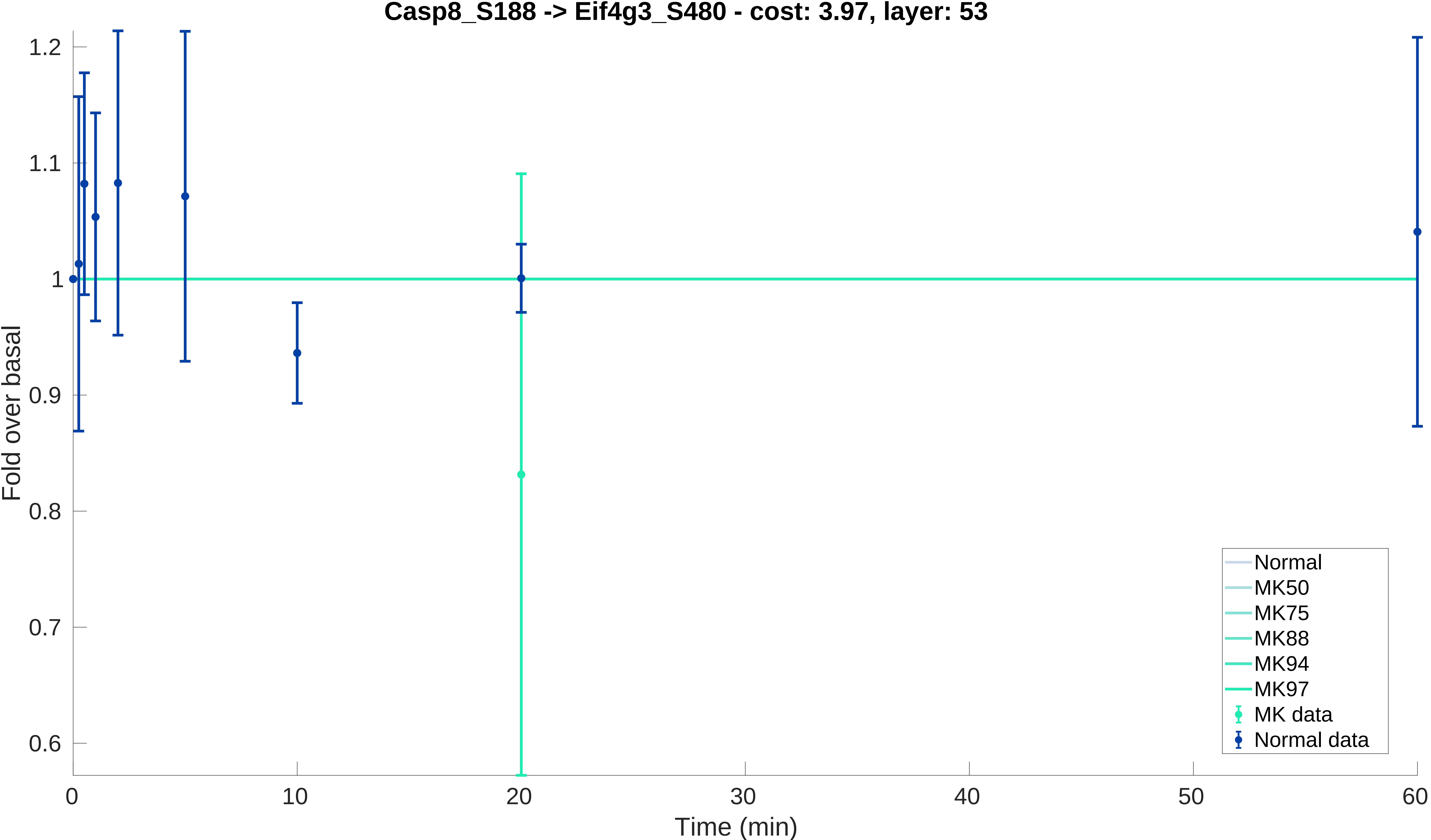

Casp8_S188 -> Eif4g3_S480 - cost: 3.97, layer: 53
Fold over basal
1.2
1.1
1
0.9
0.8
0.7
0.6
0
10
20
30
40
50
60
Time (min)
Normal
MK50
MK75
MK88
MK94
MK97
MK data
Normal data

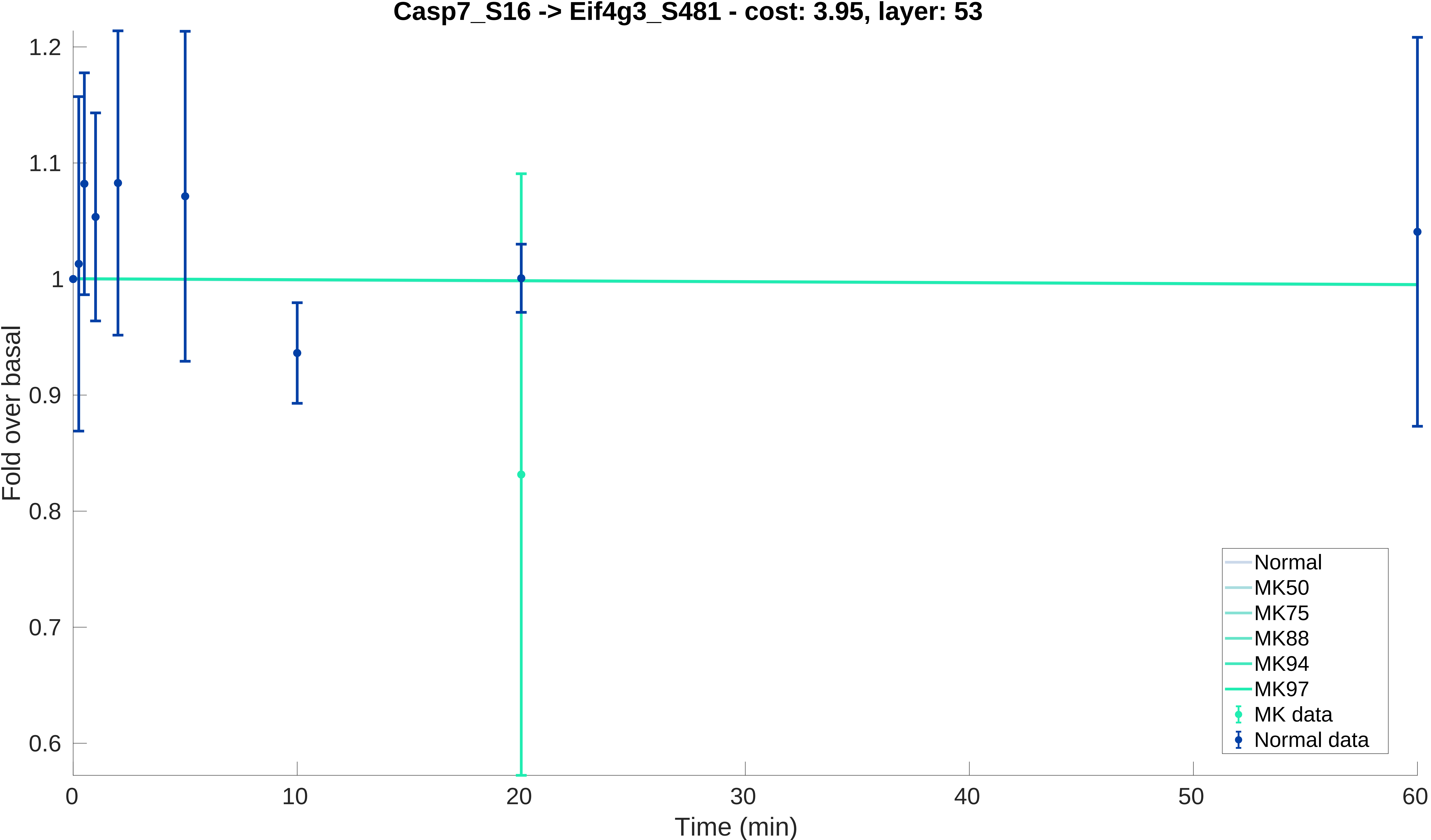
Casp7_S16 -> Eif4g3_S481 - cost: 3.95, layer: 53
Fold over basal
1.2
1.1
1
0.9
0.8
0.7
0.6
0
10
20
30
40
50
60
Time (min)
Normal
MK50
MK75
MK88
MK94
MK97
MK data
Normal data

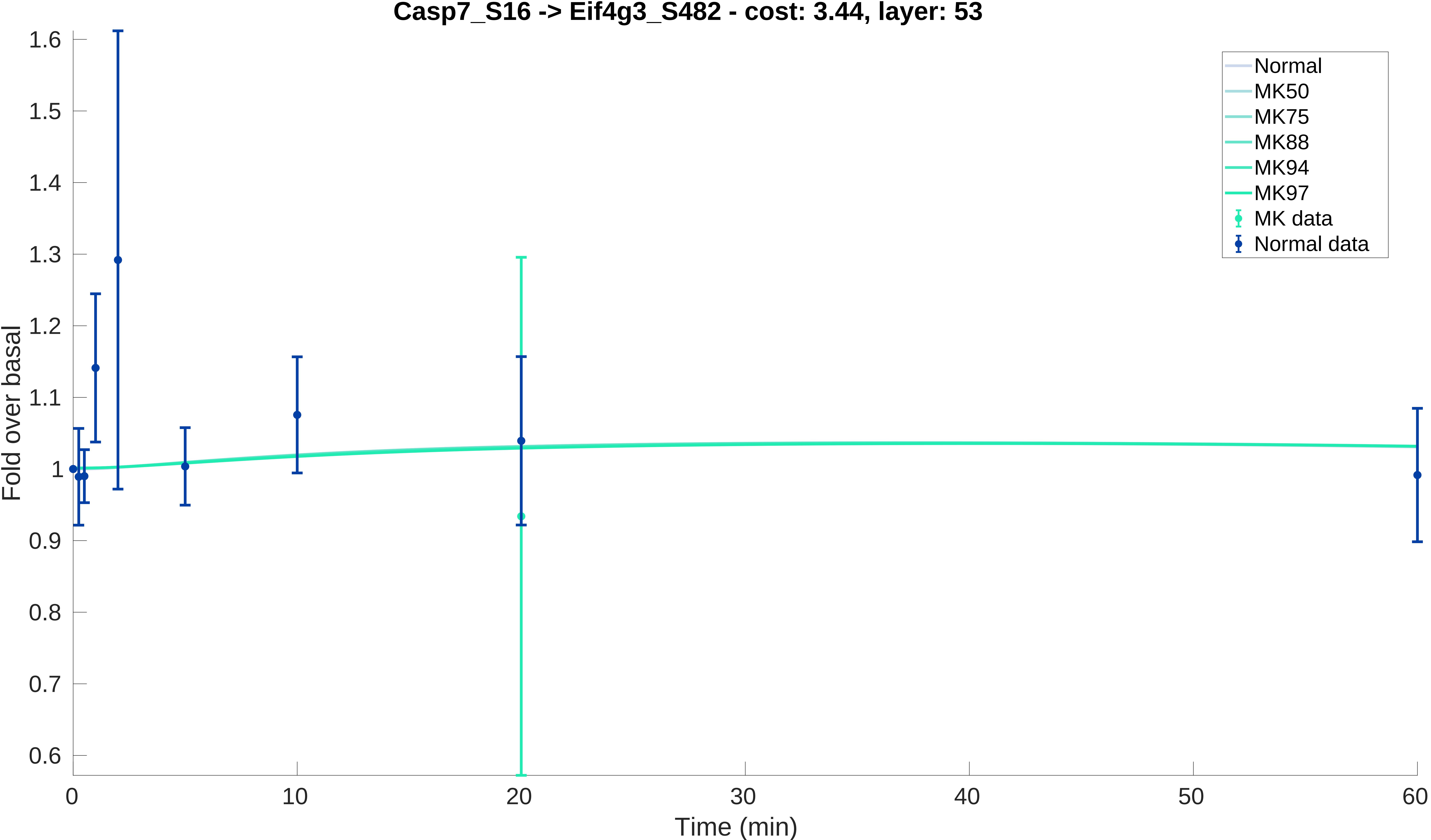

Casp7_S16 -> Eif4g3_S482 - cost: 3.44, layer: 53
Fold over basal
Time (min)
0.6
0.7
0.8
0.9
1
1.1
1.2
1.3
1.4
1.5
1.6
0
10
20
30
40
50
60
Normal
MK50
MK75
MK88
MK94
MK97
MK data
Normal data

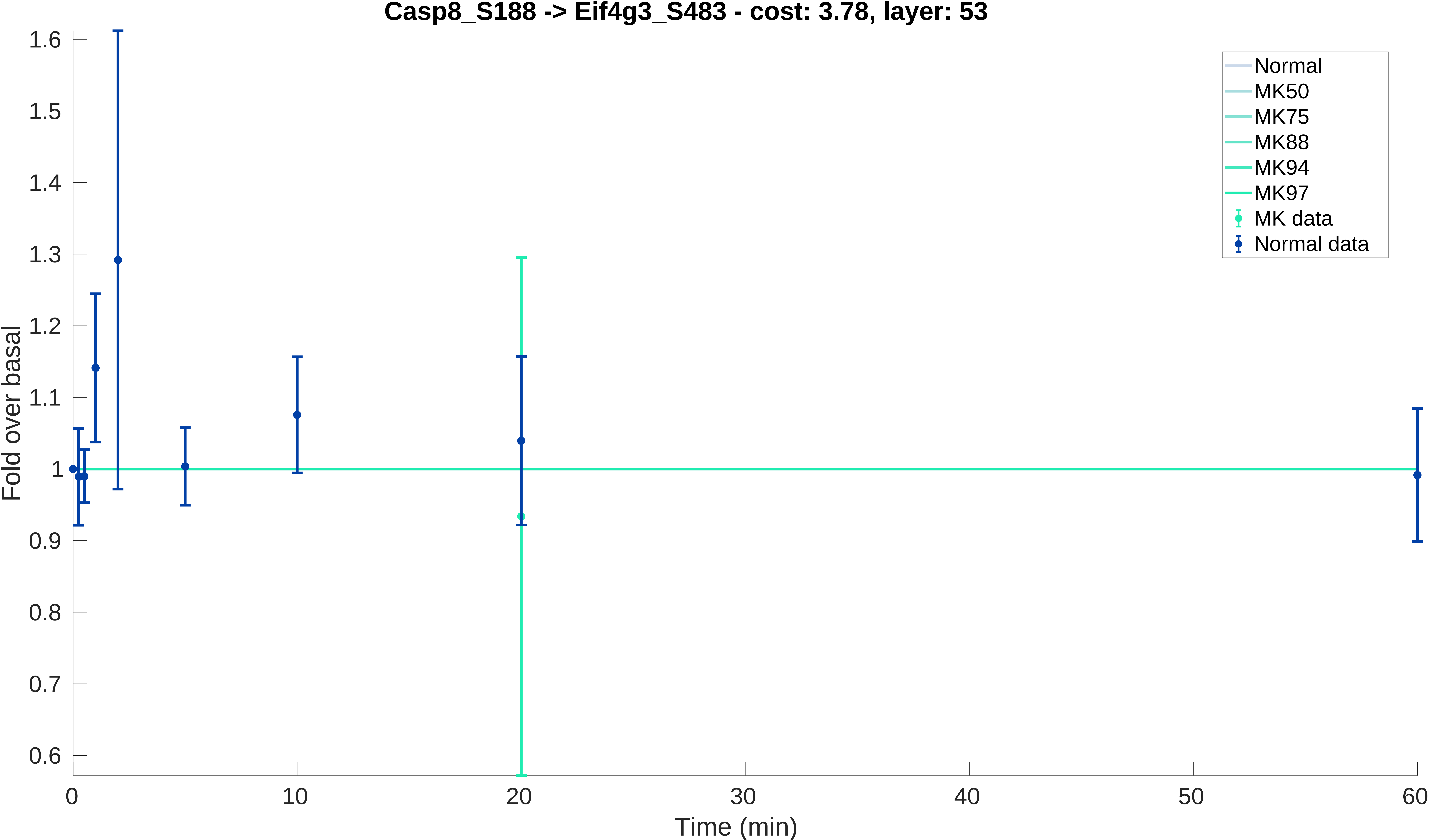

Casp8_S188 -> Eif4g3_S483 - cost: 3.78, layer: 53
Fold over basal
Time (min)
Normal
MK50
MK75
MK88
MK94
MK97
MK data
Normal data
1.6
1.5
1.4
1.3
1.2
1.1
1
0.9
0.8
0.7
0.6
0
10
20
30
40
50
60

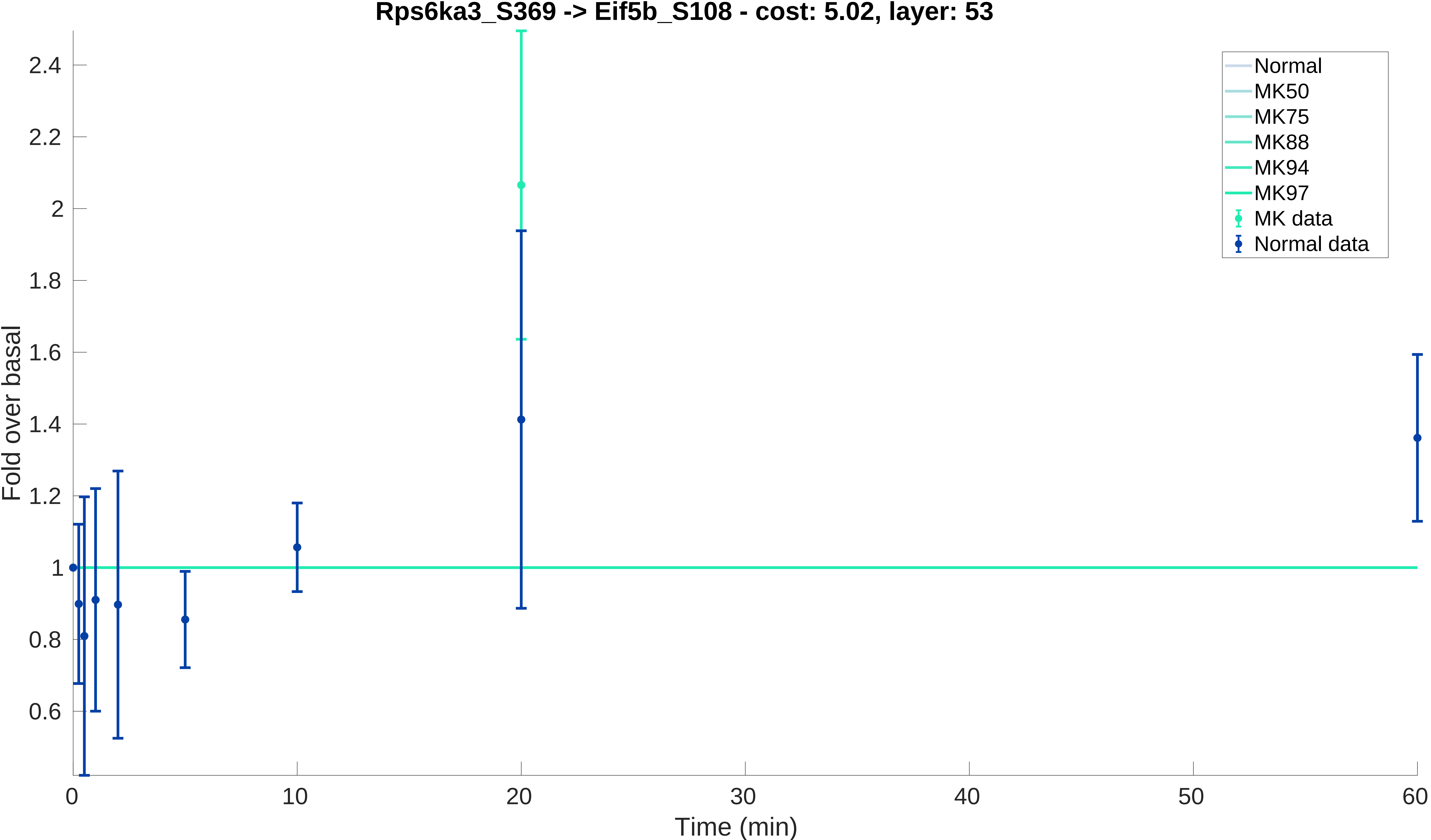

Rps6ka3_S369 -> Eif5b_S108 - cost: 5.02, layer: 53
Fold over basal
Time (min)
Normal
MK50
MK75
MK88
MK94
MK97
MK data
Normal data
2.4
2.2
2
1.8
1.6
1.4
1.2
1
0.8
0.6
0
10
20
30
40
50
60

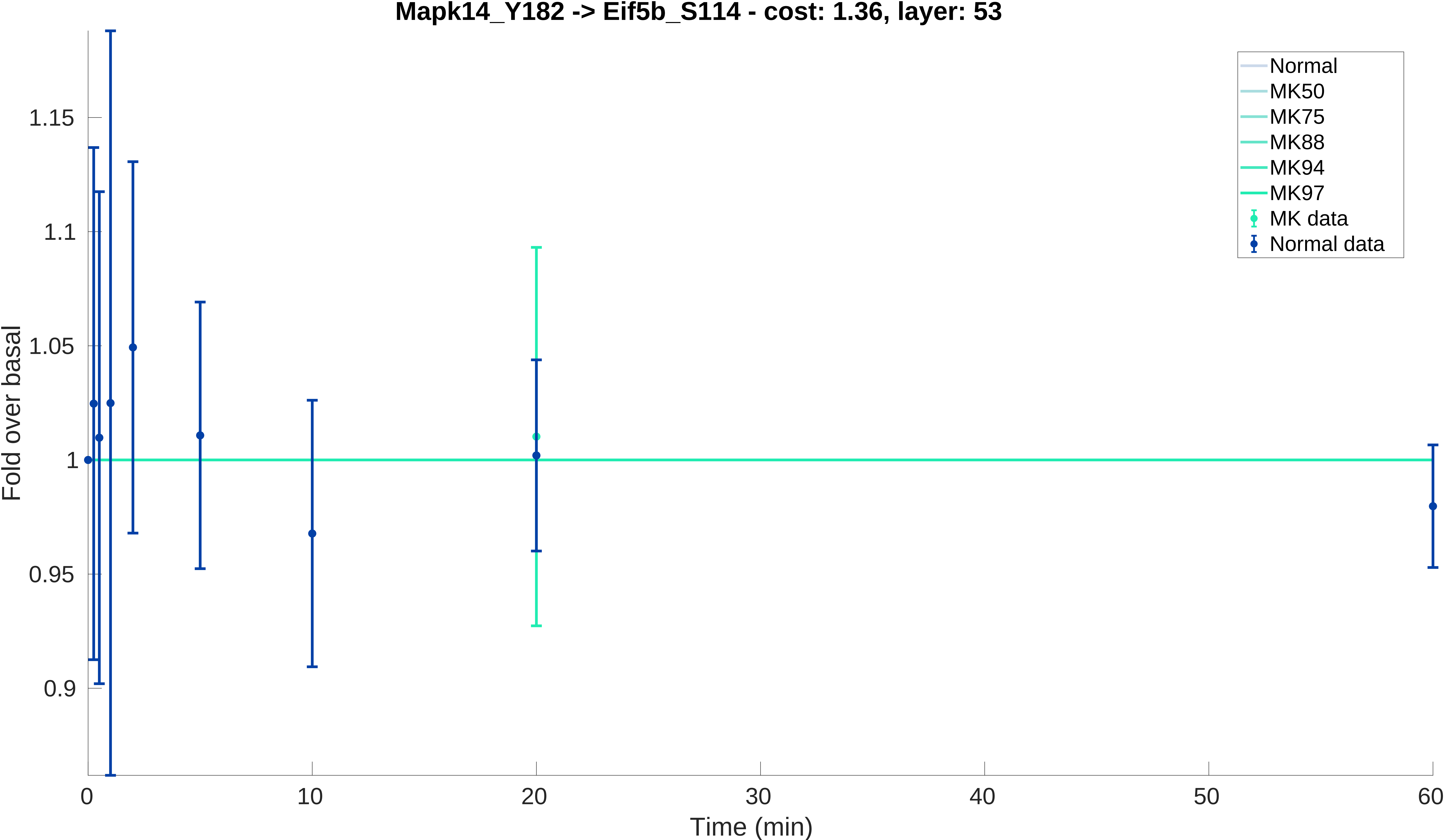

Mapk14_Y182 -> Eif5b_S114 - cost: 1.36, layer: 53
Normal
MK50
MK75
MK88
MK94
MK97
MK data
Normal data
Fold over basal
1.15
1.1
1.05
1
0.95
0.9
0
10
20
30
40
50
60
Time (min)

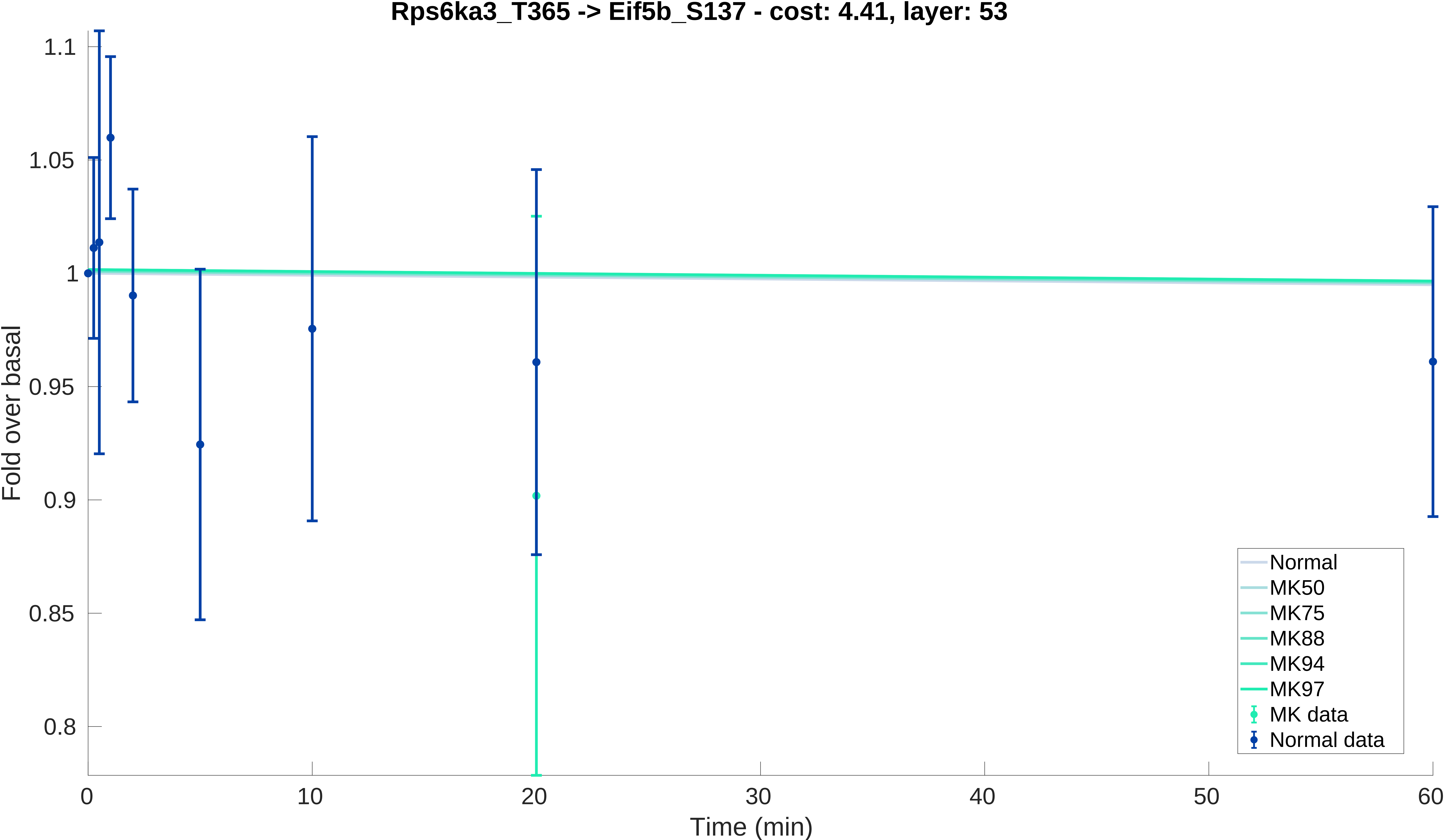
Rps6ka3_T365 -> Eif5b_S137 - cost: 4.41, layer: 53
Fold over basal
1.1
1.05
1
0.95
0.9
0.85
0.8
0
10
20
30
40
50
60
Time (min)
Normal
MK50
MK75
MK88
MK94
MK97
MK data
Normal data

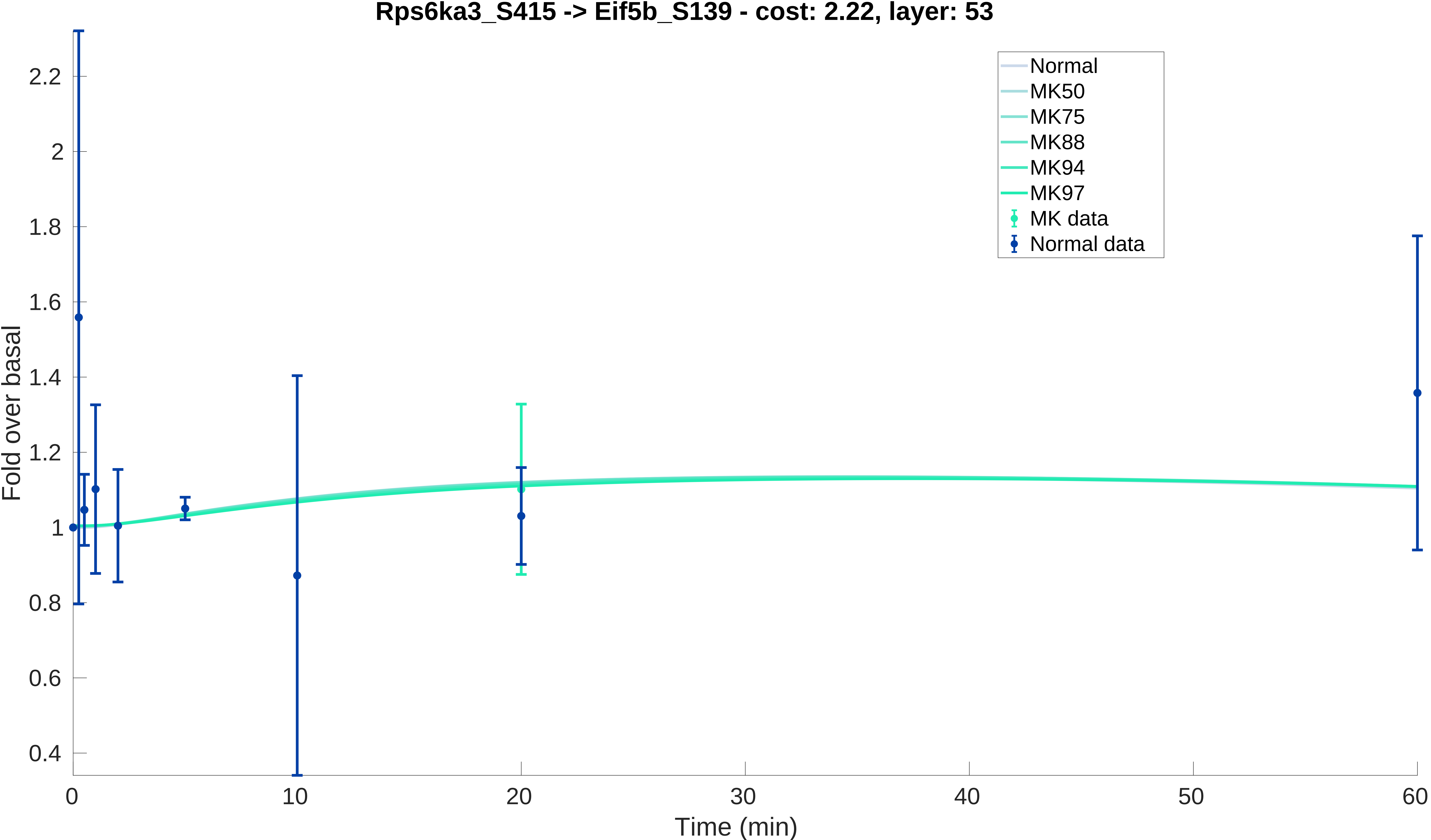

Rps6ka3_S415 -> Eif5b_S139 - cost: 2.22, layer: 53
Fold over basal
Time (min)
Normal
MK50
MK75
MK88
MK94
MK97
MK data
Normal data
0.4
0.6
0.8
1
1.2
1.4
1.6
1.8
2
2.2
0
10
20
30
40
50
60

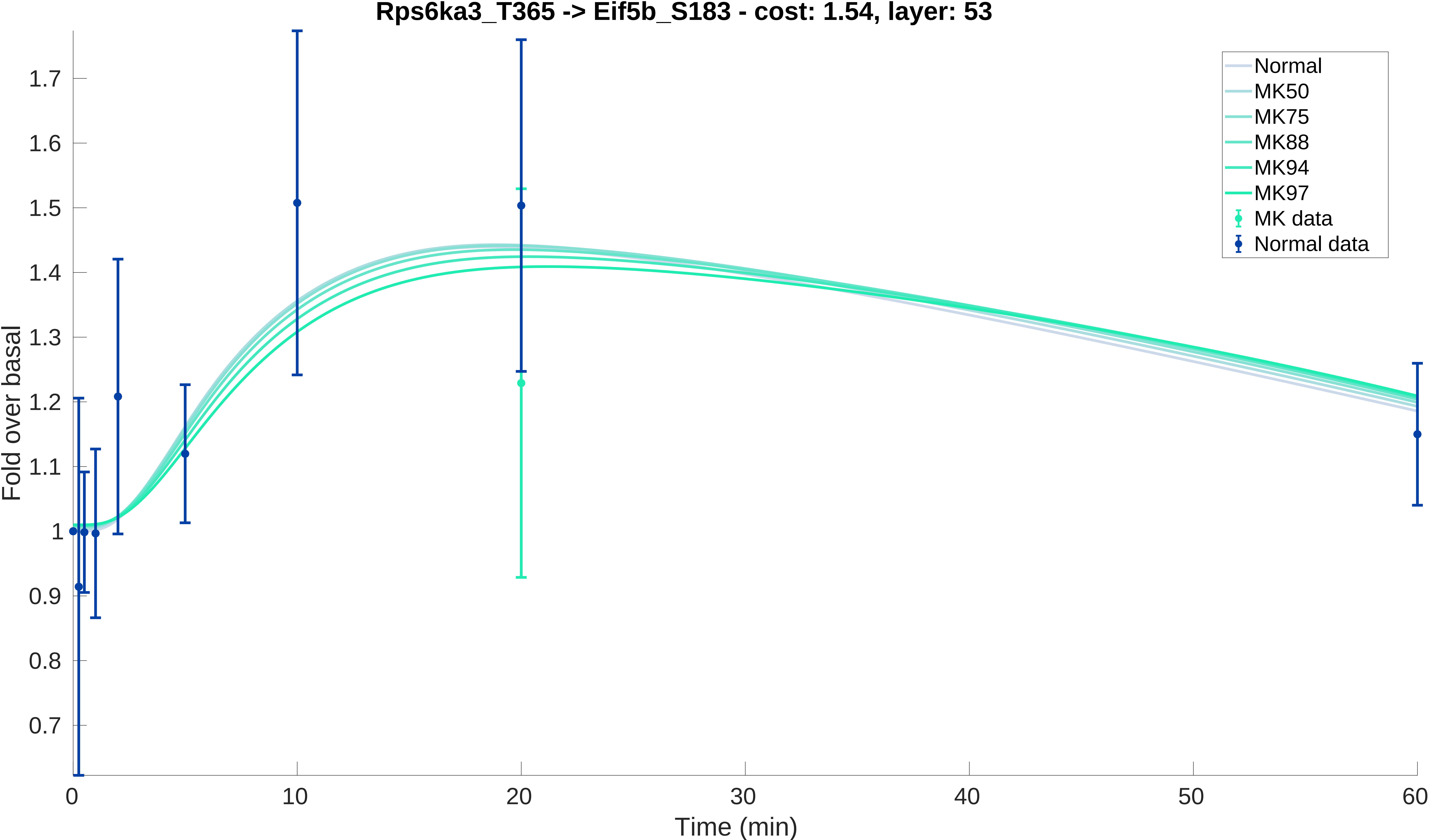
Rps6ka3_T365 -> Eif5b_S183 - cost: 1.54, layer: 53
Normal
MK50
MK75
MK88
MK94
MK97
MK data
Normal data
Fold over basal
1.7
1.6
1.5
1.4
1.3
1.2
1.1
1
0.9
0.8
0.7
0
10
20
30
40
50
60
Time (min)

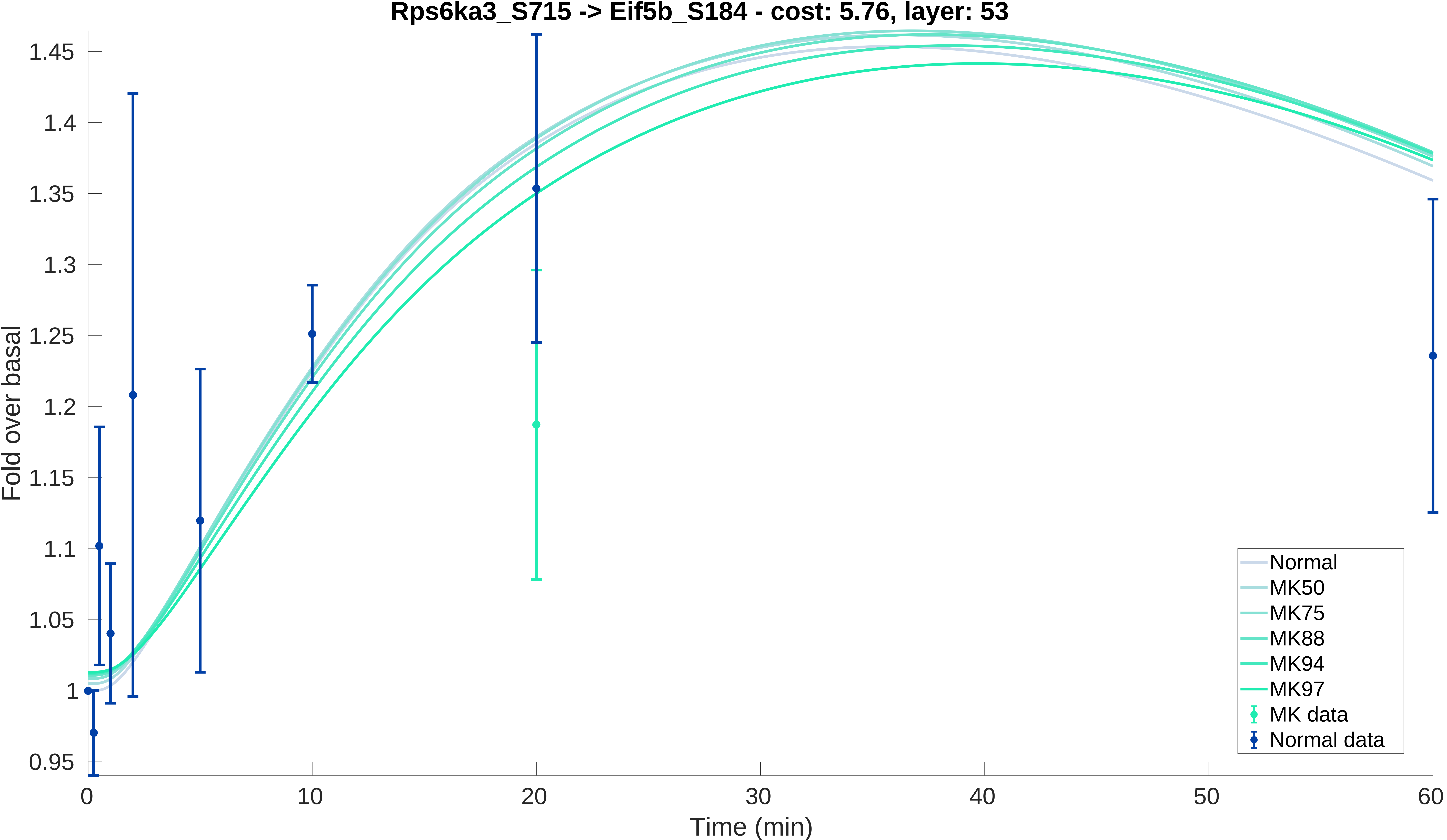

Rps6ka3_S715 -> Eif5b_S184 - cost: 5.76, layer: 53
Fold over basal
1.45
1.4
1.35
1.3
1.25
1.2
1.15
1.1
1.05
1
0.95
0
10
20
30
40
50
60
Time (min)
Normal
MK50
MK75
MK88
MK94
MK97
MK data
Normal data

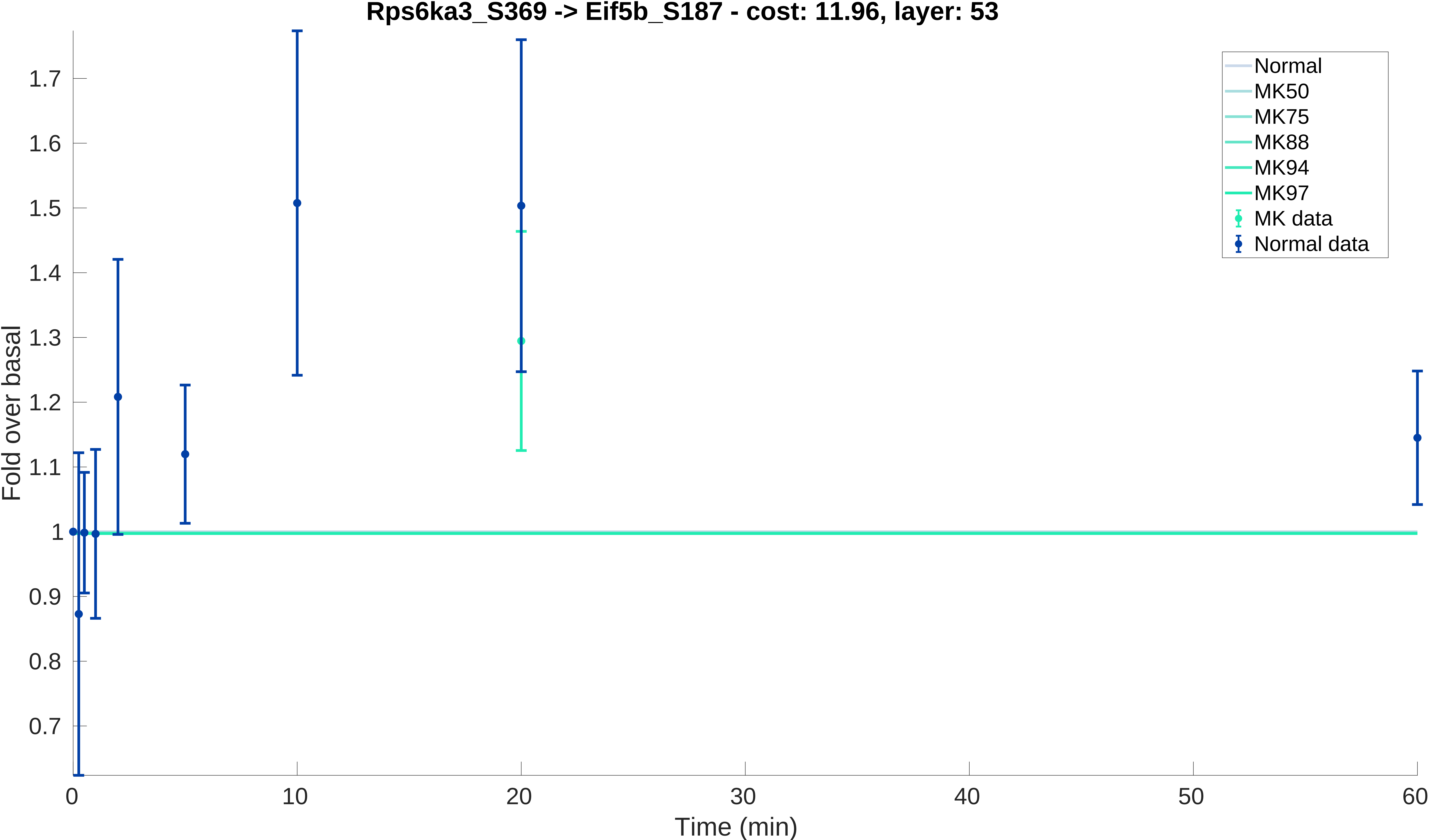

Rps6ka3_S369 -> Eif5b_S187 - cost: 11.96, layer: 53
Fold over basal
Time (min)
0.7
0.8
0.9
1
1.1
1.2
1.3
1.4
1.5
1.6
1.7
0
10
20
30
40
50
60
Normal
MK50
MK75
MK88
MK94
MK97
MK data
Normal data

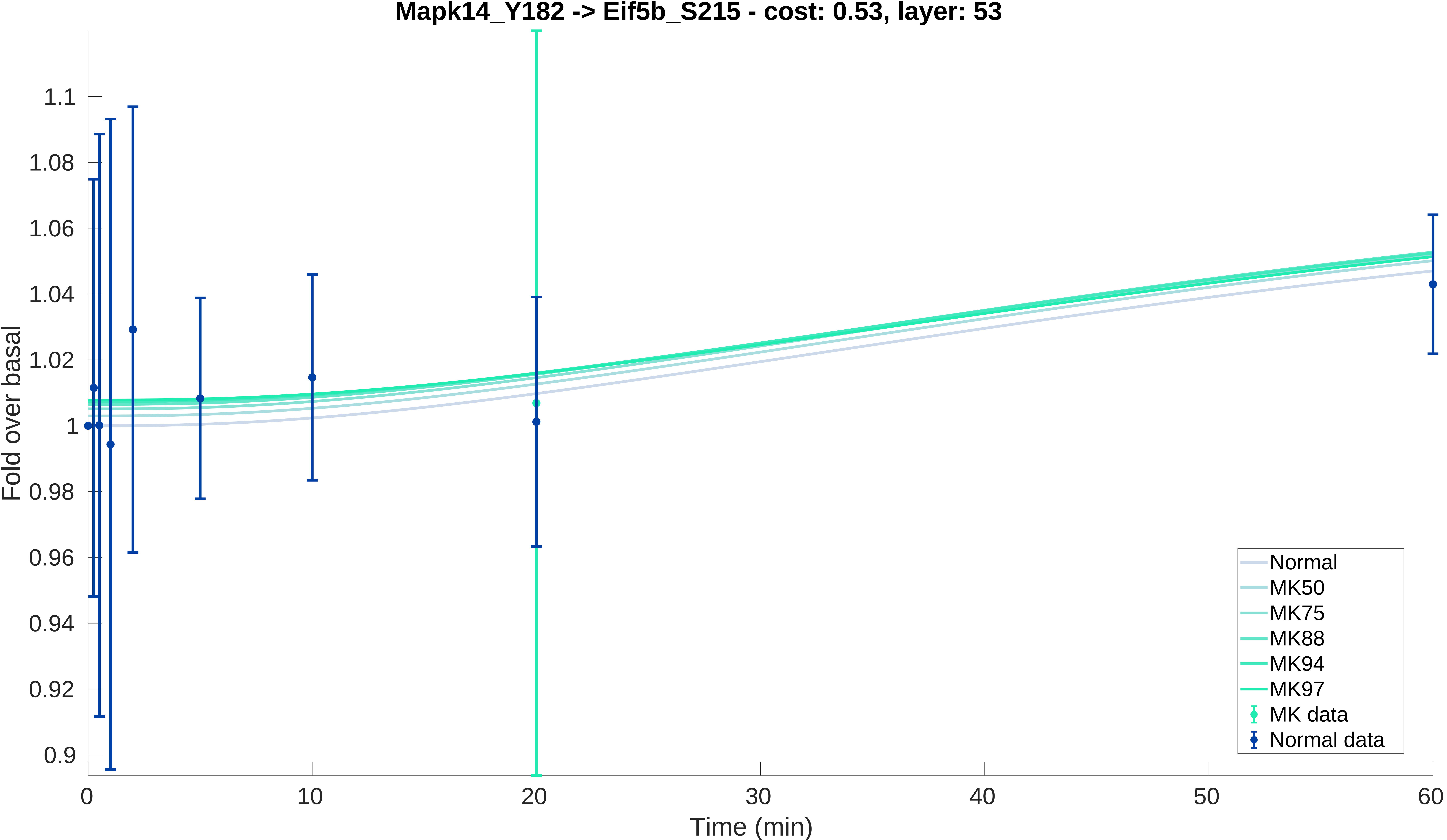

Mapk14_Y182 -> Eif5b_S215 - cost: 0.53, layer: 53
Fold over basal
1.1
1.08
1.06
1.04
1.02
1
0.98
0.96
0.94
0.92
0.9
0
10
20
30
40
50
60
Time (min)
Normal
MK50
MK75
MK88
MK94
MK97
MK data
Normal data

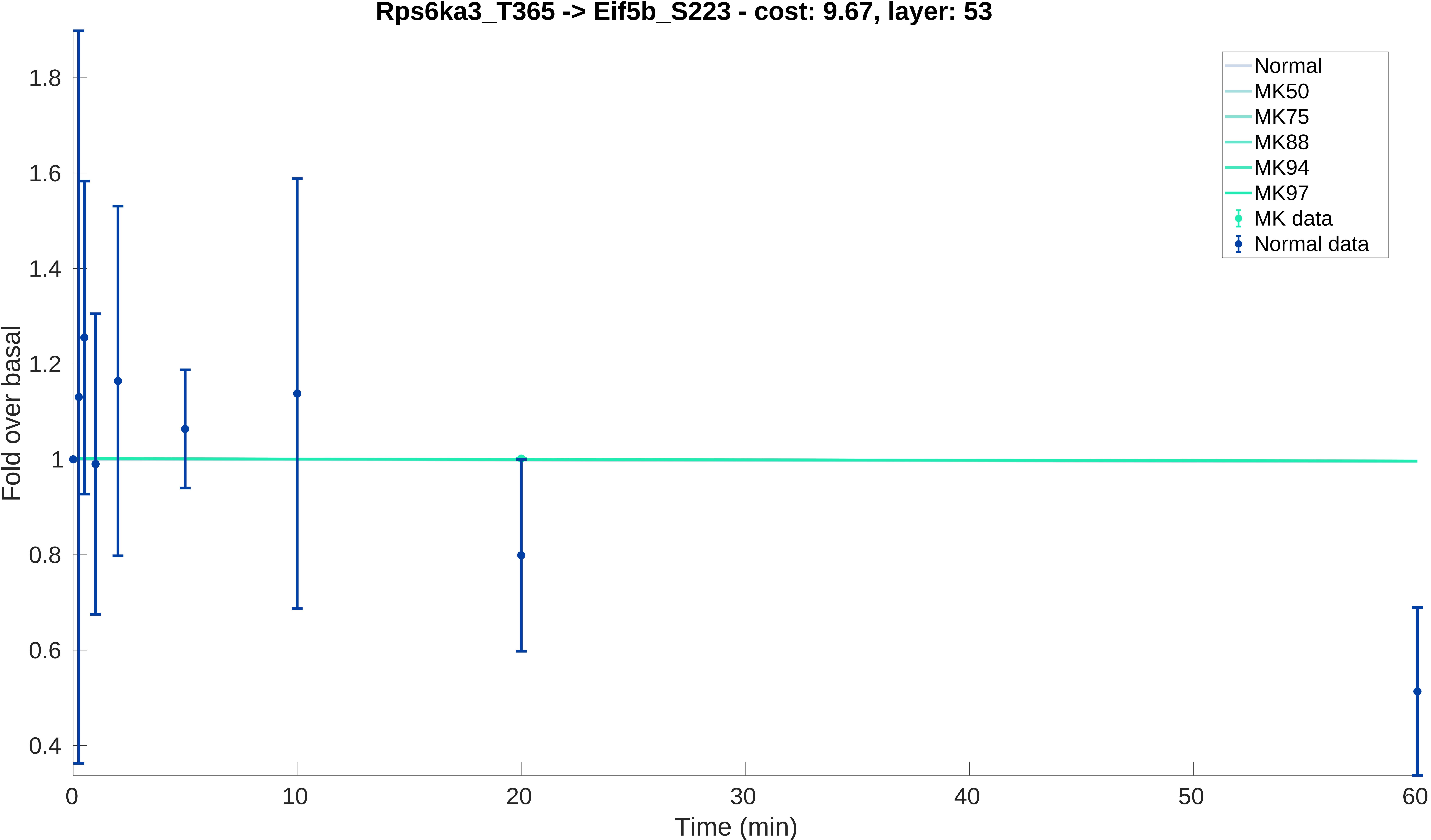

Rps6ka3_T365 -> Eif5b_S223 - cost: 9.67, layer: 53
Fold over basal
Time (min)
1.8
1.6
1.4
1.2
1
0.8
0.6
0.4
0
10
20
30
40
50
60
Normal
MK50
MK75
MK88
MK94
MK97
MK data
Normal data

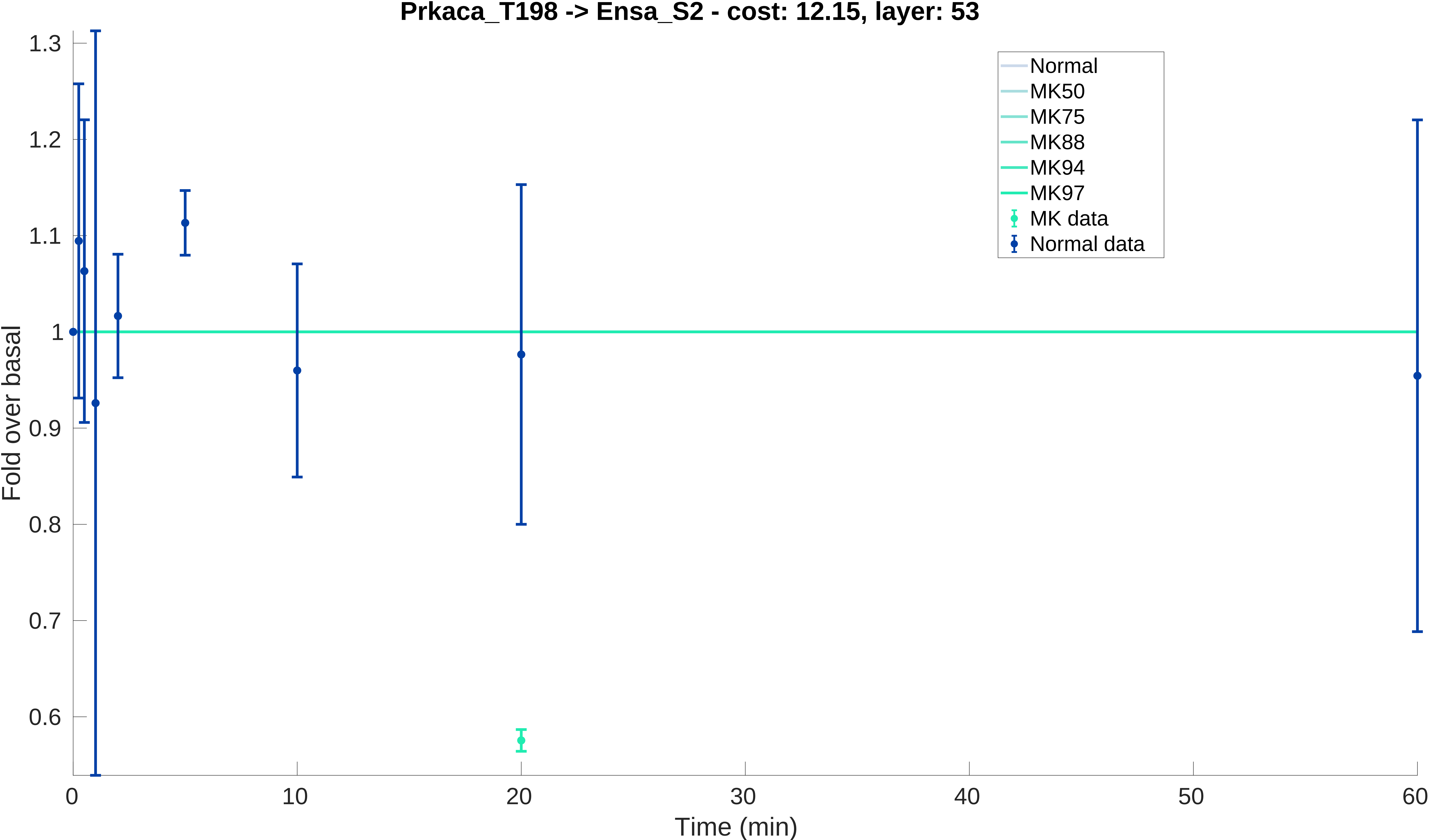

Prkaca_T198 -> Ensa_S2 - cost: 12.15, layer: 53
Fold over basal
Time (min)
Normal
MK50
MK75
MK88
MK94
MK97
MK data
Normal data
1.3
1.2
1.1
1
0.9
0.8
0.7
0.6
0
10
20
30
40
50
60

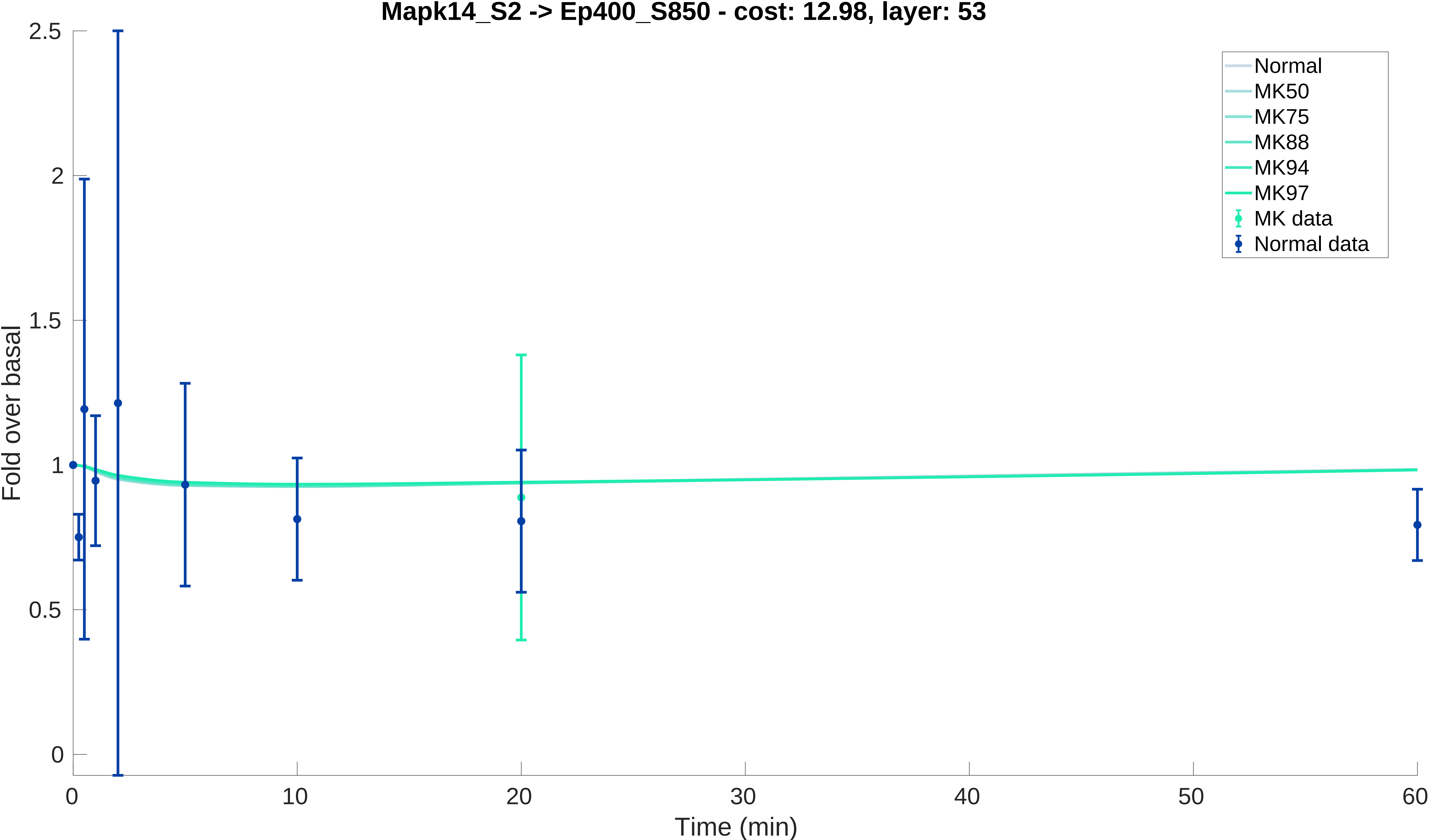
Mapk14_S2 -> Ep400_S850 - cost: 12.98, layer: 53
Fold over basal
Time (min)
0
0.5
1
1.5
2
2.5
0
10
20
30
40
50
60
Normal
MK50
MK75
MK88
MK94
MK97
MK data
Normal data

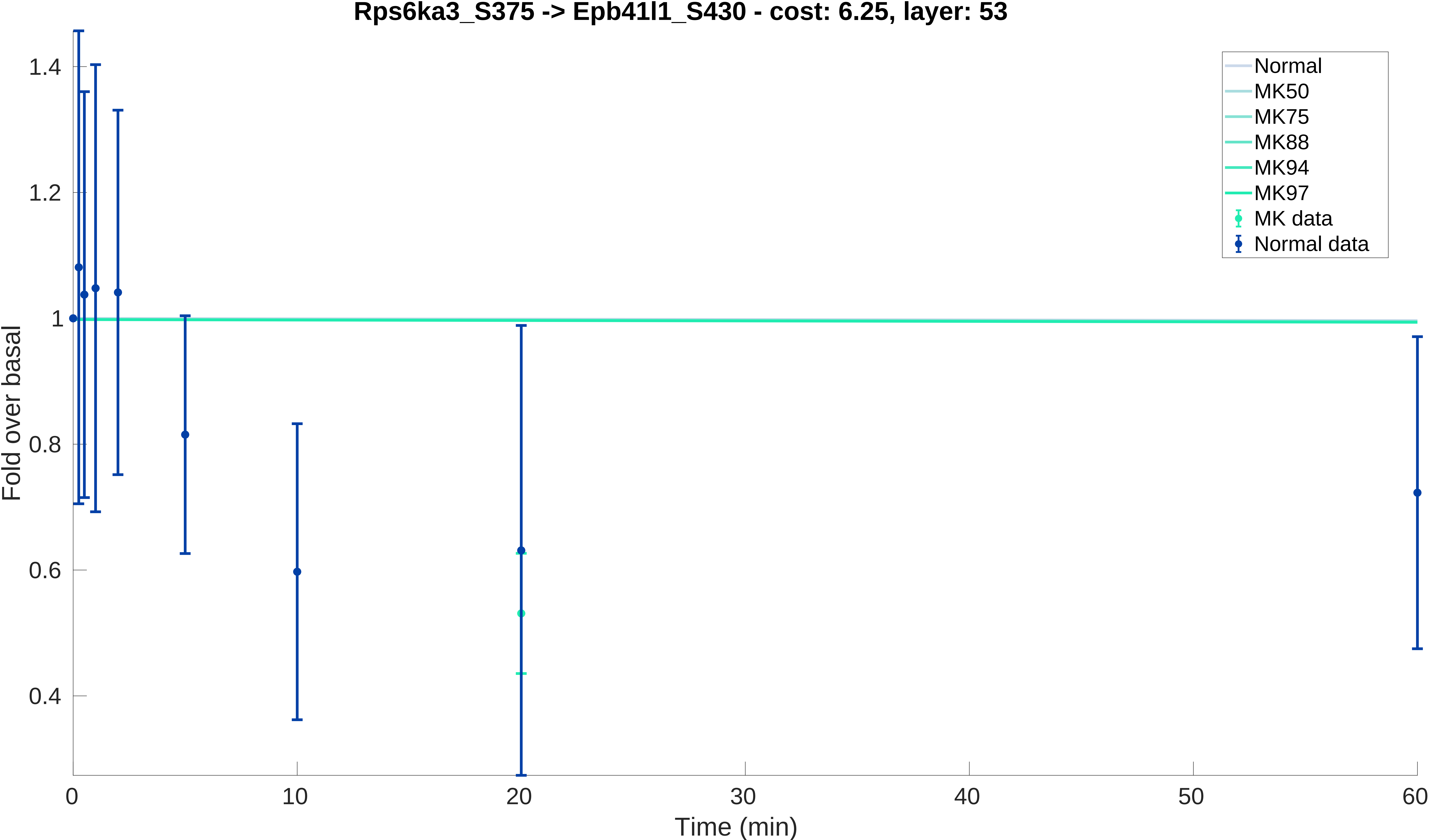

Rps6ka3_S375 -> Epb41l1_S430 - cost: 6.25, layer: 53
Fold over basal
Time (min)
1.4
1.2
1
0.8
0.6
0.4
0
10
20
30
40
50
60
Normal
MK50
MK75
MK88
MK94
MK97
MK data
Normal data

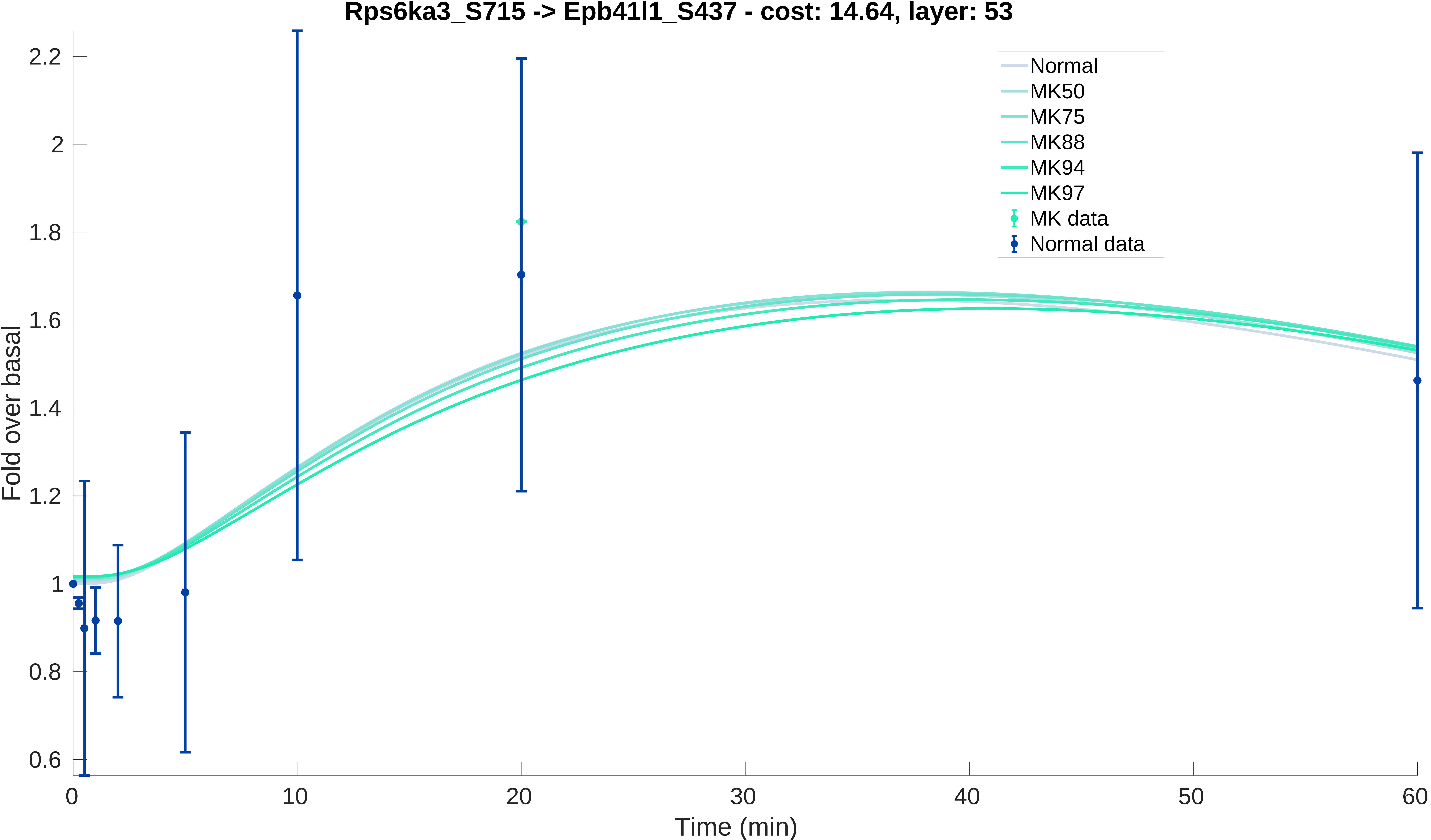
Rps6ka3_S715 -> Epb41l1_S437 - cost: 14.64, layer: 53
Normal
MK50
MK75
MK88
MK94
MK97
MK data
Normal data
Fold over basal
Time (min)
0.6
0.8
1
1.2
1.4
1.6
1.8
2
2.2
0
10
20
30
40
50
60

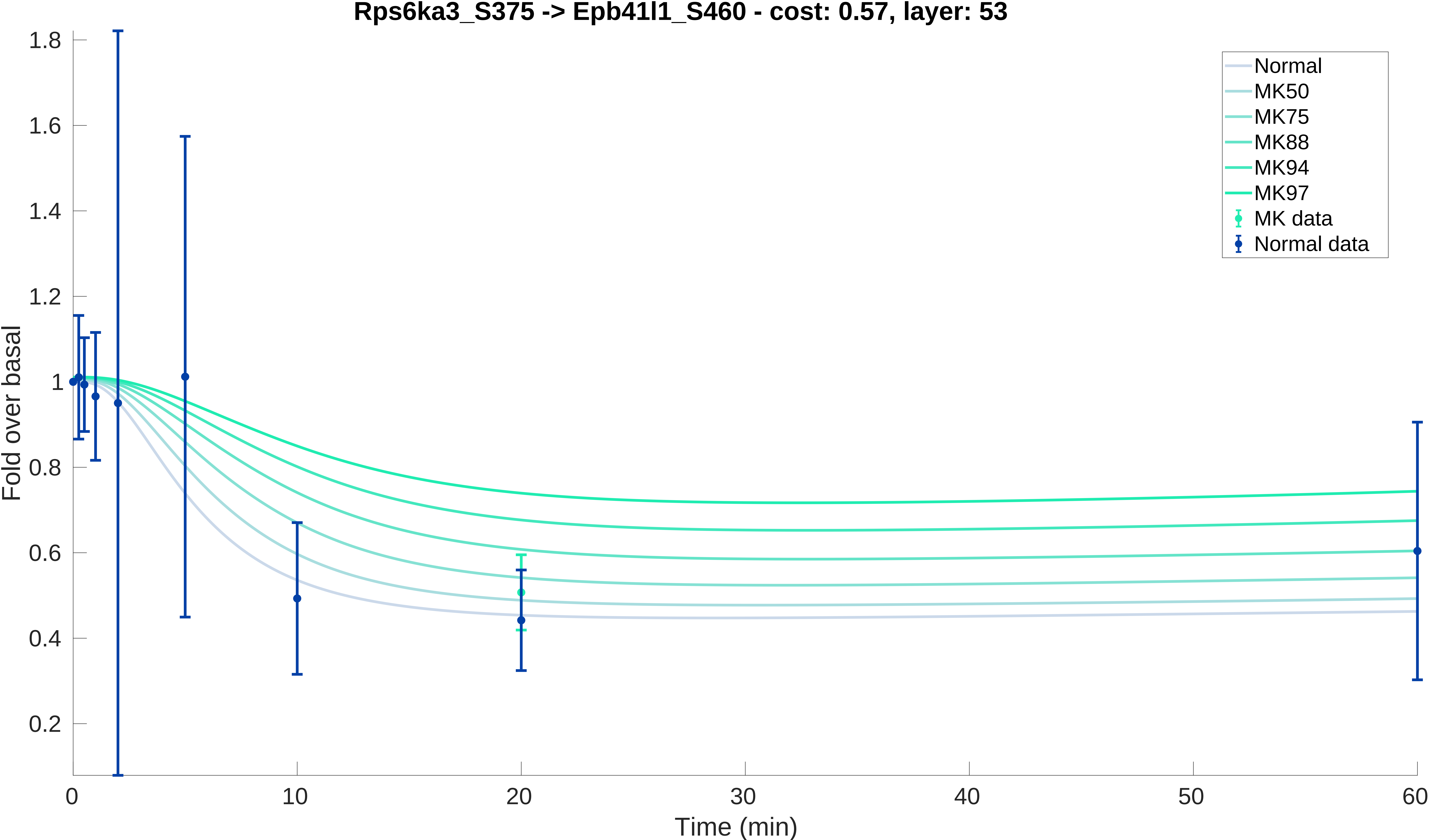
Rps6ka3_S375 -> Epb41l1_S460 - cost: 0.57, layer: 53
Fold over basal
Time (min)
0
10
20
30
40
50
60
0.2
0.4
0.6
0.8
1
1.2
1.4
1.6
1.8
Normal
MK50
MK75
MK88
MK94
MK97
MK data
Normal data

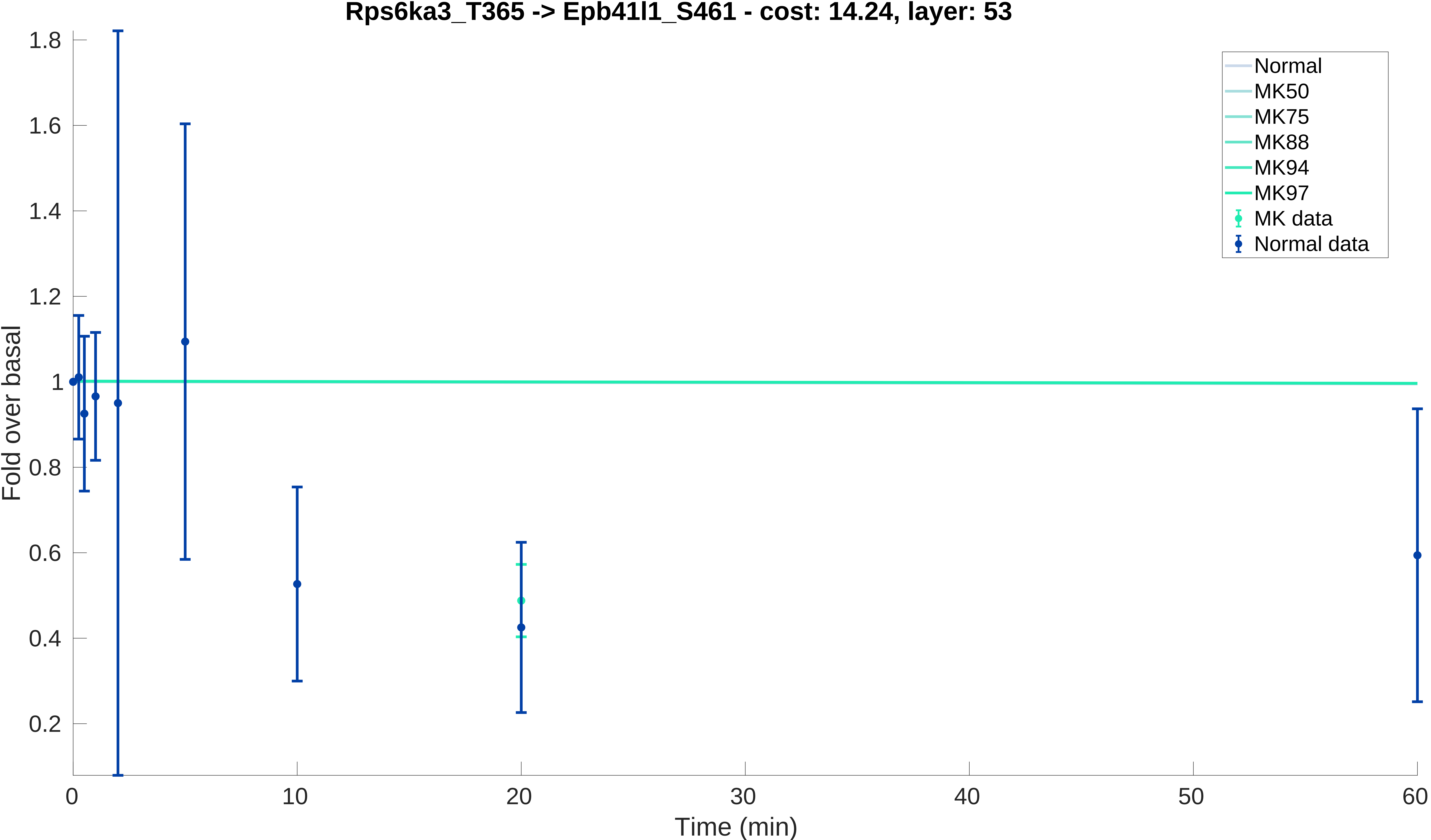

Rps6ka3_T365 -> Epb41l1_S461 - cost: 14.24, layer: 53
Fold over basal
Time (min)
Normal
MK50
MK75
MK88
MK94
MK97
MK data
Normal data
1.8
1.6
1.4
1.2
1
0.8
0.6
0.4
0.2
0
10
20
30
40
50
60

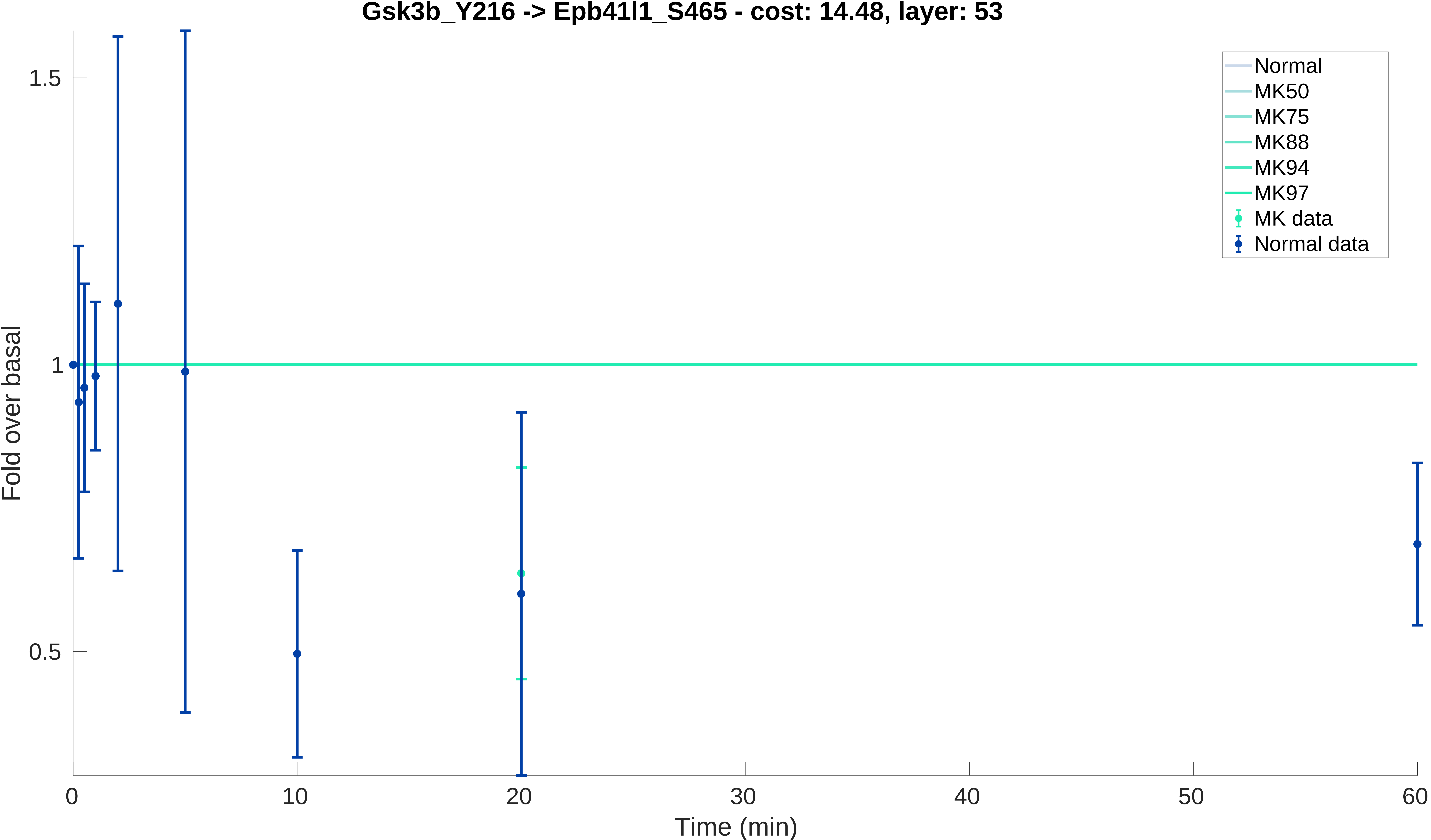

Gsk3b_Y216 -> Epb41l1_S465 - cost: 14.48, layer: 53
Fold over basal
1.5
1
0.5
Time (min)
0
10
20
30
40
50
60
Normal
MK50
MK75
MK88
MK94
MK97
MK data
Normal data

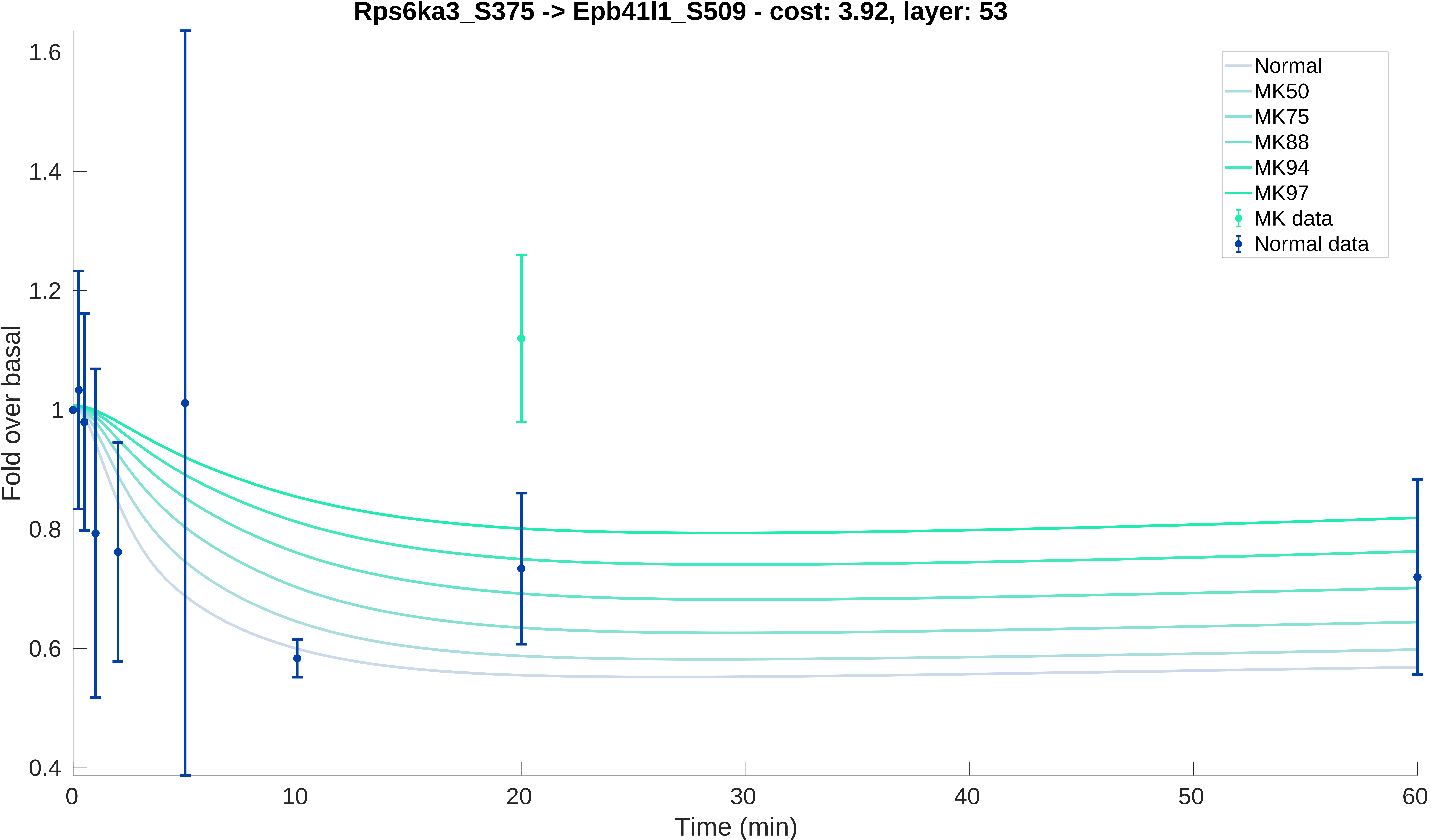

Rps6ka3_S375 -> Epb41l1_S509 - cost: 3.92, layer: 53
Normal
MK50
MK75
MK88
MK94
MK97
MK data
Normal data
Fold over basal
Time (min)
1.6
1.4
1.2
1
0.8
0.6
0.4
0
10
20
30
40
50
60

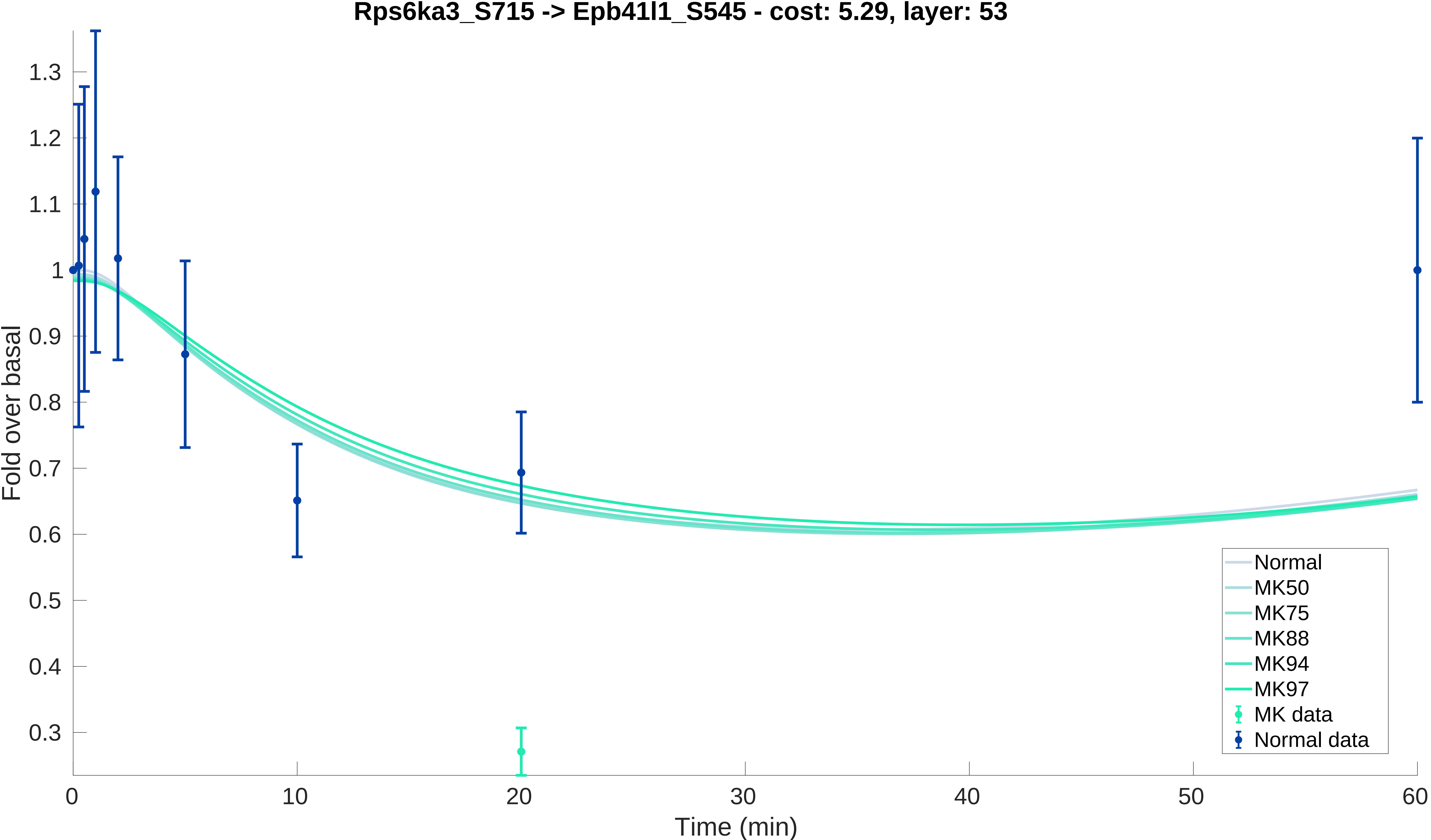

Rps6ka3_S715 -> Epb41l1_S545 - cost: 5.29, layer: 53
Fold over basal
Time (min)
1.3
1.2
1.1
1
0.9
0.8
0.7
0.6
0.5
0.4
0.3
0
10
20
30
40
50
60
Normal
MK50
MK75
MK88
MK94
MK97
MK data
Normal data

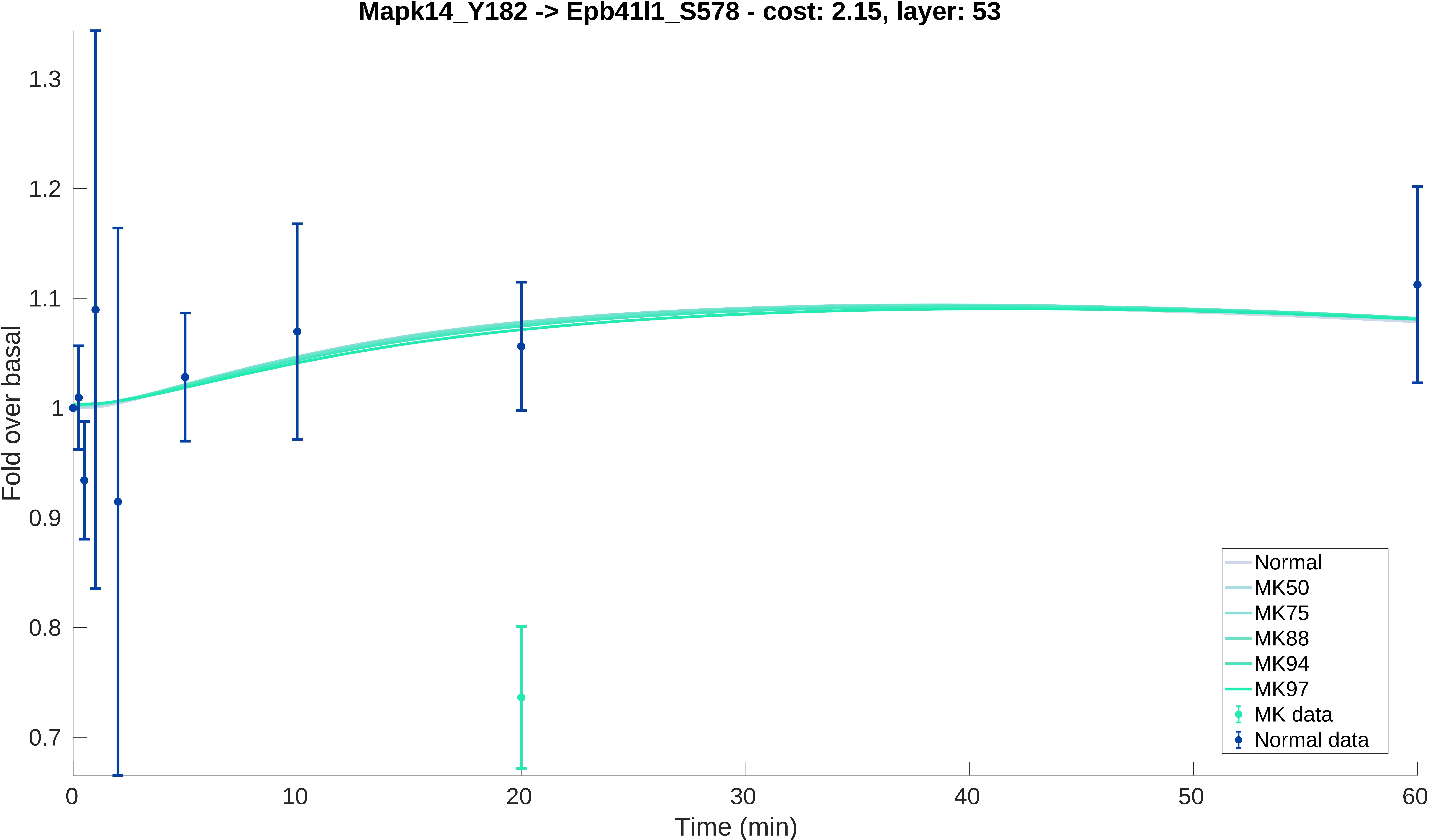

Mapk14_Y182 -> Epb41l1_S578 - cost: 2.15, layer: 53
Fold over basal
1.3
1.2
1.1
1
0.9
0.8
0.7
0
10
20
30
40
50
60
Time (min)
Normal
MK50
MK75
MK88
MK94
MK97
MK data
Normal data

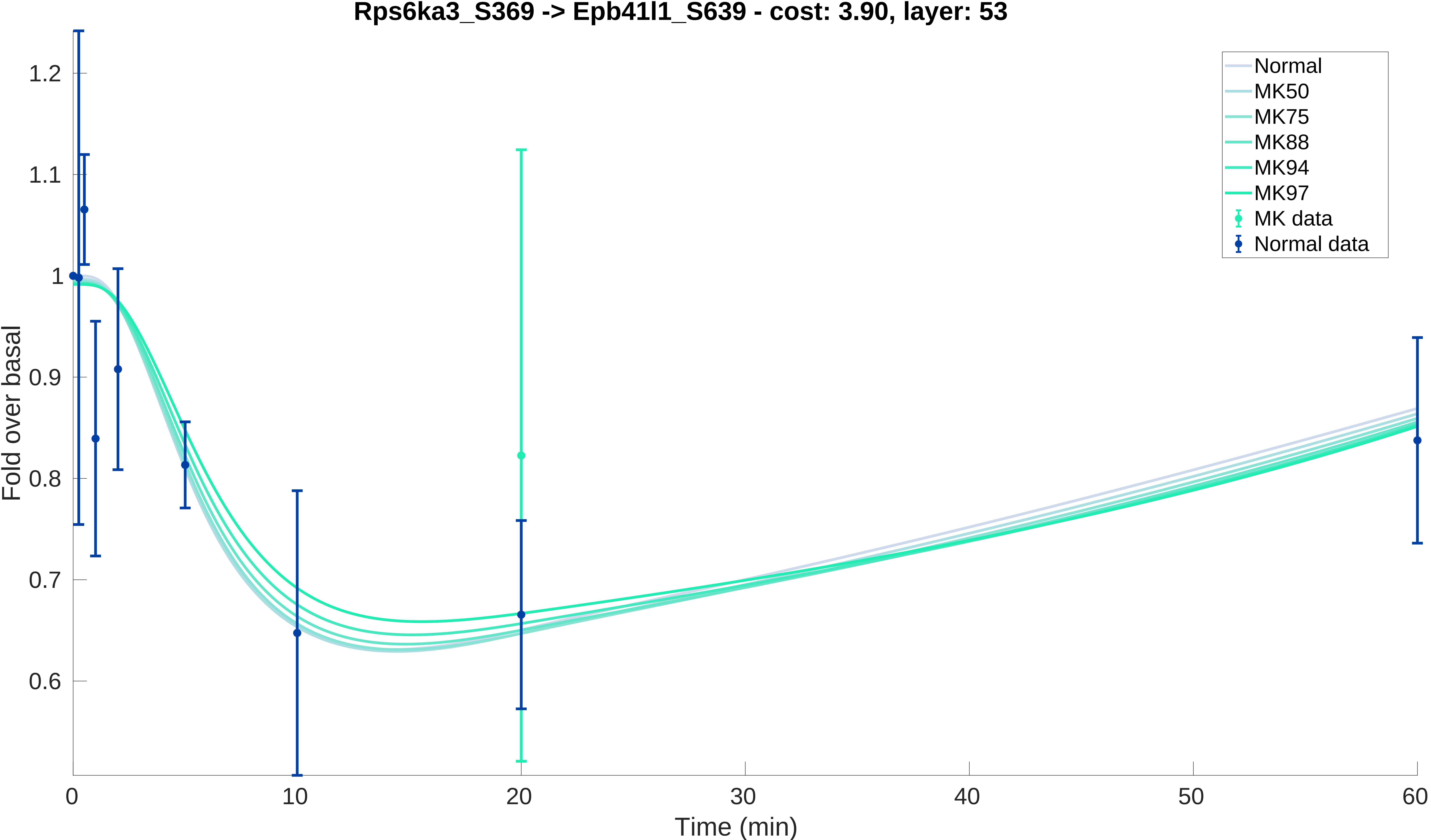
Rps6ka3_S369 -> Epb41l1_S639 - cost: 3.90, layer: 53
Fold over basal
Time (min)
0.6
0.7
0.8
0.9
1
1.1
1.2
0
10
20
30
40
50
60
Normal
MK50
MK75
MK88
MK94
MK97
MK data
Normal data

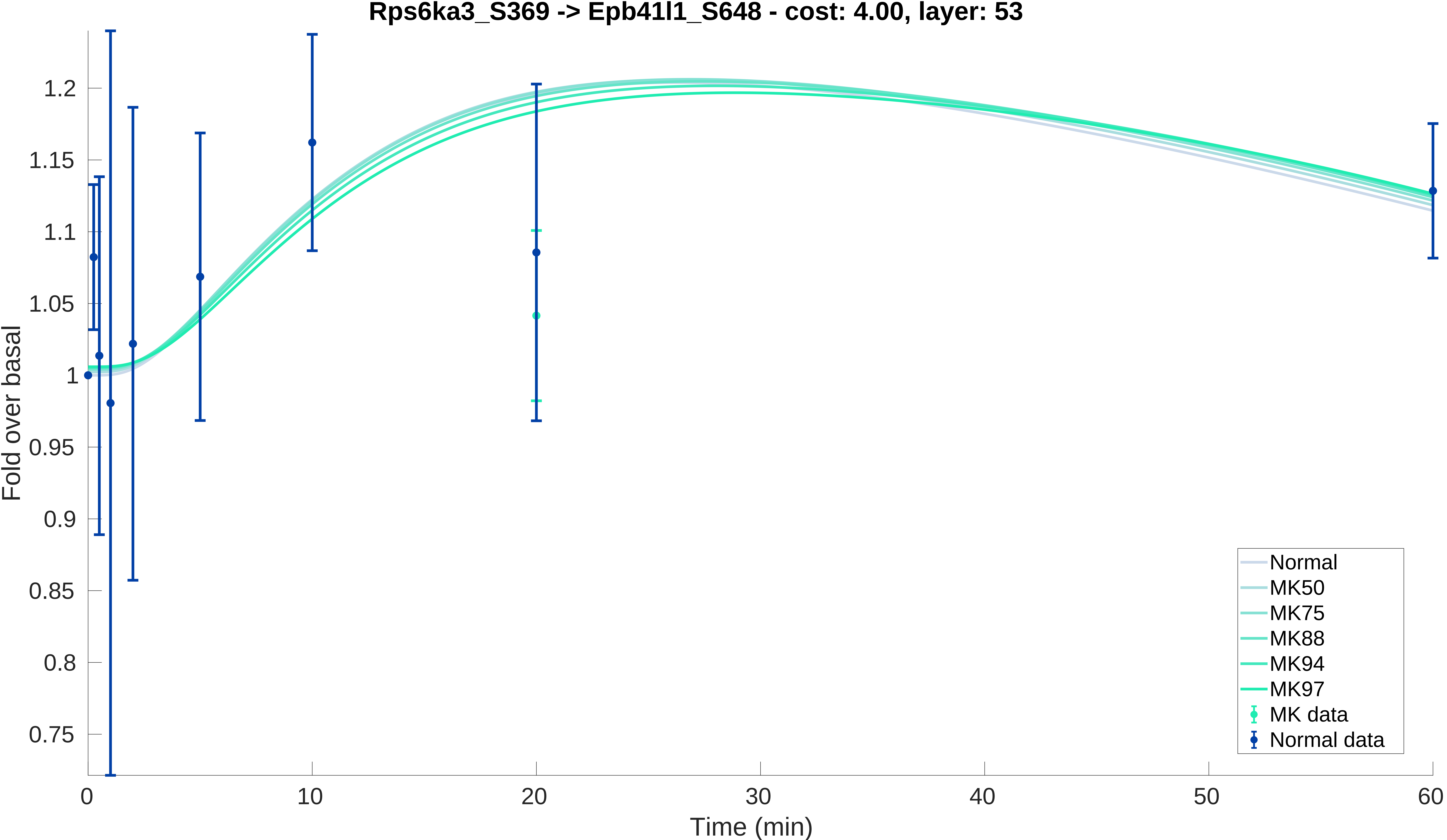

Rps6ka3_S369 -> Epb41l1_S648 - cost: 4.00, layer: 53
Fold over basal
1.2
1.15
1.1
1.05
1
0.95
0.9
0.85
0.8
0.75
0
10
20
30
40
50
60
Time (min)
Normal
MK50
MK75
MK88
MK94
MK97
MK data
Normal data

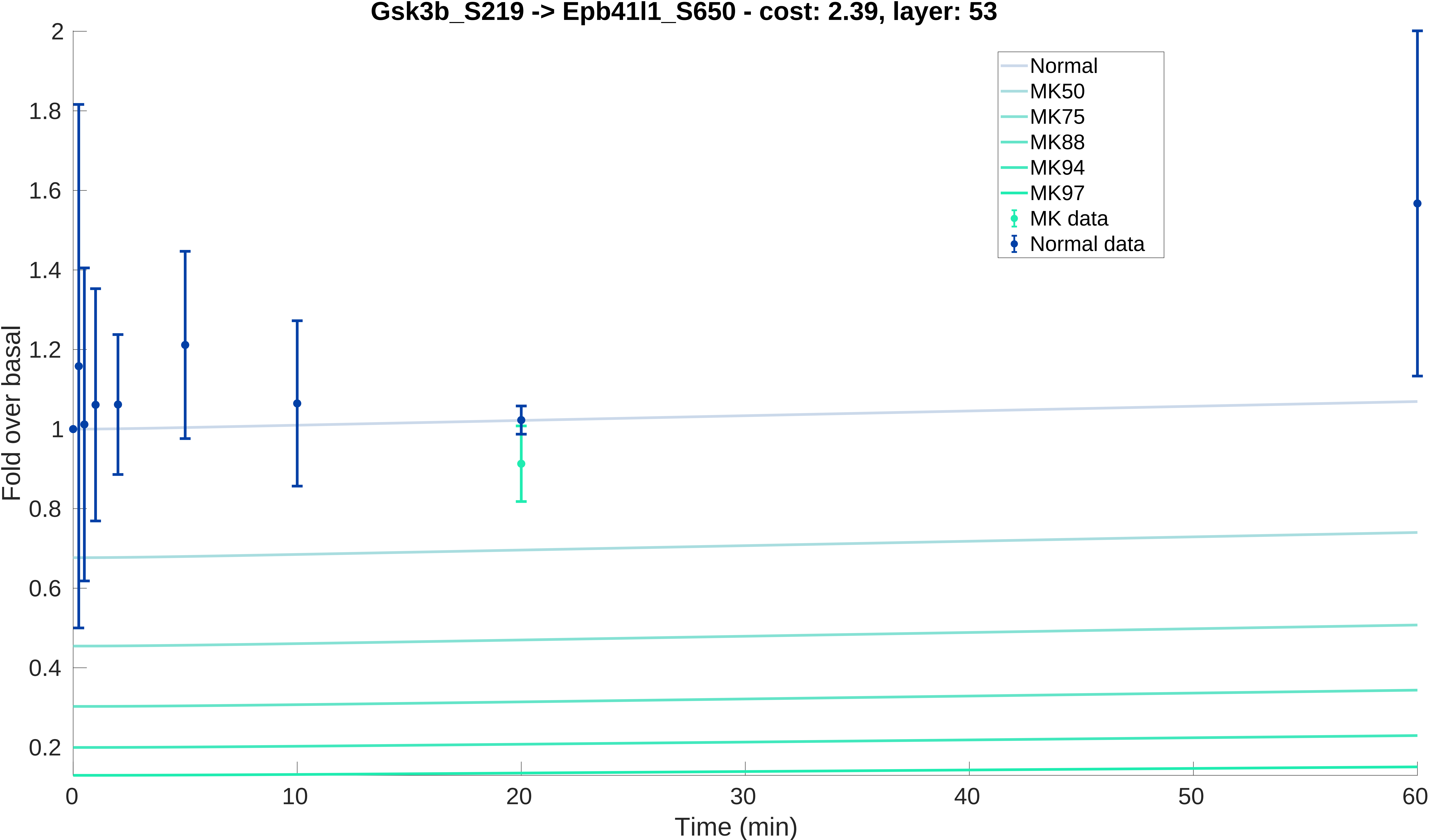

Gsk3b_S219 -> Epb41l1_S650 - cost: 2.39, layer: 53
Fold over basal
Time (min)
2
1.8
1.6
1.4
1.2
1
0.8
0.6
0.4
0.2
0
10
20
30
40
50
60
Normal
MK50
MK75
MK88
MK94
MK97
MK data
Normal data

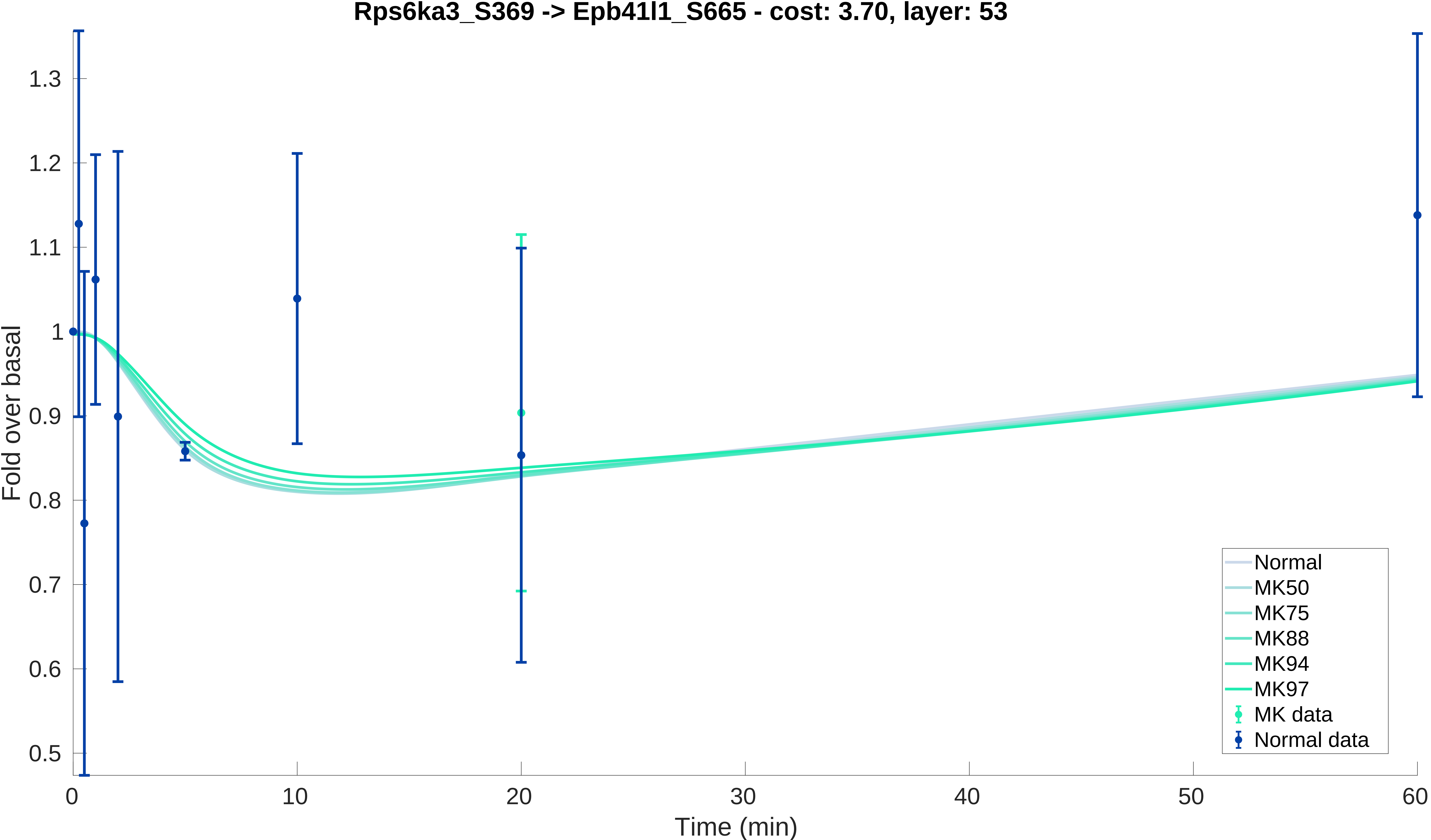

Rps6ka3_S369 -> Epb41l1_S665 - cost: 3.70, layer: 53
Fold over basal
1.3
1.2
1.1
1
0.9
0.8
0.7
0.6
0.5
0
10
20
30
40
50
60
Time (min)
Normal
MK50
MK75
MK88
MK94
MK97
MK data
Normal data

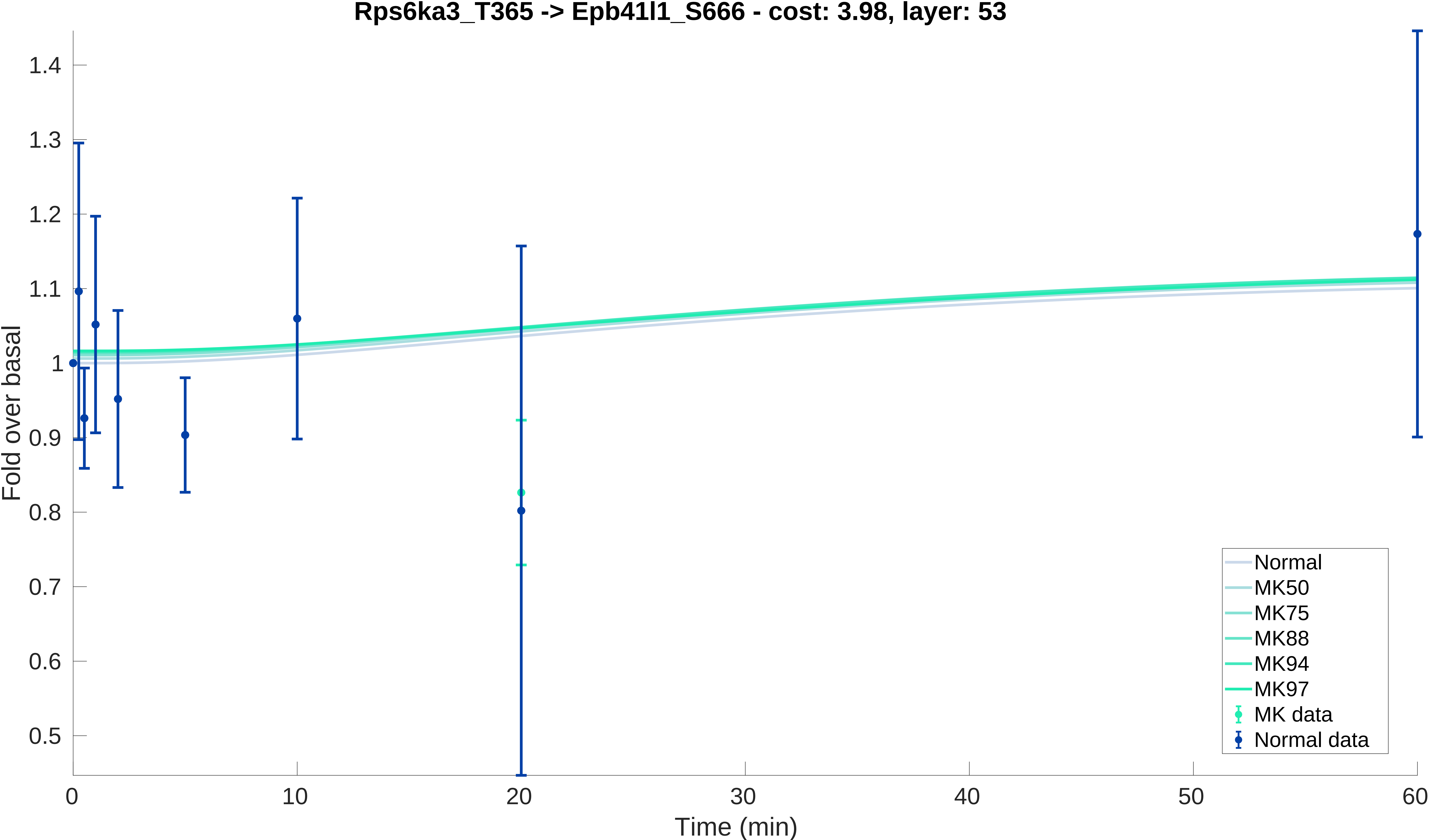

Rps6ka3_T365 -> Epb41l1_S666 - cost: 3.98, layer: 53
Fold over basal
Time (min)
1.4
1.3
1.2
1.1
1
0.9
0.8
0.7
0.6
0.5
0
10
20
30
40
50
60
Normal
MK50
MK75
MK88
MK94
MK97
MK data
Normal data

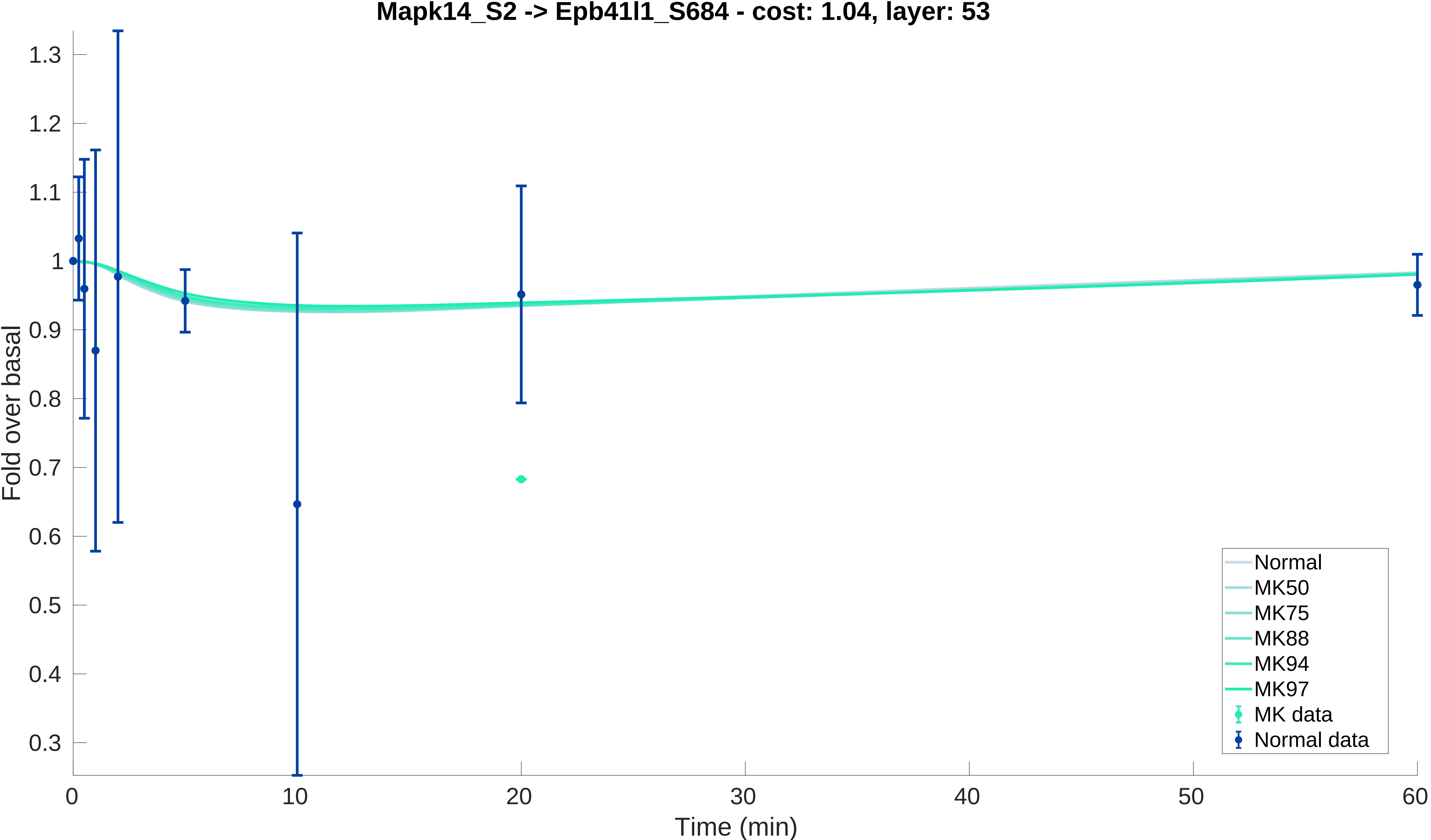

Mapk14_S2 -> Epb41l1_S684 - cost: 1.04, layer: 53
Fold over basal
1.3
1.2
1.1
1
0.9
0.8
0.7
0.6
0.5
0.4
0.3
0
10
20
30
40
50
60
Time (min)
Normal
MK50
MK75
MK88
MK94
MK97
MK data
Normal data

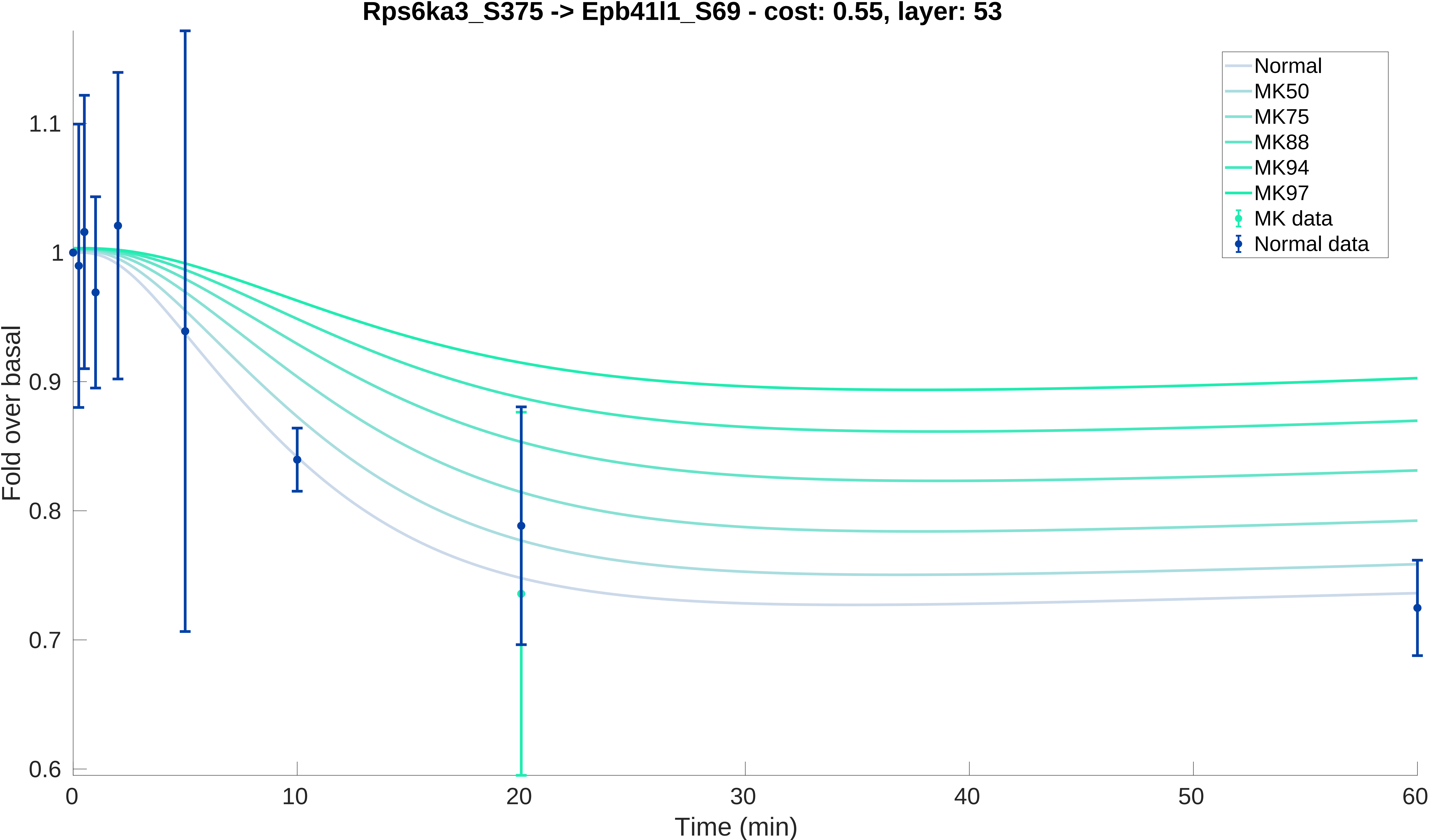

Rps6ka3_S375 -> Epb41l1_S69 - cost: 0.55, layer: 53
Fold over basal
Time (min)
1.1
1
0.9
0.8
0.7
0.6
0
10
20
30
40
50
60
Normal
MK50
MK75
MK88
MK94
MK97
MK data
Normal data

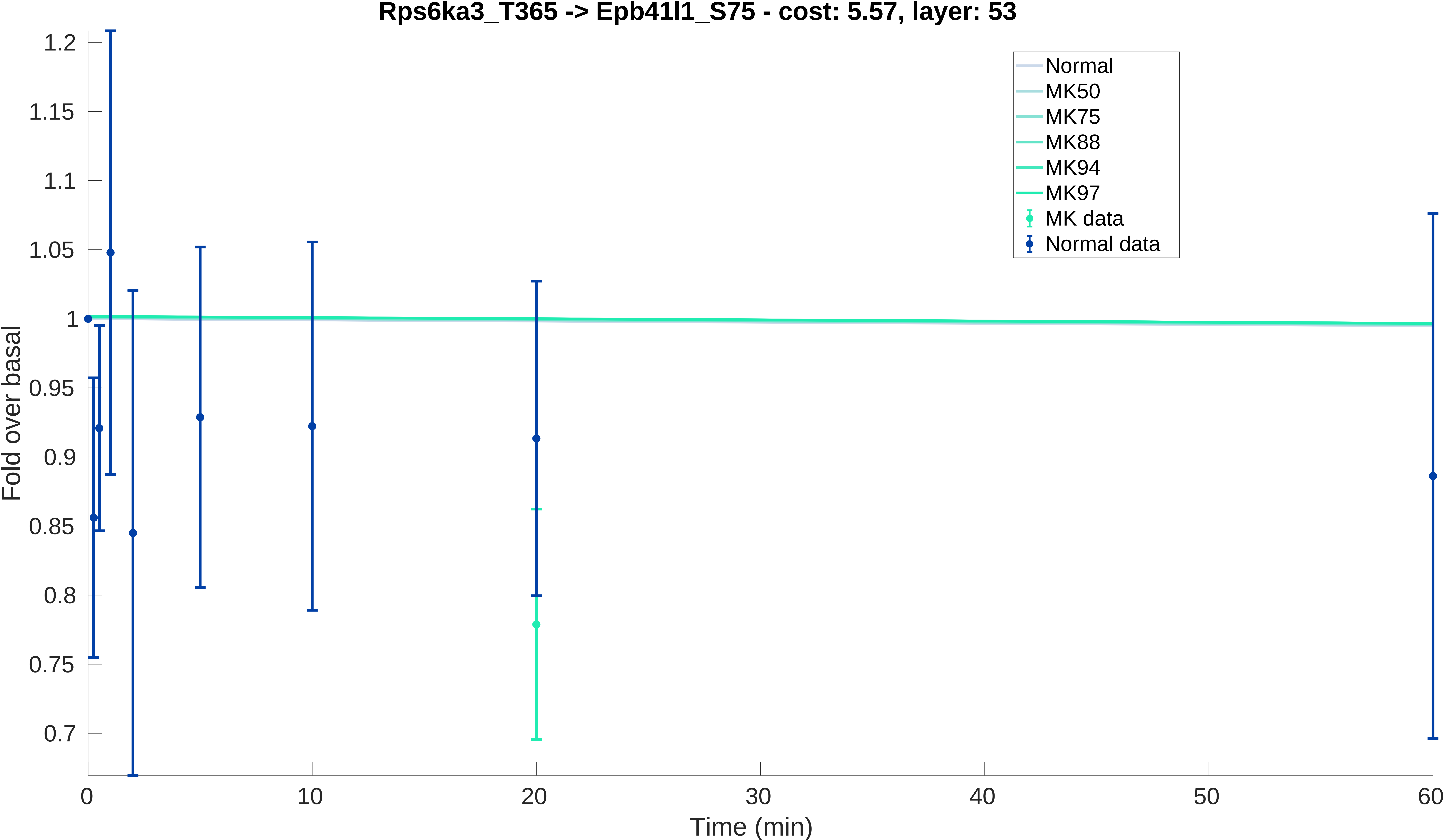

Rps6ka3_T365 -> Epb41l1_S75 - cost: 5.57, layer: 53
Fold over basal
Time (min)
1.2
1.15
1.1
1.05
1
0.95
0.9
0.85
0.8
0.75
0.7
0
10
20
30
40
50
60
Normal
MK50
MK75
MK88
MK94
MK97
MK data
Normal data

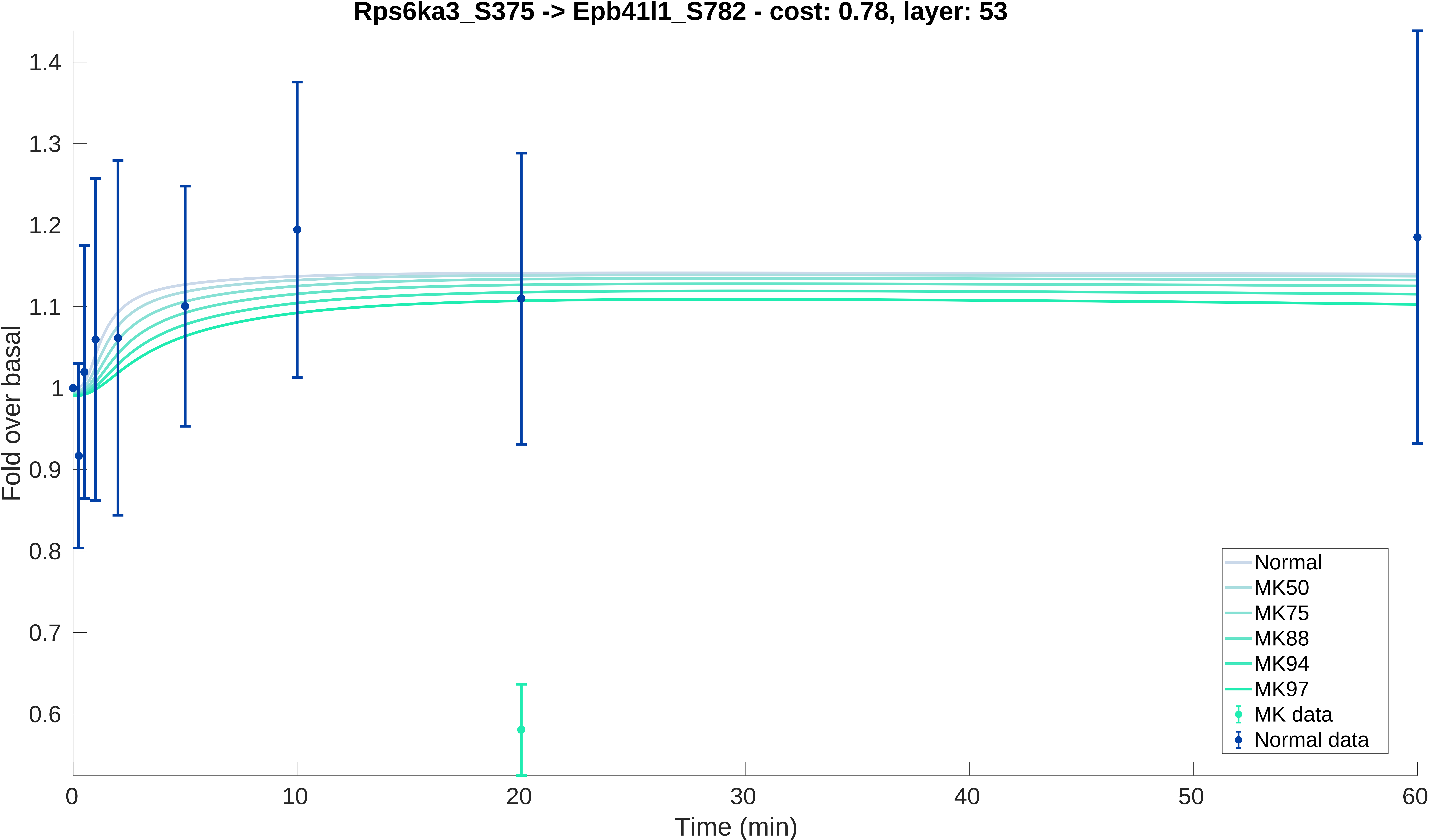
Rps6ka3_S375 -> Epb41l1_S782 - cost: 0.78, layer: 53
Fold over basal
1.4
1.3
1.2
1.1
1
0.9
0.8
0.7
0.6
0
10
20
30
40
50
60
Time (min)
Normal
MK50
MK75
MK88
MK94
MK97
MK data
Normal data

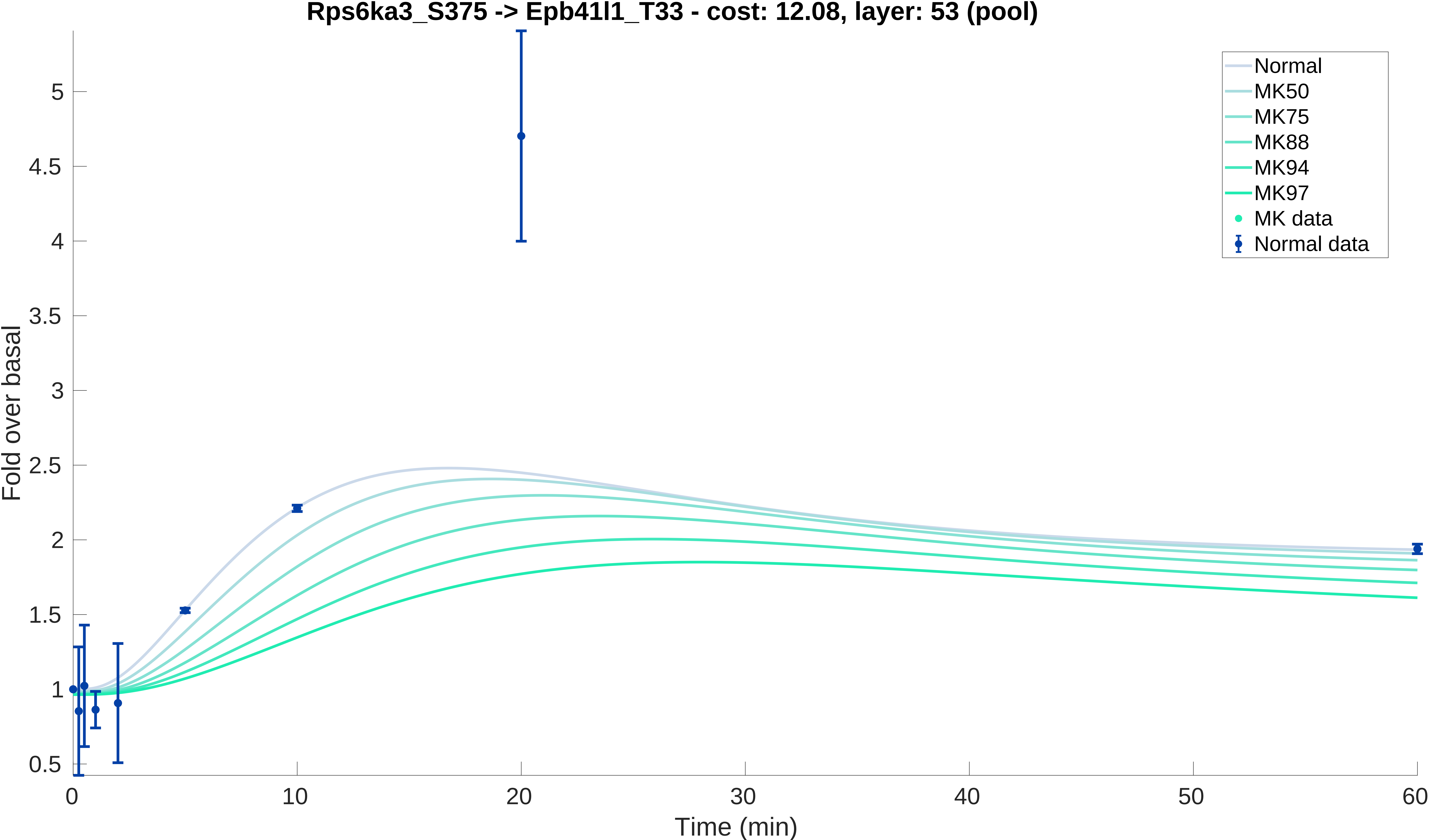

Rps6ka3_S375 -> Epb41l1_T33 - cost: 12.08, layer: 53 (pool)
Fold over basal
Time (min)
0.5
1
1.5
2
2.5
3
3.5
4
4.5
5
0
10
20
30
40
50
60
Normal
MK50
MK75
MK88
MK94
MK97
MK data
Normal data

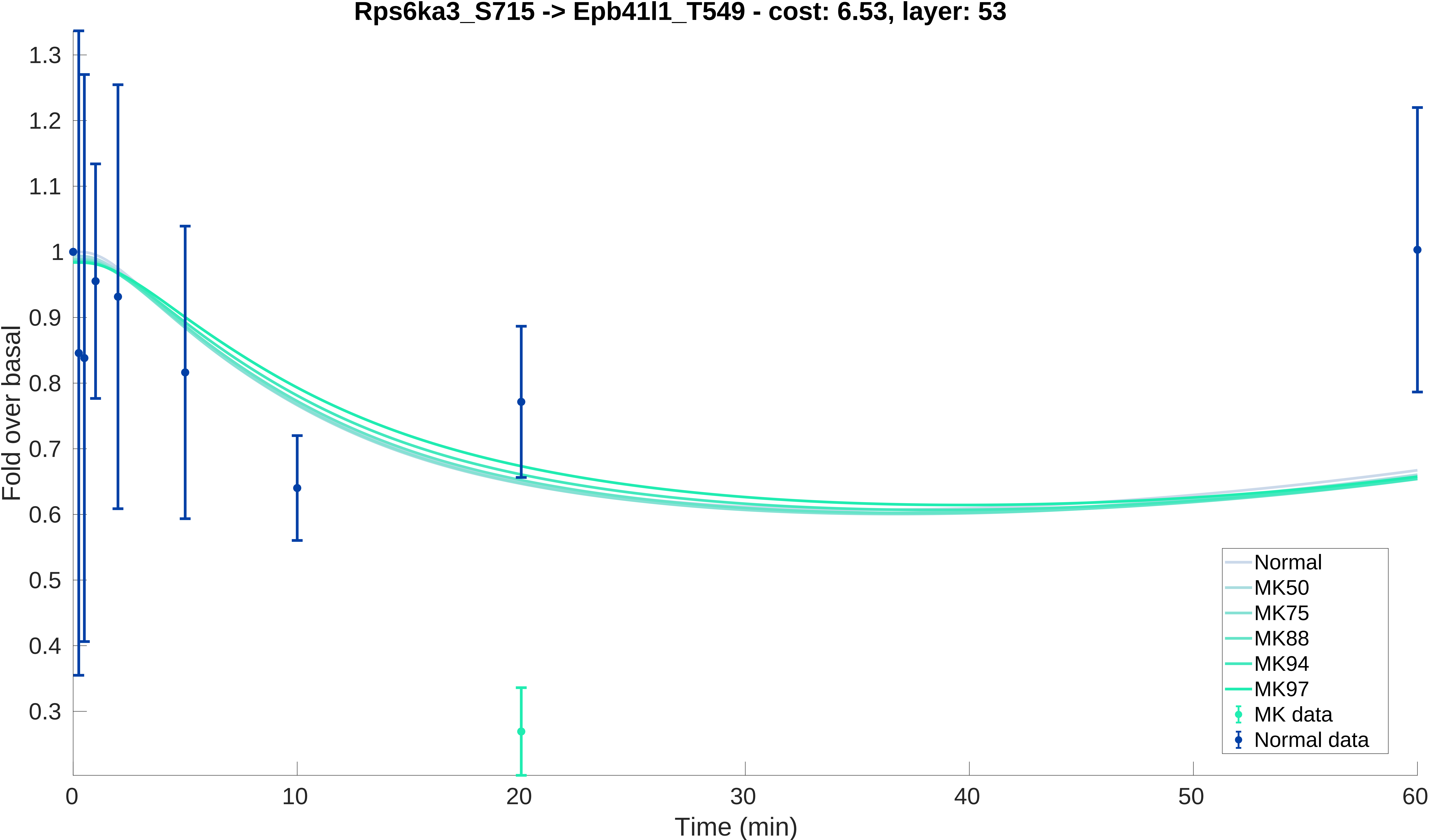

Rps6ka3_S715 -> Epb41l1_T549 - cost: 6.53, layer: 53
Fold over basal
1.3
1.2
1.1
1
0.9
0.8
0.7
0.6
0.5
0.4
0.3
0
10
20
30
40
50
60
Time (min)
Normal
MK50
MK75
MK88
MK94
MK97
MK data
Normal data

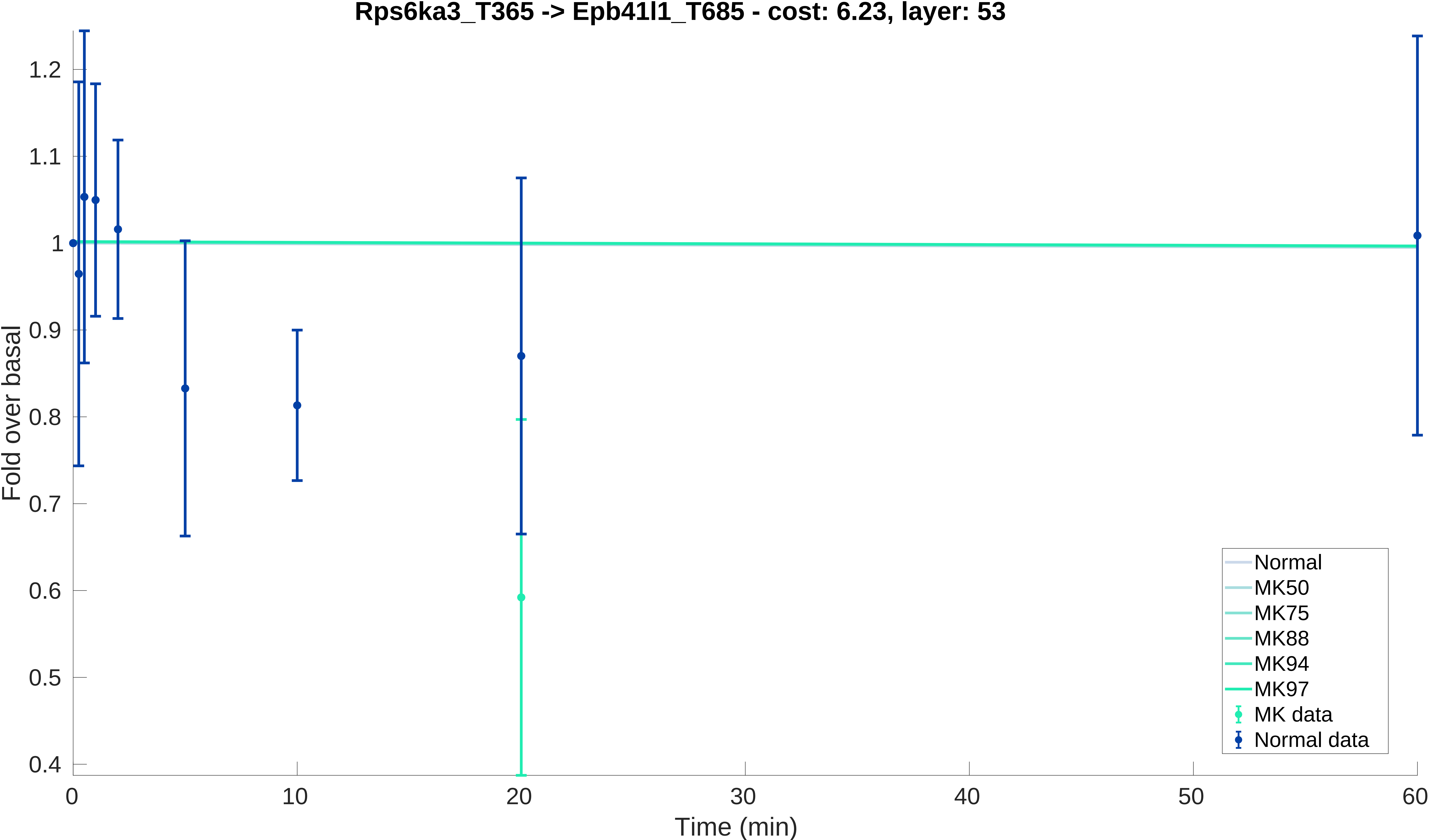

Rps6ka3_T365 -> Epb41l1_T685 - cost: 6.23, layer: 53
Fold over basal
Time (min)
0.4
0.5
0.6
0.7
0.8
0.9
1
1.1
1.2
0
10
20
30
40
50
60
Normal
MK50
MK75
MK88
MK94
MK97
MK data
Normal data

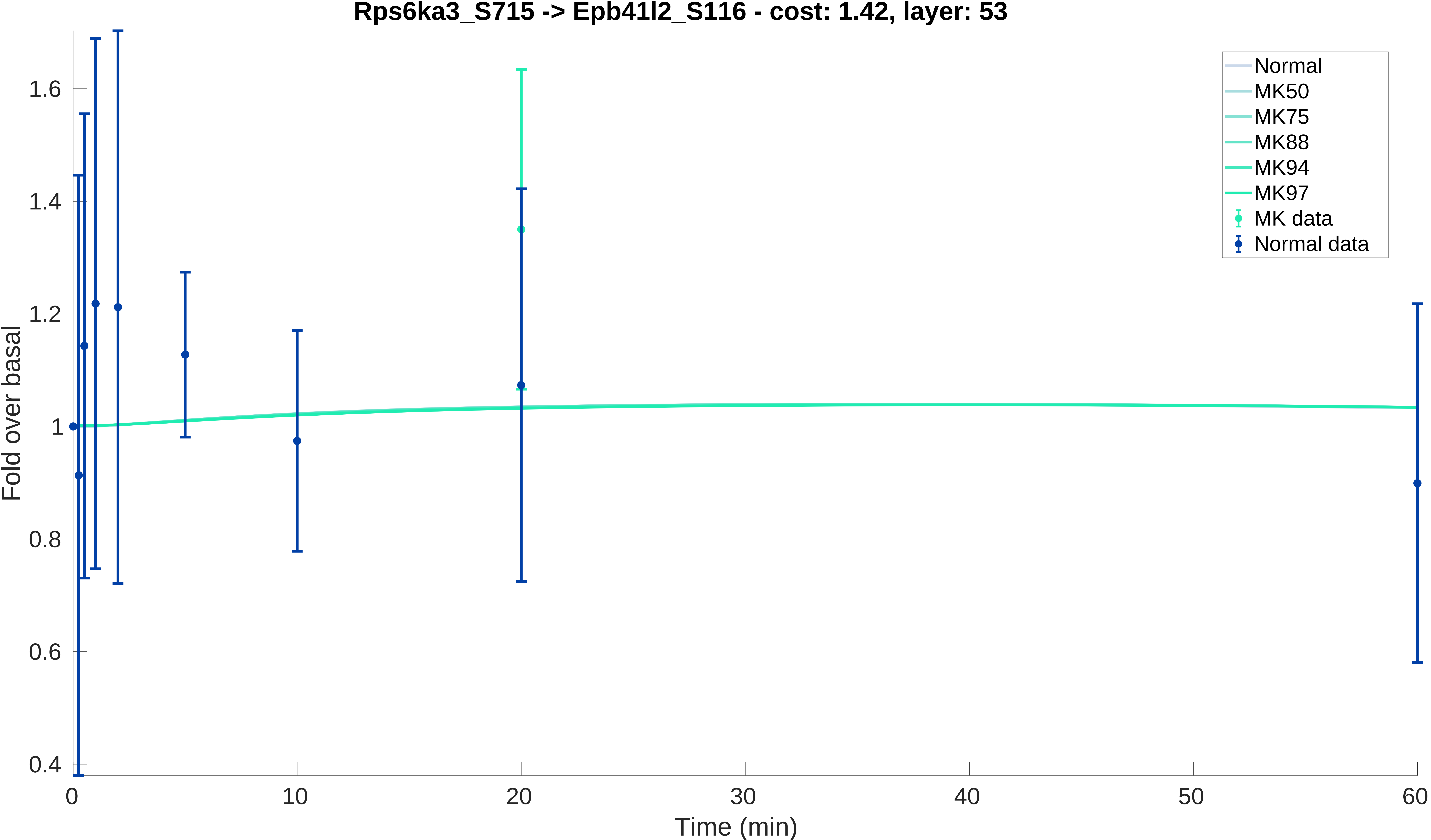
Rps6ka3_S715 -> Epb41l2_S116 - cost: 1.42, layer: 53
Fold over basal
Time (min)
0.4
0.6
0.8
1
1.2
1.4
1.6
0
10
20
30
40
50
60
Normal
MK50
MK75
MK88
MK94
MK97
MK data
Normal data

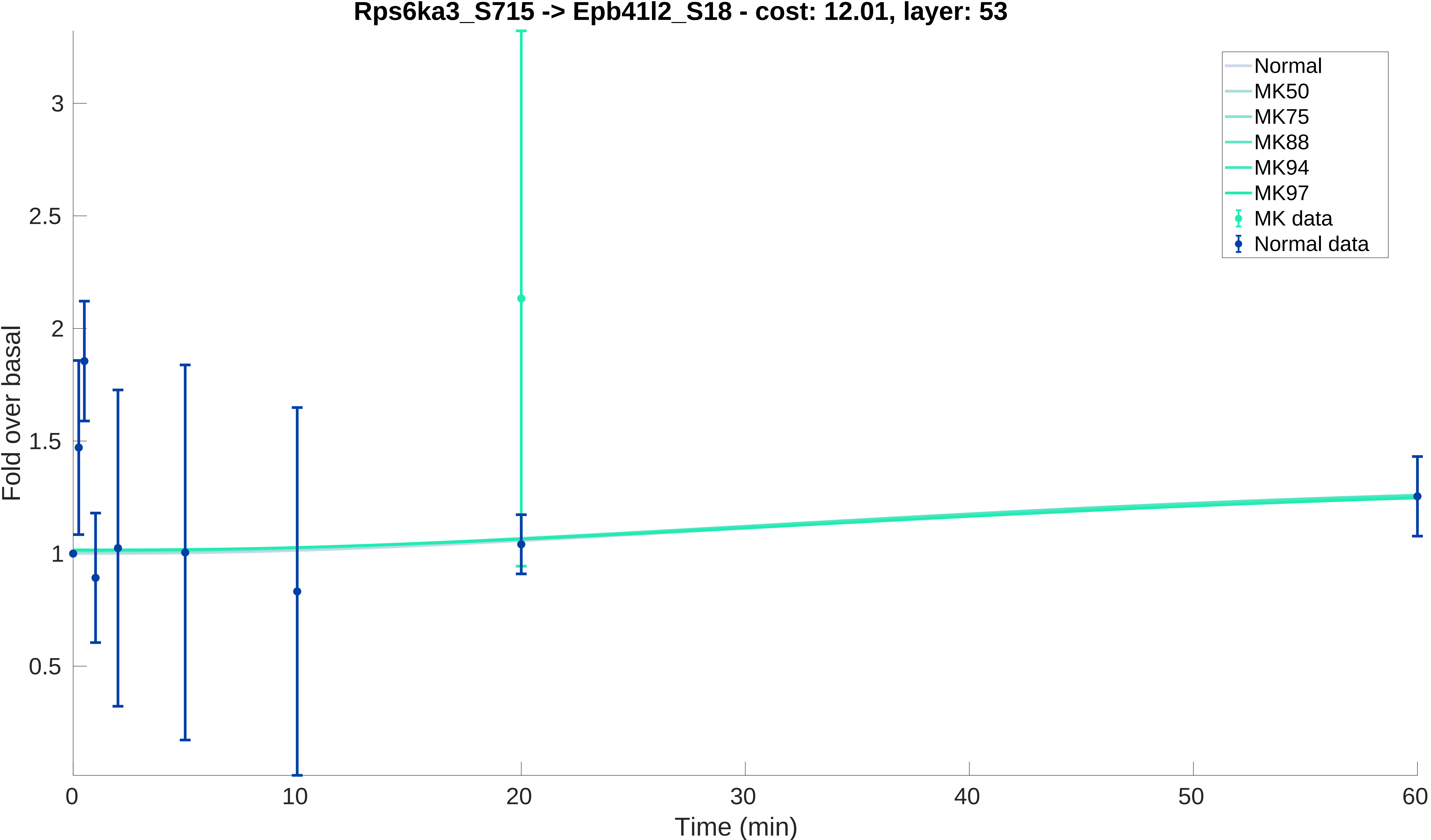

Rps6ka3_S715 -> Epb41l2_S18 - cost: 12.01, layer: 53
Fold over basal
Time (min)
0
10
20
30
40
50
60
0.5
1
1.5
2
2.5
3
Normal
MK50
MK75
MK88
MK94
MK97
MK data
Normal data

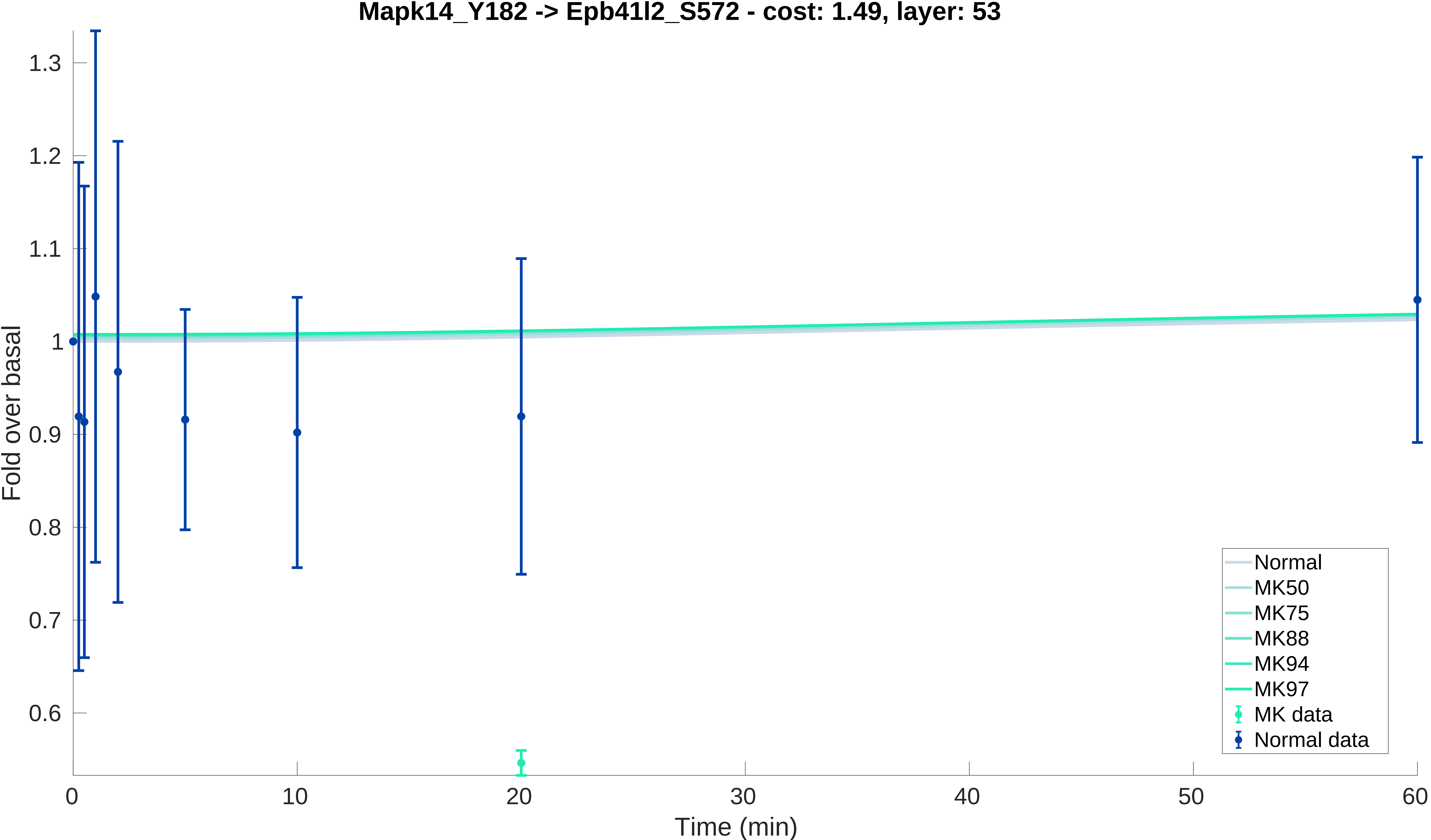

Mapk14_Y182 -> Epb41l2_S572 - cost: 1.49, layer: 53
Fold over basal
Time (min)
1.3
1.2
1.1
1
0.9
0.8
0.7
0.6
0
10
20
30
40
50
60
Normal
MK50
MK75
MK88
MK94
MK97
MK data
Normal data

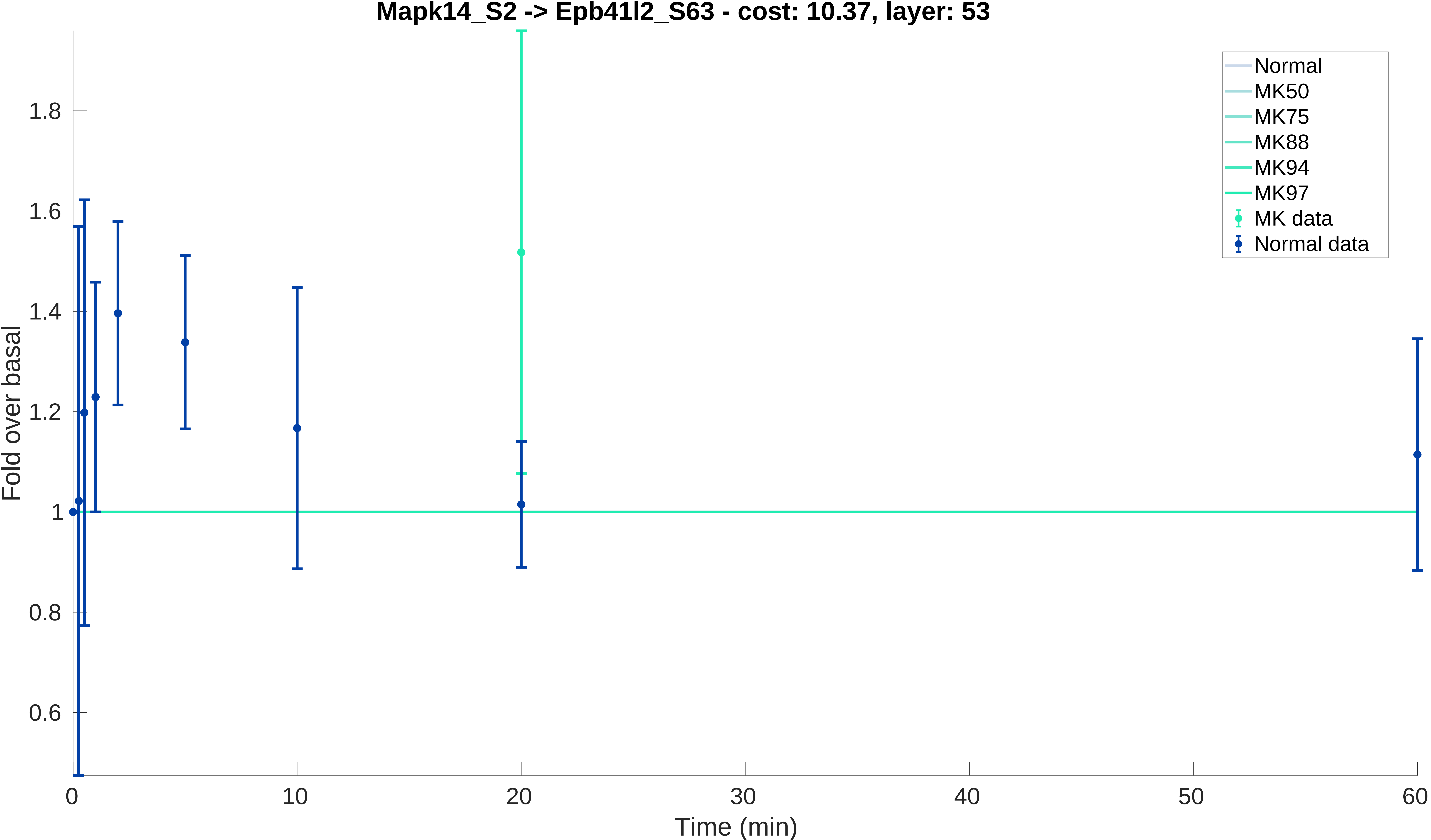

Mapk14_S2 -> Epb41l2_S63 - cost: 10.37, layer: 53
Fold over basal
Time (min)
1.8
1.6
1.4
1.2
1
0.8
0.6
0
10
20
30
40
50
60
Normal
MK50
MK75
MK88
MK94
MK97
MK data
Normal data

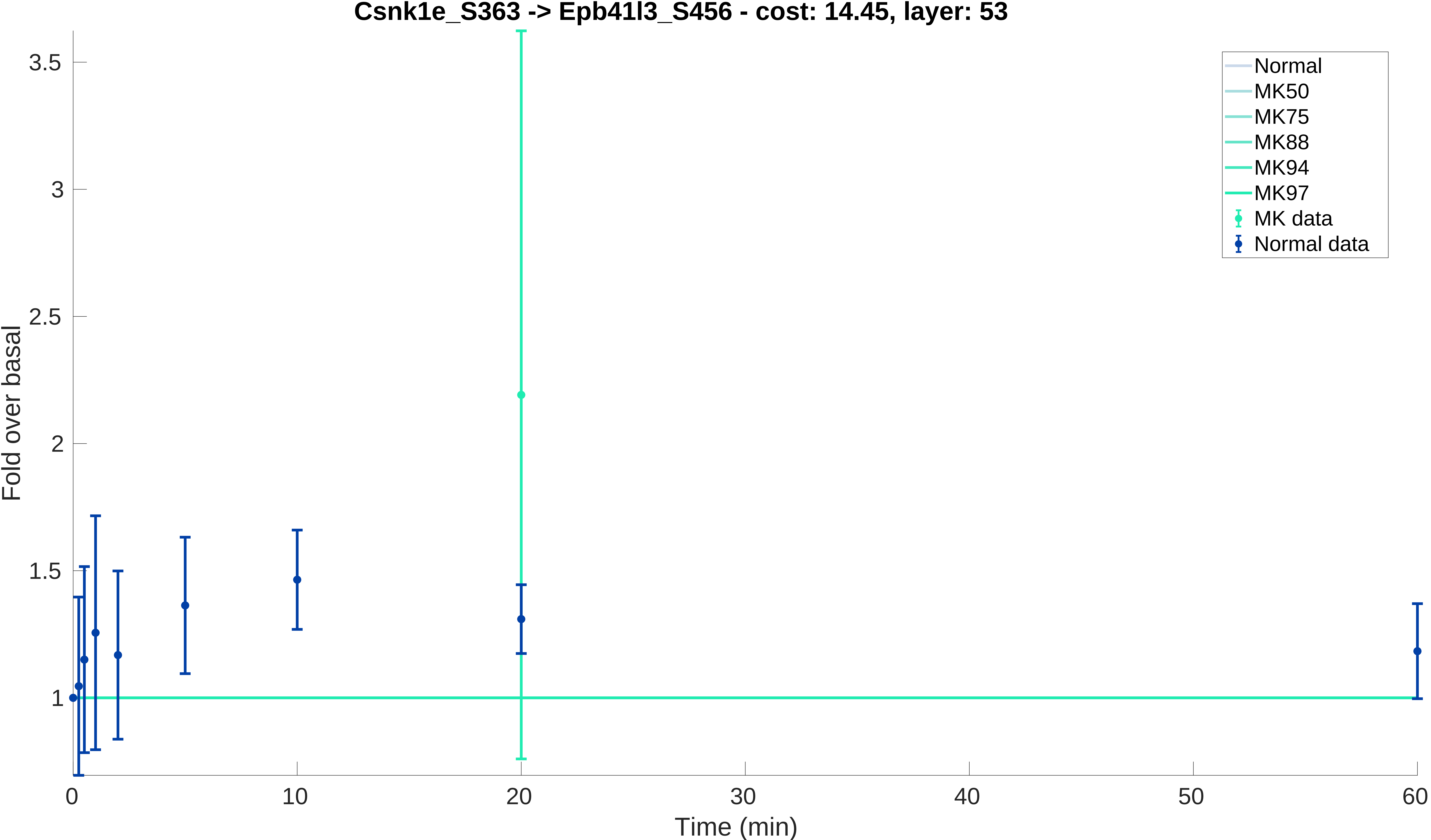
Csnk1e_S363 -> Epb41l3_S456 - cost: 14.45, layer: 53
Fold over basal
3.5
3
2.5
2
1.5
1
Time (min)
0
10
20
30
40
50
60
Normal
MK50
MK75
MK88
MK94
MK97
MK data
Normal data

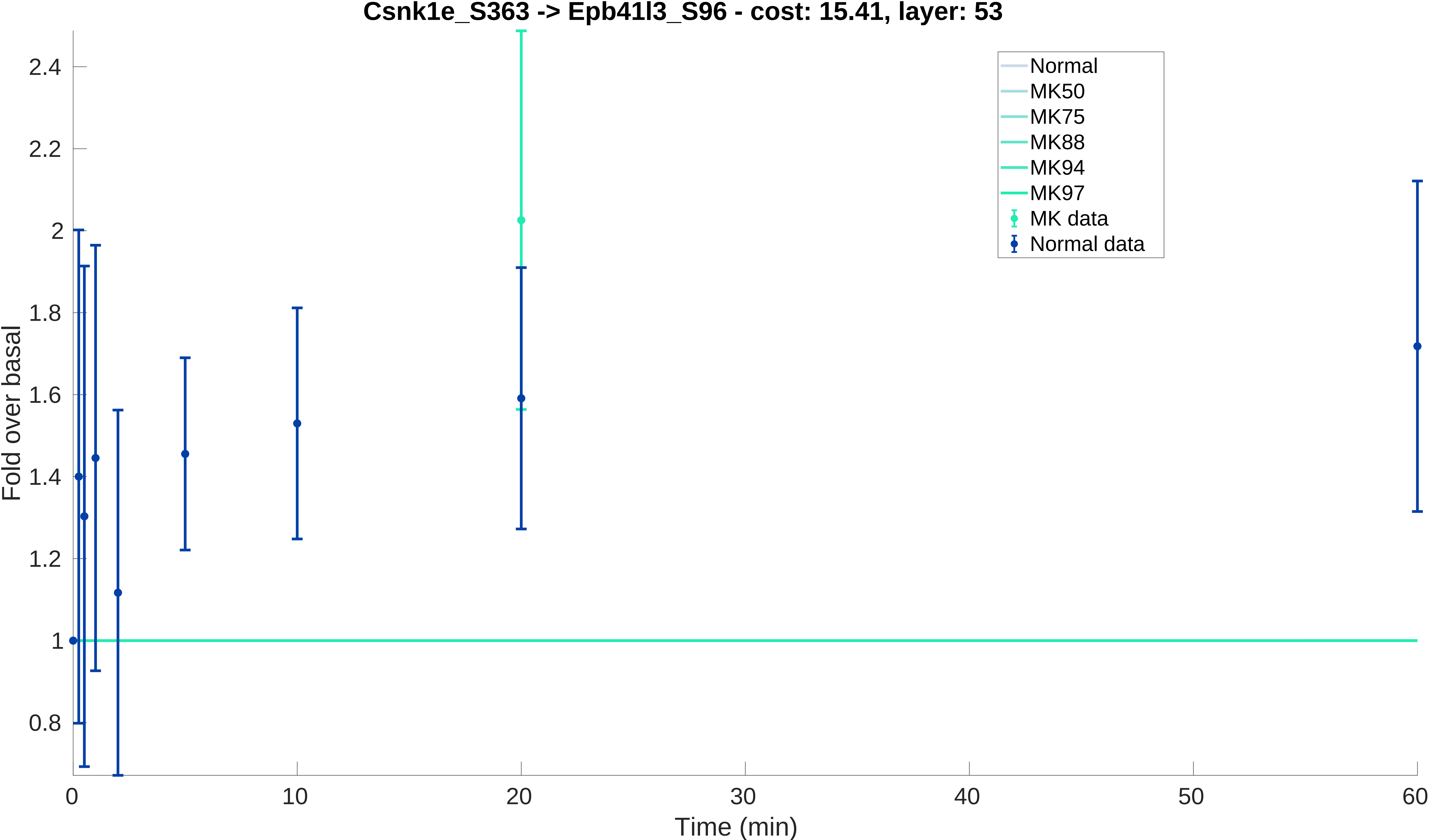

Csnk1e_S363 -> Epb41l3_S96 - cost: 15.41, layer: 53
Fold over basal
Time (min)
Normal
MK50
MK75
MK88
MK94
MK97
MK data
Normal data
2.4
2.2
2
1.8
1.6
1.4
1.2
1
0.8
0
10
20
30
40
50
60

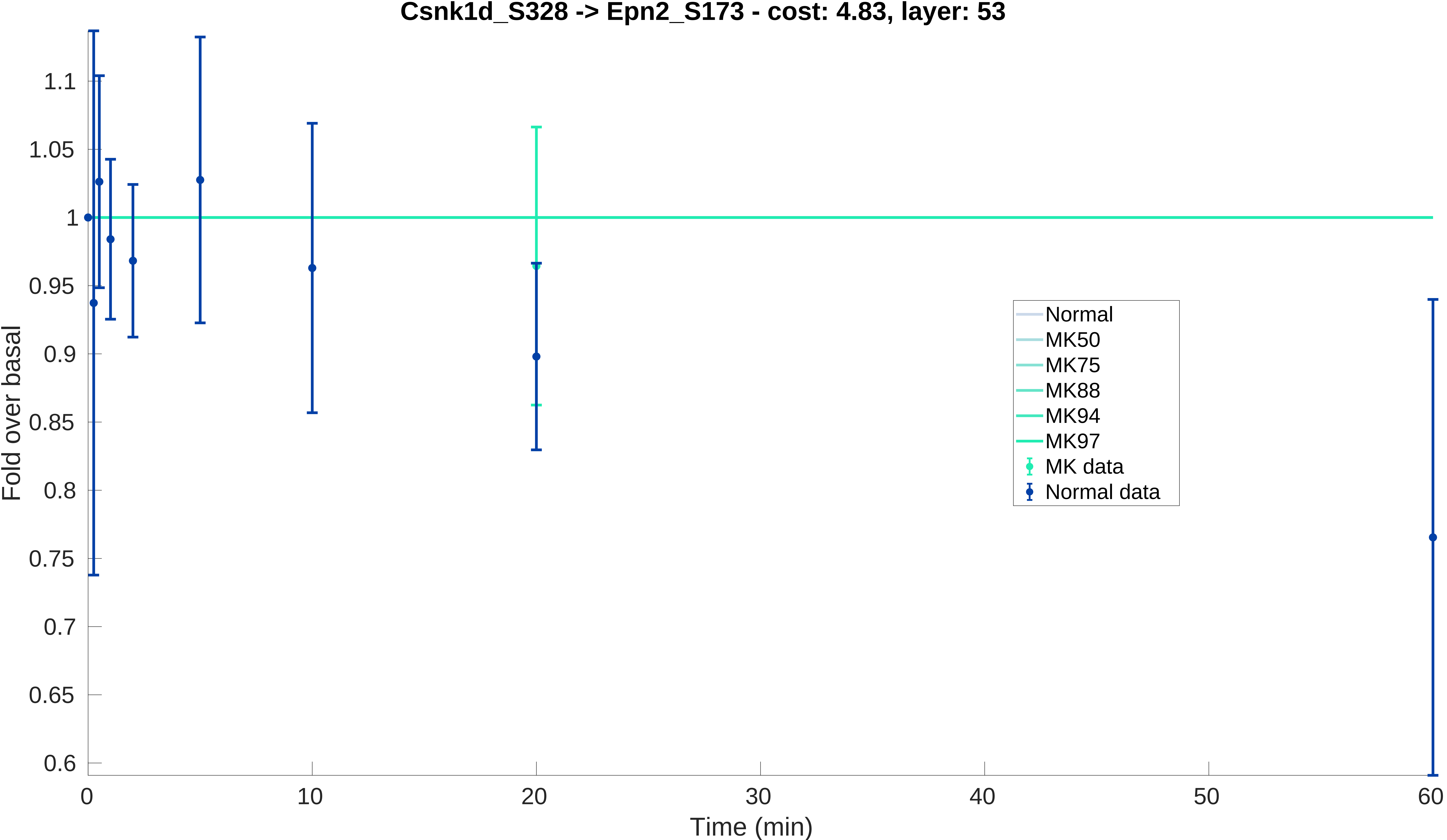

Csnk1d_S328 -> Epn2_S173 - cost: 4.83, layer: 53
Fold over basal
1.1
1.05
1
0.95
0.9
0.85
0.8
0.75
0.7
0.65
0.6
0
10
20
30
40
50
60
Time (min)
Normal
MK50
MK75
MK88
MK94
MK97
MK data
Normal data

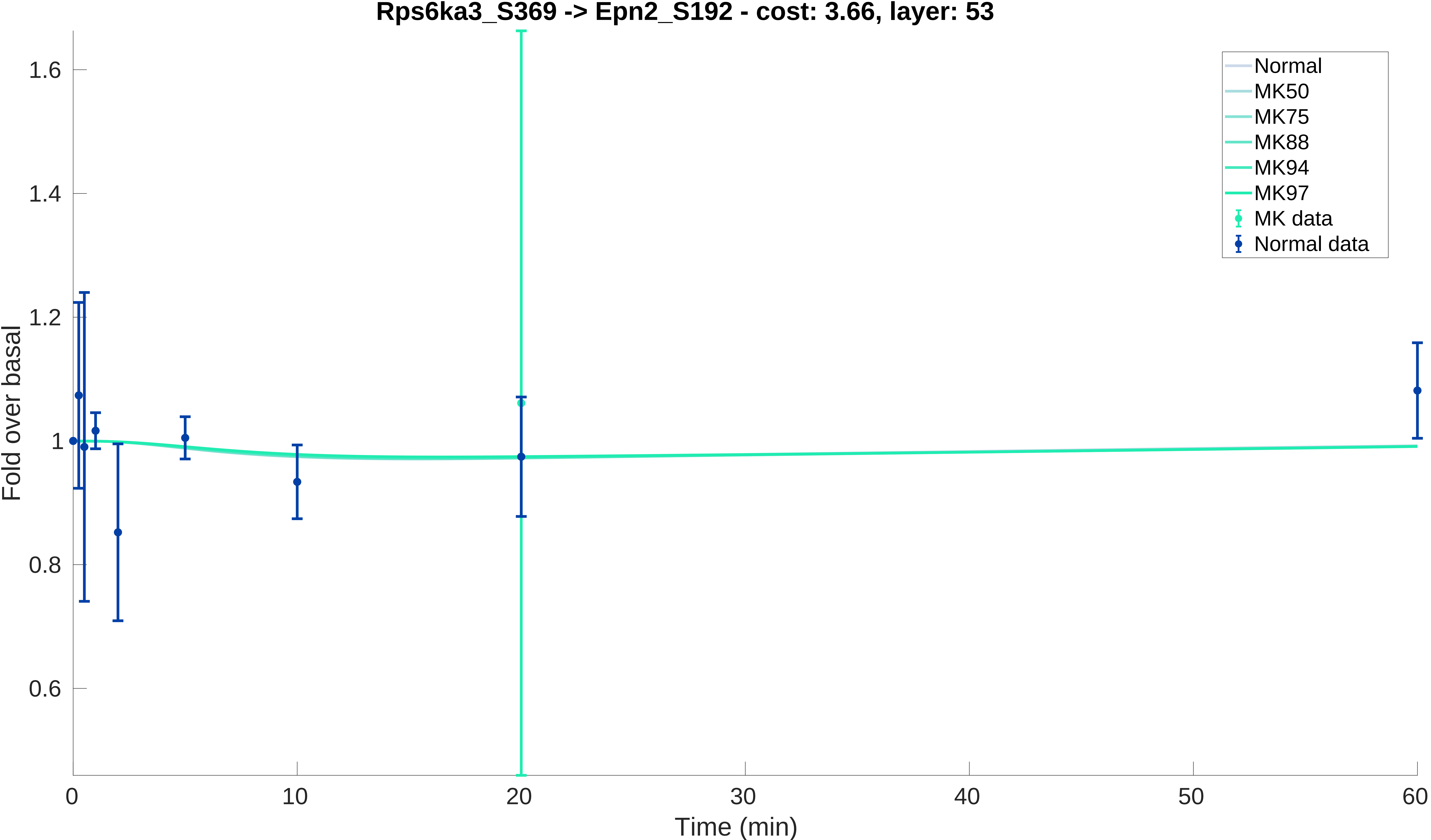

Rps6ka3_S369 -> Epn2_S192 - cost: 3.66, layer: 53
Fold over basal
Time (min)
0.6
0.8
1
1.2
1.4
1.6
0
10
20
30
40
50
60
Normal
MK50
MK75
MK88
MK94
MK97
MK data
Normal data

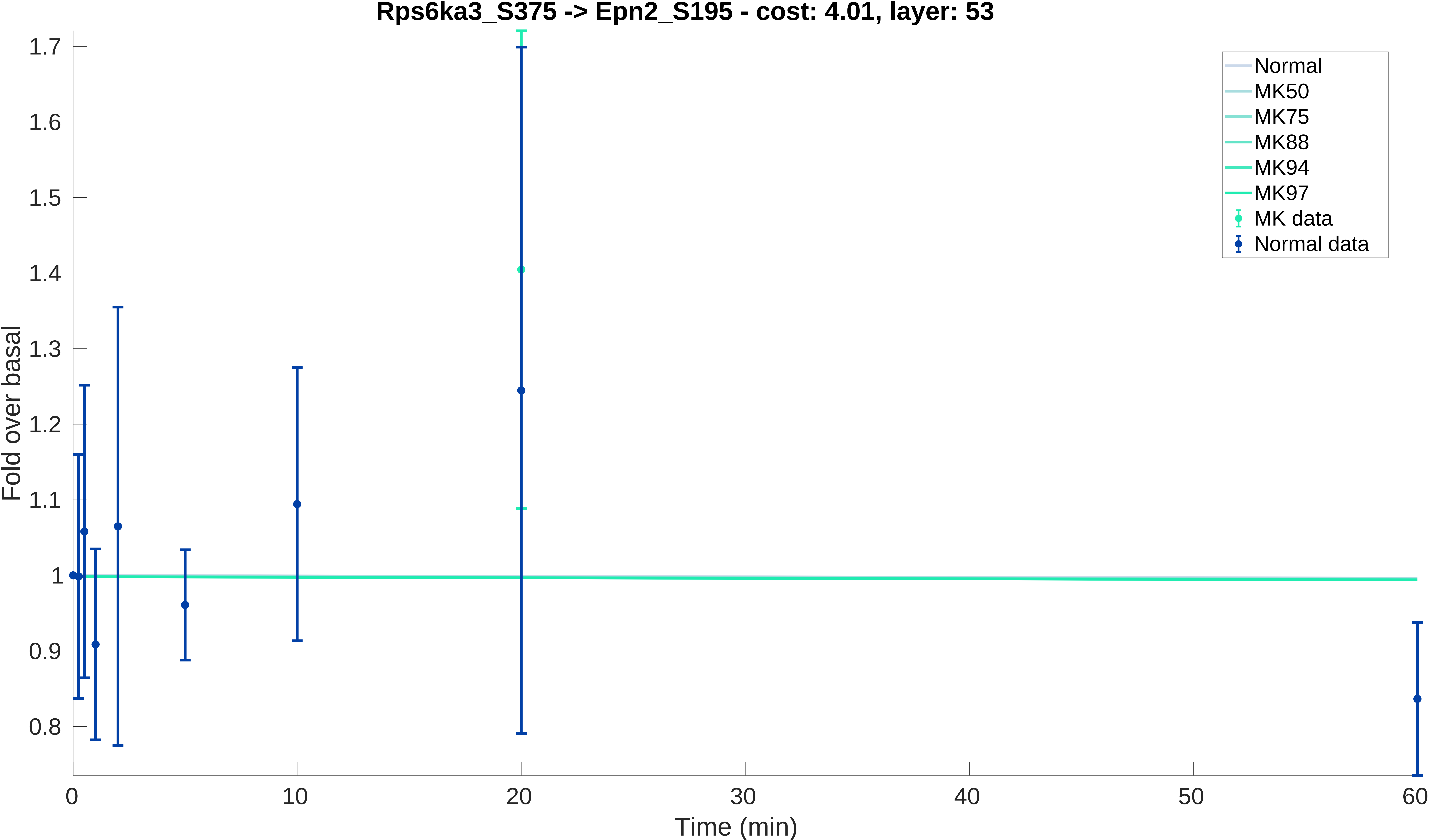

Rps6ka3_S375 -> Epn2_S195 - cost: 4.01, layer: 53
Fold over basal
Time (min)
Normal
MK50
MK75
MK88
MK94
MK97
MK data
Normal data
1.7
1.6
1.5
1.4
1.3
1.2
1.1
1
0.9
0.8
0
10
20
30
40
50
60

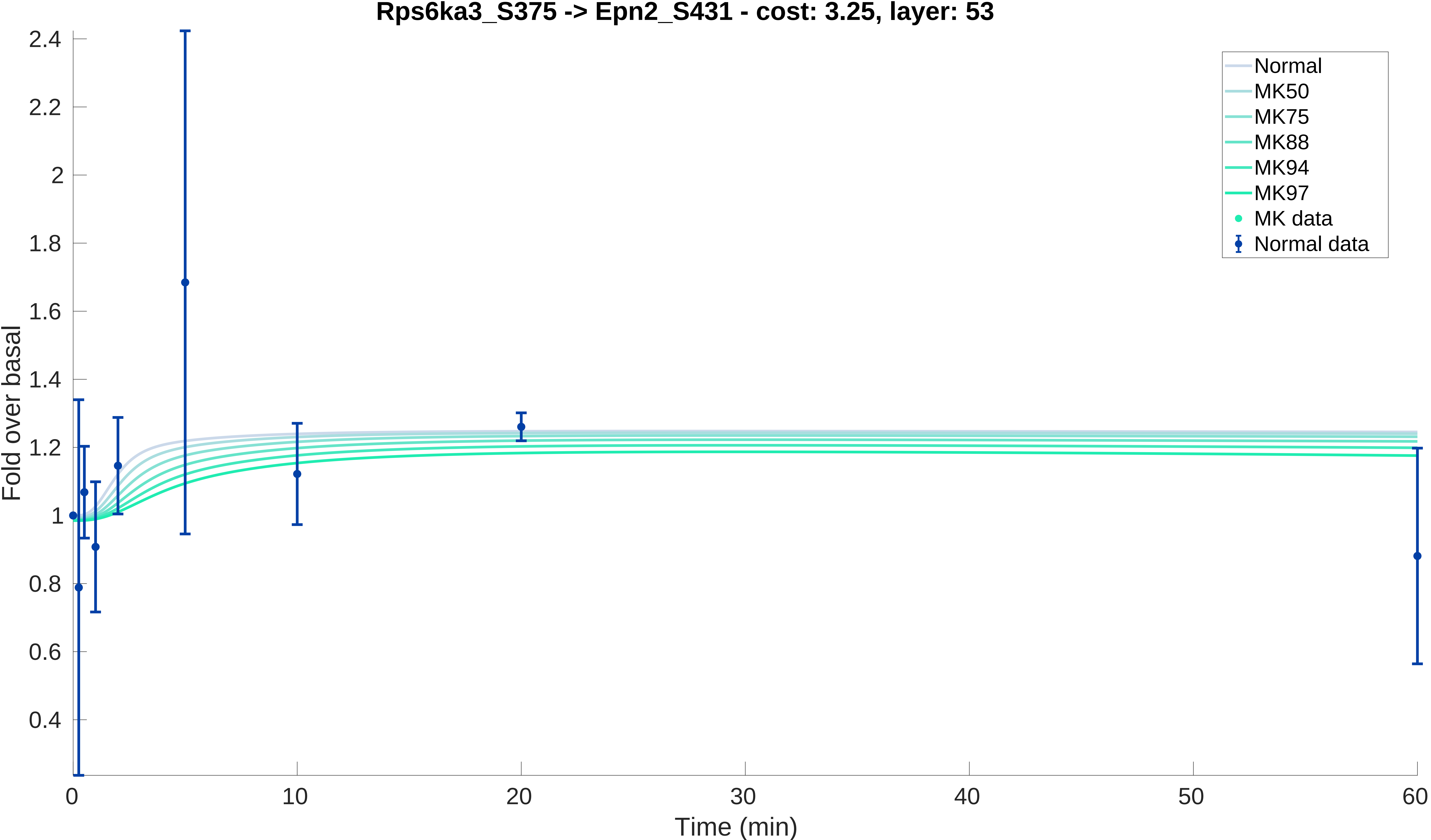

Rps6ka3_S375 -> Epn2_S431 - cost: 3.25, layer: 53
Fold over basal
Time (min)
Normal
MK50
MK75
MK88
MK94
MK97
MK data
Normal data
2.4
2.2
2
1.8
1.6
1.4
1.2
1
0.8
0.6
0.4
0
10
20
30
40
50
60

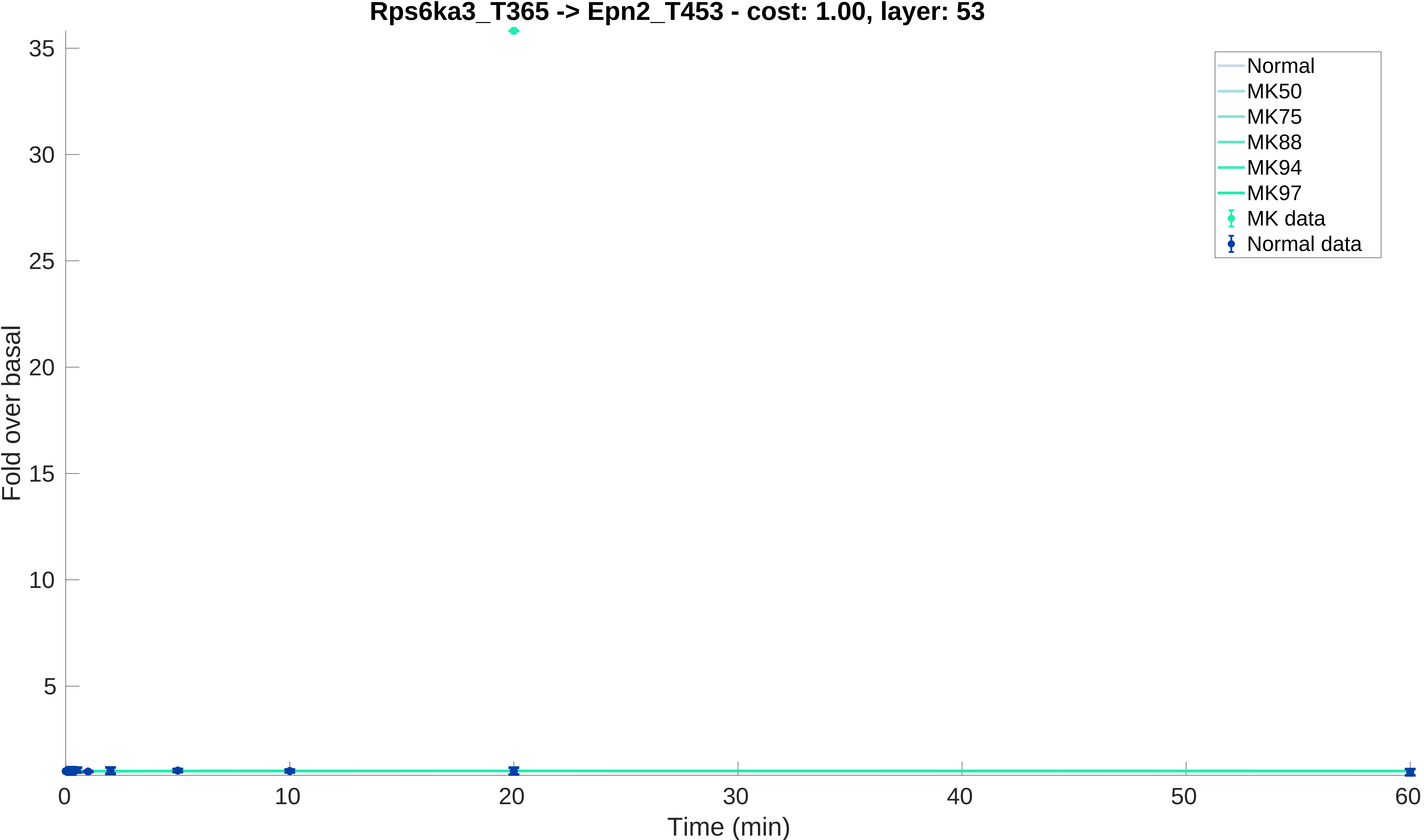

Rps6ka3_T365 -> Epn2_T453 - cost: 1.00, layer: 53
Fold over basal
Time (min)
0
10
20
30
40
50
60
5
10
15
20
25
30
35
Normal
MK50
MK75
MK88
MK94
MK97
MK data
Normal data

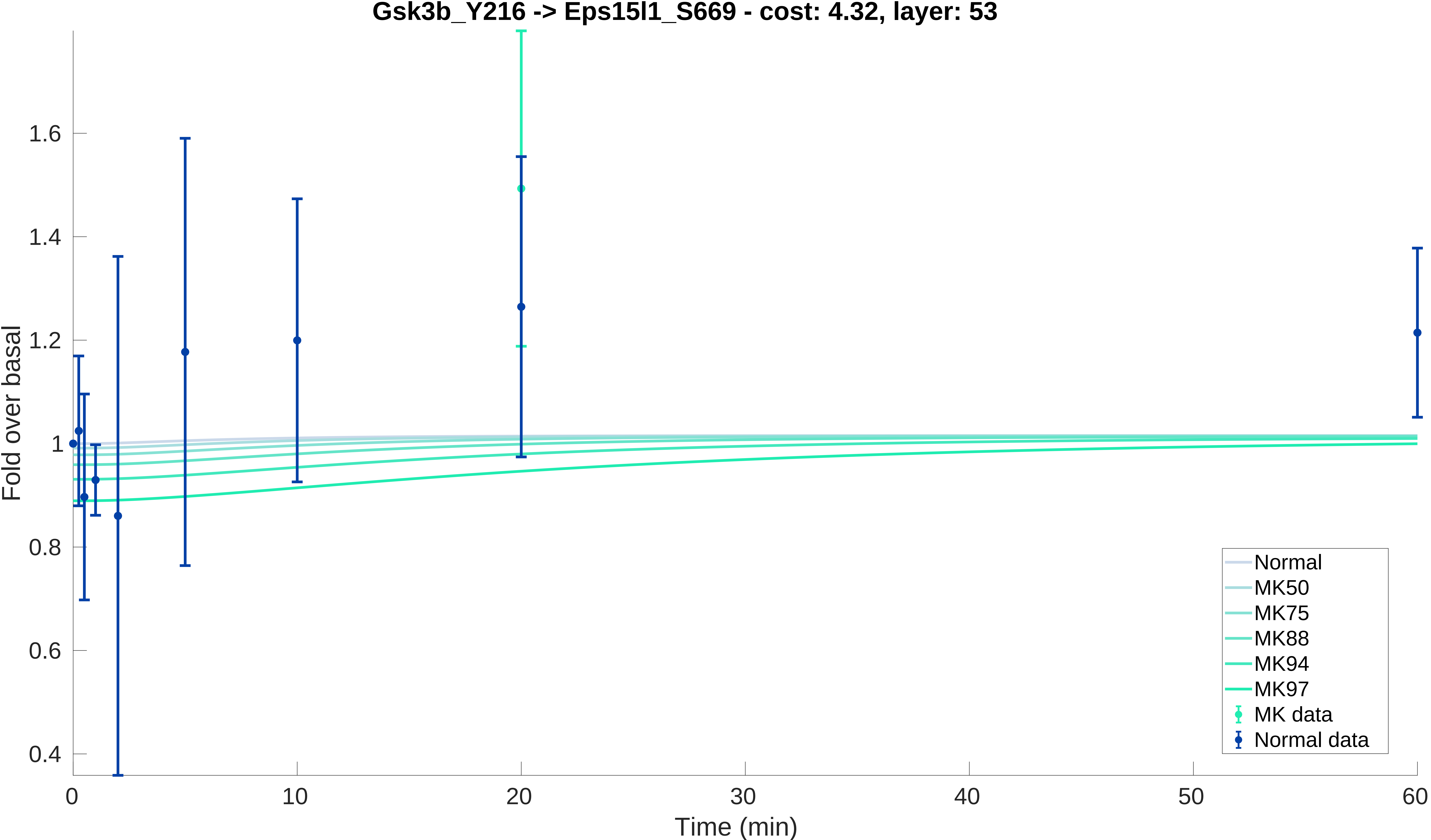

Gsk3b_Y216 -> Eps15l1_S669 - cost: 4.32, layer: 53
Fold over basal
1.6
1.4
1.2
1
0.8
0.6
0.4
0
10
20
30
40
50
60
Time (min)
Normal
MK50
MK75
MK88
MK94
MK97
MK data
Normal data

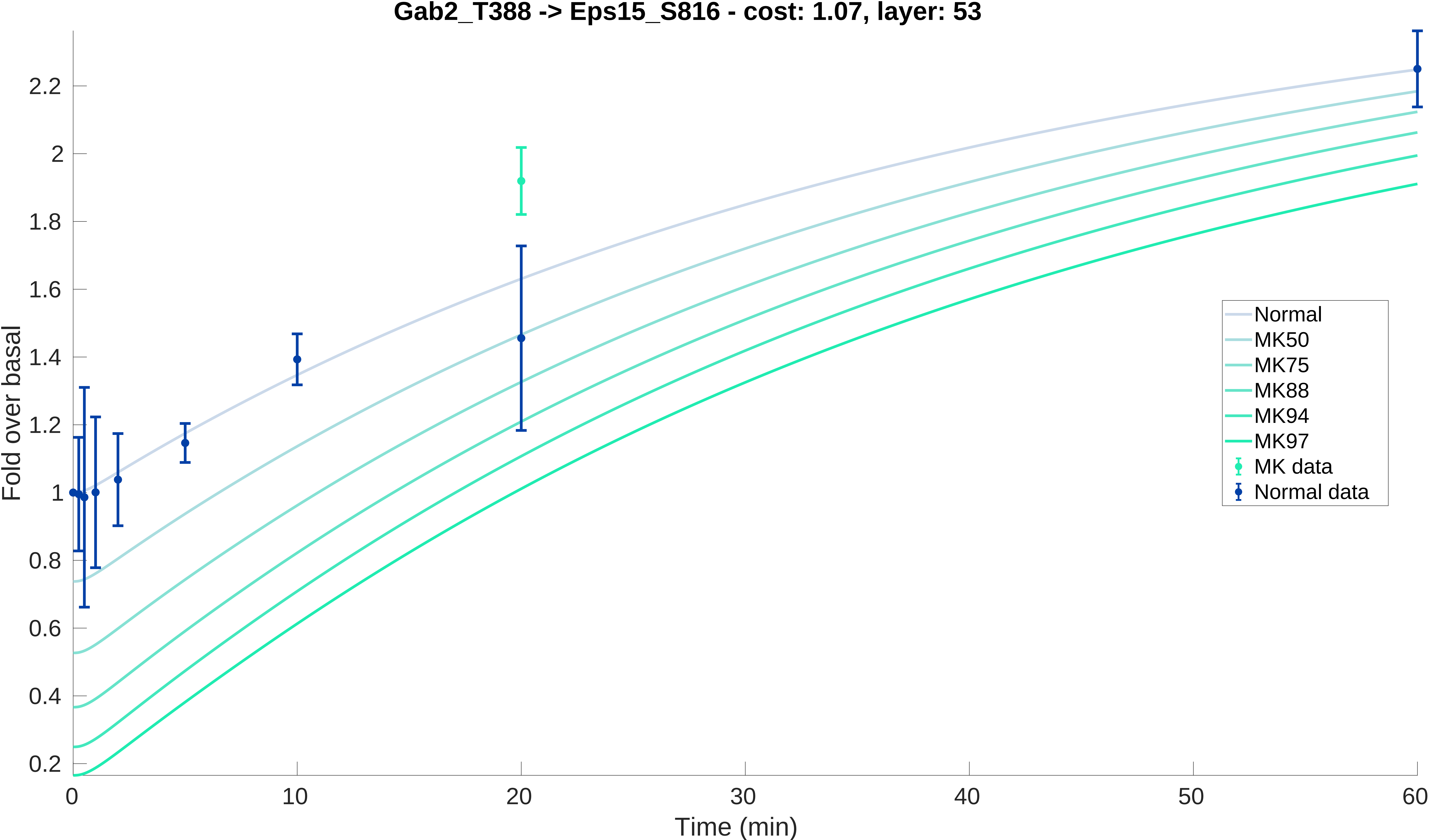

Gab2_T388 -> Eps15_S816 - cost: 1.07, layer: 53
Fold over basal
Time (min)
0
10
20
30
40
50
60
0.2
0.4
0.6
0.8
1
1.2
1.4
1.6
1.8
2
2.2
Normal
MK50
MK75
MK88
MK94
MK97
MK data
Normal data

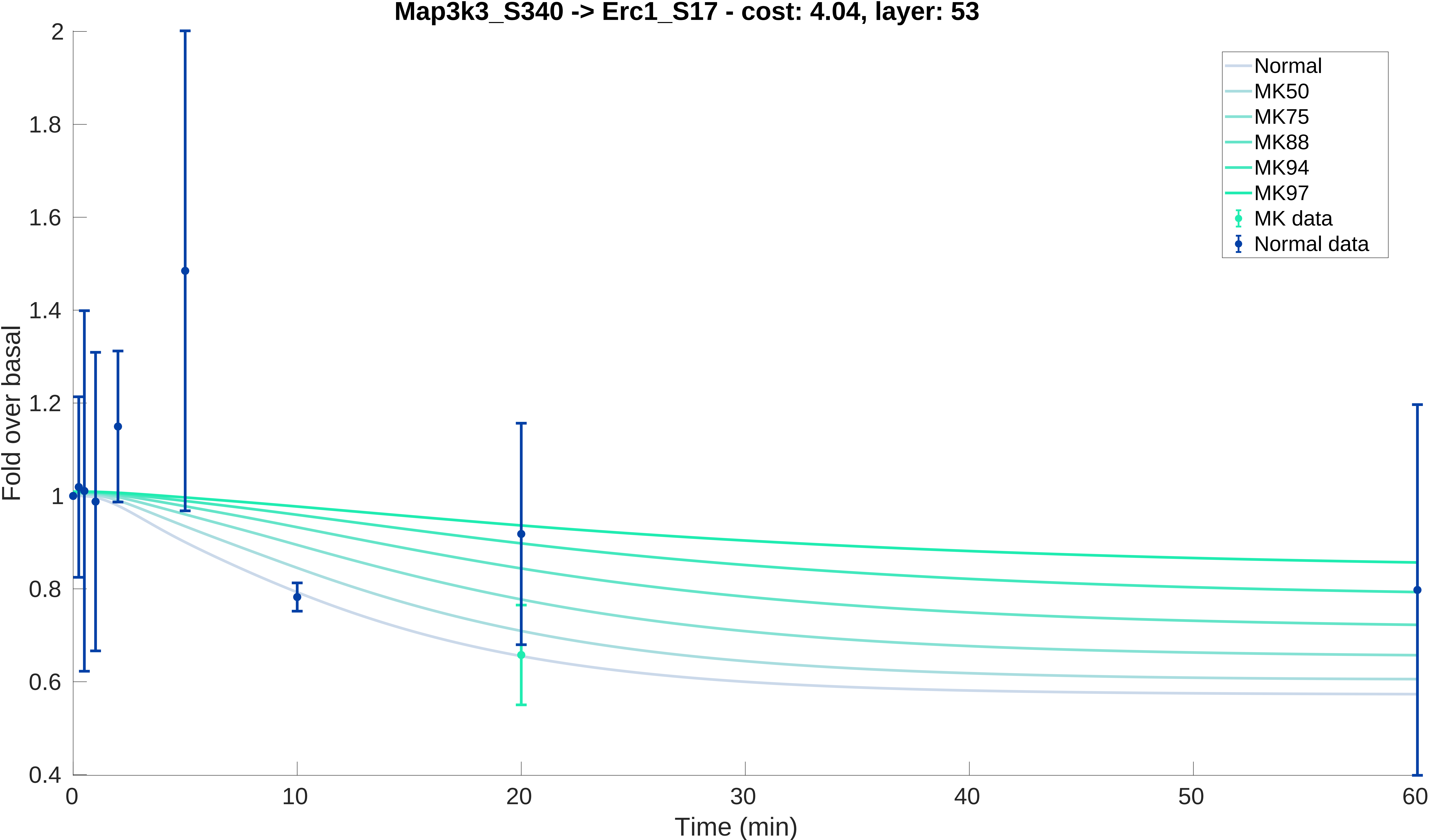

Map3k3_S340 -> Erc1_S17 - cost: 4.04, layer: 53
Fold over basal
Time (min)
0.4
0.6
0.8
1
1.2
1.4
1.6
1.8
2
0
10
20
30
40
50
60
Normal
MK50
MK75
MK88
MK94
MK97
MK data
Normal data

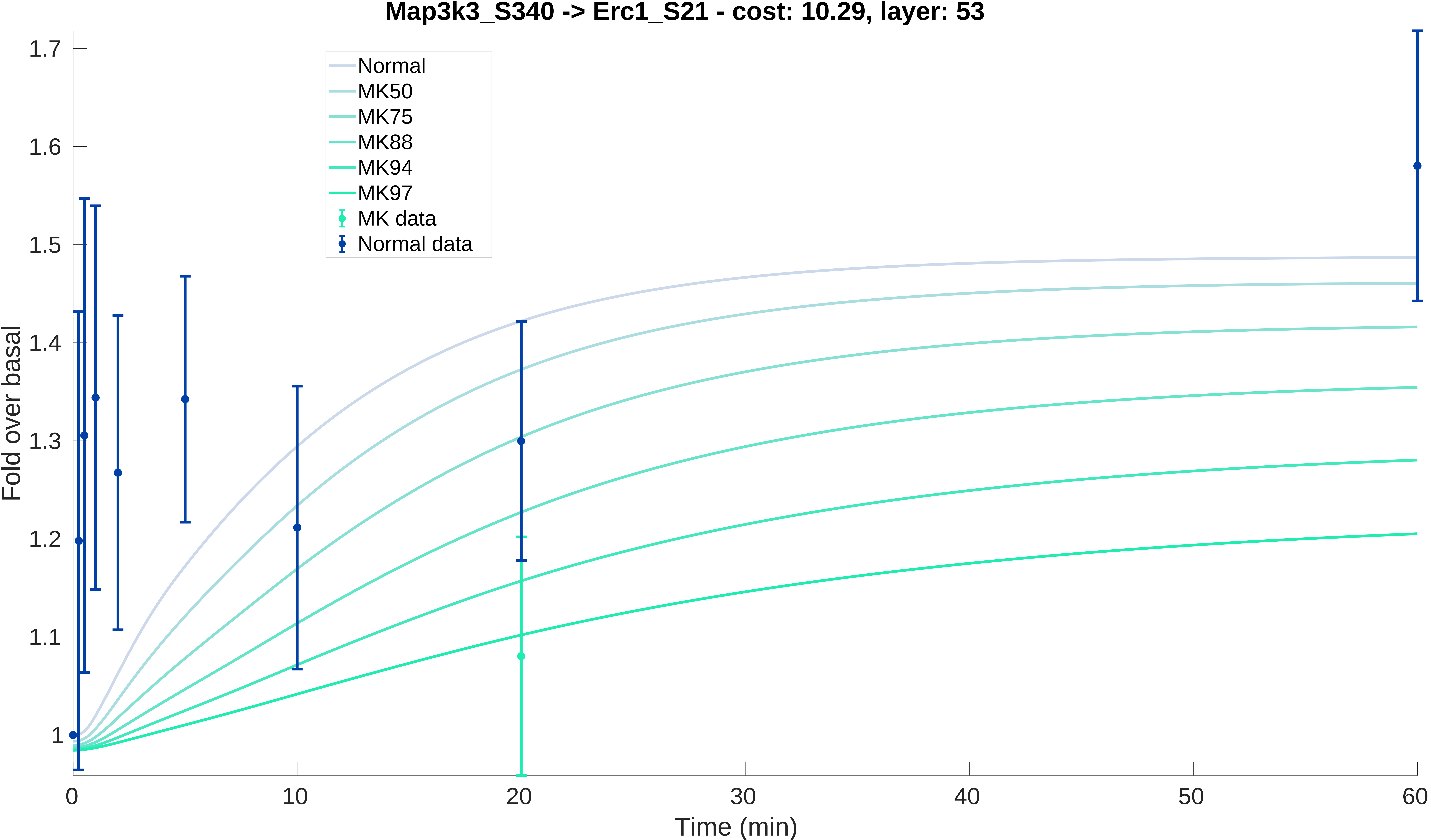

Map3k3_S340 -> Erc1_S21 - cost: 10.29, layer: 53
Normal
MK50
MK75
MK88
MK94
MK97
MK data
Normal data
Fold over basal
1.7
1.6
1.5
1.4
1.3
1.2
1.1
1
0
10
20
30
40
50
60
Time (min)

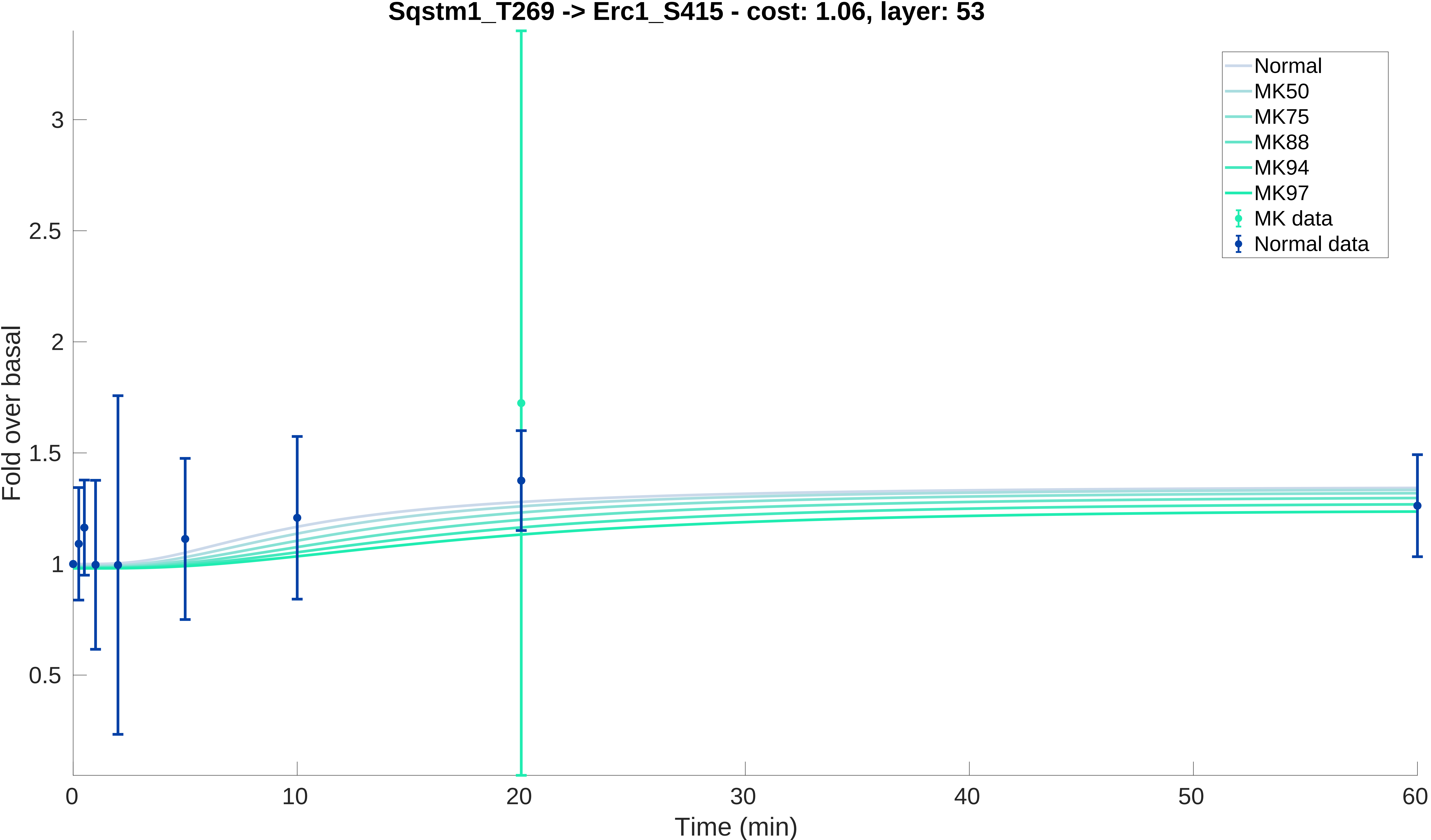

Sqstm1_T269 -> Erc1_S415 - cost: 1.06, layer: 53
Fold over basal
Time (min)
0
10
20
30
40
50
60
0.5
1
1.5
2
2.5
3
Normal
MK50
MK75
MK88
MK94
MK97
MK data
Normal data

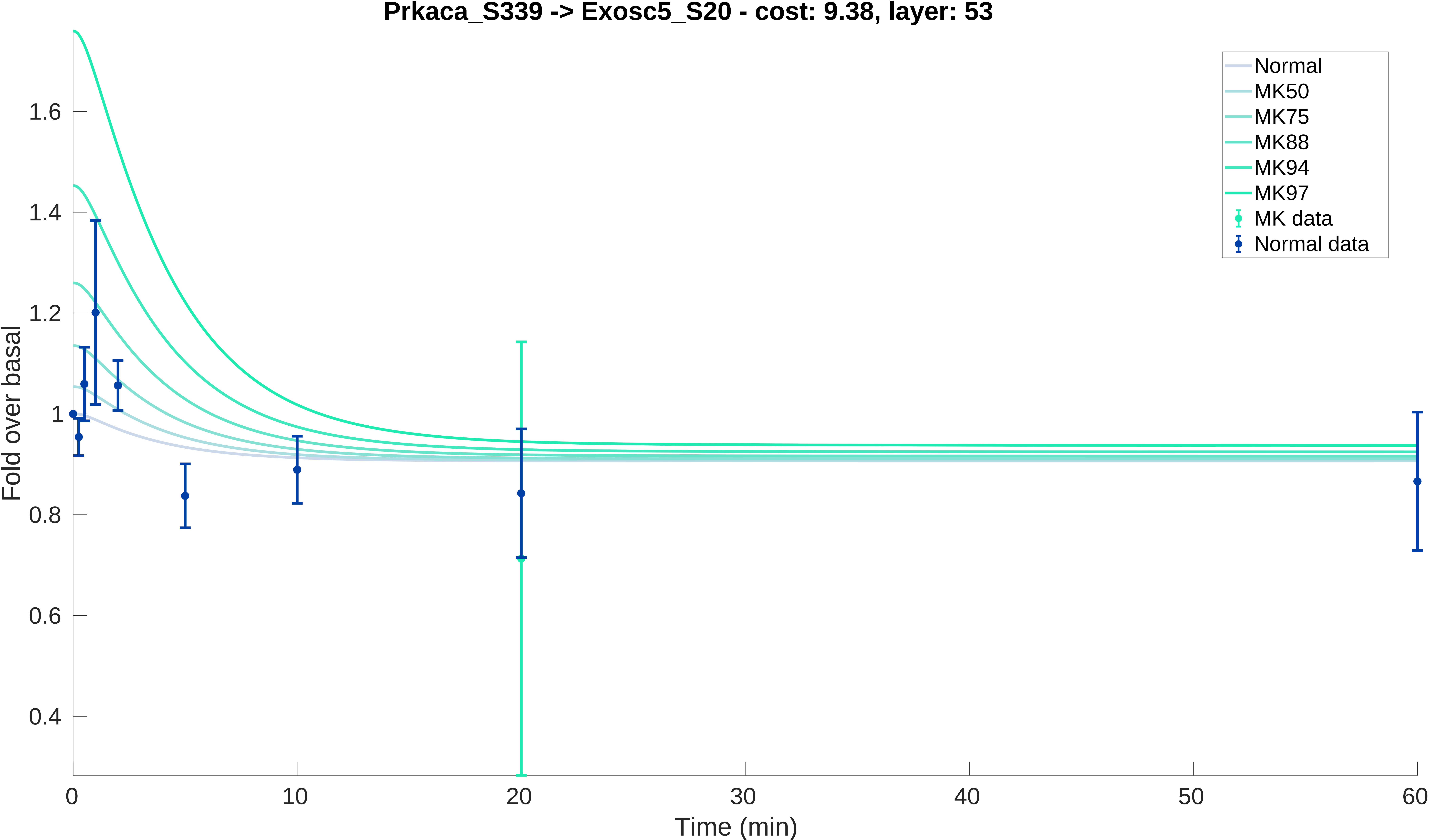

Prrkaca_S339 -> Exosc5_S20 - cost: 9.38, layer: 53
Normal
MK50
MK75
MK88
MK94
MK97
MK data
Normal data
Fold over basal
1.6
1.4
1.2
1
0.8
0.6
0.4
0
10
20
30
40
50
60
Time (min)

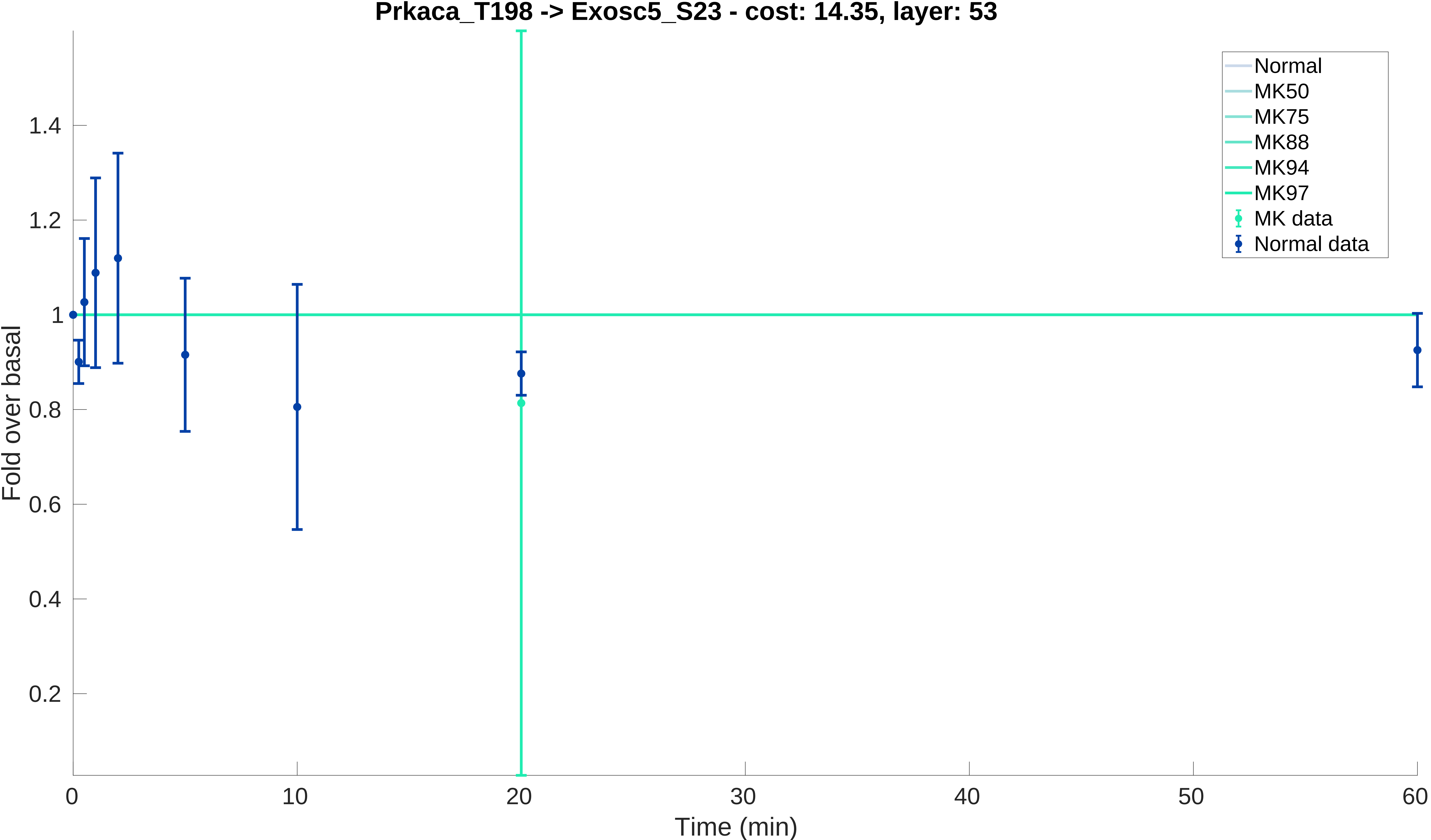

Prkaca_T198 -> Exosc5_S23 - cost: 14.35, layer: 53
Normal
MK50
MK75
MK88
MK94
MK97
MK data
Normal data
Fold over basal
Time (min)
0.2
0.4
0.6
0.8
1
1.2
1.4
0
10
20
30
40
50
60

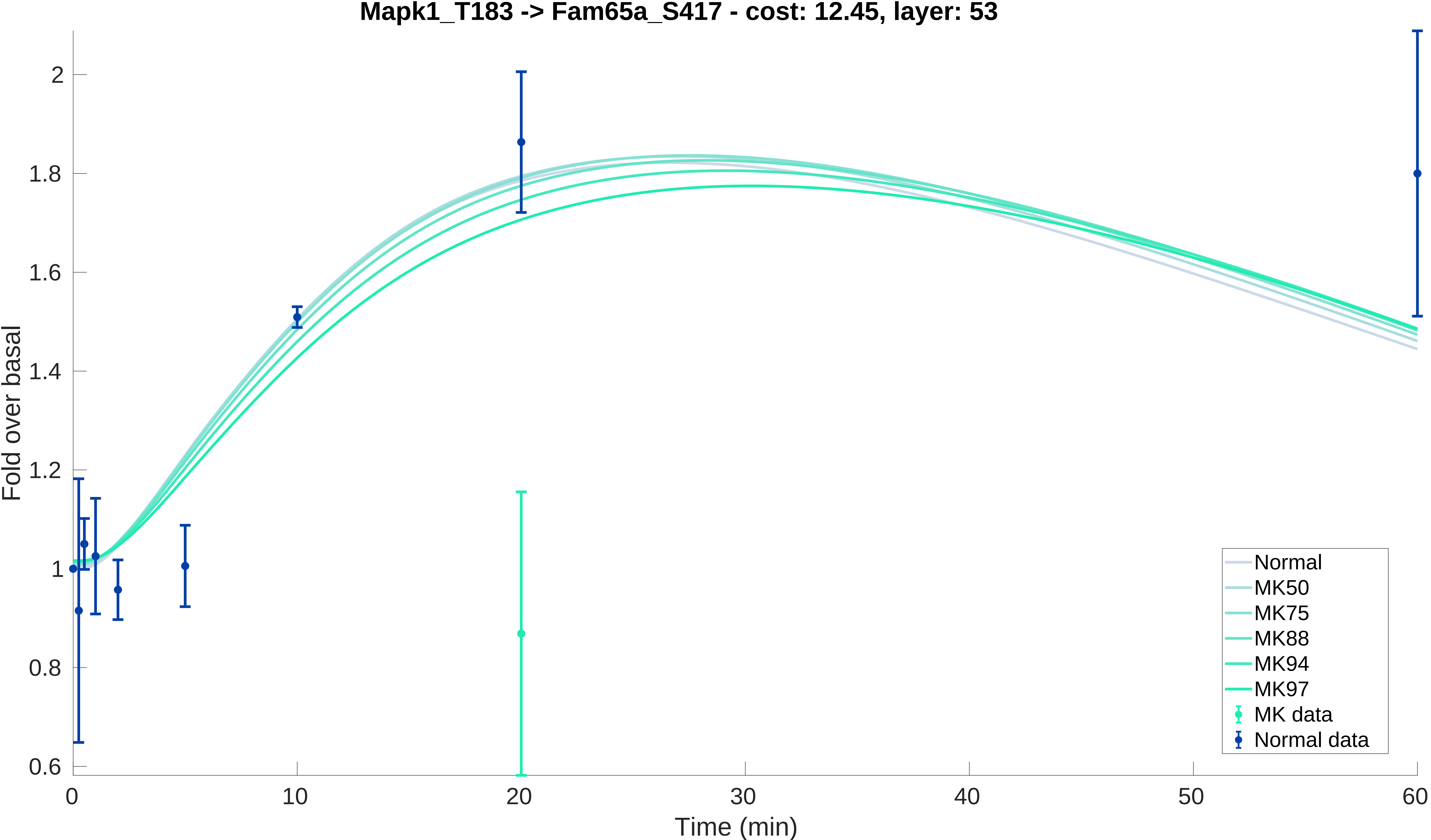
Mapk1_T183 -> Fam65a_S417 - cost: 12.45, layer: 53
Fold over basal
2
1.8
1.6
1.4
1.2
1
0.8
0.6
0
10
20
30
40
50
60
Time (min)
Normal
MK50
MK75
MK88
MK94
MK97
MK data
Normal data

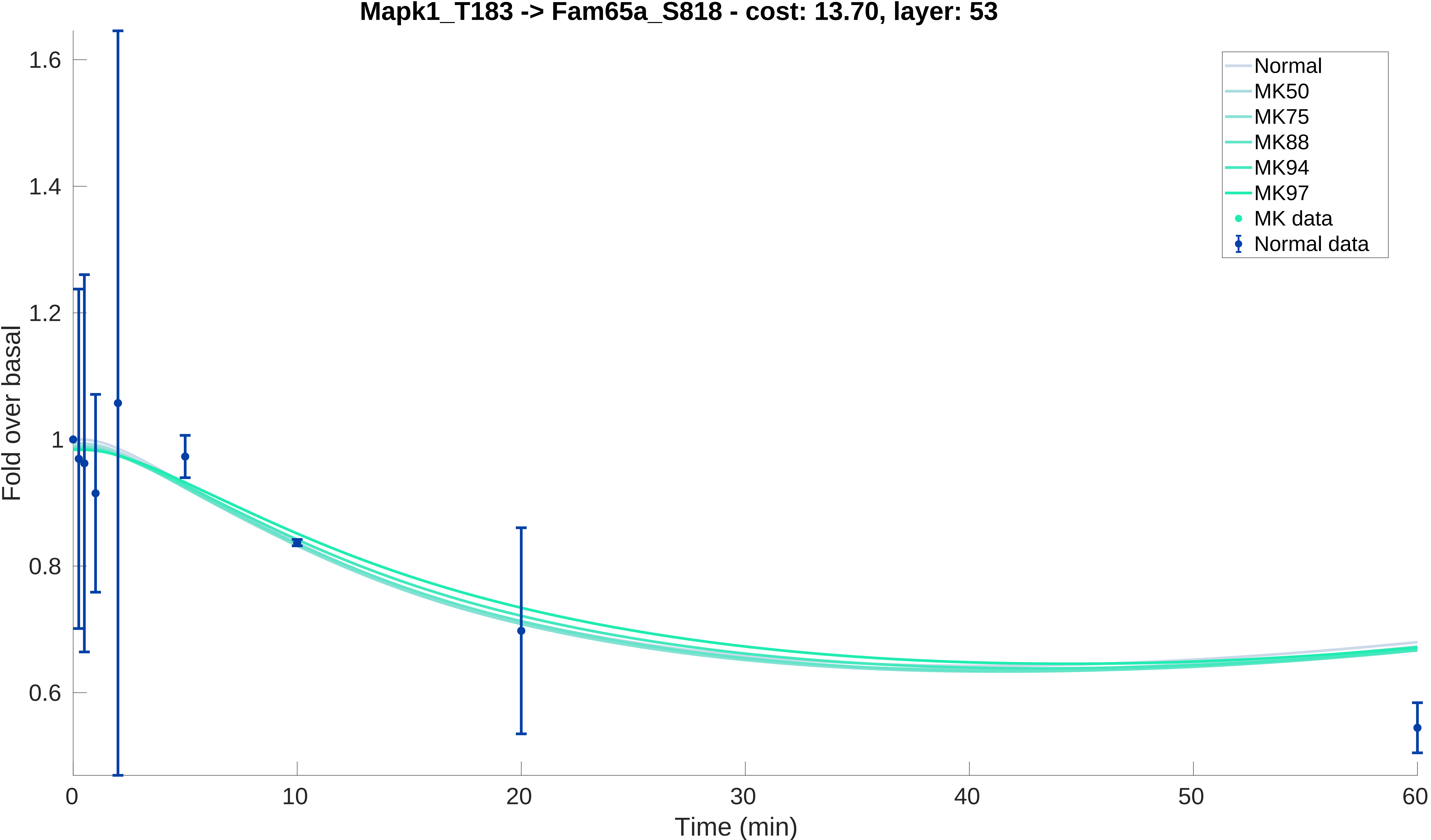
Mapk1_T183 -> Fam65a_S818 - cost: 13.70, layer: 53
Fold over basal
1.6
1.4
1.2
1
0.8
0.6
0
10
20
30
40
50
60
Time (min)
Normal
MK50
MK75
MK88
MK94
MK97
MK data
Normal data

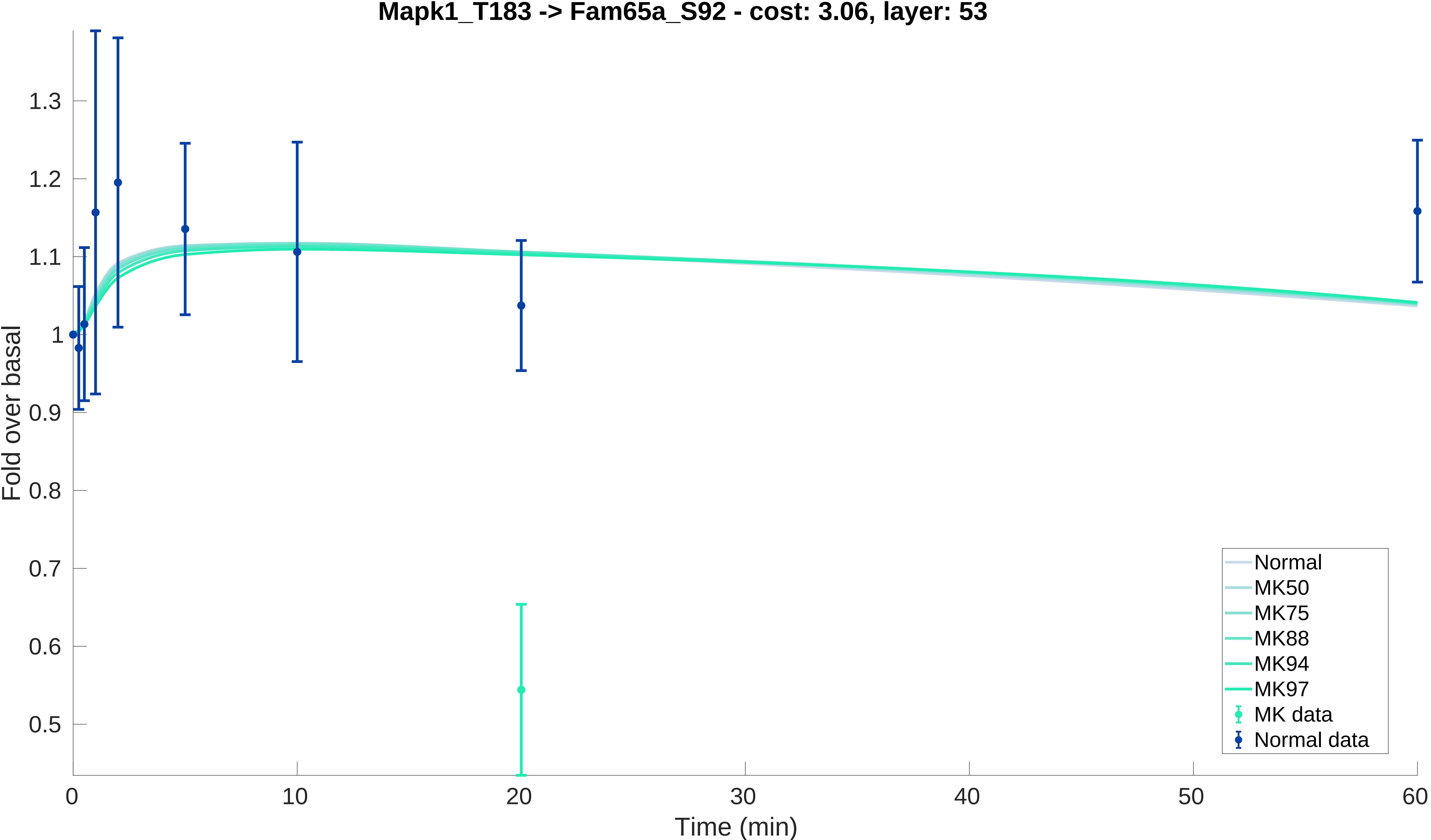

Mapk1_T183 -> Fam65a_S92 - cost: 3.06, layer: 53
1.3
1.2
1.1
1
0.9
0.8
0.7
0.6
0.5
Fold over basal
0
10
20
30
40
50
60
Time (min)
Normal
MK50
MK75
MK88
MK94
MK97
MK data
Normal data

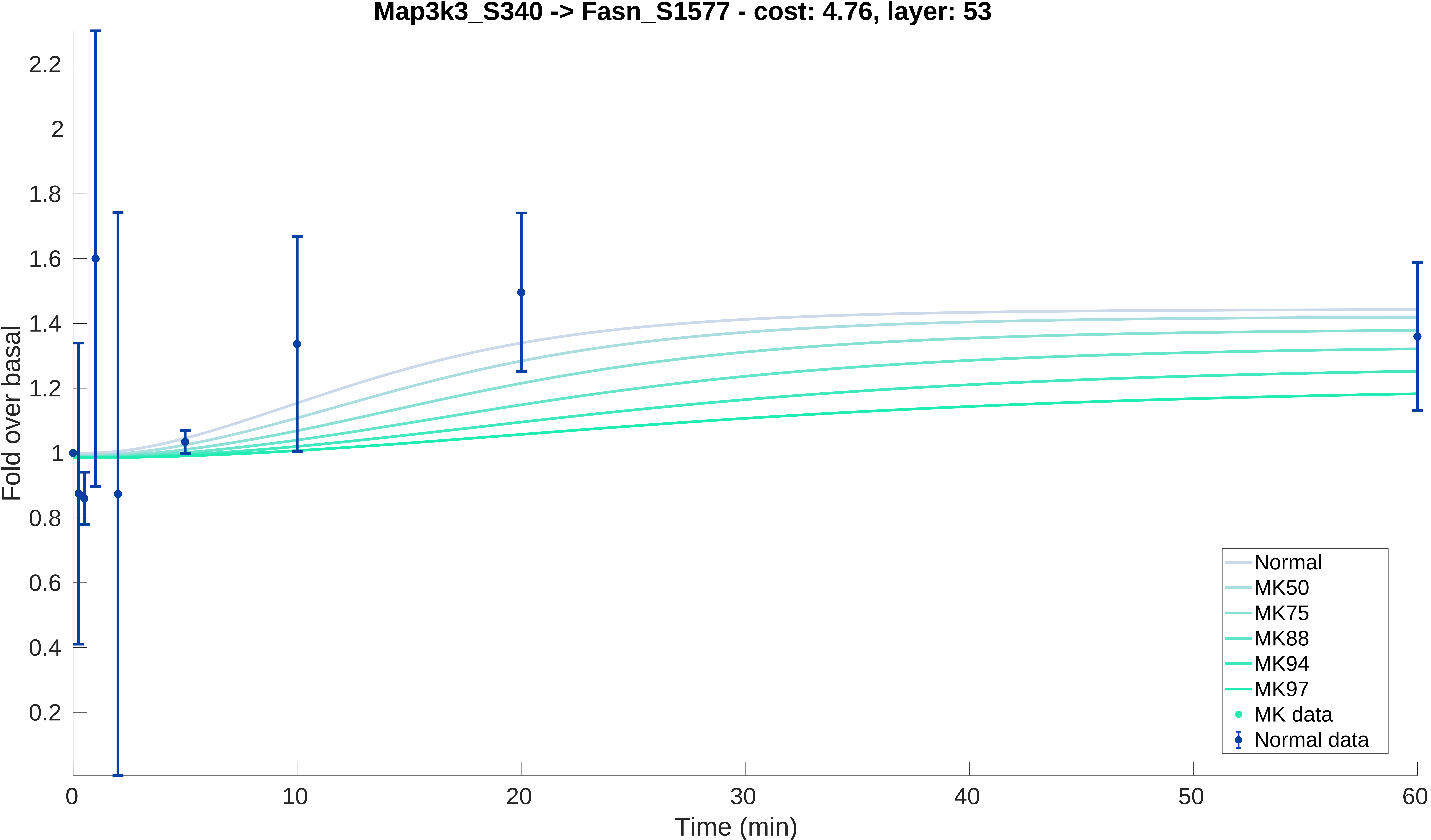
Map3k3_S340 -> Fasn_S1577 - cost: 4.76, layer: 53
Fold over basal
2.2
2
1.8
1.6
1.4
1.2
1
0.8
0.6
0.4
0.2
0
10
20
30
40
50
60
Time (min)
Normal
MK50
MK75
MK88
MK94
MK97
MK data
Normal data

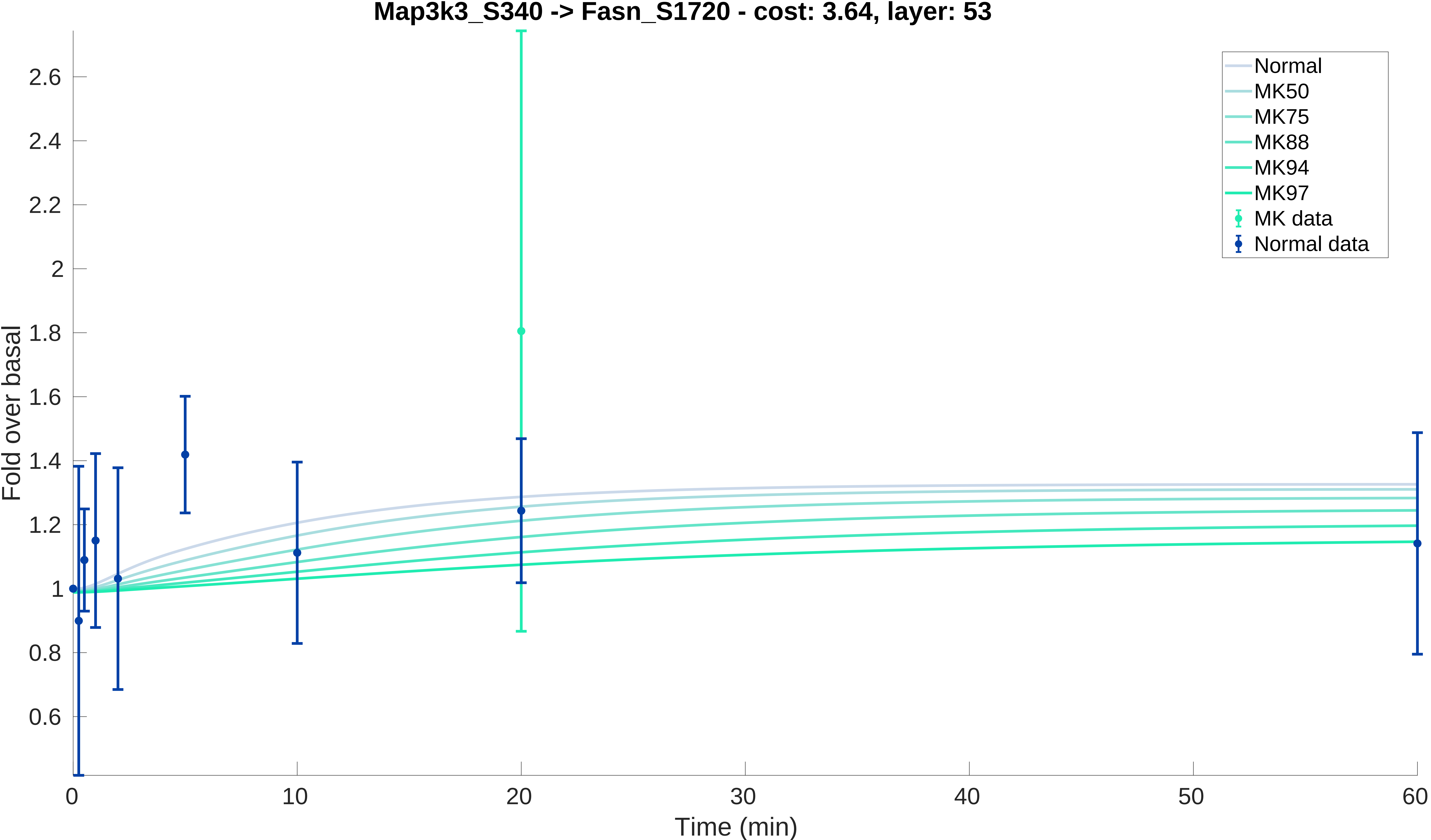

Map3k3_S340 -> Fasn_S1720 - cost: 3.64, layer: 53
Fold over basal
Time (min)
2.6
2.4
2.2
2
1.8
1.6
1.4
1.2
1
0.8
0.6
0
10
20
30
40
50
60
Normal
MK50
MK75
MK88
MK94
MK97
MK data
Normal data

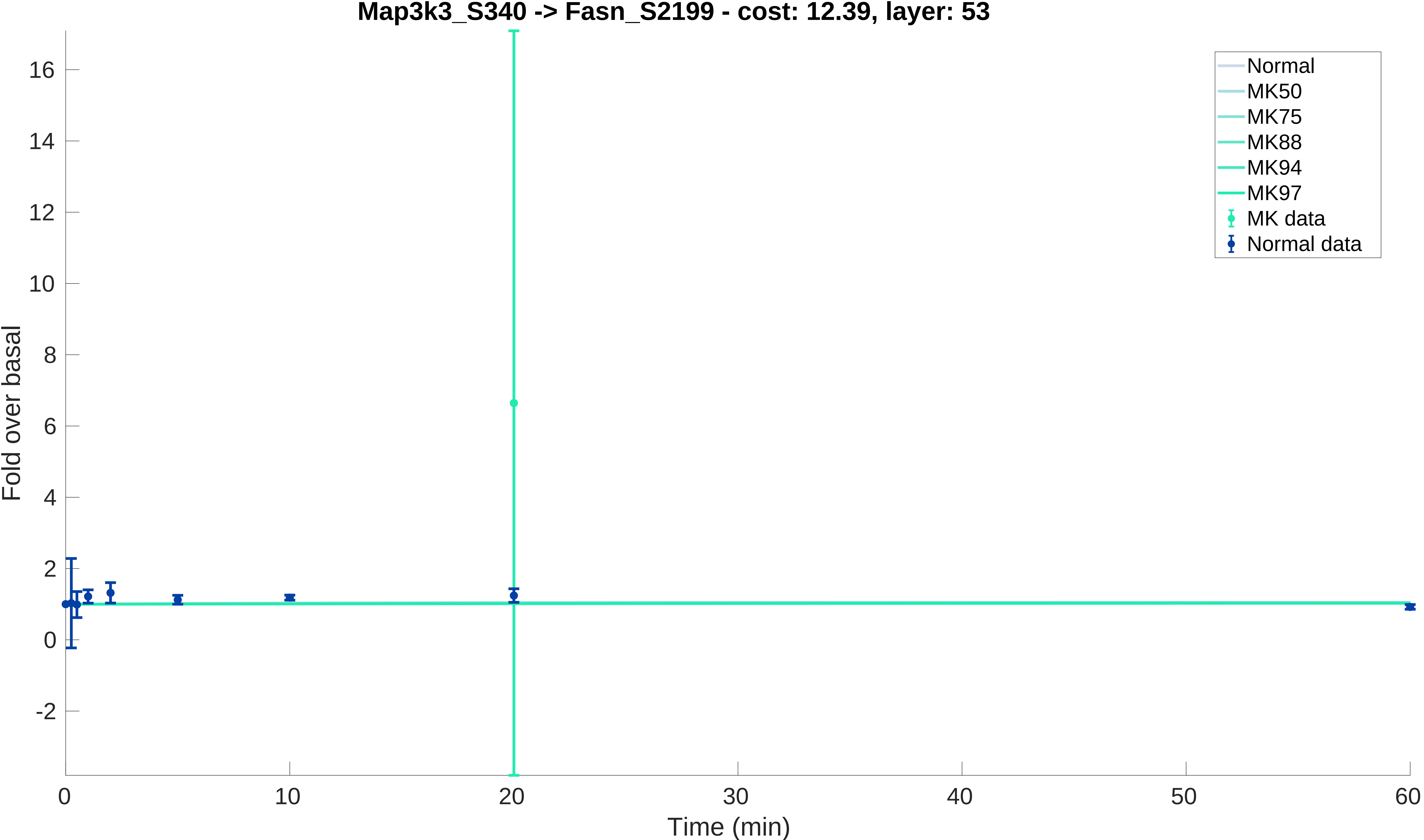

Map3k3_S340 -> Fasn_S2199 - cost: 12.39, layer: 53
Fold over basal
Time (min)
16
14
12
10
8
6
4
2
0
-2
0
10
20
30
40
50
60
Normal
MK50
MK75
MK88
MK94
MK97
MK data
Normal data

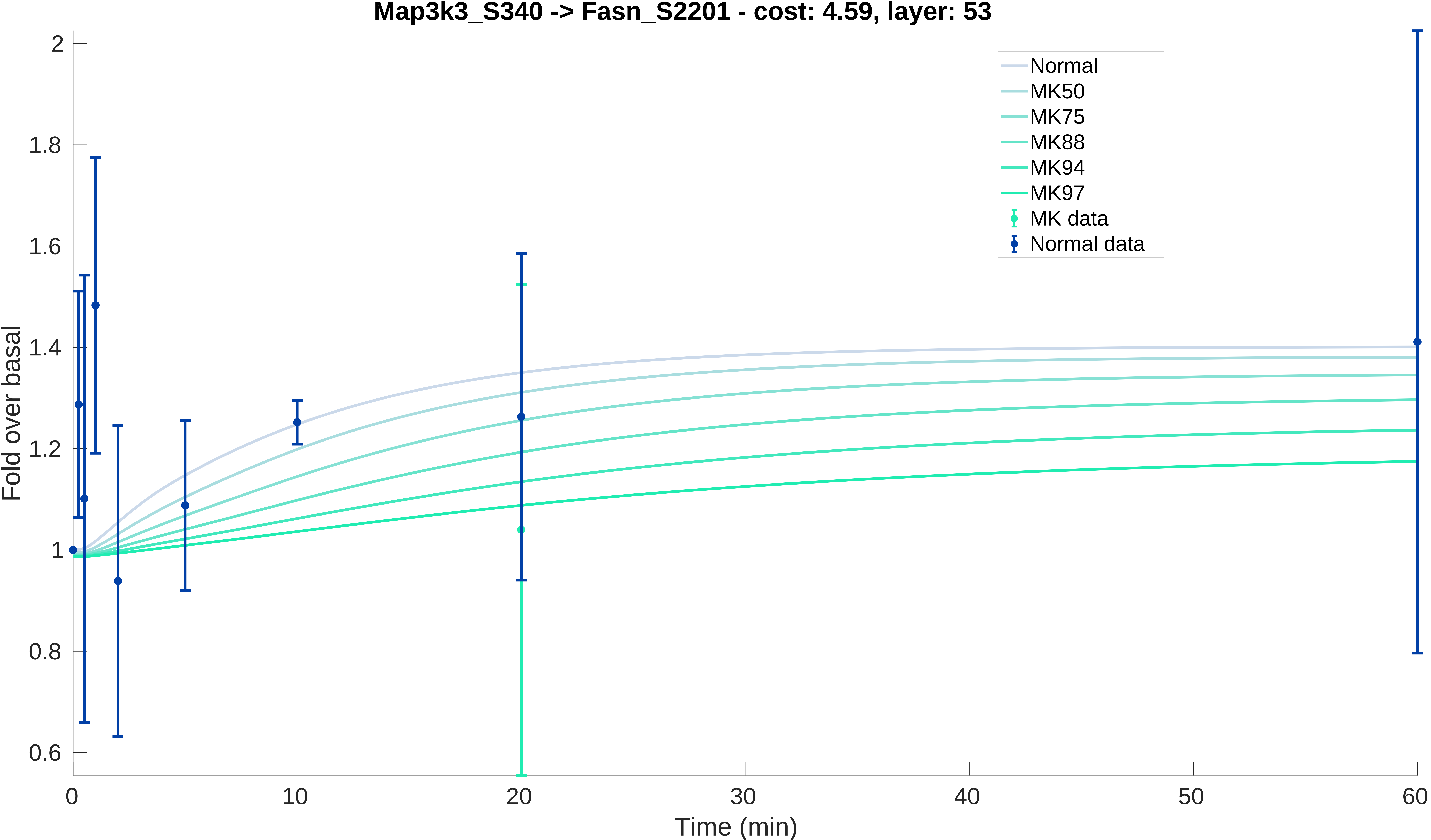
Map3k3_S340 -> Fasn_S2201 - cost: 4.59, layer: 53
Fold over basal
Time (min)
0.6
0.8
1
1.2
1.4
1.6
1.8
2
0
10
20
30
40
50
60
Normal
MK50
MK75
MK88
MK94
MK97
MK data
Normal data

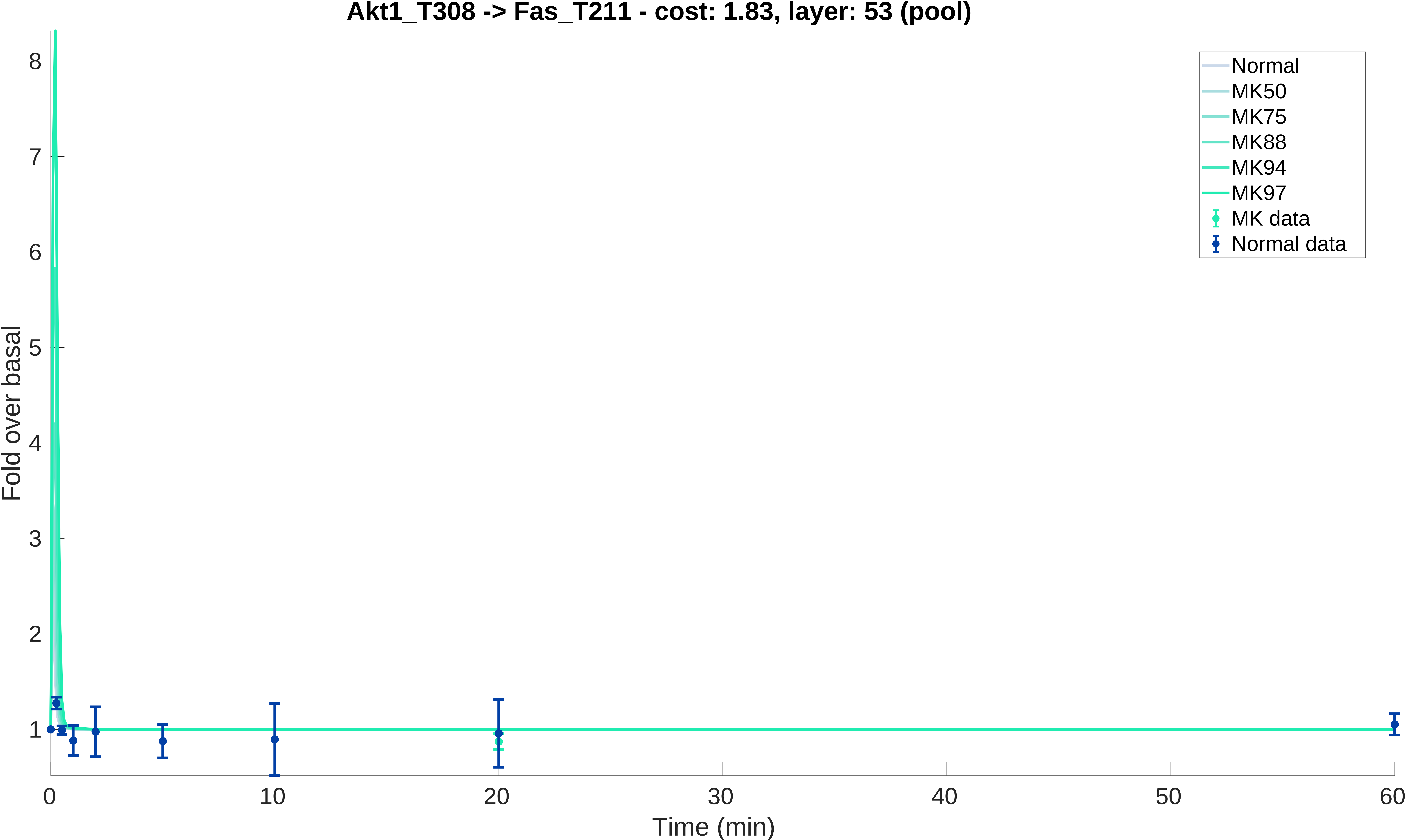

Akt1_T308 -> Fas_T211 - cost: 1.83, layer: 53 (pool)
Fold over basal
Time (min)
0
10
20
30
40
50
60
1
2
3
4
5
6
7
8
Normal
MK50
MK75
MK88
MK94
MK97
MK data
Normal data

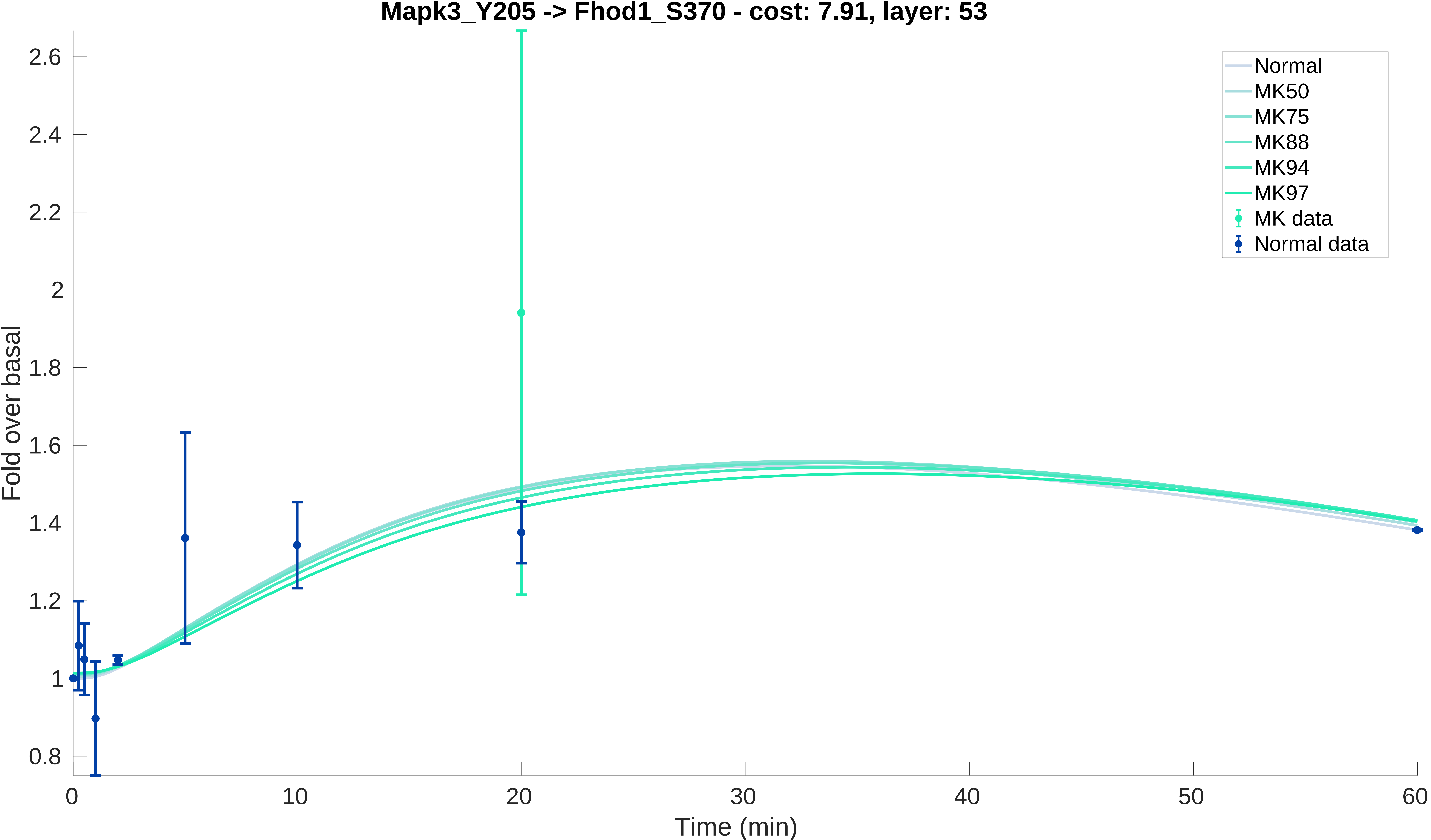

Mapk3_Y205 -> Fhod1_S370 - cost: 7.91, layer: 53
Fold over basal
2.6
2.4
2.2
2
1.8
1.6
1.4
1.2
1
0.8
0
10
20
30
40
50
60
Time (min)
Normal
MK50
MK75
MK88
MK94
MK97
MK data
Normal data

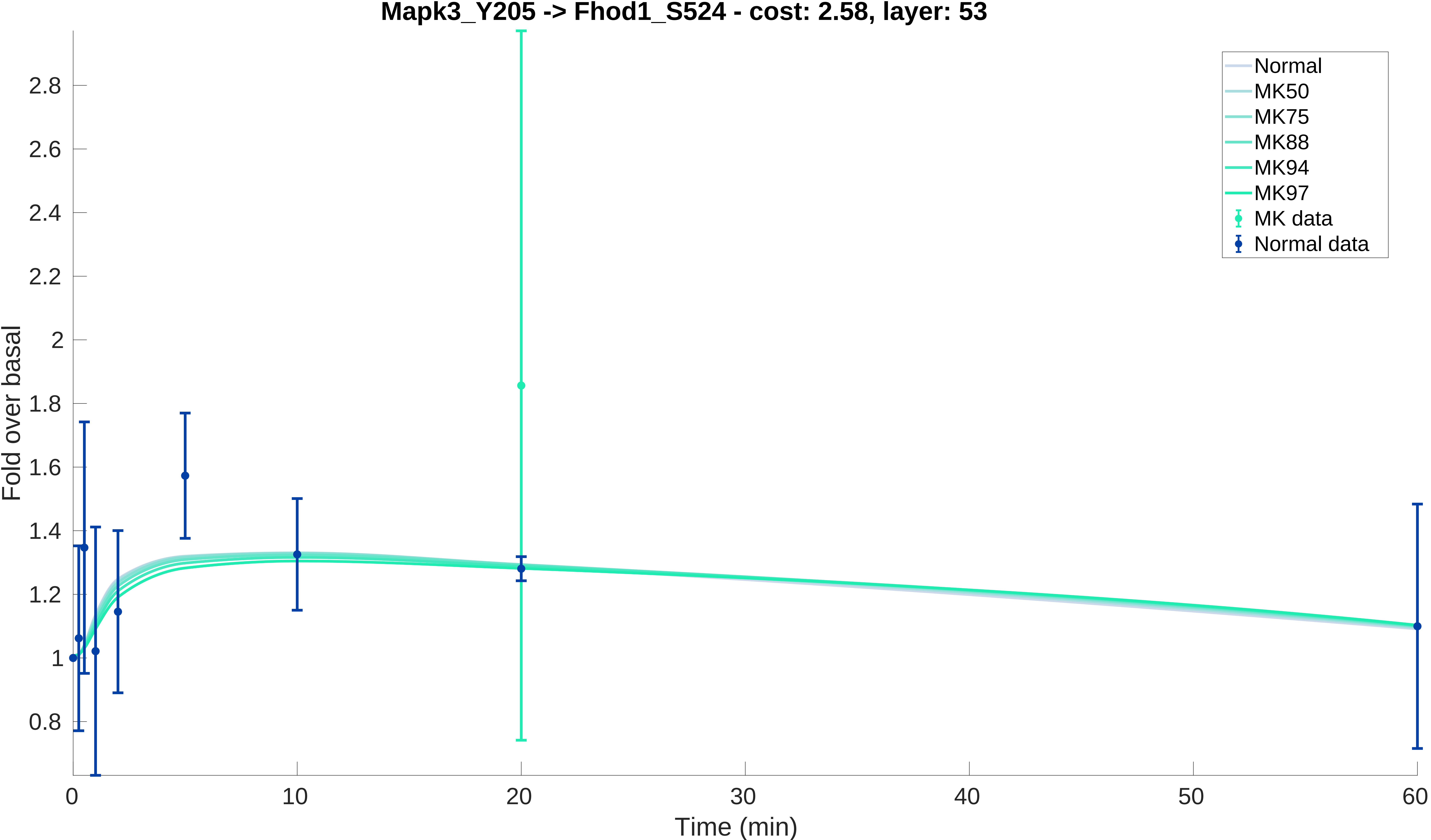
Mapk3_Y205 -> Fhod1_S524 - cost: 2.58, layer: 53
Fold over basal
Time (min)
0
10
20
30
40
50
60
0.8
1
1.2
1.4
1.6
1.8
2
2.2
2.4
2.6
2.8
Normal
MK50
MK75
MK88
MK94
MK97
MK data
Normal data

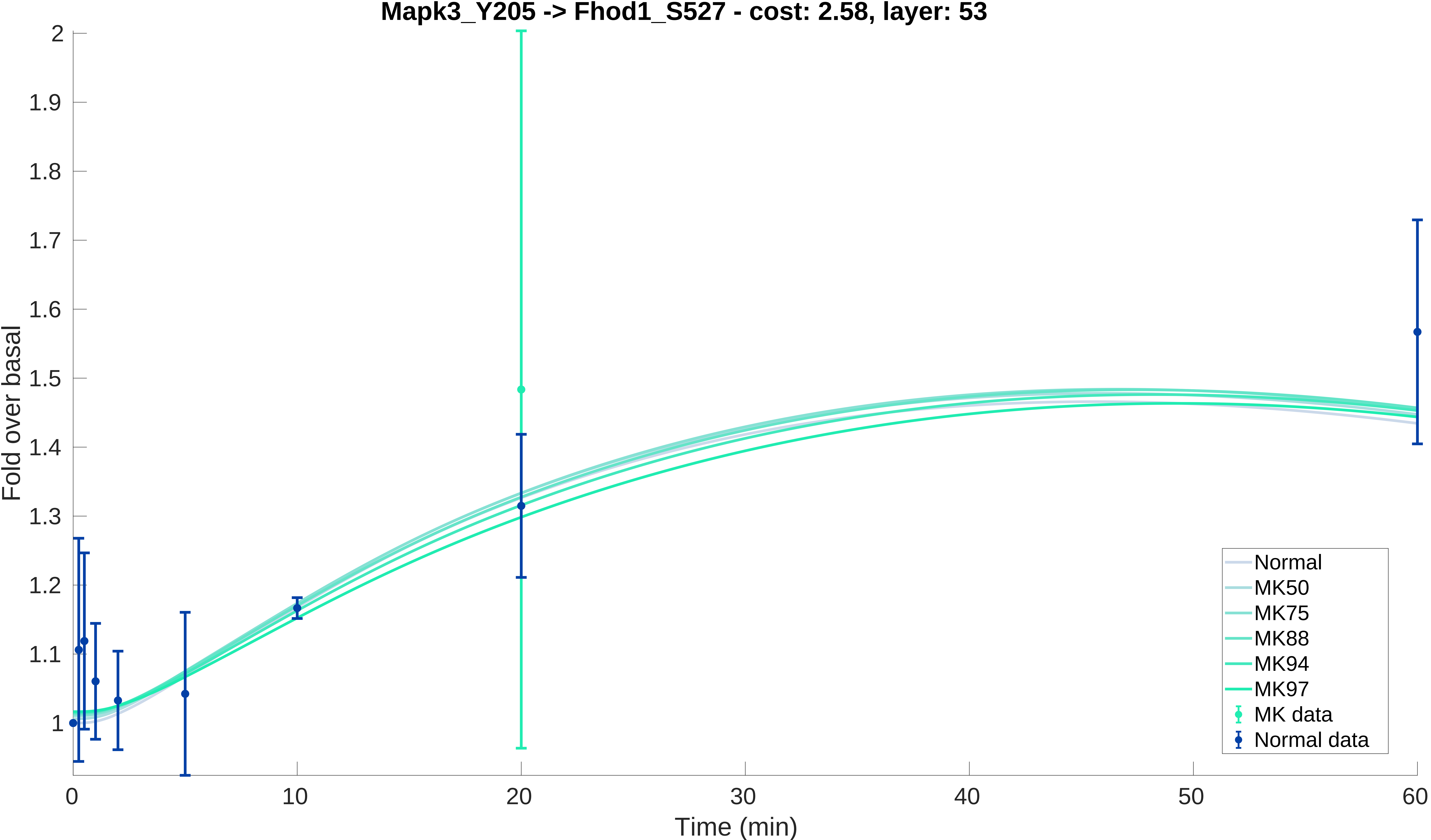
Mapk3_Y205 -> Fhod1_S527 - cost: 2.58, layer: 53
Fold over basal
2
1.9
1.8
1.7
1.6
1.5
1.4
1.3
1.2
1.1
1
0
10
20
30
40
50
60
Time (min)
Normal
MK50
MK75
MK88
MK94
MK97
MK data
Normal data

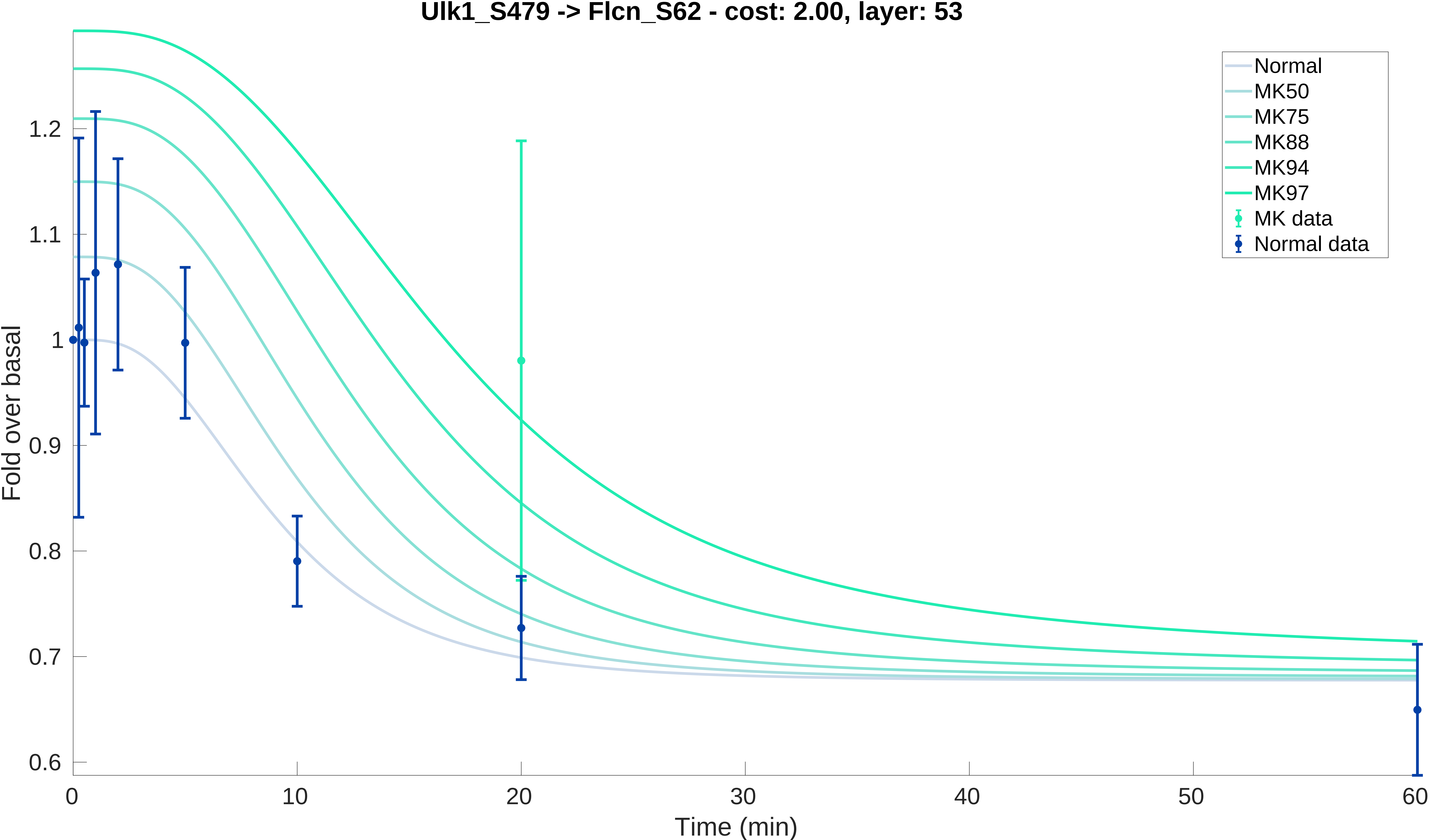

Ulk1_S479 -> Flcn_S62 - cost: 2.00, layer: 53
Fold over basal
Time (min)
0.6
0.7
0.8
0.9
1
1.1
1.2
0
10
20
30
40
50
60
Normal
MK50
MK75
MK88
MK94
MK97
MK data
Normal data

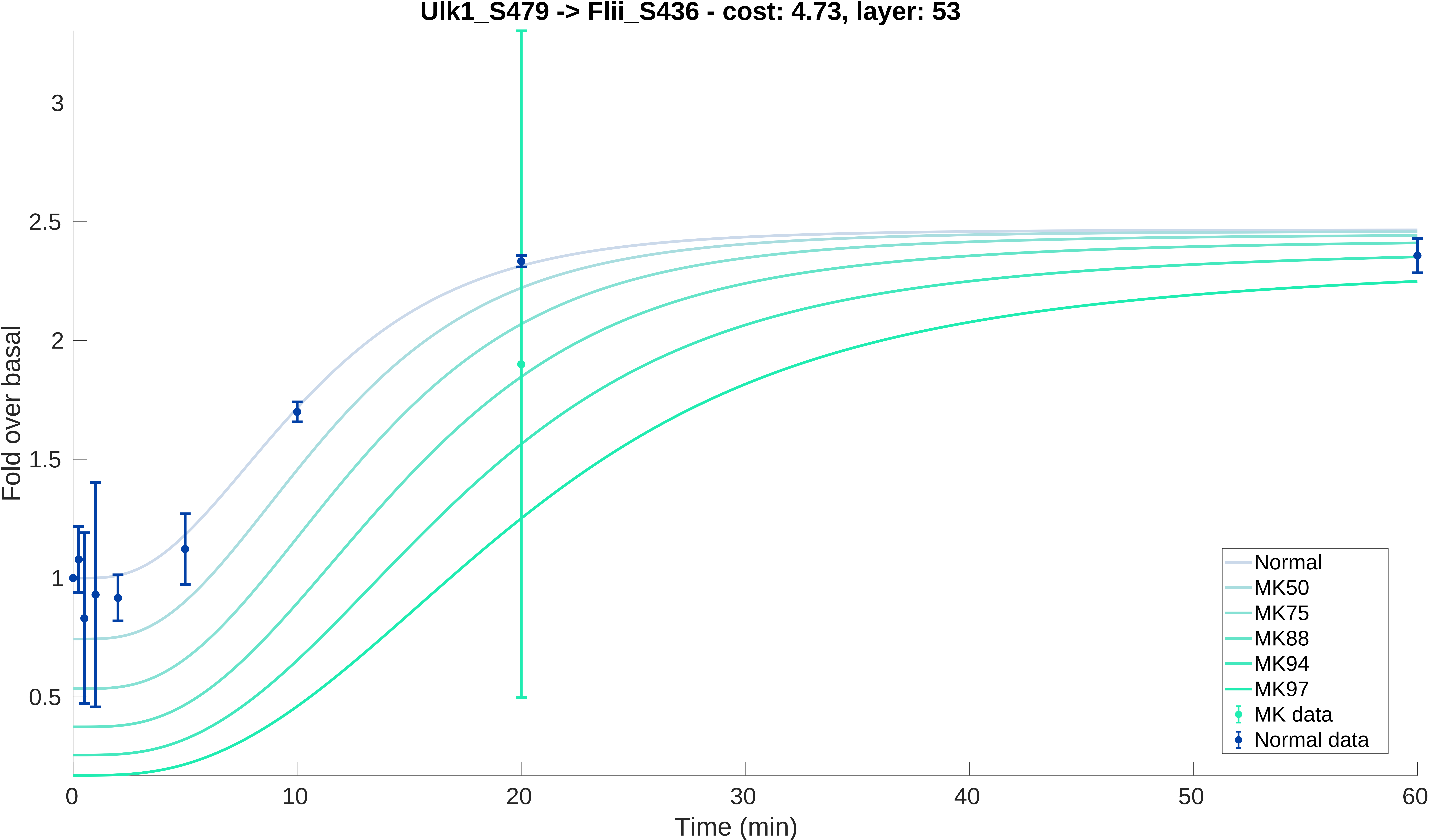

Ulk1_S479 -> Flii_S436 - cost: 4.73, layer: 53
Fold over basal
Time (min)
0
10
20
30
40
50
60
0.5
1
1.5
2
2.5
3
Normal
MK50
MK75
MK88
MK94
MK97
MK data
Normal data

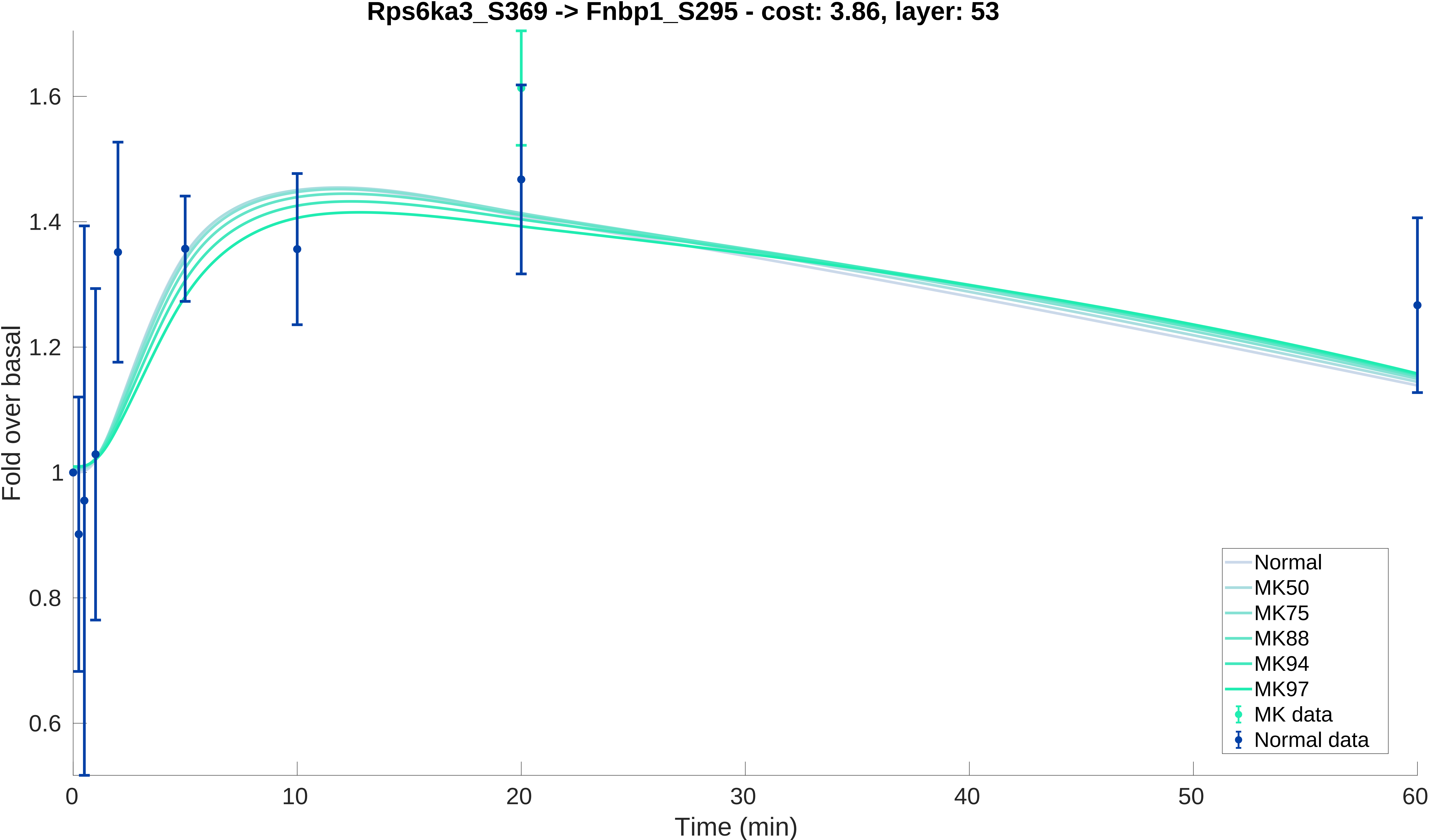

Rps6ka3_S369 -> Fnbp1_S295 - cost: 3.86, layer: 53
Fold over basal
1.6
1.4
1.2
1
0.8
0.6
0
10
20
30
40
50
60
Time (min)
Normal
MK50
MK75
MK88
MK94
MK97
MK data
Normal data

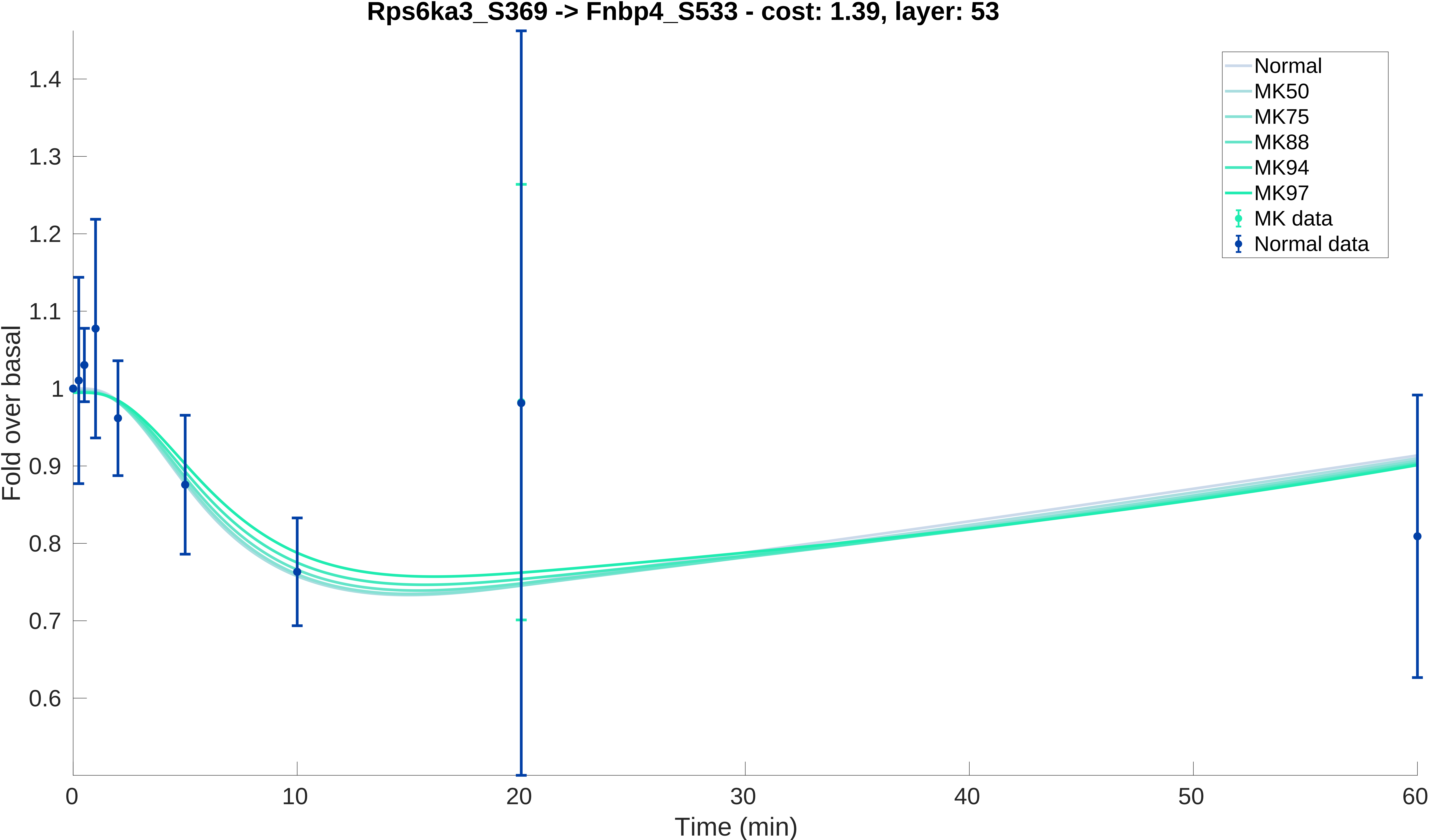
Rps6ka3_S369 -> Fnbp4_S533 - cost: 1.39, layer: 53
Fold over basal
1.4
1.3
1.2
1.1
1
0.9
0.8
0.7
0.6
0
10
20
30
40
50
60
Time (min)
Normal
MK50
MK75
MK88
MK94
MK97
MK data
Normal data

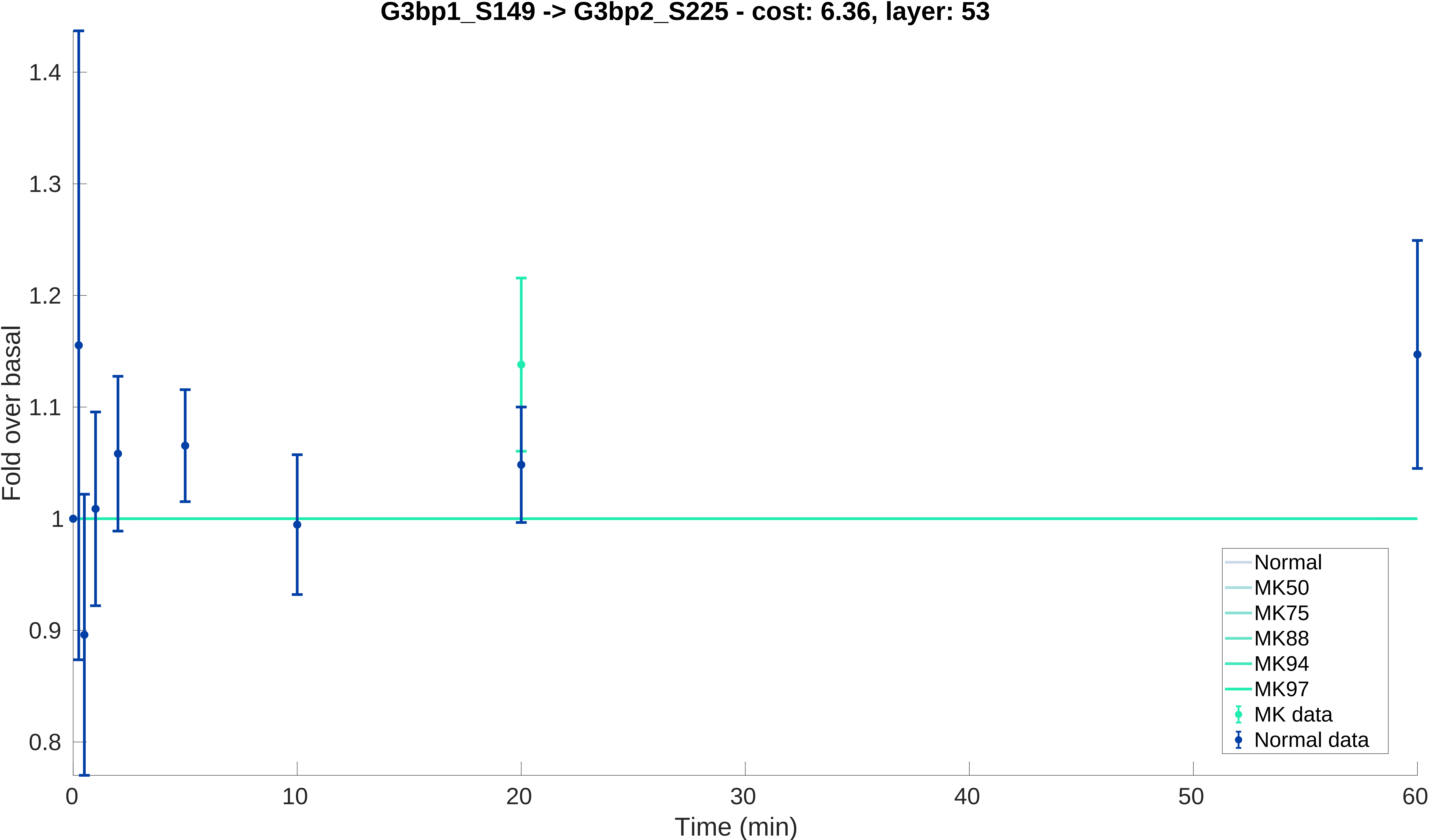

G3bp1_S149 -> G3bp2_S225 - cost: 6.36, layer: 53
Fold over basal
1.4
1.3
1.2
1.1
1
0.9
0.8
0
10
20
30
40
50
60
Time (min)
Normal
MK50
MK75
MK88
MK94
MK97
MK data
Normal data

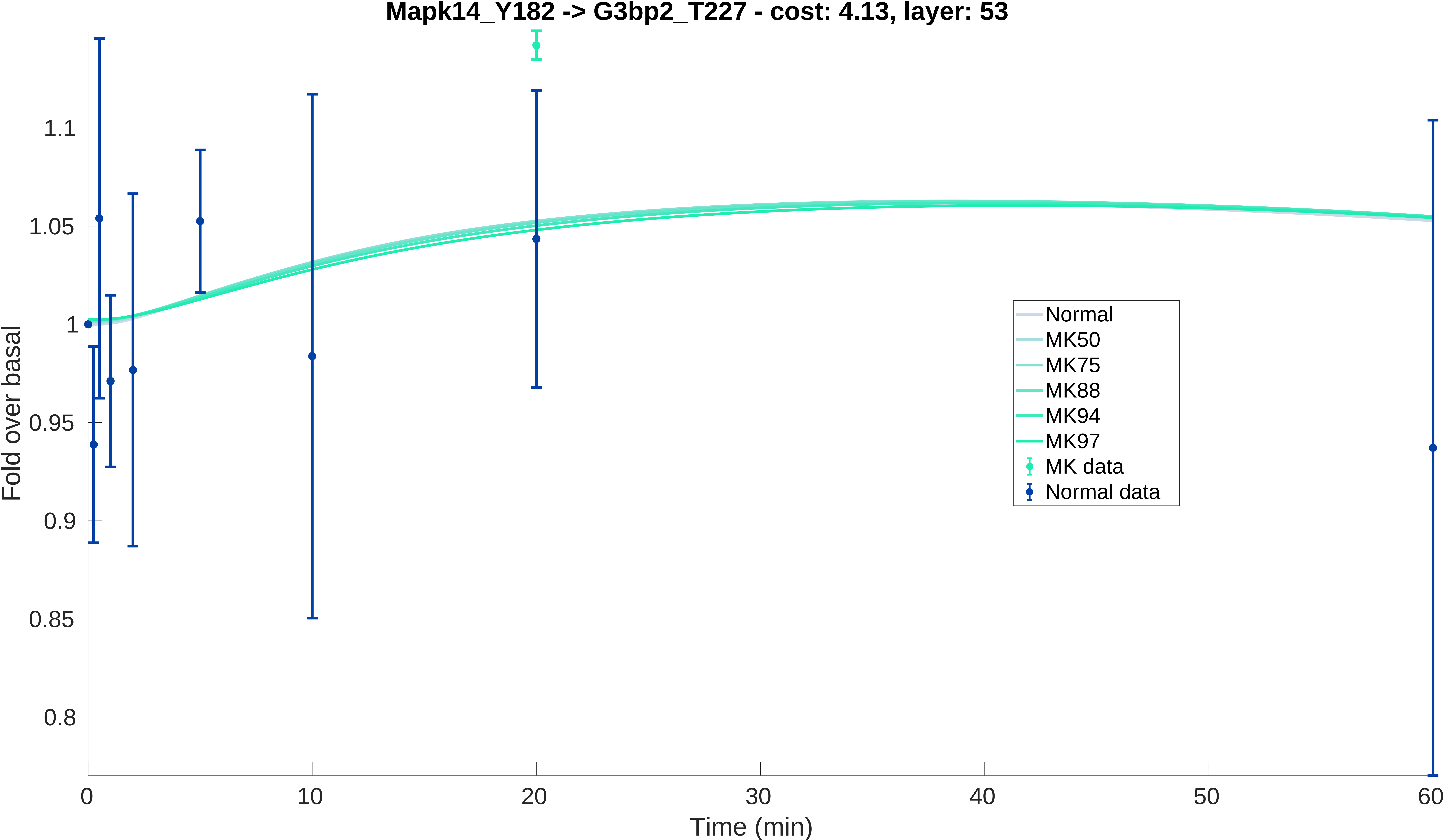

Mapk14_Y182 -> G3bp2_T227 - cost: 4.13, layer: 53
Fold over basal
Time (min)
1.1
1.05
1
0.95
0.9
0.85
0.8
0
10
20
30
40
50
60
Normal
MK50
MK75
MK88
MK94
MK97
MK data
Normal data

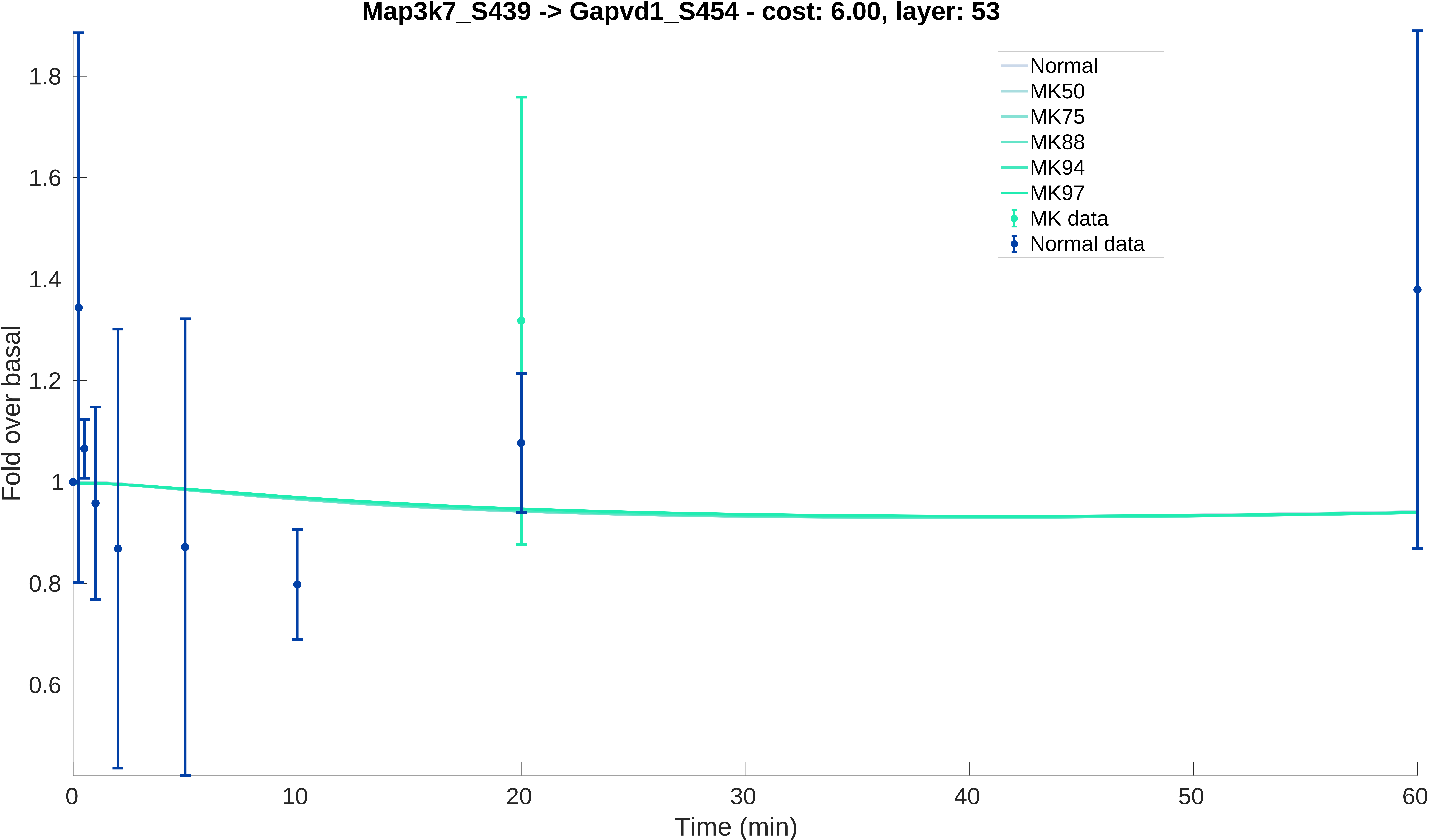

Map3k7_S439 -> Gapvd1_S454 - cost: 6.00, layer: 53
Normal
MK50
MK75
MK88
MK94
MK97
MK data
Normal data
Fold over basal
1.8
1.6
1.4
1.2
1
0.8
0.6
0
10
20
30
40
50
60
Time (min)

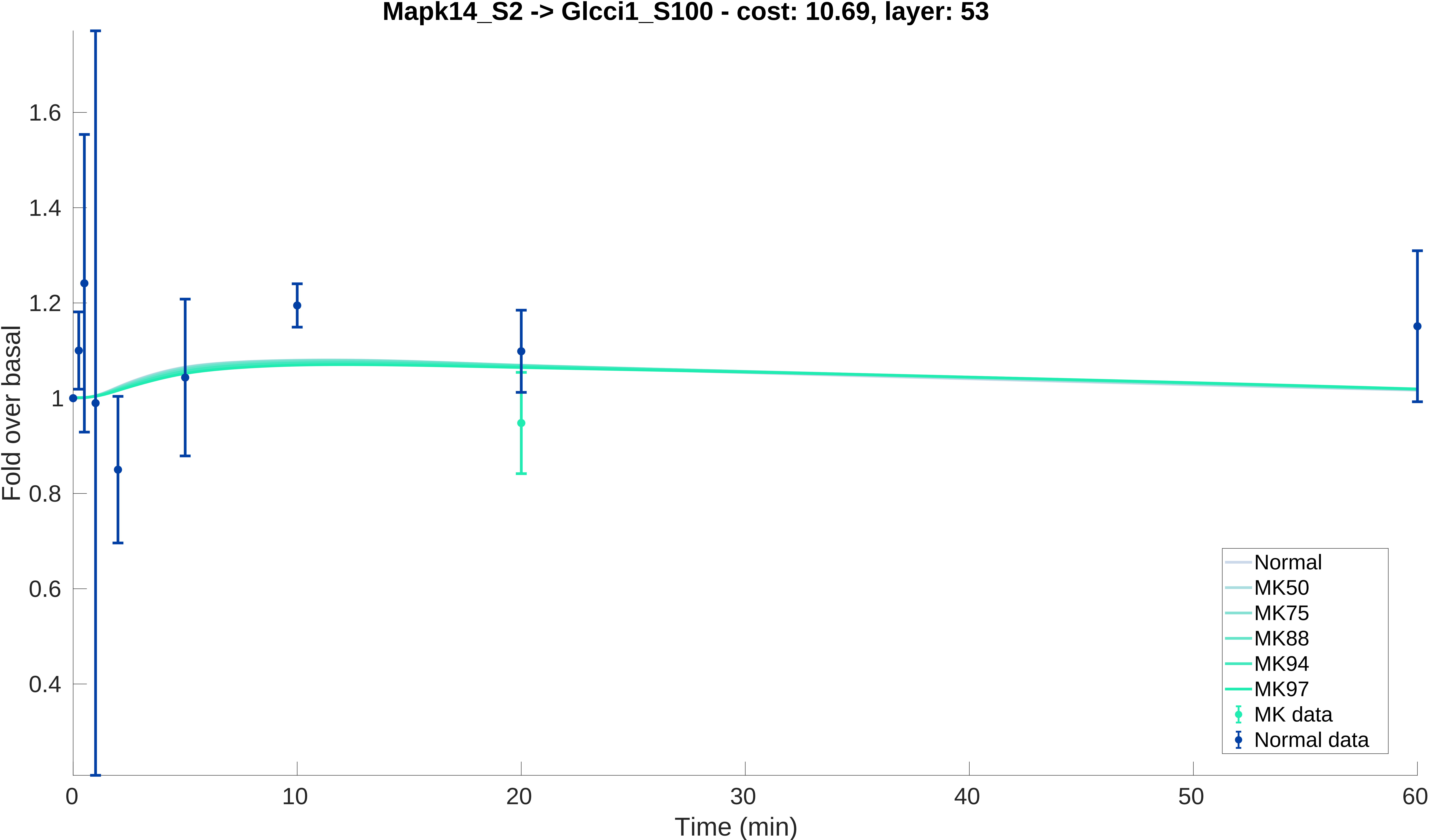

Mapk14_S2 -> Glcci1_S100 - cost: 10.69, layer: 53
Fold over basal
1.6
1.4
1.2
1
0.8
0.6
0.4
0
10
20
30
40
50
60
Time (min)
Normal
MK50
MK75
MK88
MK94
MK97
MK data
Normal data

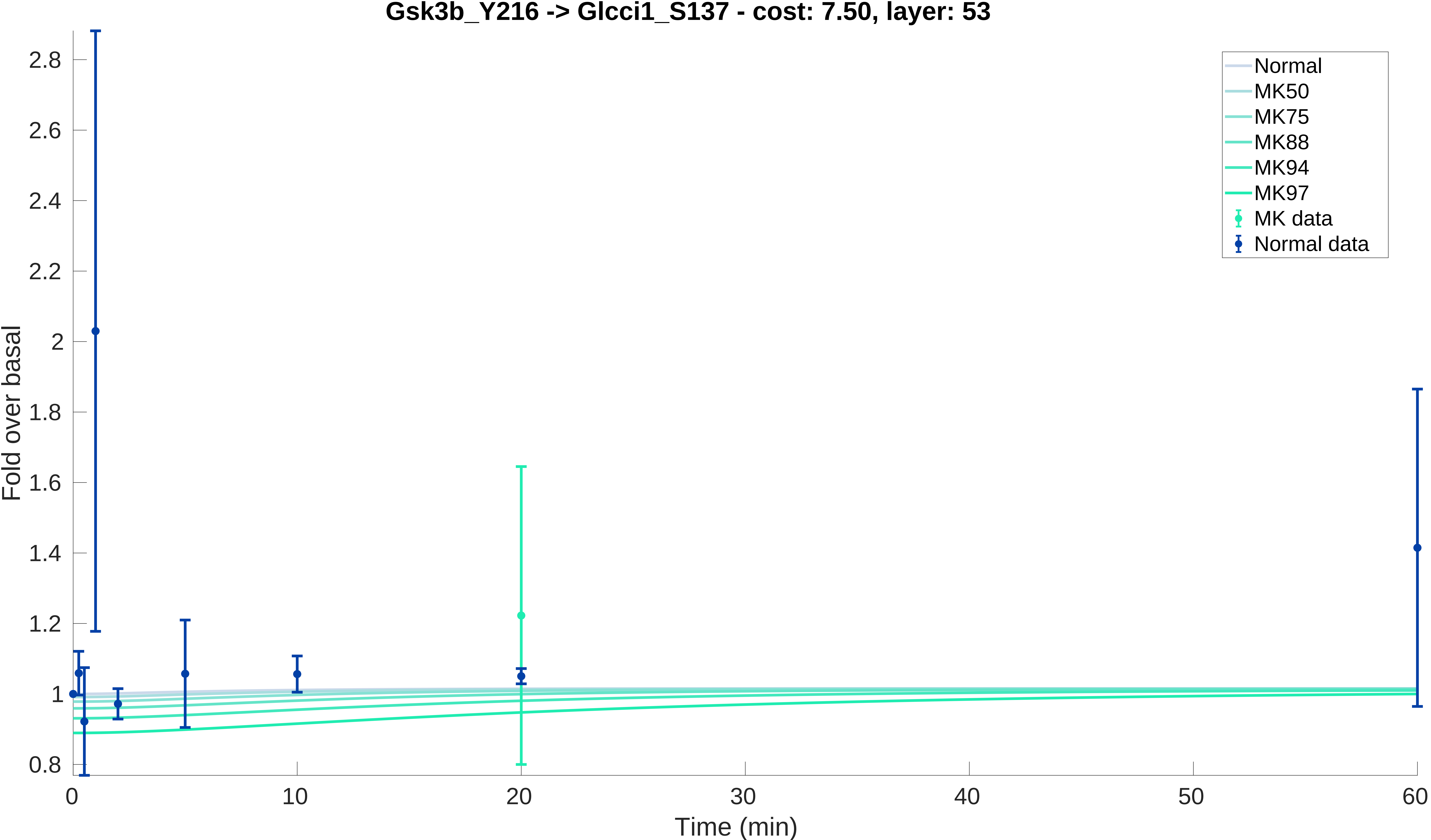

Gsk3b_Y216 -> Glcci1_S137 - cost: 7.50, layer: 53
Fold over basal
Time (min)
2.8
2.6
2.4
2.2
2
1.8
1.6
1.4
1.2
1
0.8
0
10
20
30
40
50
60
Normal
MK50
MK75
MK88
MK94
MK97
MK data
Normal data

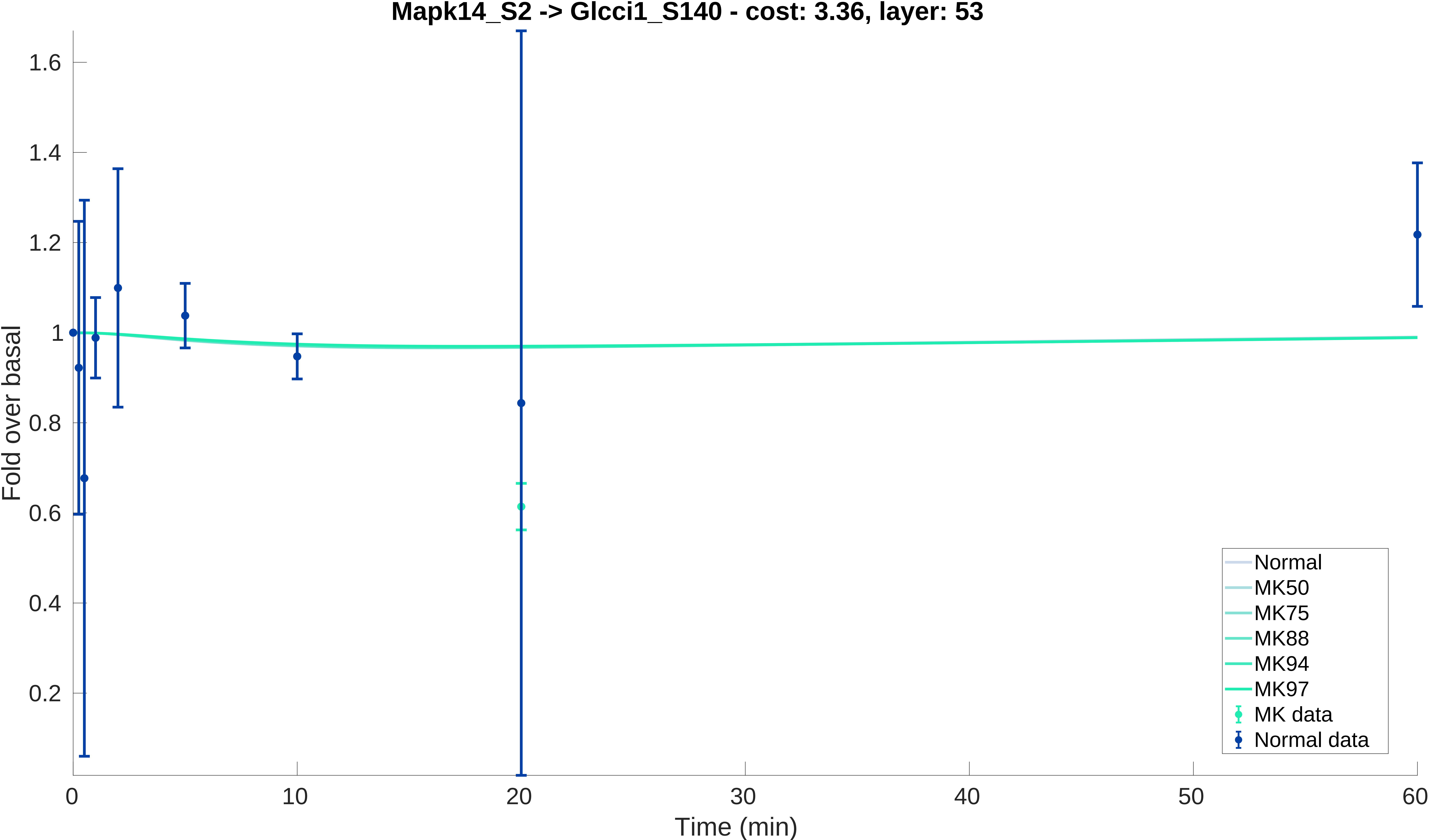

Mapk14_S2 -> Glcci1_S140 - cost: 3.36, layer: 53
Fold over basal
1.6
1.4
1.2
1
0.8
0.6
0.4
0.2
0
10
20
30
40
50
60
Time (min)
Normal
MK50
MK75
MK88
MK94
MK97
MK data
Normal data

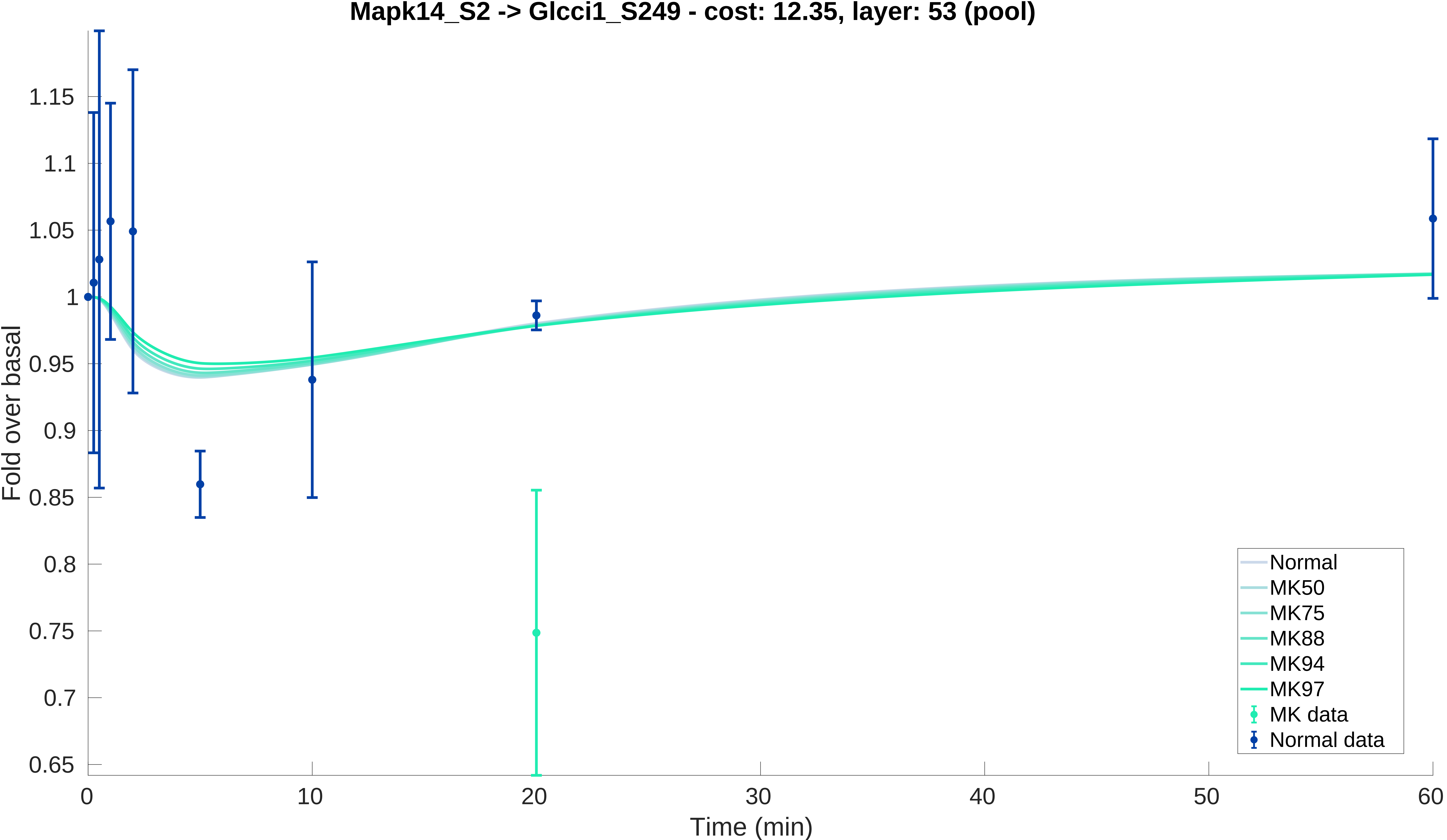

Mapk14_S2 -> Glcci1_S249 - cost: 12.35, layer: 53 (pool)
Fold over basal
1.15
1.1
1.05
1
0.95
0.9
0.85
0.8
0.75
0.7
0.65
0
10
20
30
40
50
60
Time (min)
Normal
MK50
MK75
MK88
MK94
MK97
MK data
Normal data

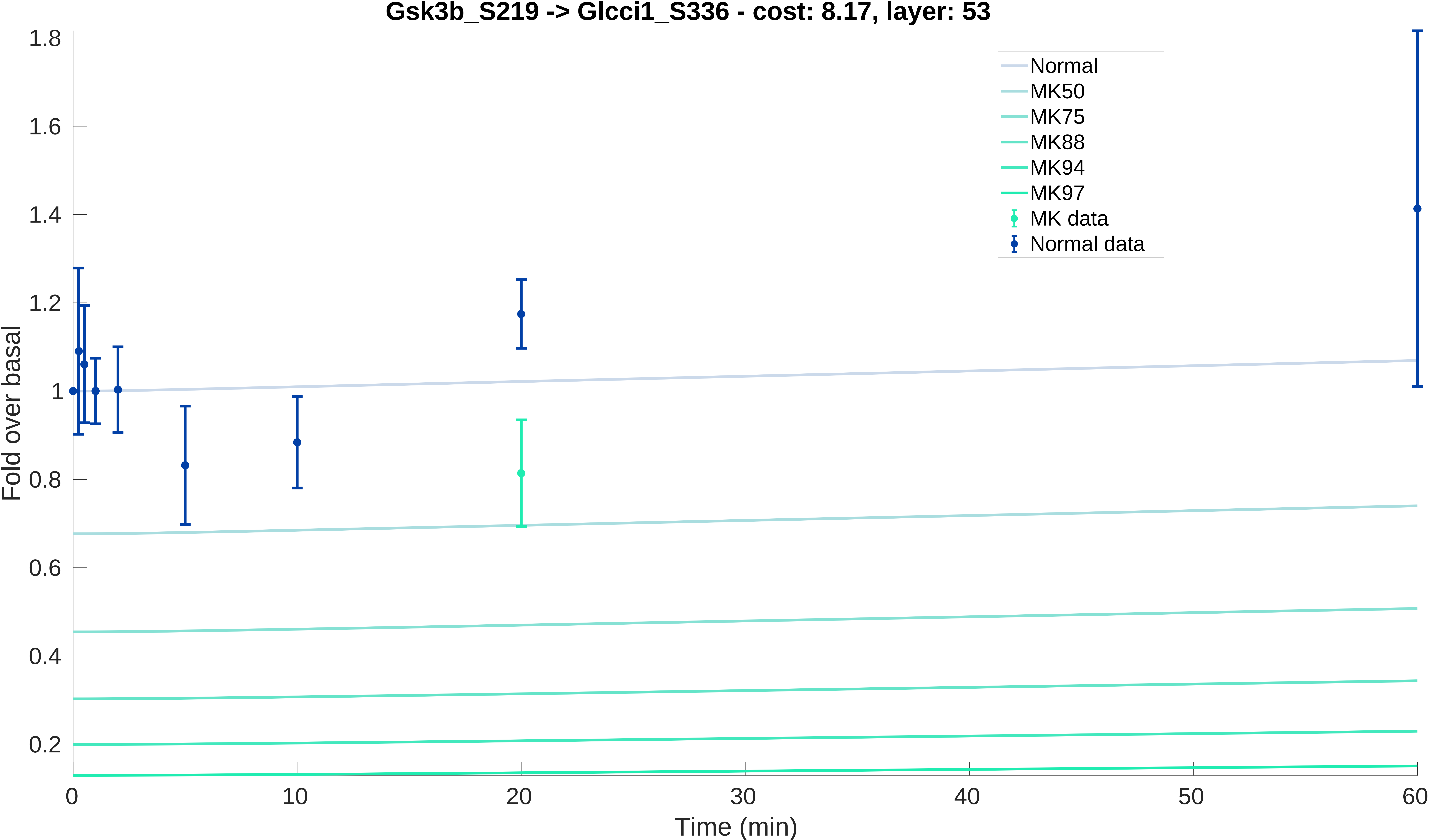
Gsk3b_S219 -> Glcci1_S336 - cost: 8.17, layer: 53
Fold over basal
Time (min)
Normal
MK50
MK75
MK88
MK94
MK97
MK data
Normal data
1.8
1.6
1.4
1.2
1
0.8
0.6
0.4
0.2
0
10
20
30
40
50
60

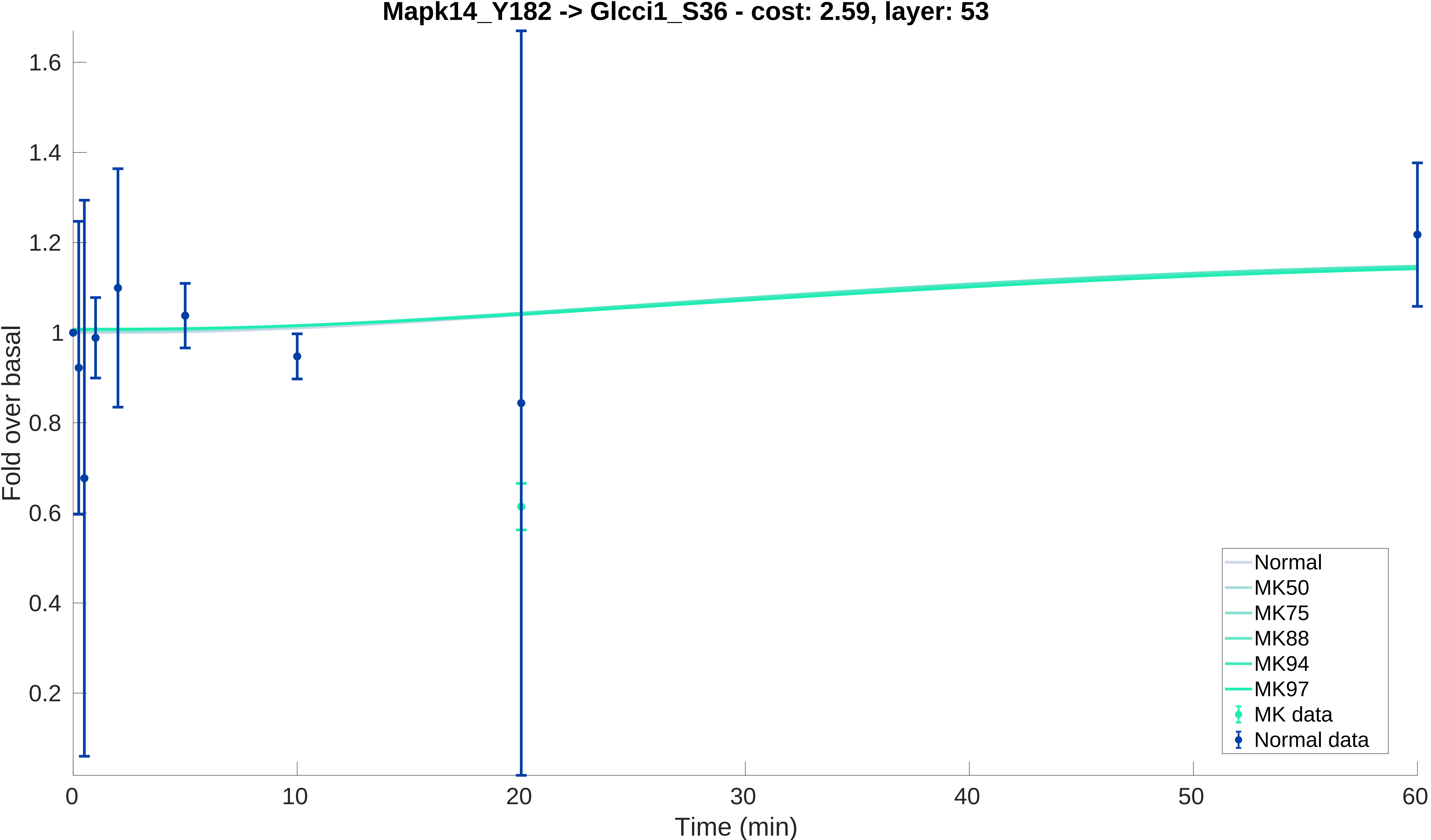

Mapk14_Y182 -> Glcci1_S36 - cost: 2.59, layer: 53
Fold over basal
1.6
1.4
1.2
1
0.8
0.6
0.4
0.2
0
10
20
30
40
50
60
Time (min)
Normal
MK50
MK75
MK88
MK94
MK97
MK data
Normal data

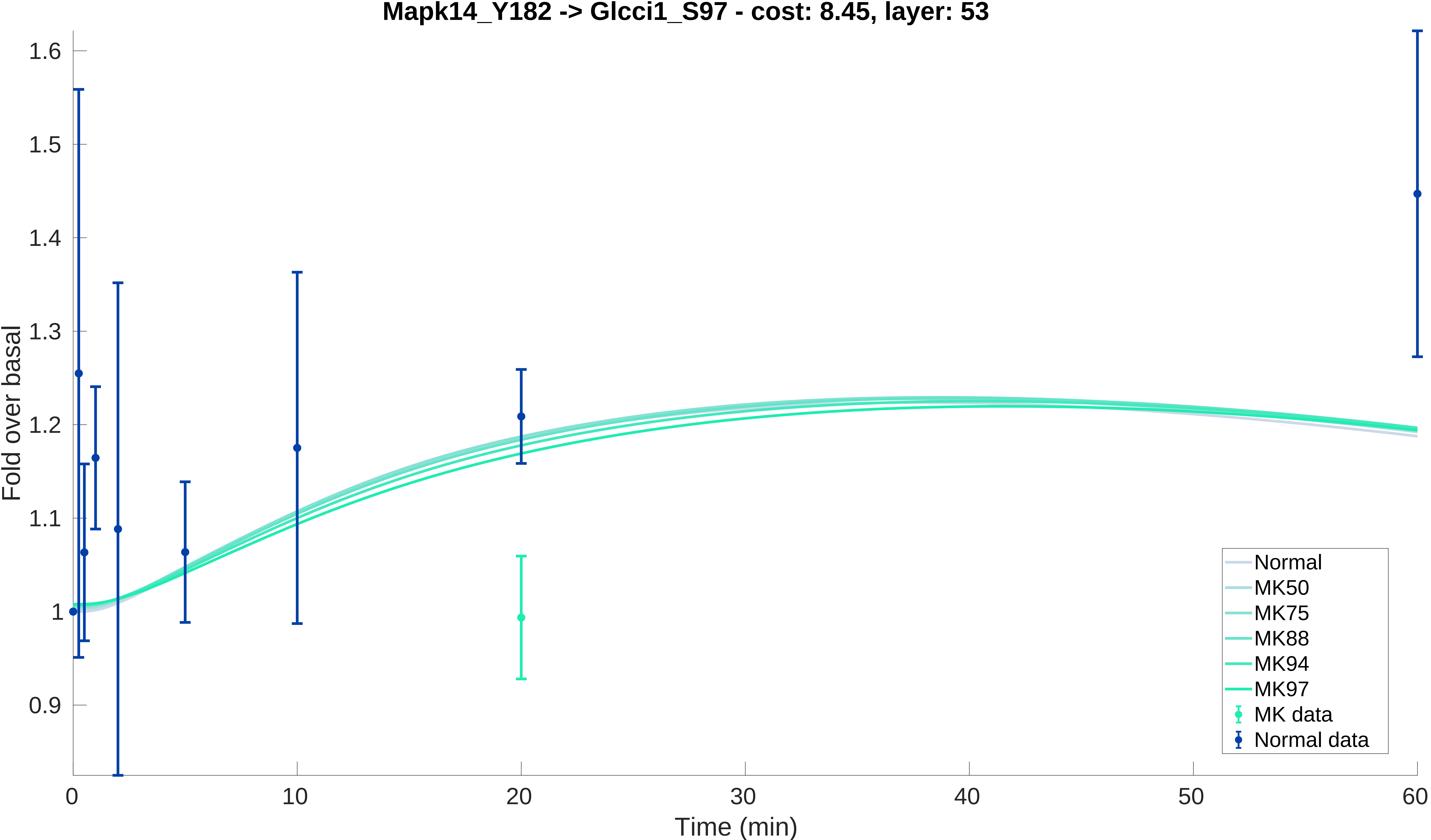
Mapk14_Y182 -> Glcci1_S97 - cost: 8.45, layer: 53
Fold over basal
Time (min)
0.9
1
1.1
1.2
1.3
1.4
1.5
1.6
0
10
20
30
40
50
60
Normal
MK50
MK75
MK88
MK94
MK97
MK data
Normal data

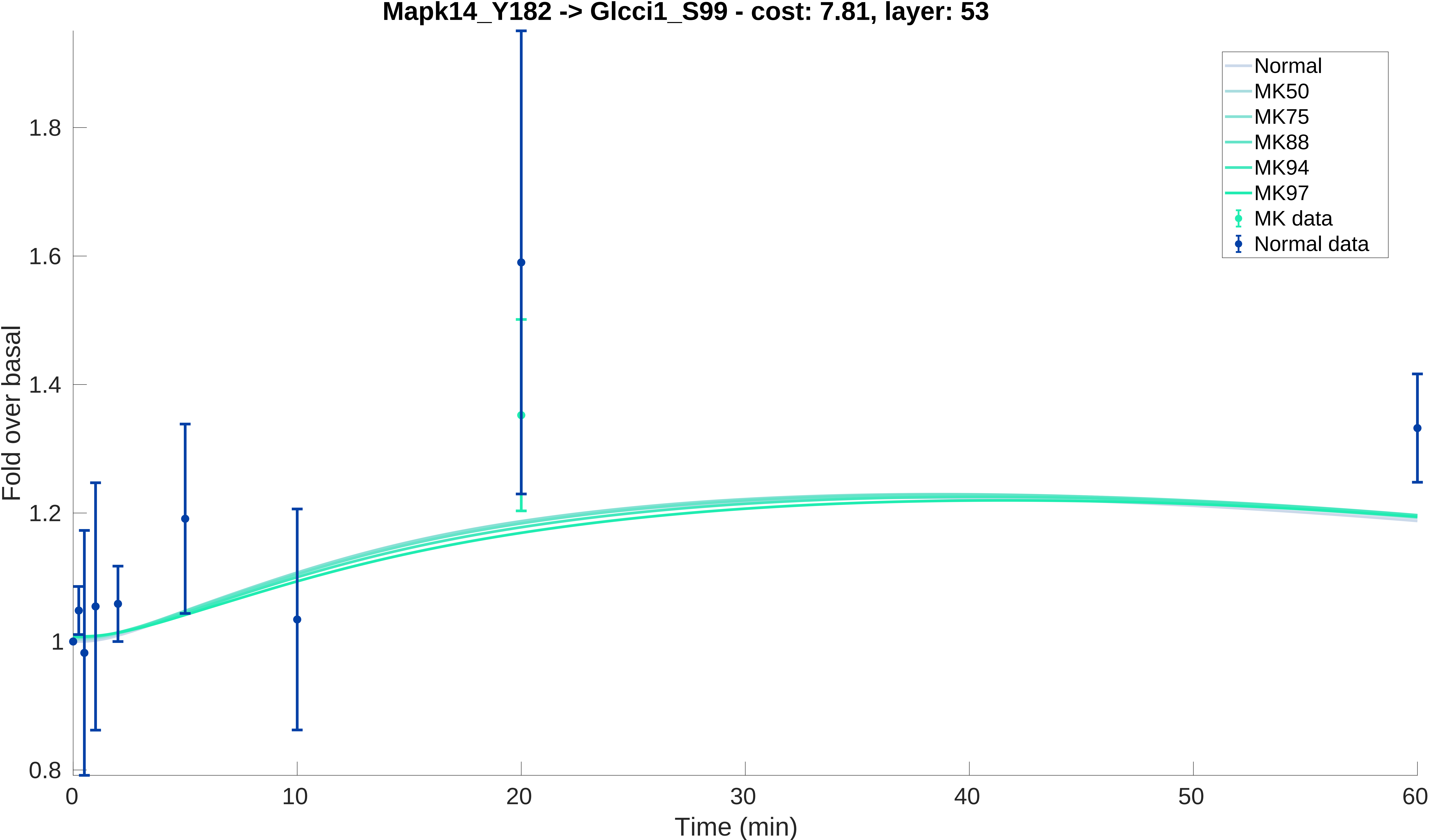

Mapk14_Y182 -> Glcci1_S99 - cost: 7.81, layer: 53
Fold over basal
Time (min)
0.8
1
1.2
1.4
1.6
1.8
0
10
20
30
40
50
60
Normal
MK50
MK75
MK88
MK94
MK97
MK data
Normal data

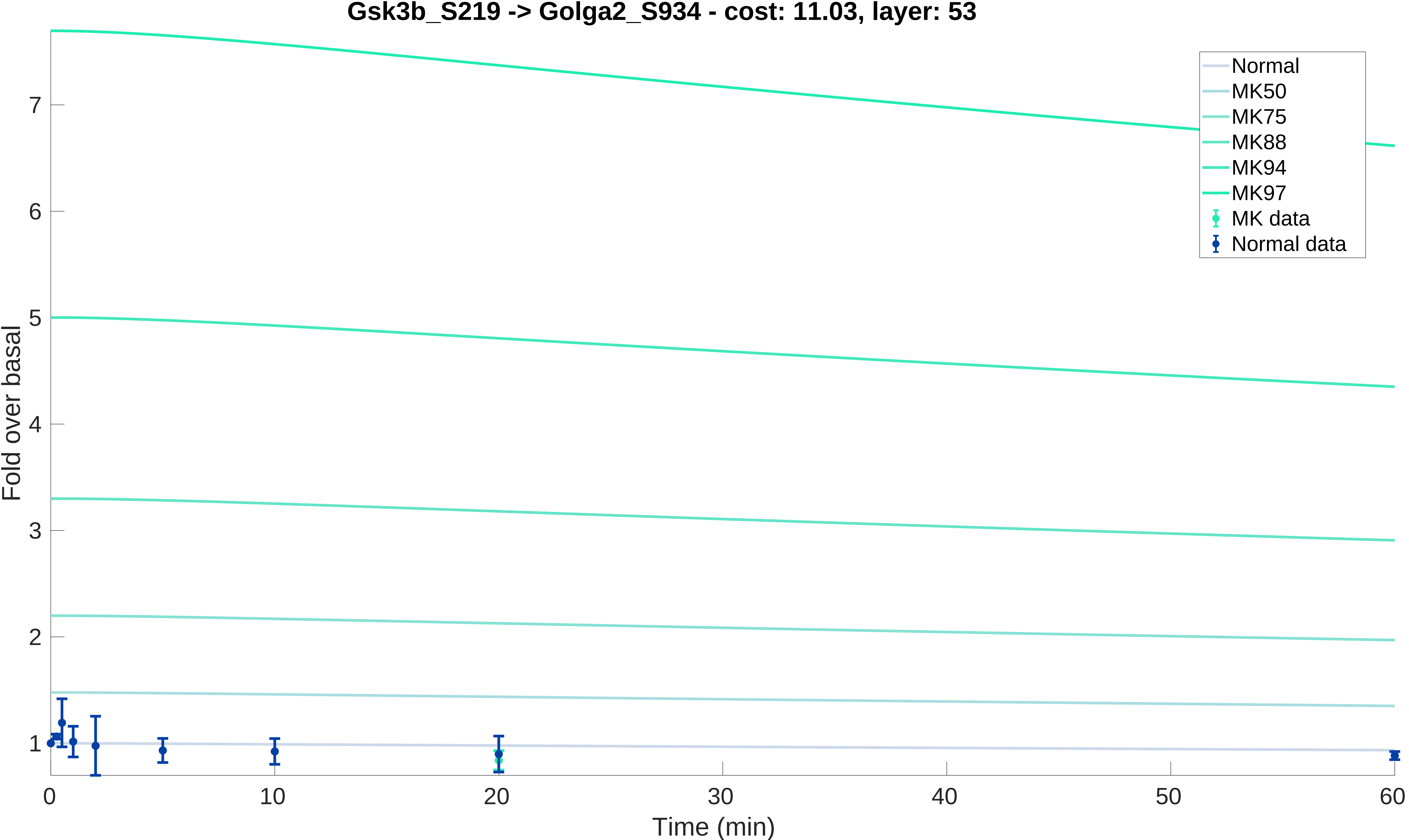

Gsk3b_S219 -> Golga2_S934 - cost: 11.03, layer: 53
Fold over basal
Time (min)
0
10
20
30
40
50
60
1
2
3
4
5
6
7
Normal
MK50
MK75
MK88
MK94
MK97
MK data
Normal data

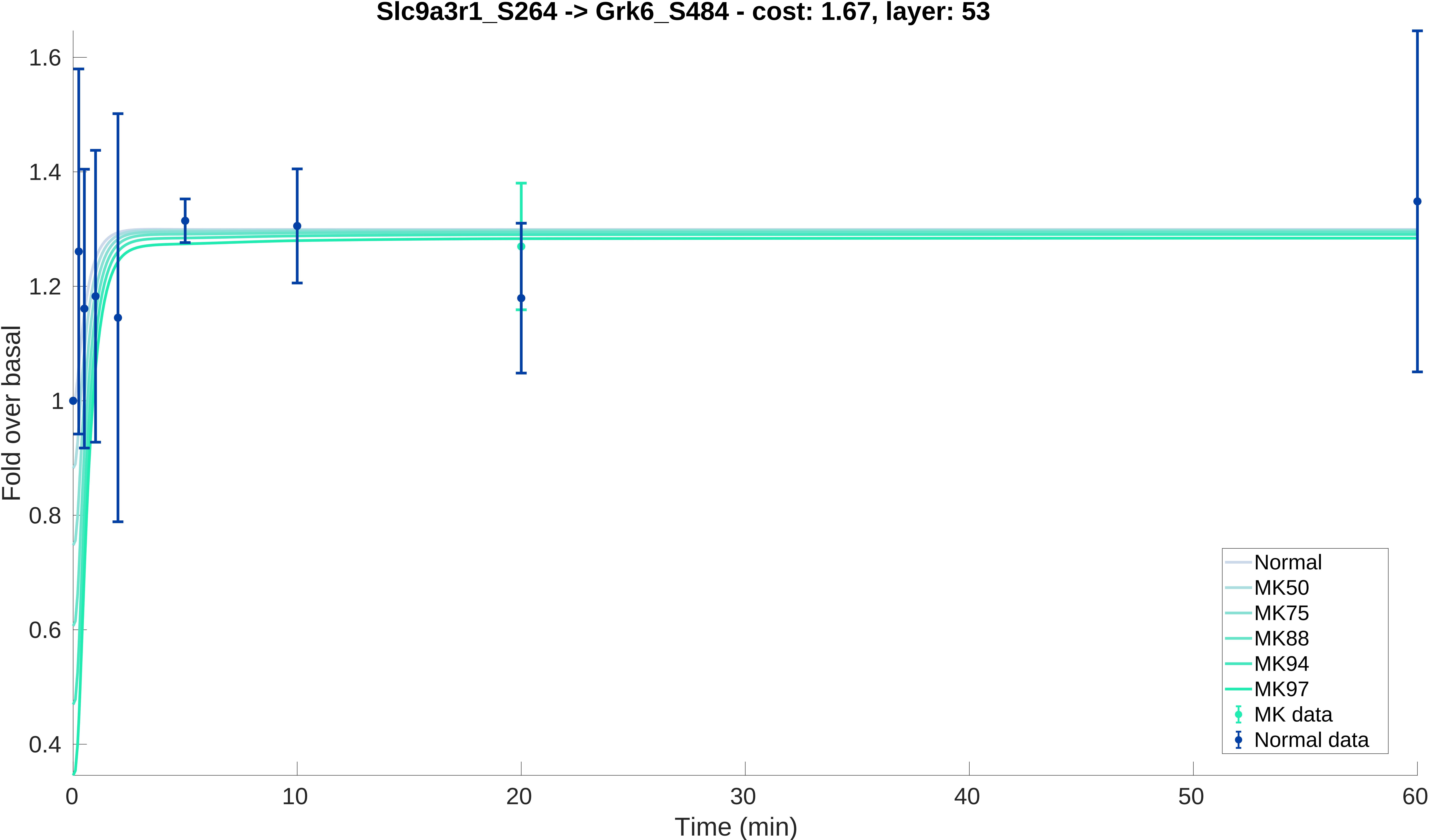

Slc9a3r1_S264 -> Grk6_S484 - cost: 1.67, layer: 53
Fold over basal
Time (min)
1.6
1.4
1.2
1
0.8
0.6
0.4
0
10
20
30
40
50
60
Normal
MK50
MK75
MK88
MK94
MK97
MK data
Normal data

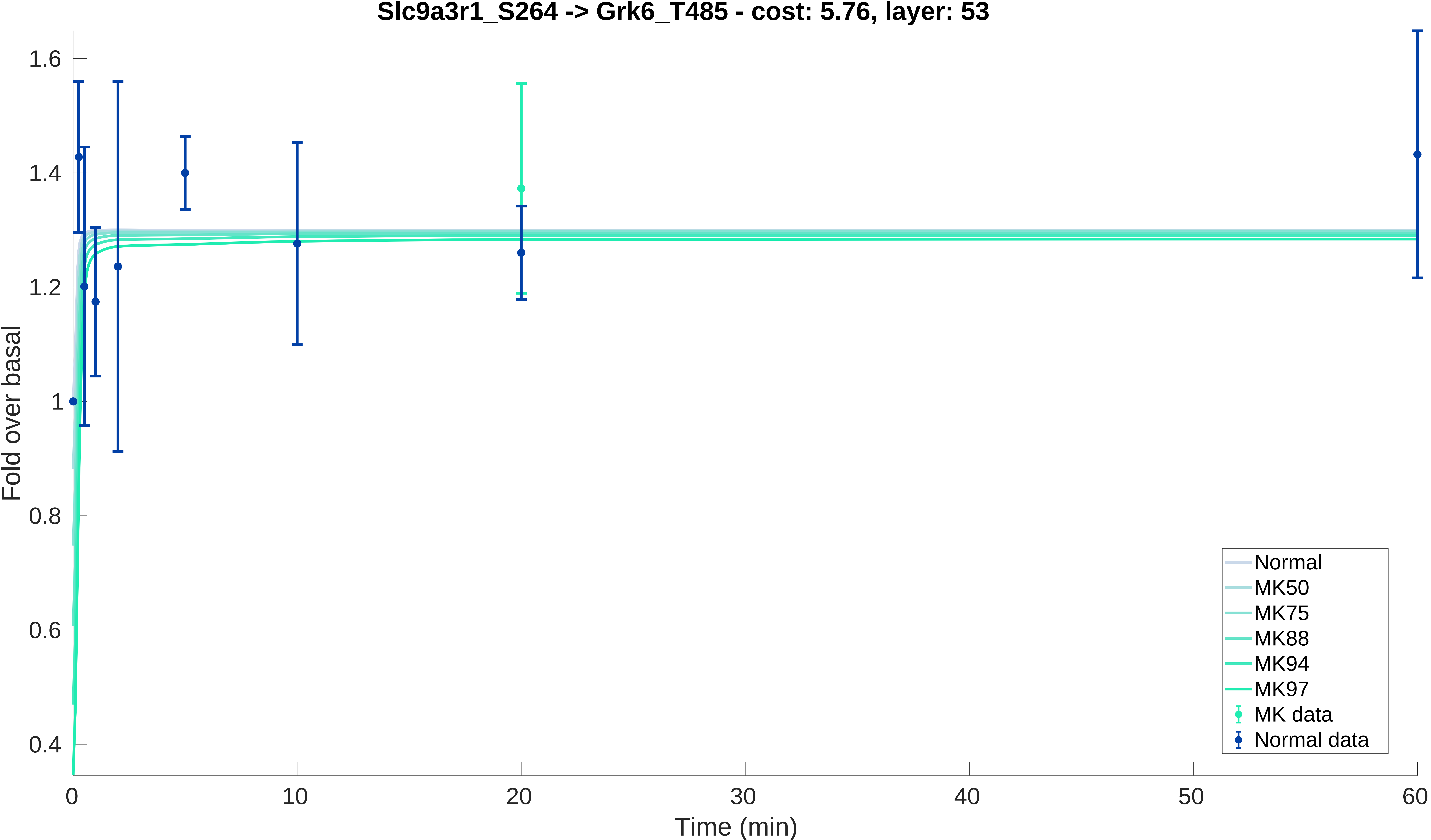
Slc9a3r1_S264 -> Grk6_T485 - cost: 5.76, layer: 53
Fold over basal
Time (min)
0.4
0.6
0.8
1
1.2
1.4
1.6
0
10
20
30
40
50
60
Normal
MK50
MK75
MK88
MK94
MK97
MK data
Normal data

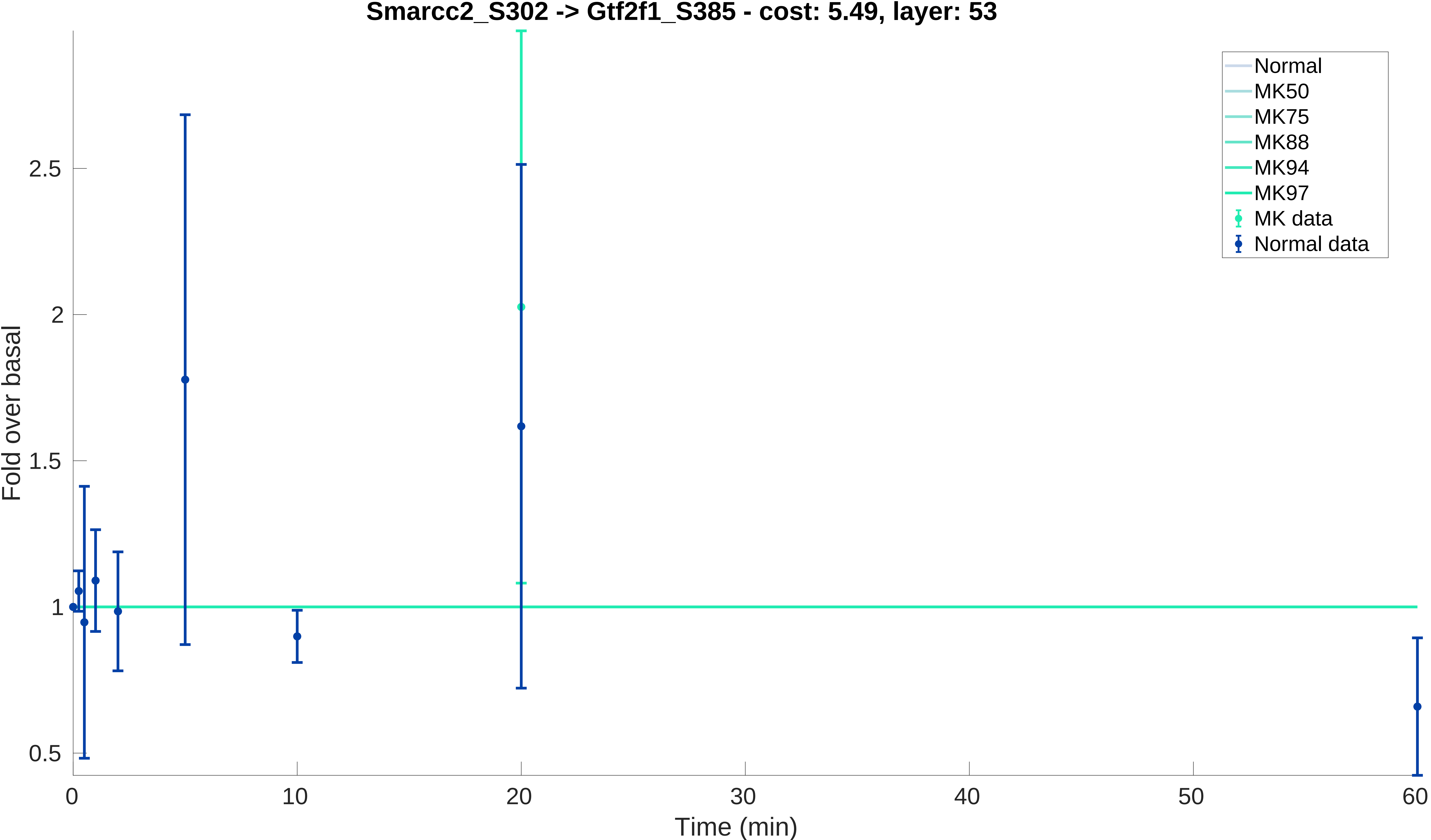
Smarcc2_S302 -> Gtf2f1_S385 - cost: 5.49, layer: 53
Fold over basal
Time (min)
0.5
1
1.5
2
2.5
0
10
20
30
40
50
60
Normal
MK50
MK75
MK88
MK94
MK97
MK data
Normal data

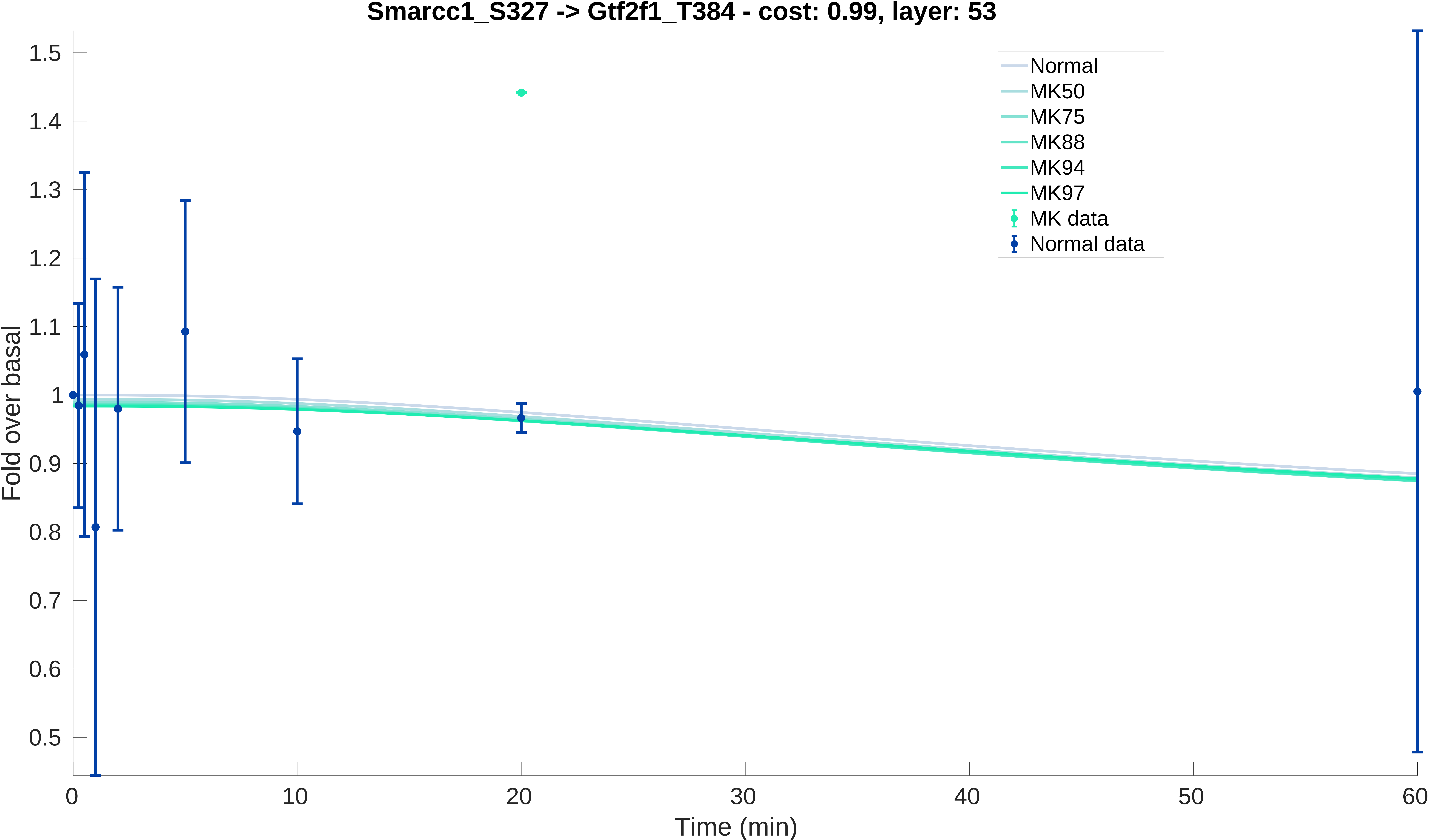

Smarcc1_S327 -> Gtf2f1_T384 - cost: 0.99, layer: 53
Fold over basal
Time (min)
1.5
1.4
1.3
1.2
1.1
1
0.9
0.8
0.7
0.6
0.5
0
10
20
30
40
50
60
Normal
MK50
MK75
MK88
MK94
MK97
MK data
Normal data

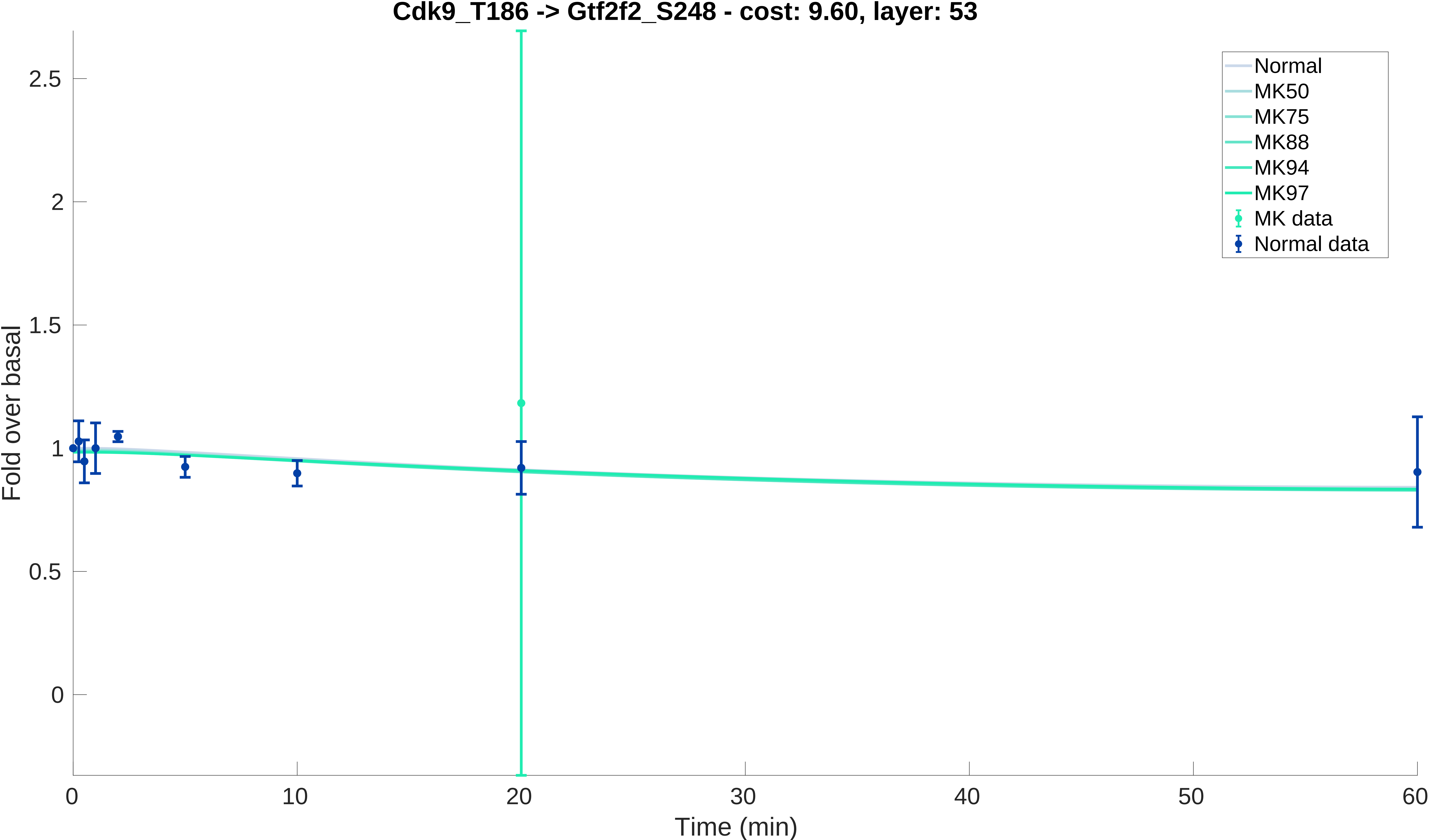

Cdk9_T186 -> Gtf2f2_S248 - cost: 9.60, layer: 53
Fold over basal
Time (min)
2.5
2
1.5
1
0.5
0
0
10
20
30
40
50
60
Normal
MK50
MK75
MK88
MK94
MK97
MK data
Normal data

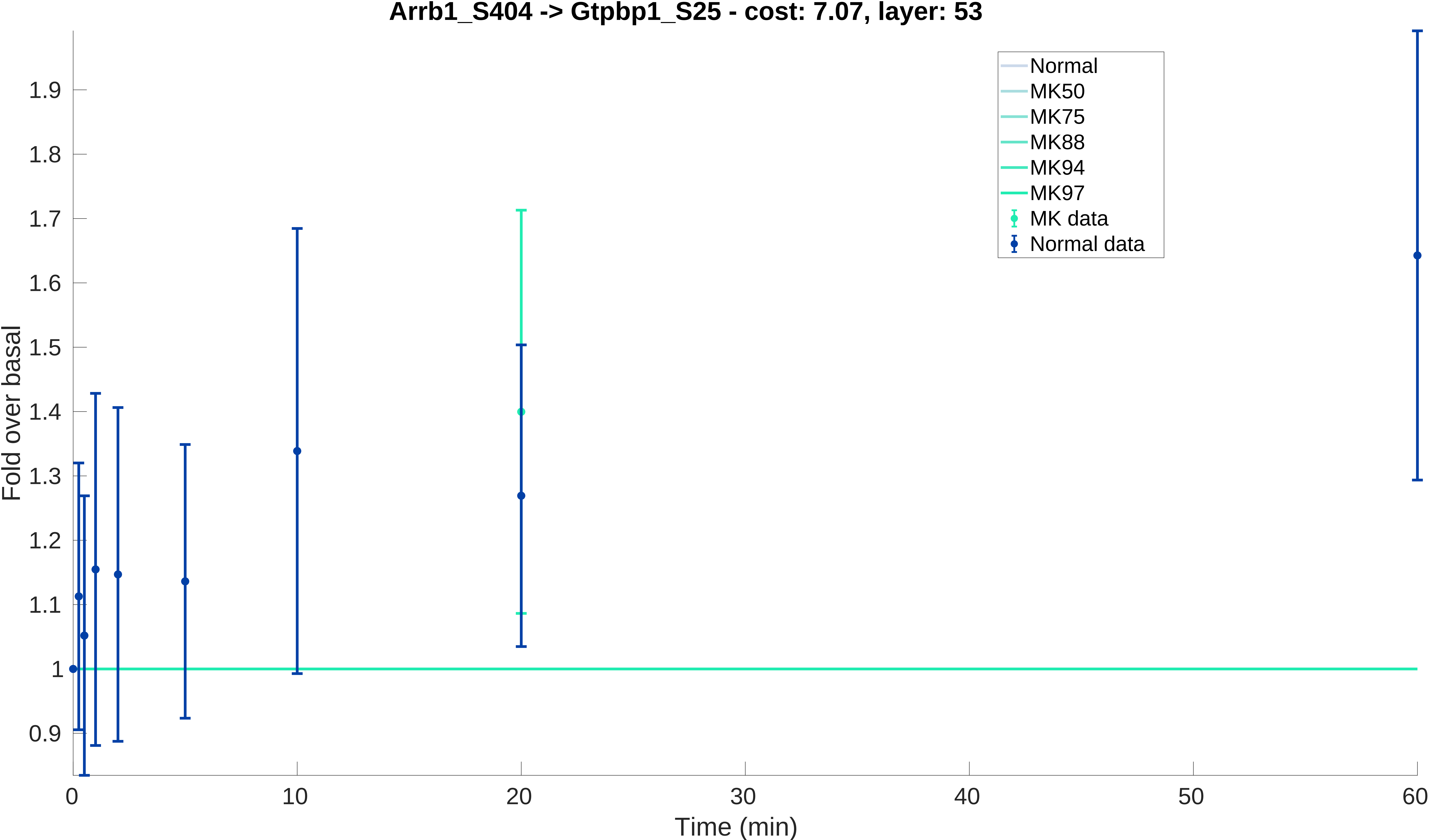

Arrb1_S404 -> Gtpbp1_S25 - cost: 7.07, layer: 53
Fold over basal
Time (min)
0.9
1
1.1
1.2
1.3
1.4
1.5
1.6
1.7
1.8
1.9
0
10
20
30
40
50
60
Normal
MK50
MK75
MK88
MK94
MK97
MK data
Normal data

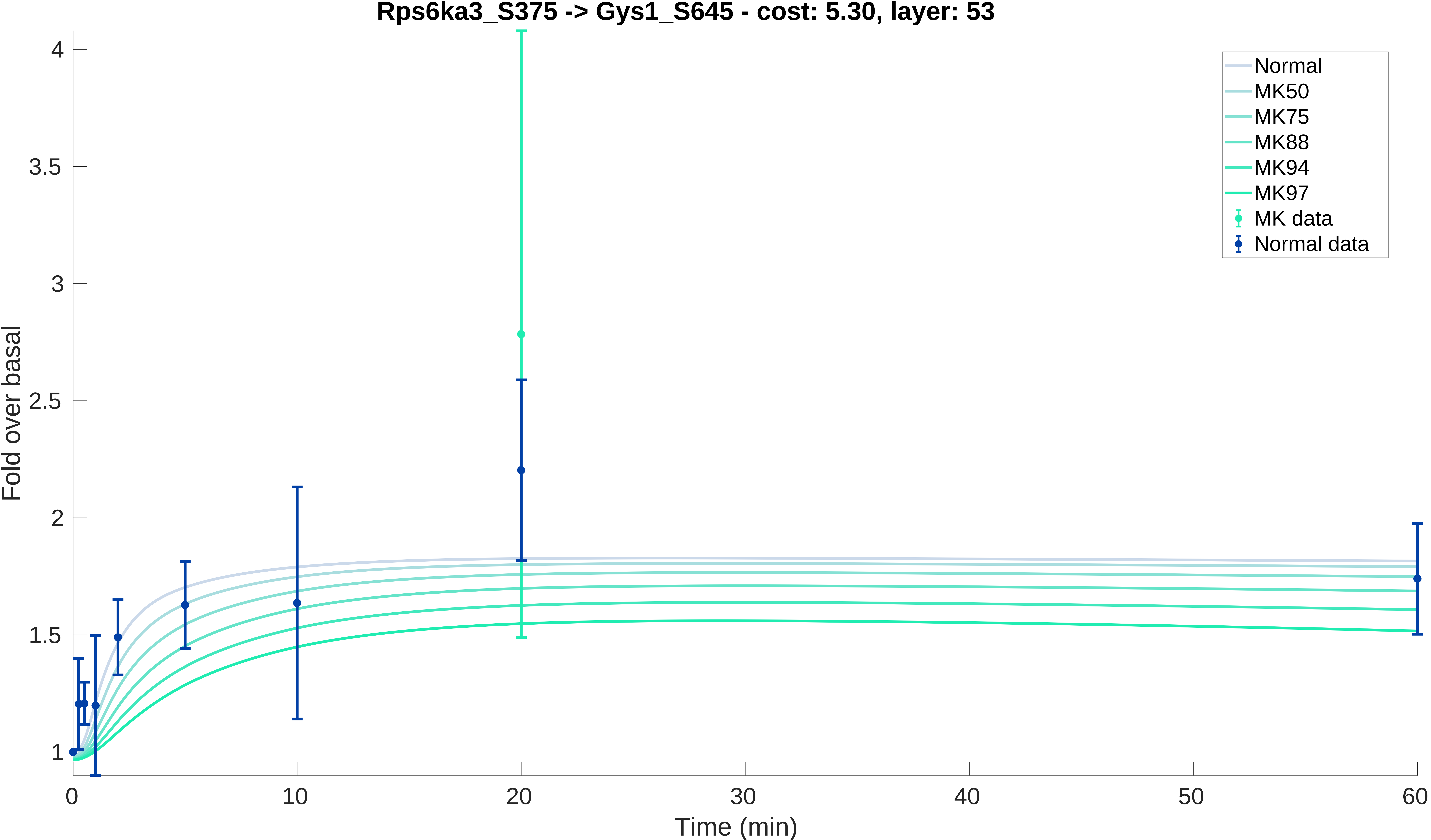

Rps6ka3_S375 -> Gys1_S645 - cost: 5.30, layer: 53
Fold over basal
Time (min)
4
3.5
3
2.5
2
1.5
1
0
10
20
30
40
50
60
Normal
MK50
MK75
MK88
MK94
MK97
MK data
Normal data

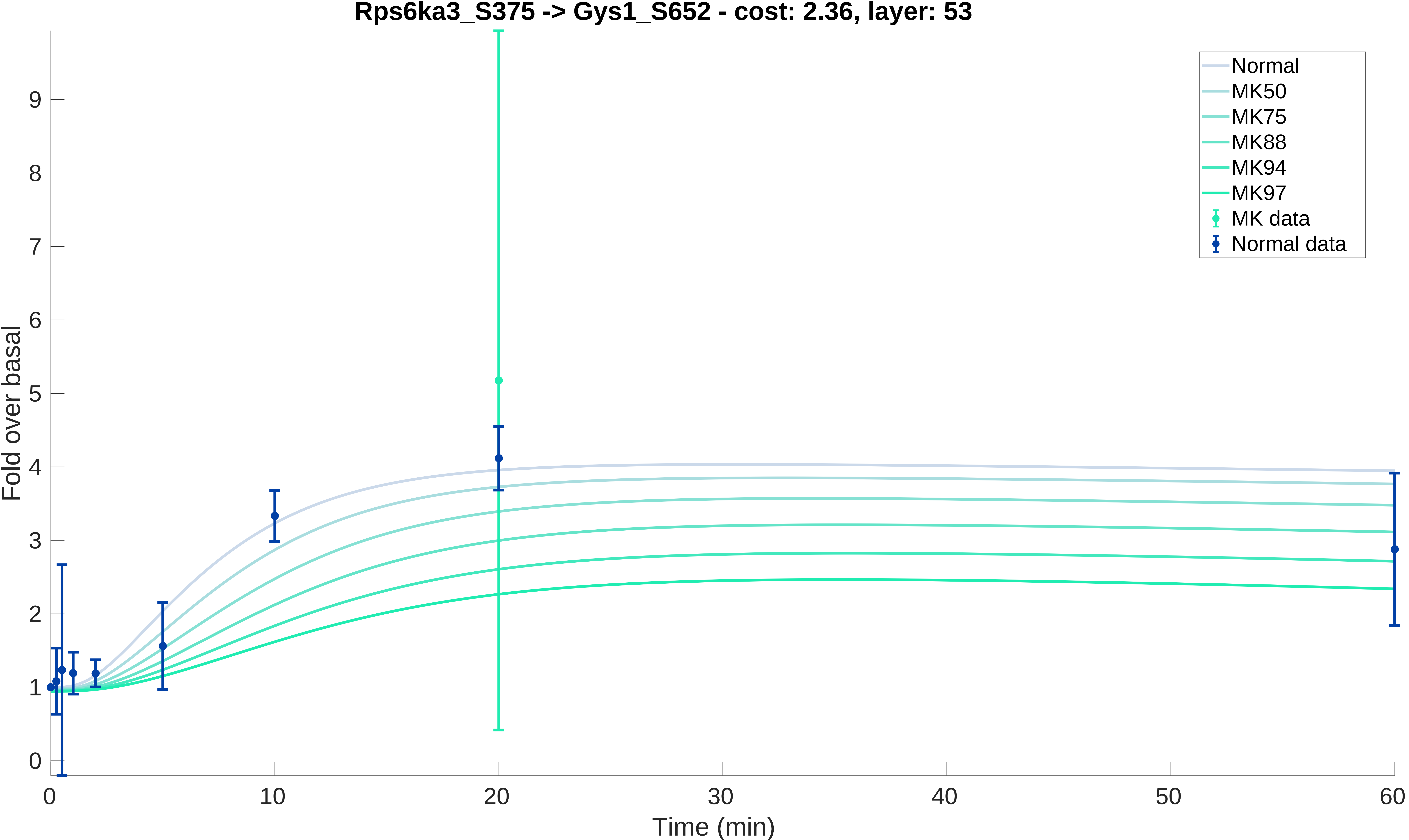

Rps6ka3_S375 -> Gys1_S652 - cost: 2.36, layer: 53
Fold over basal
0
1
2
3
4
5
6
7
8
9
0
10
20
30
40
50
60
Time (min)
Normal
MK50
MK75
MK88
MK94
MK97
MK data
Normal data

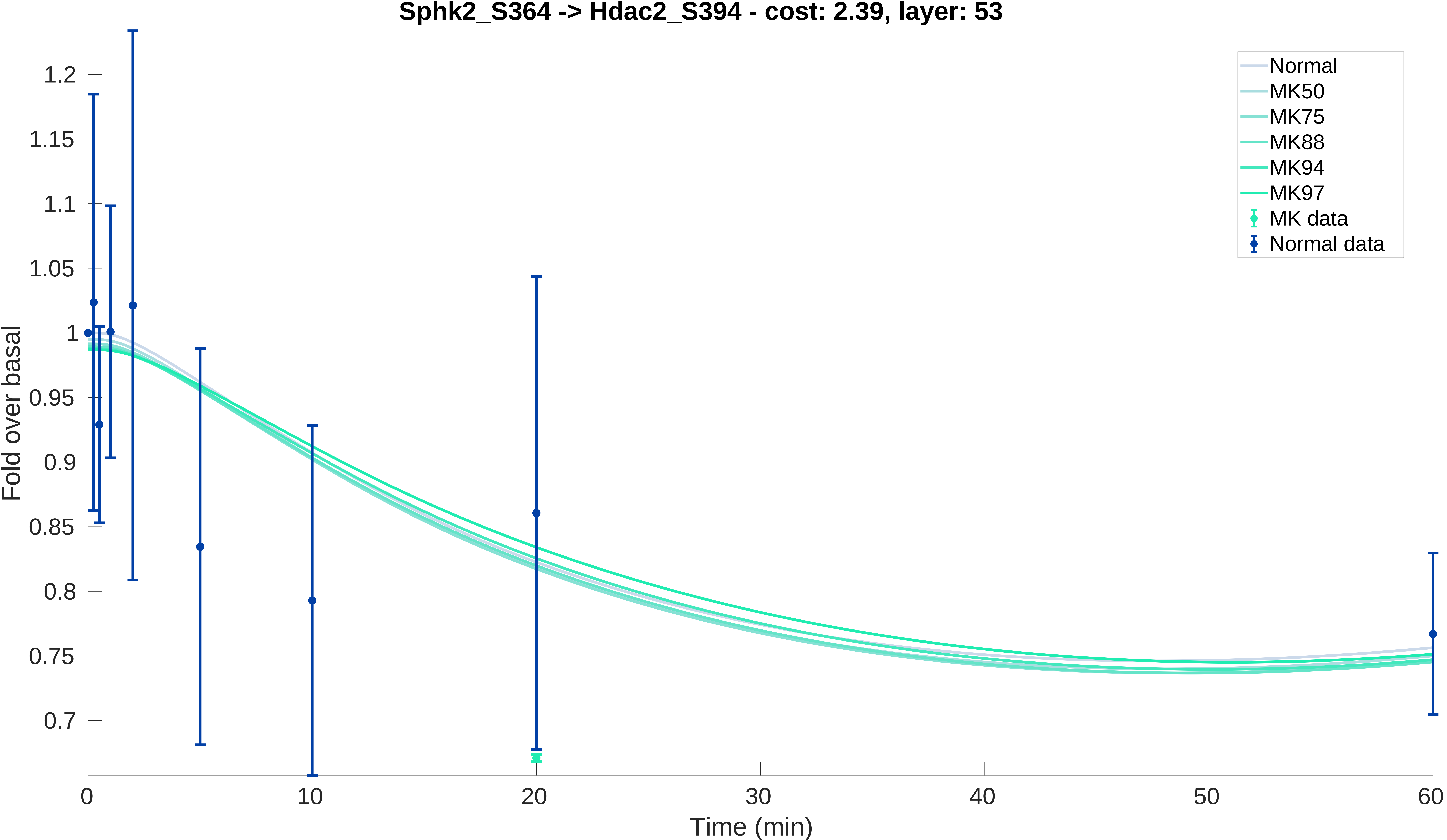

Sphk2_S364 -> Hdac2_S394 - cost: 2.39, layer: 53
Normal
MK50
MK75
MK88
MK94
MK97
MK data
Normal data
Fold over basal
1.2
1.15
1.1
1.05
1
0.95
0.9
0.85
0.8
0.75
0.7
Time (min)
0
10
20
30
40
50
60

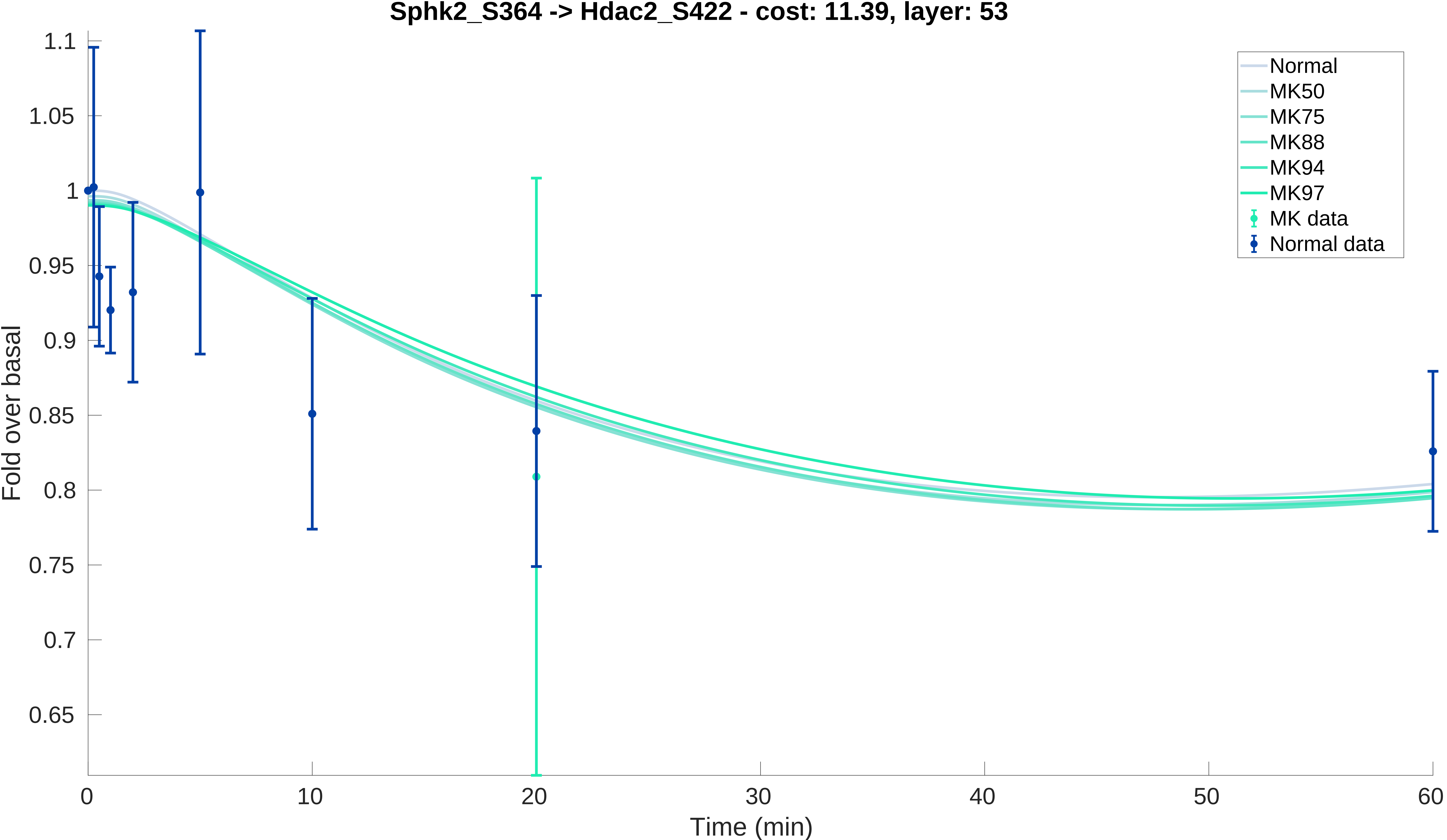

Sphk2_S364 -> Hdac2_S422 - cost: 11.39, layer: 53
Fold over basal
Time (min)
0
10
20
30
40
50
60
1.1
1.05
1
0.95
0.9
0.85
0.8
0.75
0.7
0.65
Normal
MK50
MK75
MK88
MK94
MK97
MK data
Normal data

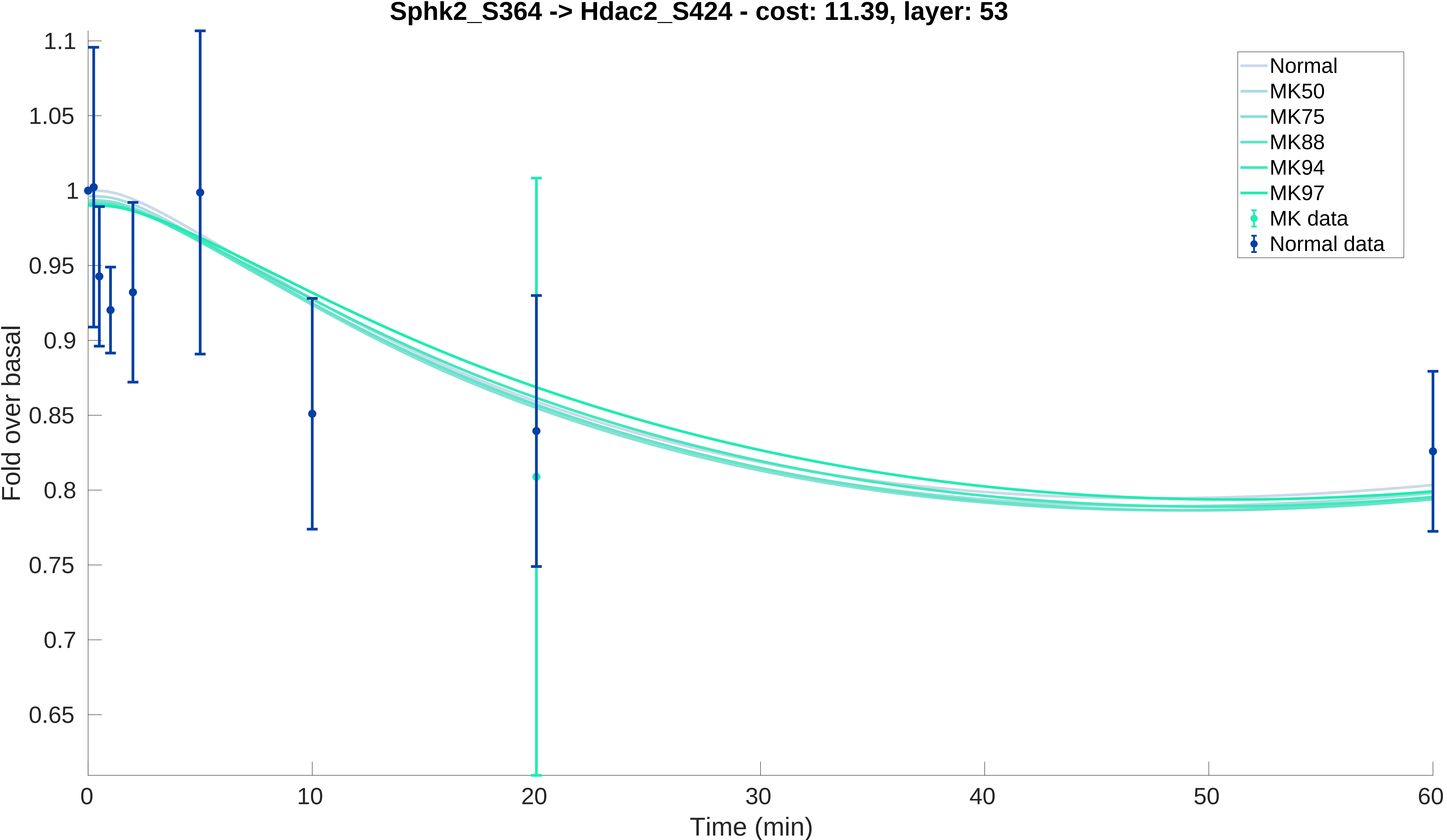

Sphk2_S364 -> Hdac2_S424 - cost: 11.39, layer: 53
Fold over basal
1.1
1.05
1
0.95
0.9
0.85
0.8
0.75
0.7
0.65
0
10
20
30
40
50
60
Time (min)
Normal
MK50
MK75
MK88
MK94
MK97
MK data
Normal data

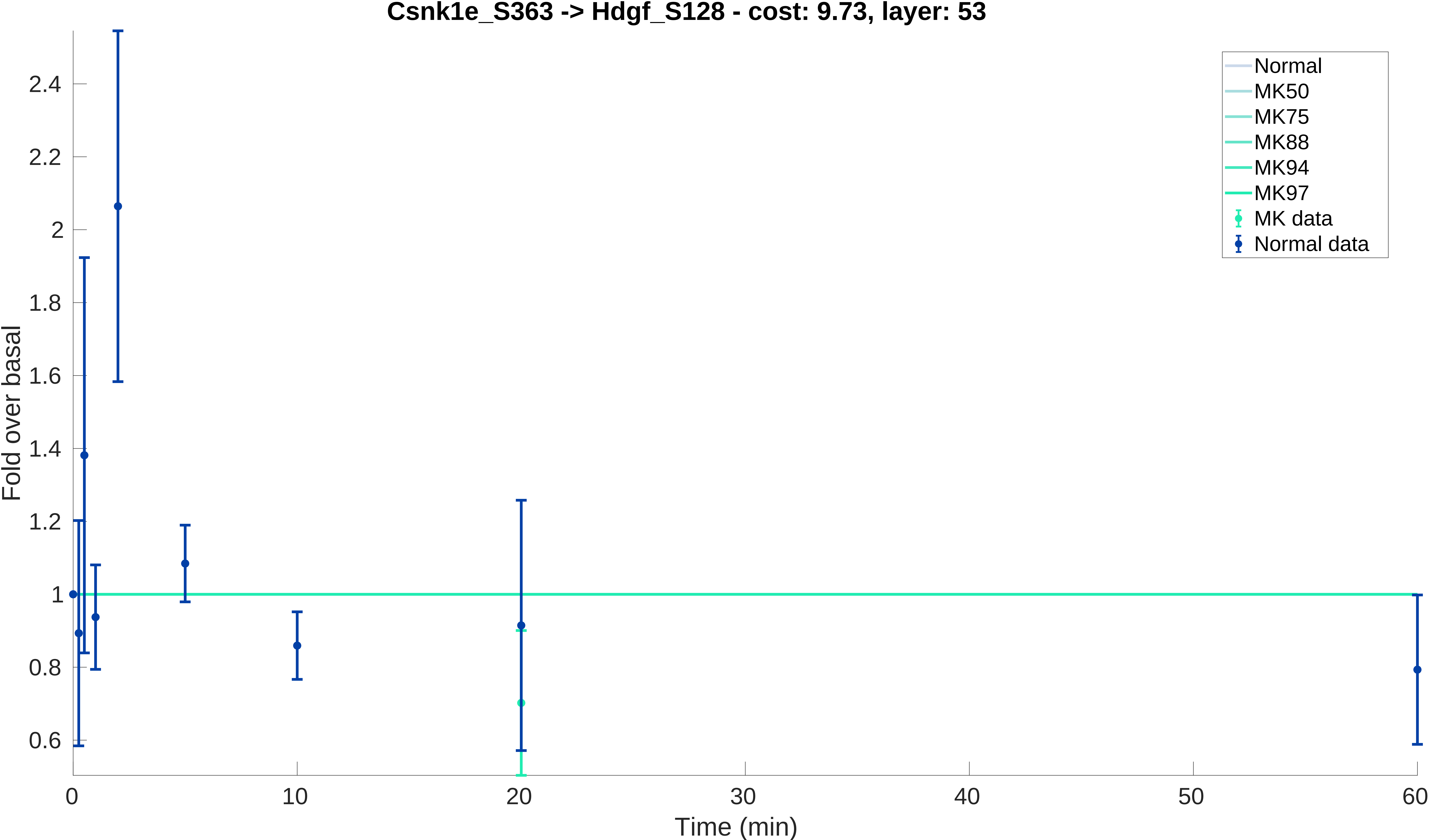
Csnk1e_S363 -> Hdgf_S128 - cost: 9.73, layer: 53
Fold over basal
Time (min)
Normal
MK50
MK75
MK88
MK94
MK97
MK data
Normal data
0.6
0.8
1
1.2
1.4
1.6
1.8
2
2.2
2.4
0
10
20
30
40
50
60

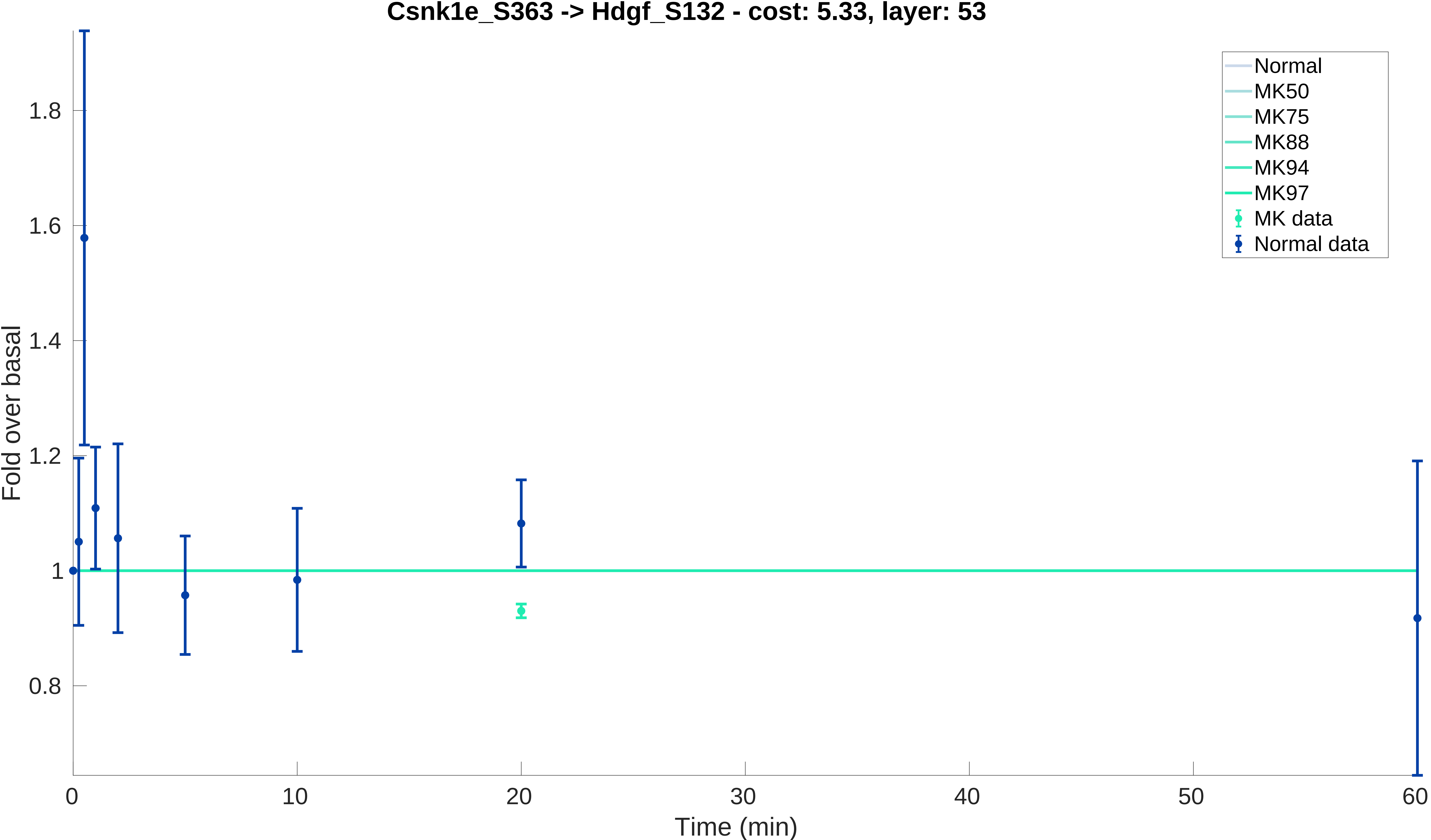

Csnk1e_S363 -> Hdgf_S132 - cost: 5.33, layer: 53
Normal
MK50
MK75
MK88
MK94
MK97
MK data
Normal data
Fold over basal
1.8
1.6
1.4
1.2
1
0.8
0
10
20
30
40
50
60
Time (min)

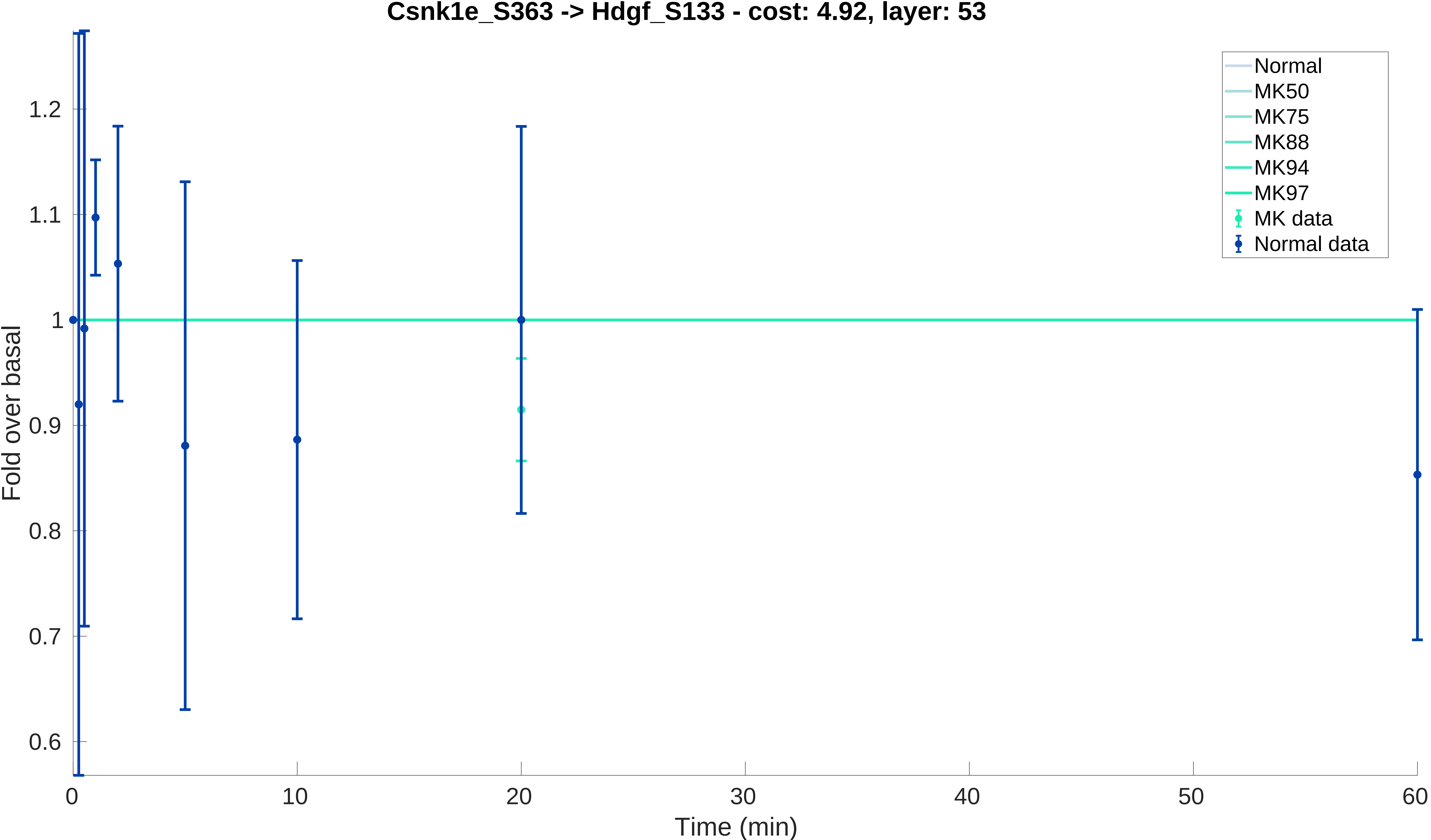

Csnk1e_S363 -> Hdgf_S133 - cost: 4.92, layer: 53
Fold over basal
Time (min)
0.6
0.7
0.8
0.9
1
1.1
1.2
0
10
20
30
40
50
60
Normal
MK50
MK75
MK88
MK94
MK97
MK data
Normal data

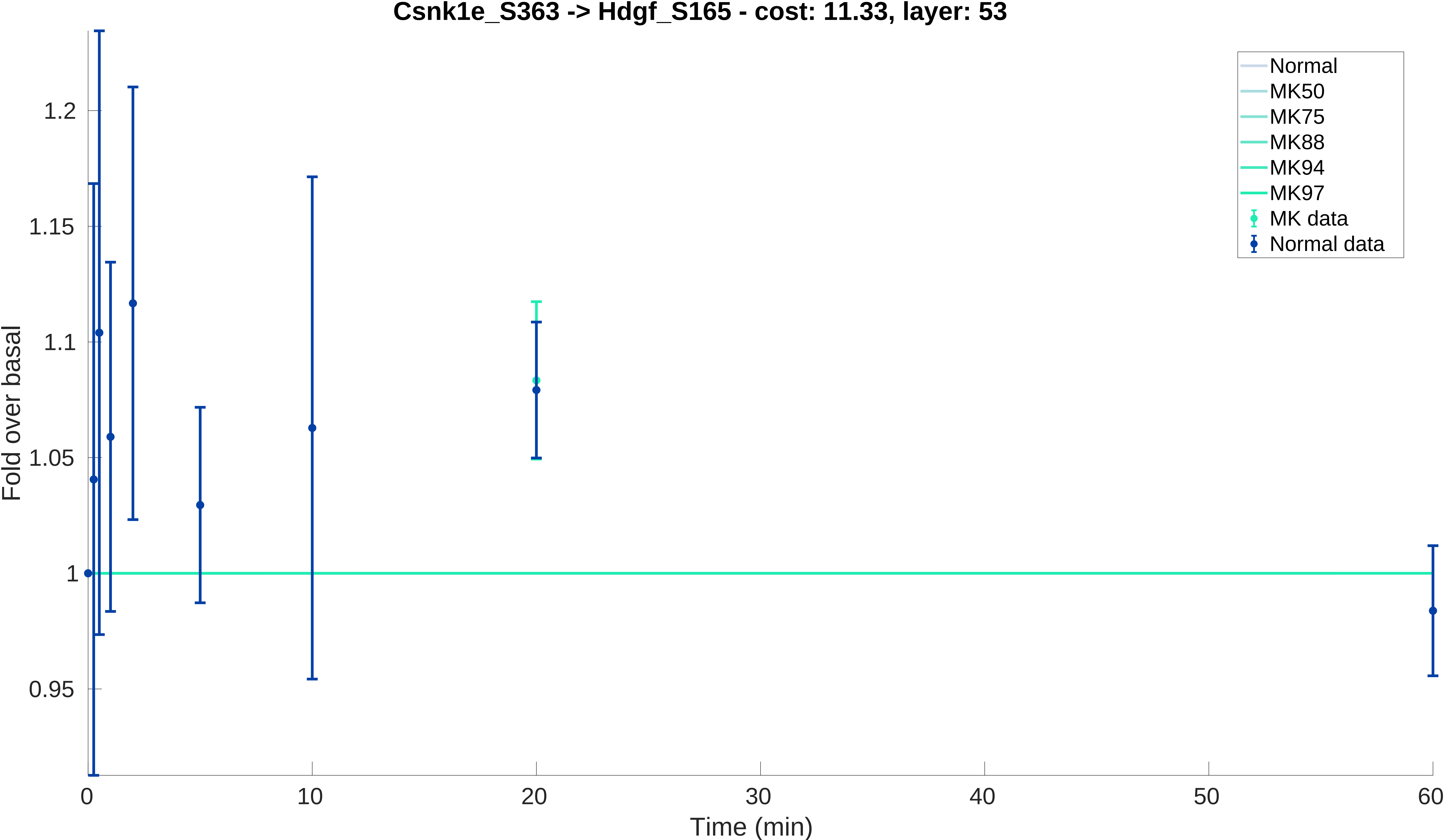

Csnk1e_S363 -> Hdgf_S165 - cost: 11.33, layer: 53
Normal
MK50
MK75
MK88
MK94
MK97
MK data
Normal data
Fold over basal
1.2
1.15
1.1
1.05
1
0.95
0
10
20
30
40
50
60
Time (min)

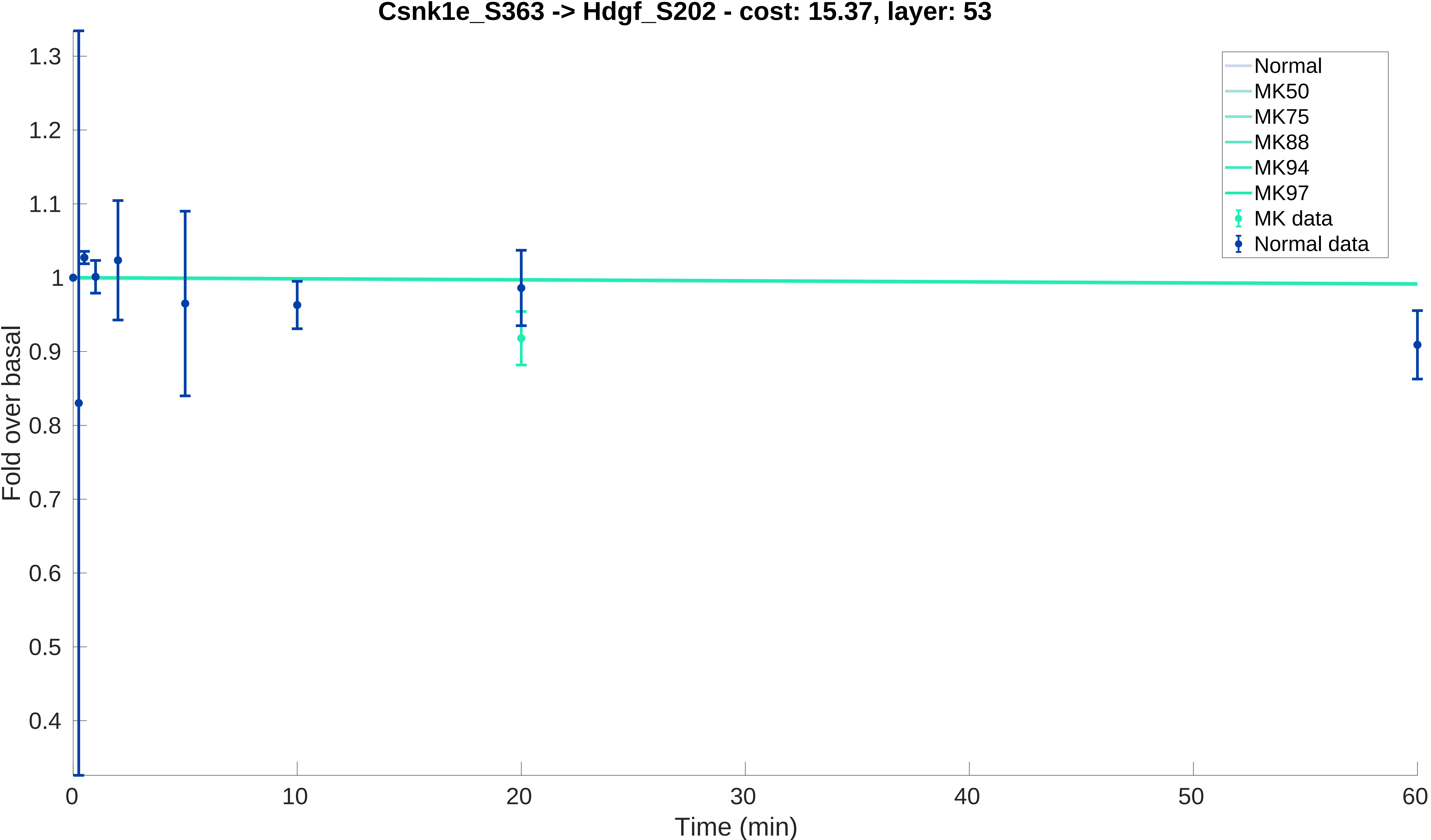

Csnk1e_S363 -> Hdgf_S202 - cost: 15.37, layer: 53
Fold over basal
Time (min)
1.3
1.2
1.1
1
0.9
0.8
0.7
0.6
0.5
0.4
0
10
20
30
40
50
60
Normal
MK50
MK75
MK88
MK94
MK97
MK data
Normal data

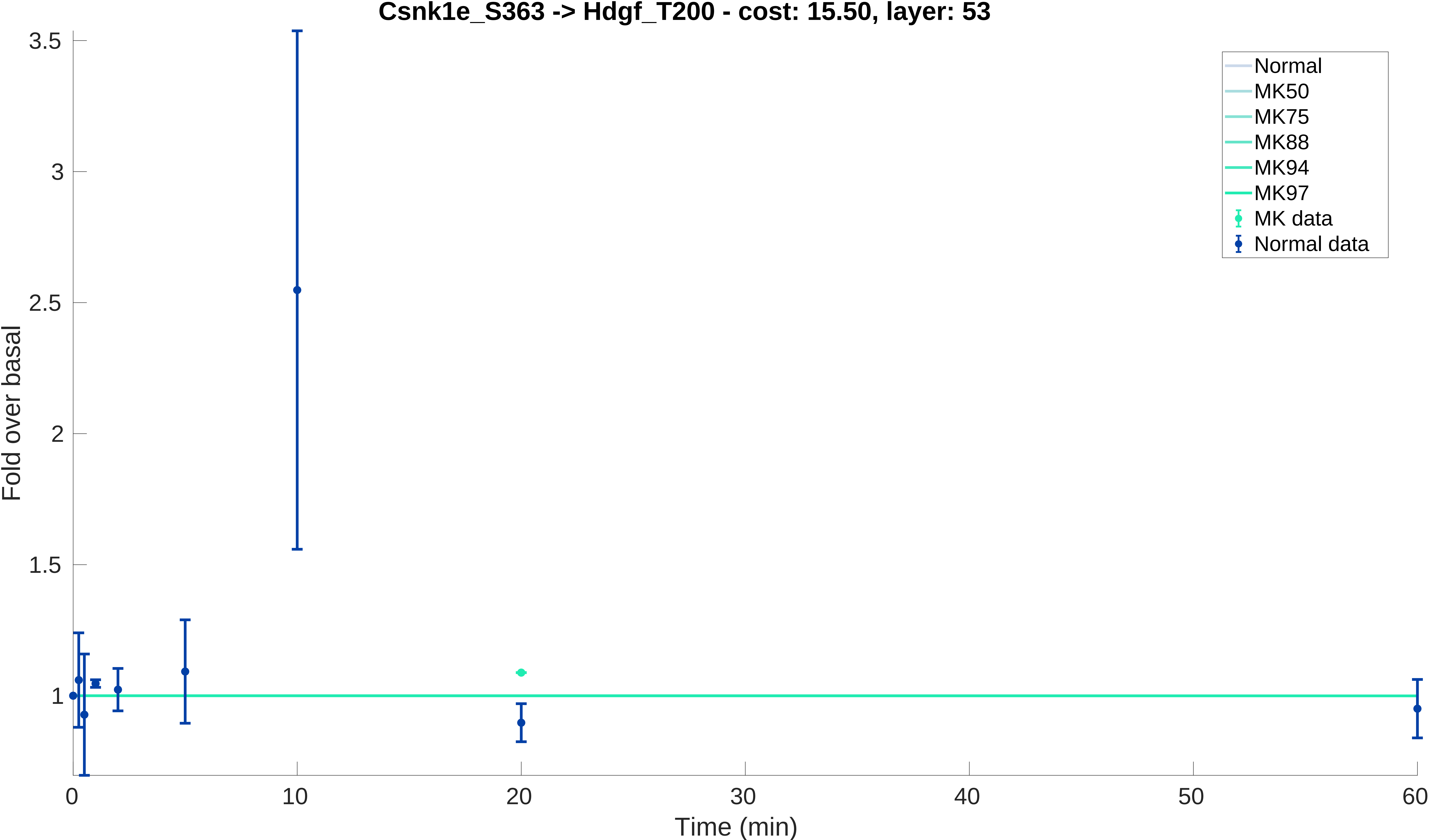

Csnk1e_S363 -> Hdgf_T200 - cost: 15.50, layer: 53
Fold over basal
3.5
3
2.5
2
1.5
1
0
10
20
30
40
50
60
Time (min)
Normal
MK50
MK75
MK88
MK94
MK97
MK data
Normal data

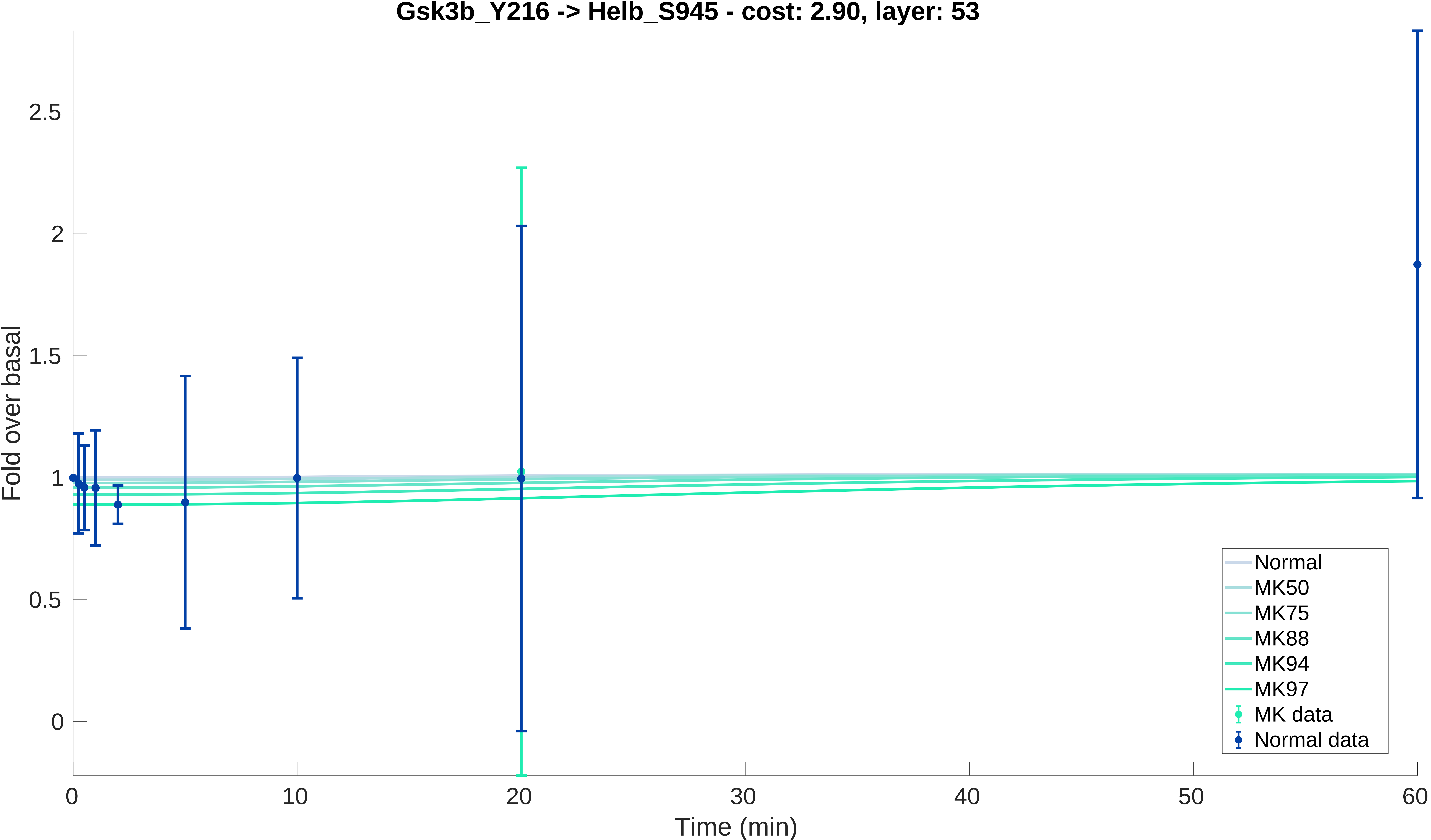

Gsk3b_Y216 -> Helb_S945 - cost: 2.90, layer: 53
Fold over basal
Time (min)
0
0.5
1
1.5
2
2.5
0
10
20
30
40
50
60
Normal
MK50
MK75
MK88
MK94
MK97
MK data
Normal data

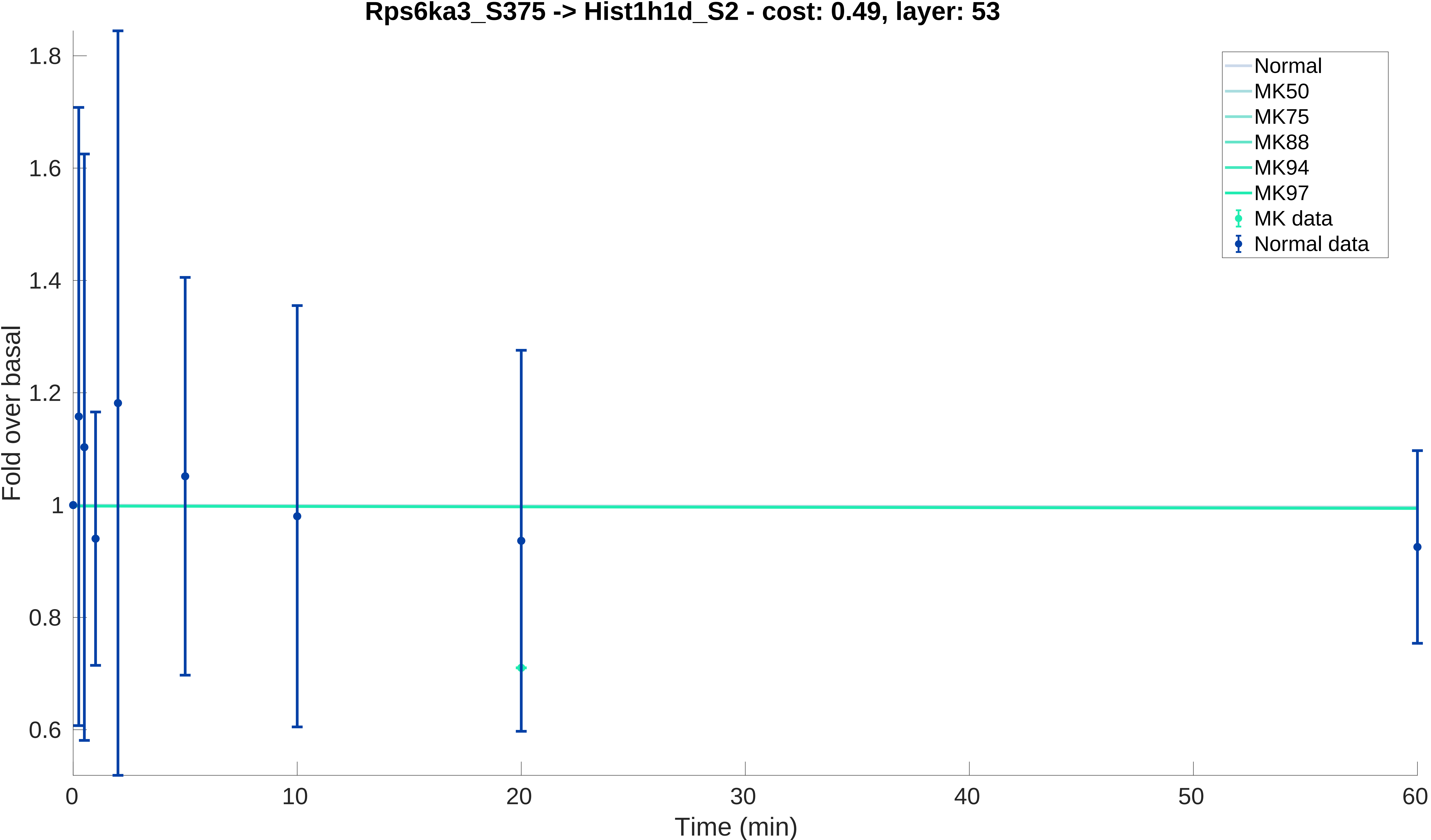
Rps6ka3_S375 -> Hist1h1d_S2 - cost: 0.49, layer: 53
Fold over basal
Time (min)
0.6
0.8
1
1.2
1.4
1.6
1.8
0
10
20
30
40
50
60
Normal
MK50
MK75
MK88
MK94
MK97
MK data
Normal data

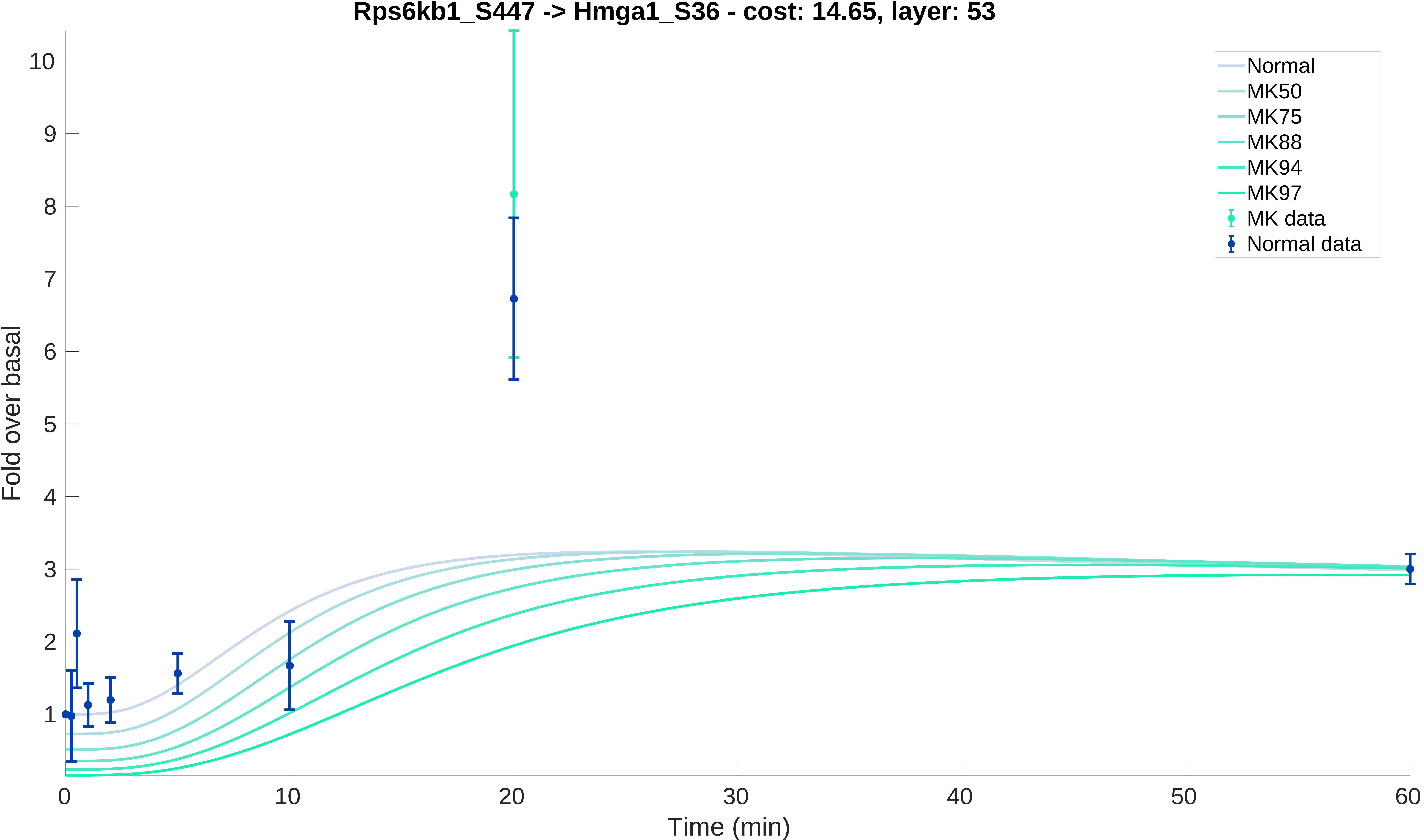

Rps6kb1_S447 -> Hmga1_S36 - cost: 14.65, layer: 53
Fold over basal
Time (min)
0
10
20
30
40
50
60
1
2
3
4
5
6
7
8
9
10
Normal
MK50
MK75
MK88
MK94
MK97
MK data
Normal data

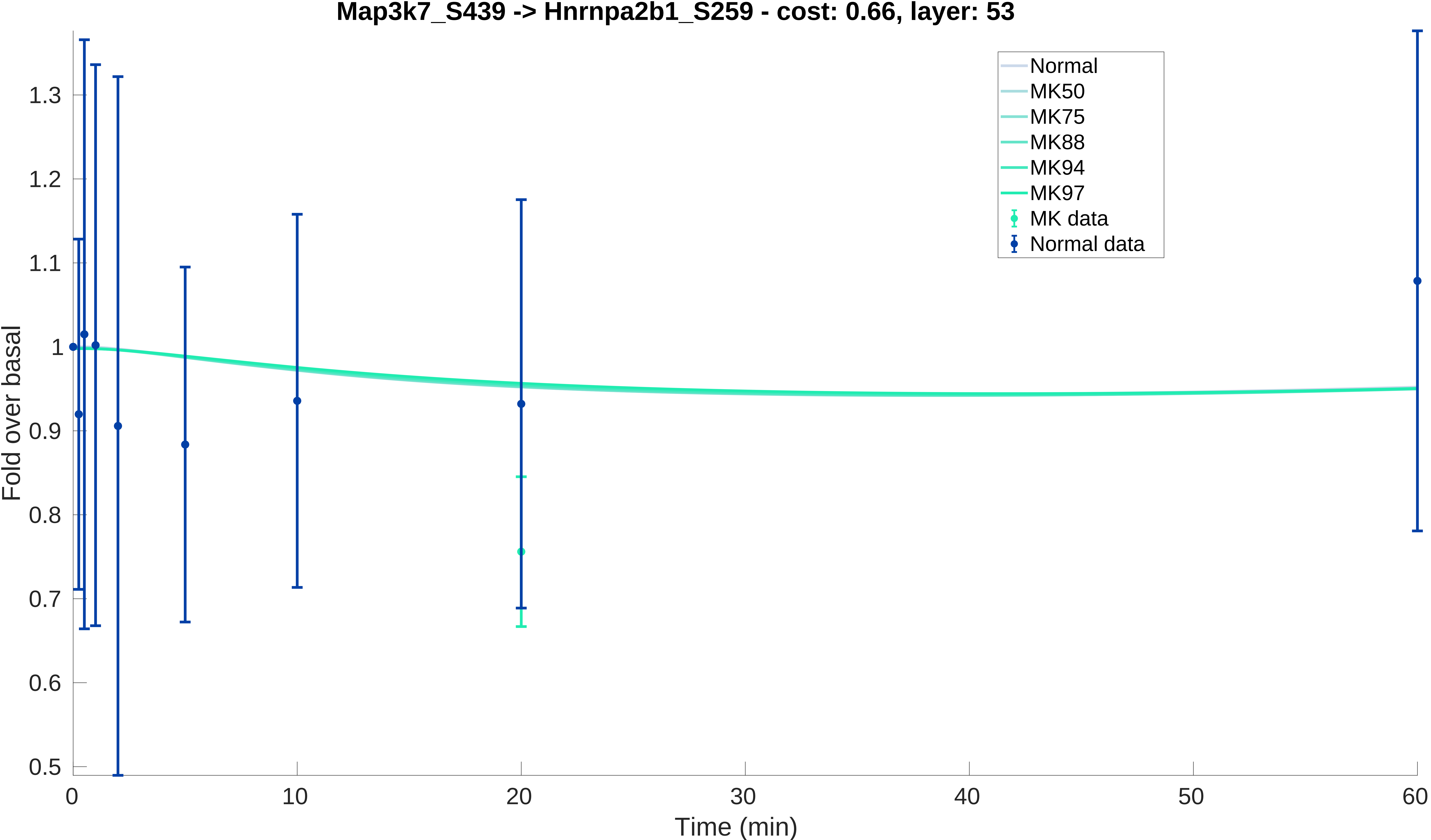
Map3k7_S439 -> Hnrnpa2b1_S259 - cost: 0.66, layer: 53
Fold over basal
Time (min)
0.5
0.6
0.7
0.8
0.9
1
1.1
1.2
1.3
0
10
20
30
40
50
60
Normal
MK50
MK75
MK88
MK94
MK97
MK data
Normal data

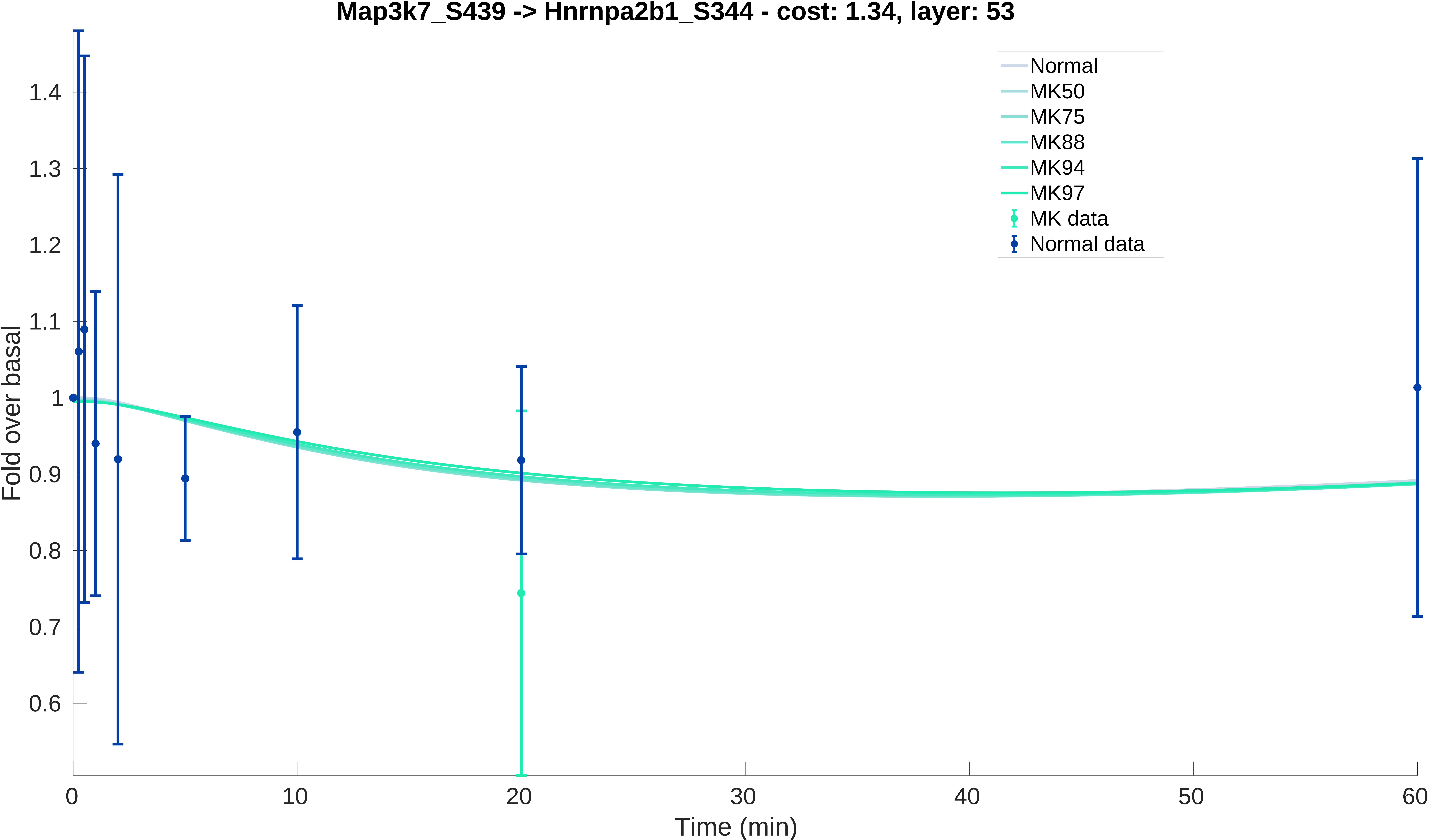

Map3k7_S439 -> Hnrnpa2b1_S344 - cost: 1.34, layer: 53
Normal
MK50
MK75
MK88
MK94
MK97
MK data
Normal data
Fold over basal
1.4
1.3
1.2
1.1
1
0.9
0.8
0.7
0.6
0
10
20
30
40
50
60
Time (min)

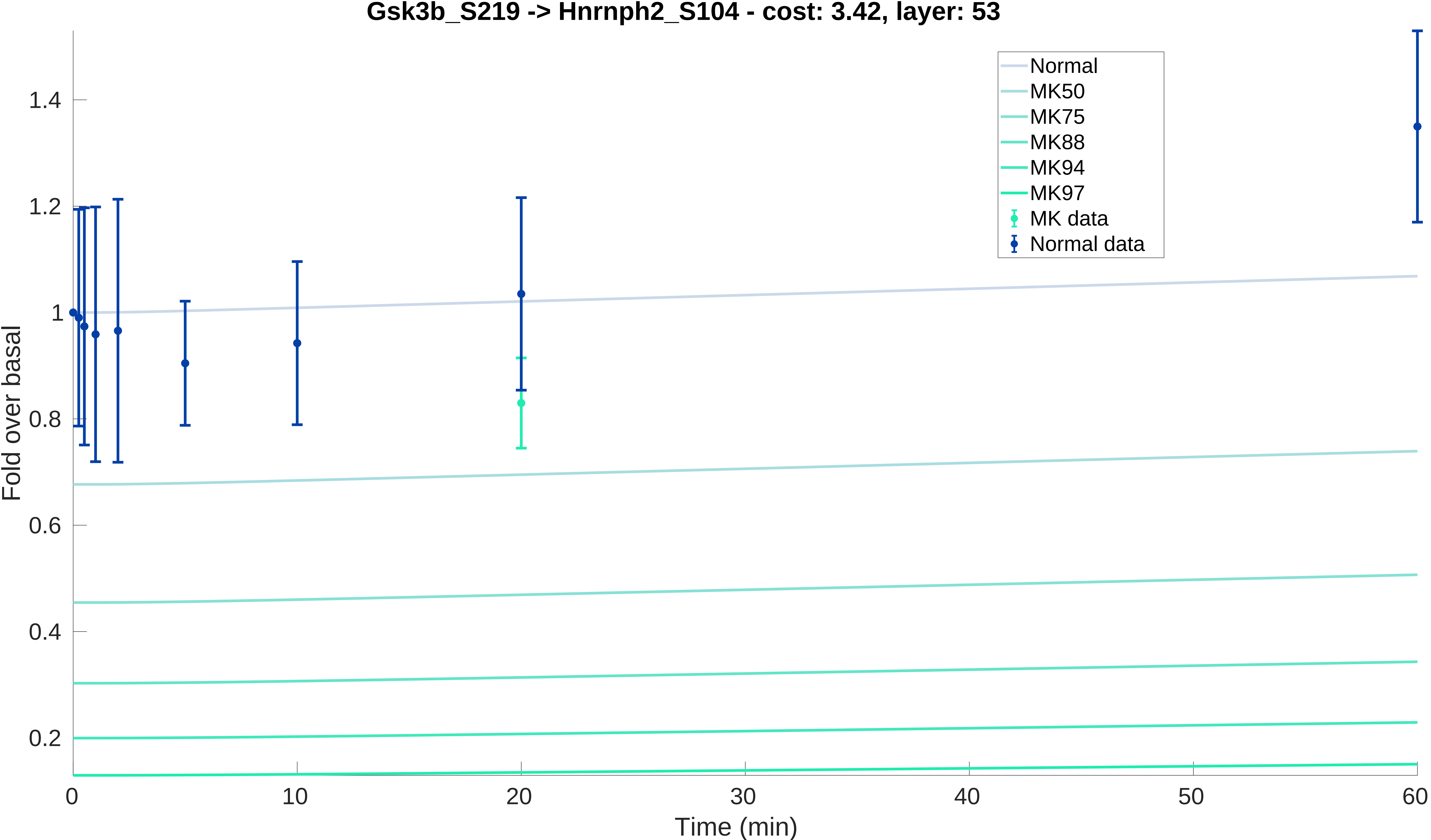

Gsk3b_S219 -> Hnrnph2_S104 - cost: 3.42, layer: 53
Fold over basal
Time (min)
1.4
1.2
1
0.8
0.6
0.4
0.2
0
10
20
30
40
50
60
Normal
MK50
MK75
MK88
MK94
MK97
MK data
Normal data

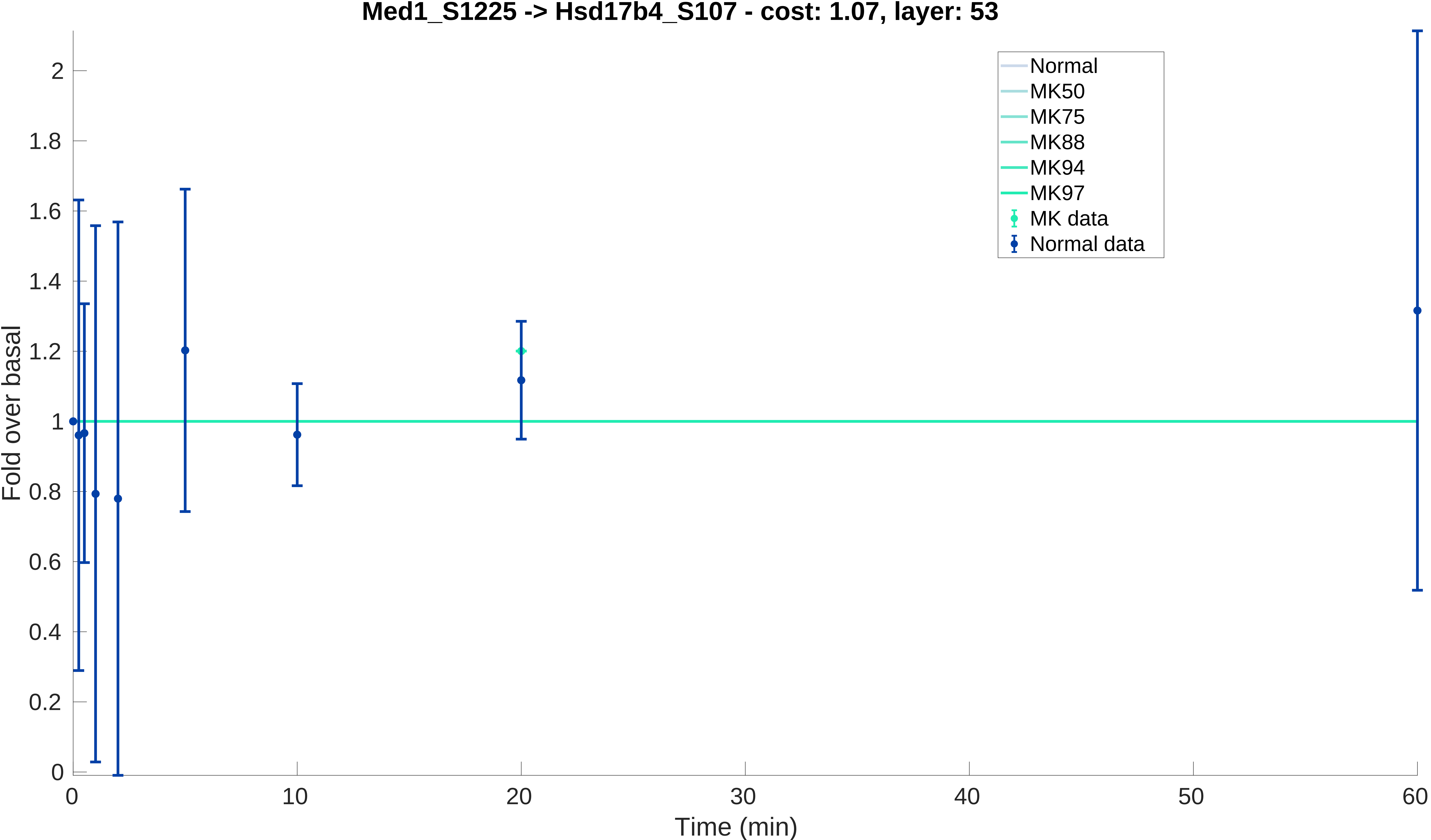

Med1_S1225 -> Hsd17b4_S107 - cost: 1.07, layer: 53
Normal
MK50
MK75
MK88
MK94
MK97
MK data
Normal data
Fold over basal
Time (min)
0
0.2
0.4
0.6
0.8
1
1.2
1.4
1.6
1.8
2
0
10
20
30
40
50
60

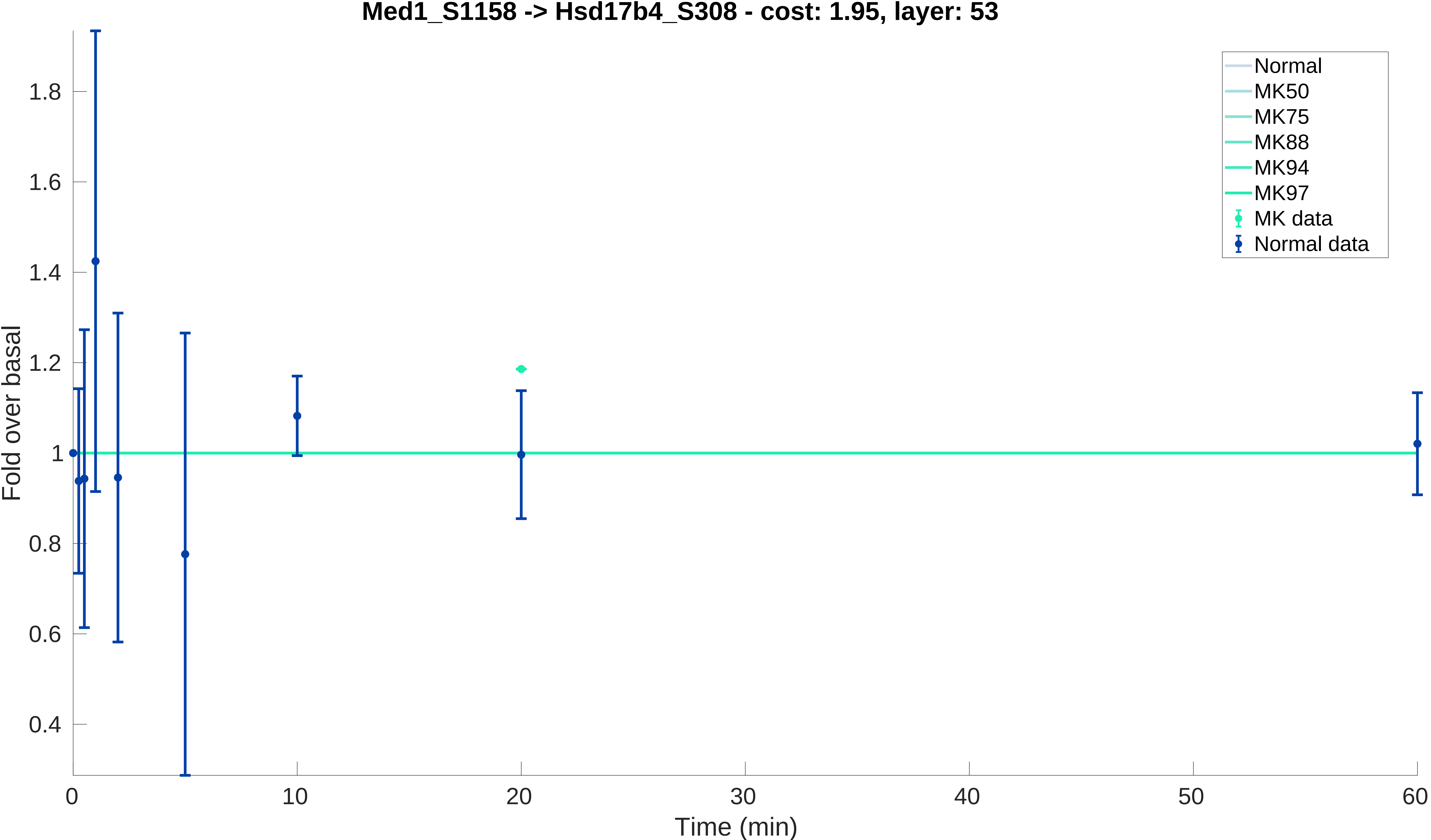
Med1_S1158 -> Hsd17b4_S308 - cost: 1.95, layer: 53
Fold over basal
0.4
0.6
0.8
1
1.2
1.4
1.6
1.8
0
10
20
30
40
50
60
Time (min)
Normal
MK50
MK75
MK88
MK94
MK97
MK data
Normal data

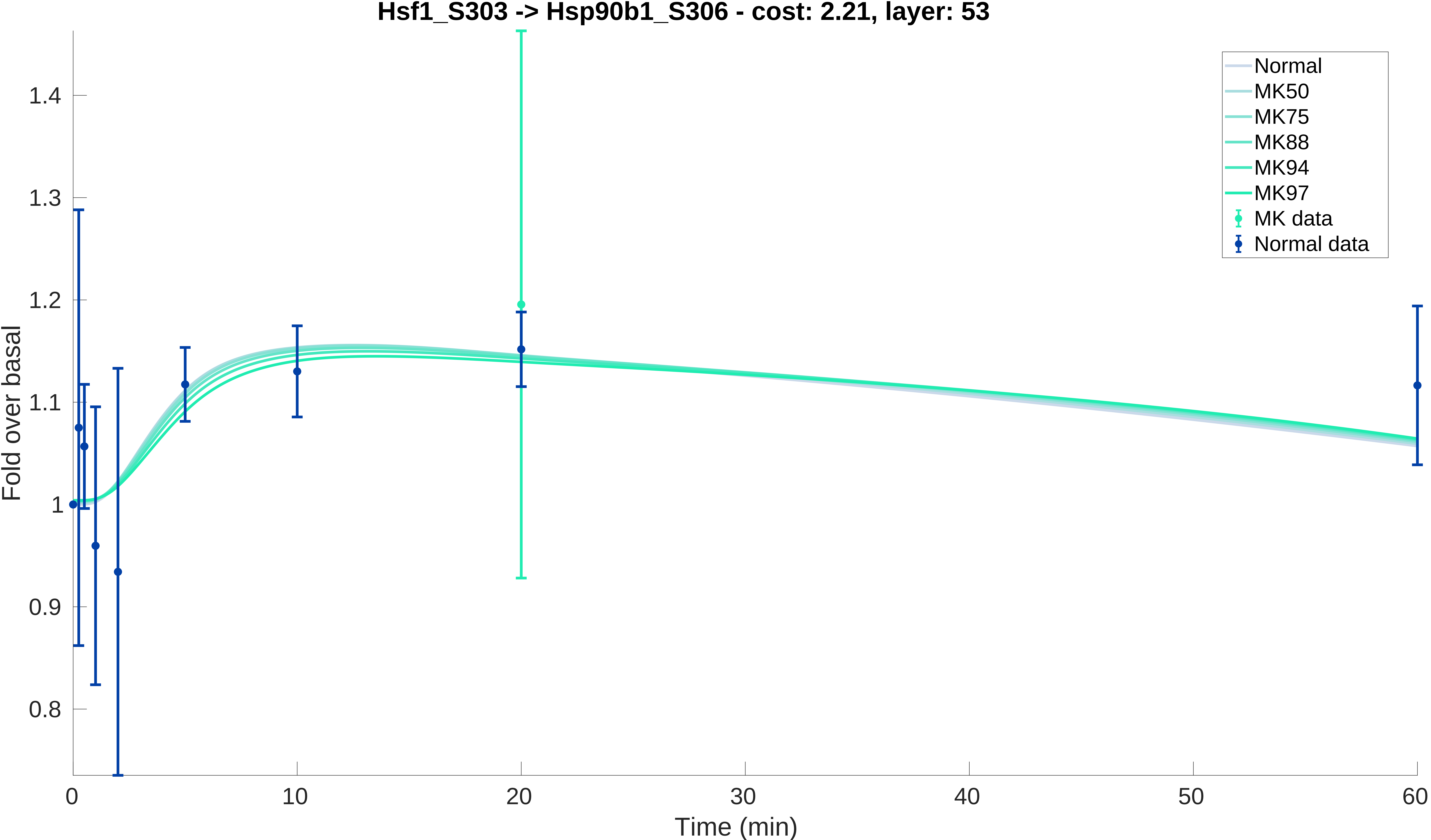

Hsf1_S303 -> Hsp90b1_S306 - cost: 2.21, layer: 53
Fold over basal
Time (min)
1.4
1.3
1.2
1.1
1
0.9
0.8
0
10
20
30
40
50
60
Normal
MK50
MK75
MK88
MK94
MK97
MK data
Normal data

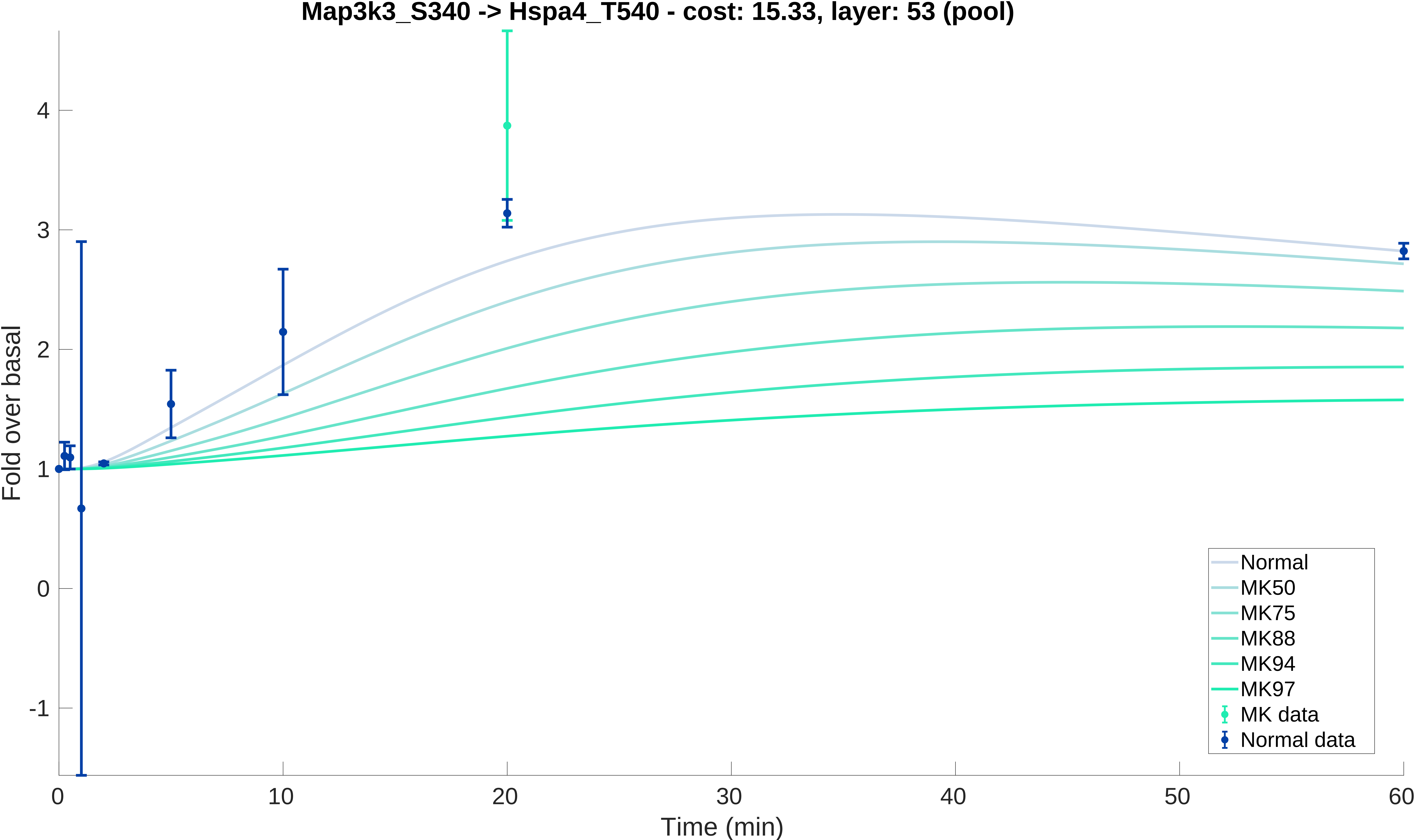

Map3k3_S340 -> Hspa4_T540 - cost: 15.33, layer: 53 (pool)
Fold over basal
4
3
2
1
0
-1
0
10
20
30
40
50
60
Time (min)
Normal
MK50
MK75
MK88
MK94
MK97
MK data
Normal data

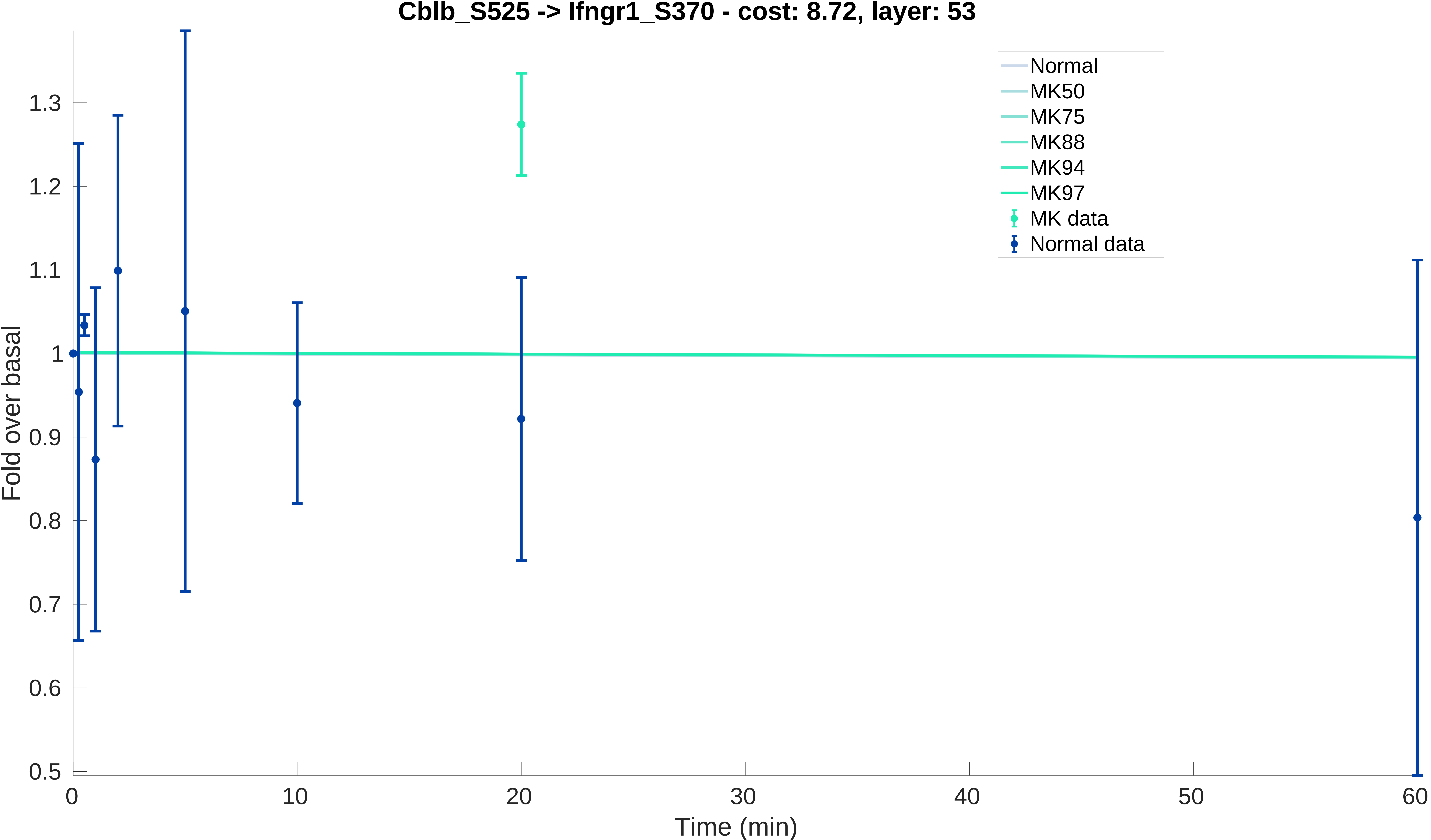

Cblb_S525 -> Ifngr1_S370 - cost: 8.72, layer: 53
Normal
MK50
MK75
MK88
MK94
MK97
MK data
Normal data
Fold over basal
1.3
1.2
1.1
1
0.9
0.8
0.7
0.6
0.5
0
10
20
30
40
50
60
Time (min)

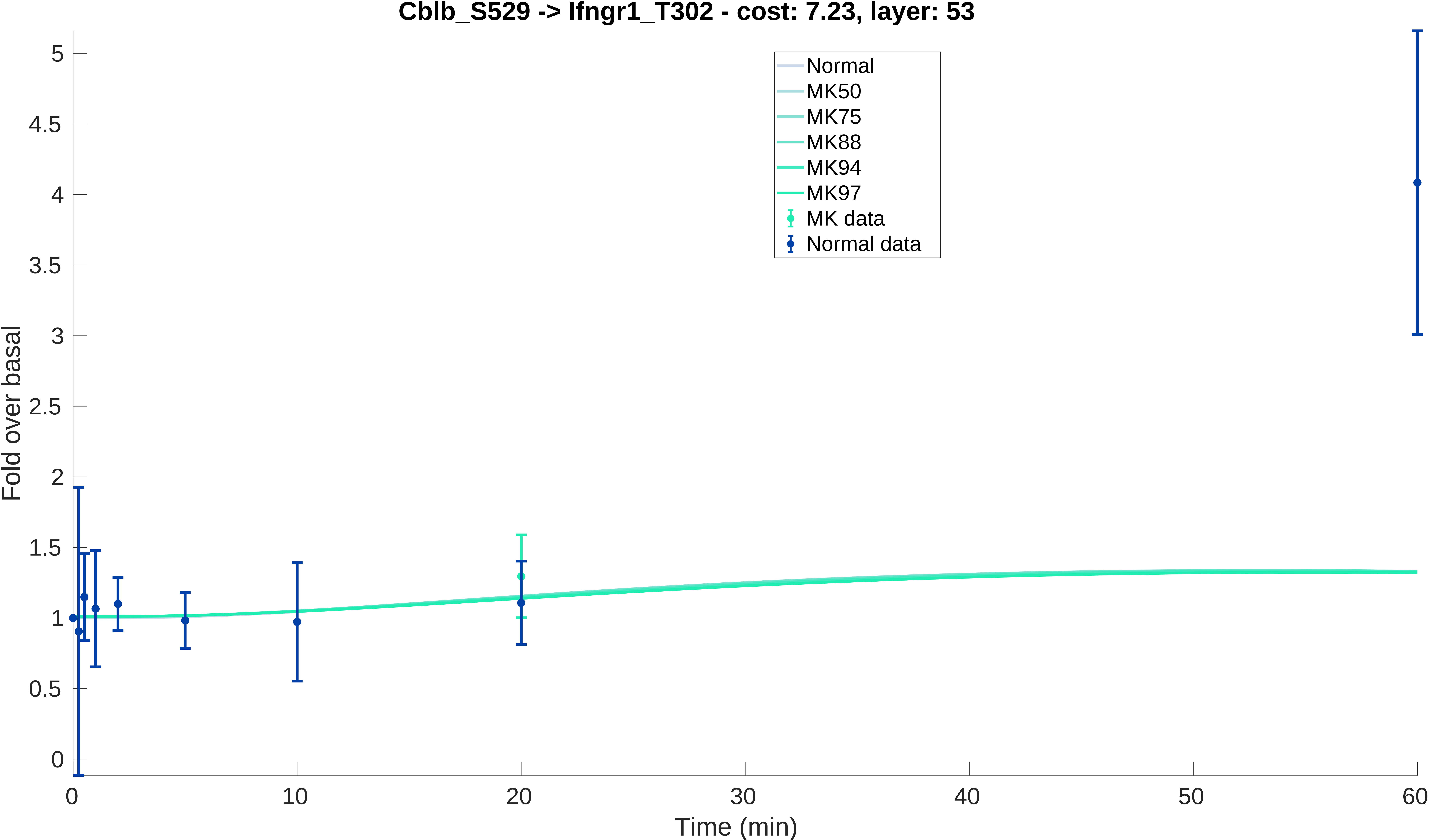

Cblb_S529 -> Ifngr1_T302 - cost: 7.23, layer: 53
Normal
MK50
MK75
MK88
MK94
MK97
MK data
Normal data
Fold over basal
Time (min)
0
0.5
1
1.5
2
2.5
3
3.5
4
4.5
5
0
10
20
30
40
50
60

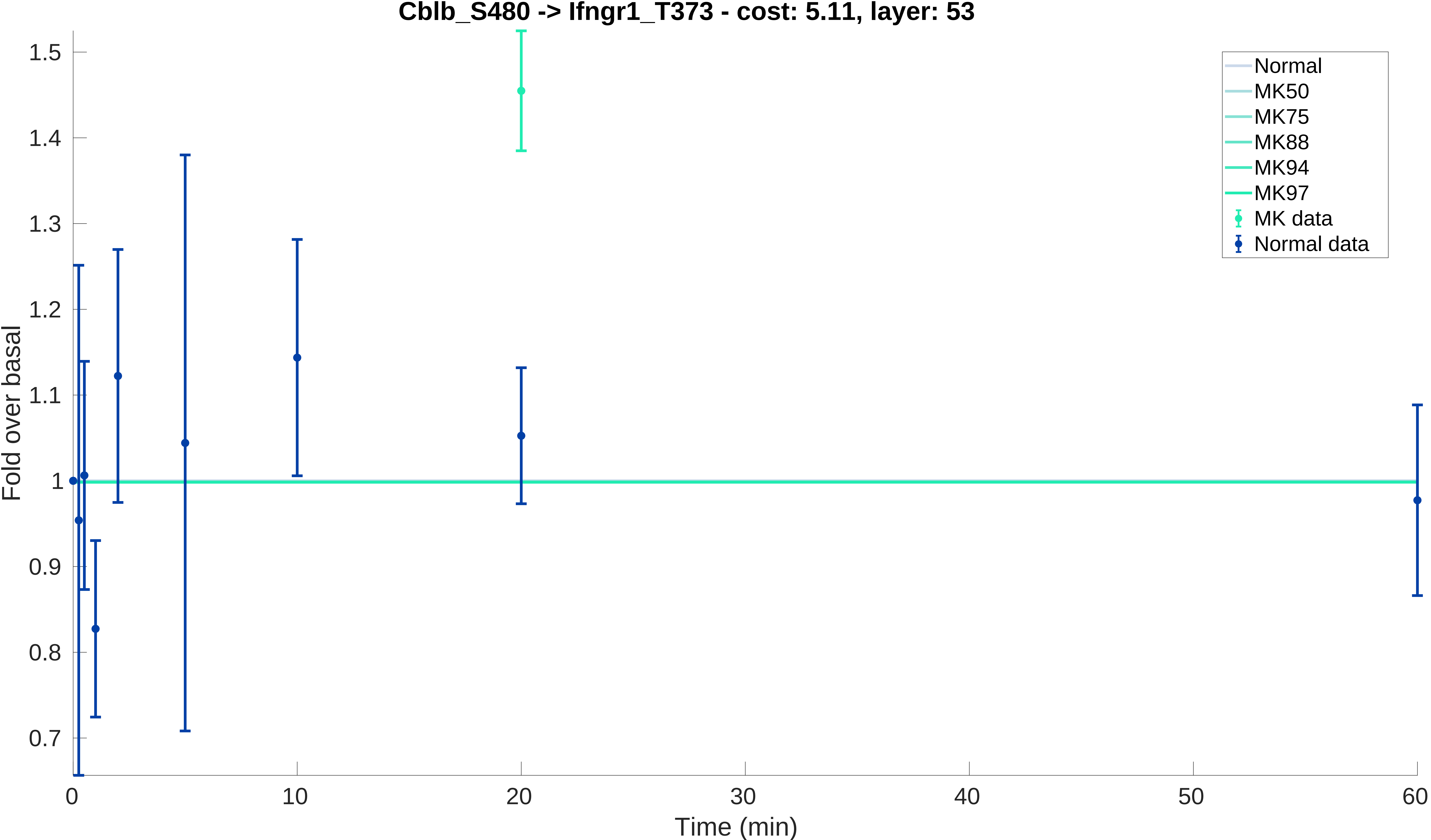

Cblb_S480 -> Ifngr1_T373 - cost: 5.11, layer: 53
Fold over basal
1.5
1.4
1.3
1.2
1.1
1
0.9
0.8
0.7
0
10
20
30
40
50
60
Time (min)
Normal
MK50
MK75
MK88
MK94
MK97
MK data
Normal data

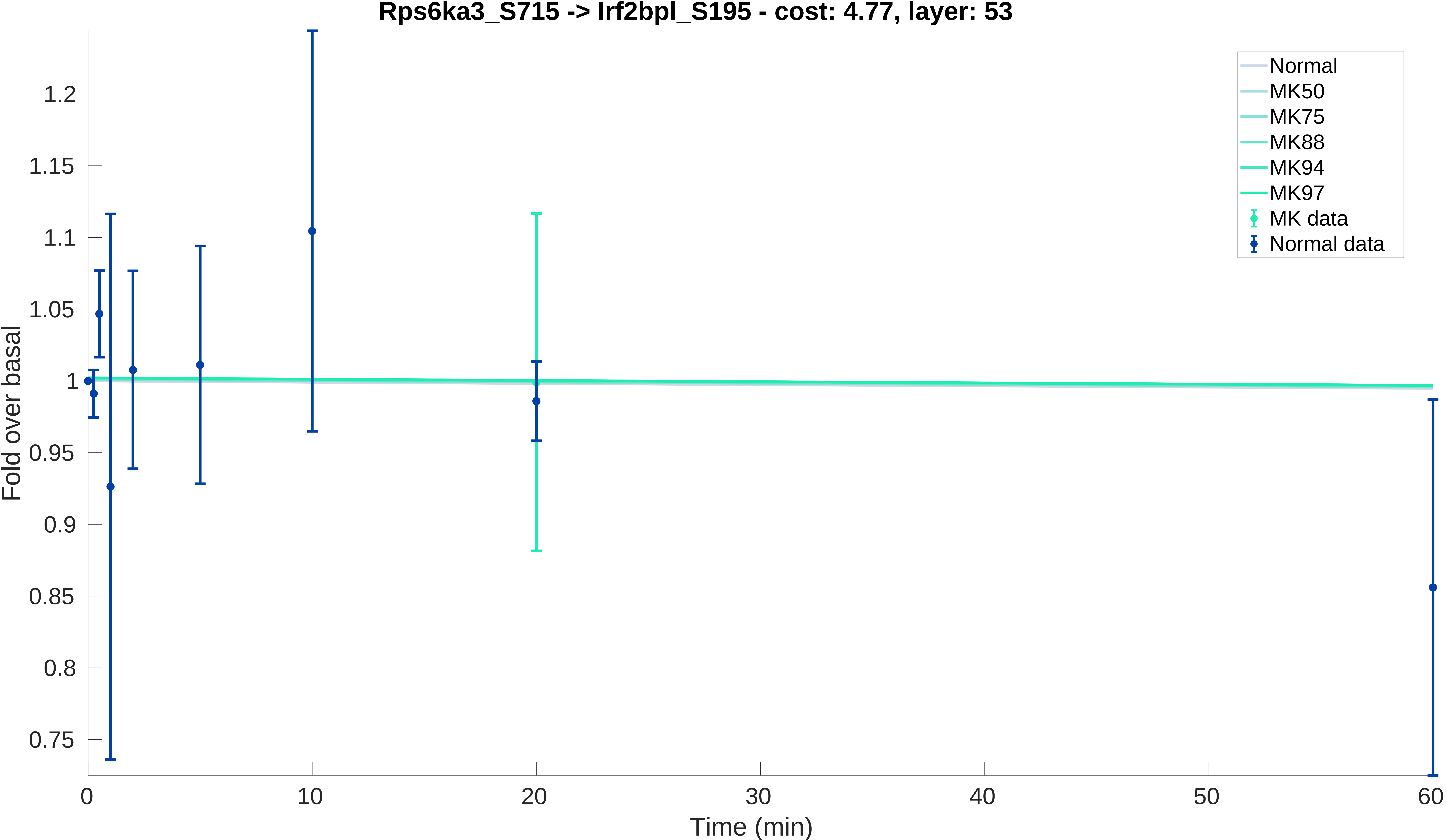

Rps6ka3_S715 -> Irf2bpl_S195 - cost: 4.77, layer: 53
Normal
MK50
MK75
MK88
MK94
MK97
MK data
Normal data
Fold over basal
1.2
1.15
1.1
1.05
1
0.95
0.9
0.85
0.8
0.75
0
10
20
30
40
50
60
Time (min)

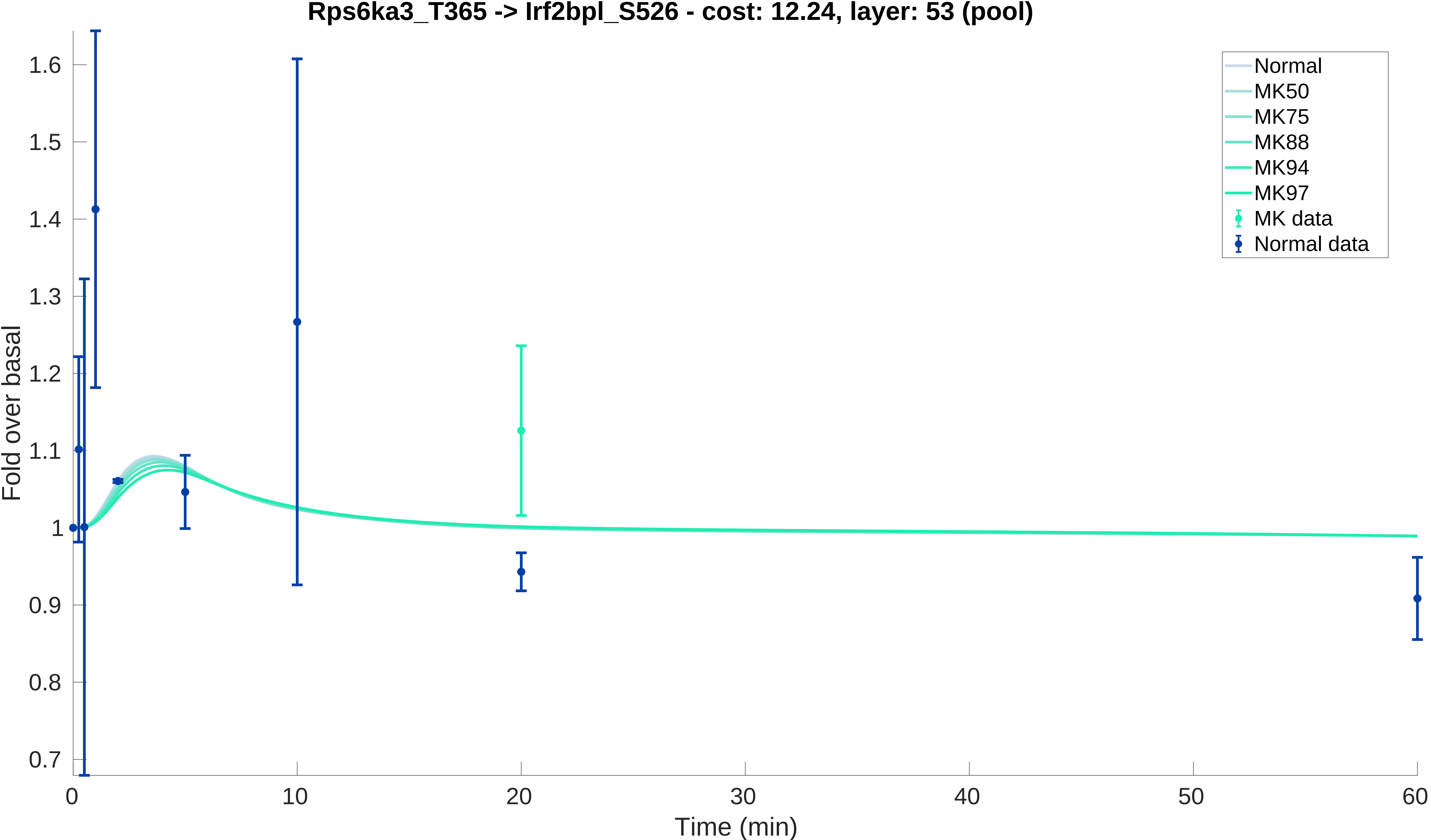

Rps6ka3_T365 -> Irf2bpl_S526 - cost: 12.24, layer: 53 (pool)
Fold over basal
Time (min)
0.7
0.8
0.9
1
1.1
1.2
1.3
1.4
1.5
1.6
0
10
20
30
40
50
60
Normal
MK50
MK75
MK88
MK94
MK97
MK data
Normal data

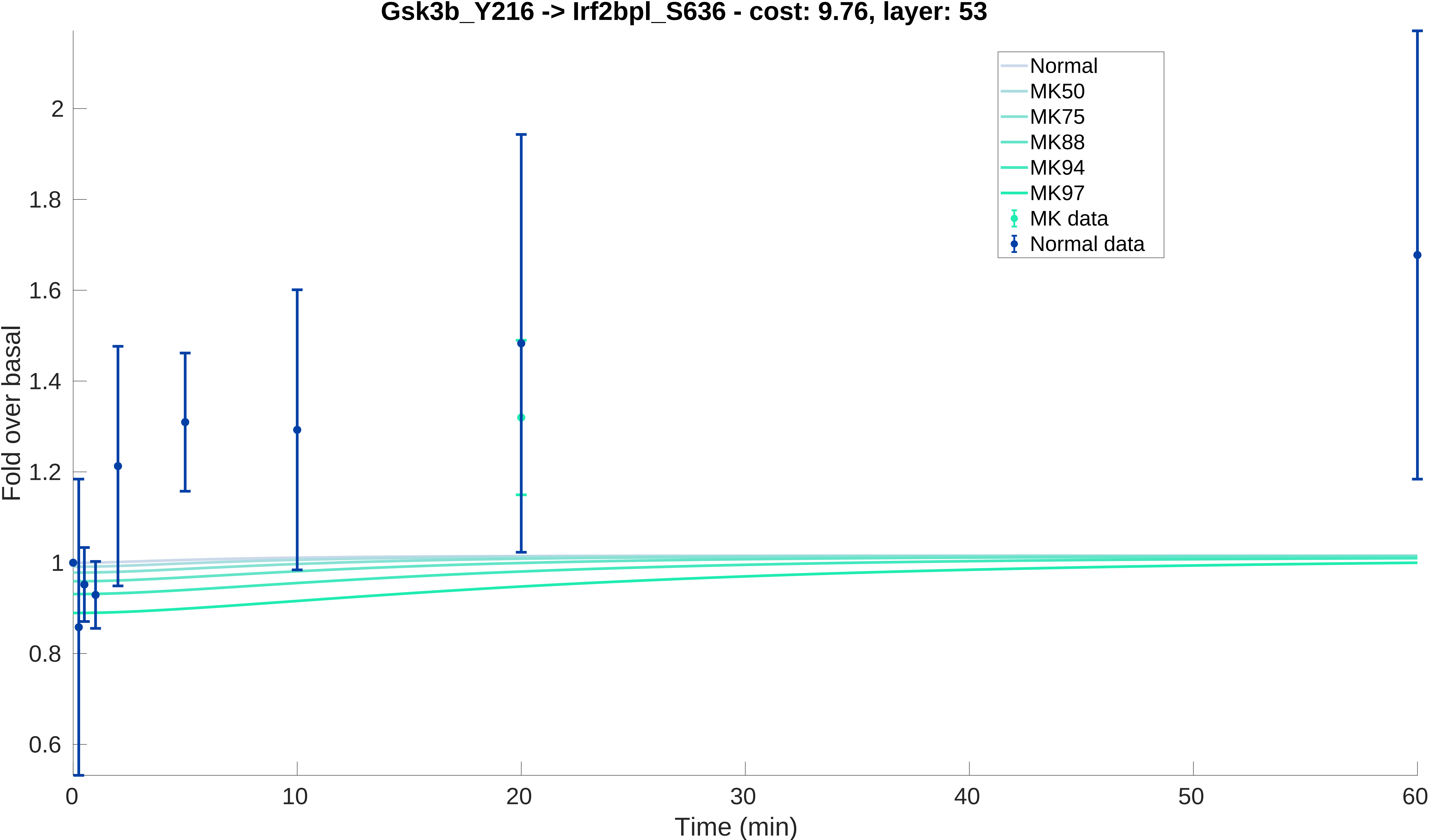
Gsk3b_Y216 -> Irf2bpl_S636 - cost: 9.76, layer: 53
Fold over basal
Time (min)
0.6
0.8
1
1.2
1.4
1.6
1.8
2
0
10
20
30
40
50
60
Normal
MK50
MK75
MK88
MK94
MK97
MK data
Normal data

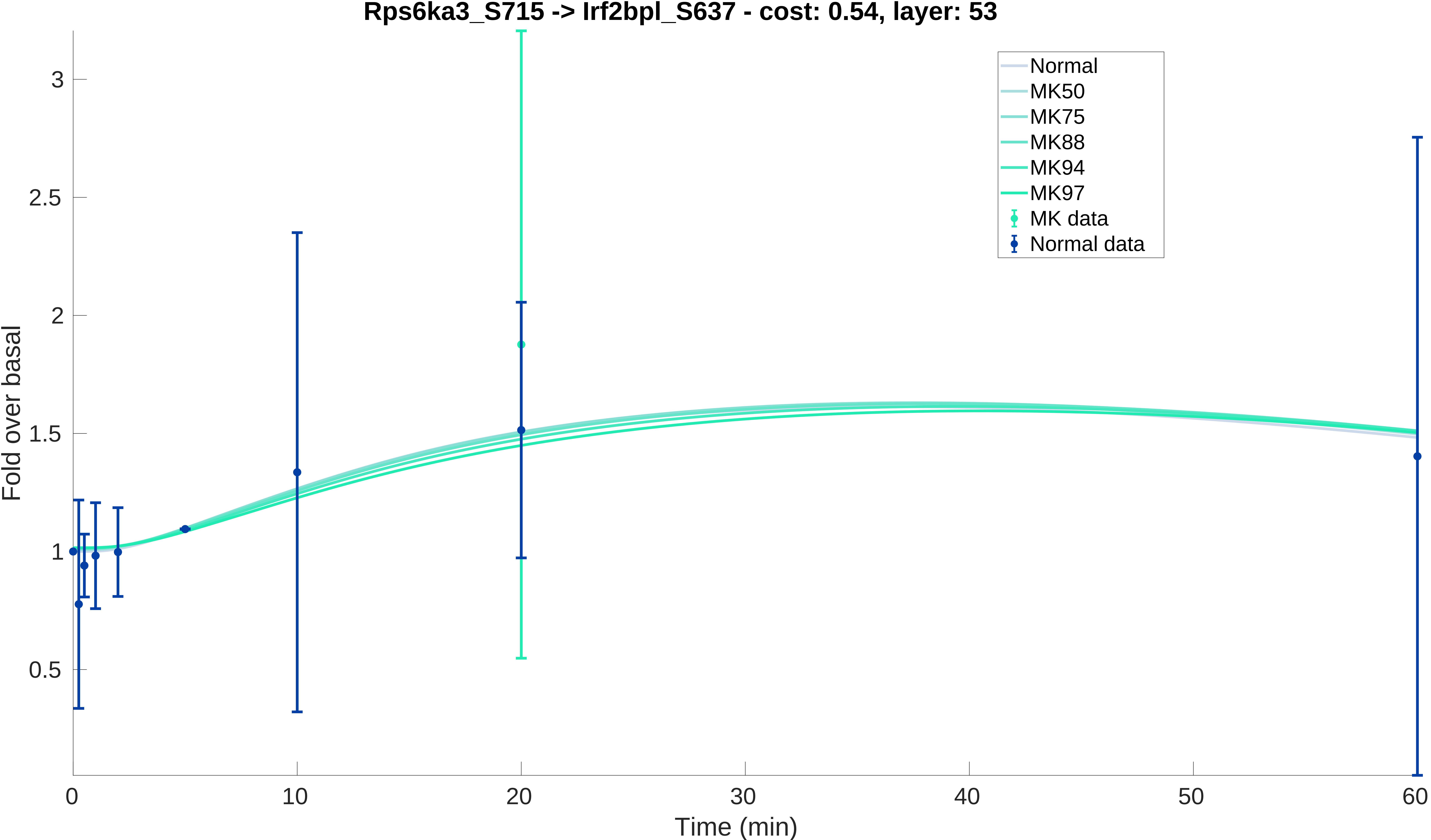

Rps6ka3_S715 -> Irf2bpl_S637 - cost: 0.54, layer: 53
Fold over basal
Time (min)
0
10
20
30
40
50
60
0.5
1
1.5
2
2.5
3
Normal
MK50
MK75
MK88
MK94
MK97
MK data
Normal data

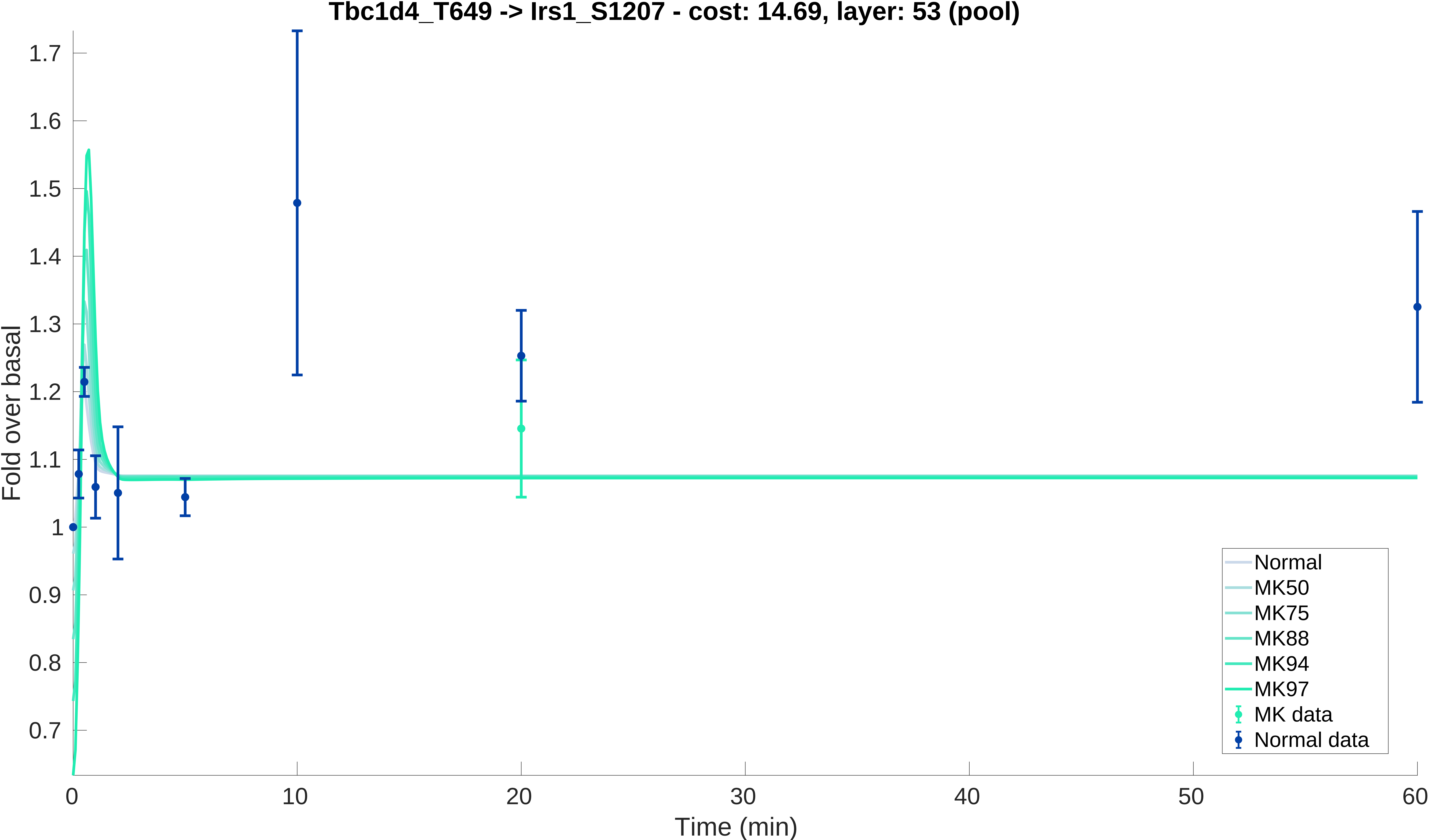

Tbc1d4_T649 -> Irs1_S1207 - cost: 14.69, layer: 53 (pool)
Fold over basal
Time (min)
1.7
1.6
1.5
1.4
1.3
1.2
1.1
1
0.9
0.8
0.7
0
10
20
30
40
50
60
Normal
MK50
MK75
MK88
MK94
MK97
MK data
Normal data

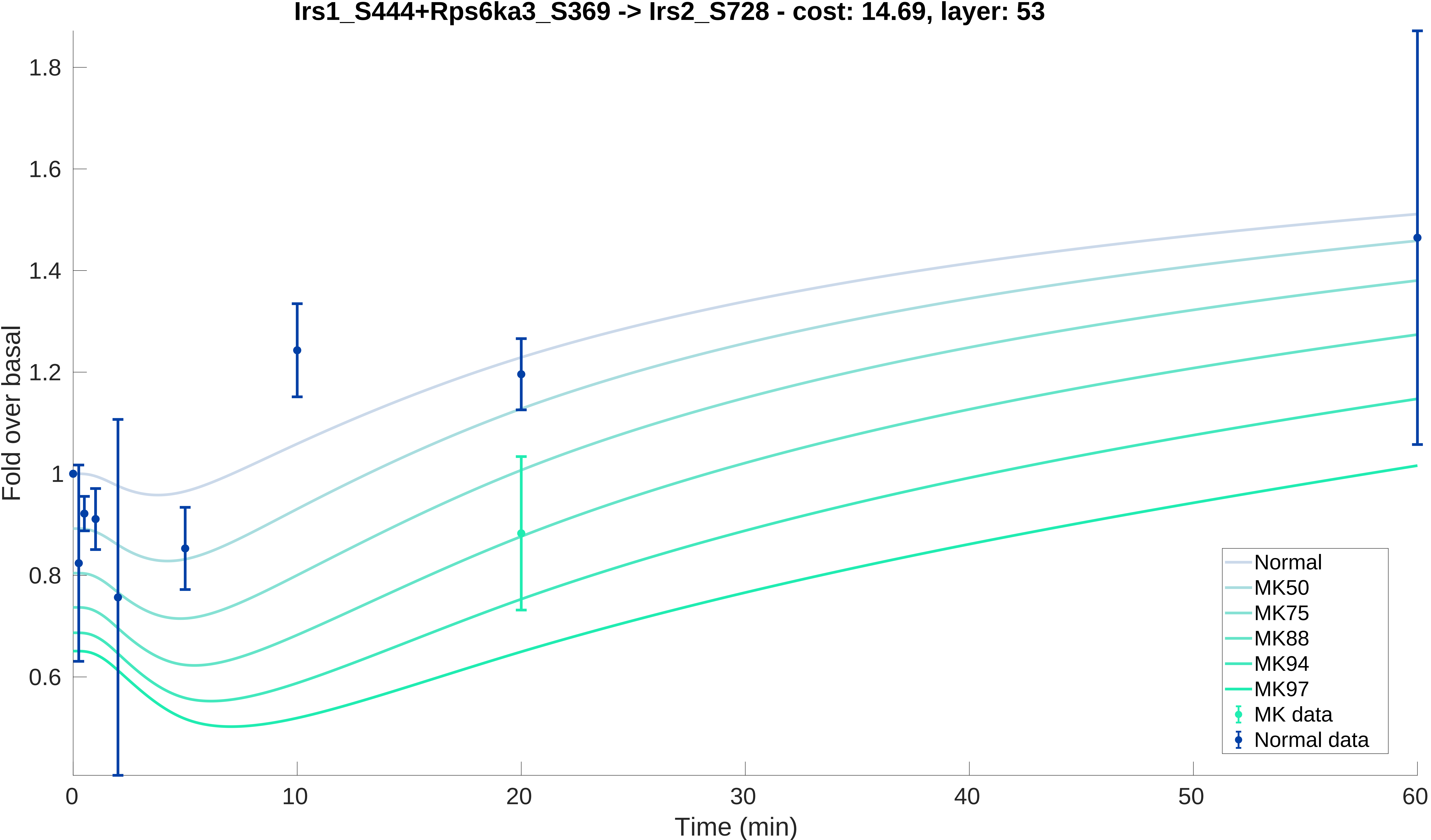
Irs1_S444+Rps6ka3_S369 -> Irs2_S728 - cost: 14.69, layer: 53
Fold over basal
Time (min)
0.6
0.8
1
1.2
1.4
1.6
1.8
0
10
20
30
40
50
60
Normal
MK50
MK75
MK88
MK94
MK97
MK data
Normal data

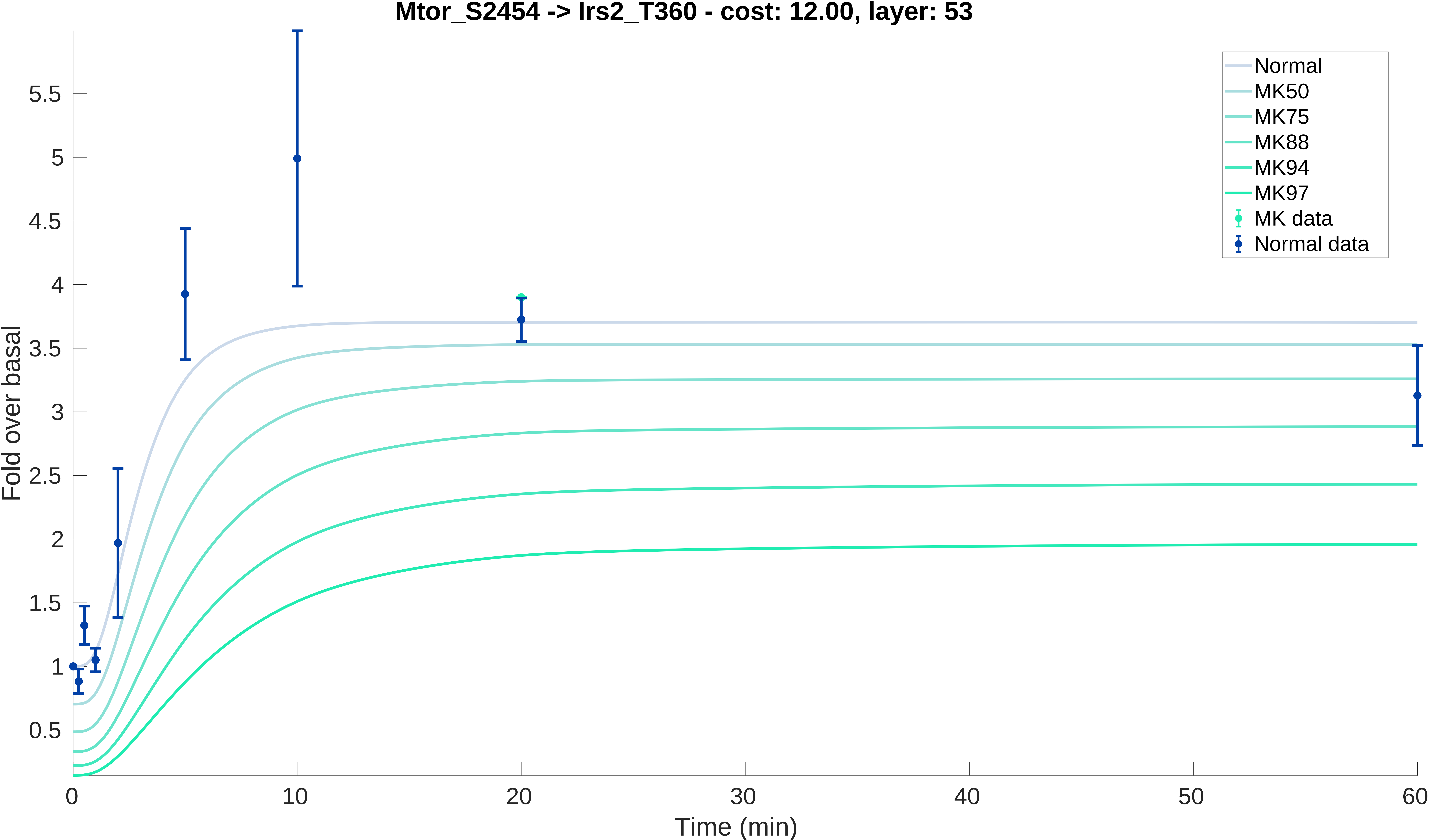

Mtor_S2454 -> Irs2_T360 - cost: 12.00, layer: 53
Fold over basal
5.5
5
4.5
4
3.5
3
2.5
2
1.5
1
0.5
0
10
20
30
40
50
60
Time (min)
Normal
MK50
MK75
MK88
MK94
MK97
MK data
Normal data

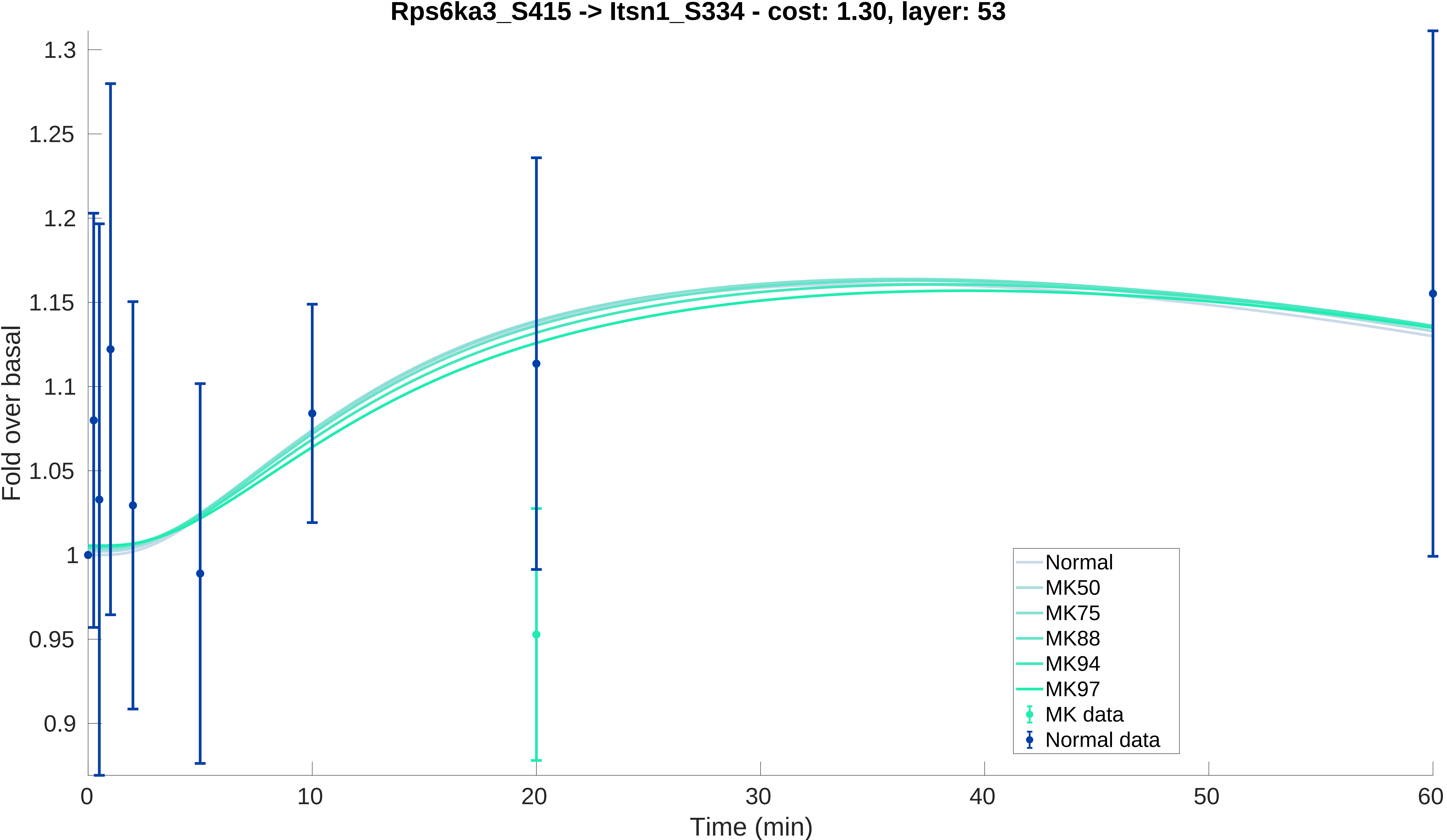
Rps6ka3_S415 -> Itsn1_S334 - cost: 1.30, layer: 53
Fold over basal
Time (min)
1.3
1.25
1.2
1.15
1.1
1.05
1
0.95
0.9
0
10
20
30
40
50
60
Normal
MK50
MK75
MK88
MK94
MK97
MK data
Normal data

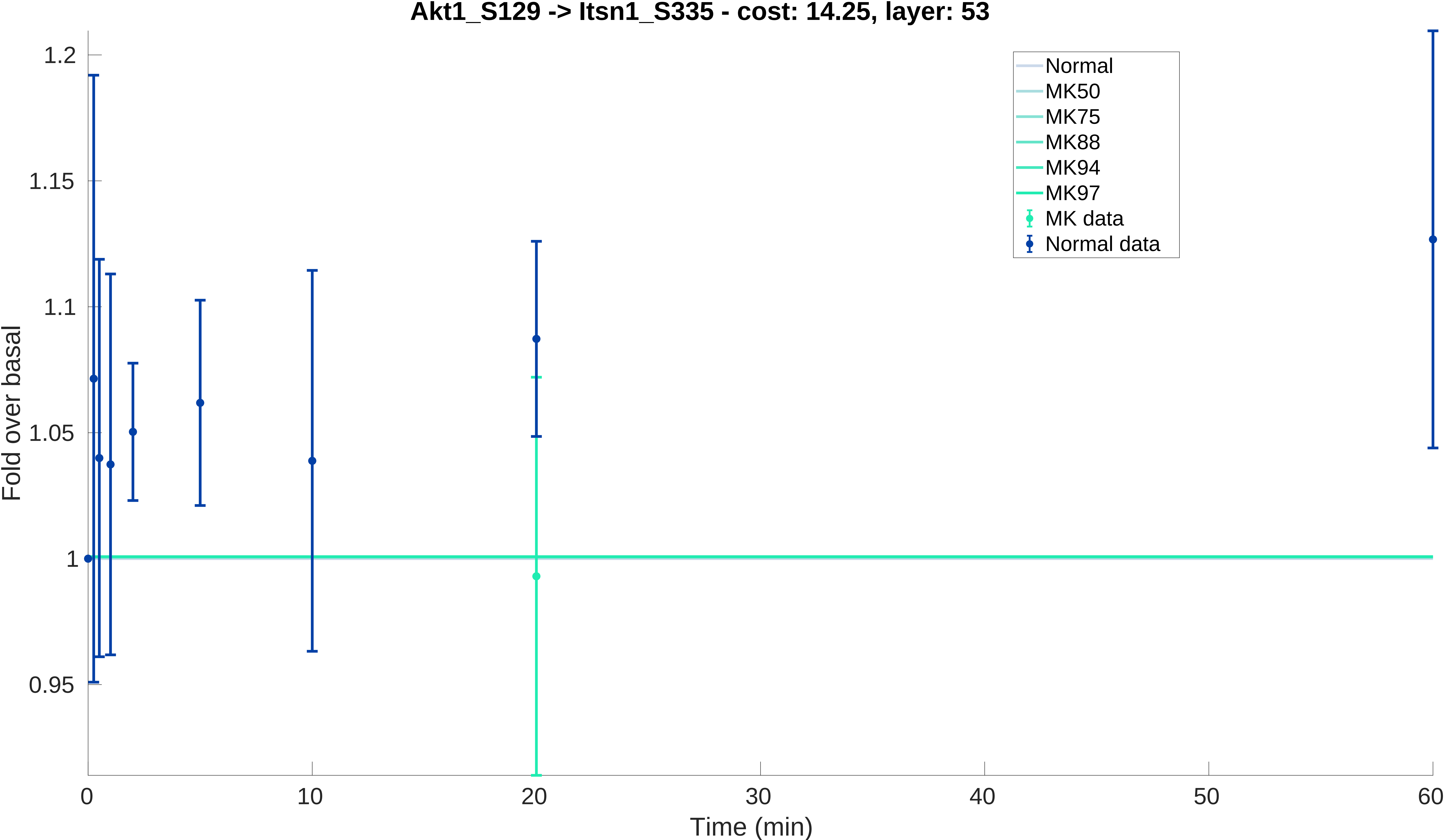

Akt1_S129 -> Itsn1_S335 - cost: 14.25, layer: 53
Normal
MK50
MK75
MK88
MK94
MK97
MK data
Normal data
Fold over basal
1.2
1.15
1.1
1.05
1
0.95
0
10
20
30
40
50
60
Time (min)

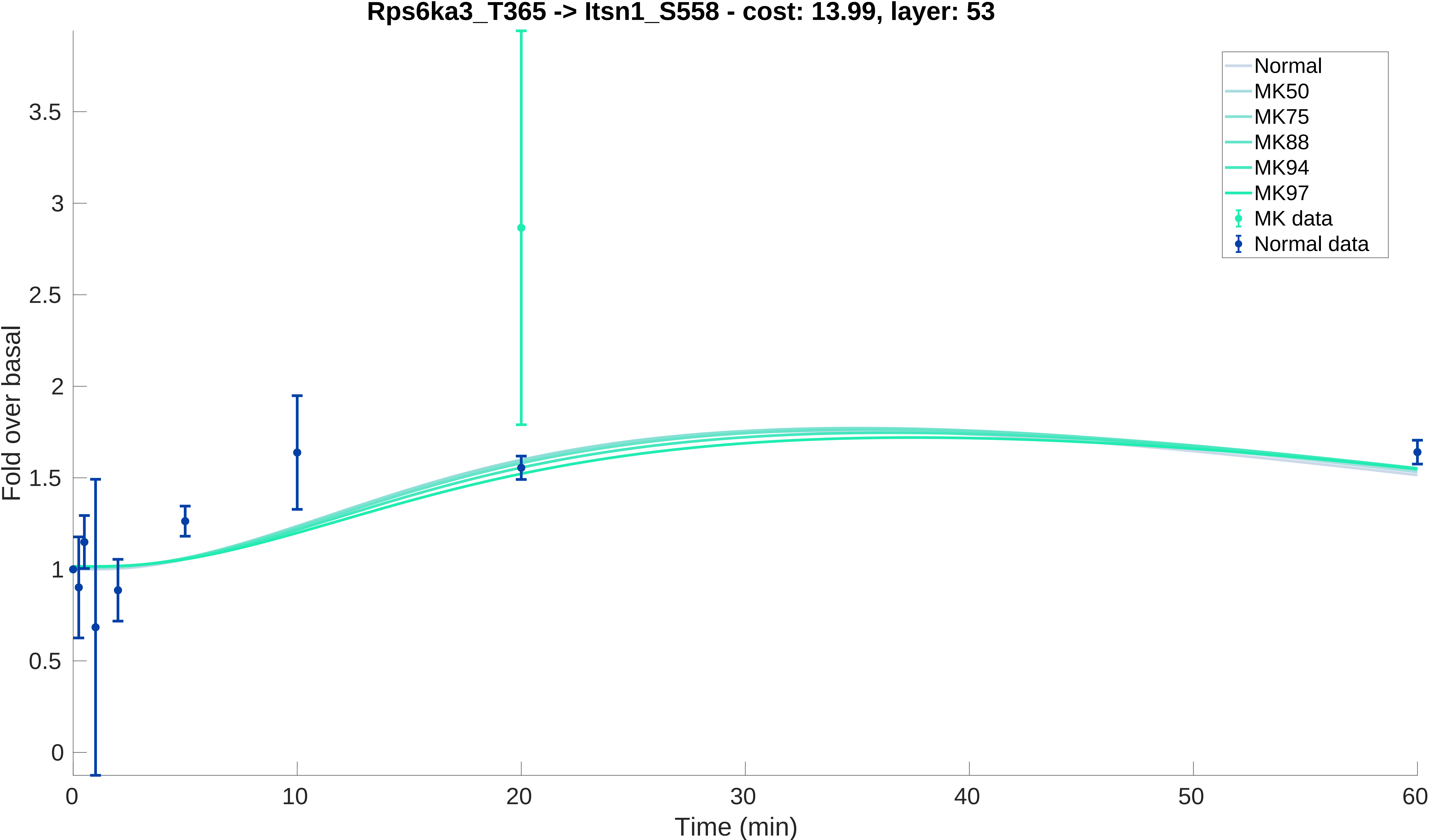

Rps6ka3_T365 -> Itsn1_S558 - cost: 13.99, layer: 53
Normal
MK50
MK75
MK88
MK94
MK97
MK data
Normal data
Fold over basal
Time (min)
0
0.5
1
1.5
2
2.5
3
3.5
0
10
20
30
40
50
60

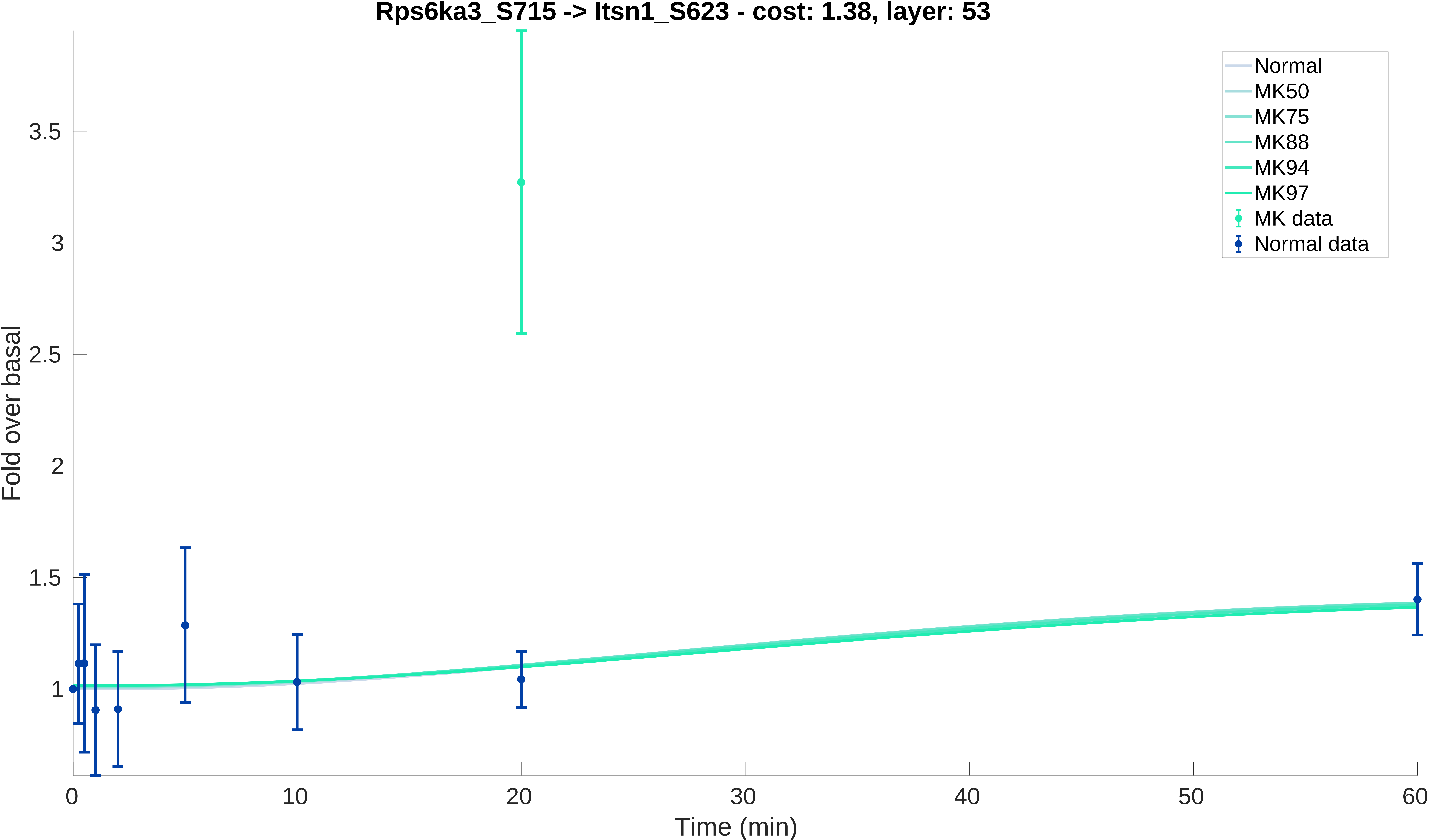

Rps6ka3_S715 -> Itsn1_S623 - cost: 1.38, layer: 53
Fold over basal
Time (min)
3.5
3
2.5
2
1.5
1
0
10
20
30
40
50
60
Normal
MK50
MK75
MK88
MK94
MK97
MK data
Normal data

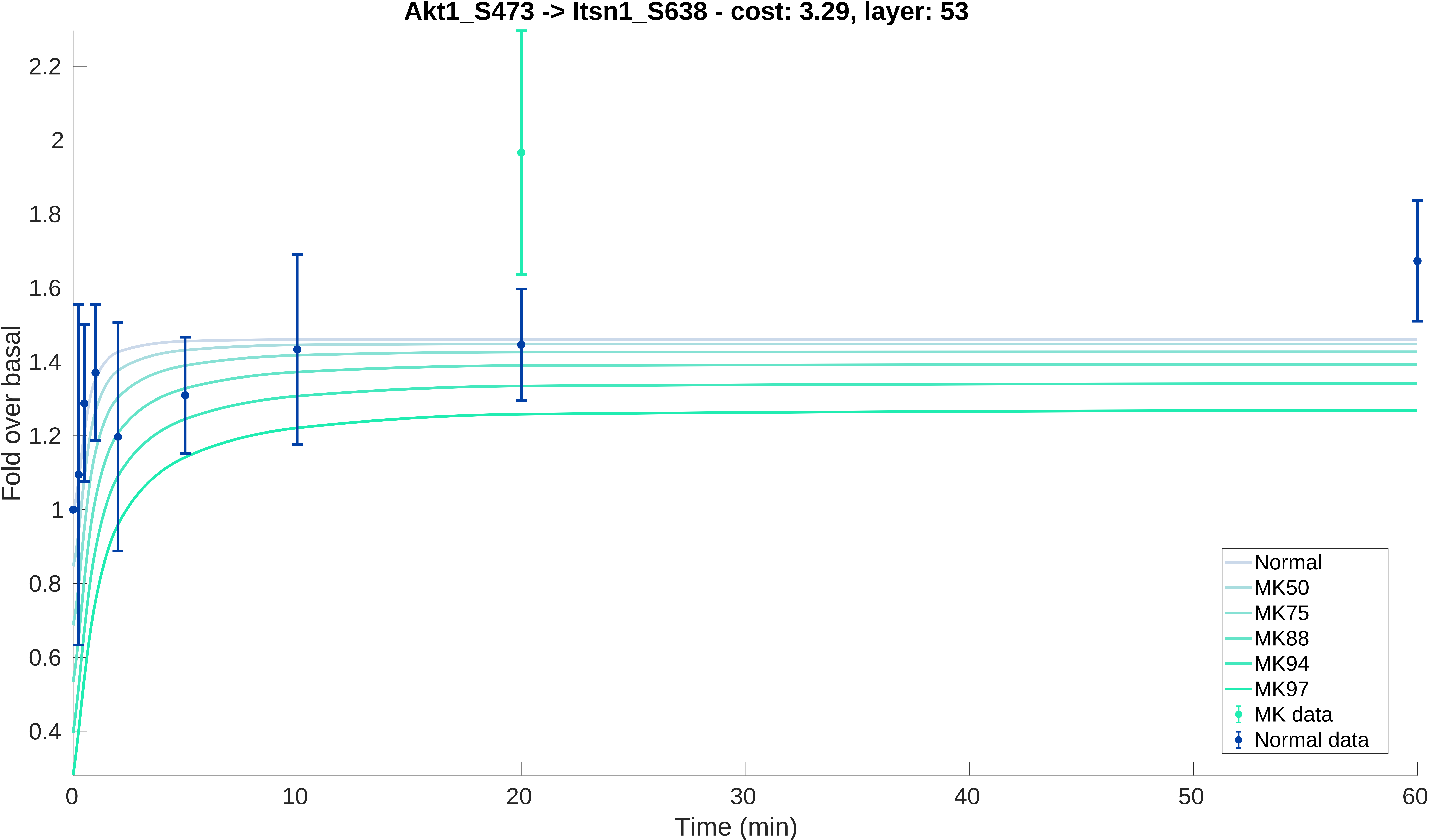
Akt1_S473 -> Itsn1_S638 - cost: 3.29, layer: 53
Fold over basal
Time (min)
2.2
2
1.8
1.6
1.4
1.2
1
0.8
0.6
0.4
0
10
20
30
40
50
60
Normal
MK50
MK75
MK88
MK94
MK97
MK data
Normal data

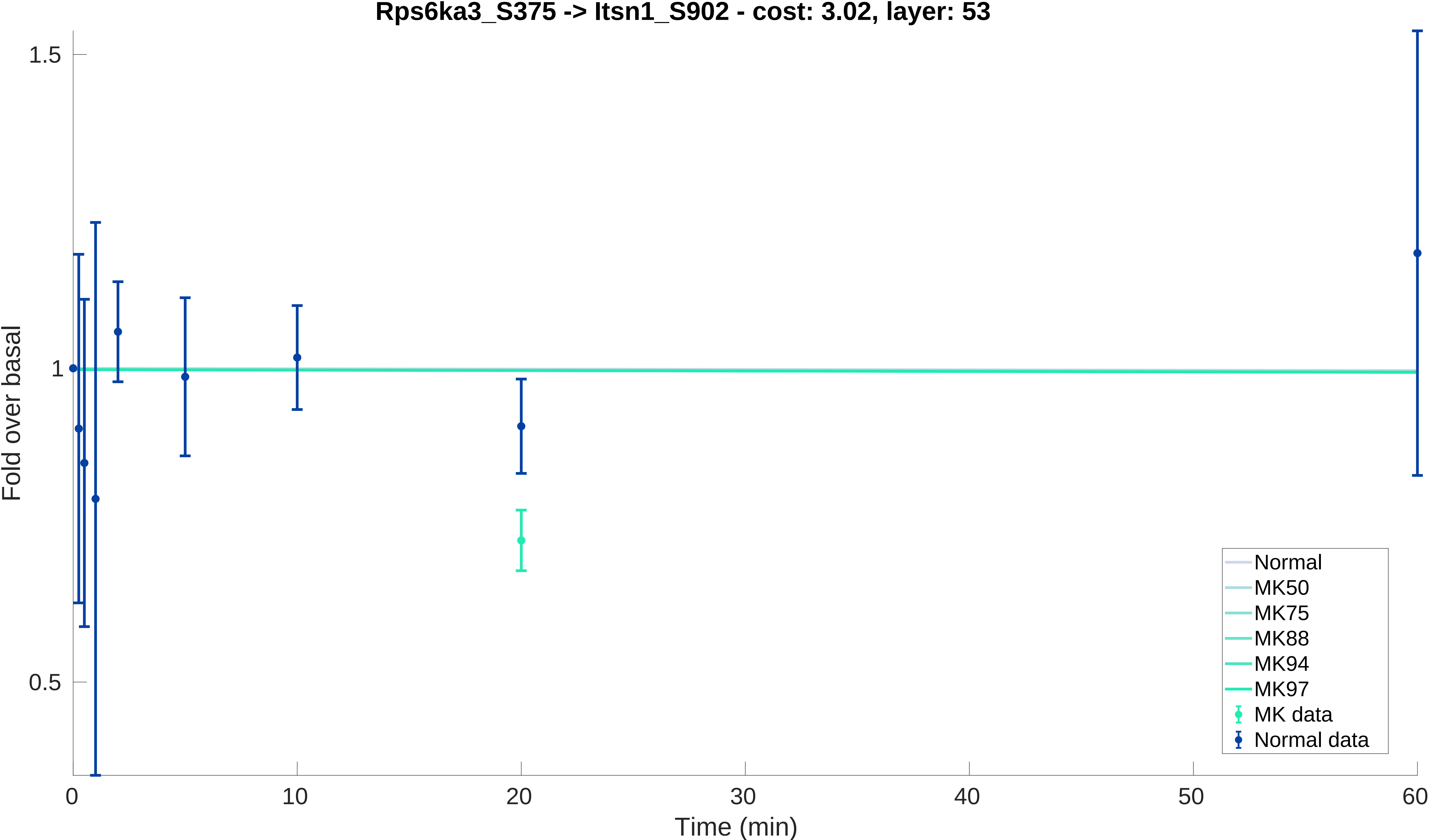
Rps6ka3_S375 -> Itsn1_S902 - cost: 3.02, layer: 53
Fold over basal
1.5
1
0.5
0
10
20
30
40
50
60
Time (min)
Normal
MK50
MK75
MK88
MK94
MK97
MK data
Normal data

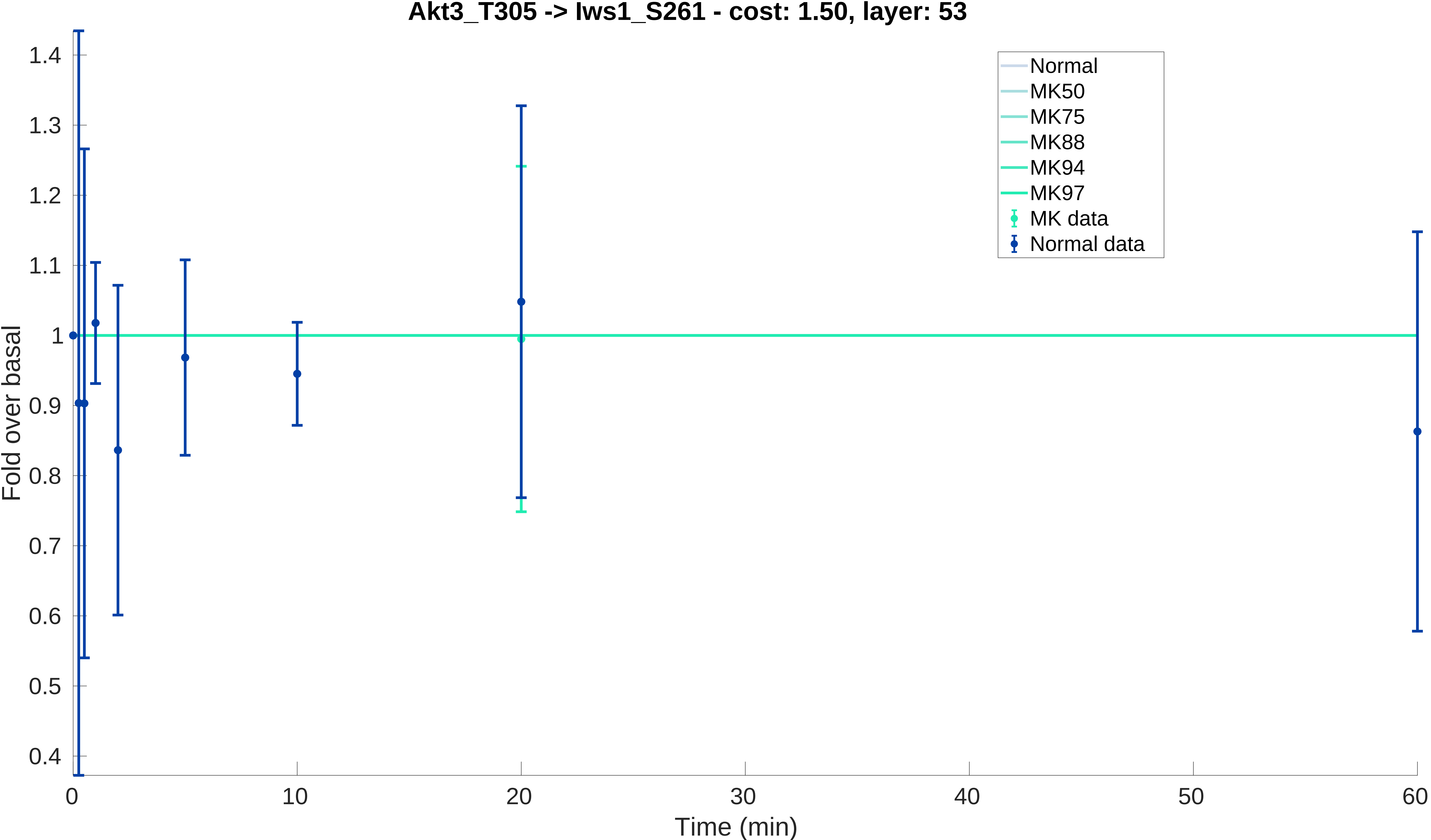
Akt3_T305 -> Iws1_S261 - cost: 1.50, layer: 53
Fold over basal
Time (min)
1.4
1.3
1.2
1.1
1
0.9
0.8
0.7
0.6
0.5
0.4
0
10
20
30
40
50
60
Normal
MK50
MK75
MK88
MK94
MK97
MK data
Normal data

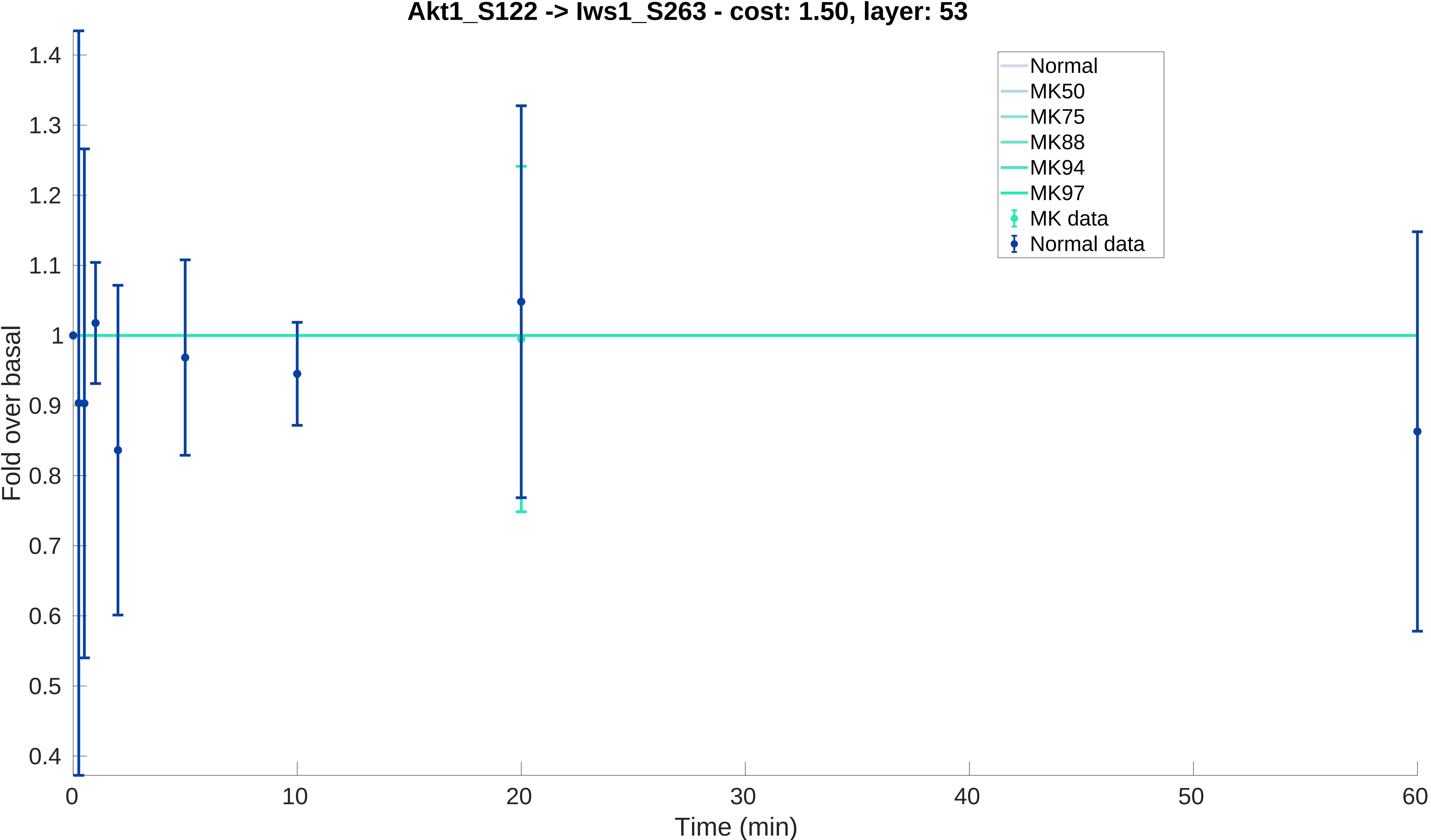
Akt1_S122 -> Iws1_S263 - cost: 1.50, layer: 53
Normal
MK50
MK75
MK88
MK94
MK97
MK data
Normal data
Fold over basal
Time (min)
1.4
1.3
1.2
1.1
1
0.9
0.8
0.7
0.6
0.5
0.4
0
10
20
30
40
50
60

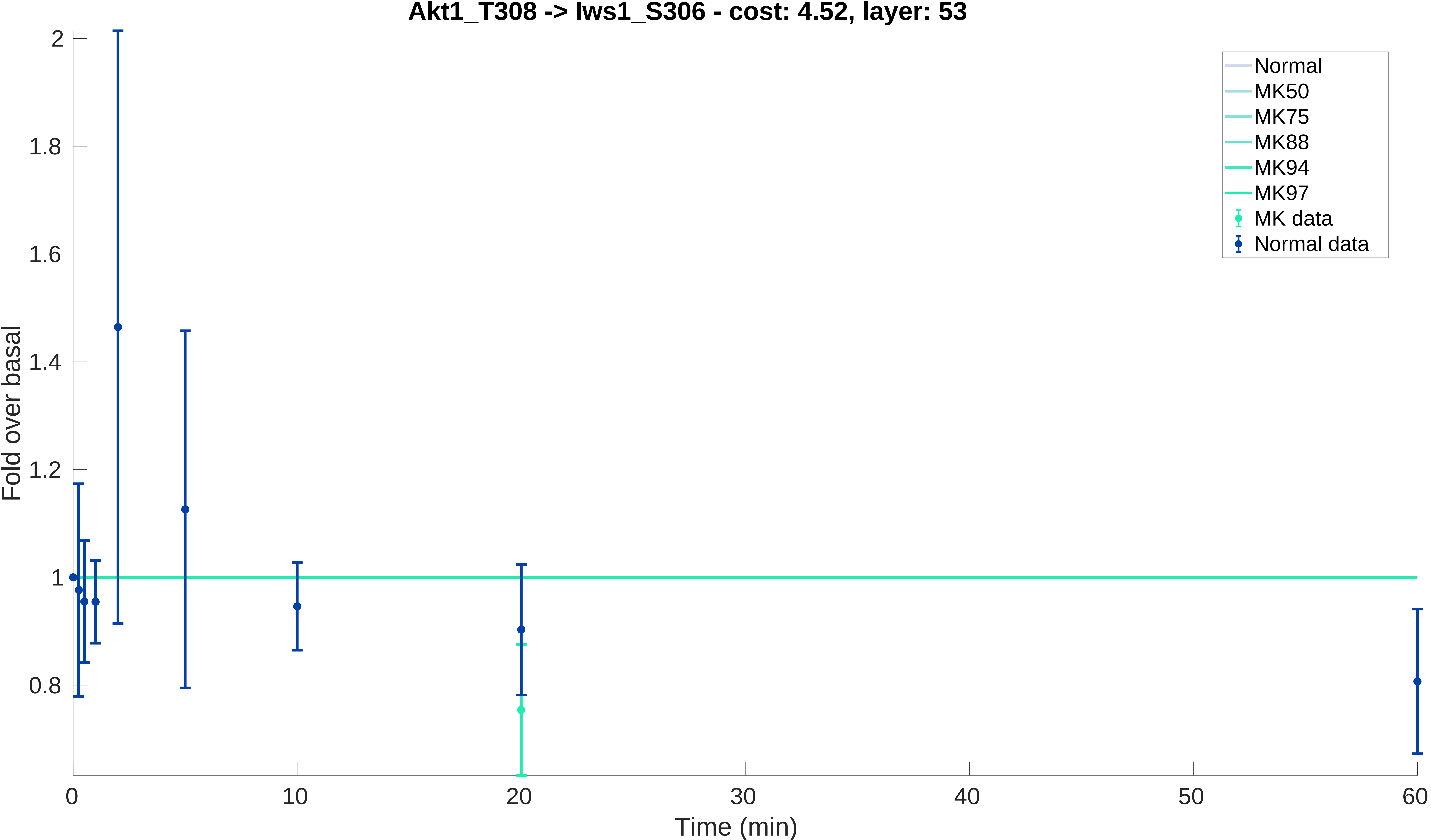
Akt1_T308 -> Iws1_S306 - cost: 4.52, layer: 53
Fold over basal
Time (min)
2
1.8
1.6
1.4
1.2
1
0.8
0
10
20
30
40
50
60
Normal
MK50
MK75
MK88
MK94
MK97
MK data
Normal data

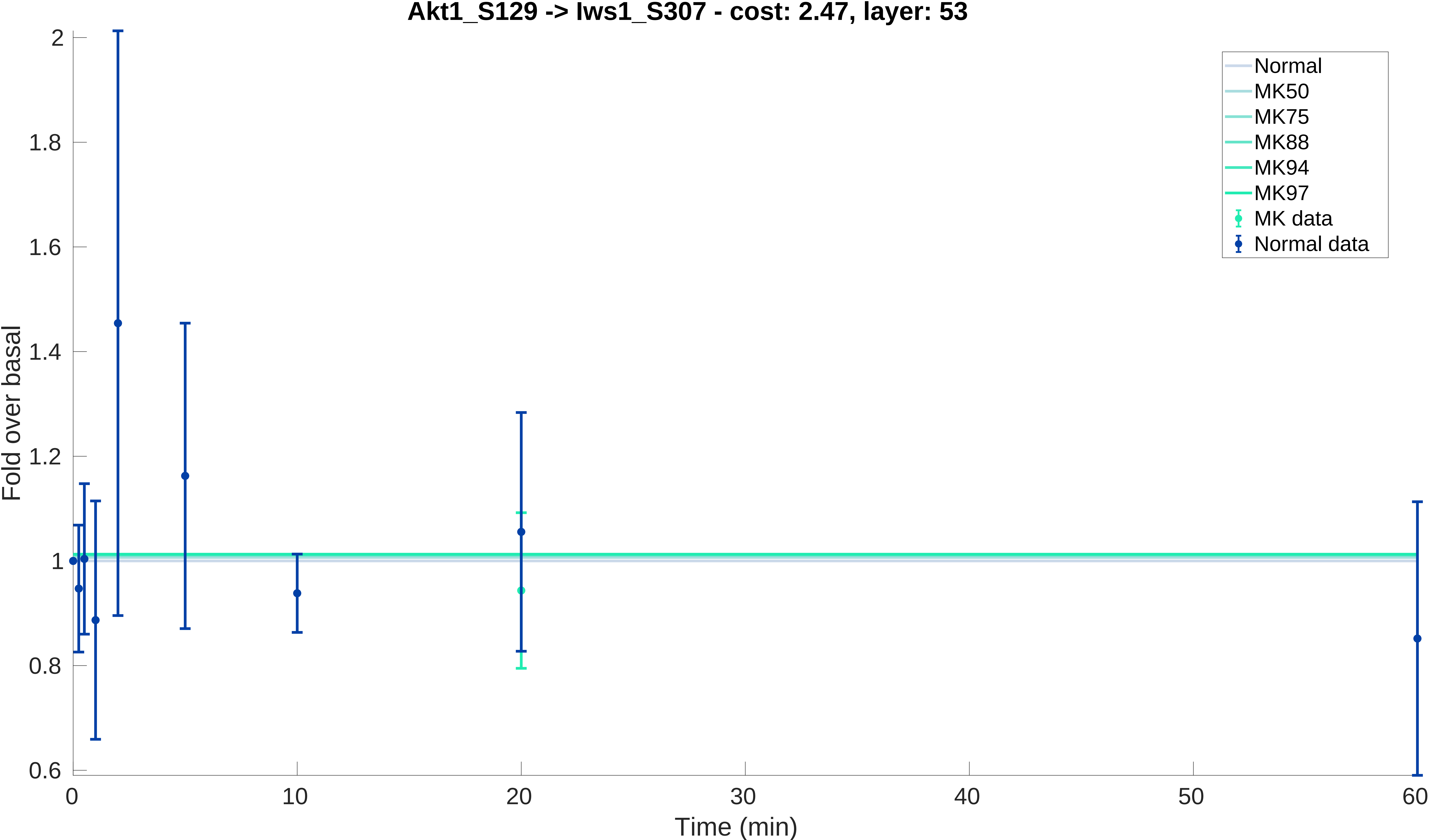

Akt1_S129 -> Iws1_S307 - cost: 2.47, layer: 53
Fold over basal
Time (min)
0
10
20
30
40
50
60
0.6
0.8
1
1.2
1.4
1.6
1.8
2
Normal
MK50
MK75
MK88
MK94
MK97
MK data
Normal data

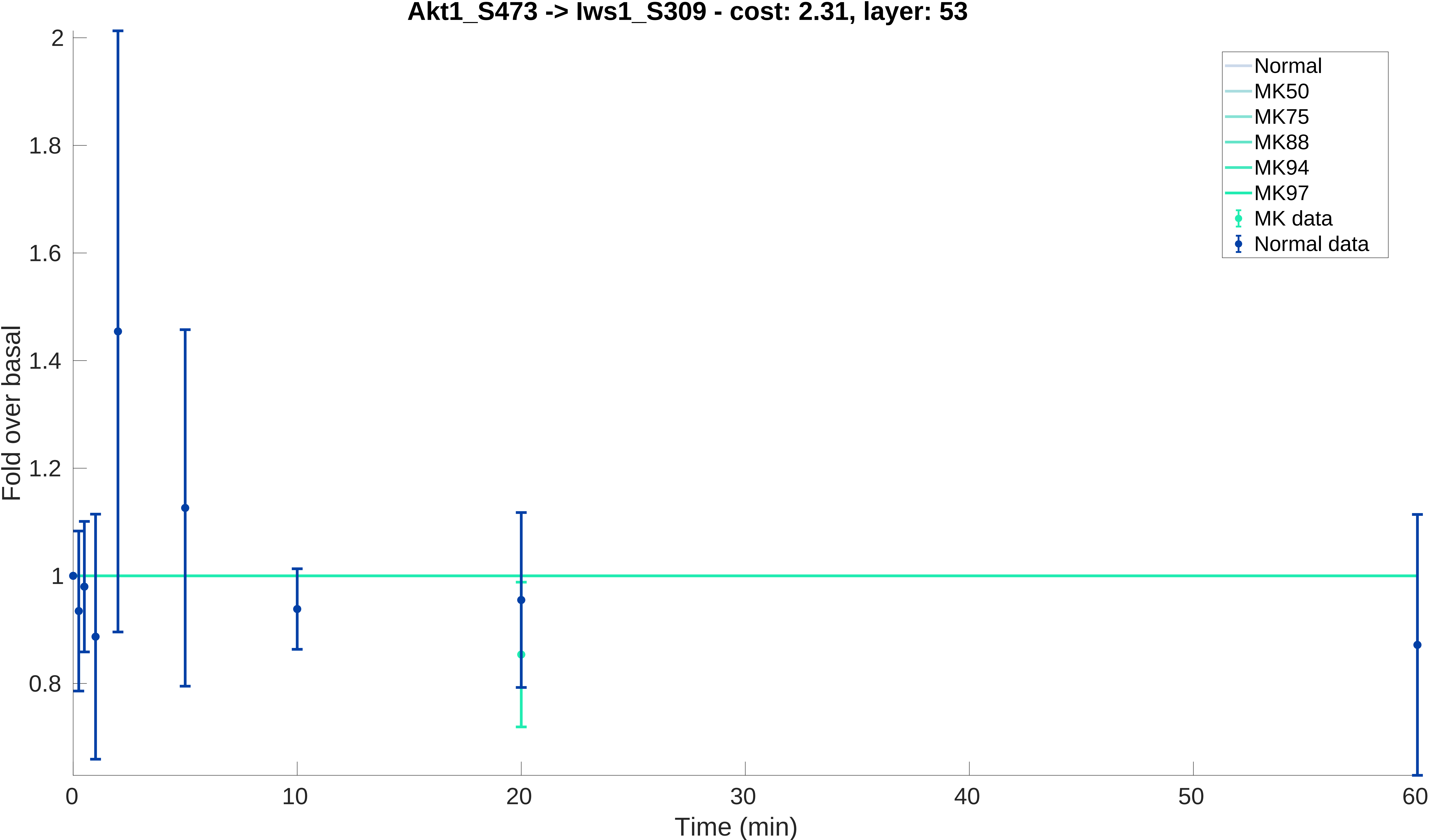

Akt1_S473 -> Iws1_S309 - cost: 2.31, layer: 53
Fold over basal
Time (min)
0.8
1
1.2
1.4
1.6
1.8
2
0
10
20
30
40
50
60
Normal
MK50
MK75
MK88
MK94
MK97
MK data
Normal data

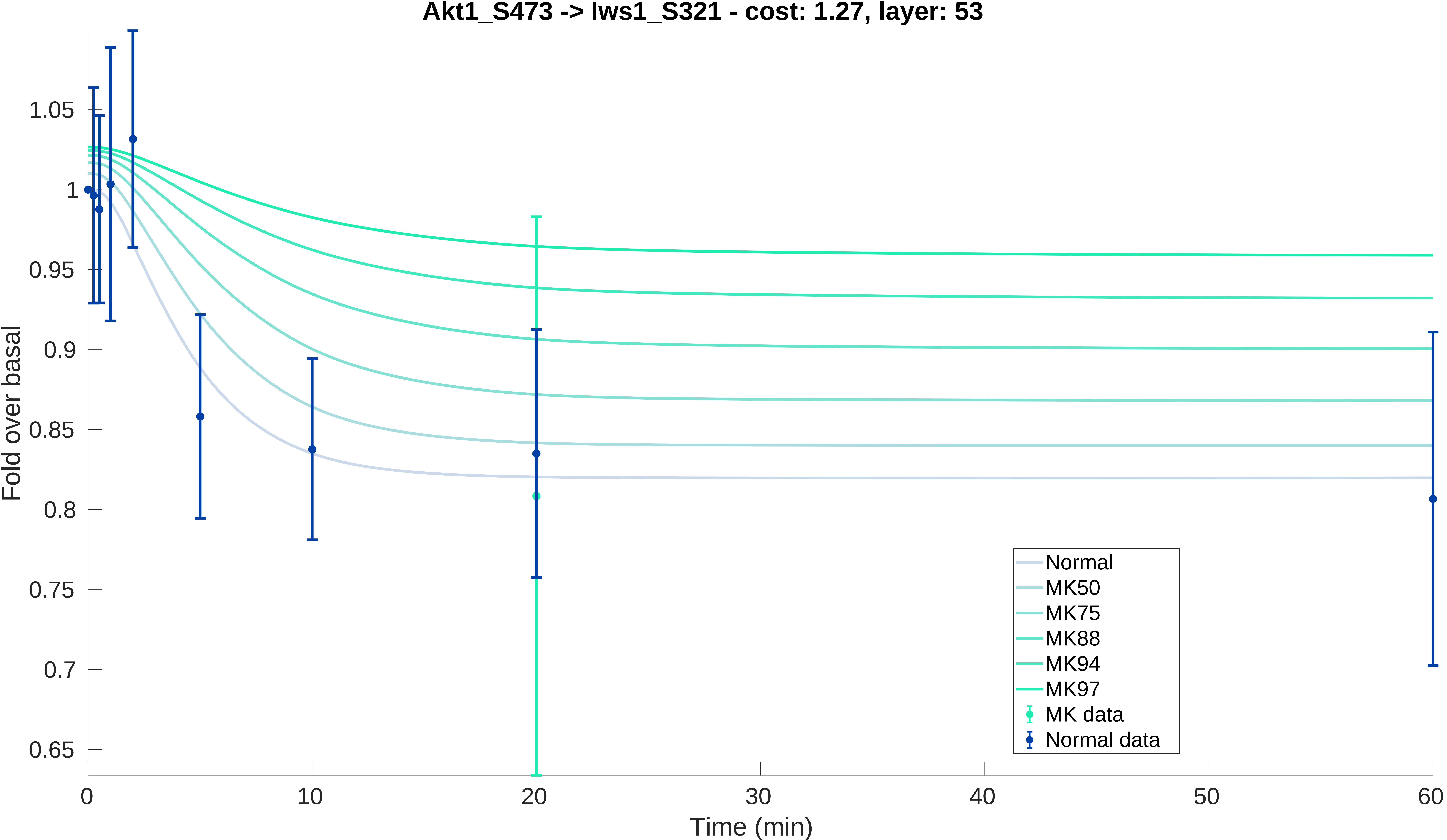

Akt1_S473 -> Iws1_S321 - cost: 1.27, layer: 53
Fold over basal
1.05
1
0.95
0.9
0.85
0.8
0.75
0.7
0.65
0
10
20
30
40
50
60
Time (min)
Normal
MK50
MK75
MK88
MK94
MK97
MK data
Normal data

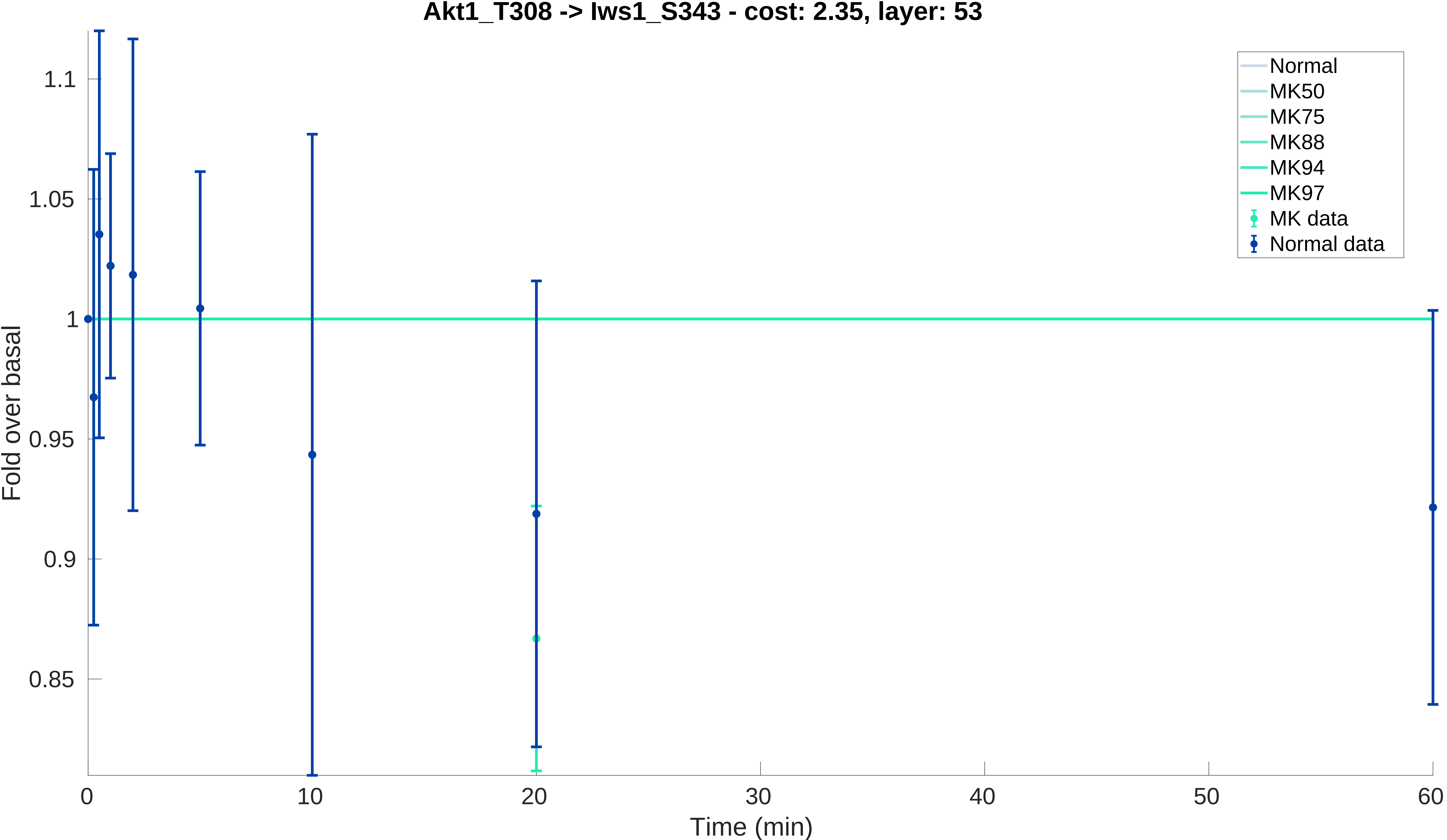
Akt1_T308 -> Iws1_S343 - cost: 2.35, layer: 53
Fold over basal
Time (min)
0
10
20
30
40
50
60
0.85
0.9
0.95
1
1.05
1.1
Normal
MK50
MK75
MK88
MK94
MK97
MK data
Normal data

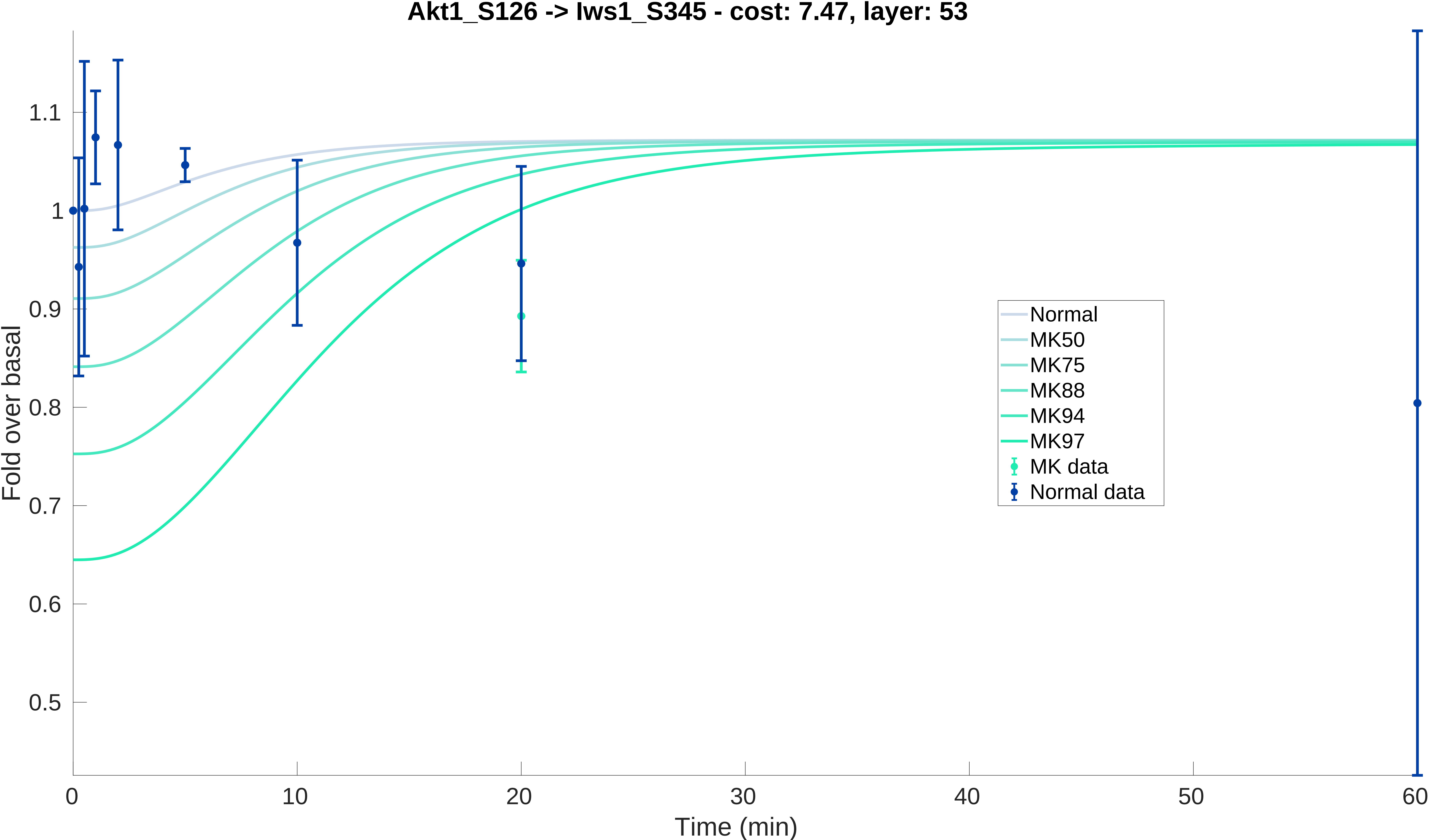
Akt1_S126 -> Iws1_S345 - cost: 7.47, layer: 53
Fold over basal
Time (min)
0
10
20
30
40
50
60
0.5
0.6
0.7
0.8
0.9
1
1.1
Normal
MK50
MK75
MK88
MK94
MK97
MK data
Normal data

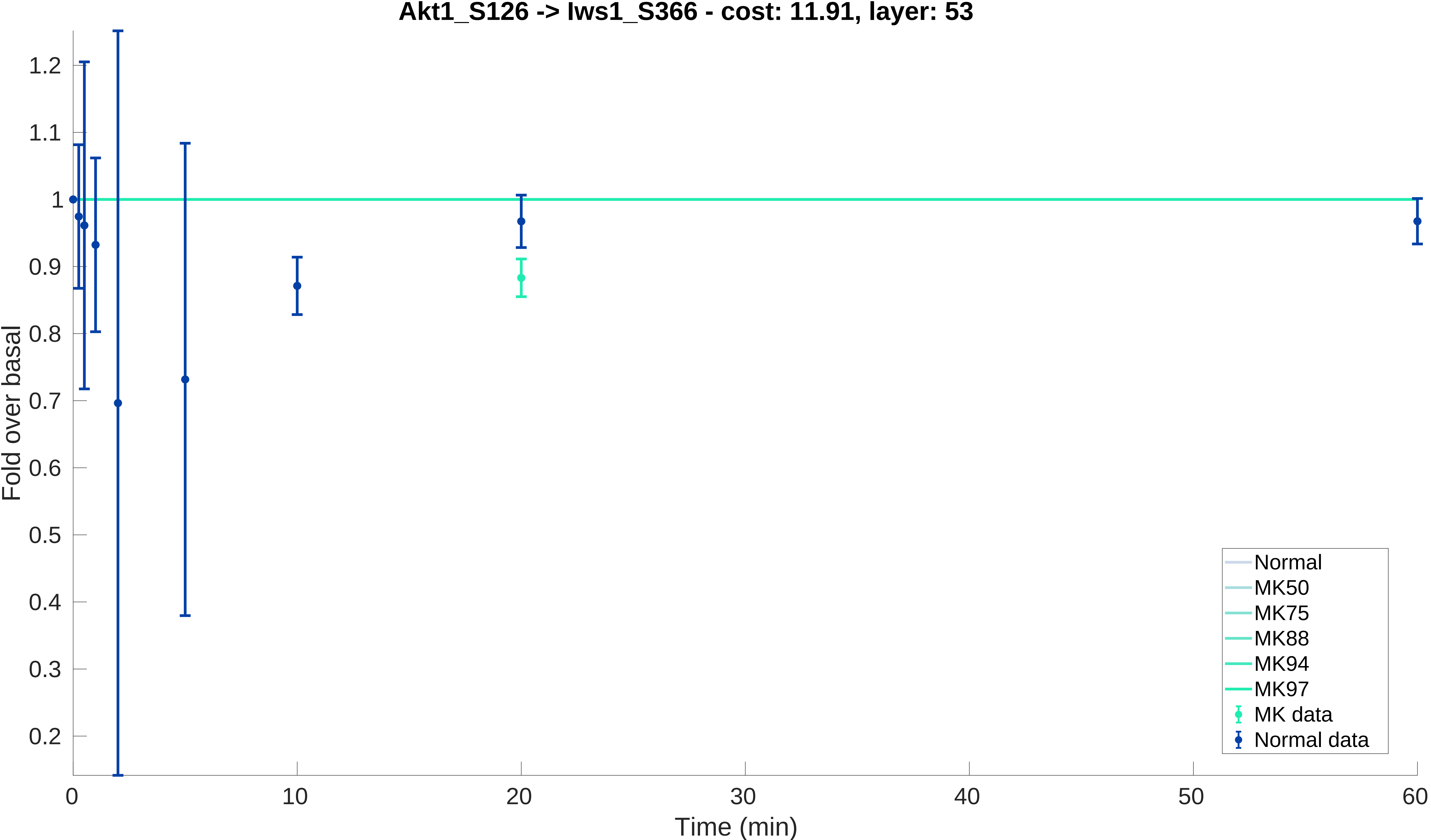

Akt1_S126 -> Iws1_S366 - cost: 11.91, layer: 53
Fold over basal
1.2
1.1
1
0.9
0.8
0.7
0.6
0.5
0.4
0.3
0.2
0
10
20
30
40
50
60
Time (min)
Normal
MK50
MK75
MK88
MK94
MK97
MK data
Normal data

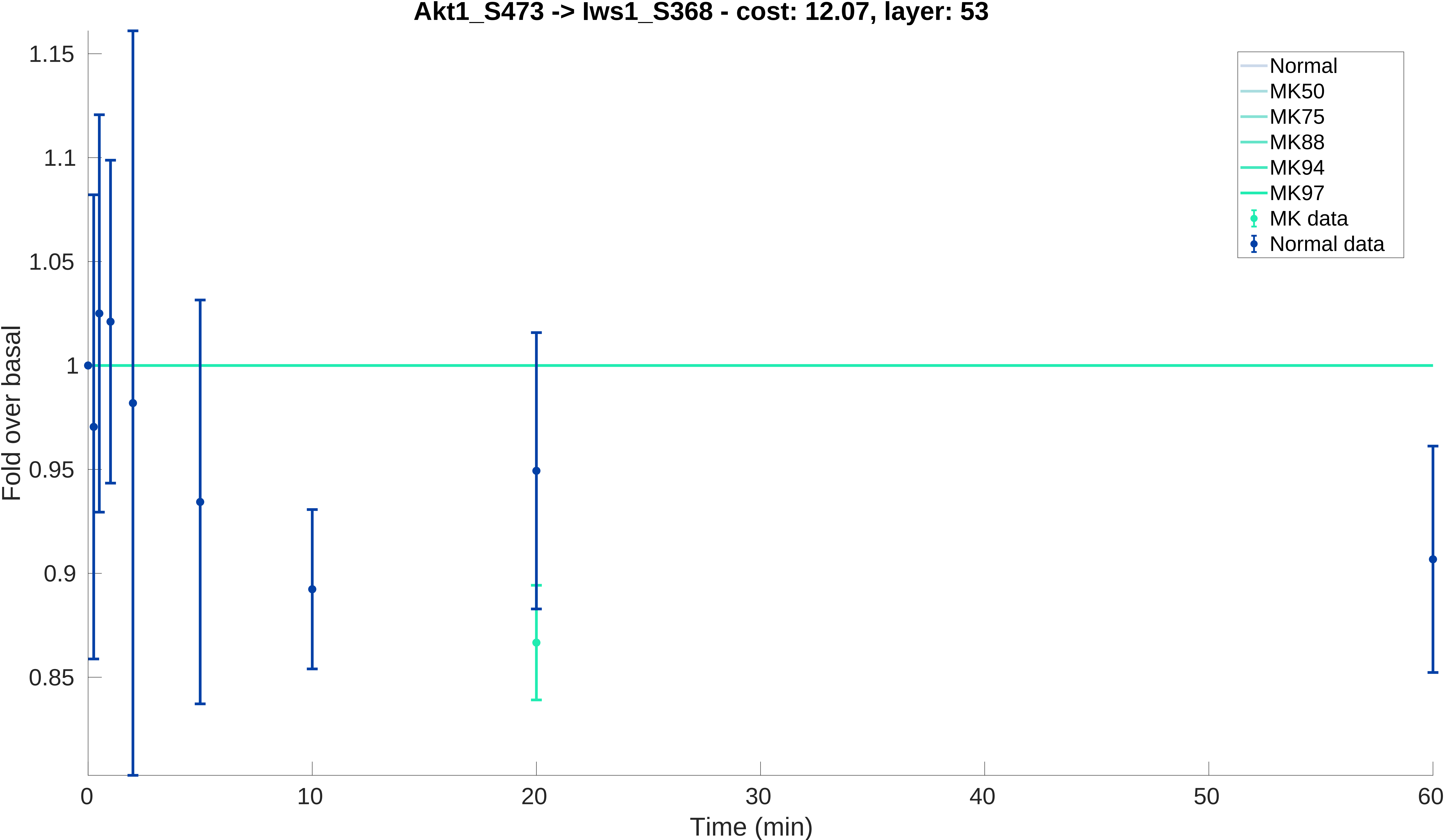

Akt1_S473 -> Iws1_S368 - cost: 12.07, layer: 53
Fold over basal
1.15
1.1
1.05
1
0.95
0.9
0.85
0
10
20
30
40
50
60
Time (min)
Normal
MK50
MK75
MK88
MK94
MK97
MK data
Normal data

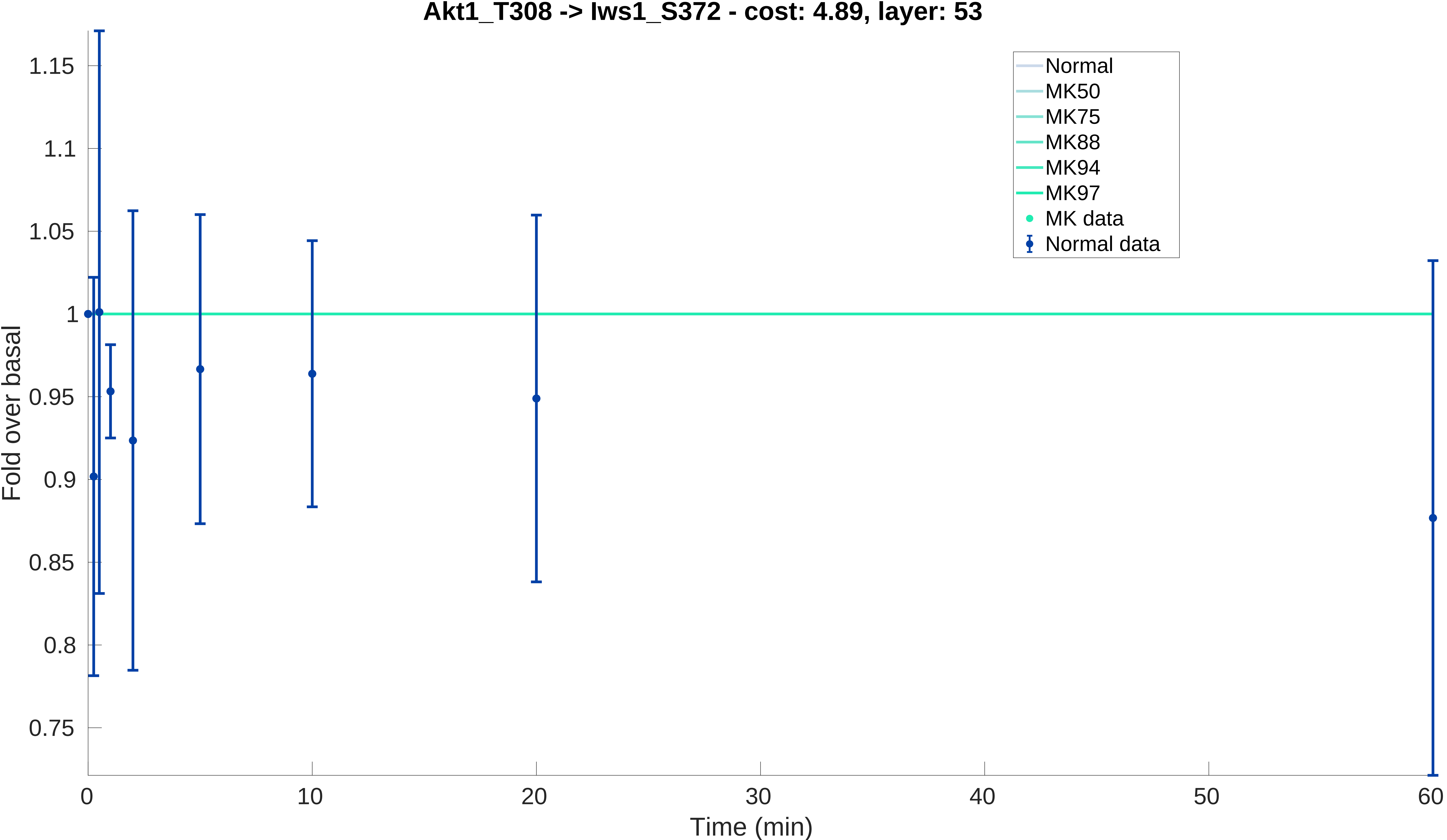

Akt1_T308 -> Iws1_S372 - cost: 4.89, layer: 53
Normal
MK50
MK75
MK88
MK94
MK97
MK data
Normal data
Fold over basal
1.15
1.1
1.05
1
0.95
0.9
0.85
0.8
0.75
0
10
20
30
40
50
60
Time (min)

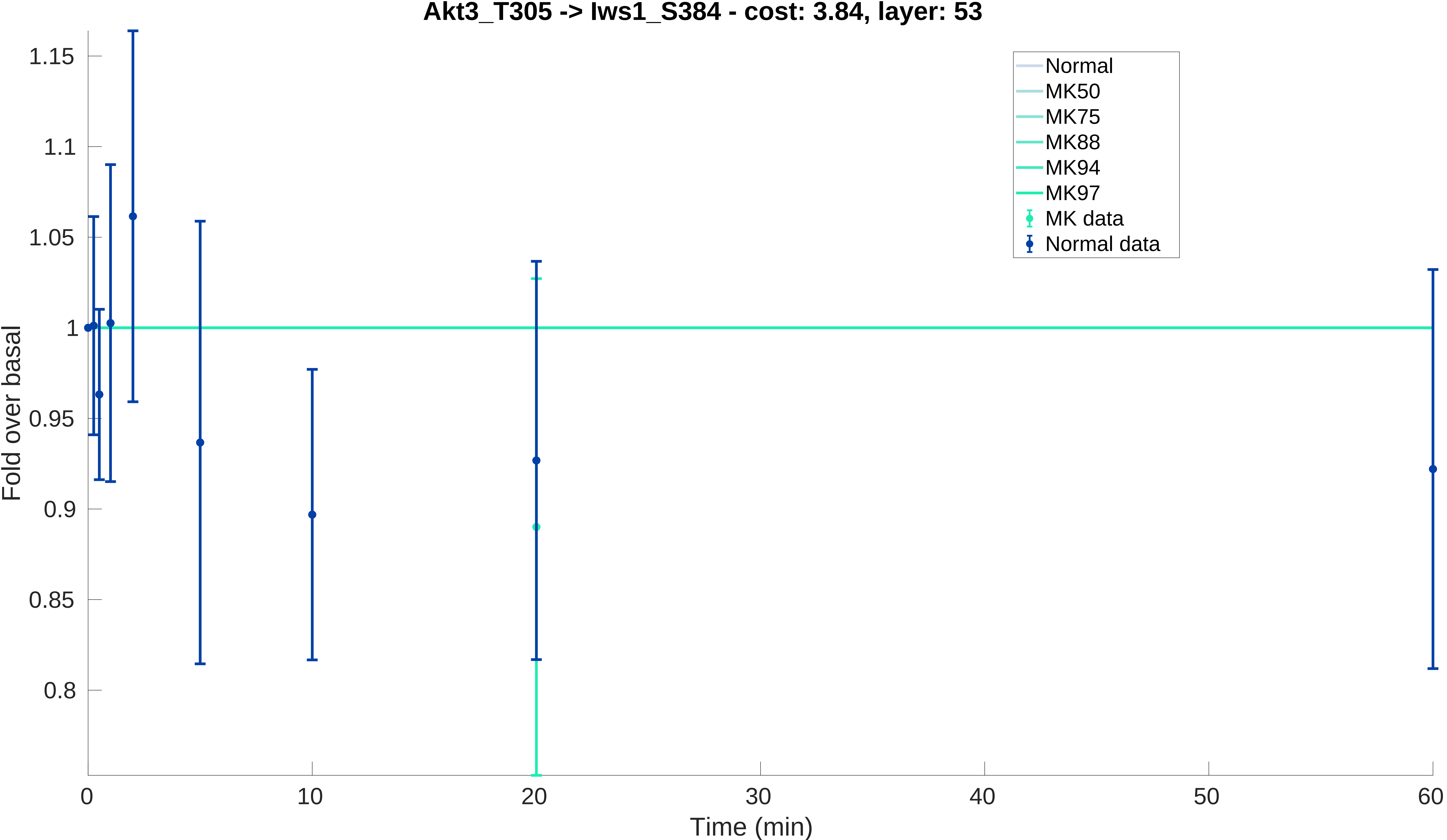

Akt3_T305 -> Iws1_S384 - cost: 3.84, layer: 53
Normal
MK50
MK75
MK88
MK94
MK97
MK data
Normal data
Fold over basal
Time (min)
1.15
1.1
1.05
1
0.95
0.9
0.85
0.8
0
10
20
30
40
50
60

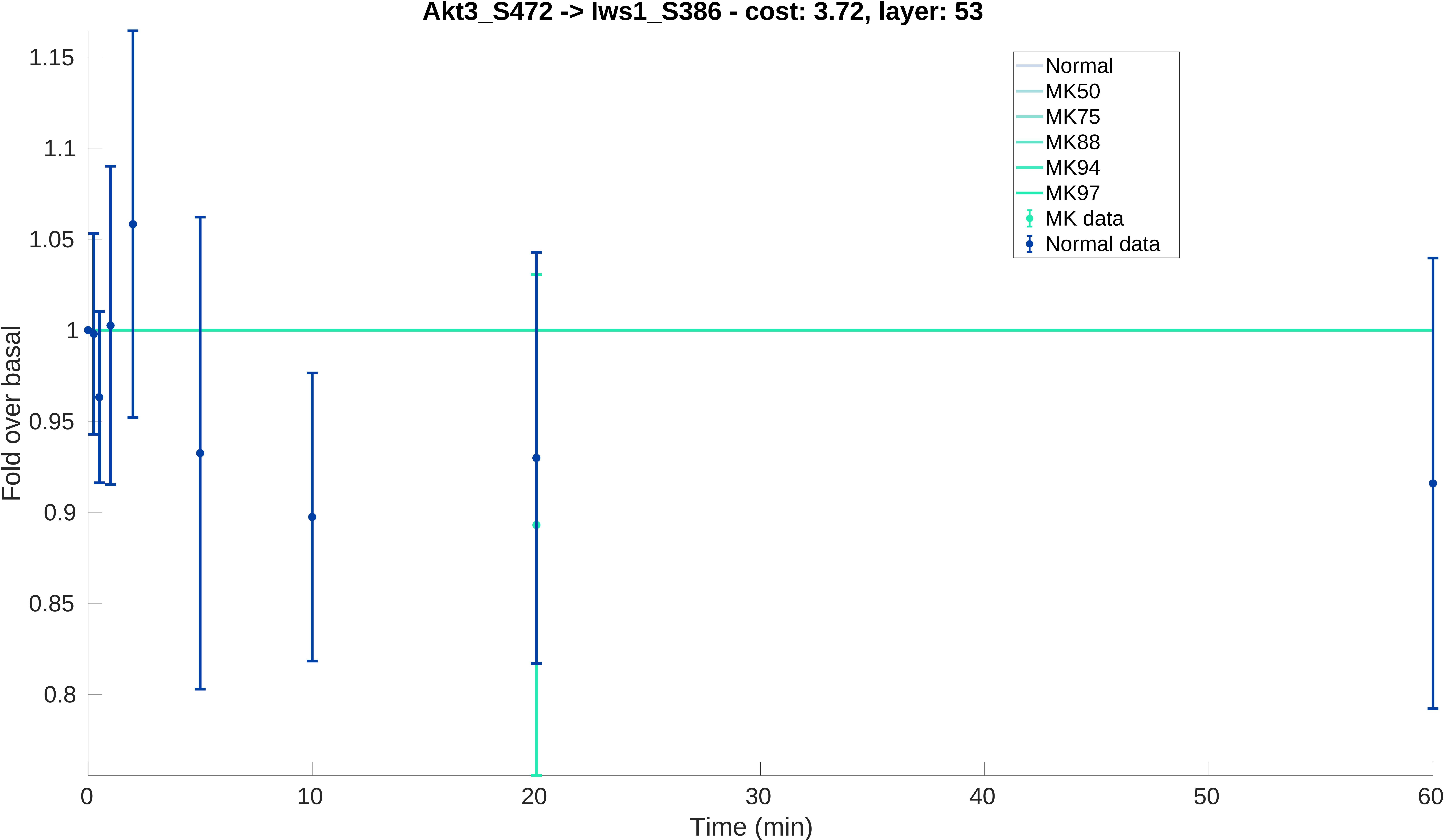
Akt3_S472 -> Iws1_S386 - cost: 3.72, layer: 53
Fold over basal
Time (min)
1.15
1.1
1.05
1
0.95
0.9
0.85
0.8
0
10
20
30
40
50
60
Normal
MK50
MK75
MK88
MK94
MK97
MK data
Normal data

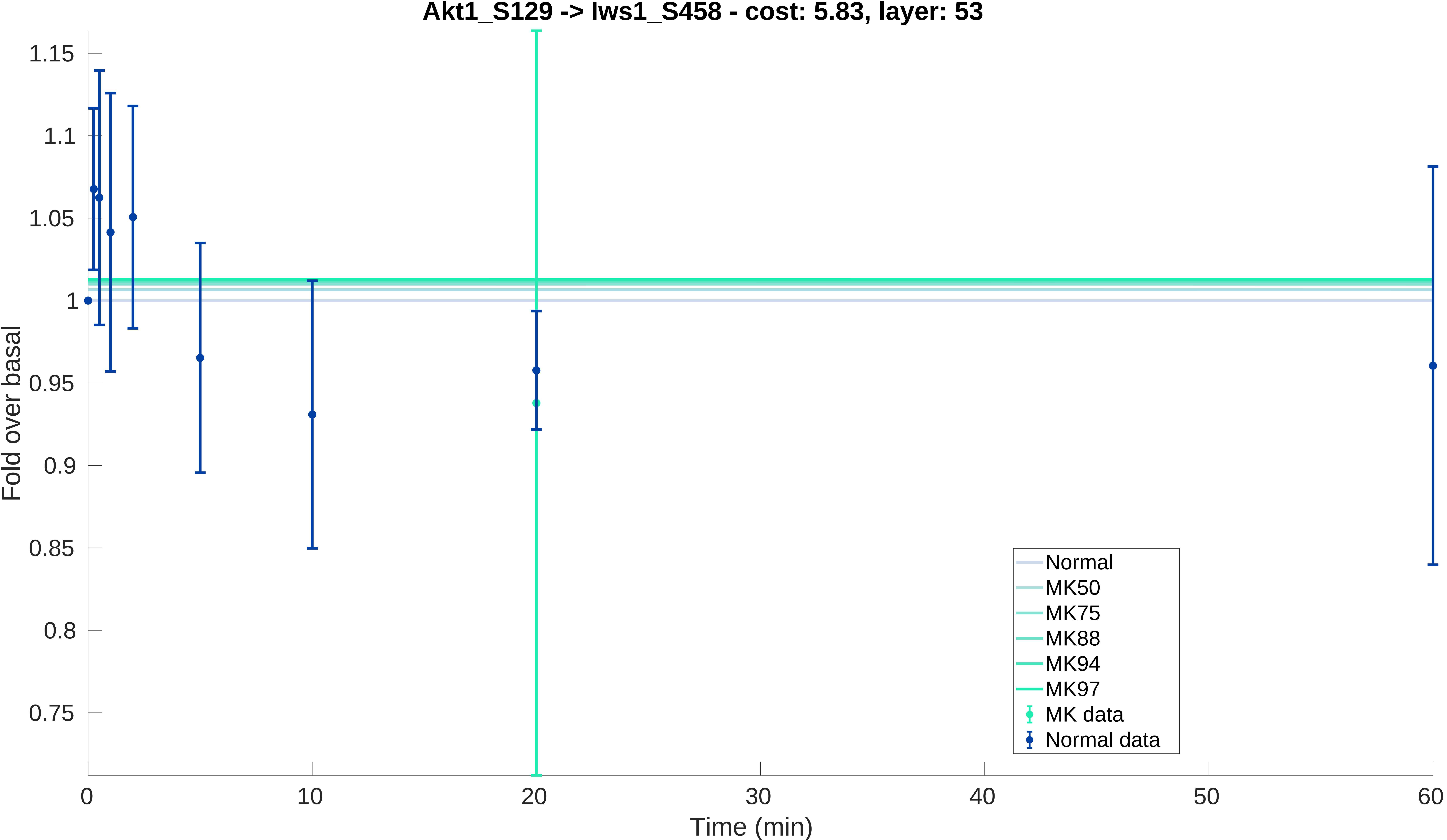

Akt1_S129 -> Iws1_S458 - cost: 5.83, layer: 53
Fold over basal
1.15
1.1
1.05
1
0.95
0.9
0.85
0.8
0.75
0
10
20
30
40
50
60
Time (min)
Normal
MK50
MK75
MK88
MK94
MK97
MK data
Normal data

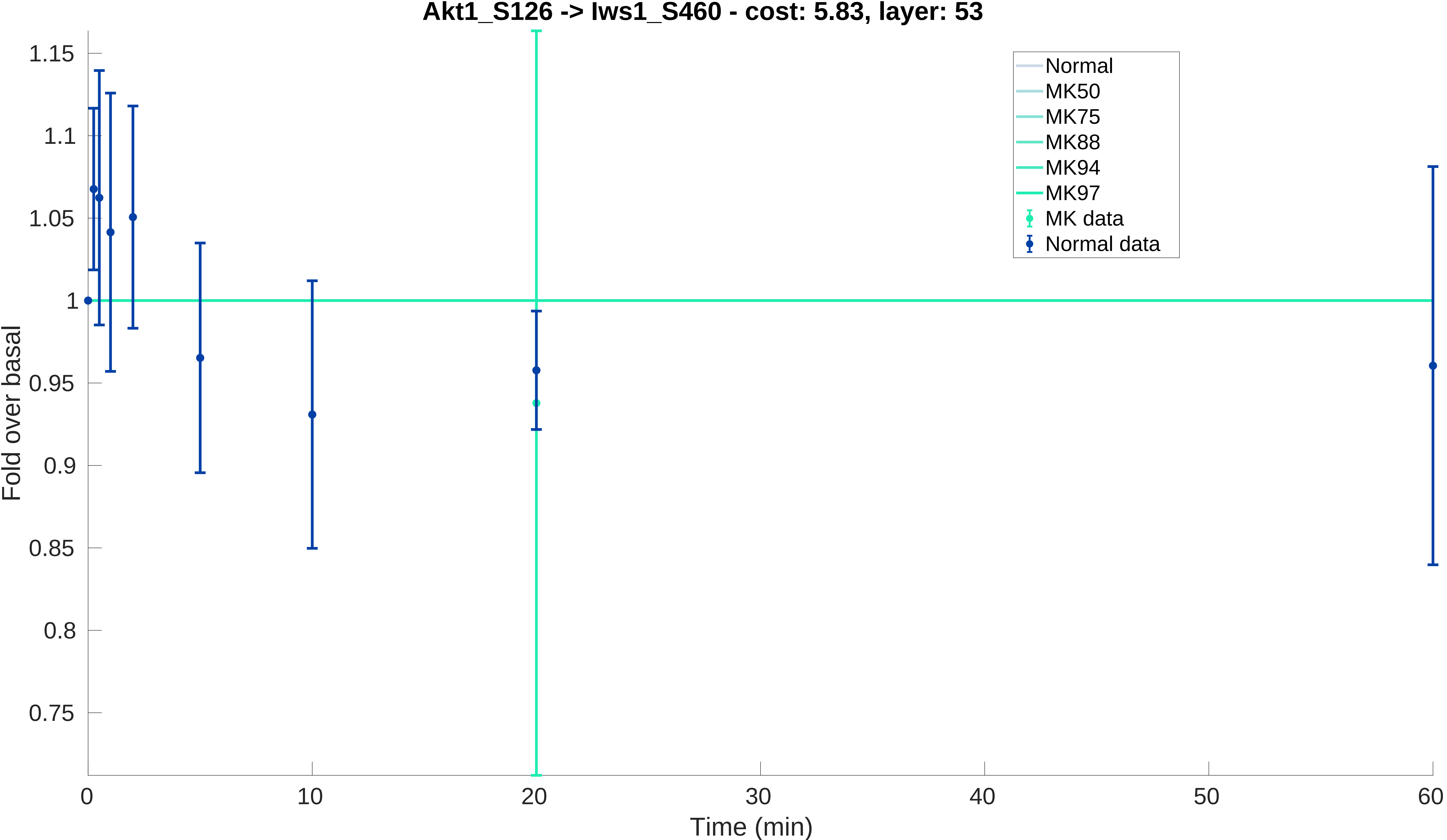

Akt1_S126 -> Iws1_S460 - cost: 5.83, layer: 53
Fold over basal
Time (min)
1.15
1.1
1.05
1
0.95
0.9
0.85
0.8
0.75
0
10
20
30
40
50
60
Normal
MK50
MK75
MK88
MK94
MK97
MK data
Normal data

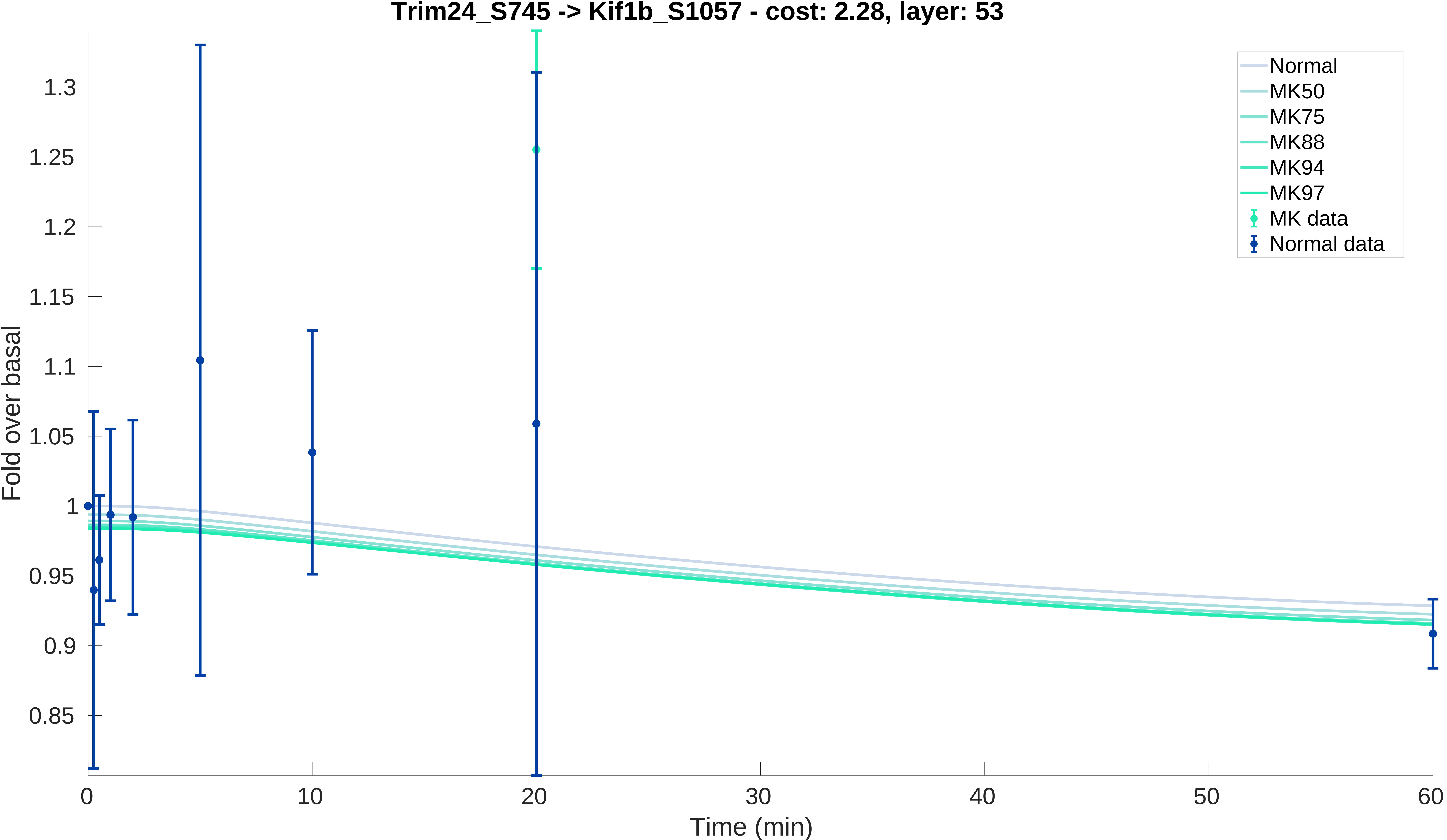

Trim24_S745 -> Kif1b_S1057 - cost: 2.28, layer: 53
Fold over basal
Time (min)
1.3
1.25
1.2
1.15
1.1
1.05
1
0.95
0.9
0.85
0
10
20
30
40
50
60
Normal
MK50
MK75
MK88
MK94
MK97
MK data
Normal data

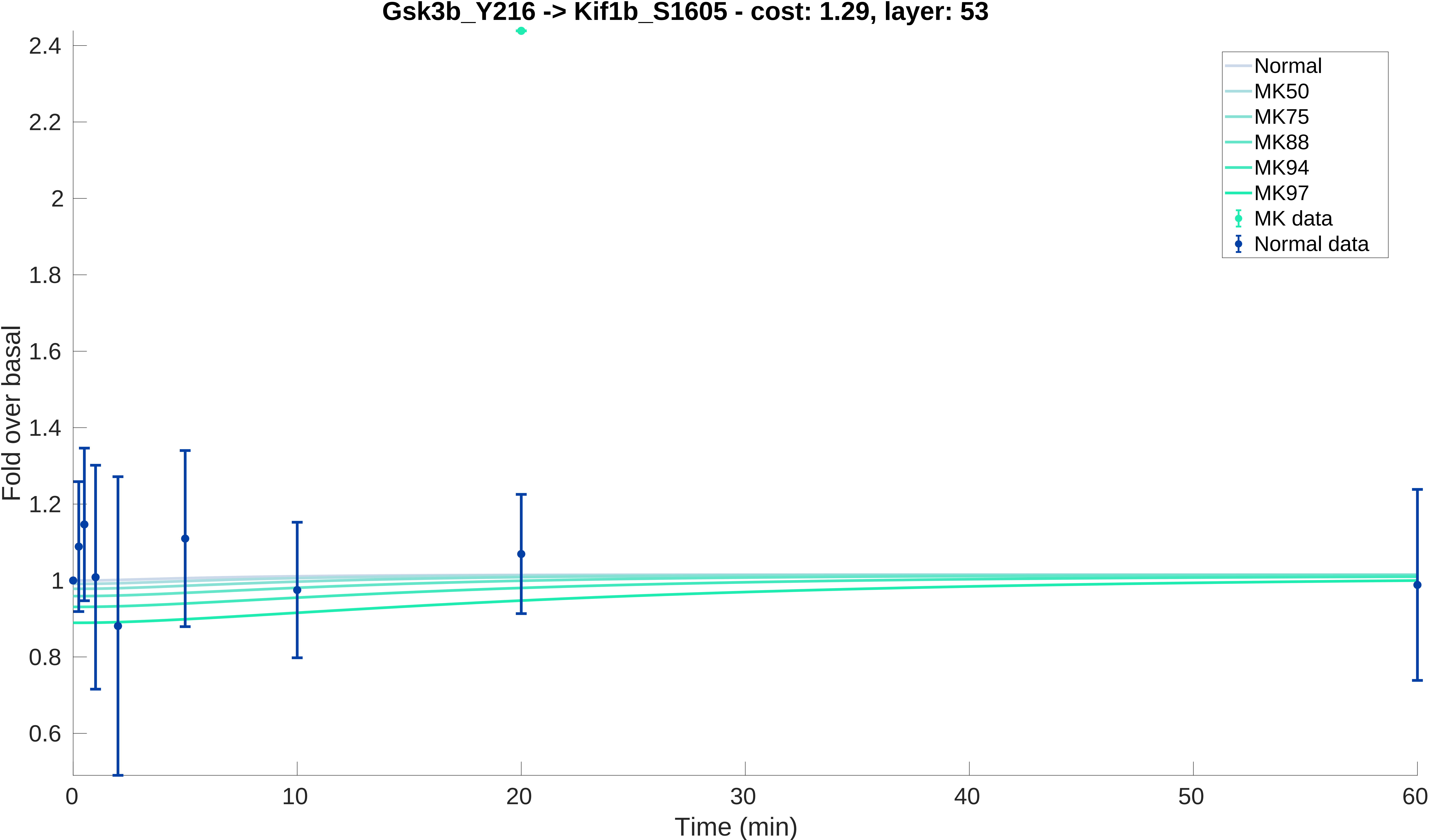

Gsk3b_Y216 -> Kif1b_S1605 - cost: 1.29, layer: 53
Fold over basal
Time (min)
2.4
2.2
2
1.8
1.6
1.4
1.2
1
0.8
0.6
0
10
20
30
40
50
60
Normal
MK50
MK75
MK88
MK94
MK97
MK data
Normal data

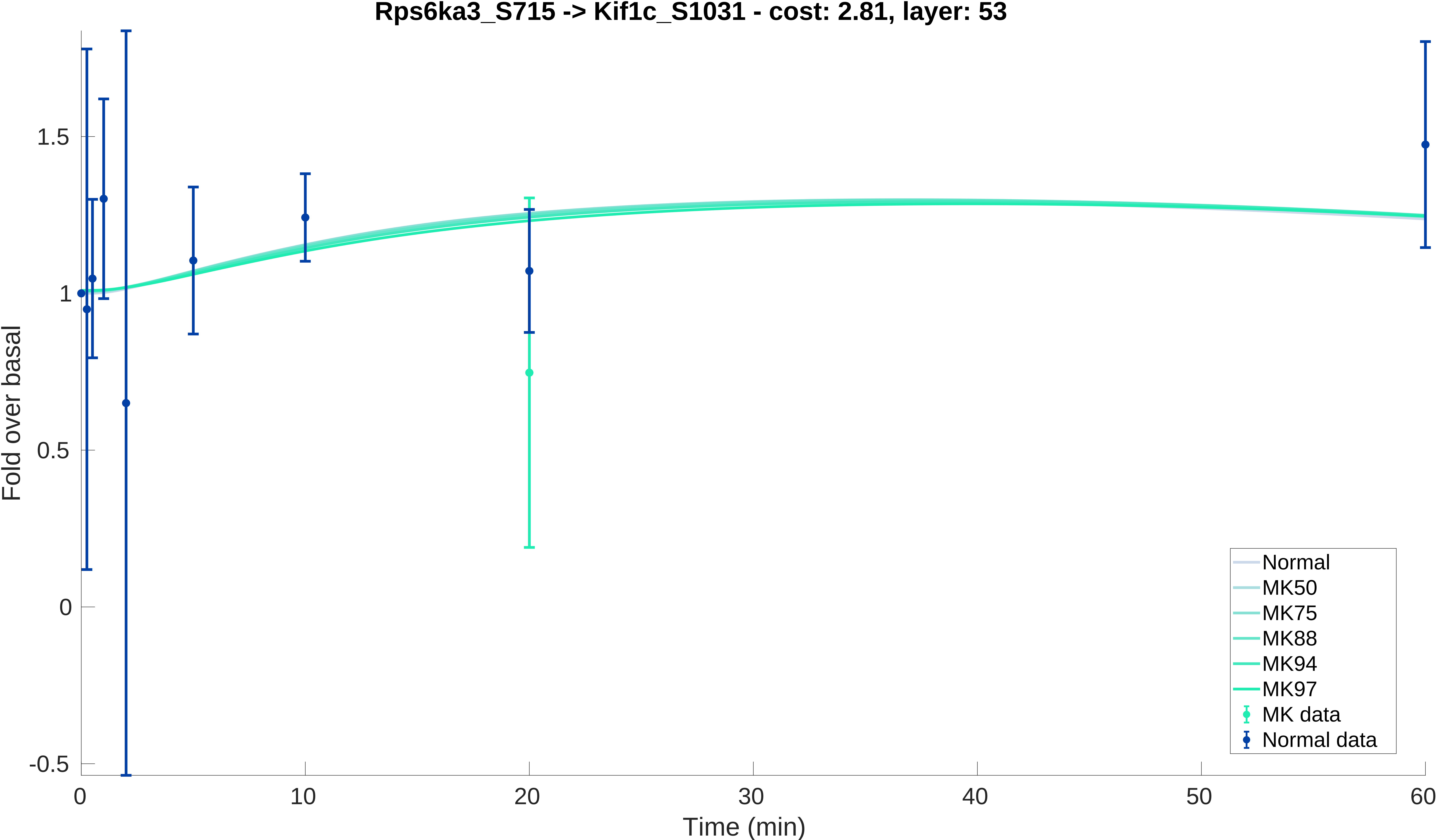

Rps6ka3_S715 -> Kif1c_S1031 - cost: 2.81, layer: 53
Fold over basal
Time (min)
1.5
1
0.5
0
-0.5
0
10
20
30
40
50
60
Normal
MK50
MK75
MK88
MK94
MK97
MK data
Normal data

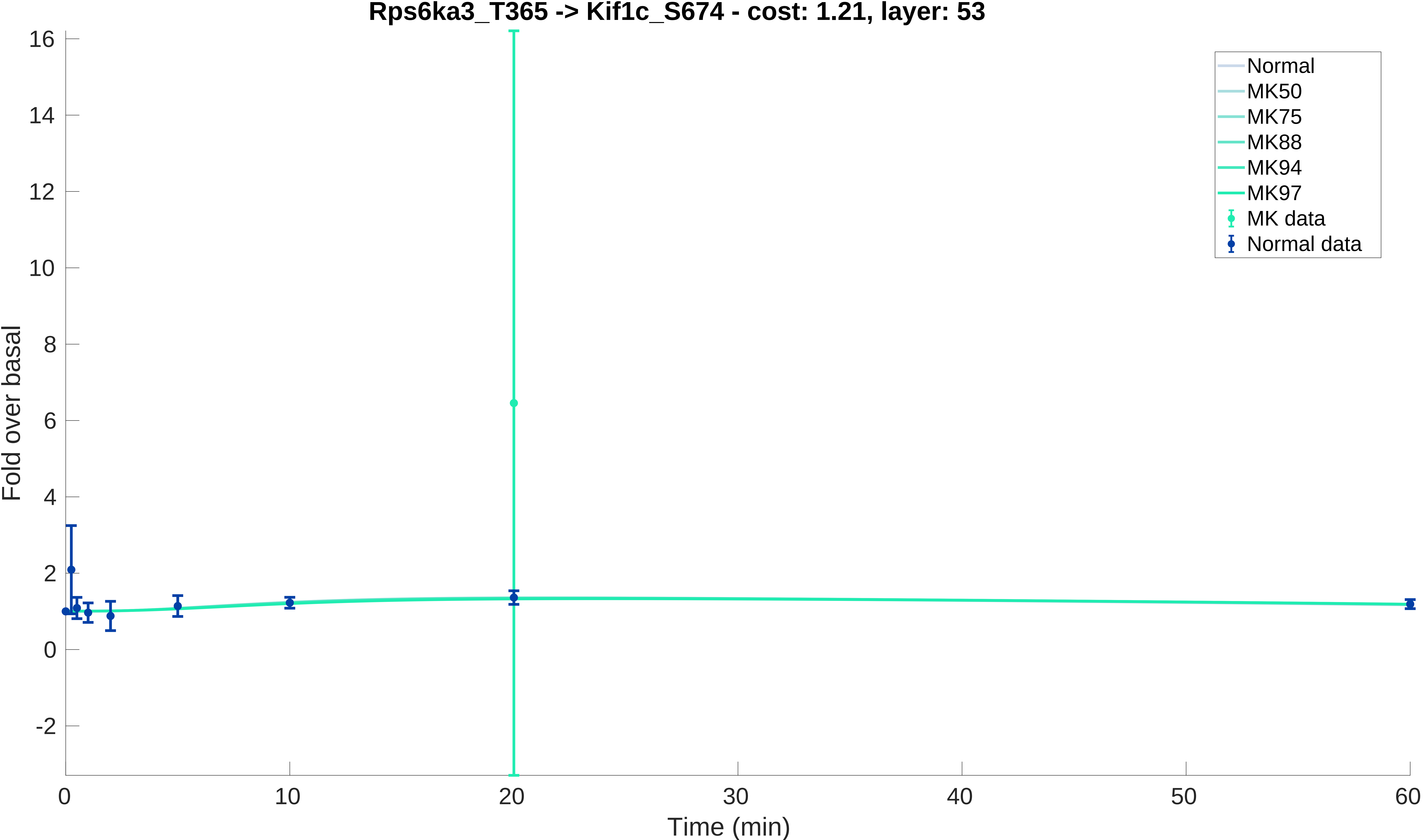

Rps6ka3_T365 -> Kif1c_S674 - cost: 1.21, layer: 53
Fold over basal
Time (min)
16
14
12
10
8
6
4
2
0
-2
0
10
20
30
40
50
60
Normal
MK50
MK75
MK88
MK94
MK97
MK data
Normal data

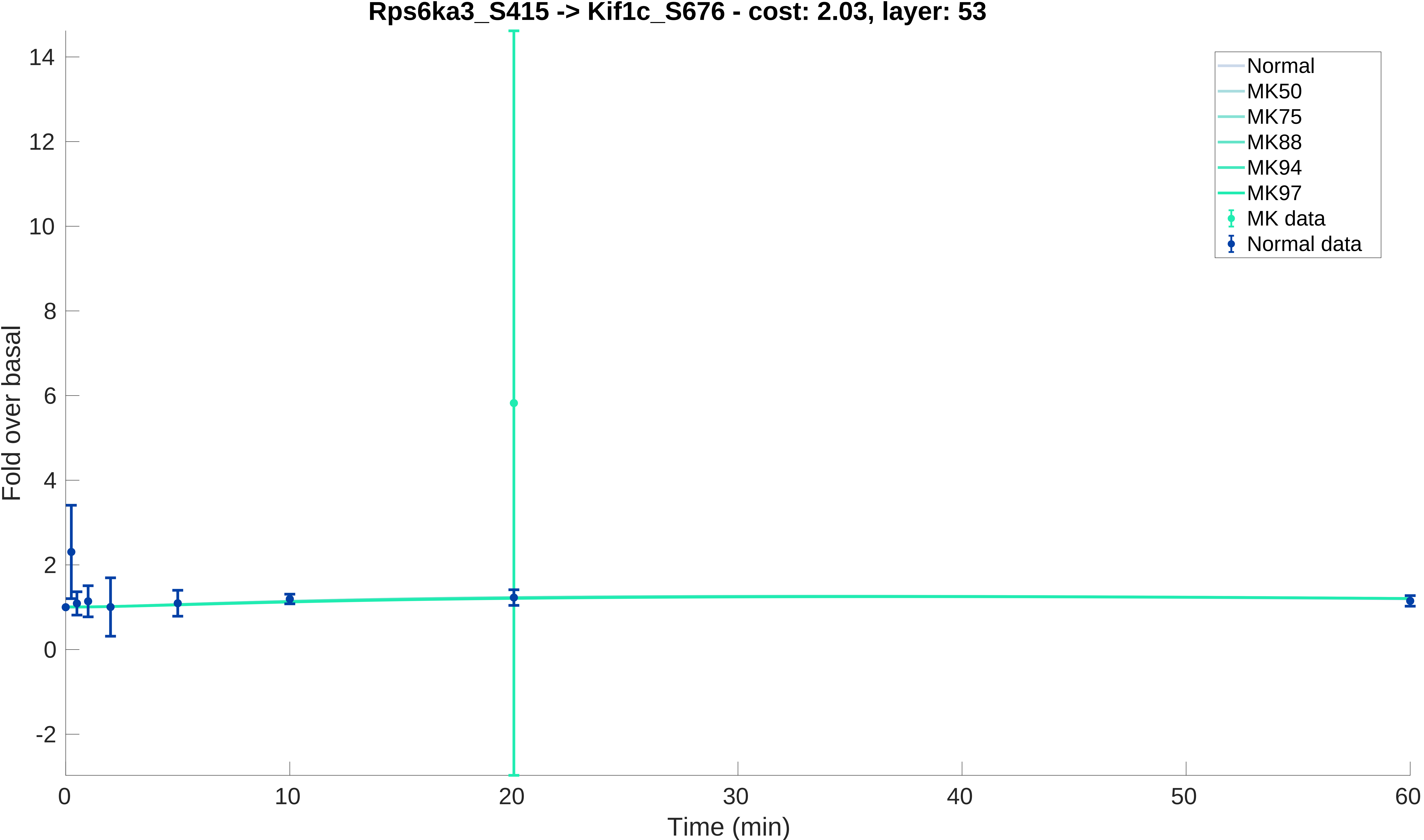

Rps6ka3_S415 -> Kif1c_S676 - cost: 2.03, layer: 53
Fold over basal
Time (min)
14
12
10
8
6
4
2
0
-2
0
10
20
30
40
50
60
Normal
MK50
MK75
MK88
MK94
MK97
MK data
Normal data

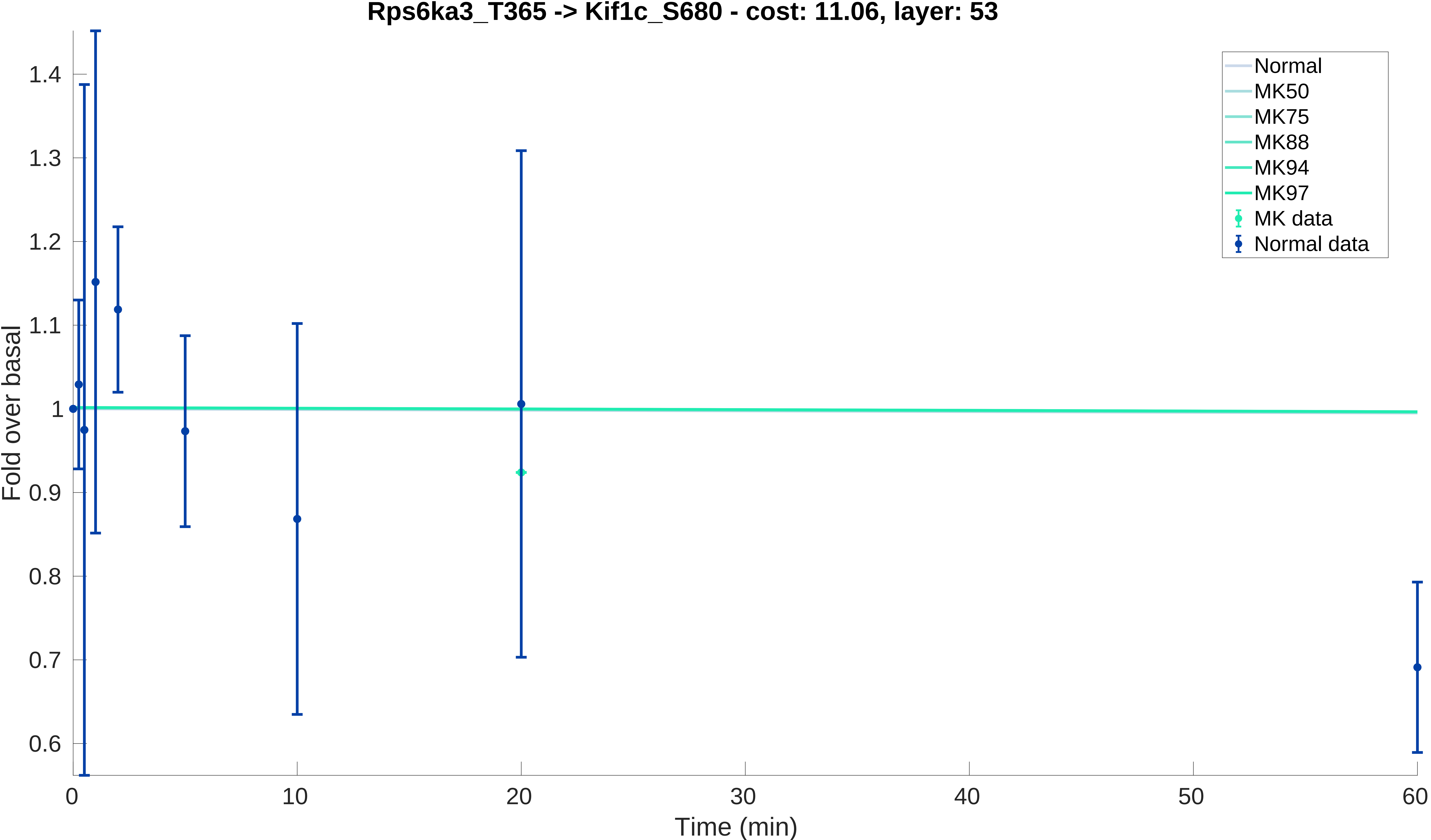

Rps6ka3_T365 -> Kif1c_S680 - cost: 11.06, layer: 53
Normal
MK50
MK75
MK88
MK94
MK97
MK data
Normal data
Fold over basal
Time (min)
1.4
1.3
1.2
1.1
1
0.9
0.8
0.7
0.6
0
10
20
30
40
50
60

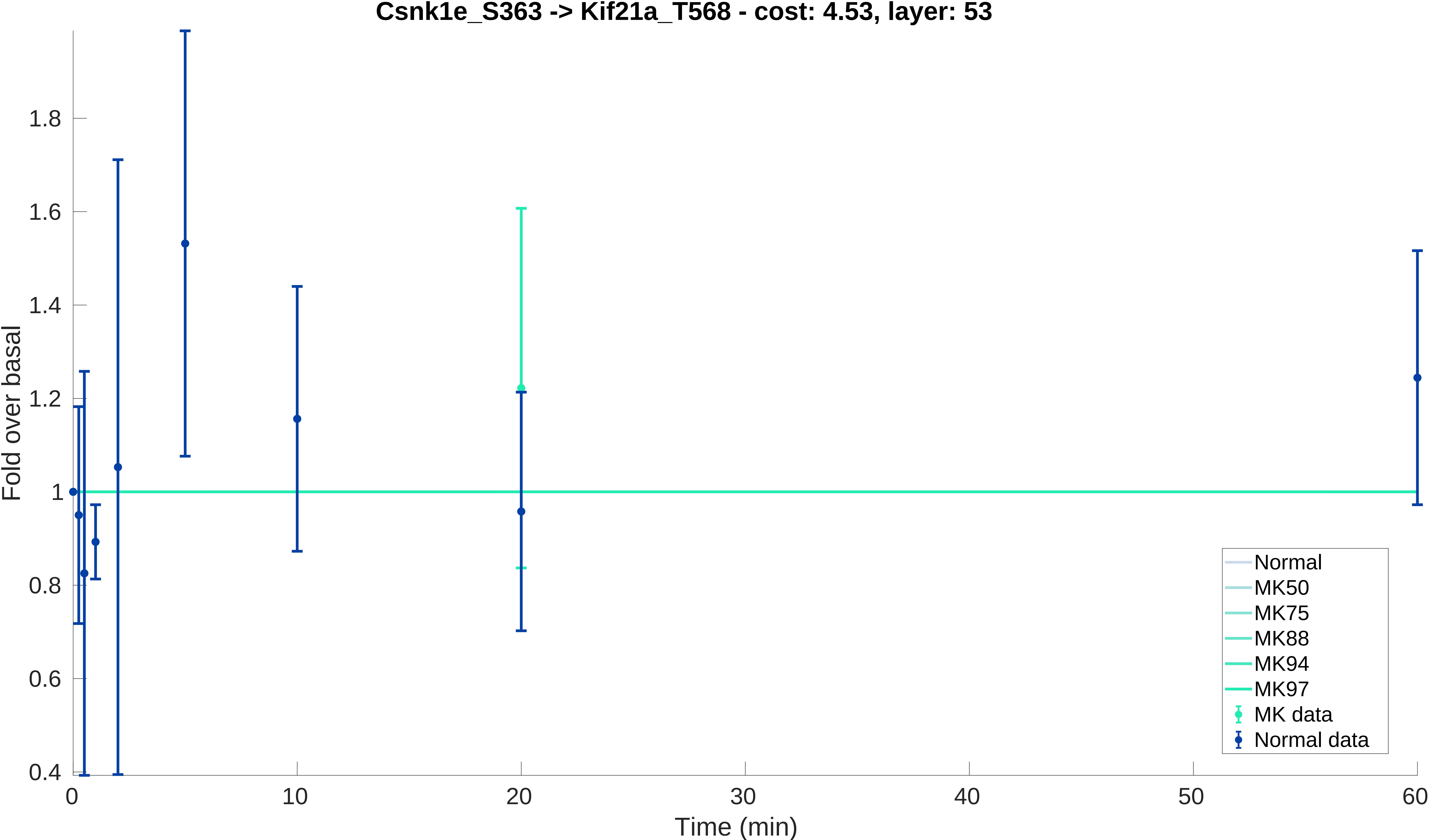

Csnk1e_S363 -> Kif21a_T568 - cost: 4.53, layer: 53
Fold over basal
1.8
1.6
1.4
1.2
1
0.8
0.6
0.4
0
10
20
30
40
50
60
Time (min)
Normal
MK50
MK75
MK88
MK94
MK97
MK data
Normal data

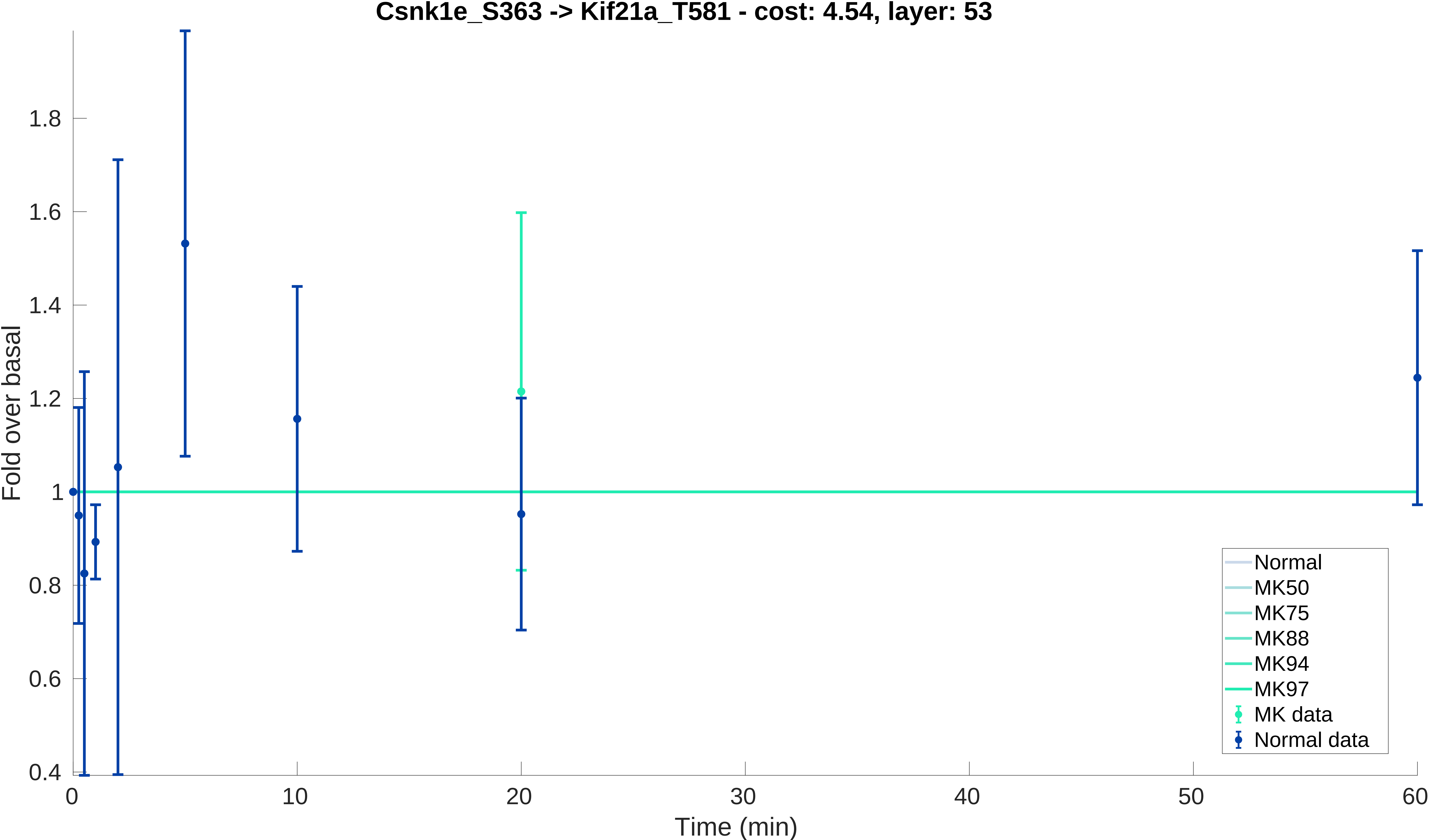

Csnk1e_S363 -> Kif21a_T581 - cost: 4.54, layer: 53
Fold over basal
Time (min)
0.4
0.6
0.8
1
1.2
1.4
1.6
1.8
0
10
20
30
40
50
60
Normal
MK50
MK75
MK88
MK94
MK97
MK data
Normal data

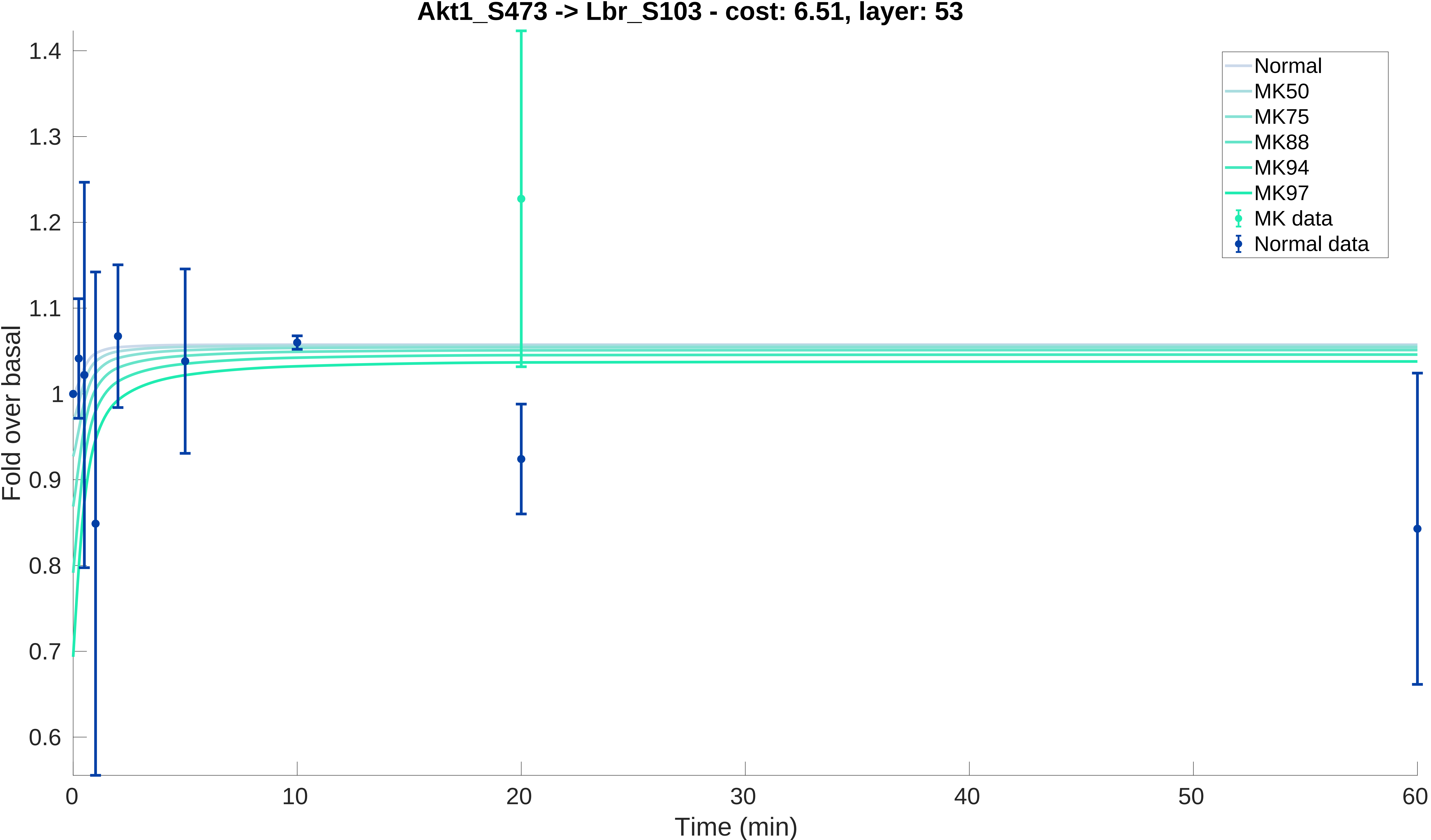
Akt1_S473 -> Lbr_S103 - cost: 6.51, layer: 53
Fold over basal
Time (min)
1.4
1.3
1.2
1.1
1
0.9
0.8
0.7
0.6
0
10
20
30
40
50
60
Normal
MK50
MK75
MK88
MK94
MK97
MK data
Normal data

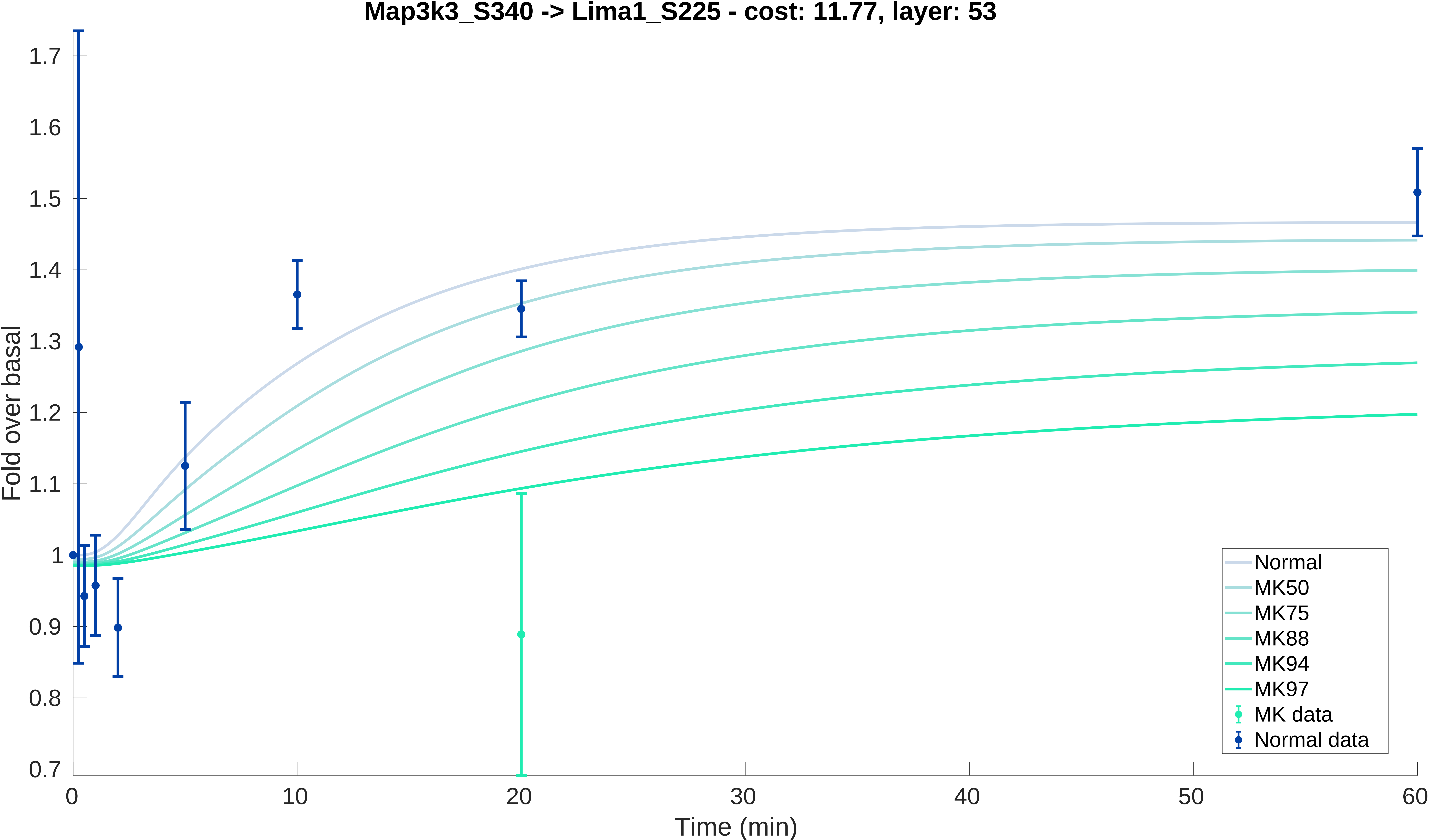

Map3k3_S340 -> Lima1_S225 - cost: 11.77, layer: 53
Fold over basal
1.7
1.6
1.5
1.4
1.3
1.2
1.1
1
0.9
0.8
0.7
0
10
20
30
40
50
60
Time (min)
Normal
MK50
MK75
MK88
MK94
MK97
MK data
Normal data

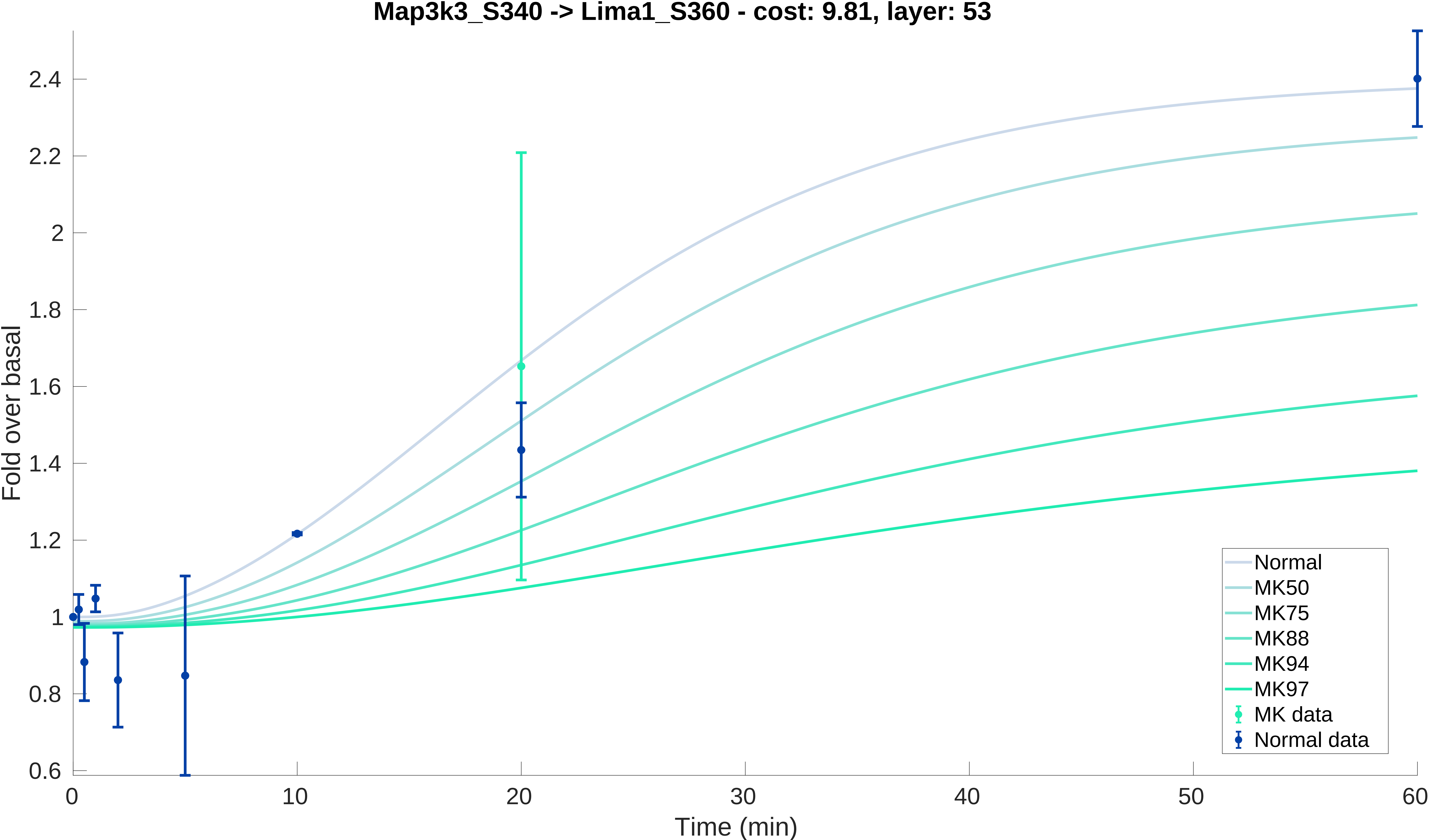
Map3k3_S340 -> Lima1_S360 - cost: 9.81, layer: 53
Fold over basal
Time (min)
0
10
20
30
40
50
60
0.6
0.8
1
1.2
1.4
1.6
1.8
2
2.2
2.4
Normal
MK50
MK75
MK88
MK94
MK97
MK data
Normal data

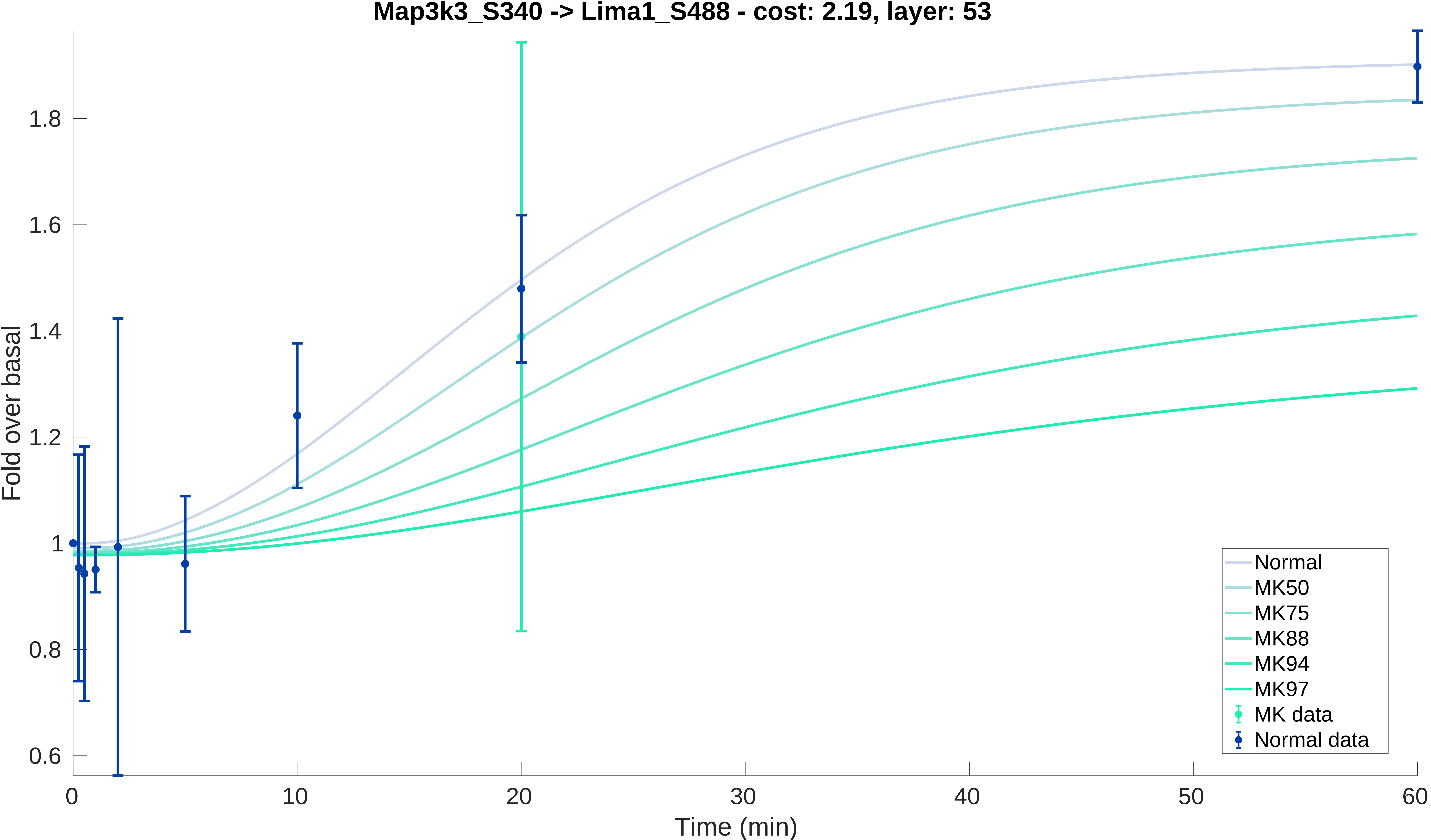
Map3k3_S340 -> Lima1_S488 - cost: 2.19, layer: 53
Fold over basal
Time (min)
0.6
0.8
1
1.2
1.4
1.6
1.8
0
10
20
30
40
50
60
Normal
MK50
MK75
MK88
MK94
MK97
MK data
Normal data

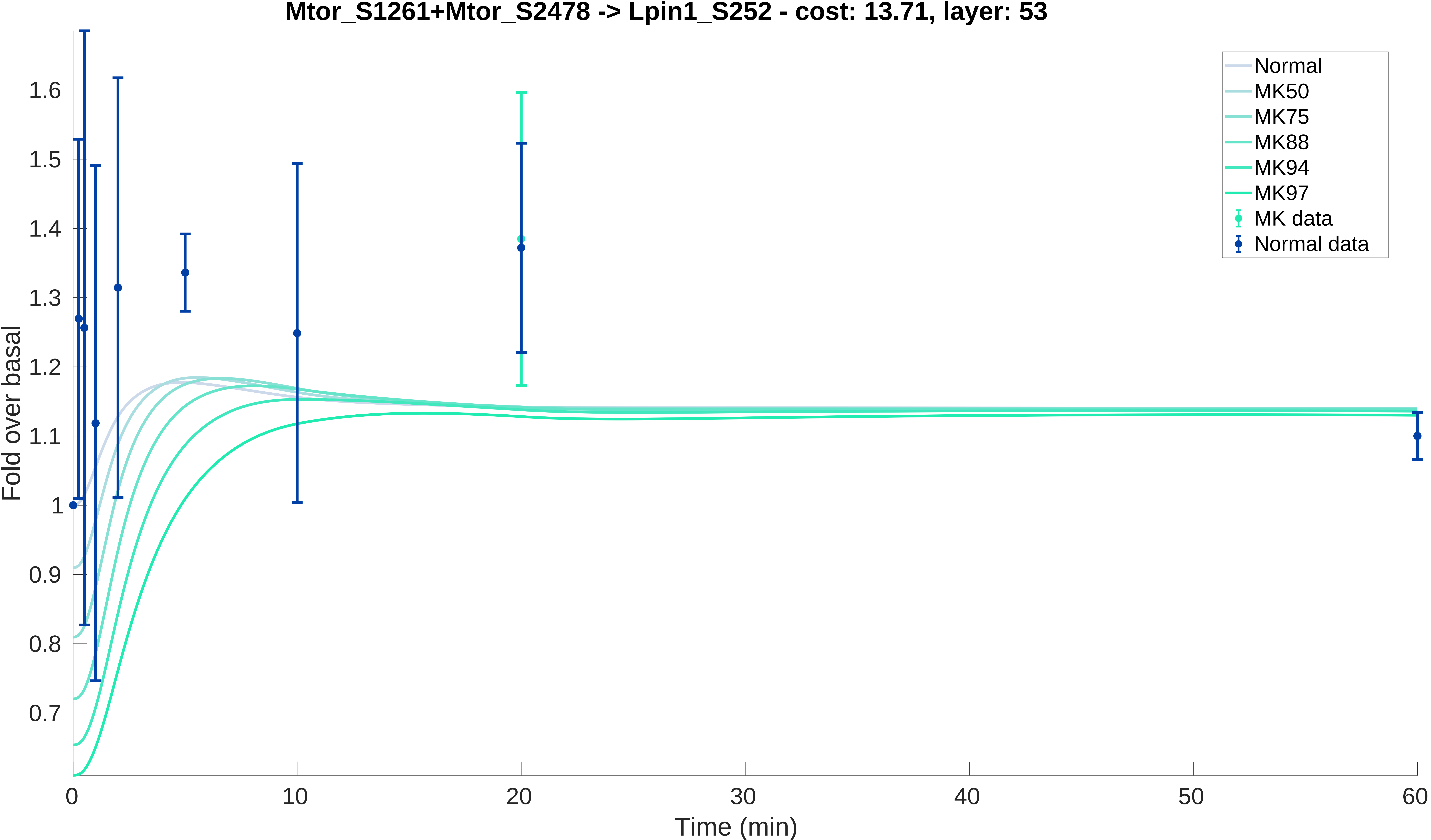

Mtor_S1261+Mtor_S2478 -> Lpin1_S252 - cost: 13.71, layer: 53
Fold over basal
Time (min)
0.7
0.8
0.9
1
1.1
1.2
1.3
1.4
1.5
1.6
0
10
20
30
40
50
60
Normal
MK50
MK75
MK88
MK94
MK97
MK data
Normal data

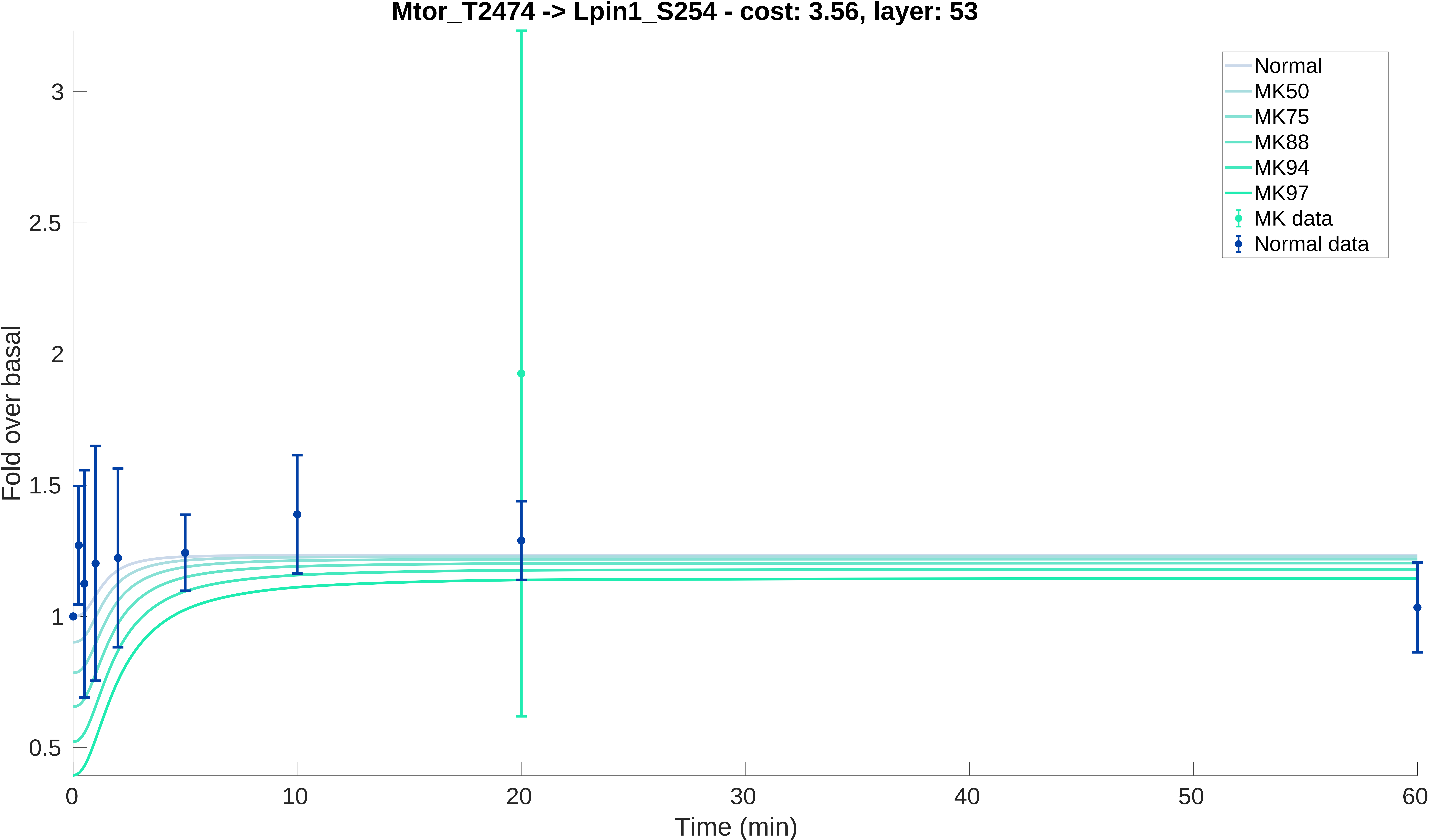
Mtor_T2474 -> Lpin1_S254 - cost: 3.56, layer: 53
Fold over basal
Time (min)
Normal
MK50
MK75
MK88
MK94
MK97
MK data
Normal data
0.5
1
1.5
2
2.5
3
0
10
20
30
40
50
60

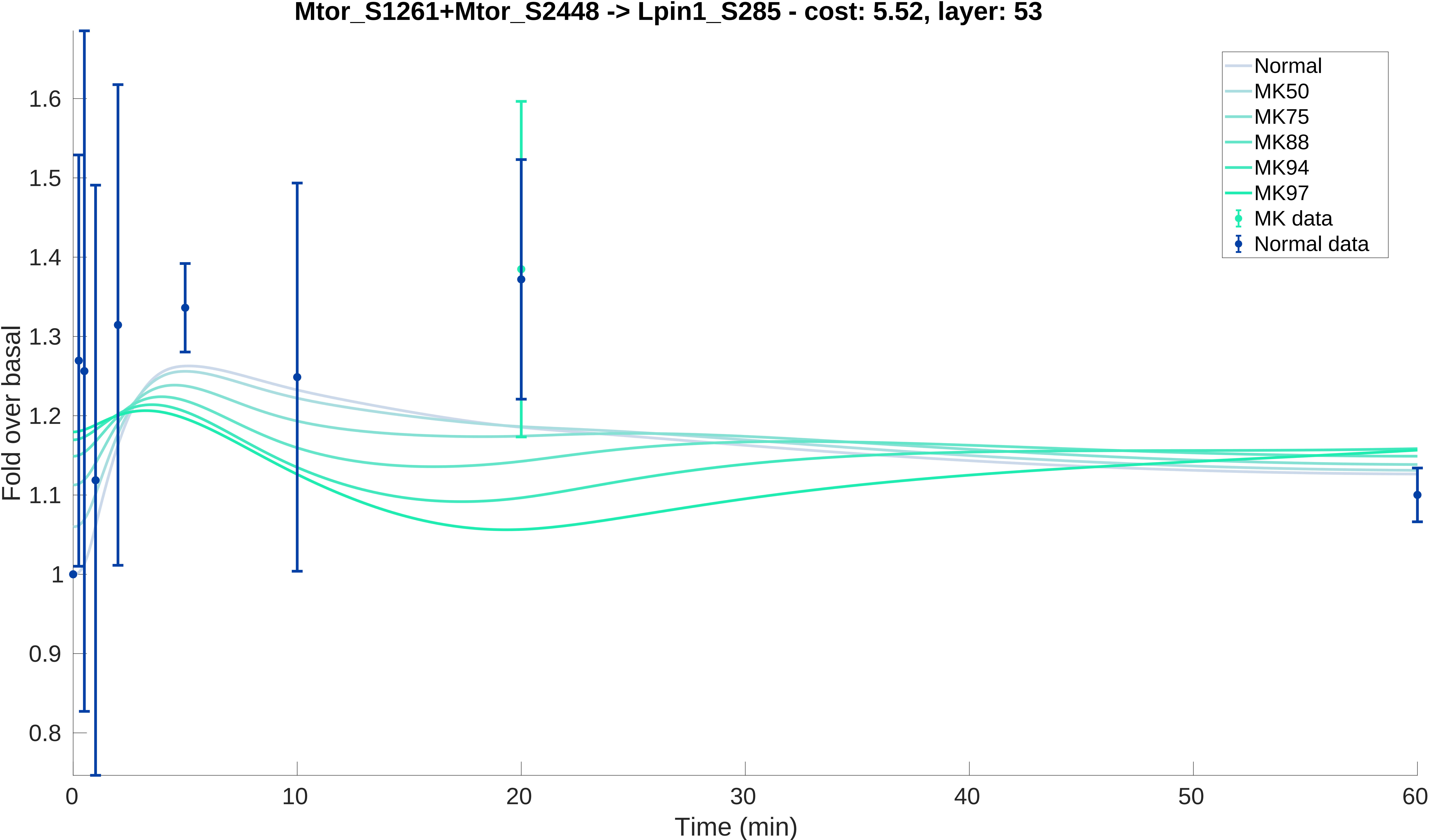

Mtor_S1261+Mtor_S2448 -> Lpin1_S285 - cost: 5.52, layer: 53
Fold over basal
Time (min)
0.8
0.9
1
1.1
1.2
1.3
1.4
1.5
1.6
0
10
20
30
40
50
60
Normal
MK50
MK75
MK88
MK94
MK97
MK data
Normal data

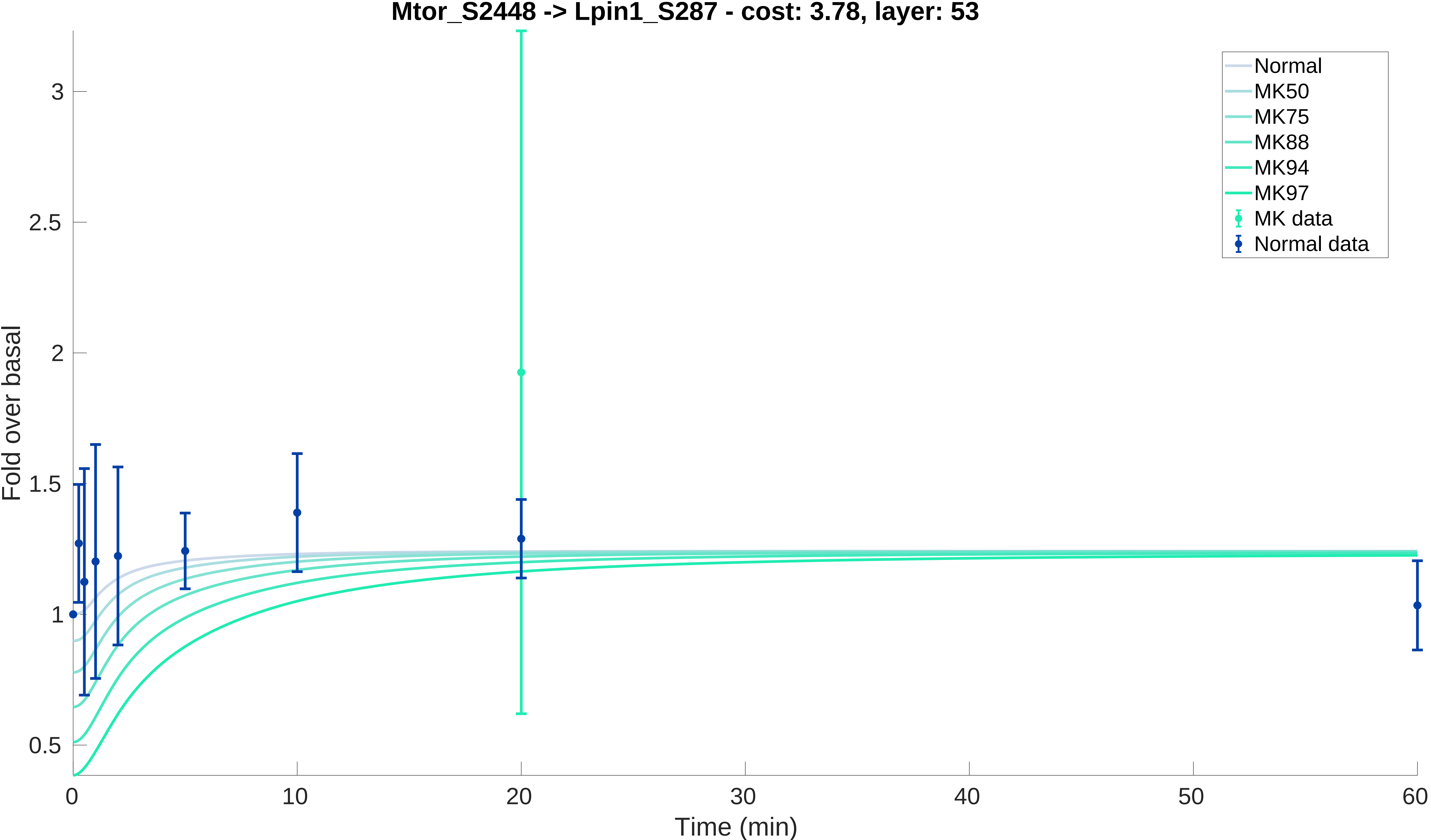

Mtor_S2448 -> Lpin1_S287 - cost: 3.78, layer: 53
Normal
MK50
MK75
MK88
MK94
MK97
MK data
Normal data
Fold over basal
Time (min)
0.5
1
1.5
2
2.5
3
0
10
20
30
40
50
60

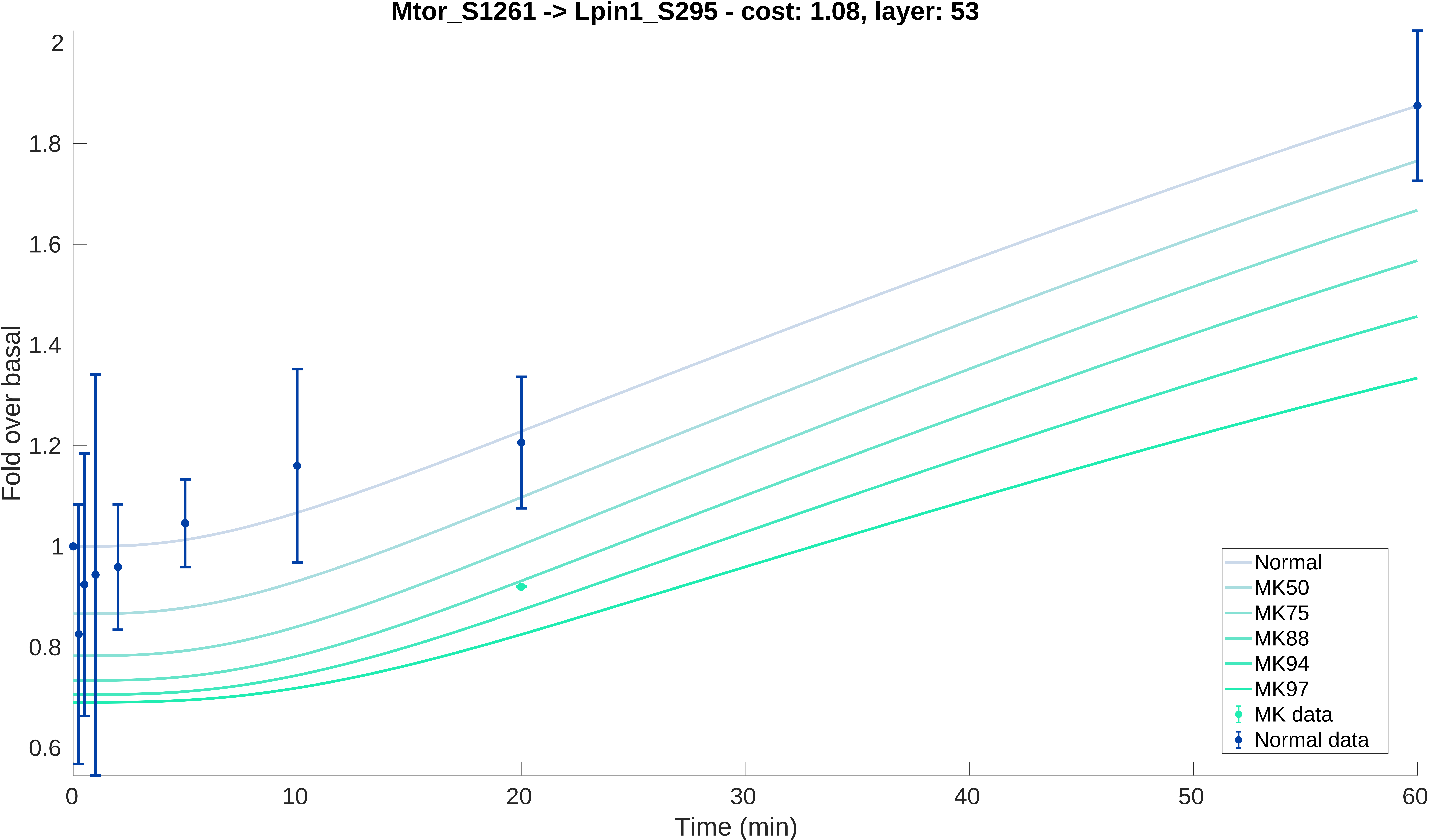

Mtor_S1261 -> Lpin1_S295 - cost: 1.08, layer: 53
Fold over basal
Time (min)
2
1.8
1.6
1.4
1.2
1
0.8
0.6
0
10
20
30
40
50
60
Normal
MK50
MK75
MK88
MK94
MK97
MK data
Normal data

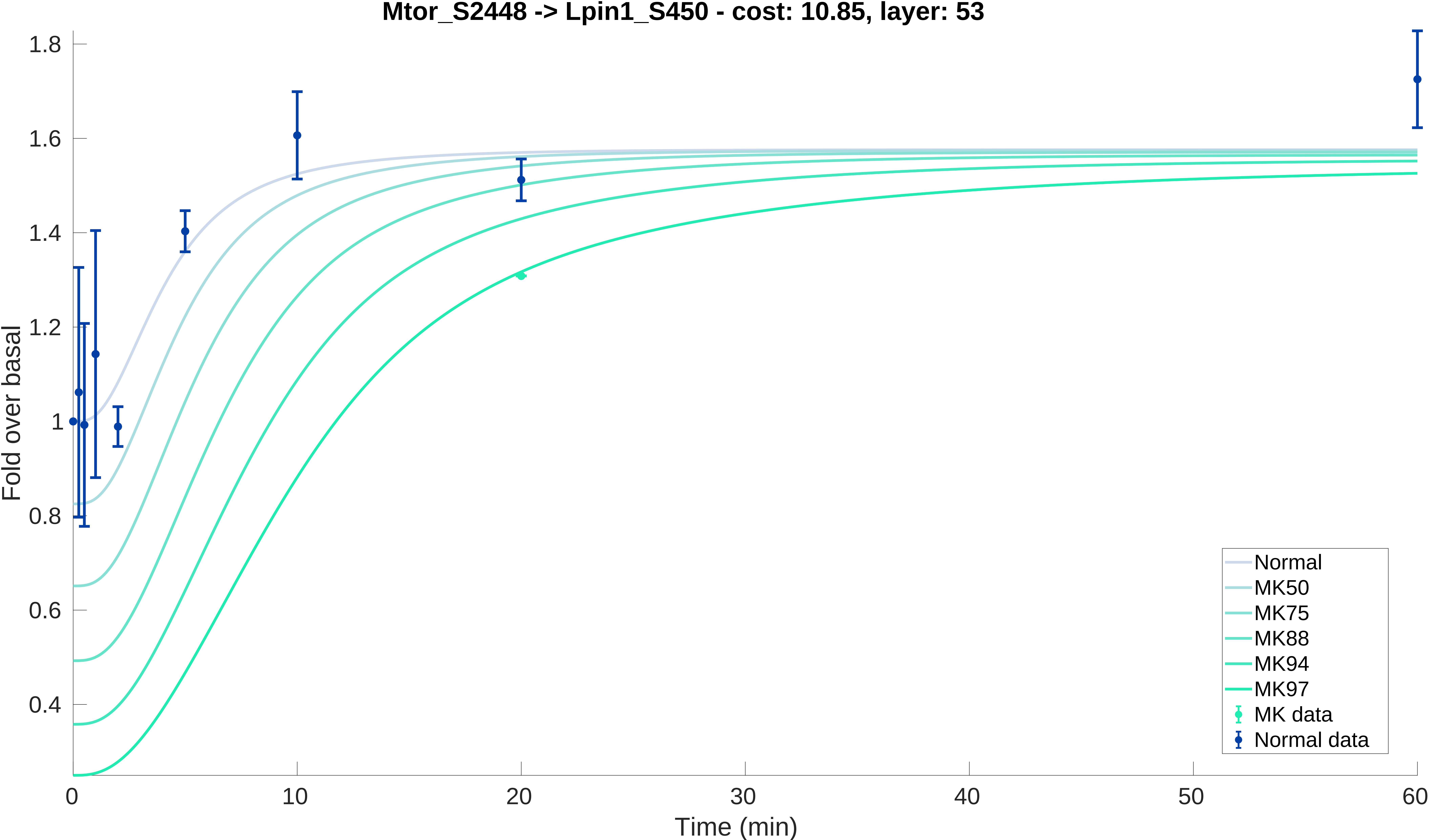
Mtor_S2448 -> Lpin1_S450 - cost: 10.85, layer: 53
Fold over basal
1.8
1.6
1.4
1.2
1
0.8
0.6
0.4
Time (min)
0
10
20
30
40
50
60
Normal
MK50
MK75
MK88
MK94
MK97
MK data
Normal data

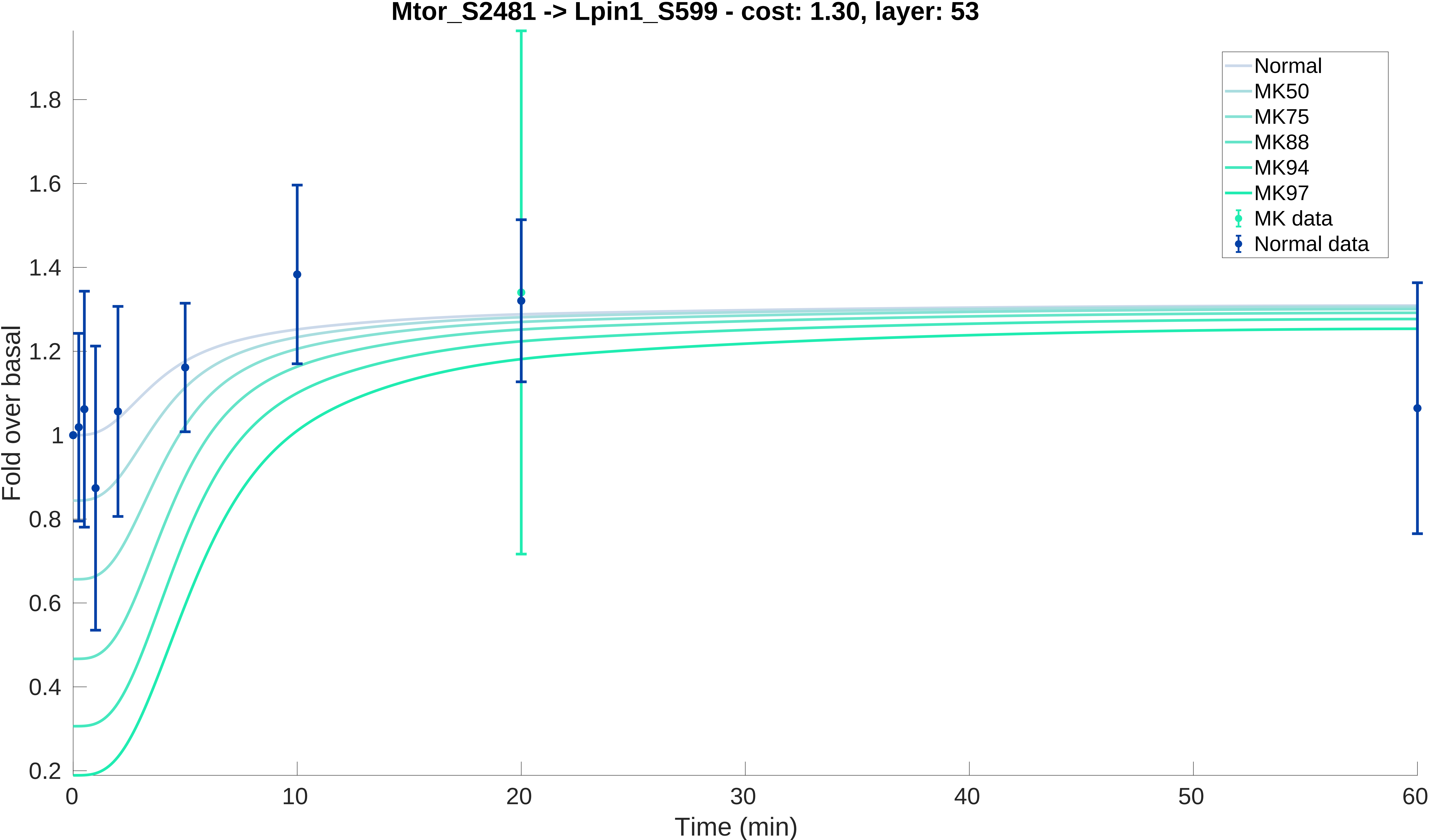
Mtor_S2481 -> Lpin1_S599 - cost: 1.30, layer: 53
Normal
MK50
MK75
MK88
MK94
MK97
MK data
Normal data
Fold over basal
Time (min)
0.2
0.4
0.6
0.8
1
1.2
1.4
1.6
1.8
0
10
20
30
40
50
60

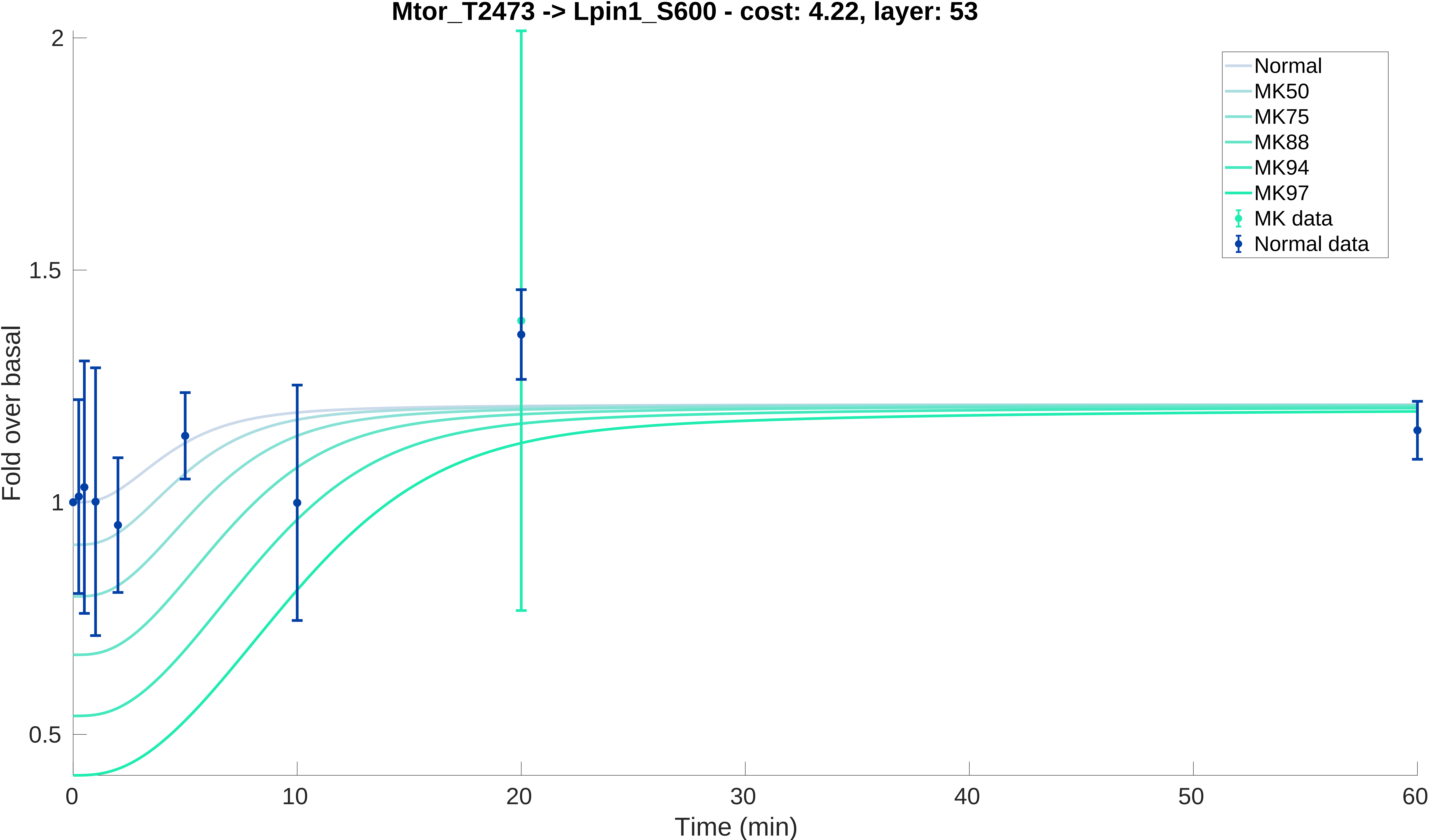

Mtor_T2473 -> Lpin1_S600 - cost: 4.22, layer: 53
Fold over basal
Time (min)
0.5
1
1.5
2
0
10
20
30
40
50
60
Normal
MK50
MK75
MK88
MK94
MK97
MK data
Normal data

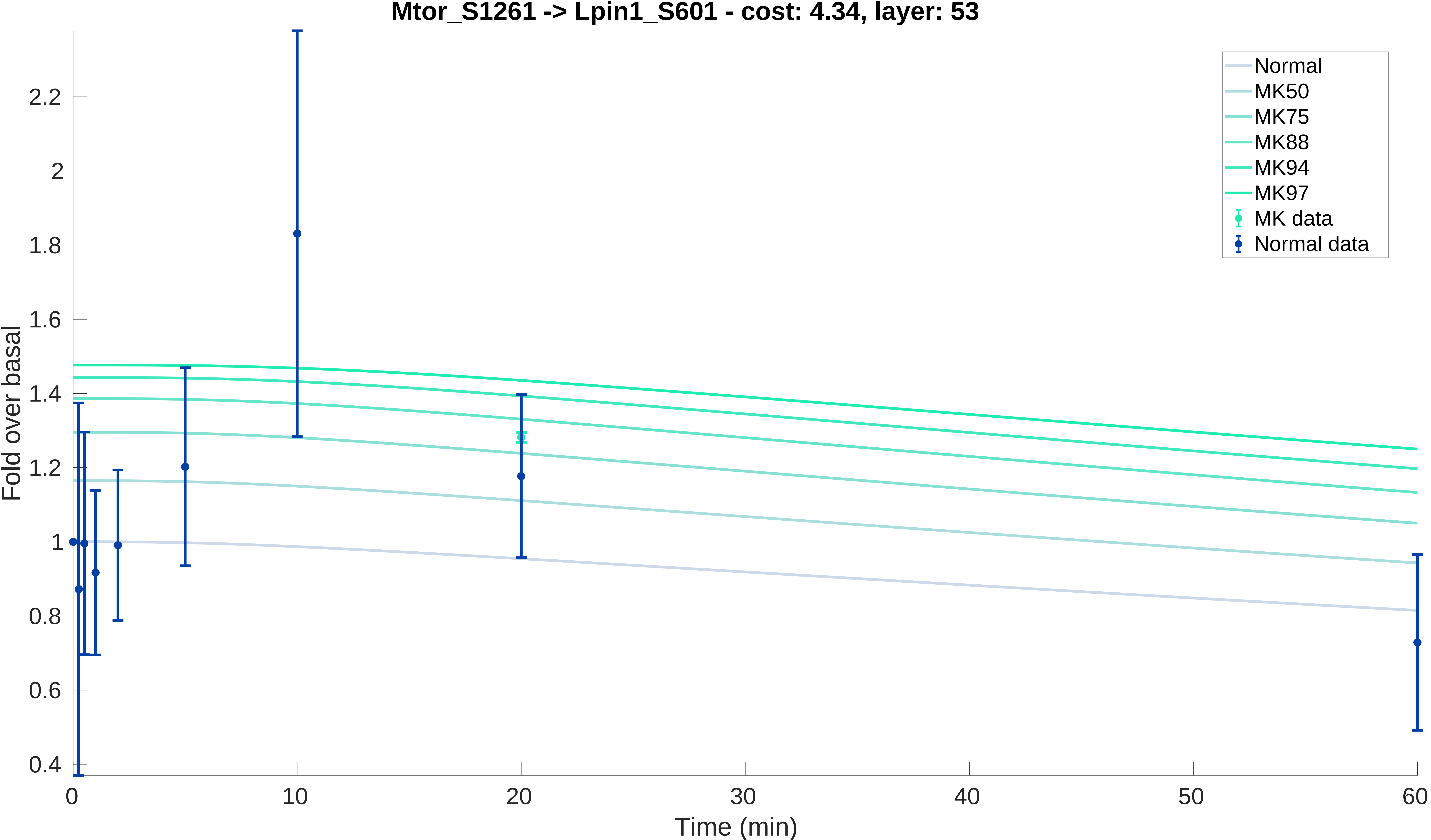

Mtor_S1261 -> Lpin1_S601 - cost: 4.34, layer: 53
Normal
MK50
MK75
MK88
MK94
MK97
MK data
Normal data
Fold over basal
2.2
2
1.8
1.6
1.4
1.2
1
0.8
0.6
0.4
0
10
20
30
40
50
60
Time (min)

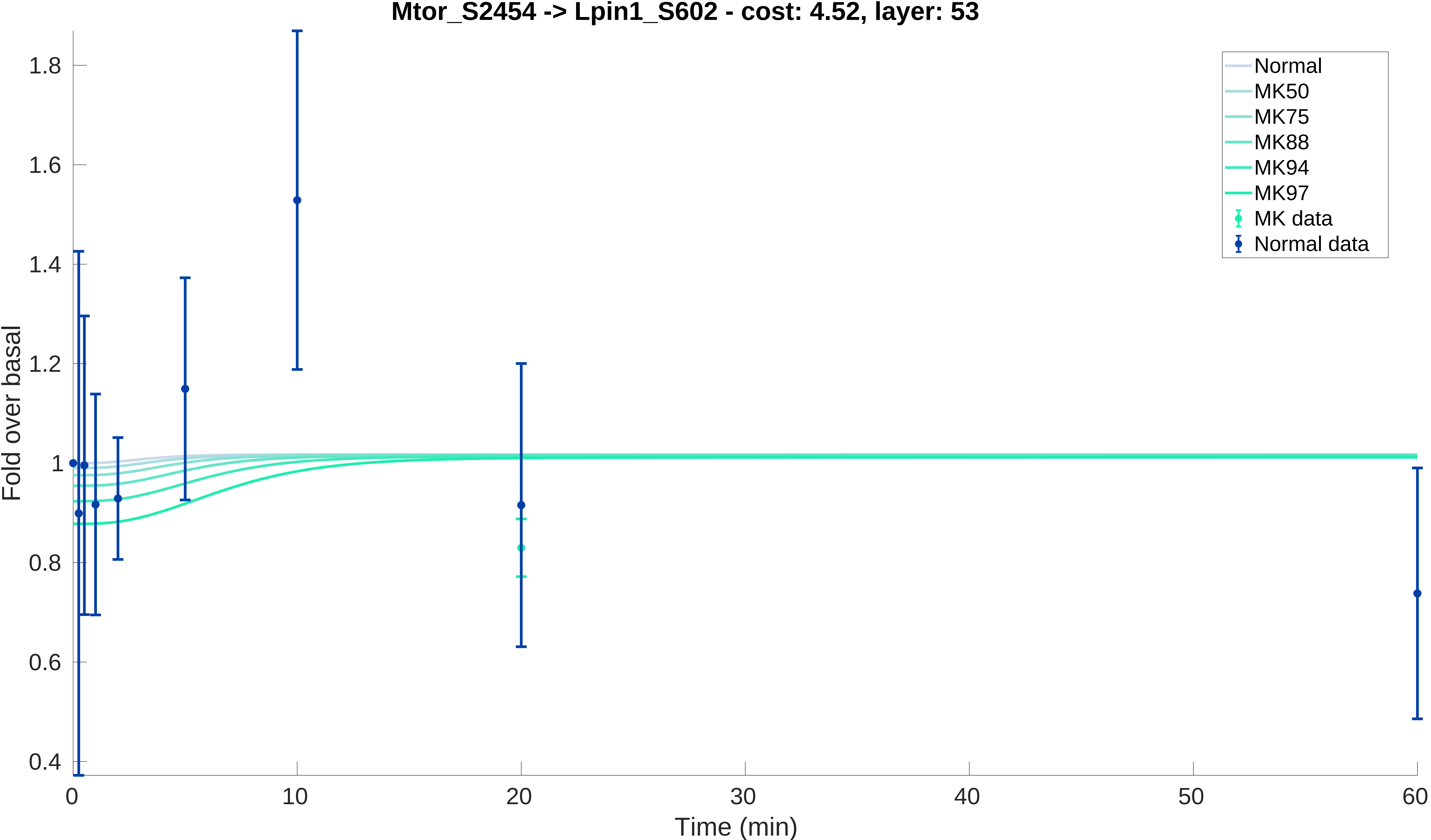

Mtor_S2454 -> Lpin1_S602 - cost: 4.52, layer: 53
Fold over basal
1.8
1.6
1.4
1.2
1
0.8
0.6
0.4
0
10
20
30
40
50
60
Time (min)
Normal
MK50
MK75
MK88
MK94
MK97
MK data
Normal data

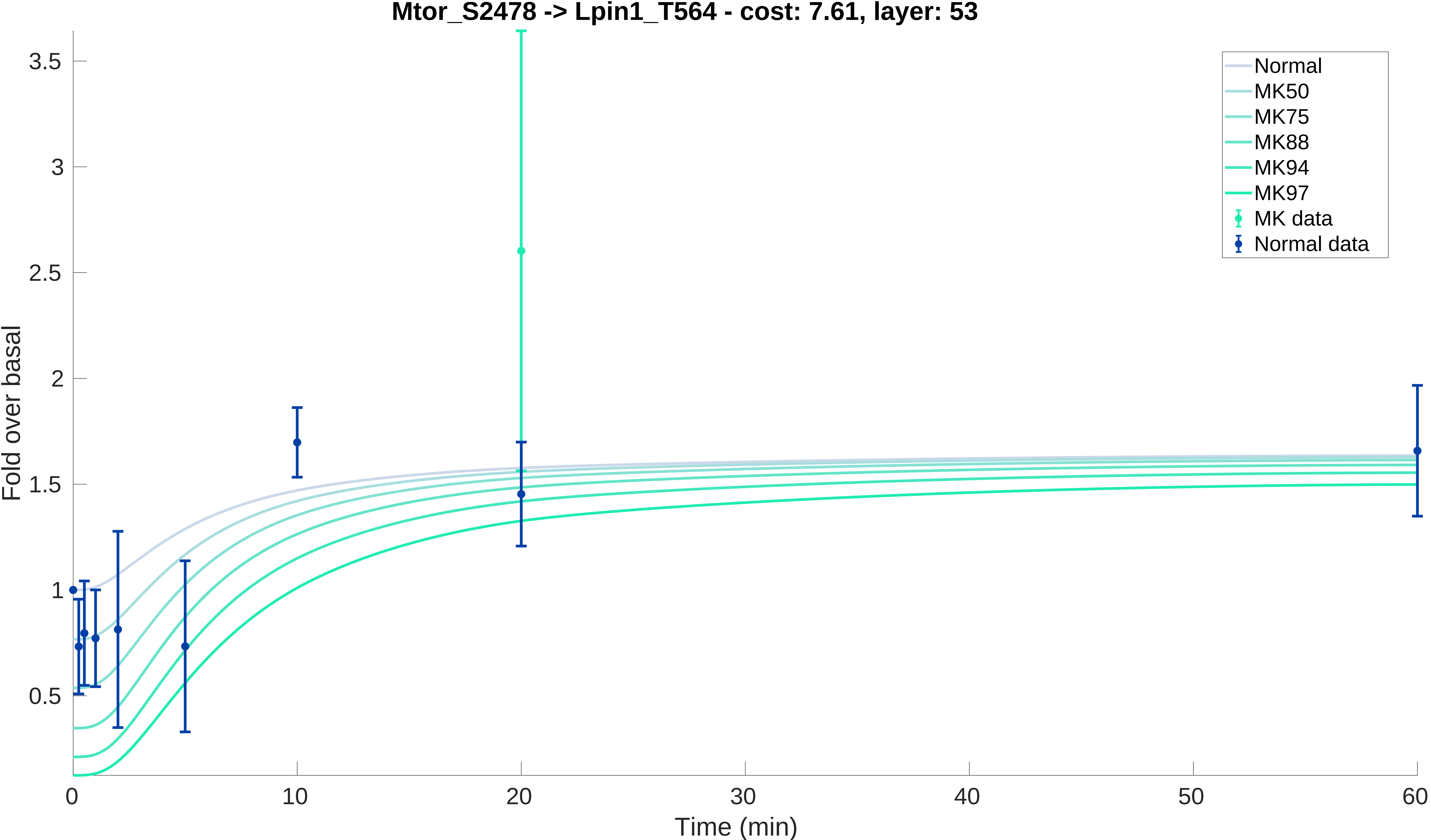

Mtor_S2478 -> Lpin1_T564 - cost: 7.61, layer: 53
Fold over basal
Time (min)
Normal
MK50
MK75
MK88
MK94
MK97
MK data
Normal data
3.5
3
2.5
2
1.5
1
0.5
0
10
20
30
40
50
60

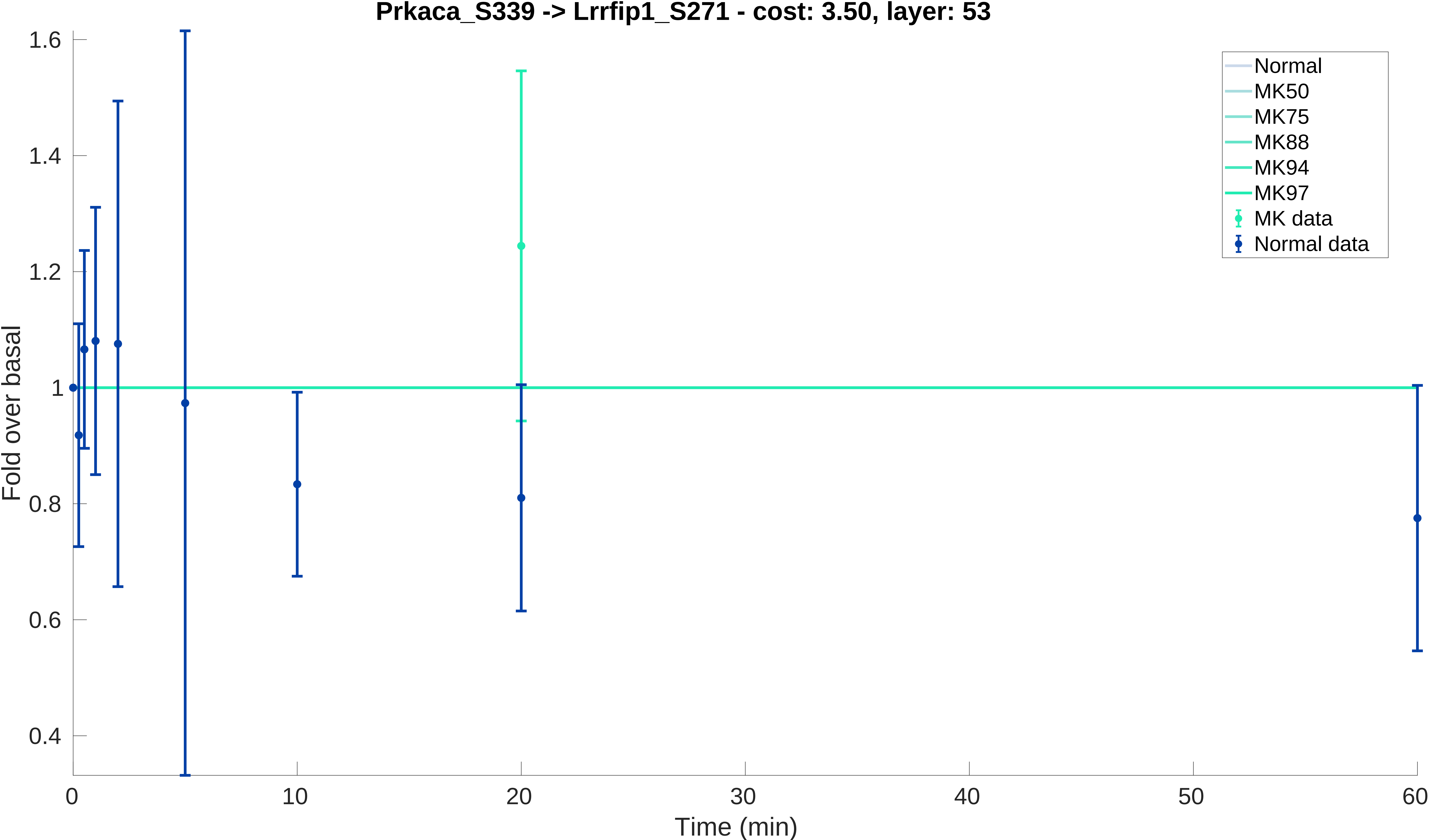

Prkaca_S339 -> Lrrfip1_S271 - cost: 3.50, layer: 53
Fold over basal
1.6
1.4
1.2
1
0.8
0.6
0.4
0
10
20
30
40
50
60
Time (min)
Normal
MK50
MK75
MK88
MK94
MK97
MK data
Normal data

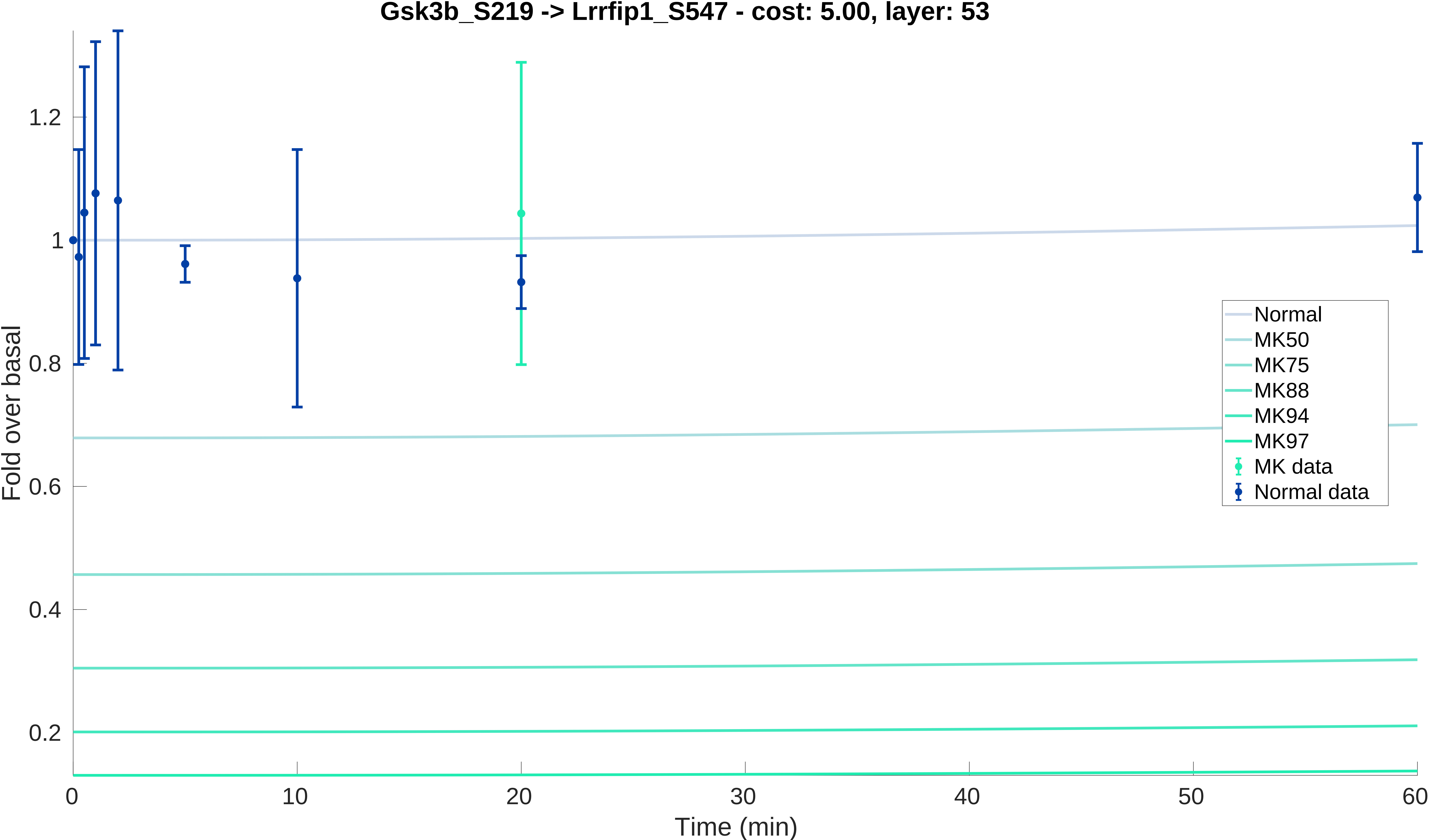

Gsk3b_S219 -> Lrrfip1_S547 - cost: 5.00, layer: 53
Fold over basal
1.2
1
0.8
0.6
0.4
0.2
0
10
20
30
40
50
60
Time (min)
Normal
MK50
MK75
MK88
MK94
MK97
MK data
Normal data

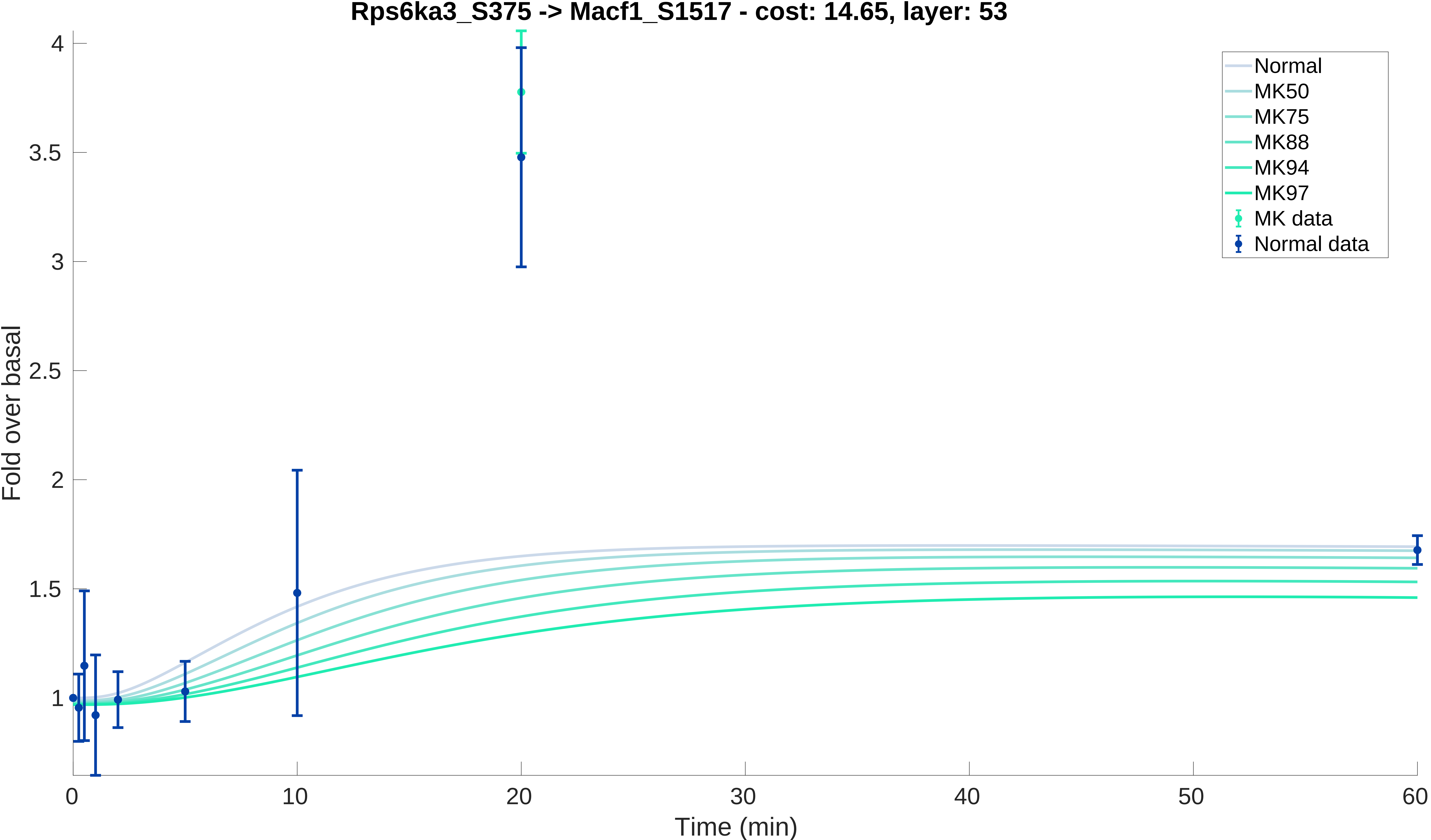

Rps6ka3_S375 -> Macf1_S1517 - cost: 14.65, layer: 53
Fold over basal
Time (min)
Normal
MK50
MK75
MK88
MK94
MK97
MK data
Normal data
4
3.5
3
2.5
2
1.5
1
0
10
20
30
40
50
60

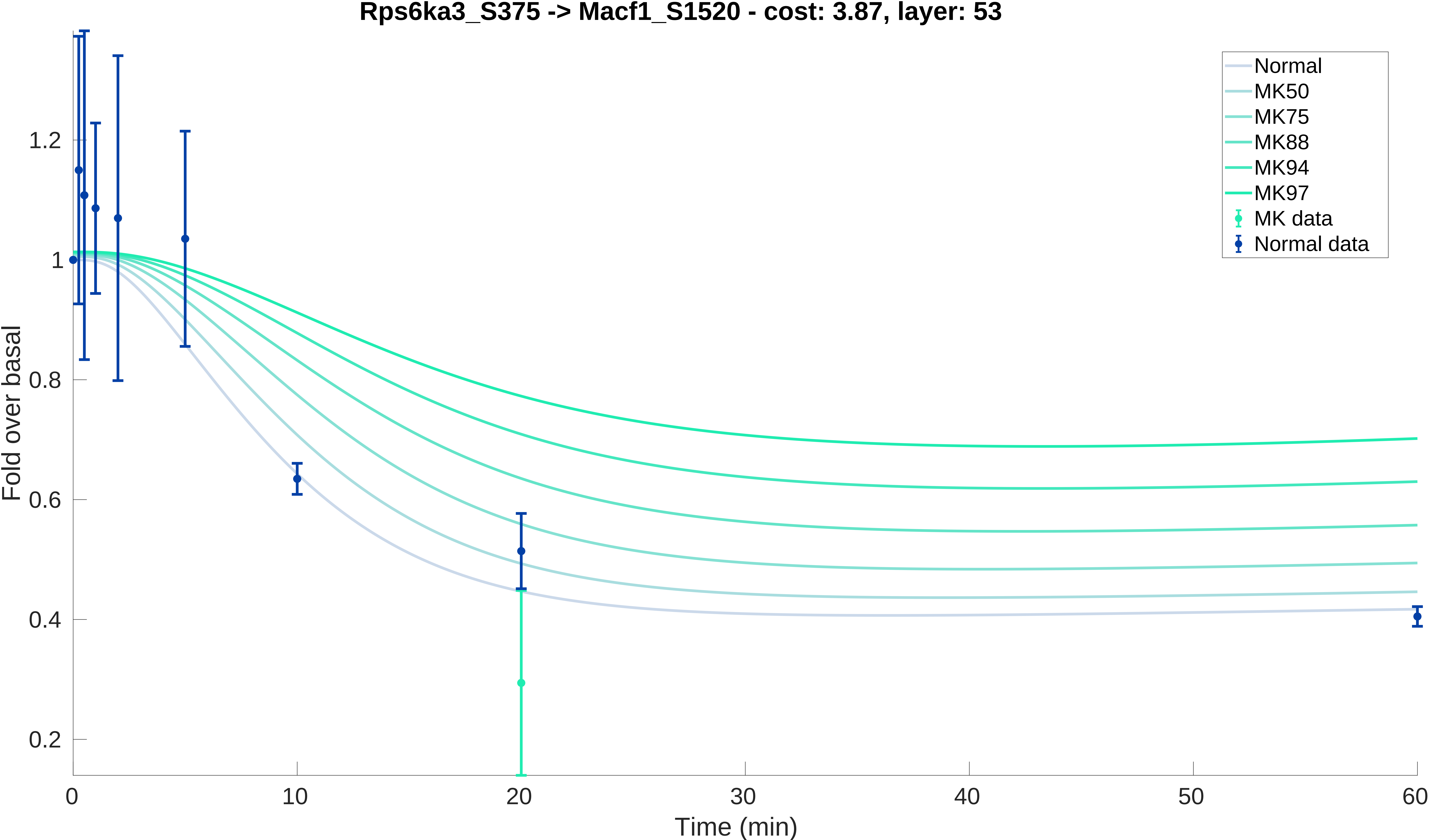

Rps6ka3_S375 -> Macf1_S1520 - cost: 3.87, layer: 53
Fold over basal
Time (min)
0.2
0.4
0.6
0.8
1
1.2
0
10
20
30
40
50
60
Normal
MK50
MK75
MK88
MK94
MK97
MK data
Normal data

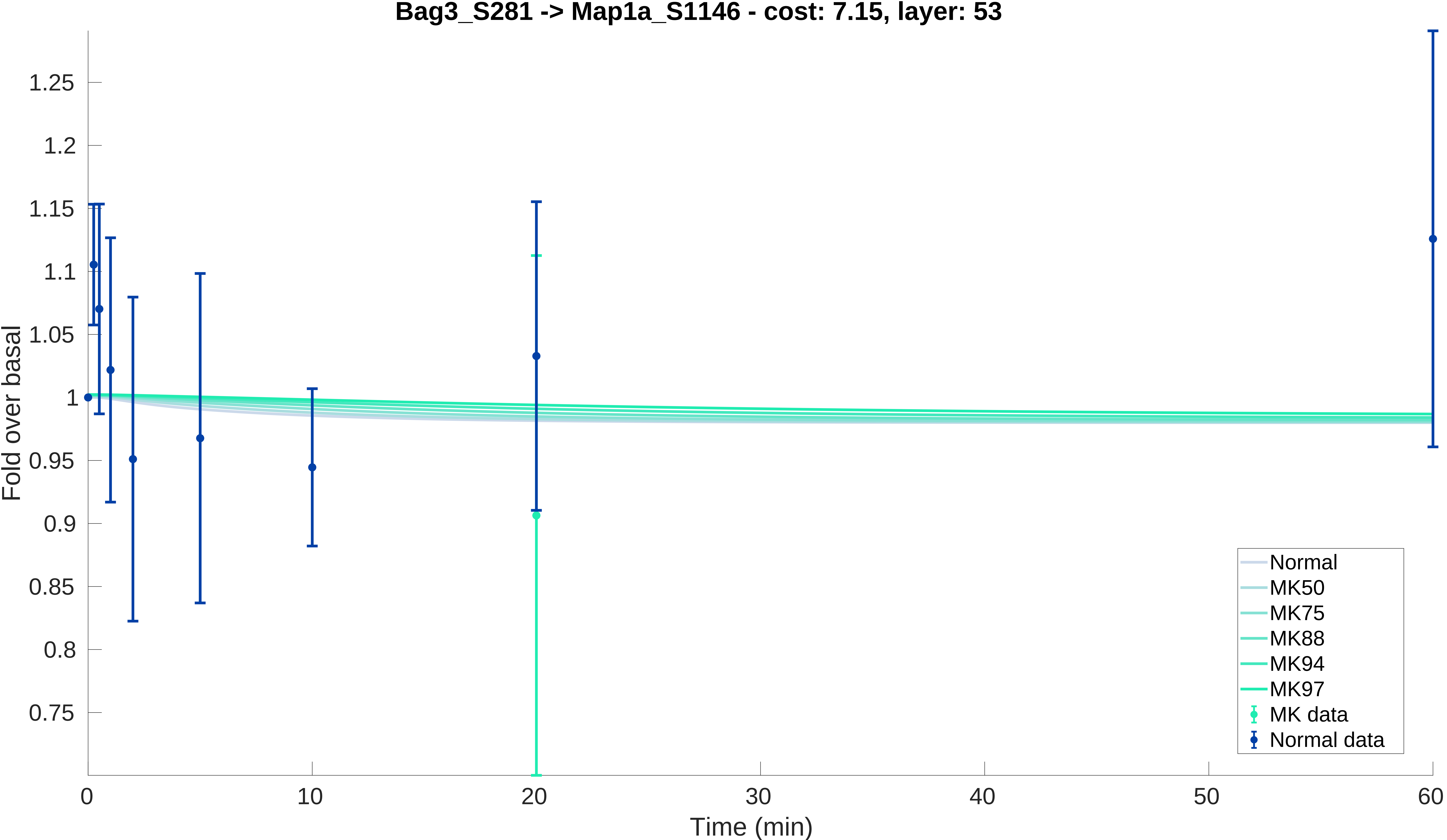

Bag3_S281 -> Map1a_S1146 - cost: 7.15, layer: 53
Fold over basal
1.25
1.2
1.15
1.1
1.05
1
0.95
0.9
0.85
0.8
0.75
0
10
20
30
40
50
60
Time (min)
Normal
MK50
MK75
MK88
MK94
MK97
MK data
Normal data

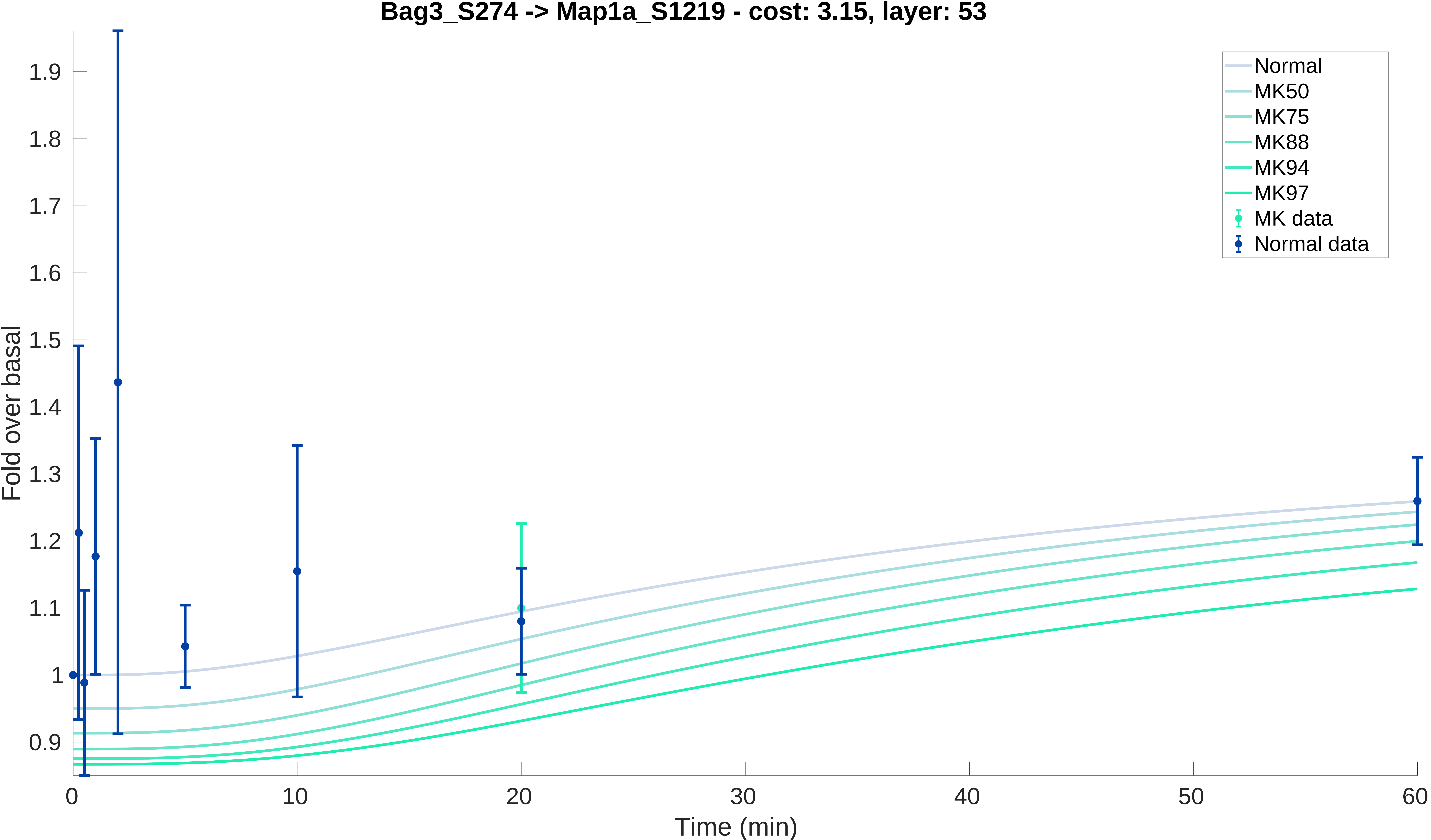

Bag3_S274 -> Map1a_S1219 - cost: 3.15, layer: 53
Fold over basal
Time (min)
0.9
1
1.1
1.2
1.3
1.4
1.5
1.6
1.7
1.8
1.9
0
10
20
30
40
50
60
Normal
MK50
MK75
MK88
MK94
MK97
MK data
Normal data

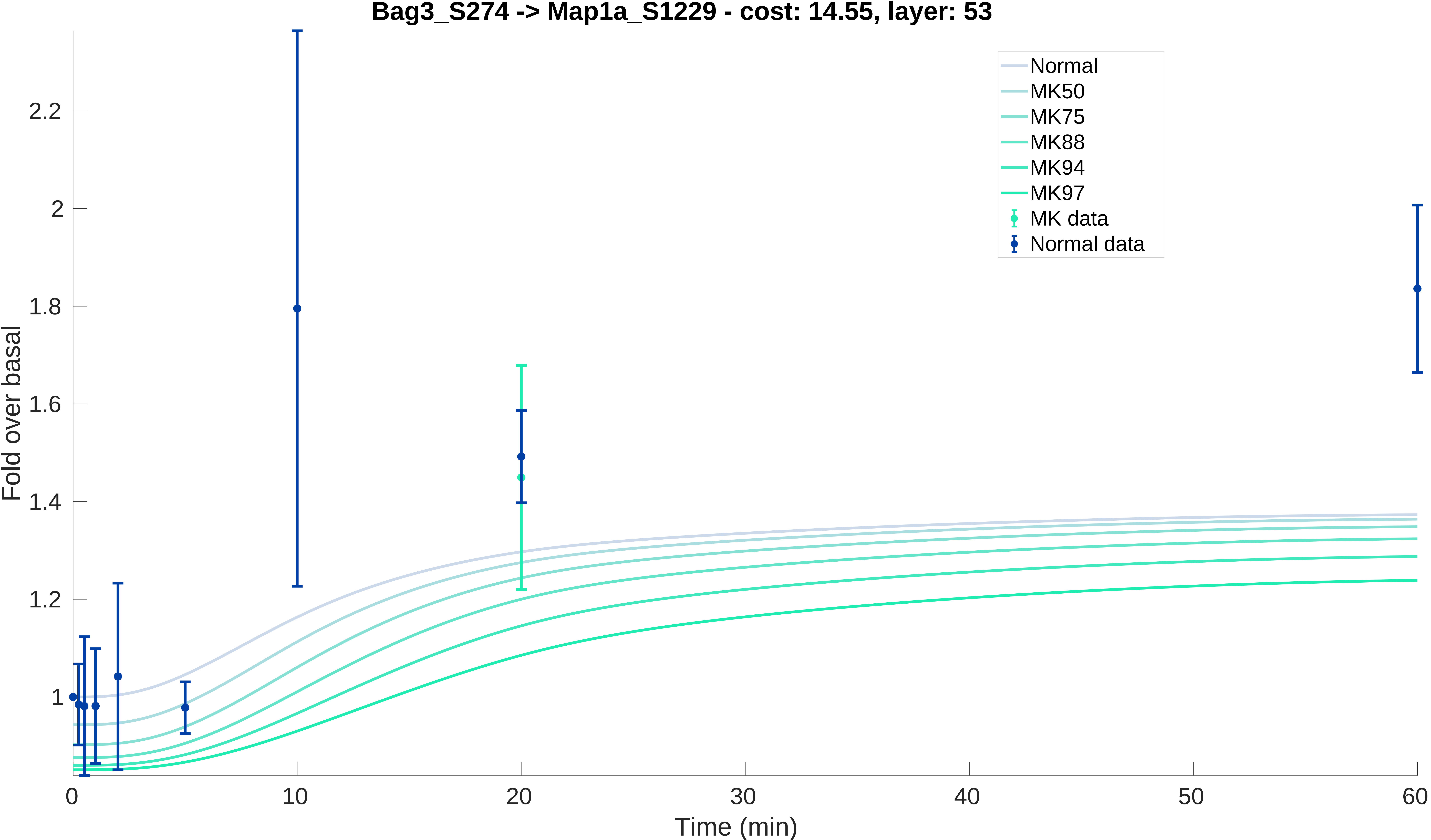

Bag3_S274 -> Map1a_S1229 - cost: 14.55, layer: 53
Normal
MK50
MK75
MK88
MK94
MK97
MK data
Normal data
Fold over basal
Time (min)
2.2
2
1.8
1.6
1.4
1.2
1
0
10
20
30
40
50
60

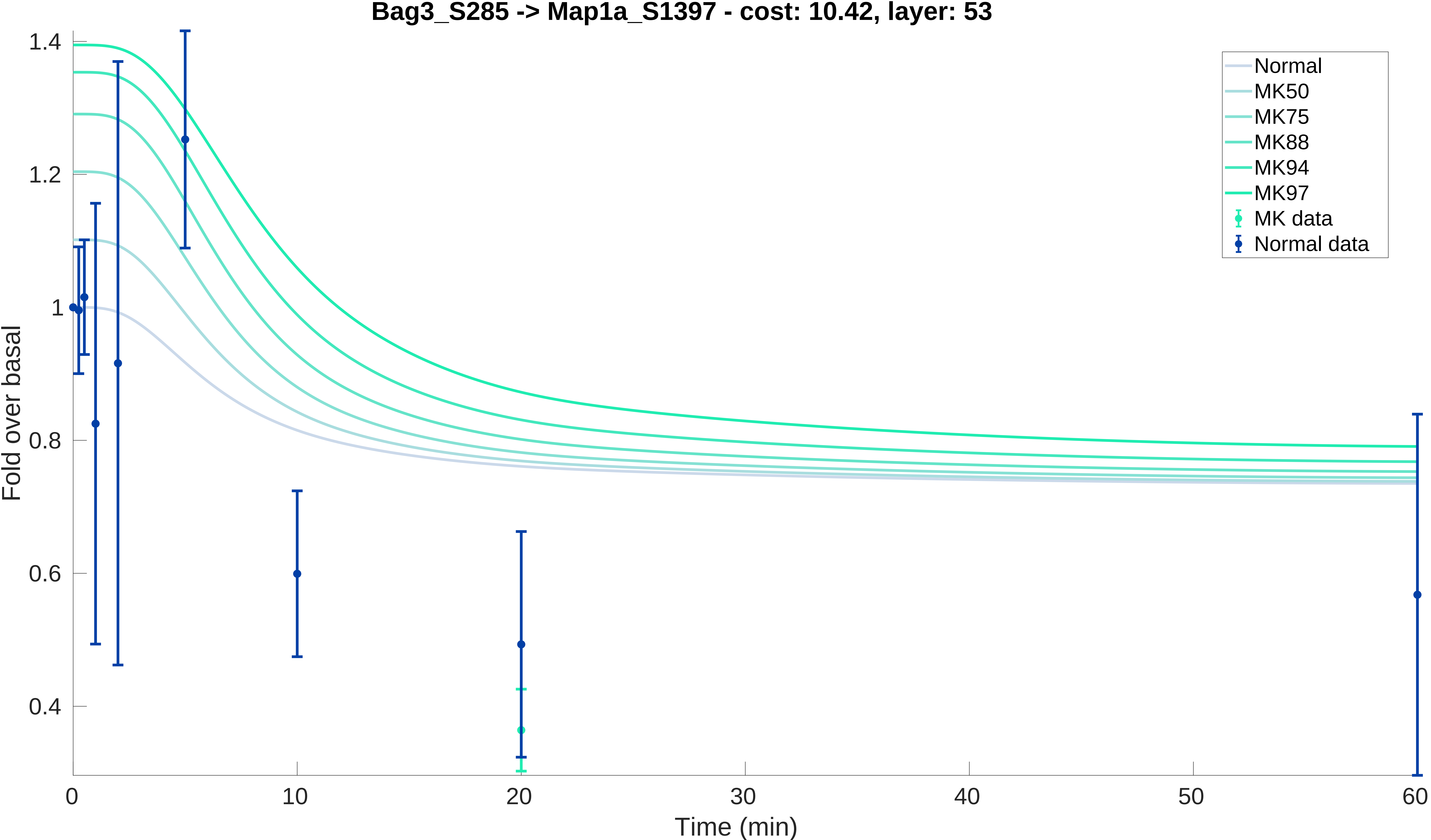
Bag3_S285 -> Map1a_S1397 - cost: 10.42, layer: 53
1.4
1.2
1
0.8
0.6
0.4
Fold over basal
0
10
20
30
40
50
60
Time (min)
Normal
MK50
MK75
MK88
MK94
MK97
MK data
Normal data

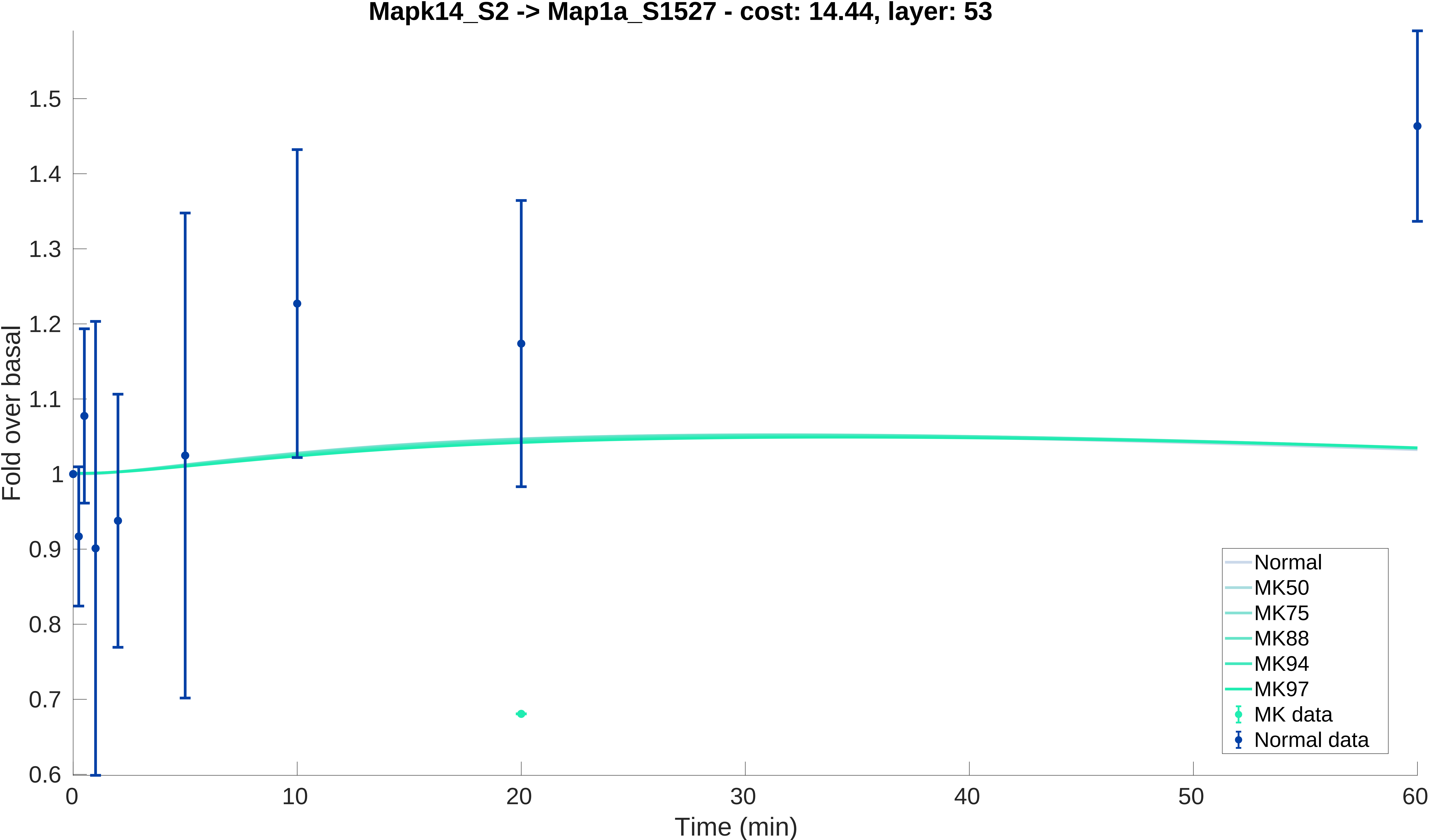
Mapk14_S2 -> Map1a_S1527 - cost: 14.44, layer: 53
Fold over basal
Time (min)
0.6
0.7
0.8
0.9
1
1.1
1.2
1.3
1.4
1.5
0
10
20
30
40
50
60
Normal
MK50
MK75
MK88
MK94
MK97
MK data
Normal data

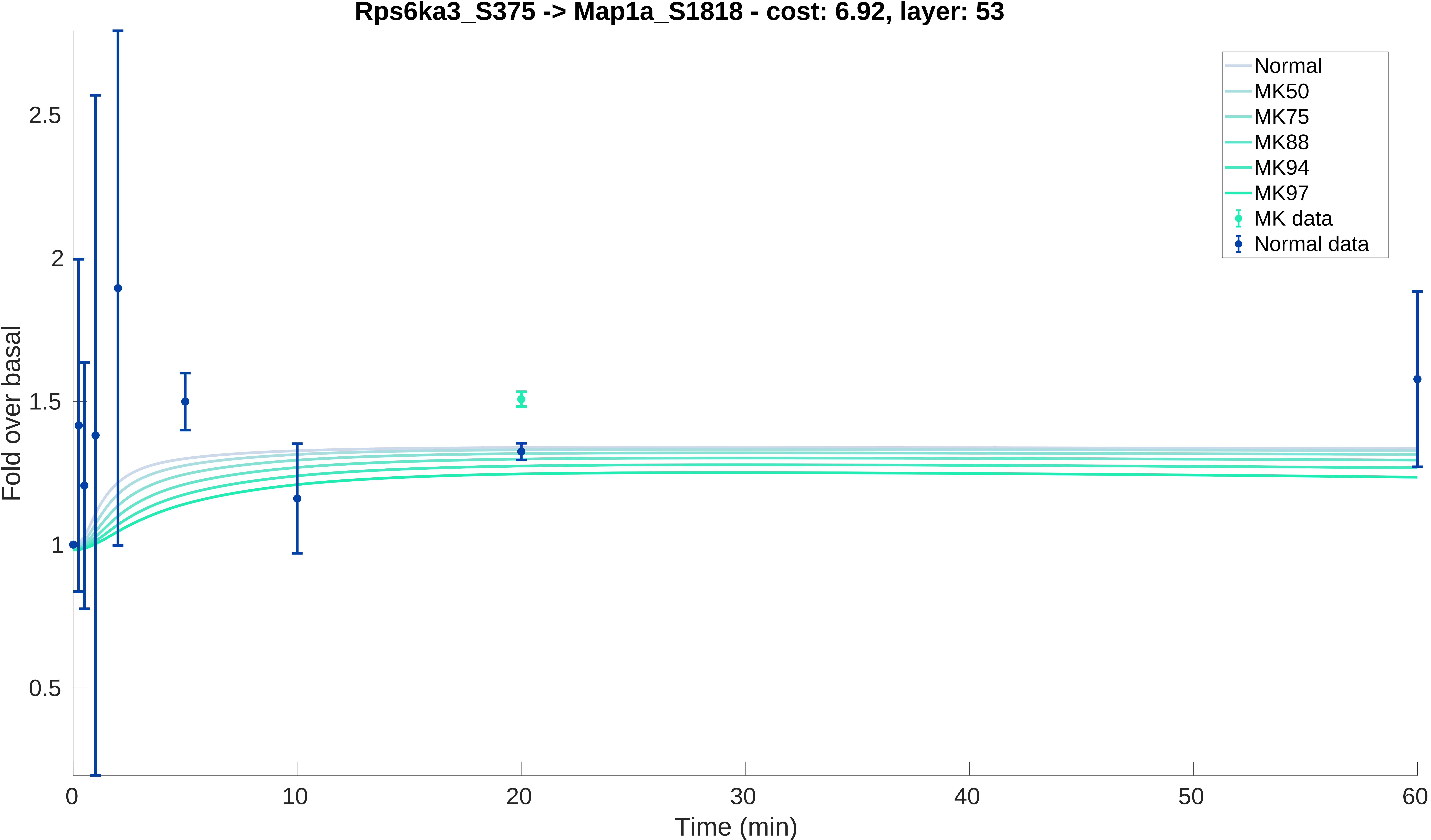

Rps6ka3_S375 -> Map1a_S1818 - cost: 6.92, layer: 53
Fold over basal
2.5
2
1.5
1
0.5
0
10
20
30
40
50
60
Time (min)
Normal
MK50
MK75
MK88
MK94
MK97
MK data
Normal data

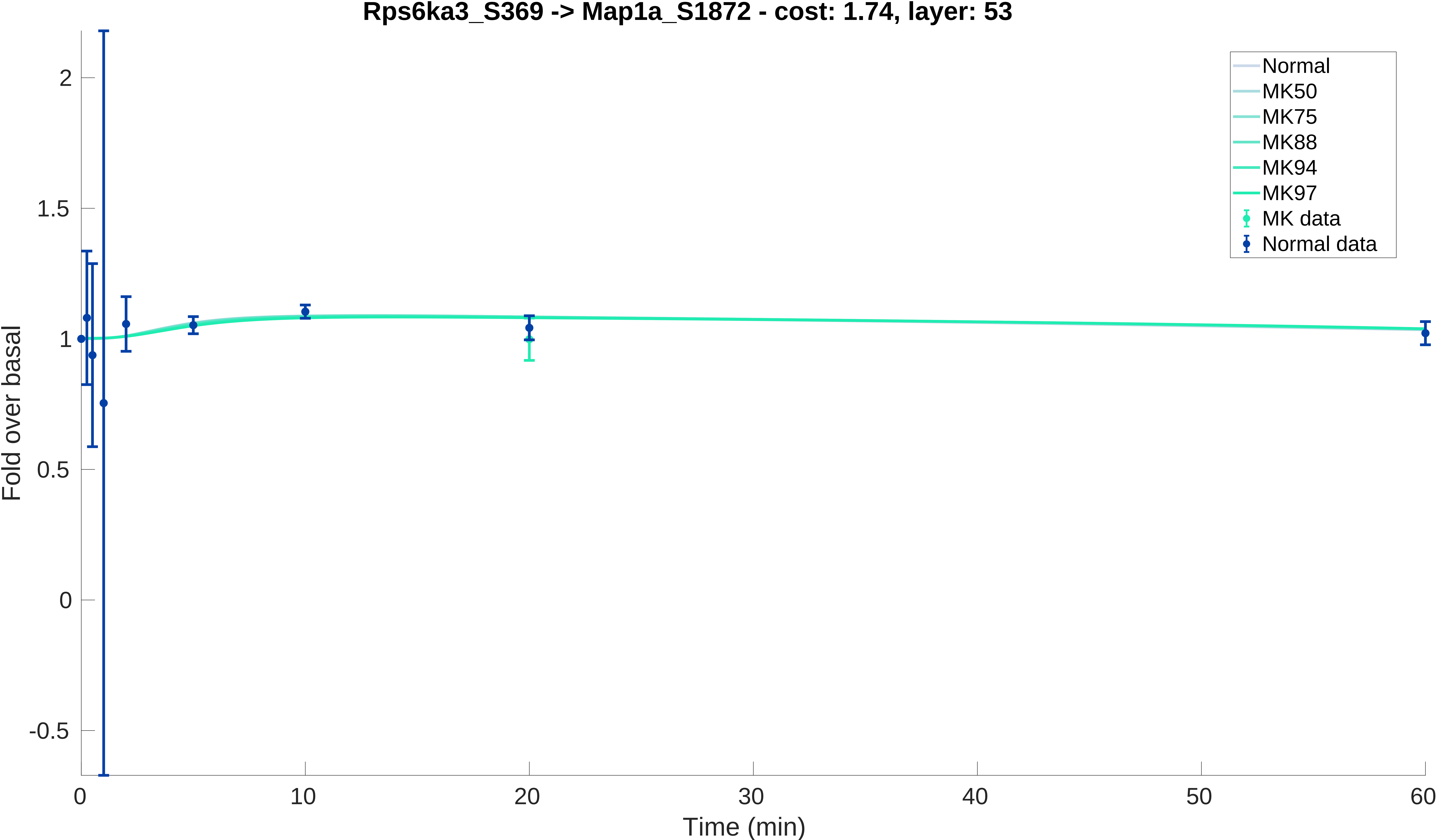
Rps6ka3_S369 -> Map1a_S1872 - cost: 1.74, layer: 53
Normal
MK50
MK75
MK88
MK94
MK97
MK data
Normal data
Fold over basal
Time (min)
2
1.5
1
0.5
0
-0.5
0
10
20
30
40
50
60

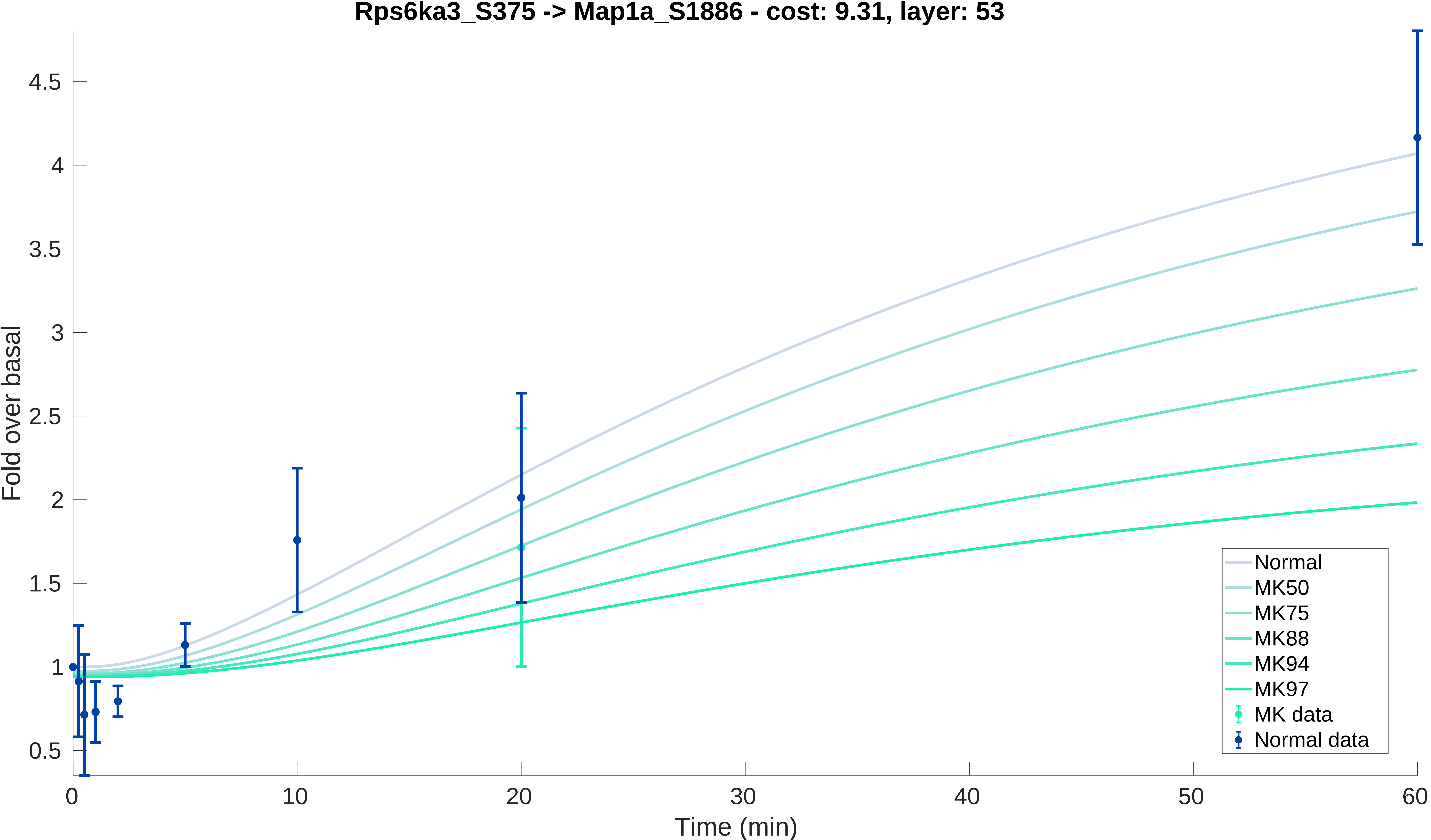

Rps6ka3_S375 -> Map1a_S1886 - cost: 9.31, layer: 53
Fold over basal
Time (min)
0.5
1
1.5
2
2.5
3
3.5
4
4.5
0
10
20
30
40
50
60
Normal
MK50
MK75
MK88
MK94
MK97
MK data
Normal data

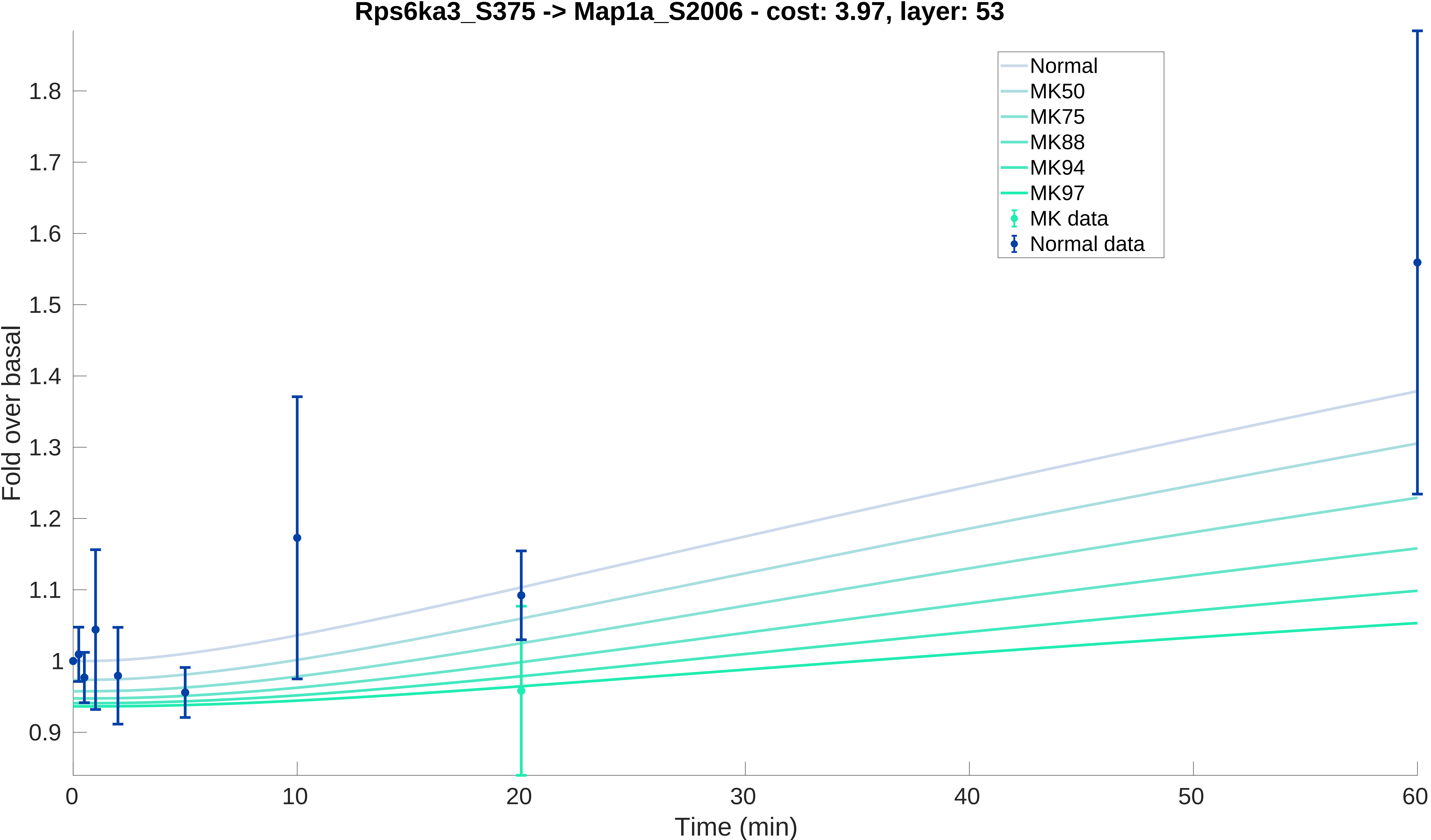
Rps6ka3_S375 -> Map1a_S2006 - cost: 3.97, layer: 53
Fold over basal
Time (min)
Normal
MK50
MK75
MK88
MK94
MK97
MK data
Normal data
0.9
1
1.1
1.2
1.3
1.4
1.5
1.6
1.7
1.8
0
10
20
30
40
50
60

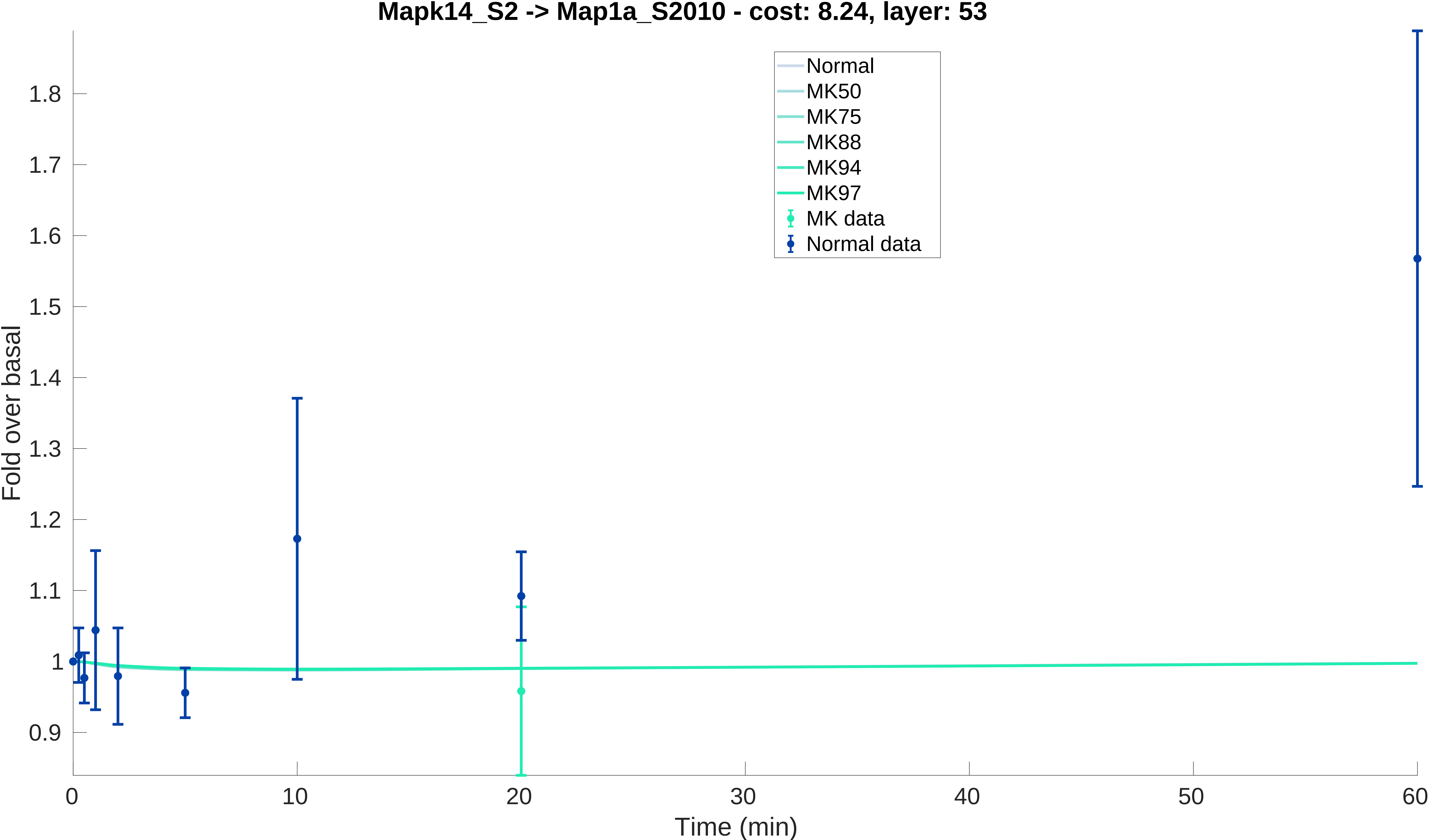
Mapk14_S2 -> Map1a_S2010 - cost: 8.24, layer: 53
Normal
MK50
MK75
MK88
MK94
MK97
MK data
Normal data
Fold over basal
1.8
1.7
1.6
1.5
1.4
1.3
1.2
1.1
1
0.9
0
10
20
30
40
50
60
Time (min)

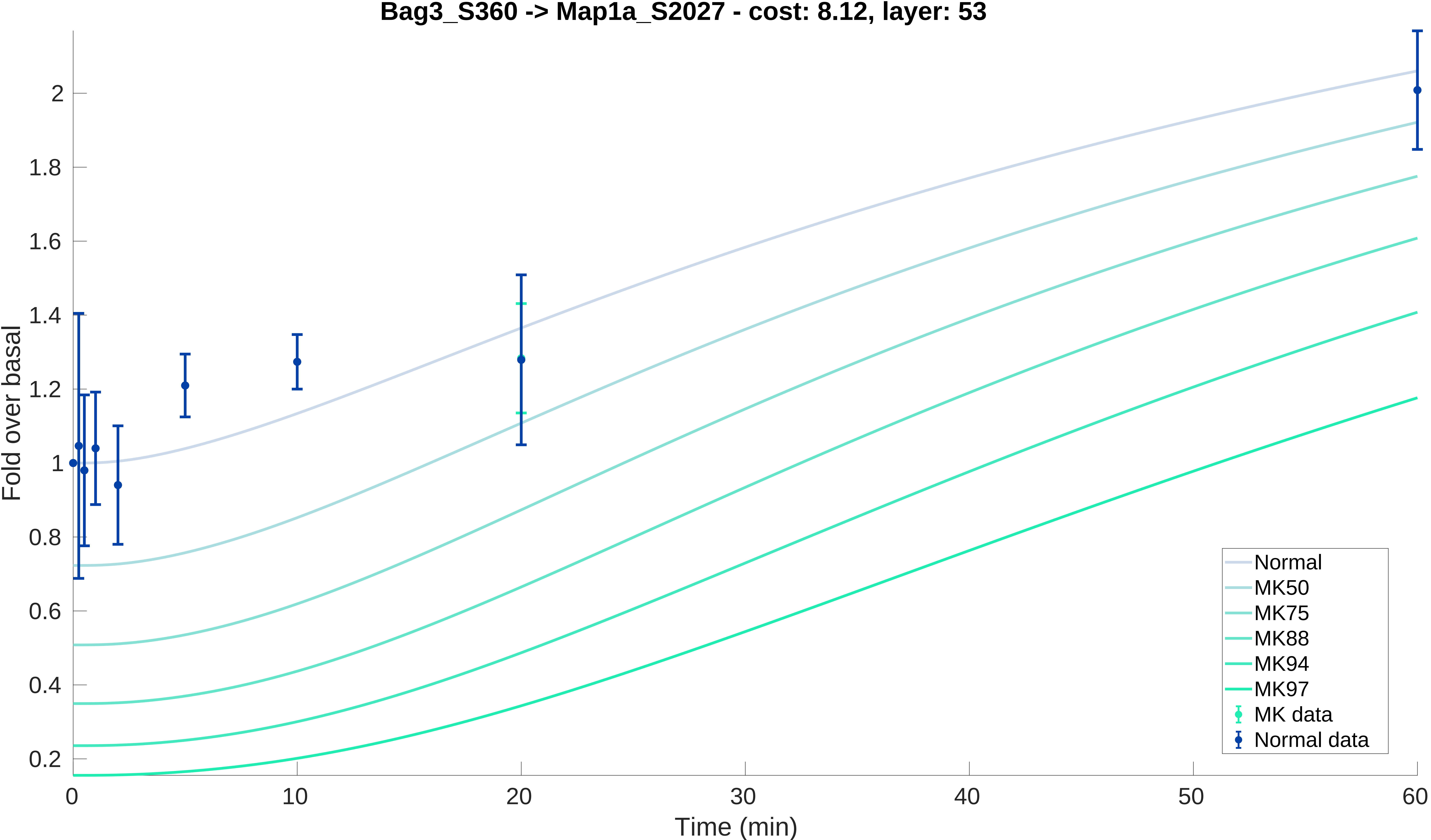

Bag3_S360 -> Map1a_S2027 - cost: 8.12, layer: 53
Fold over basal
Time (min)
0.2
0.4
0.6
0.8
1
1.2
1.4
1.6
1.8
2
0
10
20
30
40
50
60
Normal
MK50
MK75
MK88
MK94
MK97
MK data
Normal data

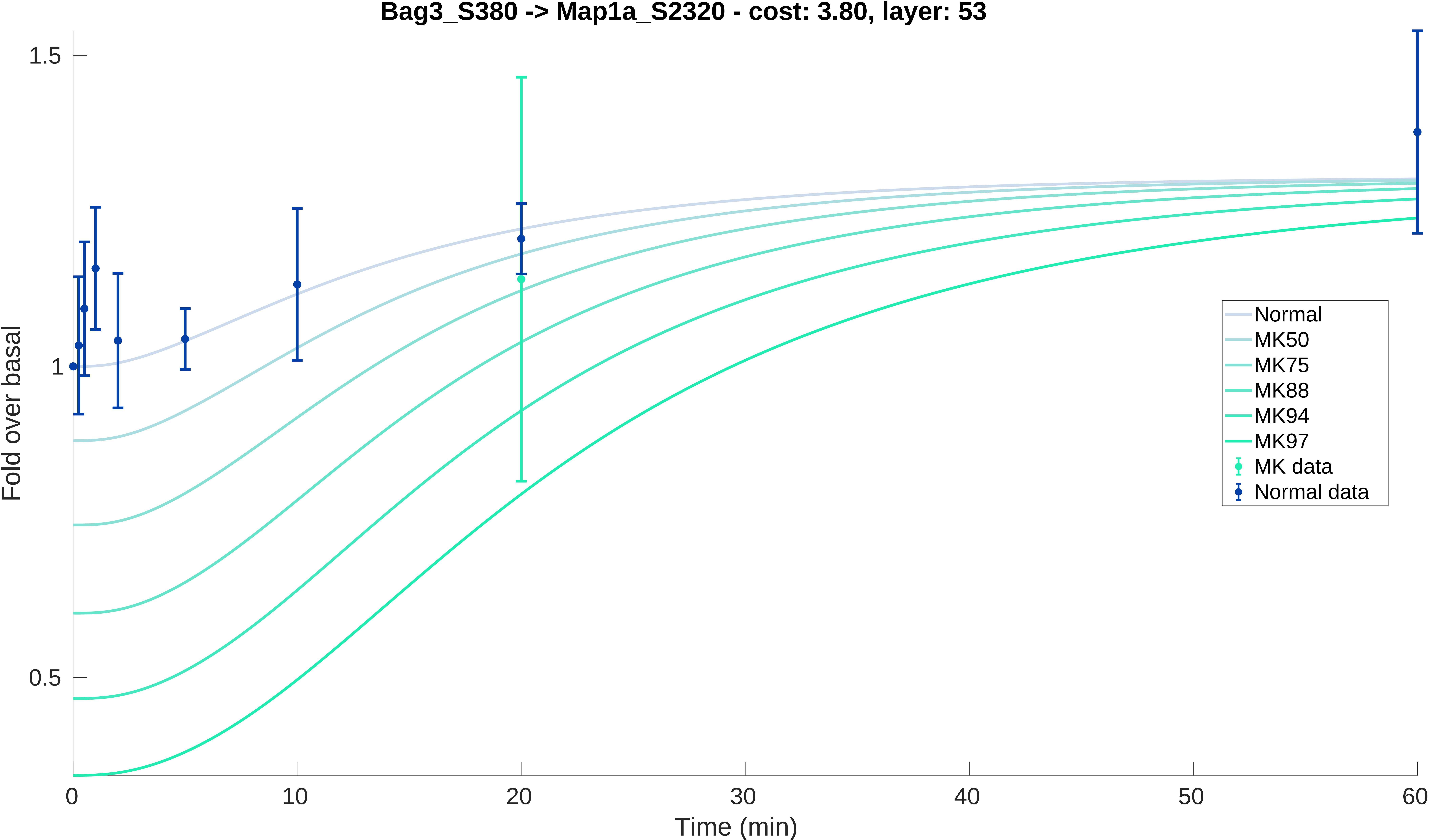

Bag3_S380 -> Map1a_S2320 - cost: 3.80, layer: 53
1.5
1
0.5
Fold over basal
0
10
20
30
40
50
60
Time (min)
Normal
MK50
MK75
MK88
MK94
MK97
MK data
Normal data

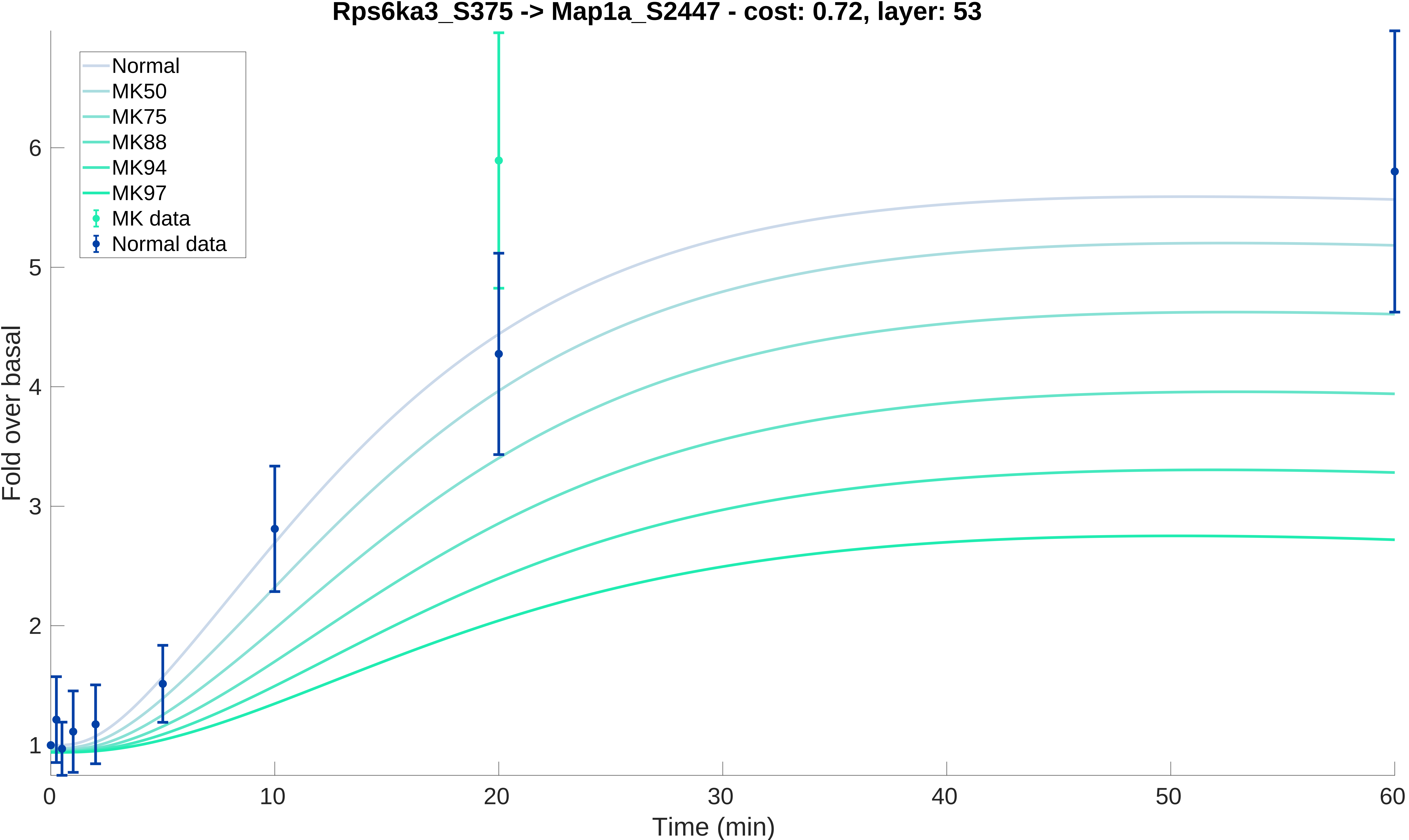
Rps6ka3_S375 -> Map1a_S2447 - cost: 0.72, layer: 53
Normal
MK50
MK75
MK88
MK94
MK97
MK data
Normal data
Fold over basal
Time (min)
0
10
20
30
40
50
60
1
2
3
4
5
6

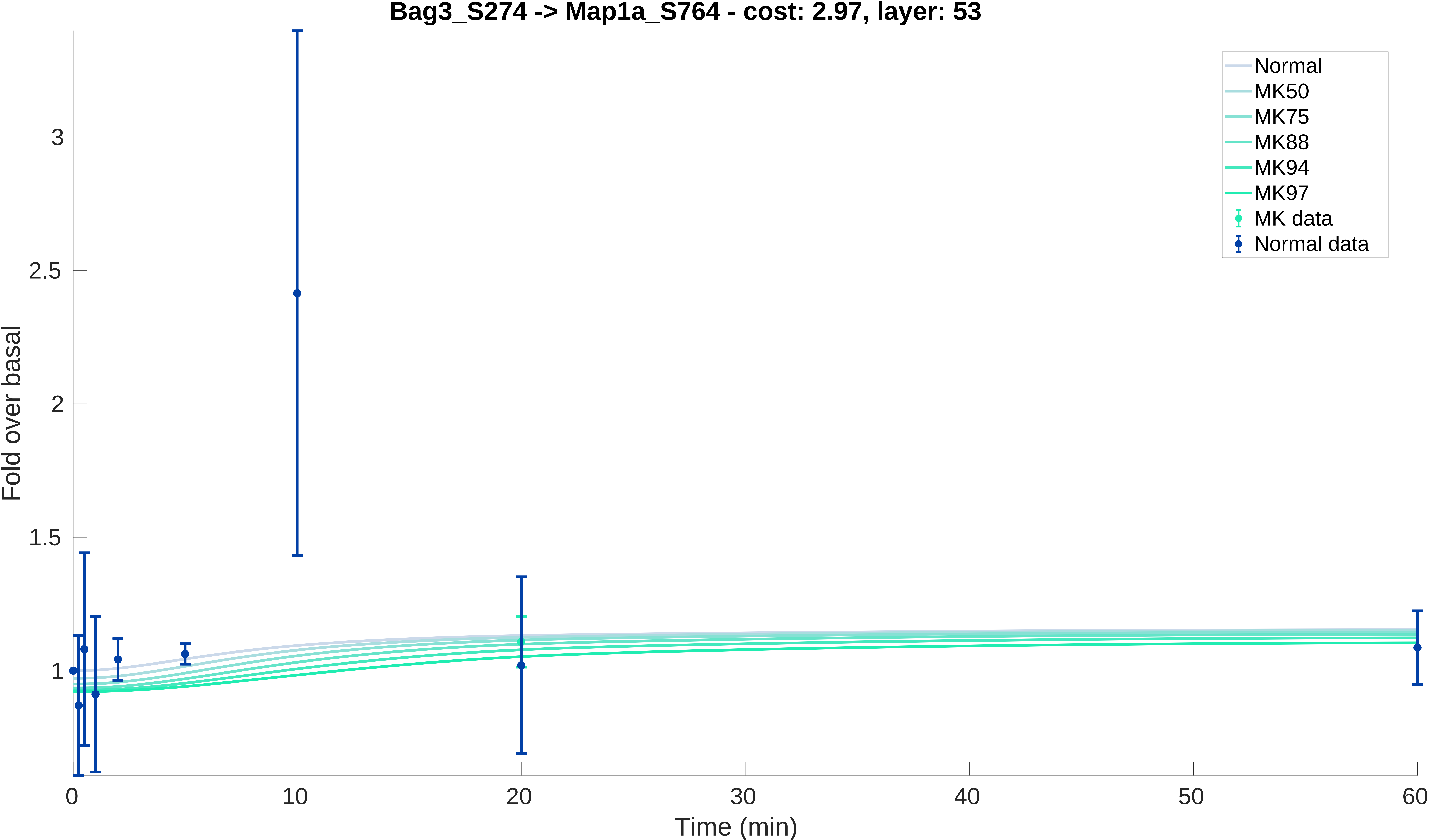

Bag3_S274 -> Map1a_S764 - cost: 2.97, layer: 53
Fold over basal
Time (min)
Normal
MK50
MK75
MK88
MK94
MK97
MK data
Normal data
0
10
20
30
40
50
60
1
1.5
2
2.5
3

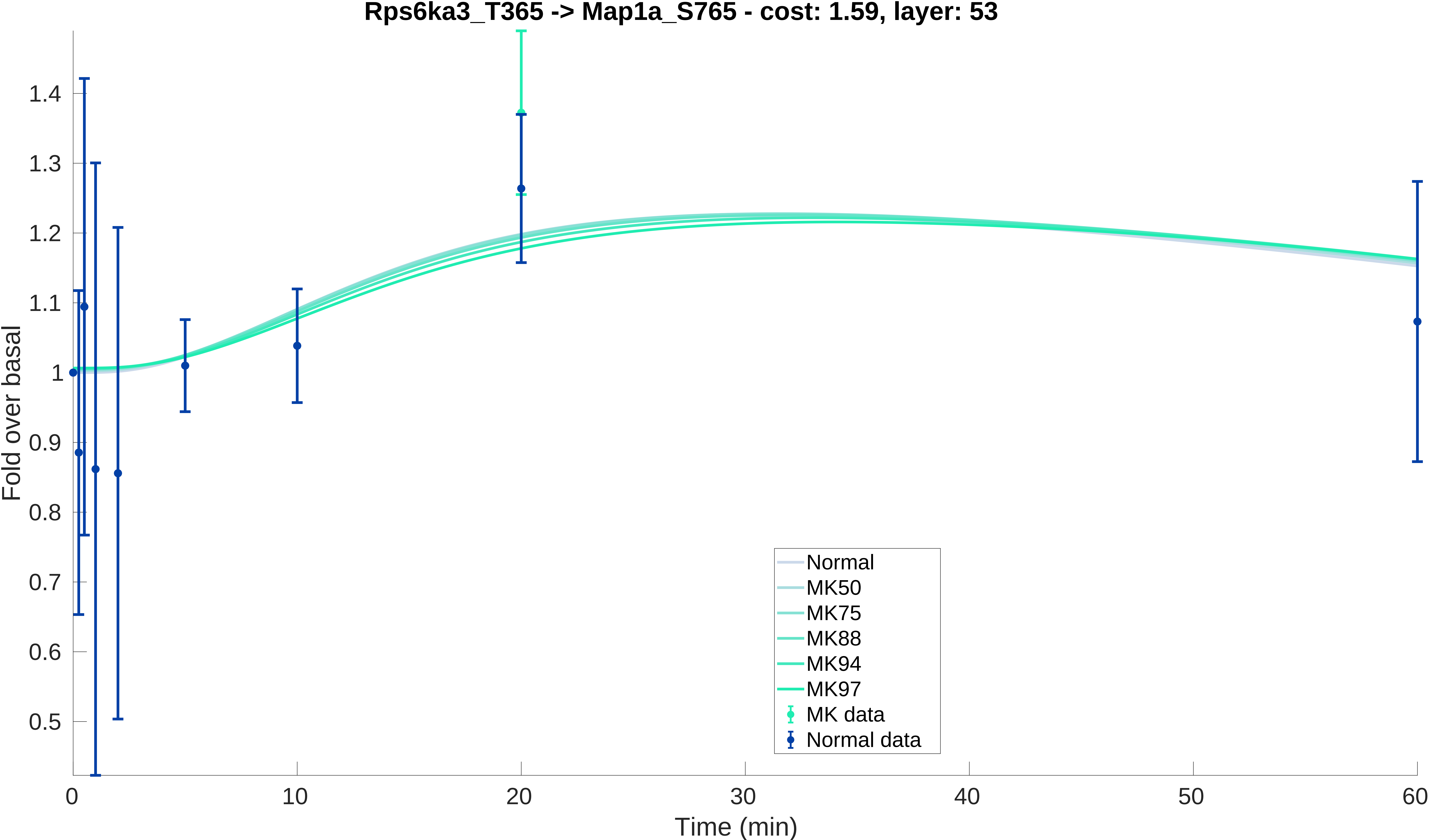
Rps6ka3_T365 -> Map1a_S765 - cost: 1.59, layer: 53
Fold over basal
1.4
1.3
1.2
1.1
1
0.9
0.8
0.7
0.6
0.5
0
10
20
30
40
50
60
Time (min)
Normal
MK50
MK75
MK88
MK94
MK97
MK data
Normal data

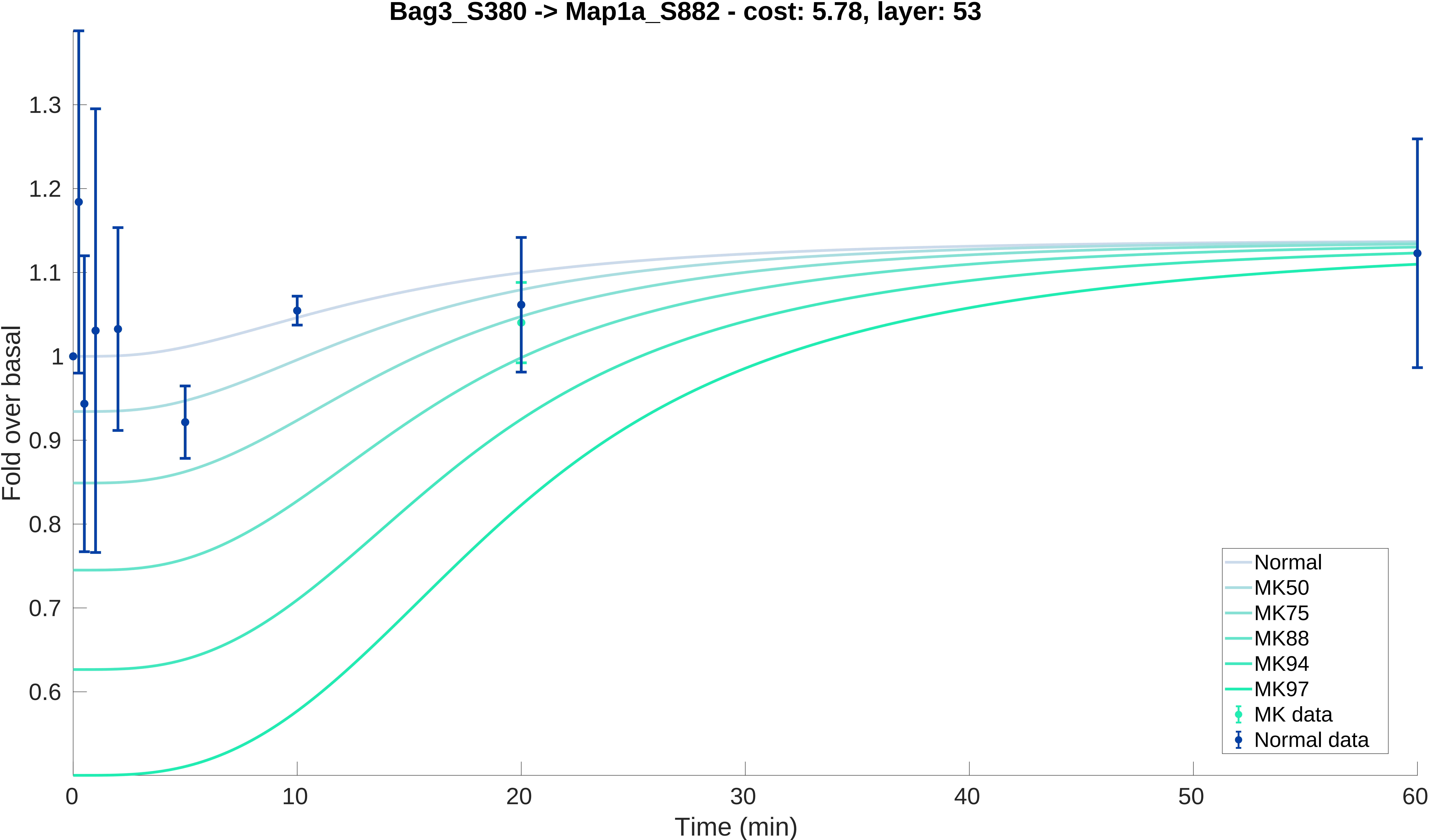
Bag3_S380 -> Map1a_S882 - cost: 5.78, layer: 53
Fold over basal
Time (min)
1.3
1.2
1.1
1
0.9
0.8
0.7
0.6
0
10
20
30
40
50
60
Normal
MK50
MK75
MK88
MK94
MK97
MK data
Normal data

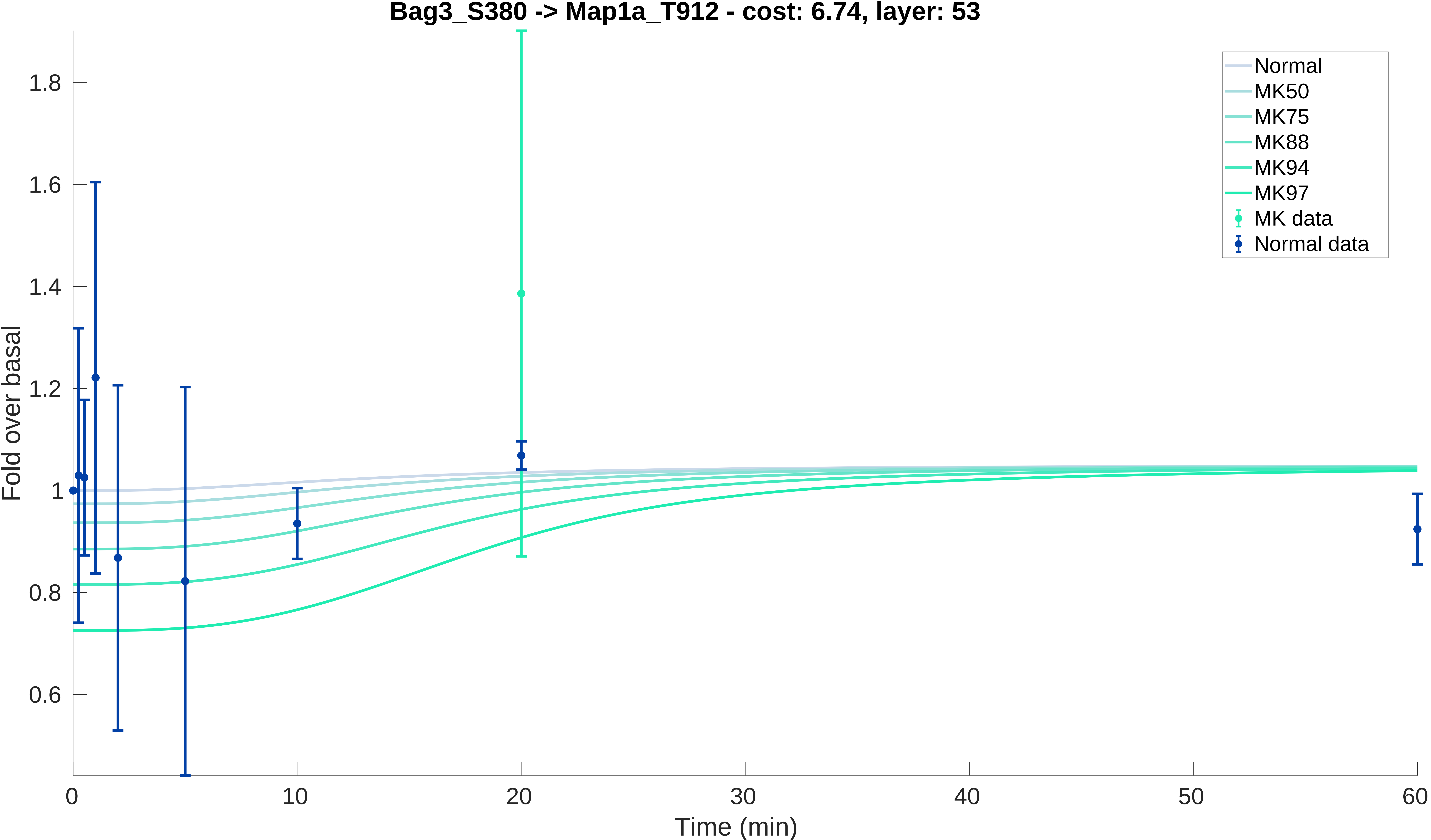

Bag3_S380 -> Map1a_T912 - cost: 6.74, layer: 53
Normal
MK50
MK75
MK88
MK94
MK97
MK data
Normal data
Fold over basal
1.8
1.6
1.4
1.2
1
0.8
0.6
0
10
20
30
40
50
60
Time (min)

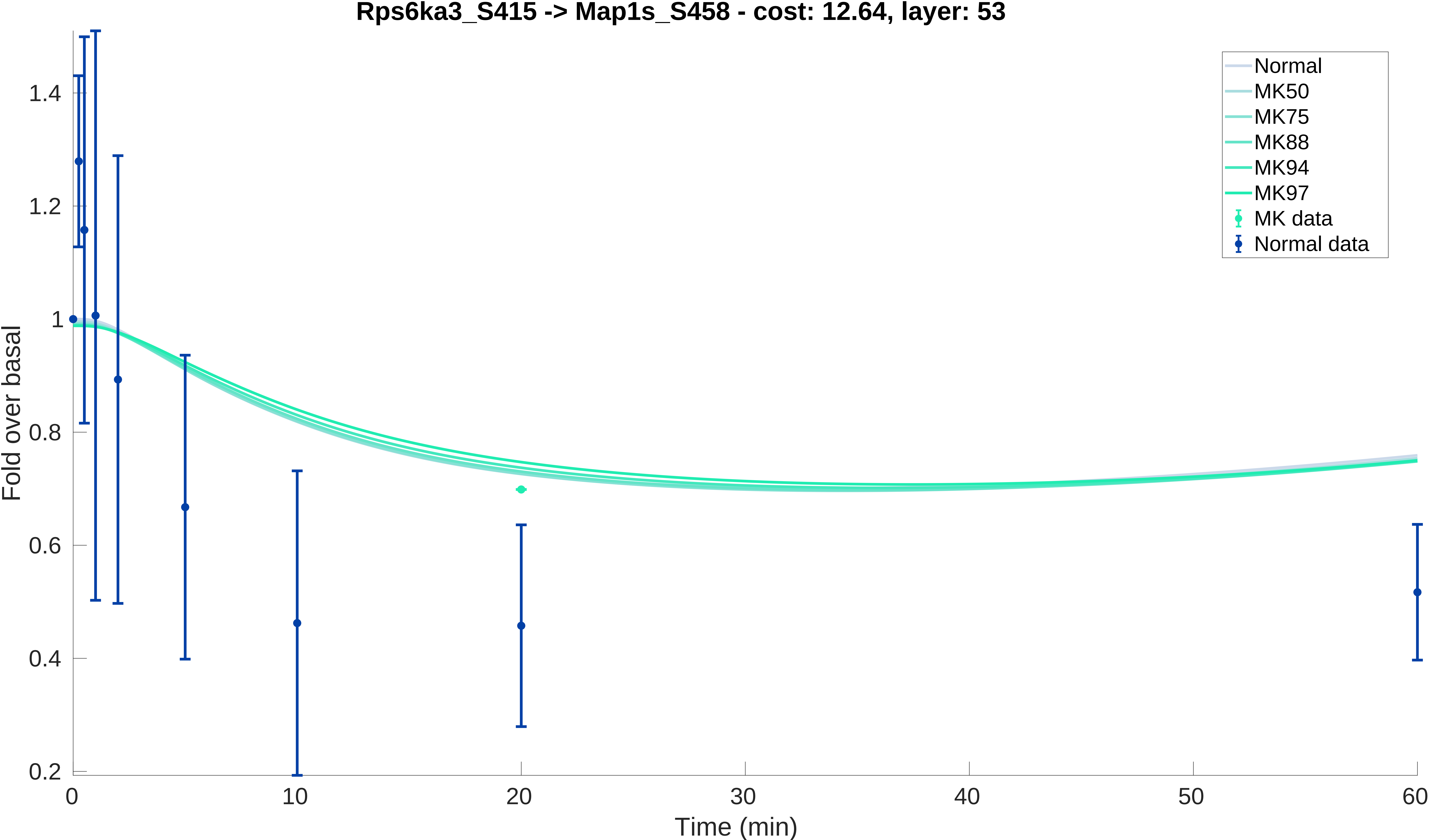

Rps6ka3_S415 -> Map1s_S458 - cost: 12.64, layer: 53
Fold over basal
Time (min)
0.2
0.4
0.6
0.8
1
1.2
1.4
0
10
20
30
40
50
60
Normal
MK50
MK75
MK88
MK94
MK97
MK data
Normal data

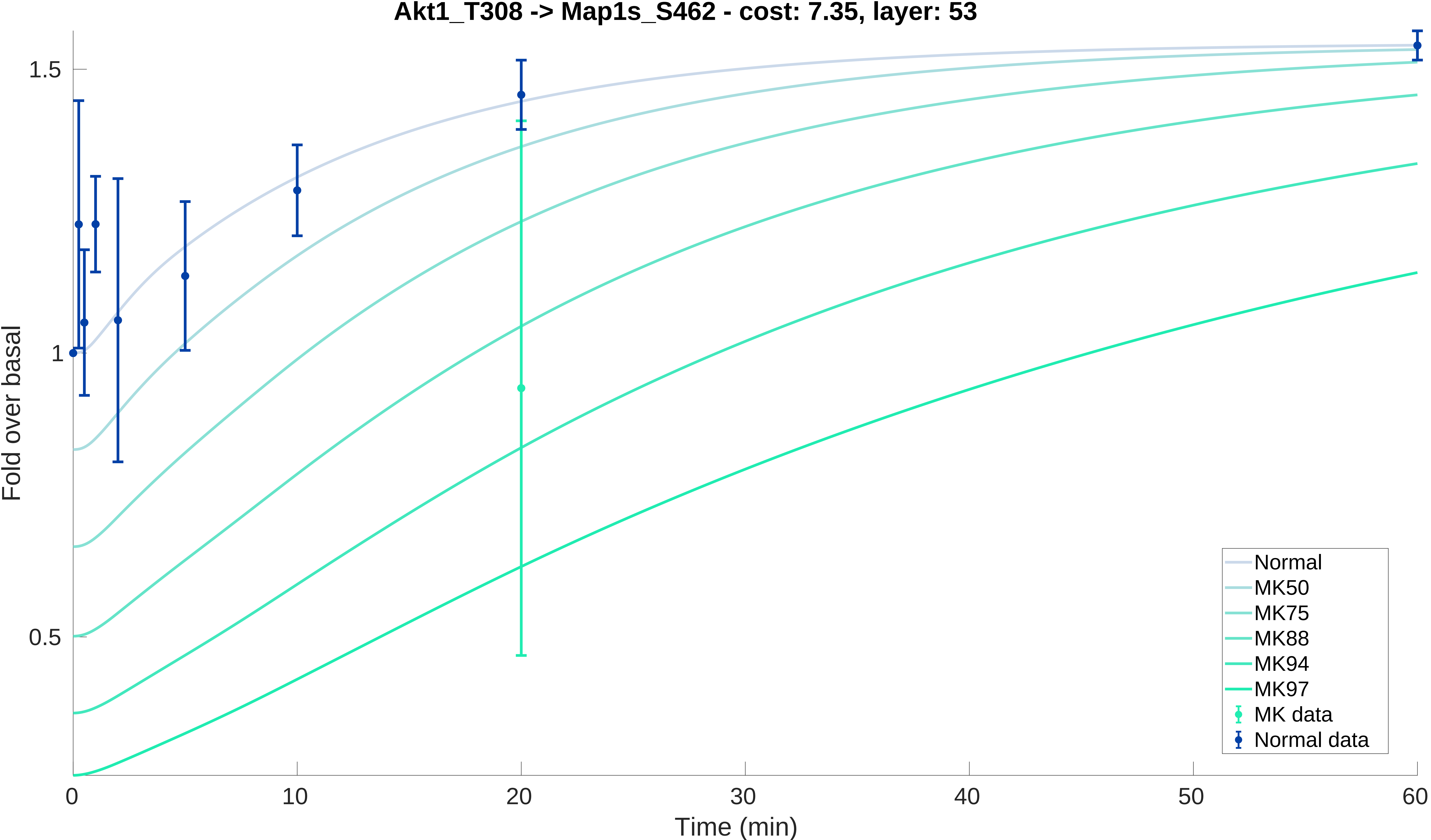

Akt1_T308 -> Map1s_S462 - cost: 7.35, layer: 53
Fold over basal
1.5
1
0.5
0
10
20
30
40
50
60
Time (min)
Normal
MK50
MK75
MK88
MK94
MK97
MK data
Normal data

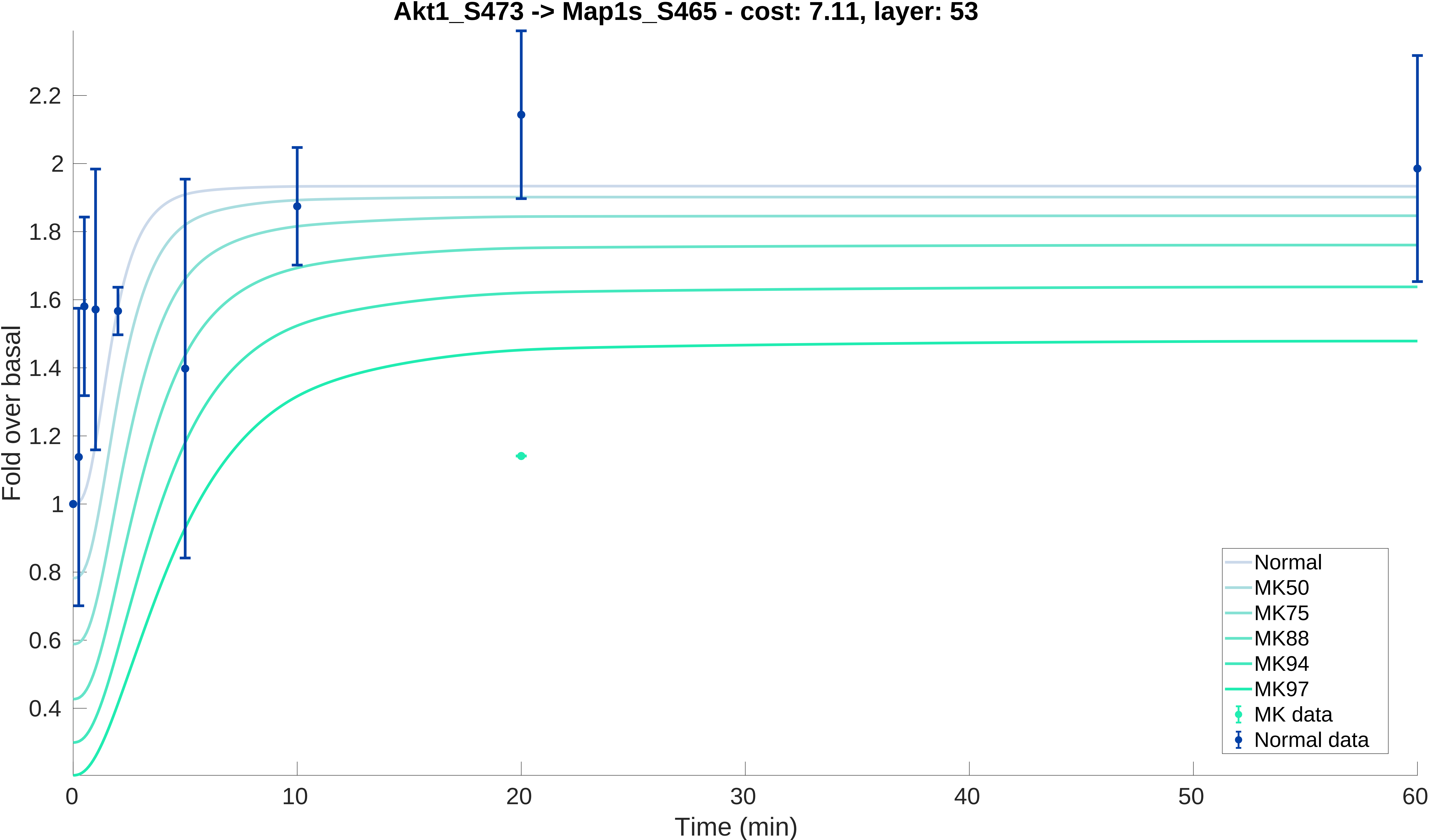

Akt1_S473 -> Map1s_S465 - cost: 7.11, layer: 53
Fold over basal
Time (min)
0
10
20
30
40
50
60
0.4
0.6
0.8
1
1.2
1.4
1.6
1.8
2
2.2
Normal
MK50
MK75
MK88
MK94
MK97
MK data
Normal data

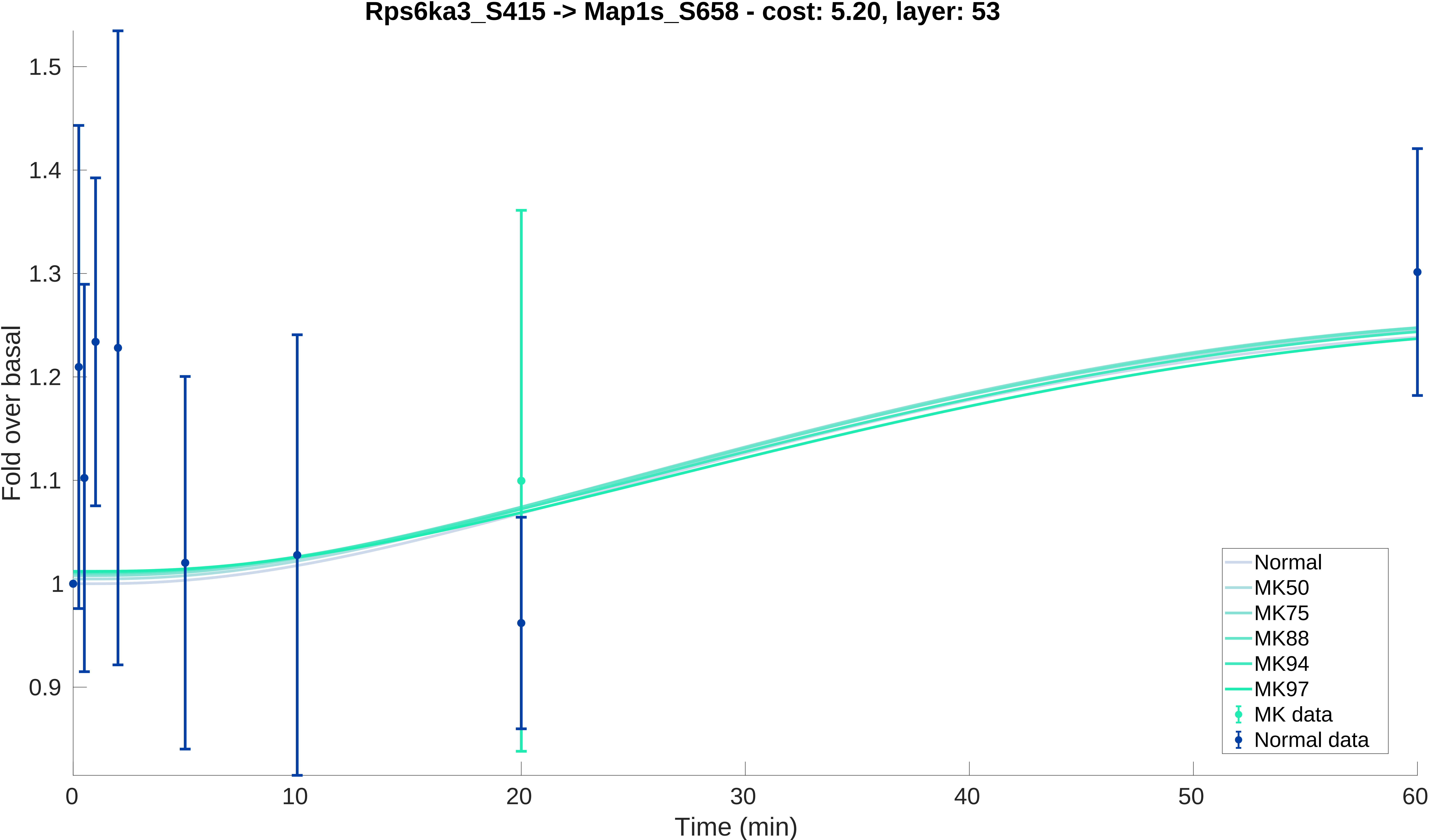

Rps6ka3_S415 -> Map1s_S658 - cost: 5.20, layer: 53
Fold over basal
Time (min)
0.9
1
1.1
1.2
1.3
1.4
1.5
0
10
20
30
40
50
60
Normal
MK50
MK75
MK88
MK94
MK97
MK data
Normal data

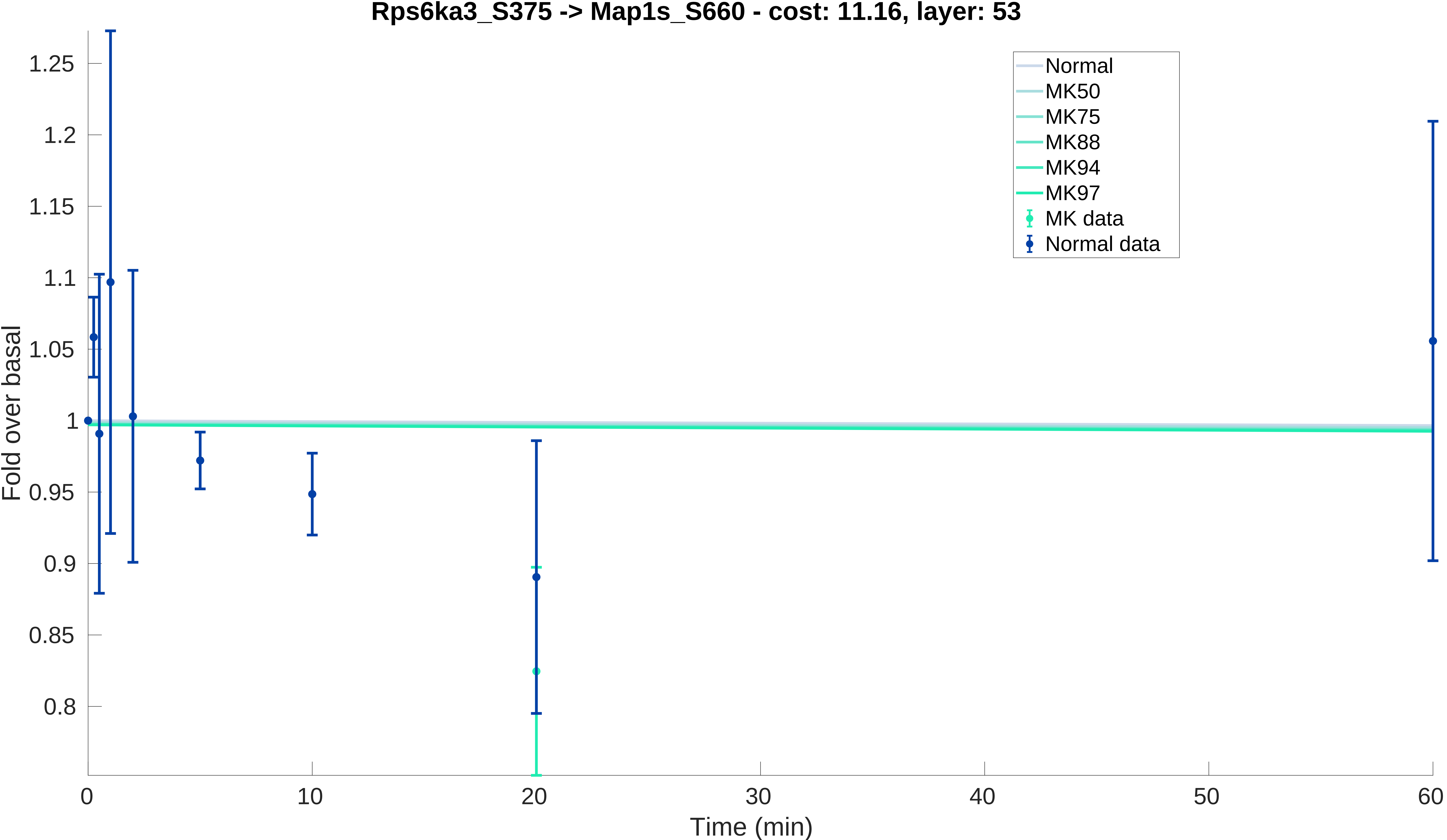

Rps6ka3_S375 -> Map1s_S660 - cost: 11.16, layer: 53
Normal
MK50
MK75
MK88
MK94
MK97
MK data
Normal data
Fold over basal
1.25
1.2
1.15
1.1
1.05
1
0.95
0.9
0.85
0.8
0
10
20
30
40
50
60
Time (min)

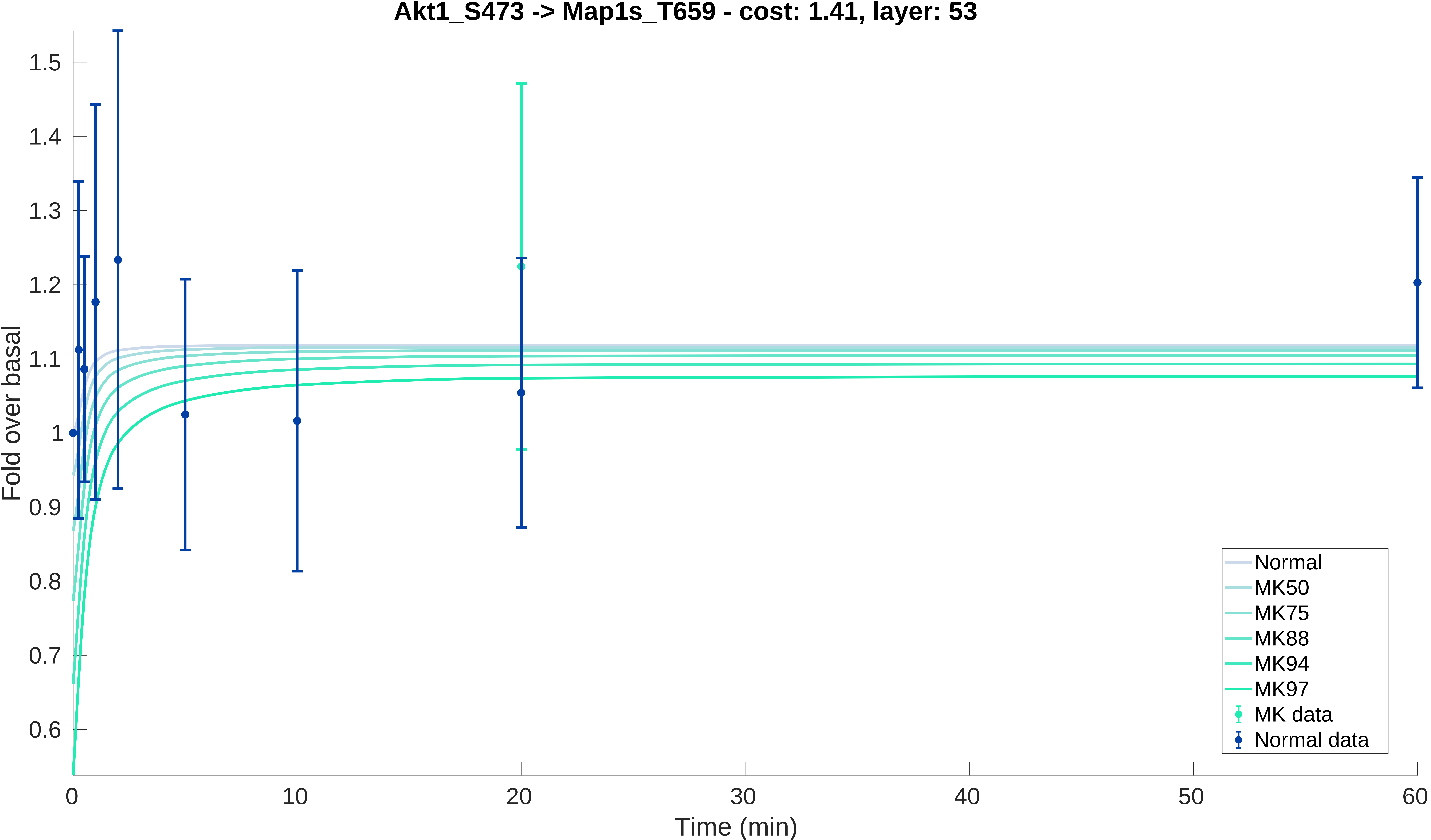

Akt1_S473 -> Map1s_T659 - cost: 1.41, layer: 53
Fold over basal
1.5
1.4
1.3
1.2
1.1
1
0.9
0.8
0.7
0.6
0
10
20
30
40
50
60
Time (min)
Normal
MK50
MK75
MK88
MK94
MK97
MK data
Normal data

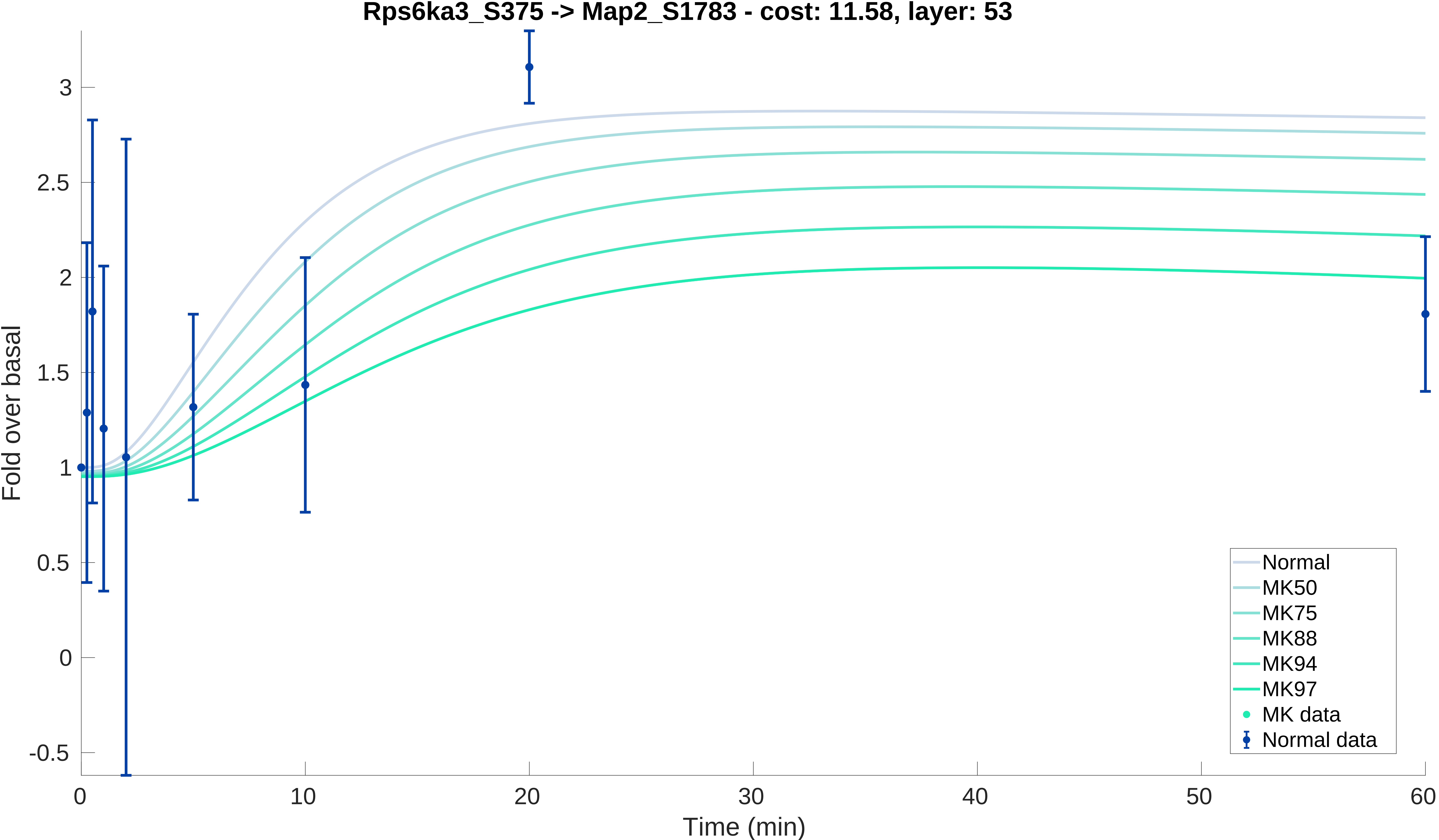

Rps6ka3_S375 -> Map2_S1783 - cost: 11.58, layer: 53
Fold over basal
Time (min)
3
2.5
2
1.5
1
0.5
0
-0.5
0
10
20
30
40
50
60
Normal
MK50
MK75
MK88
MK94
MK97
MK data
Normal data

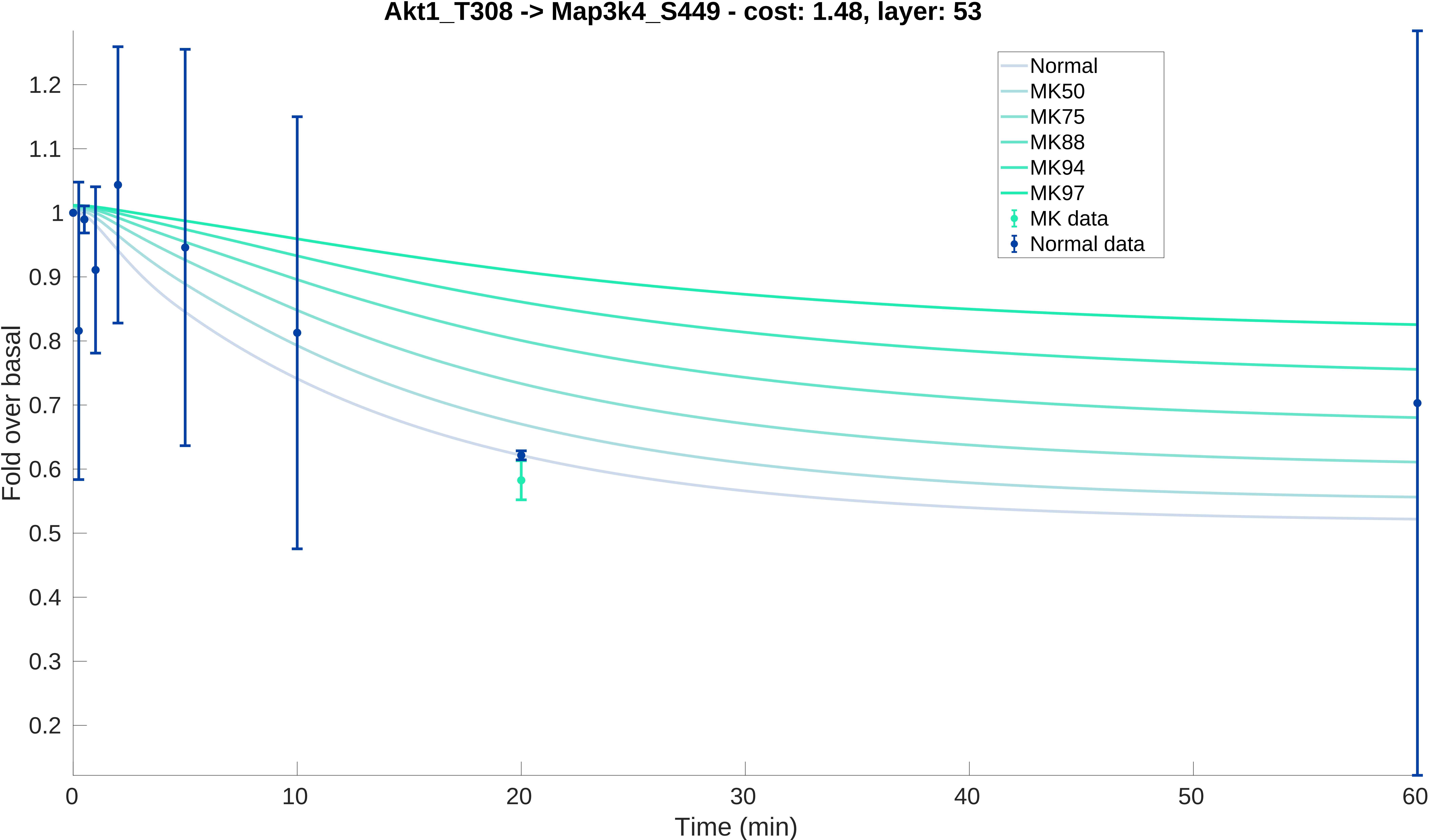

Akt1_T308 -> Map3k4_S449 - cost: 1.48, layer: 53
Fold over basal
Time (min)
0
10
20
30
40
50
60
0.2
0.3
0.4
0.5
0.6
0.7
0.8
0.9
1
1.1
1.2
Normal
MK50
MK75
MK88
MK94
MK97
MK data
Normal data

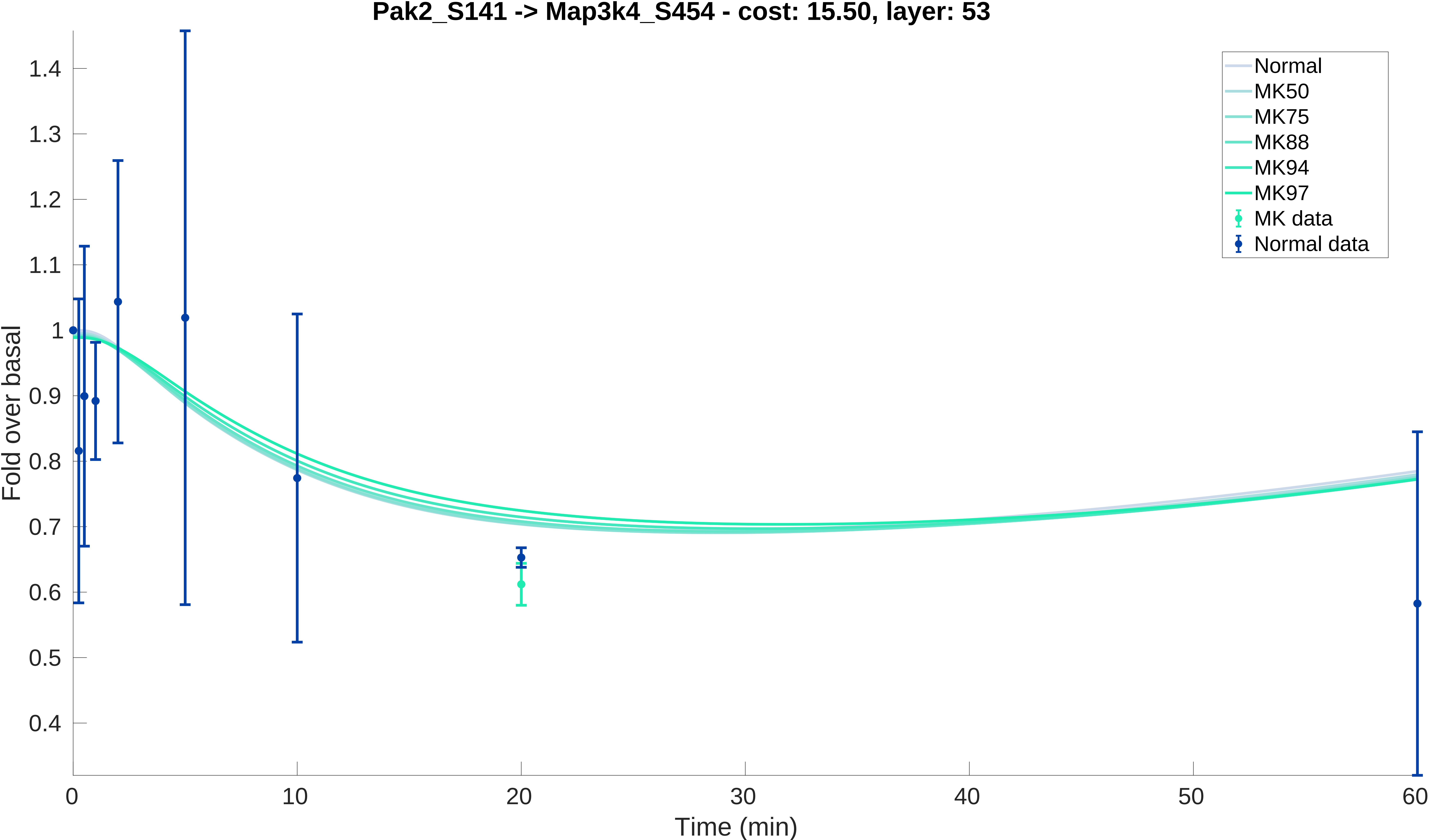

Pak2_S141 -> Map3k4_S454 - cost: 15.50, layer: 53
Fold over basal
Time (min)
0.4
0.5
0.6
0.7
0.8
0.9
1
1.1
1.2
1.3
1.4
0
10
20
30
40
50
60
Normal
MK50
MK75
MK88
MK94
MK97
MK data
Normal data

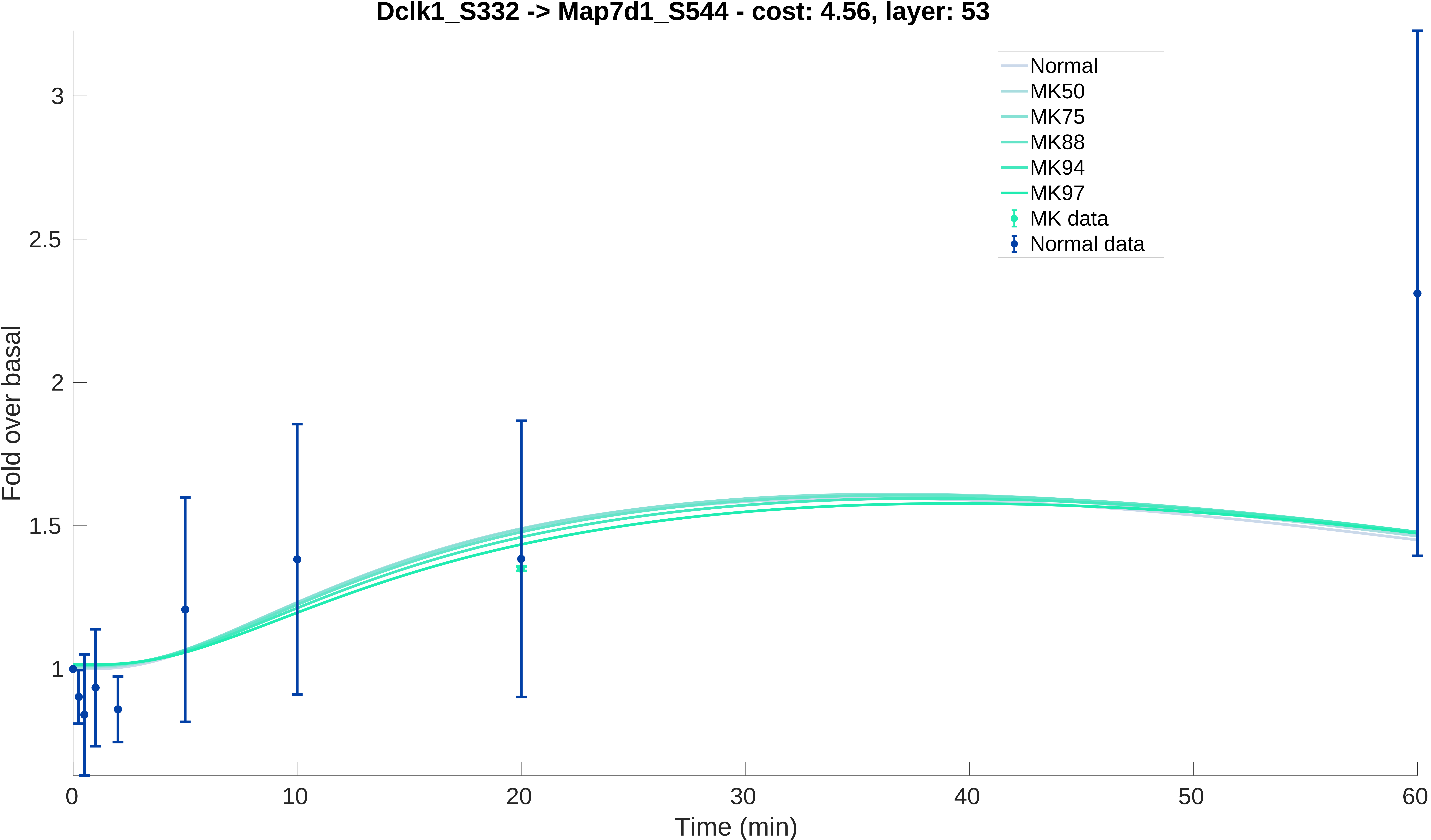
Dclk1_S332 -> Map7d1_S544 - cost: 4.56, layer: 53
Fold over basal
Time (min)
0
10
20
30
40
50
60
1
1.5
2
2.5
3
Normal
MK50
MK75
MK88
MK94
MK97
MK data
Normal data

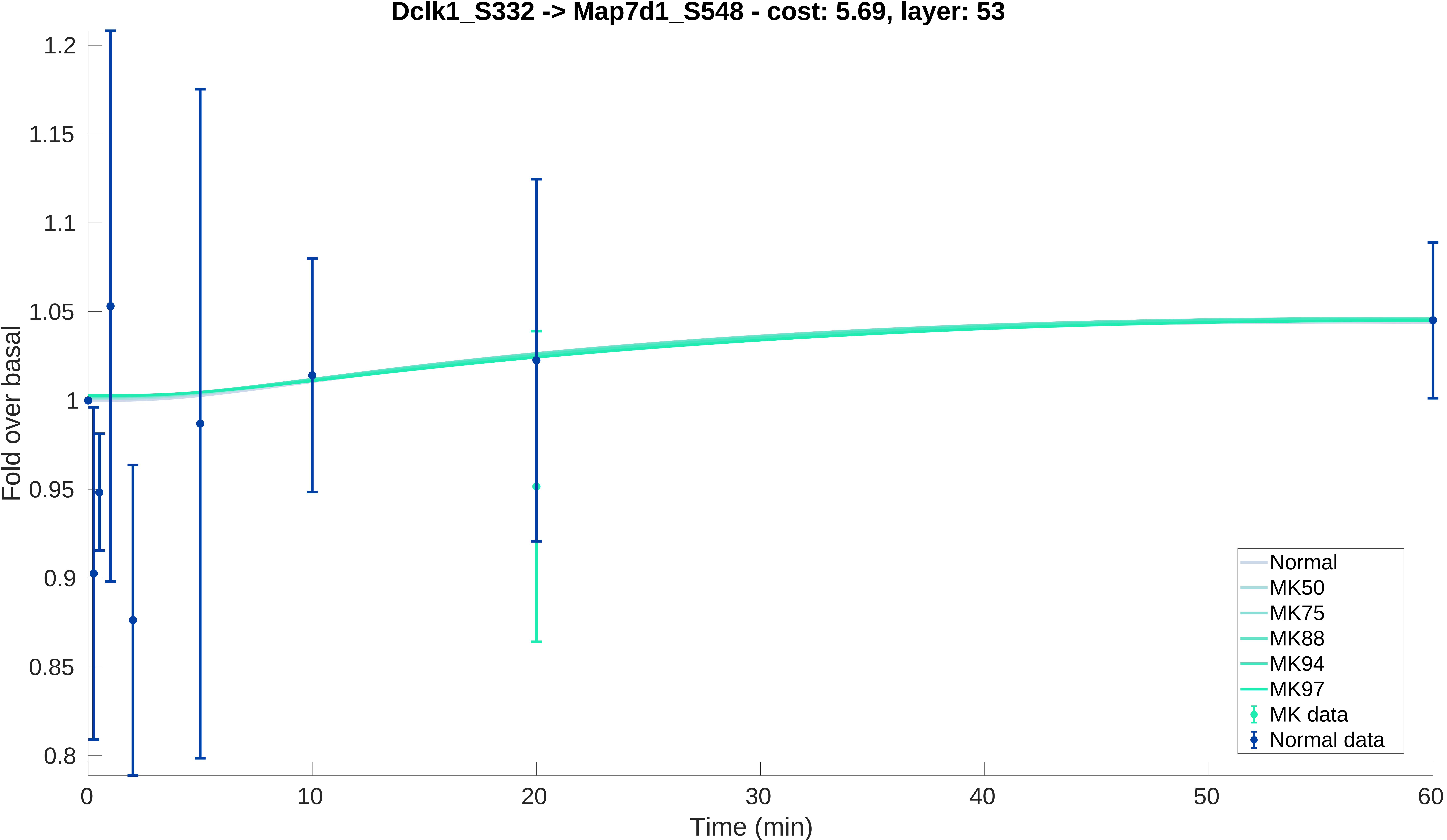

Dclk1_S332 -> Map7d1_S548 - cost: 5.69, layer: 53
Fold over basal
Time (min)
1.2
1.15
1.1
1.05
1
0.95
0.9
0.85
0.8
0
10
20
30
40
50
60
Normal
MK50
MK75
MK88
MK94
MK97
MK data
Normal data

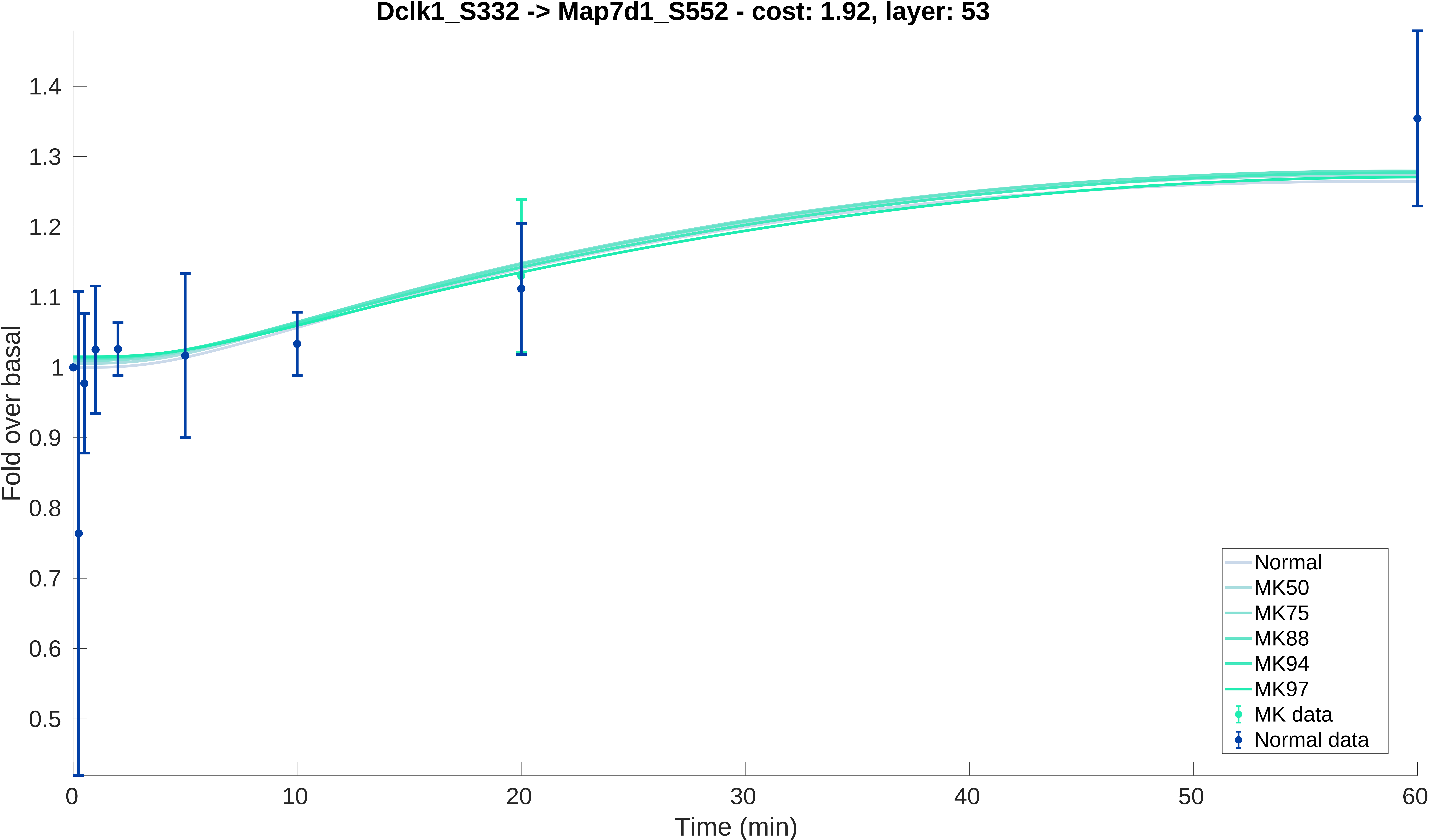

Dclk1_S332 -> Map7d1_S552 - cost: 1.92, layer: 53
Fold over basal
Time (min)
0.5
0.6
0.7
0.8
0.9
1
1.1
1.2
1.3
1.4
0
10
20
30
40
50
60
Normal
MK50
MK75
MK88
MK94
MK97
MK data
Normal data

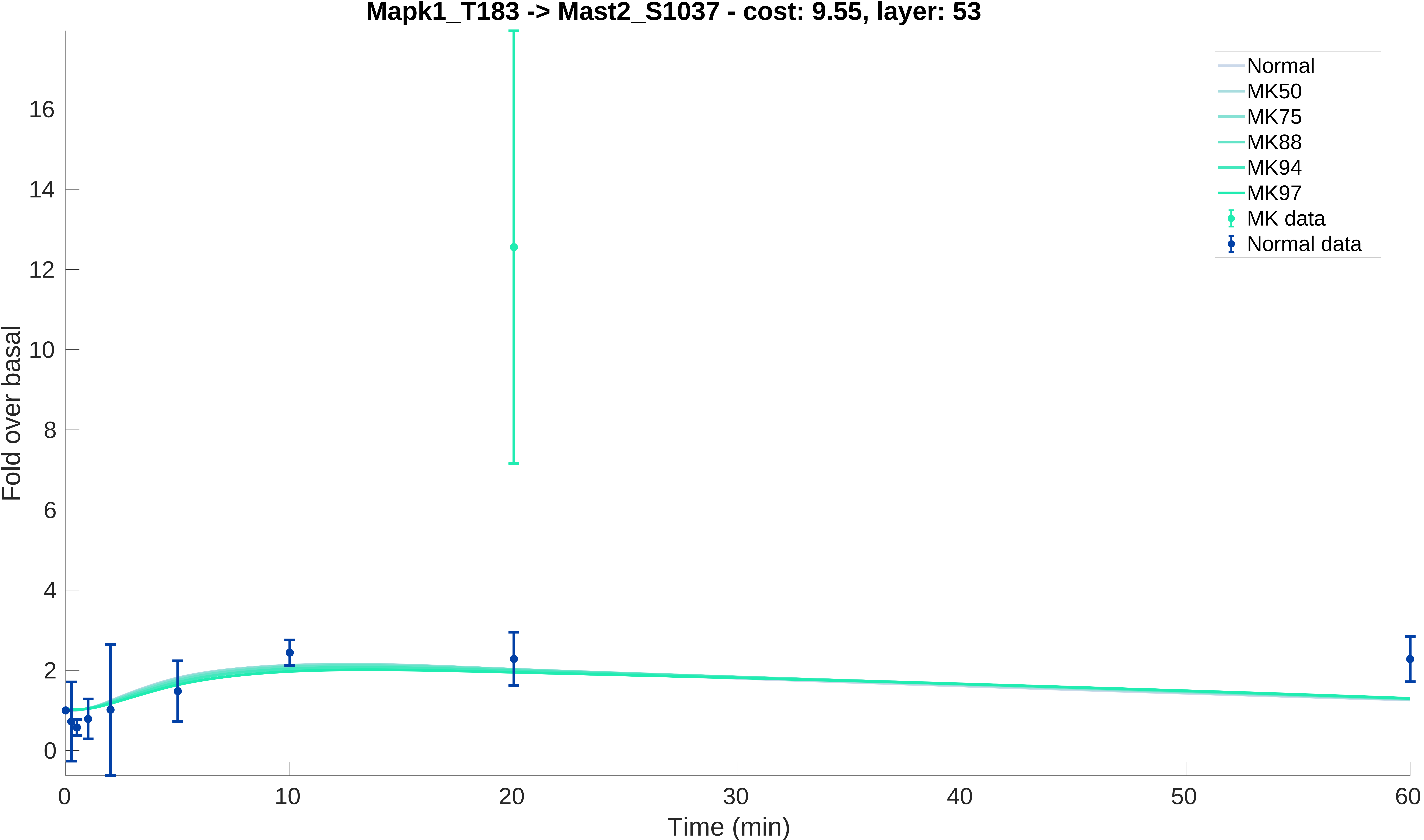
Mapk1_T183 -> Mast2_S1037 - cost: 9.55, layer: 53
Fold over basal
Time (min)
0
2
4
6
8
10
12
14
16
0
10
20
30
40
50
60
Normal
MK50
MK75
MK88
MK94
MK97
MK data
Normal data

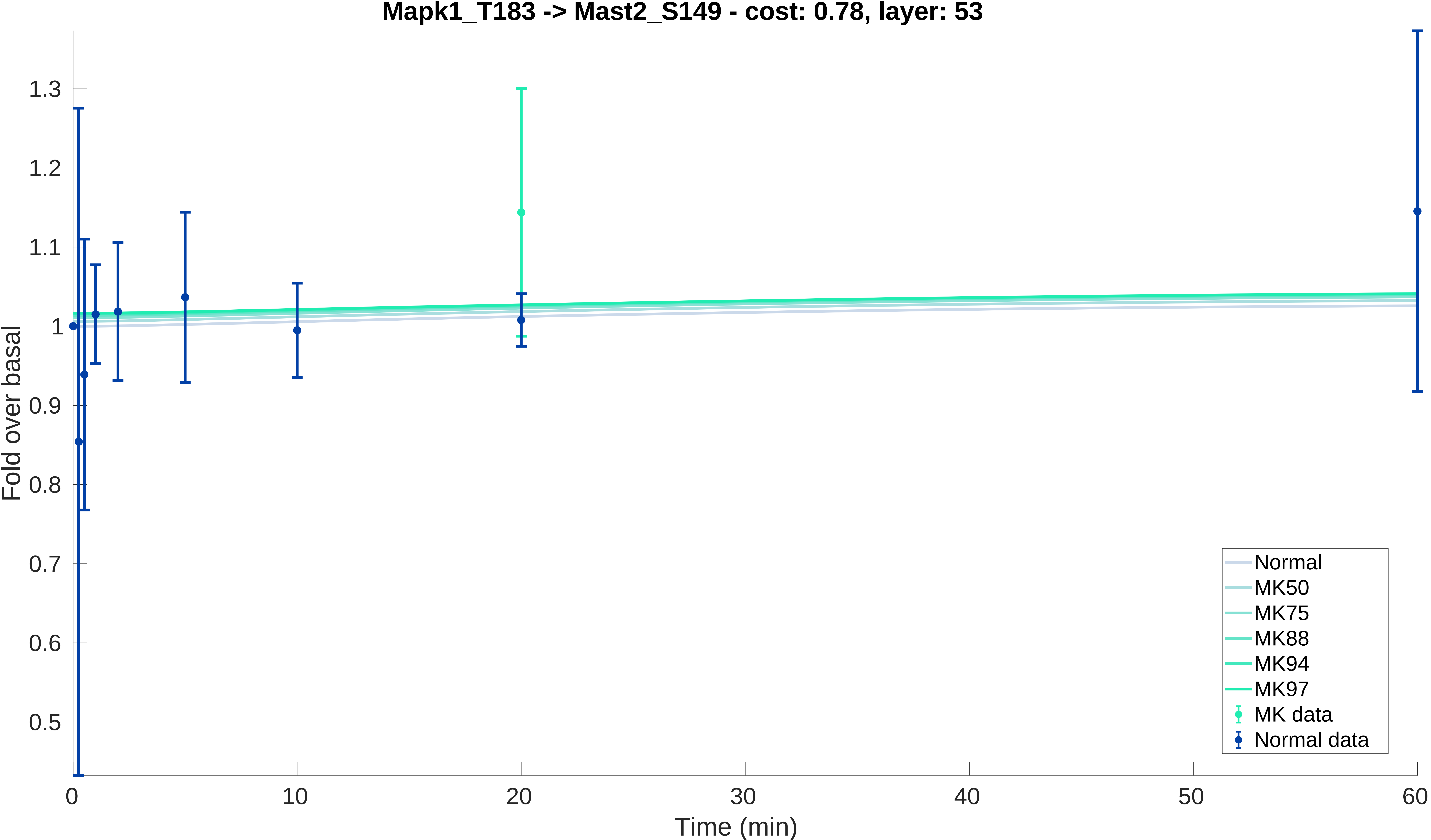

Mapk1_T183 -> Mast2_S149 - cost: 0.78, layer: 53
Fold over basal
Time (min)
1.3
1.2
1.1
1
0.9
0.8
0.7
0.6
0.5
0
10
20
30
40
50
60
Normal
MK50
MK75
MK88
MK94
MK97
MK data
Normal data

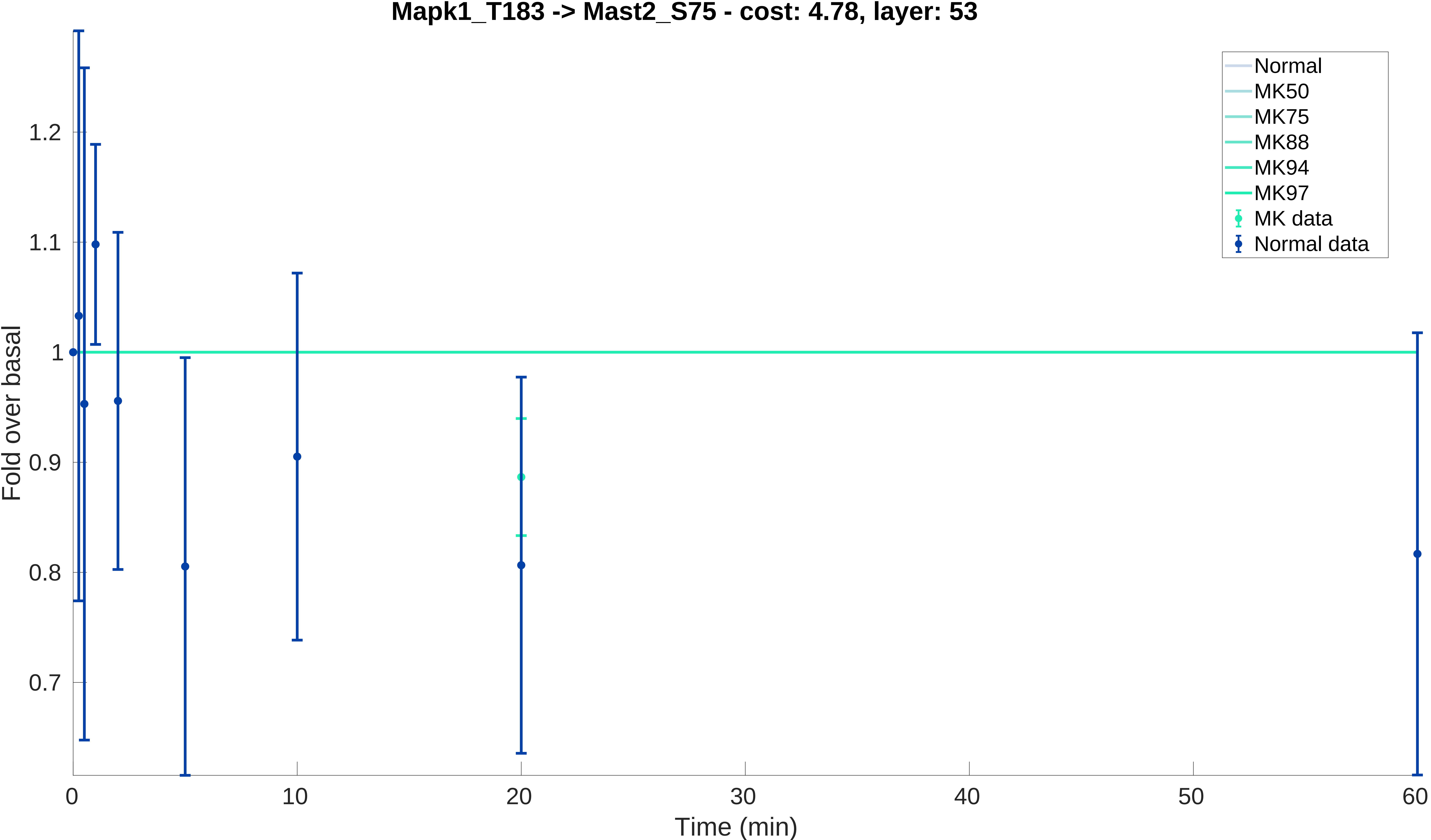
Mapk1_T183 -> Mast2_S75 - cost: 4.78, layer: 53
Normal
MK50
MK75
MK88
MK94
MK97
MK data
Normal data
Fold over basal
Time (min)
0.7
0.8
0.9
1
1.1
1.2
0
10
20
30
40
50
60

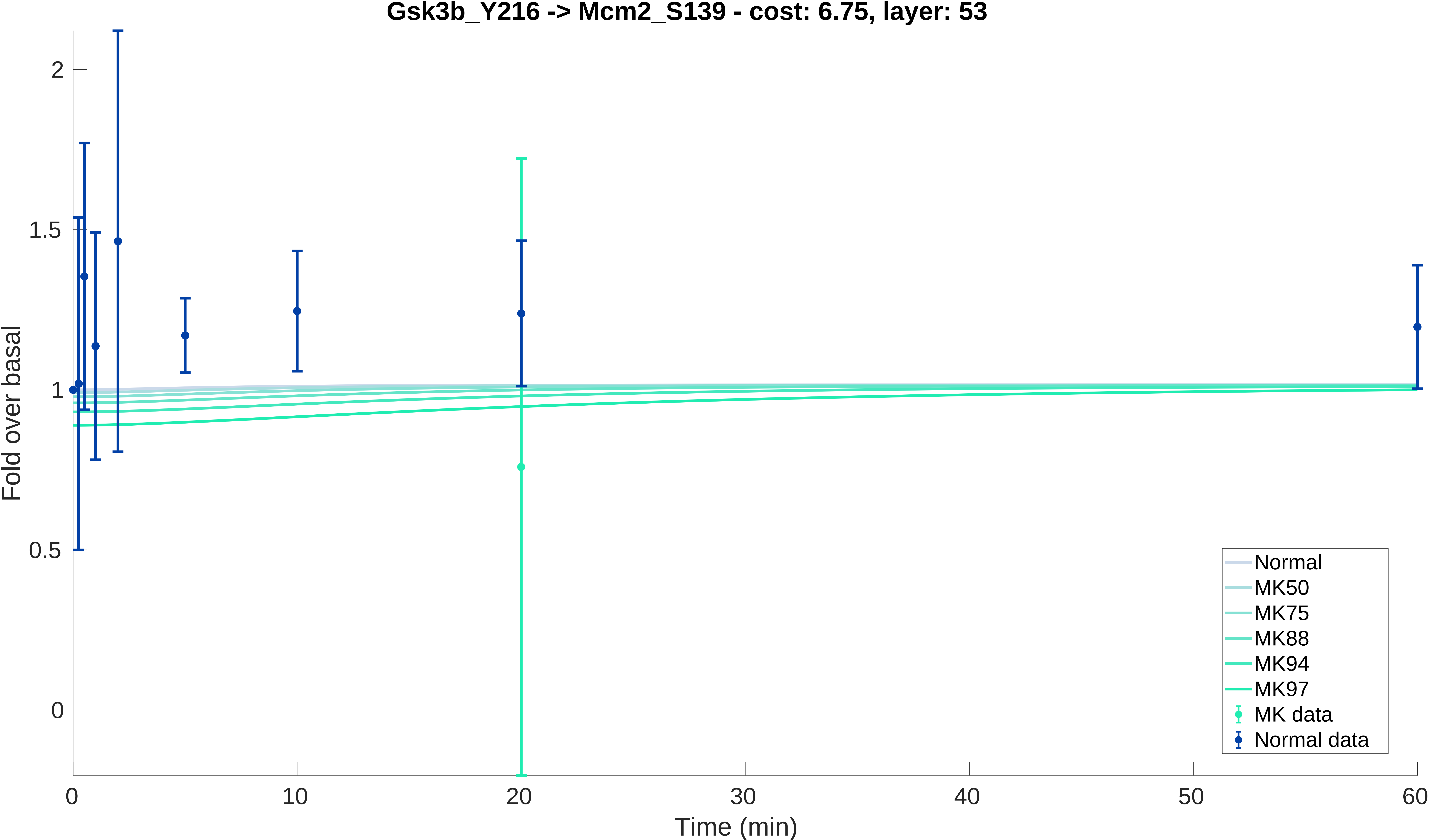

Gsk3b_Y216 -> Mcm2_S139 - cost: 6.75, layer: 53
Fold over basal
Time (min)
0
0.5
1
1.5
2
0
10
20
30
40
50
60
Normal
MK50
MK75
MK88
MK94
MK97
MK data
Normal data

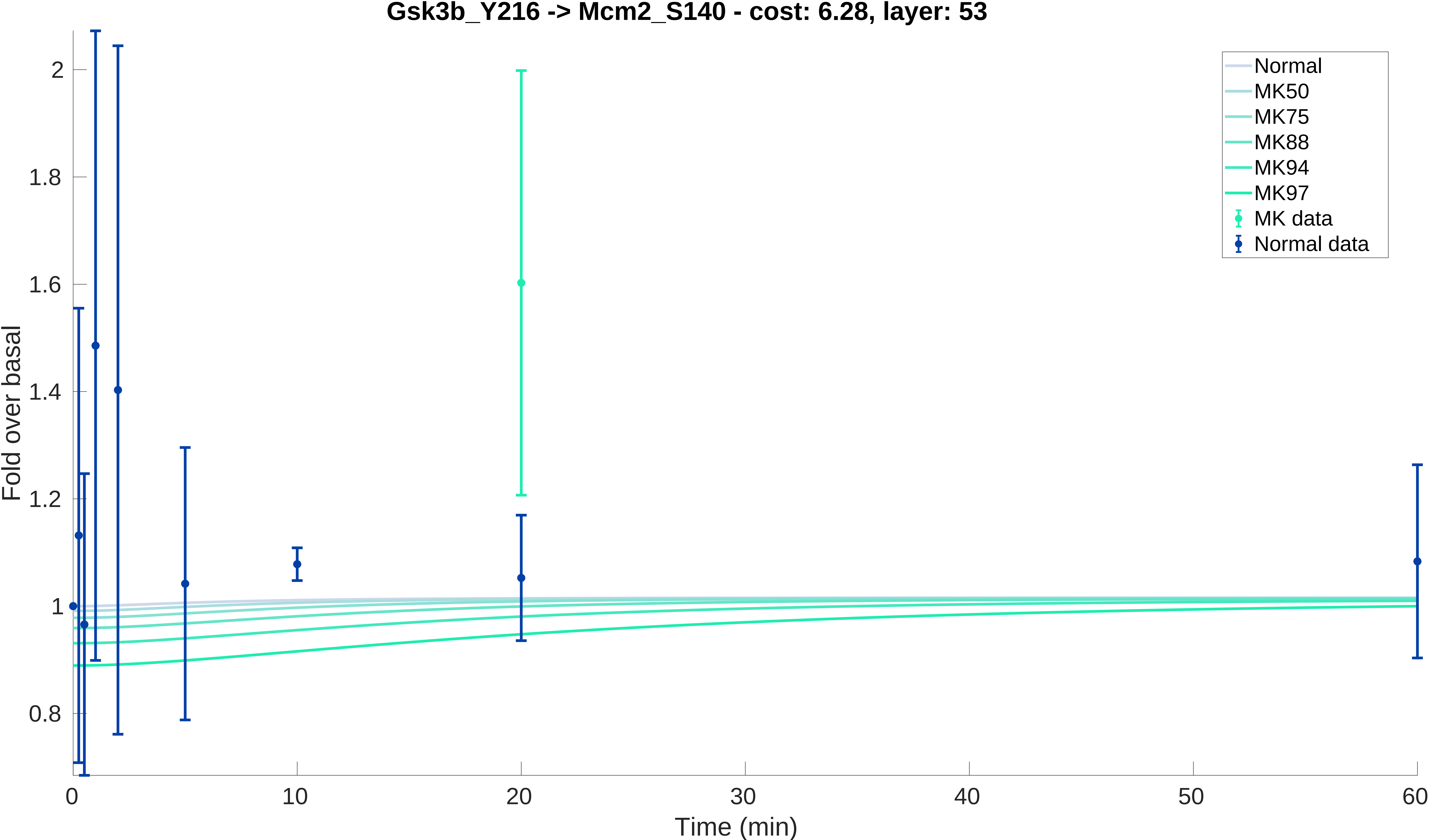

Gsk3b_Y216 -> Mcm2_S140 - cost: 6.28, layer: 53
Fold over basal
Time (min)
2
1.8
1.6
1.4
1.2
1
0.8
0
10
20
30
40
50
60
Normal
MK50
MK75
MK88
MK94
MK97
MK data
Normal data

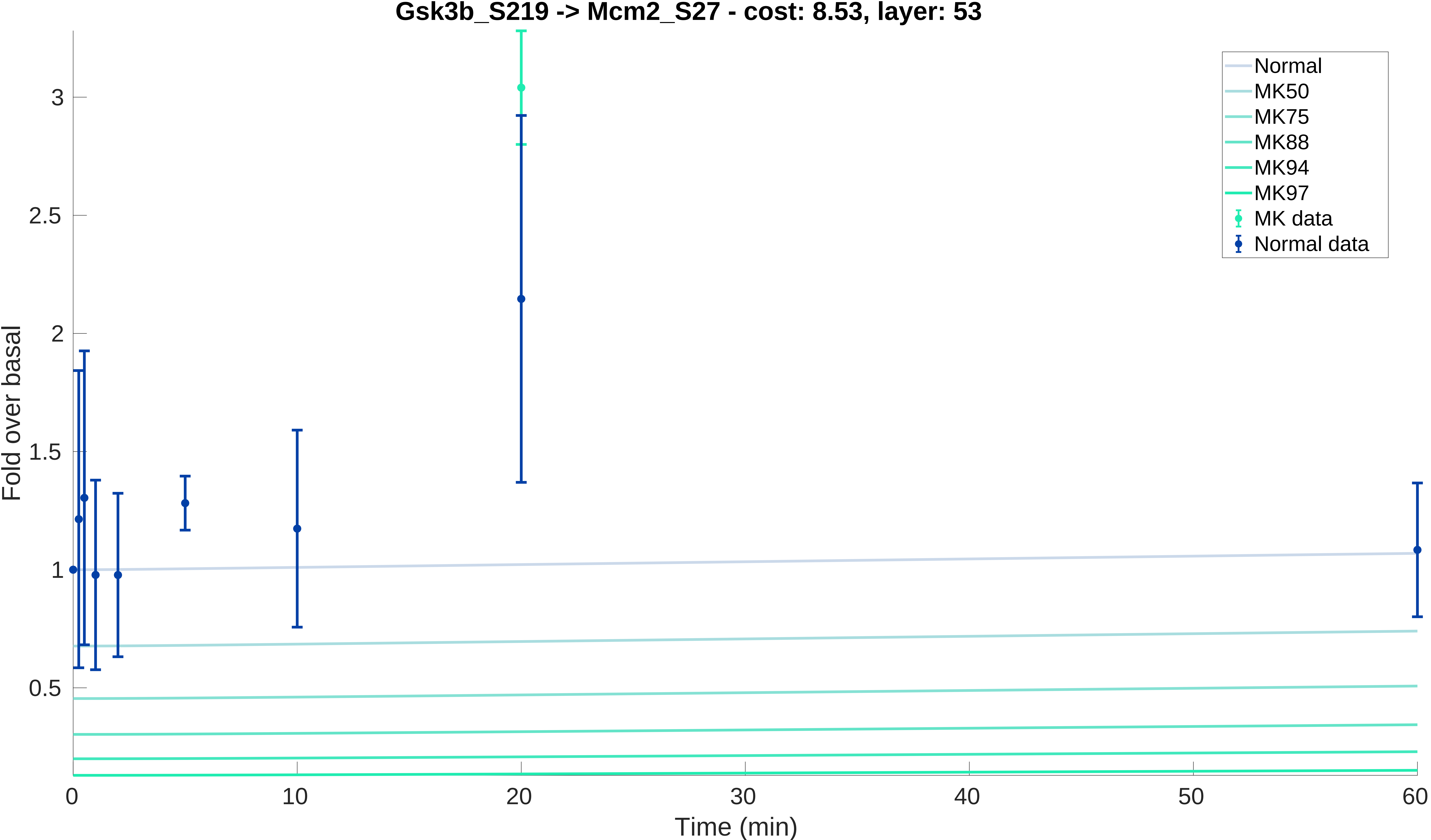
Gsk3b_S219 -> Mcm2_S27 - cost: 8.53, layer: 53
Fold over basal
0.5
1
1.5
2
2.5
3
0
10
20
30
40
50
60
Time (min)
Normal
MK50
MK75
MK88
MK94
MK97
MK data
Normal data

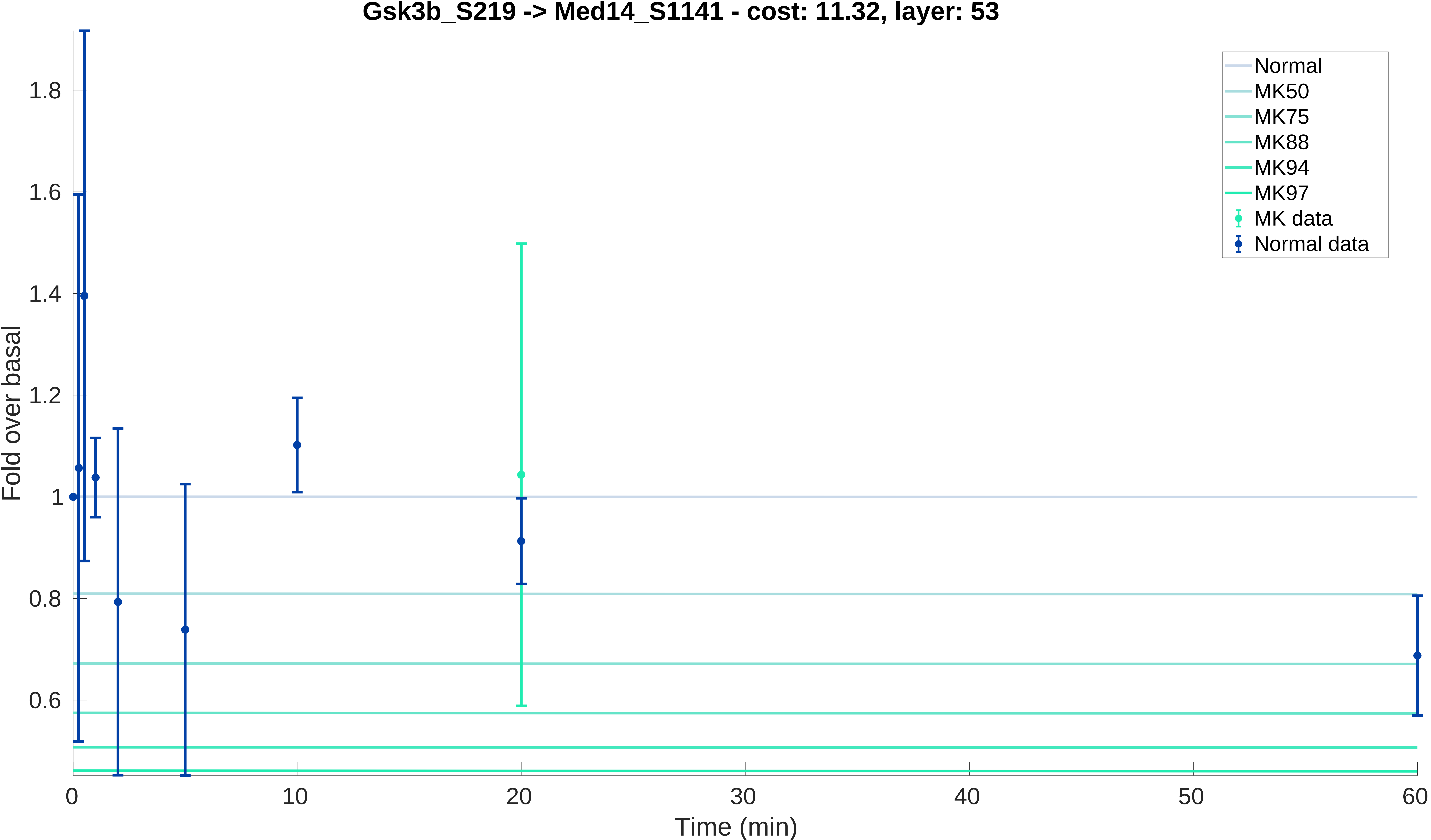

Gsk3b_S219 -> Med14_S1141 - cost: 11.32, layer: 53
Fold over basal
Time (min)
1.8
1.6
1.4
1.2
1
0.8
0.6
0
10
20
30
40
50
60
Normal
MK50
MK75
MK88
MK94
MK97
MK data
Normal data

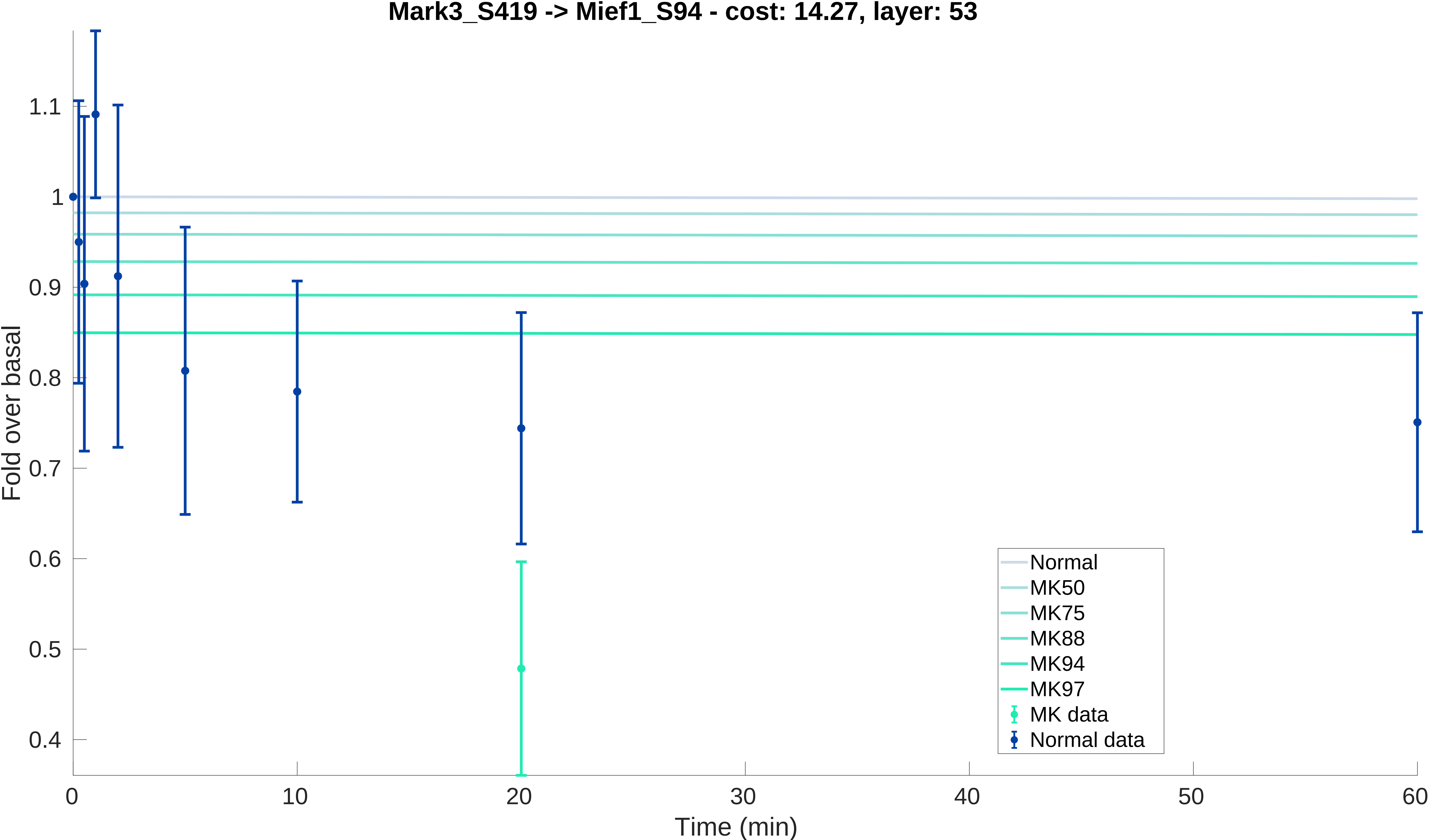
Mark3_S419 -> Mief1_S94 - cost: 14.27, layer: 53
Fold over basal
1.1
1
0.9
0.8
0.7
0.6
0.5
0.4
0
10
20
30
40
50
60
Time (min)
Normal
MK50
MK75
MK88
MK94
MK97
MK data
Normal data

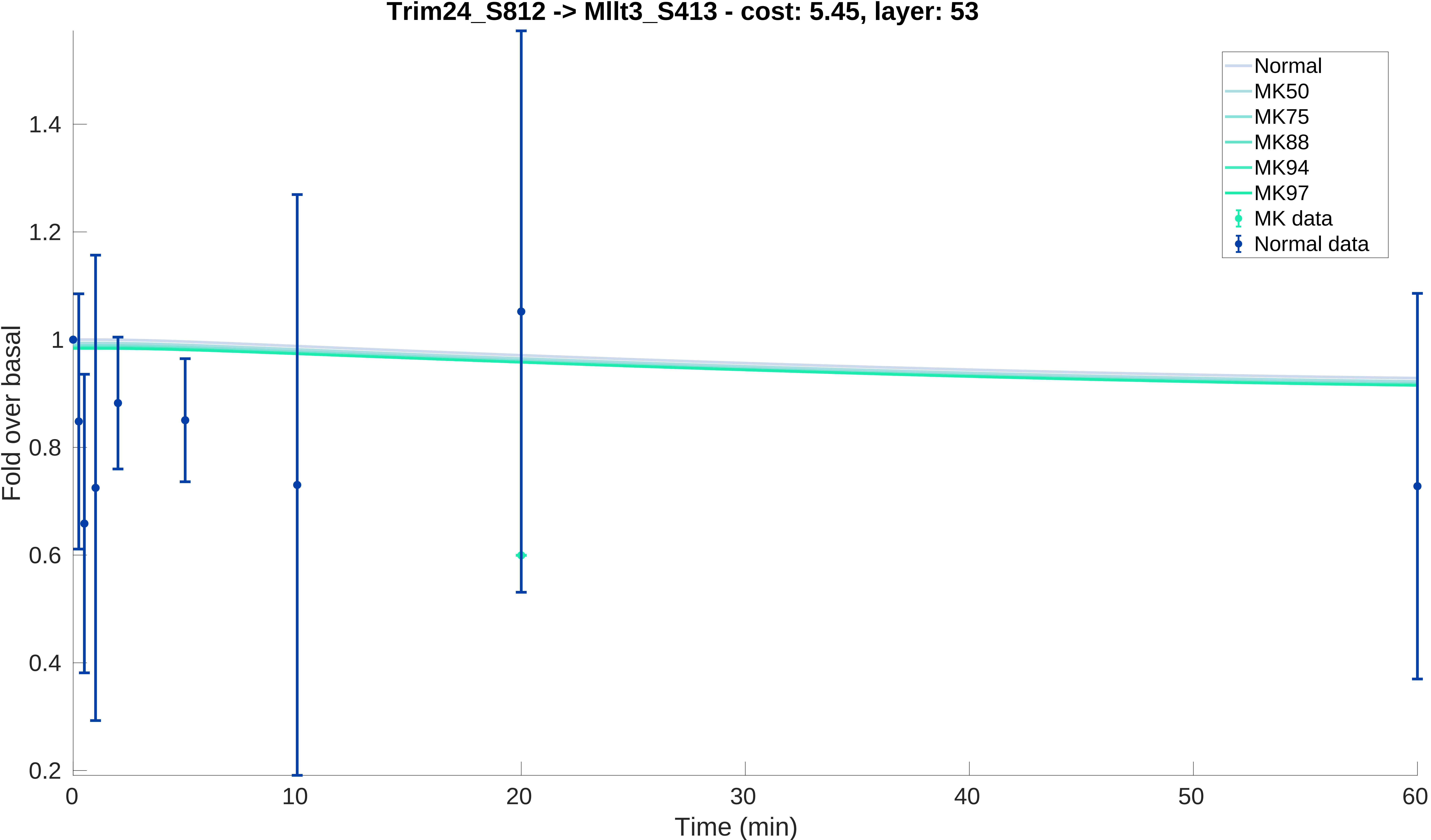
Trim24_S812 -> Mllt3_S413 - cost: 5.45, layer: 53
Normal
MK50
MK75
MK88
MK94
MK97
MK data
Normal data
Fold over basal
1.4
1.2
1
0.8
0.6
0.4
0.2
0
10
20
30
40
50
60
Time (min)

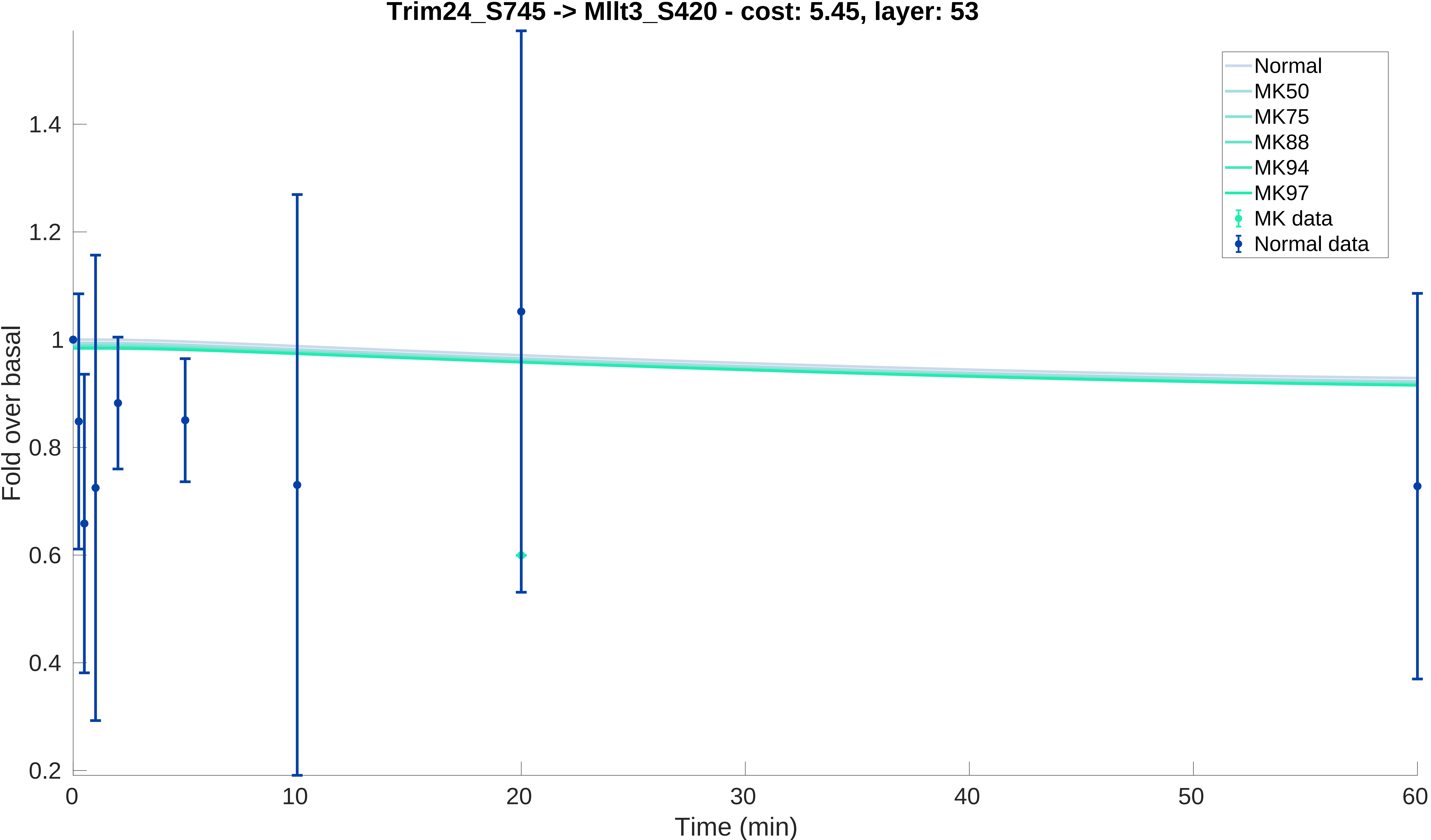

Trim24_S745 -> Mllt3_S420 - cost: 5.45, layer: 53
Normal
MK50
MK75
MK88
MK94
MK97
MK data
Normal data
Fold over basal
Time (min)
0.2
0.4
0.6
0.8
1
1.2
1.4
0
10
20
30
40
50
60

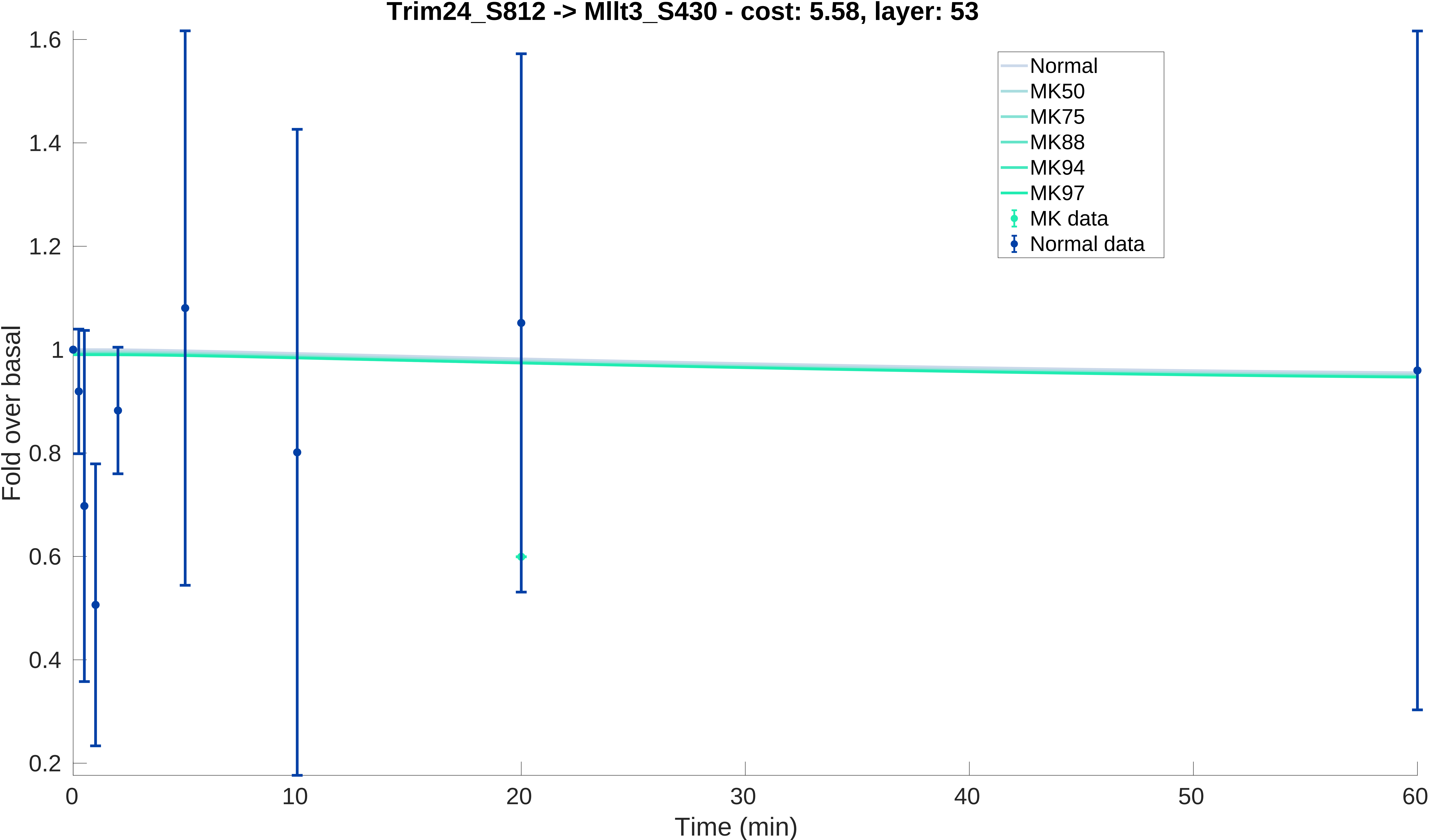

Trim24_S812 -> Mllt3_S430 - cost: 5.58, layer: 53
Fold over basal
Time (min)
0.2
0.4
0.6
0.8
1
1.2
1.4
1.6
0
10
20
30
40
50
60
Normal
MK50
MK75
MK88
MK94
MK97
MK data
Normal data

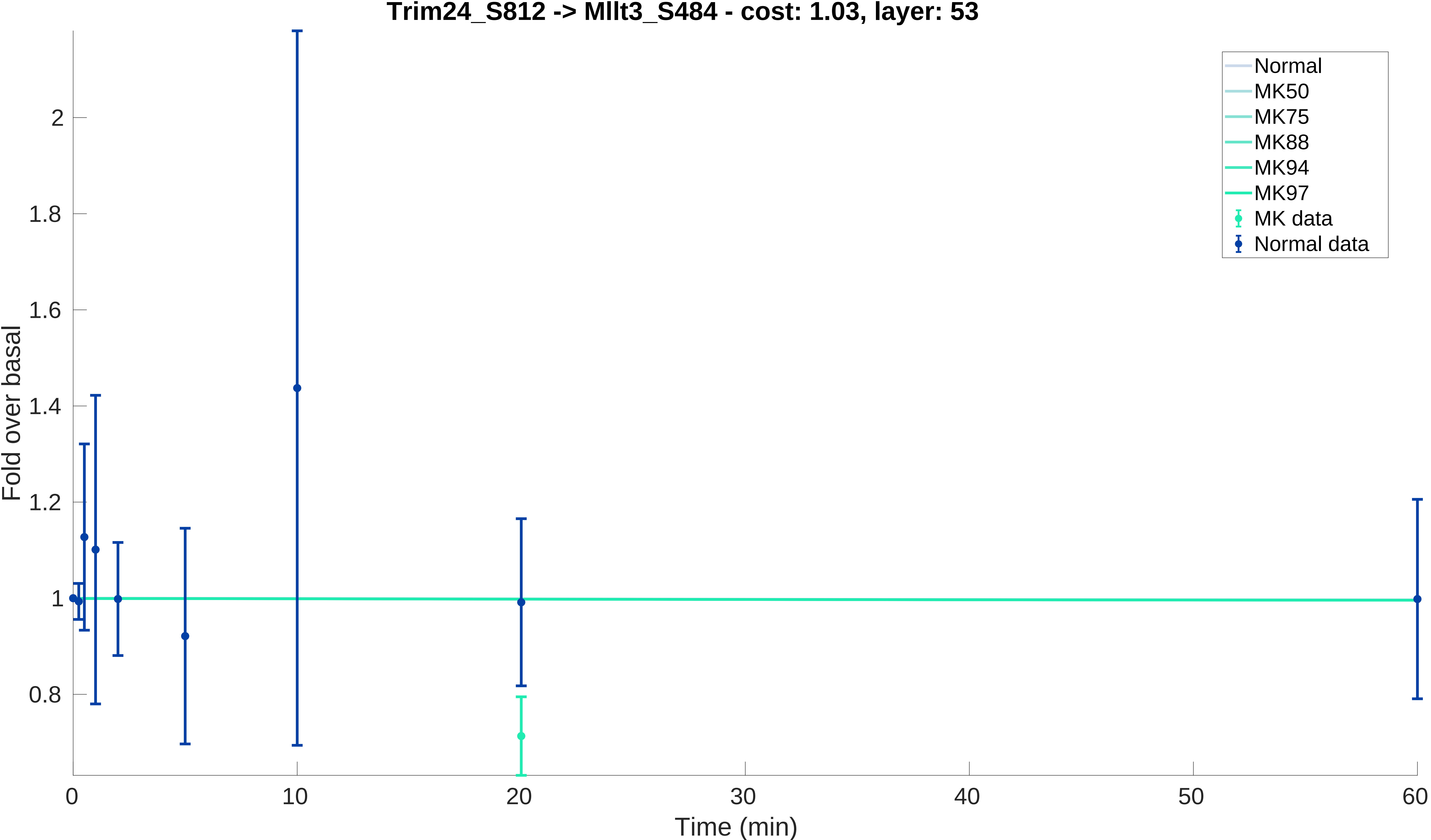

Trim24_S812 -> Mllt3_S484 - cost: 1.03, layer: 53
Fold over basal
2
1.8
1.6
1.4
1.2
1
0.8
0
10
20
30
40
50
60
Time (min)
Normal
MK50
MK75
MK88
MK94
MK97
MK data
Normal data

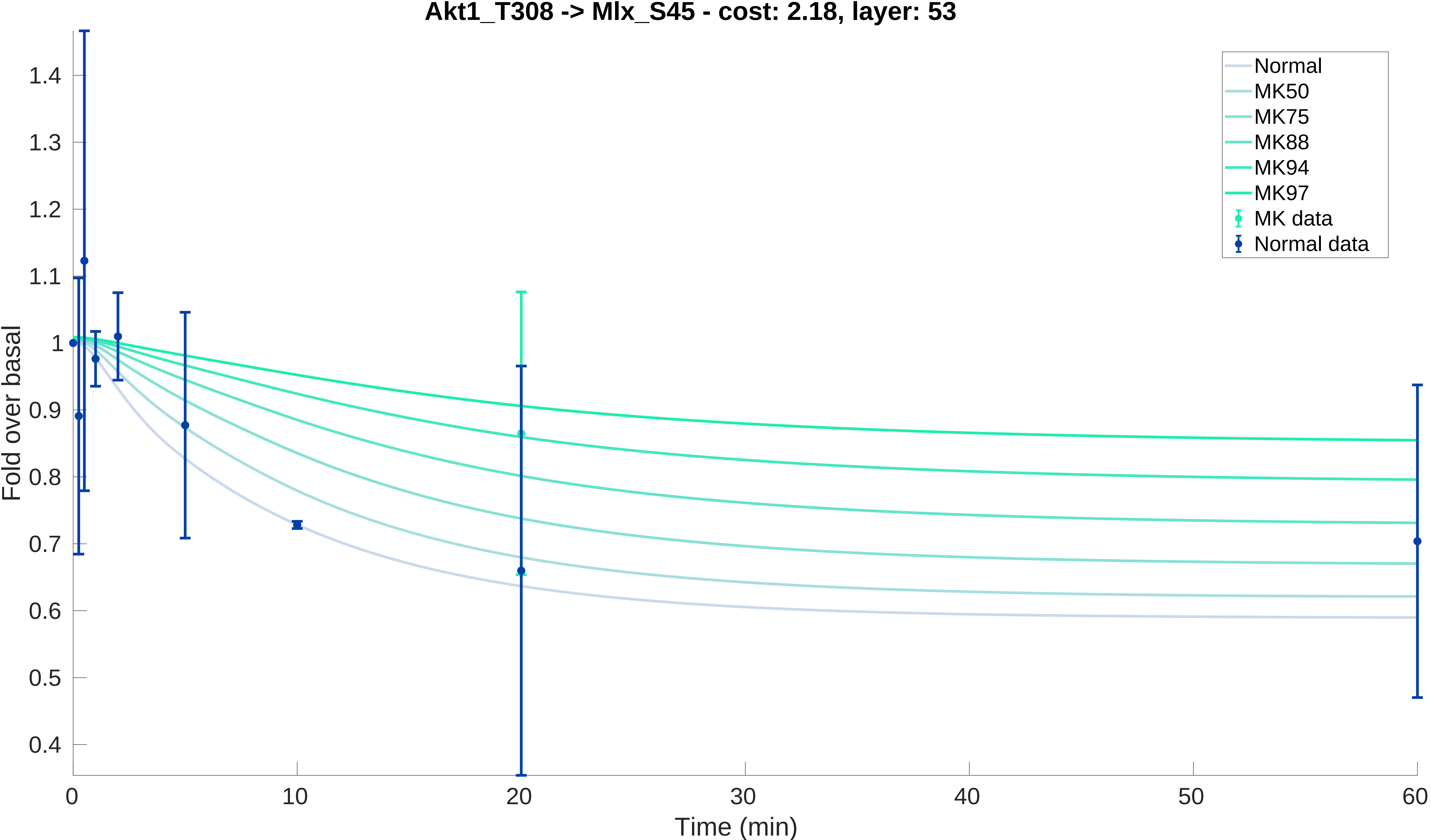

Akt1_T308 -> Mlx_S45 - cost: 2.18, layer: 53
Fold over basal
1.4
1.3
1.2
1.1
1
0.9
0.8
0.7
0.6
0.5
0.4
0
10
20
30
40
50
60
Time (min)
Normal
MK50
MK75
MK88
MK94
MK97
MK data
Normal data

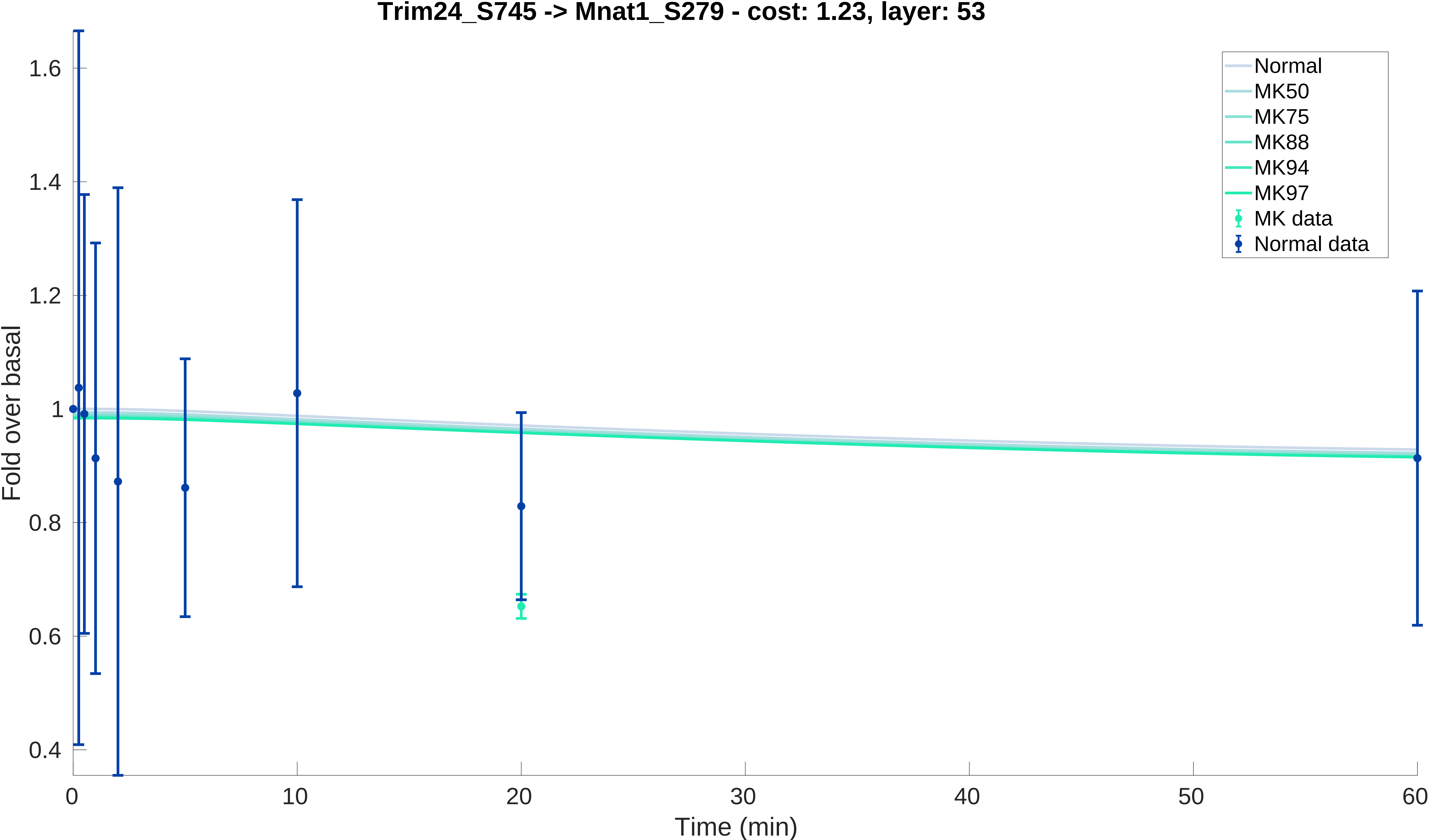

Trim24_S745 -> Mnat1_S279 - cost: 1.23, layer: 53
Normal
MK50
MK75
MK88
MK94
MK97
MK data
Normal data
Fold over basal
1.6
1.4
1.2
1
0.8
0.6
0.4
0
10
20
30
40
50
60
Time (min)

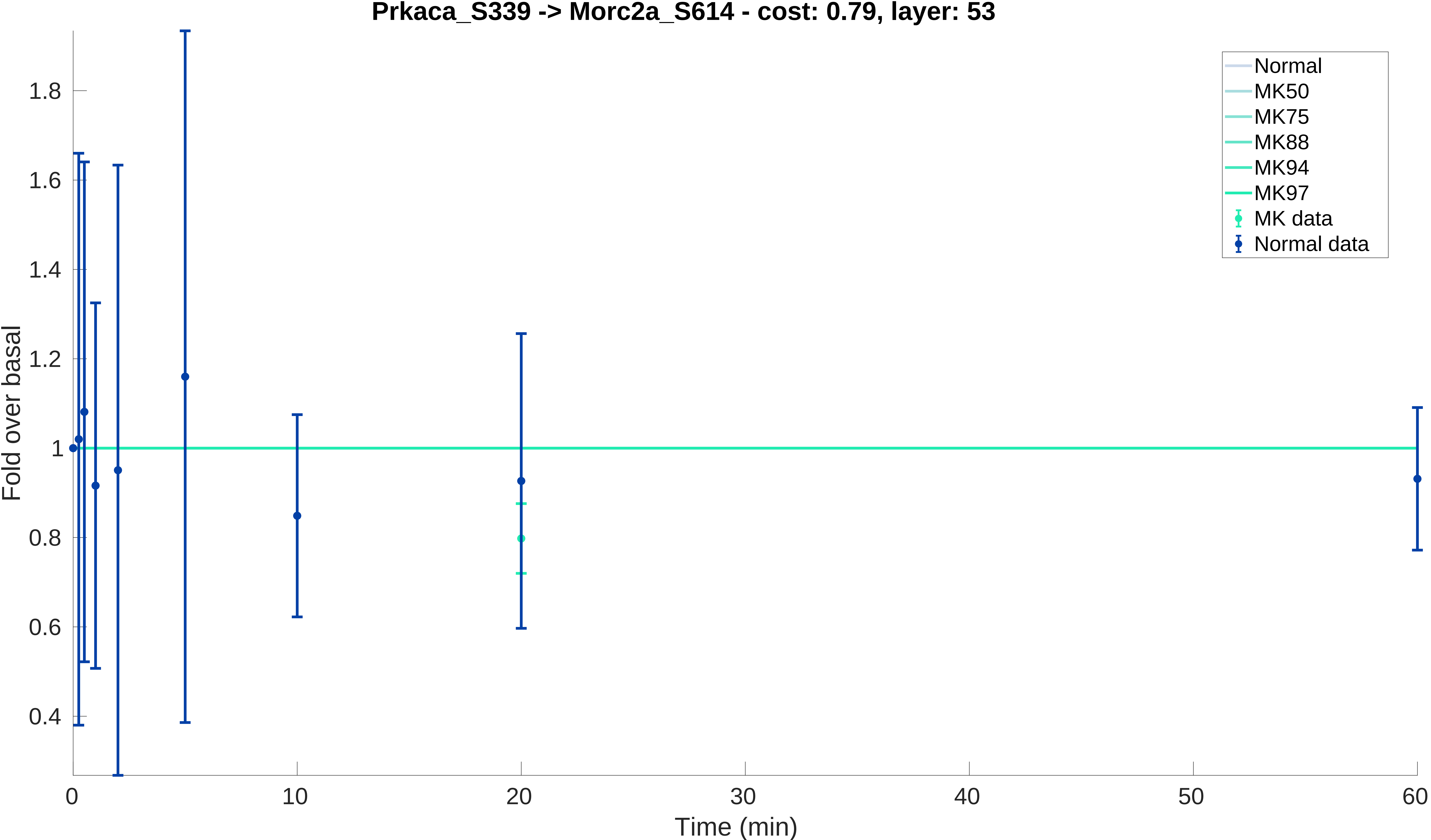

Prkaca_S339 -> Morc2a_S614 - cost: 0.79, layer: 53
Fold over basal
Time (min)
0.4
0.6
0.8
1
1.2
1.4
1.6
1.8
0
10
20
30
40
50
60
Normal
MK50
MK75
MK88
MK94
MK97
MK data
Normal data

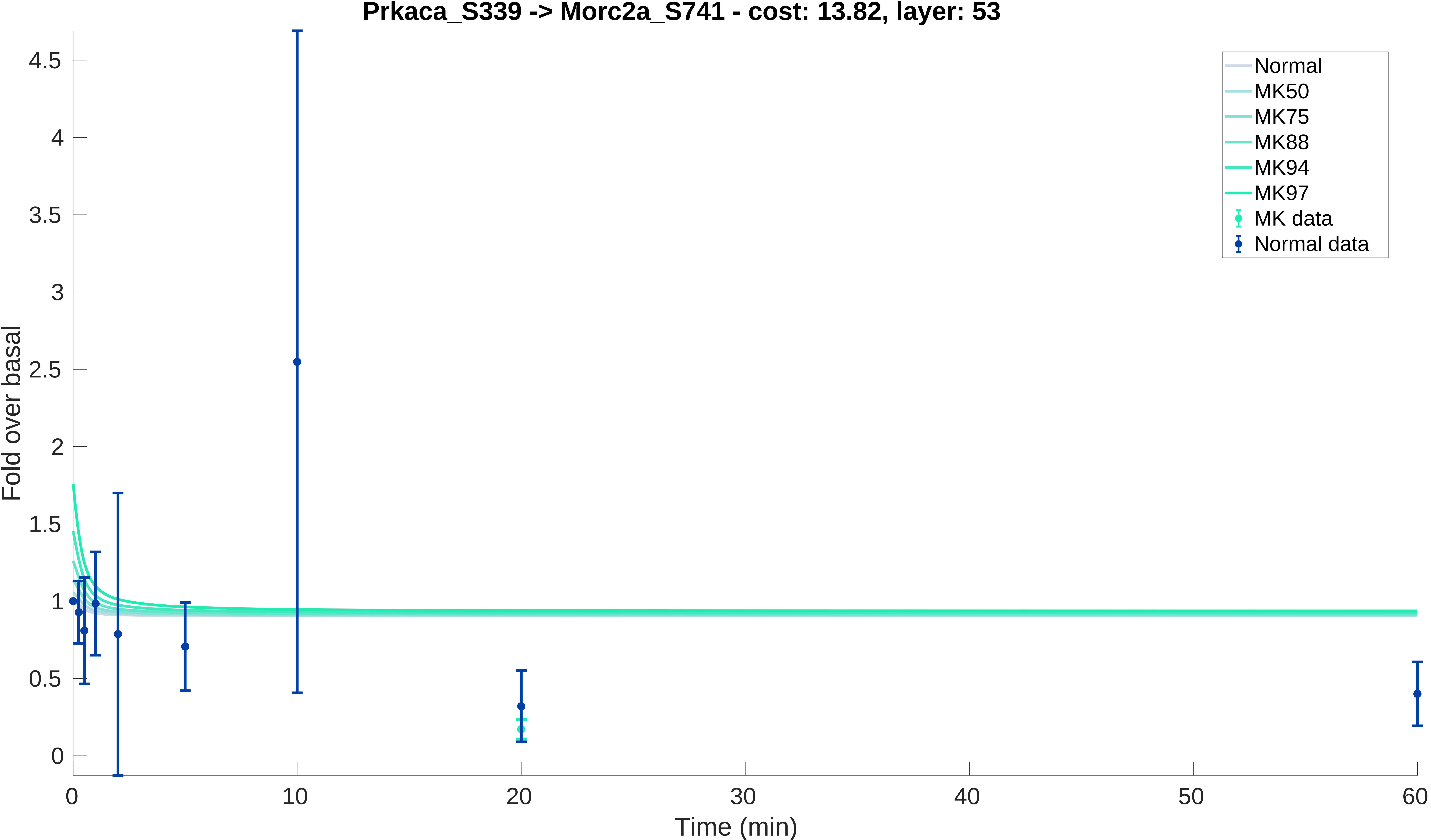

Prkaca_S339 -> Morc2a_S741 - cost: 13.82, layer: 53
Fold over basal
Time (min)
0
0.5
1
1.5
2
2.5
3
3.5
4
4.5
10
20
30
40
50
60
Normal
MK50
MK75
MK88
MK94
MK97
MK data
Normal data

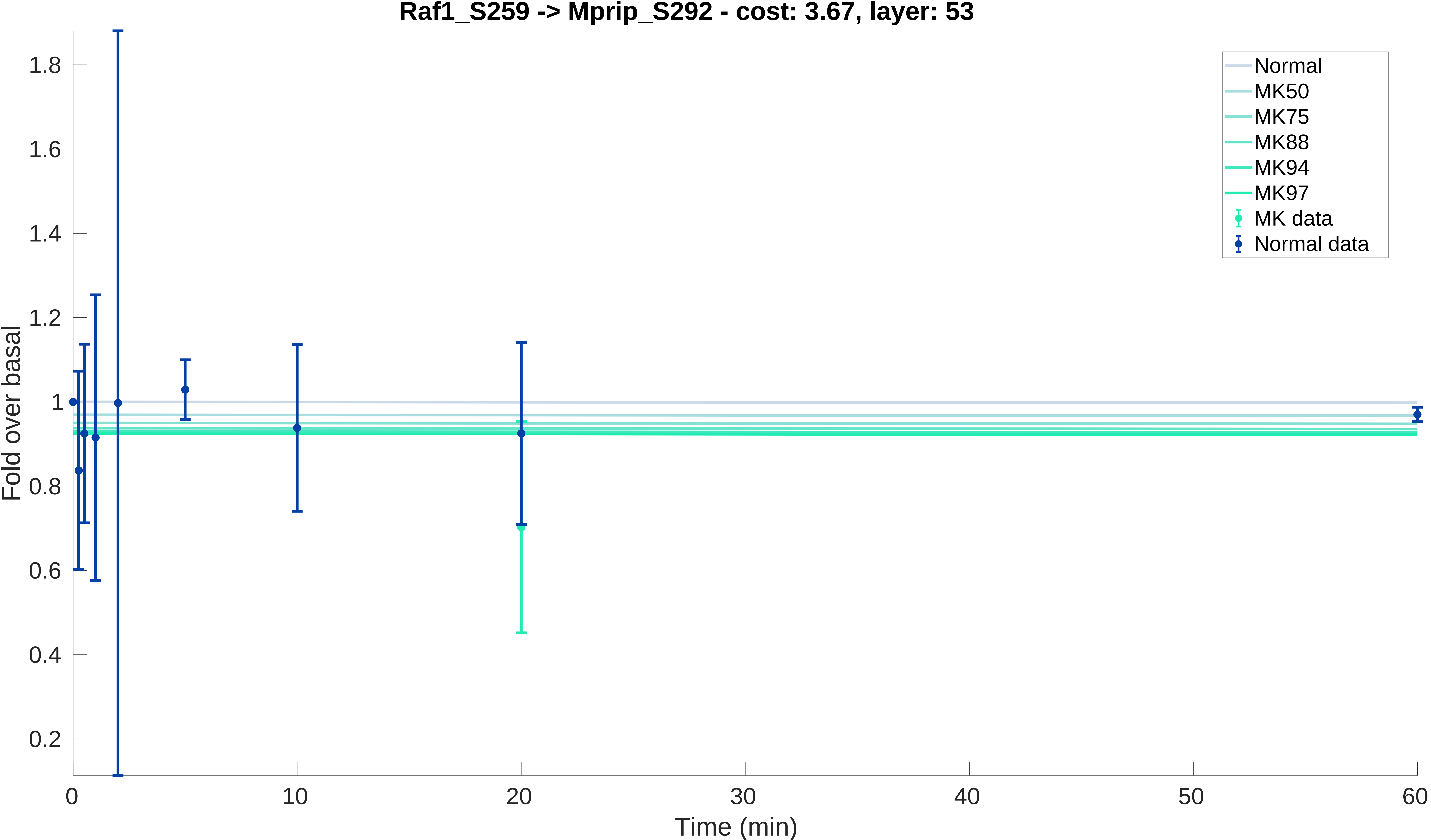

Raf1_S259 -> Mprip_S292 - cost: 3.67, layer: 53
Fold over basal
Time (min)
1.8
1.6
1.4
1.2
1
0.8
0.6
0.4
0.2
0
10
20
30
40
50
60
Normal
MK50
MK75
MK88
MK94
MK97
MK data
Normal data

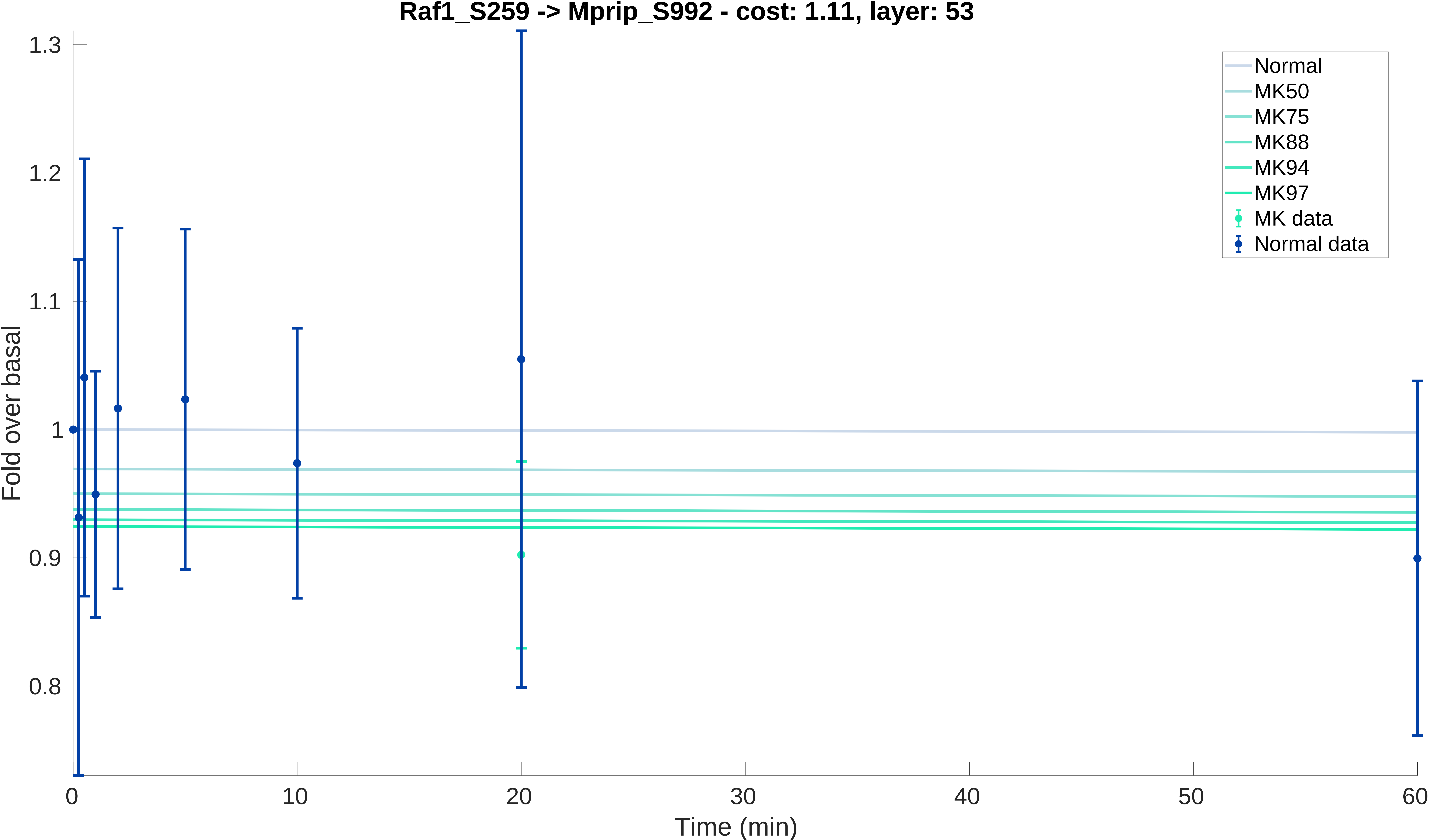

Raf1_S259 -> Mprip_S992 - cost: 1.11, layer: 53
Fold over basal
Time (min)
1.3
1.2
1.1
1
0.9
0.8
0
10
20
30
40
50
60
Normal
MK50
MK75
MK88
MK94
MK97
MK data
Normal data

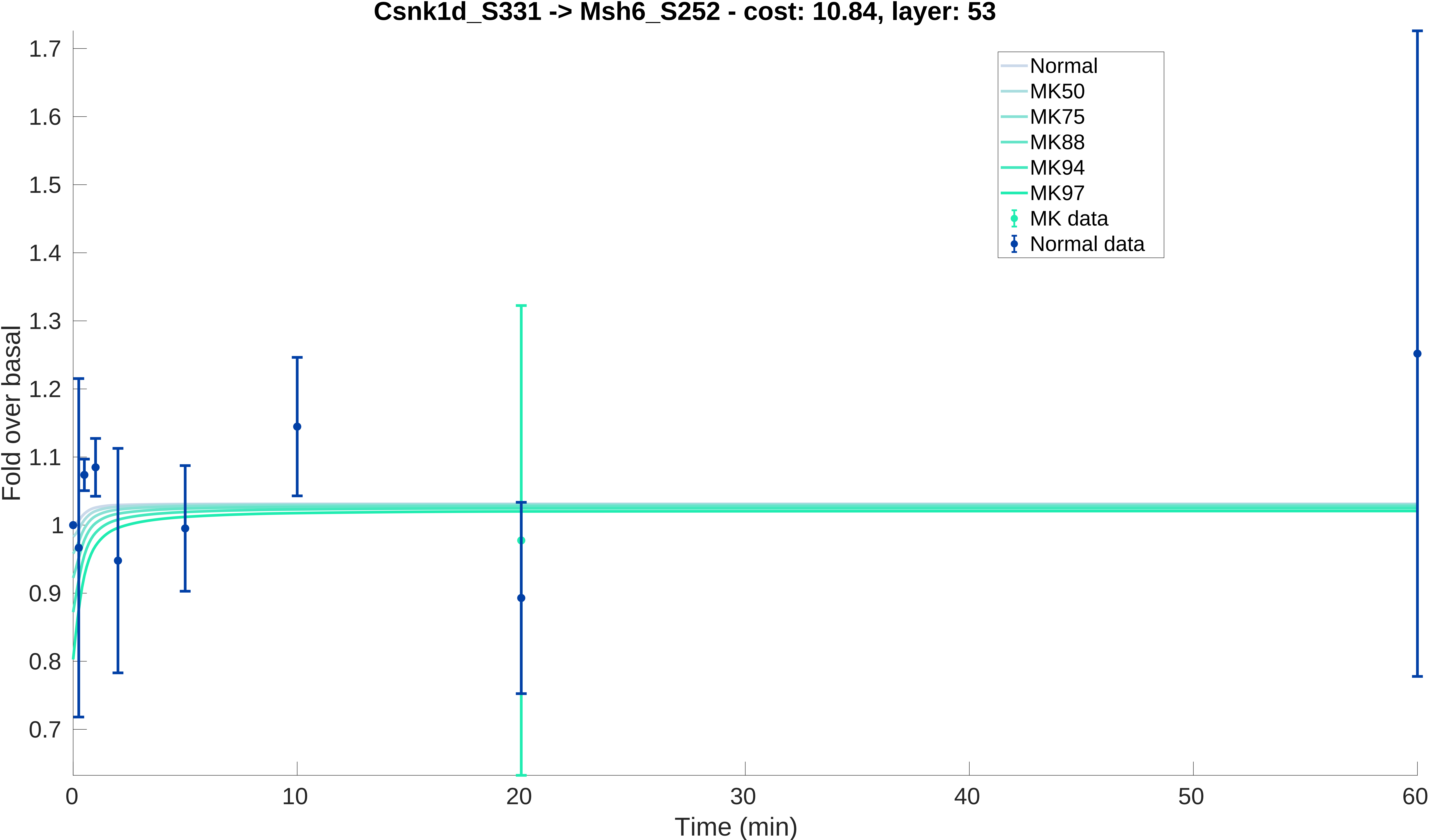

Csnk1d_S331 -> Msh6_S252 - cost: 10.84, layer: 53
Normal
MK50
MK75
MK88
MK94
MK97
MK data
Normal data
Fold over basal
Time (min)
0.7
0.8
0.9
1
1.1
1.2
1.3
1.4
1.5
1.6
1.7
0
10
20
30
40
50
60

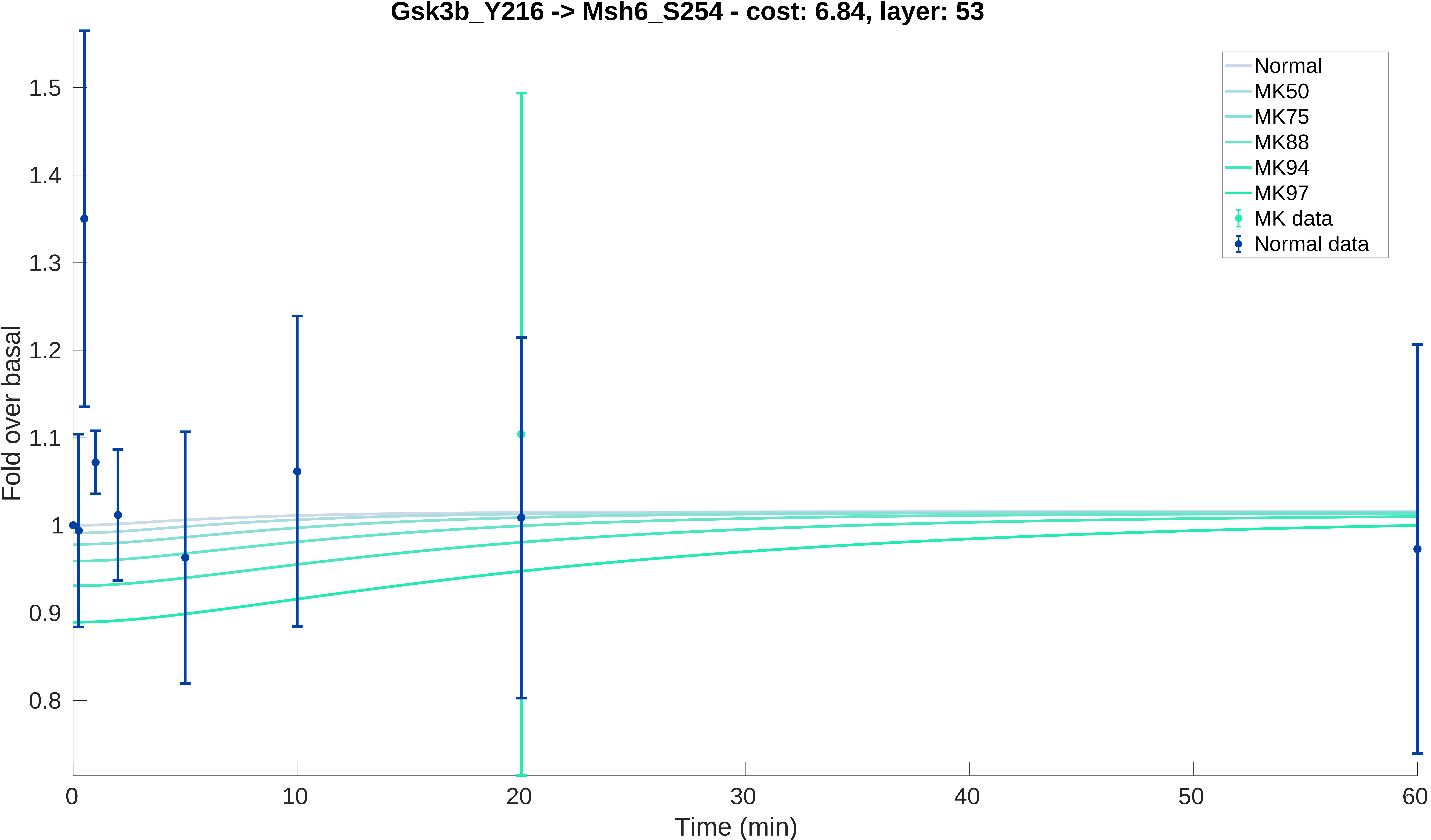
Gsk3b_Y216 -> Msh6_S254 - cost: 6.84, layer: 53
Fold over basal
1.5
1.4
1.3
1.2
1.1
1
0.9
0.8
0
10
20
30
40
50
60
Time (min)
Normal
MK50
MK75
MK88
MK94
MK97
MK data
Normal data

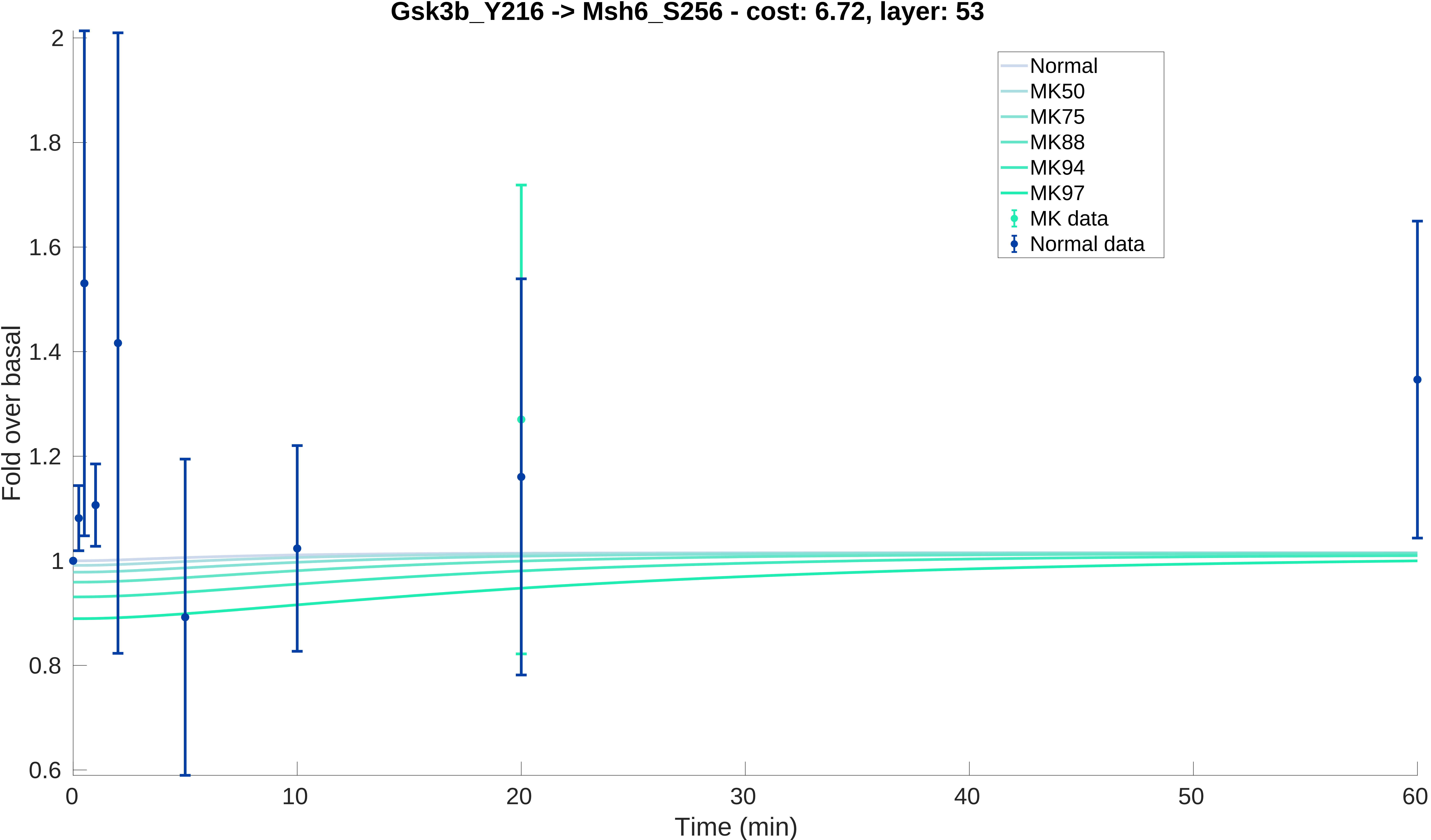

Gsk3b_Y216 -> Msh6_S256 - cost: 6.72, layer: 53
Fold over basal
Time (min)
2
1.8
1.6
1.4
1.2
1
0.8
0.6
0
10
20
30
40
50
60
Normal
MK50
MK75
MK88
MK94
MK97
MK data
Normal data

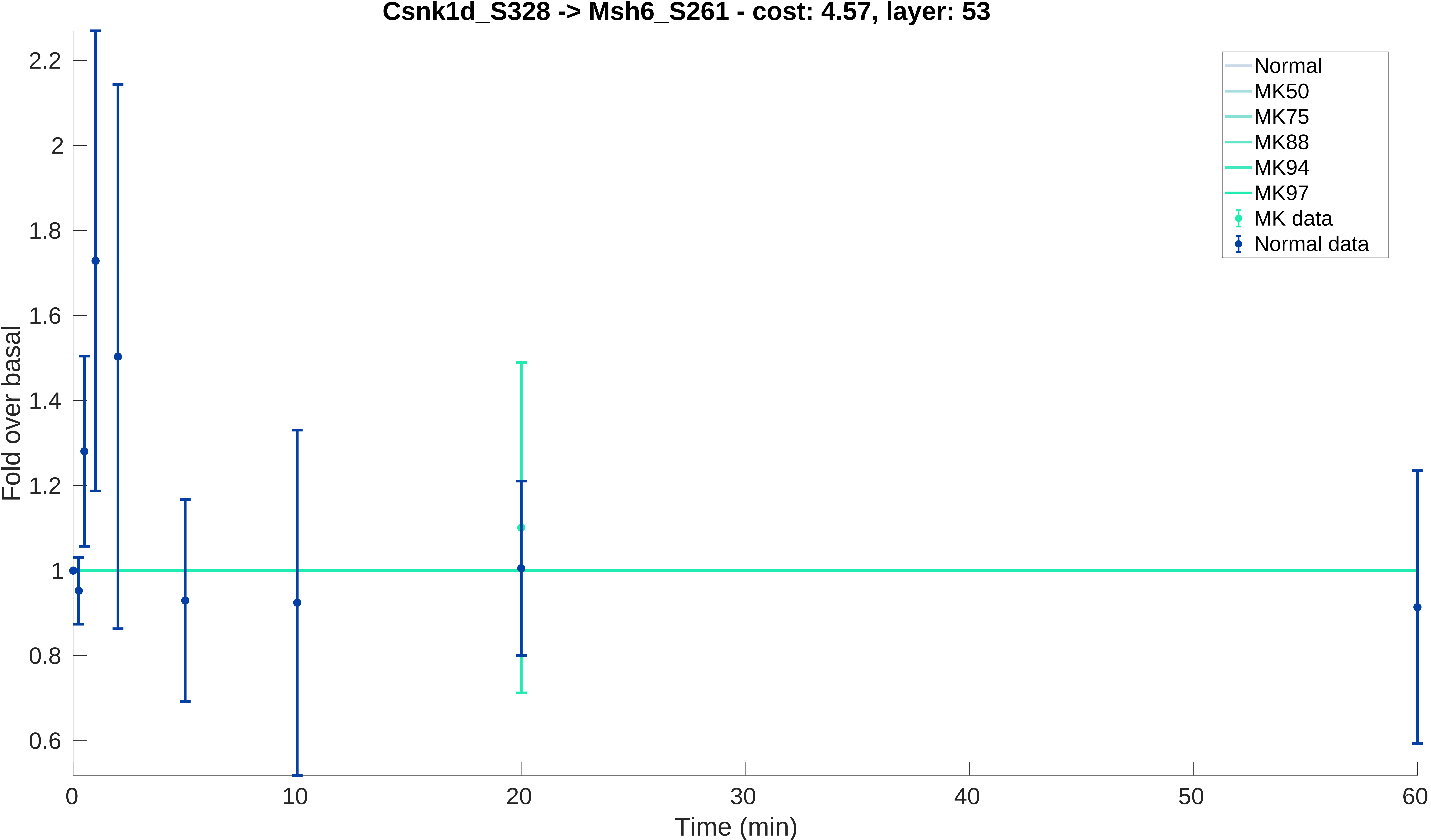

Csnk1d_S328 -> Msh6_S261 - cost: 4.57, layer: 53
Fold over basal
Time (min)
Normal
MK50
MK75
MK88
MK94
MK97
MK data
Normal data
2.2
2
1.8
1.6
1.4
1.2
1
0.8
0.6
0
10
20
30
40
50
60

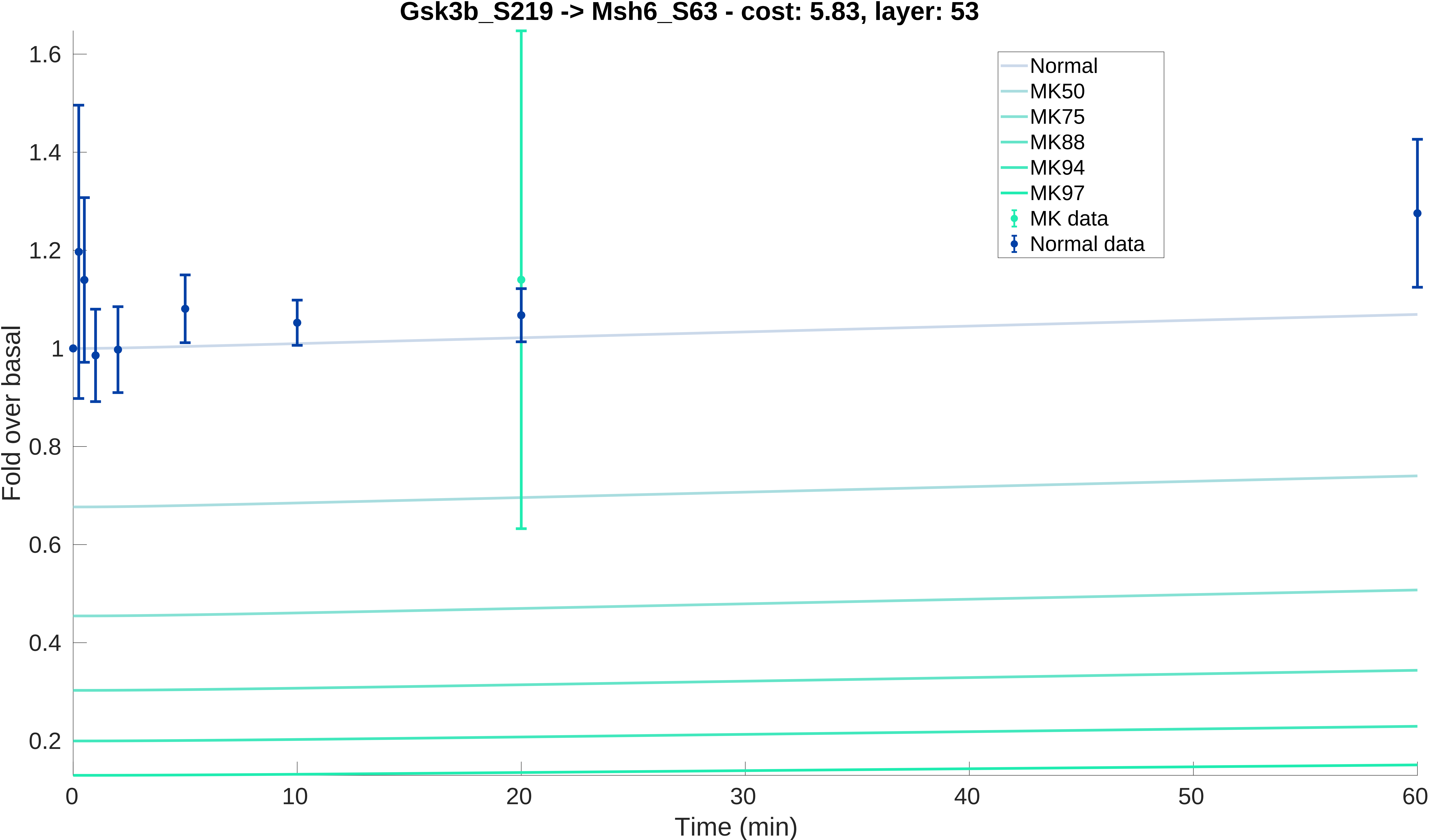

Gsk3b_S219 -> Msh6_S63 - cost: 5.83, layer: 53
Fold over basal
Time (min)
1.6
1.4
1.2
1
0.8
0.6
0.4
0.2
0
10
20
30
40
50
60
Normal
MK50
MK75
MK88
MK94
MK97
MK data
Normal data

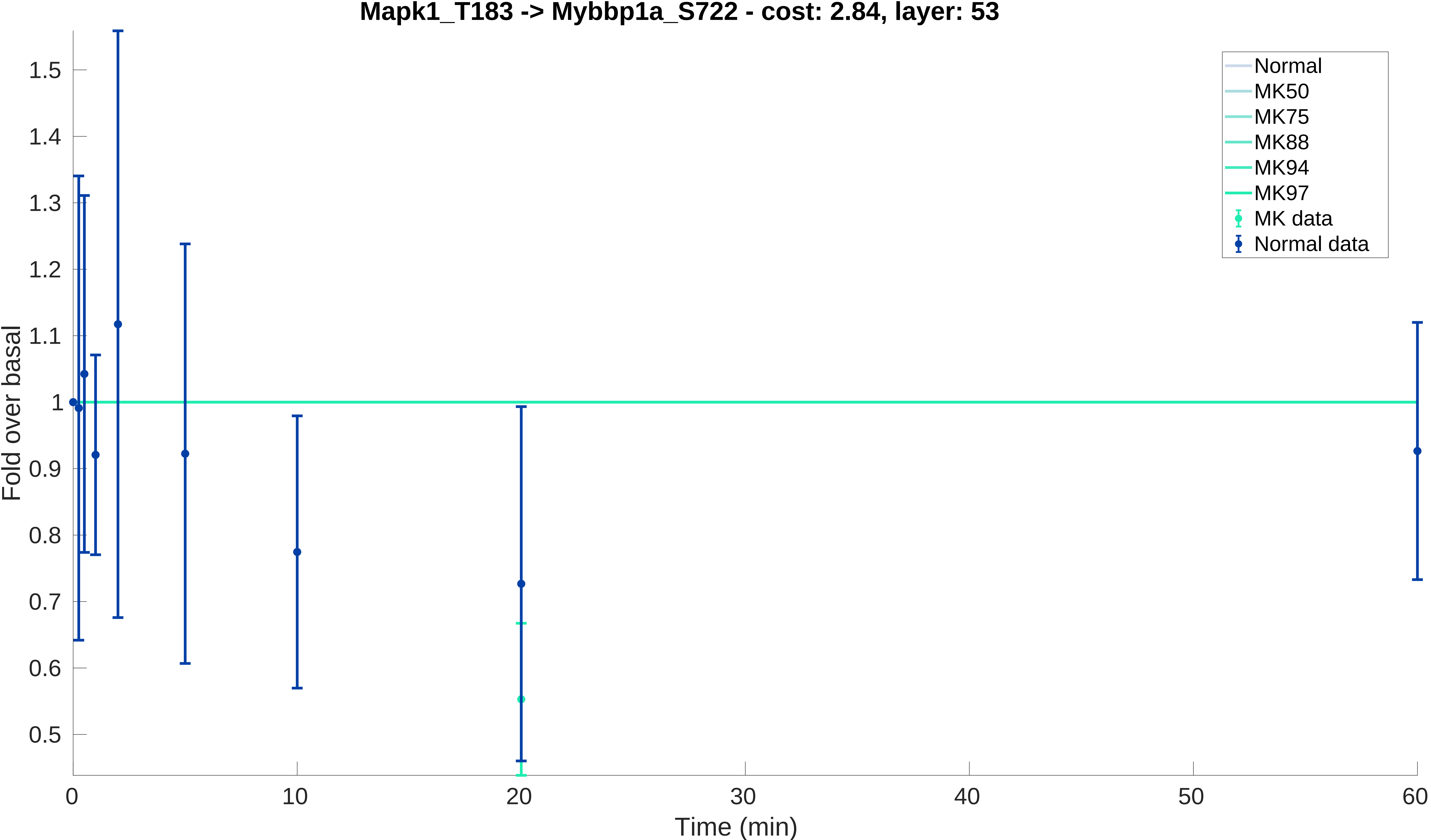

Mapk1_T183 -> Mybbp1a_S722 - cost: 2.84, layer: 53
Fold over basal
1.5
1.4
1.3
1.2
1.1
1
0.9
0.8
0.7
0.6
0.5
0
10
20
30
40
50
60
Time (min)
Normal
MK50
MK75
MK88
MK94
MK97
MK data
Normal data

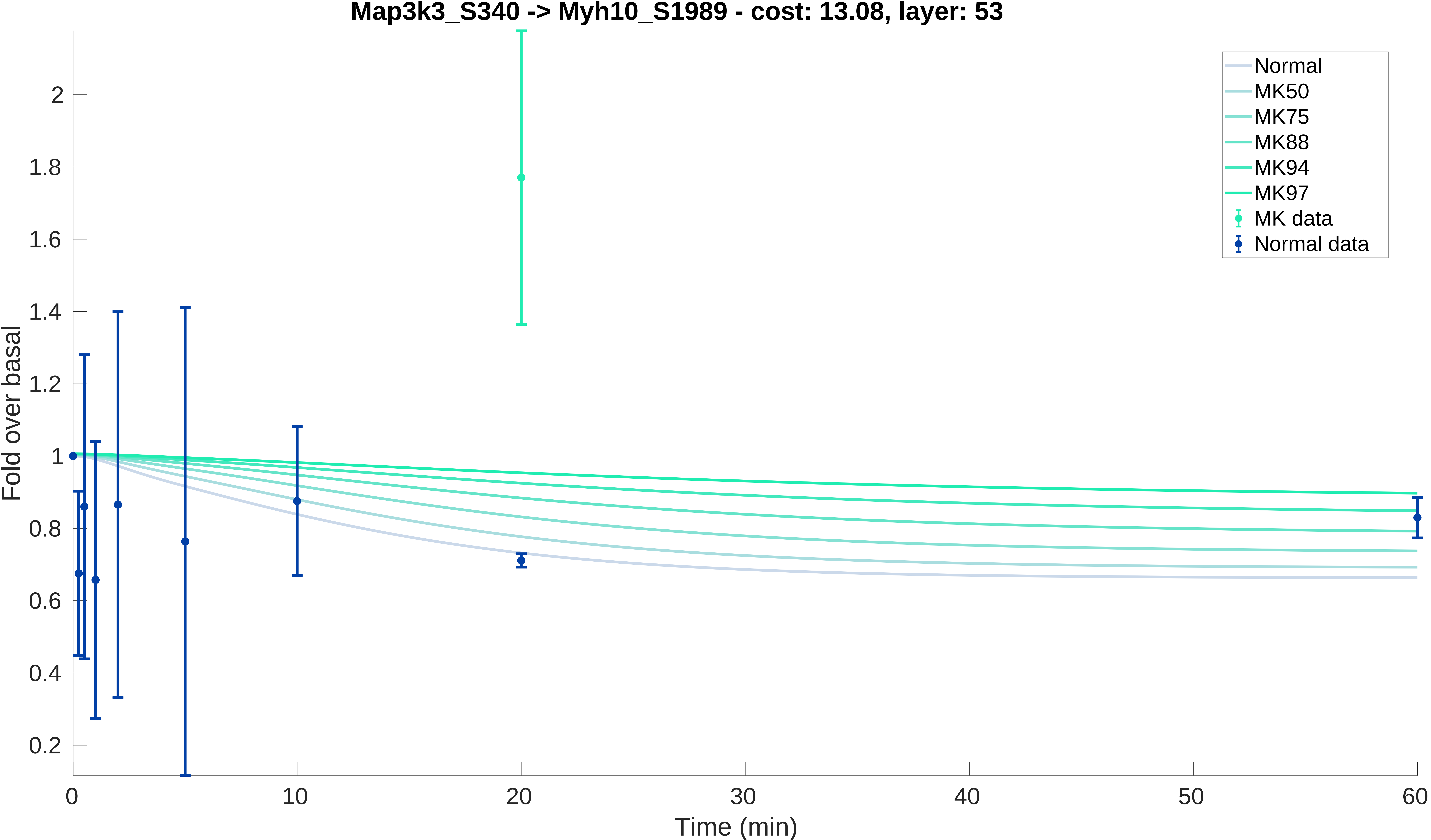
Map3k3_S340 -> Myh10_S1989 - cost: 13.08, layer: 53
Fold over basal
Time (min)
Normal
MK50
MK75
MK88
MK94
MK97
MK data
Normal data
0
10
20
30
40
50
60
0.2
0.4
0.6
0.8
1
1.2
1.4
1.6
1.8
2

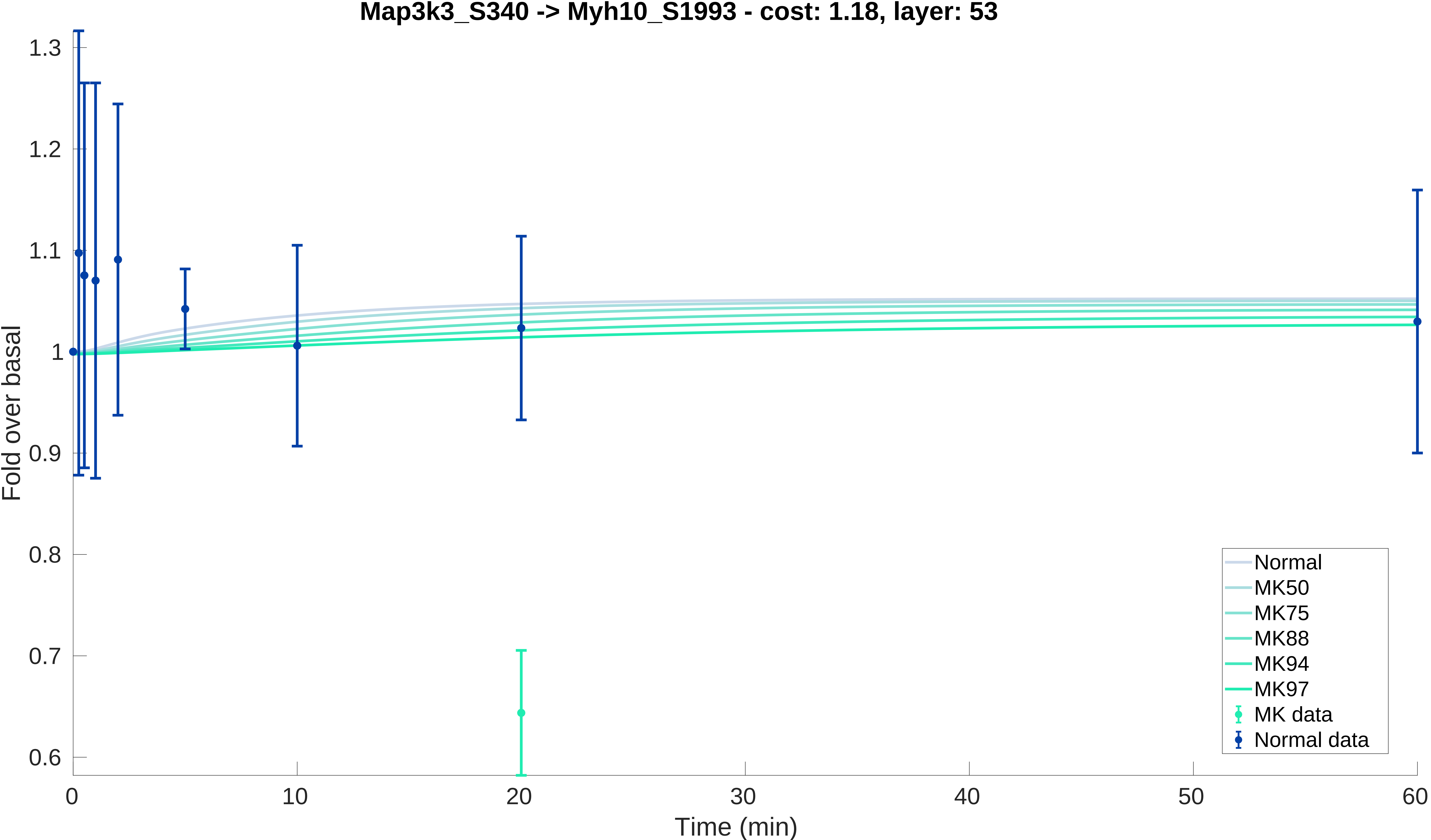

Map3k3_S340 -> Myh10_S1993 - cost: 1.18, layer: 53
Fold over basal
1.3
1.2
1.1
1
0.9
0.8
0.7
0.6
0
10
20
30
40
50
60
Time (min)
Normal
MK50
MK75
MK88
MK94
MK97
MK data
Normal data

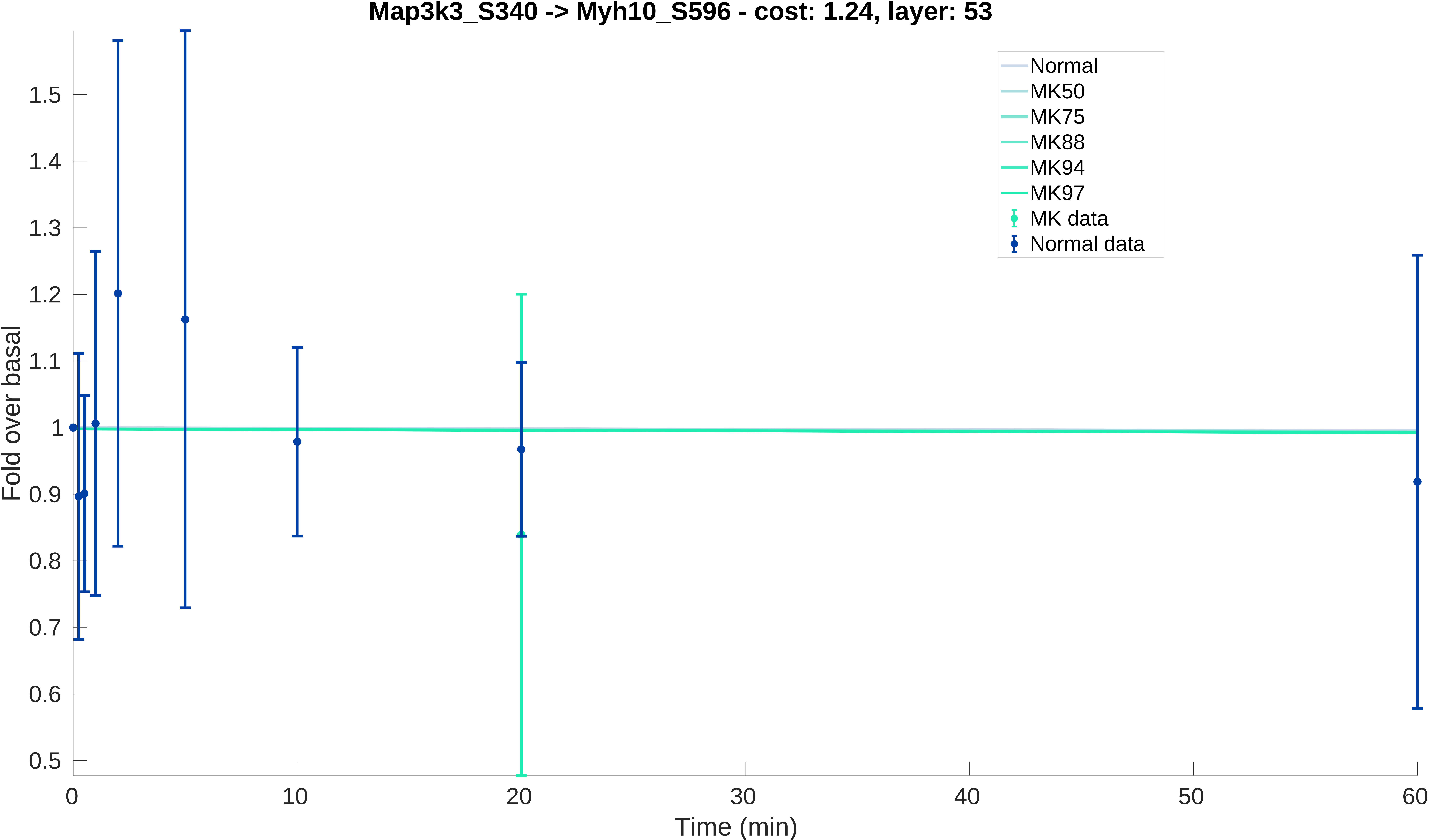

Map3k3_S340 -> Myh10_S596 - cost: 1.24, layer: 53
Normal
MK50
MK75
MK88
MK94
MK97
MK data
Normal data
Fold over basal
1.5
1.4
1.3
1.2
1.1
1
0.9
0.8
0.7
0.6
0.5
0
10
20
30
40
50
60
Time (min)

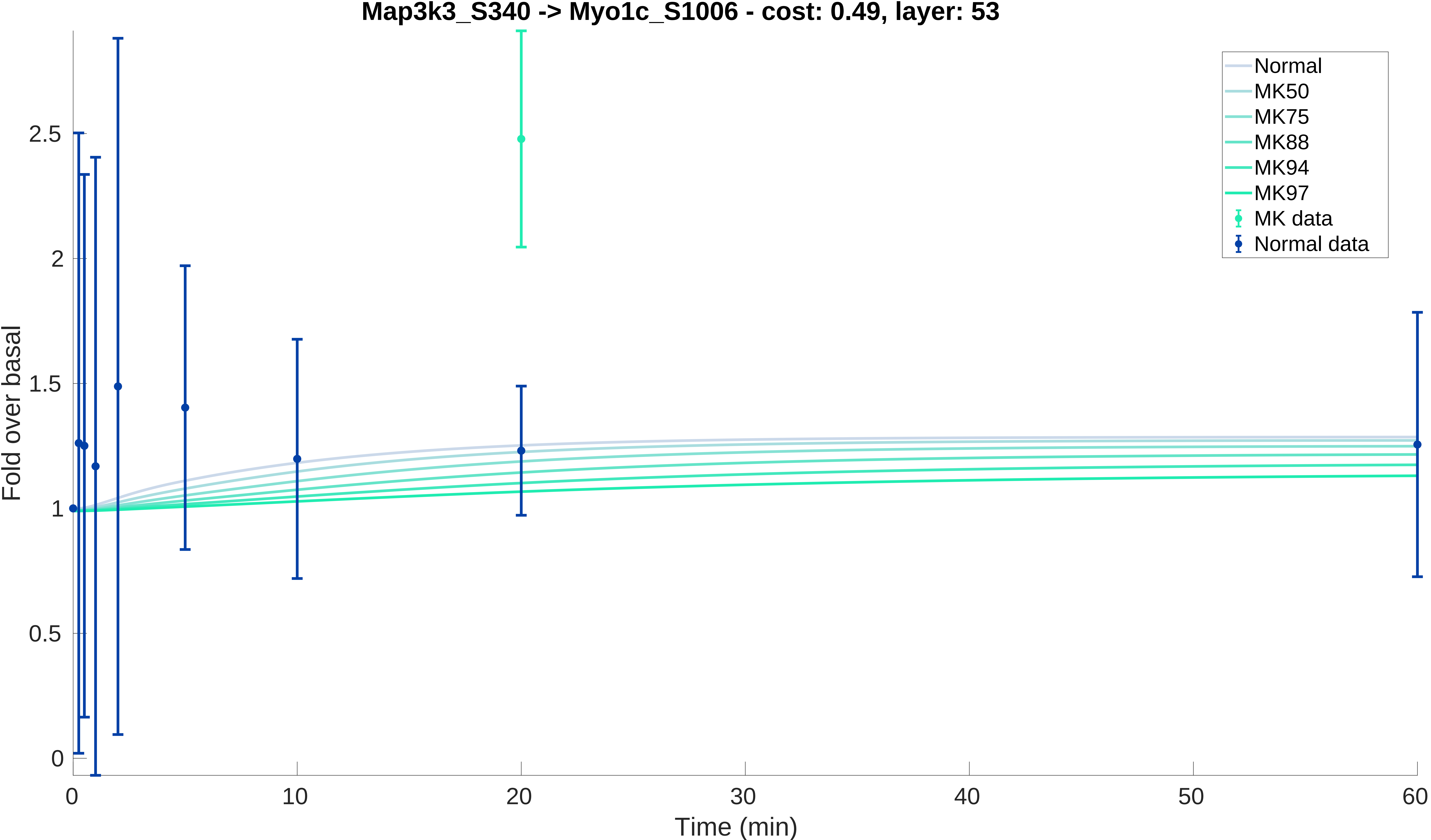

Map3k3_S340 -> Myo1c_S1006 - cost: 0.49, layer: 53
Normal
MK50
MK75
MK88
MK94
MK97
MK data
Normal data
Fold over basal
0
0.5
1
1.5
2
2.5
0
10
20
30
40
50
60
Time (min)

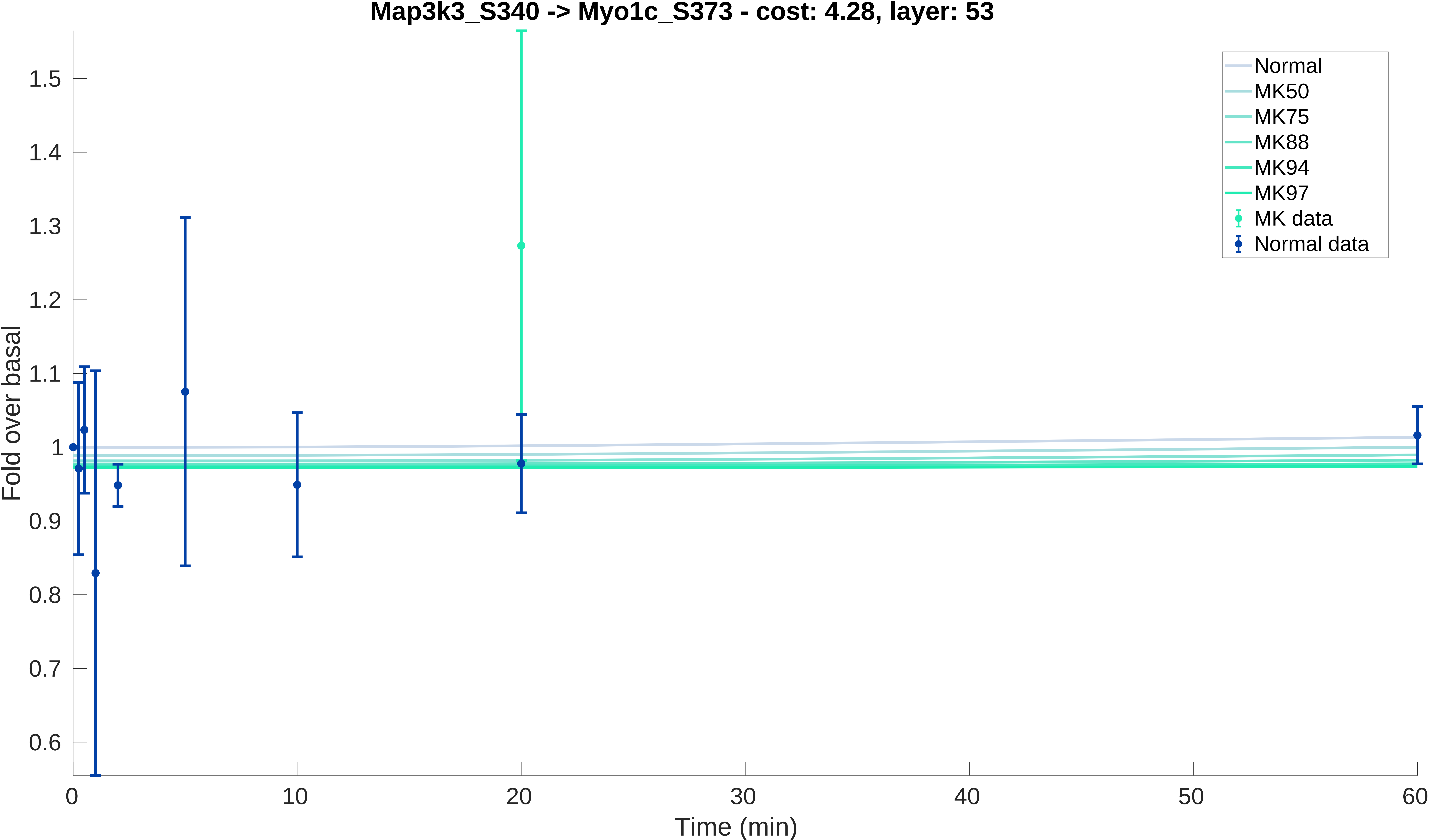

Map3k3_S340 -> Myo1c_S373 - cost: 4.28, layer: 53
Fold over basal
Time (min)
1.5
1.4
1.3
1.2
1.1
1
0.9
0.8
0.7
0.6
0
10
20
30
40
50
60
Normal
MK50
MK75
MK88
MK94
MK97
MK data
Normal data

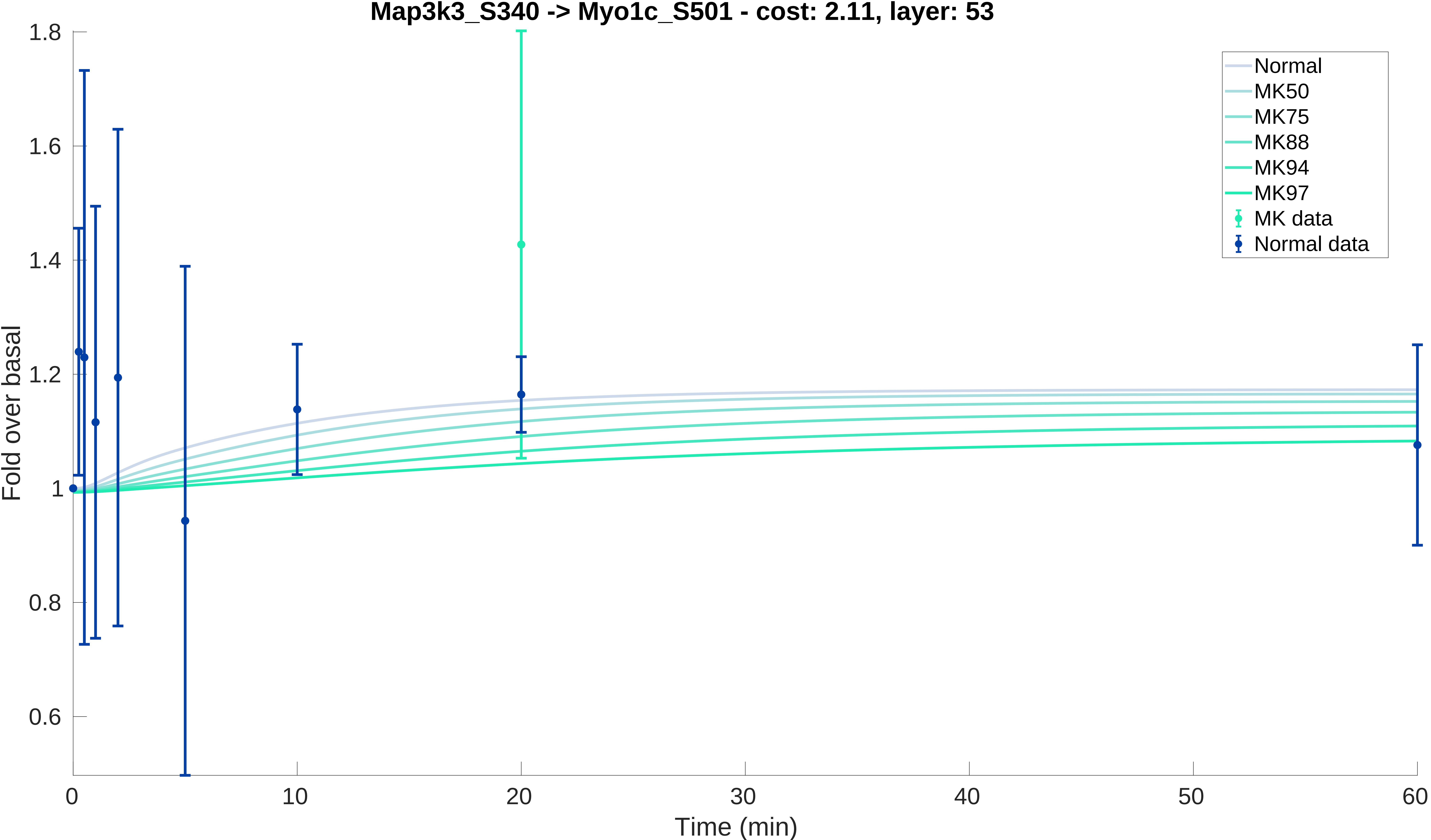

Map3k3_S340 -> Myo1c_S501 - cost: 2.11, layer: 53
Fold over basal
Time (min)
Normal
MK50
MK75
MK88
MK94
MK97
MK data
Normal data
1.8
1.6
1.4
1.2
1
0.8
0.6
0
10
20
30
40
50
60

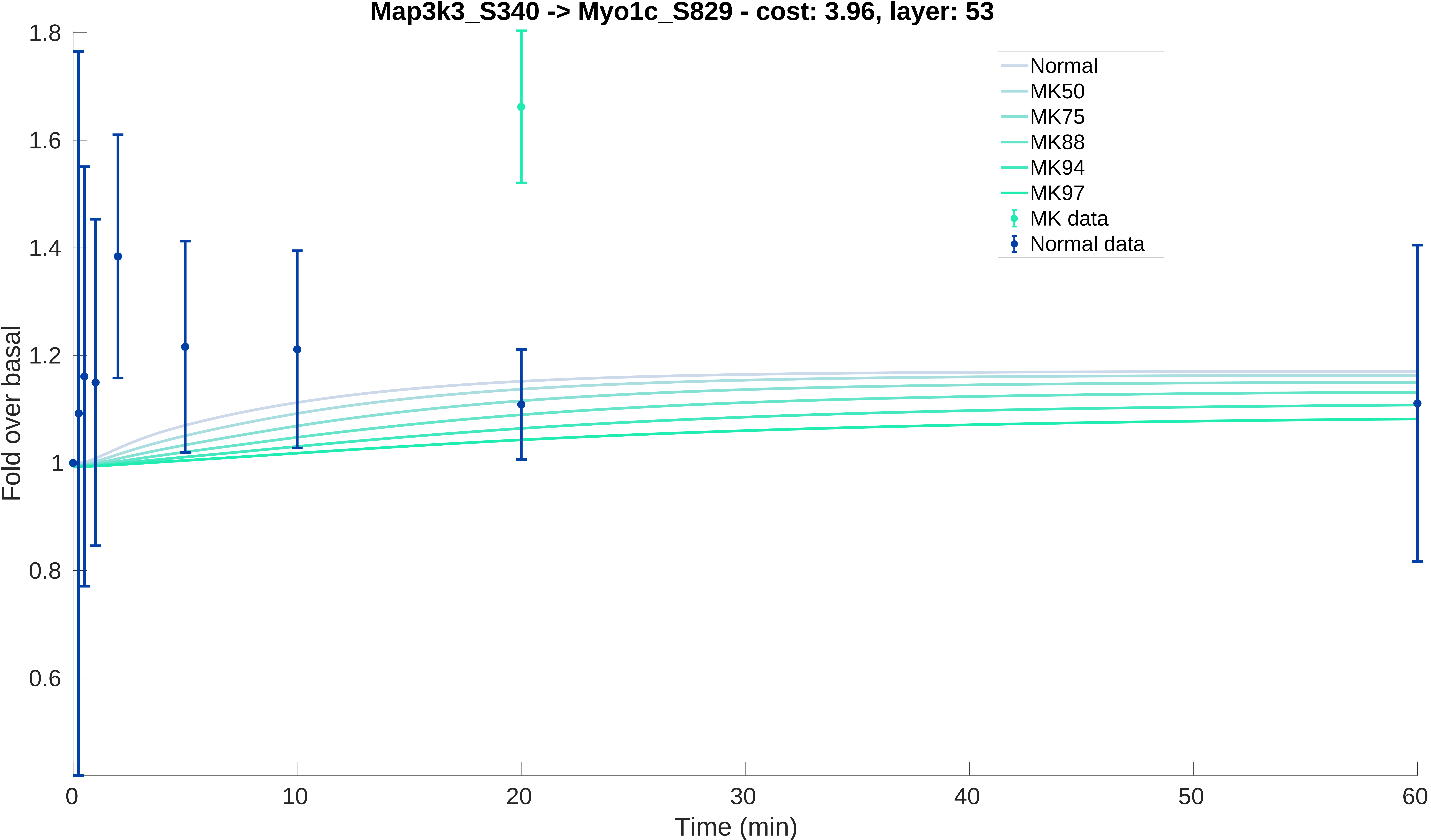
Map3k3_S340 -> Myo1c_S829 - cost: 3.96, layer: 53
Fold over basal
1.8
1.6
1.4
1.2
1
0.8
0.6
0
10
20
30
40
50
60
Time (min)
Normal
MK50
MK75
MK88
MK94
MK97
MK data
Normal data

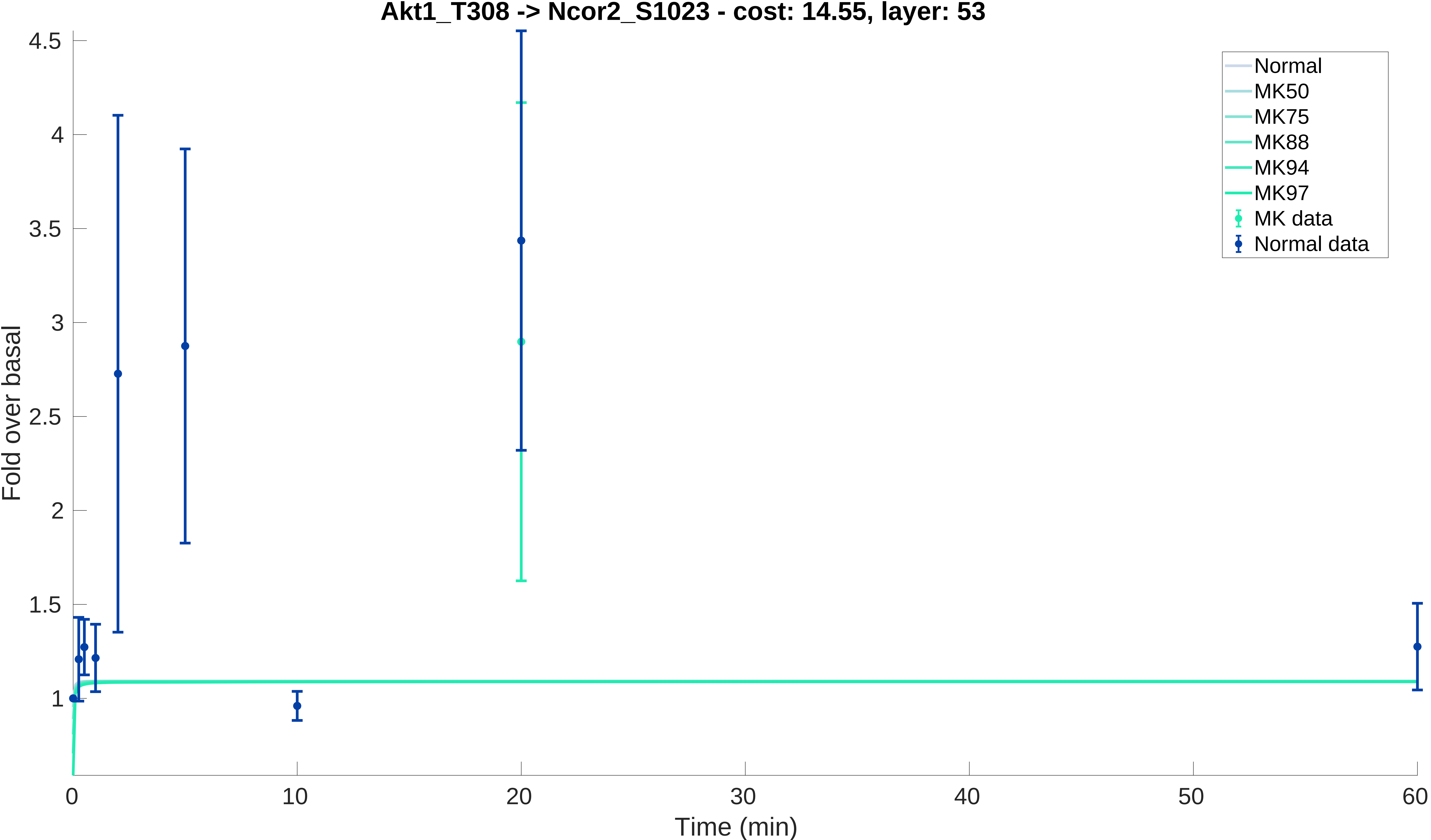

Akt1_T308 -> Ncor2_S1023 - cost: 14.55, layer: 53
Fold over basal
Time (min)
4.5
4
3.5
3
2.5
2
1.5
1
0
10
20
30
40
50
60
Normal
MK50
MK75
MK88
MK94
MK97
MK data
Normal data

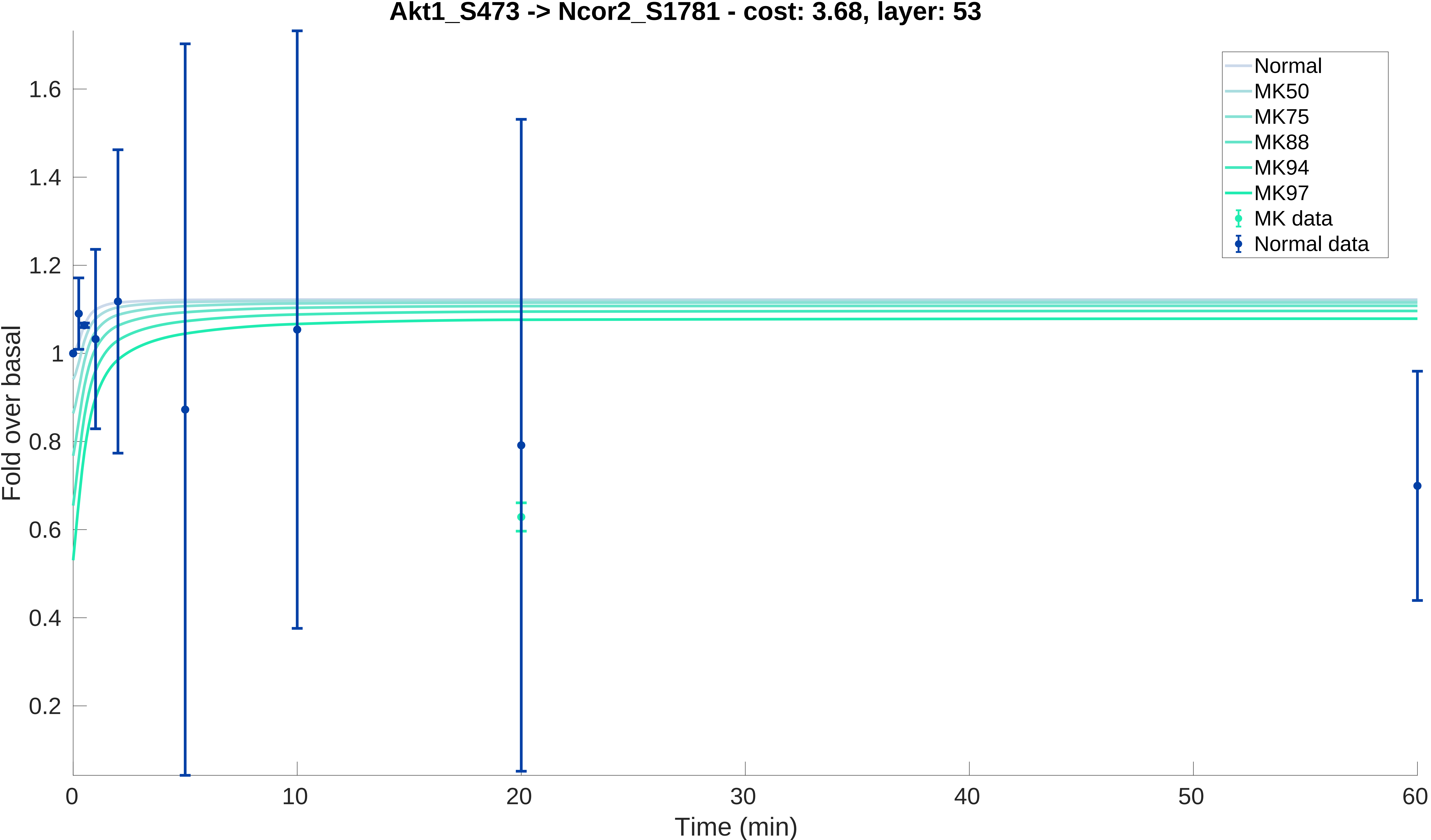
Akt1_S473 -> Ncor2_S1781 - cost: 3.68, layer: 53
Fold over basal
Time (min)
0
10
20
30
40
50
60
0.2
0.4
0.6
0.8
1
1.2
1.4
1.6
Normal
MK50
MK75
MK88
MK94
MK97
MK data
Normal data

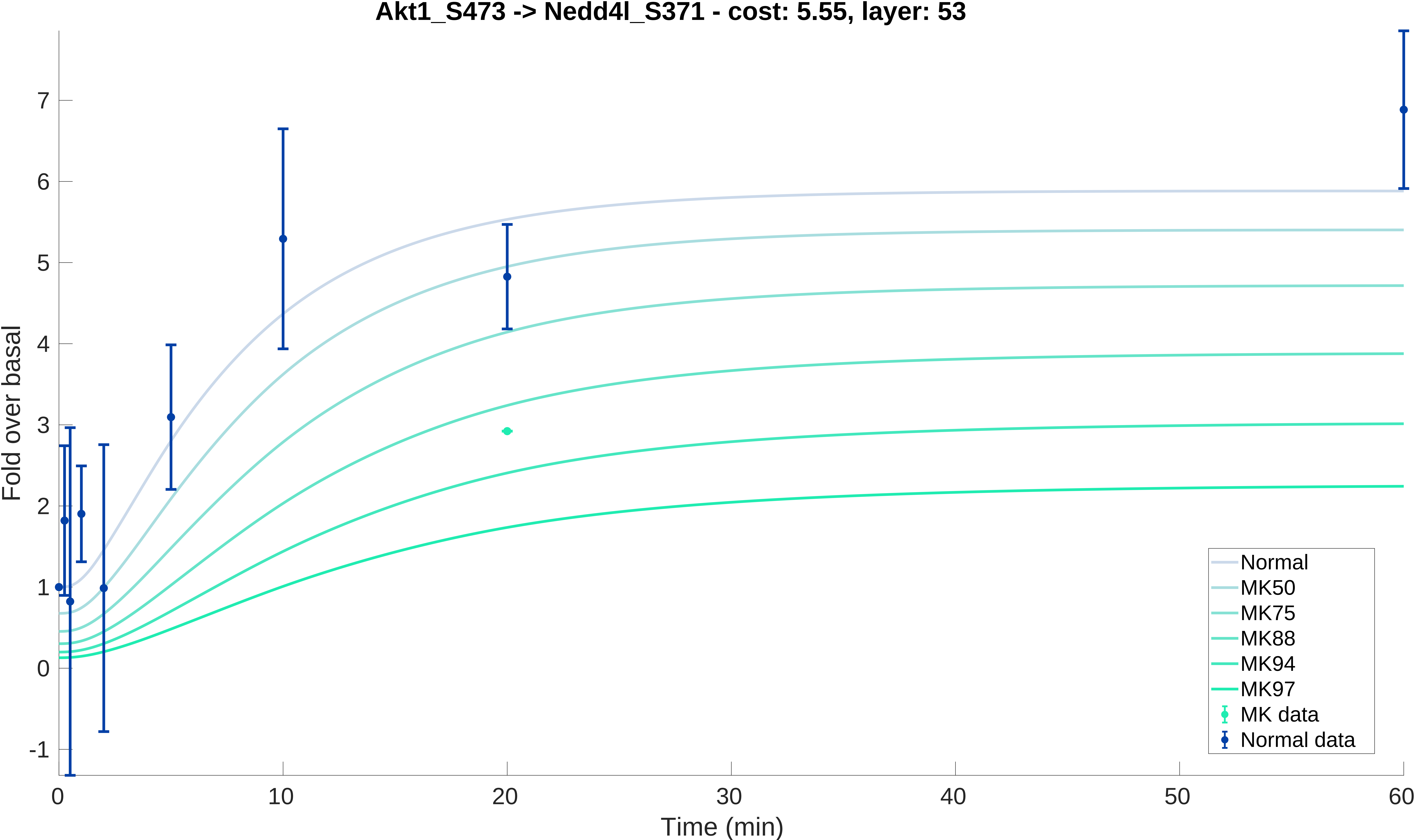
Akt1_S473 -> Nedd4l_S371 - cost: 5.55, layer: 53
Fold over basal
Time (min)
-1
0
1
2
3
4
5
6
7
0
10
20
30
40
50
60
Normal
MK50
MK75
MK88
MK94
MK97
MK data
Normal data

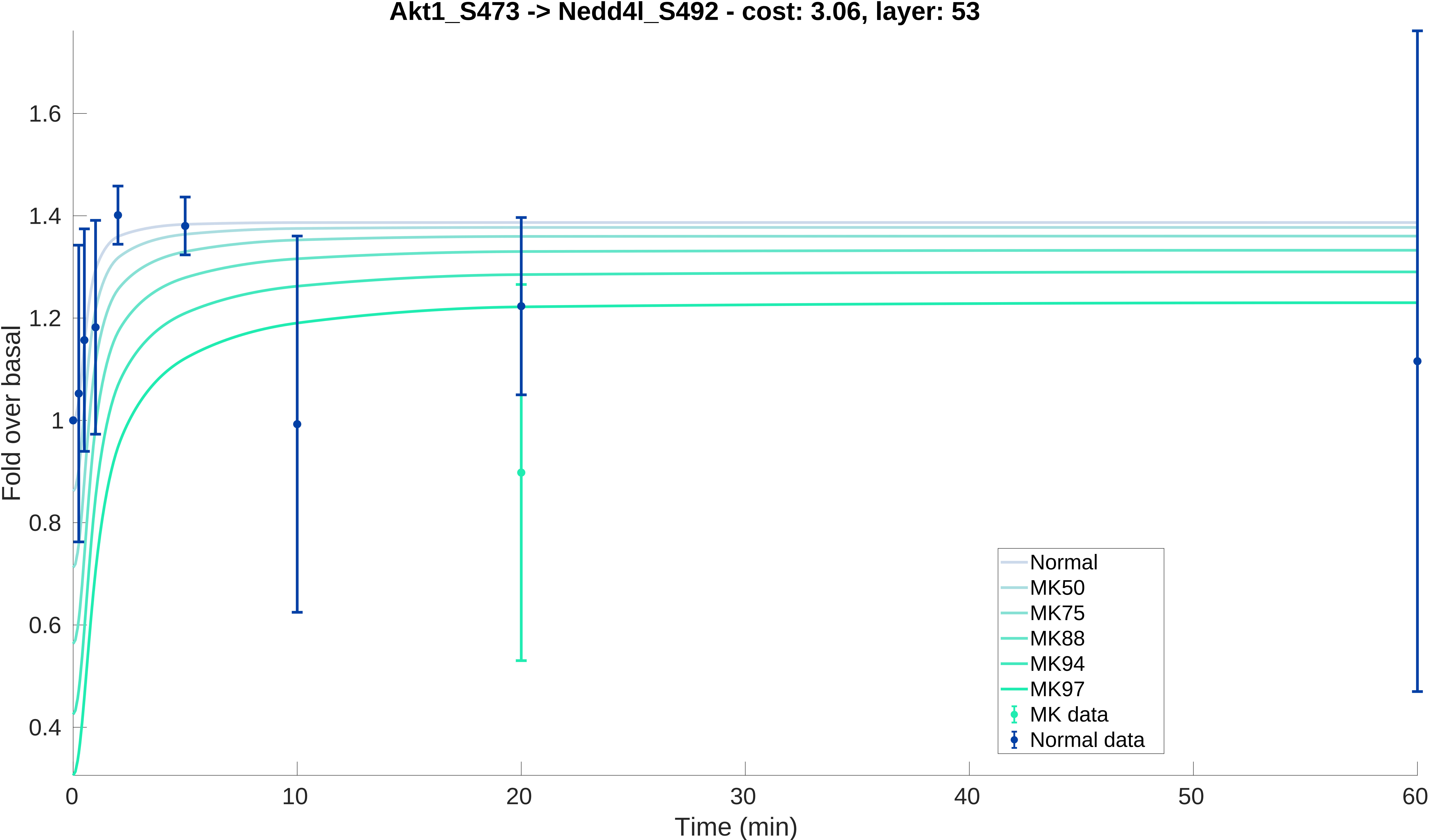
Akt1_S473 -> Nedd4l_S492 - cost: 3.06, layer: 53
Fold over basal
Time (min)
0.4
0.6
0.8
1
1.2
1.4
1.6
0
10
20
30
40
50
60
Normal
MK50
MK75
MK88
MK94
MK97
MK data
Normal data

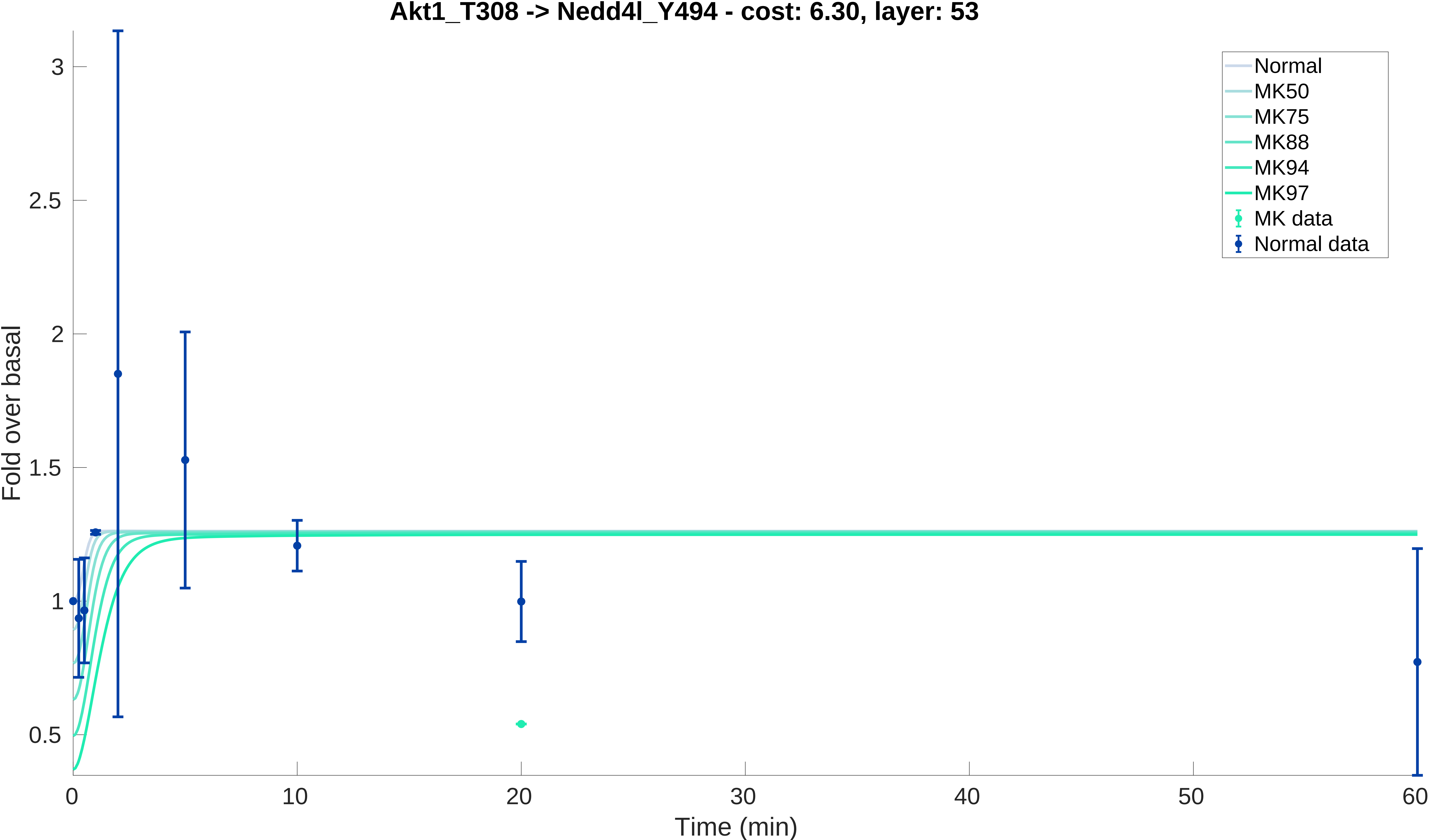

Akt1_T308 -> Nedd4l_Y494 - cost: 6.30, layer: 53
Fold over basal
3
2.5
2
1.5
1
0.5
0
10
20
30
40
50
60
Time (min)
Normal
MK50
MK75
MK88
MK94
MK97
MK data
Normal data

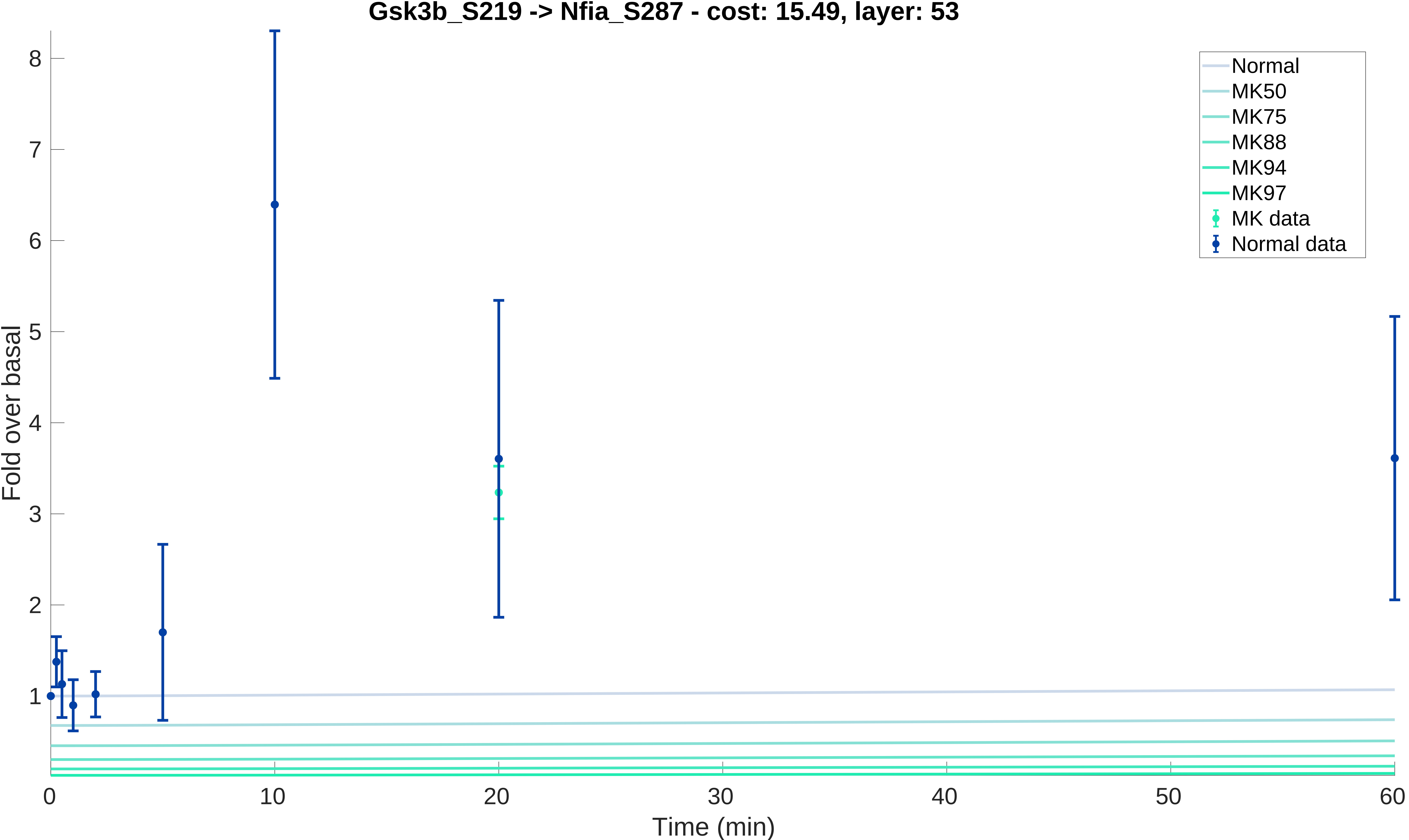

Gsk3b_S219 -> Nfia_S287 - cost: 15.49, layer: 53
Fold over basal
Time (min)
0
10
20
30
40
50
60
1
2
3
4
5
6
7
8
Normal
MK50
MK75
MK88
MK94
MK97
MK data
Normal data

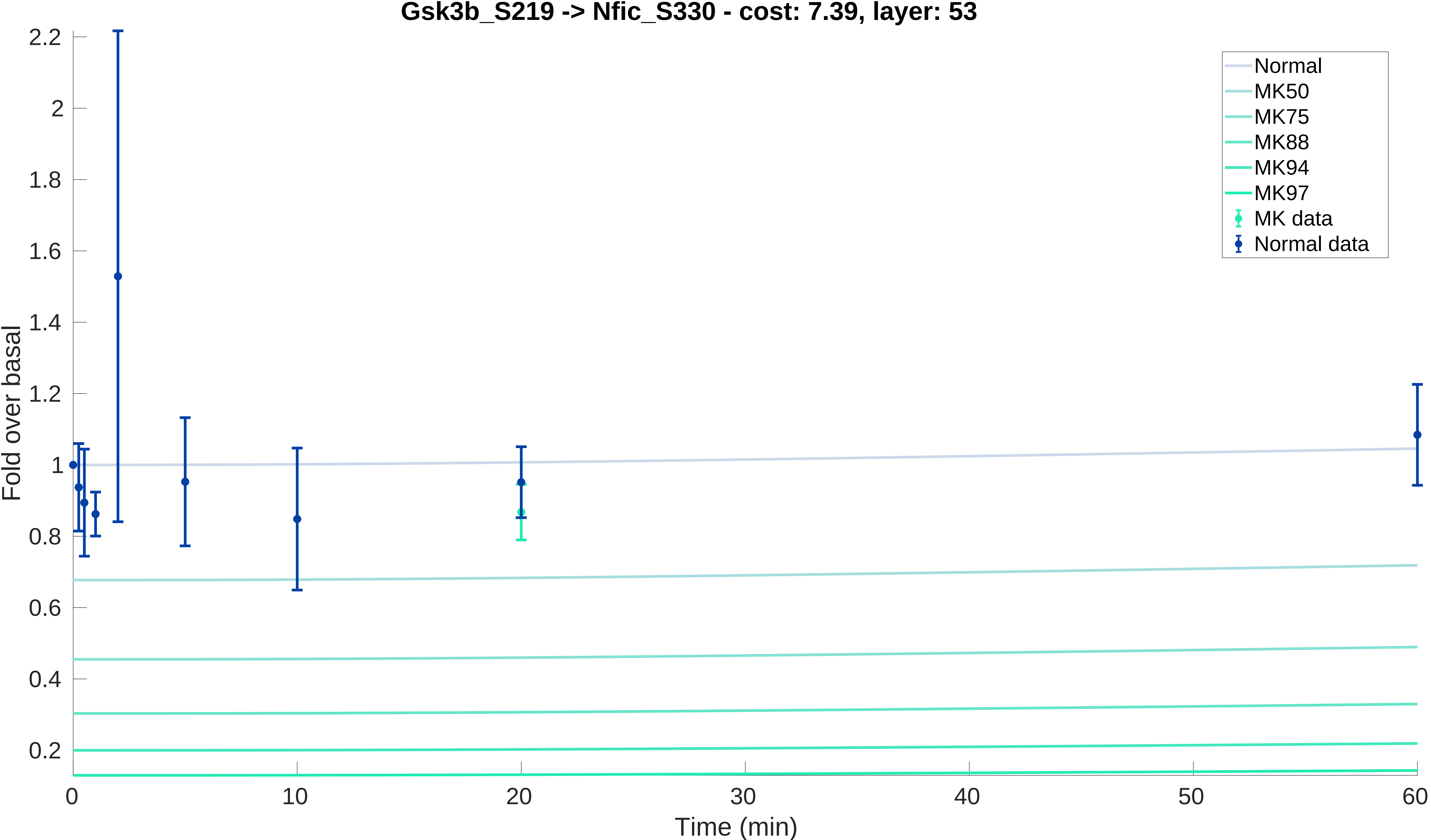

Gsk3b_S219 -> Nfic_S330 - cost: 7.39, layer: 53
Fold over basal
2.2
2
1.8
1.6
1.4
1.2
1
0.8
0.6
0.4
0.2
0
10
20
30
40
50
60
Time (min)
Normal
MK50
MK75
MK88
MK94
MK97
MK data
Normal data

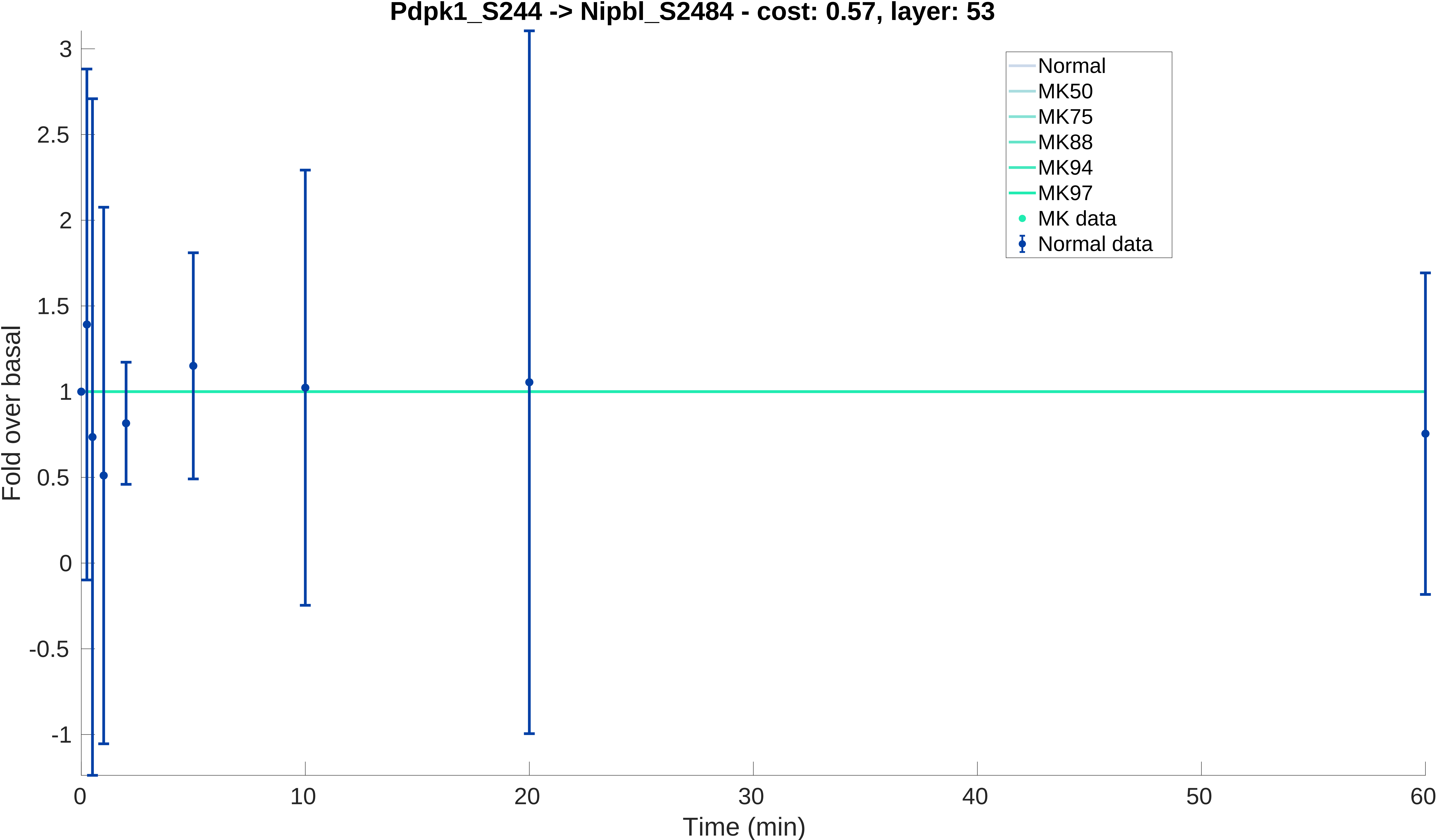

Pdpk1_S244 -> Nipbl_S2484 - cost: 0.57, layer: 53
Fold over basal
Time (min)
3
2.5
2
1.5
1
0.5
0
-0.5
-1
0
10
20
30
40
50
60
Normal
MK50
MK75
MK88
MK94
MK97
MK data
Normal data

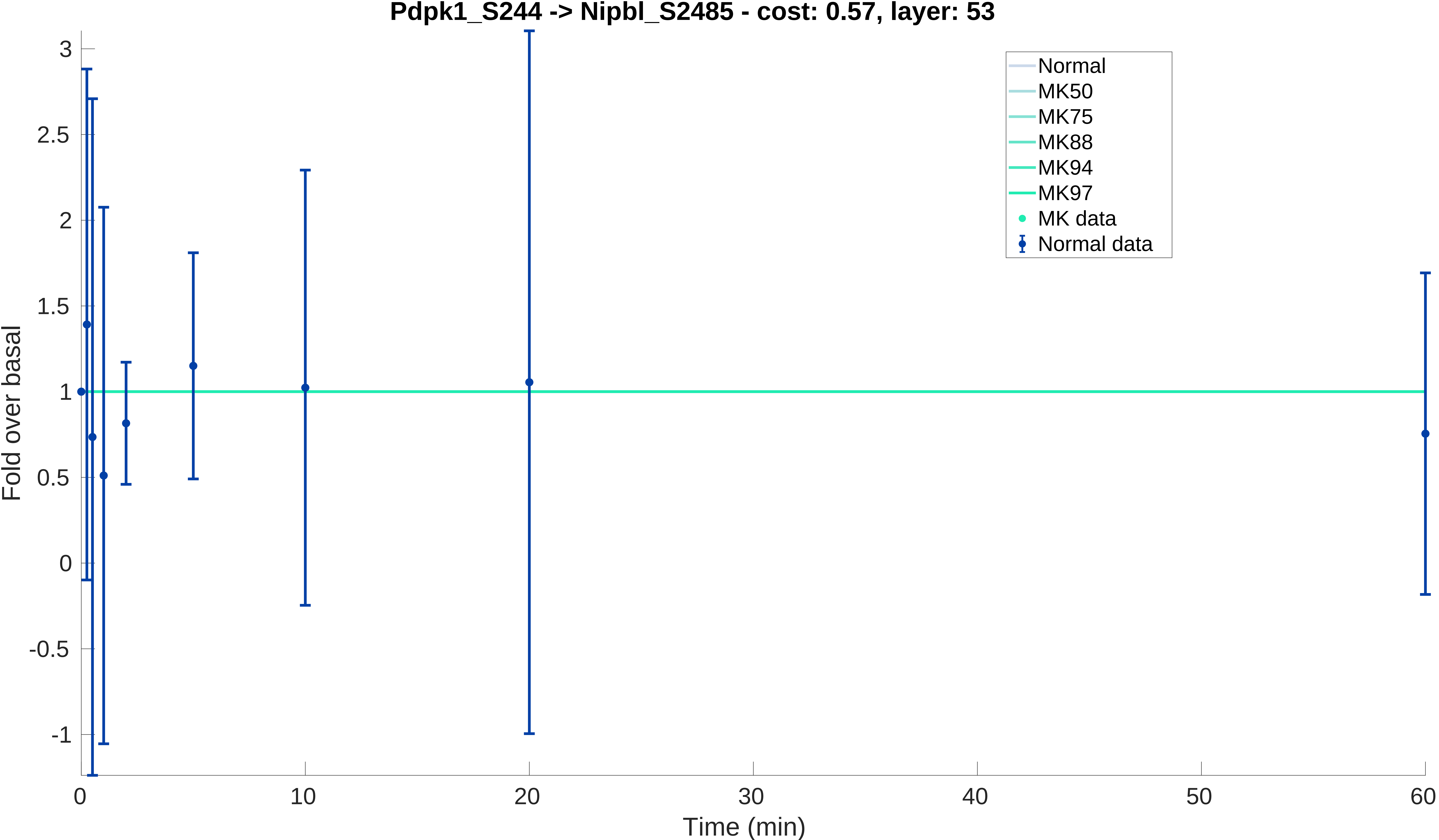

Pdpk1_S244 -> Nipbl_S2485 - cost: 0.57, layer: 53
Normal
MK50
MK75
MK88
MK94
MK97
MK data
Normal data
Fold over basal
Time (min)
3
2.5
2
1.5
1
0.5
0
-0.5
-1
0
10
20
30
40
50
60

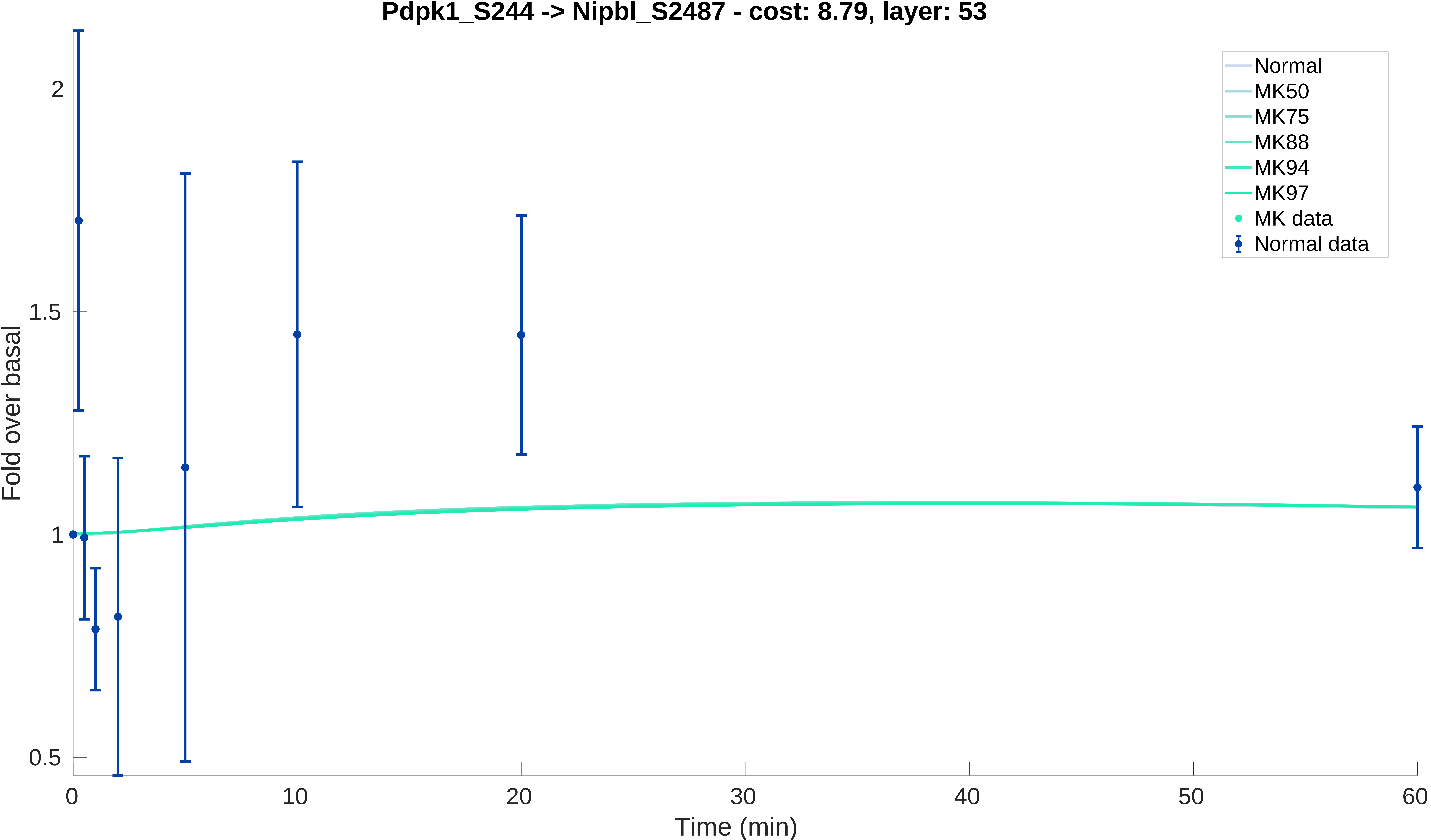

Pdpk1_S244 -> Nipbl_S2487 - cost: 8.79, layer: 53
Fold over basal
Time (min)
0.5
1
1.5
2
0
10
20
30
40
50
60
Normal
MK50
MK75
MK88
MK94
MK97
MK data
Normal data

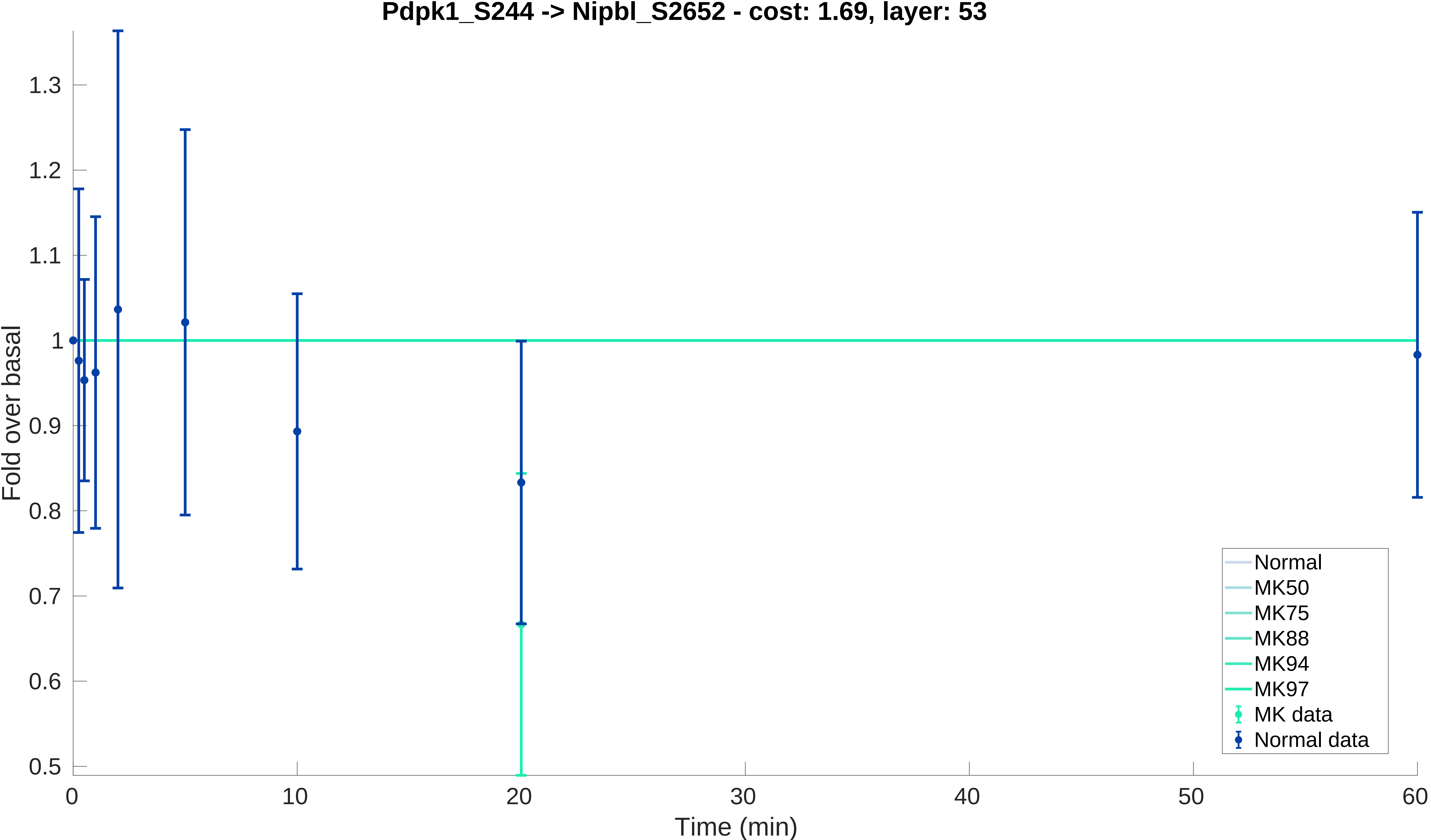
Pdpk1_S244 -> Nipbl_S2652 - cost: 1.69, layer: 53
Fold over basal
Time (min)
0
10
20
30
40
50
60
0.5
0.6
0.7
0.8
0.9
1
1.1
1.2
1.3
Normal
MK50
MK75
MK88
MK94
MK97
MK data
Normal data

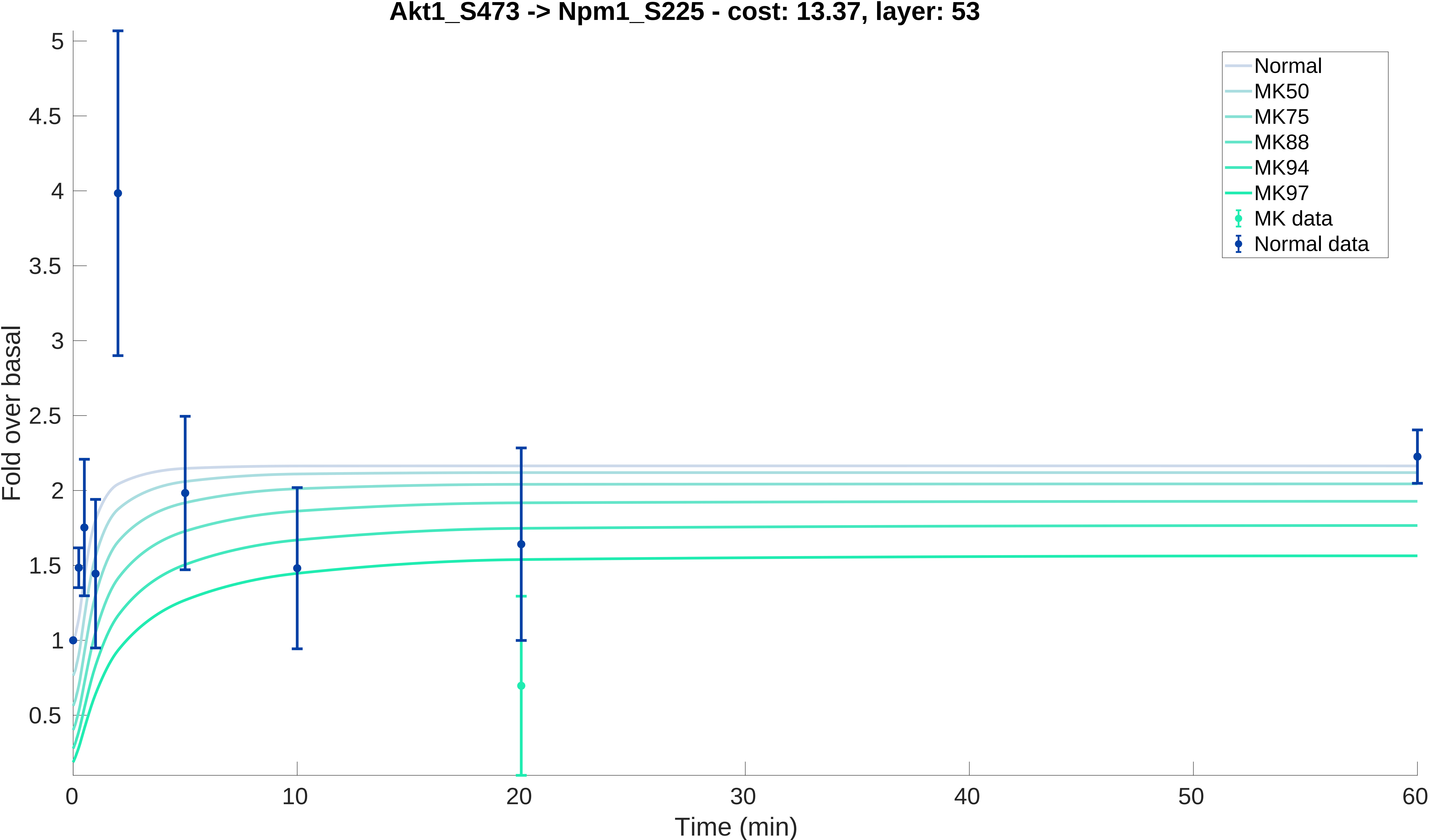

Akt1_S473 -> Npm1_S225 - cost: 13.37, layer: 53
Fold over basal
Time (min)
0
10
20
30
40
50
60
0.5
1
1.5
2
2.5
3
3.5
4
4.5
5
Normal
MK50
MK75
MK88
MK94
MK97
MK data
Normal data

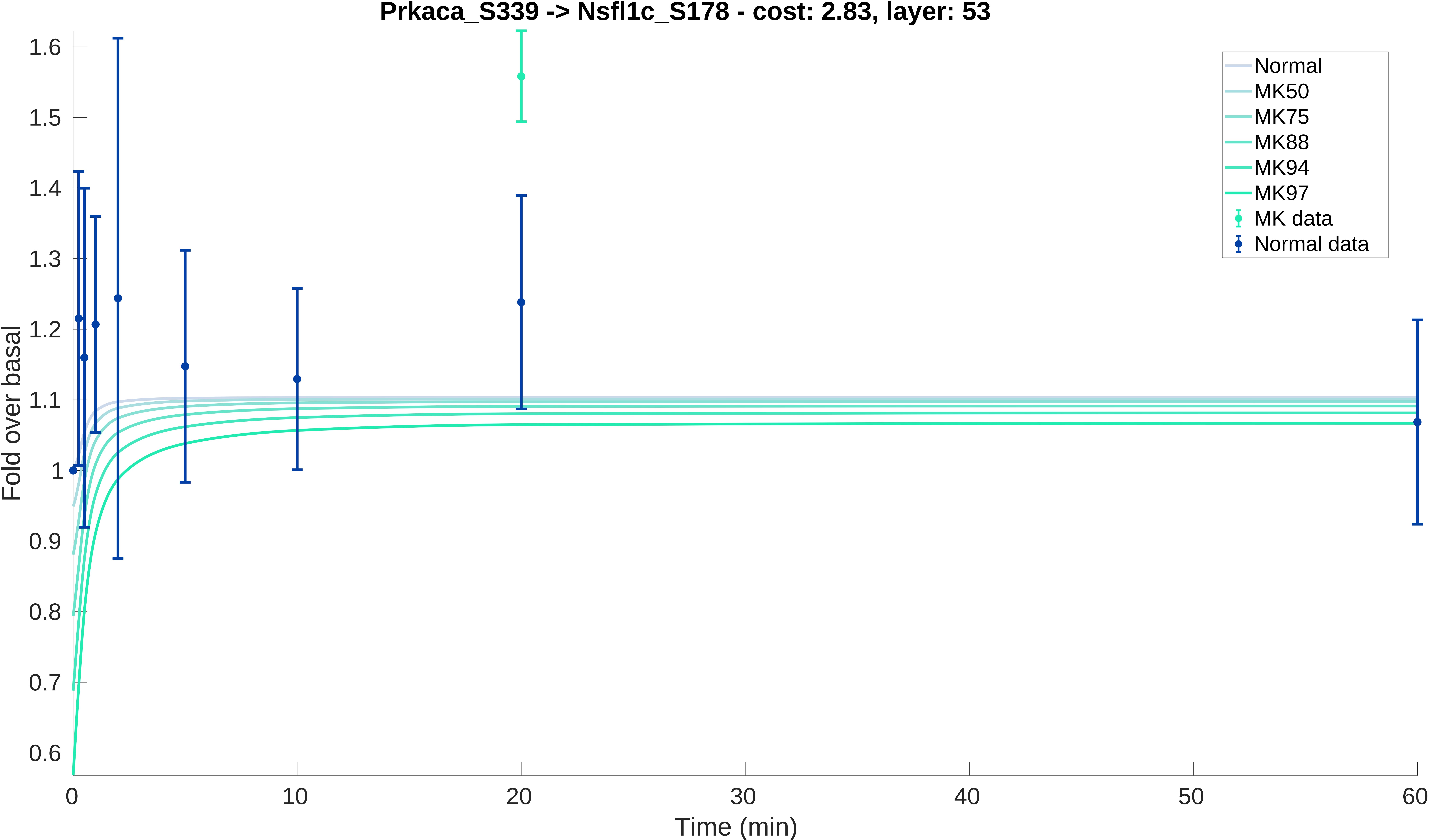
Prkaca_S339 -> Nsfl1c_S178 - cost: 2.83, layer: 53
Fold over basal
Time (min)
1.6
1.5
1.4
1.3
1.2
1.1
1
0.9
0.8
0.7
0.6
0
10
20
30
40
50
60
Normal
MK50
MK75
MK88
MK94
MK97
MK data
Normal data

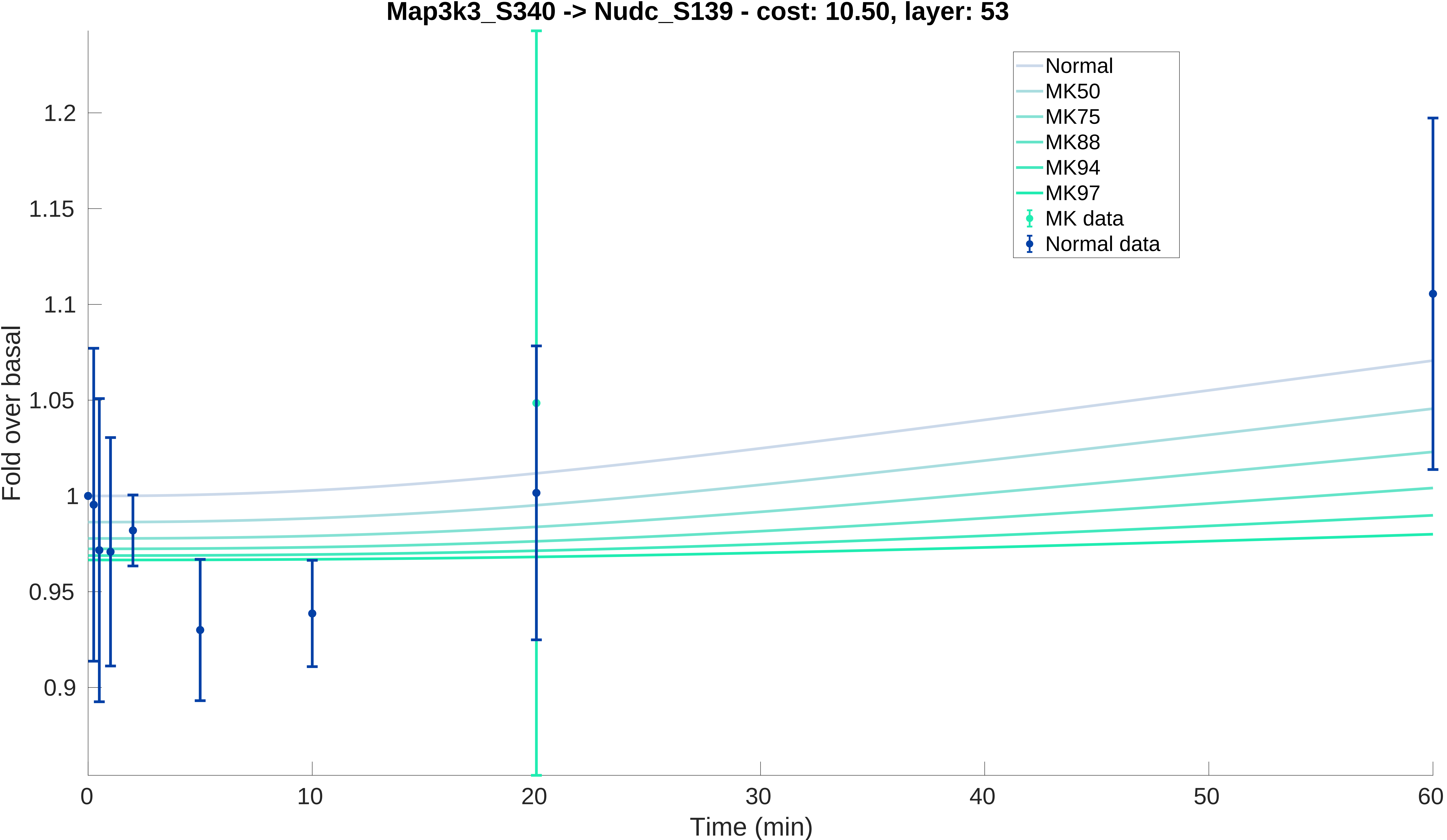
Map3k3_S340 -> Nudc_S139 - cost: 10.50, layer: 53
Normal
MK50
MK75
MK88
MK94
MK97
MK data
Normal data
Fold over basal
1.2
1.15
1.1
1.05
1
0.95
0.9
0
10
20
30
40
50
60
Time (min)

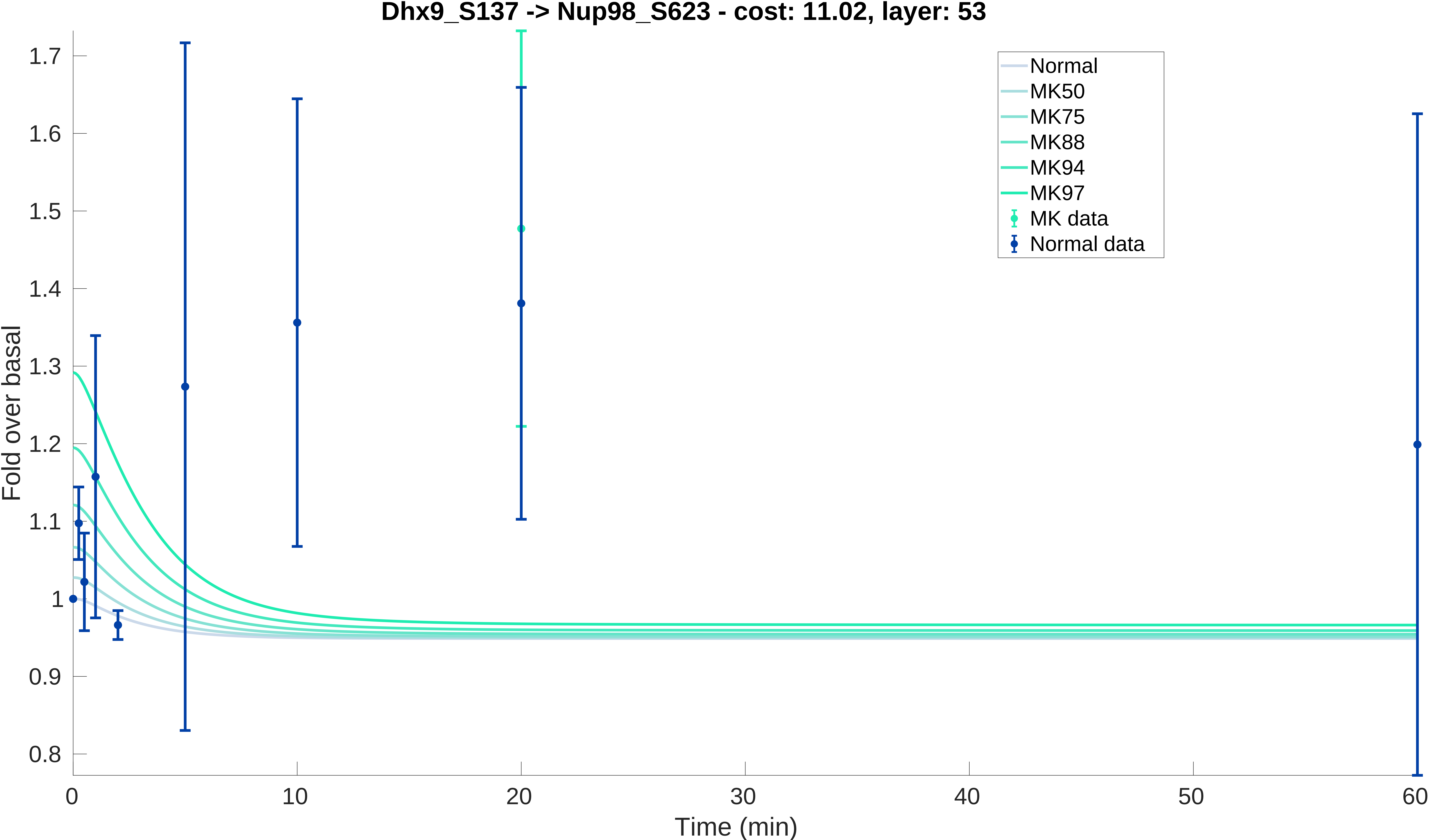

Dhx9_S137 -> Nup98_S623 - cost: 11.02, layer: 53
Fold over basal
Time (min)
0.8
0.9
1
1.1
1.2
1.3
1.4
1.5
1.6
1.7
0
10
20
30
40
50
60
Normal
MK50
MK75
MK88
MK94
MK97
MK data
Normal data

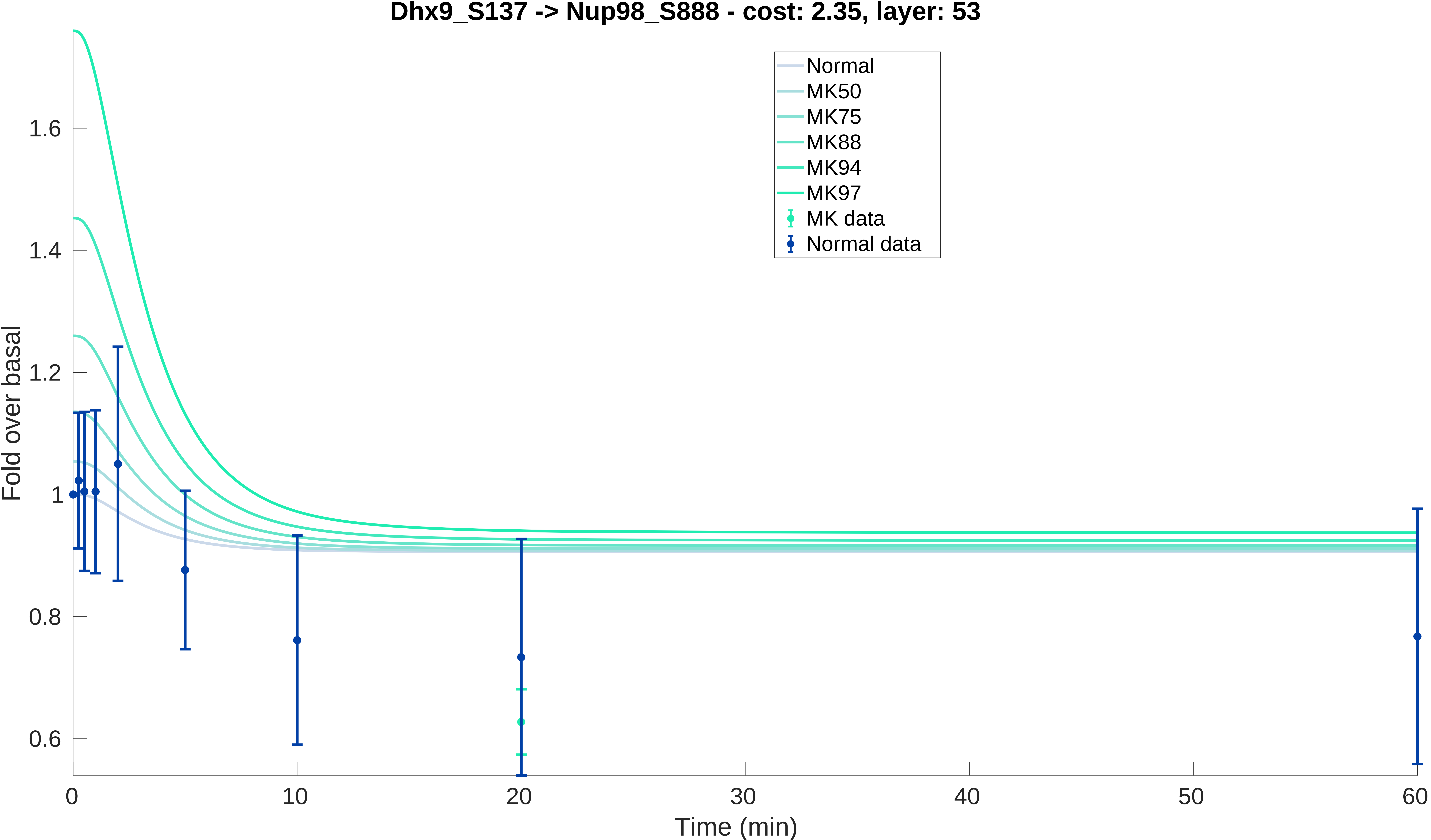

Dhx9_S137 -> Nup98_S888 - cost: 2.35, layer: 53
Normal
MK50
MK75
MK88
MK94
MK97
MK data
Normal data
1.6
1.4
1.2
1
0.8
0.6
Fold over basal
0
10
20
30
40
50
60
Time (min)

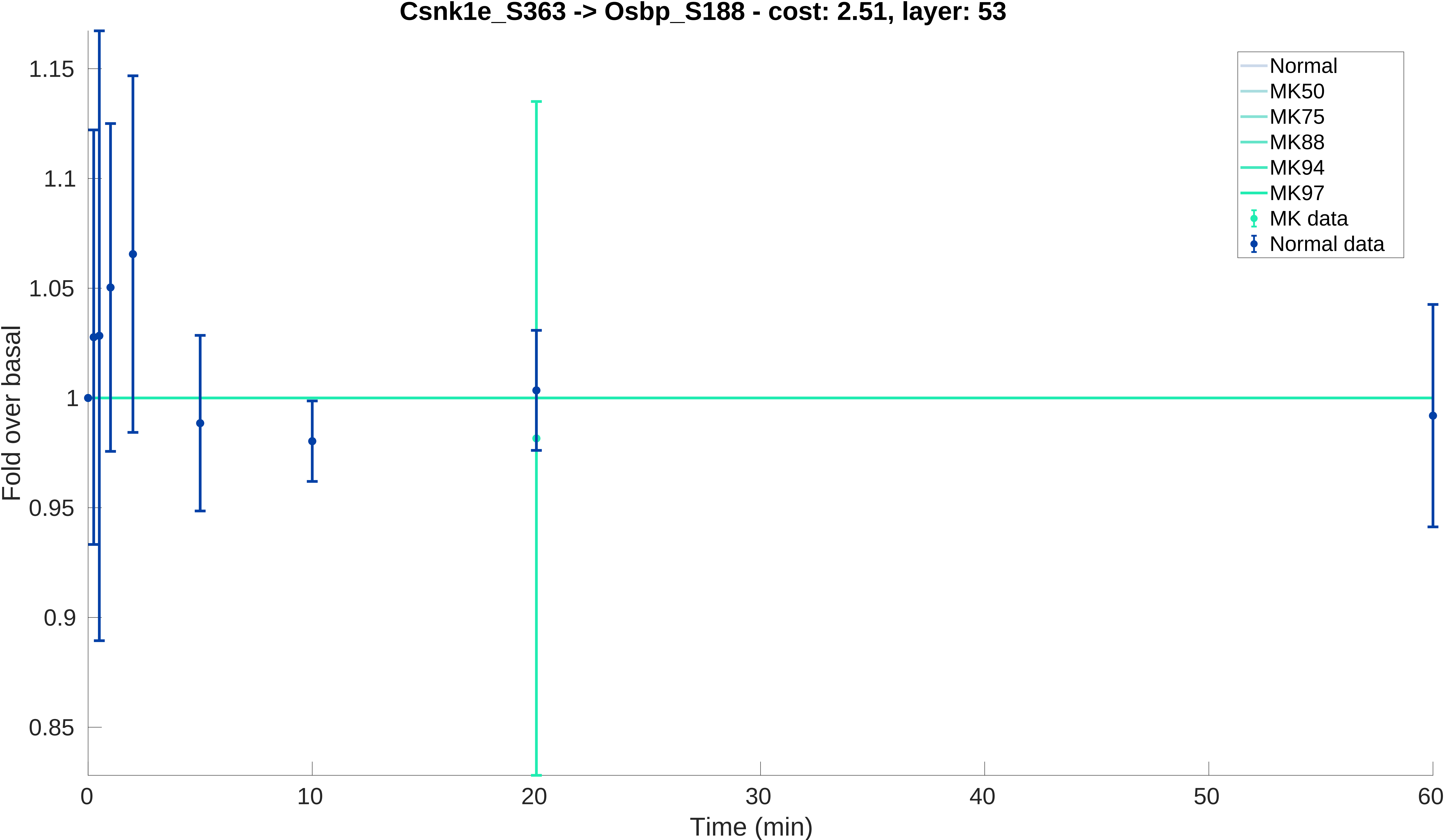

Csnk1e_S363 -> Osbp_S188 - cost: 2.51, layer: 53
Fold over basal
Time (min)
1.15
1.1
1.05
1
0.95
0.9
0.85
0
10
20
30
40
50
60
Normal
MK50
MK75
MK88
MK94
MK97
MK data
Normal data

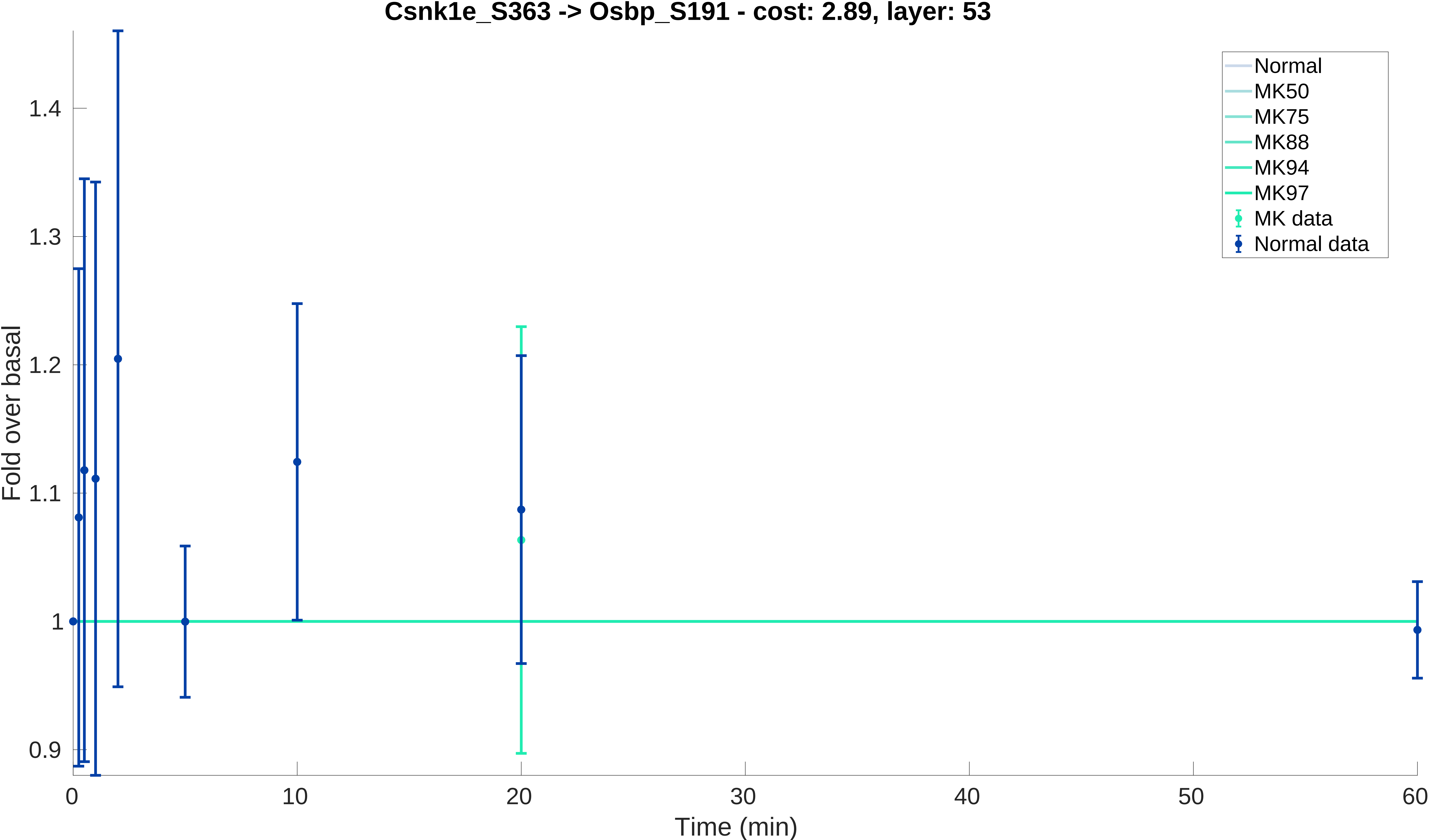

Csnk1e_S363 -> Osbp_S191 - cost: 2.89, layer: 53
Fold over basal
1.4
1.3
1.2
1.1
1
0.9
0
10
20
30
40
50
60
Time (min)
Normal
MK50
MK75
MK88
MK94
MK97
MK data
Normal data

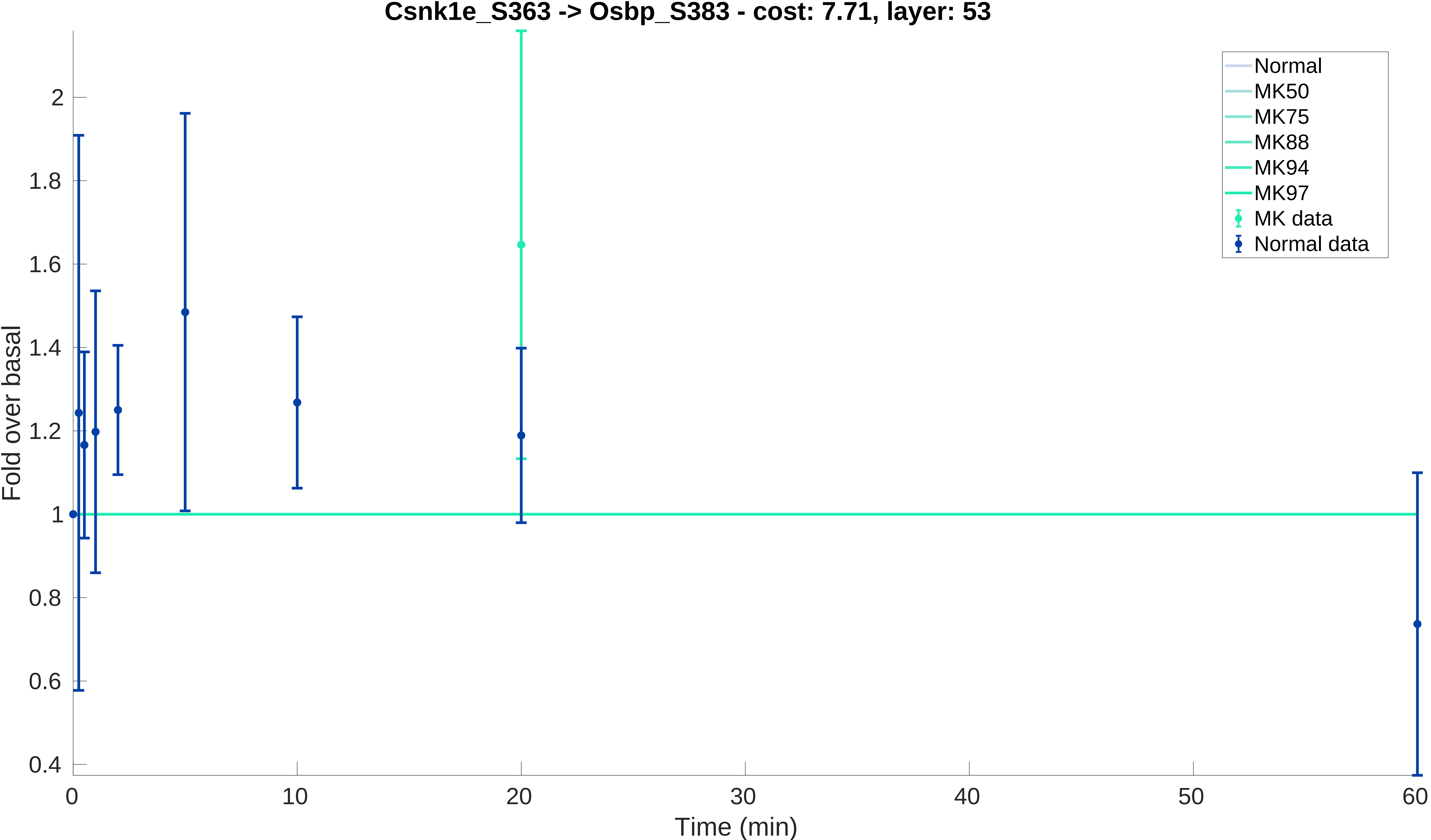

Csnk1e_S363 -> Osbp_S383 - cost: 7.71, layer: 53
Fold over basal
Time (min)
0
10
20
30
40
50
60
0.4
0.6
0.8
1
1.2
1.4
1.6
1.8
2
Normal
MK50
MK75
MK88
MK94
MK97
MK data
Normal data

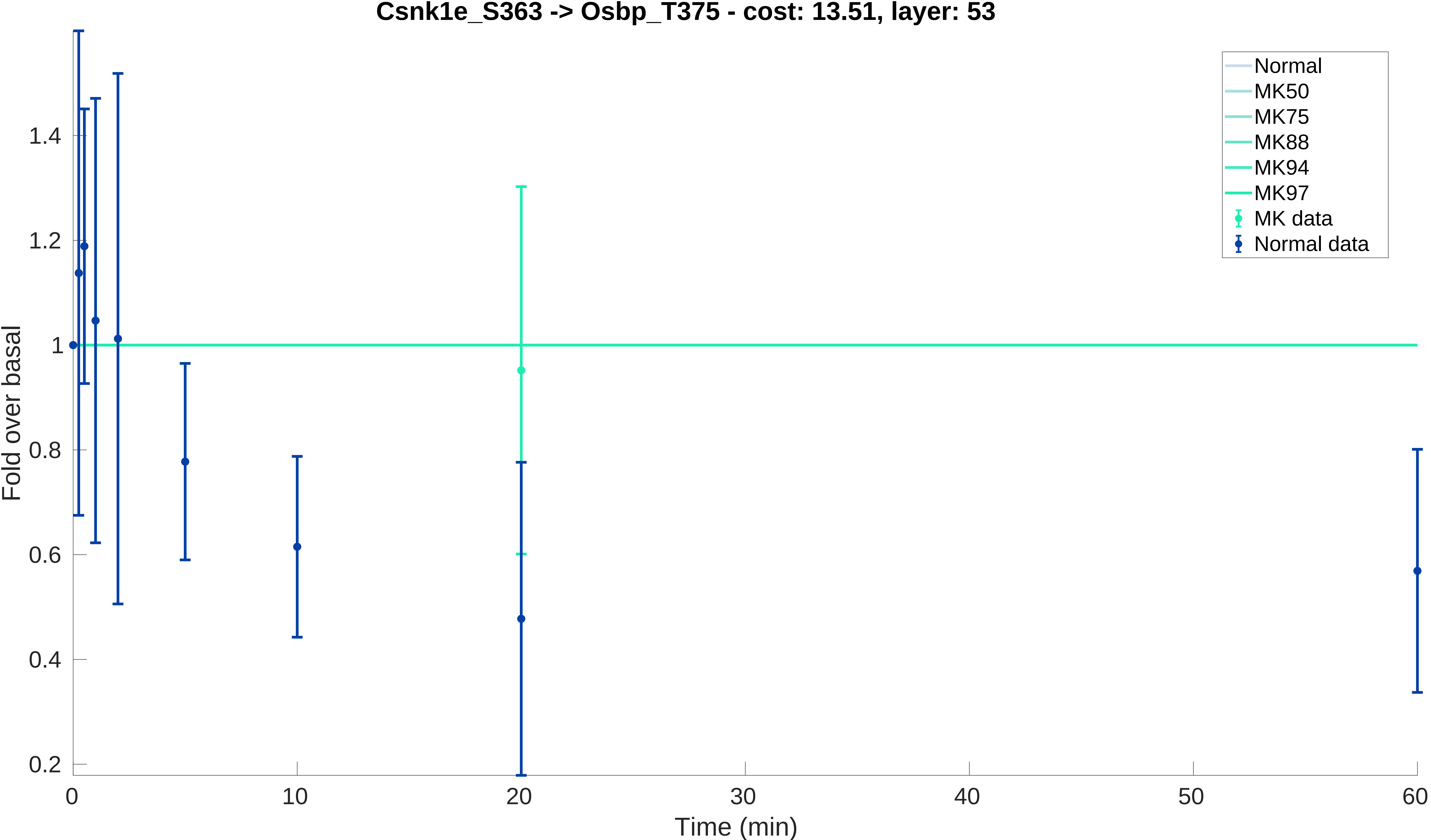

Csnk1e_S363 -> Osbp_T375 - cost: 13.51, layer: 53
Fold over basal
Time (min)
0
10
20
30
40
50
60
0.2
0.4
0.6
0.8
1
1.2
1.4
Normal
MK50
MK75
MK88
MK94
MK97
MK data
Normal data

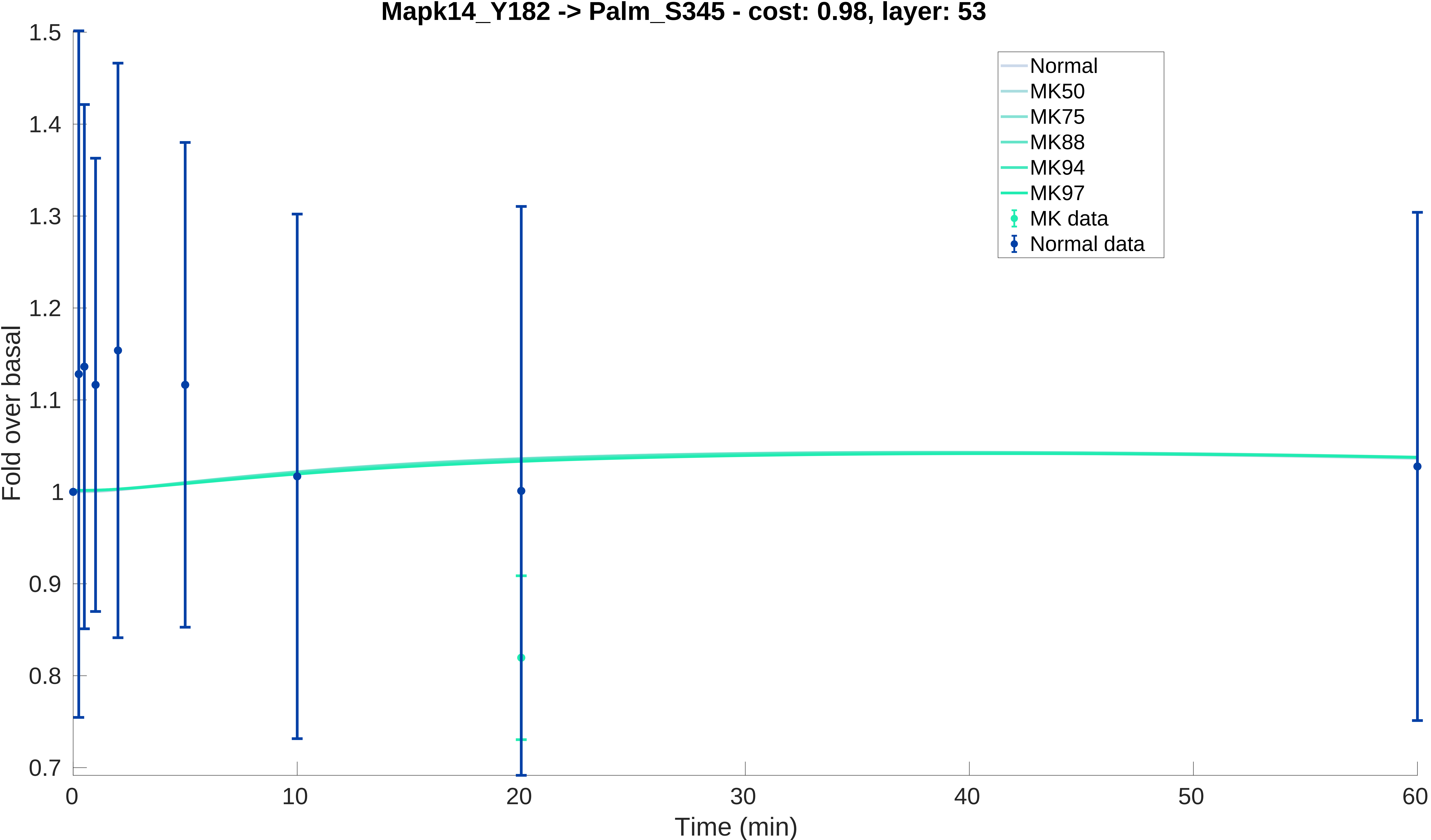
Mapk14_Y182 -> Palm_S345 - cost: 0.98, layer: 53
Fold over basal
Time (min)
1.5
1.4
1.3
1.2
1.1
1
0.9
0.8
0.7
0
10
20
30
40
50
60
Normal
MK50
MK75
MK88
MK94
MK97
MK data
Normal data

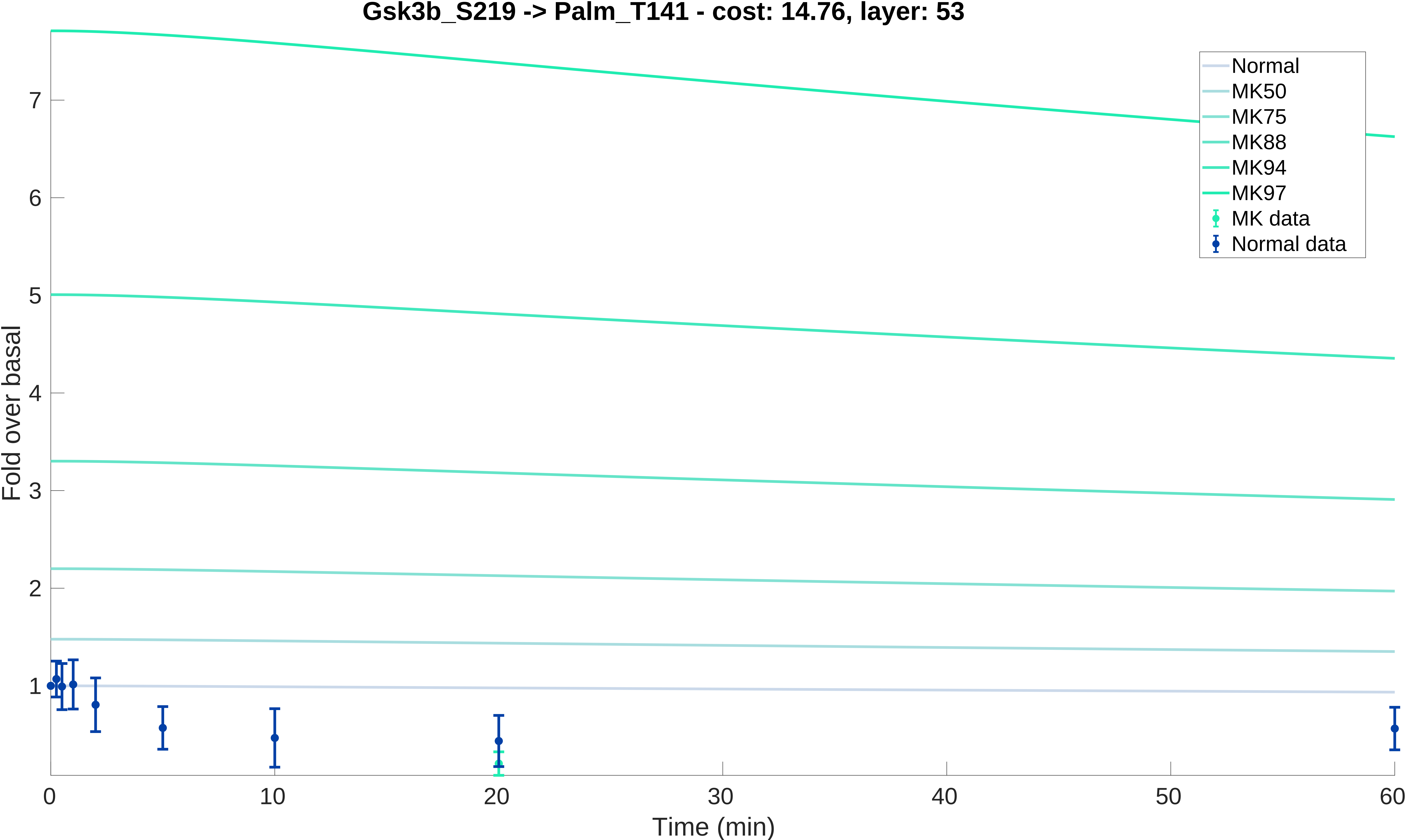

Gsk3b_S219 -> Palm_T141 - cost: 14.76, layer: 53
Normal
MK50
MK75
MK88
MK94
MK97
MK data
Normal data
Fold over basal
Time (min)
1
2
3
4
5
6
7
0
10
20
30
40
50
60

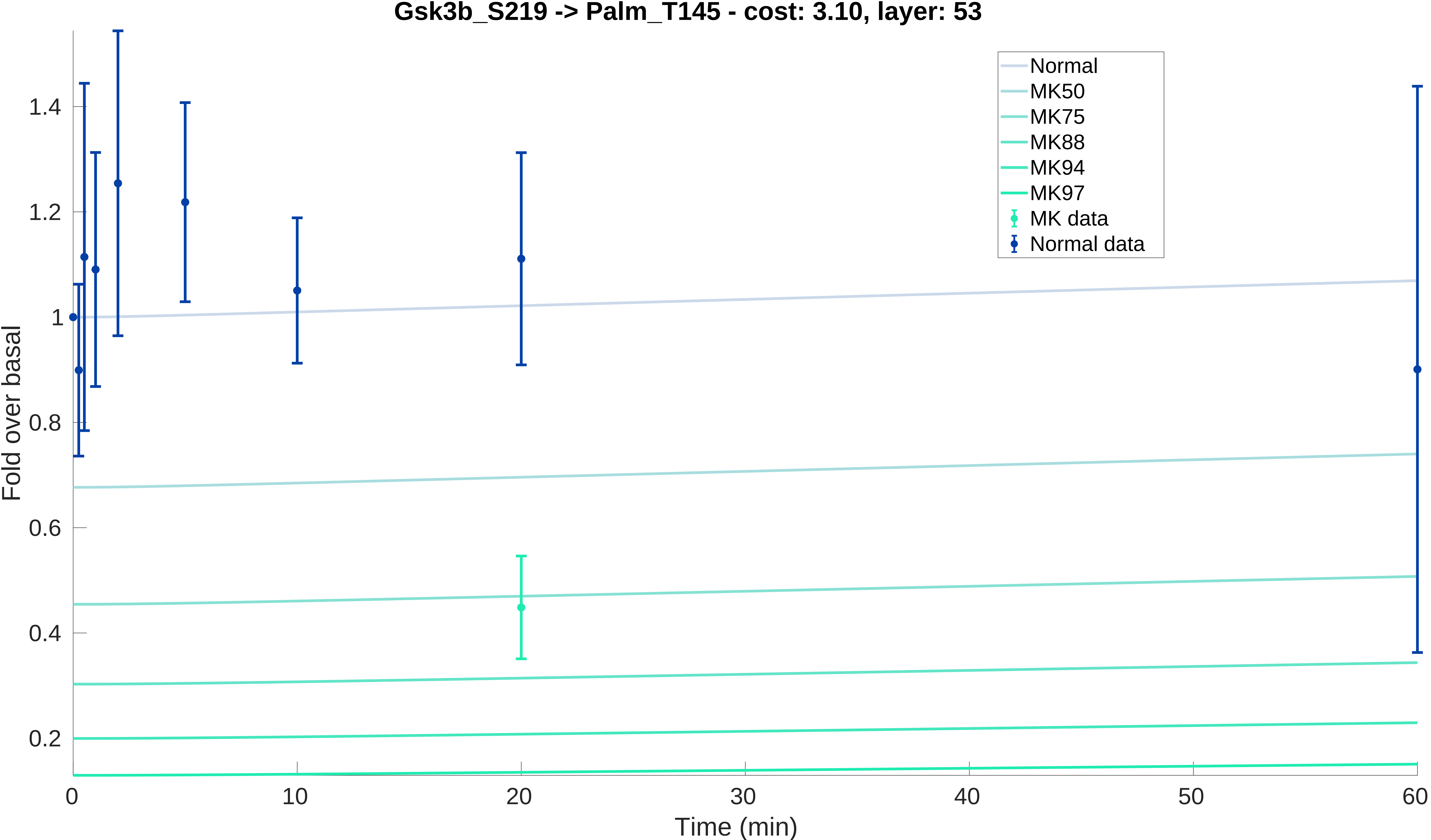

Gsk3b_S219 -> Palm_T145 - cost: 3.10, layer: 53
Fold over basal
Time (min)
0.2
0.4
0.6
0.8
1
1.2
1.4
0
10
20
30
40
50
60
Normal
MK50
MK75
MK88
MK94
MK97
MK data
Normal data

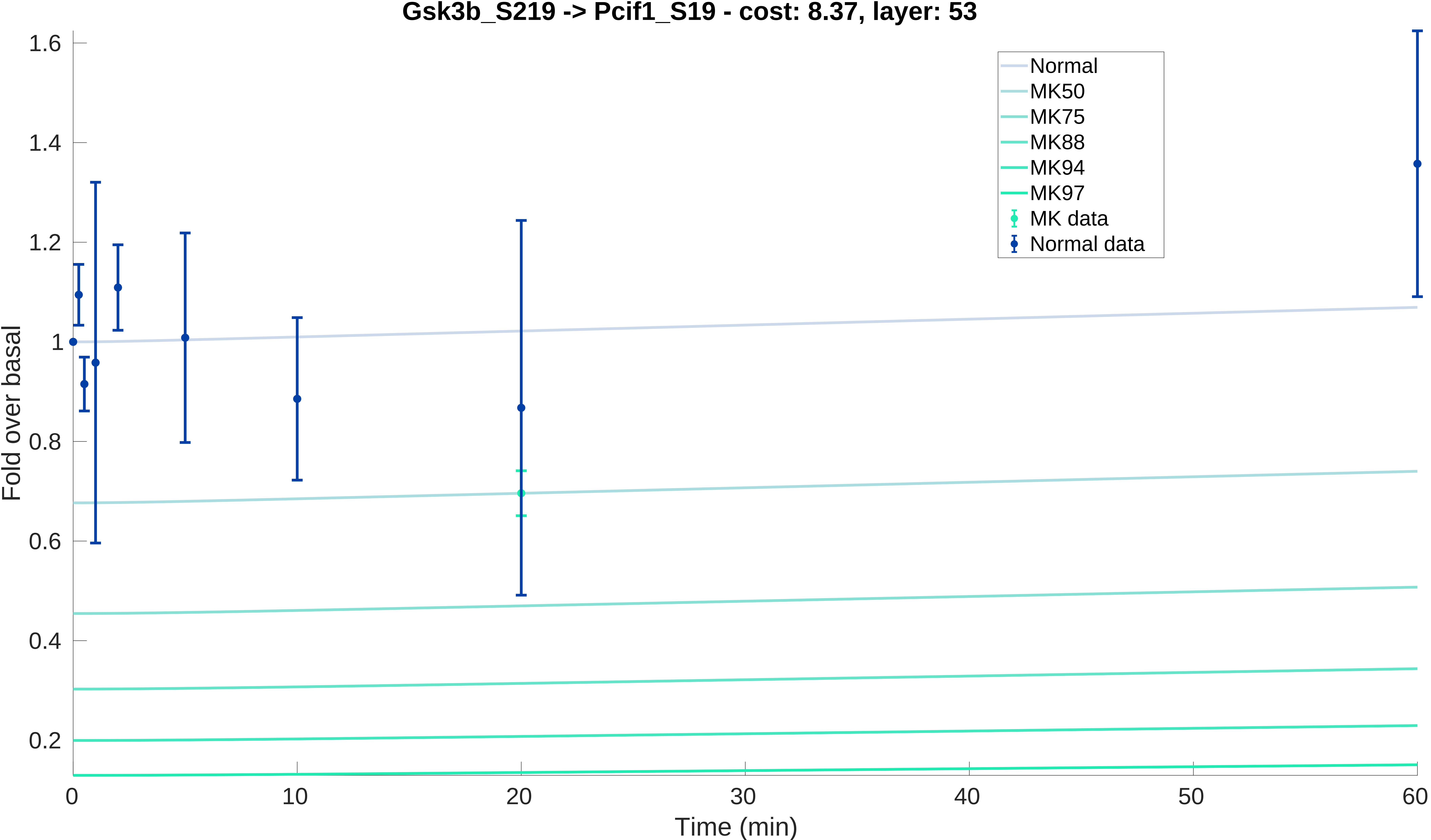

Gsk3b_S219 -> Pcif1_S19 - cost: 8.37, layer: 53
Fold over basal
Time (min)
0
10
20
30
40
50
60
0.2
0.4
0.6
0.8
1
1.2
1.4
1.6
Normal
MK50
MK75
MK88
MK94
MK97
MK data
Normal data

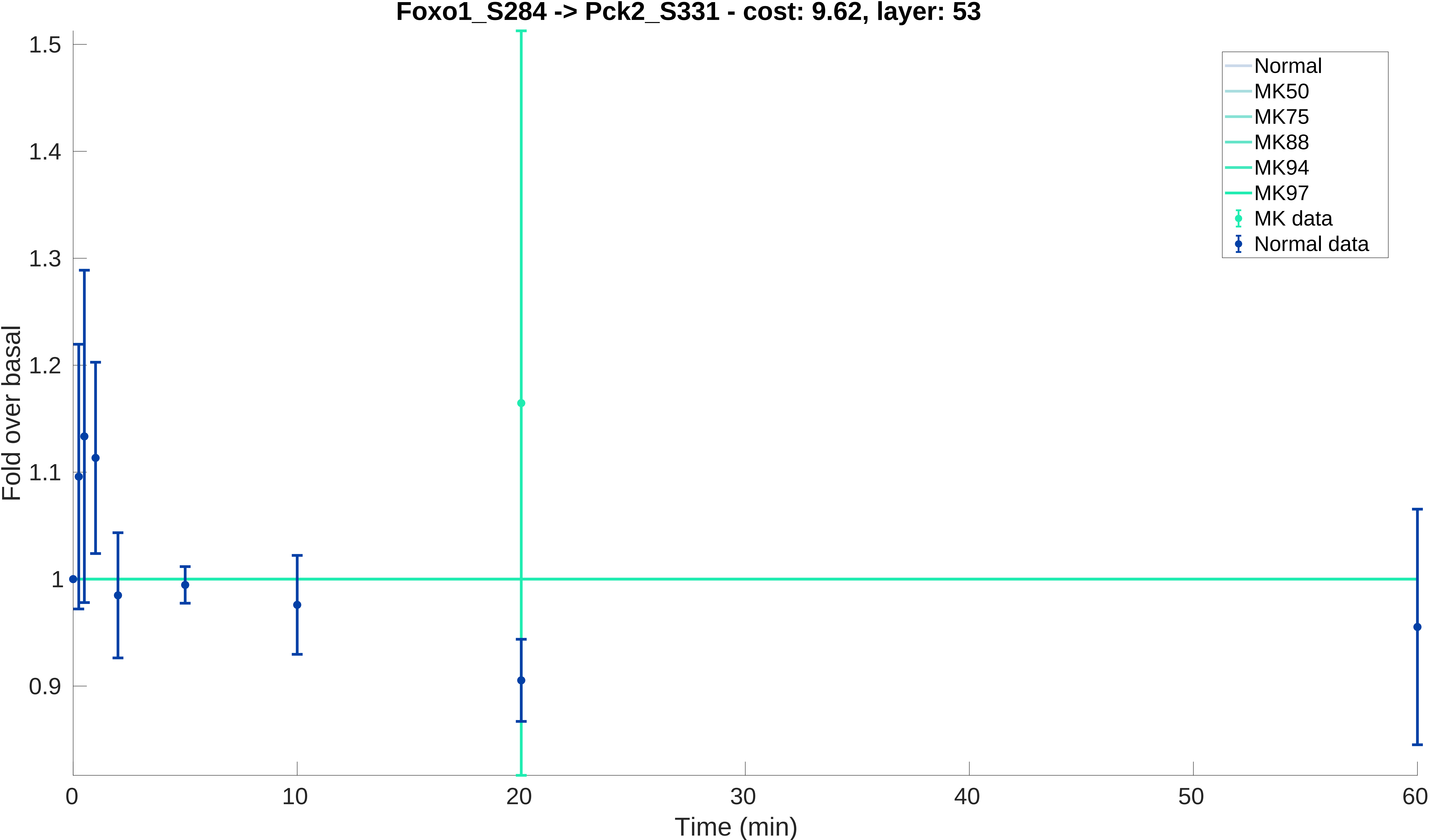

Foxo1_S284 -> Pck2_S331 - cost: 9.62, layer: 53
Fold over basal
Time (min)
1.5
1.4
1.3
1.2
1.1
1
0.9
0
10
20
30
40
50
60
Normal
MK50
MK75
MK88
MK94
MK97
MK data
Normal data

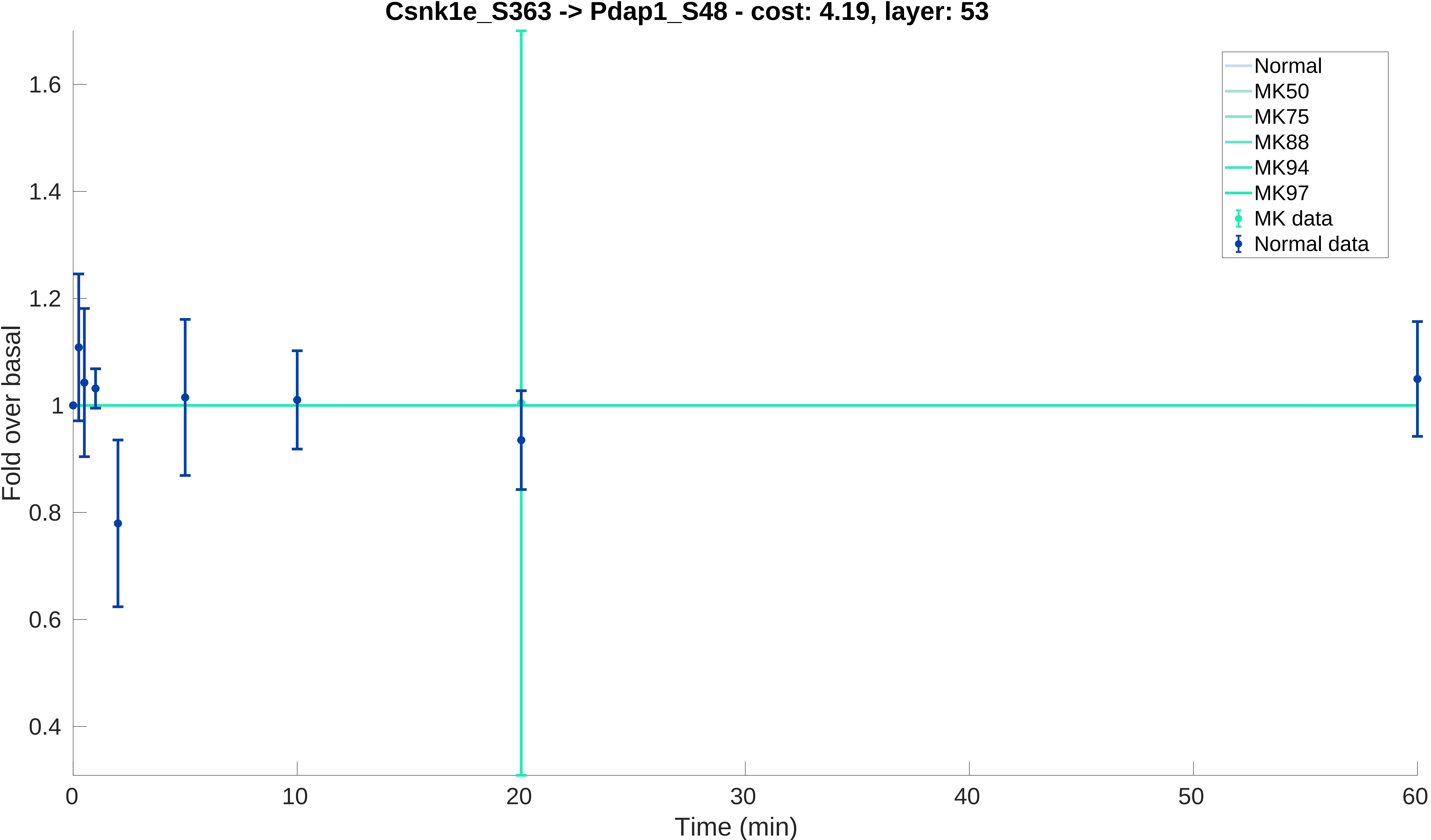

Csnk1e_S363 -> Pdap1_S48 - cost: 4.19, layer: 53
Normal
MK50
MK75
MK88
MK94
MK97
MK data
Normal data
Fold over basal
1.6
1.4
1.2
1
0.8
0.6
0.4
0
10
20
30
40
50
60
Time (min)

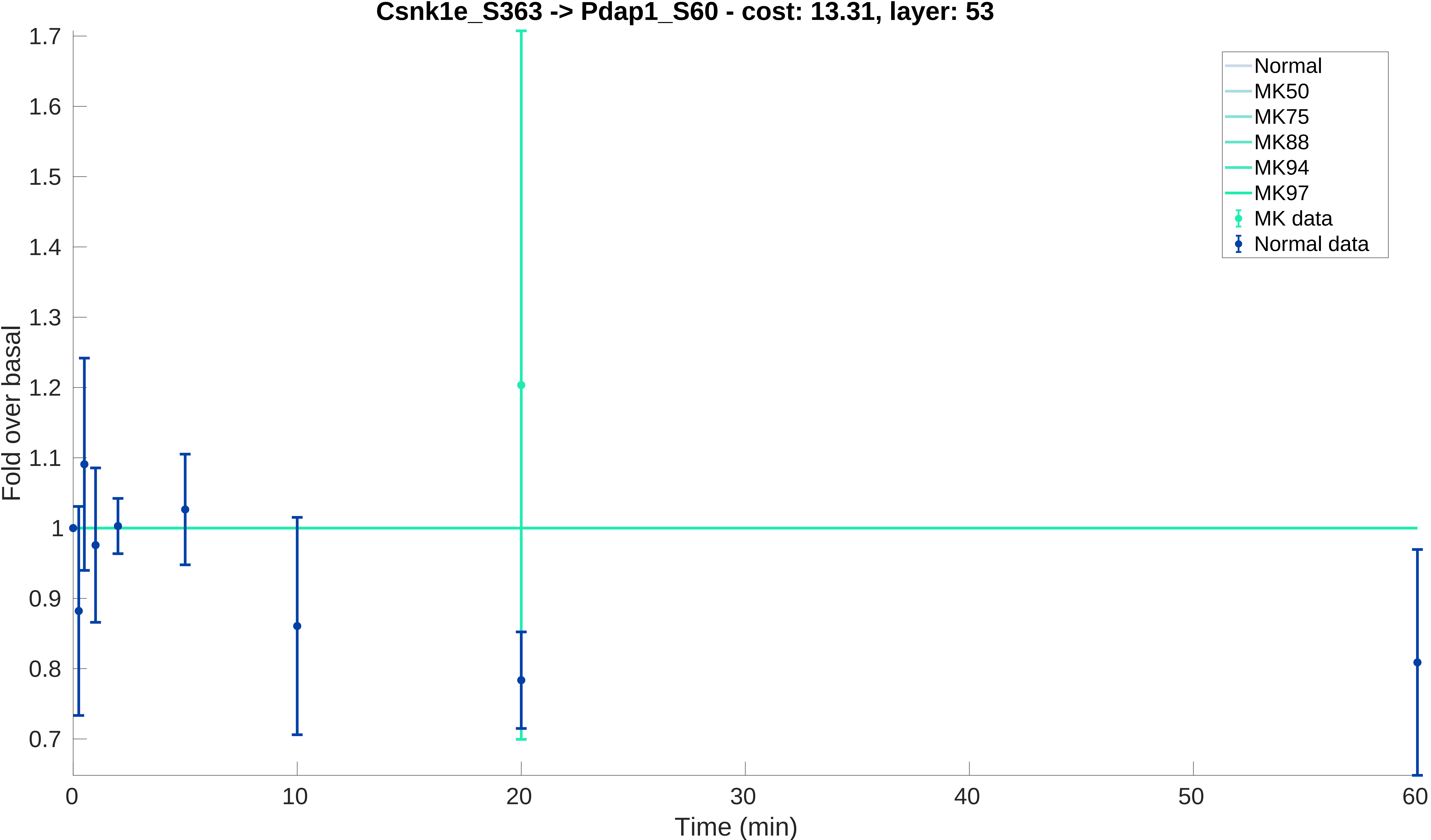

Csnk1e_S363 -> Pdap1_S60 - cost: 13.31, layer: 53
Fold over basal
Time (min)
1.7
1.6
1.5
1.4
1.3
1.2
1.1
1
0.9
0.8
0.7
0
10
20
30
40
50
60
Normal
MK50
MK75
MK88
MK94
MK97
MK data
Normal data

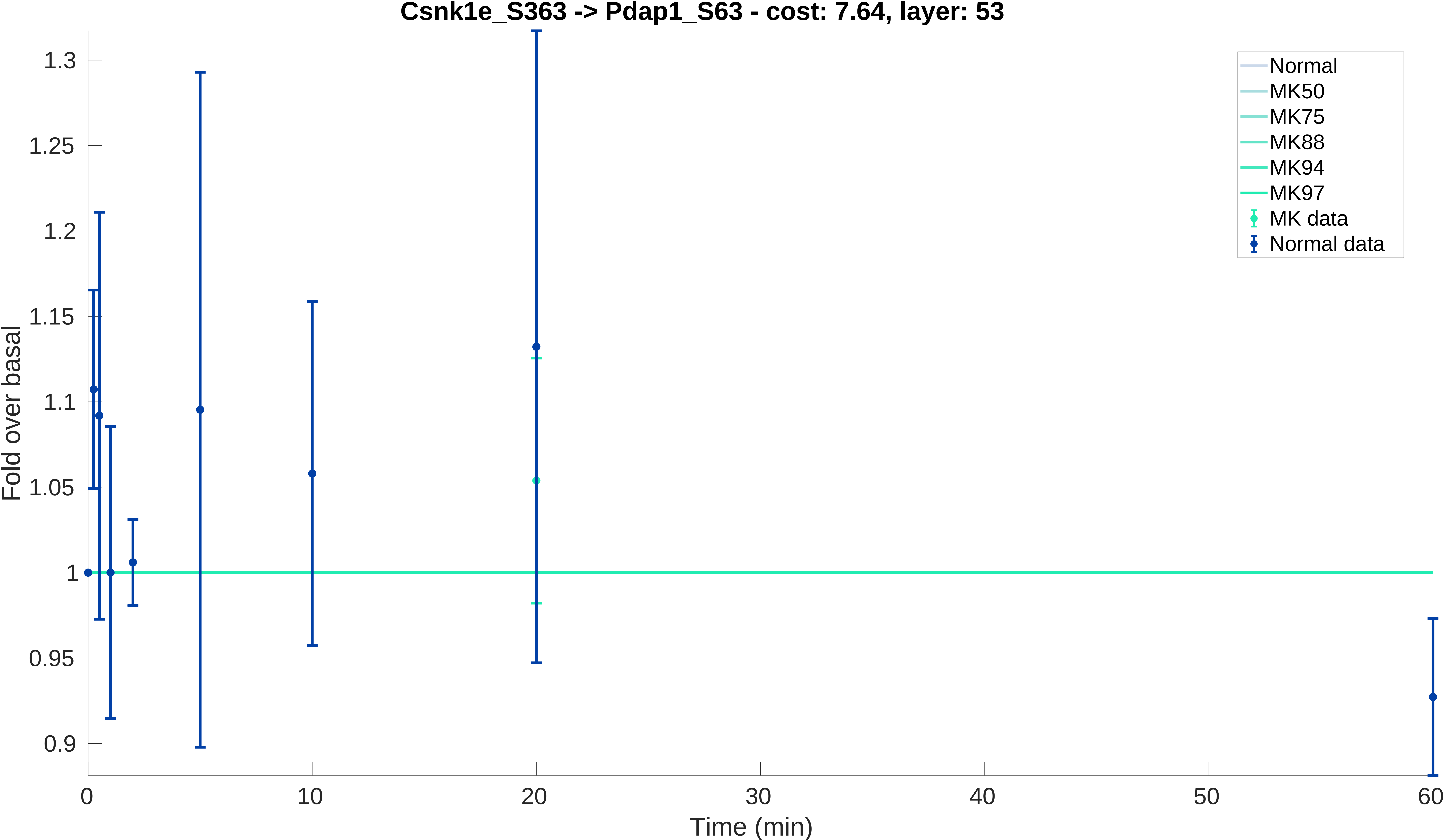

Csnk1e_S363 -> Pdap1_S63 - cost: 7.64, layer: 53
Fold over basal
Time (min)
1.3
1.25
1.2
1.15
1.1
1.05
1
0.95
0.9
0
10
20
30
40
50
60
Normal
MK50
MK75
MK88
MK94
MK97
MK data
Normal data

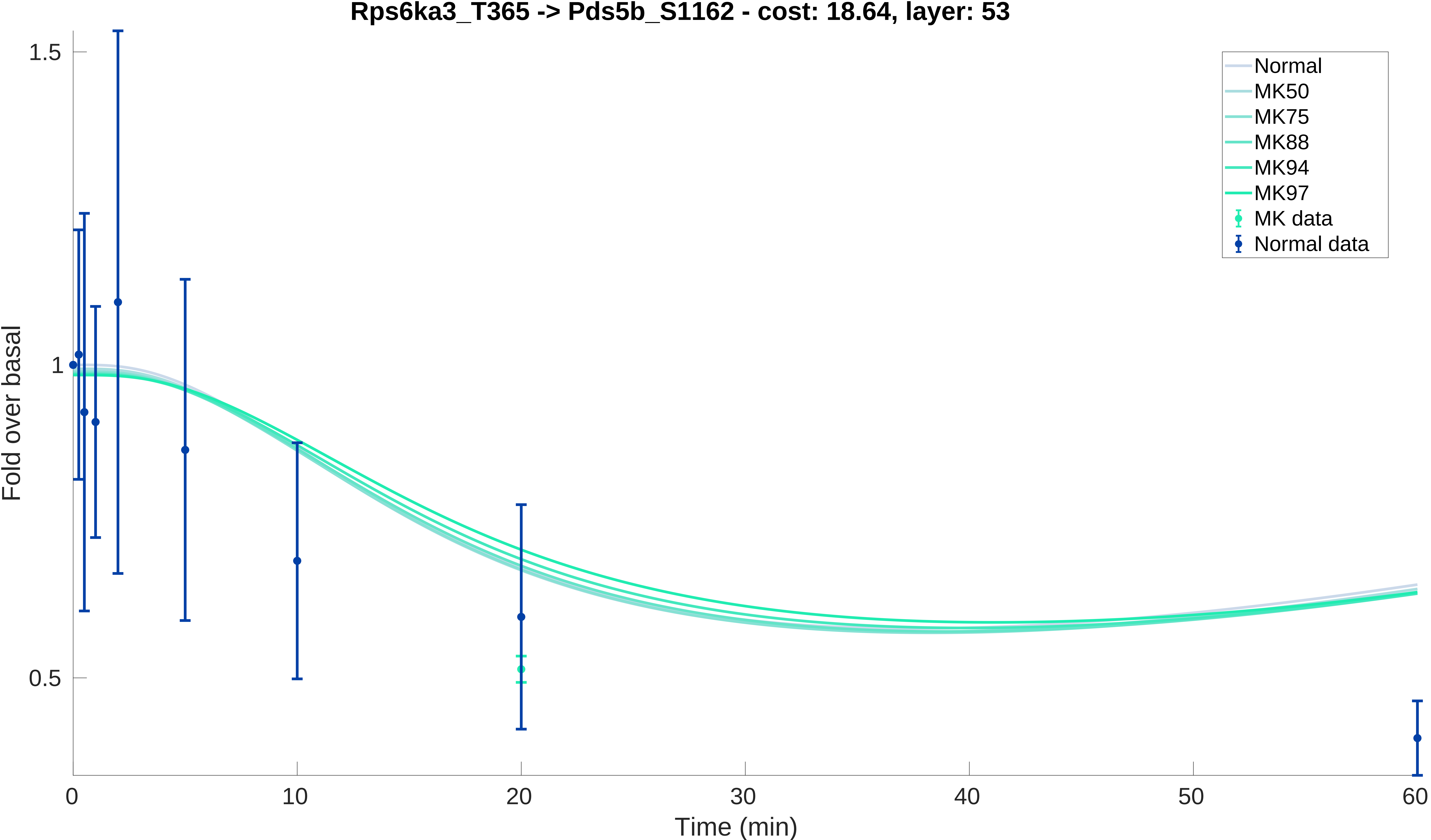

Rps6ka3_T365 -> Pds5b_S1162 - cost: 18.64, layer: 53
Fold over basal
1.5
1
0.5
0
10
20
30
40
50
60
Time (min)
Normal
MK50
MK75
MK88
MK94
MK97
MK data
Normal data

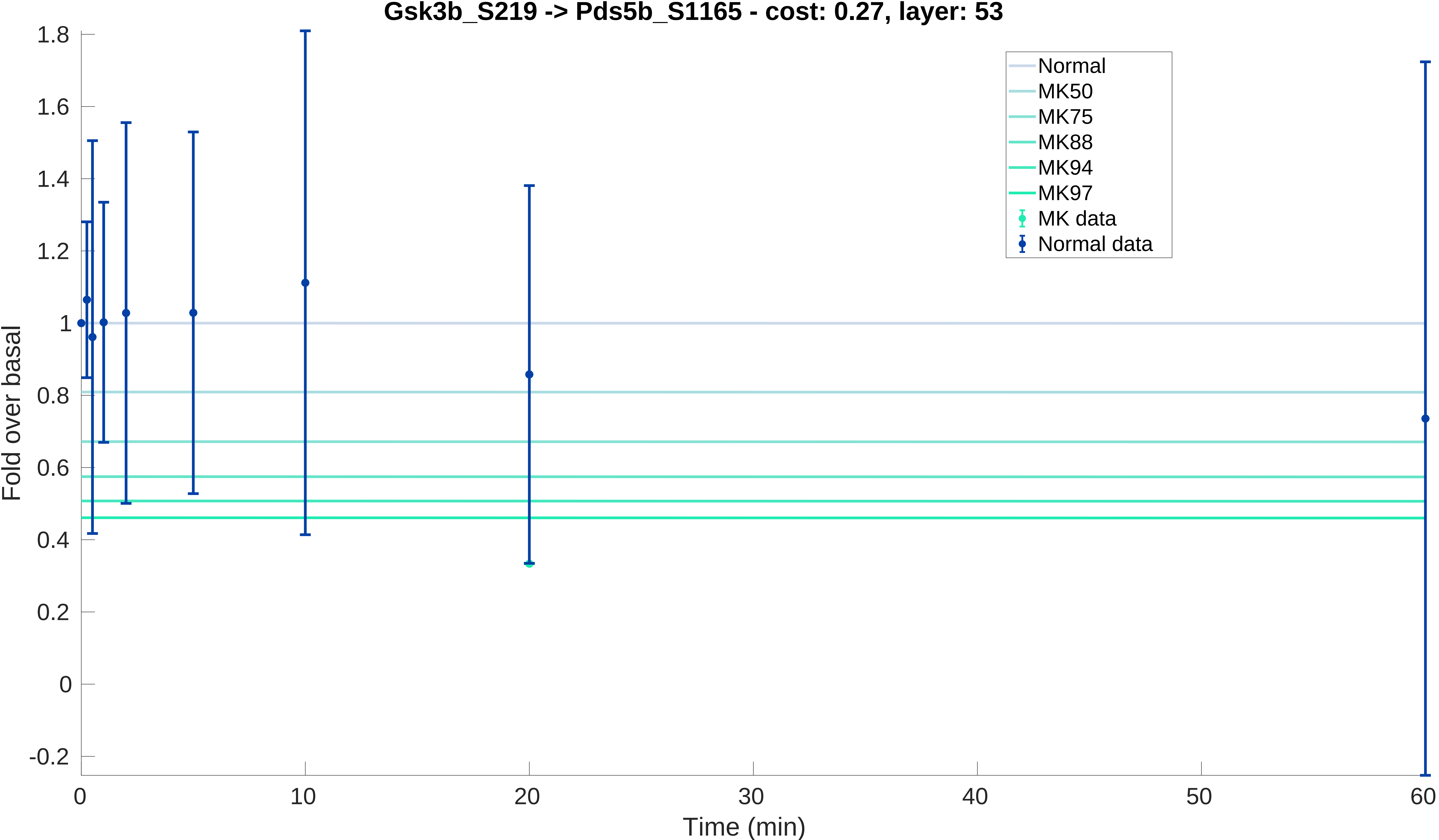

Gsk3b_S219 -> Pds5b_S1165 - cost: 0.27, layer: 53
Normal
MK50
MK75
MK88
MK94
MK97
MK data
Normal data
Fold over basal
Time (min)
1.8
1.6
1.4
1.2
1
0.8
0.6
0.4
0.2
0
-0.2
0
10
20
30
40
50
60

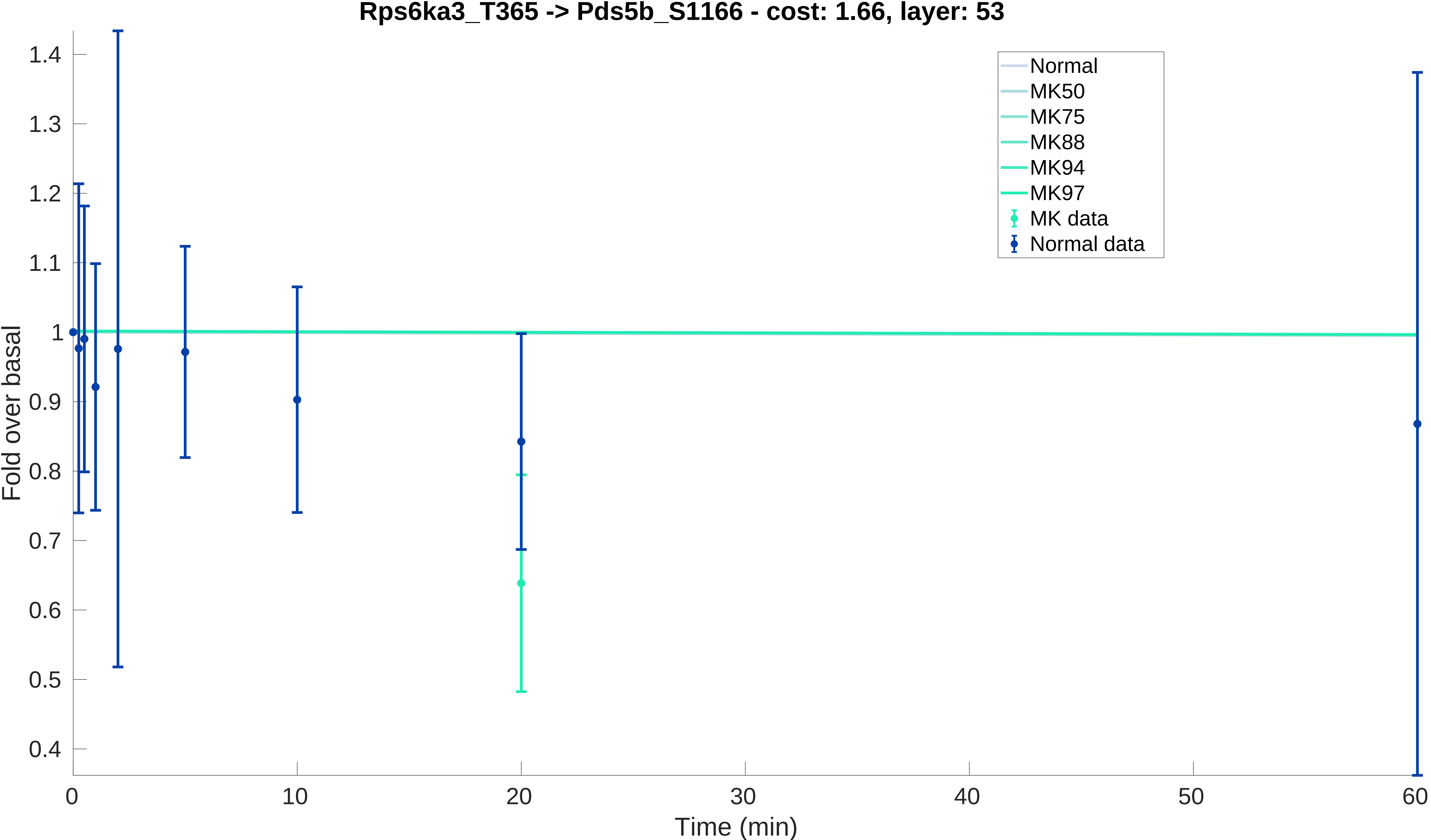

Rps6ka3_T365 -> Pds5b_S1166 - cost: 1.66, layer: 53
Fold over basal
Time (min)
Normal
MK50
MK75
MK88
MK94
MK97
MK data
Normal data
0.4
0.5
0.6
0.7
0.8
0.9
1
1.1
1.2
1.3
1.4
0
10
20
30
40
50
60

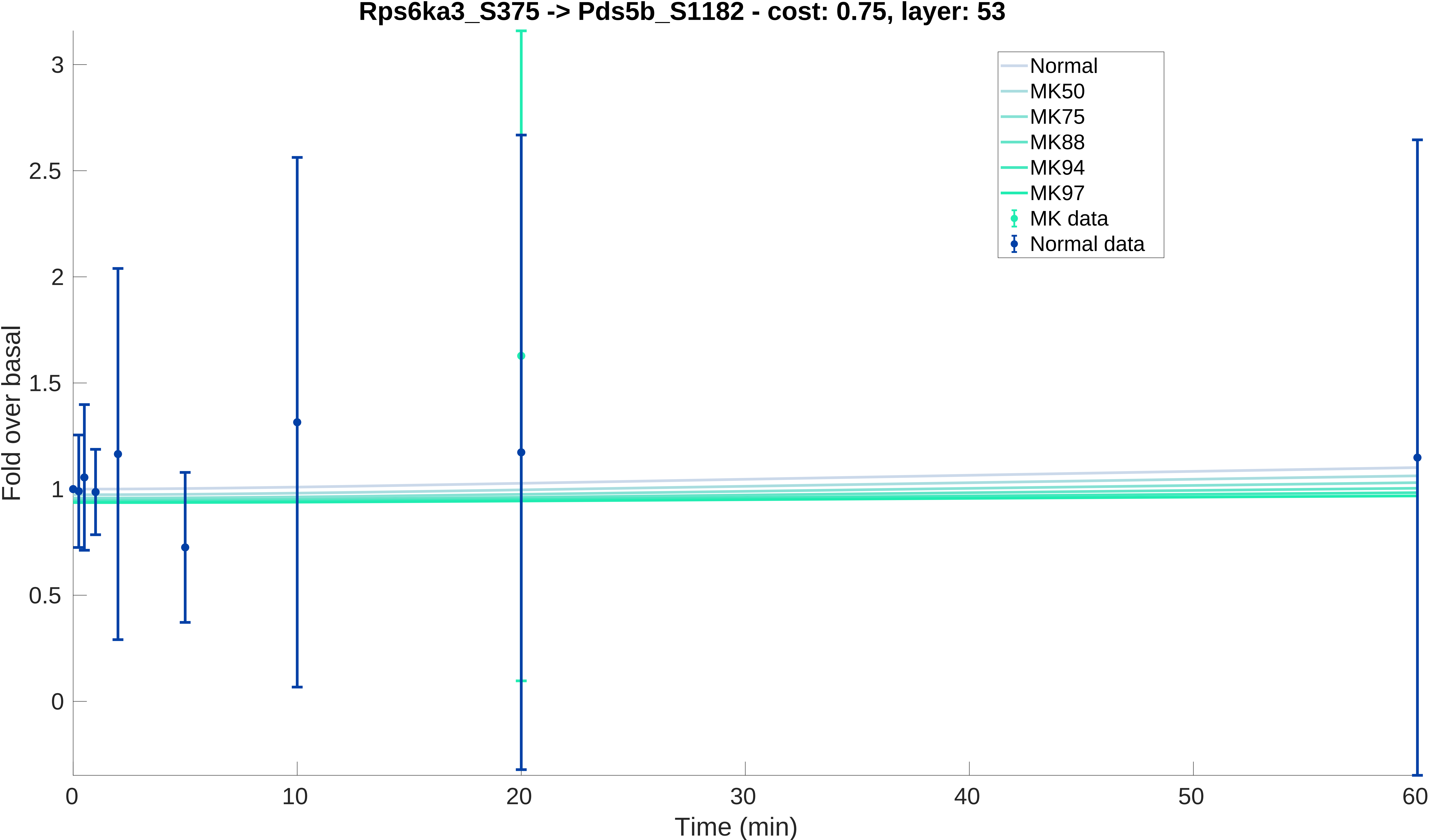

Rps6ka3_S375 -> Pds5b_S1182 - cost: 0.75, layer: 53
Fold over basal
Time (min)
0
0.5
1
1.5
2
2.5
3
0
10
20
30
40
50
60
Normal
MK50
MK75
MK88
MK94
MK97
MK data
Normal data

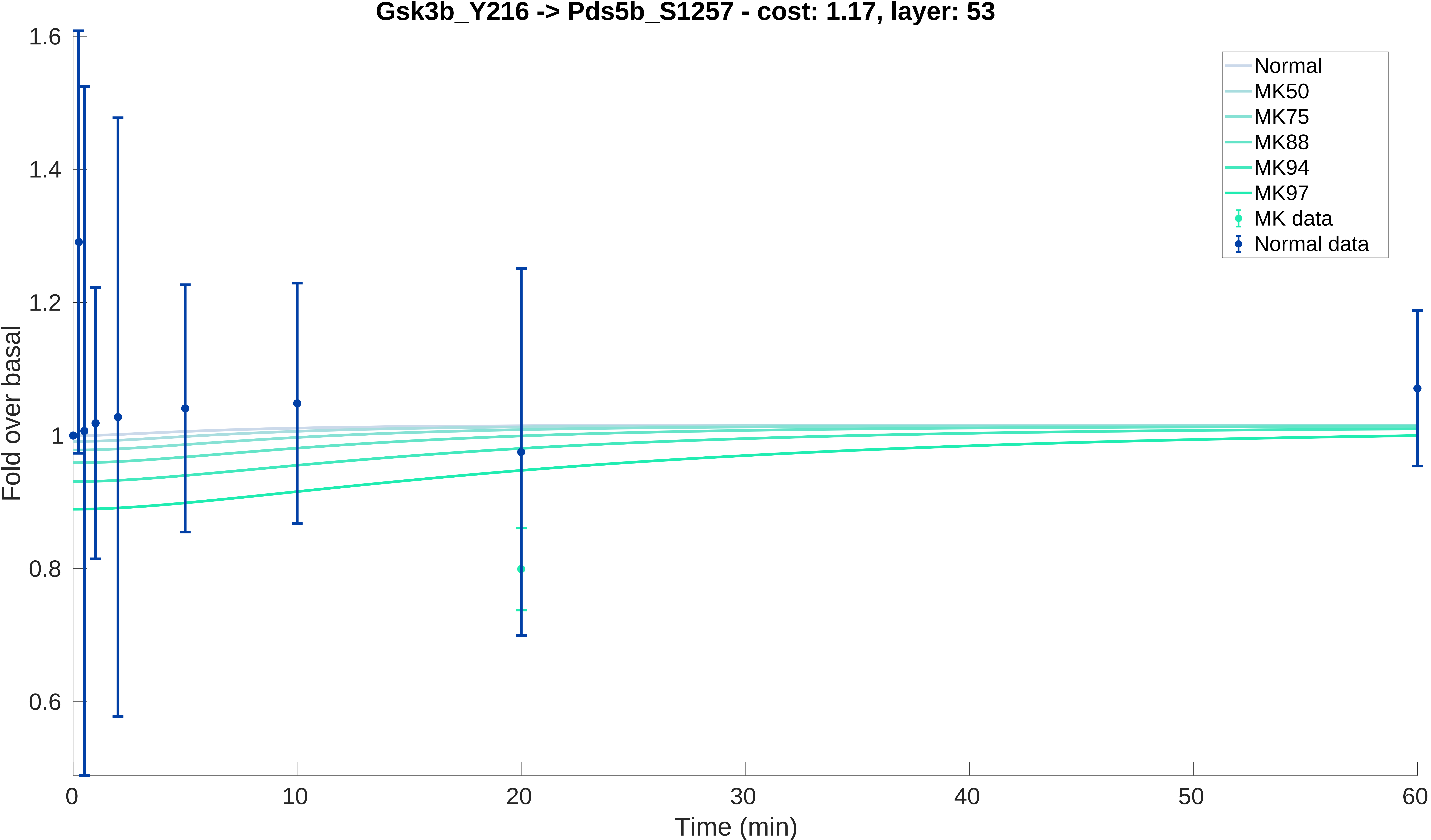

Gsk3b_Y216 -> Pds5b_S1257 - cost: 1.17, layer: 53
Fold over basal
1.6
1.4
1.2
1
0.8
0.6
Time (min)
0
10
20
30
40
50
60
Normal
MK50
MK75
MK88
MK94
MK97
MK data
Normal data

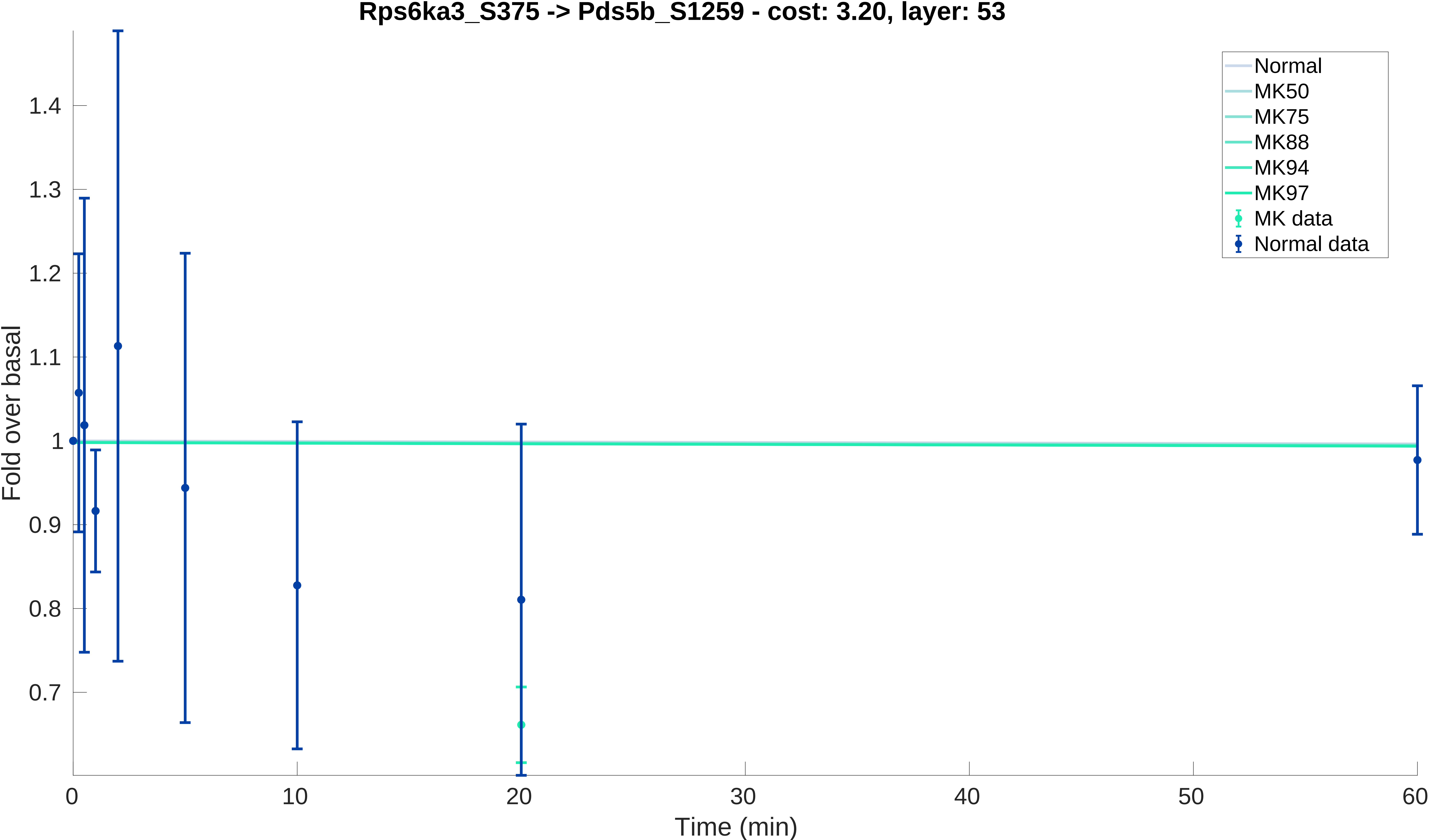

Rps6ka3_S375 -> Pds5b_S1259 - cost: 3.20, layer: 53
Fold over basal
Time (min)
1.4
1.3
1.2
1.1
1
0.9
0.8
0.7
0
10
20
30
40
50
60
Normal
MK50
MK75
MK88
MK94
MK97
MK data
Normal data

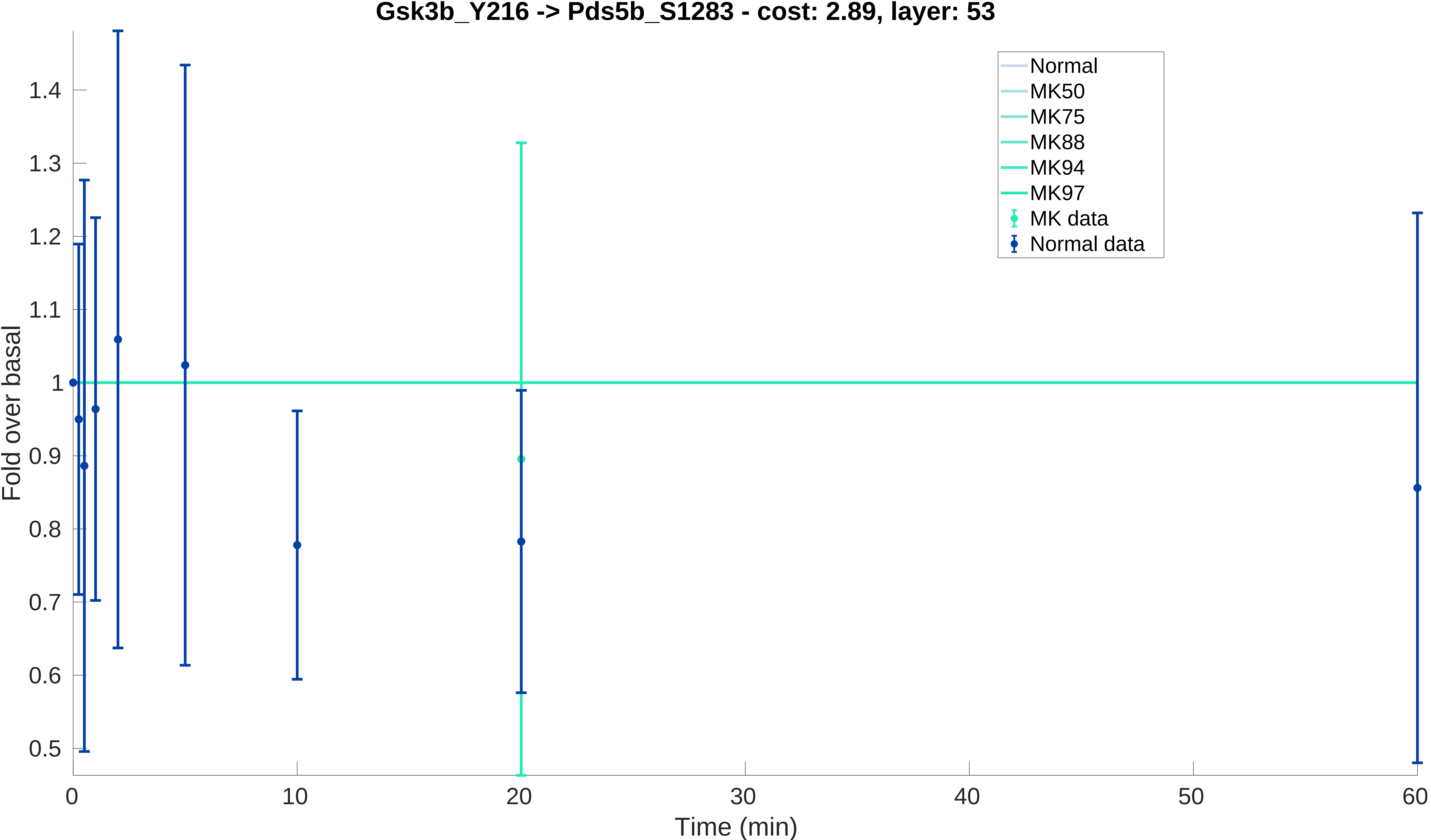

Gsk3b_Y216 -> Pds5b_S1283 - cost: 2.89, layer: 53
Fold over basal
Time (min)
Normal
MK50
MK75
MK88
MK94
MK97
MK data
Normal data
1.4
1.3
1.2
1.1
1
0.9
0.8
0.7
0.6
0.5
0
10
20
30
40
50
60

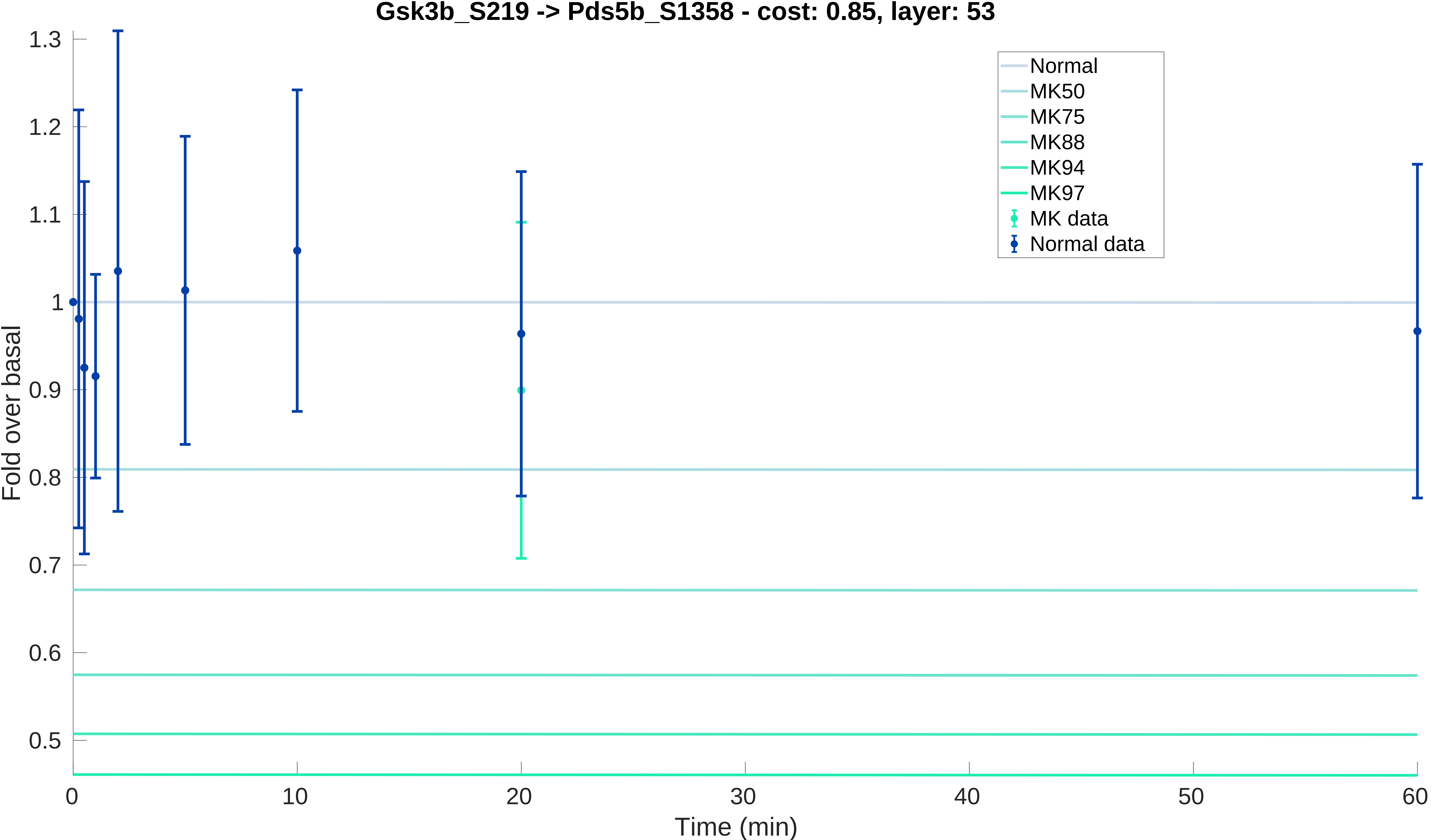

Gsk3b_S219 -> Pds5b_S1358 - cost: 0.85, layer: 53
Fold over basal
Time (min)
Normal
MK50
MK75
MK88
MK94
MK97
MK data
Normal data
0
10
20
30
40
50
60
0.5
0.6
0.7
0.8
0.9
1
1.1
1.2
1.3

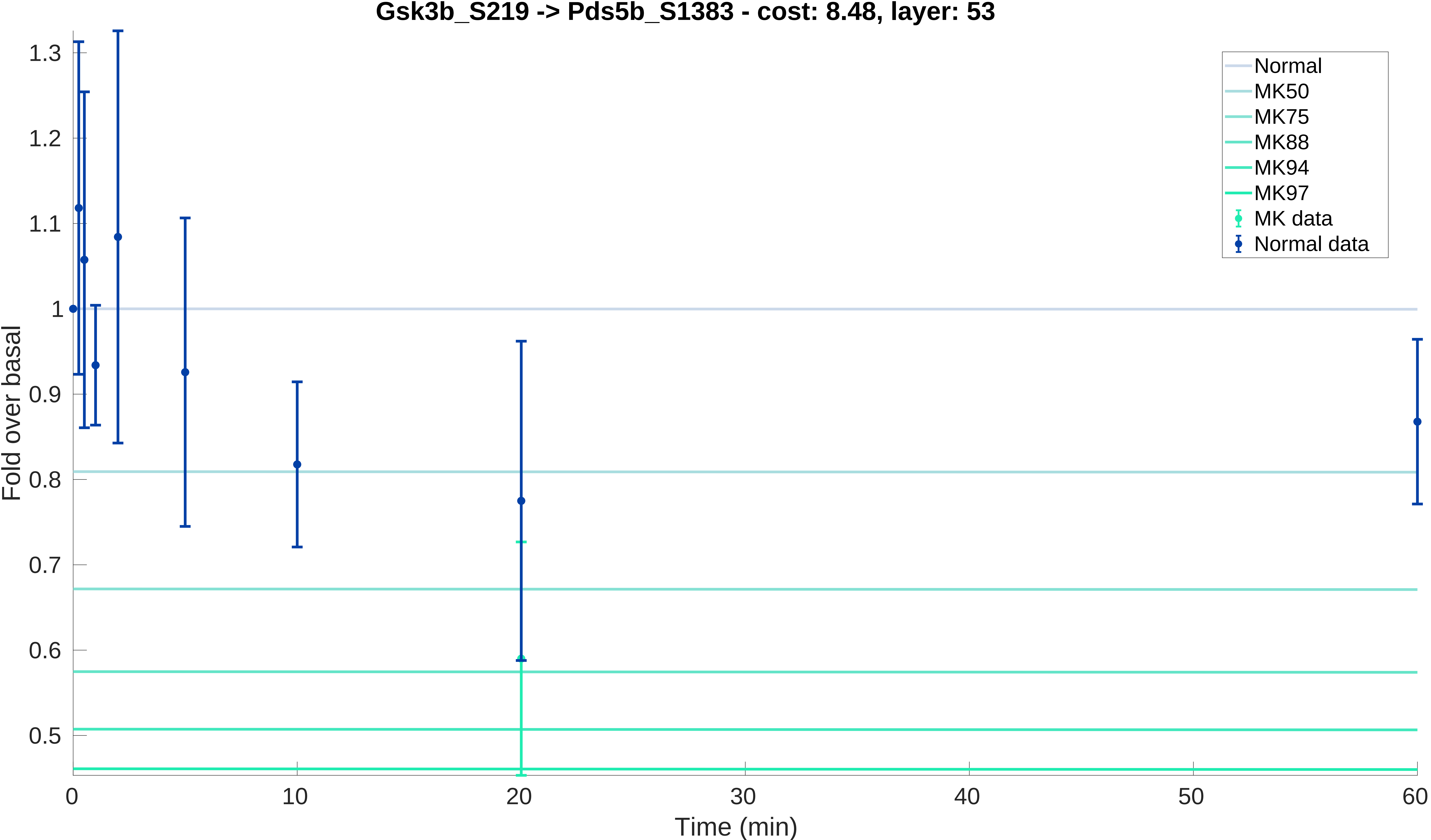

Gsk3b_S219 -> Pds5b_S1383 - cost: 8.48, layer: 53
Fold over basal
1.3
1.2
1.1
1
0.9
0.8
0.7
0.6
0.5
0
10
20
30
40
50
60
Time (min)
Normal
MK50
MK75
MK88
MK94
MK97
MK data
Normal data

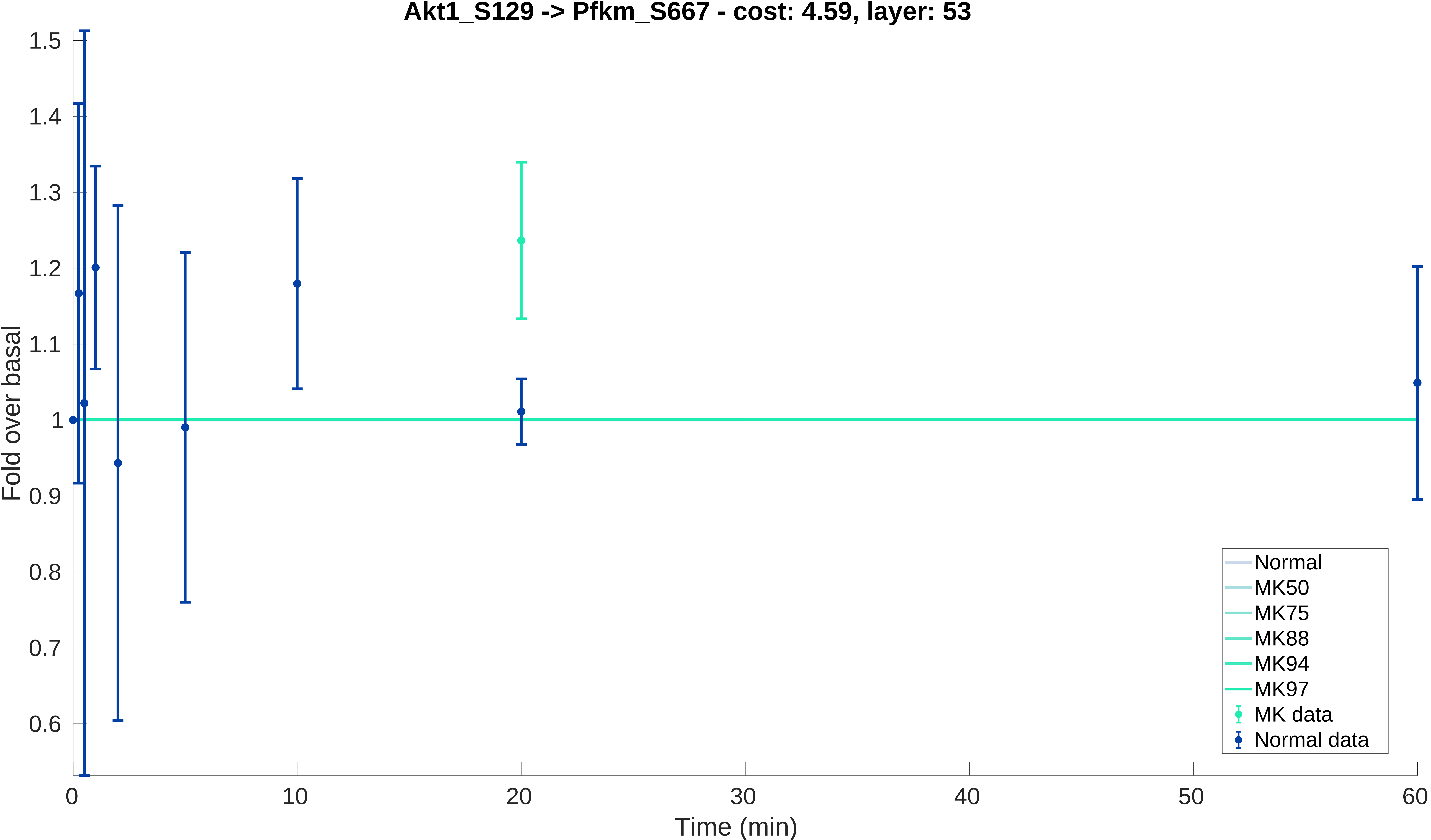
Akt1_S129 -> Pfkm_S667 - cost: 4.59, layer: 53
Fold over basal
1.5
1.4
1.3
1.2
1.1
1
0.9
0.8
0.7
0.6
0
10
20
30
40
50
60
Time (min)
Normal
MK50
MK75
MK88
MK94
MK97
MK data
Normal data

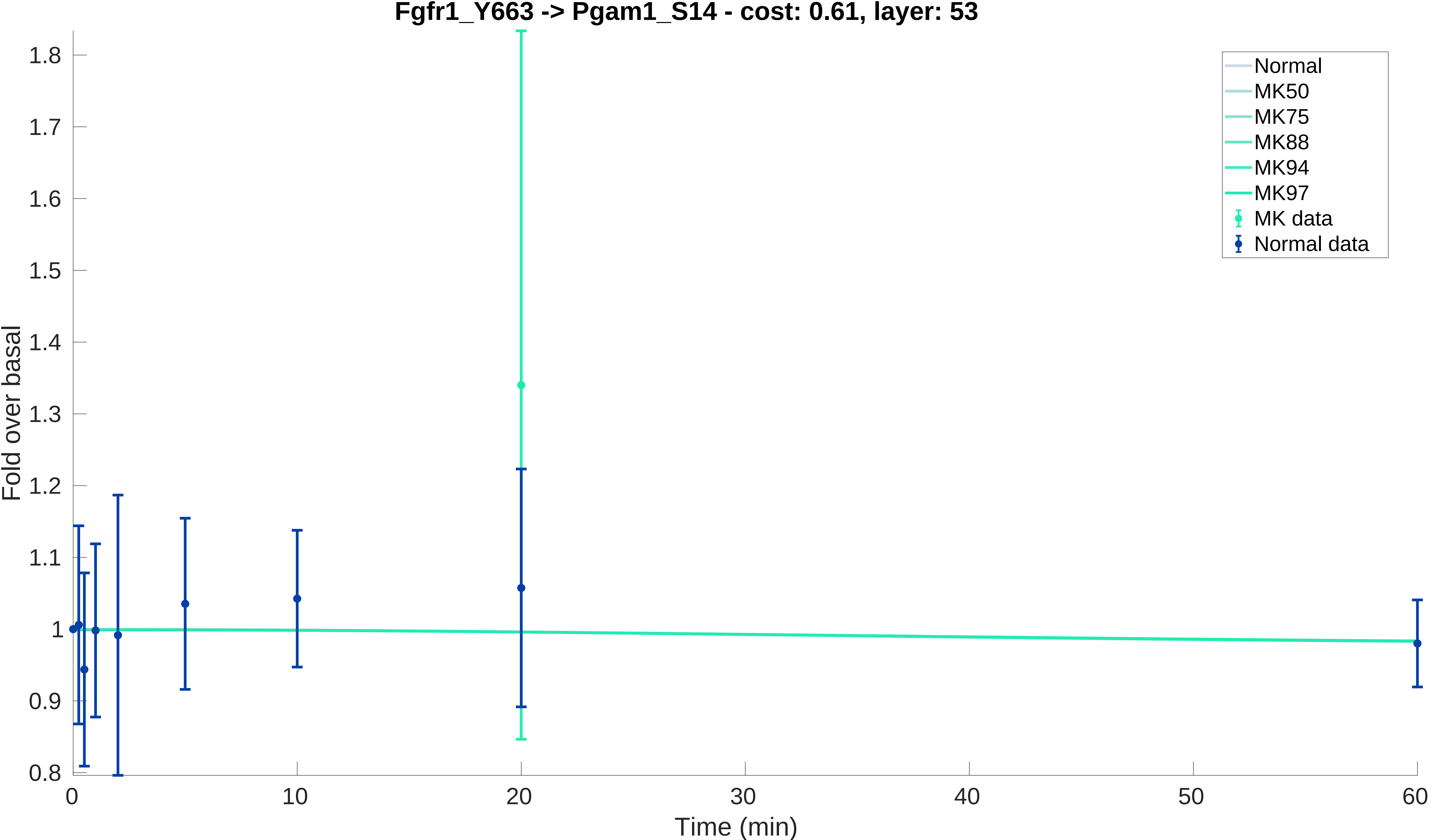

Fgfr1_Y663 -> Pgam1_S14 - cost: 0.61, layer: 53
Fold over basal
Time (min)
1.8
1.7
1.6
1.5
1.4
1.3
1.2
1.1
1
0.9
0.8
0
10
20
30
40
50
60
Normal
MK50
MK75
MK88
MK94
MK97
MK data
Normal data

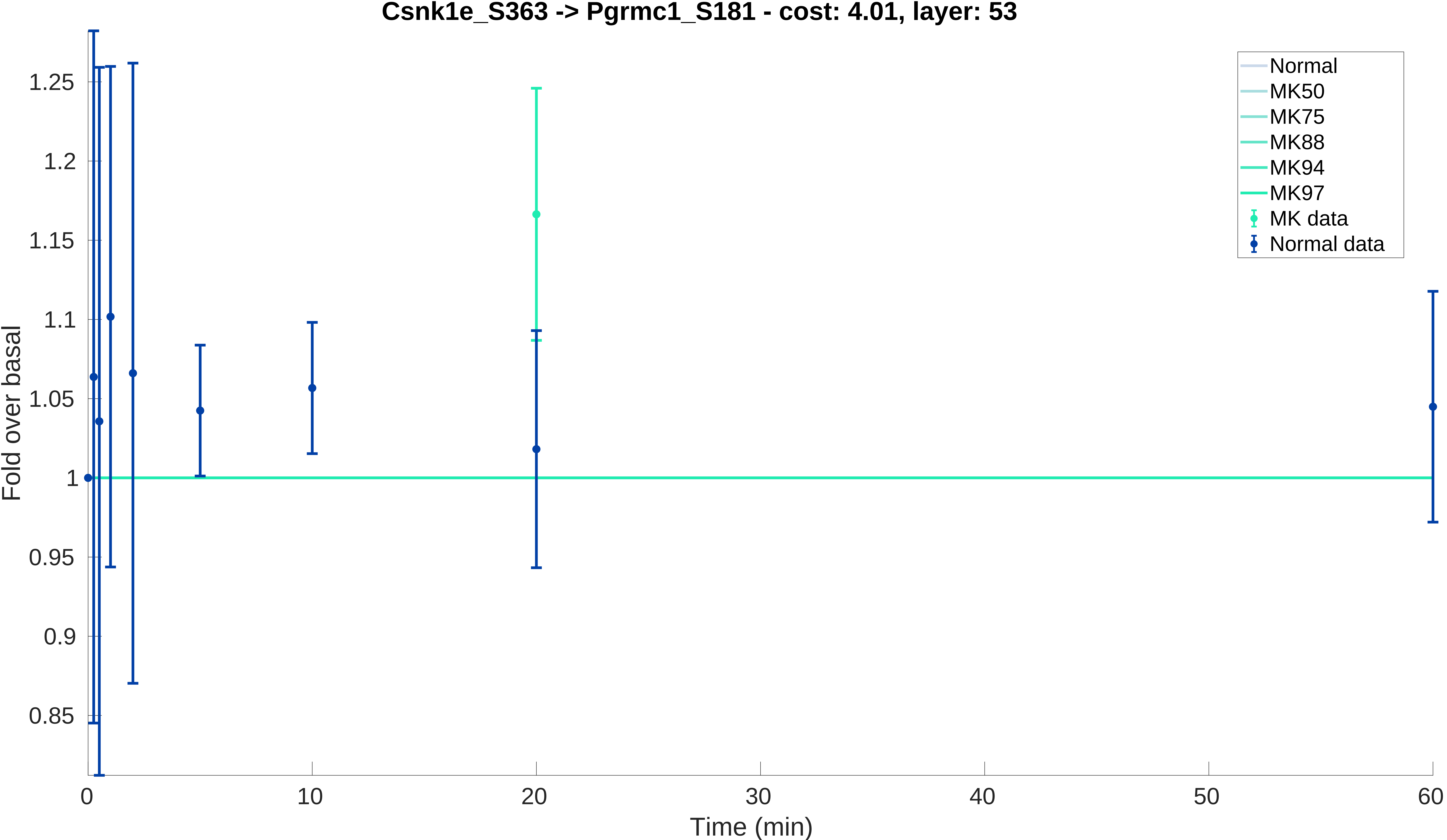

Csnk1e_S363 -> Pgrmc1_S181 - cost: 4.01, layer: 53
Normal
MK50
MK75
MK88
MK94
MK97
MK data
Normal data
Fold over basal
1.25
1.2
1.15
1.1
1.05
1
0.95
0.9
0.85
0
10
20
30
40
50
60
Time (min)

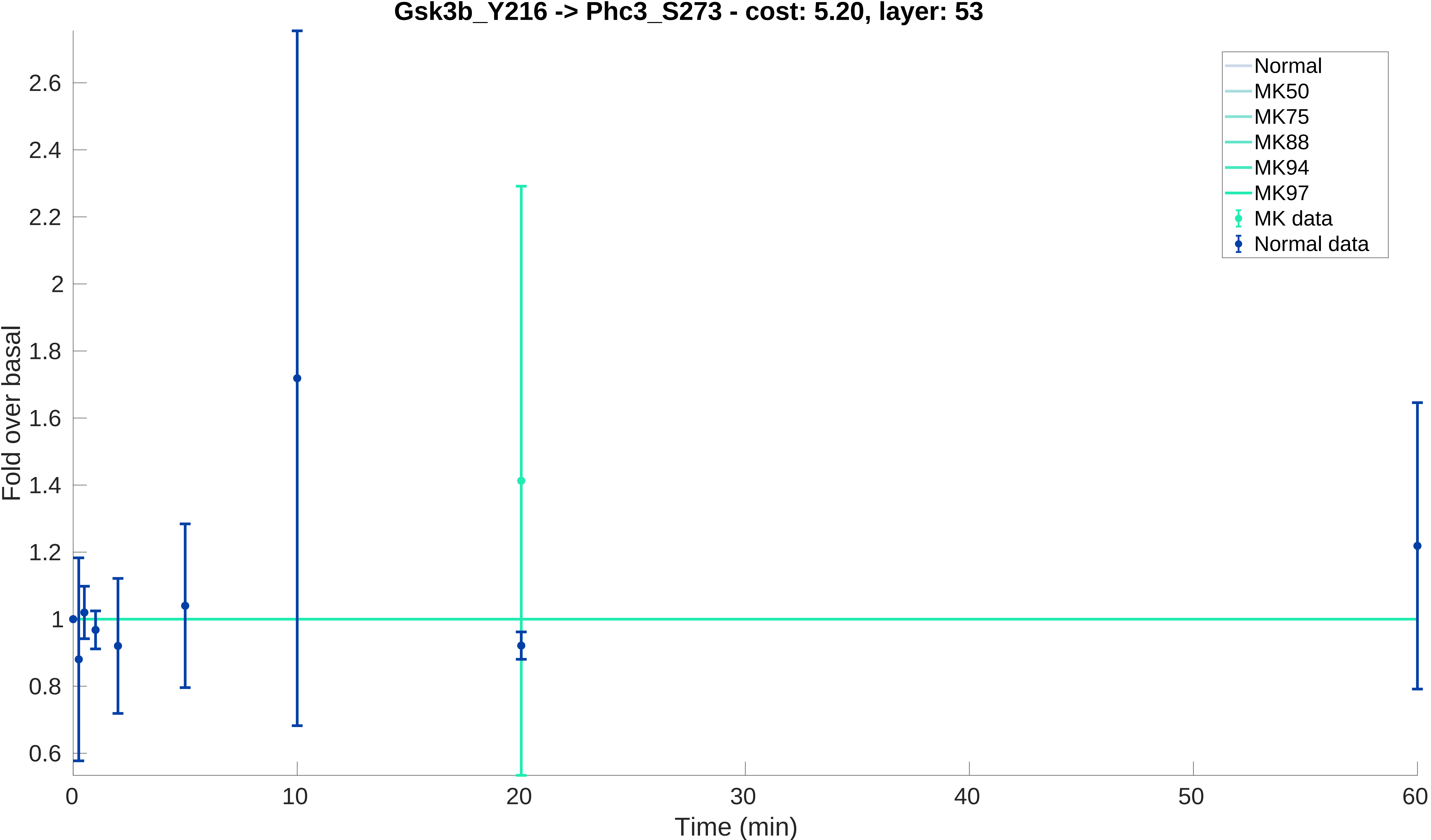

Gsk3b_Y216 -> Phc3_S273 - cost: 5.20, layer: 53
Normal
MK50
MK75
MK88
MK94
MK97
MK data
Normal data
Fold over basal
2.6
2.4
2.2
2
1.8
1.6
1.4
1.2
1
0.8
0.6
0
10
20
30
40
50
60
Time (min)

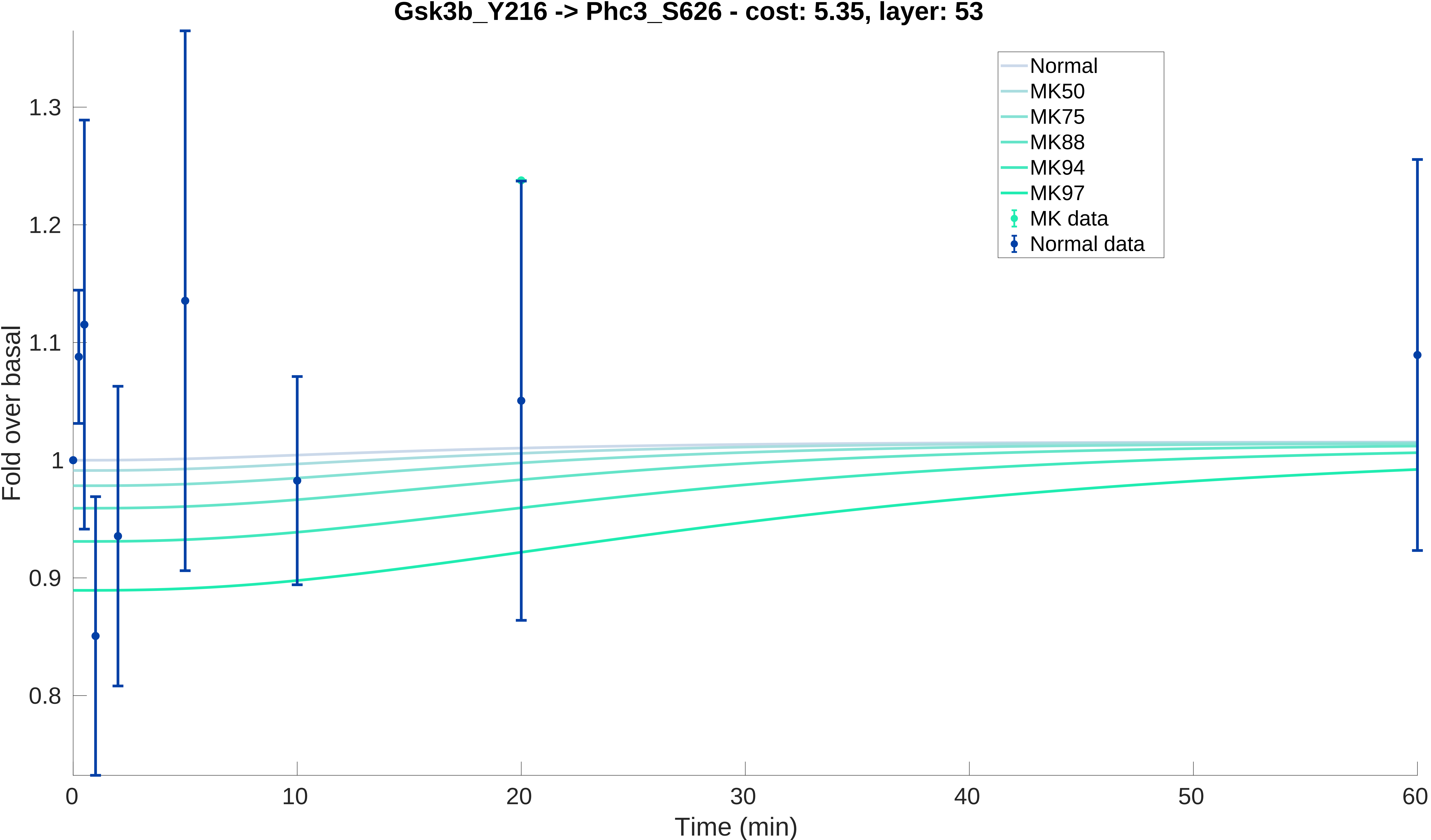

Gsk3b_Y216 -> Phc3_S626 - cost: 5.35, layer: 53
Normal
MK50
MK75
MK88
MK94
MK97
MK data
Normal data
1.3
1.2
1.1
1
0.9
0.8
Fold over basal
0
10
20
30
40
50
60
Time (min)

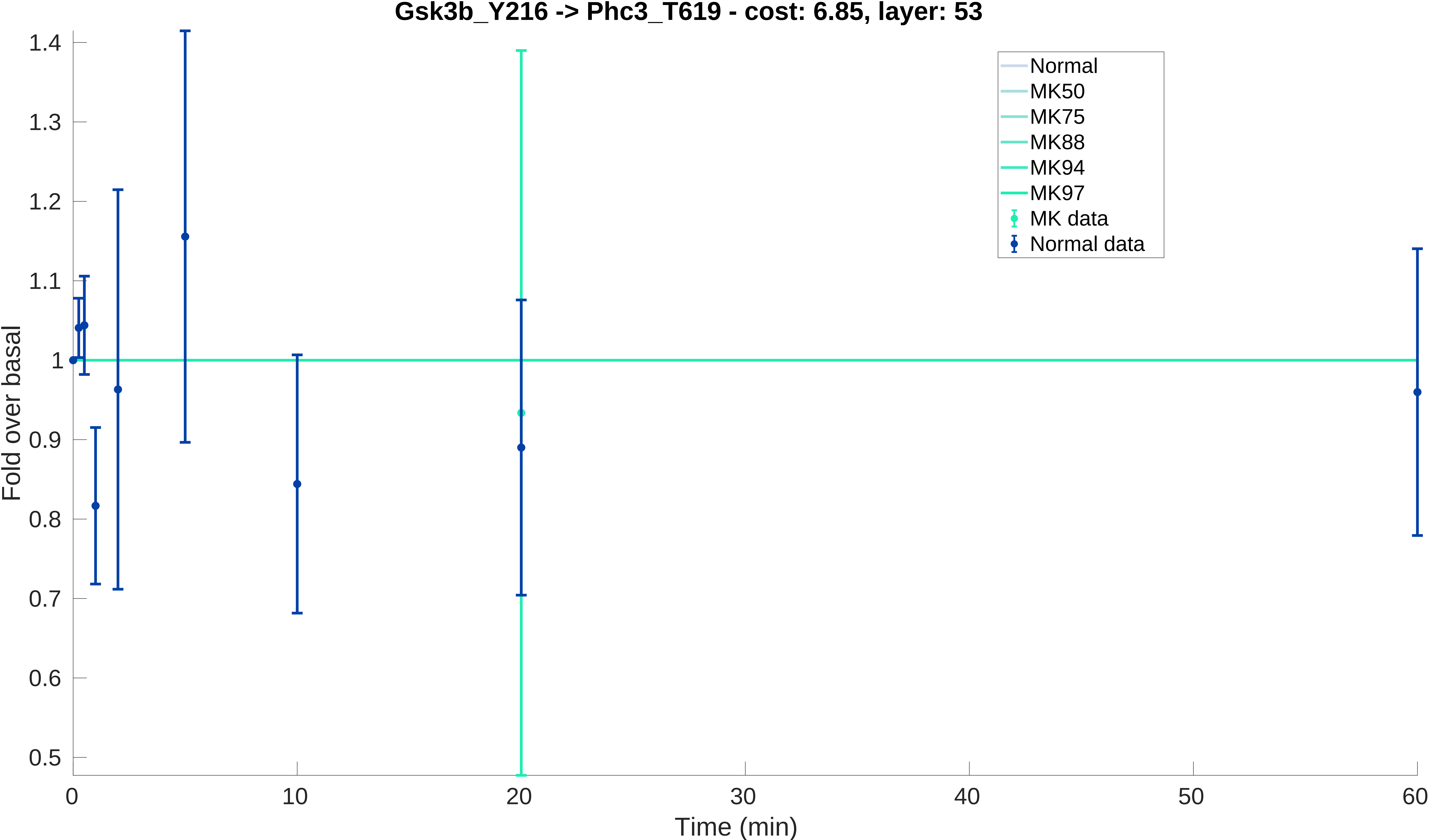

Gsk3b_Y216 -> Phc3_T619 - cost: 6.85, layer: 53
Fold over basal
Time (min)
0.5
0.6
0.7
0.8
0.9
1
1.1
1.2
1.3
1.4
0
10
20
30
40
50
60
Normal
MK50
MK75
MK88
MK94
MK97
MK data
Normal data

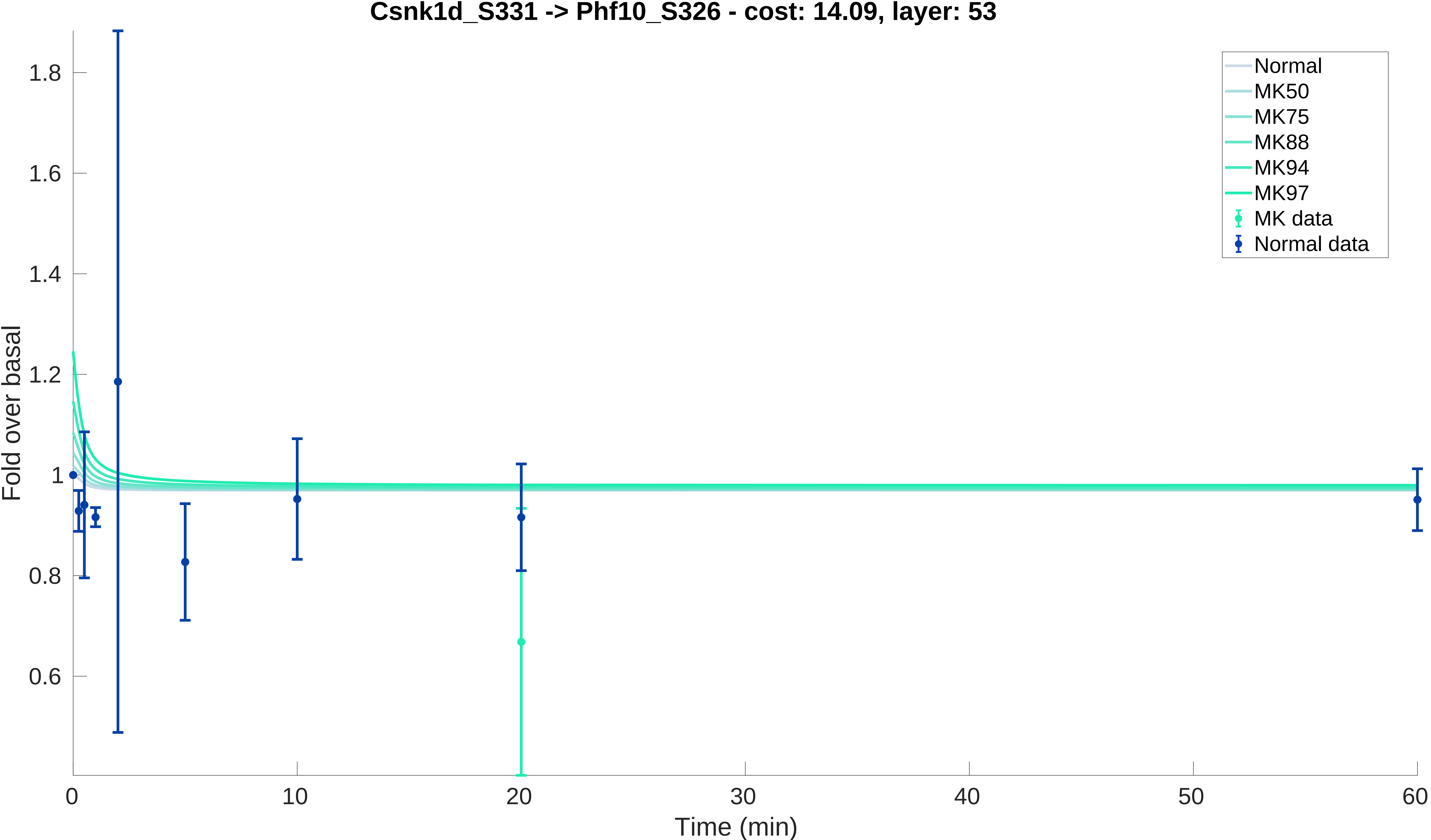

Csnk1d_S331 -> Phf10_S326 - cost: 14.09, layer: 53
Fold over basal
Time (min)
1.8
1.6
1.4
1.2
1
0.8
0.6
0
10
20
30
40
50
60
Normal
MK50
MK75
MK88
MK94
MK97
MK data
Normal data

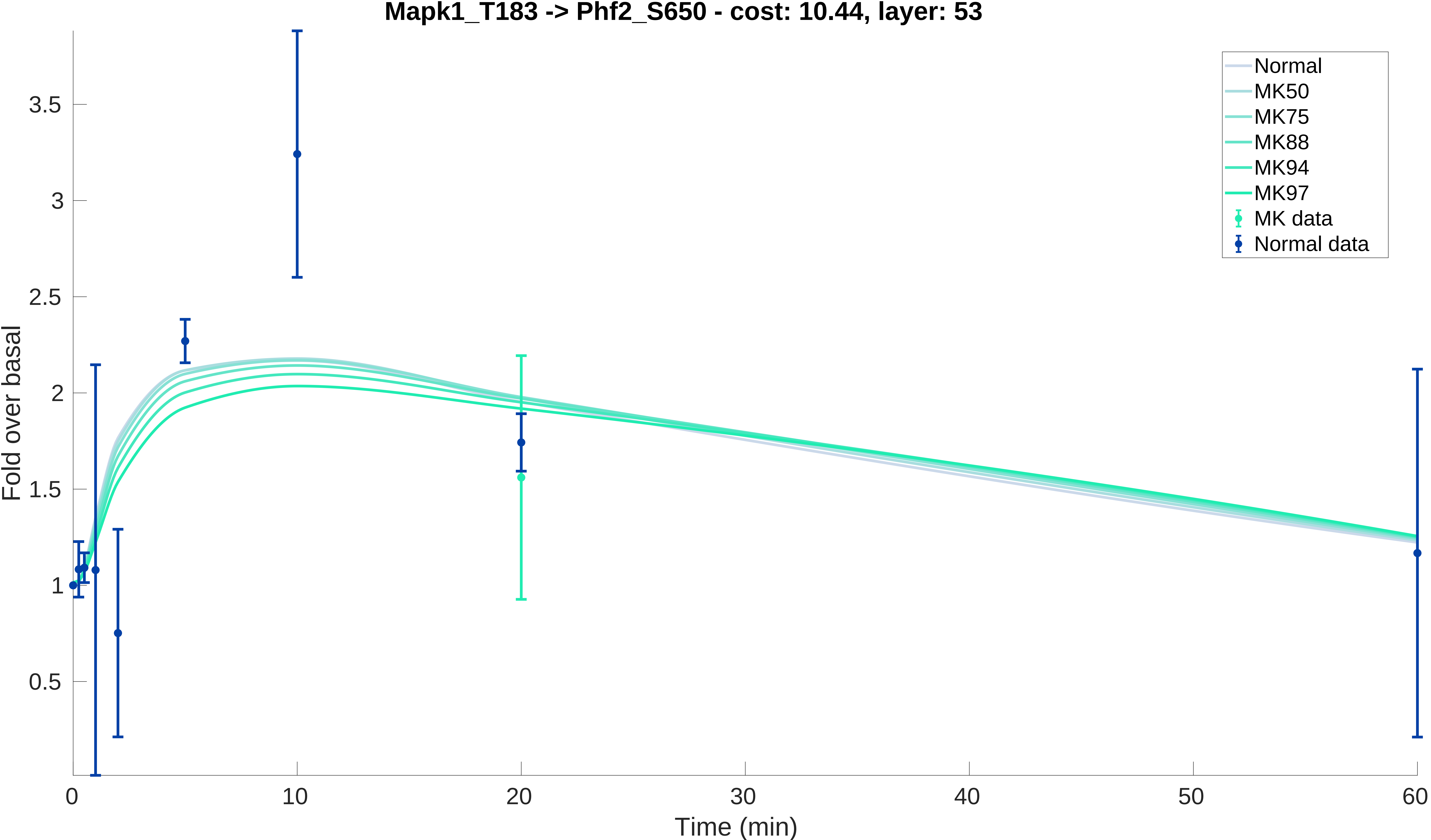

Mapk1_T183 -> Phf2_S650 - cost: 10.44, layer: 53
Fold over basal
Time (min)
0.5
1
1.5
2
2.5
3
3.5
0
10
20
30
40
50
60
Normal
MK50
MK75
MK88
MK94
MK97
MK data
Normal data

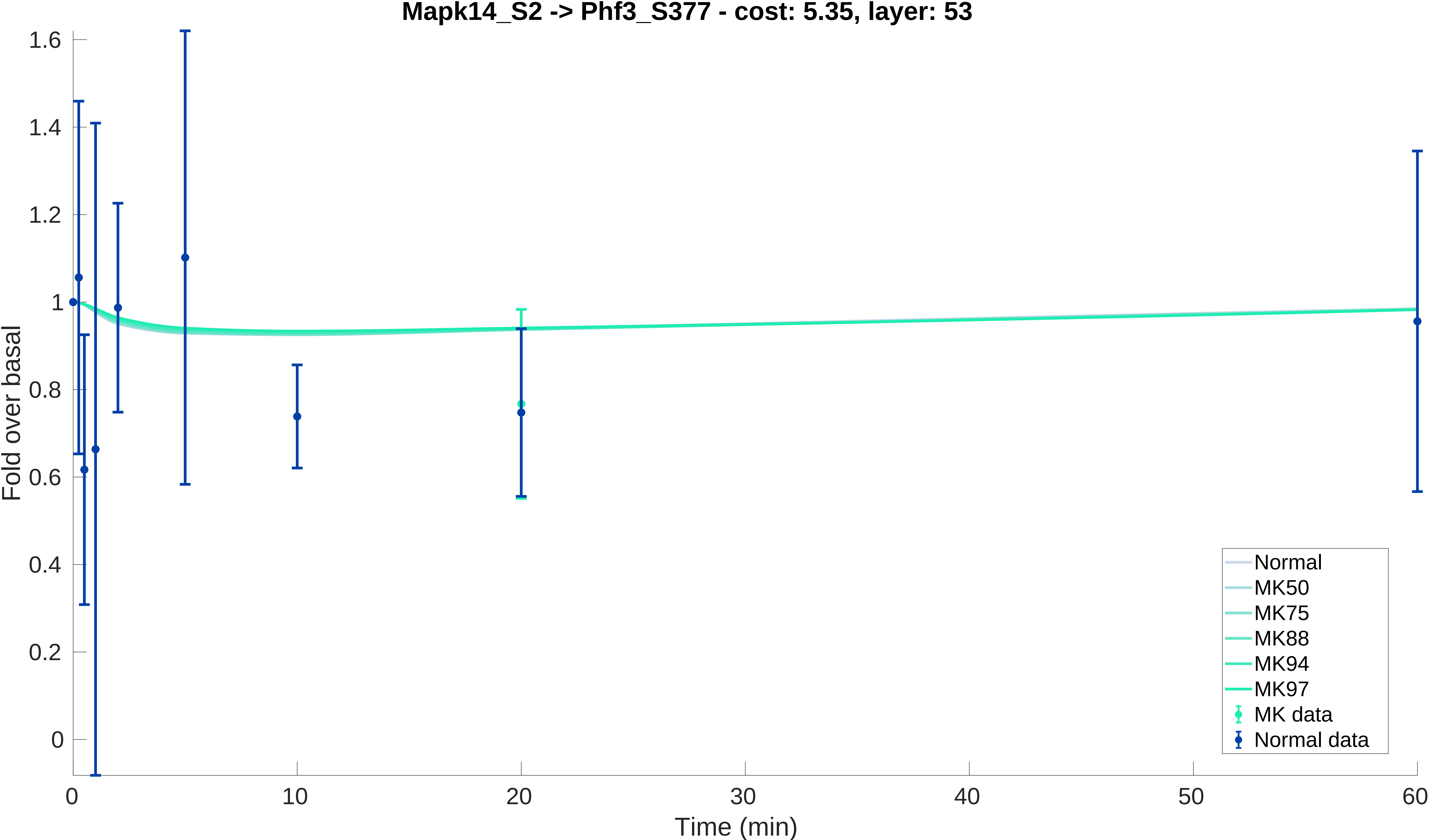

Mapk14_S2 -> Phf3_S377 - cost: 5.35, layer: 53
1.6
1.4
1.2
1
0.8
0.6
0.4
0.2
0
Fold over basal
0
10
20
30
40
50
60
Time (min)
Normal
MK50
MK75
MK88
MK94
MK97
MK data
Normal data

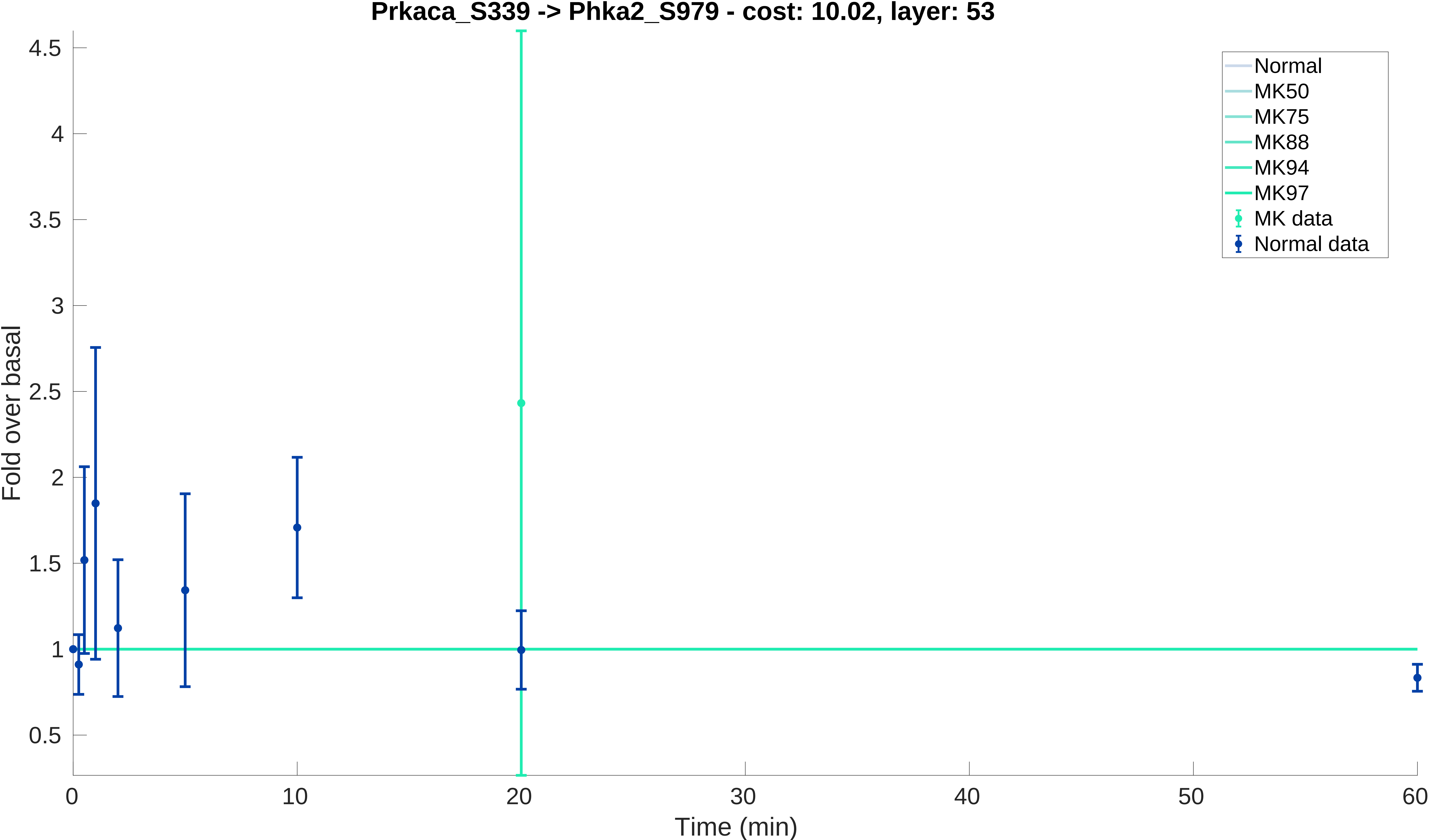

Prkaca_S339 -> Phka2_S979 - cost: 10.02, layer: 53
Fold over basal
Time (min)
0.5
1
1.5
2
2.5
3
3.5
4
4.5
0
10
20
30
40
50
60
Normal
MK50
MK75
MK88
MK94
MK97
MK data
Normal data

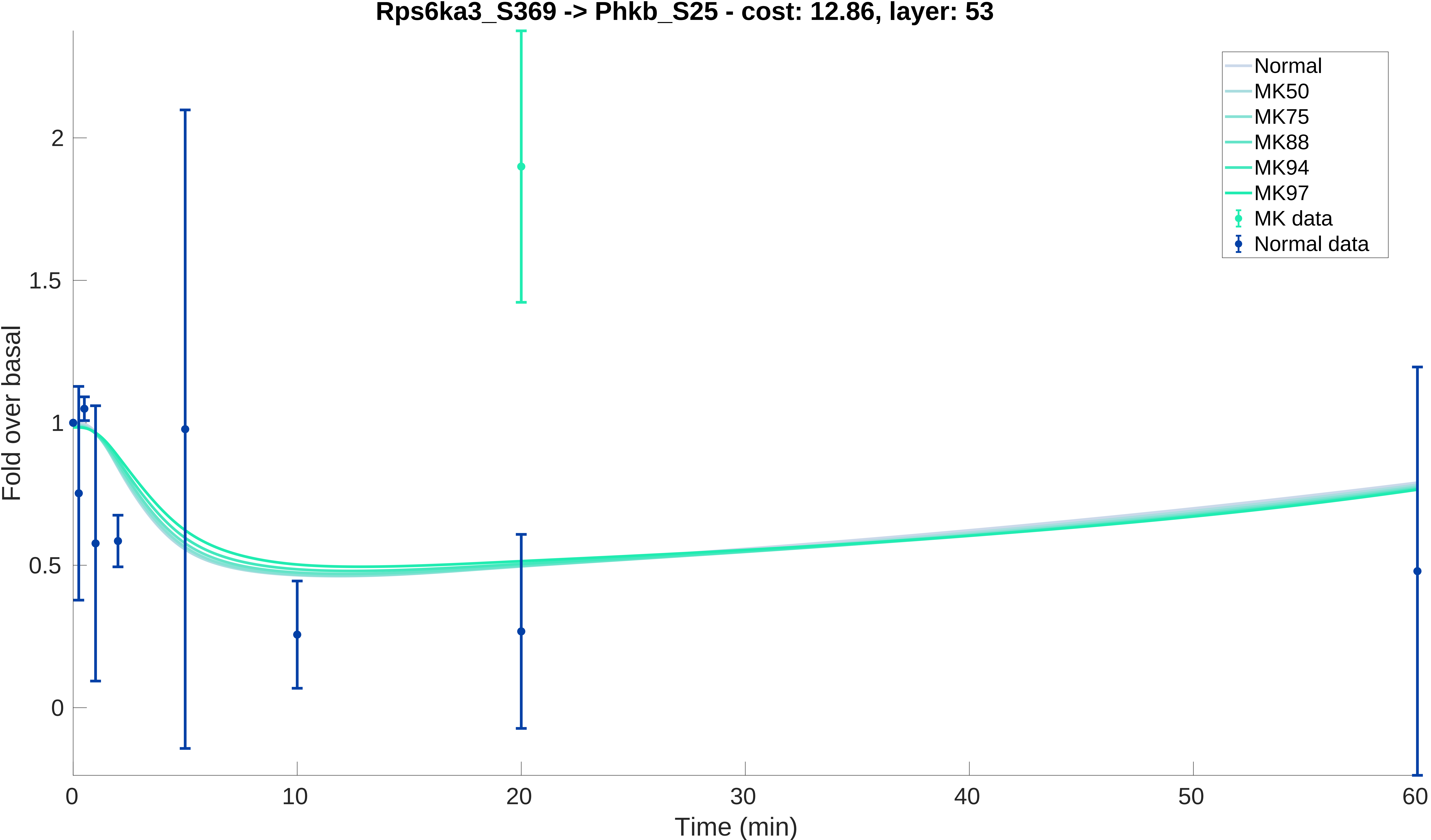
Rps6ka3_S369 -> Phkb_S25 - cost: 12.86, layer: 53
Fold over basal
Time (min)
0
0.5
1
1.5
2
0
10
20
30
40
50
60
Normal
MK50
MK75
MK88
MK94
MK97
MK data
Normal data

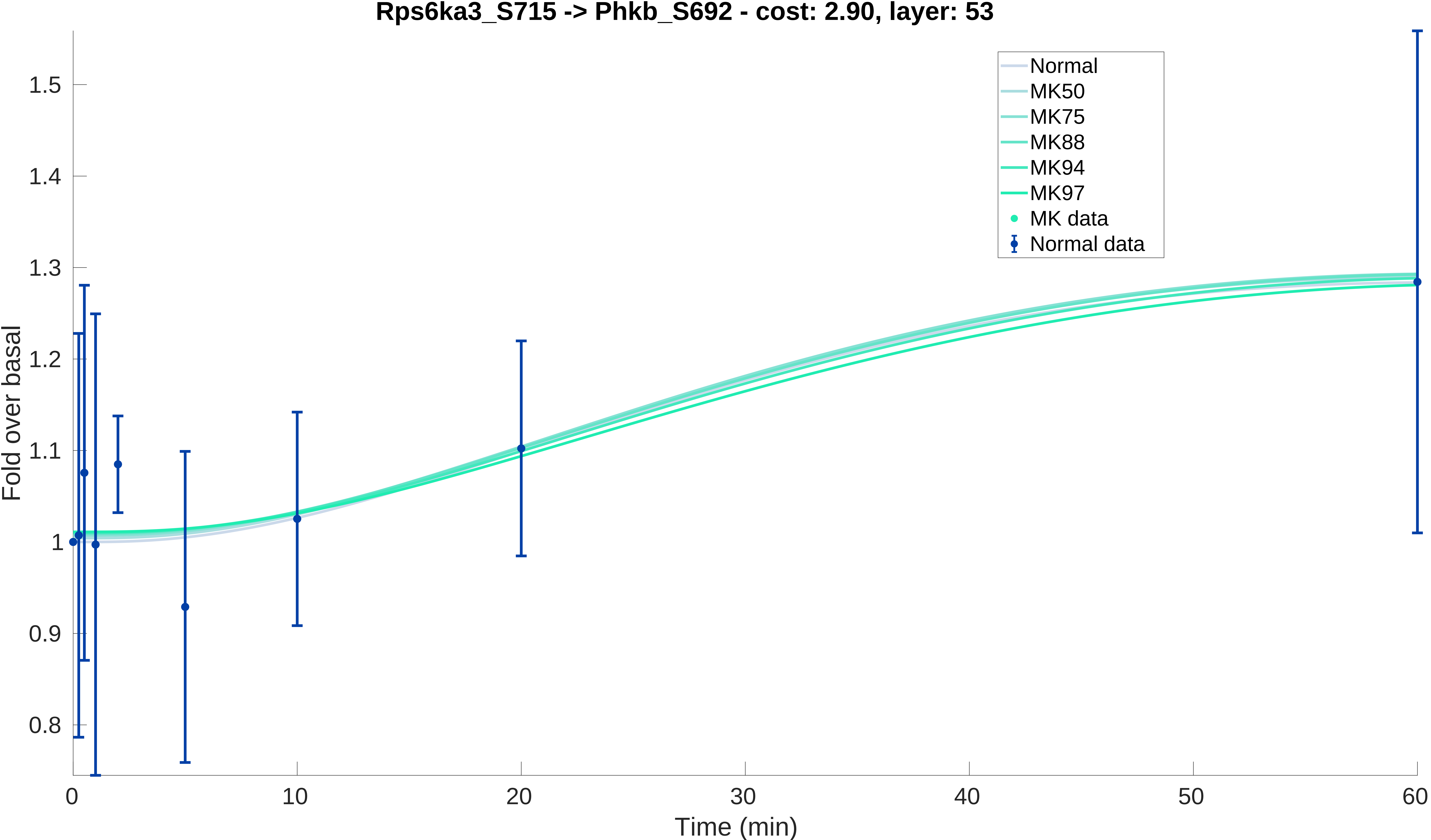

Rps6ka3_S715 -> Phkb_S692 - cost: 2.90, layer: 53
Normal
MK50
MK75
MK88
MK94
MK97
MK data
Normal data
Fold over basal
1.5
1.4
1.3
1.2
1.1
1
0.9
0.8
0
10
20
30
40
50
60
Time (min)

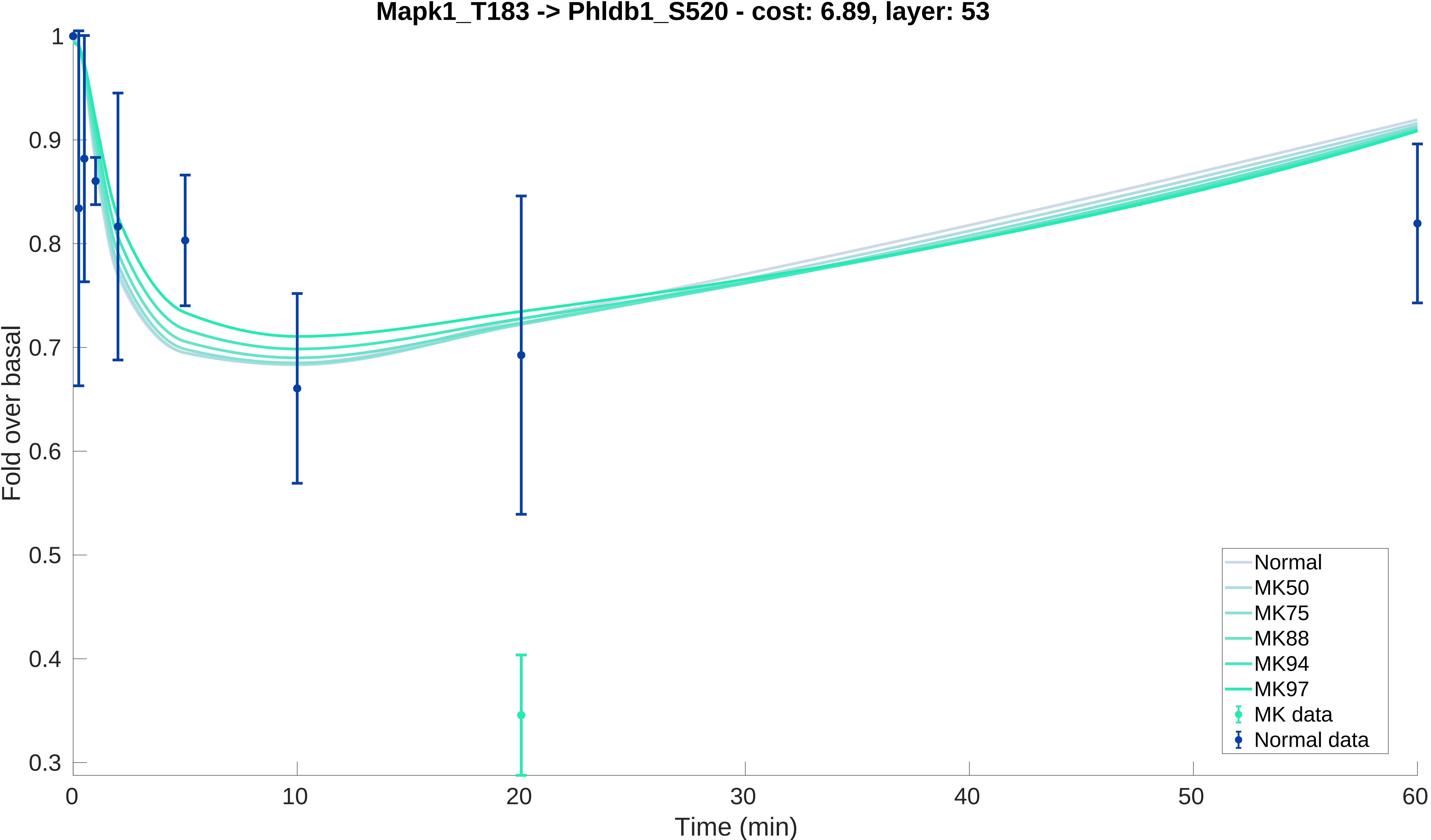

Mapk1_T183 -> Phldb1_S520 - cost: 6.89, layer: 53
Fold over basal
1
0.9
0.8
0.7
0.6
0.5
0.4
0.3
0
10
20
30
40
50
60
Time (min)
Normal
MK50
MK75
MK88
MK94
MK97
MK data
Normal data

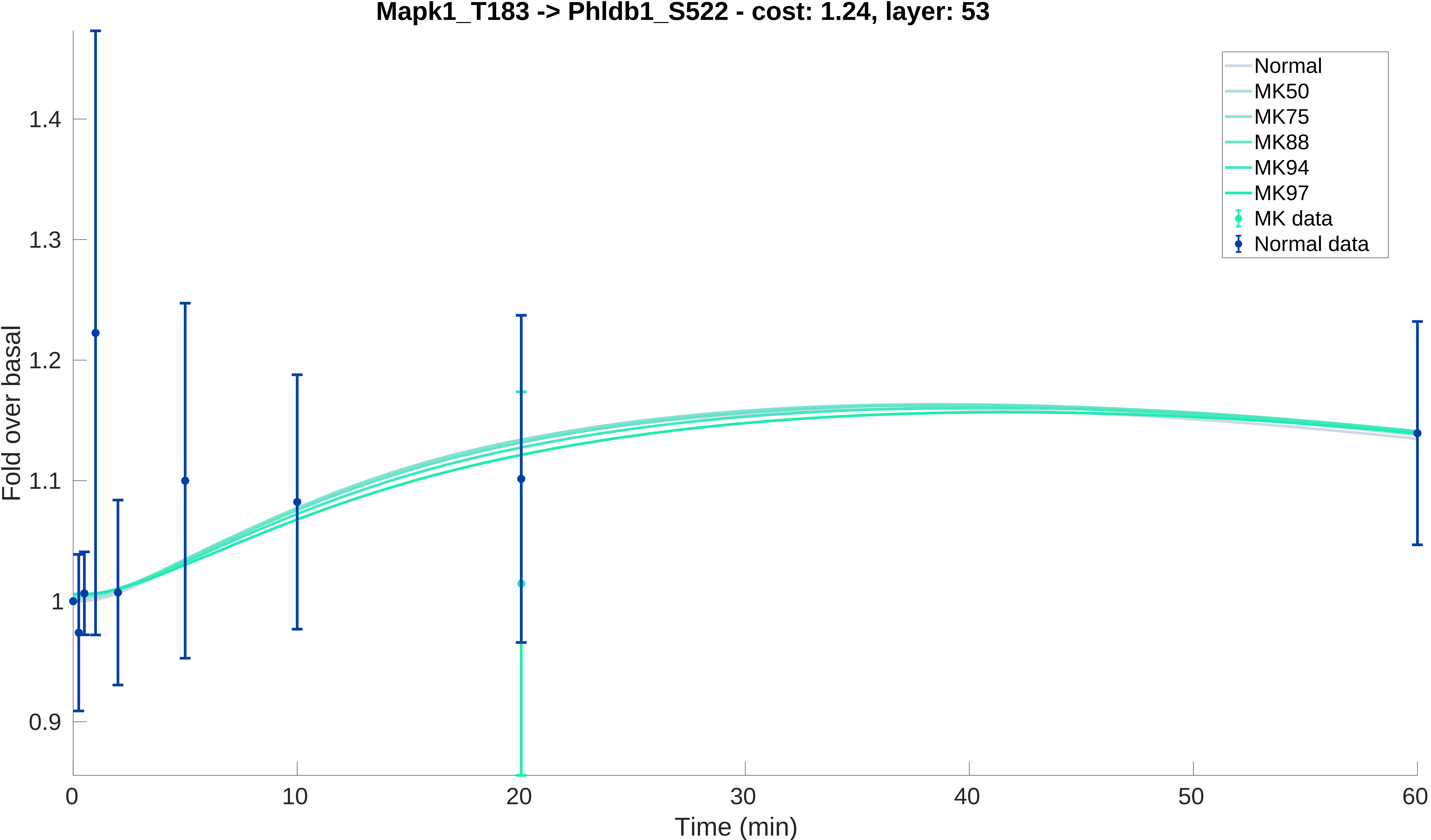

Mapk1_T183 -> Phldb1_S522 - cost: 1.24, layer: 53
Fold over basal
Time (min)
0.9
1
1.1
1.2
1.3
1.4
0
10
20
30
40
50
60
Normal
MK50
MK75
MK88
MK94
MK97
MK data
Normal data

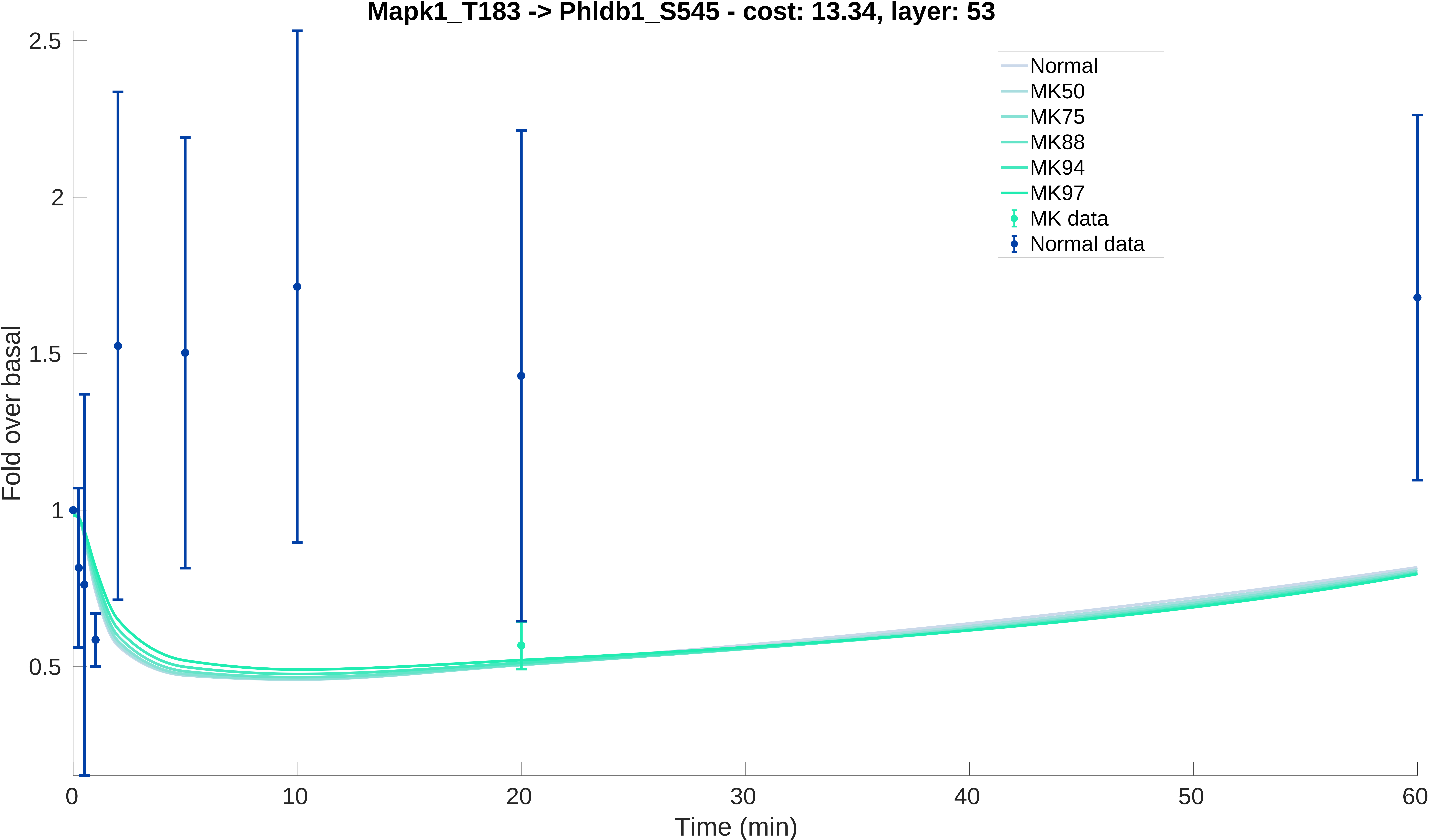

Mapk1_T183 -> Phldb1_S545 - cost: 13.34, layer: 53
Fold over basal
Time (min)
0.5
1
1.5
2
2.5
0
10
20
30
40
50
60
Normal
MK50
MK75
MK88
MK94
MK97
MK data
Normal data

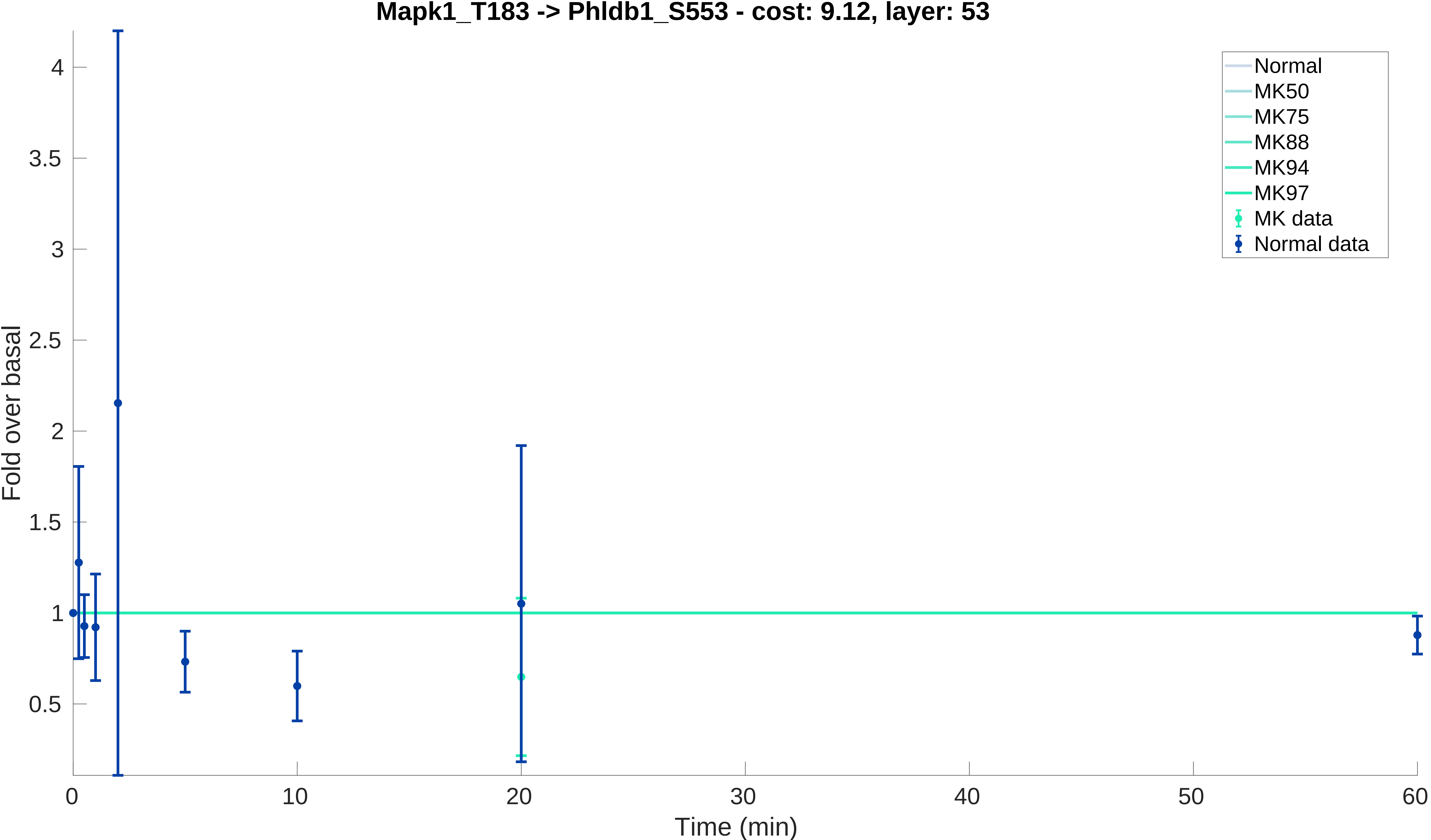
Mapk1_T183 -> Phldb1_S553 - cost: 9.12, layer: 53
Fold over basal
Time (min)
0
10
20
30
40
50
60
0.5
1
1.5
2
2.5
3
3.5
4
Normal
MK50
MK75
MK88
MK94
MK97
MK data
Normal data

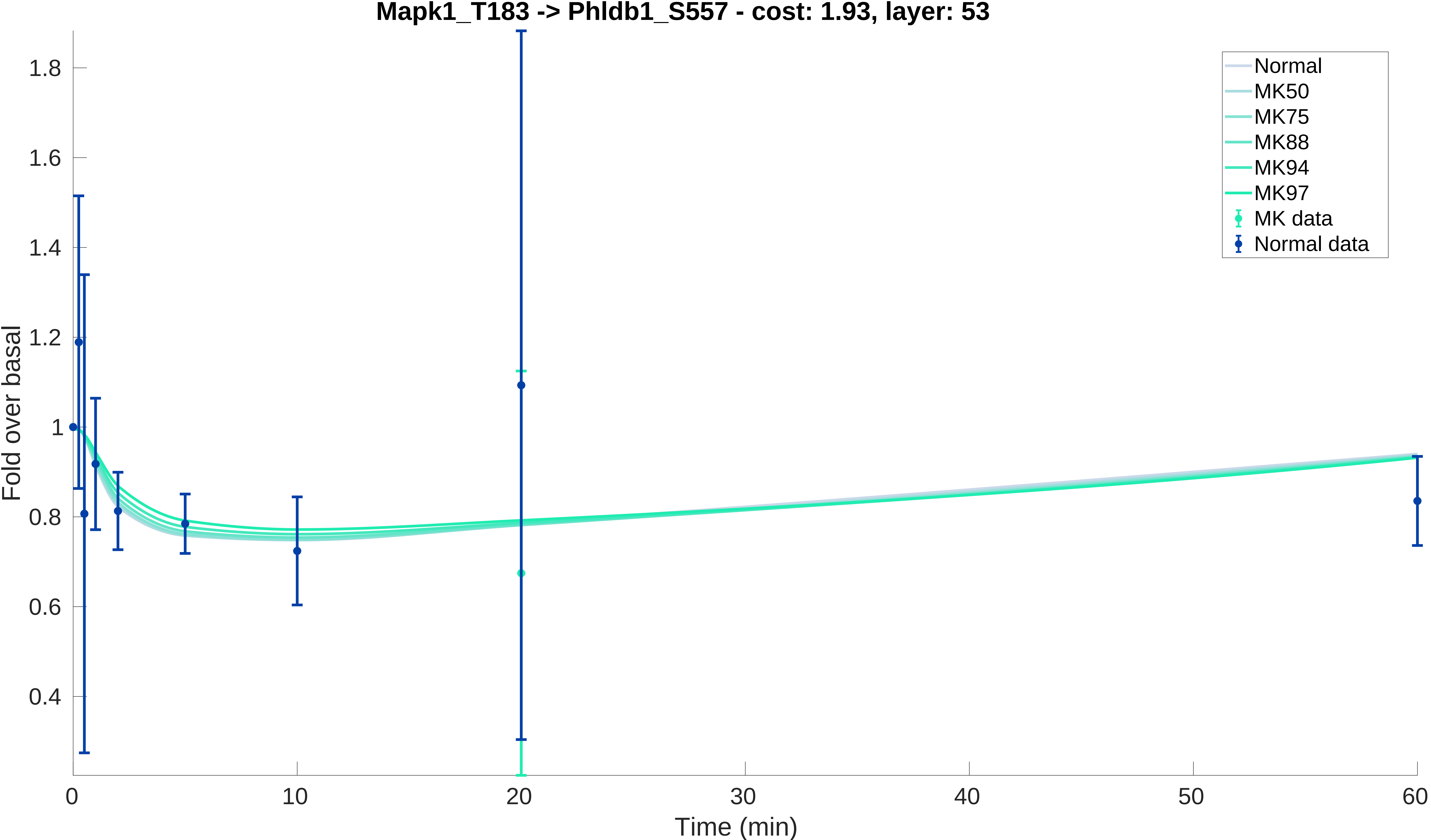

Mapk1_T183 -> Phldb1_S557 - cost: 1.93, layer: 53
Fold over basal
Time (min)
1.8
1.6
1.4
1.2
1
0.8
0.6
0.4
0
10
20
30
40
50
60
Normal
MK50
MK75
MK88
MK94
MK97
MK data
Normal data

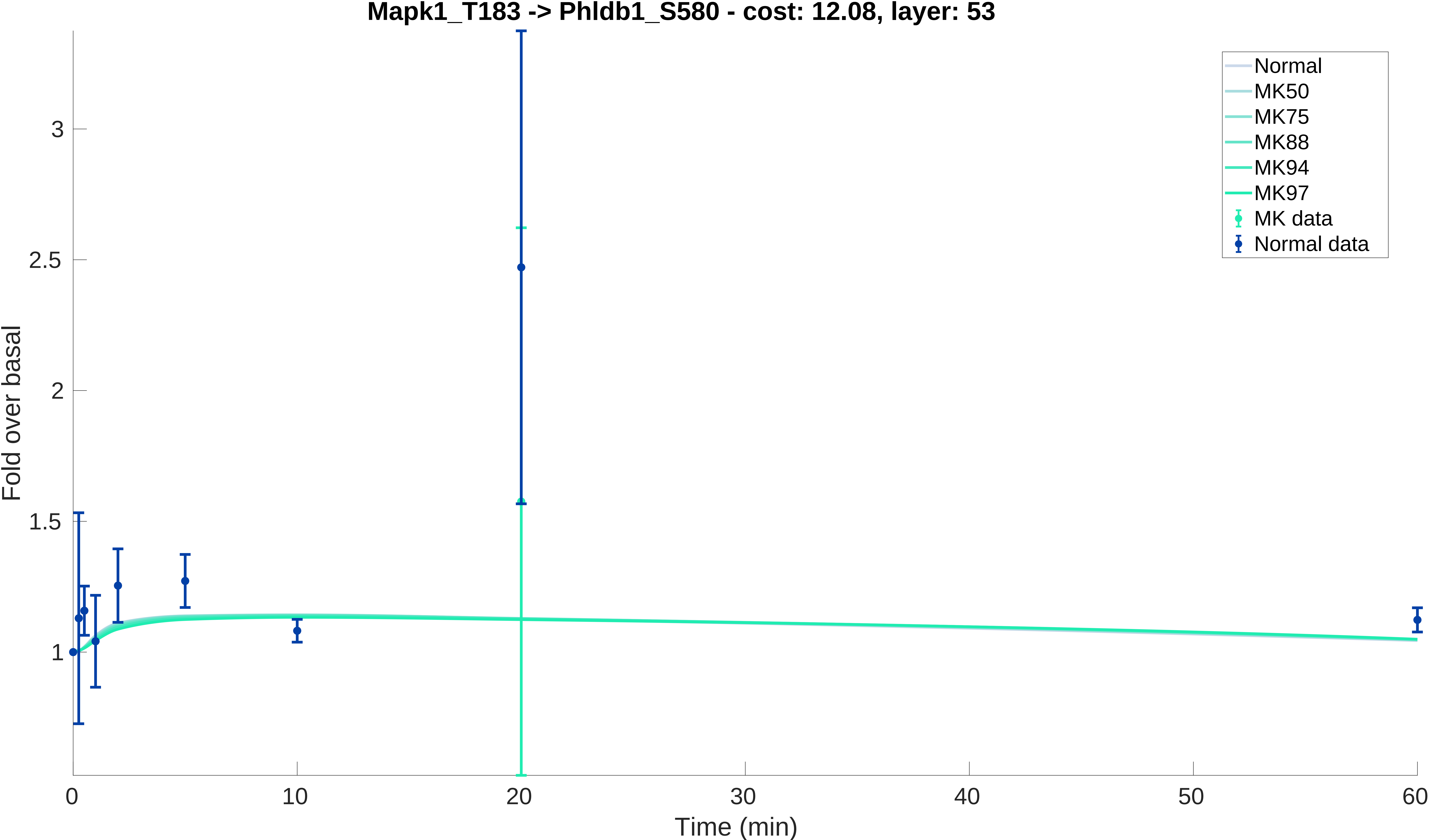

Mapk1_T183 -> Phldb1_S580 - cost: 12.08, layer: 53
Fold over basal
Time (min)
3
2.5
2
1.5
1
0
10
20
30
40
50
60
Normal
MK50
MK75
MK88
MK94
MK97
MK data
Normal data

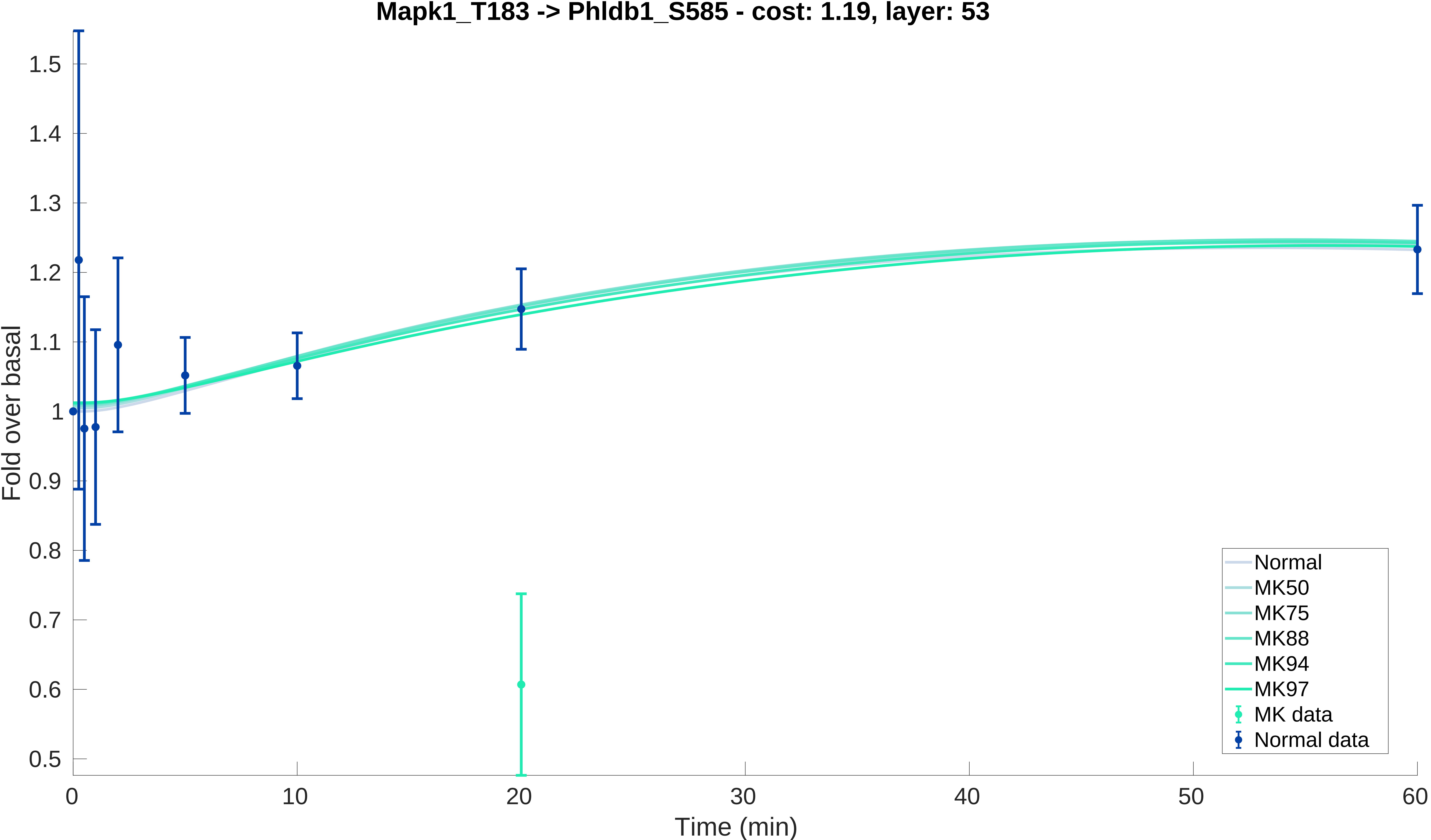

Mapk1_T183 -> Phldb1_S585 - cost: 1.19, layer: 53
Fold over basal
Time (min)
0
10
20
30
40
50
60
0.5
0.6
0.7
0.8
0.9
1
1.1
1.2
1.3
1.4
1.5
Normal
MK50
MK75
MK88
MK94
MK97
MK data
Normal data

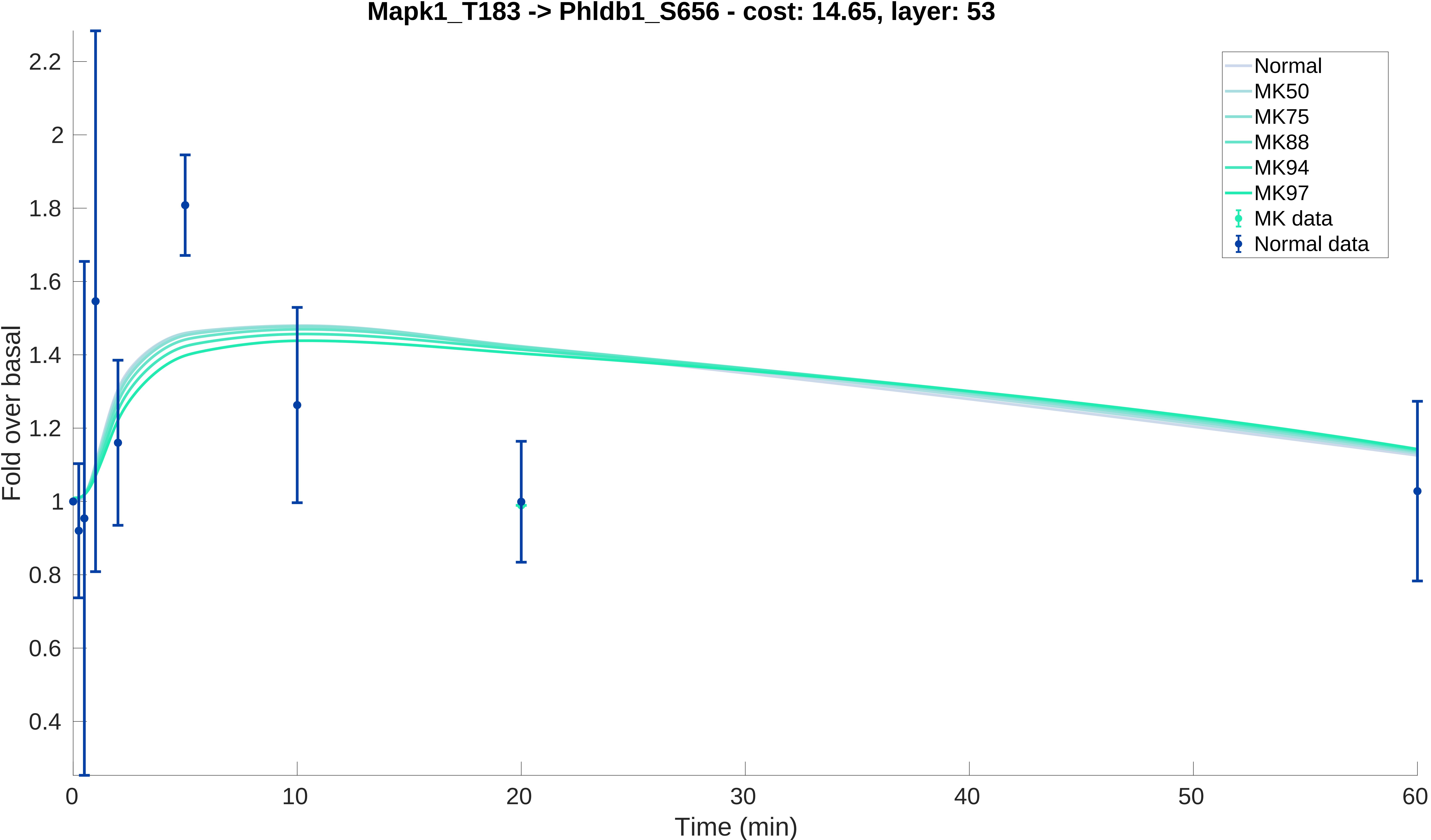
Mapk1_T183 -> Phldb1_S656 - cost: 14.65, layer: 53
Fold over basal
Time (min)
0.4
0.6
0.8
1
1.2
1.4
1.6
1.8
2
2.2
0
10
20
30
40
50
60
Normal
MK50
MK75
MK88
MK94
MK97
MK data
Normal data

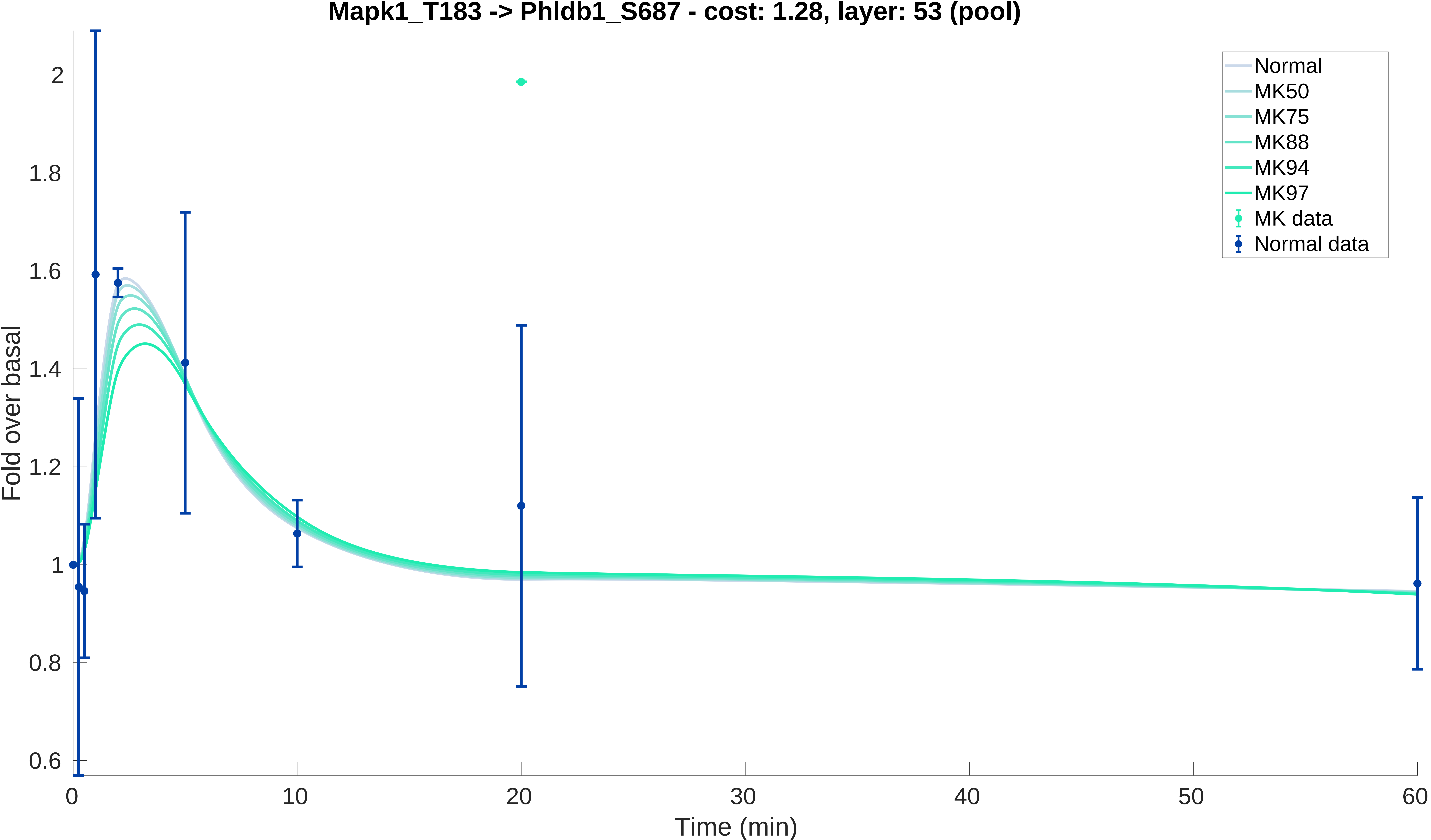

Mapk1_T183 -> Phldb1_S687 - cost: 1.28, layer: 53 (pool)
Fold over basal
Time (min)
0.6
0.8
1
1.2
1.4
1.6
1.8
2
0
10
20
30
40
50
60
Normal
MK50
MK75
MK88
MK94
MK97
MK data
Normal data

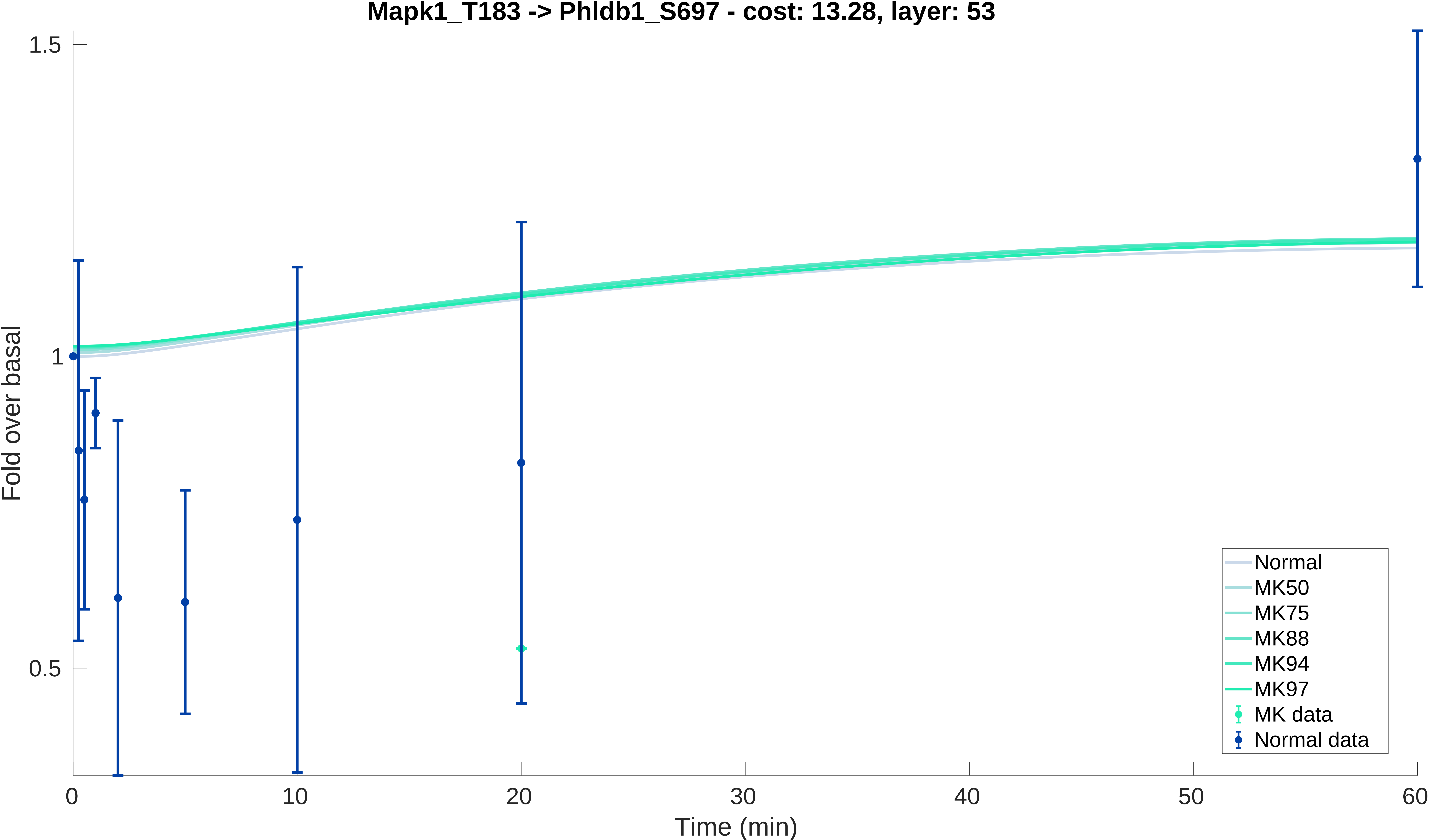
Mapk1_T183 -> Phldb1_S697 - cost: 13.28, layer: 53
Fold over basal
1.5
1
0.5
0
10
20
30
40
50
60
Time (min)
Normal
MK50
MK75
MK88
MK94
MK97
MK data
Normal data

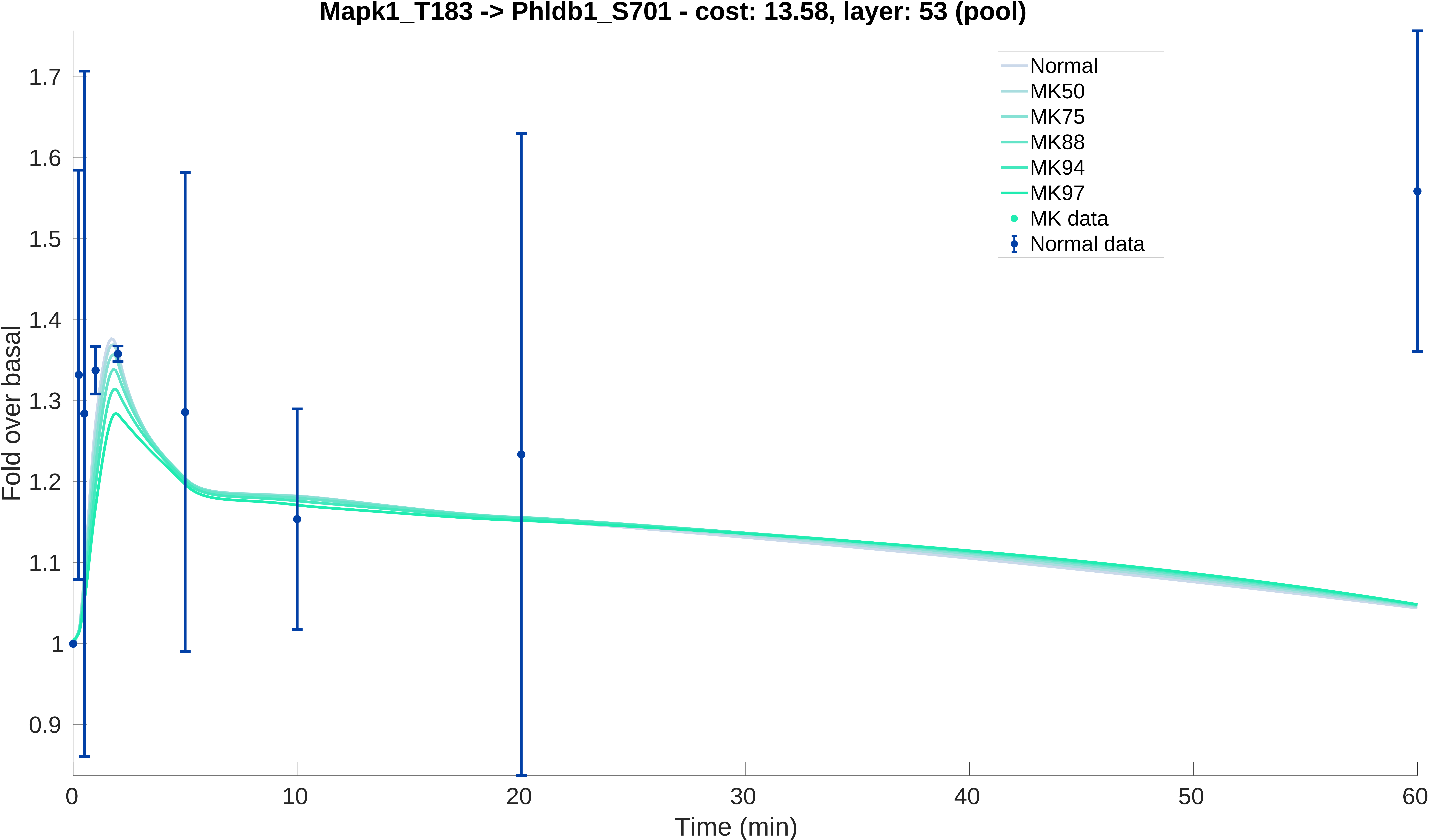

Mapk1_T183 -> Phldb1_S701 - cost: 13.58, layer: 53 (pool)
Fold over basal
Time (min)
0.9
1
1.1
1.2
1.3
1.4
1.5
1.6
1.7
0
10
20
30
40
50
60
Normal
MK50
MK75
MK88
MK94
MK97
MK data
Normal data

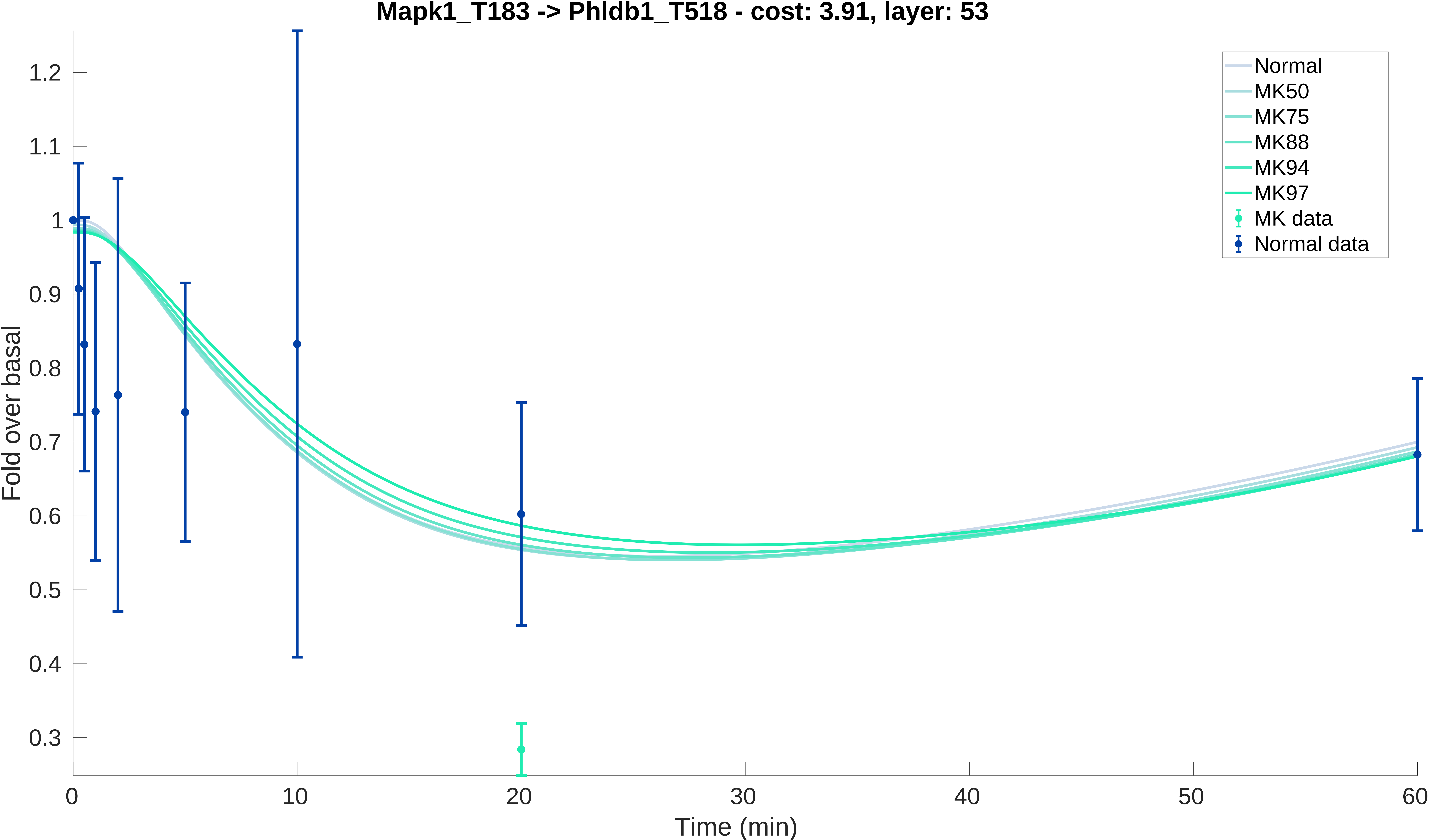

Mapk1_T183 -> Phldb1_T518 - cost: 3.91, layer: 53
Fold over basal
Time (min)
0
10
20
30
40
50
60
0.3
0.4
0.5
0.6
0.7
0.8
0.9
1
1.1
1.2
Normal
MK50
MK75
MK88
MK94
MK97
MK data
Normal data

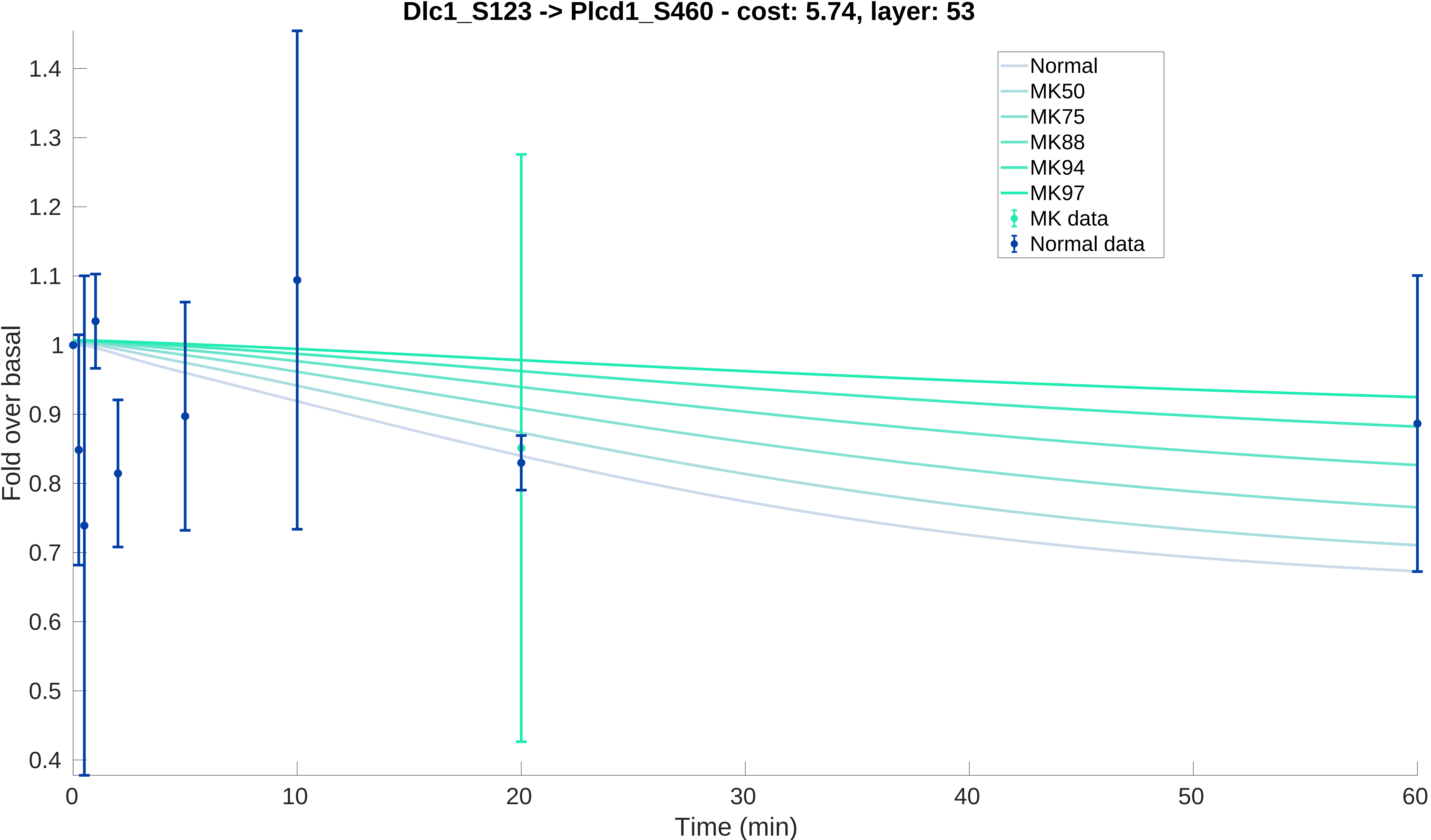

Dlc1_S123 -> Plcd1_S460 - cost: 5.74, layer: 53
Normal
MK50
MK75
MK88
MK94
MK97
MK data
Normal data
Fold over basal
Time (min)
0.4
0.5
0.6
0.7
0.8
0.9
1
1.1
1.2
1.3
1.4
0
10
20
30
40
50
60

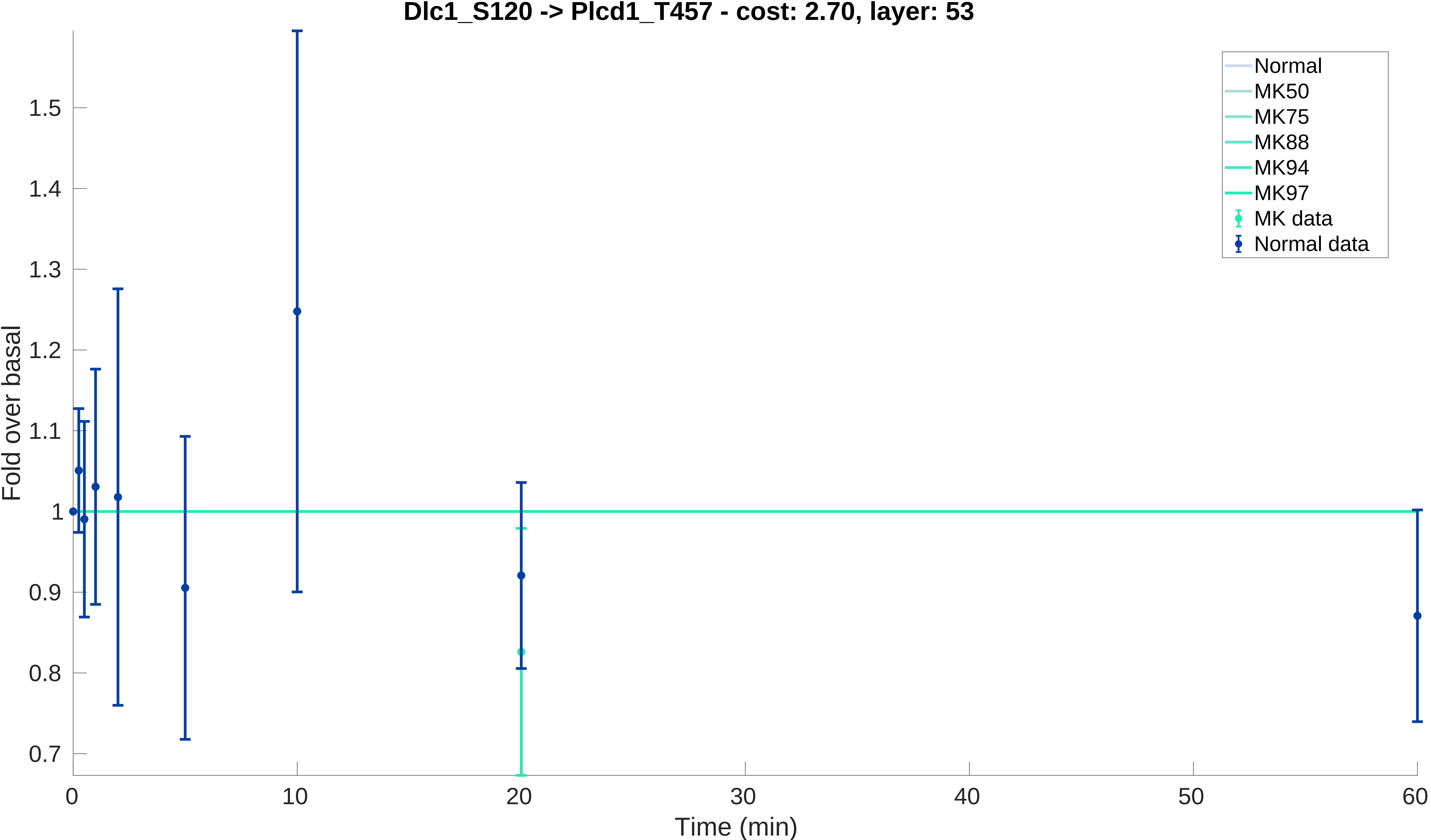

Dlc1_S120 -> Plcd1_T457 - cost: 2.70, layer: 53
Normal
MK50
MK75
MK88
MK94
MK97
MK data
Normal data
Fold over basal
1.5
1.4
1.3
1.2
1.1
1
0.9
0.8
0.7
0
10
20
30
40
50
60
Time (min)

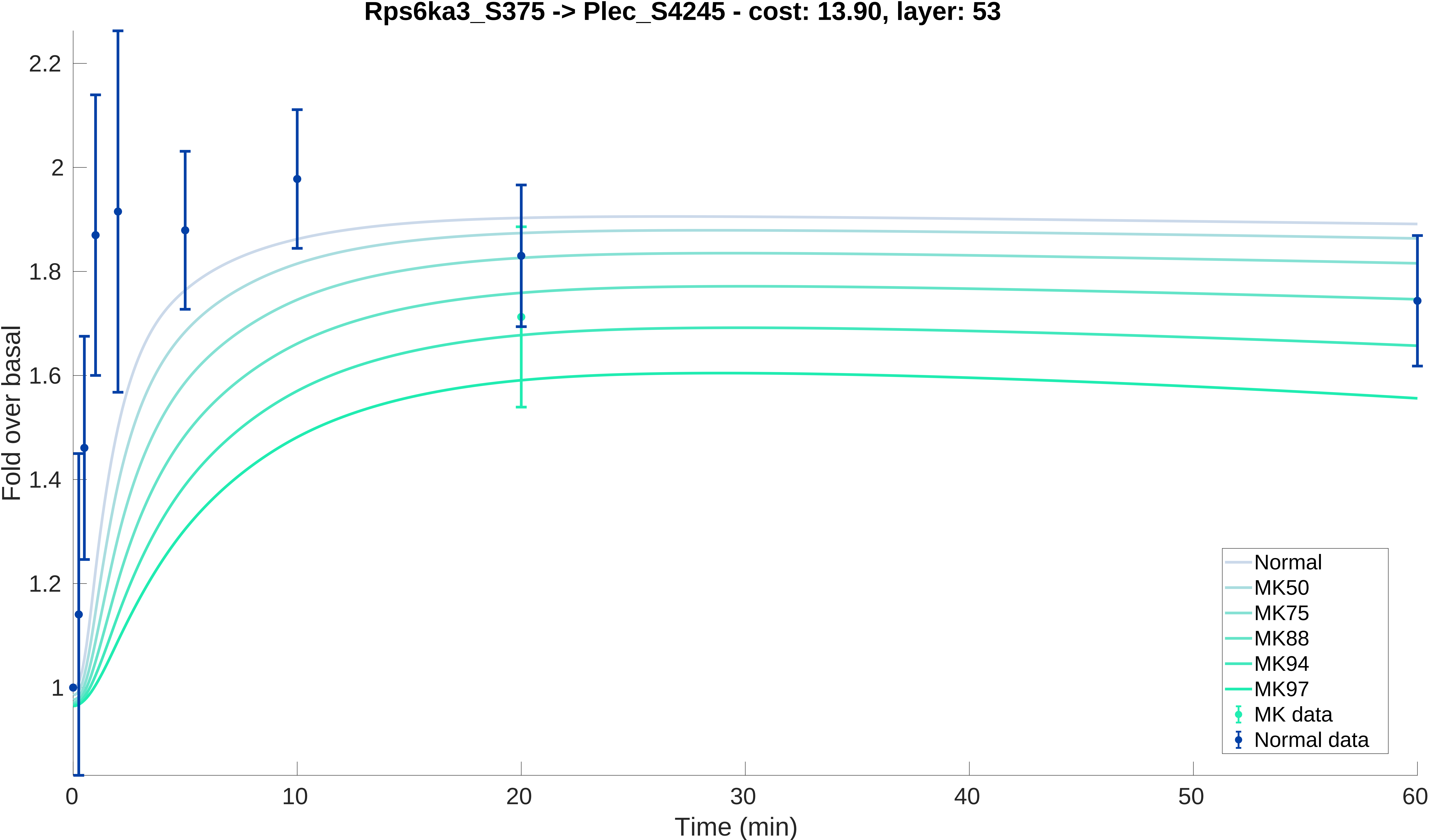
Rps6ka3_S375 -> Plec_S4245 - cost: 13.90, layer: 53
Fold over basal
Time (min)
2.2
2
1.8
1.6
1.4
1.2
1
0
10
20
30
40
50
60
Normal
MK50
MK75
MK88
MK94
MK97
MK data
Normal data

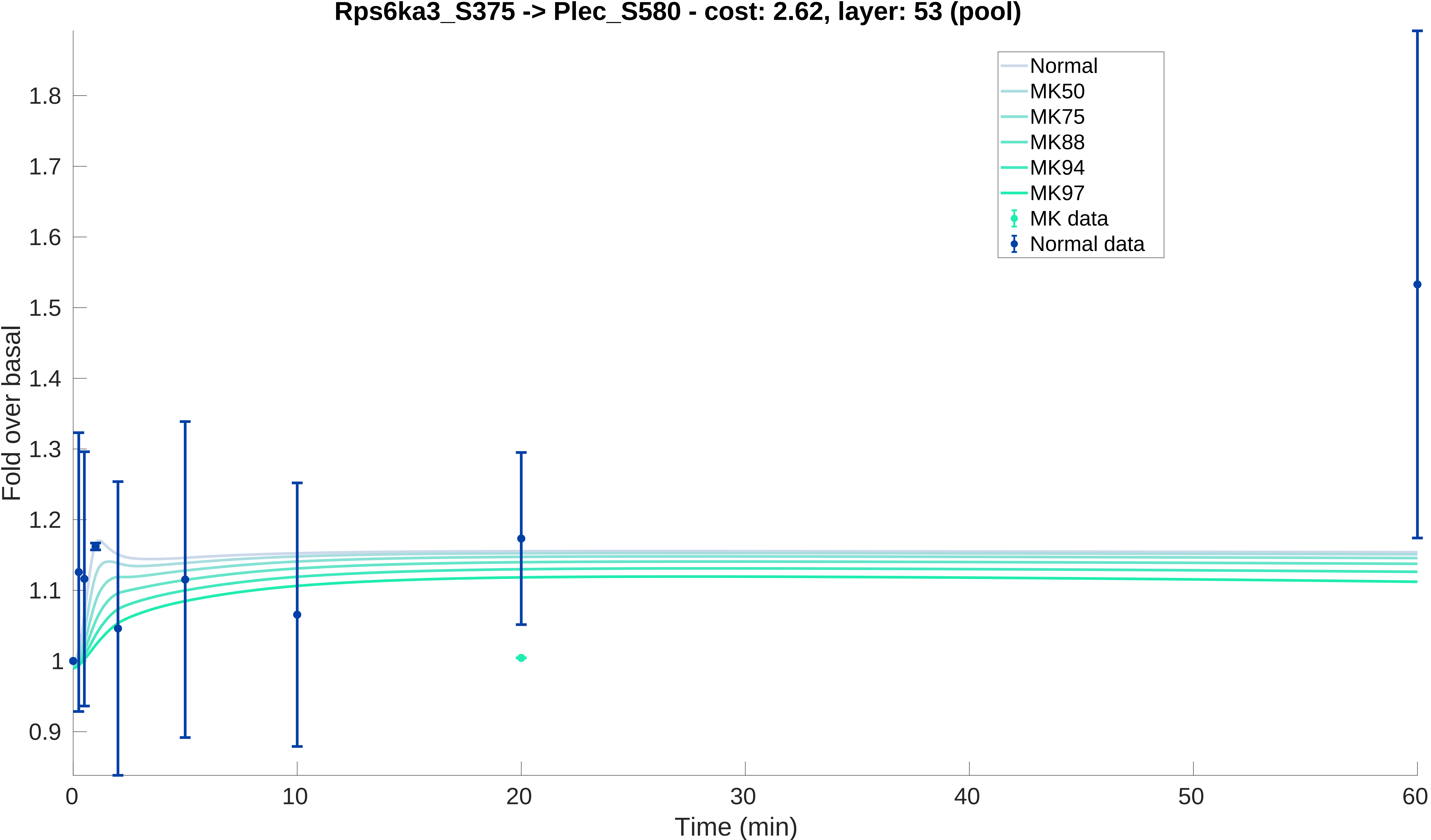
Rps6ka3_S375 -> Plec_S580 - cost: 2.62, layer: 53 (pool)
Normal
MK50
MK75
MK88
MK94
MK97
MK data
Normal data
Fold over basal
Time (min)
0.9
1
1.1
1.2
1.3
1.4
1.5
1.6
1.7
1.8
0
10
20
30
40
50
60

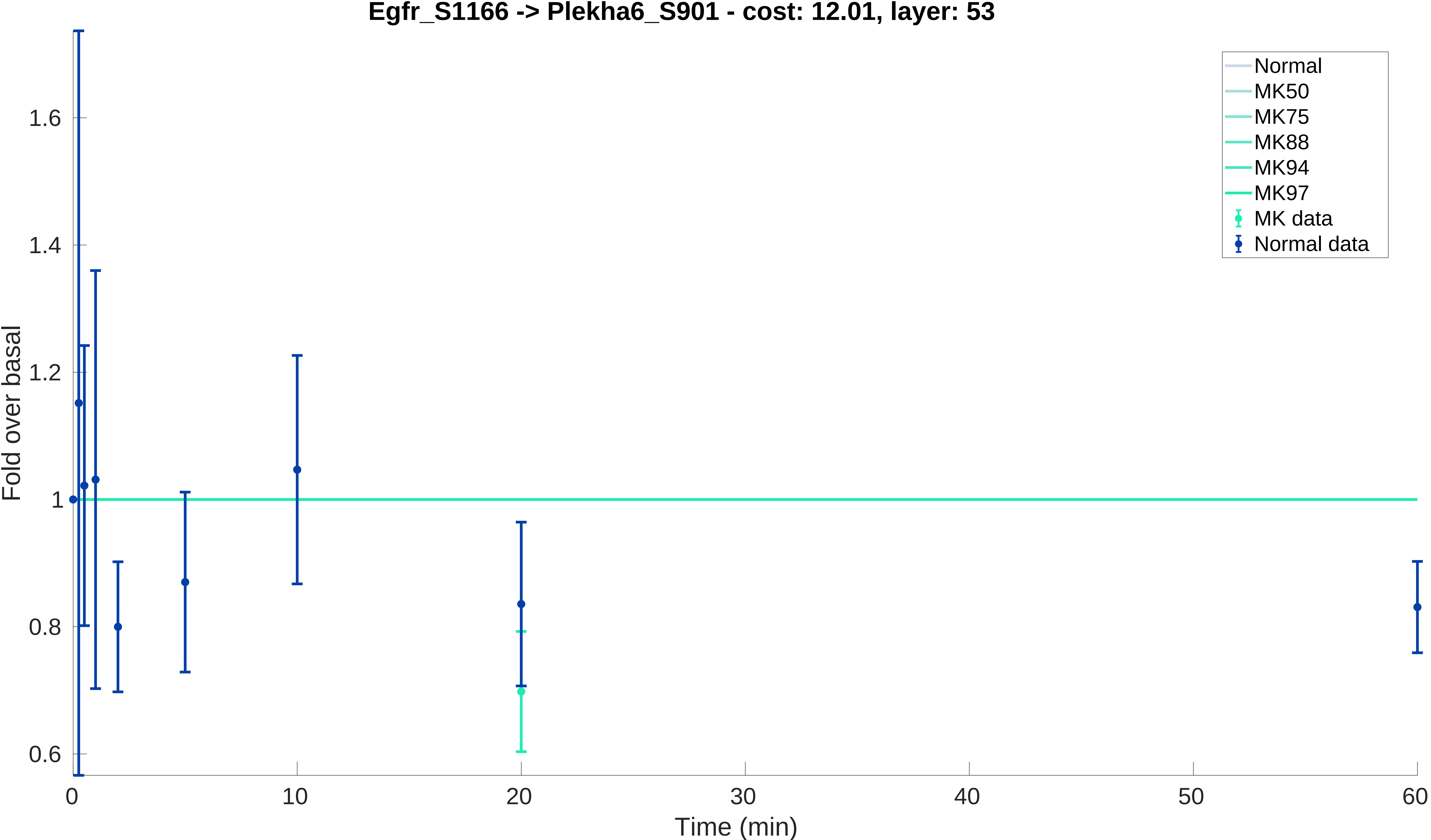

Egfr_S1166 -> Plekha6_S901 - cost: 12.01, layer: 53
Normal
MK50
MK75
MK88
MK94
MK97
MK data
Normal data
Fold over basal
1.6
1.4
1.2
1
0.8
0.6
0
10
20
30
40
50
60
Time (min)

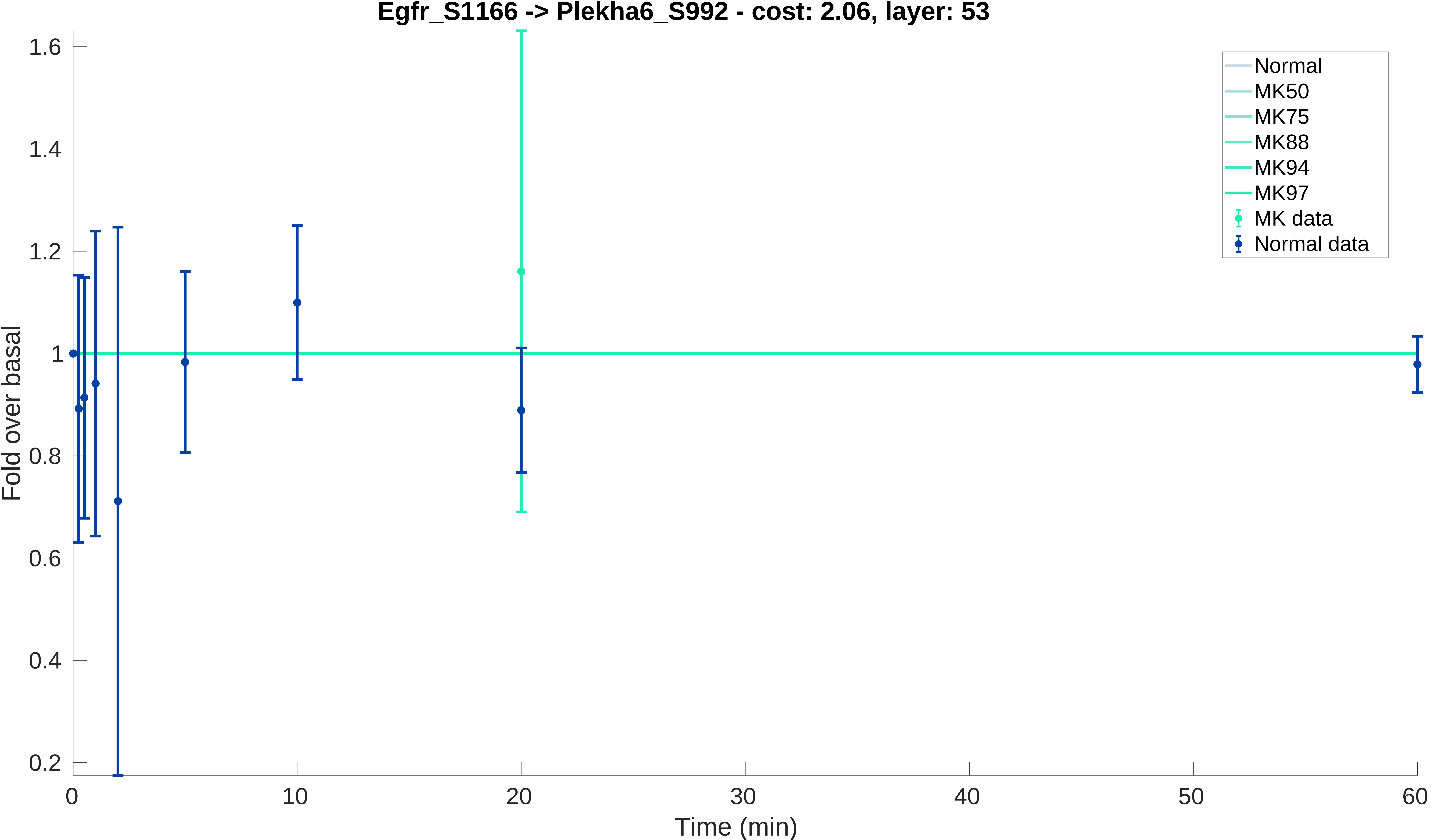
Egfr_S1166 -> Plekha6_S992 - cost: 2.06, layer: 53
Fold over basal
1.6
1.4
1.2
1
0.8
0.6
0.4
0.2
0
10
20
30
40
50
60
Time (min)
Normal
MK50
MK75
MK88
MK94
MK97
MK data
Normal data

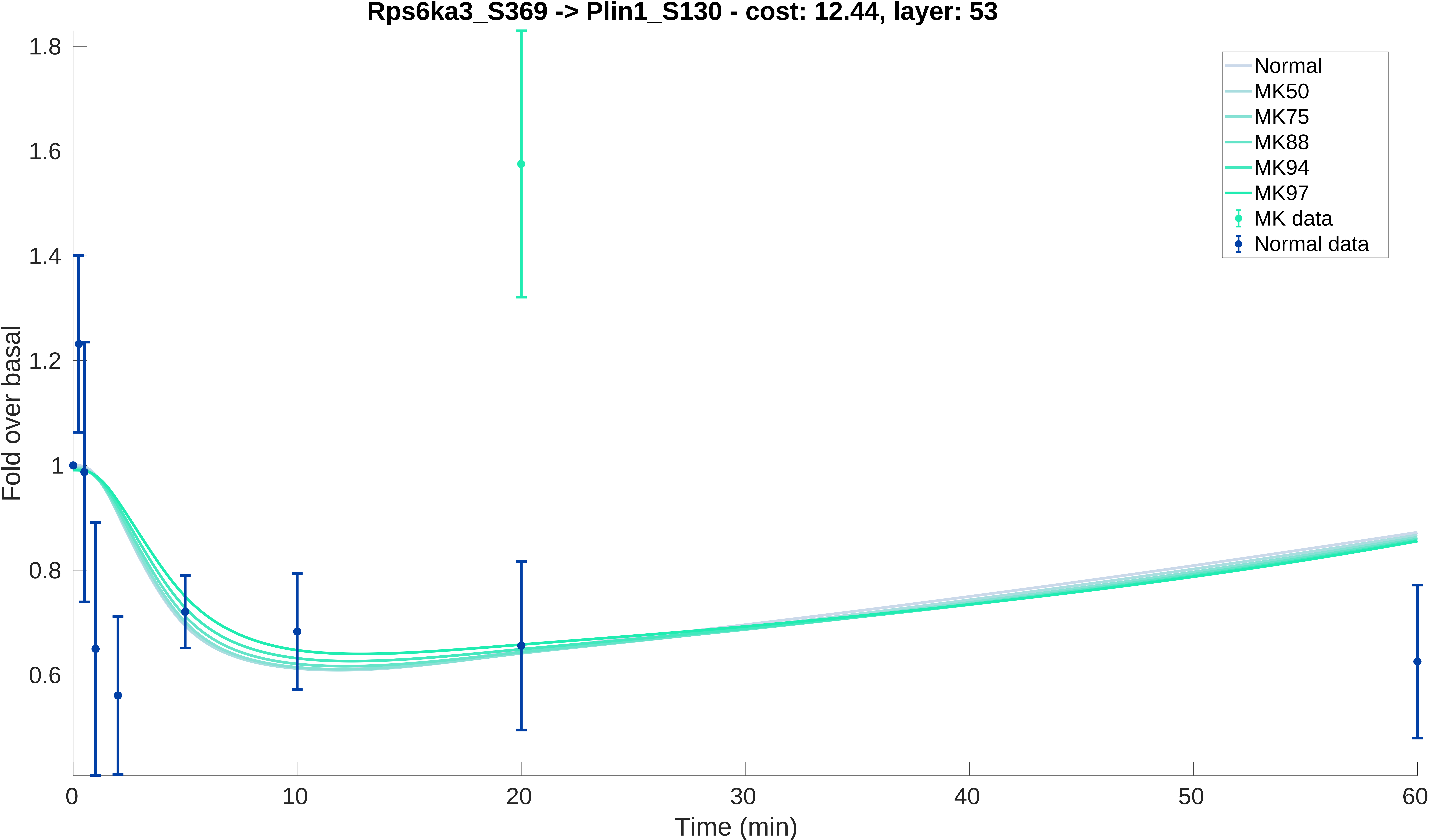
Rps6ka3_S369 -> Plin1_S130 - cost: 12.44, layer: 53
Fold over basal
Time (min)
0.6
0.8
1
1.2
1.4
1.6
1.8
0
10
20
30
40
50
60
Normal
MK50
MK75
MK88
MK94
MK97
MK data
Normal data

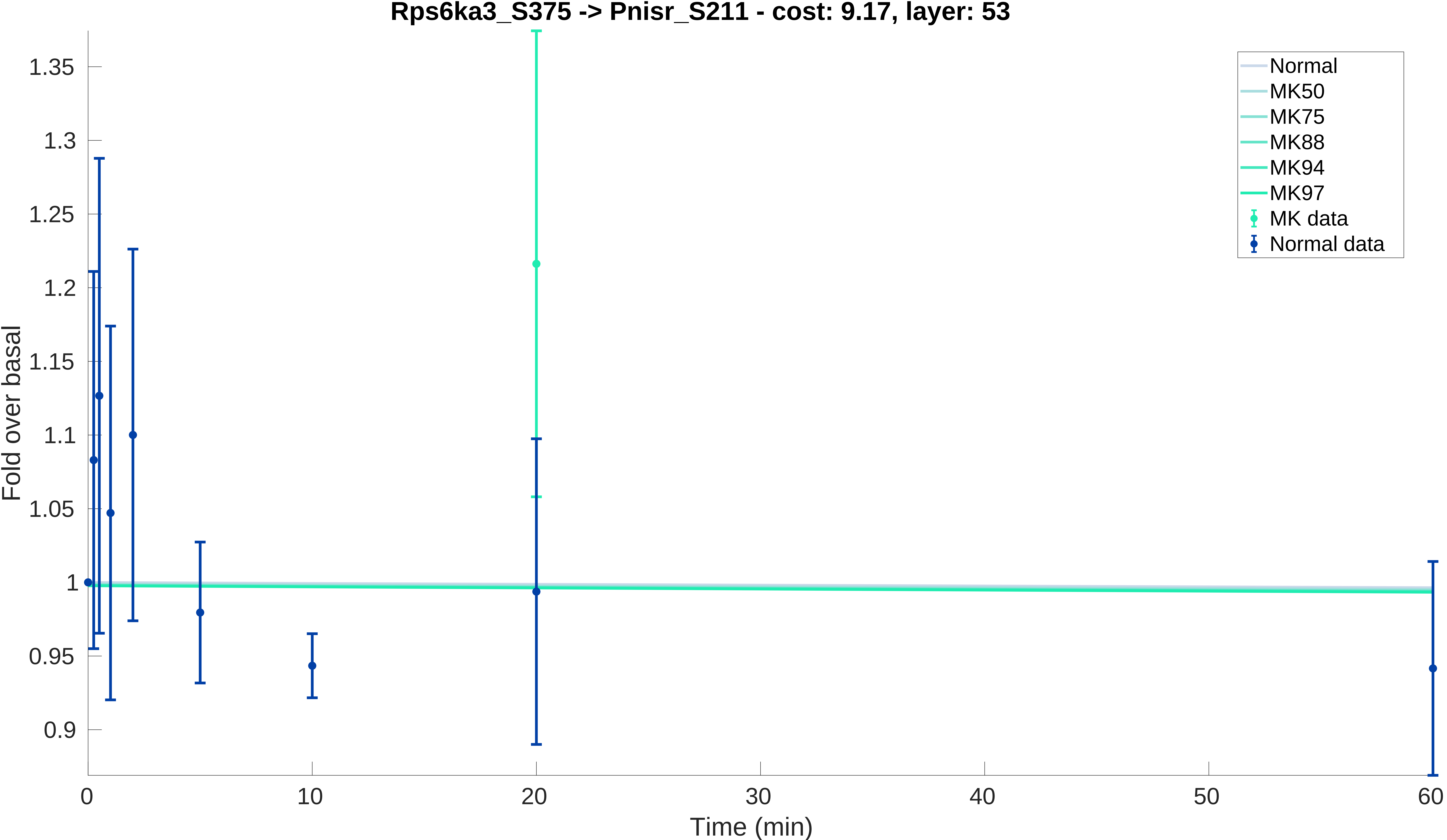

Rps6ka3_S375 -> Pnisr_S211 - cost: 9.17, layer: 53
Fold over basal
Time (min)
1.35
1.3
1.25
1.2
1.15
1.1
1.05
1
0.95
0.9
0
10
20
30
40
50
60
Normal
MK50
MK75
MK88
MK94
MK97
MK data
Normal data

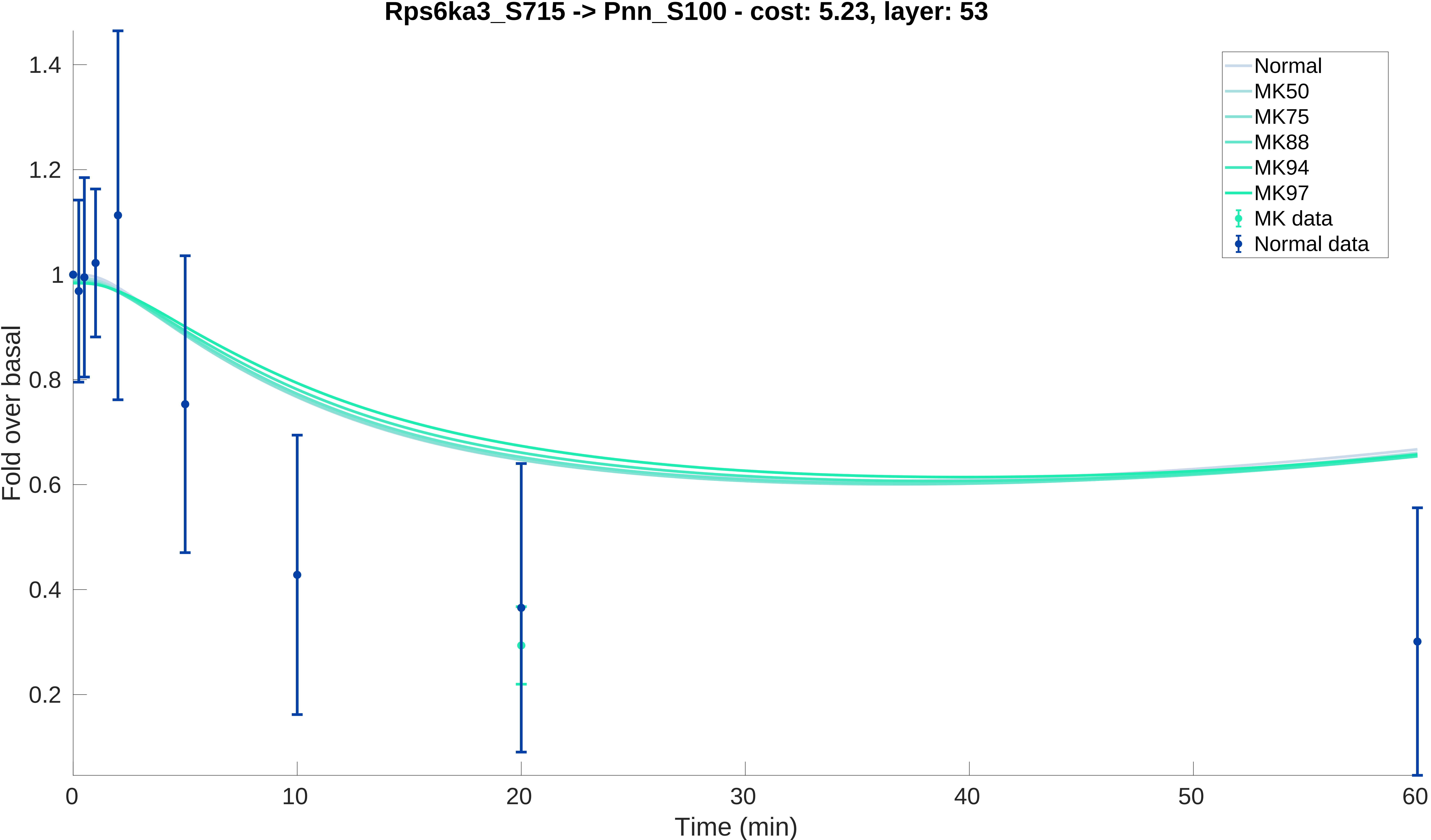
Rps6ka3_S715 -> Pnn_S100 - cost: 5.23, layer: 53
Fold over basal
Time (min)
0
10
20
30
40
50
60
0.2
0.4
0.6
0.8
1
1.2
1.4
Normal
MK50
MK75
MK88
MK94
MK97
MK data
Normal data

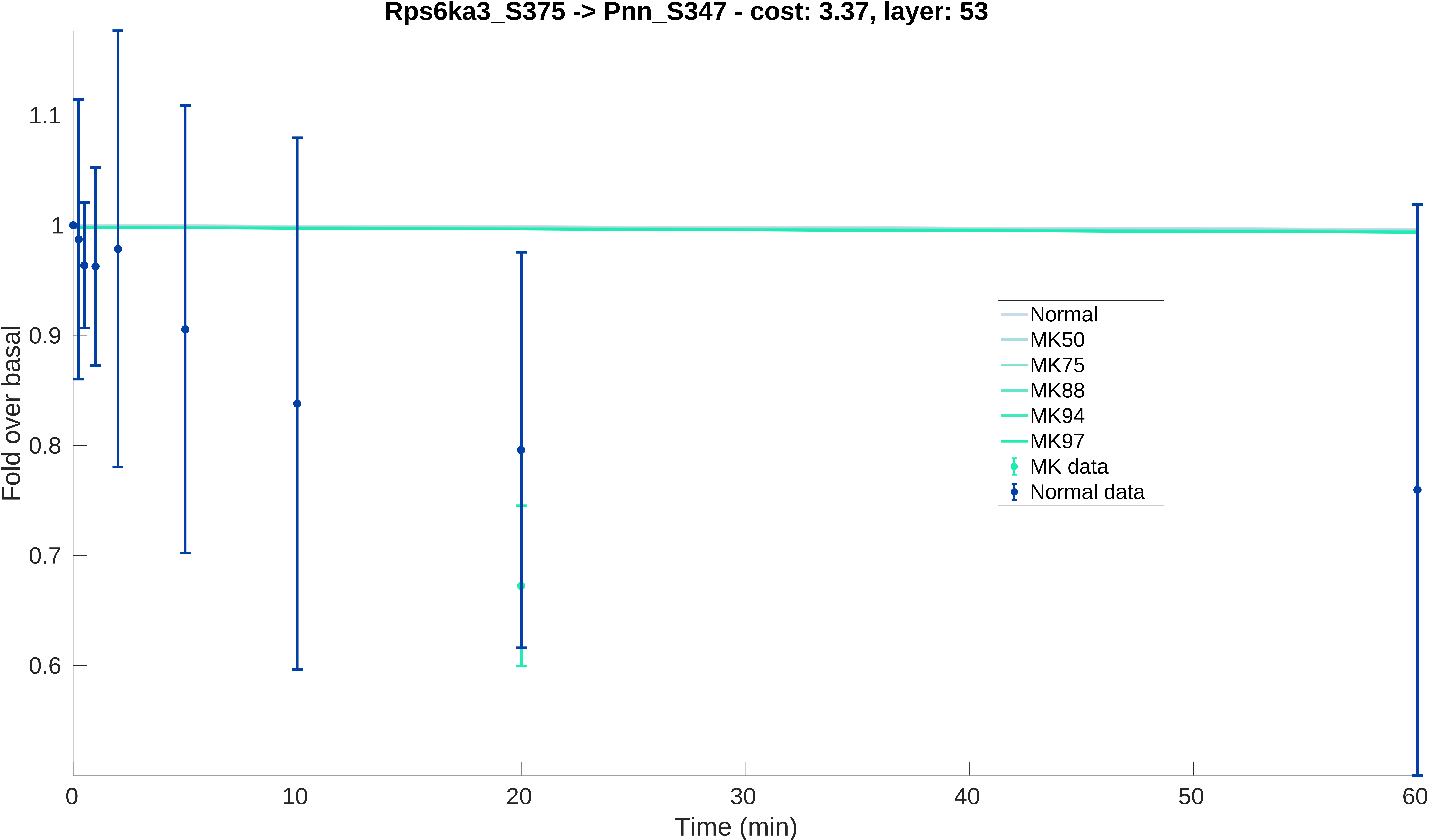
Rps6ka3_S375 -> Pnn_S347 - cost: 3.37, layer: 53
Fold over basal
1.1
1
0.9
0.8
0.7
0.6
0
10
20
30
40
50
60
Time (min)
Normal
MK50
MK75
MK88
MK94
MK97
MK data
Normal data

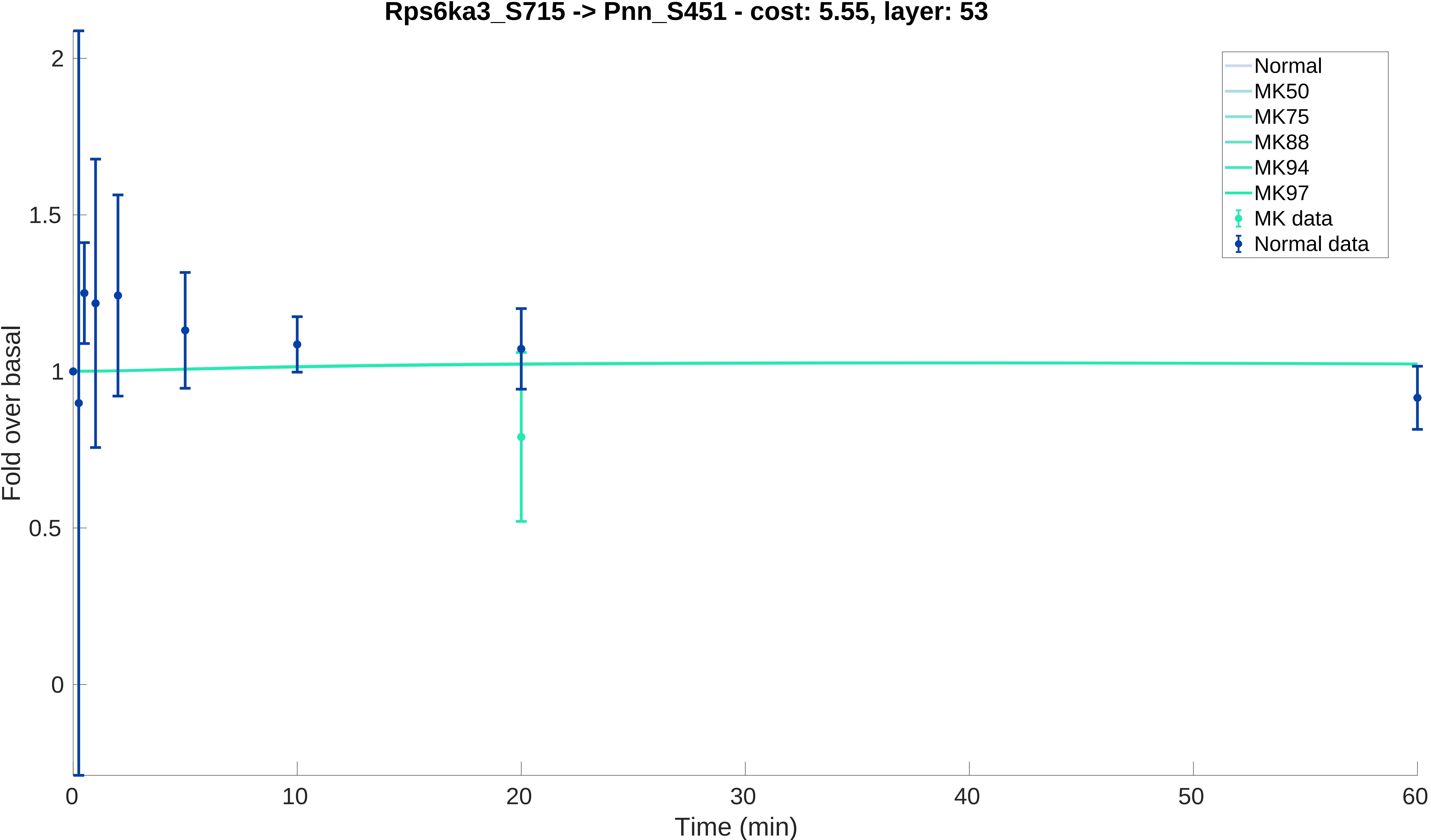

Rps6ka3_S715 -> Pnn_S451 - cost: 5.55, layer: 53
Normal
MK50
MK75
MK88
MK94
MK97
MK data
Normal data
Fold over basal
Time (min)
2
1.5
1
0.5
0
0
10
20
30
40
50
60

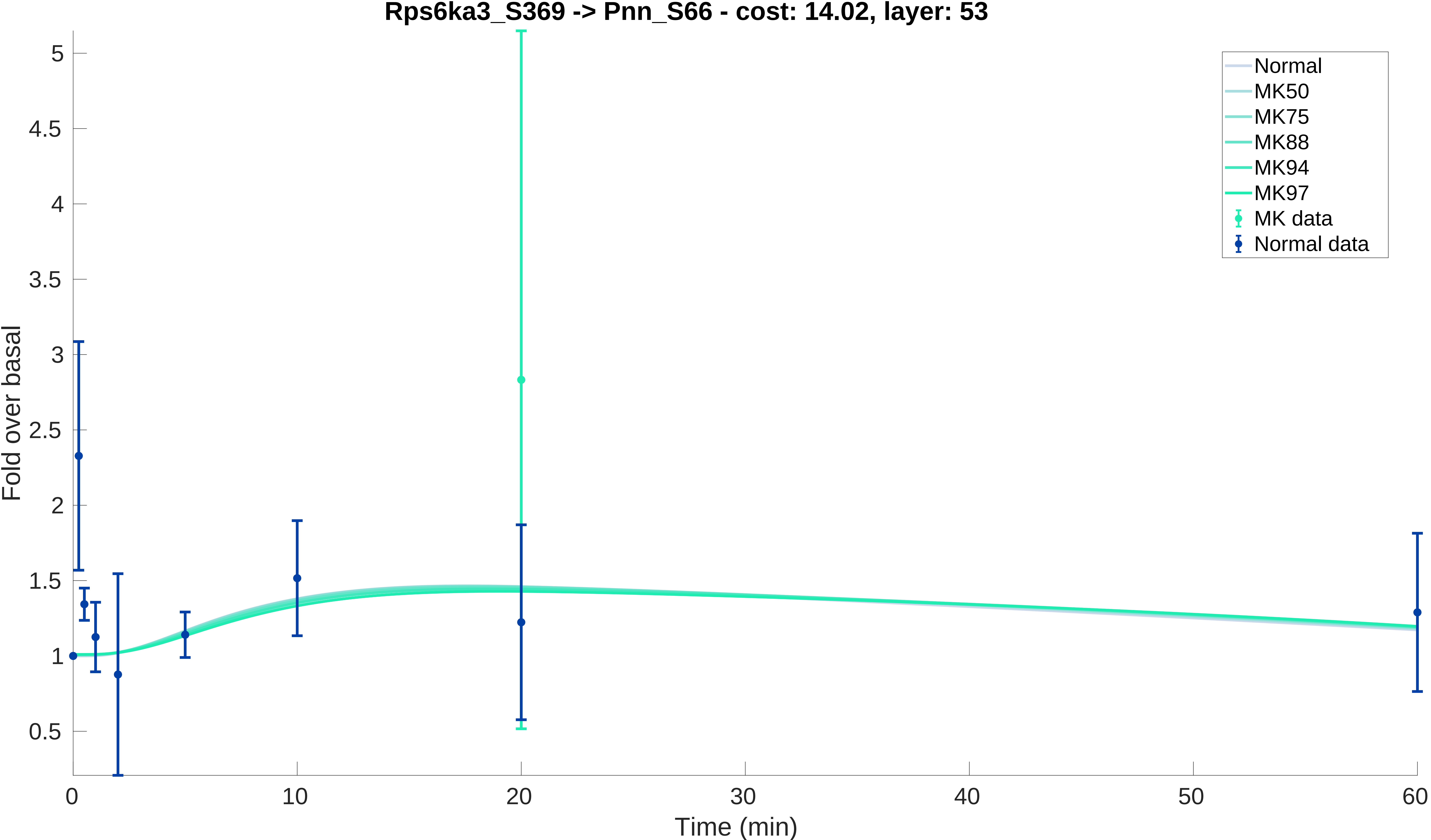
Rps6ka3_S369 -> Pnn_S66 - cost: 14.02, layer: 53
Fold over basal
Time (min)
0.5
1
1.5
2
2.5
3
3.5
4
4.5
5
0
10
20
30
40
50
60
Normal
MK50
MK75
MK88
MK94
MK97
MK data
Normal data

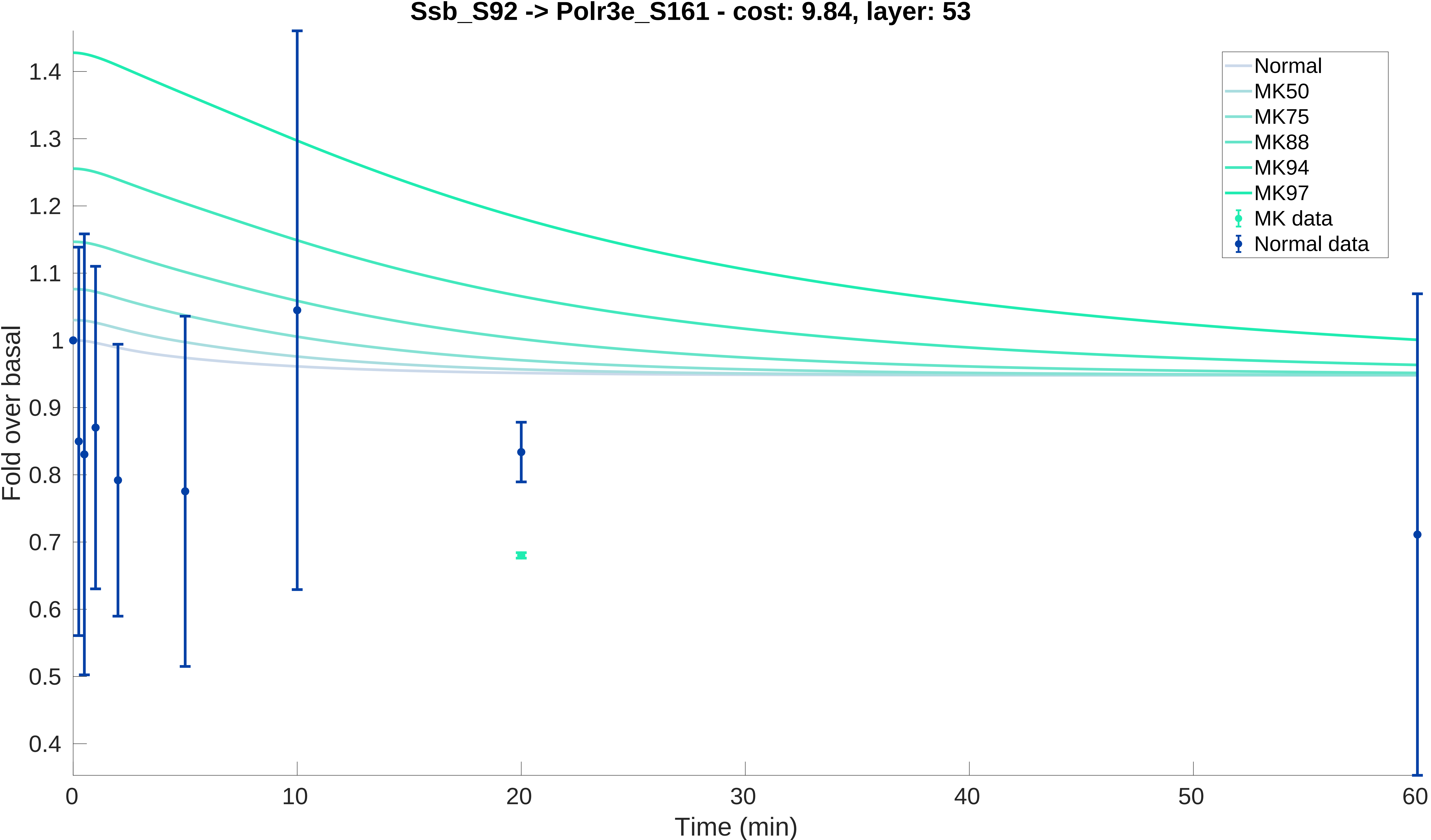

Ssb_S92 -> Polr3e_S161 - cost: 9.84, layer: 53
Fold over basal
Time (min)
0
10
20
30
40
50
60
0.4
0.5
0.6
0.7
0.8
0.9
1
1.1
1.2
1.3
1.4
Normal
MK50
MK75
MK88
MK94
MK97
MK data
Normal data

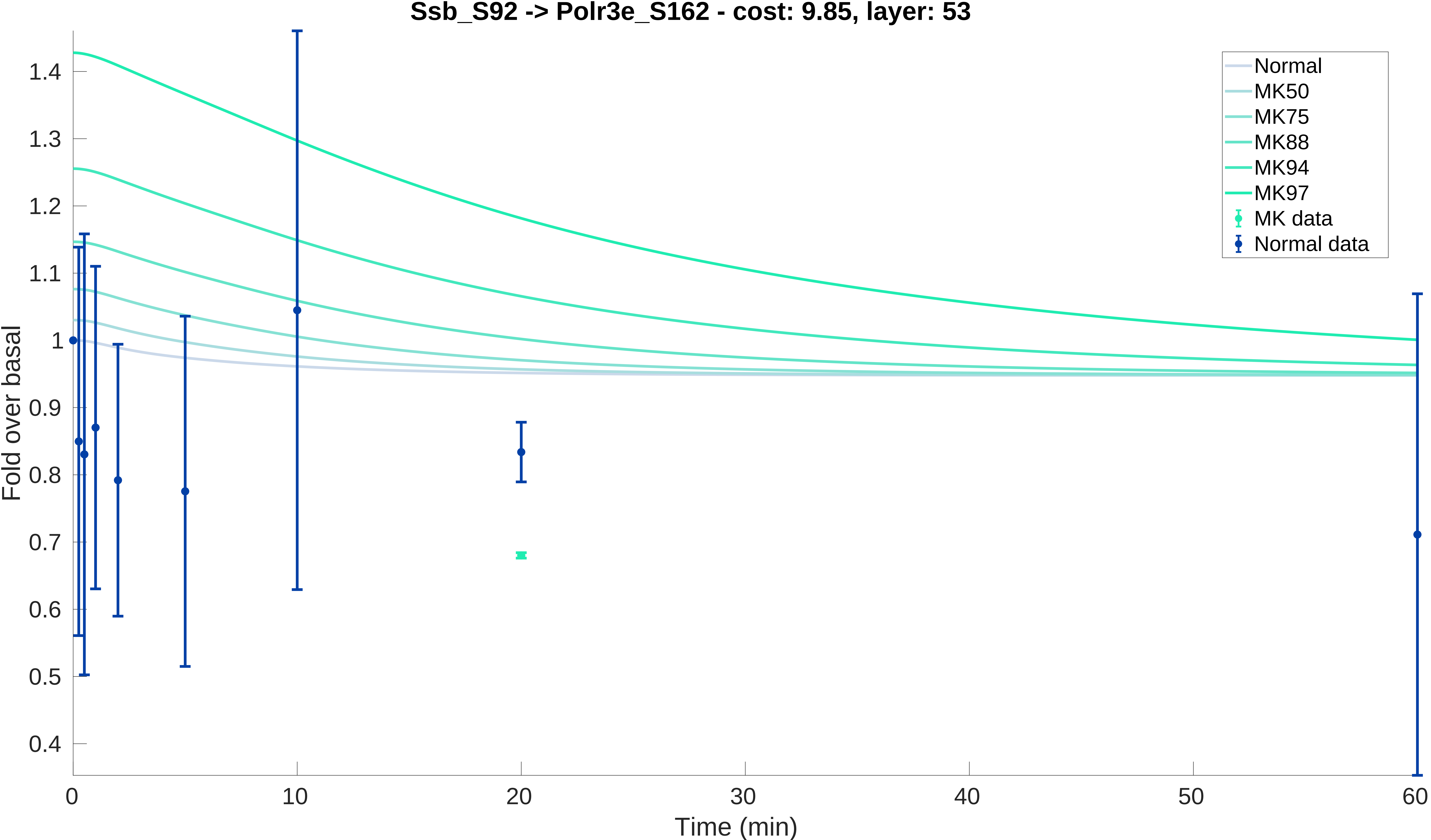

Ssb_S92 -> Polr3e_S162 - cost: 9.85, layer: 53
Fold over basal
1.4
1.3
1.2
1.1
1
0.9
0.8
0.7
0.6
0.5
0.4
0
10
20
30
40
50
60
Time (min)
Normal
MK50
MK75
MK88
MK94
MK97
MK data
Normal data

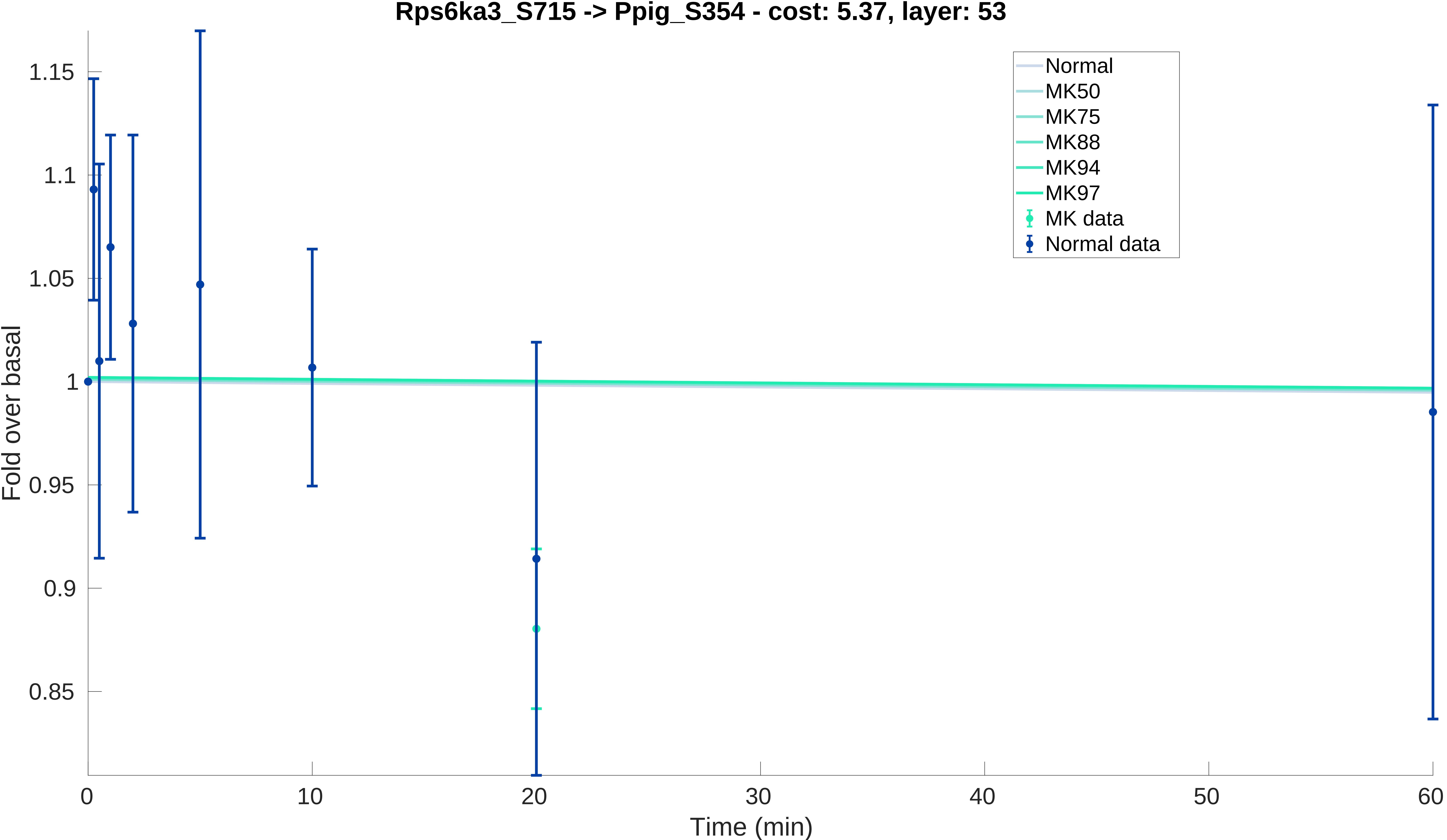

Rps6ka3_S715 -> Ppig_S354 - cost: 5.37, layer: 53
Fold over basal
1.15
1.1
1.05
1
0.95
0.9
0.85
0
10
20
30
40
50
60
Time (min)
Normal
MK50
MK75
MK88
MK94
MK97
MK data
Normal data

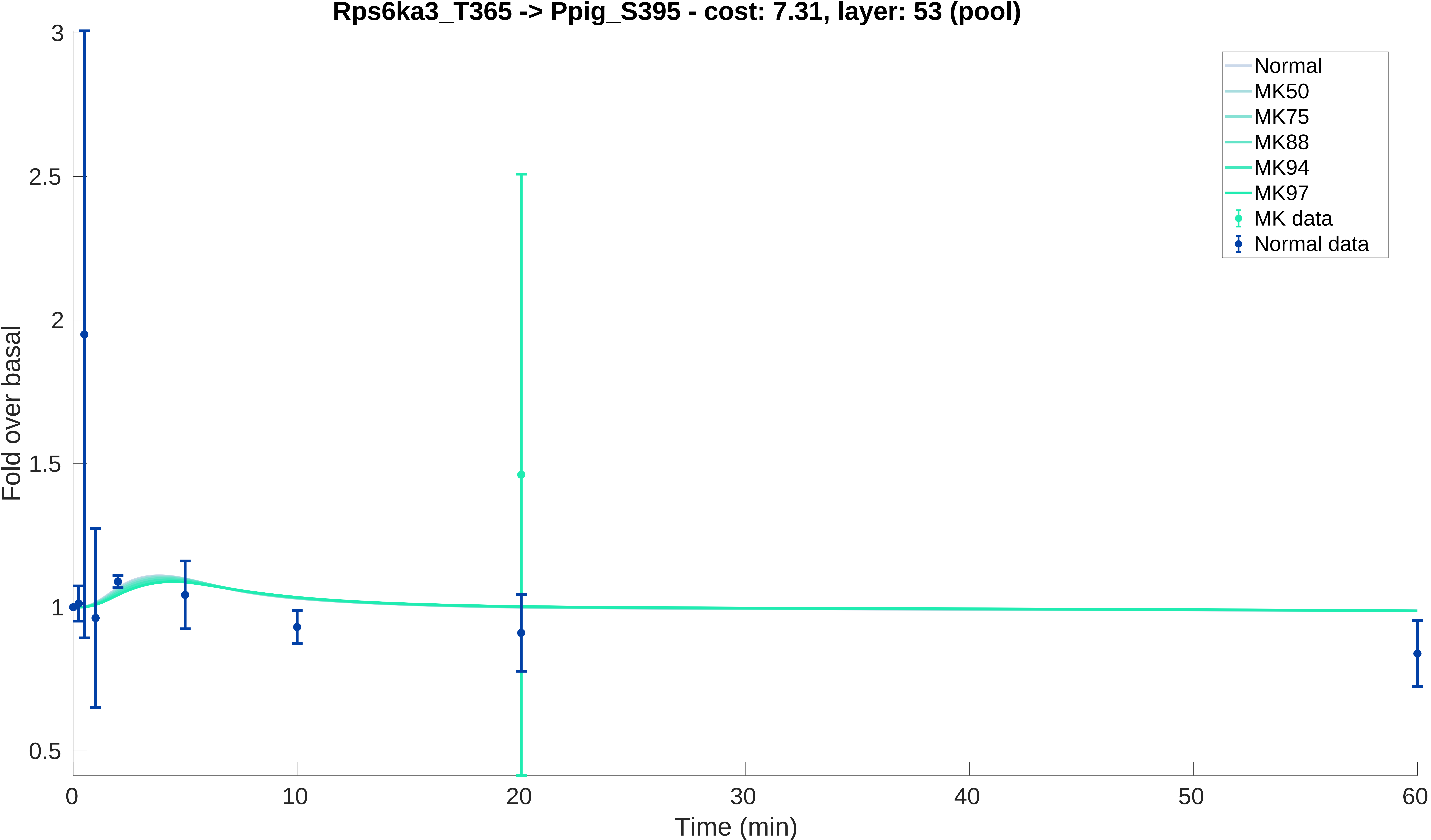

Rps6ka3_T365 -> Ppig_S395 - cost: 7.31, layer: 53 (pool)
Fold over basal
Time (min)
0.5
1
1.5
2
2.5
3
0
10
20
30
40
50
60
Normal
MK50
MK75
MK88
MK94
MK97
MK data
Normal data

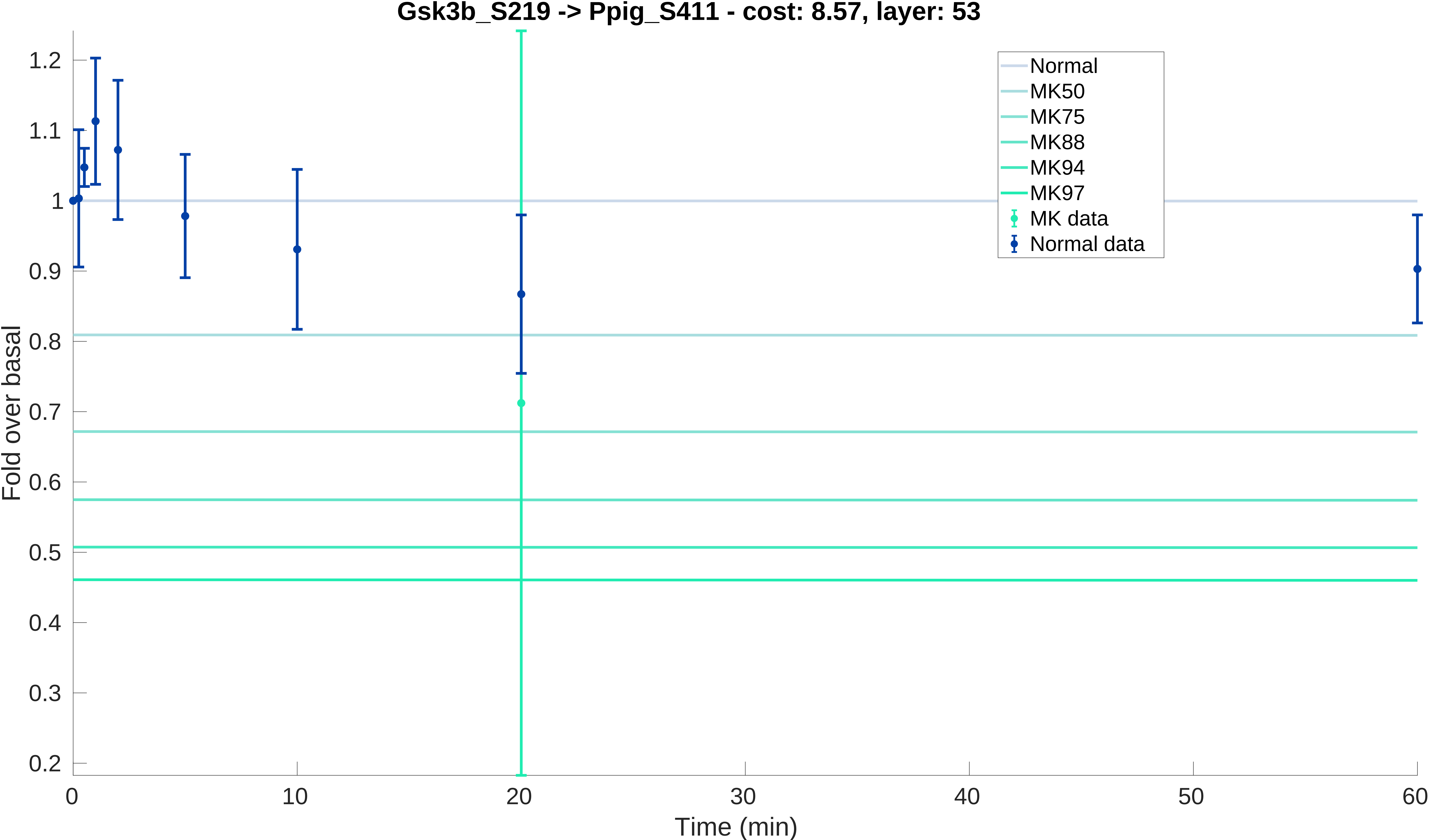

Gsk3b_S219 -> Ppig_S411 - cost: 8.57, layer: 53
Fold over basal
Time (min)
1.2
1.1
1
0.9
0.8
0.7
0.6
0.5
0.4
0.3
0.2
0
10
20
30
40
50
60
Normal
MK50
MK75
MK88
MK94
MK97
MK data
Normal data

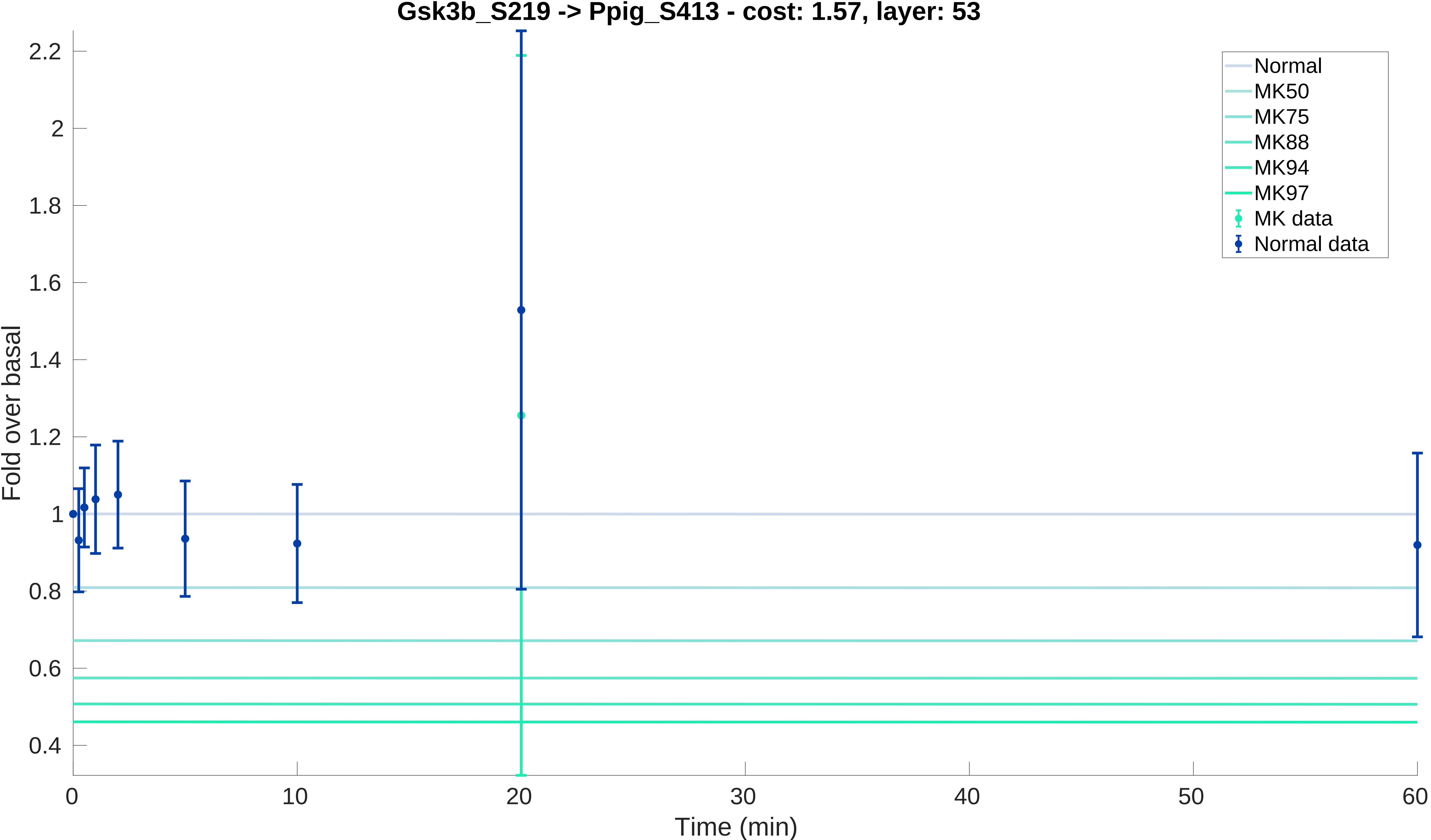

Gsk3b_S219 -> Ppig_S413 - cost: 1.57, layer: 53
Fold over basal
Time (min)
0.4
0.6
0.8
1
1.2
1.4
1.6
1.8
2
2.2
0
10
20
30
40
50
60
Normal
MK50
MK75
MK88
MK94
MK97
MK data
Normal data

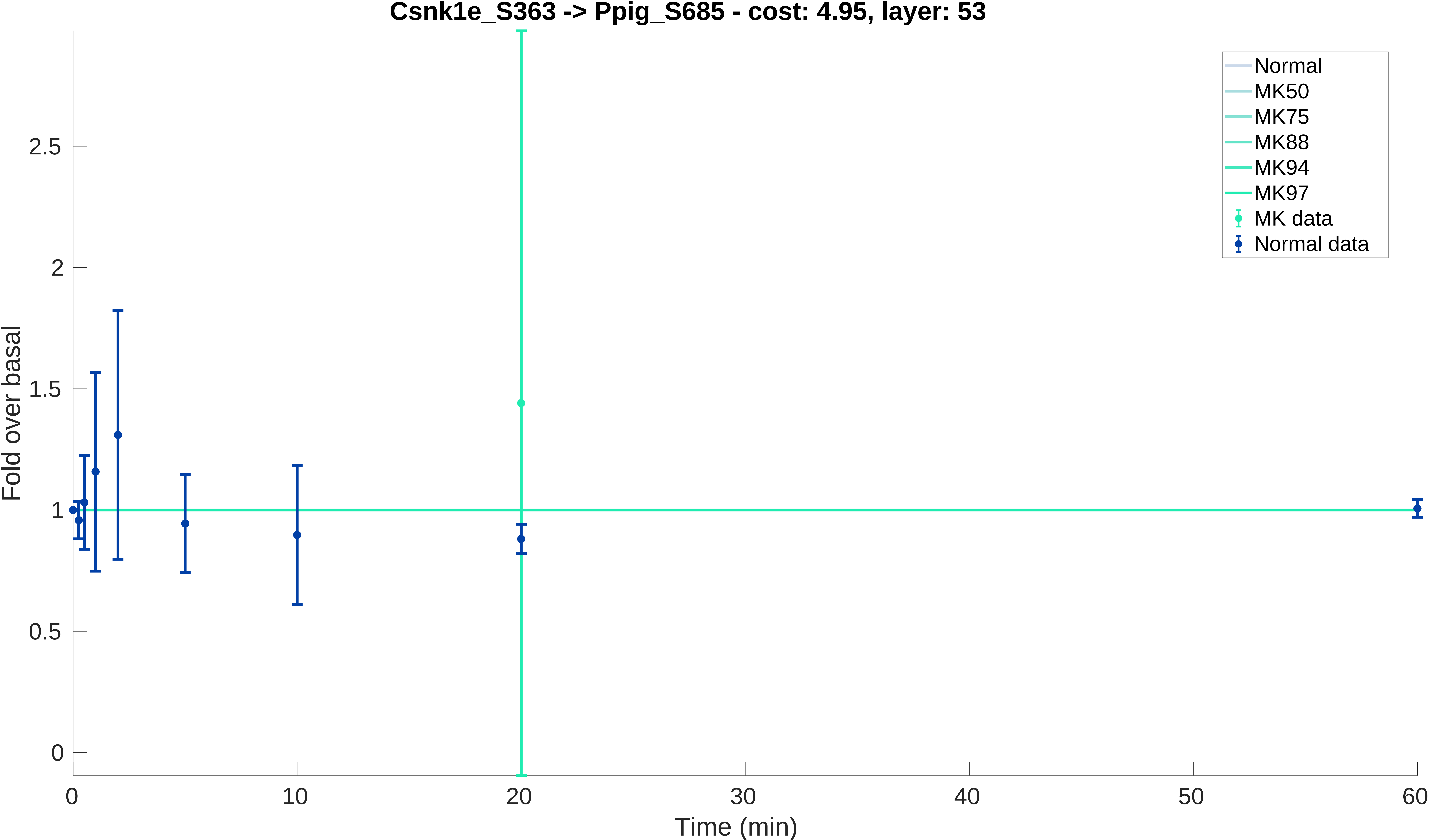

Csnk1e_S363 -> Ppig_S685 - cost: 4.95, layer: 53
Normal
MK50
MK75
MK88
MK94
MK97
MK data
Normal data
Fold over basal
0
0.5
1
1.5
2
2.5
0
10
20
30
40
50
60
Time (min)

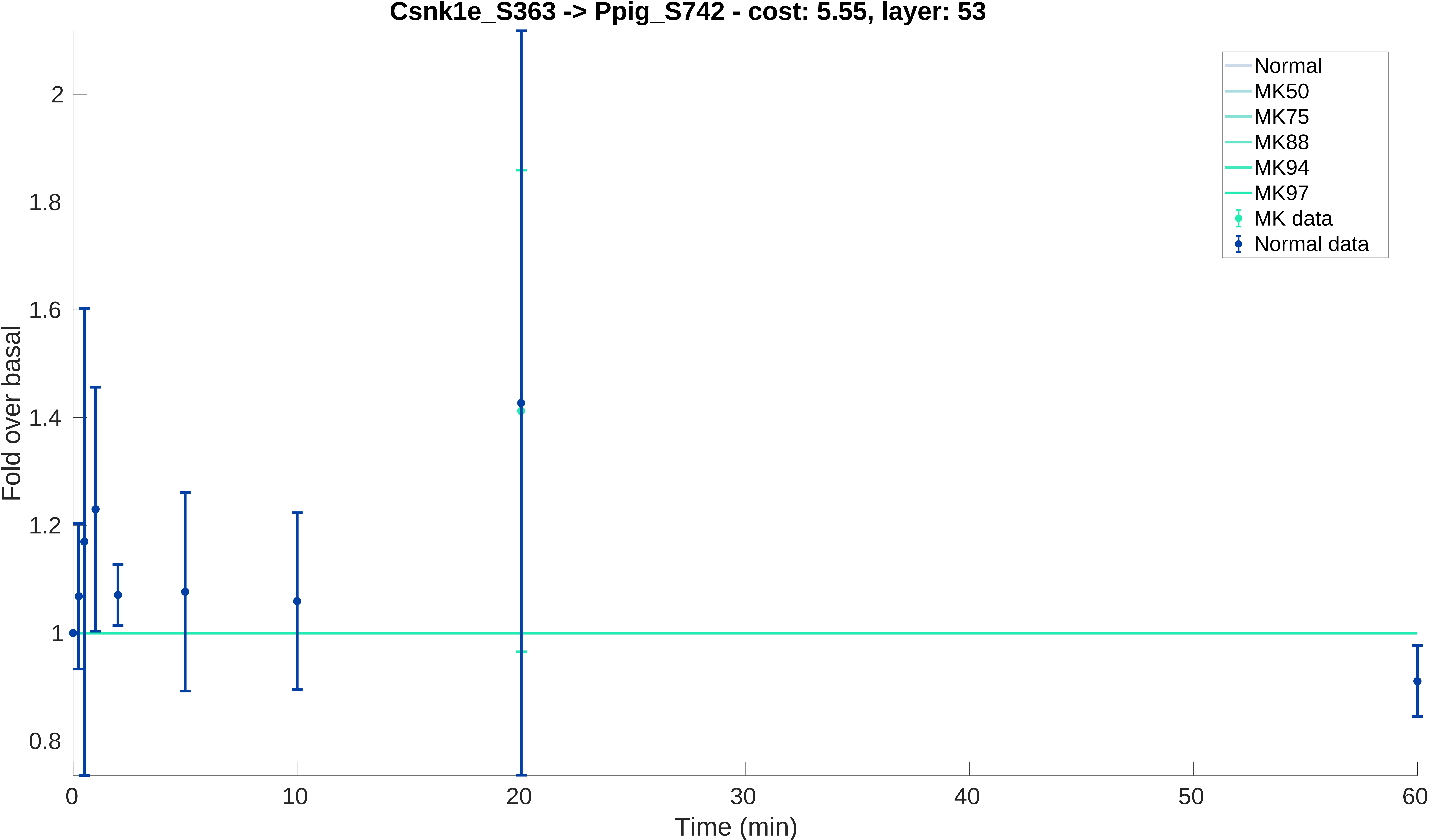

Csnk1e_S363 -> Ppig_S742 - cost: 5.55, layer: 53
Normal
MK50
MK75
MK88
MK94
MK97
MK data
Normal data
Fold over basal
Time (min)
0.8
1
1.2
1.4
1.6
1.8
2
0
10
20
30
40
50
60

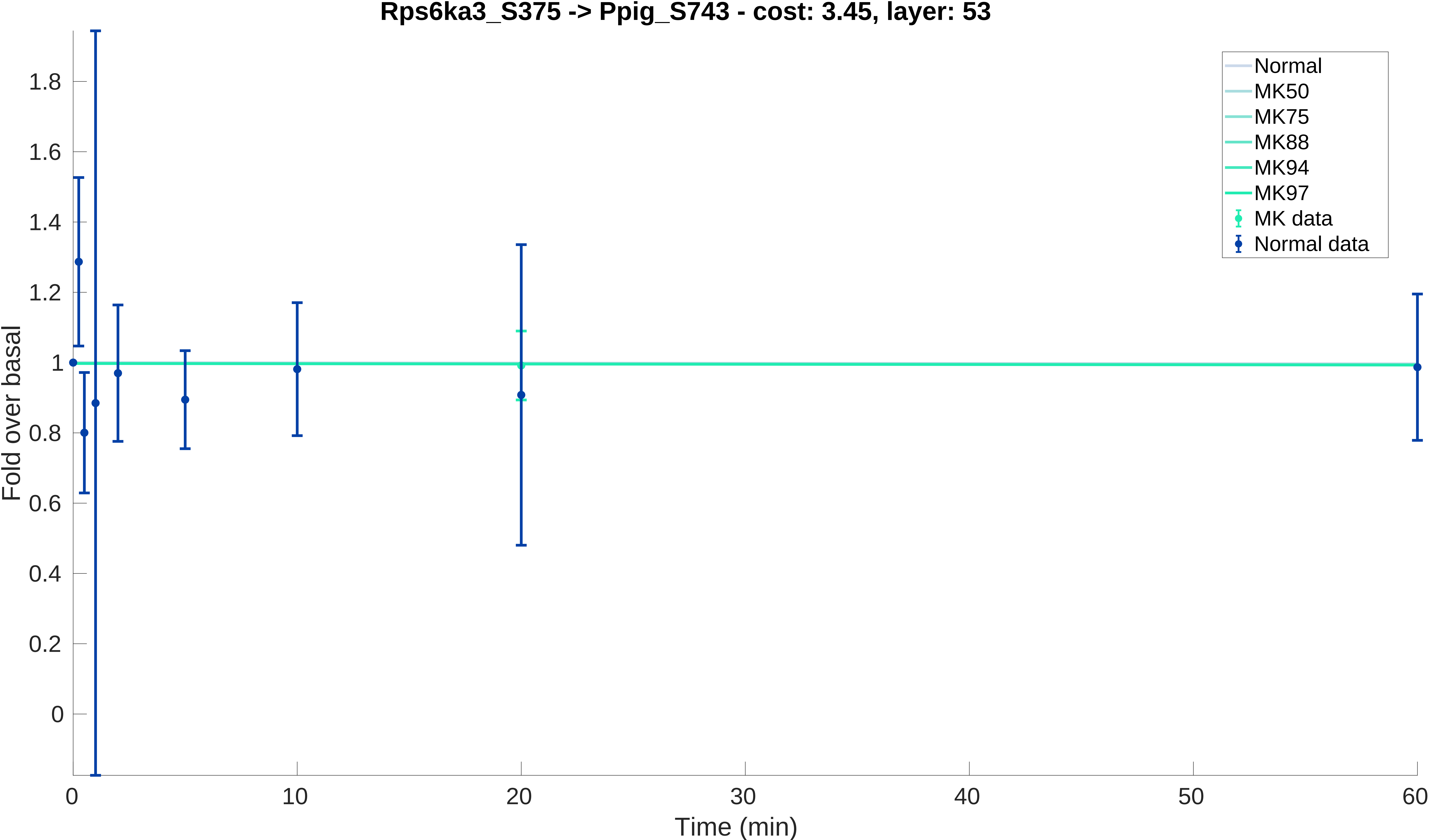

Rps6ka3_S375 -> Ppig_S743 - cost: 3.45, layer: 53
Fold over basal
Time (min)
0
0.2
0.4
0.6
0.8
1
1.2
1.4
1.6
1.8
0
10
20
30
40
50
60
Normal
MK50
MK75
MK88
MK94
MK97
MK data
Normal data

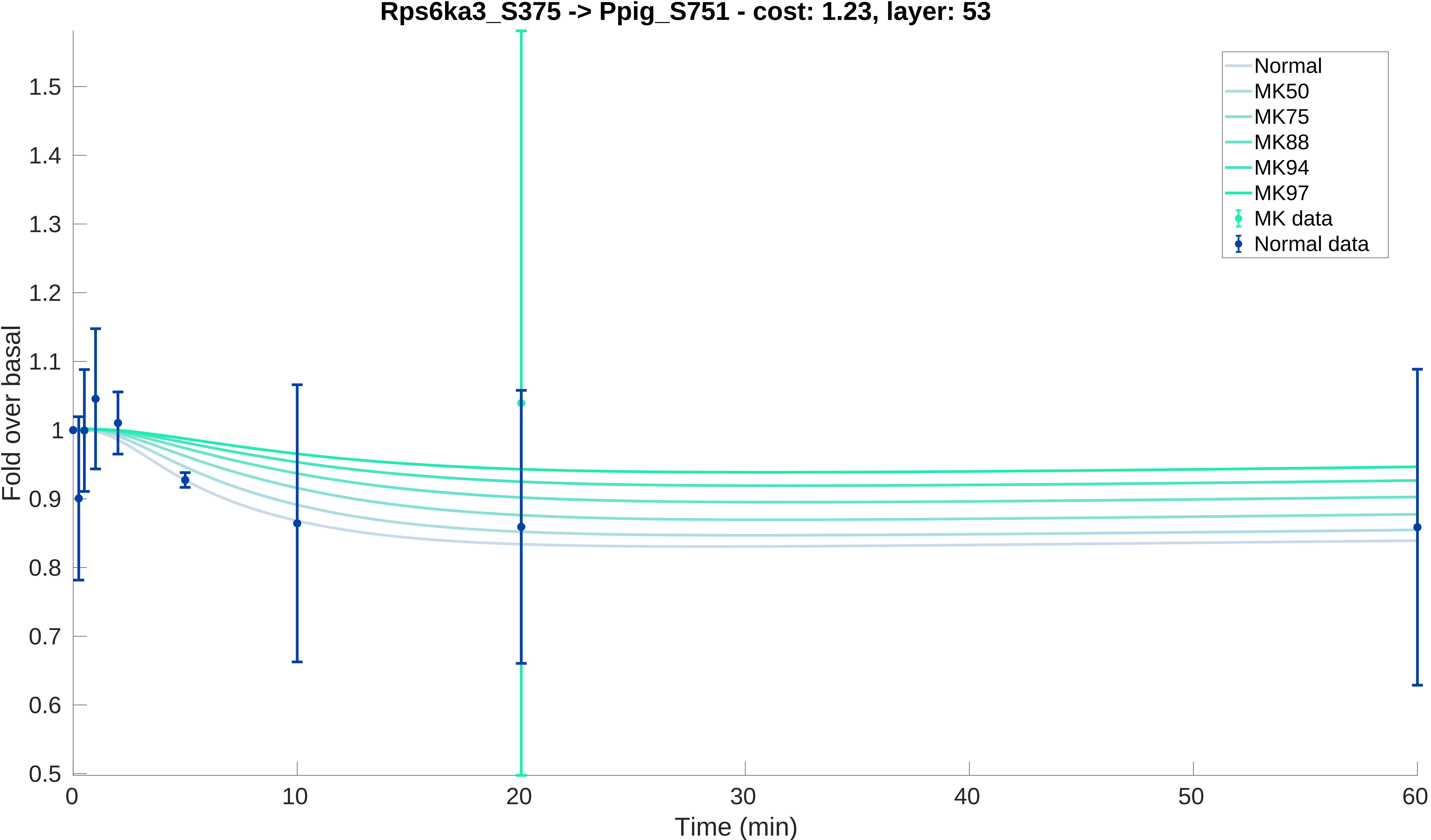

Rps6ka3_S375 -> Ppig_S751 - cost: 1.23, layer: 53
1.5
1.4
1.3
1.2
1.1
1
0.9
0.8
0.7
0.6
0.5
Fold over basal
0
10
20
30
40
50
60
Time (min)
Normal
MK50
MK75
MK88
MK94
MK97
MK data
Normal data

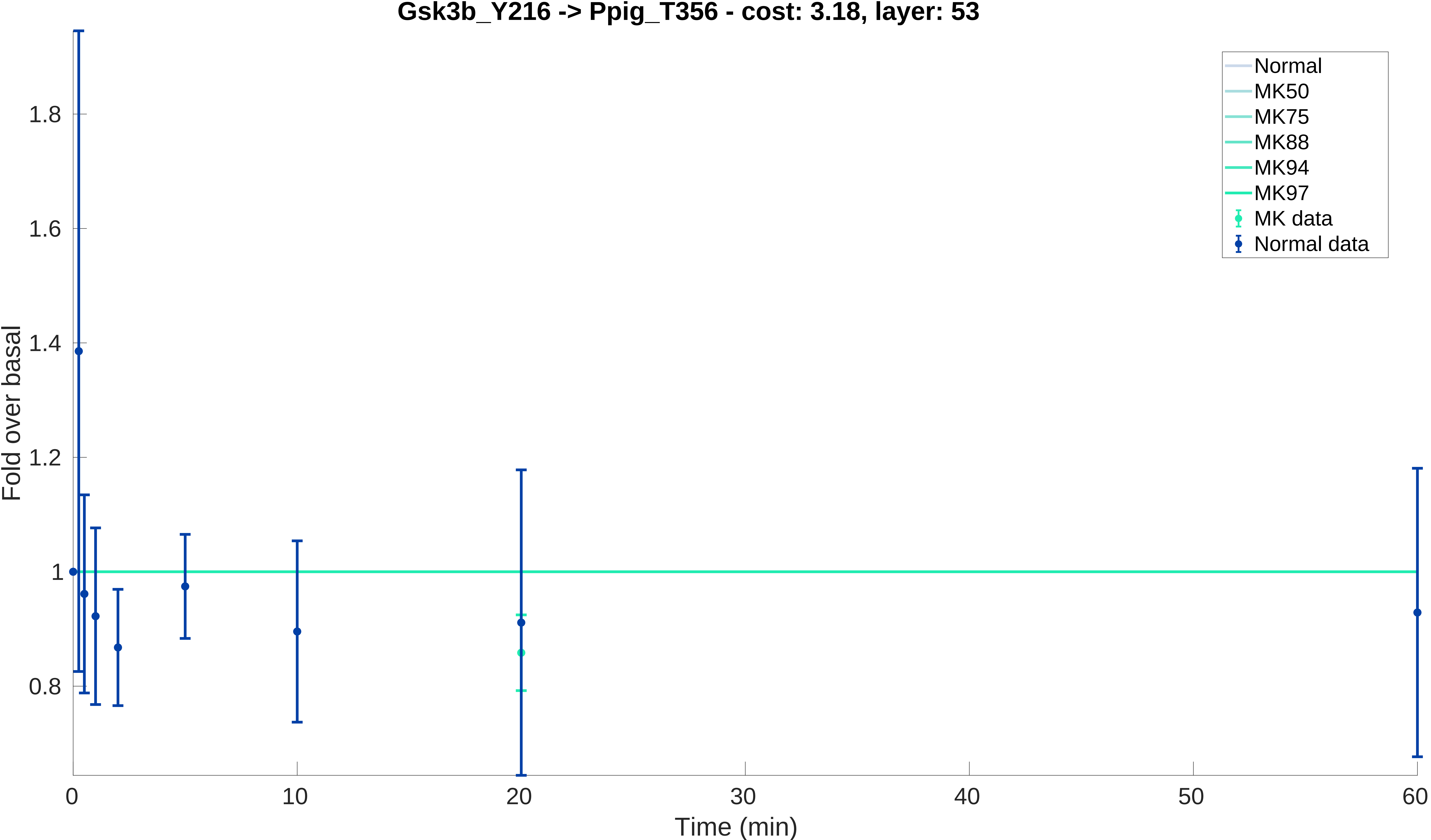

Gsk3b_Y216 -> Ppig_T356 - cost: 3.18, layer: 53
Normal
MK50
MK75
MK88
MK94
MK97
MK data
Normal data
Fold over basal
1.8
1.6
1.4
1.2
1
0.8
0
10
20
30
40
50
60
Time (min)

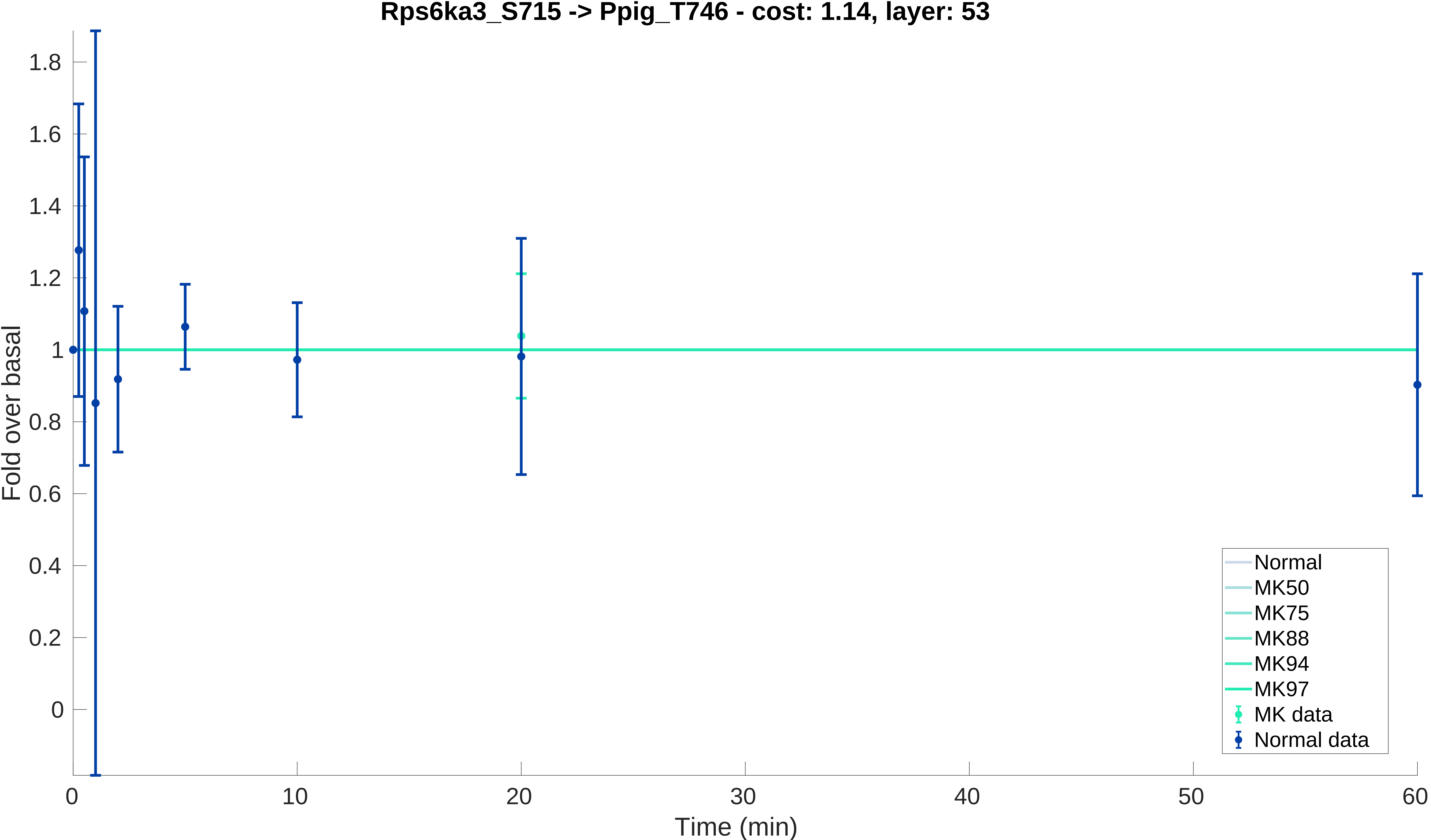

Rps6ka3_S715 -> Ppig_T746 - cost: 1.14, layer: 53
Fold over basal
Time (min)
1.8
1.6
1.4
1.2
1
0.8
0.6
0.4
0.2
0
0
10
20
30
40
50
60
Normal
MK50
MK75
MK88
MK94
MK97
MK data
Normal data

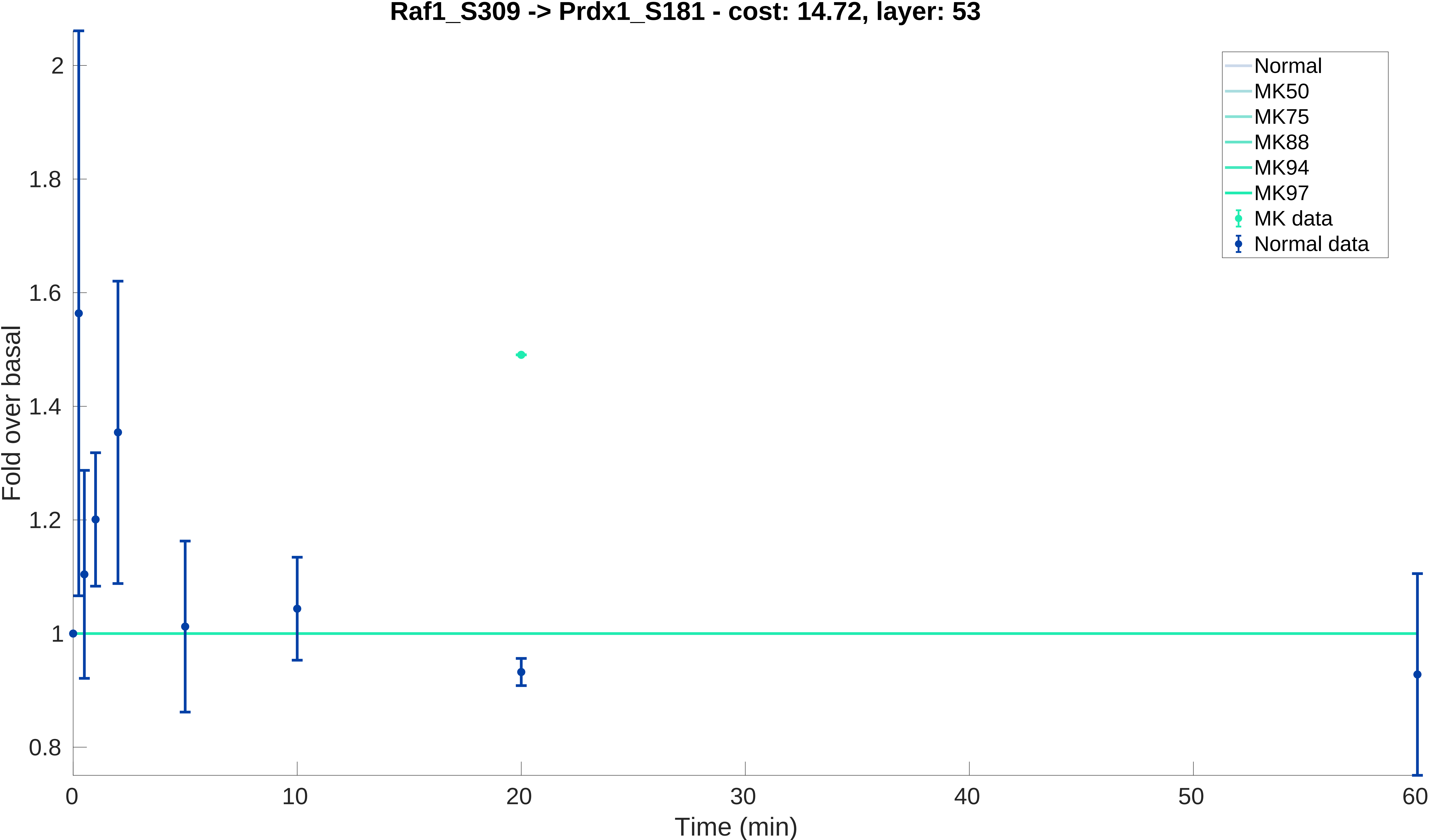

Raf1_S309 -> Prdx1_S181 - cost: 14.72, layer: 53
Normal
MK50
MK75
MK88
MK94
MK97
MK data
Normal data
Fold over basal
2
1.8
1.6
1.4
1.2
1
0.8
0
10
20
30
40
50
60
Time (min)

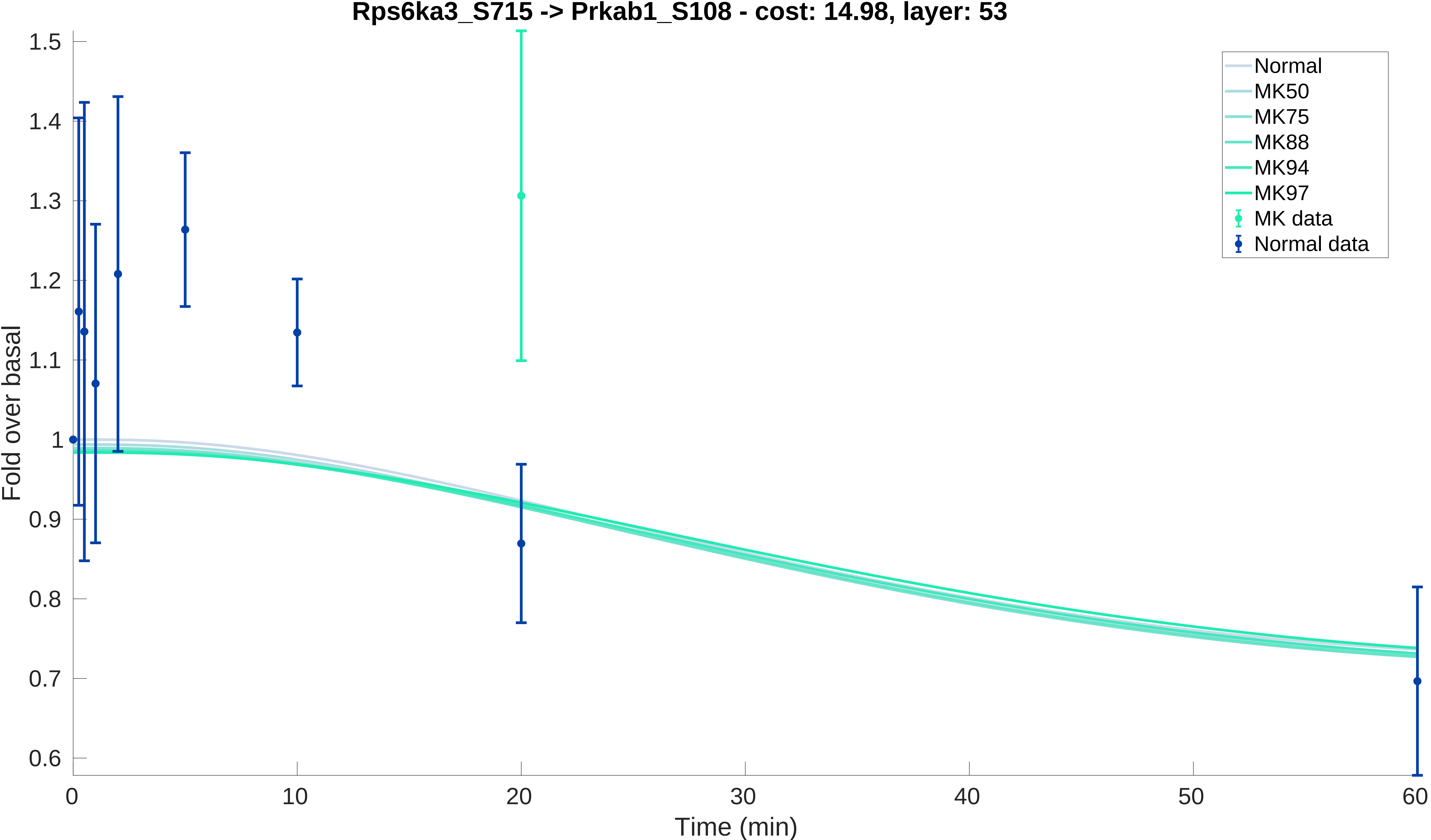

Rps6ka3_S715 -> Prkab1_S108 - cost: 14.98, layer: 53
Normal
MK50
MK75
MK88
MK94
MK97
MK data
Normal data
Fold over basal
1.5
1.4
1.3
1.2
1.1
1
0.9
0.8
0.7
0.6
0
10
20
30
40
50
60
Time (min)

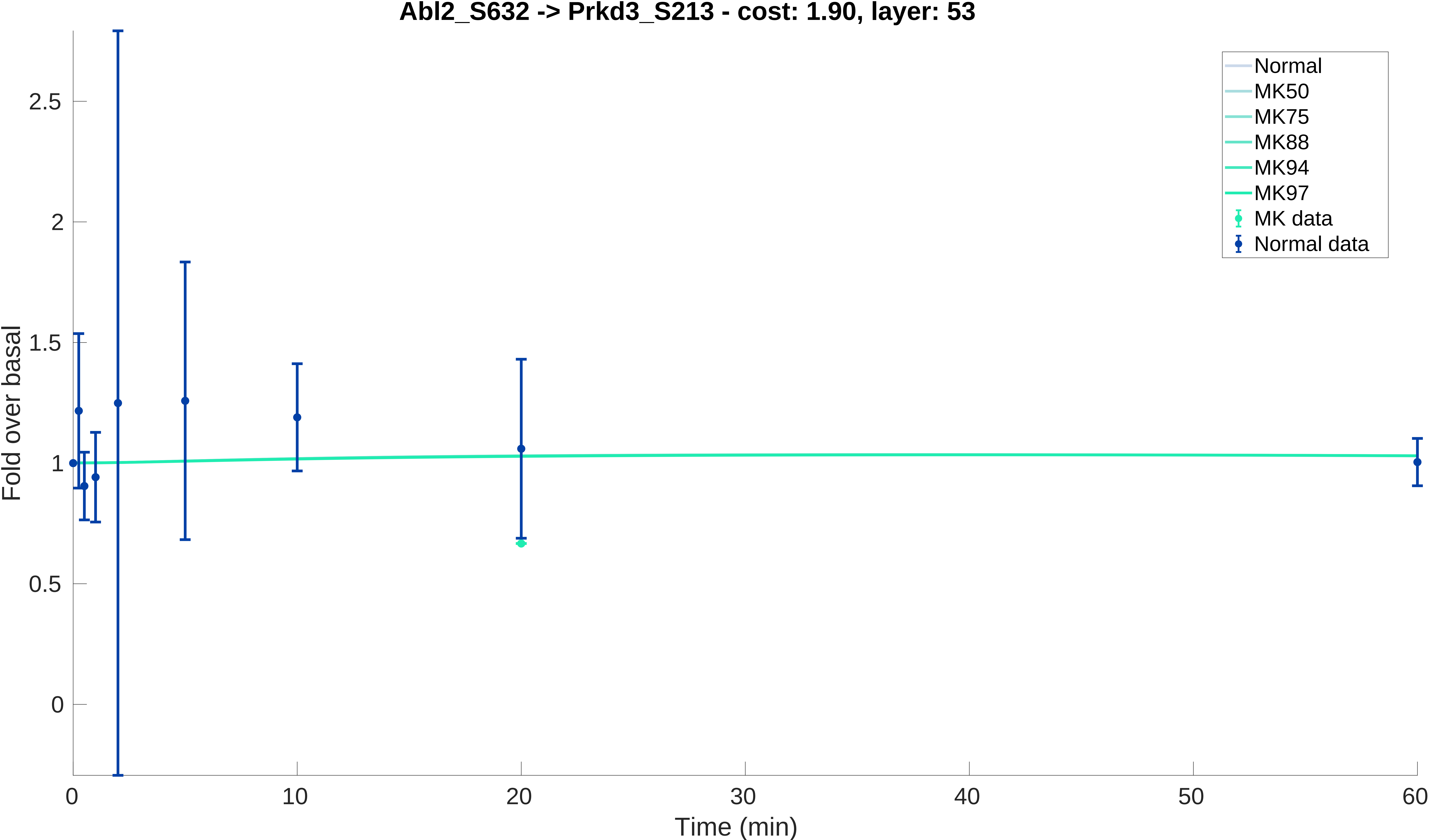

Abl2_S632 -> Prkd3_S213 - cost: 1.90, layer: 53
Fold over basal
Time (min)
Normal
MK50
MK75
MK88
MK94
MK97
MK data
Normal data
0
0.5
1
1.5
2
2.5
0
10
20
30
40
50
60

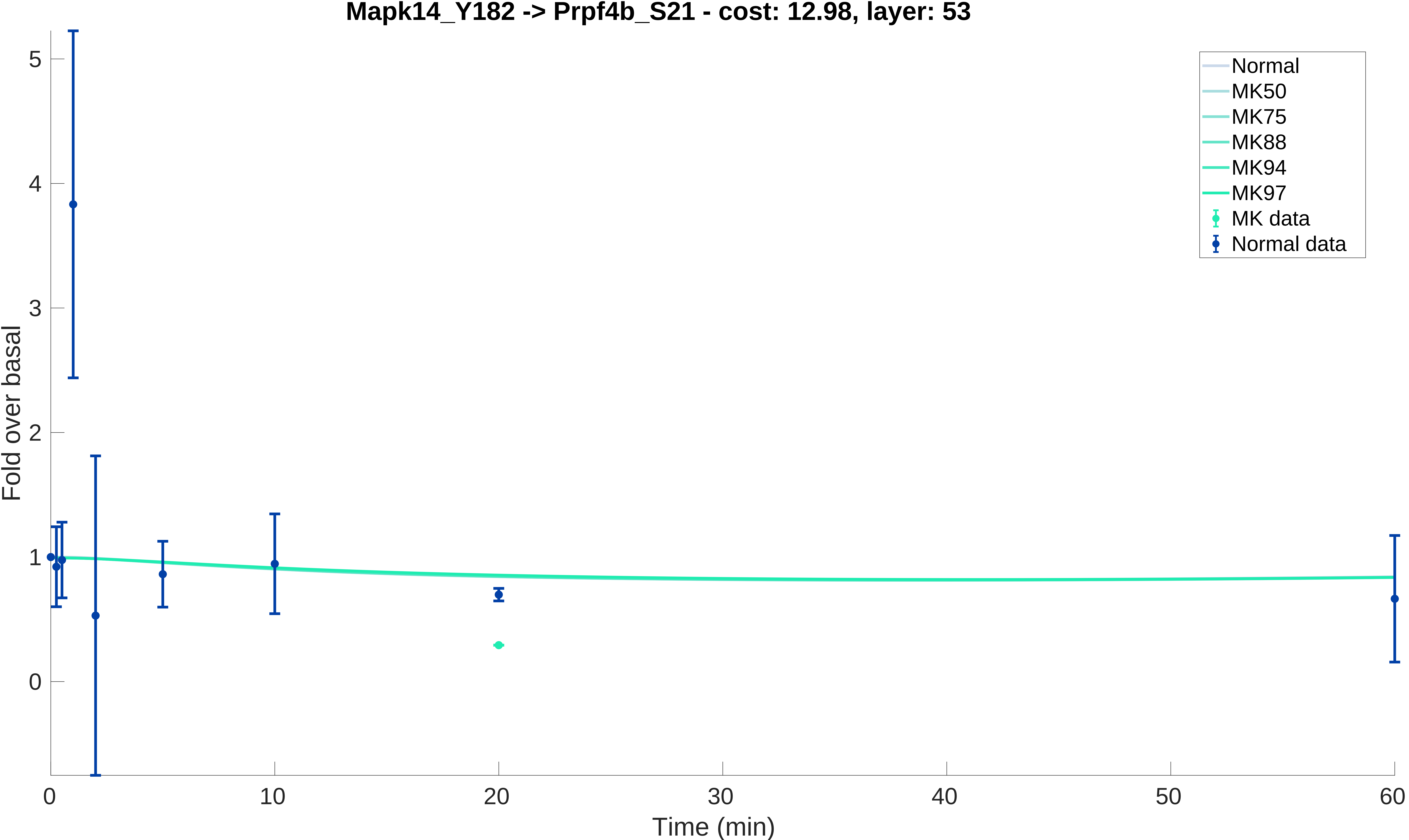
Mapk14_Y182 -> Prpf4b_S21 - cost: 12.98, layer: 53
Fold over basal
Time (min)
0
1
2
3
4
5
0
10
20
30
40
50
60
Normal
MK50
MK75
MK88
MK94
MK97
MK data
Normal data

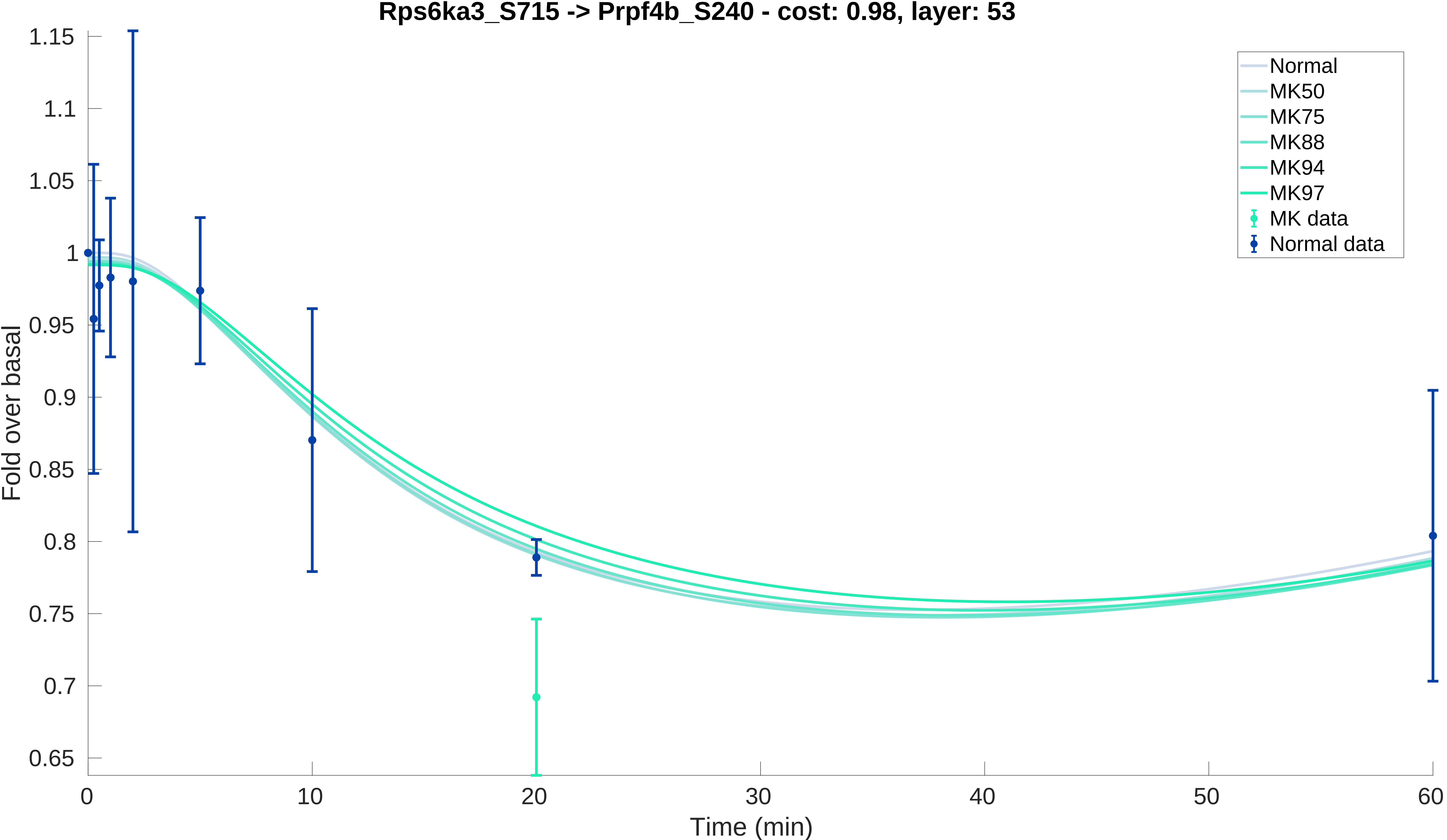
Rps6ka3_S715 -> Prpf4b_S240 - cost: 0.98, layer: 53
Fold over basal
Time (min)
1.15
1.1
1.05
1
0.95
0.9
0.85
0.8
0.75
0.7
0.65
0
10
20
30
40
50
60
Normal
MK50
MK75
MK88
MK94
MK97
MK data
Normal data

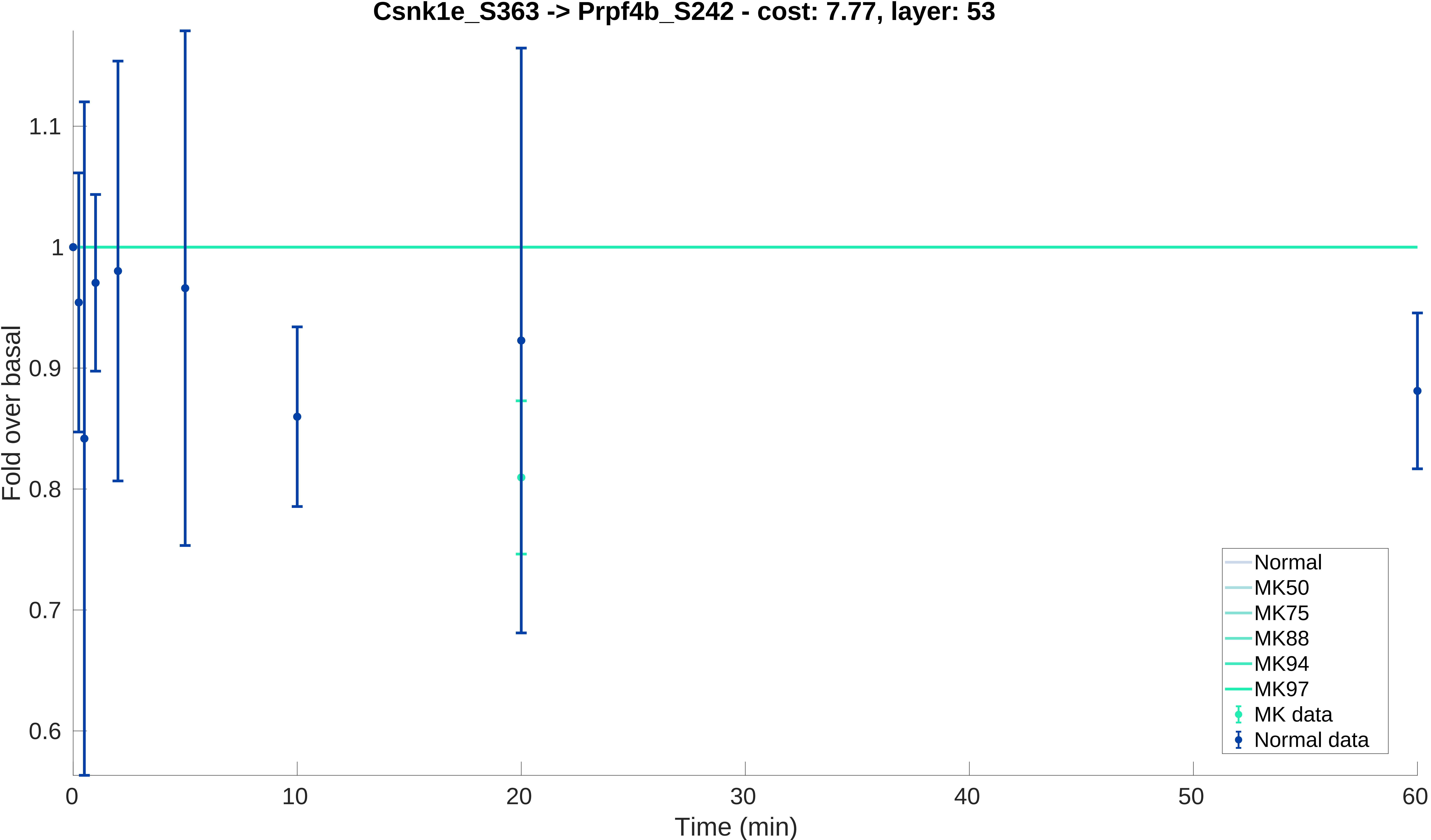
Csnk1e_S363 -> Prpf4b_S242 - cost: 7.77, layer: 53
Fold over basal
1.1
1
0.9
0.8
0.7
0.6
0
10
20
30
40
50
60
Time (min)
Normal
MK50
MK75
MK88
MK94
MK97
MK data
Normal data

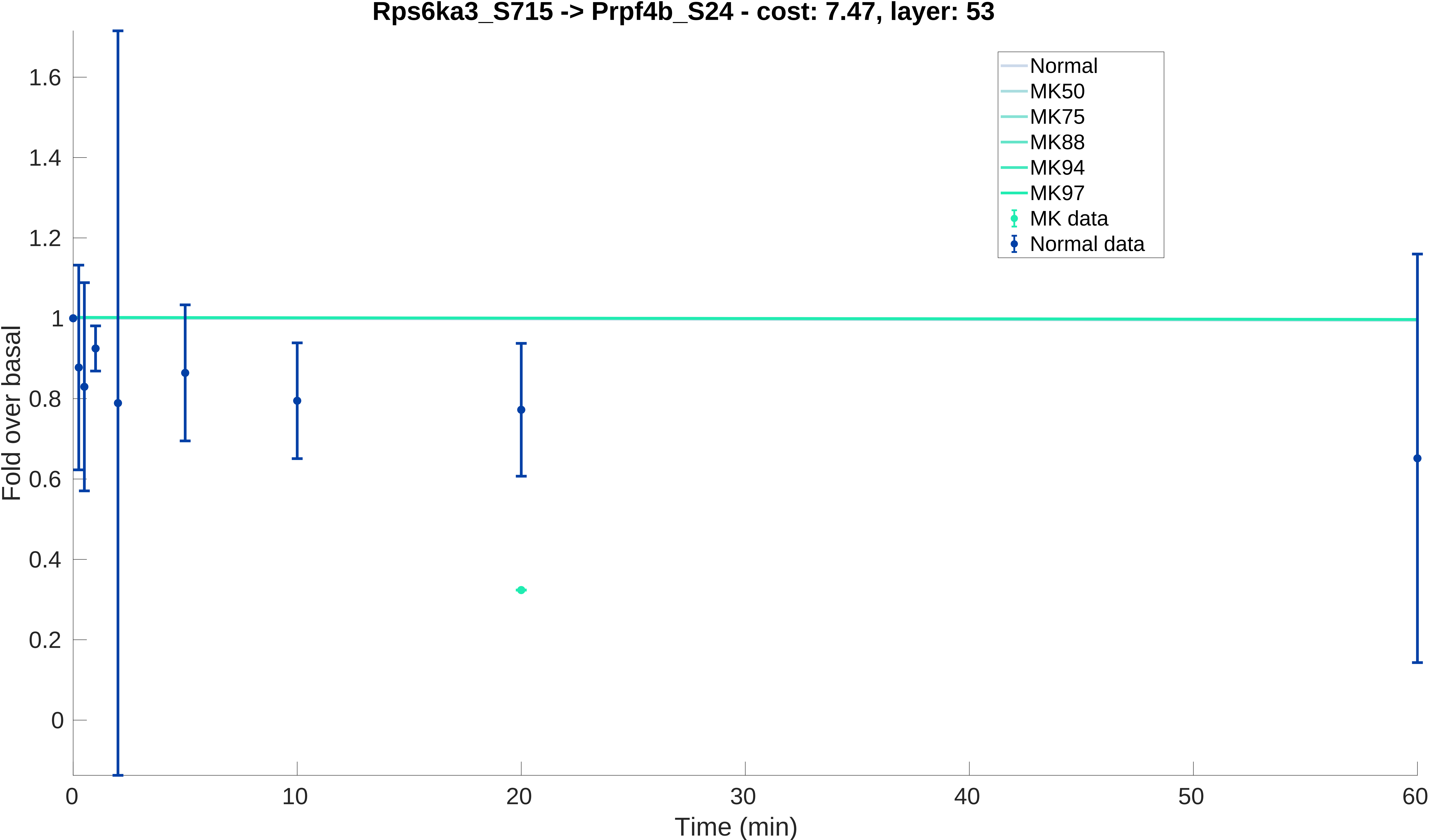
Rps6ka3_S715 -> Prpf4b_S24 - cost: 7.47, layer: 53
Fold over basal
Time (min)
0
0.2
0.4
0.6
0.8
1
1.2
1.4
1.6
0
10
20
30
40
50
60
Normal
MK50
MK75
MK88
MK94
MK97
MK data
Normal data

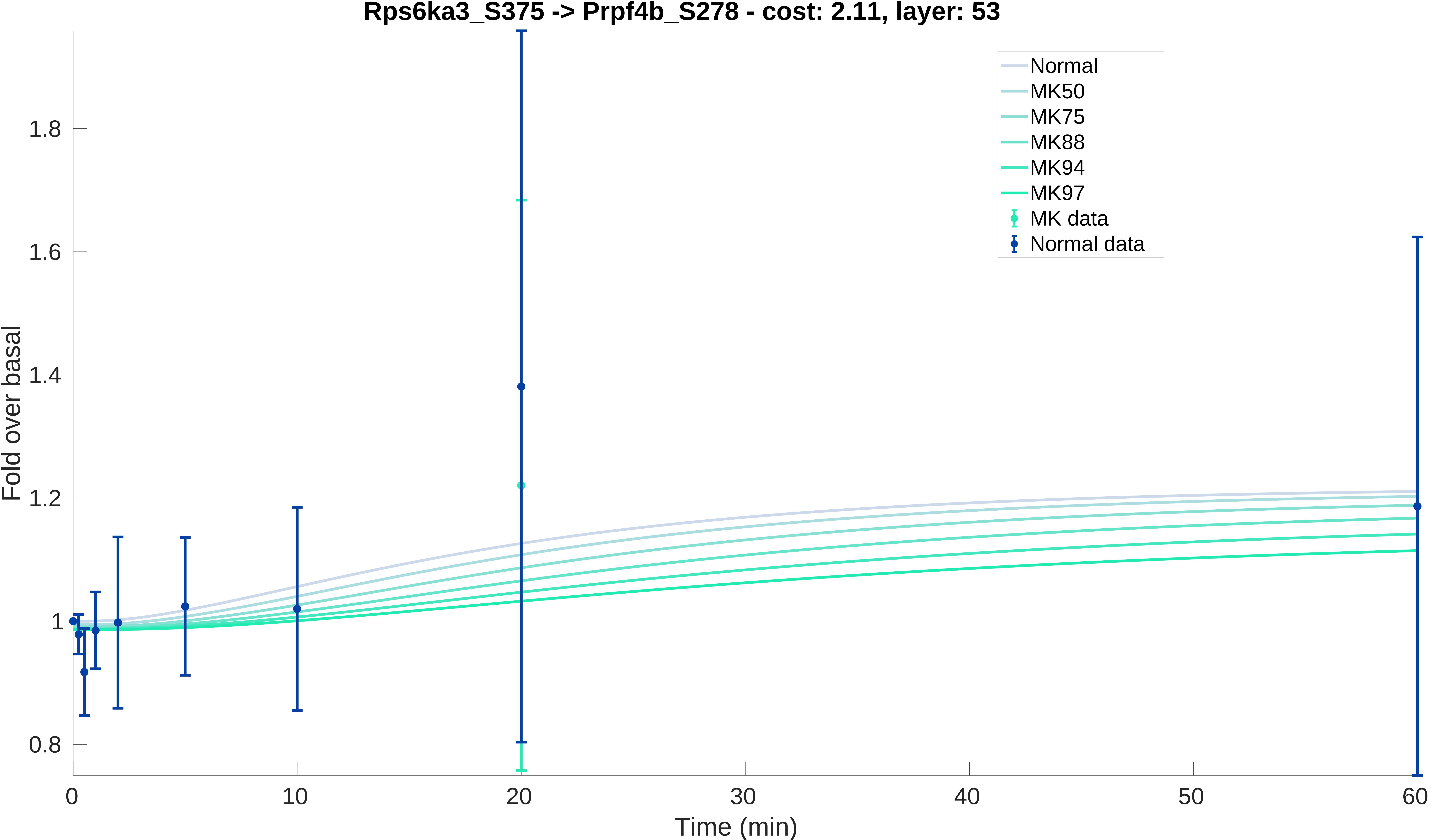

Rps6ka3_S375 -> Prpf4b_S278 - cost: 2.11, layer: 53
Normal
MK50
MK75
MK88
MK94
MK97
MK data
Normal data
Fold over basal
Time (min)
0.8
1
1.2
1.4
1.6
1.8
0
10
20
30
40
50
60

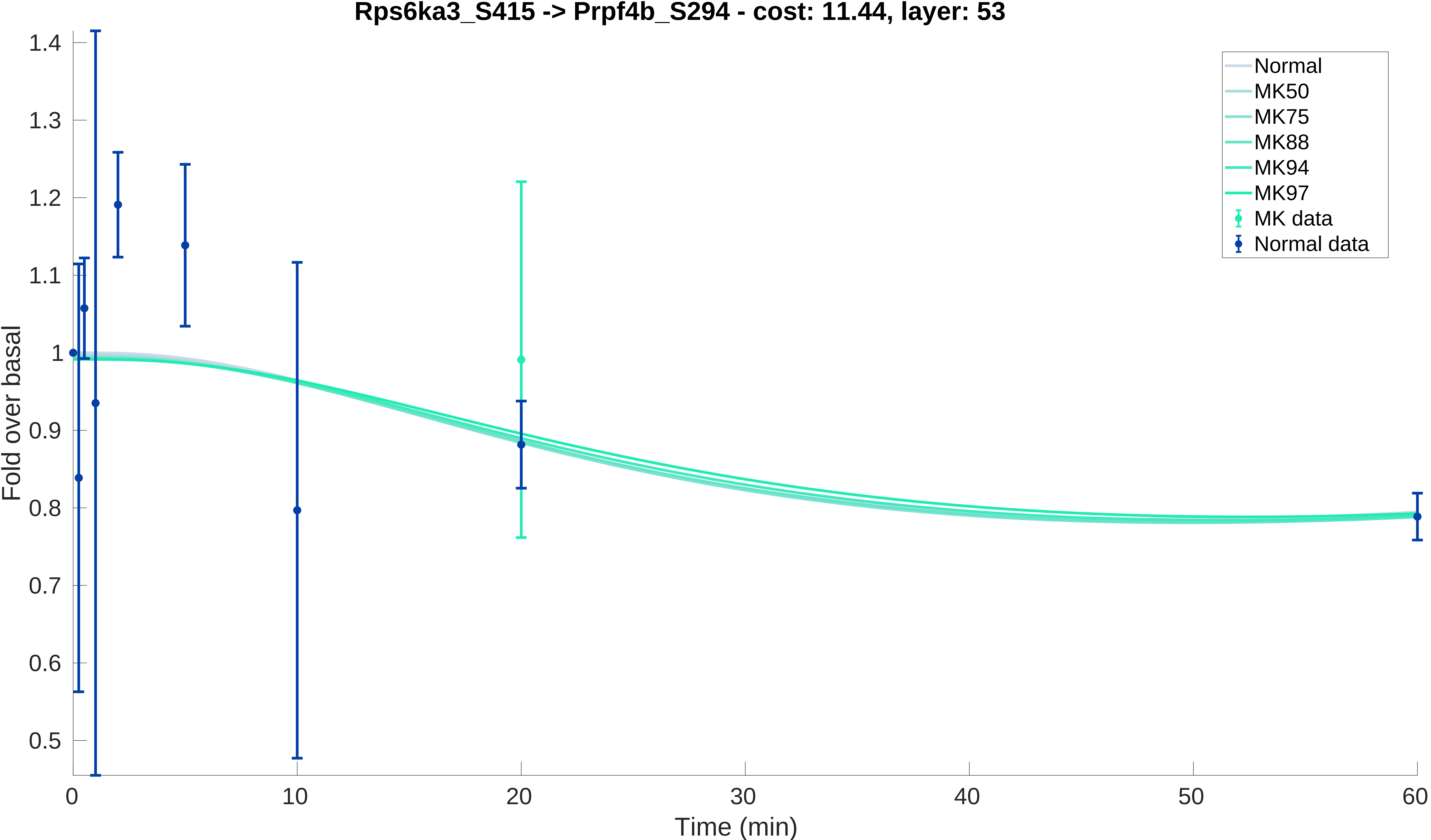
Rps6ka3_S415 -> Prpf4b_S294 - cost: 11.44, layer: 53
Fold over basal
Time (min)
Normal
MK50
MK75
MK88
MK94
MK97
MK data
Normal data
0
10
20
30
40
50
60
0.5
0.6
0.7
0.8
0.9
1
1.1
1.2
1.3
1.4

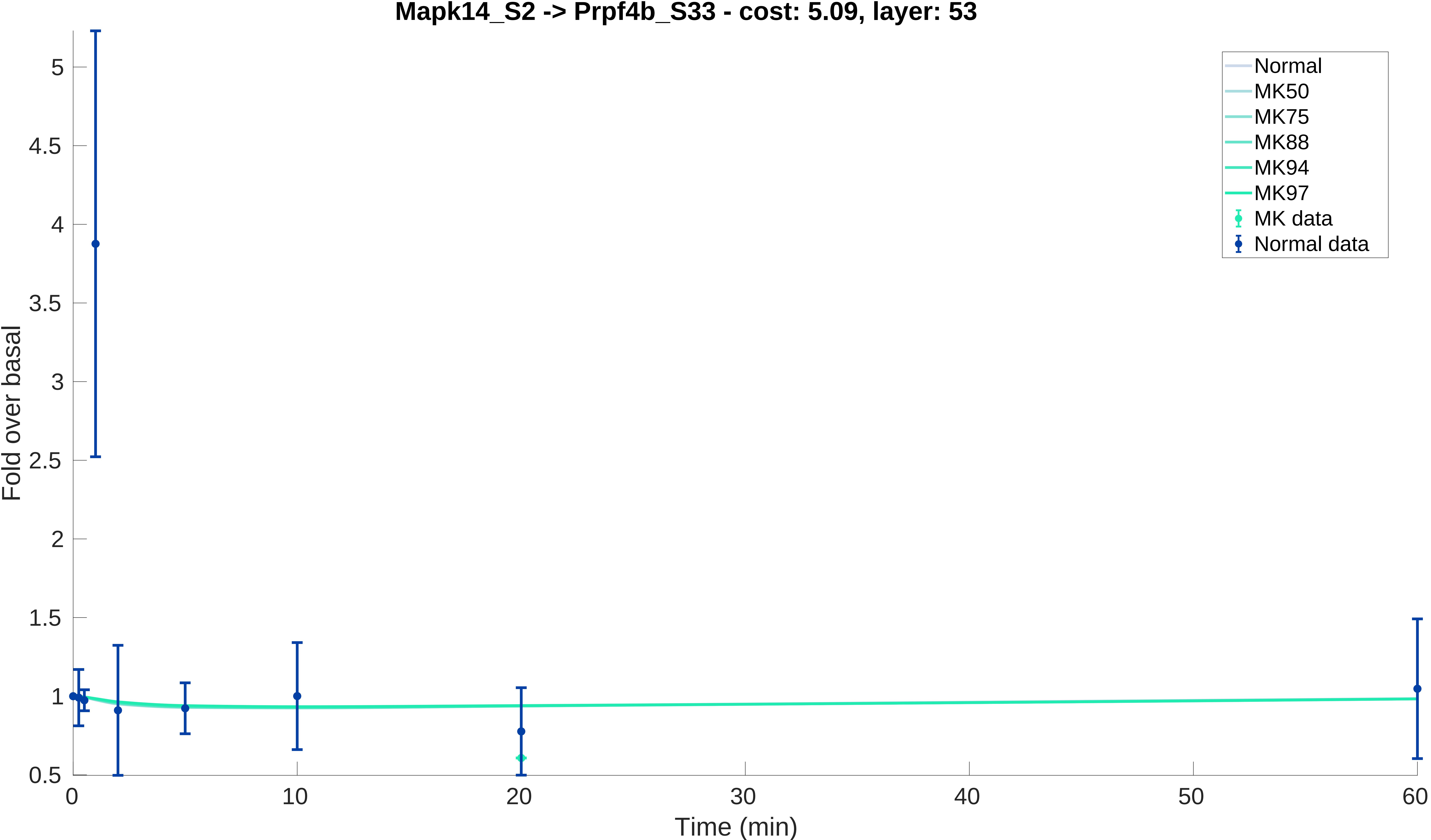
Mapk14_S2 -> Prpf4b_S33 - cost: 5.09, layer: 53
Fold over basal
Time (min)
0.5
1
1.5
2
2.5
3
3.5
4
4.5
5
0
10
20
30
40
50
60
Normal
MK50
MK75
MK88
MK94
MK97
MK data
Normal data

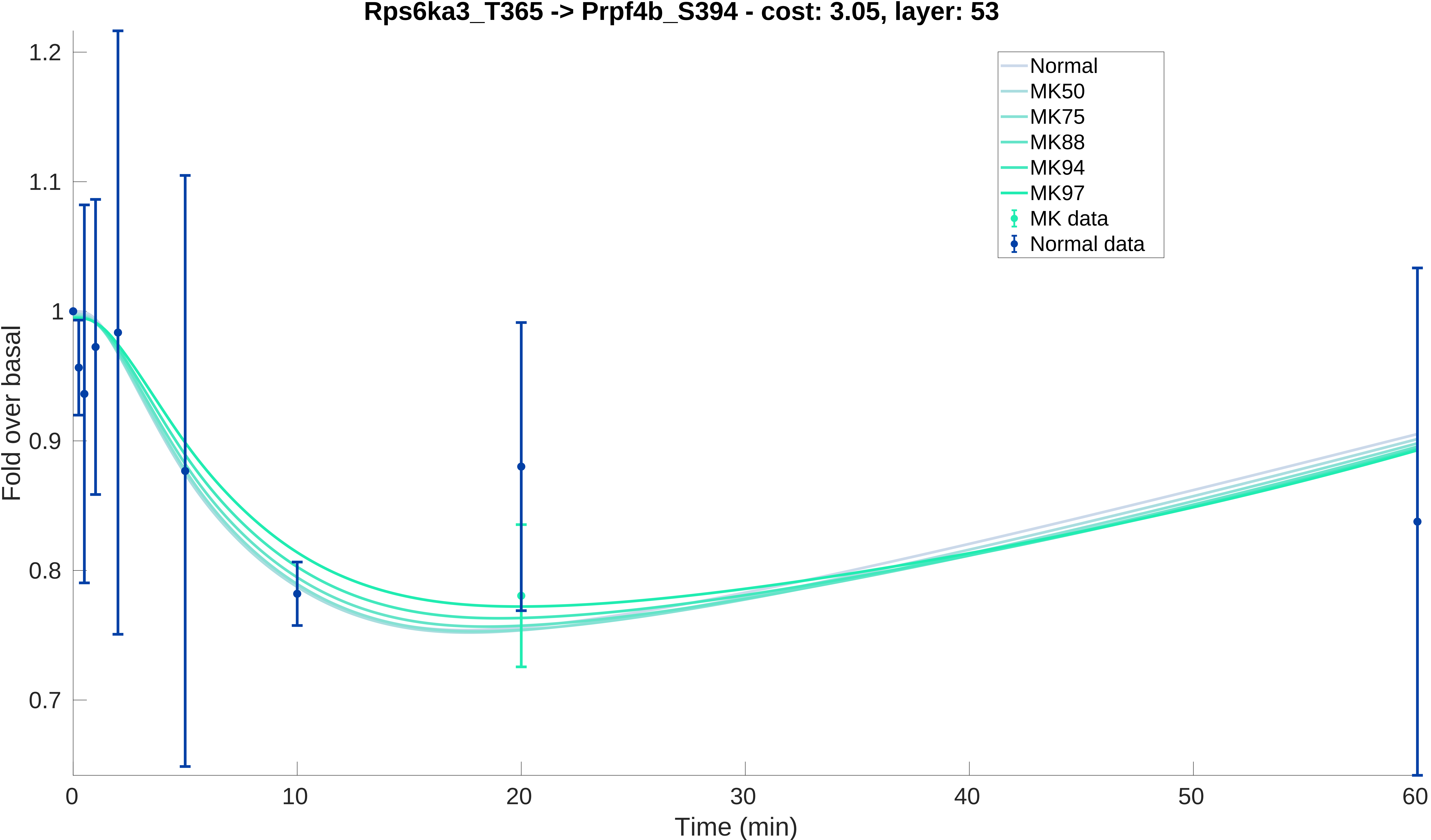
Rps6ka3_T365 -> Prpf4b_S394 - cost: 3.05, layer: 53
Fold over basal
Time (min)
0.7
0.8
0.9
1
1.1
1.2
0
10
20
30
40
50
60
Normal
MK50
MK75
MK88
MK94
MK97
MK data
Normal data

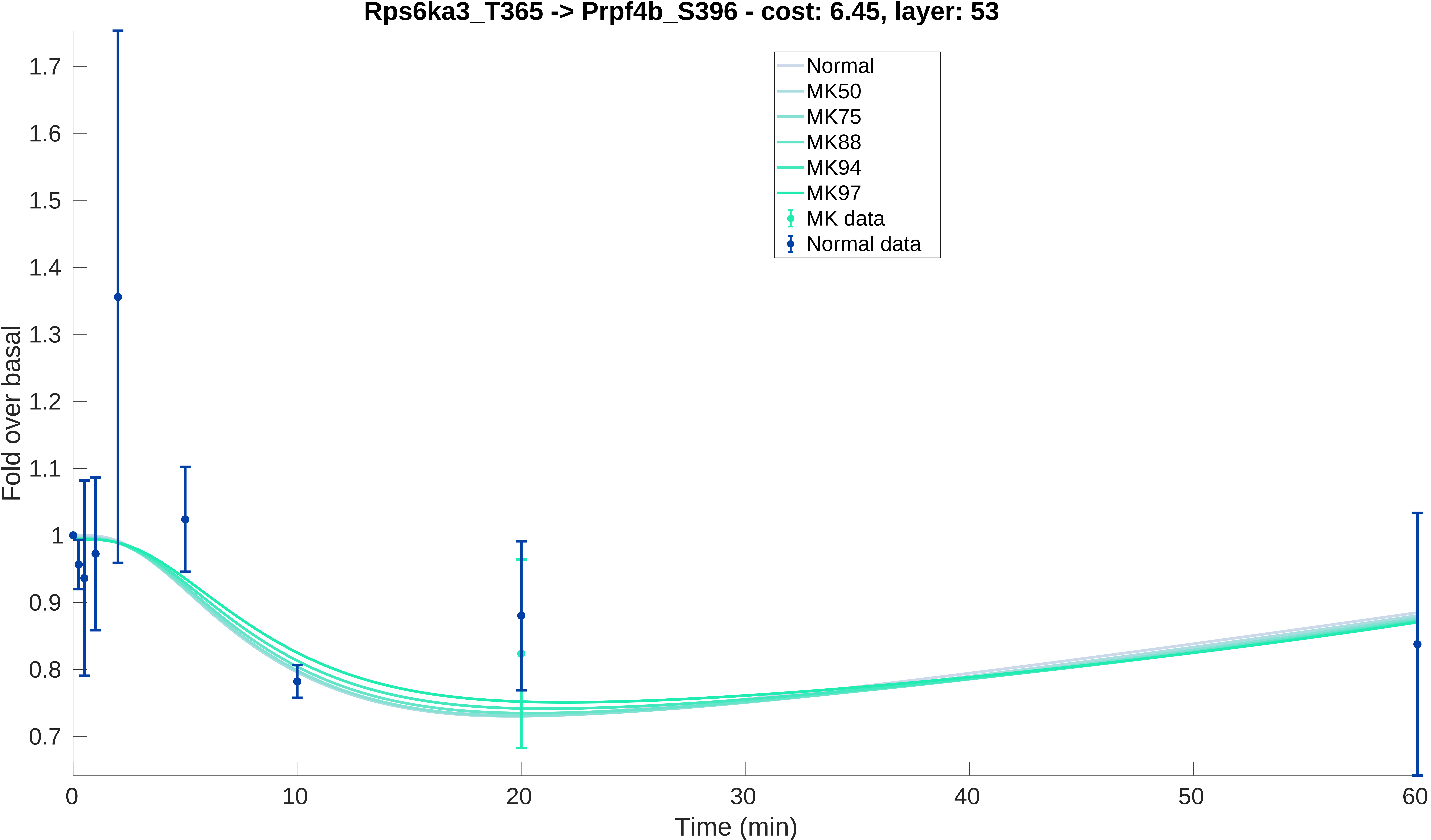
Rps6ka3_T365 -> Prpf4b_S396 - cost: 6.45, layer: 53
Fold over basal
Time (min)
Normal
MK50
MK75
MK88
MK94
MK97
MK data
Normal data
0.7
0.8
0.9
1
1.1
1.2
1.3
1.4
1.5
1.6
1.7
0
10
20
30
40
50
60

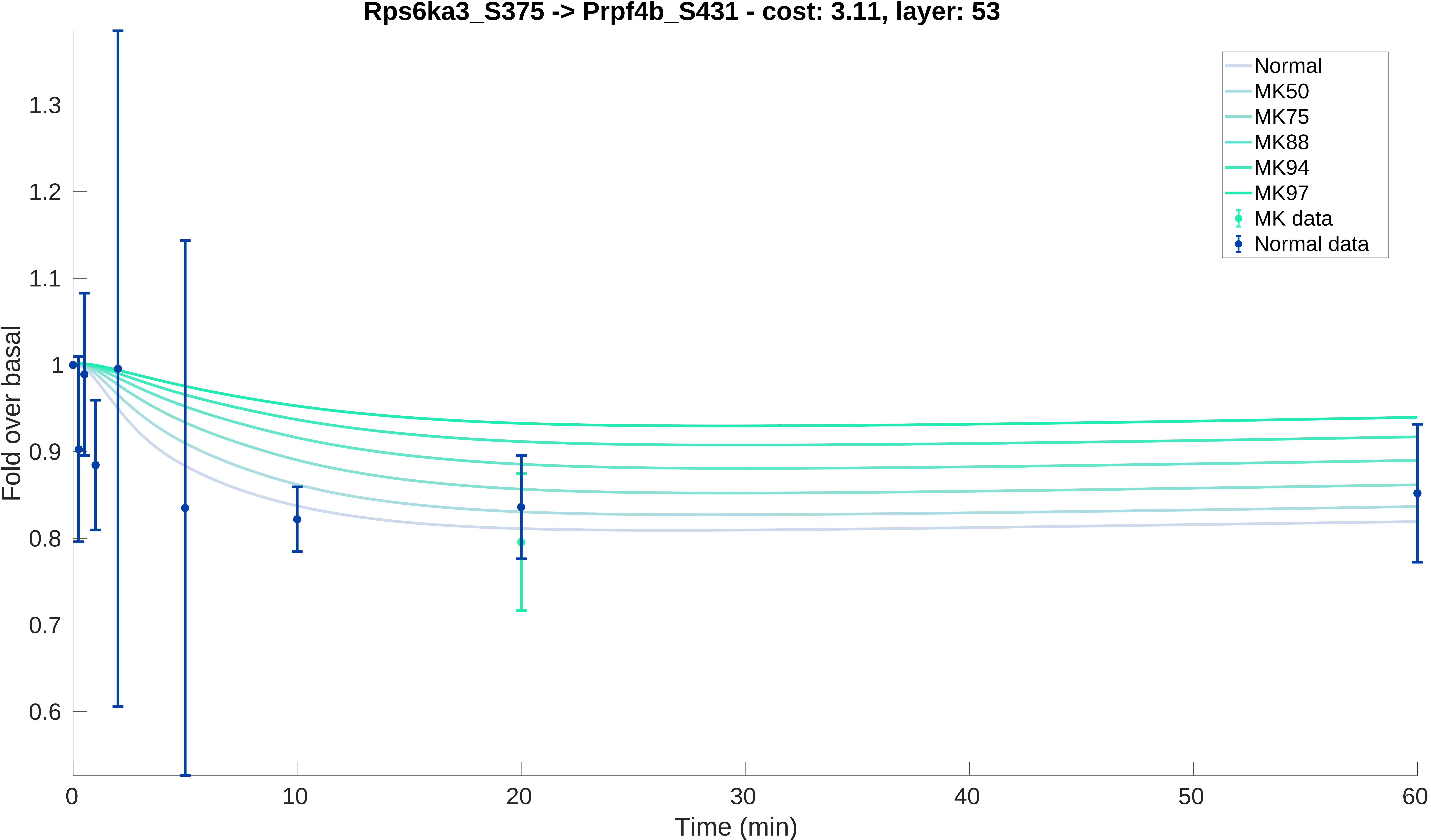
Rps6ka3_S375 -> Prpf4b_S431 - cost: 3.11, layer: 53
Normal
MK50
MK75
MK88
MK94
MK97
MK data
Normal data
Fold over basal
Time (min)
1.3
1.2
1.1
1
0.9
0.8
0.7
0.6
0
10
20
30
40
50
60

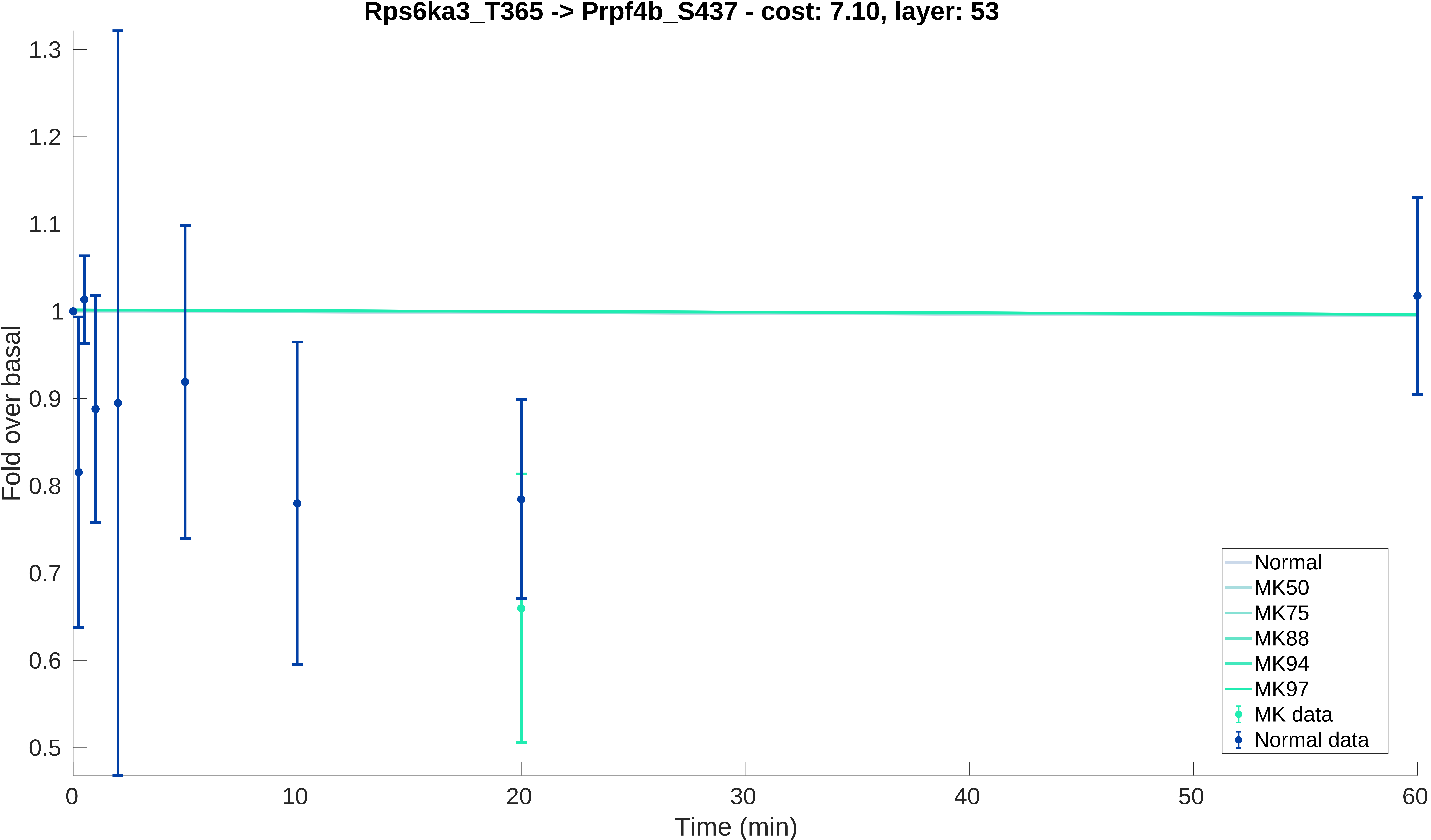

Rps6ka3_T365 -> Prpf4b_S437 - cost: 7.10, layer: 53
1.3
1.2
1.1
1
0.9
0.8
0.7
0.6
0.5
Fold over basal
0
10
20
30
40
50
60
Time (min)
Normal
MK50
MK75
MK88
MK94
MK97
MK data
Normal data

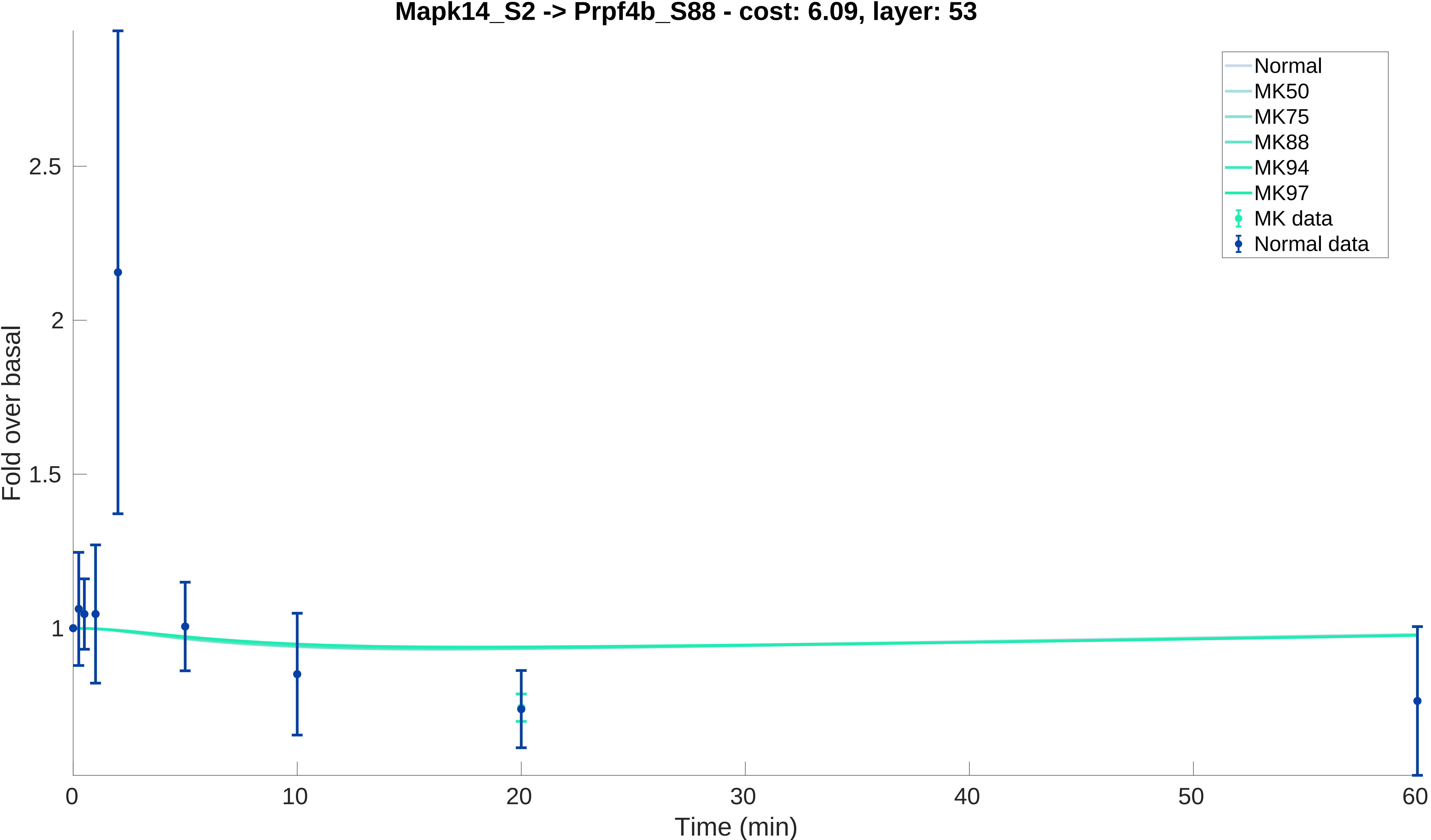

Mapk14_S2 -> Prpf4b_S88 - cost: 6.09, layer: 53
Fold over basal
2.5
2
1.5
1
0
10
20
30
40
50
60
Time (min)
Normal
MK50
MK75
MK88
MK94
MK97
MK data
Normal data

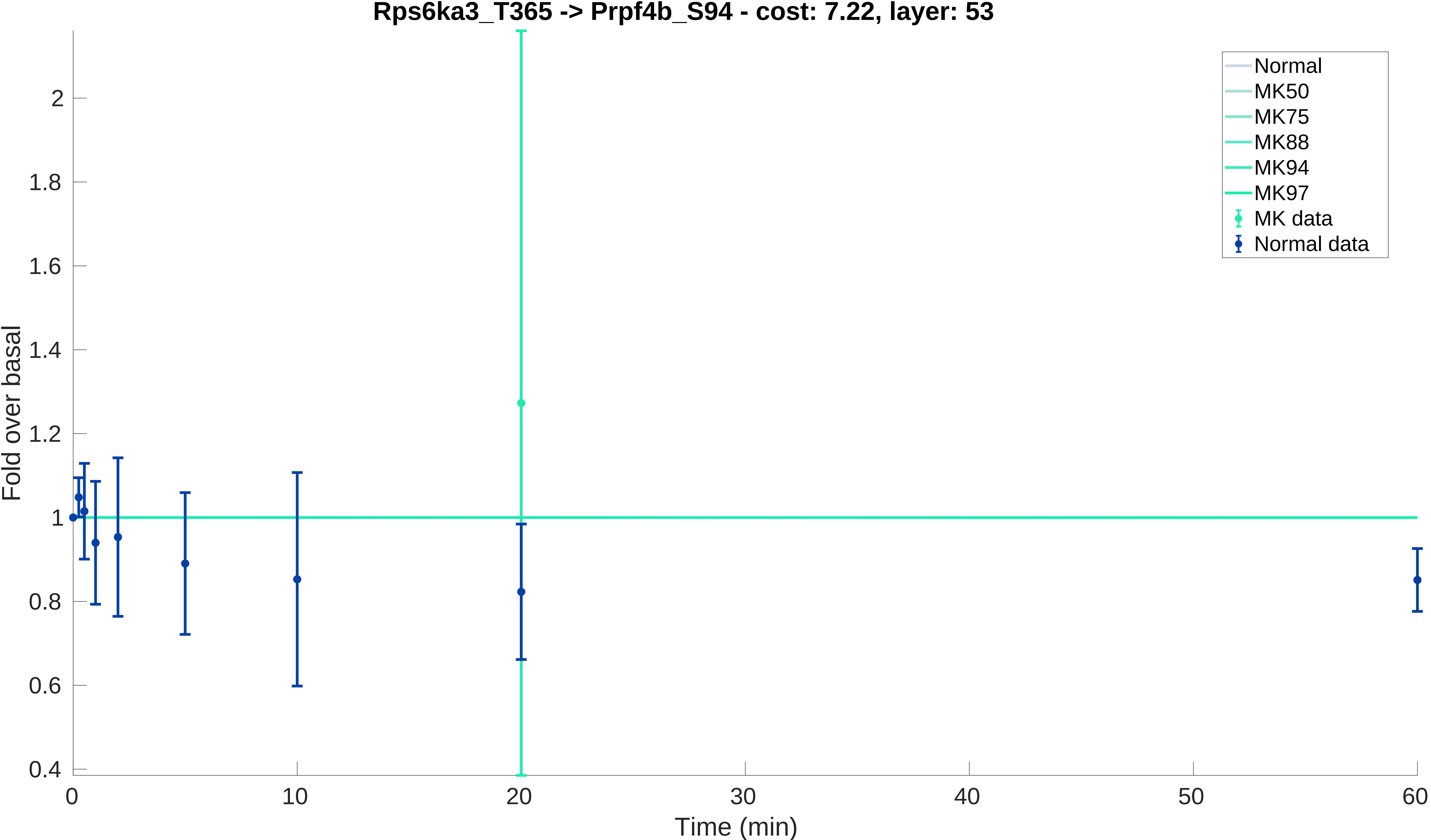

Rps6ka3_T365 -> Prpf4b_S94 - cost: 7.22, layer: 53
Normal
MK50
MK75
MK88
MK94
MK97
MK data
Normal data
Fold over basal
Time (min)
0
10
20
30
40
50
60
0.4
0.6
0.8
1
1.2
1.4
1.6
1.8
2

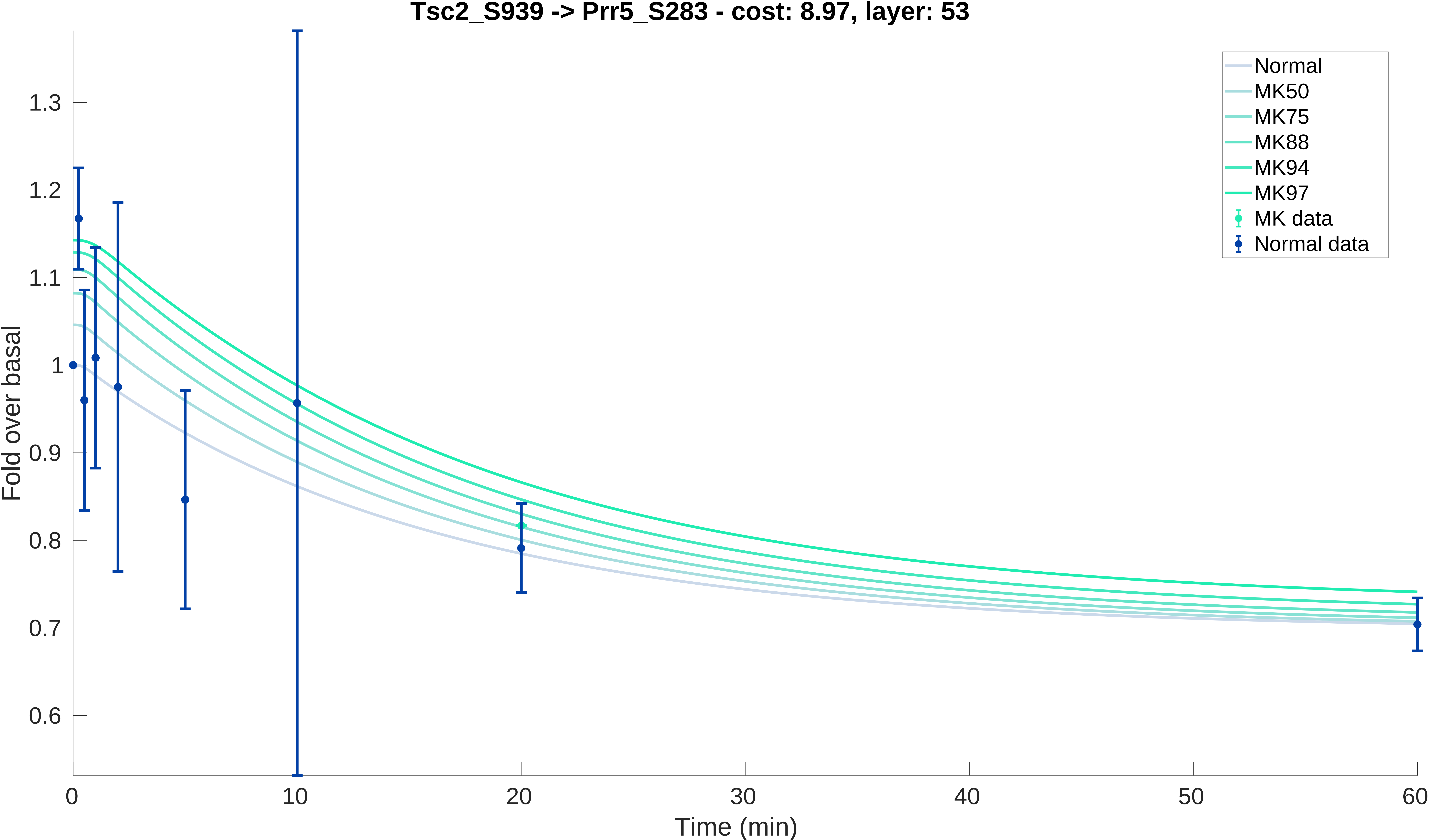

Tsc2_S939 -> Prr5_S283 - cost: 8.97, layer: 53
Fold over basal
1.3
1.2
1.1
1
0.9
0.8
0.7
0.6
0
10
20
30
40
50
60
Time (min)
Normal
MK50
MK75
MK88
MK94
MK97
MK data
Normal data

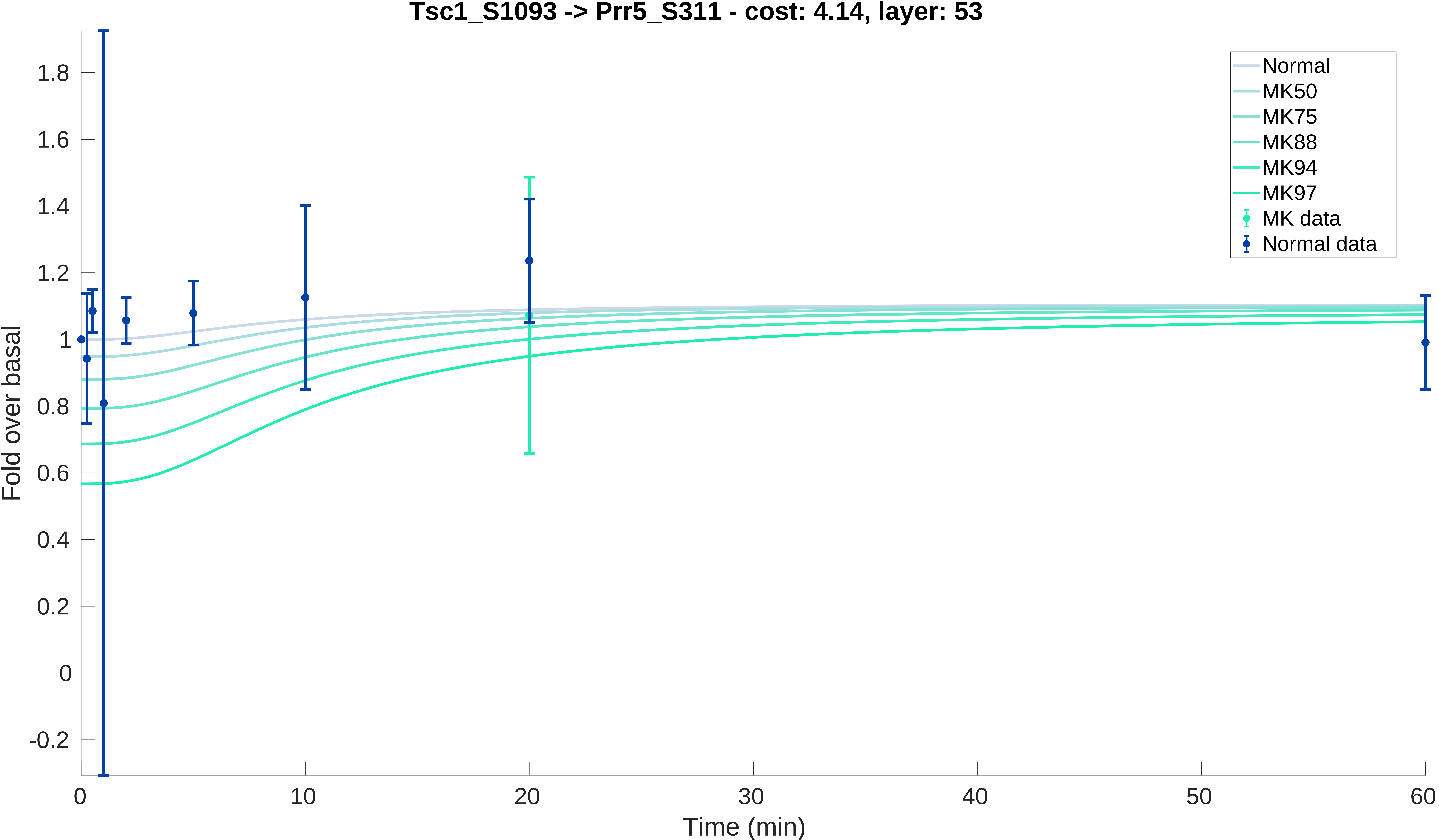

Tsc1_S1093 -> Prr5_S311 - cost: 4.14, layer: 53
Normal
MK50
MK75
MK88
MK94
MK97
MK data
Normal data
Fold over basal
Time (min)
1.8
1.6
1.4
1.2
1
0.8
0.6
0.4
0.2
0
-0.2
0
10
20
30
40
50
60

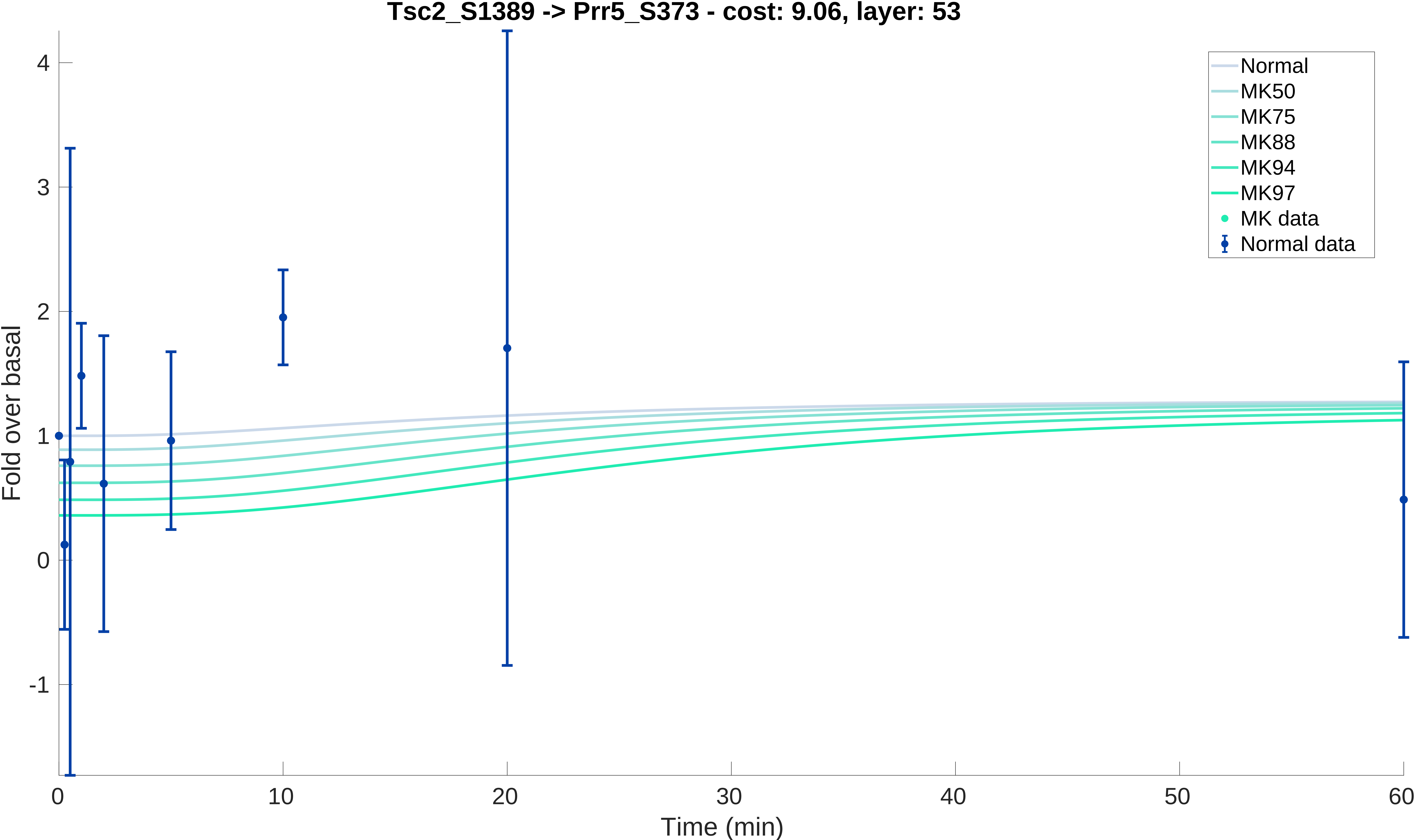

Tsc2_S1389 -> Prr5_S373 - cost: 9.06, layer: 53
Fold over basal
Time (min)
4
3
2
1
0
-1
0
10
20
30
40
50
60
Normal
MK50
MK75
MK88
MK94
MK97
MK data
Normal data

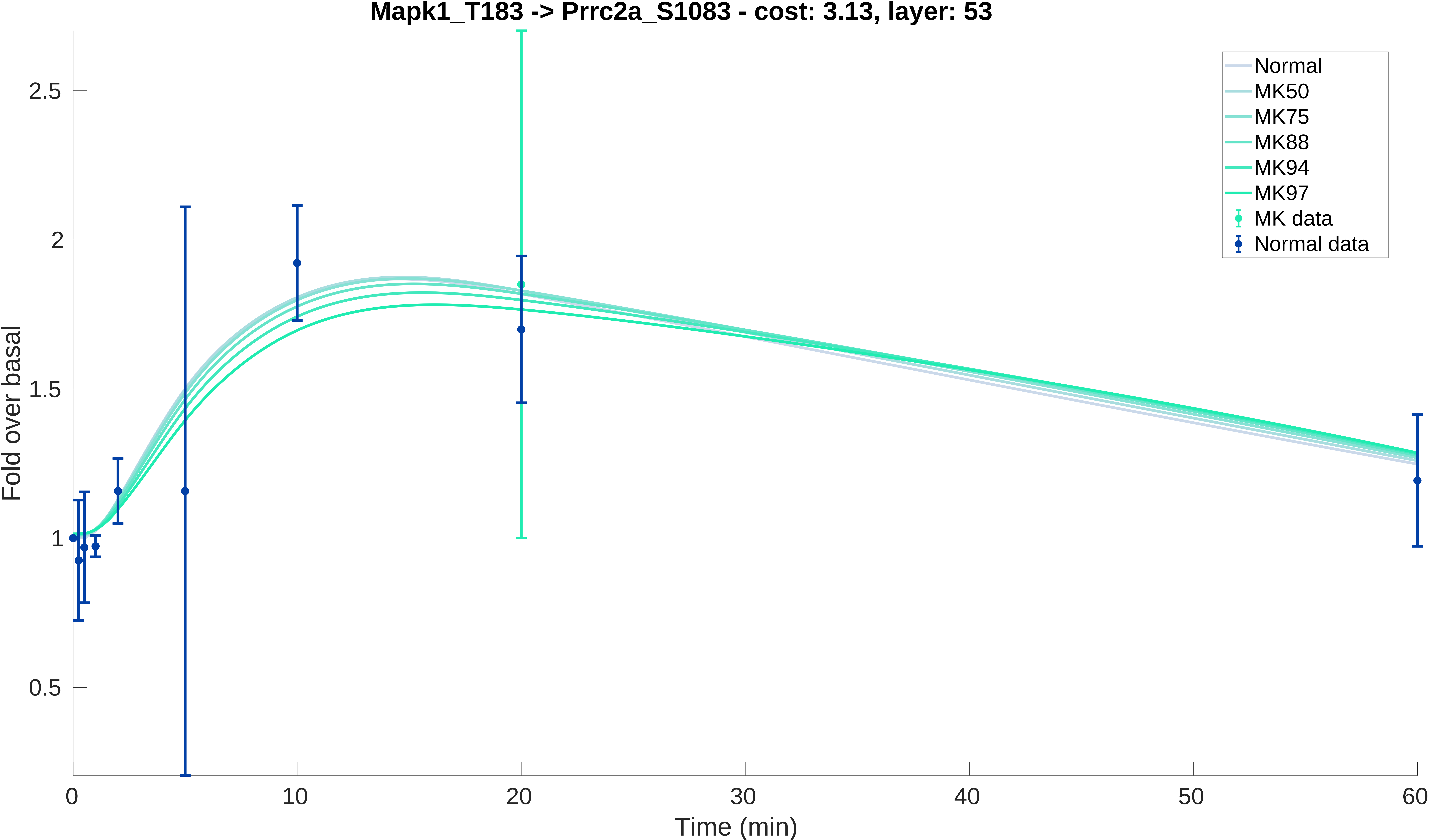

Mapk1_T183 -> Prrc2a_S1083 - cost: 3.13, layer: 53
Normal
MK50
MK75
MK88
MK94
MK97
MK data
Normal data
Fold over basal
2.5
2
1.5
1
0.5
0
10
20
30
40
50
60
Time (min)

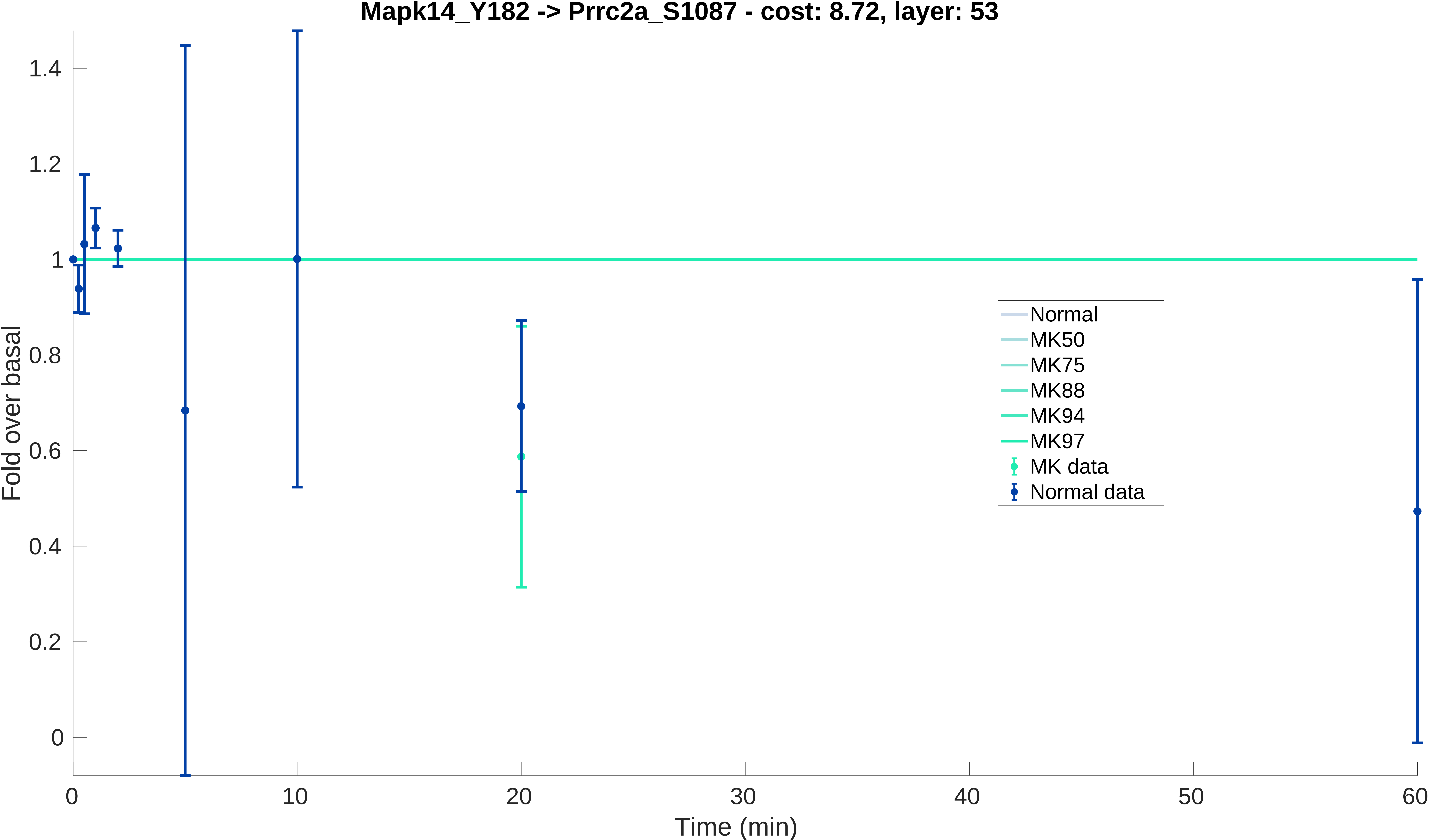

Mapk14_Y182 -> Prrc2a_S1087 - cost: 8.72, layer: 53
Fold over basal
Time (min)
0
0.2
0.4
0.6
0.8
1
1.2
1.4
0
10
20
30
40
50
60
Normal
MK50
MK75
MK88
MK94
MK97
MK data
Normal data

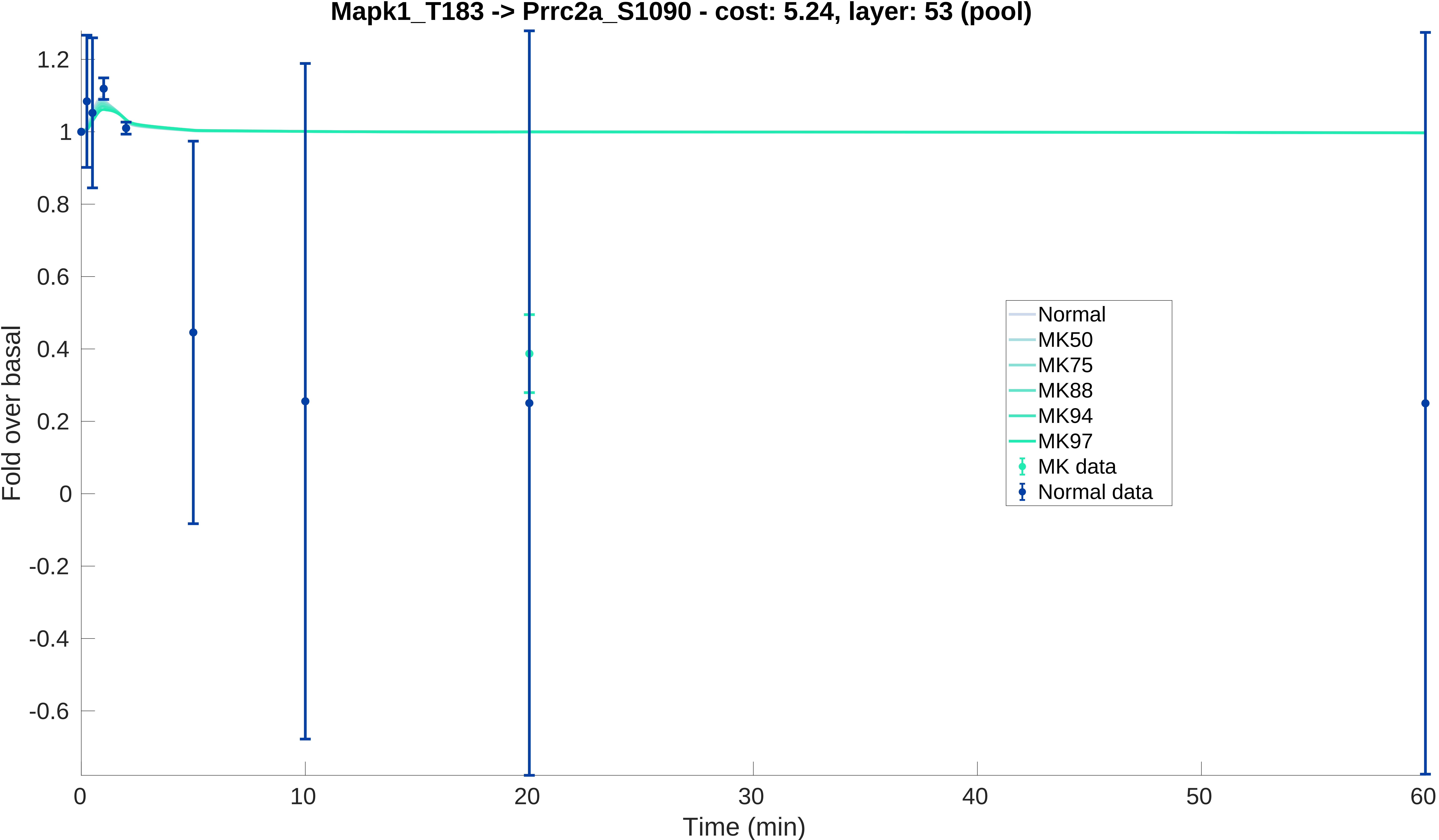

Mapk1_T183 -> Prrc2a_S1090 - cost: 5.24, layer: 53 (pool)
Fold over basal
1.2
1
0.8
0.6
0.4
0.2
0
-0.2
-0.4
-0.6
0
10
20
30
40
50
60
Time (min)
Normal
MK50
MK75
MK88
MK94
MK97
MK data
Normal data

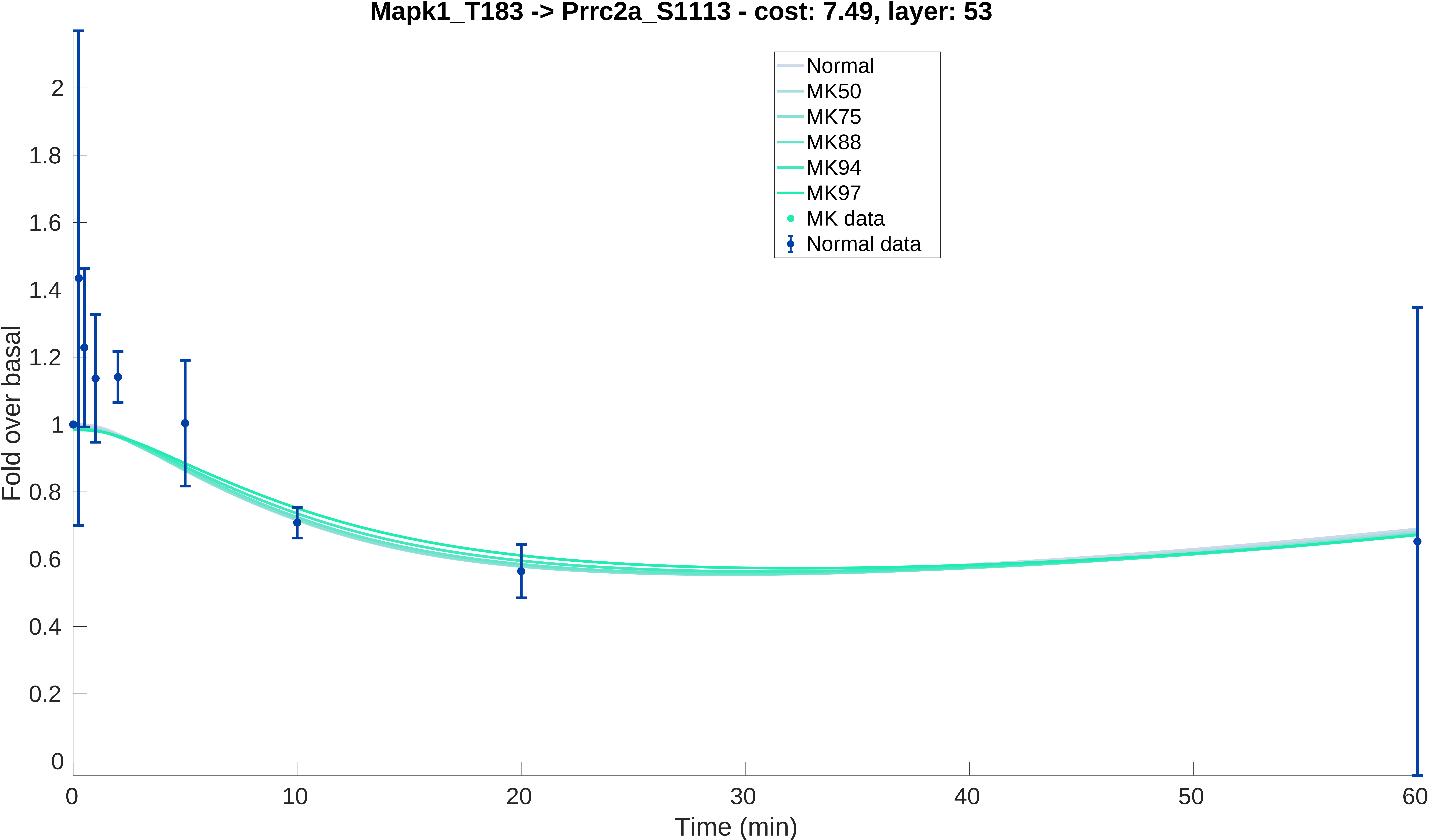

Mapk1_T183 -> Prrc2a_S1113 - cost: 7.49, layer: 53
Normal
MK50
MK75
MK88
MK94
MK97
MK data
Normal data
Fold over basal
0
0.2
0.4
0.6
0.8
1
1.2
1.4
1.6
1.8
2
Time (min)
0
10
20
30
40
50
60

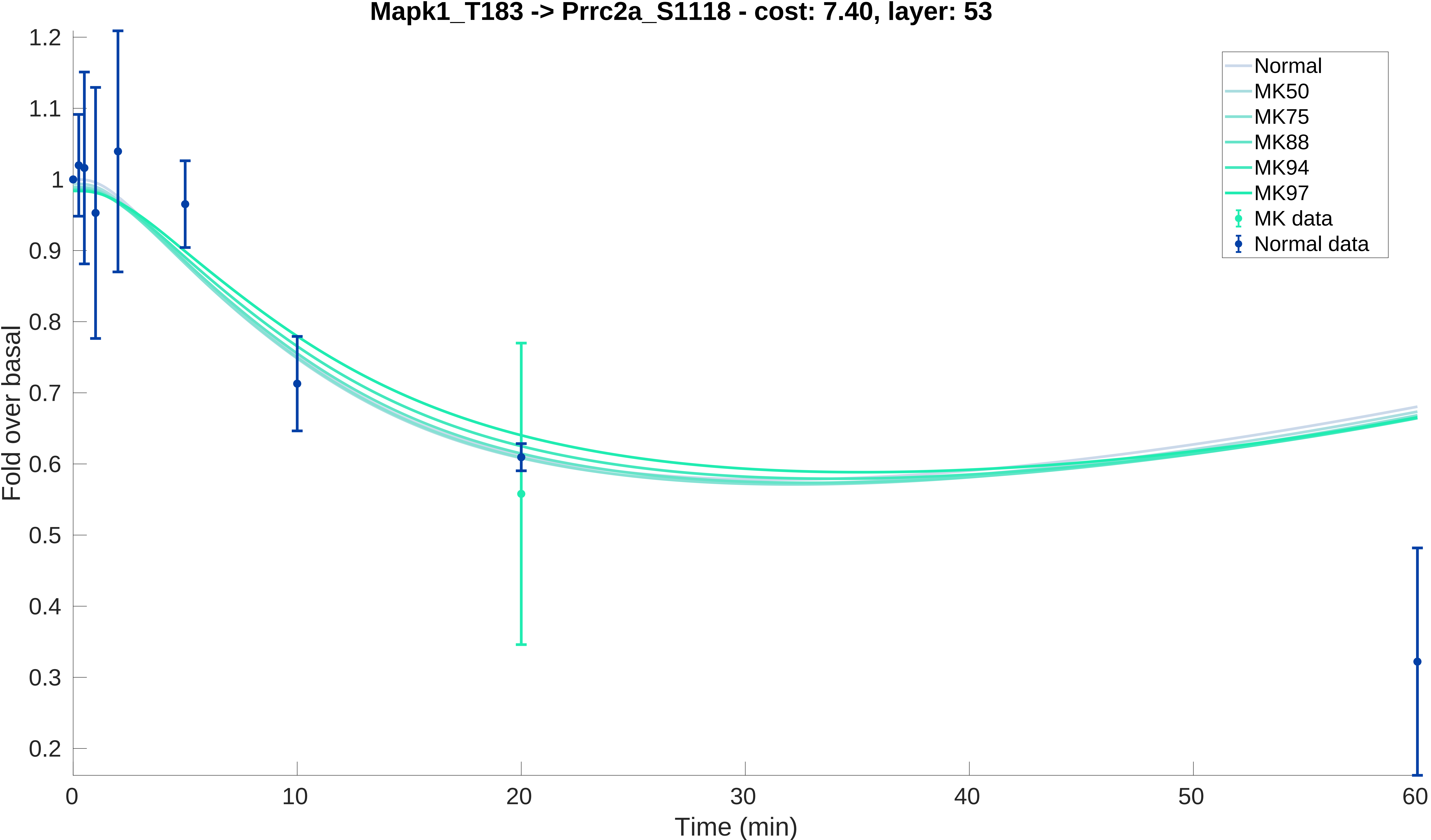

Mapk1_T183 -> Prrc2a_S1118 - cost: 7.40, layer: 53
Fold over basal
Time (min)
1.2
1.1
1
0.9
0.8
0.7
0.6
0.5
0.4
0.3
0.2
0
10
20
30
40
50
60
Normal
MK50
MK75
MK88
MK94
MK97
MK data
Normal data

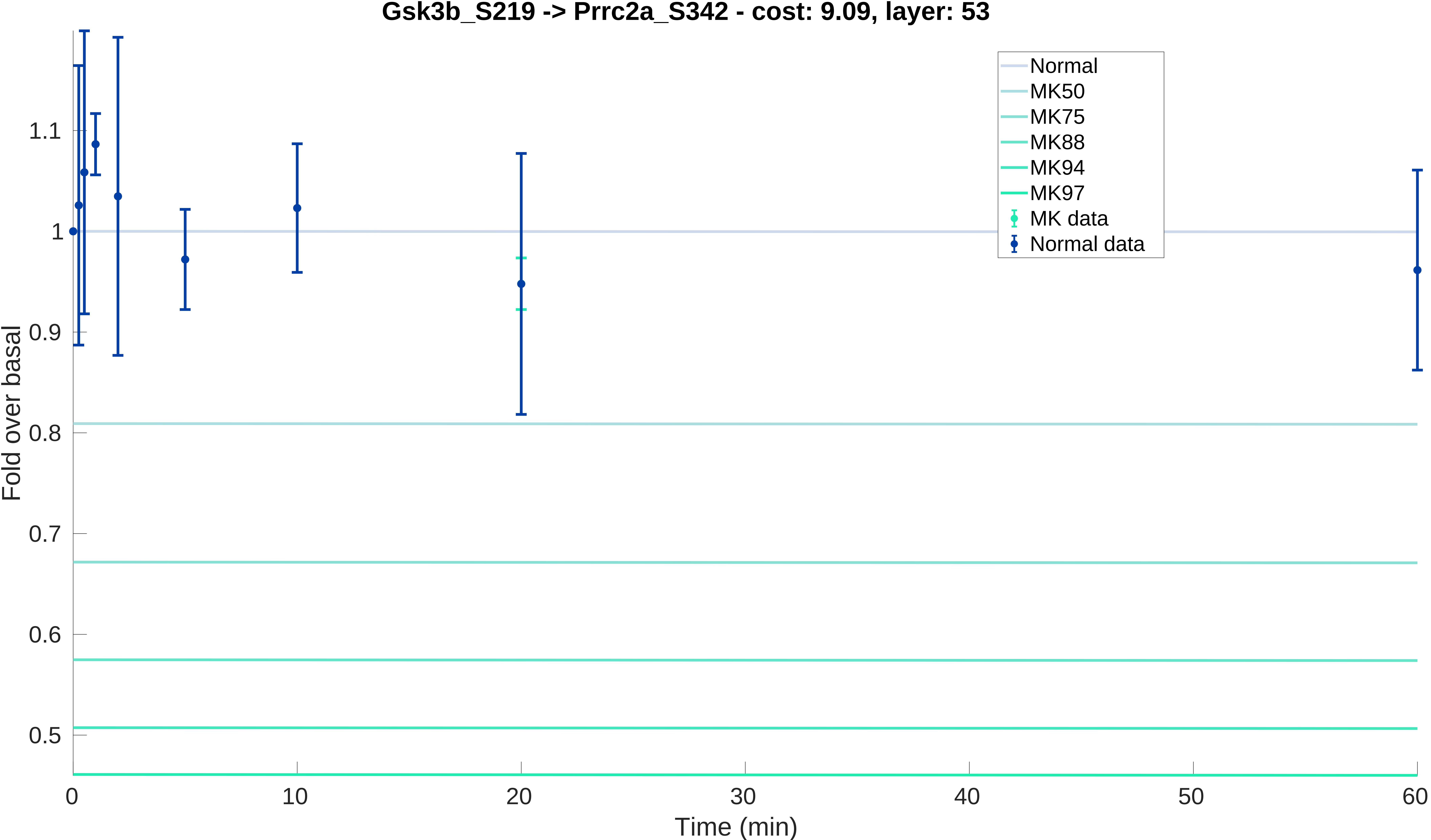

Gsk3b_S219 -> Prrc2a_S342 - cost: 9.09, layer: 53
Fold over basal
Time (min)
1.1
1
0.9
0.8
0.7
0.6
0.5
0
10
20
30
40
50
60
Normal
MK50
MK75
MK88
MK94
MK97
MK data
Normal data

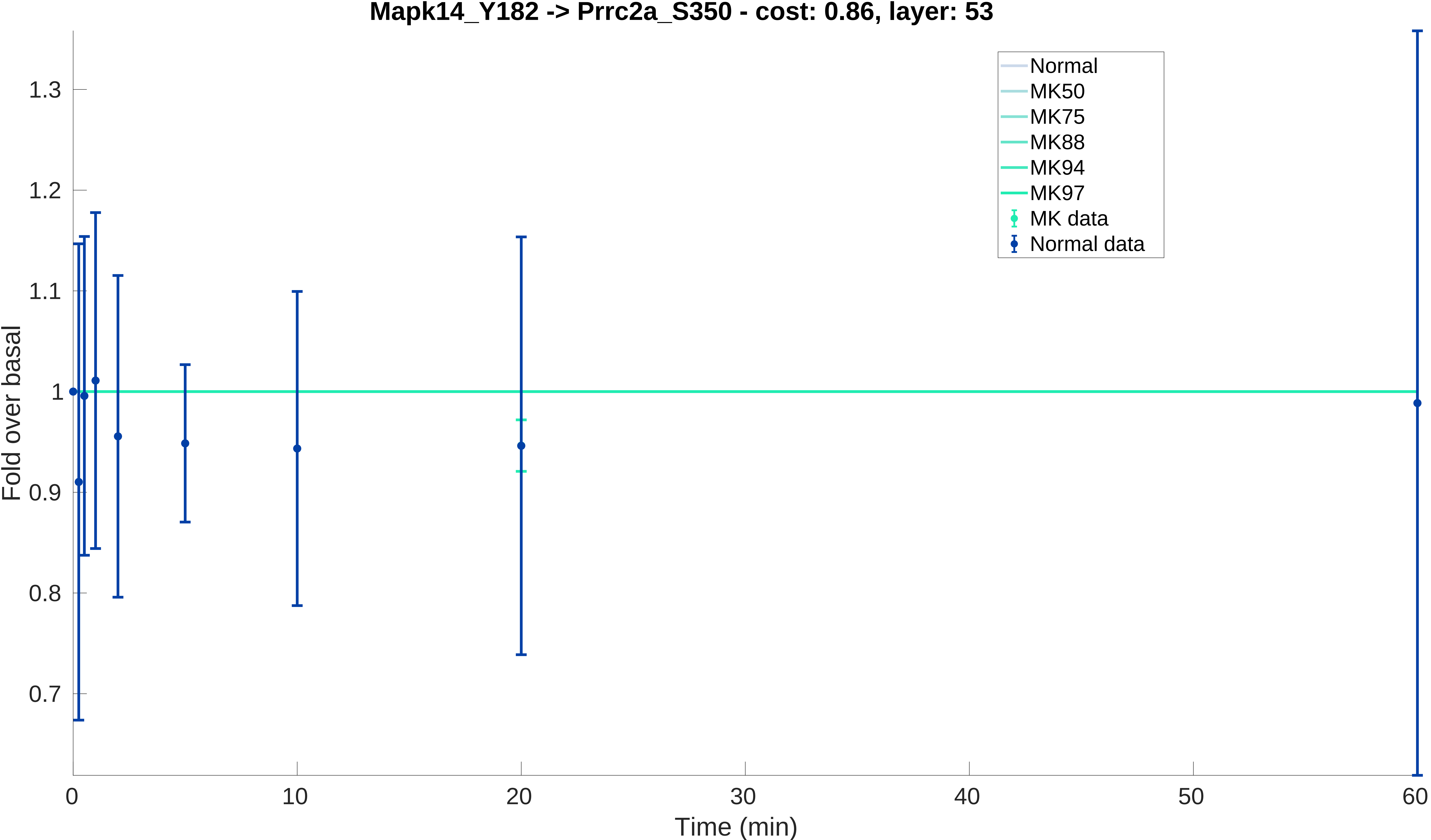

Mapk14_Y182 -> Prrc2a_S350 - cost: 0.86, layer: 53
Normal
MK50
MK75
MK88
MK94
MK97
MK data
Normal data
Fold over basal
1.3
1.2
1.1
1
0.9
0.8
0.7
0
10
20
30
40
50
60
Time (min)

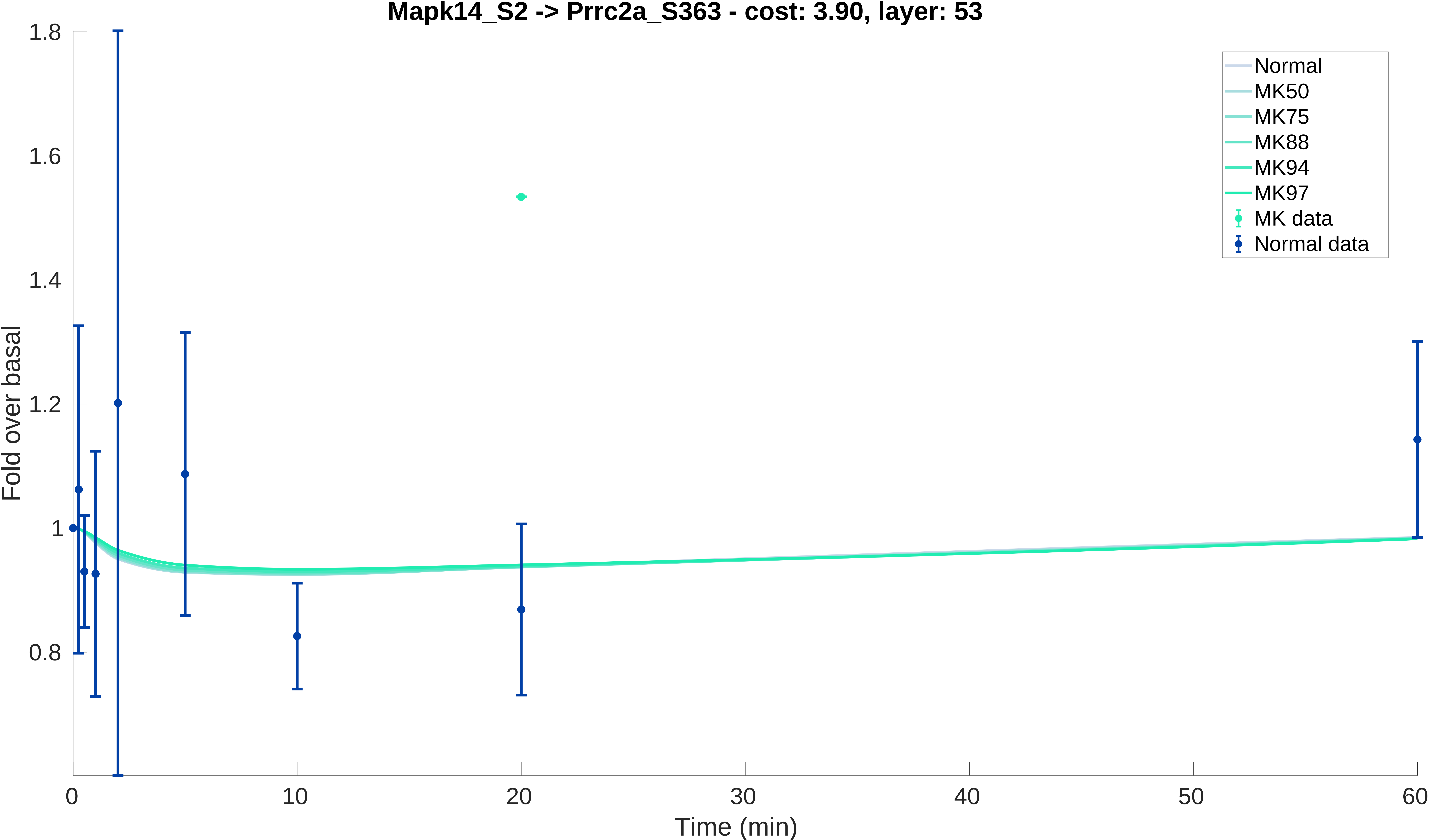

Mapk14_S2 -> Prrc2a_S363 - cost: 3.90, layer: 53
Fold over basal
Time (min)
1.8
1.6
1.4
1.2
1
0.8
0
10
20
30
40
50
60
Normal
MK50
MK75
MK88
MK94
MK97
MK data
Normal data

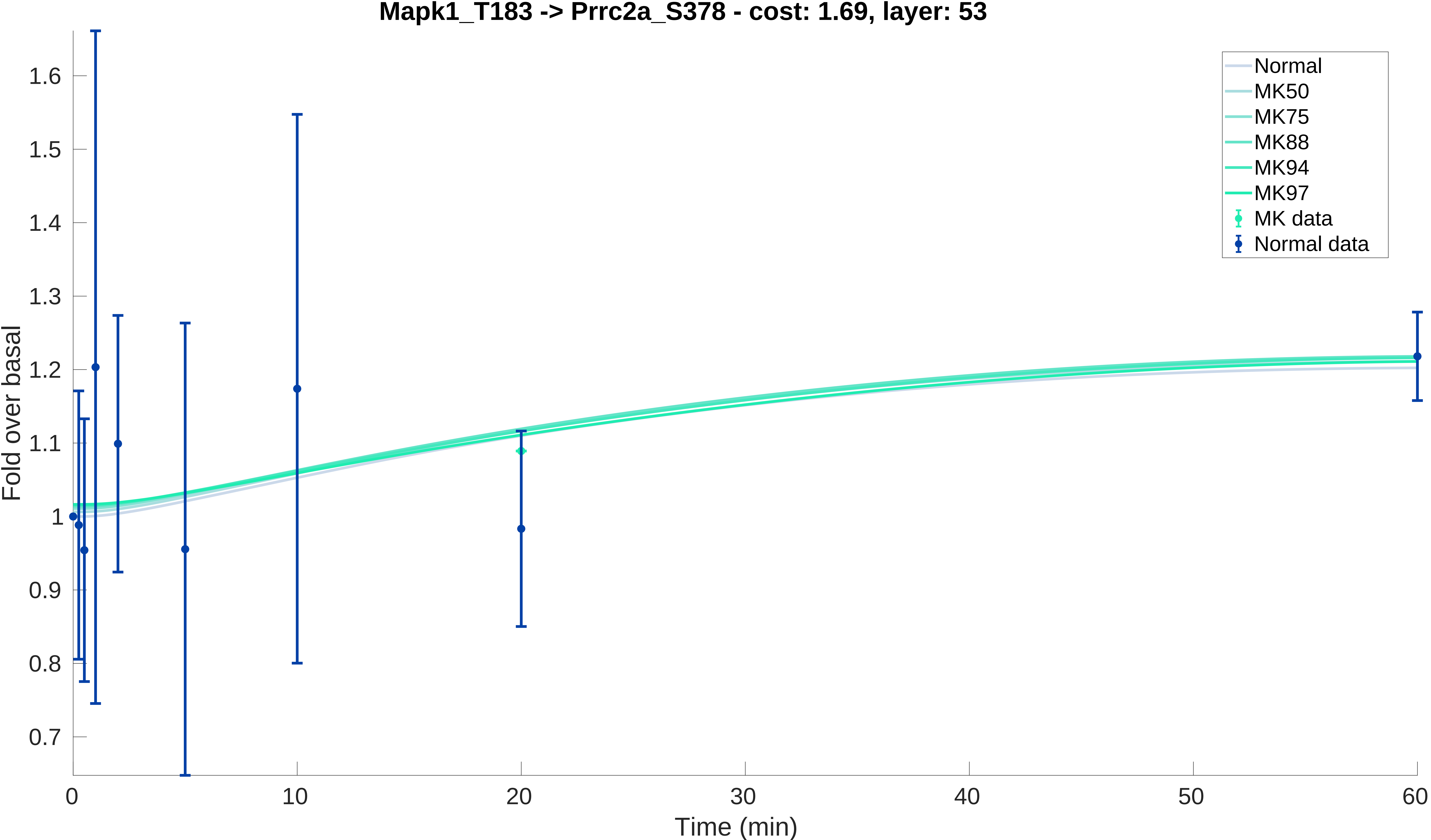

Mapk1_T183 -> Prrc2a_S378 - cost: 1.69, layer: 53
Fold over basal
Time (min)
0.7
0.8
0.9
1
1.1
1.2
1.3
1.4
1.5
1.6
0
10
20
30
40
50
60
Normal
MK50
MK75
MK88
MK94
MK97
MK data
Normal data

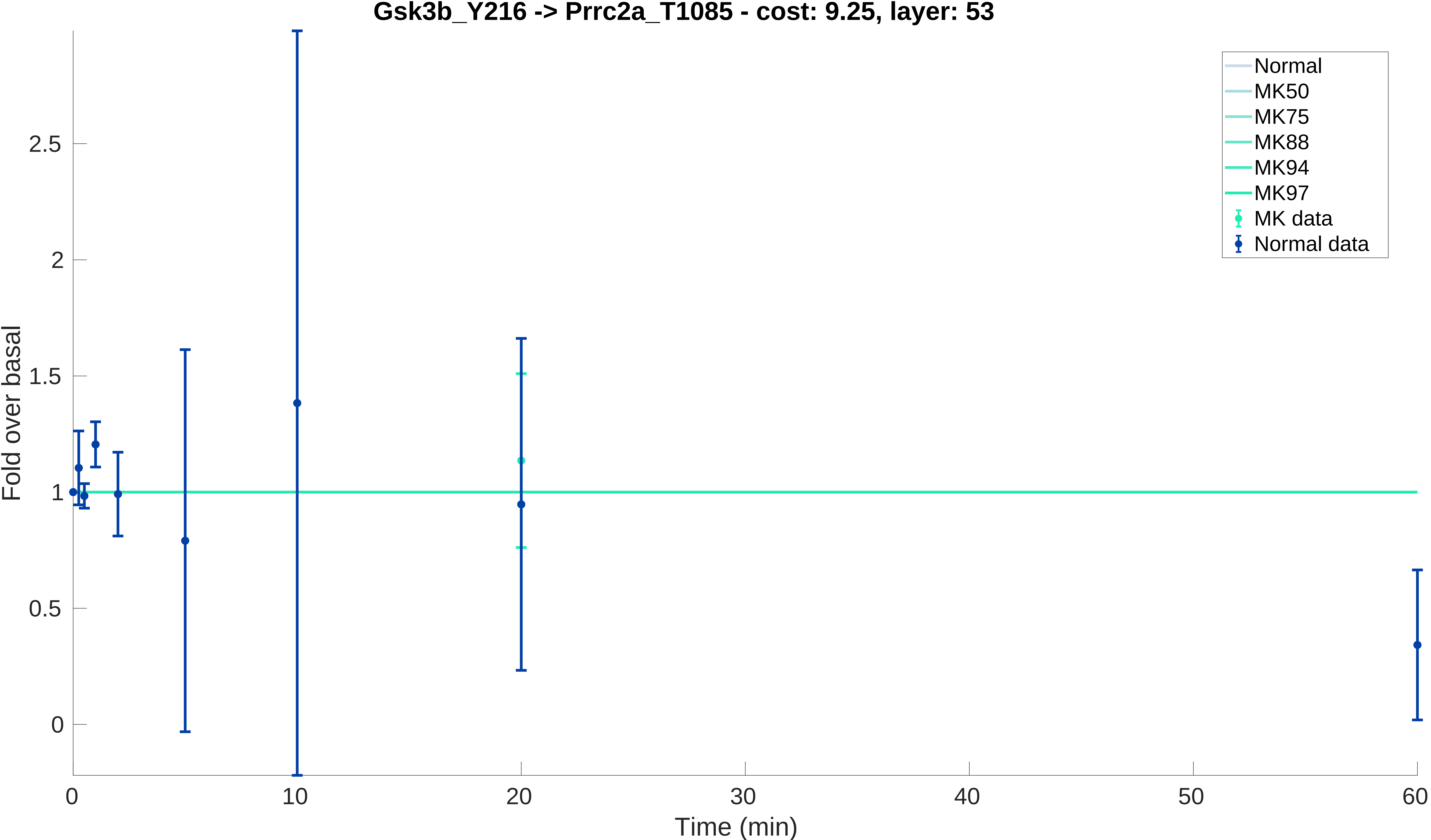

Gsk3b_Y216 -> Prrc2a_T1085 - cost: 9.25, layer: 53
Normal
MK50
MK75
MK88
MK94
MK97
MK data
Normal data
Fold over basal
Time (min)
0
0.5
1
1.5
2
2.5
0
10
20
30
40
50
60

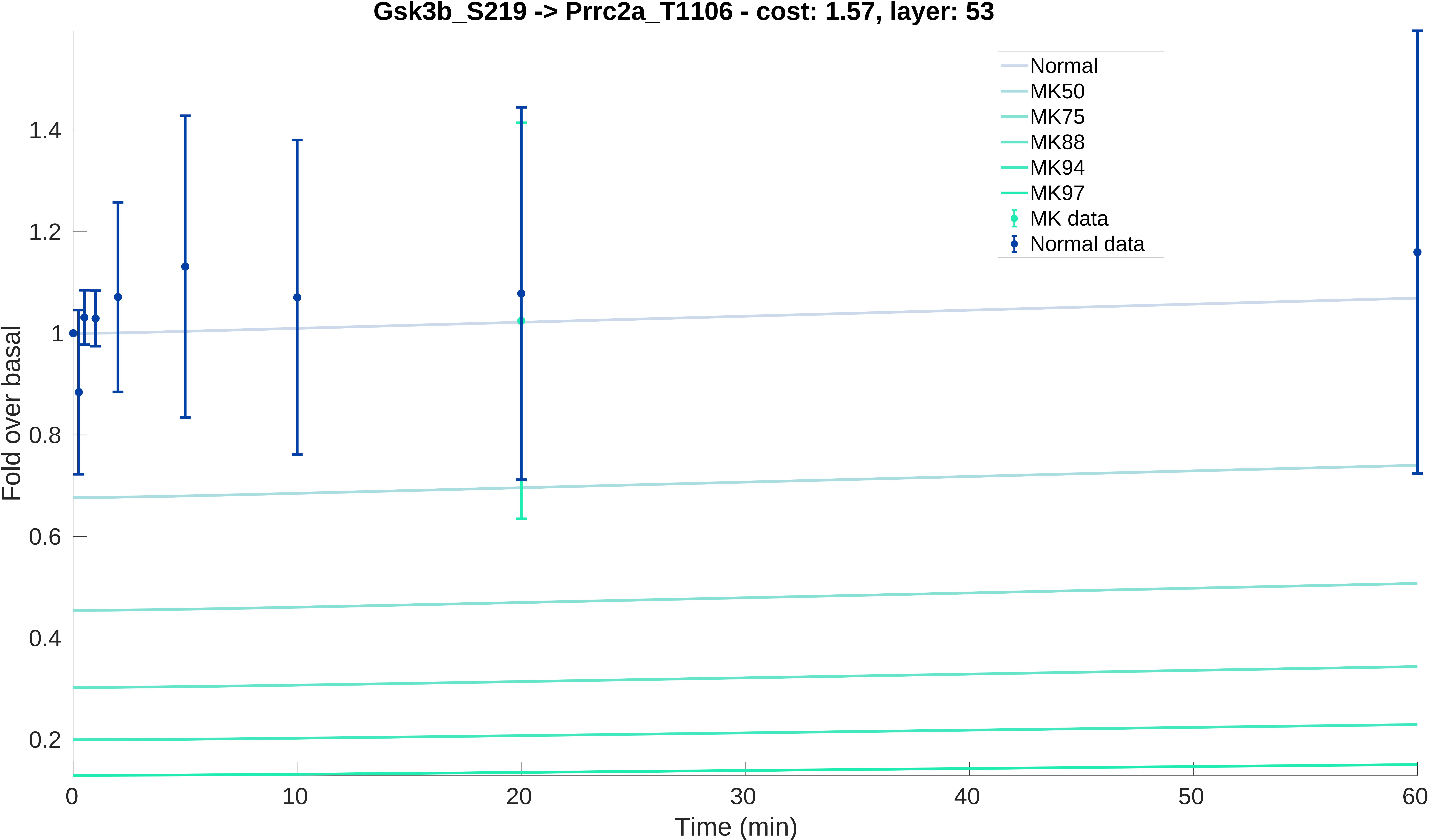

Gsk3b_S219 -> Prrc2a_T1106 - cost: 1.57, layer: 53
Normal
MK50
MK75
MK88
MK94
MK97
MK data
Normal data
Fold over basal
Time (min)
0
10
20
30
40
50
60
0.2
0.4
0.6
0.8
1
1.2
1.4

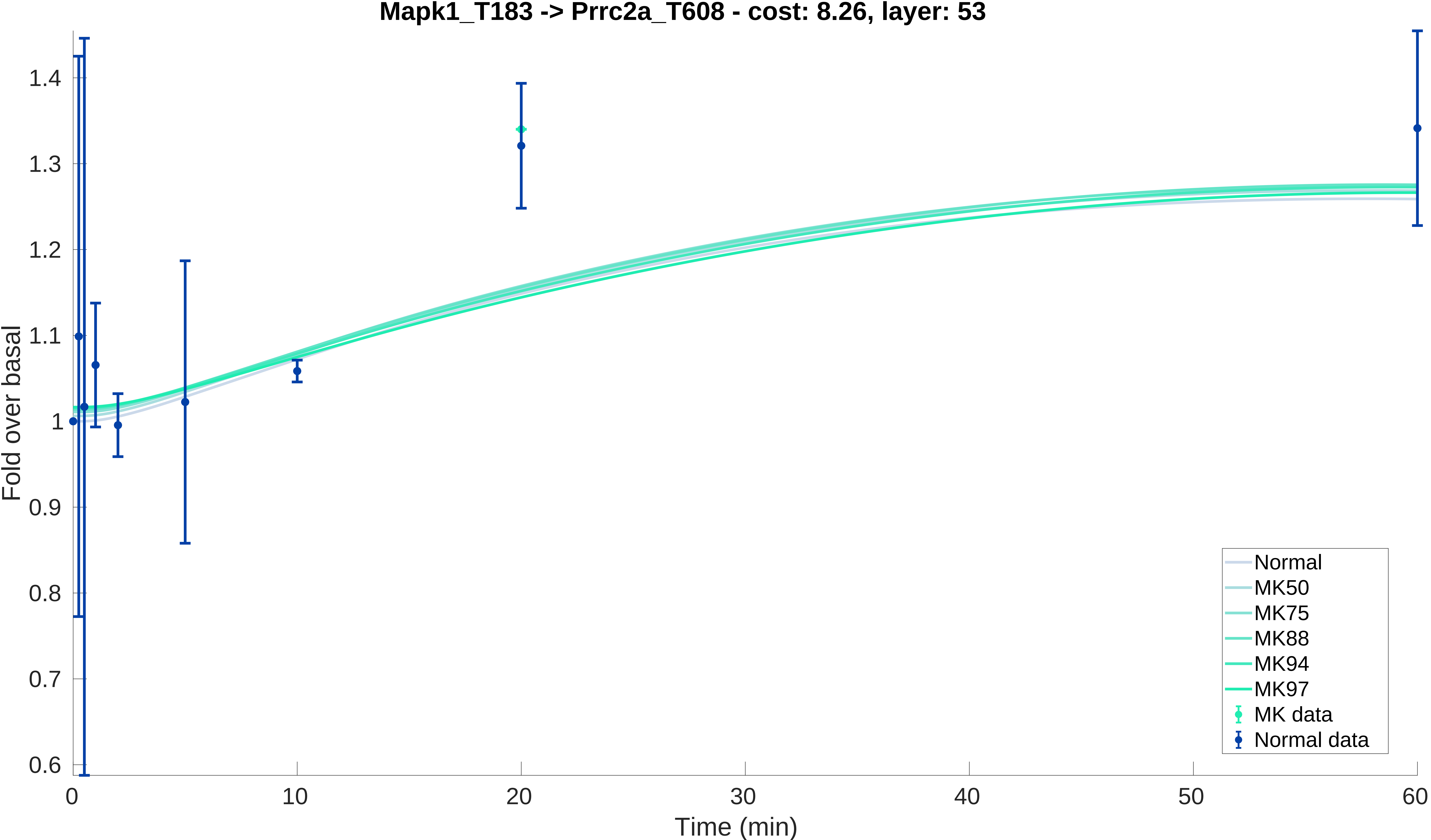

Mapk1_T183 -> Prrc2a_T608 - cost: 8.26, layer: 53
Fold over basal
1.4
1.3
1.2
1.1
1
0.9
0.8
0.7
0.6
0
10
20
30
40
50
60
Time (min)
Normal
MK50
MK75
MK88
MK94
MK97
MK data
Normal data

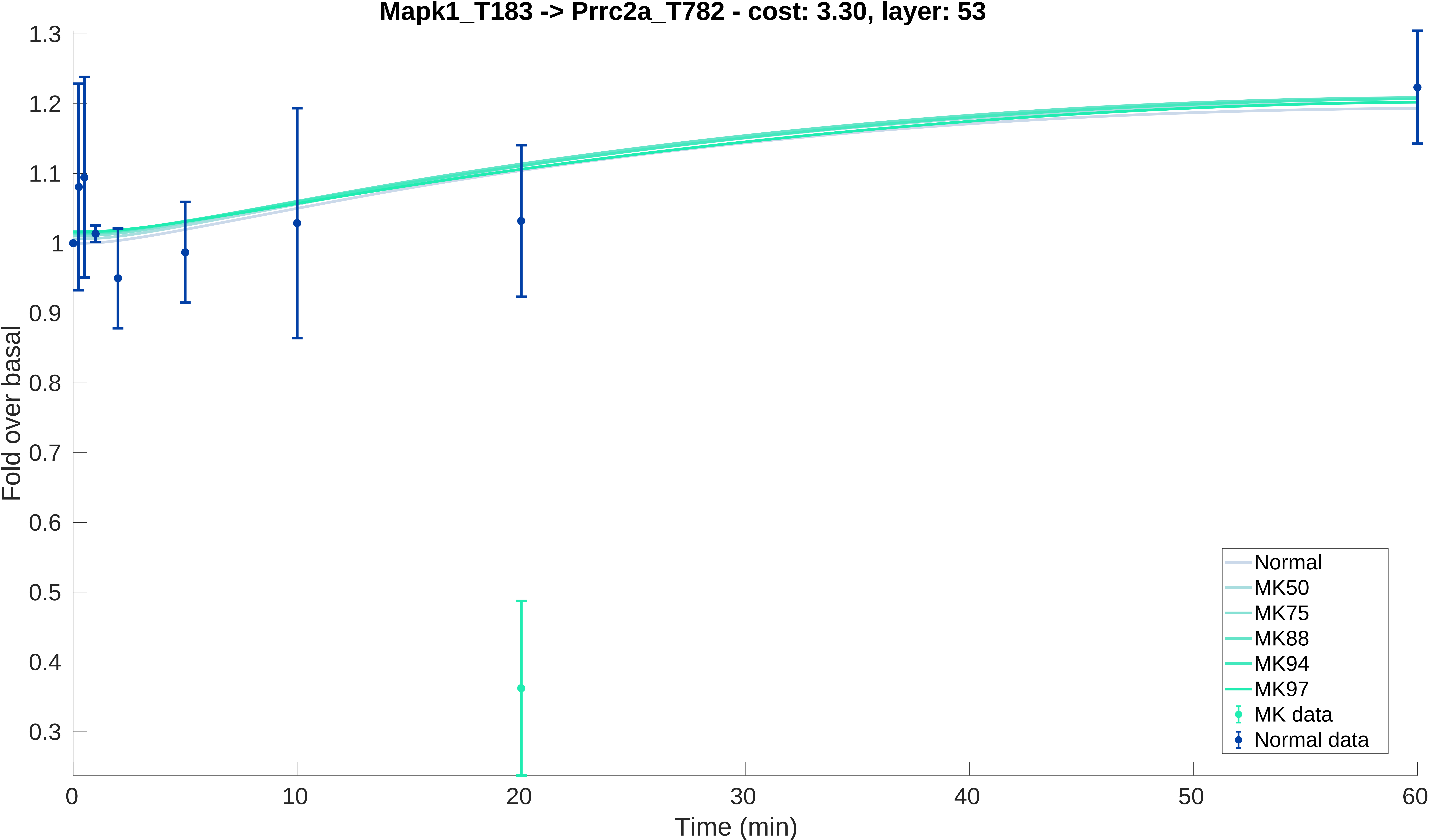
Mapk1_T183 -> Prrc2a_T782 - cost: 3.30, layer: 53
Fold over basal
Time (min)
1.3
1.2
1.1
1
0.9
0.8
0.7
0.6
0.5
0.4
0.3
0
10
20
30
40
50
60
Normal
MK50
MK75
MK88
MK94
MK97
MK data
Normal data

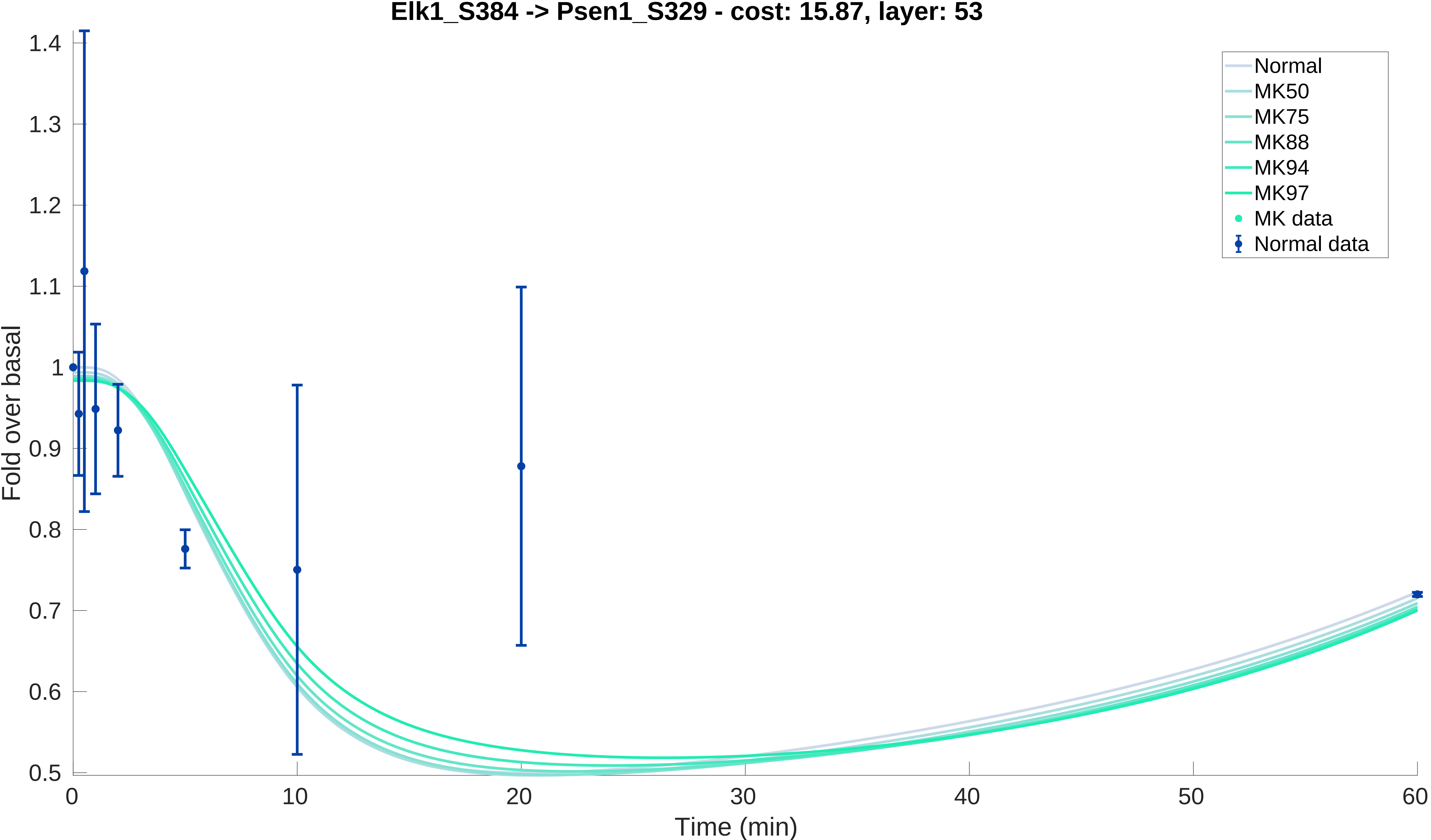

Elk1_S384 -> Psen1_S329 - cost: 15.87, layer: 53
Fold over basal
1.4
1.3
1.2
1.1
1
0.9
0.8
0.7
0.6
0.5
0
10
20
30
40
50
60
Time (min)
Normal
MK50
MK75
MK88
MK94
MK97
MK data
Normal data

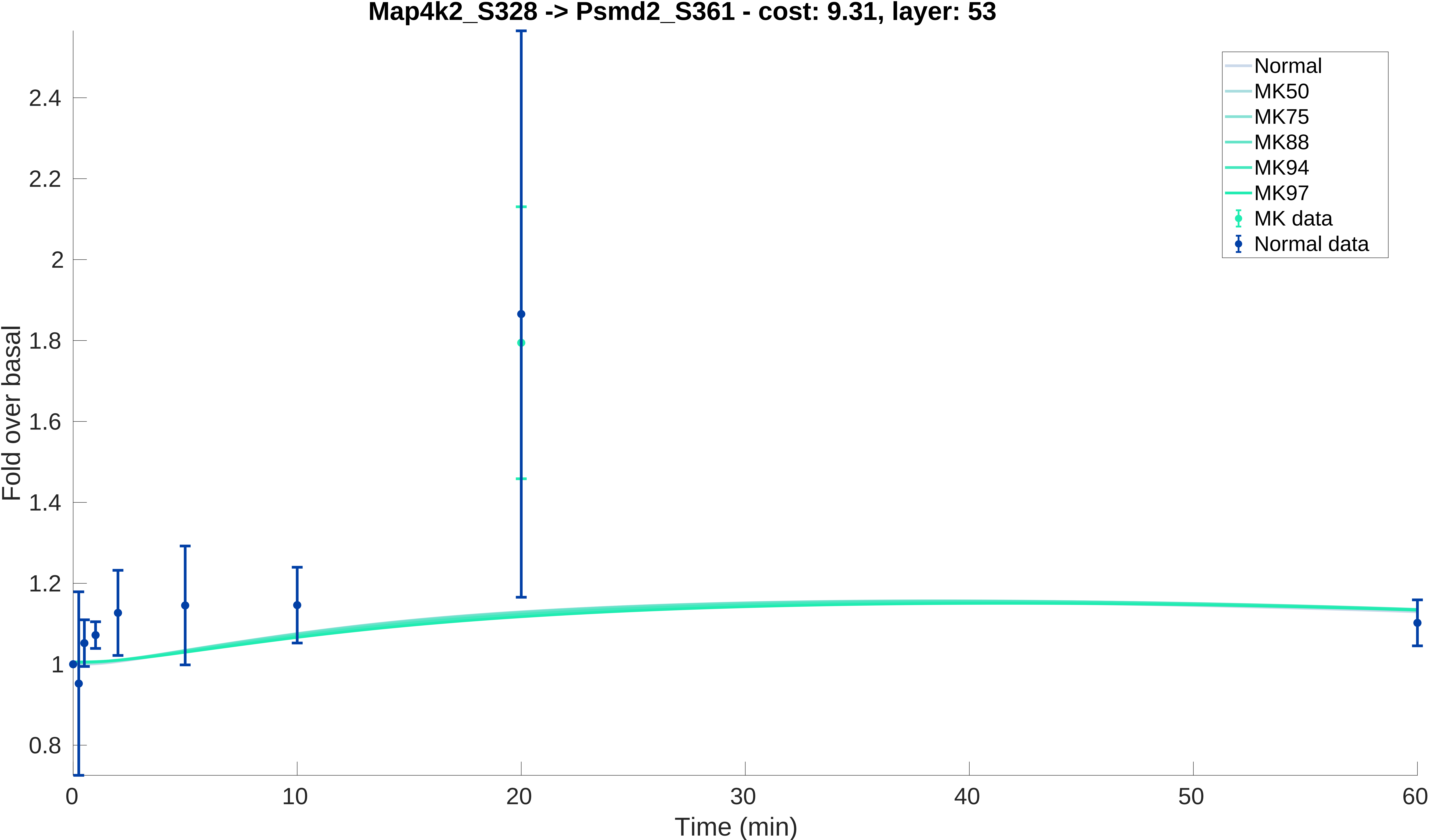

Map4k2_S328 -> Psmd2_S361 - cost: 9.31, layer: 53
Fold over basal
2.4
2.2
2
1.8
1.6
1.4
1.2
1
0.8
0
10
20
30
40
50
60
Time (min)
Normal
MK50
MK75
MK88
MK94
MK97
MK data
Normal data

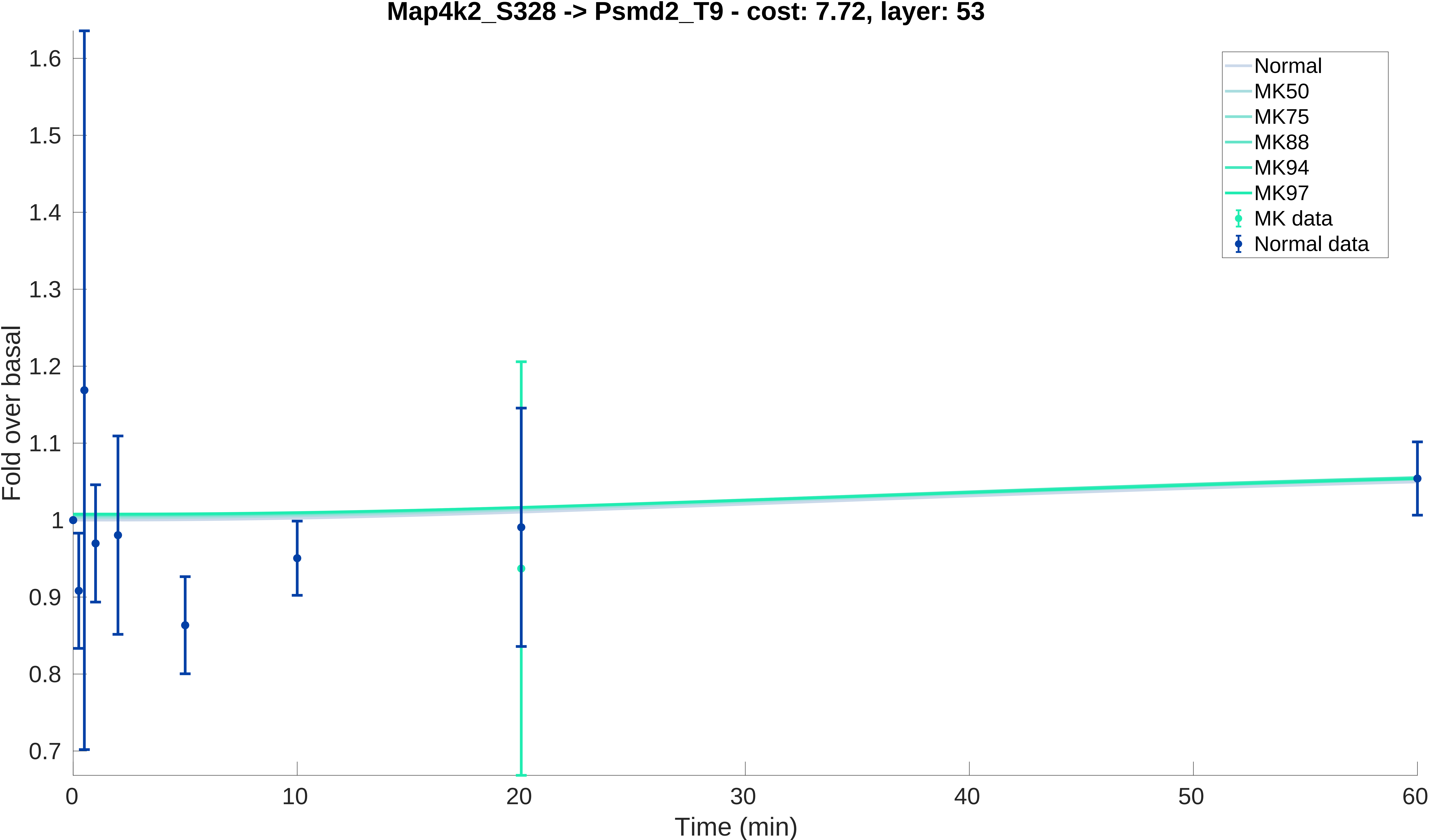

Map4k2_S328 -> Psmd2_T9 - cost: 7.72, layer: 53
Normal
MK50
MK75
MK88
MK94
MK97
MK data
Normal data
Fold over basal
Time (min)
1.6
1.5
1.4
1.3
1.2
1.1
1
0.9
0.8
0.7
0
10
20
30
40
50
60

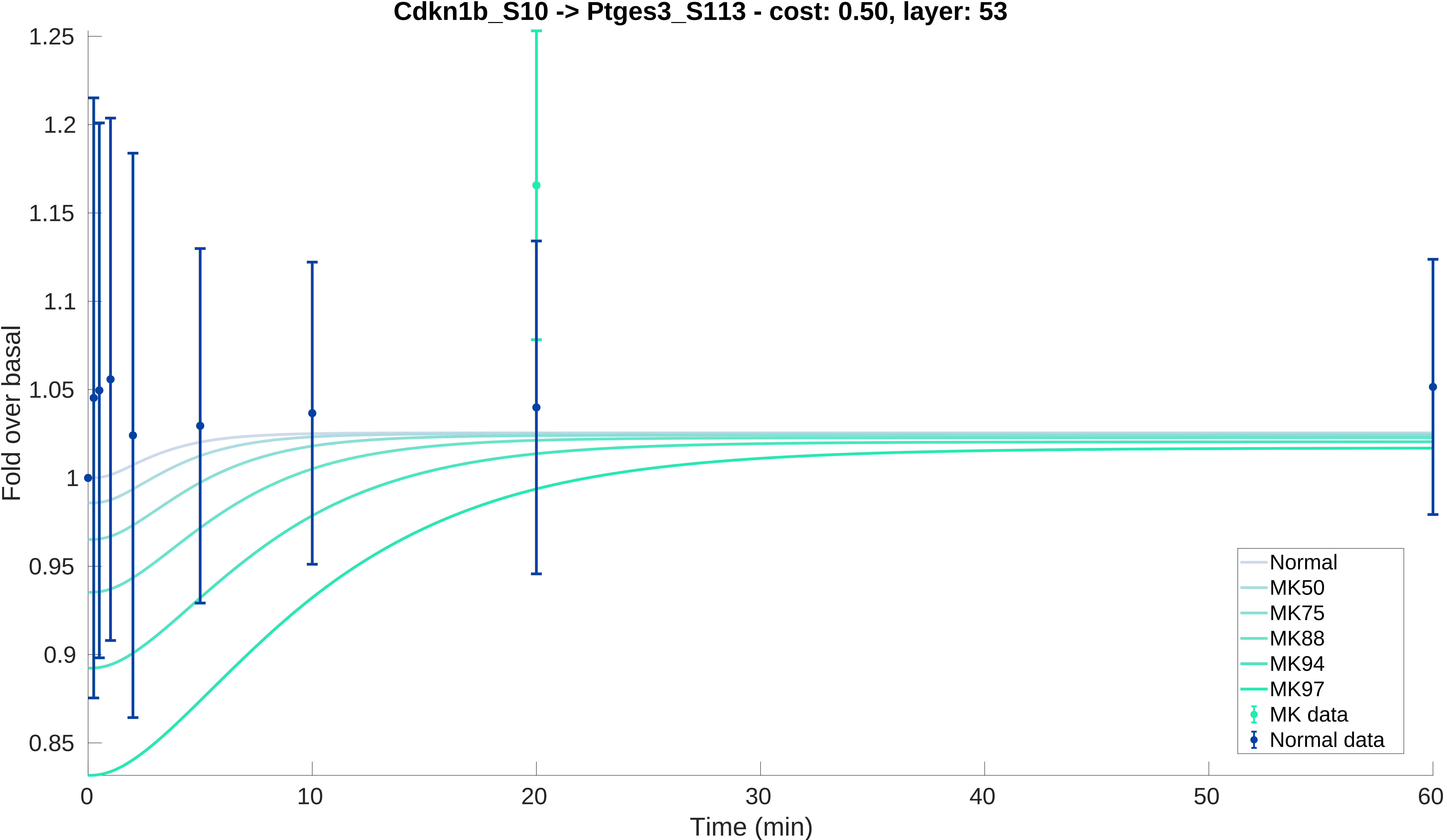

Cdkn1b_S10 -> Ptges3_S113 - cost: 0.50, layer: 53
1.25
1.2
1.15
1.1
1.05
1
0.95
0.9
0.85
Fold over basal
0
10
20
30
40
50
60
Time (min)
Normal
MK50
MK75
MK88
MK94
MK97
MK data
Normal data

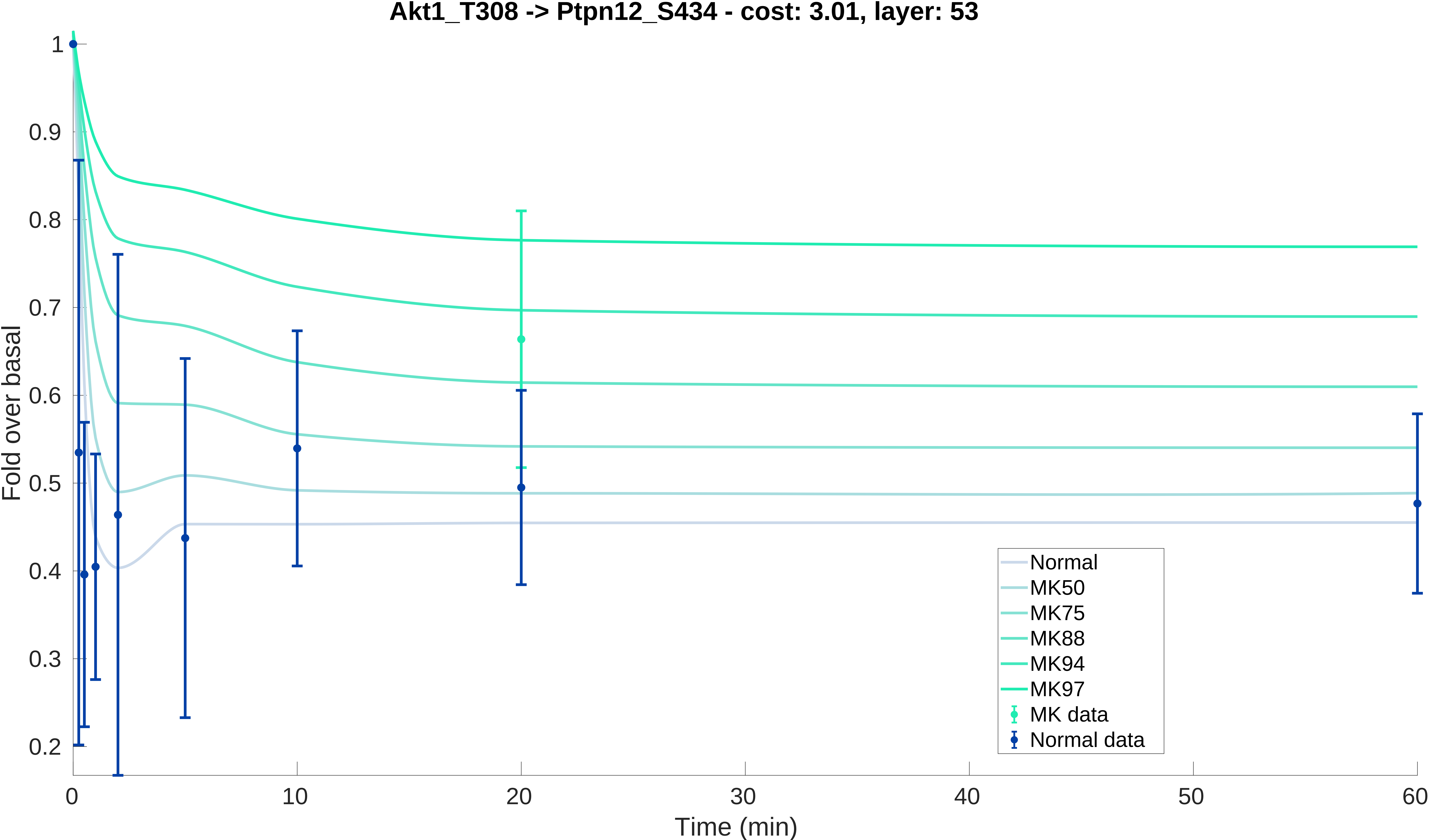
Akt1_T308 -> Ptpn12_S434 - cost: 3.01, layer: 53
Fold over basal
1
0.9
0.8
0.7
0.6
0.5
0.4
0.3
0.2
0
10
20
30
40
50
60
Time (min)
Normal
MK50
MK75
MK88
MK94
MK97
MK data
Normal data

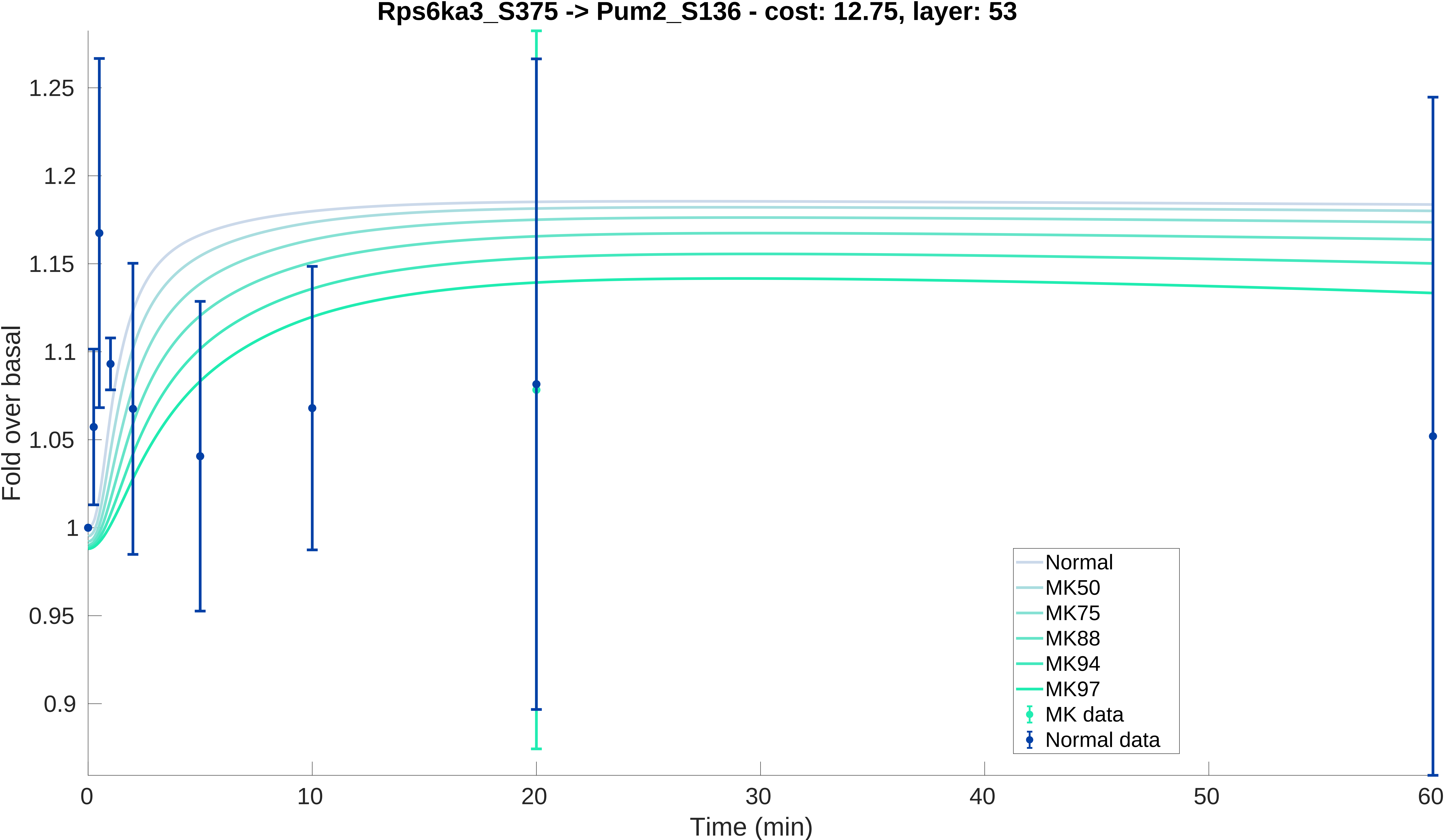
Rps6ka3_S375 -> Pum2_S136 - cost: 12.75, layer: 53
Fold over basal
1.25
1.2
1.15
1.1
1.05
1
0.95
0.9
0
10
20
30
40
50
60
Time (min)
Normal
MK50
MK75
MK88
MK94
MK97
MK data
Normal data

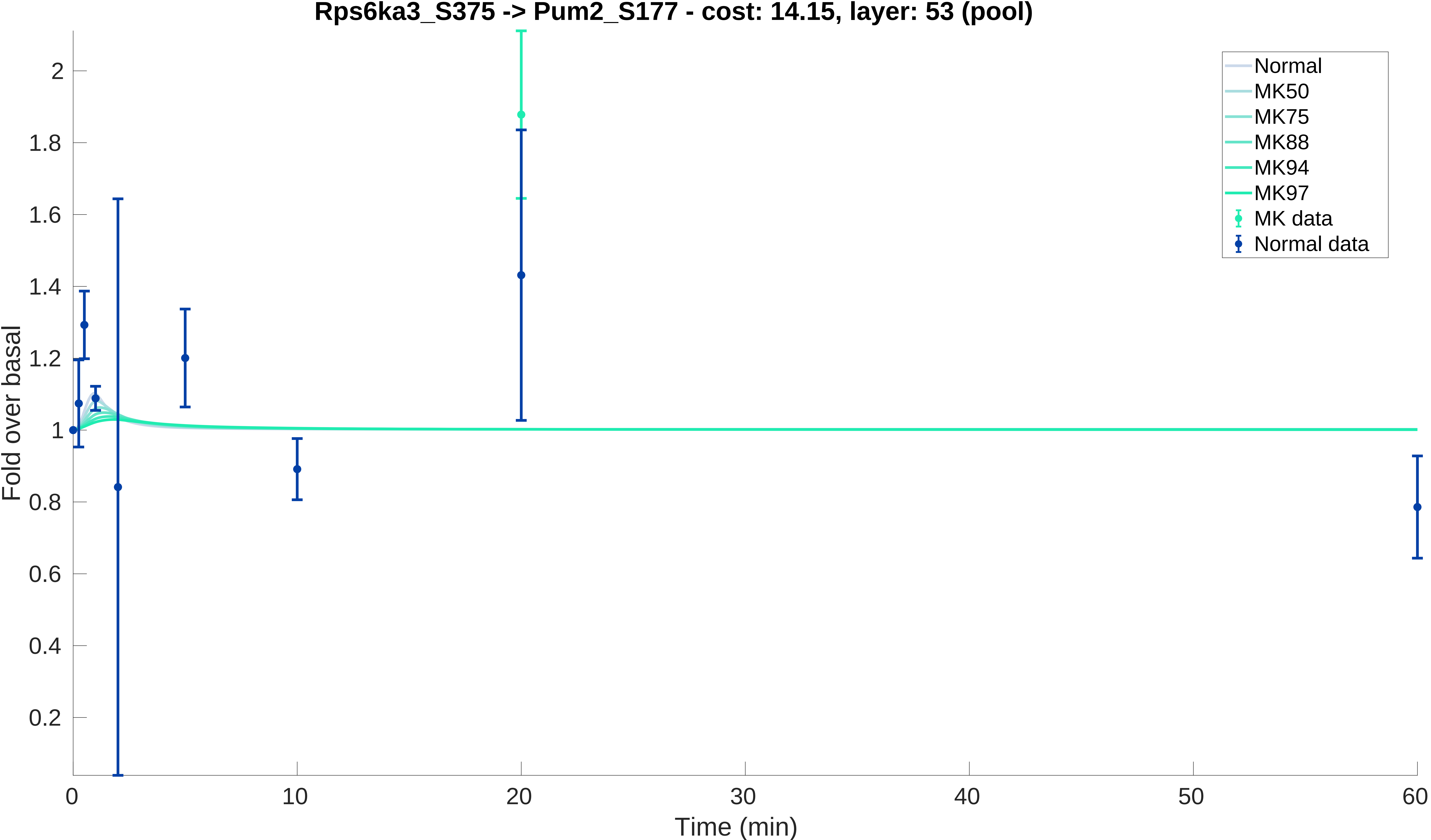
Rps6ka3_S375 -> Pum2_S177 - cost: 14.15, layer: 53 (pool)
Fold over basal
Time (min)
0
10
20
30
40
50
60
0.2
0.4
0.6
0.8
1
1.2
1.4
1.6
1.8
2
Normal
MK50
MK75
MK88
MK94
MK97
MK data
Normal data

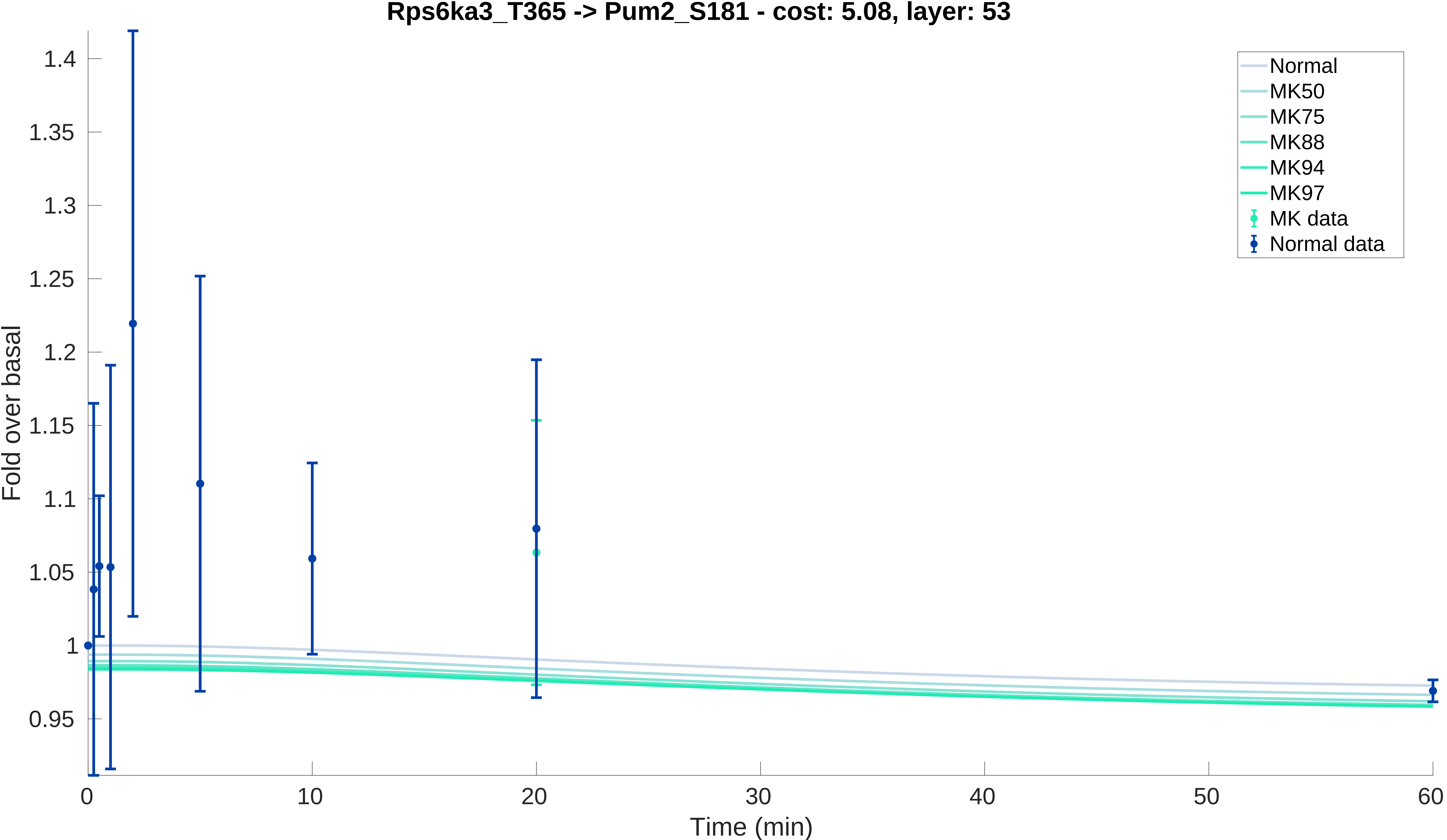

Rps6ka3_T365 -> Pum2_S181 - cost: 5.08, layer: 53
Normal
MK50
MK75
MK88
MK94
MK97
MK data
Normal data
Fold over basal
1.4
1.35
1.3
1.25
1.2
1.15
1.1
1.05
1
0.95
0
10
20
30
40
50
60
Time (min)

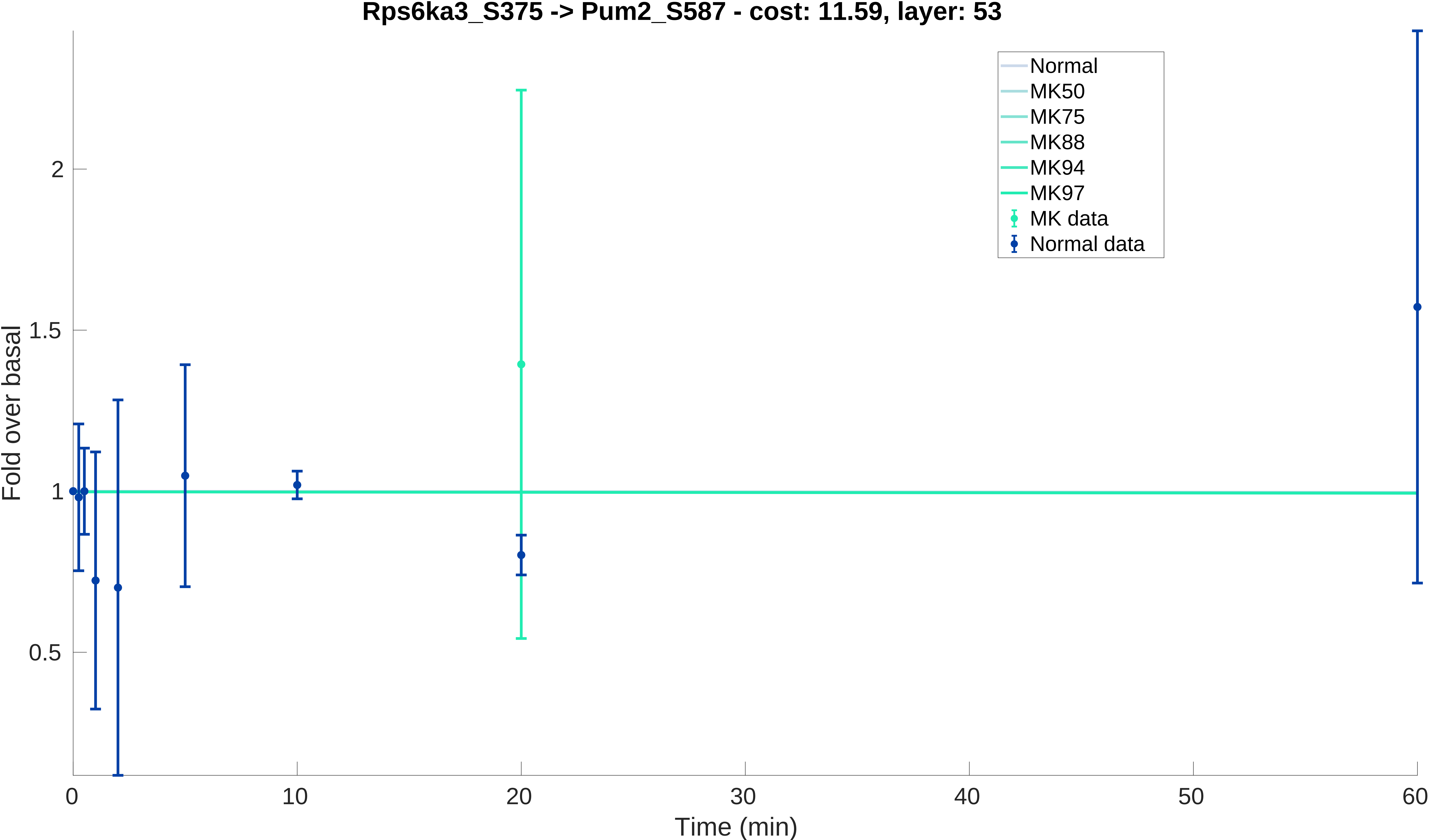
Rps6ka3_S375 -> Pum2_S587 - cost: 11.59, layer: 53
Fold over basal
Time (min)
0.5
1
1.5
2
0
10
20
30
40
50
60
Normal
MK50
MK75
MK88
MK94
MK97
MK data
Normal data

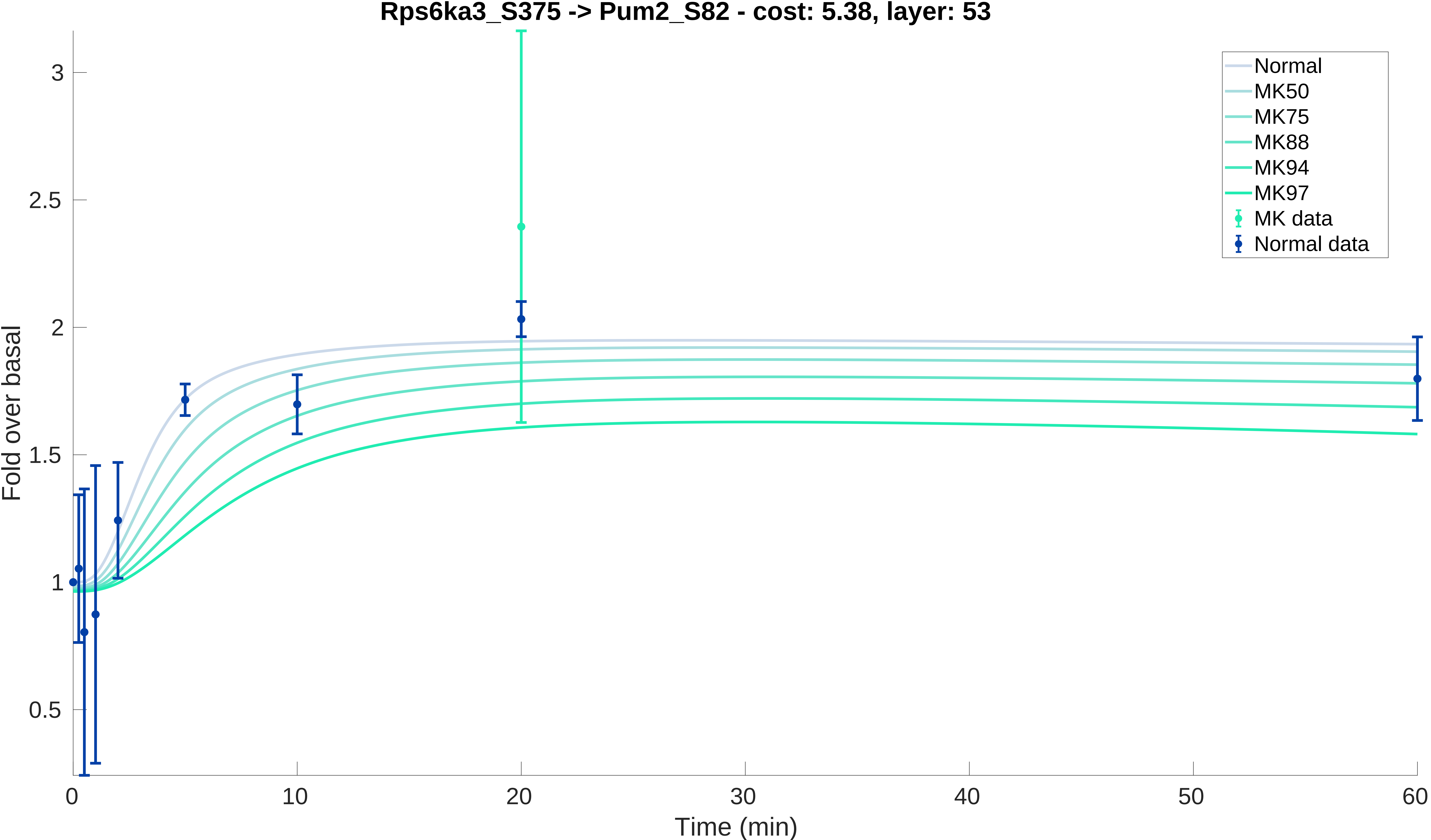
Rps6ka3_S375 -> Pum2_S82 - cost: 5.38, layer: 53
Fold over basal
Time (min)
0
10
20
30
40
50
60
0.5
1
1.5
2
2.5
3
Normal
MK50
MK75
MK88
MK94
MK97
MK data
Normal data

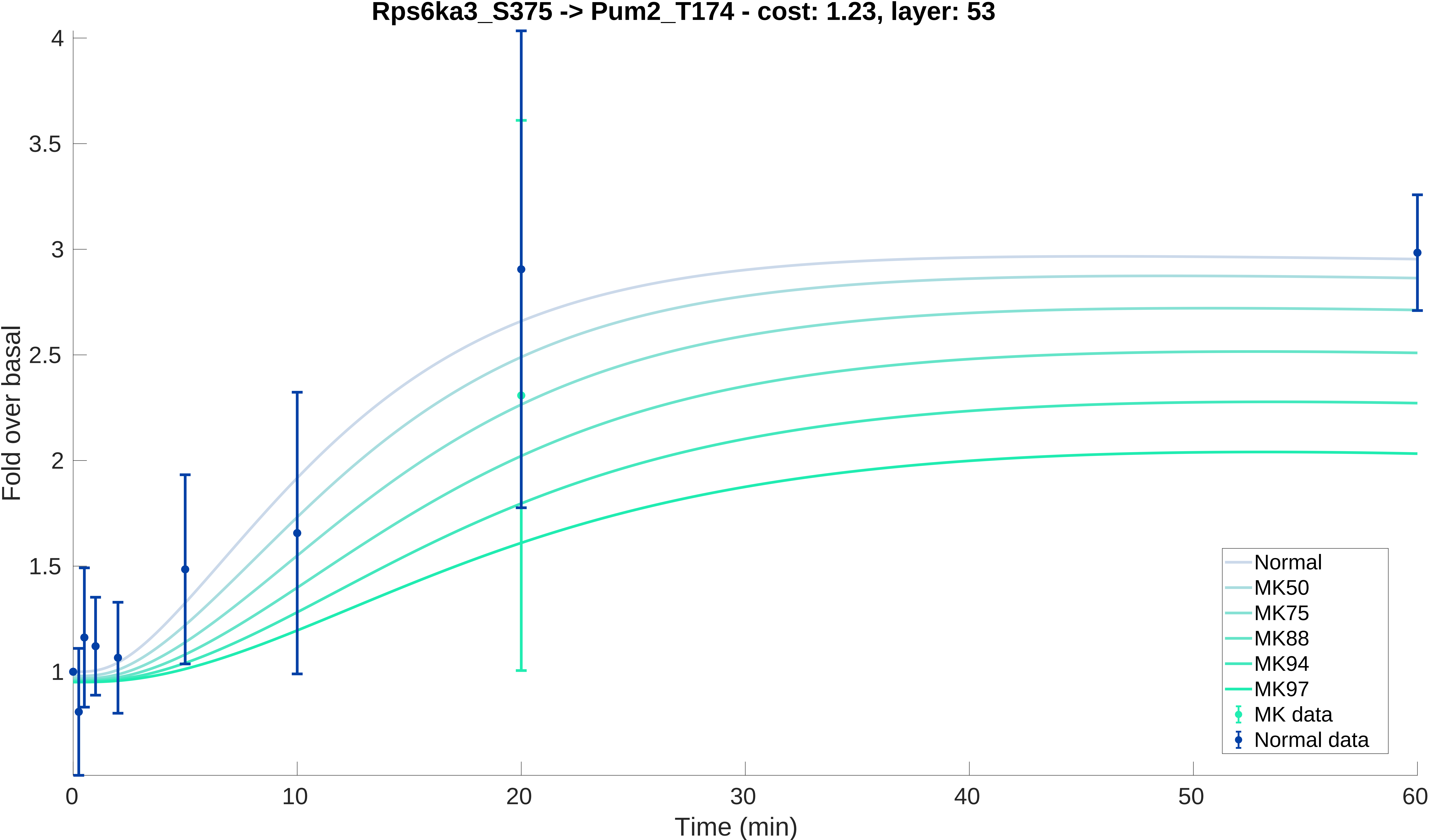
Rps6ka3_S375 -> Pum2_T174 - cost: 1.23, layer: 53
Fold over basal
Time (min)
0
10
20
30
40
50
60
1
1.5
2
2.5
3
3.5
4
Normal
MK50
MK75
MK88
MK94
MK97
MK data
Normal data

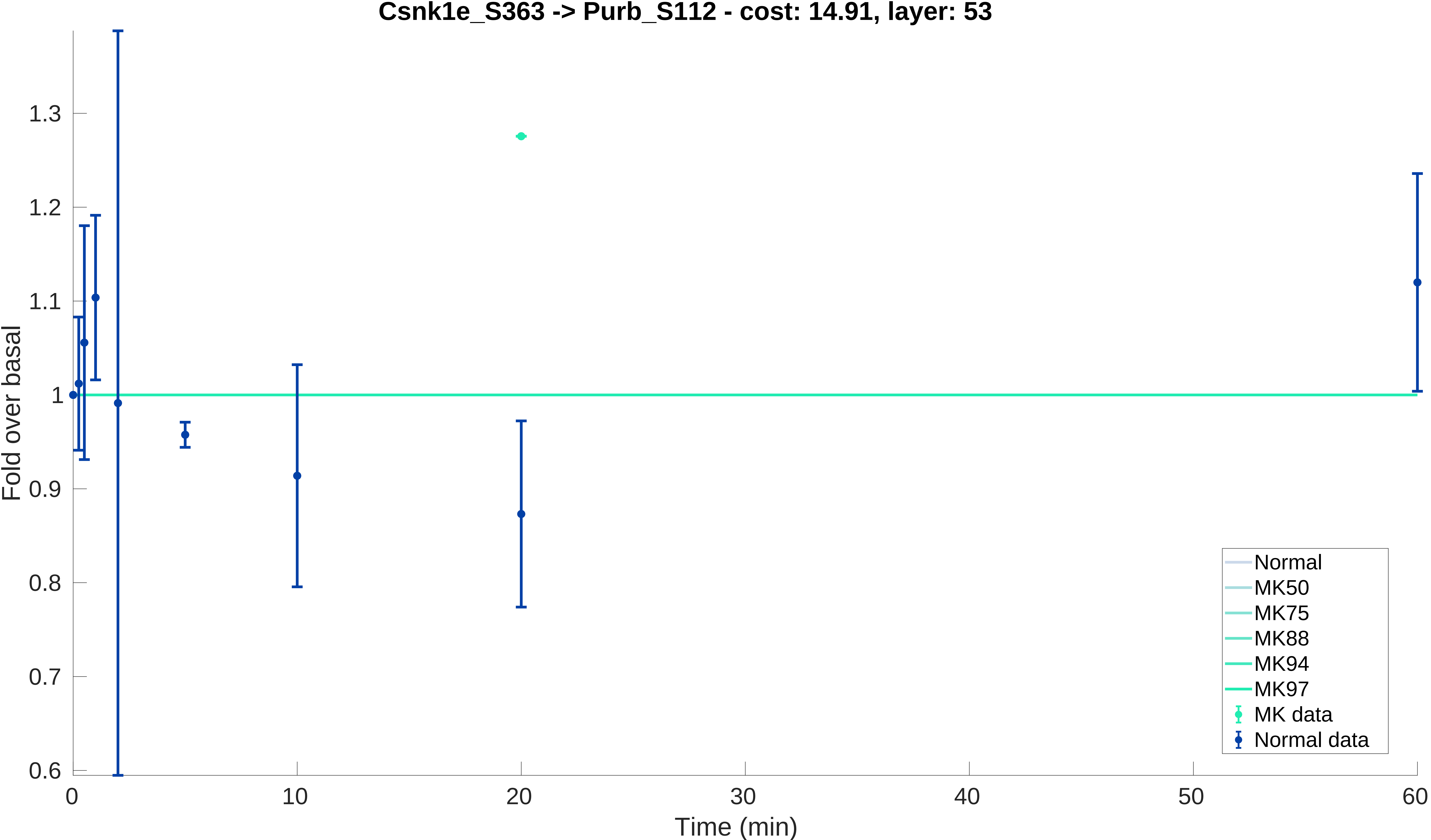

Csnk1e_S363 -> Purb_S112 - cost: 14.91, layer: 53
Fold over basal
1.3
1.2
1.1
1
0.9
0.8
0.7
0.6
0
10
20
30
40
50
60
Time (min)
Normal
MK50
MK75
MK88
MK94
MK97
MK data
Normal data

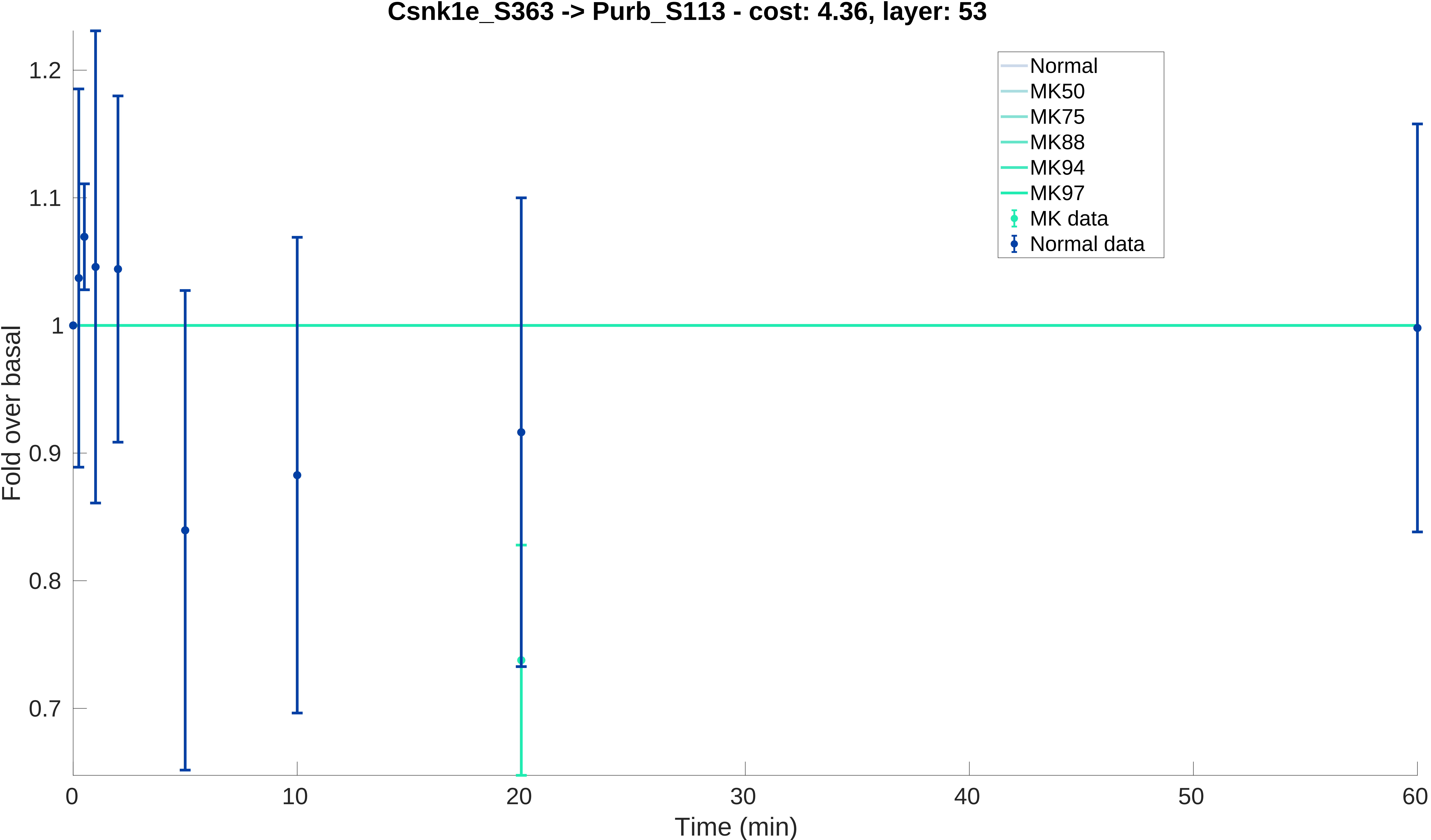
Csnk1e_S363 -> Purb_S113 - cost: 4.36, layer: 53
Fold over basal
Time (min)
0.7
0.8
0.9
1
1.1
1.2
0
10
20
30
40
50
60
Normal
MK50
MK75
MK88
MK94
MK97
MK data
Normal data

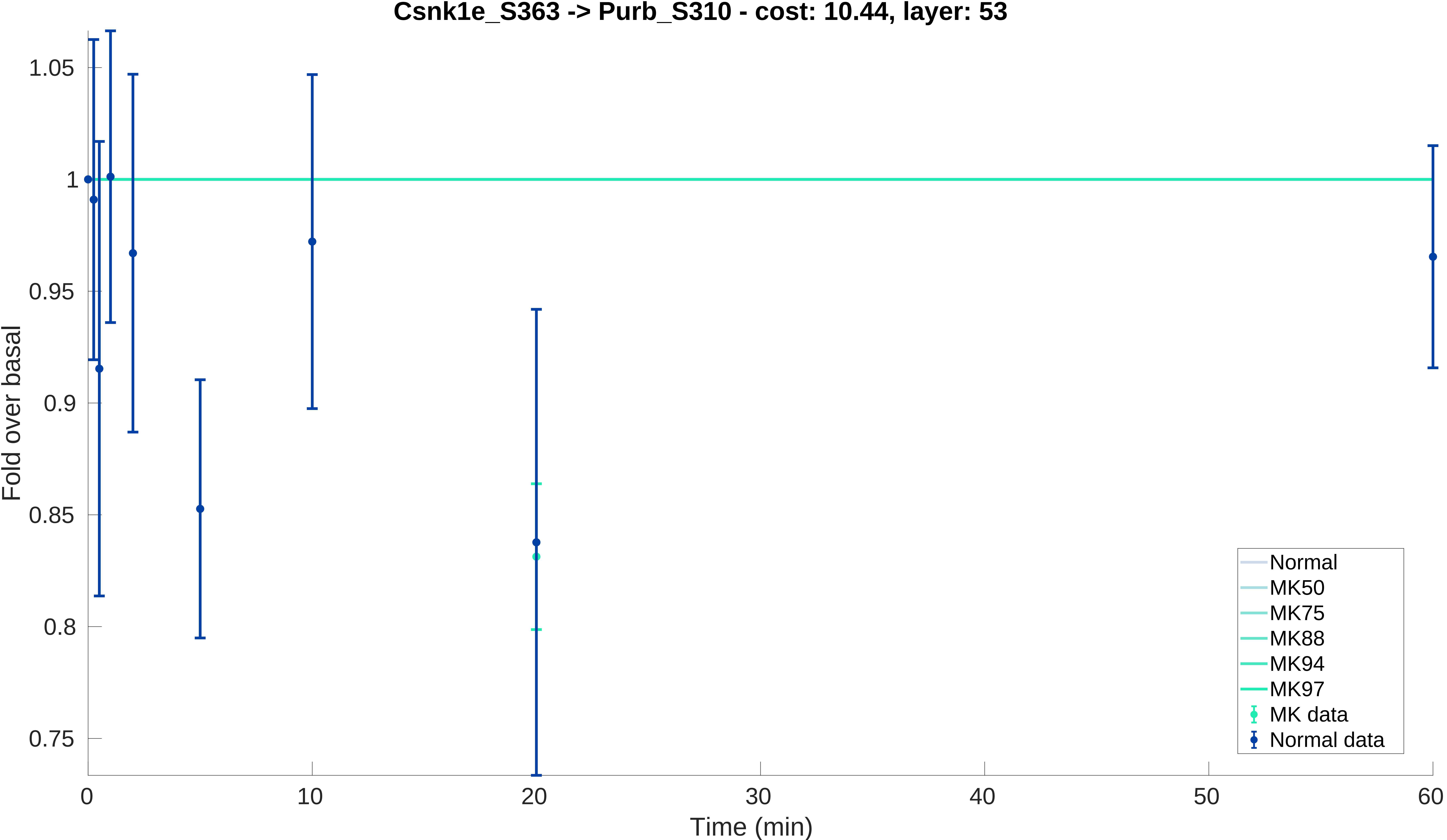

Csnk1e_S363 -> Purb_S310 - cost: 10.44, layer: 53
Fold over basal
1.05
1
0.95
0.9
0.85
0.8
0.75
0
10
20
30
40
50
60
Time (min)
Normal
MK50
MK75
MK88
MK94
MK97
MK data
Normal data

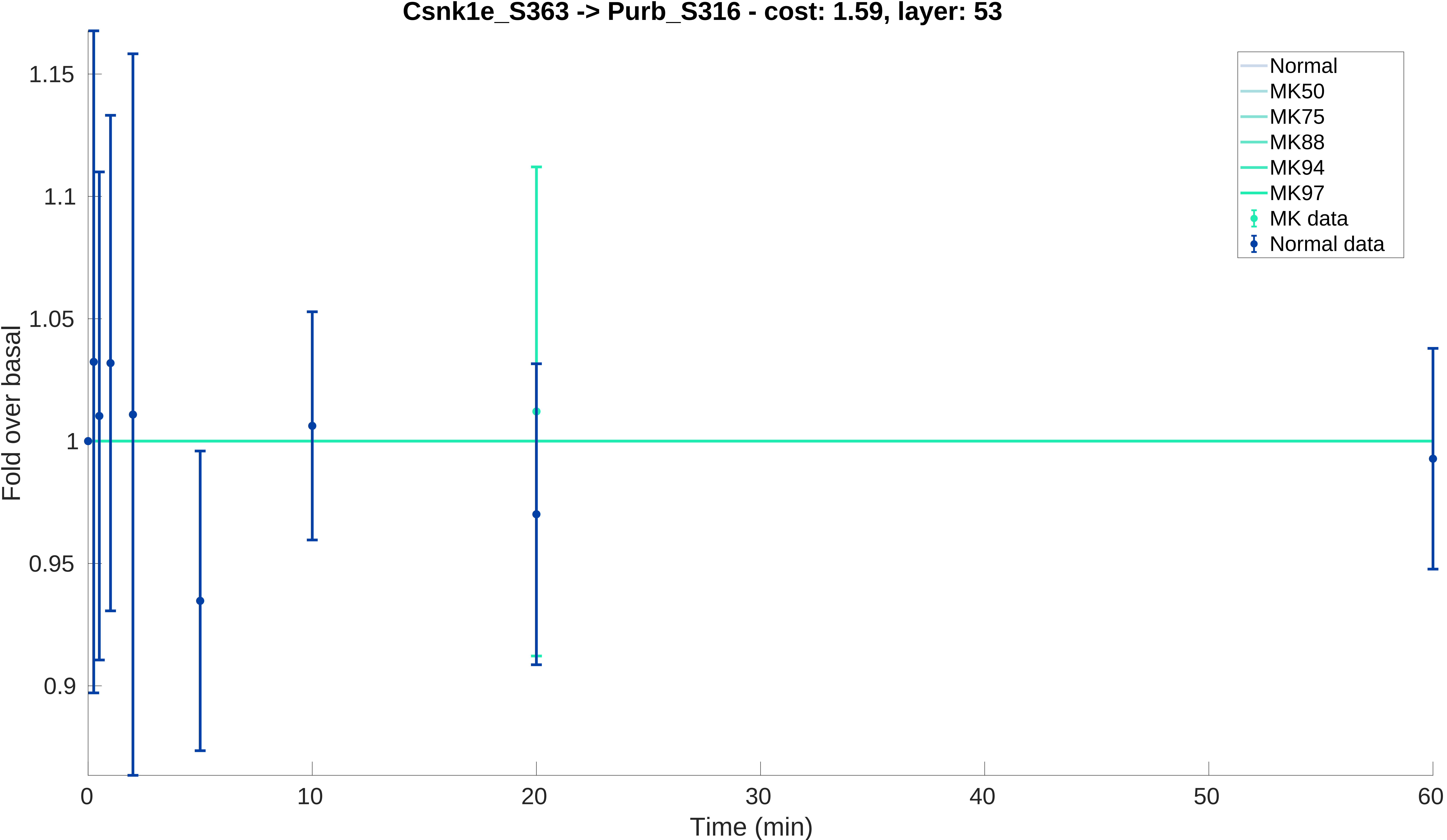

Csnk1e_S363 -> Purb_S316 - cost: 1.59, layer: 53
Fold over basal
Time (min)
1.15
1.1
1.05
1
0.95
0.9
0
10
20
30
40
50
60
Normal
MK50
MK75
MK88
MK94
MK97
MK data
Normal data

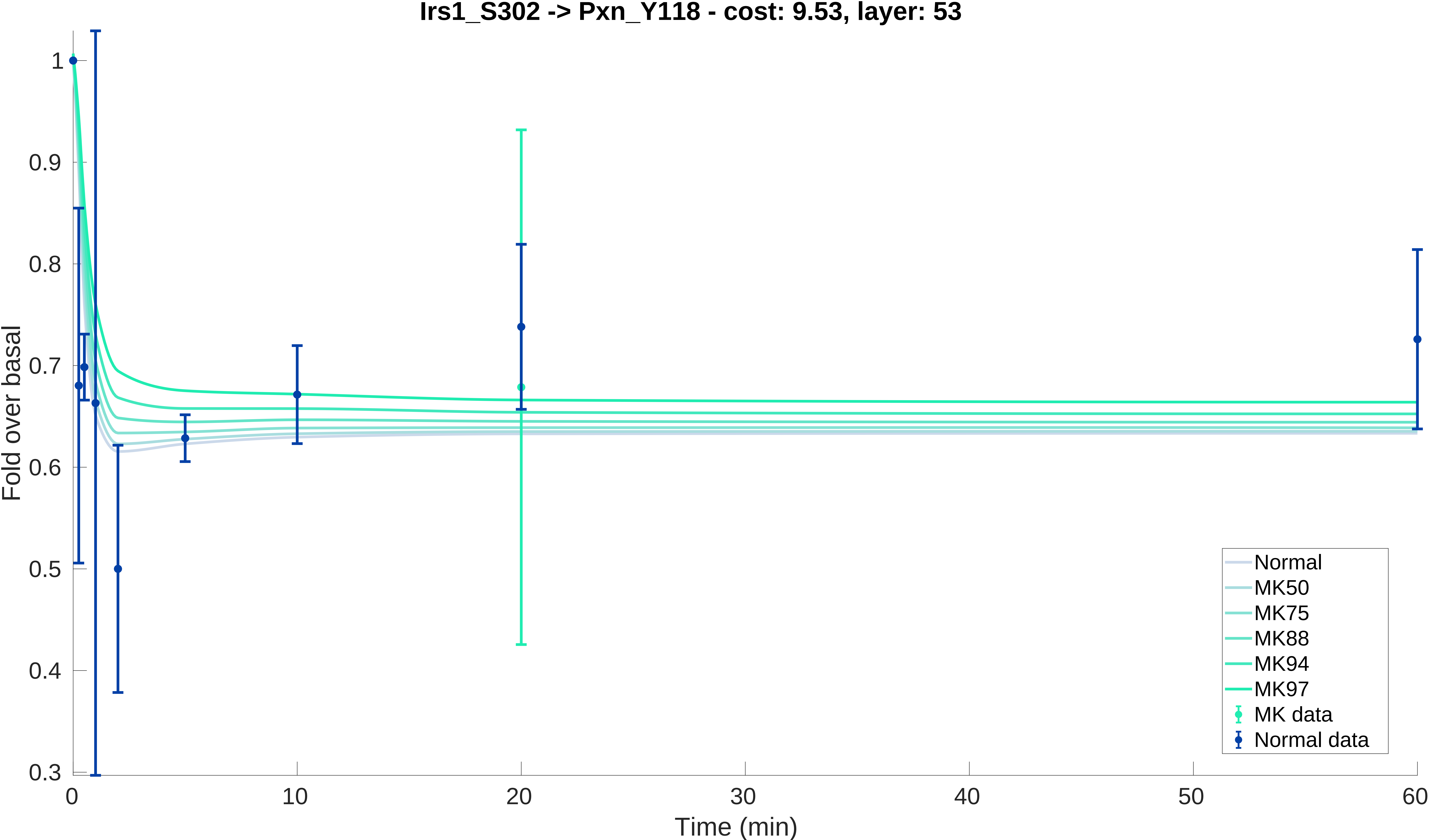

Irs1_S302 -> Pxn_Y118 - cost: 9.53, layer: 53
Fold over basal
Time (min)
1
0.9
0.8
0.7
0.6
0.5
0.4
0.3
0
10
20
30
40
50
60
Normal
MK50
MK75
MK88
MK94
MK97
MK data
Normal data

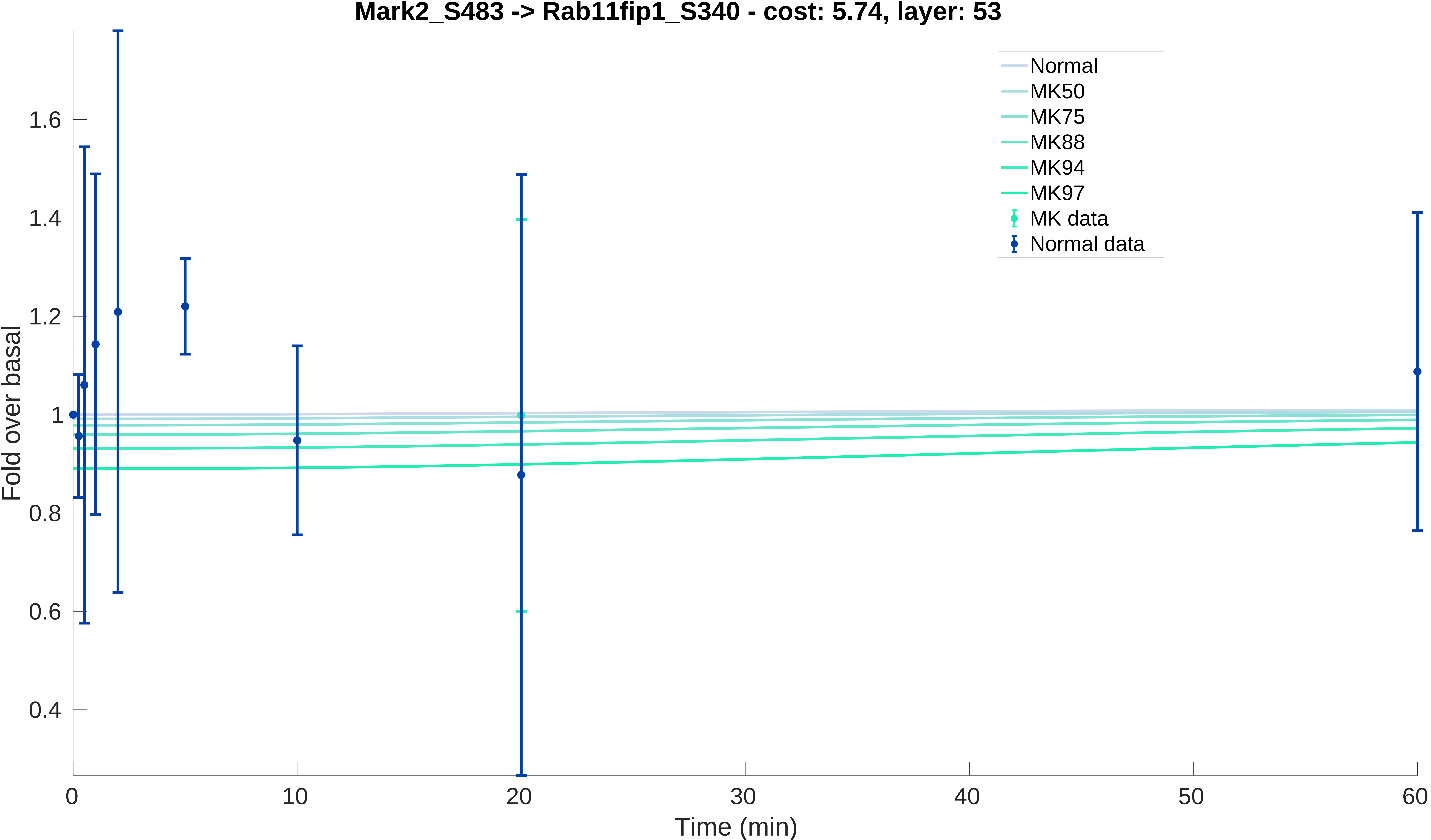
Mark2_S483 -> Rab11fip1_S340 - cost: 5.74, layer: 53
Fold over basal
1.6
1.4
1.2
1
0.8
0.6
0.4
0
10
20
30
40
50
60
Time (min)
Normal
MK50
MK75
MK88
MK94
MK97
MK data
Normal data

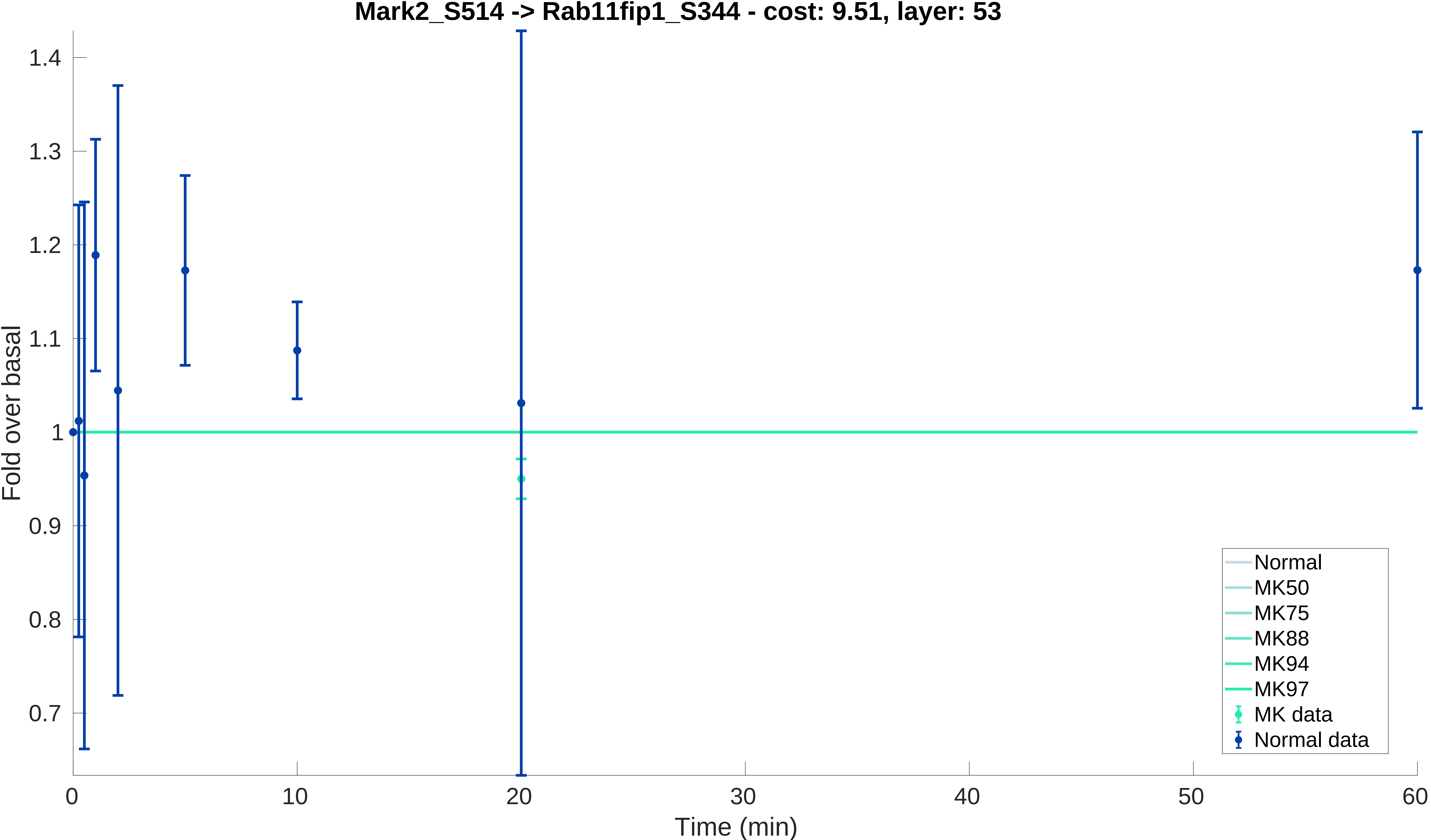
Mark2_S514 -> Rab11fip1_S344 - cost: 9.51, layer: 53
Fold over basal
1.4
1.3
1.2
1.1
1
0.9
0.8
0.7
Time (min)
0
10
20
30
40
50
60
Normal
MK50
MK75
MK88
MK94
MK97
MK data
Normal data

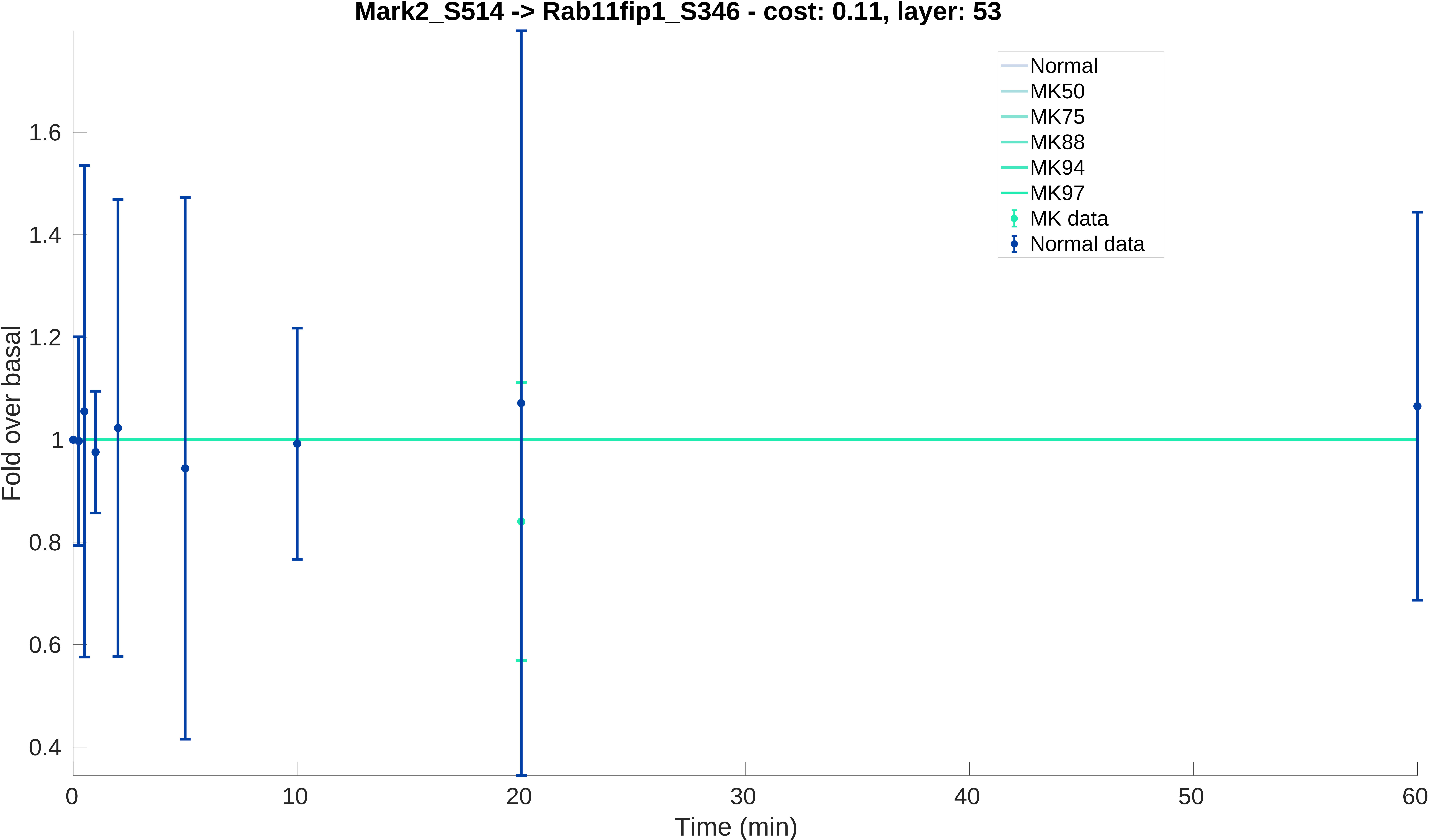
Mark2_S514 -> Rab11fip1_S346 - cost: 0.11, layer: 53
Fold over basal
Time (min)
0.4
0.6
0.8
1
1.2
1.4
1.6
0
10
20
30
40
50
60
Normal
MK50
MK75
MK88
MK94
MK97
MK data
Normal data

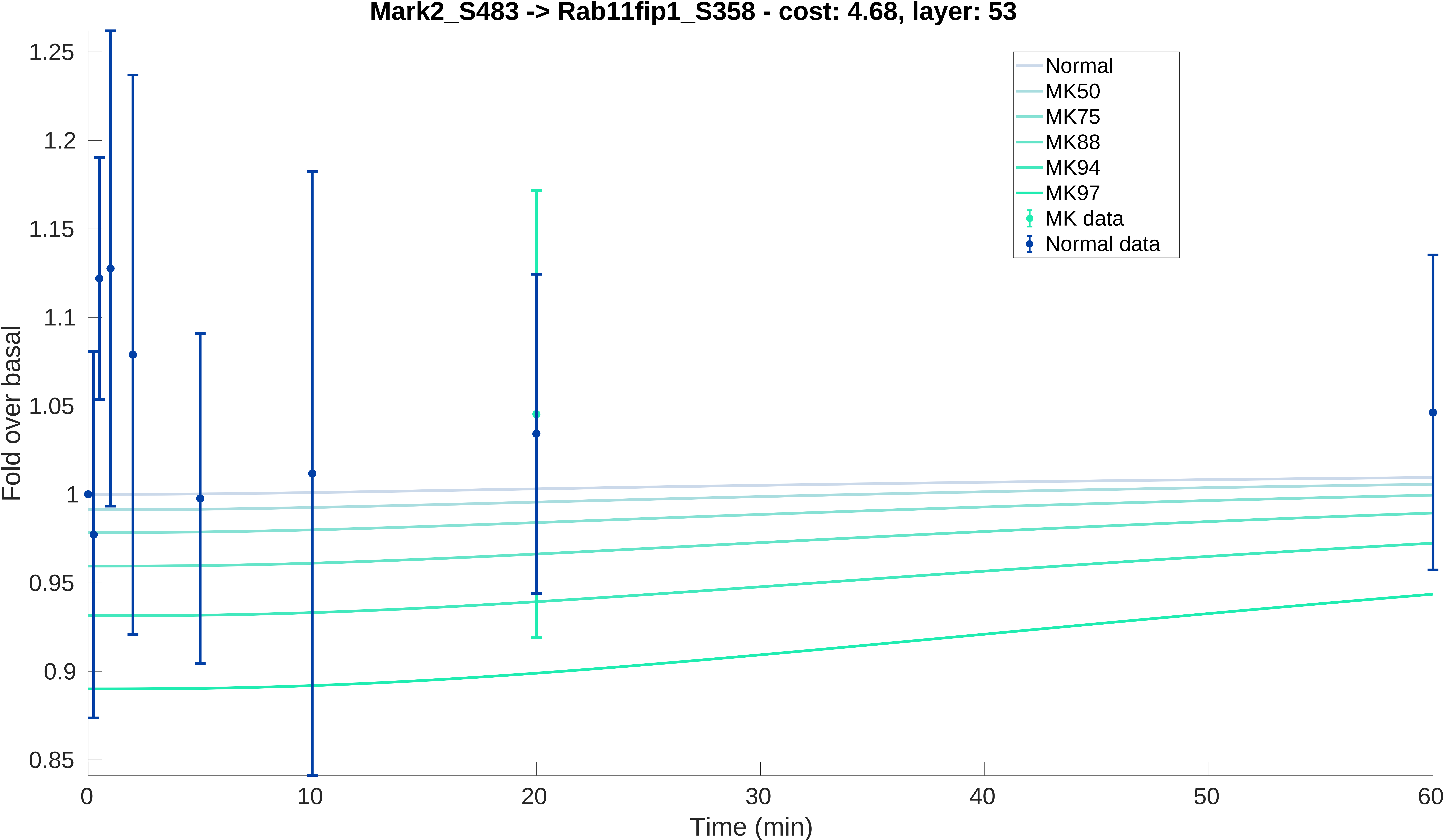
Mark2_S483 -> Rab11fip1_S358 - cost: 4.68, layer: 53
Fold over basal
Time (min)
1.25
1.2
1.15
1.1
1.05
1
0.95
0.9
0.85
0
10
20
30
40
50
60
Normal
MK50
MK75
MK88
MK94
MK97
MK data
Normal data

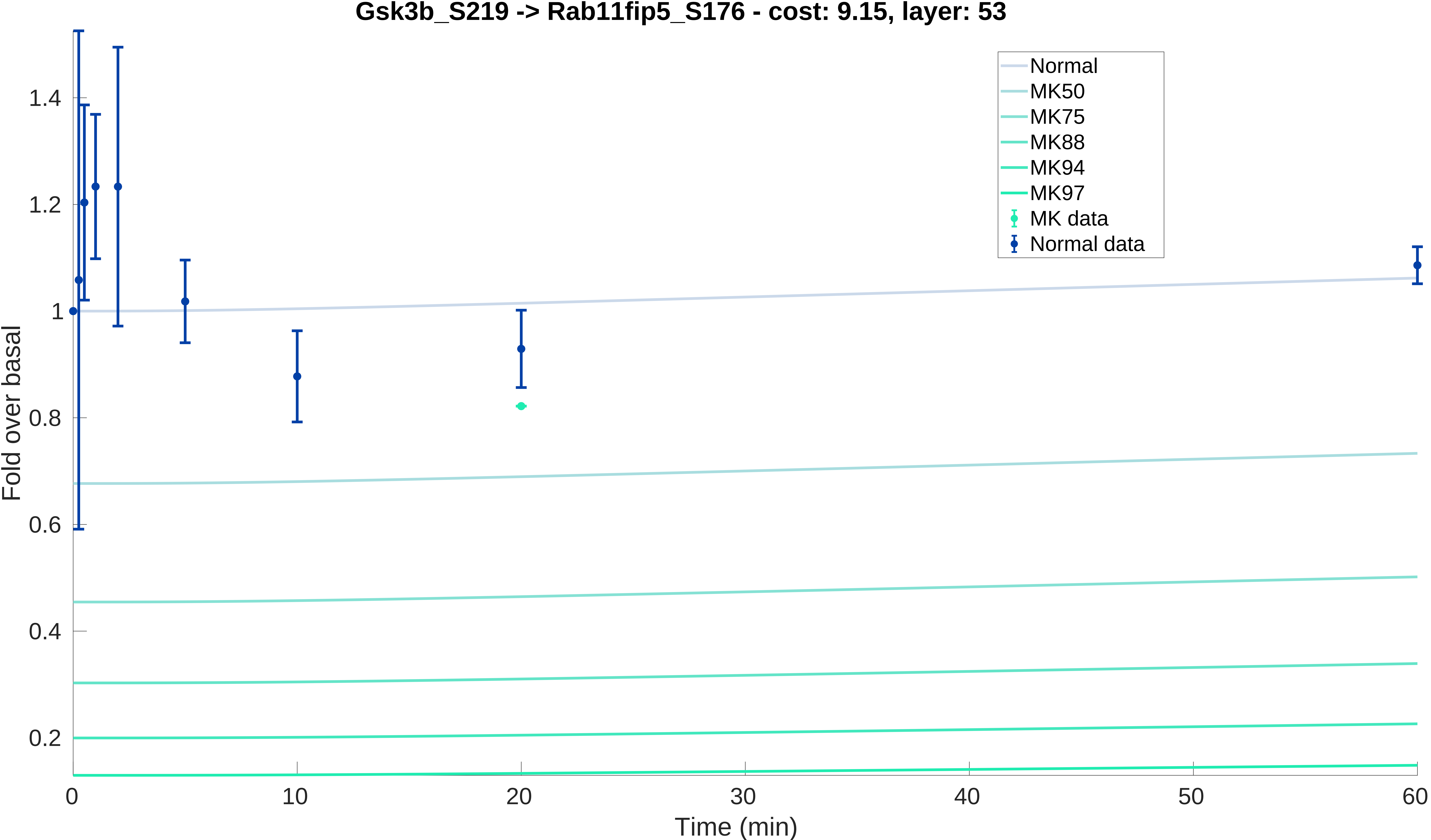
Gsk3b_S219 -> Rab11fip5_S176 - cost: 9.15, layer: 53
Normal
MK50
MK75
MK88
MK94
MK97
MK data
Normal data
Fold over basal
1.4
1.2
1
0.8
0.6
0.4
0.2
0
10
20
30
40
50
60
Time (min)

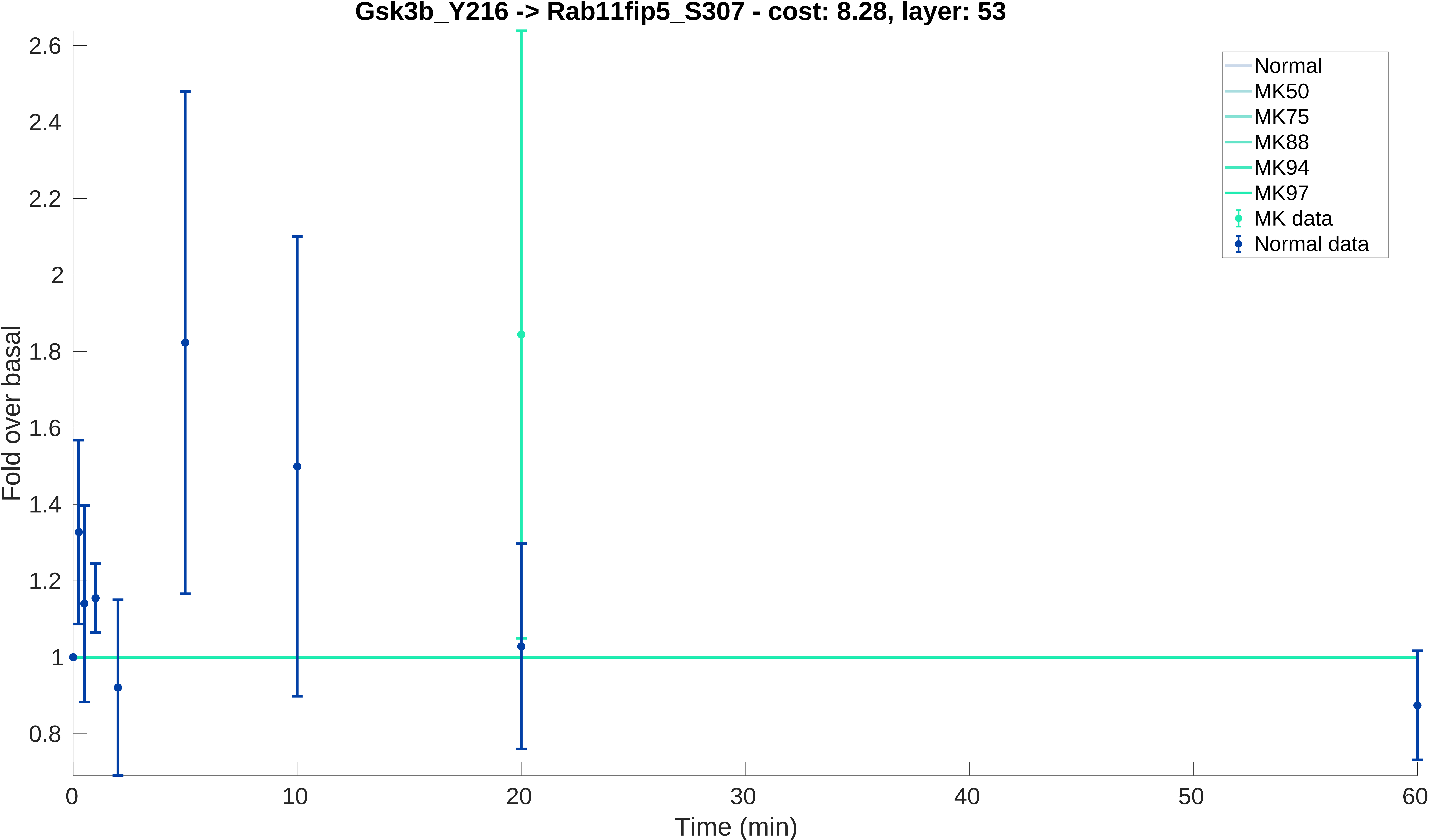
Gsk3b_Y216 -> Rab11fip5_S307 - cost: 8.28, layer: 53
Fold over basal
Time (min)
Normal
MK50
MK75
MK88
MK94
MK97
MK data
Normal data
2.6
2.4
2.2
2
1.8
1.6
1.4
1.2
1
0.8
0
10
20
30
40
50
60

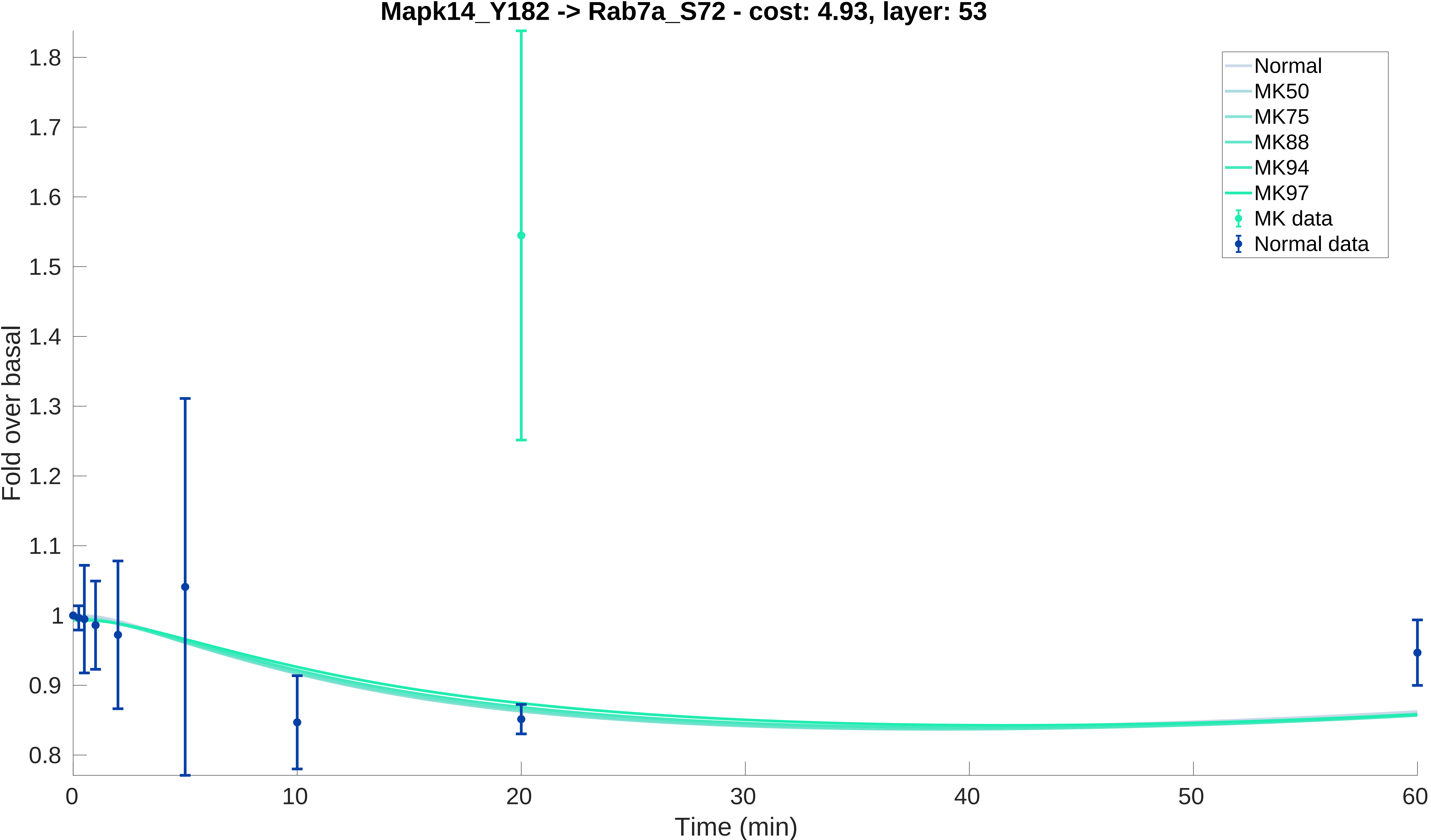

Mapk14_Y182 -> Rab7a_S72 - cost: 4.93, layer: 53
Fold over basal
Time (min)
0.8
0.9
1
1.1
1.2
1.3
1.4
1.5
1.6
1.7
1.8
0
10
20
30
40
50
60
Normal
MK50
MK75
MK88
MK94
MK97
MK data
Normal data

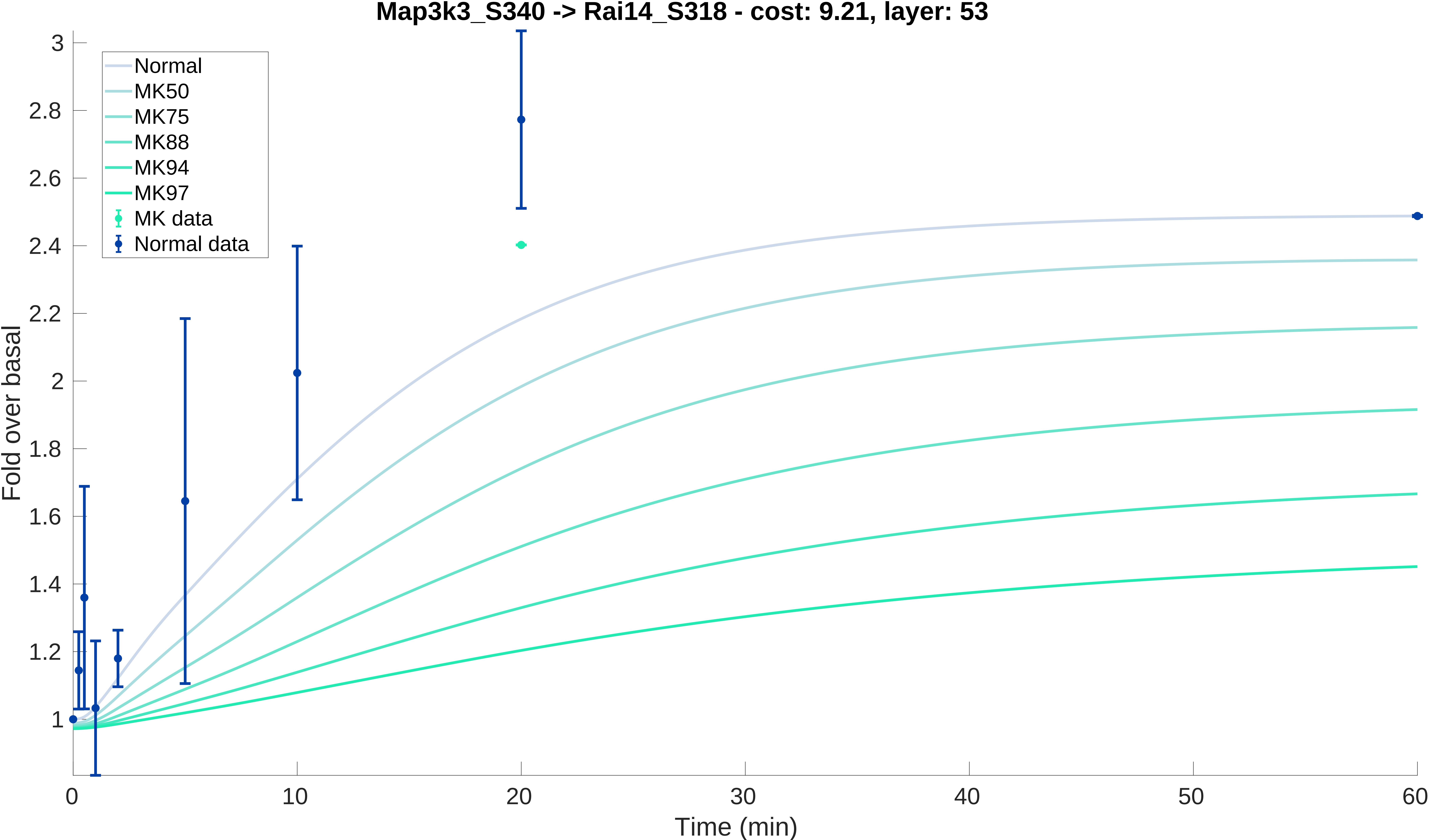

Map3k3_S340 -> Rai14_S318 - cost: 9.21, layer: 53
Normal
MK50
MK75
MK88
MK94
MK97
MK data
Normal data
Fold over basal
Time (min)
3
2.8
2.6
2.4
2.2
2
1.8
1.6
1.4
1.2
1
0
10
20
30
40
50
60

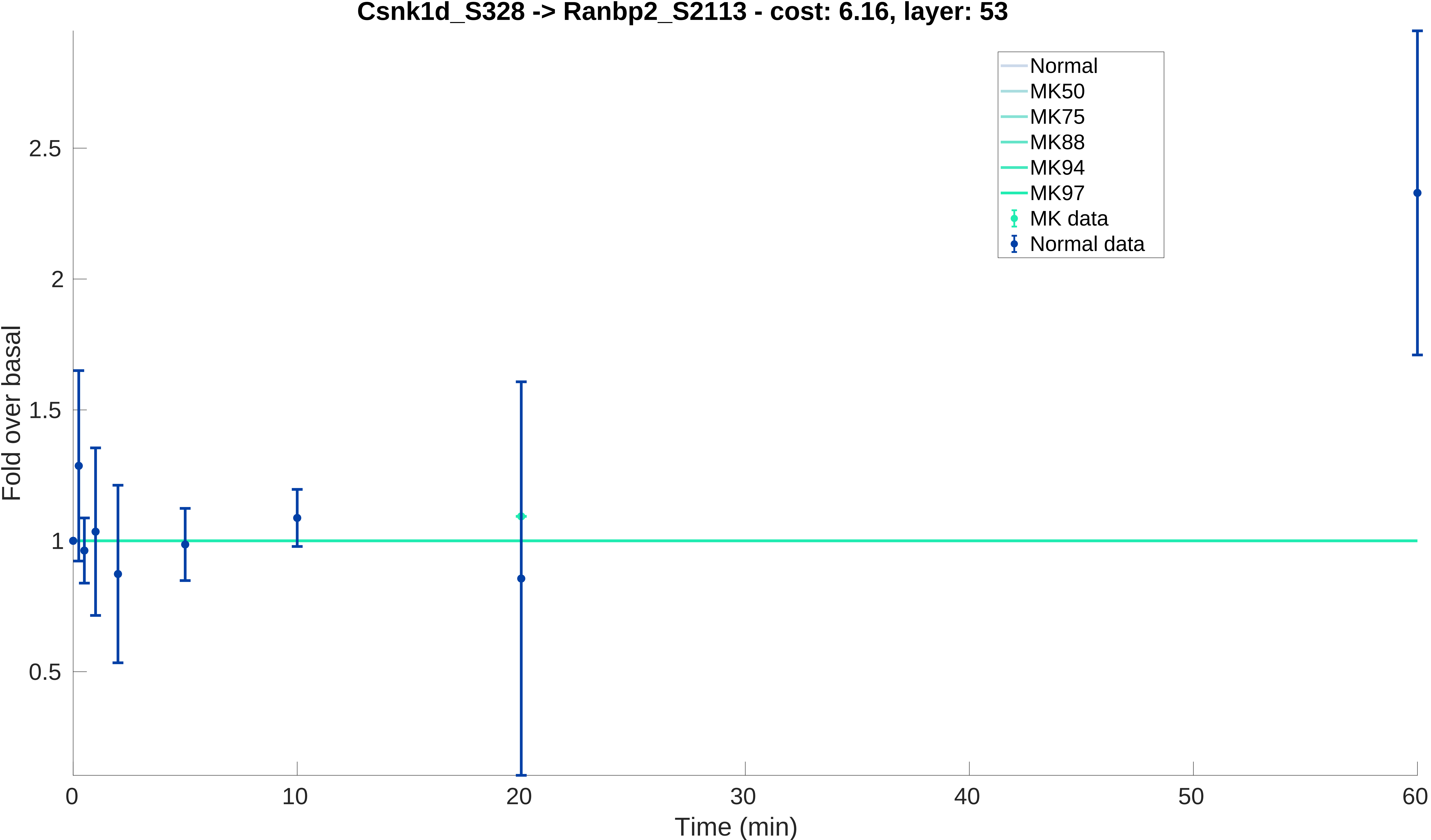

Csnk1d_S328 -> Ranbp2_S2113 - cost: 6.16, layer: 53
Normal
MK50
MK75
MK88
MK94
MK97
MK data
Normal data
Fold over basal
0.5
1
1.5
2
2.5
0
10
20
30
40
50
60
Time (min)

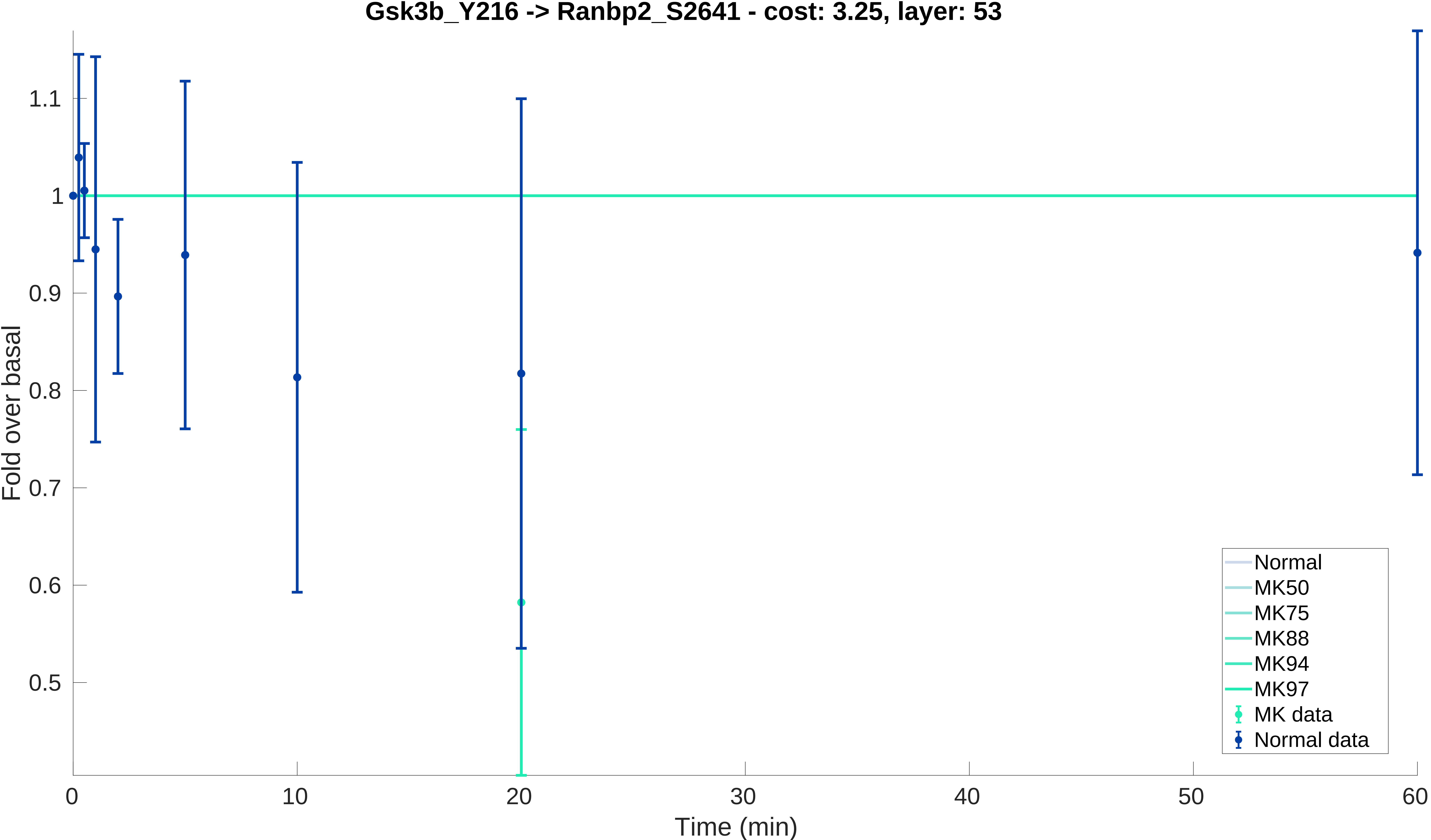

Gsk3b_Y216 -> Ranbp2_S2641 - cost: 3.25, layer: 53
1.1
1
0.9
0.8
0.7
0.6
0.5
Fold over basal
0
10
20
30
40
50
60
Time (min)
Normal
MK50
MK75
MK88
MK94
MK97
MK data
Normal data

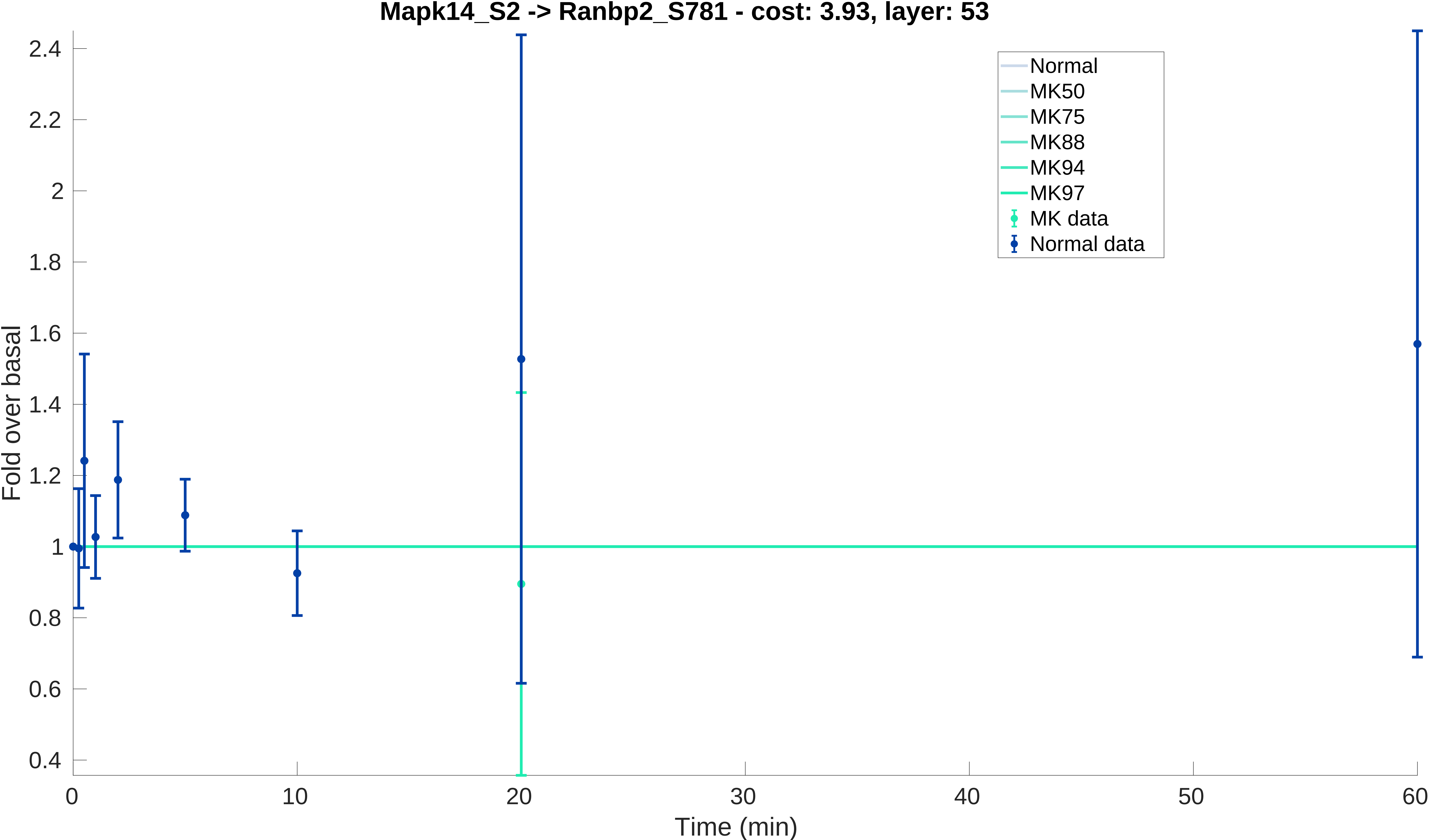

Mapk14_S2 -> Ranbp2_S781 - cost: 3.93, layer: 53
Fold over basal
Time (min)
2.4
2.2
2
1.8
1.6
1.4
1.2
1
0.8
0.6
0.4
0
10
20
30
40
50
60
Normal
MK50
MK75
MK88
MK94
MK97
MK data
Normal data

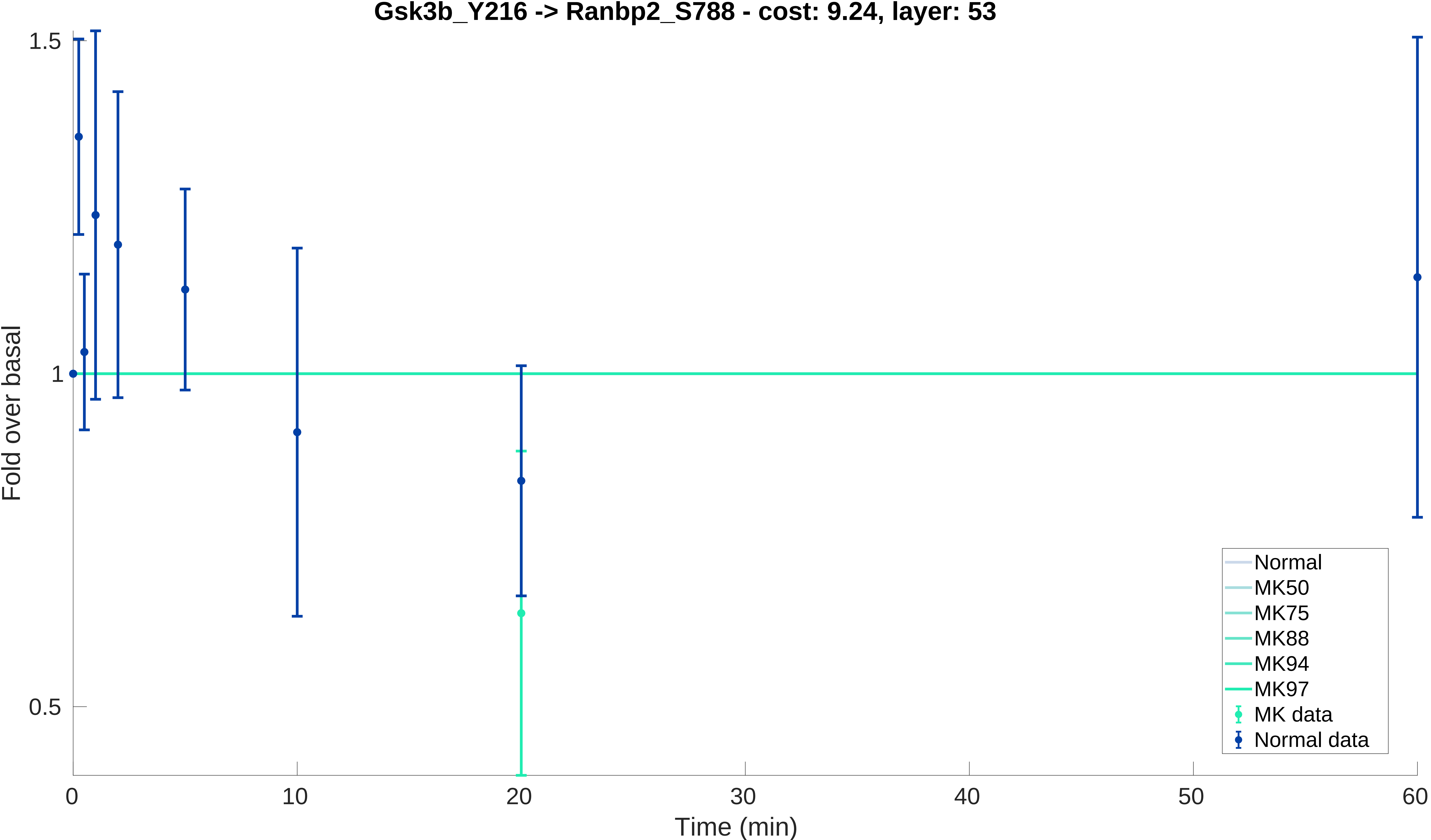
Gsk3b_Y216 -> Ranbp2_S788 - cost: 9.24, layer: 53
Fold over basal
1.5
1
0.5
0
10
20
30
40
50
60
Time (min)
Normal
MK50
MK75
MK88
MK94
MK97
MK data
Normal data

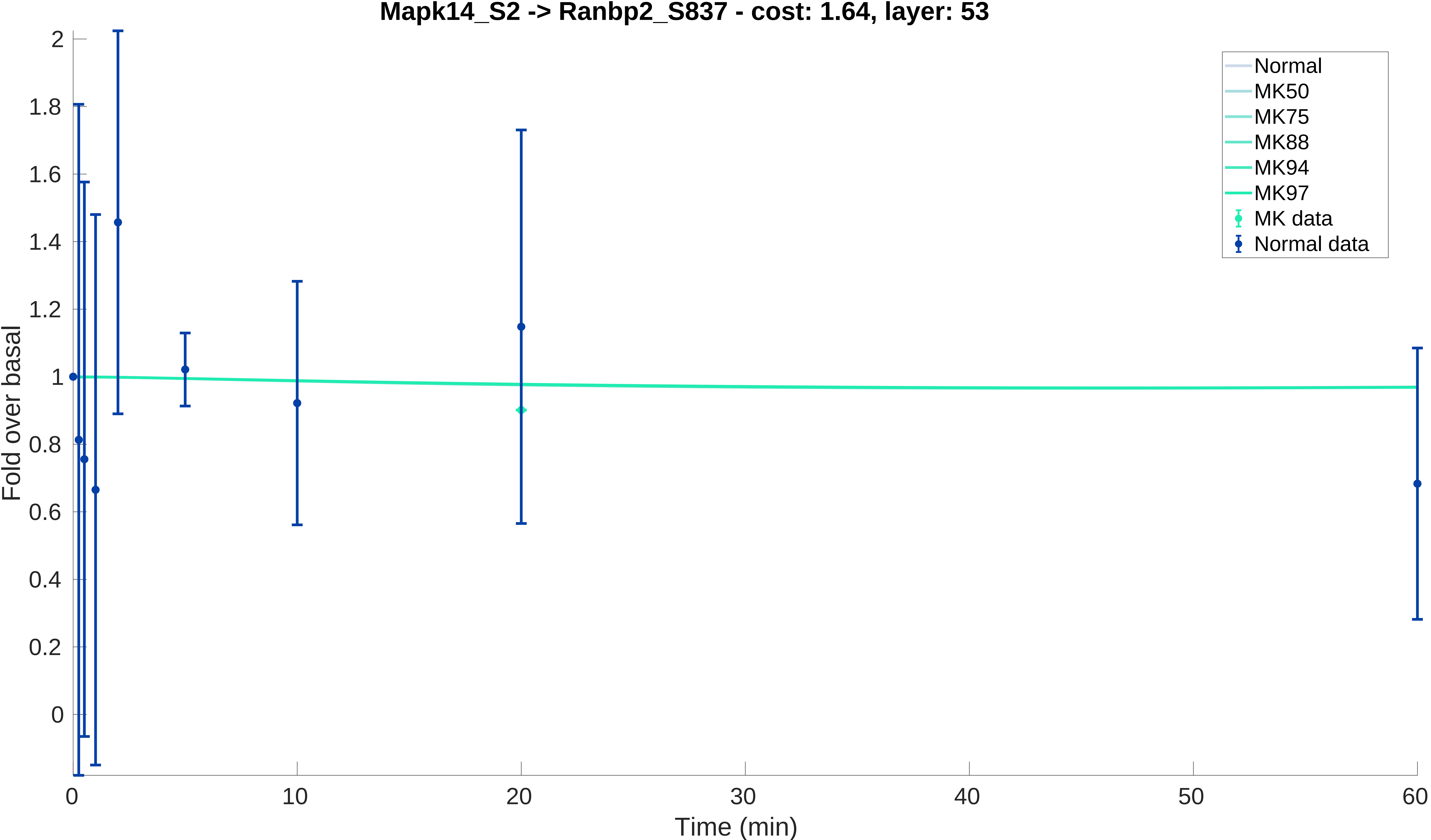
Mapk14_S2 -> Ranbp2_S837 - cost: 1.64, layer: 53
Fold over basal
Time (min)
2
1.8
1.6
1.4
1.2
1
0.8
0.6
0.4
0.2
0
0
10
20
30
40
50
60
Normal
MK50
MK75
MK88
MK94
MK97
MK data
Normal data

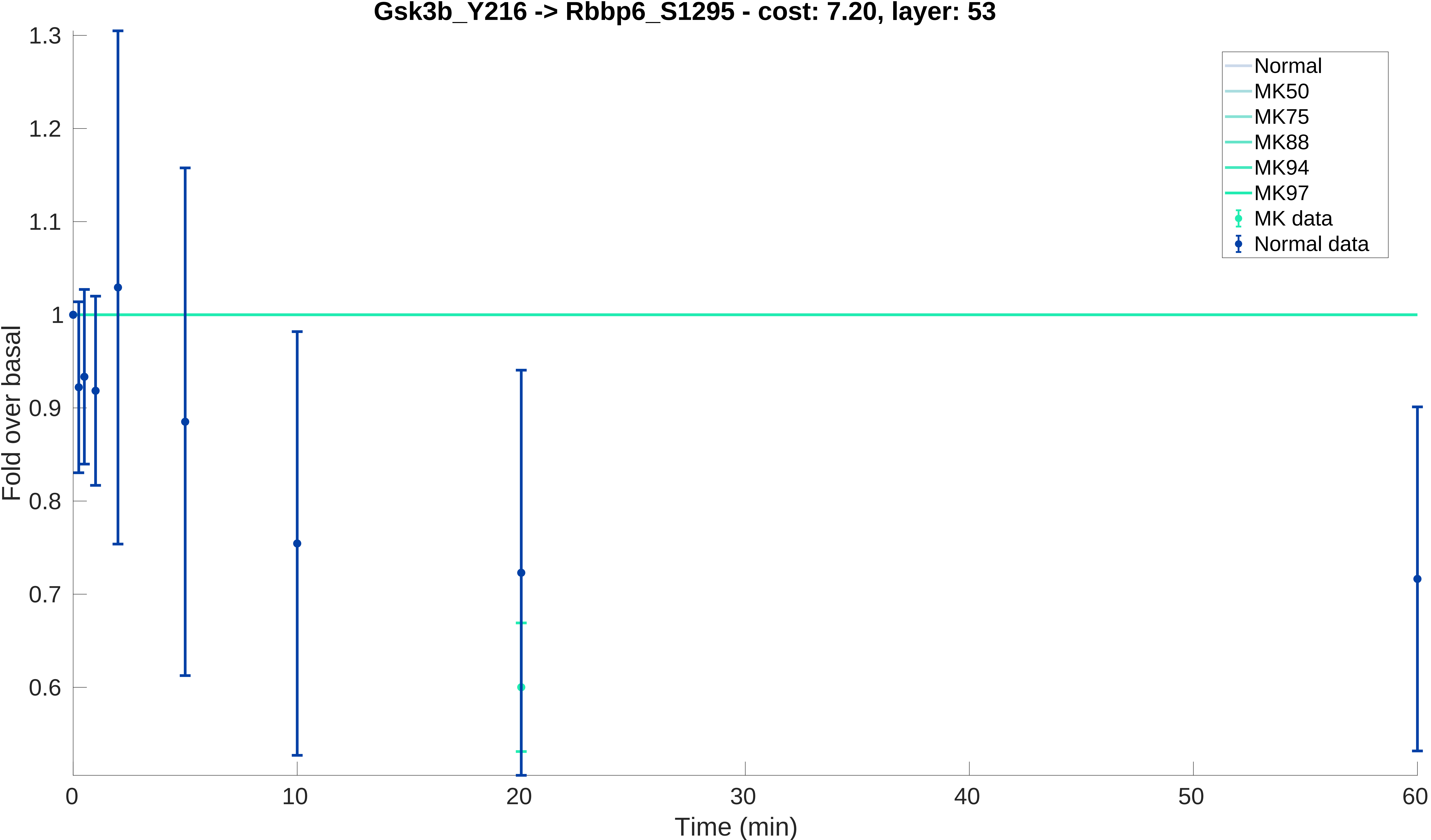
Gsk3b_Y216 -> Rbbp6_S1295 - cost: 7.20, layer: 53
Fold over basal
Time (min)
1.3
1.2
1.1
1
0.9
0.8
0.7
0.6
0
10
20
30
40
50
60
Normal
MK50
MK75
MK88
MK94
MK97
MK data
Normal data

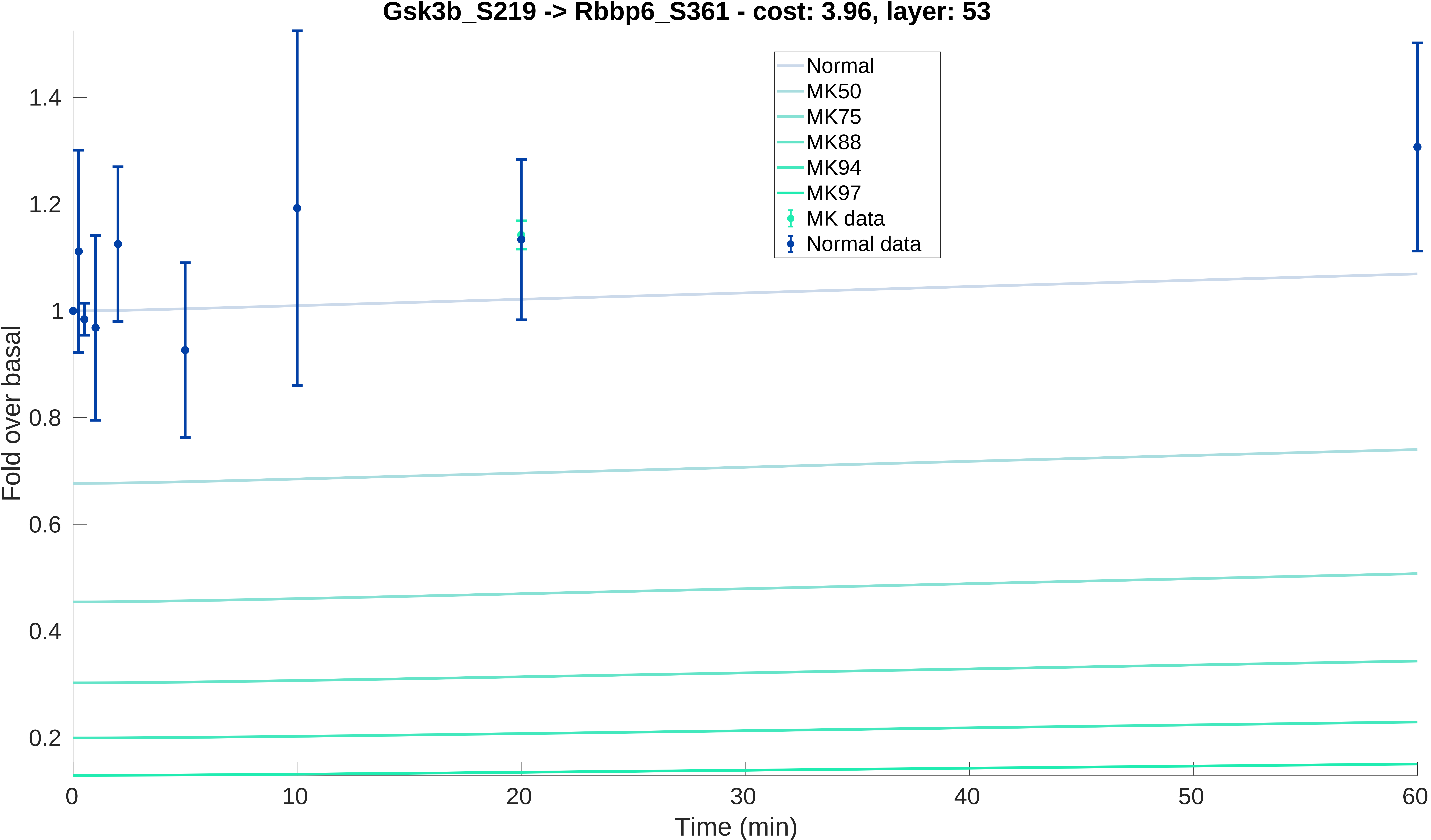

Gsk3b_S219 -> Rbbp6_S361 - cost: 3.96, layer: 53
Normal
MK50
MK75
MK88
MK94
MK97
MK data
Normal data
Fold over basal
1.4
1.2
1
0.8
0.6
0.4
0.2
0
10
20
30
40
50
60
Time (min)

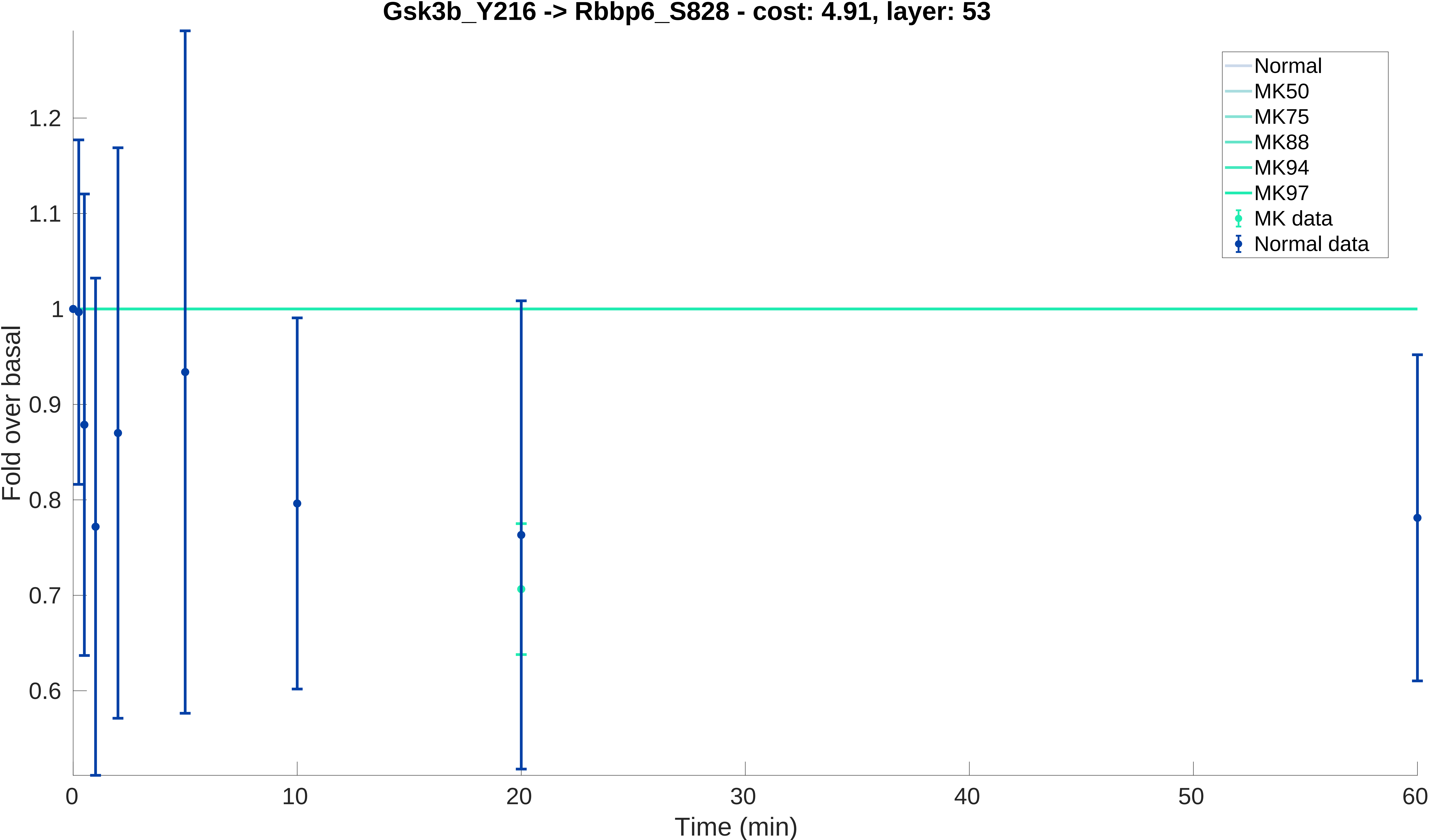

Gsk3b_Y216 -> Rbbp6_S828 - cost: 4.91, layer: 53
Fold over basal
Time (min)
1.2
1.1
1
0.9
0.8
0.7
0.6
0
10
20
30
40
50
60
Normal
MK50
MK75
MK88
MK94
MK97
MK data
Normal data

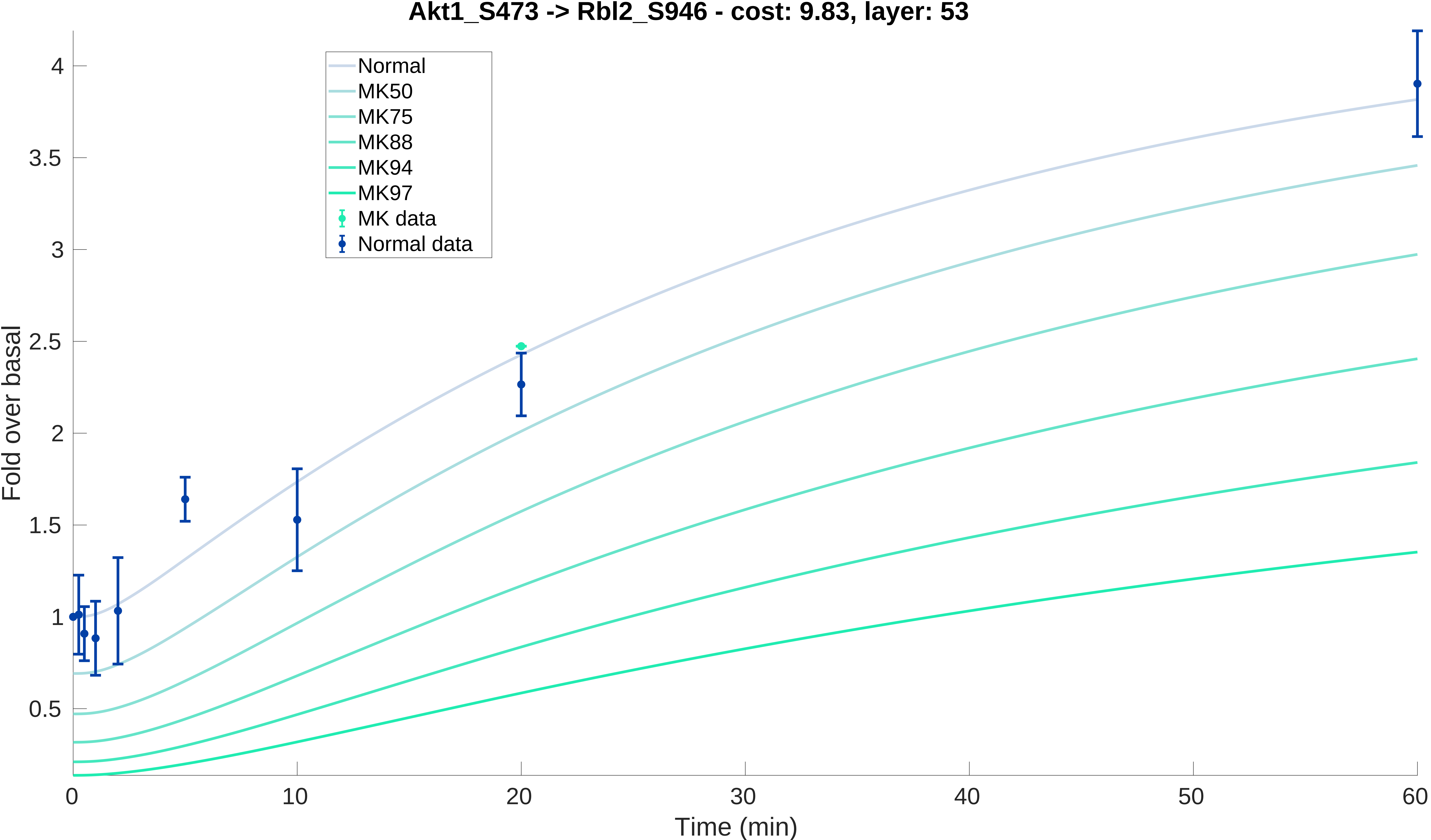

Akt1_S473 -> Rbl2_S946 - cost: 9.83, layer: 53
Normal
MK50
MK75
MK88
MK94
MK97
MK data
Normal data
Fold over basal
Time (min)
0.5
1
1.5
2
2.5
3
3.5
4
0
10
20
30
40
50
60

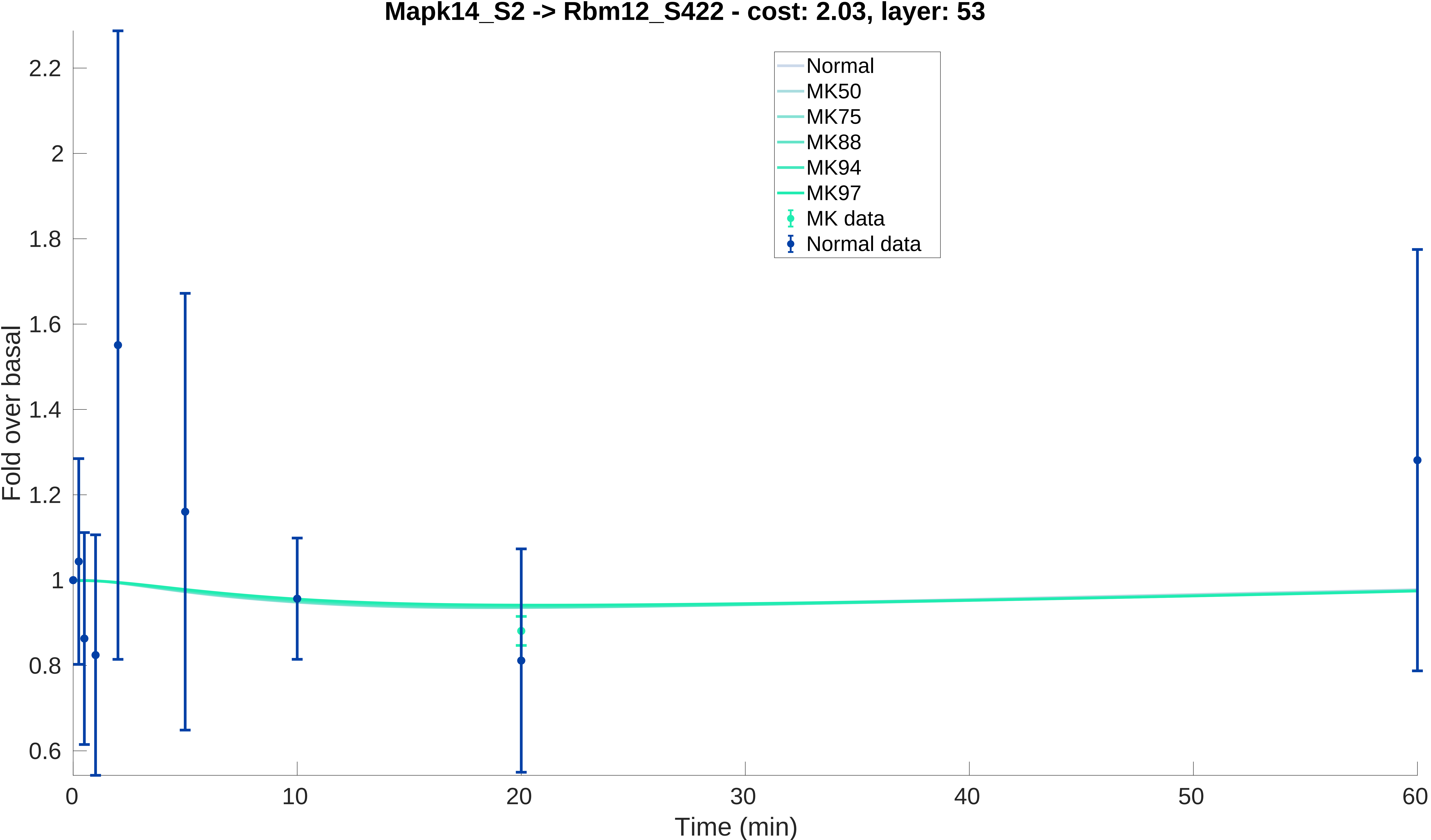

Mapk14_S2 -> Rbm12_S422 - cost: 2.03, layer: 53
Normal
MK50
MK75
MK88
MK94
MK97
MK data
Normal data
Fold over basal
2.2
2
1.8
1.6
1.4
1.2
1
0.8
0.6
0
10
20
30
40
50
60
Time (min)

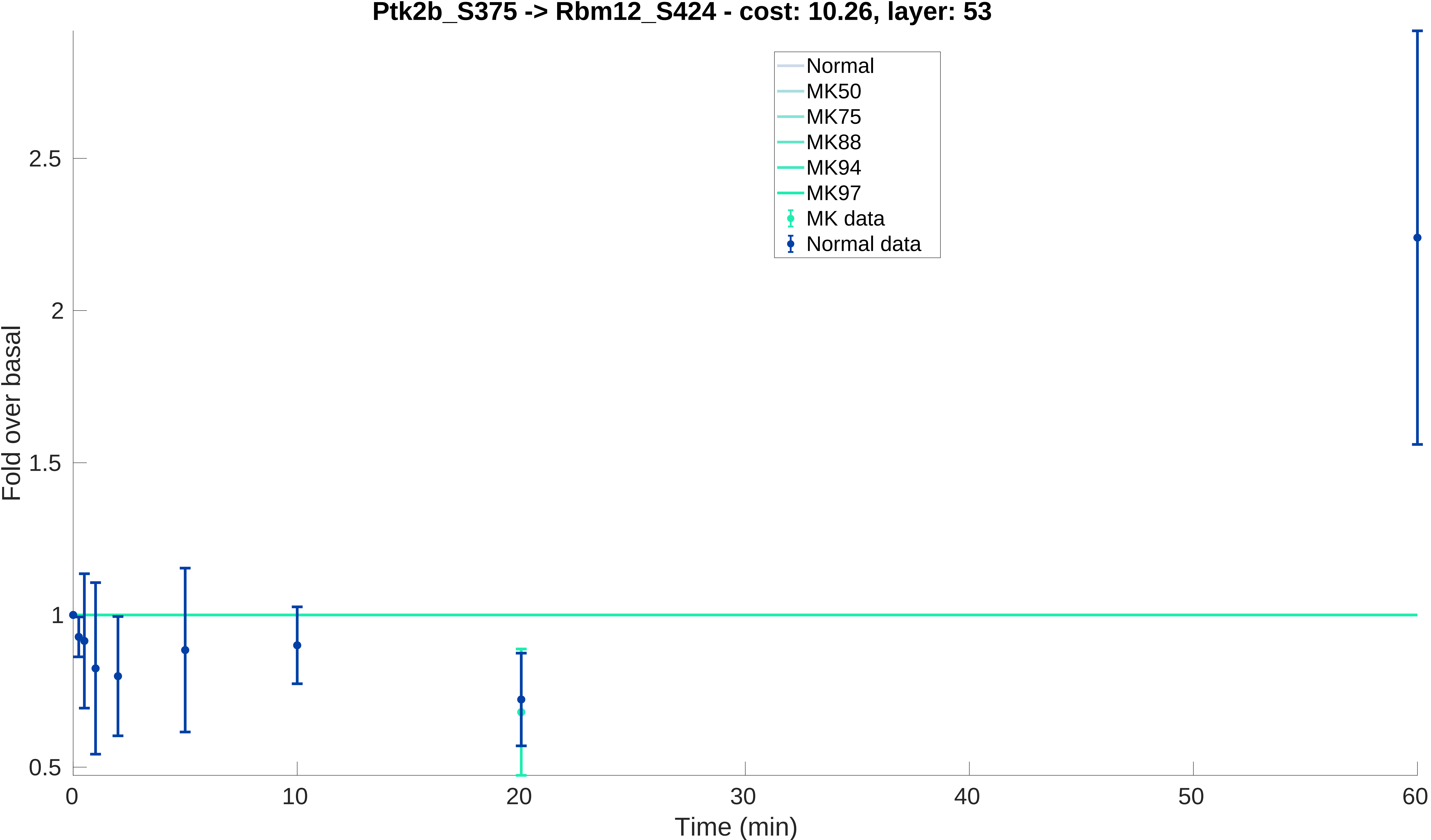
Ptk2b_S375 -> Rbm12_S424 - cost: 10.26, layer: 53
Normal
MK50
MK75
MK88
MK94
MK97
MK data
Normal data
Fold over basal
0.5
1
1.5
2
2.5
Time (min)
0
10
20
30
40
50
60

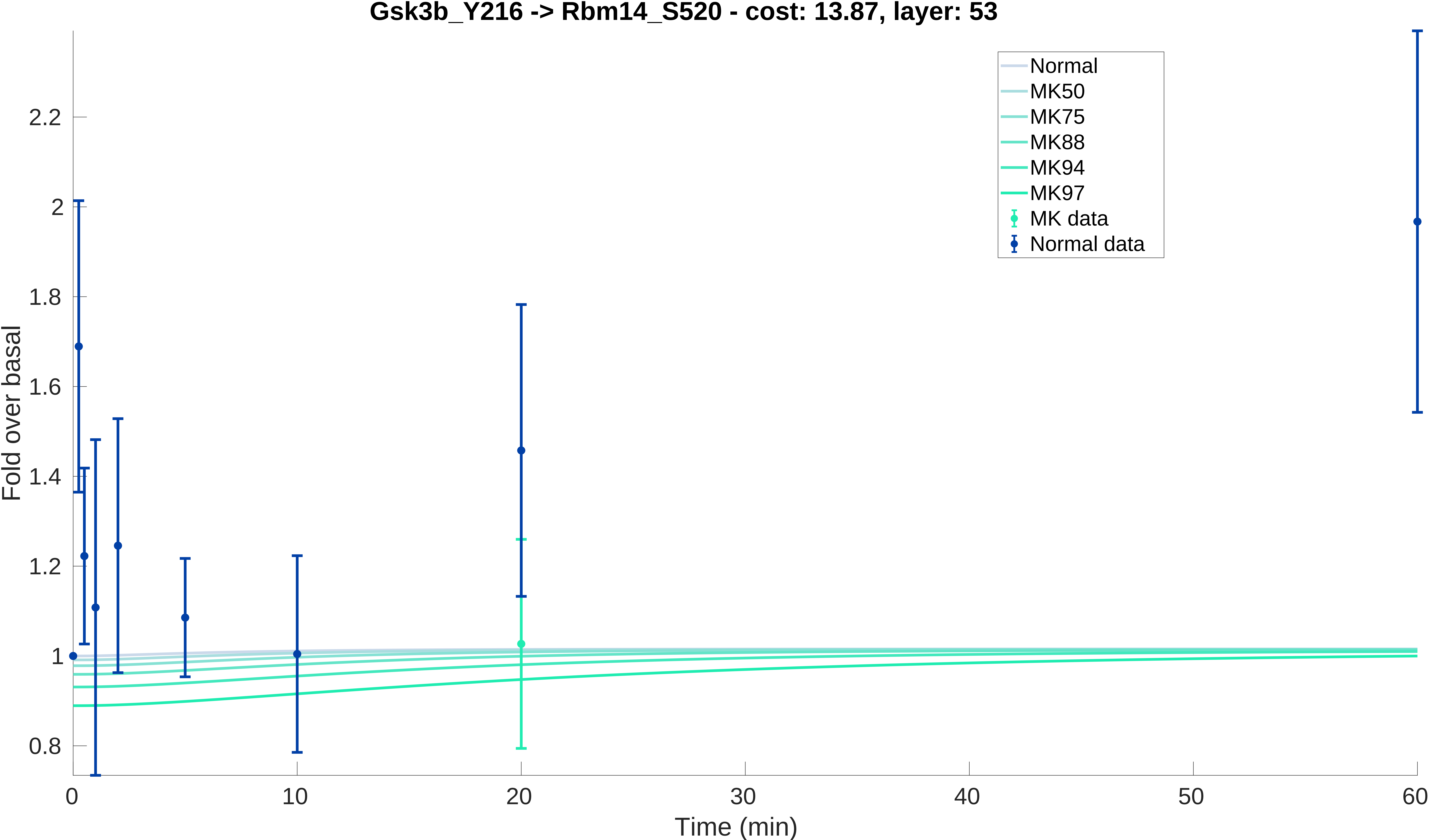

Gsk3b_Y216 -> Rbm14_S520 - cost: 13.87, layer: 53
Normal
MK50
MK75
MK88
MK94
MK97
MK data
Normal data
Fold over basal
Time (min)
0.8
1
1.2
1.4
1.6
1.8
2
2.2
0
10
20
30
40
50
60

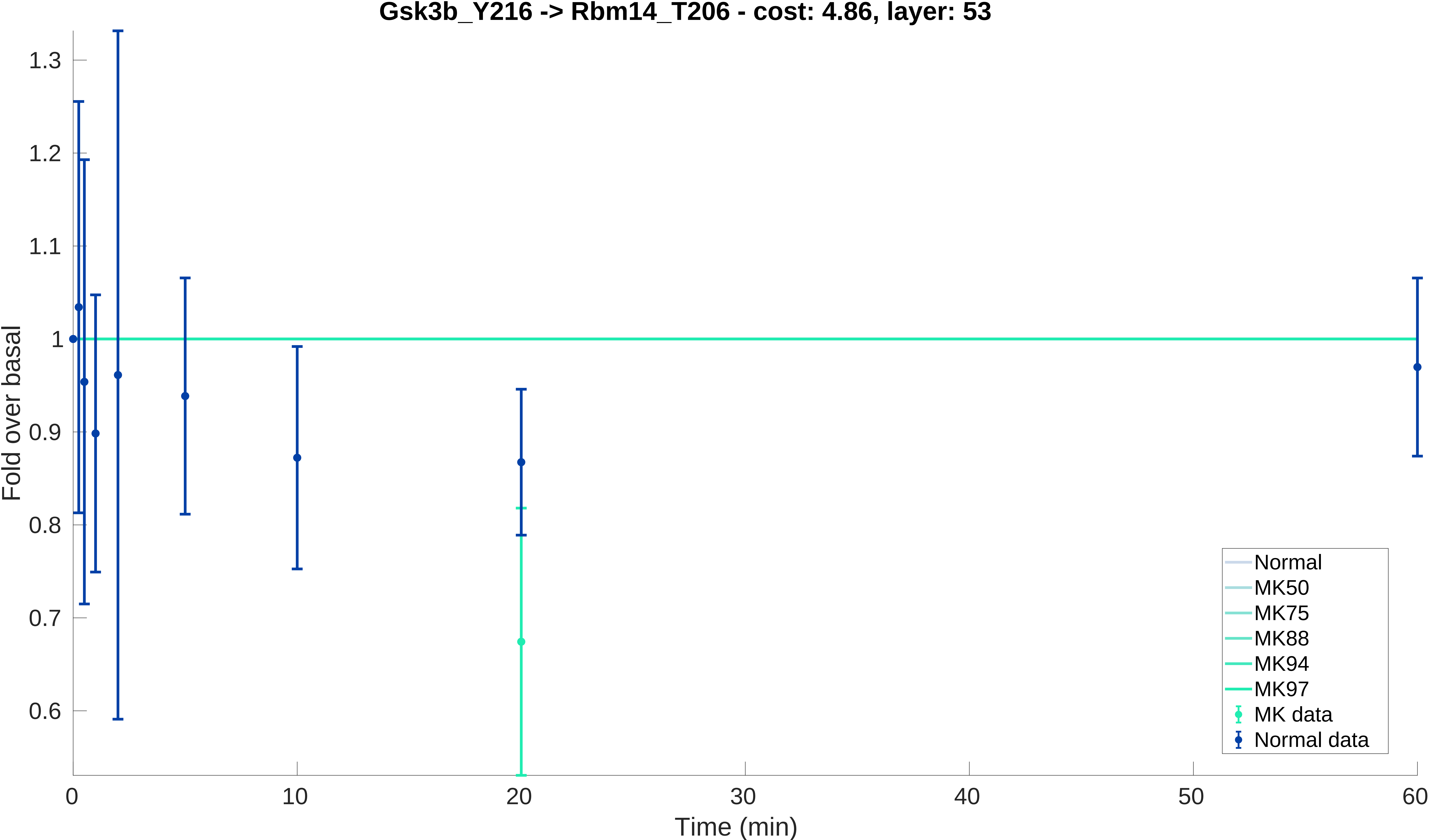
Gsk3b_Y216 -> Rbm14_T206 - cost: 4.86, layer: 53
Fold over basal
1.3
1.2
1.1
1
0.9
0.8
0.7
0.6
0
10
20
30
40
50
60
Time (min)
Normal
MK50
MK75
MK88
MK94
MK97
MK data
Normal data

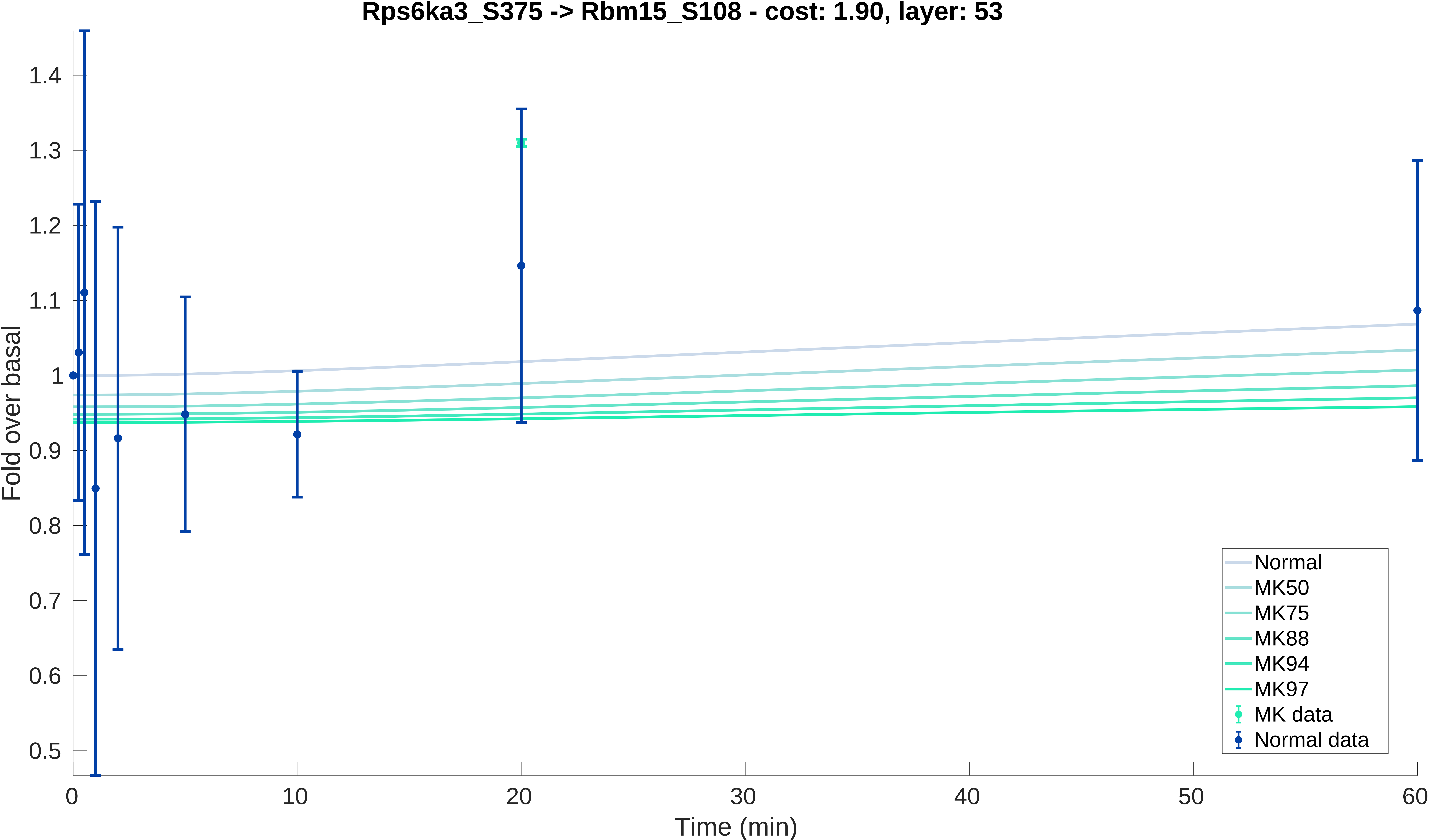

Rps6ka3_S375 -> Rbm15_S108 - cost: 1.90, layer: 53
Fold over basal
Time (min)
1.4
1.3
1.2
1.1
1
0.9
0.8
0.7
0.6
0.5
0
10
20
30
40
50
60
Normal
MK50
MK75
MK88
MK94
MK97
MK data
Normal data

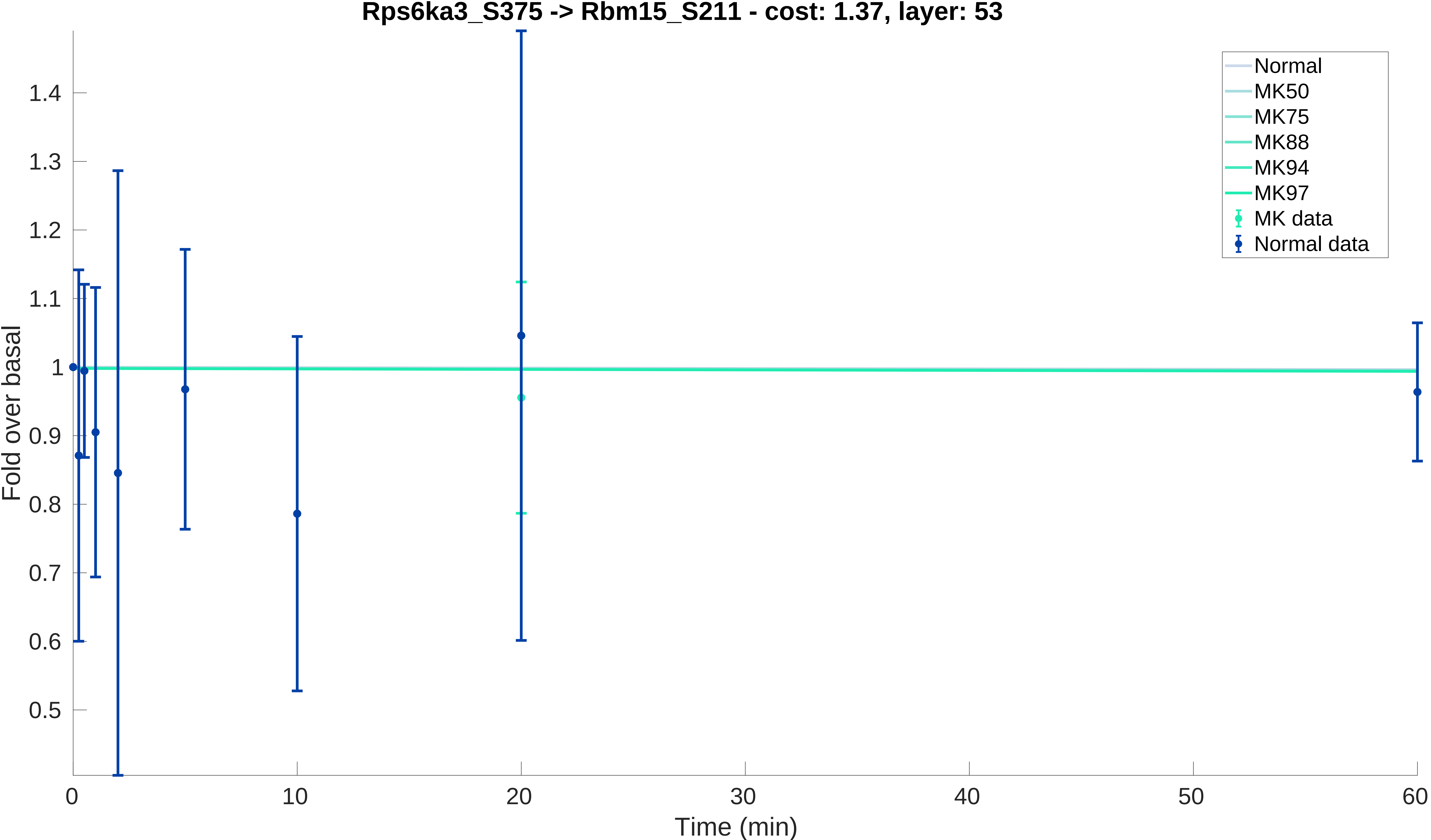

Rps6ka3_S375 -> Rbm15_S211 - cost: 1.37, layer: 53
Fold over basal
Time (min)
1.4
1.3
1.2
1.1
1
0.9
0.8
0.7
0.6
0.5
0
10
20
30
40
50
60
Normal
MK50
MK75
MK88
MK94
MK97
MK data
Normal data

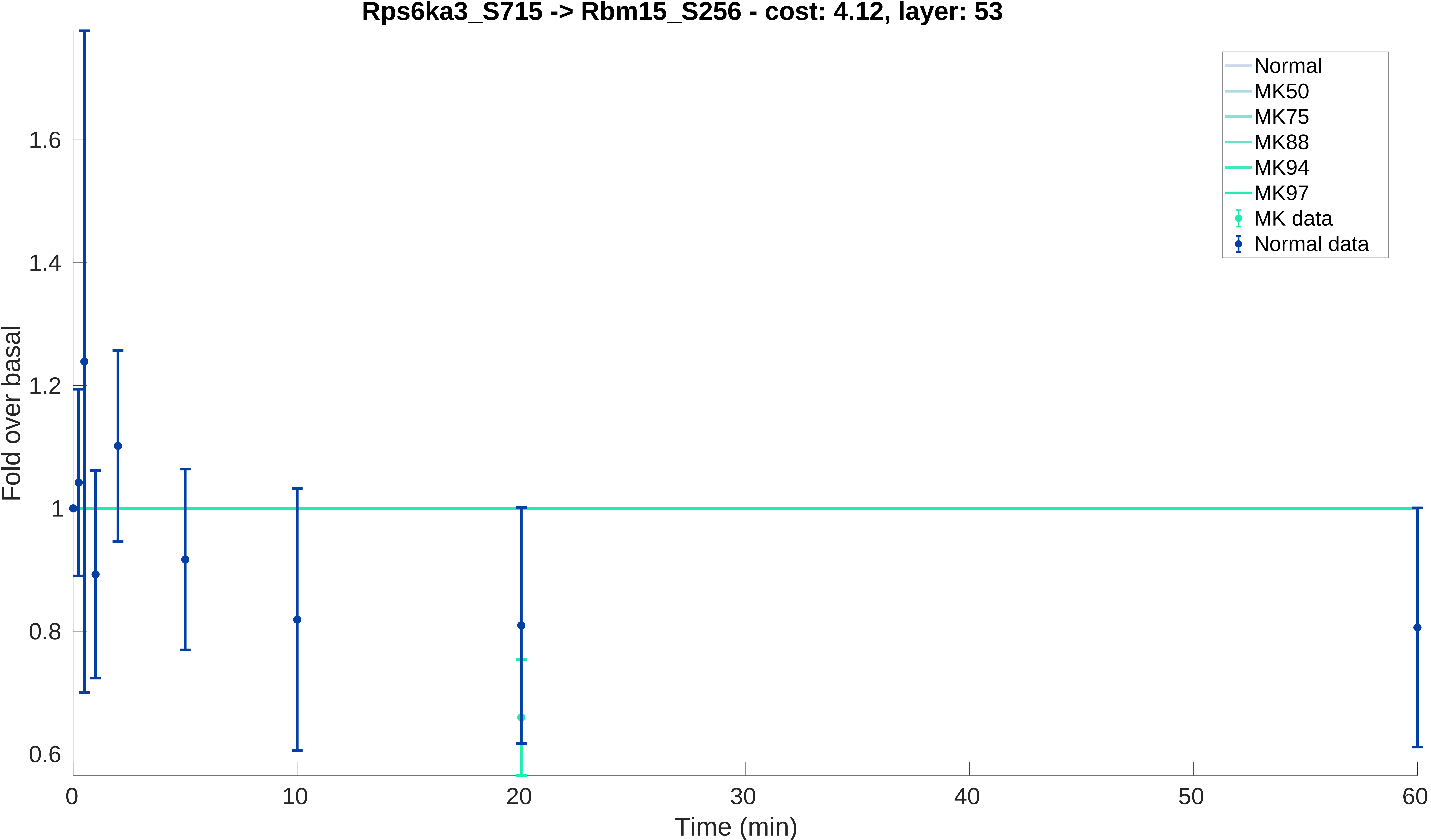

Rps6ka3_S715 -> Rbm15_S256 - cost: 4.12, layer: 53
Normal
MK50
MK75
MK88
MK94
MK97
MK data
Normal data
Fold over basal
1.6
1.4
1.2
1
0.8
0.6
0
10
20
30
40
50
60
Time (min)

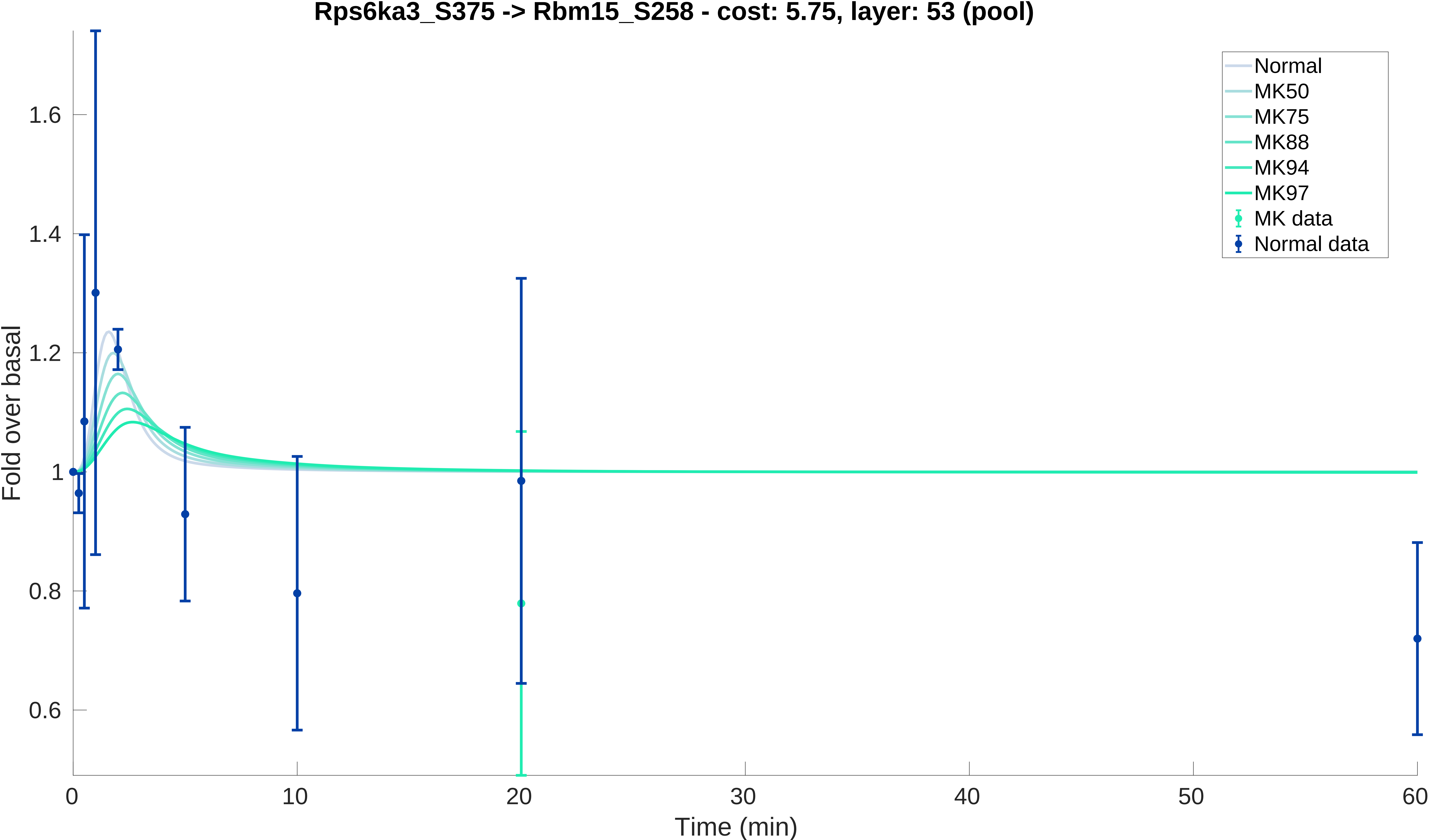
Rps6ka3_S375 -> Rbm15_S258 - cost: 5.75, layer: 53 (pool)
Fold over basal
Time (min)
0.6
0.8
1
1.2
1.4
1.6
0
10
20
30
40
50
60
Normal
MK50
MK75
MK88
MK94
MK97
MK data
Normal data

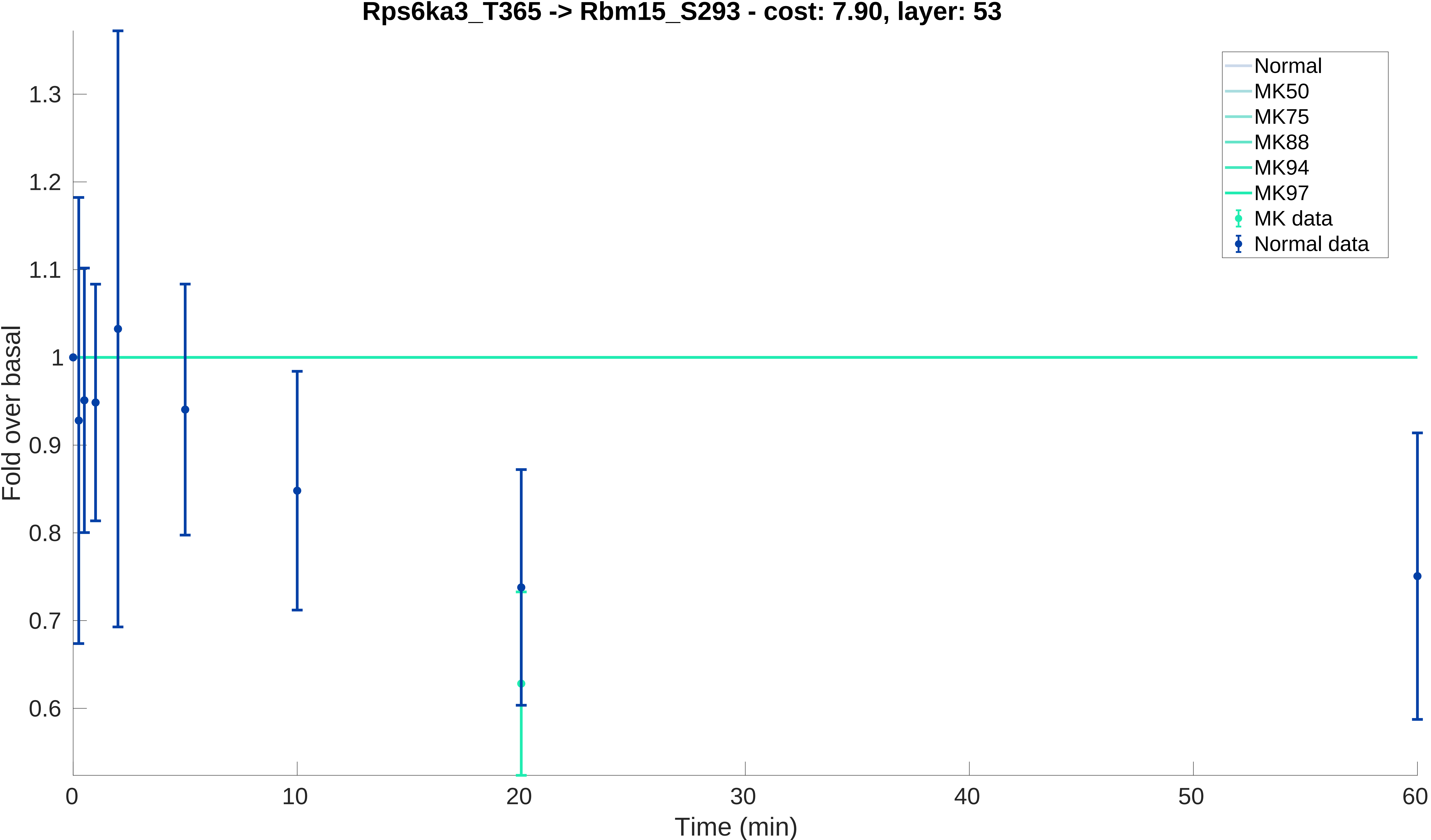

Rps6ka3_T365 -> Rbm15_S293 - cost: 7.90, layer: 53
Fold over basal
1.3
1.2
1.1
1
0.9
0.8
0.7
0.6
0
10
20
30
40
50
60
Time (min)
Normal
MK50
MK75
MK88
MK94
MK97
MK data
Normal data

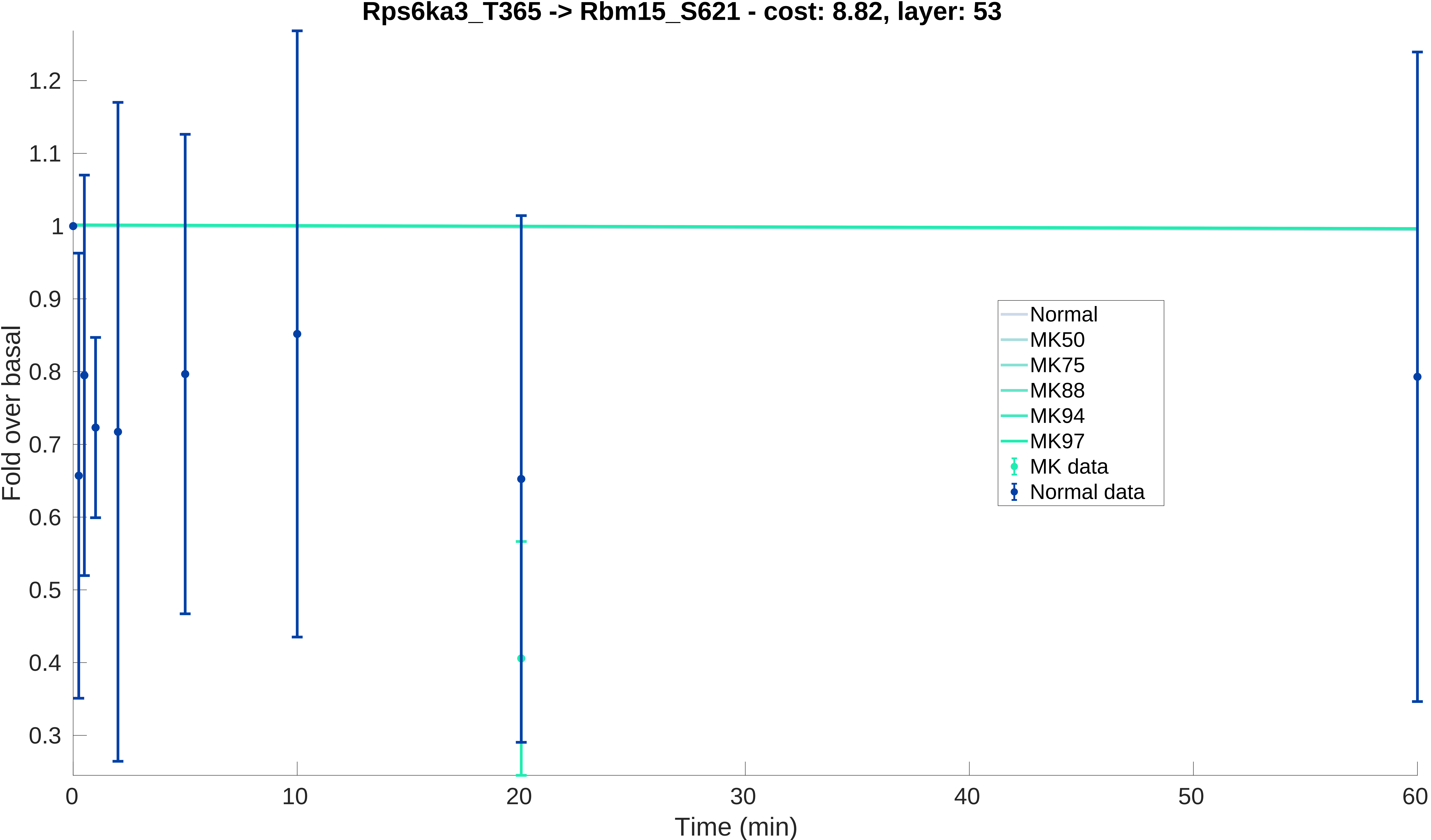

Rps6ka3_T365 -> Rbm15_S621 - cost: 8.82, layer: 53
Fold over basal
Time (min)
1.2
1.1
1
0.9
0.8
0.7
0.6
0.5
0.4
0.3
0
10
20
30
40
50
60
Normal
MK50
MK75
MK88
MK94
MK97
MK data
Normal data

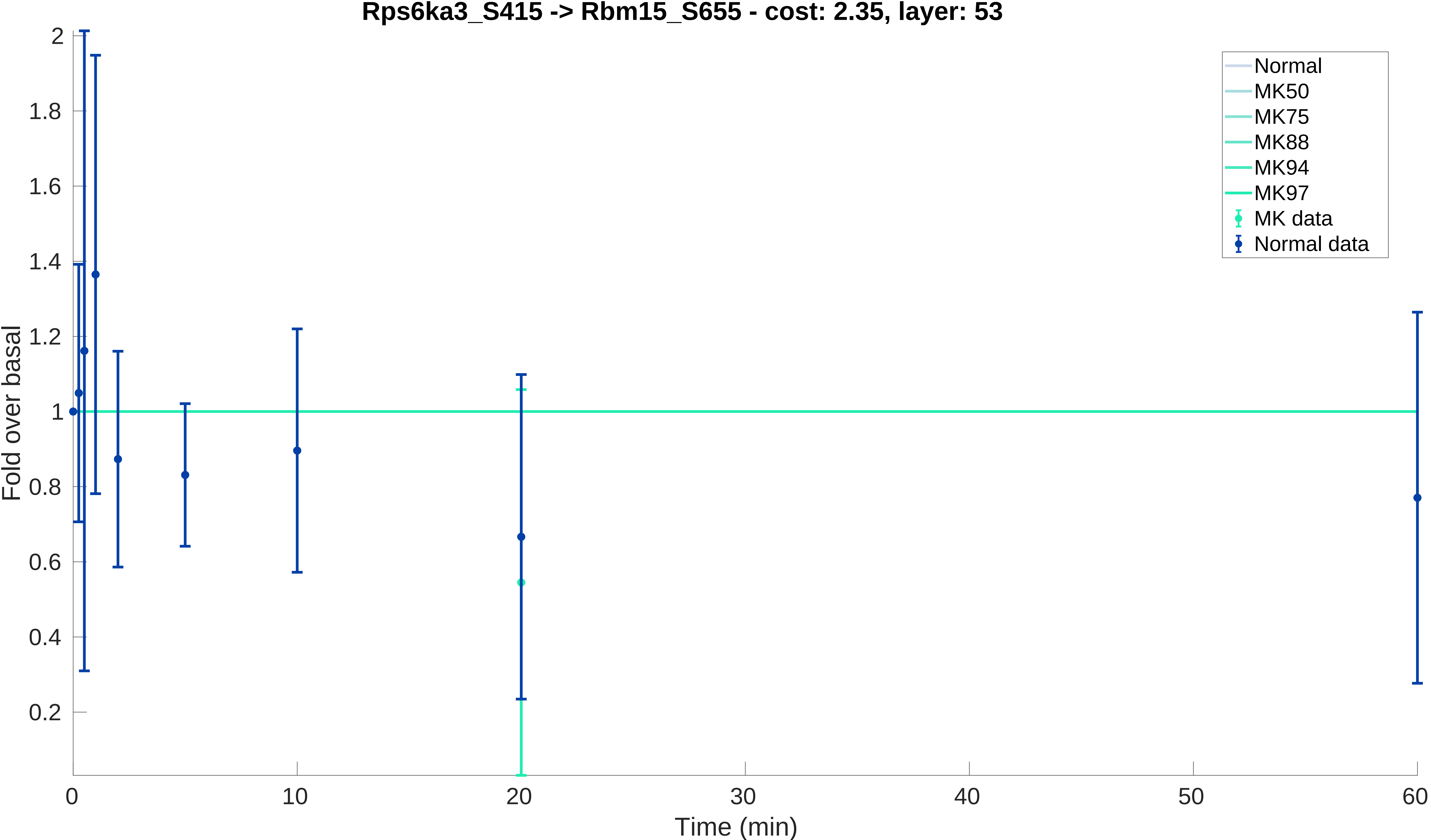

Rps6ka3_S415 -> Rbm15_S655 - cost: 2.35, layer: 53
Fold over basal
Time (min)
0
10
20
30
40
50
60
0.2
0.4
0.6
0.8
1
1.2
1.4
1.6
1.8
2
Normal
MK50
MK75
MK88
MK94
MK97
MK data
Normal data

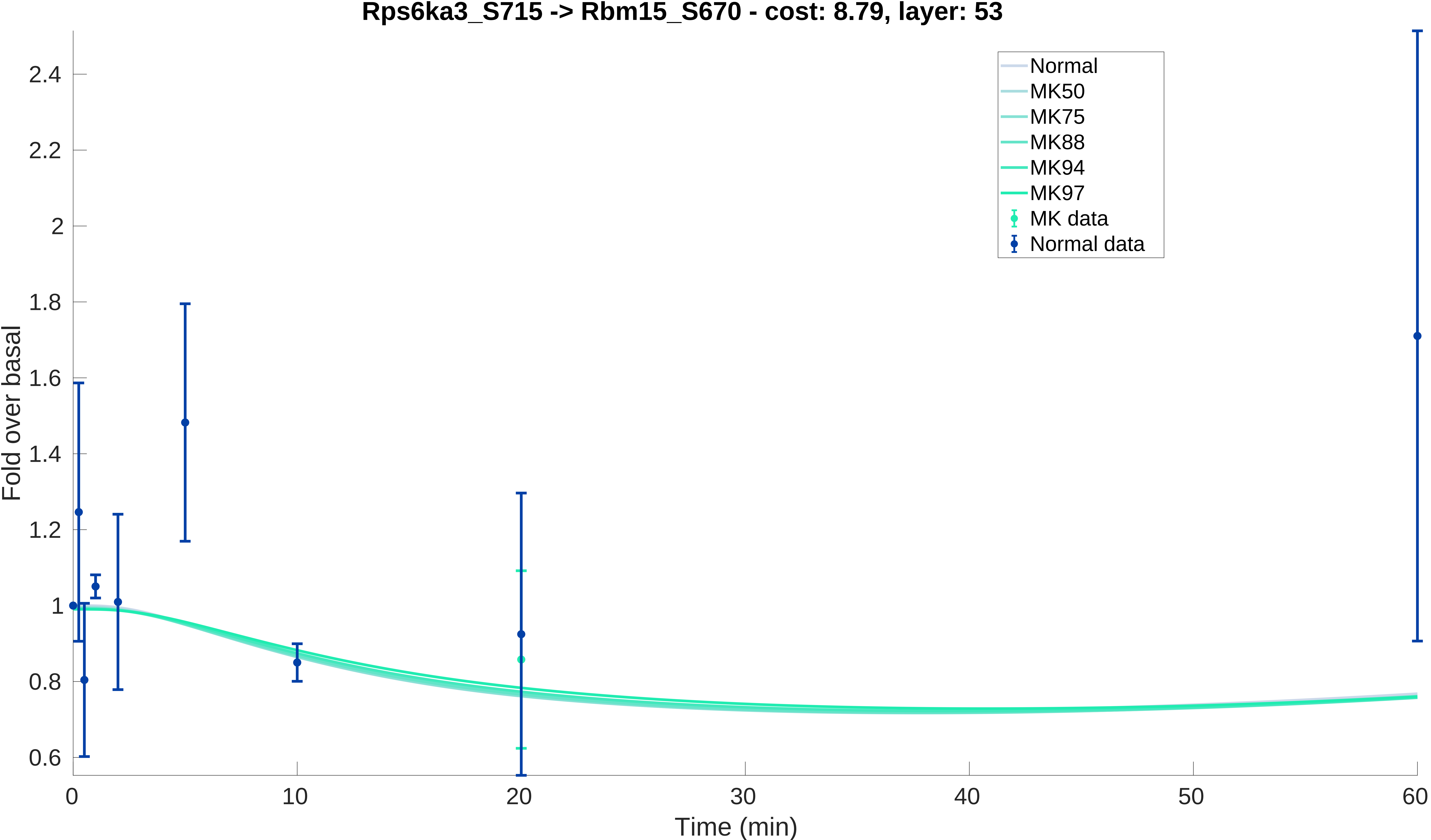

Rps6ka3_S715 -> Rbm15_S670 - cost: 8.79, layer: 53
Fold over basal
Time (min)
0.6
0.8
1
1.2
1.4
1.6
1.8
2
2.2
2.4
0
10
20
30
40
50
60
Normal
MK50
MK75
MK88
MK94
MK97
MK data
Normal data

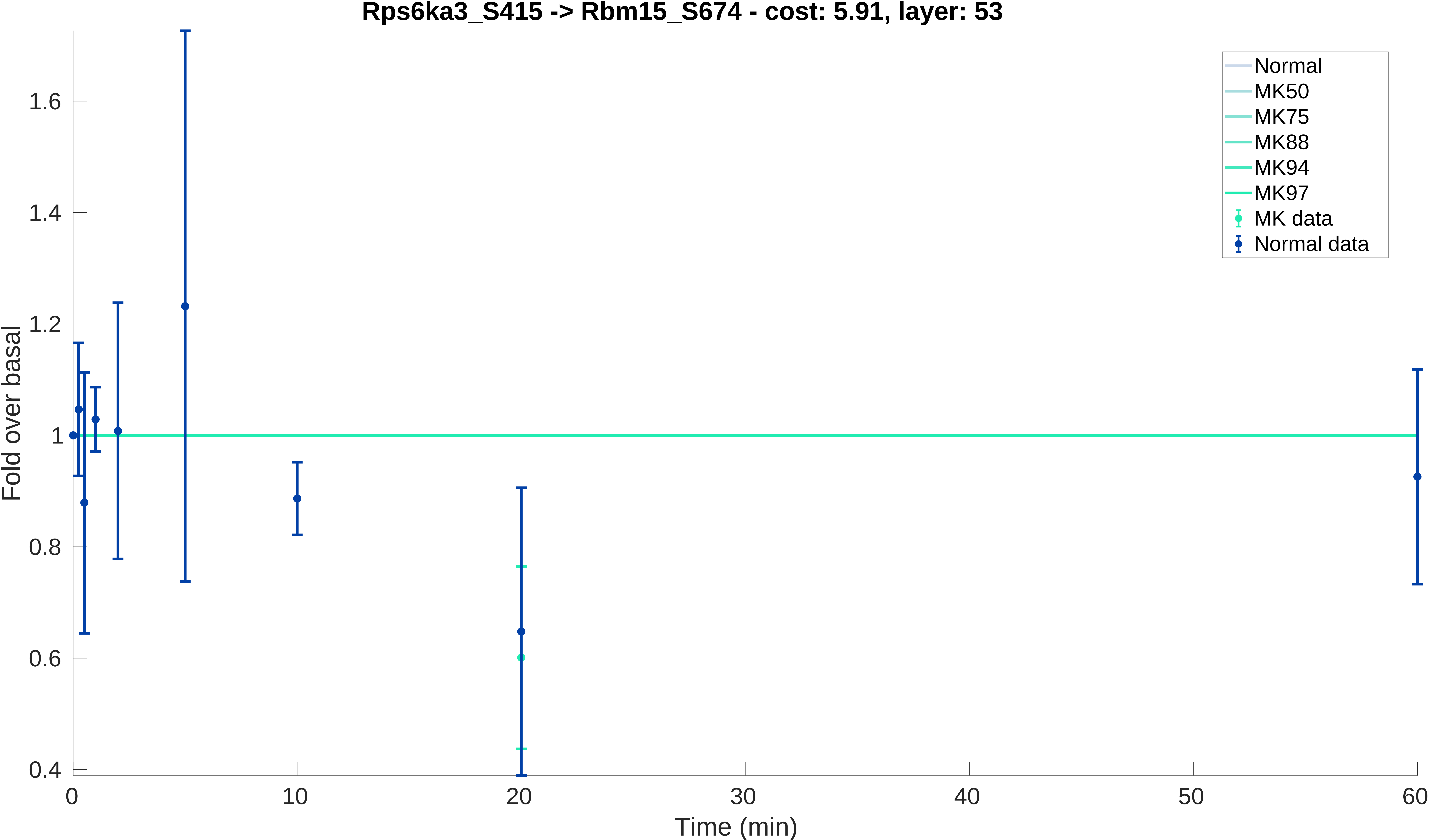

Rps6ka3_S415 -> Rbm15_S674 - cost: 5.91, layer: 53
Fold over basal
1.6
1.4
1.2
1
0.8
0.6
0.4
0
10
20
30
40
50
60
Time (min)
Normal
MK50
MK75
MK88
MK94
MK97
MK data
Normal data

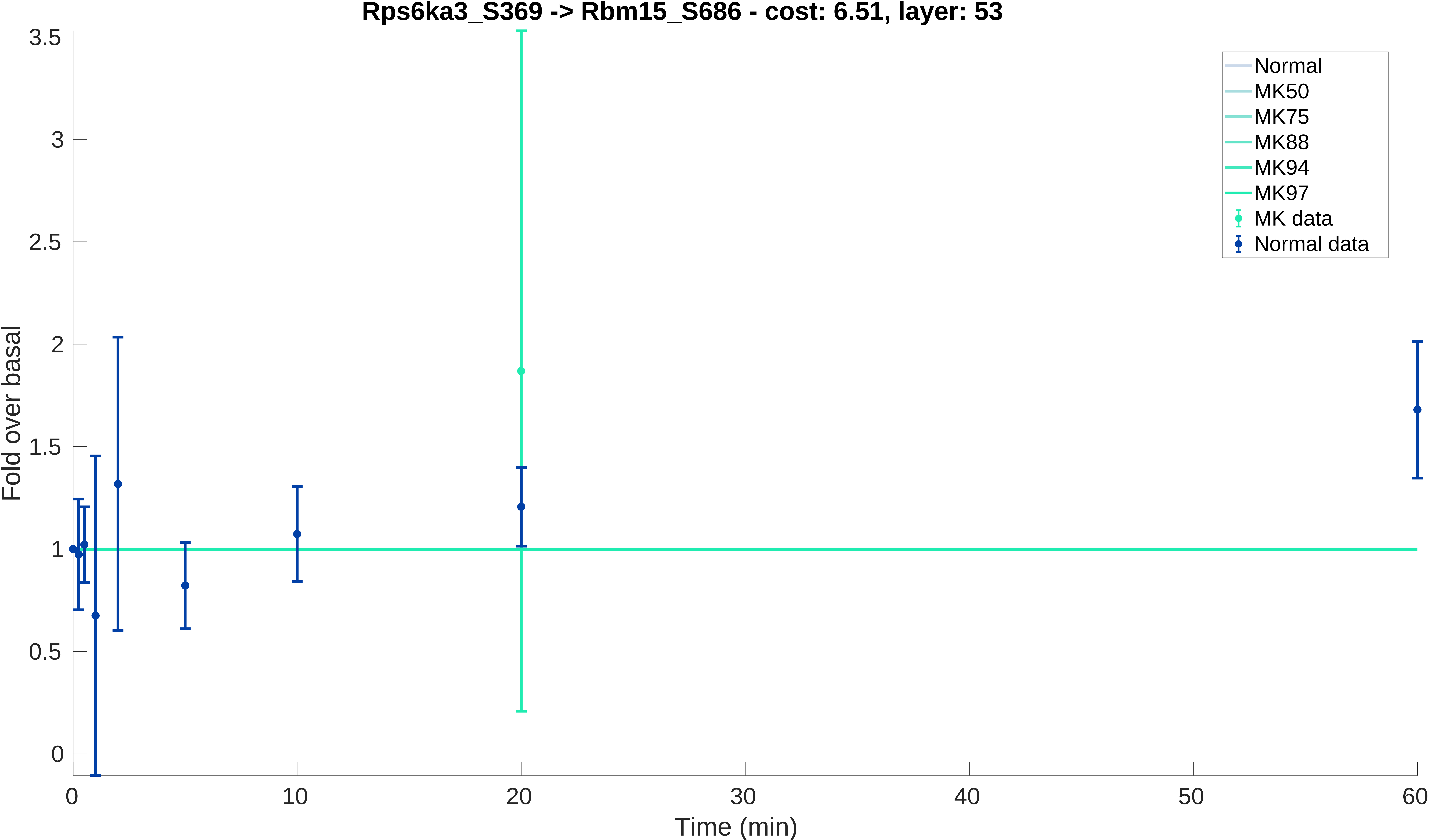

Rps6ka3_S369 -> Rbm15_S686 - cost: 6.51, layer: 53
Fold over basal
Time (min)
0
0.5
1
1.5
2
2.5
3
3.5
0
10
20
30
40
50
60
Normal
MK50
MK75
MK88
MK94
MK97
MK data
Normal data

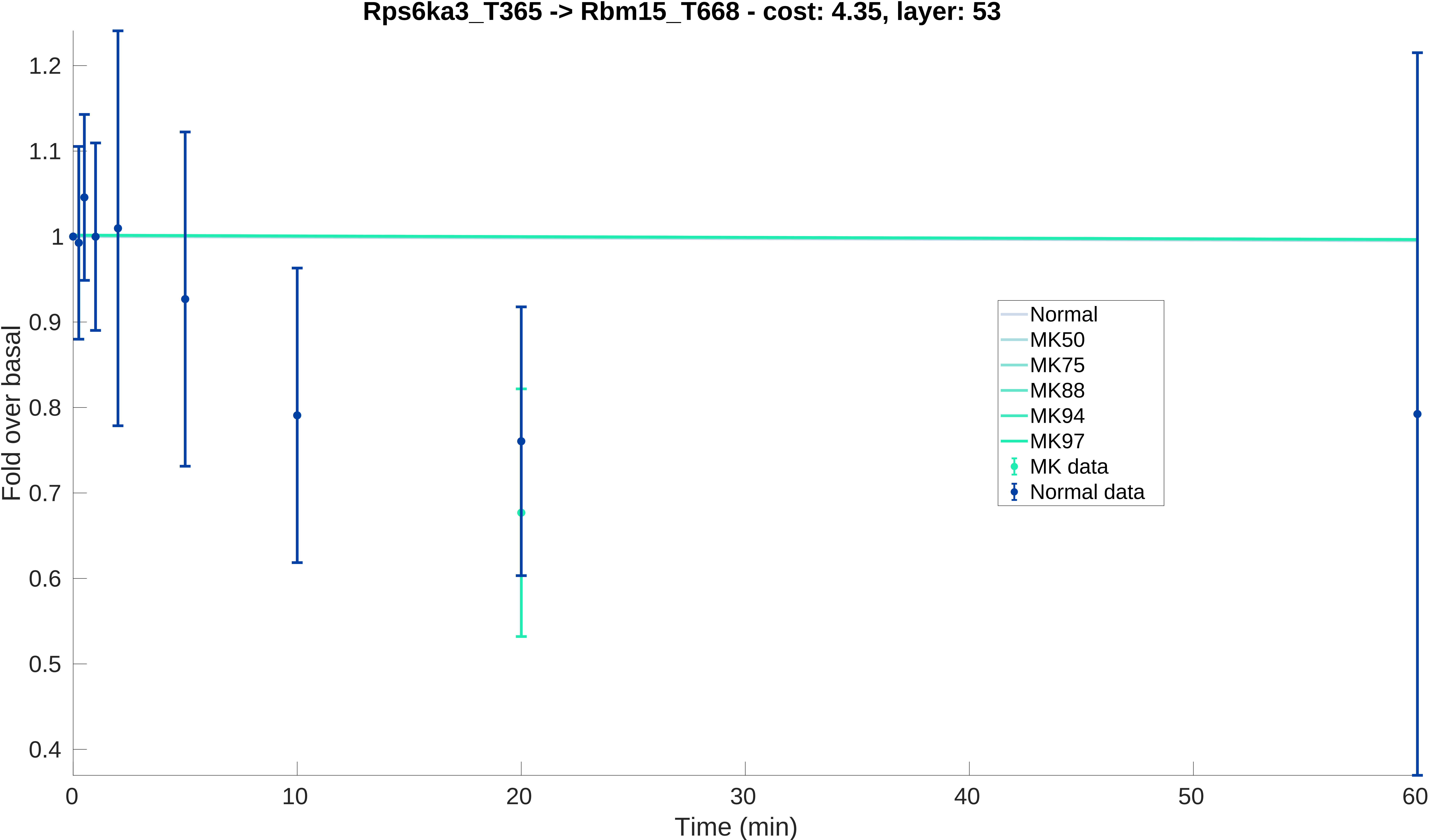

Rps6ka3_T365 -> Rbm15_T668 - cost: 4.35, layer: 53
Fold over basal
1.2
1.1
1
0.9
0.8
0.7
0.6
0.5
0.4
0
10
20
30
40
50
60
Time (min)
Normal
MK50
MK75
MK88
MK94
MK97
MK data
Normal data

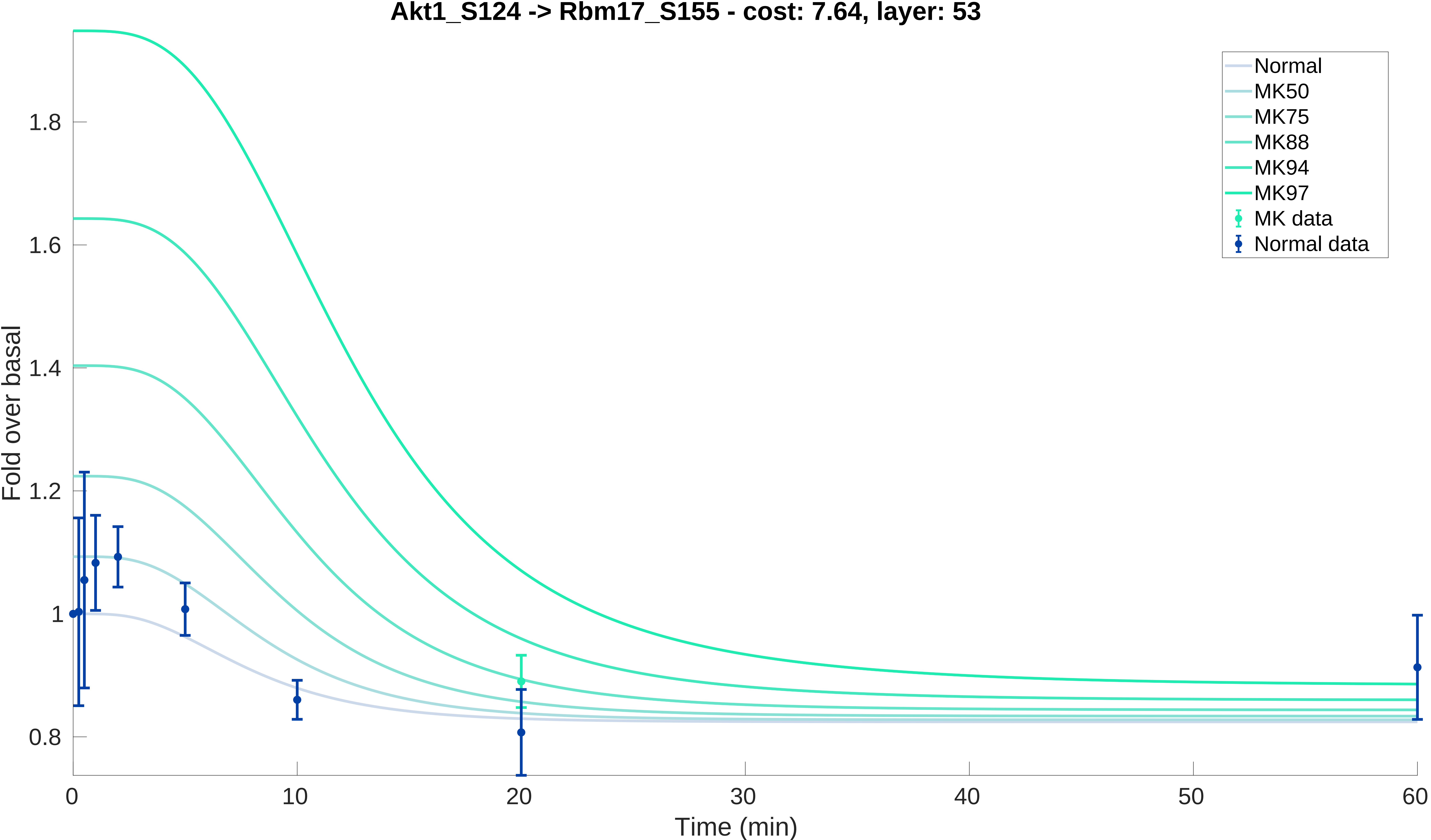

Akt1_S124 -> Rbm17_S155 - cost: 7.64, layer: 53
Normal
MK50
MK75
MK88
MK94
MK97
MK data
Normal data
Fold over basal
1.8
1.6
1.4
1.2
1
0.8
0
10
20
30
40
50
60
Time (min)

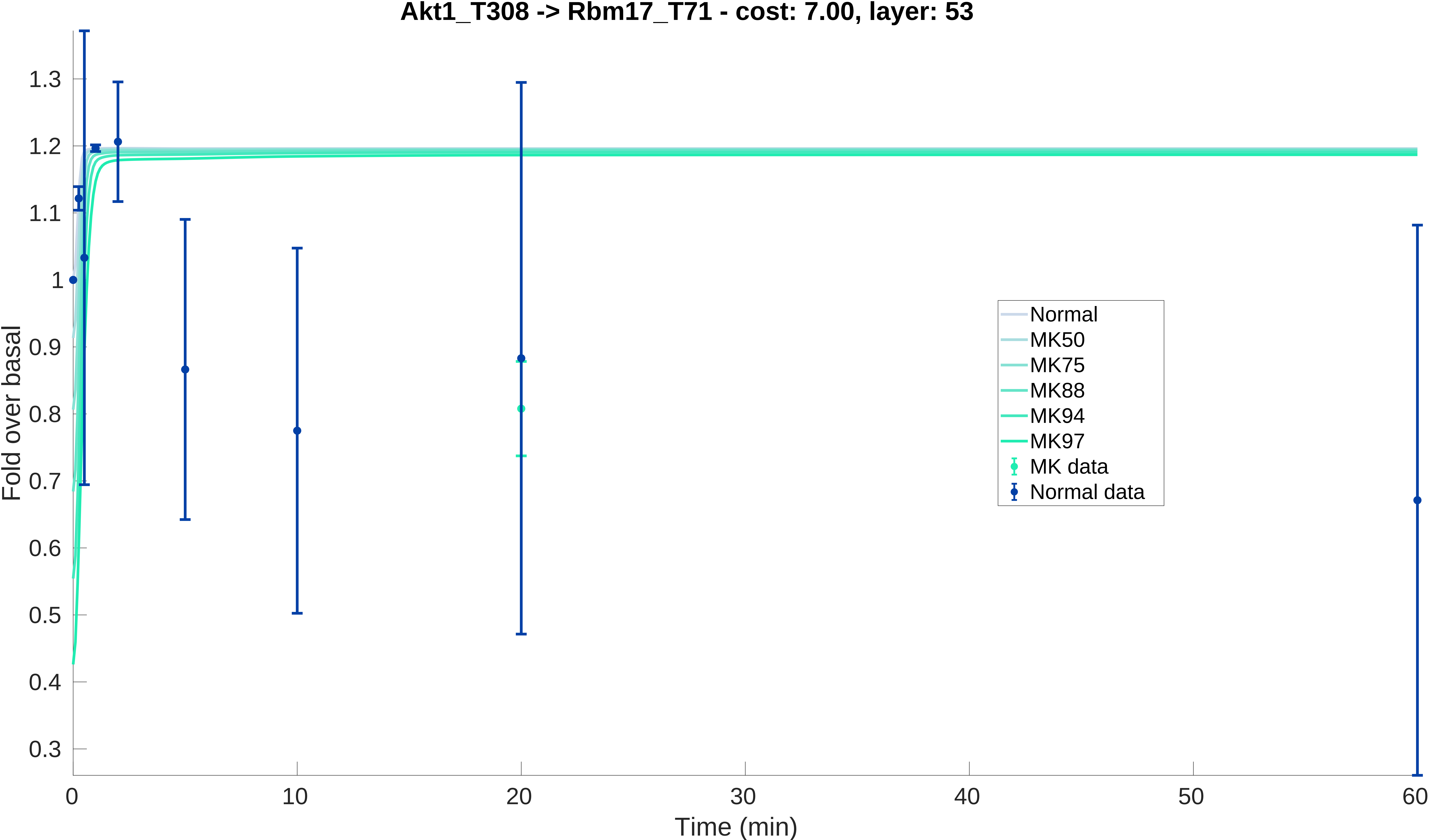

Akt1_T308 -> Rbm17_T71 - cost: 7.00, layer: 53
Fold over basal
Time (min)
1.3
1.2
1.1
1
0.9
0.8
0.7
0.6
0.5
0.4
0.3
0
10
20
30
40
50
60
Normal
MK50
MK75
MK88
MK94
MK97
MK data
Normal data

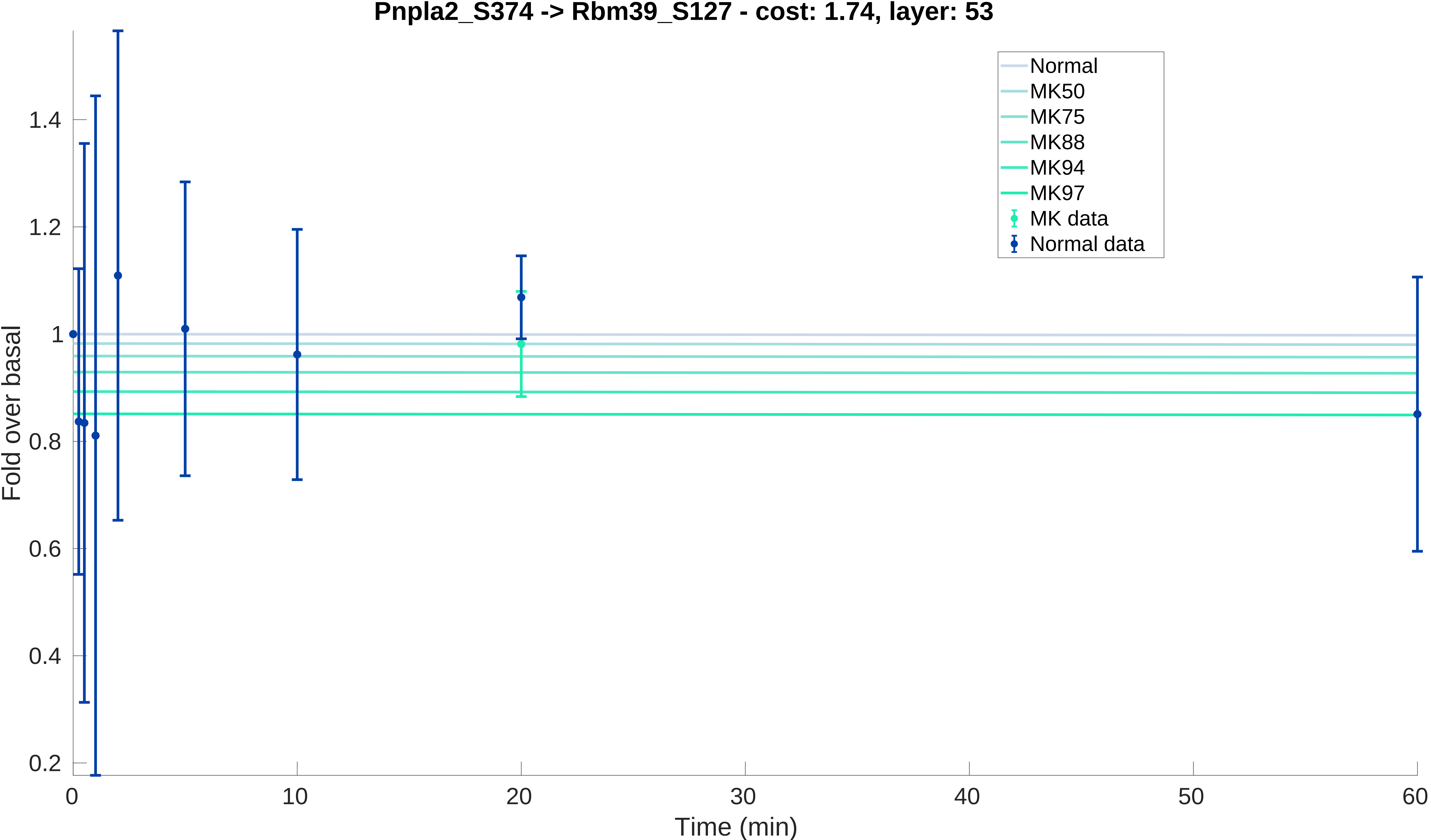

Pnpla2_S374 -> Rbm39_S127 - cost: 1.74, layer: 53
Normal
MK50
MK75
MK88
MK94
MK97
MK data
Normal data
Fold over basal
Time (min)
0.2
0.4
0.6
0.8
1
1.2
1.4
0
10
20
30
40
50
60

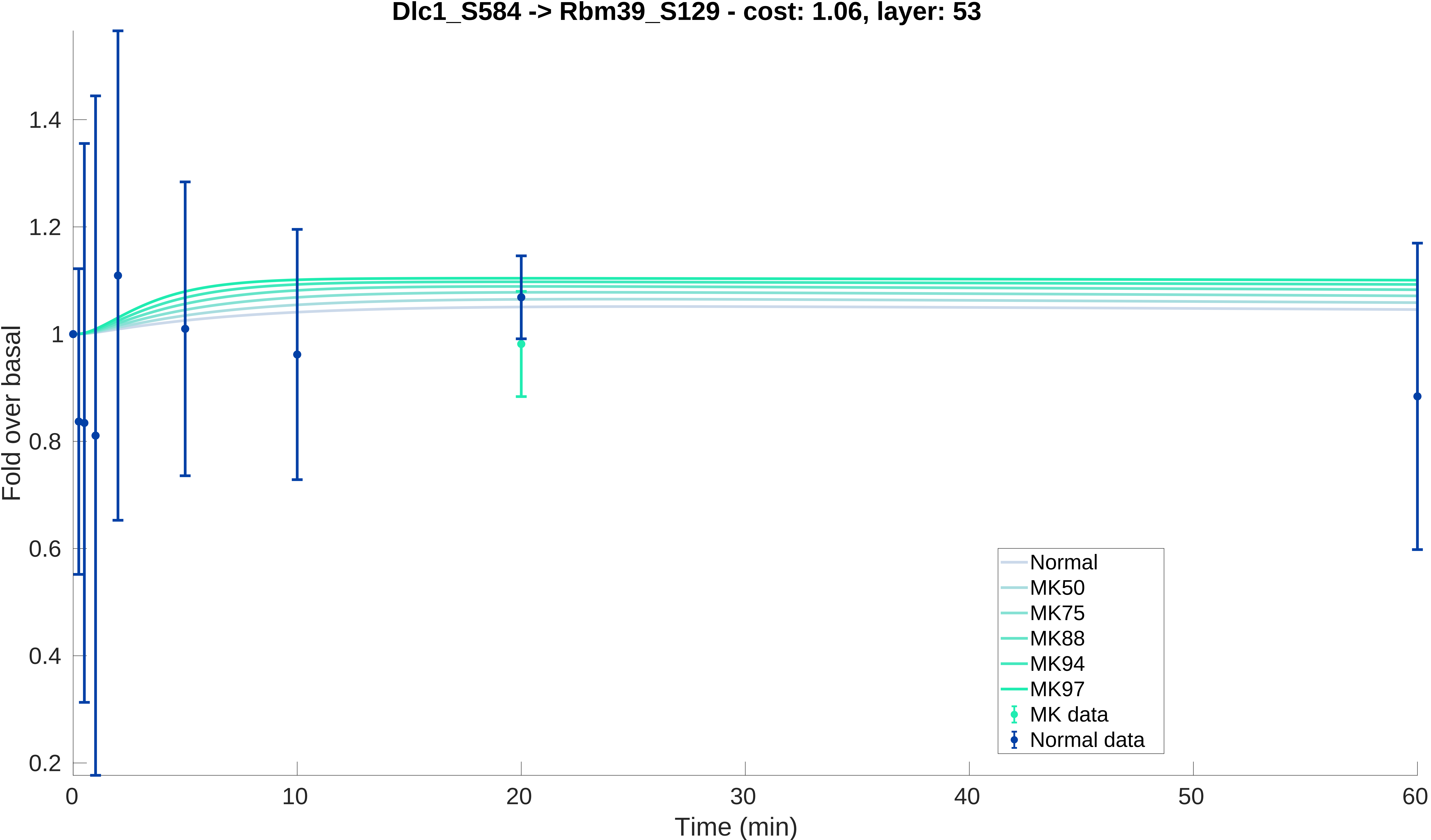
Dlc1_S584 -> Rbm39_S129 - cost: 1.06, layer: 53
Fold over basal
Time (min)
0
10
20
30
40
50
60
0.2
0.4
0.6
0.8
1
1.2
1.4
Normal
MK50
MK75
MK88
MK94
MK97
MK data
Normal data

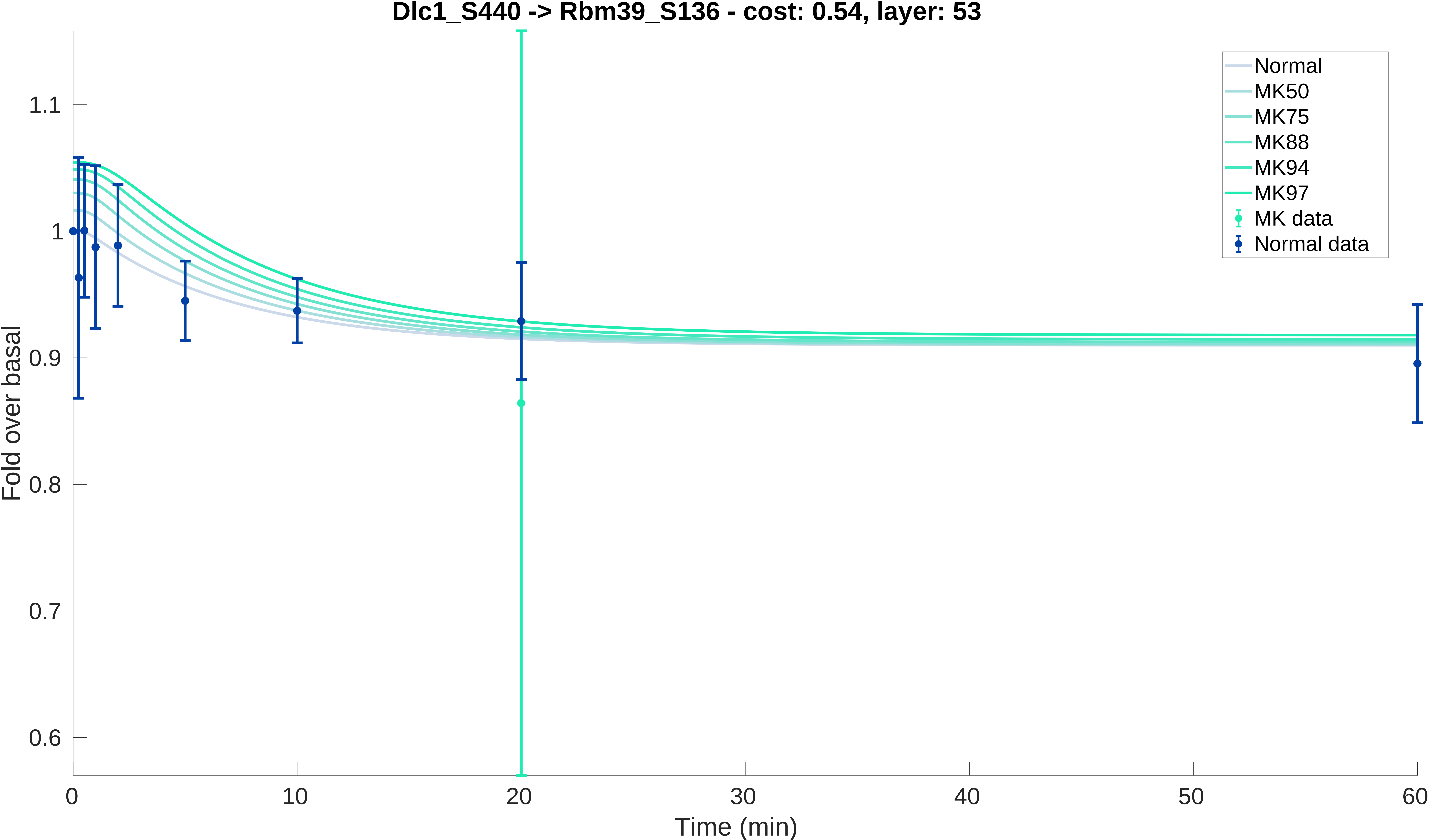

Dlc1_S440 -> Rbm39_S136 - cost: 0.54, layer: 53
Fold over basal
Time (min)
1.1
1
0.9
0.8
0.7
0.6
0
10
20
30
40
50
60
Normal
MK50
MK75
MK88
MK94
MK97
MK data
Normal data

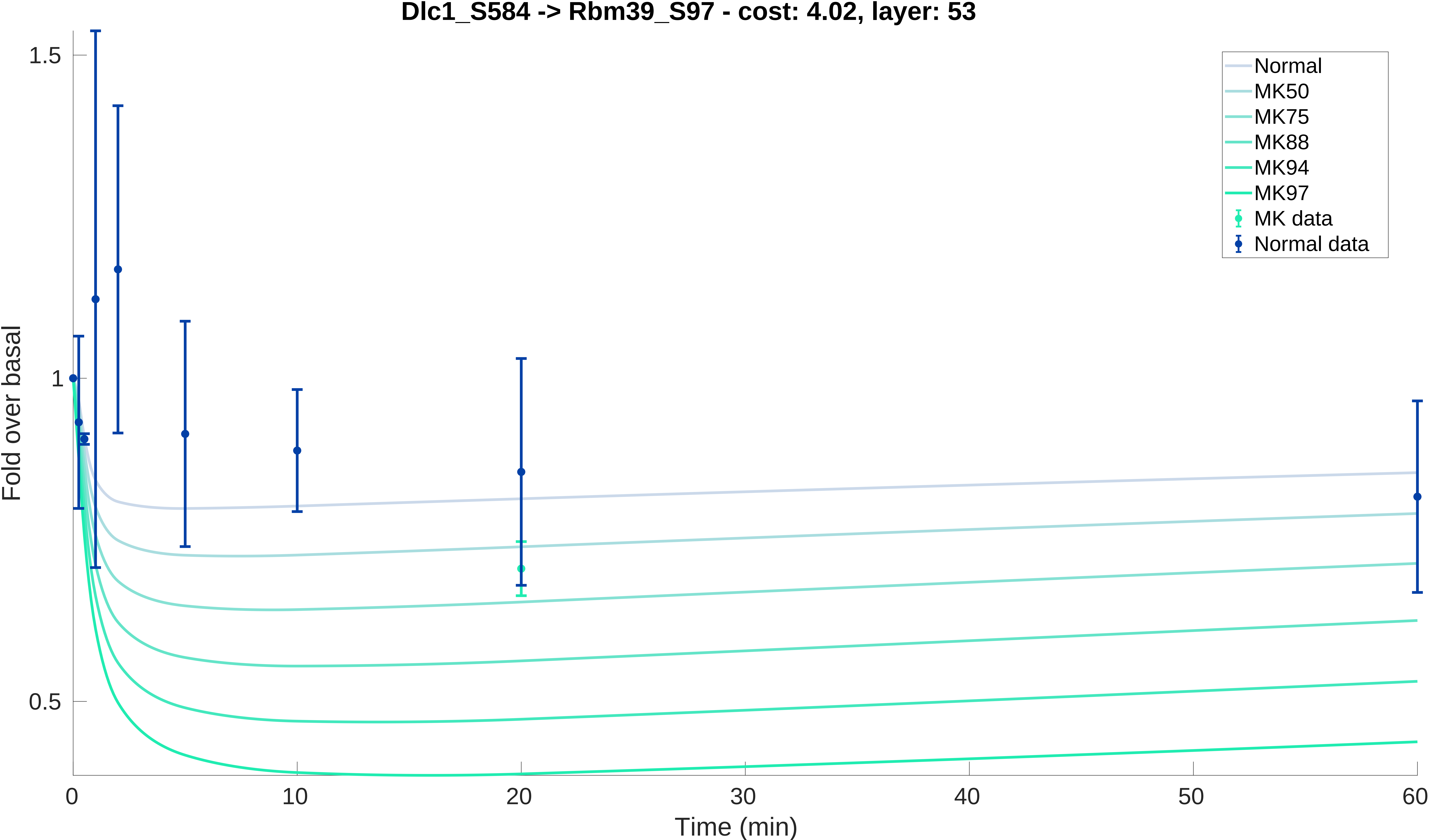

Dlc1_S584 -> Rbm39_S97 - cost: 4.02, layer: 53
Fold over basal
1.5
1
0.5
0
10
20
30
40
50
60
Time (min)
Normal
MK50
MK75
MK88
MK94
MK97
MK data
Normal data

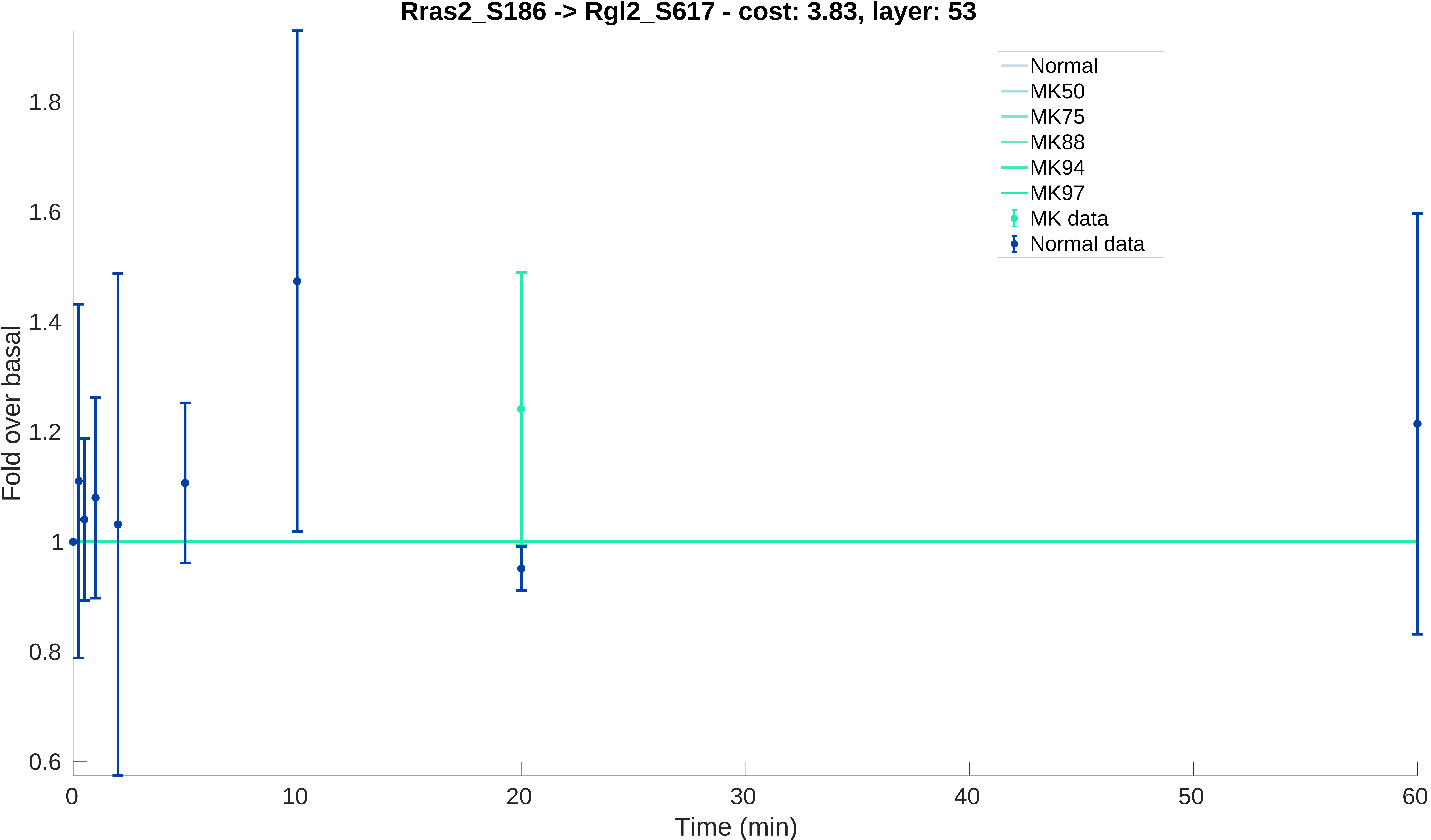

Rras2_S186 -> Rgl2_S617 - cost: 3.83, layer: 53
Normal
MK50
MK75
MK88
MK94
MK97
MK data
Normal data
Fold over basal
Time (min)
0.6
0.8
1
1.2
1.4
1.6
1.8
0
10
20
30
40
50
60

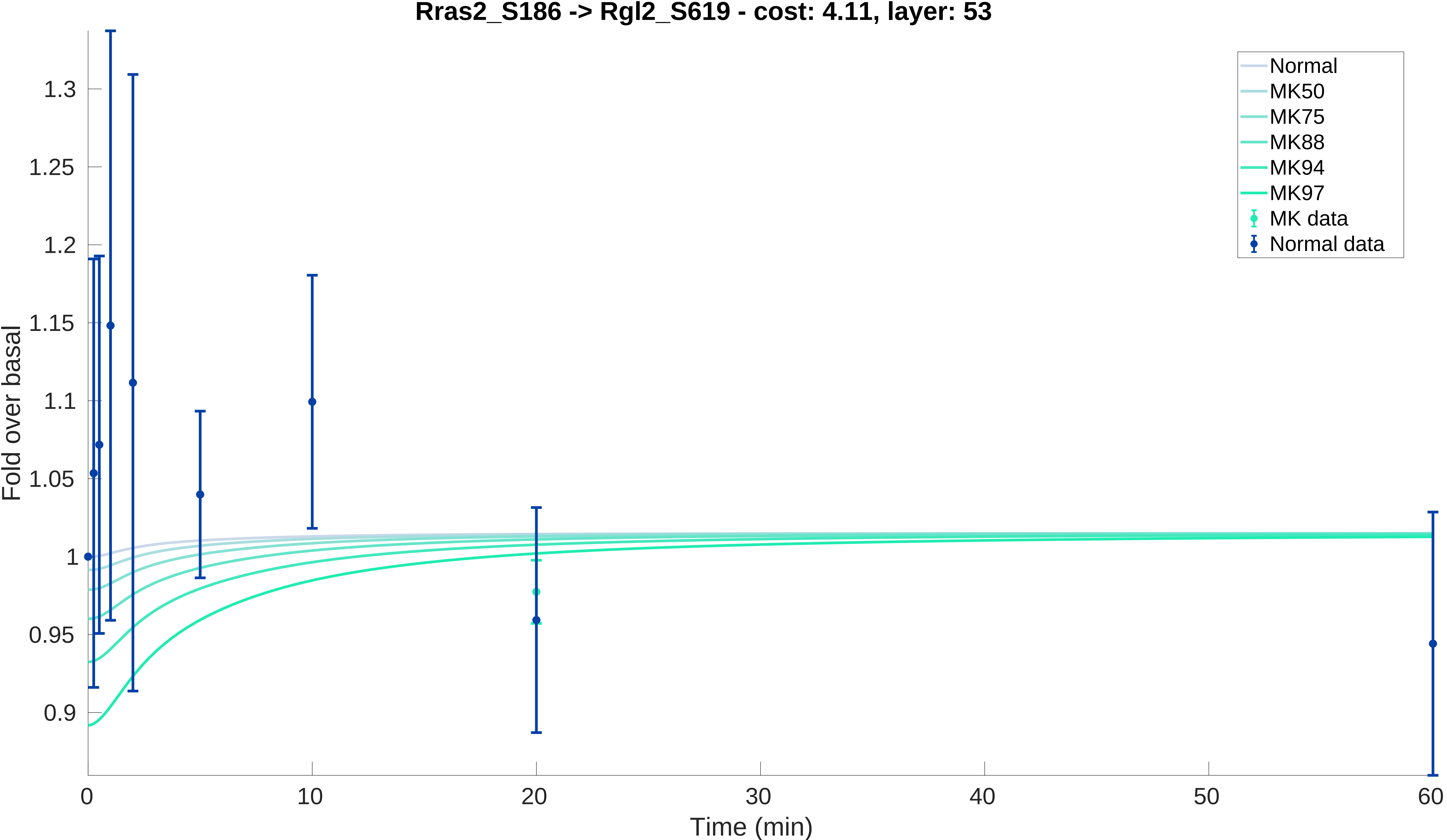
Rras2_S186 -> Rgl2_S619 - cost: 4.11, layer: 53
Fold over basal
1.3
1.25
1.2
1.15
1.1
1.05
1
0.95
0.9
0
10
20
30
40
50
60
Time (min)
Normal
MK50
MK75
MK88
MK94
MK97
MK data
Normal data

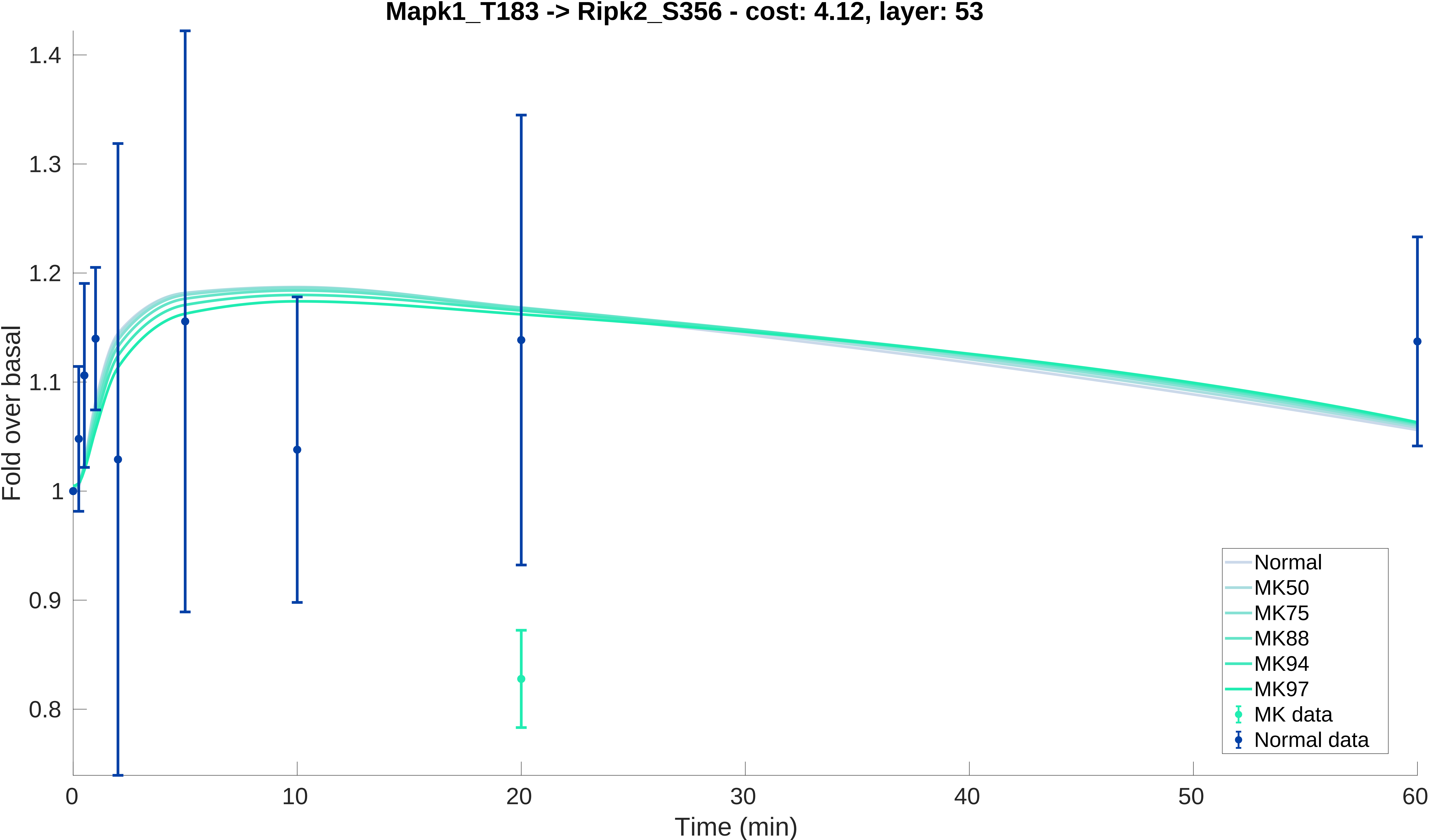

Mapk1_T183 -> Ripk2_S356 - cost: 4.12, layer: 53
Fold over basal
1.4
1.3
1.2
1.1
1
0.9
0.8
0
10
20
30
40
50
60
Time (min)
Normal
MK50
MK75
MK88
MK94
MK97
MK data
Normal data

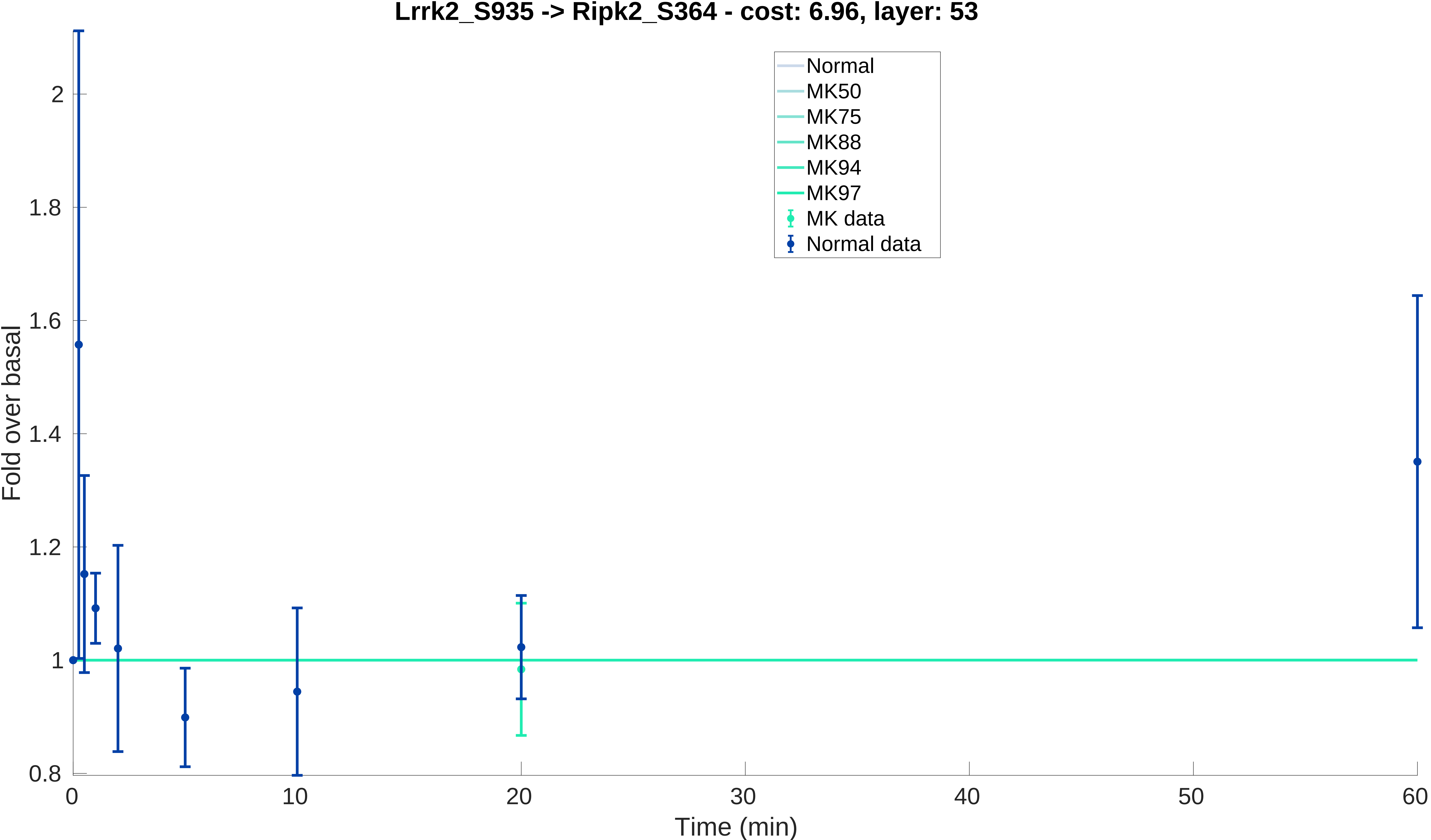

Lrrk2_S935 -> Ripk2_S364 - cost: 6.96, layer: 53
Normal
MK50
MK75
MK88
MK94
MK97
MK data
Normal data
Fold over basal
0.8
1
1.2
1.4
1.6
1.8
2
0
10
20
30
40
50
60
Time (min)

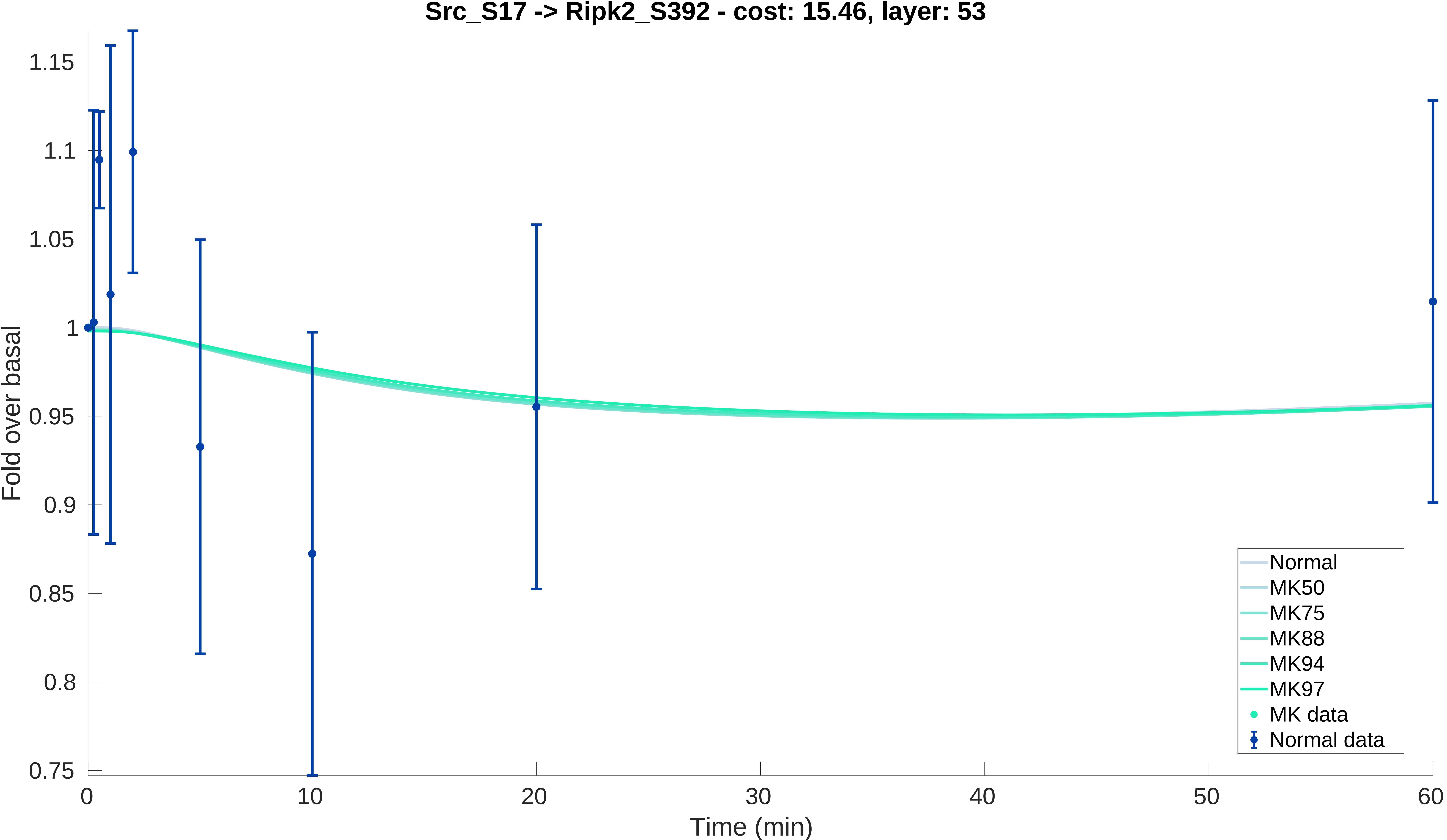

Src_S17 -> Ripk2_S392 - cost: 15.46, layer: 53
Fold over basal
1.15
1.1
1.05
1
0.95
0.9
0.85
0.8
0.75
0
10
20
30
40
50
60
Time (min)
Normal
MK50
MK75
MK88
MK94
MK97
MK data
Normal data

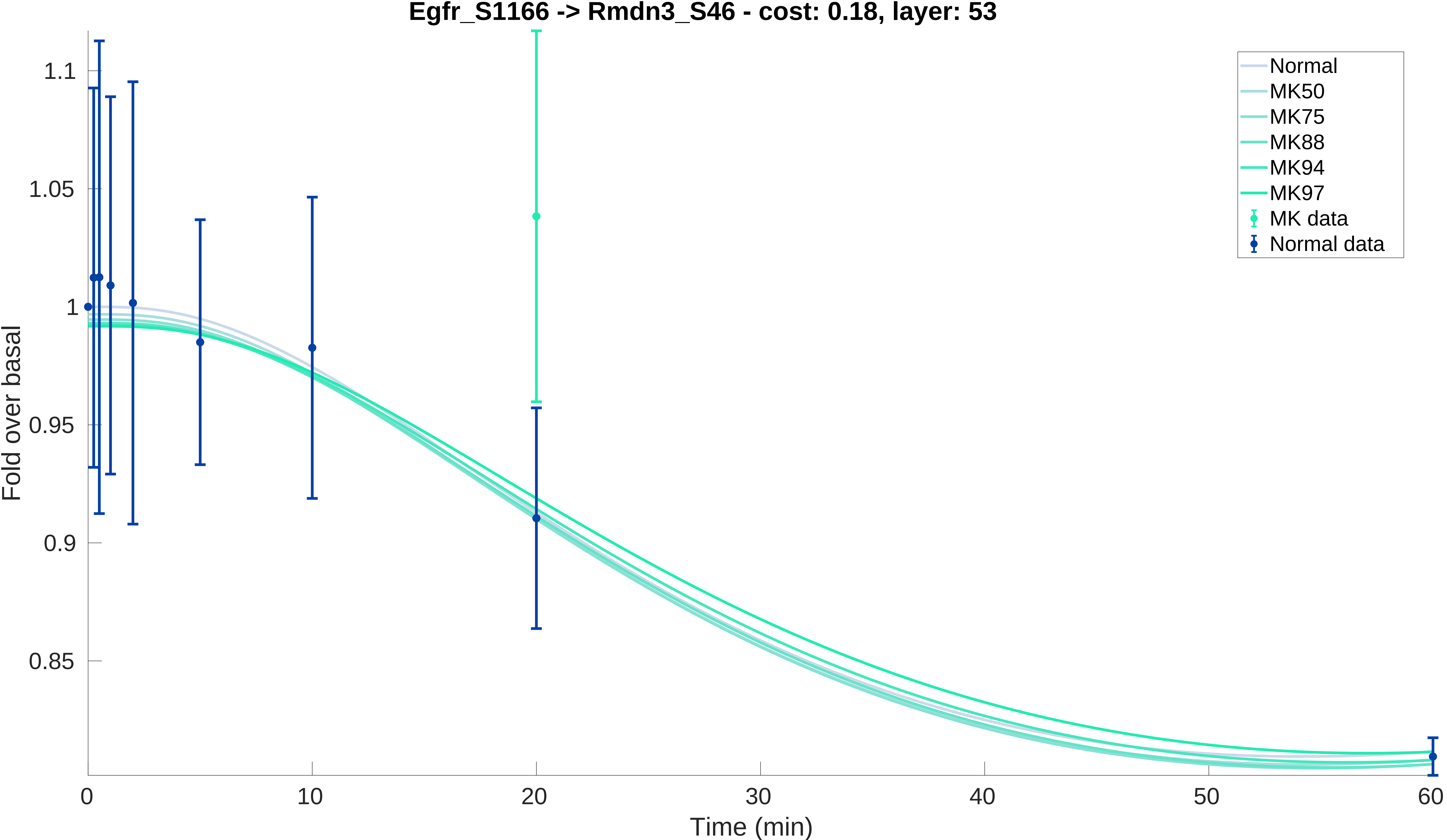

Egfr_S1166 -> Rmdn3_S46 - cost: 0.18, layer: 53
Fold over basal
1.1
1.05
1
0.95
0.9
0.85
0
10
20
30
40
50
60
Time (min)
Normal
MK50
MK75
MK88
MK94
MK97
MK data
Normal data

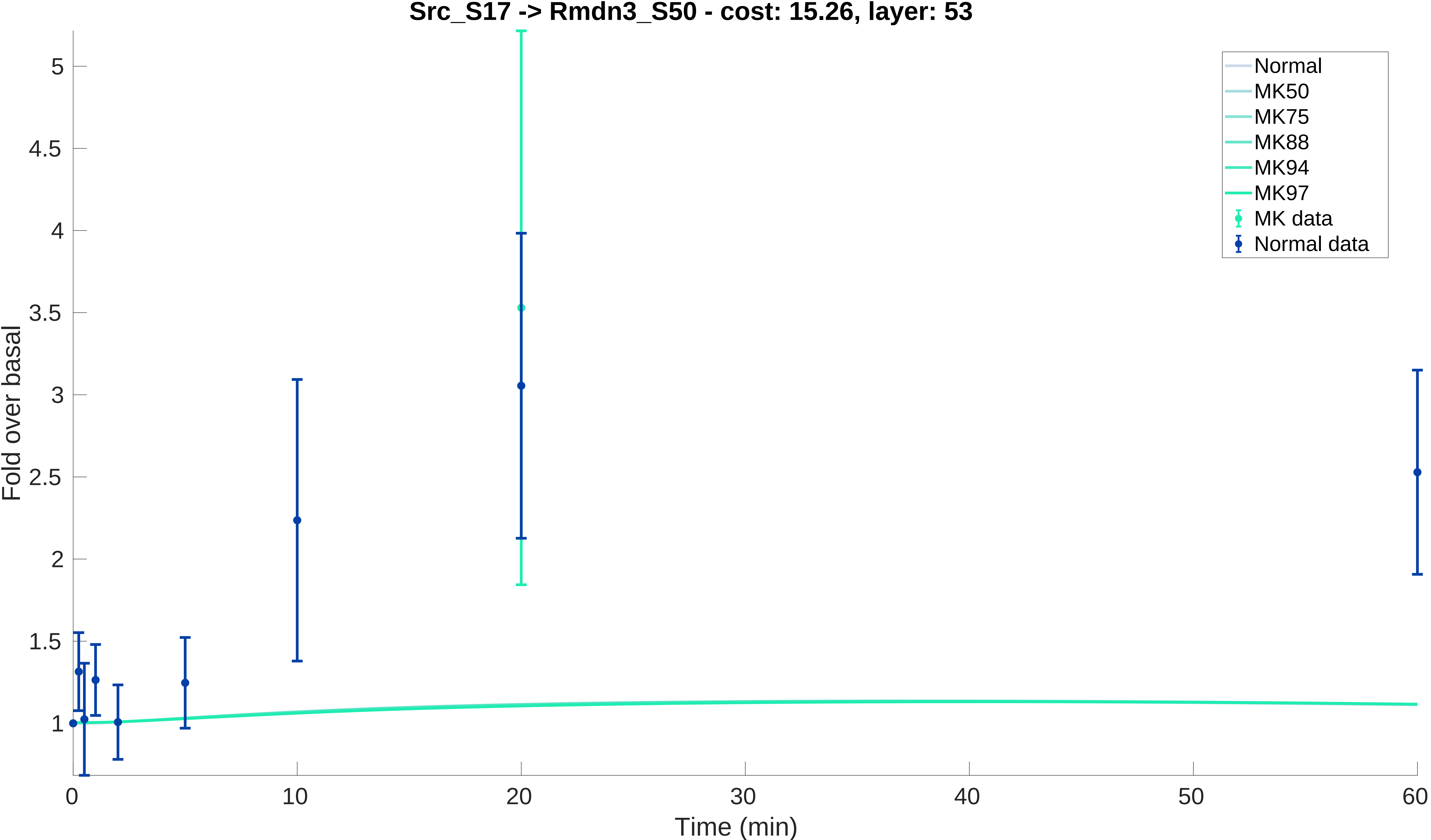

Src_S17 -> Rmdn3_S50 - cost: 15.26, layer: 53
Fold over basal
Time (min)
5
4.5
4
3.5
3
2.5
2
1.5
1
0
10
20
30
40
50
60
Normal
MK50
MK75
MK88
MK94
MK97
MK data
Normal data

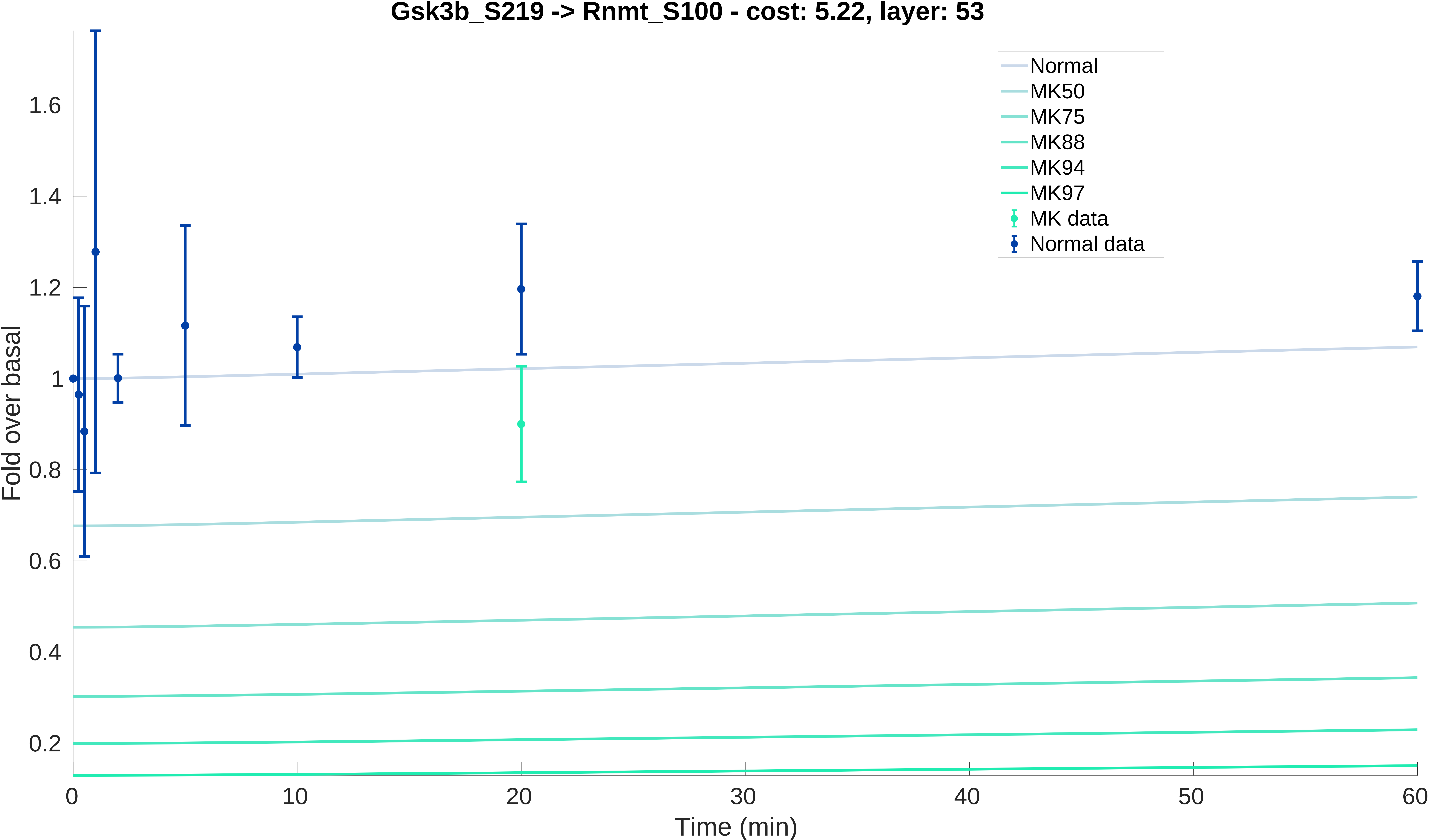

Gsk3b_S219 -> Rnmt_S100 - cost: 5.22, layer: 53
Normal
MK50
MK75
MK88
MK94
MK97
MK data
Normal data
Fold over basal
Time (min)
0
10
20
30
40
50
60
0.2
0.4
0.6
0.8
1
1.2
1.4
1.6

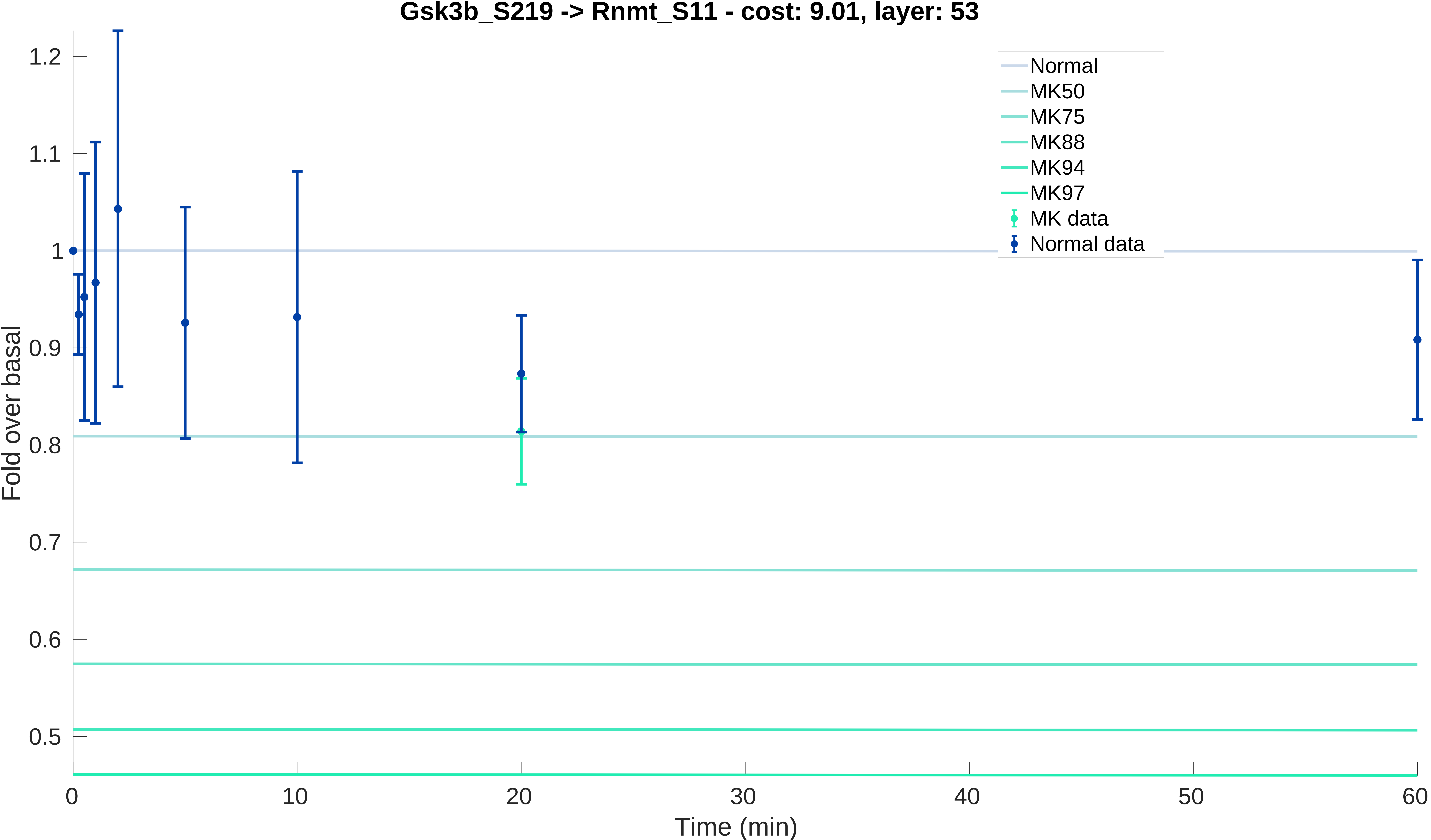

Gsk3b_S219 -> Rnmt_S11 - cost: 9.01, layer: 53
Fold over basal
Time (min)
1.2
1.1
1
0.9
0.8
0.7
0.6
0.5
0
10
20
30
40
50
60
Normal
MK50
MK75
MK88
MK94
MK97
MK data
Normal data

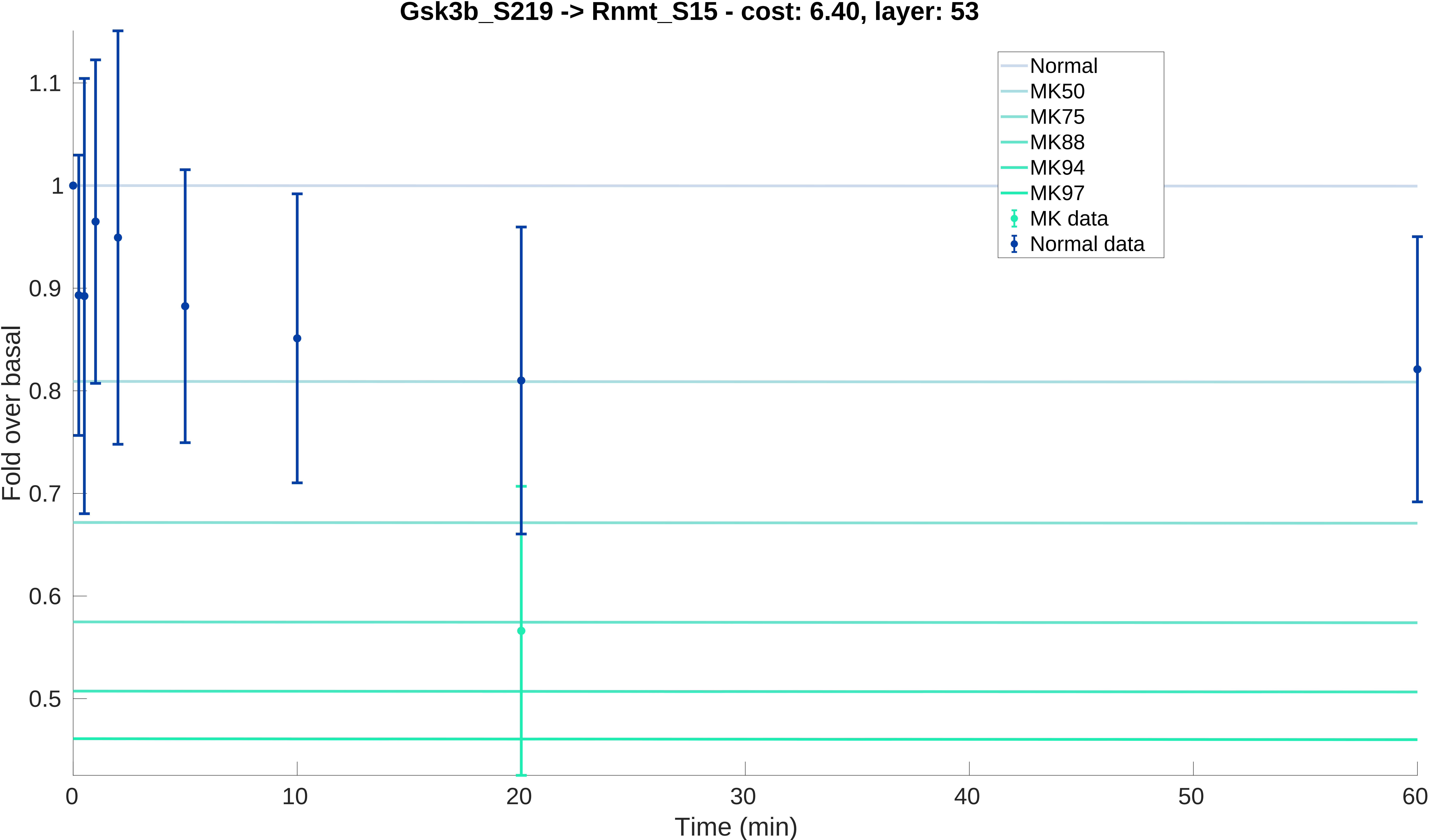

Gsk3b_S219 -> Rnmt_S15 - cost: 6.40, layer: 53
Fold over basal
Time (min)
1.1
1
0.9
0.8
0.7
0.6
0.5
0
10
20
30
40
50
60
Normal
MK50
MK75
MK88
MK94
MK97
MK data
Normal data

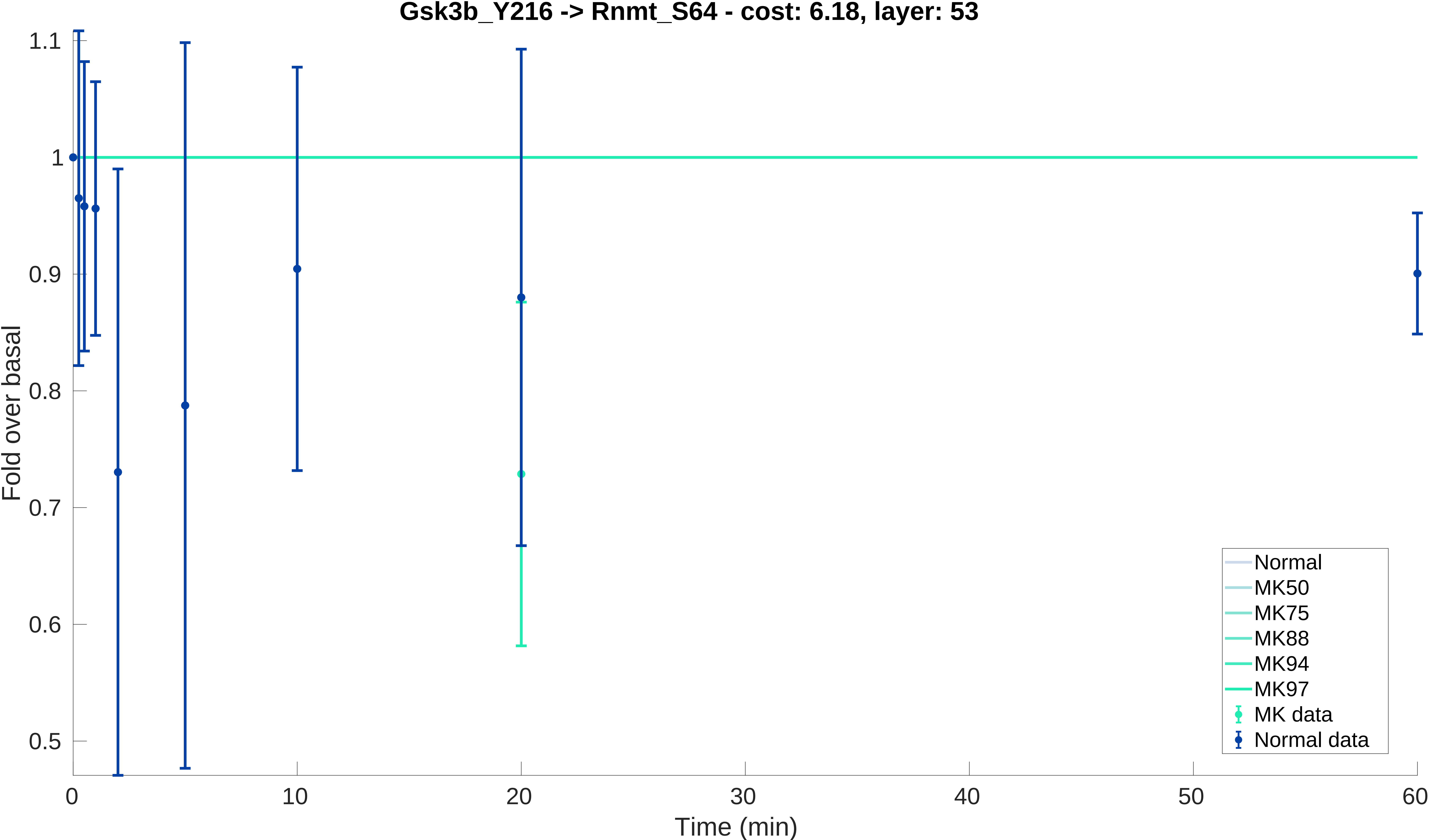

Gsk3b_Y216 -> Rnmt_S64 - cost: 6.18, layer: 53
Fold over basal
Time (min)
1.1
1
0.9
0.8
0.7
0.6
0.5
0
10
20
30
40
50
60
Normal
MK50
MK75
MK88
MK94
MK97
MK data
Normal data

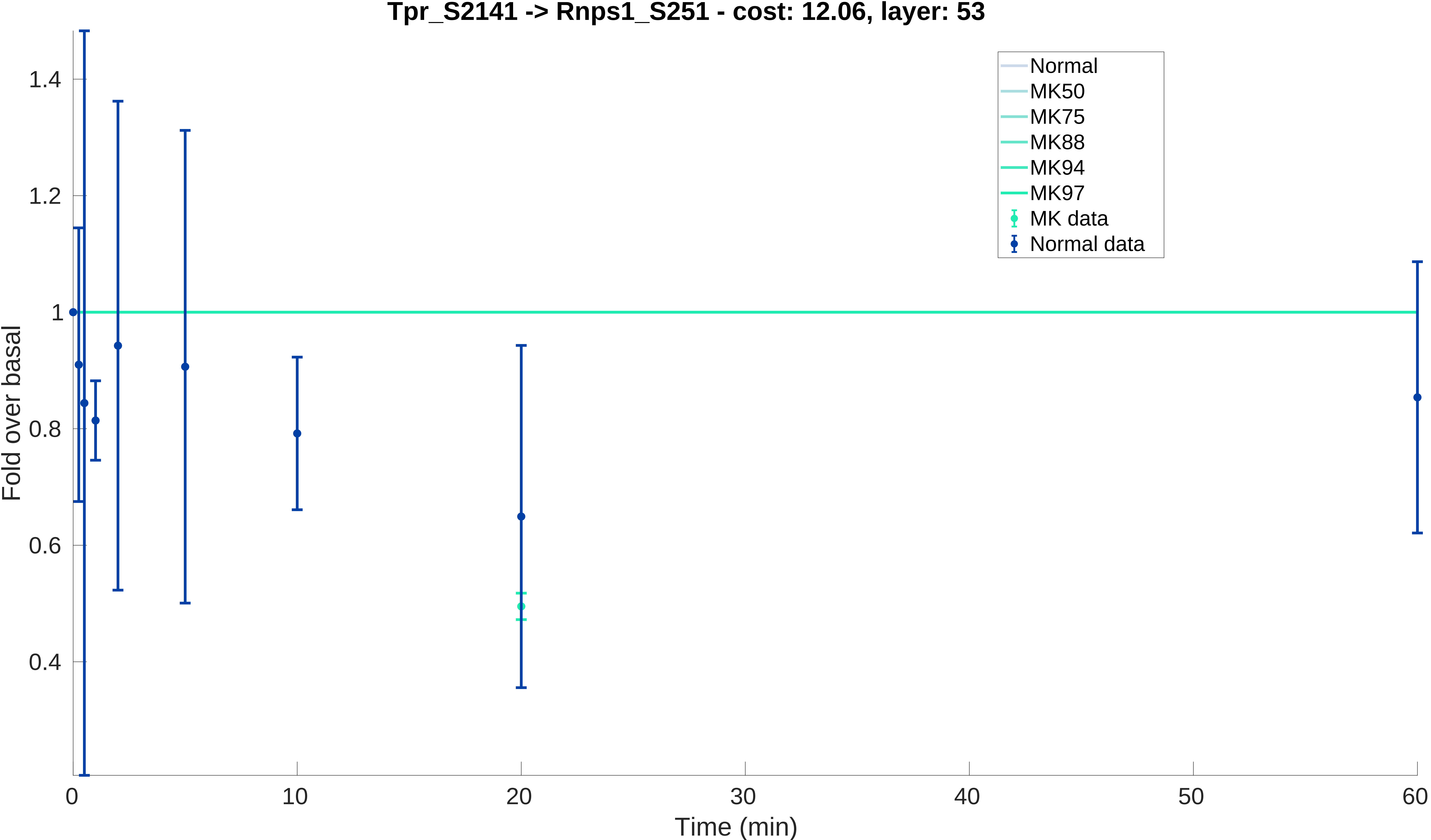

Tpr_S2141 -> Rnps1_S251 - cost: 12.06, layer: 53
Normal
MK50
MK75
MK88
MK94
MK97
MK data
Normal data
Fold over basal
1.4
1.2
1
0.8
0.6
0.4
0
10
20
30
40
50
60
Time (min)

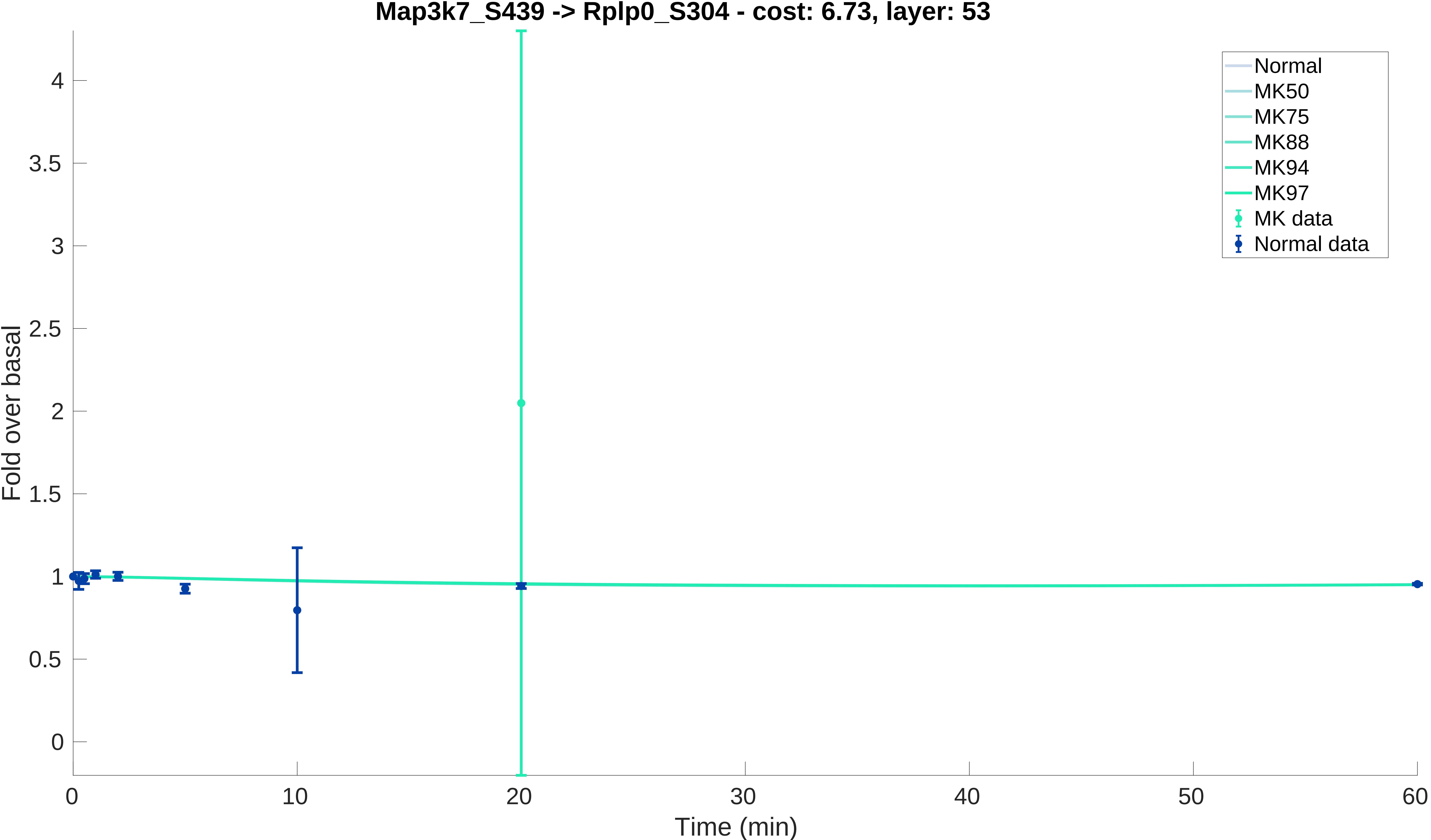

Map3k7_S439 -> Rplp0_S304 - cost: 6.73, layer: 53
Fold over basal
Time (min)
0
0.5
1
1.5
2
2.5
3
3.5
4
0
10
20
30
40
50
60
Normal
MK50
MK75
MK88
MK94
MK97
MK data
Normal data

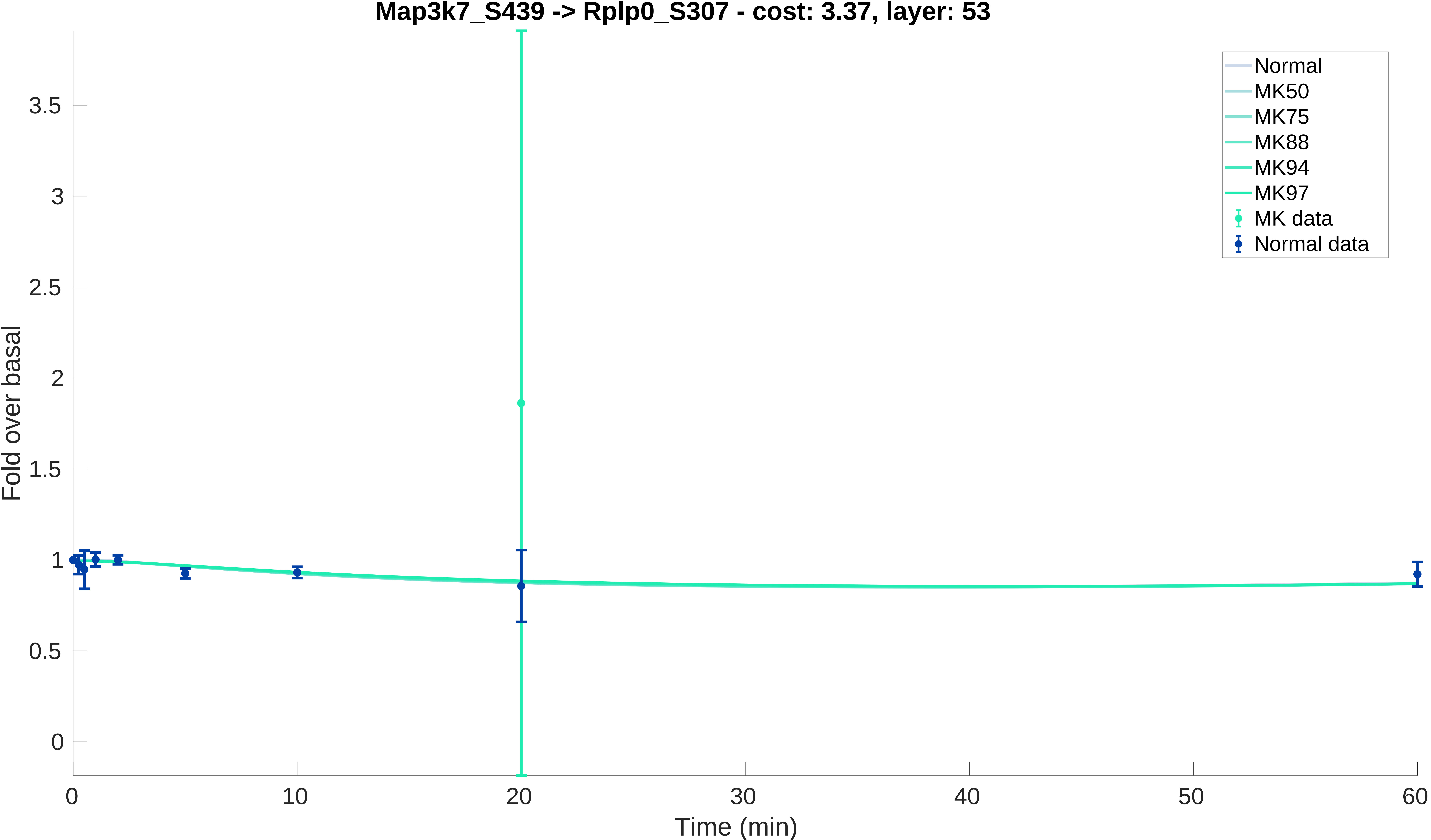

Map3k7_S439 -> Rplp0_S307 - cost: 3.37, layer: 53
Fold over basal
Time (min)
0
0.5
1
1.5
2
2.5
3
3.5
0
10
20
30
40
50
60
Normal
MK50
MK75
MK88
MK94
MK97
MK data
Normal data

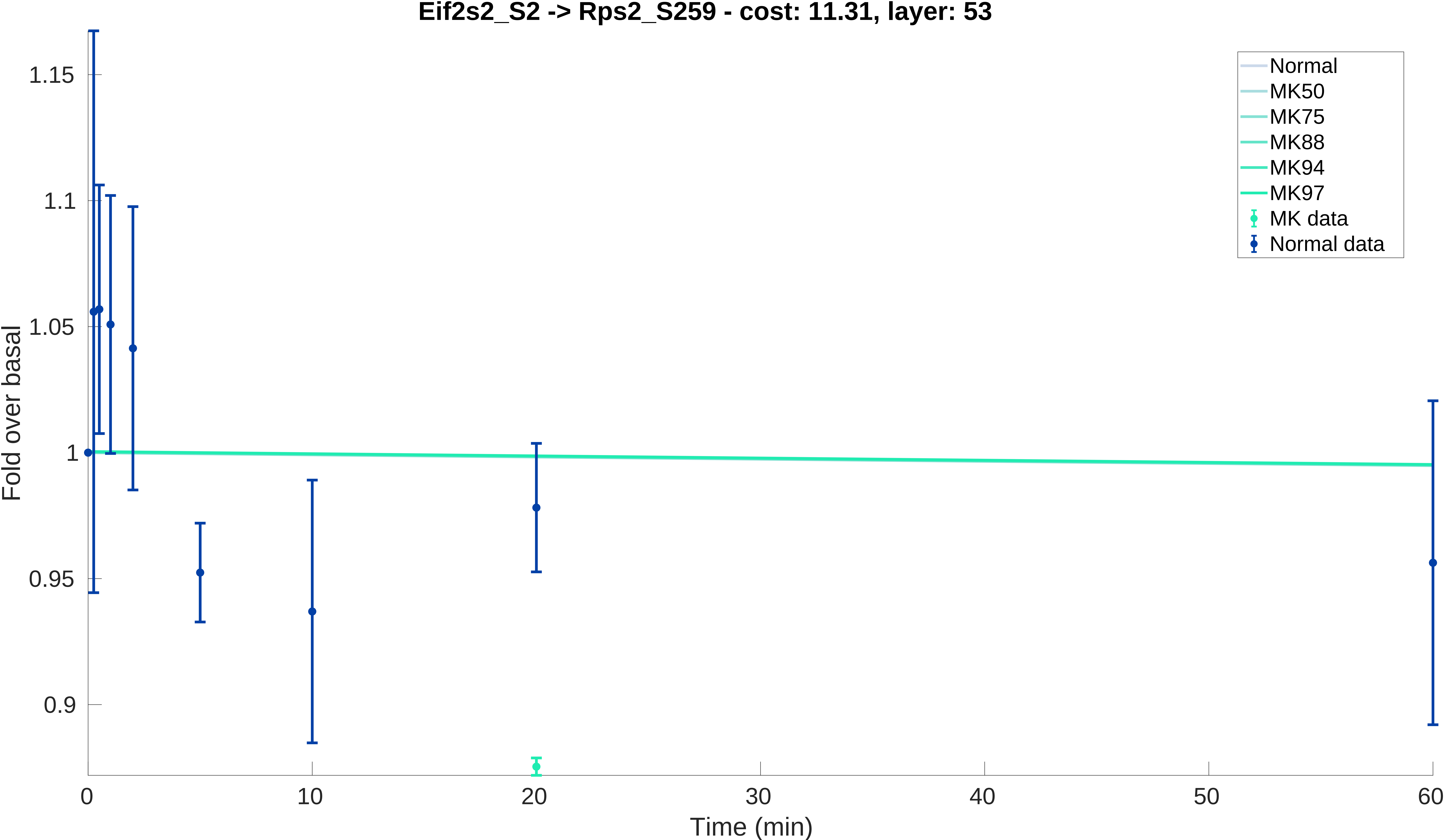

Eif2s2_S2 -> Rps2_S259 - cost: 11.31, layer: 53
Normal
MK50
MK75
MK88
MK94
MK97
MK data
Normal data
Fold over basal
1.15
1.1
1.05
1
0.95
0.9
0
10
20
30
40
50
60
Time (min)

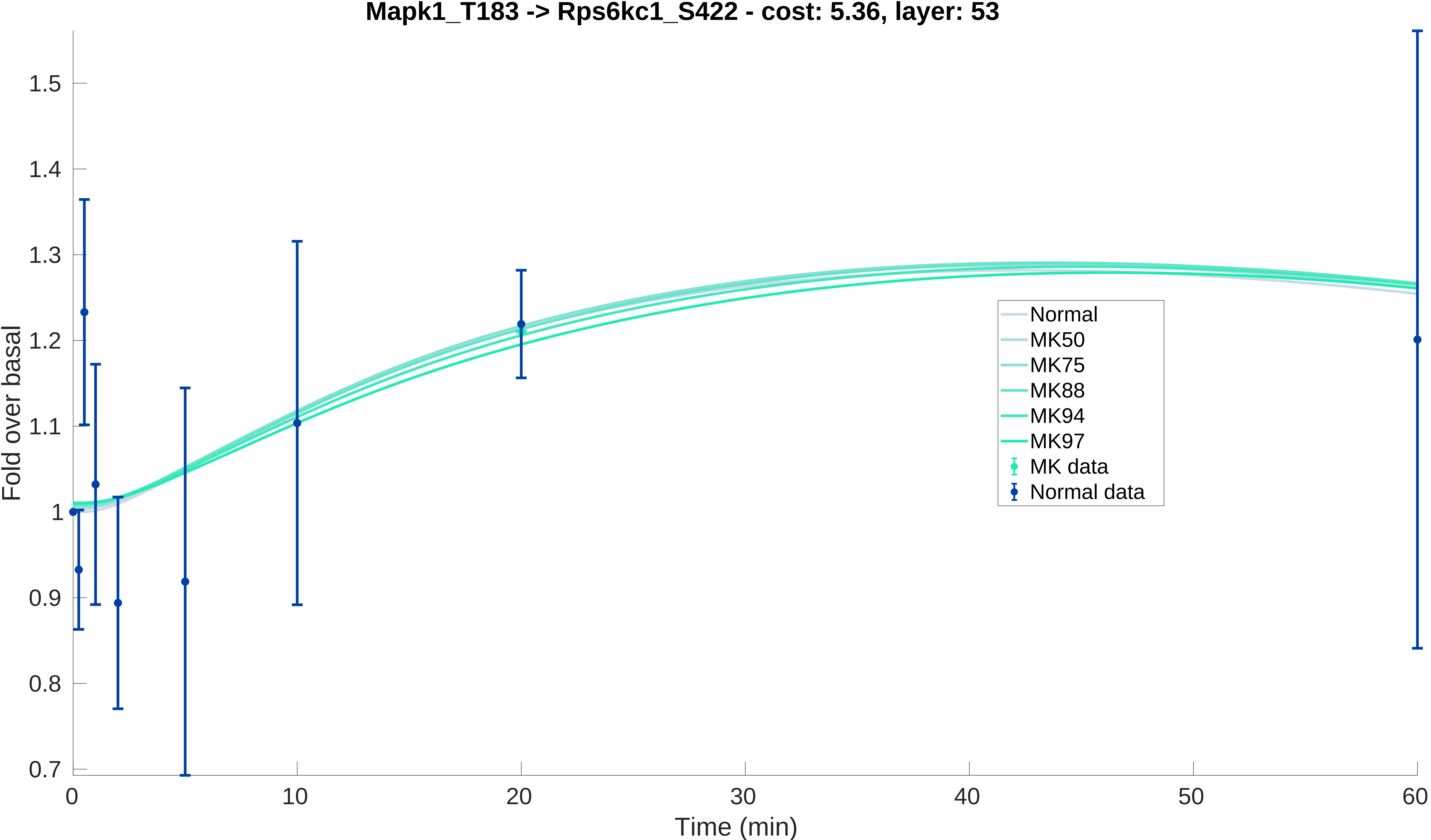
Mapk1_T183 -> Rps6kc1_S422 - cost: 5.36, layer: 53
Fold over basal
Time (min)
1.5
1.4
1.3
1.2
1.1
1
0.9
0.8
0.7
0
10
20
30
40
50
60
Normal
MK50
MK75
MK88
MK94
MK97
MK data
Normal data

Mapk1_T183 -> Rps6kc1_S426 - cost: 1.37, layer: 53

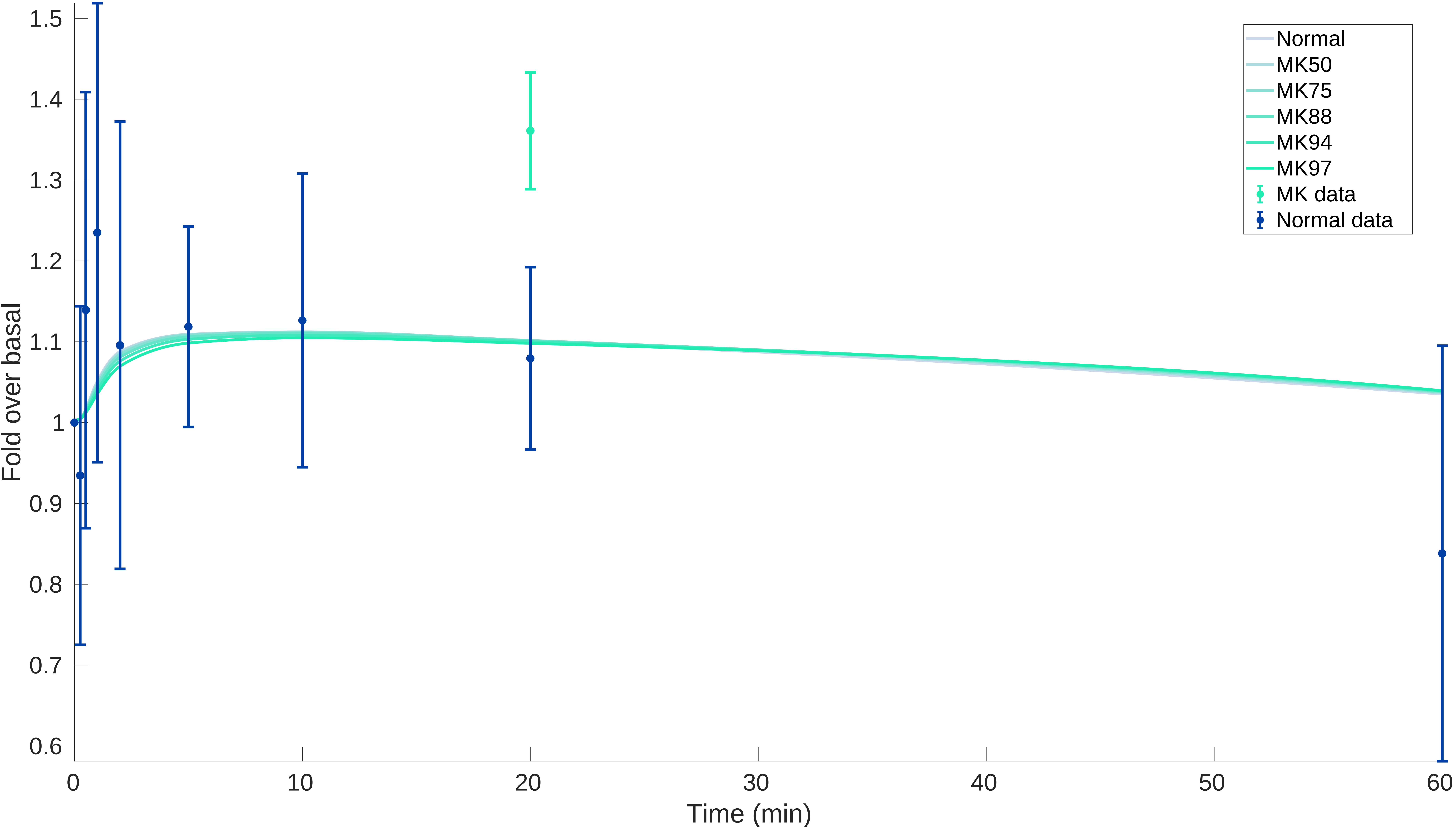

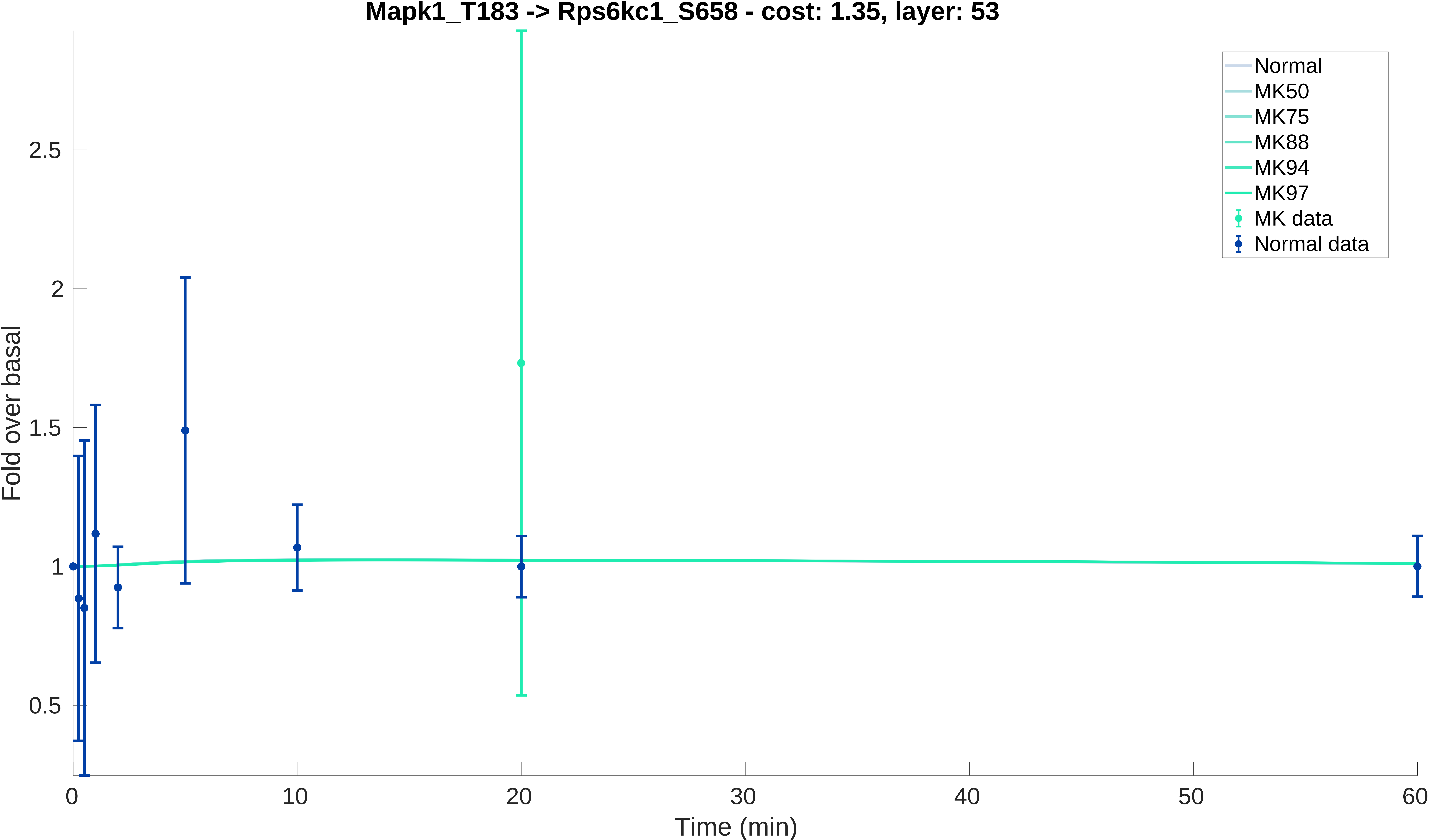

Mapk1_T183 -> Rps6kc1_S658 - cost: 1.35, layer: 53
Fold over basal
0.5
1
1.5
2
2.5
0
10
20
30
40
50
60
Time (min)
Normal
MK50
MK75
MK88
MK94
MK97
MK data
Normal data

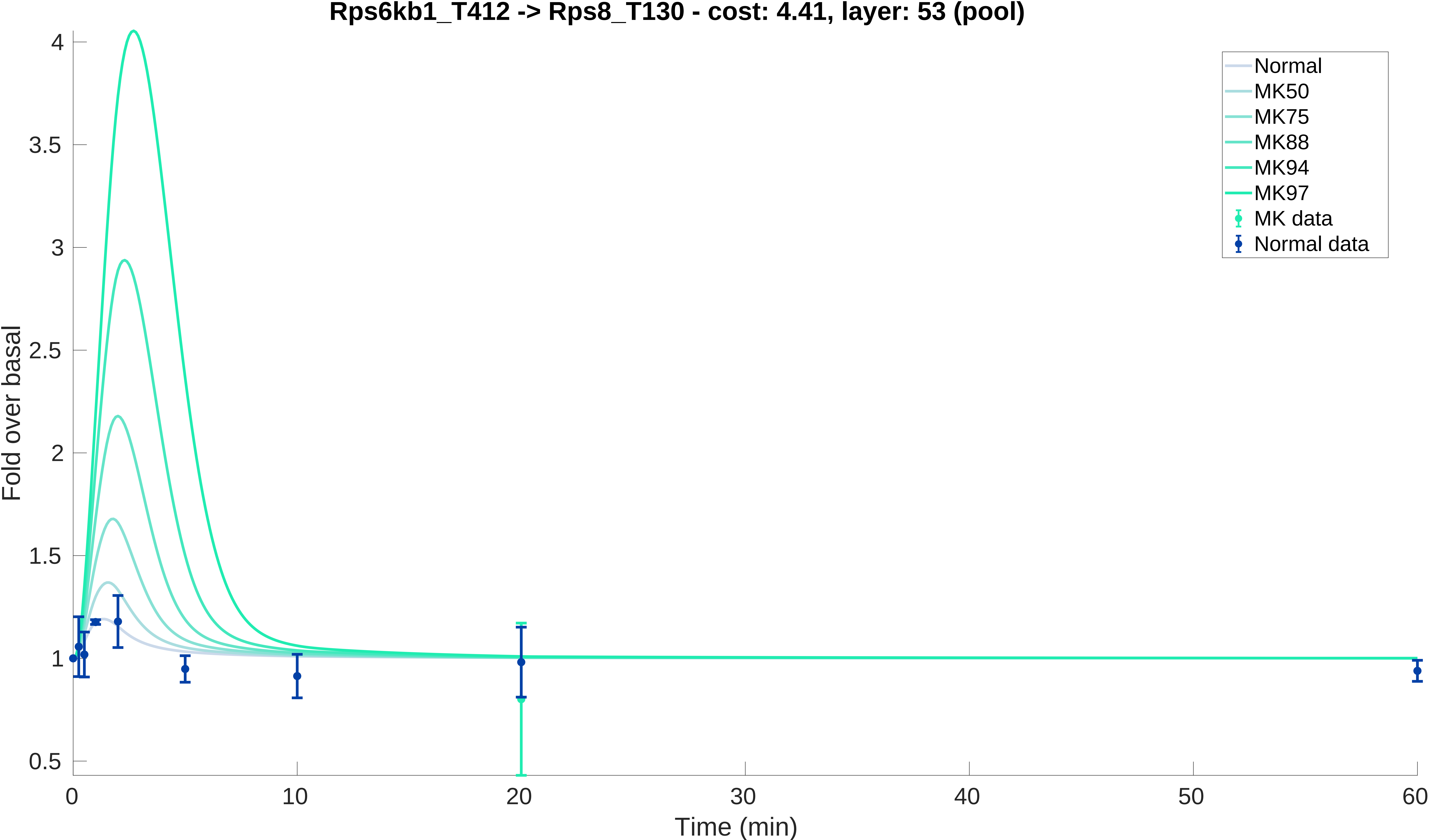
Rps6kb1_T412 -> Rps8_T130 - cost: 4.41, layer: 53 (pool)
Fold over basal
Time (min)
Normal
MK50
MK75
MK88
MK94
MK97
MK data
Normal data
0.5
1
1.5
2
2.5
3
3.5
4
0
10
20
30
40
50
60

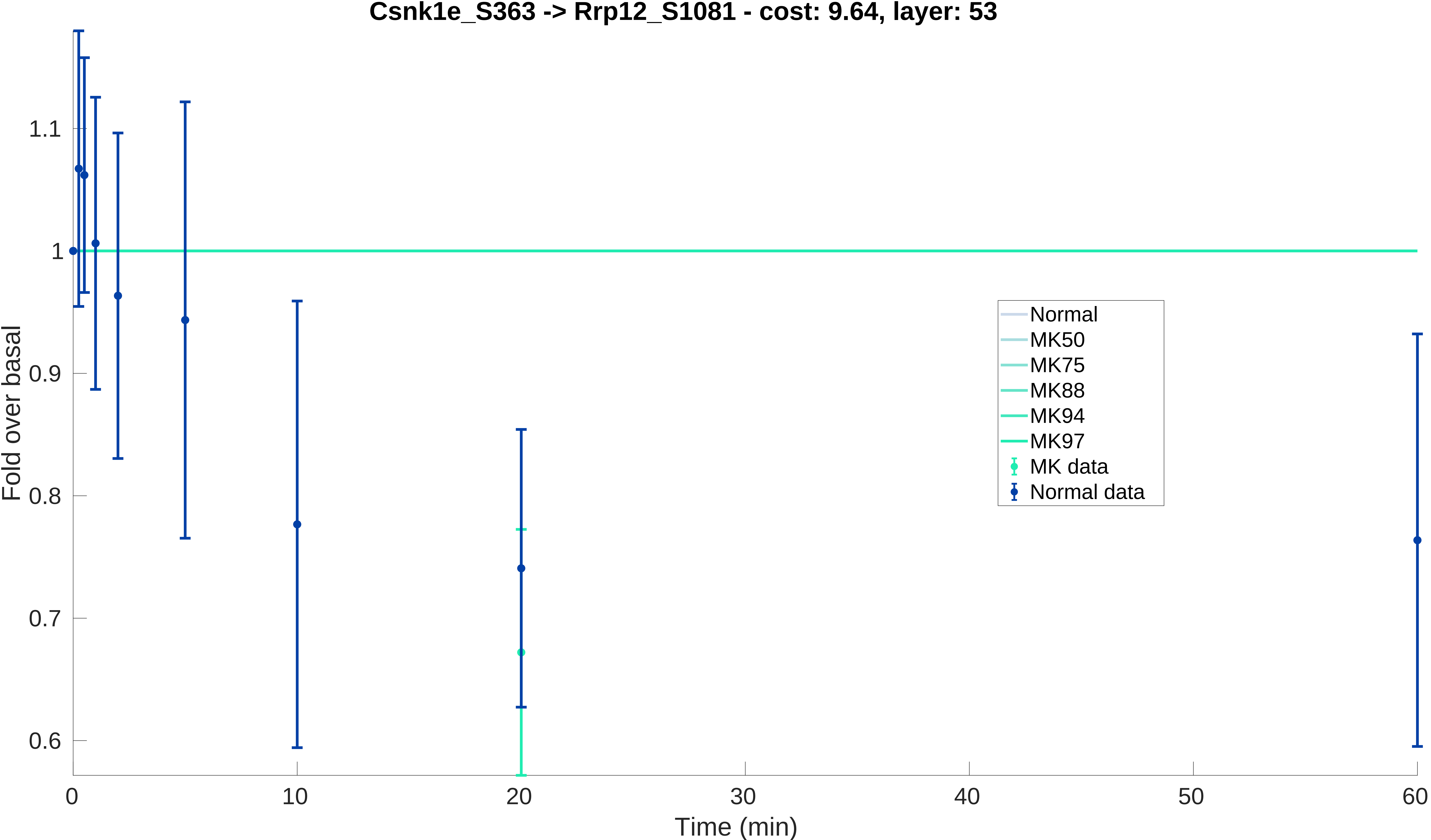

Csnk1e_S363 -> Rrp12_S1081 - cost: 9.64, layer: 53
Fold over basal
1.1
1
0.9
0.8
0.7
0.6
0
10
20
30
40
50
60
Time (min)
Normal
MK50
MK75
MK88
MK94
MK97
MK data
Normal data

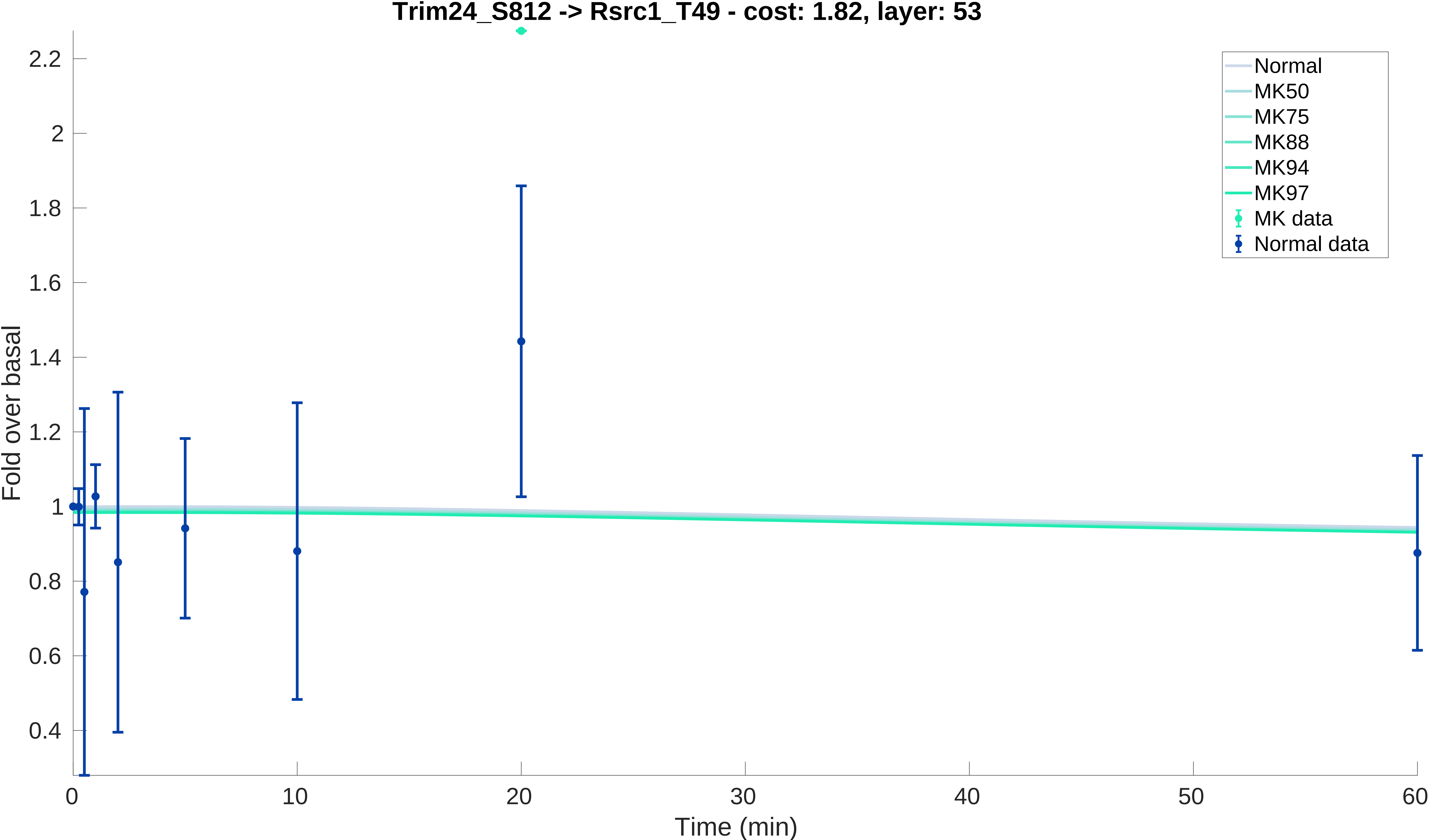

Trim24_S812 -> Rsrc1_T49 - cost: 1.82, layer: 53
Fold over basal
Time (min)
0
10
20
30
40
50
60
0.4
0.6
0.8
1
1.2
1.4
1.6
1.8
2
2.2
Normal
MK50
MK75
MK88
MK94
MK97
MK data
Normal data

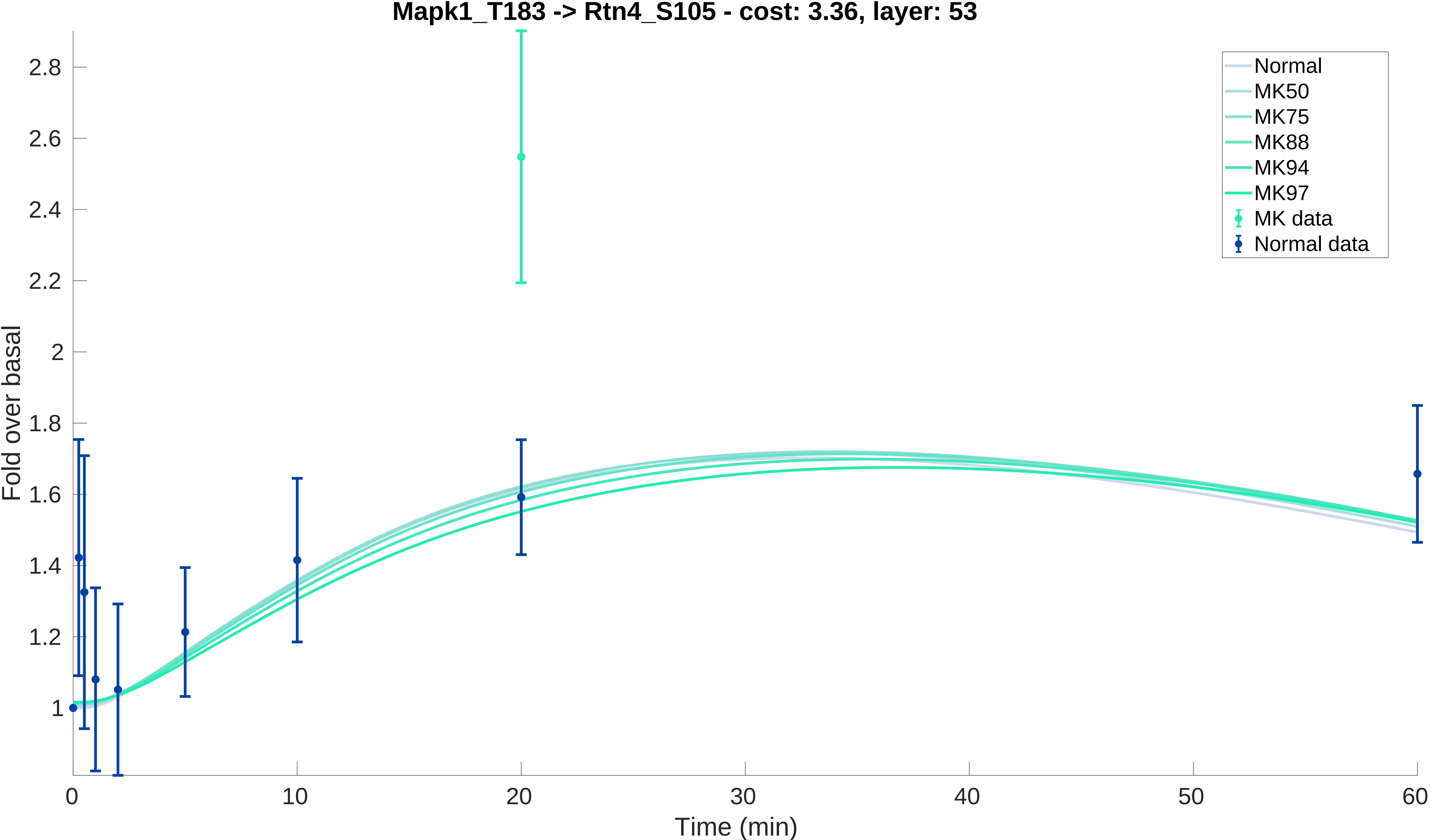

Mapk1_T183 -> Rtn4_S105 - cost: 3.36, layer: 53
Fold over basal
Time (min)
2.8
2.6
2.4
2.2
2
1.8
1.6
1.4
1.2
1
0
10
20
30
40
50
60
Normal
MK50
MK75
MK88
MK94
MK97
MK data
Normal data

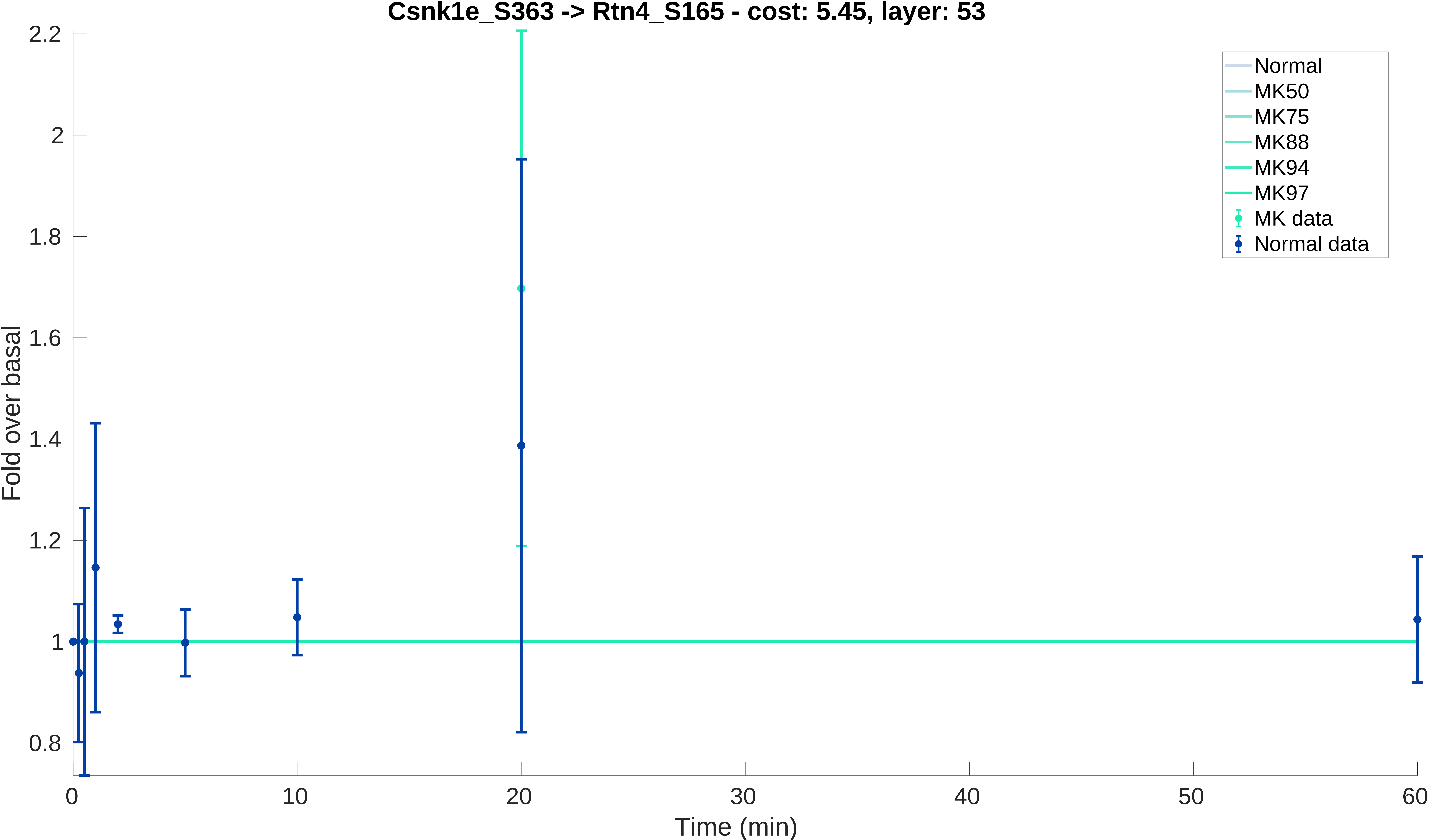

Csnk1e_S363 -> Rtn4_S165 - cost: 5.45, layer: 53
Fold over basal
Time (min)
2.2
2
1.8
1.6
1.4
1.2
1
0.8
0
10
20
30
40
50
60
Normal
MK50
MK75
MK88
MK94
MK97
MK data
Normal data

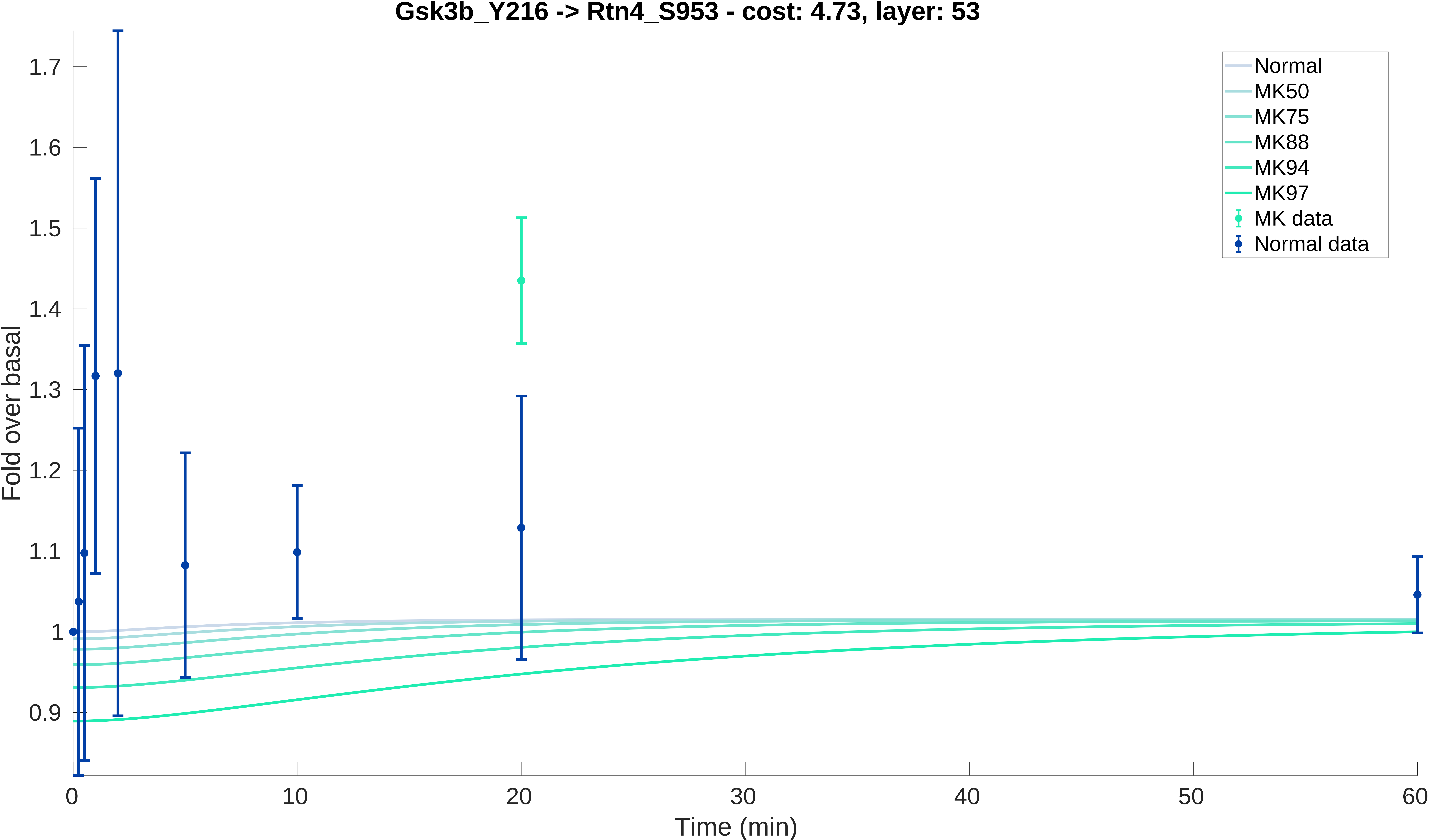

Gsk3b_Y216 -> Rtn4_S953 - cost: 4.73, layer: 53
Fold over basal
Time (min)
0.9
1
1.1
1.2
1.3
1.4
1.5
1.6
1.7
0
10
20
30
40
50
60
Normal
MK50
MK75
MK88
MK94
MK97
MK data
Normal data

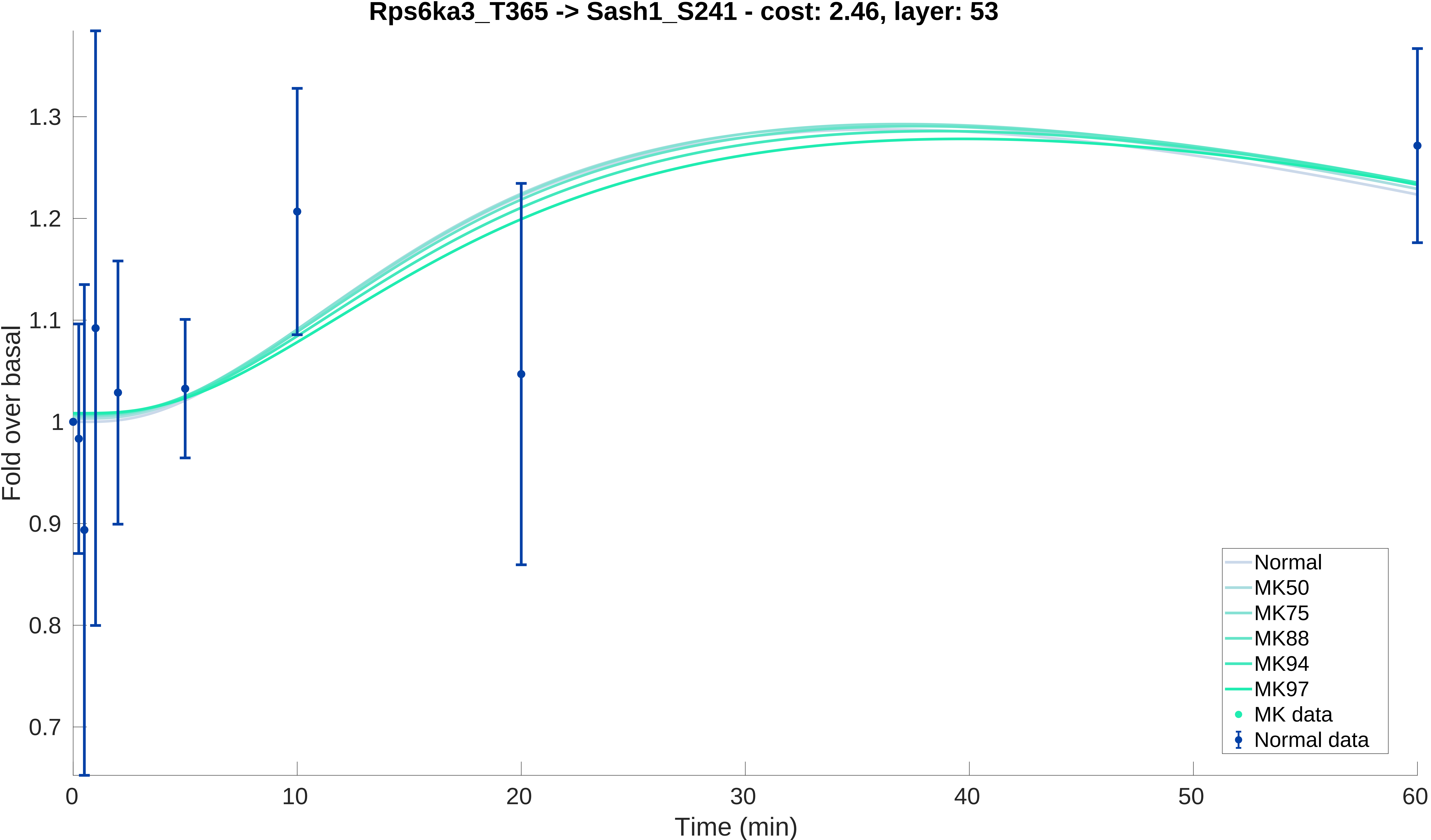
Rps6ka3_T365 -> Sash1_S241 - cost: 2.46, layer: 53
Fold over basal
1.3
1.2
1.1
1
0.9
0.8
0.7
0
10
20
30
40
50
60
Time (min)
Normal
MK50
MK75
MK88
MK94
MK97
MK data
Normal data

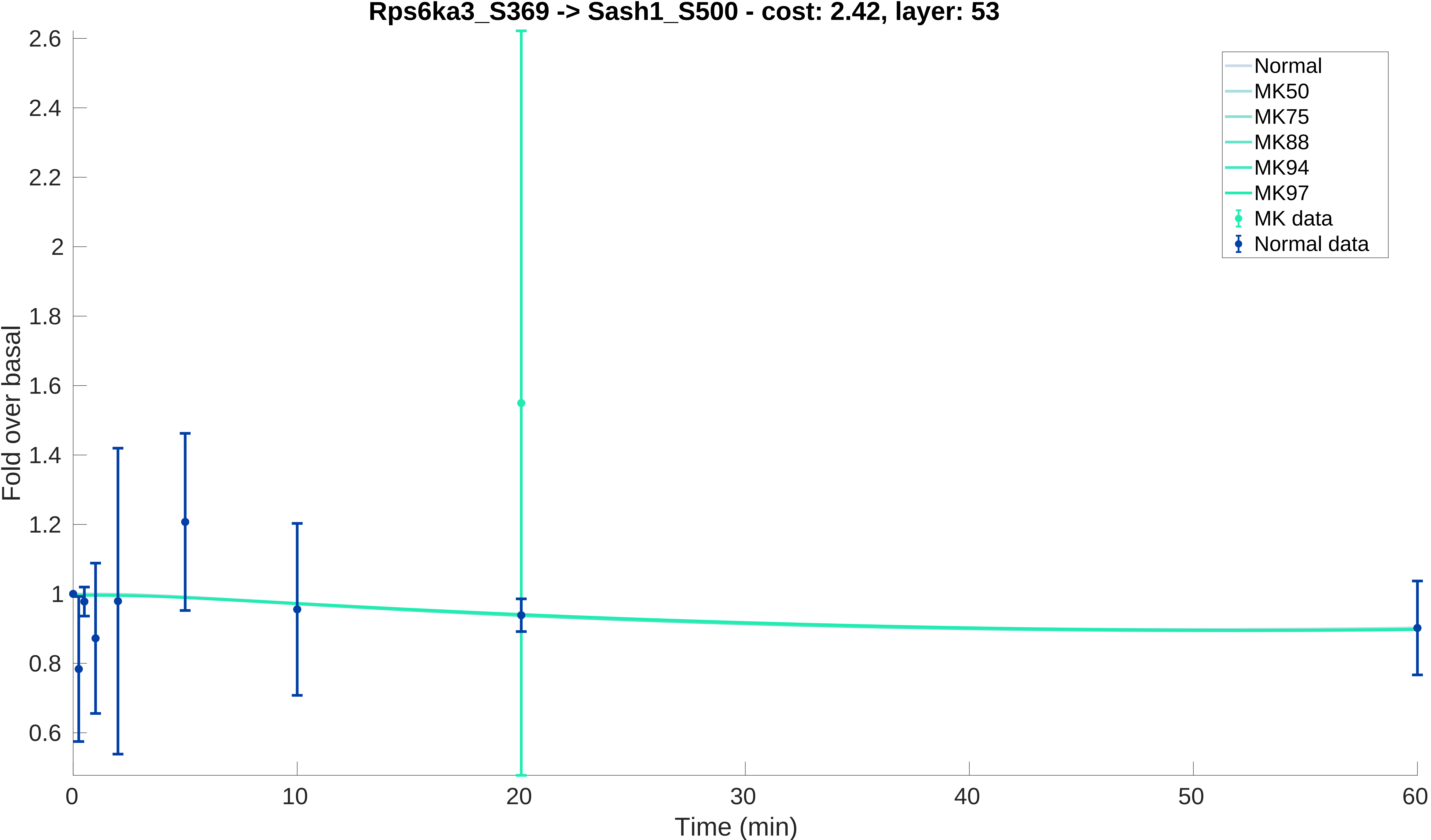

Rps6ka3_S369 -> Sash1_S500 - cost: 2.42, layer: 53
Fold over basal
Time (min)
Normal
MK50
MK75
MK88
MK94
MK97
MK data
Normal data
2.6
2.4
2.2
2
1.8
1.6
1.4
1.2
1
0.8
0.6
0
10
20
30
40
50
60

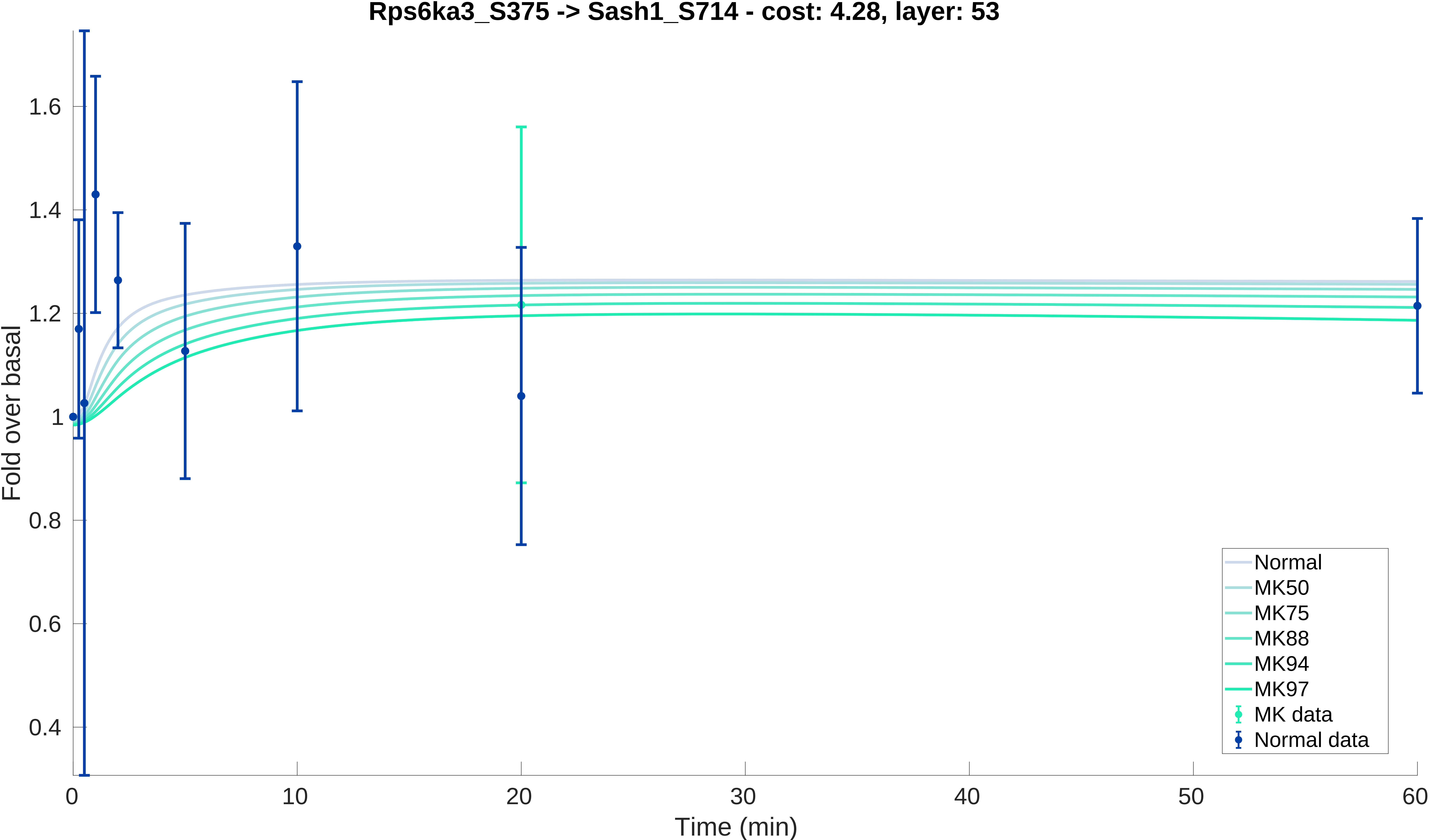

Rps6ka3_S375 -> Sash1_S714 - cost: 4.28, layer: 53
Fold over basal
1.6
1.4
1.2
1
0.8
0.6
0.4
0
10
20
30
40
50
60
Time (min)
Normal
MK50
MK75
MK88
MK94
MK97
MK data
Normal data

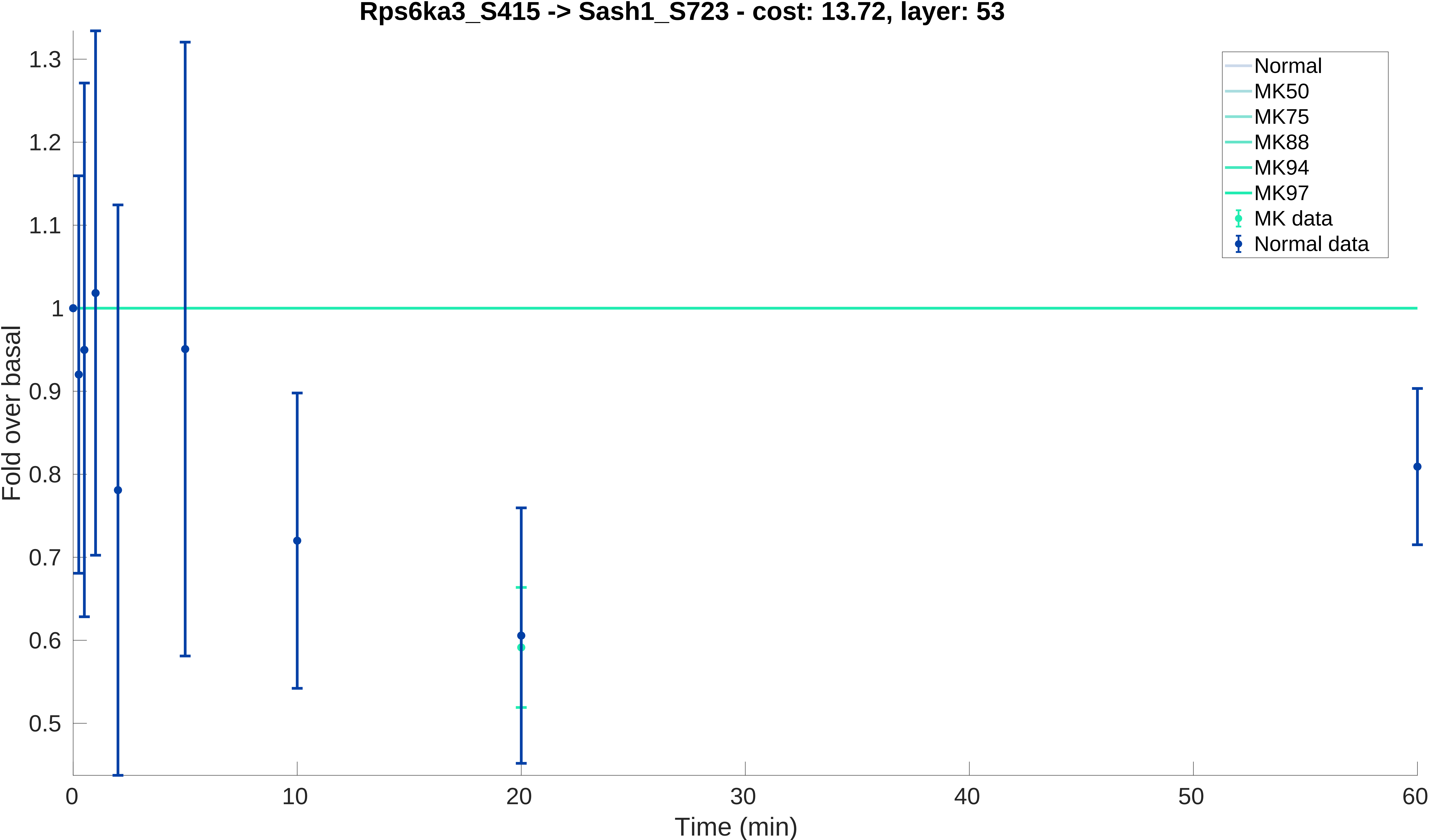

Rps6ka3_S415 -> Sash1_S723 - cost: 13.72, layer: 53
Fold over basal
Time (min)
0.5
0.6
0.7
0.8
0.9
1
1.1
1.2
1.3
0
10
20
30
40
50
60
Normal
MK50
MK75
MK88
MK94
MK97
MK data
Normal data

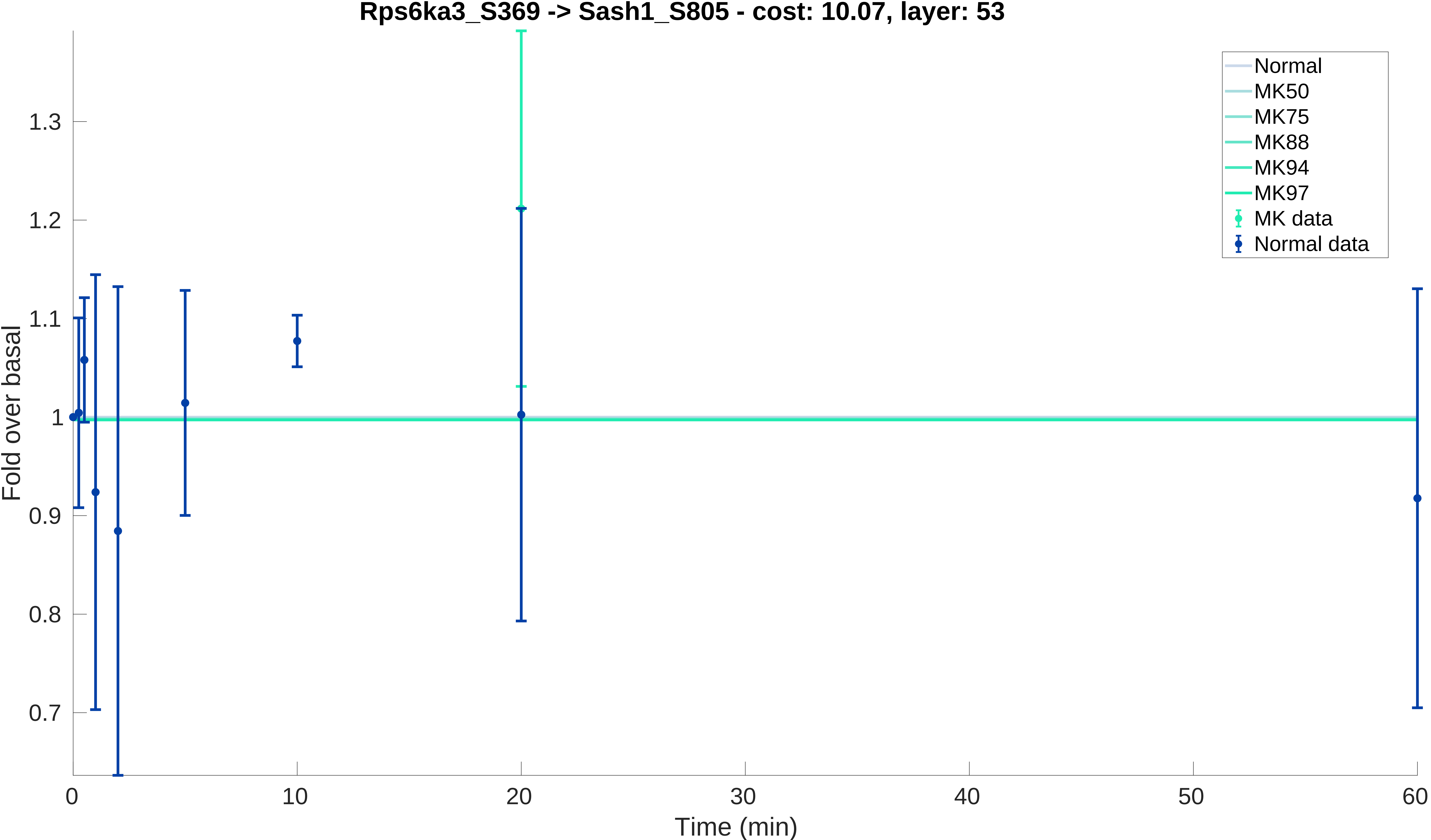
Rps6ka3_S369 -> Sash1_S805 - cost: 10.07, layer: 53
Fold over basal
Time (min)
1.3
1.2
1.1
1
0.9
0.8
0.7
0
10
20
30
40
50
60
Normal
MK50
MK75
MK88
MK94
MK97
MK data
Normal data

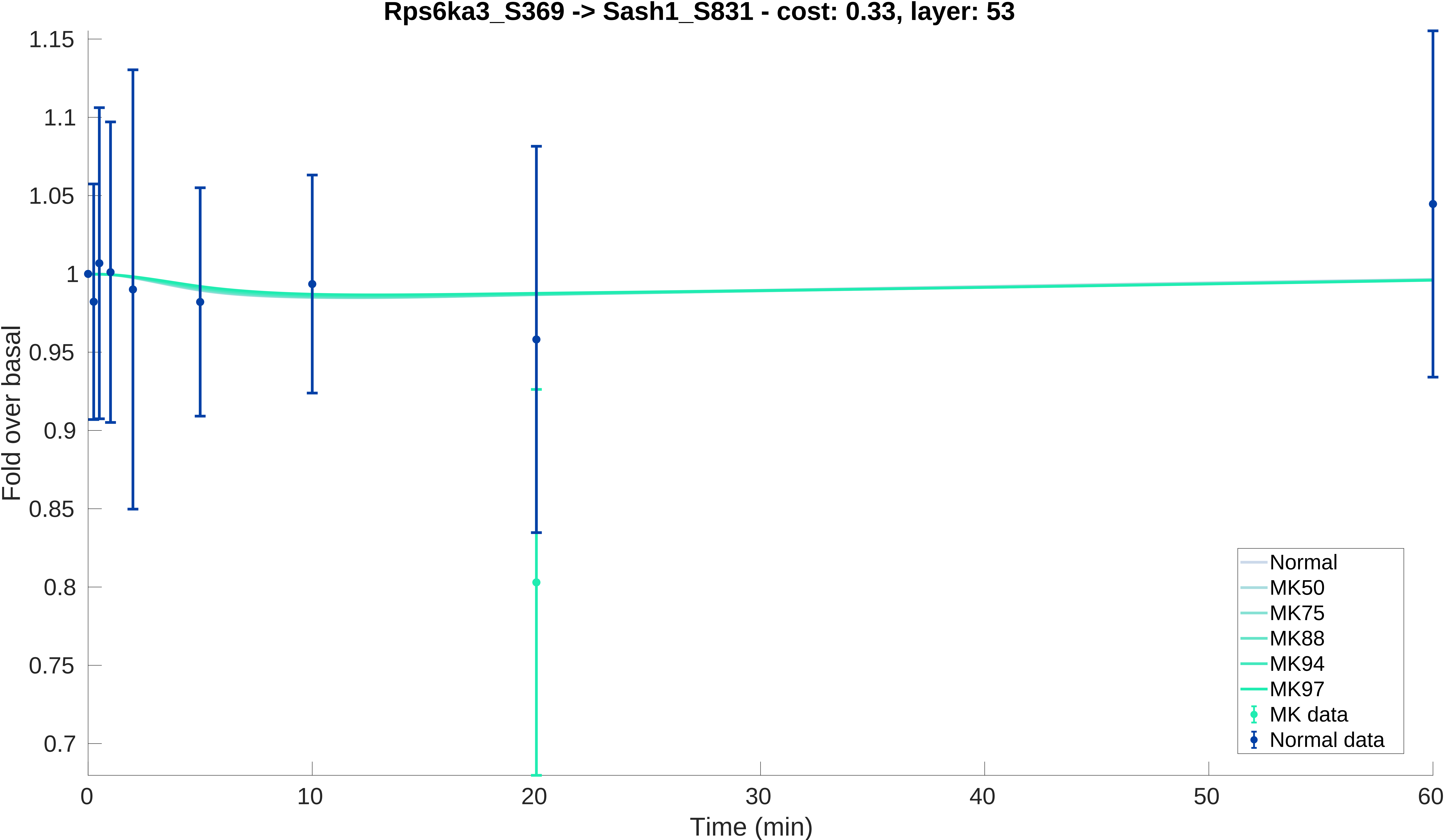

Rps6ka3_S369 -> Sash1_S831 - cost: 0.33, layer: 53
Fold over basal
1.15
1.1
1.05
1
0.95
0.9
0.85
0.8
0.75
0.7
0
10
20
30
40
50
60
Time (min)
Normal
MK50
MK75
MK88
MK94
MK97
MK data
Normal data

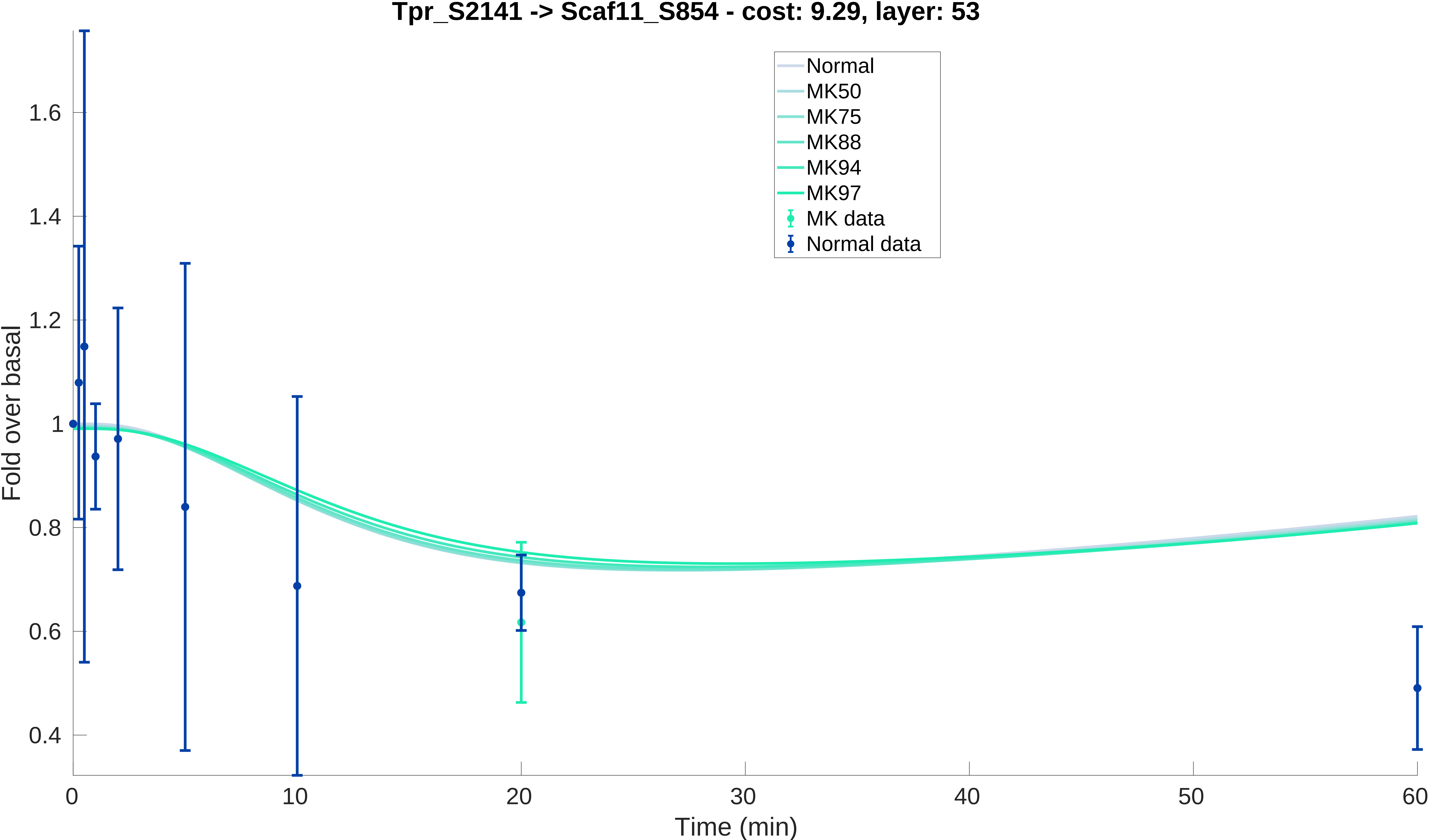

Tpr_S2141 -> Scaf11_S854 - cost: 9.29, layer: 53
Normal
MK50
MK75
MK88
MK94
MK97
MK data
Normal data
Fold over basal
0.4
0.6
0.8
1
1.2
1.4
1.6
0
10
20
30
40
50
60
Time (min)

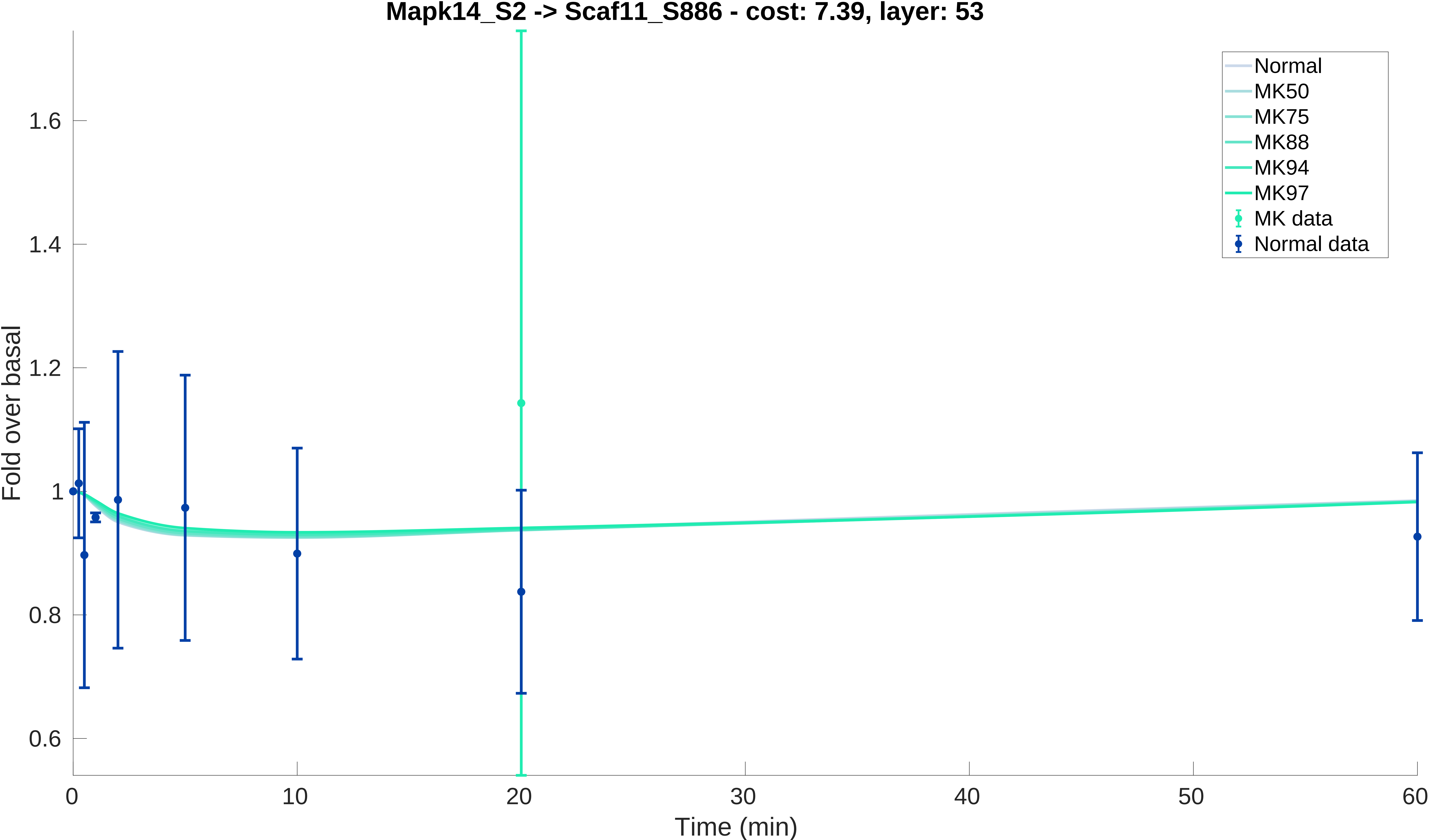
Mapk14_S2 -> Scaf11_S886 - cost: 7.39, layer: 53
Fold over basal
0.6
0.8
1
1.2
1.4
1.6
0
10
20
30
40
50
60
Time (min)
Normal
MK50
MK75
MK88
MK94
MK97
MK data
Normal data

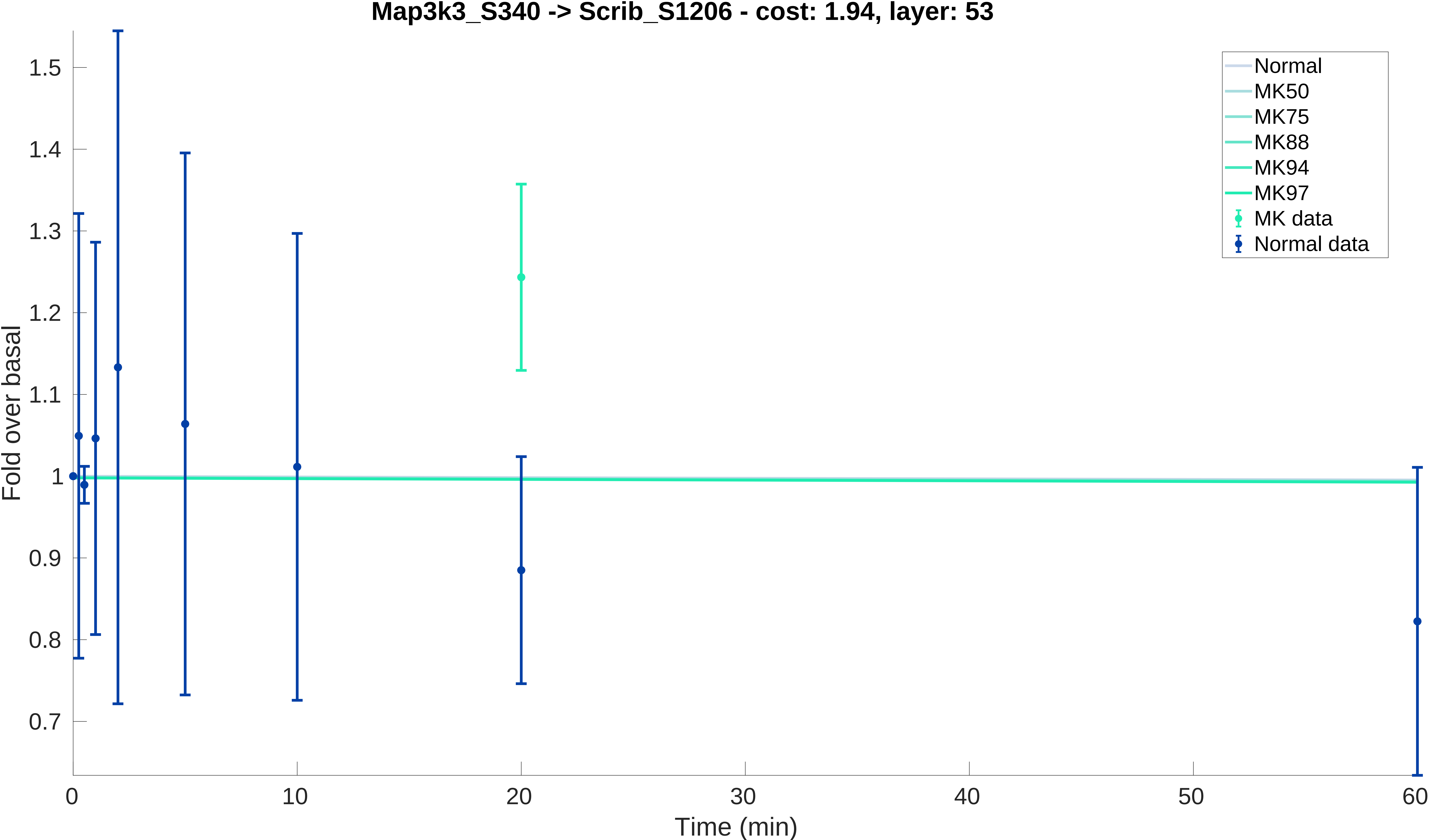

Map3k3_S340 -> Scrib_S1206 - cost: 1.94, layer: 53
Fold over basal
Time (min)
Normal
MK50
MK75
MK88
MK94
MK97
MK data
Normal data
1.5
1.4
1.3
1.2
1.1
1
0.9
0.8
0.7
0
10
20
30
40
50
60

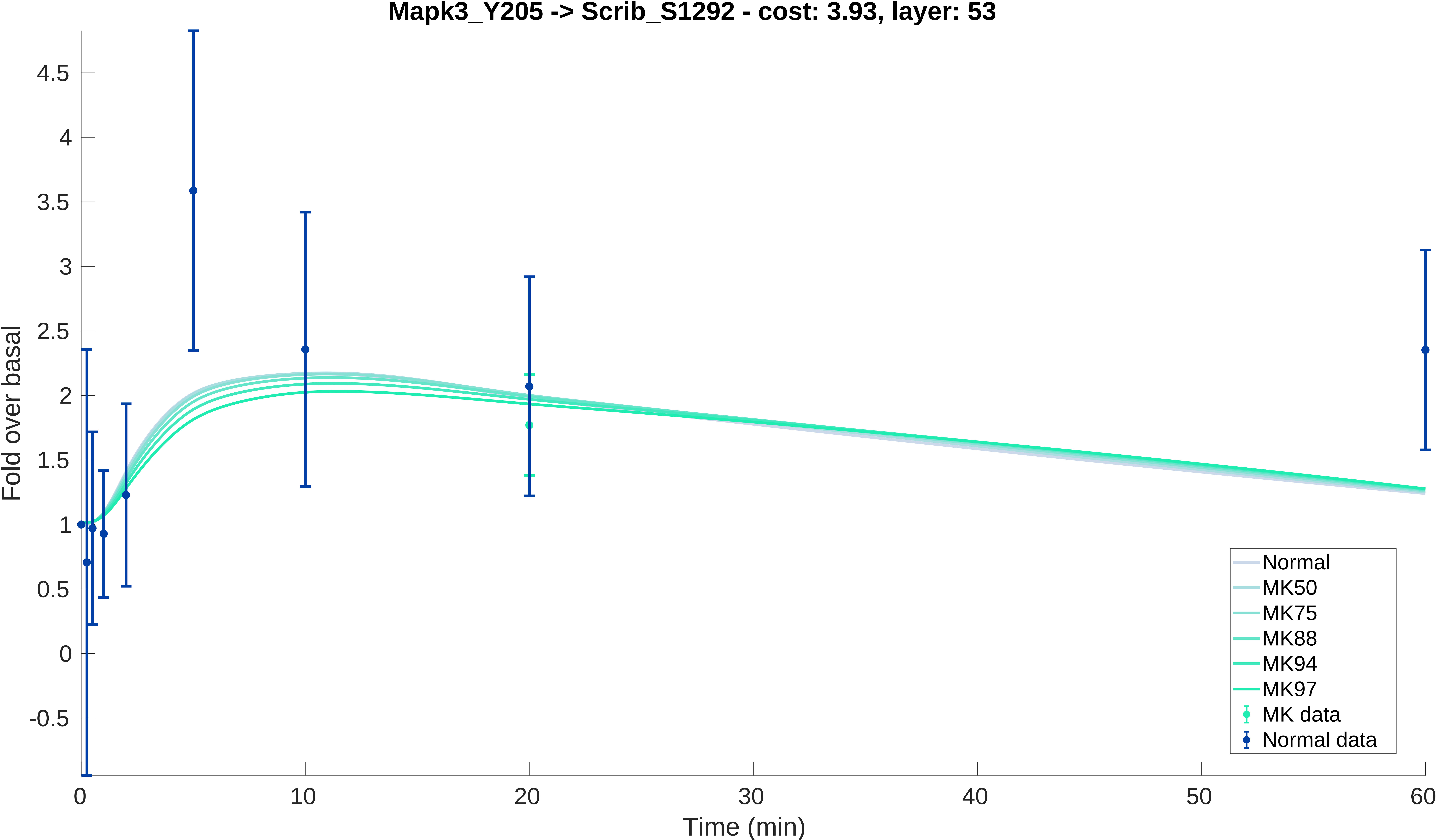

Mapk3_Y205 -> Scrib_S1292 - cost: 3.93, layer: 53
Fold over basal
Time (min)
4.5
4
3.5
3
2.5
2
1.5
1
0.5
0
-0.5
0
10
20
30
40
50
60
Normal
MK50
MK75
MK88
MK94
MK97
MK data
Normal data

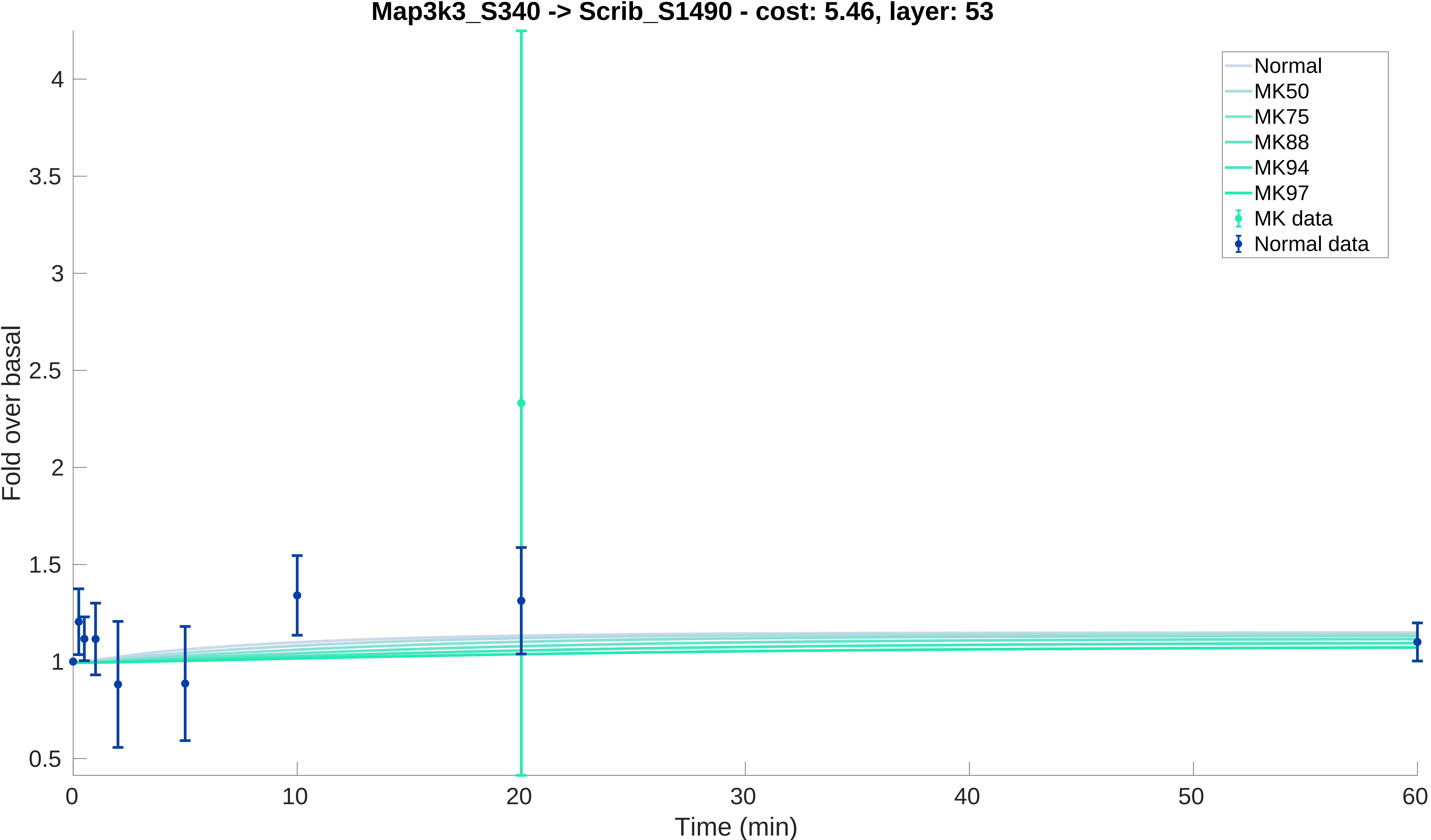

Map3k3_S340 -> Scrib_S1490 - cost: 5.46, layer: 53
Fold over basal
Time (min)
0.5
1
1.5
2
2.5
3
3.5
4
0
10
20
30
40
50
60
Normal
MK50
MK75
MK88
MK94
MK97
MK data
Normal data

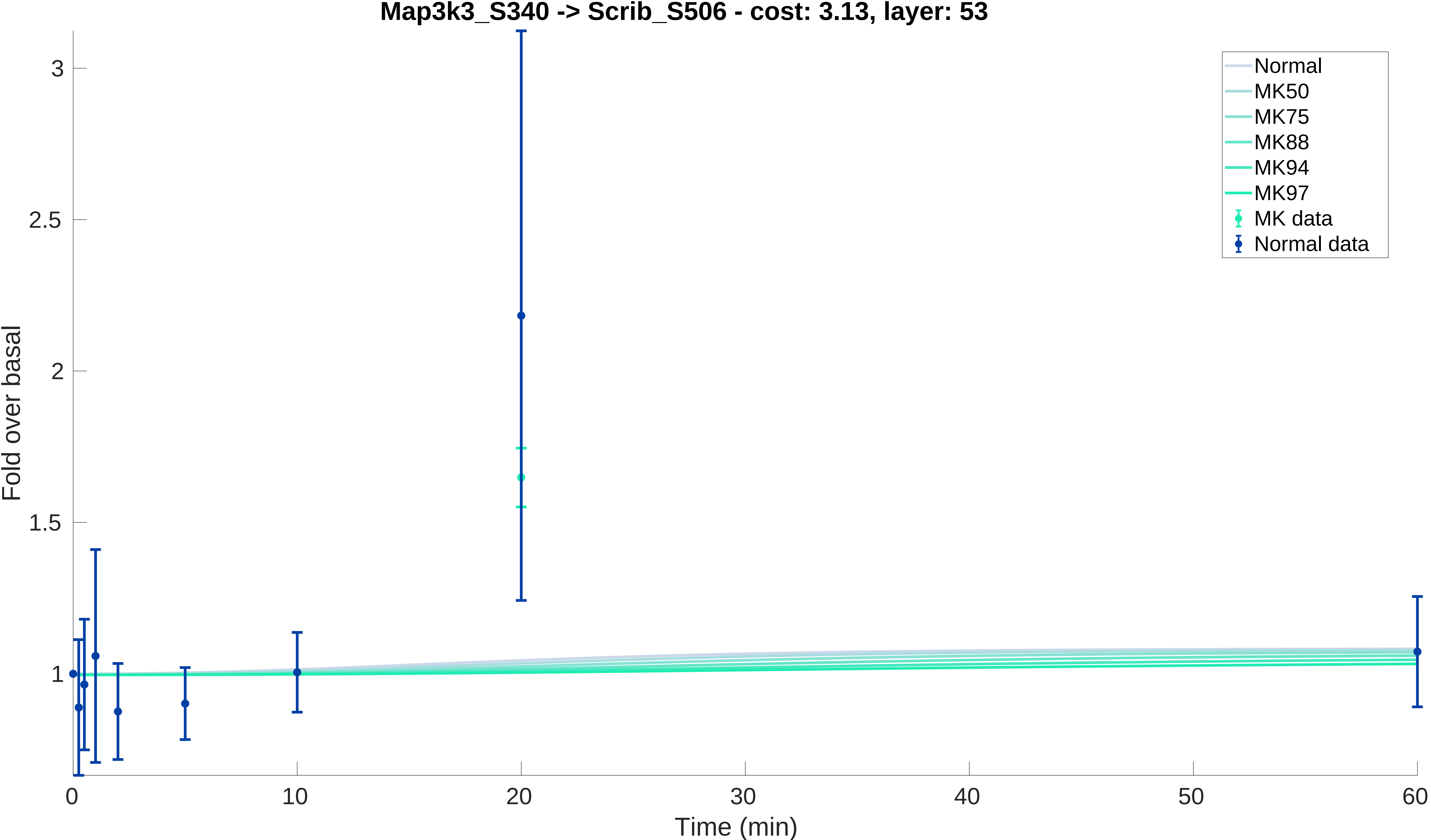
Map3k3_S340 -> Scrib_S506 - cost: 3.13, layer: 53
Fold over basal
Time (min)
0
10
20
30
40
50
60
1
1.5
2
2.5
3
Normal
MK50
MK75
MK88
MK94
MK97
MK data
Normal data

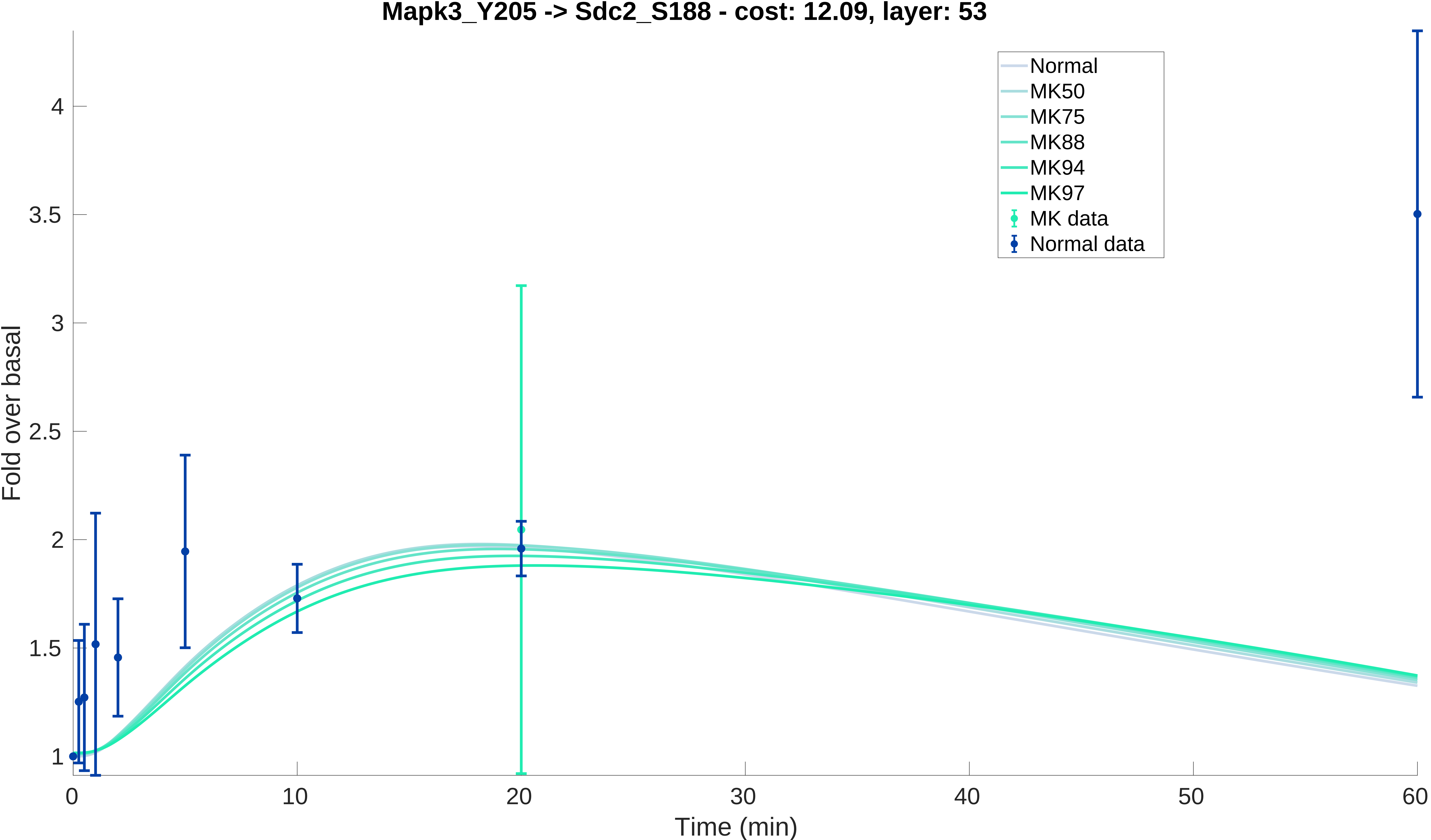

Mapk3_Y205 -> Sdc2_S188 - cost: 12.09, layer: 53
Normal
MK50
MK75
MK88
MK94
MK97
MK data
Normal data
Fold over basal
Time (min)
4
3.5
3
2.5
2
1.5
1
0
10
20
30
40
50
60

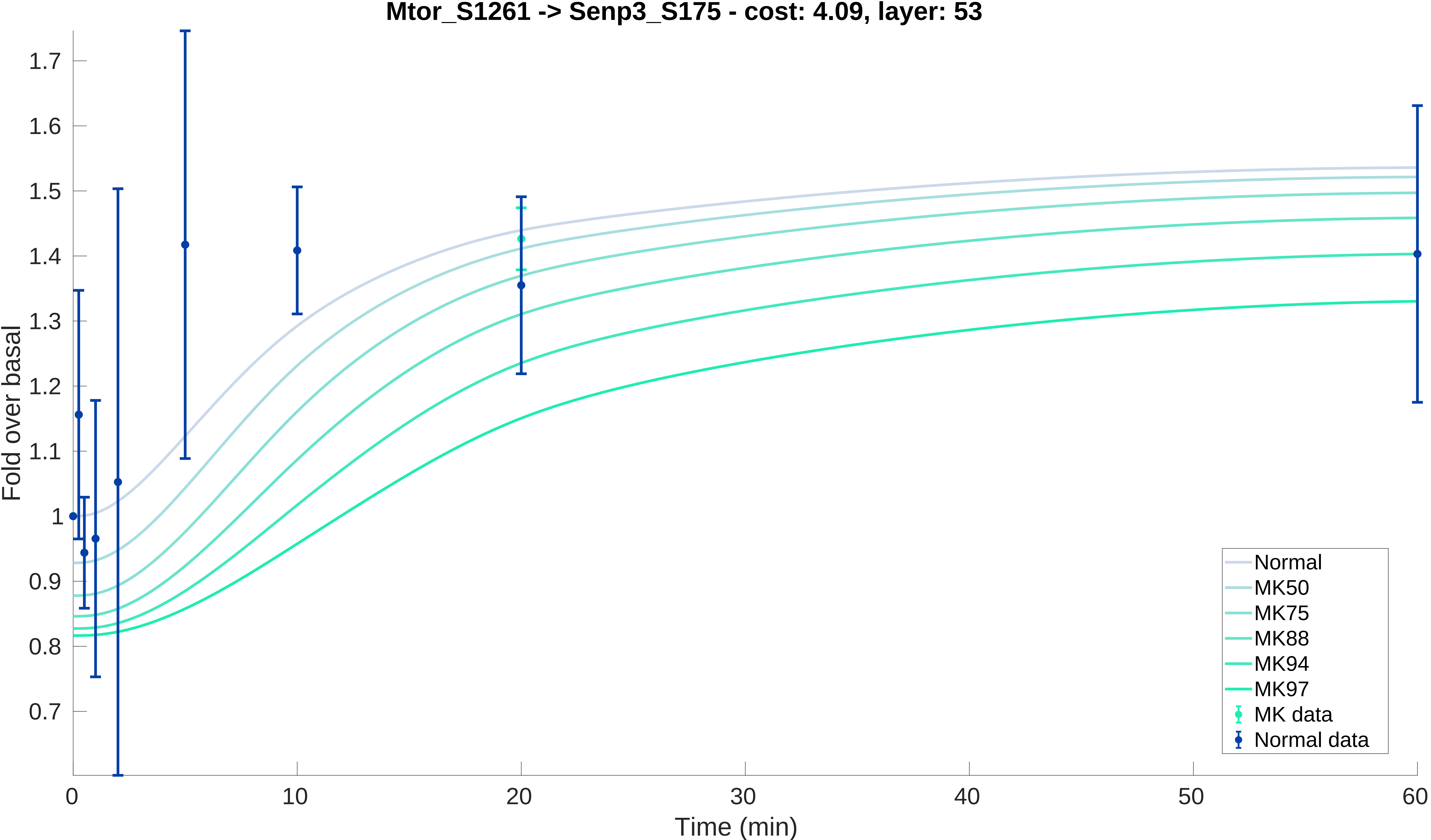

Mtor_S1261 -> Senp3_S175 - cost: 4.09, layer: 53
Fold over basal
1.7
1.6
1.5
1.4
1.3
1.2
1.1
1
0.9
0.8
0.7
0
10
20
30
40
50
60
Time (min)
Normal
MK50
MK75
MK88
MK94
MK97
MK data
Normal data

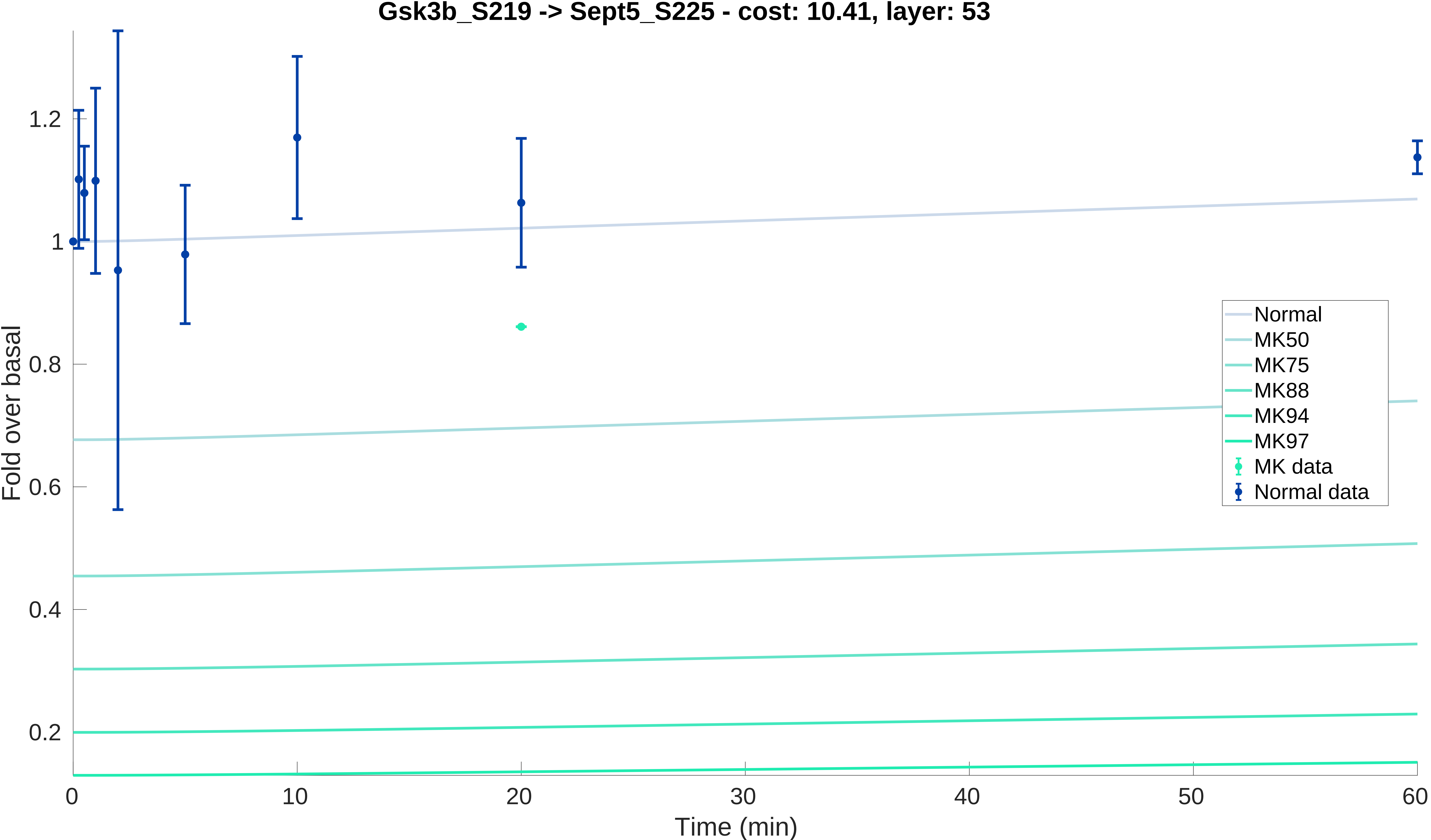

Gsk3b_S219 -> Sept5_S225 - cost: 10.41, layer: 53
Fold over basal
1.2
1
0.8
0.6
0.4
0.2
0
10
20
30
40
50
60
Time (min)
Normal
MK50
MK75
MK88
MK94
MK97
MK data
Normal data

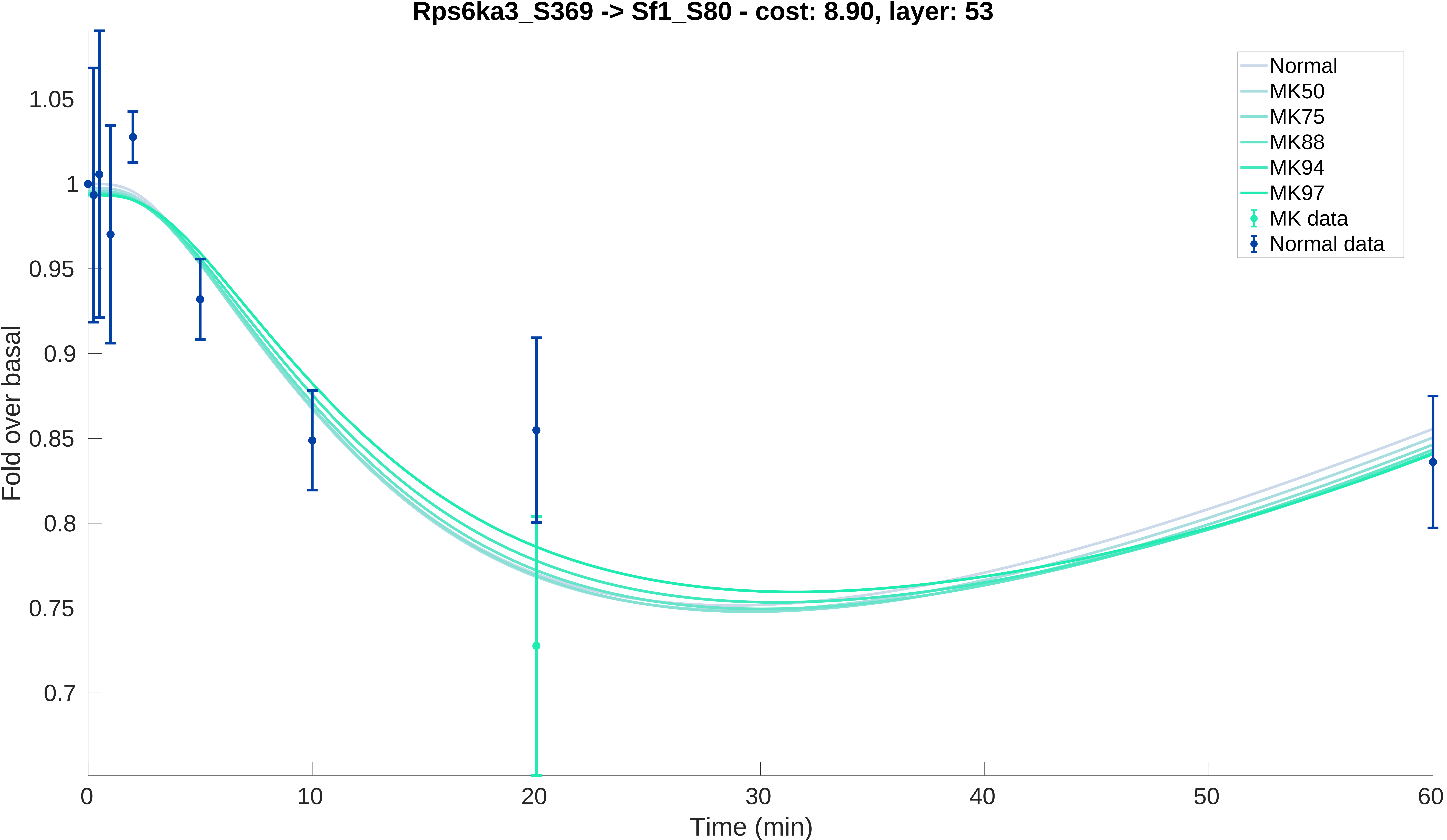

Rps6ka3_S369 -> Sf1_S80 - cost: 8.90, layer: 53
1.05
1
0.95
0.9
0.85
0.8
0.75
0.7
Fold over basal
0
10
20
30
40
50
60
Time (min)
Normal
MK50
MK75
MK88
MK94
MK97
MK data
Normal data

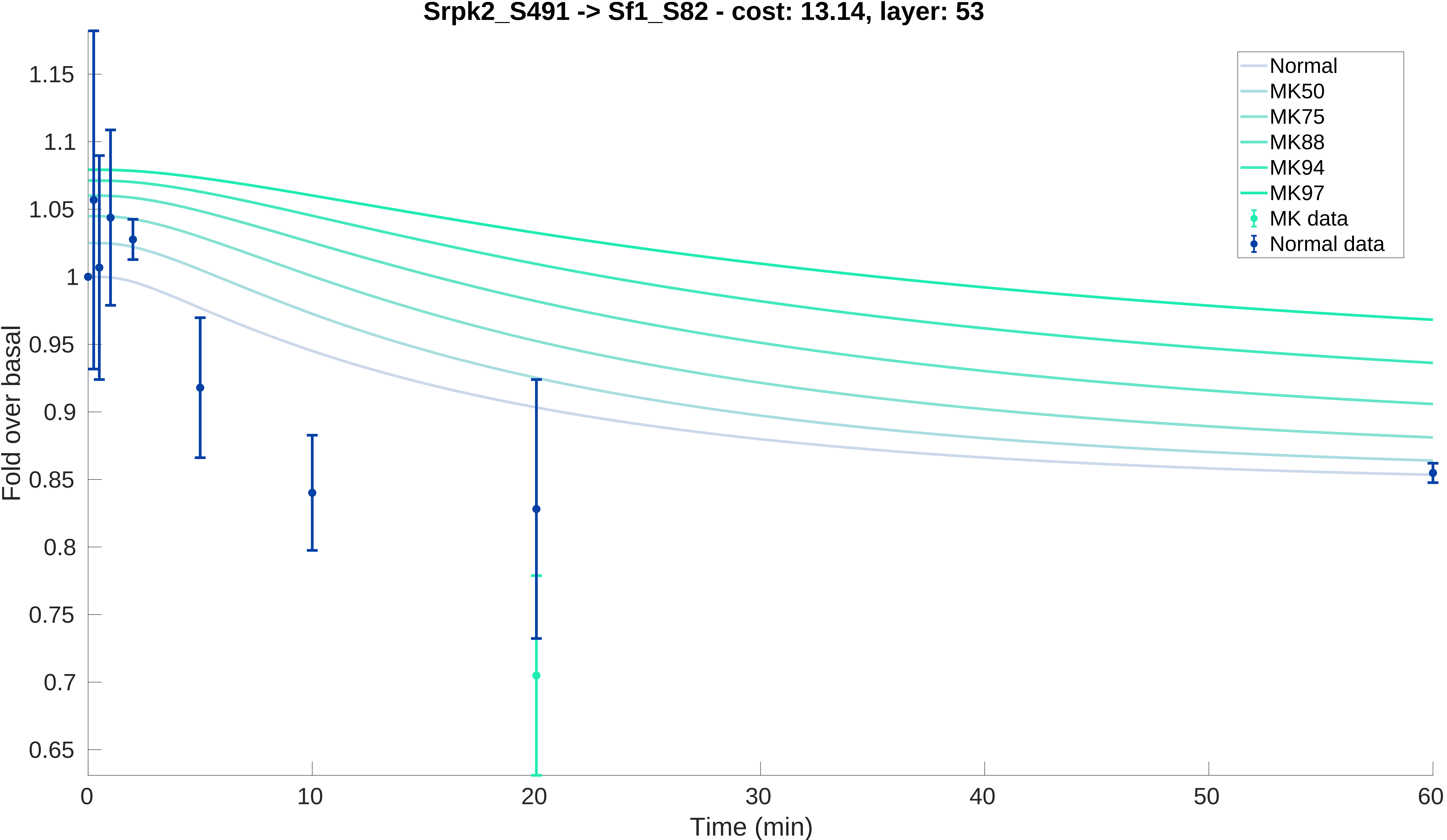

Srpk2_S491 -> Sf1_S82 - cost: 13.14, layer: 53
Fold over basal
Time (min)
1.15
1.1
1.05
1
0.95
0.9
0.85
0.8
0.75
0.7
0.65
0
10
20
30
40
50
60
Normal
MK50
MK75
MK88
MK94
MK97
MK data
Normal data

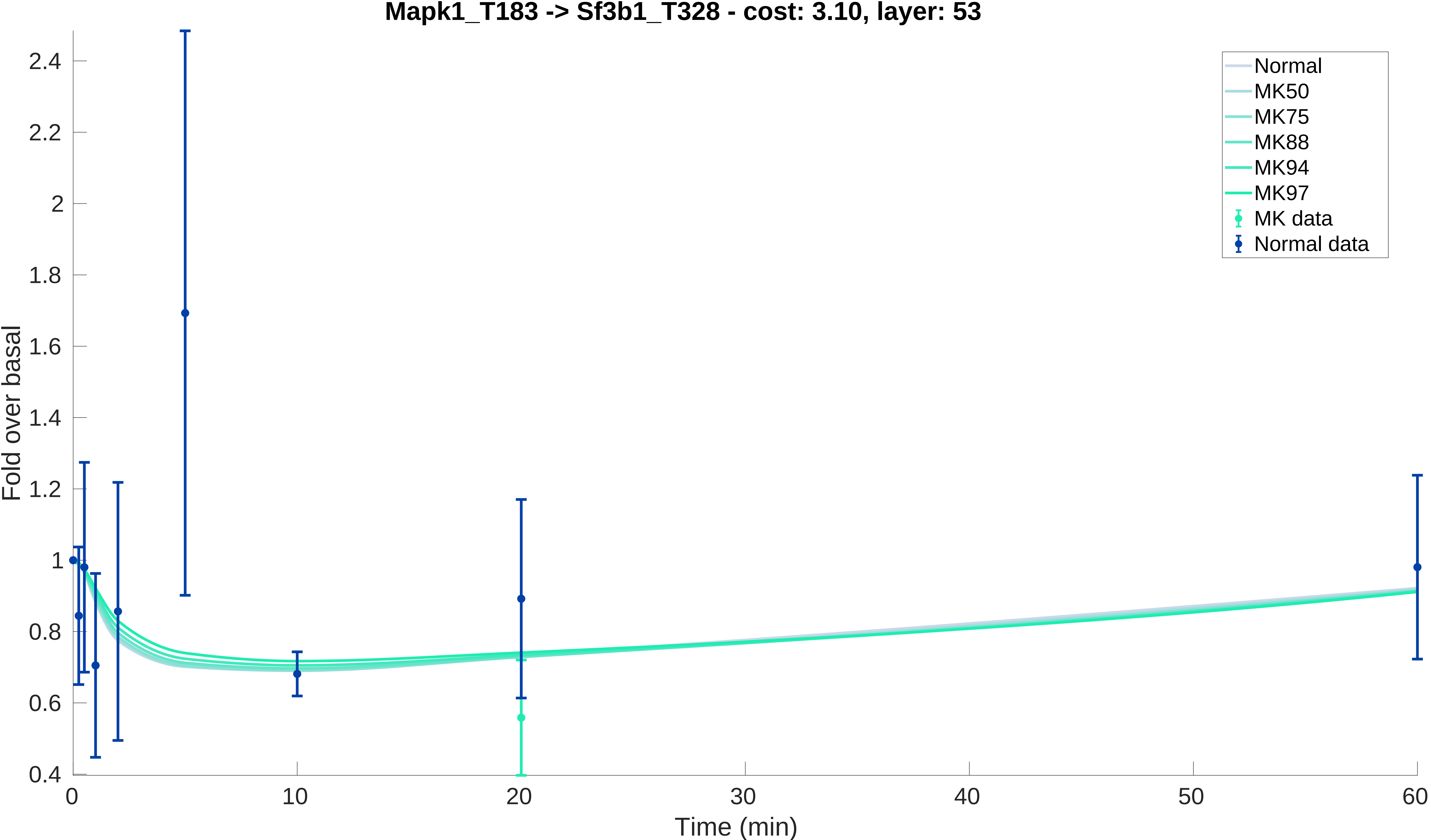

Mapk1_T183 -> Sf3b1_T328 - cost: 3.10, layer: 53
Normal
MK50
MK75
MK88
MK94
MK97
MK data
Normal data
Fold over basal
Time (min)
2.4
2.2
2
1.8
1.6
1.4
1.2
1
0.8
0.6
0.4
0
10
20
30
40
50
60

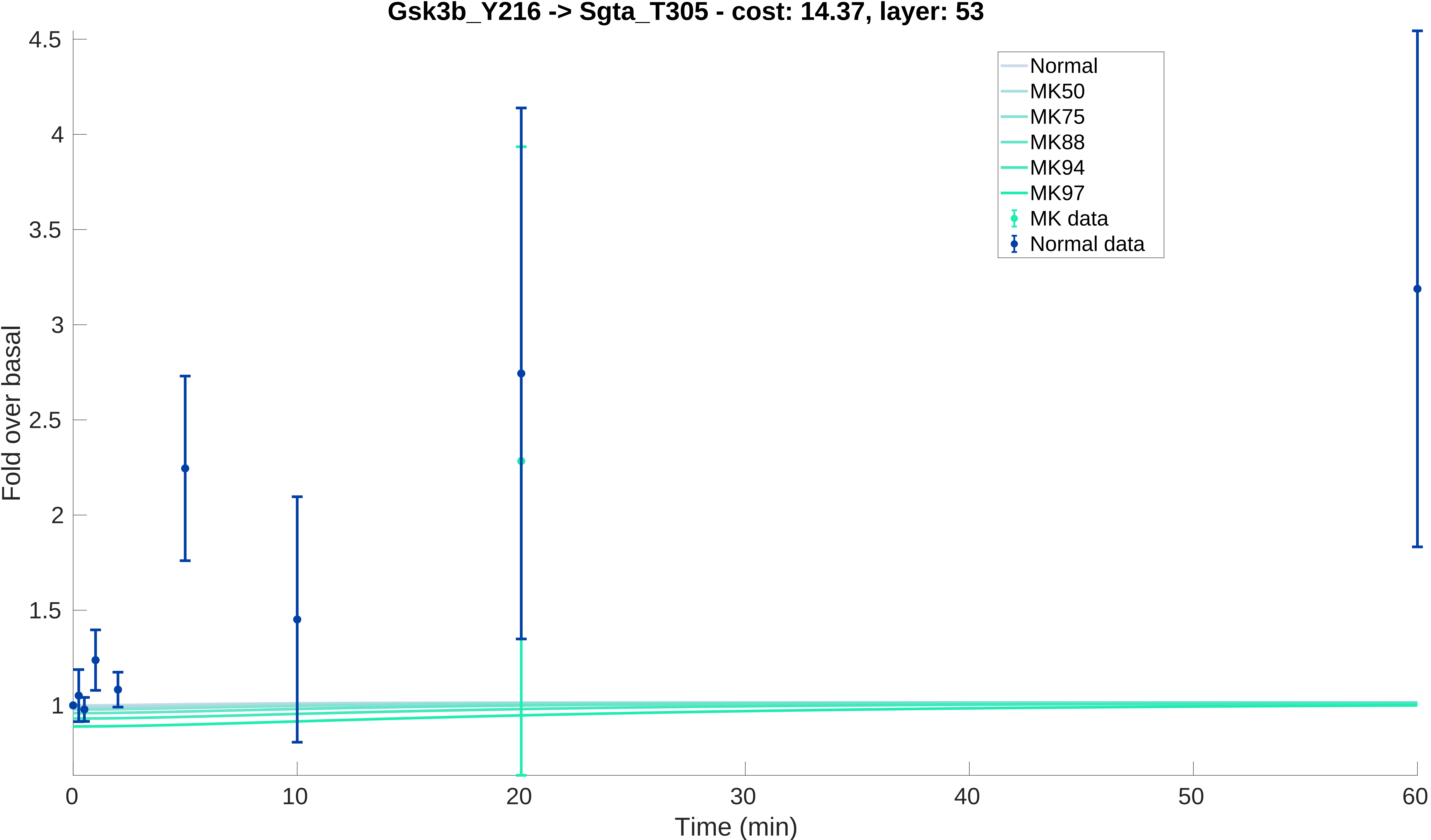

Gsk3b_Y216 -> Sgta_T305 - cost: 14.37, layer: 53
Fold over basal
Time (min)
Normal
MK50
MK75
MK88
MK94
MK97
MK data
Normal data
4.5
4
3.5
3
2.5
2
1.5
1
0
10
20
30
40
50
60

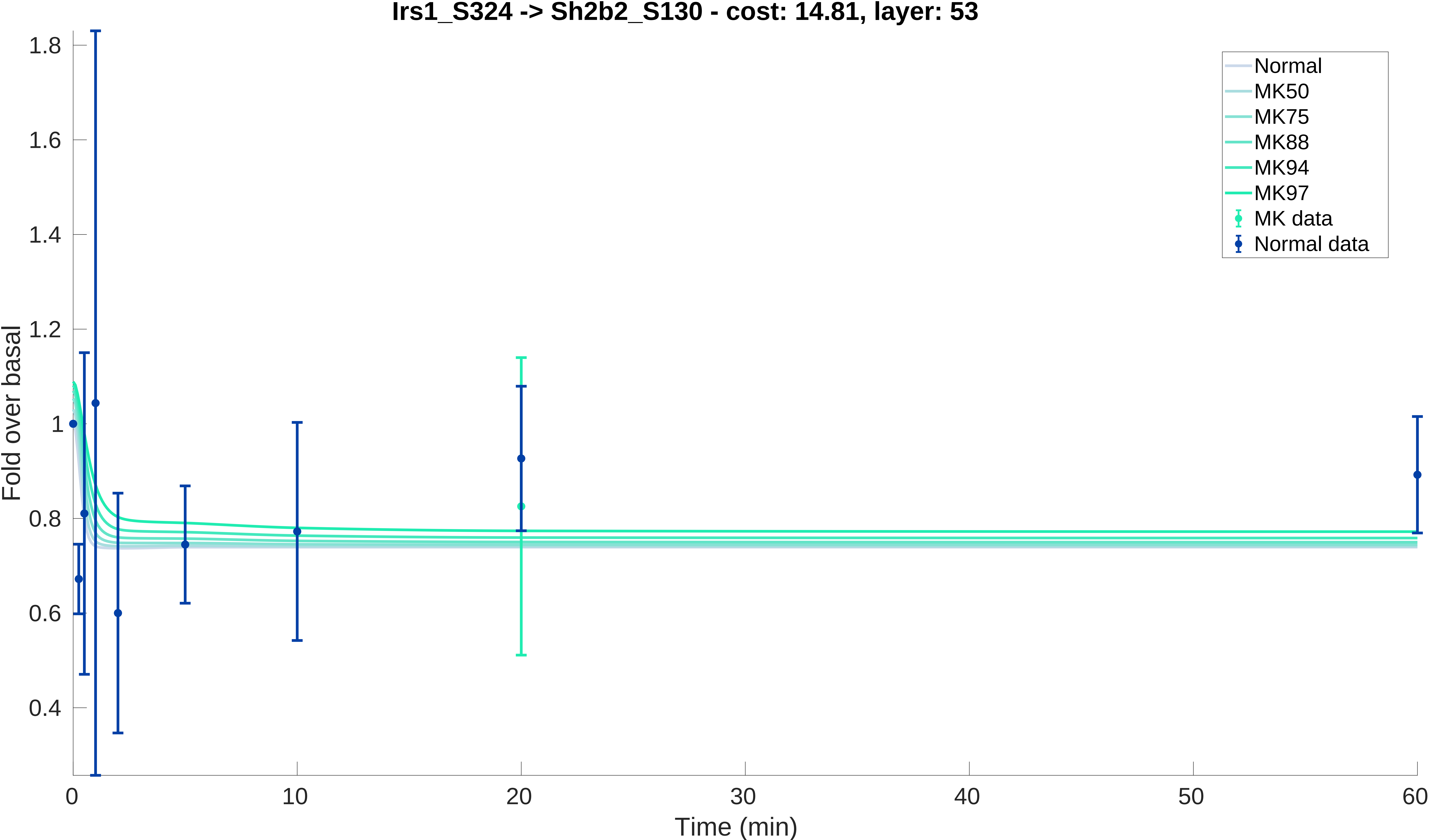

Irs1_S324 -> Sh2b2_S130 - cost: 14.81, layer: 53
Fold over basal
1.8
1.6
1.4
1.2
1
0.8
0.6
0.4
0
10
20
30
40
50
60
Time (min)
Normal
MK50
MK75
MK88
MK94
MK97
MK data
Normal data

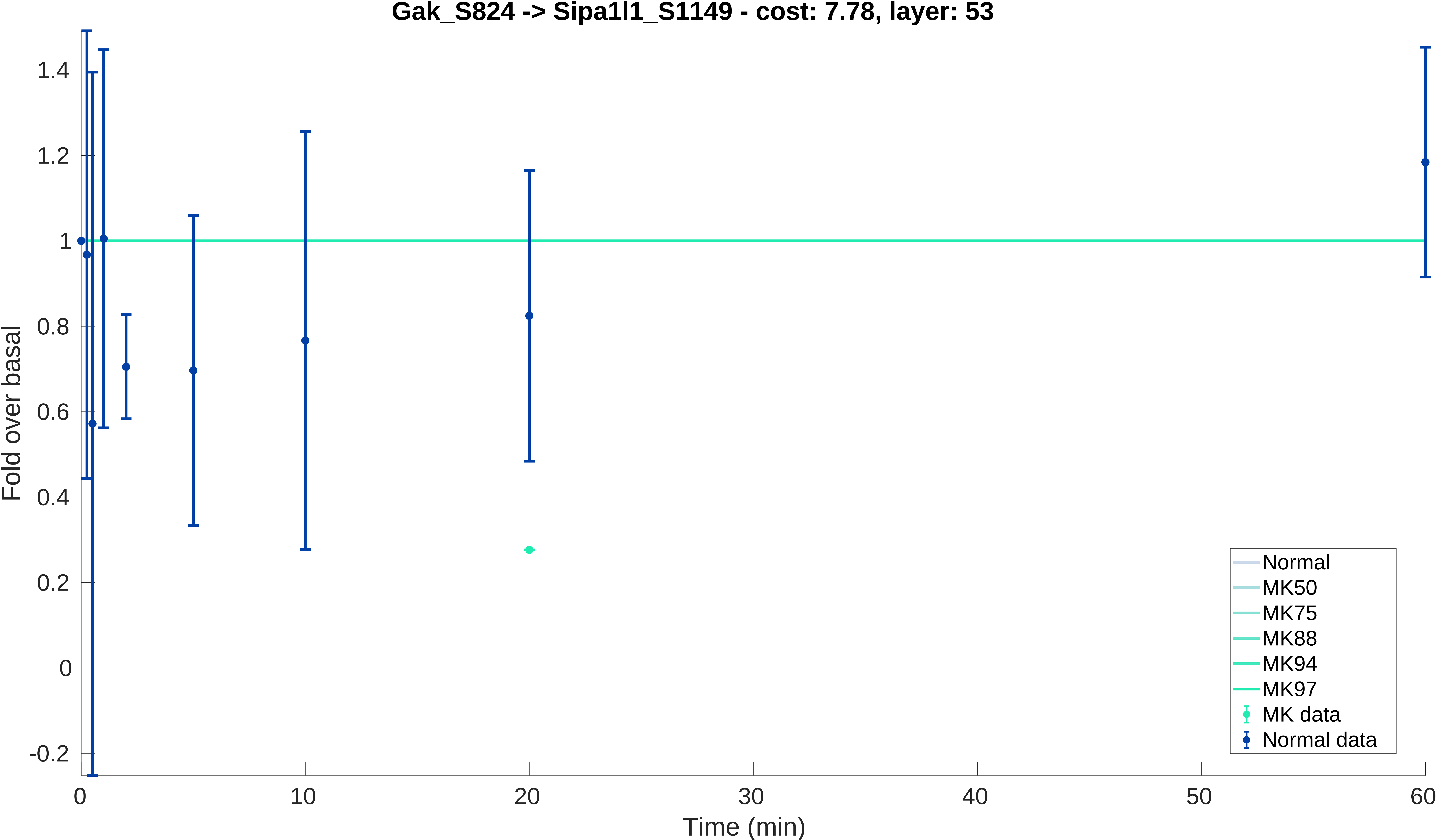

Gak_S824 -> Sipa1l1_S1149 - cost: 7.78, layer: 53
Fold over basal
1.4
1.2
1
0.8
0.6
0.4
0.2
0
-0.2
0
10
20
30
40
50
60
Time (min)
Normal
MK50
MK75
MK88
MK94
MK97
MK data
Normal data

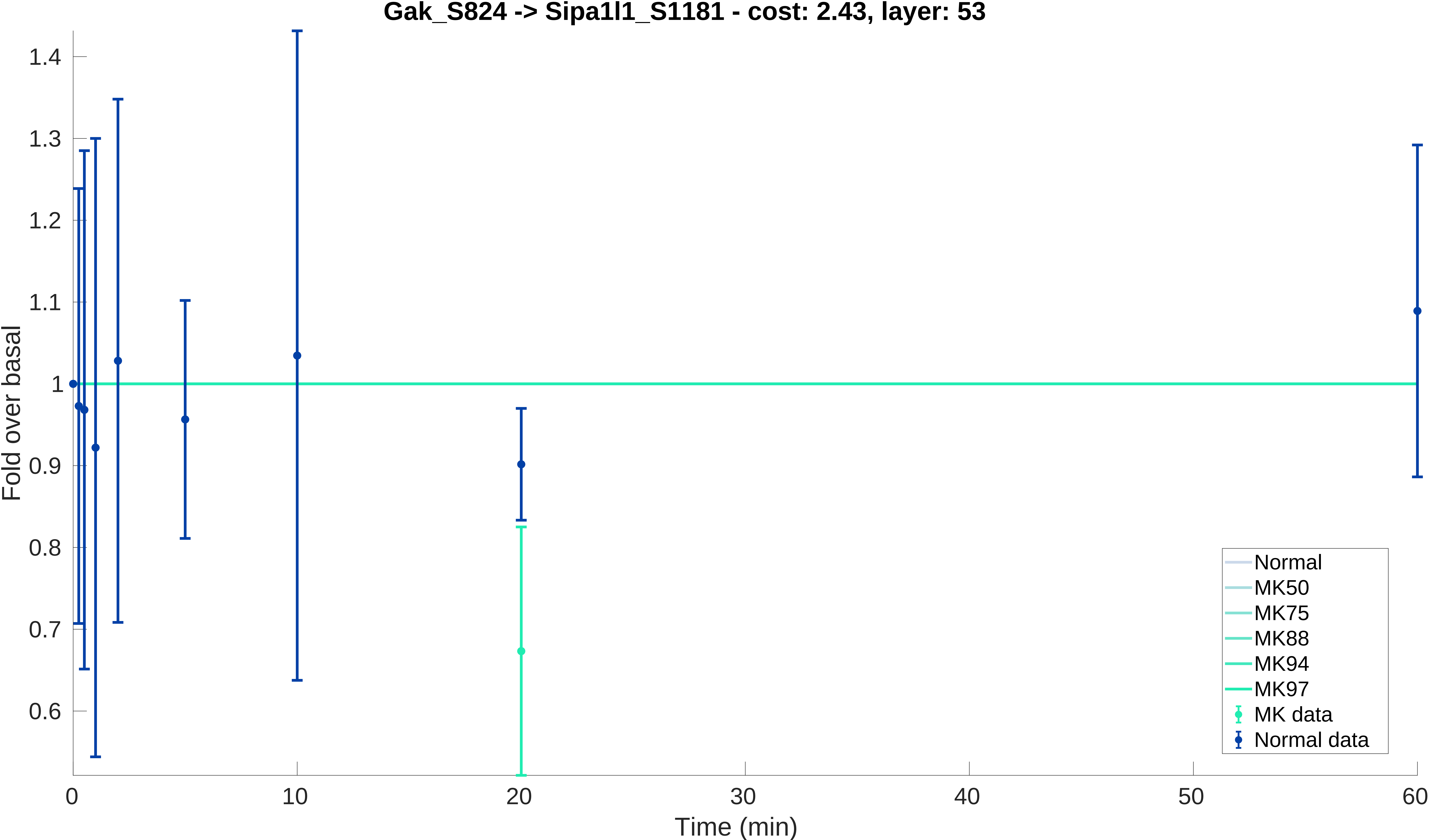

Gak_S824 -> Sipa1l1_S1181 - cost: 2.43, layer: 53
Fold over basal
Time (min)
0.6
0.7
0.8
0.9
1
1.1
1.2
1.3
1.4
0
10
20
30
40
50
60
Normal
MK50
MK75
MK88
MK94
MK97
MK data
Normal data

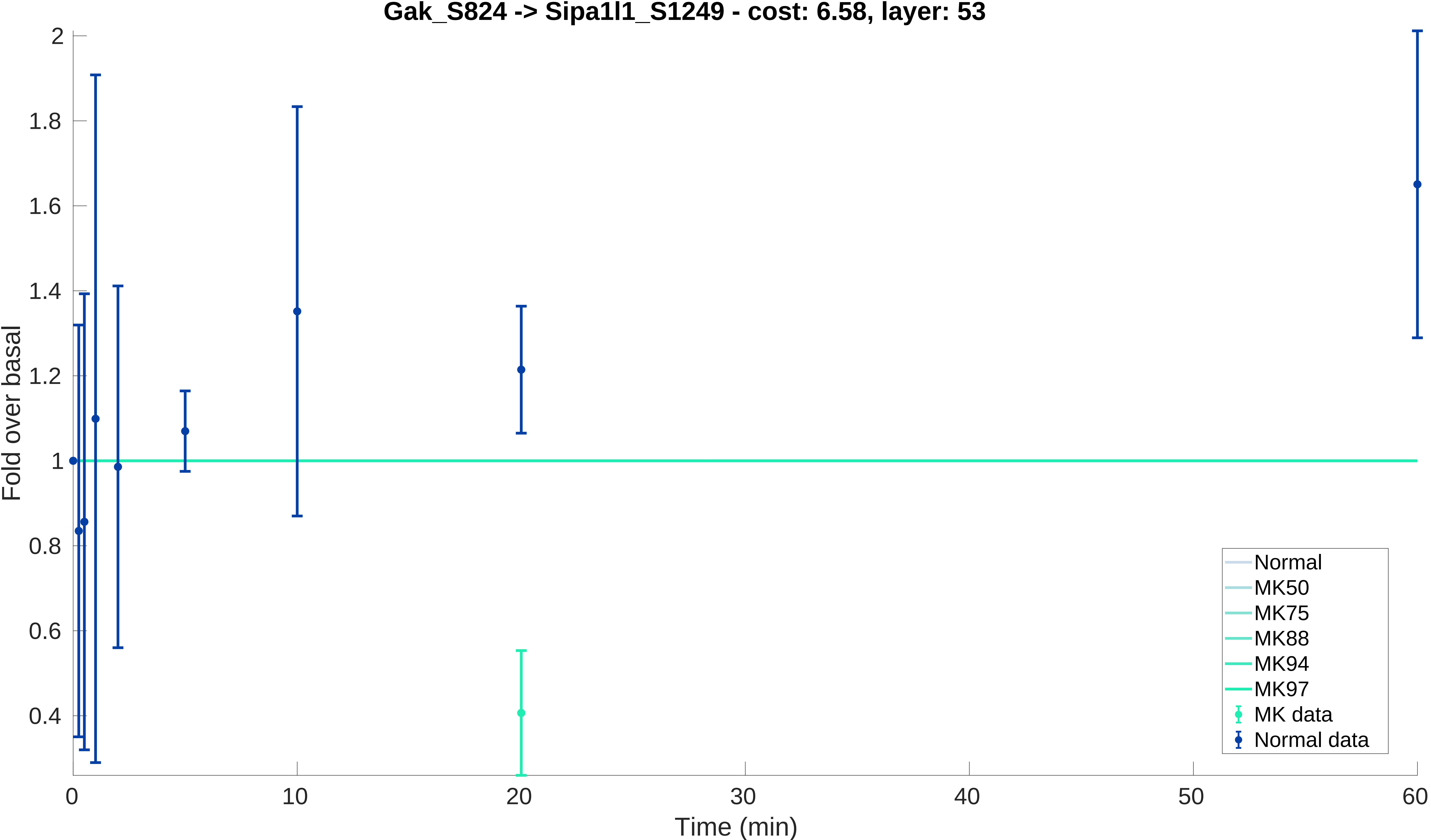
Gak_S824 -> Sipa1l1_S1249 - cost: 6.58, layer: 53
Fold over basal
Time (min)
0
10
20
30
40
50
60
0.4
0.6
0.8
1
1.2
1.4
1.6
1.8
2
Normal
MK50
MK75
MK88
MK94
MK97
MK data
Normal data

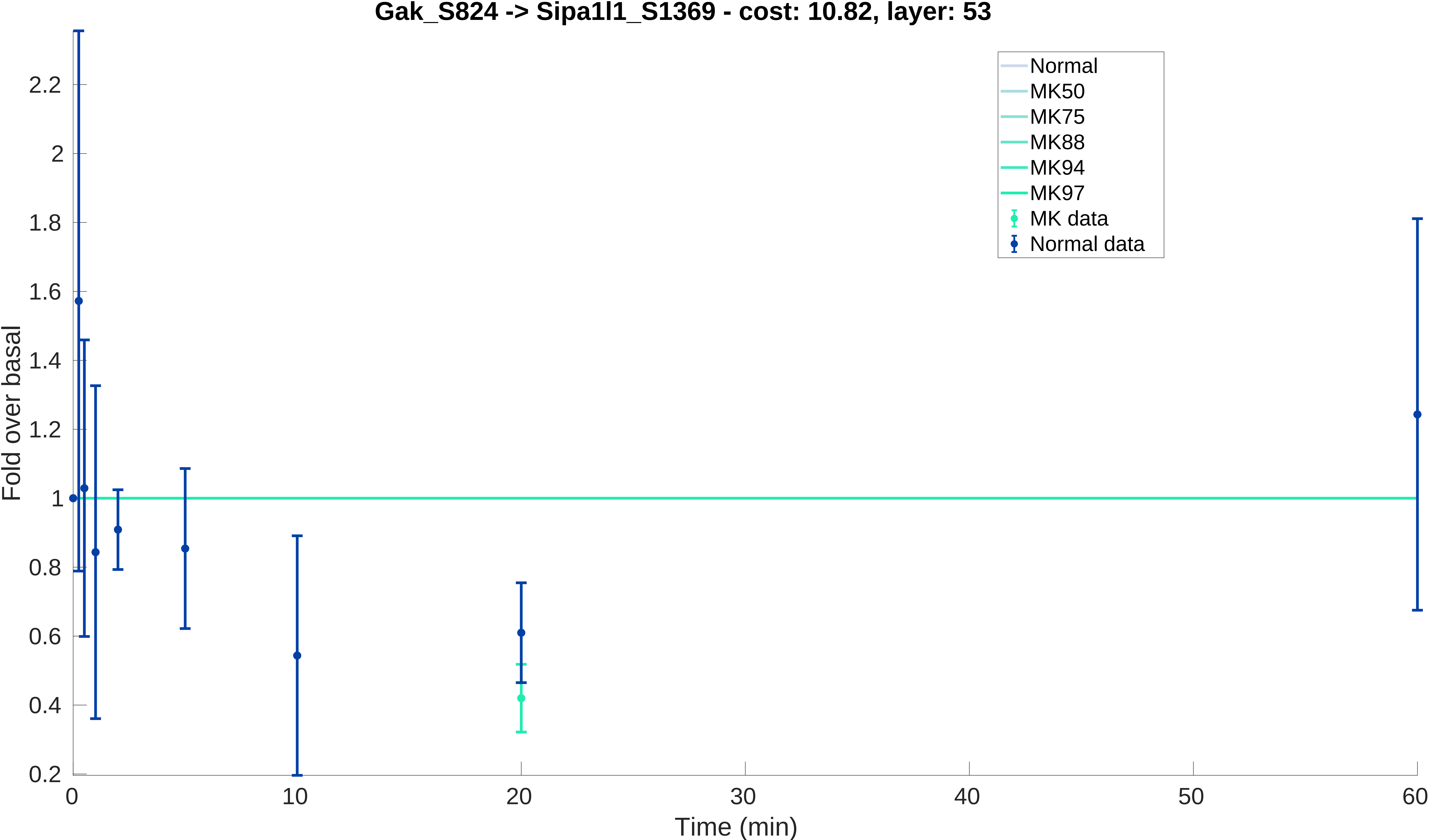

Gak_S824 -> Sipa1l1_S1369 - cost: 10.82, layer: 53
Fold over basal
Time (min)
Normal
MK50
MK75
MK88
MK94
MK97
MK data
Normal data
0.2
0.4
0.6
0.8
1
1.2
1.4
1.6
1.8
2
2.2
0
10
20
30
40
50
60

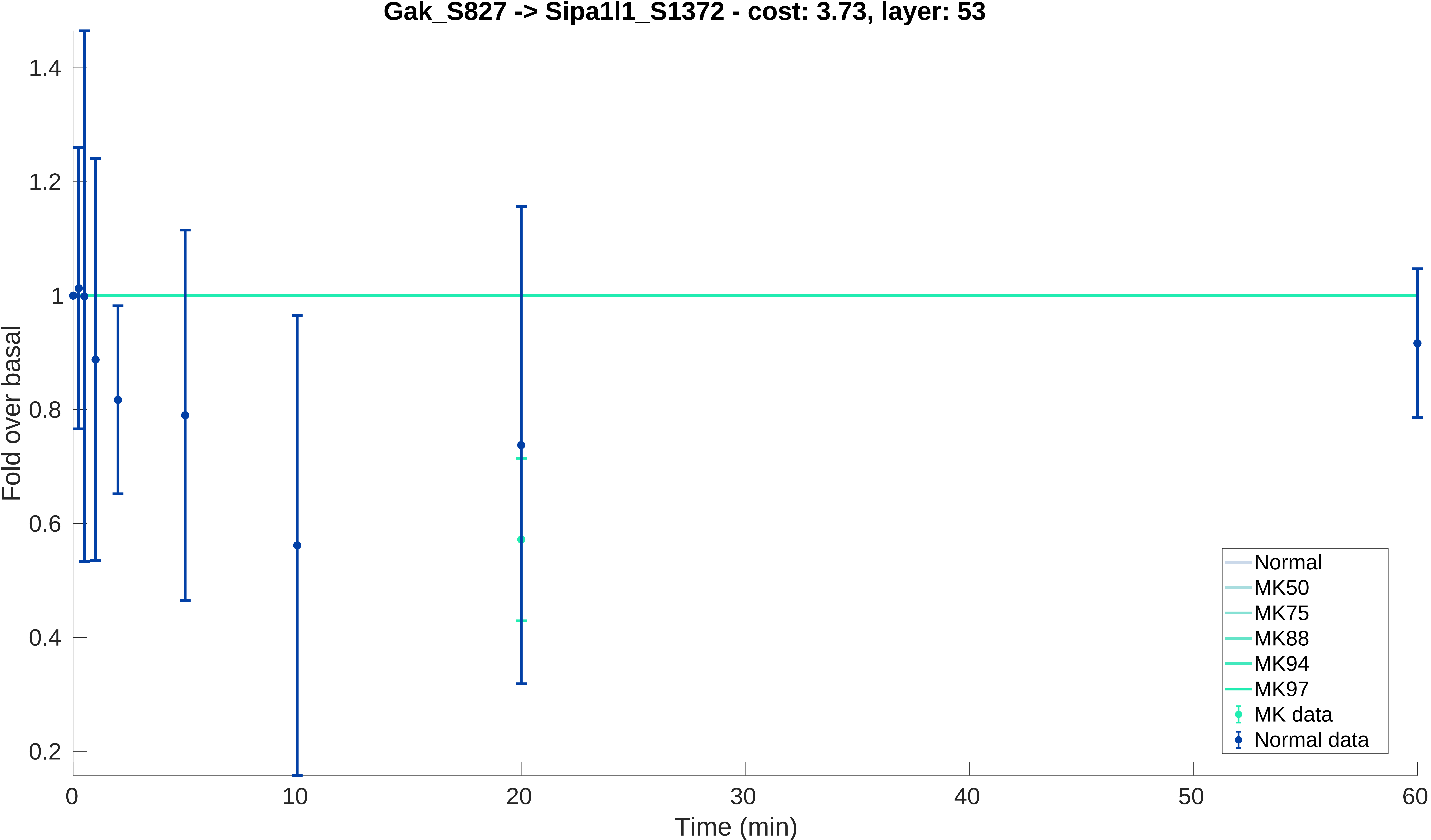

Gak_S827 -> Sipa1l1_S1372 - cost: 3.73, layer: 53
Fold over basal
Time (min)
0
10
20
30
40
50
60
0.2
0.4
0.6
0.8
1
1.2
1.4
Normal
MK50
MK75
MK88
MK94
MK97
MK data
Normal data

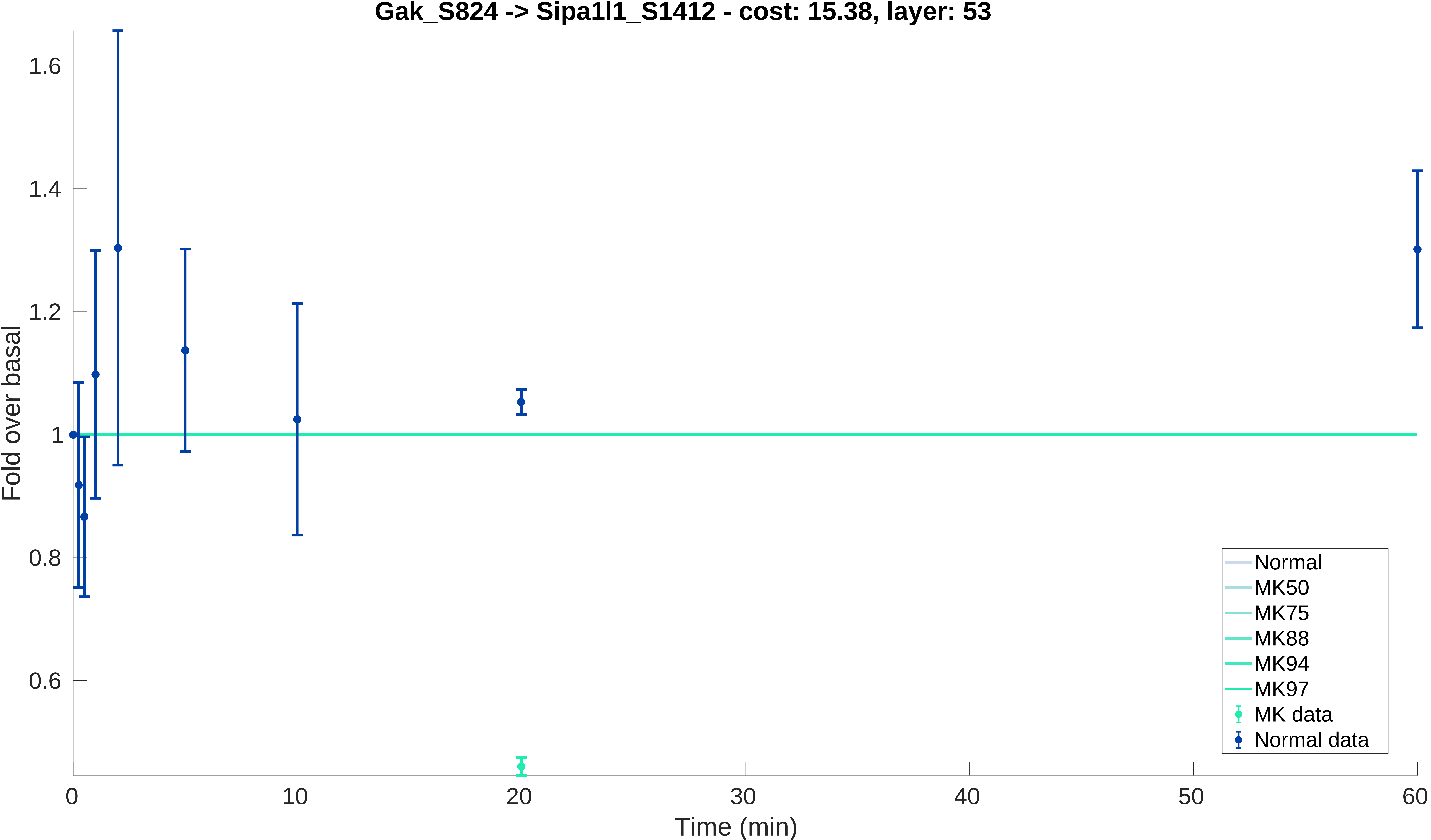

Gak_S824 -> Sipa1l1_S1412 - cost: 15.38, layer: 53
Fold over basal
1.6
1.4
1.2
1
0.8
0.6
0
10
20
30
40
50
60
Time (min)
Normal
MK50
MK75
MK88
MK94
MK97
MK data
Normal data

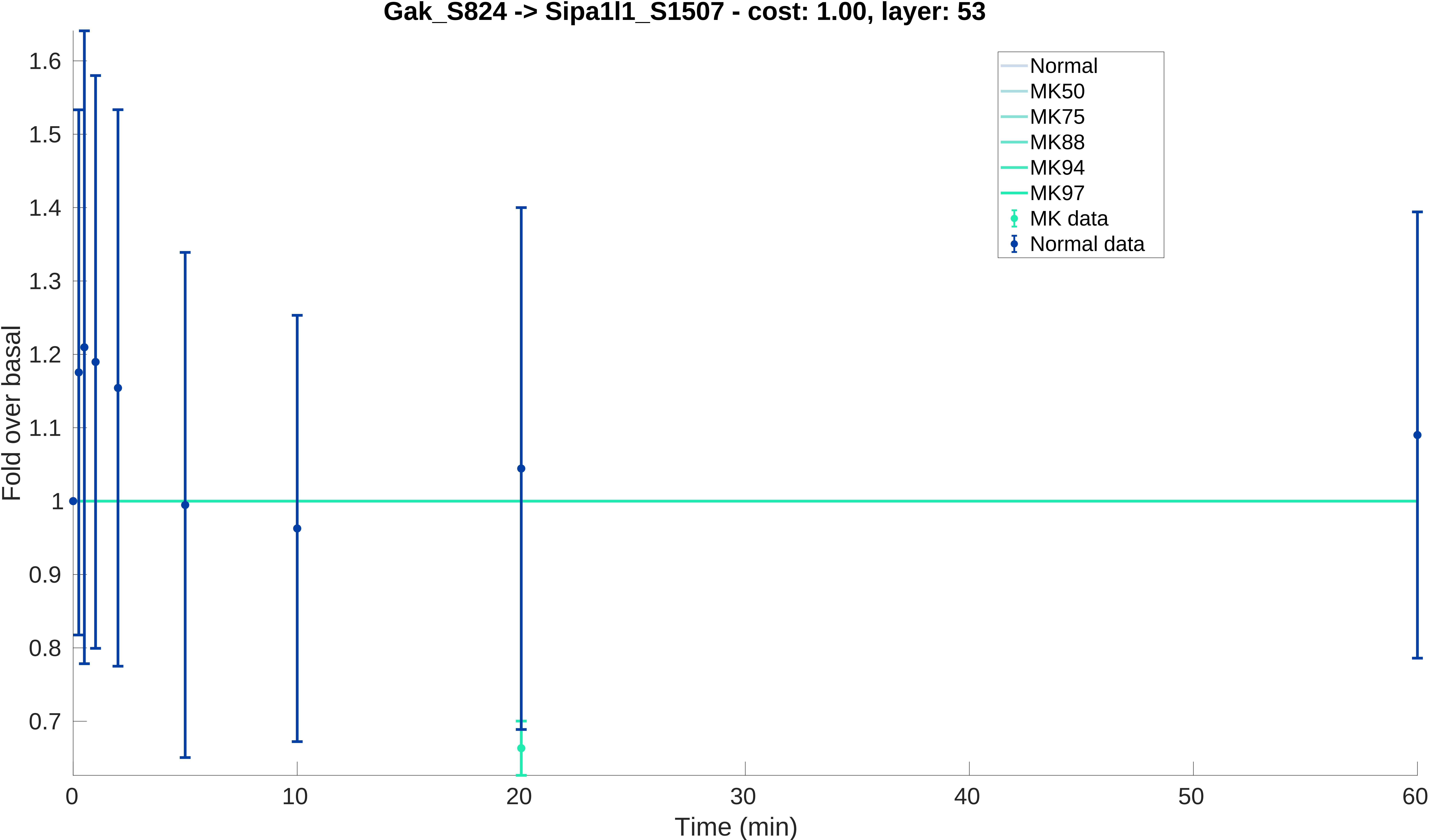

Gak_S824 -> Sipa1l1_S1507 - cost: 1.00, layer: 53
Normal
MK50
MK75
MK88
MK94
MK97
MK data
Normal data
Fold over basal
Time (min)
1.6
1.5
1.4
1.3
1.2
1.1
1
0.9
0.8
0.7
0
10
20
30
40
50
60

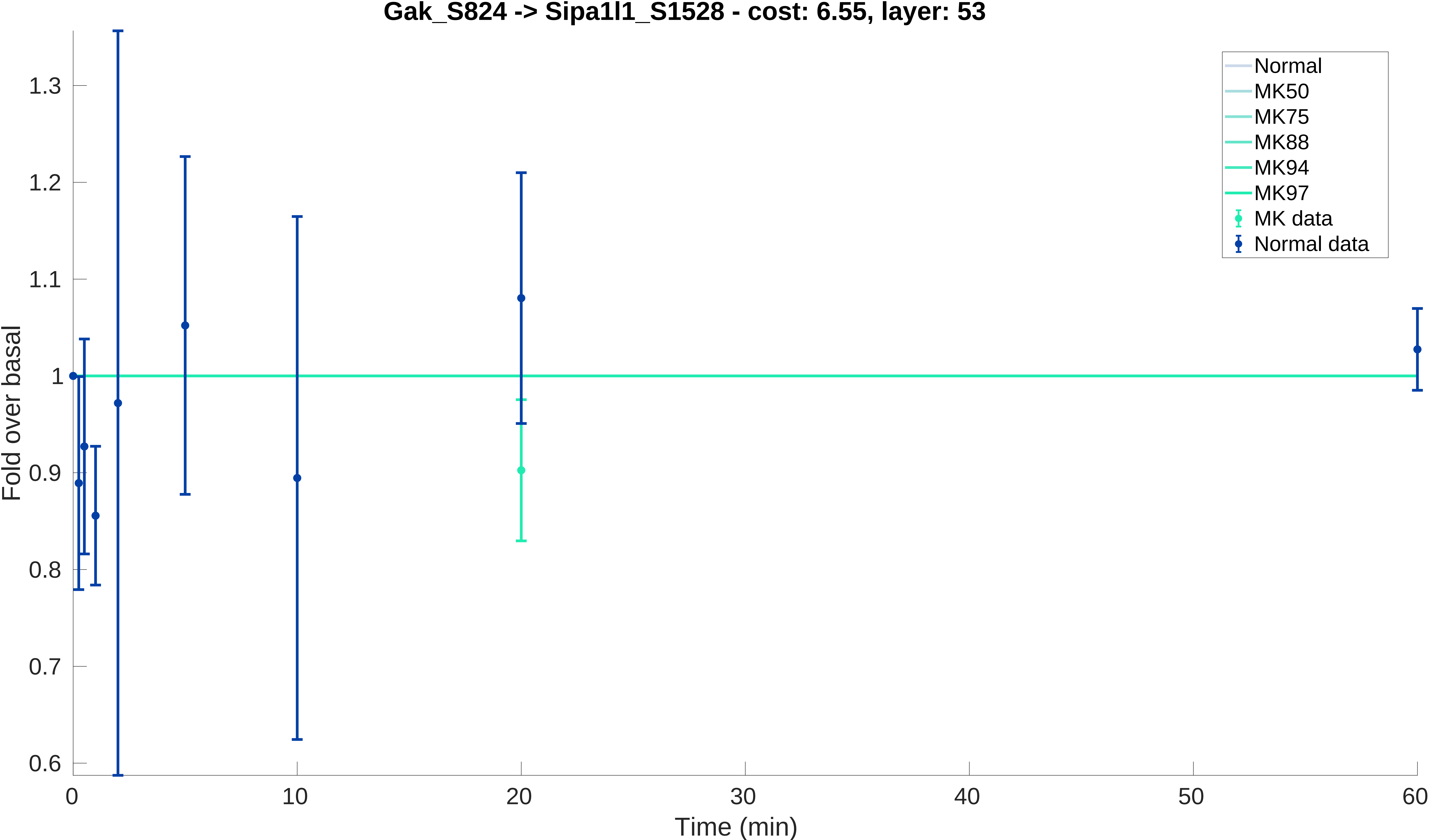

Gak_S824 -> Sipa1l1_S1528 - cost: 6.55, layer: 53
Fold over basal
1.3
1.2
1.1
1
0.9
0.8
0.7
0.6
0
10
20
30
40
50
60
Time (min)
Normal
MK50
MK75
MK88
MK94
MK97
MK data
Normal data

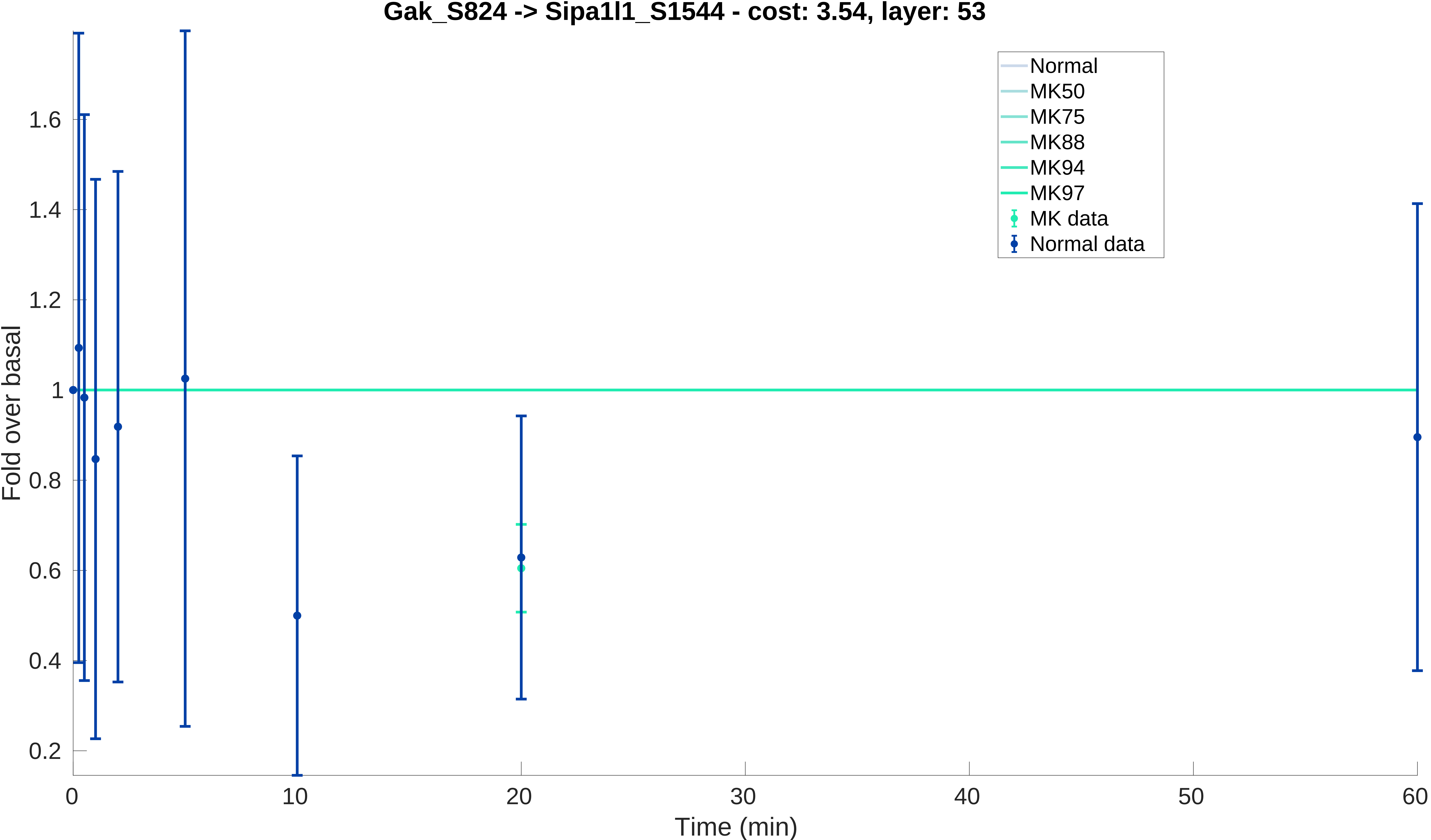
Gak_S824 -> Sipa1l1_S1544 - cost: 3.54, layer: 53
Normal
MK50
MK75
MK88
MK94
MK97
MK data
Normal data
Fold over basal
0.2
0.4
0.6
0.8
1
1.2
1.4
1.6
0
10
20
30
40
50
60
Time (min)

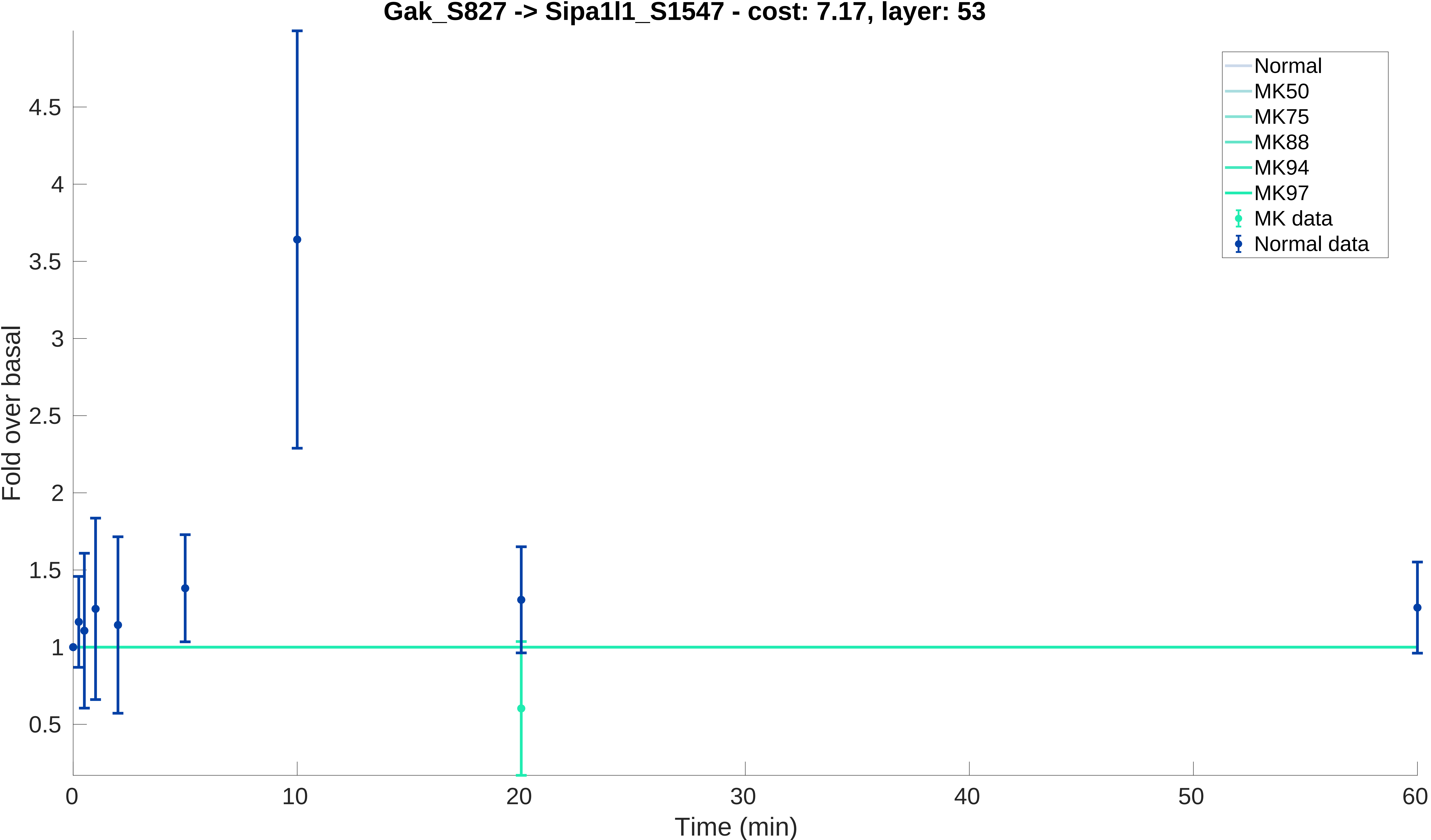
Gak_S827 -> Sipa1l1_S1547 - cost: 7.17, layer: 53
Fold over basal
Time (min)
0.5
1
1.5
2
2.5
3
3.5
4
4.5
0
10
20
30
40
50
60
Normal
MK50
MK75
MK88
MK94
MK97
MK data
Normal data

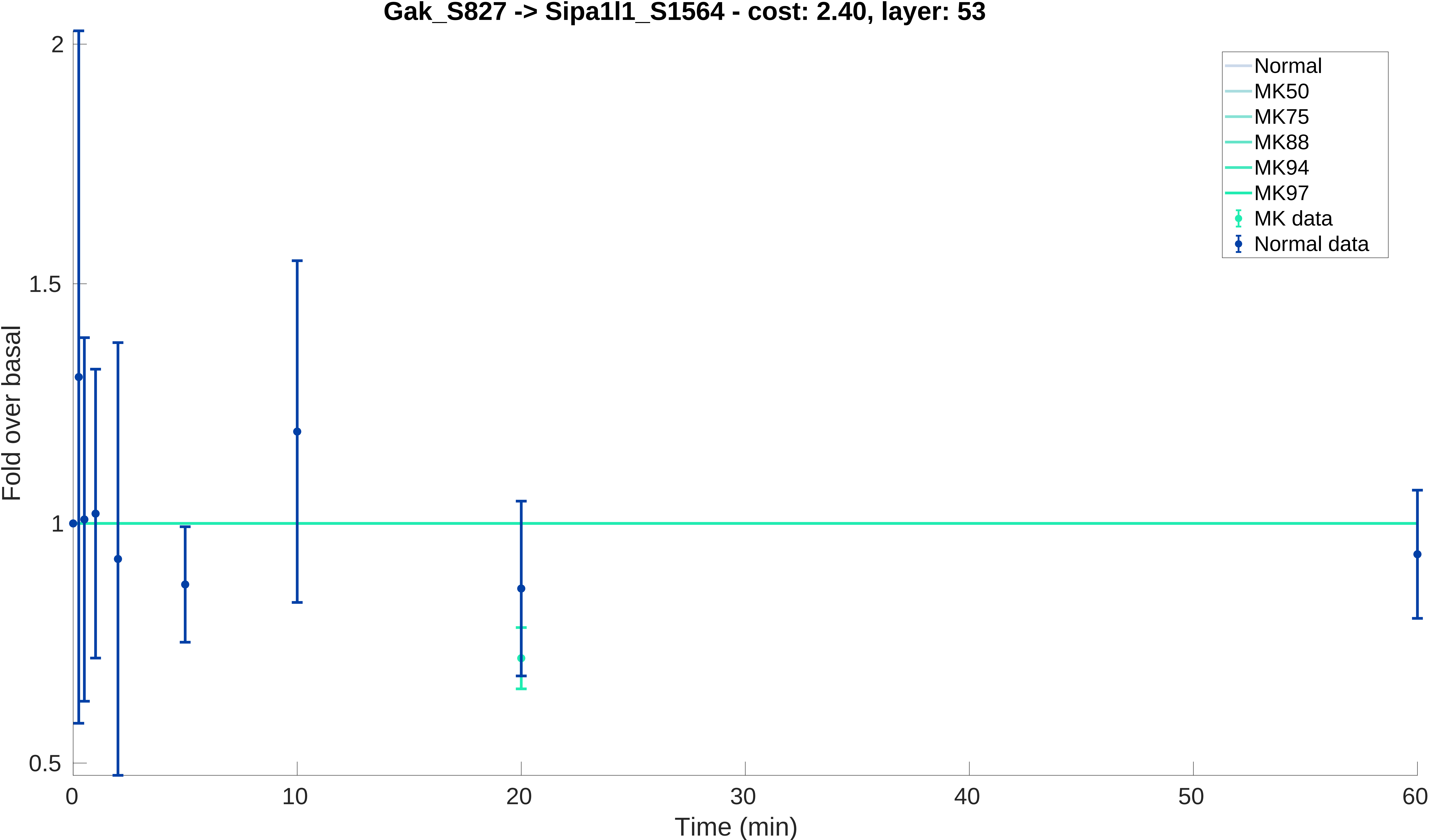

Gak_S827 -> Sipa1l1_S1564 - cost: 2.40, layer: 53
Normal
MK50
MK75
MK88
MK94
MK97
MK data
Normal data
Fold over basal
Time (min)
2
1.5
1
0.5
0
10
20
30
40
50
60

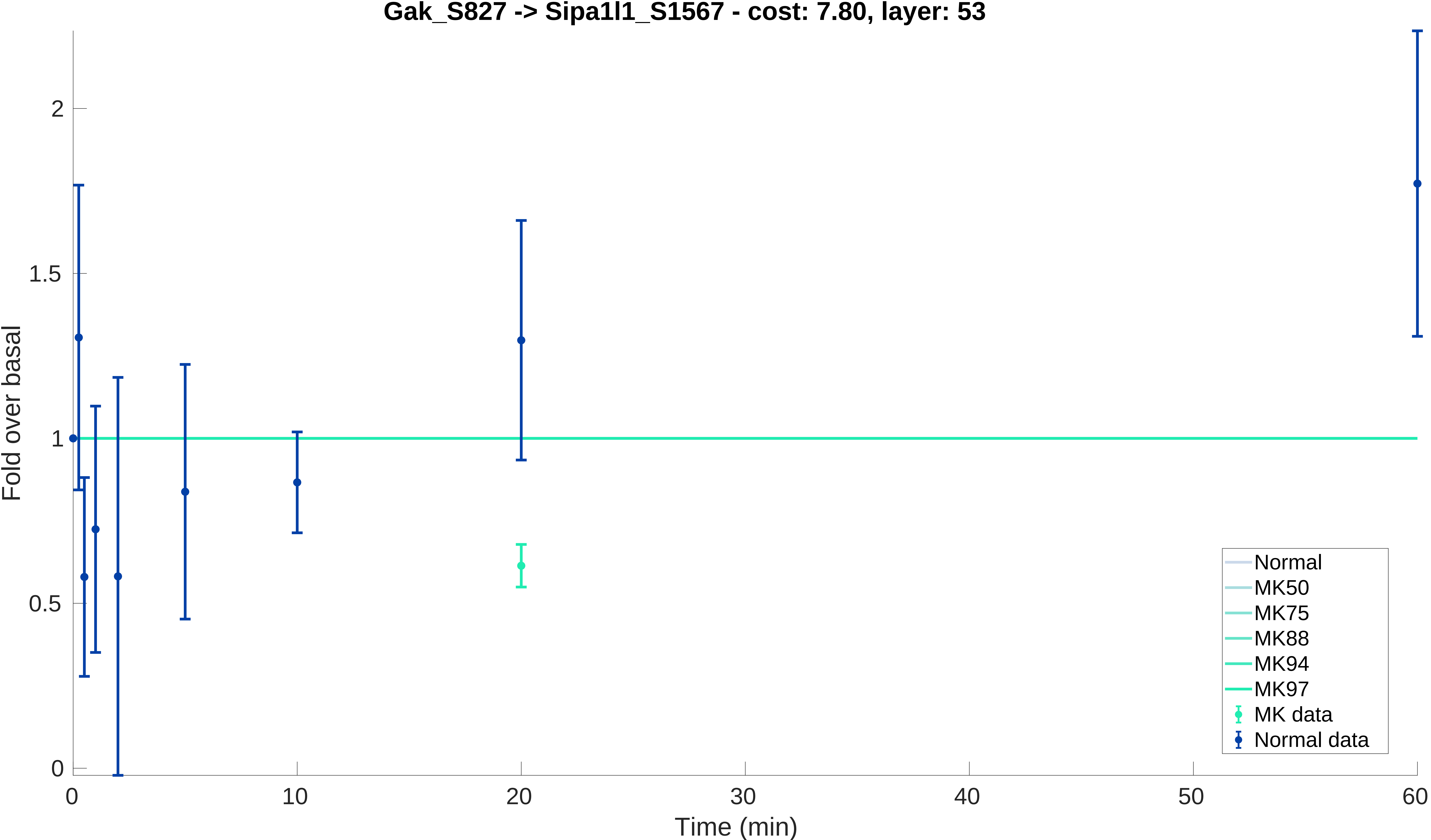

Gak_S827 -> Sipa1l1_S1567 - cost: 7.80, layer: 53
Fold over basal
Time (min)
0
0.5
1
1.5
2
0
10
20
30
40
50
60
Normal
MK50
MK75
MK88
MK94
MK97
MK data
Normal data

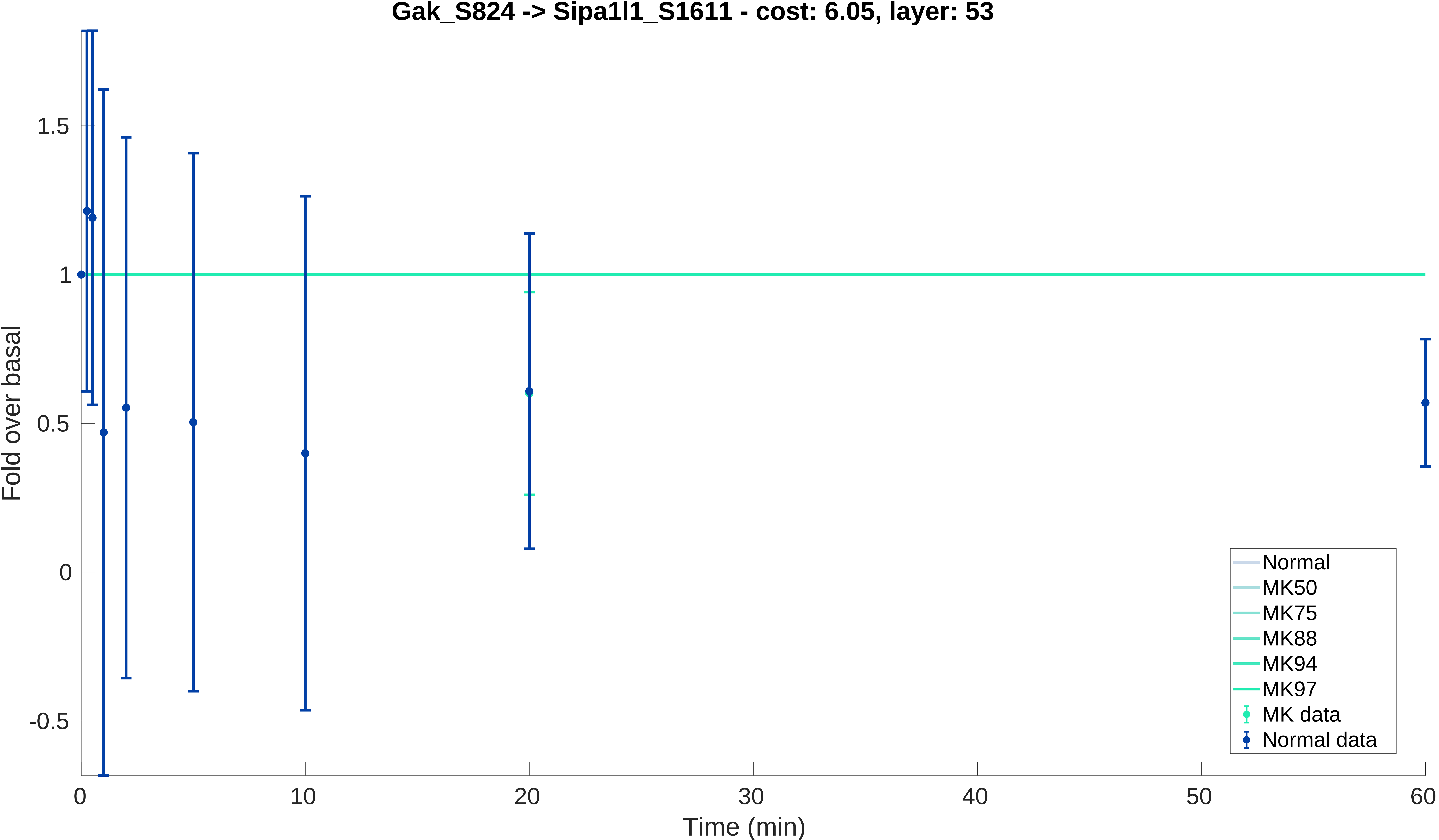

Gak_S824 -> Sipa1l1_S1611 - cost: 6.05, layer: 53
Fold over basal
1.5
1
0.5
0
-0.5
0
10
20
30
40
50
60
Time (min)
Normal
MK50
MK75
MK88
MK94
MK97
MK data
Normal data

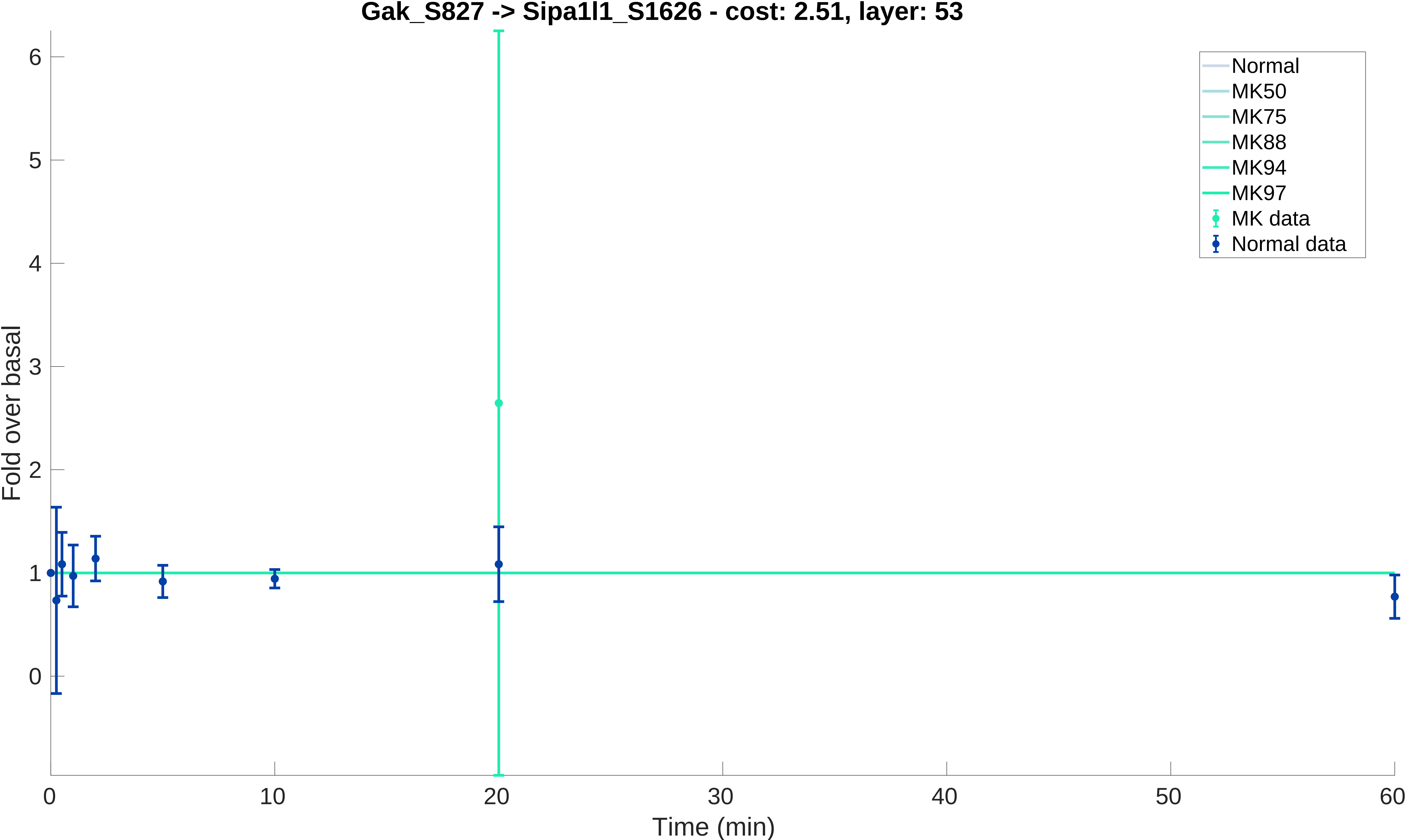

Gak_S827 -> Sipa1l1_S1626 - cost: 2.51, layer: 53
Fold over basal
Time (min)
Normal
MK50
MK75
MK88
MK94
MK97
MK data
Normal data
0
1
2
3
4
5
6
0
10
20
30
40
50
60

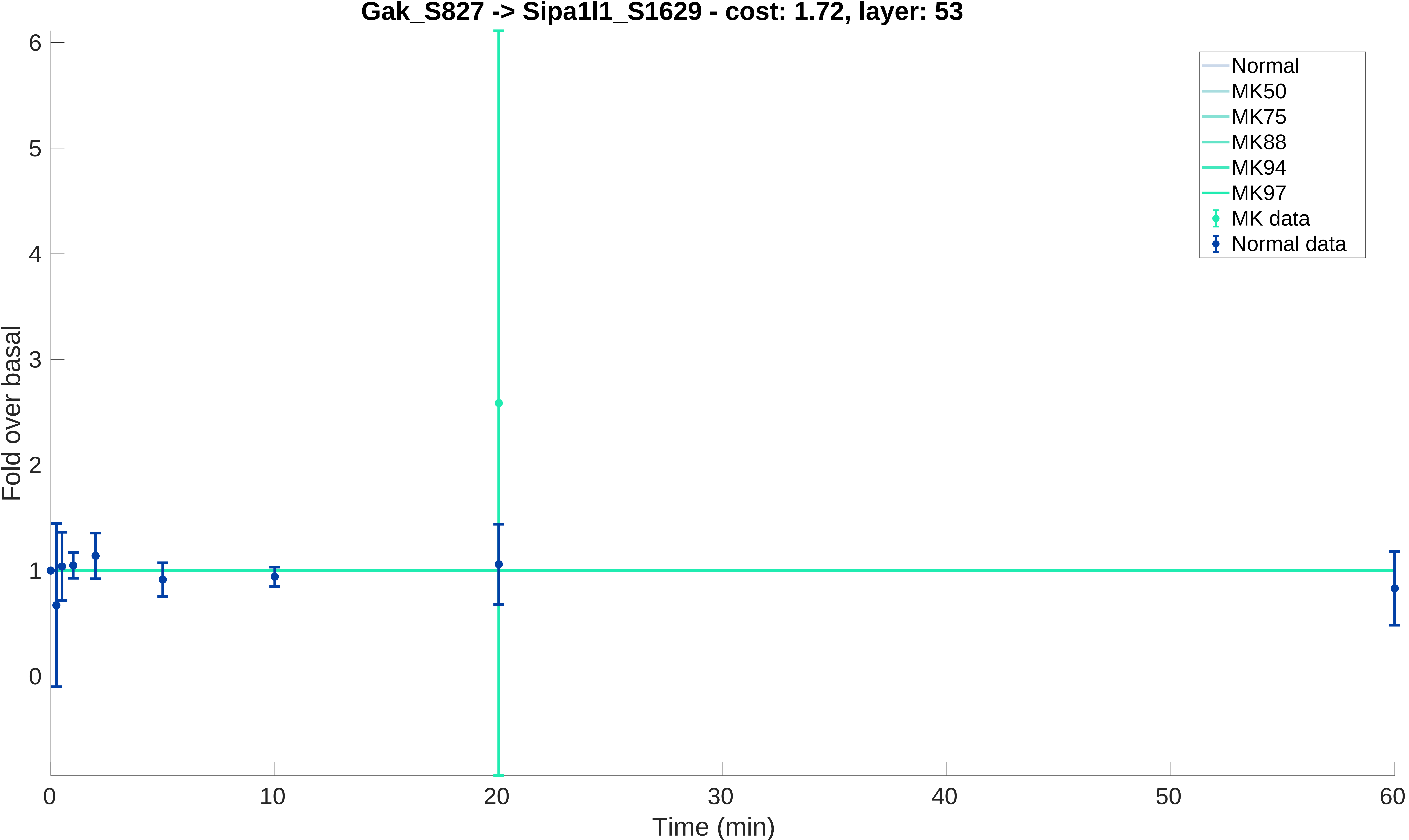

Gak_S827 -> Sipa1l1_S1629 - cost: 1.72, layer: 53
Fold over basal
Time (min)
Normal
MK50
MK75
MK88
MK94
MK97
MK data
Normal data

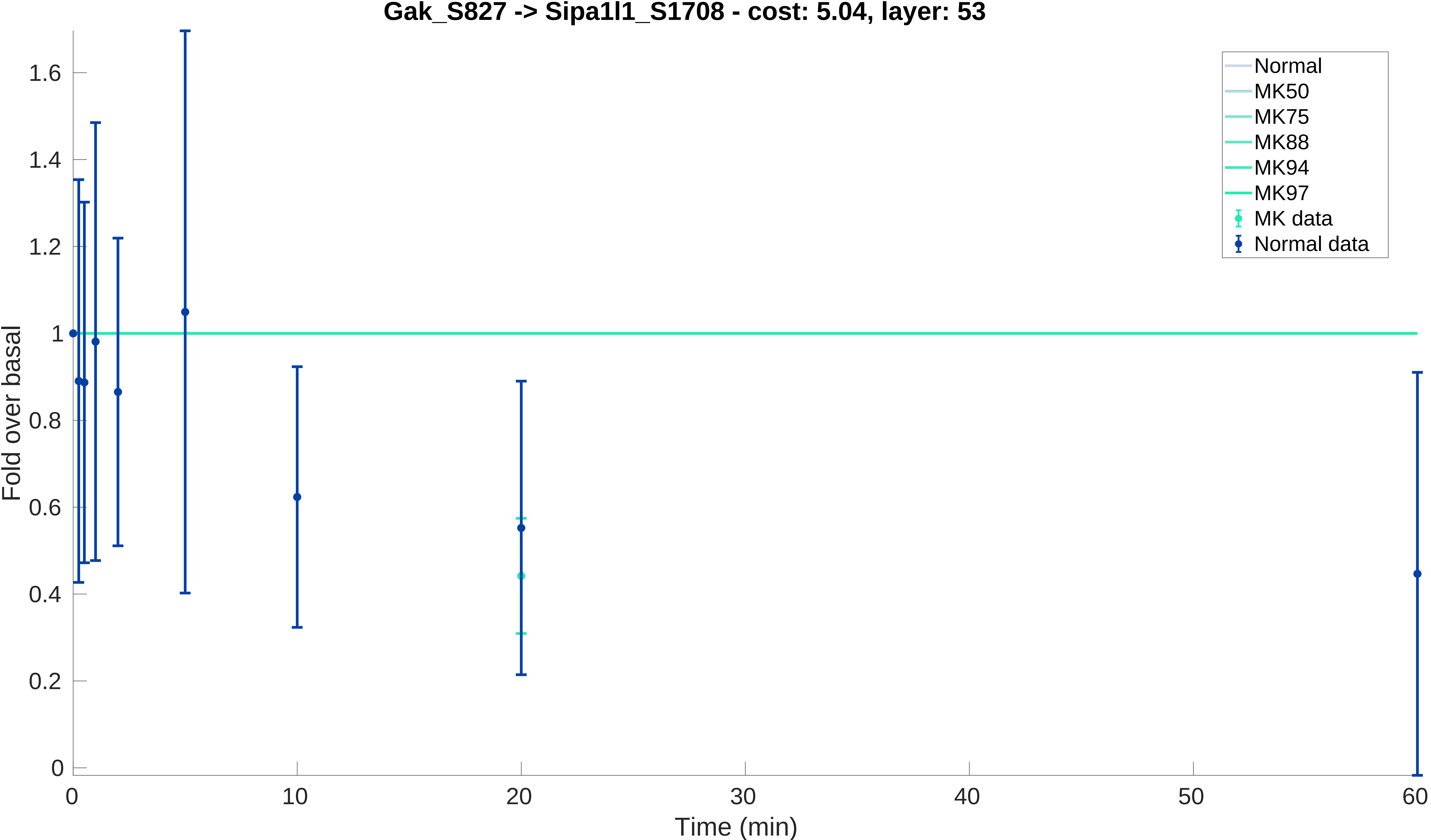

Gak_S827 -> Sipa1l1_S1708 - cost: 5.04, layer: 53
Fold over basal
Time (min)
0
0.2
0.4
0.6
0.8
1
1.2
1.4
1.6
0
10
20
30
40
50
60
Normal
MK50
MK75
MK88
MK94
MK97
MK data
Normal data

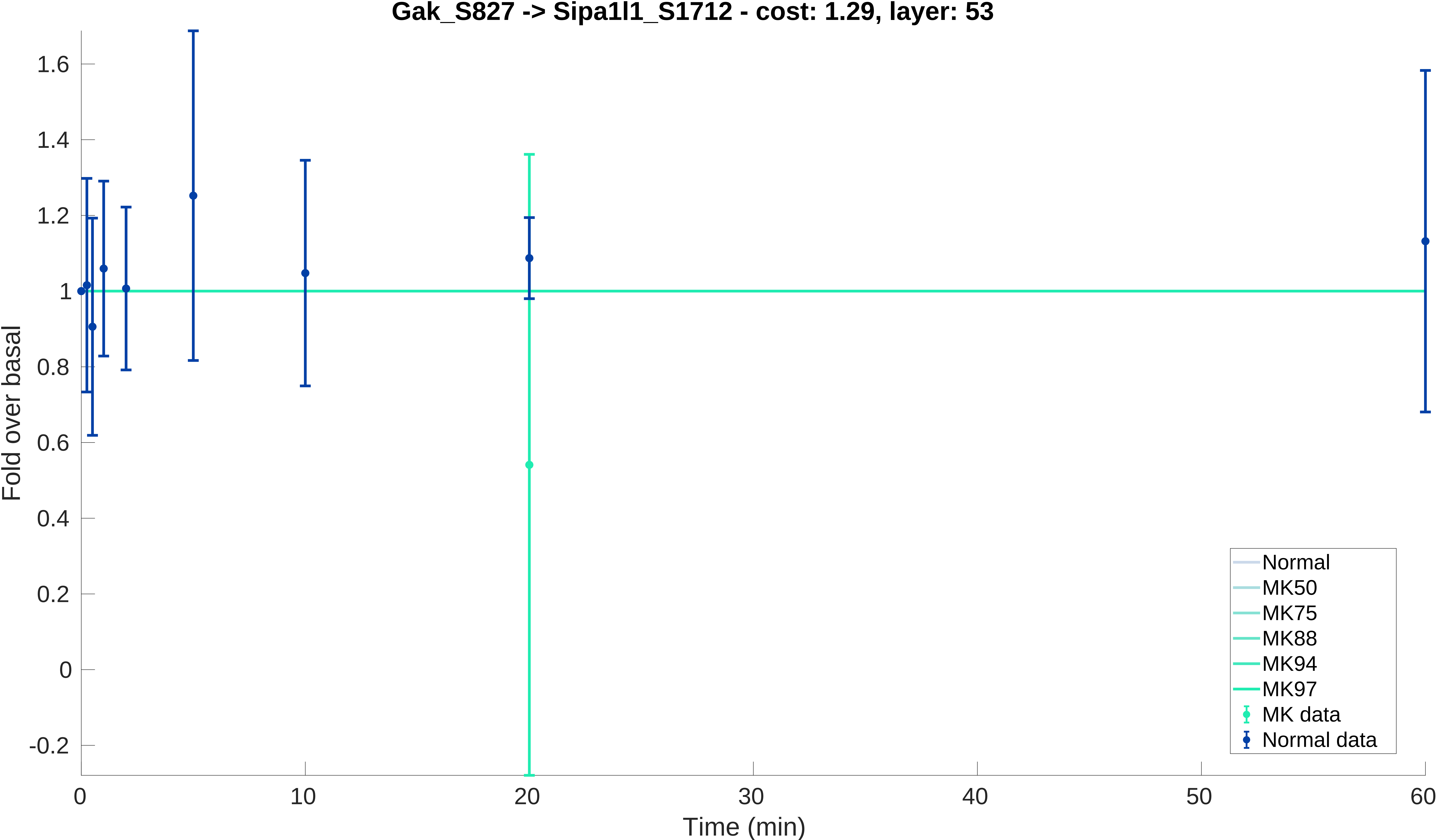

Gak_S827 -> Sipa1l1_S1712 - cost: 1.29, layer: 53
Fold over basal
1.6
1.4
1.2
1
0.8
0.6
0.4
0.2
0
-0.2
0
10
20
30
40
50
60
Time (min)
Normal
MK50
MK75
MK88
MK94
MK97
MK data
Normal data

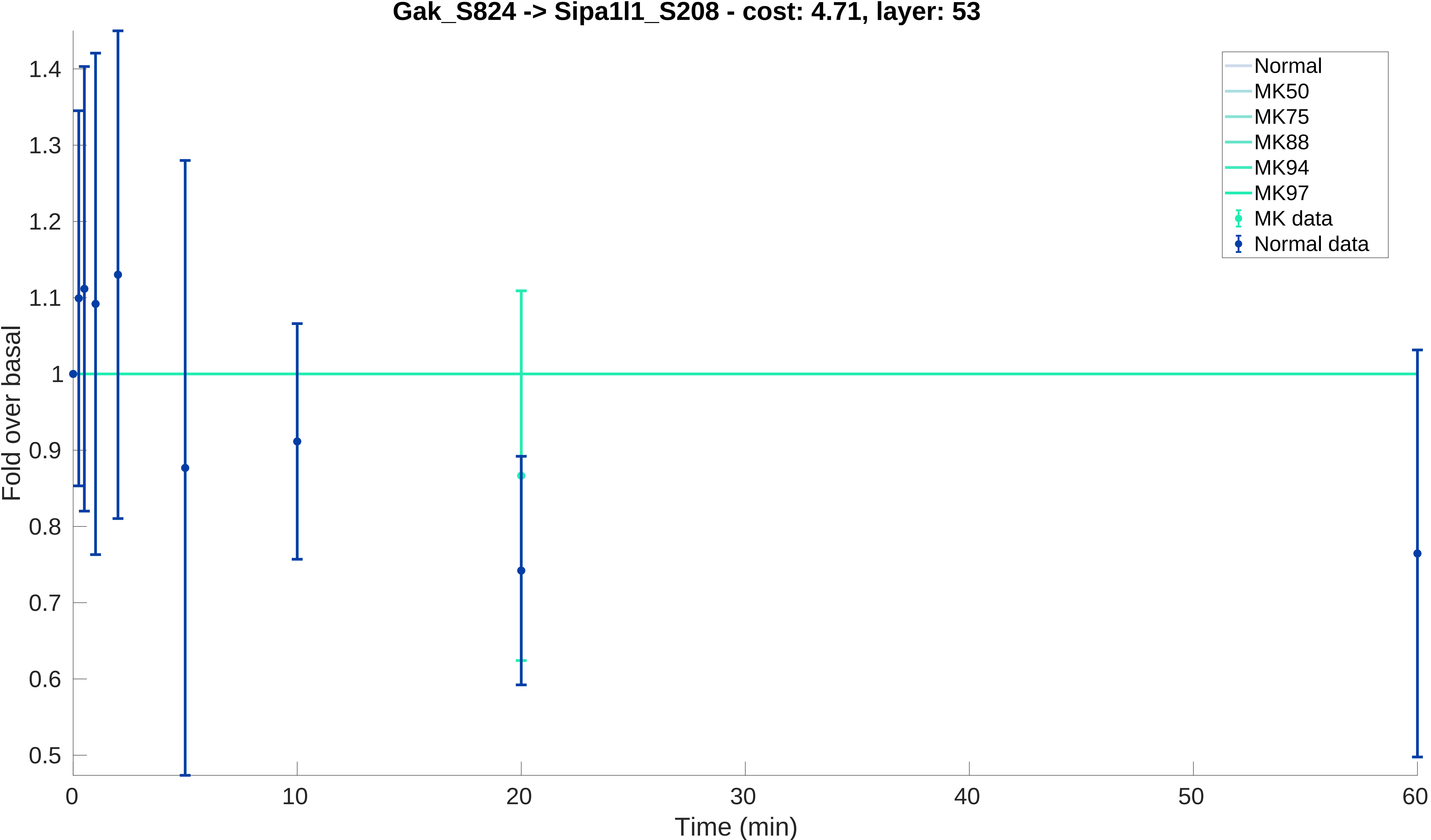

Gak_S824 -> Sipa1l1_S208 - cost: 4.71, layer: 53
Normal
MK50
MK75
MK88
MK94
MK97
MK data
Normal data
Fold over basal
1.4
1.3
1.2
1.1
1
0.9
0.8
0.7
0.6
0.5
0
10
20
30
40
50
60
Time (min)

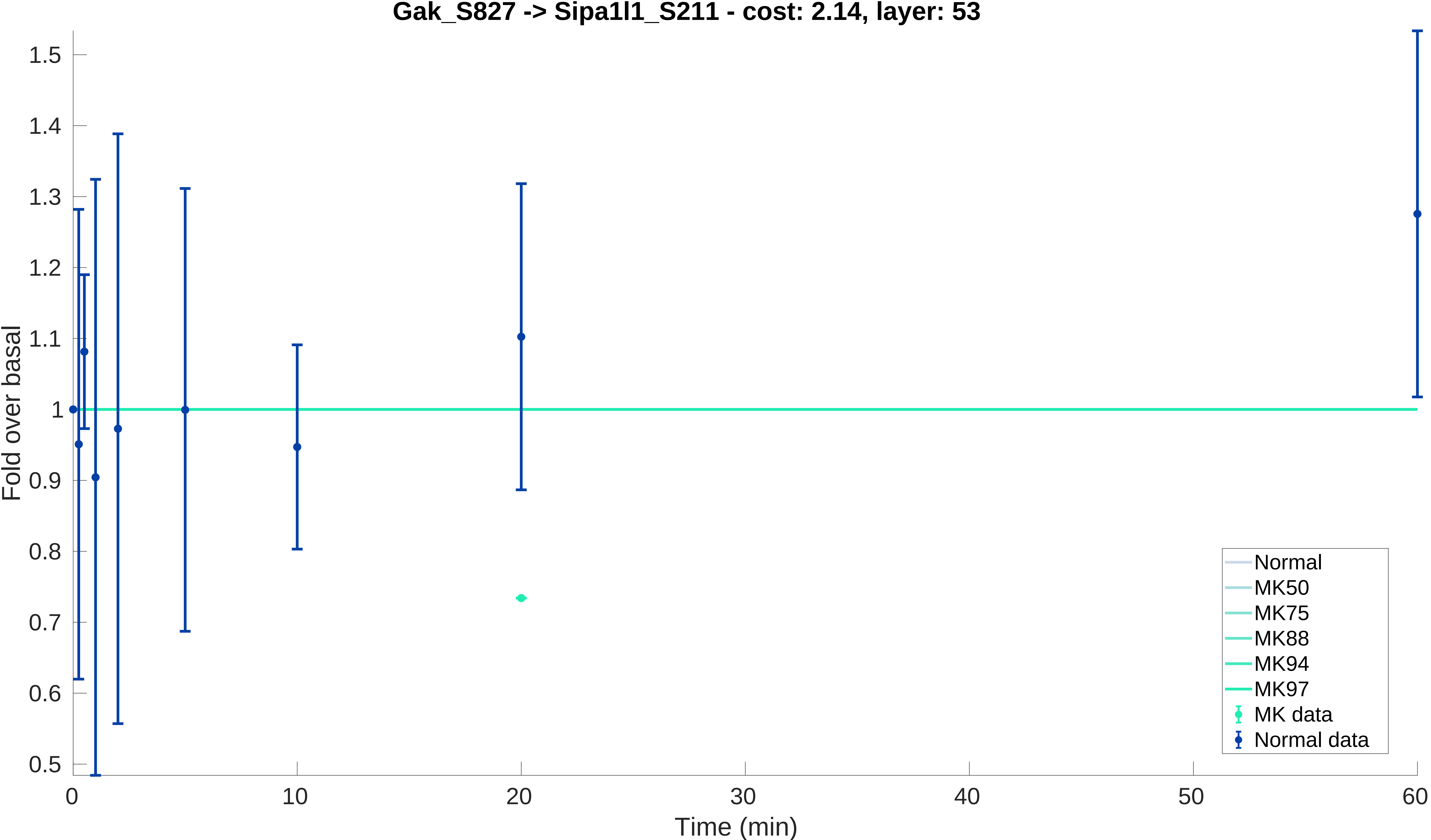

Gak_S827 -> Sipa1l1_S211 - cost: 2.14, layer: 53
Fold over basal
1.5
1.4
1.3
1.2
1.1
1
0.9
0.8
0.7
0.6
0.5
0
10
20
30
40
50
60
Time (min)
Normal
MK50
MK75
MK88
MK94
MK97
MK data
Normal data

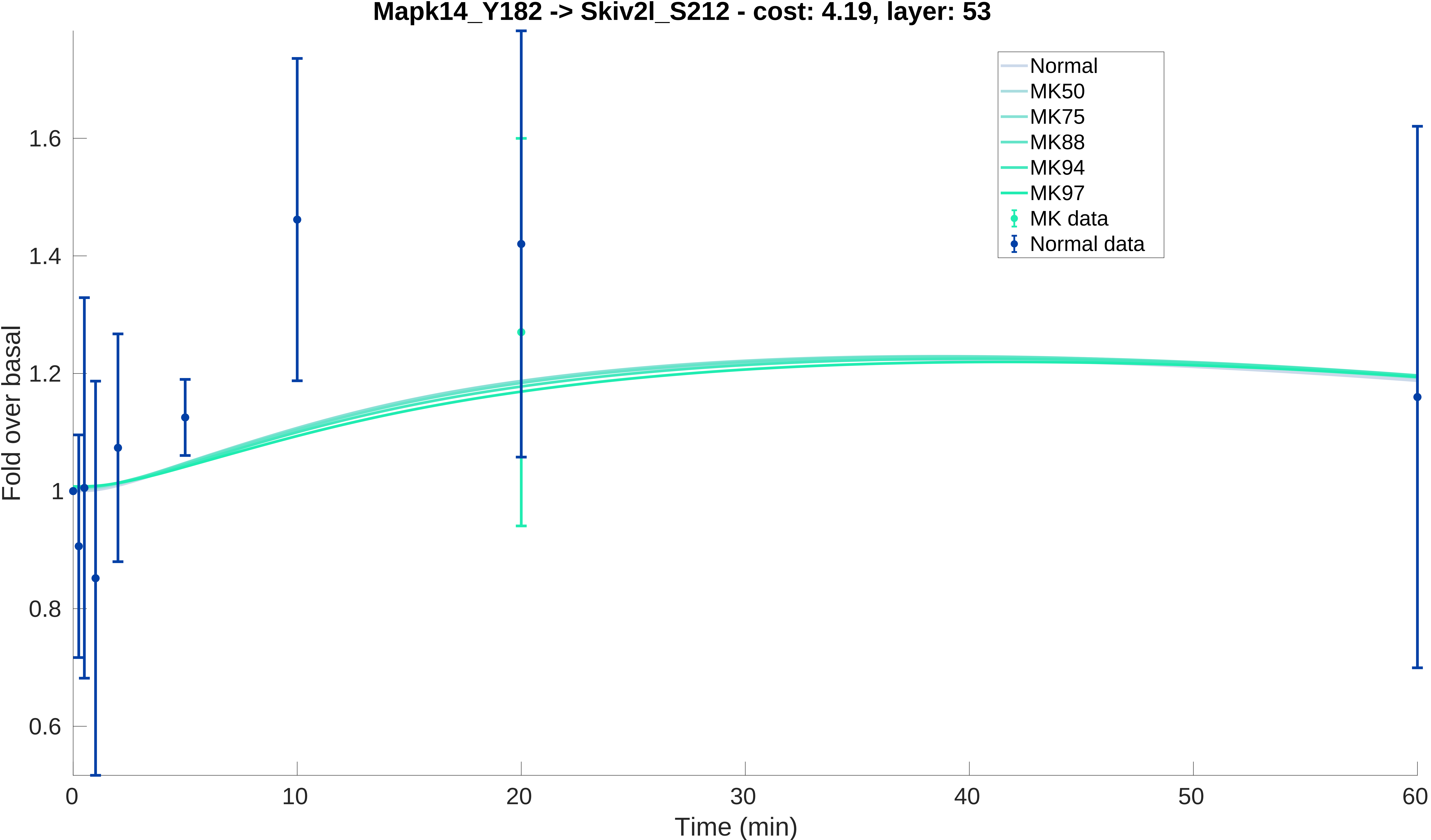

Mapk14_Y182 -> Skiv2l_S212 - cost: 4.19, layer: 53
Fold over basal
Time (min)
1.6
1.4
1.2
1
0.8
0.6
0
10
20
30
40
50
60
Normal
MK50
MK75
MK88
MK94
MK97
MK data
Normal data

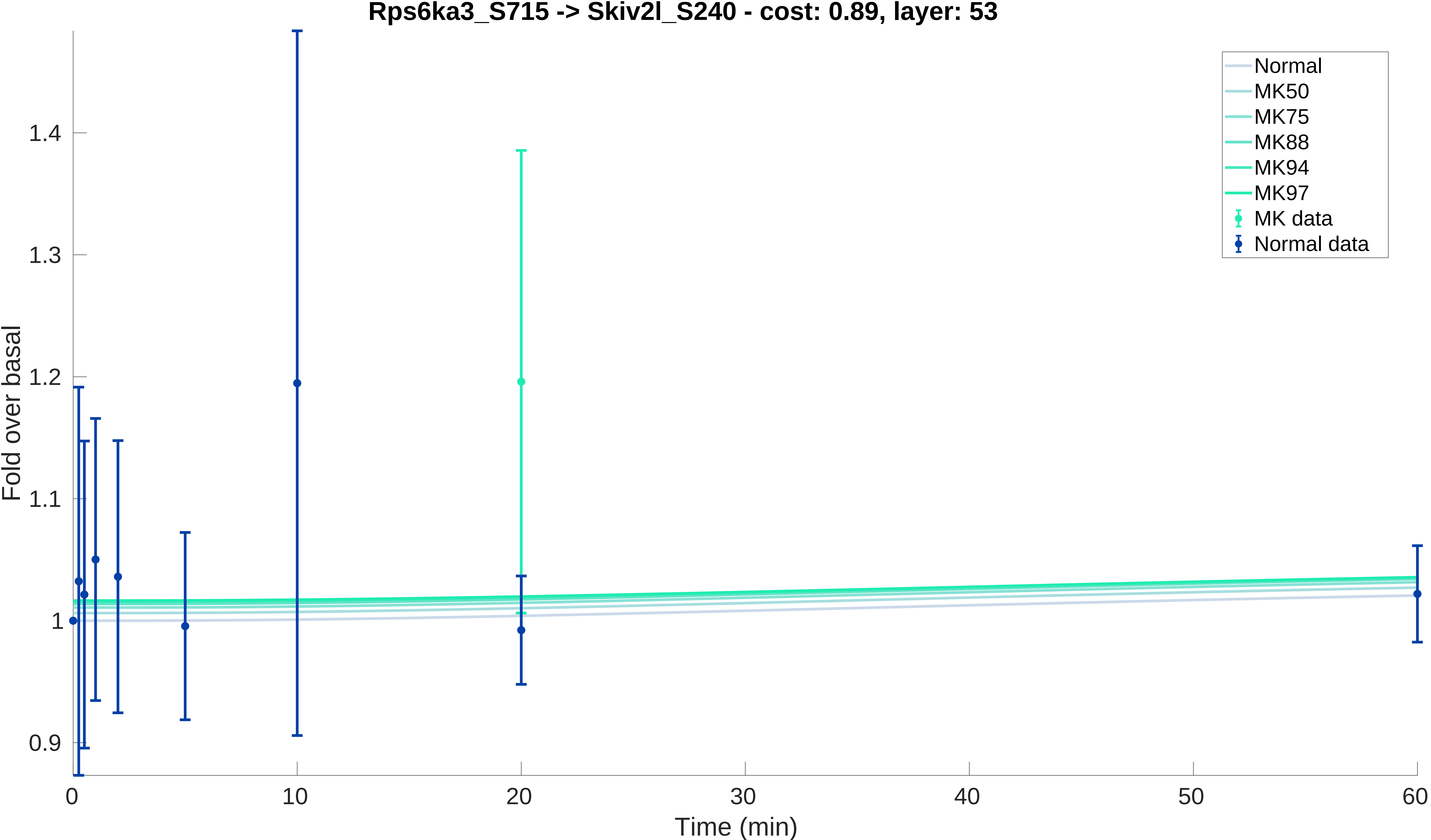

Rps6ka3_S715 -> Skiv2l_S240 - cost: 0.89, layer: 53
Fold over basal
Time (min)
1.4
1.3
1.2
1.1
1
0.9
0
10
20
30
40
50
60
Normal
MK50
MK75
MK88
MK94
MK97
MK data
Normal data

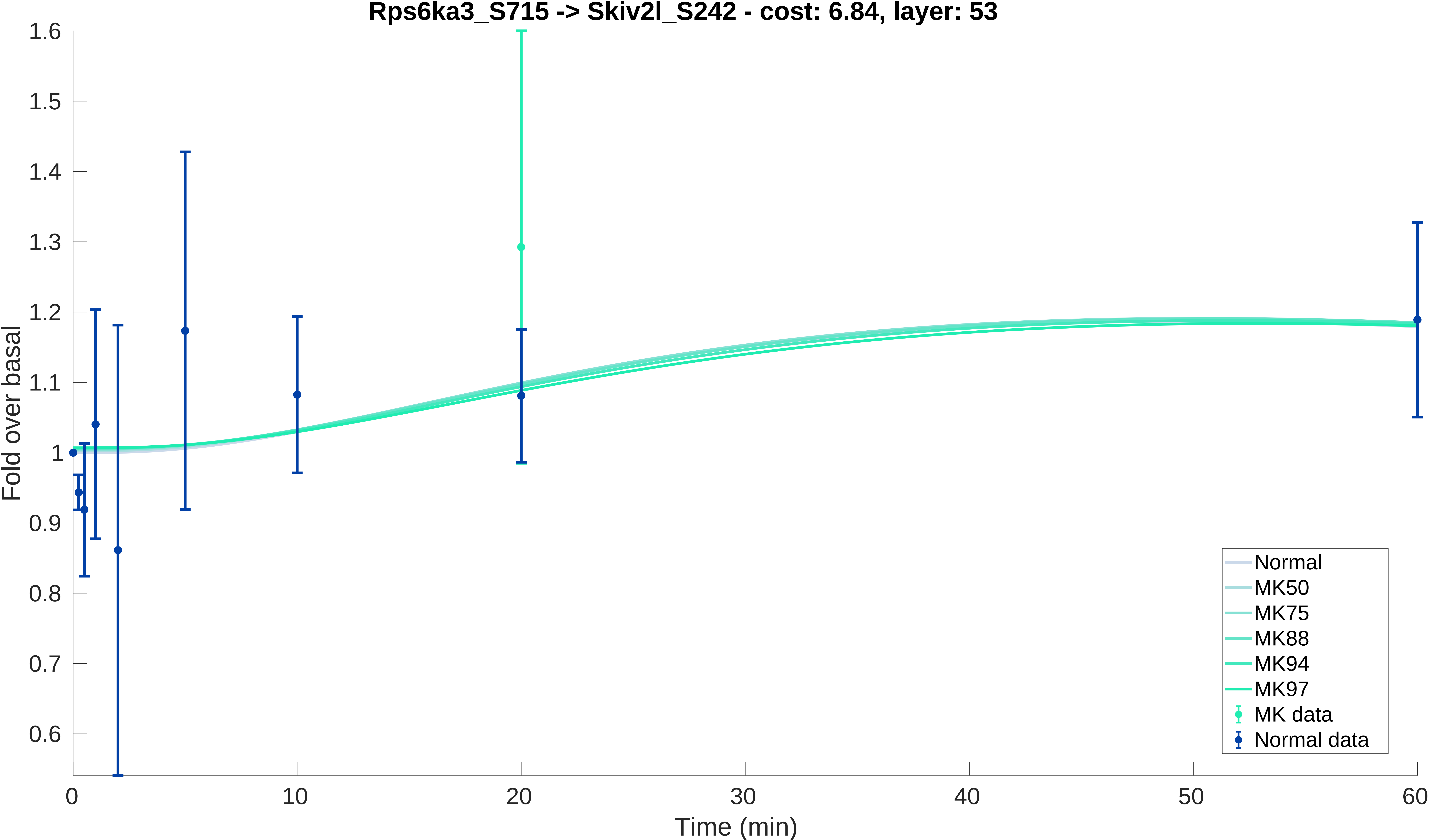

Rps6ka3_S715 -> Skiv2l_S242 - cost: 6.84, layer: 53
Fold over basal
Time (min)
1.6
1.5
1.4
1.3
1.2
1.1
1
0.9
0.8
0.7
0.6
0
10
20
30
40
50
60
Normal
MK50
MK75
MK88
MK94
MK97
MK data
Normal data

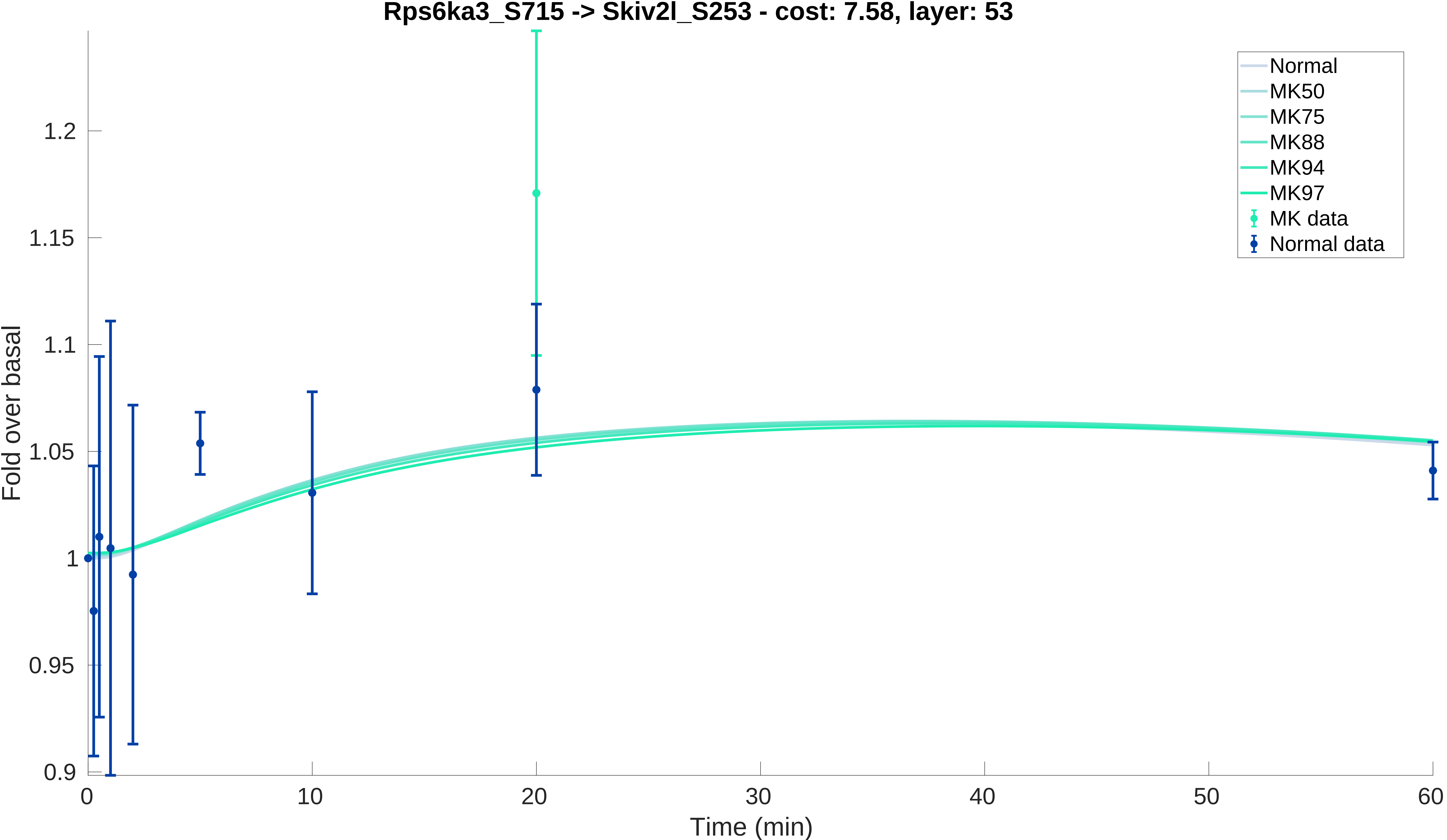
Rps6ka3_S715 -> Skiv2l_S253 - cost: 7.58, layer: 53
Fold over basal
1.2
1.15
1.1
1.05
1
0.95
0.9
0
10
20
30
40
50
60
Time (min)
Normal
MK50
MK75
MK88
MK94
MK97
MK data
Normal data

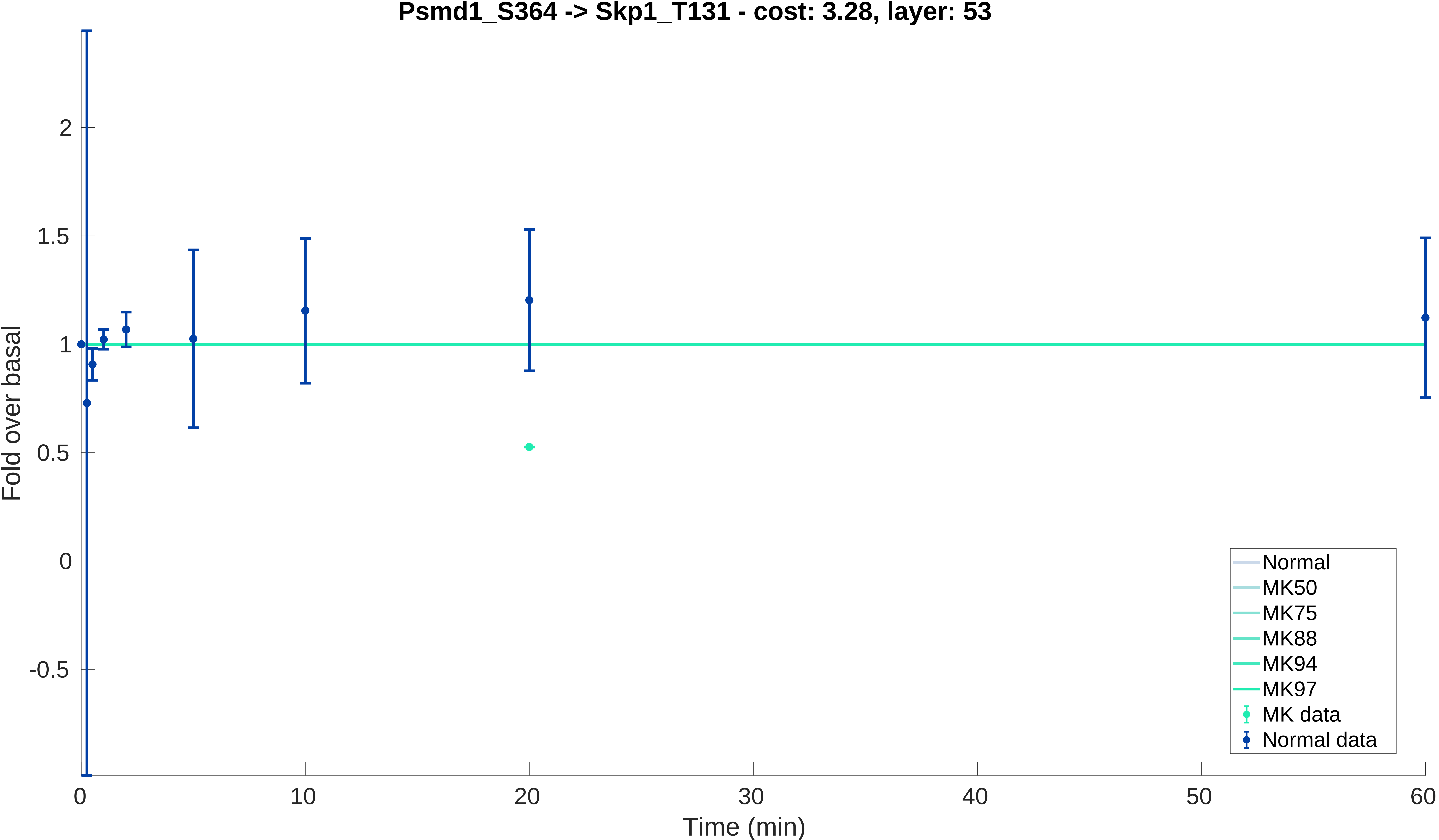

Psmd1_S364 -> Skp1_T131 - cost: 3.28, layer: 53
Fold over basal
2
1.5
1
0.5
0
-0.5
0
10
20
30
40
50
60
Time (min)
Normal
MK50
MK75
MK88
MK94
MK97
MK data
Normal data

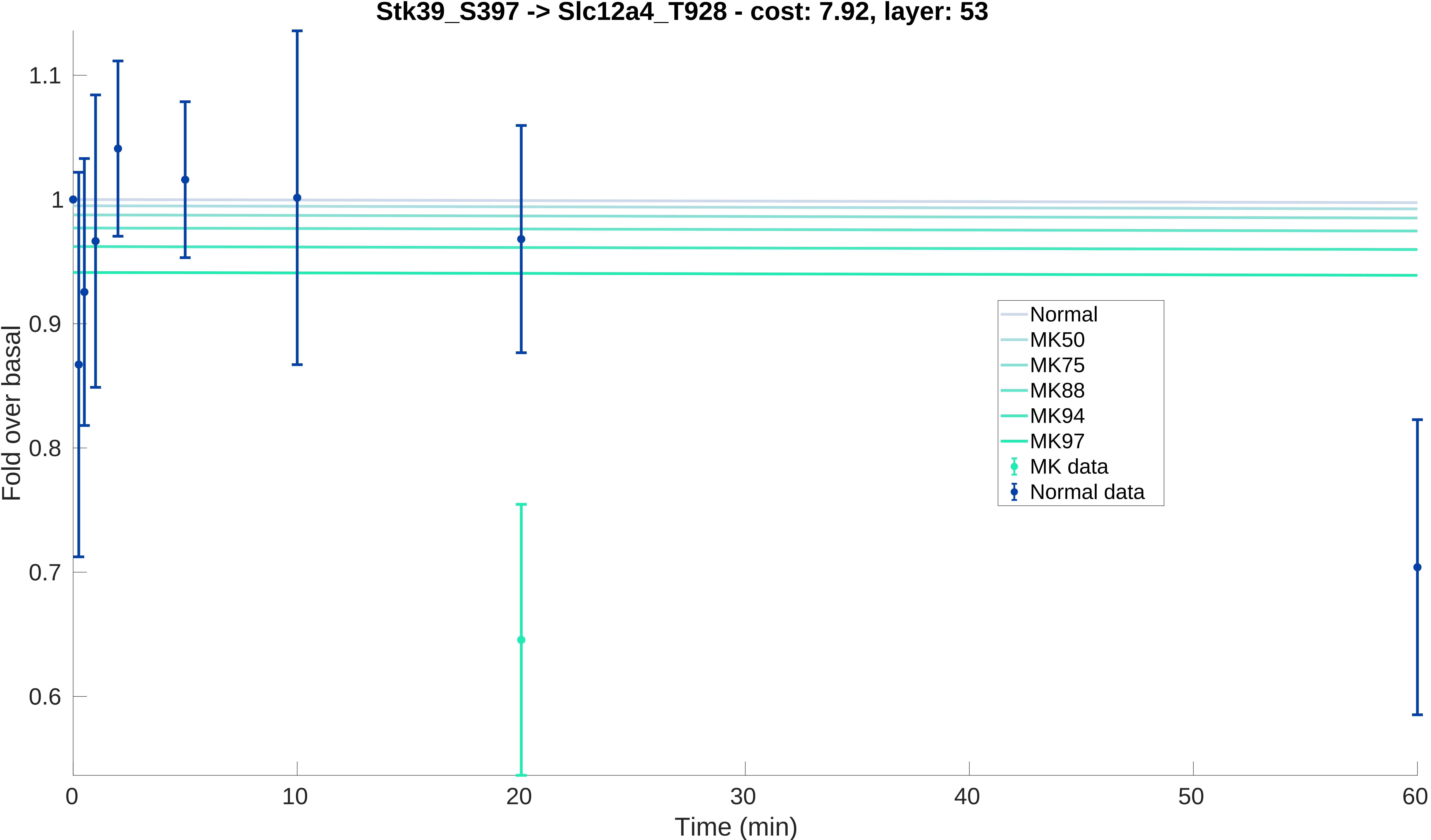

Stk39_S397 -> Slc12a4_T928 - cost: 7.92, layer: 53
Fold over basal
1.1
1
0.9
0.8
0.7
0.6
0
10
20
30
40
50
60
Time (min)
Normal
MK50
MK75
MK88
MK94
MK97
MK data
Normal data

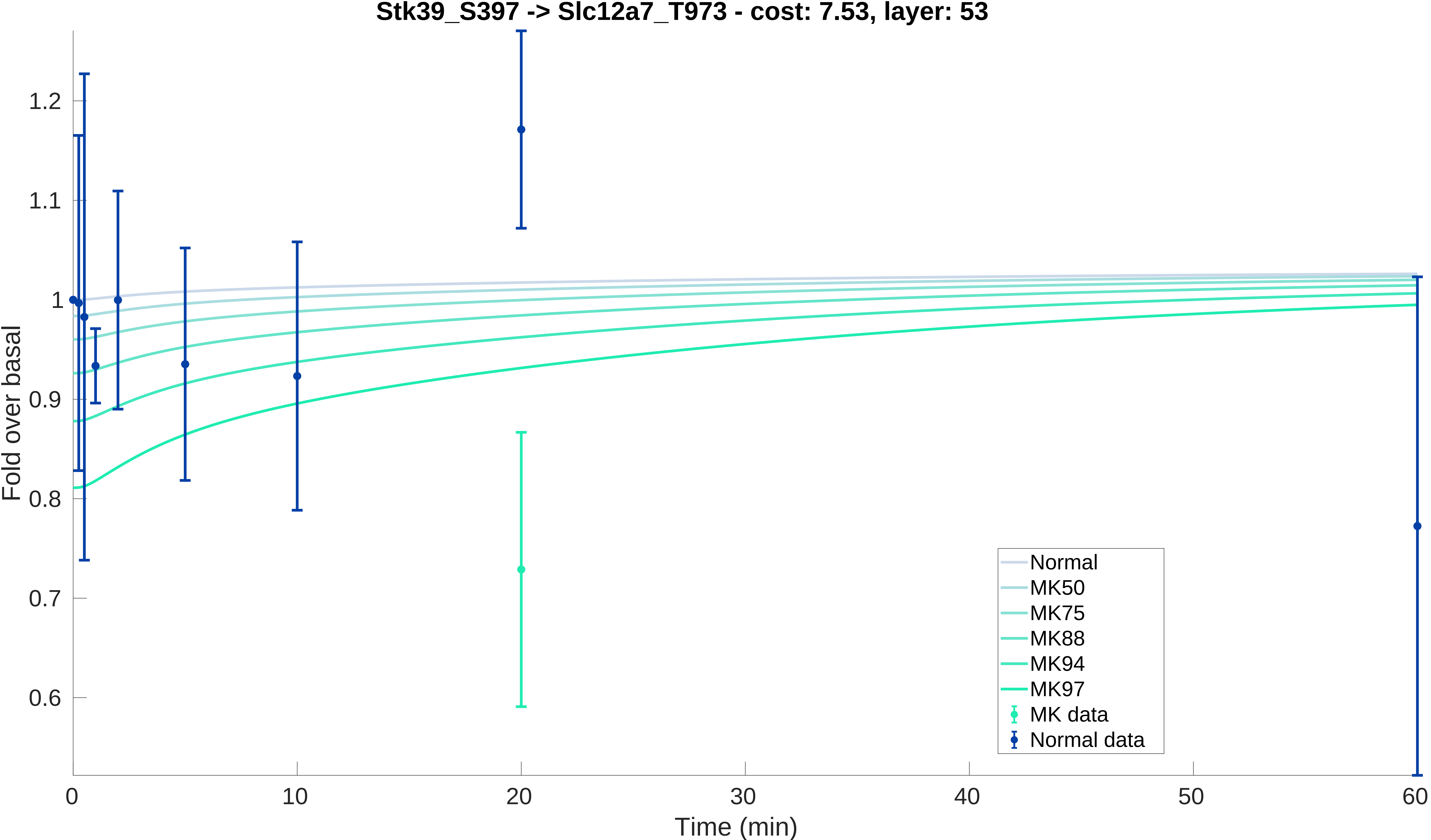

Stk39_S397 -> Slc12a7_T973 - cost: 7.53, layer: 53
Fold over basal
Time (min)
0.6
0.7
0.8
0.9
1
1.1
1.2
0
10
20
30
40
50
60
Normal
MK50
MK75
MK88
MK94
MK97
MK data
Normal data

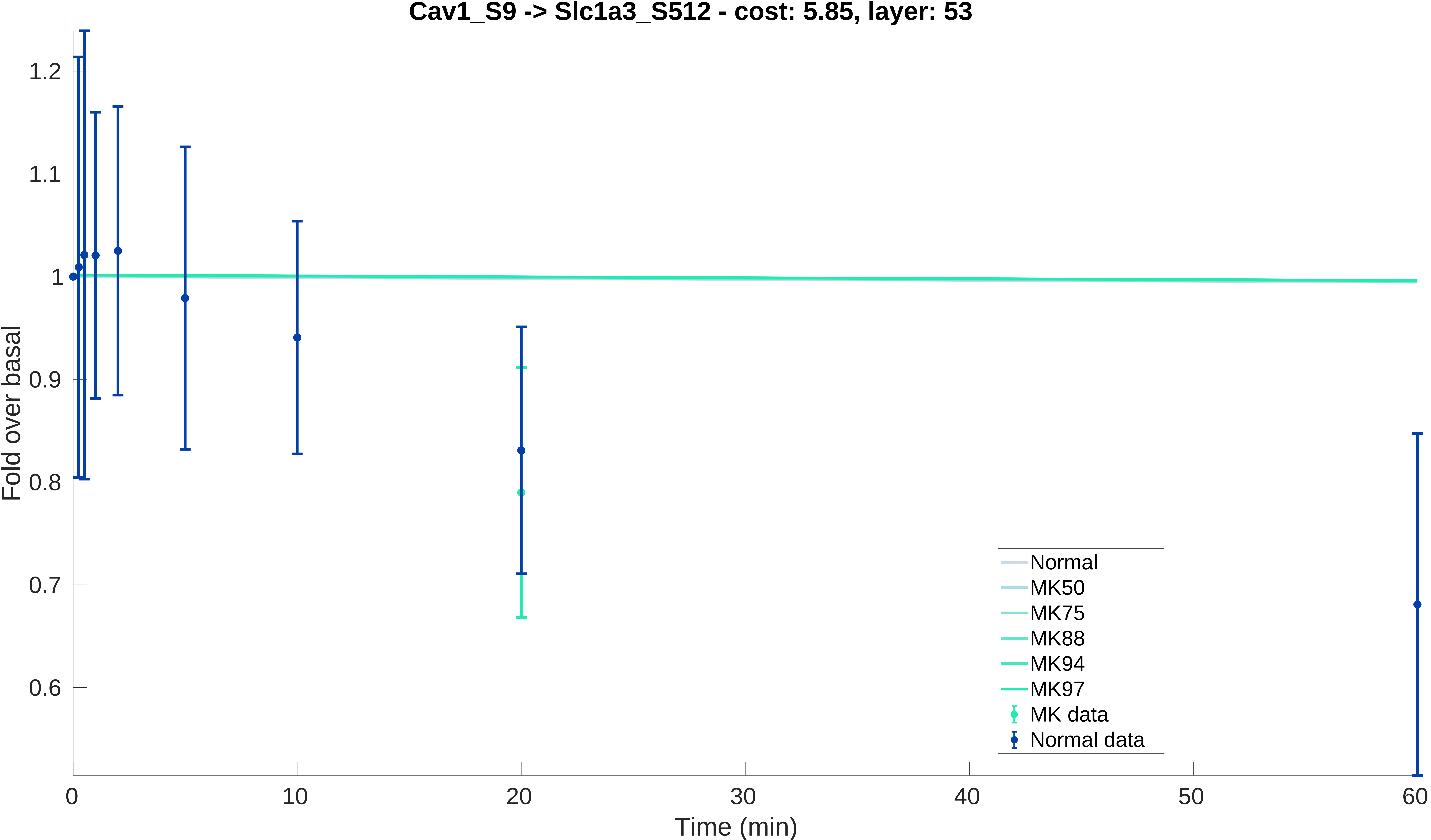

Cav1_S9 -> Slc1a3_S512 - cost: 5.85, layer: 53
Fold over basal
1.2
1.1
1
0.9
0.8
0.7
0.6
0
10
20
30
40
50
60
Time (min)
Normal
MK50
MK75
MK88
MK94
MK97
MK data
Normal data

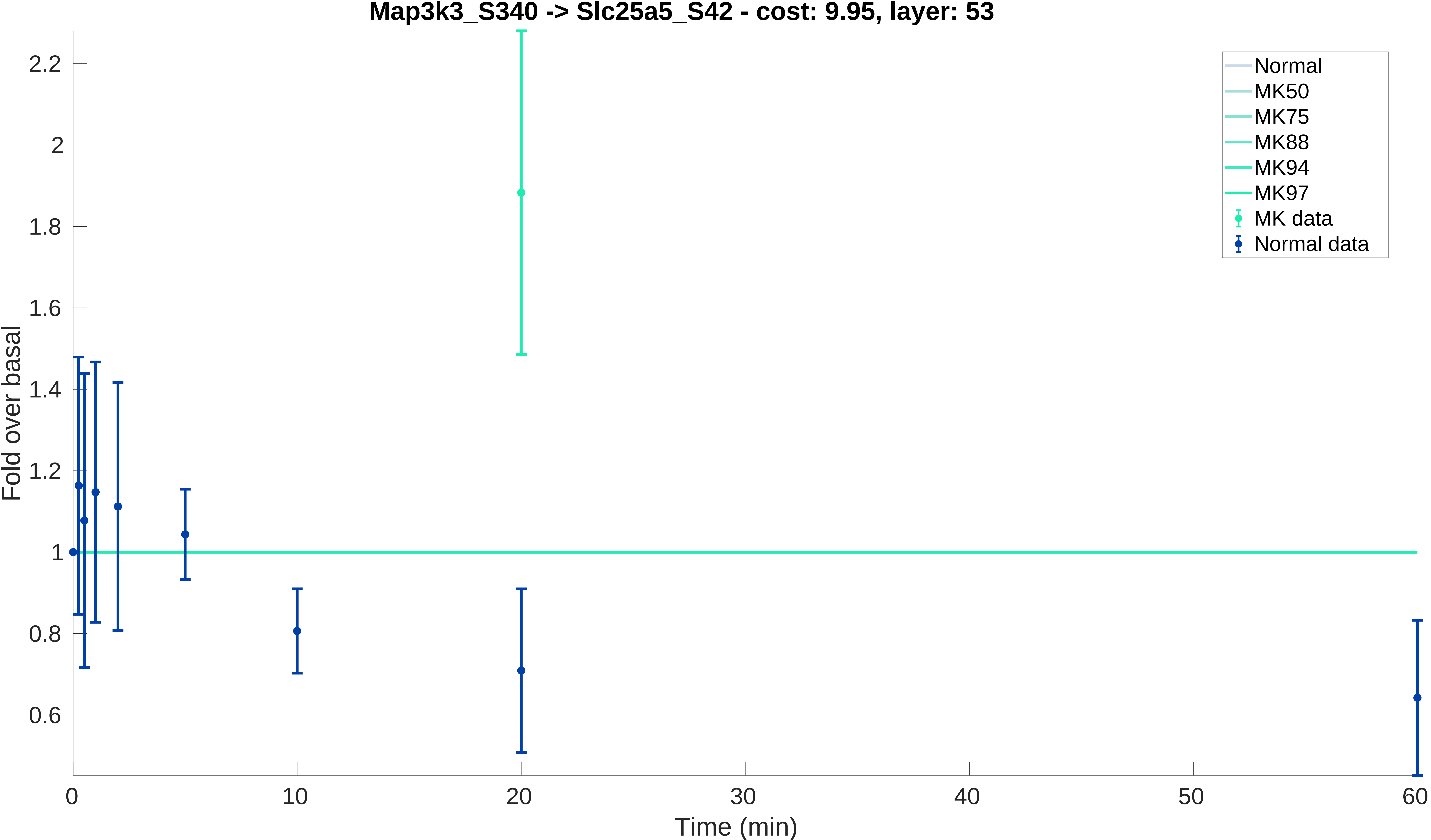

Map3k3_S340 -> Slc25a5_S42 - cost: 9.95, layer: 53
Fold over basal
Time (min)
0.6
0.8
1
1.2
1.4
1.6
1.8
2
2.2
0
10
20
30
40
50
60
Normal
MK50
MK75
MK88
MK94
MK97
MK data
Normal data

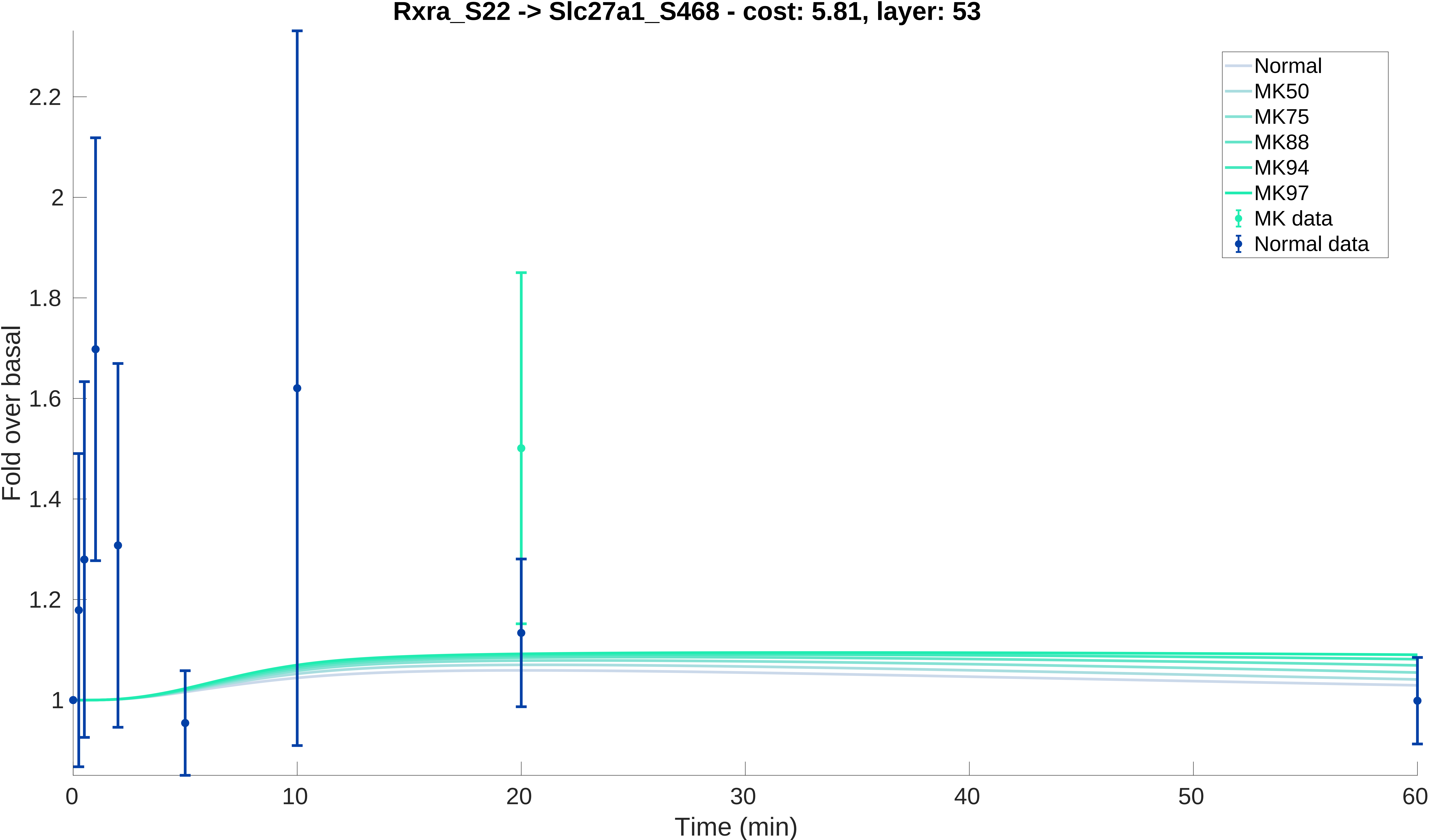

Rxra_S22 -> Slc27a1_S468 - cost: 5.81, layer: 53
Fold over basal
Time (min)
2.2
2
1.8
1.6
1.4
1.2
1
0
10
20
30
40
50
60
Normal
MK50
MK75
MK88
MK94
MK97
MK data
Normal data

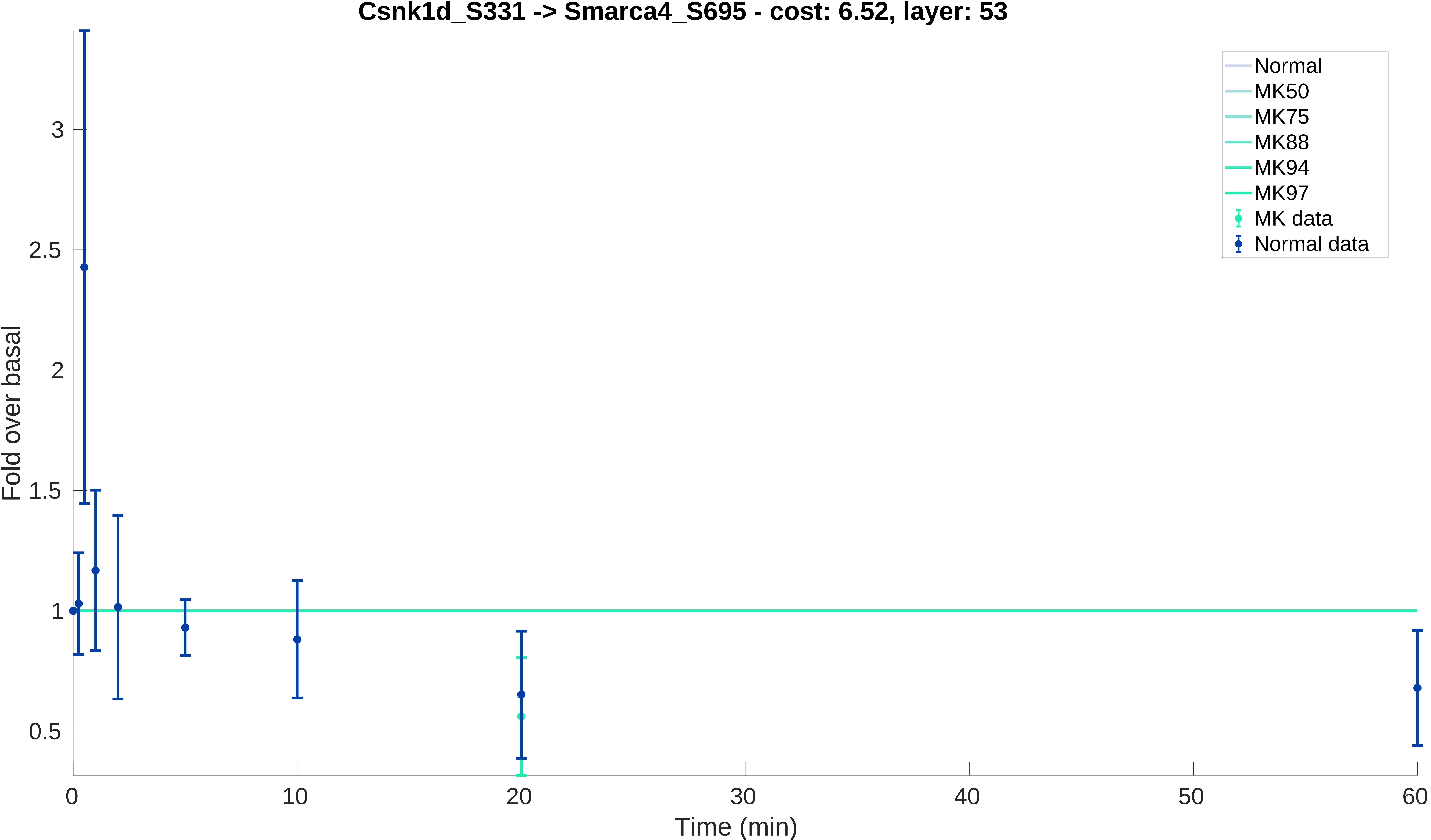

Csnk1d_S331 -> Smarca4_S695 - cost: 6.52, layer: 53
Fold over basal
Time (min)
0.5
1
1.5
2
2.5
3
0
10
20
30
40
50
60
Normal
MK50
MK75
MK88
MK94
MK97
MK data
Normal data

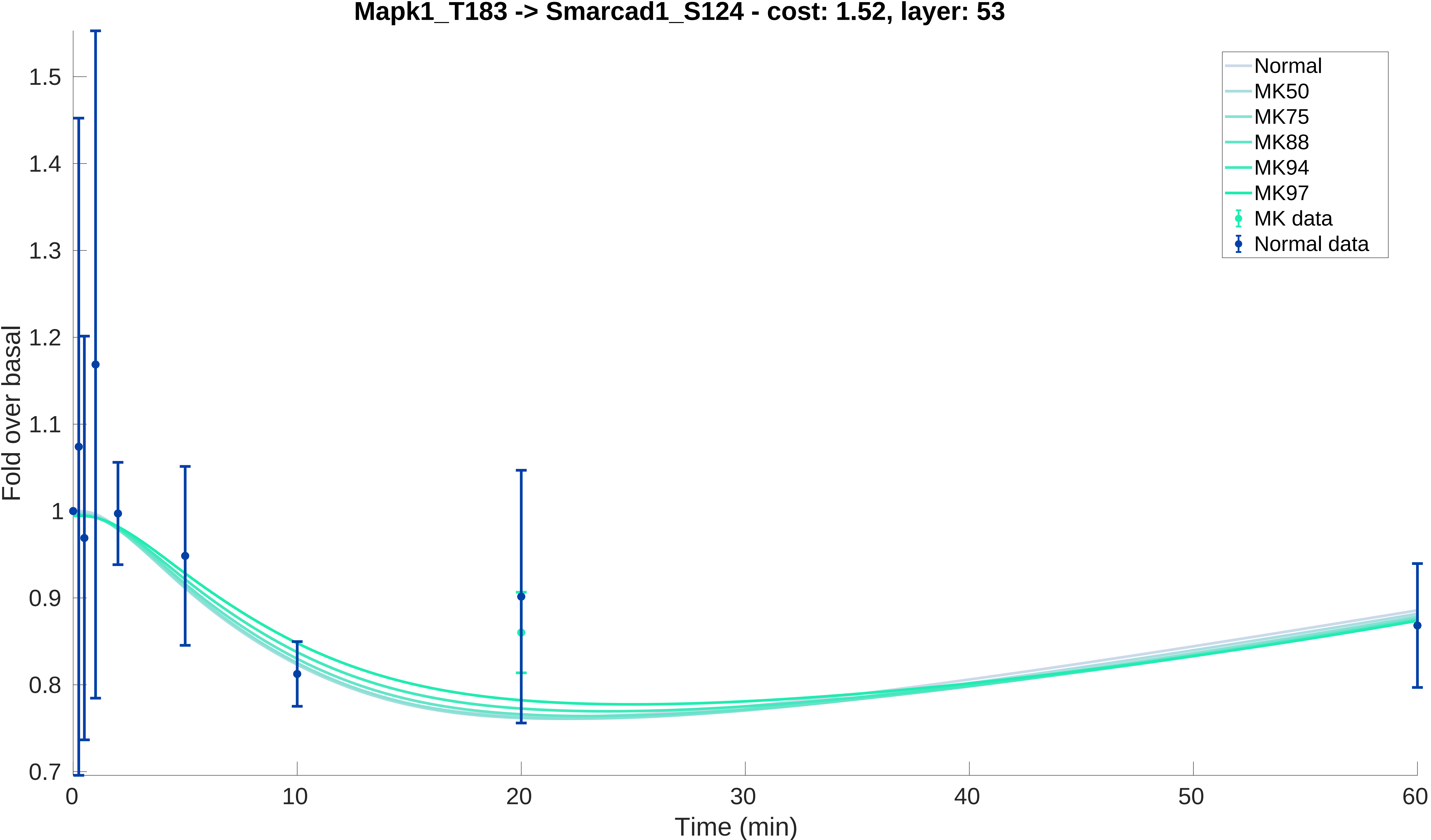
Mapk1_T183 -> Smarcad1_S124 - cost: 1.52, layer: 53
Fold over basal
Time (min)
0.7
0.8
0.9
1
1.1
1.2
1.3
1.4
1.5
0
10
20
30
40
50
60
Normal
MK50
MK75
MK88
MK94
MK97
MK data
Normal data

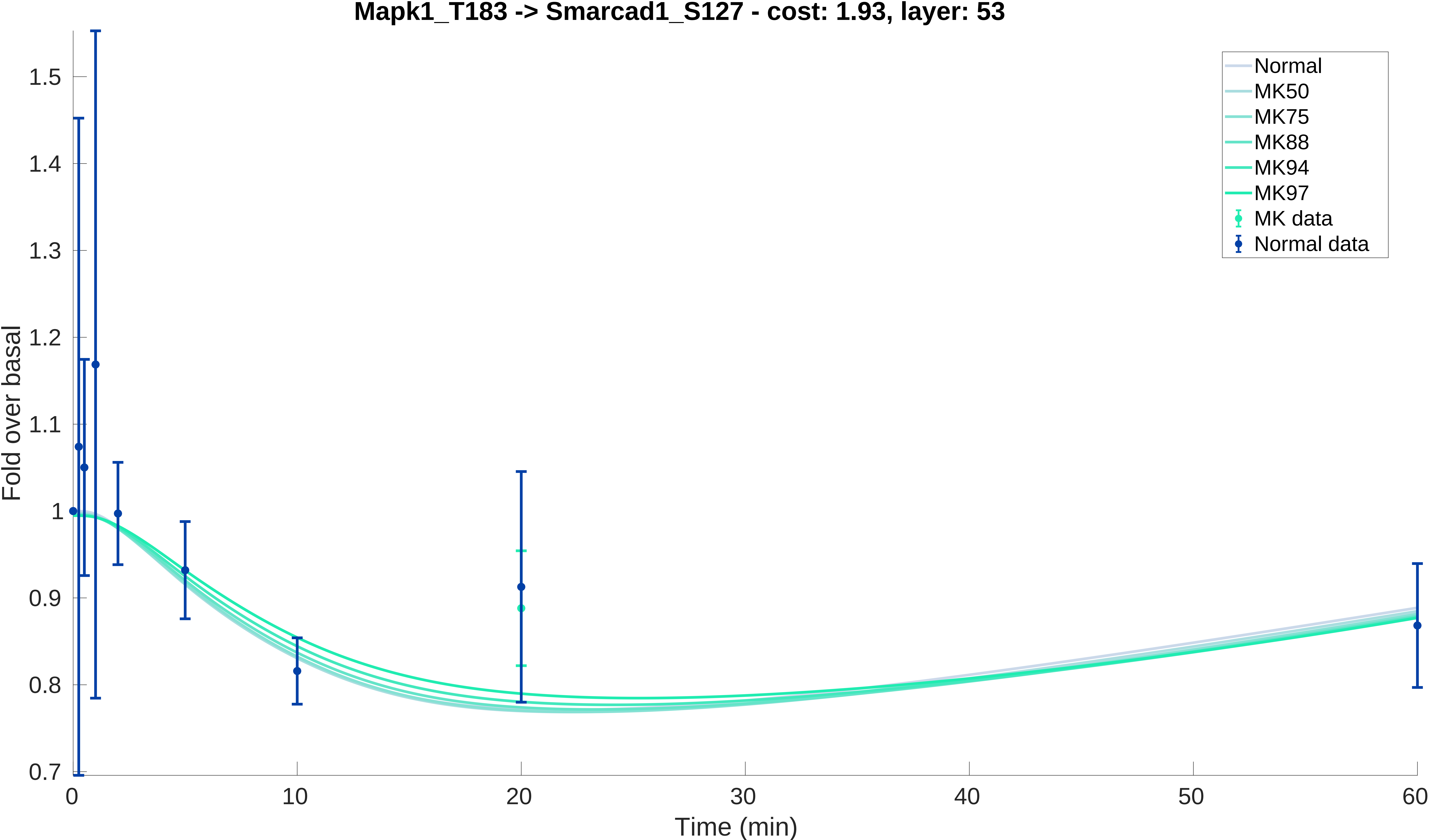
Mapk1_T183 -> Smarcad1_S127 - cost: 1.93, layer: 53
Fold over basal
Time (min)
0.7
0.8
0.9
1
1.1
1.2
1.3
1.4
1.5
0
10
20
30
40
50
60
Normal
MK50
MK75
MK88
MK94
MK97
MK data
Normal data

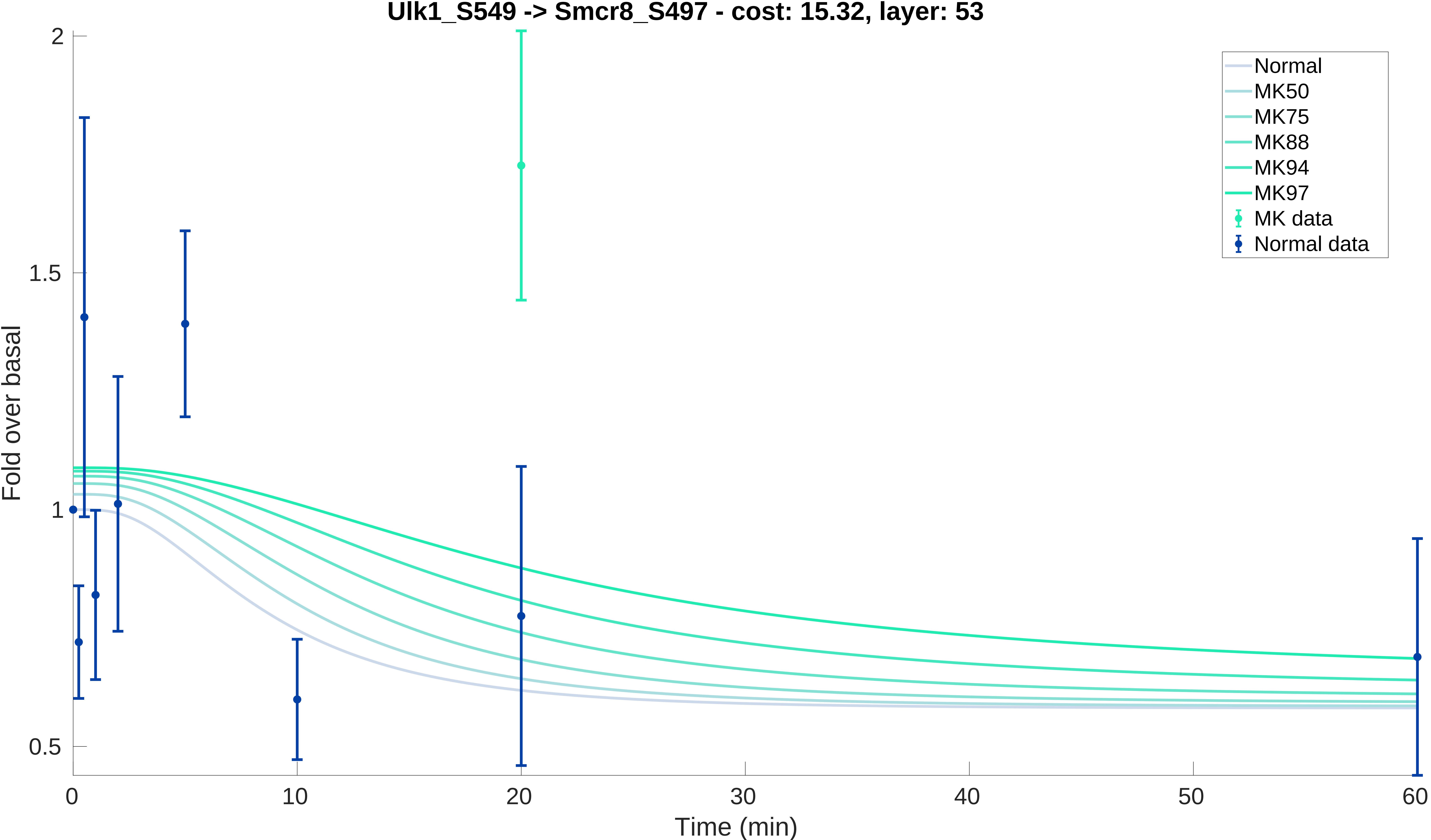

Ulk1_S549 -> Smcr8_S497 - cost: 15.32, layer: 53
Fold over basal
Time (min)
2
1.5
1
0.5
0
10
20
30
40
50
60
Normal
MK50
MK75
MK88
MK94
MK97
MK data
Normal data

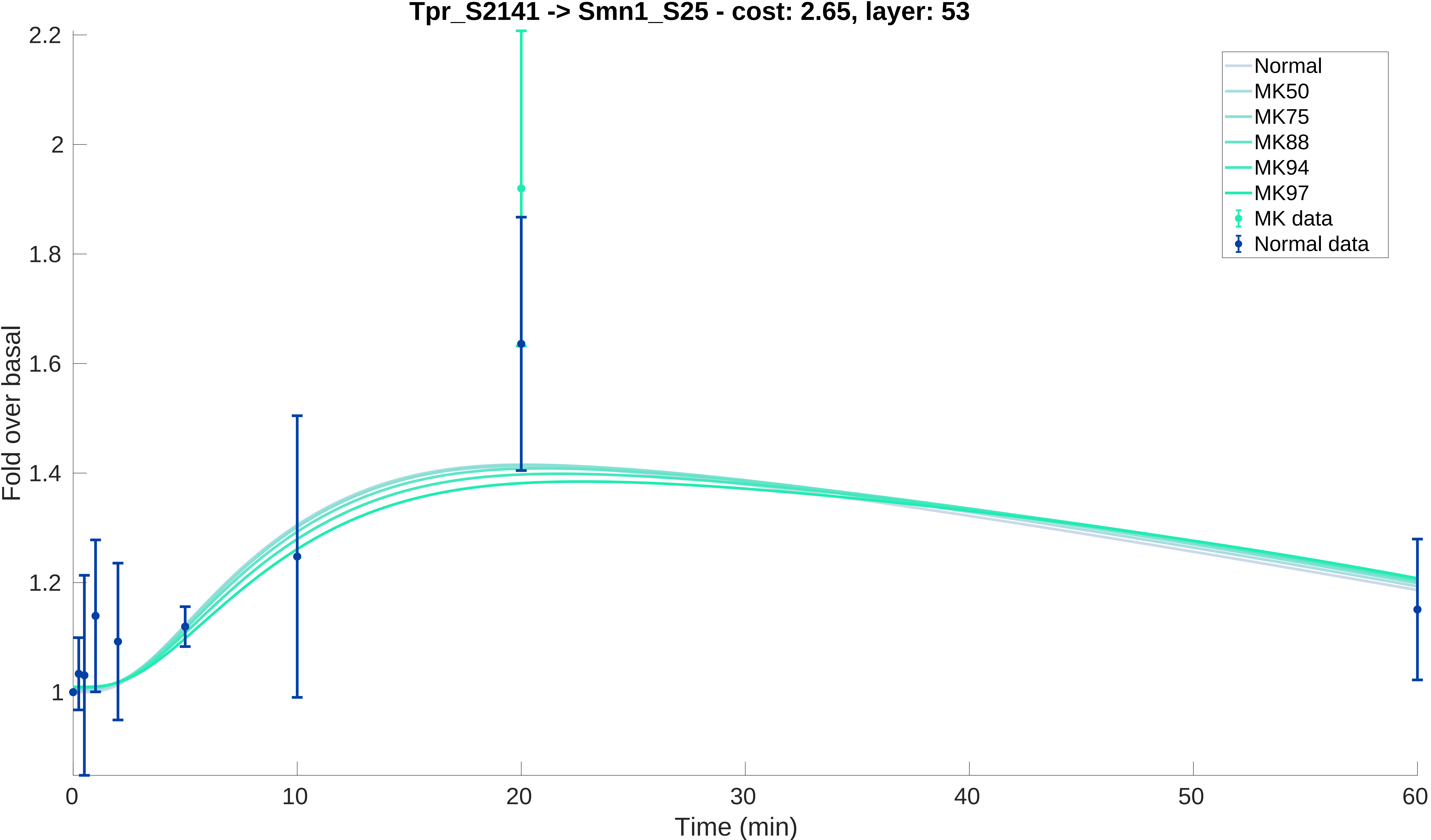
Tpr_S2141 -> Smn1_S25 - cost: 2.65, layer: 53
Fold over basal
Time (min)
2.2
2
1.8
1.6
1.4
1.2
1
0
10
20
30
40
50
60
Normal
MK50
MK75
MK88
MK94
MK97
MK data
Normal data

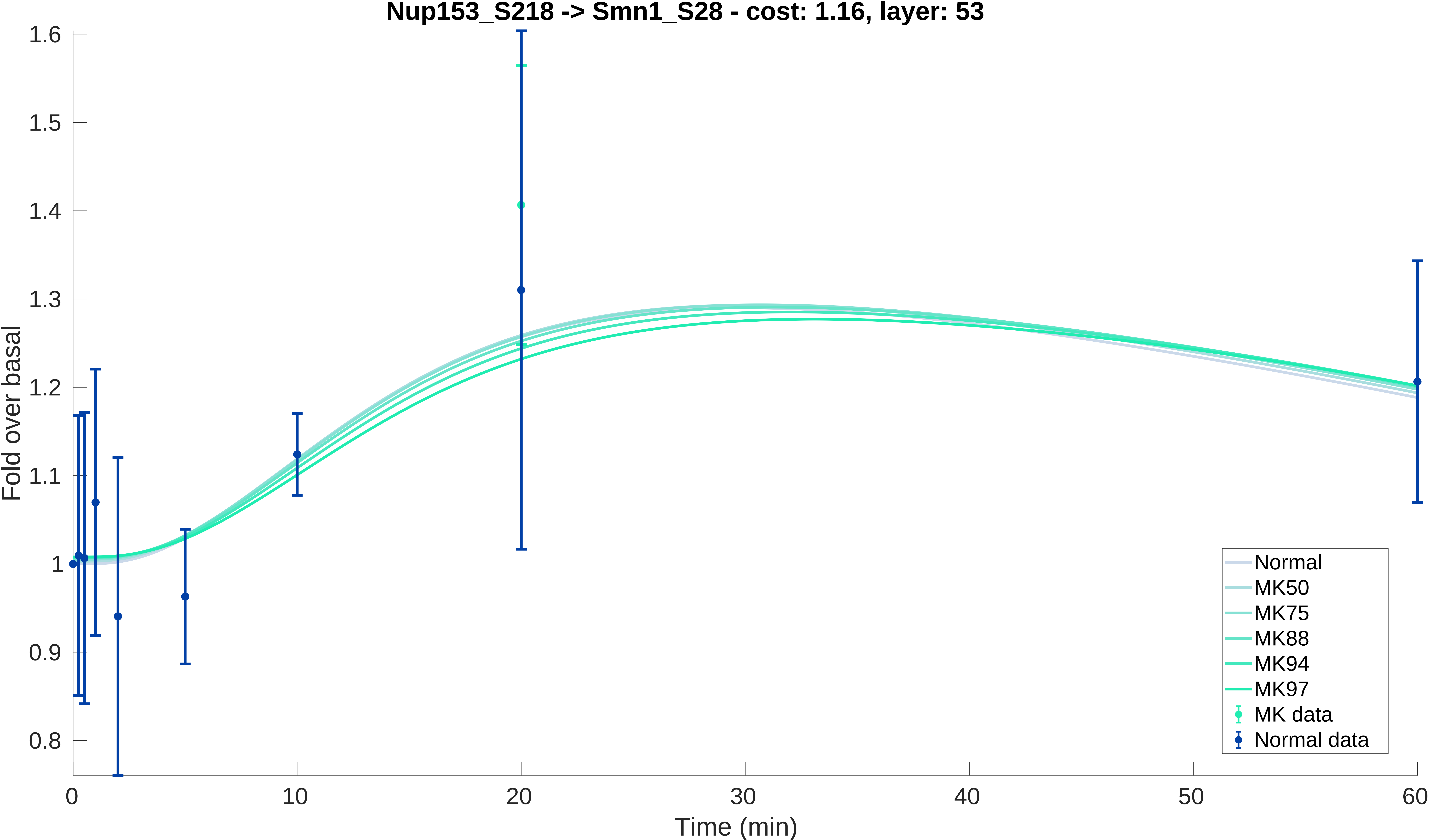

Nup153_S218 -> Smn1_S28 - cost: 1.16, layer: 53
Fold over basal
Time (min)
0.8
0.9
1
1.1
1.2
1.3
1.4
1.5
1.6
0
10
20
30
40
50
60
Normal
MK50
MK75
MK88
MK94
MK97
MK data
Normal data

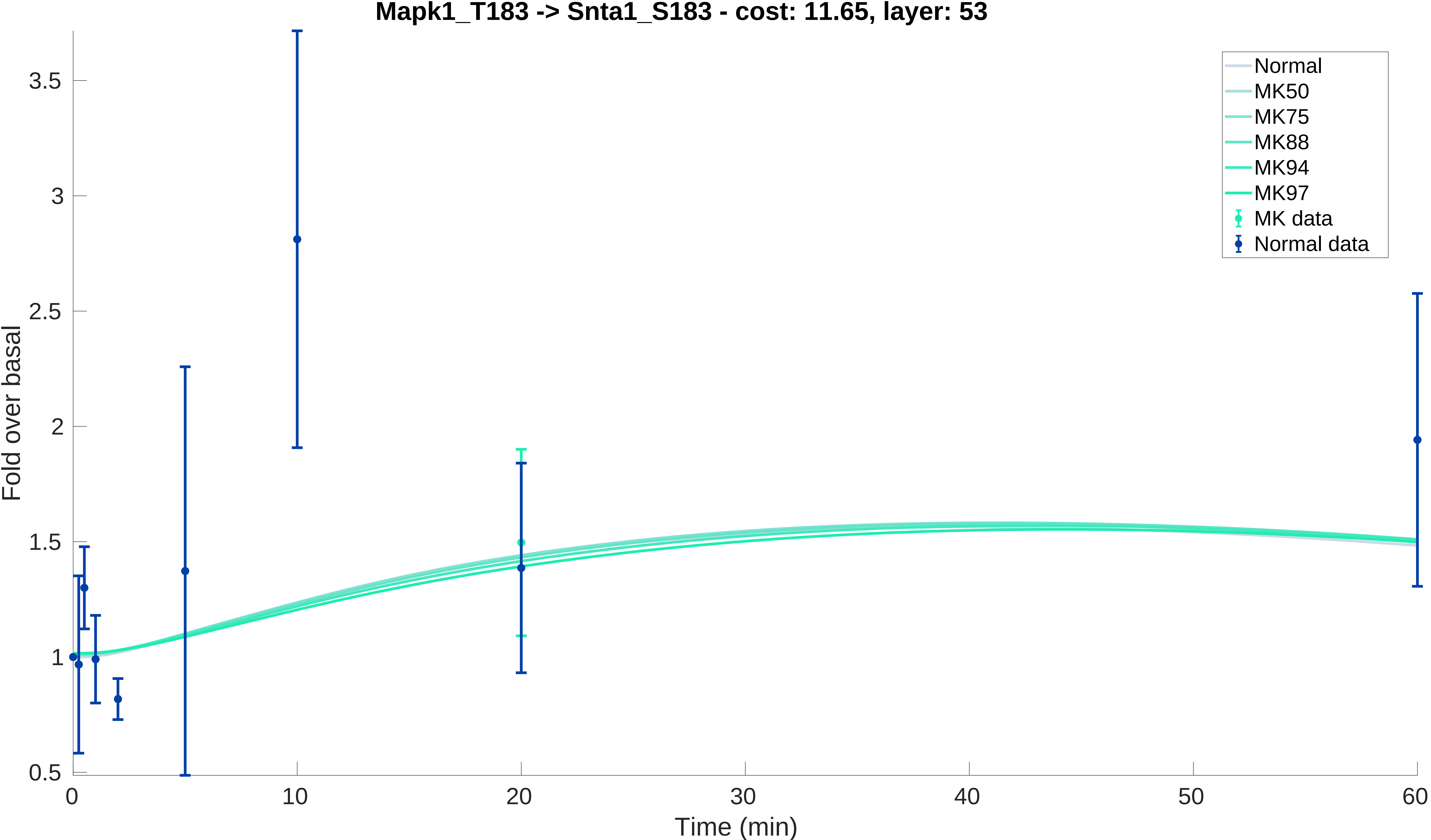
Mapk1_T183 -> Snta1_S183 - cost: 11.65, layer: 53
Fold over basal
Time (min)
0.5
1
1.5
2
2.5
3
3.5
0
10
20
30
40
50
60
Normal
MK50
MK75
MK88
MK94
MK97
MK data
Normal data

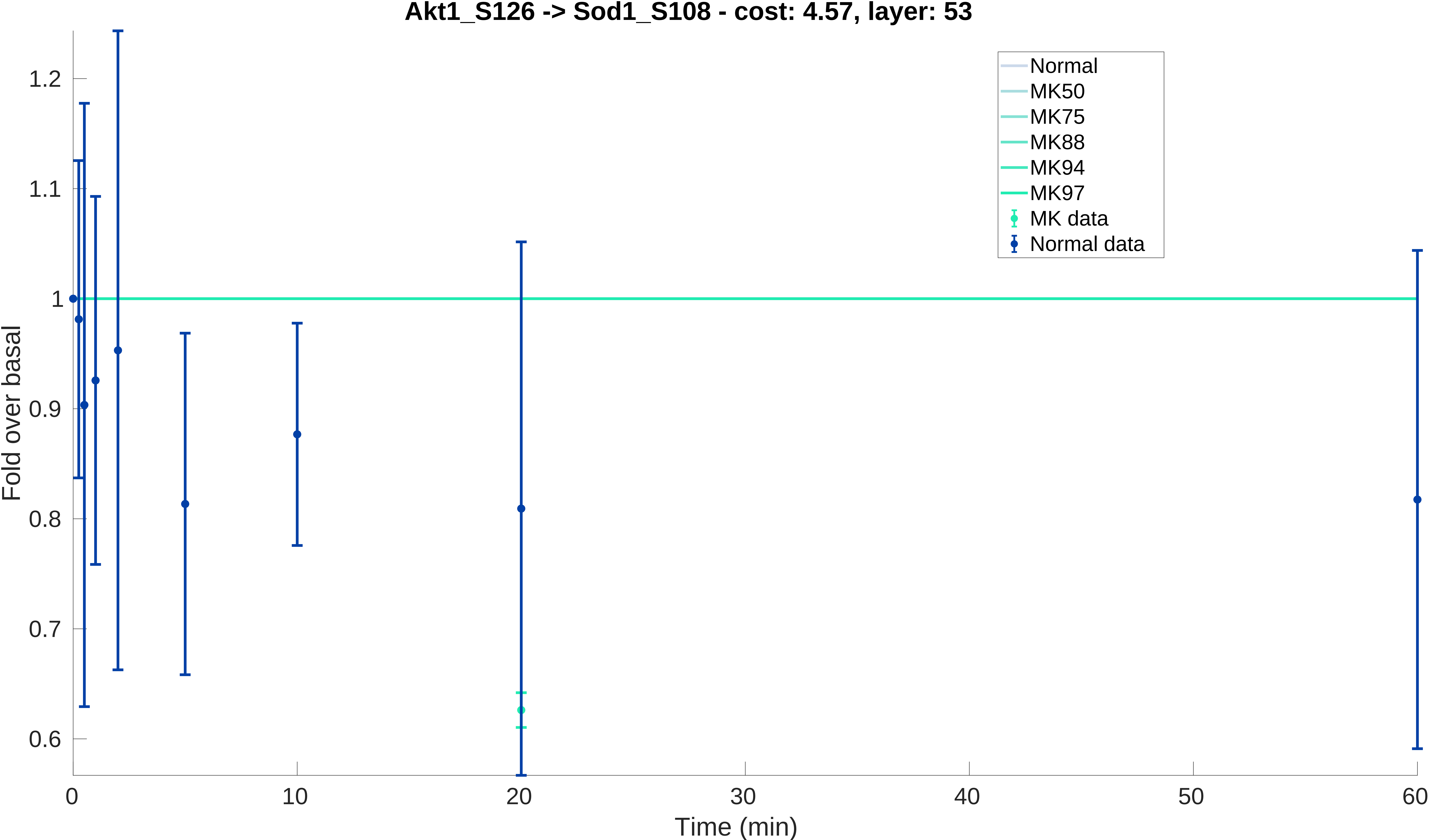

Akt1_S126 -> Sod1_S108 - cost: 4.57, layer: 53
Normal
MK50
MK75
MK88
MK94
MK97
MK data
Normal data
Fold over basal
1.2
1.1
1
0.9
0.8
0.7
0.6
0
10
20
30
40
50
60
Time (min)

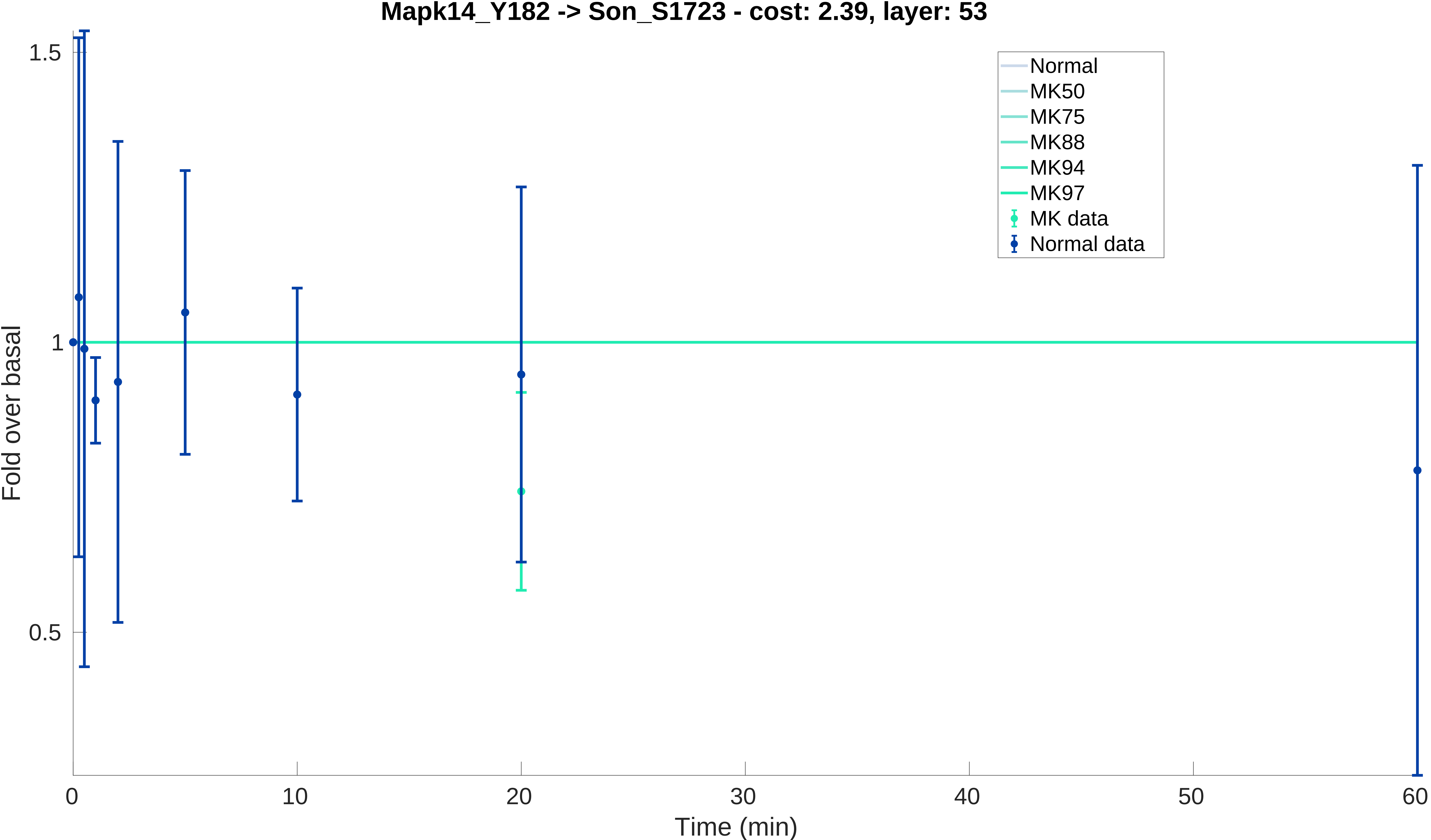

Mapk14_Y182 -> Son_S1723 - cost: 2.39, layer: 53
Fold over basal
1.5
1
0.5
0
10
20
30
40
50
60
Time (min)
Normal
MK50
MK75
MK88
MK94
MK97
MK data
Normal data

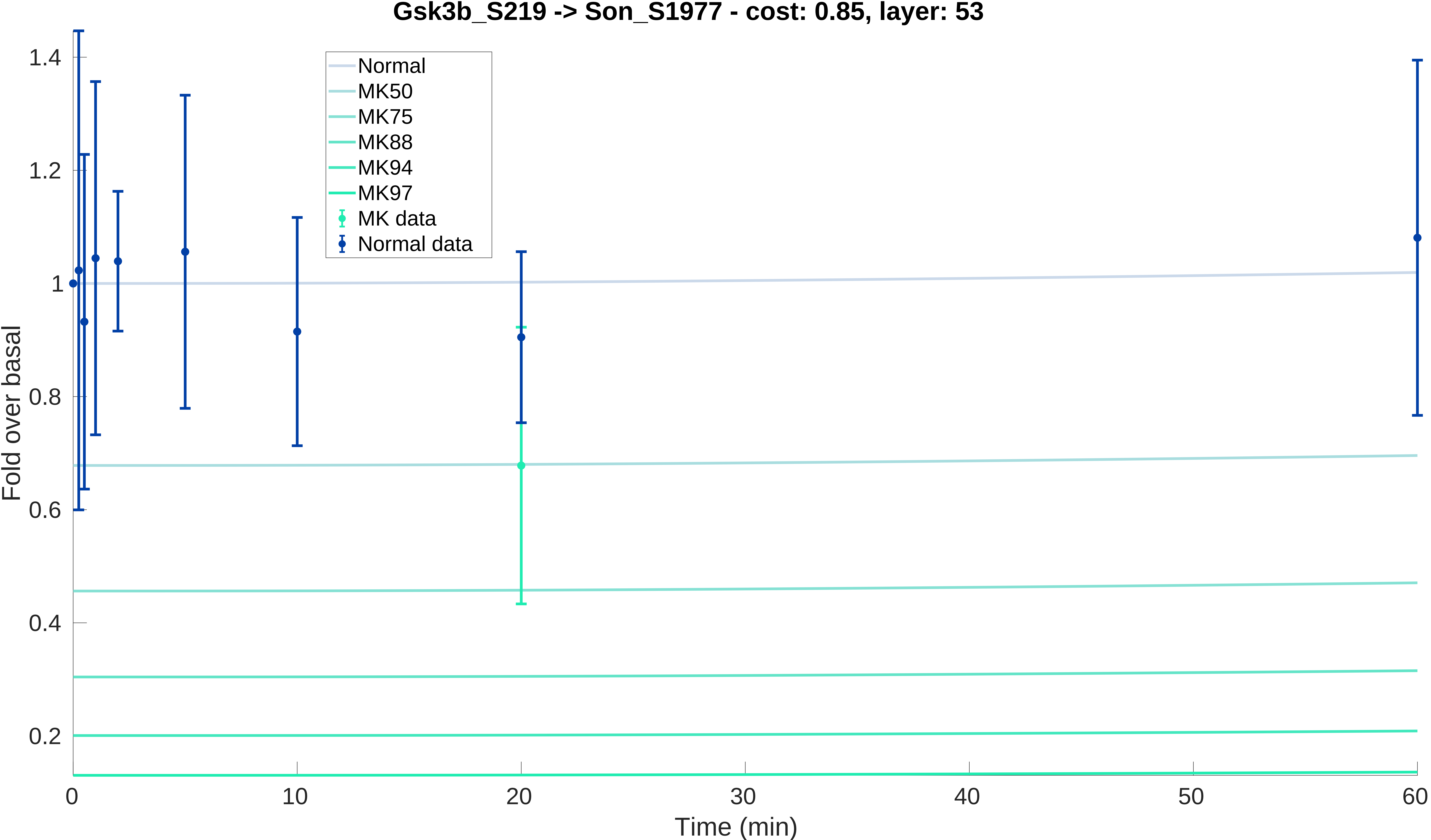

Gsk3b_S219 -> Son_S1977 - cost: 0.85, layer: 53
Normal
MK50
MK75
MK88
MK94
MK97
MK data
Normal data
Fold over basal
1.4
1.2
1
0.8
0.6
0.4
0.2
0
10
20
30
40
50
60
Time (min)

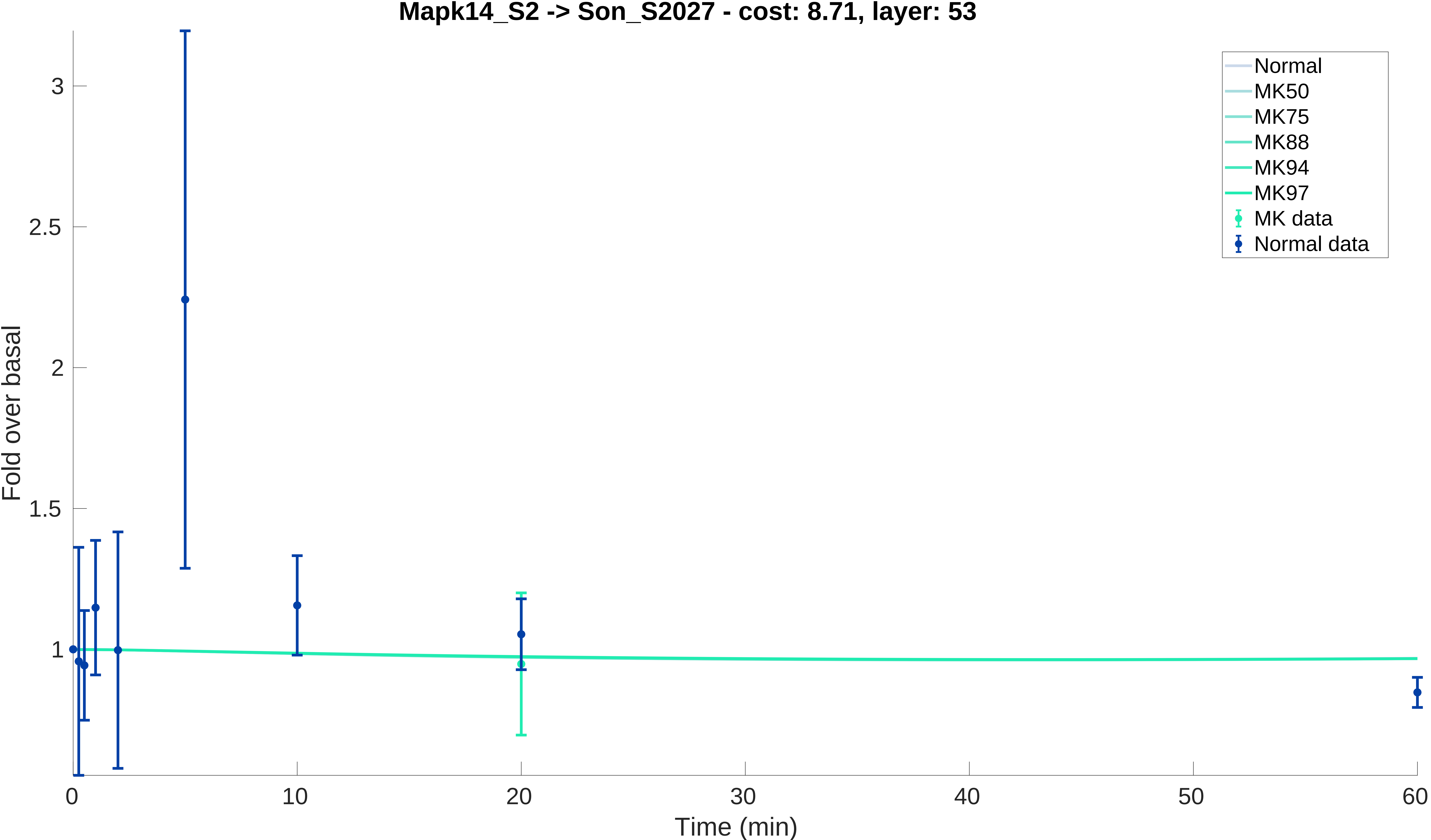

Mapk14_S2 -> Son_S2027 - cost: 8.71, layer: 53
Normal
MK50
MK75
MK88
MK94
MK97
MK data
Normal data
Fold over basal
3
2.5
2
1.5
1
0
10
20
30
40
50
60
Time (min)

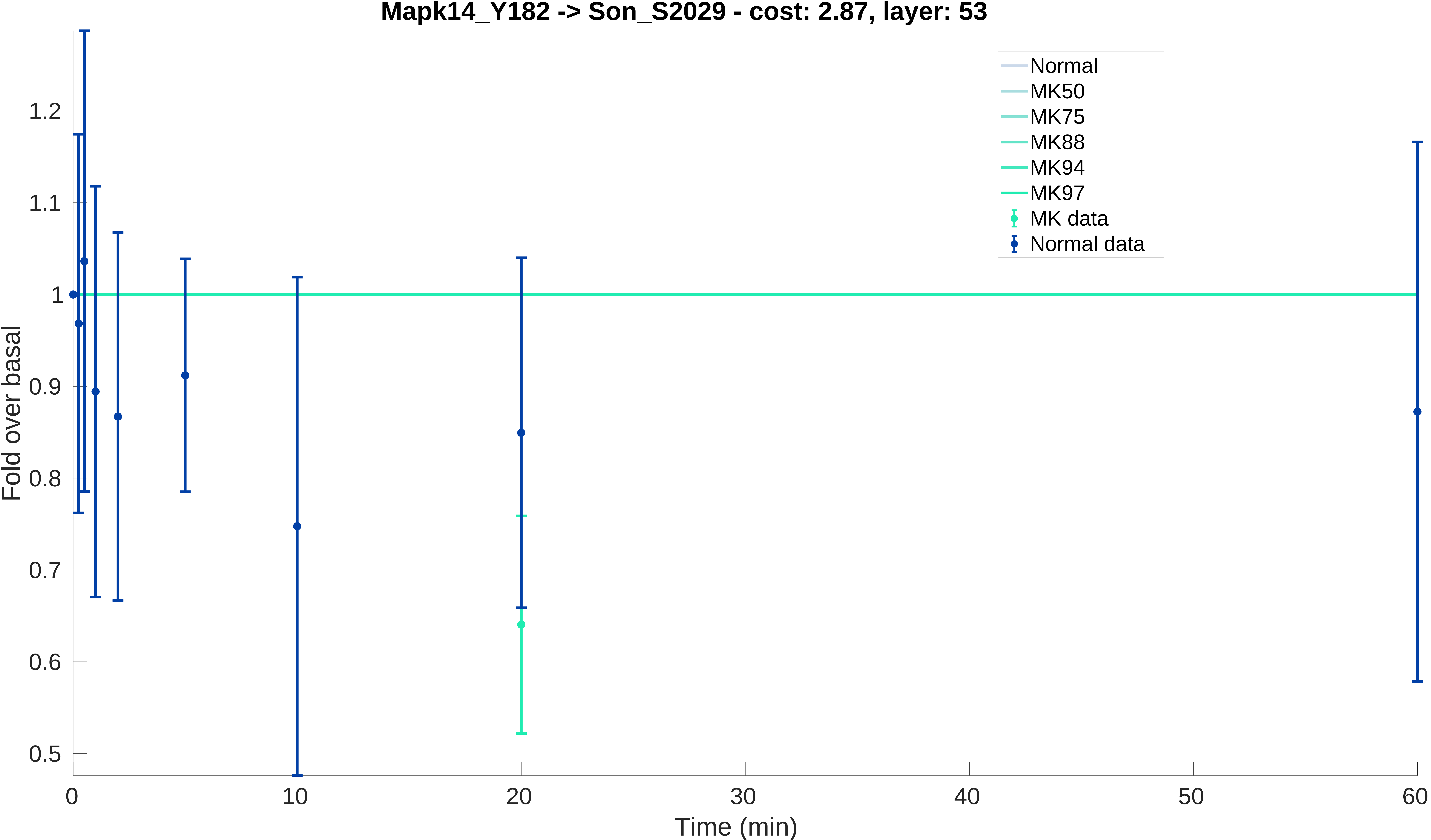

Mapk14_Y182 -> Son_S2029 - cost: 2.87, layer: 53
Normal
MK50
MK75
MK88
MK94
MK97
MK data
Normal data
Fold over basal
Time (min)
0.5
0.6
0.7
0.8
0.9
1
1.1
1.2
0
10
20
30
40
50
60

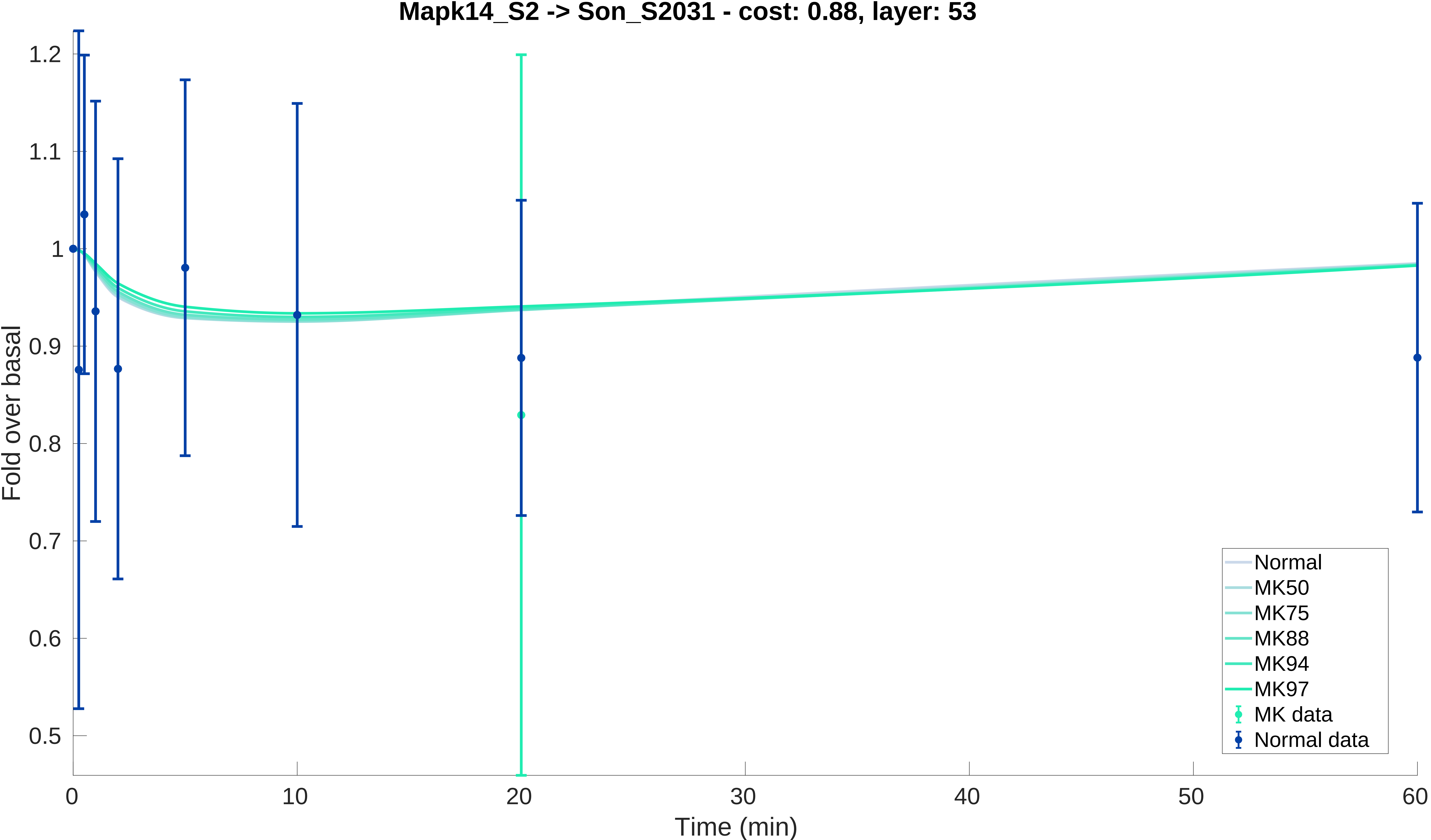
Mapk14_S2 -> Son_S2031 - cost: 0.88, layer: 53
Fold over basal
1.2
1.1
1
0.9
0.8
0.7
0.6
0.5
0
10
20
30
40
50
60
Time (min)
Normal
MK50
MK75
MK88
MK94
MK97
MK data
Normal data

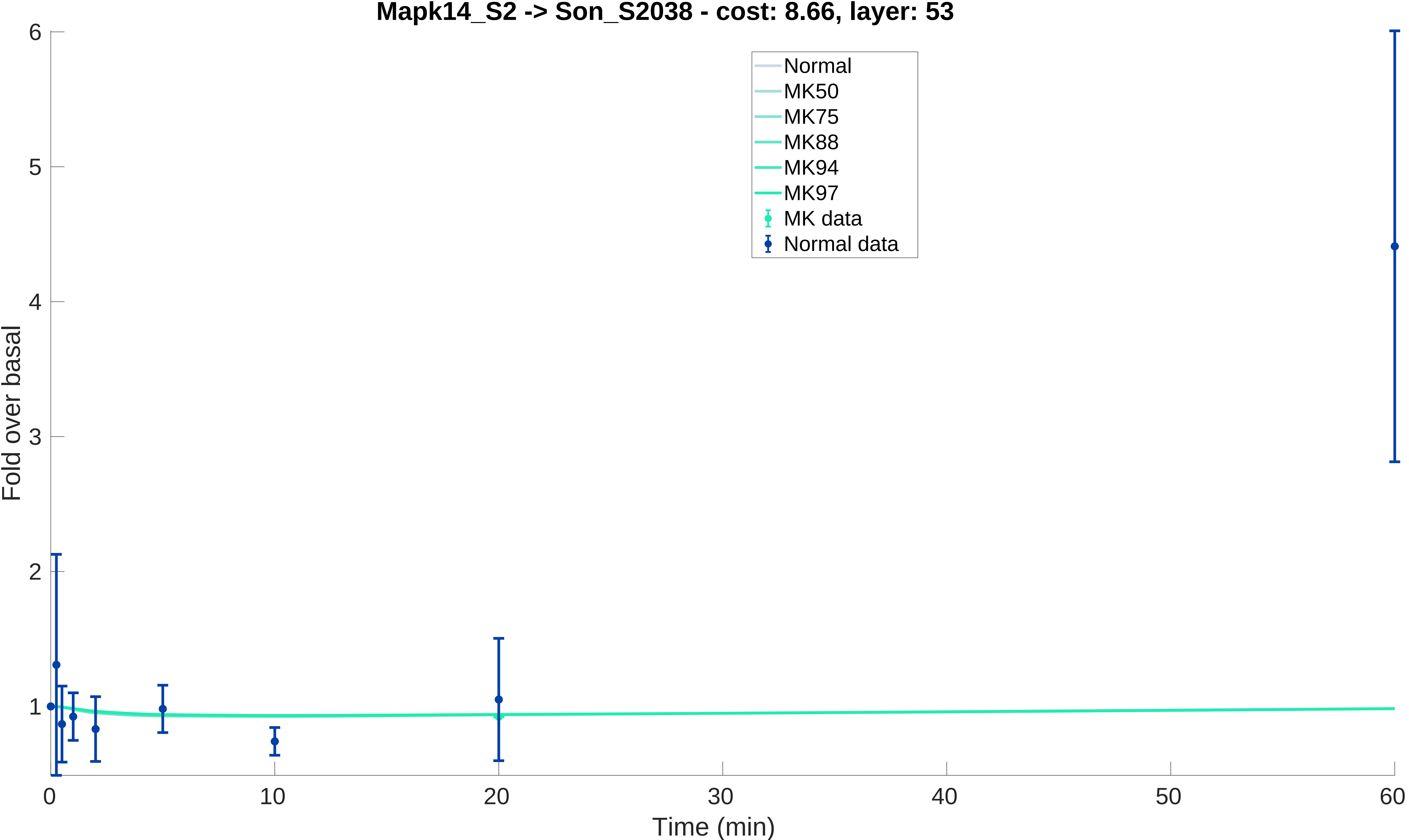

Mapk14_S2 -> Son_S2038 - cost: 8.66, layer: 53
Fold over basal
Time (min)
0
10
20
30
40
50
60
1
2
3
4
5
6
Normal
MK50
MK75
MK88
MK94
MK97
MK data
Normal data

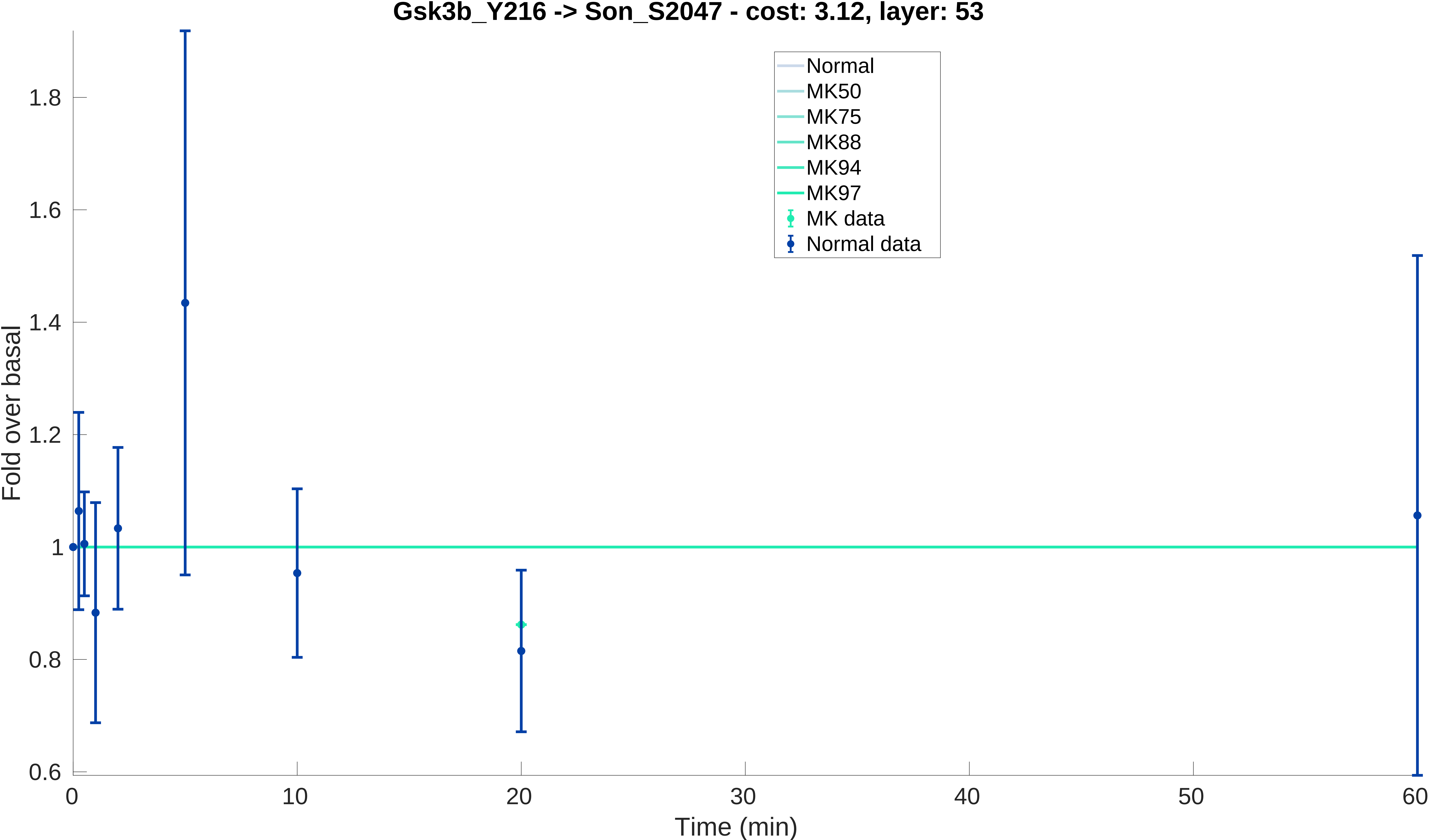
Gsk3b_Y216 -> Son_S2047 - cost: 3.12, layer: 53
Normal
MK50
MK75
MK88
MK94
MK97
MK data
Normal data
Fold over basal
0.6
0.8
1
1.2
1.4
1.6
1.8
0
10
20
30
40
50
60
Time (min)

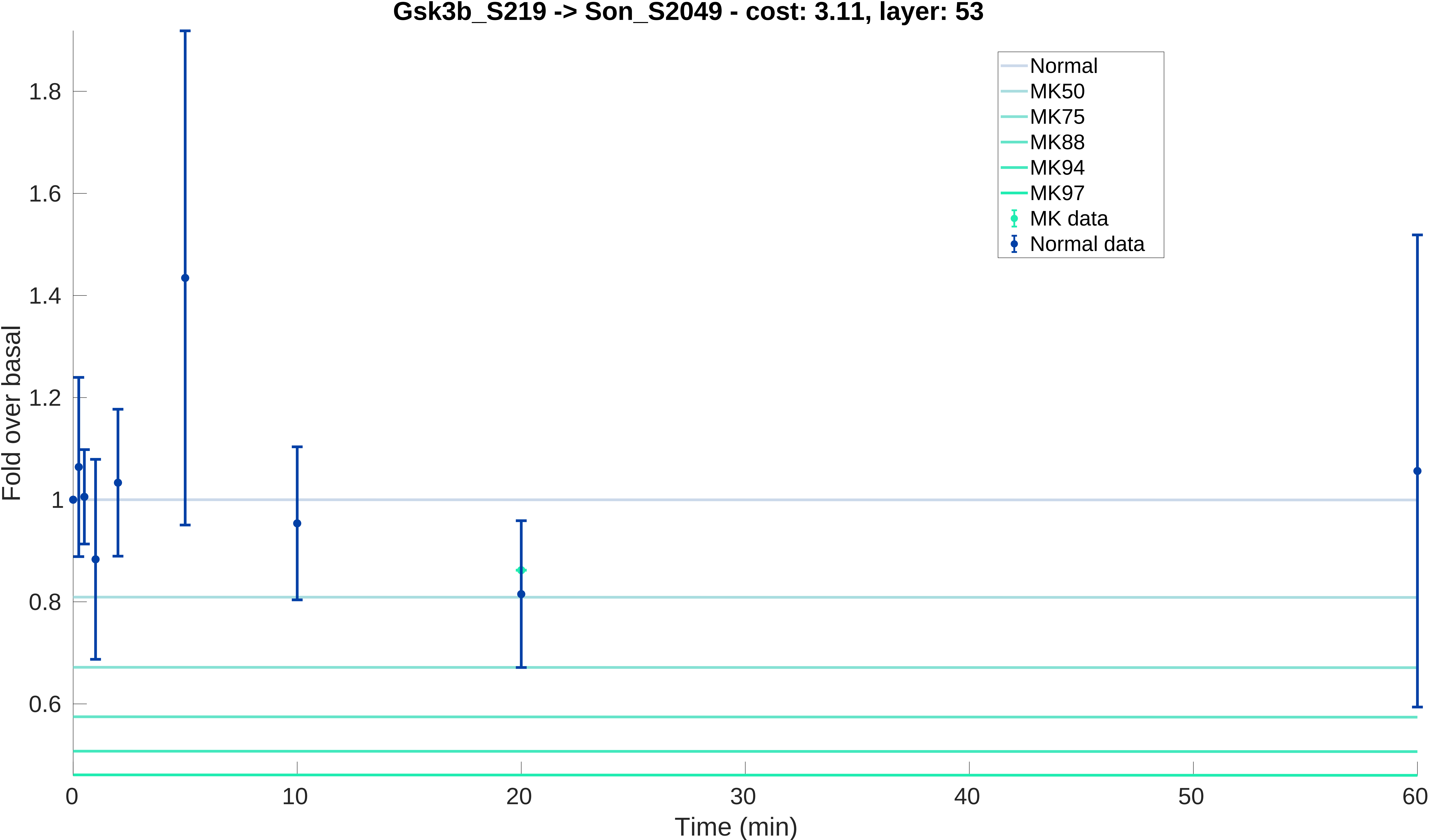

Gsk3b_S219 -> Son_S2049 - cost: 3.11, layer: 53
Normal
MK50
MK75
MK88
MK94
MK97
MK data
Normal data
Fold over basal
1.8
1.6
1.4
1.2
1
0.8
0.6
0
10
20
30
40
50
60
Time (min)

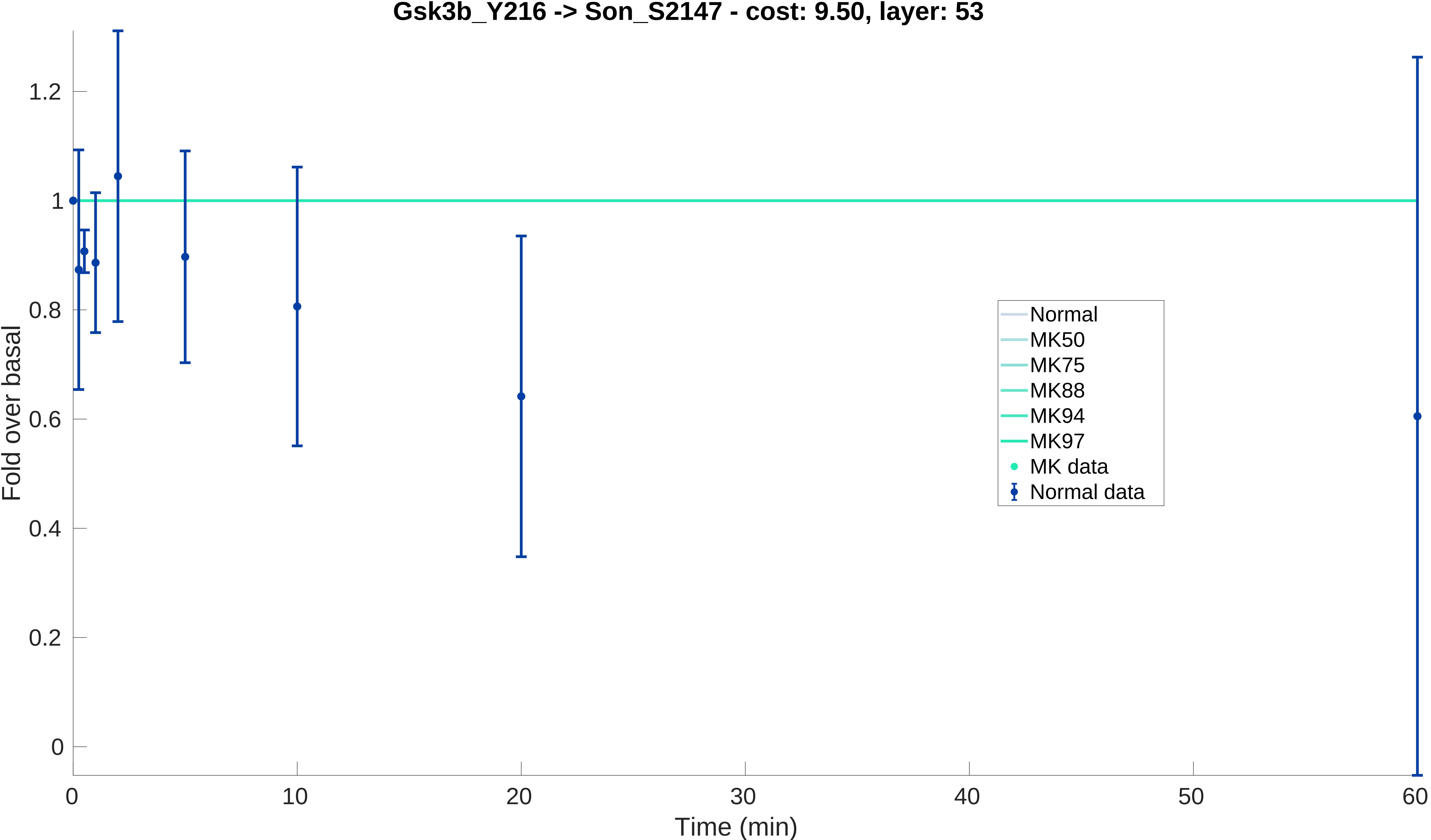
Gsk3b_Y216 -> Son_S2147 - cost: 9.50, layer: 53
Fold over basal
Time (min)
0
0.2
0.4
0.6
0.8
1
1.2
0
10
20
30
40
50
60
Normal
MK50
MK75
MK88
MK94
MK97
MK data
Normal data

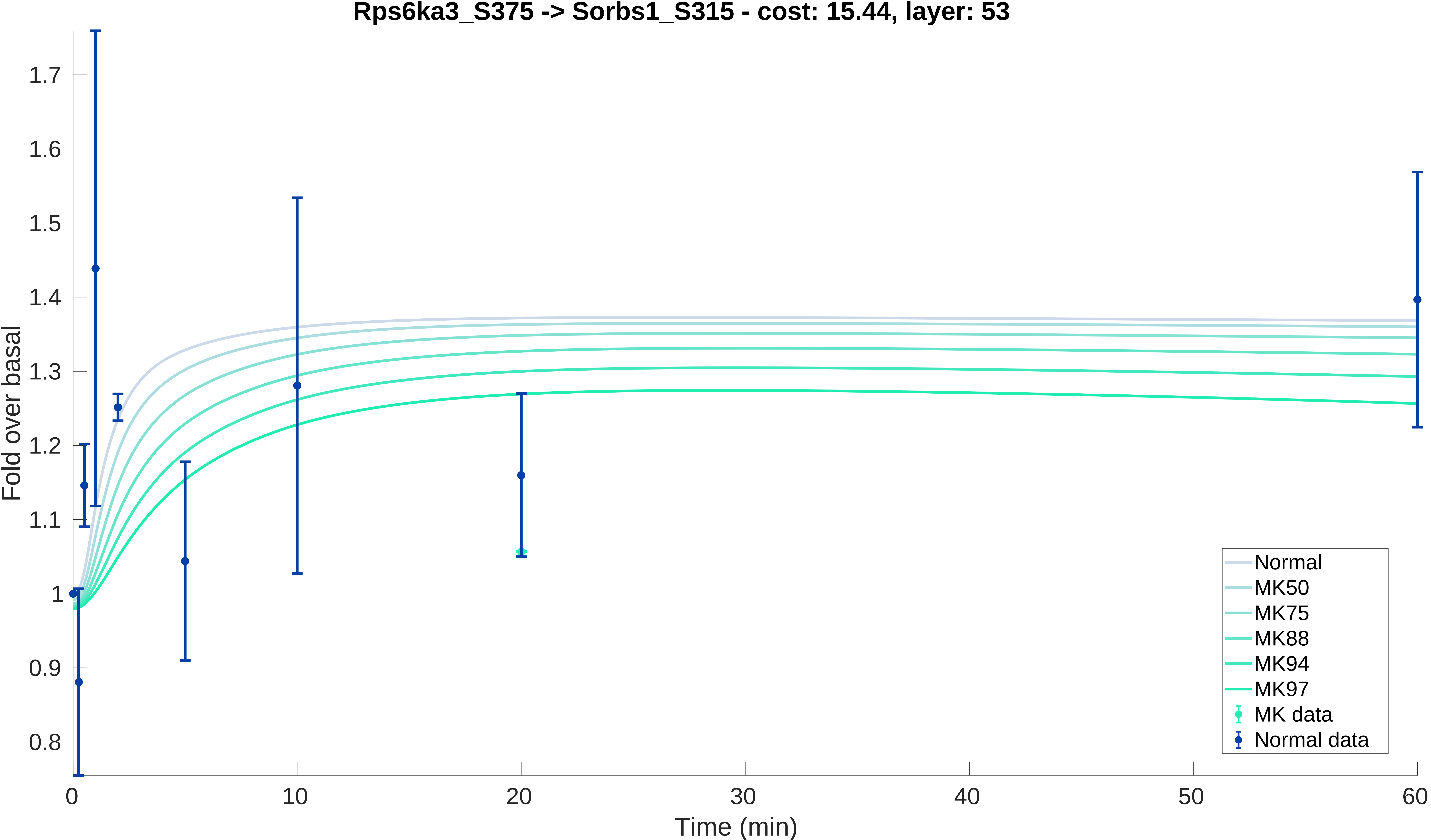
Rps6ka3_S375 -> Sorbs1_S315 - cost: 15.44, layer: 53
Fold over basal
Time (min)
0.8
0.9
1
1.1
1.2
1.3
1.4
1.5
1.6
1.7
0
10
20
30
40
50
60
Normal
MK50
MK75
MK88
MK94
MK97
MK data
Normal data

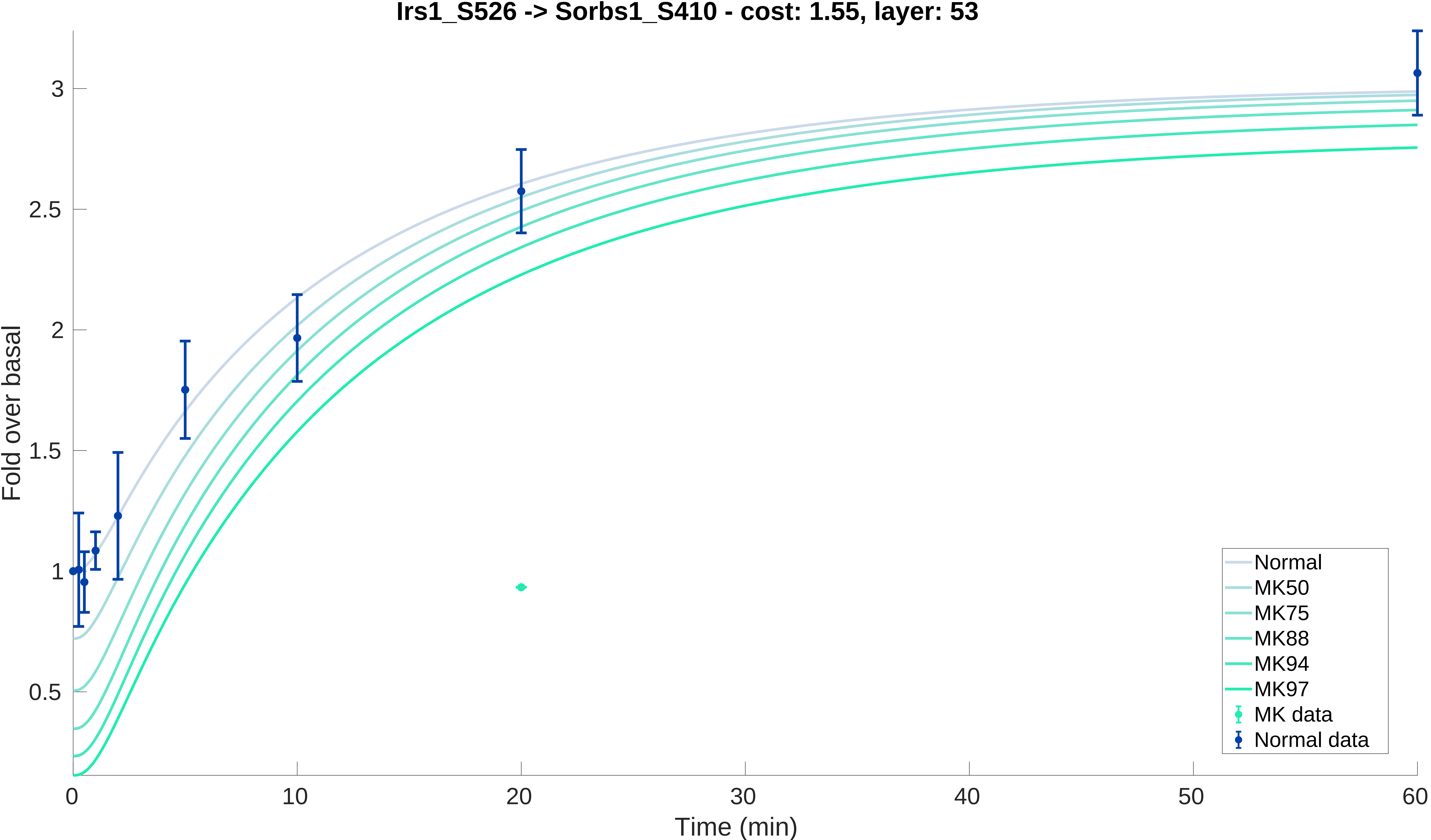

Irs1_S526 -> Sorbs1_S410 - cost: 1.55, layer: 53
Fold over basal
Time (min)
0
10
20
30
40
50
60
0.5
1
1.5
2
2.5
3
Normal
MK50
MK75
MK88
MK94
MK97
MK data
Normal data

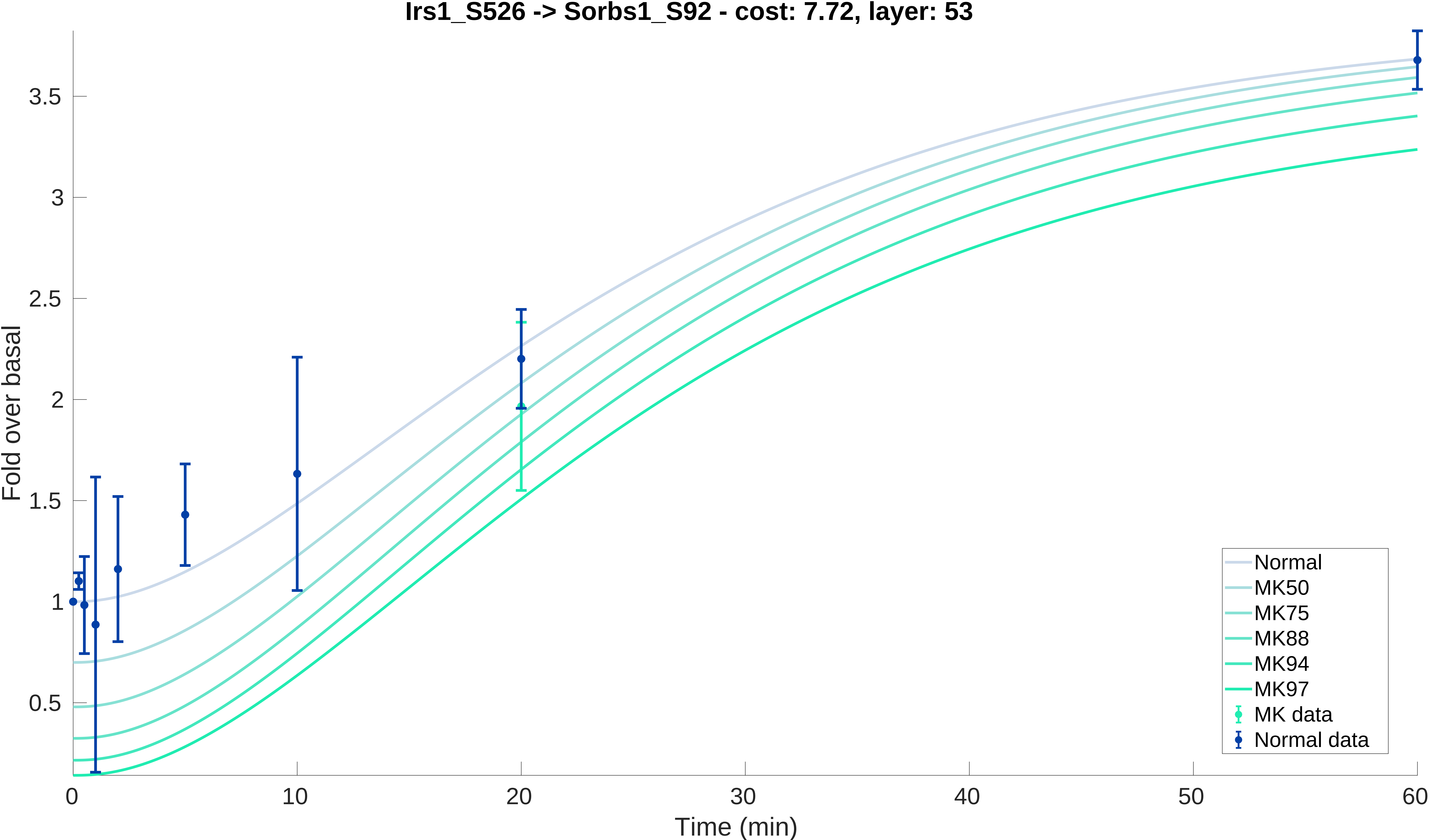

Irs1_S526 -> Sorbs1_S92 - cost: 7.72, layer: 53
Fold over basal
Time (min)
0.5
1
1.5
2
2.5
3
3.5
0
10
20
30
40
50
60
Normal
MK50
MK75
MK88
MK94
MK97
MK data
Normal data

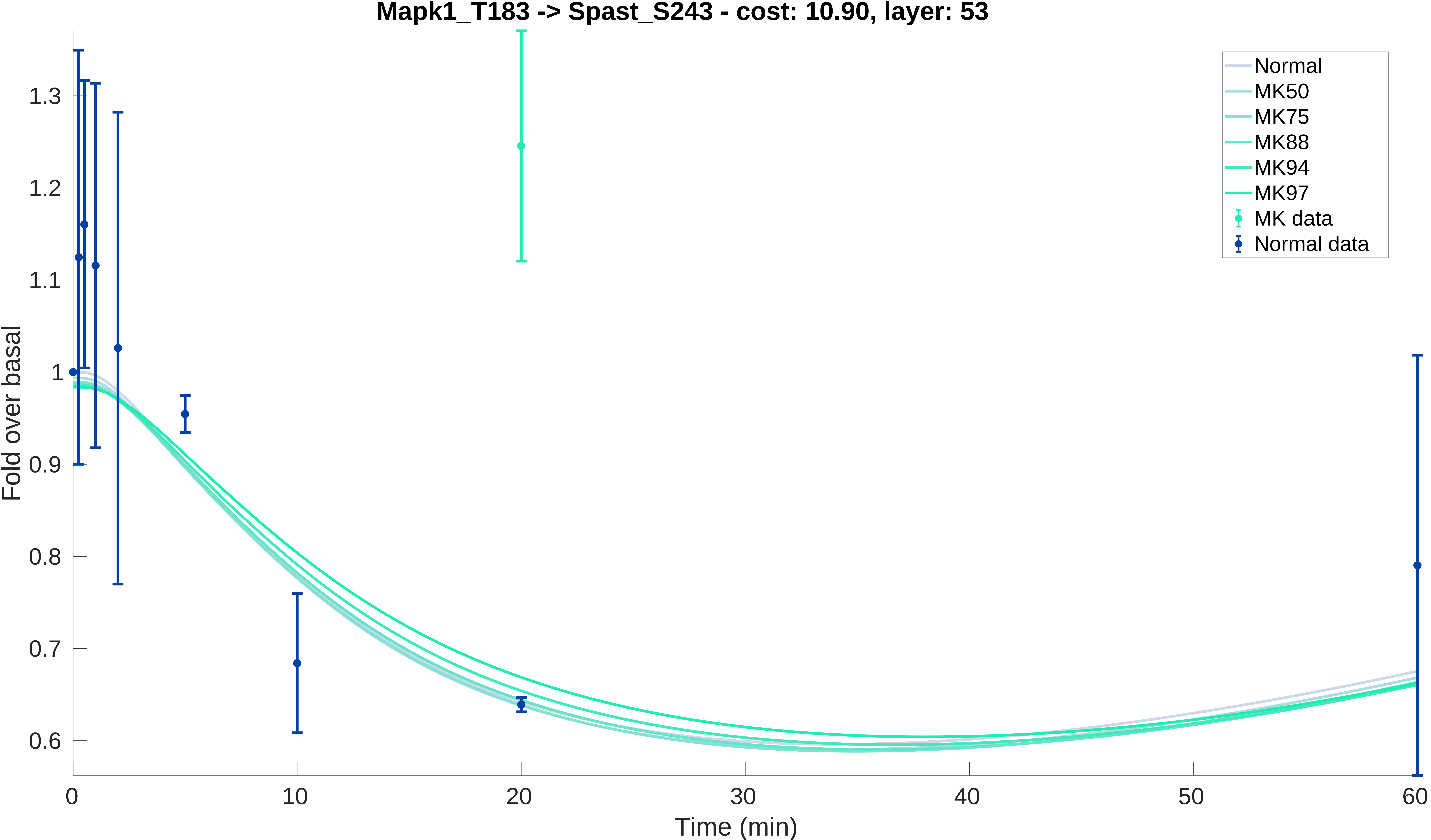

Mapk1_T183 -> Spast_S243 - cost: 10.90, layer: 53
Normal
MK50
MK75
MK88
MK94
MK97
MK data
Normal data
Fold over basal
Time (min)
0
10
20
30
40
50
60
0.6
0.7
0.8
0.9
1
1.1
1.2
1.3

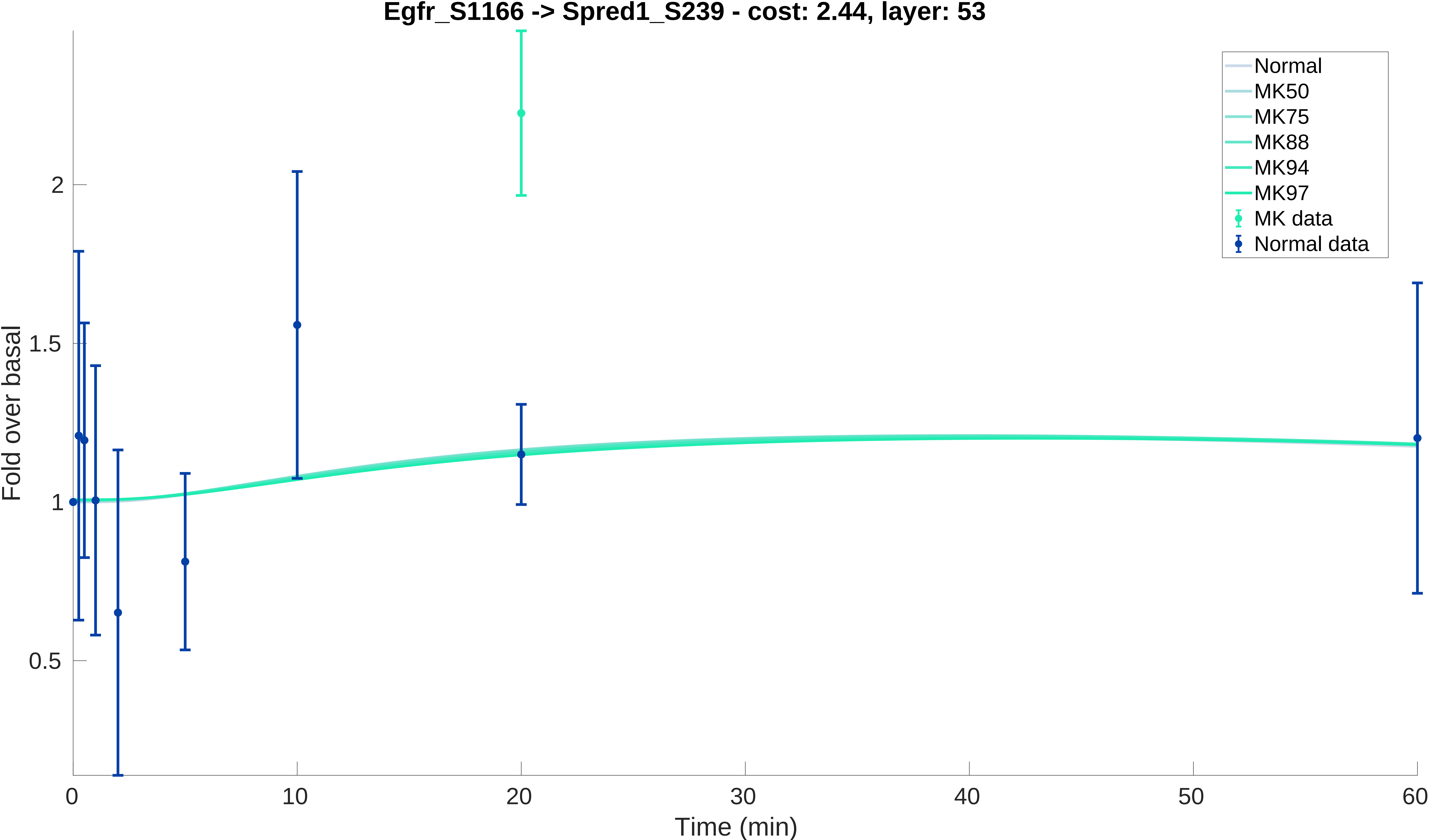

Egfr_S1166 -> Spred1_S239 - cost: 2.44, layer: 53
Fold over basal
Time (min)
0.5
1
1.5
2
0
10
20
30
40
50
60
Normal
MK50
MK75
MK88
MK94
MK97
MK data
Normal data

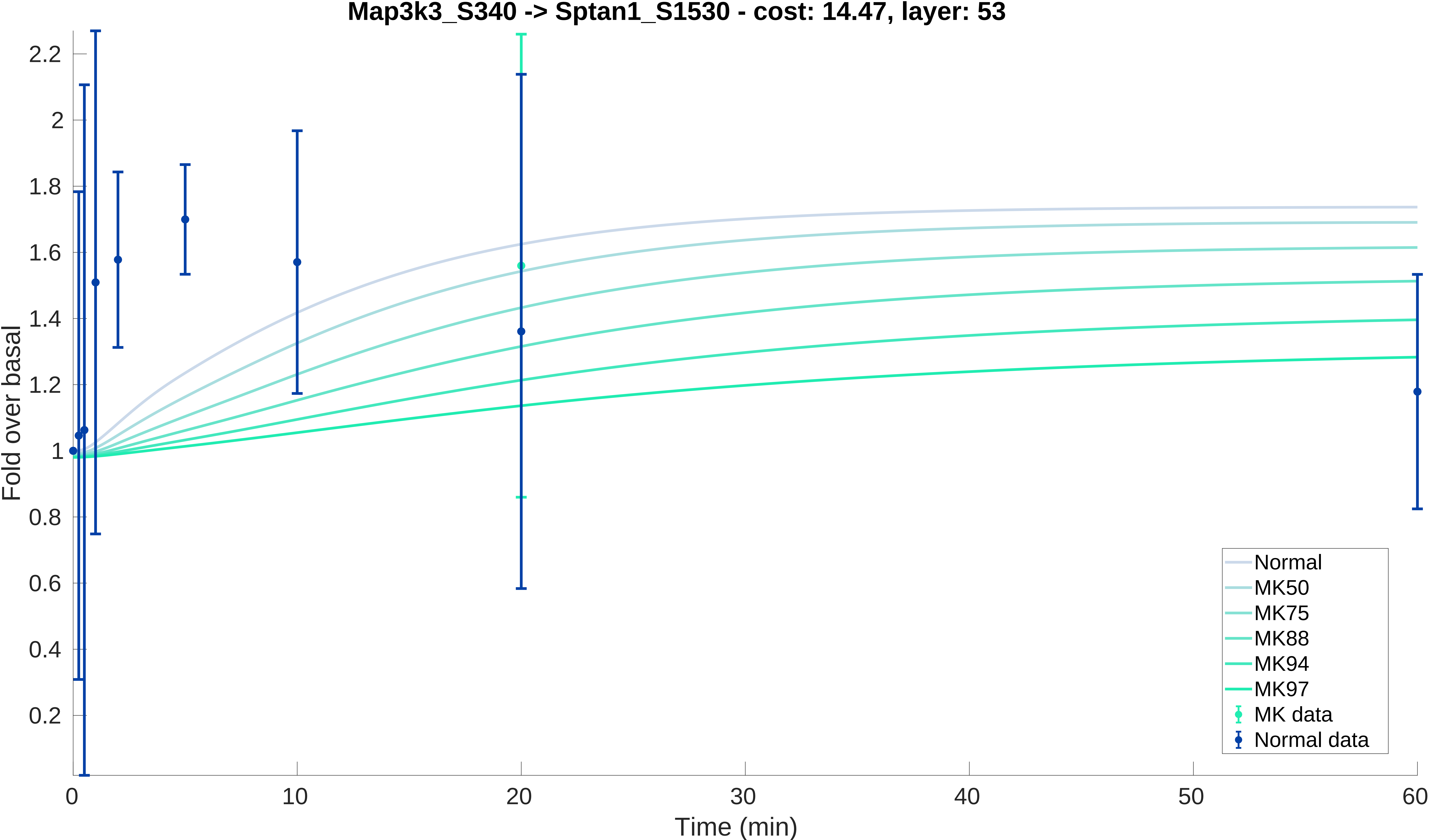

Map3k3_S340 -> Sptan1_S1530 - cost: 14.47, layer: 53
Fold over basal
Time (min)
0.2
0.4
0.6
0.8
1
1.2
1.4
1.6
1.8
2
2.2
0
10
20
30
40
50
60
Normal
MK50
MK75
MK88
MK94
MK97
MK data
Normal data

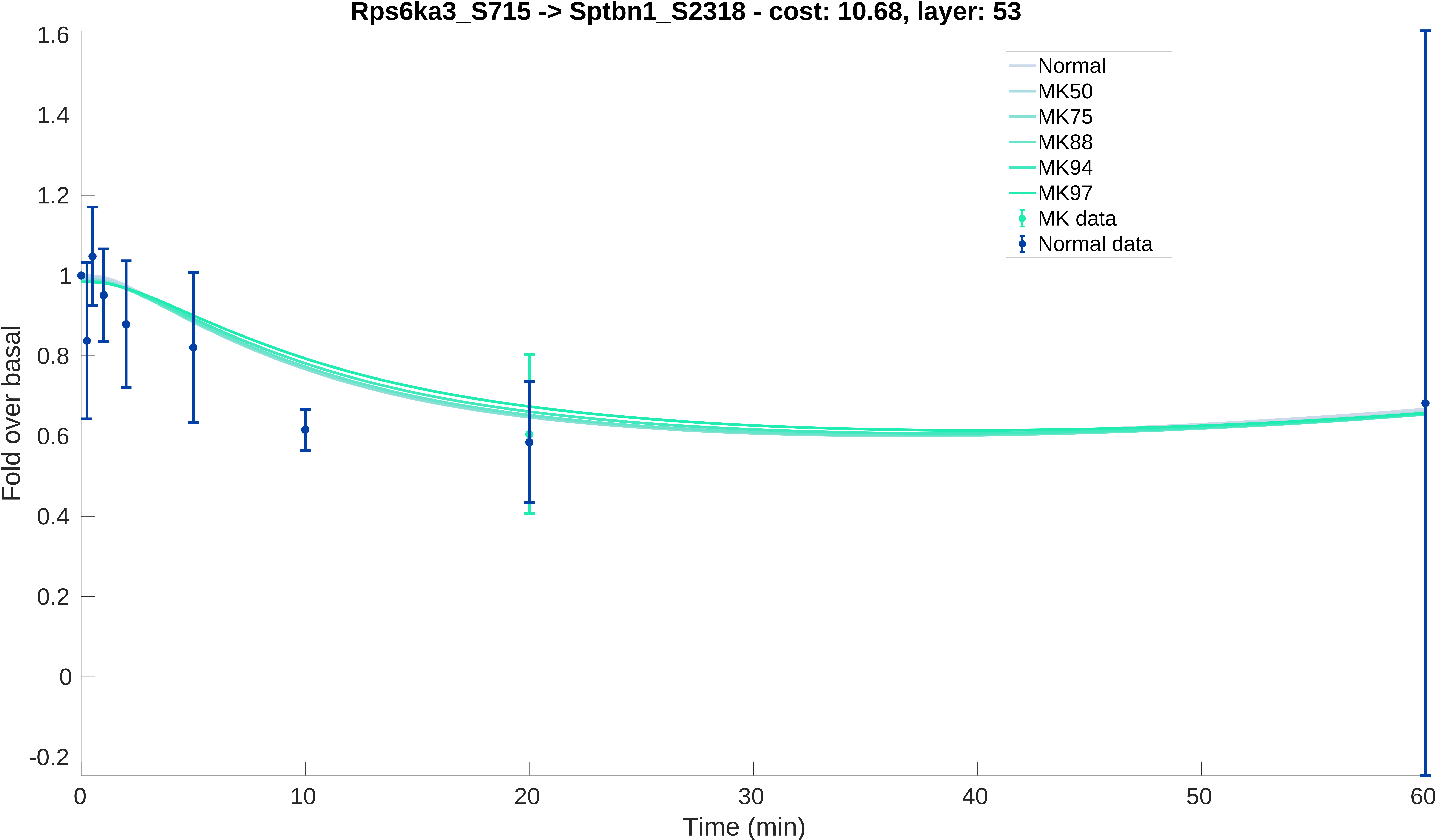

Rps6ka3_S715 -> Sptbn1_S2318 - cost: 10.68, layer: 53
Fold over basal
Time (min)
1.6
1.4
1.2
1
0.8
0.6
0.4
0.2
0
-0.2
0
10
20
30
40
50
60
Normal
MK50
MK75
MK88
MK94
MK97
MK data
Normal data

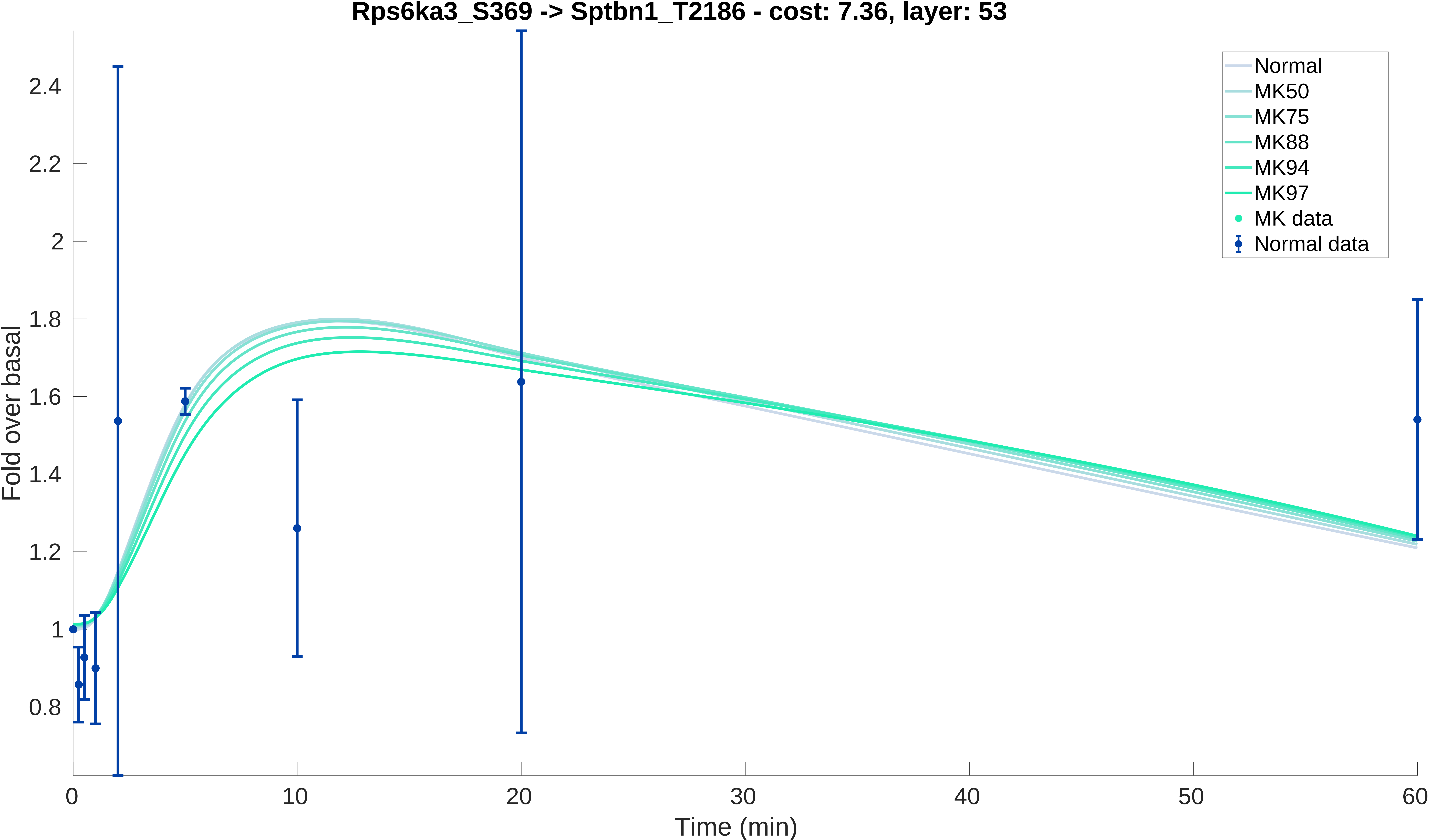

Rps6ka3_S369 -> Sptbn1_T2186 - cost: 7.36, layer: 53
Fold over basal
Time (min)
2.4
2.2
2
1.8
1.6
1.4
1.2
1
0.8
0
10
20
30
40
50
60
Normal
MK50
MK75
MK88
MK94
MK97
MK data
Normal data

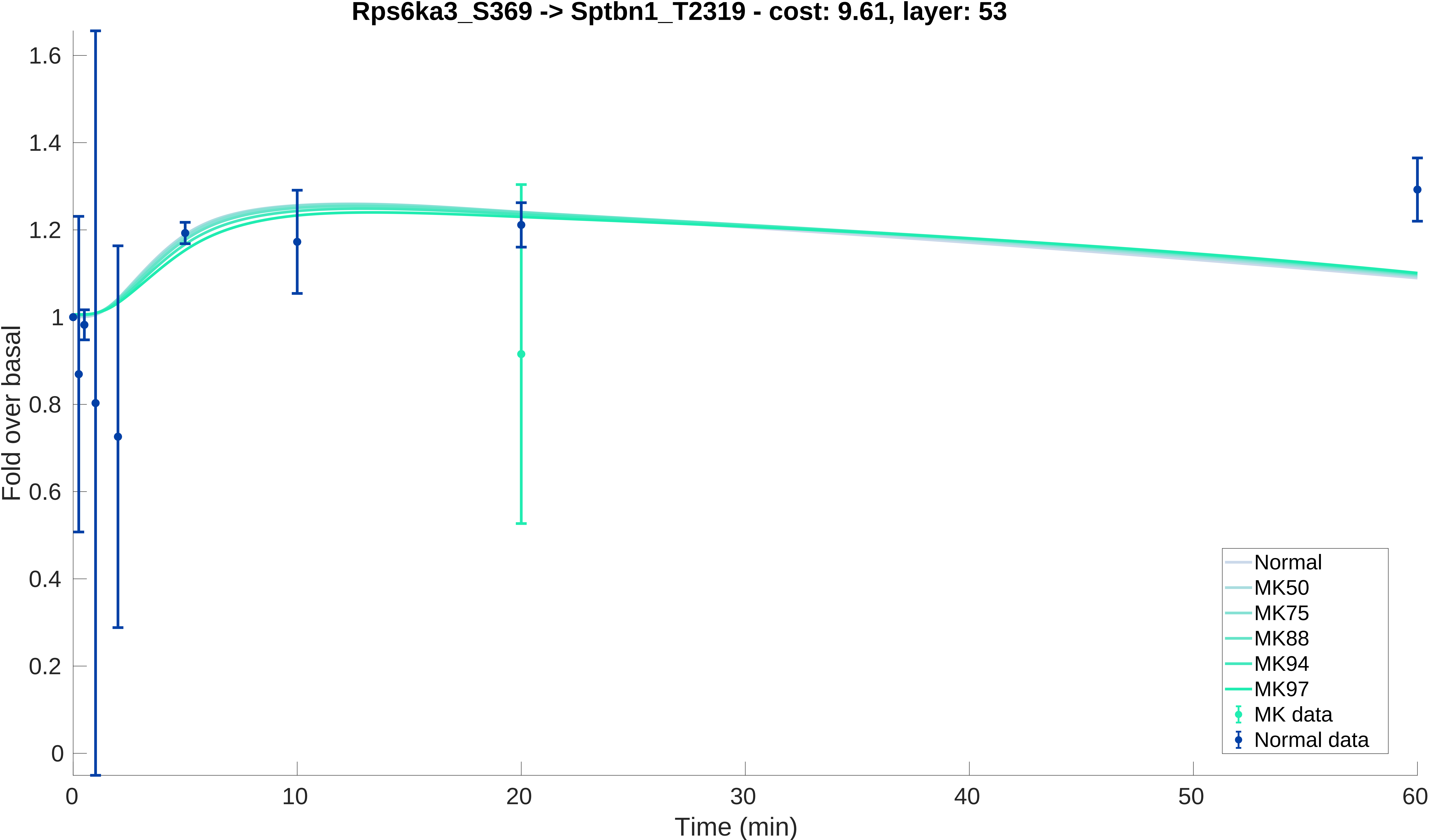

Rps6ka3_S369 -> Sptbn1_T2319 - cost: 9.61, layer: 53
Fold over basal
Time (min)
0
0.2
0.4
0.6
0.8
1
1.2
1.4
1.6
0
10
20
30
40
50
60
Normal
MK50
MK75
MK88
MK94
MK97
MK data
Normal data

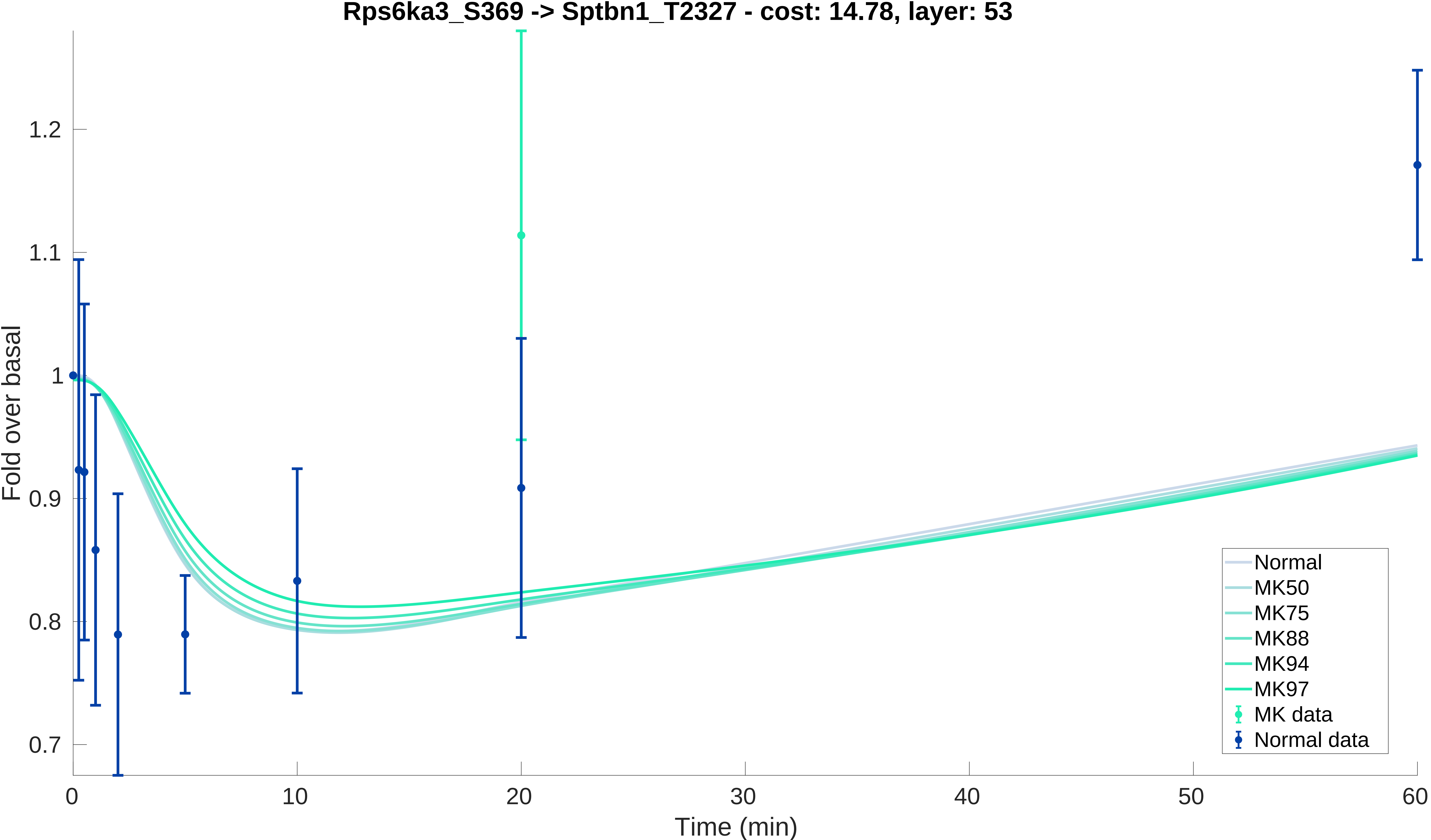

Rps6ka3_S369 -> Sptbn1_T2327 - cost: 14.78, layer: 53
Fold over basal
Time (min)
1.2
1.1
1
0.9
0.8
0.7
0
10
20
30
40
50
60
Normal
MK50
MK75
MK88
MK94
MK97
MK data
Normal data

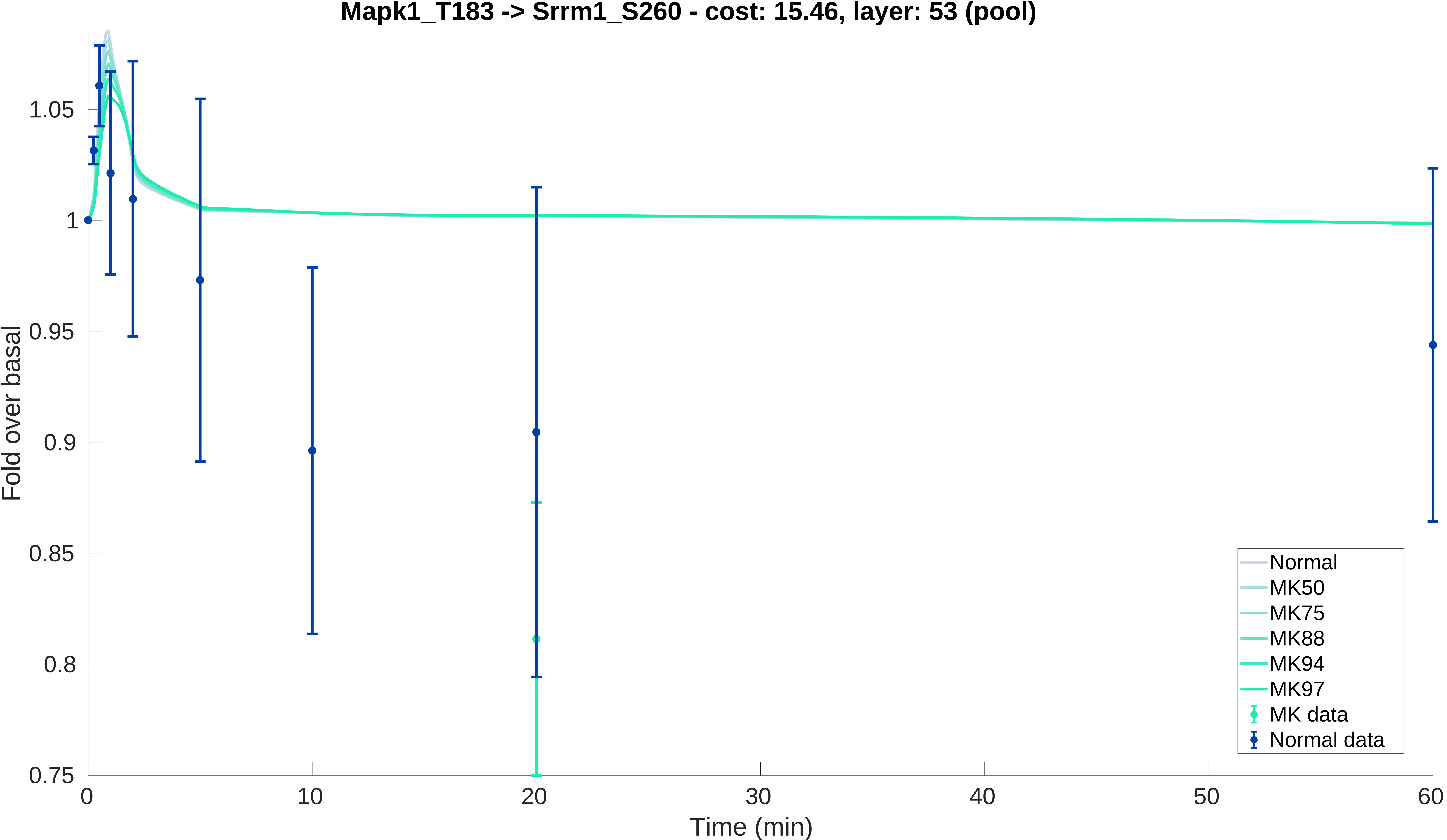

Mapk1_T183 -> Srrm1_S260 - cost: 15.46, layer: 53 (pool)
Fold over basal
1.05
1
0.95
0.9
0.85
0.8
0.75
0
10
20
30
40
50
60
Time (min)
Normal
MK50
MK75
MK88
MK94
MK97
MK data
Normal data

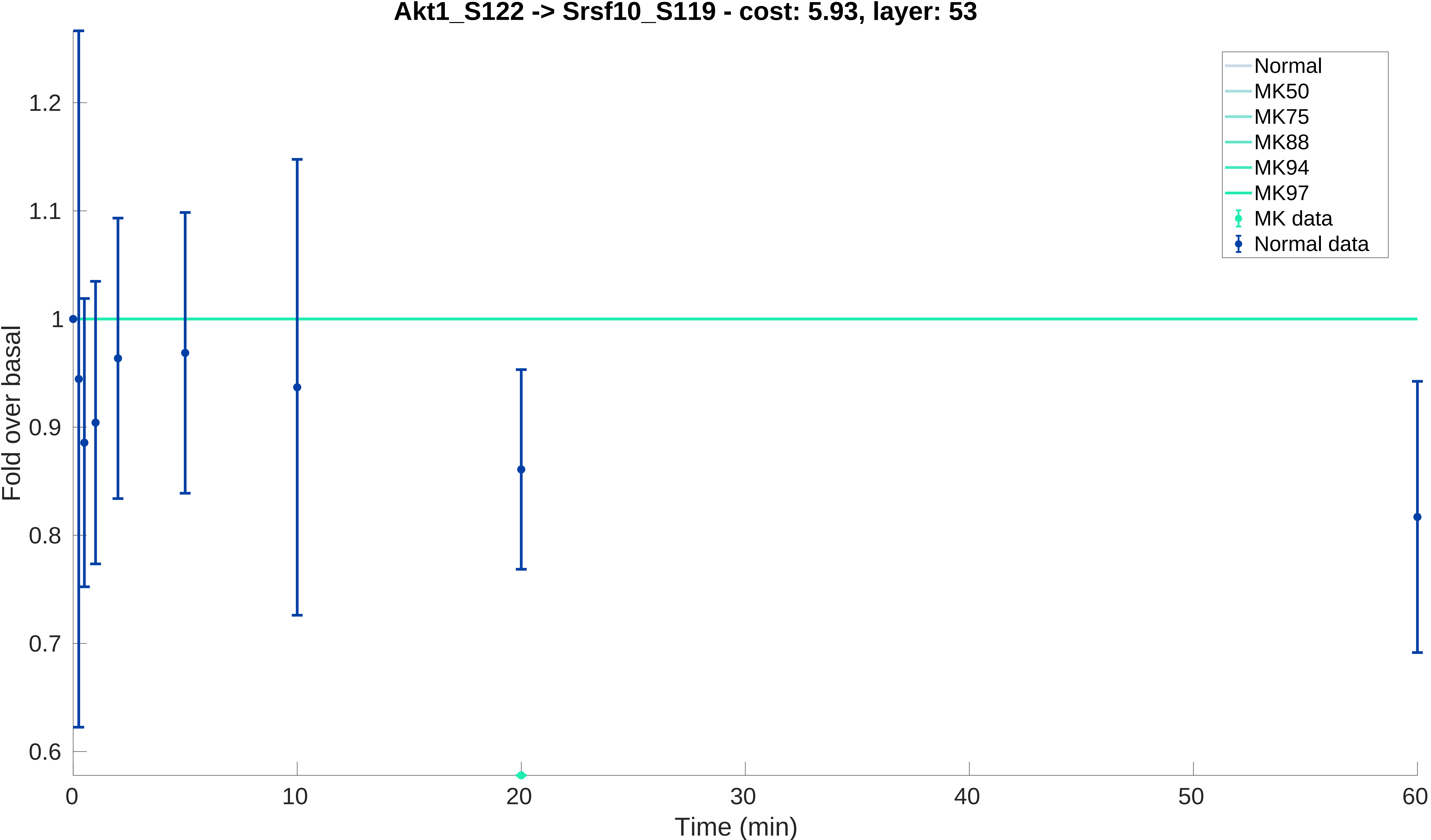

Akt1_S122 -> Srsf10_S119 - cost: 5.93, layer: 53
Fold over basal
Time (min)
0.6
0.7
0.8
0.9
1
1.1
1.2
0
10
20
30
40
50
60
Normal
MK50
MK75
MK88
MK94
MK97
MK data
Normal data

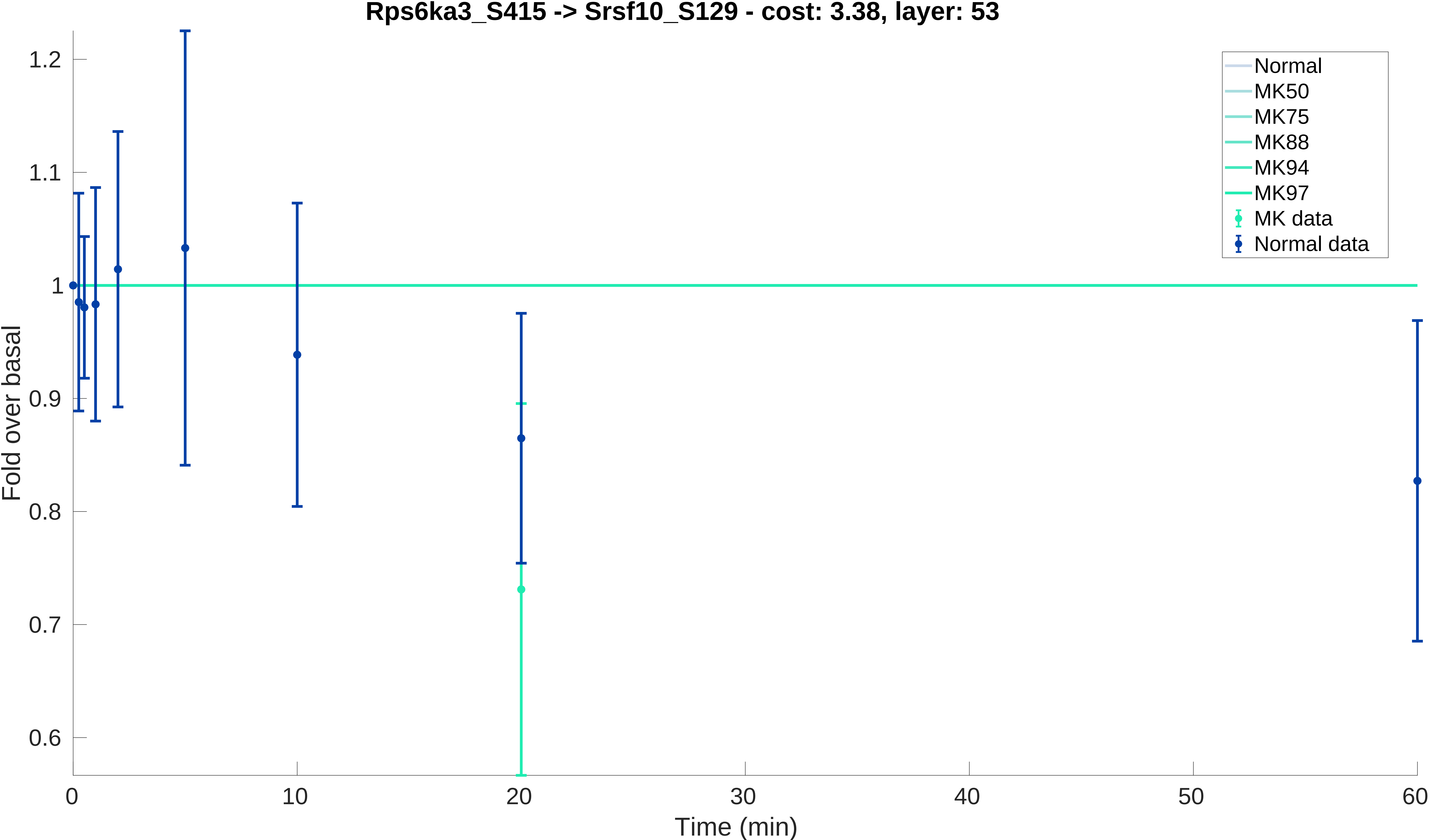

Rps6ka3_S415 -> Srsf10_S129 - cost: 3.38, layer: 53
Normal
MK50
MK75
MK88
MK94
MK97
MK data
Normal data
Fold over basal
Time (min)
0.6
0.7
0.8
0.9
1
1.1
1.2
0
10
20
30
40
50
60

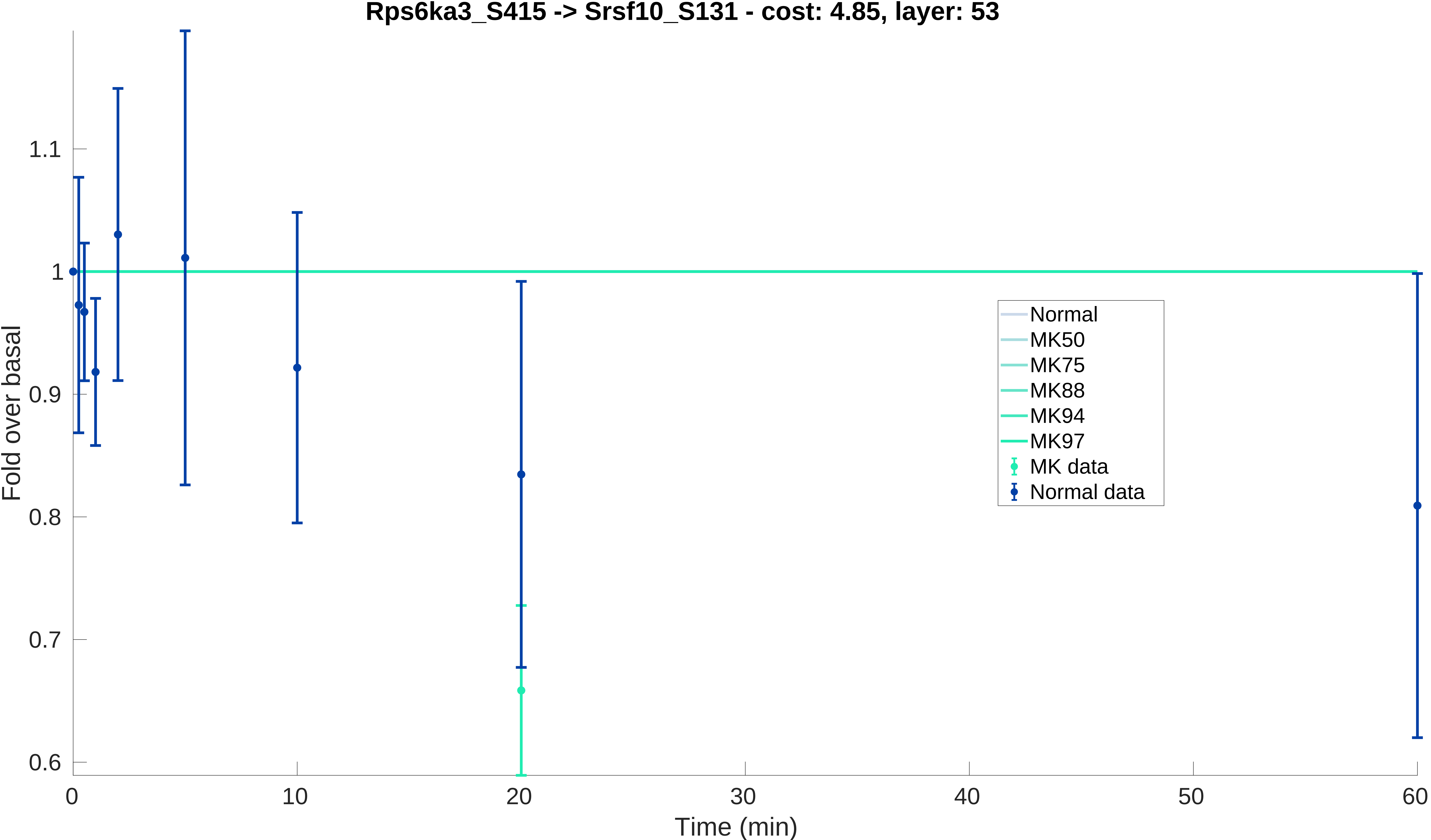
Rps6ka3_S415 -> Srsf10_S131 - cost: 4.85, layer: 53
Fold over basal
Time (min)
0.6
0.7
0.8
0.9
1
1.1
0
10
20
30
40
50
60
Normal
MK50
MK75
MK88
MK94
MK97
MK data
Normal data

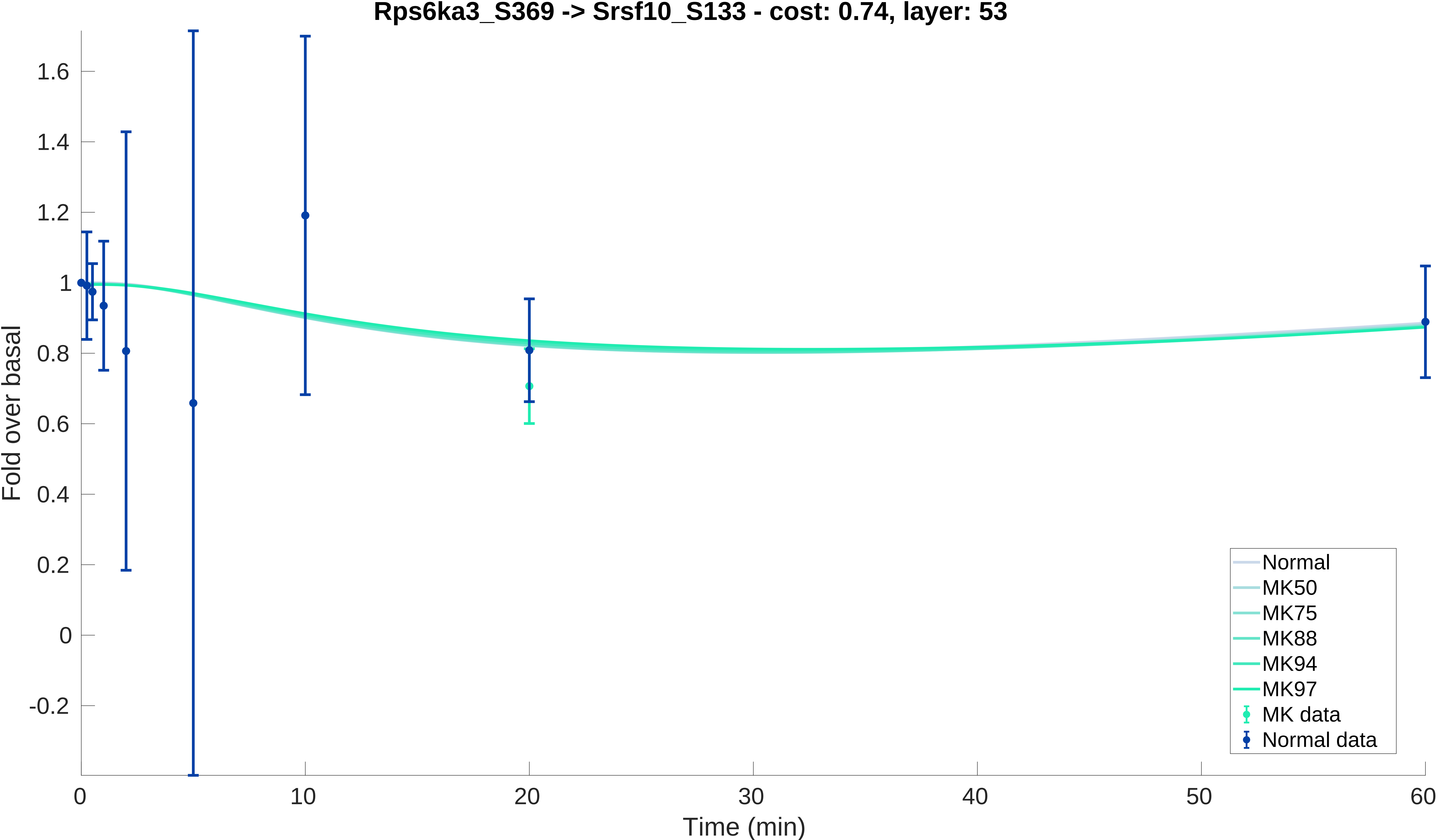

Rps6ka3_S369 -> Srsf10_S133 - cost: 0.74, layer: 53
Fold over basal
1.6
1.4
1.2
1
0.8
0.6
0.4
0.2
0
-0.2
0
10
20
30
40
50
60
Time (min)
Normal
MK50
MK75
MK88
MK94
MK97
MK data
Normal data

Akt1_S122 -> Srsf7_S192 - cost: 2.06, layer: 53

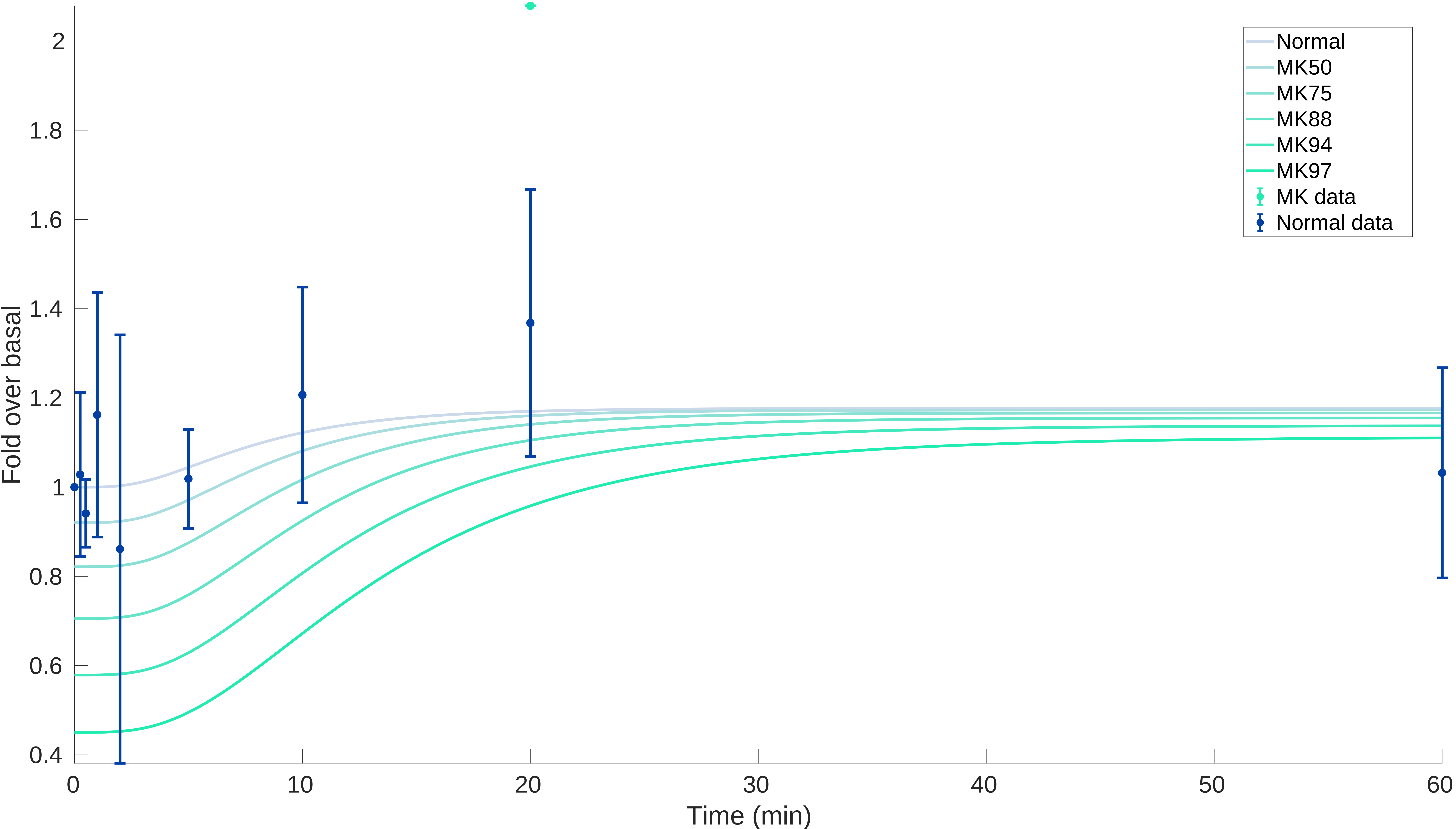

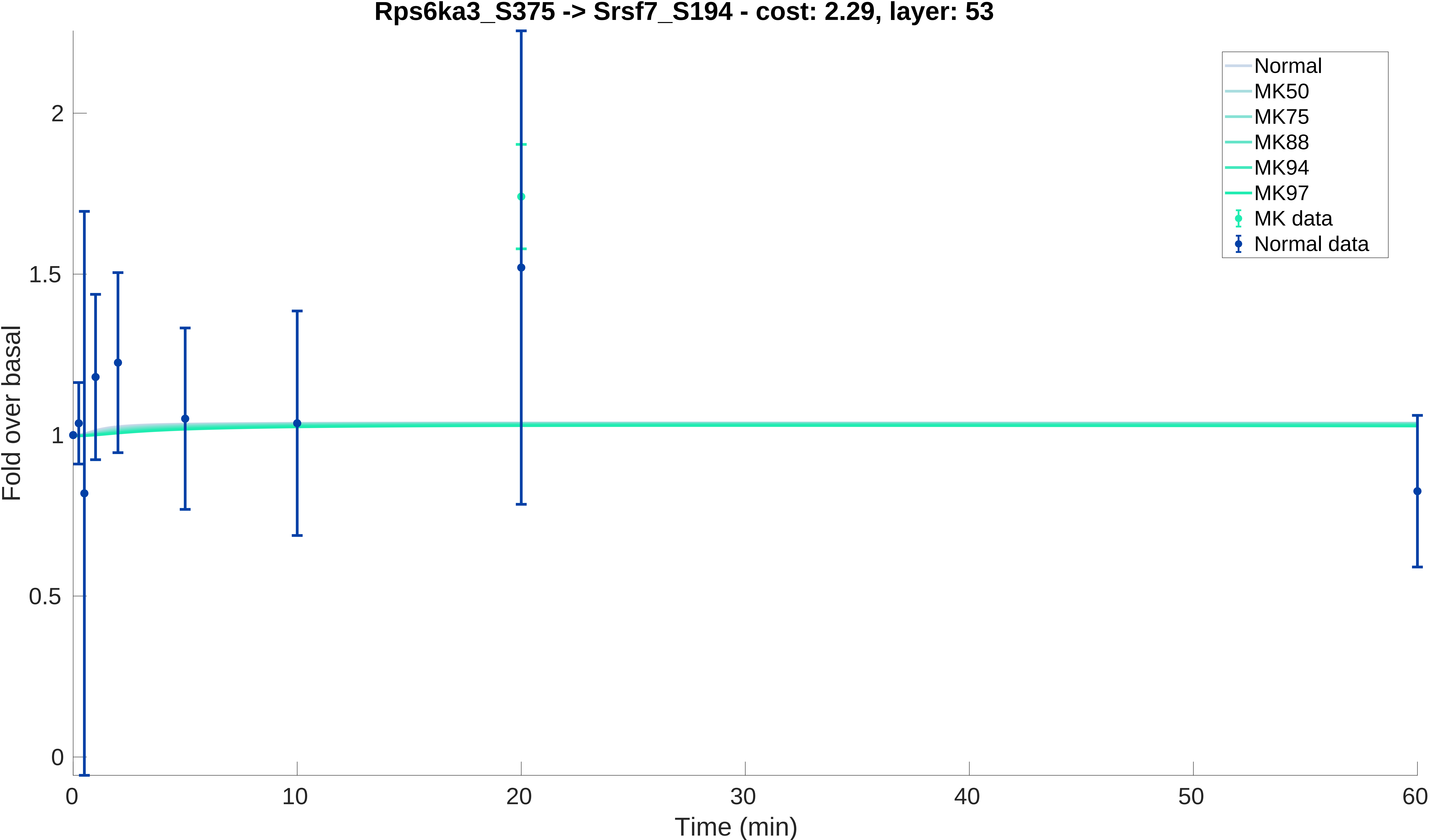

Rps6ka3_S375 -> Srsf7_S194 - cost: 2.29, layer: 53
Fold over basal
Time (min)
0
0.5
1
1.5
2
0
10
20
30
40
50
60
Normal
MK50
MK75
MK88
MK94
MK97
MK data
Normal data

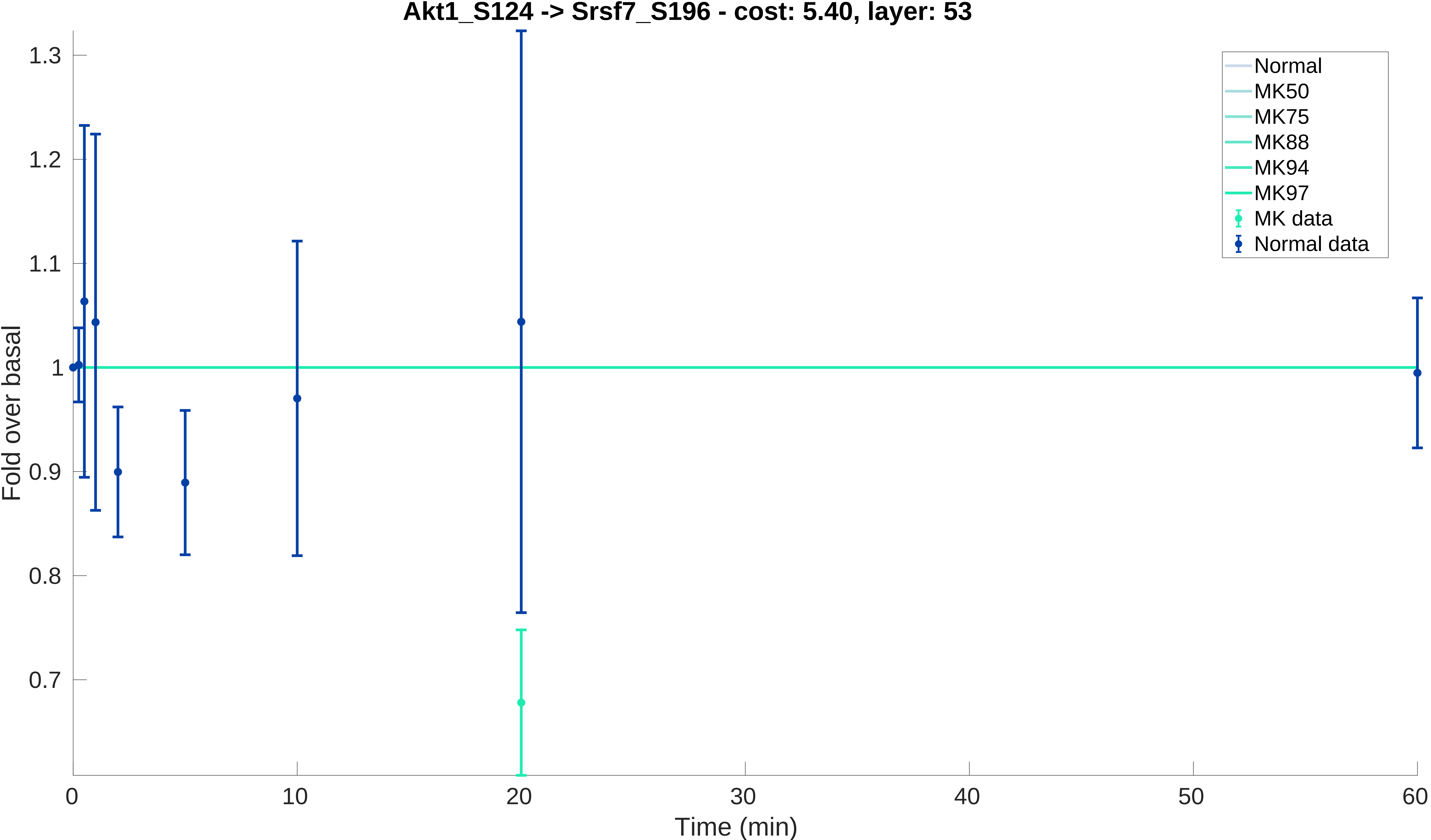
Akt1_S124 -> Srsf7_S196 - cost: 5.40, layer: 53
Fold over basal
Time (min)
1.3
1.2
1.1
1
0.9
0.8
0.7
0
10
20
30
40
50
60
Normal
MK50
MK75
MK88
MK94
MK97
MK data
Normal data

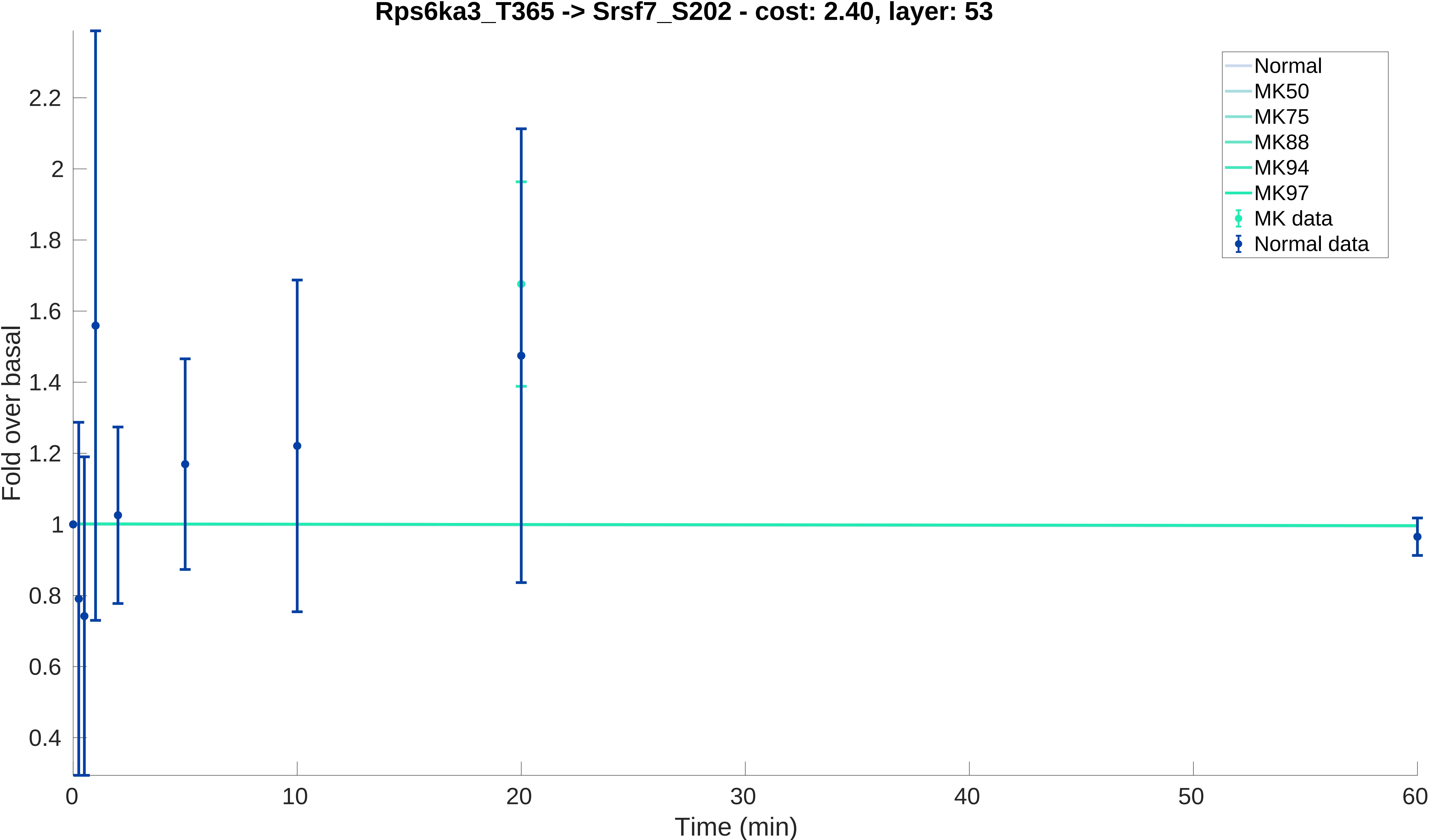

Rps6ka3_T365 -> Srsf7_S202 - cost: 2.40, layer: 53
Normal
MK50
MK75
MK88
MK94
MK97
MK data
Normal data
Fold over basal
2.2
2
1.8
1.6
1.4
1.2
1
0.8
0.6
0.4
0
10
20
30
40
50
60
Time (min)

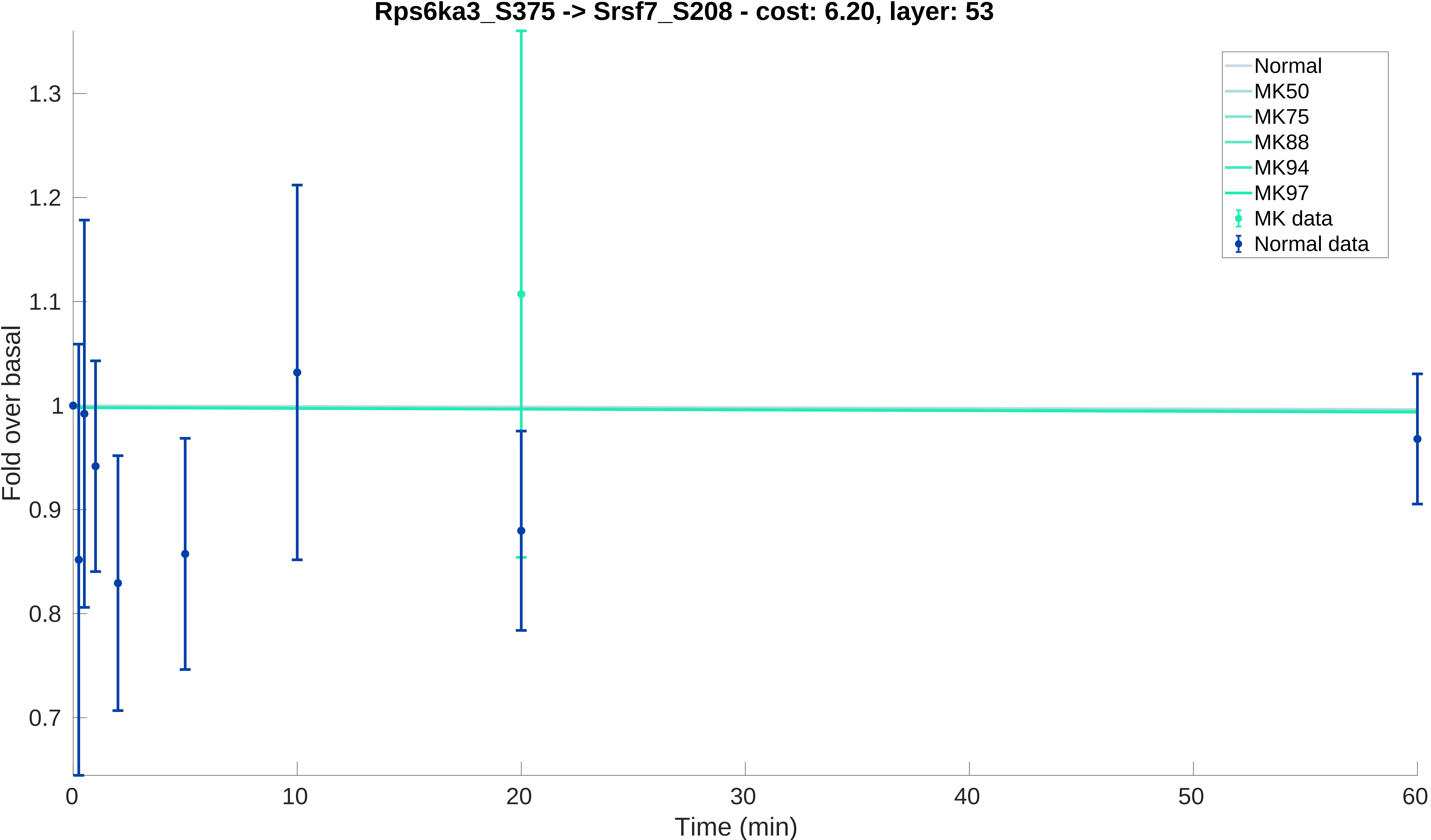

Rps6ka3_S375 -> Srsf7_S208 - cost: 6.20, layer: 53
Normal
MK50
MK75
MK88
MK94
MK97
MK data
Normal data
Fold over basal
1.3
1.2
1.1
1
0.9
0.8
0.7
0
10
20
30
40
50
60
Time (min)

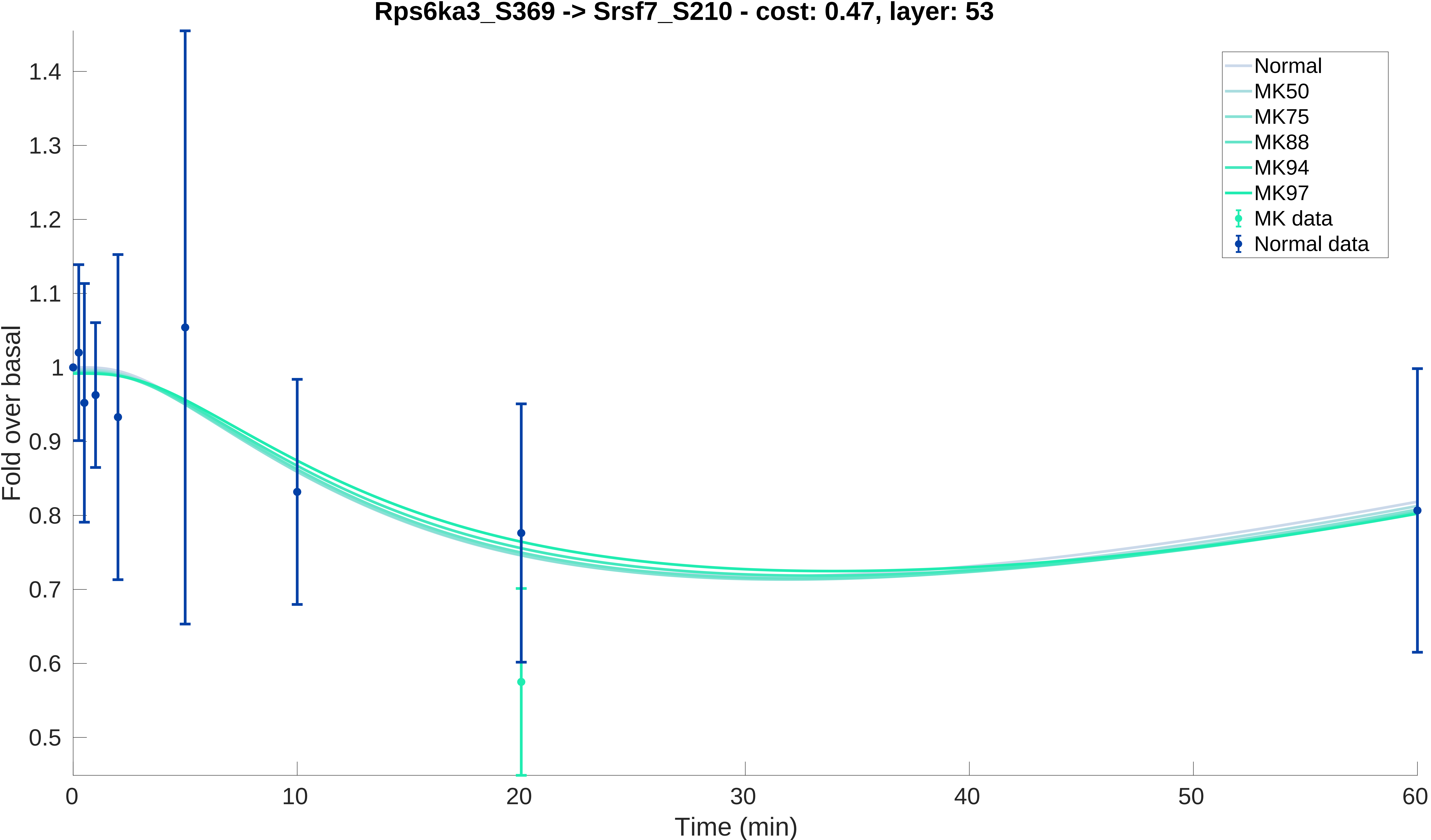
Rps6ka3_S369 -> Srsf7_S210 - cost: 0.47, layer: 53
Fold over basal
1.4
1.3
1.2
1.1
1
0.9
0.8
0.7
0.6
0.5
0
10
20
30
40
50
60
Time (min)
Normal
MK50
MK75
MK88
MK94
MK97
MK data
Normal data

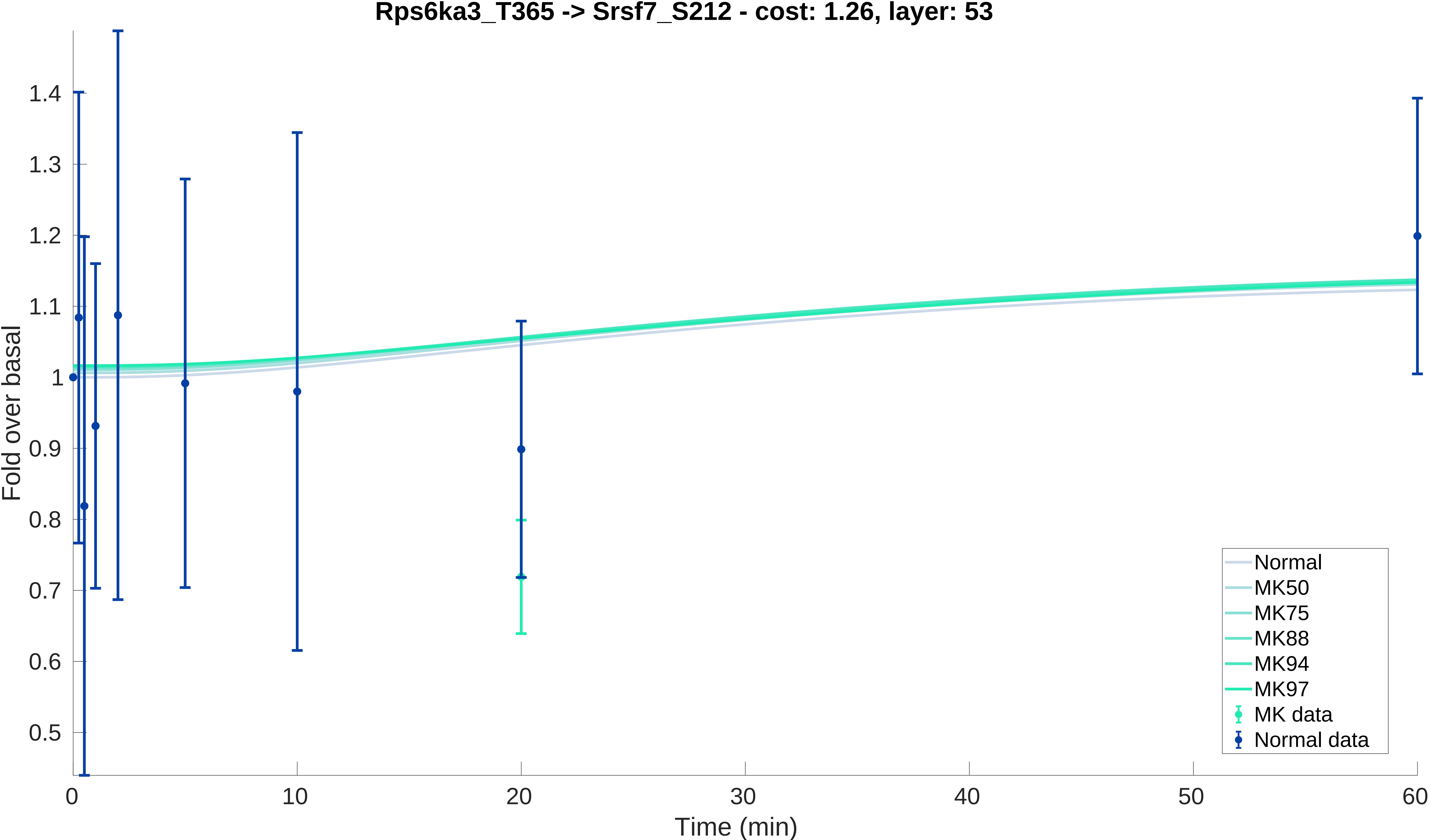

Rps6ka3_T365 -> Srsf7_S212 - cost: 1.26, layer: 53
Fold over basal
Time (min)
0.5
0.6
0.7
0.8
0.9
1
1.1
1.2
1.3
1.4
0
10
20
30
40
50
60
Normal
MK50
MK75
MK88
MK94
MK97
MK data
Normal data

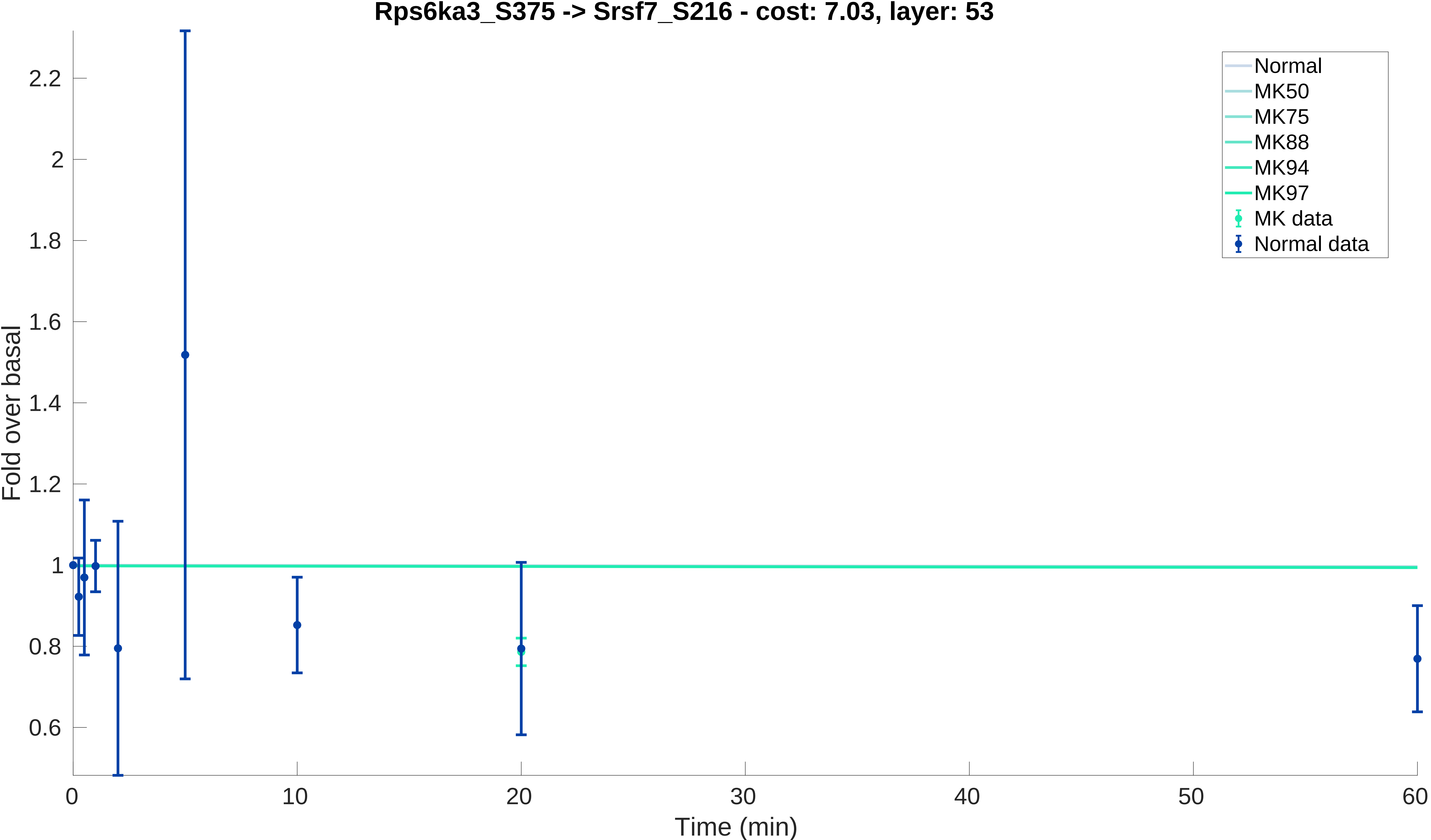

Rps6ka3_S375 -> Srsf7_S216 - cost: 7.03, layer: 53
Fold over basal
Time (min)
0.6
0.8
1
1.2
1.4
1.6
1.8
2
2.2
0
10
20
30
40
50
60
Normal
MK50
MK75
MK88
MK94
MK97
MK data
Normal data

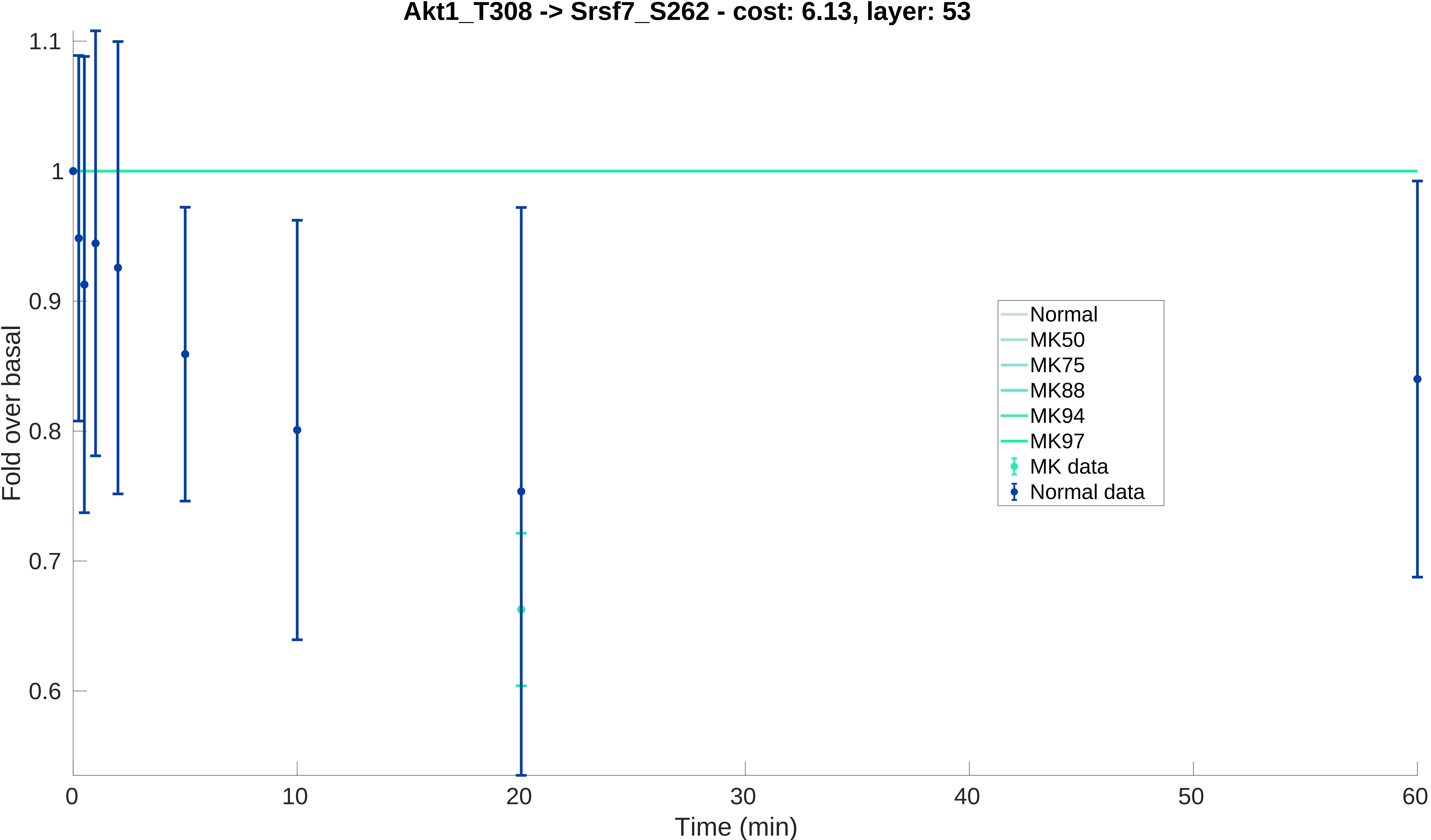

Akt1_T308 -> Srsf7_S262 - cost: 6.13, layer: 53
Fold over basal
Time (min)
Normal
MK50
MK75
MK88
MK94
MK97
MK data
Normal data
1.1
1
0.9
0.8
0.7
0.6
0
10
20
30
40
50
60

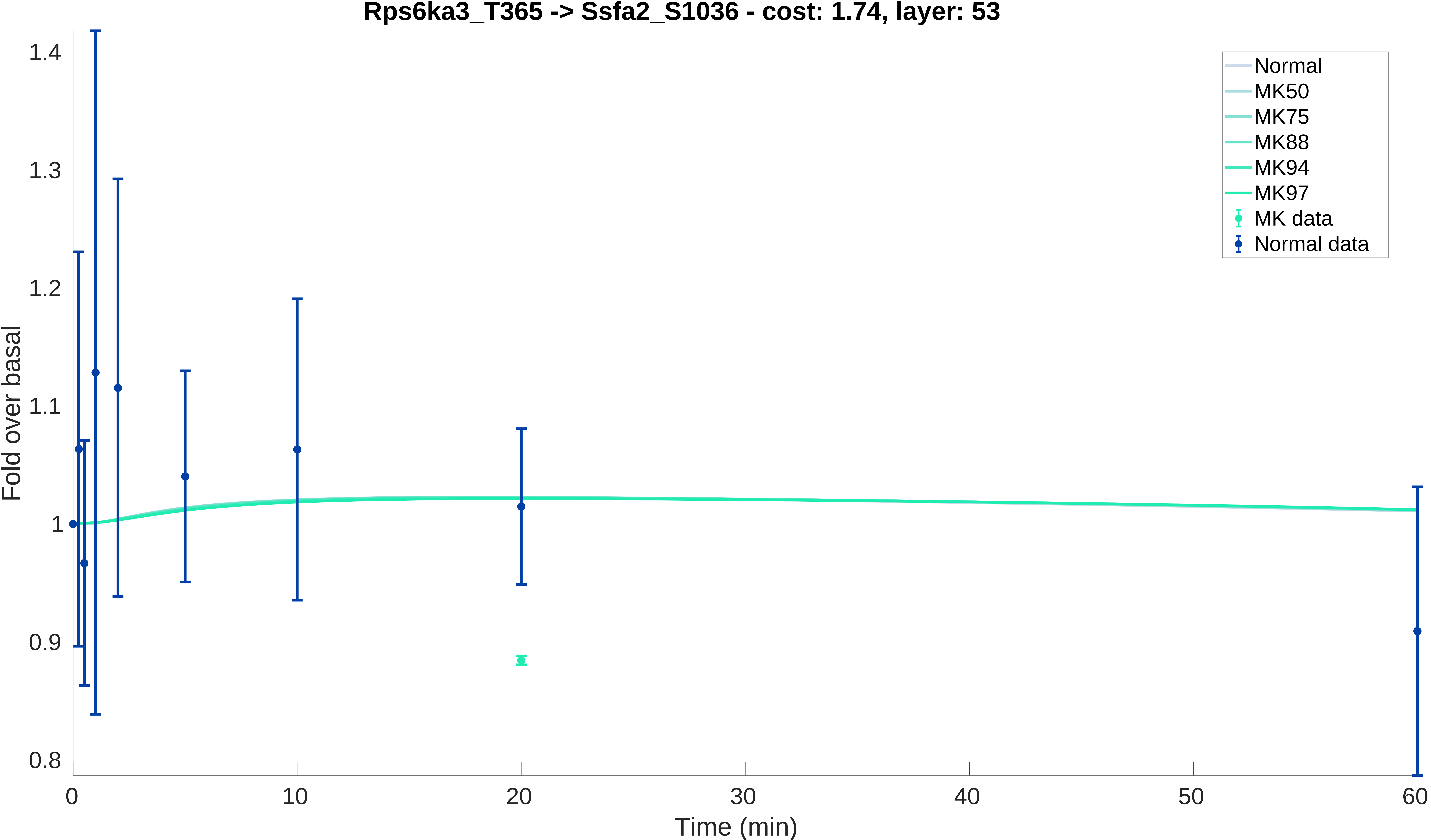

Rps6ka3_T365 -> Ssfa2_S1036 - cost: 1.74, layer: 53
Fold over basal
Time (min)
1.4
1.3
1.2
1.1
1
0.9
0.8
0
10
20
30
40
50
60
Normal
MK50
MK75
MK88
MK94
MK97
MK data
Normal data

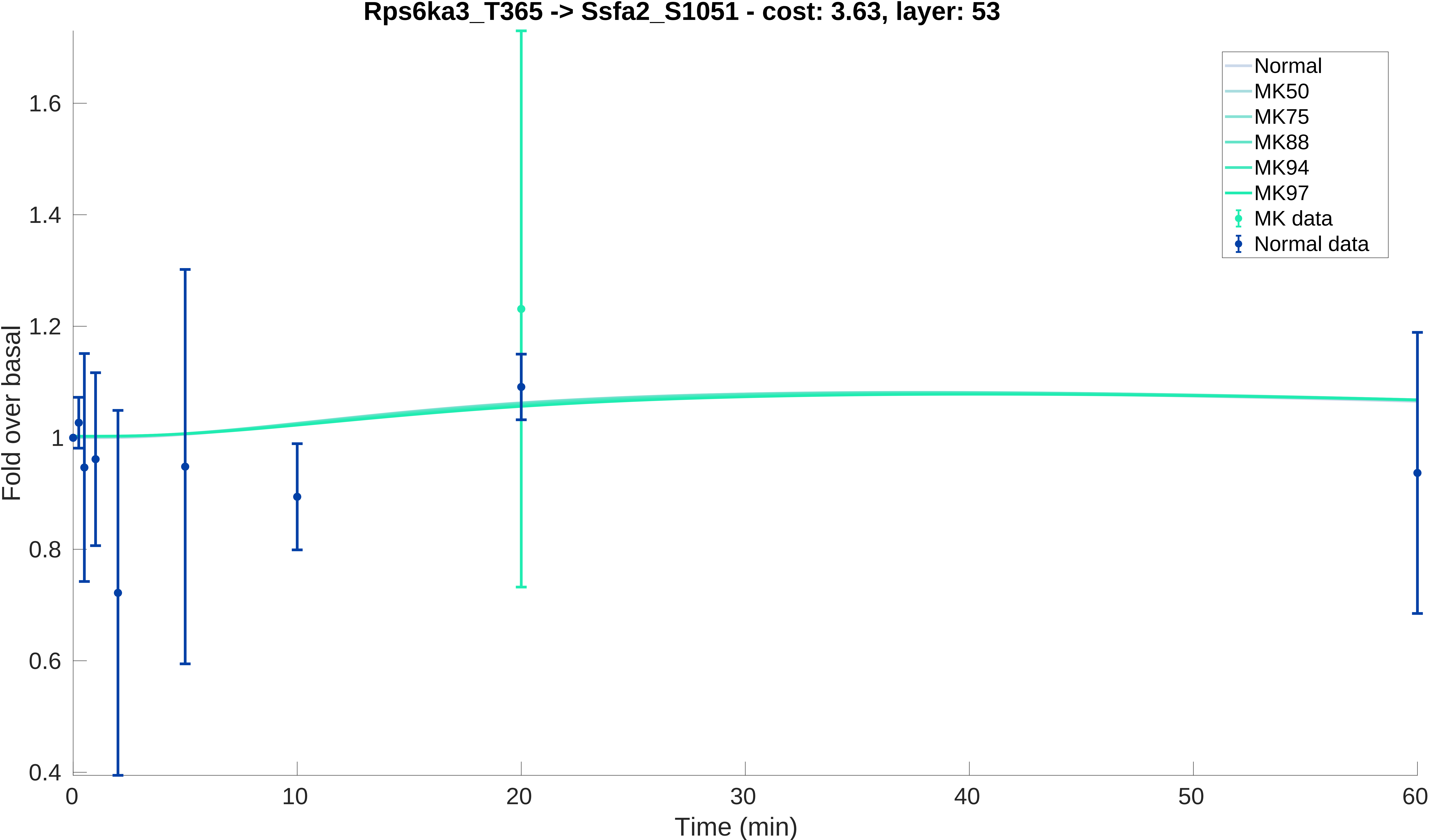
Rps6ka3_T365 -> Ssfa2_S1051 - cost: 3.63, layer: 53
Fold over basal
Time (min)
0.4
0.6
0.8
1
1.2
1.4
1.6
0
10
20
30
40
50
60
Normal
MK50
MK75
MK88
MK94
MK97
MK data
Normal data

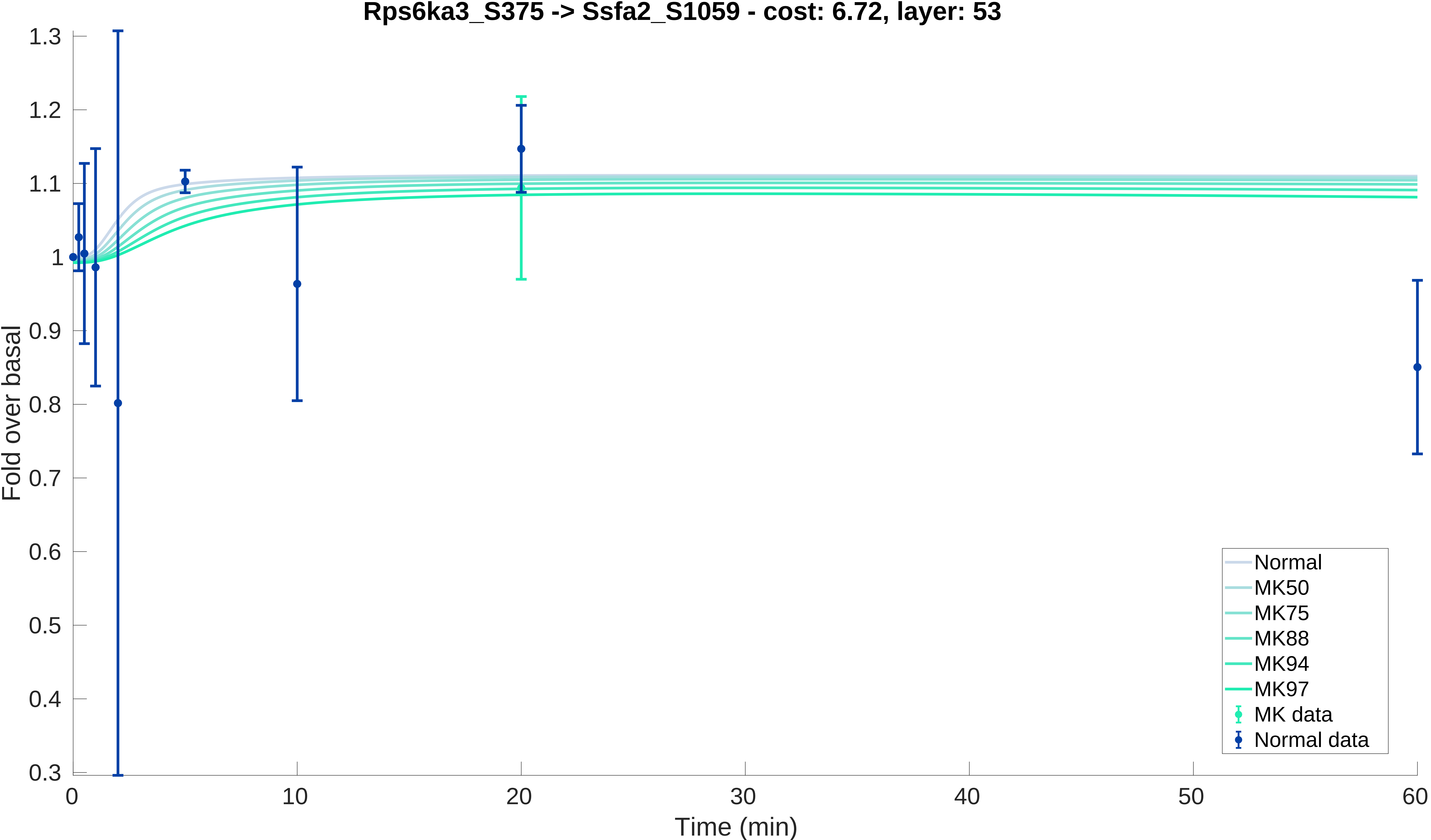

Rps6ka3_S375 -> Ssfa2_S1059 - cost: 6.72, layer: 53
Fold over basal
Time (min)
1.3
1.2
1.1
1
0.9
0.8
0.7
0.6
0.5
0.4
0.3
0
10
20
30
40
50
60
Normal
MK50
MK75
MK88
MK94
MK97
MK data
Normal data

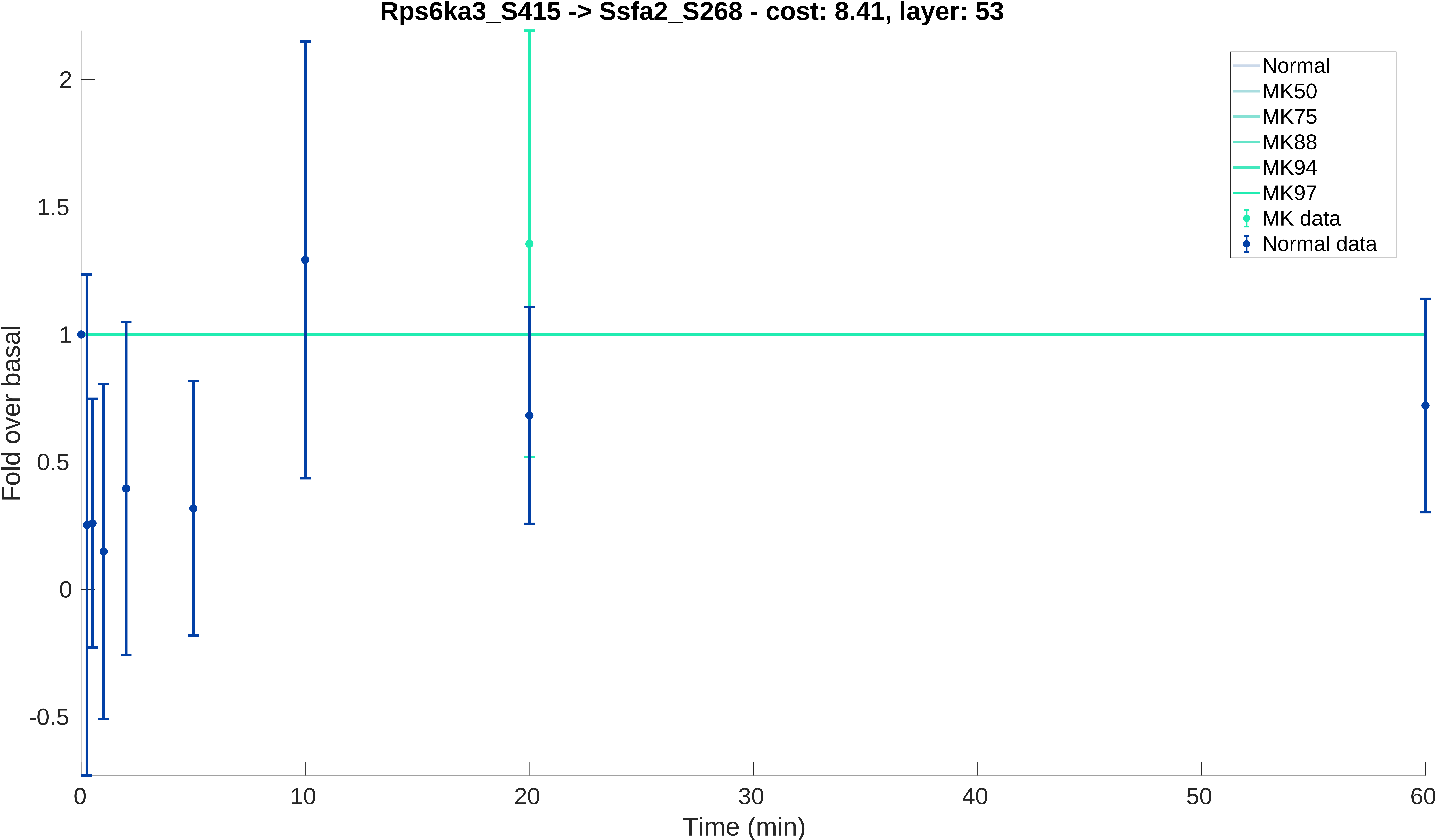
Rps6ka3_S415 -> Ssfa2_S268 - cost: 8.41, layer: 53
Fold over basal
Time (min)
Normal
MK50
MK75
MK88
MK94
MK97
MK data
Normal data
2
1.5
1
0.5
0
-0.5
0
10
20
30
40
50
60

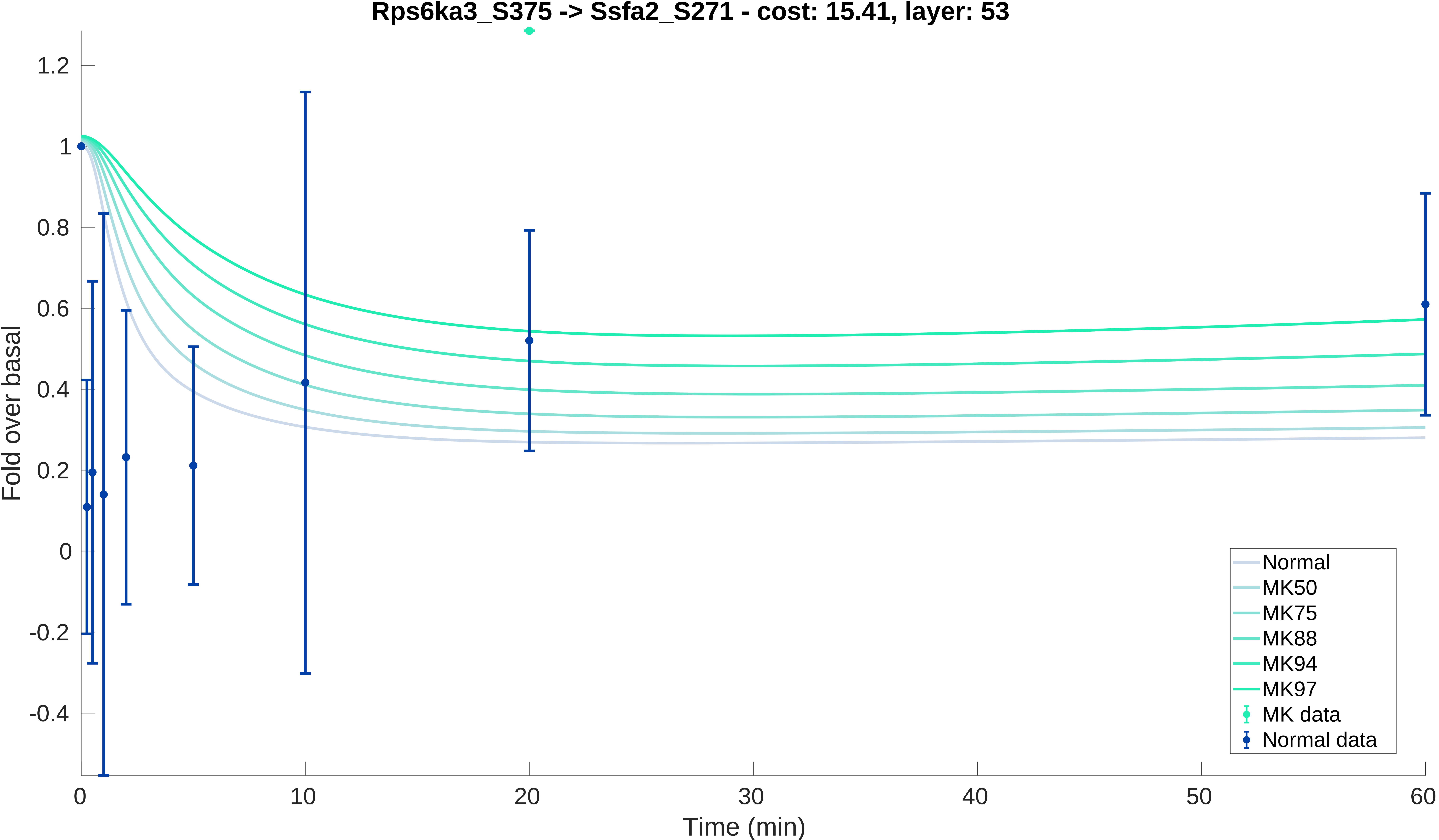

Rps6ka3_S375 -> Ssfa2_S271 - cost: 15.41, layer: 53
Fold over basal
Time (min)
1.2
1
0.8
0.6
0.4
0.2
0
-0.2
-0.4
0
10
20
30
40
50
60
Normal
MK50
MK75
MK88
MK94
MK97
MK data
Normal data

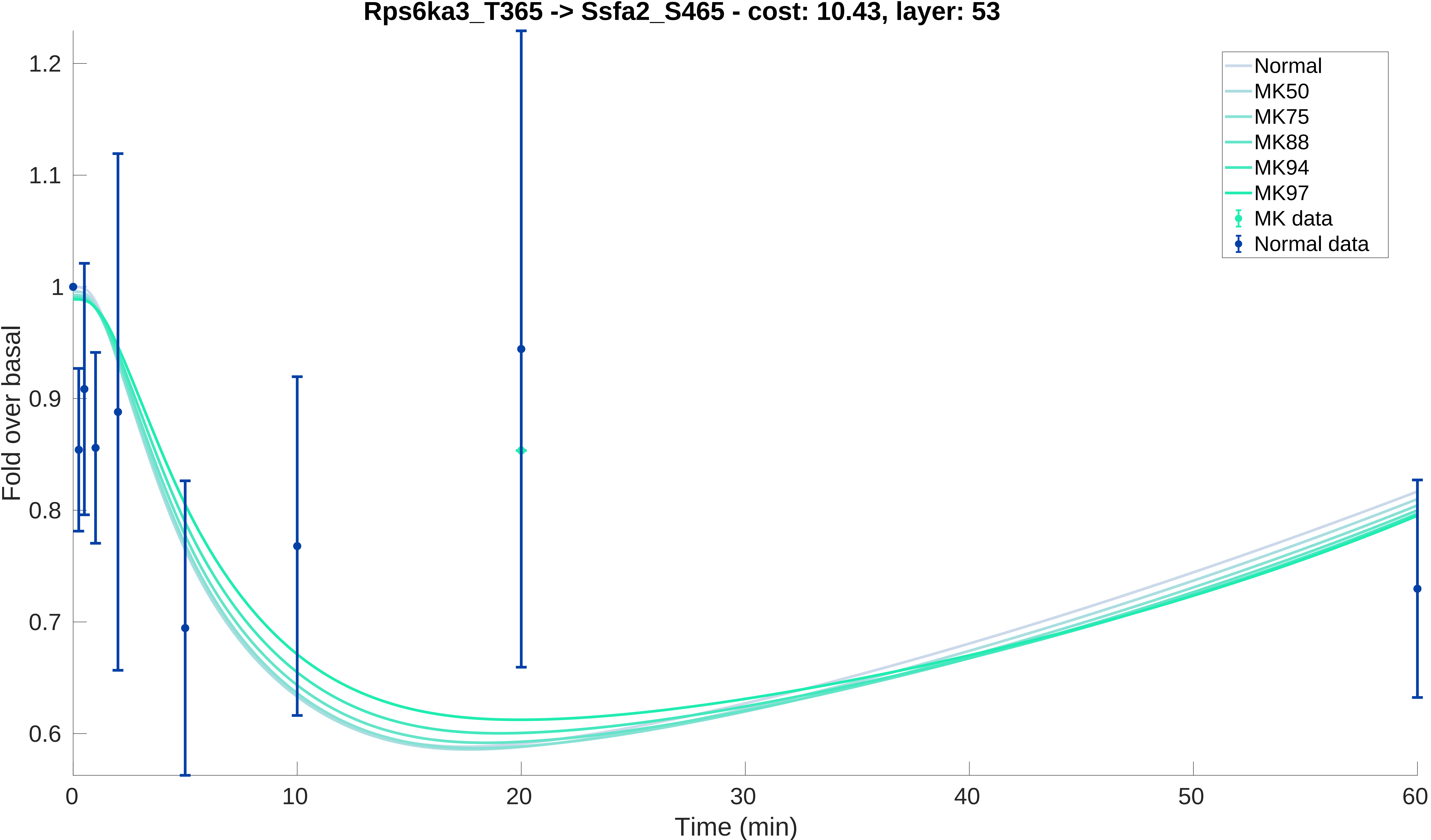

Rps6ka3_T365 -> Ssfa2_S465 - cost: 10.43, layer: 53
Fold over basal
Time (min)
Normal
MK50
MK75
MK88
MK94
MK97
MK data
Normal data
0.6
0.7
0.8
0.9
1
1.1
1.2
0
10
20
30
40
50
60

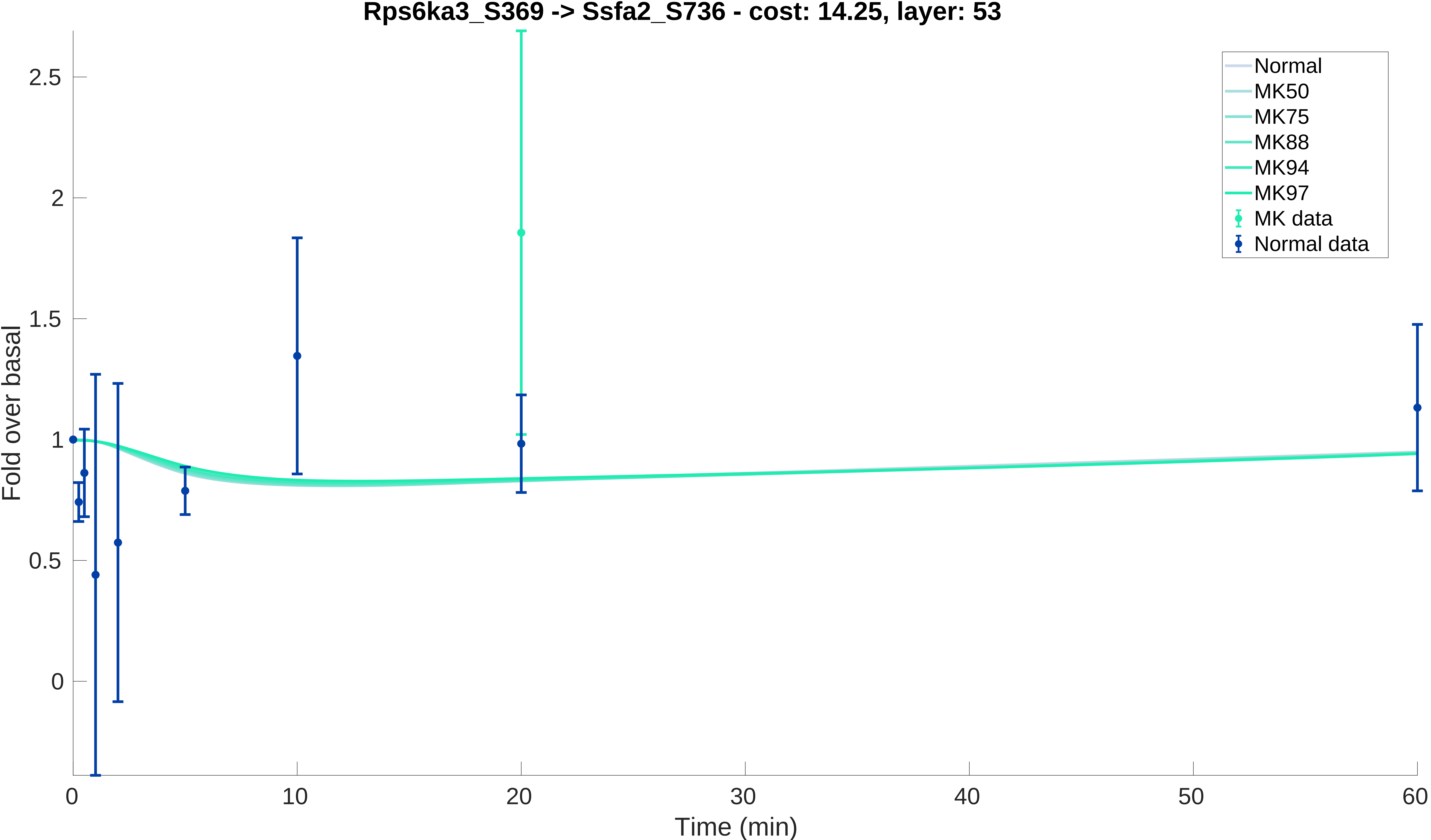

Rps6ka3_S369 -> Ssfa2_S736 - cost: 14.25, layer: 53
Fold over basal
Time (min)
0
0.5
1
1.5
2
2.5
0
10
20
30
40
50
60
Normal
MK50
MK75
MK88
MK94
MK97
MK data
Normal data

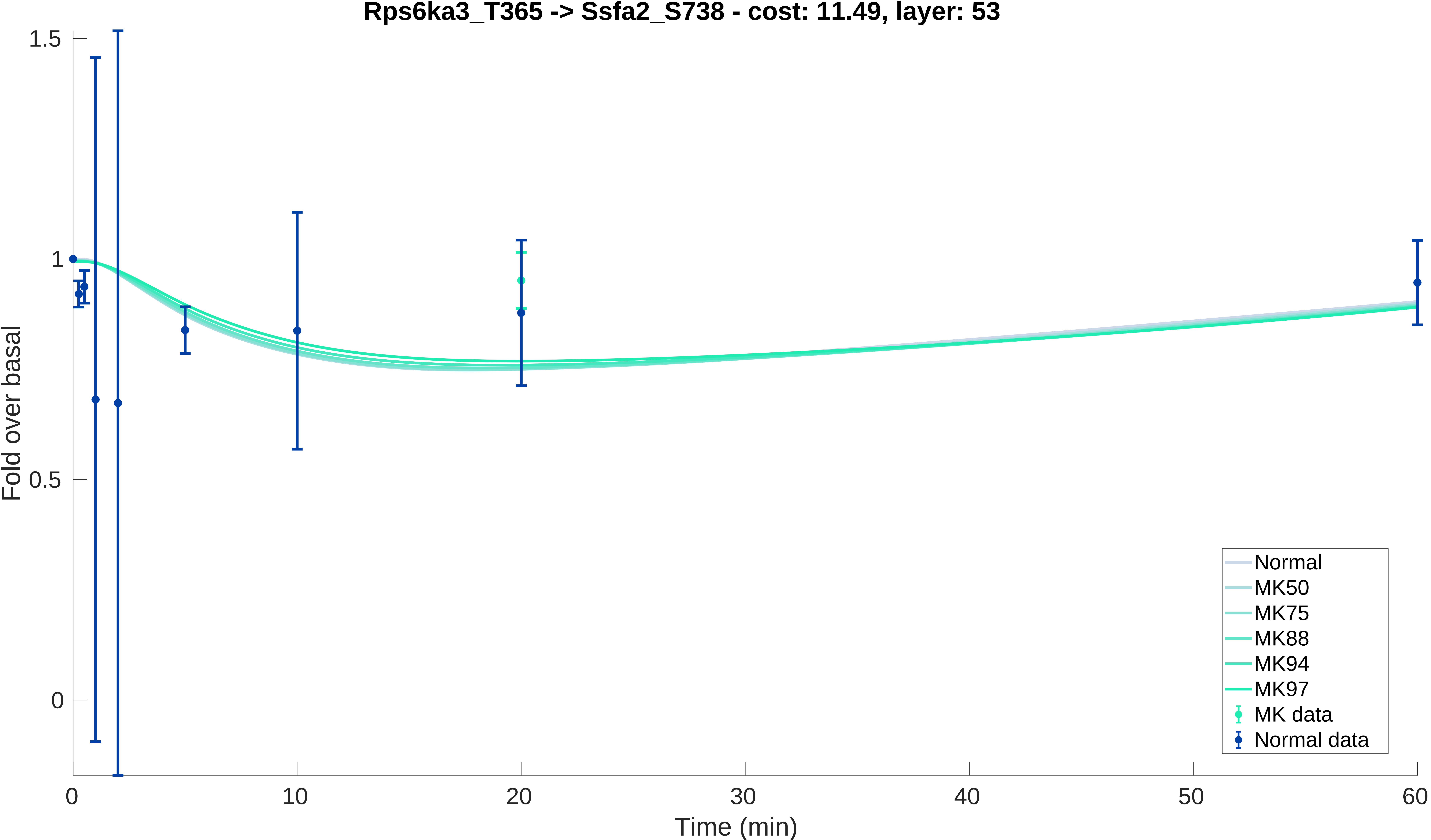

Rps6ka3_T365 -> Ssfa2_S738 - cost: 11.49, layer: 53
1.5
1
0.5
0
Fold over basal
0
10
20
30
40
50
60
Time (min)
Normal
MK50
MK75
MK88
MK94
MK97
MK data
Normal data

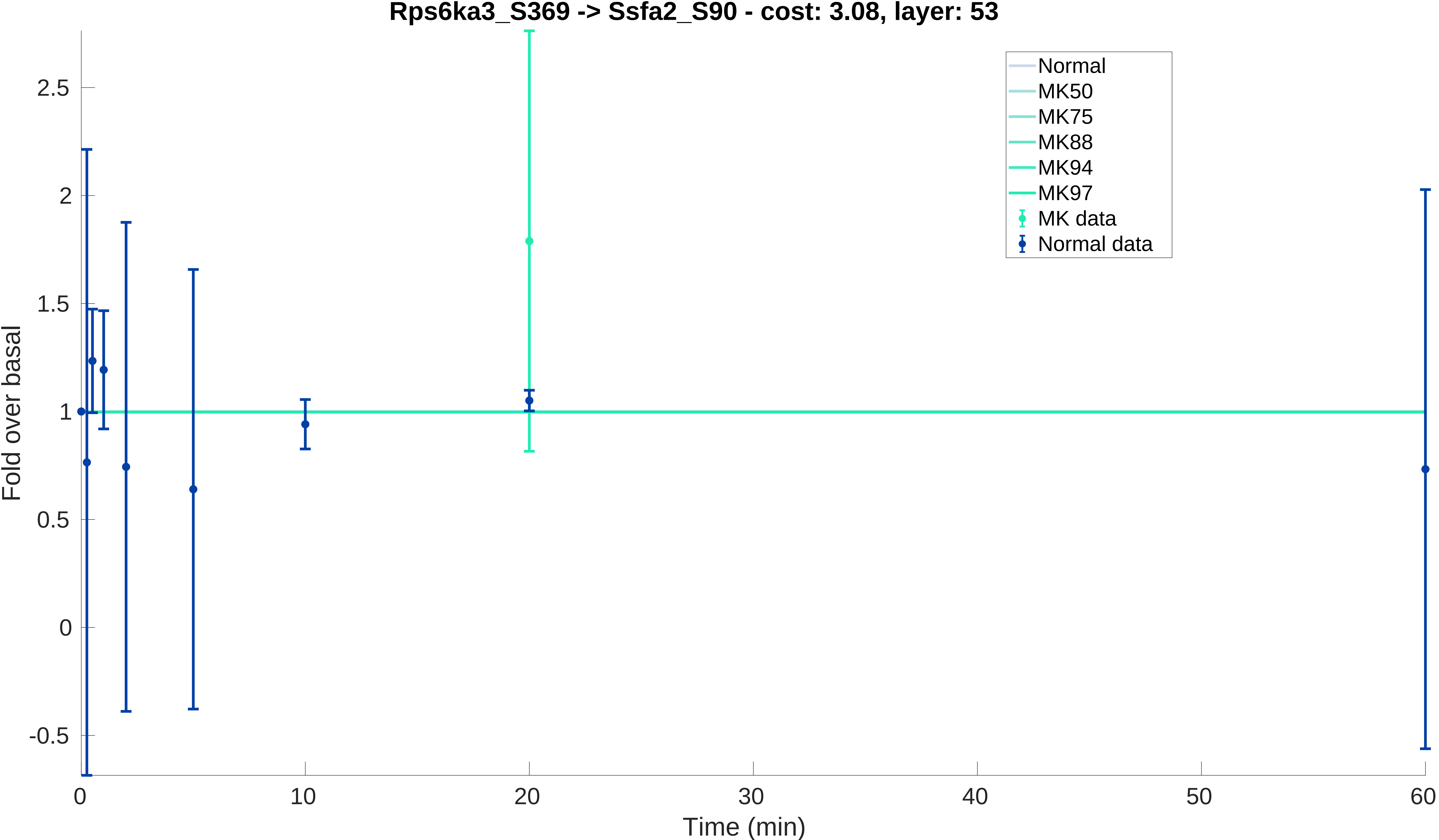

Rps6ka3_S369 -> Ssfa2_S90 - cost: 3.08, layer: 53
Normal
MK50
MK75
MK88
MK94
MK97
MK data
Normal data
Fold over basal
Time (min)
2.5
2
1.5
1
0.5
0
-0.5
0
10
20
30
40
50
60

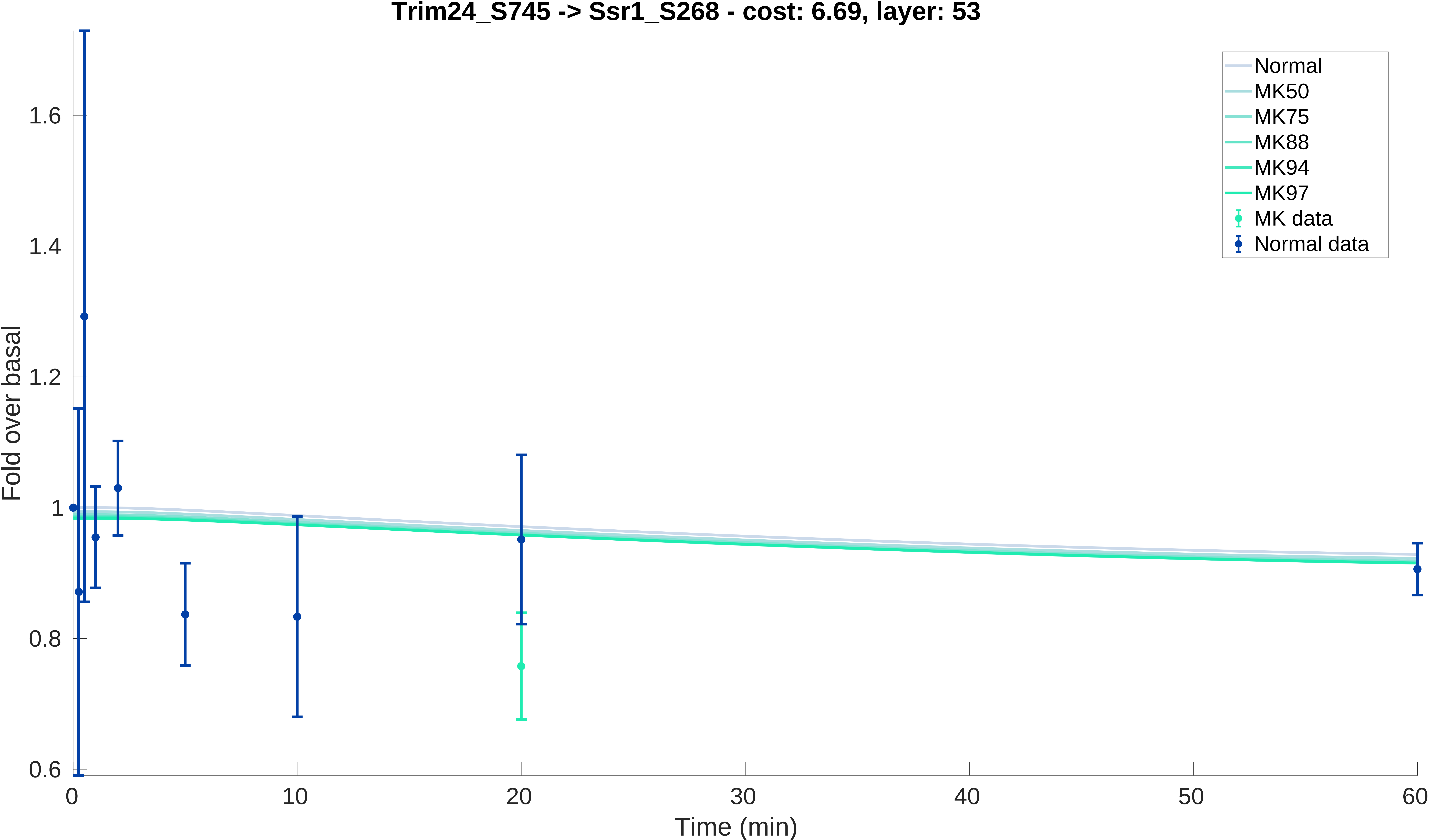

Trim24_S745 -> Ssr1_S268 - cost: 6.69, layer: 53
Fold over basal
Time (min)
1.6
1.4
1.2
1
0.8
0.6
0
10
20
30
40
50
60
Normal
MK50
MK75
MK88
MK94
MK97
MK data
Normal data

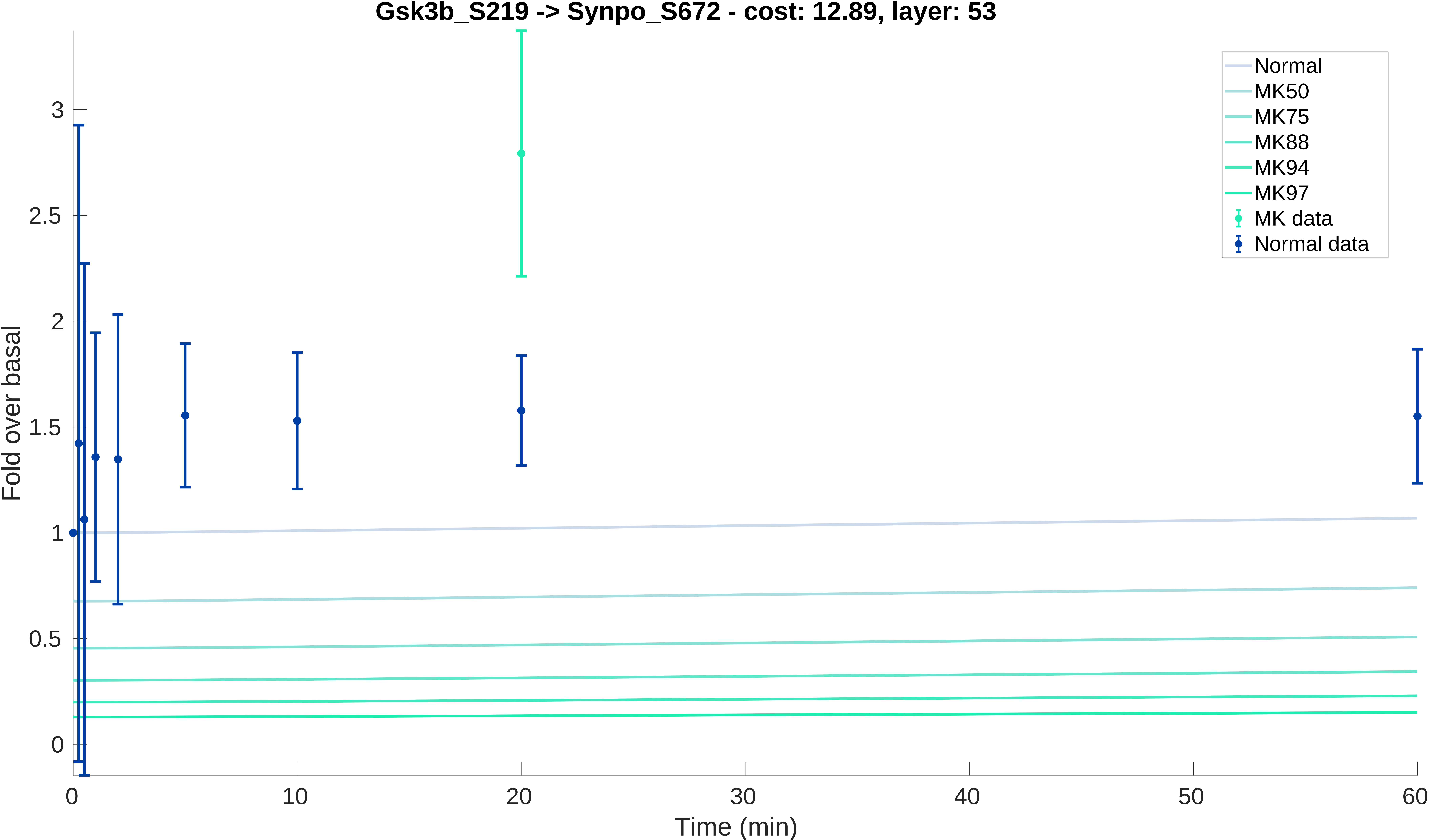

Gsk3b_S219 -> Synpo_S672 - cost: 12.89, layer: 53
Fold over basal
Time (min)
Normal
MK50
MK75
MK88
MK94
MK97
MK data
Normal data
0
0.5
1
1.5
2
2.5
3
0
10
20
30
40
50
60

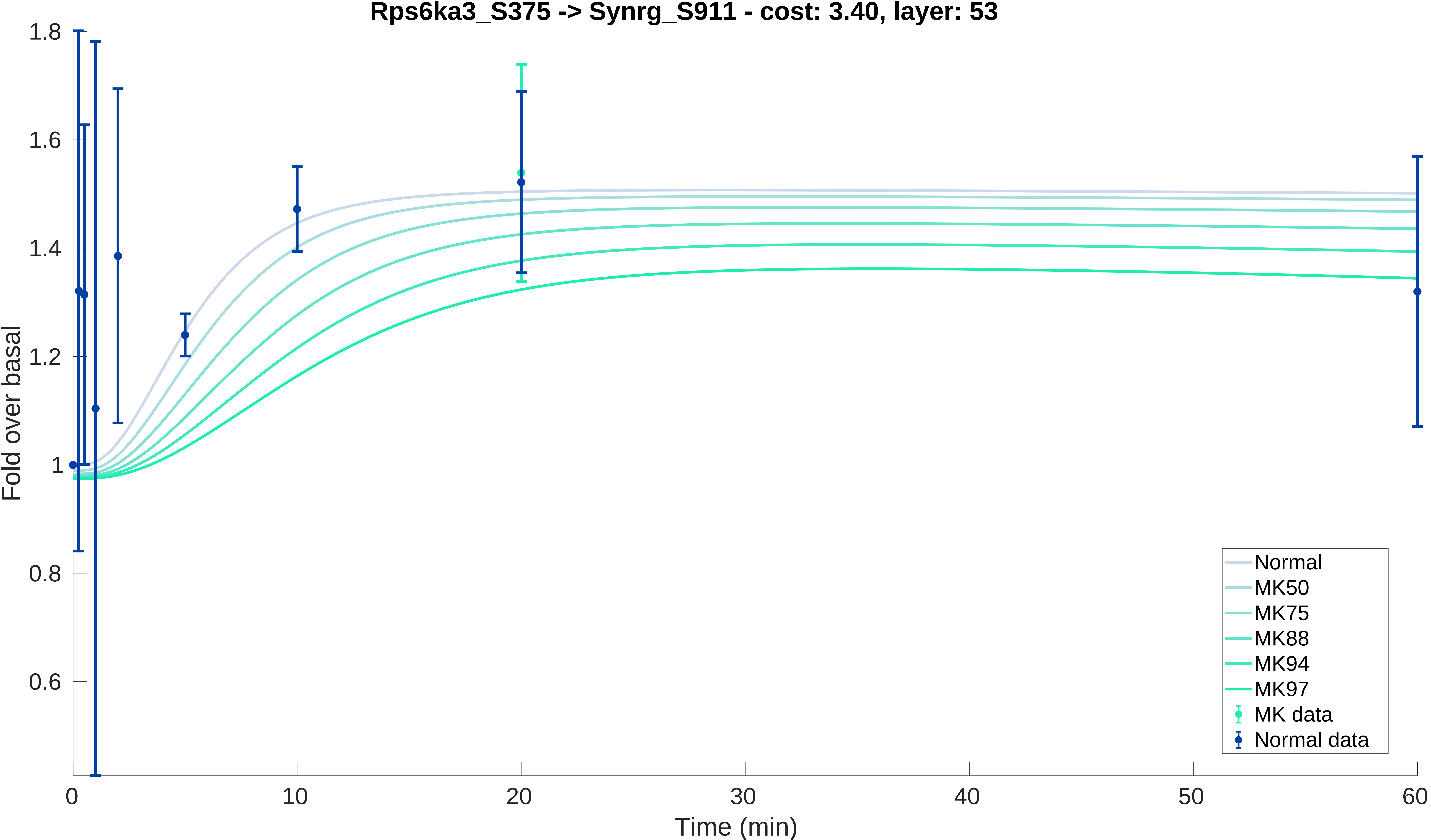
Rps6ka3_S375 -> Synrg_S911 - cost: 3.40, layer: 53
Fold over basal
1.8
1.6
1.4
1.2
1
0.8
0.6
0
10
20
30
40
50
60
Time (min)
Normal
MK50
MK75
MK88
MK94
MK97
MK data
Normal data

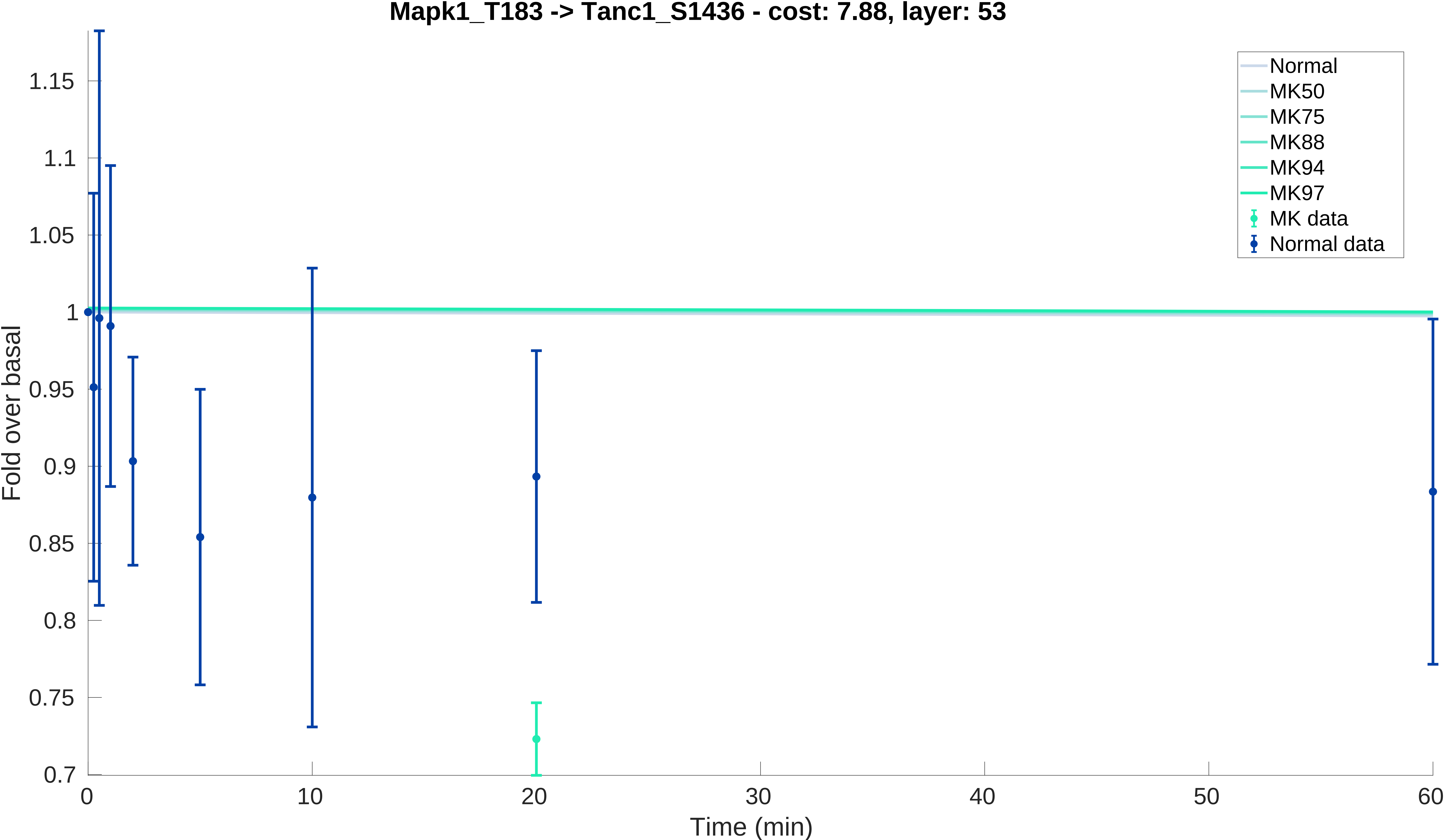

Mapk1_T183 -> Tanc1_S1436 - cost: 7.88, layer: 53
Normal
MK50
MK75
MK88
MK94
MK97
MK data
Normal data
Fold over basal
1.15
1.1
1.05
1
0.95
0.9
0.85
0.8
0.75
0.7
0
10
20
30
40
50
60
Time (min)

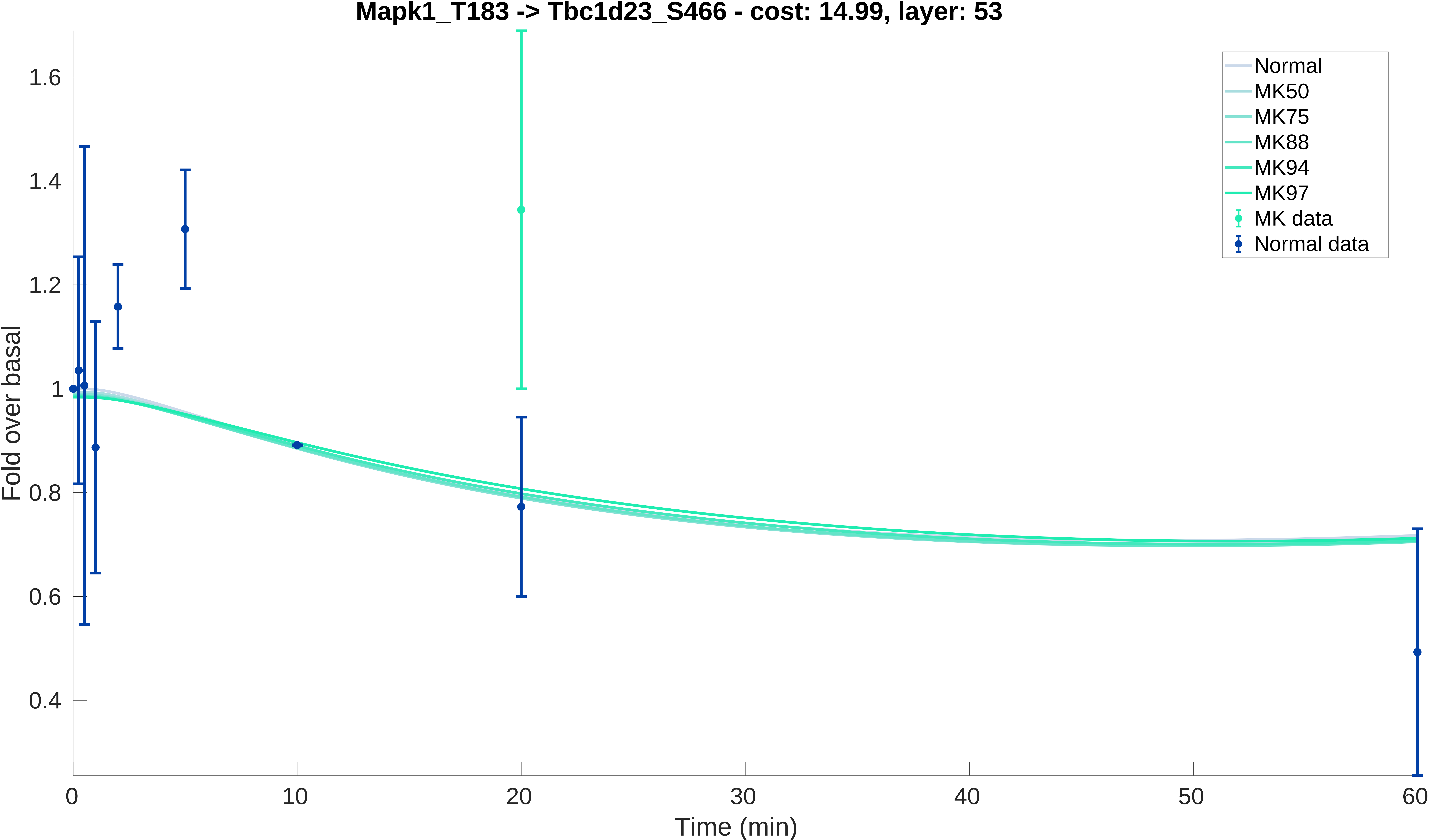

Mapk1_T183 -> Tbc1d23_S466 - cost: 14.99, layer: 53
Fold over basal
Time (min)
0
10
20
30
40
50
60
0.4
0.6
0.8
1
1.2
1.4
1.6
Normal
MK50
MK75
MK88
MK94
MK97
MK data
Normal data

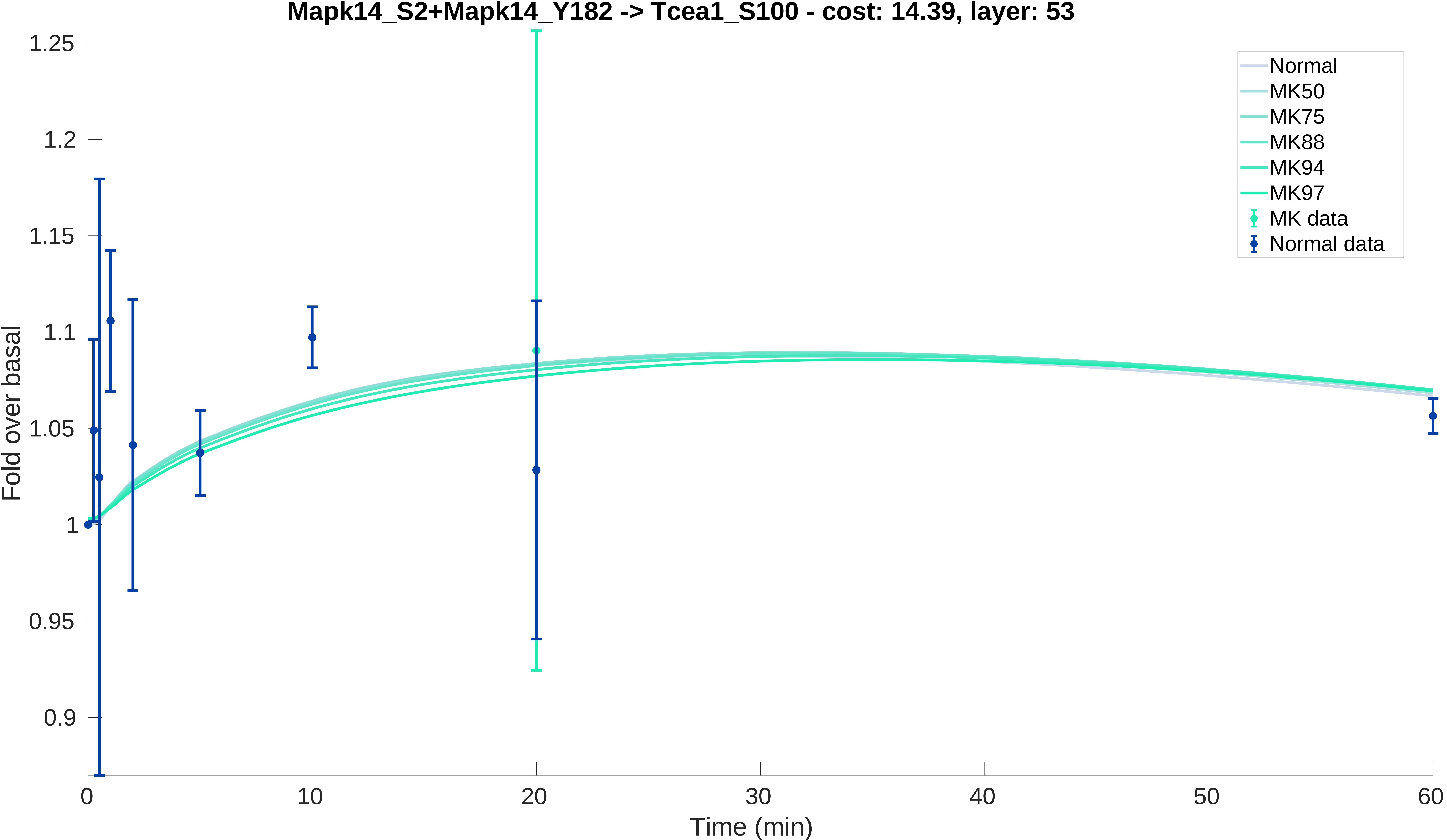

Mapk14_S2+Mapk14_Y182 -> Tcea1_S100 - cost: 14.39, layer: 53
Fold over basal
1.25
1.2
1.15
1.1
1.05
1
0.95
0.9
0
10
20
30
40
50
60
Time (min)
Normal
MK50
MK75
MK88
MK94
MK97
MK data
Normal data

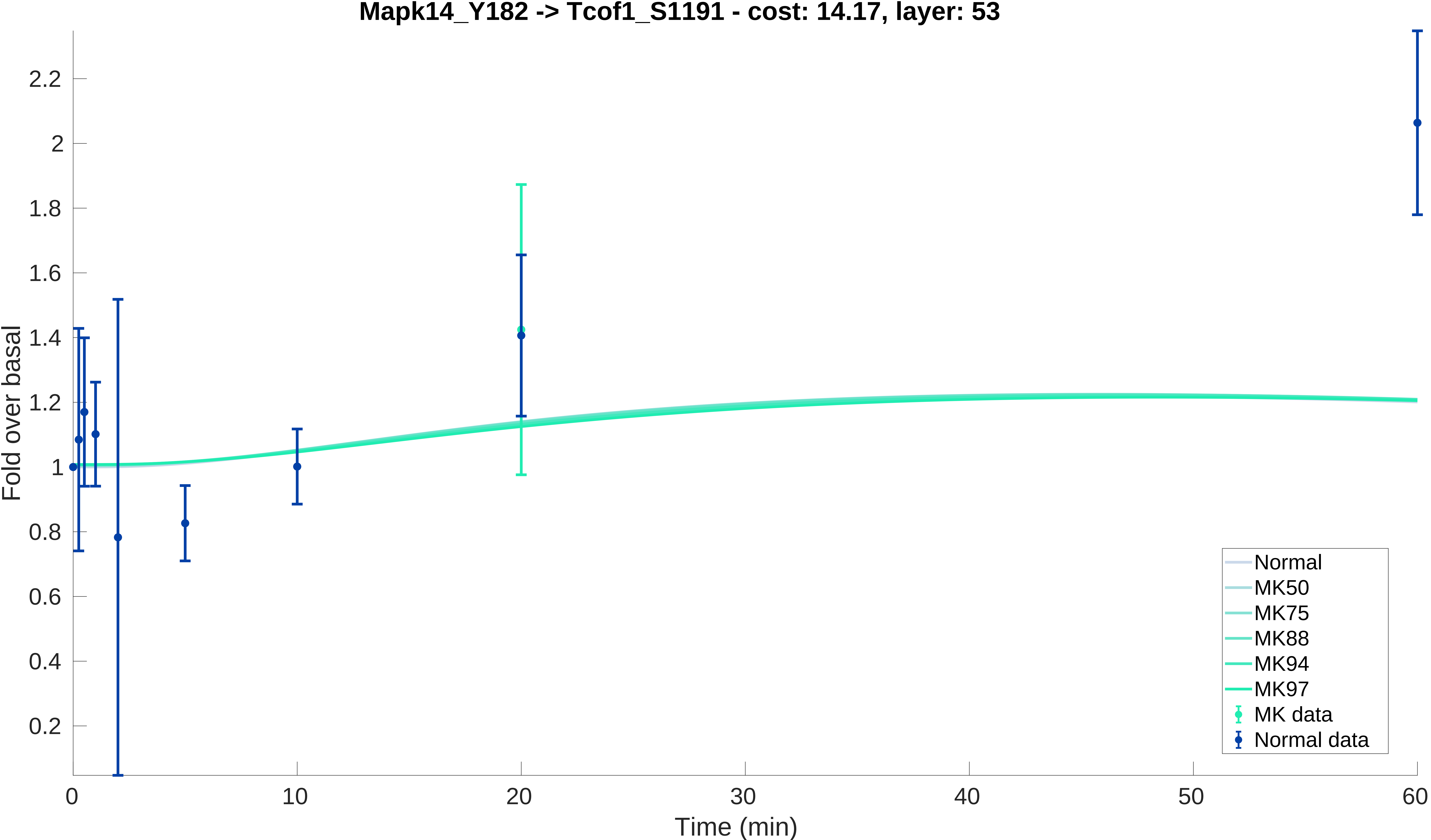
Mapk14_Y182 -> Tcof1_S1191 - cost: 14.17, layer: 53
Fold over basal
2.2
2
1.8
1.6
1.4
1.2
1
0.8
0.6
0.4
0.2
0
10
20
30
40
50
60
Time (min)
Normal
MK50
MK75
MK88
MK94
MK97
MK data
Normal data

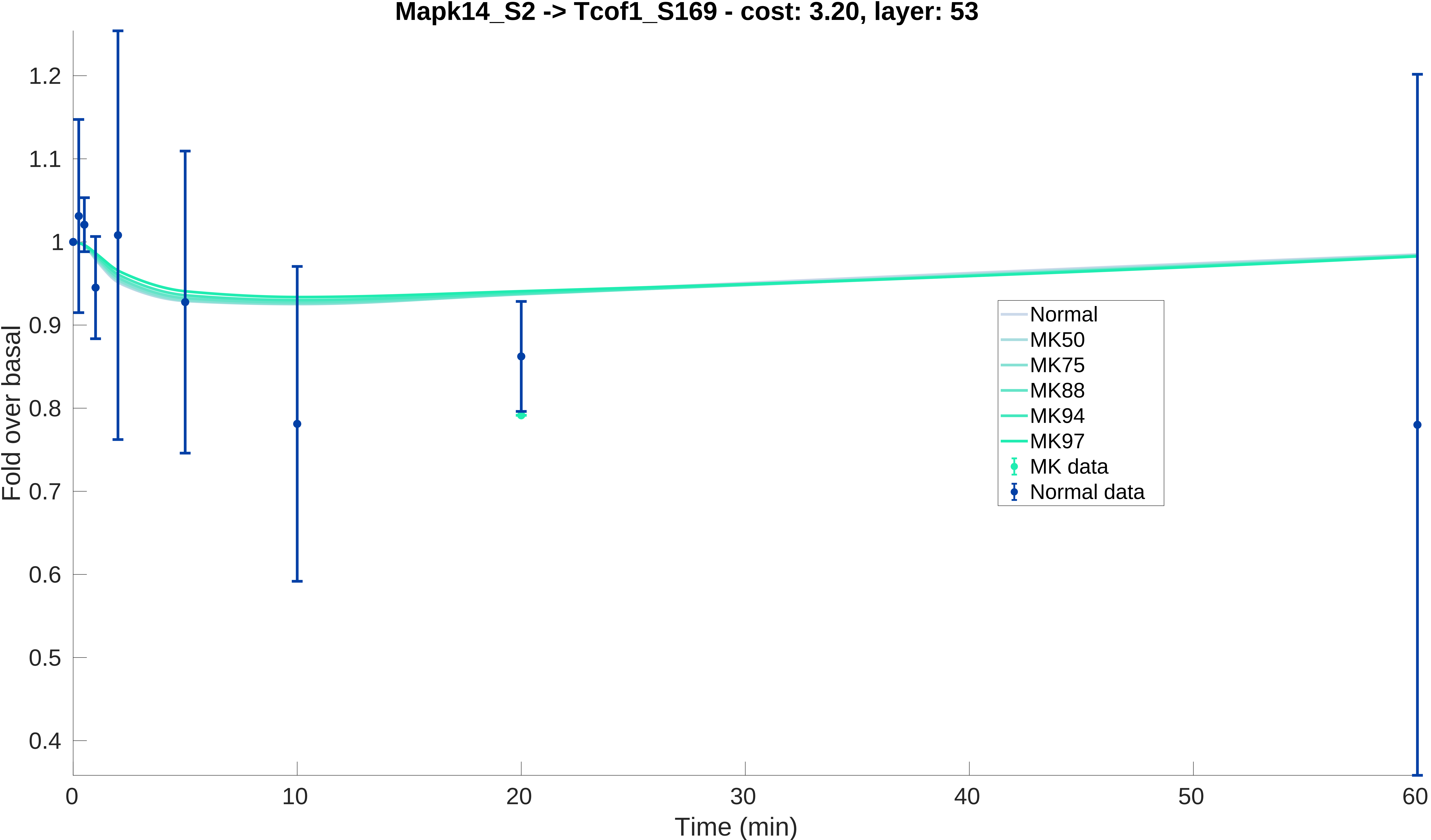

Mapk14_S2 -> Tcof1_S169 - cost: 3.20, layer: 53
Fold over basal
Time (min)
0
10
20
30
40
50
60
0.4
0.5
0.6
0.7
0.8
0.9
1
1.1
1.2
Normal
MK50
MK75
MK88
MK94
MK97
MK data
Normal data

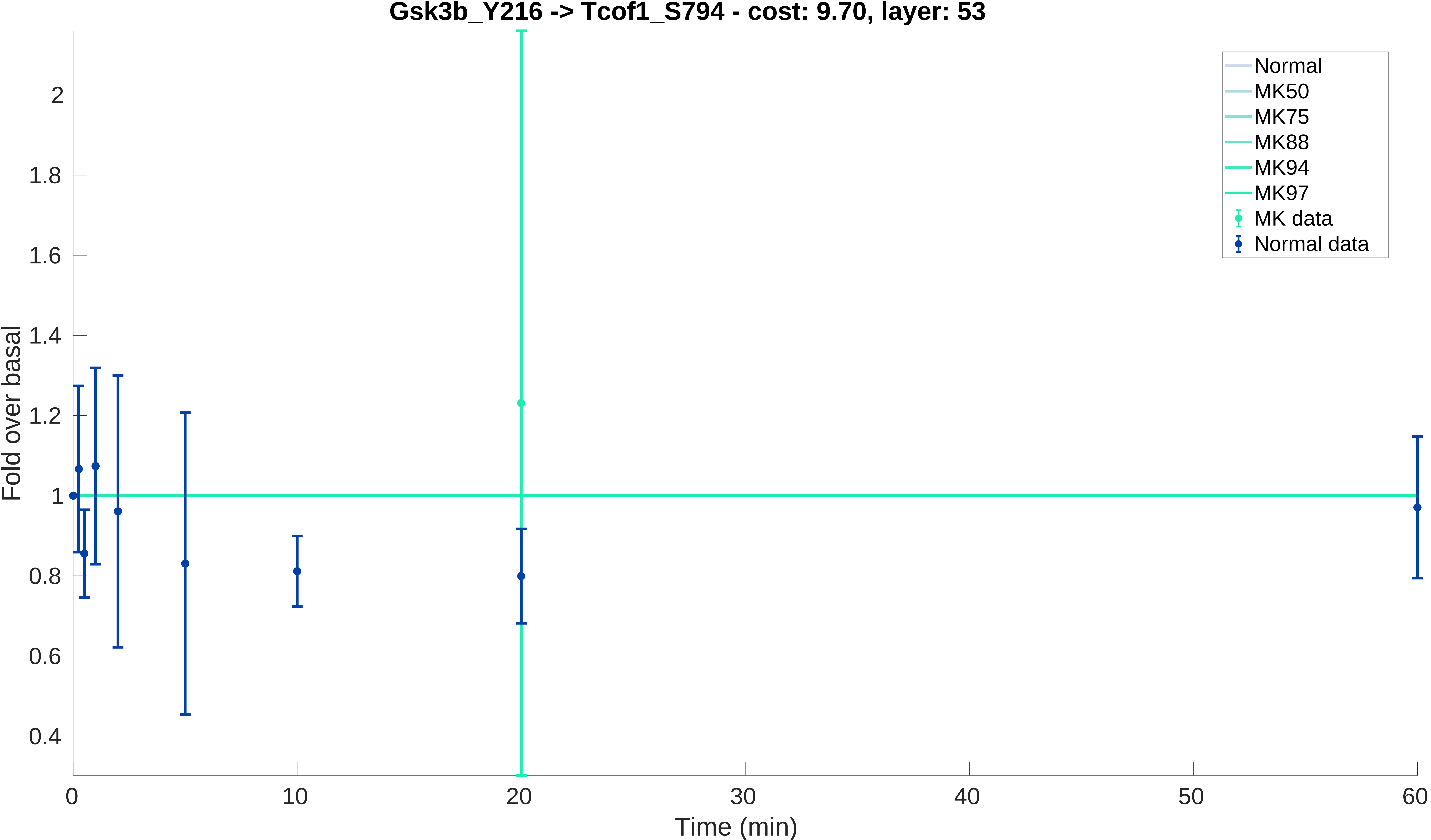

Gsk3b_Y216 -> Tcof1_S794 - cost: 9.70, layer: 53
Fold over basal
Time (min)
Normal
MK50
MK75
MK88
MK94
MK97
MK data
Normal data
0.4
0.6
0.8
1
1.2
1.4
1.6
1.8
2
0
10
20
30
40
50
60

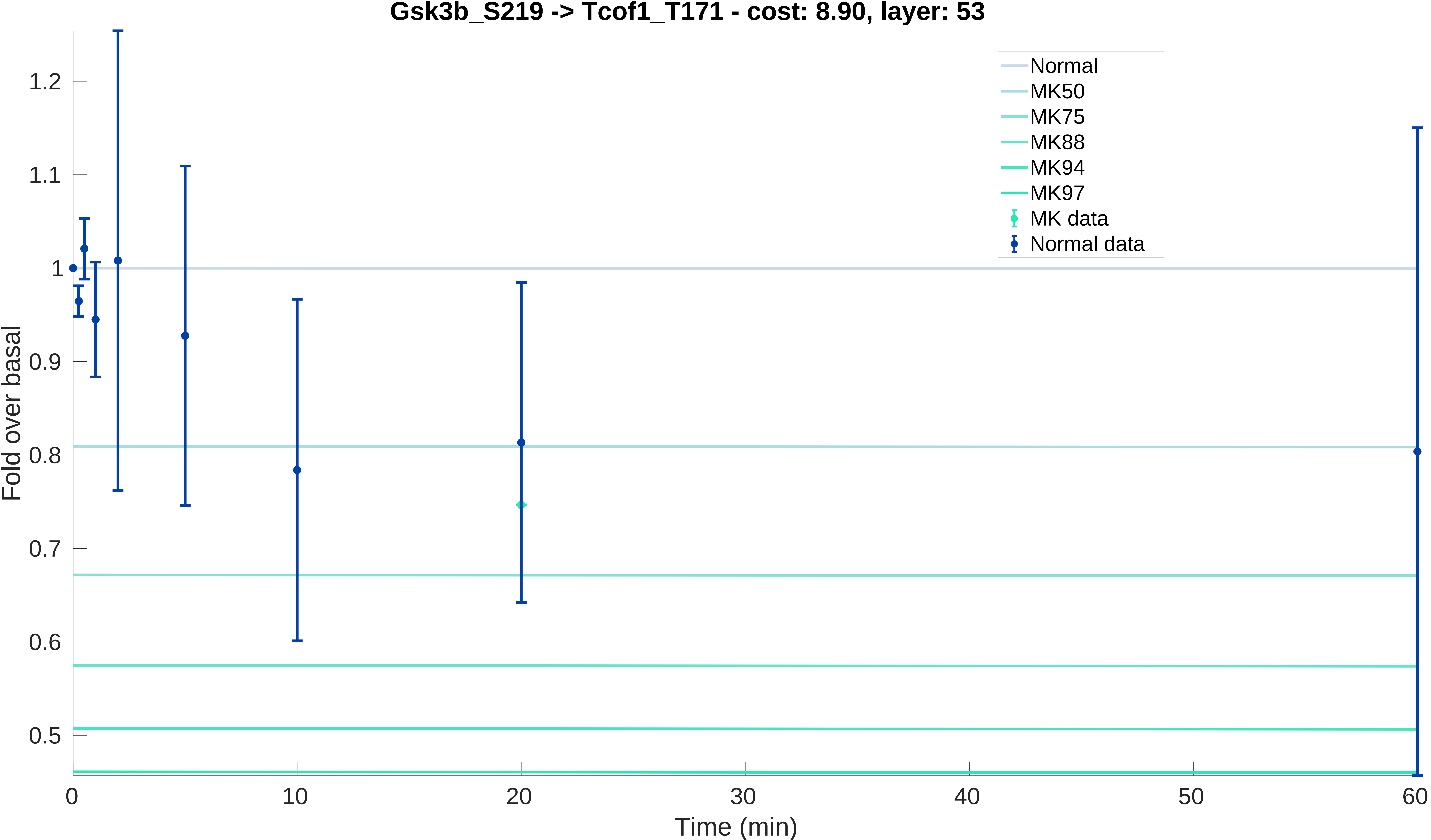

Gsk3b_S219 -> Tcof1_T171 - cost: 8.90, layer: 53
Fold over basal
Time (min)
0.5
0.6
0.7
0.8
0.9
1
1.1
1.2
0
10
20
30
40
50
60
Normal
MK50
MK75
MK88
MK94
MK97
MK data
Normal data

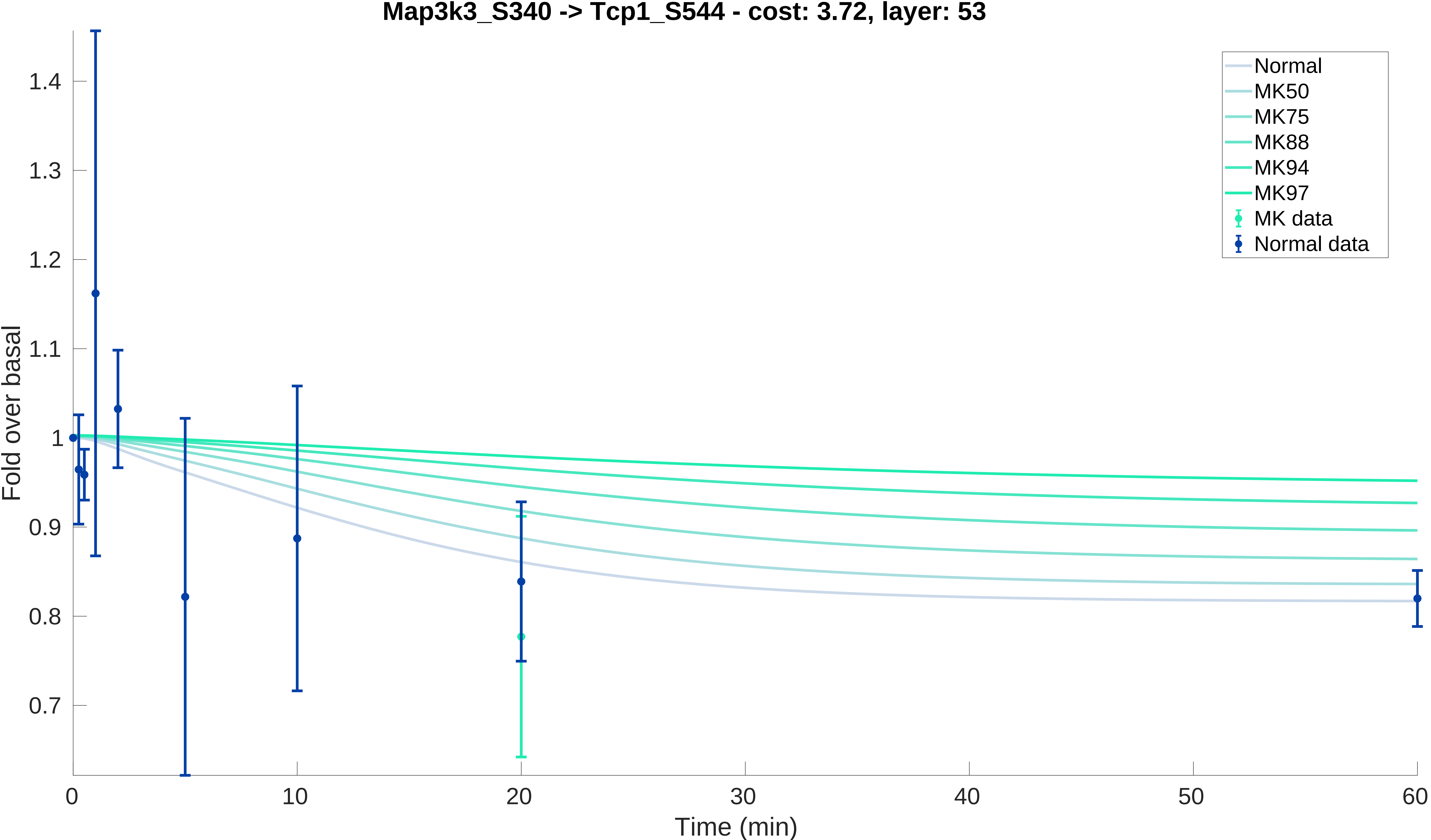

Map3k3_S340 -> Tcp1_S544 - cost: 3.72, layer: 53
Normal
MK50
MK75
MK88
MK94
MK97
MK data
Normal data
Fold over basal
1.4
1.3
1.2
1.1
1
0.9
0.8
0.7
0
10
20
30
40
50
60
Time (min)

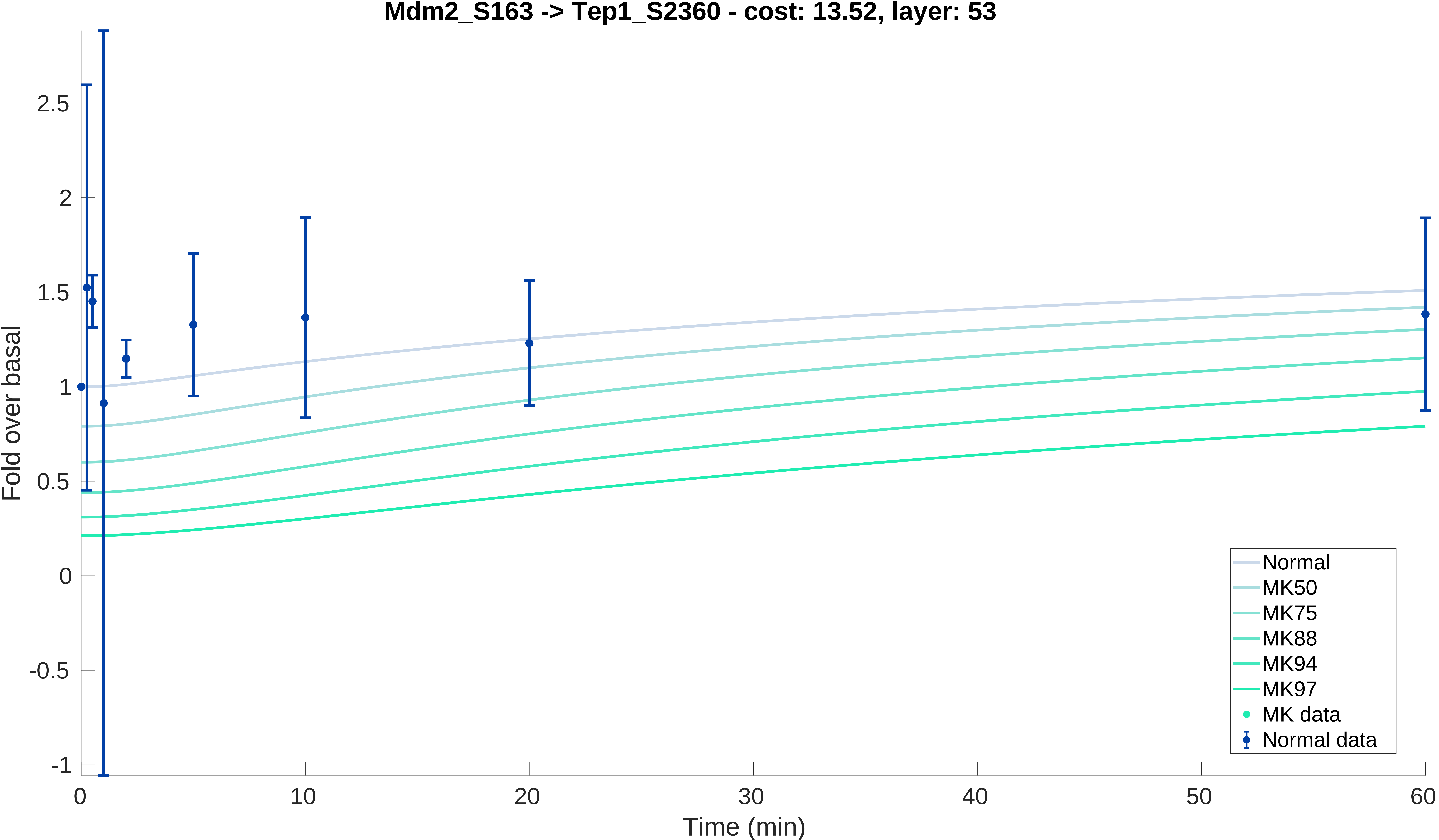

Mdm2_S163 -> Tep1_S2360 - cost: 13.52, layer: 53
Fold over basal
2.5
2
1.5
1
0.5
0
-0.5
-1
0
10
20
30
40
50
60
Time (min)
Normal
MK50
MK75
MK88
MK94
MK97
MK data
Normal data

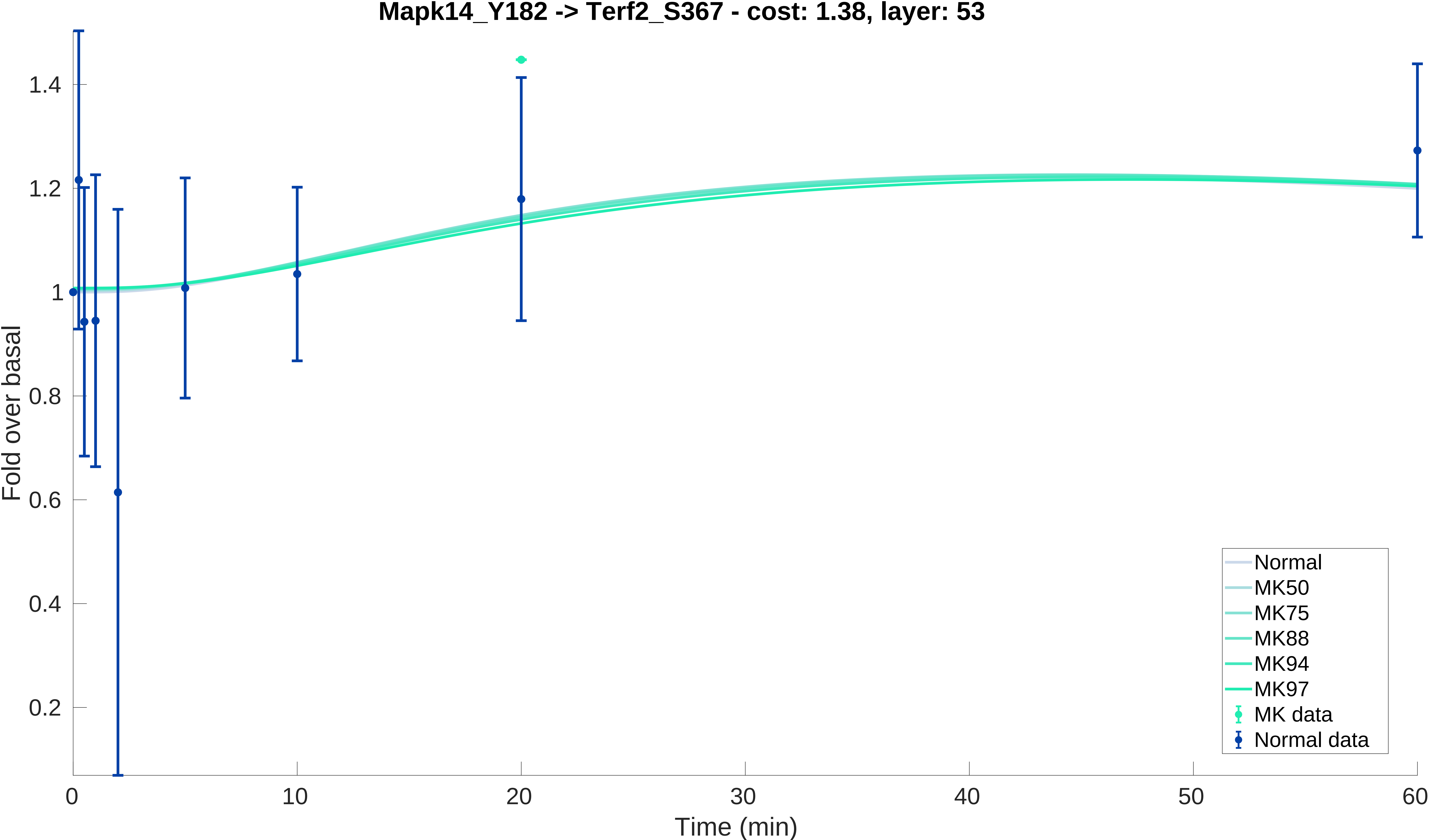

Mapk14_Y182 -> Terf2_S367 - cost: 1.38, layer: 53
Fold over basal
1.4
1.2
1
0.8
0.6
0.4
0.2
0
10
20
30
40
50
60
Time (min)
Normal
MK50
MK75
MK88
MK94
MK97
MK data
Normal data

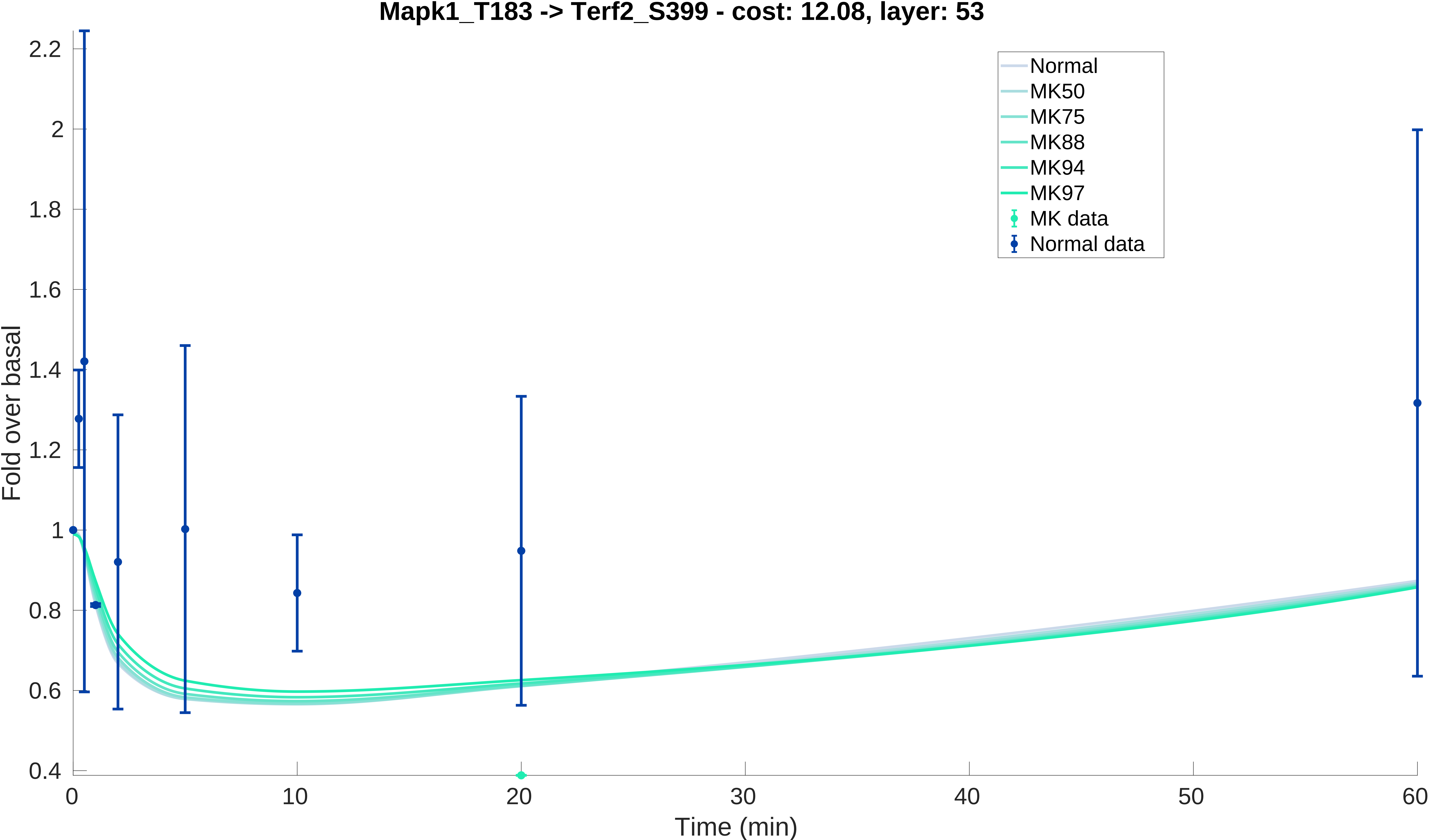
Mapk1_T183 -> Terf2_S399 - cost: 12.08, layer: 53
Fold over basal
Time (min)
0.4
0.6
0.8
1
1.2
1.4
1.6
1.8
2
2.2
0
10
20
30
40
50
60
Normal
MK50
MK75
MK88
MK94
MK97
MK data
Normal data

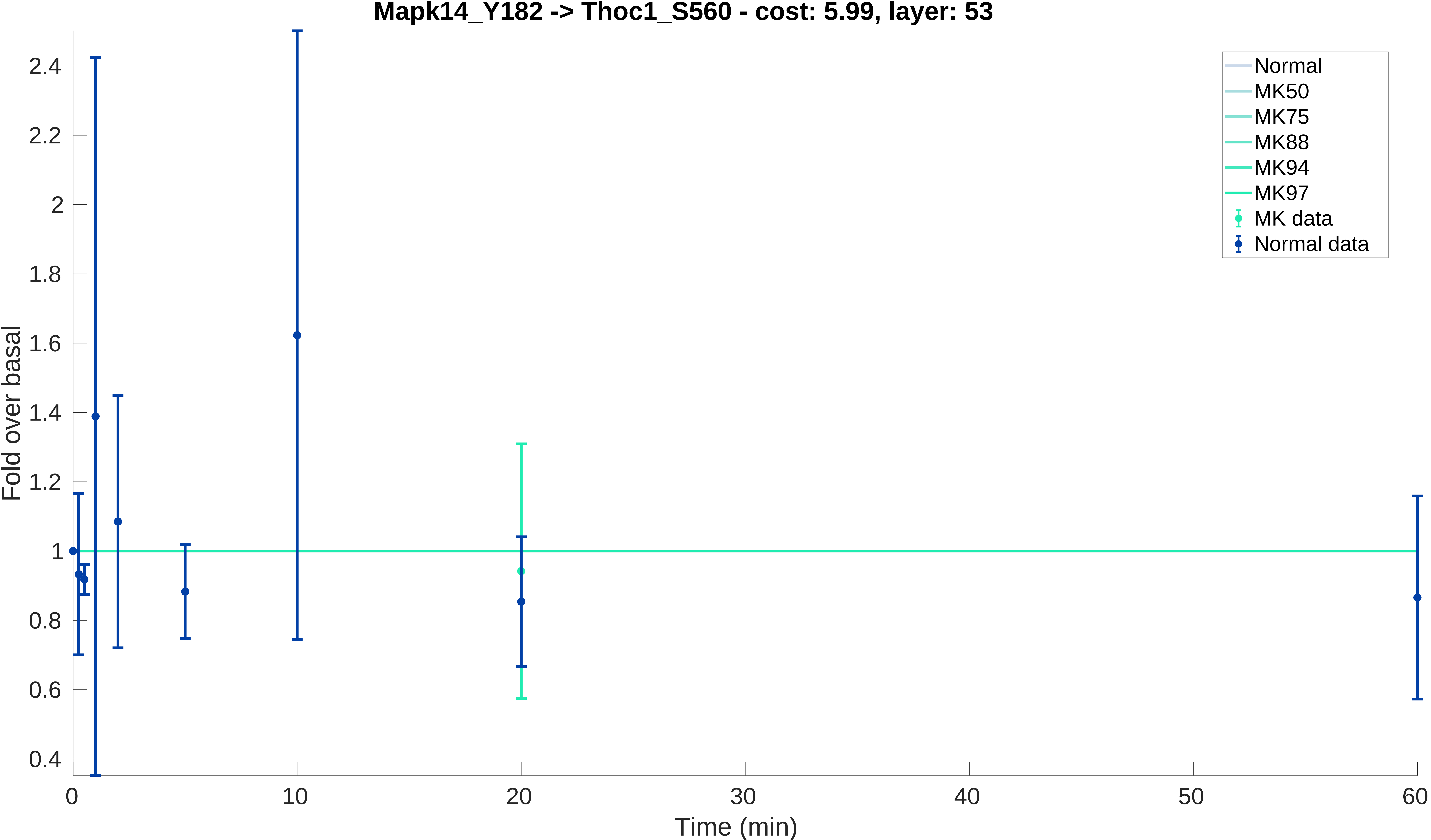

Mapk14_Y182 -> Thoc1_S560 - cost: 5.99, layer: 53
Fold over basal
Time (min)
0.4
0.6
0.8
1
1.2
1.4
1.6
1.8
2
2.2
2.4
0
10
20
30
40
50
60
Normal
MK50
MK75
MK88
MK94
MK97
MK data
Normal data

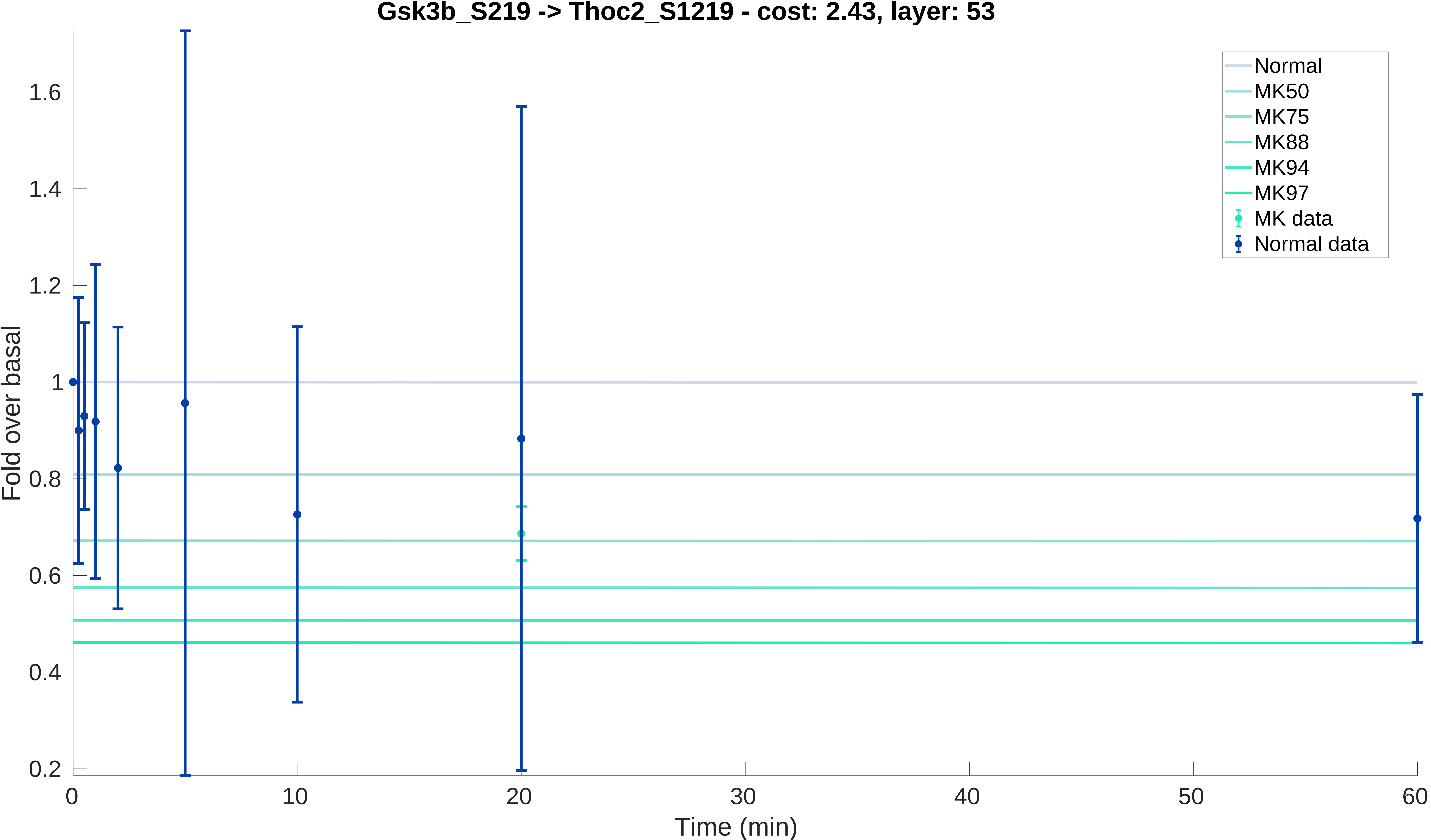

Gsk3b_S219 -> Thoc2_S1219 - cost: 2.43, layer: 53
Fold over basal
Time (min)
0.2
0.4
0.6
0.8
1
1.2
1.4
1.6
0
10
20
30
40
50
60
Normal
MK50
MK75
MK88
MK94
MK97
MK data
Normal data

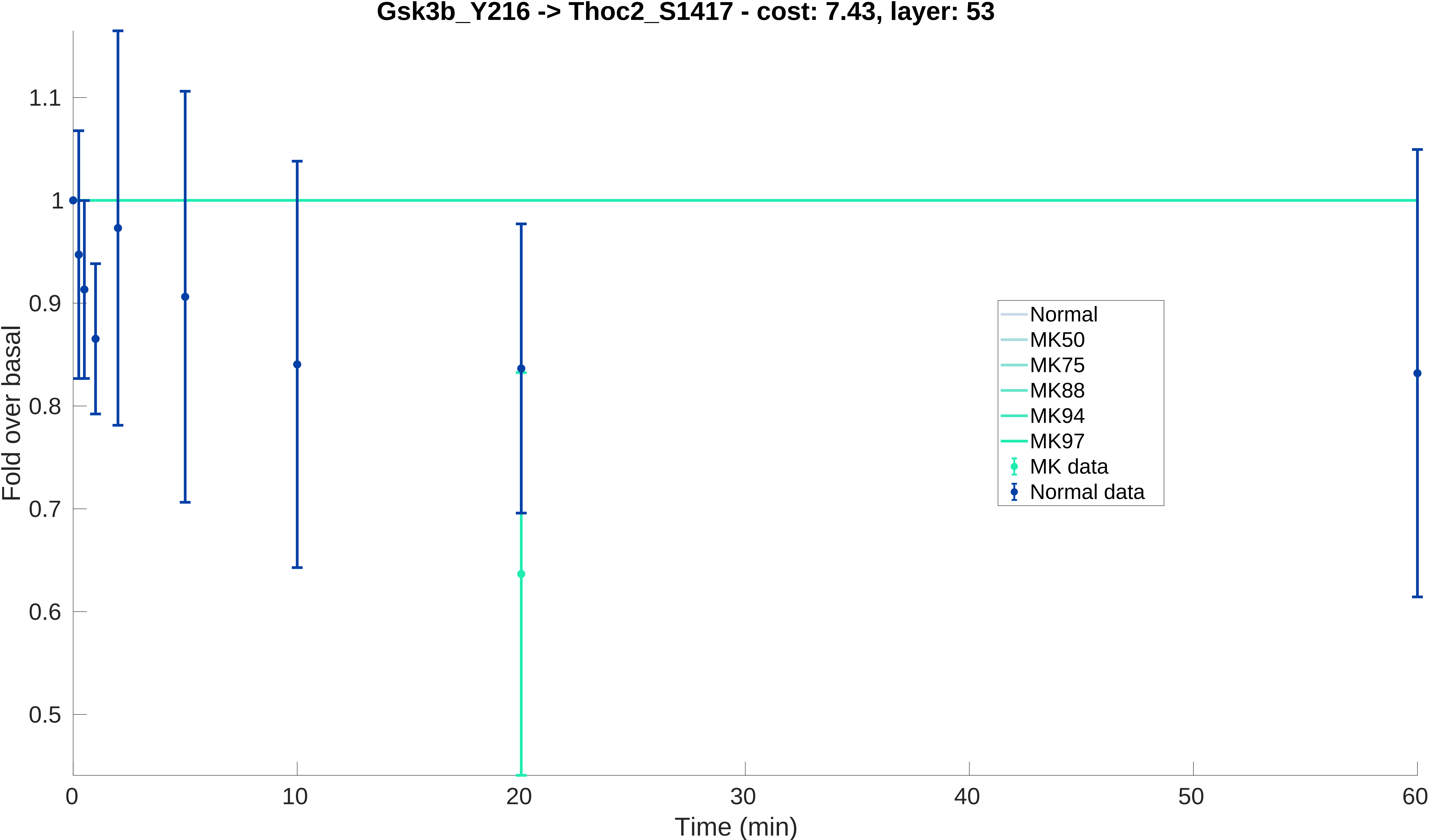

Gsk3b_Y216 -> Thoc2_S1417 - cost: 7.43, layer: 53
Fold over basal
1.1
1
0.9
0.8
0.7
0.6
0.5
0
10
20
30
40
50
60
Time (min)
Normal
MK50
MK75
MK88
MK94
MK97
MK data
Normal data

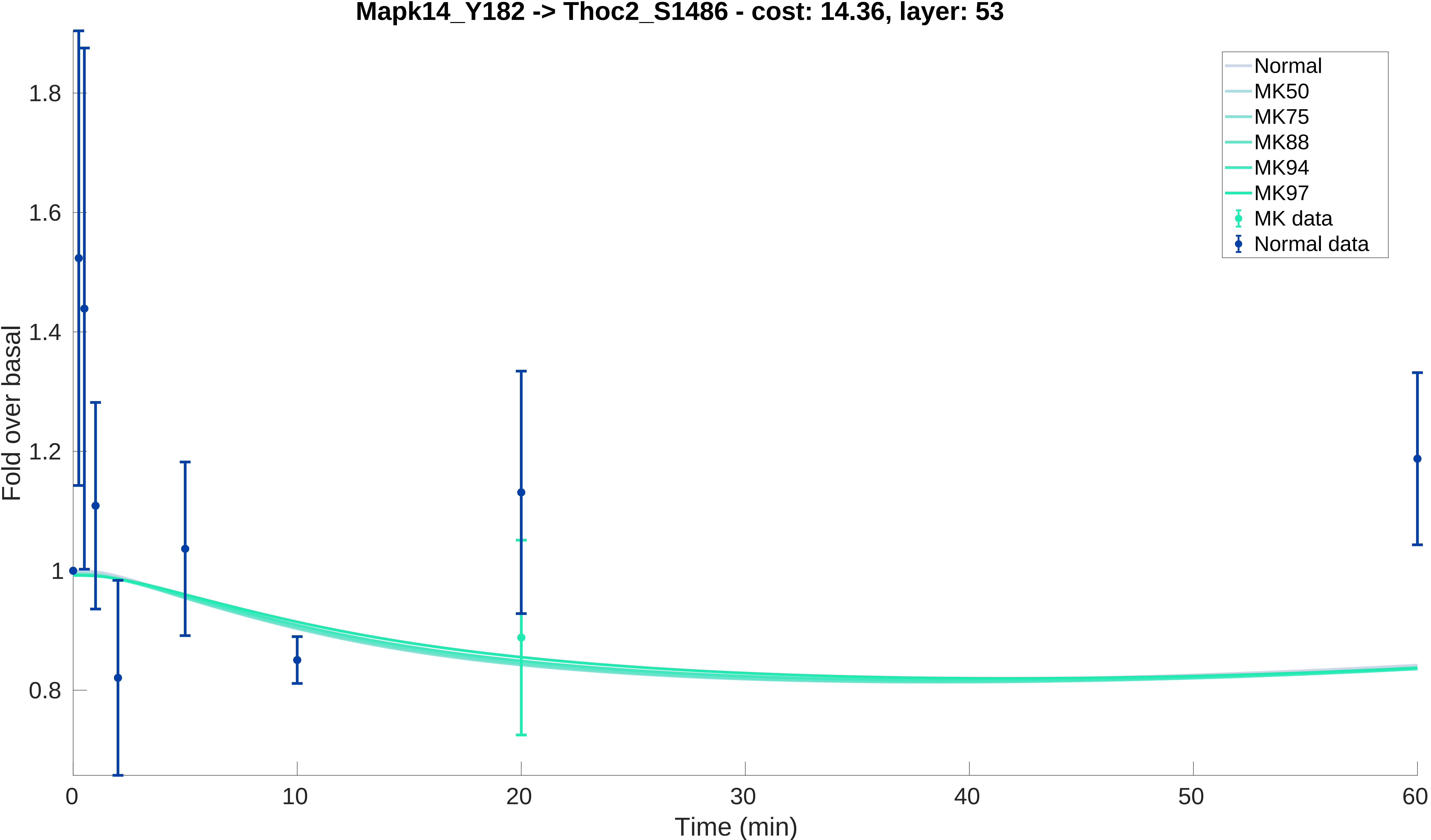

Mapk14_Y182 -> Thoc2_S1486 - cost: 14.36, layer: 53
Fold over basal
1.8
1.6
1.4
1.2
1
0.8
0
10
20
30
40
50
60
Time (min)
Normal
MK50
MK75
MK88
MK94
MK97
MK data
Normal data

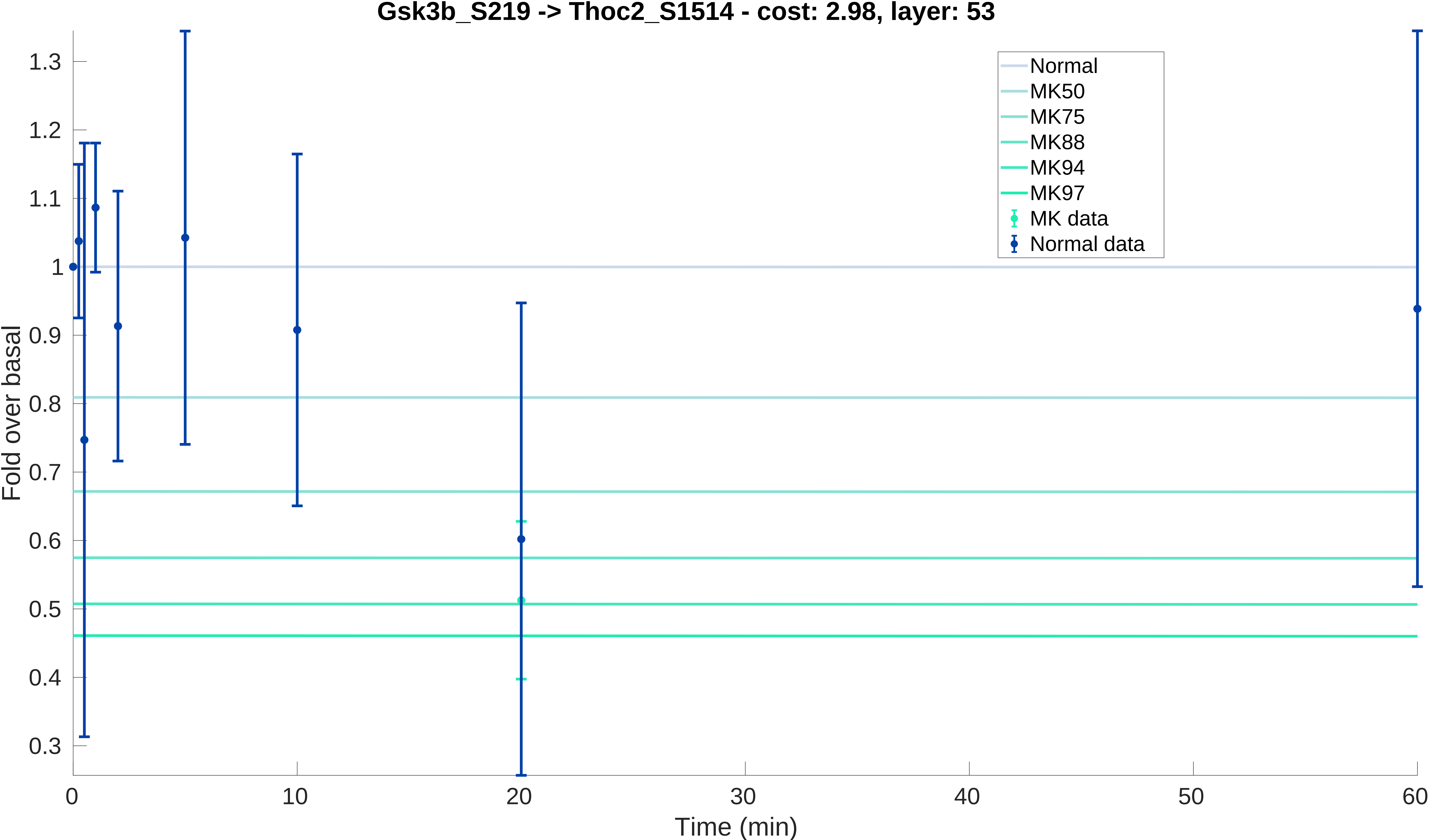

Gsk3b_S219 -> Thoc2_S1514 - cost: 2.98, layer: 53
Normal
MK50
MK75
MK88
MK94
MK97
MK data
Normal data
Fold over basal
Time (min)
1.3
1.2
1.1
1
0.9
0.8
0.7
0.6
0.5
0.4
0.3
0
10
20
30
40
50
60

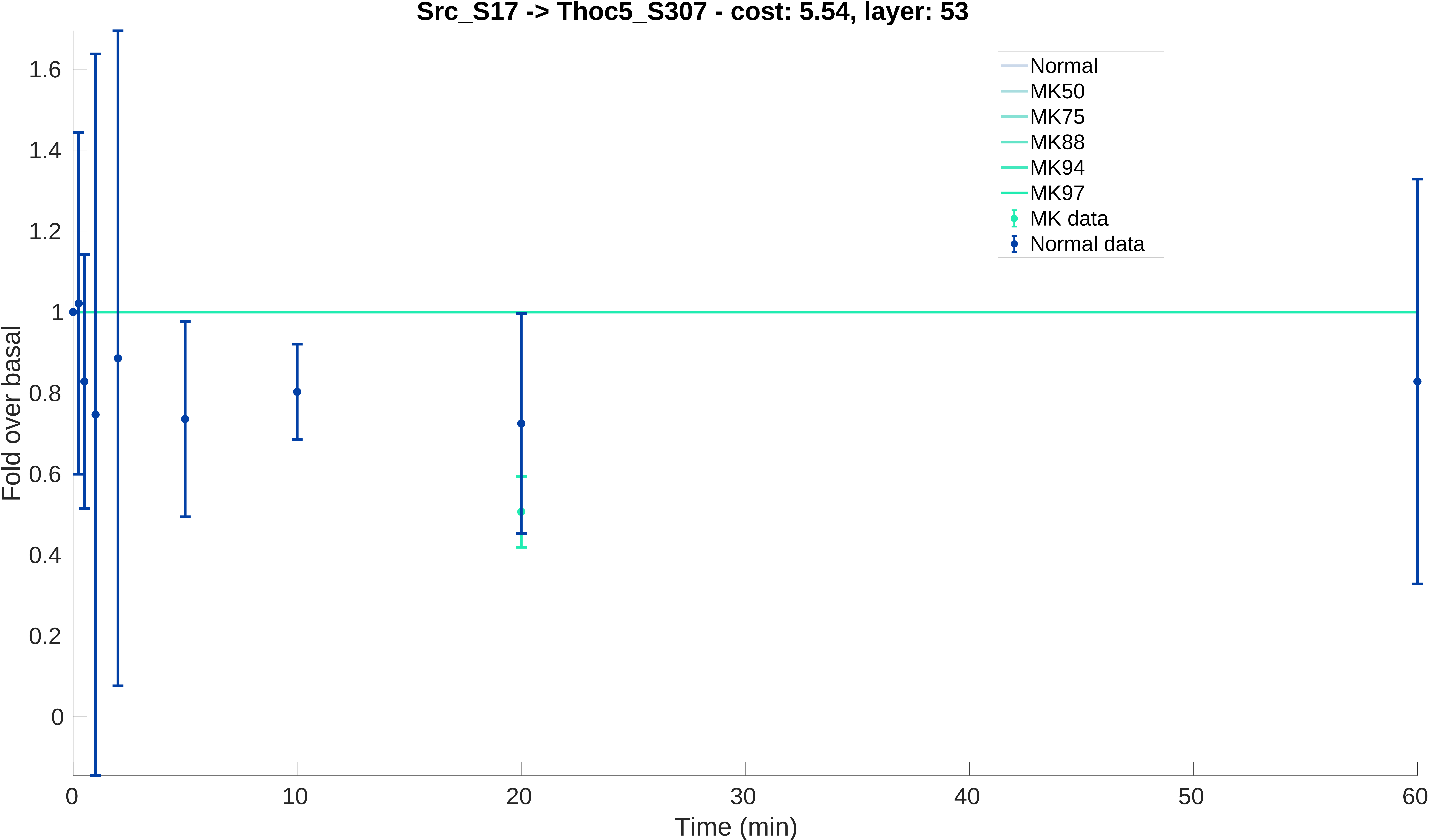

Src_S17 -> Thoc5_S307 - cost: 5.54, layer: 53
Fold over basal
Time (min)
Normal
MK50
MK75
MK88
MK94
MK97
MK data
Normal data
0
0.2
0.4
0.6
0.8
1
1.2
1.4
1.6
0
10
20
30
40
50
60

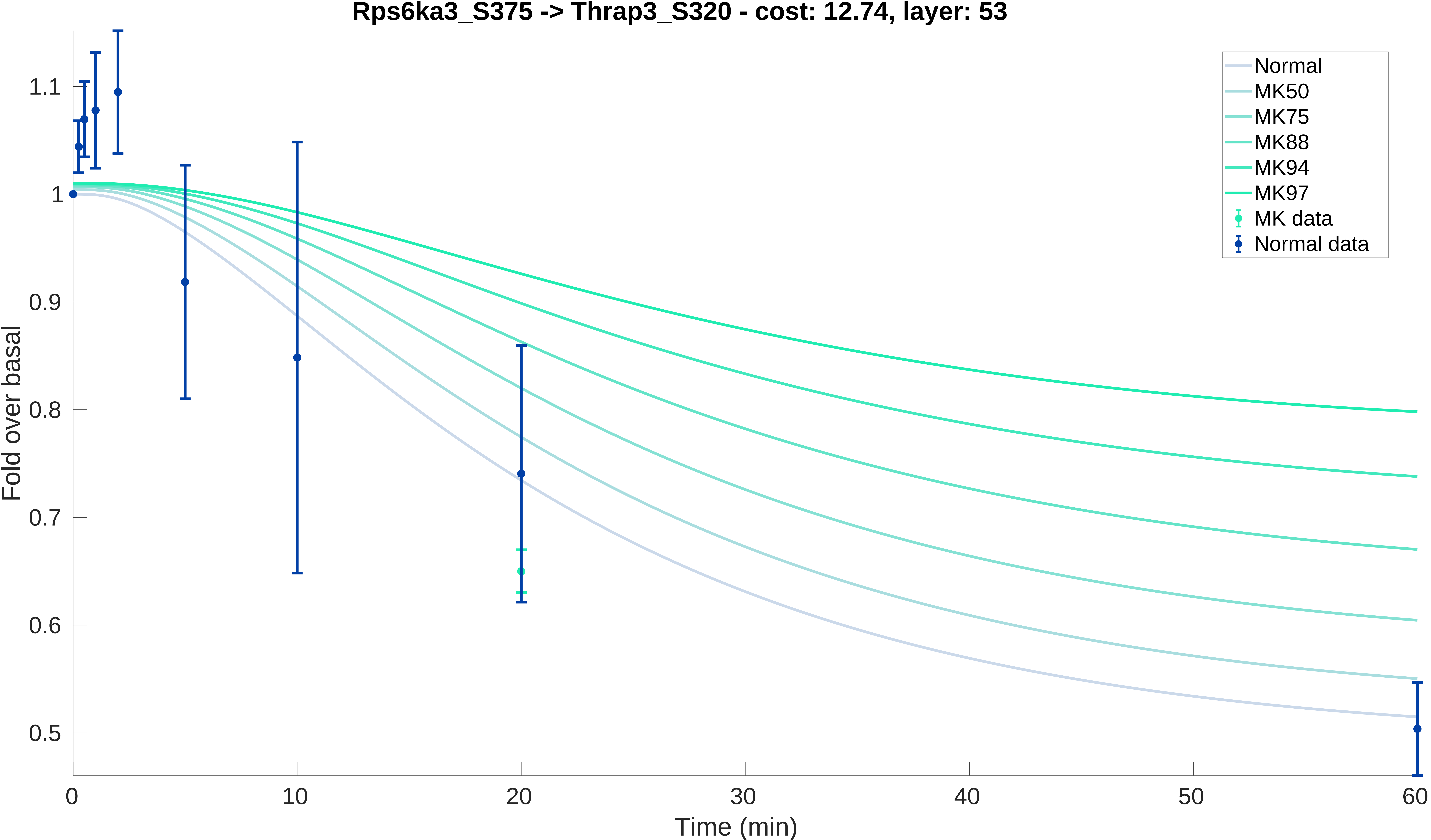

Rps6ka3_S375 -> Thrap3_S320 - cost: 12.74, layer: 53
Fold over basal
Time (min)
0.5
0.6
0.7
0.8
0.9
1
1.1
0
10
20
30
40
50
60
Normal
MK50
MK75
MK88
MK94
MK97
MK data
Normal data

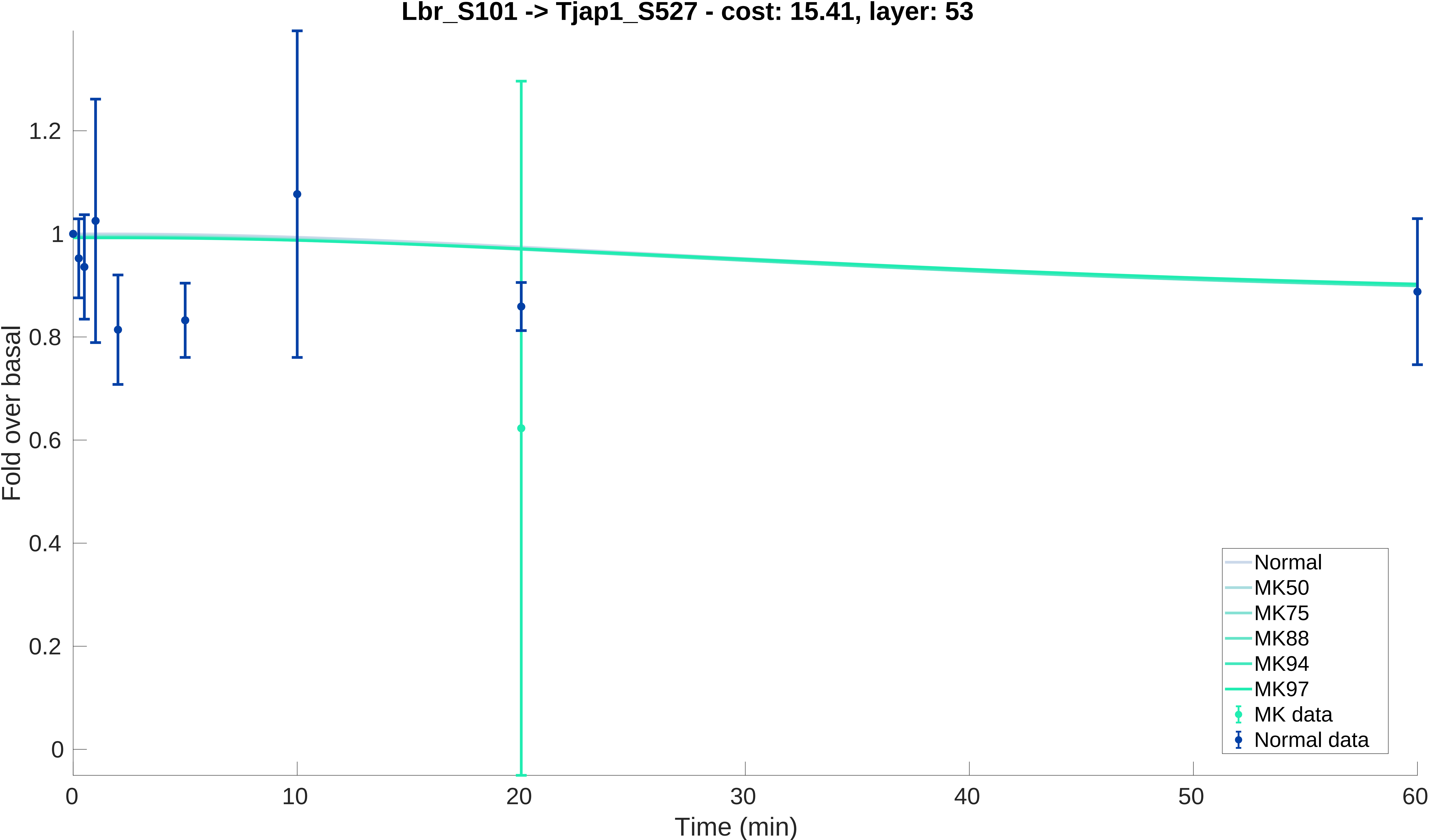

Lbr_S101 -> Tjap1_S527 - cost: 15.41, layer: 53
Fold over basal
1.2
1
0.8
0.6
0.4
0.2
0
0
10
20
30
40
50
60
Time (min)
Normal
MK50
MK75
MK88
MK94
MK97
MK data
Normal data

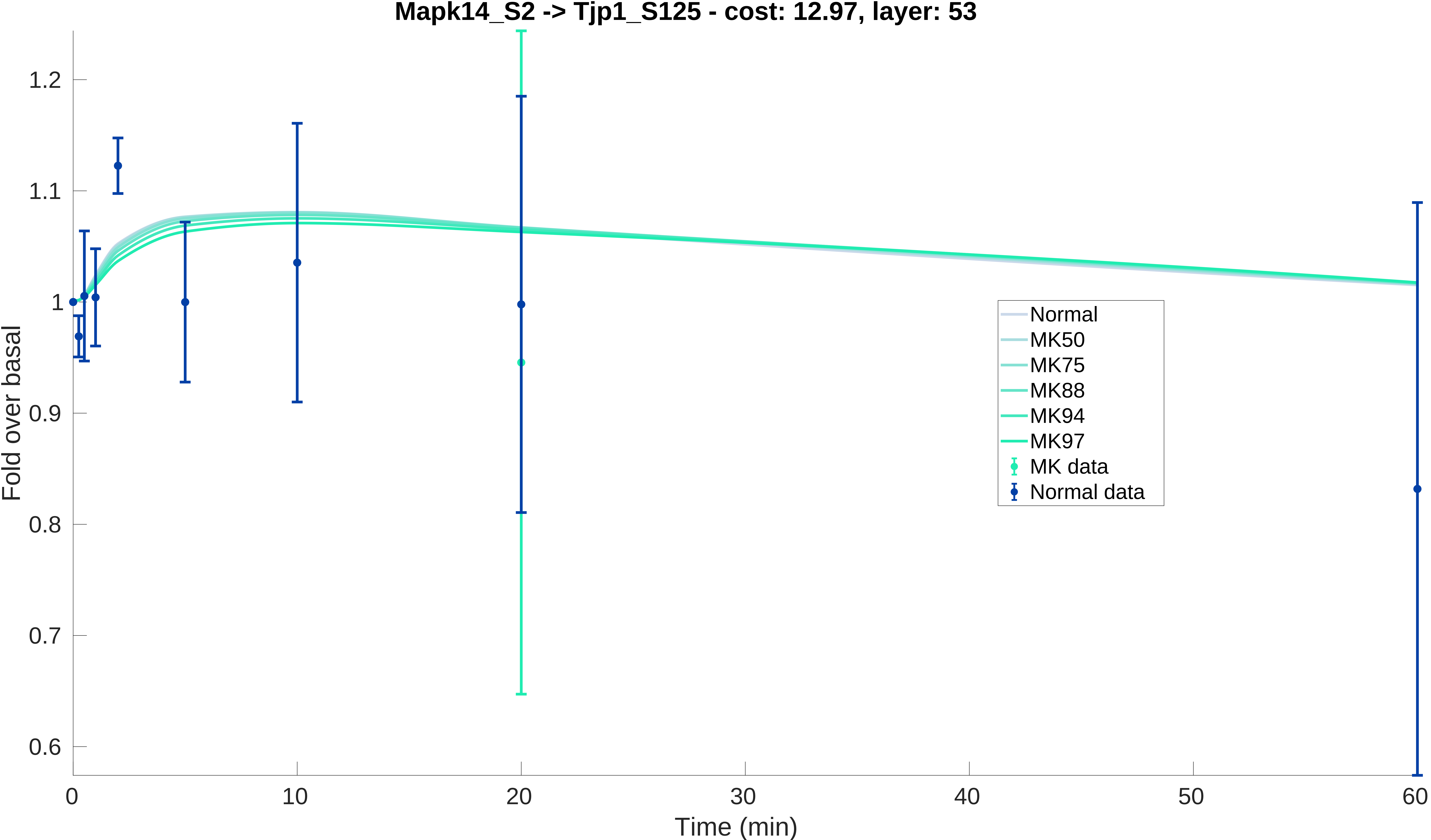

Mapk14_S2 -> Tjp1_S125 - cost: 12.97, layer: 53
Fold over basal
Time (min)
0.6
0.7
0.8
0.9
1
1.1
1.2
0
10
20
30
40
50
60
Normal
MK50
MK75
MK88
MK94
MK97
MK data
Normal data

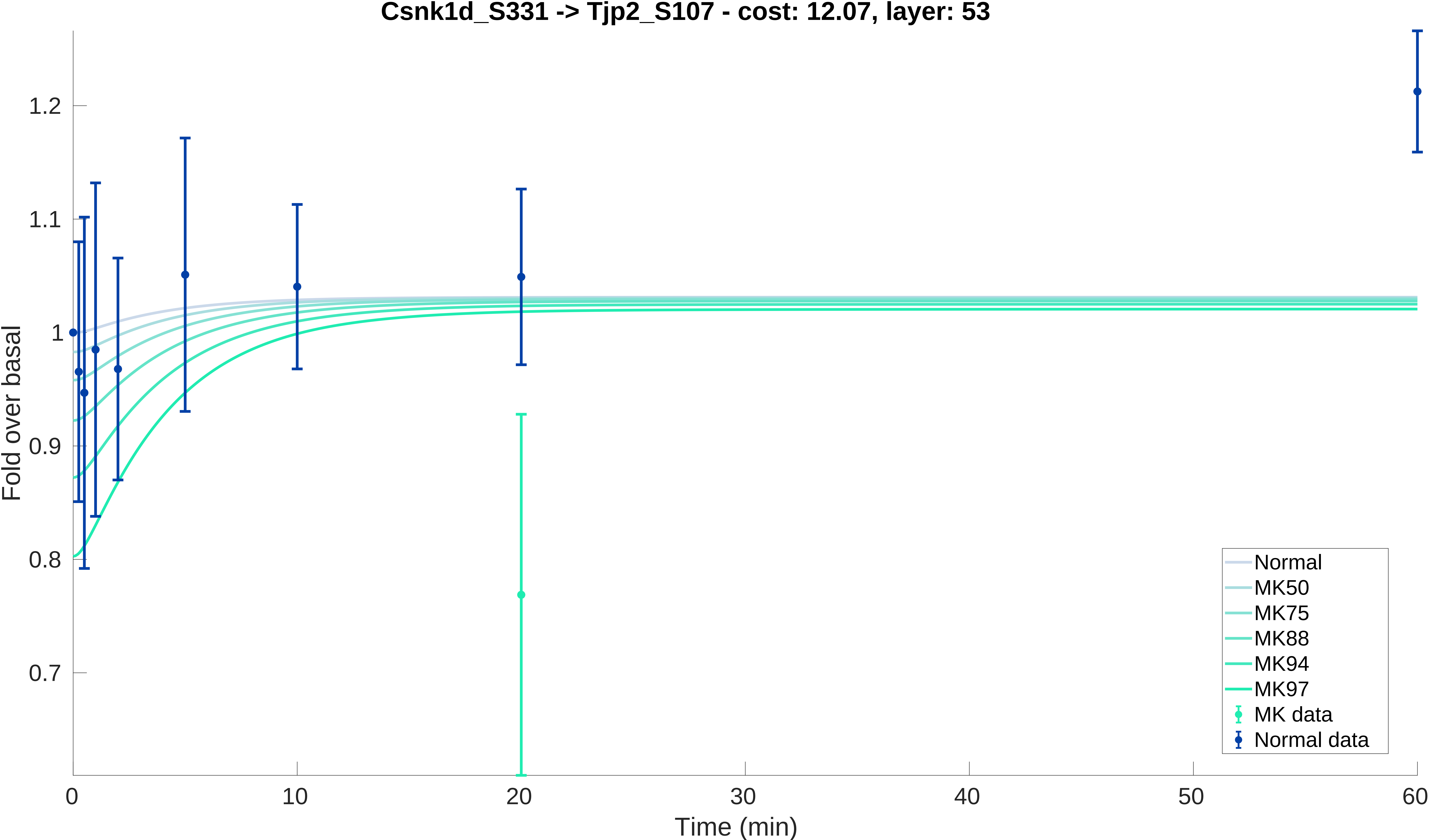
Csnk1d_S331 -> Tjp2_S107 - cost: 12.07, layer: 53
Fold over basal
1.2
1.1
1
0.9
0.8
0.7
0
10
20
30
40
50
60
Time (min)
Normal
MK50
MK75
MK88
MK94
MK97
MK data
Normal data

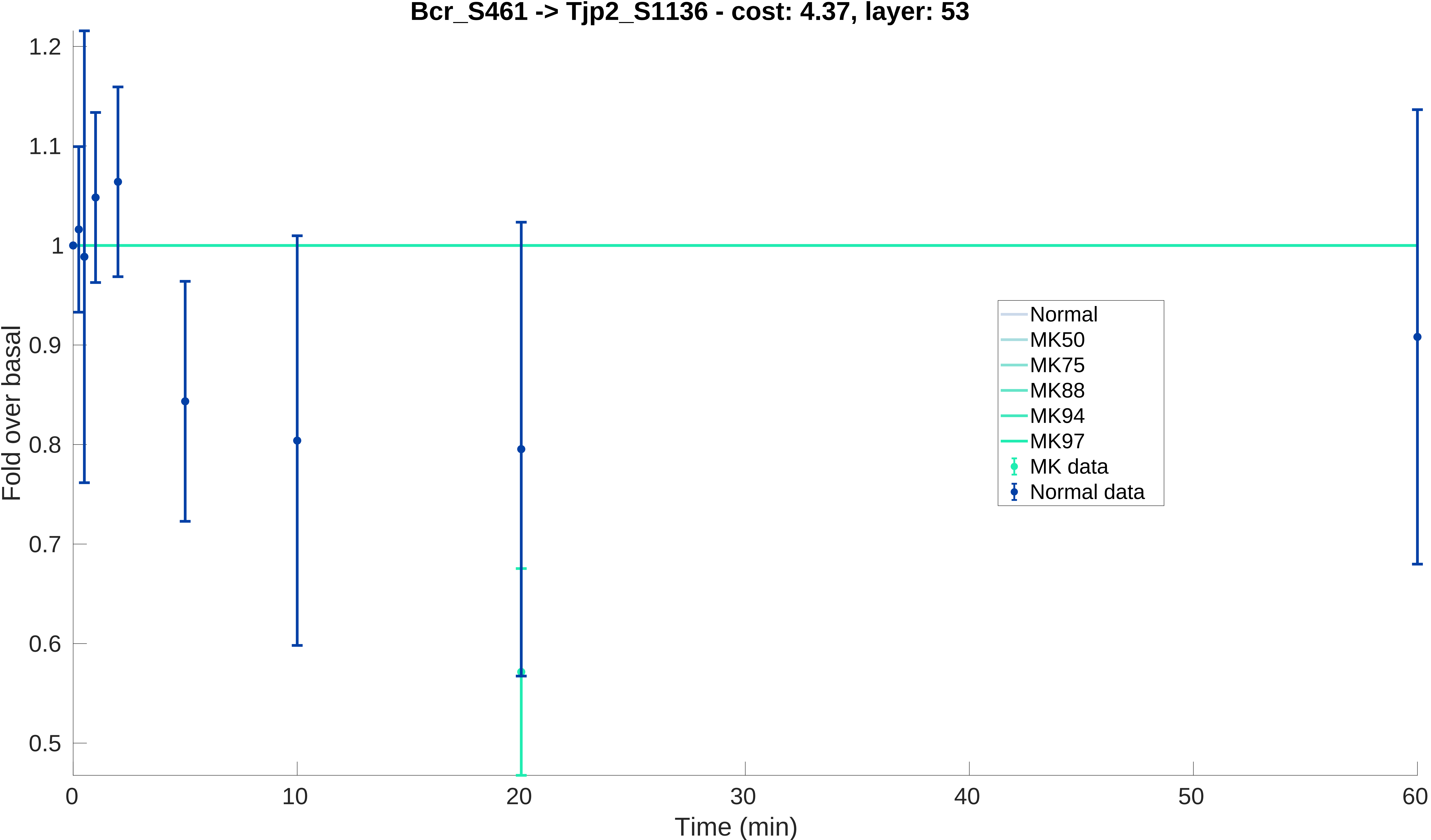

Bcr_S461 -> Tjp2_S1136 - cost: 4.37, layer: 53
Fold over basal
Time (min)
0.5
0.6
0.7
0.8
0.9
1
1.1
1.2
0
10
20
30
40
50
60
Normal
MK50
MK75
MK88
MK94
MK97
MK data
Normal data

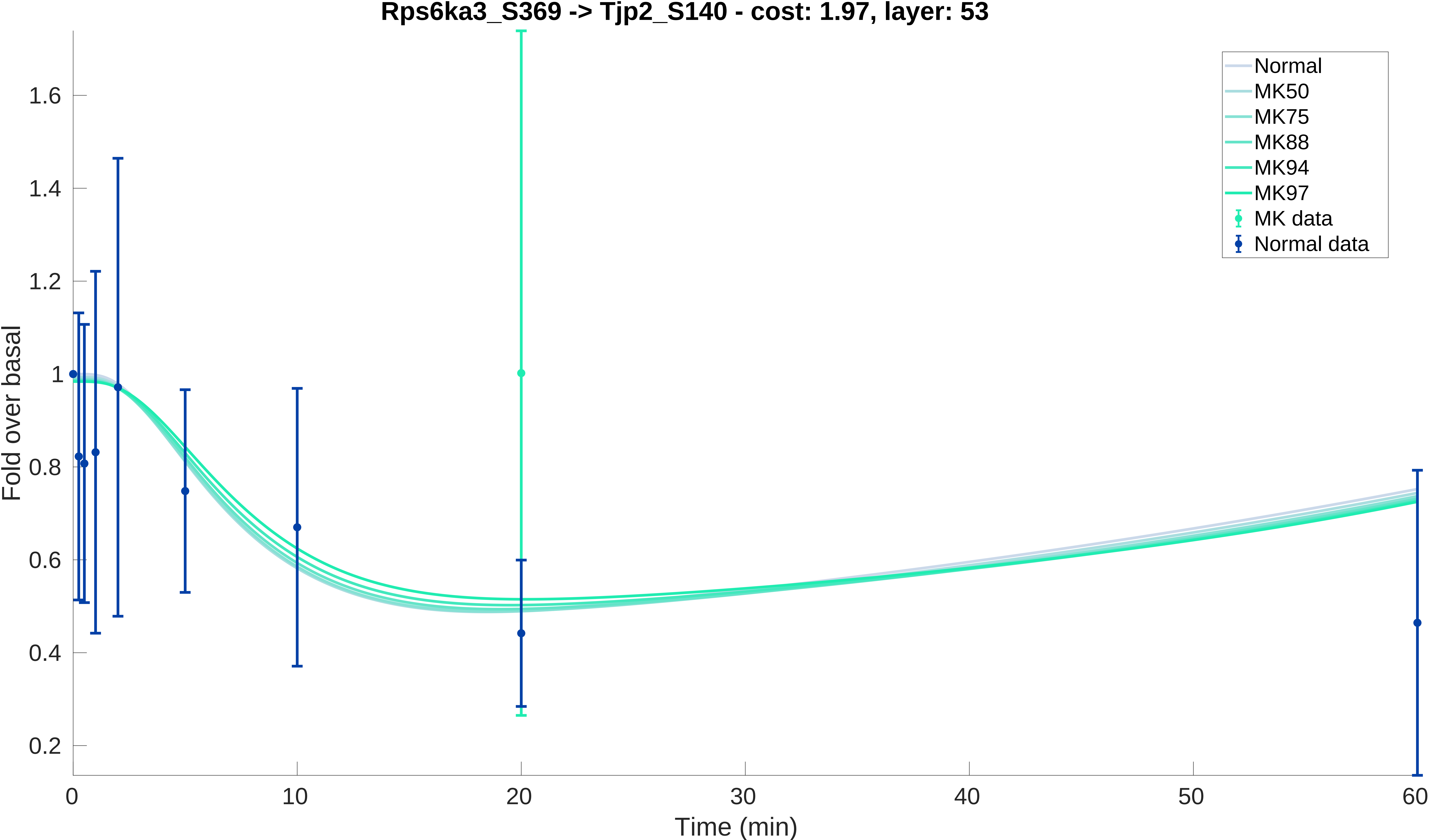

Rps6ka3_S369 -> Tjp2_S140 - cost: 1.97, layer: 53
Fold over basal
Time (min)
0.2
0.4
0.6
0.8
1
1.2
1.4
1.6
0
10
20
30
40
50
60
Normal
MK50
MK75
MK88
MK94
MK97
MK data
Normal data

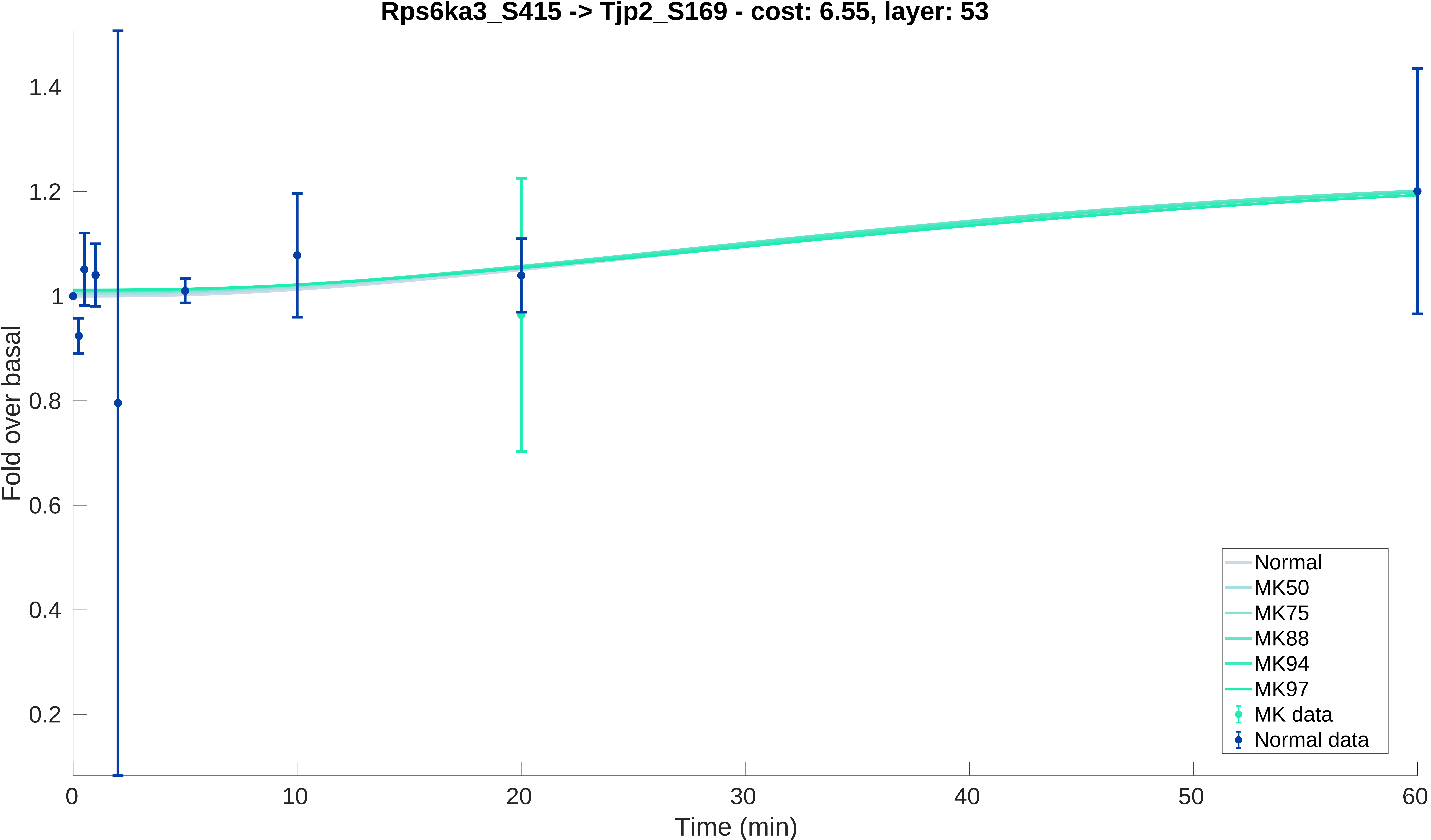

Rps6ka3_S415 -> Tjp2_S169 - cost: 6.55, layer: 53
Fold over basal
1.4
1.2
1
0.8
0.6
0.4
0.2
0
10
20
30
40
50
60
Time (min)
Normal
MK50
MK75
MK88
MK94
MK97
MK data
Normal data

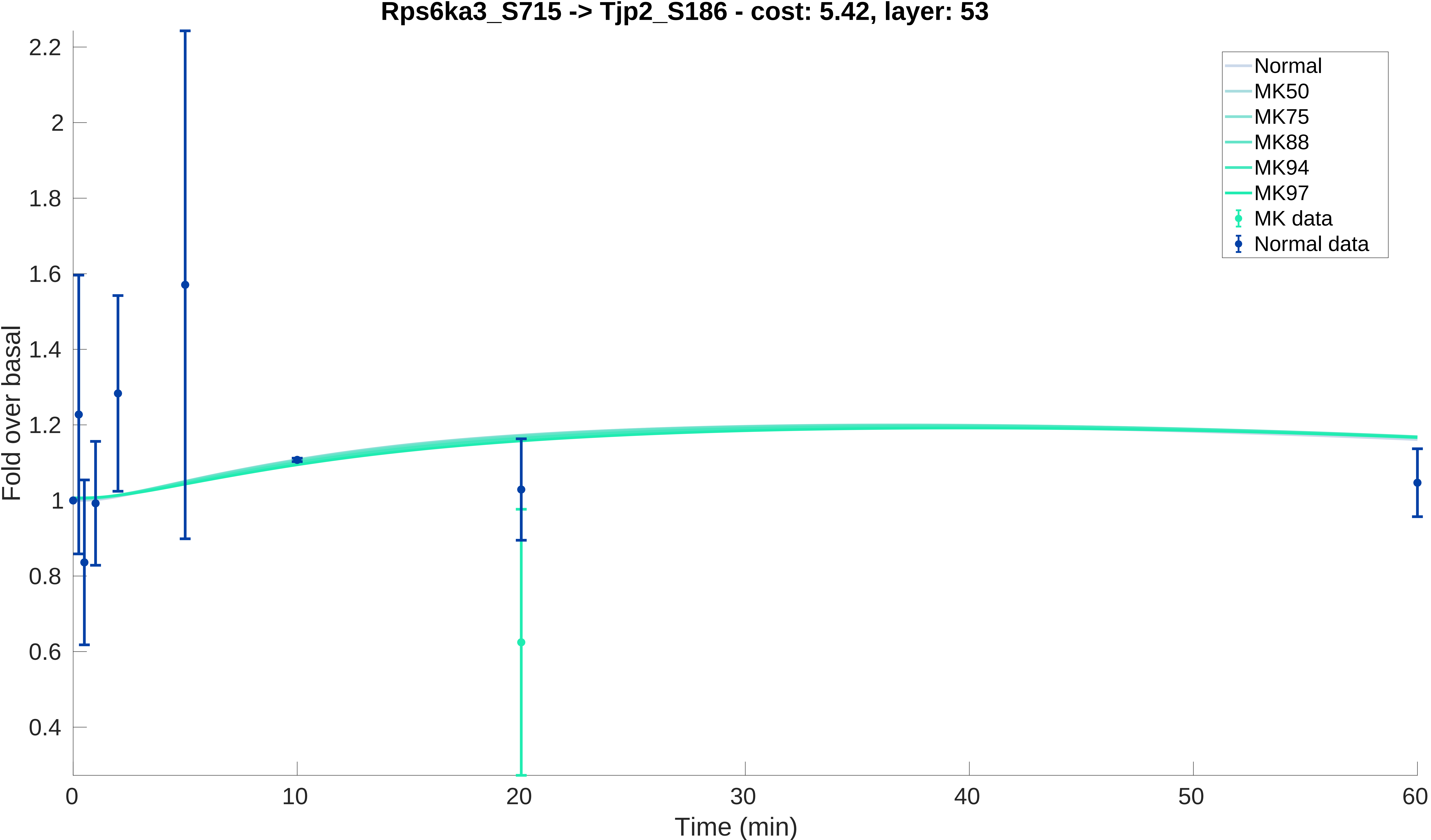

Rps6ka3_S715 -> Tjp2_S186 - cost: 5.42, layer: 53
Fold over basal
Time (min)
Normal
MK50
MK75
MK88
MK94
MK97
MK data
Normal data
0.4
0.6
0.8
1
1.2
1.4
1.6
1.8
2
2.2
0
10
20
30
40
50
60

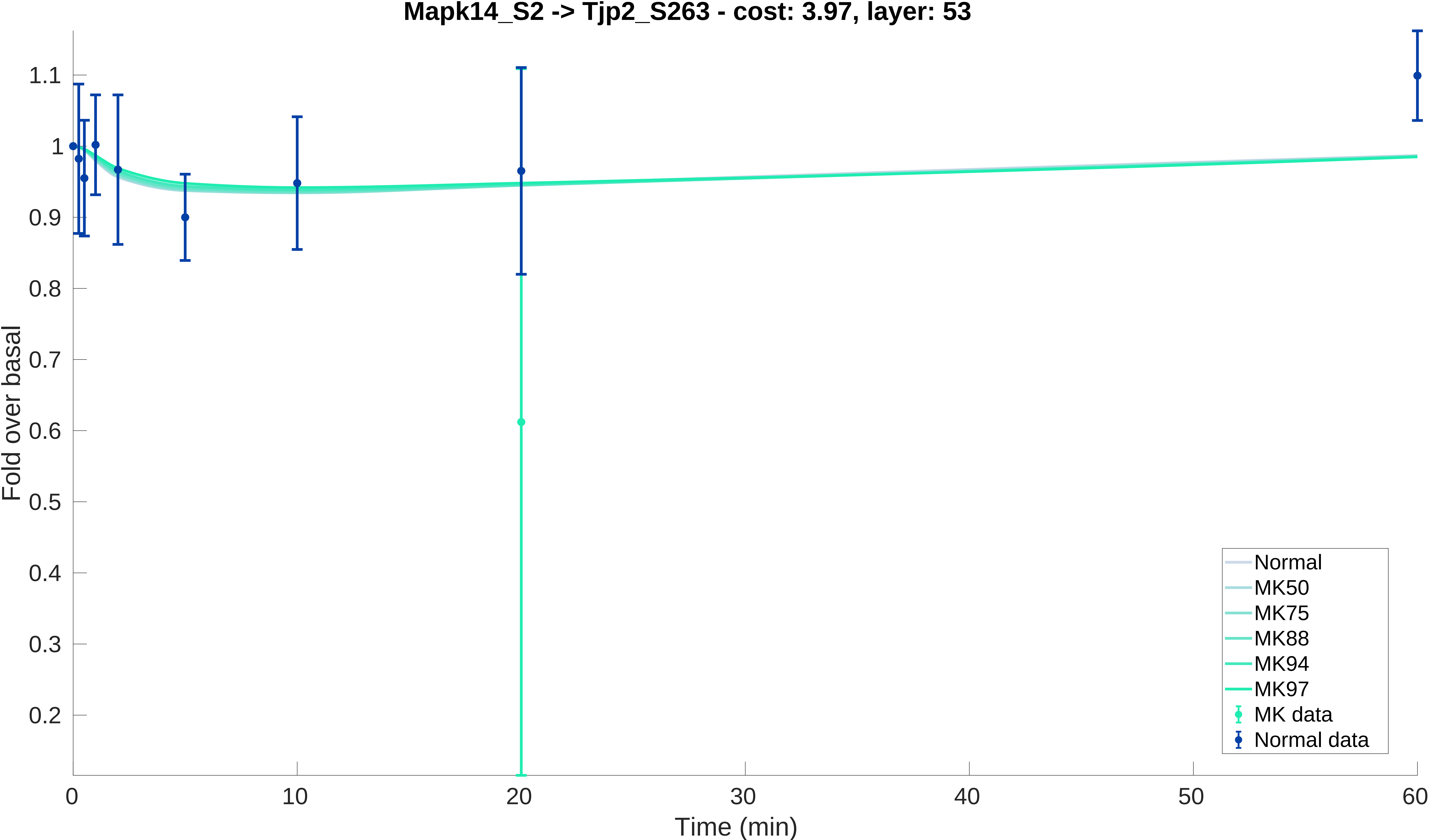

Mapk14_S2 -> Tjp2_S263 - cost: 3.97, layer: 53
Fold over basal
Time (min)
1.1
1
0.9
0.8
0.7
0.6
0.5
0.4
0.3
0.2
0
10
20
30
40
50
60
Normal
MK50
MK75
MK88
MK94
MK97
MK data
Normal data

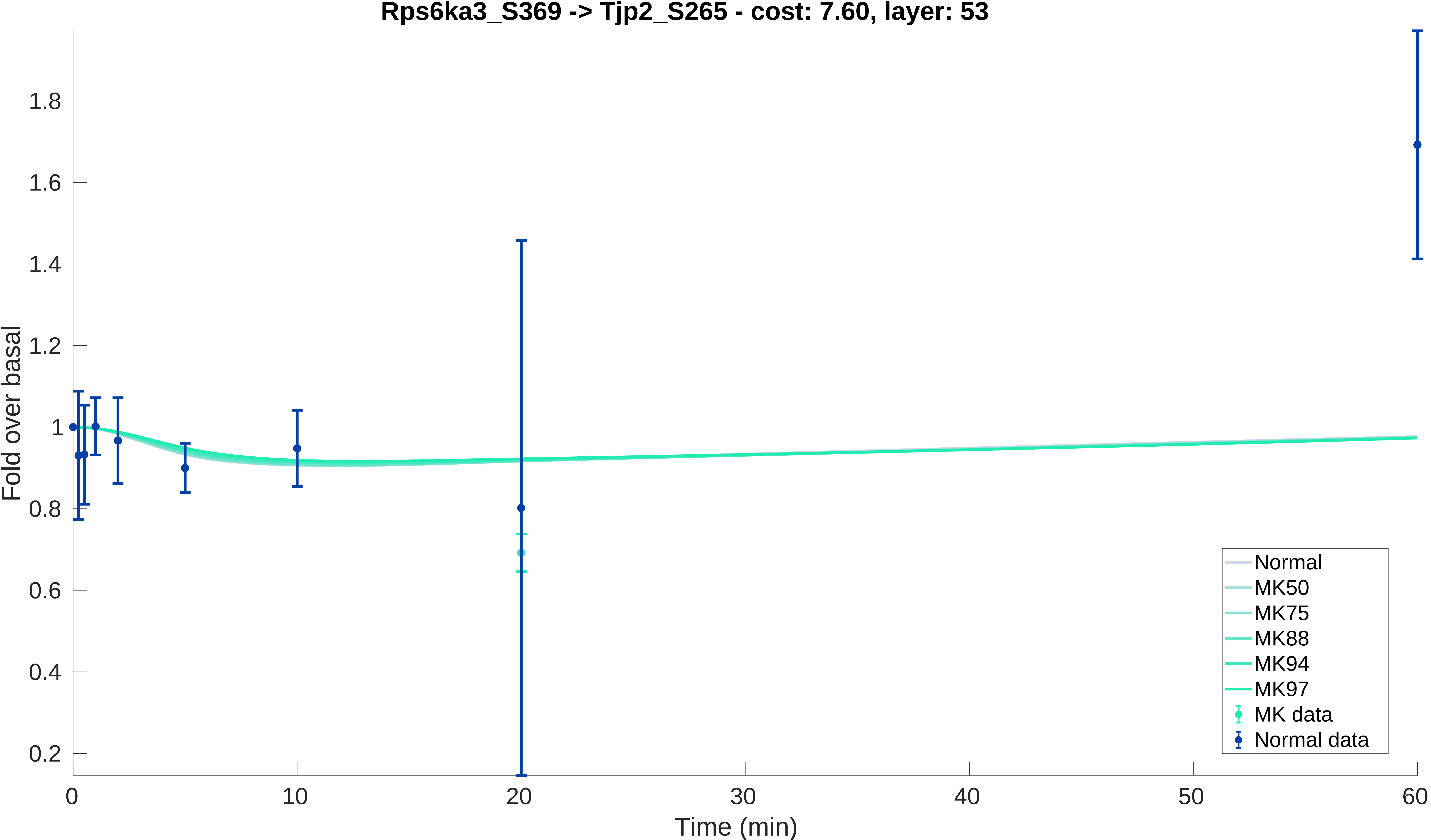

Rps6ka3_S369 -> Tjp2_S265 - cost: 7.60, layer: 53
Fold over basal
1.8
1.6
1.4
1.2
1
0.8
0.6
0.4
0.2
0
10
20
30
40
50
60
Time (min)
Normal
MK50
MK75
MK88
MK94
MK97
MK data
Normal data

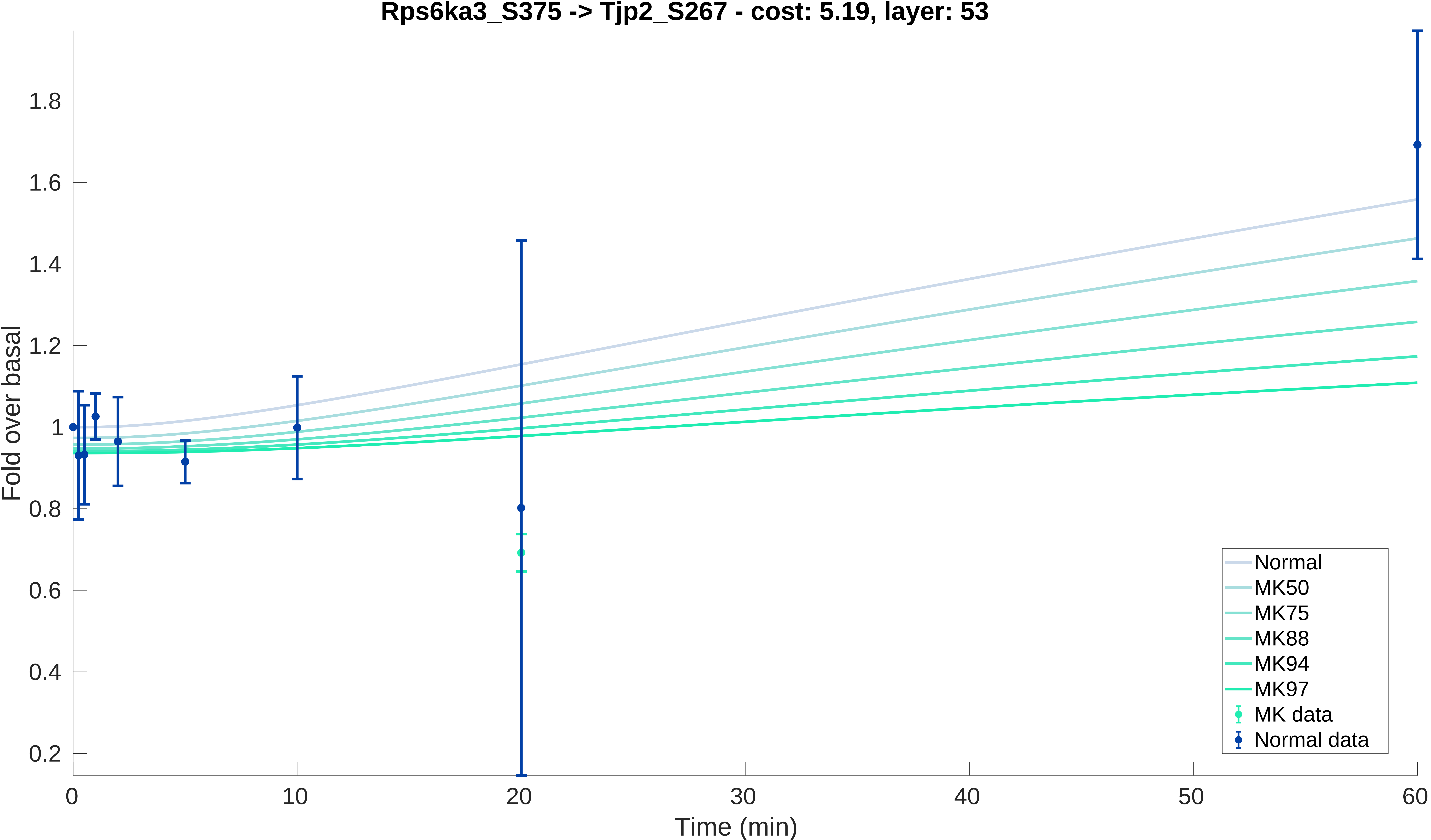

Rps6ka3_S375 -> Tjp2_S267 - cost: 5.19, layer: 53
Fold over basal
1.8
1.6
1.4
1.2
1
0.8
0.6
0.4
0.2
0
10
20
30
40
50
60
Time (min)
Normal
MK50
MK75
MK88
MK94
MK97
MK data
Normal data

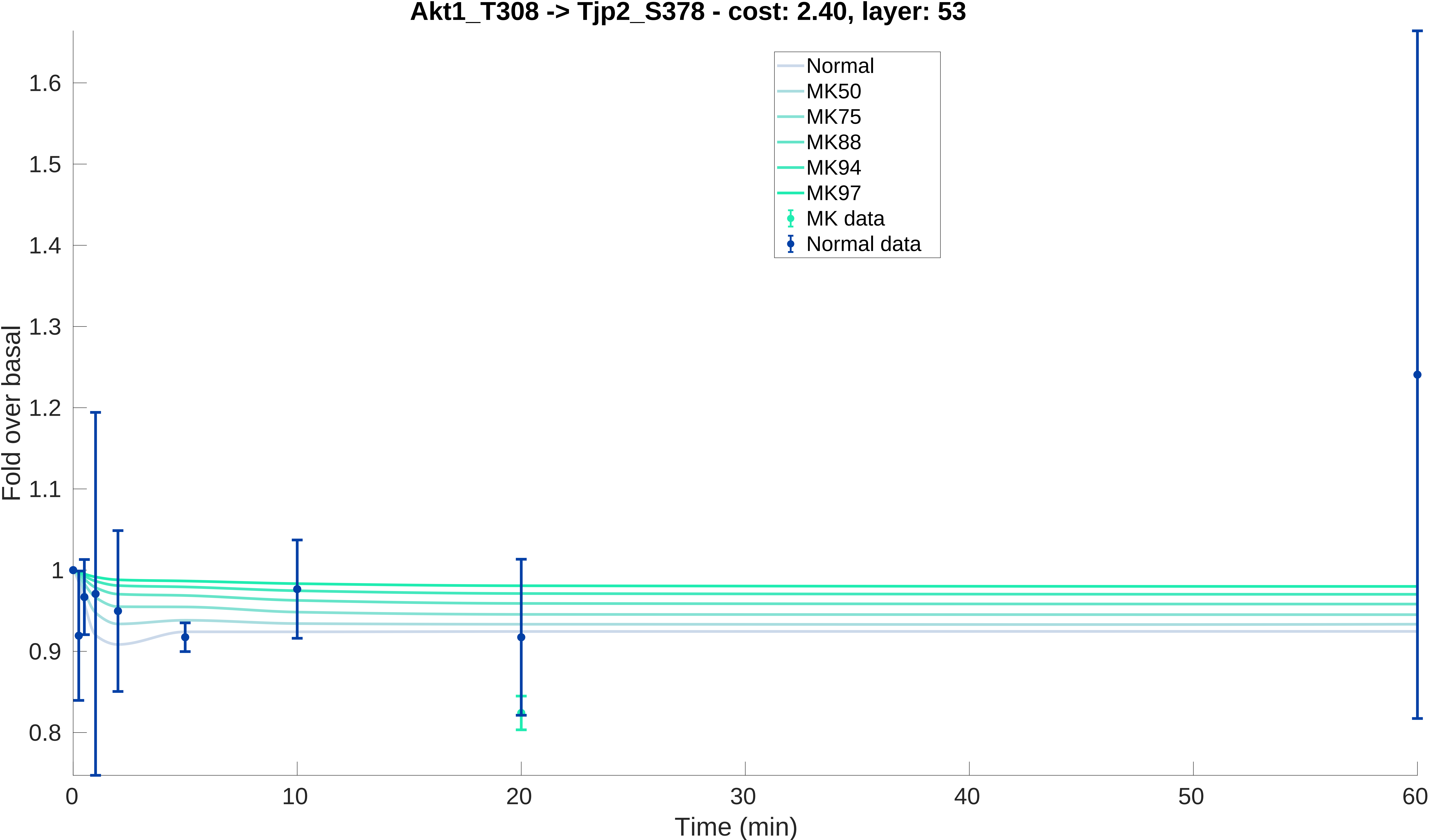

Akt1_T308 -> Tjp2_S378 - cost: 2.40, layer: 53
Fold over basal
Time (min)
Normal
MK50
MK75
MK88
MK94
MK97
MK data
Normal data

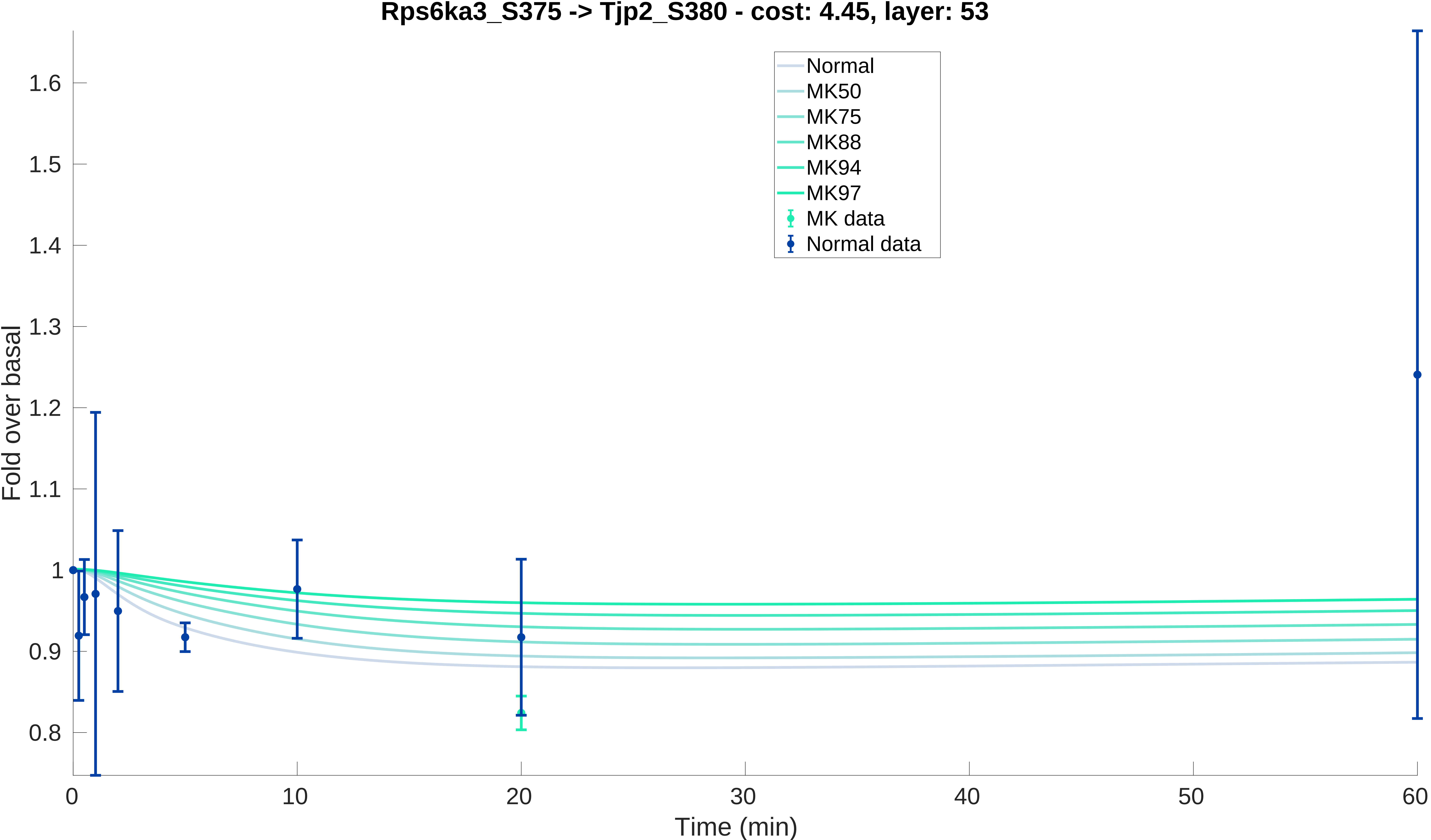
Rps6ka3_S375 -> Tjp2_S380 - cost: 4.45, layer: 53
Fold over basal
Time (min)
0.8
0.9
1
1.1
1.2
1.3
1.4
1.5
1.6
0
10
20
30
40
50
60
Normal
MK50
MK75
MK88
MK94
MK97
MK data
Normal data

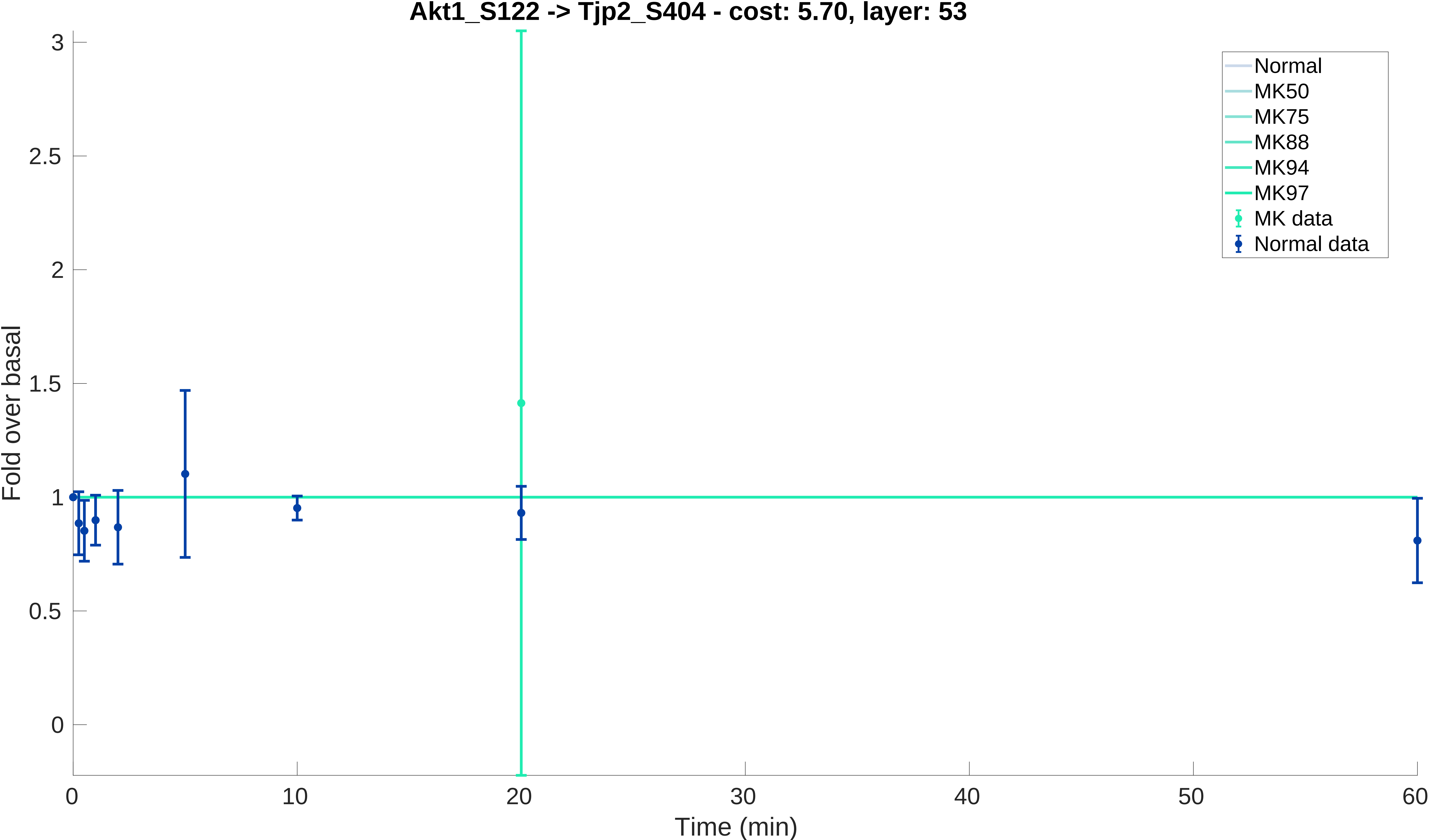

Akt1_S122 -> Tjp2_S404 - cost: 5.70, layer: 53
Fold over basal
Time (min)
0
0.5
1
1.5
2
2.5
3
0
10
20
30
40
50
60
Normal
MK50
MK75
MK88
MK94
MK97
MK data
Normal data

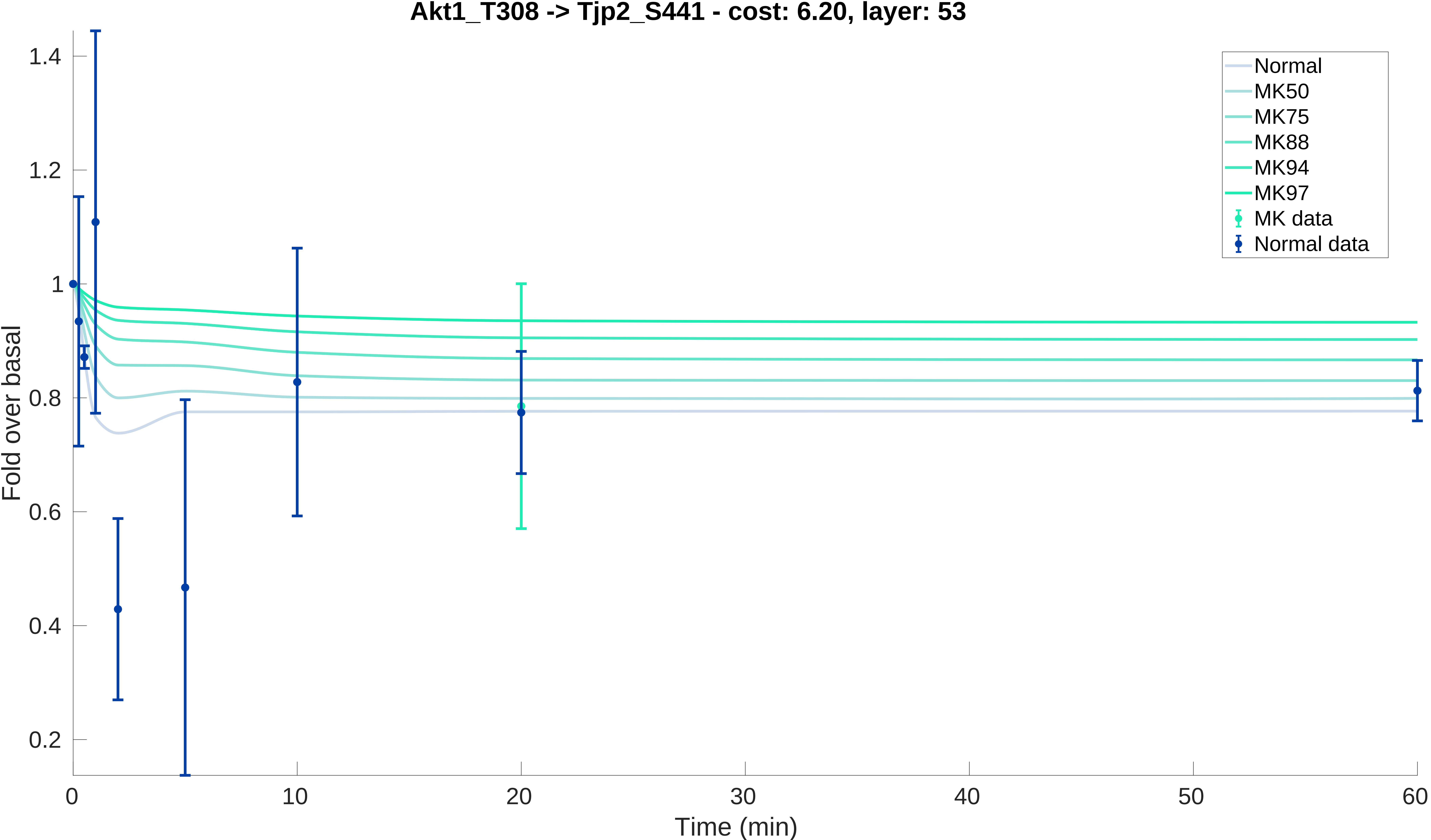

Akt1_T308 -> Tjp2_S441 - cost: 6.20, layer: 53
Normal
MK50
MK75
MK88
MK94
MK97
MK data
Normal data
Fold over basal
1.4
1.2
1
0.8
0.6
0.4
0.2
0
10
20
30
40
50
60
Time (min)

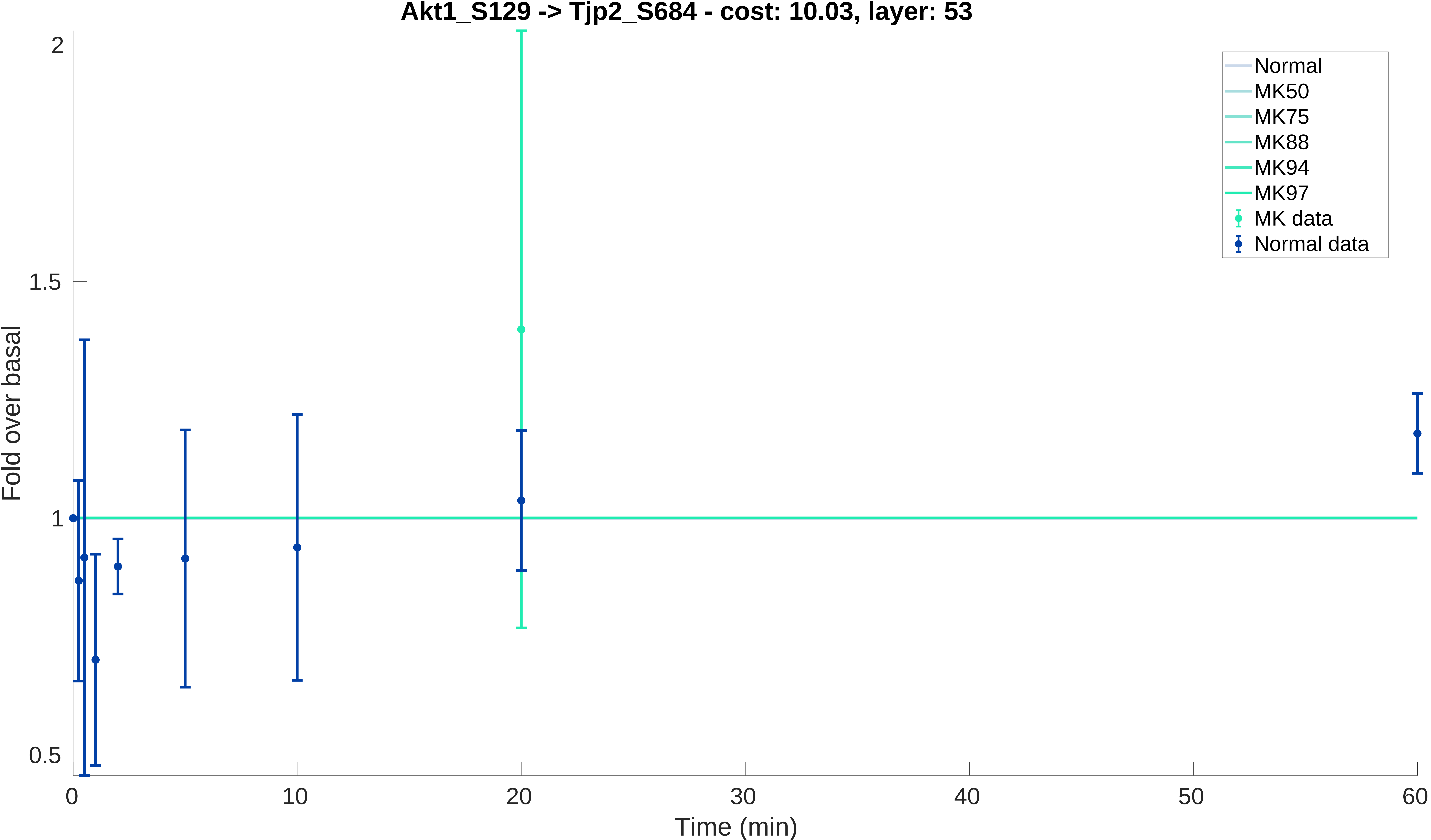

Akt1_S129 -> Tjp2_S684 - cost: 10.03, layer: 53
Normal
MK50
MK75
MK88
MK94
MK97
MK data
Normal data
Fold over basal
2
1.5
1
0.5
0
10
20
30
40
50
60
Time (min)

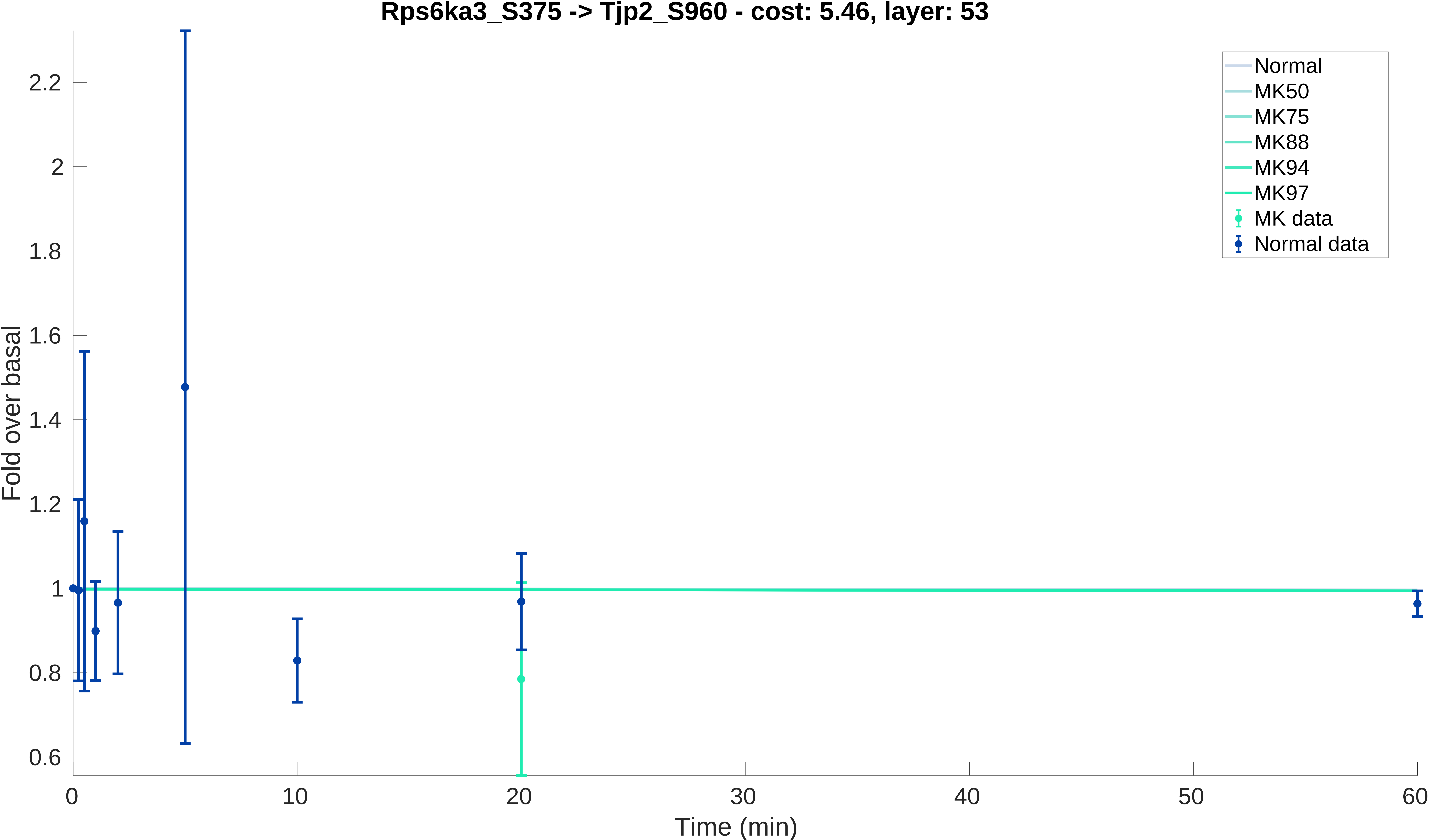

Rps6ka3_S375 -> Tjp2_S960 - cost: 5.46, layer: 53
Fold over basal
Time (min)
2.2
2
1.8
1.6
1.4
1.2
1
0.8
0.6
0
10
20
30
40
50
60
Normal
MK50
MK75
MK88
MK94
MK97
MK data
Normal data

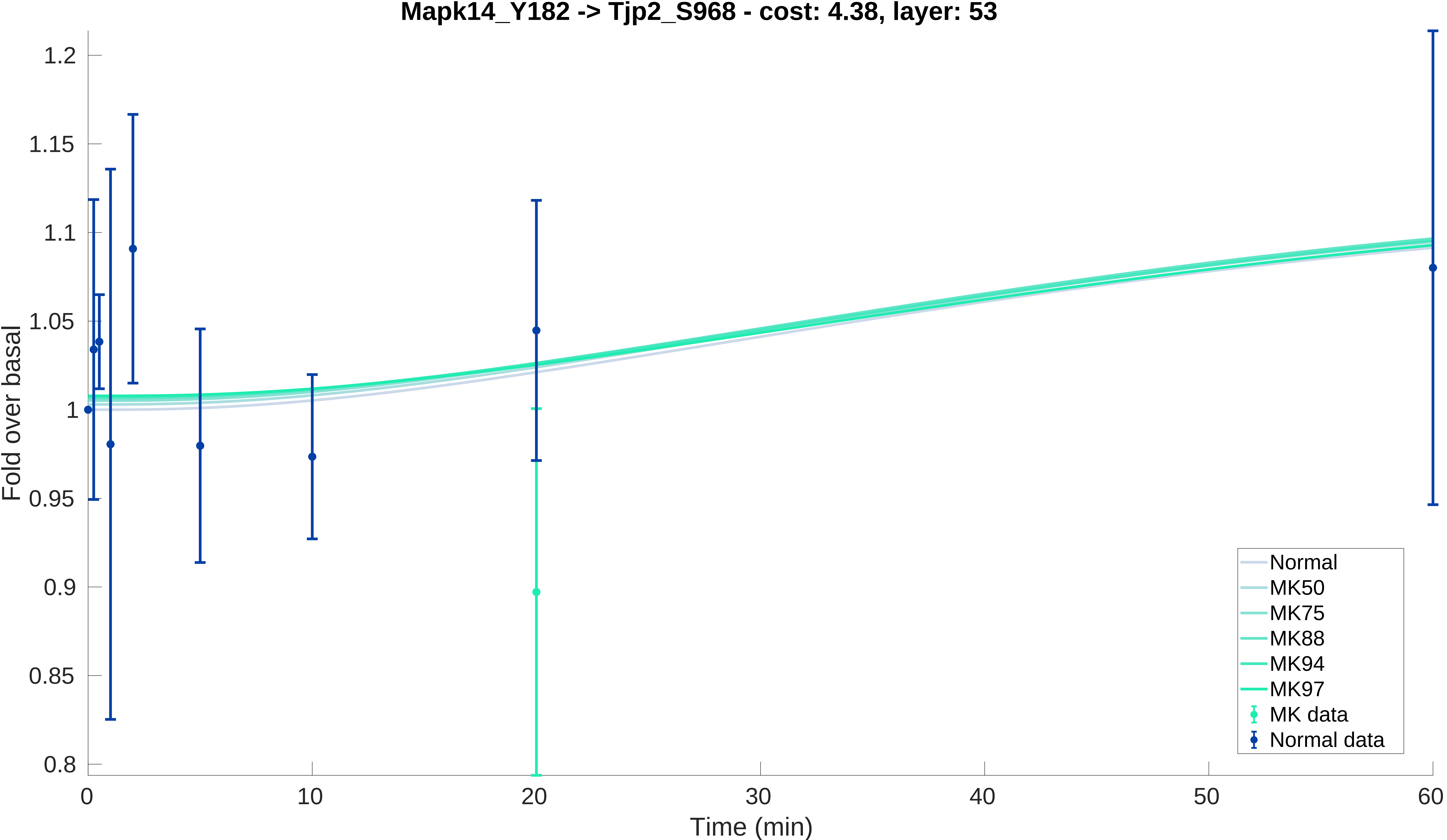
Mapk14_Y182 -> Tjp2_S968 - cost: 4.38, layer: 53
Fold over basal
Time (min)
1.2
1.15
1.1
1.05
1
0.95
0.9
0.85
0.8
0
10
20
30
40
50
60
Normal
MK50
MK75
MK88
MK94
MK97
MK data
Normal data

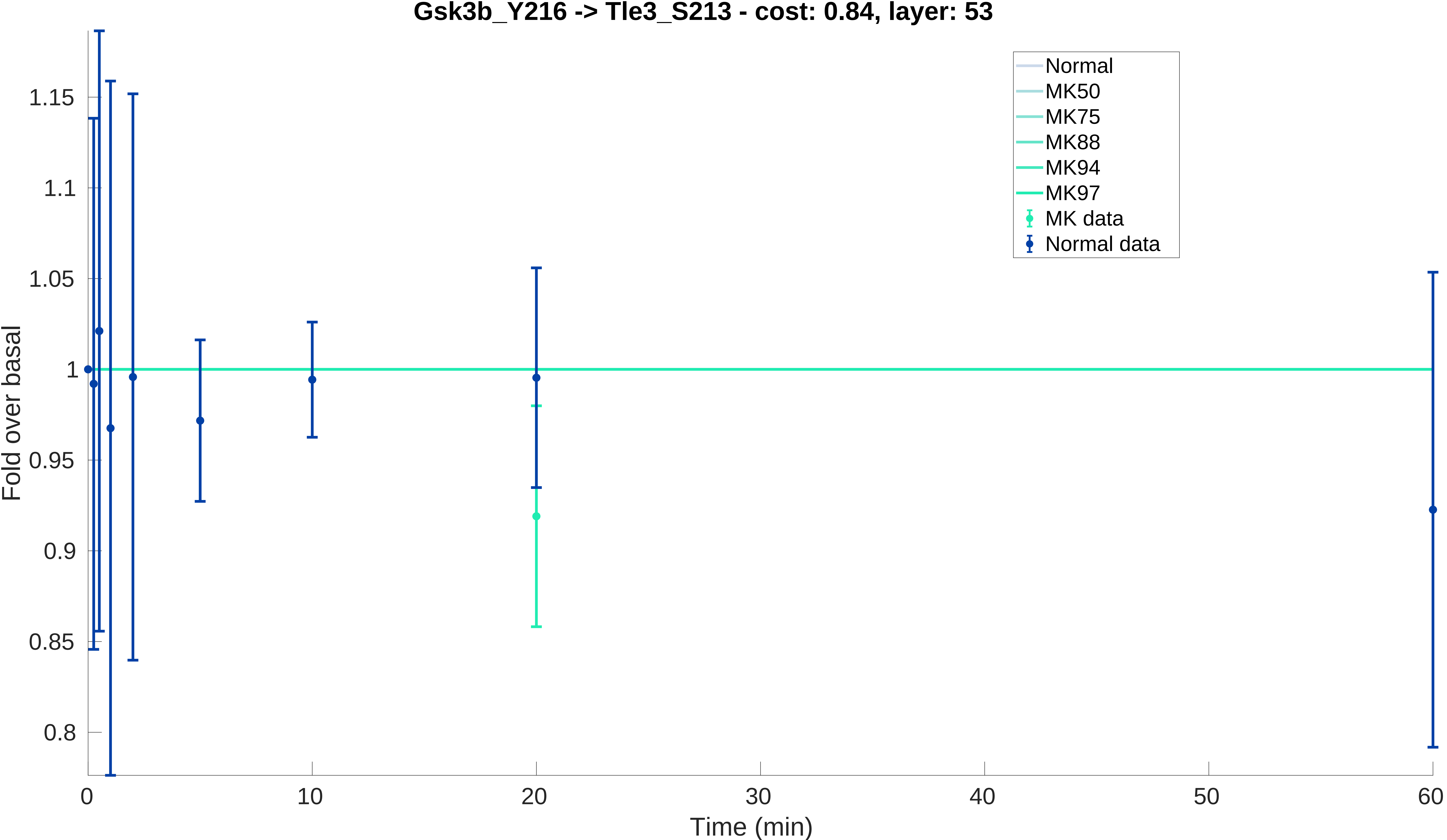
Gsk3b_Y216 -> Tle3_S213 - cost: 0.84, layer: 53
Fold over basal
Time (min)
0.8
0.85
0.9
0.95
1
1.05
1.1
1.15
0
10
20
30
40
50
60
Normal
MK50
MK75
MK88
MK94
MK97
MK data
Normal data

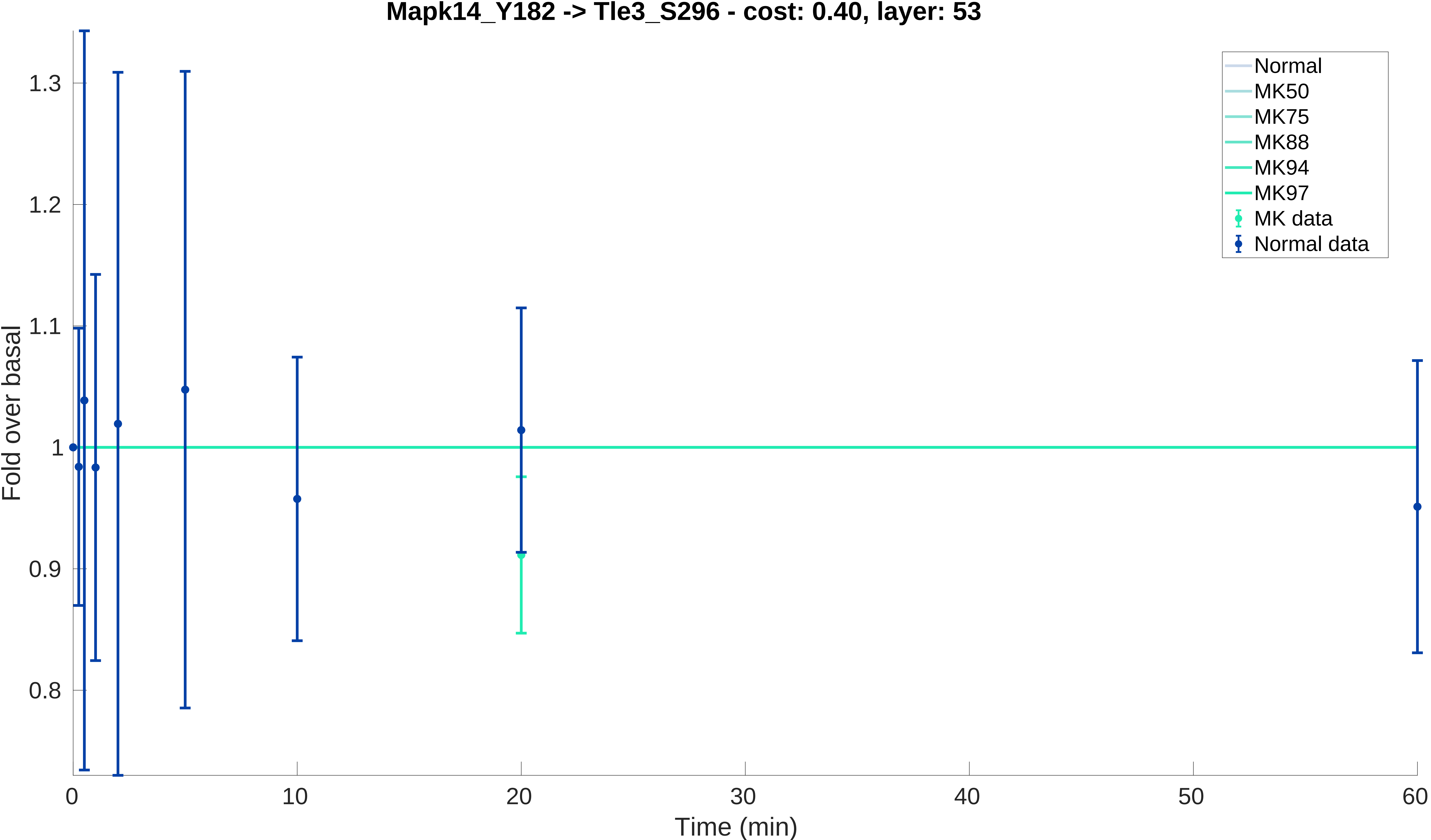

Mapk14_Y182 -> Tle3_S296 - cost: 0.40, layer: 53
Fold over basal
Time (min)
0.8
0.9
1
1.1
1.2
1.3
0
10
20
30
40
50
60
Normal
MK50
MK75
MK88
MK94
MK97
MK data
Normal data

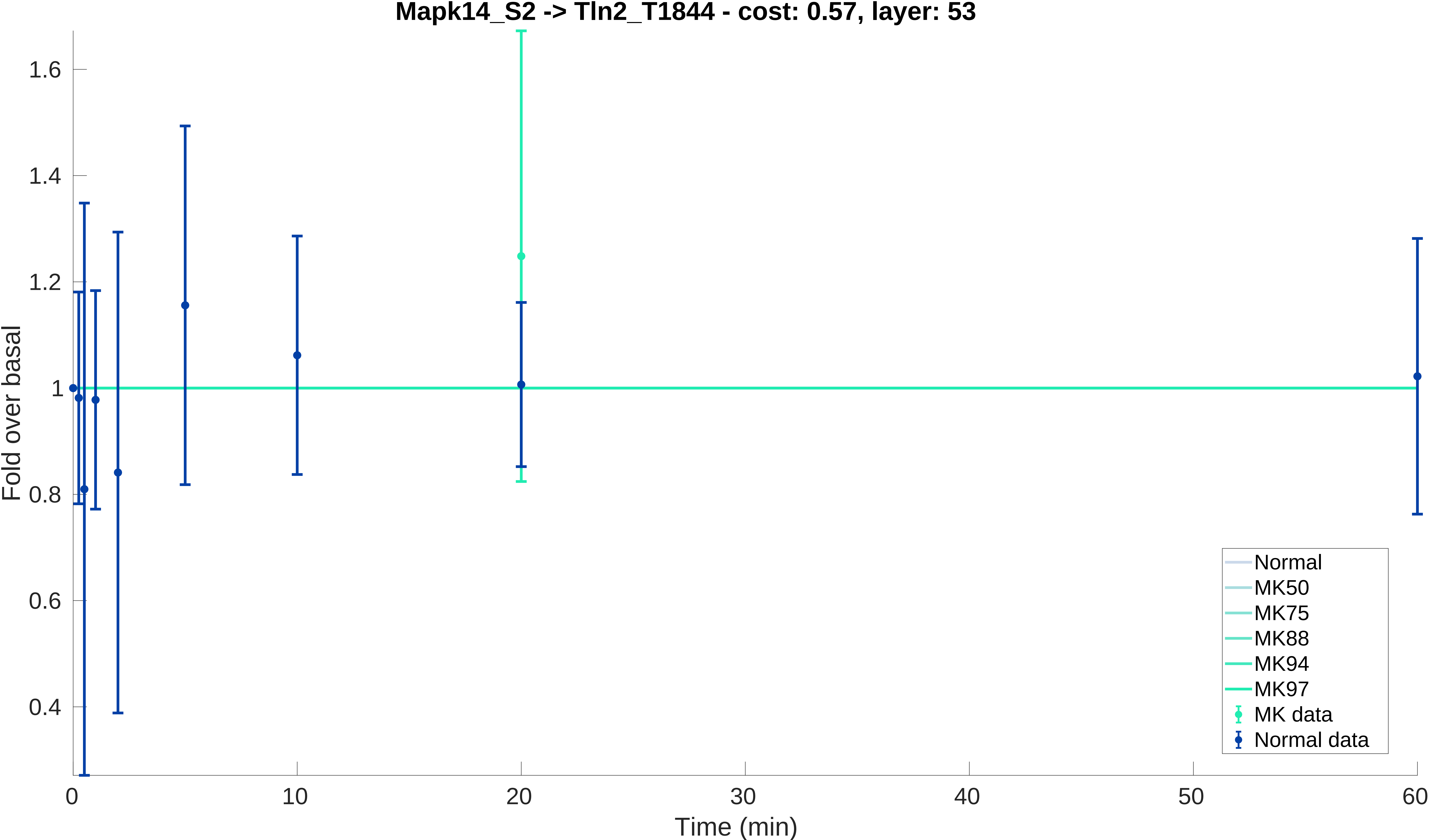

Mapk14_S2 -> Tln2_T1844 - cost: 0.57, layer: 53
Fold over basal
Time (min)
0.4
0.6
0.8
1
1.2
1.4
1.6
0
10
20
30
40
50
60
Normal
MK50
MK75
MK88
MK94
MK97
MK data
Normal data

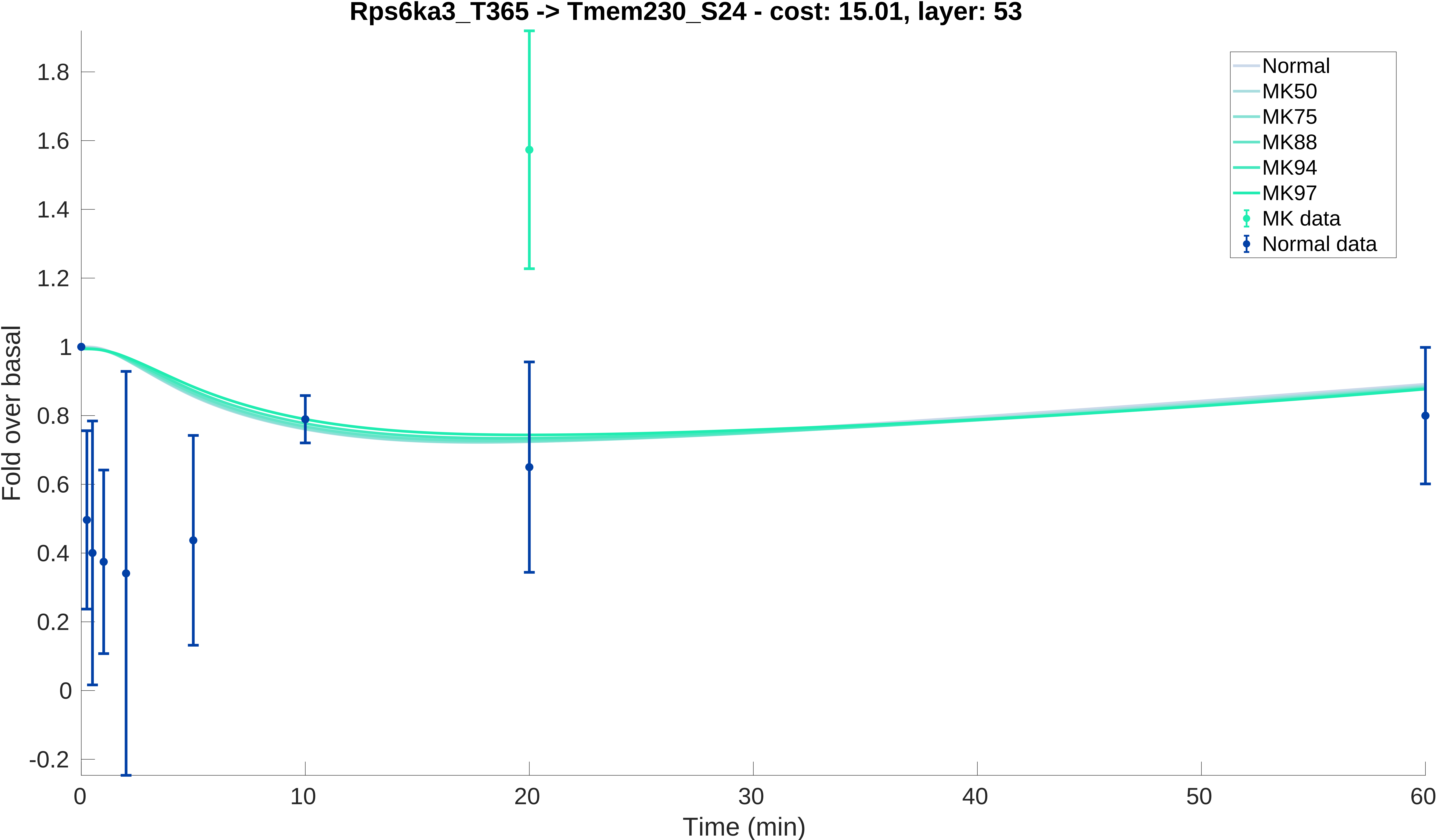

Rps6ka3_T365 -> Tmem230_S24 - cost: 15.01, layer: 53
Fold over basal
Time (min)
1.8
1.6
1.4
1.2
1
0.8
0.6
0.4
0.2
0
-0.2
0
10
20
30
40
50
60
Normal
MK50
MK75
MK88
MK94
MK97
MK data
Normal data

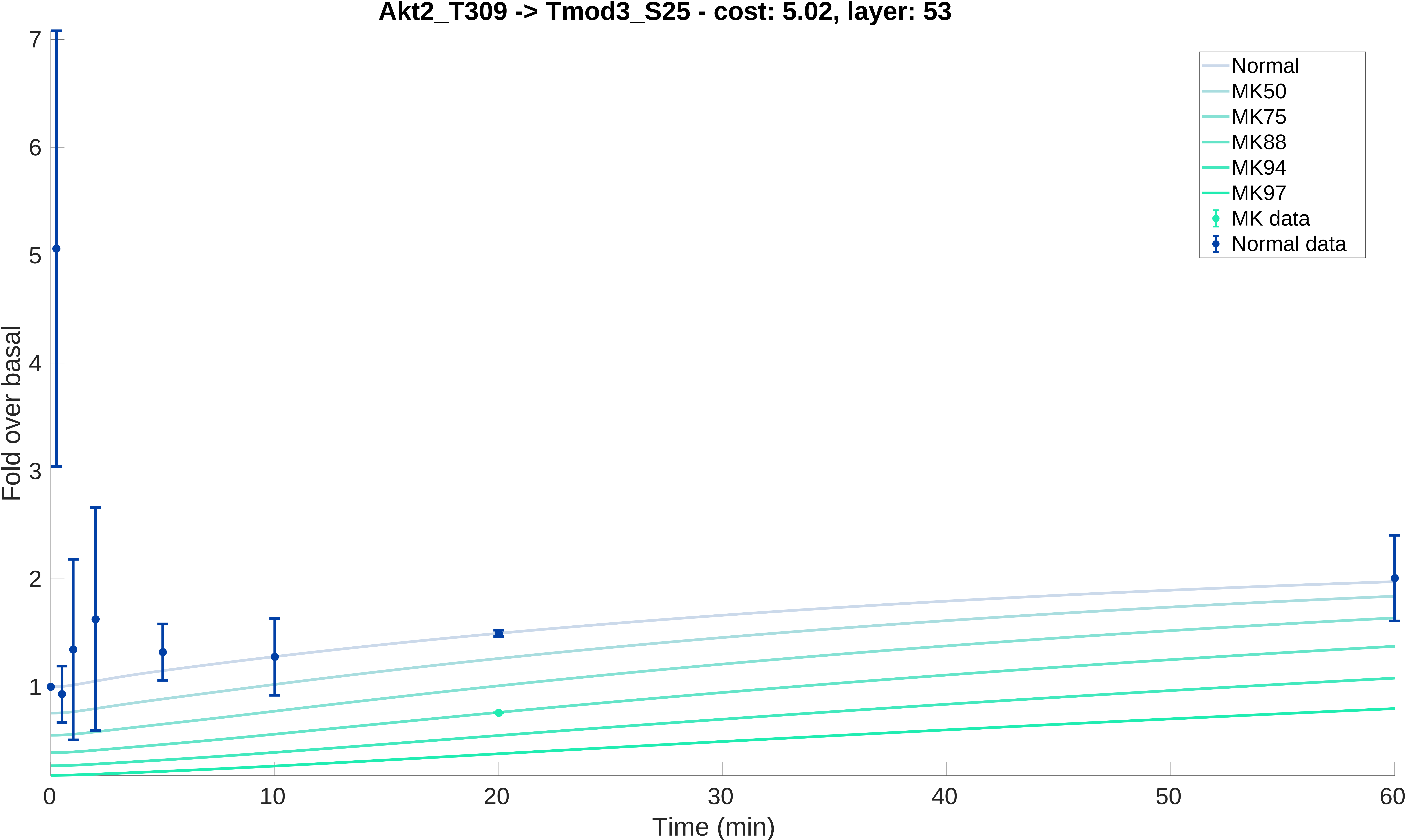
Akt2_T309 -> Tmod3_S25 - cost: 5.02, layer: 53
Fold over basal
Time (min)
0
10
20
30
40
50
60
1
2
3
4
5
6
7
Normal
MK50
MK75
MK88
MK94
MK97
MK data
Normal data

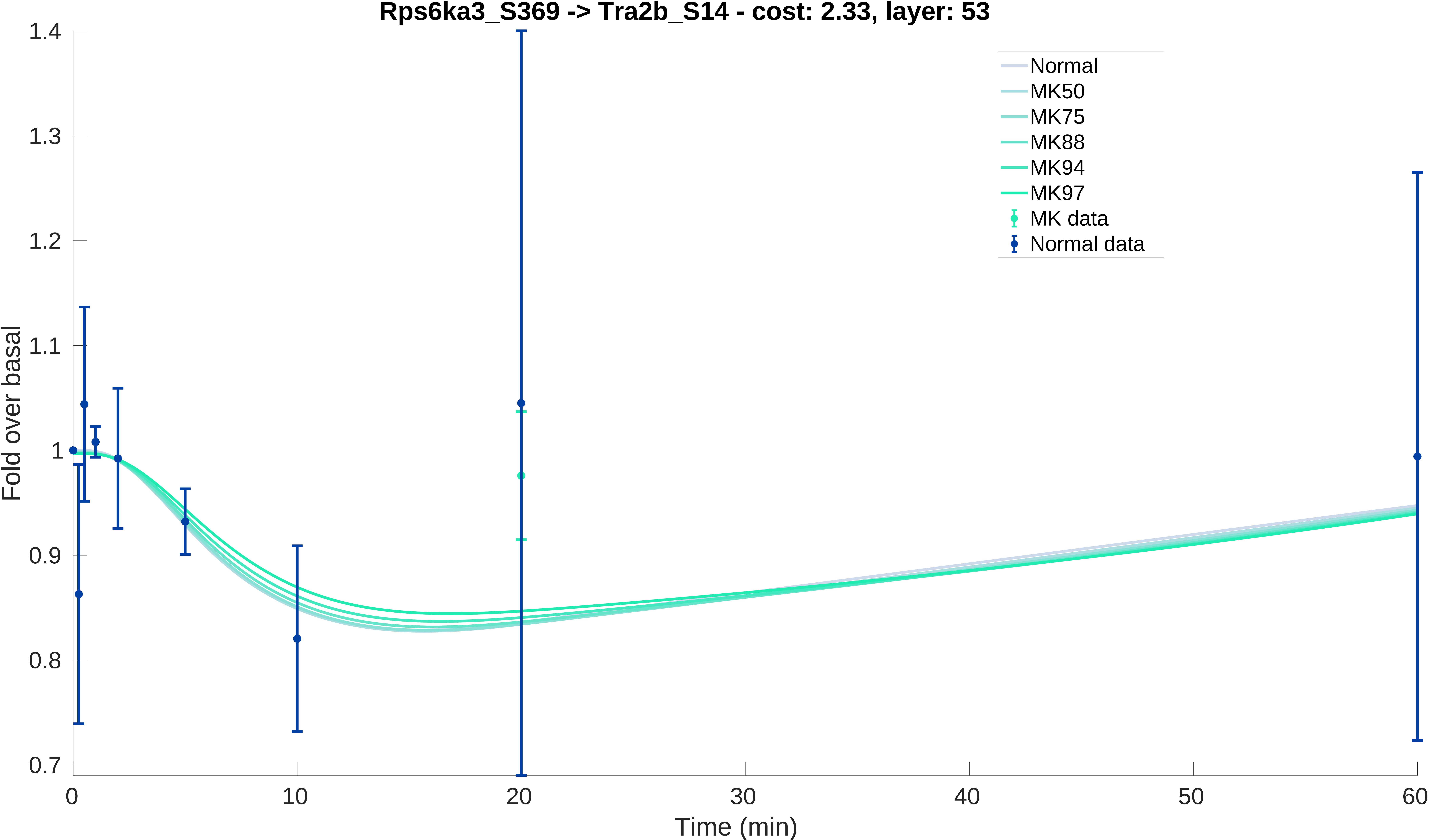

Rps6ka3_S369 -> Tra2b_S14 - cost: 2.33, layer: 53
Fold over basal
Time (min)
1.4
1.3
1.2
1.1
1
0.9
0.8
0.7
0
10
20
30
40
50
60
Normal
MK50
MK75
MK88
MK94
MK97
MK data
Normal data

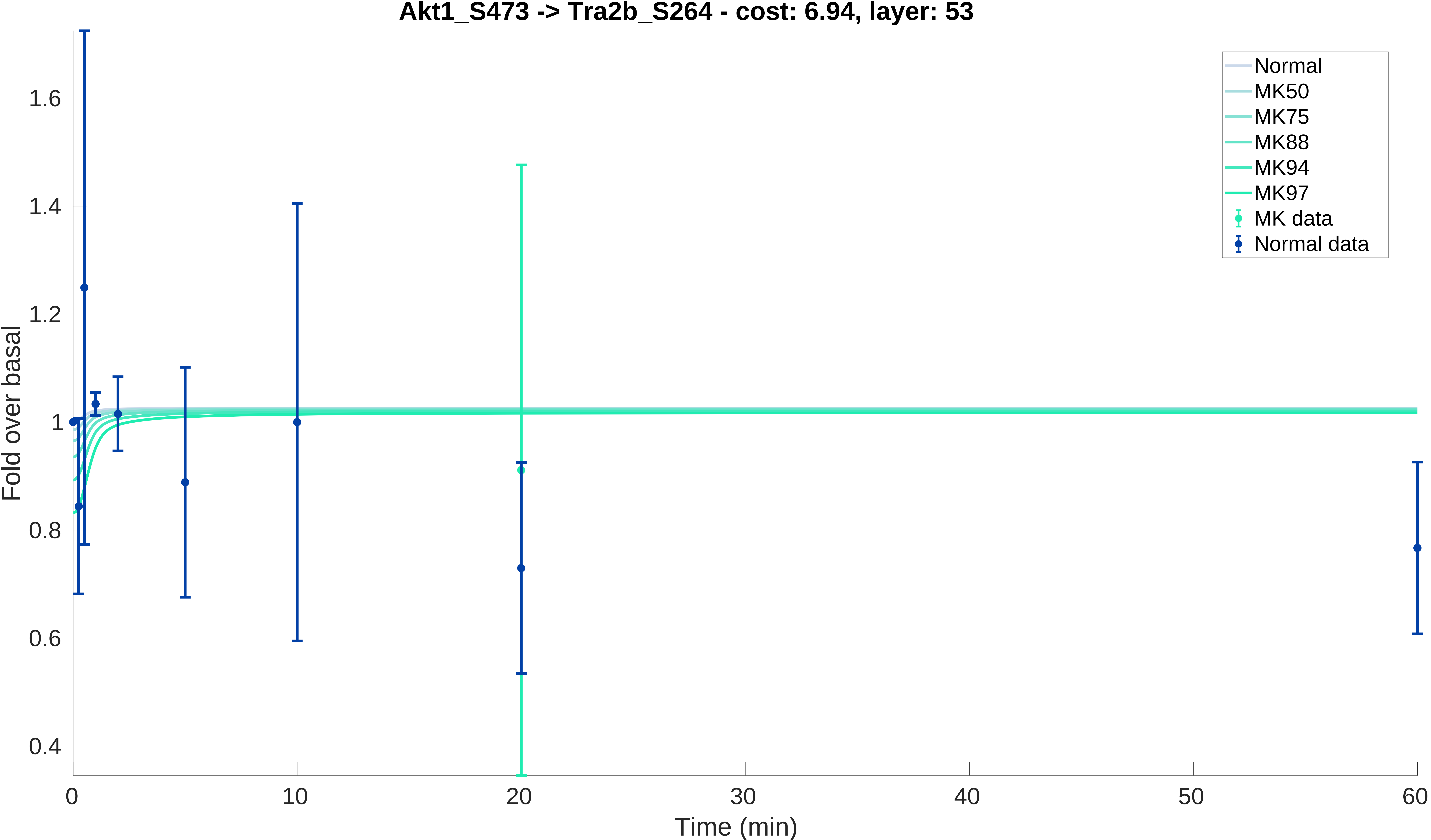

Akt1_S473 -> Tra2b_S264 - cost: 6.94, layer: 53
Fold over basal
Time (min)
1.6
1.4
1.2
1
0.8
0.6
0.4
0
10
20
30
40
50
60
Normal
MK50
MK75
MK88
MK94
MK97
MK data
Normal data

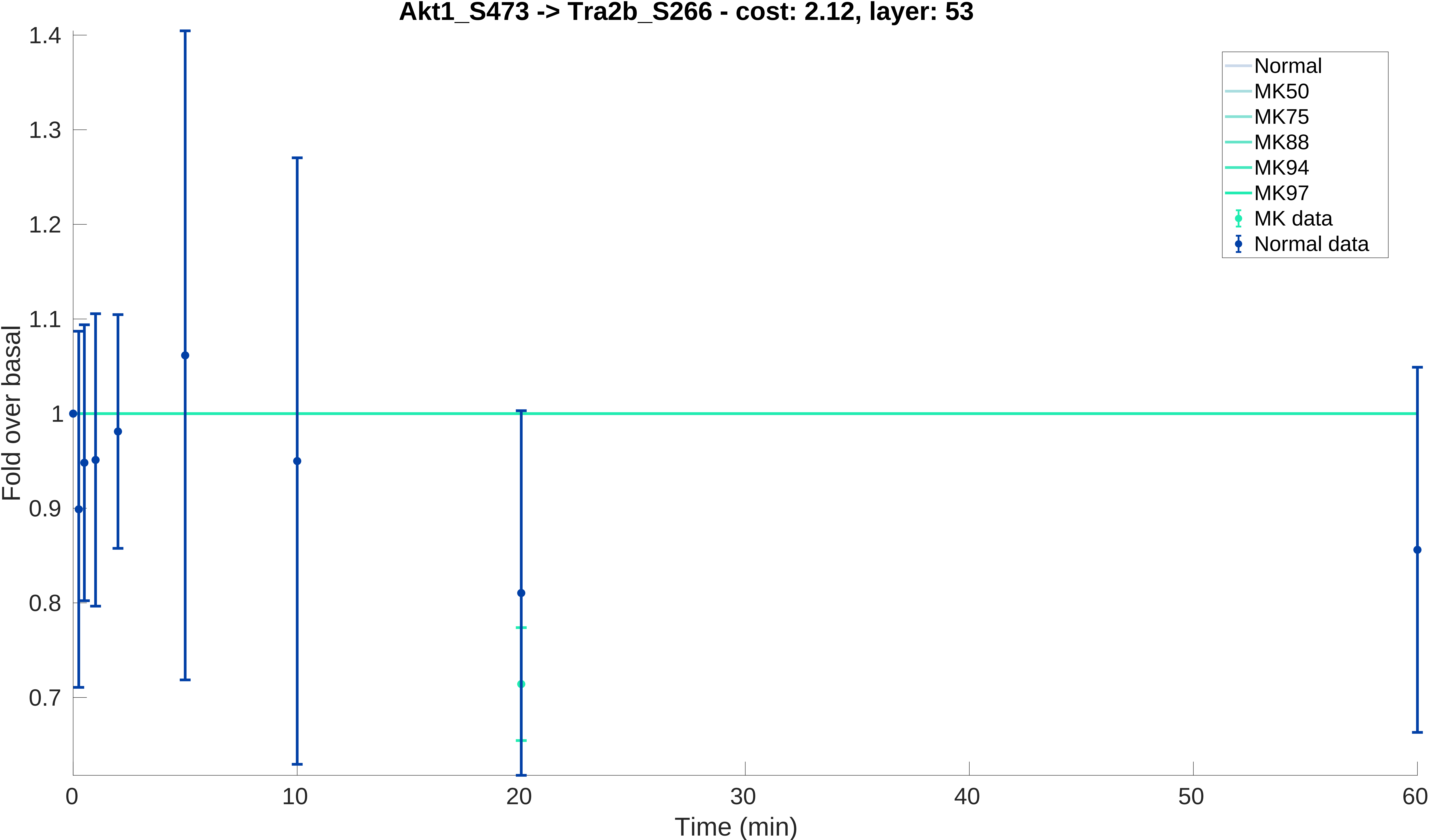

Akt1_S473 -> Tra2b_S266 - cost: 2.12, layer: 53
Fold over basal
Time (min)
1.4
1.3
1.2
1.1
1
0.9
0.8
0.7
0
10
20
30
40
50
60
Normal
MK50
MK75
MK88
MK94
MK97
MK data
Normal data

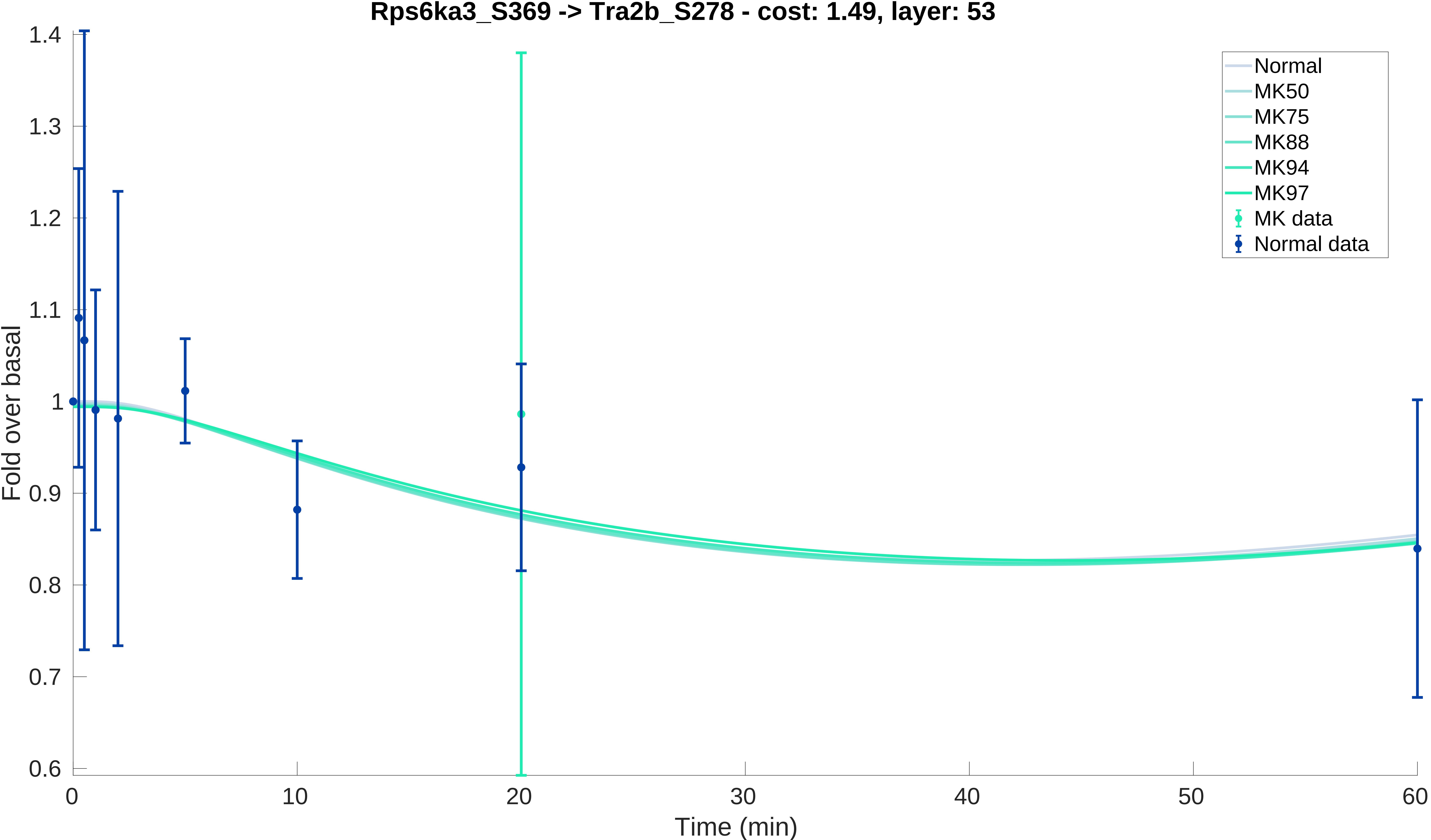

Rps6ka3_S369 -> Tra2b_S278 - cost: 1.49, layer: 53
Fold over basal
Time (min)
1.4
1.3
1.2
1.1
1
0.9
0.8
0.7
0.6
0
10
20
30
40
50
60
Normal
MK50
MK75
MK88
MK94
MK97
MK data
Normal data

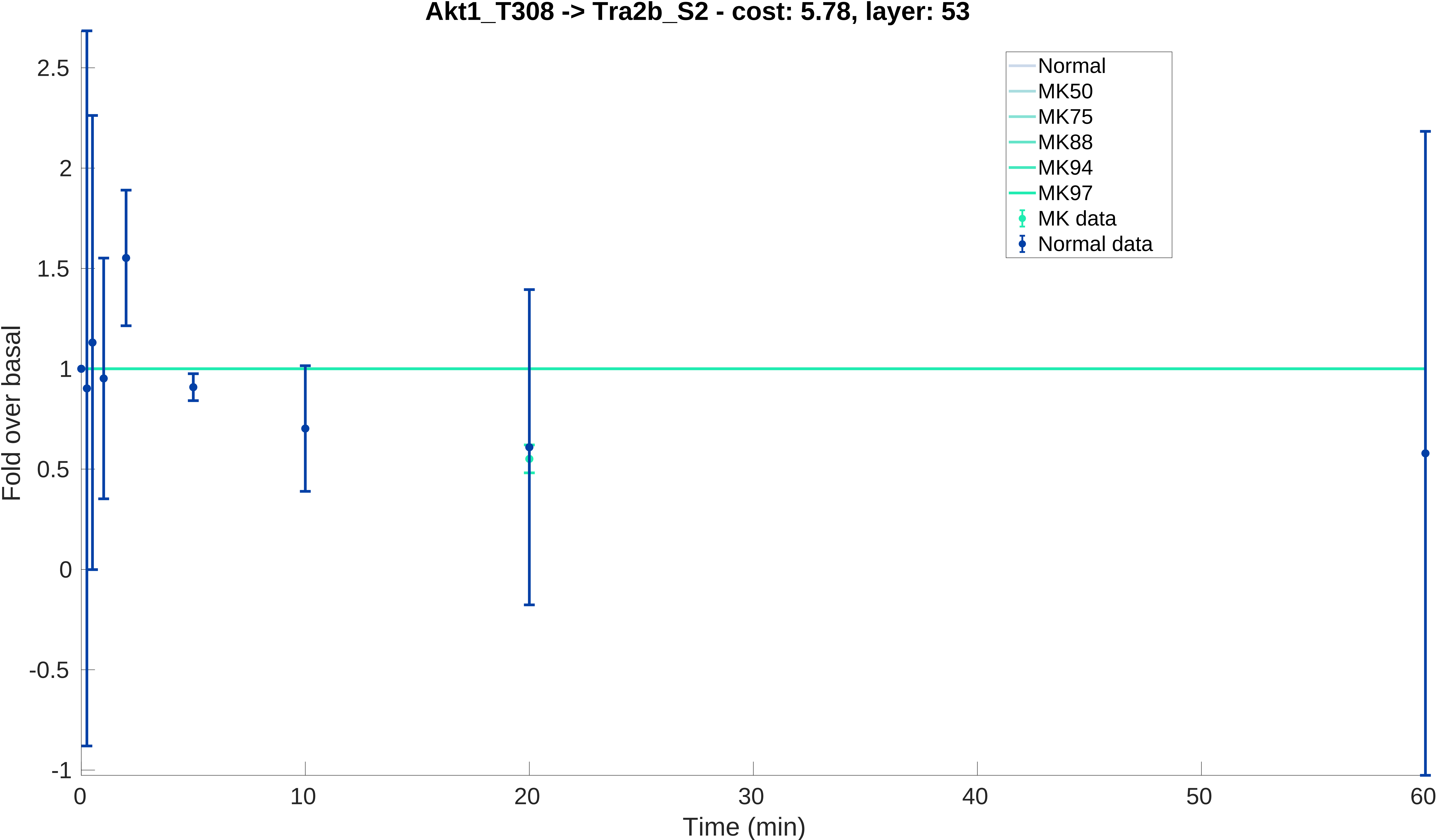

Akt1_T308 -> Tra2b_S2 - cost: 5.78, layer: 53
Normal
MK50
MK75
MK88
MK94
MK97
MK data
Normal data
Fold over basal
Time (min)
2.5
2
1.5
1
0.5
0
-0.5
-1
0
10
20
30
40
50
60

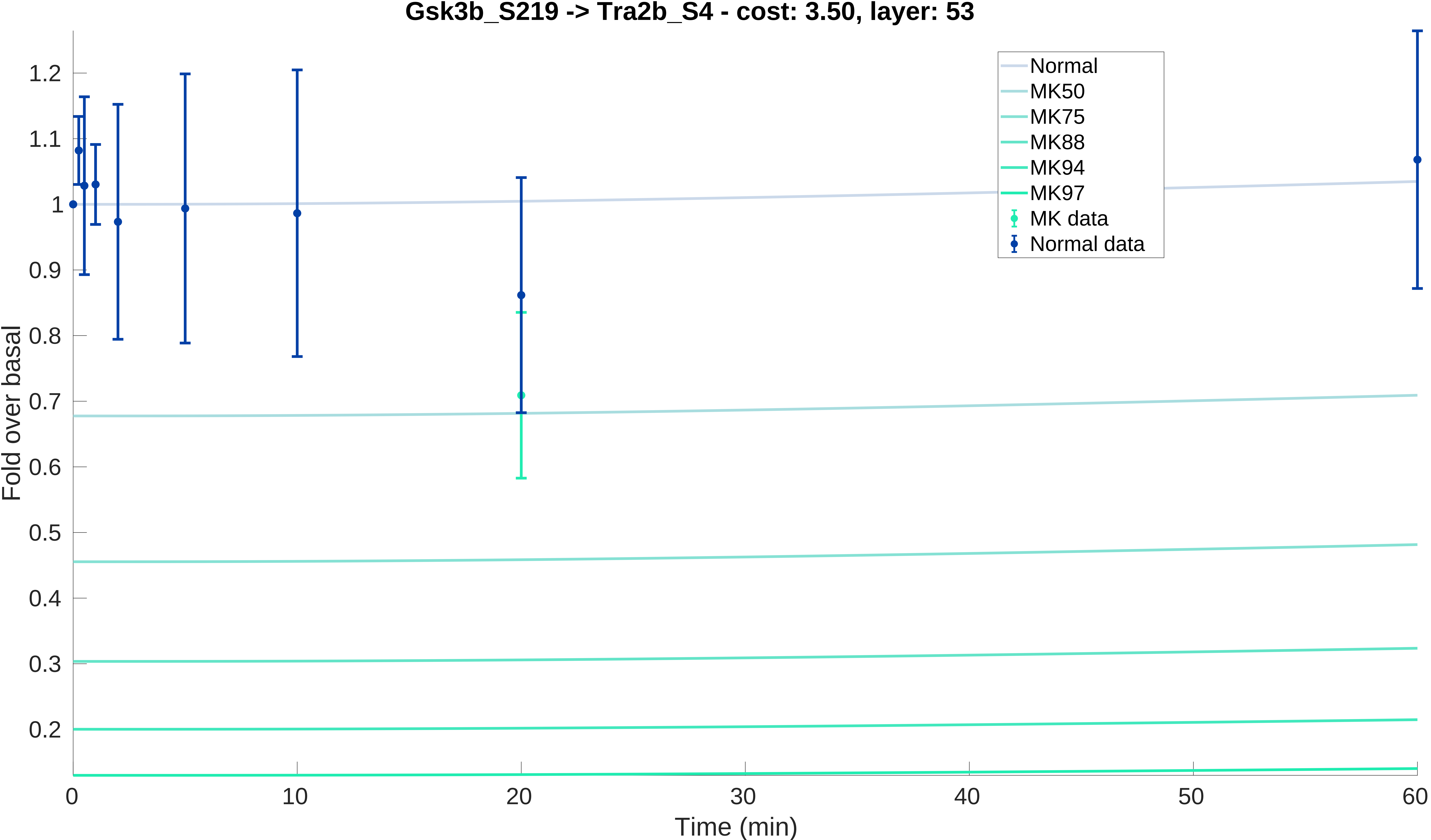

Gsk3b_S219 -> Tra2b_S4 - cost: 3.50, layer: 53
Normal
MK50
MK75
MK88
MK94
MK97
MK data
Normal data
Fold over basal
Time (min)
0
10
20
30
40
50
60
0.2
0.3
0.4
0.5
0.6
0.7
0.8
0.9
1
1.1
1.2

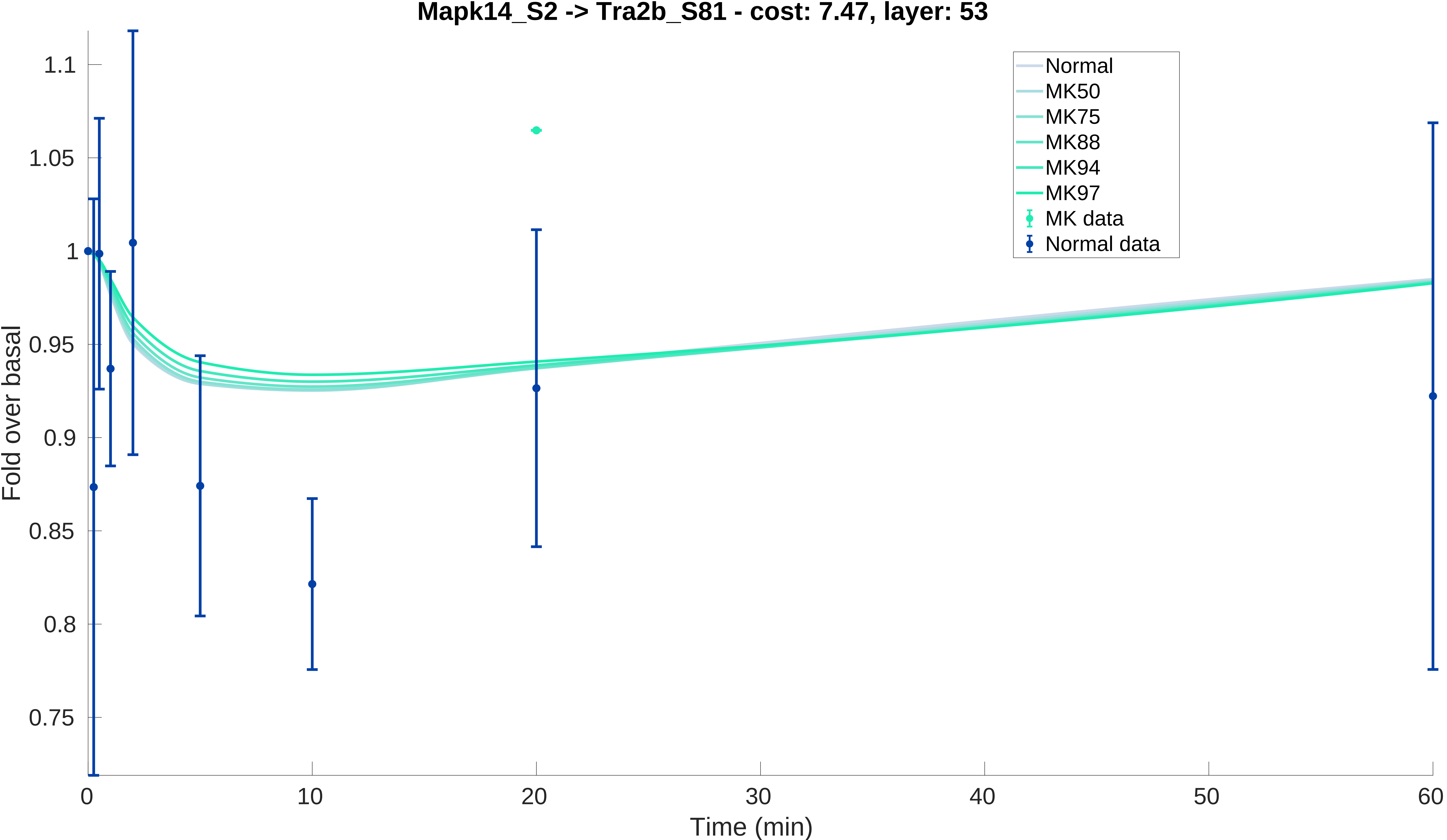
Mapk14_S2 -> Tra2b_S81 - cost: 7.47, layer: 53
Fold over basal
1.1
1.05
1
0.95
0.9
0.85
0.8
0.75
Time (min)
0
10
20
30
40
50
60
Normal
MK50
MK75
MK88
MK94
MK97
MK data
Normal data

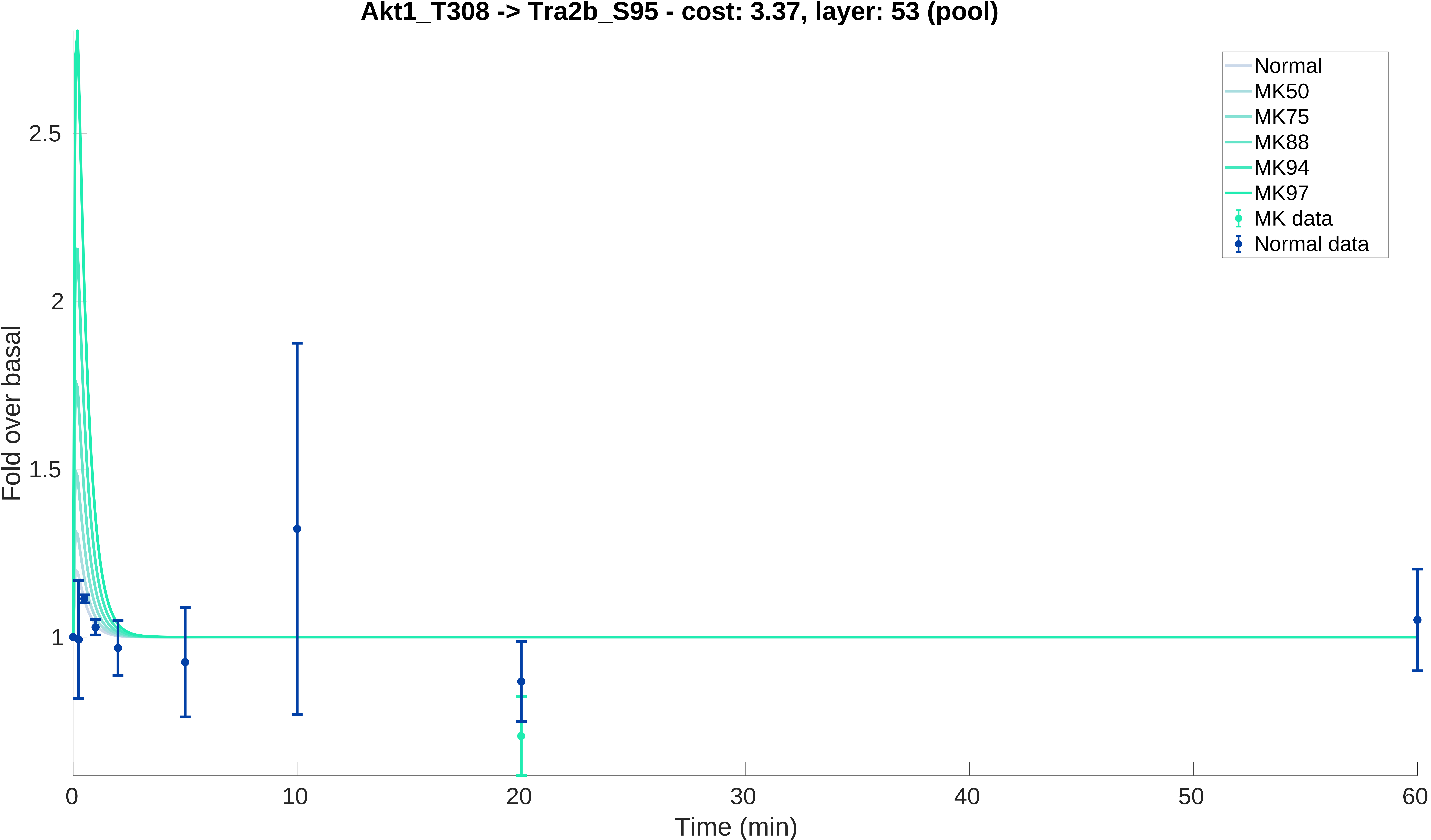
Akt1_T308 -> Tra2b_S95 - cost: 3.37, layer: 53 (pool)
Fold over basal
Time (min)
0
10
20
30
40
50
60
1
1.5
2
2.5
Normal
MK50
MK75
MK88
MK94
MK97
MK data
Normal data

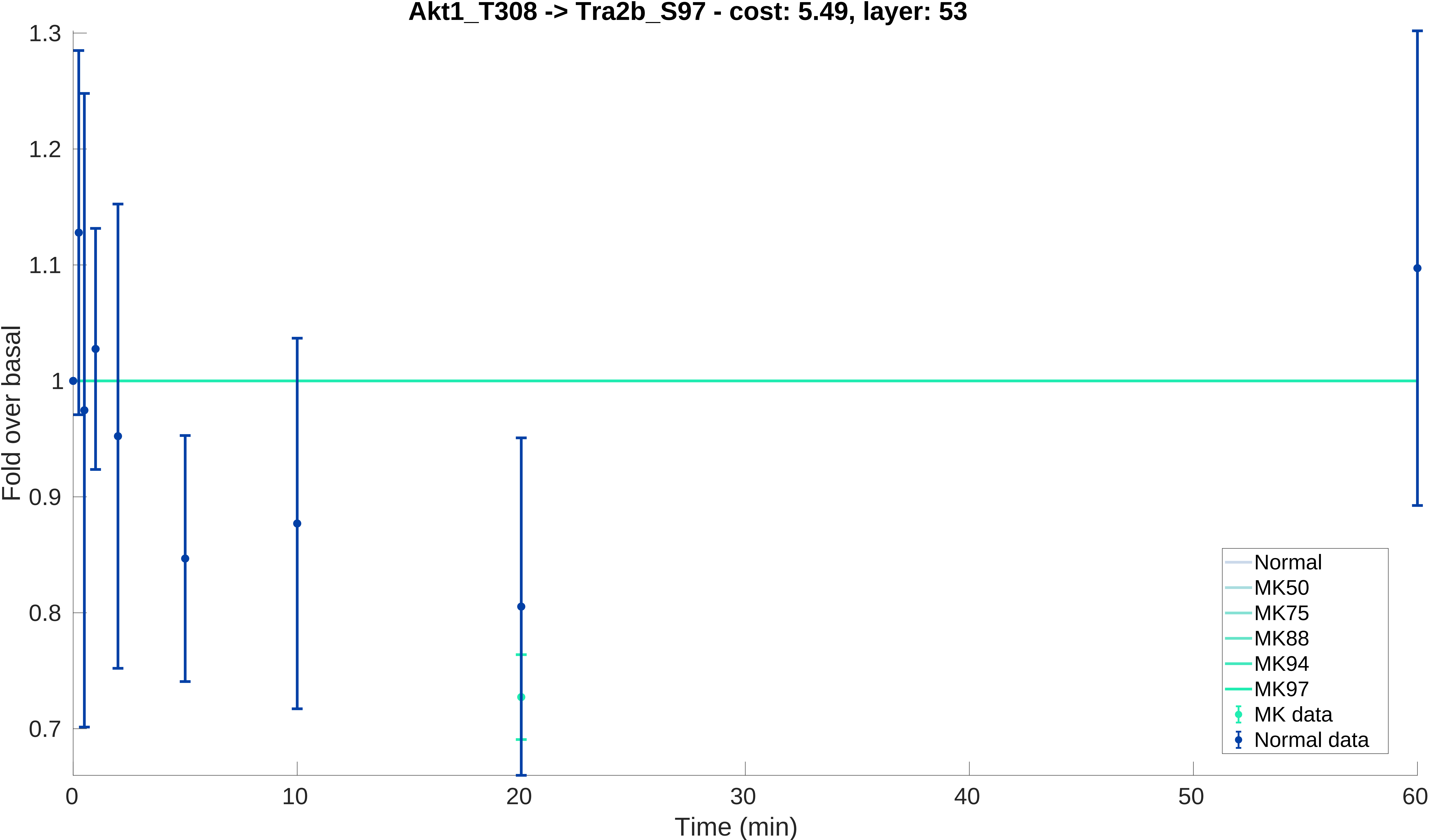

Akt1_T308 -> Tra2b_S97 - cost: 5.49, layer: 53
Fold over basal
1.3
1.2
1.1
1
0.9
0.8
0.7
0
10
20
30
40
50
60
Time (min)
Normal
MK50
MK75
MK88
MK94
MK97
MK data
Normal data

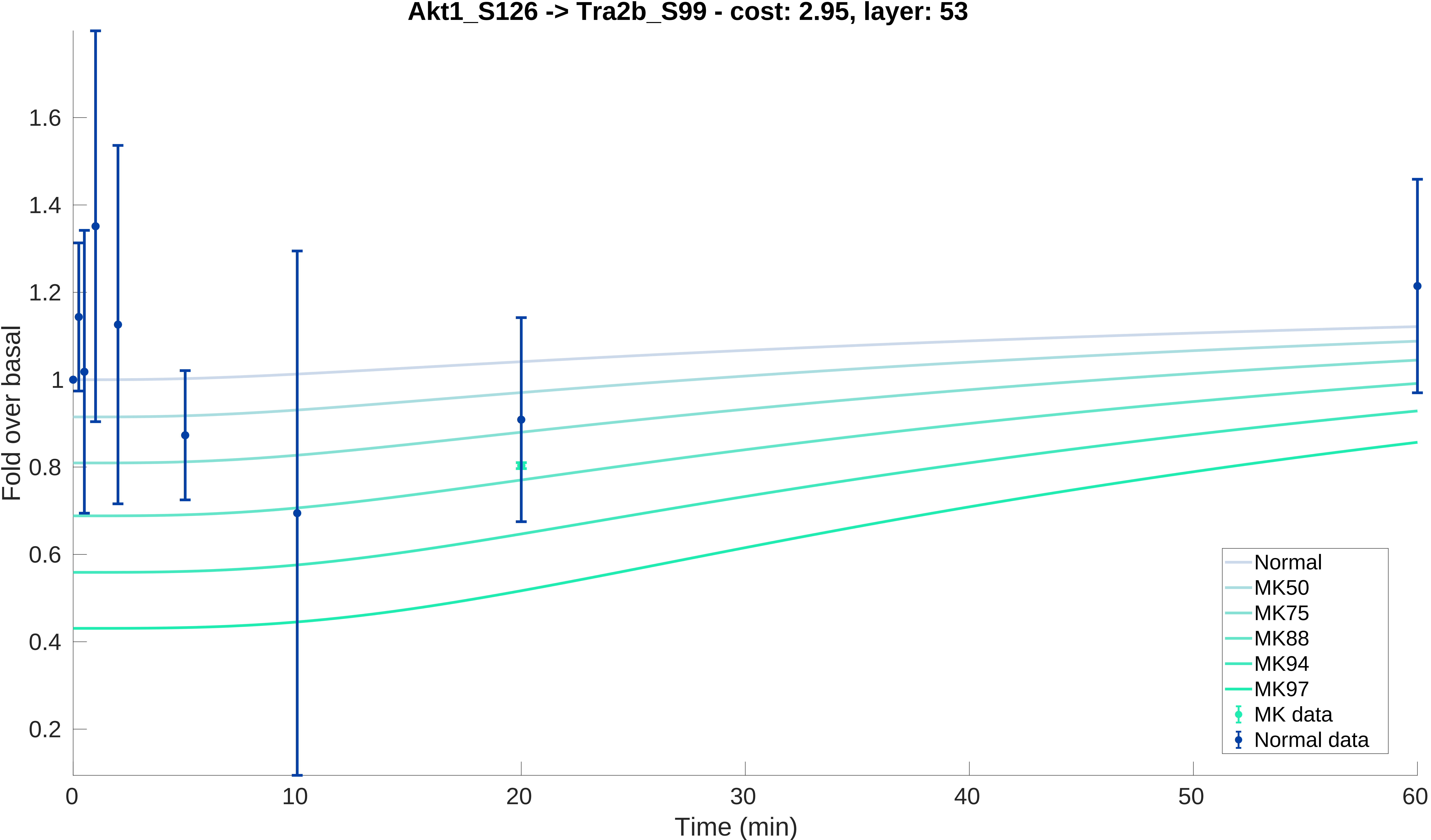

Akt1_S126 -> Tra2b_S99 - cost: 2.95, layer: 53
Fold over basal
Time (min)
0.2
0.4
0.6
0.8
1
1.2
1.4
1.6
0
10
20
30
40
50
60
Normal
MK50
MK75
MK88
MK94
MK97
MK data
Normal data

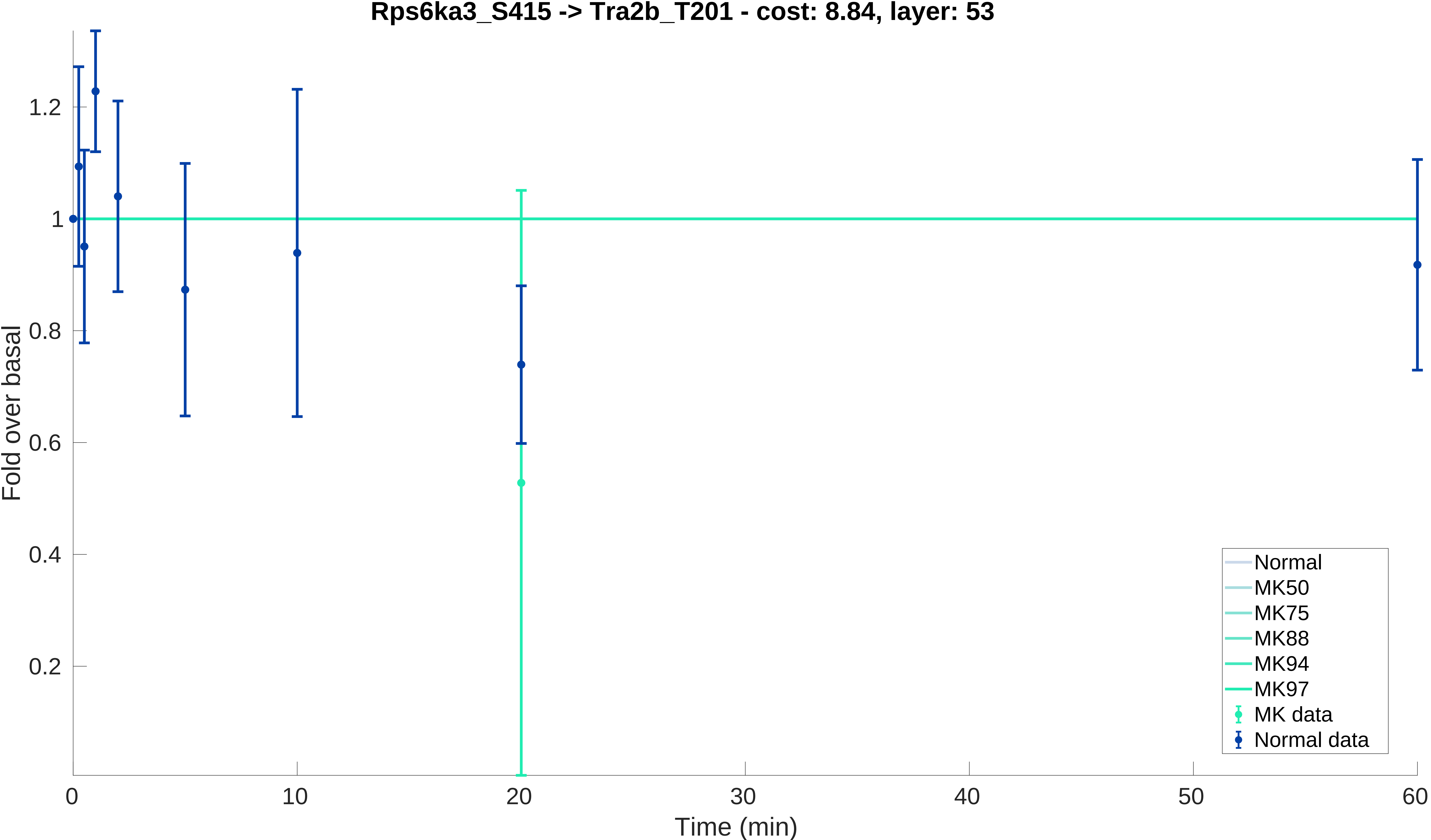
Rps6ka3_S415 -> Tra2b_T201 - cost: 8.84, layer: 53
Fold over basal
1.2
1
0.8
0.6
0.4
0.2
0
10
20
30
40
50
60
Time (min)
Normal
MK50
MK75
MK88
MK94
MK97
MK data
Normal data

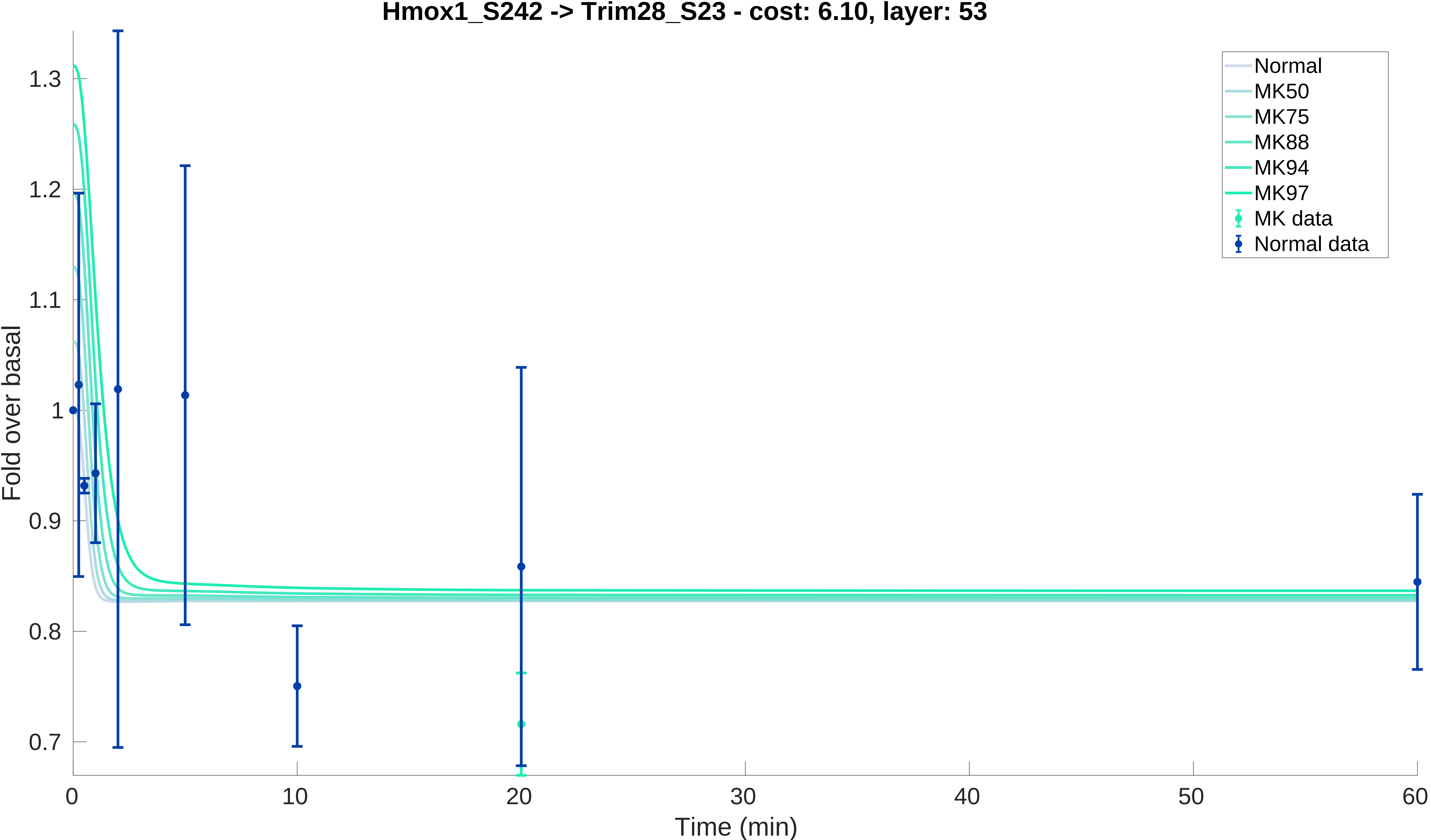
Hmox1_S242 -> Trim28_S23 - cost: 6.10, layer: 53
Fold over basal
1.3
1.2
1.1
1
0.9
0.8
0.7
0
10
20
30
40
50
60
Time (min)
Normal
MK50
MK75
MK88
MK94
MK97
MK data
Normal data

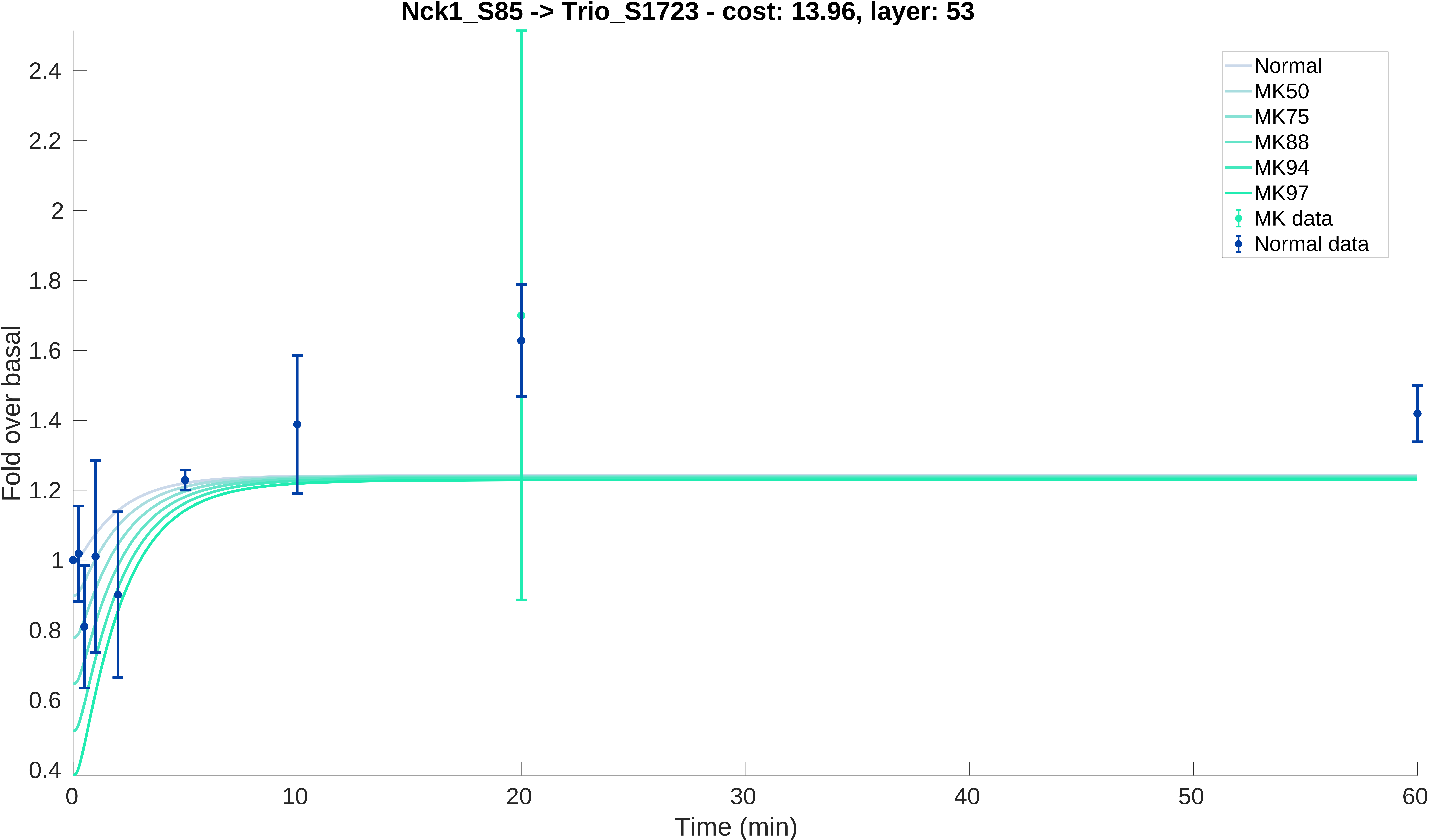
Nck1_S85 -> Trio_S1723 - cost: 13.96, layer: 53
Fold over basal
Time (min)
0.4
0.6
0.8
1
1.2
1.4
1.6
1.8
2
2.2
2.4
0
10
20
30
40
50
60
Normal
MK50
MK75
MK88
MK94
MK97
MK data
Normal data

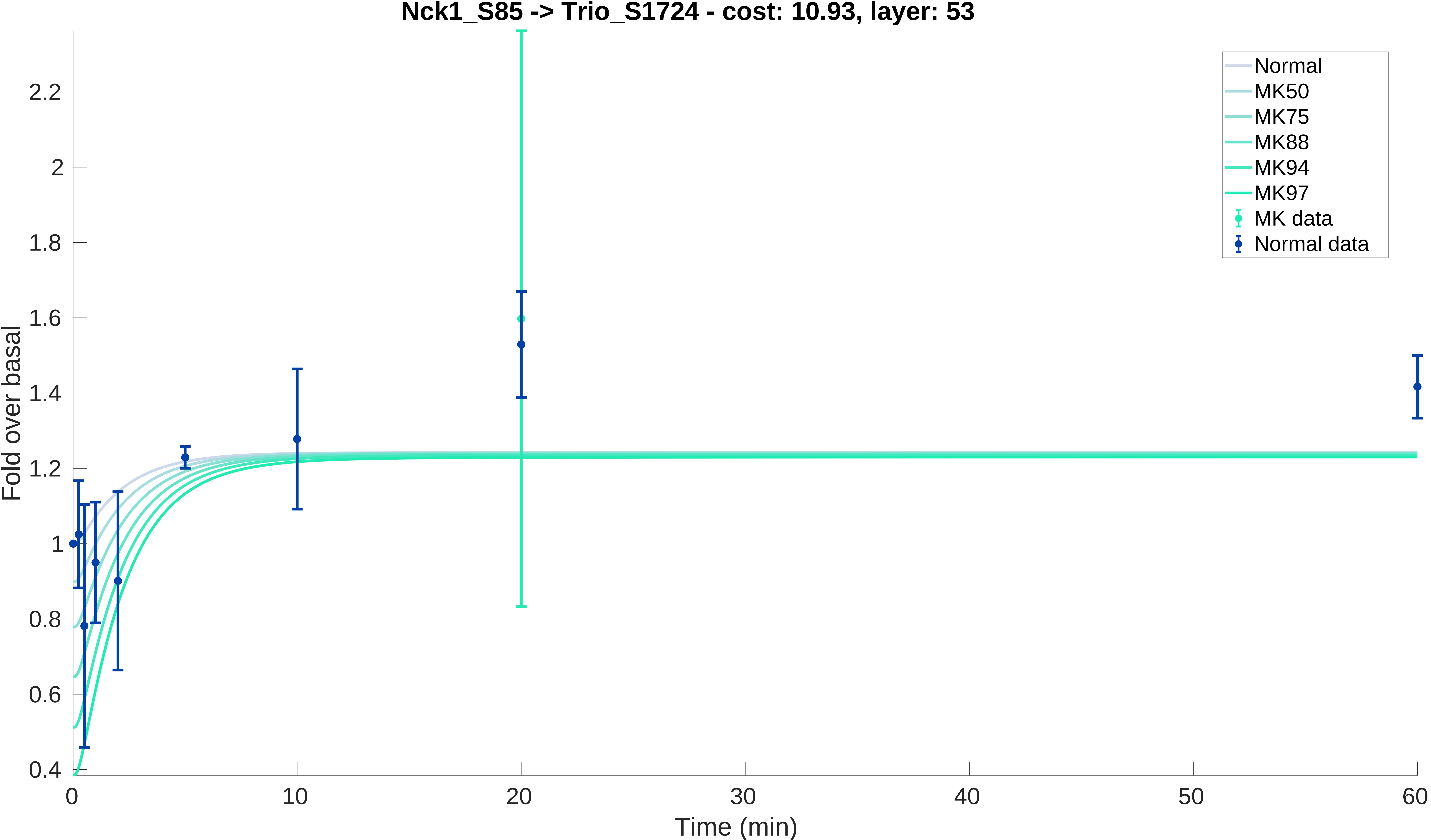
Nck1_S85 -> Trio_S1724 - cost: 10.93, layer: 53
Normal
MK50
MK75
MK88
MK94
MK97
MK data
Normal data
Fold over basal
Time (min)
0.4
0.6
0.8
1
1.2
1.4
1.6
1.8
2
2.2
0
10
20
30
40
50
60

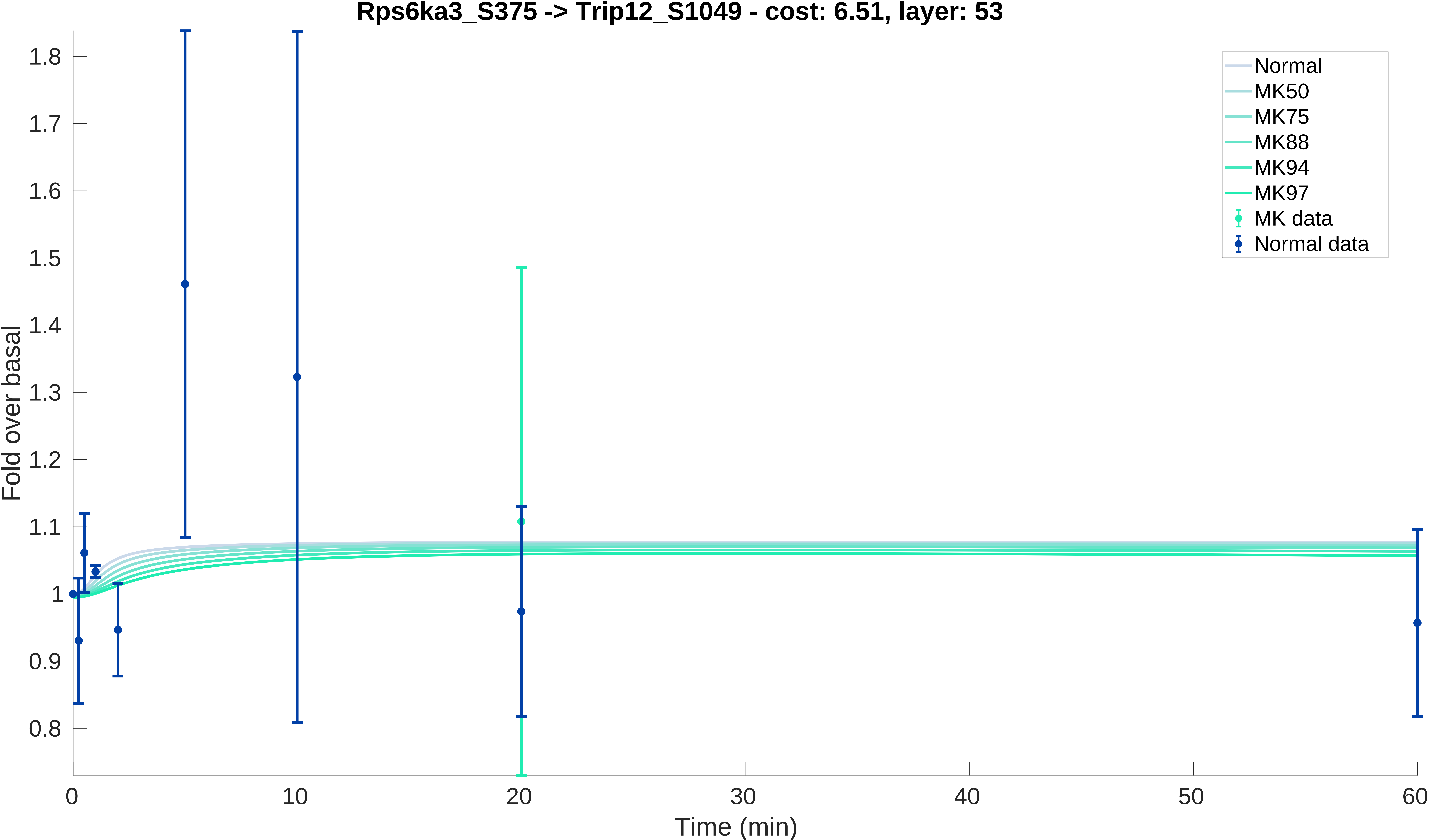

Rps6ka3_S375 -> Trip12_S1049 - cost: 6.51, layer: 53
Fold over basal
Time (min)
Normal
MK50
MK75
MK88
MK94
MK97
MK data
Normal data
0.8
0.9
1
1.1
1.2
1.3
1.4
1.5
1.6
1.7
1.8
0
10
20
30
40
50
60

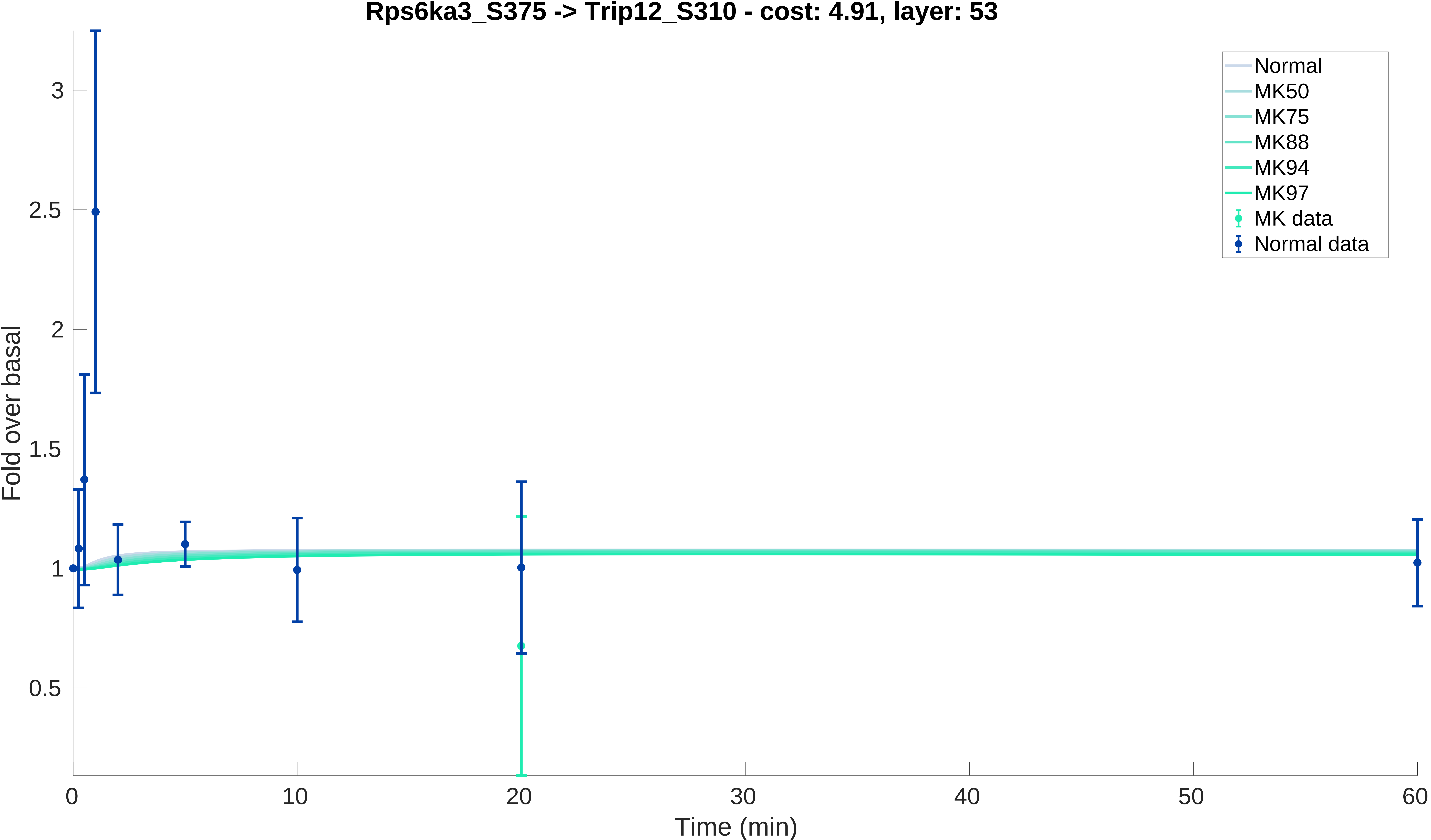
Rps6ka3_S375 -> Trip12_S310 - cost: 4.91, layer: 53
Fold over basal
Time (min)
0.5
1
1.5
2
2.5
3
0
10
20
30
40
50
60
Normal
MK50
MK75
MK88
MK94
MK97
MK data
Normal data

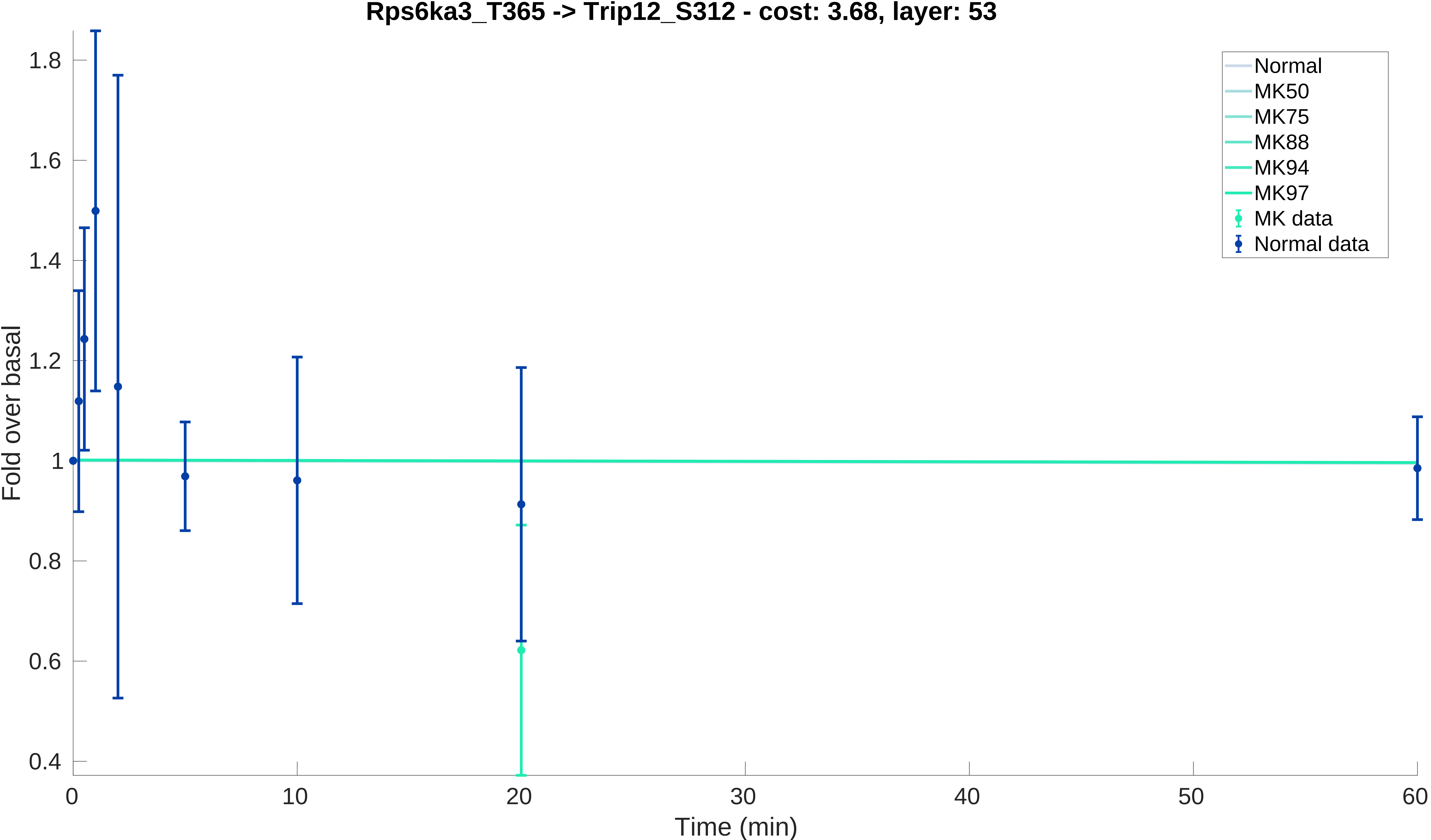

Rps6ka3_T365 -> Trip12_S312 - cost: 3.68, layer: 53
Fold over basal
Time (min)
0.4
0.6
0.8
1
1.2
1.4
1.6
1.8
0
10
20
30
40
50
60
Normal
MK50
MK75
MK88
MK94
MK97
MK data
Normal data

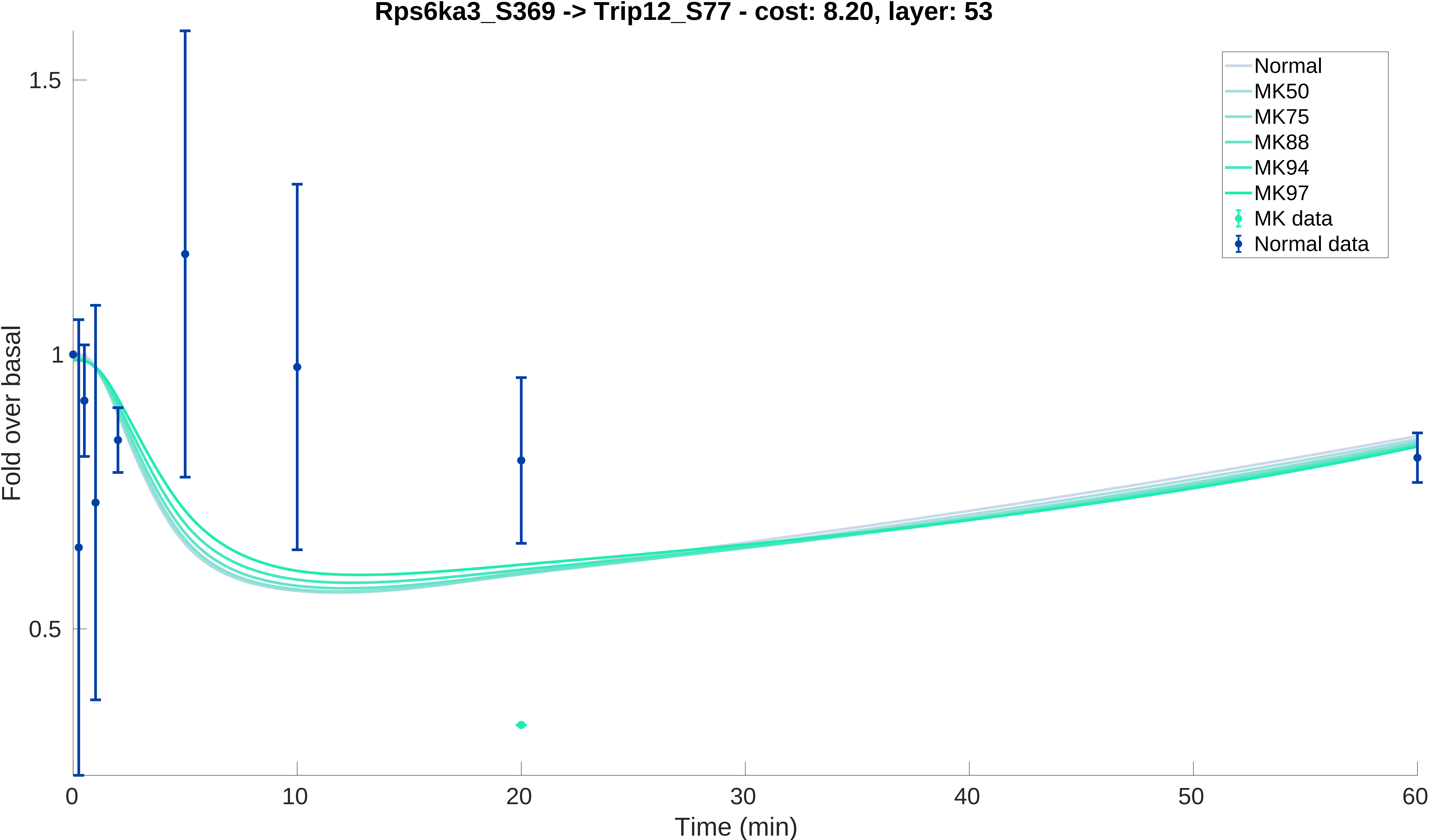

Rps6ka3_S369 -> Trip12_S77 - cost: 8.20, layer: 53
Fold over basal
Time (min)
1.5
1
0.5
0
10
20
30
40
50
60
Normal
MK50
MK75
MK88
MK94
MK97
MK data
Normal data

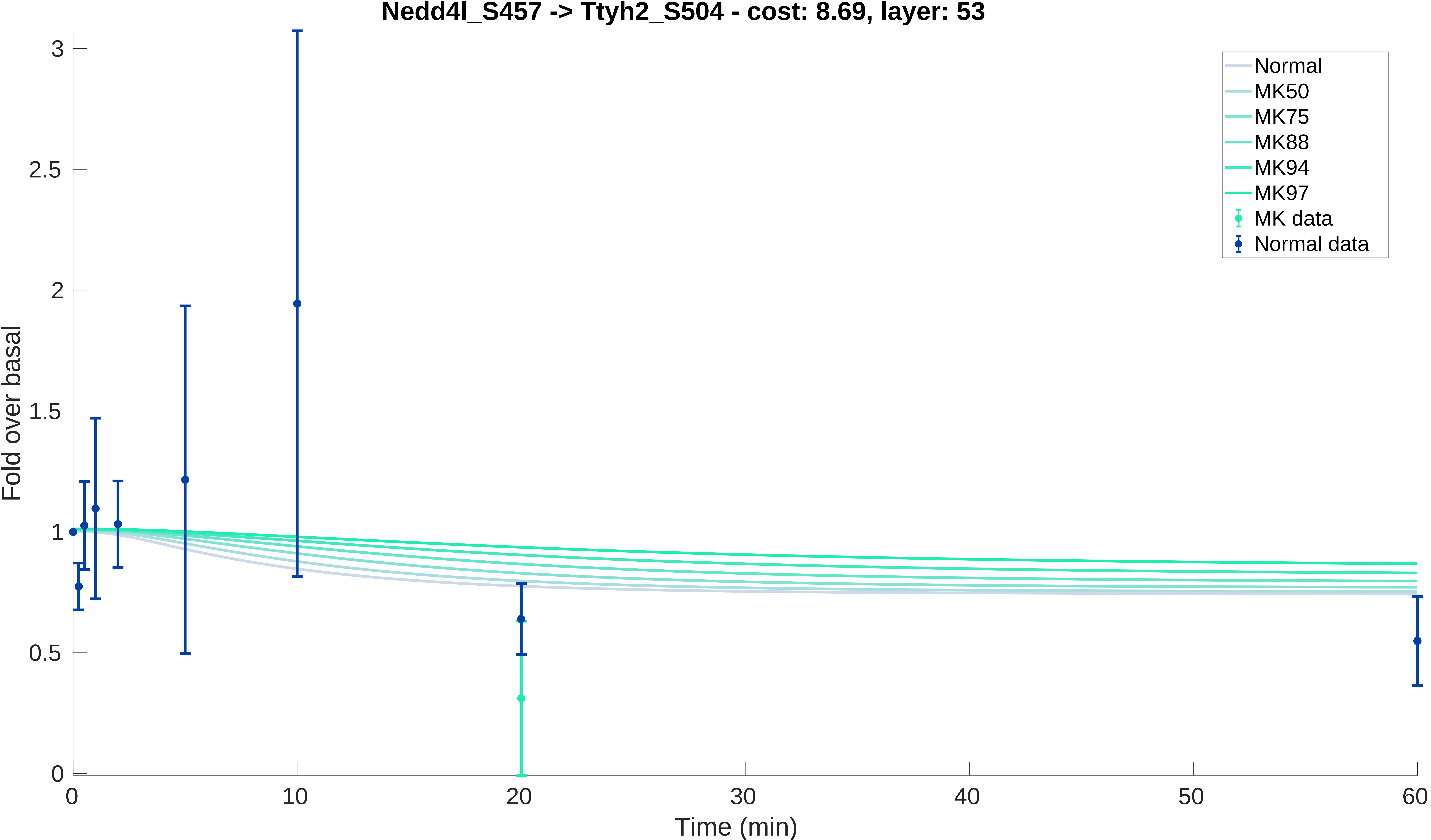

Nedd4l_S457 -> Ttyh2_S504 - cost: 8.69, layer: 53
Fold over basal
Time (min)
0
0.5
1
1.5
2
2.5
3
0
10
20
30
40
50
60
Normal
MK50
MK75
MK88
MK94
MK97
MK data
Normal data

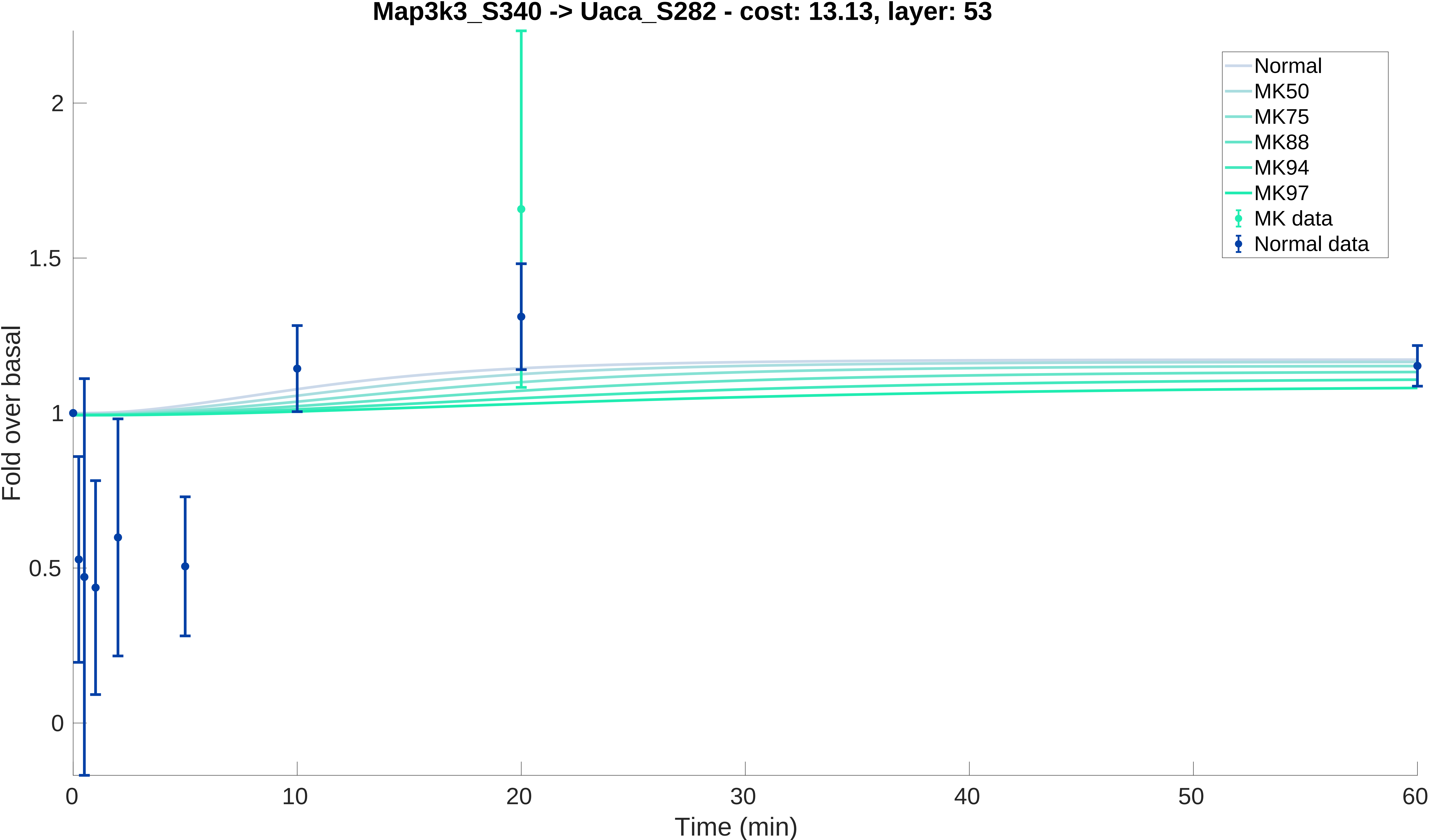

Map3k3_S340 -> Uaca_S282 - cost: 13.13, layer: 53
Normal
MK50
MK75
MK88
MK94
MK97
MK data
Normal data
Fold over basal
Time (min)
0
0.5
1
1.5
2
0
10
20
30
40
50
60

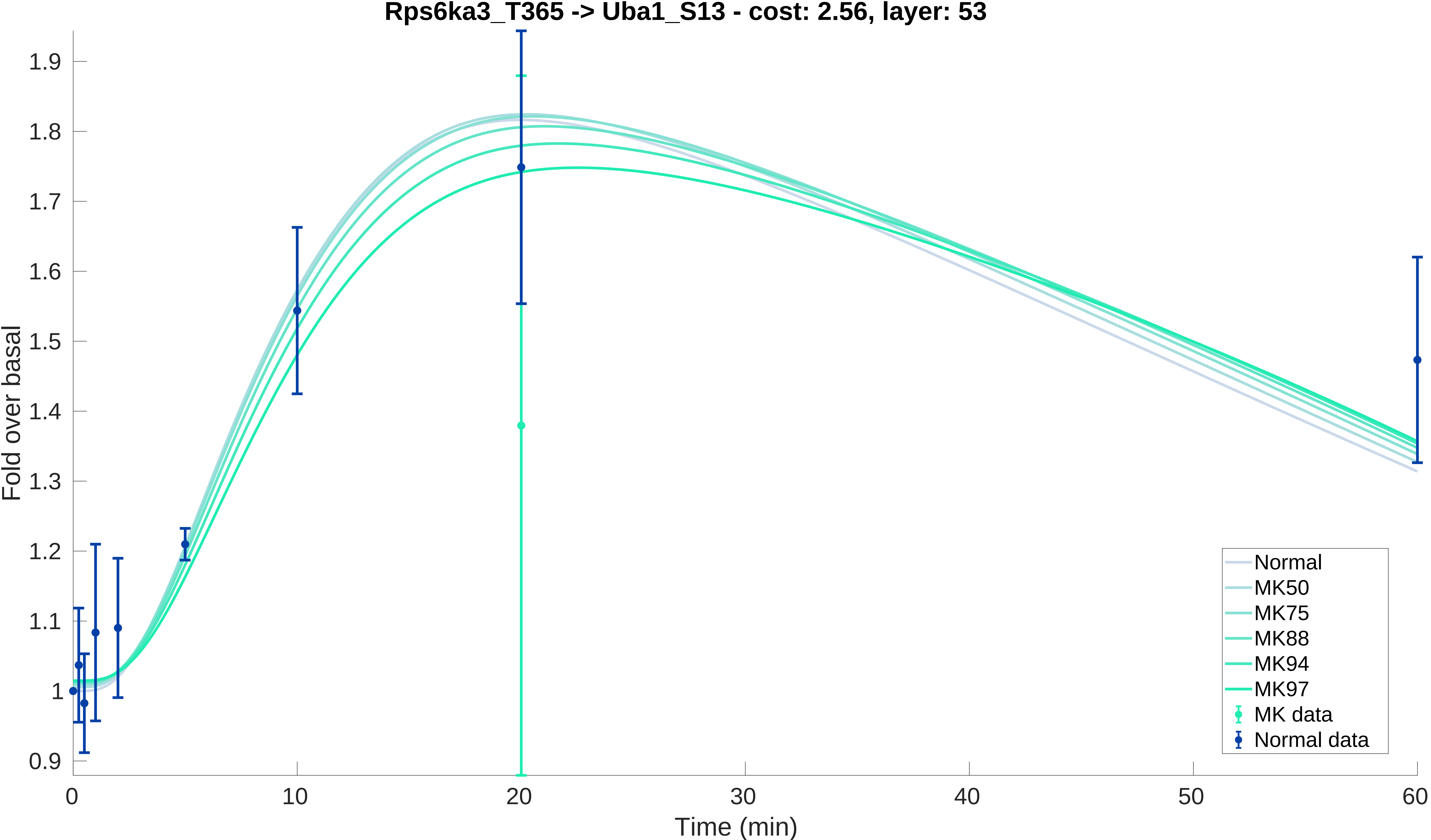

Rps6ka3_T365 -> Uba1_S13 - cost: 2.56, layer: 53
Fold over basal
1.9
1.8
1.7
1.6
1.5
1.4
1.3
1.2
1.1
1
0.9
0
10
20
30
40
50
60
Time (min)
Normal
MK50
MK75
MK88
MK94
MK97
MK data
Normal data

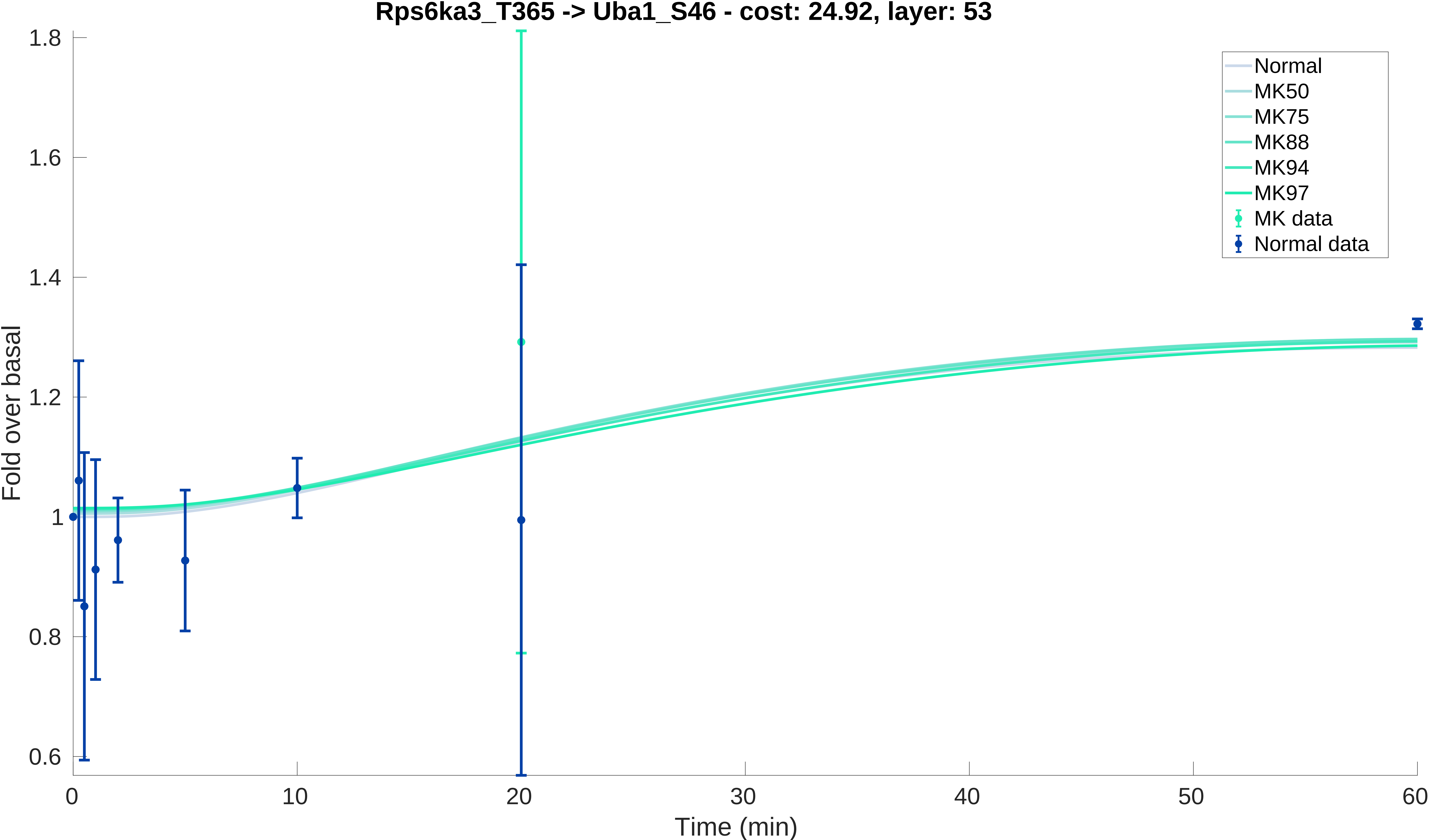

Rps6ka3_T365 -> Uba1_S46 - cost: 24.92, layer: 53
Fold over basal
Time (min)
0.6
0.8
1
1.2
1.4
1.6
1.8
0
10
20
30
40
50
60
Normal
MK50
MK75
MK88
MK94
MK97
MK data
Normal data

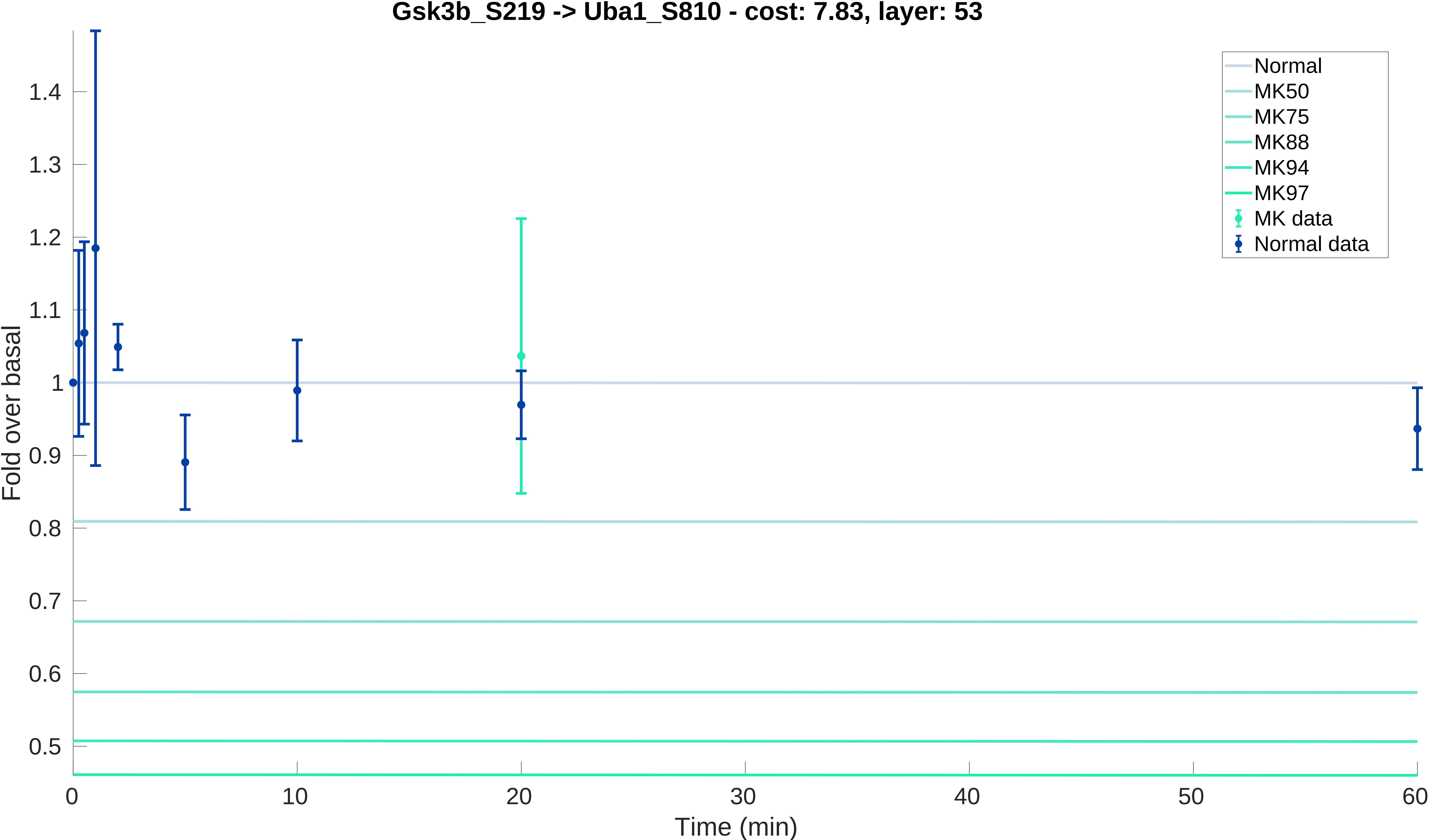

Gsk3b_S219 -> Uba1_S810 - cost: 7.83, layer: 53
Normal
MK50
MK75
MK88
MK94
MK97
MK data
Normal data
Fold over basal
1.4
1.3
1.2
1.1
1
0.9
0.8
0.7
0.6
0.5
0
10
20
30
40
50
60
Time (min)

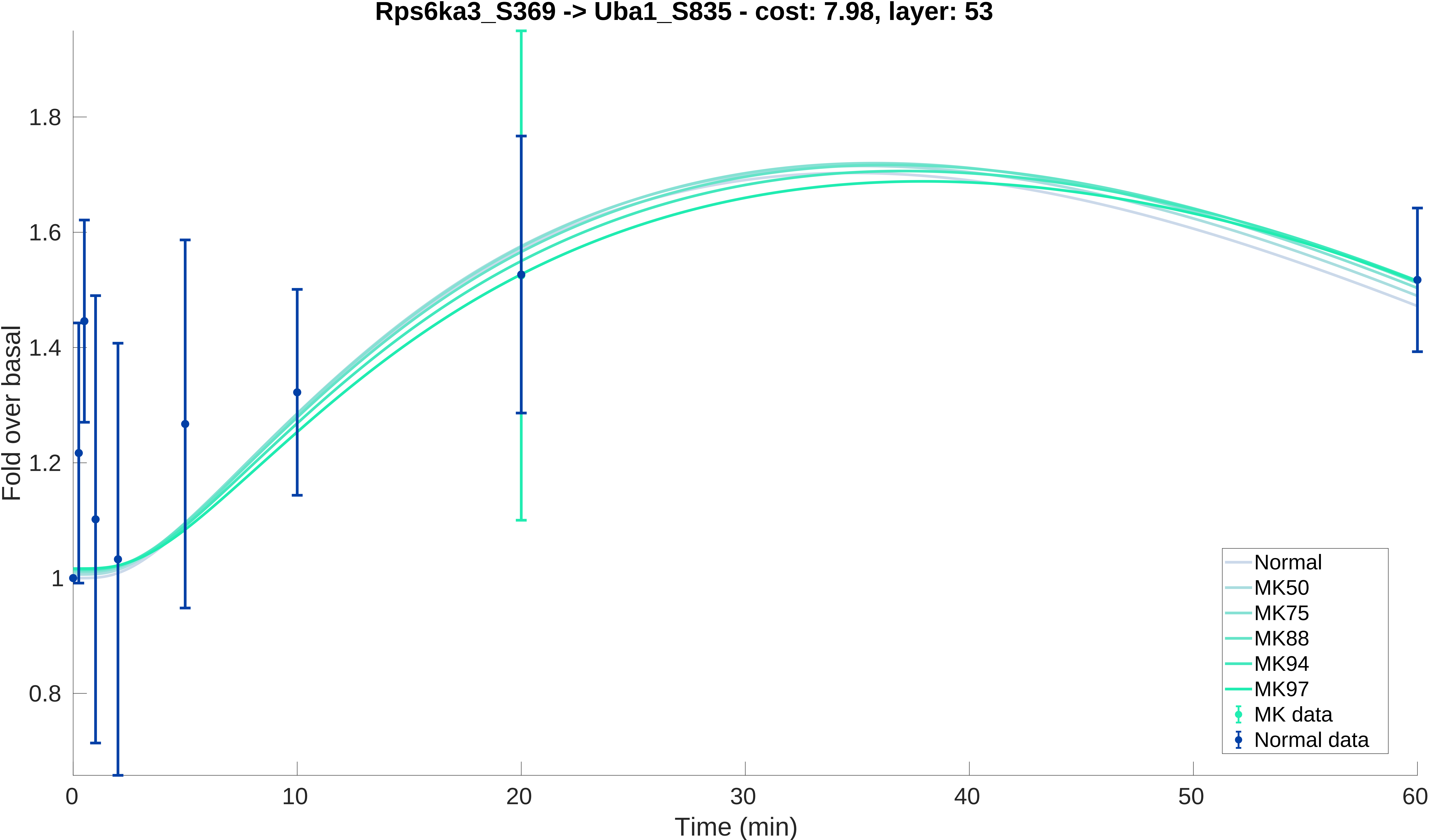

Rps6ka3_S369 -> Uba1_S835 - cost: 7.98, layer: 53
Fold over basal
Time (min)
1.8
1.6
1.4
1.2
1
0.8
0
10
20
30
40
50
60
Normal
MK50
MK75
MK88
MK94
MK97
MK data
Normal data

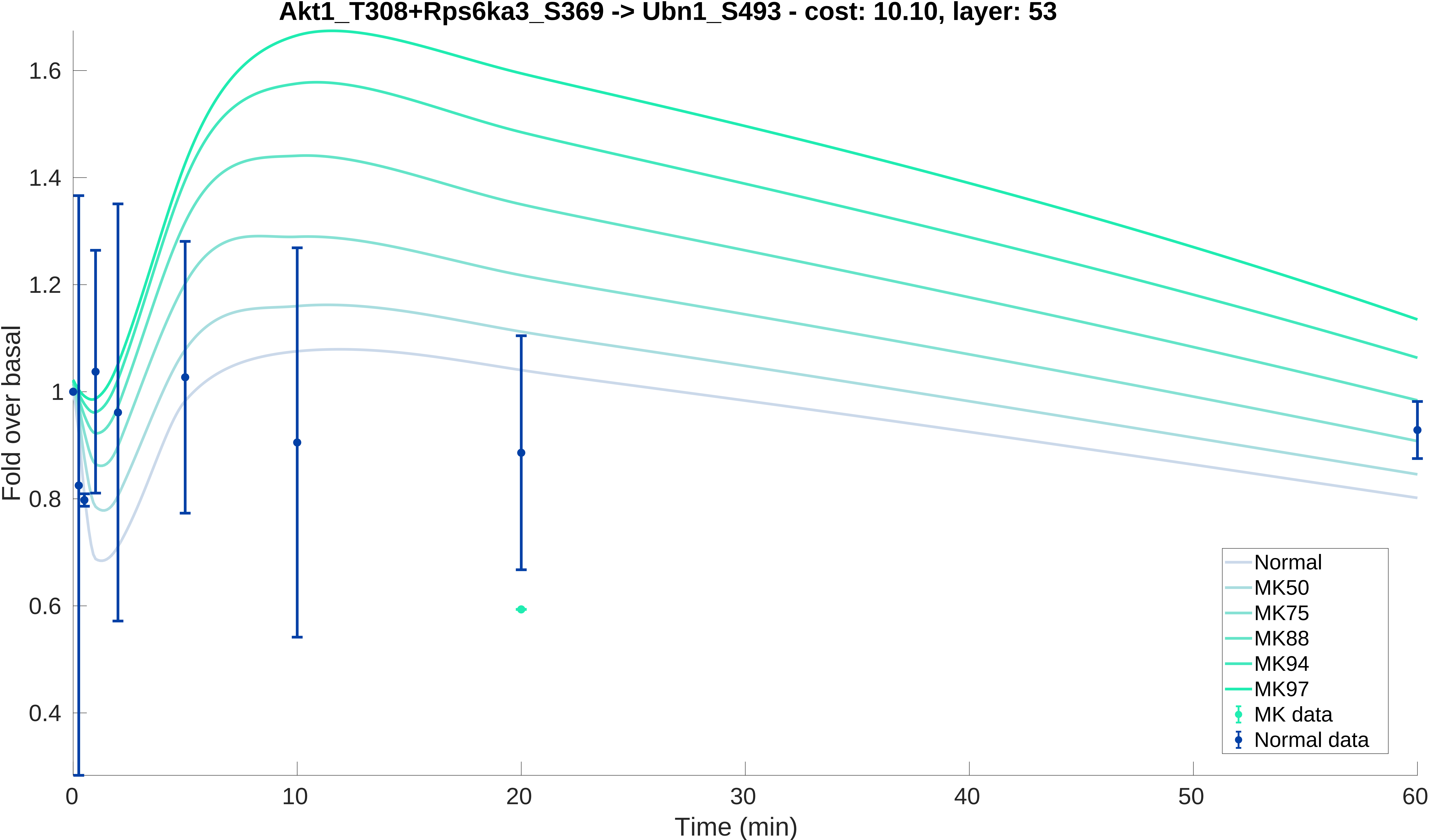

Akt1_T308+Rps6ka3_S369 -> Ubn1_S493 - cost: 10.10, layer: 53
Fold over basal
Time (min)
0.4
0.6
0.8
1
1.2
1.4
1.6
0
10
20
30
40
50
60
Normal
MK50
MK75
MK88
MK94
MK97
MK data
Normal data

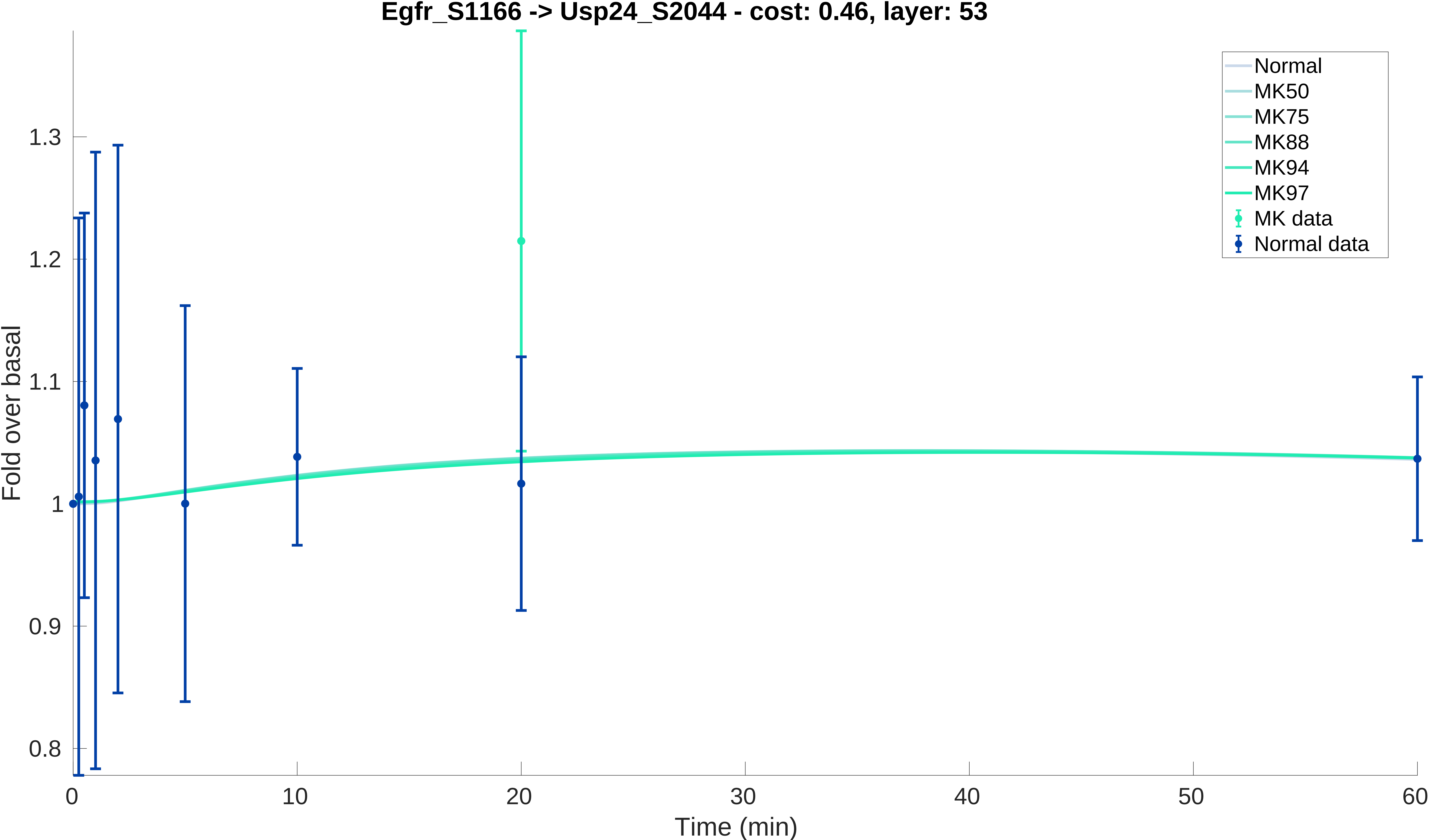

Egfr_S1166 -> Usp24_S2044 - cost: 0.46, layer: 53
Fold over basal
Time (min)
0.8
0.9
1
1.1
1.2
1.3
0
10
20
30
40
50
60
Normal
MK50
MK75
MK88
MK94
MK97
MK data
Normal data

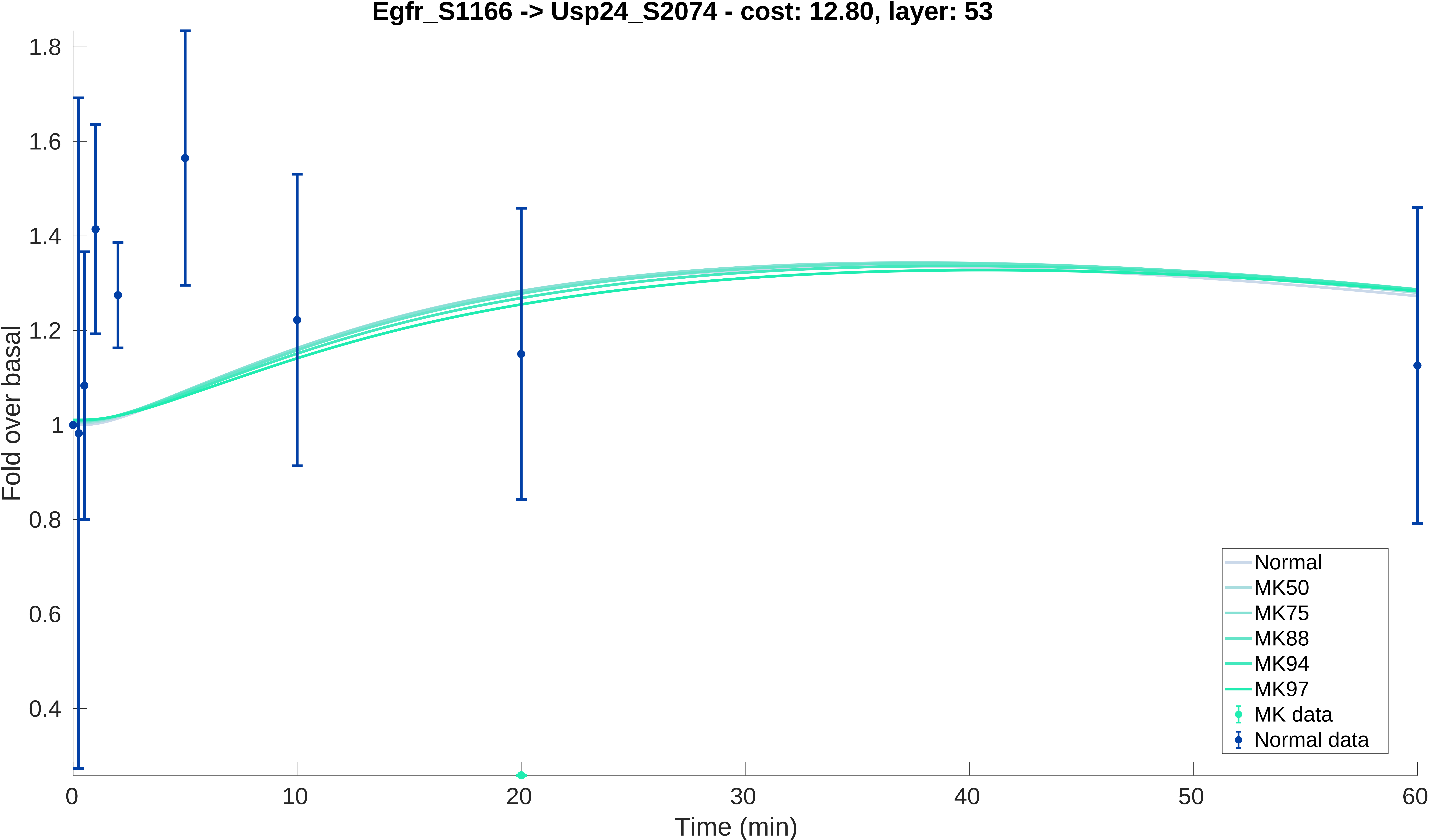
Egfr_S1166 -> Usp24_S2074 - cost: 12.80, layer: 53
Fold over basal
Time (min)
0.4
0.6
0.8
1
1.2
1.4
1.6
1.8
0
10
20
30
40
50
60
Normal
MK50
MK75
MK88
MK94
MK97
MK data
Normal data

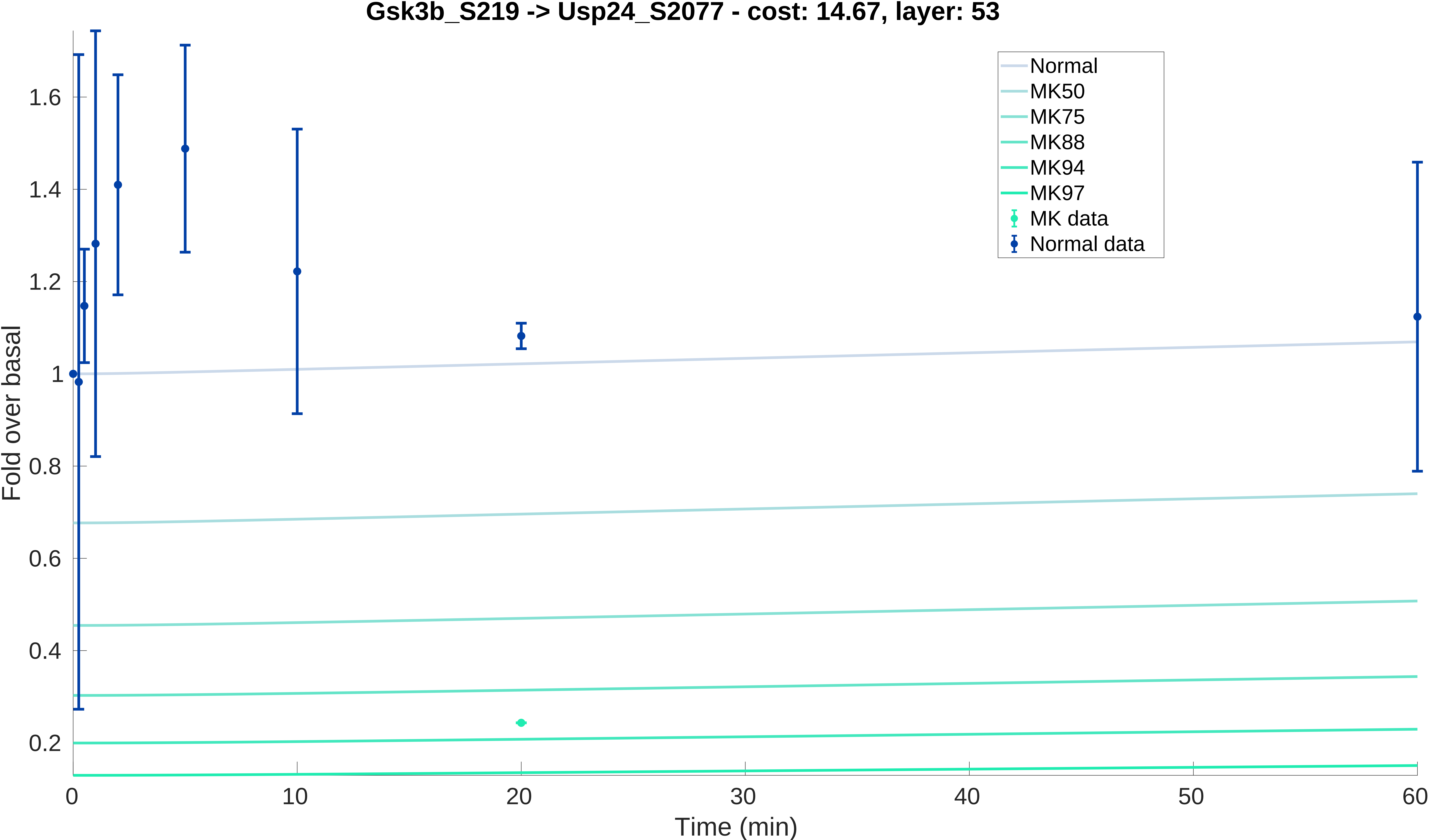
Gsk3b_S219 -> Usp24_S2077 - cost: 14.67, layer: 53
Fold over basal
Time (min)
0
10
20
30
40
50
60
0.2
0.4
0.6
0.8
1
1.2
1.4
1.6
Normal
MK50
MK75
MK88
MK94
MK97
MK data
Normal data

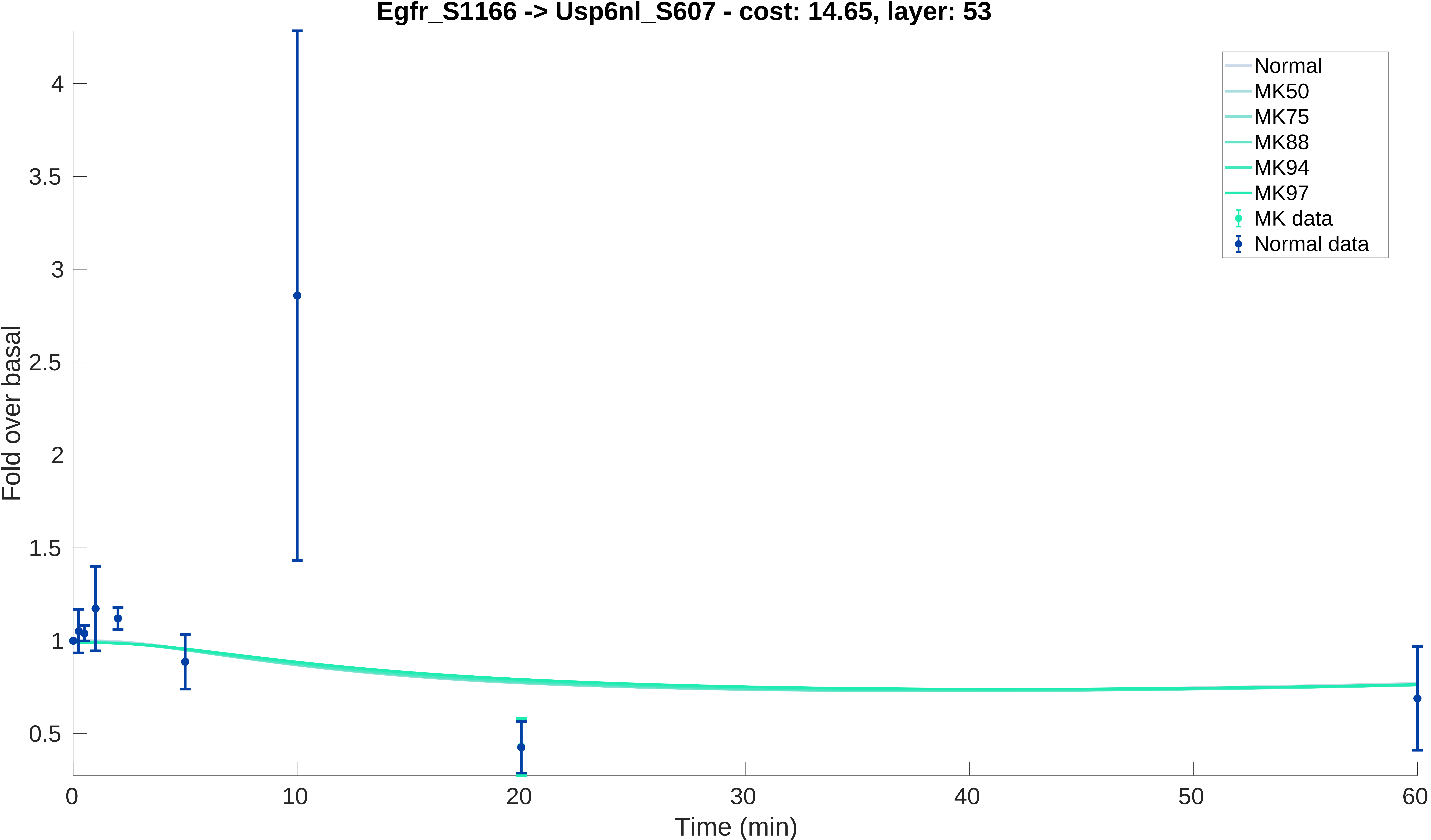

Egfr_S1166 -> Usp6nl_S607 - cost: 14.65, layer: 53
Fold over basal
Time (min)
4
3.5
3
2.5
2
1.5
1
0.5
0
10
20
30
40
50
60
Normal
MK50
MK75
MK88
MK94
MK97
MK data
Normal data

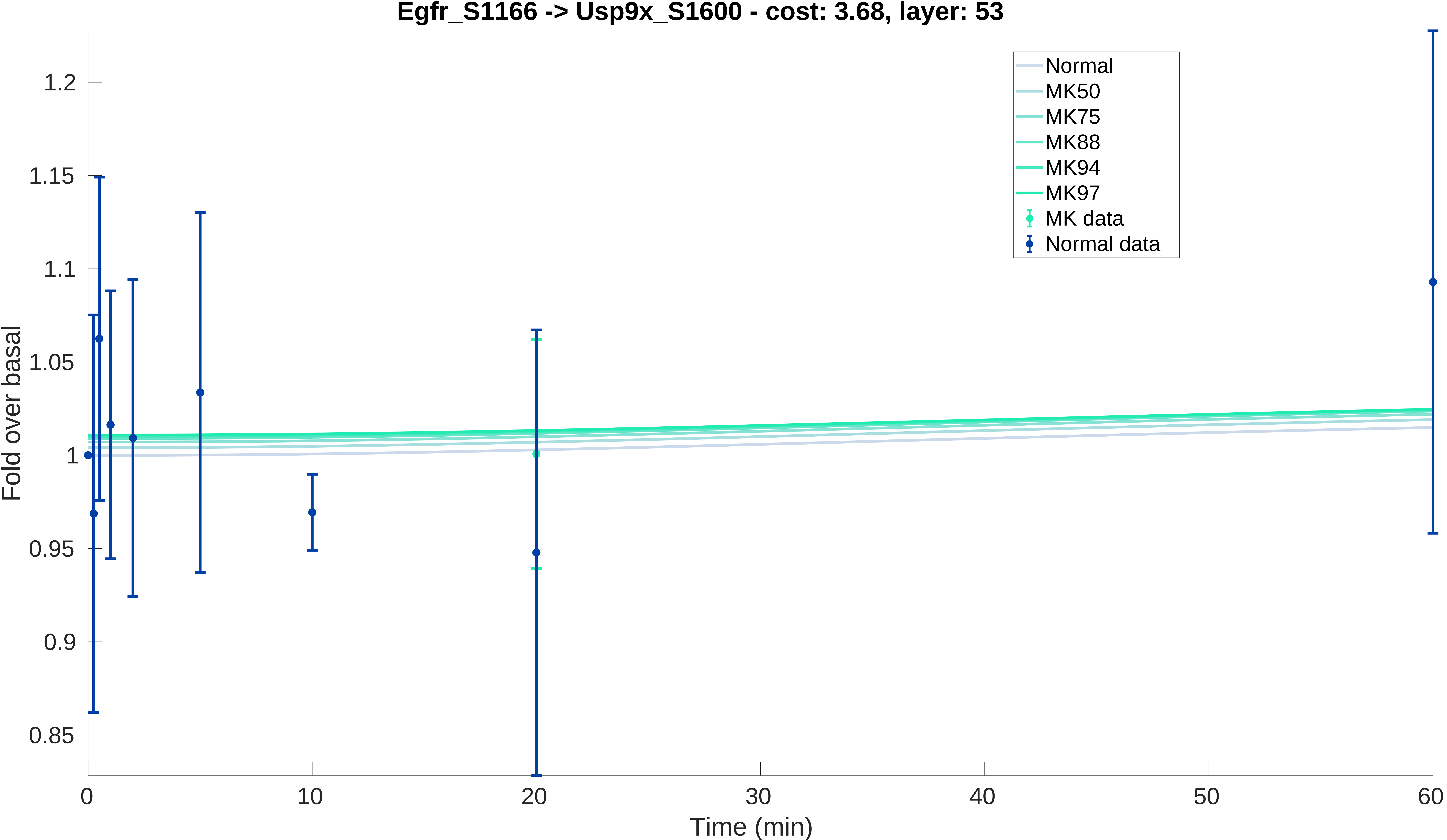

Egfr_S1166 -> Usp9x_S1600 - cost: 3.68, layer: 53
Normal
MK50
MK75
MK88
MK94
MK97
MK data
Normal data
Fold over basal
1.2
1.15
1.1
1.05
1
0.95
0.9
0.85
0
10
20
30
40
50
60
Time (min)

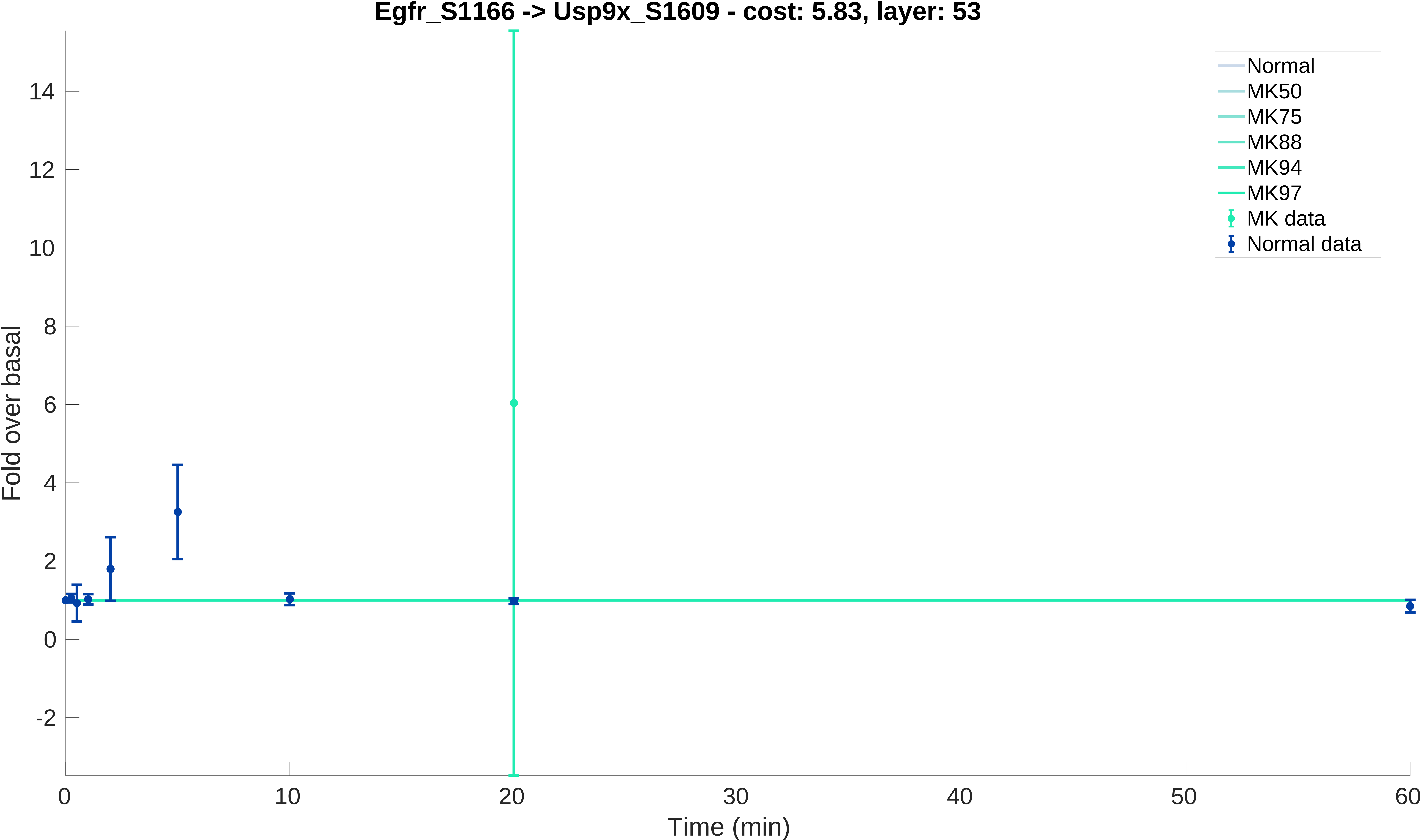

Egfr_S1166 -> Usp9x_S1609 - cost: 5.83, layer: 53
Normal
MK50
MK75
MK88
MK94
MK97
MK data
Normal data
Fold over basal
14
12
10
8
6
4
2
0
-2
0
10
20
30
40
50
60
Time (min)

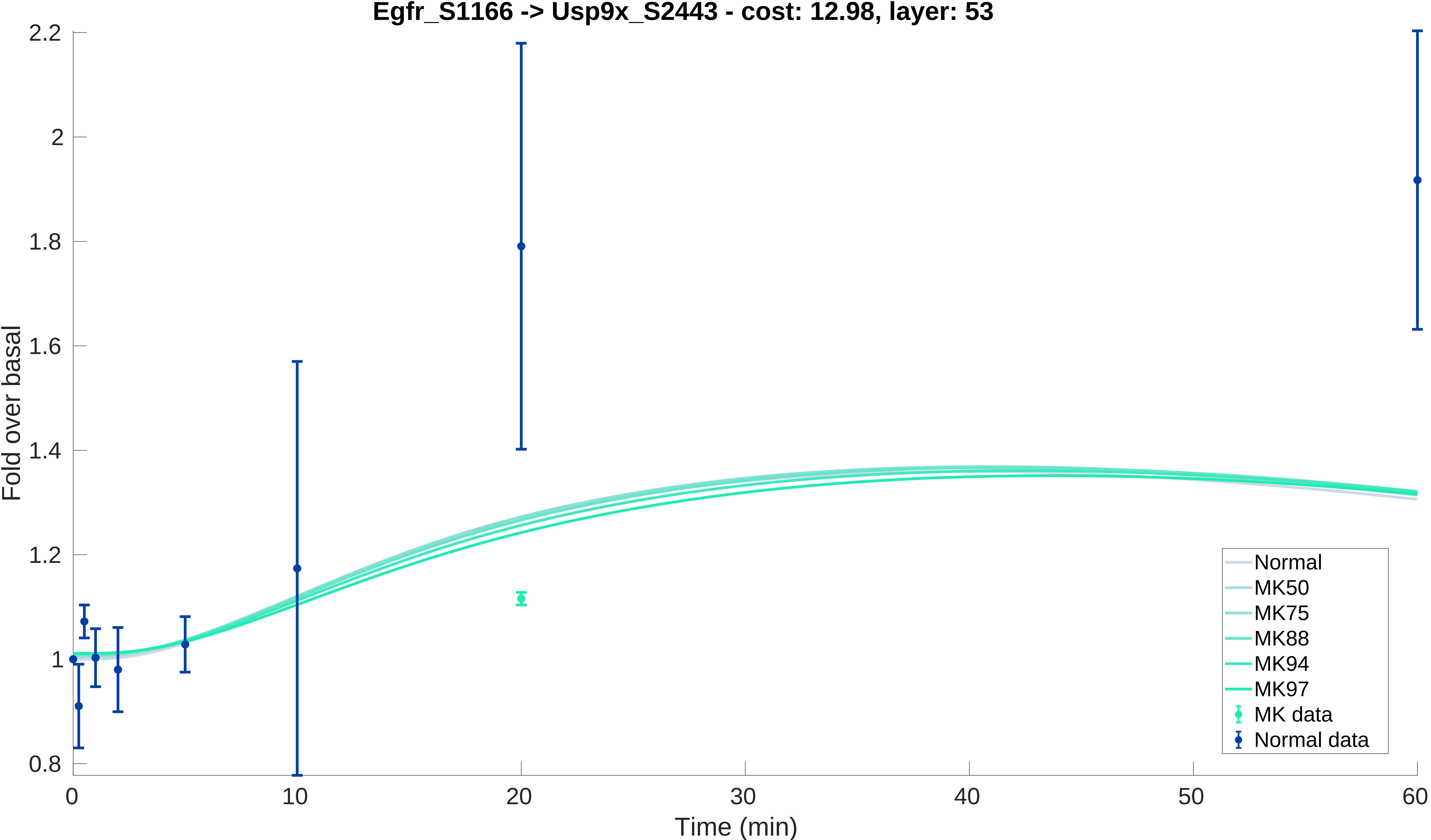

Egfr_S1166 -> Usp9x_S2443 - cost: 12.98, layer: 53
Fold over basal
Time (min)
0.8
1
1.2
1.4
1.6
1.8
2
2.2
0
10
20
30
40
50
60
Normal
MK50
MK75
MK88
MK94
MK97
MK data
Normal data

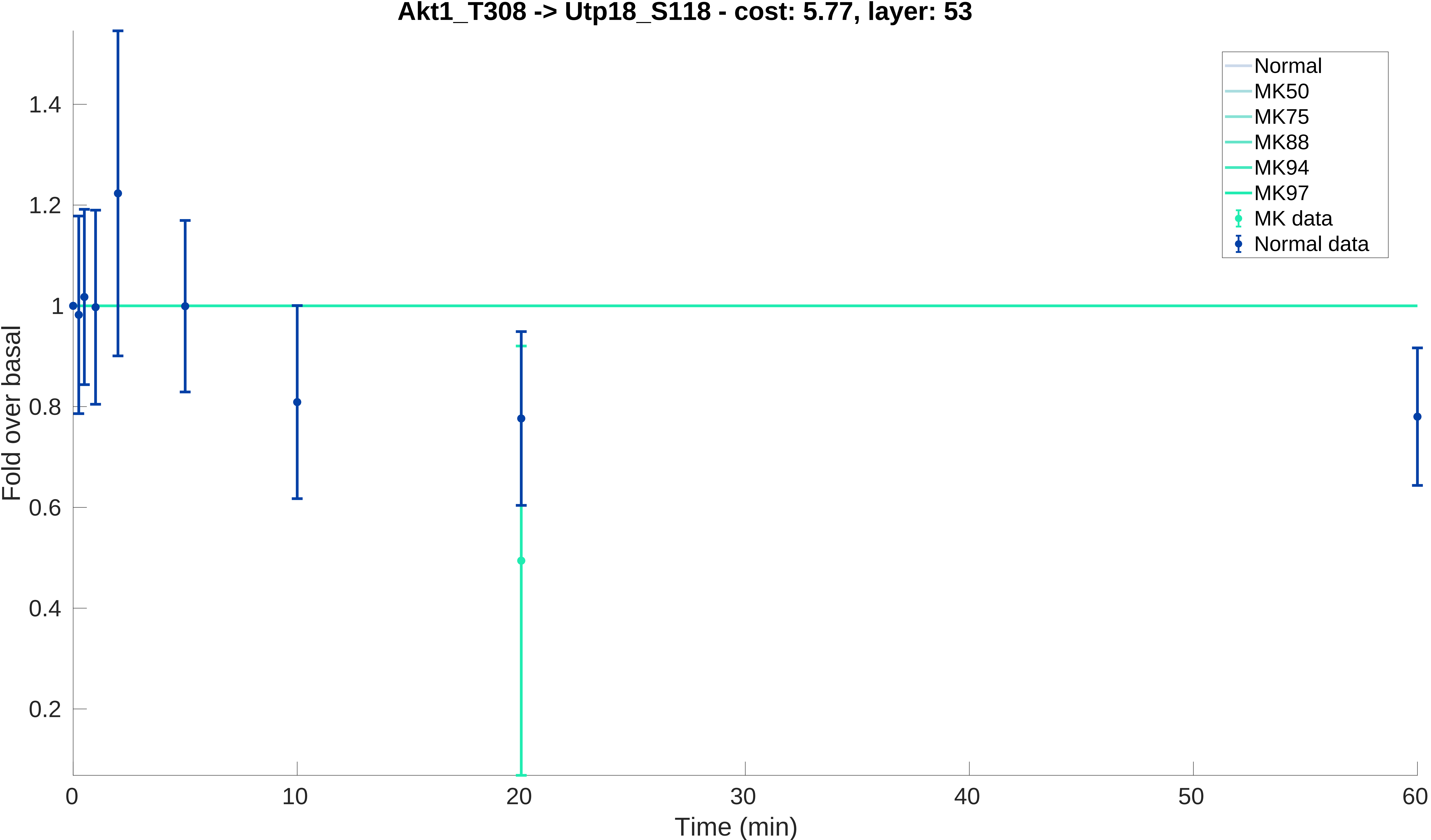

Akt1_T308 -> Utp18_S118 - cost: 5.77, layer: 53
Fold over basal
Time (min)
0
10
20
30
40
50
60
0.2
0.4
0.6
0.8
1
1.2
1.4
Normal
MK50
MK75
MK88
MK94
MK97
MK data
Normal data

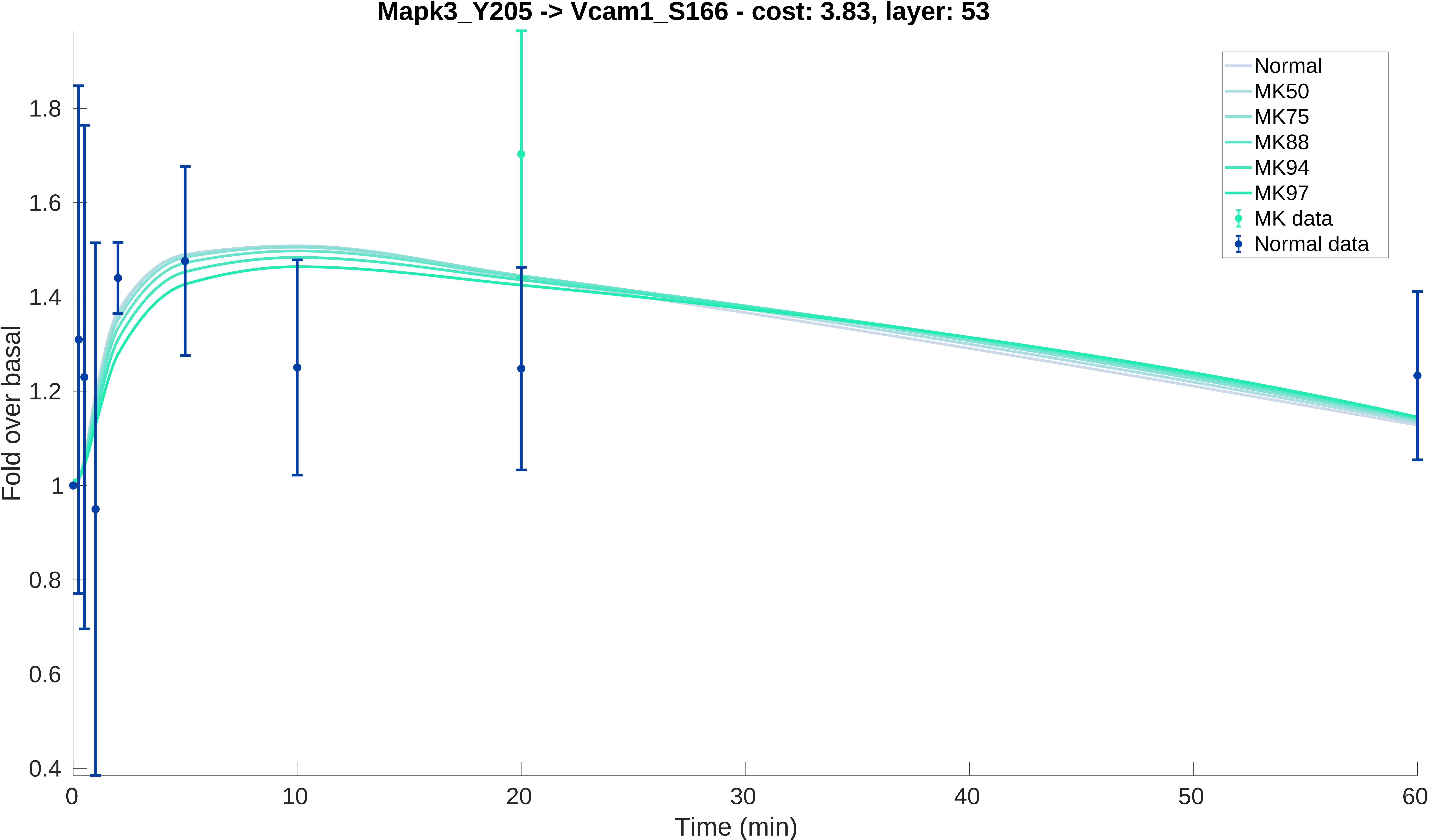

Mapk3_Y205 -> Vcam1_S166 - cost: 3.83, layer: 53
Fold over basal
Time (min)
0.4
0.6
0.8
1
1.2
1.4
1.6
1.8
0
10
20
30
40
50
60
Normal
MK50
MK75
MK88
MK94
MK97
MK data
Normal data

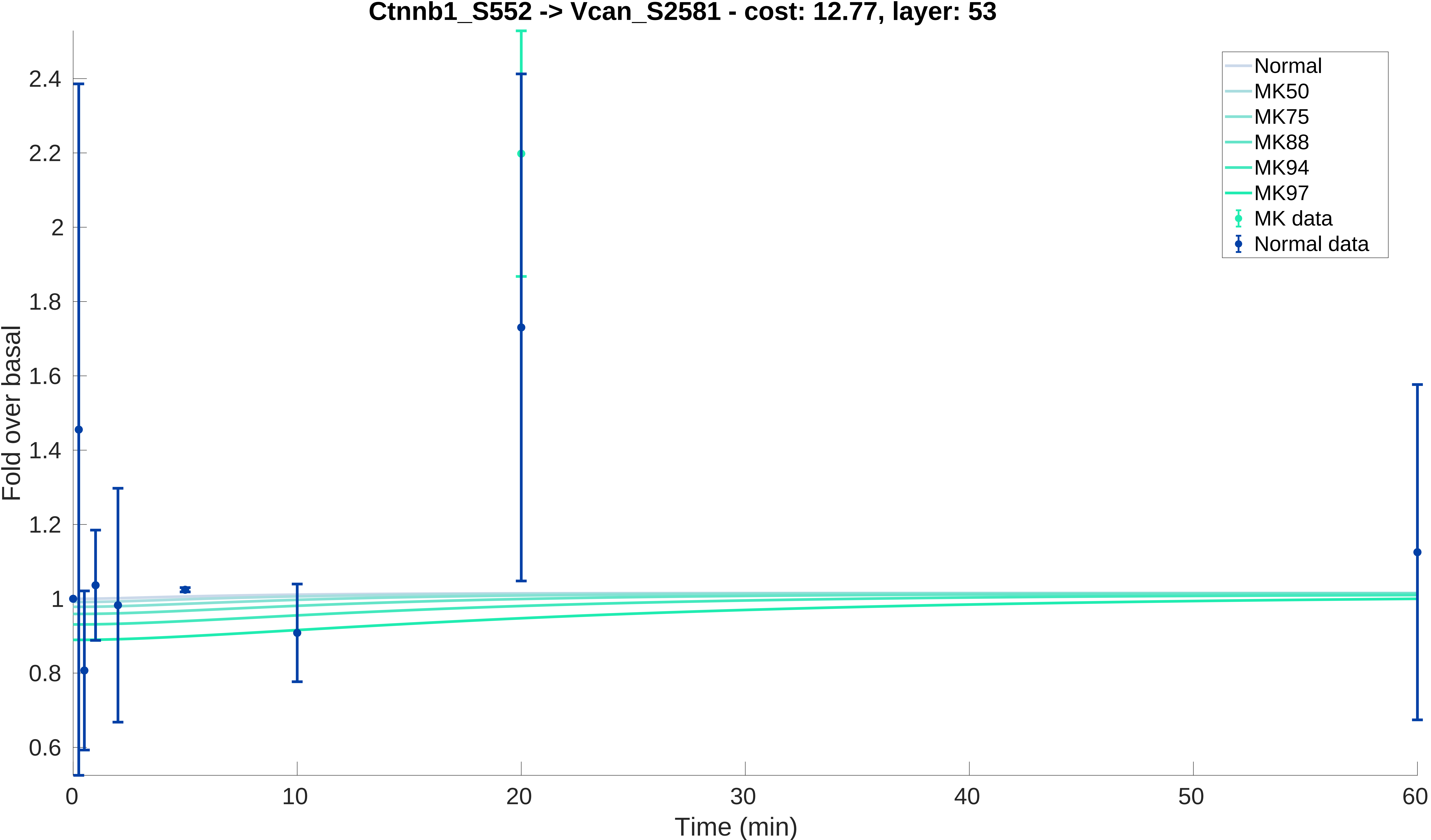

Ctnnb1_S552 -> Vcan_S2581 - cost: 12.77, layer: 53
Fold over basal
Time (min)
0.6
0.8
1
1.2
1.4
1.6
1.8
2
2.2
2.4
0
10
20
30
40
50
60
Normal
MK50
MK75
MK88
MK94
MK97
MK data
Normal data

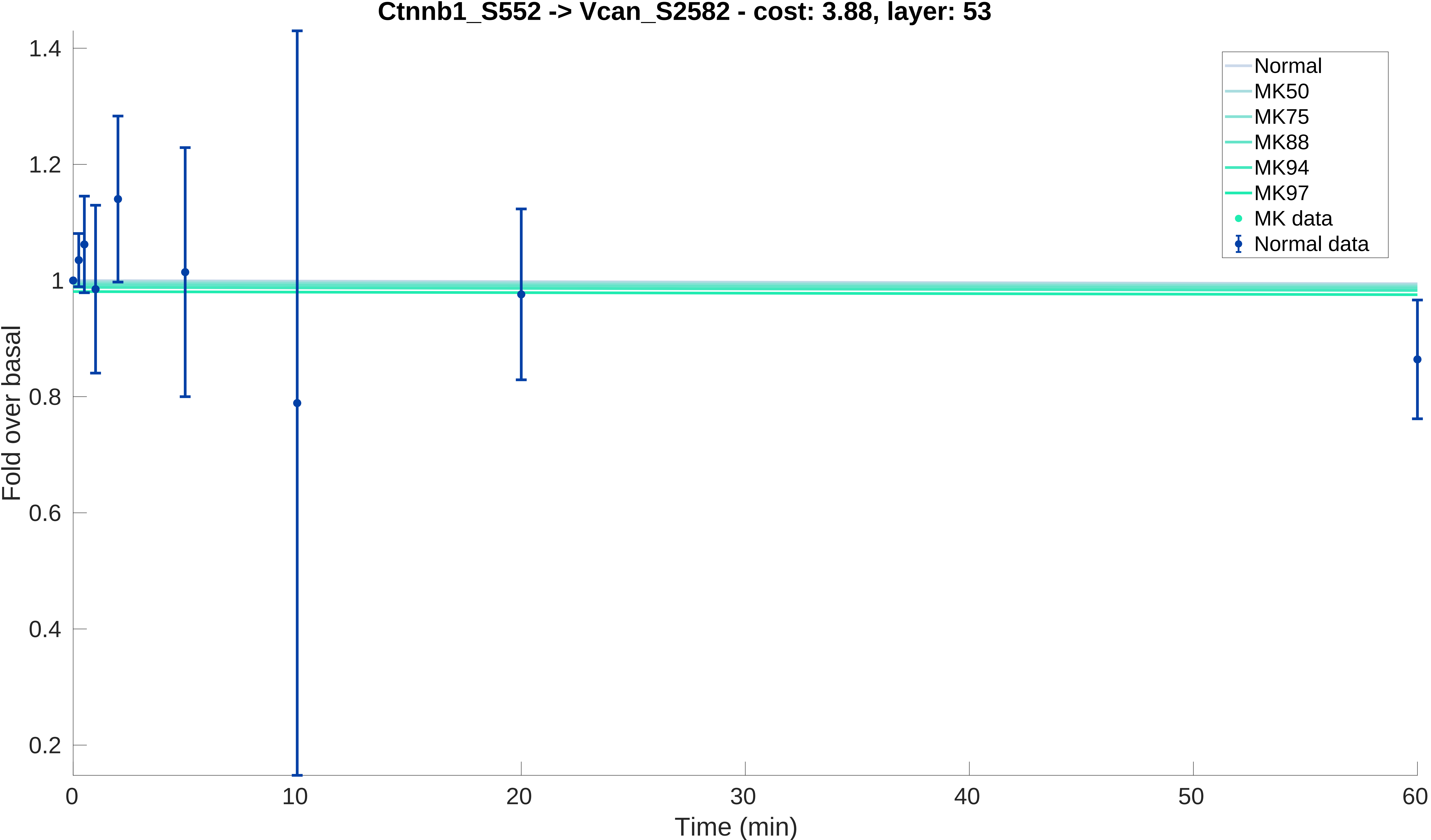

Ctnnb1_S552 -> Vcan_S2582 - cost: 3.88, layer: 53
Fold over basal
Time (min)
1.4
1.2
1
0.8
0.6
0.4
0.2
0
10
20
30
40
50
60
Normal
MK50
MK75
MK88
MK94
MK97
MK data
Normal data

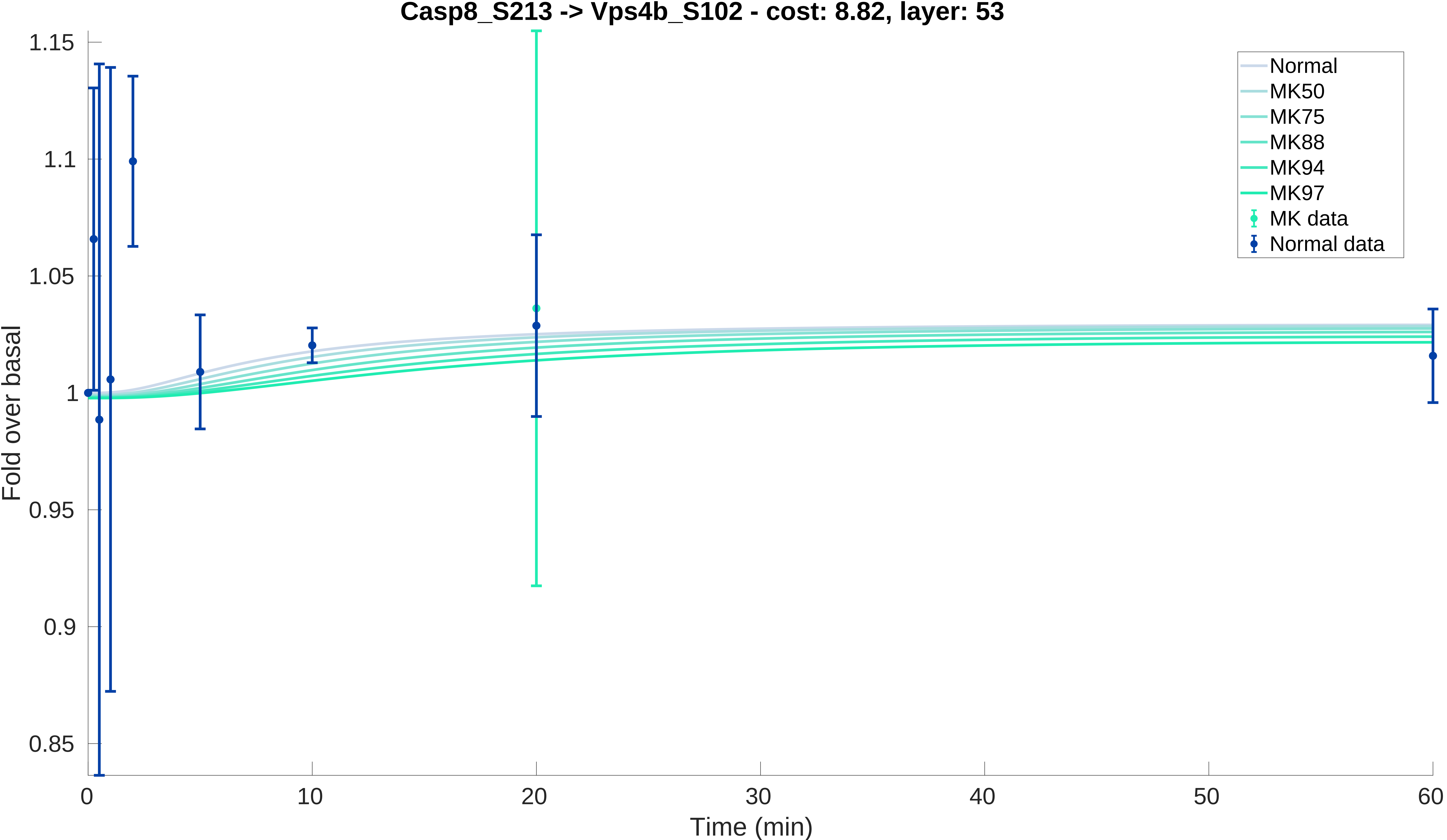

Casp8_S213 -> Vps4b_S102 - cost: 8.82, layer: 53
1.15
1.1
1.05
1
0.95
0.9
0.85
Fold over basal
0
10
20
30
40
50
60
Time (min)
Normal
MK50
MK75
MK88
MK94
MK97
MK data
Normal data

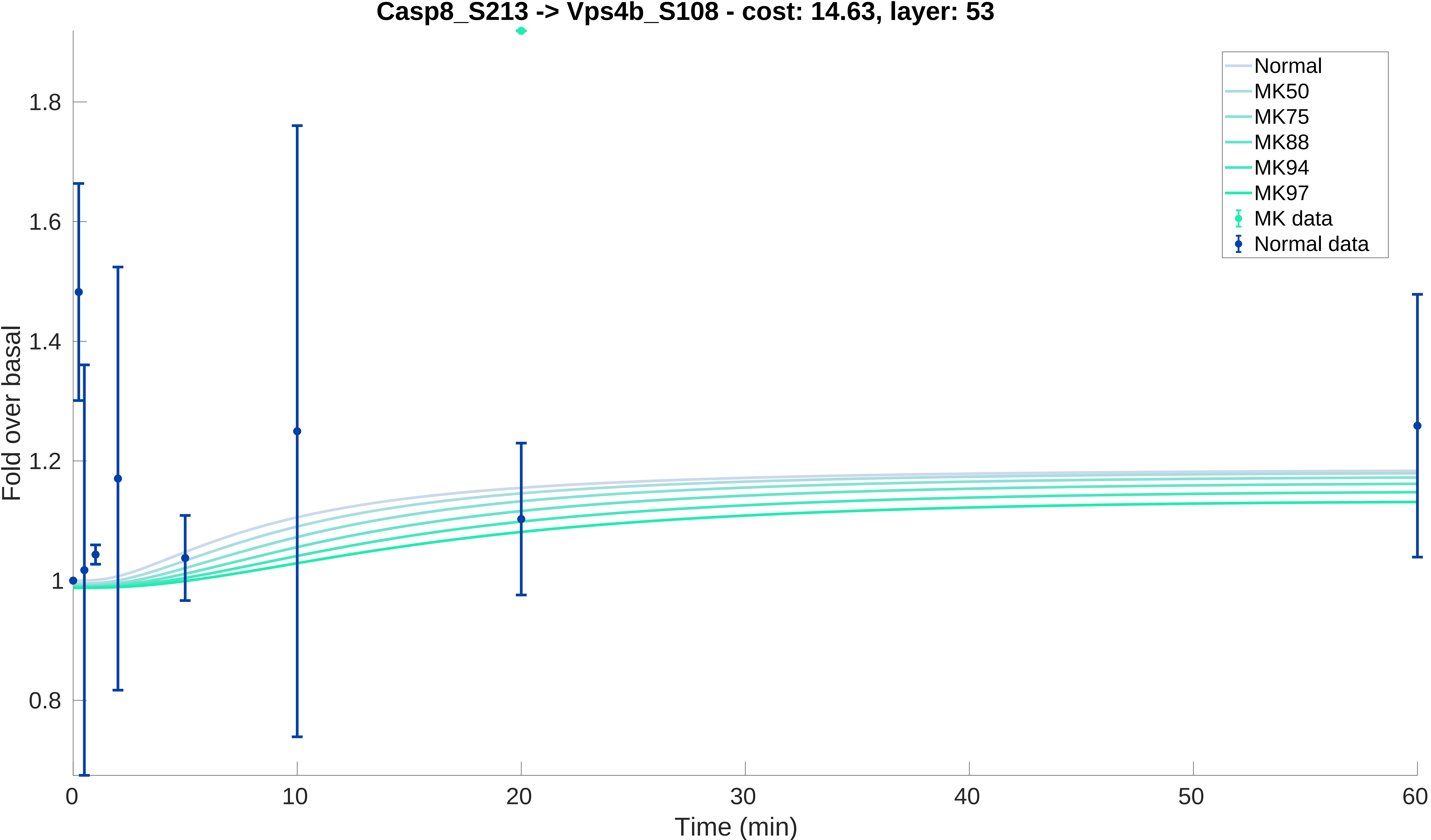

Casp8_S213 -> Vps4b_S108 - cost: 14.63, layer: 53
Normal
MK50
MK75
MK88
MK94
MK97
MK data
Normal data
Fold over basal
1.8
1.6
1.4
1.2
1
0.8
0
10
20
30
40
50
60
Time (min)

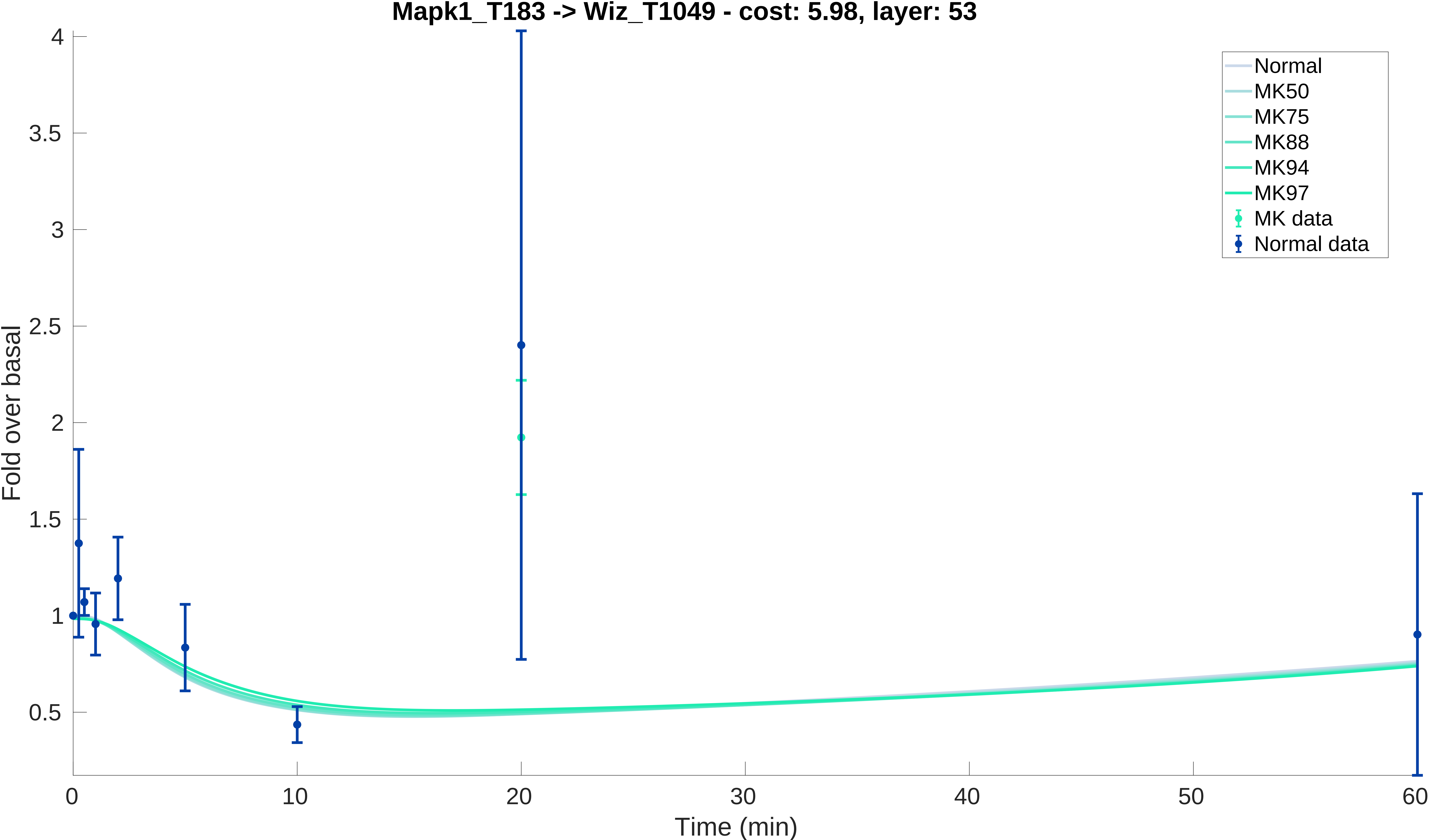

Mapk1_T183 -> Wiz_T1049 - cost: 5.98, layer: 53
Fold over basal
Time (min)
4
3.5
3
2.5
2
1.5
1
0.5
0
10
20
30
40
50
60
Normal
MK50
MK75
MK88
MK94
MK97
MK data
Normal data

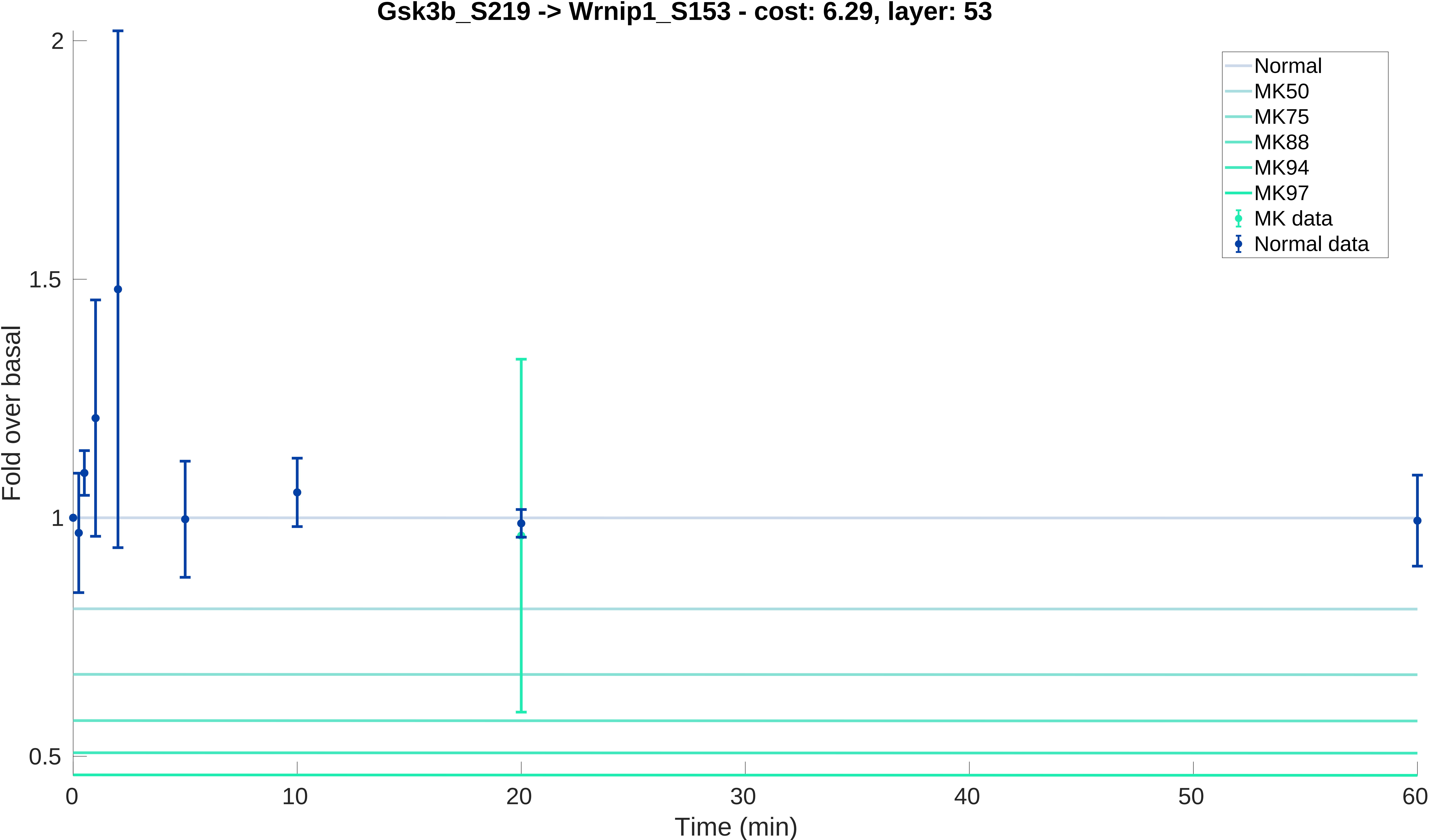

Gsk3b_S219 -> Wrnip1_S153 - cost: 6.29, layer: 53
Fold over basal
Time (min)
0.5
1
1.5
2
0
10
20
30
40
50
60
Normal
MK50
MK75
MK88
MK94
MK97
MK data
Normal data

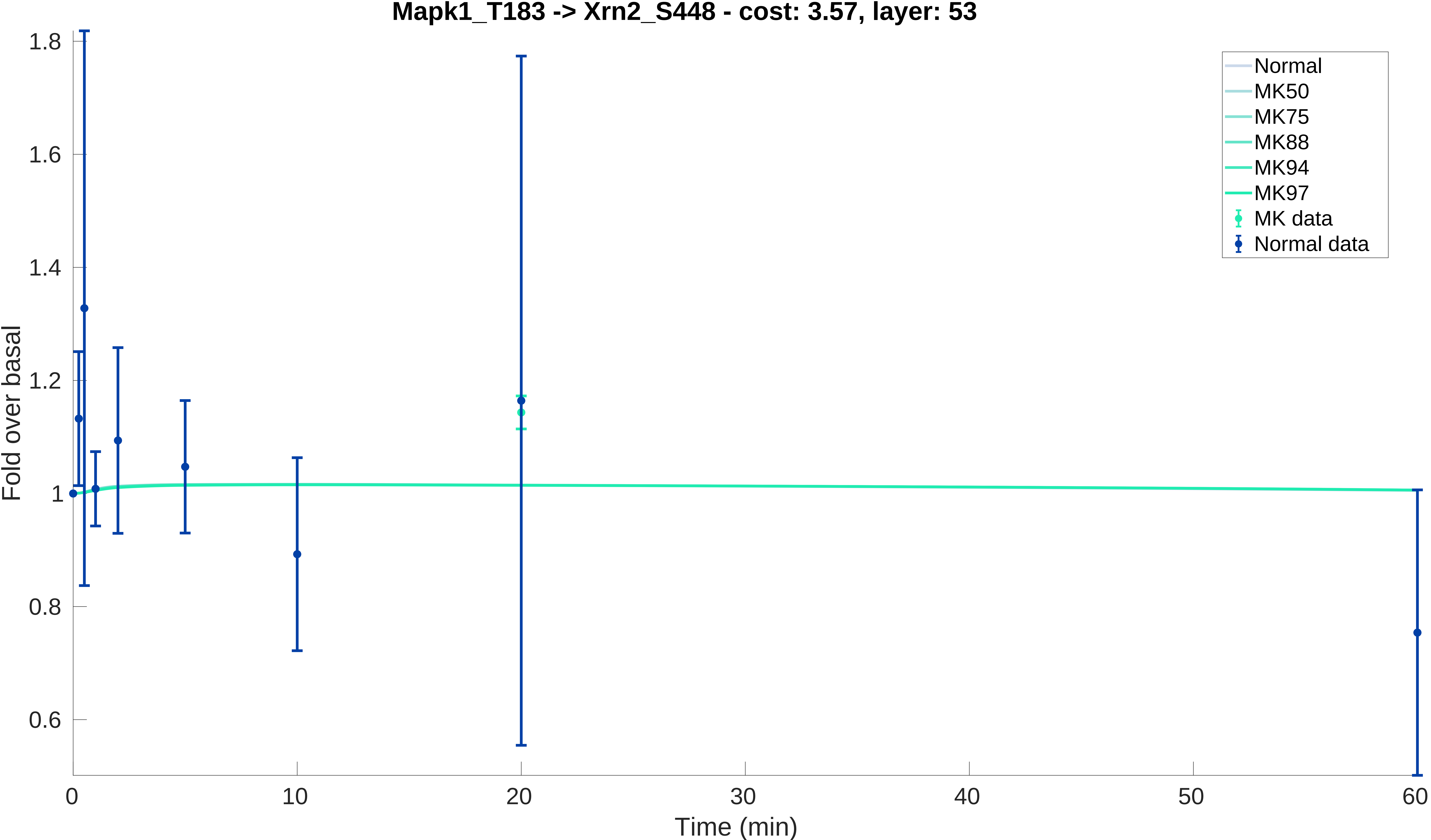

Mapk1_T183 -> Xrn2_S448 - cost: 3.57, layer: 53
Fold over basal
Time (min)
0.6
0.8
1
1.2
1.4
1.6
1.8
0
10
20
30
40
50
60
Normal
MK50
MK75
MK88
MK94
MK97
MK data
Normal data

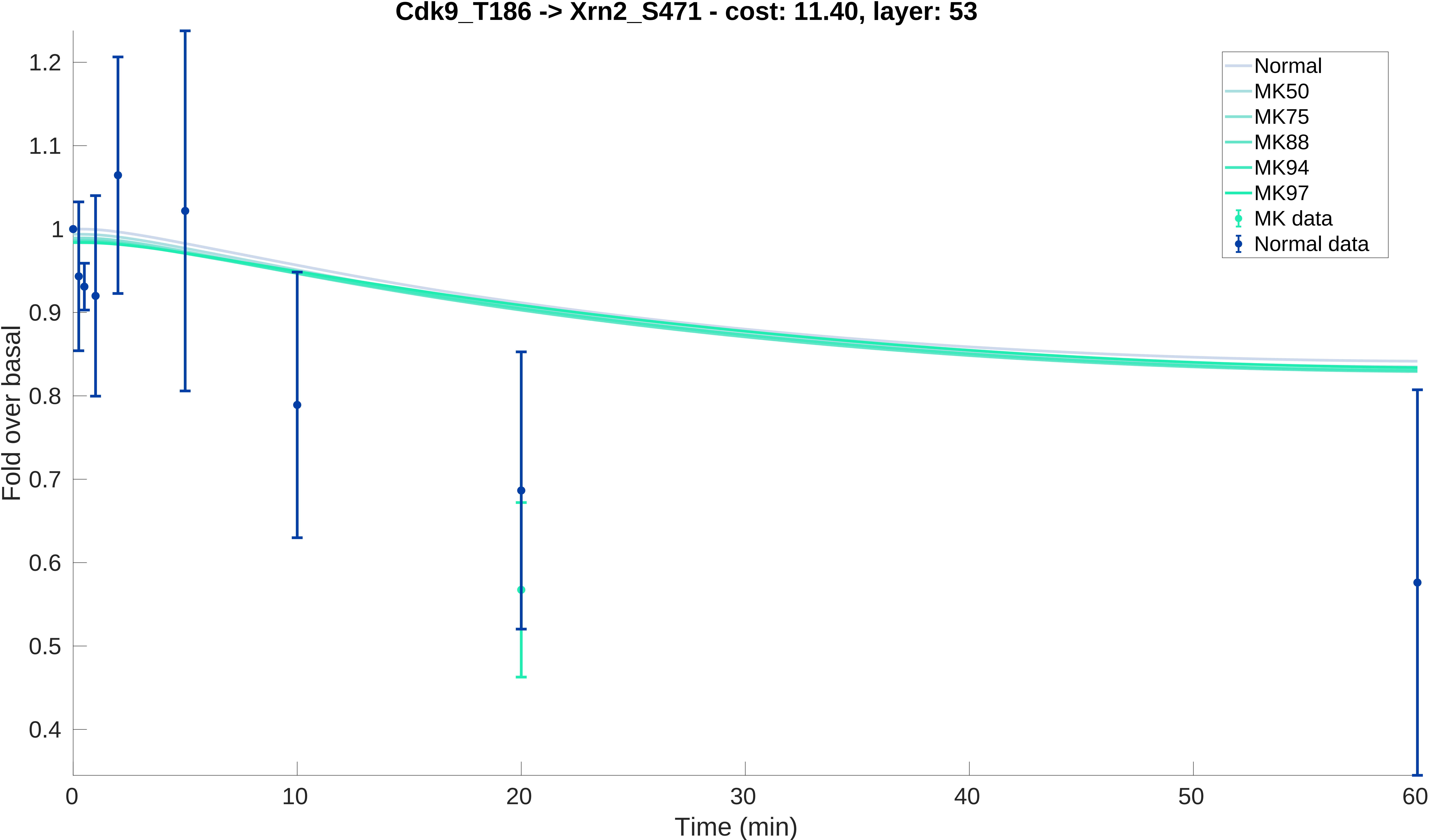
Cdk9_T186 -> Xrn2_S471 - cost: 11.40, layer: 53
Fold over basal
1.2
1.1
1
0.9
0.8
0.7
0.6
0.5
0.4
Time (min)
0
10
20
30
40
50
60
Normal
MK50
MK75
MK88
MK94
MK97
MK data
Normal data

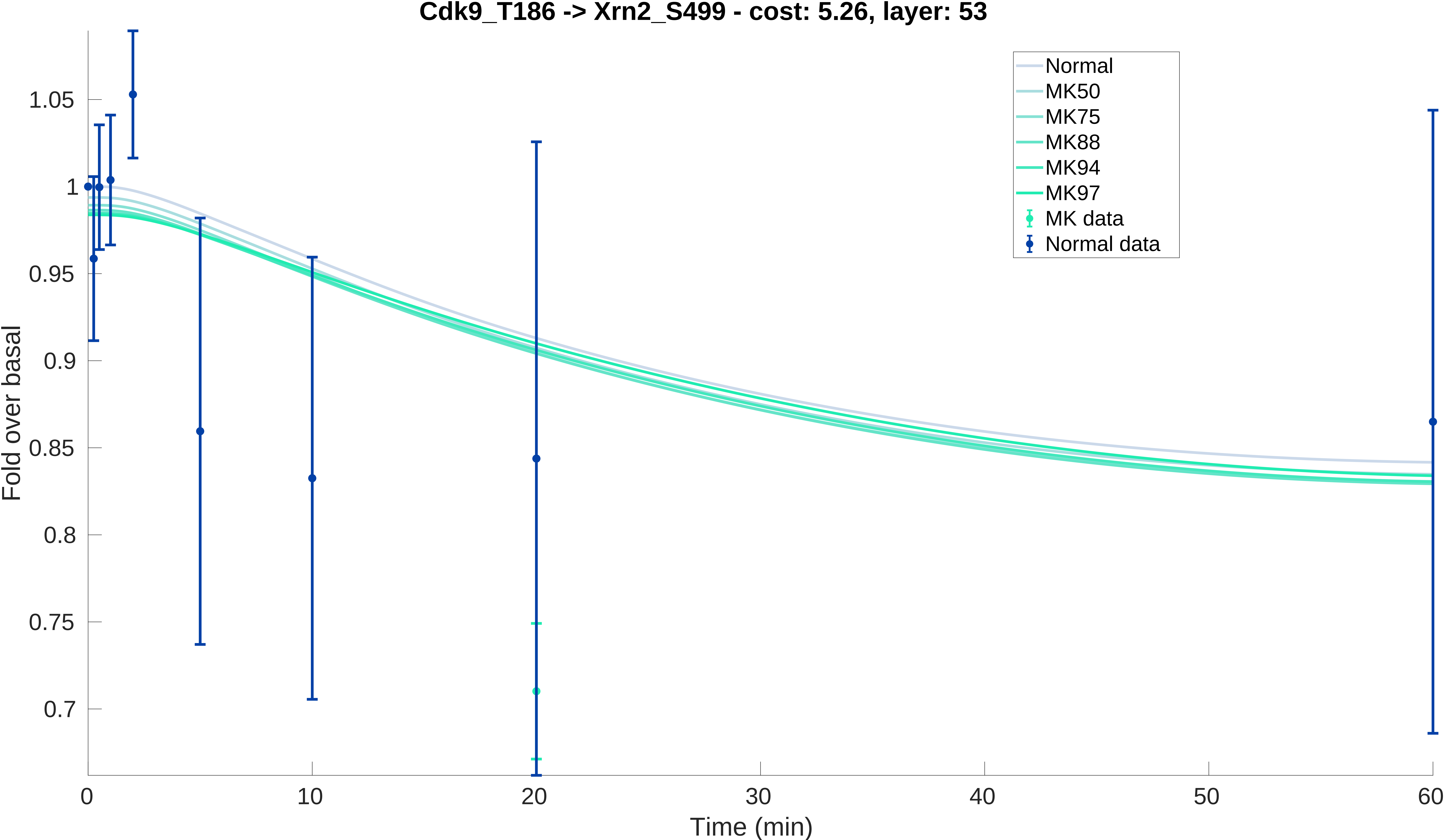
Cdk9_T186 -> Xrn2_S499 - cost: 5.26, layer: 53
Normal
MK50
MK75
MK88
MK94
MK97
MK data
Normal data
Fold over basal
Time (min)
1.05
1
0.95
0.9
0.85
0.8
0.75
0.7
0
10
20
30
40
50
60

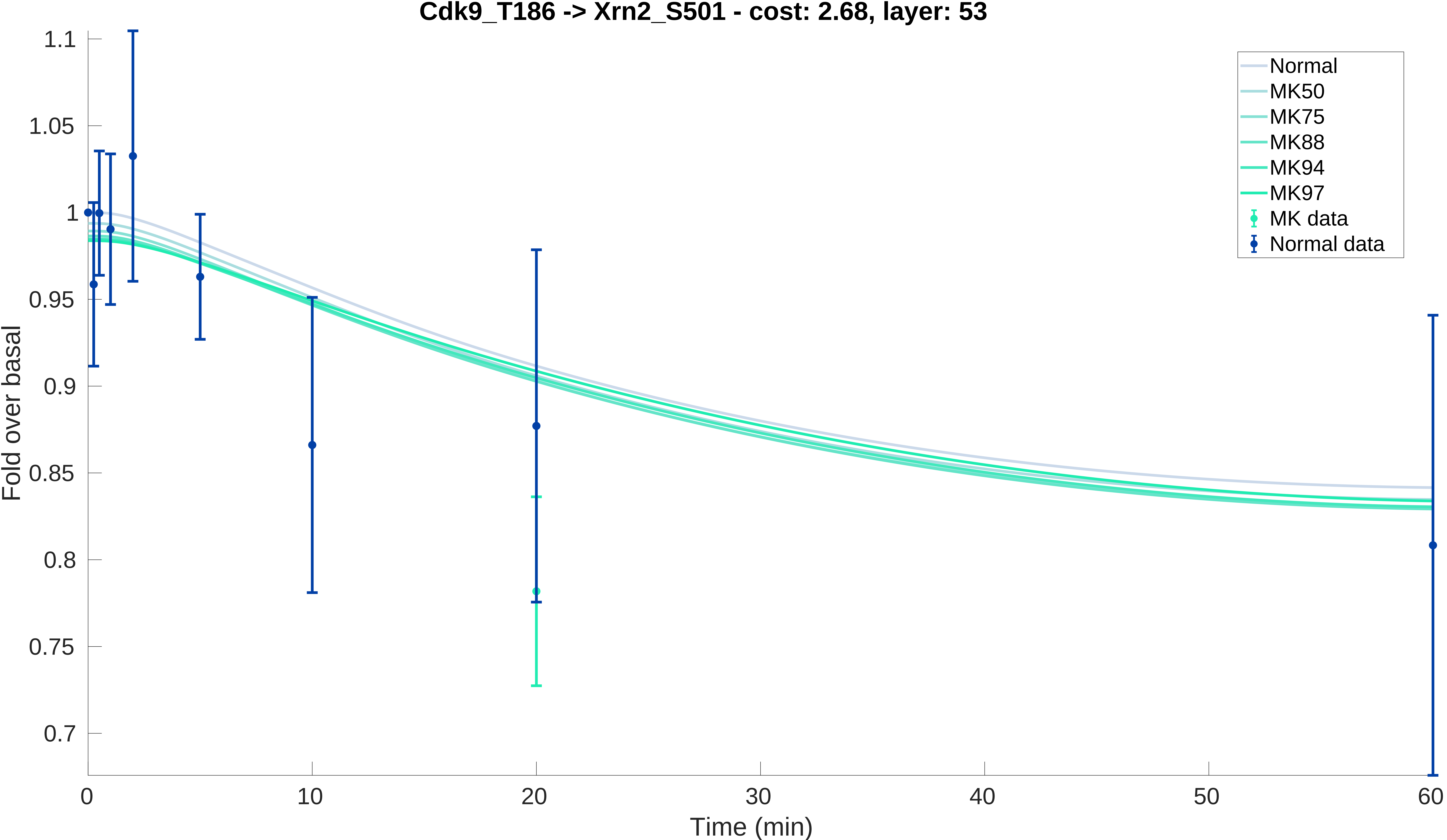

Cdk9_T186 -> Xrn2_S501 - cost: 2.68, layer: 53
1.1
1.05
1
0.95
0.9
0.85
0.8
0.75
0.7
Fold over basal
0
10
20
30
40
50
60
Time (min)
Normal
MK50
MK75
MK88
MK94
MK97
MK data
Normal data

#### Gsk3b_Y216 -> Zc3hav1_S269 - cost: 4.71, layer: 53

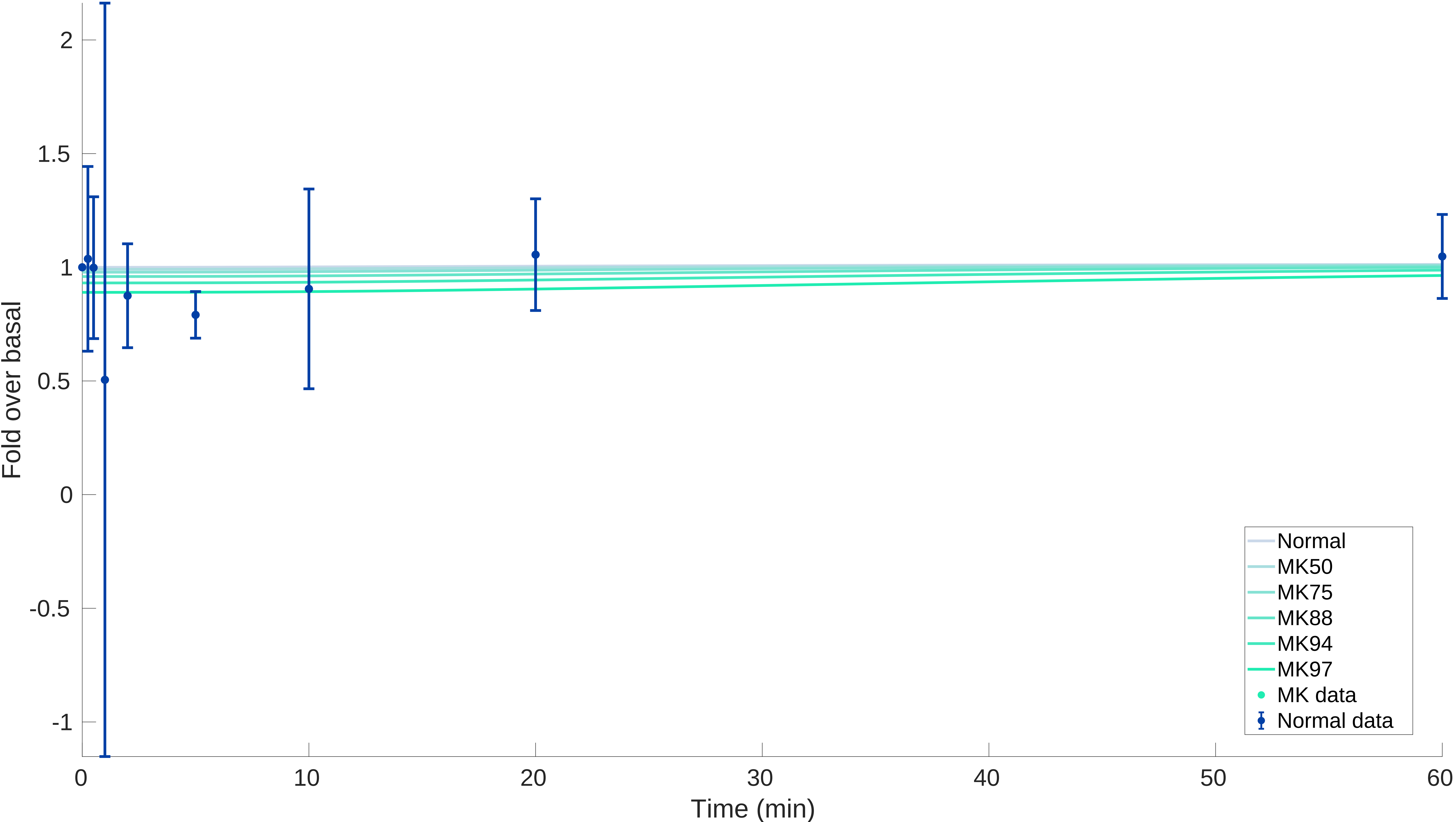

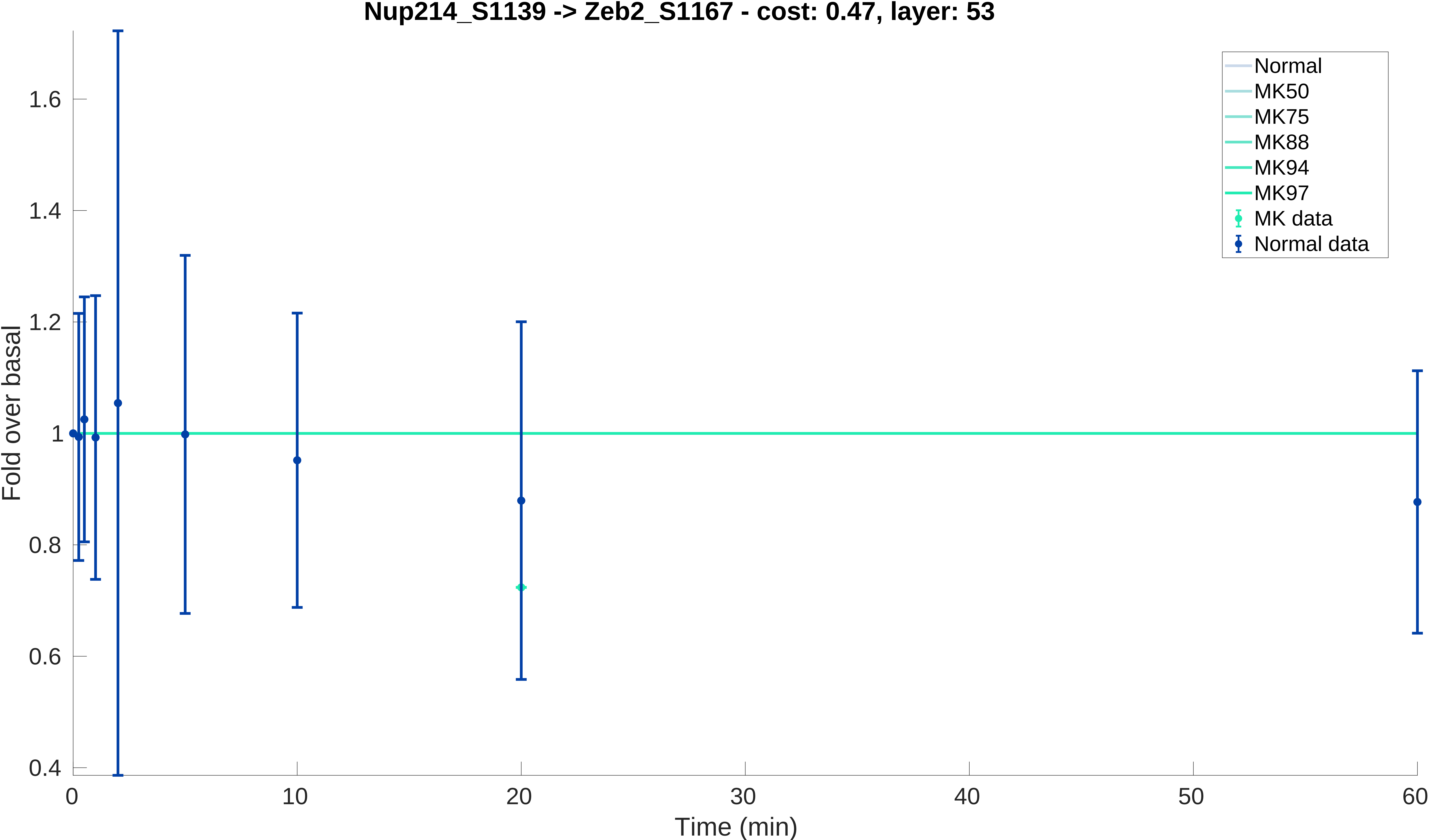

Nup214_S1139 -> Zeb2_S1167 - cost: 0.47, layer: 53
Normal
MK50
MK75
MK88
MK94
MK97
MK data
Normal data
Fold over basal
Time (min)
0.4
0.6
0.8
1
1.2
1.4
1.6
0
10
20
30
40
50
60

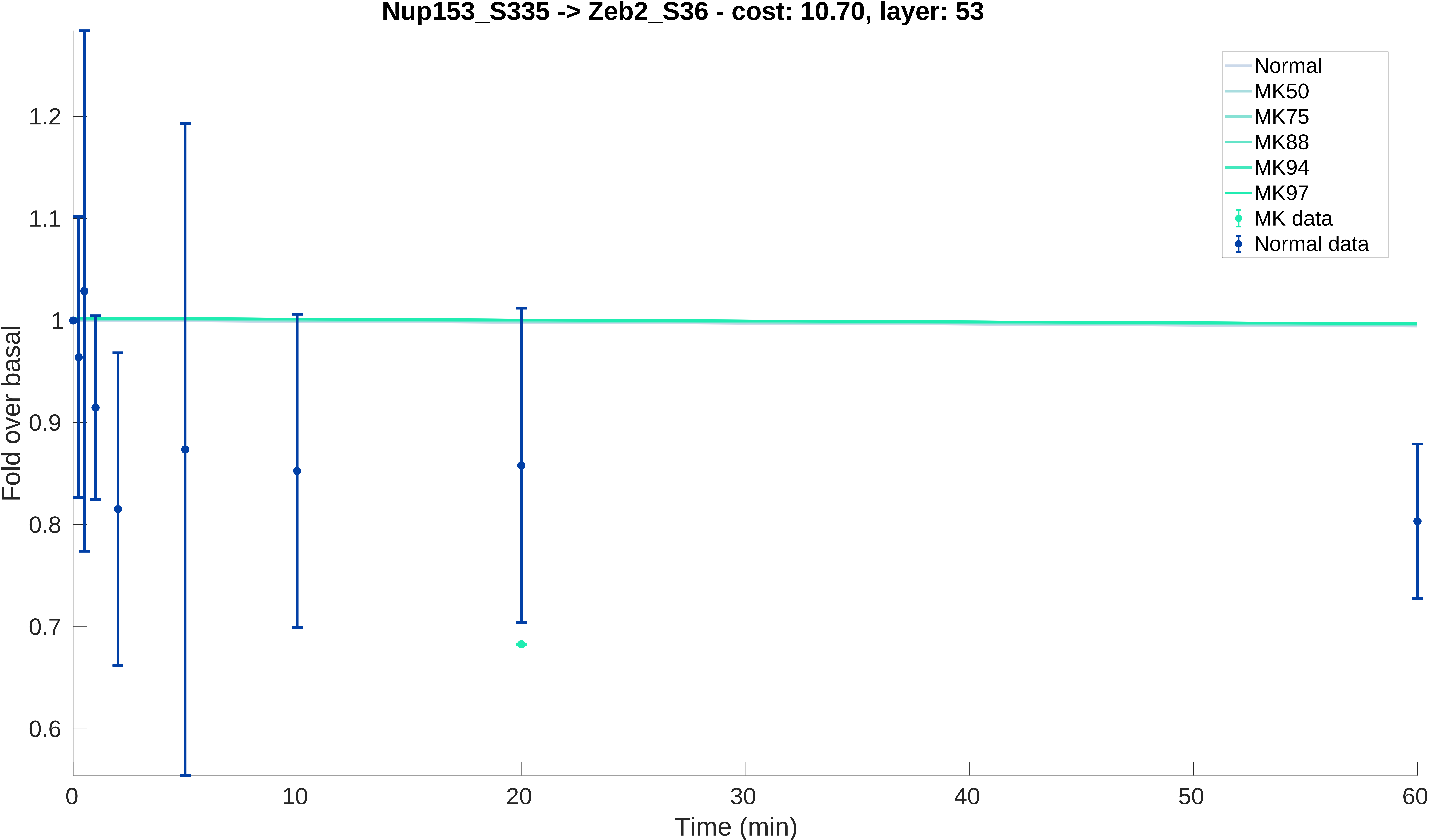

Nup153_S335 -> Zeb2_S36 - cost: 10.70, layer: 53
Fold over basal
Time (min)
0.6
0.7
0.8
0.9
1
1.1
1.2
0
10
20
30
40
50
60
Normal
MK50
MK75
MK88
MK94
MK97
MK data
Normal data

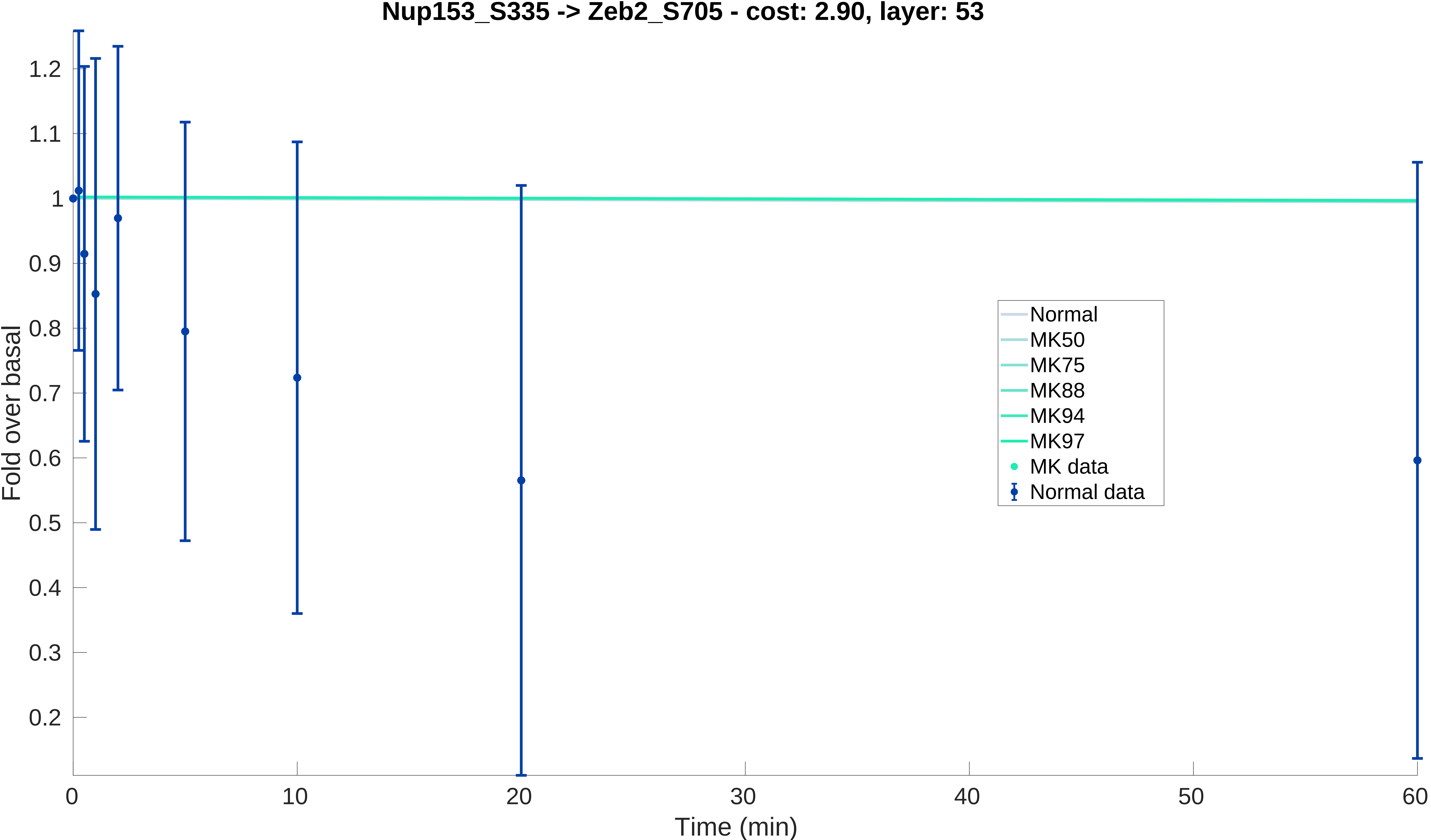

Nup153_S335 -> Zeb2_S705 - cost: 2.90, layer: 53
Fold over basal
1.2
1.1
1
0.9
0.8
0.7
0.6
0.5
0.4
0.3
0.2
0
10
20
30
40
50
60
Time (min)
Normal
MK50
MK75
MK88
MK94
MK97
MK data
Normal data

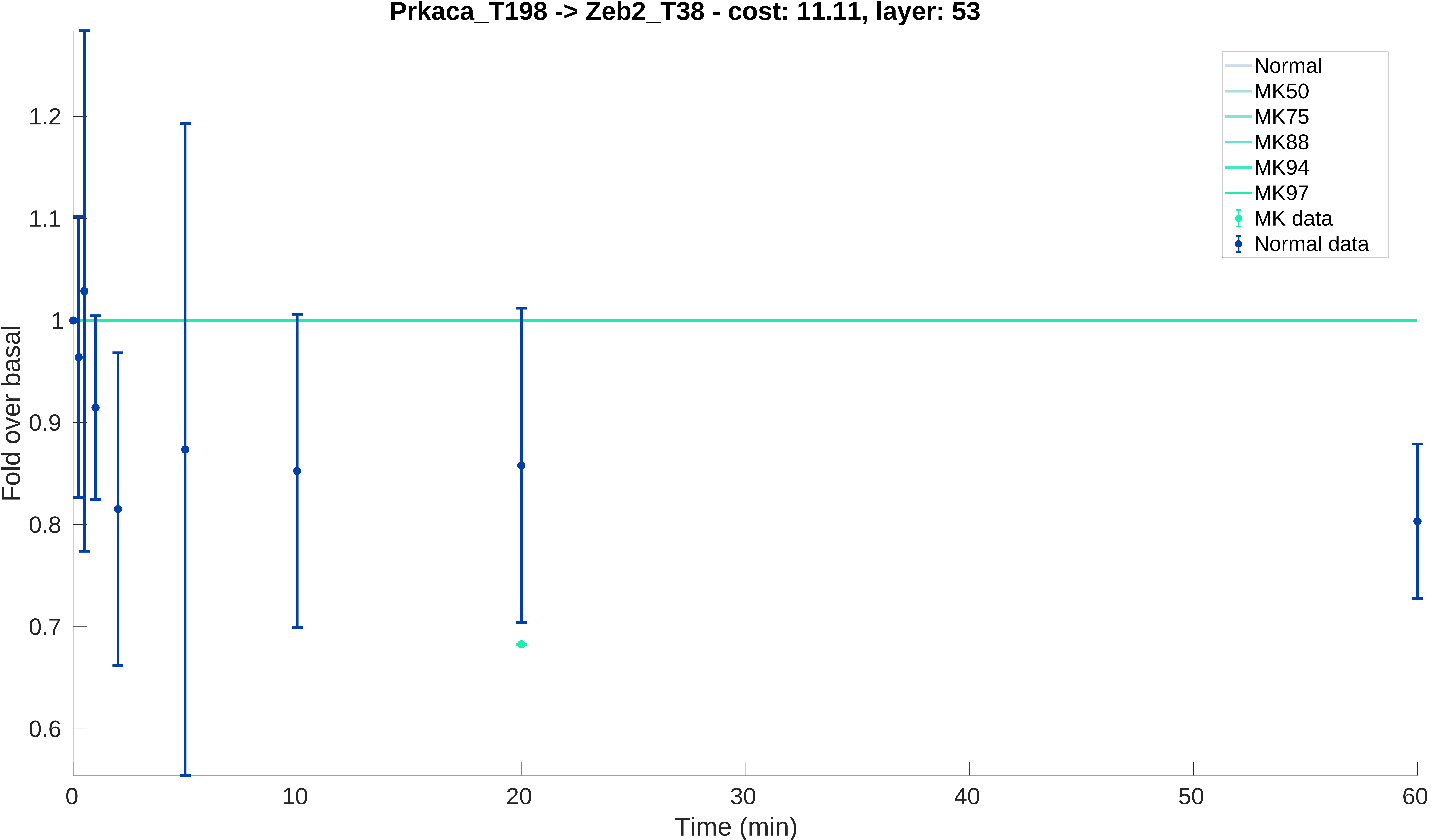
Prrkaca_T198 -> Zeb2_T38 - cost: 11.11, layer: 53
Normal
MK50
MK75
MK88
MK94
MK97
MK data
Normal data
Fold over basal
1.2
1.1
1
0.9
0.8
0.7
0.6
0
10
20
30
40
50
60
Time (min)

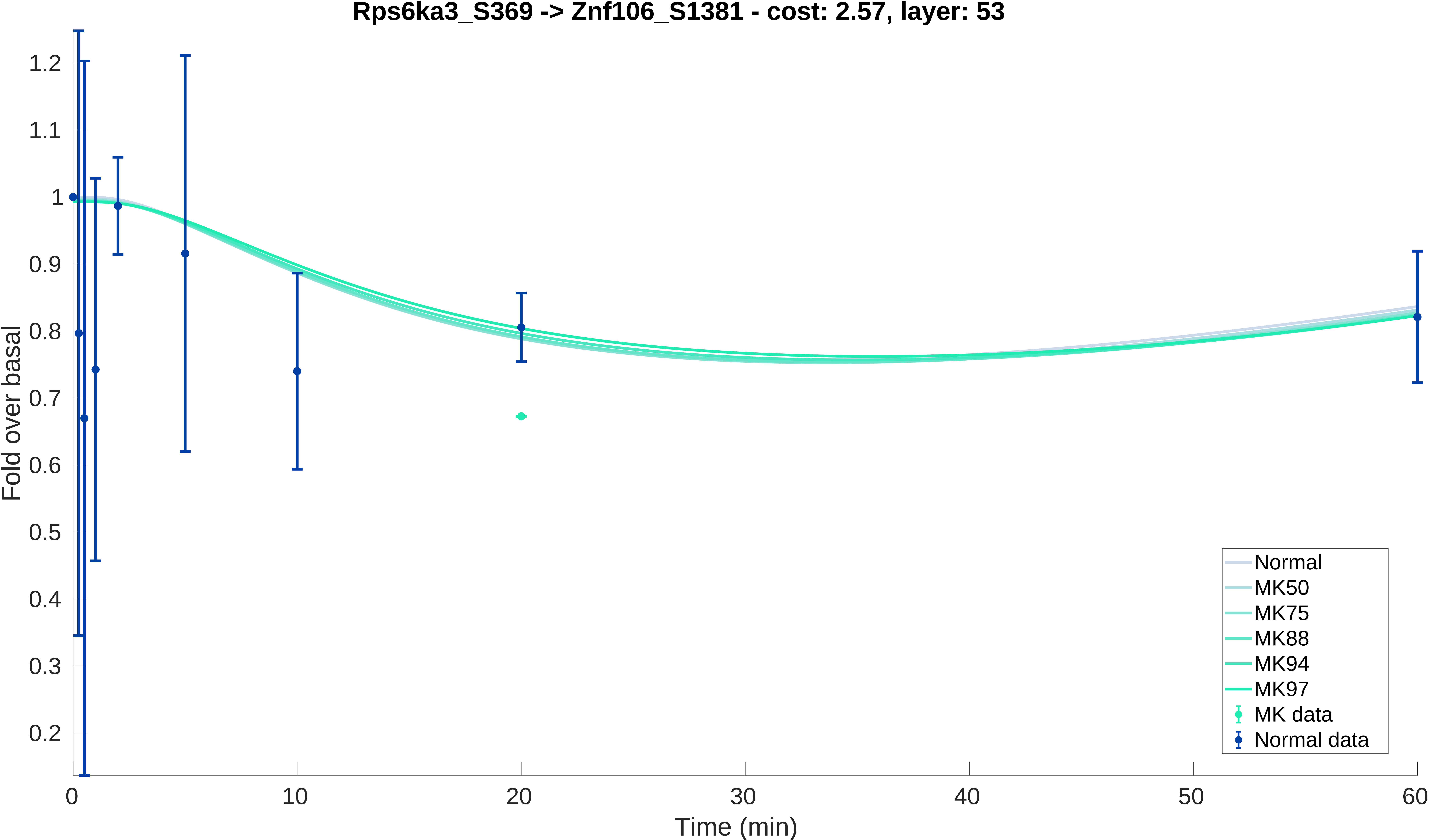

Rps6ka3_S369 -> Znf106_S1381 - cost: 2.57, layer: 53
Fold over basal
Time (min)
0.2
0.3
0.4
0.5
0.6
0.7
0.8
0.9
1
1.1
1.2
0
10
20
30
40
50
60
Normal
MK50
MK75
MK88
MK94
MK97
MK data
Normal data

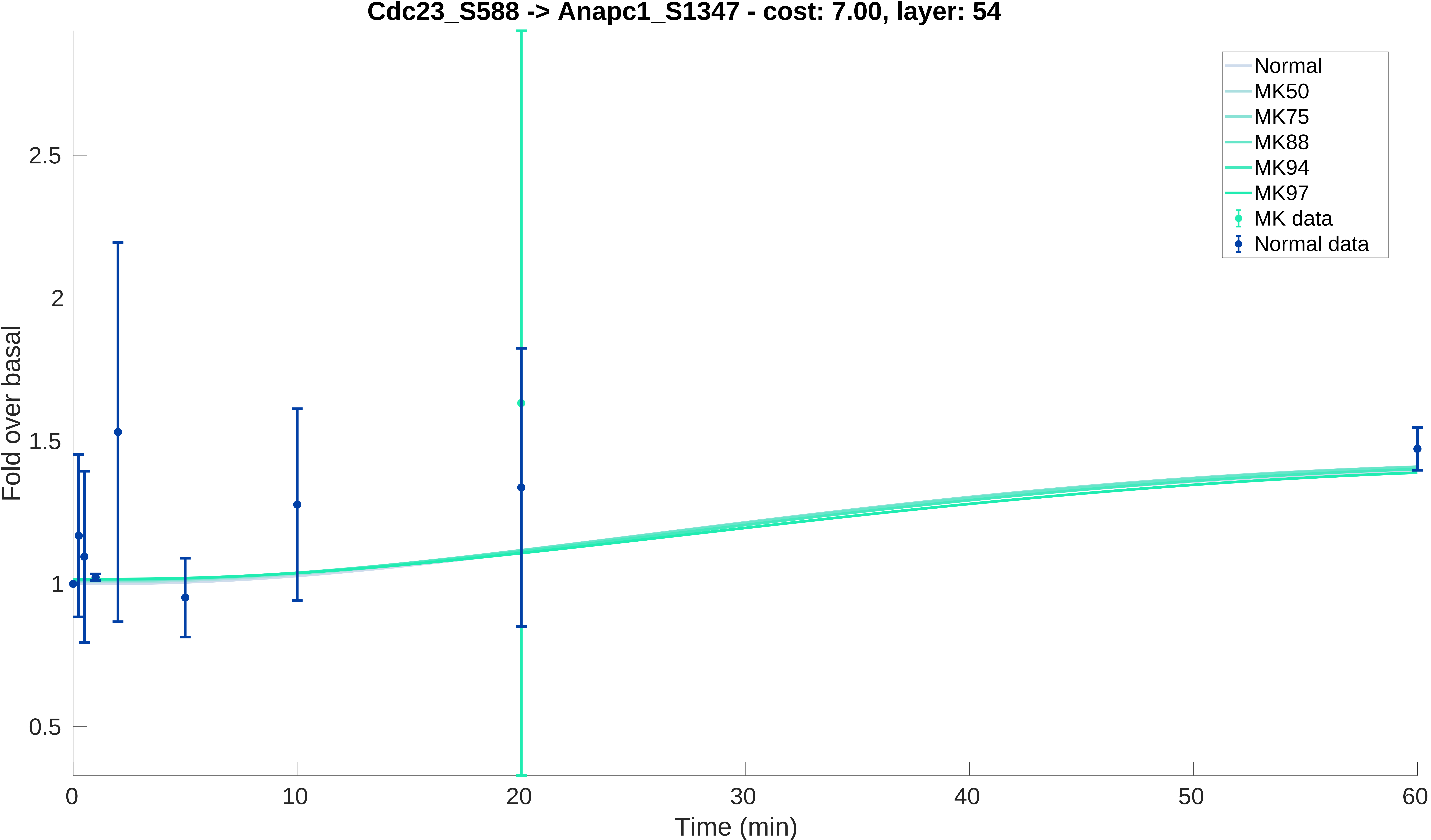

Cdc23_S588 -> Anapc1_S1347 - cost: 7.00, layer: 54
Normal
MK50
MK75
MK88
MK94
MK97
MK data
Normal data
Fold over basal
0.5
1
1.5
2
2.5
Time (min)
0
10
20
30
40
50
60

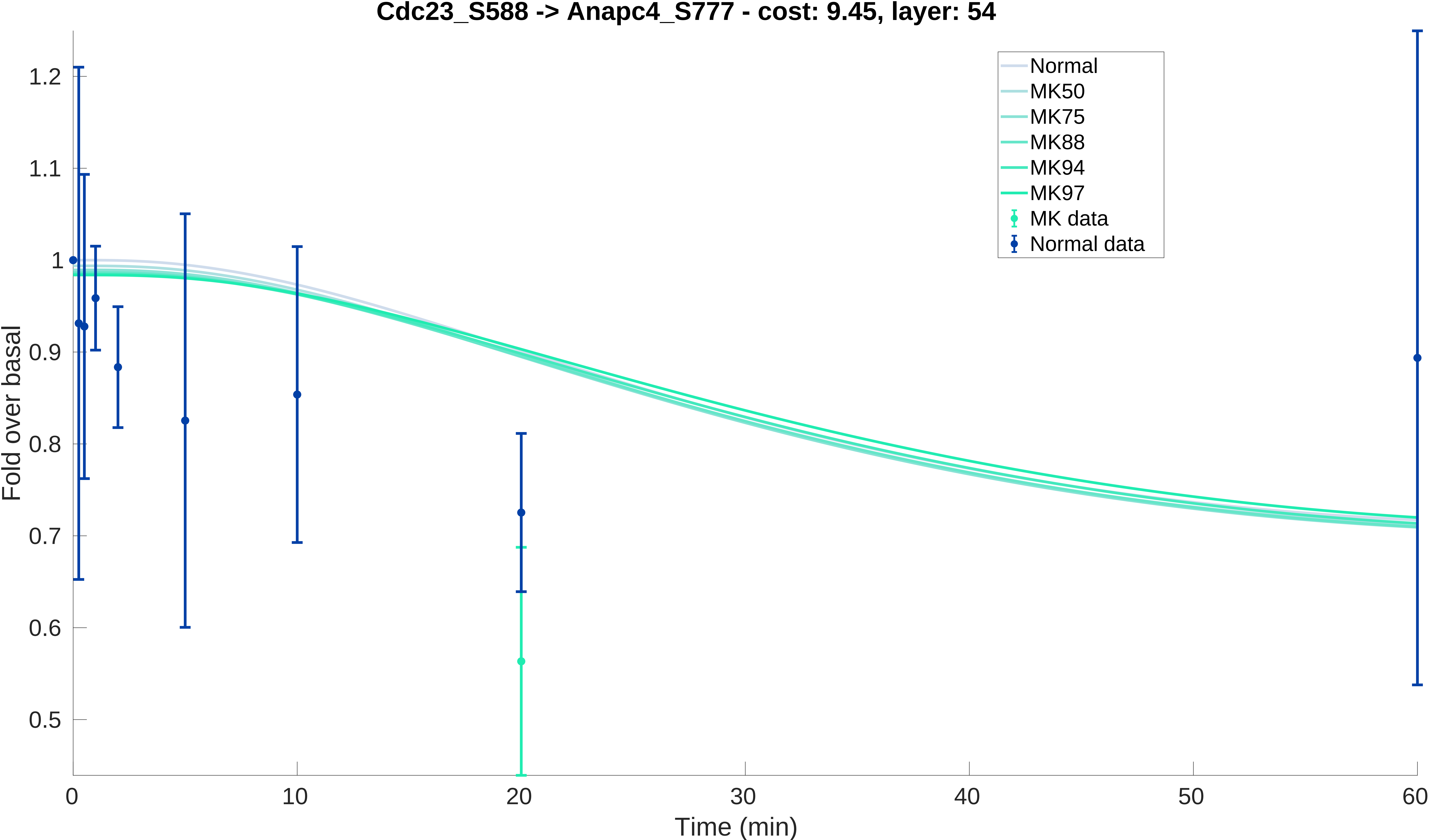

Cdc23_S588 -> Anapc4_S777 - cost: 9.45, layer: 54
Normal
MK50
MK75
MK88
MK94
MK97
MK data
Normal data
Fold over basal
Time (min)
1.2
1.1
1
0.9
0.8
0.7
0.6
0.5
0
10
20
30
40
50
60

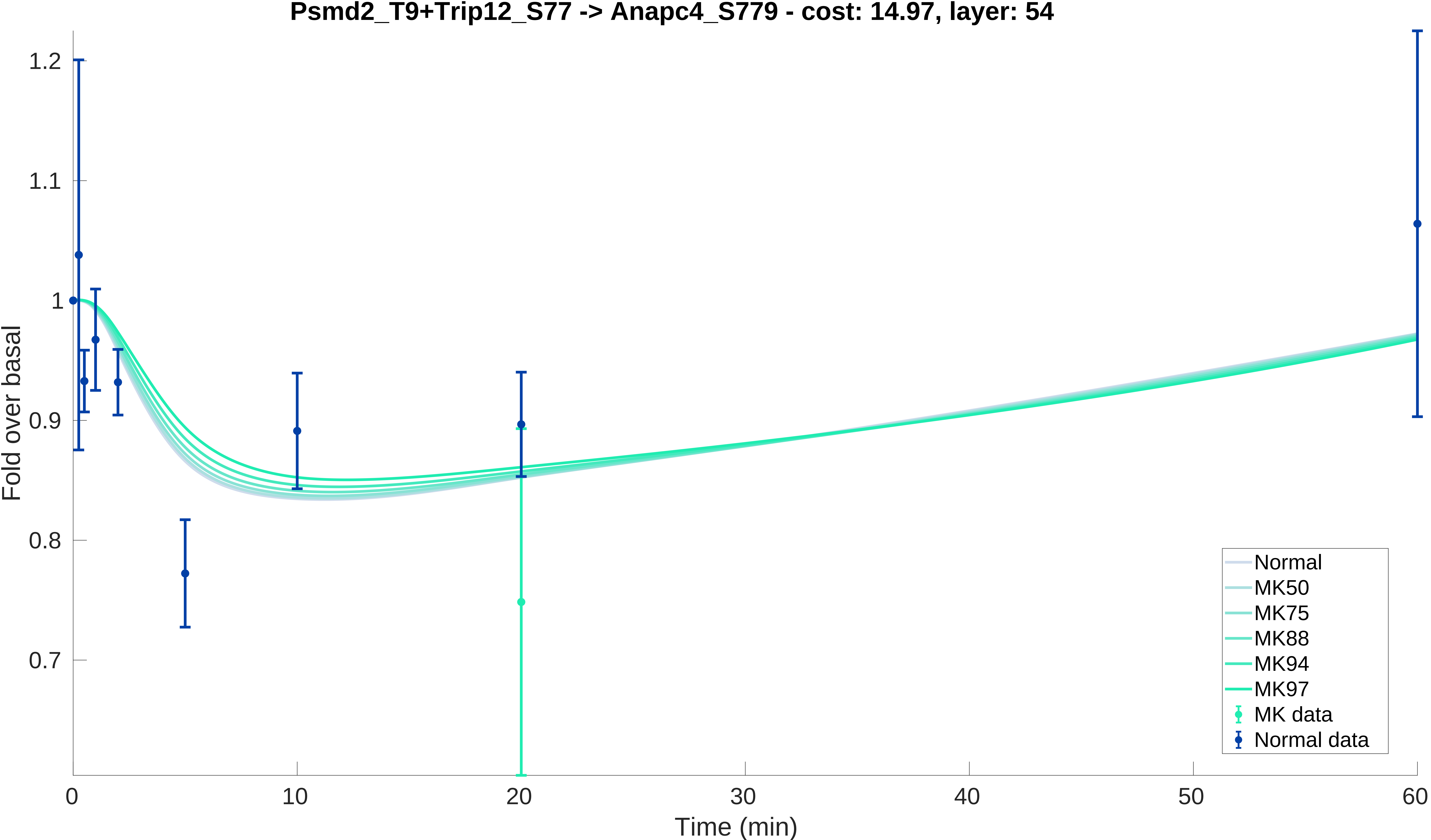

Psmd2_T9+Trip12_S77 -> Anapc4_S779 - cost: 14.97, layer: 54
Fold over basal
1.2
1.1
1
0.9
0.8
0.7
0
10
20
30
40
50
60
Time (min)
Normal
MK50
MK75
MK88
MK94
MK97
MK data
Normal data

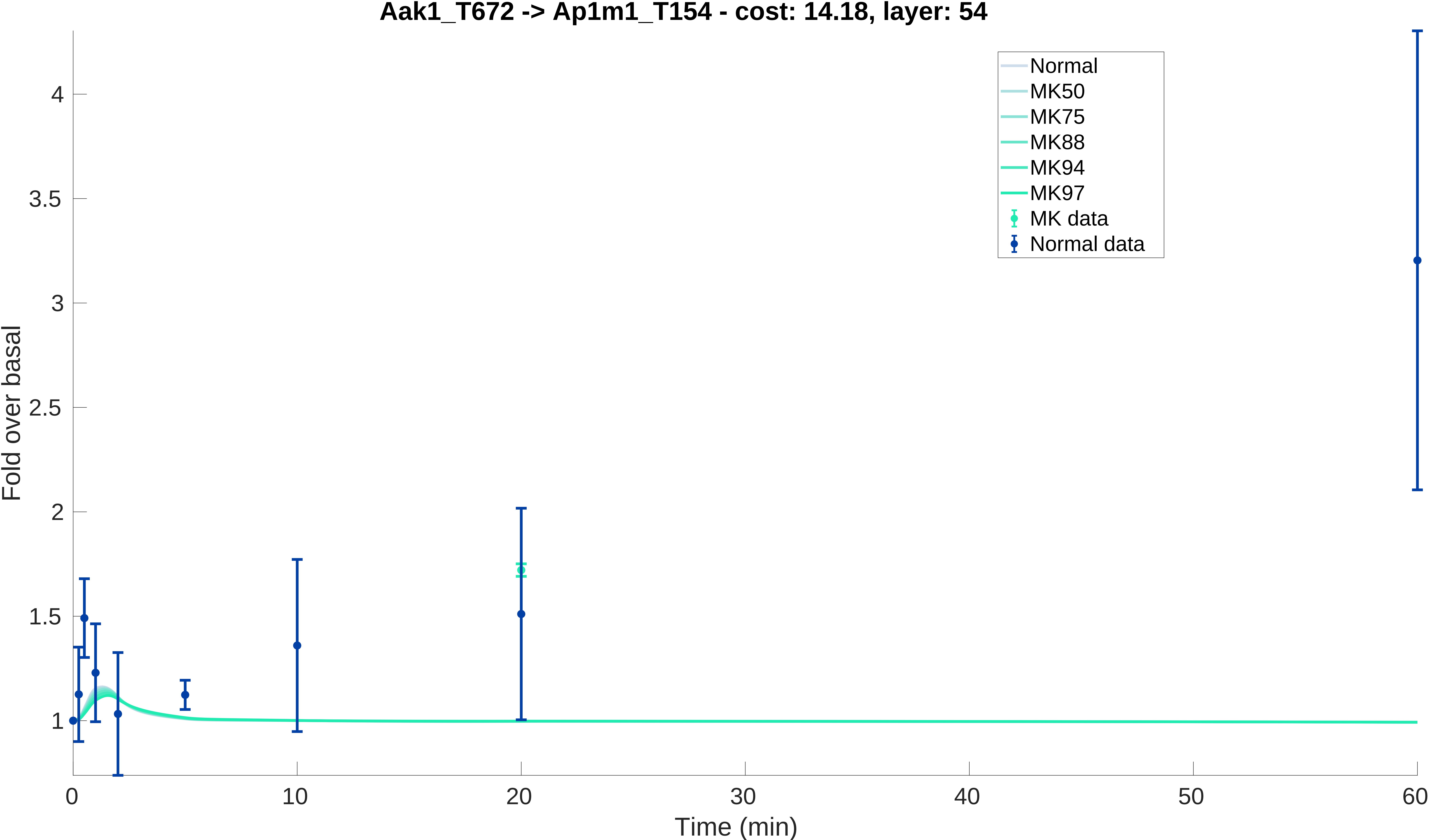
Aak1_T672 -> Ap1m1_T154 - cost: 14.18, layer: 54
Normal
MK50
MK75
MK88
MK94
MK97
MK data
Normal data
Fold over basal
Time (min)
4
3.5
3
2.5
2
1.5
1
0
10
20
30
40
50
60

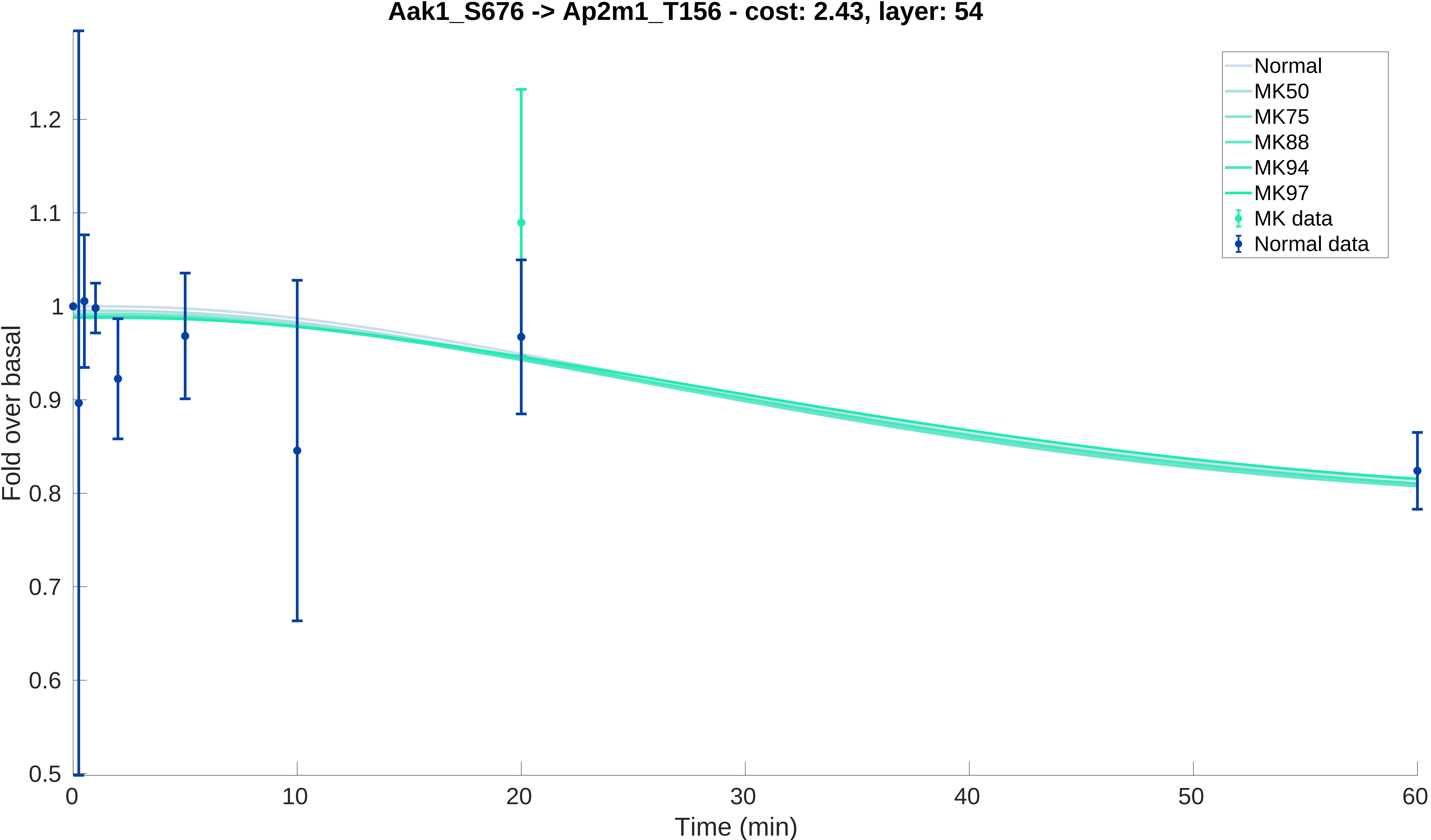

Aak1_S676 -> Ap2m1_T156 - cost: 2.43, layer: 54
Normal
MK50
MK75
MK88
MK94
MK97
MK data
Normal data
Fold over basal
Time (min)
0.5
0.6
0.7
0.8
0.9
1
1.1
1.2
0
10
20
30
40
50
60

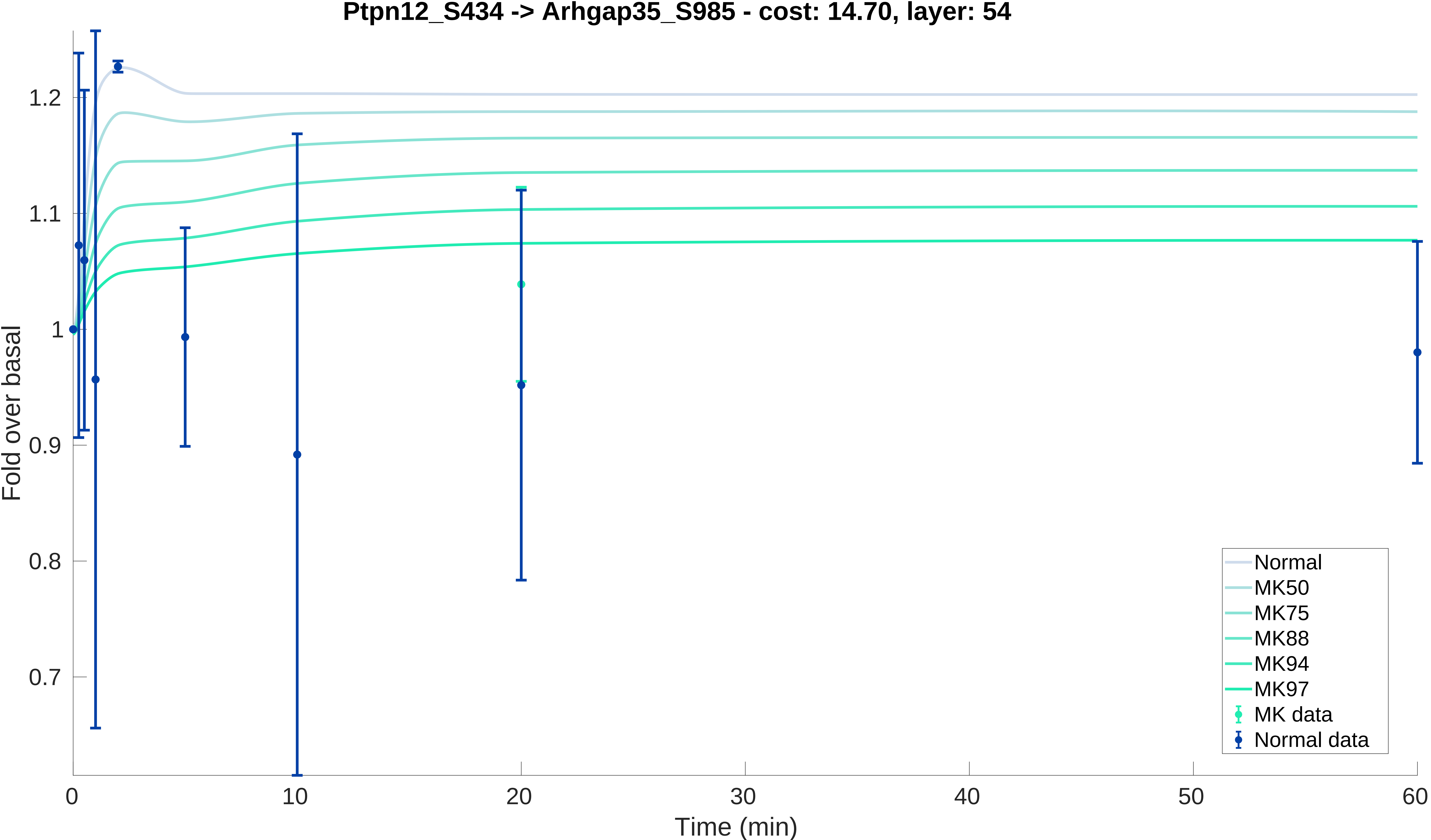

Ptpn12_S434 -> Arhgap35_S985 - cost: 14.70, layer: 54
Fold over basal
Time (min)
1.2
1.1
1
0.9
0.8
0.7
0
10
20
30
40
50
60
Normal
MK50
MK75
MK88
MK94
MK97
MK data
Normal data

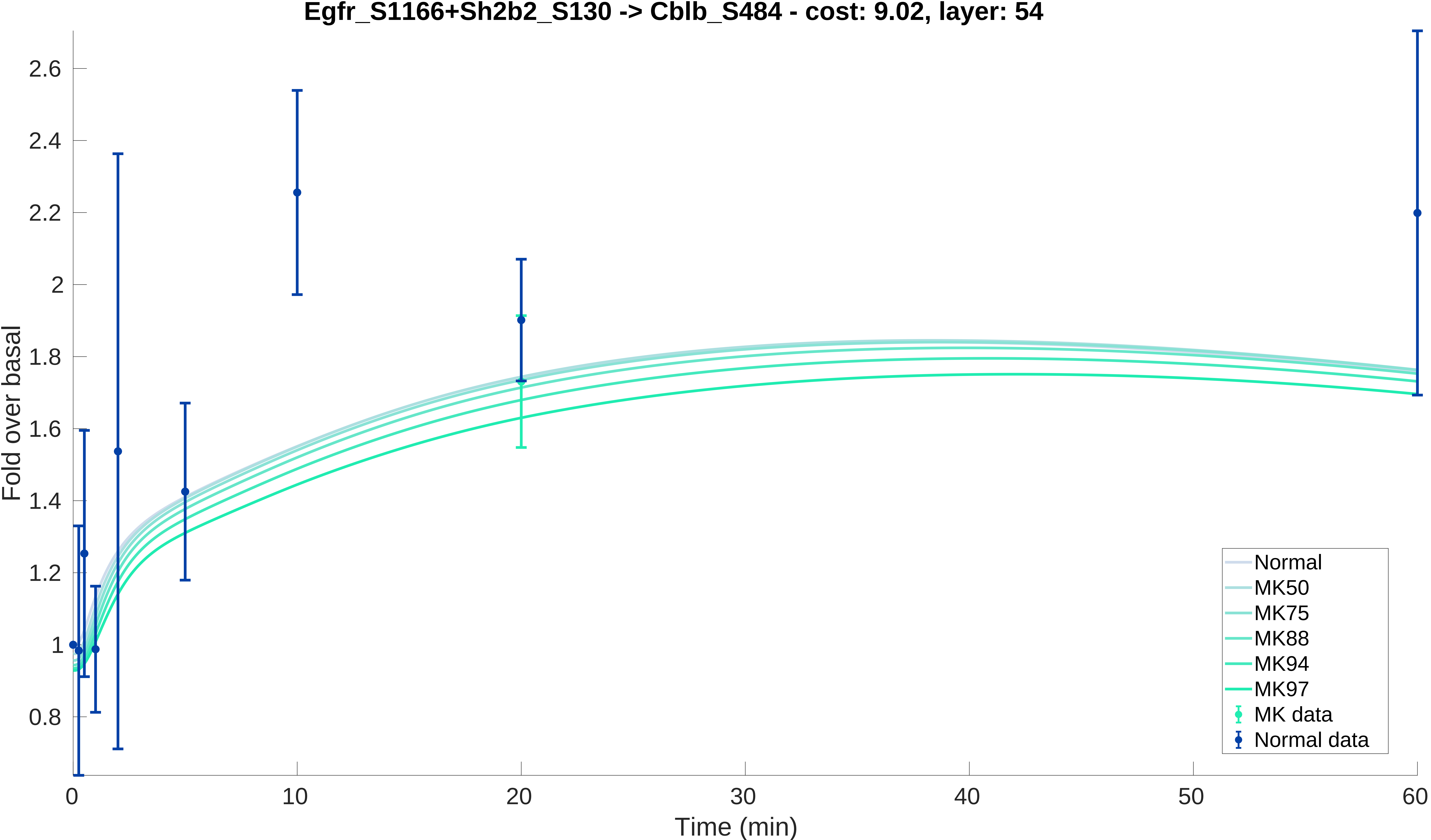

Egfr_S1166+Sh2b2_S130 -> Cblb_S484 - cost: 9.02, layer: 54
Fold over basal
2.6
2.4
2.2
2
1.8
1.6
1.4
1.2
1
0.8
0
10
20
30
40
50
60
Time (min)
Normal
MK50
MK75
MK88
MK94
MK97
MK data
Normal data

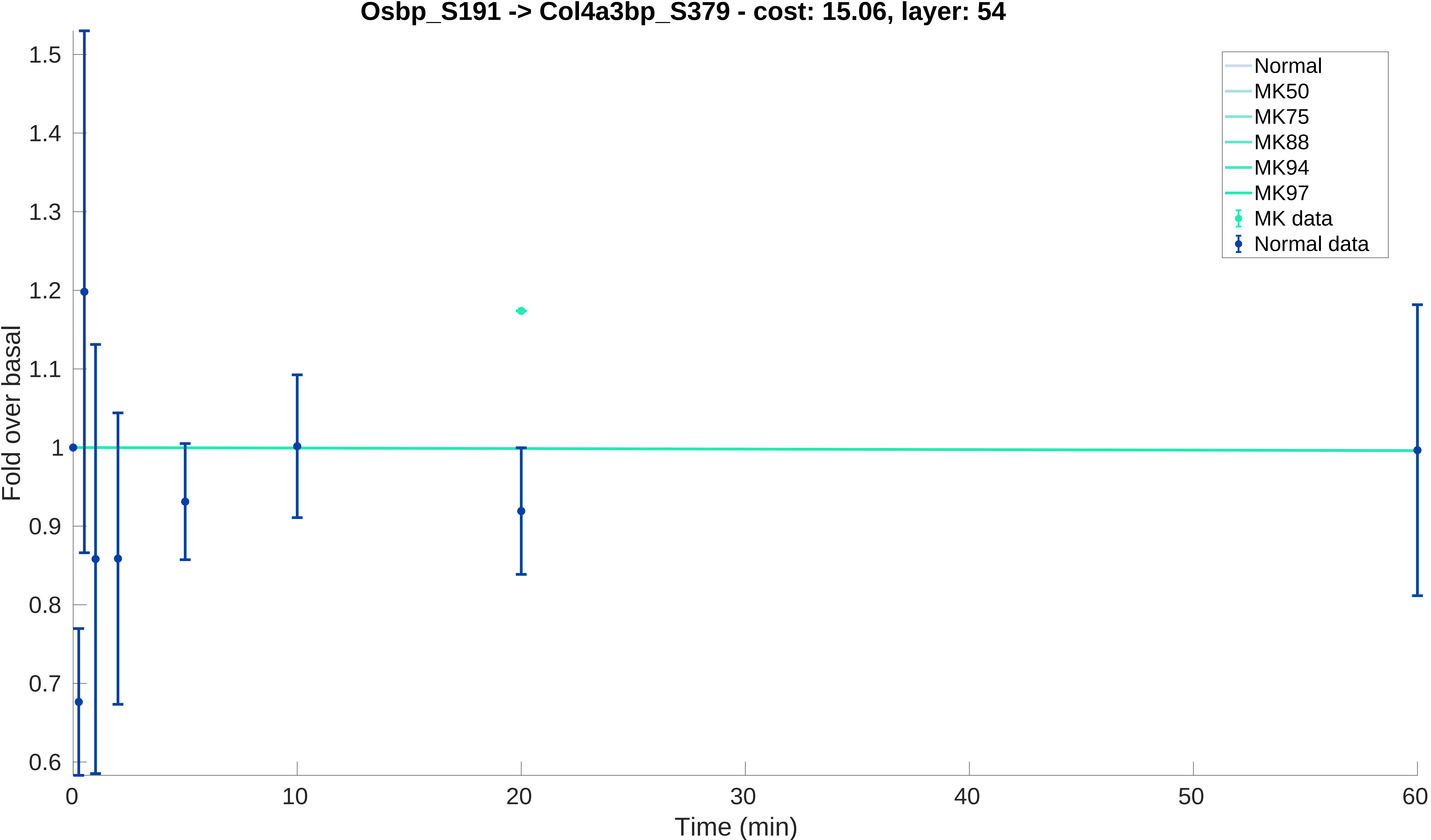

Osbp_S191 -> Col4a3bp_S379 - cost: 15.06, layer: 54
Normal
MK50
MK75
MK88
MK94
MK97
MK data
Normal data
Fold over basal
Time (min)
1.5
1.4
1.3
1.2
1.1
1
0.9
0.8
0.7
0.6
0
10
20
30
40
50
60

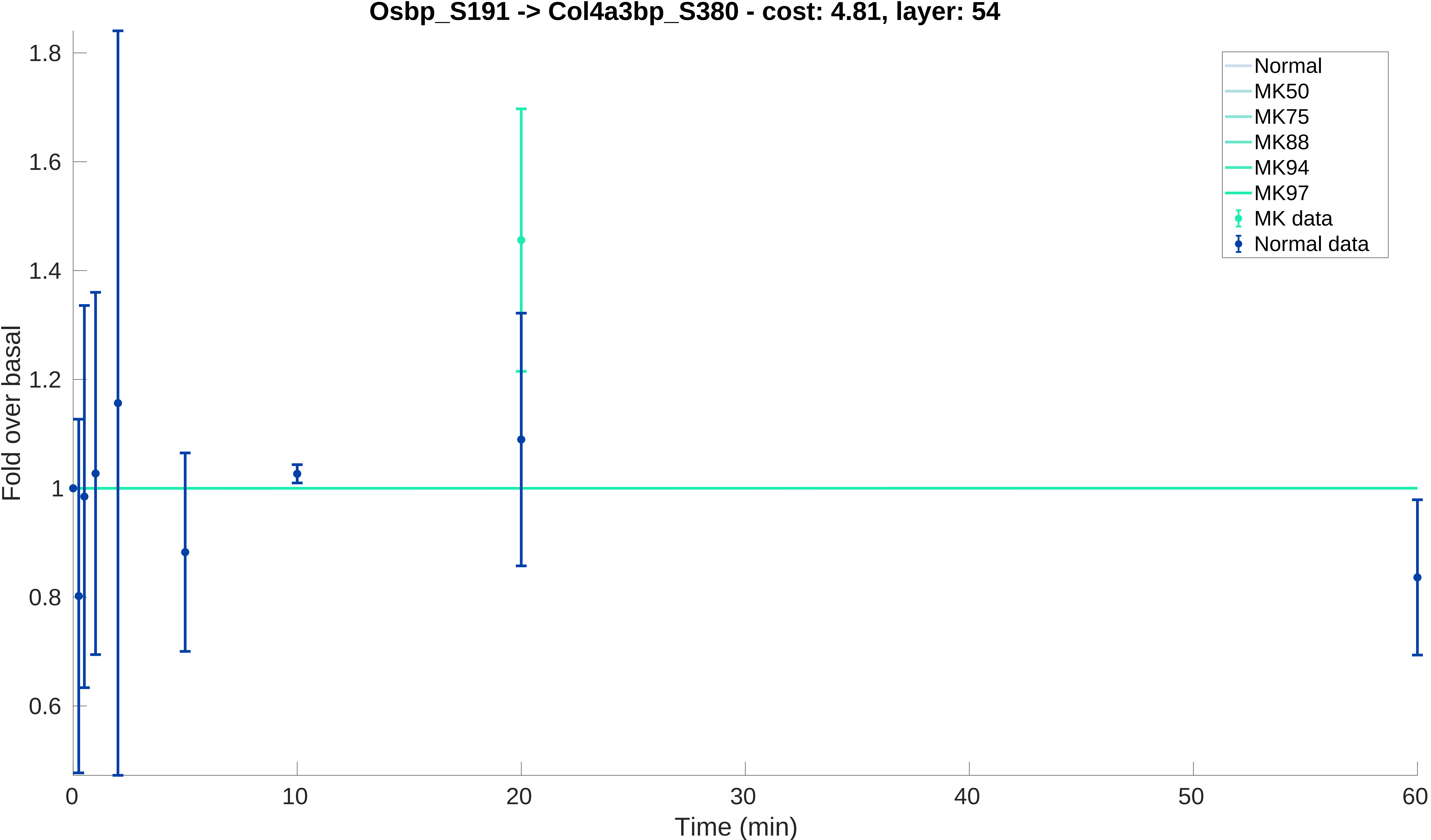

Osbp_S191 -> Col4a3bp_S380 - cost: 4.81, layer: 54
Fold over basal
Time (min)
1.8
1.6
1.4
1.2
1
0.8
0.6
0
10
20
30
40
50
60
Normal
MK50
MK75
MK88
MK94
MK97
MK data
Normal data

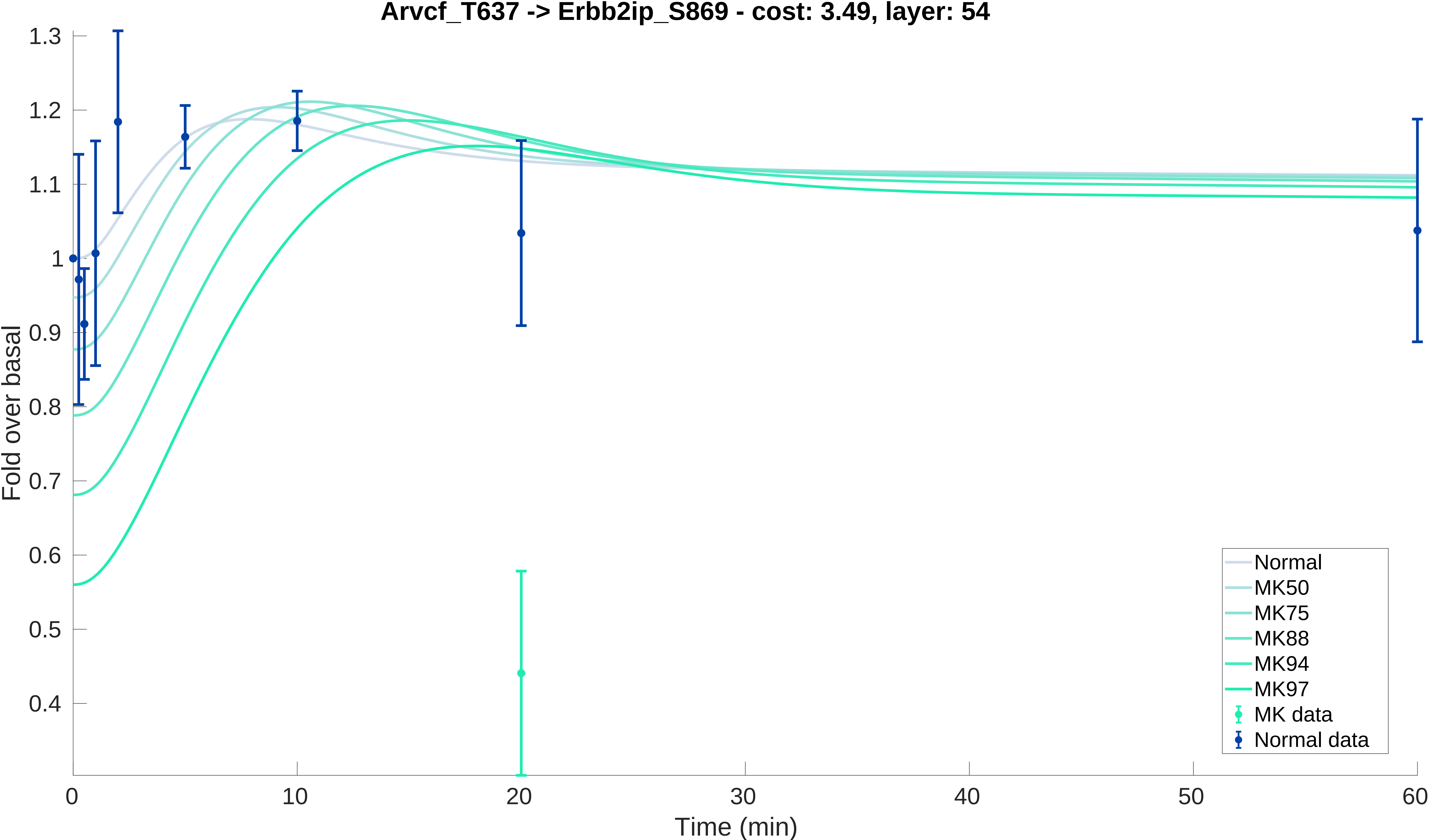

Arvcf_T637 -> Erbb2ip_S869 - cost: 3.49, layer: 54
Fold over basal
Time (min)
1.3
1.2
1.1
1
0.9
0.8
0.7
0.6
0.5
0.4
0
10
20
30
40
50
60
Normal
MK50
MK75
MK88
MK94
MK97
MK data
Normal data

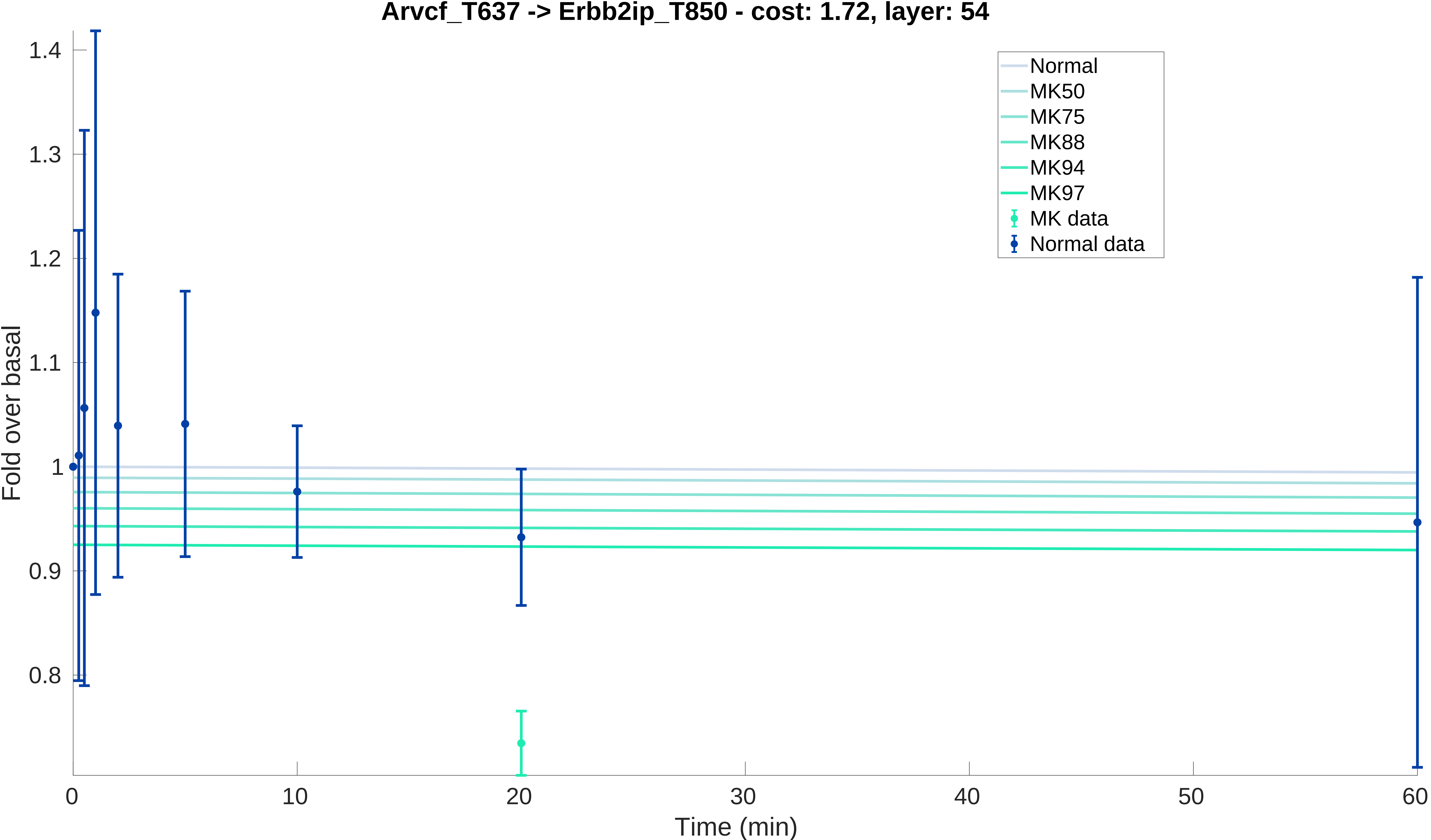

Arvcf_T637 -> Erbb2ip_T850 - cost: 1.72, layer: 54
Normal
MK50
MK75
MK88
MK94
MK97
MK data
Normal data
Fold over basal
1.4
1.3
1.2
1.1
1
0.9
0.8
0
10
20
30
40
50
60
Time (min)

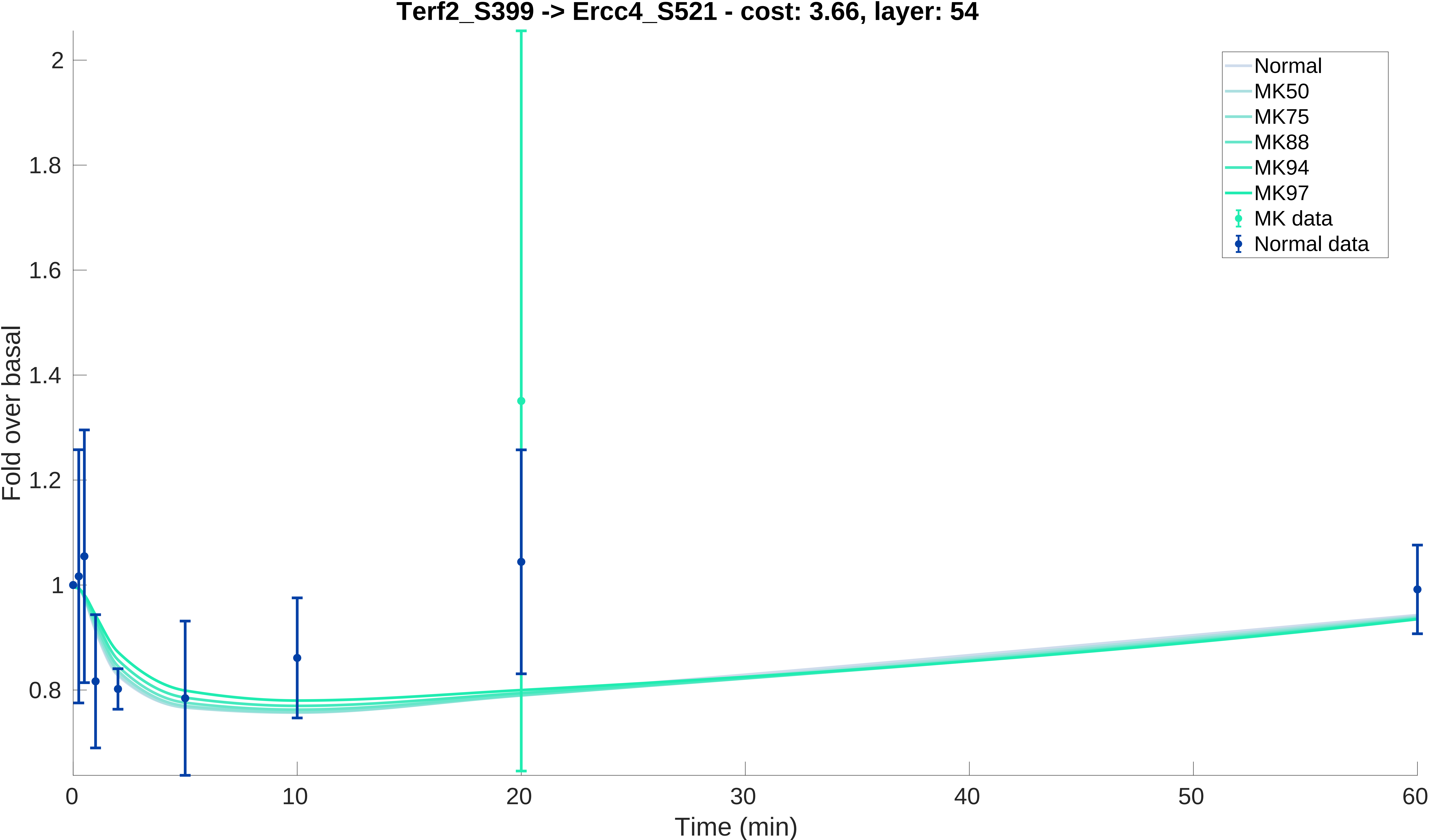

Terf2_S399 -> Ercc4_S521 - cost: 3.66, layer: 54
Normal
MK50
MK75
MK88
MK94
MK97
MK data
Normal data
Fold over basal
Time (min)
0
10
20
30
40
50
60
0.8
1
1.2
1.4
1.6
1.8
2

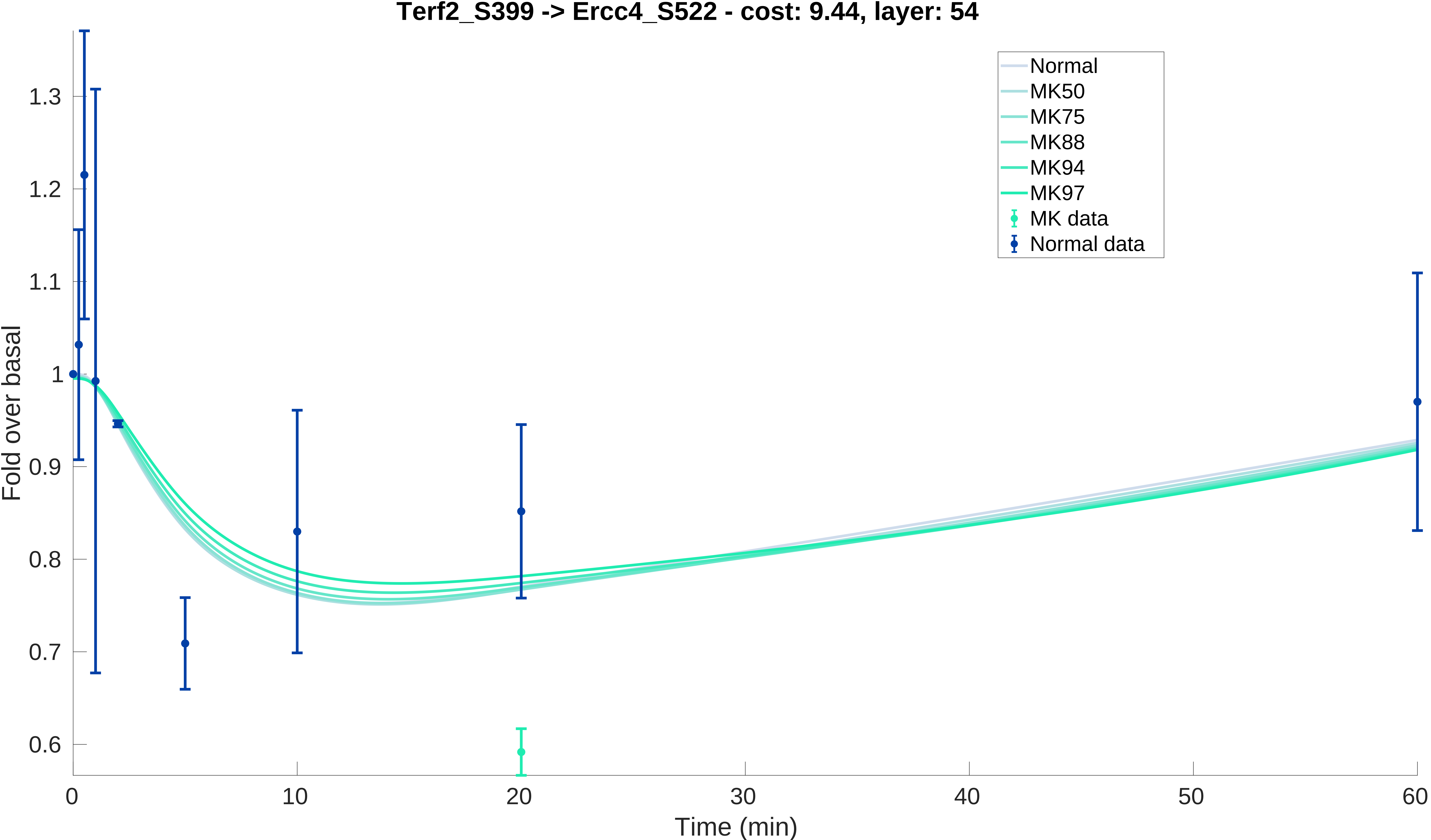

Terf2_S399 -> Ercc4_S522 - cost: 9.44, layer: 54
Fold over basal
Time (min)
1.3
1.2
1.1
1
0.9
0.8
0.7
0.6
0
10
20
30
40
50
60
Normal
MK50
MK75
MK88
MK94
MK97
MK data
Normal data

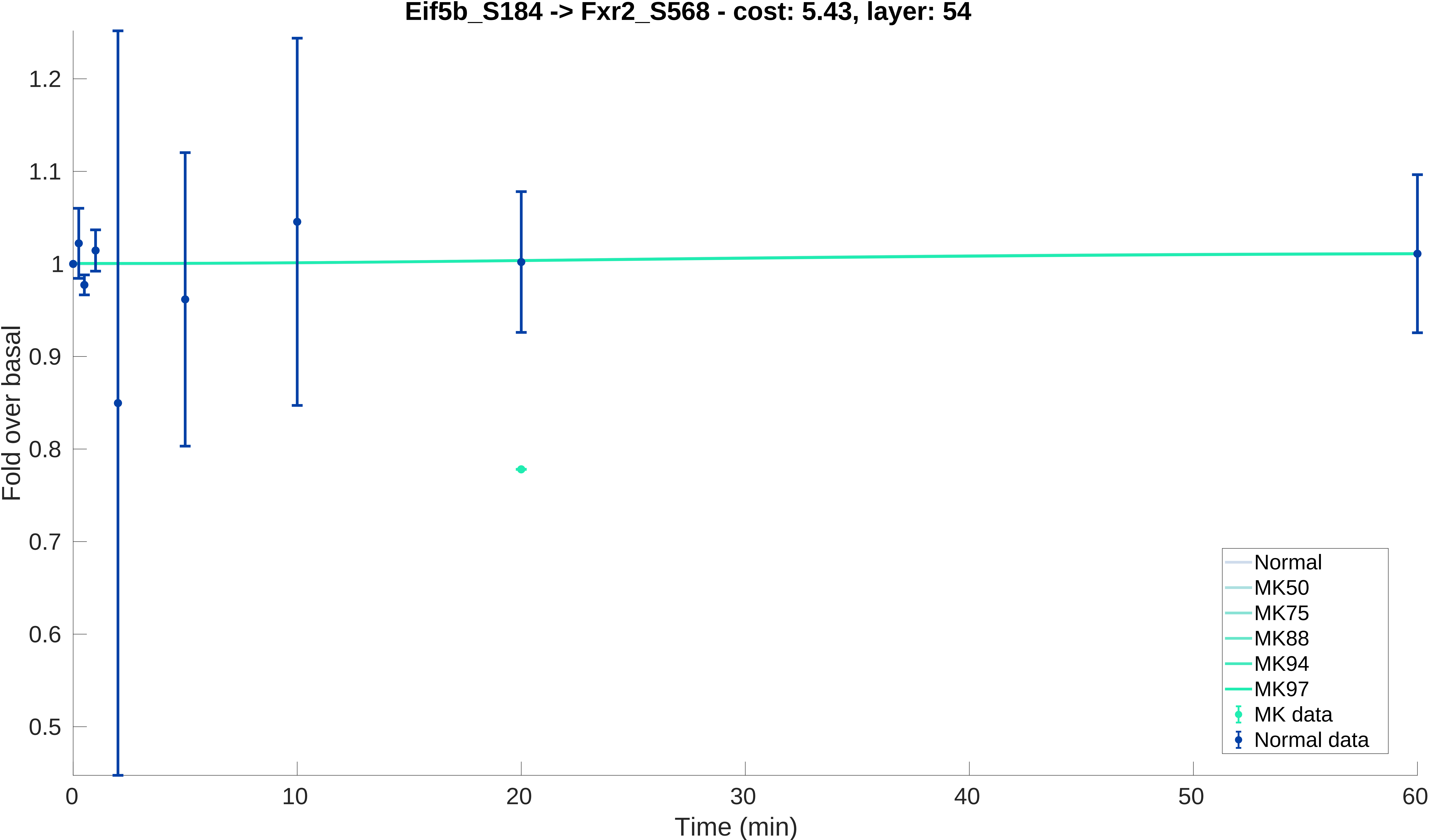
Eif5b_S184 -> Fxr2_S568 - cost: 5.43, layer: 54
Fold over basal
Time (min)
0.5
0.6
0.7
0.8
0.9
1
1.1
1.2
0
10
20
30
40
50
60
Normal
MK50
MK75
MK88
MK94
MK97
MK data
Normal data

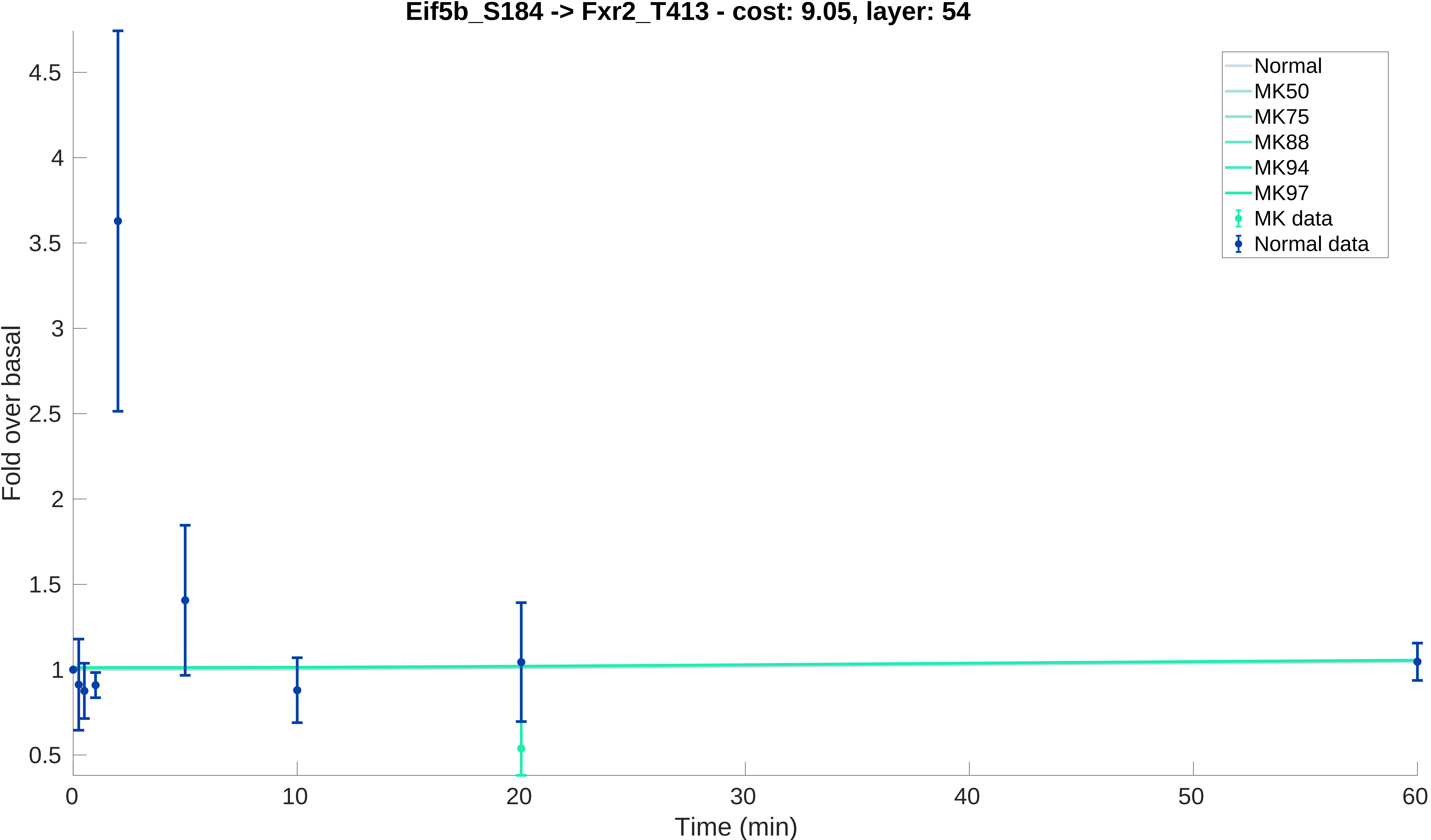

Eif5b_S184 -> Fxr2_T413 - cost: 9.05, layer: 54
Fold over basal
4.5
4
3.5
3
2.5
2
1.5
1
0.5
0
10
20
30
40
50
60
Time (min)
Normal
MK50
MK75
MK88
MK94
MK97
MK data
Normal data

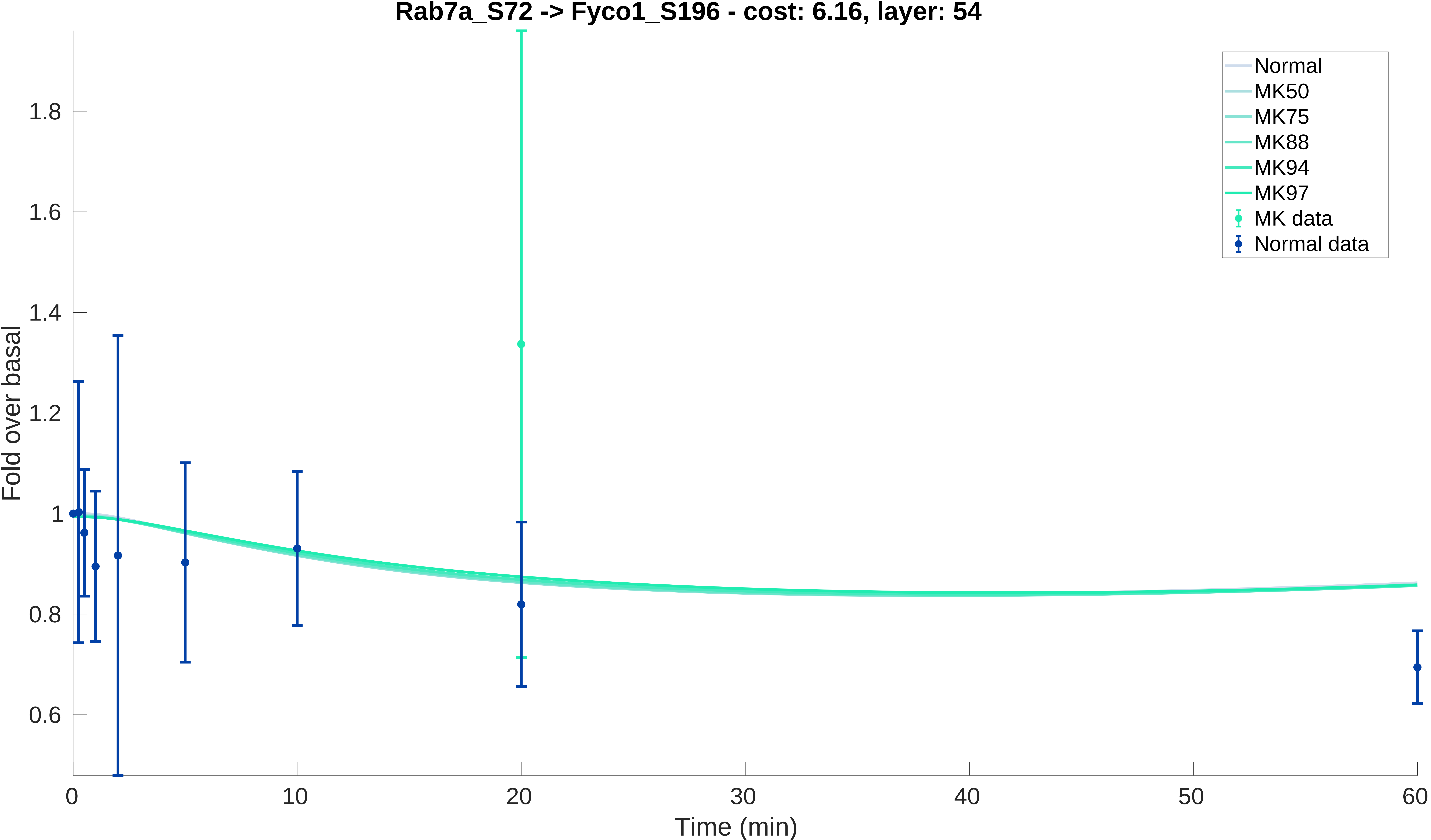

Rab7a_S72 -> Fyco1_S196 - cost: 6.16, layer: 54
Fold over basal
Time (min)
0.6
0.8
1
1.2
1.4
1.6
1.8
0
10
20
30
40
50
60
Normal
MK50
MK75
MK88
MK94
MK97
MK data
Normal data

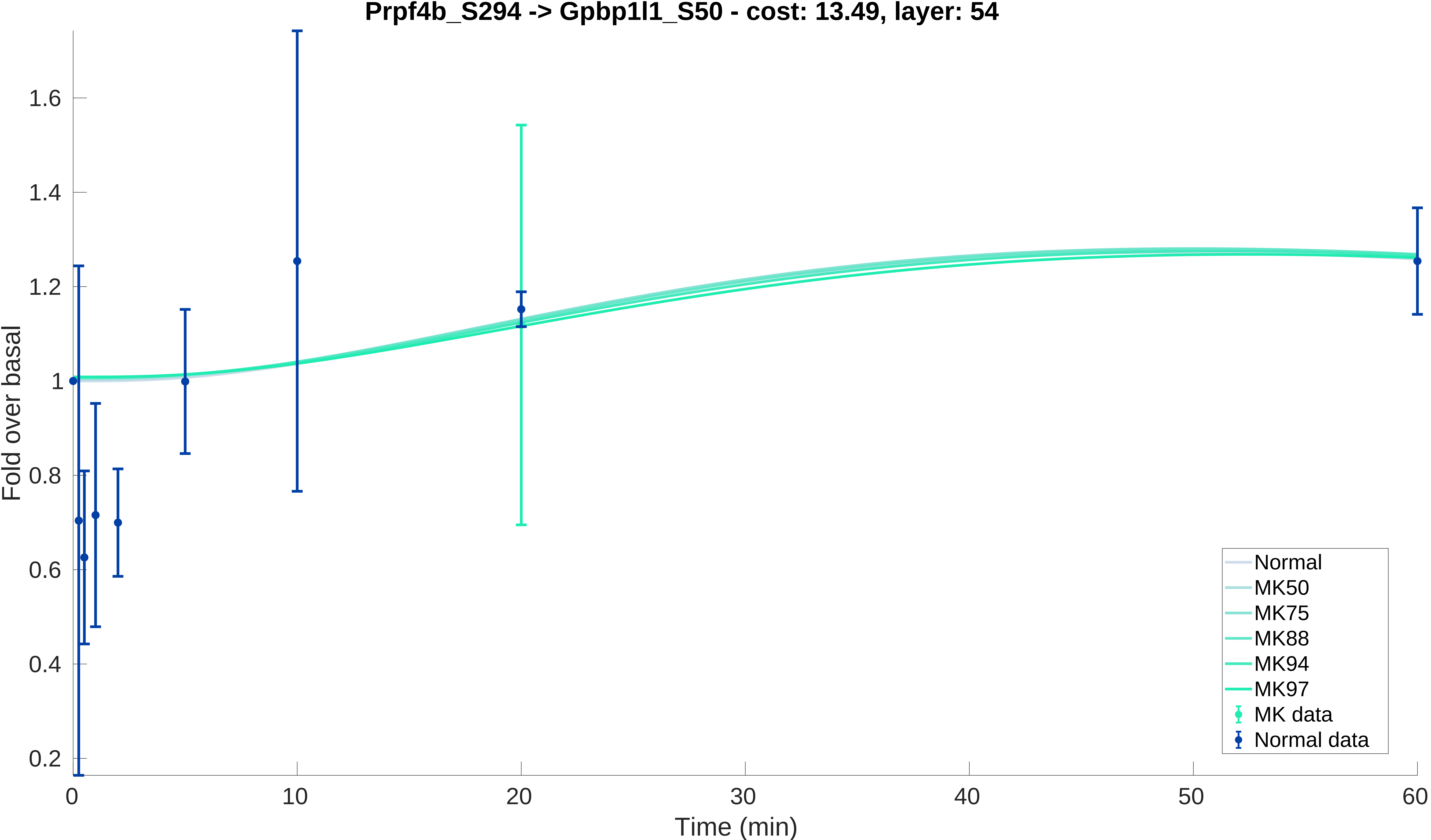

Prpf4b_S294 -> Gpbp1l1_S50 - cost: 13.49, layer: 54
Fold over basal
1.6
1.4
1.2
1
0.8
0.6
0.4
0.2
0
10
20
30
40
50
60
Time (min)
Normal
MK50
MK75
MK88
MK94
MK97
MK data
Normal data

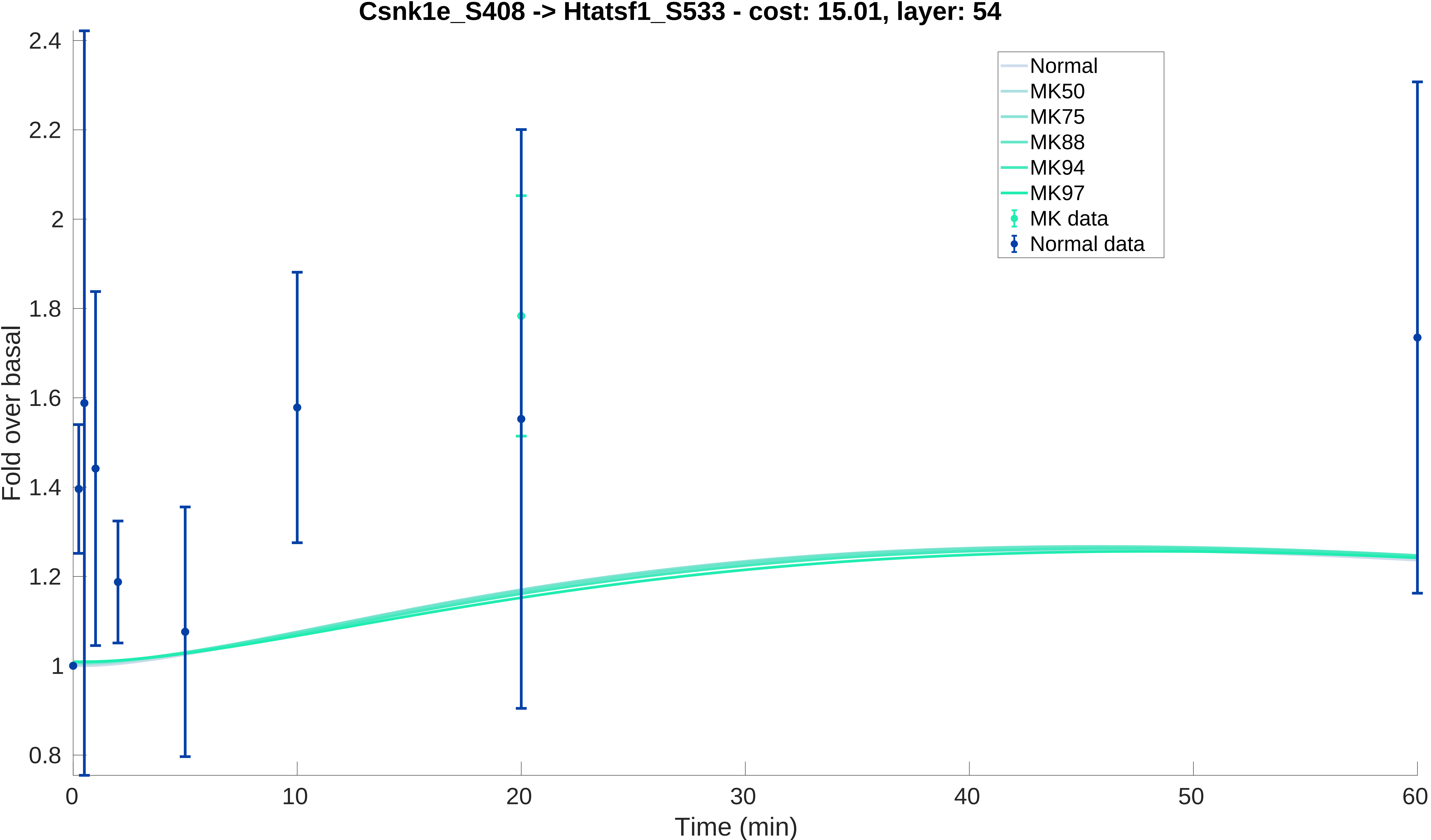
Csnk1e_S408 -> Htatsf1_S533 - cost: 15.01, layer: 54
Fold over basal
Time (min)
Normal
MK50
MK75
MK88
MK94
MK97
MK data
Normal data
2.4
2.2
2
1.8
1.6
1.4
1.2
1
0.8
0
10
20
30
40
50
60

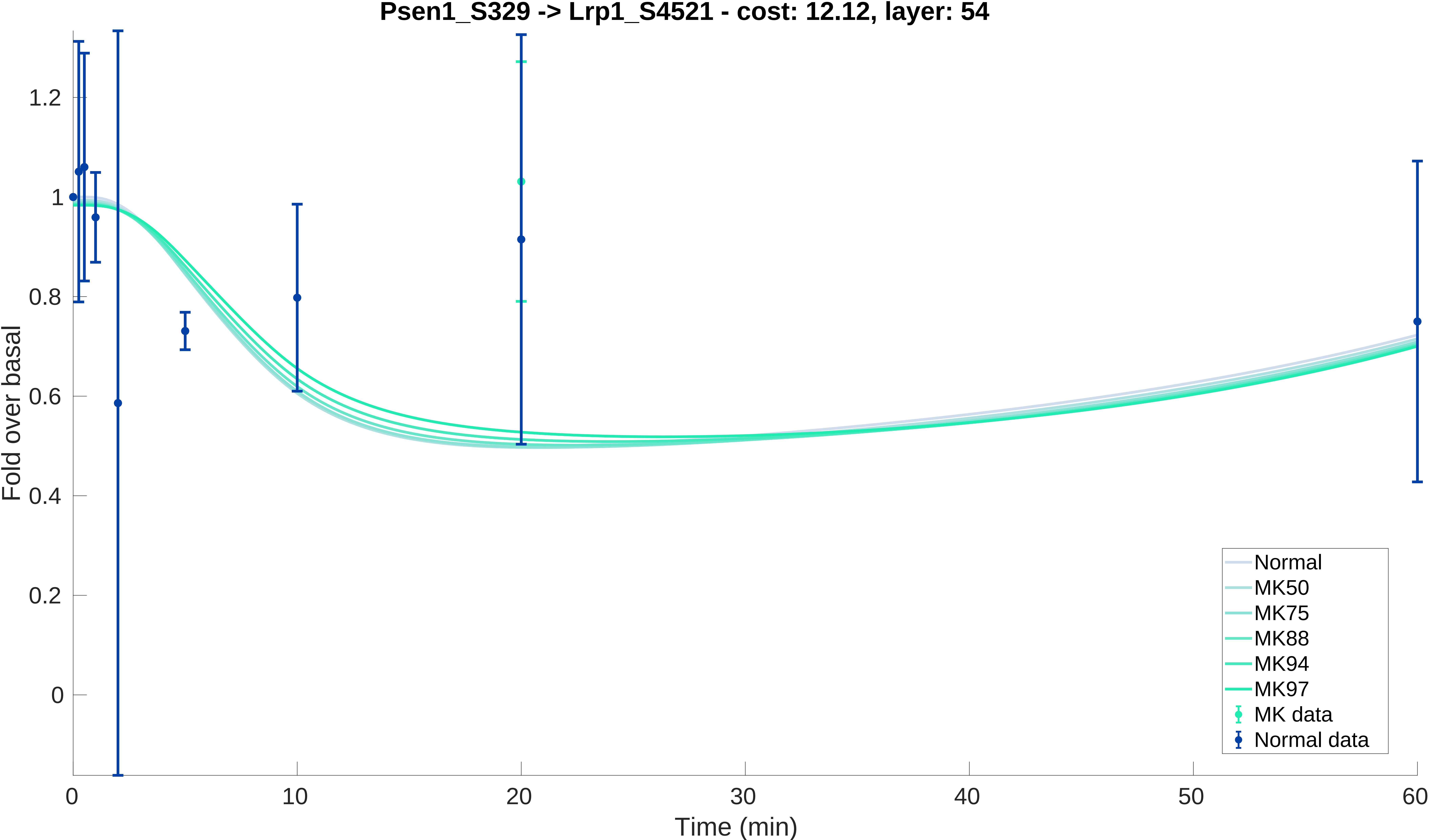

Psen1_S329 -> Lrp1_S4521 - cost: 12.12, layer: 54
Fold over basal
1.2
1
0.8
0.6
0.4
0.2
0
0
10
20
30
40
50
60
Time (min)
Normal
MK50
MK75
MK88
MK94
MK97
MK data
Normal data

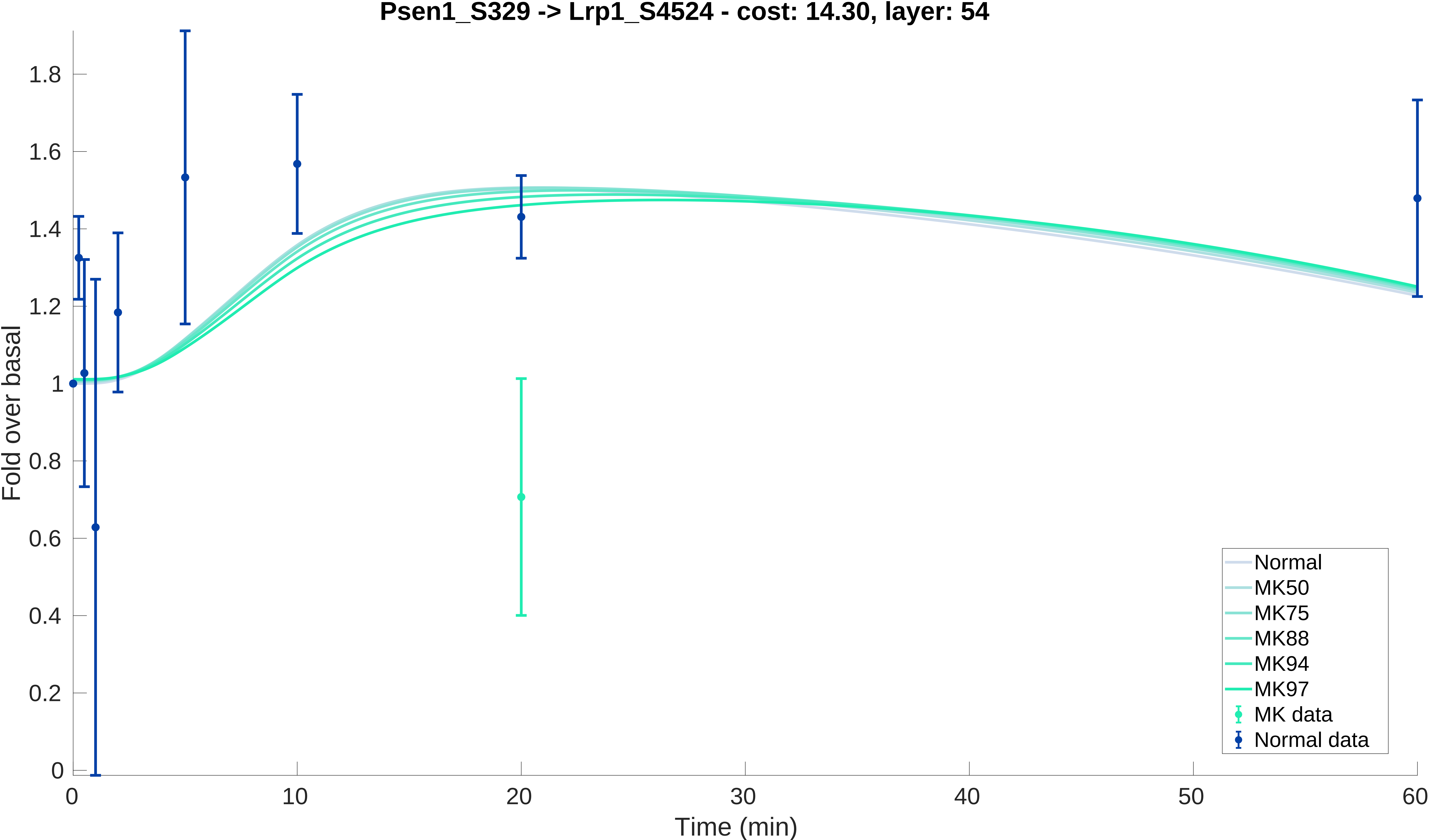

Psen1_S329 -> Lrp1_S4524 - cost: 14.30, layer: 54
Fold over basal
0
0.2
0.4
0.6
0.8
1
1.2
1.4
1.6
1.8
Time (min)
0
10
20
30
40
50
60
Normal
MK50
MK75
MK88
MK94
MK97
MK data
Normal data

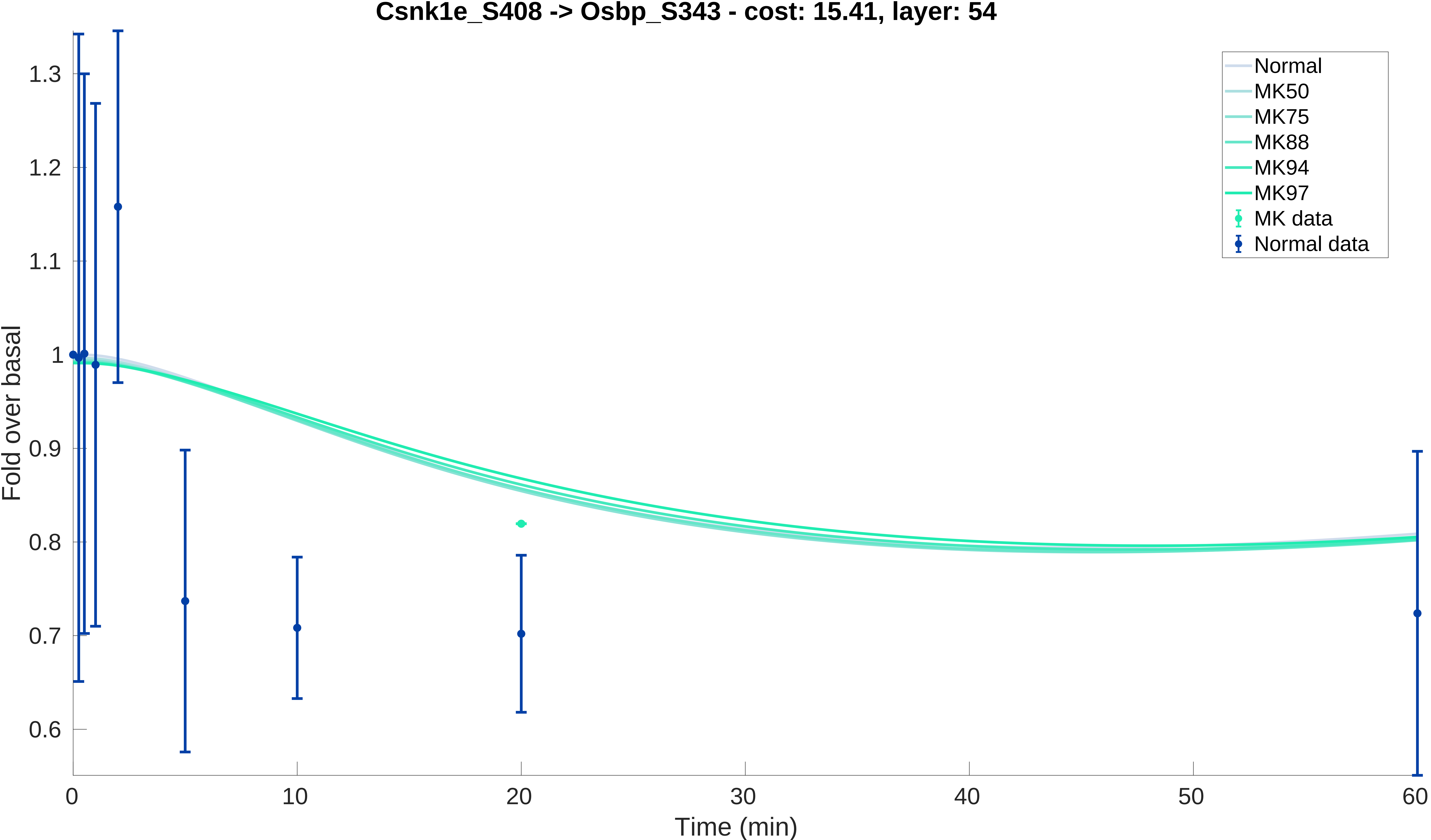

Csnk1e_S408 -> Osbp_S343 - cost: 15.41, layer: 54
Fold over basal
Time (min)
1.3
1.2
1.1
1
0.9
0.8
0.7
0.6
0
10
20
30
40
50
60
Normal
MK50
MK75
MK88
MK94
MK97
MK data
Normal data

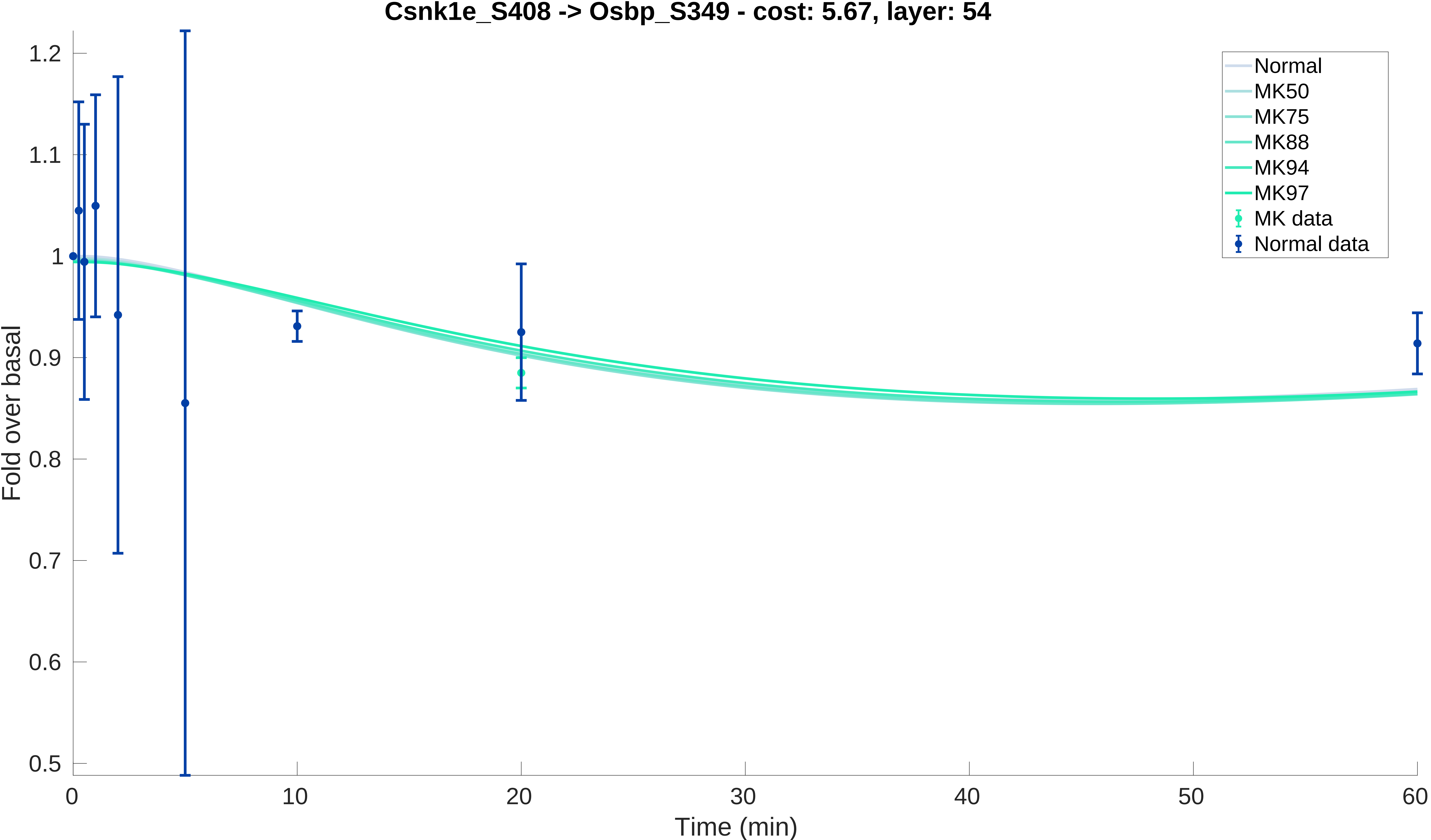

Csnk1e_S408 -> Osbp_S349 - cost: 5.67, layer: 54
Fold over basal
Time (min)
1.2
1.1
1
0.9
0.8
0.7
0.6
0.5
0
10
20
30
40
50
60
Normal
MK50
MK75
MK88
MK94
MK97
MK data
Normal data

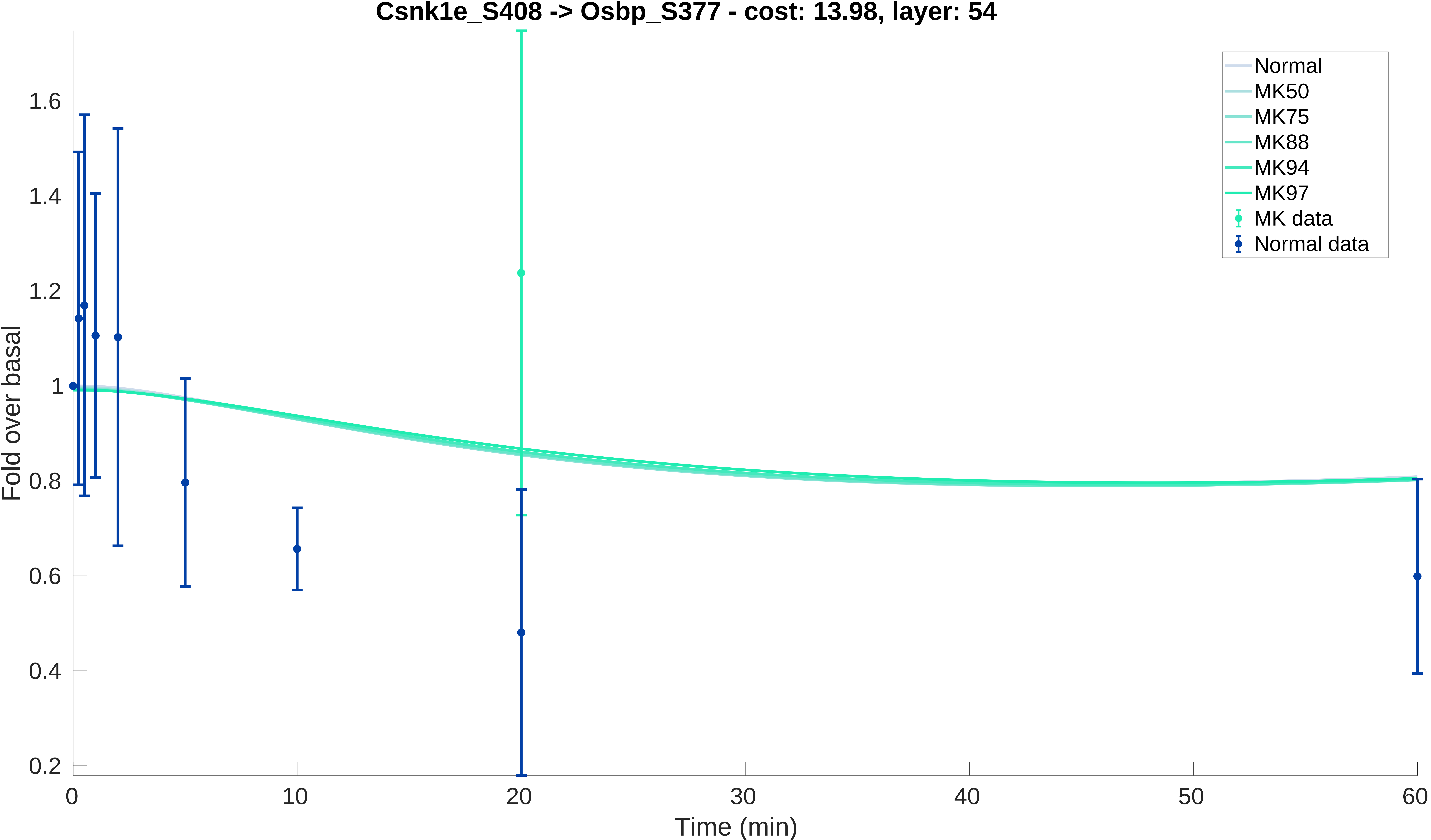

Csnk1e_S408 -> Osbp_S377 - cost: 13.98, layer: 54
Fold over basal
Time (min)
0.2
0.4
0.6
0.8
1
1.2
1.4
1.6
0
10
20
30
40
50
60
Normal
MK50
MK75
MK88
MK94
MK97
MK data
Normal data

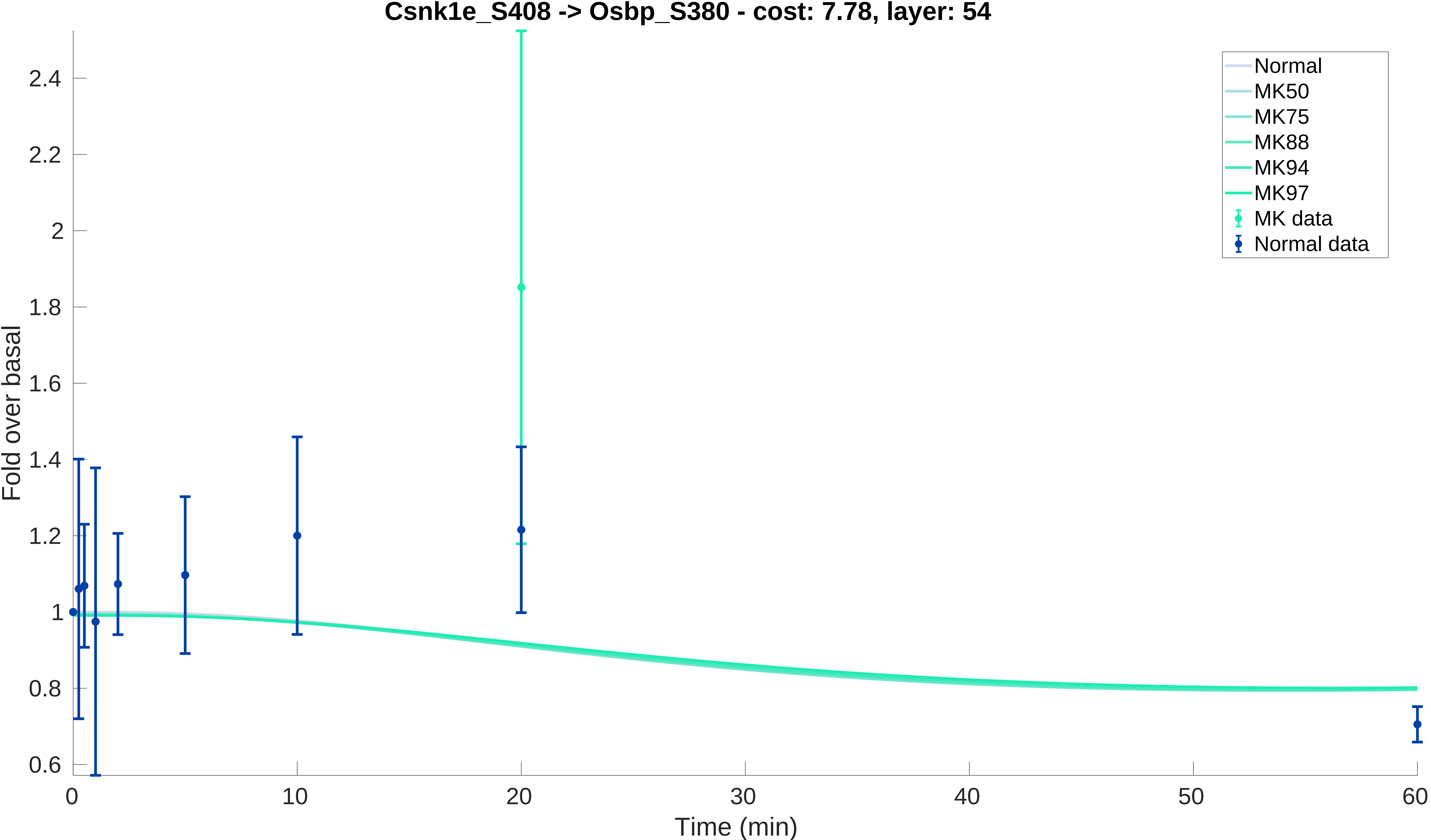

Csnk1e_S408 -> Osbp_S380 - cost: 7.78, layer: 54
Fold over basal
Time (min)
Normal
MK50
MK75
MK88
MK94
MK97
MK data
Normal data
2.4
2.2
2
1.8
1.6
1.4
1.2
1
0.8
0.6
0
10
20
30
40
50
60

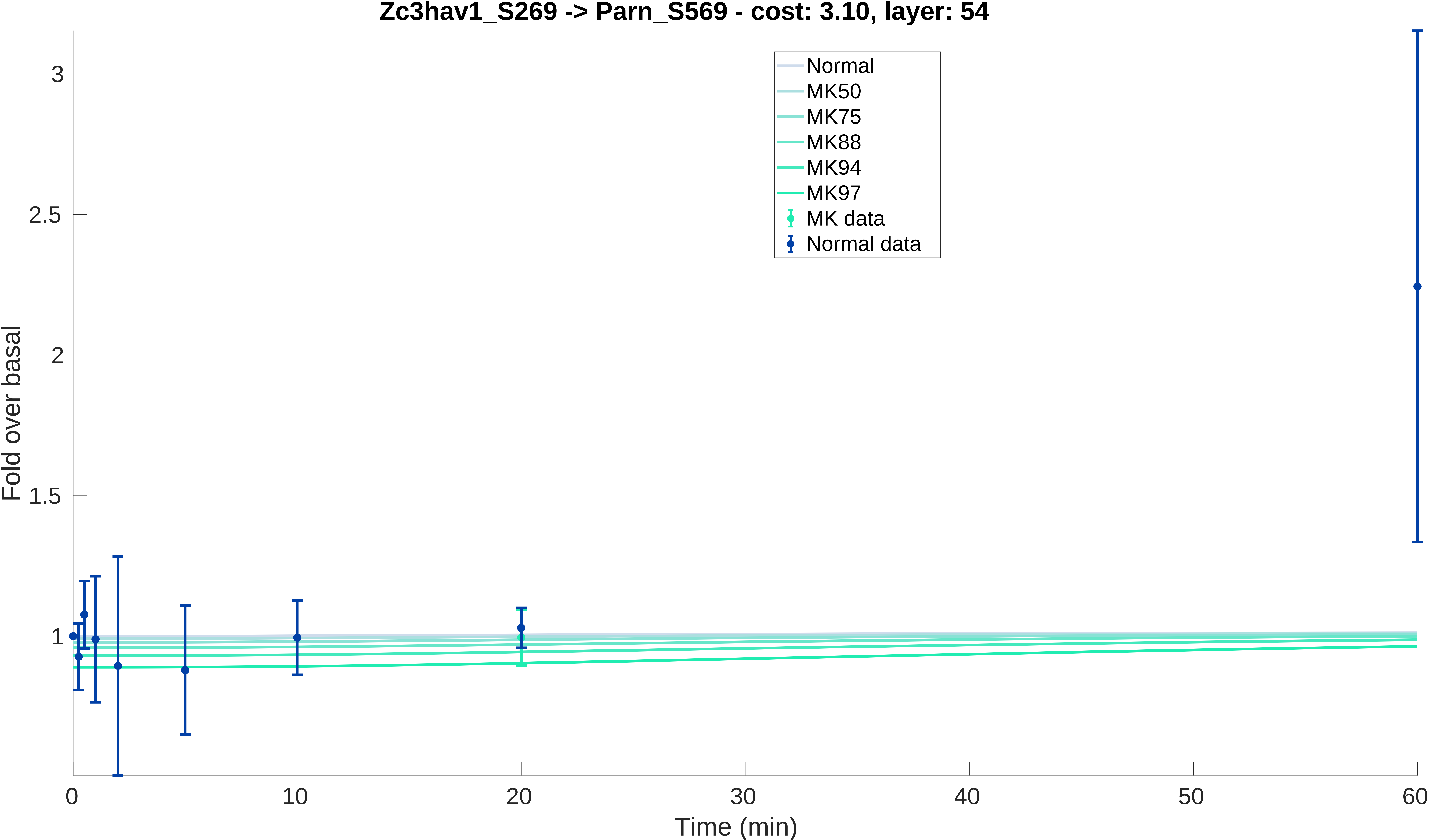

Zc3hav1_S269 -> Parn_S569 - cost: 3.10, layer: 54
Fold over basal
Time (min)
0
10
20
30
40
50
60
1
1.5
2
2.5
3
Normal
MK50
MK75
MK88
MK94
MK97
MK data
Normal data

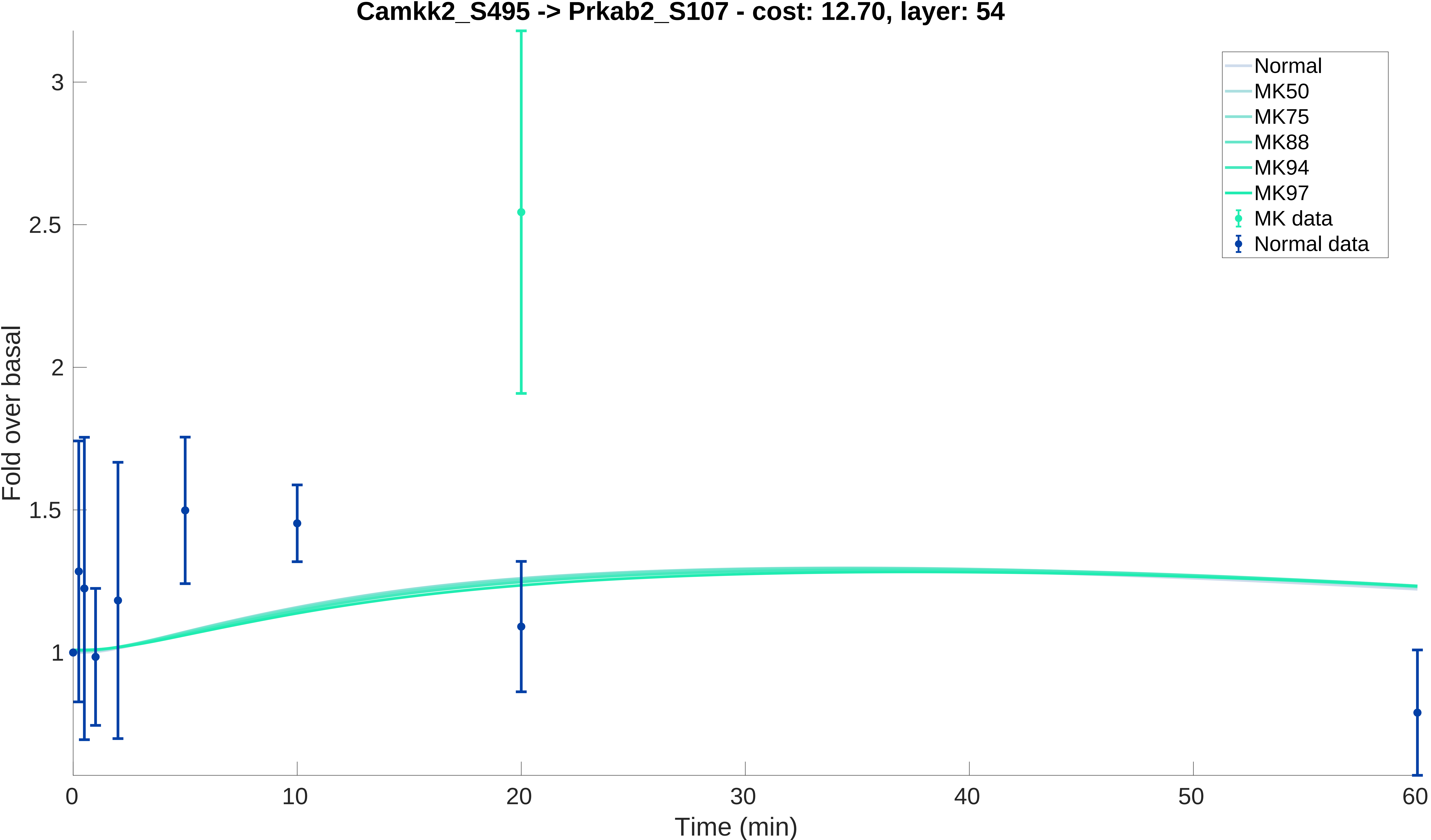

Camkk2_S495 -> Prkab2_S107 - cost: 12.70, layer: 54
Fold over basal
Time (min)
3
2.5
2
1.5
1
0
10
20
30
40
50
60
Normal
MK50
MK75
MK88
MK94
MK97
MK data
Normal data

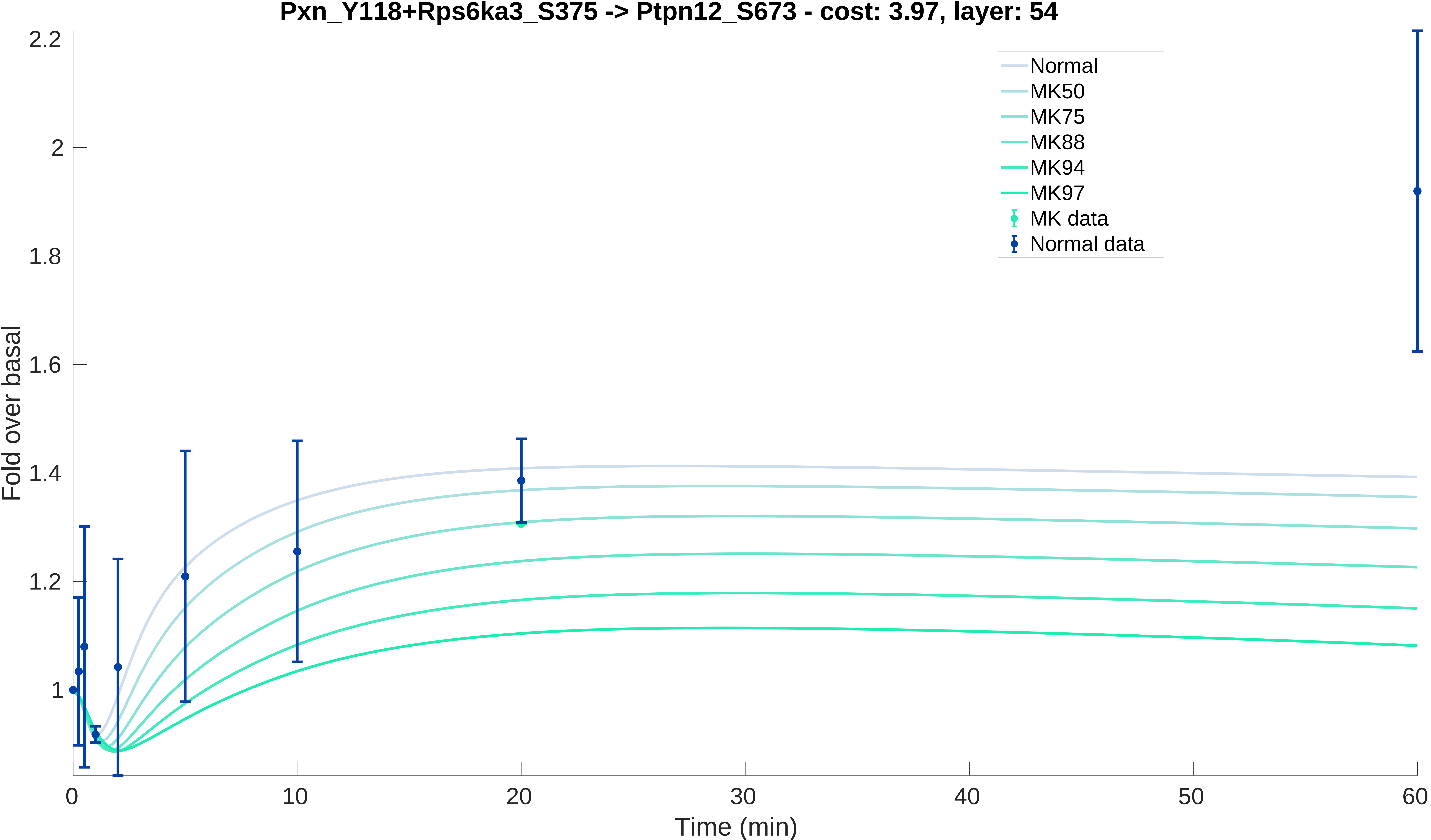

Pxn_Y118+Rps6ka3_S375 -> Ptpn12_S673 - cost: 3.97, layer: 54
Normal
MK50
MK75
MK88
MK94
MK97
MK data
Normal data
Fold over basal
Time (min)
2.2
2
1.8
1.6
1.4
1.2
1
0
10
20
30
40
50
60

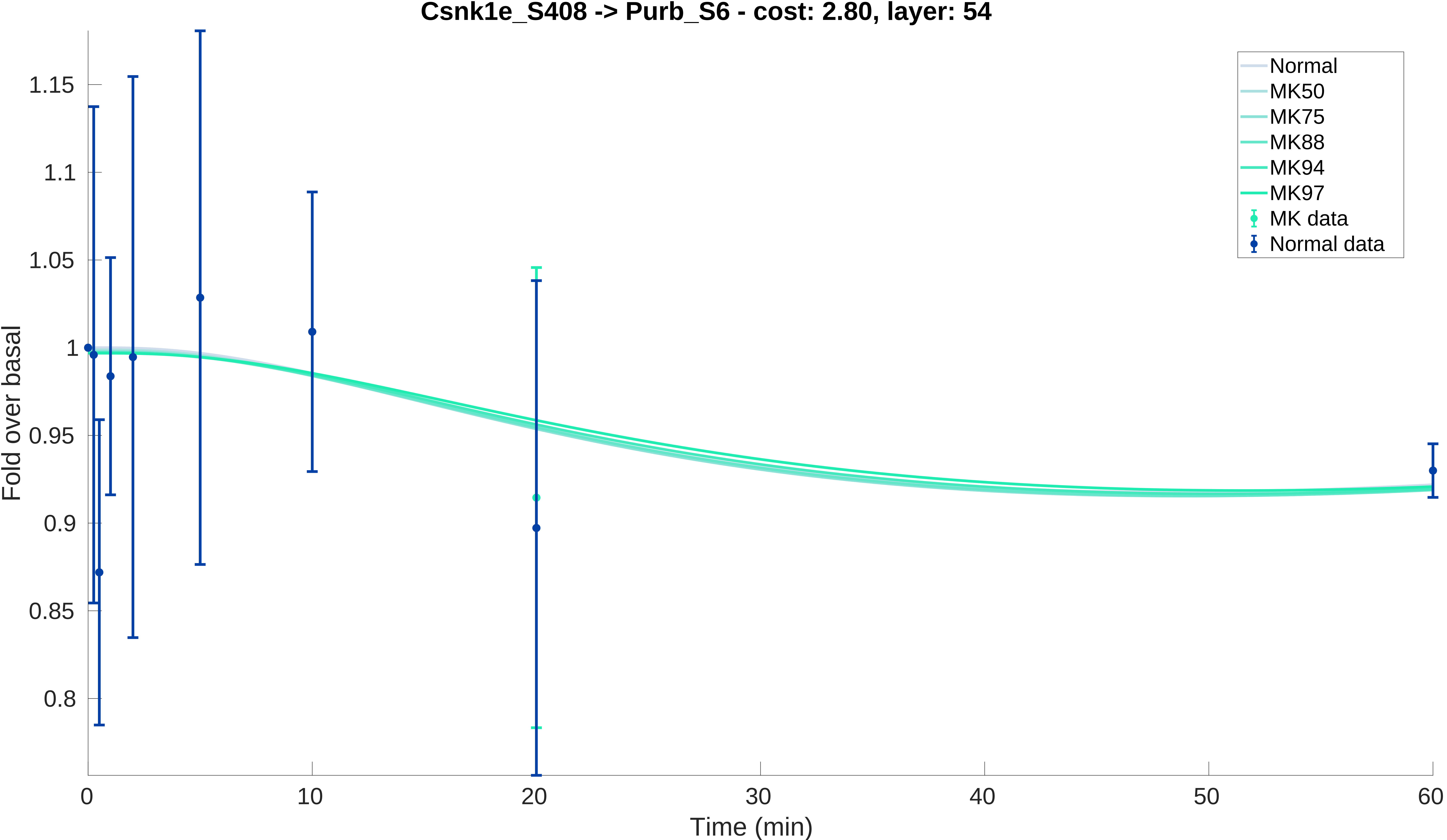
Csnk1e_S408 -> Purb_S6 - cost: 2.80, layer: 54
Fold over basal
1.15
1.1
1.05
1
0.95
0.9
0.85
0.8
0
10
20
30
40
50
60
Time (min)
Normal
MK50
MK75
MK88
MK94
MK97
MK data
Normal data

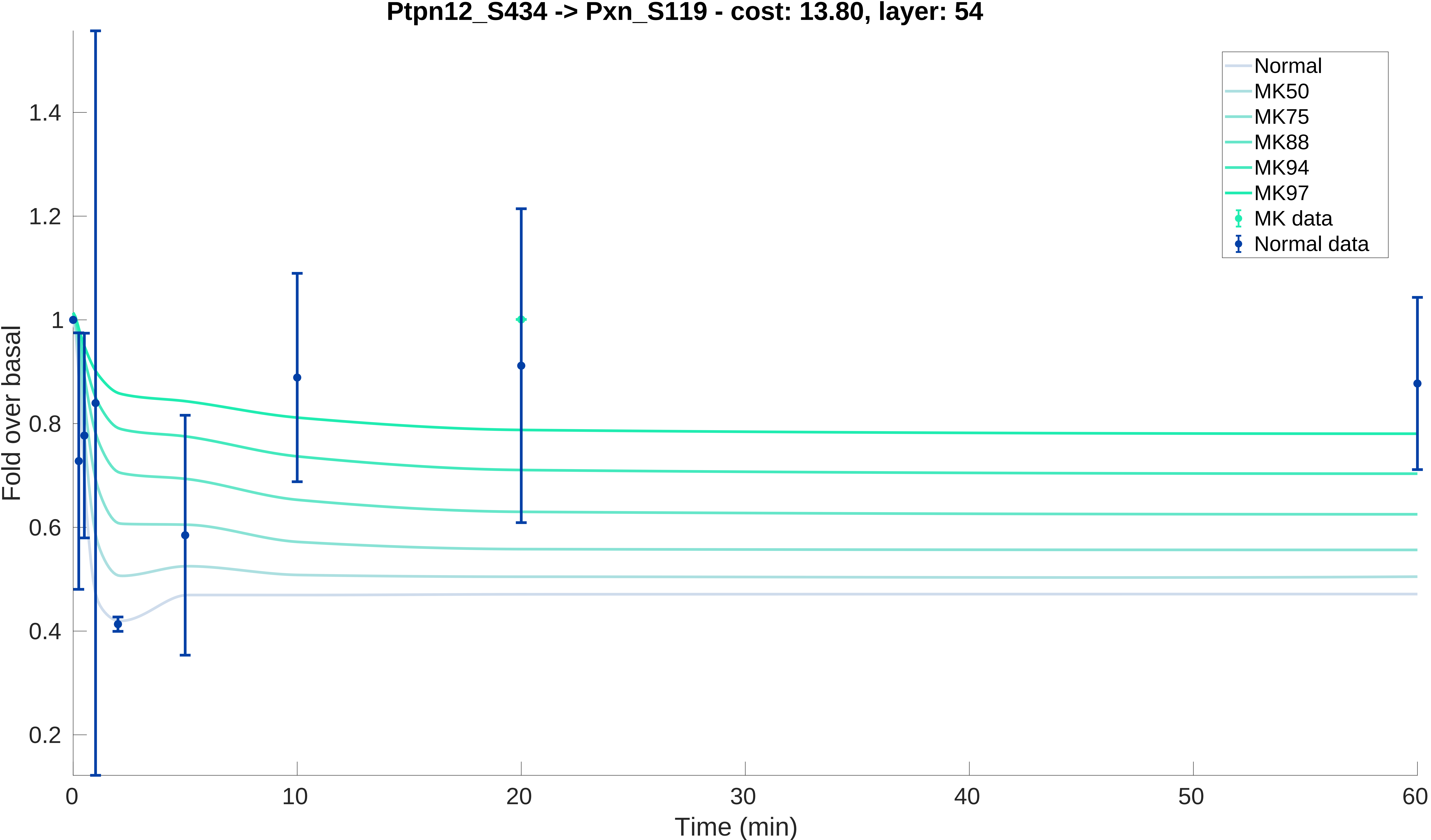

Ptpn12_S434 -> Pxn_S119 - cost: 13.80, layer: 54
Normal
MK50
MK75
MK88
MK94
MK97
MK data
Normal data
Fold over basal
Time (min)
0.2
0.4
0.6
0.8
1
1.2
1.4
0
10
20
30
40
50
60

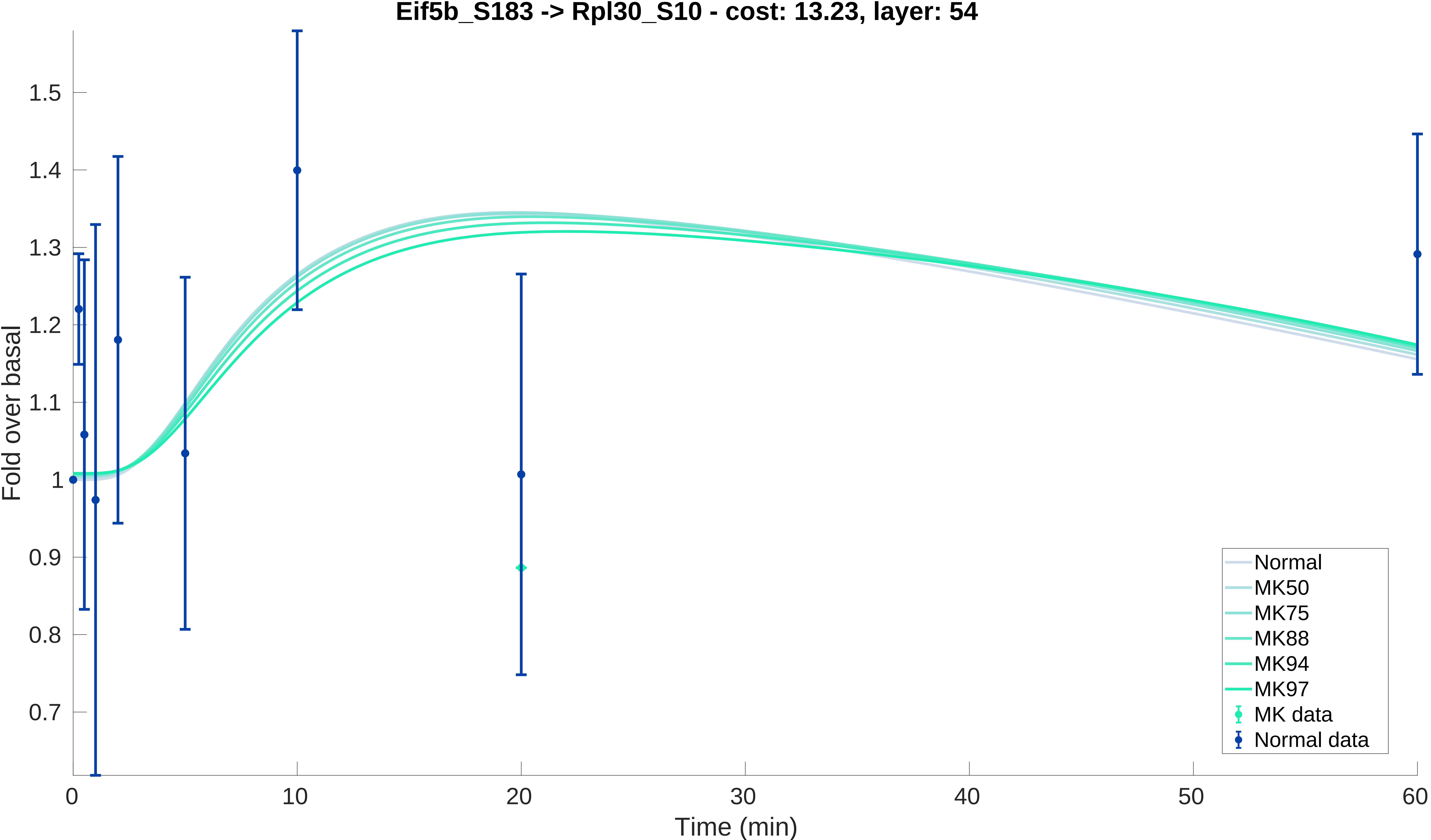

Eif5b_S183 -> Rpl30_S10 - cost: 13.23, layer: 54
Fold over basal
Time (min)
1.5
1.4
1.3
1.2
1.1
1
0.9
0.8
0.7
0
10
20
30
40
50
60
Normal
MK50
MK75
MK88
MK94
MK97
MK data
Normal data

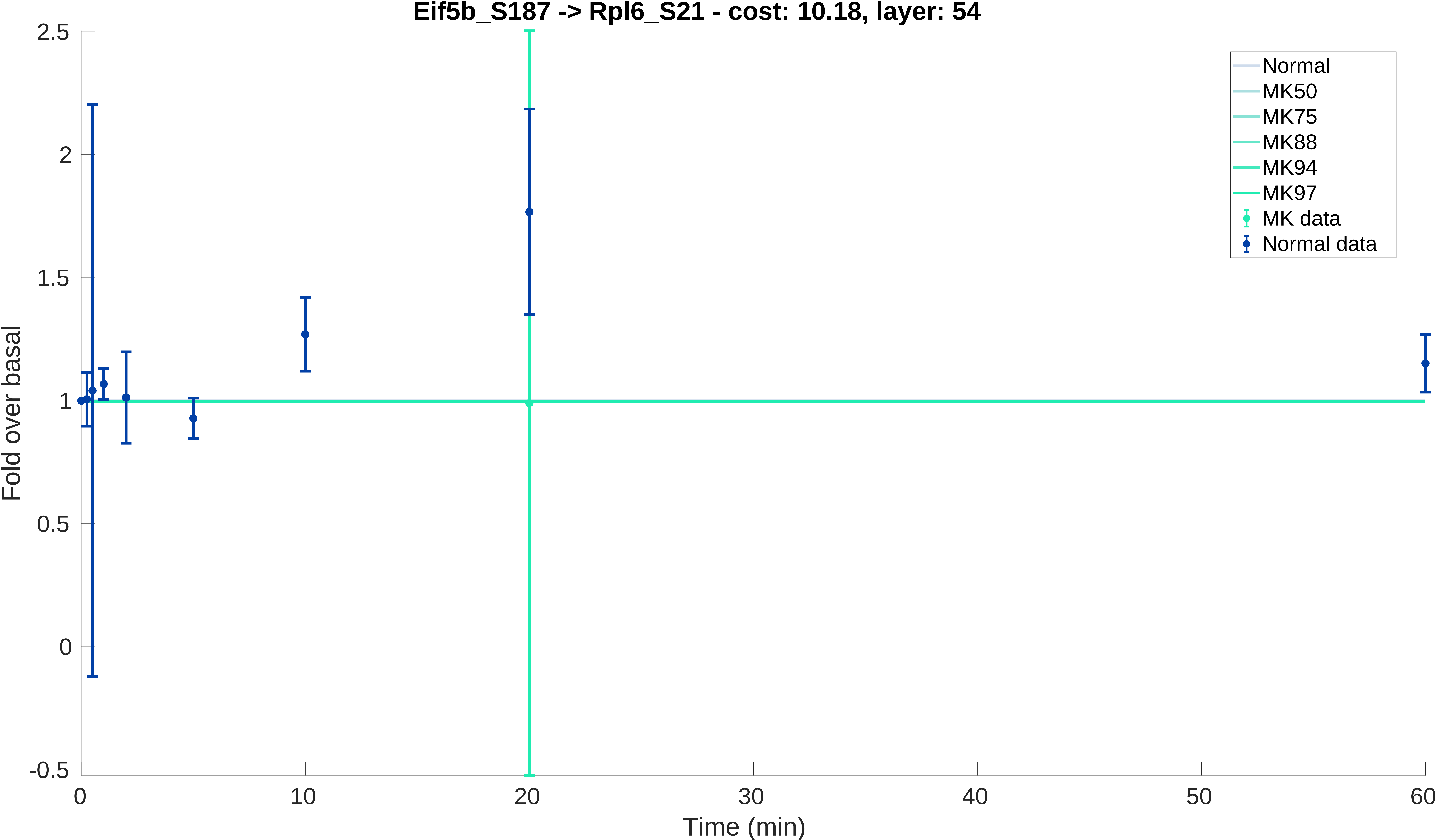

Eif5b_S187 -> Rpl6_S21 - cost: 10.18, layer: 54
Fold over basal
Time (min)
2.5
2
1.5
1
0.5
0
-0.5
0
10
20
30
40
50
60
Normal
MK50
MK75
MK88
MK94
MK97
MK data
Normal data

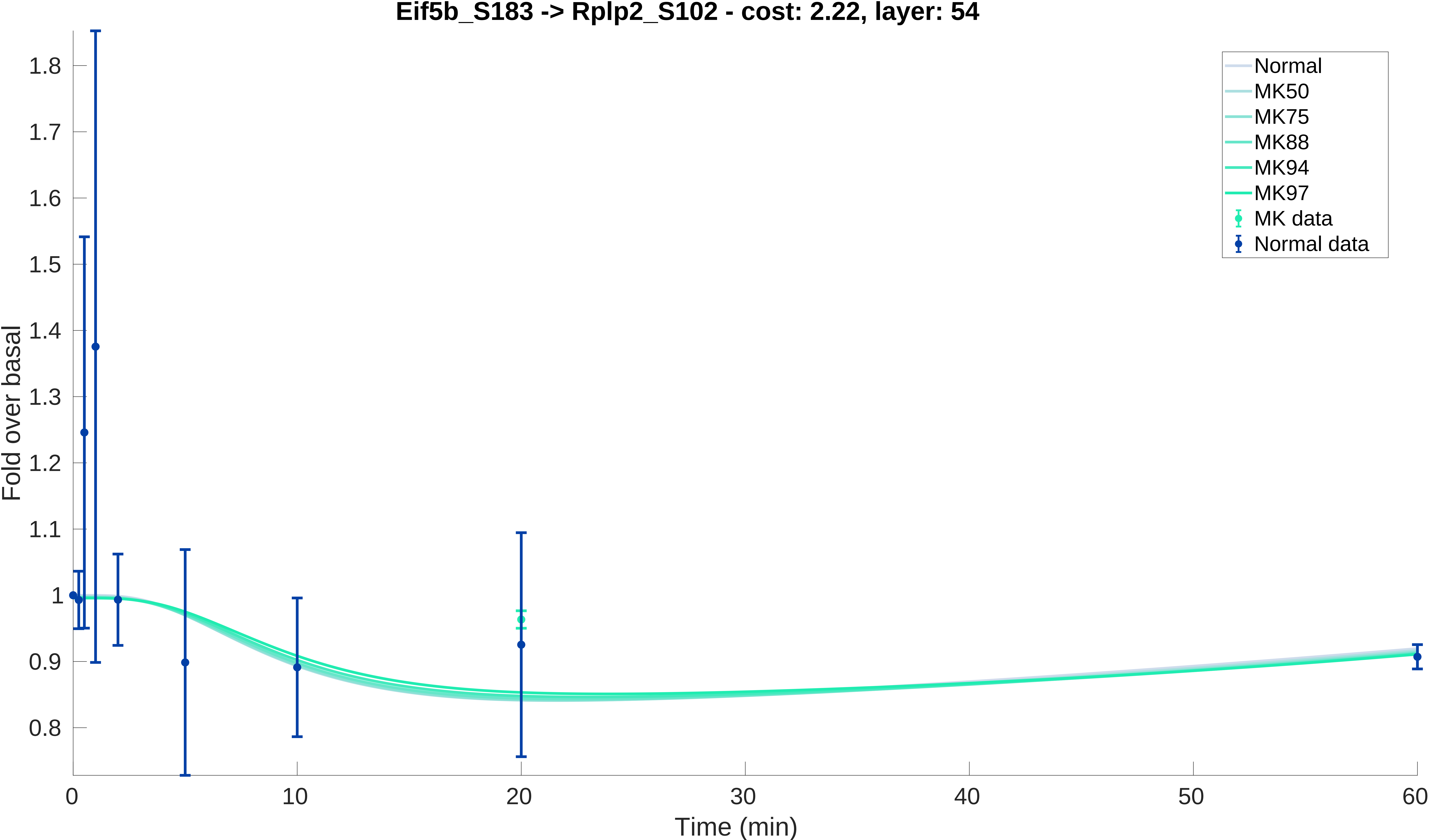

Eif5b_S183 -> Rplp2_S102 - cost: 2.22, layer: 54
Normal
MK50
MK75
MK88
MK94
MK97
MK data
Normal data
Fold over basal
Time (min)
0
10
20
30
40
50
60
0.8
0.9
1
1.1
1.2
1.3
1.4
1.5
1.6
1.7
1.8

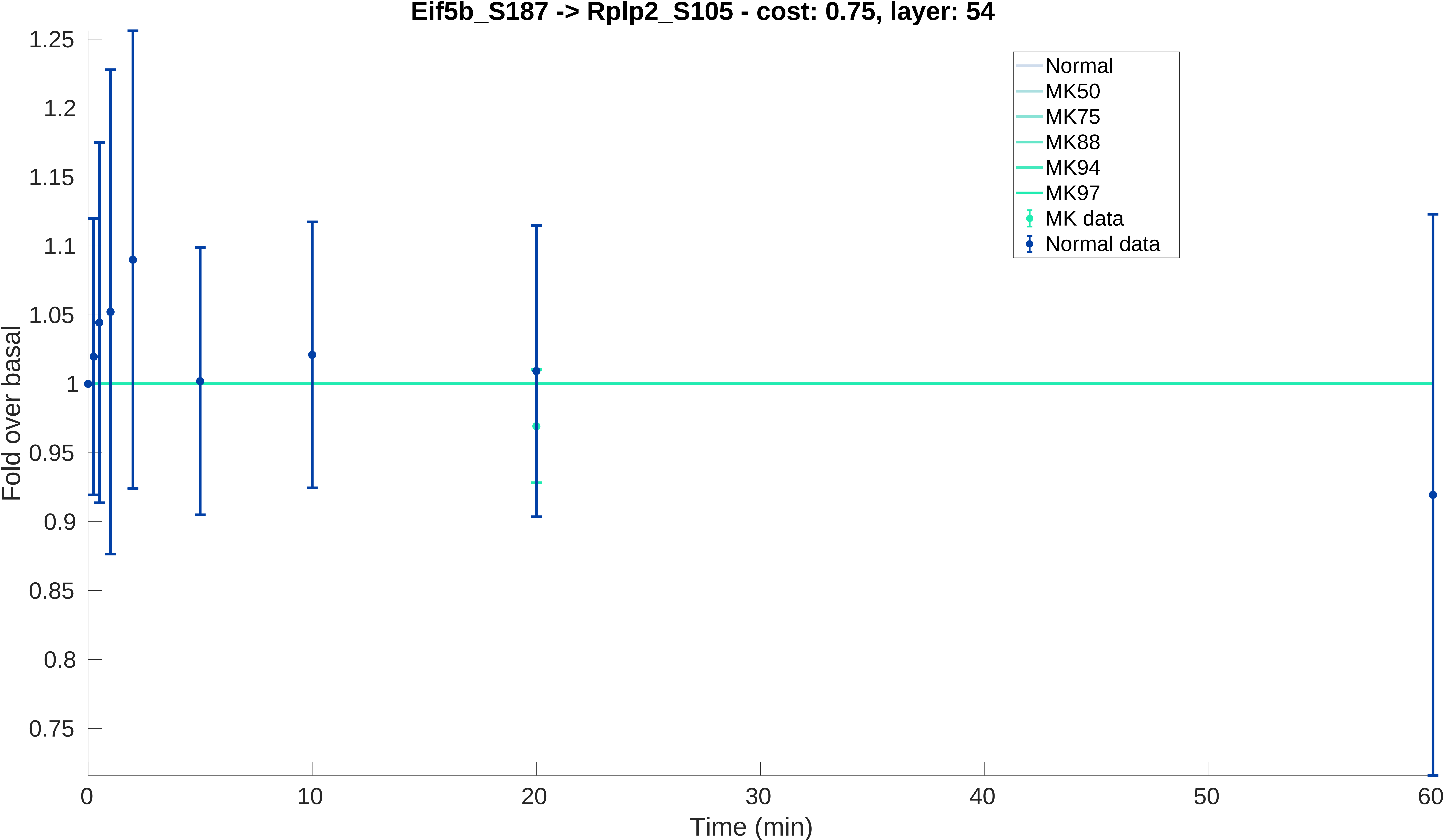

Eif5b_S187 -> Rplp2_S105 - cost: 0.75, layer: 54
Normal
MK50
MK75
MK88
MK94
MK97
MK data
Normal data
Fold over basal
Time (min)
1.25
1.2
1.15
1.1
1.05
1
0.95
0.9
0.85
0.8
0.75
0
10
20
30
40
50
60

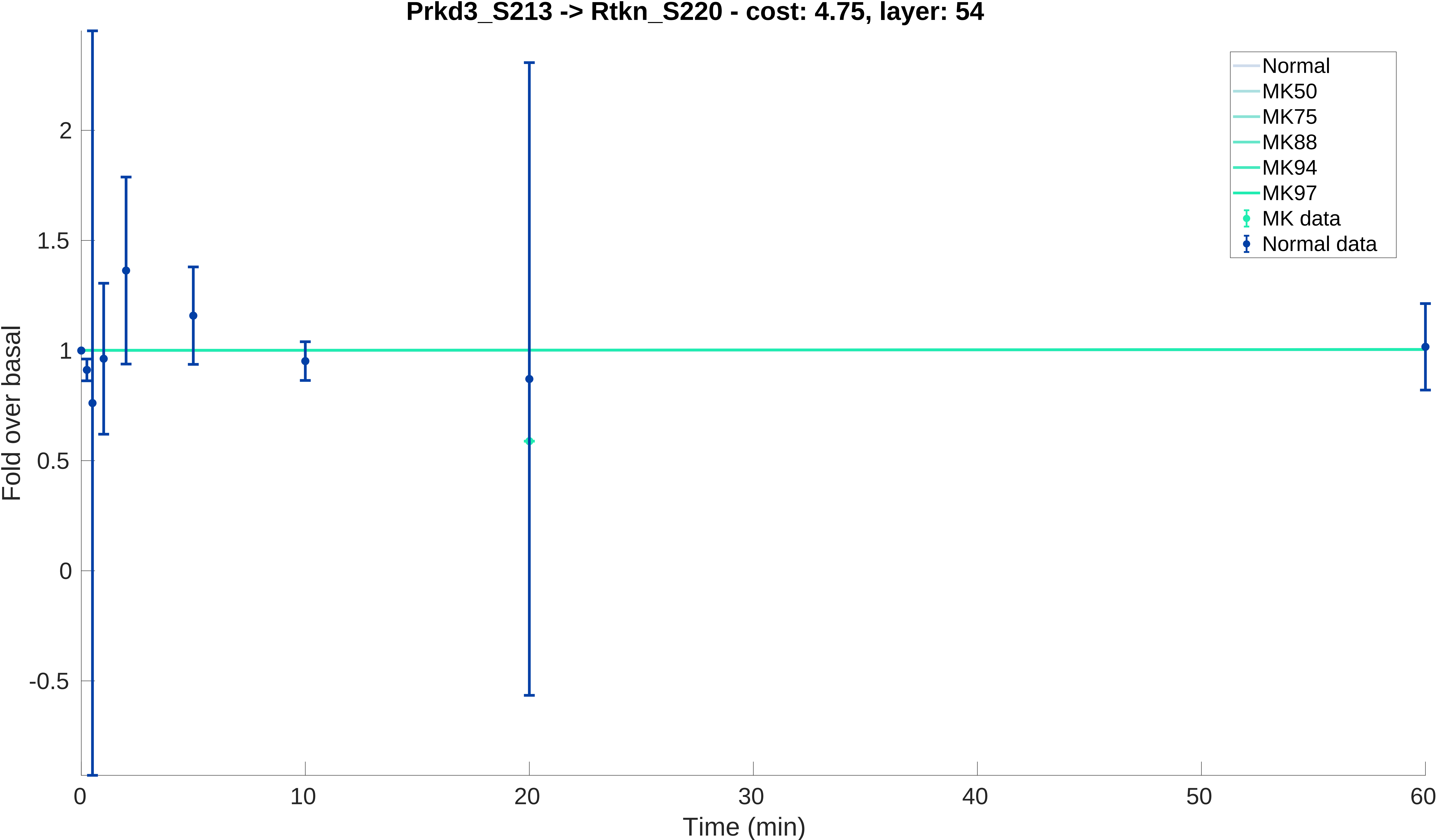

Prkd3_S213 -> Rtkn_S220 - cost: 4.75, layer: 54
Normal
MK50
MK75
MK88
MK94
MK97
MK data
Normal data
Fold over basal
Time (min)
2
1.5
1
0.5
0
-0.5
0
10
20
30
40
50
60

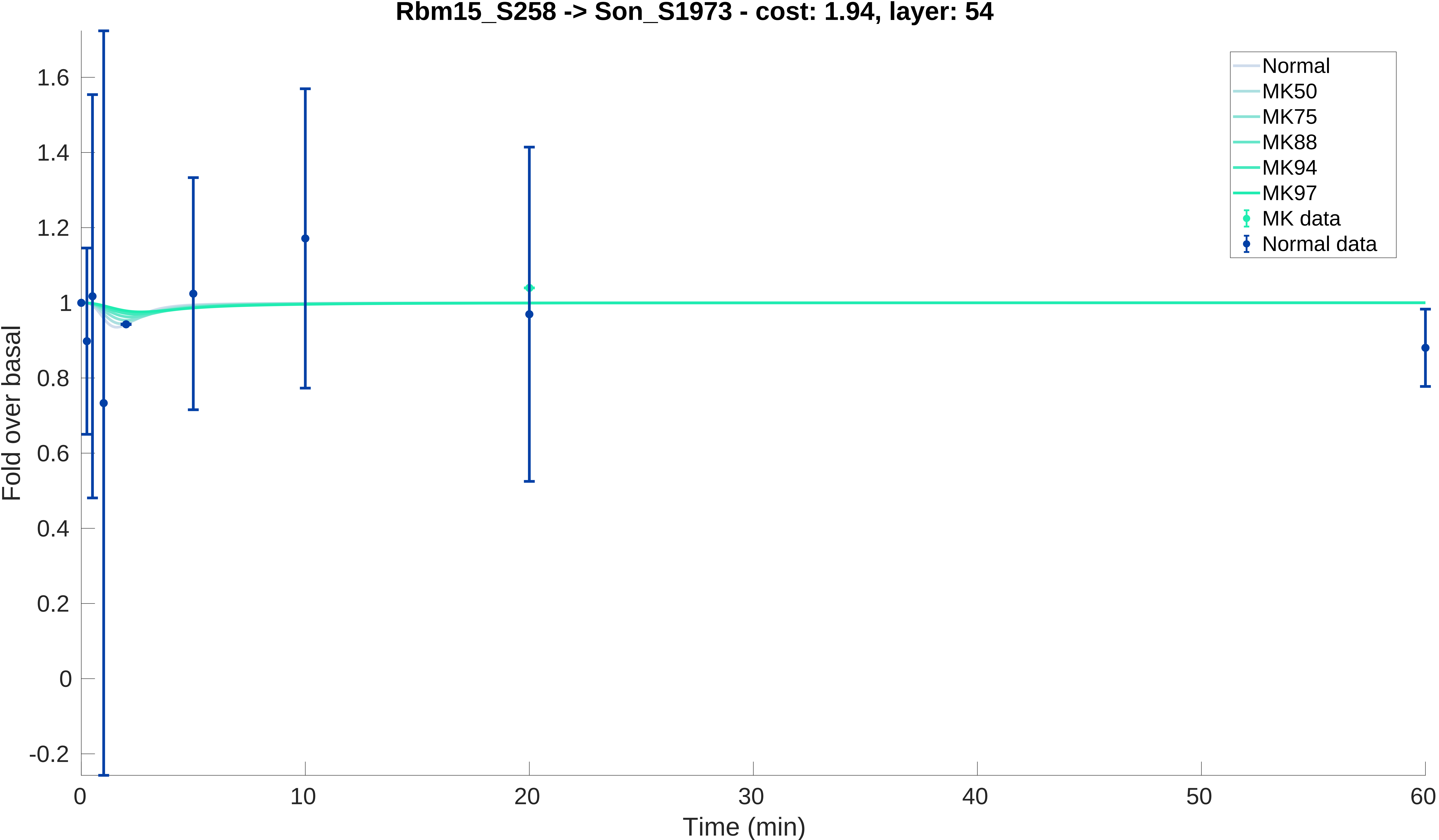

Rbm15_S258 -> Son_S1973 - cost: 1.94, layer: 54
Normal
MK50
MK75
MK88
MK94
MK97
MK data
Normal data
Fold over basal
Time (min)
1.6
1.4
1.2
1
0.8
0.6
0.4
0.2
0
-0.2
0
10
20
30
40
50
60

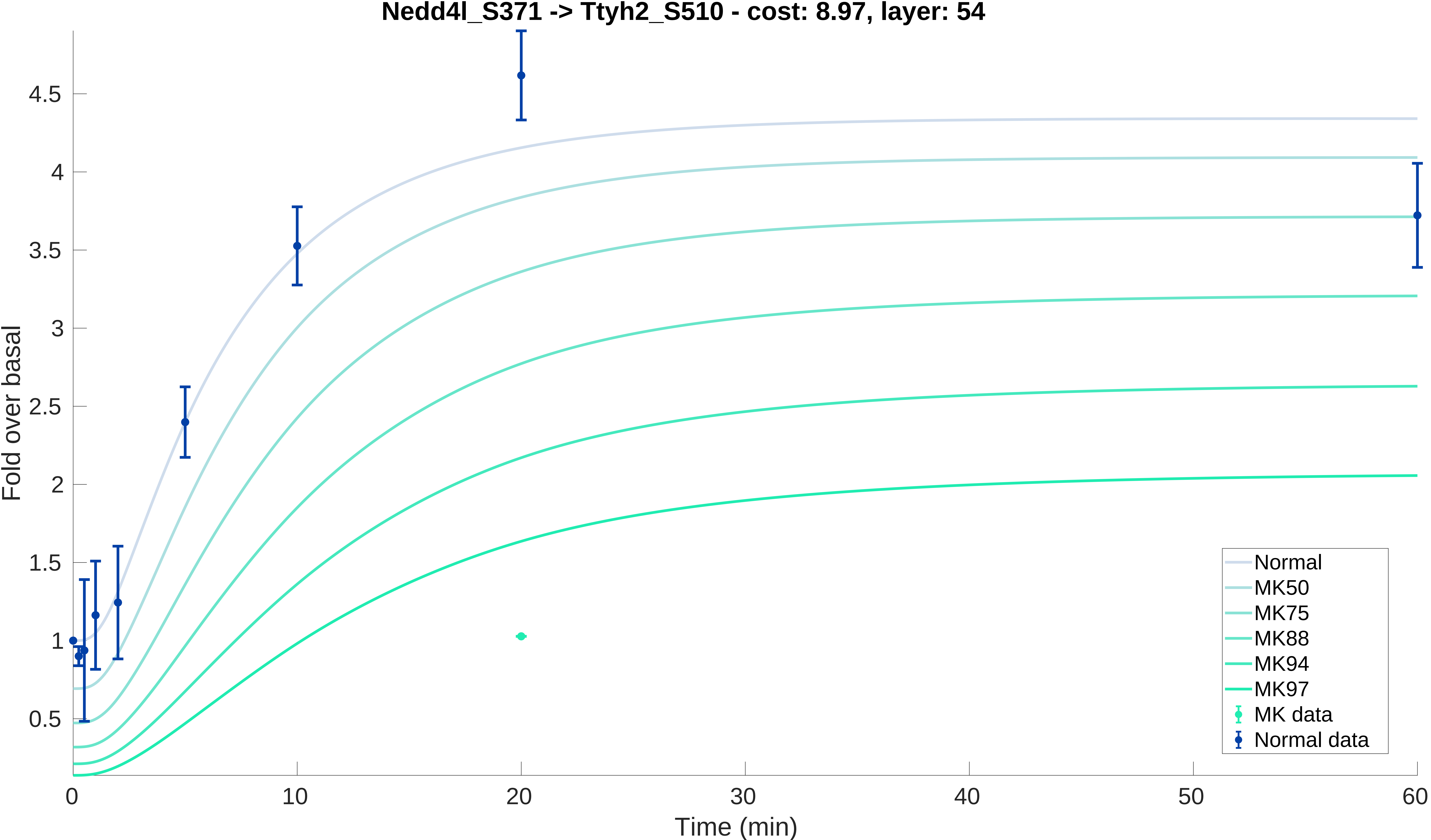

Nedd4l_S371 -> Ttyh2_S510 - cost: 8.97, layer: 54
Fold over basal
Time (min)
0.5
1
1.5
2
2.5
3
3.5
4
4.5
0
10
20
30
40
50
60
Normal
MK50
MK75
MK88
MK94
MK97
MK data
Normal data

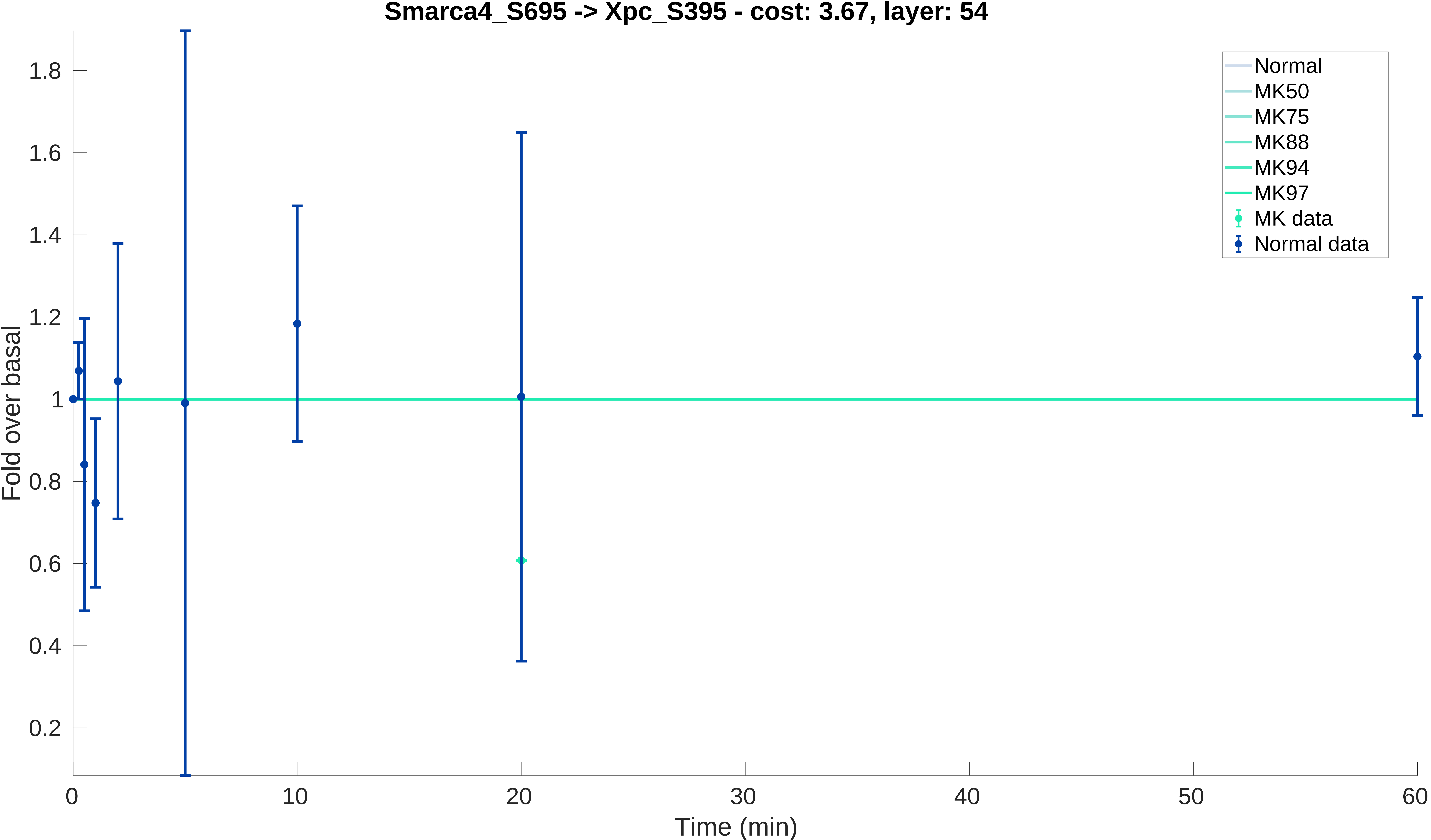

Smarca4_S695 -> Xpc_S395 - cost: 3.67, layer: 54
Fold over basal
Time (min)
0.2
0.4
0.6
0.8
1
1.2
1.4
1.6
1.8
0
10
20
30
40
50
60
Normal
MK50
MK75
MK88
MK94
MK97
MK data
Normal data

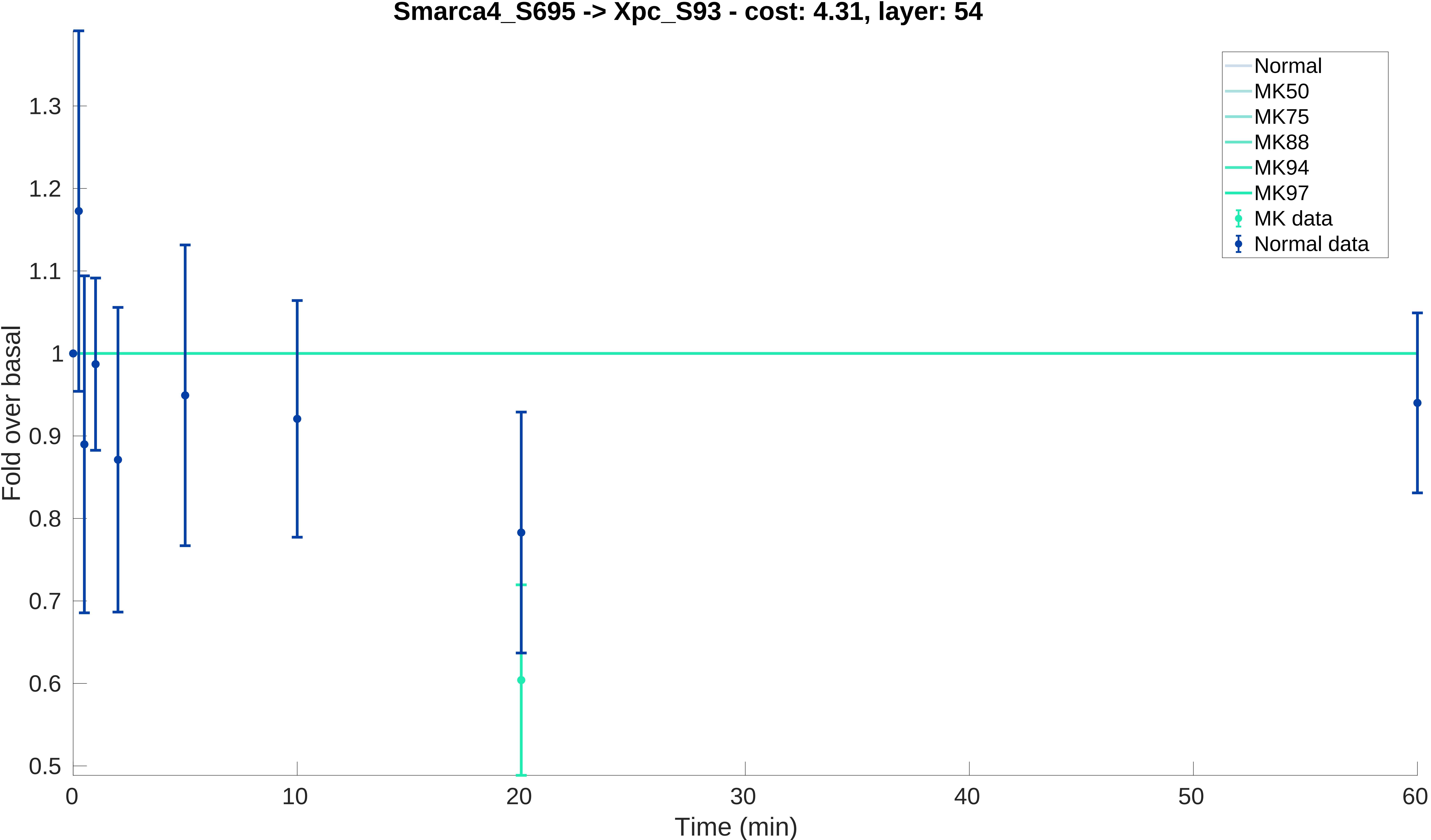

Smarca4_S695 -> Xpc_S93 - cost: 4.31, layer: 54
Normal
MK50
MK75
MK88
MK94
MK97
MK data
Normal data
Fold over basal
1.3
1.2
1.1
1
0.9
0.8
0.7
0.6
0.5
0
10
20
30
40
50
60
Time (min)

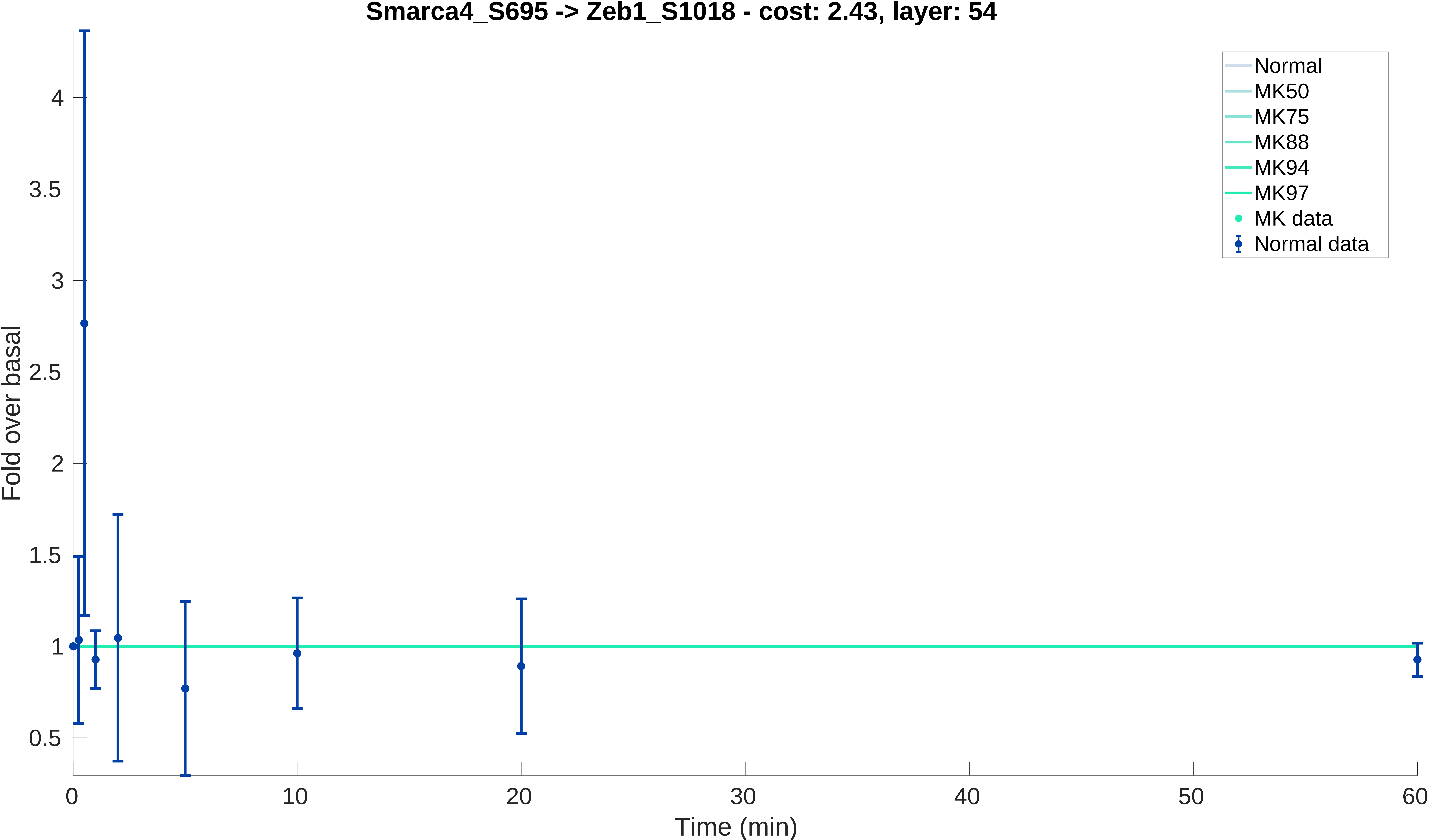

Smarca4_S695 -> Zeb1_S1018 - cost: 2.43, layer: 54
Fold over basal
Time (min)
0
10
20
30
40
50
60
0.5
1
1.5
2
2.5
3
3.5
4
Normal
MK50
MK75
MK88
MK94
MK97
MK data
Normal data

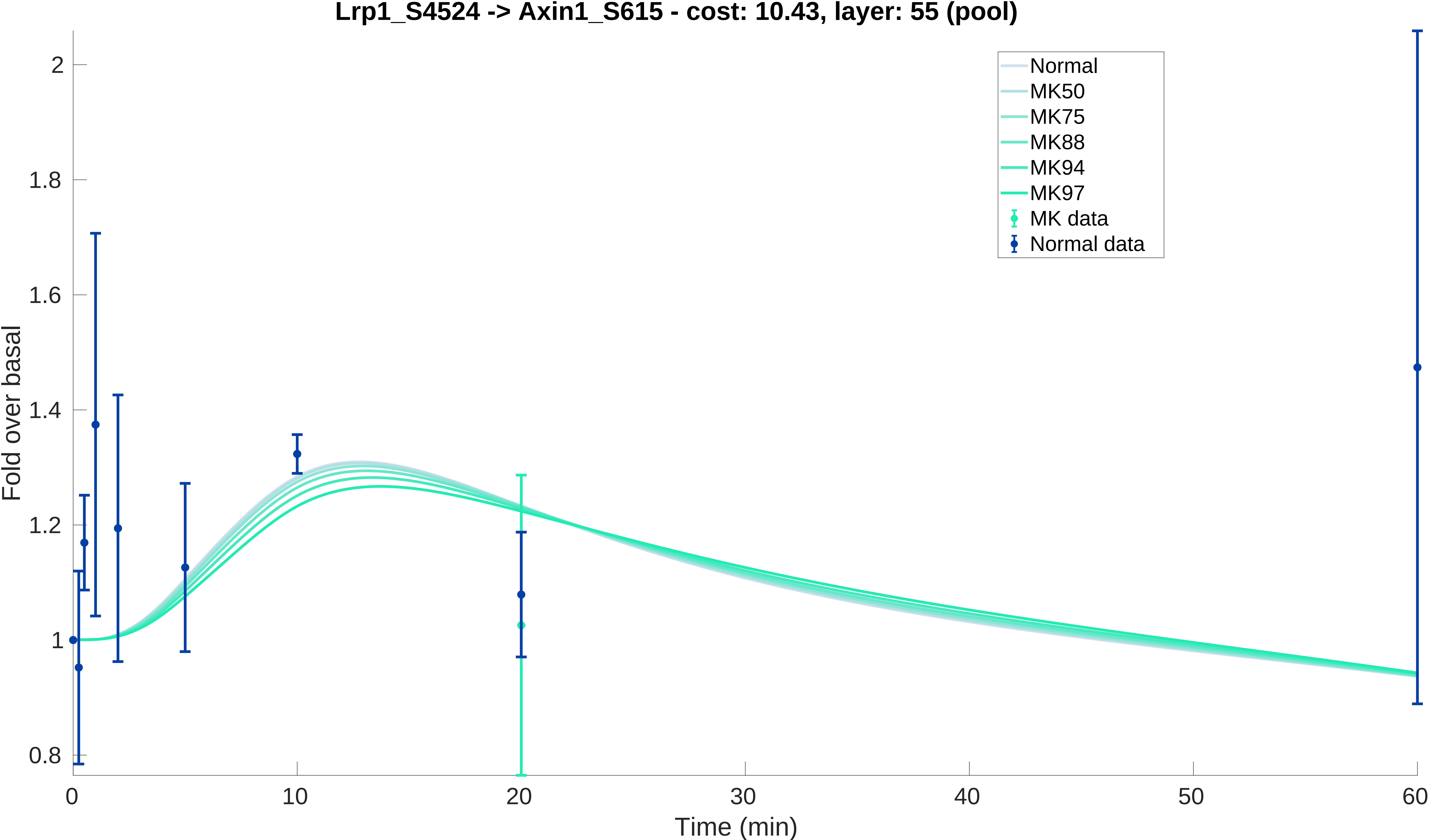

Lrp1_S4524 -> Axin1_S615 - cost: 10.43, layer: 55 (pool)
Fold over basal
Time (min)
0.8
1
1.2
1.4
1.6
1.8
2
0
10
20
30
40
50
60
Normal
MK50
MK75
MK88
MK94
MK97
MK data
Normal data

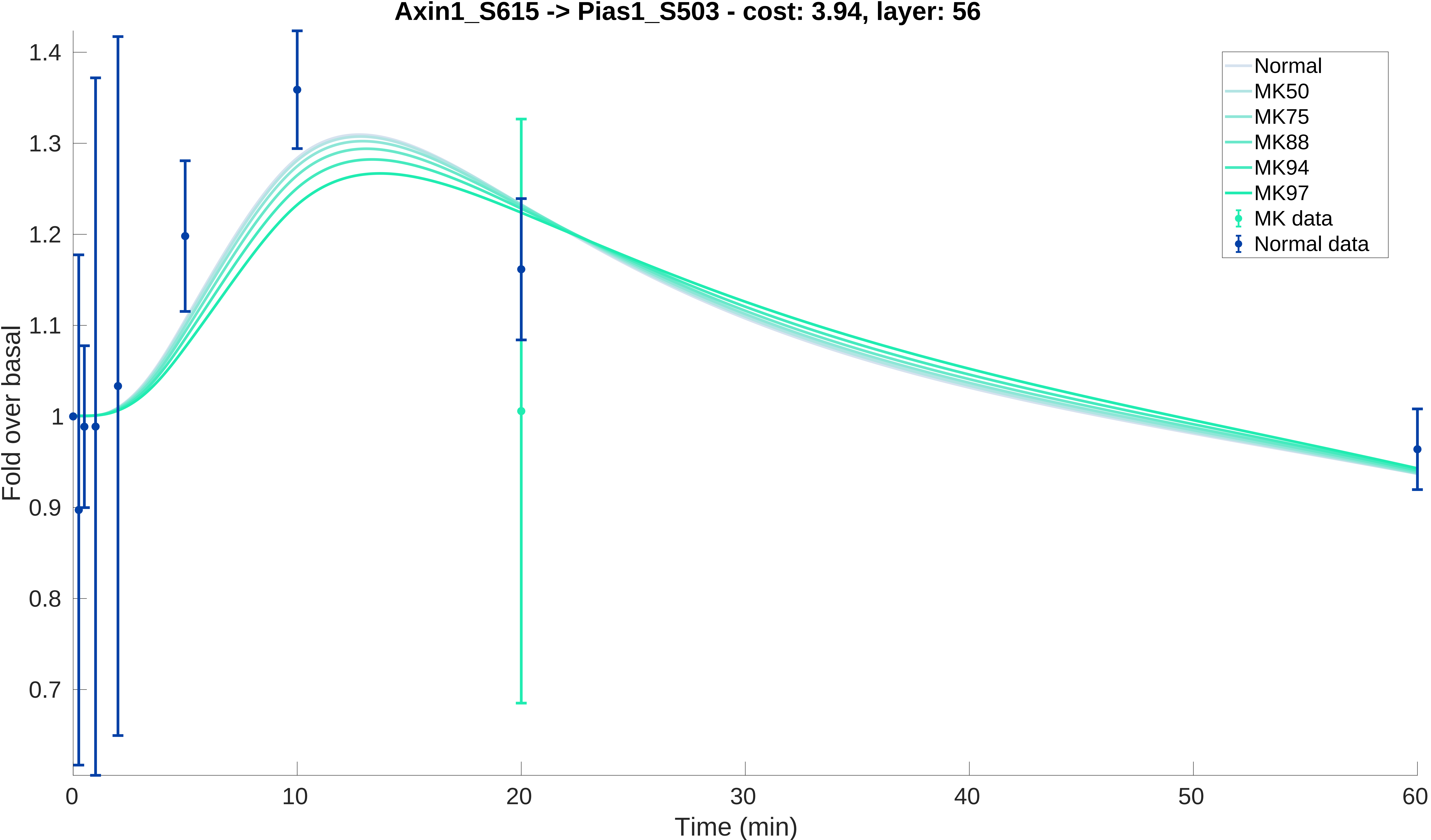

Axin1_S615 -> Pias1_S503 - cost: 3.94, layer: 56
Fold over basal
Time (min)
1.4
1.3
1.2
1.1
1
0.9
0.8
0.7
0
10
20
30
40
50
60
Normal
MK50
MK75
MK88
MK94
MK97
MK data
Normal data

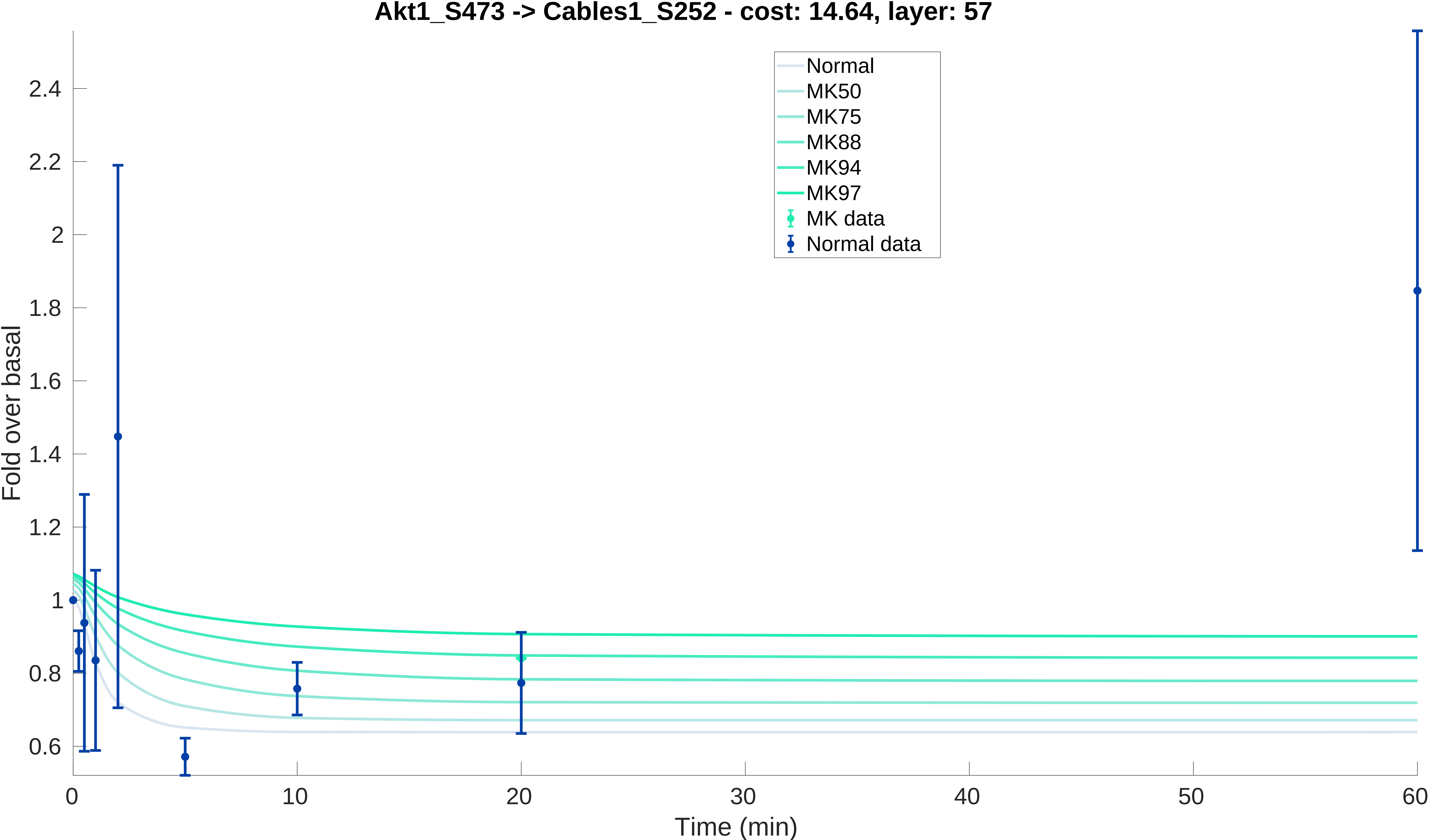

Akt1_S473 -> Cables1_S252 - cost: 14.64, layer: 57
Normal
MK50
MK75
MK88
MK94
MK97
MK data
Normal data
Fold over basal
2.4
2.2
2
1.8
1.6
1.4
1.2
1
0.8
0.6
0
10
20
30
40
50
60
Time (min)

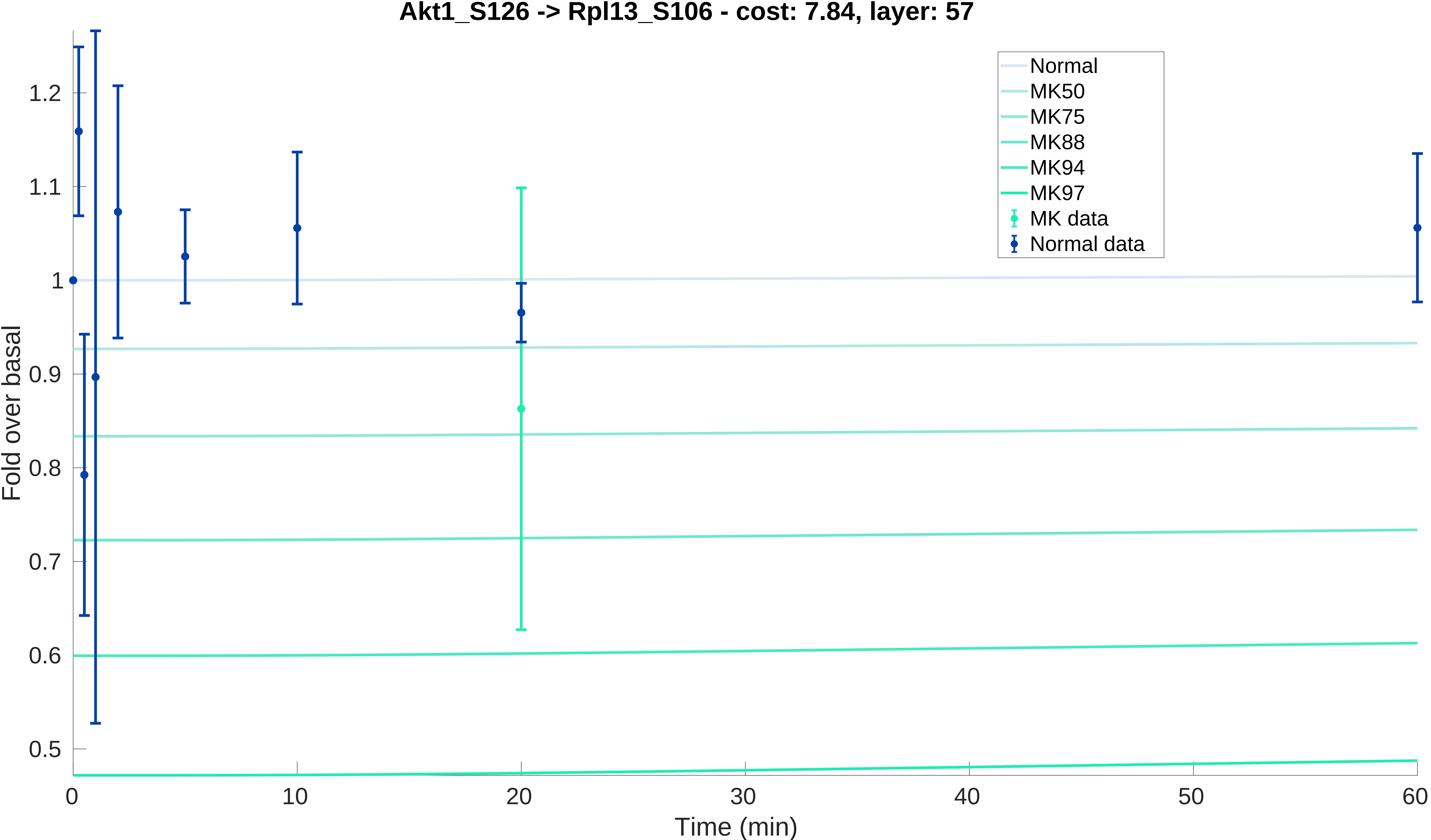
Akt1_S126 -> Rpl13_S106 - cost: 7.84, layer: 57
Normal
MK50
MK75
MK88
MK94
MK97
MK data
Normal data
Fold over basal
1.2
1.1
1
0.9
0.8
0.7
0.6
0.5
0
10
20
30
40
50
60
Time (min)

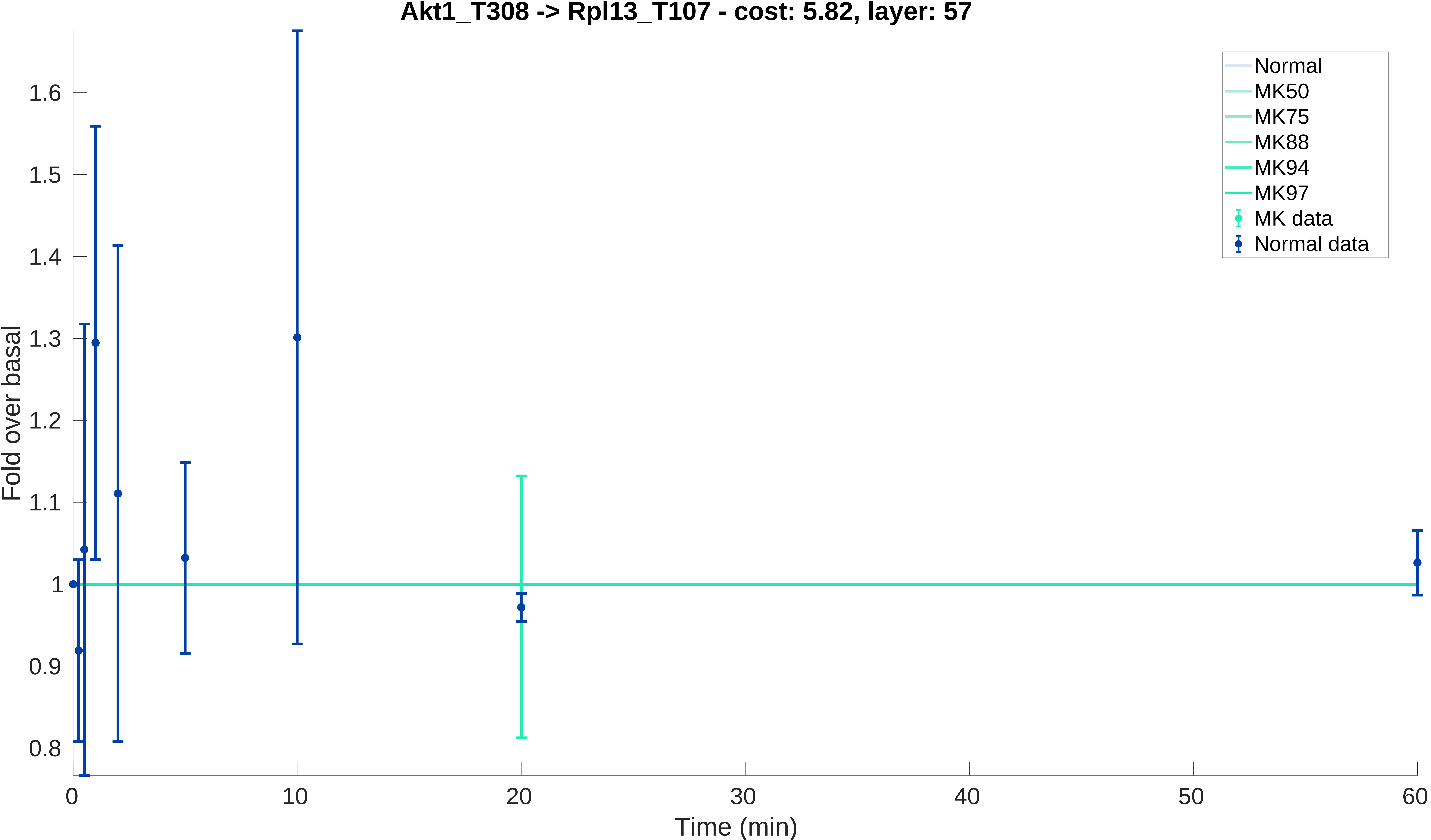

Akt1_T308 -> Rpl13_T107 - cost: 5.82, layer: 57
Fold over basal
Time (min)
0.8
0.9
1
1.1
1.2
1.3
1.4
1.5
1.6
0
10
20
30
40
50
60
Normal
MK50
MK75
MK88
MK94
MK97
MK data
Normal data

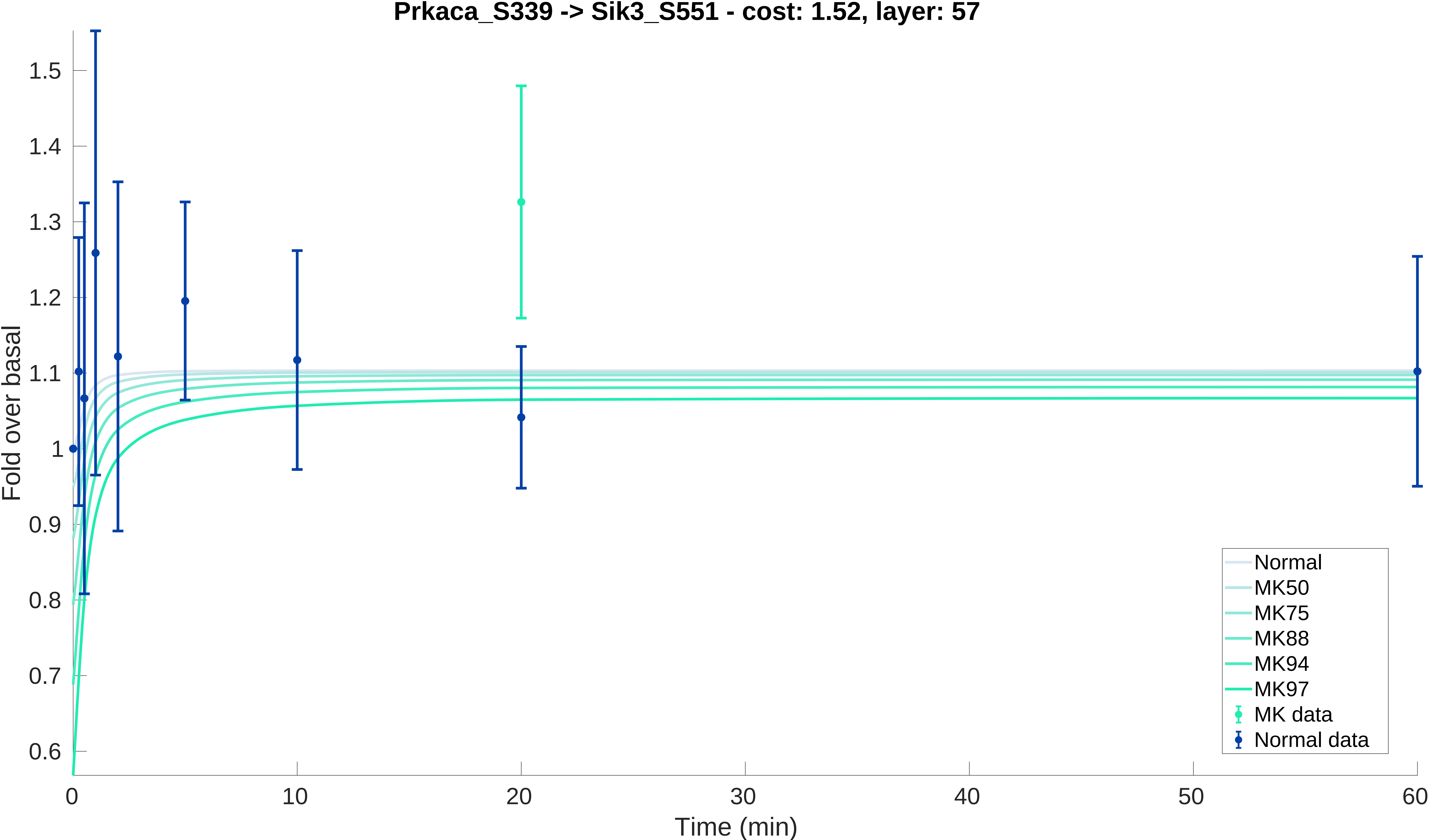

Prkaca_S339 -> Sik3_S551 - cost: 1.52, layer: 57
Fold over basal
Time (min)
0.6
0.7
0.8
0.9
1
1.1
1.2
1.3
1.4
1.5
0
10
20
30
40
50
60
Normal
MK50
MK75
MK88
MK94
MK97
MK data
Normal data

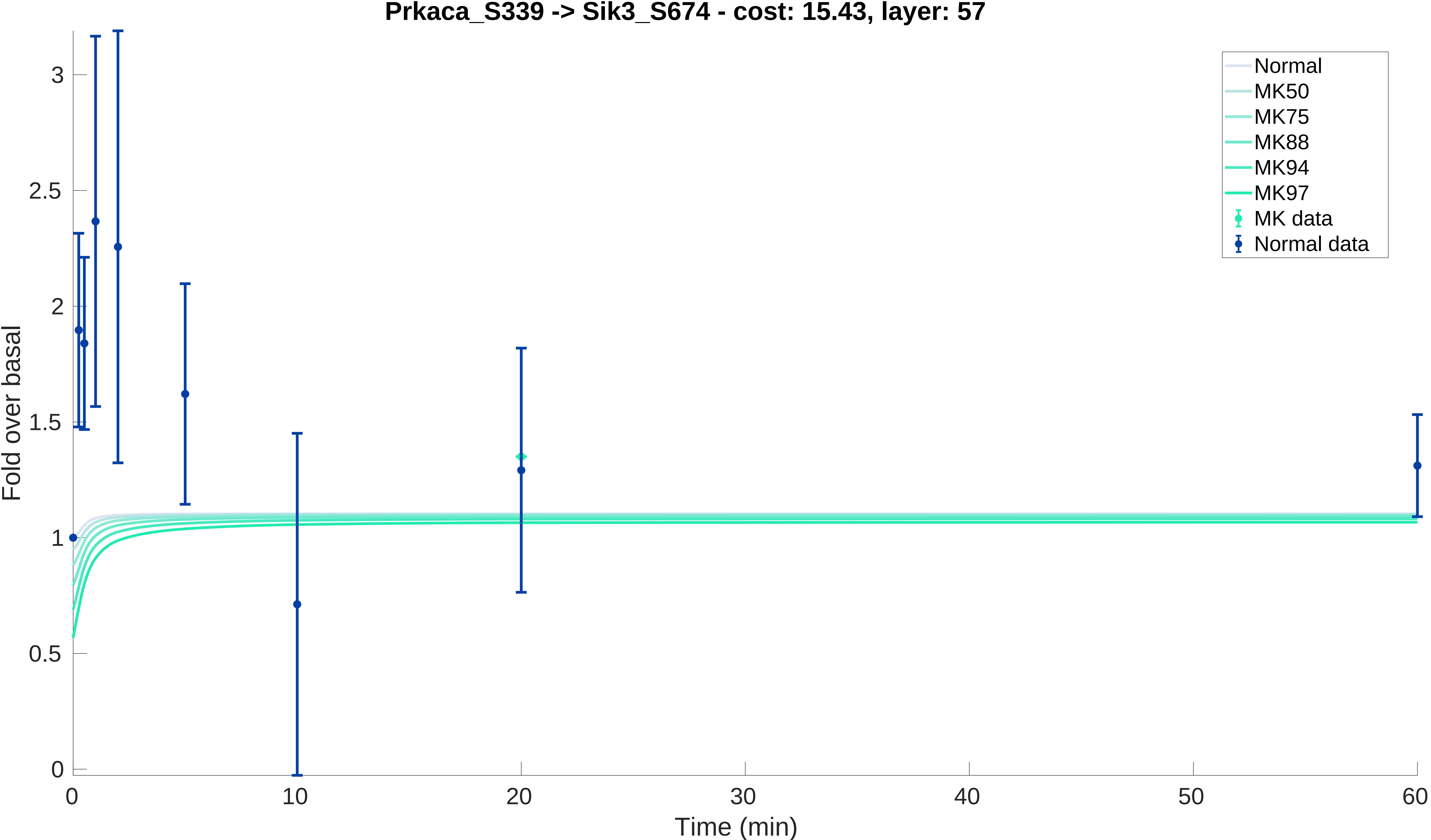

Prkaca_S339 -> Sik3_S674 - cost: 15.43, layer: 57
Fold over basal
Time (min)
0
0.5
1
1.5
2
2.5
3
0
10
20
30
40
50
60
Normal
MK50
MK75
MK88
MK94
MK97
MK data
Normal data

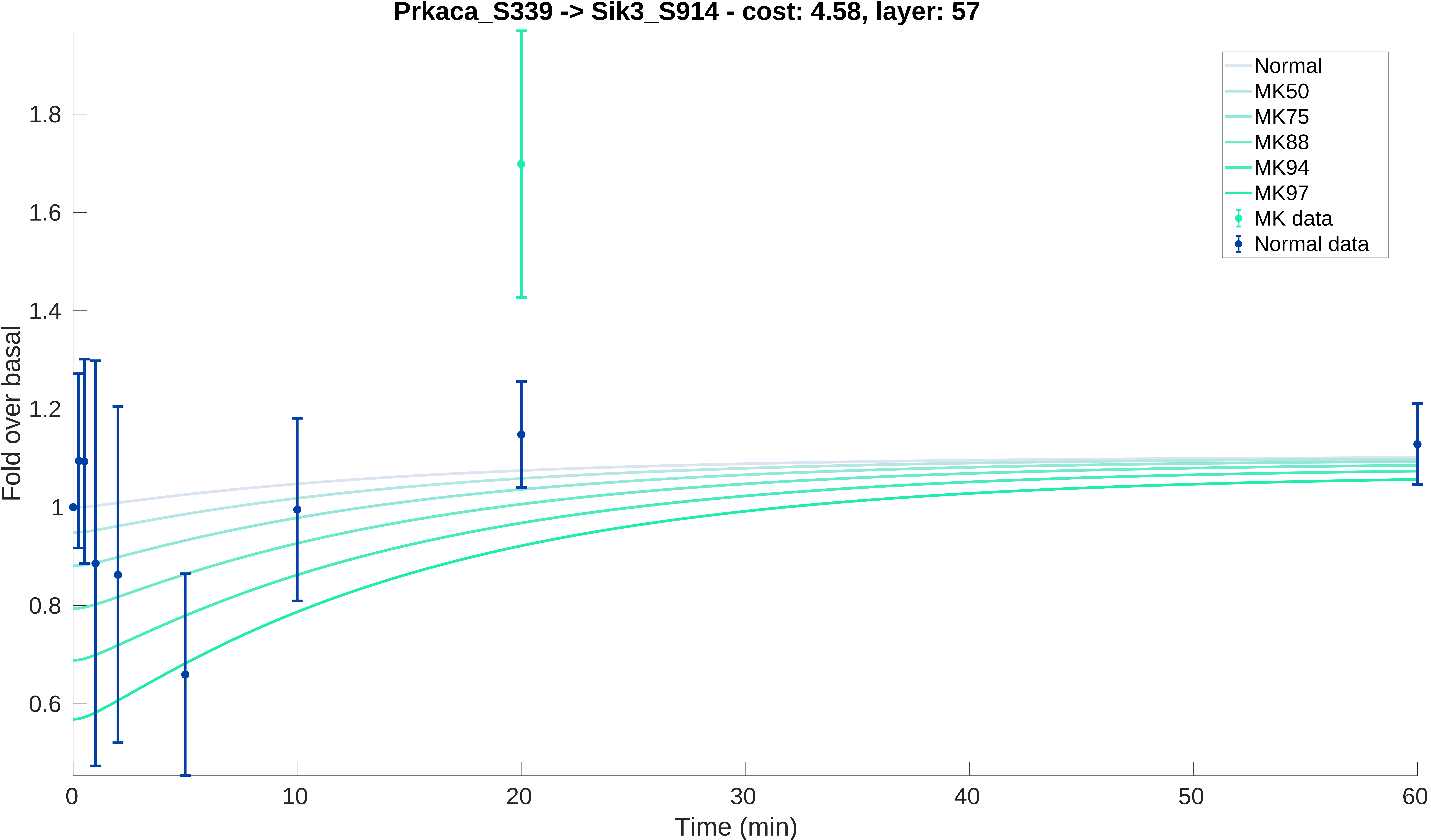

Prkaca_S339 -> Sik3_S914 - cost: 4.58, layer: 57
Fold over basal
Time (min)
0.6
0.8
1
1.2
1.4
1.6
1.8
0
10
20
30
40
50
60
Normal
MK50
MK75
MK88
MK94
MK97
MK data
Normal data

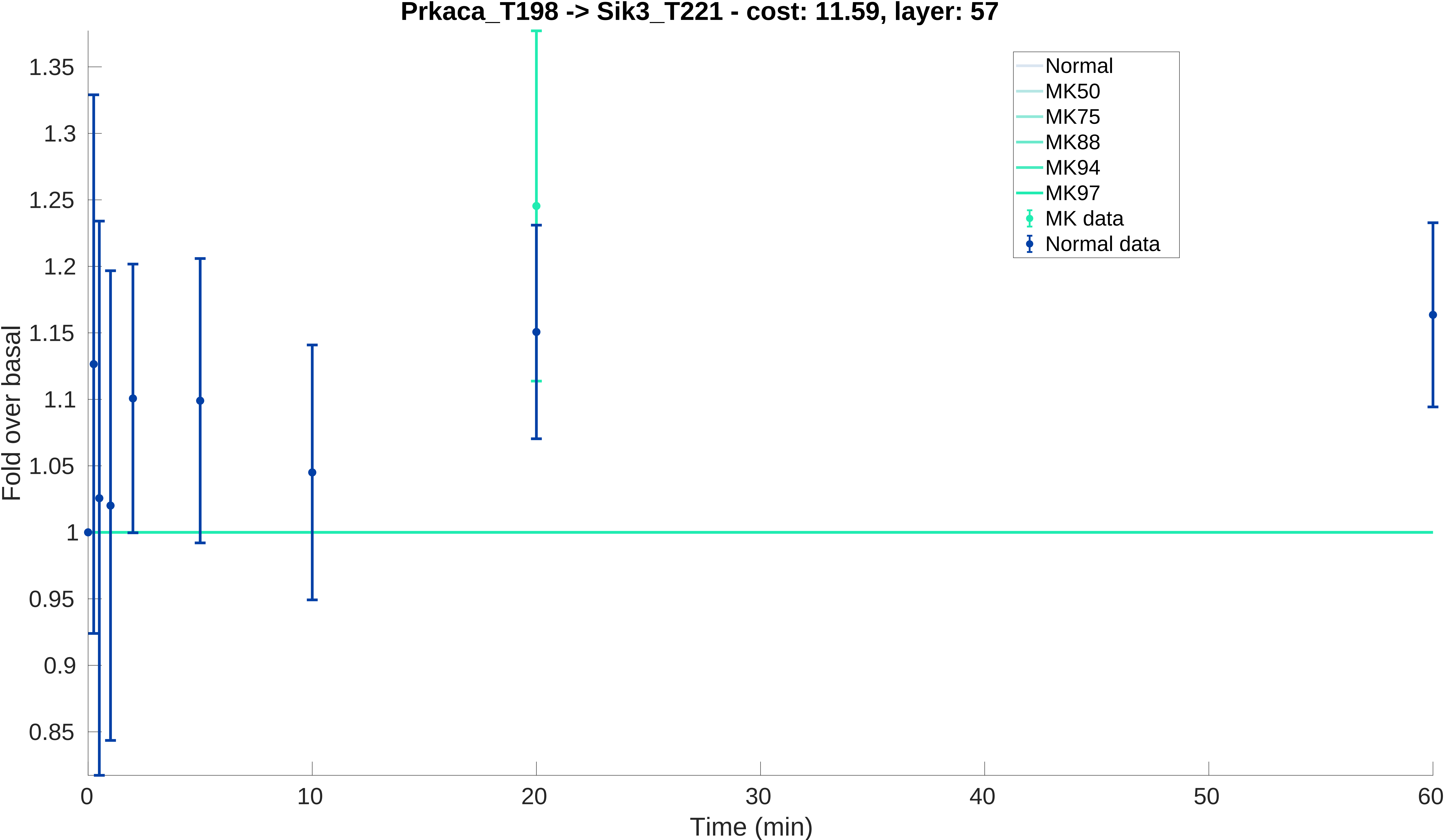

Prkaca_T198 -> Sik3_T221 - cost: 11.59, layer: 57
Fold over basal
Time (min)
0
10
20
30
40
50
60
1.35
1.3
1.25
1.2
1.15
1.1
1.05
1
0.95
0.9
0.85
Normal
MK50
MK75
MK88
MK94
MK97
MK data
Normal data

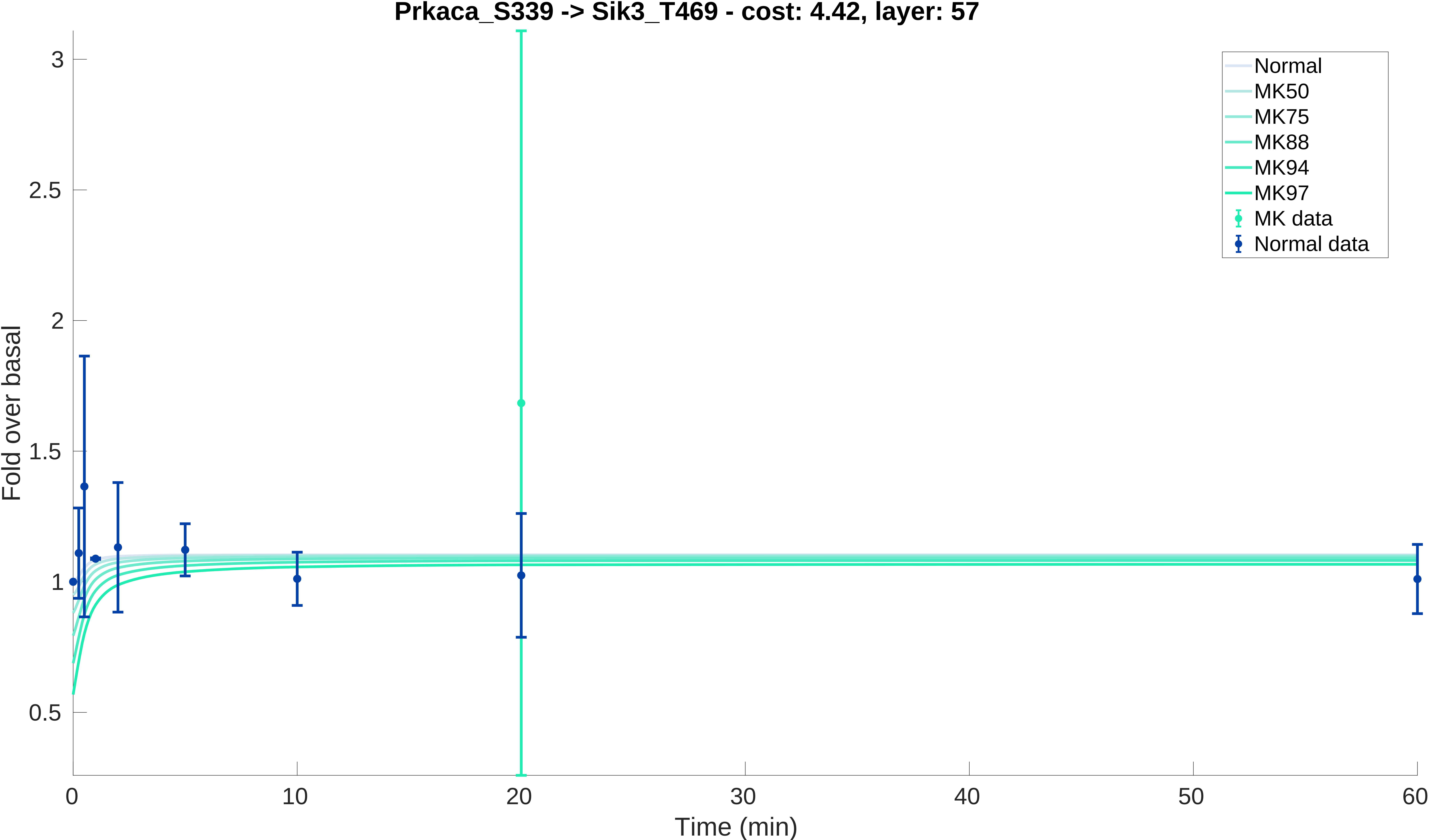

Prkaca_S339 -> Sik3_T469 - cost: 4.42, layer: 57
Fold over basal
Time (min)
0.5
1
1.5
2
2.5
3
0
10
20
30
40
50
60
Normal
MK50
MK75
MK88
MK94
MK97
MK data
Normal data

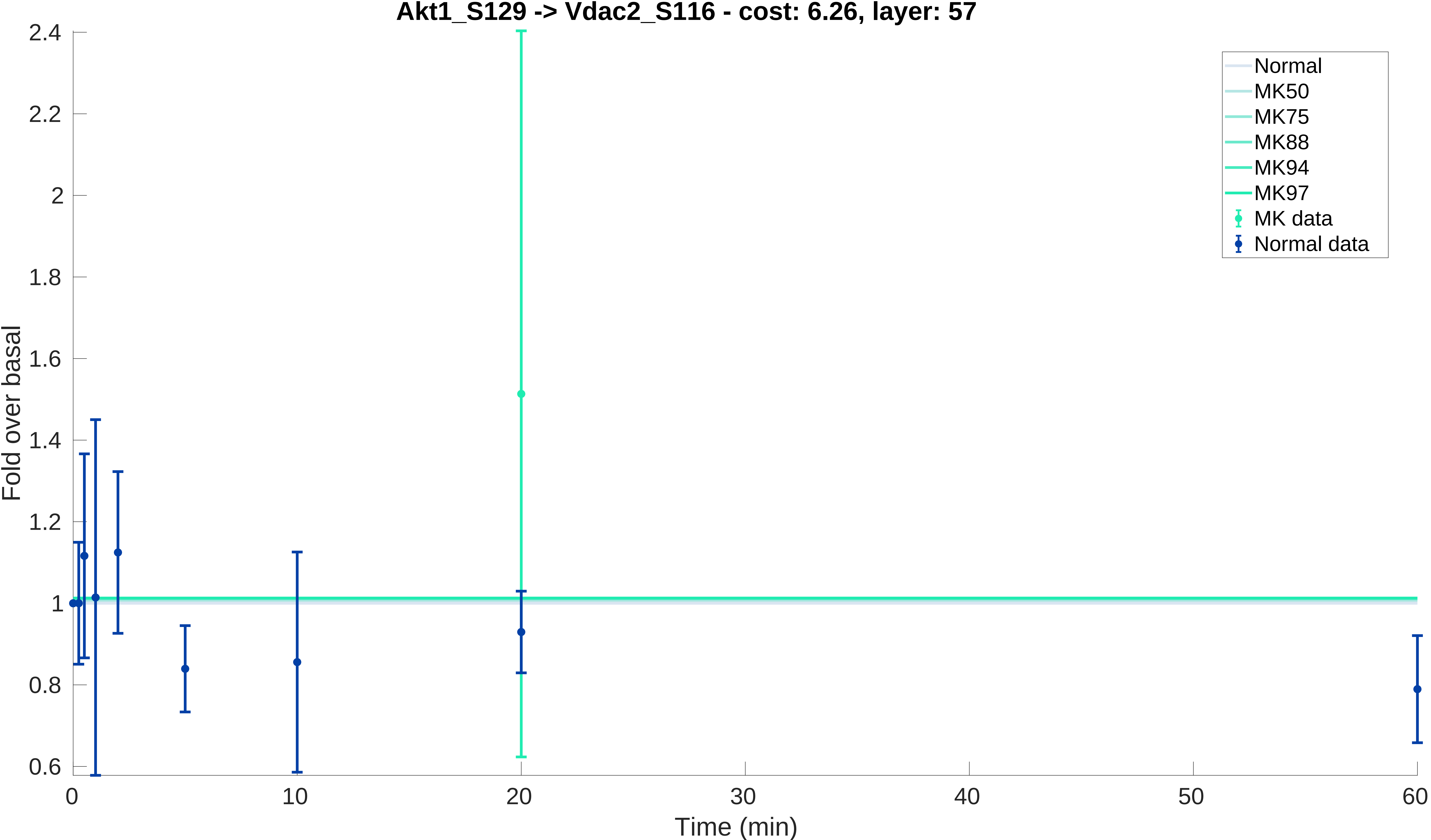

Akt1_S129 -> Vdac2_S116 - cost: 6.26, layer: 57
Fold over basal
Time (min)
0.6
0.8
1
1.2
1.4
1.6
1.8
2
2.2
2.4
0
10
20
30
40
50
60
Normal
MK50
MK75
MK88
MK94
MK97
MK data
Normal data

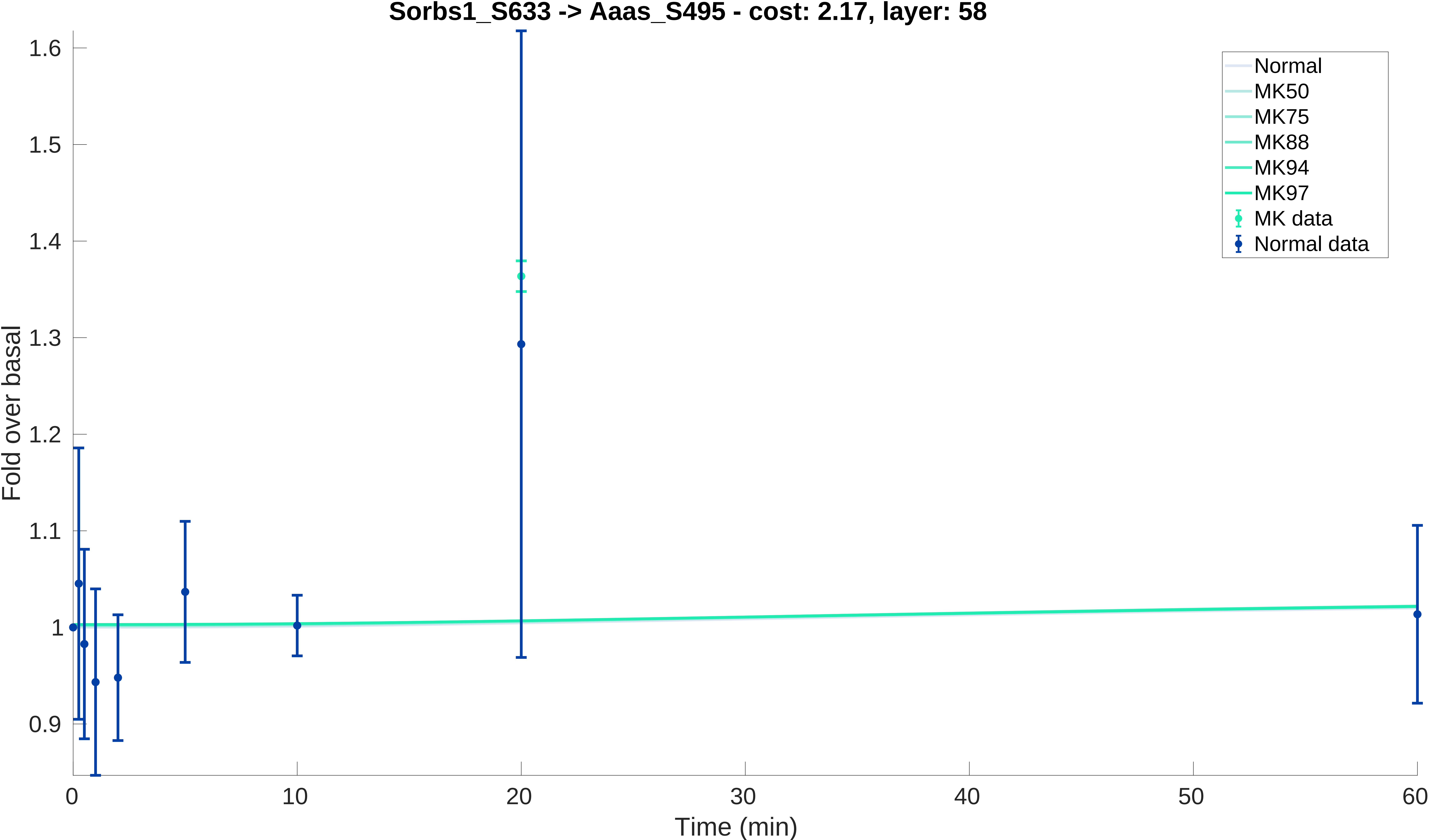

Sorbs1_S633 -> Aaas_S495 - cost: 2.17, layer: 58
Fold over basal
1.6
1.5
1.4
1.3
1.2
1.1
1
0.9
0
10
20
30
40
50
60
Time (min)
Normal
MK50
MK75
MK88
MK94
MK97
MK data
Normal data

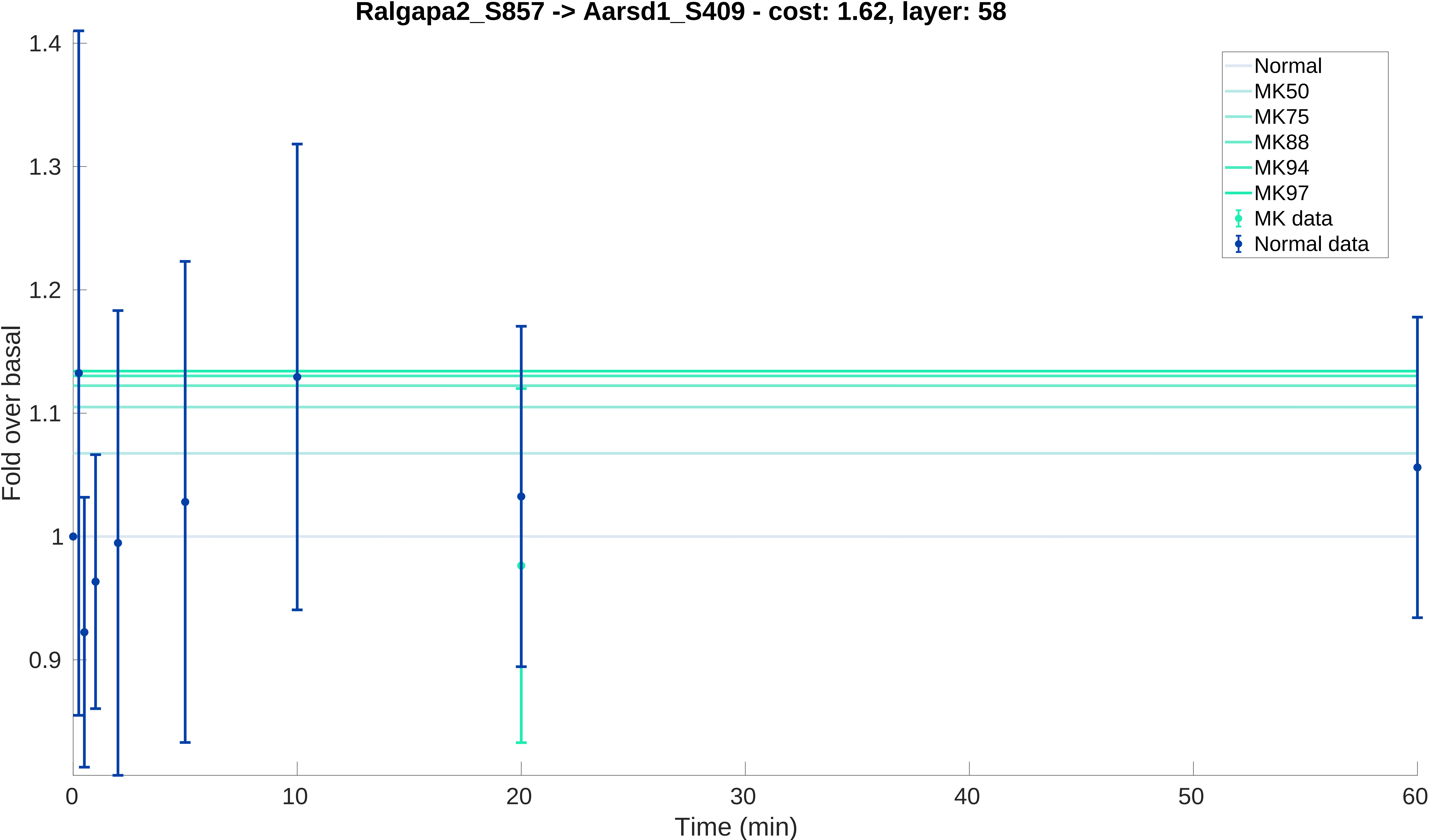

Ralgapa2_S857 -> Aarsd1_S409 - cost: 1.62, layer: 58
Fold over basal
Time (min)
Normal
MK50
MK75
MK88
MK94
MK97
MK data
Normal data
1.4
1.3
1.2
1.1
1
0.9
0
10
20
30
40
50
60

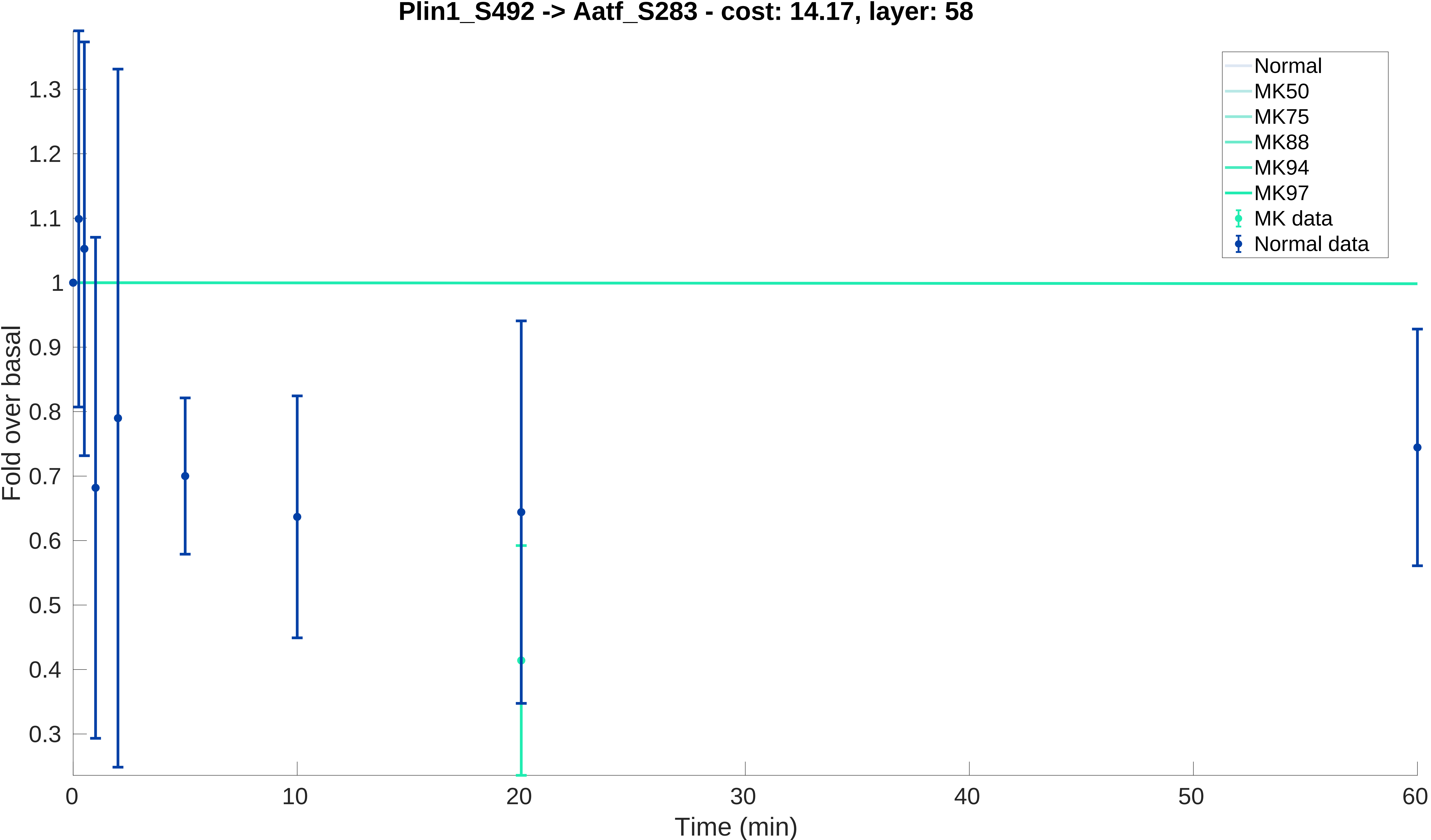
Plin1_S492 -> Aatf_S283 - cost: 14.17, layer: 58
Fold over basal
Time (min)
0
10
20
30
40
50
60
0.3
0.4
0.5
0.6
0.7
0.8
0.9
1
1.1
1.2
1.3
Normal
MK50
MK75
MK88
MK94
MK97
MK data
Normal data

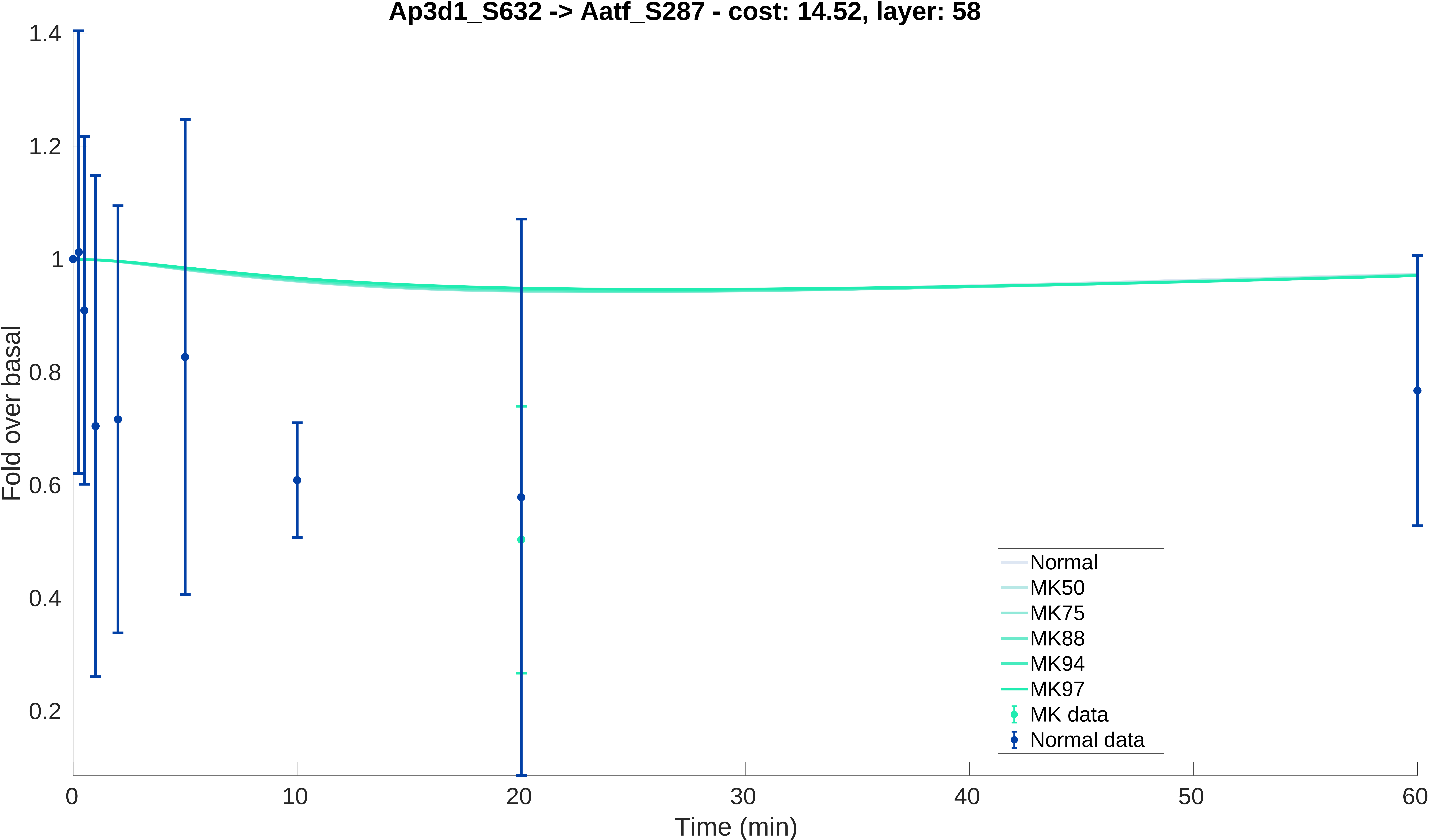

Ap3d1_S632 -> Aatf_S287 - cost: 14.52, layer: 58
Fold over basal
Time (min)
0.2
0.4
0.6
0.8
1
1.2
1.4
0
10
20
30
40
50
60
Normal
MK50
MK75
MK88
MK94
MK97
MK data
Normal data

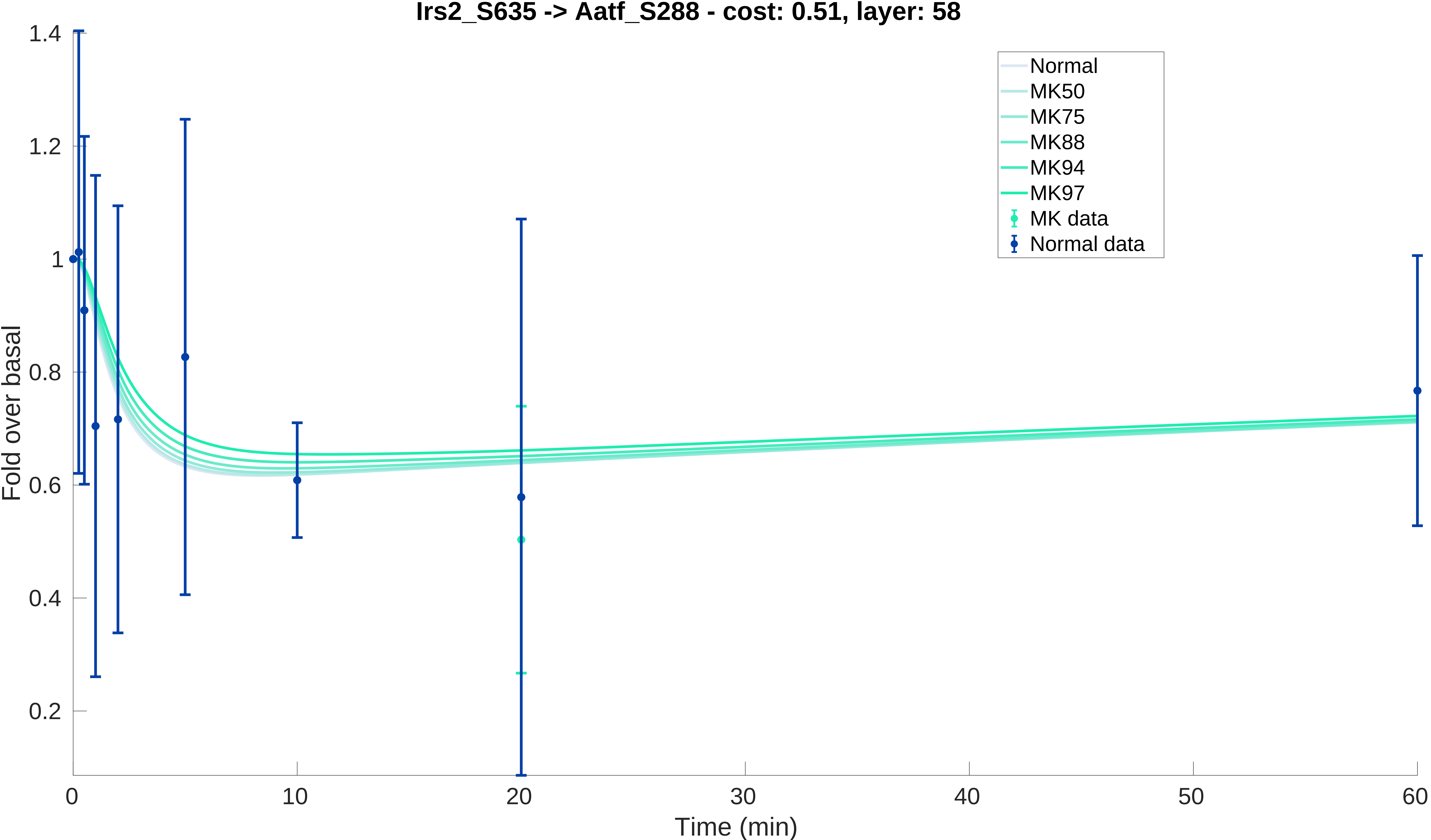

Irs2_S635 -> Aatf_S288 - cost: 0.51, layer: 58
Fold over basal
Time (min)
0.2
0.4
0.6
0.8
1
1.2
1.4
0
10
20
30
40
50
60
Normal
MK50
MK75
MK88
MK94
MK97
MK data
Normal data

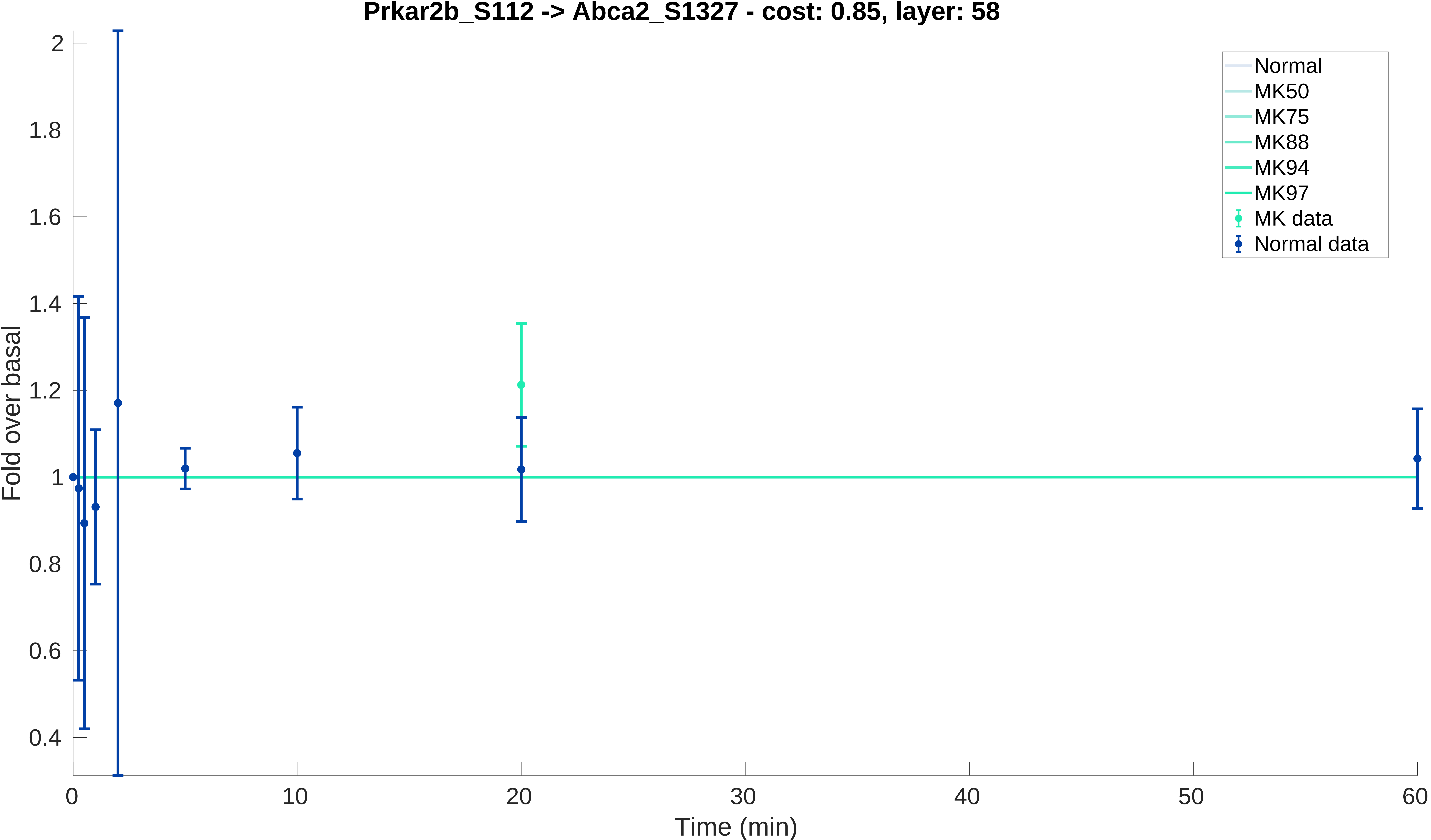

Prkar2b_S112 -> Abca2_S1327 - cost: 0.85, layer: 58
Normal
MK50
MK75
MK88
MK94
MK97
MK data
Normal data
Fold over basal
Time (min)
0
10
20
30
40
50
60
0.4
0.6
0.8
1
1.2
1.4
1.6
1.8
2

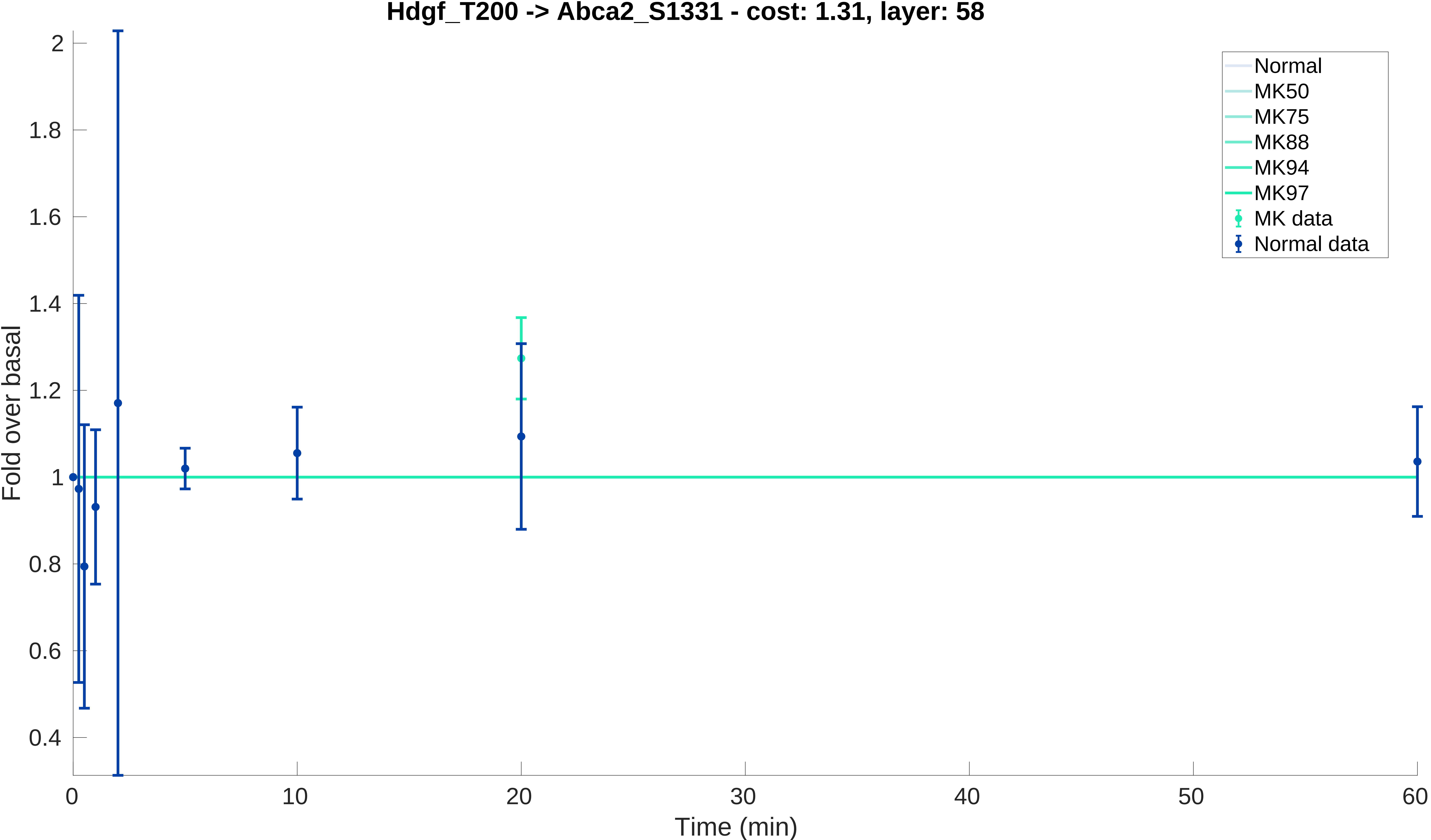

Hdgf_T200 -> Abca2_S1331 - cost: 1.31, layer: 58
Normal
MK50
MK75
MK88
MK94
MK97
MK data
Normal data
Fold over basal
Time (min)
0
10
20
30
40
50
60
0.4
0.6
0.8
1
1.2
1.4
1.6
1.8
2

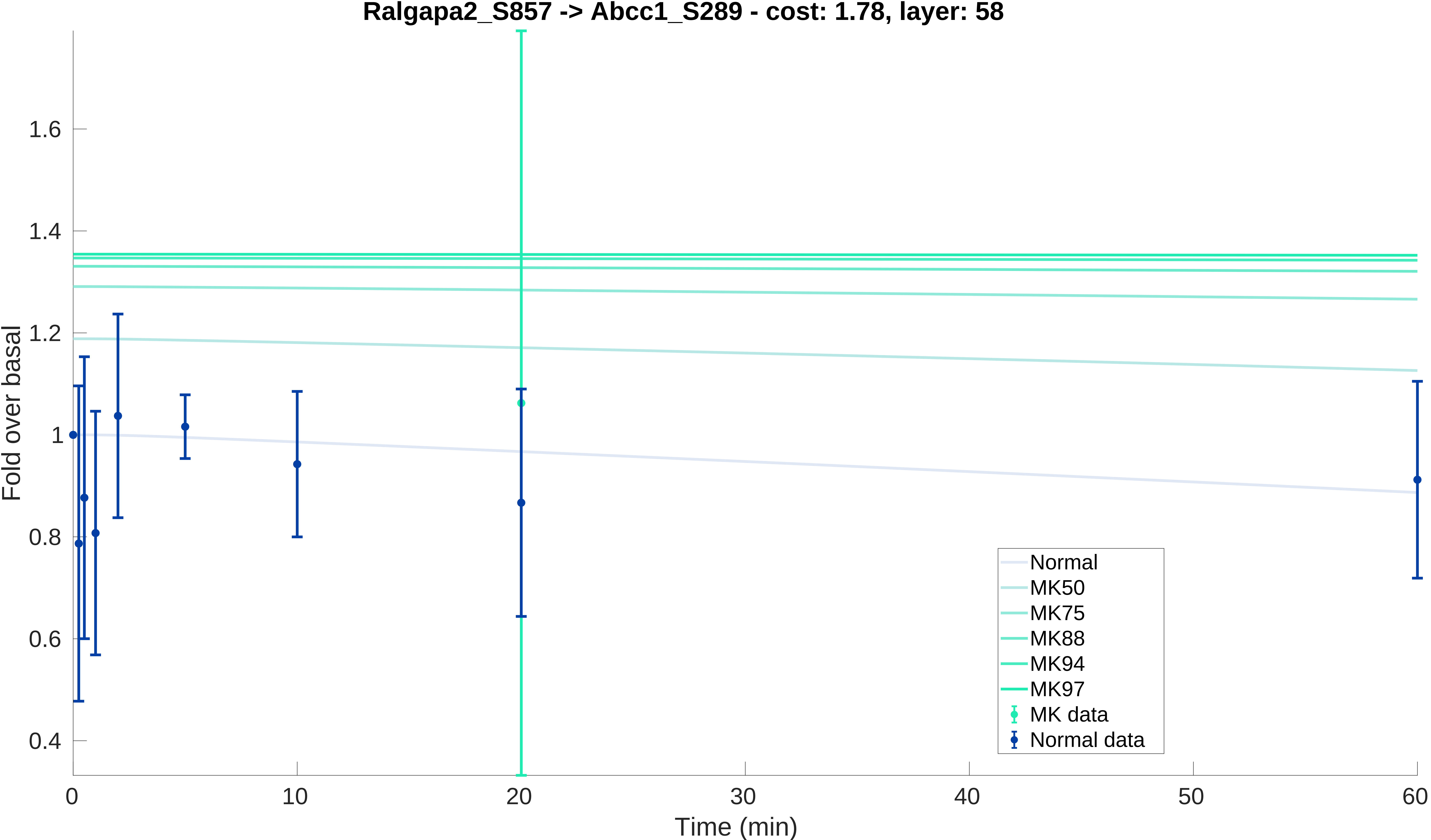

Ralgapa2_S857 -> Abcc1_S289 - cost: 1.78, layer: 58
Fold over basal
1.6
1.4
1.2
1
0.8
0.6
0.4
0
10
20
30
40
50
60
Time (min)
Normal
MK50
MK75
MK88
MK94
MK97
MK data
Normal data

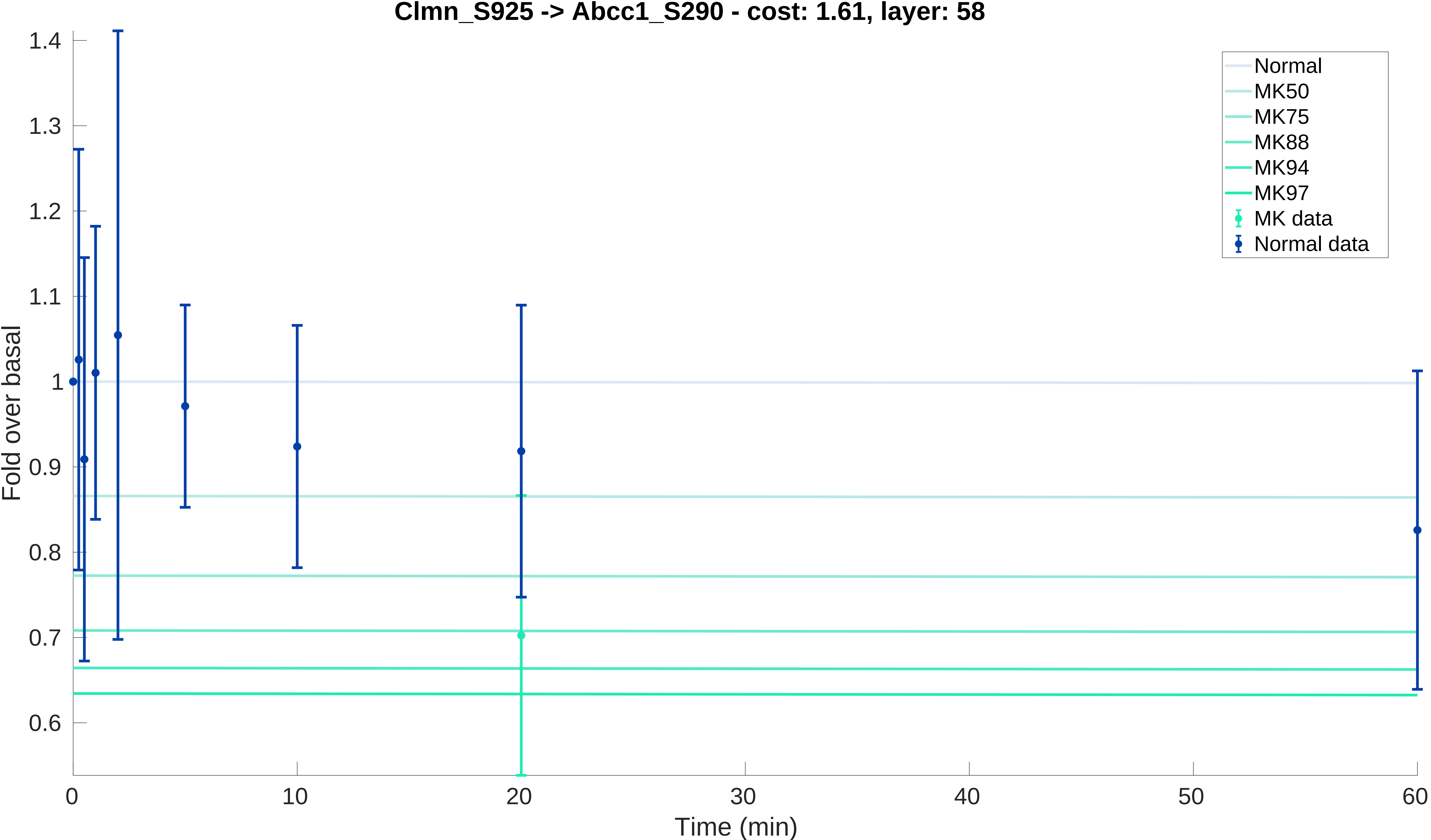
Clmn_S925 -> Abcc1_S290 - cost: 1.61, layer: 58
Fold over basal
Time (min)
1.4
1.3
1.2
1.1
1
0.9
0.8
0.7
0.6
0
10
20
30
40
50
60
Normal
MK50
MK75
MK88
MK94
MK97
MK data
Normal data

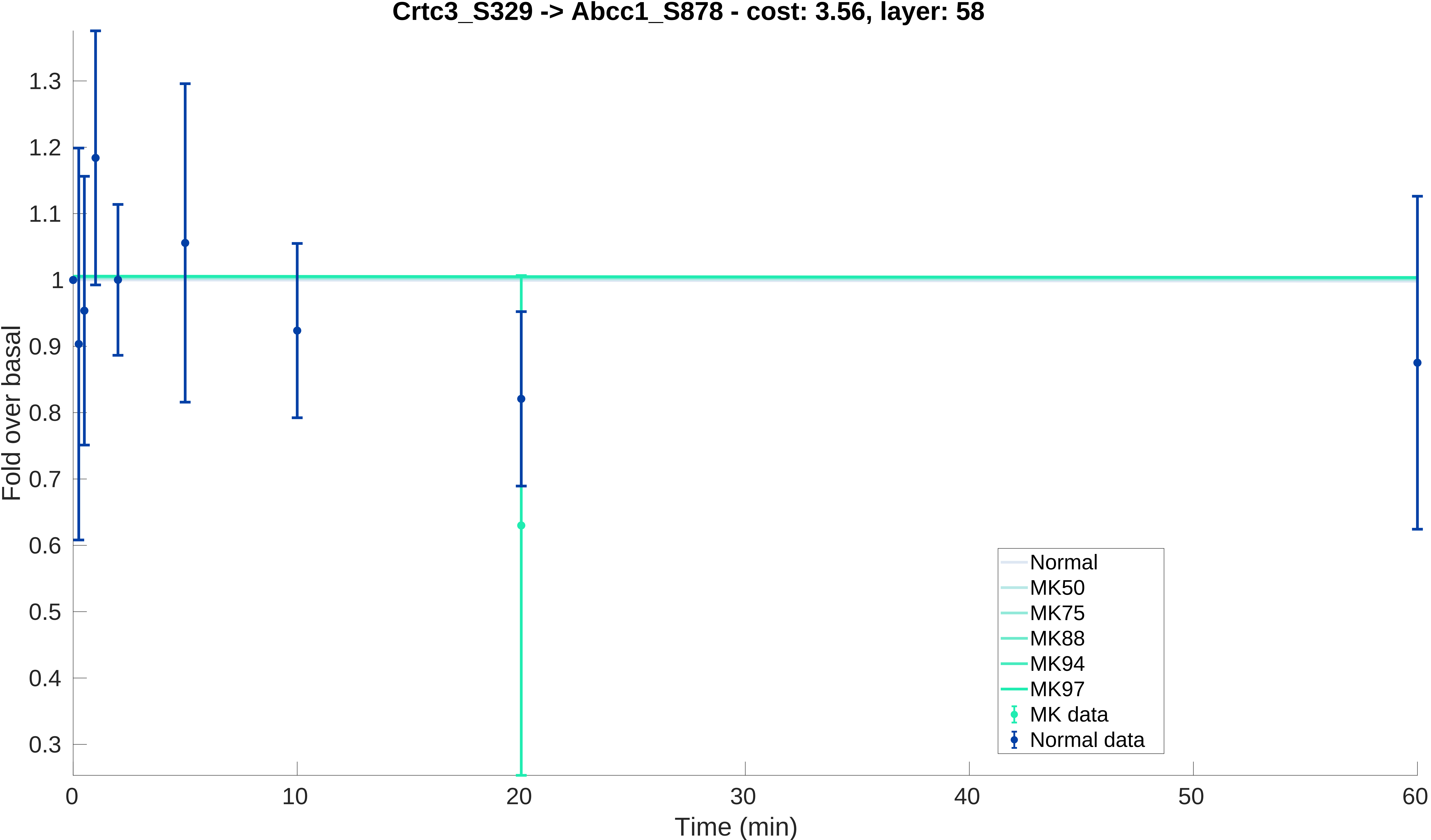

Crtc3_S329 -> Abcc1_S878 - cost: 3.56, layer: 58
Fold over basal
1.3
1.2
1.1
1
0.9
0.8
0.7
0.6
0.5
0.4
0.3
0
10
20
30
40
50
60
Time (min)
Normal
MK50
MK75
MK88
MK94
MK97
MK data
Normal data

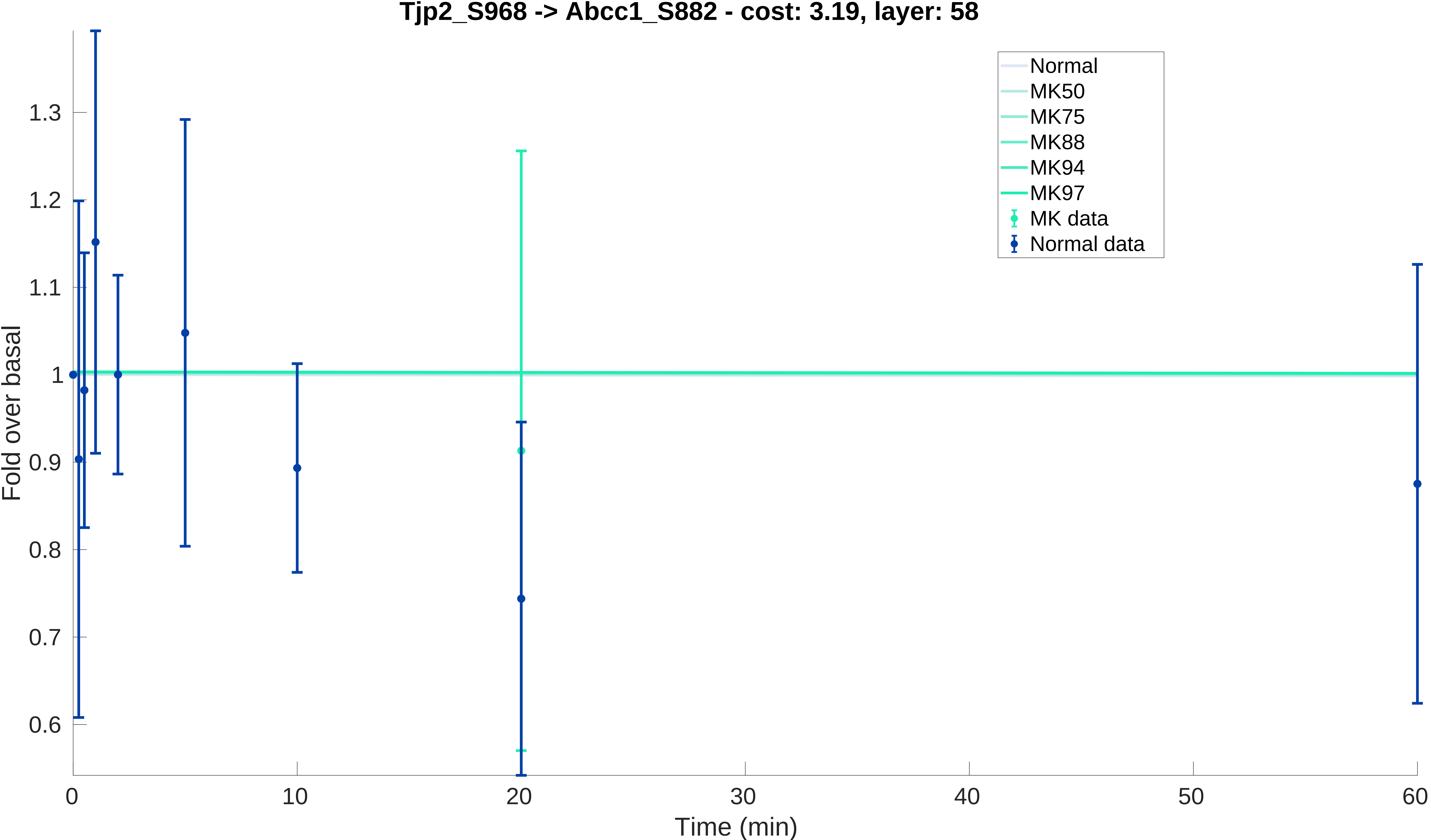
Tjp2_S968 -> Abcc1_S882 - cost: 3.19, layer: 58
Normal
MK50
MK75
MK88
MK94
MK97
MK data
Normal data
Fold over basal
Time (min)
0.6
0.7
0.8
0.9
1
1.1
1.2
1.3
0
10
20
30
40
50
60

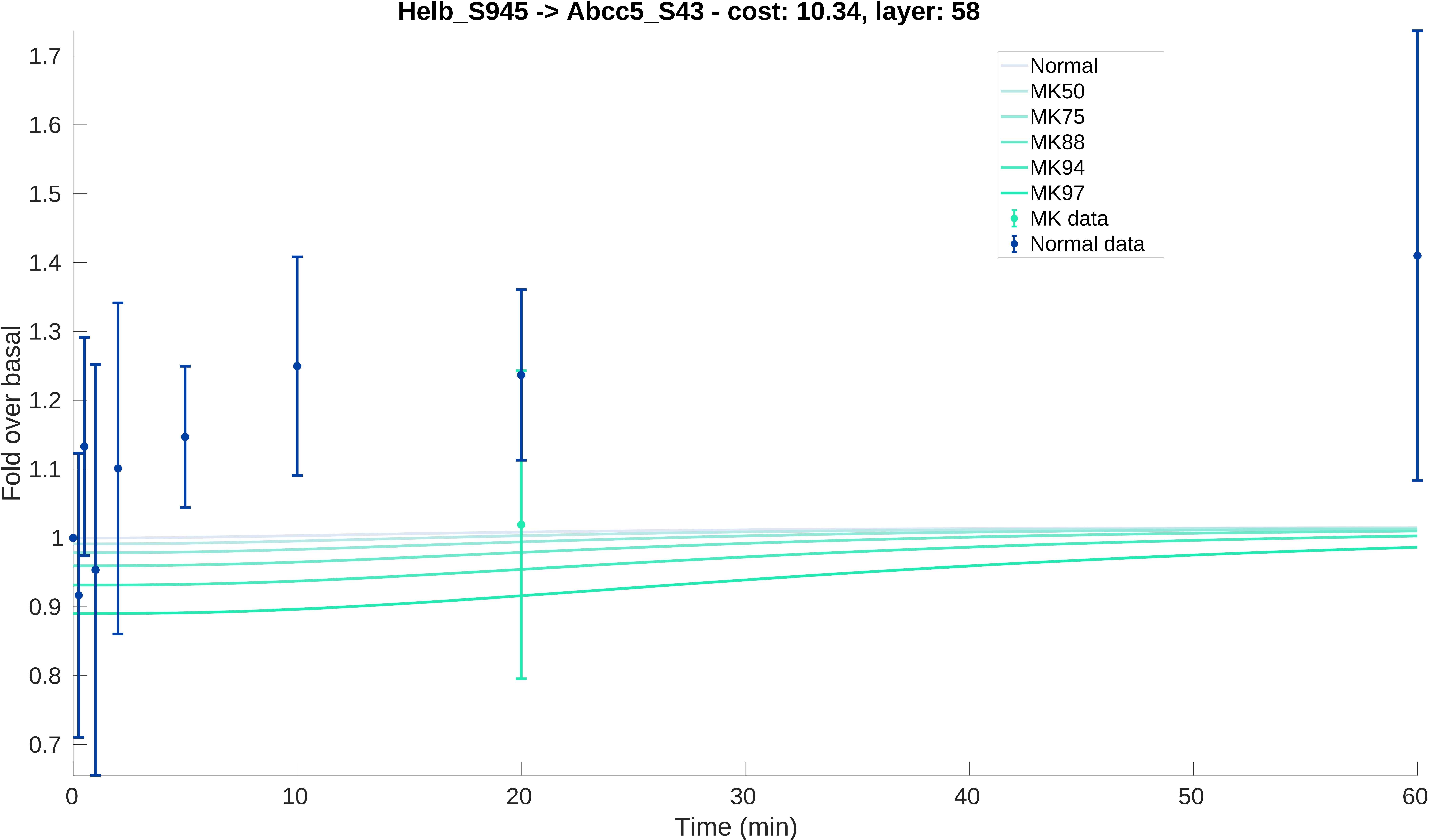

Helb_S945 -> Abcc5_S43 - cost: 10.34, layer: 58
Normal
MK50
MK75
MK88
MK94
MK97
MK data
Normal data
Fold over basal
Time (min)
1.7
1.6
1.5
1.4
1.3
1.2
1.1
1
0.9
0.8
0.7
0
10
20
30
40
50
60

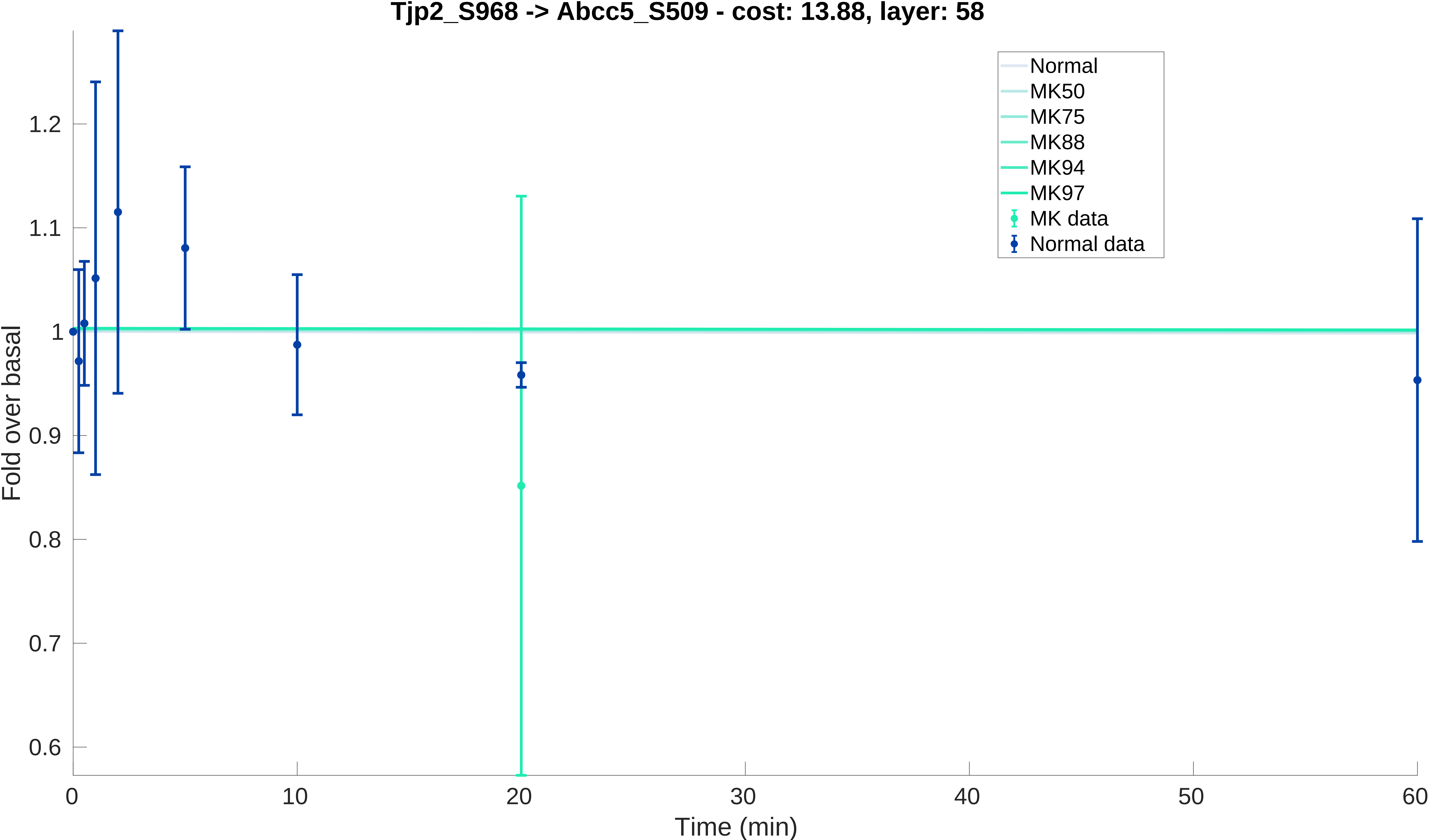

Tjp2_S968 -> Abcc5_S509 - cost: 13.88, layer: 58
Normal
MK50
MK75
MK88
MK94
MK97
MK data
Normal data
Fold over basal
1.2
1.1
1
0.9
0.8
0.7
0.6
0
10
20
30
40
50
60
Time (min)

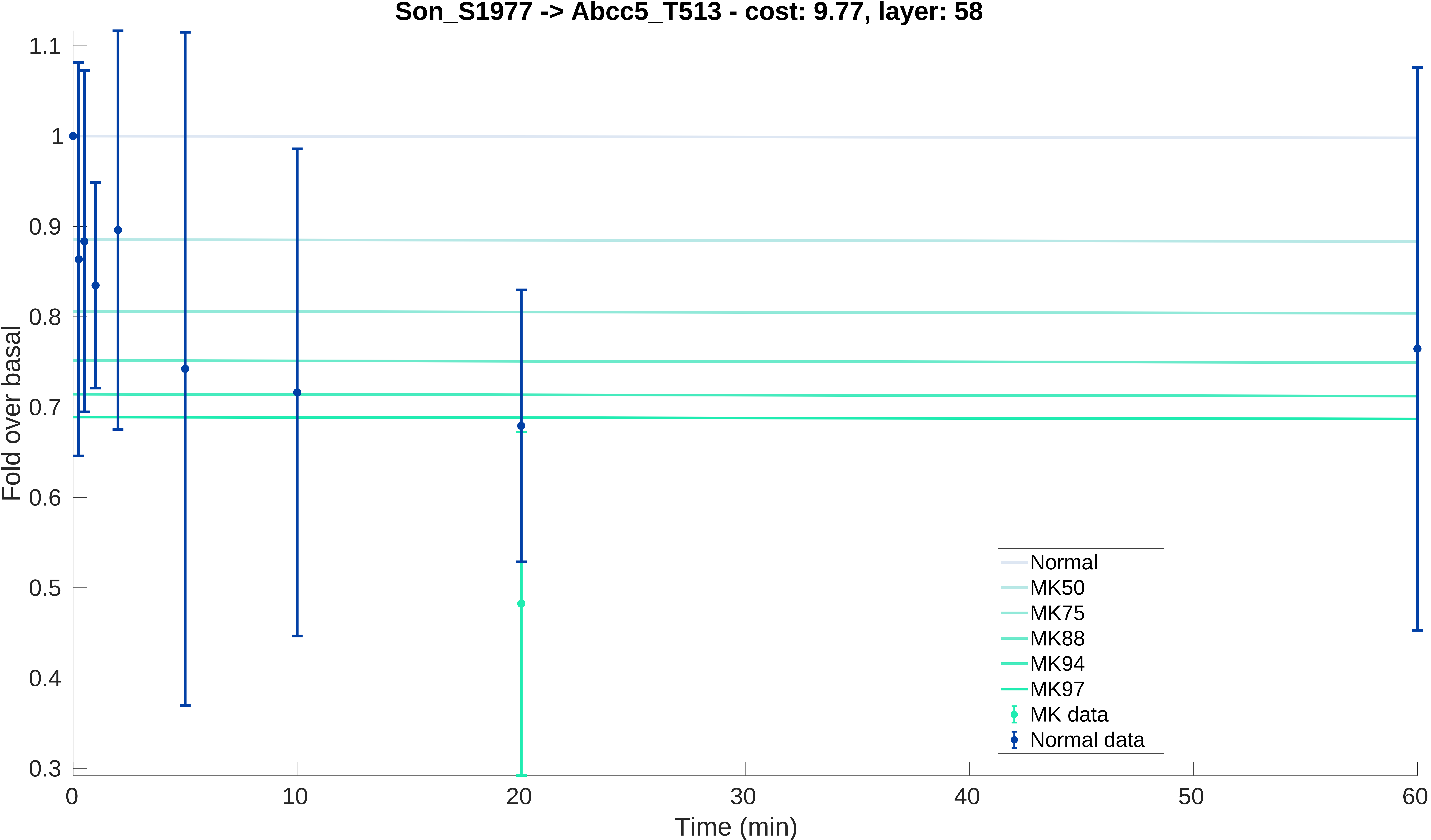

Son_S1977 -> Abcc5_T513 - cost: 9.77, layer: 58
Fold over basal
Time (min)
1.1
1
0.9
0.8
0.7
0.6
0.5
0.4
0.3
0
10
20
30
40
50
60
Normal
MK50
MK75
MK88
MK94
MK97
MK data
Normal data

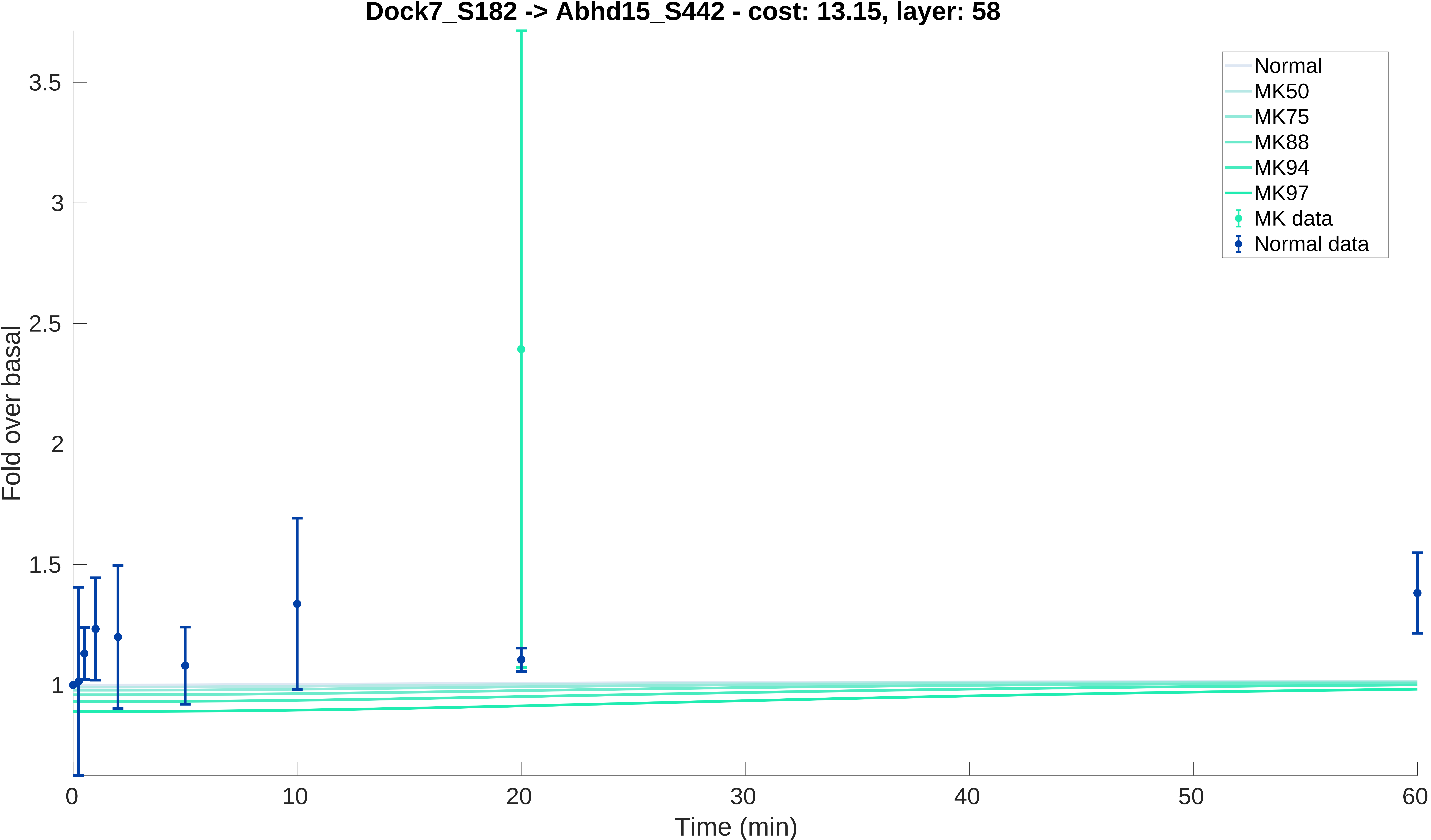

Dock7_S182 -> Abhd15_S442 - cost: 13.15, layer: 58
Fold over basal
Time (min)
3.5
3
2.5
2
1.5
1
0
10
20
30
40
50
60
Normal
MK50
MK75
MK88
MK94
MK97
MK data
Normal data

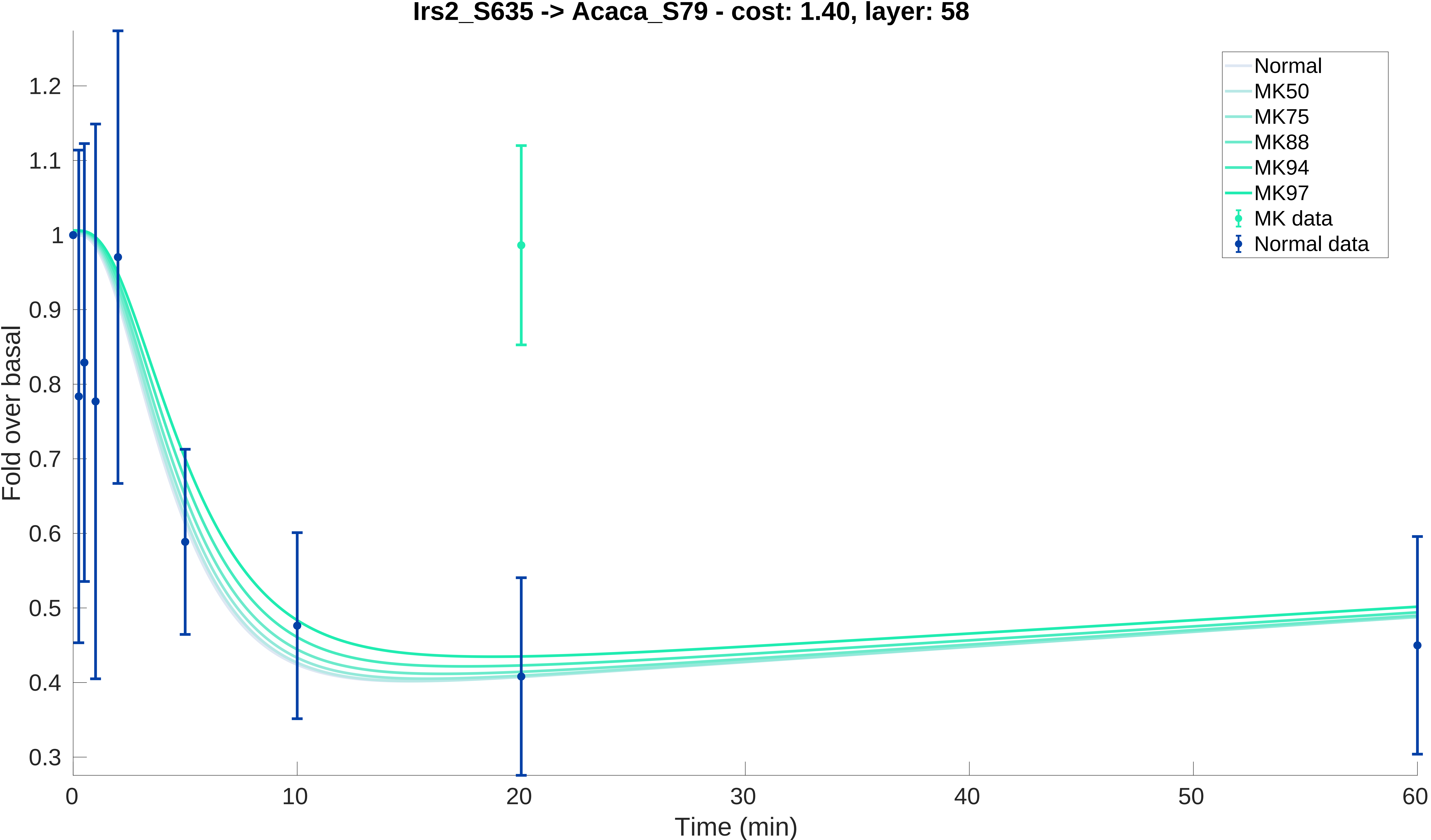
Irs2_S635 -> Acaca_S79 - cost: 1.40, layer: 58
Fold over basal
1.2
1.1
1
0.9
0.8
0.7
0.6
0.5
0.4
0.3
0
10
20
30
40
50
60
Time (min)
Normal
MK50
MK75
MK88
MK94
MK97
MK data
Normal data

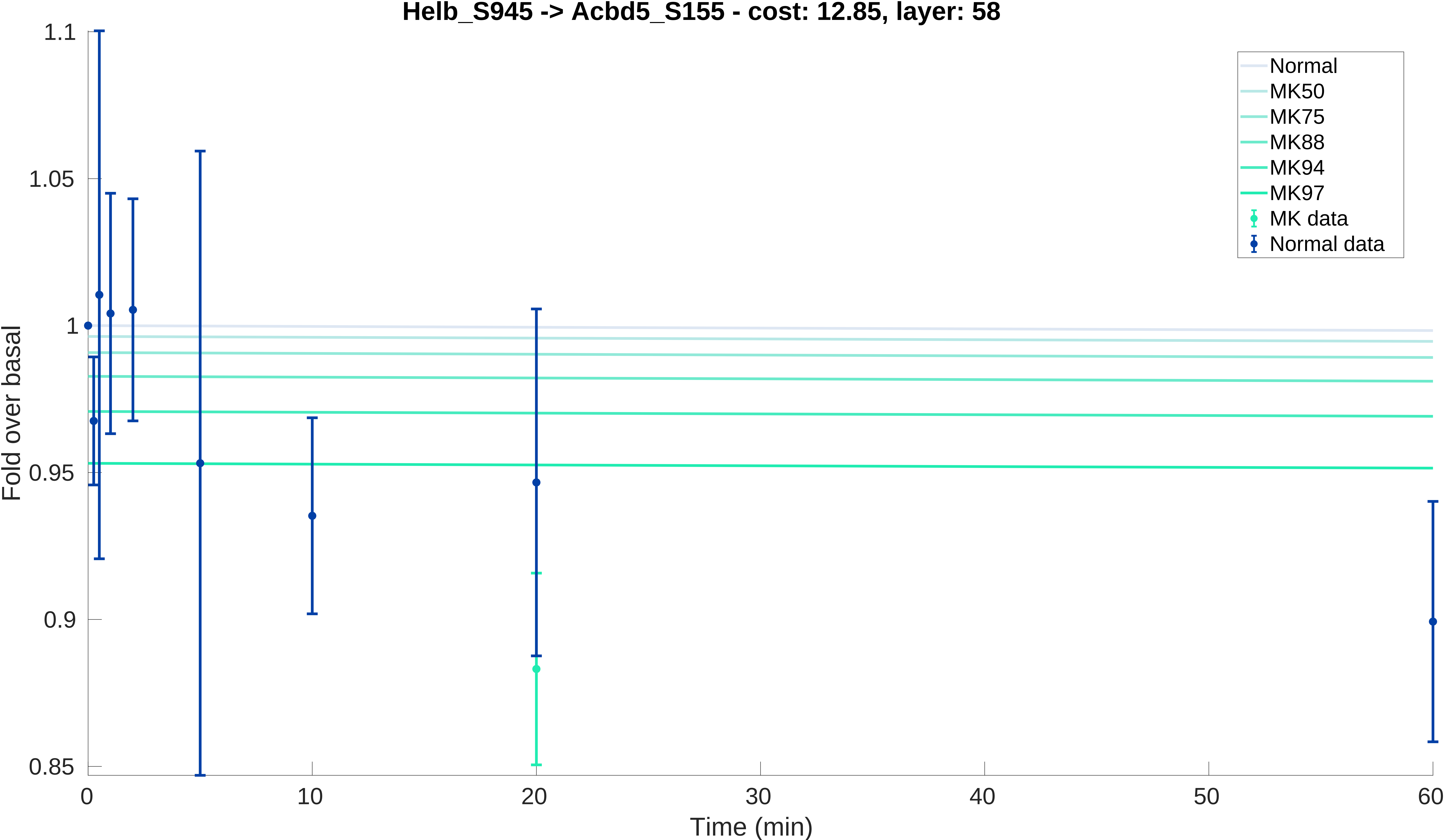
Helb_S945 -> Acbd5_S155 - cost: 12.85, layer: 58
Fold over basal
Time (min)
1.1
1.05
1
0.95
0.9
0.85
0
10
20
30
40
50
60
Normal
MK50
MK75
MK88
MK94
MK97
MK data
Normal data

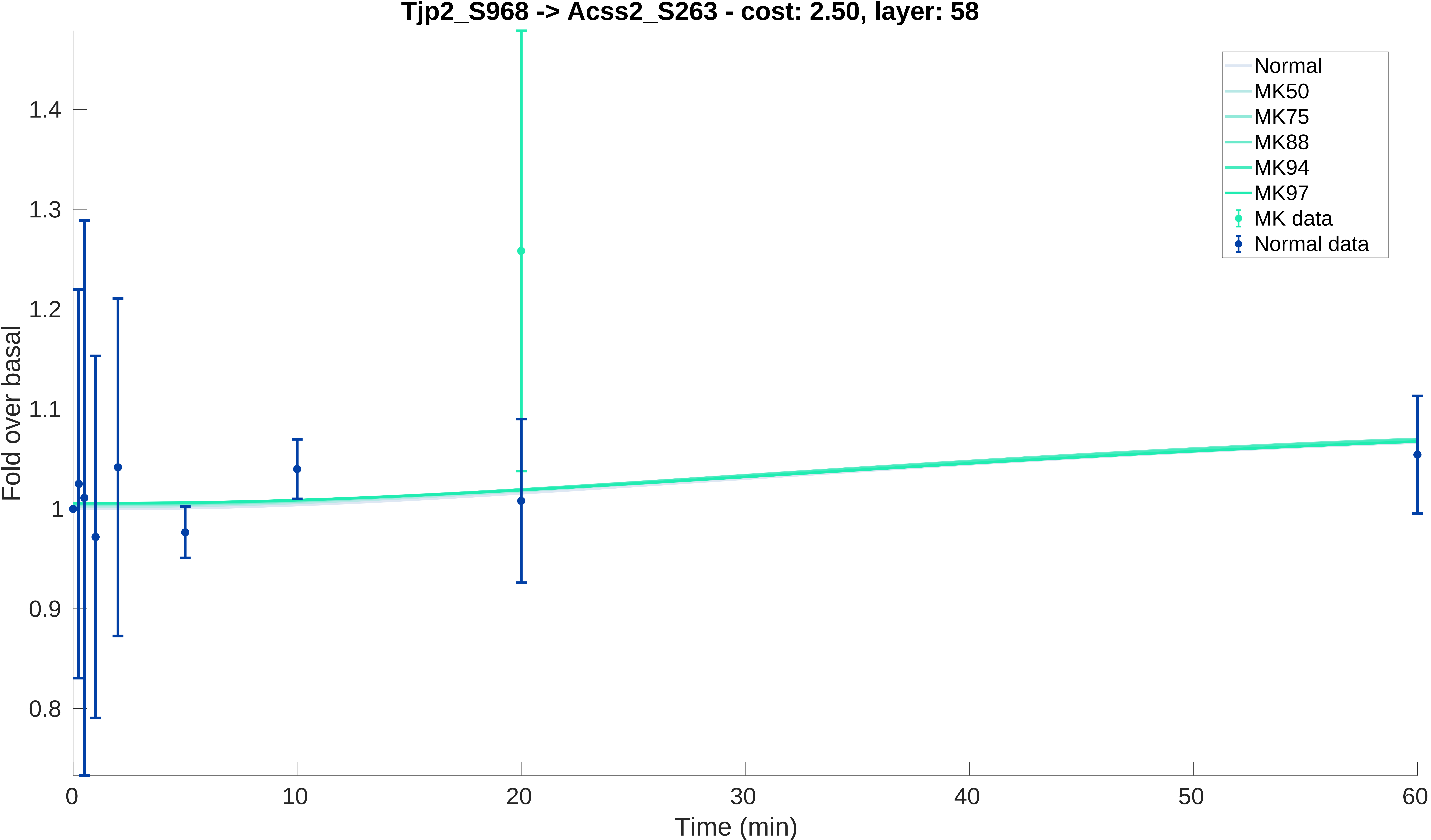

Tjp2_S968 -> Acss2_S263 - cost: 2.50, layer: 58
Fold over basal
Time (min)
Normal
MK50
MK75
MK88
MK94
MK97
MK data
Normal data
1.4
1.3
1.2
1.1
1
0.9
0.8
0
10
20
30
40
50
60

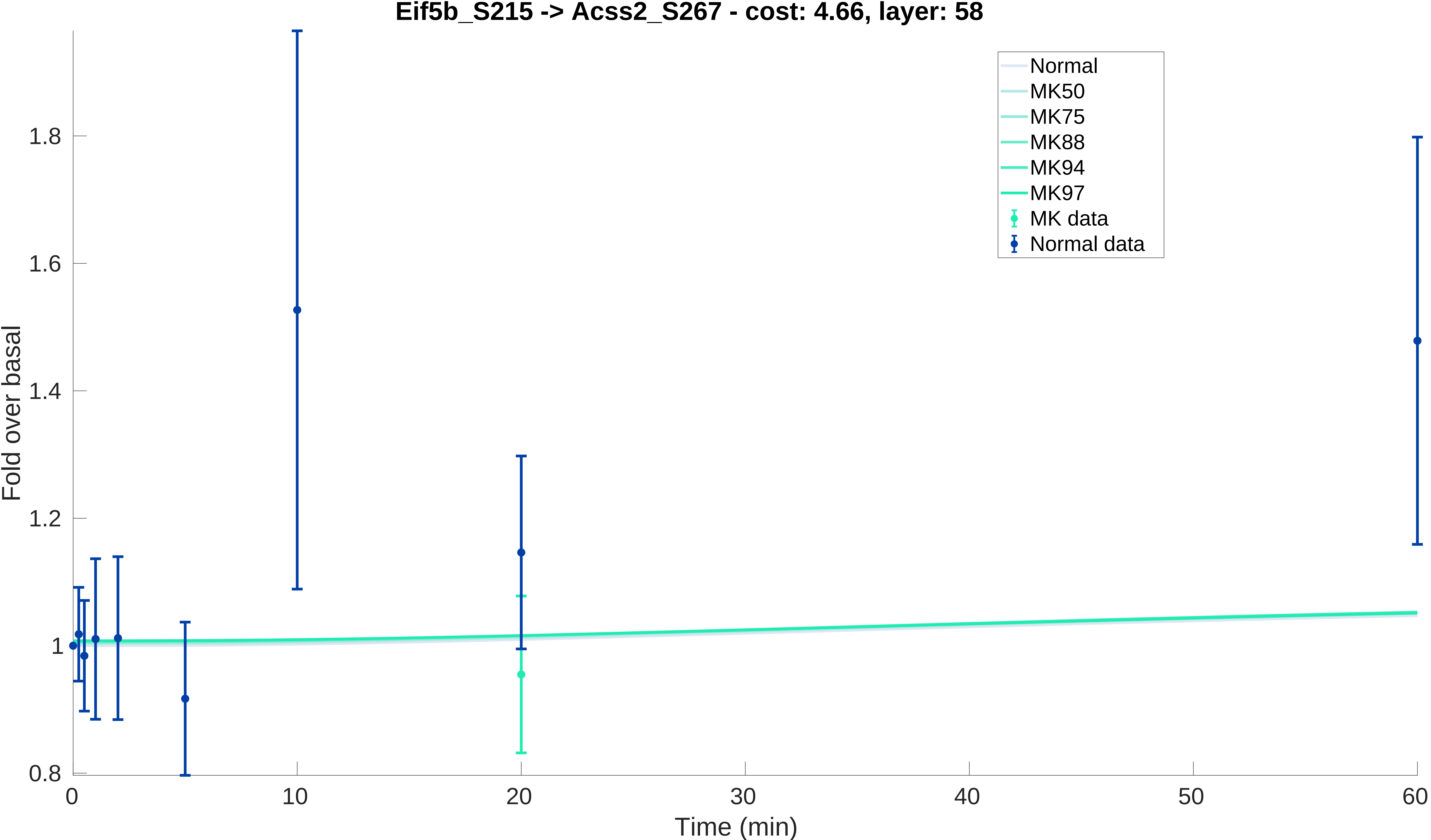

Eif5b_S215 -> Acss2_S267 - cost: 4.66, layer: 58
Normal
MK50
MK75
MK88
MK94
MK97
MK data
Normal data
Fold over basal
0.8
1
1.2
1.4
1.6
1.8
0
10
20
30
40
50
60
Time (min)

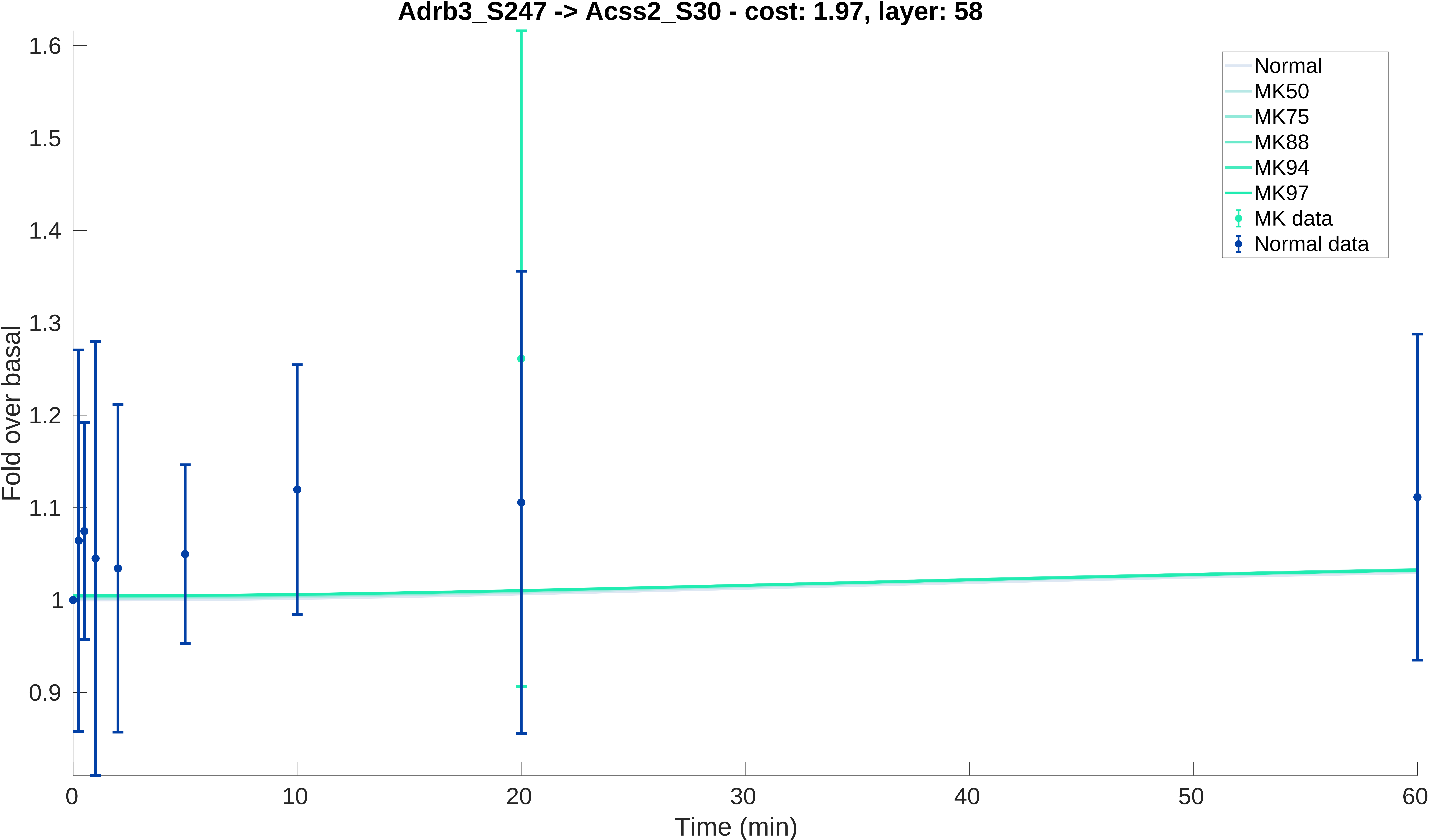
Adrb3_S247 -> Acss2_S30 - cost: 1.97, layer: 58
Fold over basal
1.6
1.5
1.4
1.3
1.2
1.1
1
0.9
0
10
20
30
40
50
60
Time (min)
Normal
MK50
MK75
MK88
MK94
MK97
MK data
Normal data

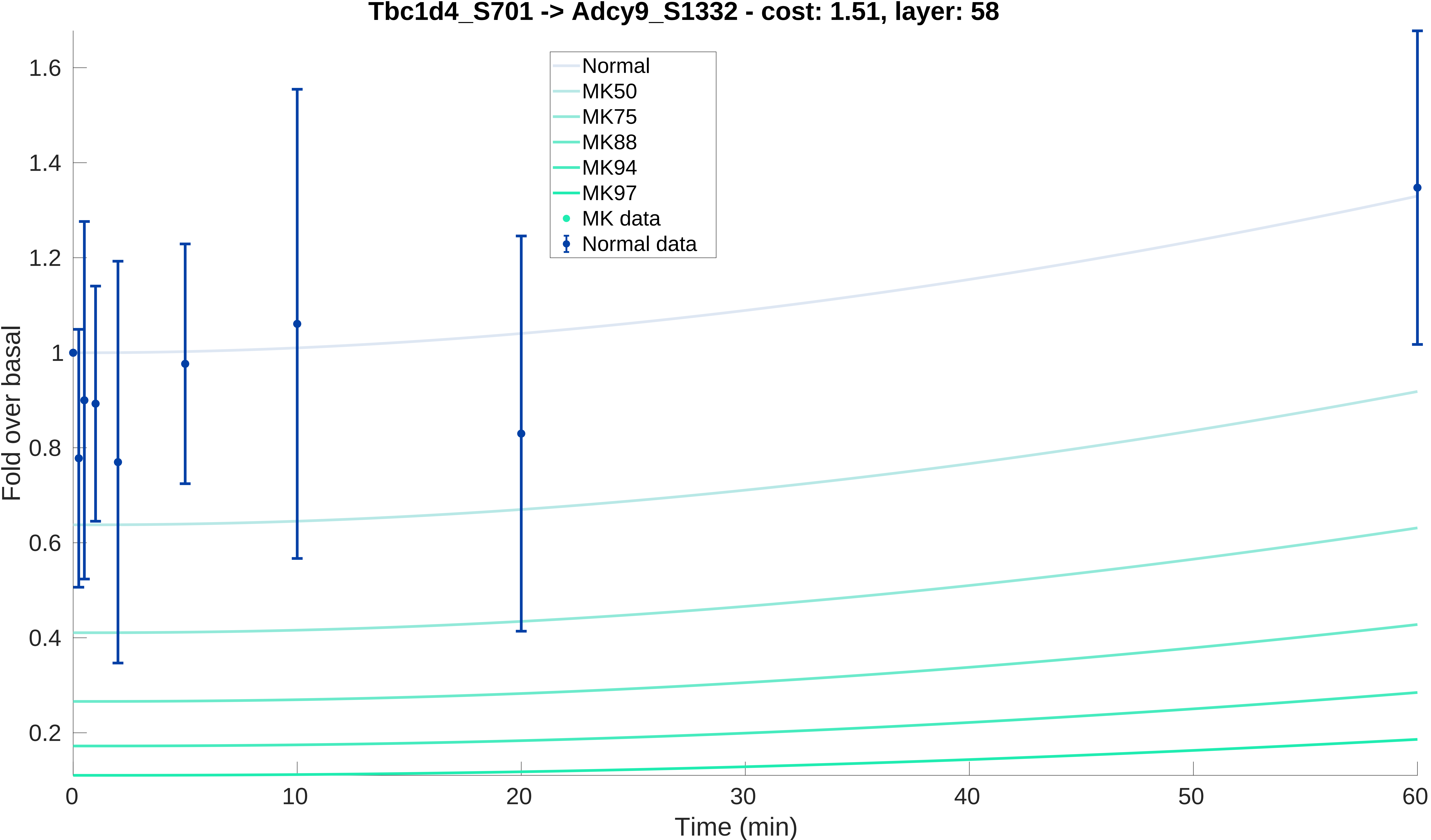

Tbc1d4_S701 -> Adcy9_S1332 - cost: 1.51, layer: 58
Normal
MK50
MK75
MK88
MK94
MK97
MK data
Normal data
Fold over basal
1.6
1.4
1.2
1
0.8
0.6
0.4
0.2
0
10
20
30
40
50
60
Time (min)

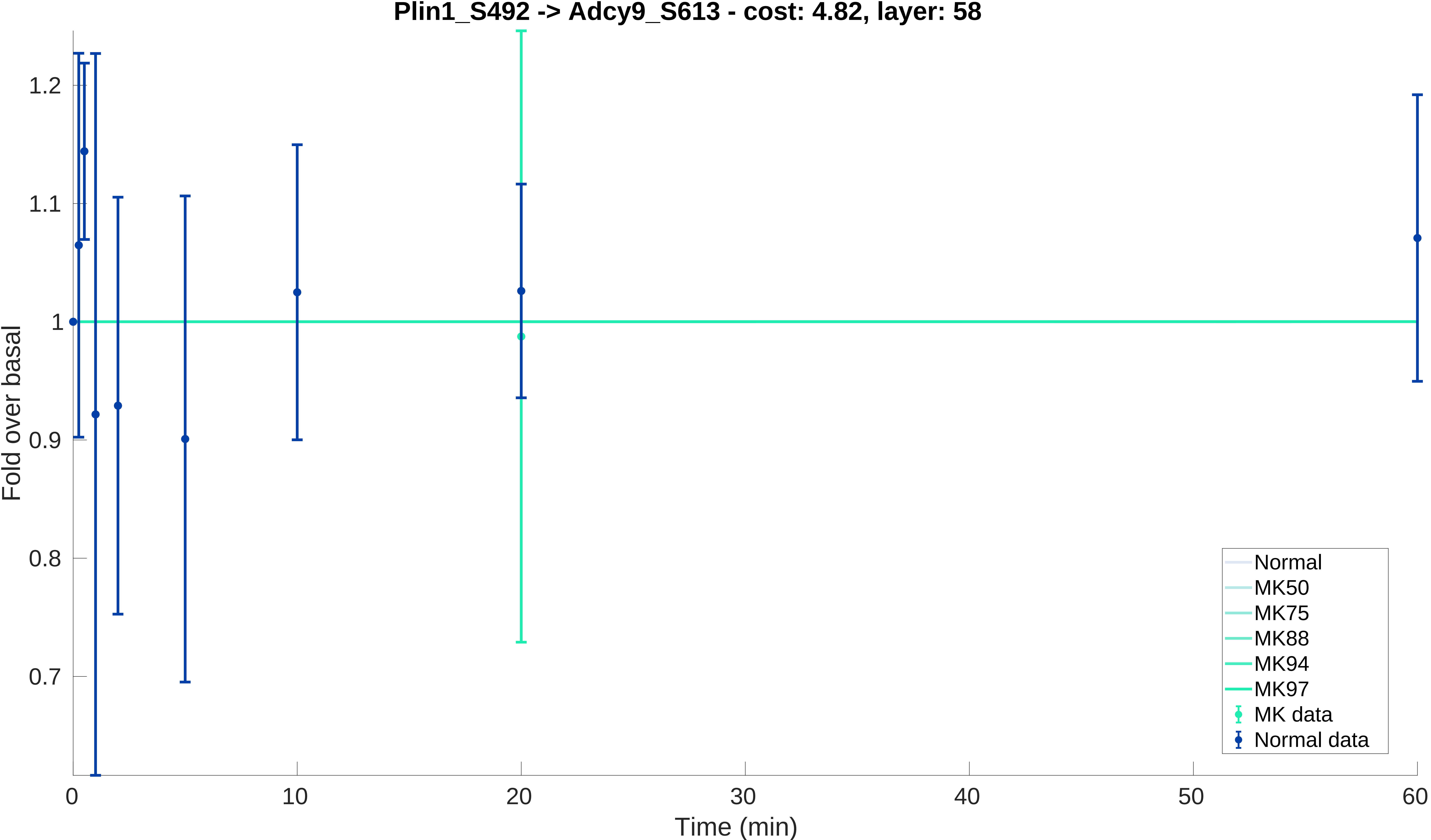

Plin1_S492 -> Adcy9_S613 - cost: 4.82, layer: 58
Fold over basal
1.2
1.1
1
0.9
0.8
0.7
0
10
20
30
40
50
60
Time (min)
Normal
MK50
MK75
MK88
MK94
MK97
MK data
Normal data

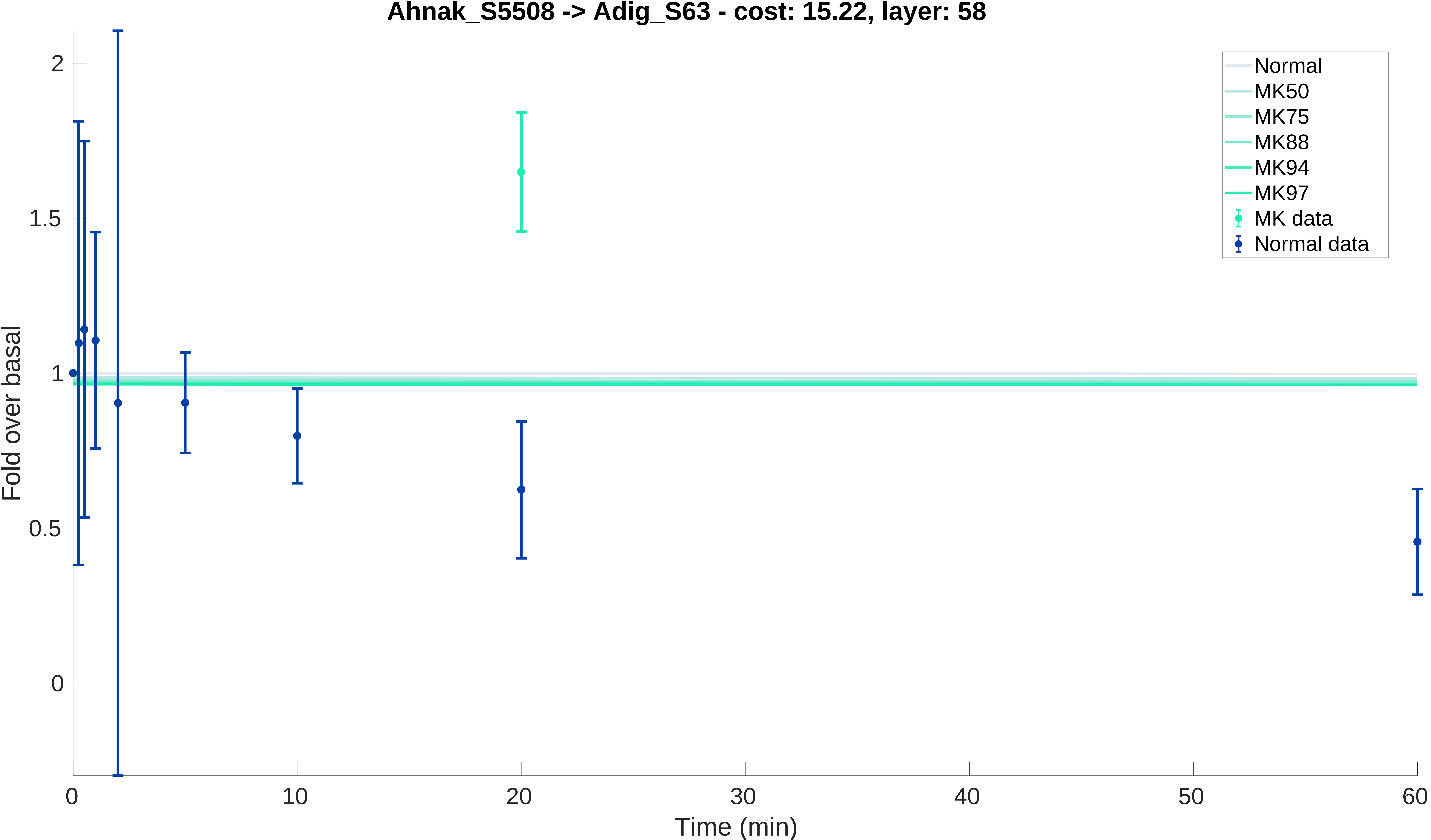

Ahnak_S5508 -> Adig_S63 - cost: 15.22, layer: 58
Normal
MK50
MK75
MK88
MK94
MK97
MK data
Normal data
Fold over basal
Time (min)
2
1.5
1
0.5
0
0
10
20
30
40
50
60

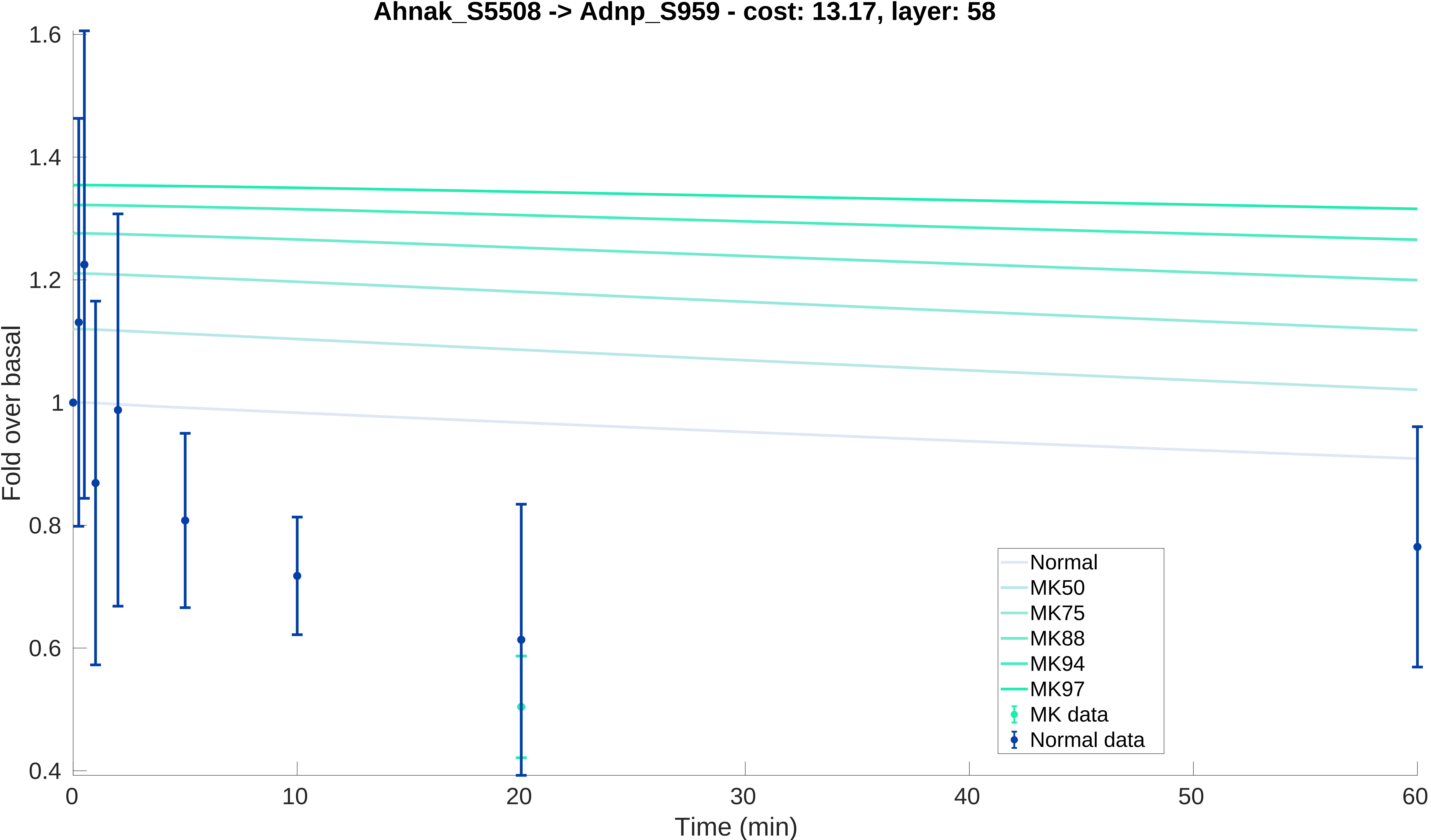

Ahnak_S5508 -> Adnp_S959 - cost: 13.17, layer: 58
Fold over basal
Time (min)
1.6
1.4
1.2
1
0.8
0.6
0.4
0
10
20
30
40
50
60
Normal
MK50
MK75
MK88
MK94
MK97
MK data
Normal data

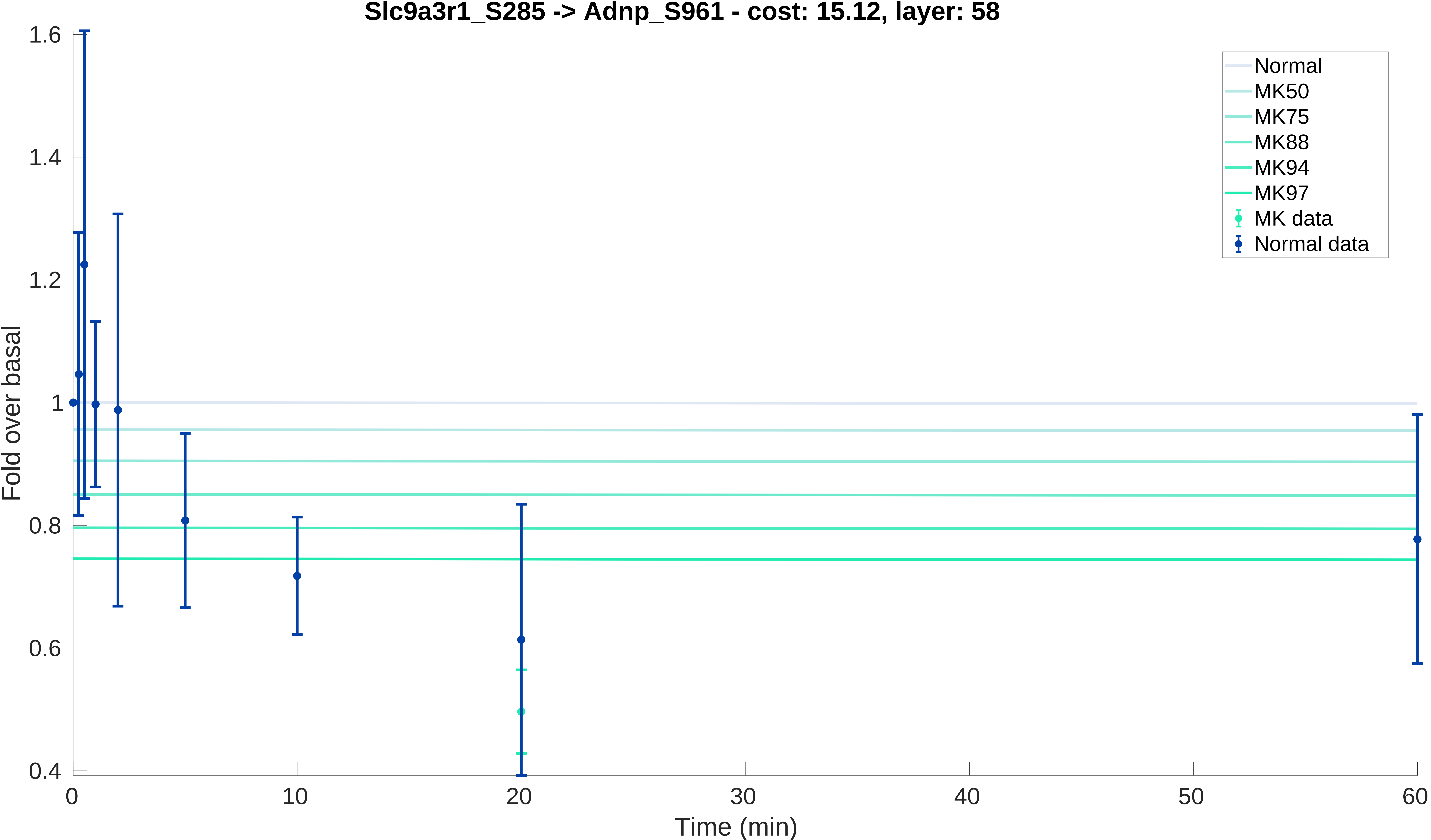

Slc9a3r1_S285 -> Adnp_S961 - cost: 15.12, layer: 58
Fold over basal
1.6
1.4
1.2
1
0.8
0.6
0.4
0
10
20
30
40
50
60
Time (min)
Normal
MK50
MK75
MK88
MK94
MK97
MK data
Normal data

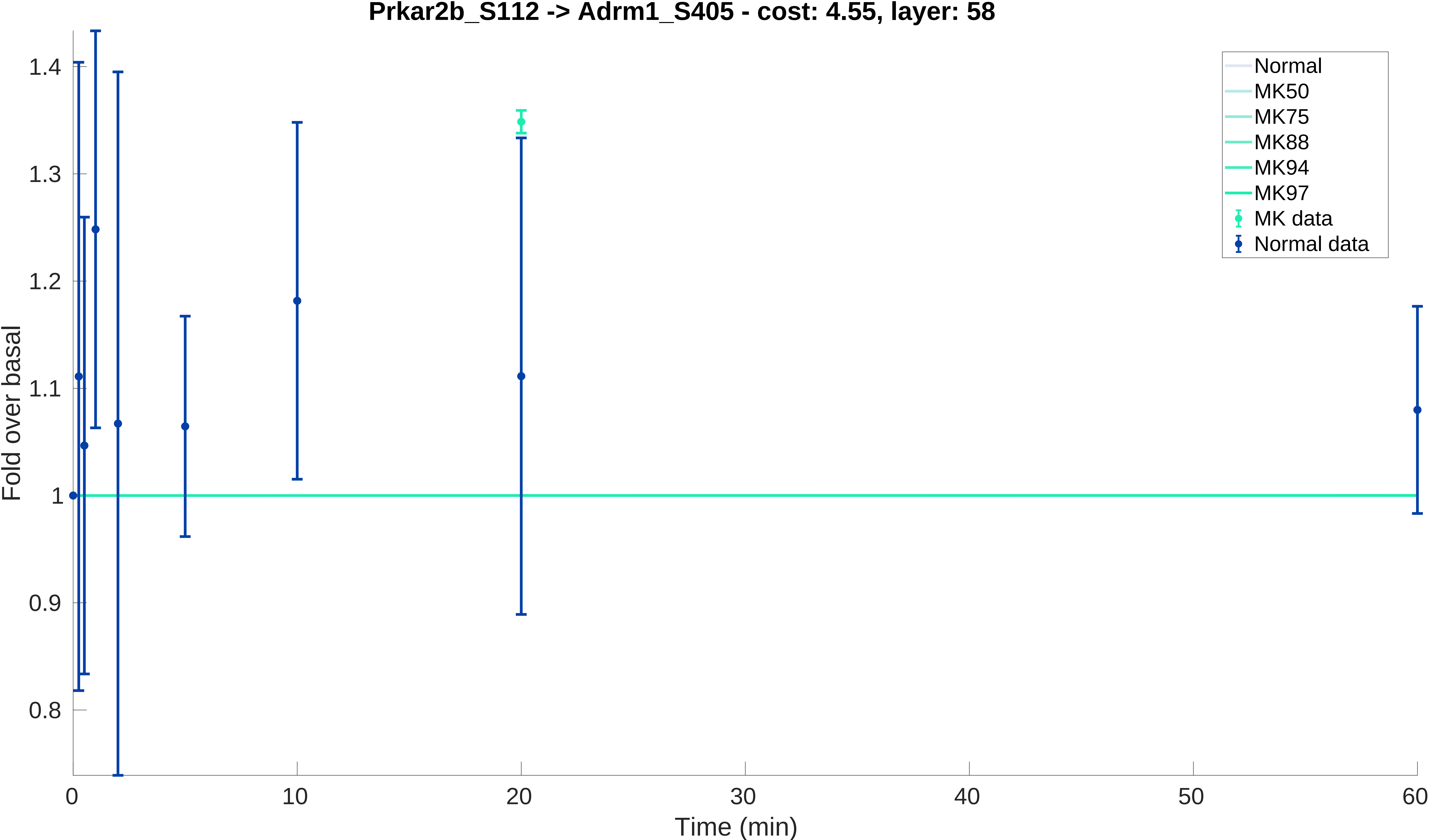

Prkar2b_S112 -> Adrm1_S405 - cost: 4.55, layer: 58
Fold over basal
1.4
1.3
1.2
1.1
1
0.9
0.8
Time (min)
0
10
20
30
40
50
60
Normal
MK50
MK75
MK88
MK94
MK97
MK data
Normal data

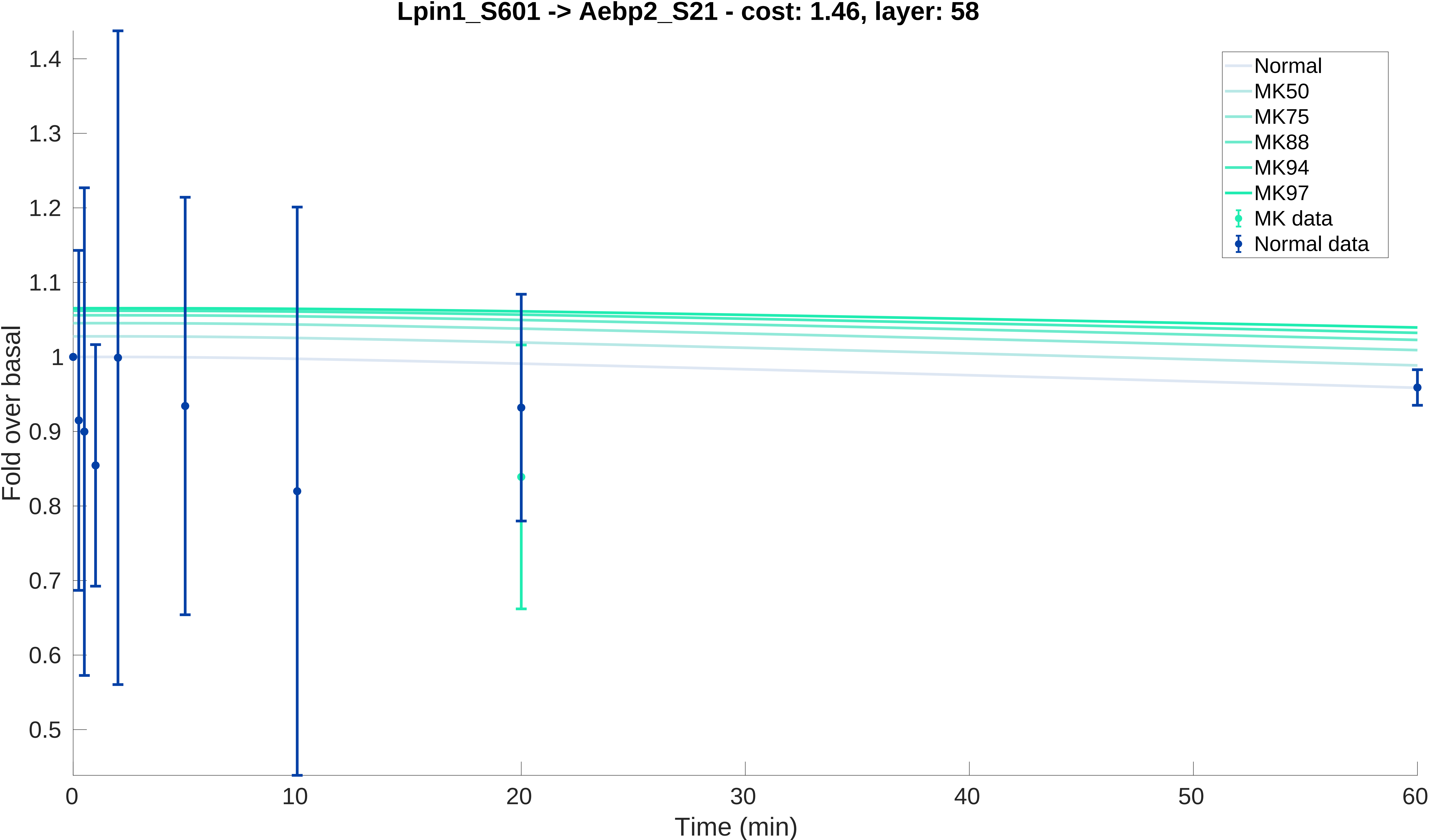

Lpin1_S601 -> Aebp2_S21 - cost: 1.46, layer: 58
Fold over basal
1.4
1.3
1.2
1.1
1
0.9
0.8
0.7
0.6
0.5
0
10
20
30
40
50
60
Time (min)
Normal
MK50
MK75
MK88
MK94
MK97
MK data
Normal data

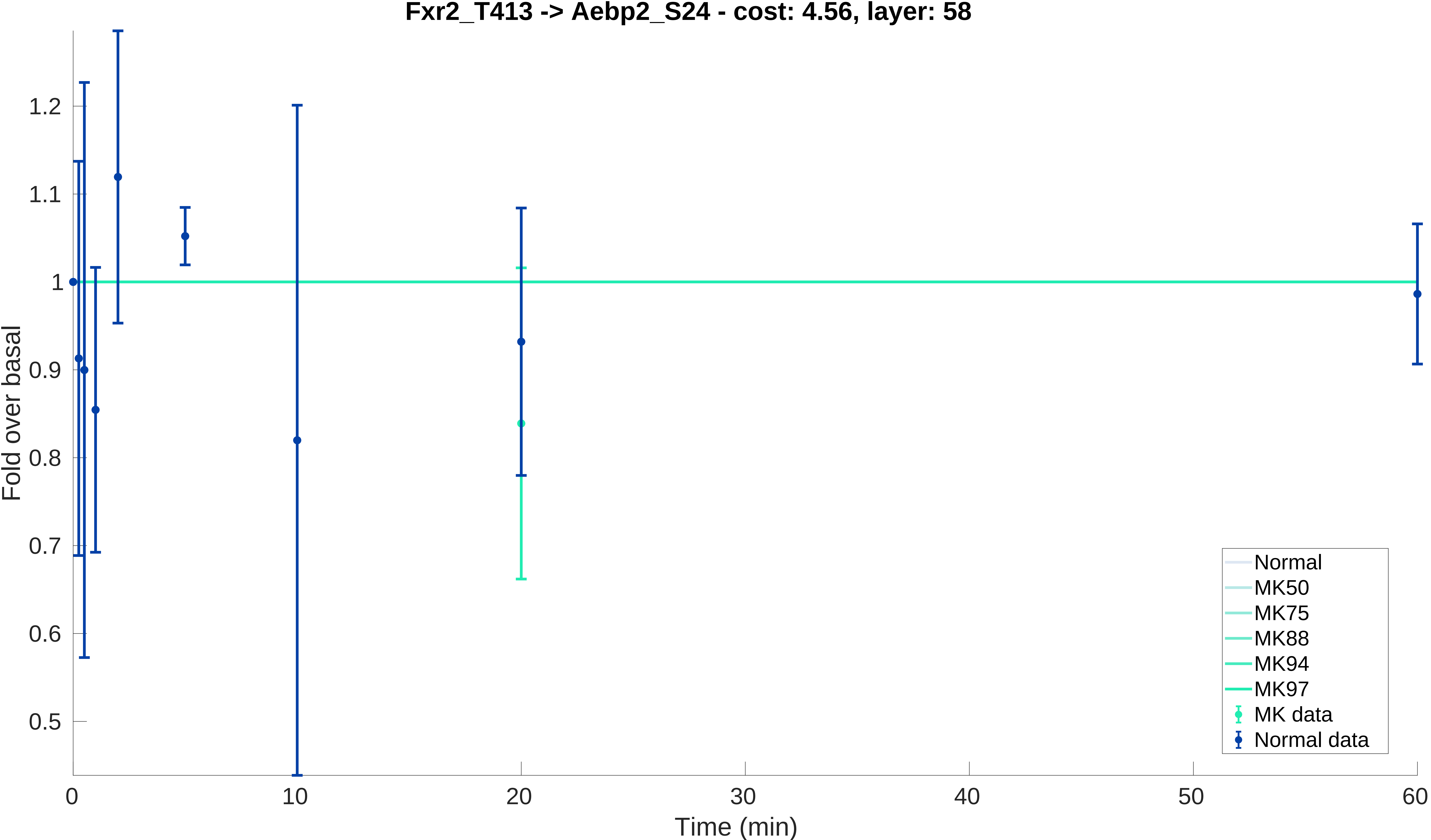

Fxr2_T413 -> Aebp2_S24 - cost: 4.56, layer: 58
Fold over basal
Time (min)
0.5
0.6
0.7
0.8
0.9
1
1.1
1.2
0
10
20
30
40
50
60
Normal
MK50
MK75
MK88
MK94
MK97
MK data
Normal data

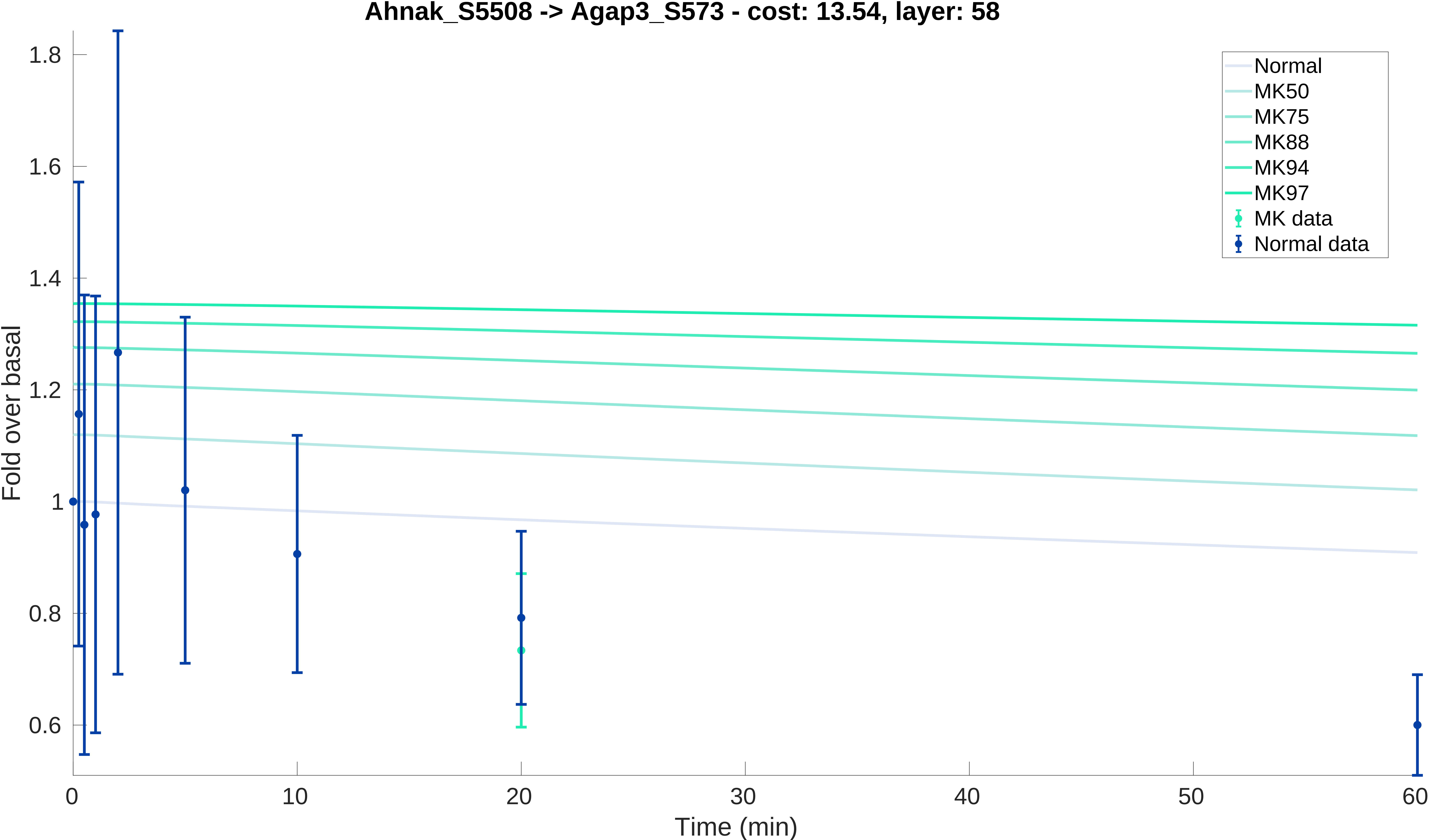

Ahnak_S5508 -> Agap3_S573 - cost: 13.54, layer: 58
Fold over basal
1.8
1.6
1.4
1.2
1
0.8
0.6
0
10
20
30
40
50
60
Time (min)
Normal
MK50
MK75
MK88
MK94
MK97
MK data
Normal data

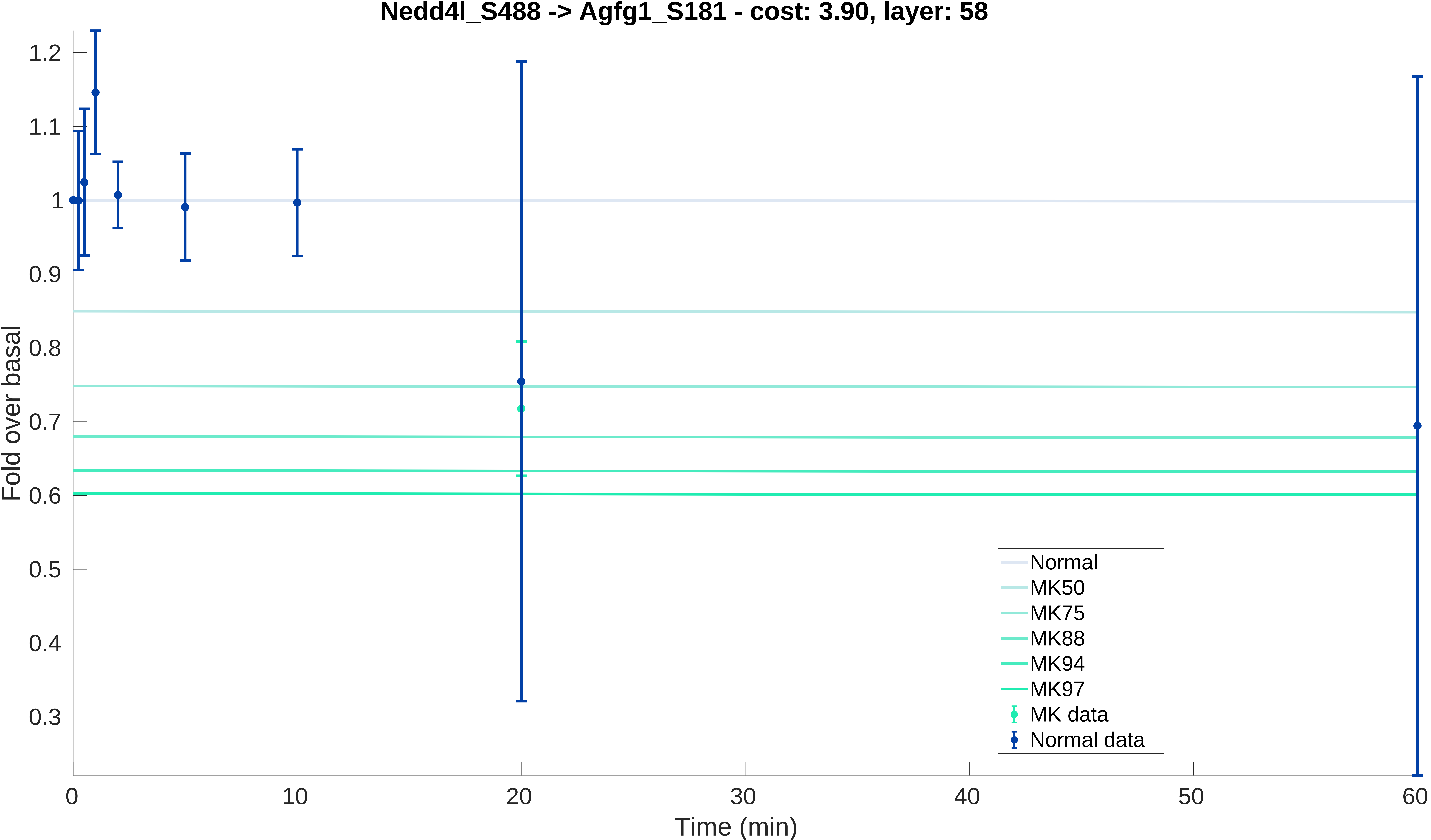

Nedd4l_S488 -> Agfg1_S181 - cost: 3.90, layer: 58
Fold over basal
Time (min)
1.2
1.1
1
0.9
0.8
0.7
0.6
0.5
0.4
0.3
0
10
20
30
40
50
60
Normal
MK50
MK75
MK88
MK94
MK97
MK data
Normal data

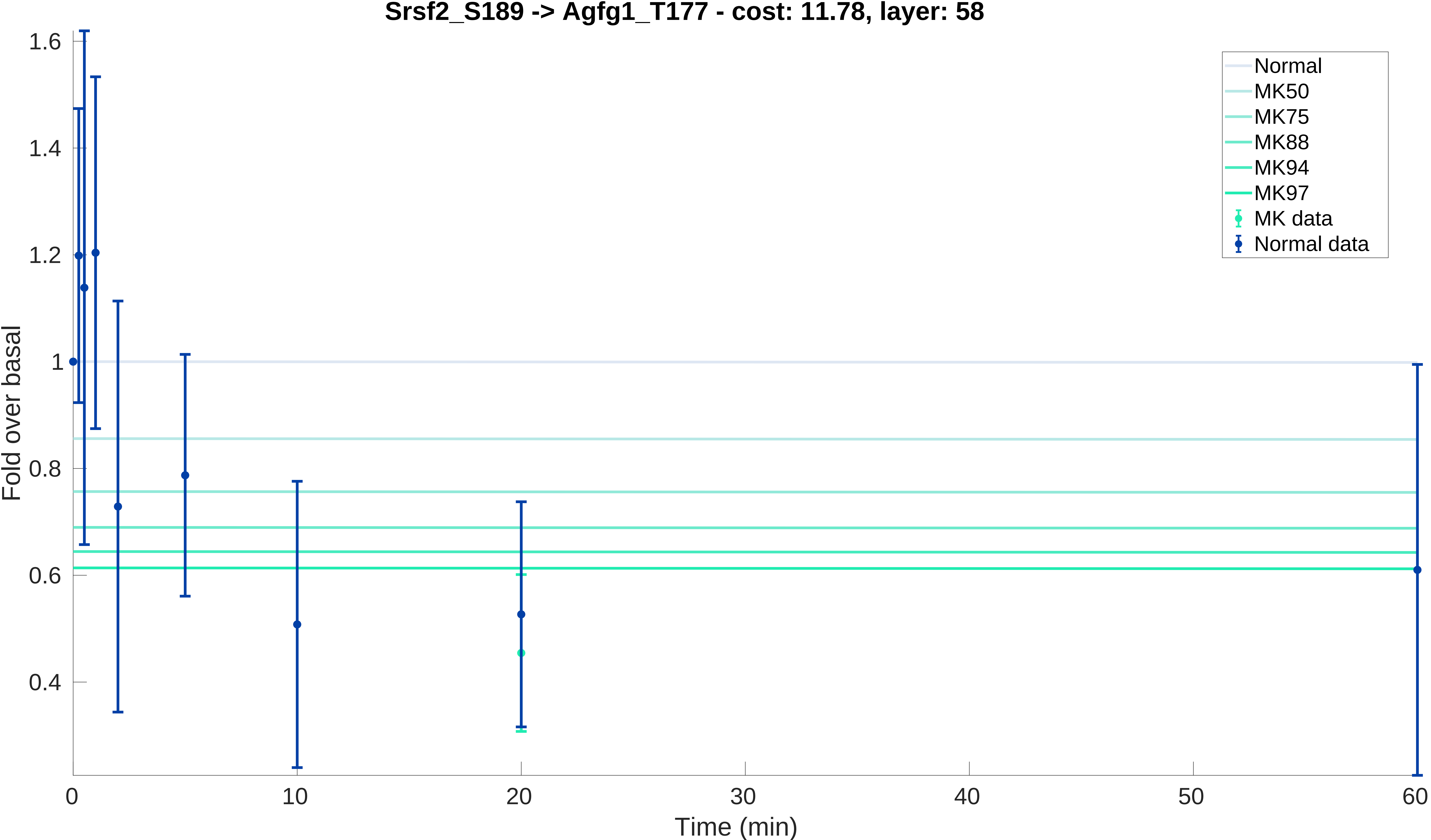

Srsf2_S189 -> Agfg1_T177 - cost: 11.78, layer: 58
Fold over basal
Time (min)
0.4
0.6
0.8
1
1.2
1.4
1.6
0
10
20
30
40
50
60
Normal
MK50
MK75
MK88
MK94
MK97
MK data
Normal data

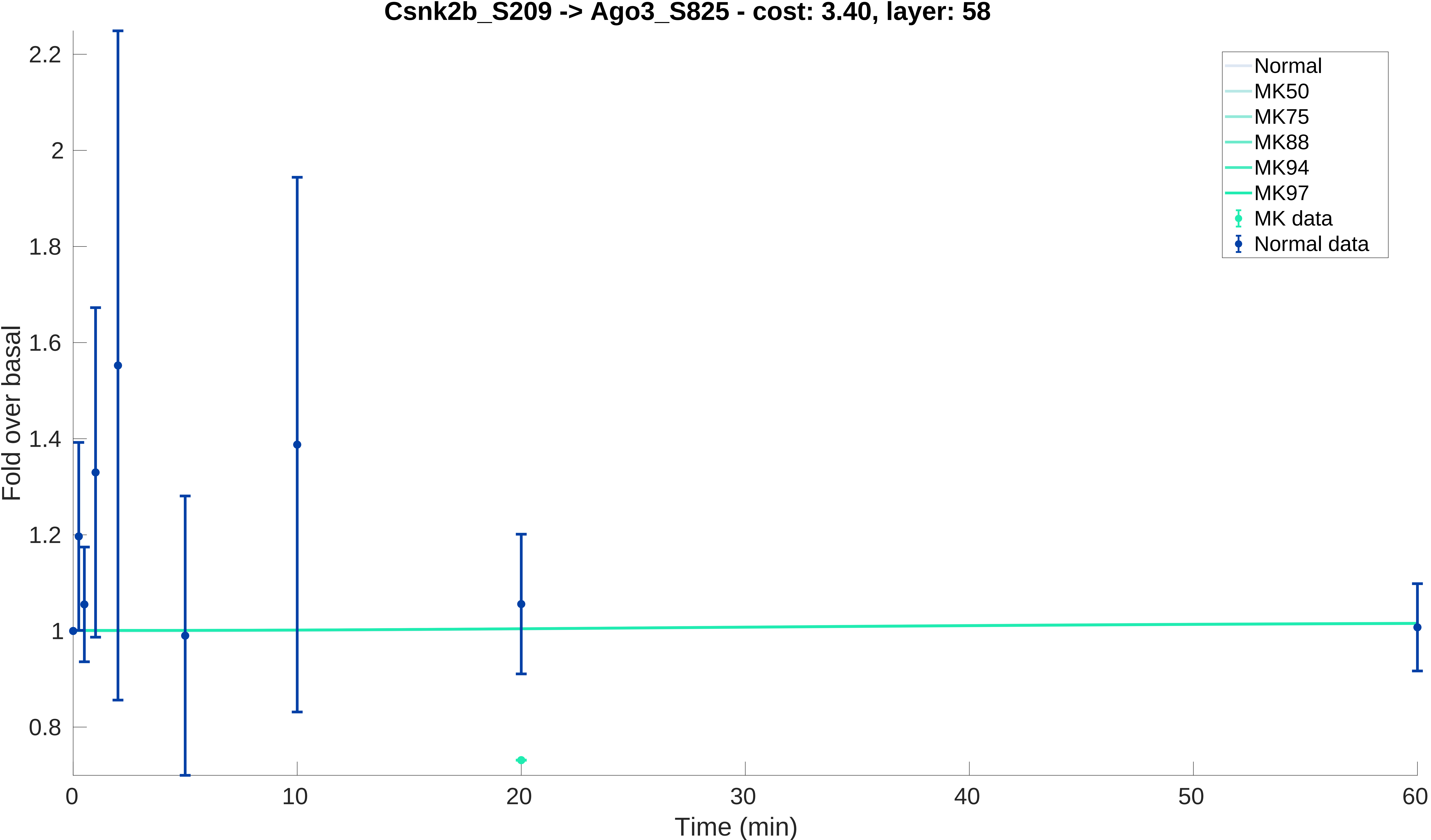

Csnk2b_S209 -> Ago3_S825 - cost: 3.40, layer: 58
Fold over basal
Time (min)
0
10
20
30
40
50
60
0.8
1
1.2
1.4
1.6
1.8
2
2.2
Normal
MK50
MK75
MK88
MK94
MK97
MK data
Normal data

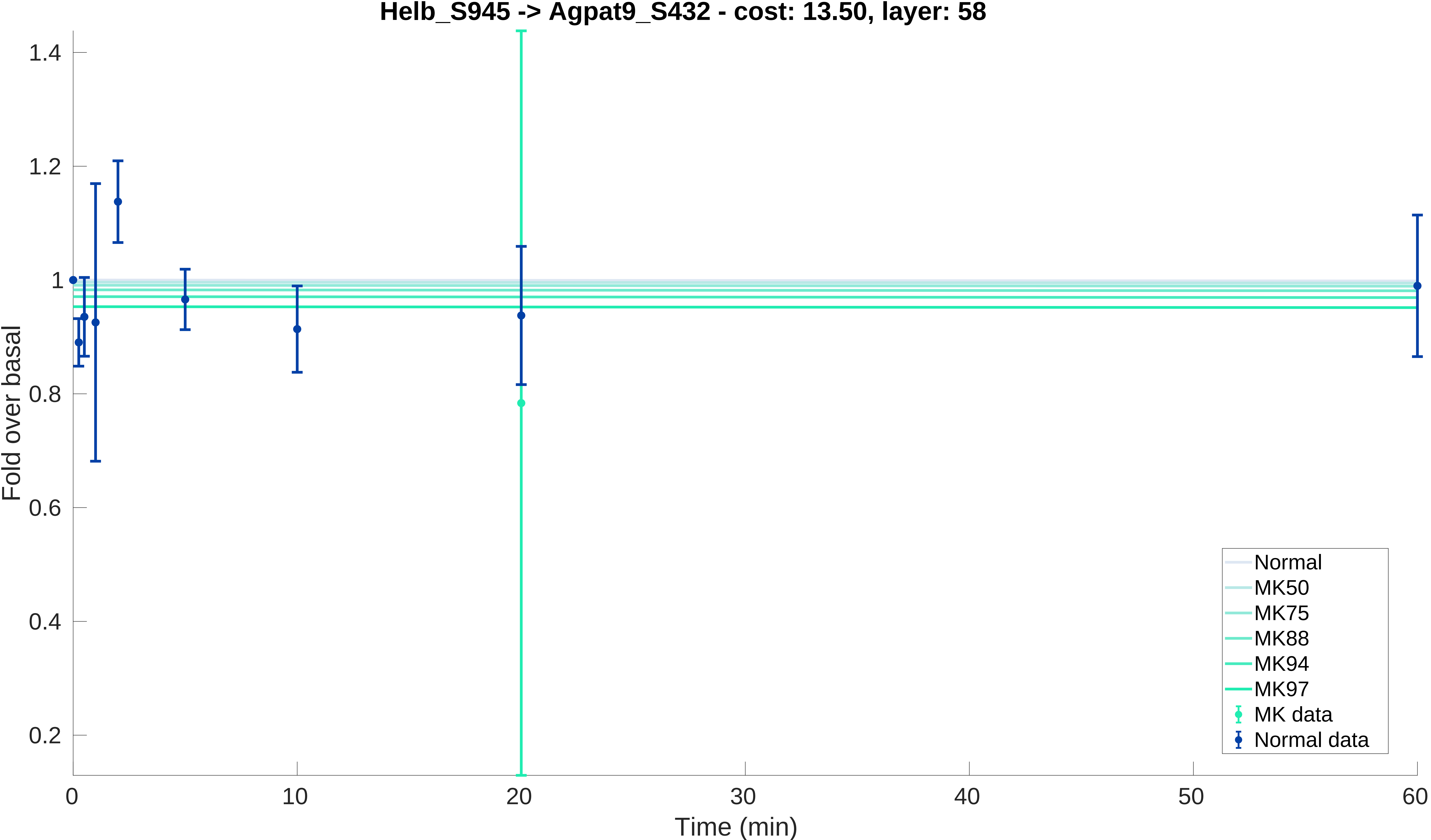

Helb_S945 -> Agpat9_S432 - cost: 13.50, layer: 58
Fold over basal
Time (min)
0
10
20
30
40
50
60
0.2
0.4
0.6
0.8
1
1.2
1.4
Normal
MK50
MK75
MK88
MK94
MK97
MK data
Normal data

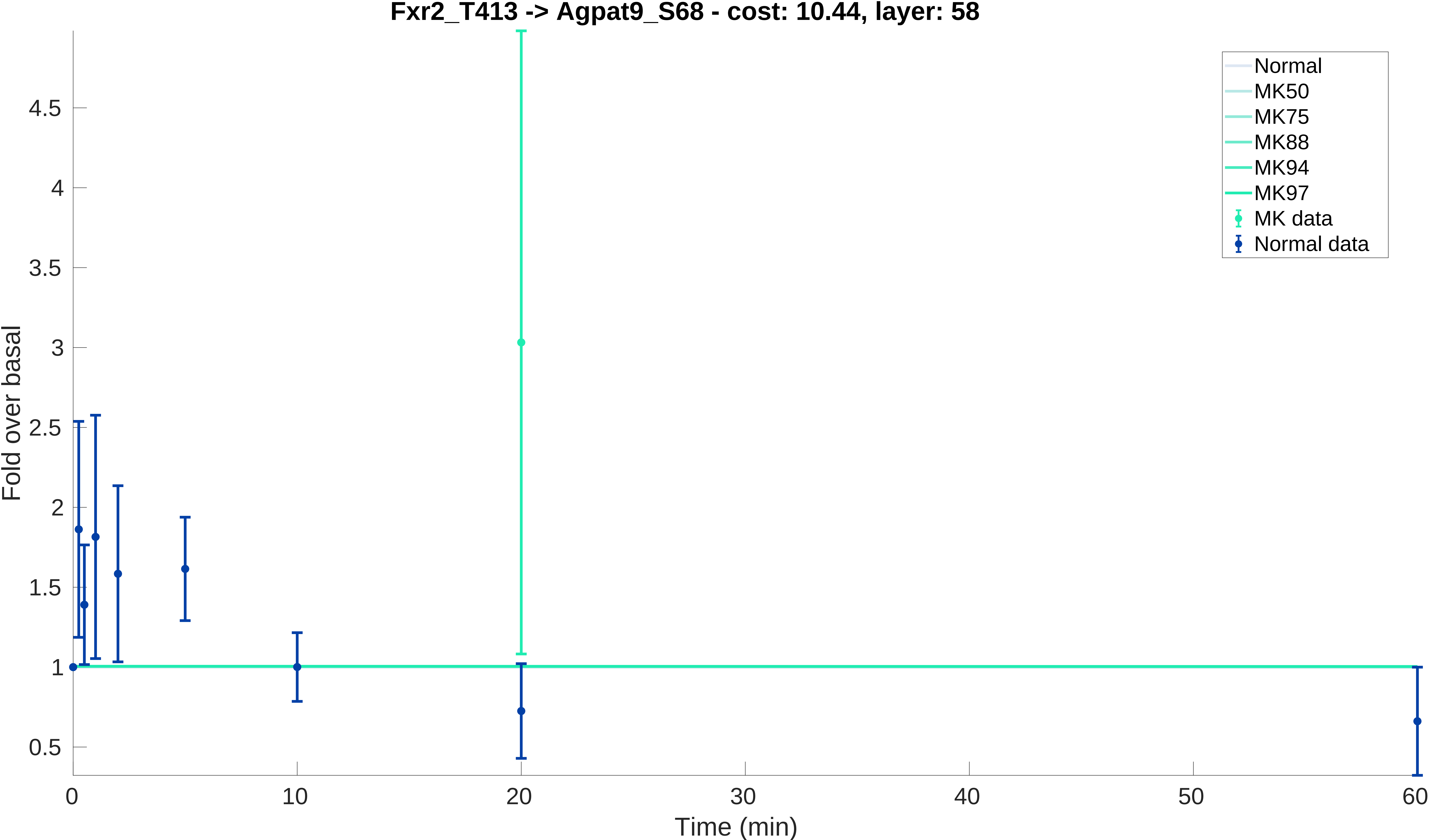

Fxr2_T413 -> Agpat9_S68 - cost: 10.44, layer: 58
Fold over basal
Time (min)
0.5
1
1.5
2
2.5
3
3.5
4
4.5
0
10
20
30
40
50
60
Normal
MK50
MK75
MK88
MK94
MK97
MK data
Normal data

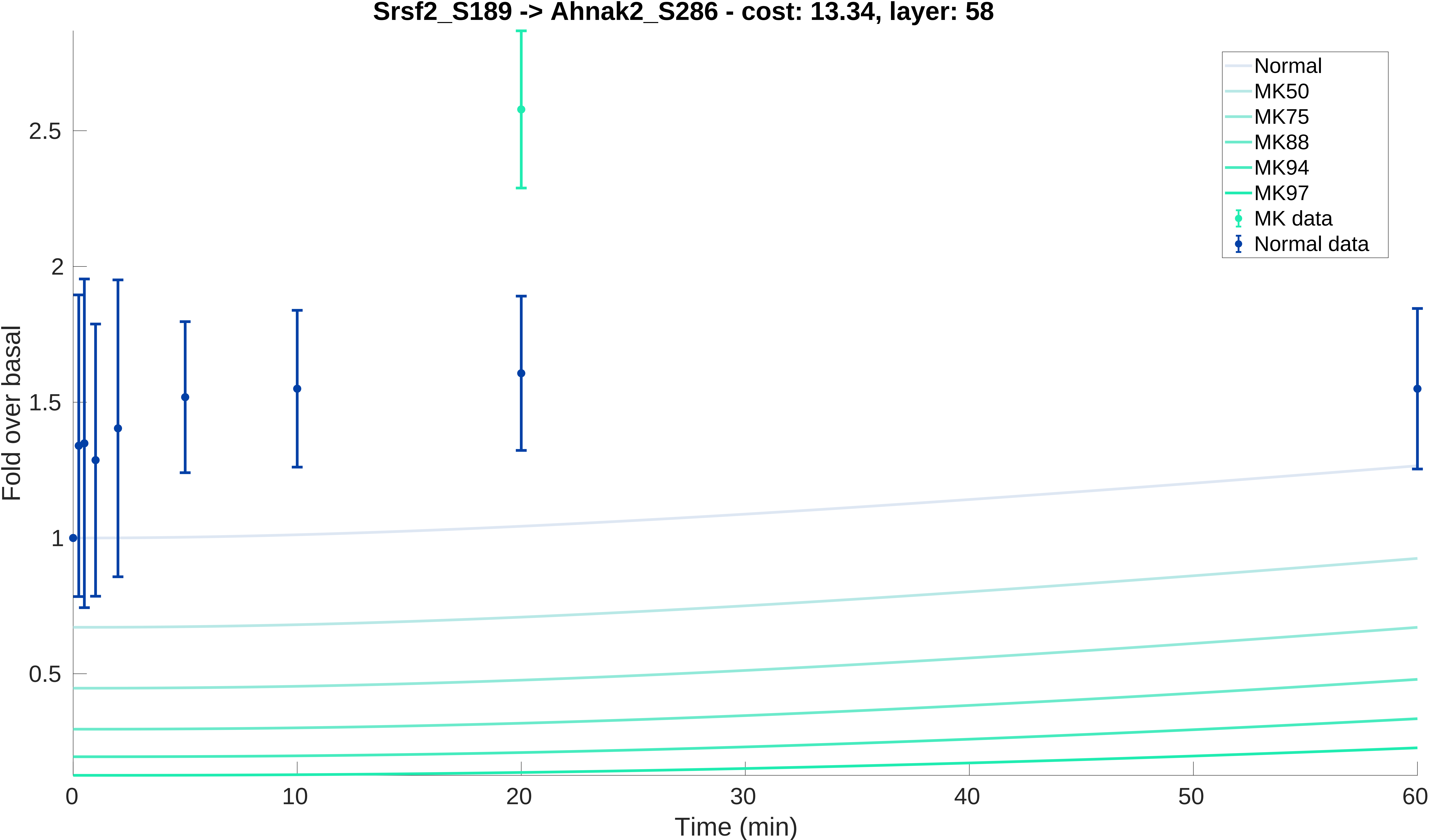

Srsf2_S189 -> Ahnak2_S286 - cost: 13.34, layer: 58
Fold over basal
Time (min)
0.5
1
1.5
2
2.5
0
10
20
30
40
50
60
Normal
MK50
MK75
MK88
MK94
MK97
MK data
Normal data

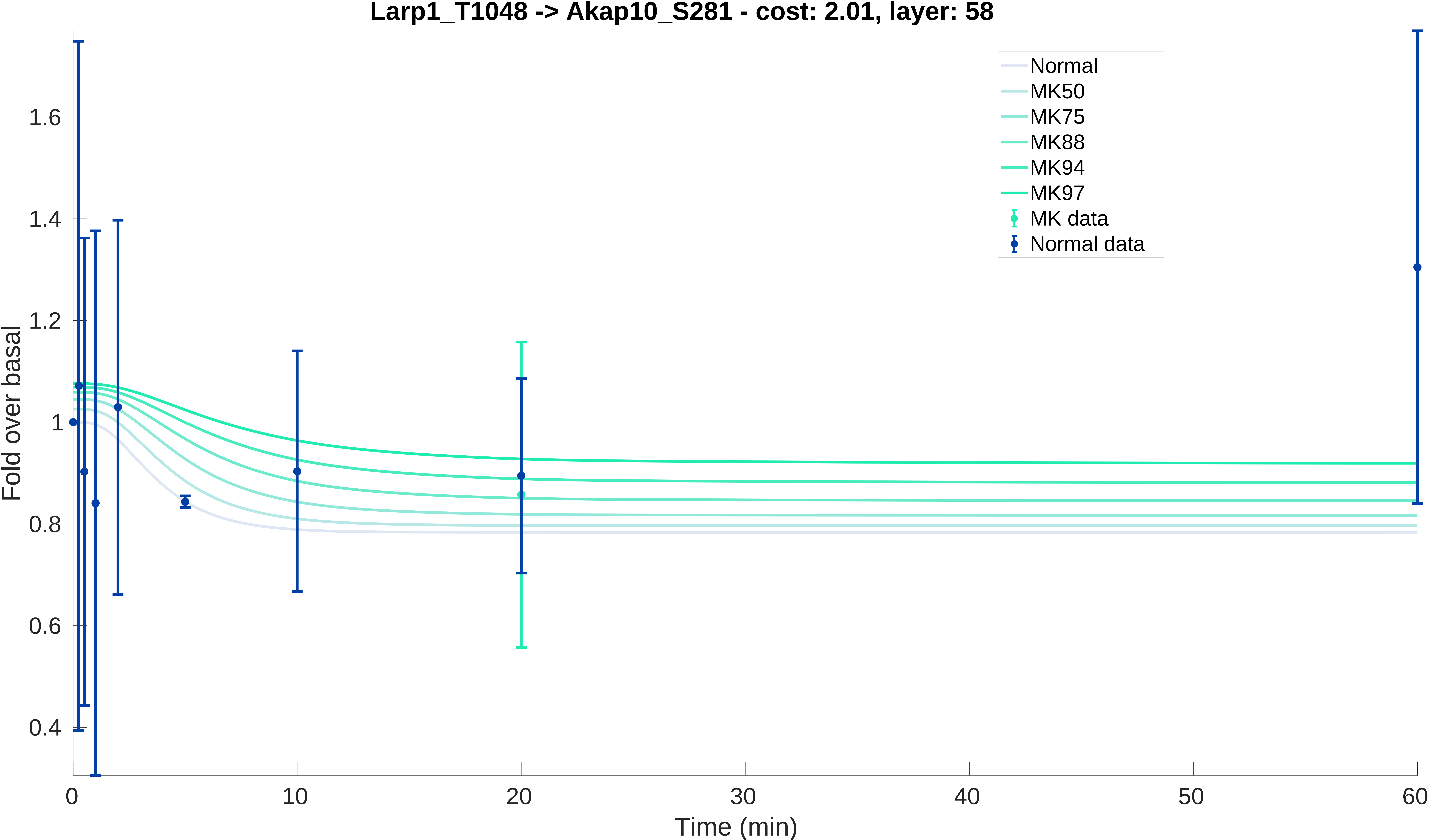
Larp1_T1048 -> Akap10_S281 - cost: 2.01, layer: 58
Normal
MK50
MK75
MK88
MK94
MK97
MK data
Normal data
Fold over basal
1.6
1.4
1.2
1
0.8
0.6
0.4
0
10
20
30
40
50
60
Time (min)

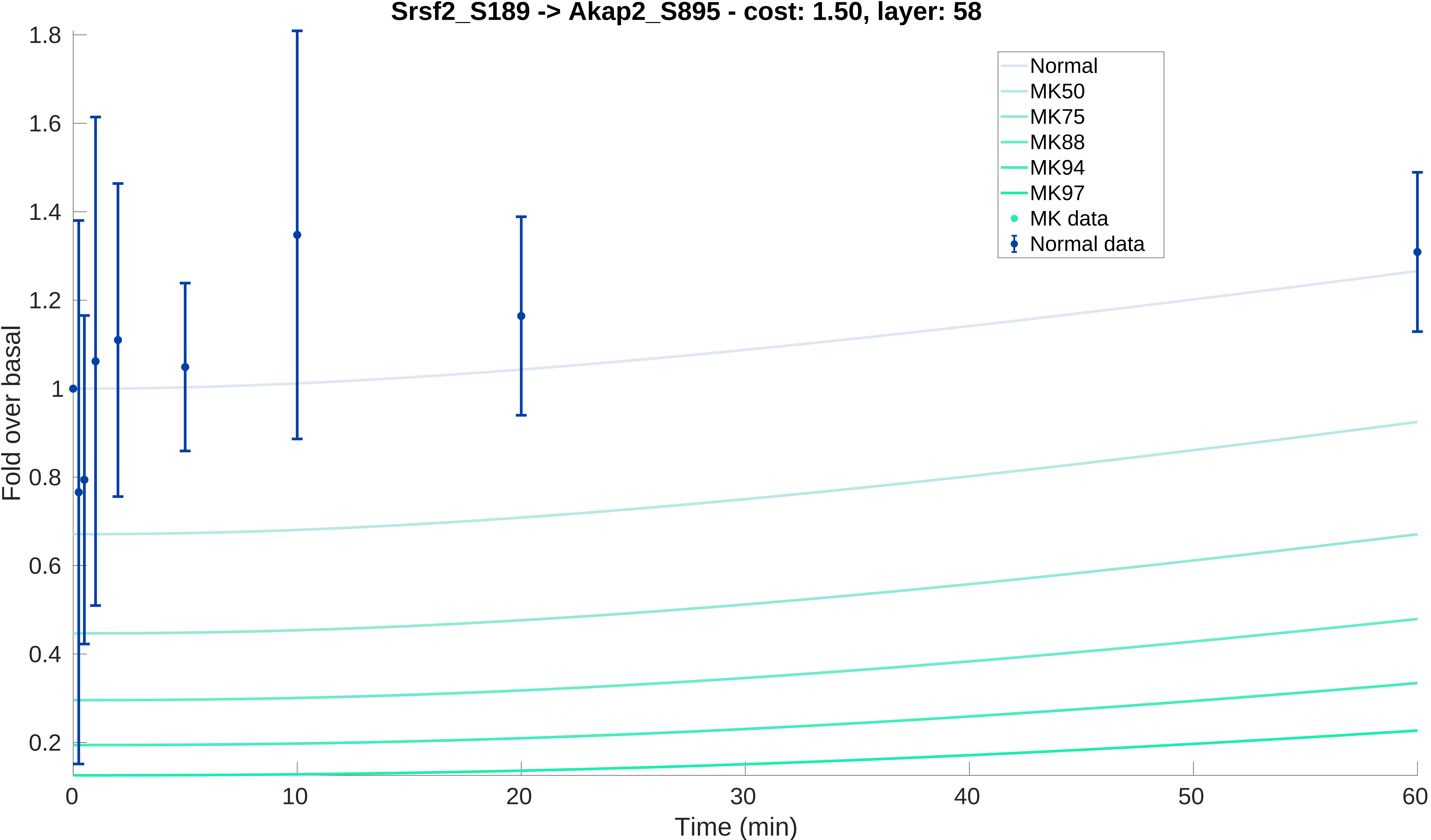
Srsf2_S189 -> Akap2_S895 - cost: 1.50, layer: 58
Fold over basal
Time (min)
1.8
1.6
1.4
1.2
1
0.8
0.6
0.4
0.2
0
10
20
30
40
50
60
Normal
MK50
MK75
MK88
MK94
MK97
MK data
Normal data

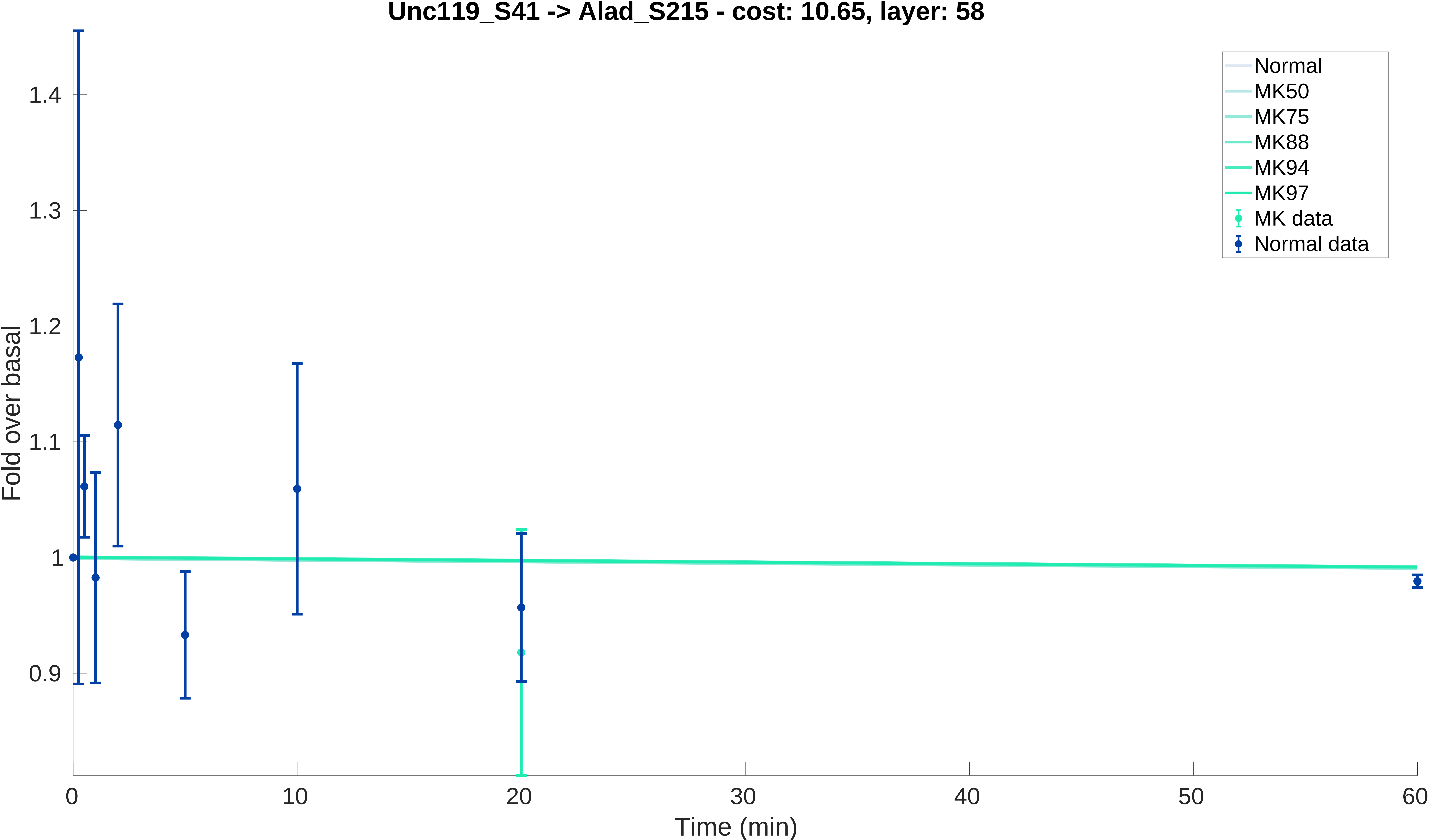
Unc119_S41 -> Alad_S215 - cost: 10.65, layer: 58
Normal
MK50
MK75
MK88
MK94
MK97
MK data
Normal data
Fold over basal
1.4
1.3
1.2
1.1
1
0.9
0
10
20
30
40
50
60
Time (min)

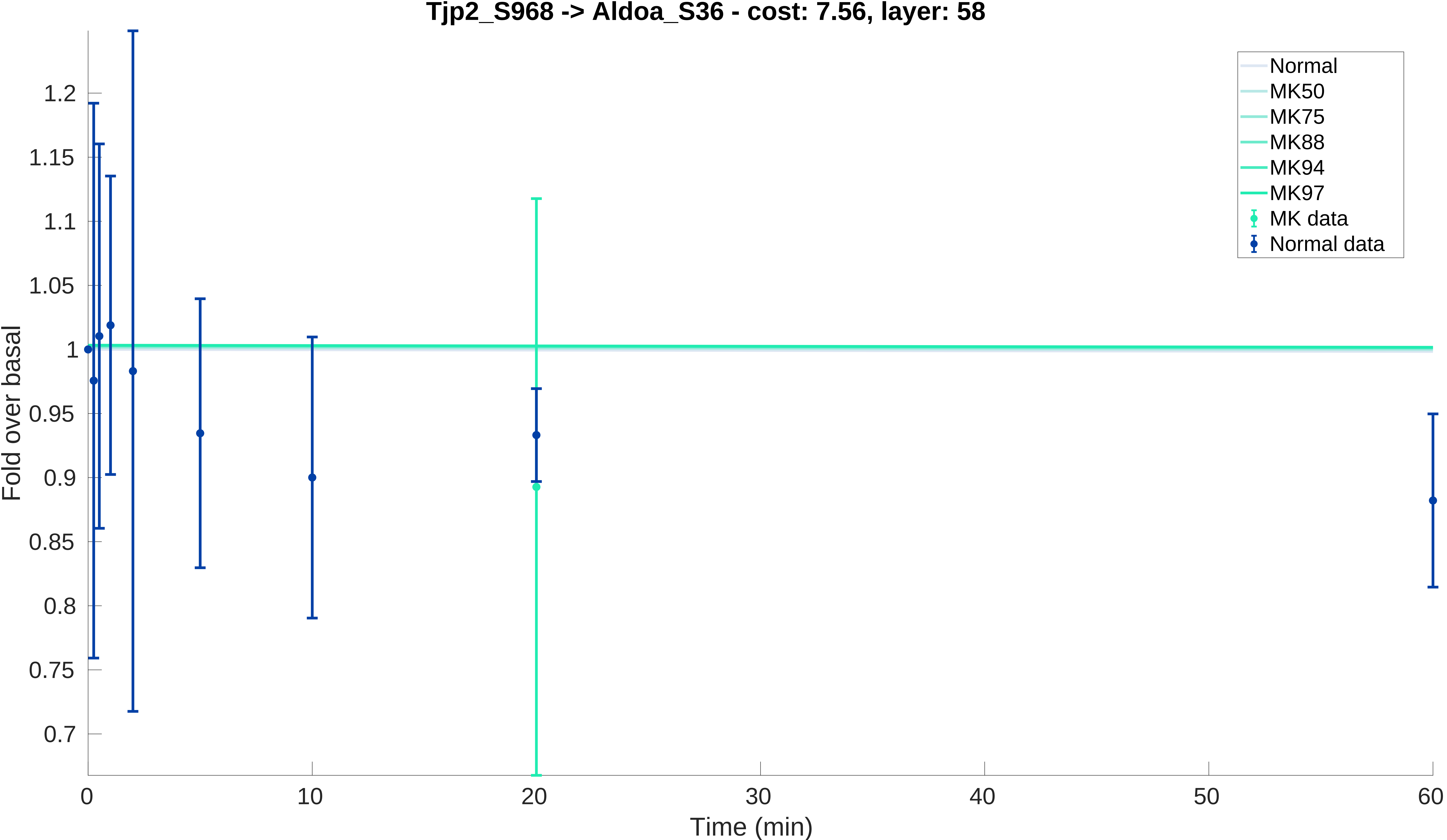
Tjp2_S968 -> Aldoa_S36 - cost: 7.56, layer: 58
Fold over basal
Time (min)
0
10
20
30
40
50
60
0.7
0.75
0.8
0.85
0.9
0.95
1
1.05
1.1
1.15
1.2
Normal
MK50
MK75
MK88
MK94
MK97
MK data
Normal data

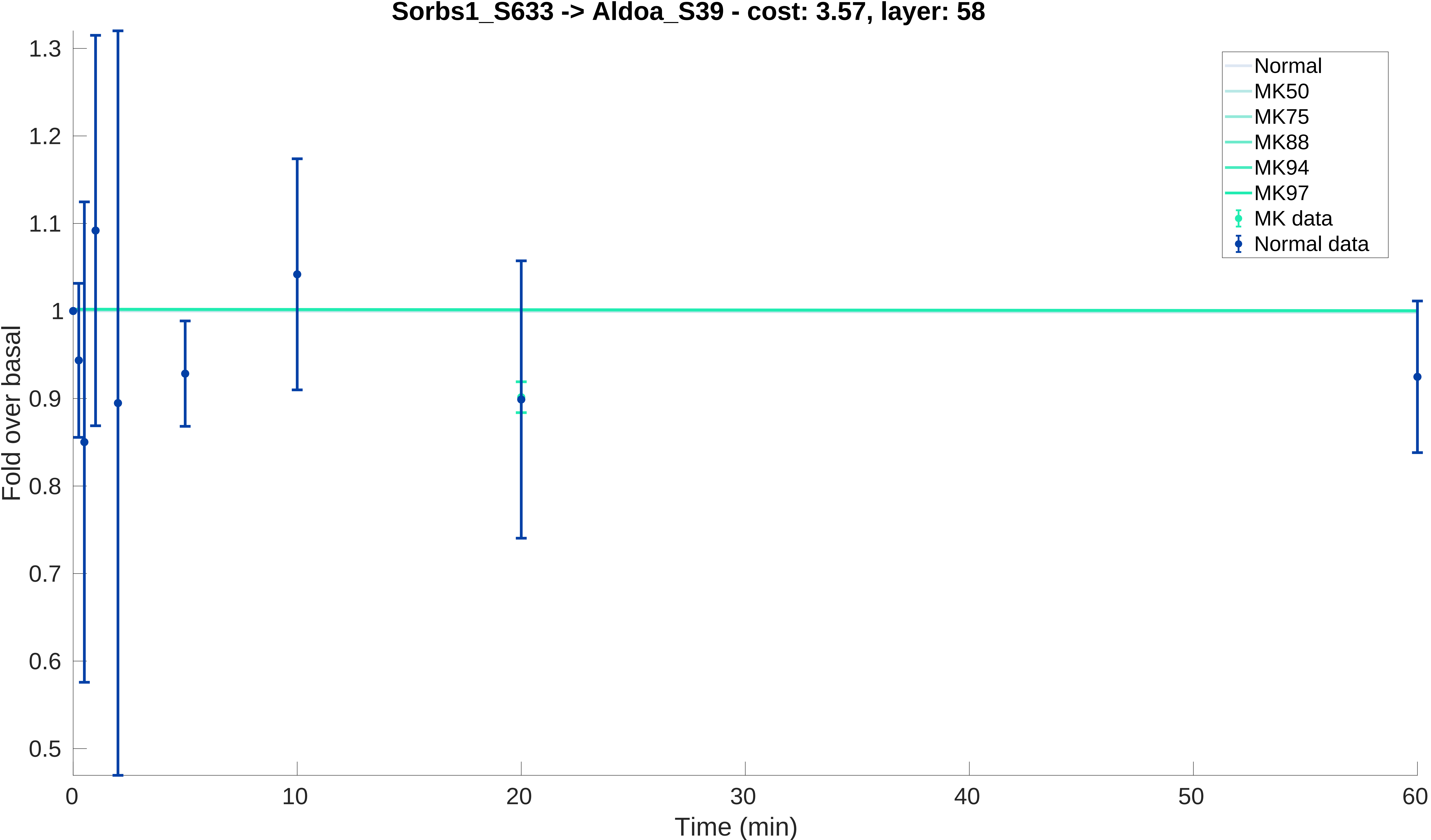

Sorbs1_S633 -> Aldoa_S39 - cost: 3.57, layer: 58
Fold over basal
Time (min)
Normal
MK50
MK75
MK88
MK94
MK97
MK data
Normal data
0.5
0.6
0.7
0.8
0.9
1
1.1
1.2
1.3
0
10
20
30
40
50
60

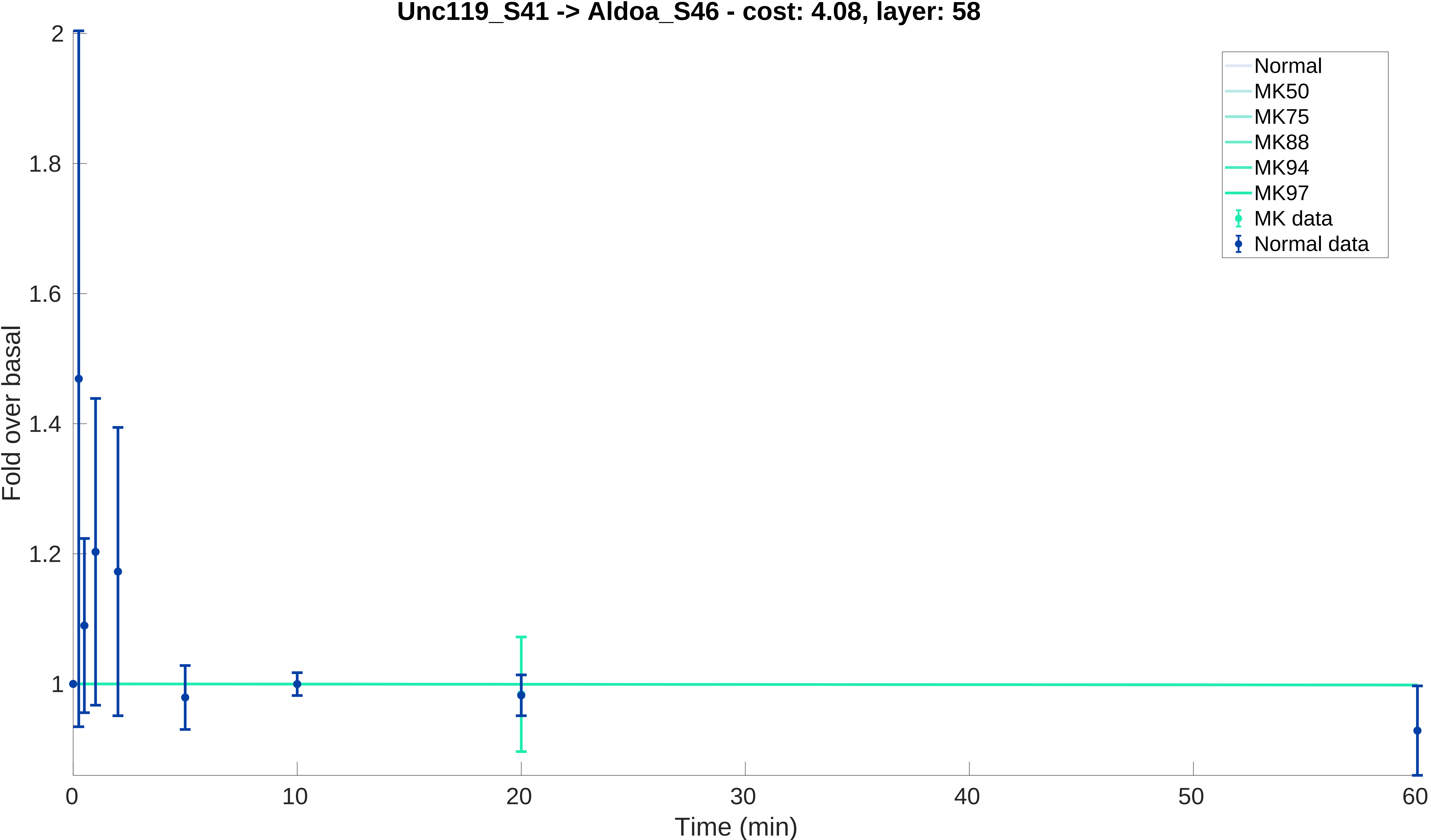

Unc119_S41 -> Aldoa_S46 - cost: 4.08, layer: 58
Fold over basal
Time (min)
2
1.8
1.6
1.4
1.2
1
0
10
20
30
40
50
60
Normal
MK50
MK75
MK88
MK94
MK97
MK data
Normal data

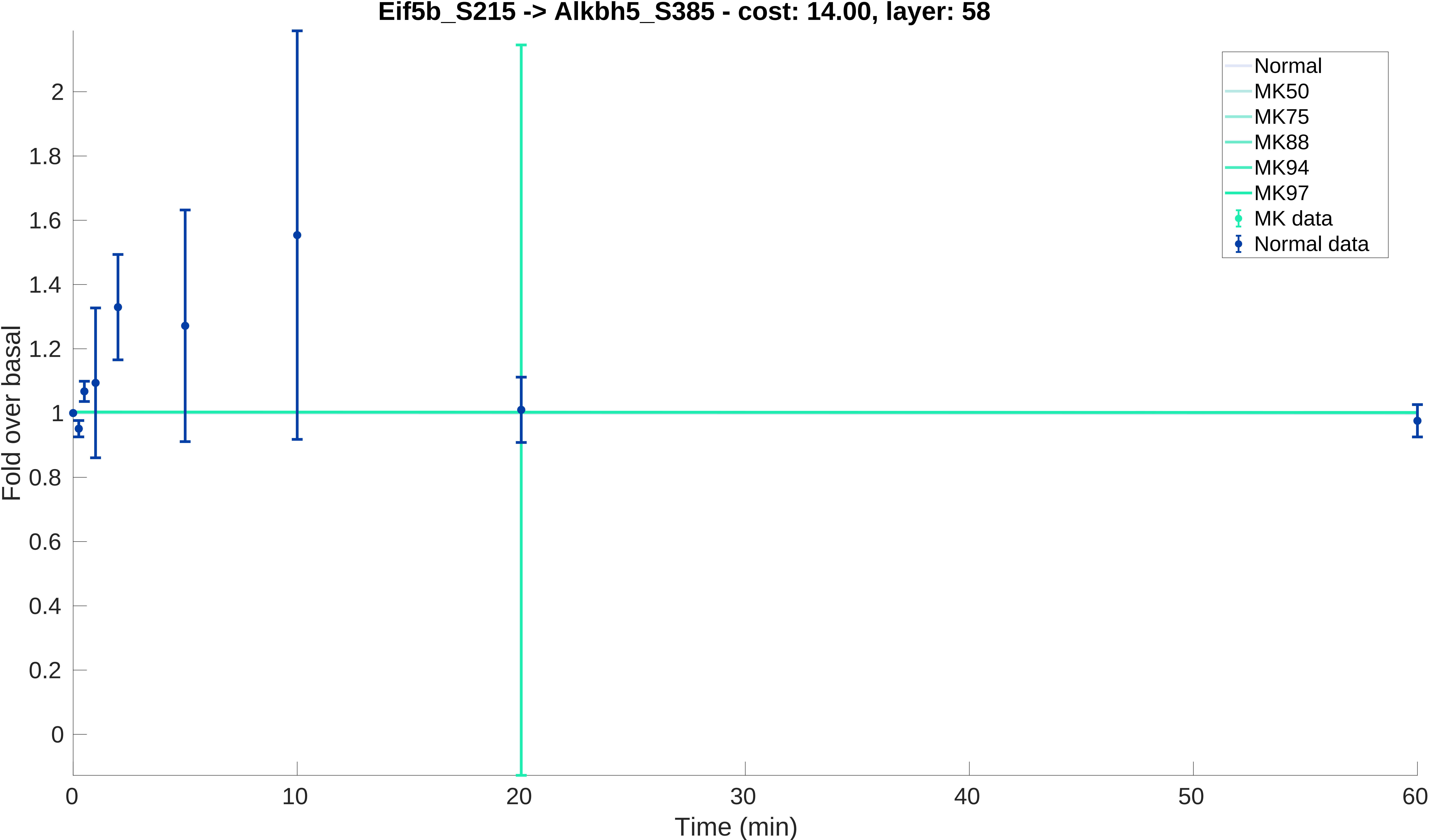

Eif5b_S215 -> Alkbh5_S385 - cost: 14.00, layer: 58
Fold over basal
Time (min)
0
0.2
0.4
0.6
0.8
1
1.2
1.4
1.6
1.8
2
0
10
20
30
40
50
60
Normal
MK50
MK75
MK88
MK94
MK97
MK data
Normal data

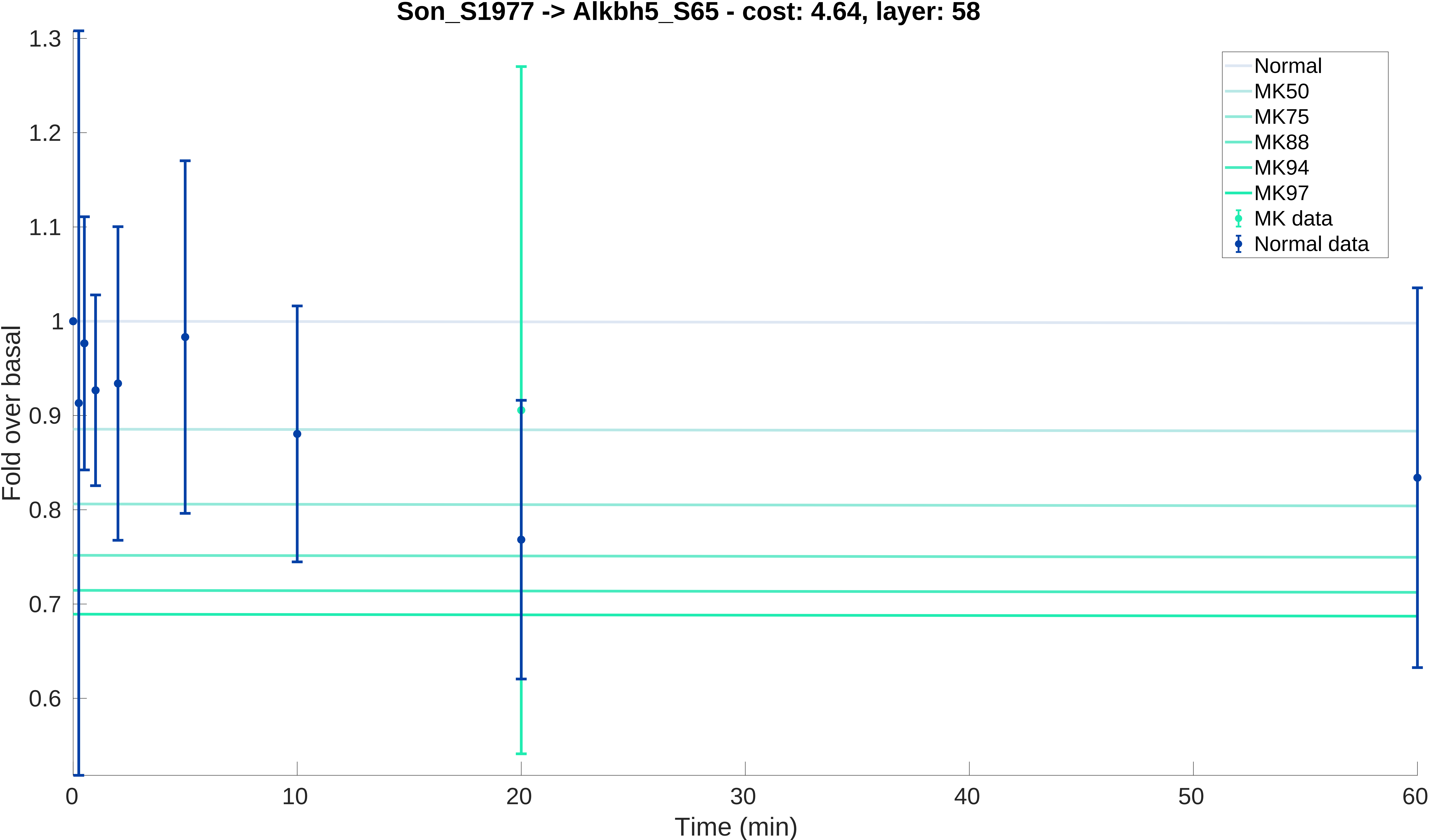

Son_S1977 -> Alkbh5_S65 - cost: 4.64, layer: 58
Fold over basal
1.3
1.2
1.1
1
0.9
0.8
0.7
0.6
0
10
20
30
40
50
60
Time (min)
Normal
MK50
MK75
MK88
MK94
MK97
MK data
Normal data

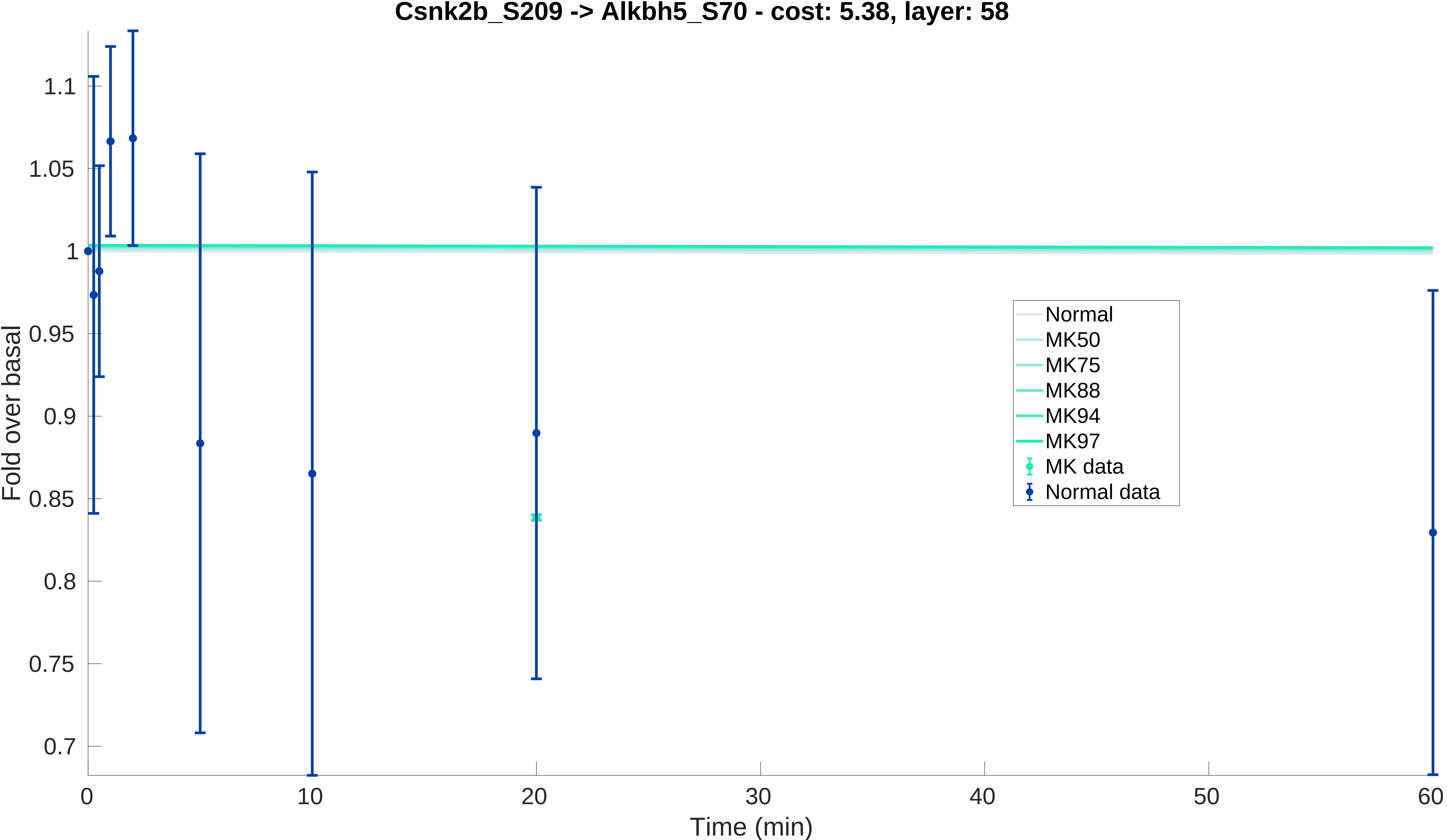
Csnk2b_S209 -> Alkbh5_S70 - cost: 5.38, layer: 58
Fold over basal
Time (min)
1.1
1.05
1
0.95
0.9
0.85
0.8
0.75
0.7
0
10
20
30
40
50
60
Normal
MK50
MK75
MK88
MK94
MK97
MK data
Normal data

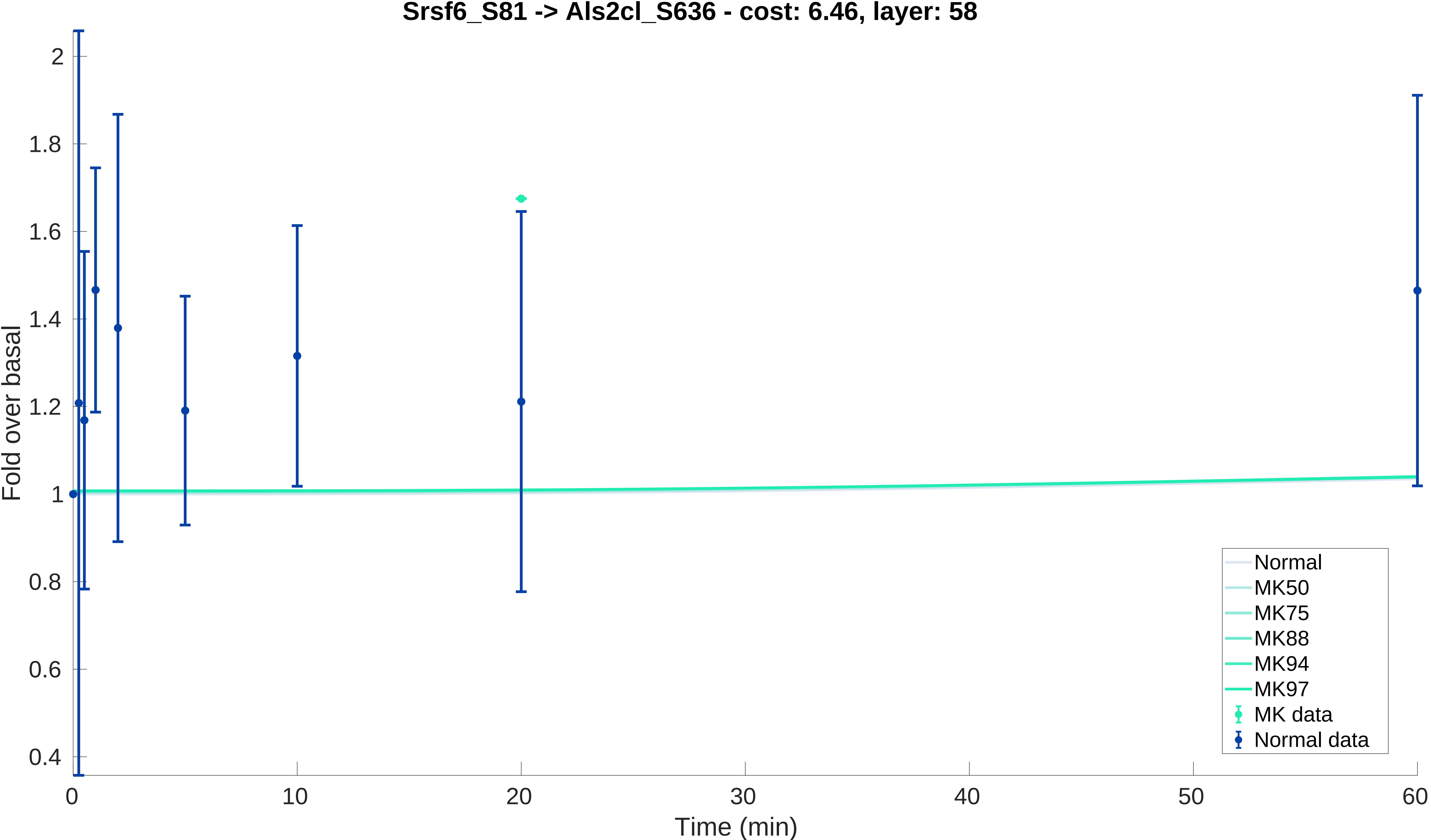
Srsf6_S81 -> Als2cl_S636 - cost: 6.46, layer: 58
Fold over basal
Time (min)
0.4
0.6
0.8
1
1.2
1.4
1.6
1.8
2
0
10
20
30
40
50
60
Normal
MK50
MK75
MK88
MK94
MK97
MK data
Normal data

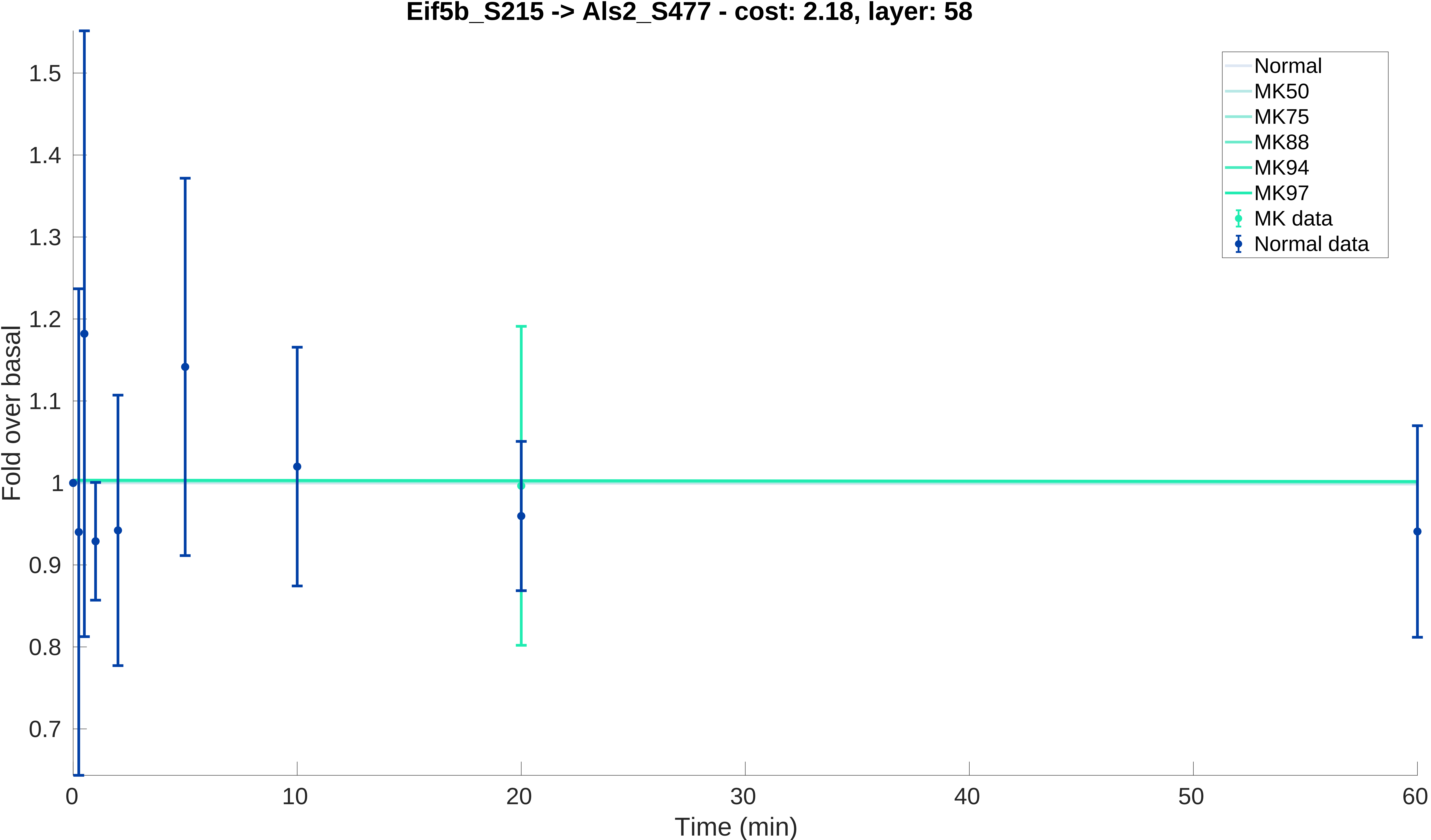

Eif5b_S215 -> Als2_S477 - cost: 2.18, layer: 58
Fold over basal
Time (min)
Normal
MK50
MK75
MK88
MK94
MK97
MK data
Normal data
1.5
1.4
1.3
1.2
1.1
1
0.9
0.8
0.7
0
10
20
30
40
50
60

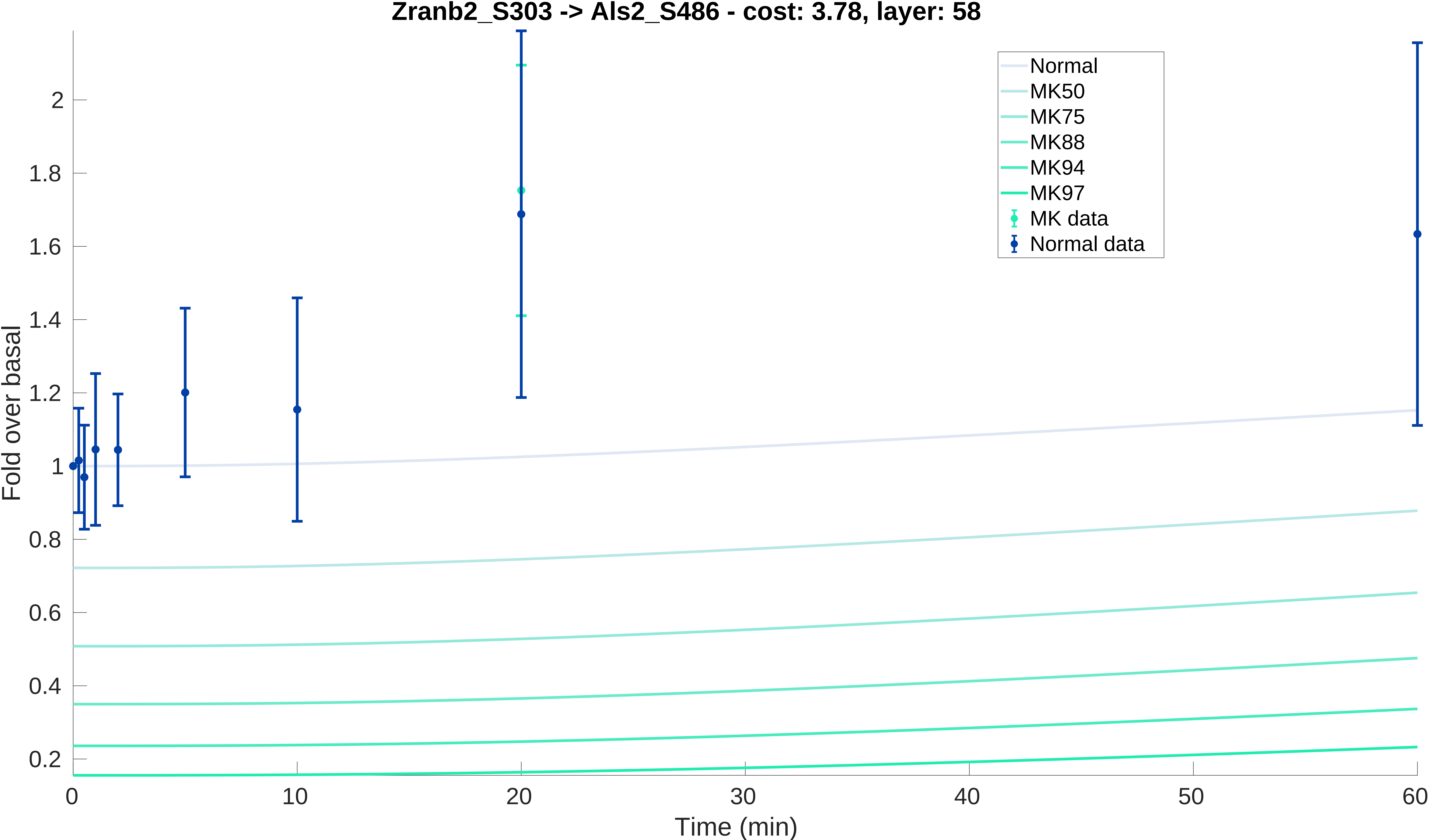
Zranb2_S303 -> Als2_S486 - cost: 3.78, layer: 58
Fold over basal
2
1.8
1.6
1.4
1.2
1
0.8
0.6
0.4
0.2
0
10
20
30
40
50
60
Time (min)
Normal
MK50
MK75
MK88
MK94
MK97
MK data
Normal data

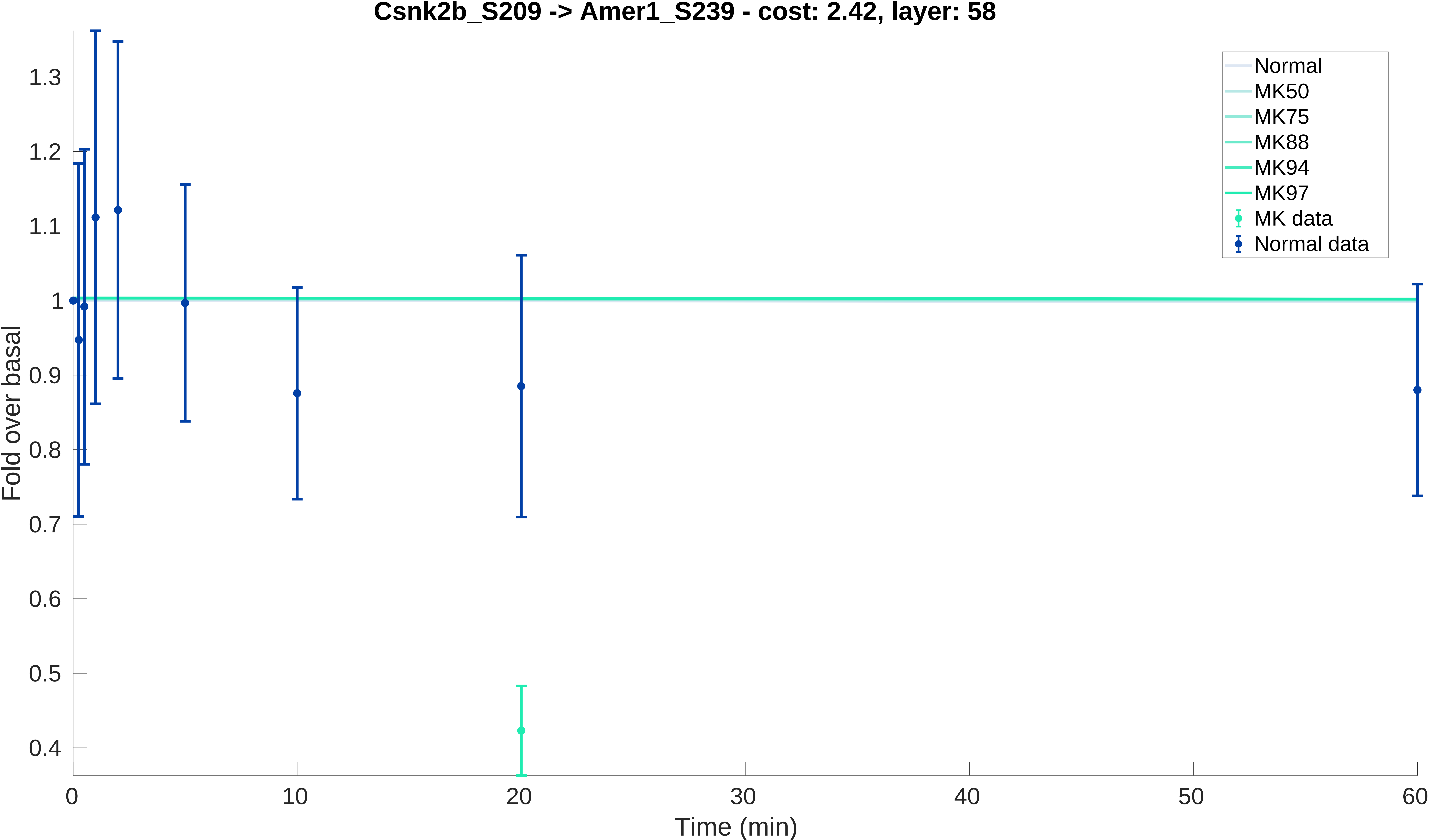

Csnk2b_S209 -> Amer1_S239 - cost: 2.42, layer: 58
Fold over basal
1.3
1.2
1.1
1
0.9
0.8
0.7
0.6
0.5
0.4
0
10
20
30
40
50
60
Time (min)
Normal
MK50
MK75
MK88
MK94
MK97
MK data
Normal data

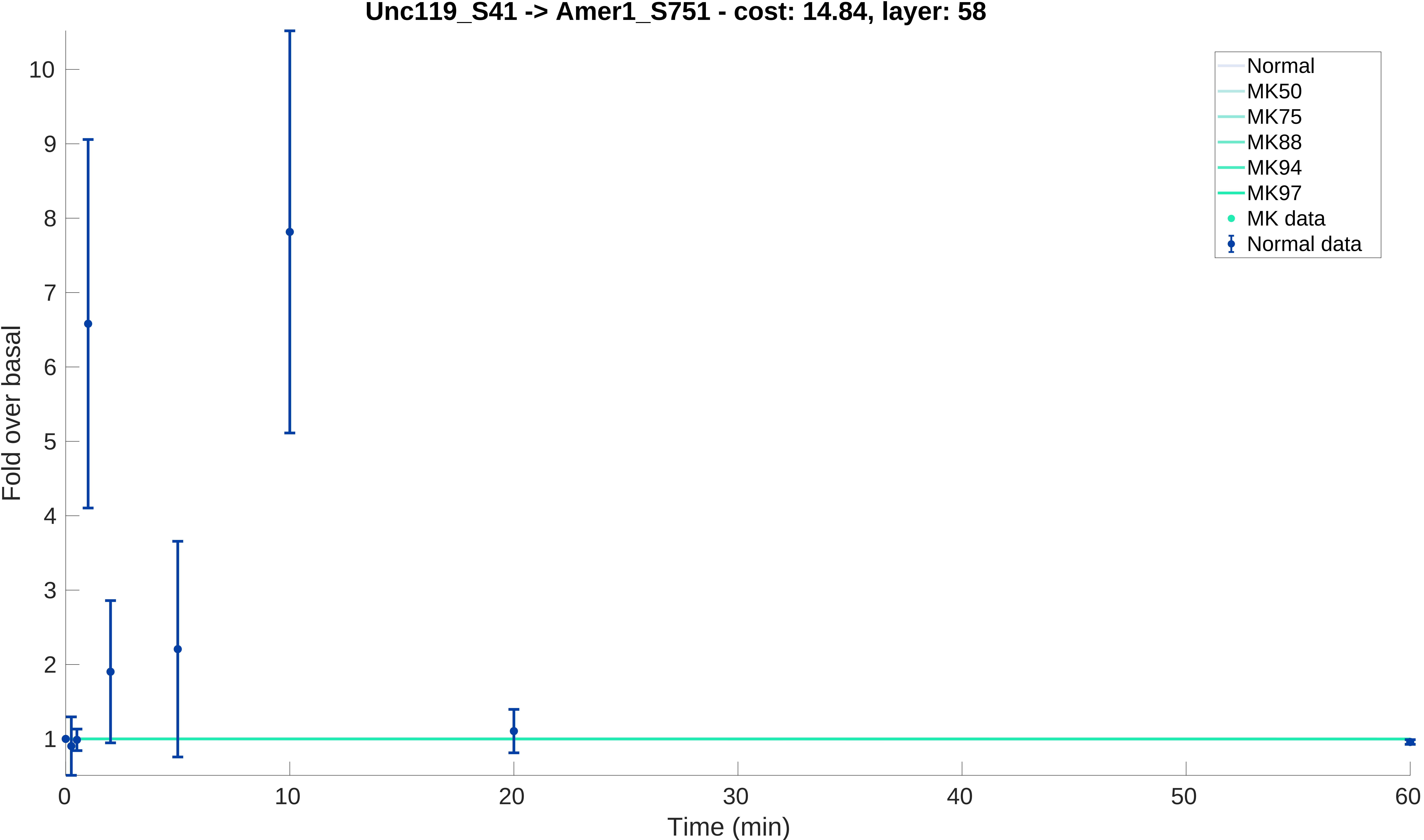
Unc119_S41 -> Amer1_S751 - cost: 14.84, layer: 58
Fold over basal
Time (min)
0
10
20
30
40
50
60
1
2
3
4
5
6
7
8
9
10
Normal
MK50
MK75
MK88
MK94
MK97
MK data
Normal data

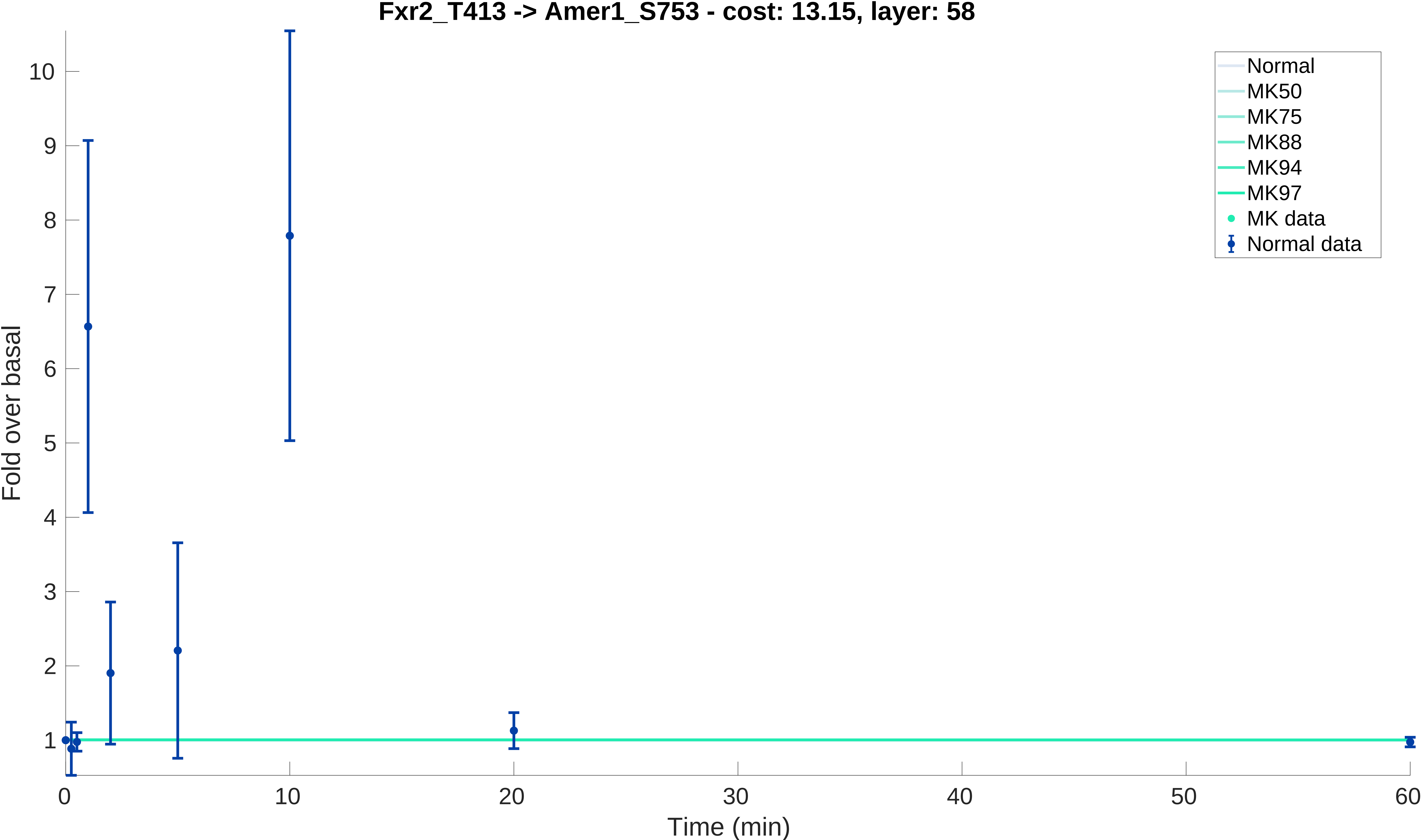
Fxr2_T413 -> Amer1_S753 - cost: 13.15, layer: 58
Fold over basal
Time (min)
0
10
20
30
40
50
60
1
2
3
4
5
6
7
8
9
10
Normal
MK50
MK75
MK88
MK94
MK97
MK data
Normal data

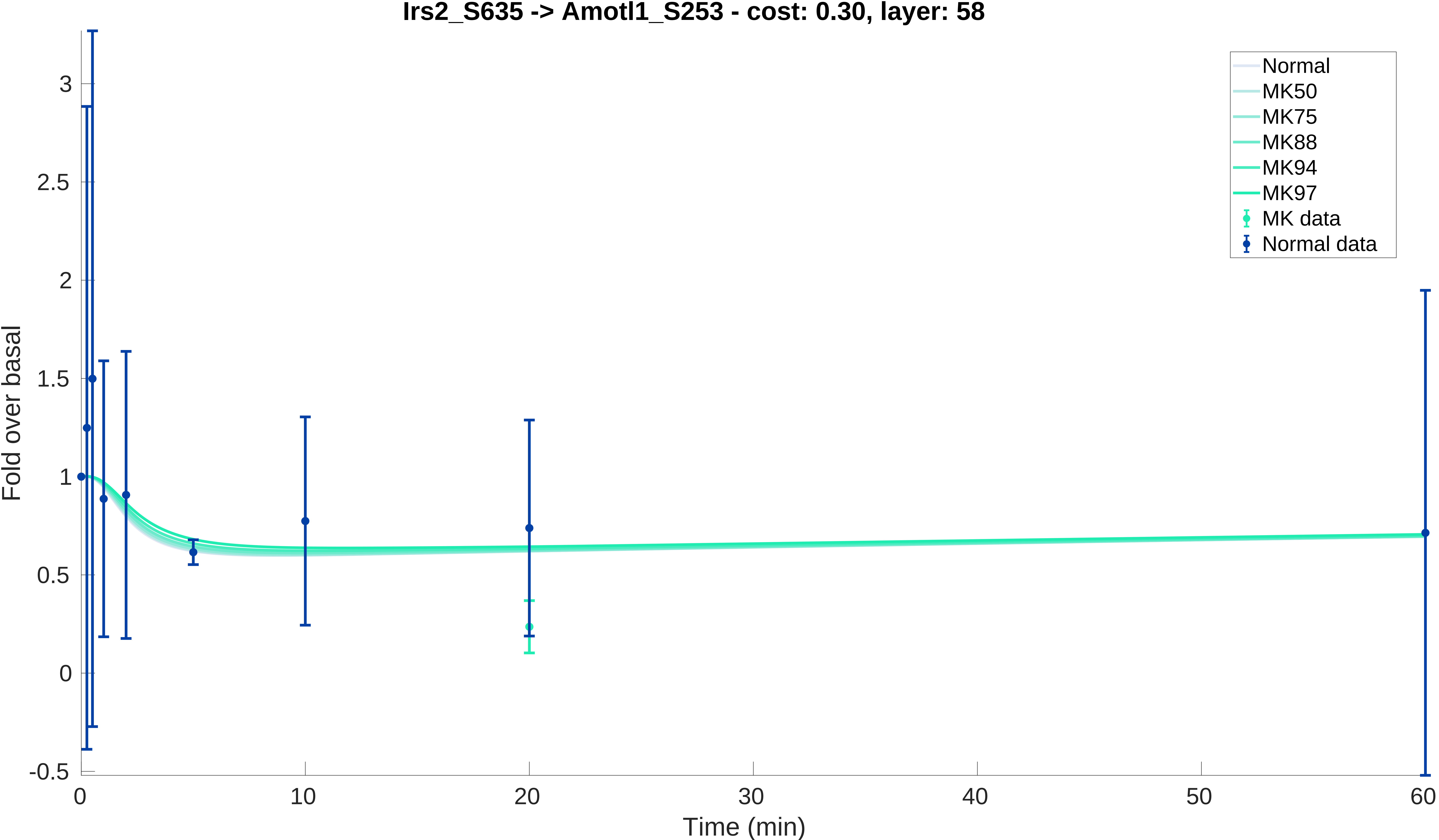

Irs2_S635 -> Amotl1_S253 - cost: 0.30, layer: 58
Normal
MK50
MK75
MK88
MK94
MK97
MK data
Normal data
Fold over basal
Time (min)
3
2.5
2
1.5
1
0.5
0
-0.5
0
10
20
30
40
50
60

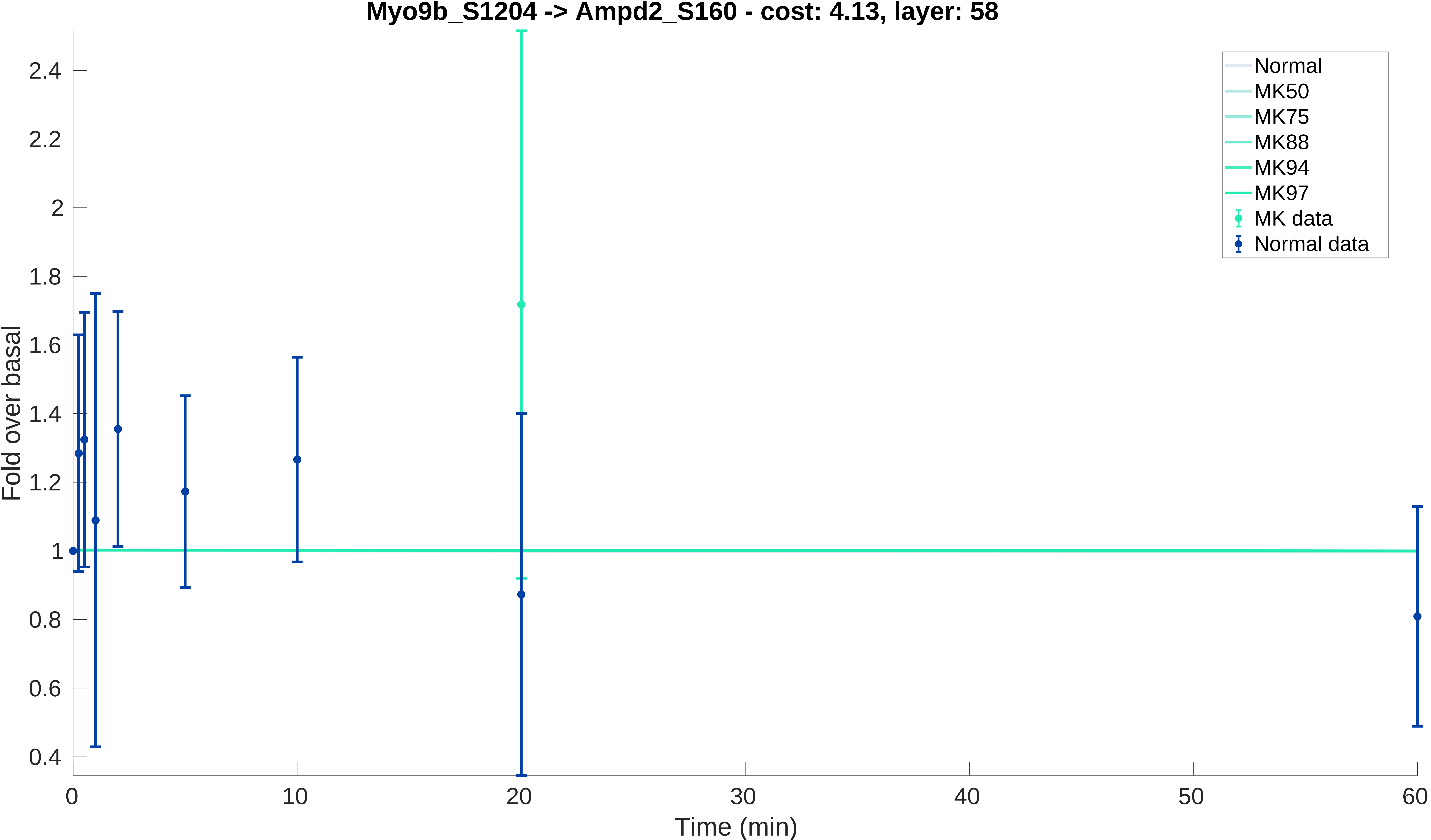

Myo9b_S1204 -> Ampd2_S160 - cost: 4.13, layer: 58
Fold over basal
Time (min)
Normal
MK50
MK75
MK88
MK94
MK97
MK data
Normal data
2.4
2.2
2
1.8
1.6
1.4
1.2
1
0.8
0.6
0.4
0
10
20
30
40
50
60

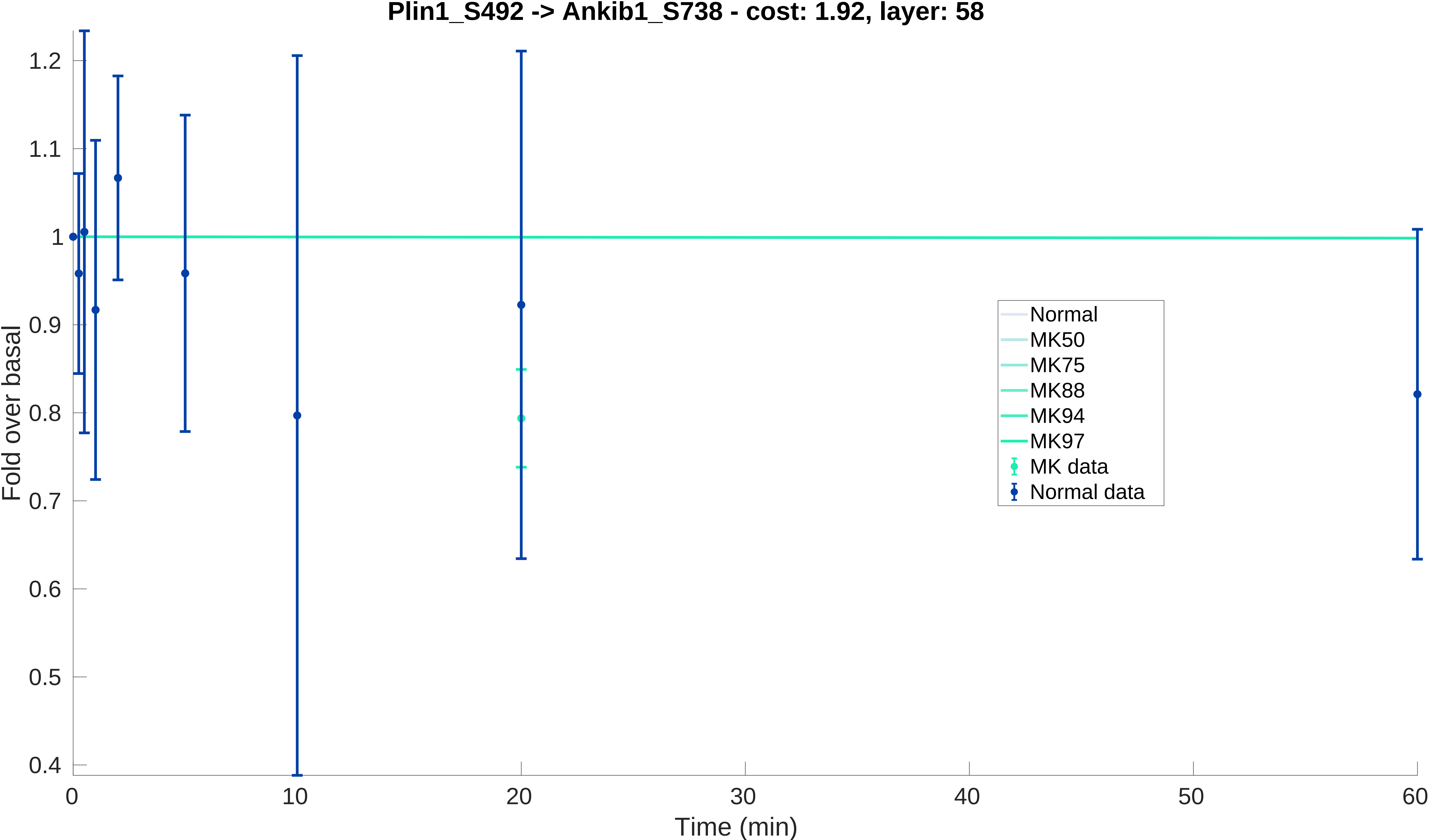

Plin1_S492 -> Ankib1_S738 - cost: 1.92, layer: 58
Fold over basal
Time (min)
0.4
0.5
0.6
0.7
0.8
0.9
1
1.1
1.2
0
10
20
30
40
50
60
Normal
MK50
MK75
MK88
MK94
MK97
MK data
Normal data

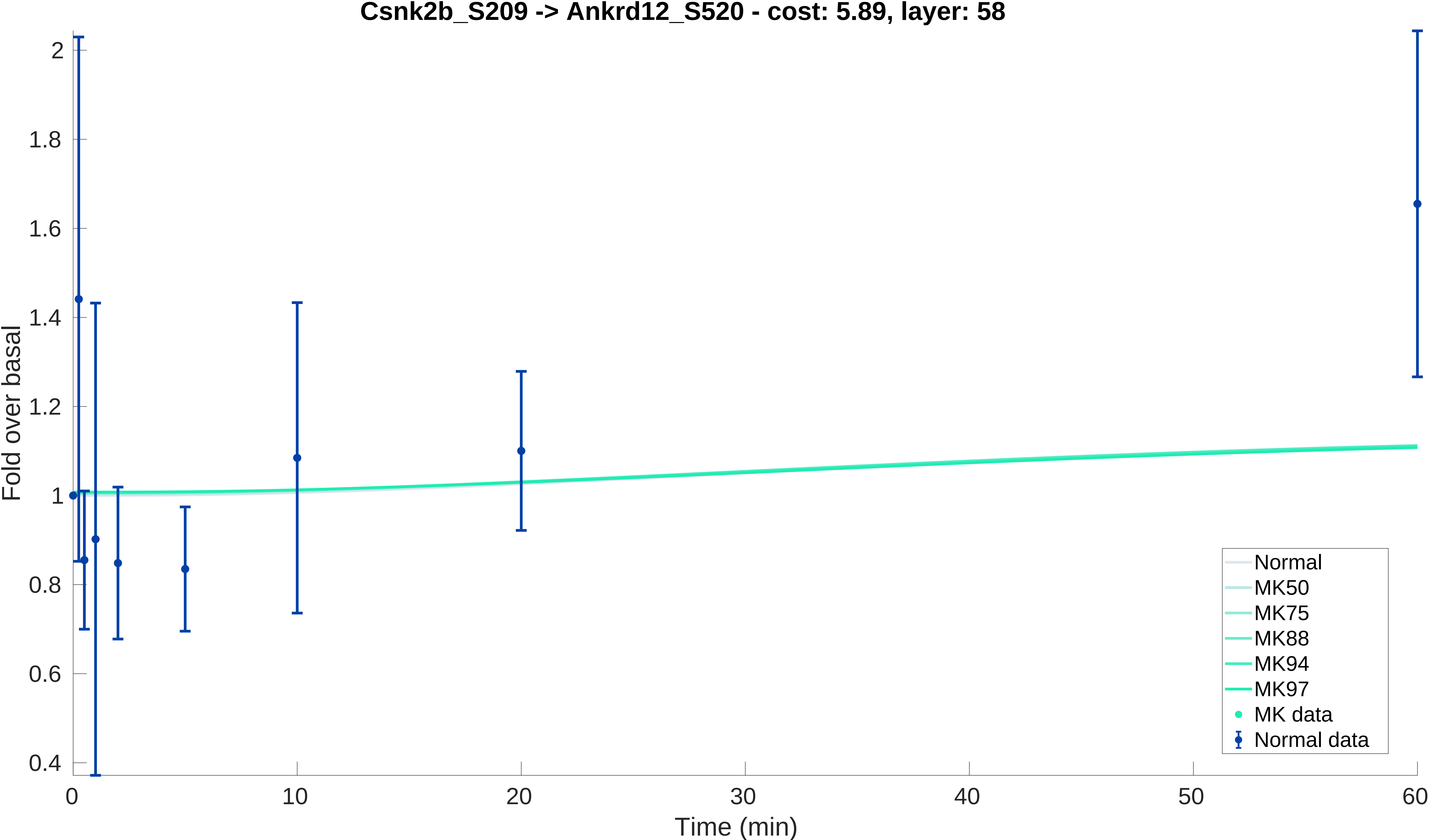

Csnk2b_S209 -> Ankrd12_S520 - cost: 5.89, layer: 58
Fold over basal
Time (min)
0.4
0.6
0.8
1
1.2
1.4
1.6
1.8
2
0
10
20
30
40
50
60
Normal
MK50
MK75
MK88
MK94
MK97
MK data
Normal data

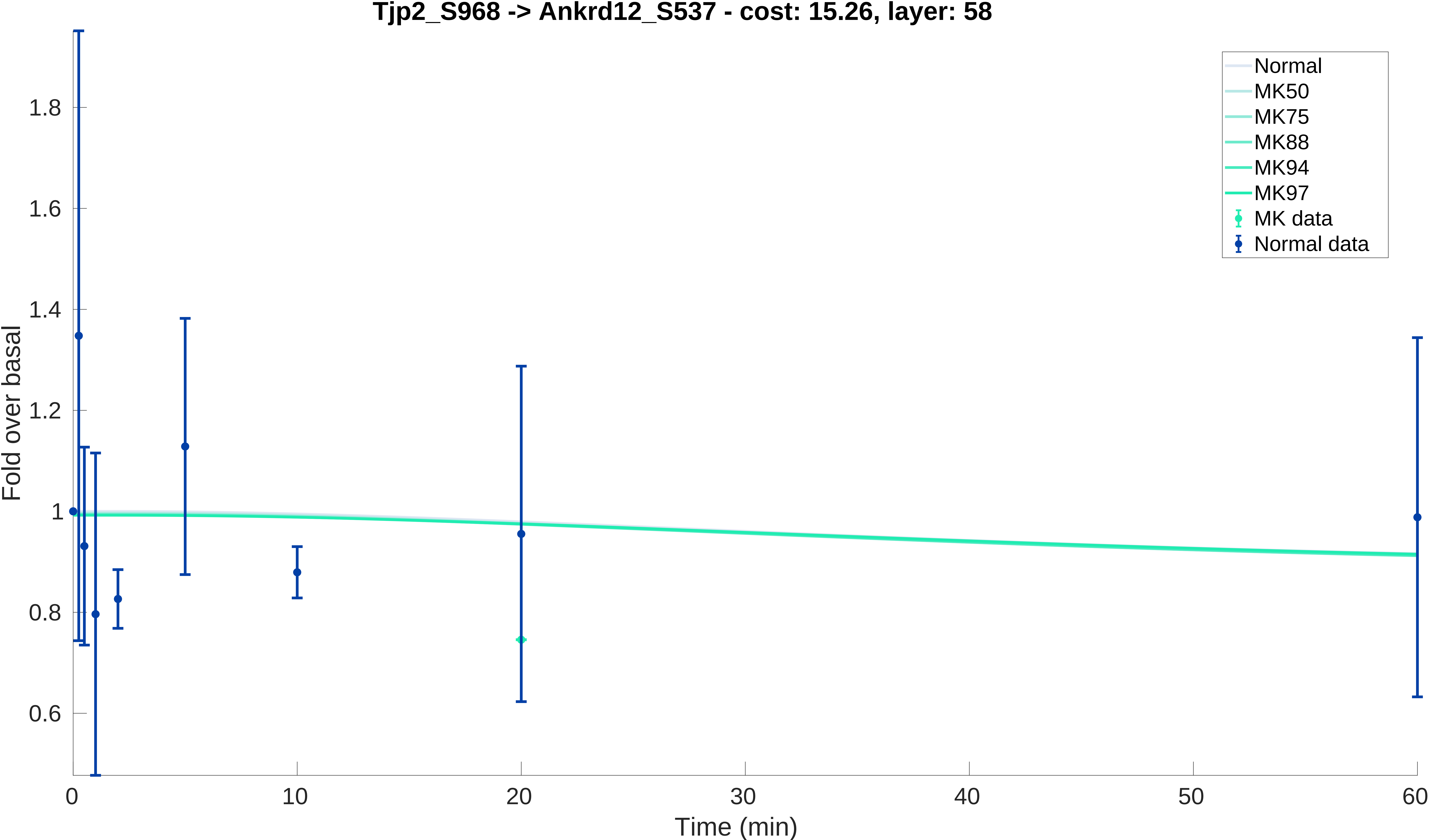

Tjp2_S968 -> Ankrd12_S537 - cost: 15.26, layer: 58
Normal
MK50
MK75
MK88
MK94
MK97
MK data
Normal data
Fold over basal
1.8
1.6
1.4
1.2
1
0.8
0.6
0
10
20
30
40
50
60
Time (min)

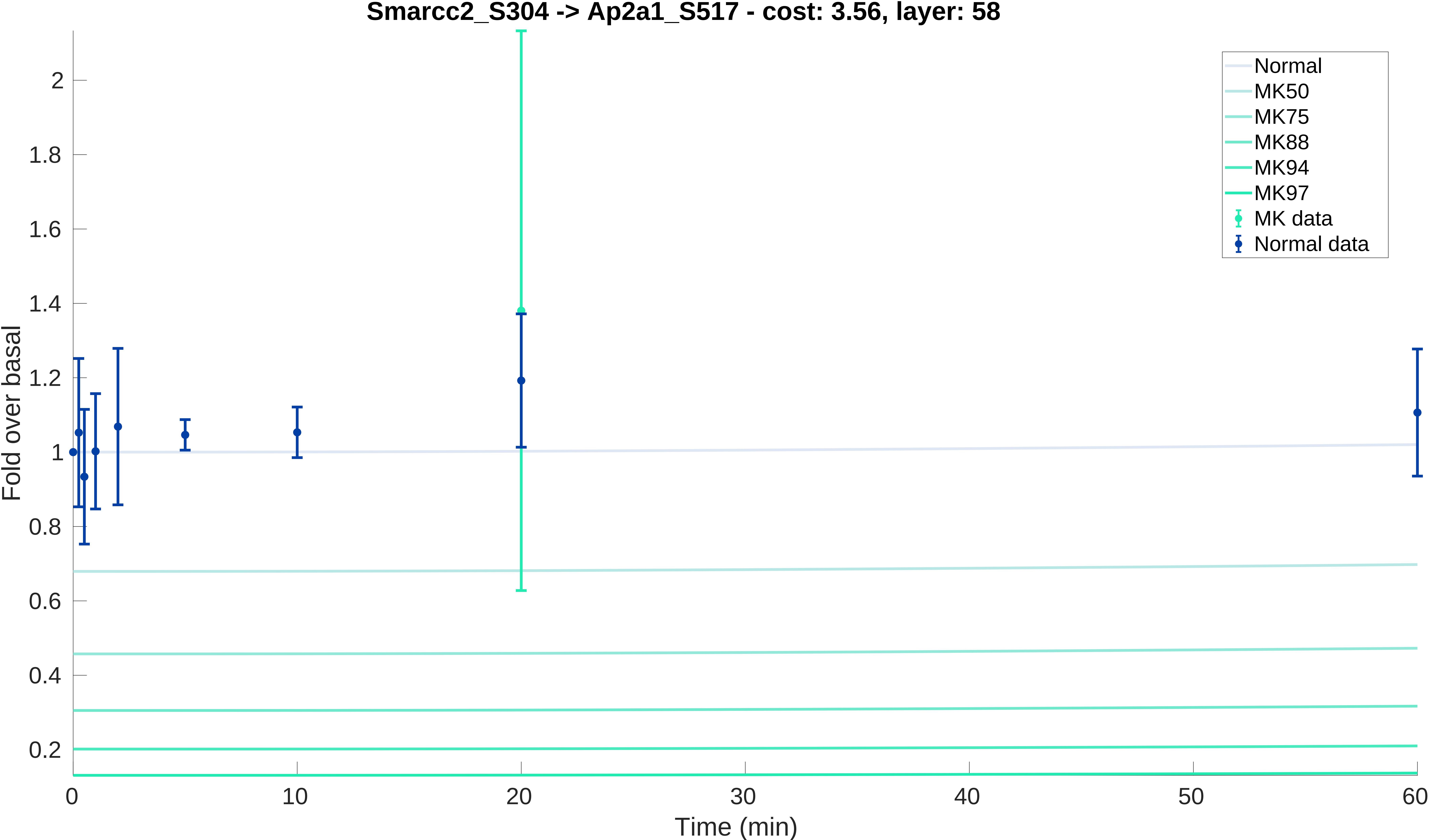
Smarcc2_S304 -> Ap2a1_S517 - cost: 3.56, layer: 58
Fold over basal
Time (min)
0
10
20
30
40
50
60
0.2
0.4
0.6
0.8
1
1.2
1.4
1.6
1.8
2
Normal
MK50
MK75
MK88
MK94
MK97
MK data
Normal data

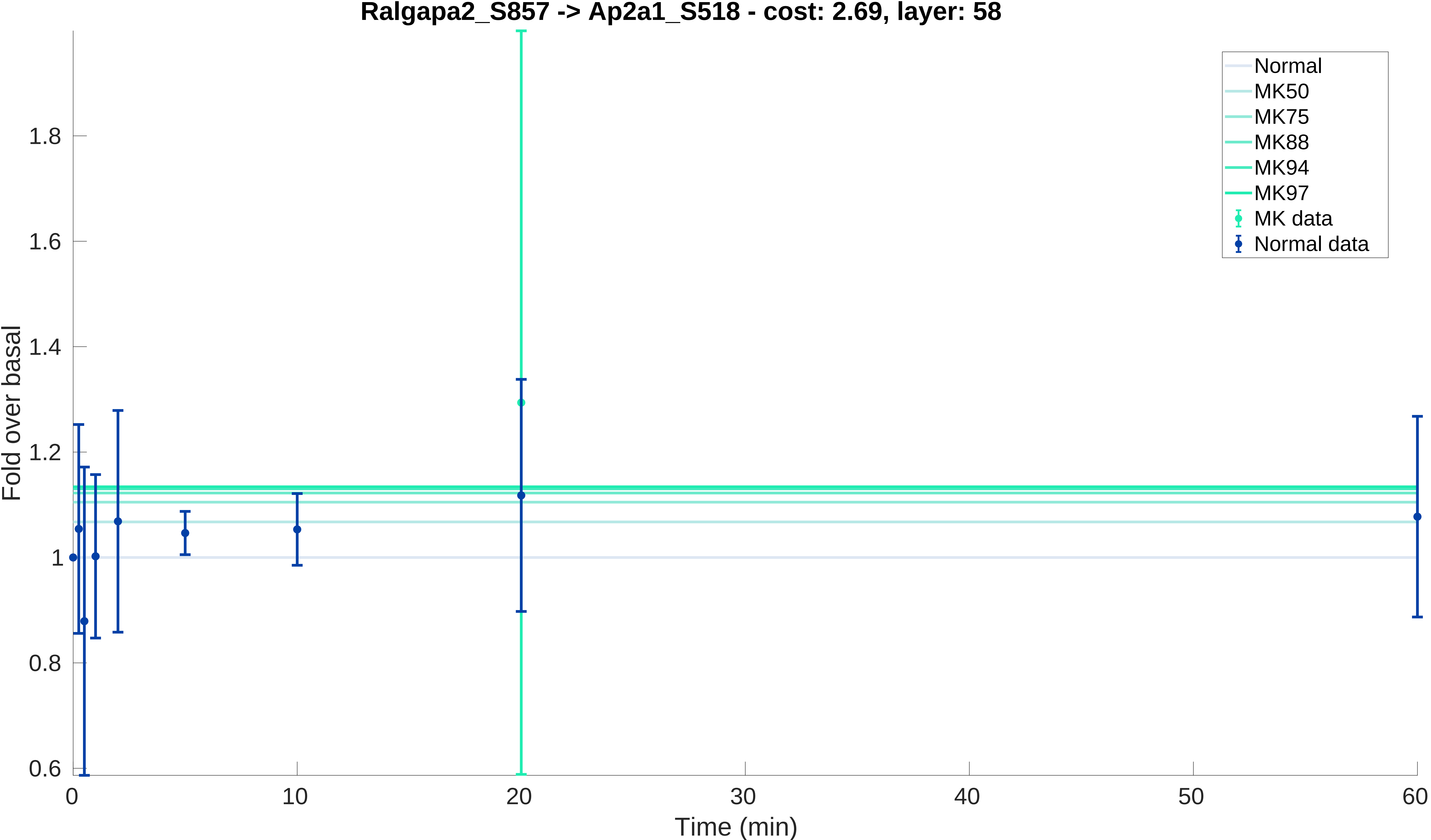

Ralgapa2_S857 -> Ap2a1_S518 - cost: 2.69, layer: 58
Fold over basal
Time (min)
0.6
0.8
1
1.2
1.4
1.6
1.8
0
10
20
30
40
50
60
Normal
MK50
MK75
MK88
MK94
MK97
MK data
Normal data

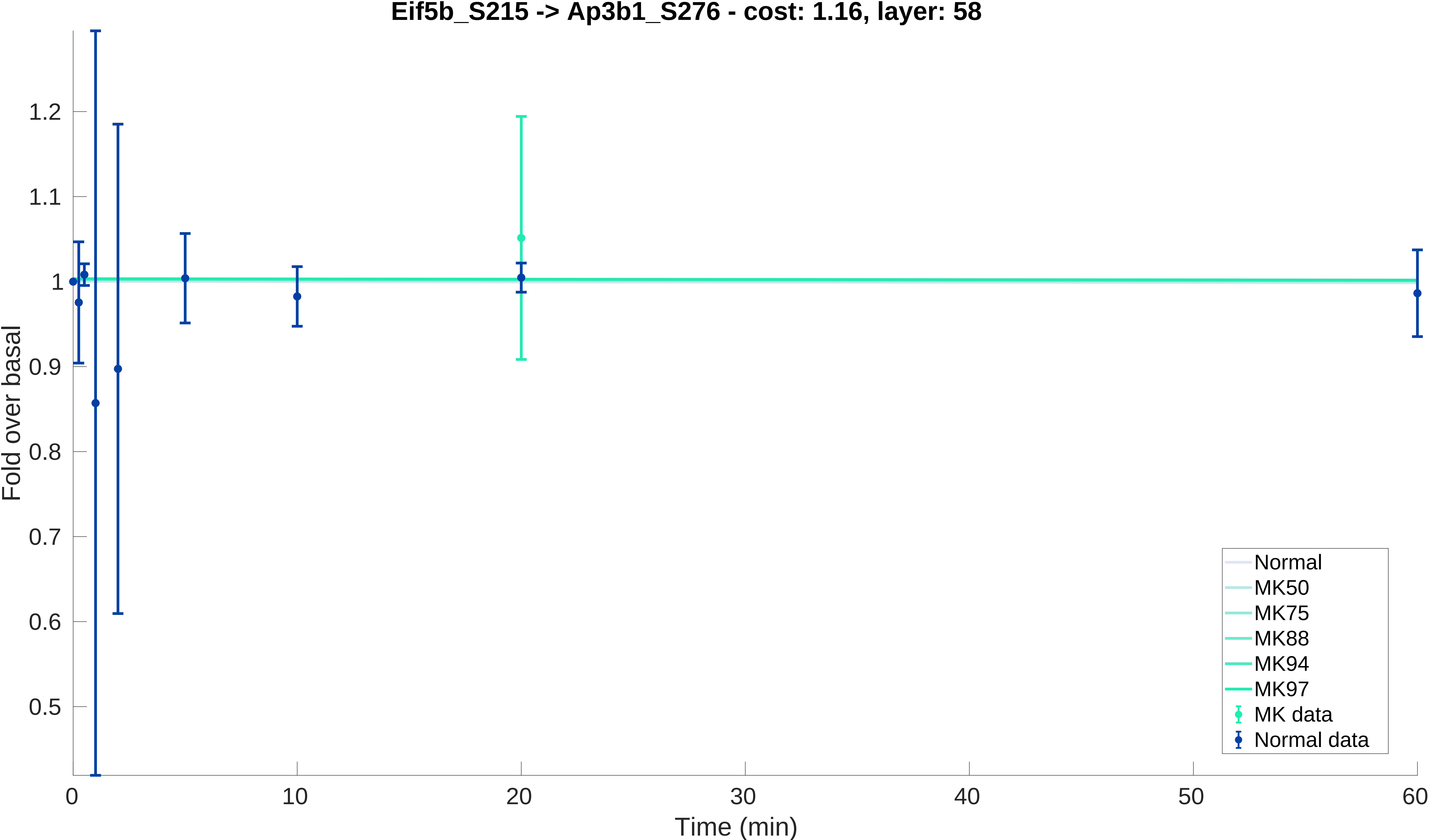

Eif5b_S215 -> Ap3b1_S276 - cost: 1.16, layer: 58
Fold over basal
Time (min)
1.2
1.1
1
0.9
0.8
0.7
0.6
0.5
0
10
20
30
40
50
60
Normal
MK50
MK75
MK88
MK94
MK97
MK data
Normal data

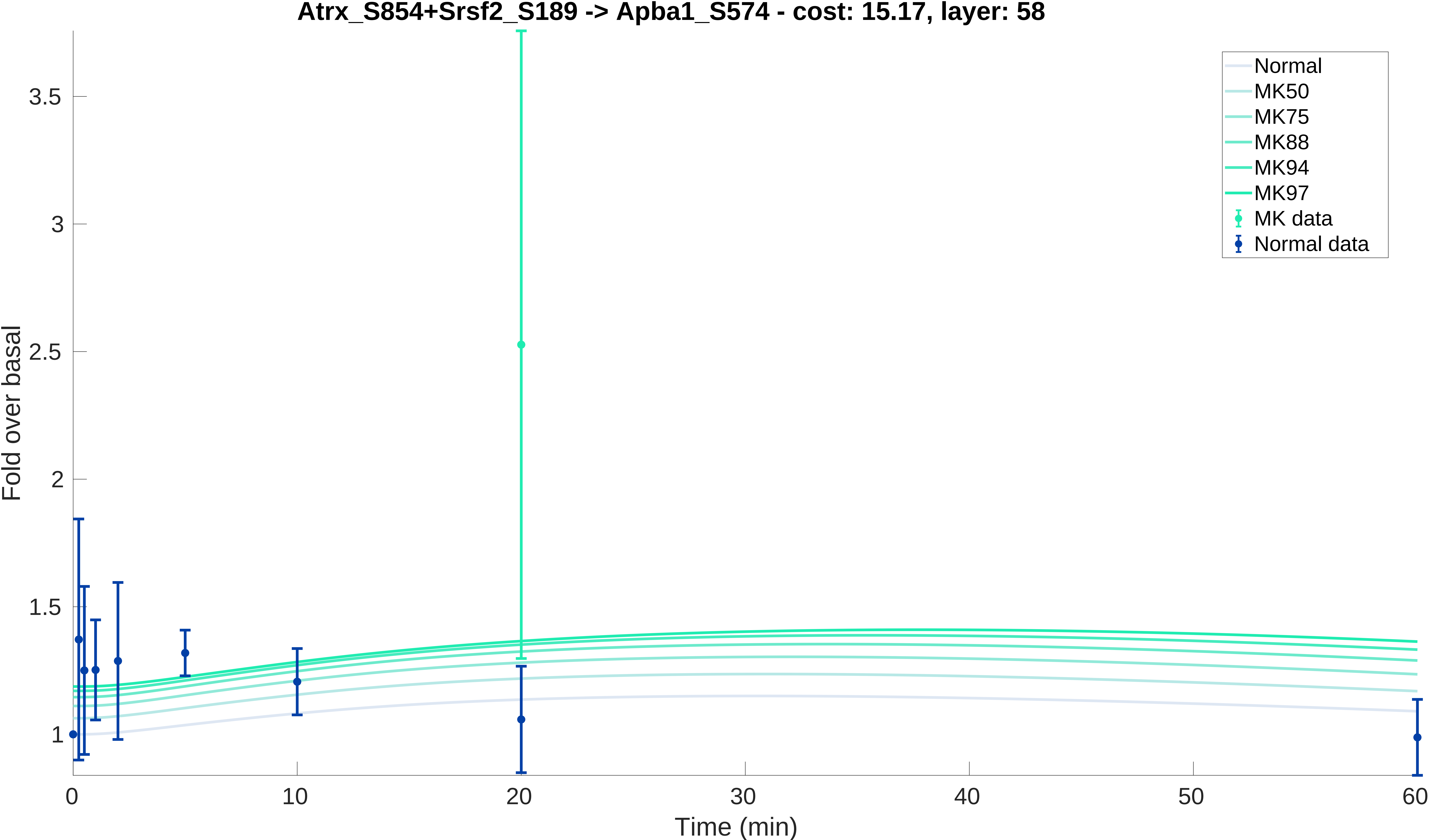
Atrx_S854+Srsf2_S189 -> Apba1_S574 - cost: 15.17, layer: 58
Fold over basal
3.5
3
2.5
2
1.5
1
0
10
20
30
40
50
60
Time (min)
Normal
MK50
MK75
MK88
MK94
MK97
MK data
Normal data

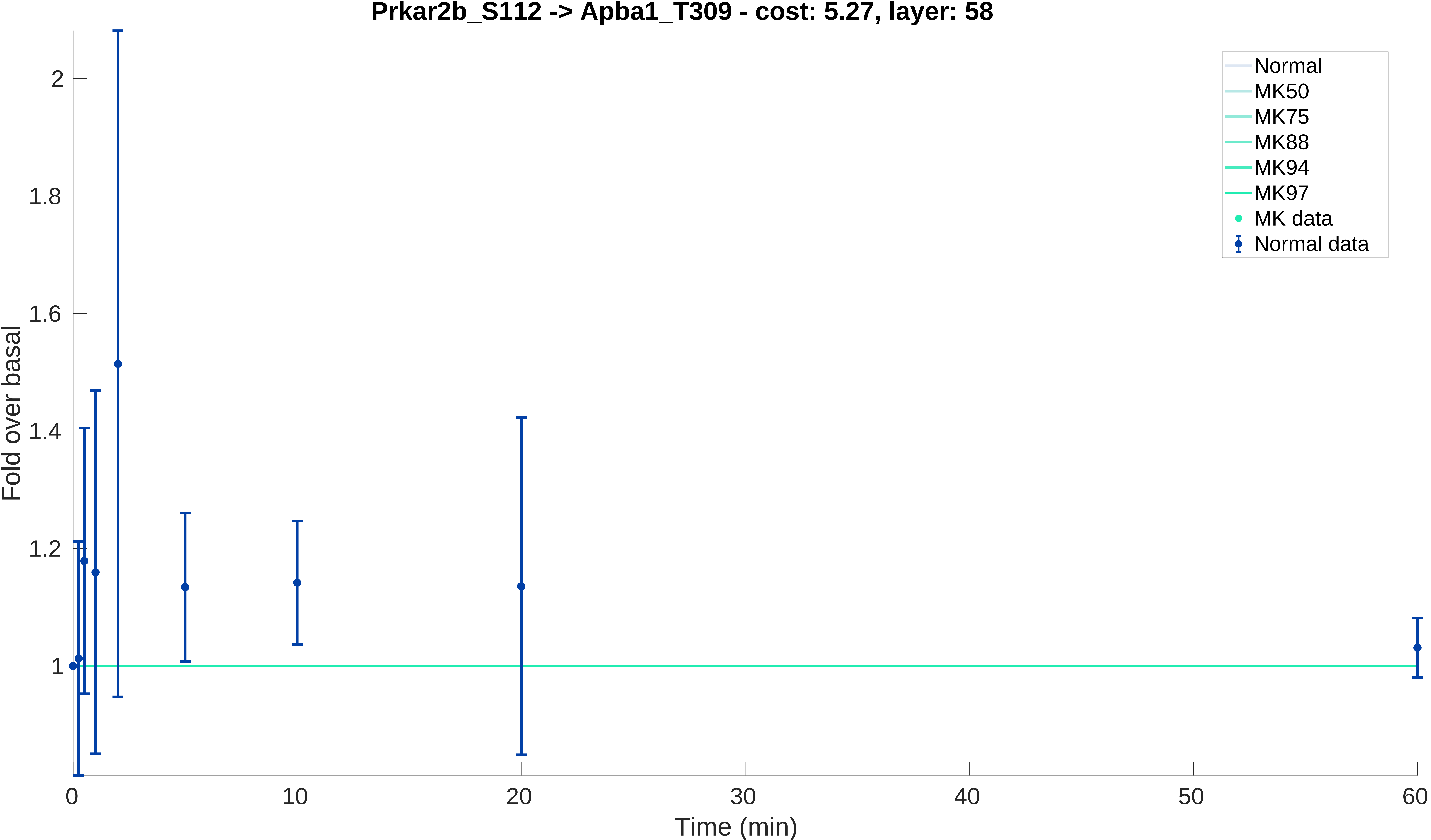

Prrkar2b_S112 -> Apba1_T309 - cost: 5.27, layer: 58
Fold over basal
Time (min)
2
1.8
1.6
1.4
1.2
1
0
10
20
30
40
50
60
Normal
MK50
MK75
MK88
MK94
MK97
MK data
Normal data

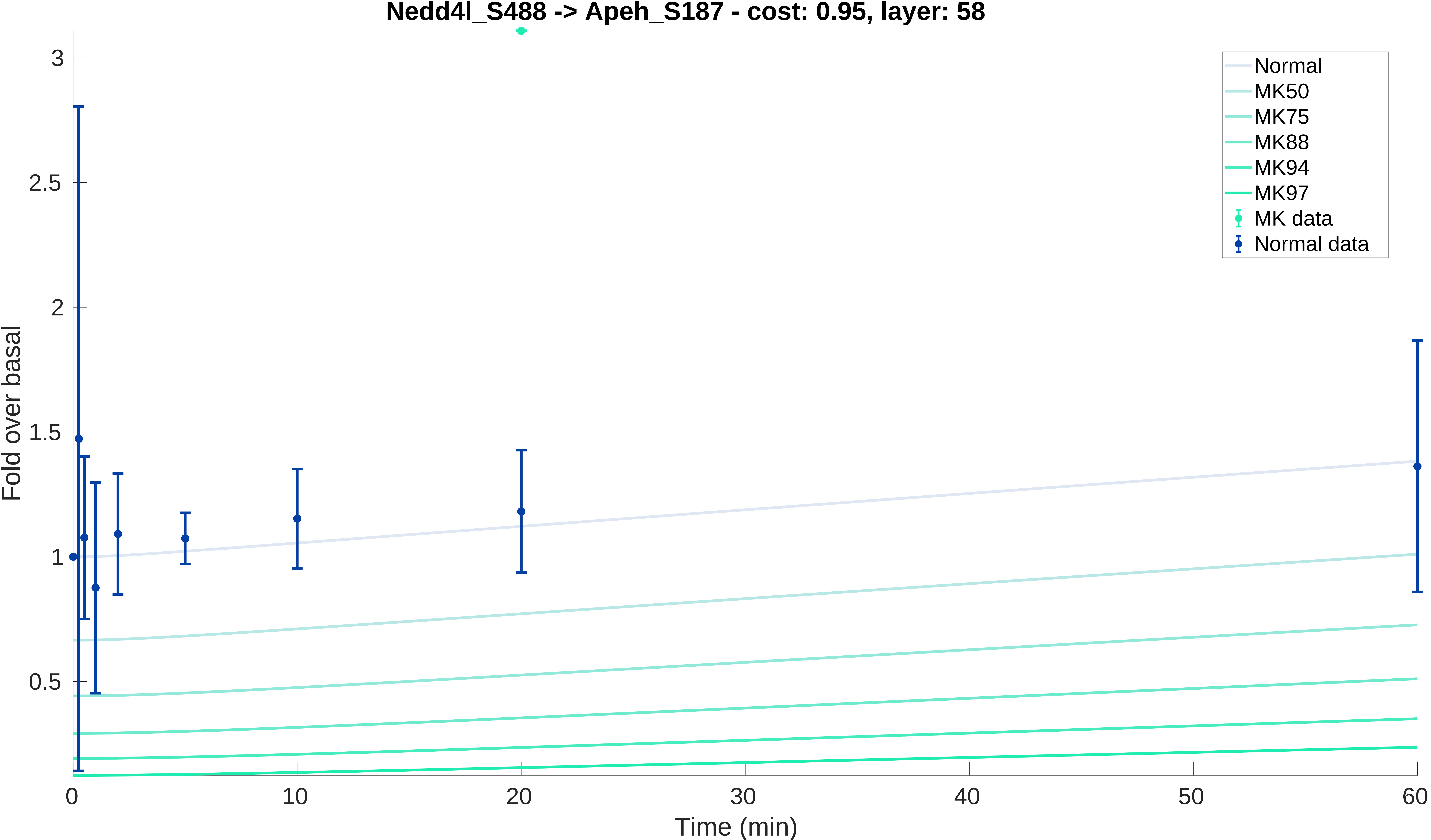
Nedd4l_S488 -> Apeh_S187 - cost: 0.95, layer: 58
Fold over basal
Time (min)
0
10
20
30
40
50
60
0.5
1
1.5
2
2.5
3
Normal
MK50
MK75
MK88
MK94
MK97
MK data
Normal data

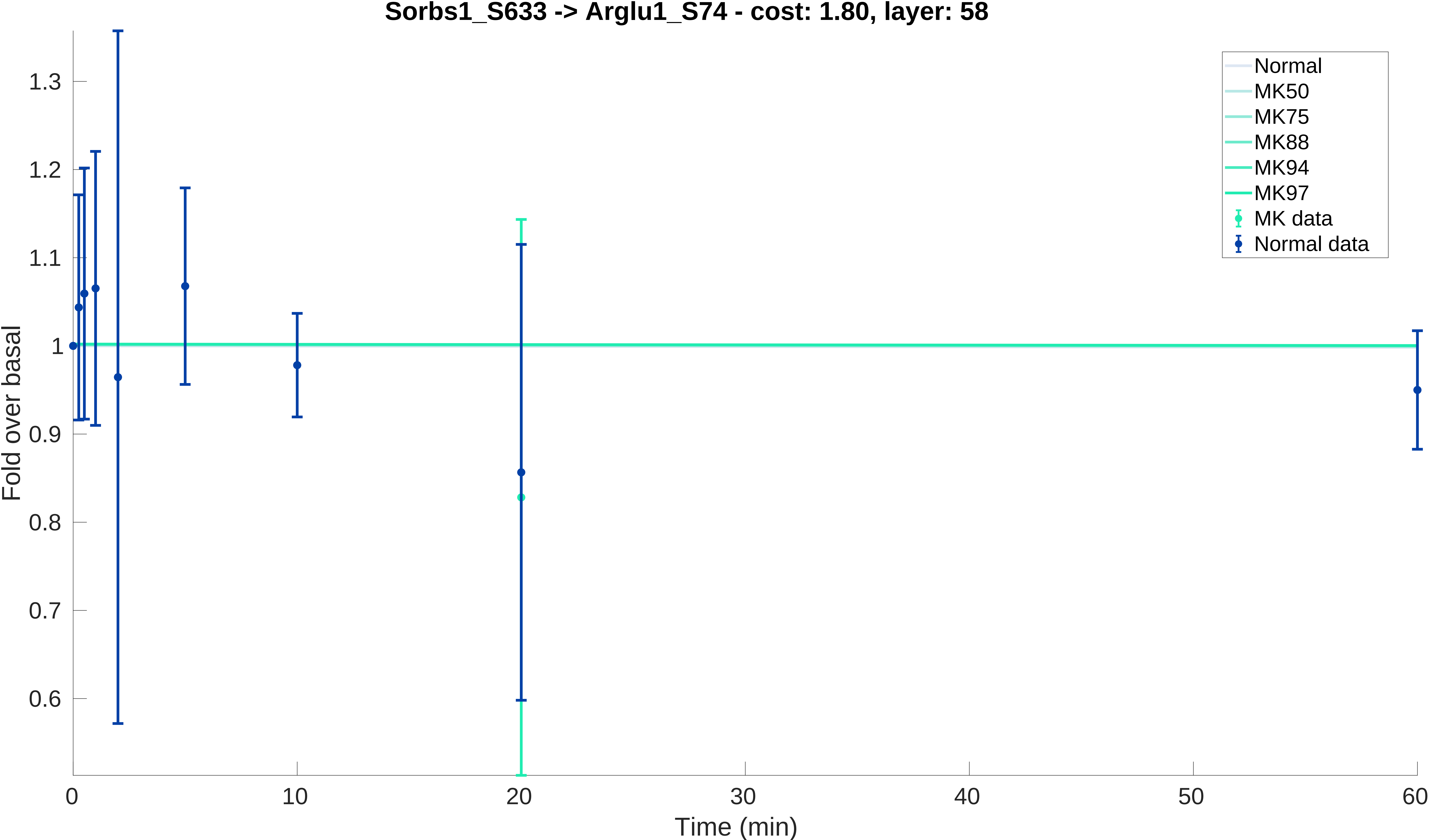

Sorbs1_S633 -> Arglu1_S74 - cost: 1.80, layer: 58
Fold over basal
Time (min)
1.3
1.2
1.1
1
0.9
0.8
0.7
0.6
0
10
20
30
40
50
60
Normal
MK50
MK75
MK88
MK94
MK97
MK data
Normal data

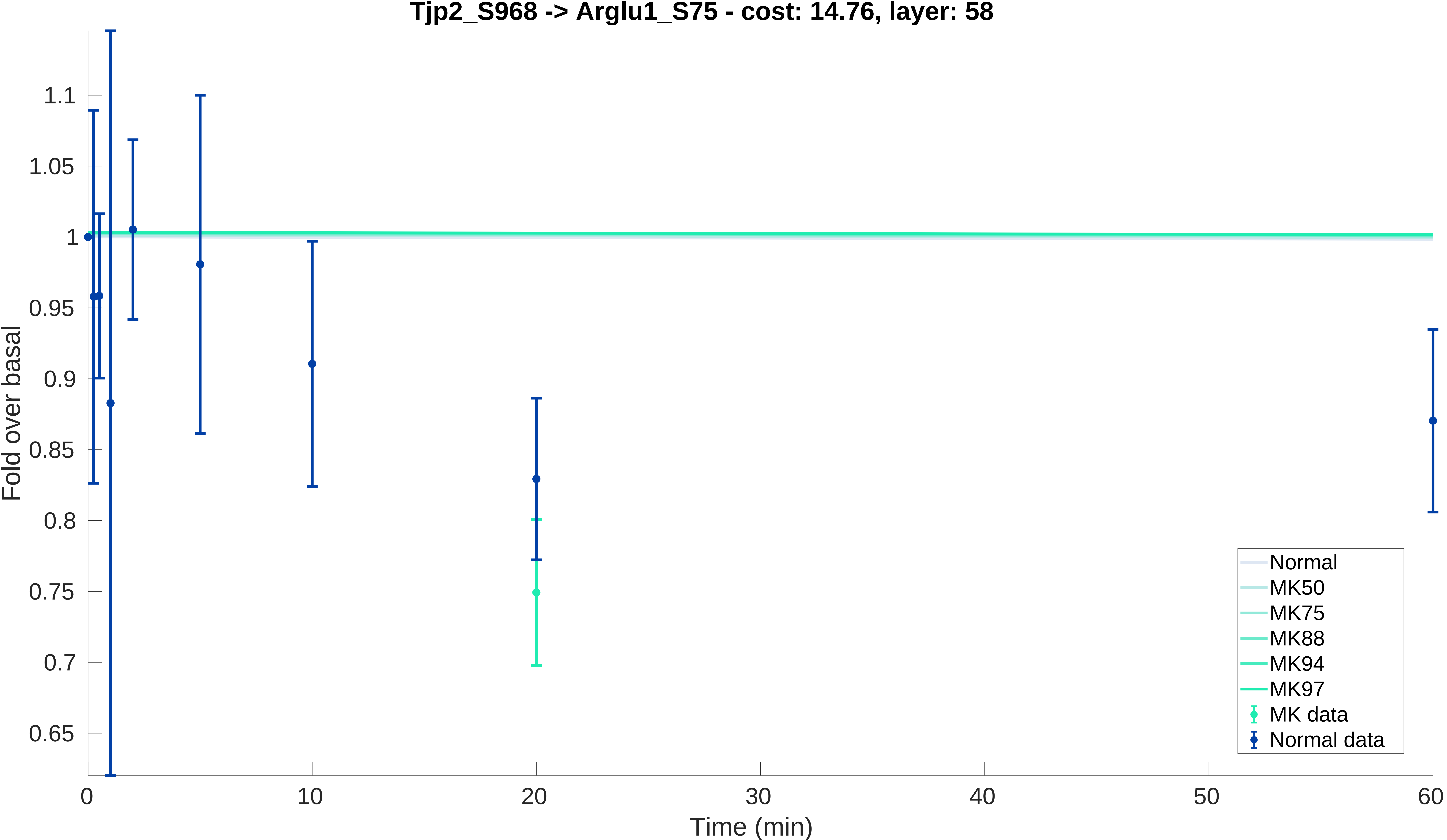
Tjp2_S968 -> Arglu1_S75 - cost: 14.76, layer: 58
Fold over basal
1.1
1.05
1
0.95
0.9
0.85
0.8
0.75
0.7
0.65
0
10
20
30
40
50
60
Time (min)
Normal
MK50
MK75
MK88
MK94
MK97
MK data
Normal data

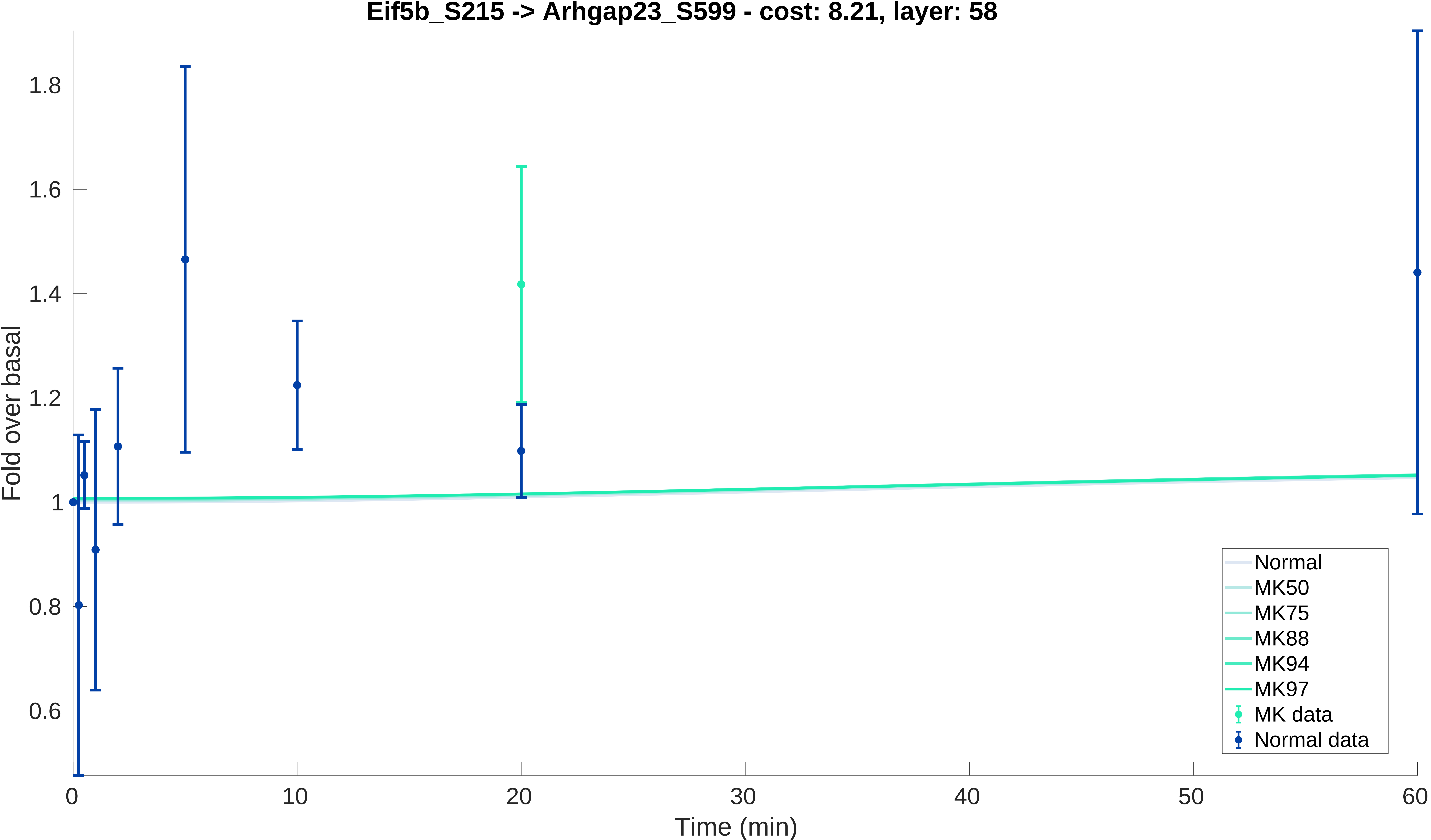
Eif5b_S215 -> Arhgap23_S599 - cost: 8.21, layer: 58
Fold over basal
Time (min)
0.6
0.8
1
1.2
1.4
1.6
1.8
0
10
20
30
40
50
60
Normal
MK50
MK75
MK88
MK94
MK97
MK data
Normal data

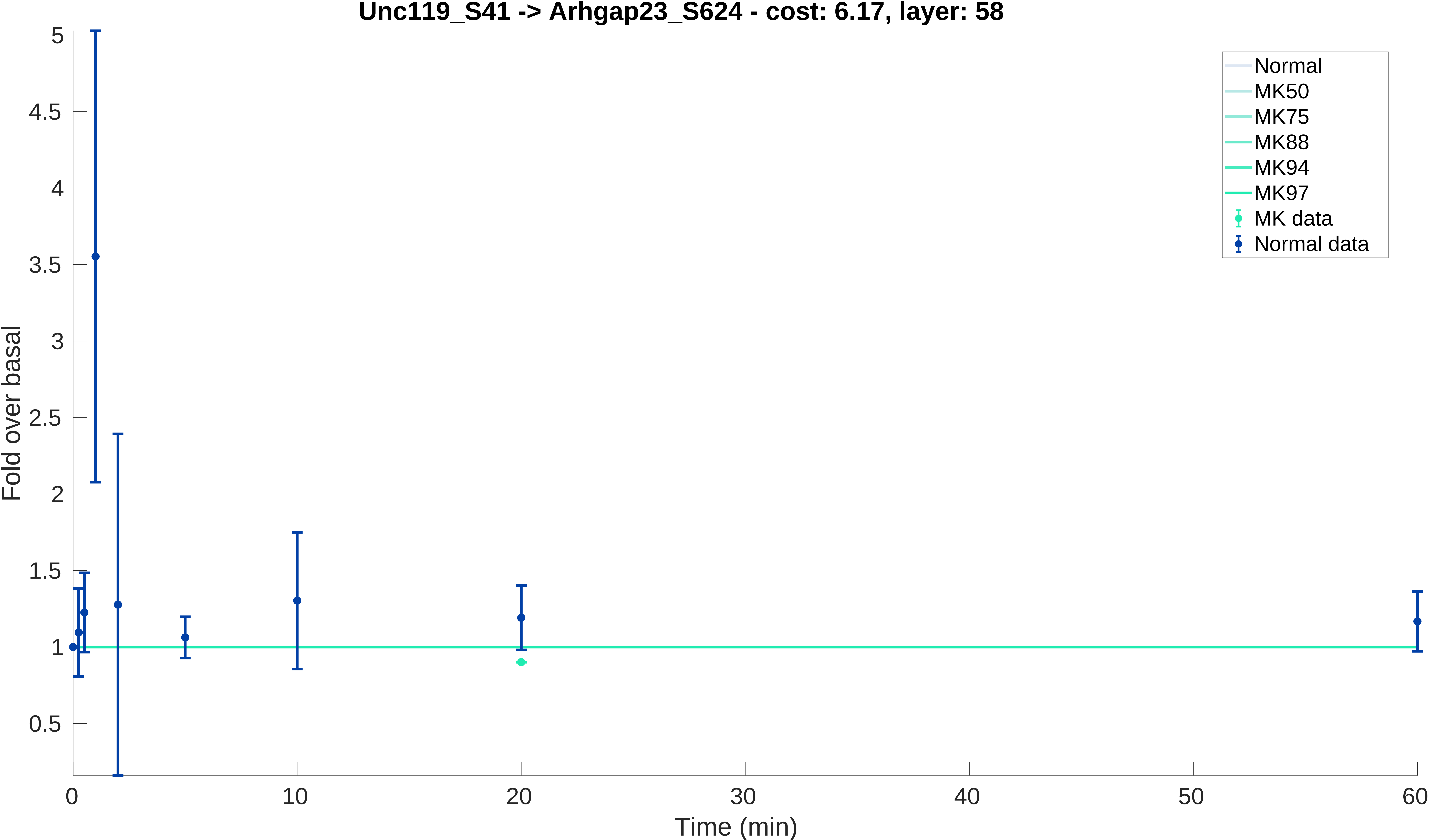

Unc119_S41 -> Arhgap23_S624 - cost: 6.17, layer: 58
Fold over basal
Time (min)
Normal
MK50
MK75
MK88
MK94
MK97
MK data
Normal data

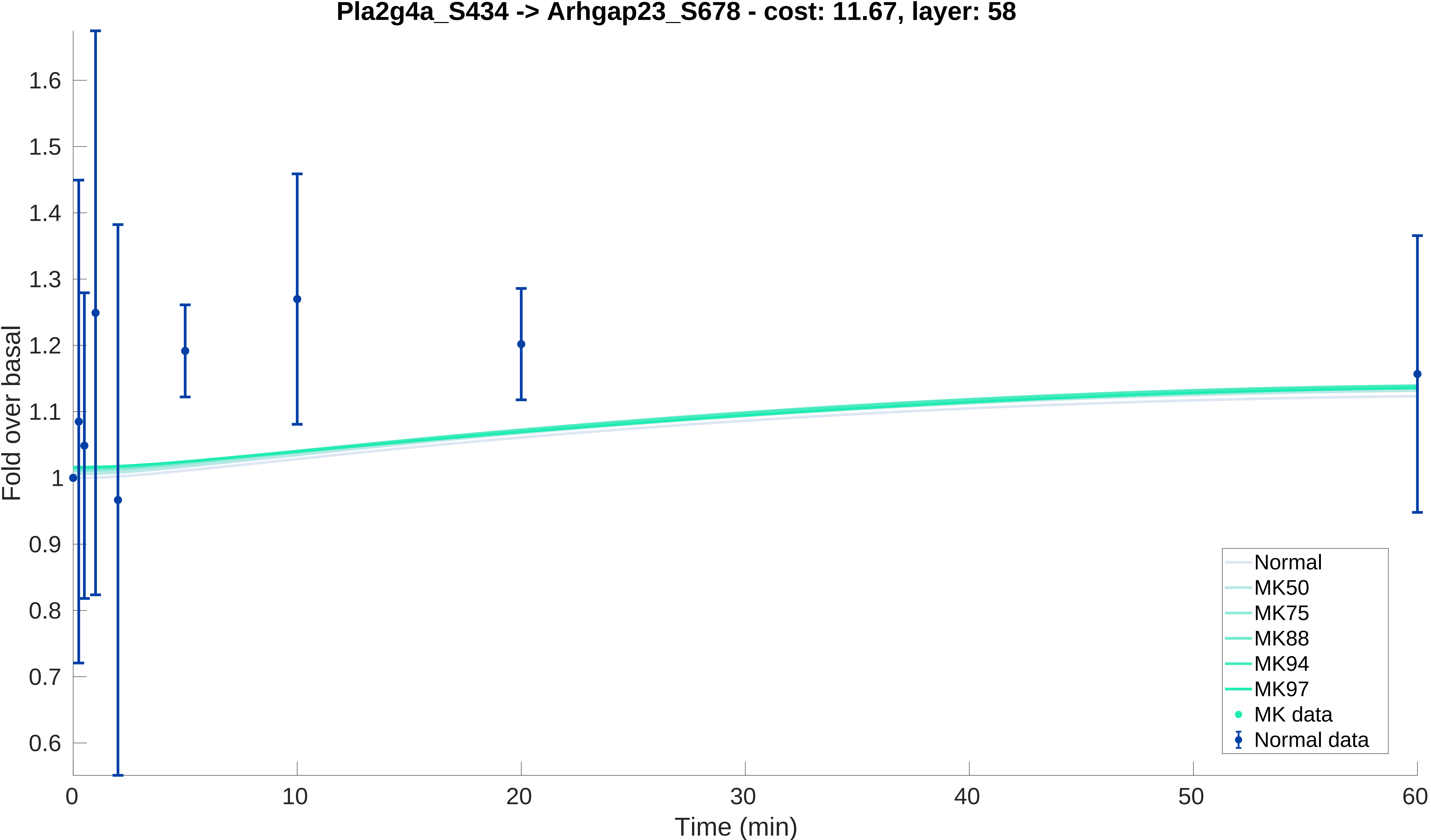

Pla2g4a_S434 -> Arhgap23_S678 - cost: 11.67, layer: 58
Fold over basal
1.6
1.5
1.4
1.3
1.2
1.1
1
0.9
0.8
0.7
0.6
0
10
20
30
40
50
60
Time (min)
Normal
MK50
MK75
MK88
MK94
MK97
MK data
Normal data

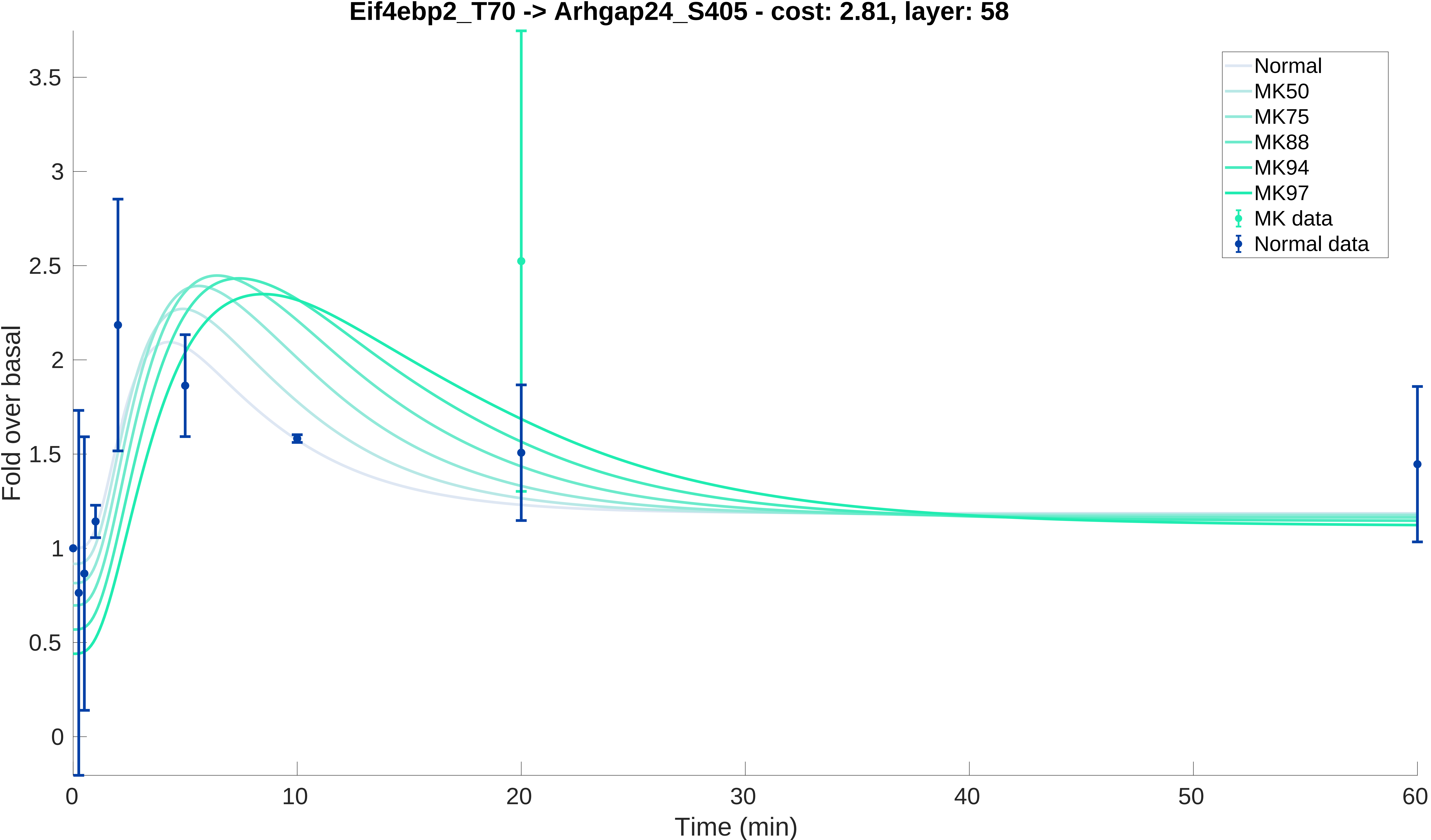

Eif4ebp2_T70 -> Arhgap24_S405 - cost: 2.81, layer: 58
Fold over basal
Time (min)
0
0.5
1
1.5
2
2.5
3
3.5
0
10
20
30
40
50
60
Normal
MK50
MK75
MK88
MK94
MK97
MK data
Normal data

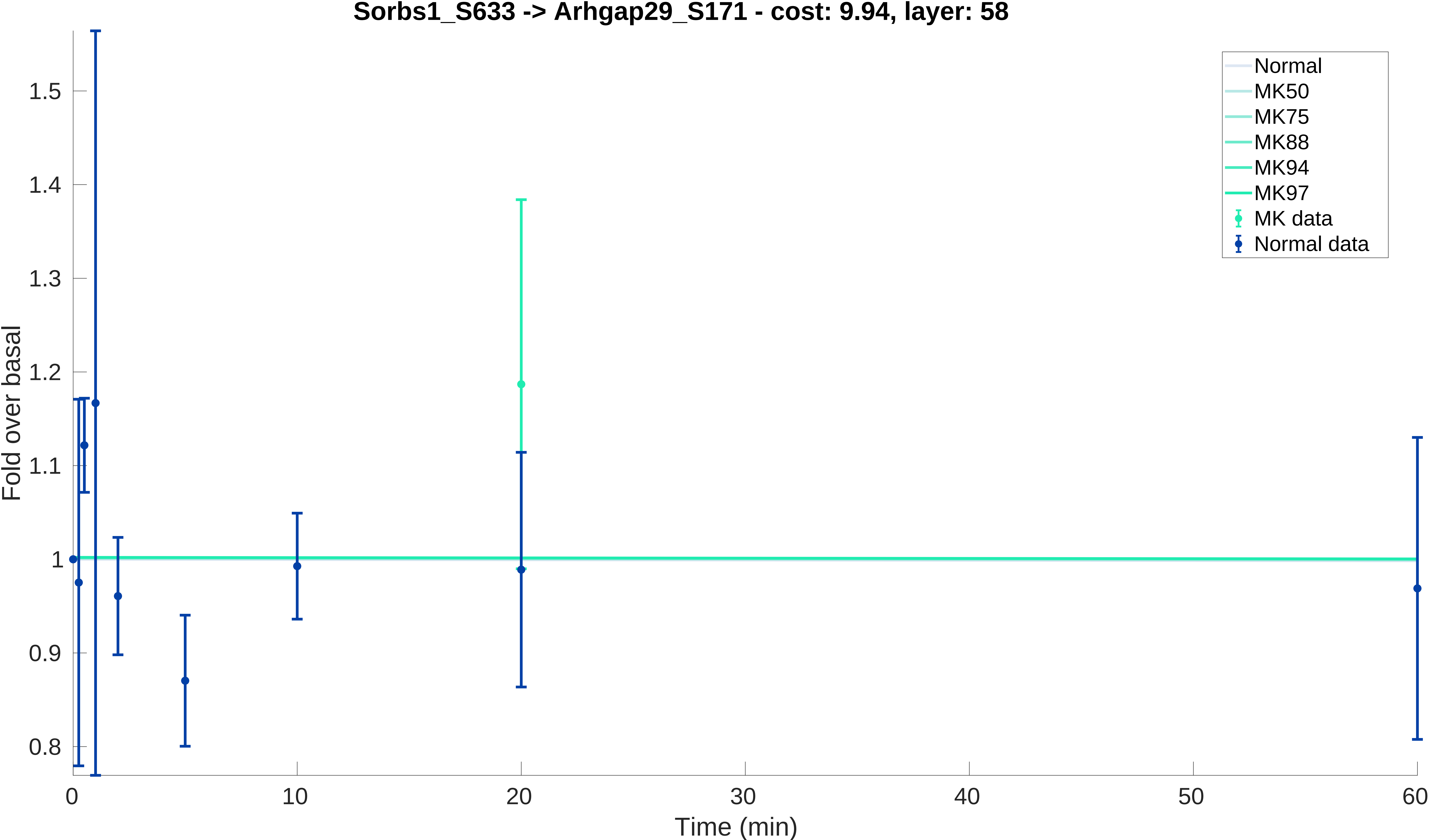

Sorbs1_S633 -> Arhgap29_S171 - cost: 9.94, layer: 58
Fold over basal
Time (min)
0.8
0.9
1
1.1
1.2
1.3
1.4
1.5
0
10
20
30
40
50
60
Normal
MK50
MK75
MK88
MK94
MK97
MK data
Normal data

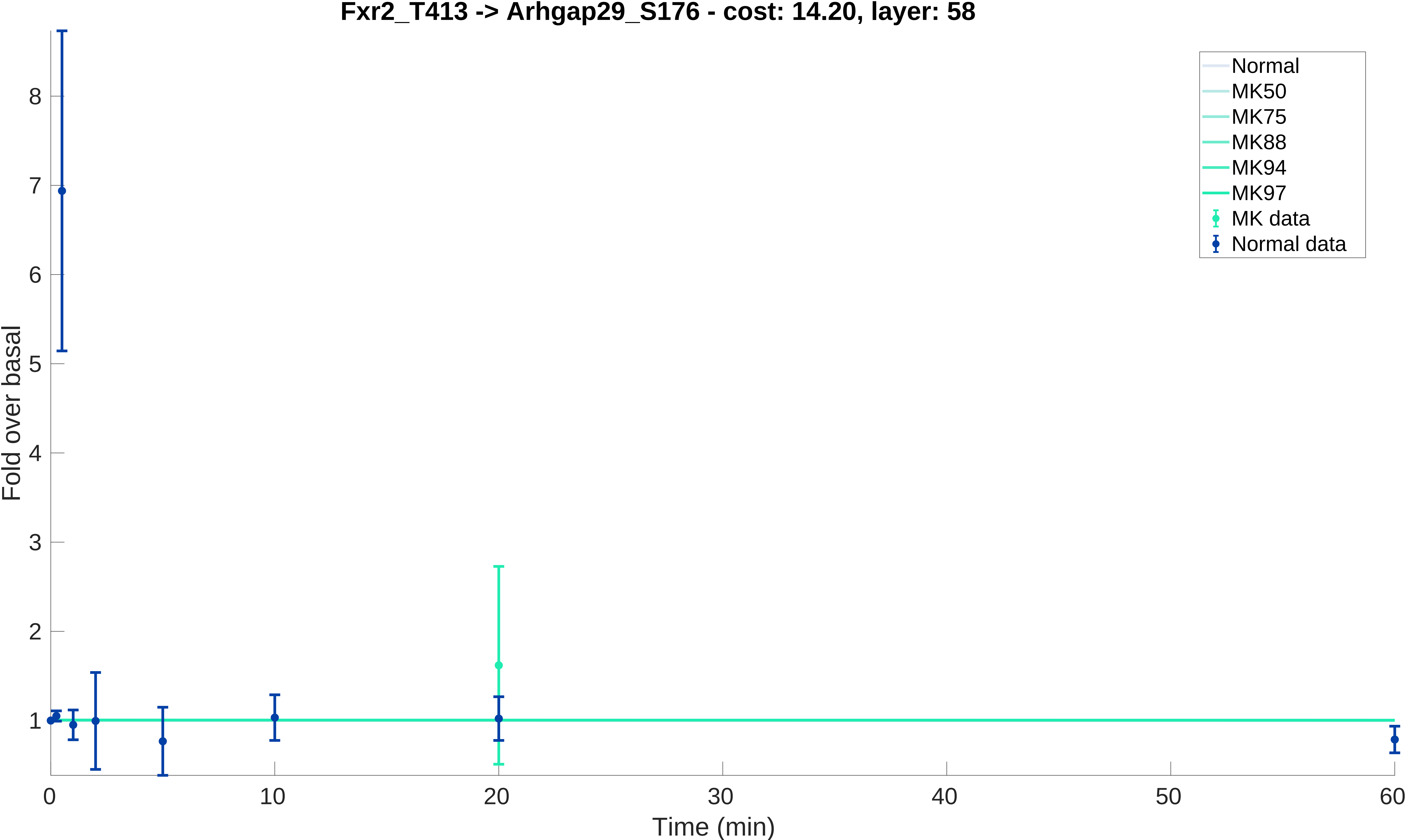

Fxr2_T413 -> Arhgap29_S176 - cost: 14.20, layer: 58
Fold over basal
Time (min)
0
10
20
30
40
50
60
1
2
3
4
5
6
7
8
Normal
MK50
MK75
MK88
MK94
MK97
MK data
Normal data

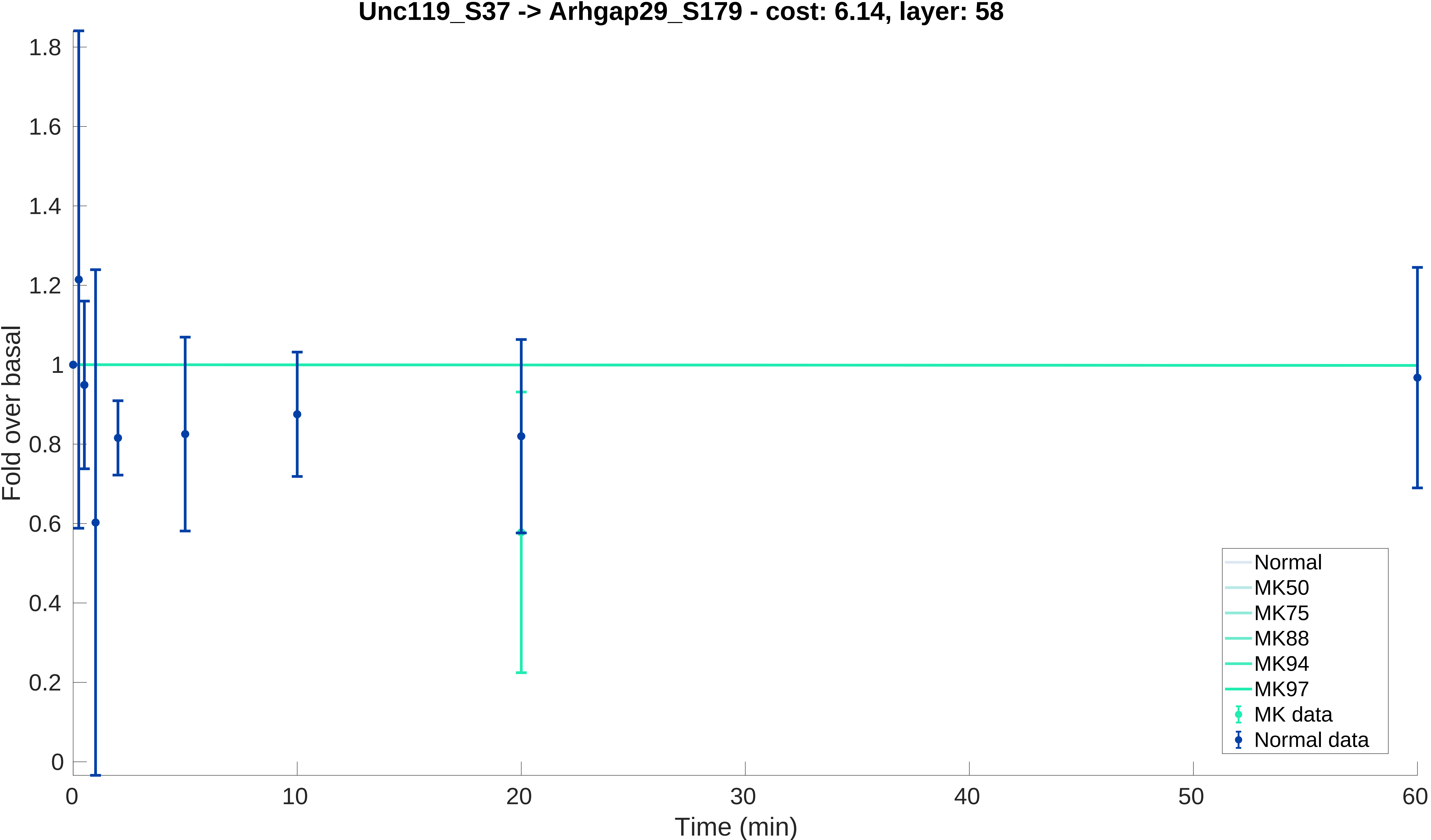

Unc119_S37 -> Arhgap29_S179 - cost: 6.14, layer: 58
Fold over basal
1.8
1.6
1.4
1.2
1
0.8
0.6
0.4
0.2
0
0
10
20
30
40
50
60
Time (min)
Normal
MK50
MK75
MK88
MK94
MK97
MK data
Normal data

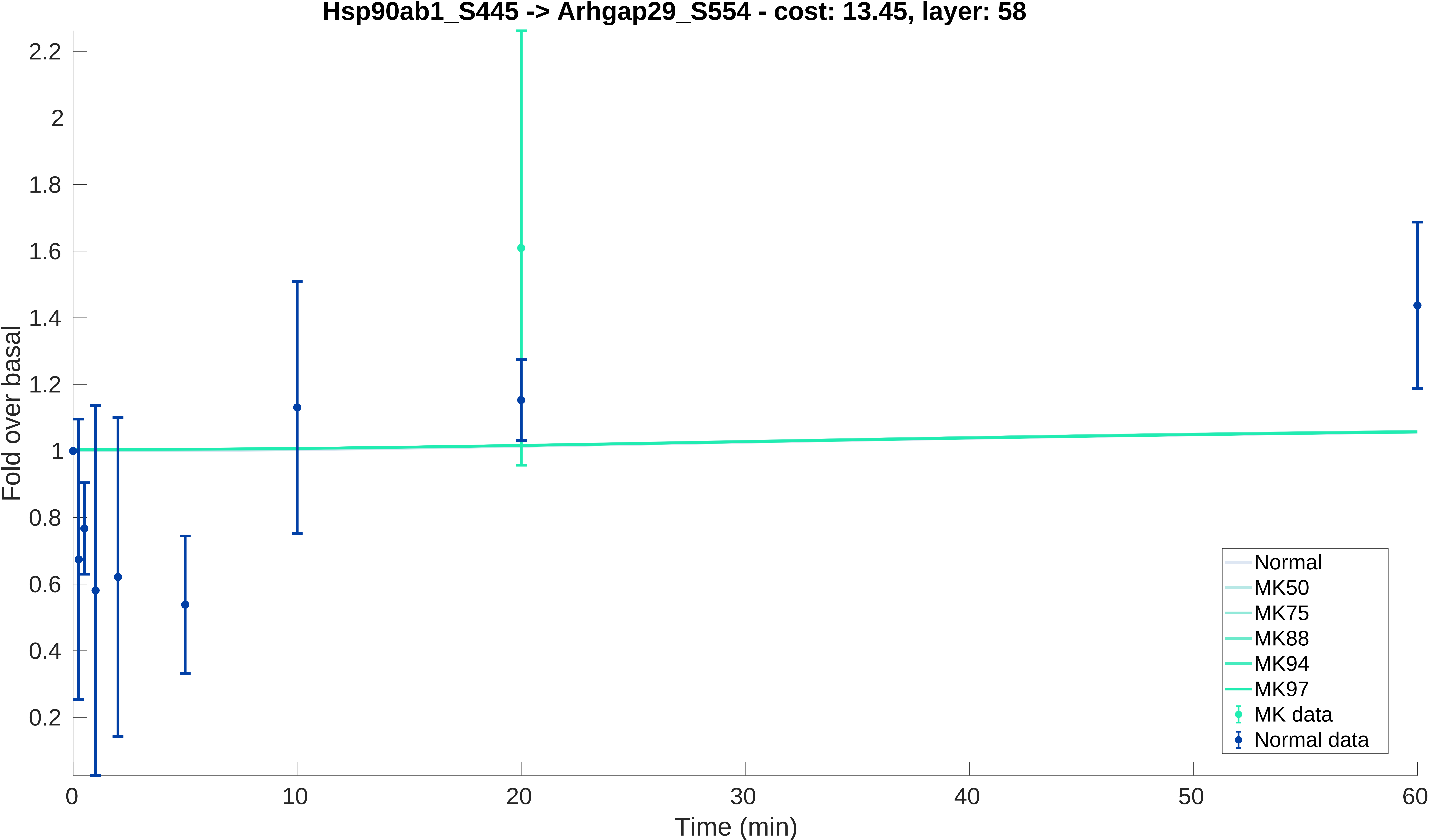

Hsp90ab1_S445 -> Arhgap29_S554 - cost: 13.45, layer: 58
Fold over basal
Time (min)
0.2
0.4
0.6
0.8
1
1.2
1.4
1.6
1.8
2
2.2
0
10
20
30
40
50
60
Normal
MK50
MK75
MK88
MK94
MK97
MK data
Normal data

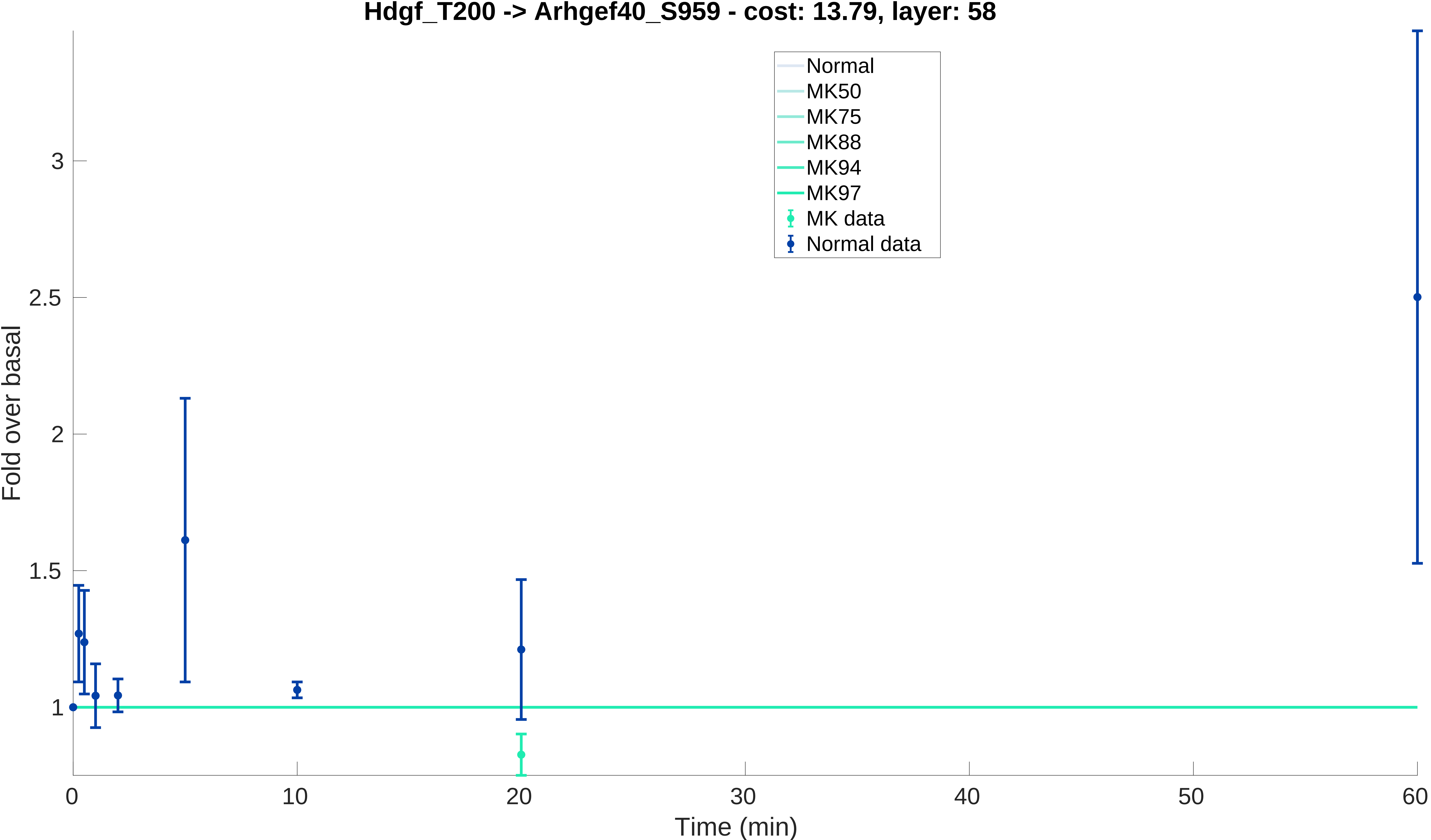

Hdgf_T200 -> Arhgef40_S959 - cost: 13.79, layer: 58
Normal
MK50
MK75
MK88
MK94
MK97
MK data
Normal data
Fold over basal
3
2.5
2
1.5
1
0
10
20
30
40
50
60
Time (min)

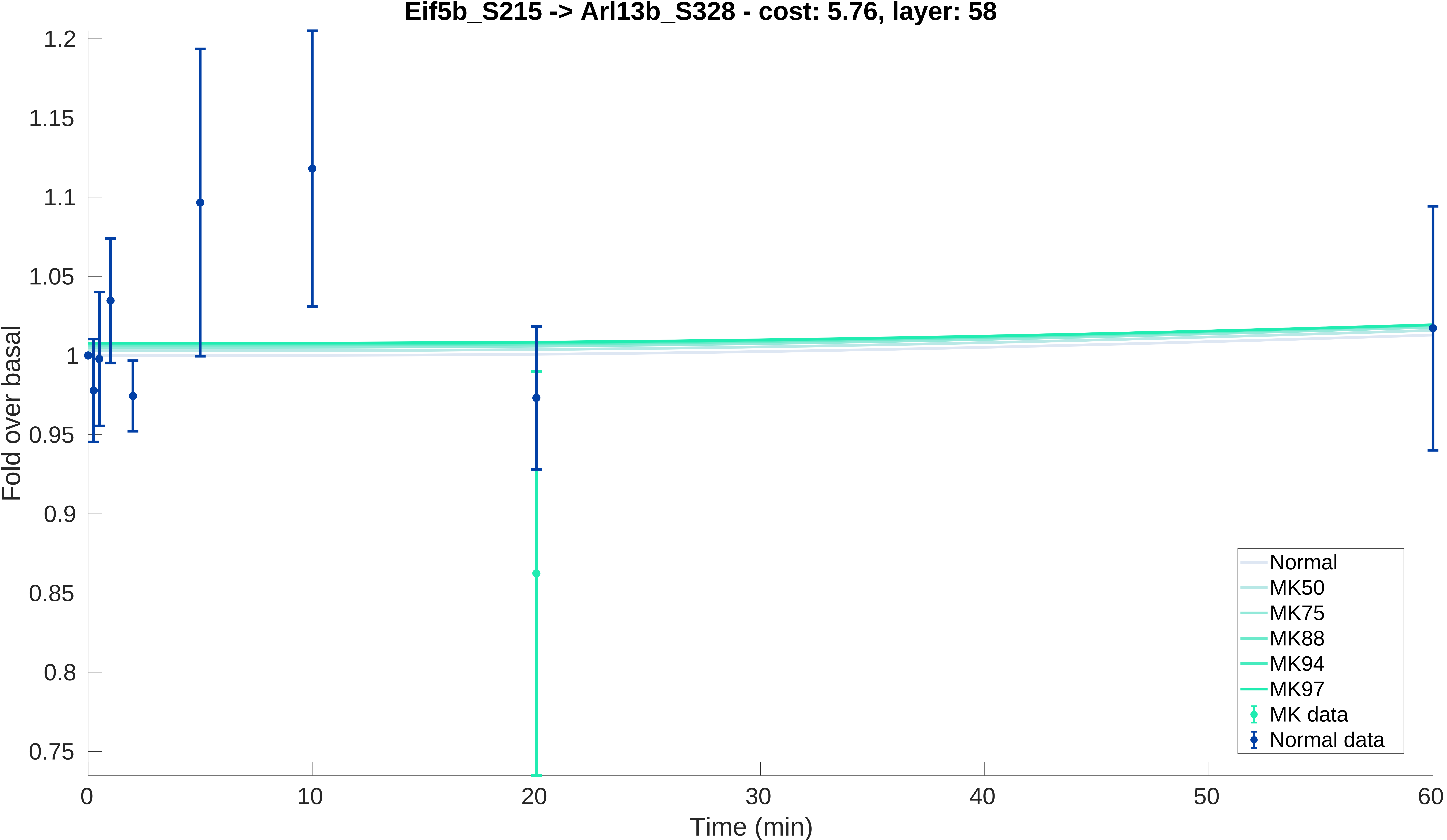

Eif5b_S215 -> Arl13b_S328 - cost: 5.76, layer: 58
Fold over basal
1.2
1.15
1.1
1.05
1
0.95
0.9
0.85
0.8
0.75
0
10
20
30
40
50
60
Time (min)
Normal
MK50
MK75
MK88
MK94
MK97
MK data
Normal data

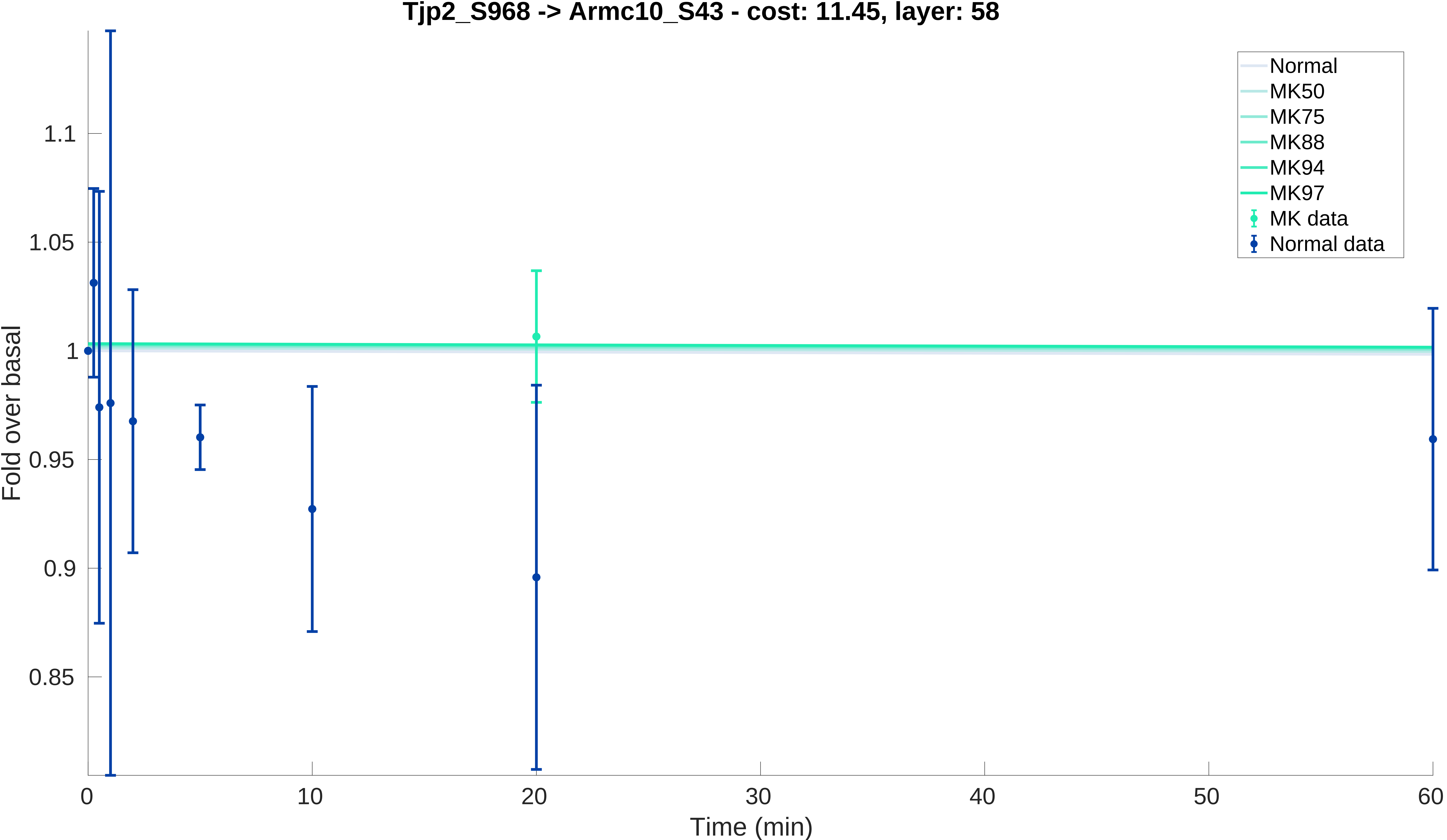

Tjp2_S968 -> Armc10_S43 - cost: 11.45, layer: 58
Normal
MK50
MK75
MK88
MK94
MK97
MK data
Normal data
Fold over basal
1.1
1.05
1
0.95
0.9
0.85
0
10
20
30
40
50
60
Time (min)

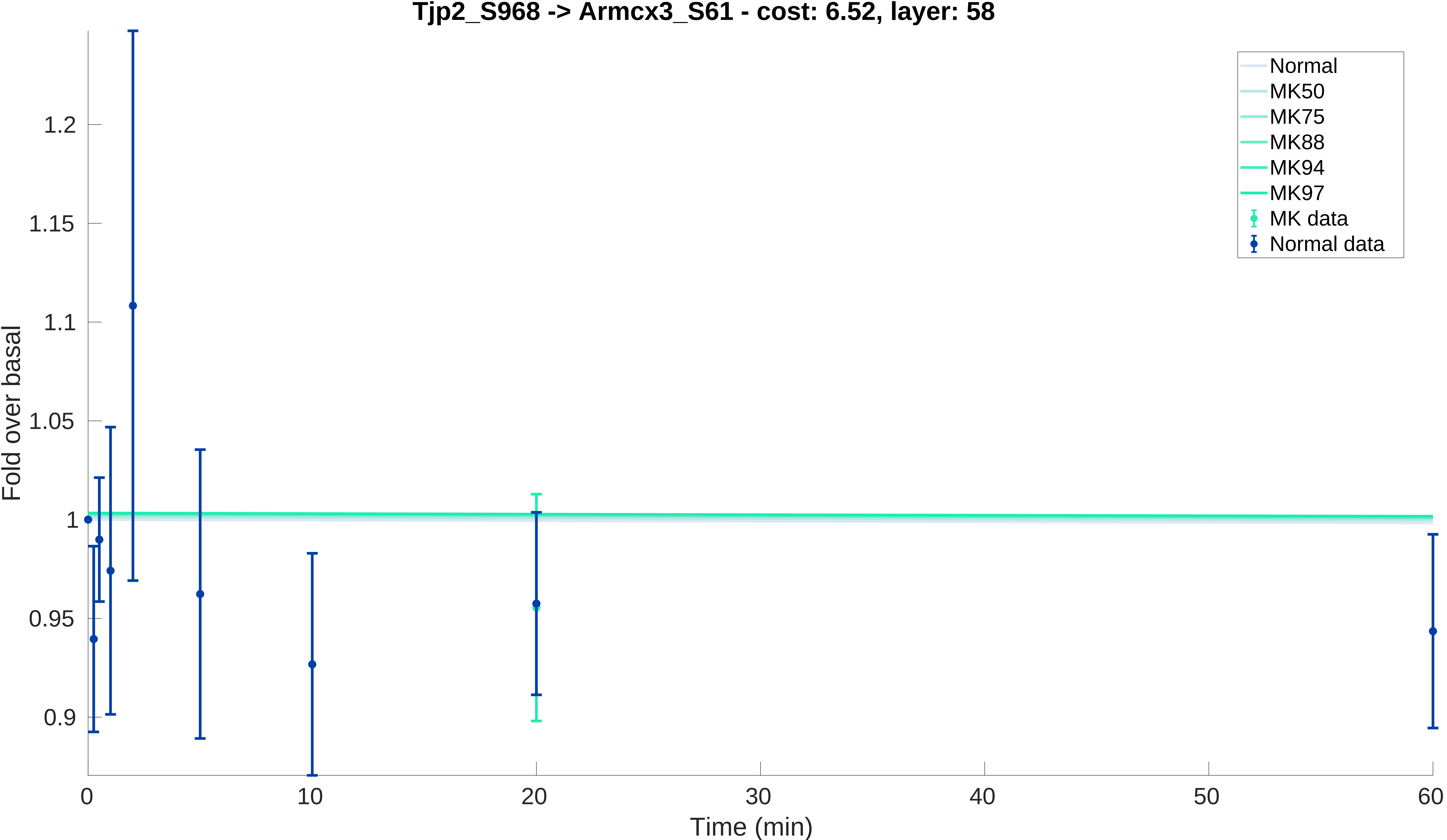

Tjp2_S968 -> Armcx3_S61 - cost: 6.52, layer: 58
Fold over basal
Time (min)
1.2
1.15
1.1
1.05
1
0.95
0.9
0
10
20
30
40
50
60
Normal
MK50
MK75
MK88
MK94
MK97
MK data
Normal data

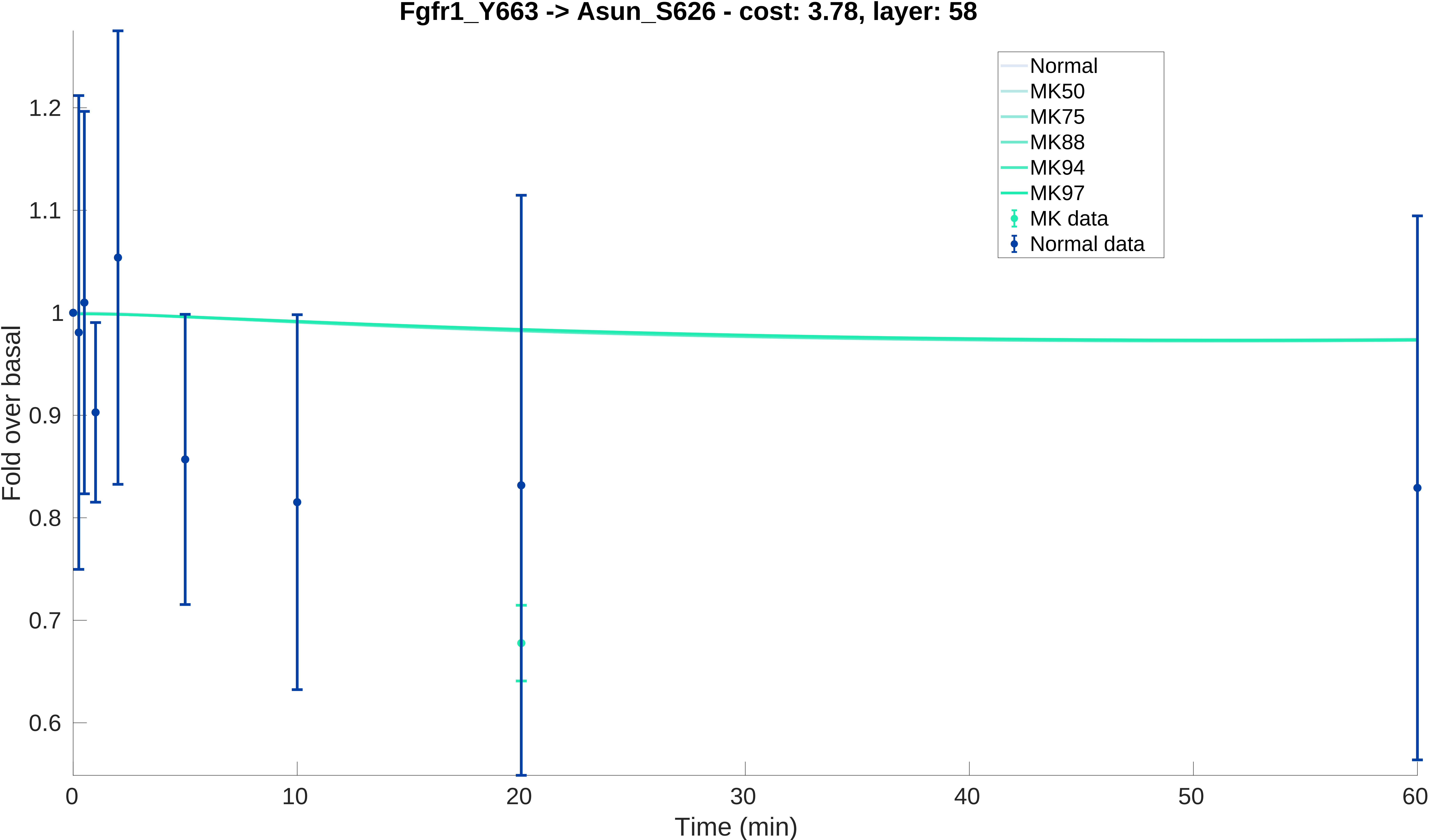

Fgfr1_Y663 -> Asun_S626 - cost: 3.78, layer: 58
Normal
MK50
MK75
MK88
MK94
MK97
MK data
Normal data
Fold over basal
1.2
1.1
1
0.9
0.8
0.7
0.6
0
10
20
30
40
50
60
Time (min)

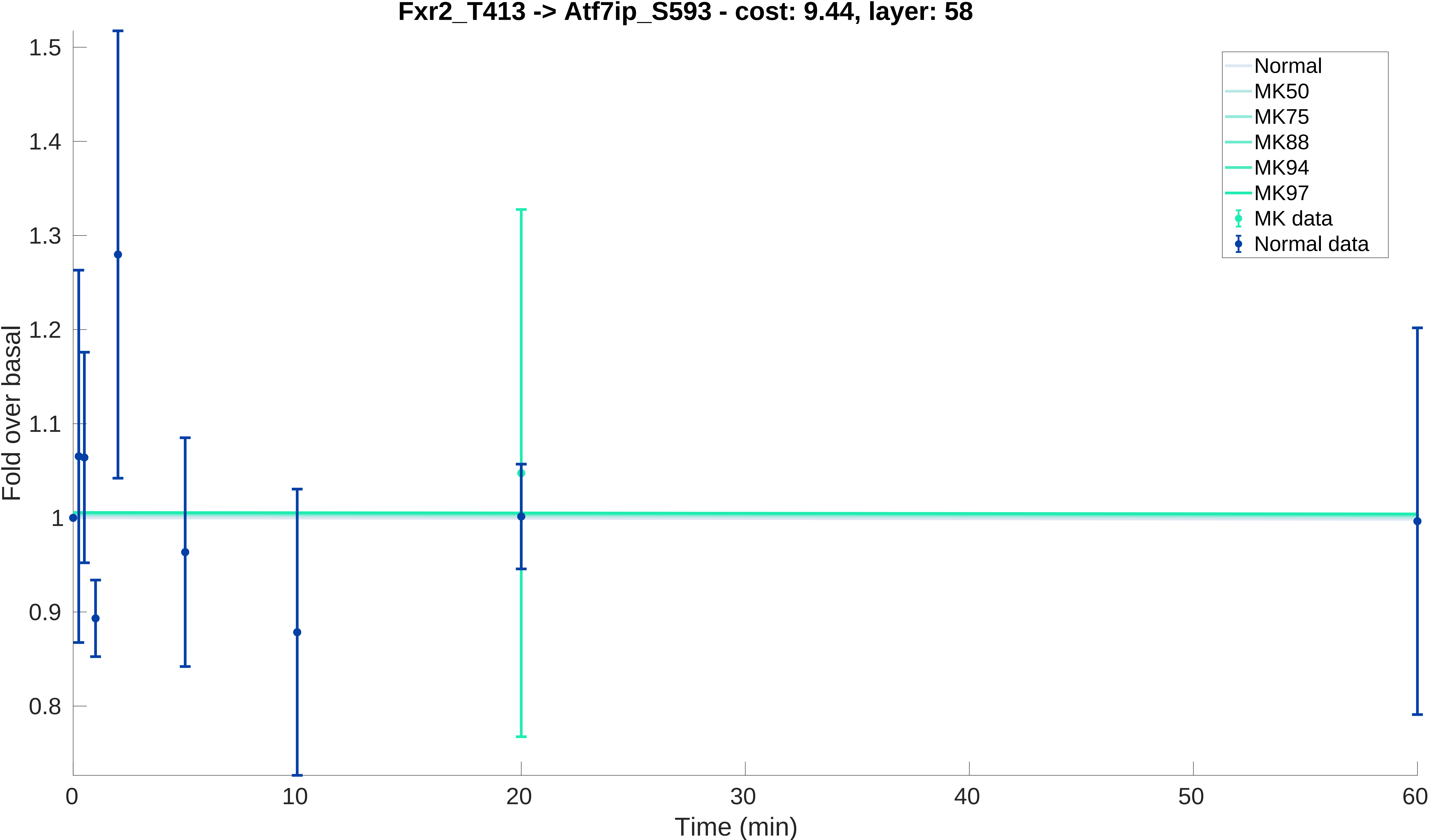

Fxr2_T413 -> Atf7ip_S593 - cost: 9.44, layer: 58
Fold over basal
Time (min)
1.5
1.4
1.3
1.2
1.1
1
0.9
0.8
0
10
20
30
40
50
60
Normal
MK50
MK75
MK88
MK94
MK97
MK data
Normal data

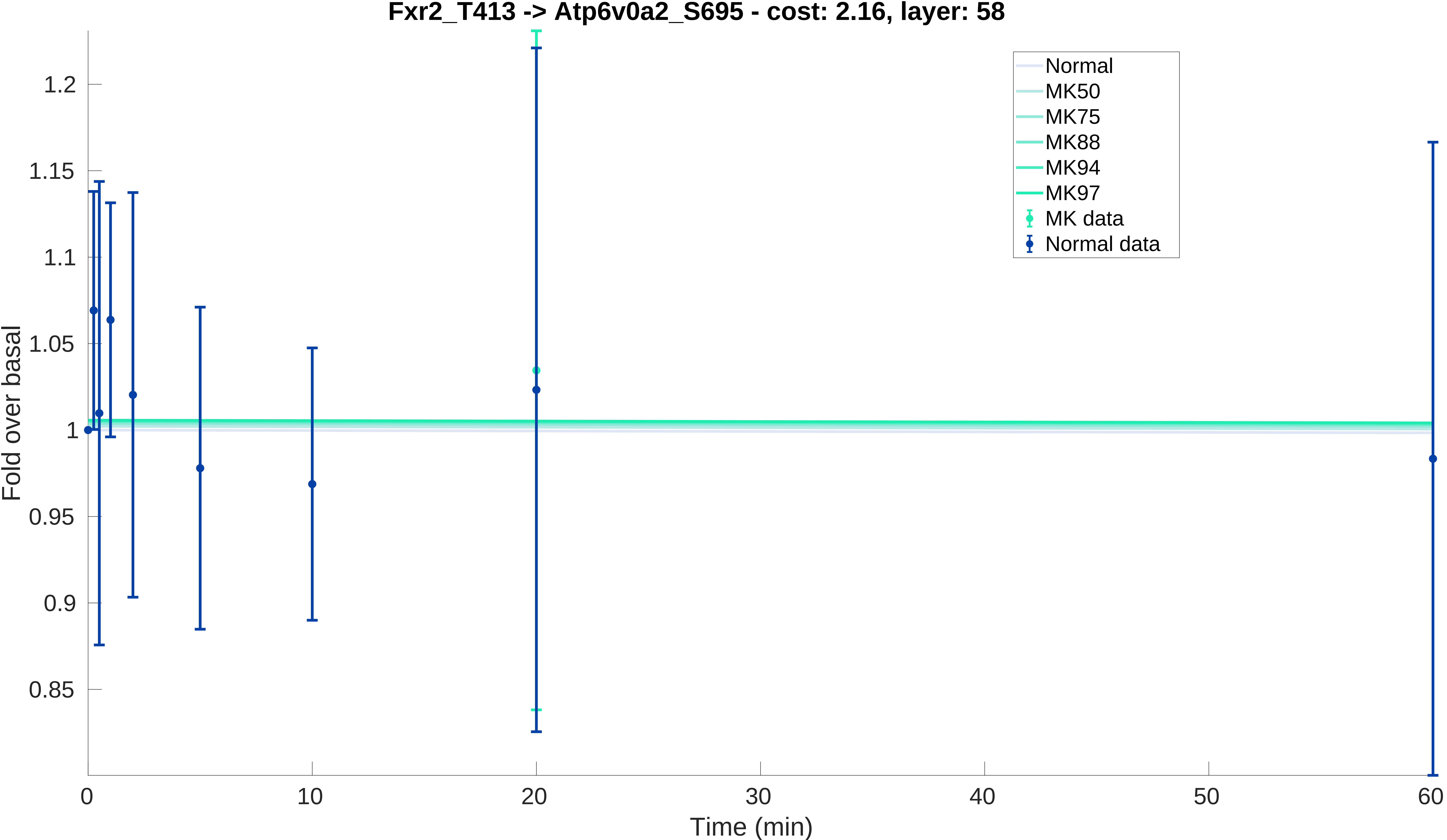

Fxr2_T413 -> Atp6v0a2_S695 - cost: 2.16, layer: 58
Normal
MK50
MK75
MK88
MK94
MK97
MK data
Normal data
1.2
1.15
1.1
1.05
1
0.95
0.9
0.85
Fold over basal
0
10
20
30
40
50
60
Time (min)

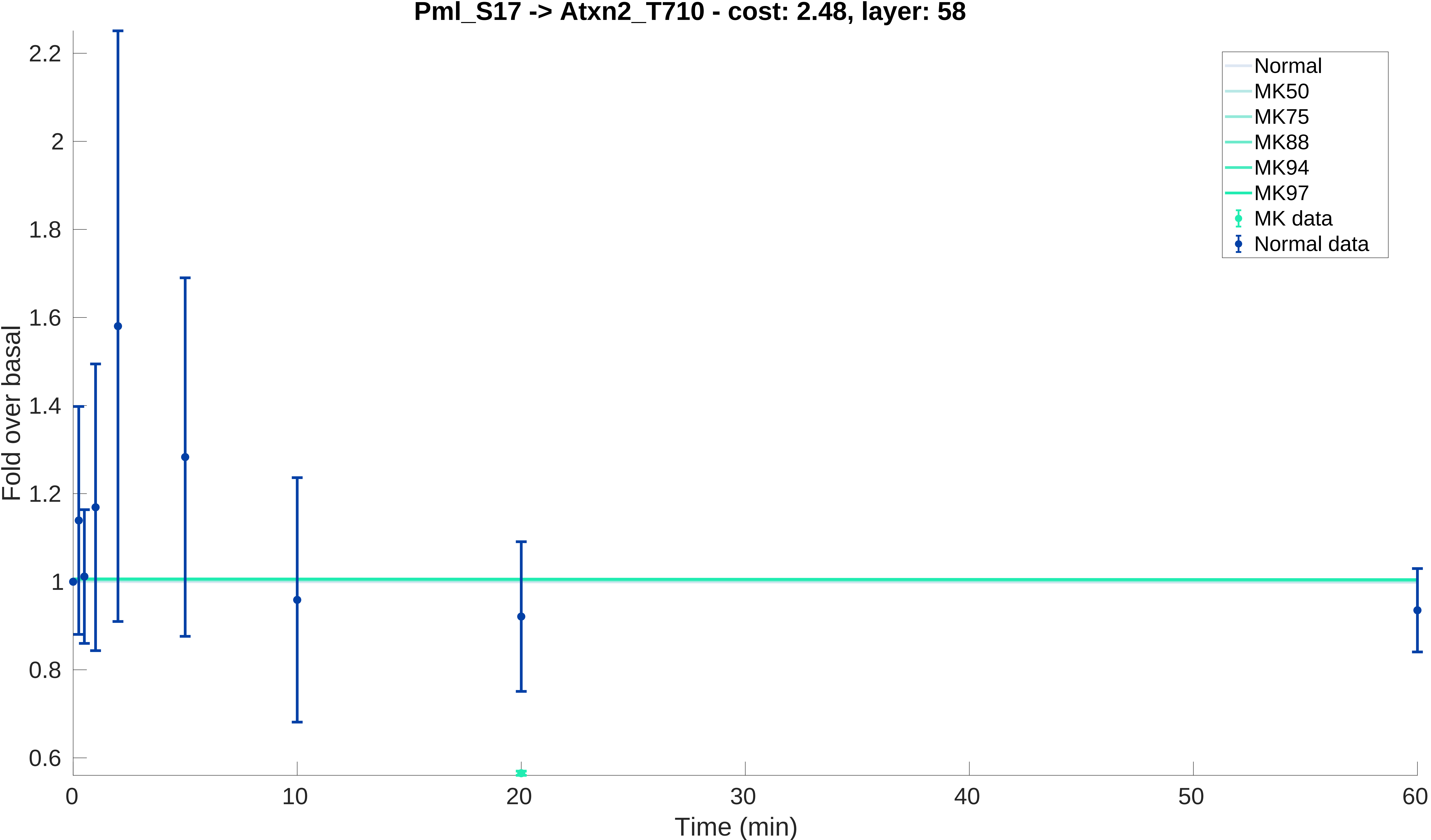
Pml_S17 -> Atxn2_T710 - cost: 2.48, layer: 58
Fold over basal
Time (min)
0.6
0.8
1
1.2
1.4
1.6
1.8
2
2.2
0
10
20
30
40
50
60
Normal
MK50
MK75
MK88
MK94
MK97
MK data
Normal data

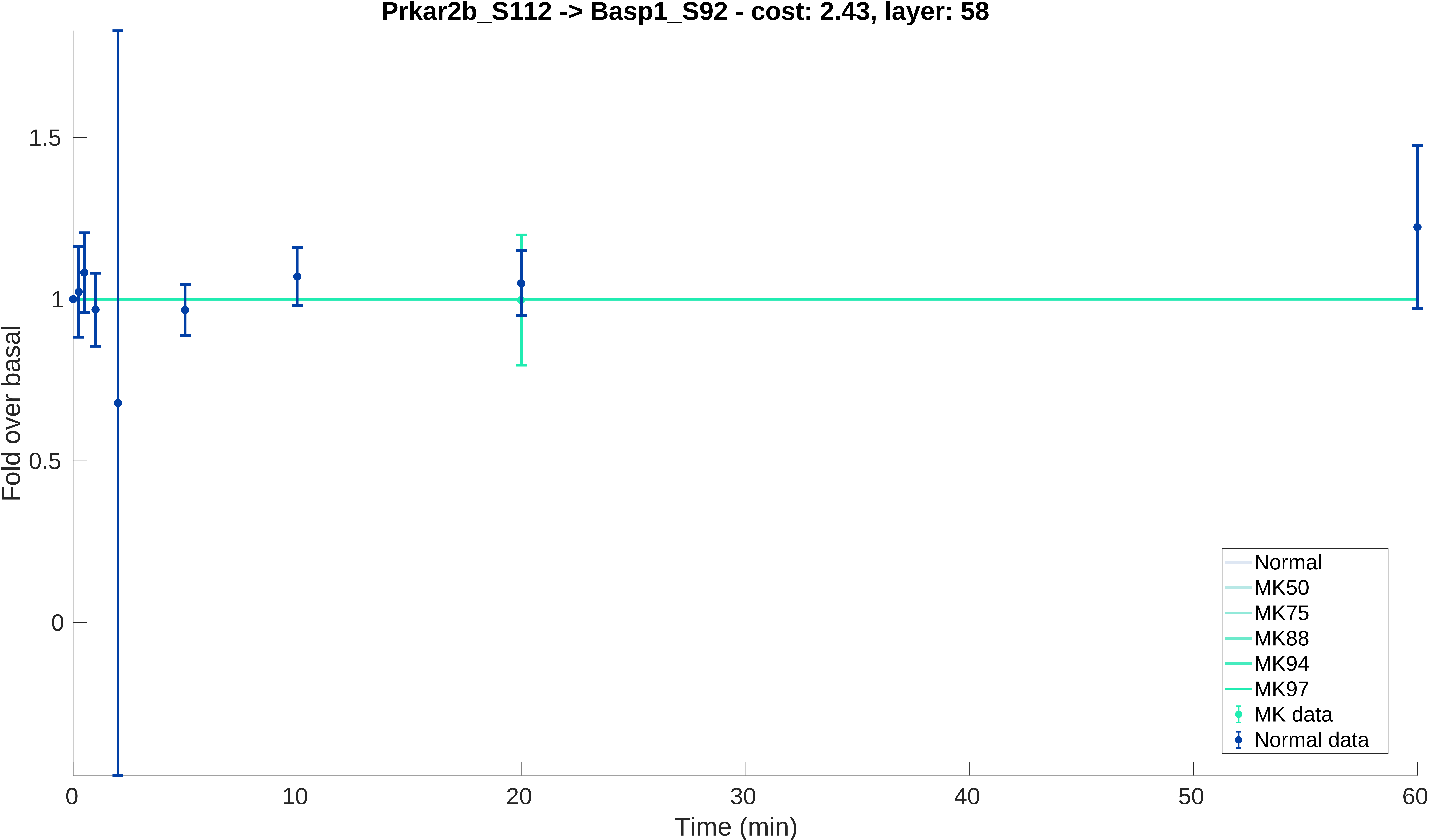

Prkar2b_S112 -> Basp1_S92 - cost: 2.43, layer: 58
Fold over basal
1.5
1
0.5
0
Time (min)
0
10
20
30
40
50
60
Normal
MK50
MK75
MK88
MK94
MK97
MK data
Normal data

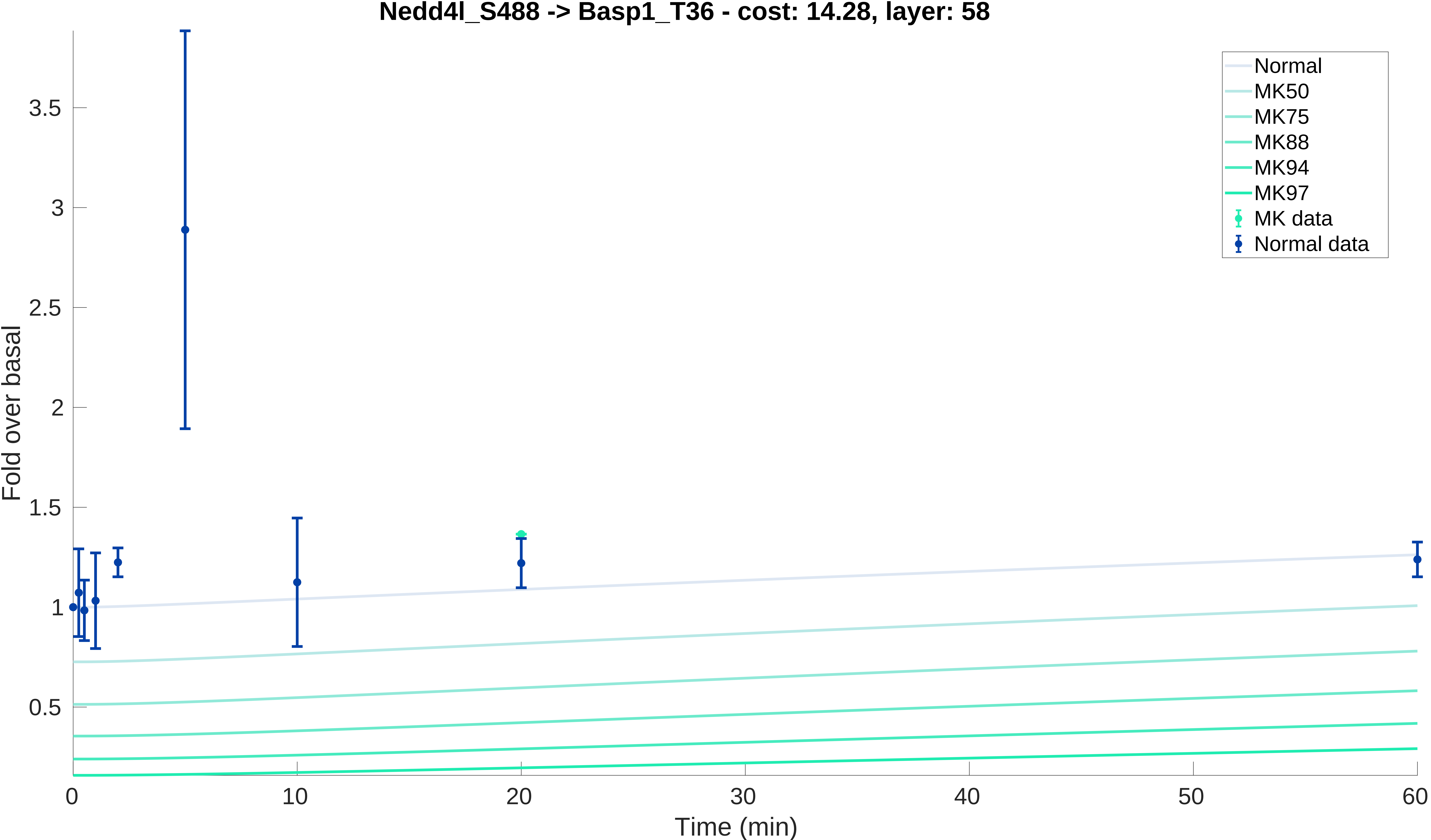
Nedd4l_S488 -> Basp1_T36 - cost: 14.28, layer: 58
Fold over basal
Time (min)
0.5
1
1.5
2
2.5
3
3.5
0
10
20
30
40
50
60
Normal
MK50
MK75
MK88
MK94
MK97
MK data
Normal data

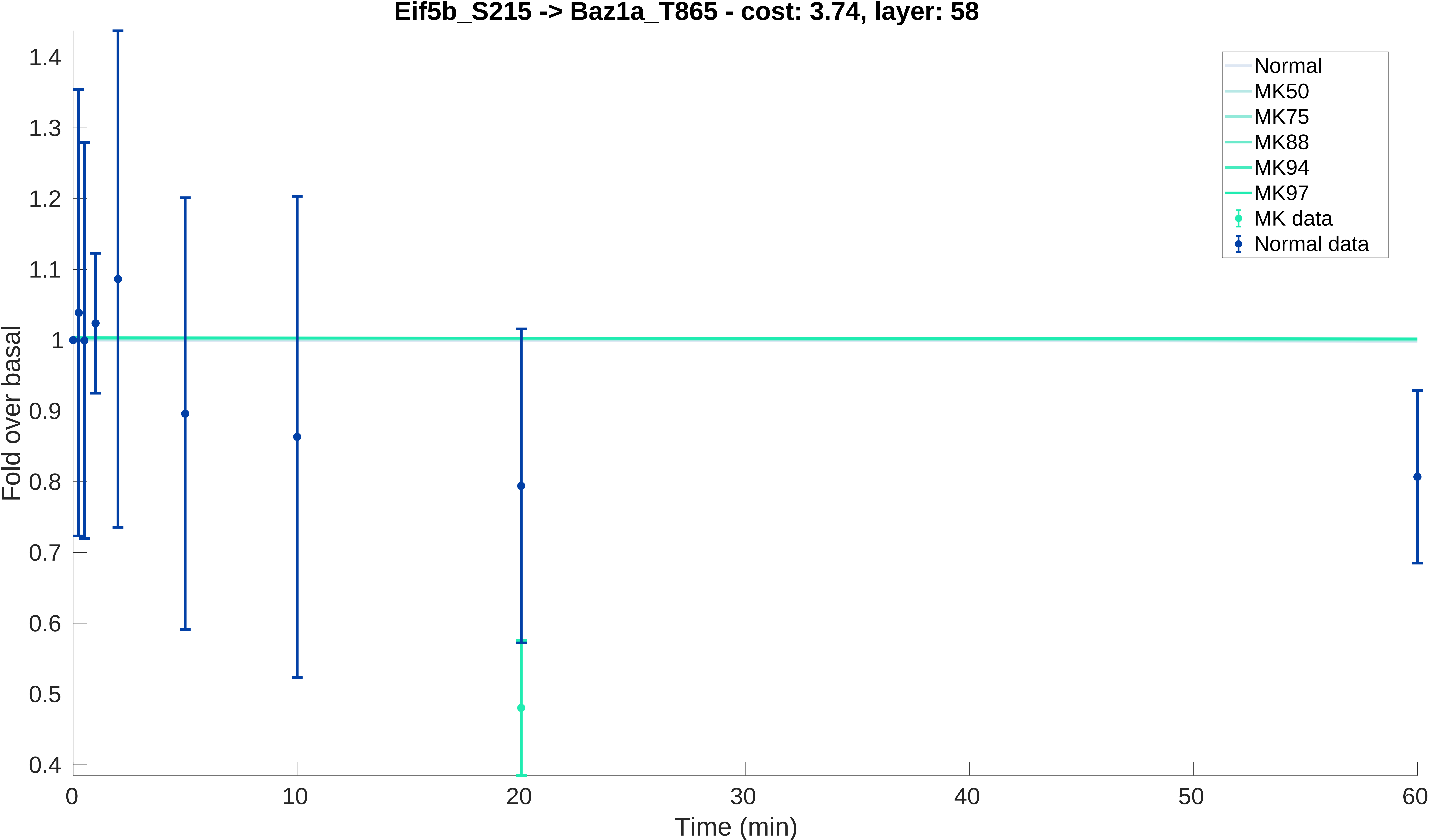

Eif5b_S215 -> Baz1a_T865 - cost: 3.74, layer: 58
Fold over basal
Time (min)
0
10
20
30
40
50
60
0.4
0.5
0.6
0.7
0.8
0.9
1
1.1
1.2
1.3
1.4
Normal
MK50
MK75
MK88
MK94
MK97
MK data
Normal data

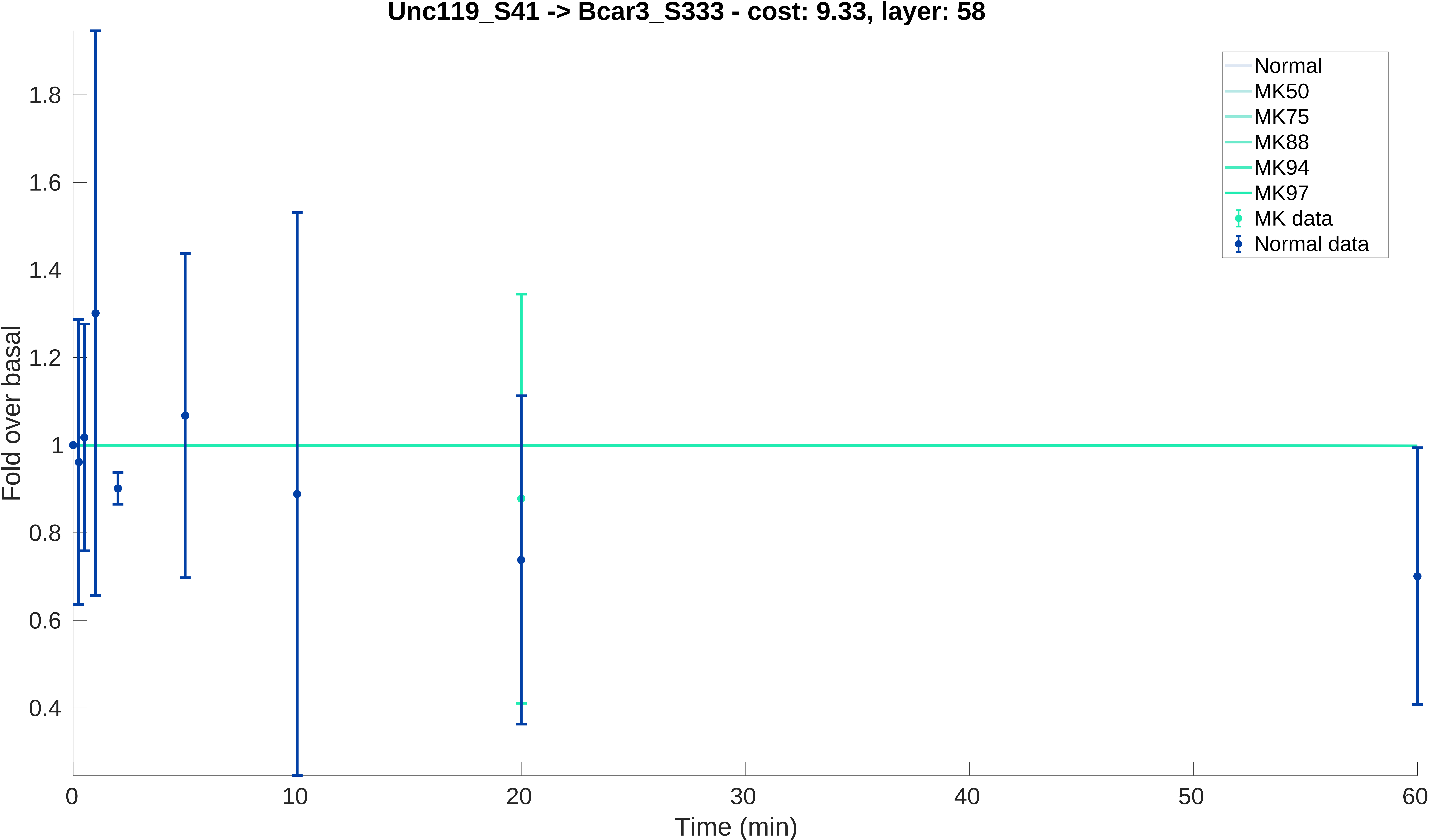

Unc119_S41 -> Bcar3_S333 - cost: 9.33, layer: 58
Fold over basal
Time (min)
0.4
0.6
0.8
1
1.2
1.4
1.6
1.8
0
10
20
30
40
50
60
Normal
MK50
MK75
MK88
MK94
MK97
MK data
Normal data

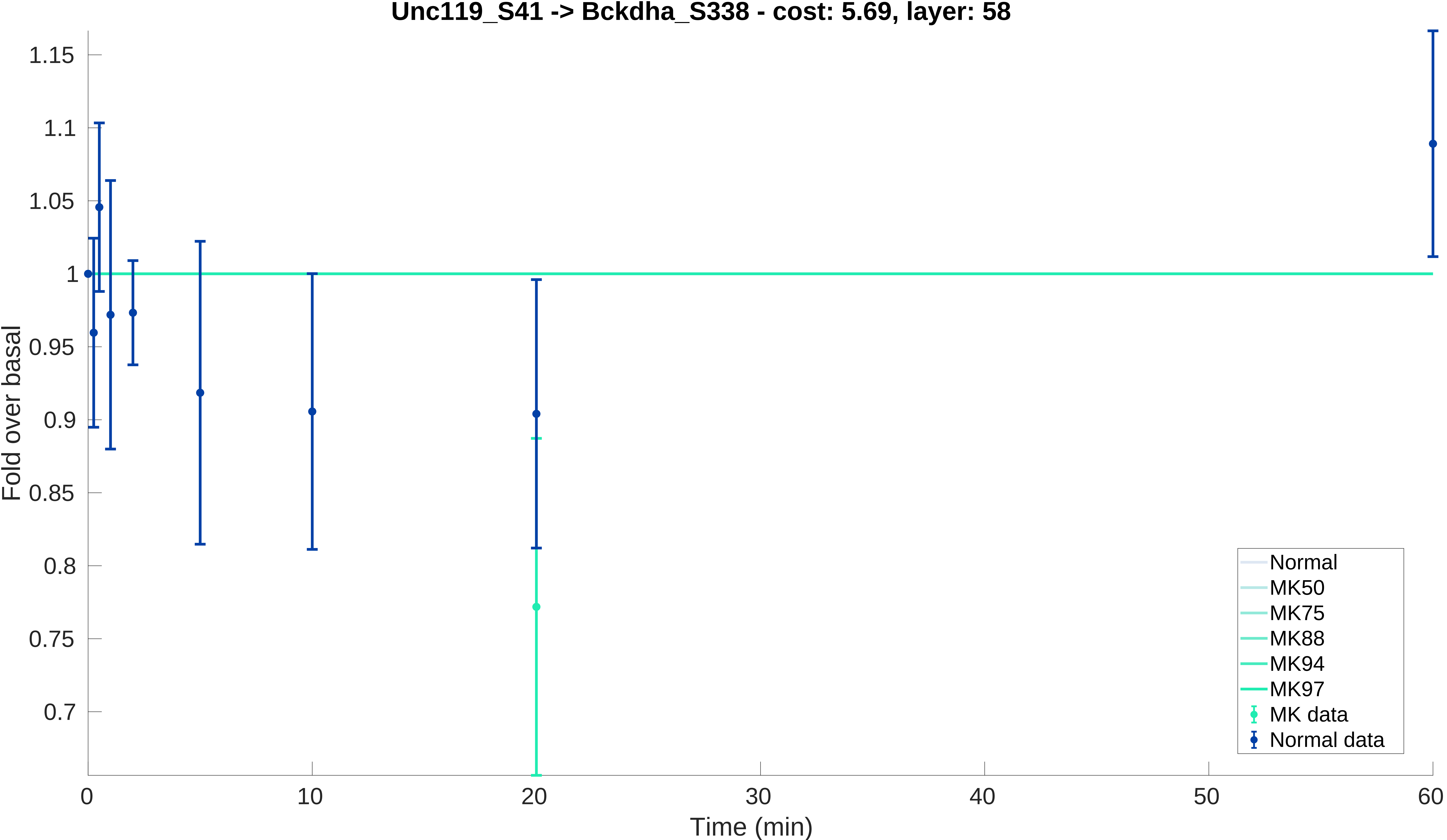
Unc119_S41 -> Bckdha_S338 - cost: 5.69, layer: 58
Fold over basal
1.15
1.1
1.05
1
0.95
0.9
0.85
0.8
0.75
0.7
0
10
20
30
40
50
60
Time (min)
Normal
MK50
MK75
MK88
MK94
MK97
MK data
Normal data

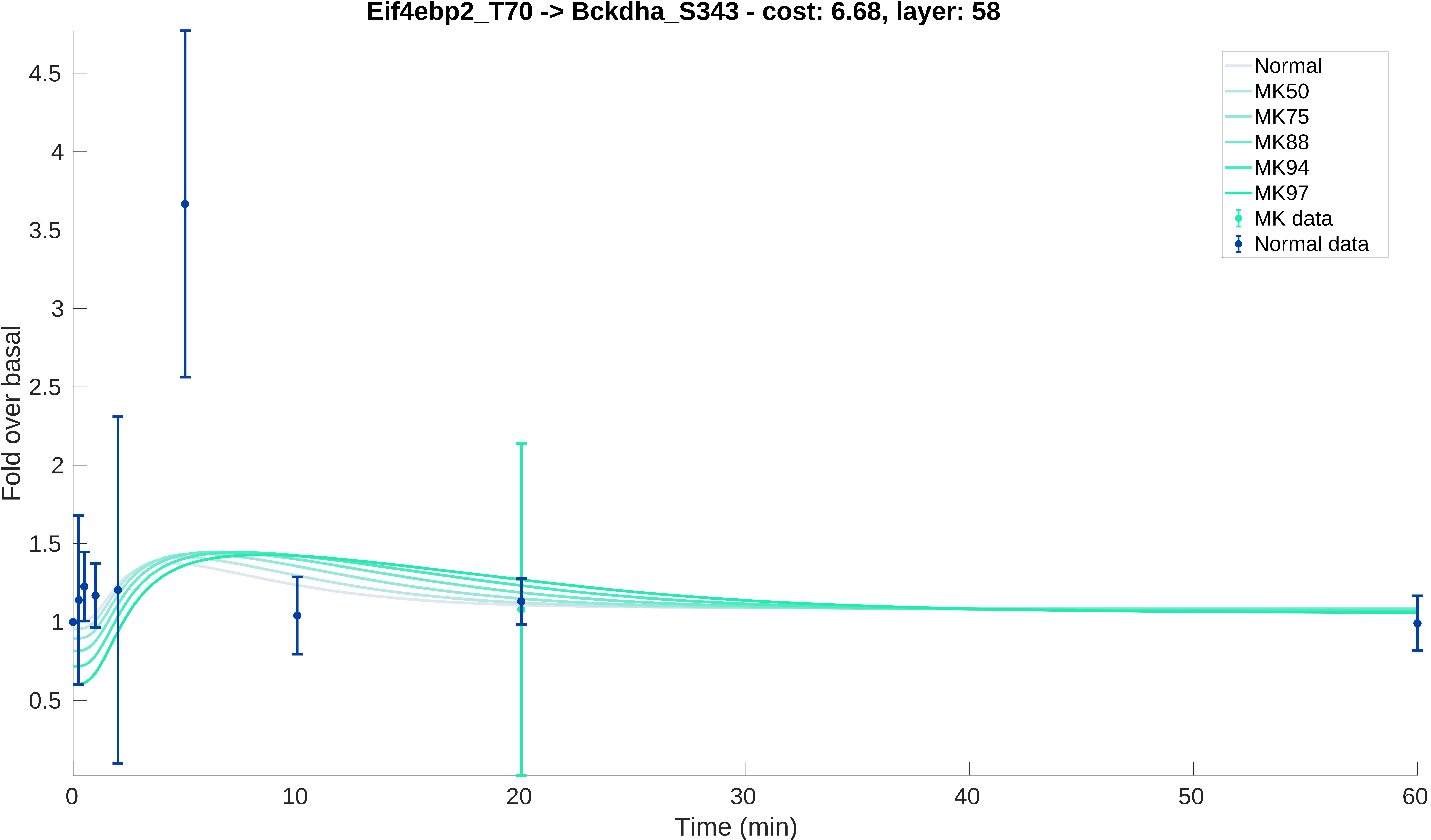

Eif4ebp2_T70 -> Bckdha_S343 - cost: 6.68, layer: 58
Fold over basal
4.5
4
3.5
3
2.5
2
1.5
1
0.5
0
10
20
30
40
50
60
Time (min)
Normal
MK50
MK75
MK88
MK94
MK97
MK data
Normal data

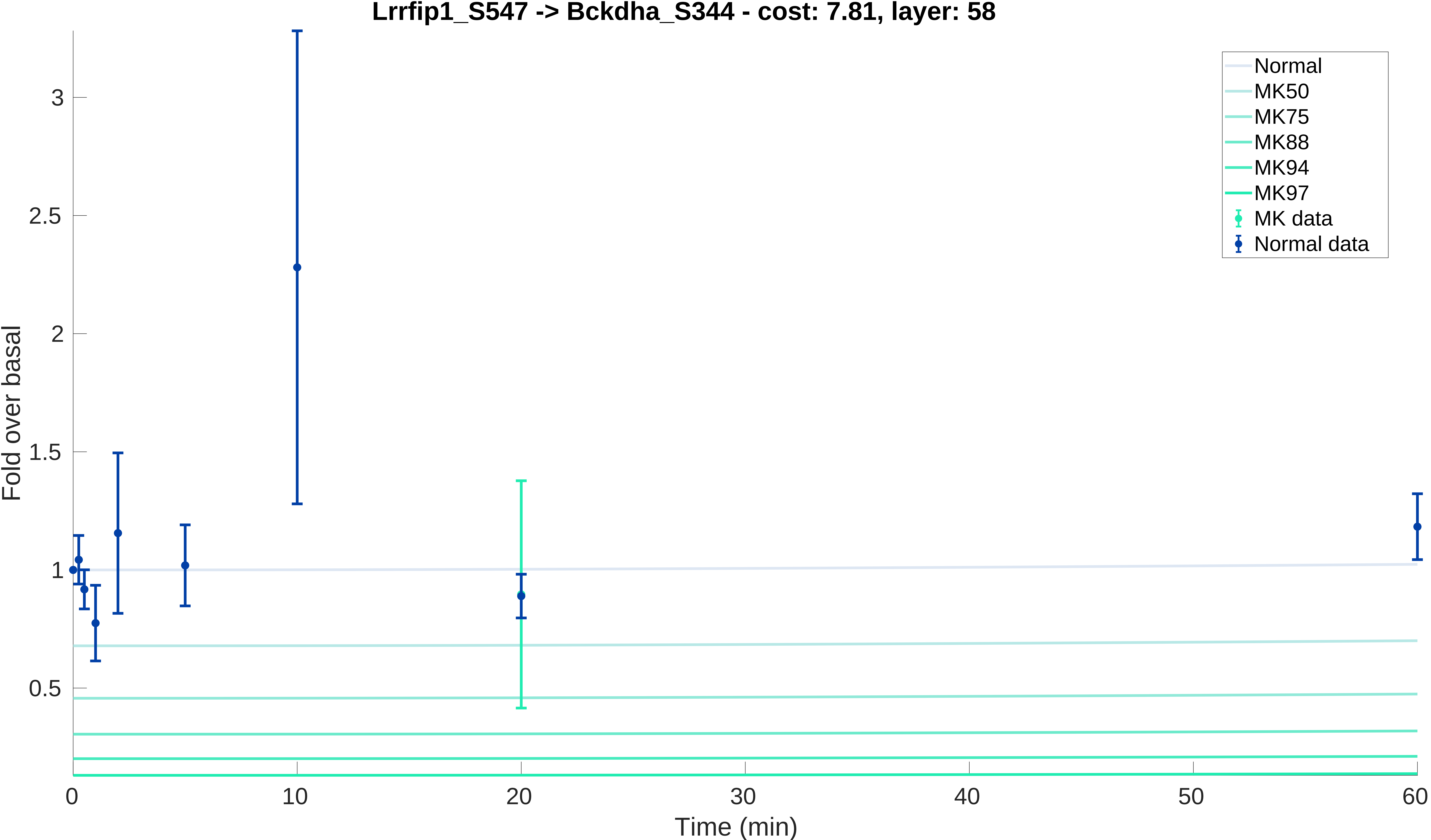
Lrrfip1_S547 -> Bckdha_S344 - cost: 7.81, layer: 58
Fold over basal
Time (min)
0.5
1
1.5
2
2.5
3
0
10
20
30
40
50
60
Normal
MK50
MK75
MK88
MK94
MK97
MK data
Normal data

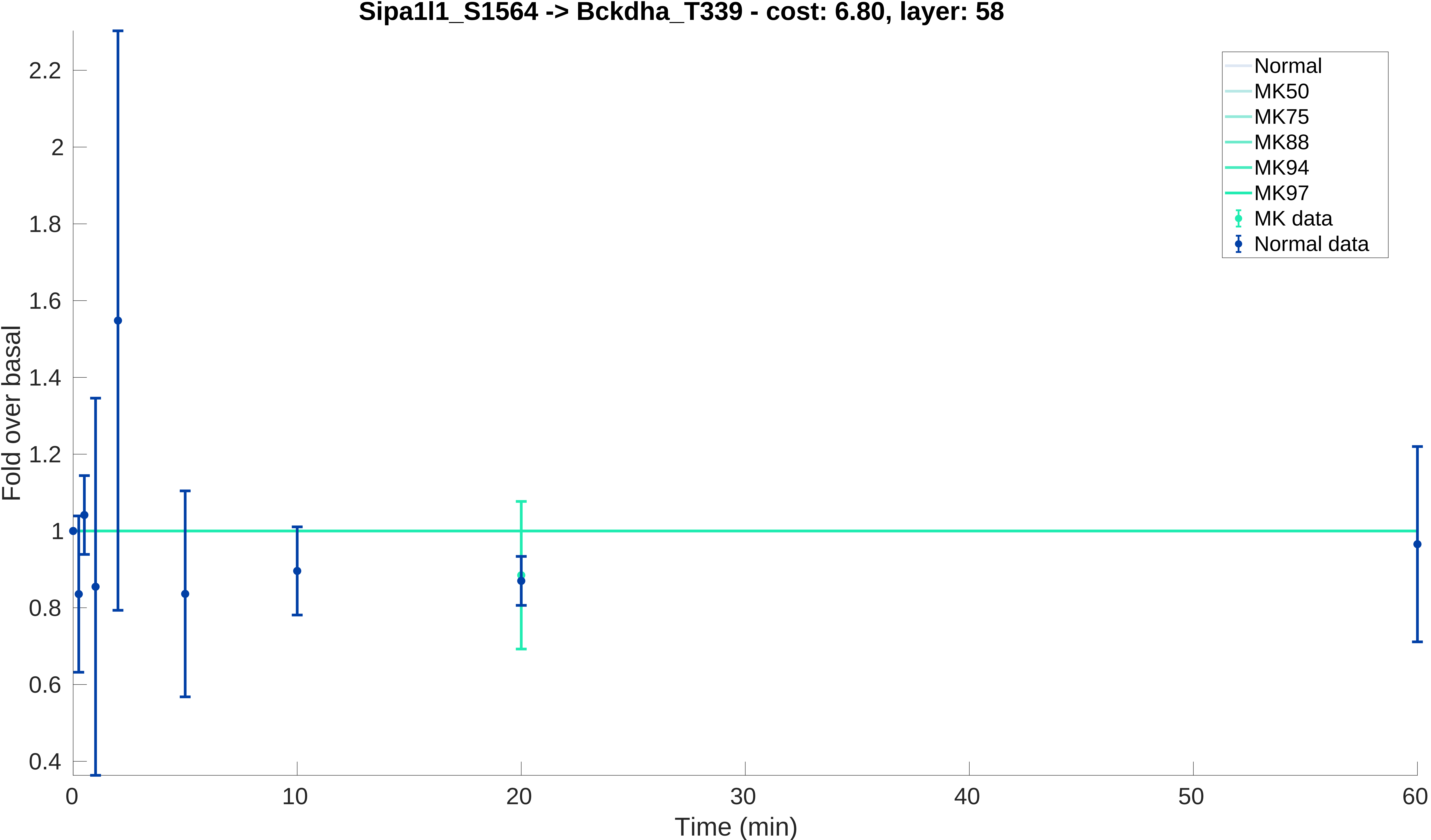

Sipa1l1_S1564 -> Bckdha_T339 - cost: 6.80, layer: 58
Fold over basal
Time (min)
Normal
MK50
MK75
MK88
MK94
MK97
MK data
Normal data
0.4
0.6
0.8
1
1.2
1.4
1.6
1.8
2
2.2
0
10
20
30
40
50
60

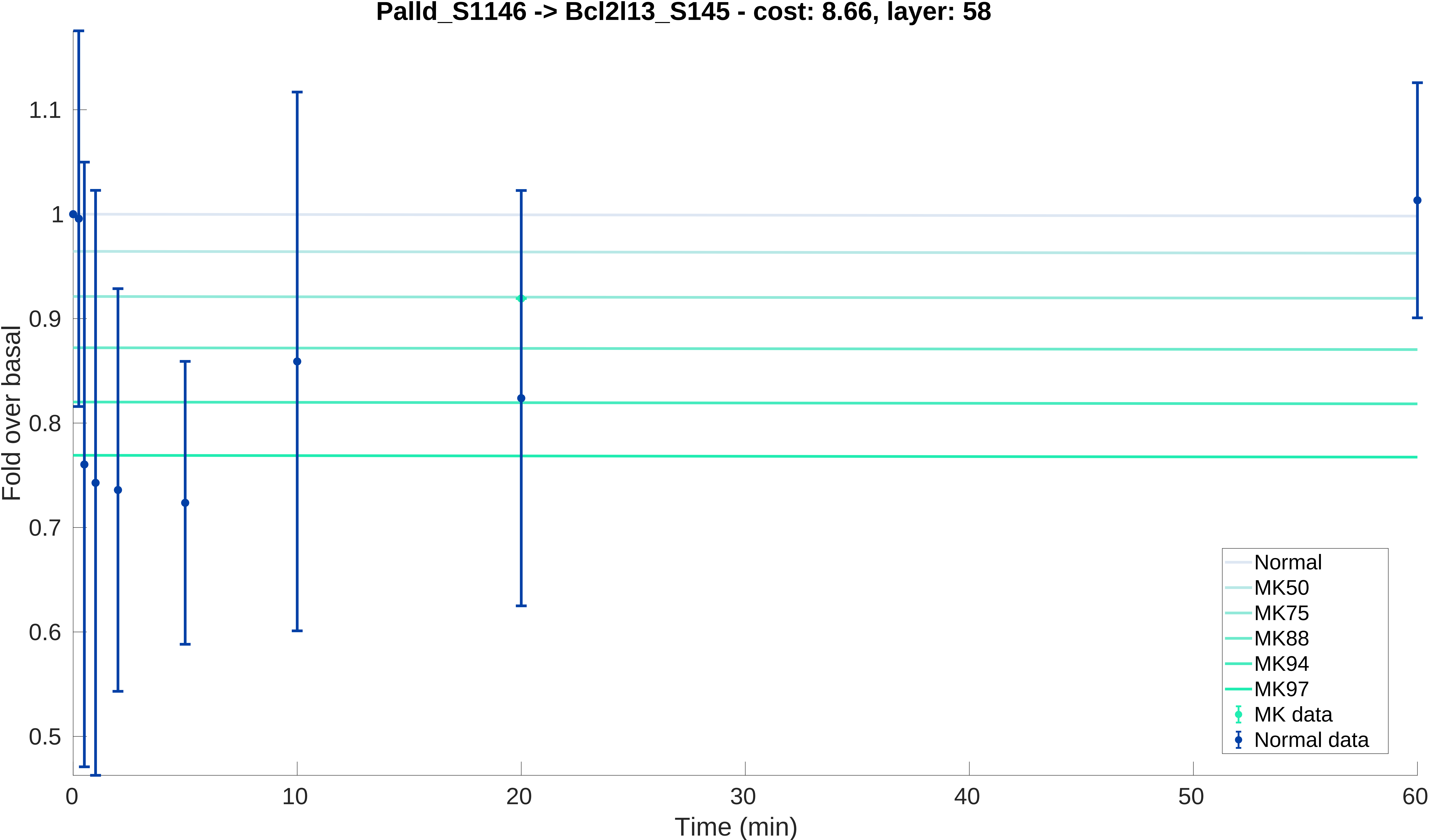

Palld_S1146 -> Bcl2l13_S145 - cost: 8.66, layer: 58
Fold over basal
Time (min)
0.5
0.6
0.7
0.8
0.9
1
1.1
0
10
20
30
40
50
60
Normal
MK50
MK75
MK88
MK94
MK97
MK data
Normal data

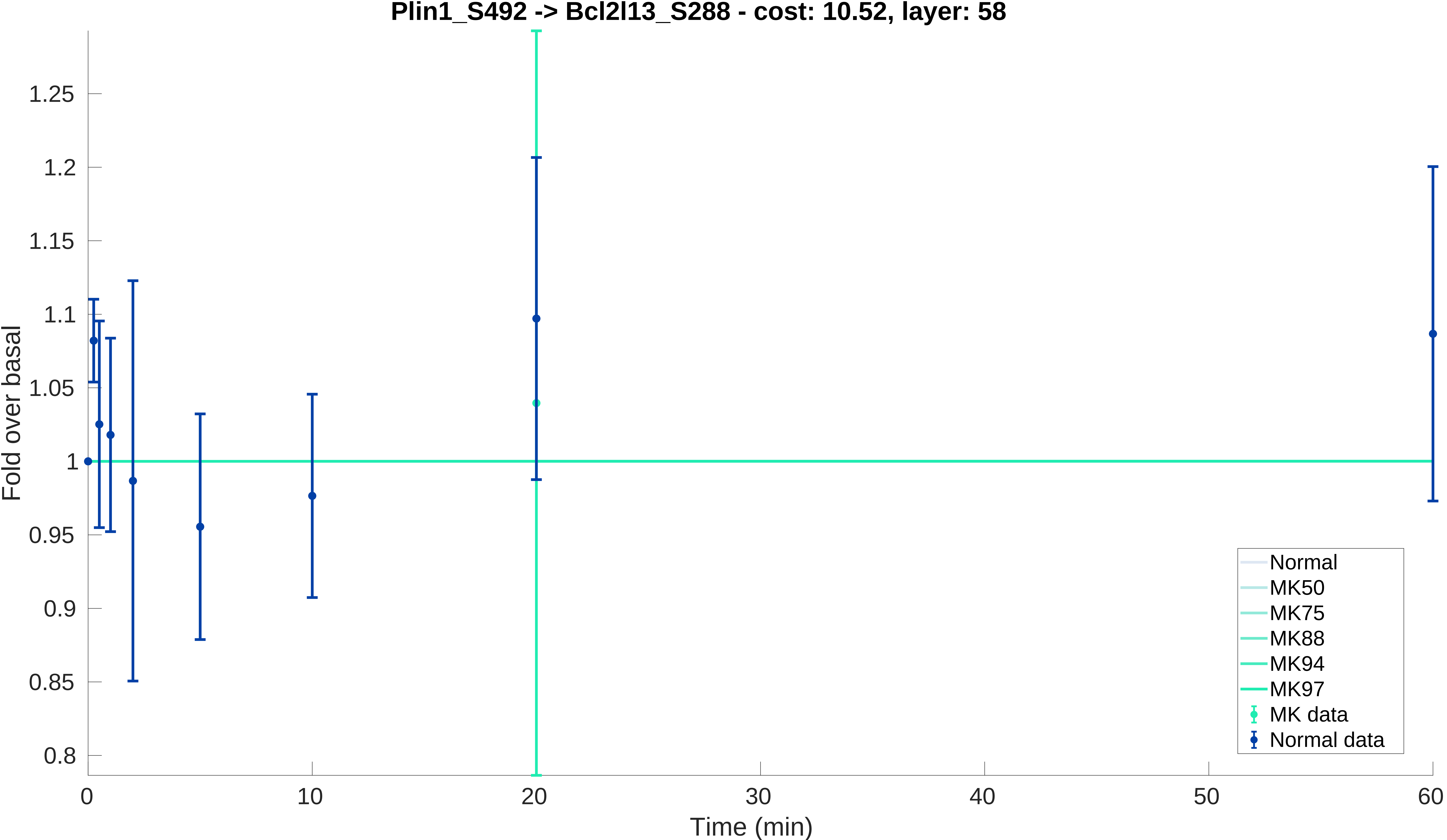

Plin1_S492 -> Bcl2l13_S288 - cost: 10.52, layer: 58
Fold over basal
1.25
1.2
1.15
1.1
1.05
1
0.95
0.9
0.85
0.8
0
10
20
30
40
50
60
Time (min)
Normal
MK50
MK75
MK88
MK94
MK97
MK data
Normal data

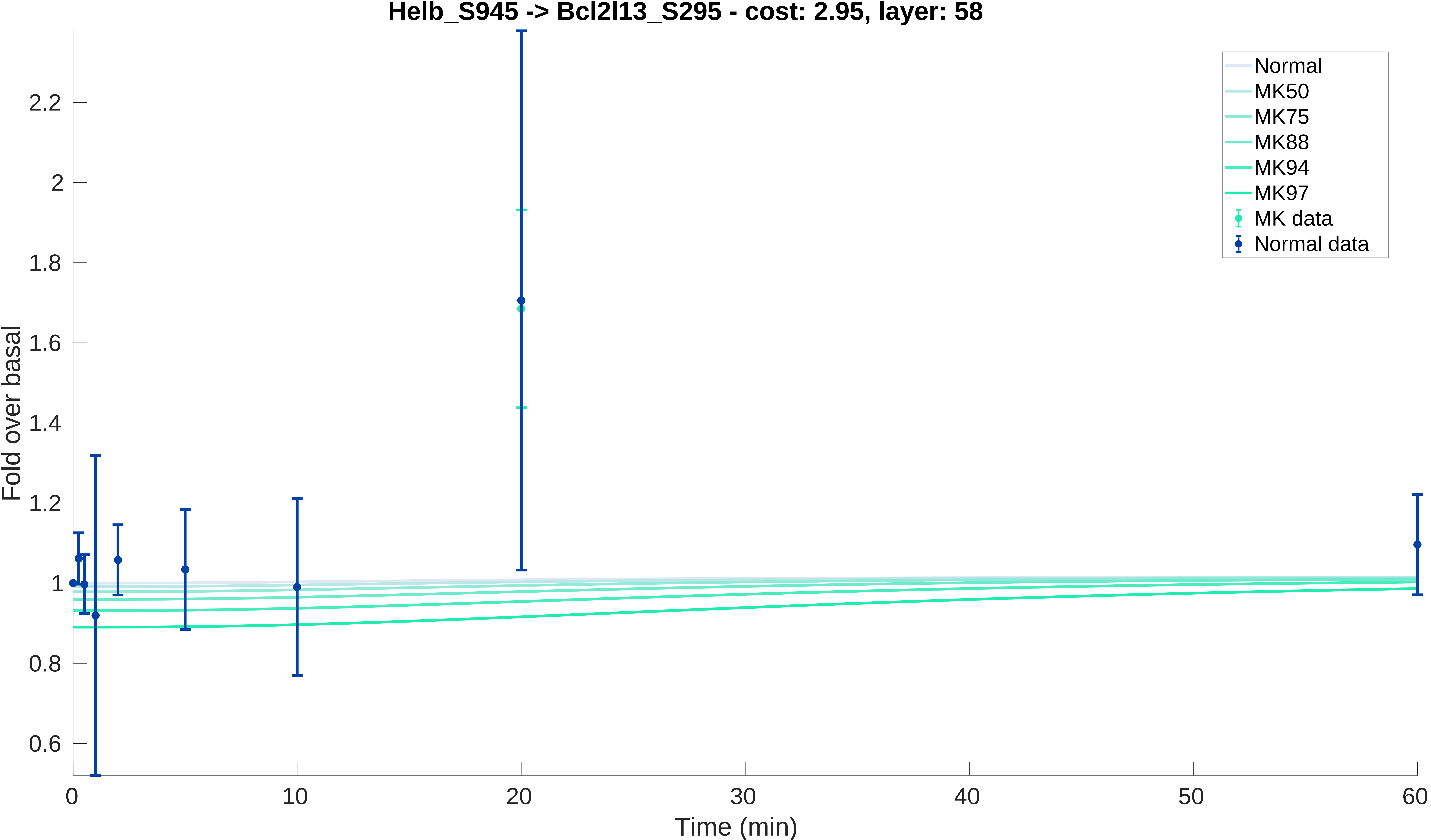

Helb_S945 -> Bcl2l13_S295 - cost: 2.95, layer: 58
Fold over basal
Time (min)
0.6
0.8
1
1.2
1.4
1.6
1.8
2
2.2
0
10
20
30
40
50
60
Normal
MK50
MK75
MK88
MK94
MK97
MK data
Normal data

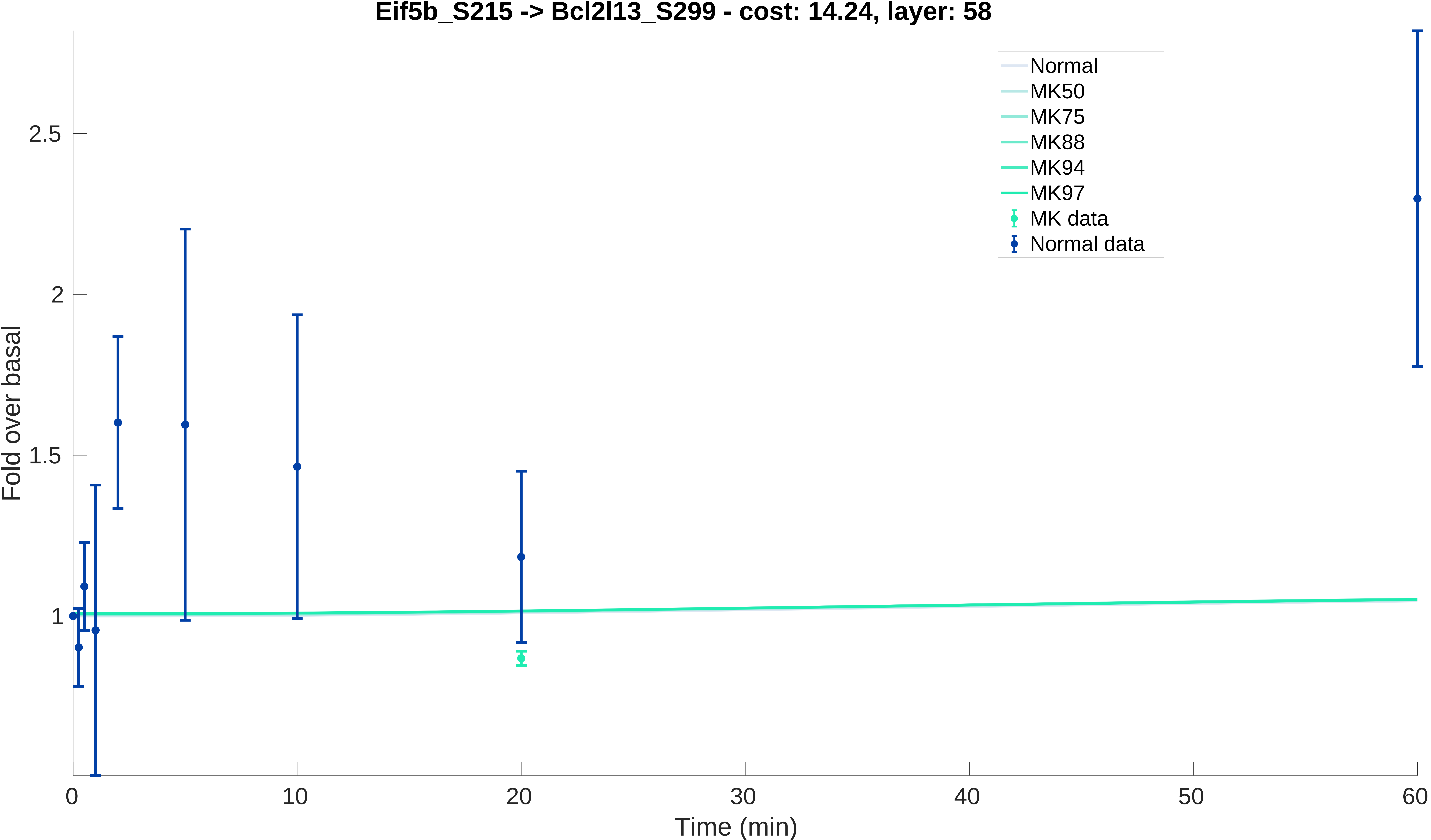

Eif5b_S215 -> Bcl2l13_S299 - cost: 14.24, layer: 58
Fold over basal
Time (min)
2.5
2
1.5
1
0
10
20
30
40
50
60
Normal
MK50
MK75
MK88
MK94
MK97
MK data
Normal data

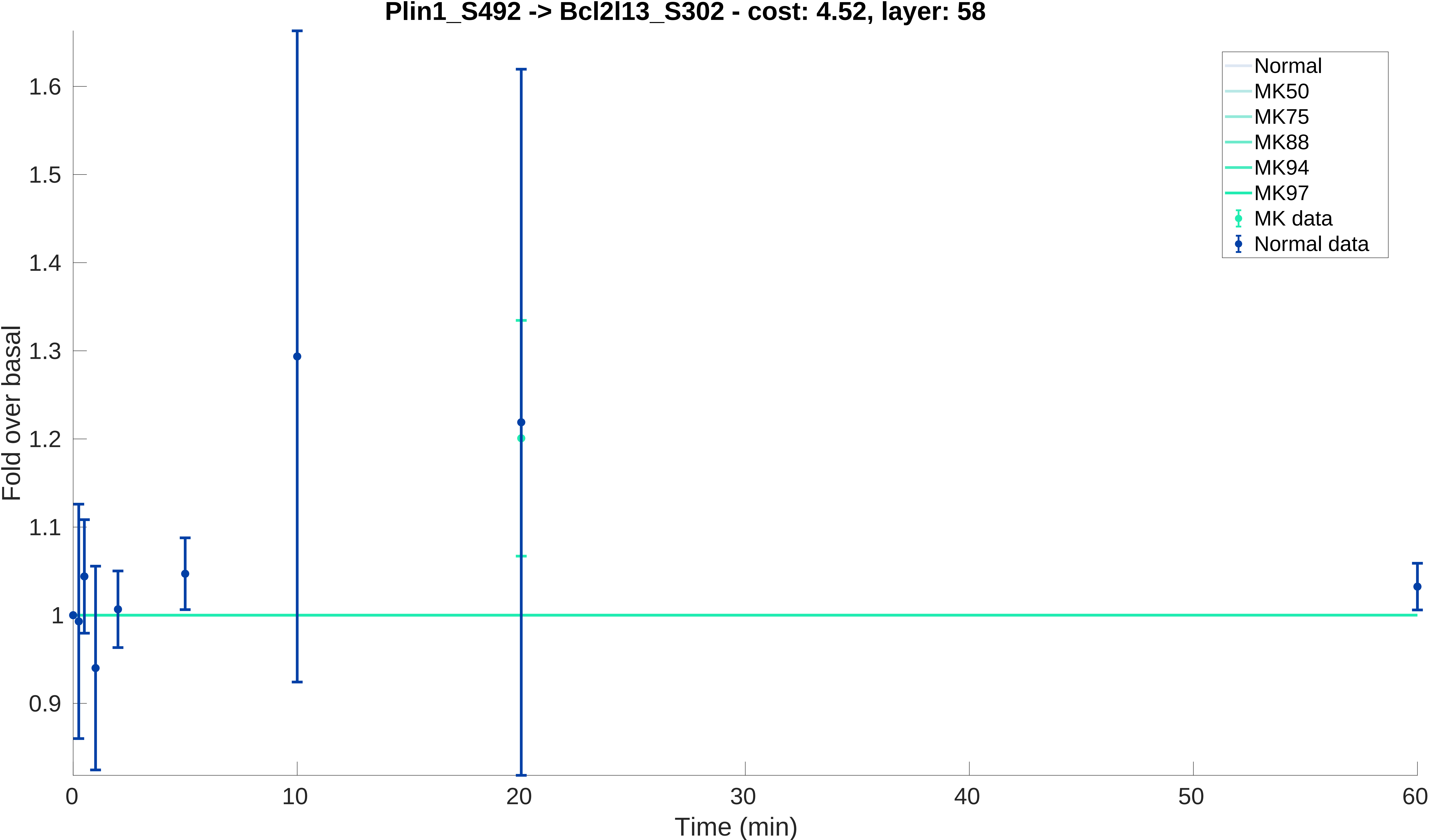

Plin1_S492 -> Bcl2l13_S302 - cost: 4.52, layer: 58
Fold over basal
Time (min)
Normal
MK50
MK75
MK88
MK94
MK97
MK data
Normal data
0.9
1
1.1
1.2
1.3
1.4
1.5
1.6
0
10
20
30
40
50
60

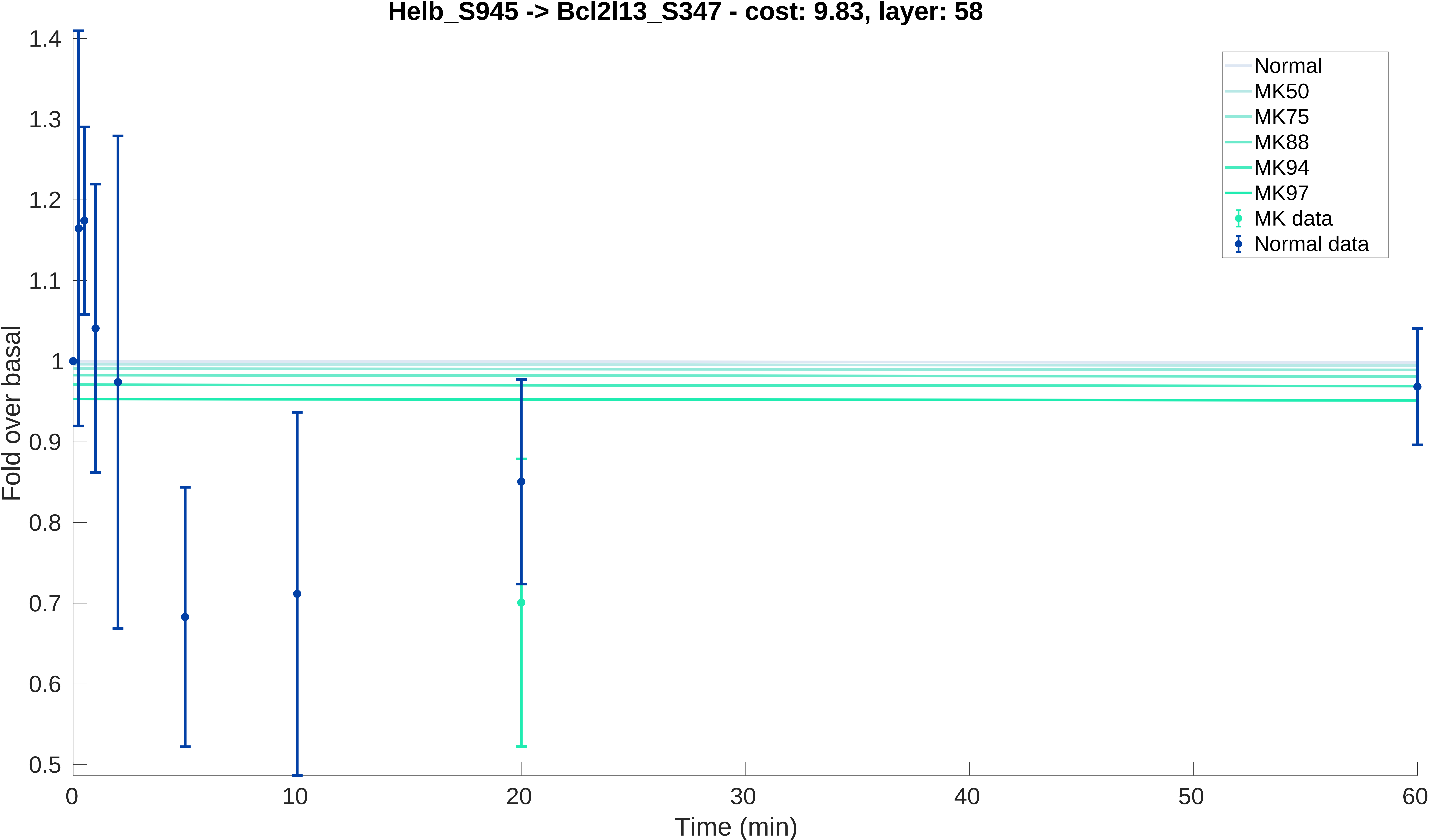

Helb_S945 -> Bcl2l13_S347 - cost: 9.83, layer: 58
Fold over basal
Time (min)
1.4
1.3
1.2
1.1
1
0.9
0.8
0.7
0.6
0.5
0
10
20
30
40
50
60
Normal
MK50
MK75
MK88
MK94
MK97
MK data
Normal data

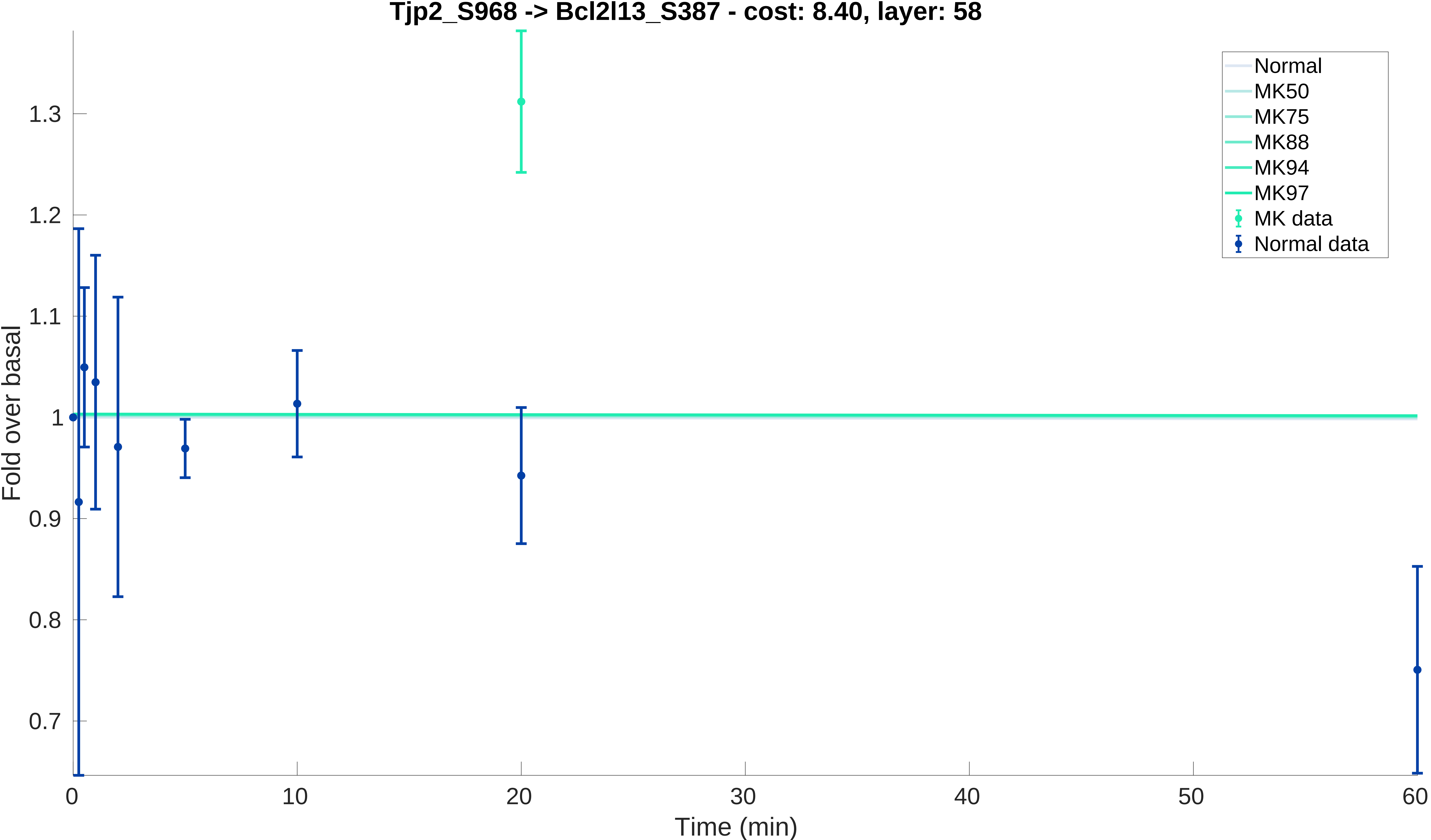

Tjp2_S968 -> Bcl2l13_S387 - cost: 8.40, layer: 58
Fold over basal
Time (min)
0.7
0.8
0.9
1
1.1
1.2
1.3
0
10
20
30
40
50
60
Normal
MK50
MK75
MK88
MK94
MK97
MK data
Normal data

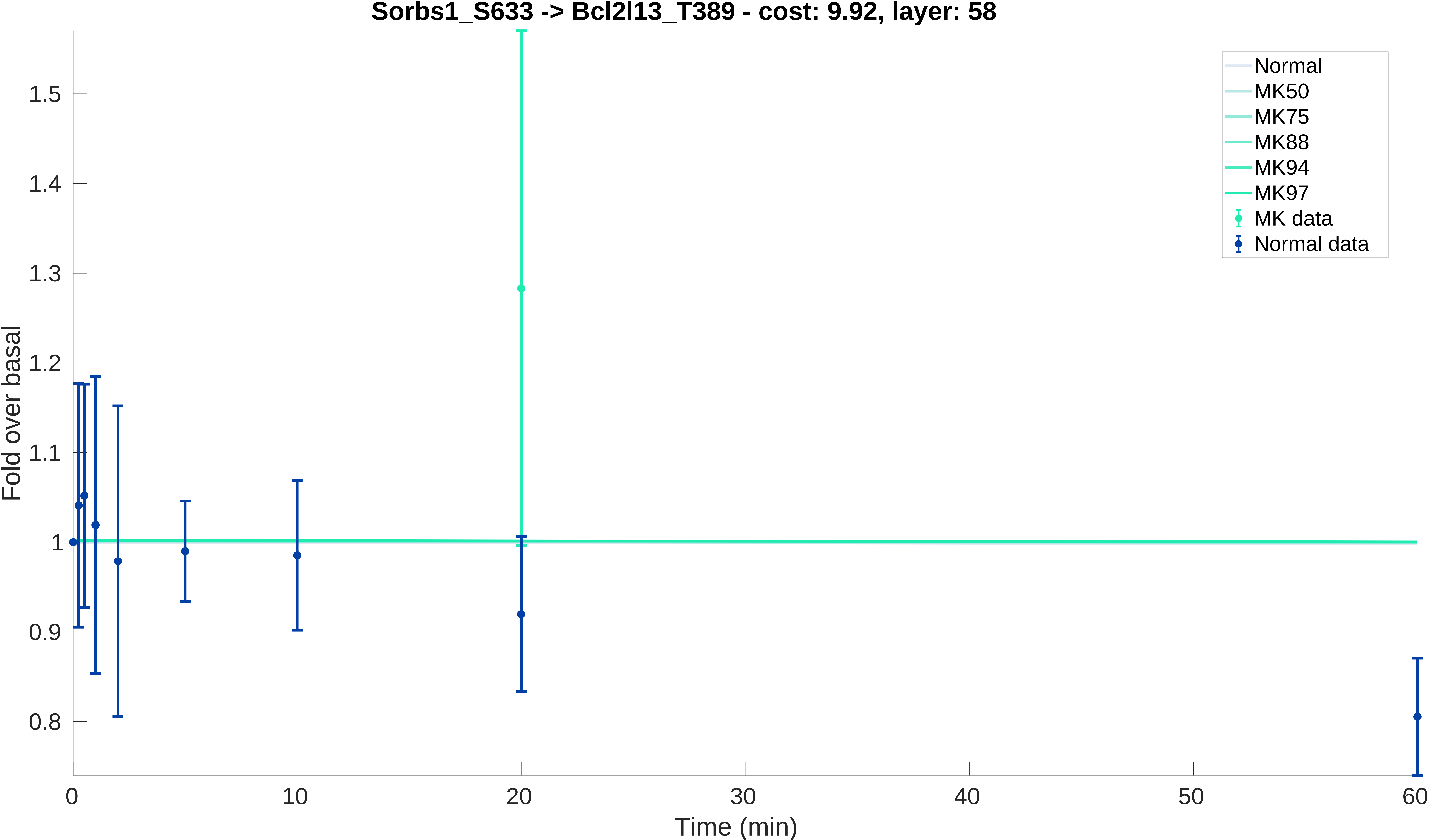

Sorbs1_S633 -> Bcl2l13_T389 - cost: 9.92, layer: 58
Fold over basal
1.5
1.4
1.3
1.2
1.1
1
0.9
0.8
0
10
20
30
40
50
60
Time (min)
Normal
MK50
MK75
MK88
MK94
MK97
MK data
Normal data

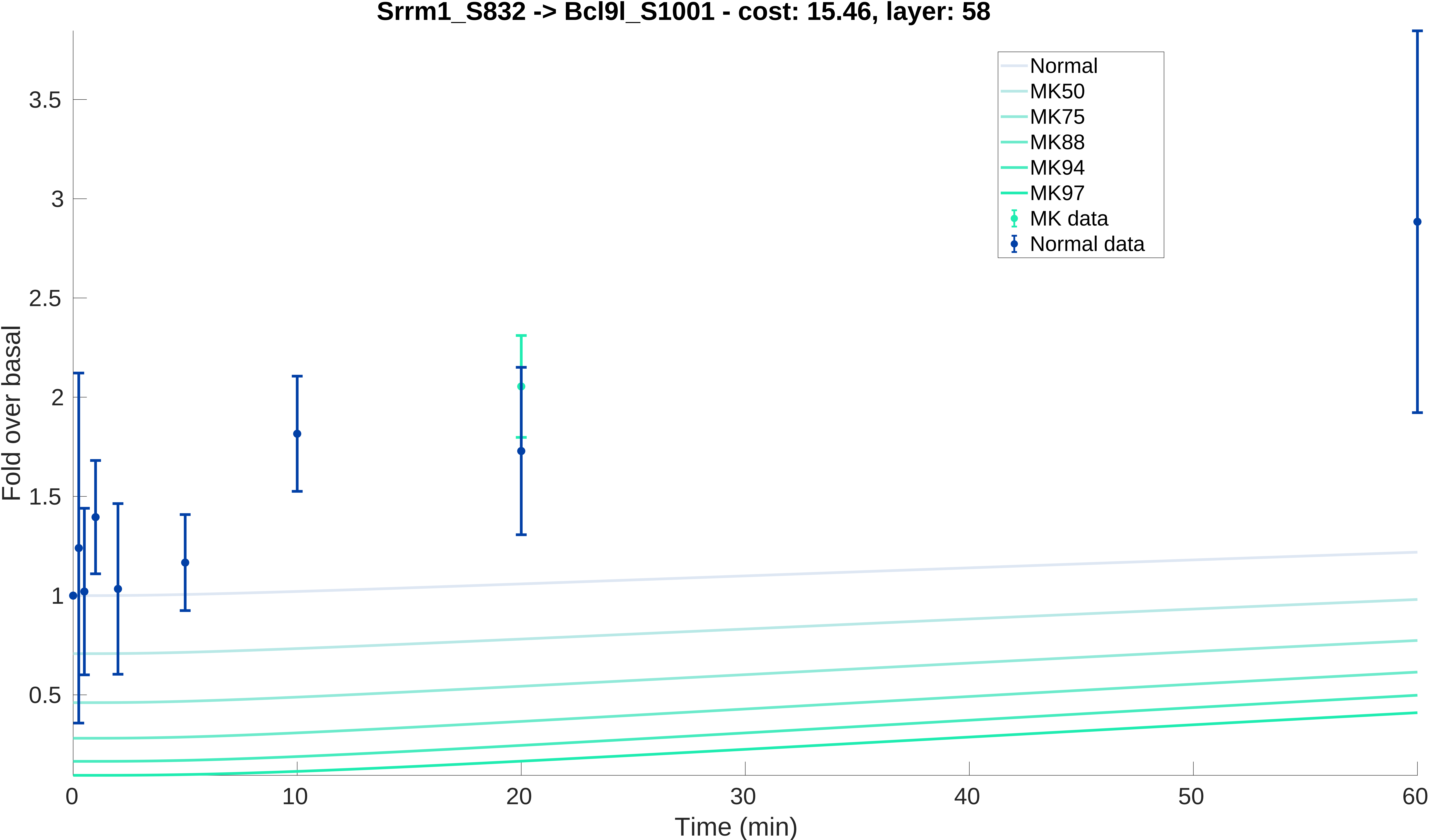

Srrm1_S832 -> Bcl9l_S1001 - cost: 15.46, layer: 58
Normal
MK50
MK75
MK88
MK94
MK97
MK data
Normal data
Fold over basal
3.5
3
2.5
2
1.5
1
0.5
0
10
20
30
40
50
60
Time (min)

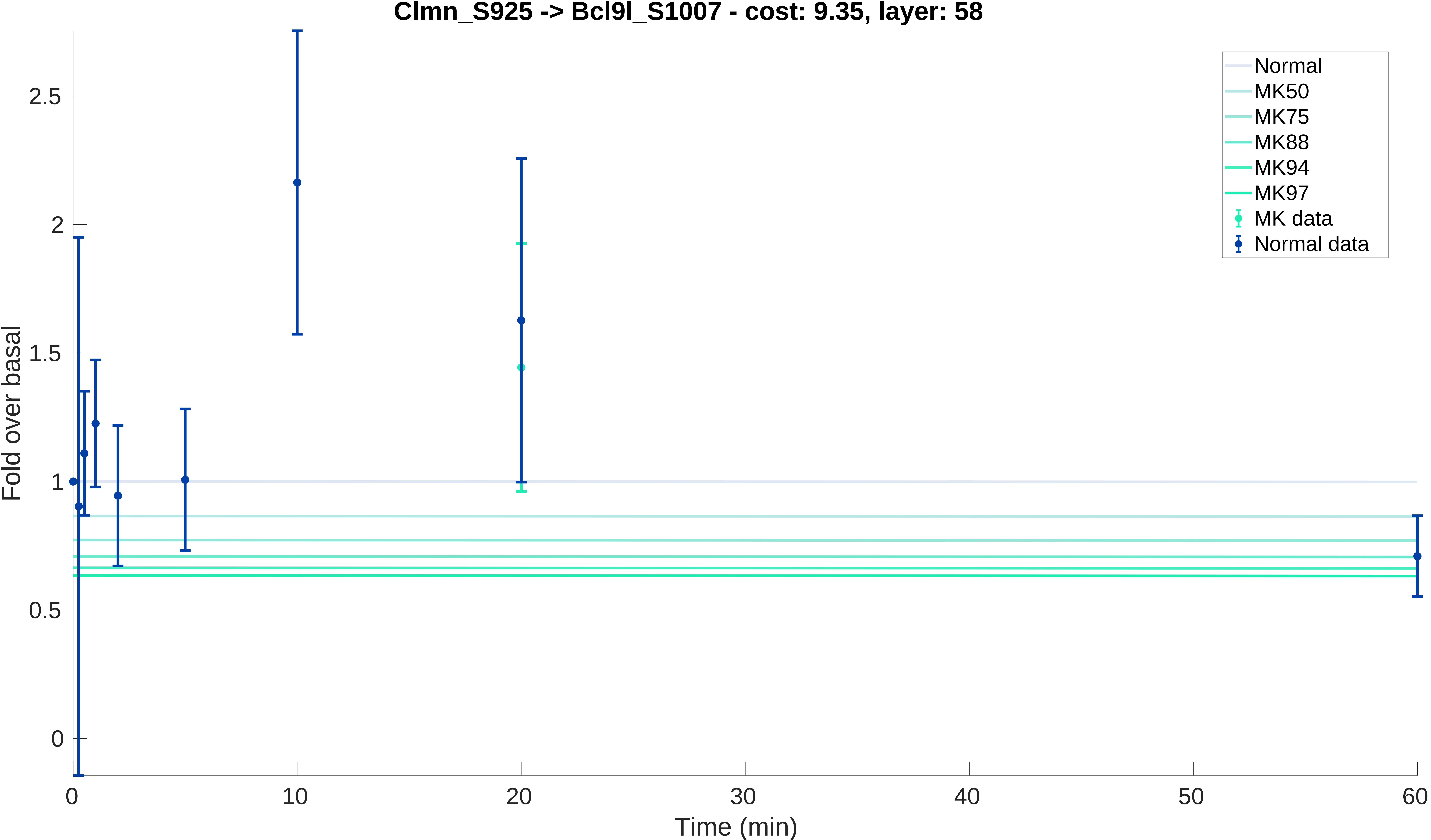

Clmn_S925 -> Bcl9l_S1007 - cost: 9.35, layer: 58
Fold over basal
Time (min)
0
0.5
1
1.5
2
2.5
0
10
20
30
40
50
60
Normal
MK50
MK75
MK88
MK94
MK97
MK data
Normal data

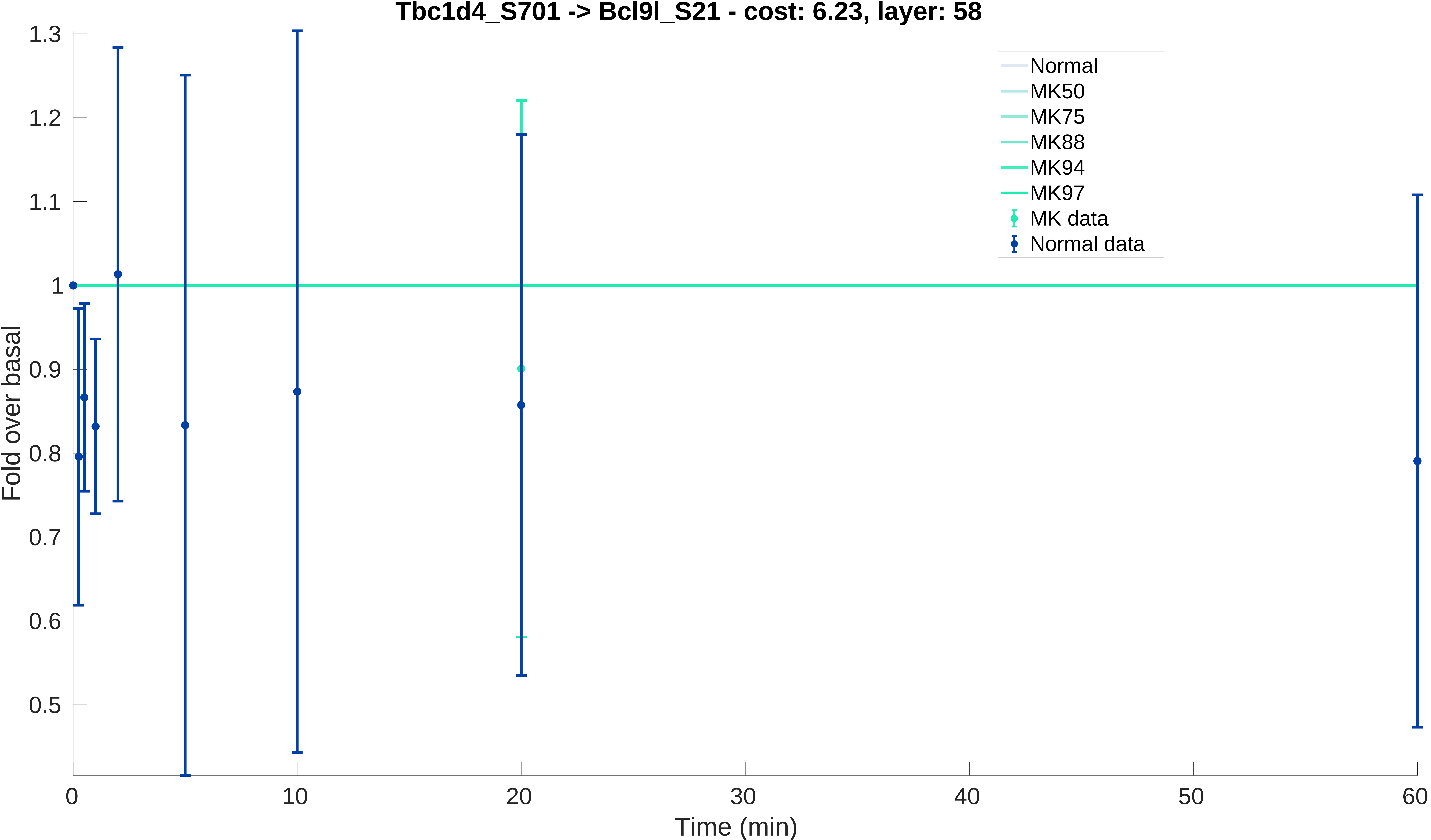
Tbc1d4_S701 -> Bcl9l_S21 - cost: 6.23, layer: 58
Fold over basal
Time (min)
1.3
1.2
1.1
1
0.9
0.8
0.7
0.6
0.5
0
10
20
30
40
50
60
Normal
MK50
MK75
MK88
MK94
MK97
MK data
Normal data

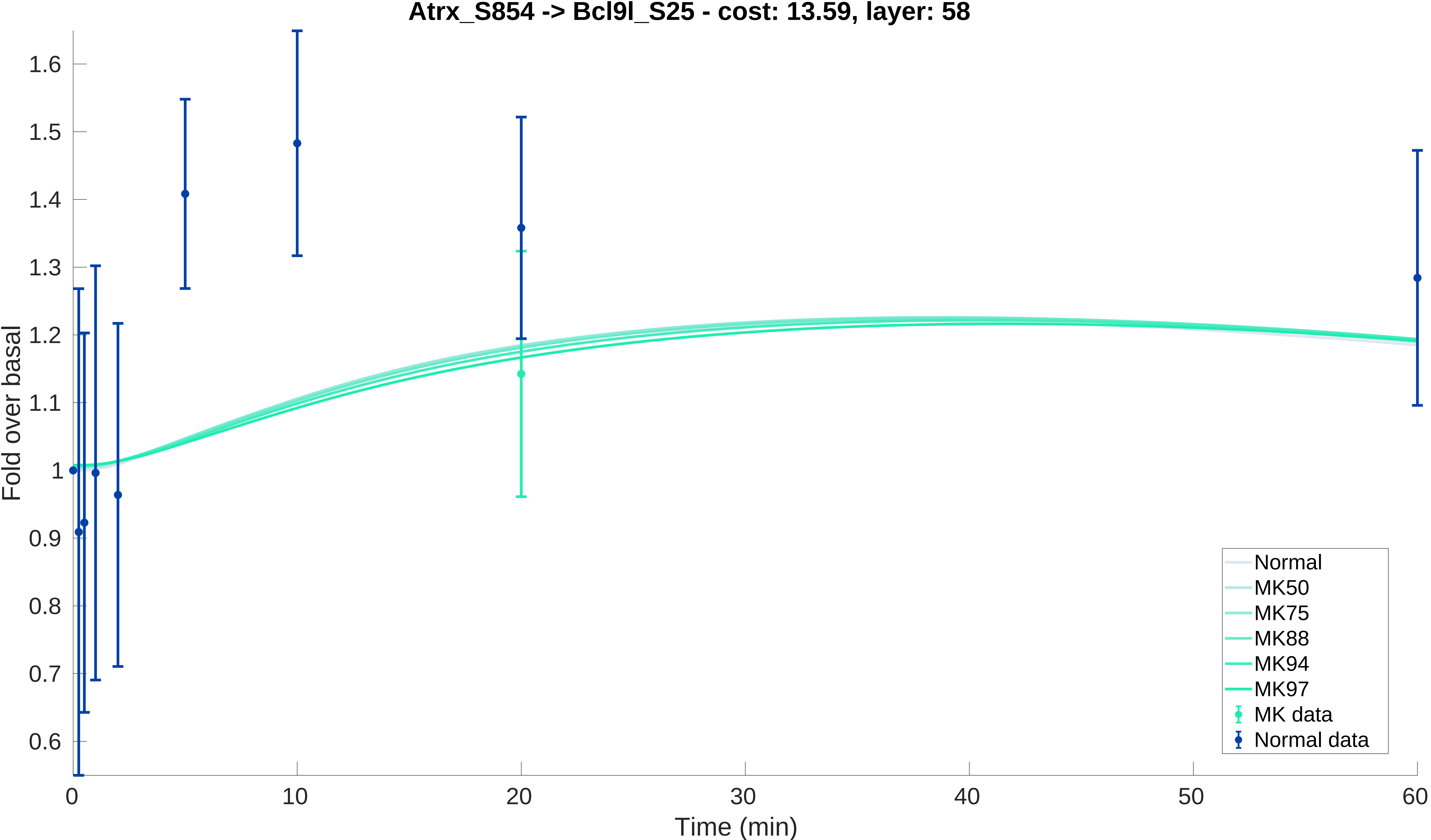

Atrx_S854 -> Bcl9l_S25 - cost: 13.59, layer: 58
Fold over basal
1.6
1.5
1.4
1.3
1.2
1.1
1
0.9
0.8
0.7
0.6
0
10
20
30
40
50
60
Time (min)
Normal
MK50
MK75
MK88
MK94
MK97
MK data
Normal data

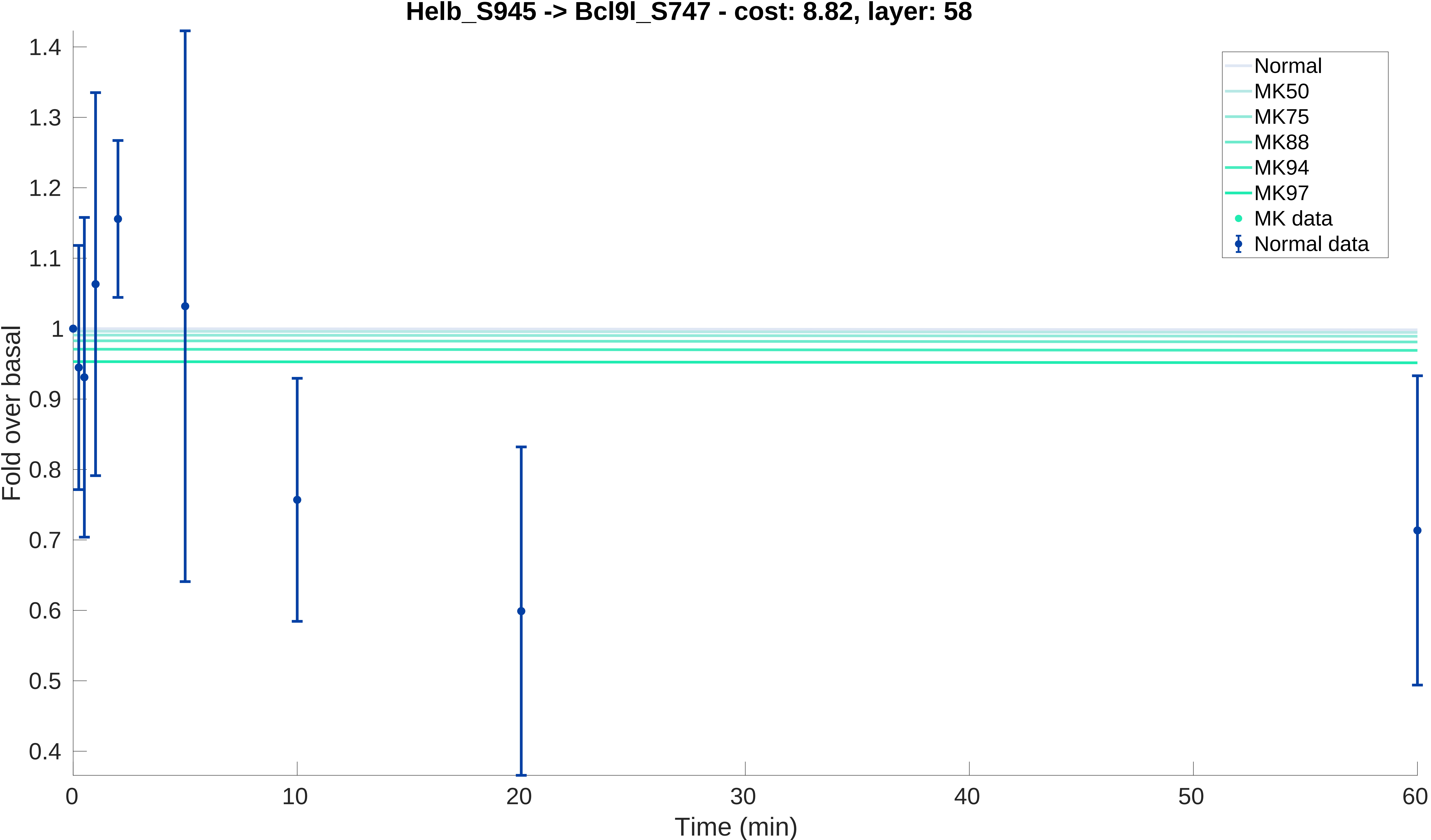

Helb_S945 -> Bcl9l_S747 - cost: 8.82, layer: 58
Fold over basal
1.4
1.3
1.2
1.1
1
0.9
0.8
0.7
0.6
0.5
0.4
0
10
20
30
40
50
60
Time (min)
Normal
MK50
MK75
MK88
MK94
MK97
MK data
Normal data

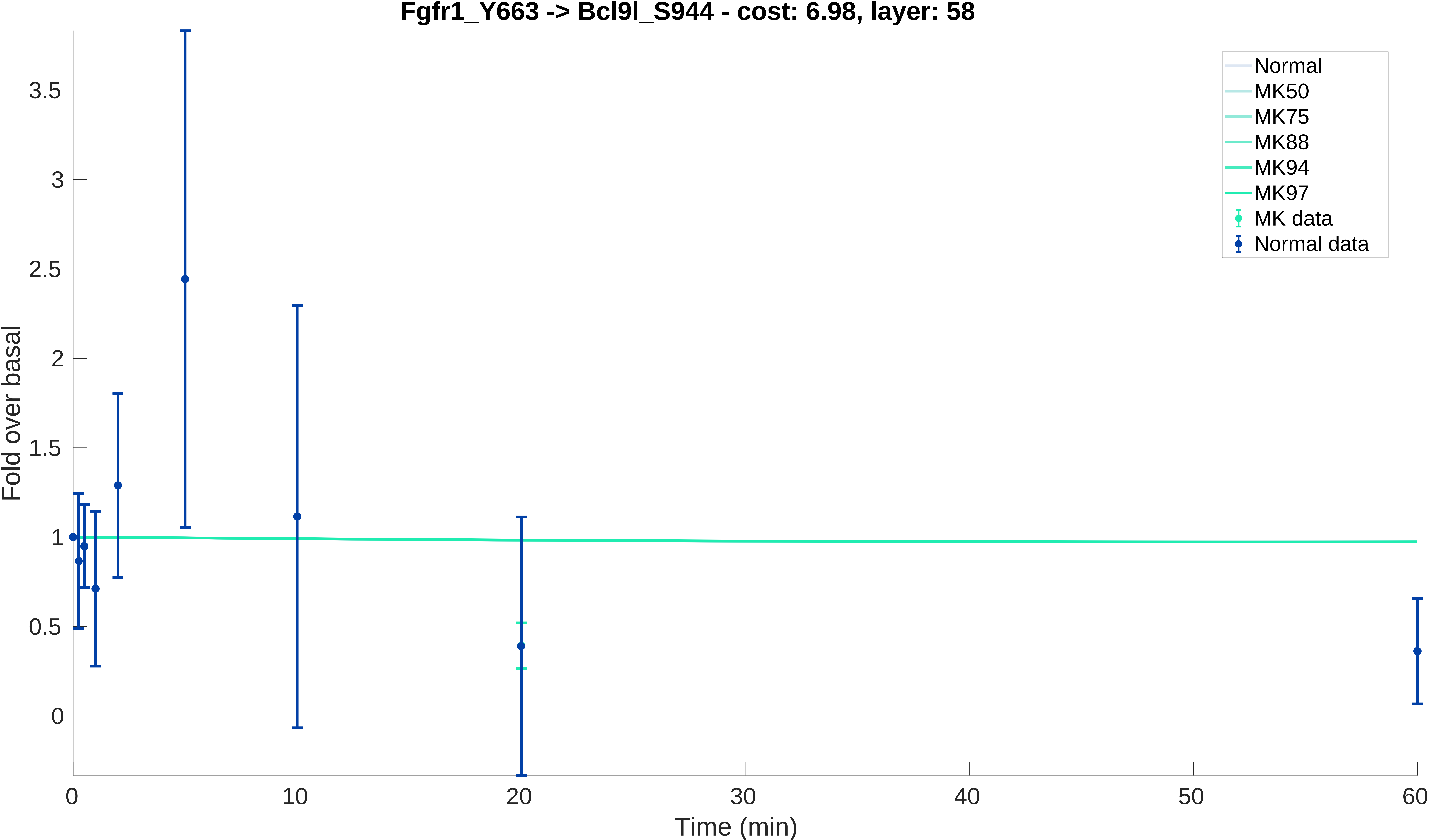

Fgfr1_Y663 -> Bcl9l_S944 - cost: 6.98, layer: 58
Fold over basal
Time (min)
0
0.5
1
1.5
2
2.5
3
3.5
0
10
20
30
40
50
60
Normal
MK50
MK75
MK88
MK94
MK97
MK data
Normal data

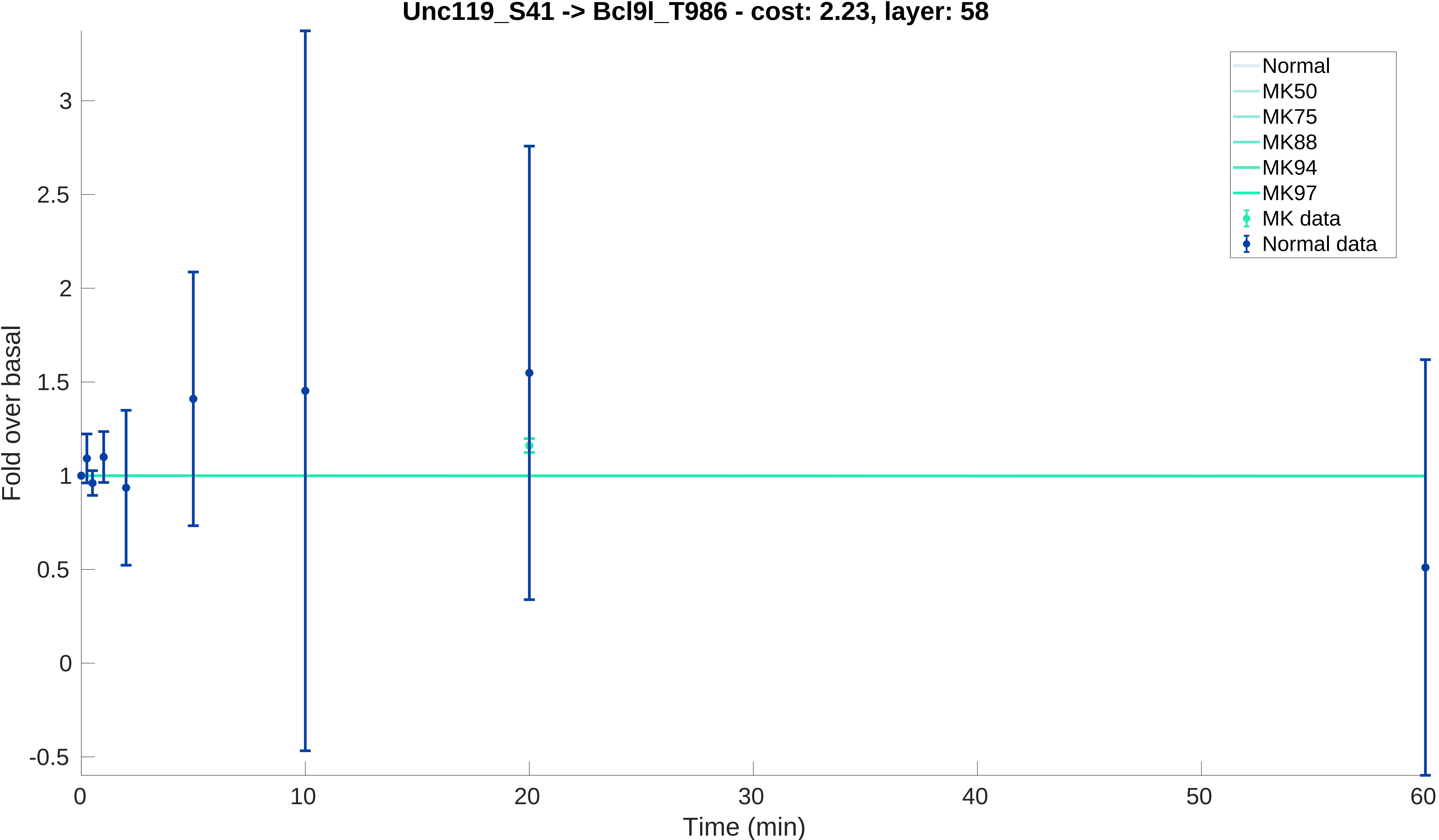

Unc119_S41 -> Bcl9l_T986 - cost: 2.23, layer: 58
Normal
MK50
MK75
MK88
MK94
MK97
MK data
Normal data
Fold over basal
Time (min)
3
2.5
2
1.5
1
0.5
0
-0.5
0
10
20
30
40
50
60

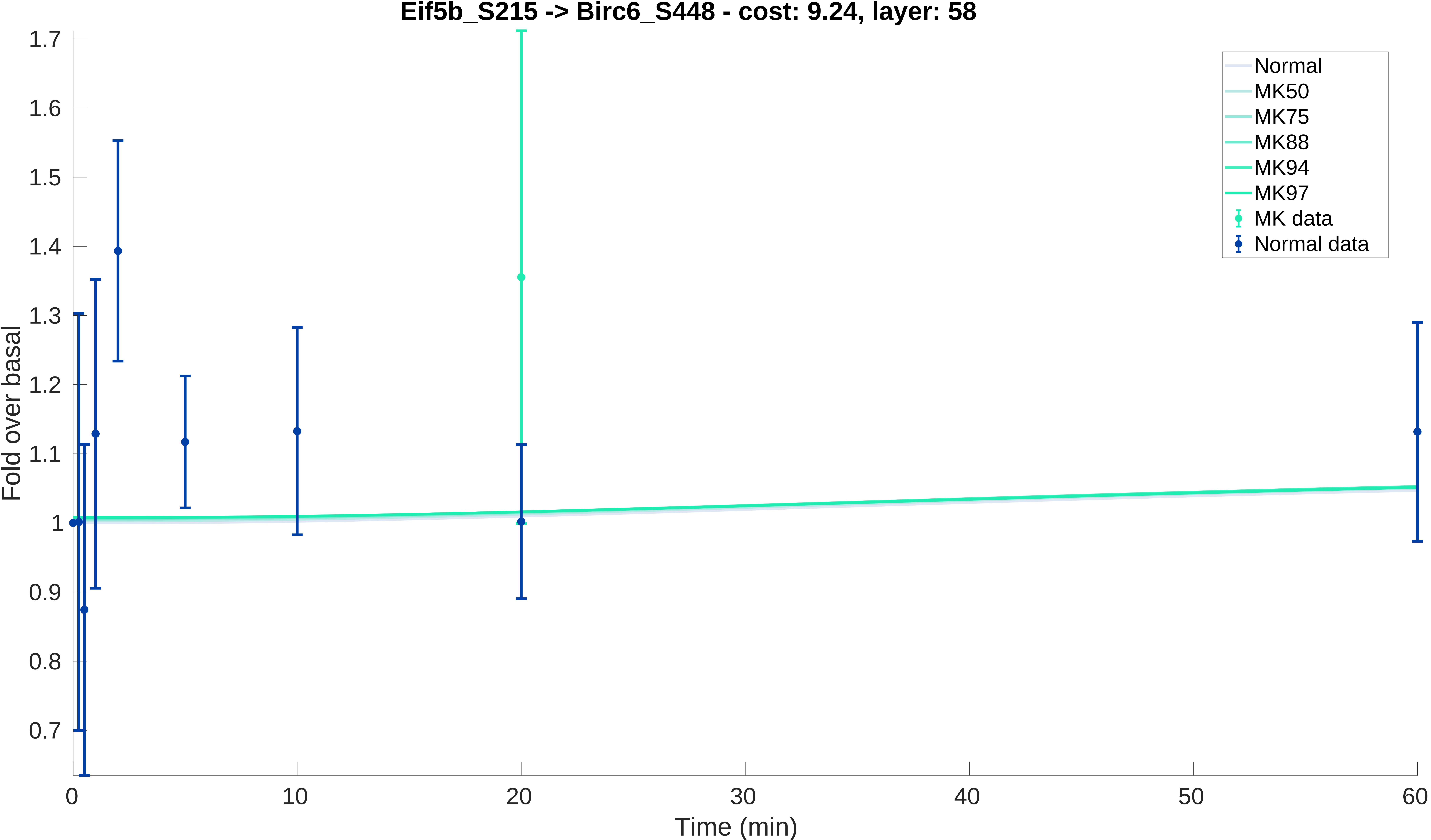

Eif5b_S215 -> Birc6_S448 - cost: 9.24, layer: 58
Fold over basal
Time (min)
Normal
MK50
MK75
MK88
MK94
MK97
MK data
Normal data
0.7
0.8
0.9
1
1.1
1.2
1.3
1.4
1.5
1.6
1.7
0
10
20
30
40
50
60

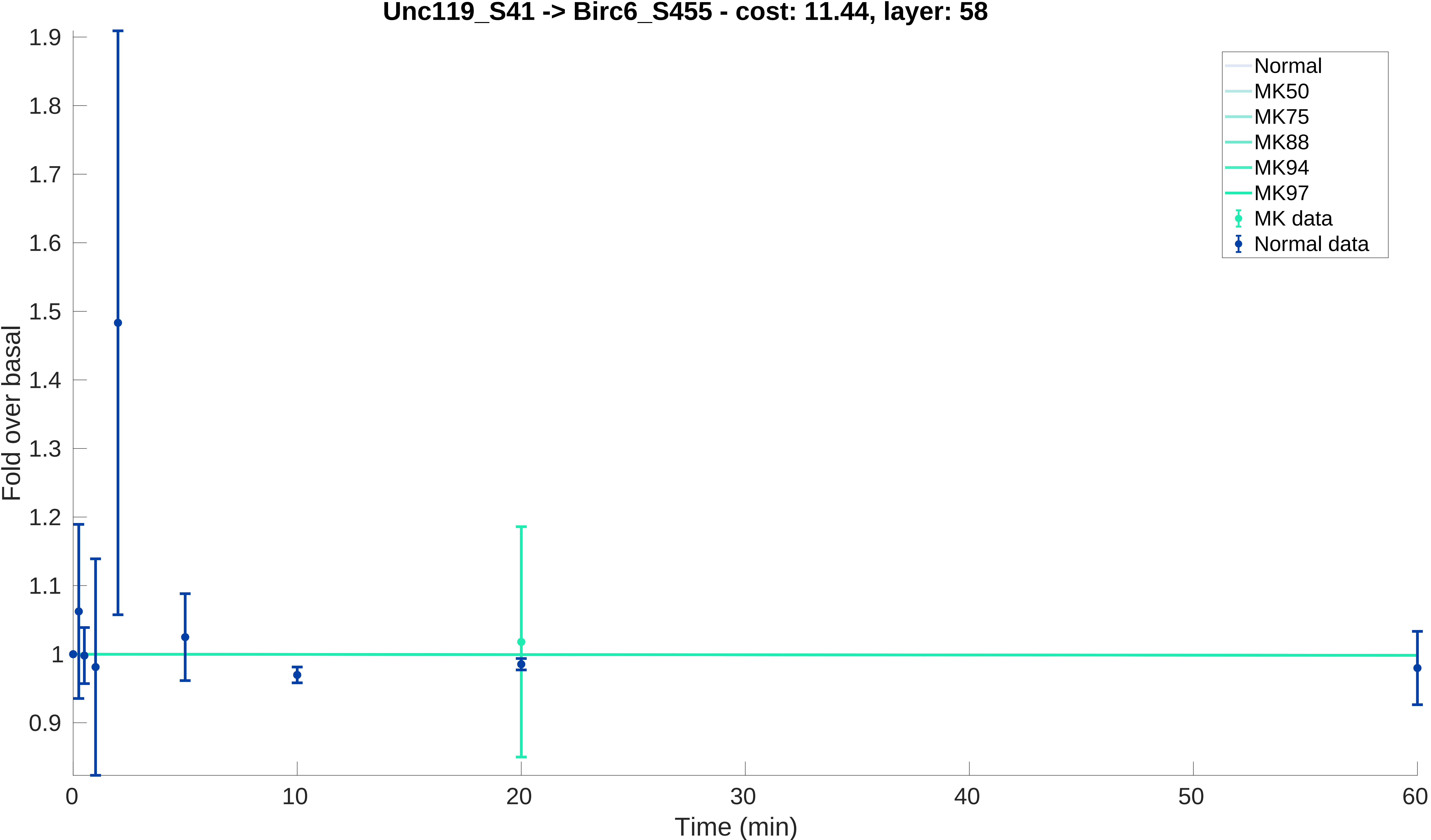
Unc119_S41 -> Birc6_S455 - cost: 11.44, layer: 58
Fold over basal
Time (min)
0.9
1
1.1
1.2
1.3
1.4
1.5
1.6
1.7
1.8
1.9
0
10
20
30
40
50
60
Normal
MK50
MK75
MK88
MK94
MK97
MK data
Normal data

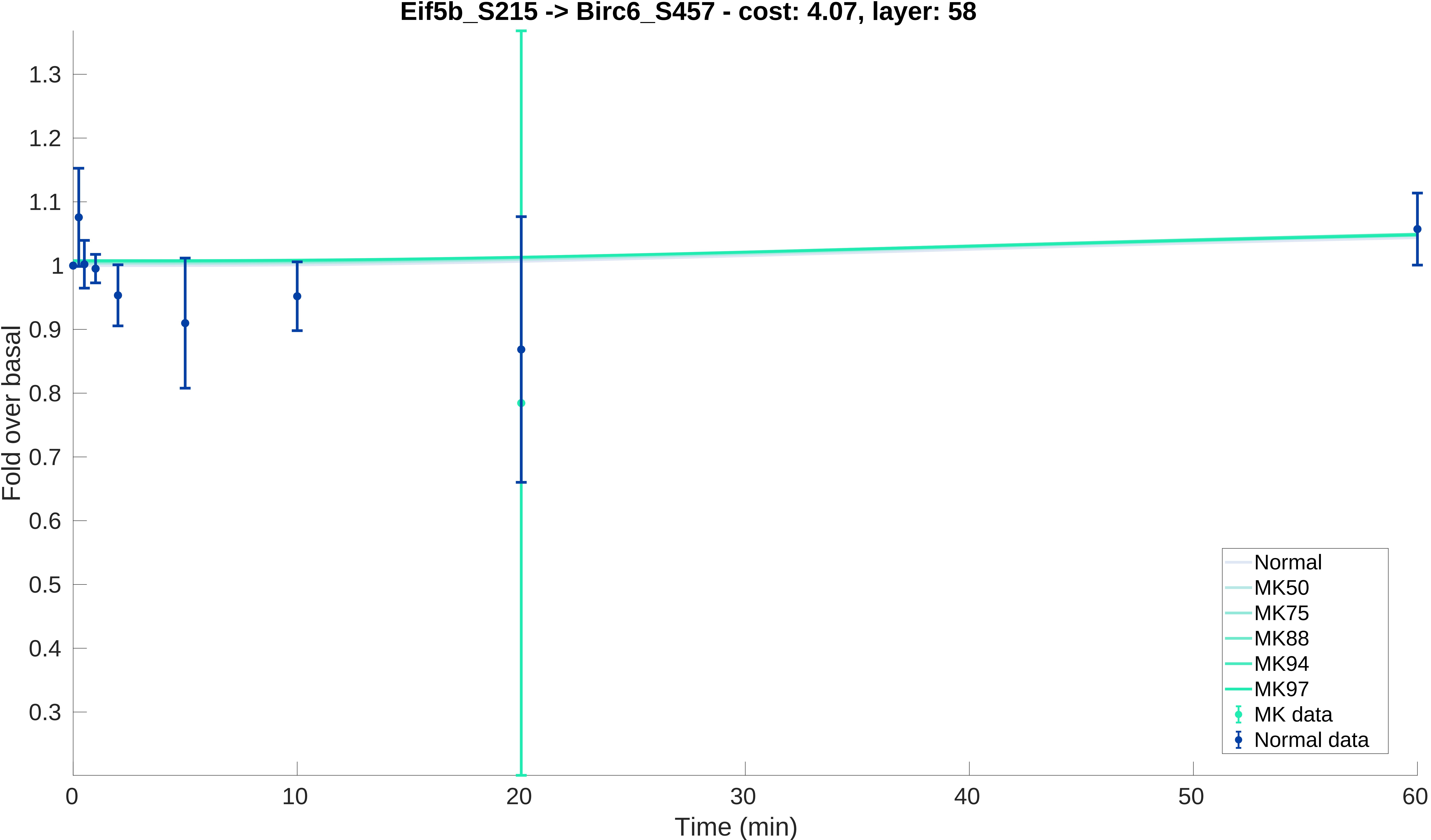
Eif5b_S215 -> Birc6_S457 - cost: 4.07, layer: 58
Fold over basal
Time (min)
1.3
1.2
1.1
1
0.9
0.8
0.7
0.6
0.5
0.4
0.3
0
10
20
30
40
50
60
Normal
MK50
MK75
MK88
MK94
MK97
MK data
Normal data

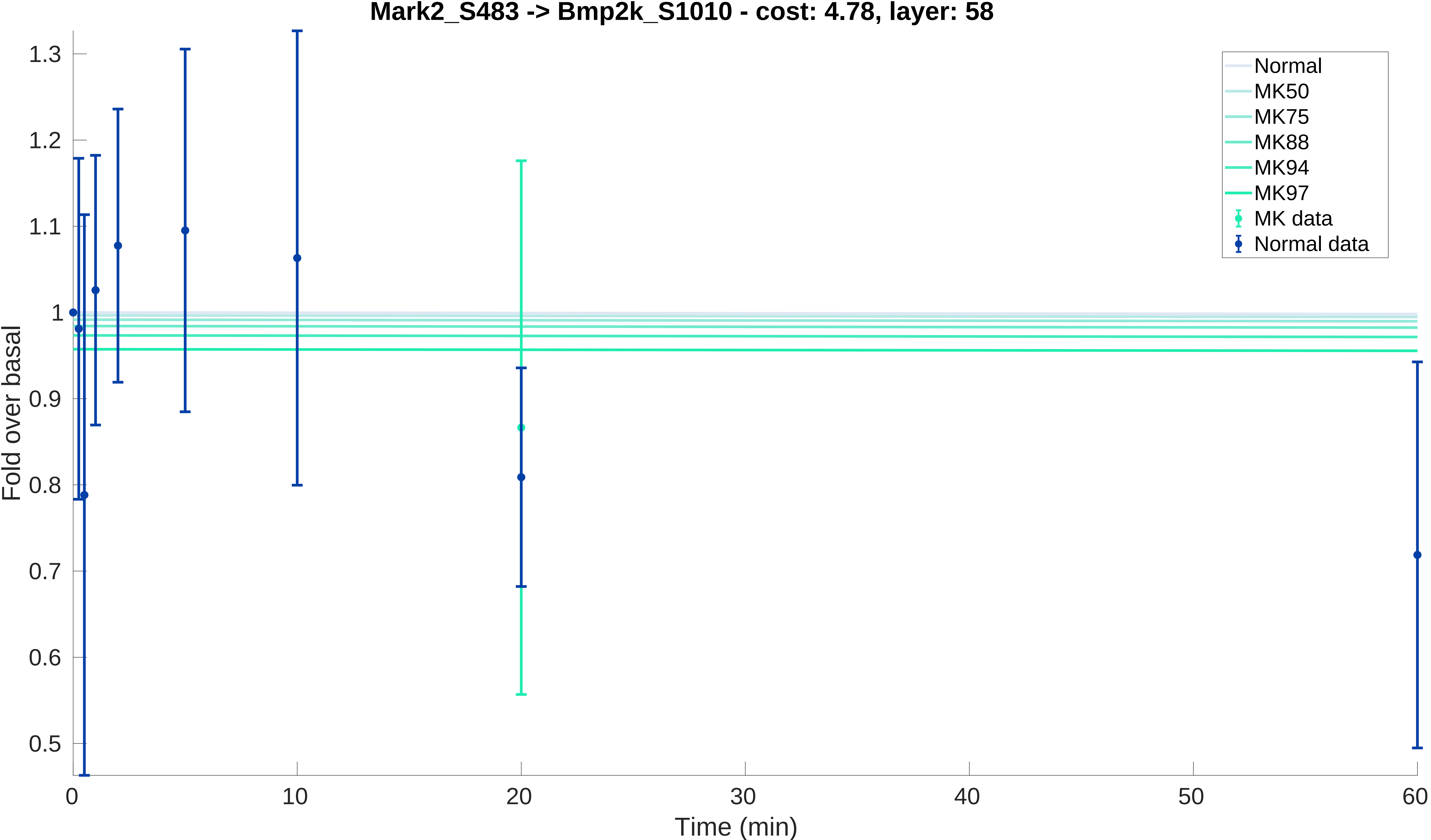

Mark2_S483 -> Bmp2k_S1010 - cost: 4.78, layer: 58
Fold over basal
Time (min)
1.3
1.2
1.1
1
0.9
0.8
0.7
0.6
0.5
0
10
20
30
40
50
60
Normal
MK50
MK75
MK88
MK94
MK97
MK data
Normal data

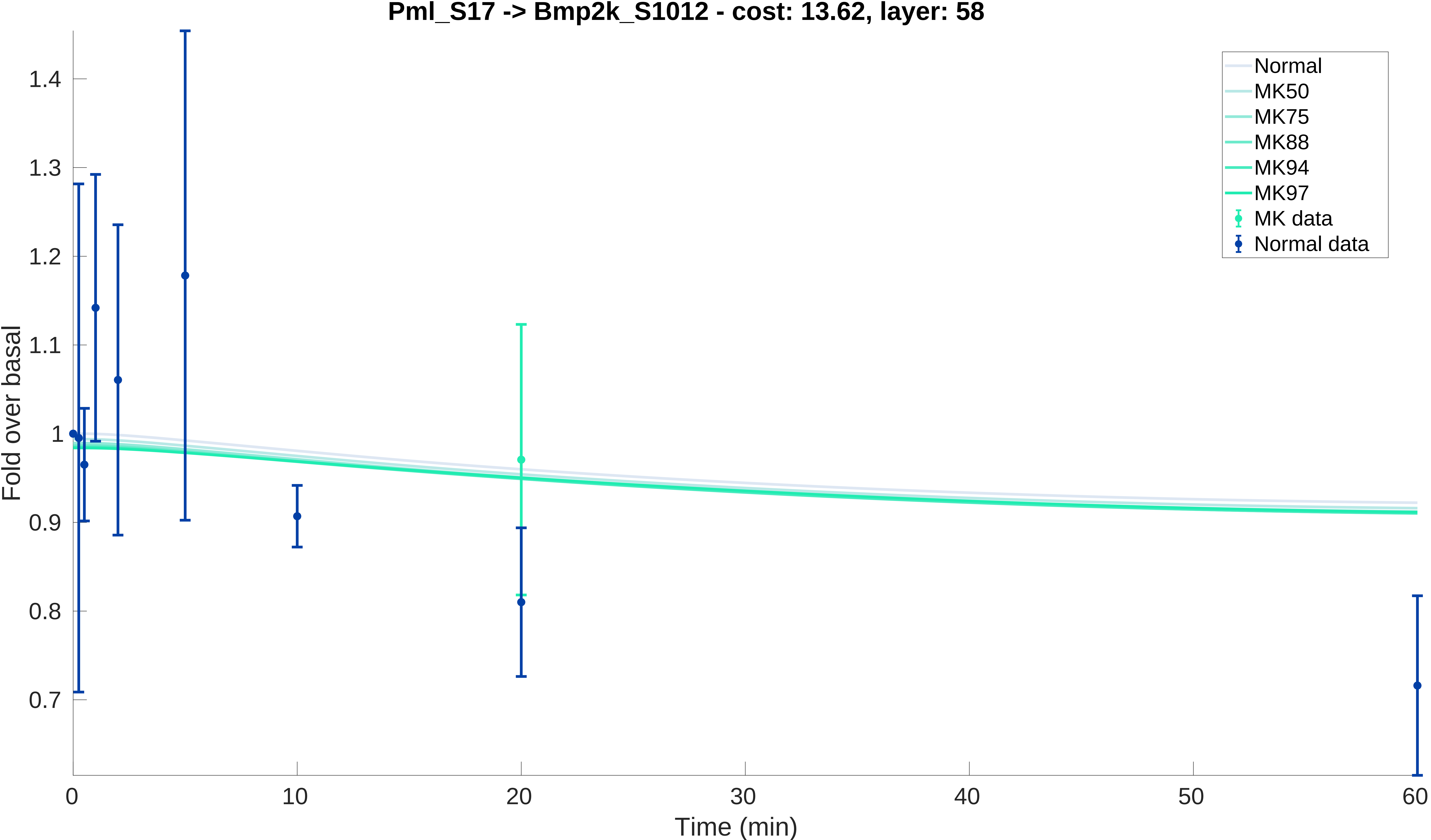

Pml_S17 -> Bmp2k_S1012 - cost: 13.62, layer: 58
Fold over basal
Time (min)
Normal
MK50
MK75
MK88
MK94
MK97
MK data
Normal data
1.4
1.3
1.2
1.1
1
0.9
0.8
0.7
0
10
20
30
40
50
60

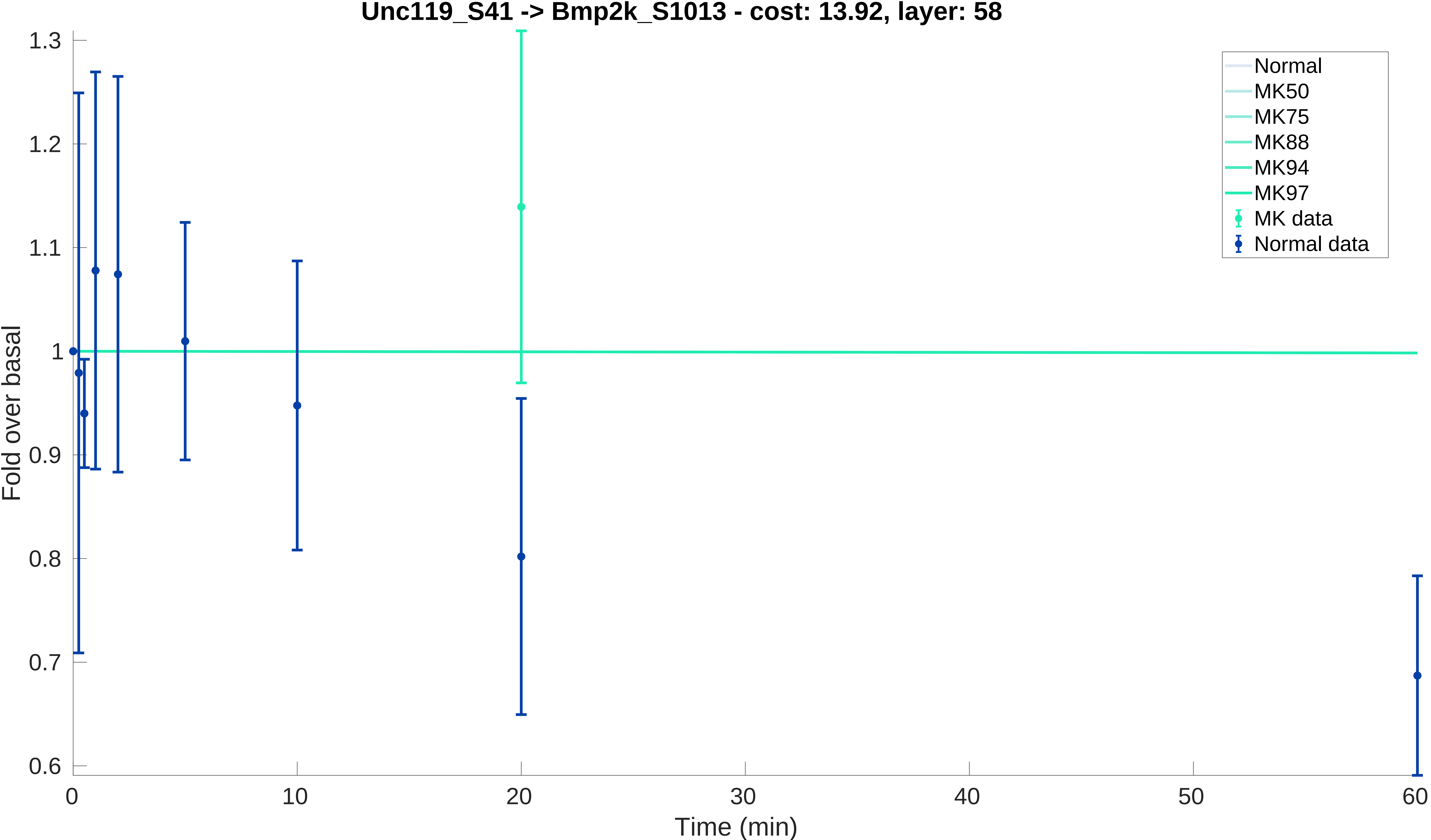

Unc119_S41 -> Bmp2k_S1013 - cost: 13.92, layer: 58
Fold over basal
Time (min)
0
10
20
30
40
50
60
0.6
0.7
0.8
0.9
1
1.1
1.2
1.3
Normal
MK50
MK75
MK88
MK94
MK97
MK data
Normal data

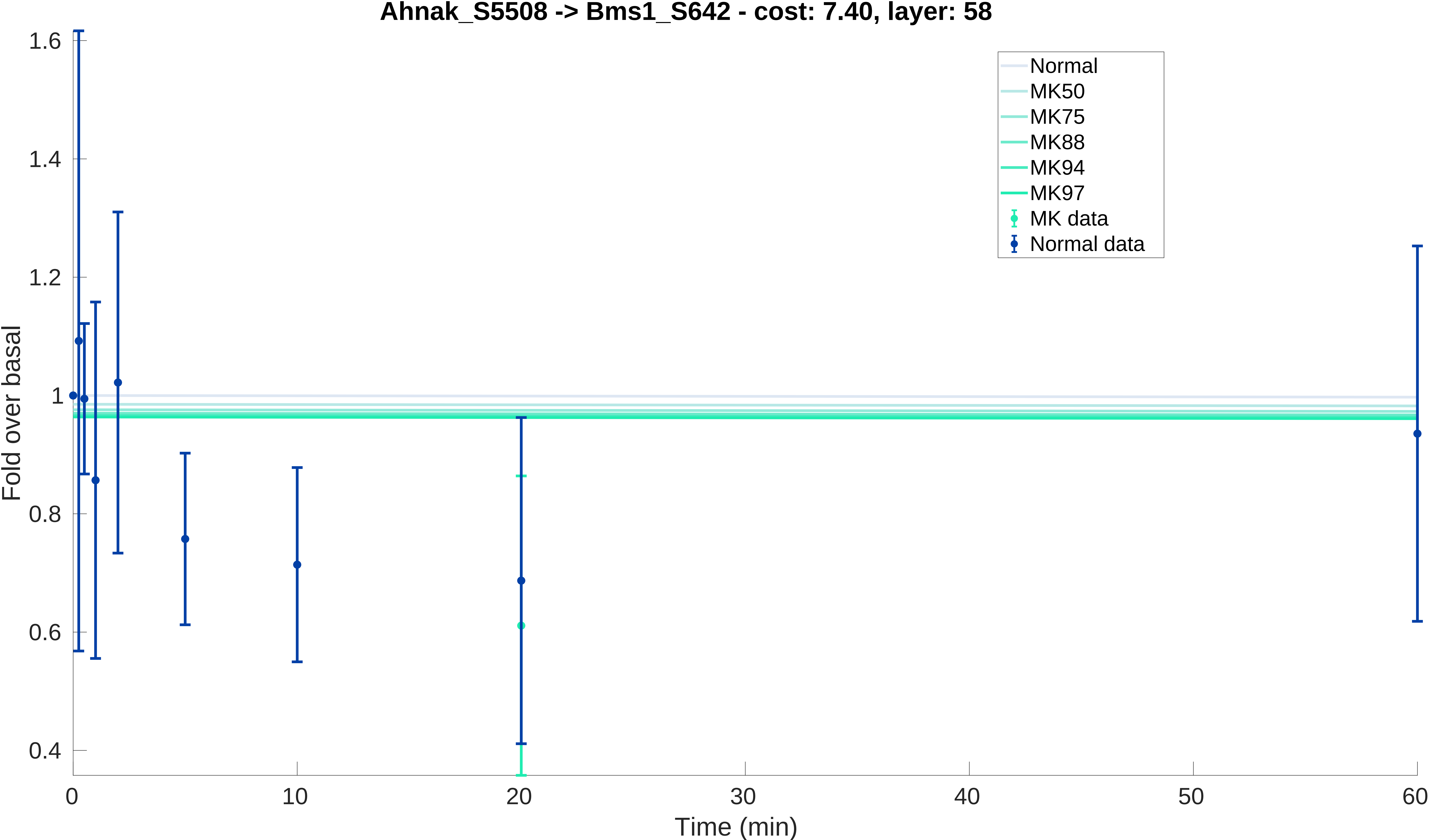

Ahnak_S5508 -> Bms1_S642 - cost: 7.40, layer: 58
Fold over basal
Time (min)
Normal
MK50
MK75
MK88
MK94
MK97
MK data
Normal data
1.6
1.4
1.2
1
0.8
0.6
0.4
0
10
20
30
40
50
60

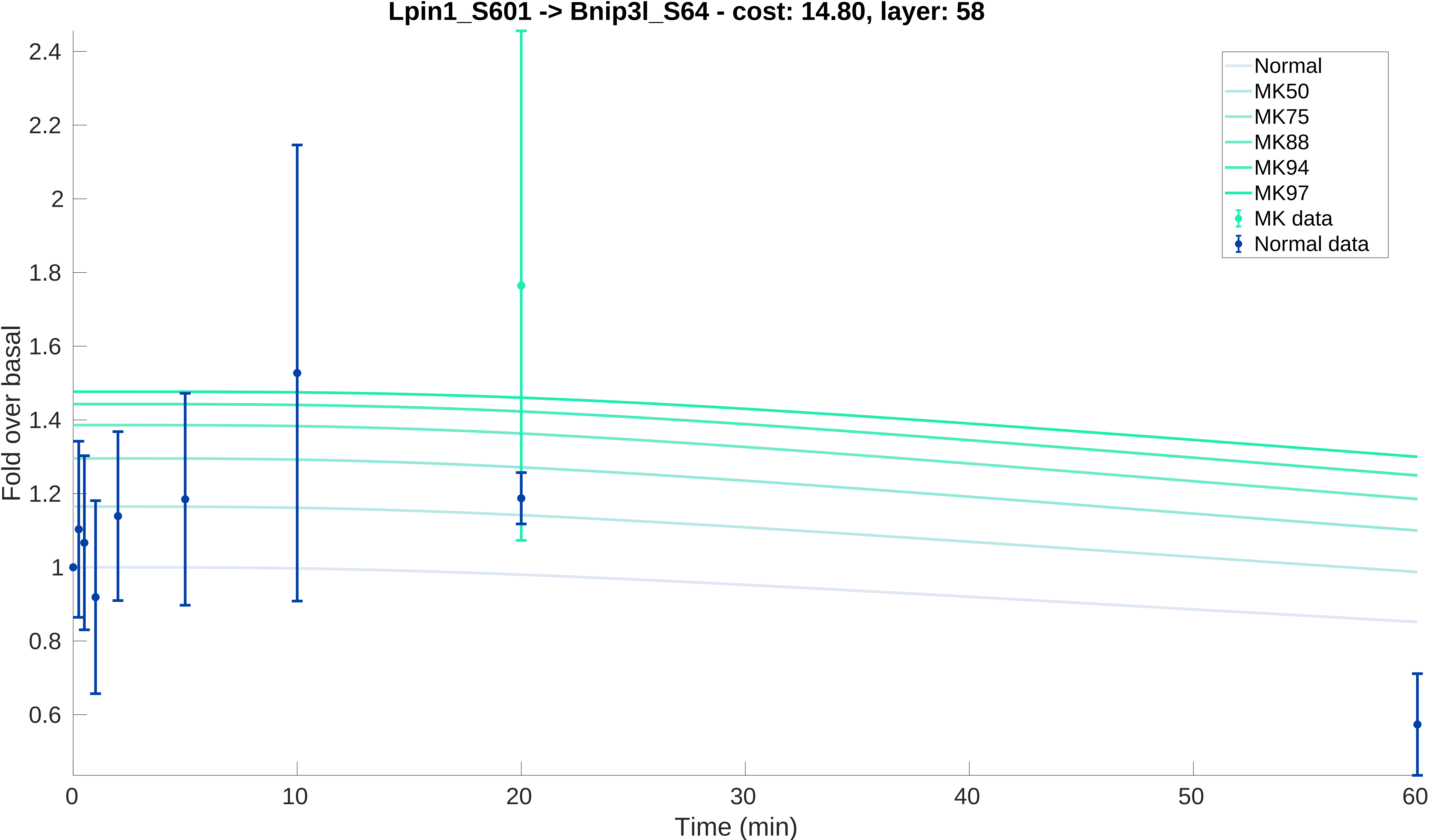
Lpin1_S601 -> Bnip3l_S64 - cost: 14.80, layer: 58
Fold over basal
Time (min)
Normal
MK50
MK75
MK88
MK94
MK97
MK data
Normal data
0.6
0.8
1
1.2
1.4
1.6
1.8
2
2.2
2.4
0
10
20
30
40
50
60

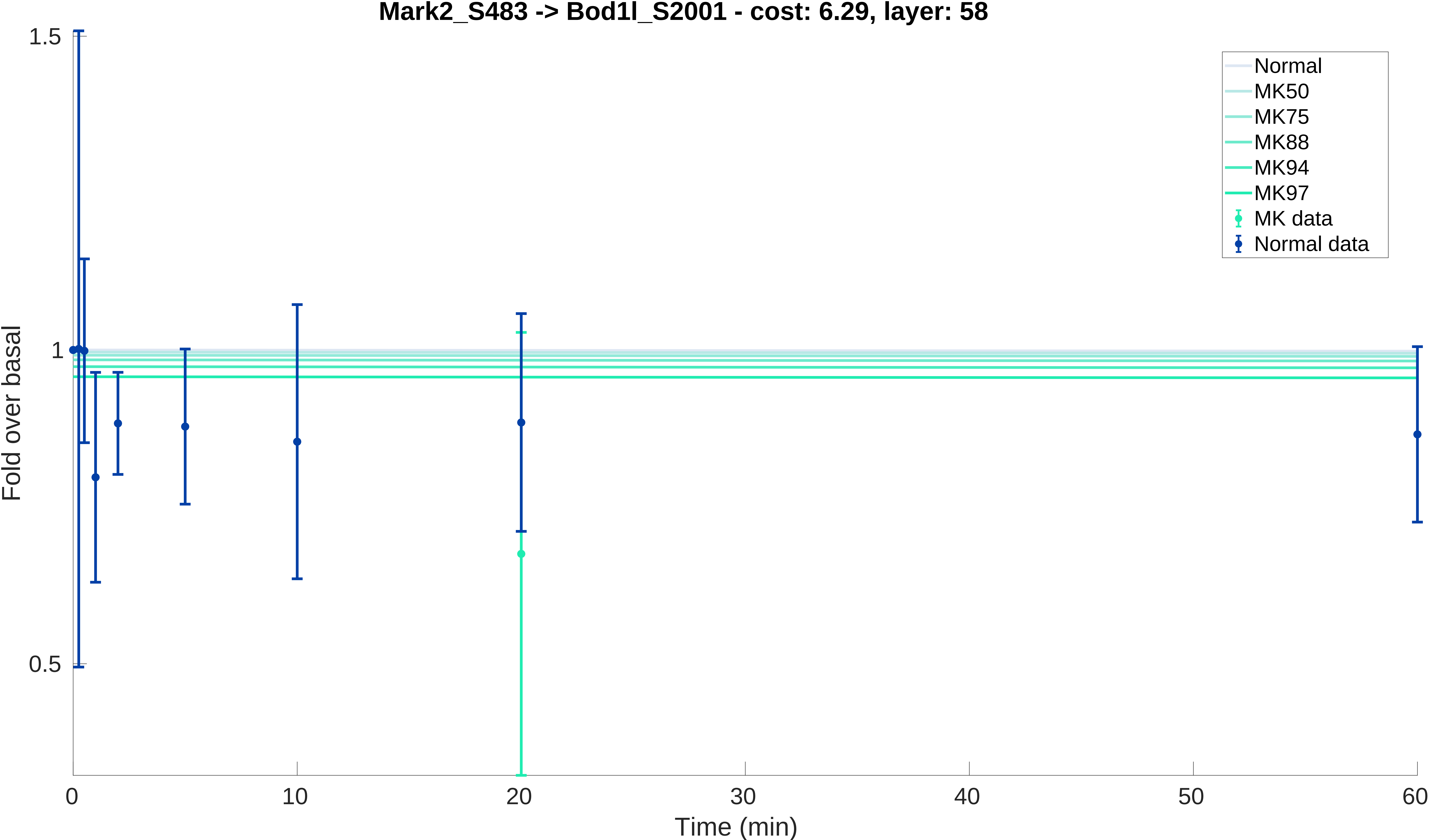

Mark2_S483 -> Bod1l_S2001 - cost: 6.29, layer: 58
Fold over basal
1.5
1
0.5
0
10
20
30
40
50
60
Time (min)
Normal
MK50
MK75
MK88
MK94
MK97
MK data
Normal data

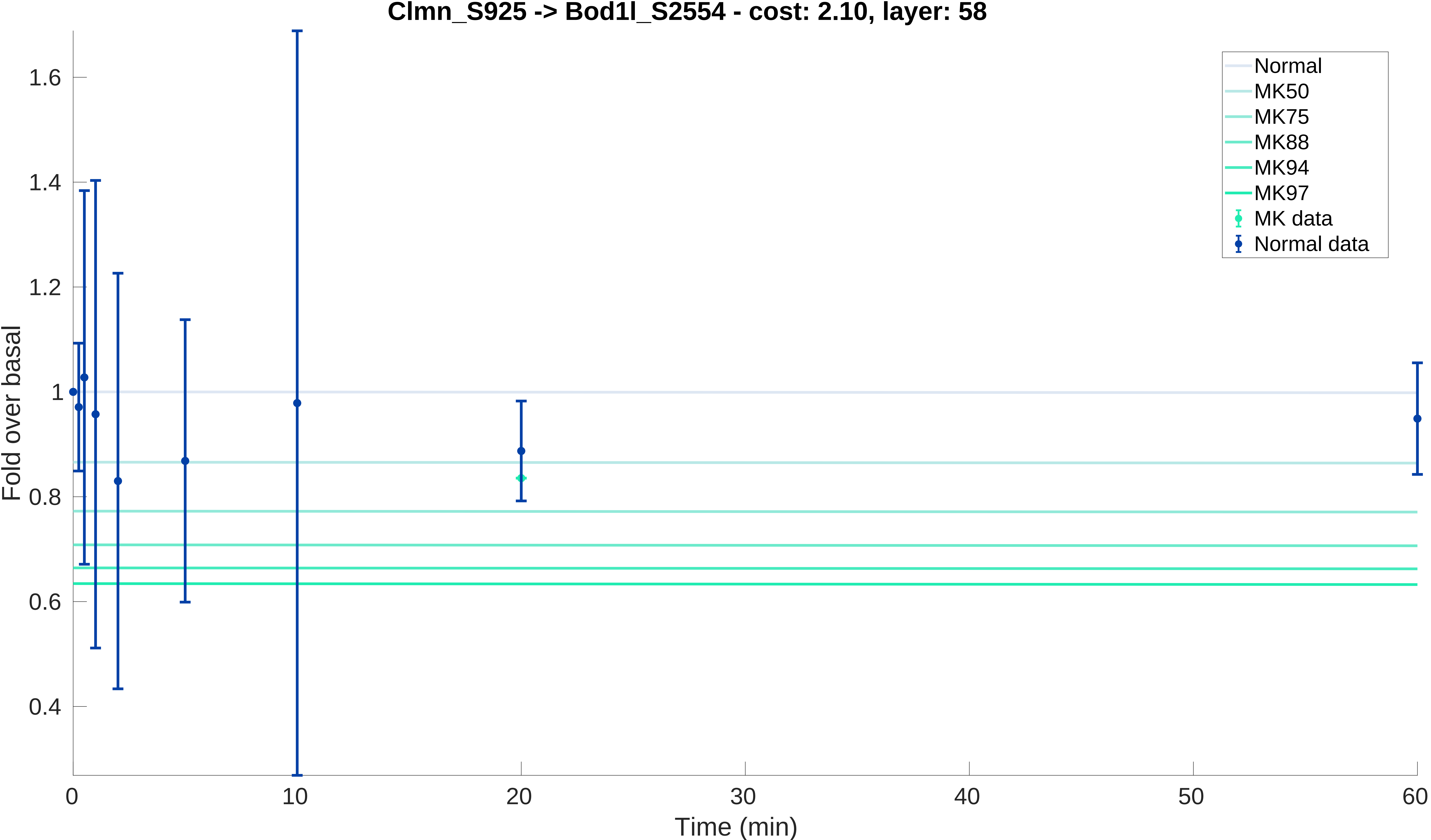
Clmn_S925 -> Bod1l_S2554 - cost: 2.10, layer: 58
Fold over basal
Time (min)
0.4
0.6
0.8
1
1.2
1.4
1.6
0
10
20
30
40
50
60
Normal
MK50
MK75
MK88
MK94
MK97
MK data
Normal data

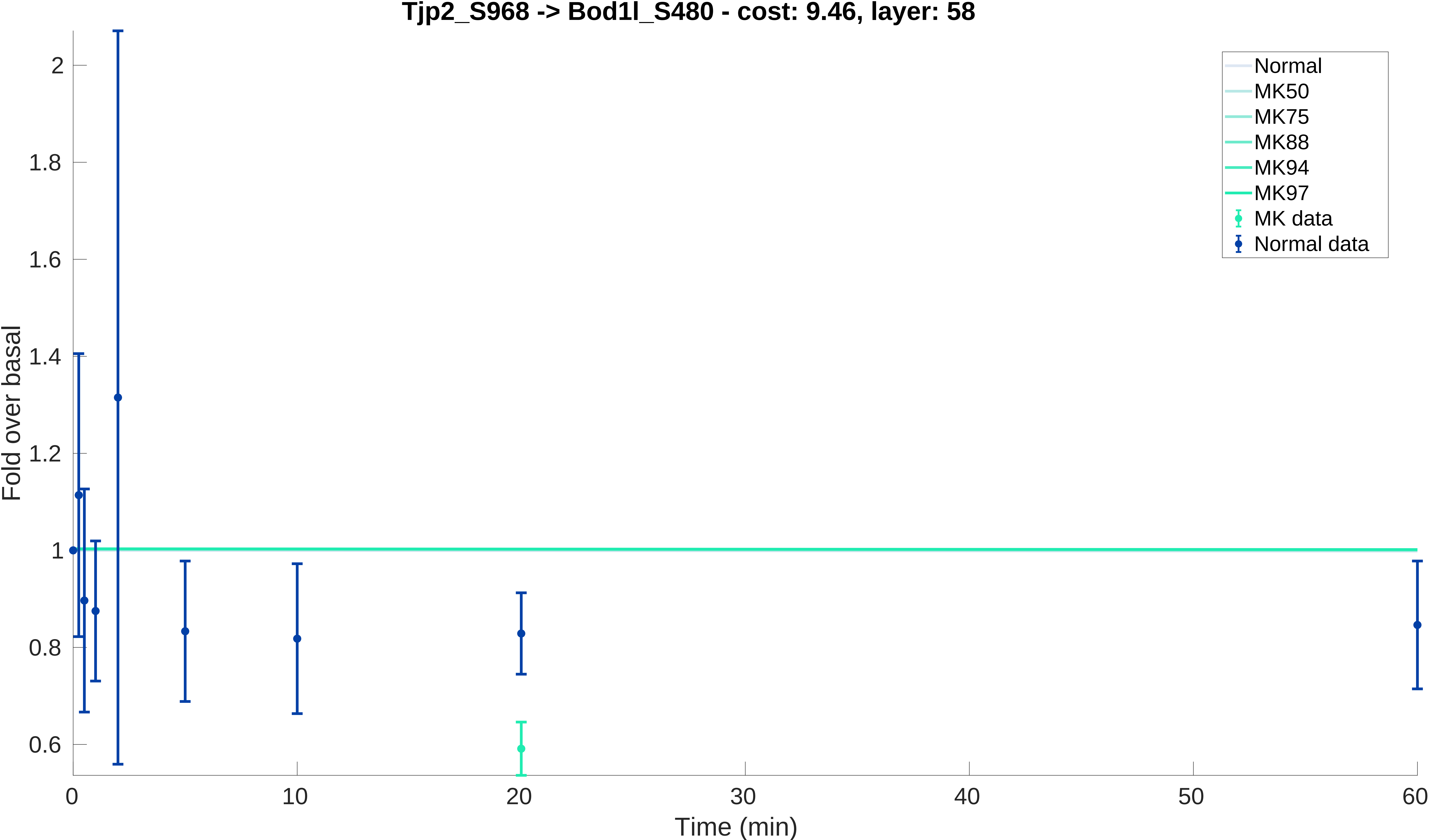
Tjp2_S968 -> Bod1l_S480 - cost: 9.46, layer: 58
Fold over basal
Time (min)
0.6
0.8
1
1.2
1.4
1.6
1.8
2
0
10
20
30
40
50
60
Normal
MK50
MK75
MK88
MK94
MK97
MK data
Normal data

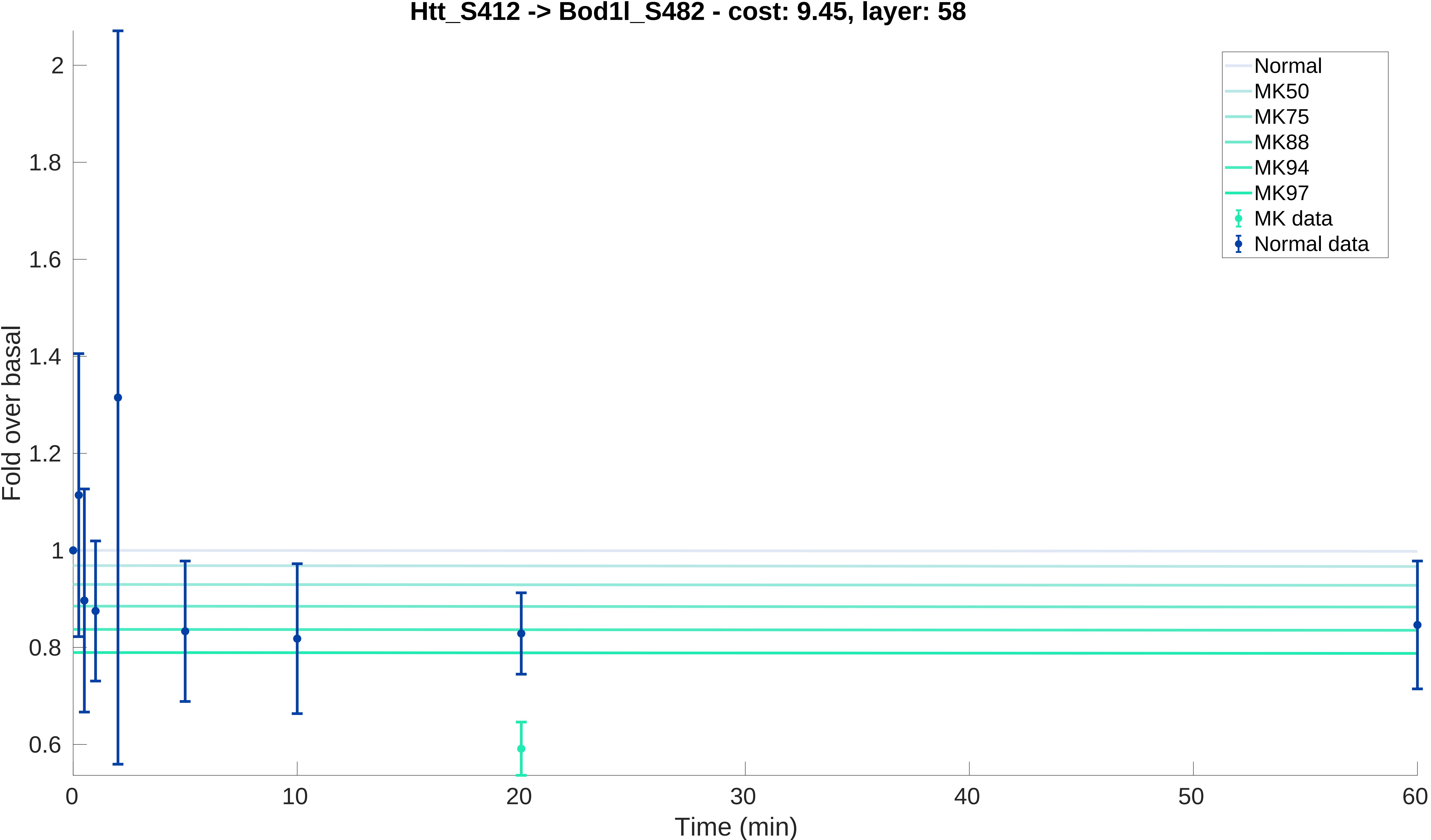

Htt_S412 -> Bod1l_S482 - cost: 9.45, layer: 58
Fold over basal
Time (min)
0.6
0.8
1
1.2
1.4
1.6
1.8
2
0
10
20
30
40
50
60
Normal
MK50
MK75
MK88
MK94
MK97
MK data
Normal data

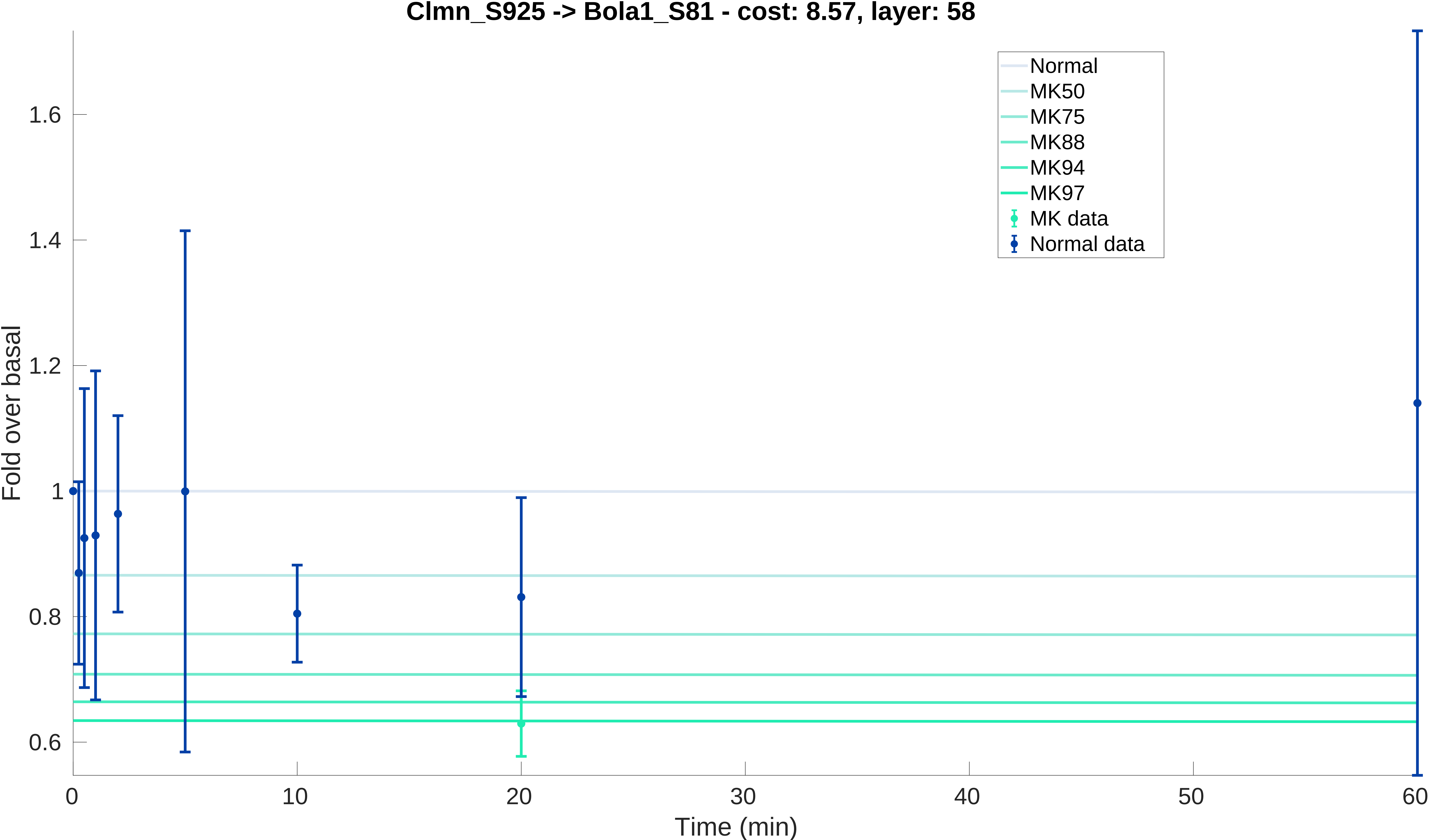

Clmn_S925 -> Bola1_S81 - cost: 8.57, layer: 58
Normal
MK50
MK75
MK88
MK94
MK97
MK data
Normal data
Fold over basal
1.6
1.4
1.2
1
0.8
0.6
0
10
20
30
40
50
60
Time (min)

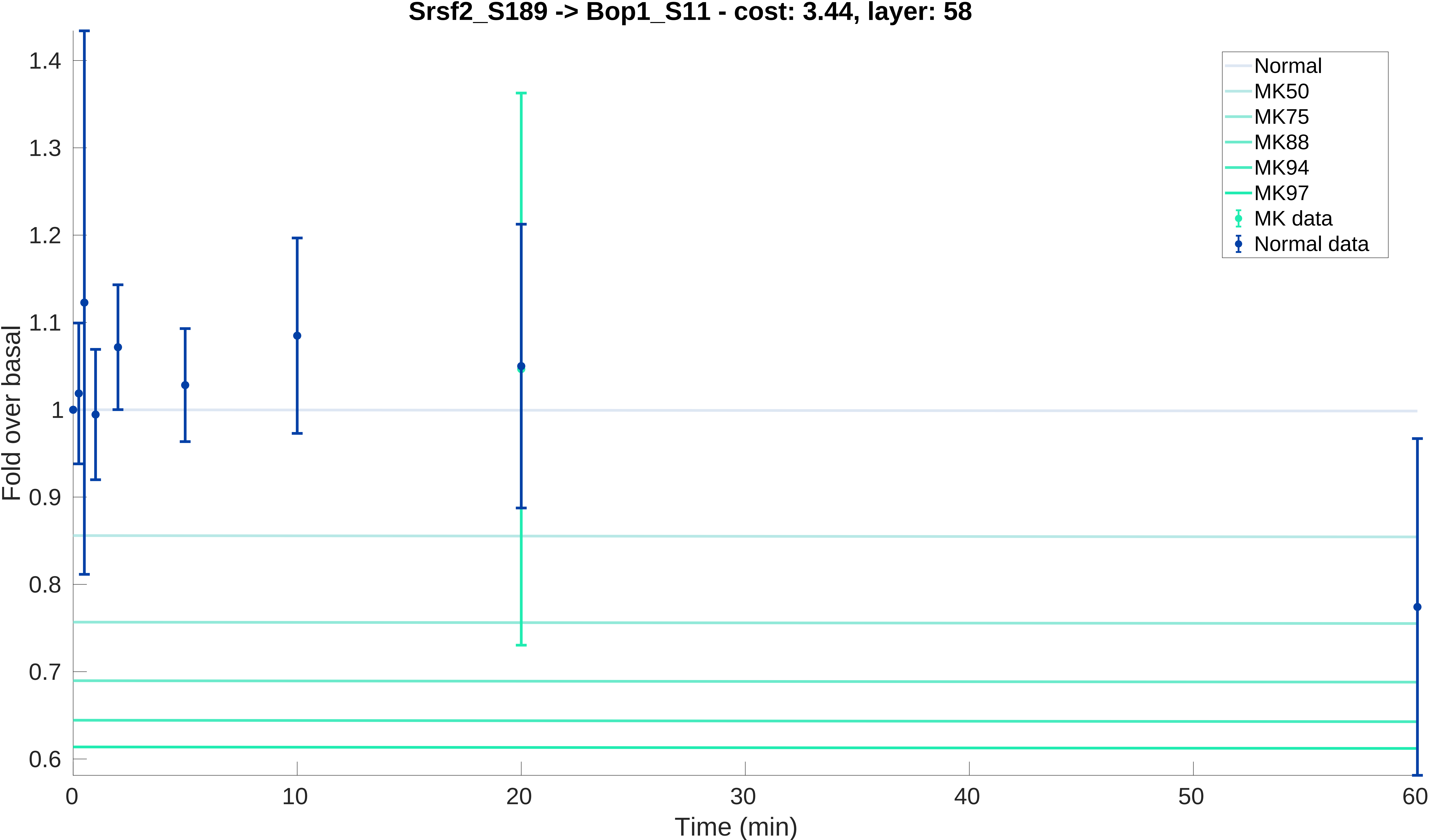

Srsf2_S189 -> Bop1_S11 - cost: 3.44, layer: 58
Fold over basal
Time (min)
1.4
1.3
1.2
1.1
1
0.9
0.8
0.7
0.6
0
10
20
30
40
50
60
Normal
MK50
MK75
MK88
MK94
MK97
MK data
Normal data

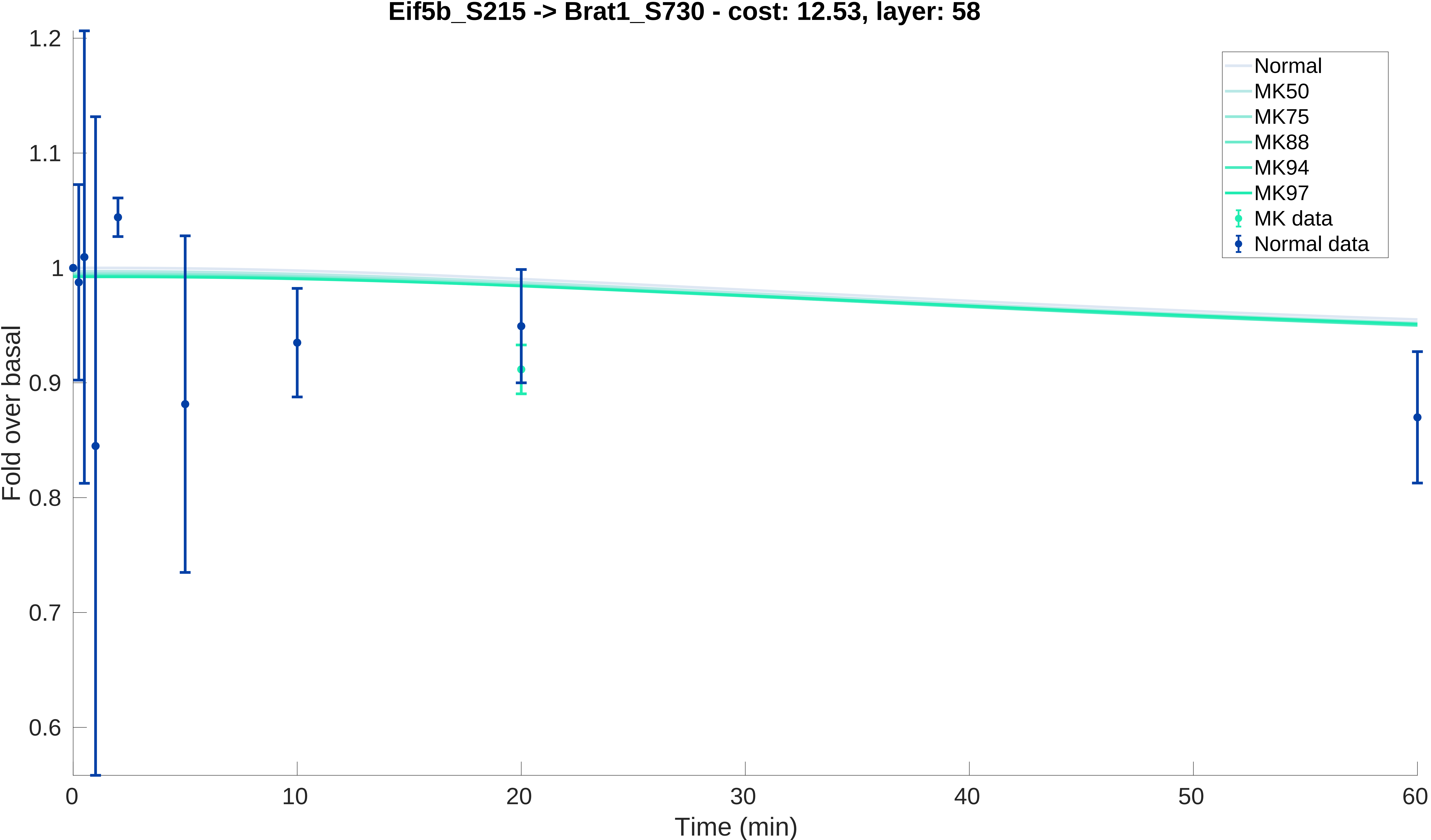

Eif5b_S215 -> Brat1_S730 - cost: 12.53, layer: 58
Fold over basal
Time (min)
Normal
MK50
MK75
MK88
MK94
MK97
MK data
Normal data
1.2
1.1
1
0.9
0.8
0.7
0.6
0
10
20
30
40
50
60

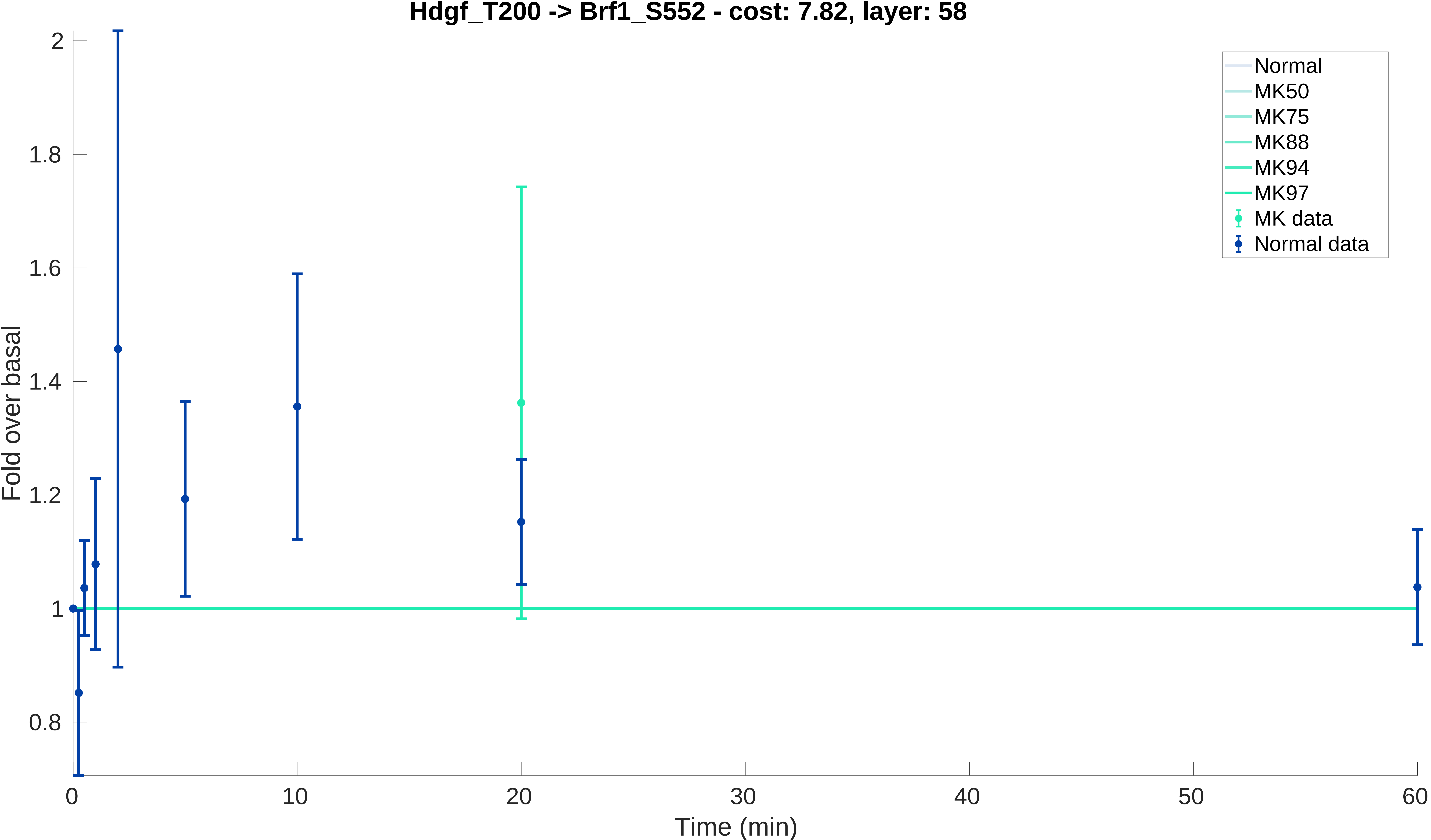

Hdgf_T200 -> Brf1_S552 - cost: 7.82, layer: 58
Fold over basal
Time (min)
0
10
20
30
40
50
60
0.8
1
1.2
1.4
1.6
1.8
2
Normal
MK50
MK75
MK88
MK94
MK97
MK data
Normal data

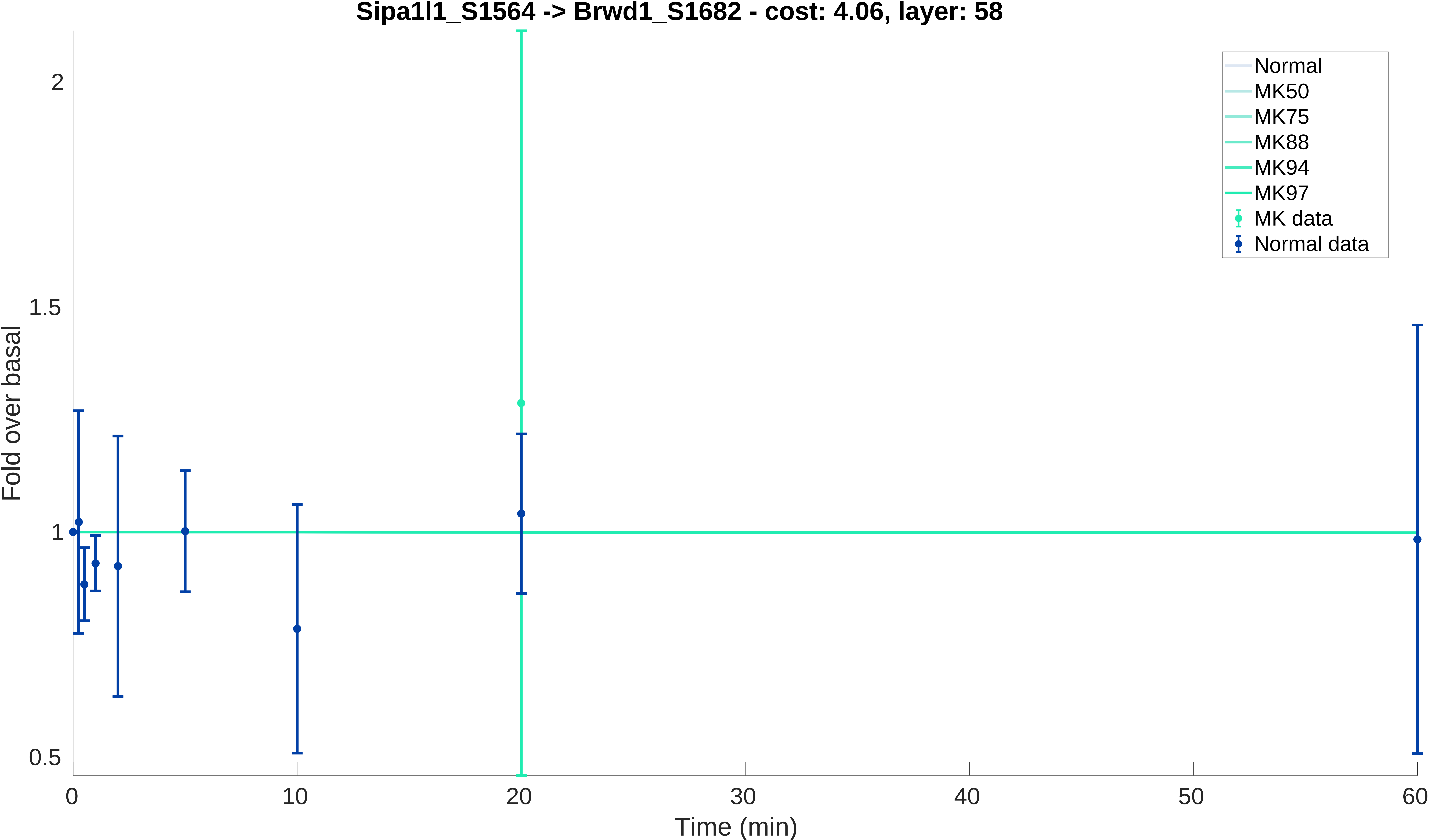

Sipa1l1_S1564 -> Brwd1_S1682 - cost: 4.06, layer: 58
Fold over basal
Time (min)
0.5
1
1.5
2
0
10
20
30
40
50
60
Normal
MK50
MK75
MK88
MK94
MK97
MK data
Normal data

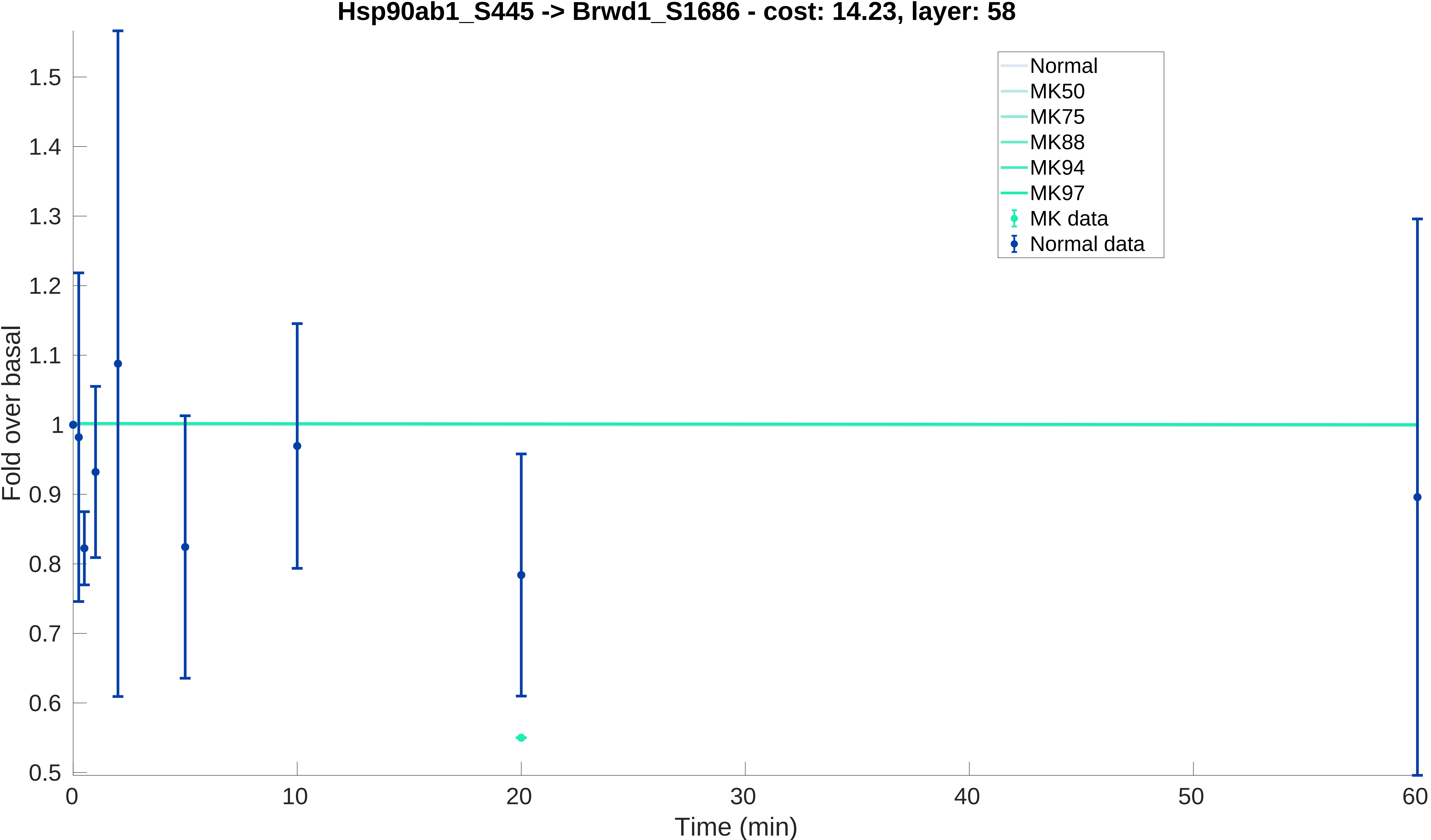
Hsp90ab1_S445 -> Brwd1_S1686 - cost: 14.23, layer: 58
Normal
MK50
MK75
MK88
MK94
MK97
MK data
Normal data
Fold over basal
Time (min)
1.5
1.4
1.3
1.2
1.1
1
0.9
0.8
0.7
0.6
0.5
0
10
20
30
40
50
60

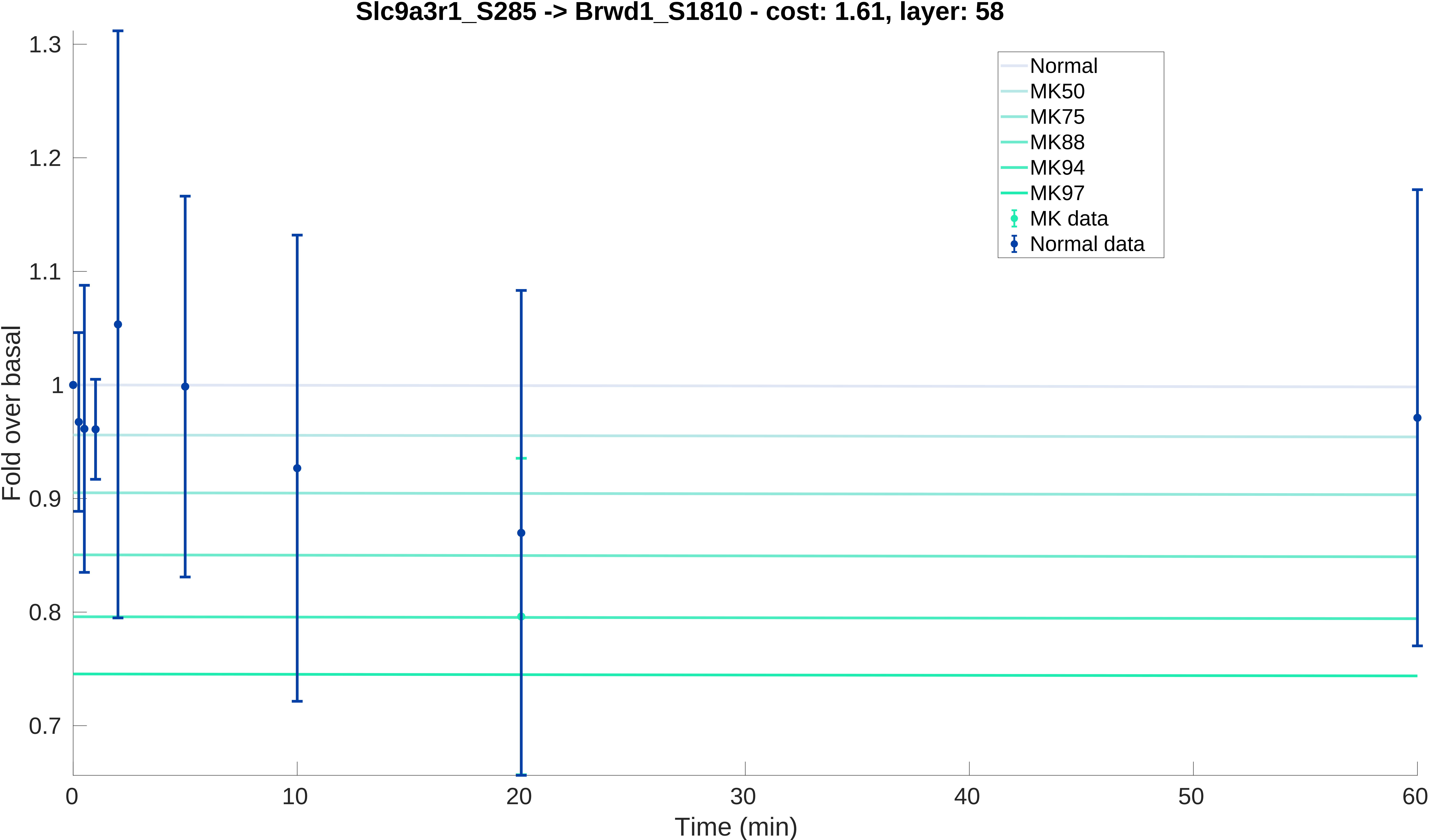

Slc9a3r1_S285 -> Brwd1_S1810 - cost: 1.61, layer: 58
Fold over basal
Time (min)
1.3
1.2
1.1
1
0.9
0.8
0.7
0
10
20
30
40
50
60
Normal
MK50
MK75
MK88
MK94
MK97
MK data
Normal data

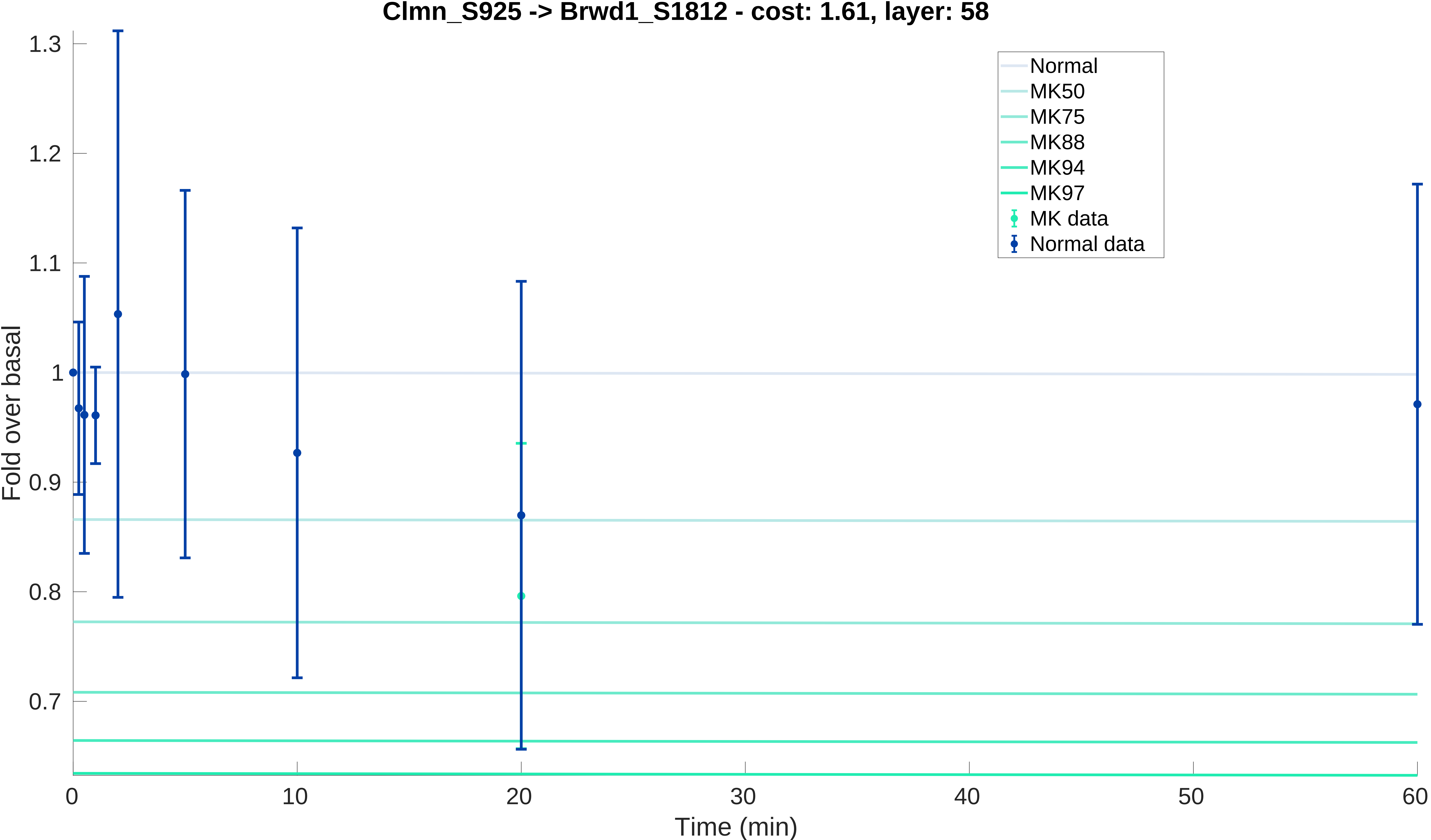

Clmn_S925 -> Brwd1_S1812 - cost: 1.61, layer: 58
Fold over basal
Time (min)
1.3
1.2
1.1
1
0.9
0.8
0.7
0
10
20
30
40
50
60
Normal
MK50
MK75
MK88
MK94
MK97
MK data
Normal data

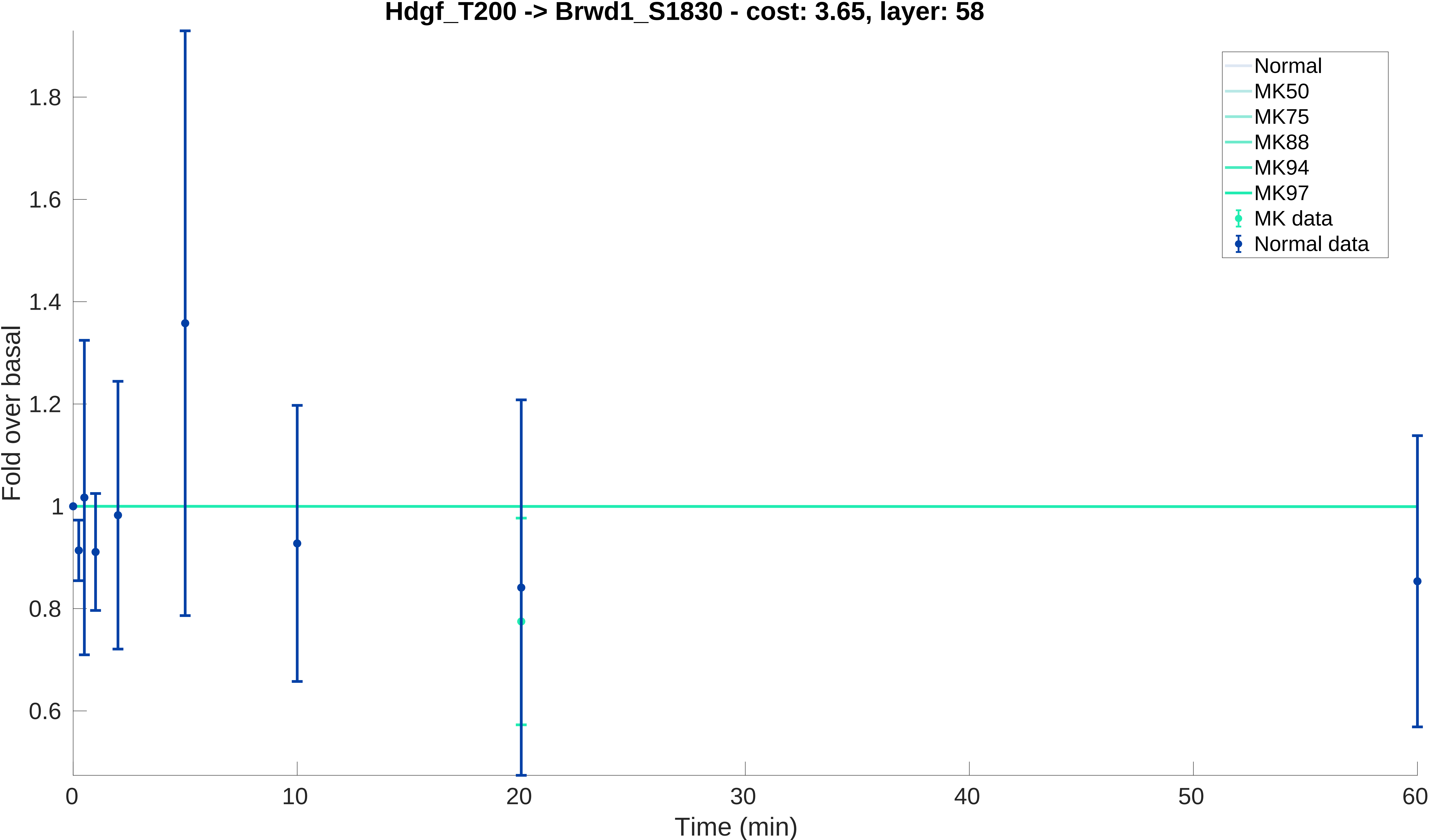

Hdgf_T200 -> Brwd1_S1830 - cost: 3.65, layer: 58
Normal
MK50
MK75
MK88
MK94
MK97
MK data
Normal data
Fold over basal
1.8
1.6
1.4
1.2
1
0.8
0.6
0
10
20
30
40
50
60
Time (min)

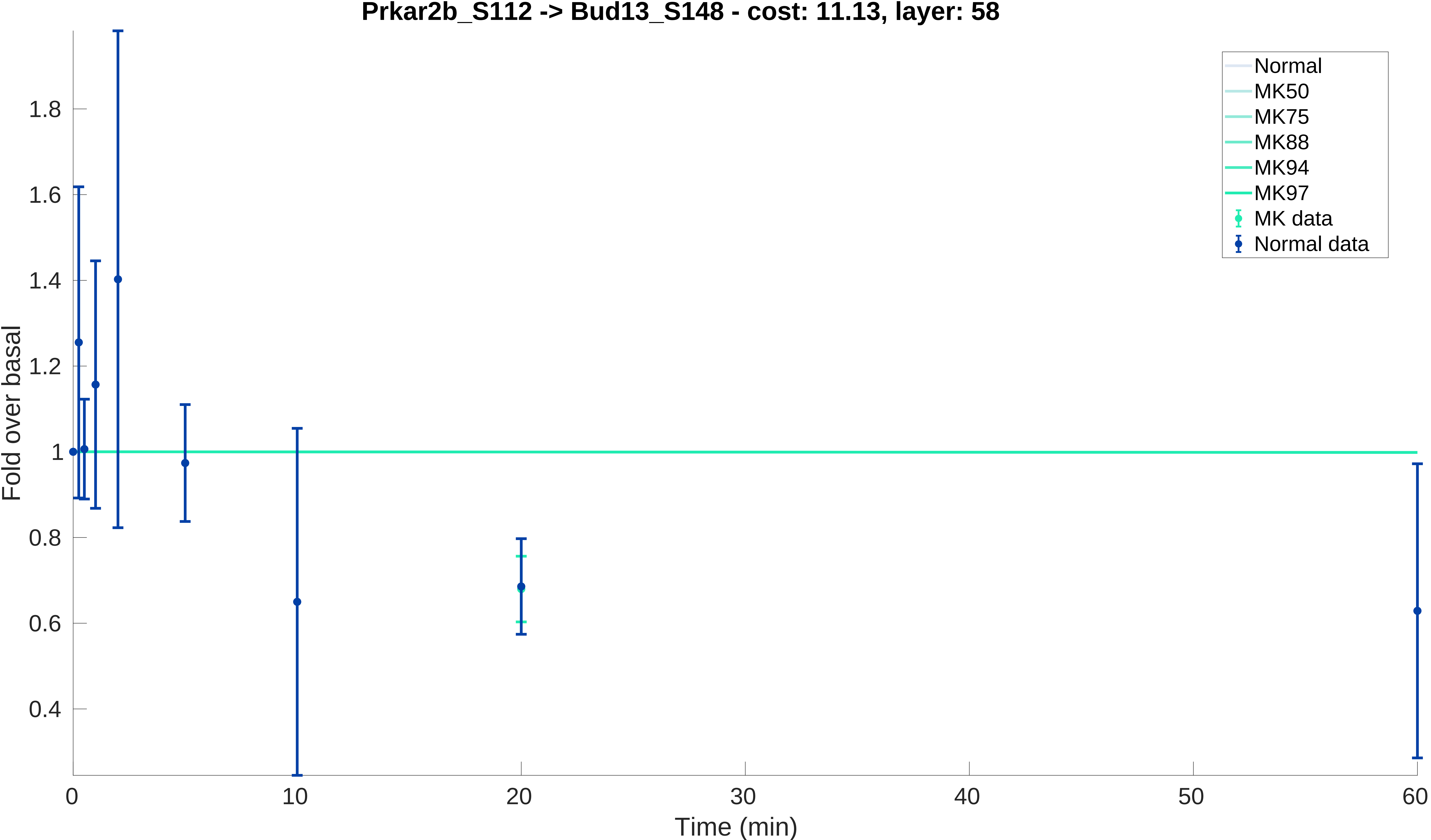
Prkar2b_S112 -> Bud13_S148 - cost: 11.13, layer: 58
Fold over basal
Time (min)
0.4
0.6
0.8
1
1.2
1.4
1.6
1.8
0
10
20
30
40
50
60
Normal
MK50
MK75
MK88
MK94
MK97
MK data
Normal data

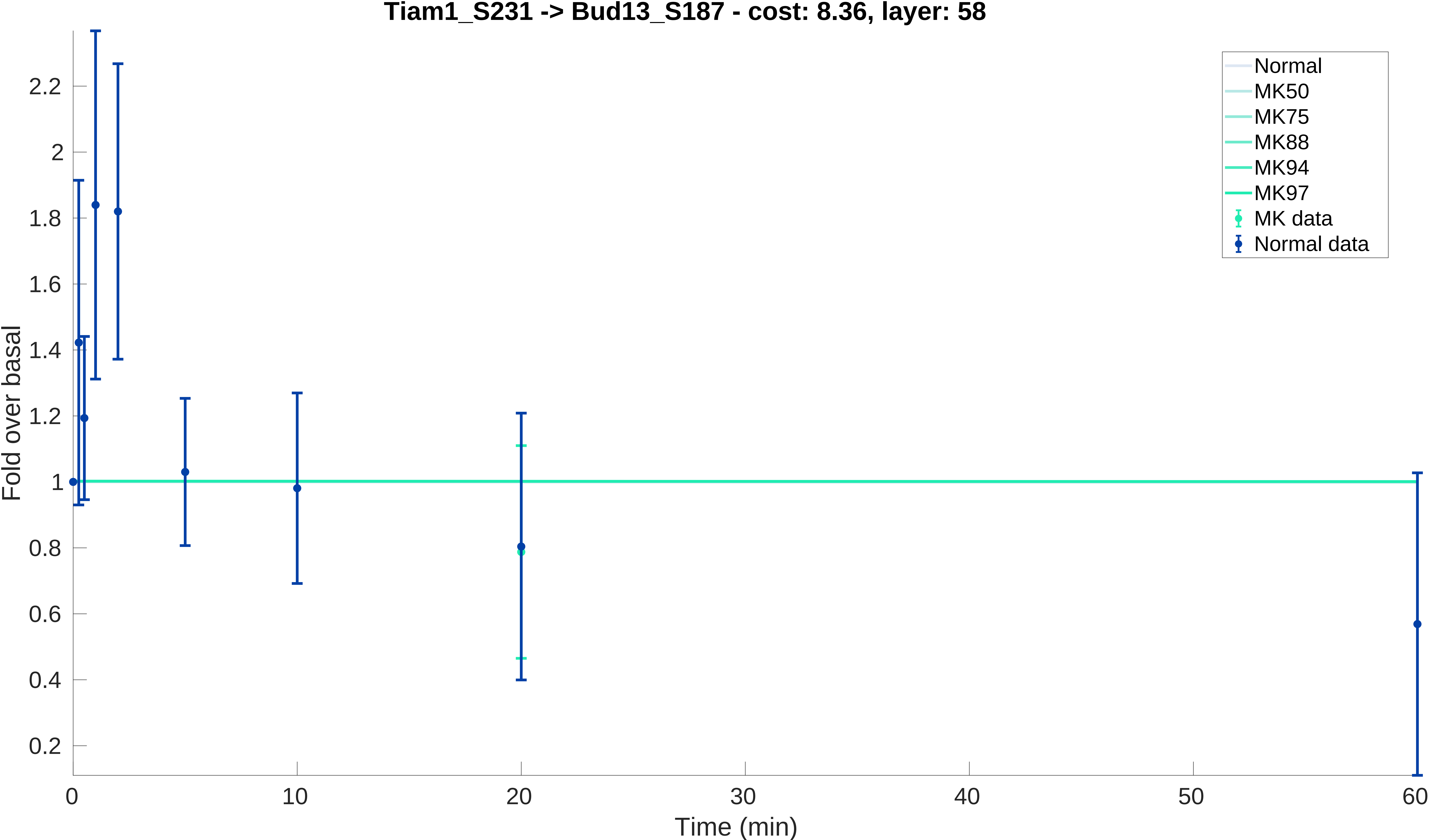

Tiam1_S231 -> Bud13_S187 - cost: 8.36, layer: 58
Fold over basal
Time (min)
0
10
20
30
40
50
60
0.2
0.4
0.6
0.8
1
1.2
1.4
1.6
1.8
2
2.2
Normal
MK50
MK75
MK88
MK94
MK97
MK data
Normal data

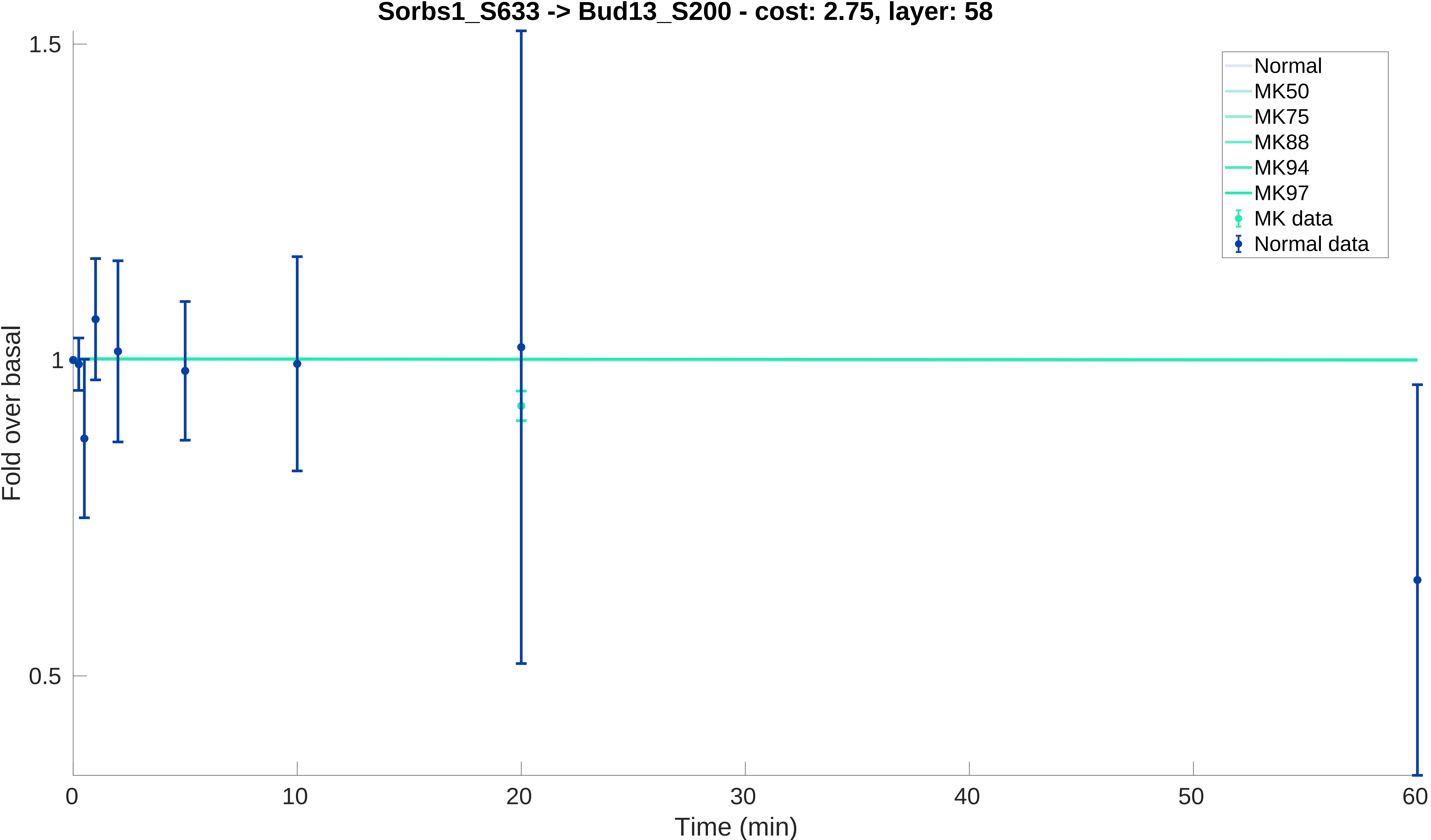

Sorbs1_S633 -> Bud13_S200 - cost: 2.75, layer: 58
Fold over basal
1.5
1
0.5
0
10
20
30
40
50
60
Time (min)
Normal
MK50
MK75
MK88
MK94
MK97
MK data
Normal data

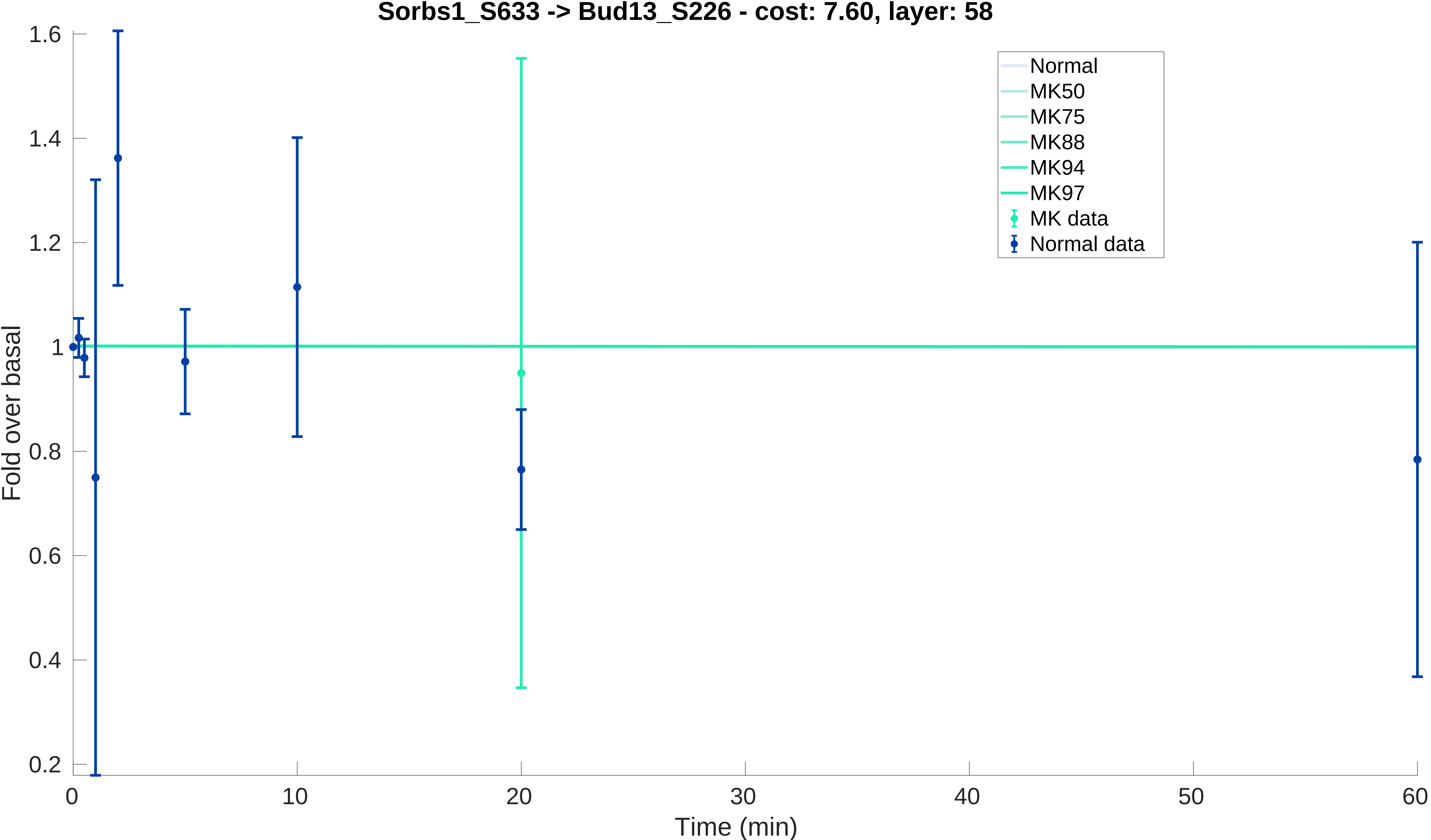

Sorbs1_S633 -> Bud13_S226 - cost: 7.60, layer: 58
Fold over basal
Time (min)
Normal
MK50
MK75
MK88
MK94
MK97
MK data
Normal data
1.6
1.4
1.2
1
0.8
0.6
0.4
0.2
0
10
20
30
40
50
60

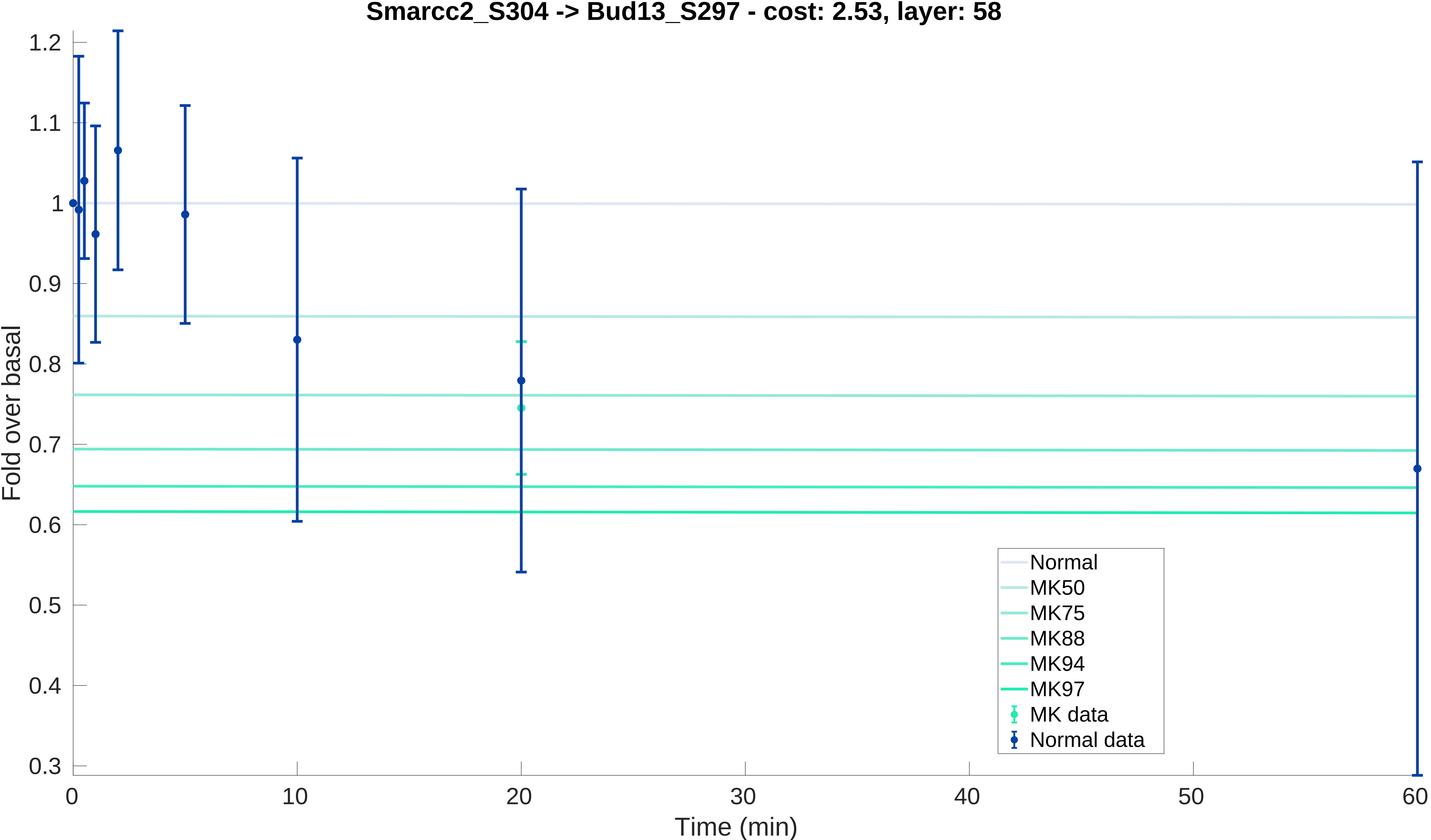

Smarcc2_S304 -> Bud13_S297 - cost: 2.53, layer: 58
Fold over basal
Time (min)
1.2
1.1
1
0.9
0.8
0.7
0.6
0.5
0.4
0.3
0
10
20
30
40
50
60
Normal
MK50
MK75
MK88
MK94
MK97
MK data
Normal data

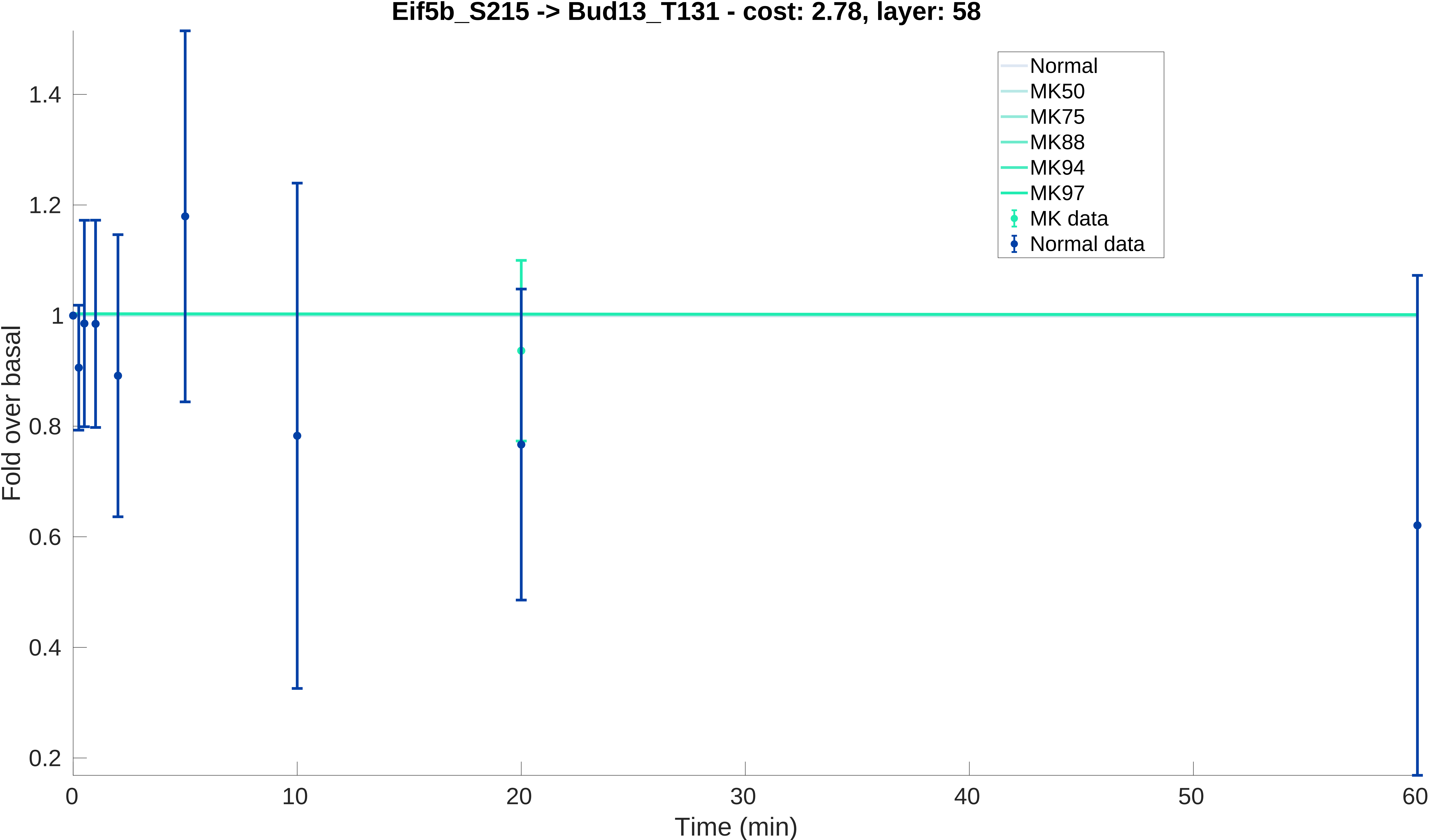
Eif5b_S215 -> Bud13_T131 - cost: 2.78, layer: 58
Normal
MK50
MK75
MK88
MK94
MK97
MK data
Normal data
Fold over basal
1.4
1.2
1
0.8
0.6
0.4
0.2
0
10
20
30
40
50
60
Time (min)

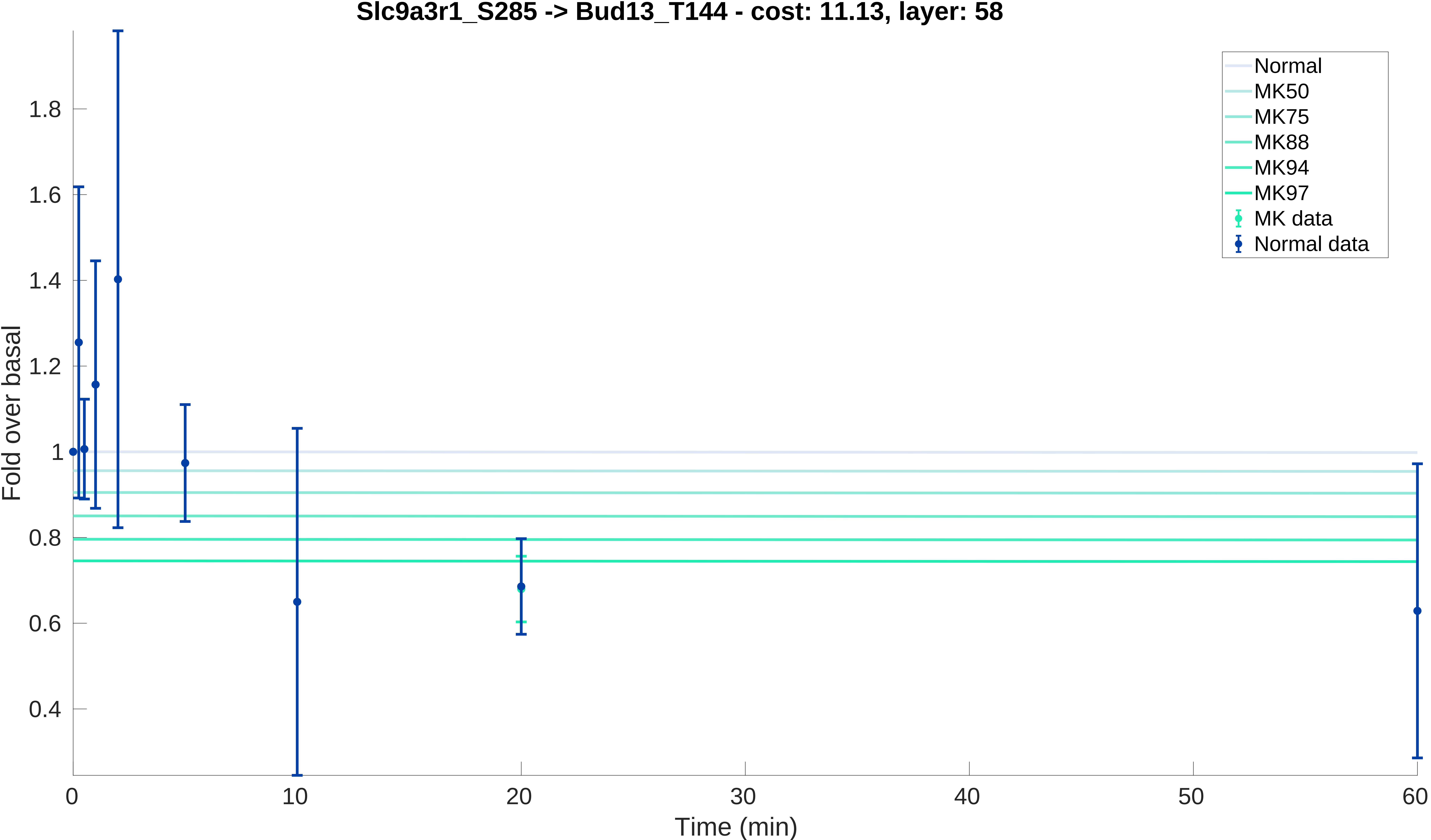

Slc9a3r1_S285 -> Bud13_T144 - cost: 11.13, layer: 58
Normal
MK50
MK75
MK88
MK94
MK97
MK data
Normal data
1.8
1.6
1.4
1.2
1
0.8
0.6
0.4
Fold over basal
0
10
20
30
40
50
60
Time (min)

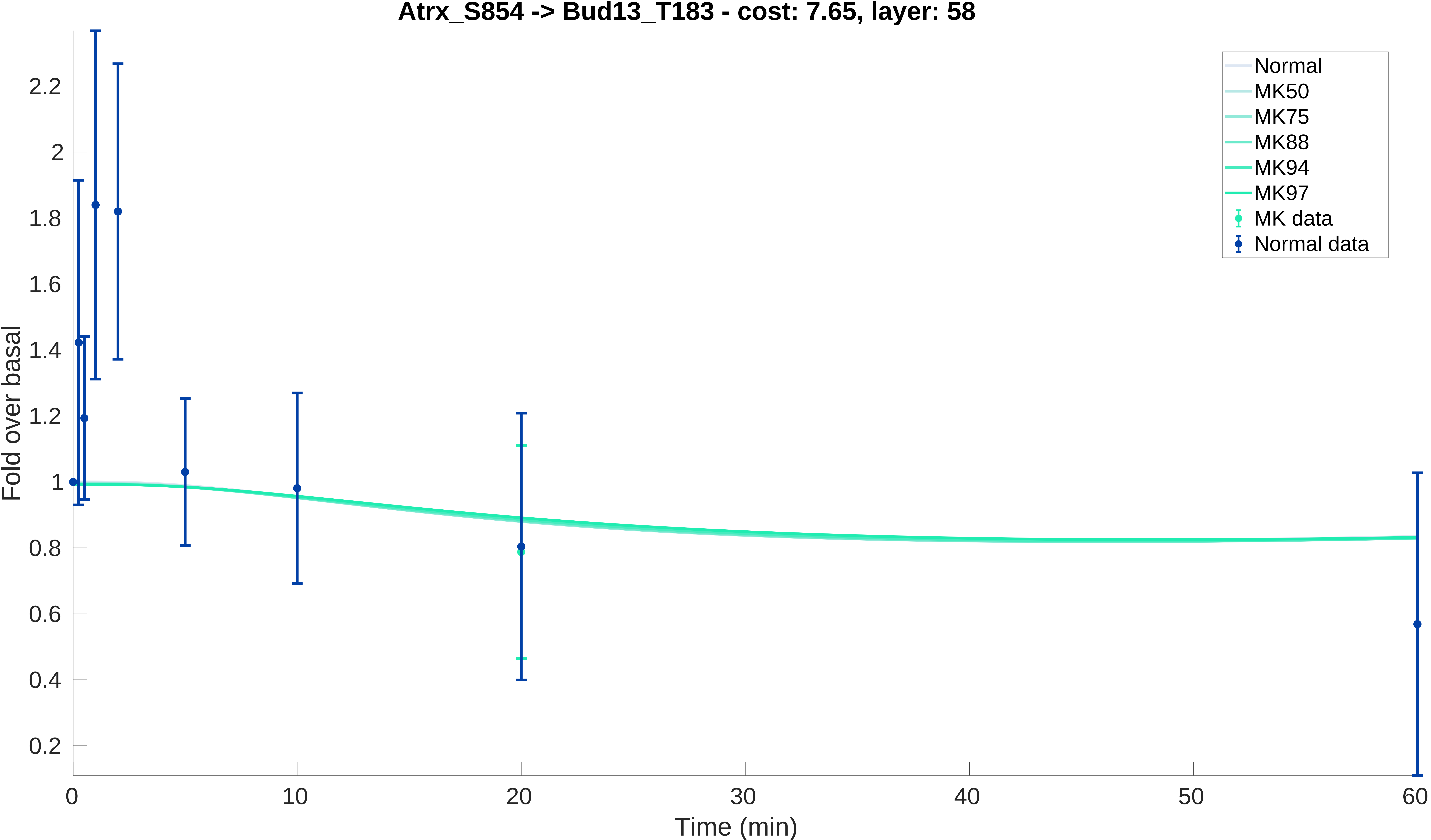

Atrx_S854 -> Bud13_T183 - cost: 7.65, layer: 58
Fold over basal
Time (min)
0.2
0.4
0.6
0.8
1
1.2
1.4
1.6
1.8
2
2.2
0
10
20
30
40
50
60
Normal
MK50
MK75
MK88
MK94
MK97
MK data
Normal data

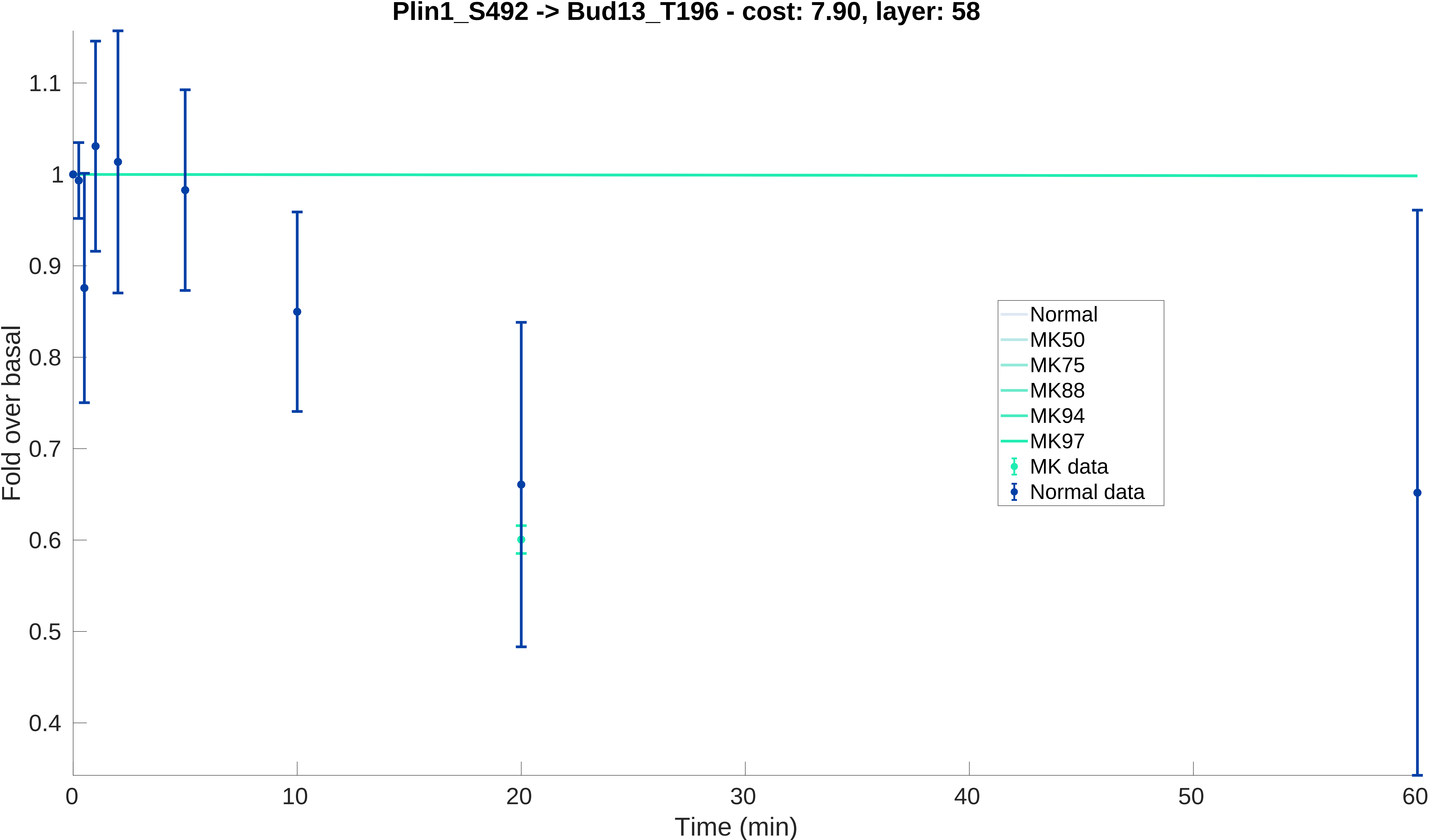
Plin1_S492 -> Bud13_T196 - cost: 7.90, layer: 58
Fold over basal
Time (min)
1.1
1
0.9
0.8
0.7
0.6
0.5
0.4
0
10
20
30
40
50
60
Normal
MK50
MK75
MK88
MK94
MK97
MK data
Normal data

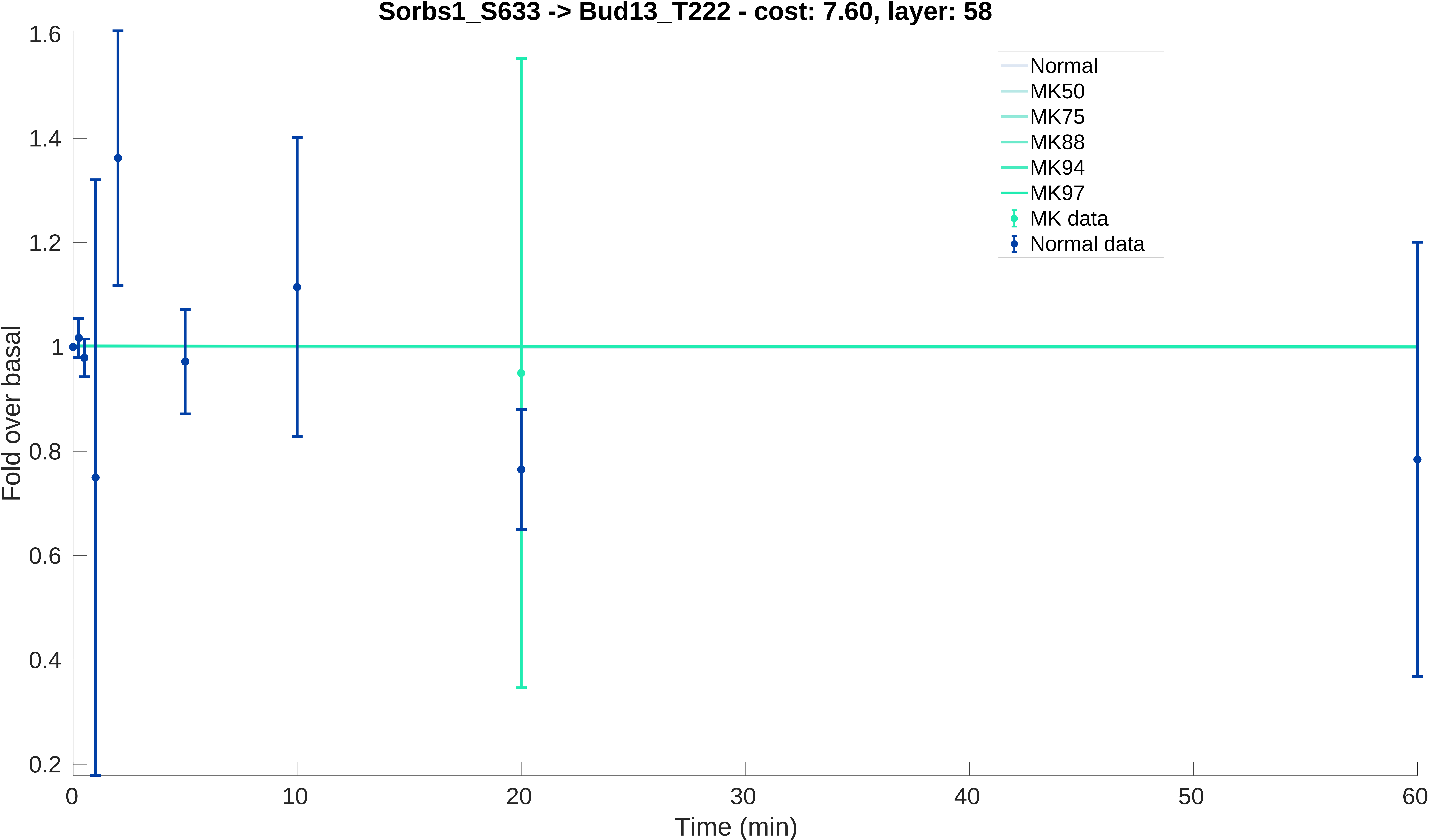

Sorbs1_S633 -> Bud13_T222 - cost: 7.60, layer: 58
Fold over basal
Time (min)
Normal
MK50
MK75
MK88
MK94
MK97
MK data
Normal data
0
10
20
30
40
50
60
0.2
0.4
0.6
0.8
1
1.2
1.4
1.6

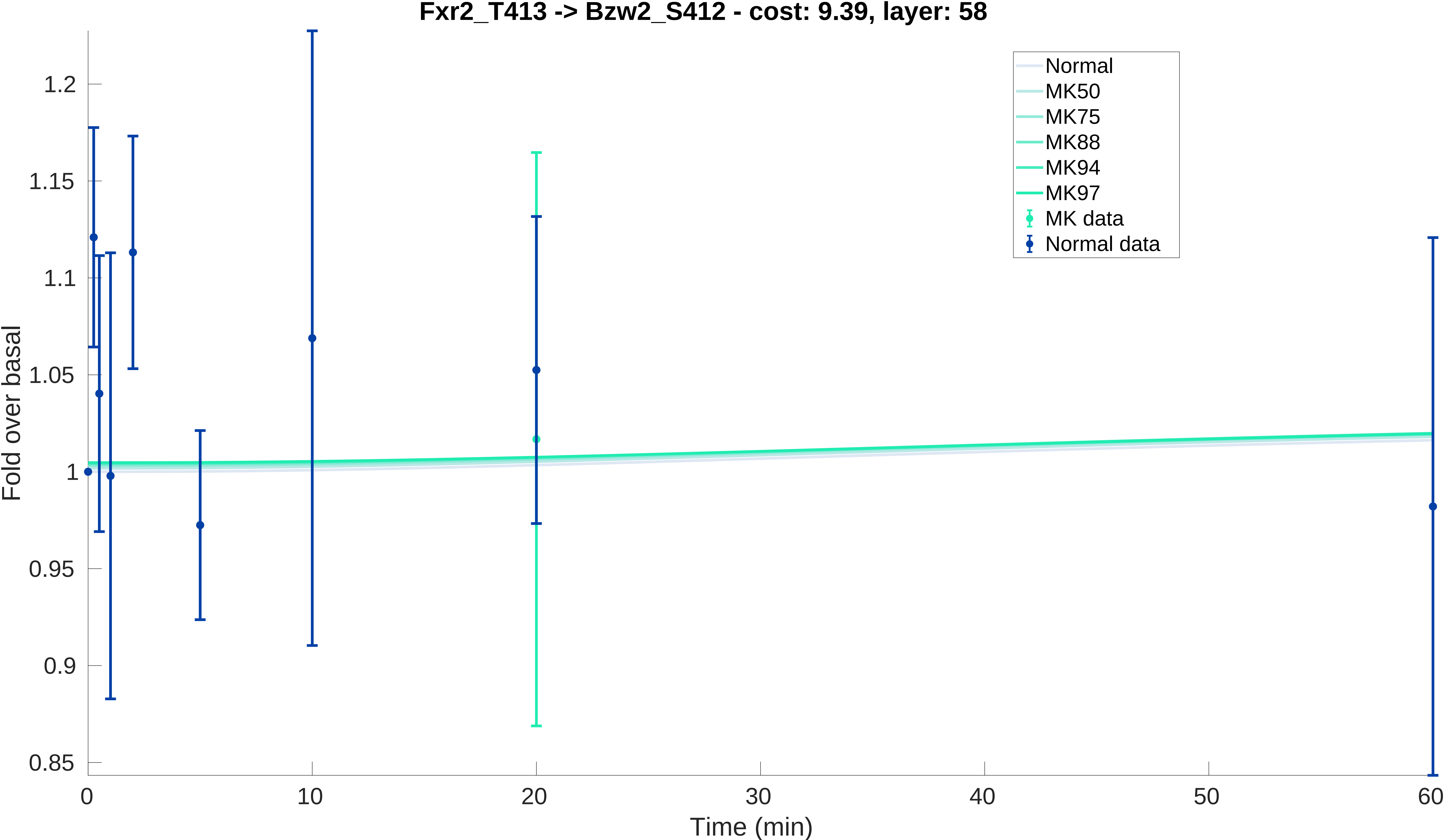

Fxr2_T413 -> Bzw2_S412 - cost: 9.39, layer: 58
Normal
MK50
MK75
MK88
MK94
MK97
MK data
Normal data
Fold over basal
1.2
1.15
1.1
1.05
1
0.95
0.9
0.85
0
10
20
30
40
50
60
Time (min)

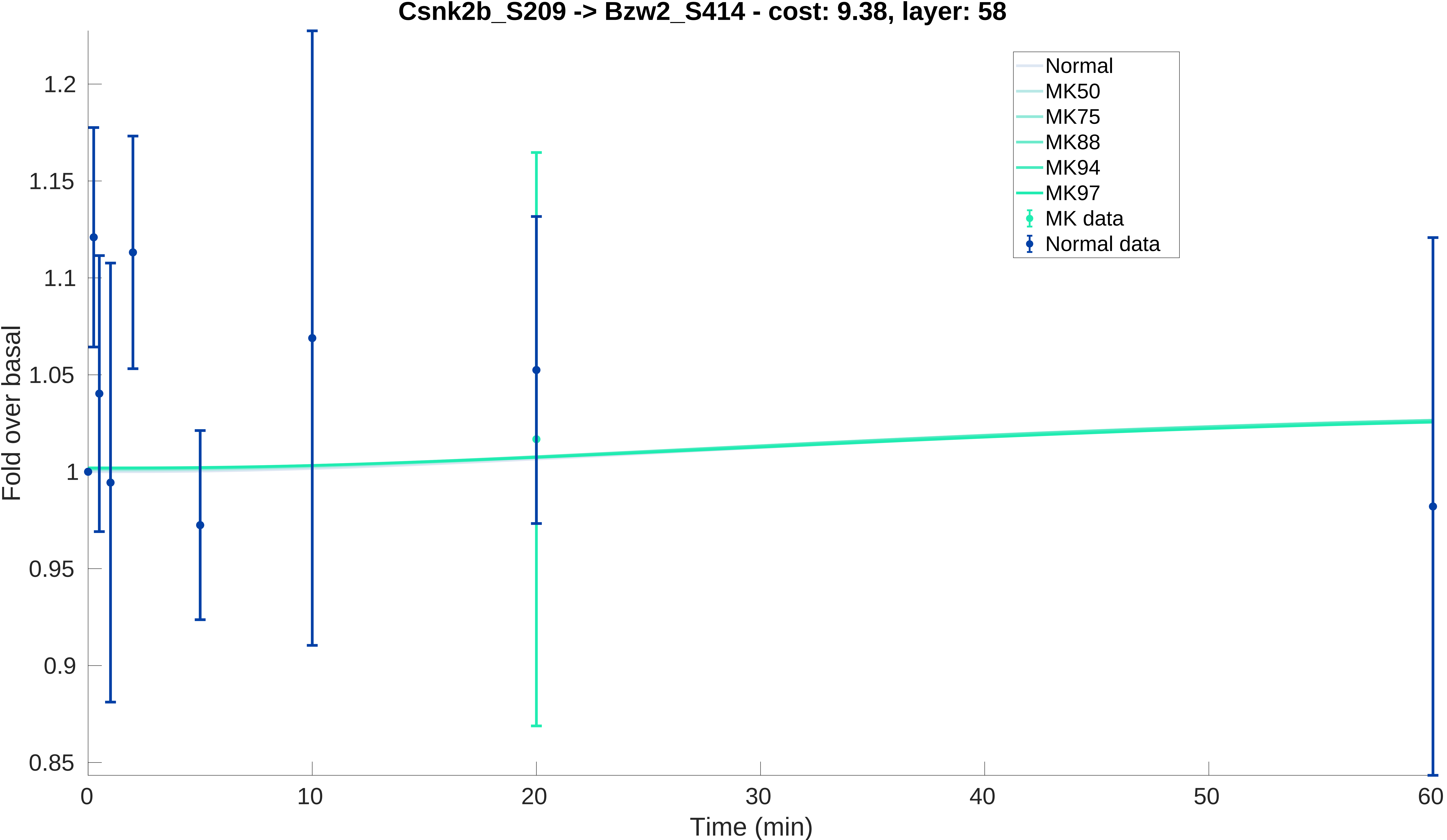

Csnk2b_S209 -> Bzw2_S414 - cost: 9.38, layer: 58
Normal
MK50
MK75
MK88
MK94
MK97
MK data
Normal data
Fold over basal
1.2
1.15
1.1
1.05
1
0.95
0.9
0.85
0
10
20
30
40
50
60
Time (min)

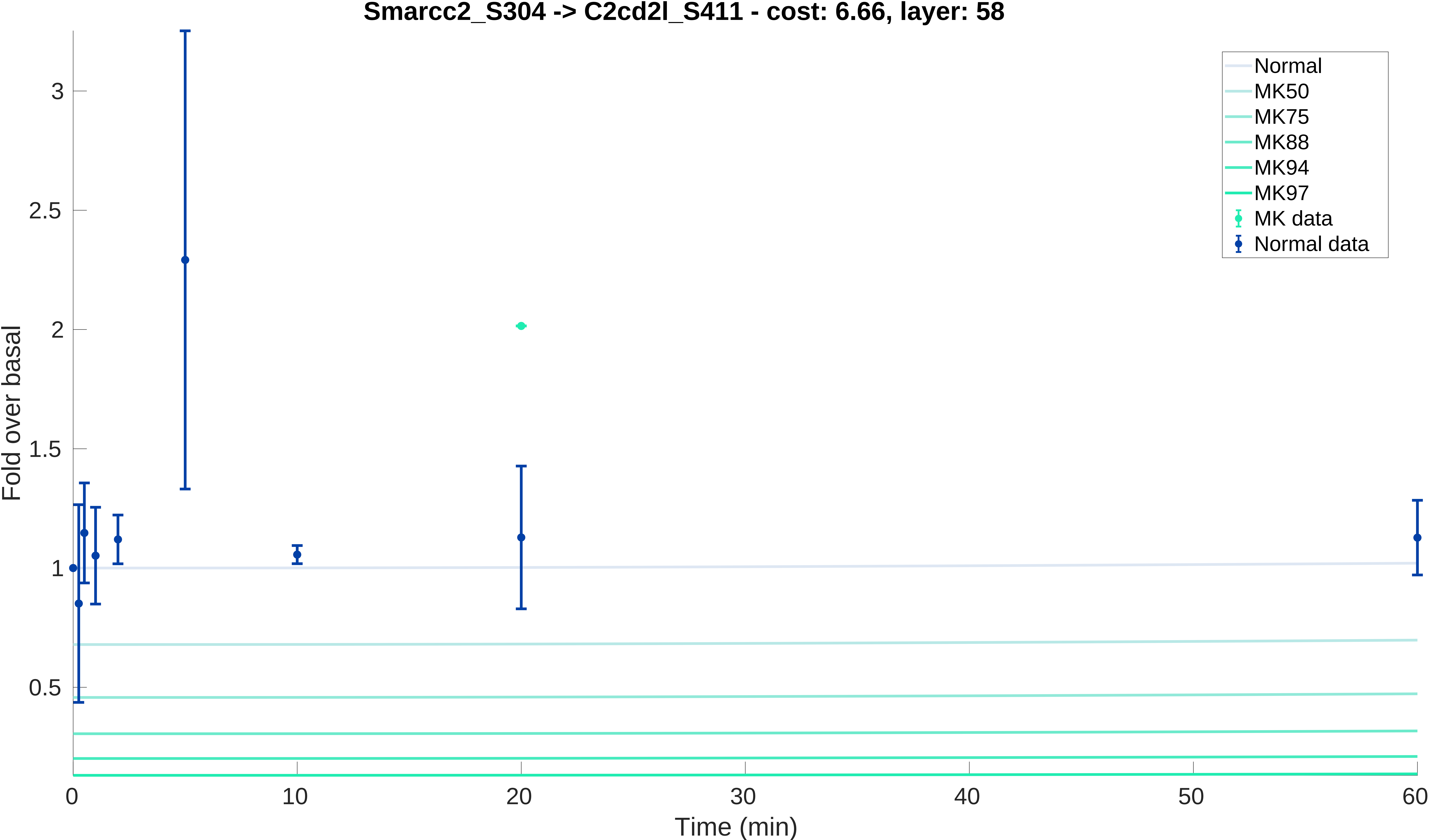

Smarcc2_S304 -> C2cd2l_S411 - cost: 6.66, layer: 58
Fold over basal
Time (min)
0.5
1
1.5
2
2.5
3
0
10
20
30
40
50
60
Normal
MK50
MK75
MK88
MK94
MK97
MK data
Normal data

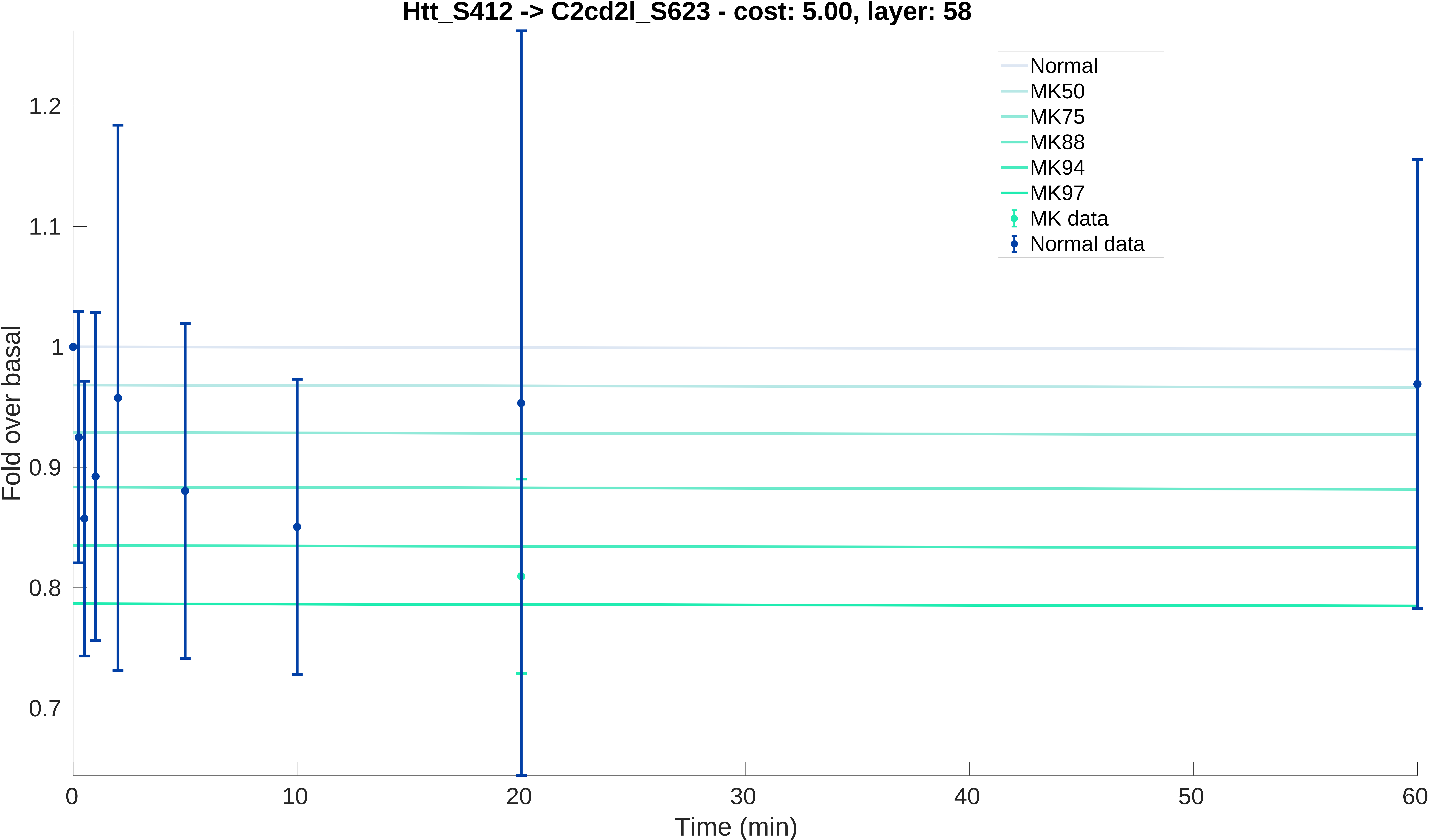

Htt_S412 -> C2cd2l_S623 - cost: 5.00, layer: 58
Normal
MK50
MK75
MK88
MK94
MK97
MK data
Normal data
Fold over basal
1.2
1.1
1
0.9
0.8
0.7
0
10
20
30
40
50
60
Time (min)

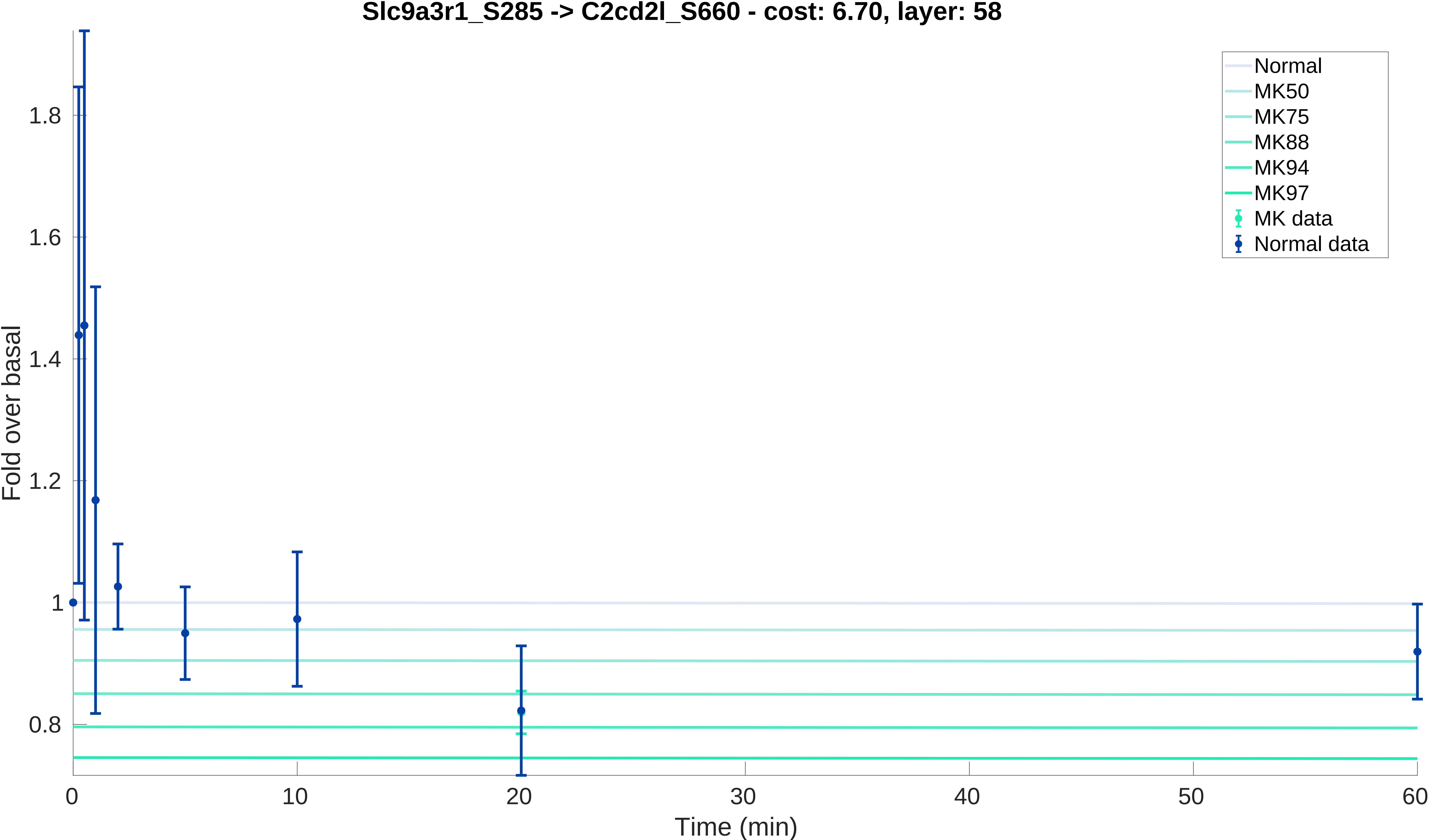
Slc9a3r1_S285 -> C2cd2l_S660 - cost: 6.70, layer: 58
Fold over basal
Time (min)
0
10
20
30
40
50
60
0.8
1
1.2
1.4
1.6
1.8
Normal
MK50
MK75
MK88
MK94
MK97
MK data
Normal data

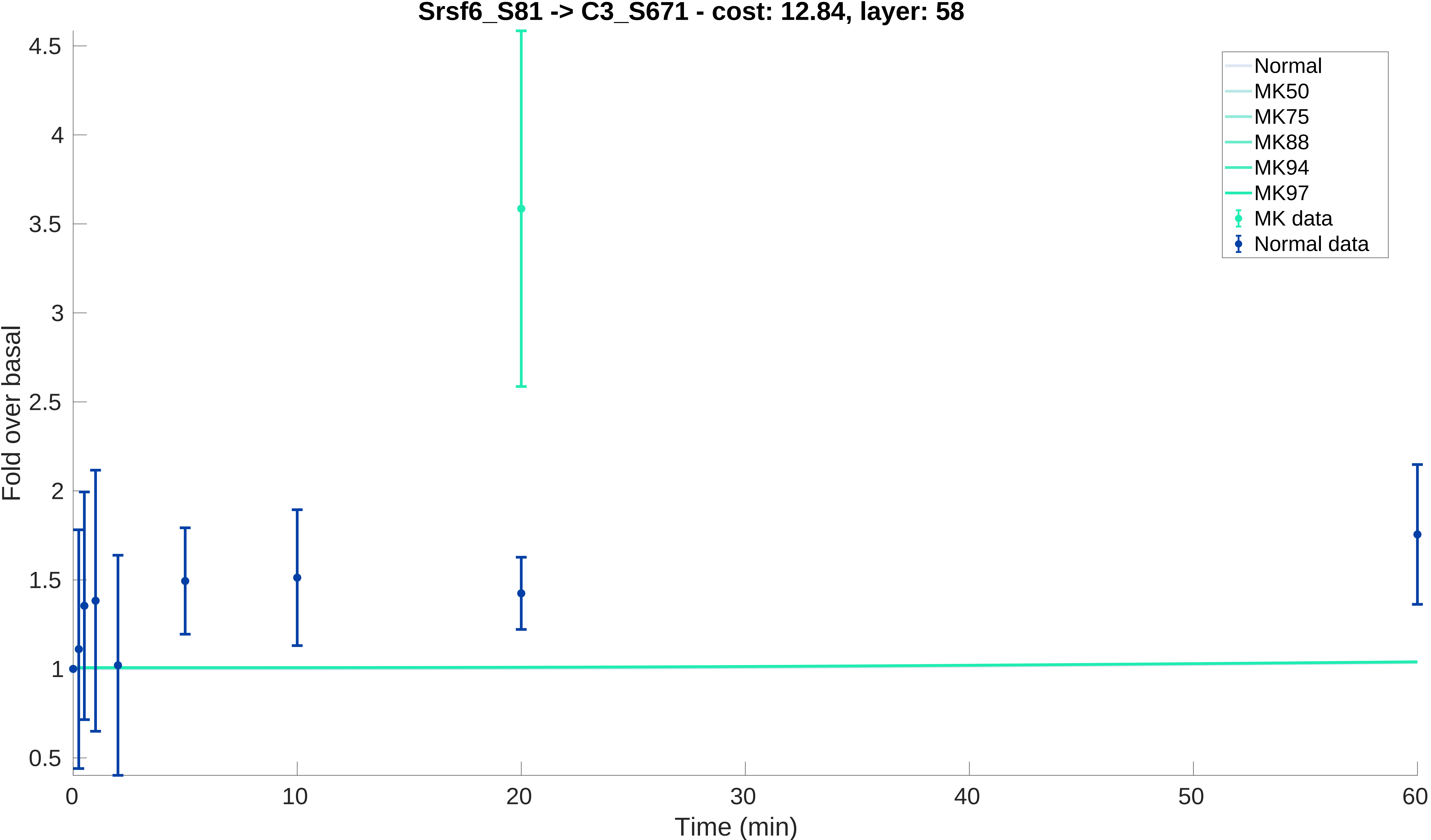

Srsf6_S81 -> C3_S671 - cost: 12.84, layer: 58
Fold over basal
Time (min)
4.5
4
3.5
3
2.5
2
1.5
1
0.5
0
10
20
30
40
50
60
Normal
MK50
MK75
MK88
MK94
MK97
MK data
Normal data

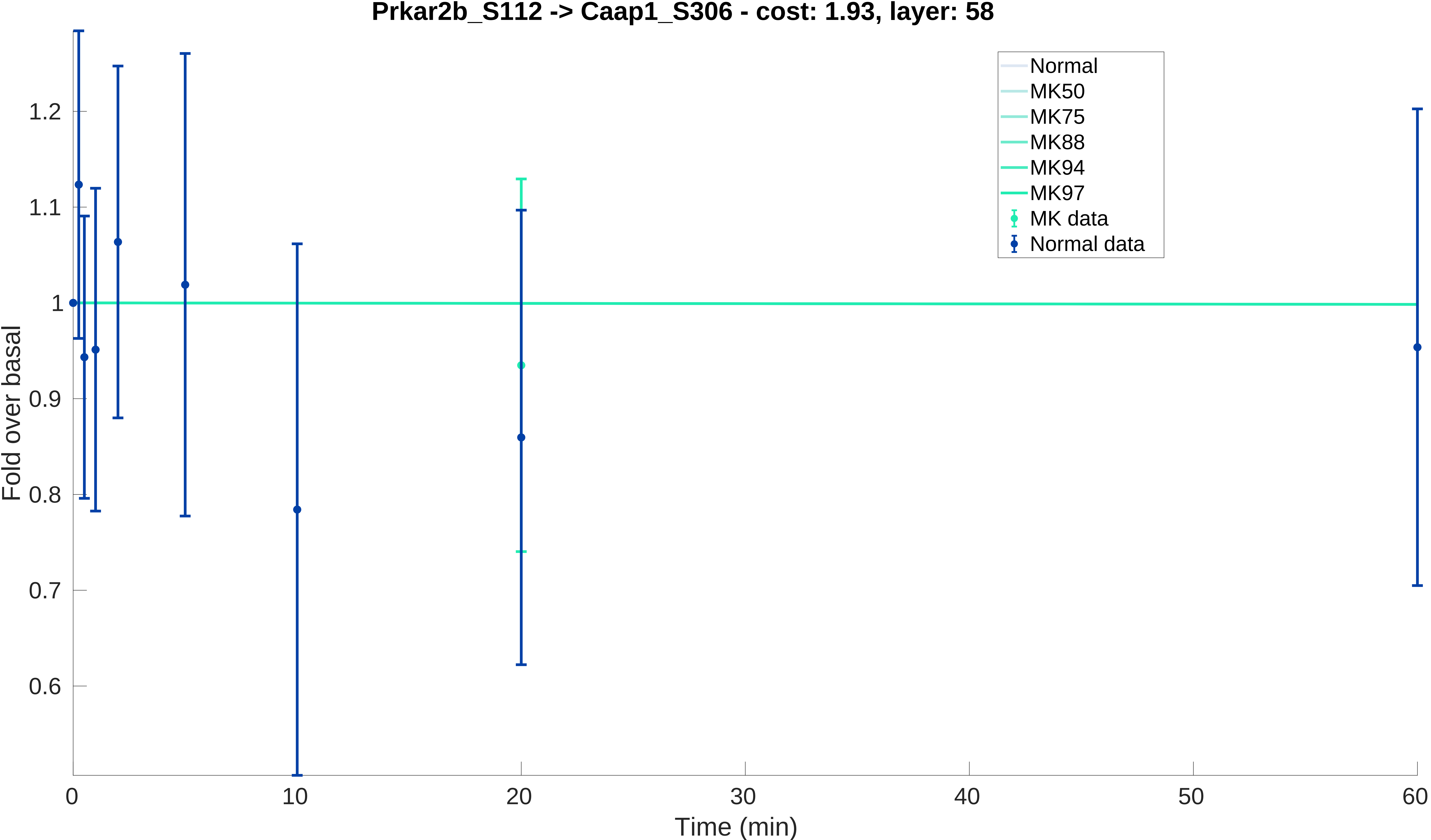
Prrkar2b_S112 -> Caap1_S306 - cost: 1.93, layer: 58
Fold over basal
Time (min)
0.6
0.7
0.8
0.9
1
1.1
1.2
0
10
20
30
40
50
60
Normal
MK50
MK75
MK88
MK94
MK97
MK data
Normal data

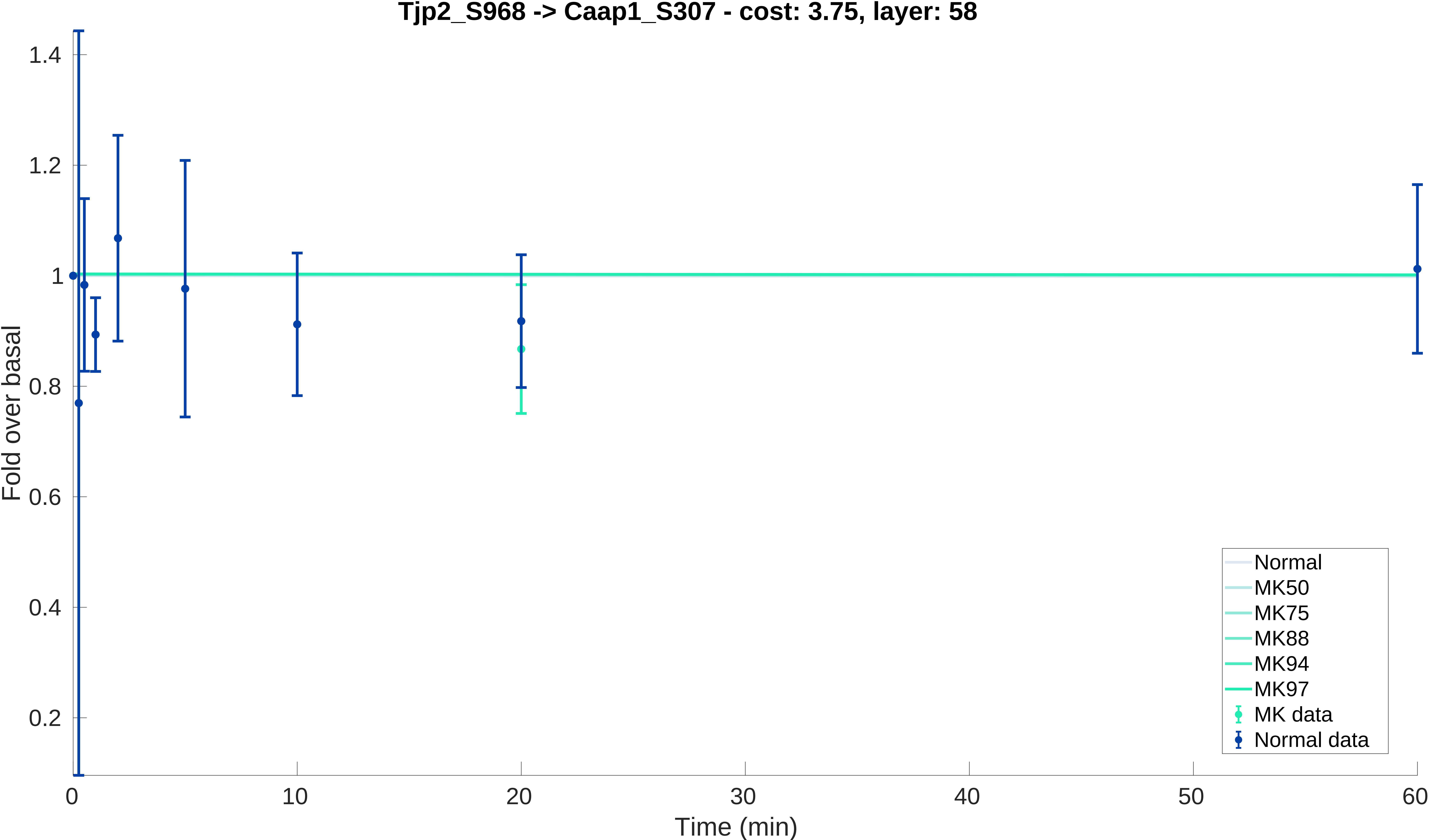

Tjp2_S968 -> Caap1_S307 - cost: 3.75, layer: 58
Fold over basal
1.4
1.2
1
0.8
0.6
0.4
0.2
0
10
20
30
40
50
60
Time (min)
Normal
MK50
MK75
MK88
MK94
MK97
MK data
Normal data

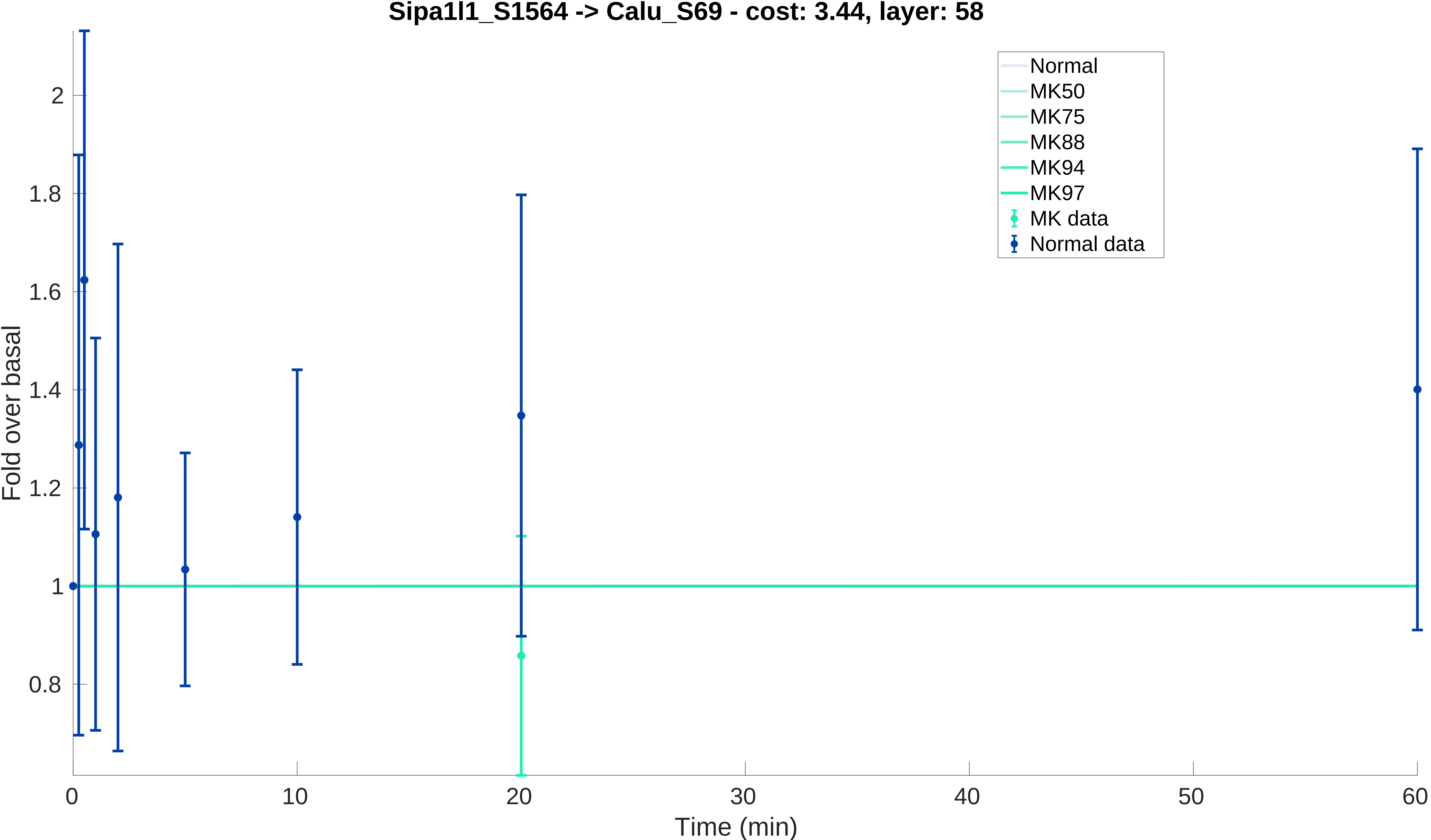

Sipa1l1_S1564 -> Calu_S69 - cost: 3.44, layer: 58
Normal
MK50
MK75
MK88
MK94
MK97
MK data
Normal data
Fold over basal
Time (min)
0.8
1
1.2
1.4
1.6
1.8
2
0
10
20
30
40
50
60

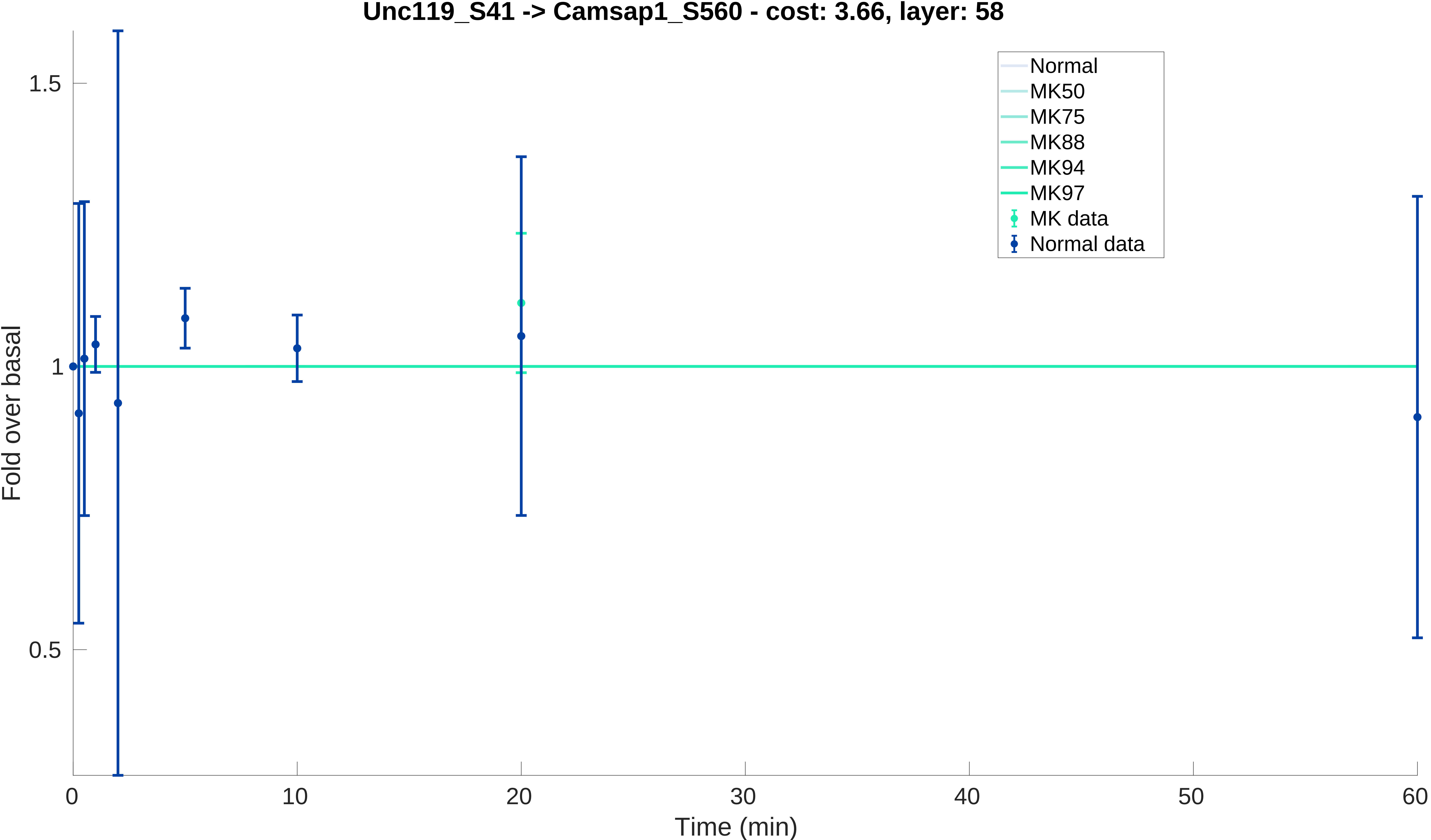

Unc119_S41 -> Camsap1_S560 - cost: 3.66, layer: 58
Normal
MK50
MK75
MK88
MK94
MK97
MK data
Normal data
Fold over basal
1.5
1
0.5
0
10
20
30
40
50
60
Time (min)

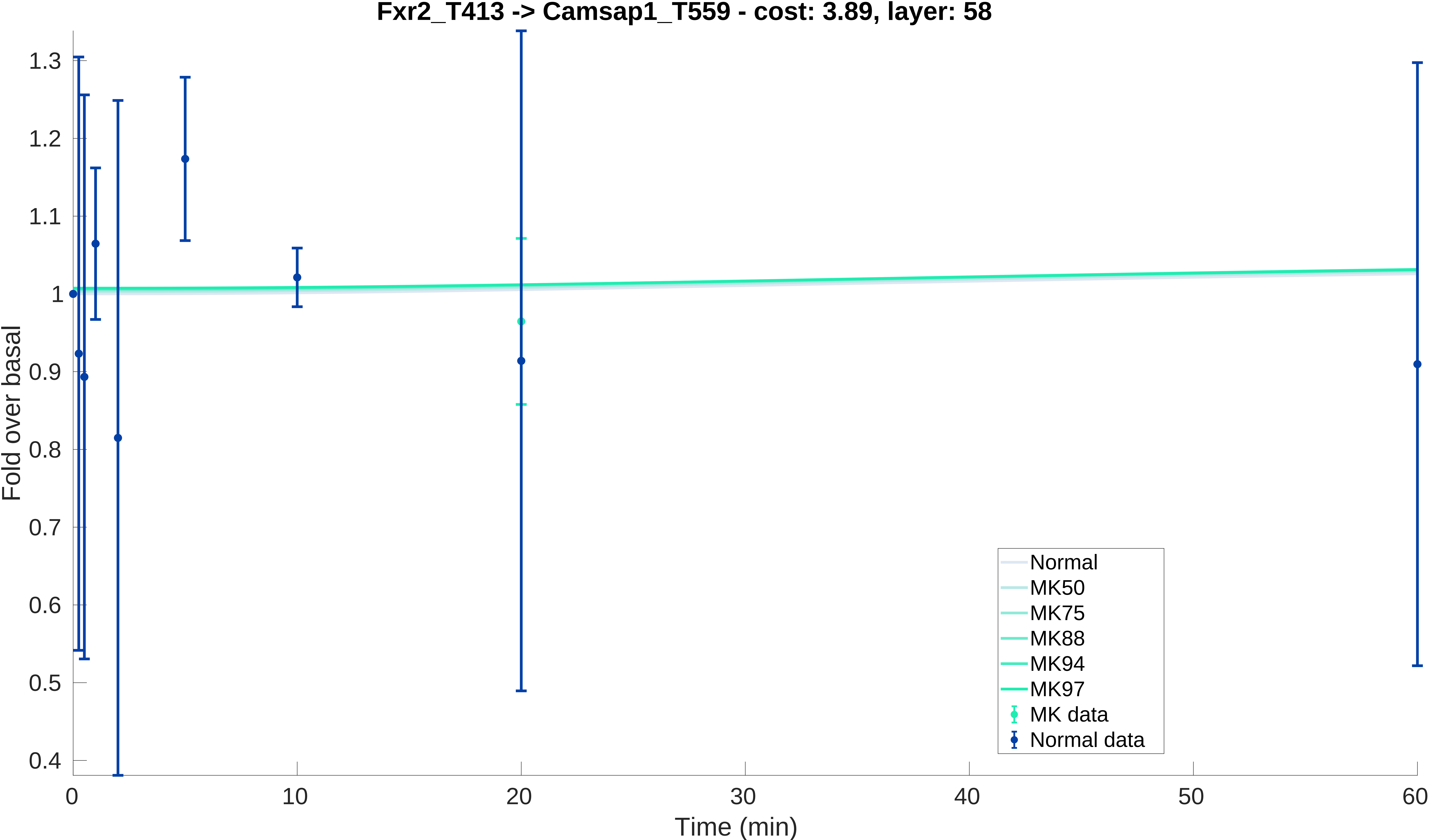

Fxr2_T413 -> Camsap1_T559 - cost: 3.89, layer: 58
Fold over basal
Time (min)
1.3
1.2
1.1
1
0.9
0.8
0.7
0.6
0.5
0.4
0
10
20
30
40
50
60
Normal
MK50
MK75
MK88
MK94
MK97
MK data
Normal data

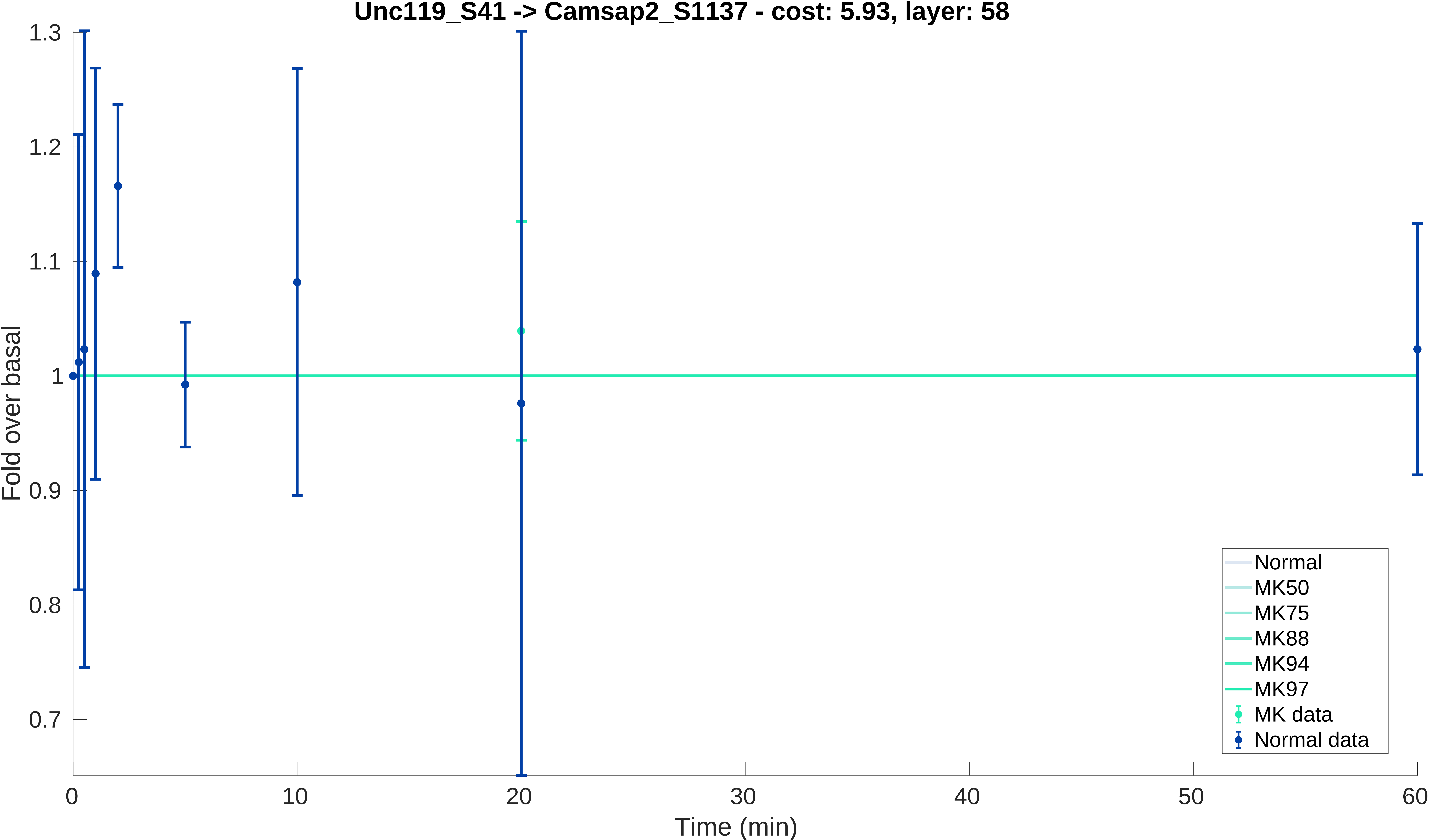

Unc119_S41 -> Camsap2_S1137 - cost: 5.93, layer: 58
Fold over basal
Time (min)
1.3
1.2
1.1
1
0.9
0.8
0.7
0
10
20
30
40
50
60
Normal
MK50
MK75
MK88
MK94
MK97
MK data
Normal data

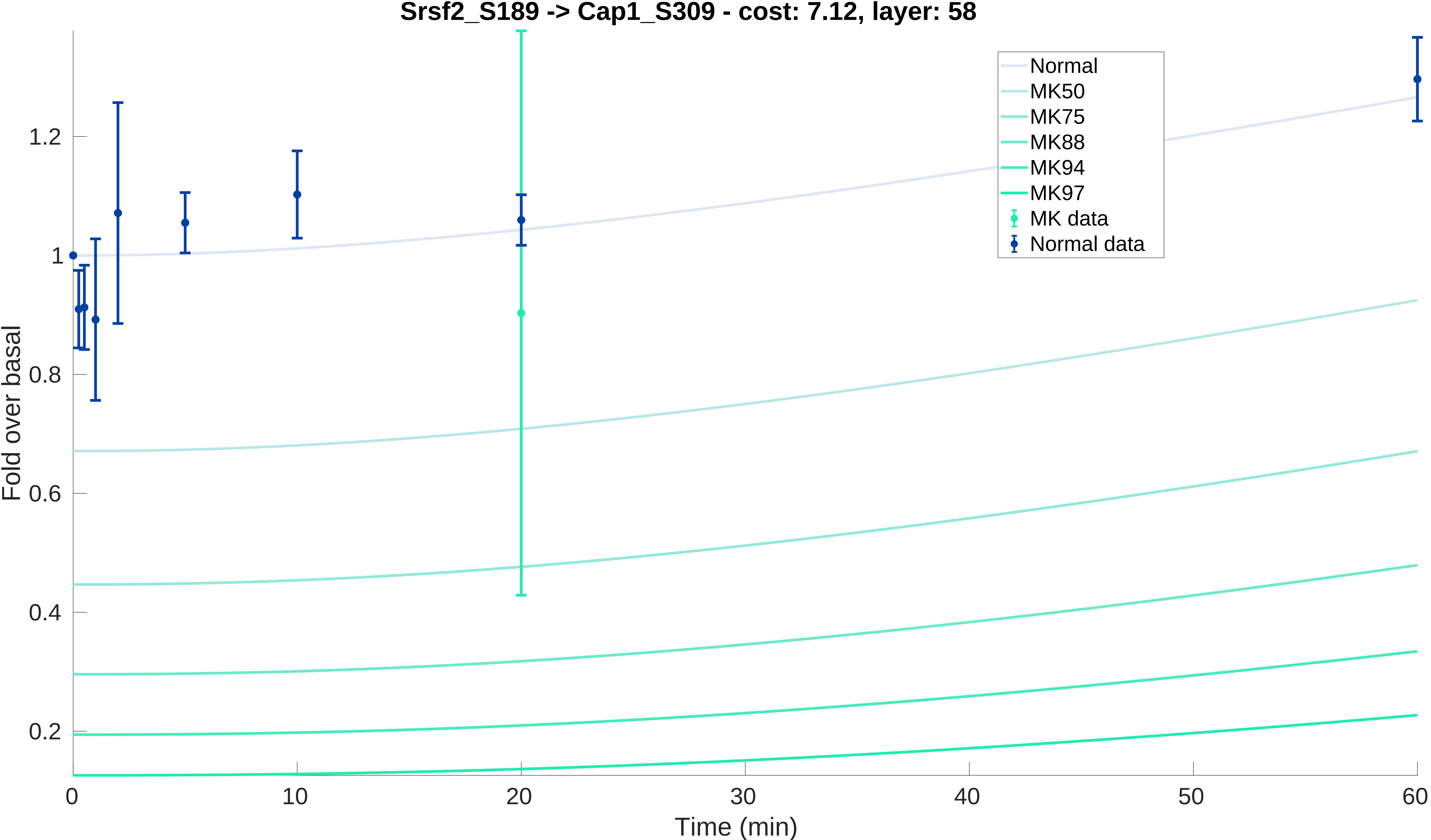

Srsf2_S189 -> Cap1_S309 - cost: 7.12, layer: 58
Normal
MK50
MK75
MK88
MK94
MK97
MK data
Normal data
Fold over basal
0.2
0.4
0.6
0.8
1
1.2
0
10
20
30
40
50
60
Time (min)

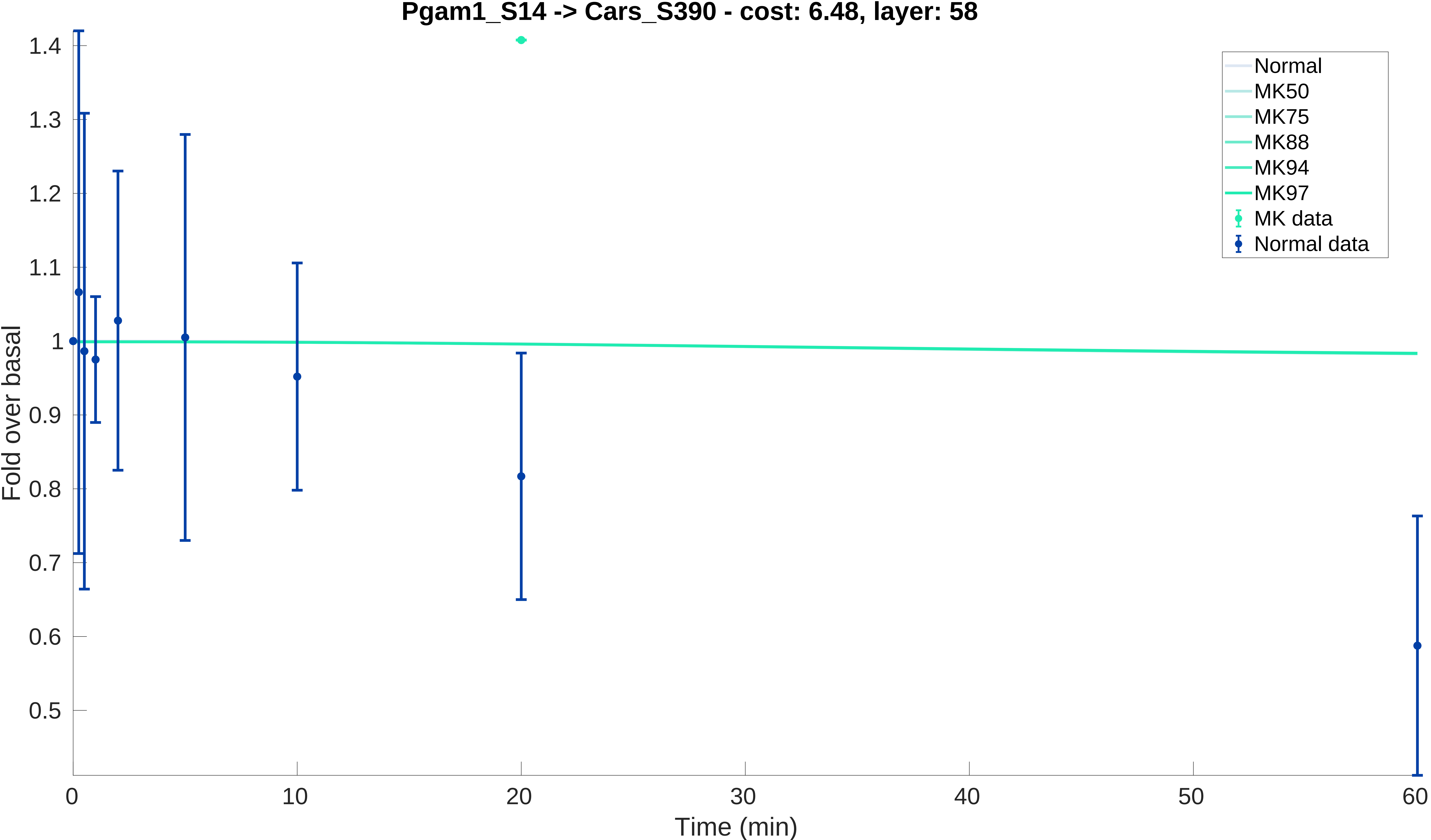
Pgam1_S14 -> Cars_S390 - cost: 6.48, layer: 58
Fold over basal
Time (min)
0.5
0.6
0.7
0.8
0.9
1
1.1
1.2
1.3
1.4
0
10
20
30
40
50
60
Normal
MK50
MK75
MK88
MK94
MK97
MK data
Normal data

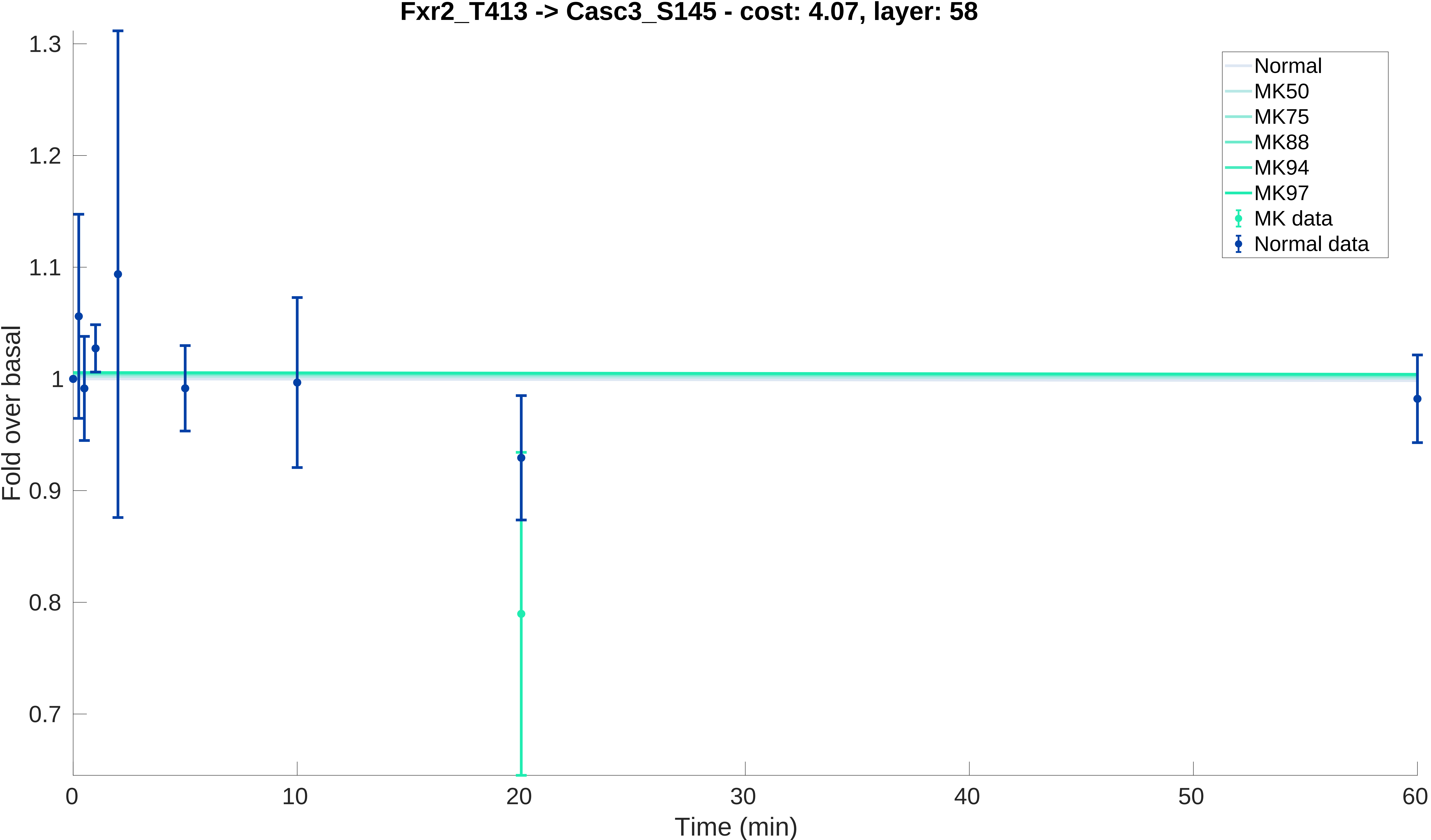

Fxr2_T413 -> Casc3_S145 - cost: 4.07, layer: 58
Fold over basal
Time (min)
1.3
1.2
1.1
1
0.9
0.8
0.7
0
10
20
30
40
50
60
Normal
MK50
MK75
MK88
MK94
MK97
MK data
Normal data

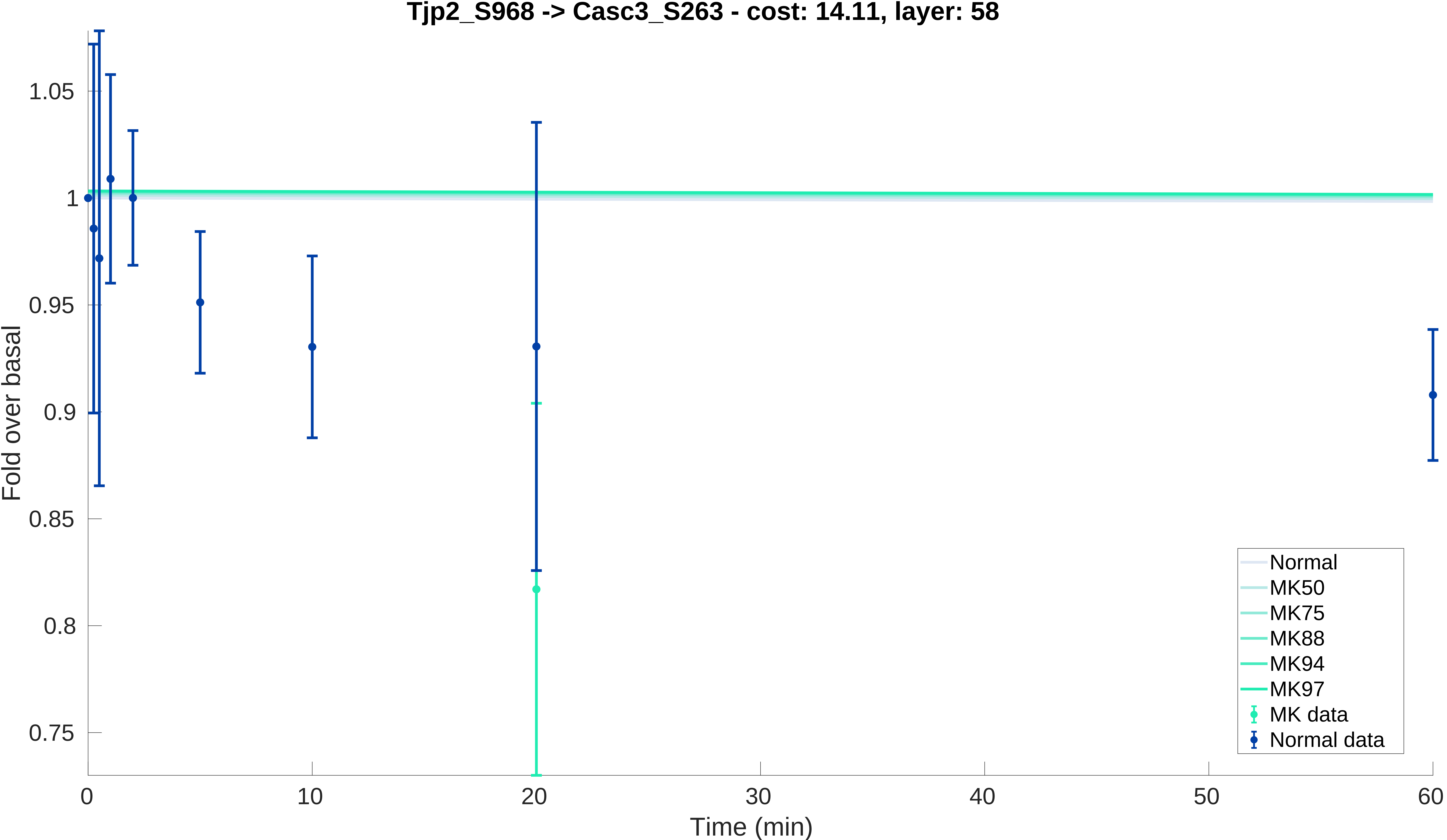

Tjp2_S968 -> Casc3_S263 - cost: 14.11, layer: 58
Fold over basal
1.05
1
0.95
0.9
0.85
0.8
0.75
0
10
20
30
40
50
60
Time (min)
Normal
MK50
MK75
MK88
MK94
MK97
MK data
Normal data

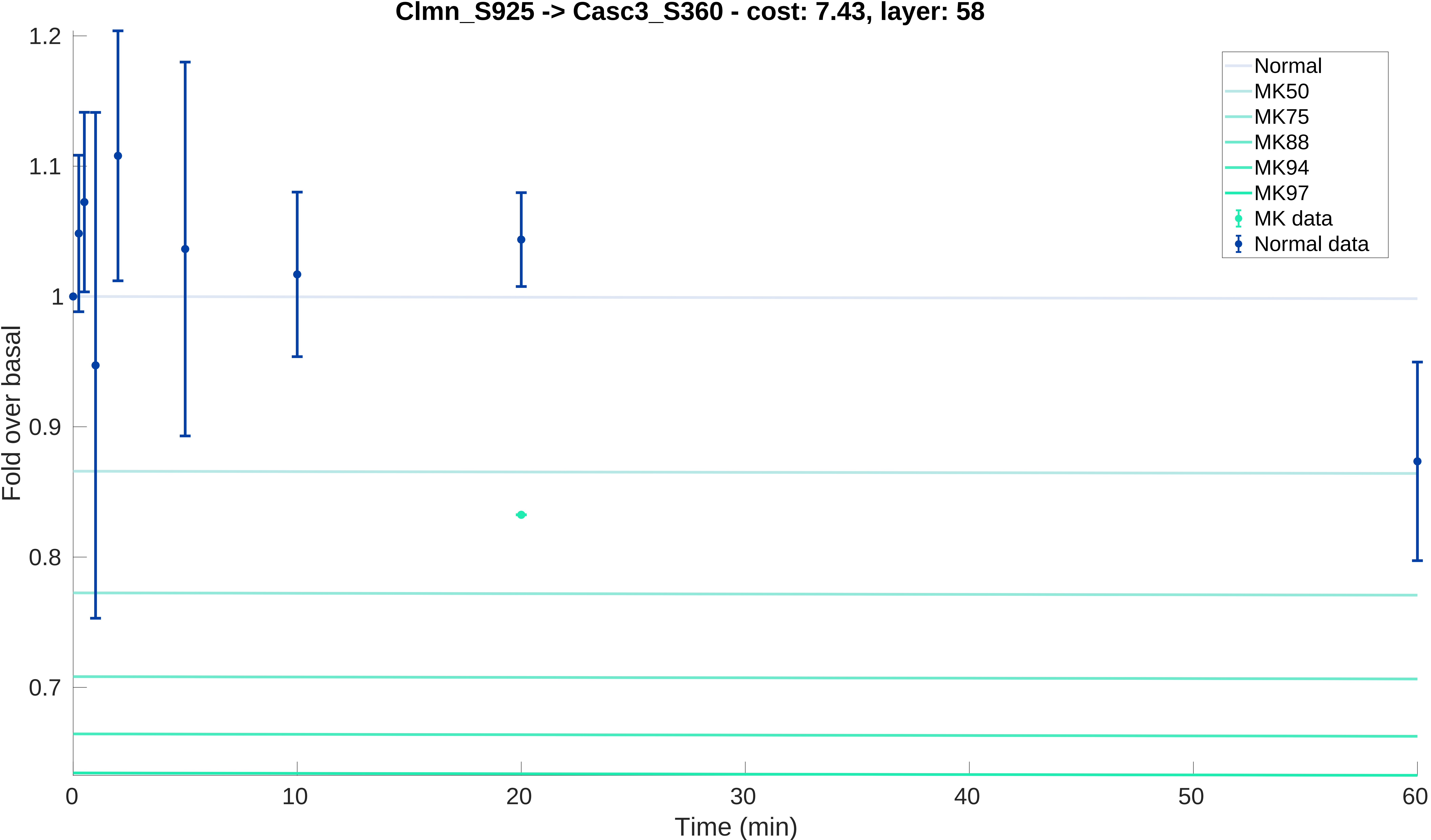
Clmn_S925 -> Casc3_S360 - cost: 7.43, layer: 58
Fold over basal
Time (min)
1.2
1.1
1
0.9
0.8
0.7
0
10
20
30
40
50
60
Normal
MK50
MK75
MK88
MK94
MK97
MK data
Normal data

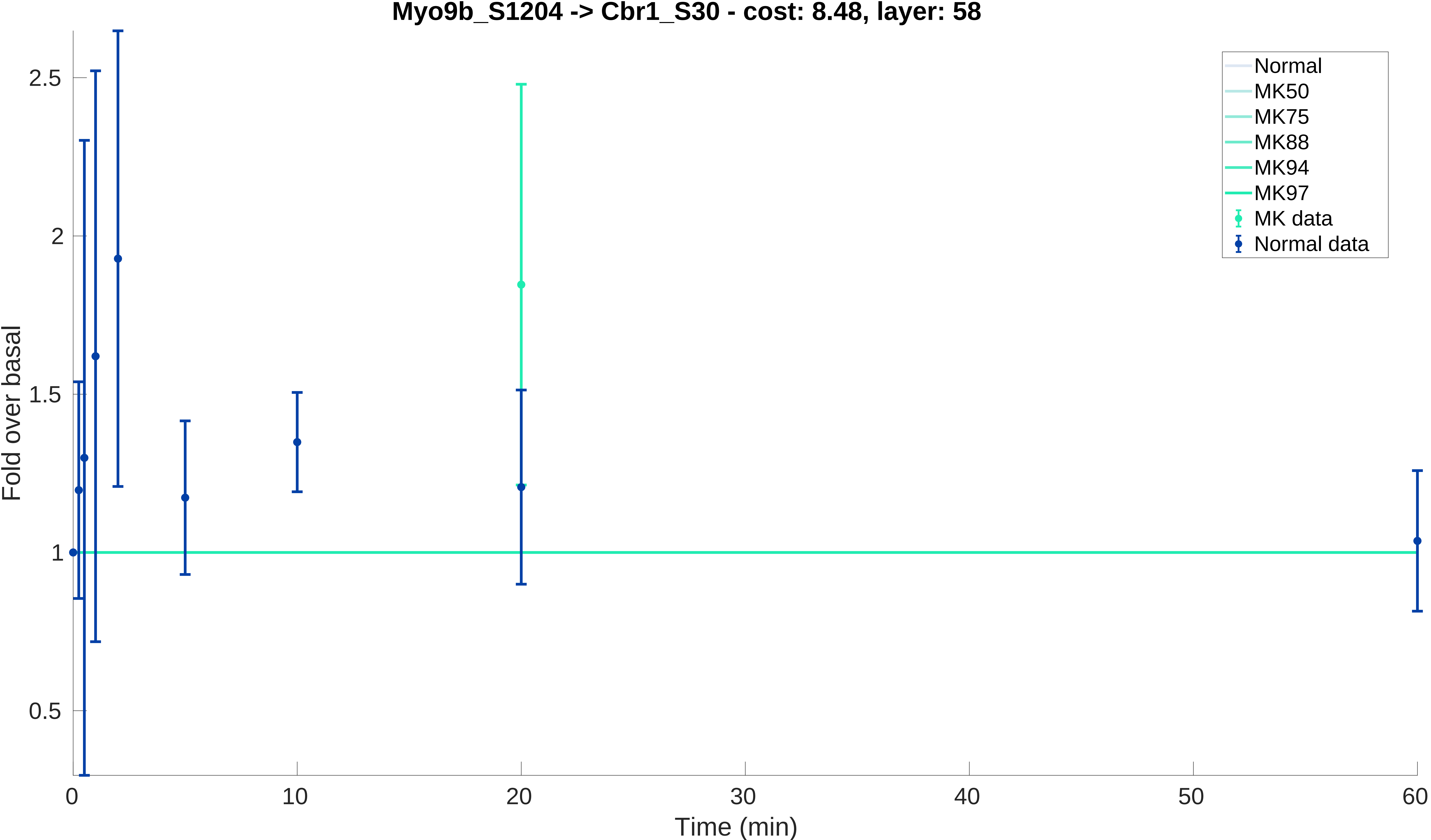

Myo9b_S1204 -> Cbr1_S30 - cost: 8.48, layer: 58
Normal
MK50
MK75
MK88
MK94
MK97
MK data
Normal data
Fold over basal
0.5
1
1.5
2
2.5
0
10
20
30
40
50
60
Time (min)

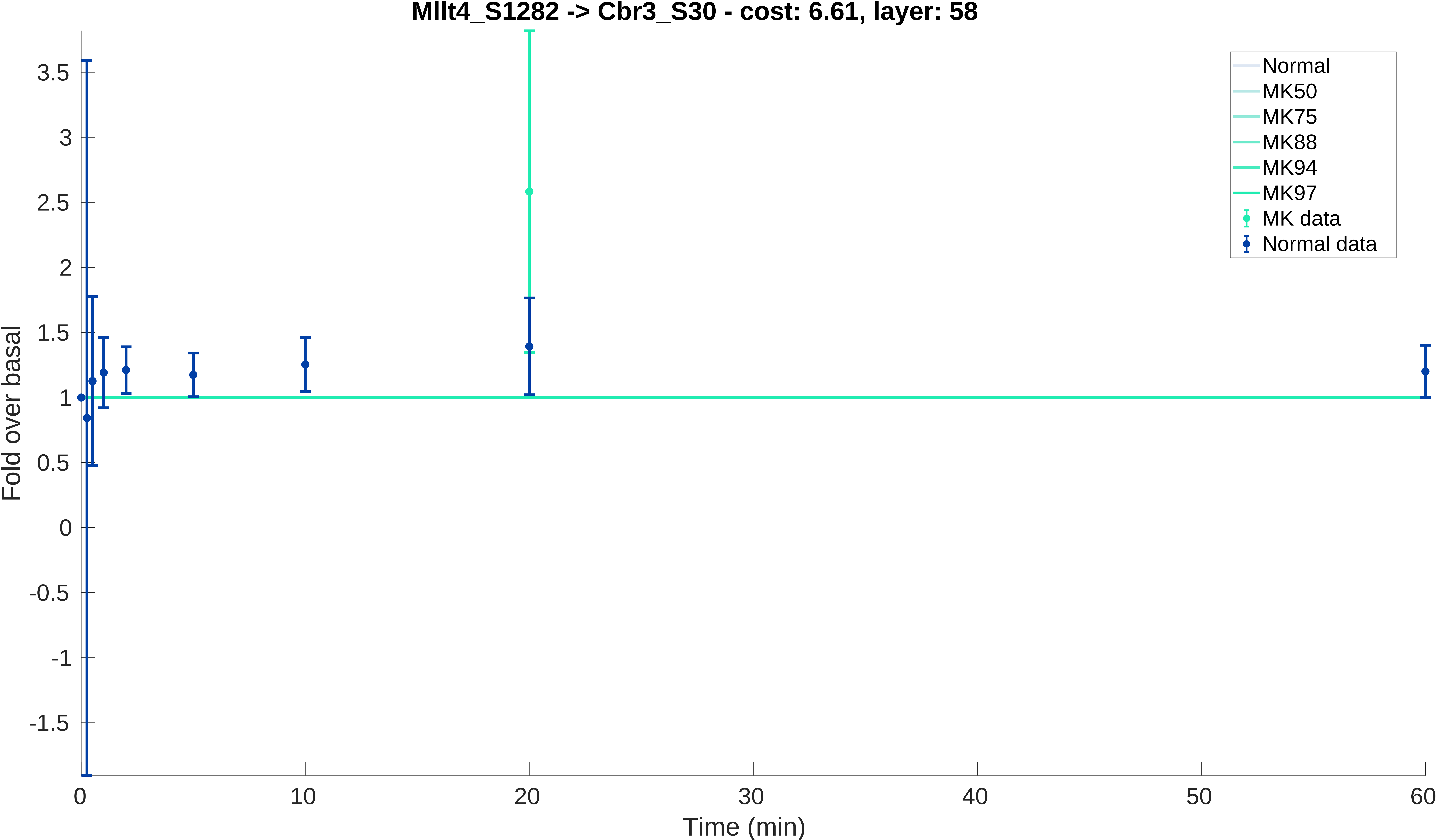

Mllt4_S1282 -> Cbr3_S30 - cost: 6.61, layer: 58
Fold over basal
3.5
3
2.5
2
1.5
1
0.5
0
-0.5
-1
-1.5
0
10
20
30
40
50
60
Time (min)
Normal
MK50
MK75
MK88
MK94
MK97
MK data
Normal data

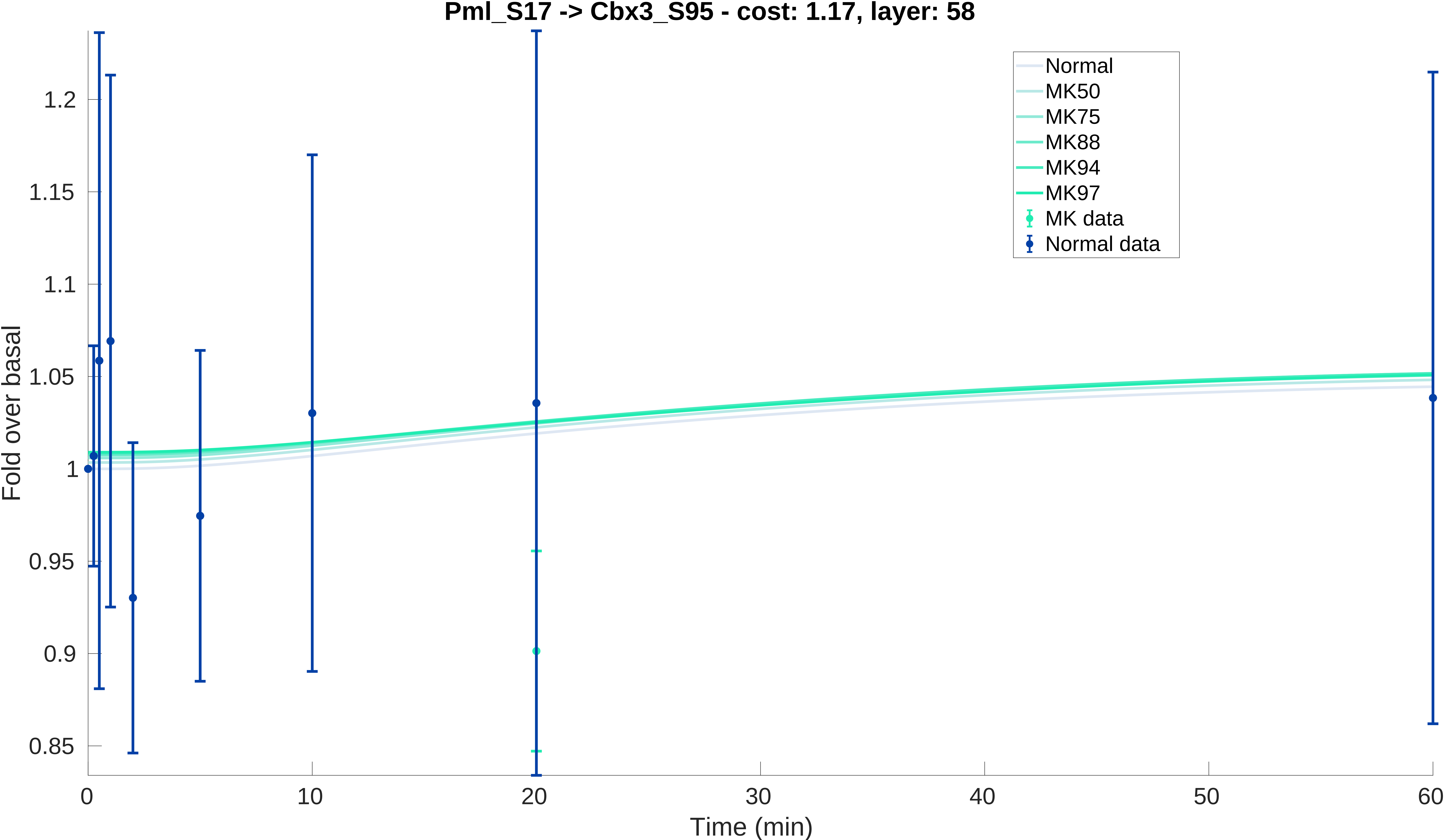

Pml_S17 -> Cbx3_S95 - cost: 1.17, layer: 58
Normal
MK50
MK75
MK88
MK94
MK97
MK data
Normal data
Fold over basal
1.2
1.15
1.1
1.05
1
0.95
0.9
0.85
0
10
20
30
40
50
60
Time (min)

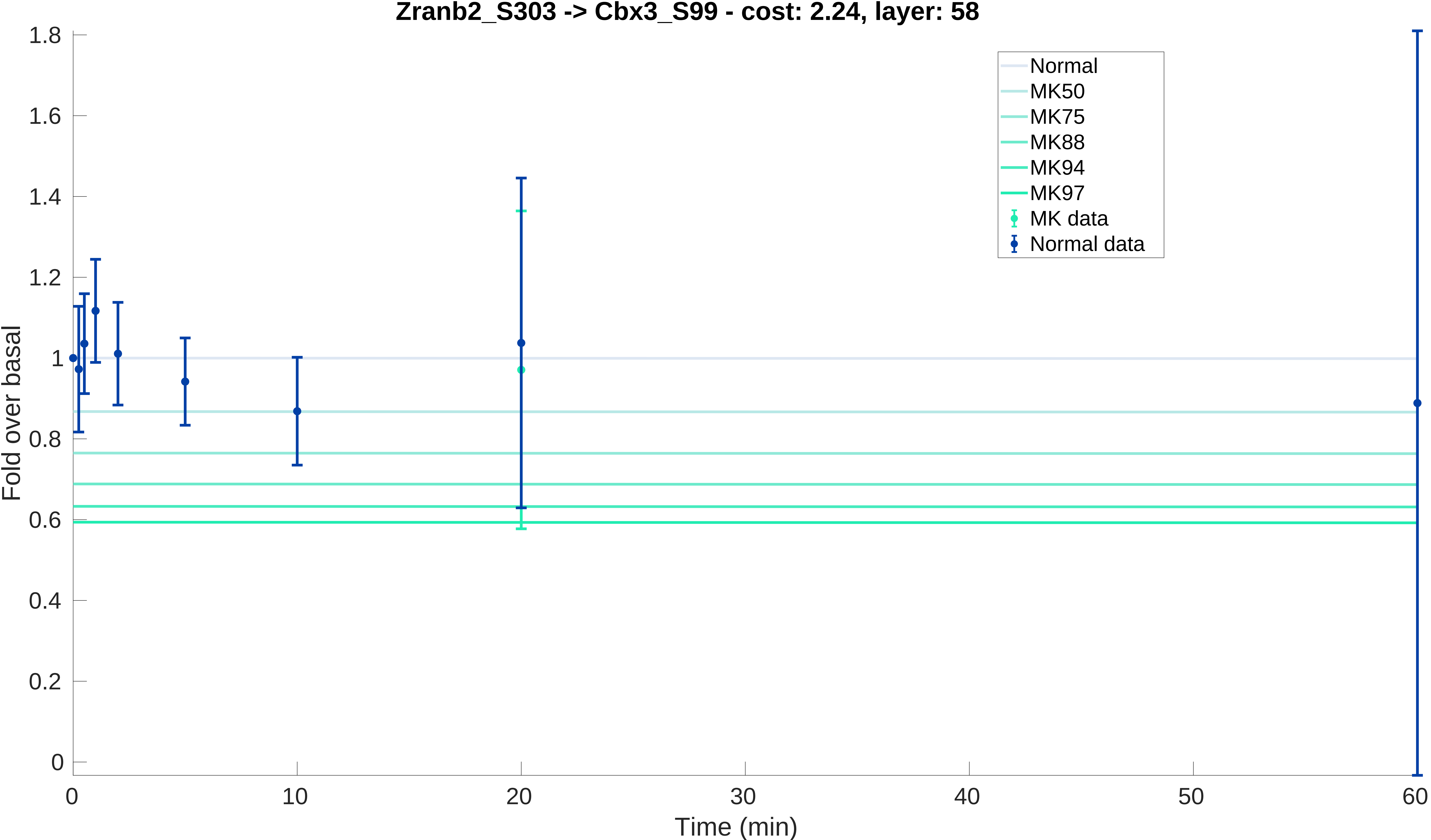

Zranb2_S303 -> Cbx3_S99 - cost: 2.24, layer: 58
Fold over basal
Time (min)
0
0.2
0.4
0.6
0.8
1
1.2
1.4
1.6
1.8
0
10
20
30
40
50
60
Normal
MK50
MK75
MK88
MK94
MK97
MK data
Normal data

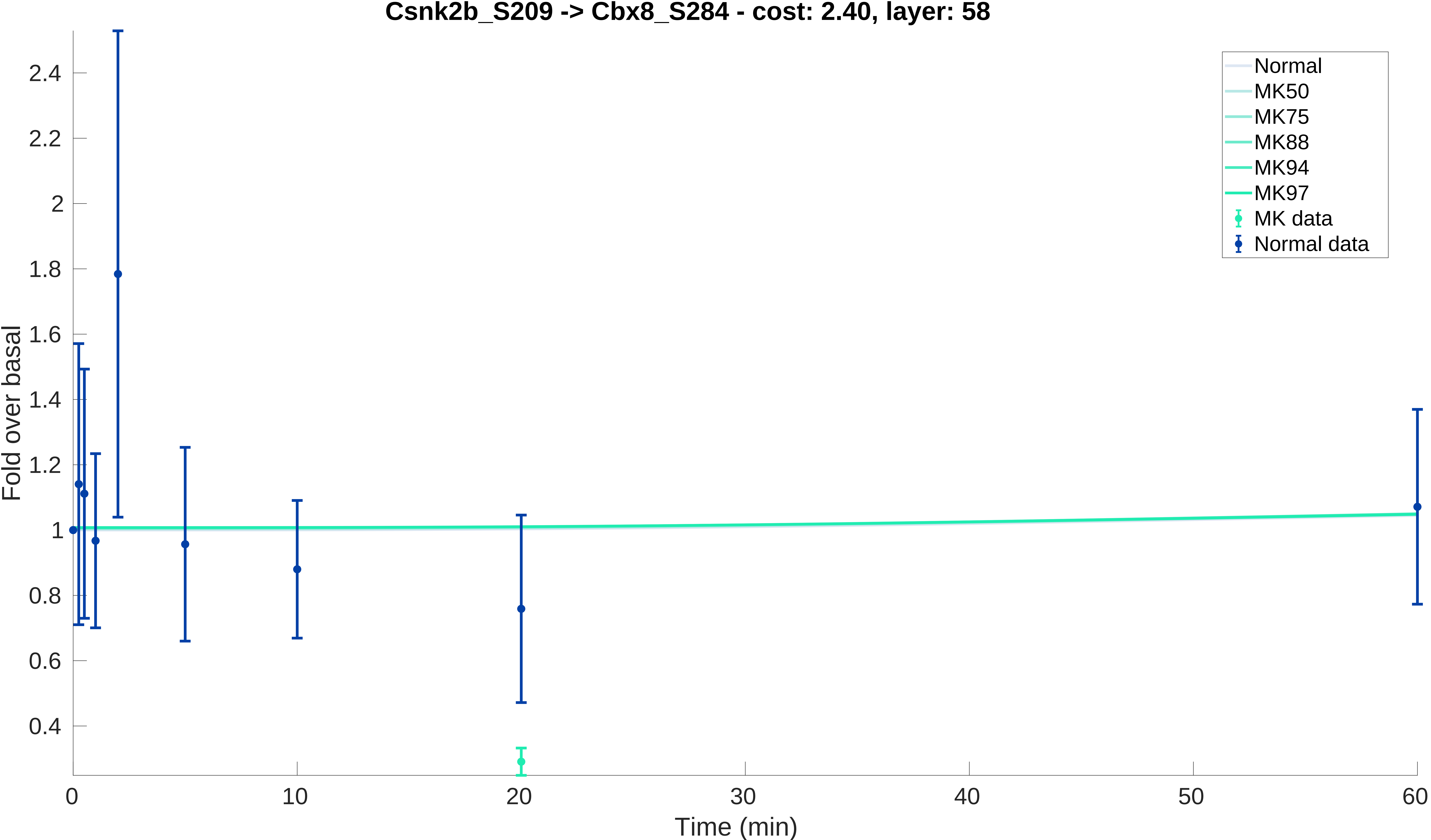
Csnk2b_S209 -> Cbx8_S284 - cost: 2.40, layer: 58
Fold over basal
Time (min)
0.4
0.6
0.8
1
1.2
1.4
1.6
1.8
2
2.2
2.4
0
10
20
30
40
50
60
Normal
MK50
MK75
MK88
MK94
MK97
MK data
Normal data

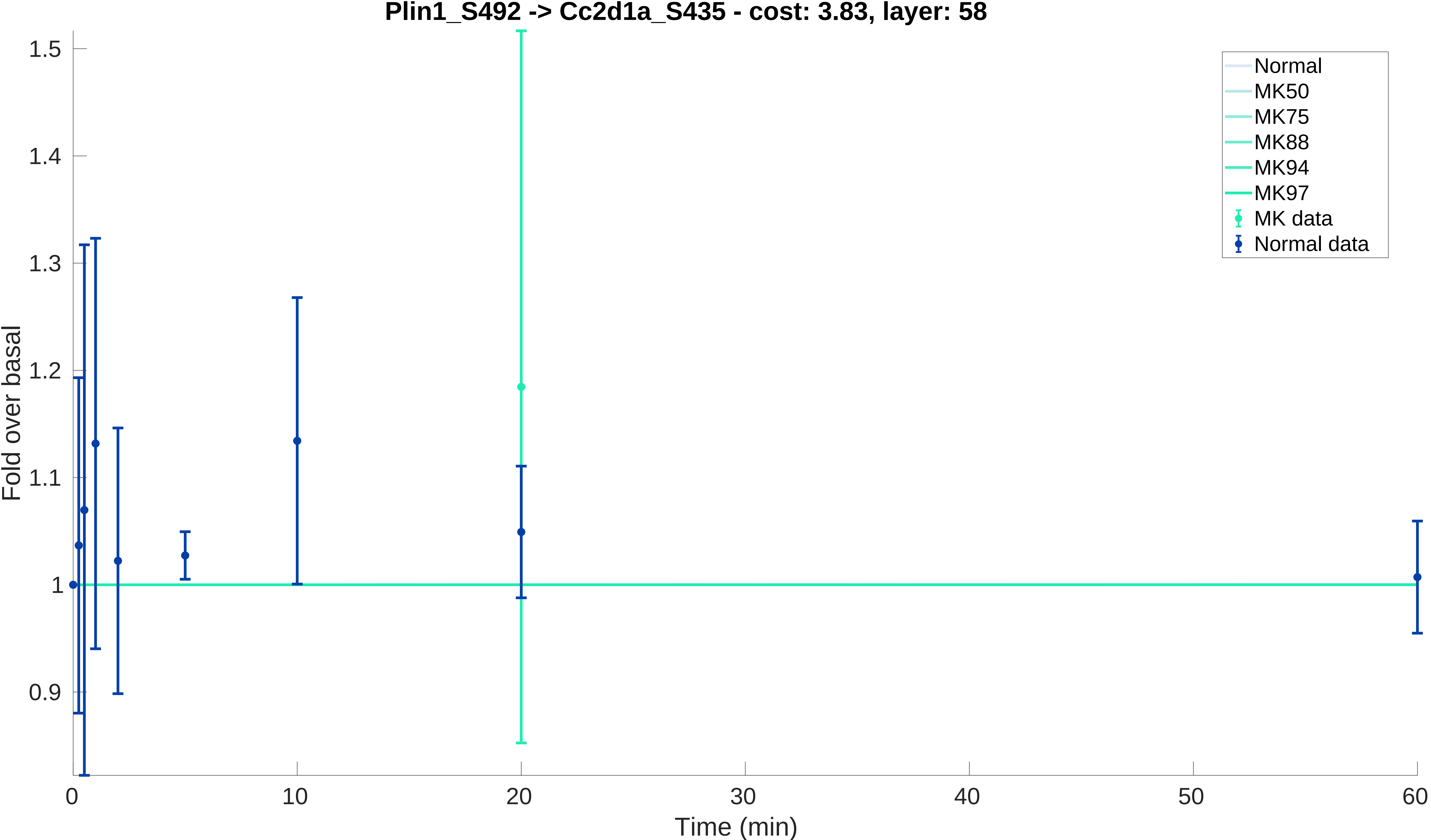

Plin1_S492 -> Cc2d1a_S435 - cost: 3.83, layer: 58
Fold over basal
1.5
1.4
1.3
1.2
1.1
1
0.9
0
10
20
30
40
50
60
Time (min)
Normal
MK50
MK75
MK88
MK94
MK97
MK data
Normal data

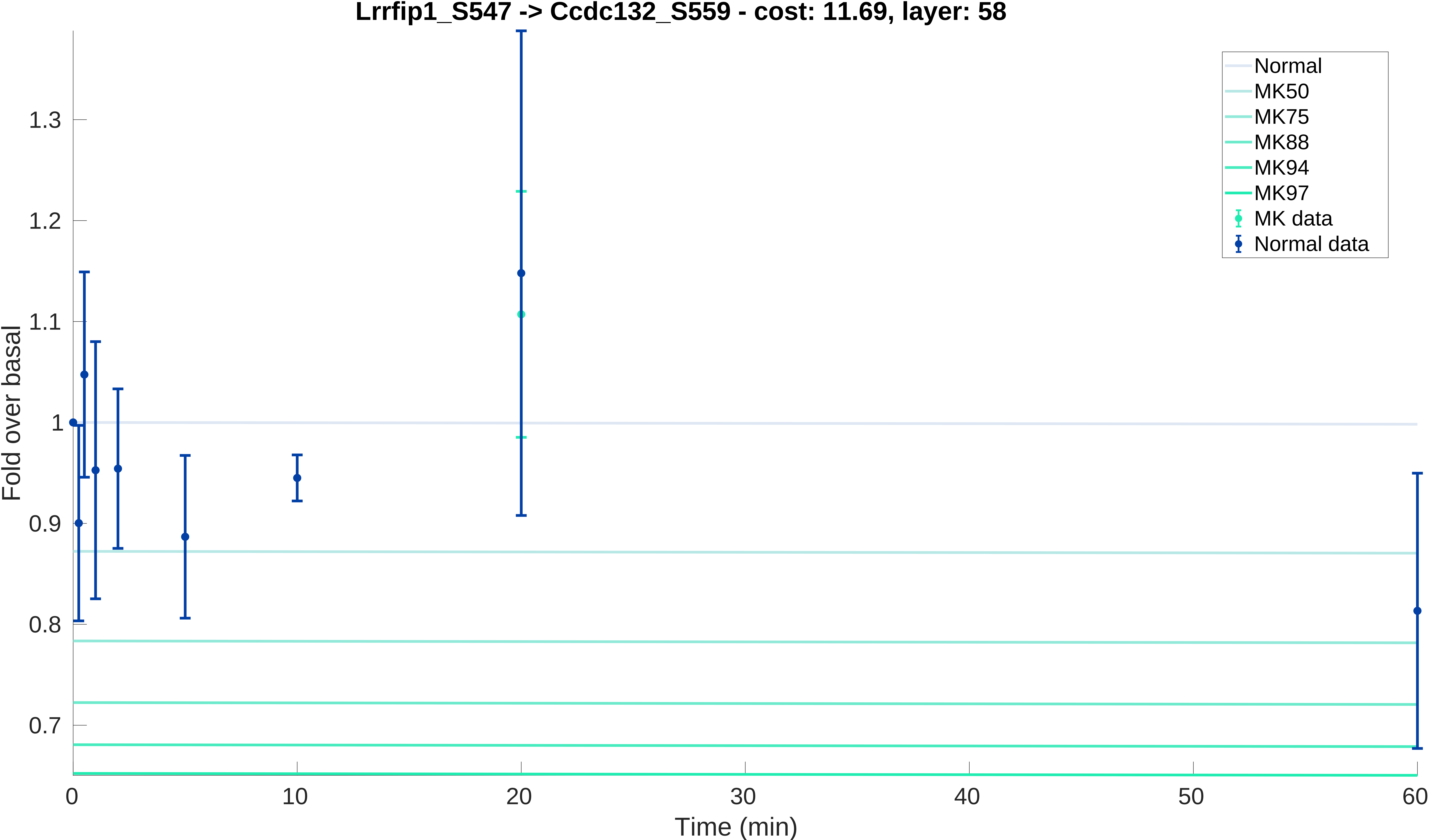

Lrrfip1_S547 -> Ccdc132_S559 - cost: 11.69, layer: 58
Fold over basal
Time (min)
0.7
0.8
0.9
1
1.1
1.2
1.3
0
10
20
30
40
50
60
Normal
MK50
MK75
MK88
MK94
MK97
MK data
Normal data

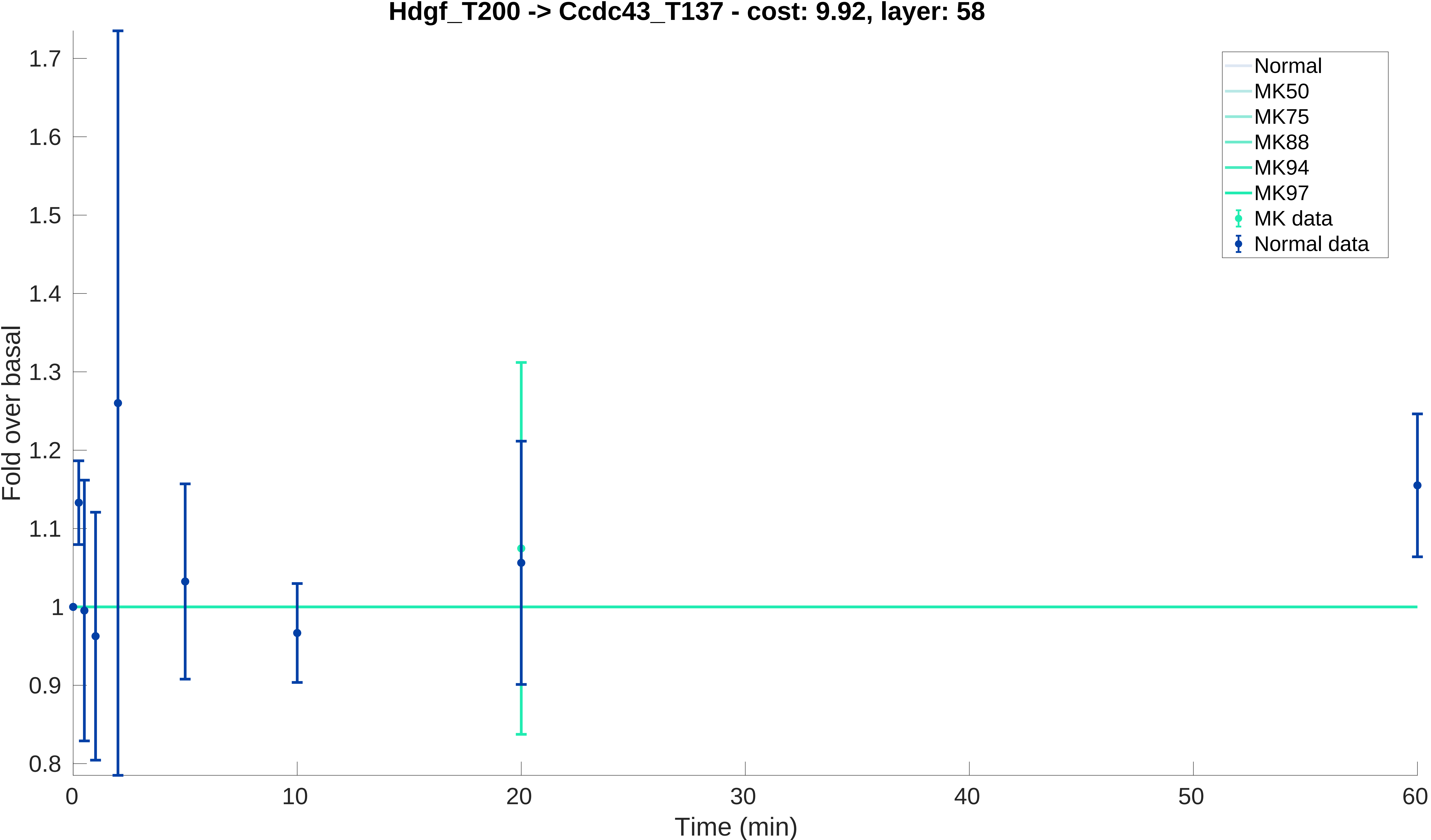

Hdgf_T200 -> Ccdc43_T137 - cost: 9.92, layer: 58
Fold over basal
Time (min)
0.8
0.9
1
1.1
1.2
1.3
1.4
1.5
1.6
1.7
0
10
20
30
40
50
60
Normal
MK50
MK75
MK88
MK94
MK97
MK data
Normal data

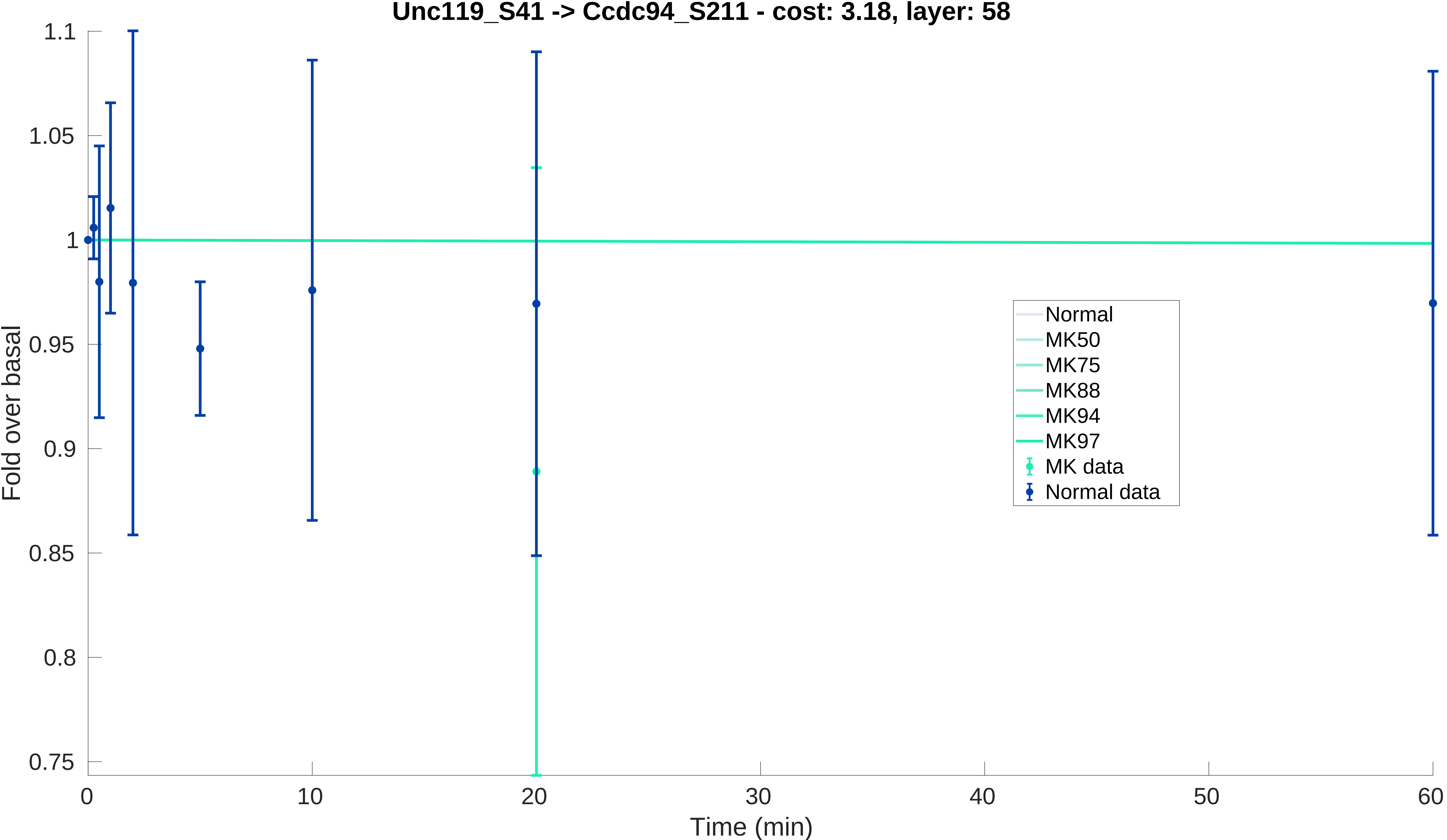

Unc119_S41 -> Ccdc94_S211 - cost: 3.18, layer: 58
Fold over basal
1.1
1.05
1
0.95
0.9
0.85
0.8
0.75
0
10
20
30
40
50
60
Time (min)
Normal
MK50
MK75
MK88
MK94
MK97
MK data
Normal data

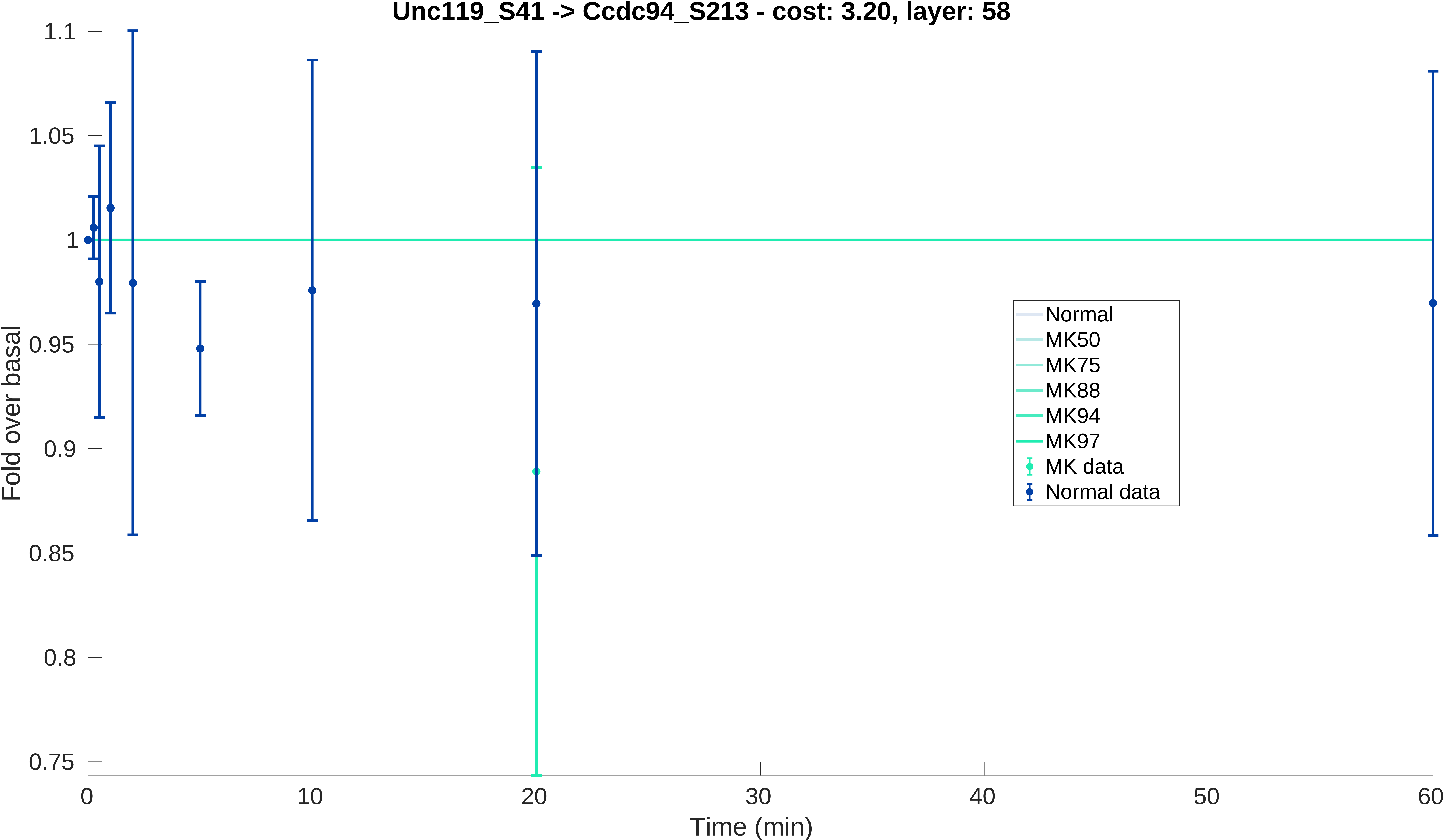

Unc119_S41 -> Ccdc94_S213 - cost: 3.20, layer: 58
Fold over basal
1.1
1.05
1
0.95
0.9
0.85
0.8
0.75
0
10
20
30
40
50
60
Time (min)
Normal
MK50
MK75
MK88
MK94
MK97
MK data
Normal data

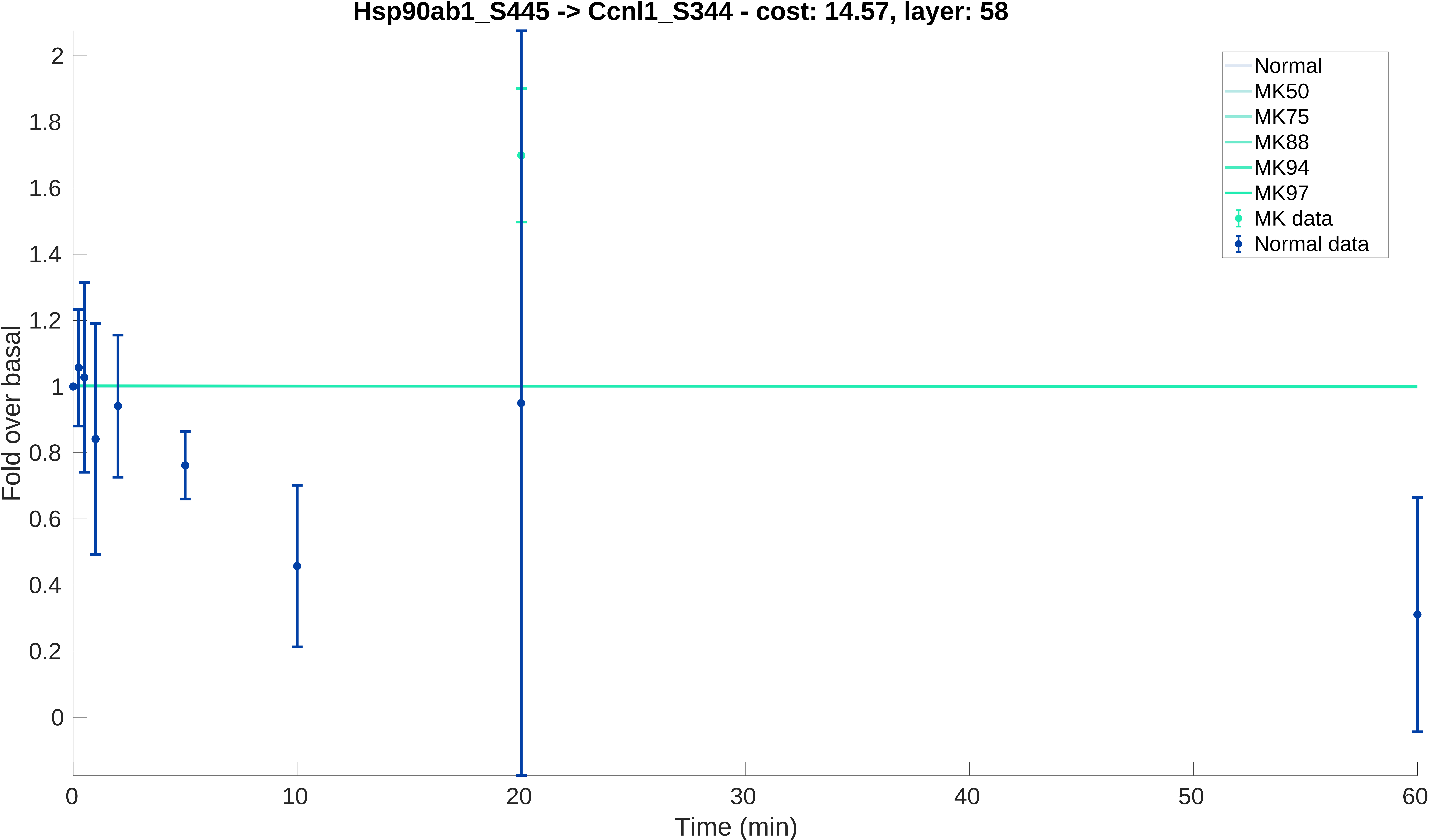
Hsp90ab1_S445 -> Ccnl1_S344 - cost: 14.57, layer: 58
Fold over basal
Time (min)
0
0.2
0.4
0.6
0.8
1
1.2
1.4
1.6
1.8
2
0
10
20
30
40
50
60
Normal
MK50
MK75
MK88
MK94
MK97
MK data
Normal data

#### Zranb2_S303 -> Ccnl1_S348 - cost: 1.61, layer: 58

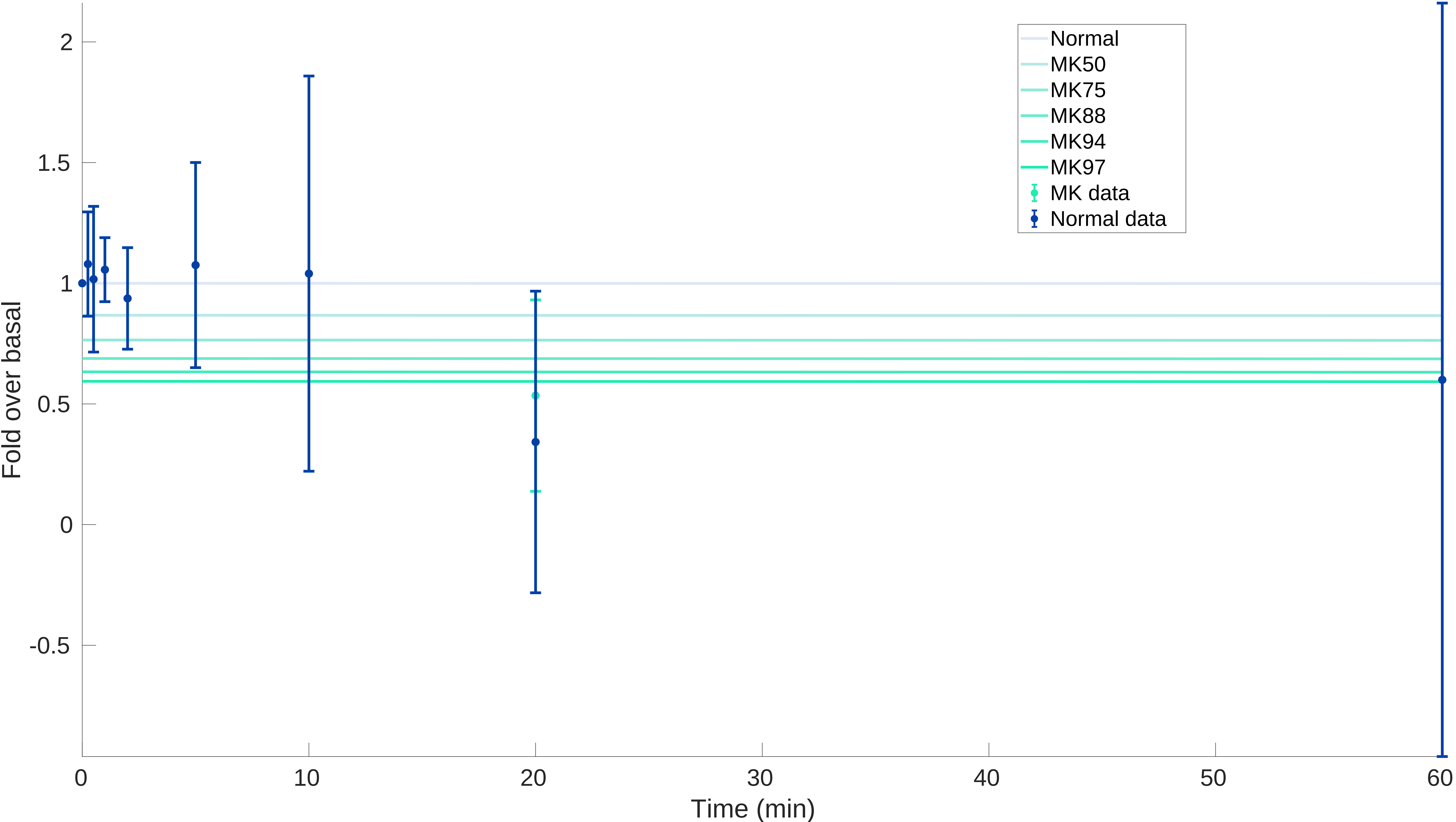

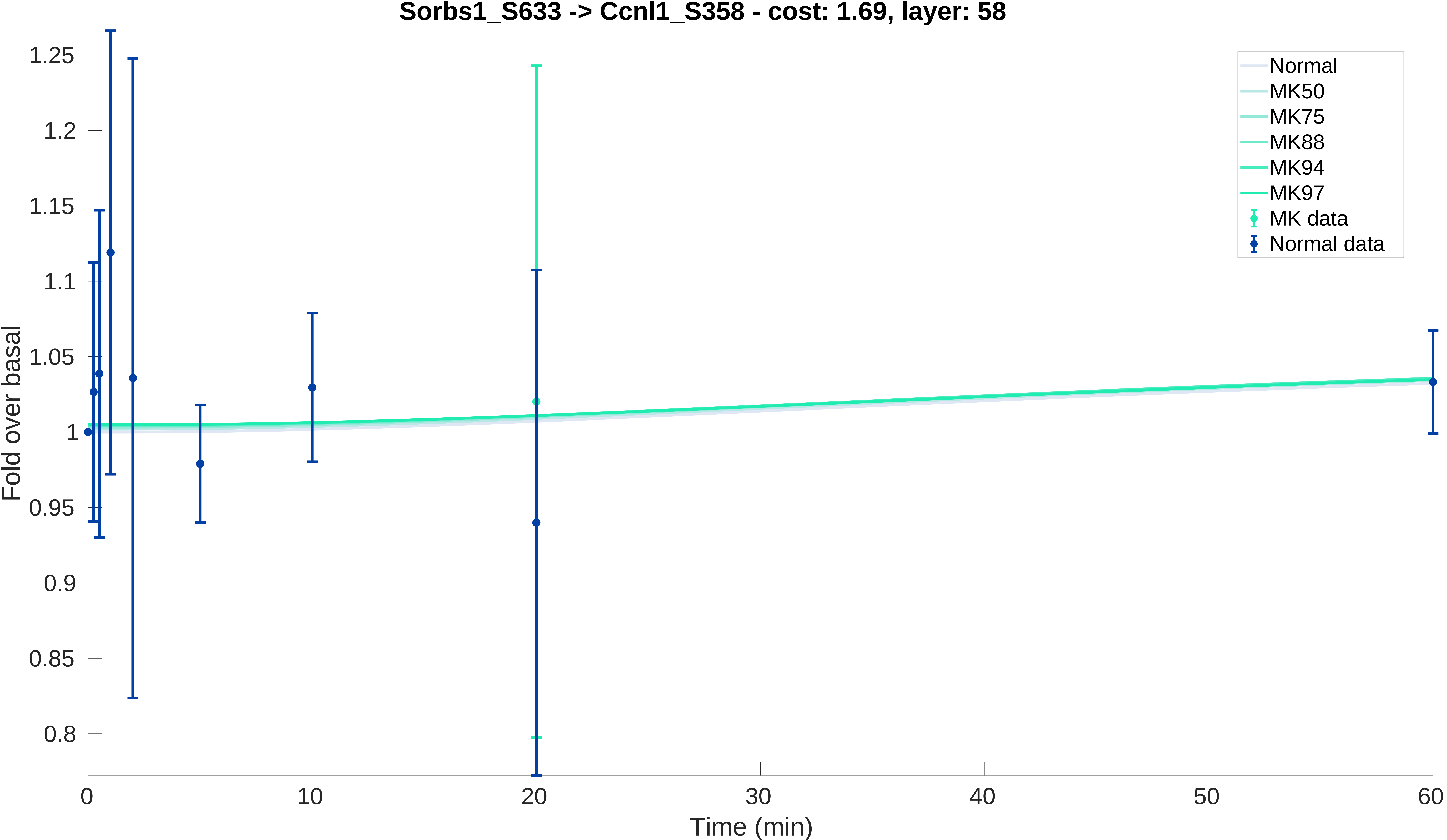

Sorbs1_S633 -> Ccnl1_S358 - cost: 1.69, layer: 58
Fold over basal
Time (min)
1.25
1.2
1.15
1.1
1.05
1
0.95
0.9
0.85
0.8
0
10
20
30
40
50
60
Normal
MK50
MK75
MK88
MK94
MK97
MK data
Normal data

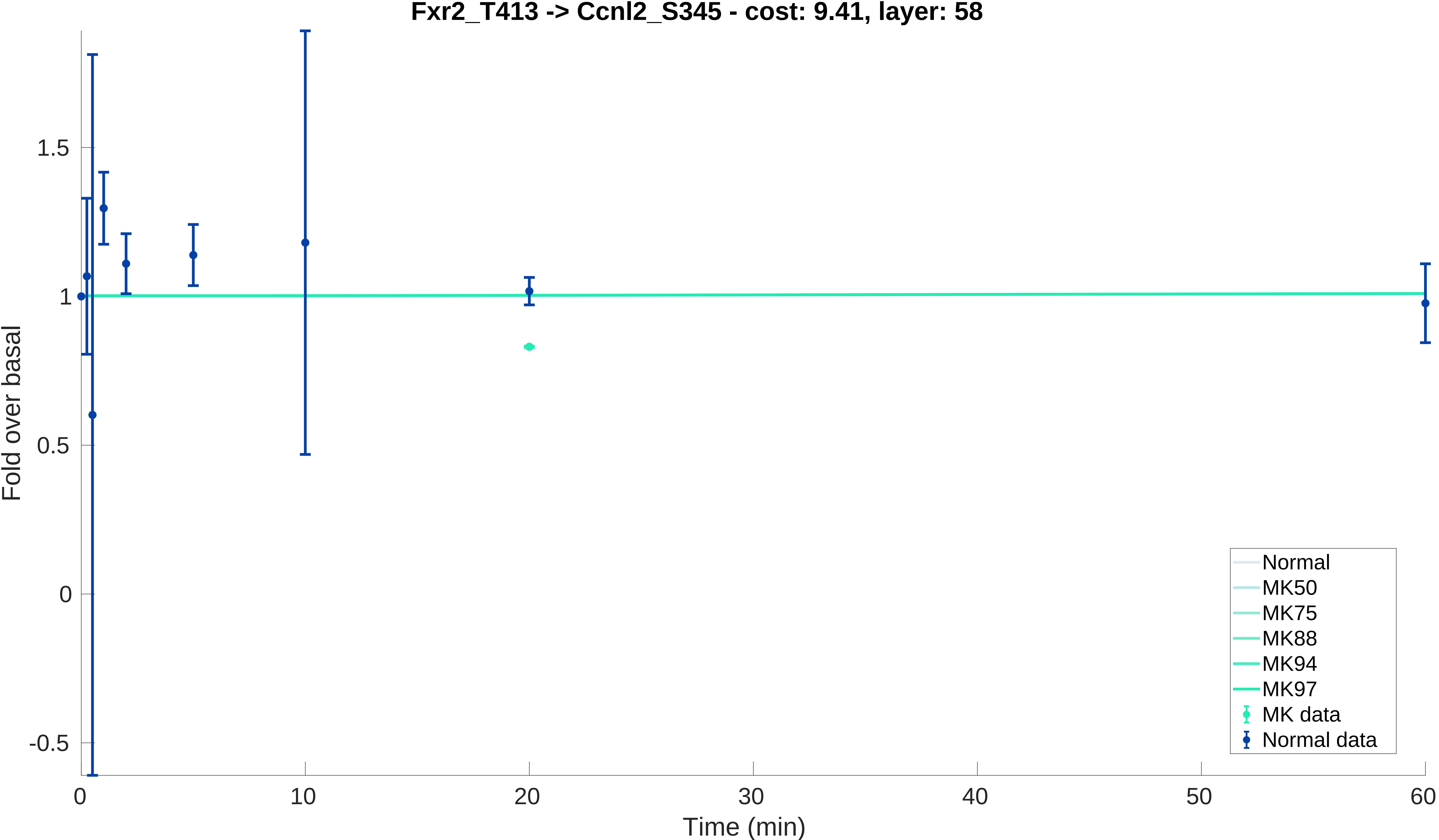

Fxr2_T413 -> Ccnl2_S345 - cost: 9.41, layer: 58
Fold over basal
Time (min)
1.5
1
0.5
0
-0.5
0
10
20
30
40
50
60
Normal
MK50
MK75
MK88
MK94
MK97
MK data
Normal data

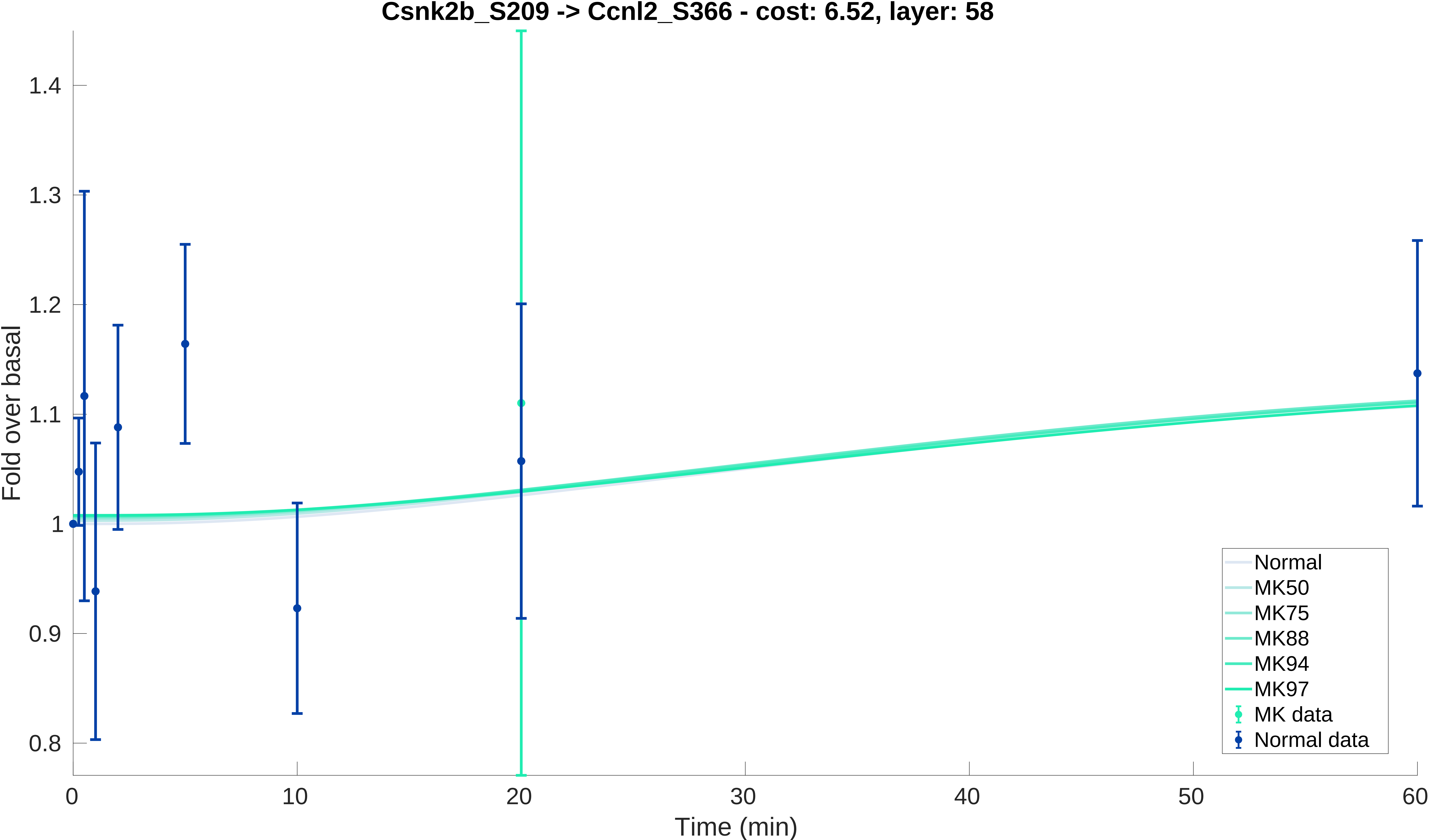

Csnk2b_S209 -> Ccnl2_S366 - cost: 6.52, layer: 58
Fold over basal
1.4
1.3
1.2
1.1
1
0.9
0.8
0
10
20
30
40
50
60
Time (min)
Normal
MK50
MK75
MK88
MK94
MK97
MK data
Normal data

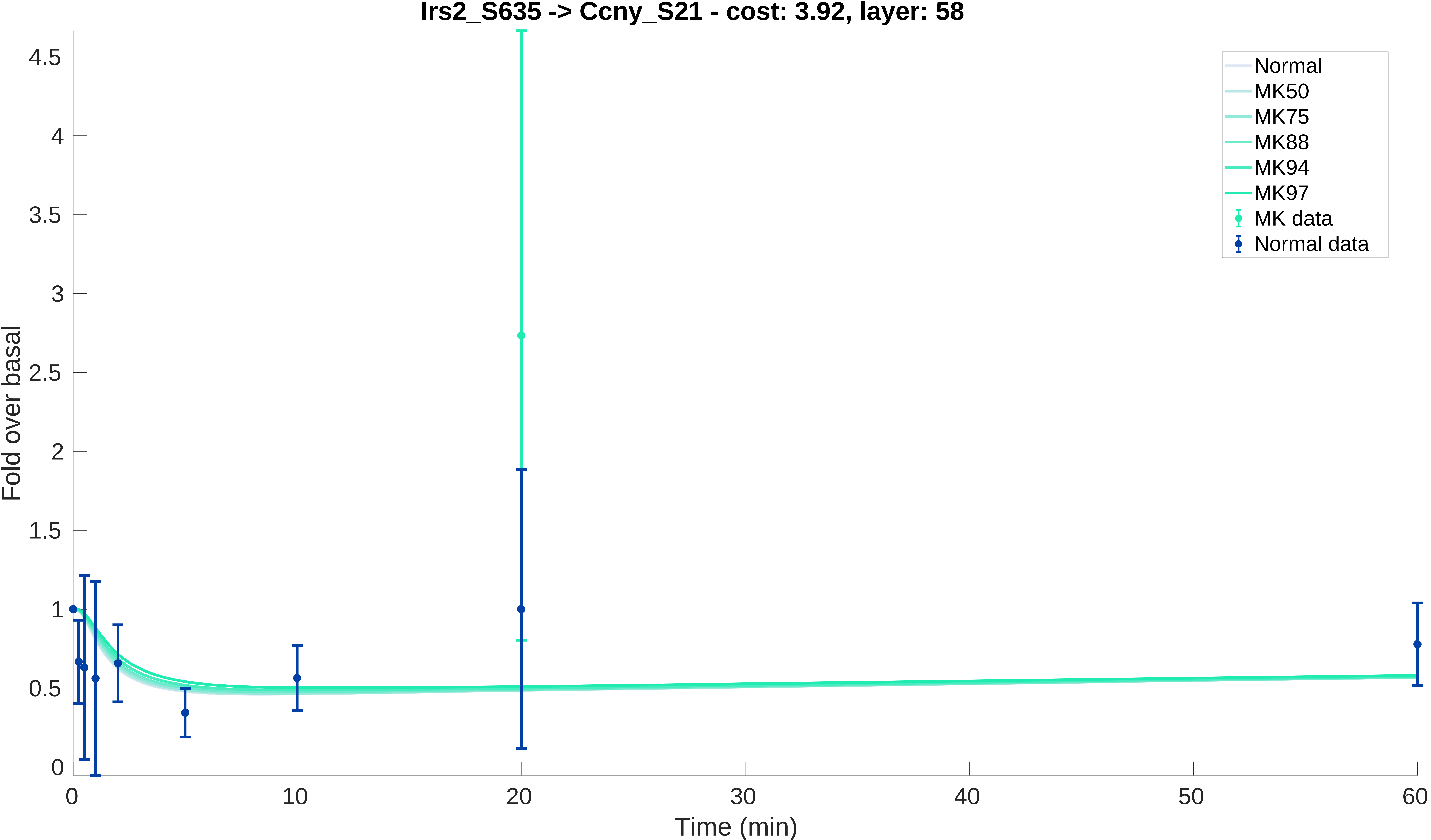

Irs2_S635 -> Ccny_S21 - cost: 3.92, layer: 58
Normal
MK50
MK75
MK88
MK94
MK97
MK data
Normal data
Fold over basal
Time (min)
0
0.5
1
1.5
2
2.5
3
3.5
4
4.5
0
10
20
30
40
50
60

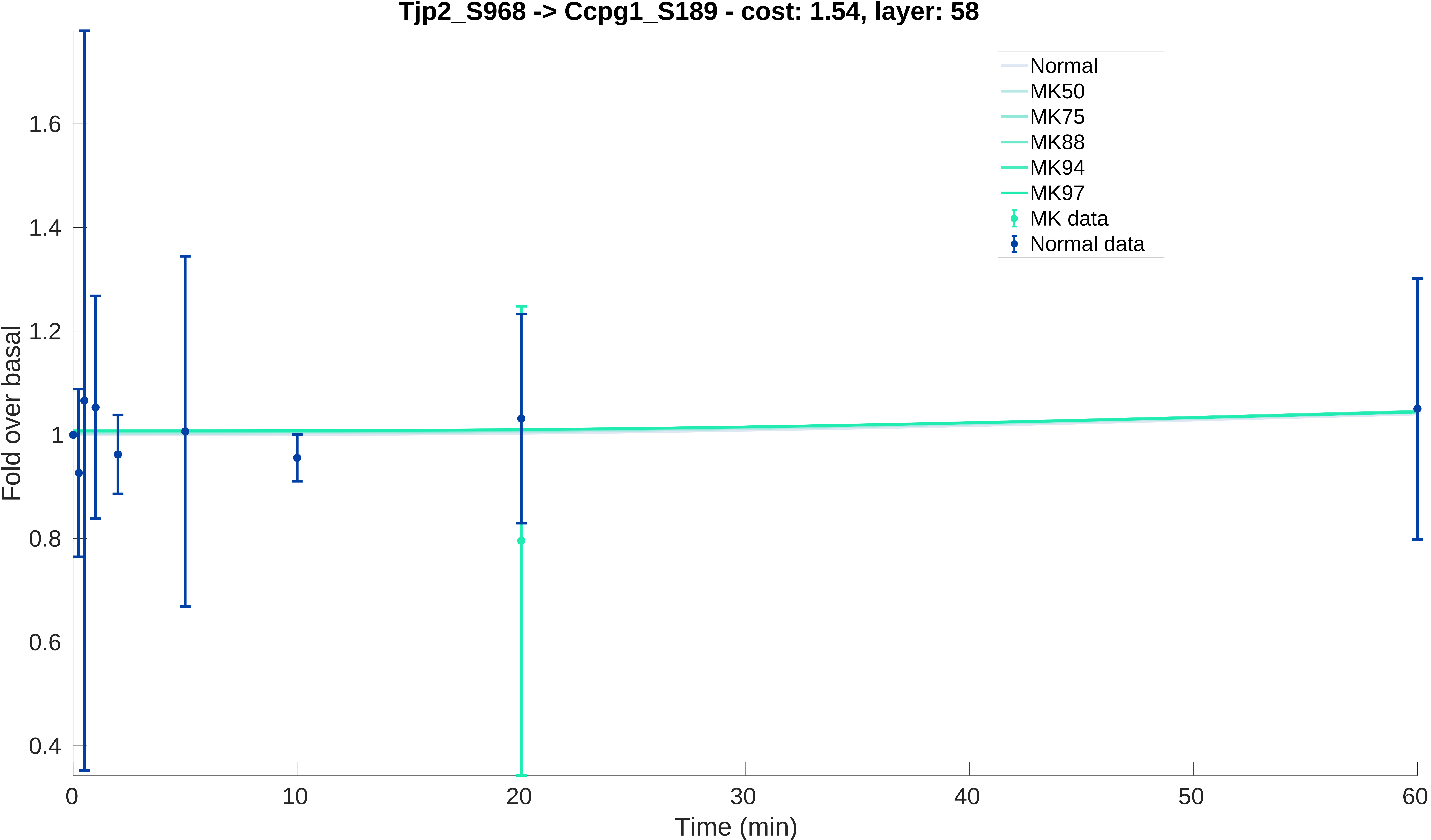
Tjp2_S968 -> Ccpg1_S189 - cost: 1.54, layer: 58
Fold over basal
Time (min)
0
10
20
30
40
50
60
0.4
0.6
0.8
1
1.2
1.4
1.6
Normal
MK50
MK75
MK88
MK94
MK97
MK data
Normal data

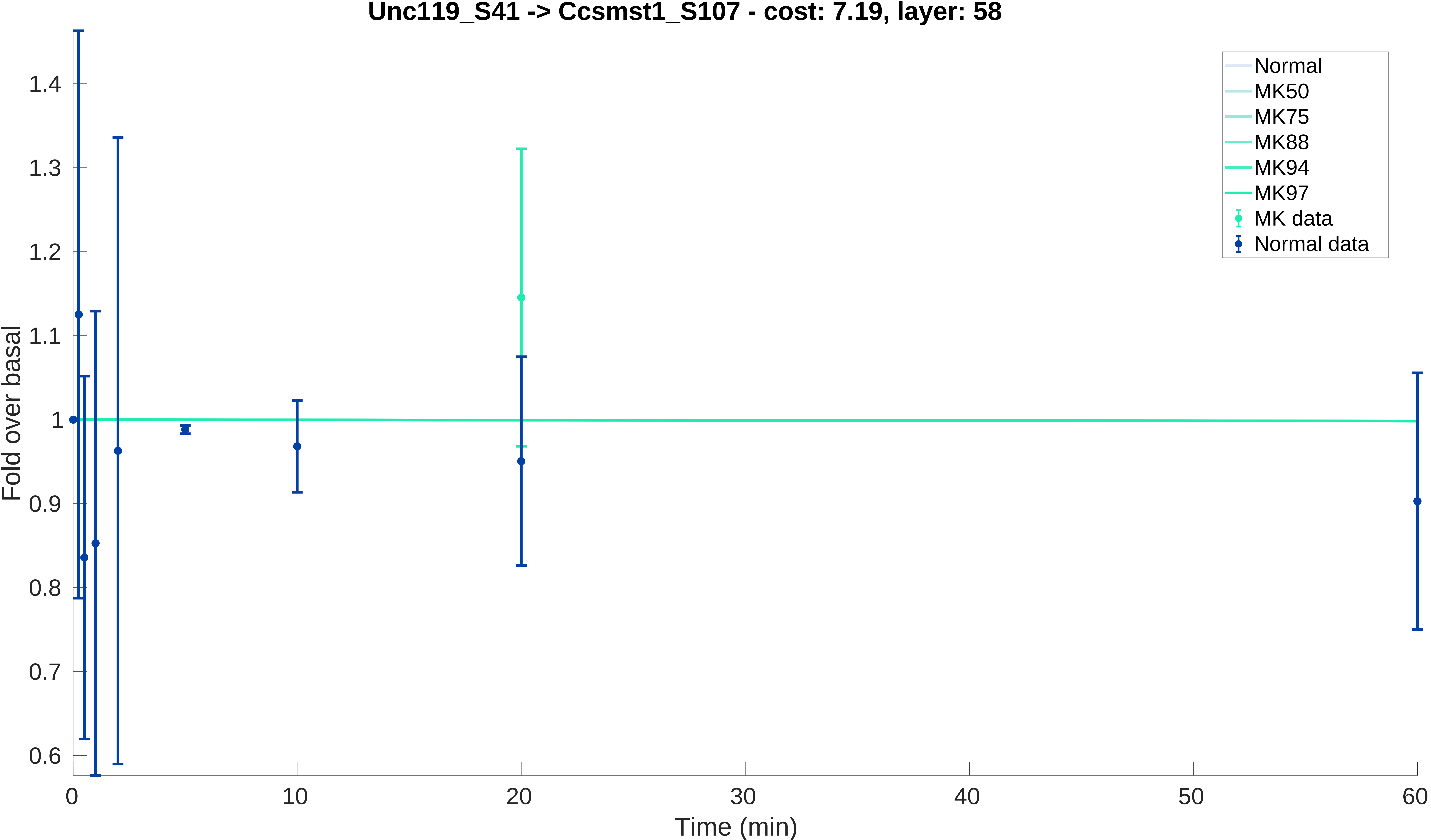

Unc119_S41 -> Ccsmst1_S107 - cost: 7.19, layer: 58
Fold over basal
Time (min)
0.6
0.7
0.8
0.9
1
1.1
1.2
1.3
1.4
0
10
20
30
40
50
60
Normal
MK50
MK75
MK88
MK94
MK97
MK data
Normal data

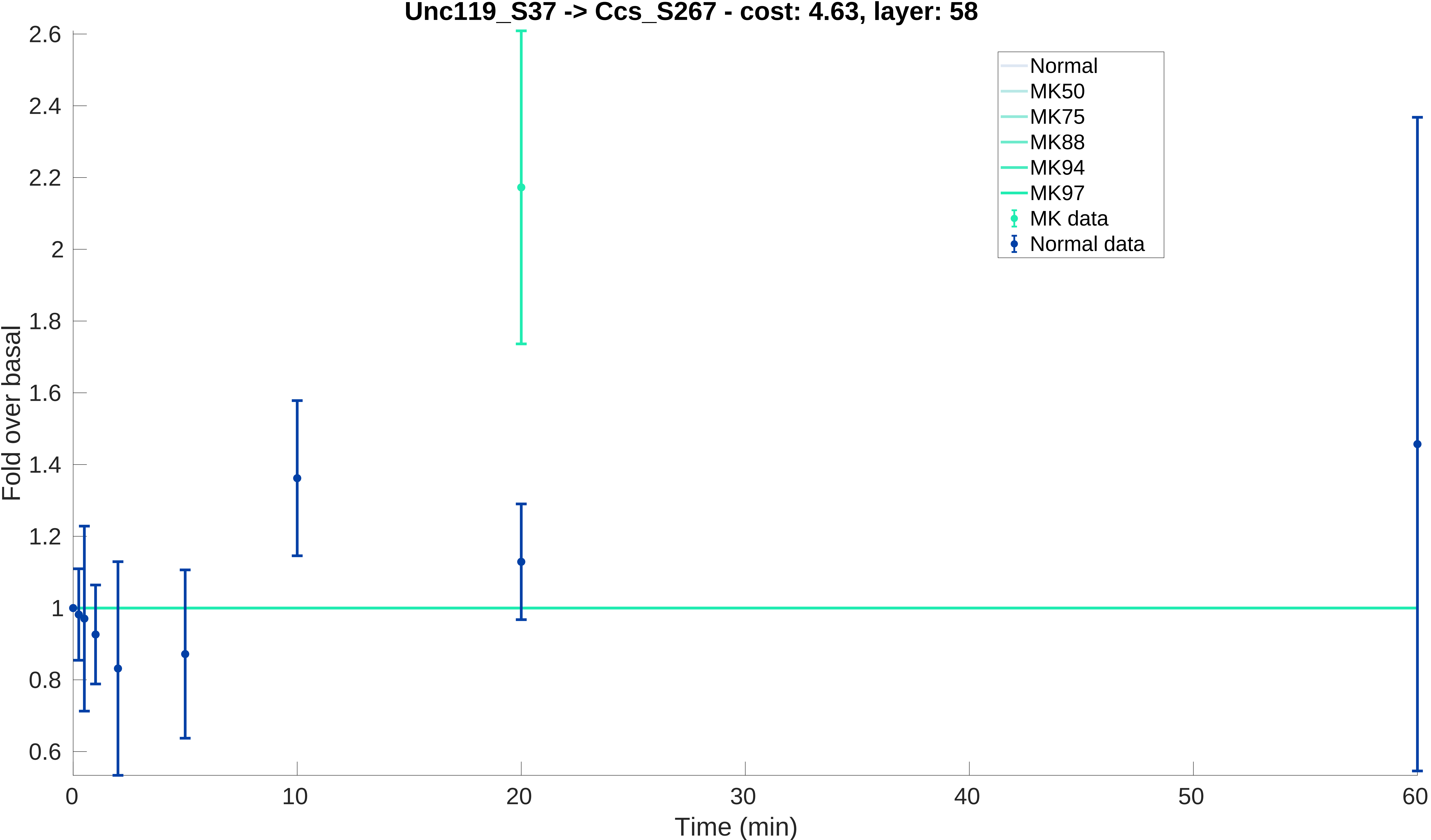
Unc119_S37 -> Ccs_S267 - cost: 4.63, layer: 58
Fold over basal
2.6
2.4
2.2
2
1.8
1.6
1.4
1.2
1
0.8
0.6
0
10
20
30
40
50
60
Time (min)
Normal
MK50
MK75
MK88
MK94
MK97
MK data
Normal data

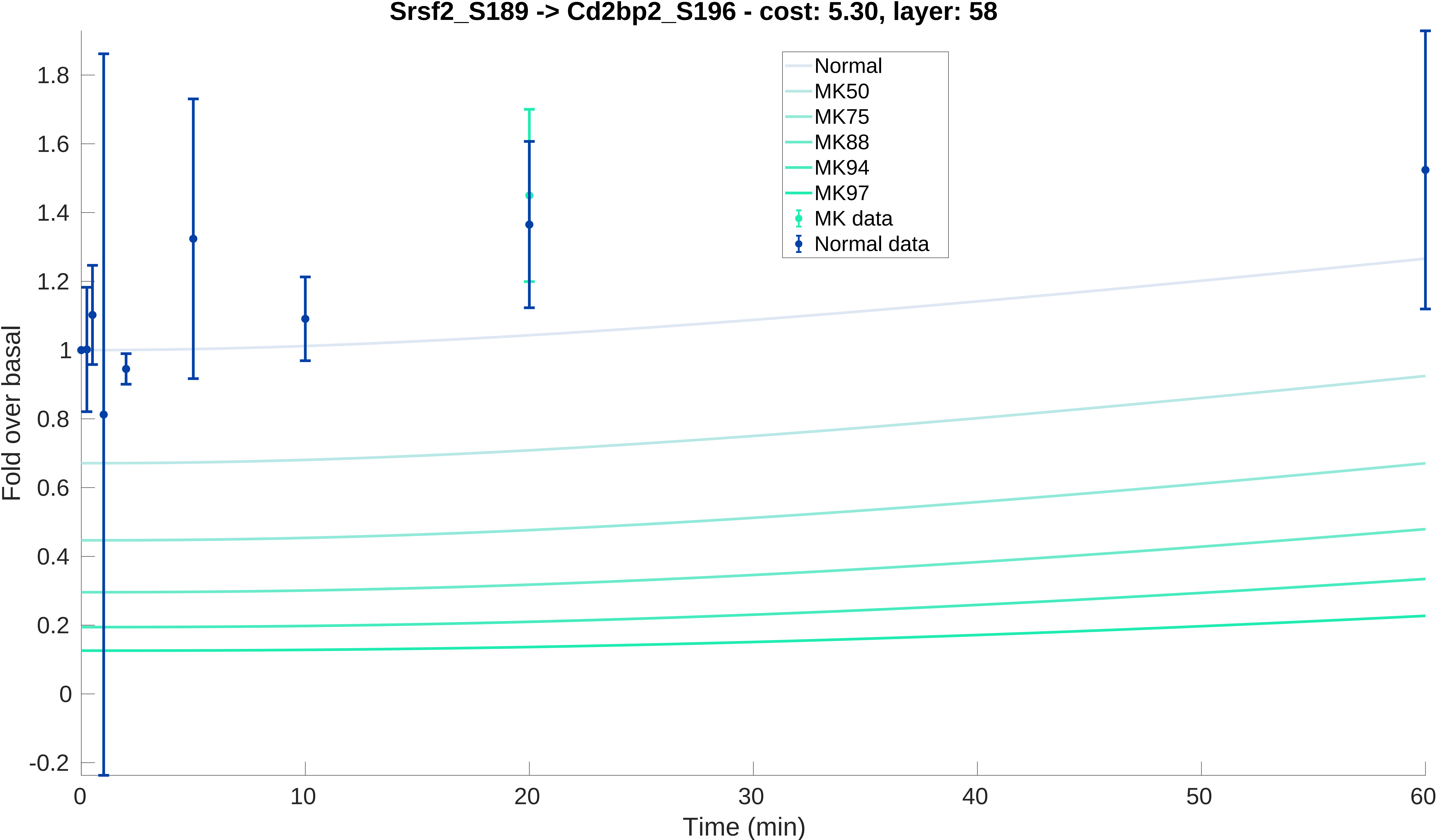

Srsf2_S189 -> Cd2bp2_S196 - cost: 5.30, layer: 58
Normal
MK50
MK75
MK88
MK94
MK97
MK data
Normal data
Fold over basal
Time (min)
-0.2
0
0.2
0.4
0.6
0.8
1
1.2
1.4
1.6
1.8
0
10
20
30
40
50
60

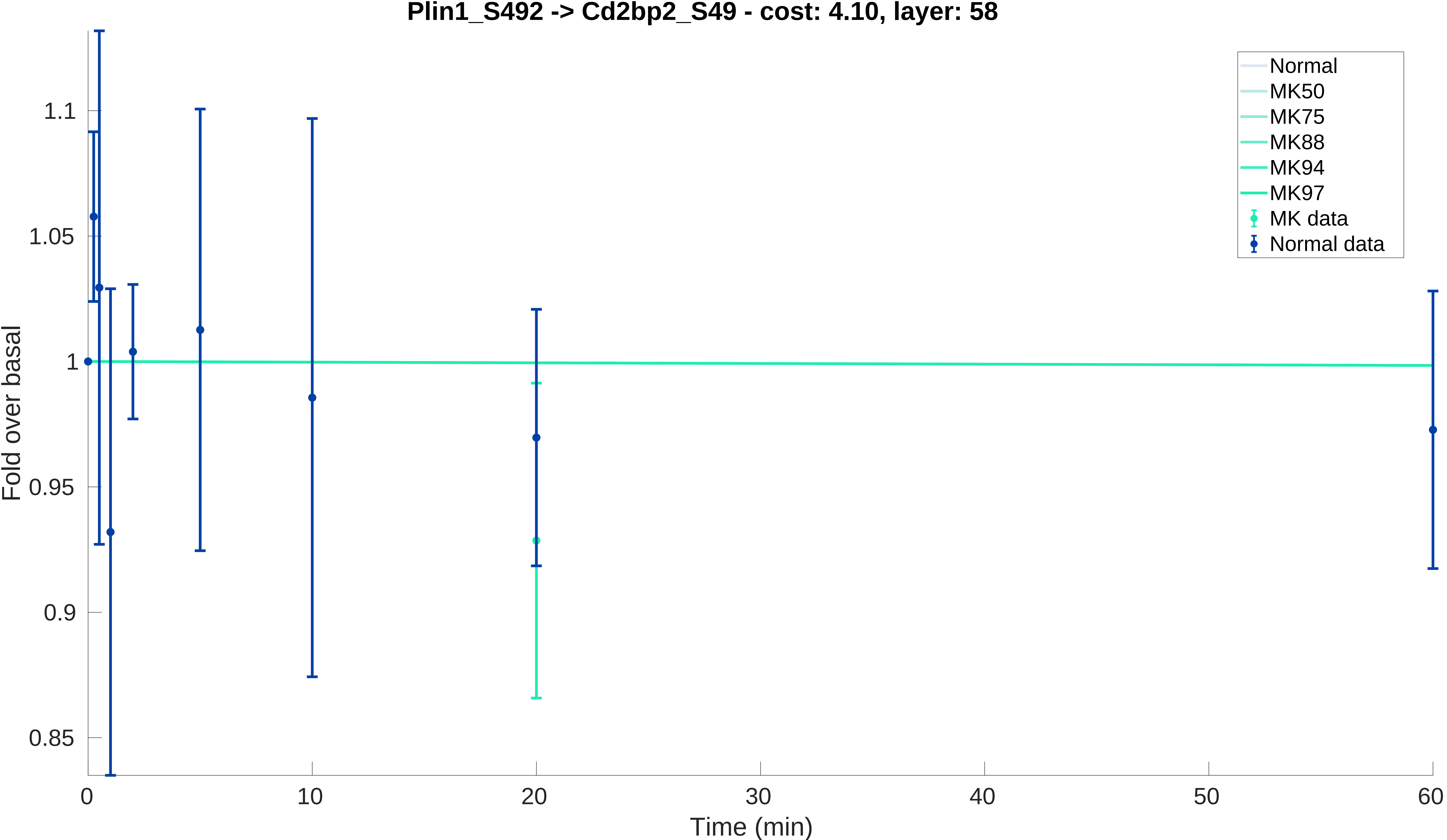

Plin1_S492 -> Cd2bp2_S49 - cost: 4.10, layer: 58
Fold over basal
1.1
1.05
1
0.95
0.9
0.85
0
10
20
30
40
50
60
Time (min)
Normal
MK50
MK75
MK88
MK94
MK97
MK data
Normal data

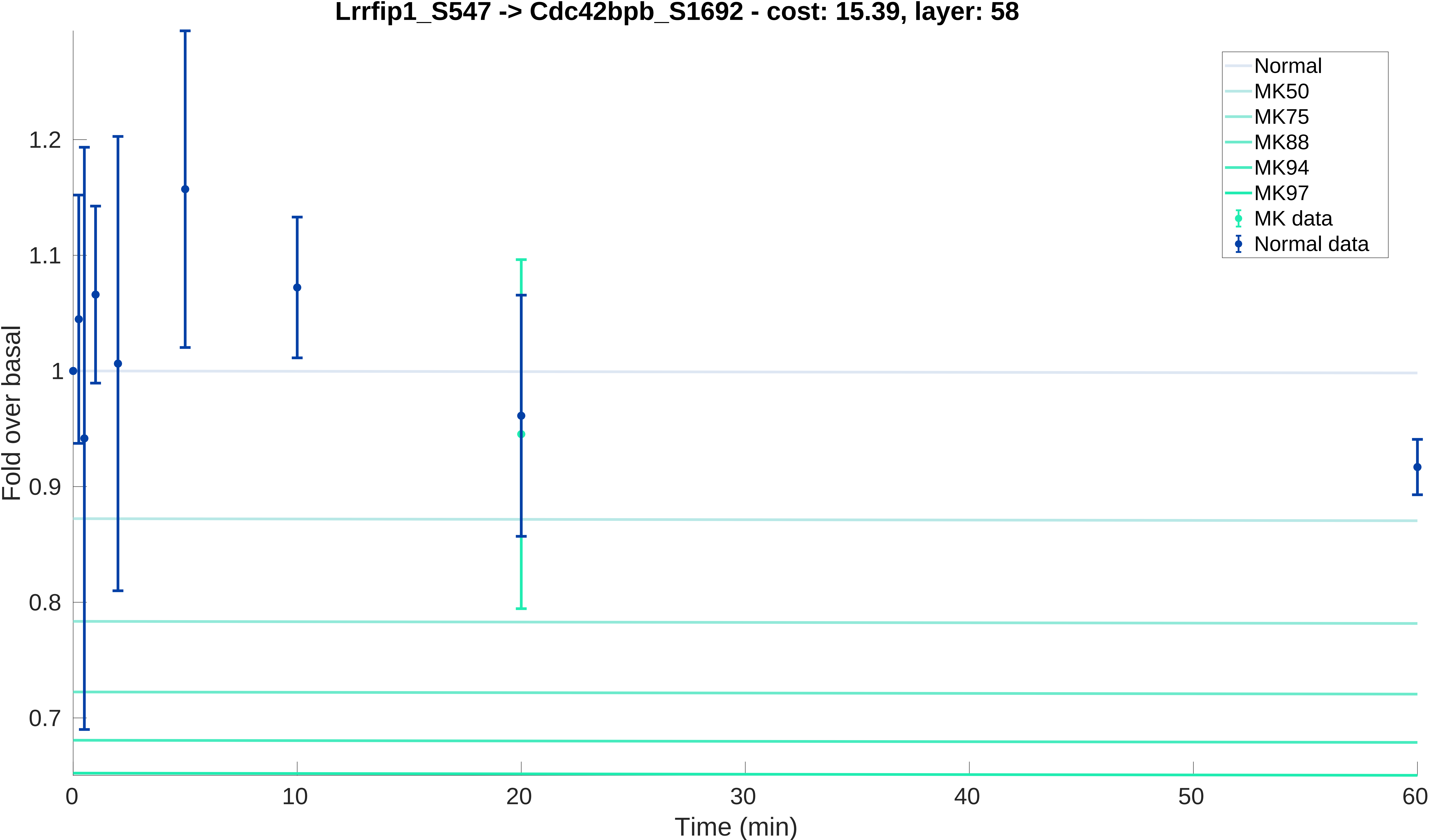
Lrrfip1_S547 -> Cdc42bpb_S1692 - cost: 15.39, layer: 58
Normal
MK50
MK75
MK88
MK94
MK97
MK data
Normal data
Fold over basal
1.2
1.1
1
0.9
0.8
0.7
Time (min)
0
10
20
30
40
50
60

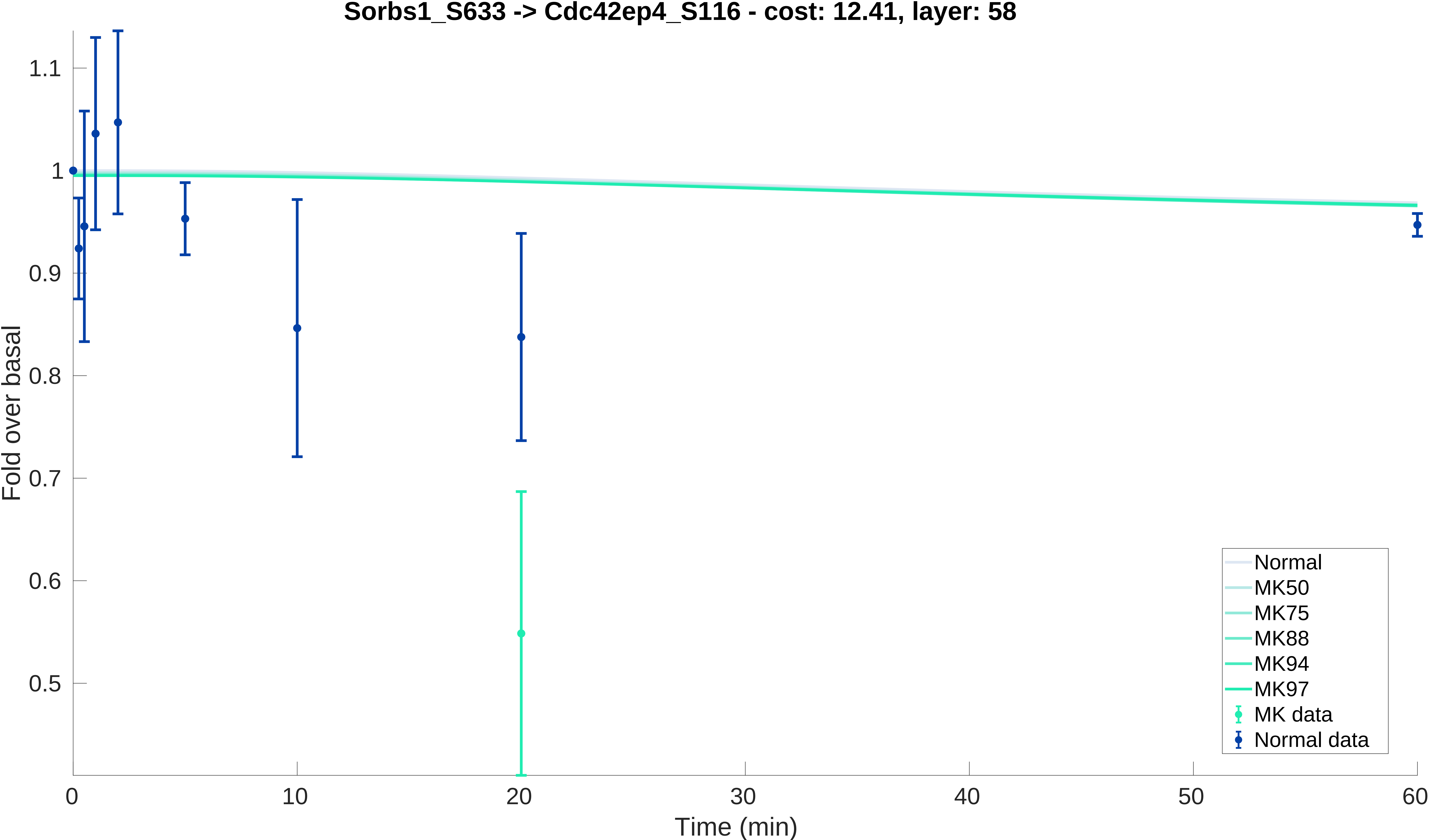

Sorbs1_S633 -> Cdc42ep4_S116 - cost: 12.41, layer: 58
Fold over basal
Time (min)
1.1
1
0.9
0.8
0.7
0.6
0.5
0
10
20
30
40
50
60
Normal
MK50
MK75
MK88
MK94
MK97
MK data
Normal data

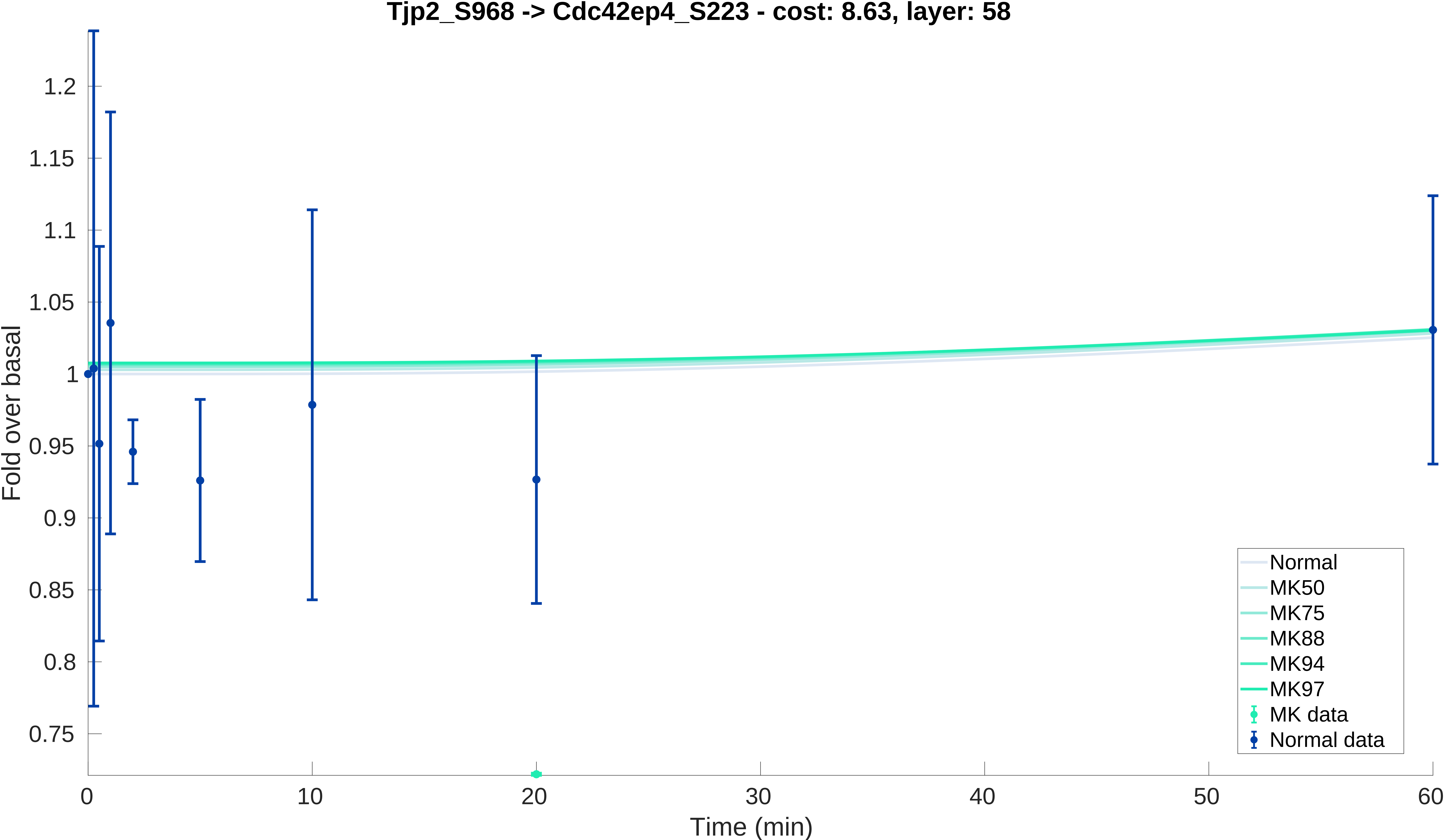
Tjp2_S968 -> Cdc42ep4_S223 - cost: 8.63, layer: 58
Fold over basal
1.2
1.15
1.1
1.05
1
0.95
0.9
0.85
0.8
0.75
0
10
20
30
40
50
60
Time (min)
Normal
MK50
MK75
MK88
MK94
MK97
MK data
Normal data

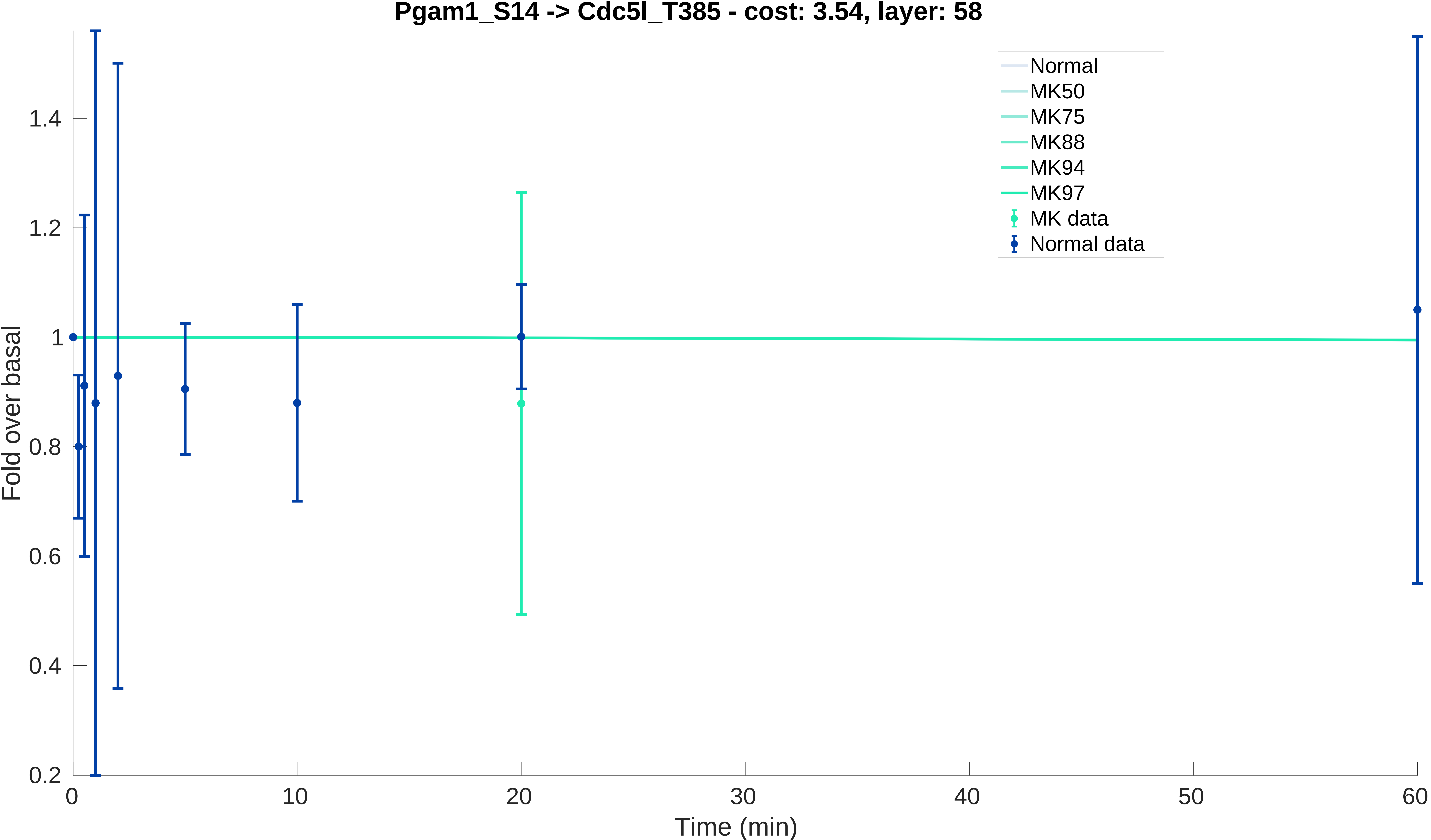

Pgam1_S14 -> Cdc5l_T385 - cost: 3.54, layer: 58
Fold over basal
Time (min)
0.2
0.4
0.6
0.8
1
1.2
1.4
0
10
20
30
40
50
60
Normal
MK50
MK75
MK88
MK94
MK97
MK data
Normal data

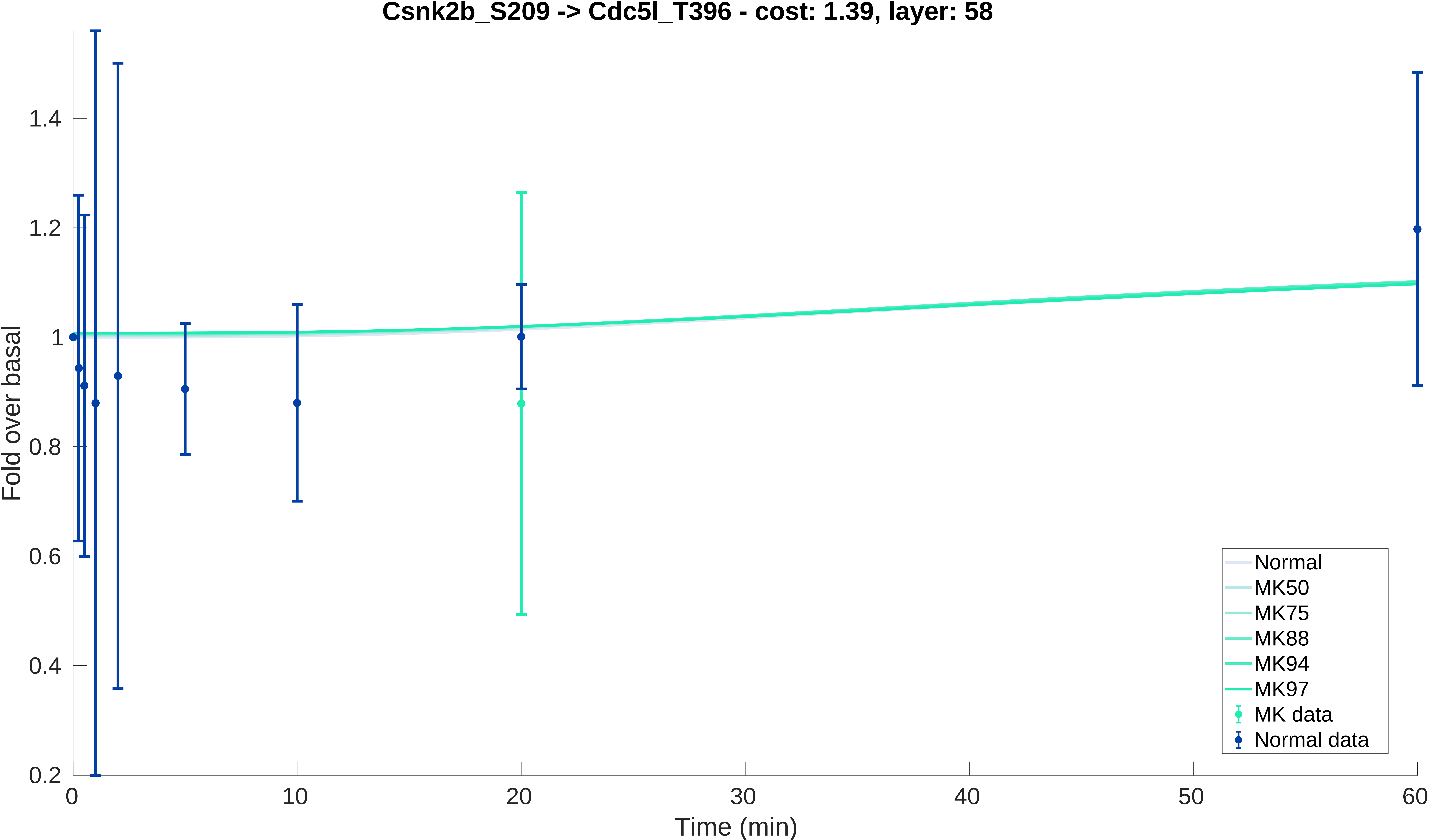

Csnk2b_S209 -> Cdc5l_T396 - cost: 1.39, layer: 58
Fold over basal
Time (min)
0.2
0.4
0.6
0.8
1
1.2
1.4
0
10
20
30
40
50
60
Normal
MK50
MK75
MK88
MK94
MK97
MK data
Normal data

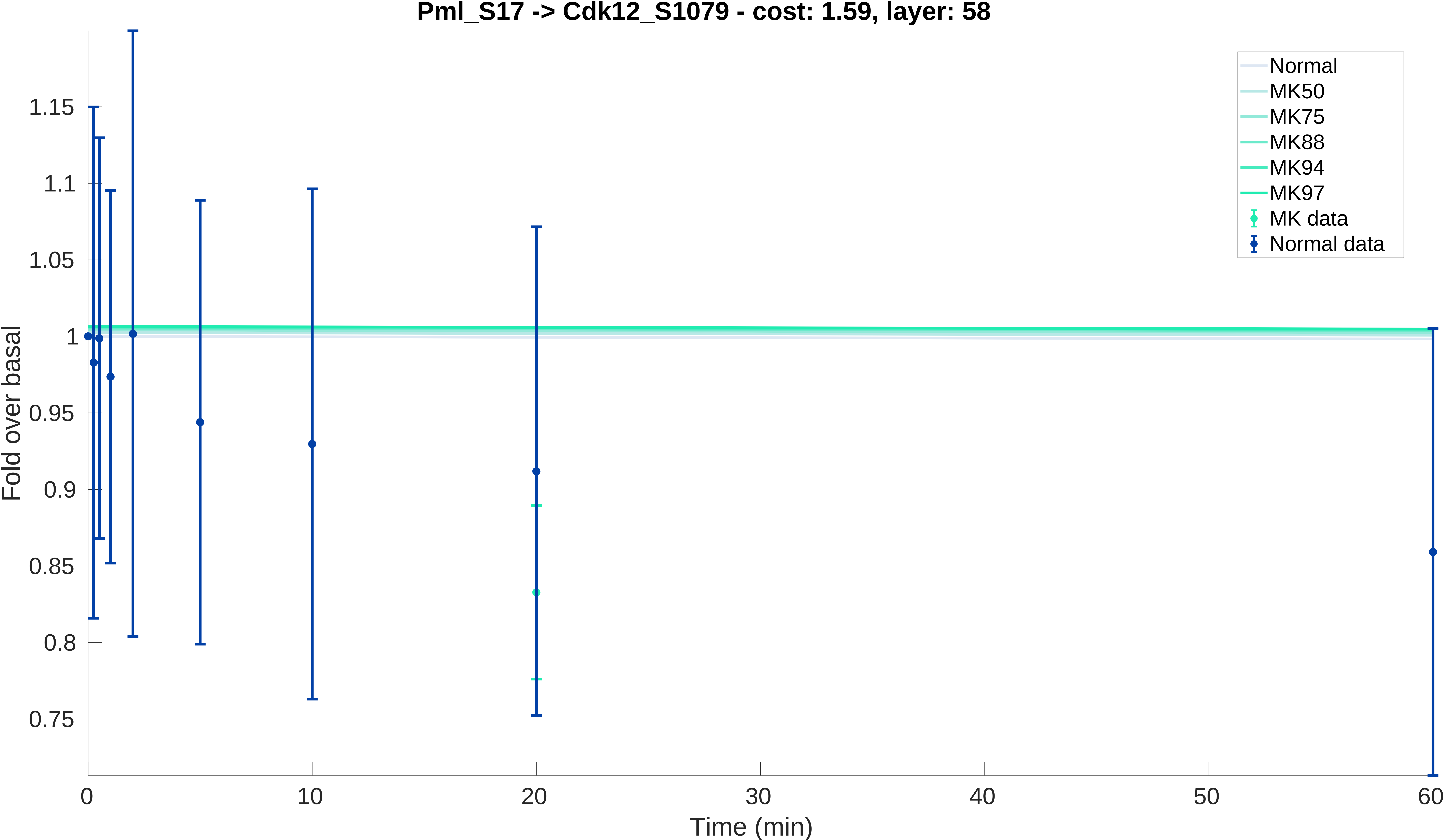

Pml_S17 -> Cdk12_S1079 - cost: 1.59, layer: 58
Fold over basal
1.15
1.1
1.05
1
0.95
0.9
0.85
0.8
0.75
0
10
20
30
40
50
60
Time (min)
Normal
MK50
MK75
MK88
MK94
MK97
MK data
Normal data

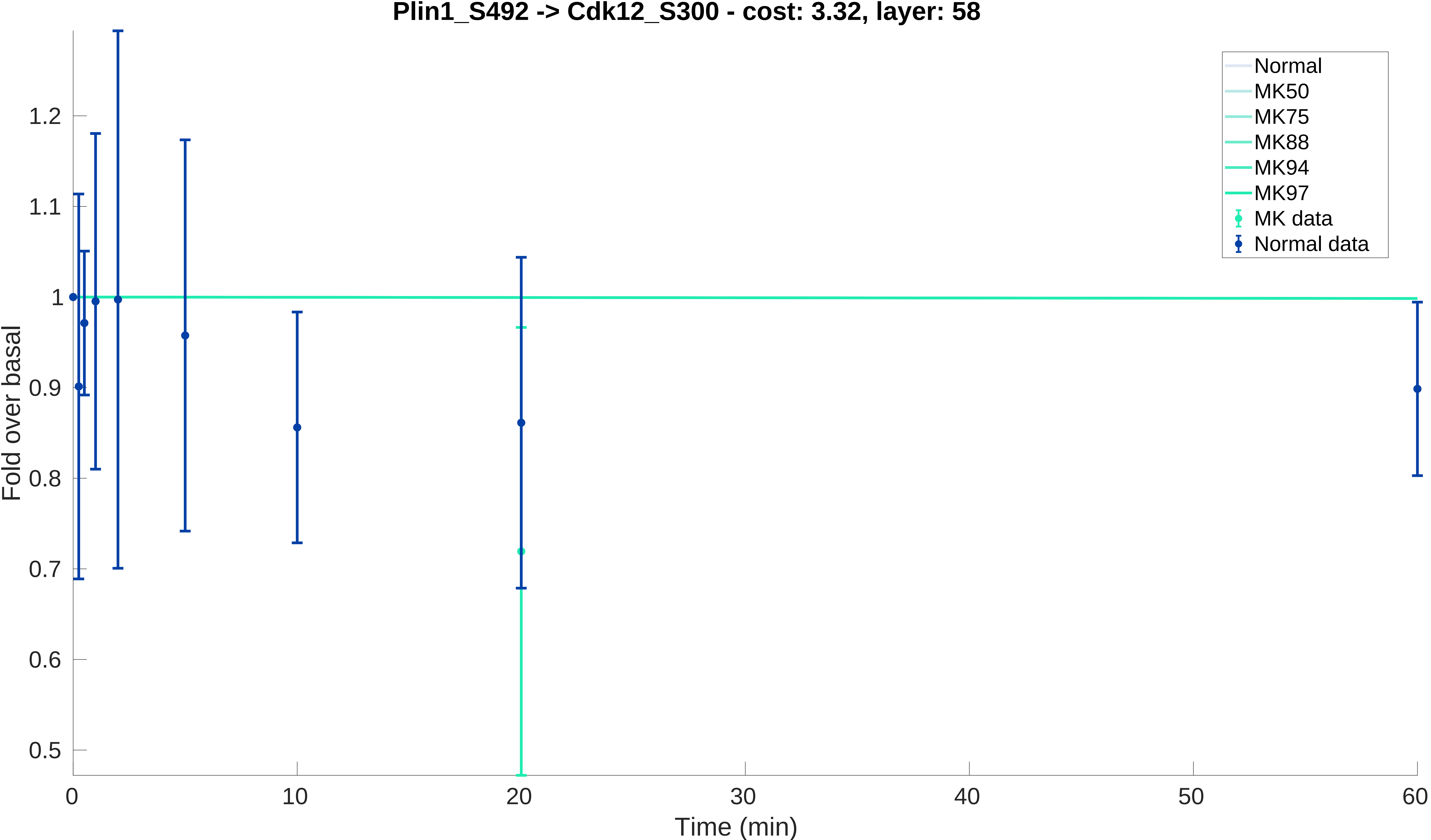

Plin1_S492 -> Cdk12_S300 - cost: 3.32, layer: 58
Normal
MK50
MK75
MK88
MK94
MK97
MK data
Normal data
Fold over basal
1.2
1.1
1
0.9
0.8
0.7
0.6
0.5
0
10
20
30
40
50
60
Time (min)

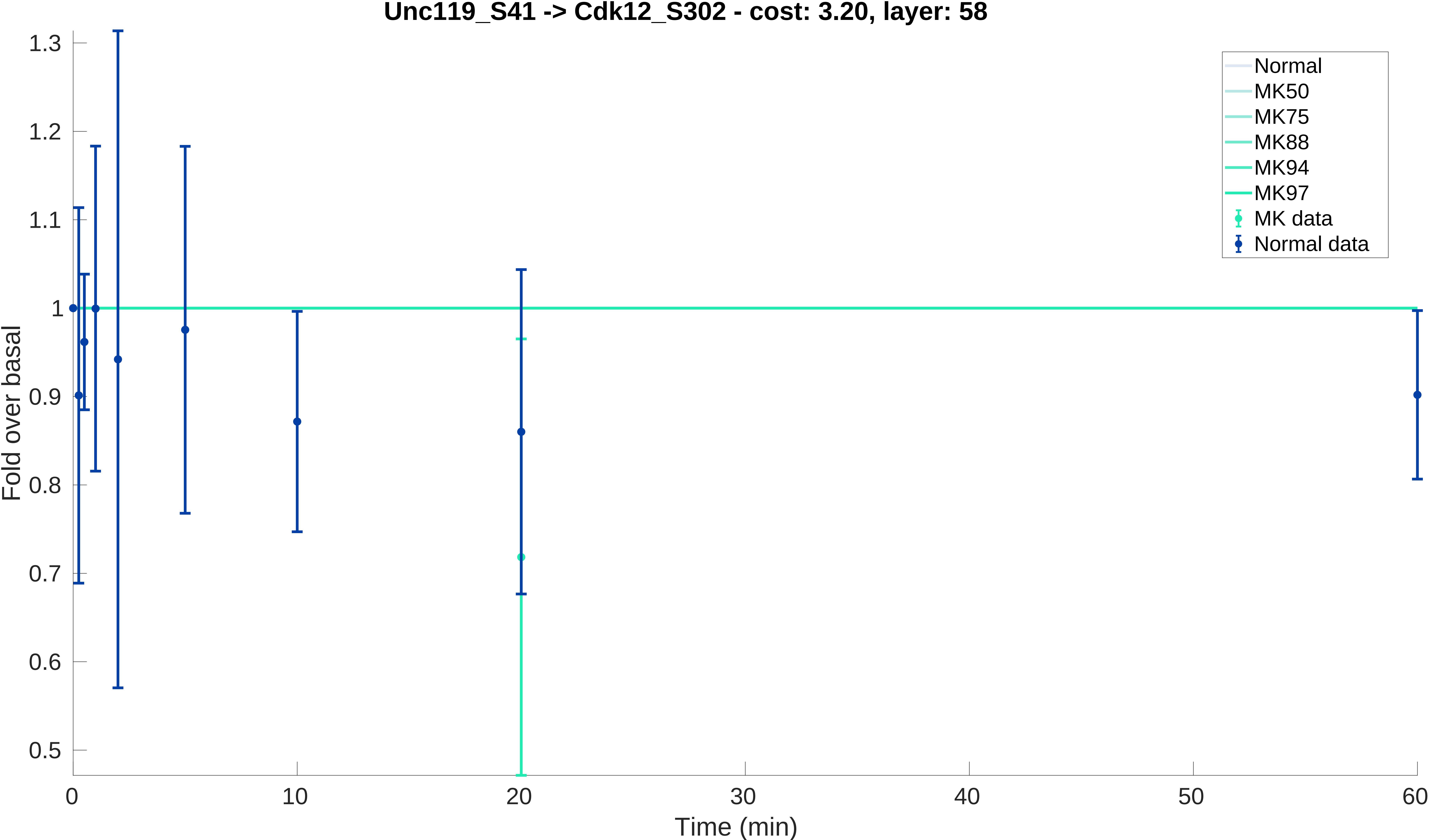

Unc119_S41 -> Cdk12_S302 - cost: 3.20, layer: 58
Fold over basal
Time (min)
1.3
1.2
1.1
1
0.9
0.8
0.7
0.6
0.5
0
10
20
30
40
50
60
Normal
MK50
MK75
MK88
MK94
MK97
MK data
Normal data

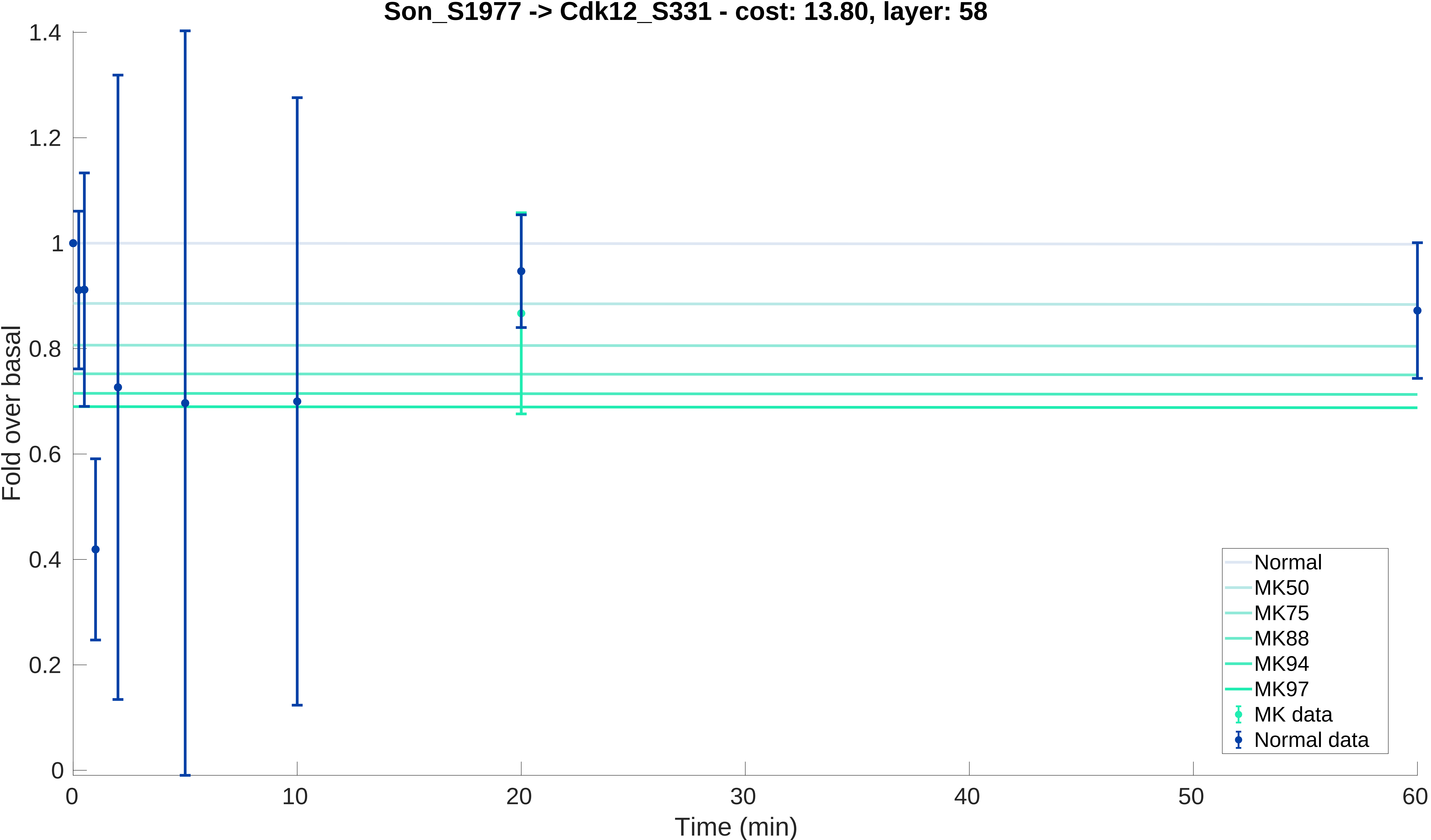

Son_S1977 -> Cdk12_S331 - cost: 13.80, layer: 58
Fold over basal
Time (min)
0
0.2
0.4
0.6
0.8
1
1.2
1.4
0
10
20
30
40
50
60
Normal
MK50
MK75
MK88
MK94
MK97
MK data
Normal data

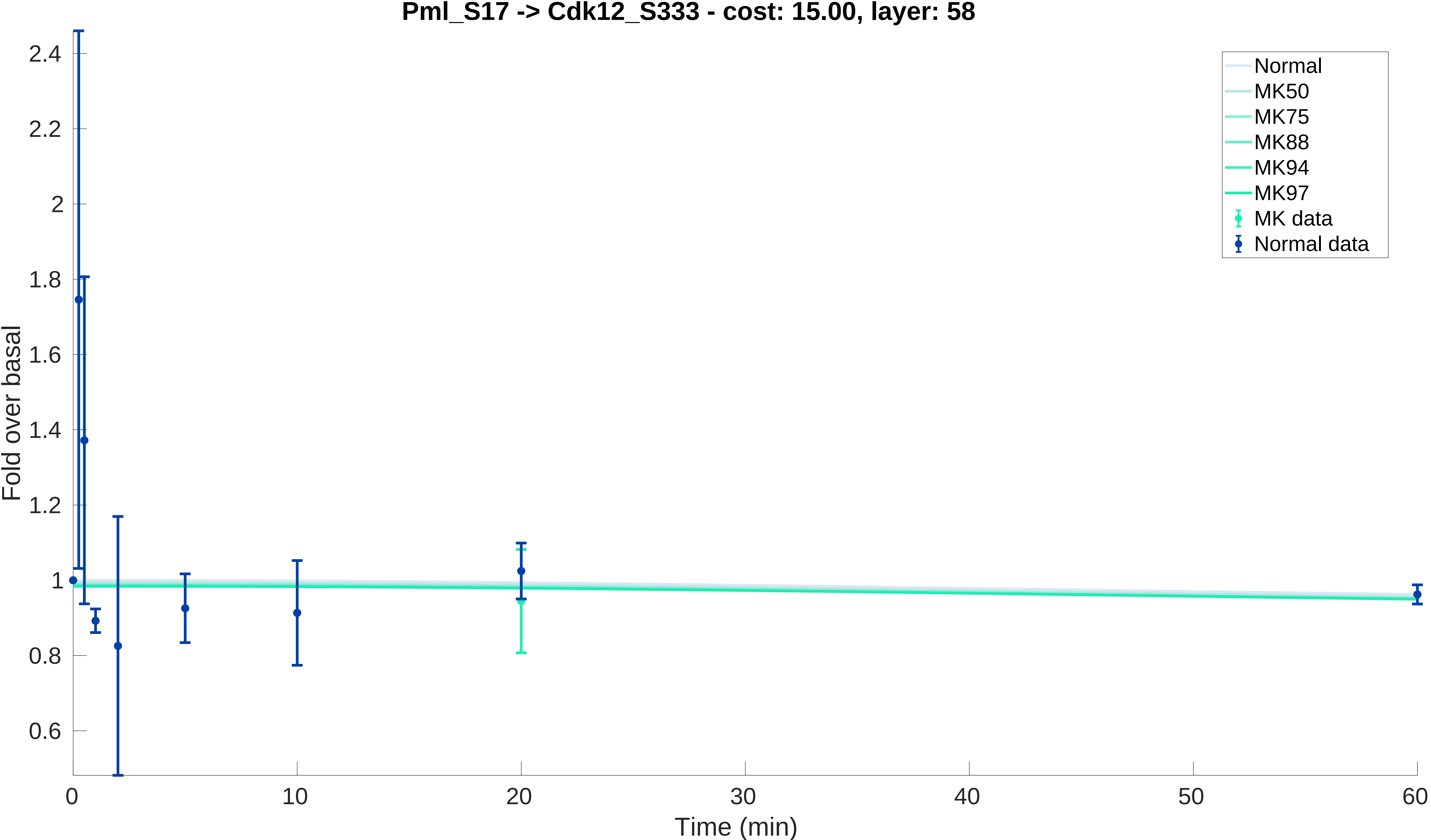
Pml_S17 -> Cdk12_S333 - cost: 15.00, layer: 58
Fold over basal
Time (min)
2.4
2.2
2
1.8
1.6
1.4
1.2
1
0.8
0.6
0
10
20
30
40
50
60
Normal
MK50
MK75
MK88
MK94
MK97
MK data
Normal data

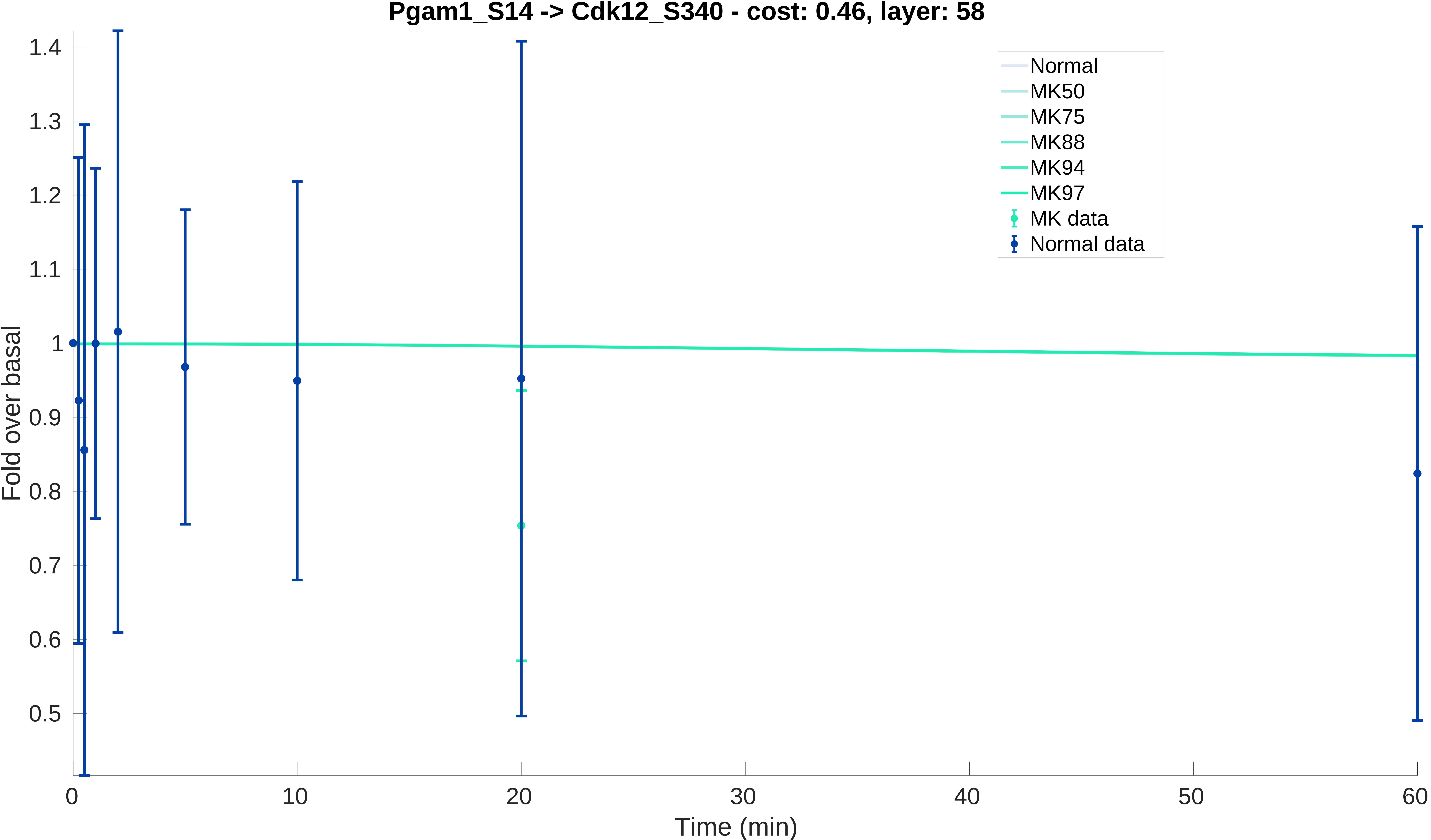
Pgam1_S14 -> Cdk12_S340 - cost: 0.46, layer: 58
Fold over basal
Time (min)
1.4
1.3
1.2
1.1
1
0.9
0.8
0.7
0.6
0.5
0
10
20
30
40
50
60
Normal
MK50
MK75
MK88
MK94
MK97
MK data
Normal data

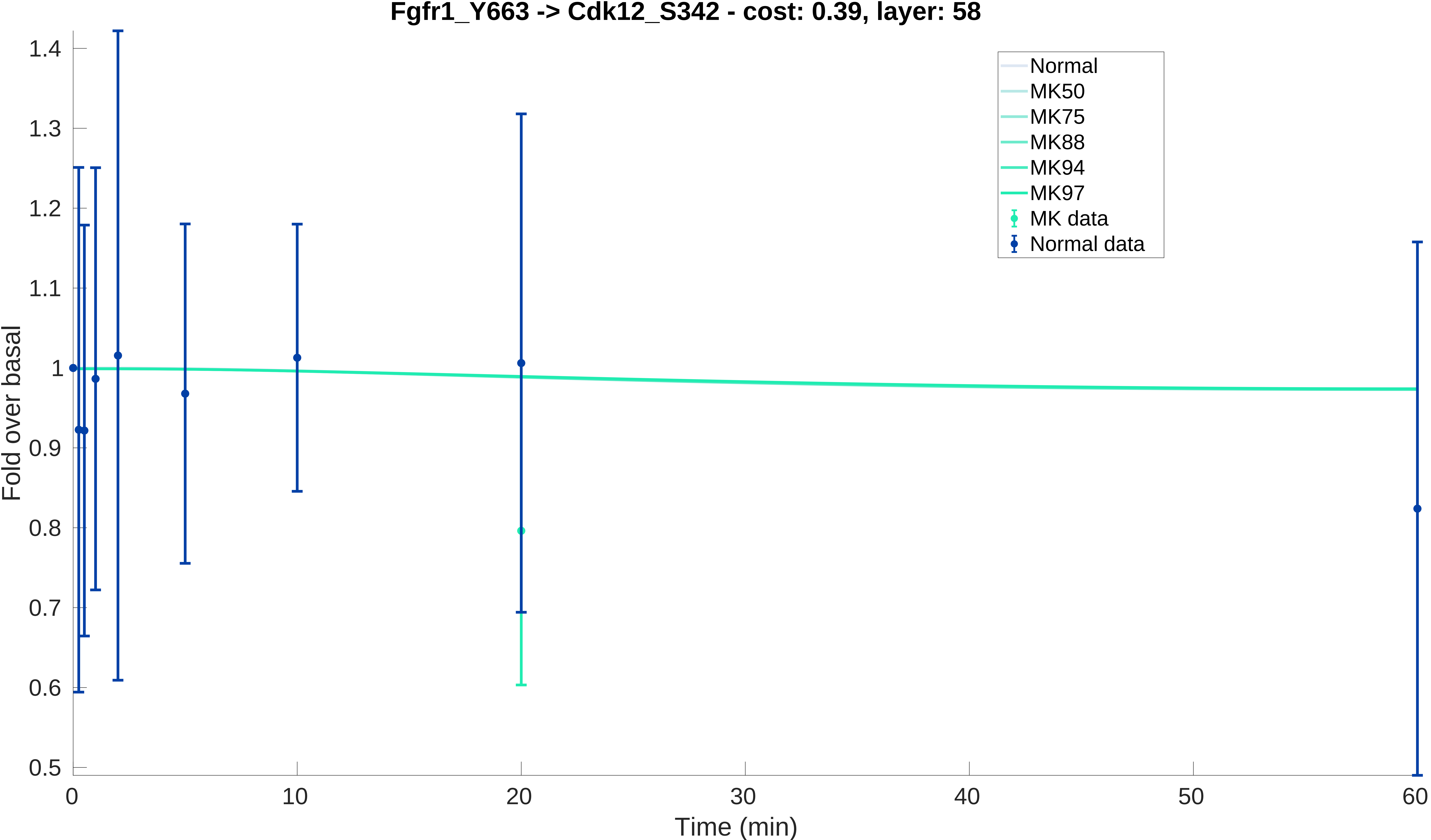

Fgfr1_Y663 -> Cdk12_S342 - cost: 0.39, layer: 58
Fold over basal
Time (min)
1.4
1.3
1.2
1.1
1
0.9
0.8
0.7
0.6
0.5
0
10
20
30
40
50
60
Normal
MK50
MK75
MK88
MK94
MK97
MK data
Normal data

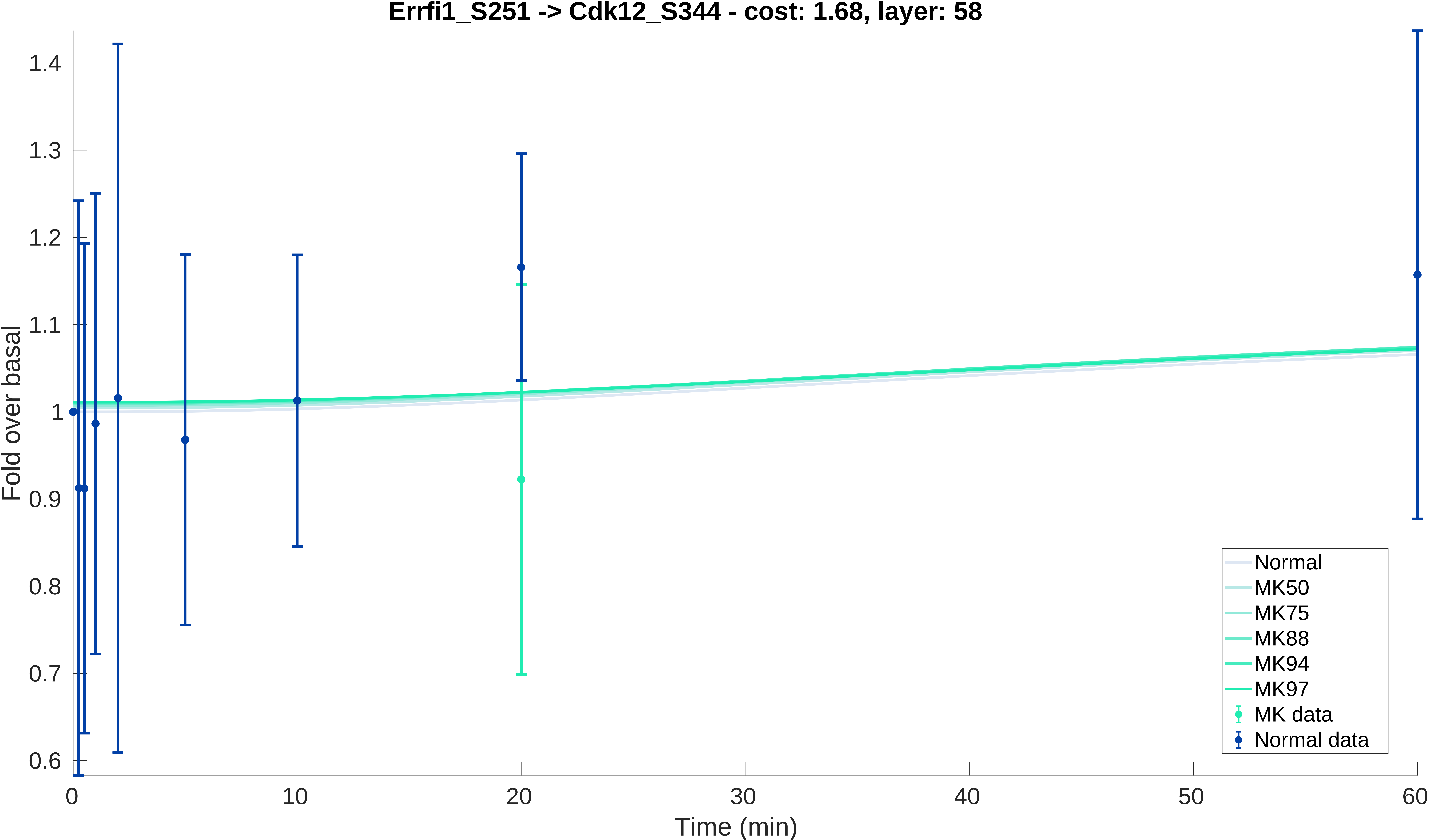

Errfi1_S251 -> Cdk12_S344 - cost: 1.68, layer: 58
Fold over basal
1.4
1.3
1.2
1.1
1
0.9
0.8
0.7
0.6
0
10
20
30
40
50
60
Time (min)
Normal
MK50
MK75
MK88
MK94
MK97
MK data
Normal data

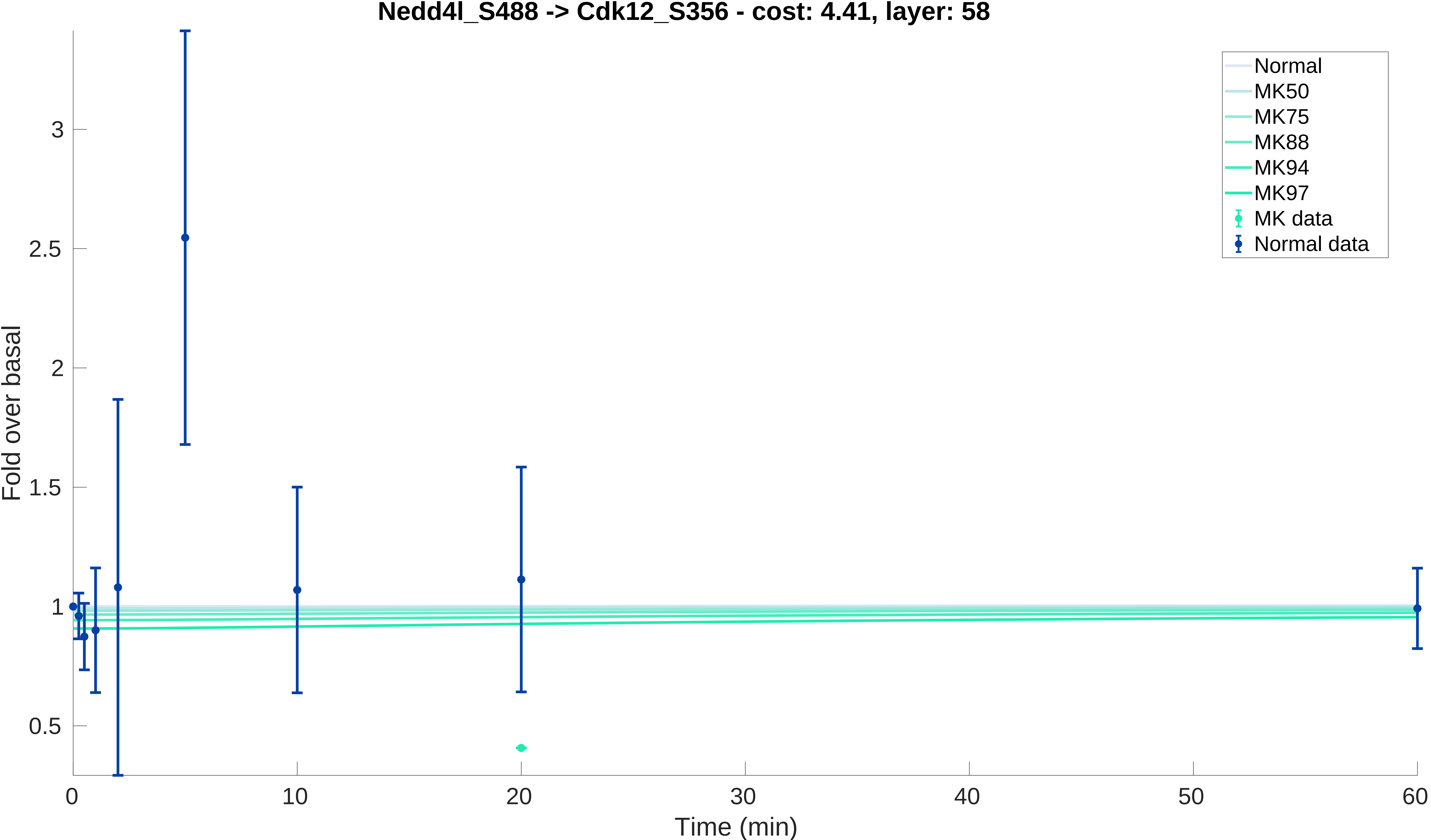

Nedd4l_S488 -> Cdk12_S356 - cost: 4.41, layer: 58
Fold over basal
Time (min)
0.5
1
1.5
2
2.5
3
0
10
20
30
40
50
60
Normal
MK50
MK75
MK88
MK94
MK97
MK data
Normal data

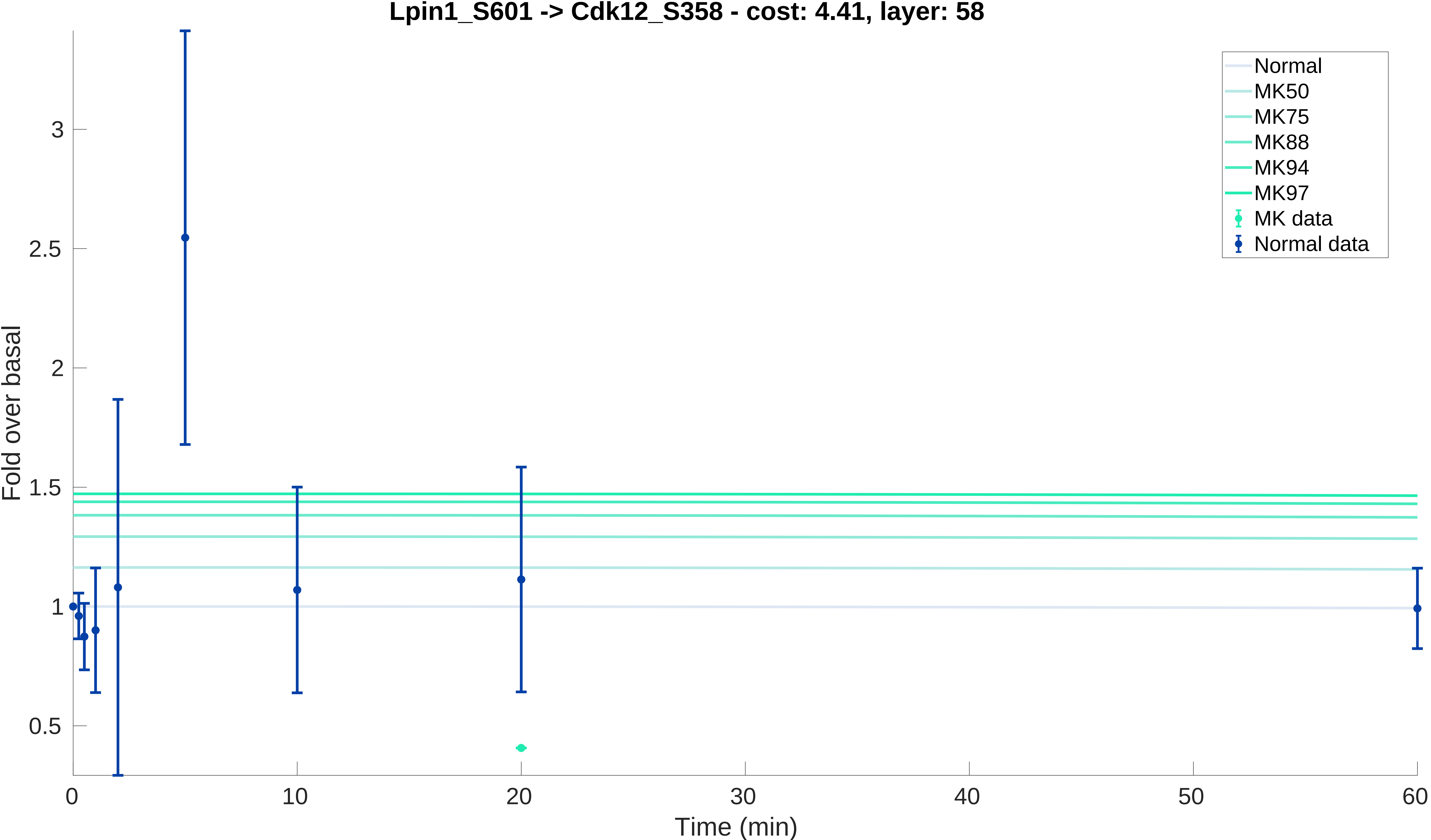

Lpin1_S601 -> Cdk12_S358 - cost: 4.41, layer: 58
Fold over basal
Time (min)
0
10
20
30
40
50
60
0.5
1
1.5
2
2.5
3
Normal
MK50
MK75
MK88
MK94
MK97
MK data
Normal data

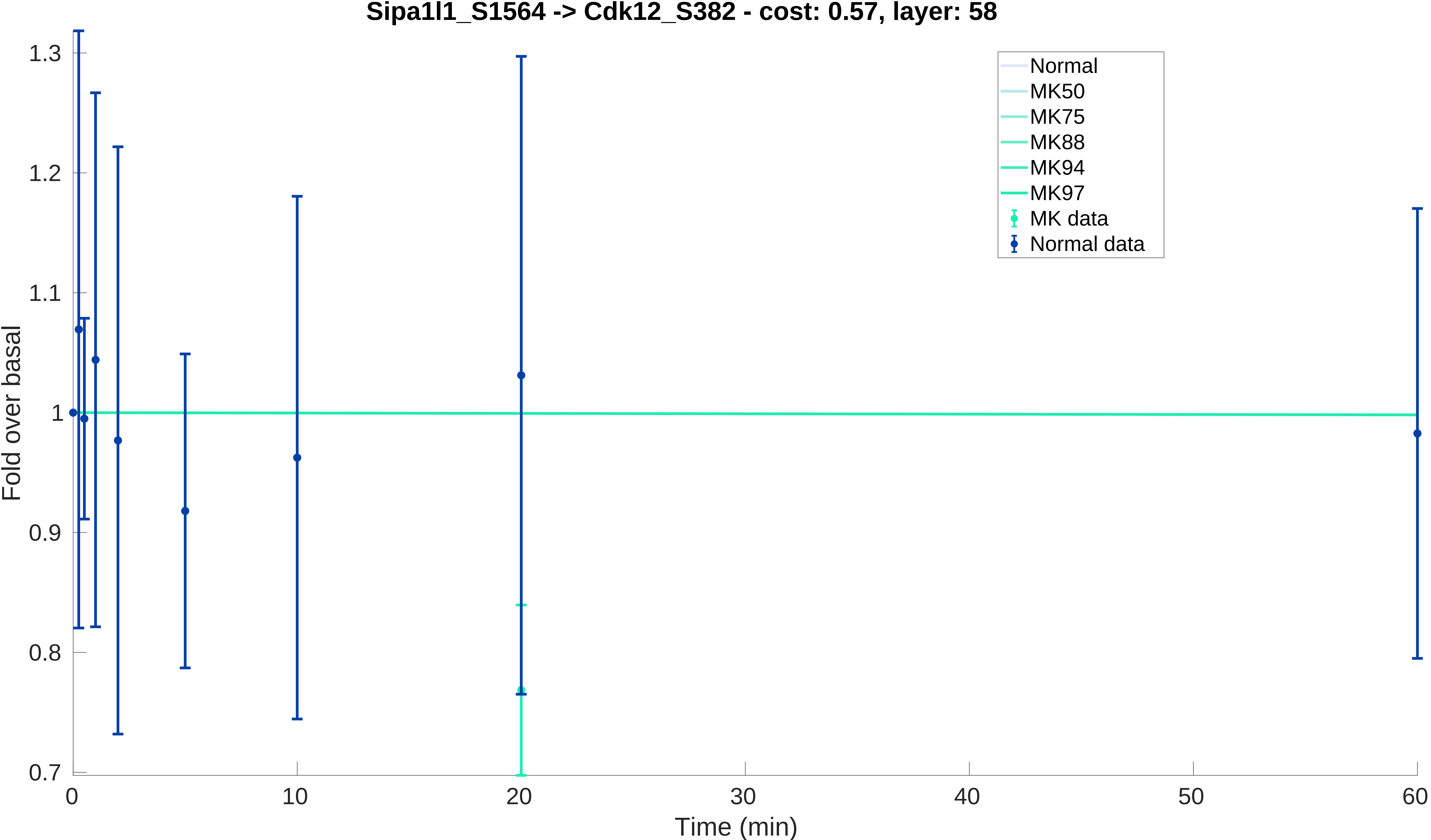
Sipa1l1_S1564 -> Cdk12_S382 - cost: 0.57, layer: 58
Fold over basal
Time (min)
0
10
20
30
40
50
60
0.7
0.8
0.9
1
1.1
1.2
1.3
Normal
MK50
MK75
MK88
MK94
MK97
MK data
Normal data

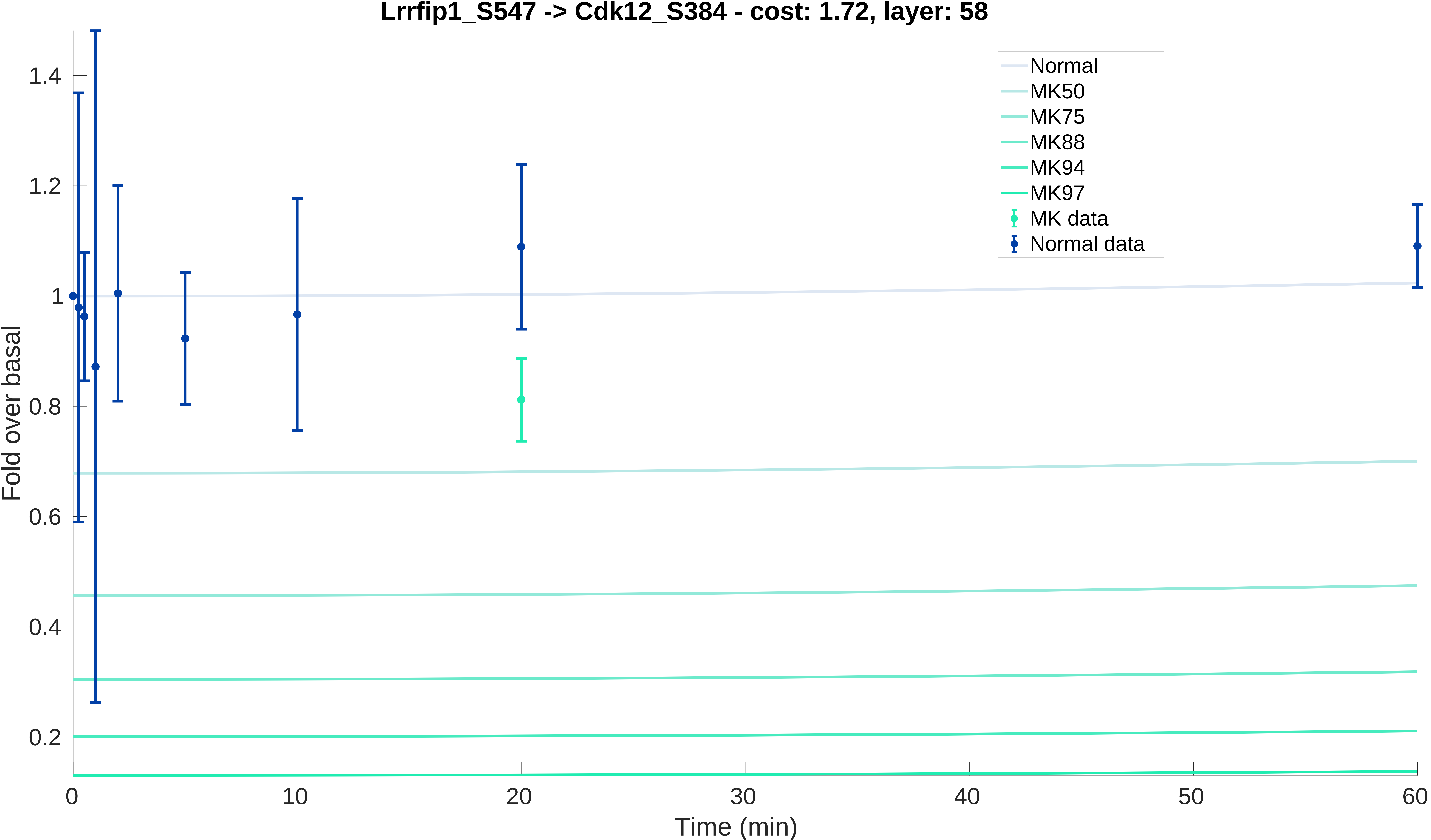

Lrrfip1_S547 -> Cdk12_S384 - cost: 1.72, layer: 58
Normal
MK50
MK75
MK88
MK94
MK97
MK data
Normal data
Fold over basal
1.4
1.2
1
0.8
0.6
0.4
0.2
0
10
20
30
40
50
60
Time (min)

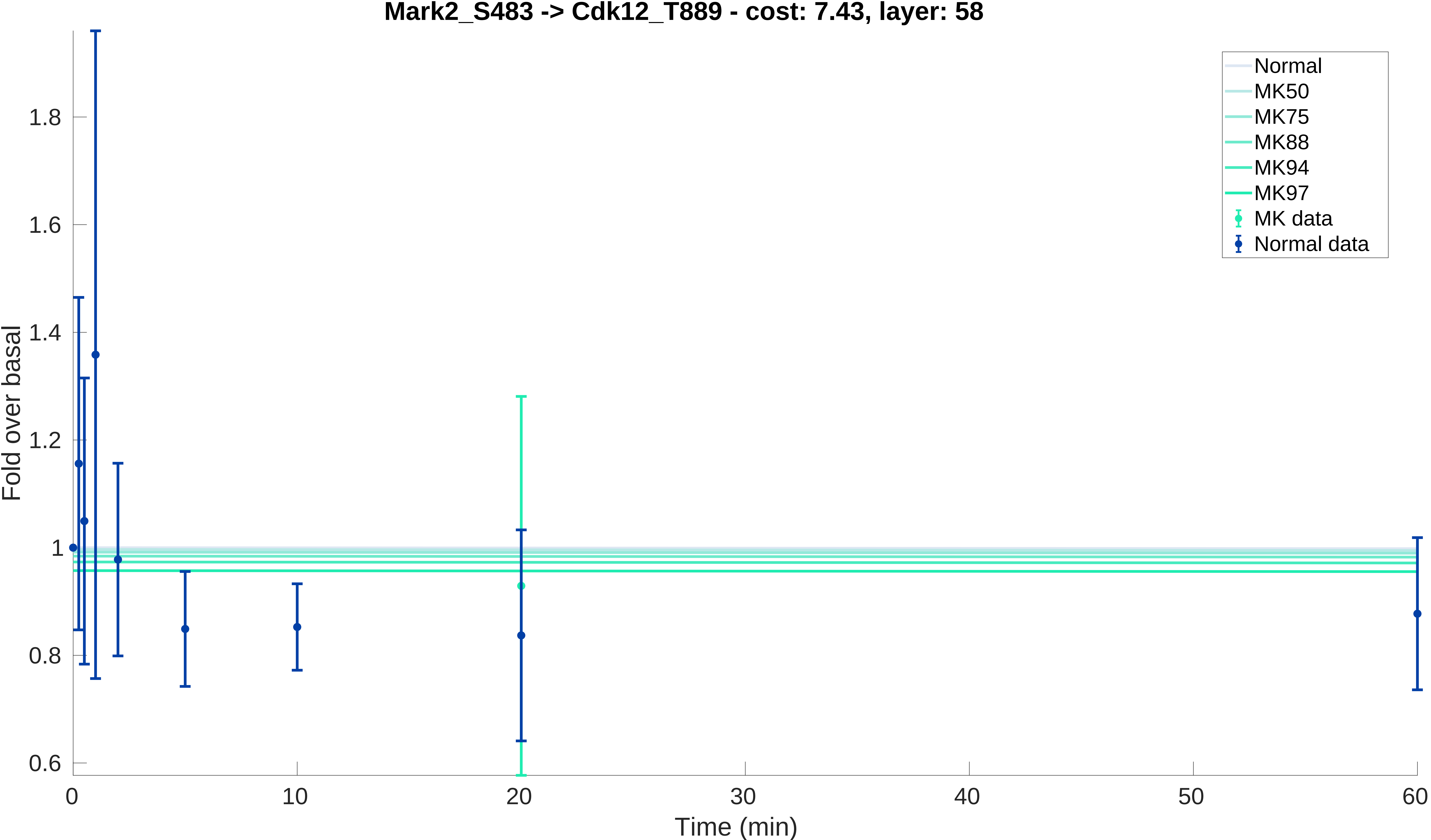

Mark2_S483 -> Cdk12_T889 - cost: 7.43, layer: 58
Fold over basal
1.8
1.6
1.4
1.2
1
0.8
0.6
0
10
20
30
40
50
60
Time (min)
Normal
MK50
MK75
MK88
MK94
MK97
MK data
Normal data

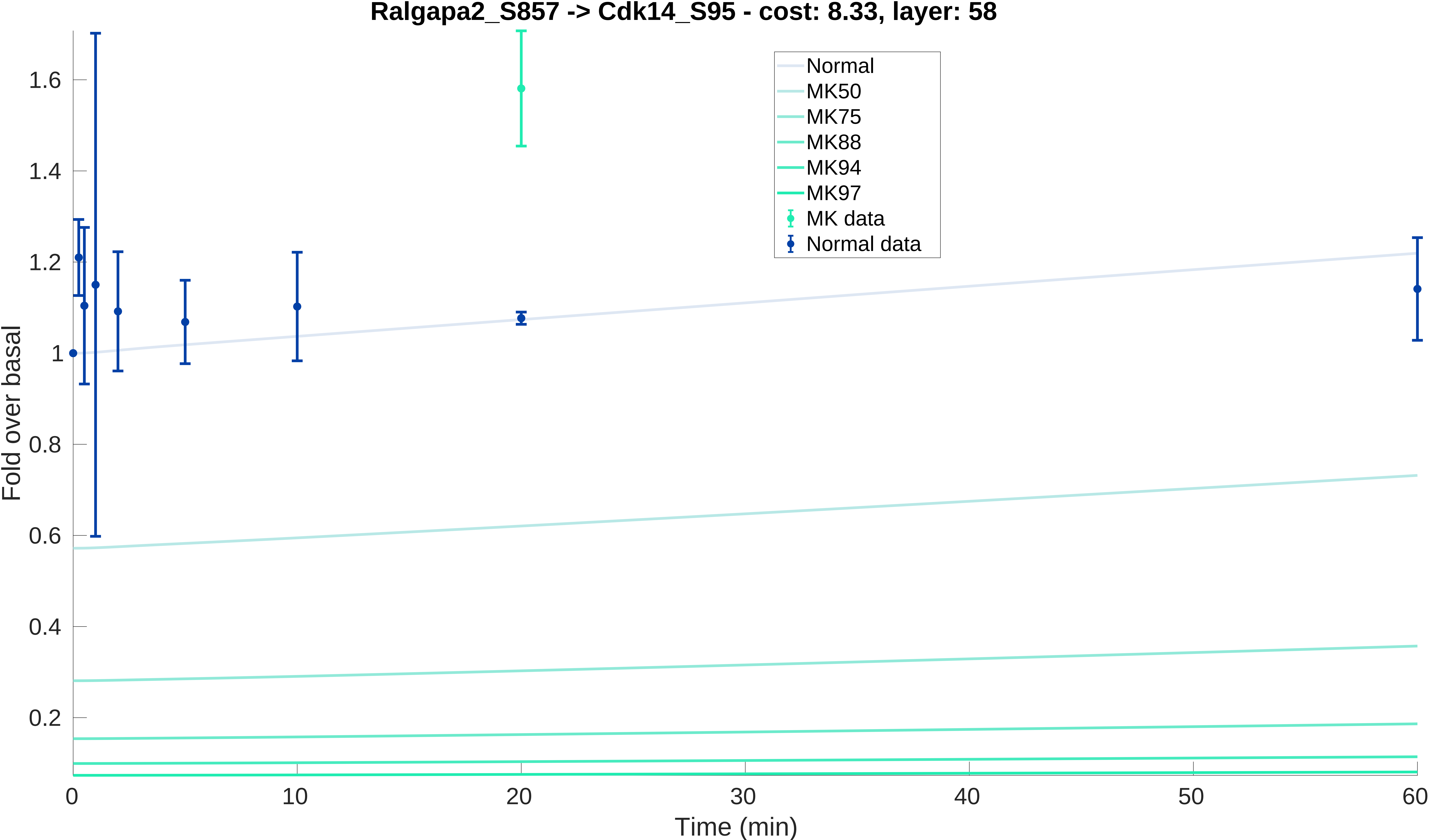
Ralgapa2_S857 -> Cdk14_S95 - cost: 8.33, layer: 58
Fold over basal
Time (min)
1.6
1.4
1.2
1
0.8
0.6
0.4
0.2
0
10
20
30
40
50
60
Normal
MK50
MK75
MK88
MK94
MK97
MK data
Normal data

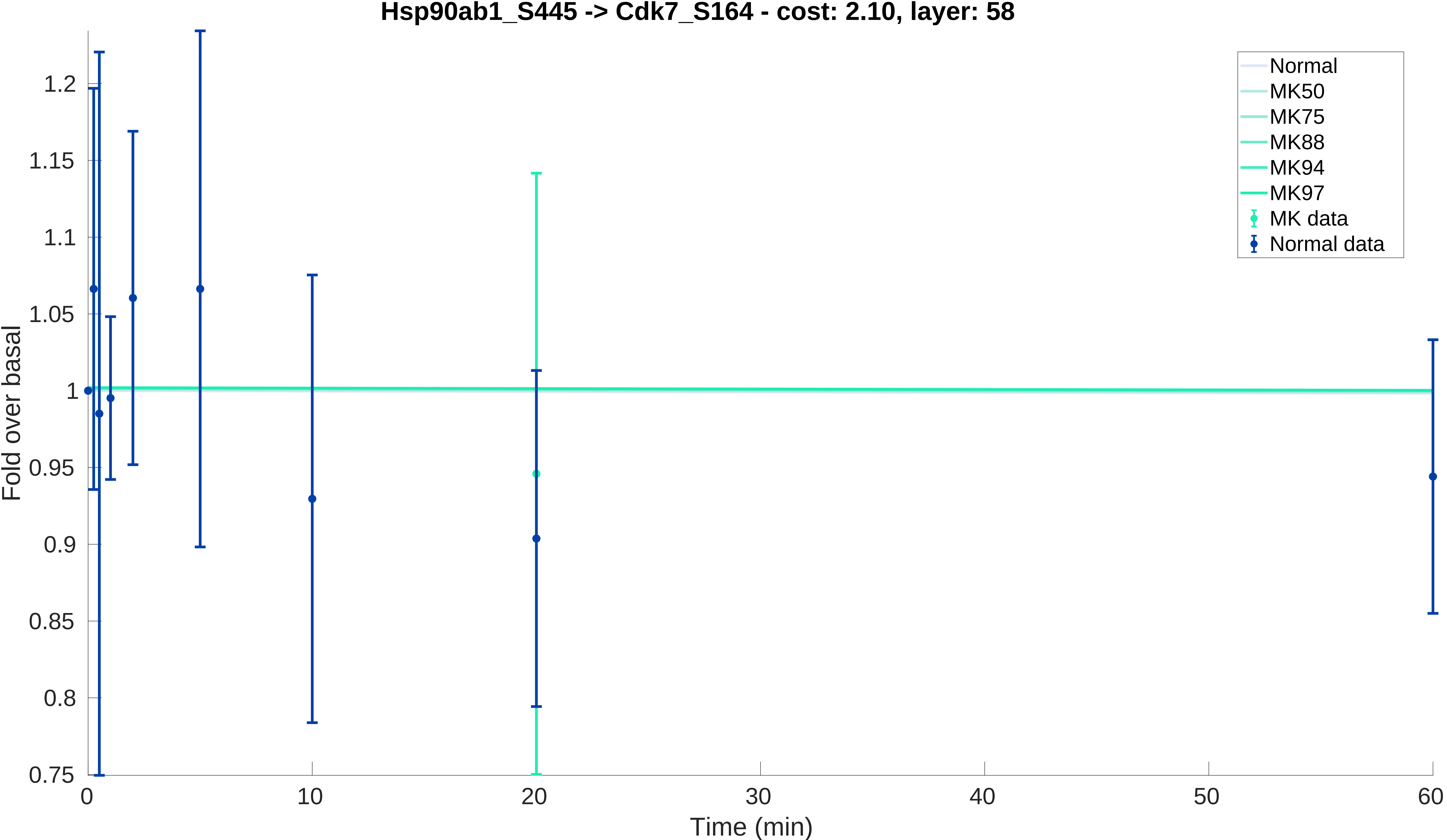

Hsp90ab1_S445 -> Cdk7_S164 - cost: 2.10, layer: 58
Fold over basal
Time (min)
0.75
0.8
0.85
0.9
0.95
1
1.05
1.1
1.15
1.2
0
10
20
30
40
50
60
Normal
MK50
MK75
MK88
MK94
MK97
MK data
Normal data

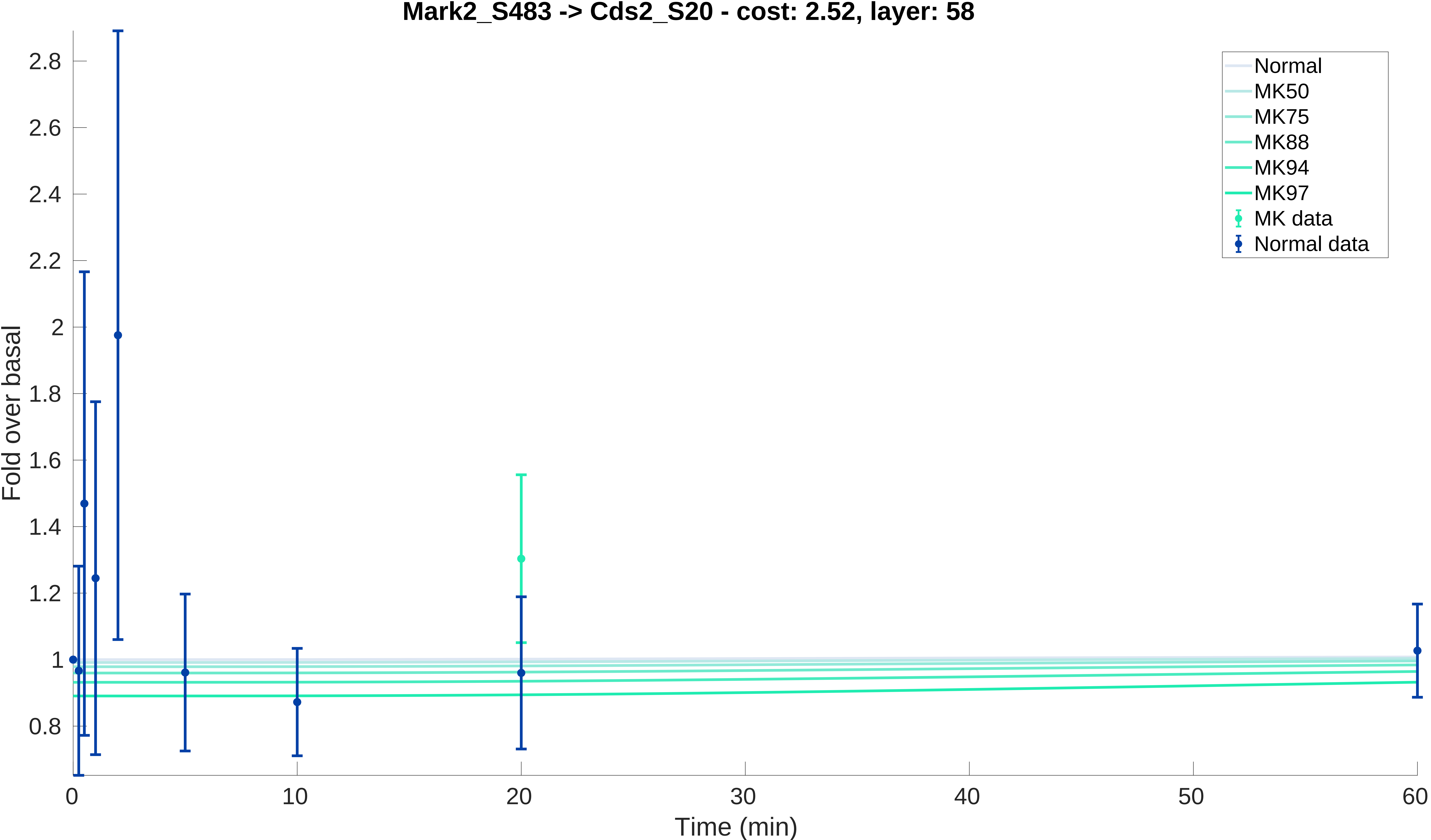
Mark2_S483 -> Cds2_S20 - cost: 2.52, layer: 58
Fold over basal
Time (min)
0.8
1
1.2
1.4
1.6
1.8
2
2.2
2.4
2.6
2.8
0
10
20
30
40
50
60
Normal
MK50
MK75
MK88
MK94
MK97
MK data
Normal data

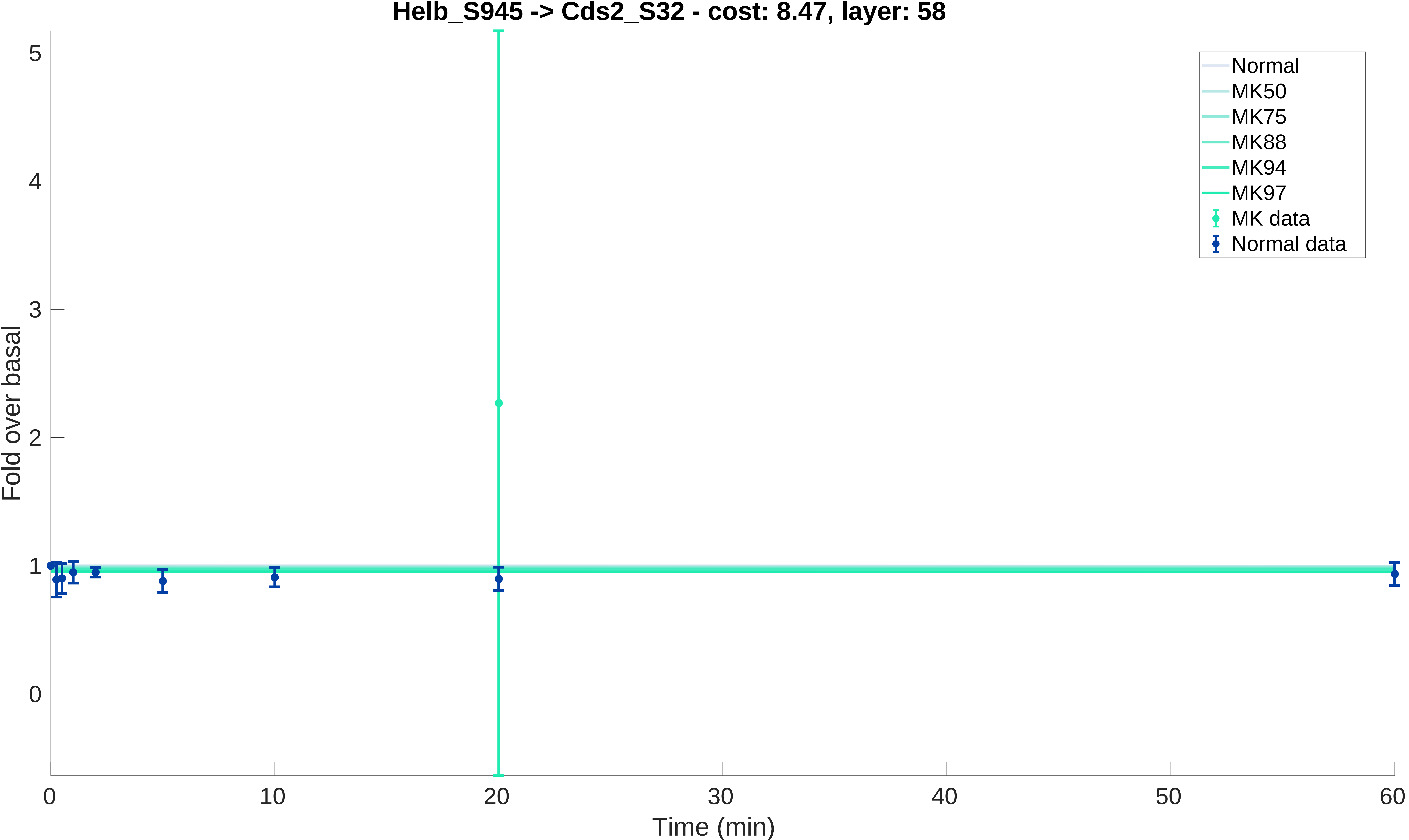

Helb_S945 -> Cds2_S32 - cost: 8.47, layer: 58
Fold over basal
Time (min)
0
10
20
30
40
50
60
5
4
3
2
1
0
Normal
MK50
MK75
MK88
MK94
MK97
MK data
Normal data

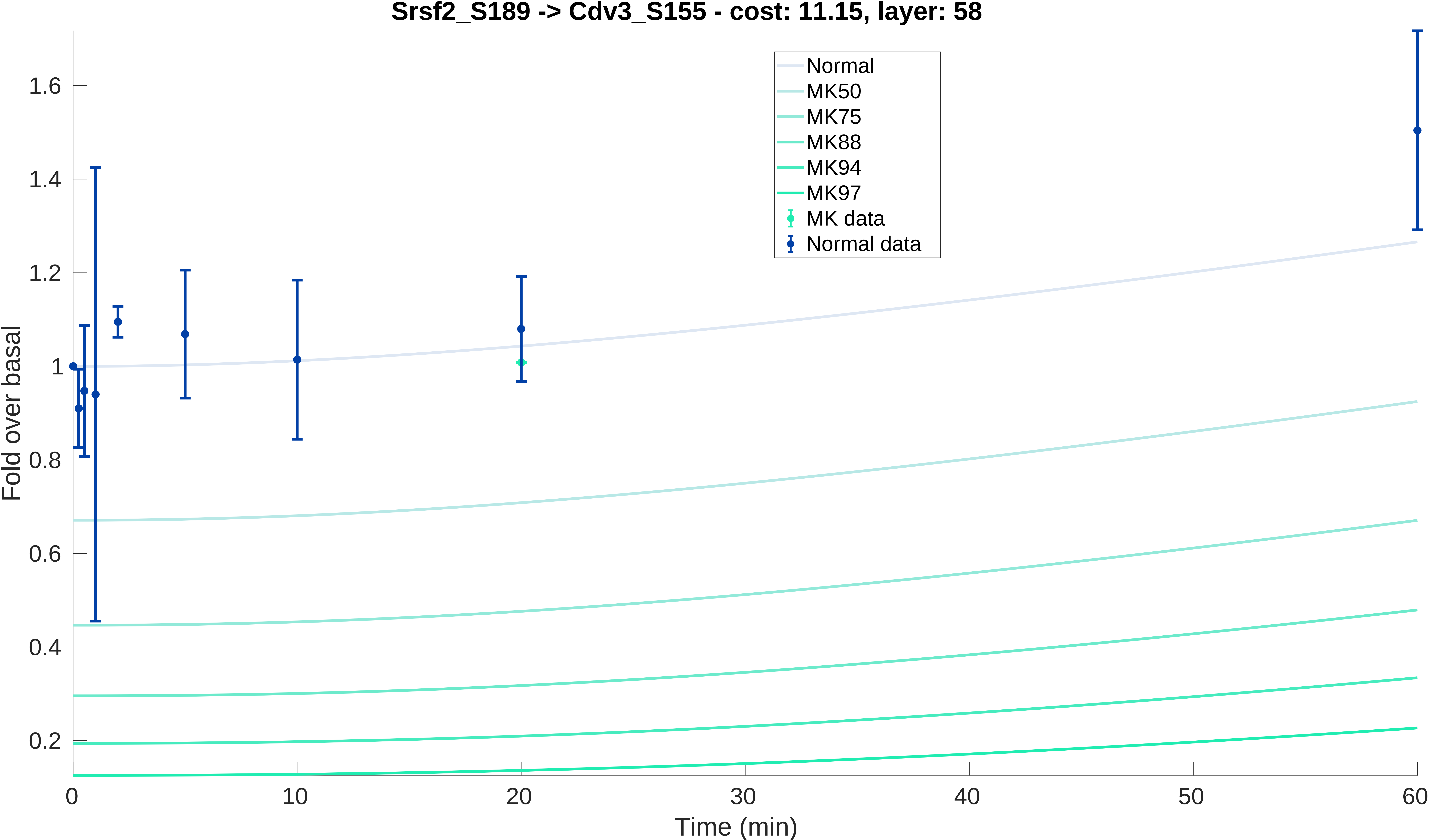

Srsf2_S189 -> Cdv3_S155 - cost: 11.15, layer: 58
Normal
MK50
MK75
MK88
MK94
MK97
MK data
Normal data
Fold over basal
1.6
1.4
1.2
1
0.8
0.6
0.4
0.2
0
10
20
30
40
50
60
Time (min)

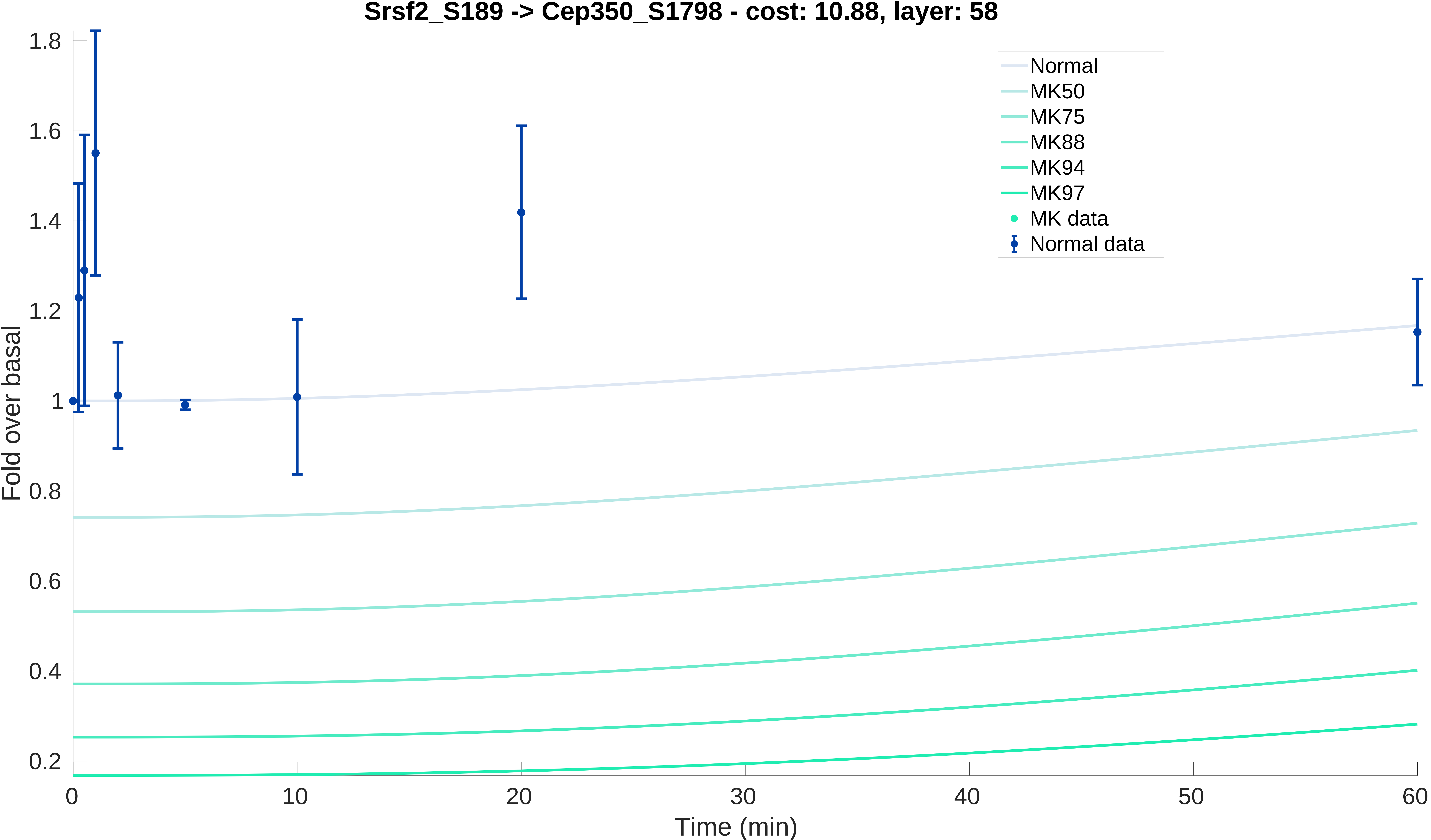

Srsf2_S189 -> Cep350_S1798 - cost: 10.88, layer: 58
Fold over basal
Time (min)
0.2
0.4
0.6
0.8
1
1.2
1.4
1.6
1.8
0
10
20
30
40
50
60
Normal
MK50
MK75
MK88
MK94
MK97
MK data
Normal data

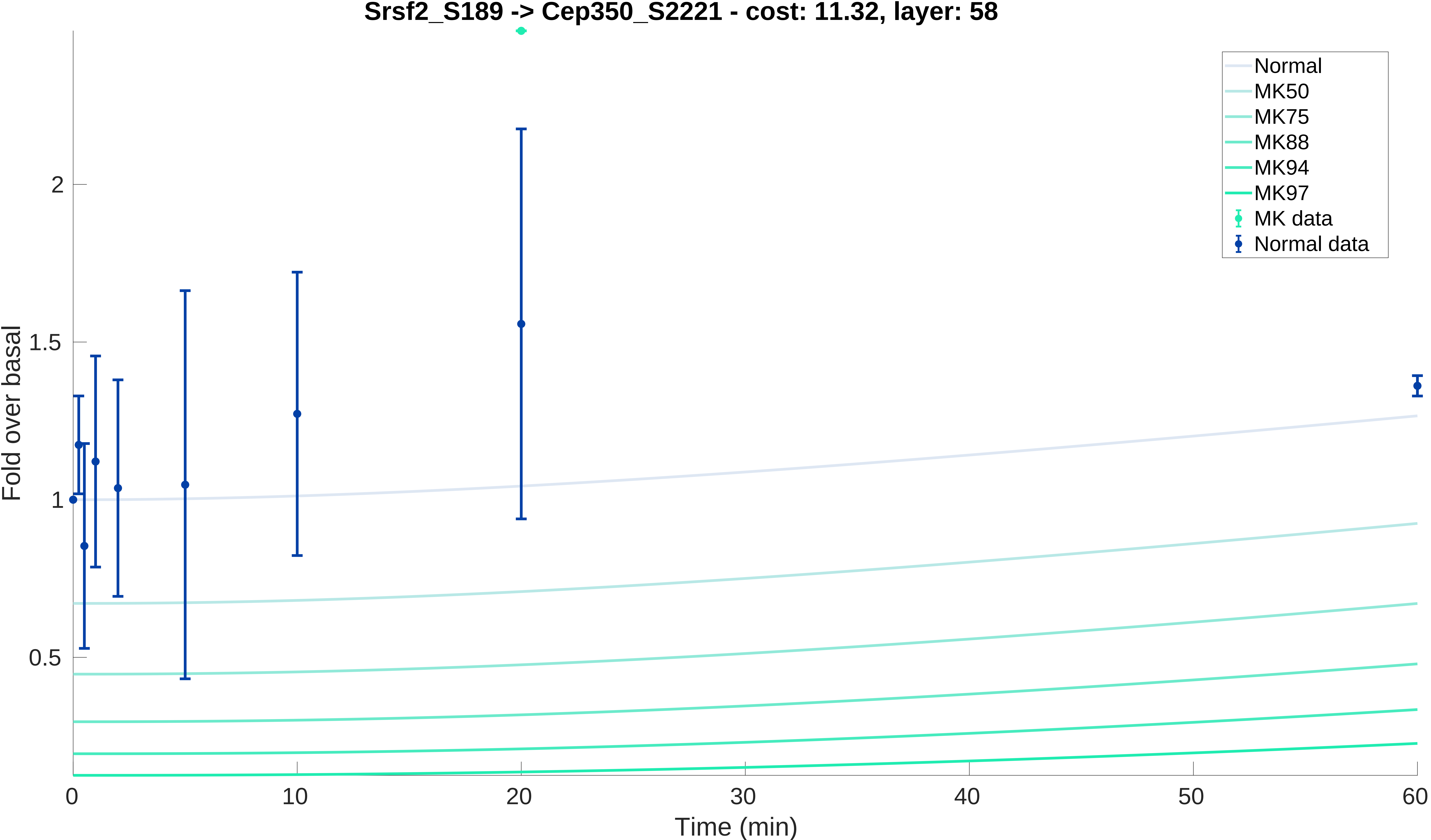

Srsf2_S189 -> Cep350_S2221 - cost: 11.32, layer: 58
Normal
MK50
MK75
MK88
MK94
MK97
MK data
Normal data
Fold over basal
2
1.5
1
0.5
0
10
20
30
40
50
60
Time (min)

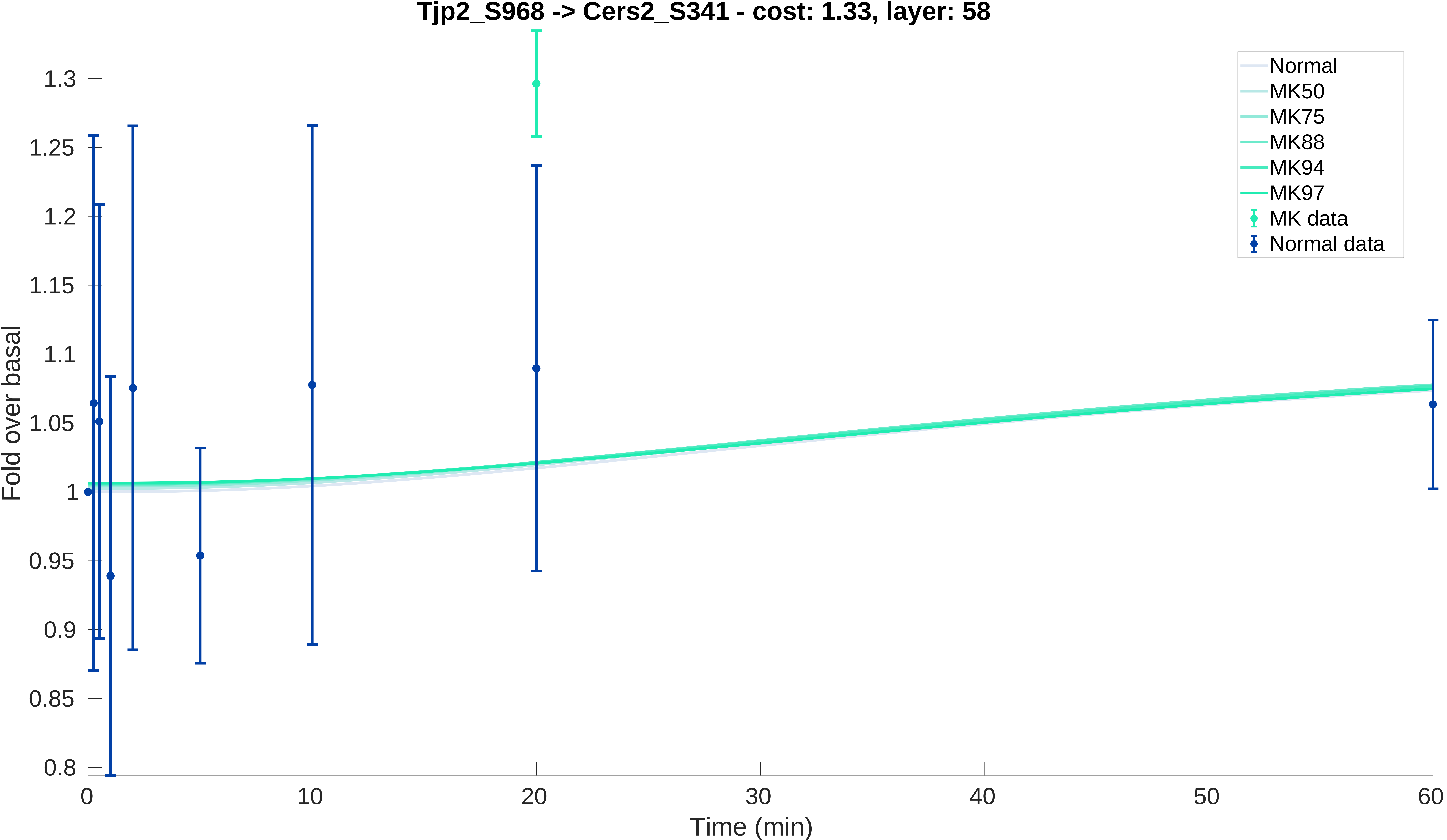

Tjp2_S968 -> Cers2_S341 - cost: 1.33, layer: 58
Fold over basal
Time (min)
1.3
1.25
1.2
1.15
1.1
1.05
1
0.95
0.9
0.85
0.8
0
10
20
30
40
50
60
Normal
MK50
MK75
MK88
MK94
MK97
MK data
Normal data

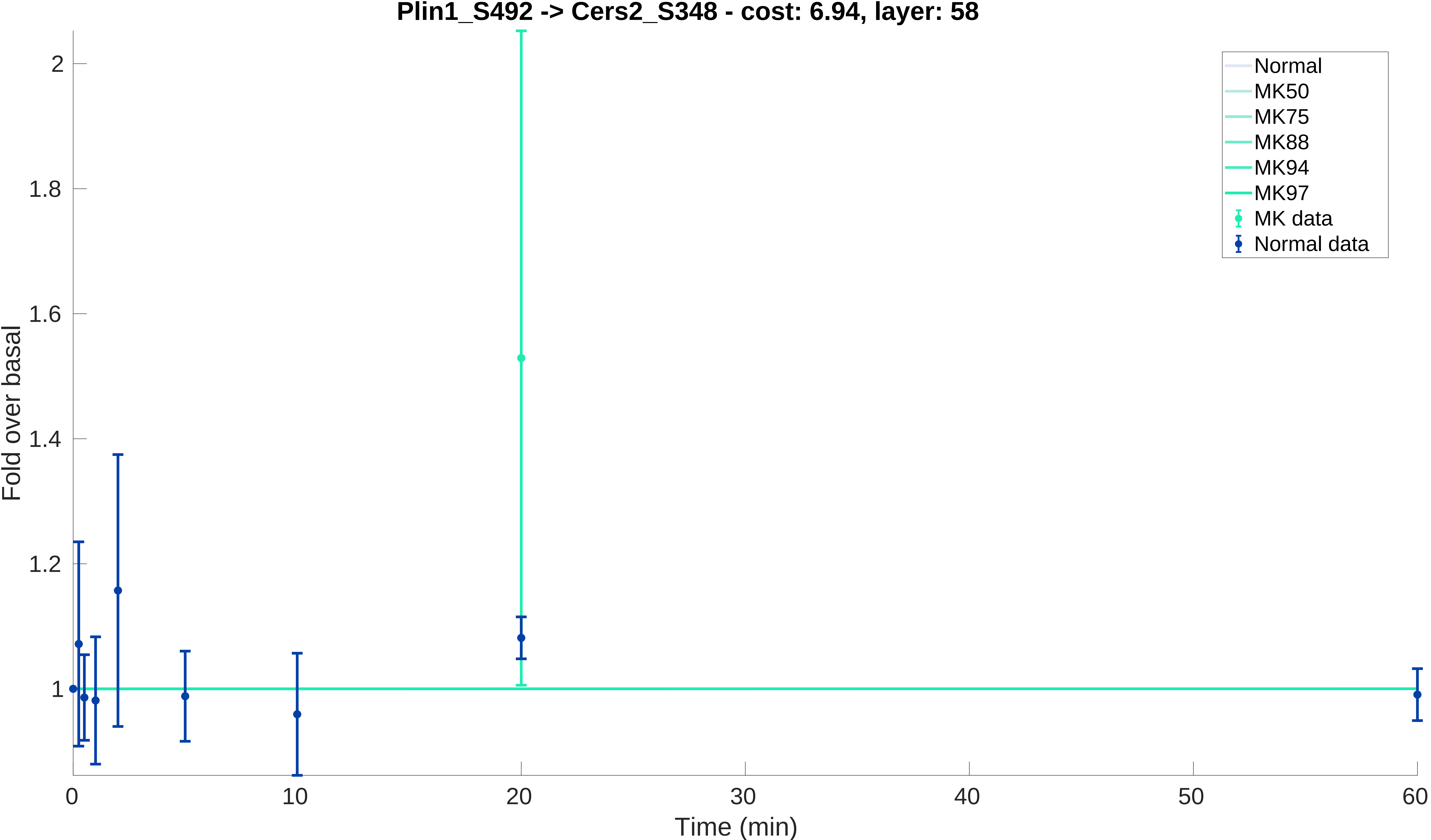

Plin1_S492 -> Cers2_S348 - cost: 6.94, layer: 58
Fold over basal
Time (min)
0
10
20
30
40
50
60
1
1.2
1.4
1.6
1.8
2
Normal
MK50
MK75
MK88
MK94
MK97
MK data
Normal data

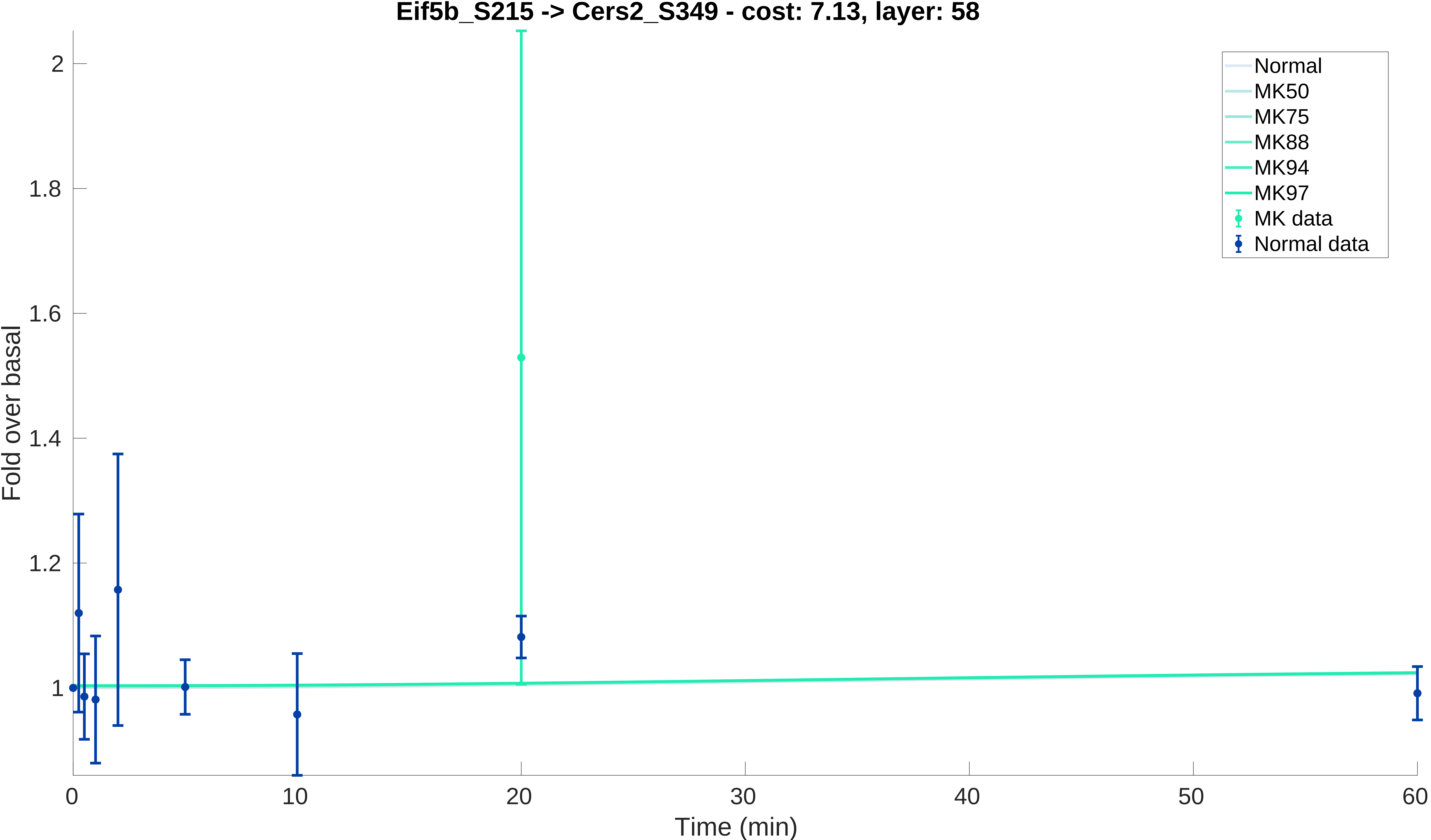

Eif5b_S215 -> Cers2_S349 - cost: 7.13, layer: 58
Fold over basal
2
1.8
1.6
1.4
1.2
1
0
10
20
30
40
50
60
Time (min)
Normal
MK50
MK75
MK88
MK94
MK97
MK data
Normal data

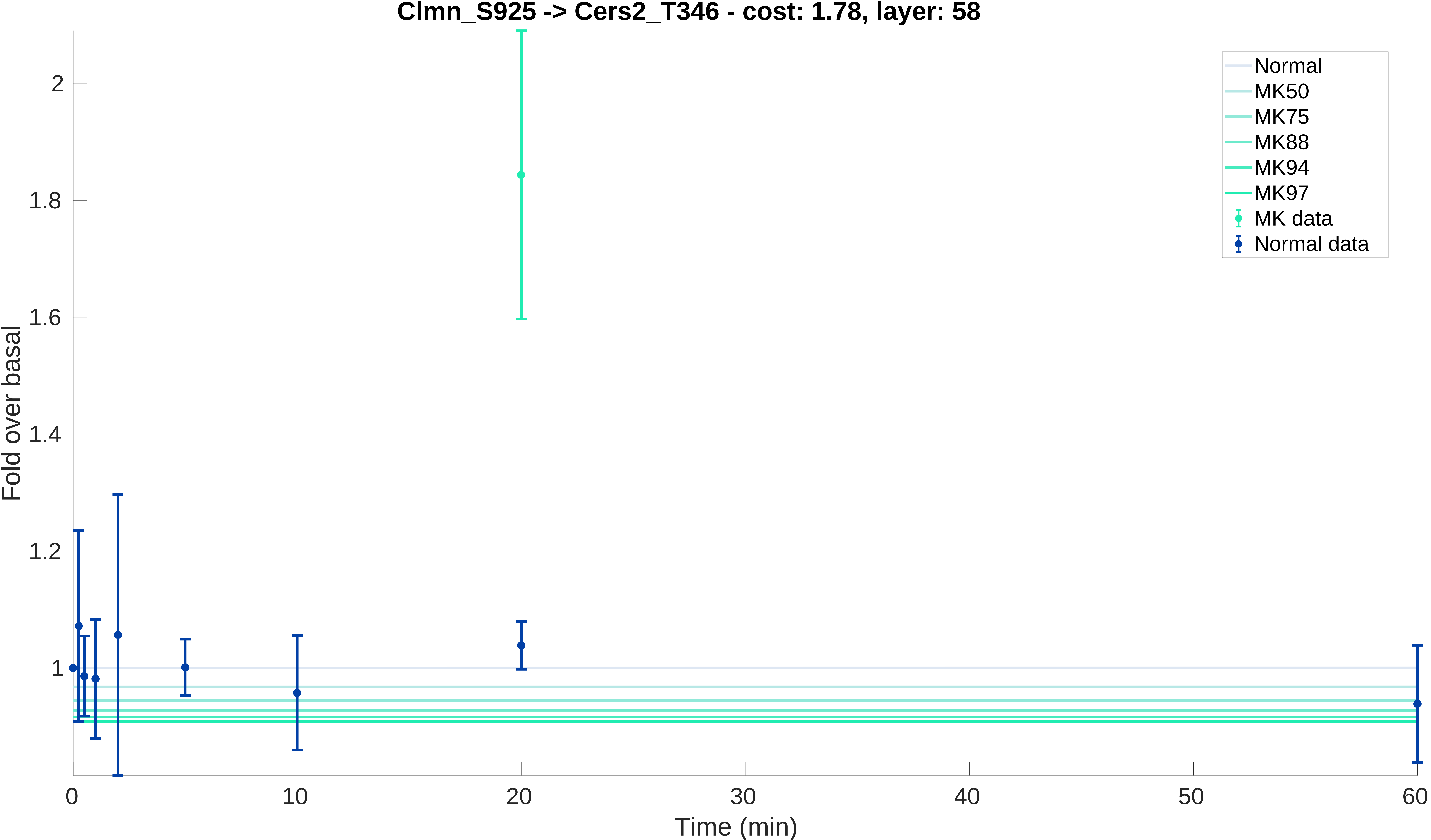

Clmn_S925 -> Cers2_T346 - cost: 1.78, layer: 58
Fold over basal
2
1.8
1.6
1.4
1.2
1
0
10
20
30
40
50
60
Time (min)
Normal
MK50
MK75
MK88
MK94
MK97
MK data
Normal data

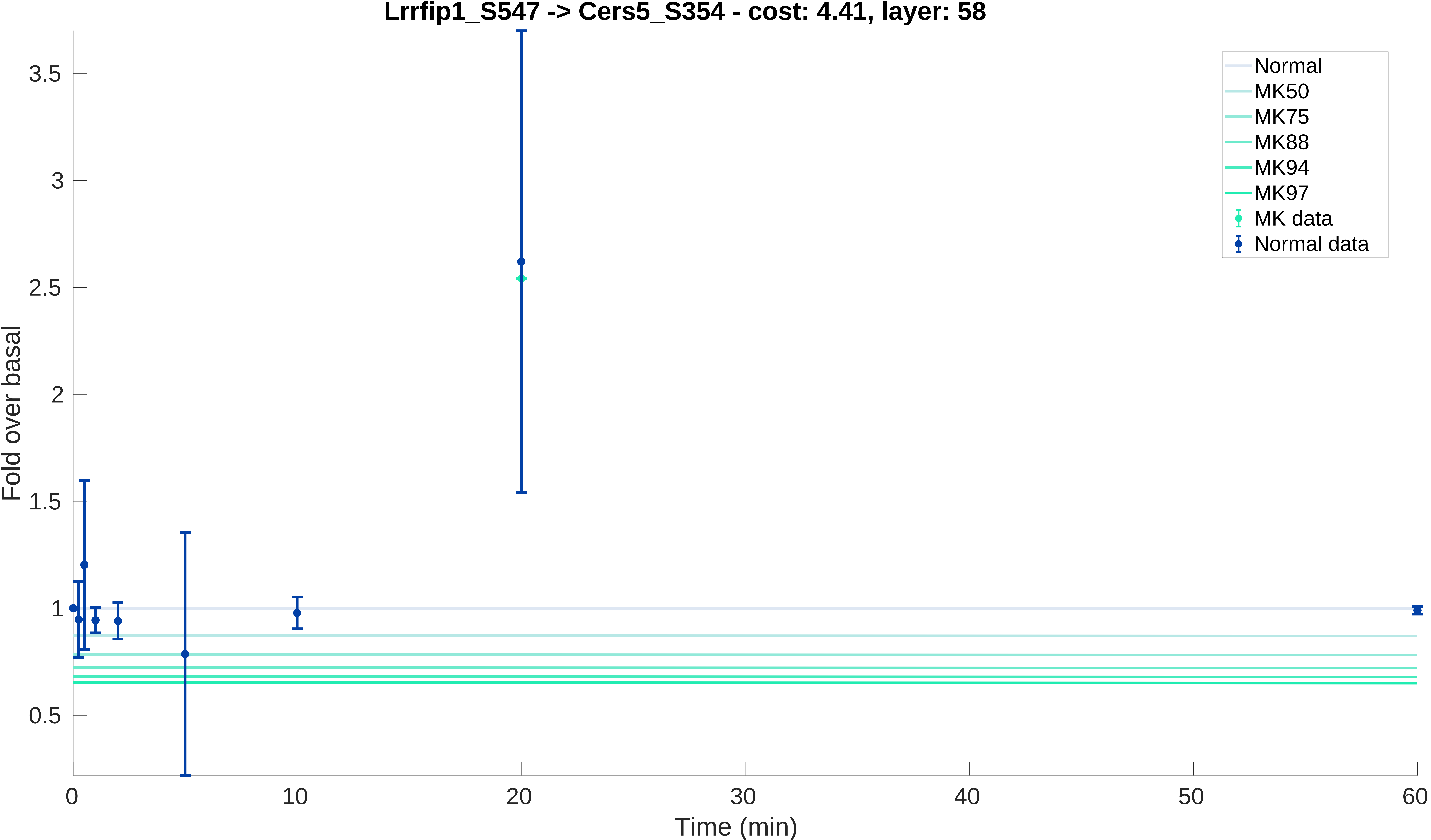

Lrrfip1_S547 -> Cers5_S354 - cost: 4.41, layer: 58
Fold over basal
Time (min)
0.5
1
1.5
2
2.5
3
3.5
0
10
20
30
40
50
60
Normal
MK50
MK75
MK88
MK94
MK97
MK data
Normal data

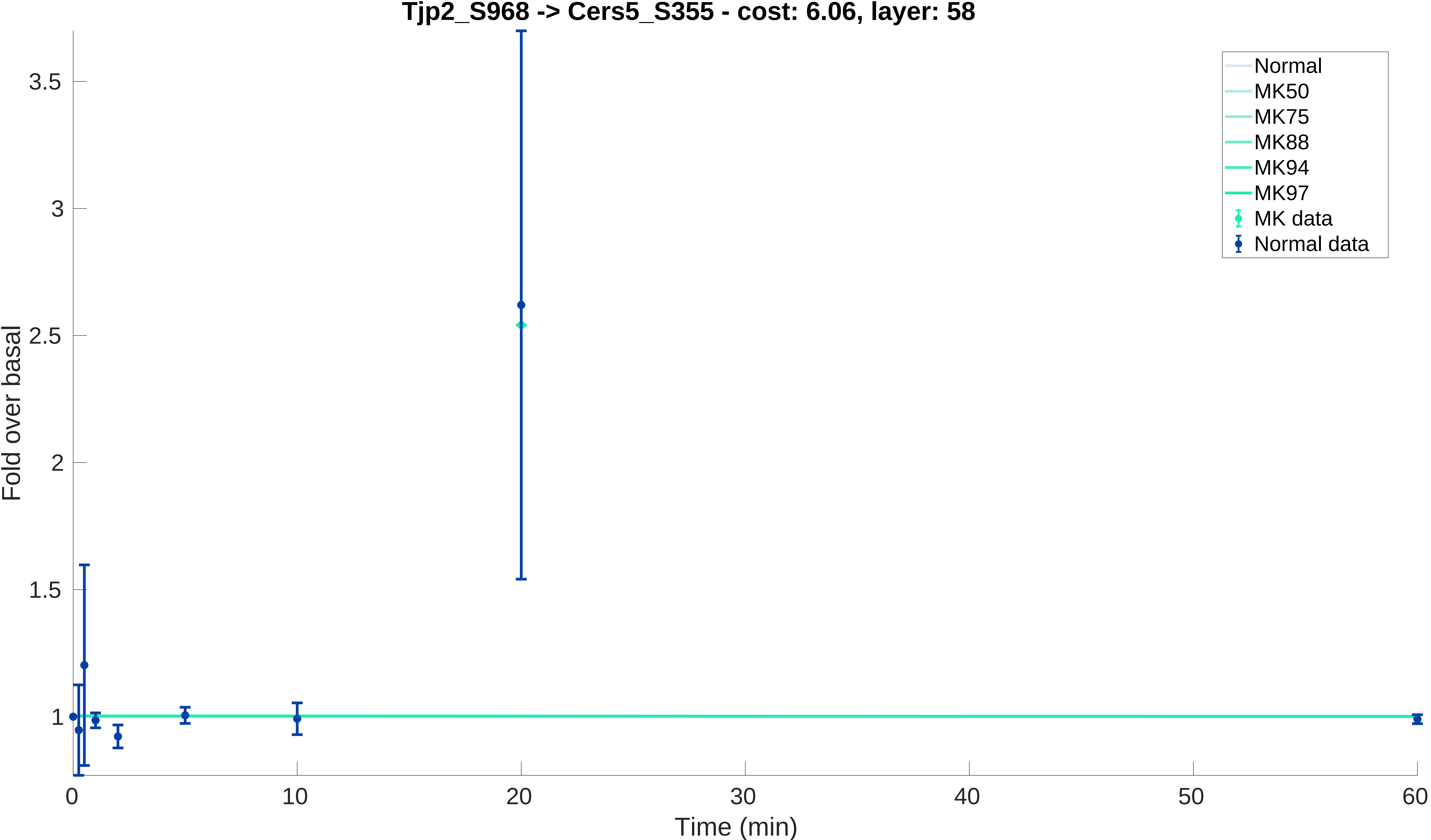

Tjp2_S968 -> Cers5_S355 - cost: 6.06, layer: 58
Fold over basal
Time (min)
3.5
3
2.5
2
1.5
1
0
10
20
30
40
50
60
Normal
MK50
MK75
MK88
MK94
MK97
MK data
Normal data

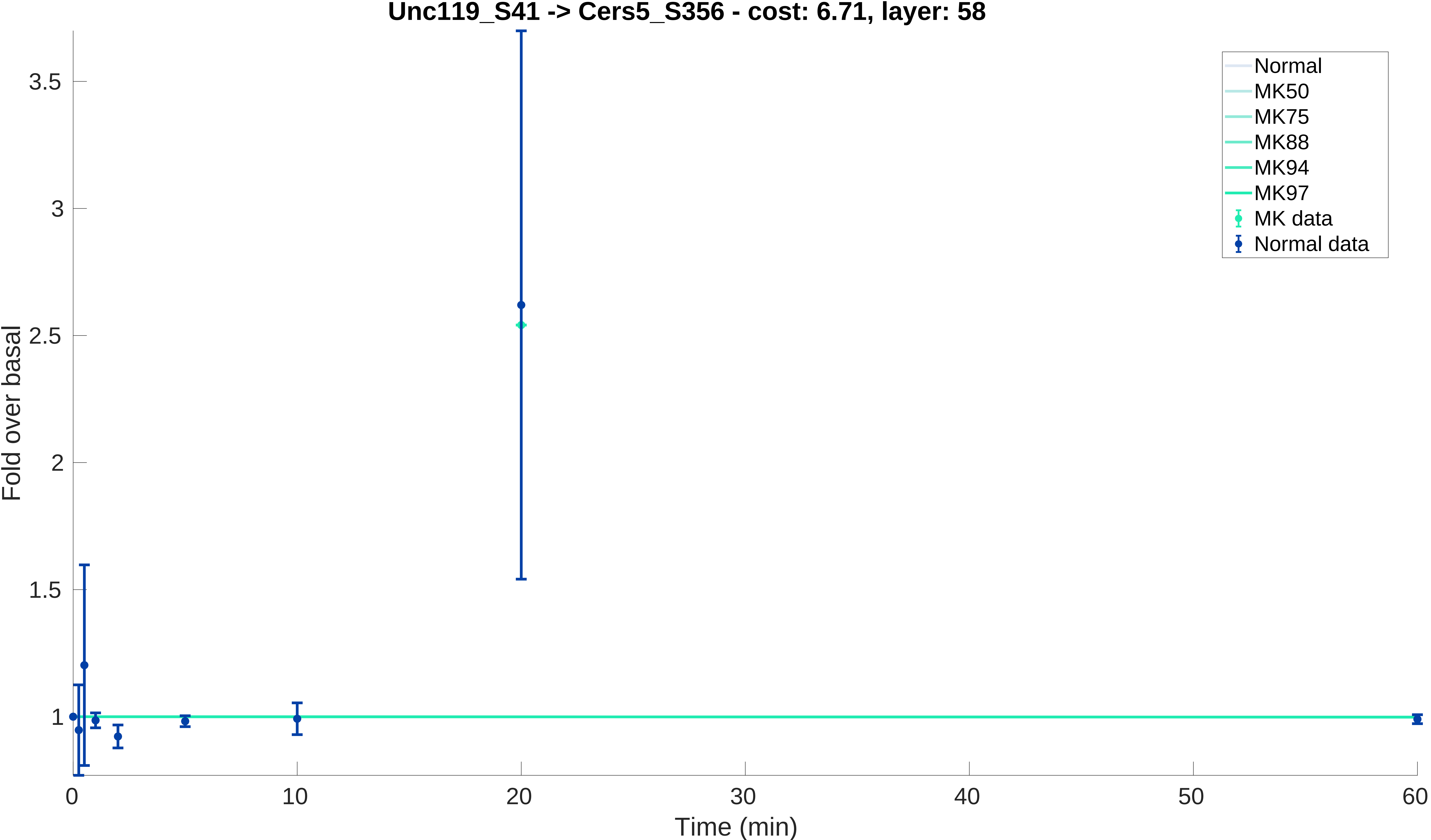

Unc119_S41 -> Cers5_S356 - cost: 6.71, layer: 58
Fold over basal
3.5
3
2.5
2
1.5
1
0
10
20
30
40
50
60
Time (min)
Normal
MK50
MK75
MK88
MK94
MK97
MK data
Normal data

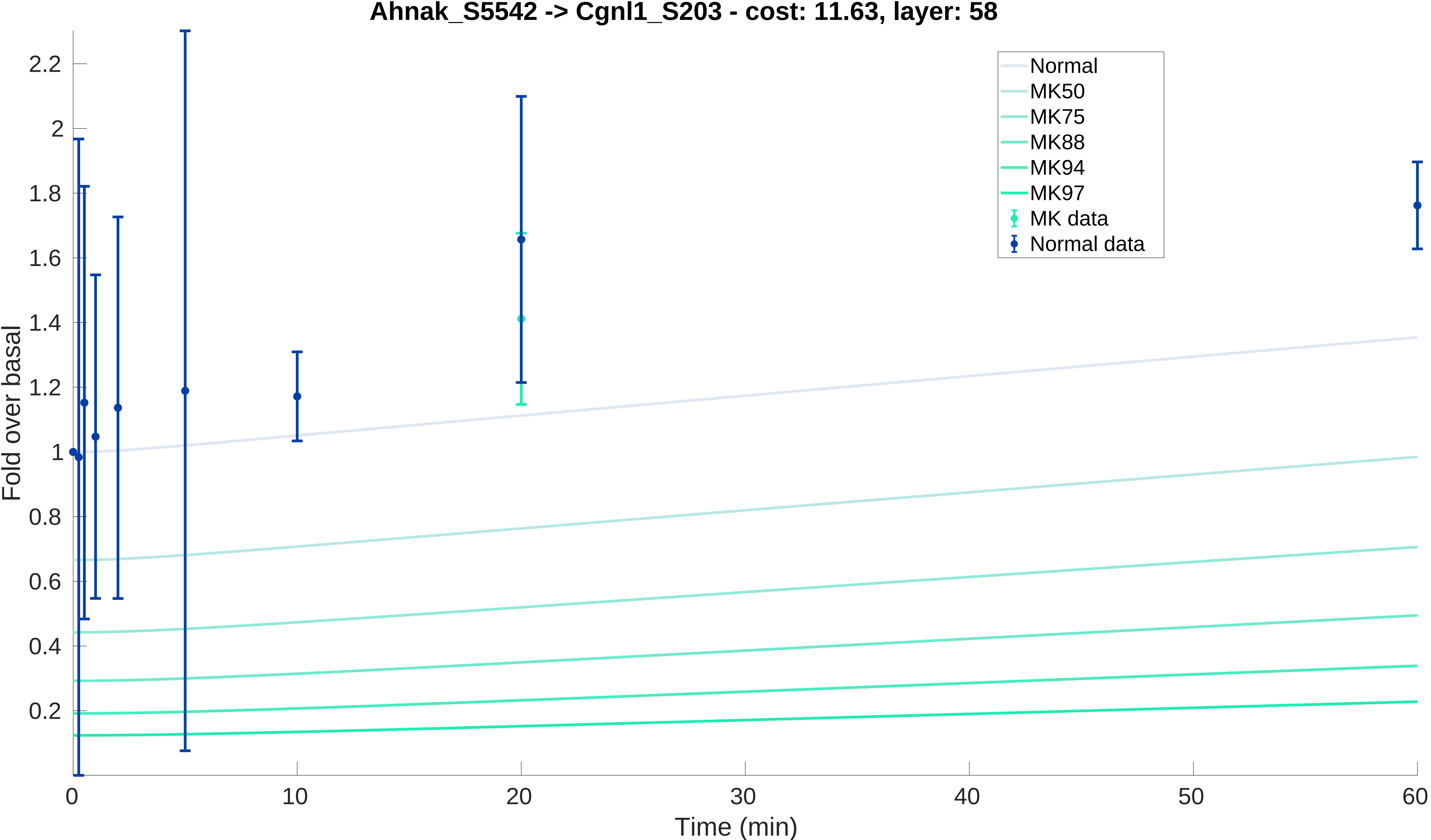

Ahnak_S5542 -> Cgnl1_S203 - cost: 11.63, layer: 58
Normal
MK50
MK75
MK88
MK94
MK97
MK data
Normal data
Fold over basal
Time (min)
2.2
2
1.8
1.6
1.4
1.2
1
0.8
0.6
0.4
0.2
0
10
20
30
40
50
60

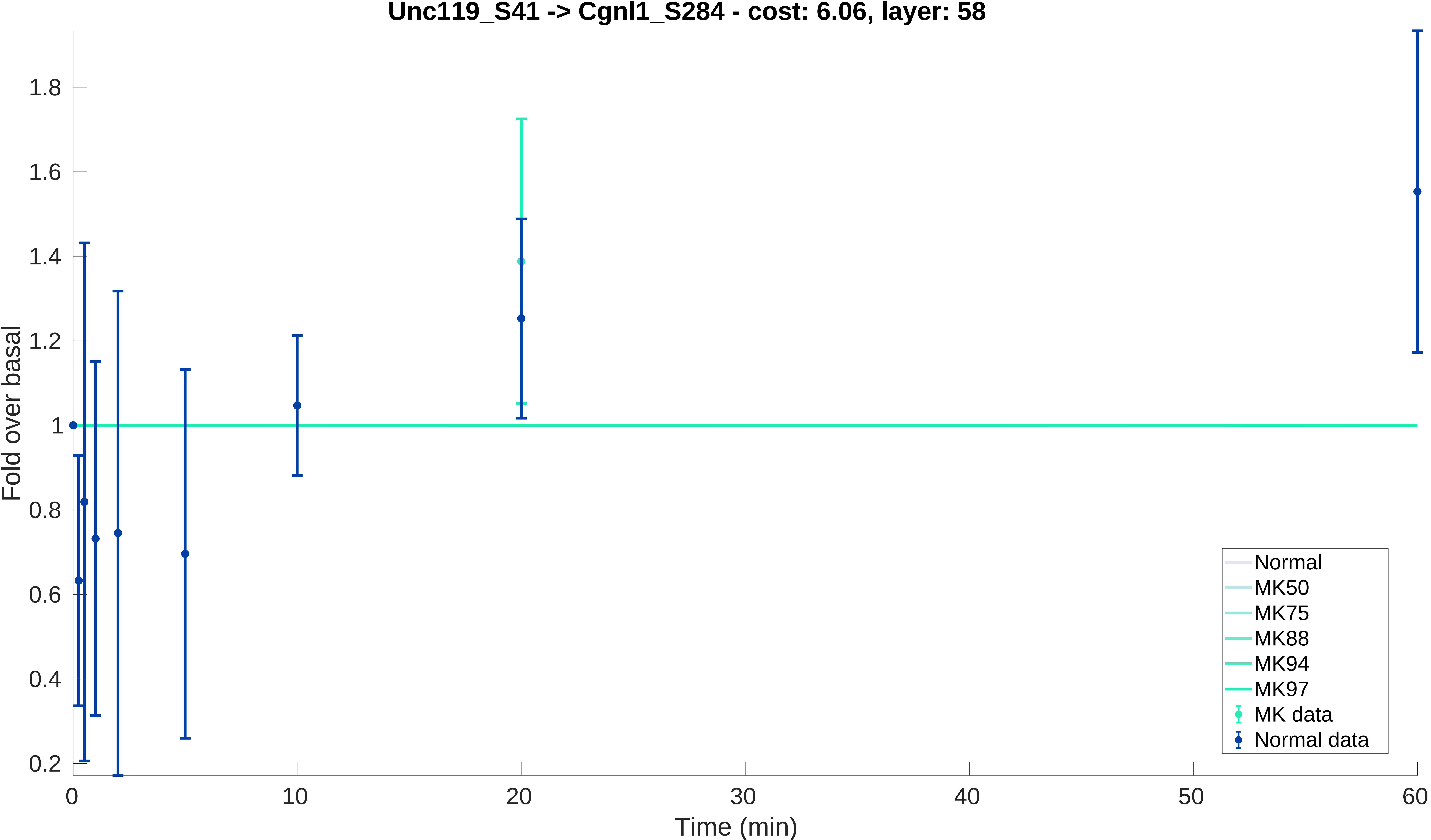

Unc119_S41 -> Cgnl1_S284 - cost: 6.06, layer: 58
Fold over basal
1.8
1.6
1.4
1.2
1
0.8
0.6
0.4
0.2
0
10
20
30
40
50
60
Time (min)
Normal
MK50
MK75
MK88
MK94
MK97
MK data
Normal data

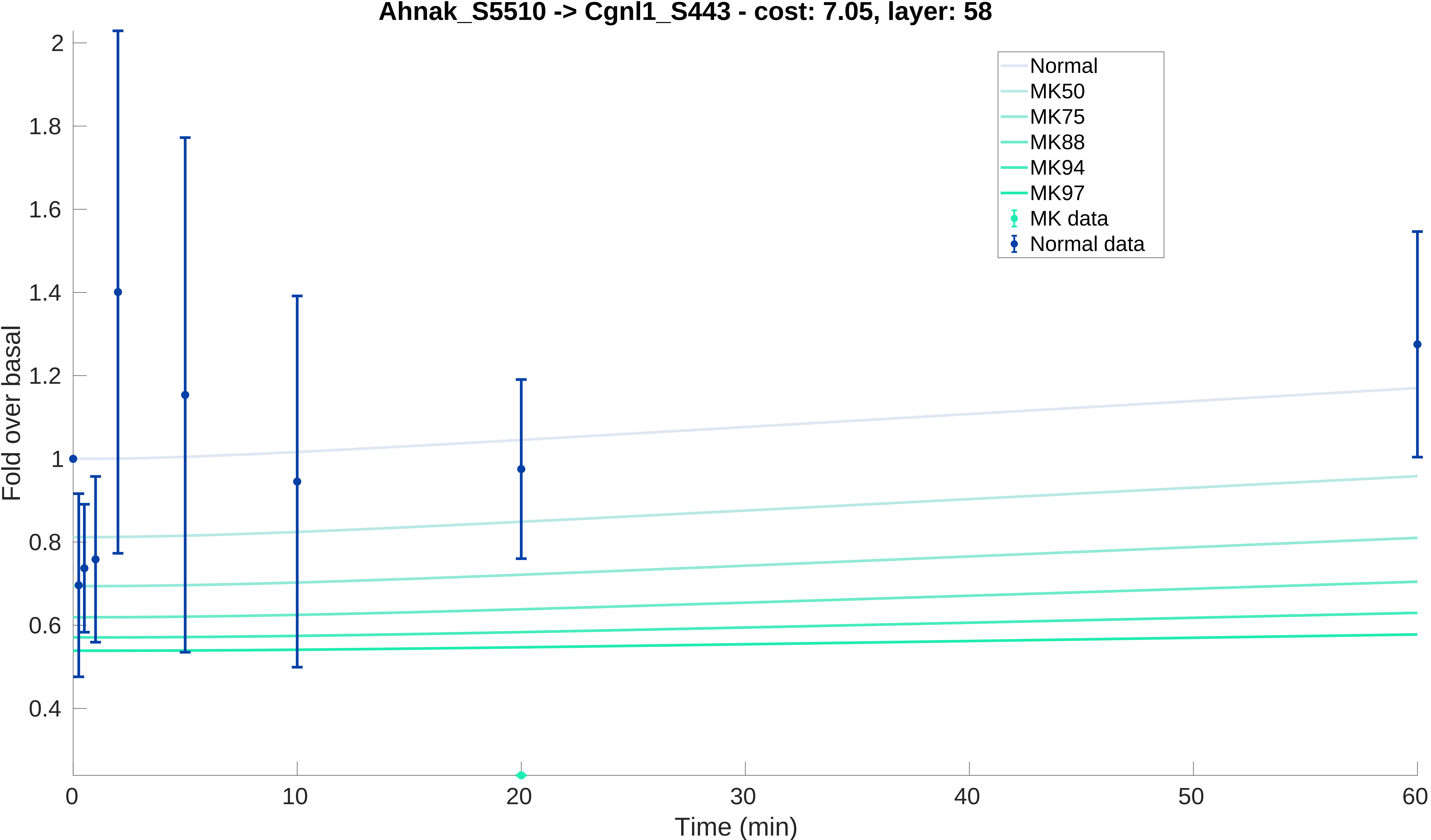

Ahnak_S5510 -> Cgnl1_S443 - cost: 7.05, layer: 58
Normal
MK50
MK75
MK88
MK94
MK97
MK data
Normal data
Fold over basal
Time (min)
0
10
20
30
40
50
60
0.4
0.6
0.8
1
1.2
1.4
1.6
1.8
2

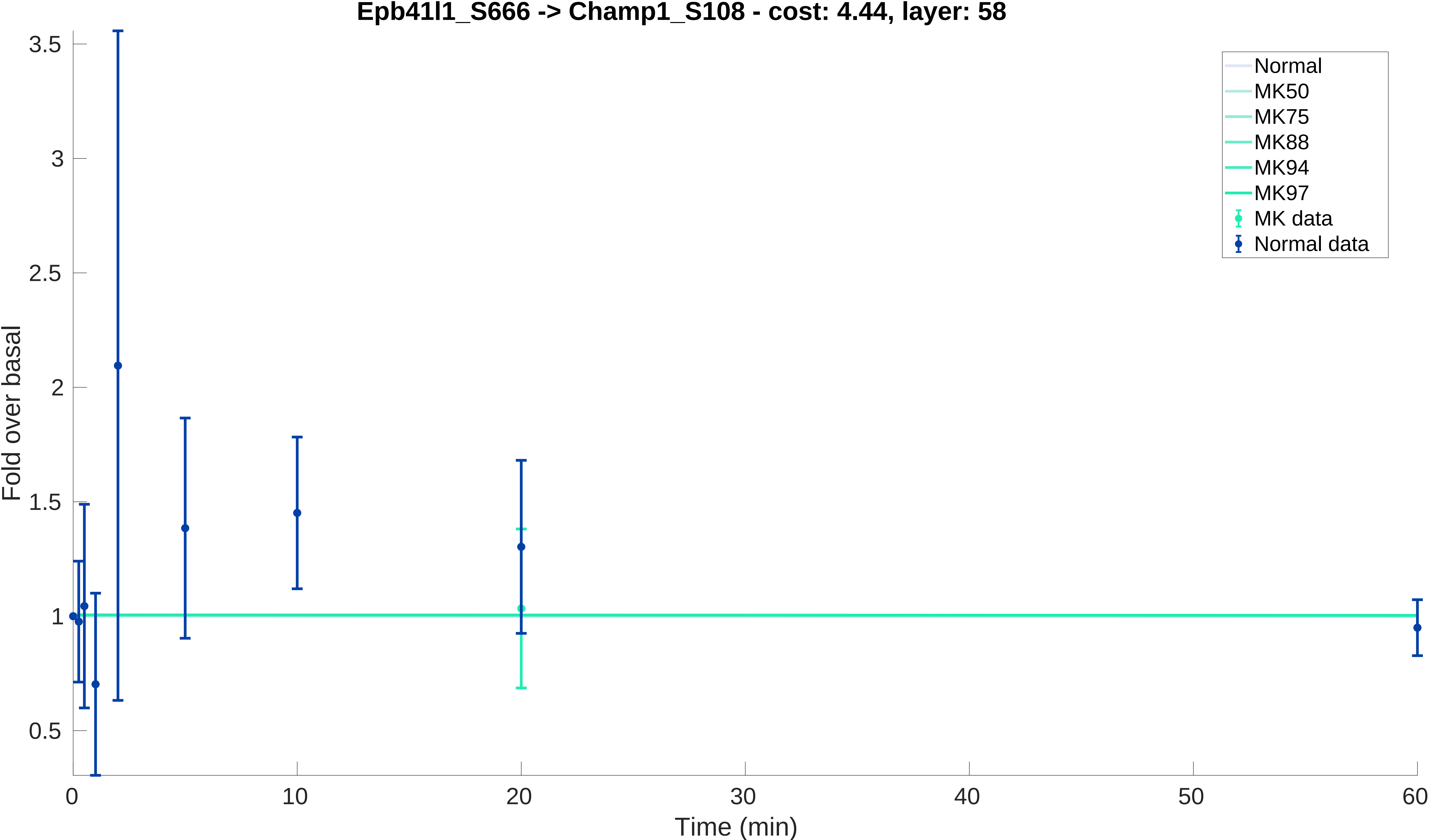

Epb41l1_S666 -> Champ1_S108 - cost: 4.44, layer: 58
Fold over basal
Time (min)
Normal
MK50
MK75
MK88
MK94
MK97
MK data
Normal data
0.5
1
1.5
2
2.5
3
3.5
0
10
20
30
40
50
60

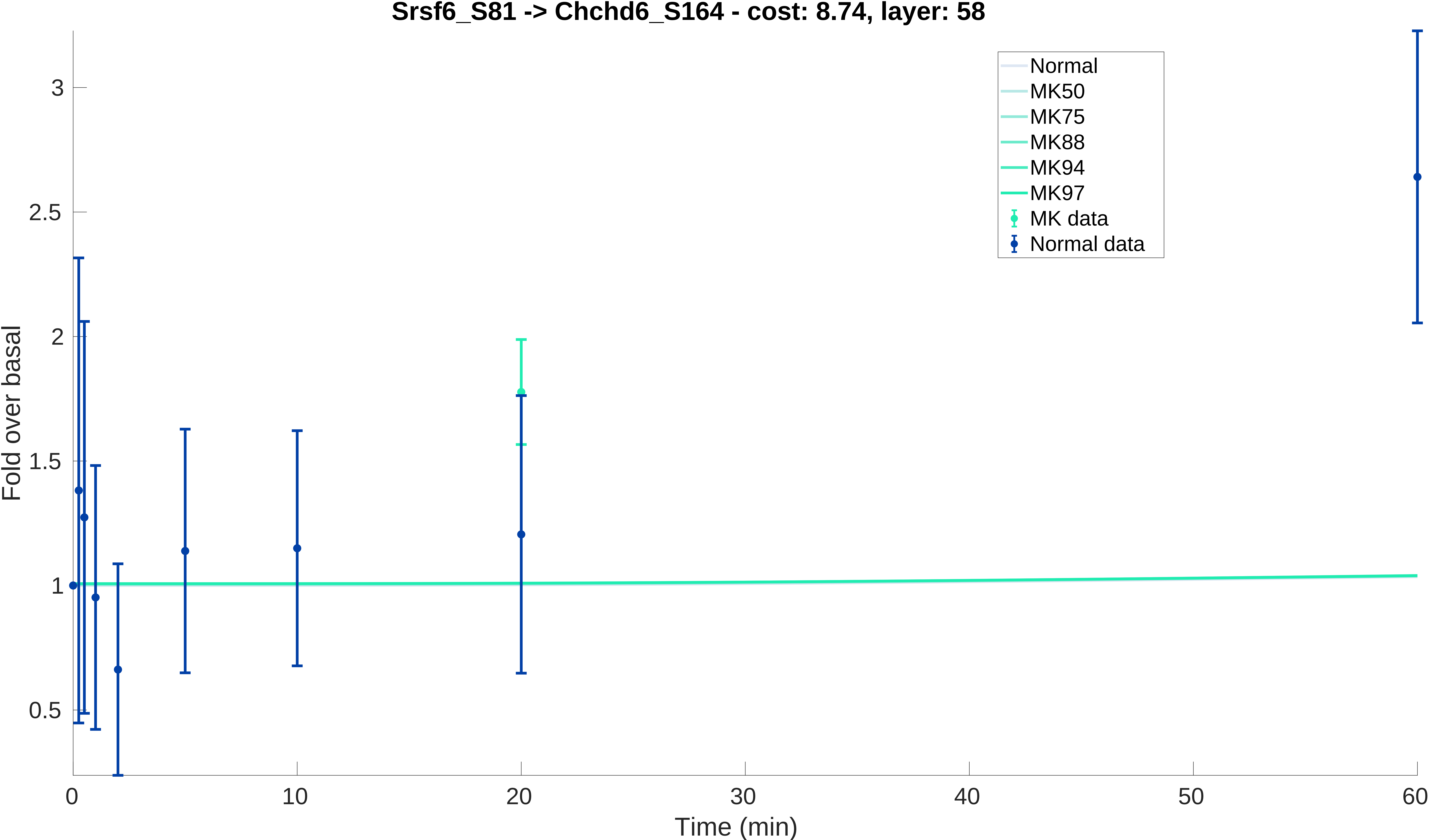

Srsf6_S81 -> Chchd6_S164 - cost: 8.74, layer: 58
Fold over basal
Time (min)
0.5
1
1.5
2
2.5
3
0
10
20
30
40
50
60
Normal
MK50
MK75
MK88
MK94
MK97
MK data
Normal data

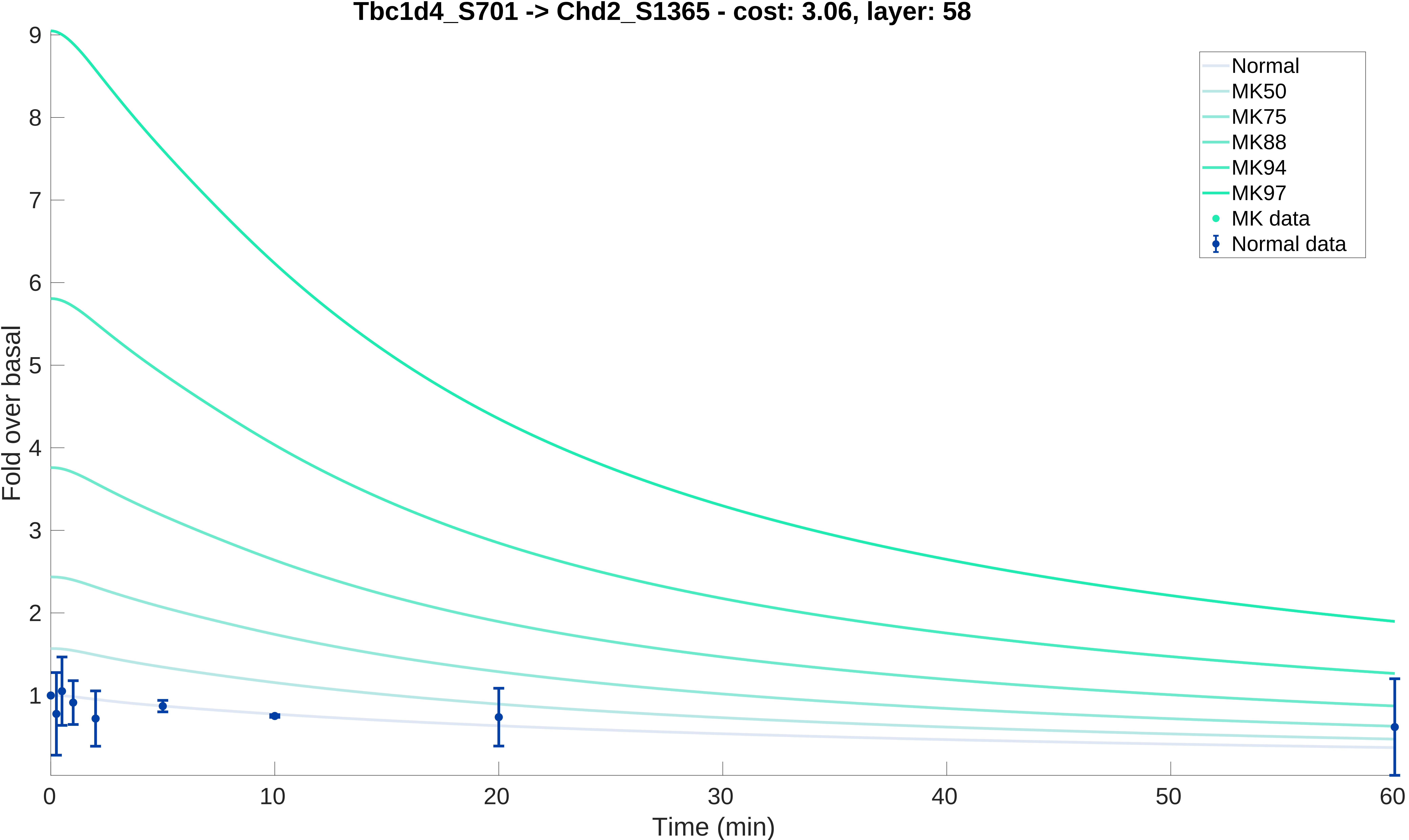
Tbc1d4_S701 -> Chd2_S1365 - cost: 3.06, layer: 58
Fold over basal
Time (min)
Normal
MK50
MK75
MK88
MK94
MK97
MK data
Normal data
0
10
20
30
40
50
60
1
2
3
4
5
6
7
8
9

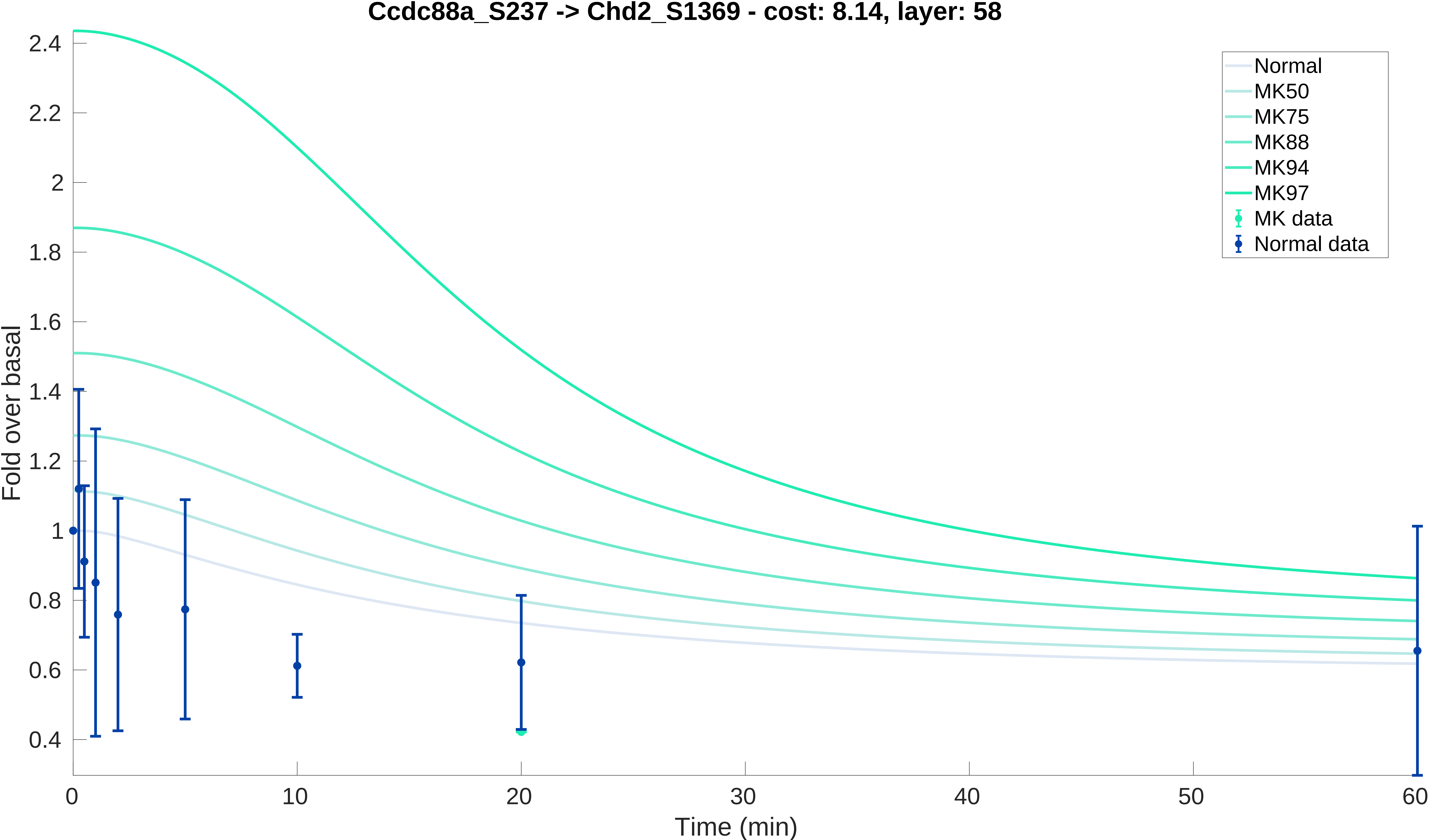

Ccdc88a_S237 -> Chd2_S1369 - cost: 8.14, layer: 58
Fold over basal
Time (min)
0
10
20
30
40
50
60
0.4
0.6
0.8
1
1.2
1.4
1.6
1.8
2
2.2
2.4
Normal
MK50
MK75
MK88
MK94
MK97
MK data
Normal data

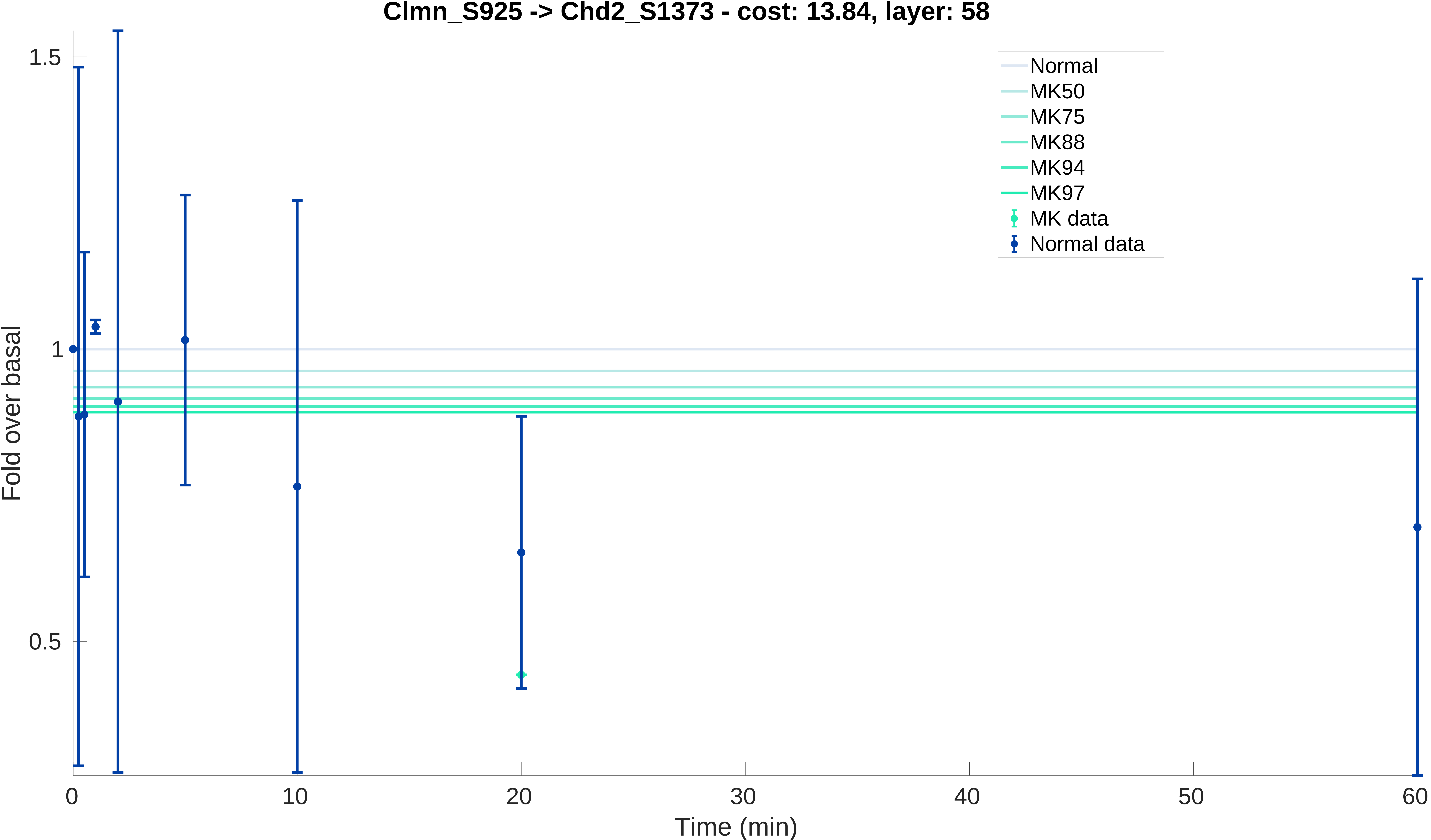

Clmn_S925 -> Chd2_S1373 - cost: 13.84, layer: 58
Normal
MK50
MK75
MK88
MK94
MK97
MK data
Normal data
Fold over basal
1.5
1
0.5
0
10
20
30
40
50
60
Time (min)

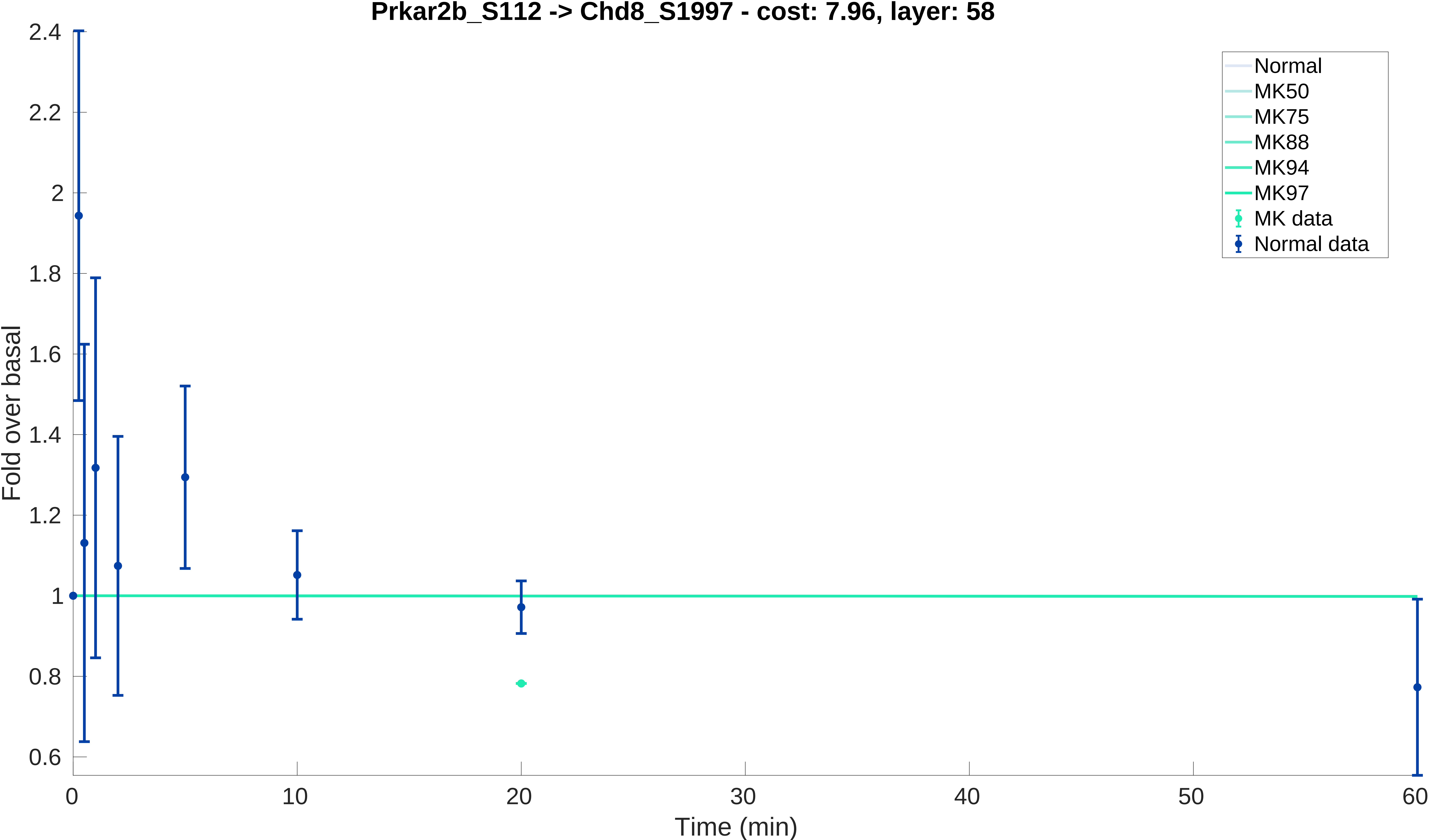
Prkar2b_S112 -> Chd8_S1997 - cost: 7.96, layer: 58
Fold over basal
2.4
2.2
2
1.8
1.6
1.4
1.2
1
0.8
0.6
0
10
20
30
40
50
60
Time (min)
Normal
MK50
MK75
MK88
MK94
MK97
MK data
Normal data

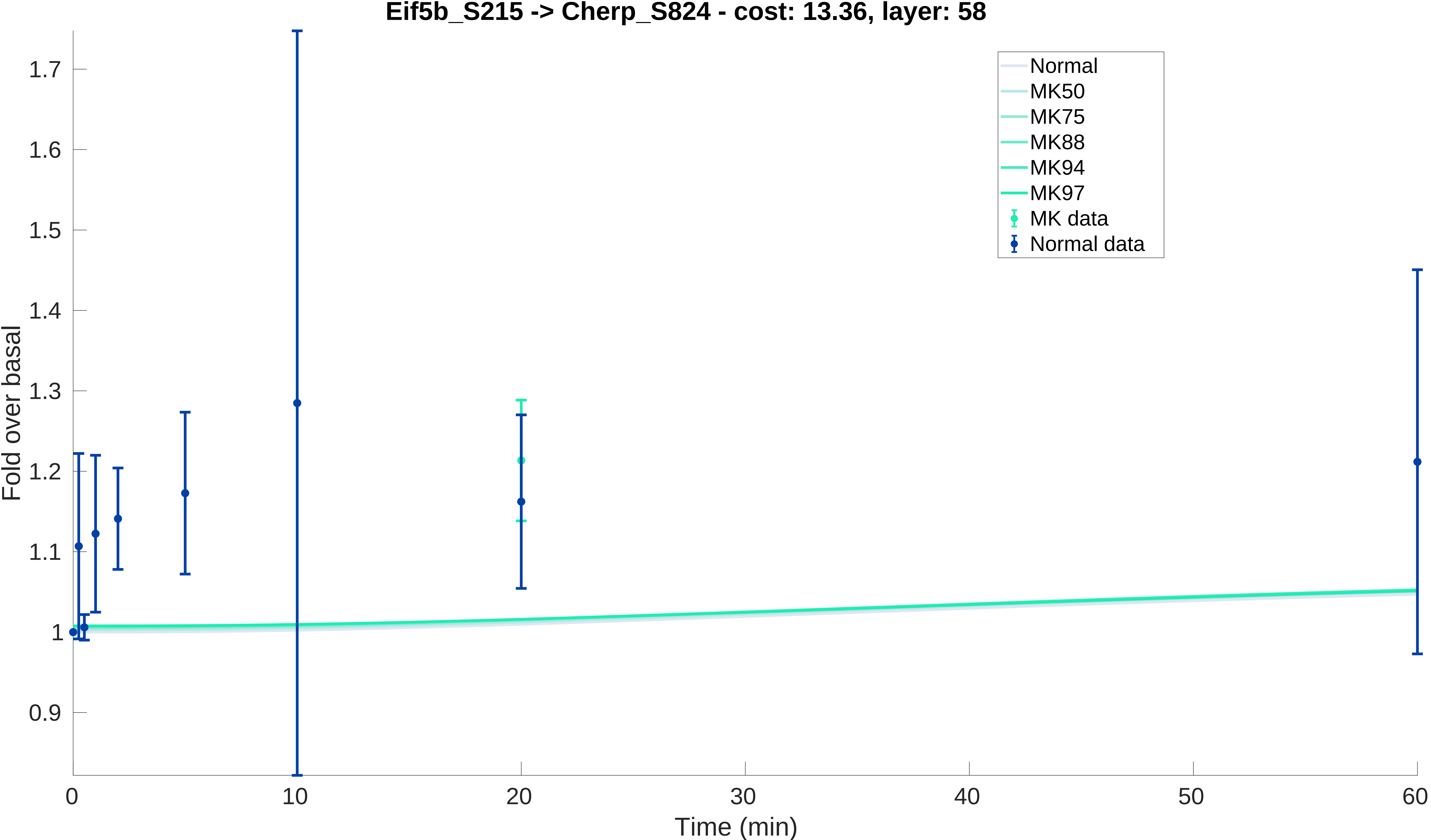

Eif5b_S215 -> Cherp_S824 - cost: 13.36, layer: 58
Normal
MK50
MK75
MK88
MK94
MK97
MK data
Normal data
Fold over basal
0.9
1
1.1
1.2
1.3
1.4
1.5
1.6
1.7
0
10
20
30
40
50
60
Time (min)

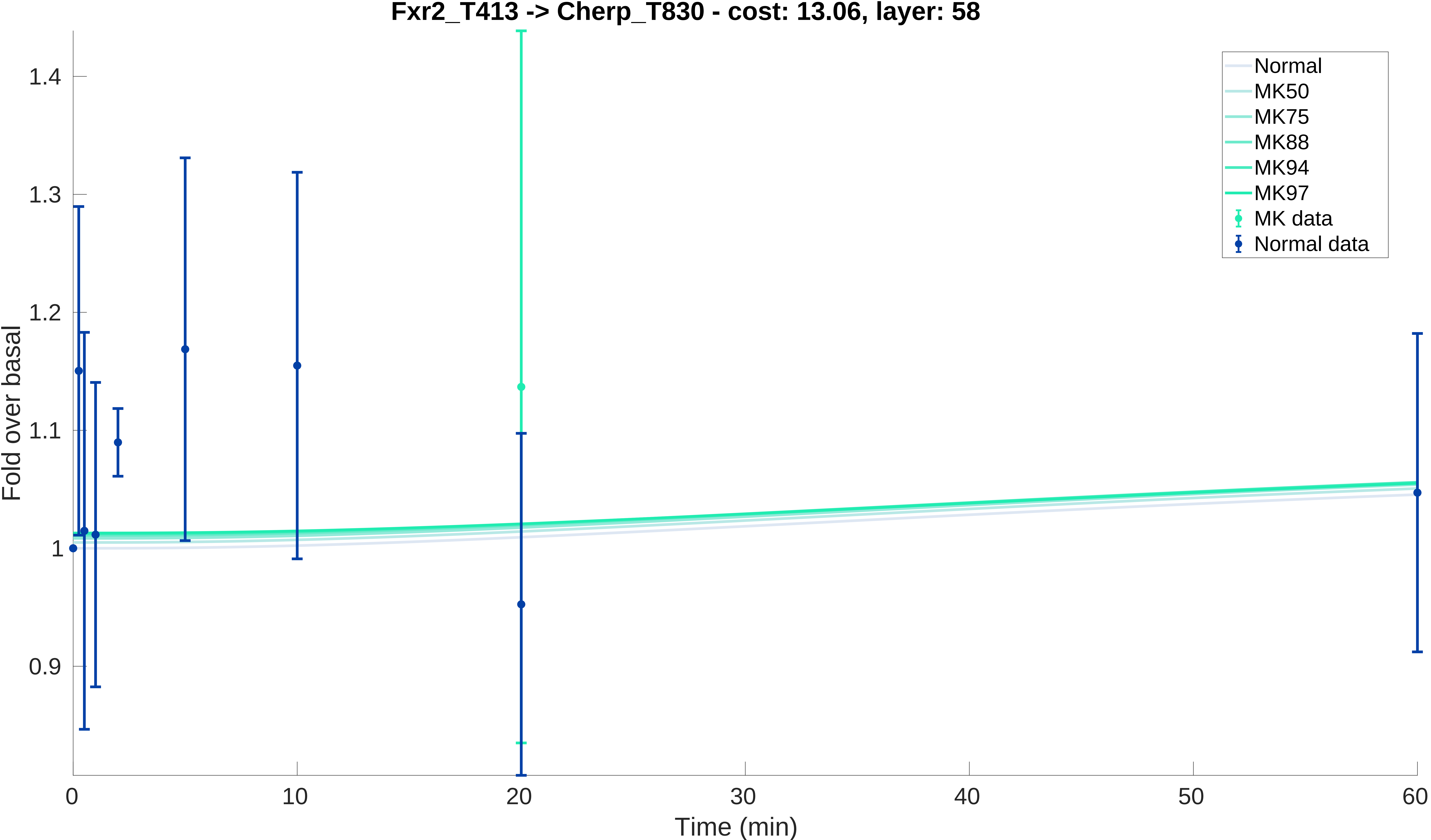
Fxr2_T413 -> Cherp_T830 - cost: 13.06, layer: 58
Fold over basal
Time (min)
1.4
1.3
1.2
1.1
1
0.9
0
10
20
30
40
50
60
Normal
MK50
MK75
MK88
MK94
MK97
MK data
Normal data

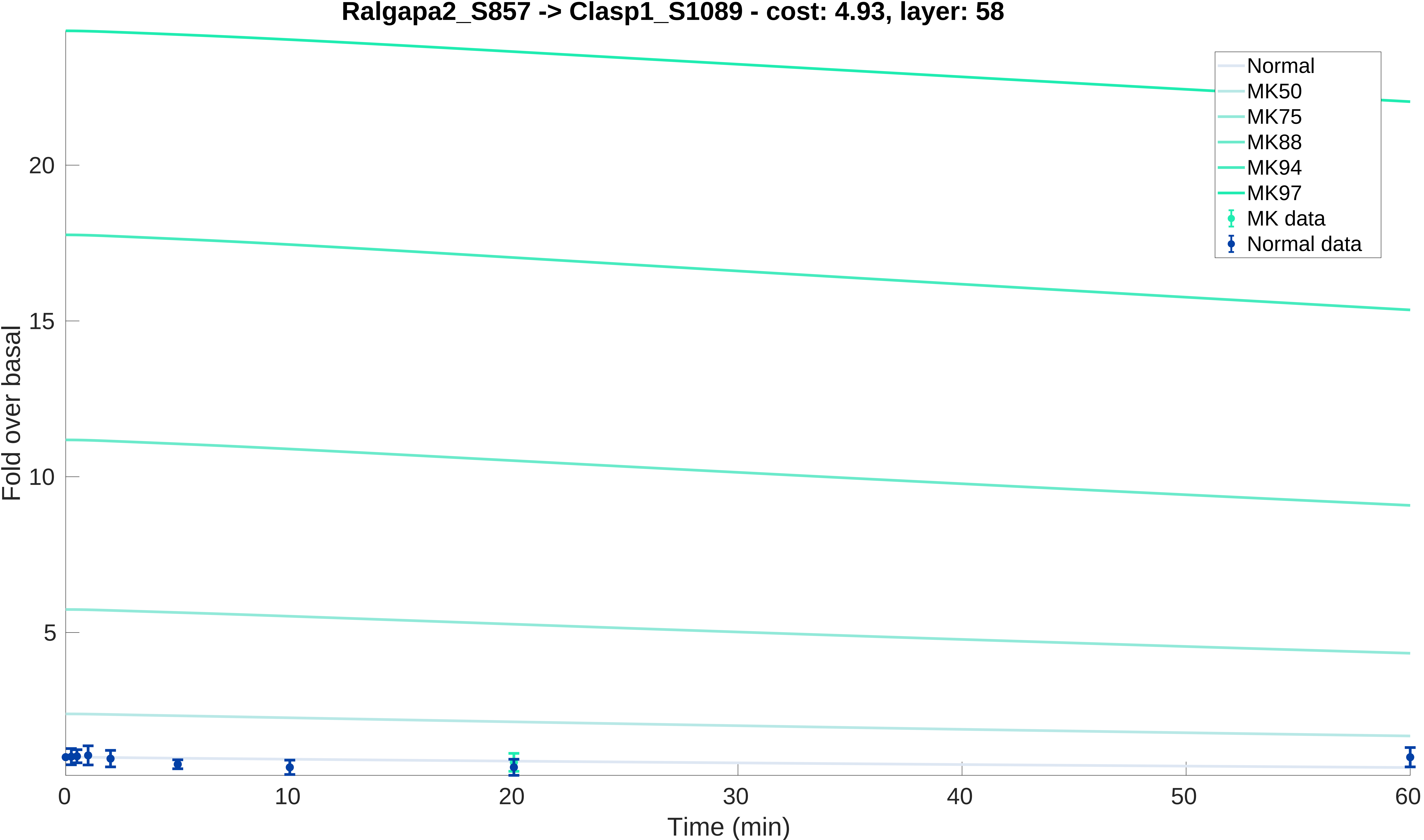

Ralgapa2_S857 -> Clasp1_S1089 - cost: 4.93, layer: 58
Normal
MK50
MK75
MK88
MK94
MK97
MK data
Normal data
Fold over basal
20
15
10
5
0
10
20
30
40
50
60
Time (min)

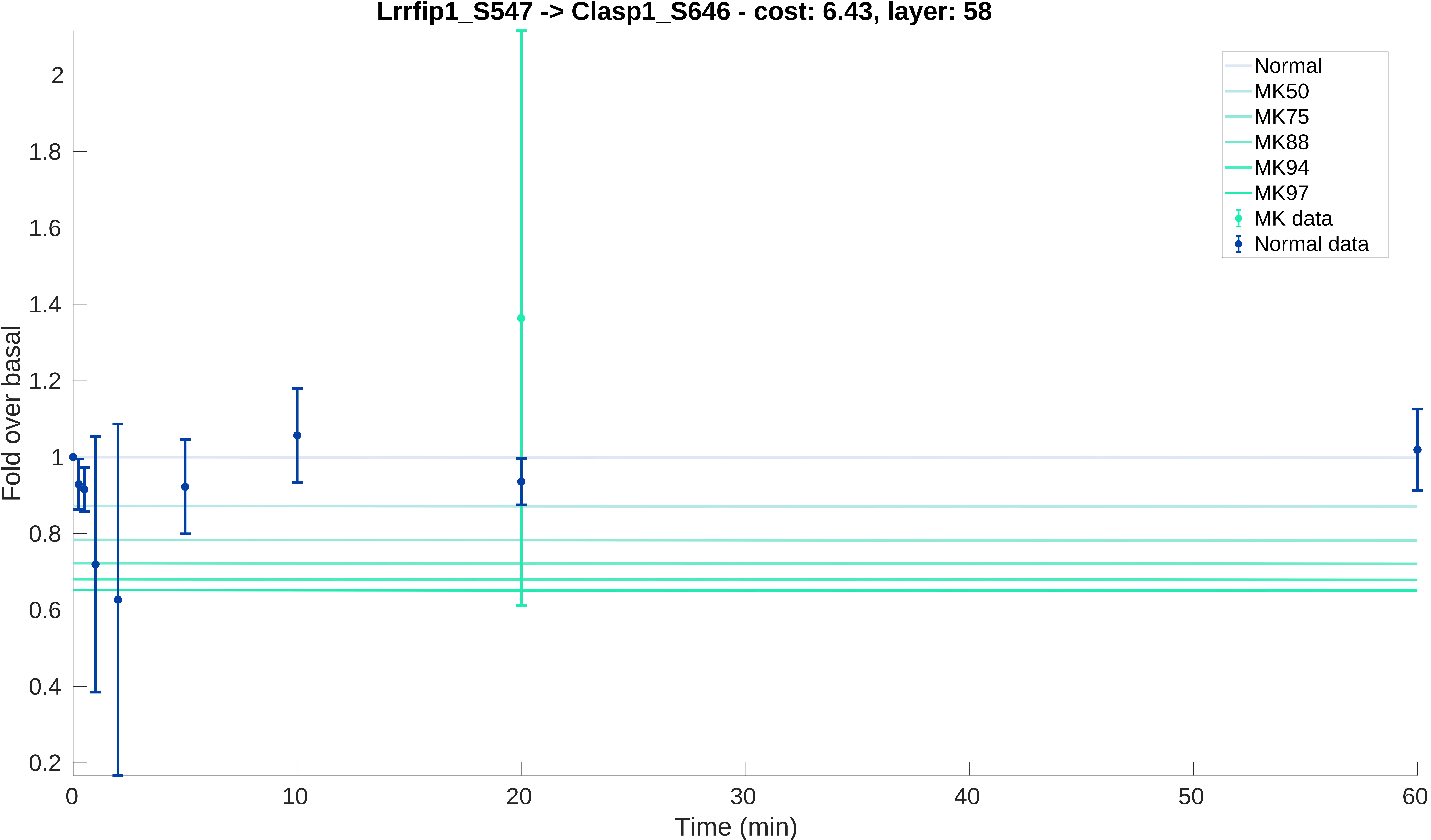

Lrrfip1_S547 -> Clasp1_S646 - cost: 6.43, layer: 58
Fold over basal
Time (min)
0.2
0.4
0.6
0.8
1
1.2
1.4
1.6
1.8
2
0
10
20
30
40
50
60
Normal
MK50
MK75
MK88
MK94
MK97
MK data
Normal data

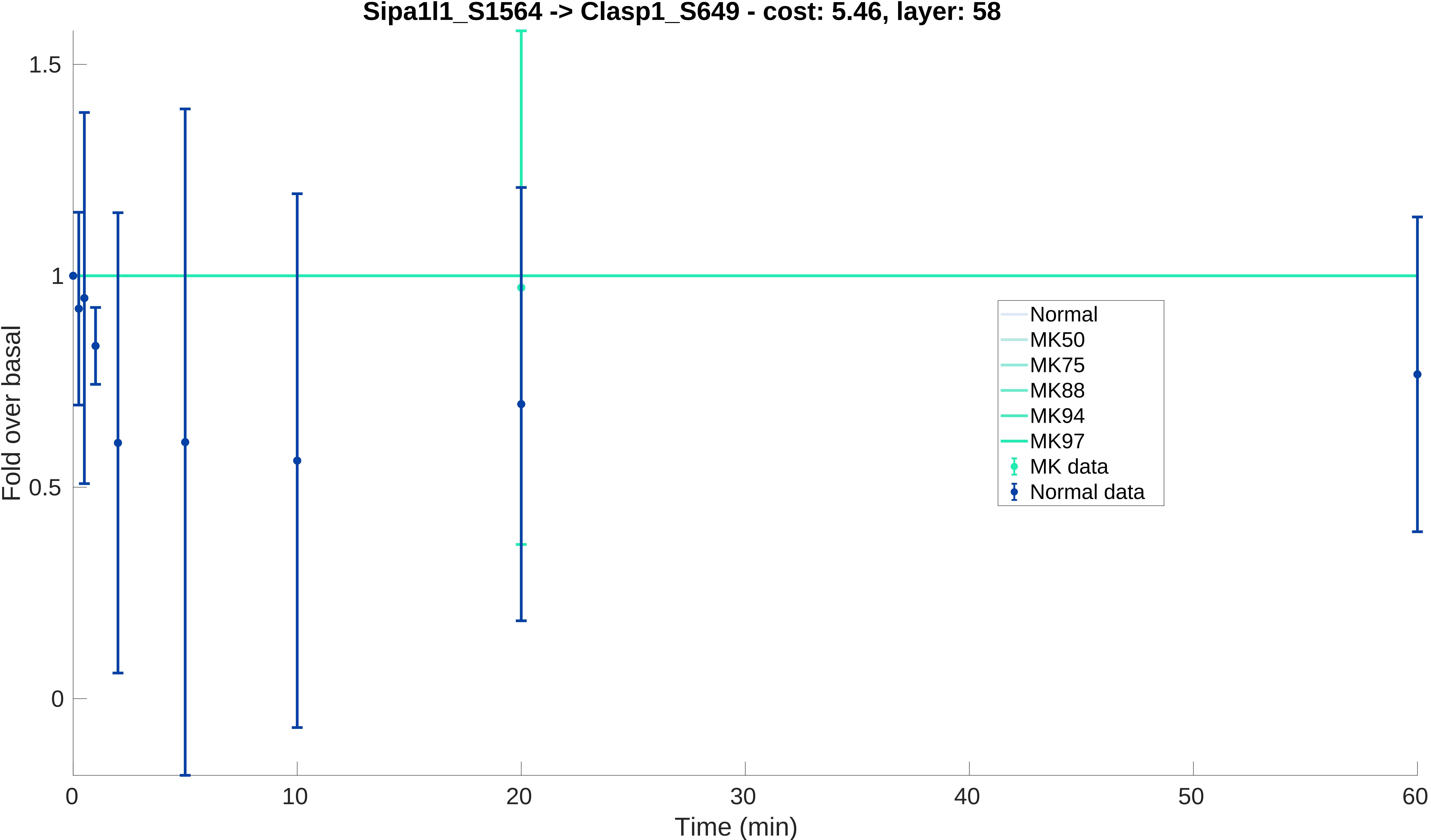

Sipa1l1_S1564 -> Clasp1_S649 - cost: 5.46, layer: 58
Fold over basal
1.5
1
0.5
0
Time (min)
0
10
20
30
40
50
60
Normal
MK50
MK75
MK88
MK94
MK97
MK data
Normal data

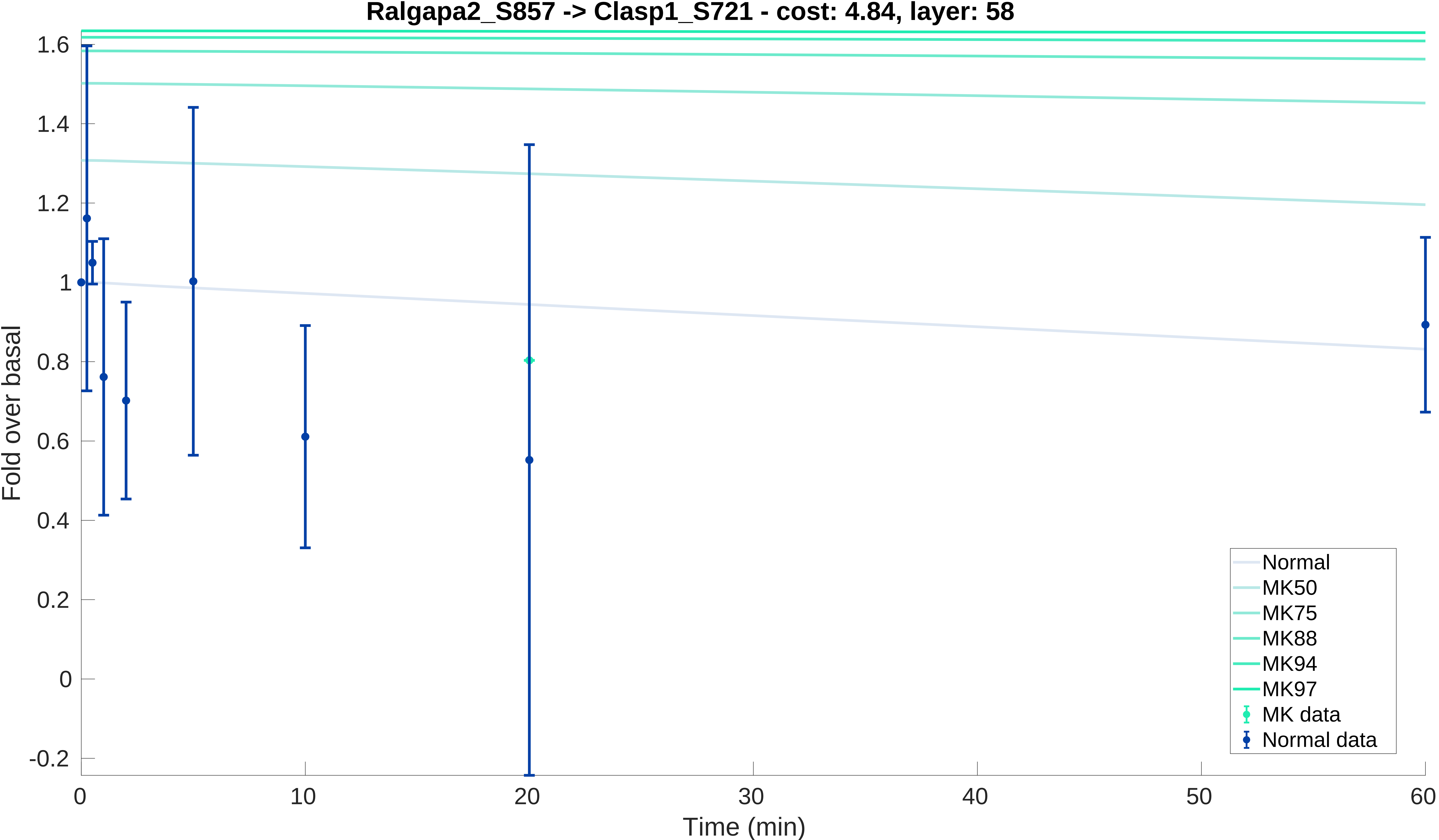

Ralgapa2_S857 -> Clasp1_S721 - cost: 4.84, layer: 58
Fold over basal
1.6
1.4
1.2
1
0.8
0.6
0.4
0.2
0
-0.2
0
10
20
30
40
50
60
Time (min)
Normal
MK50
MK75
MK88
MK94
MK97
MK data
Normal data

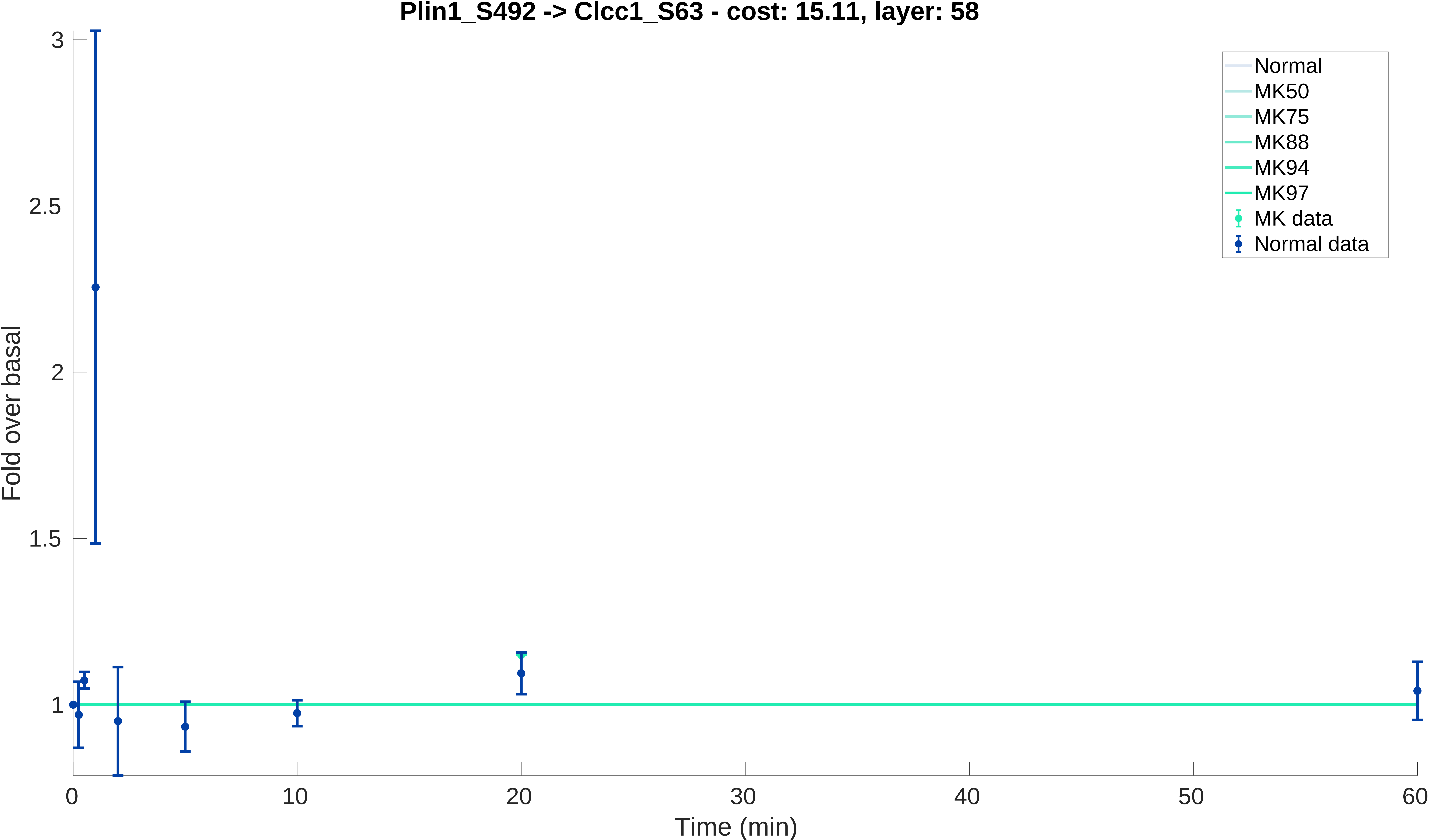
Plin1_S492 -> Clcc1_S63 - cost: 15.11, layer: 58
Normal
MK50
MK75
MK88
MK94
MK97
MK data
Normal data
Fold over basal
3
2.5
2
1.5
1
0
10
20
30
40
50
60
Time (min)

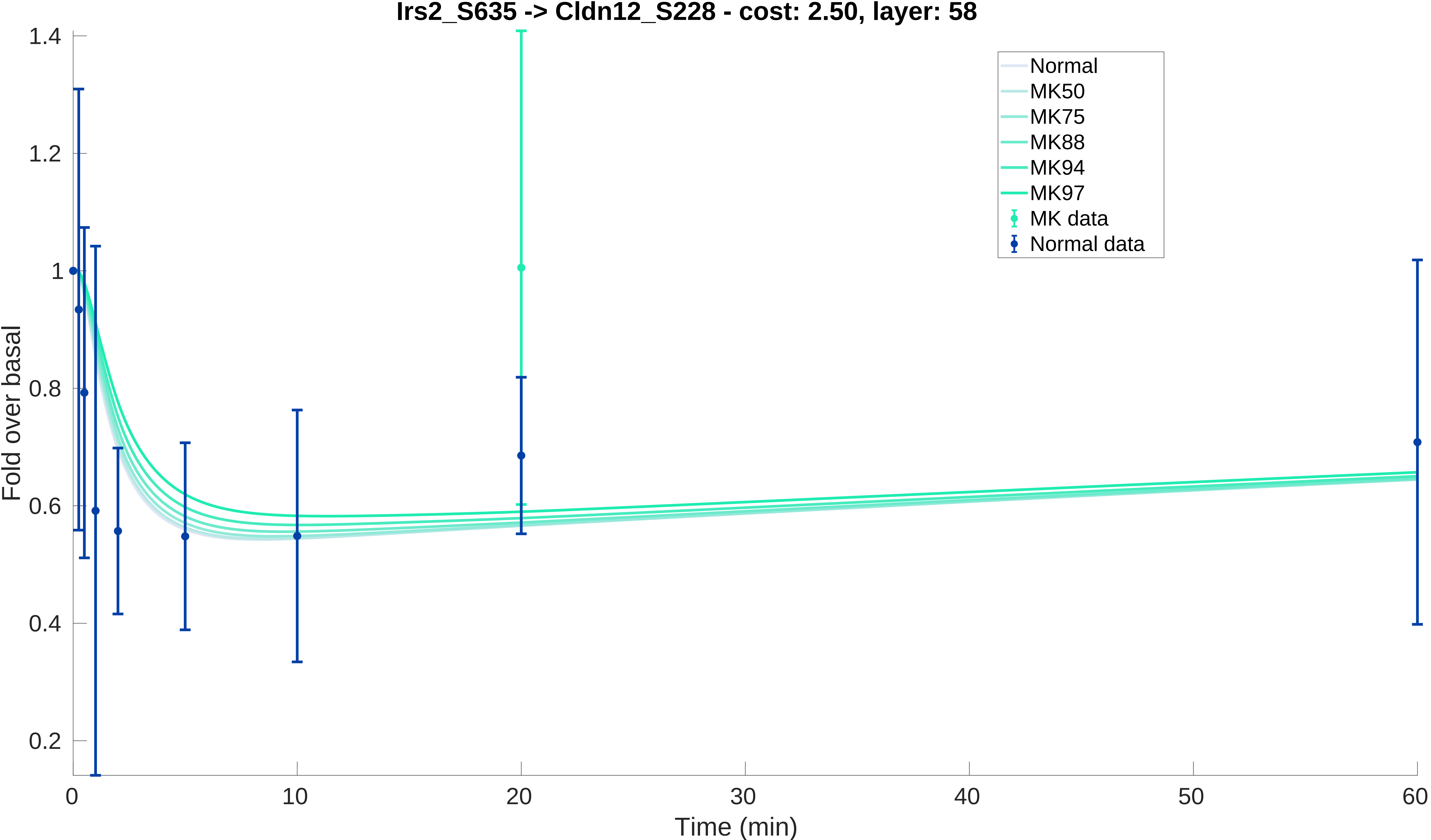

Irs2_S635 -> Cldn12_S228 - cost: 2.50, layer: 58
Fold over basal
1.4
1.2
1
0.8
0.6
0.4
0.2
0
10
20
30
40
50
60
Time (min)
Normal
MK50
MK75
MK88
MK94
MK97
MK data
Normal data

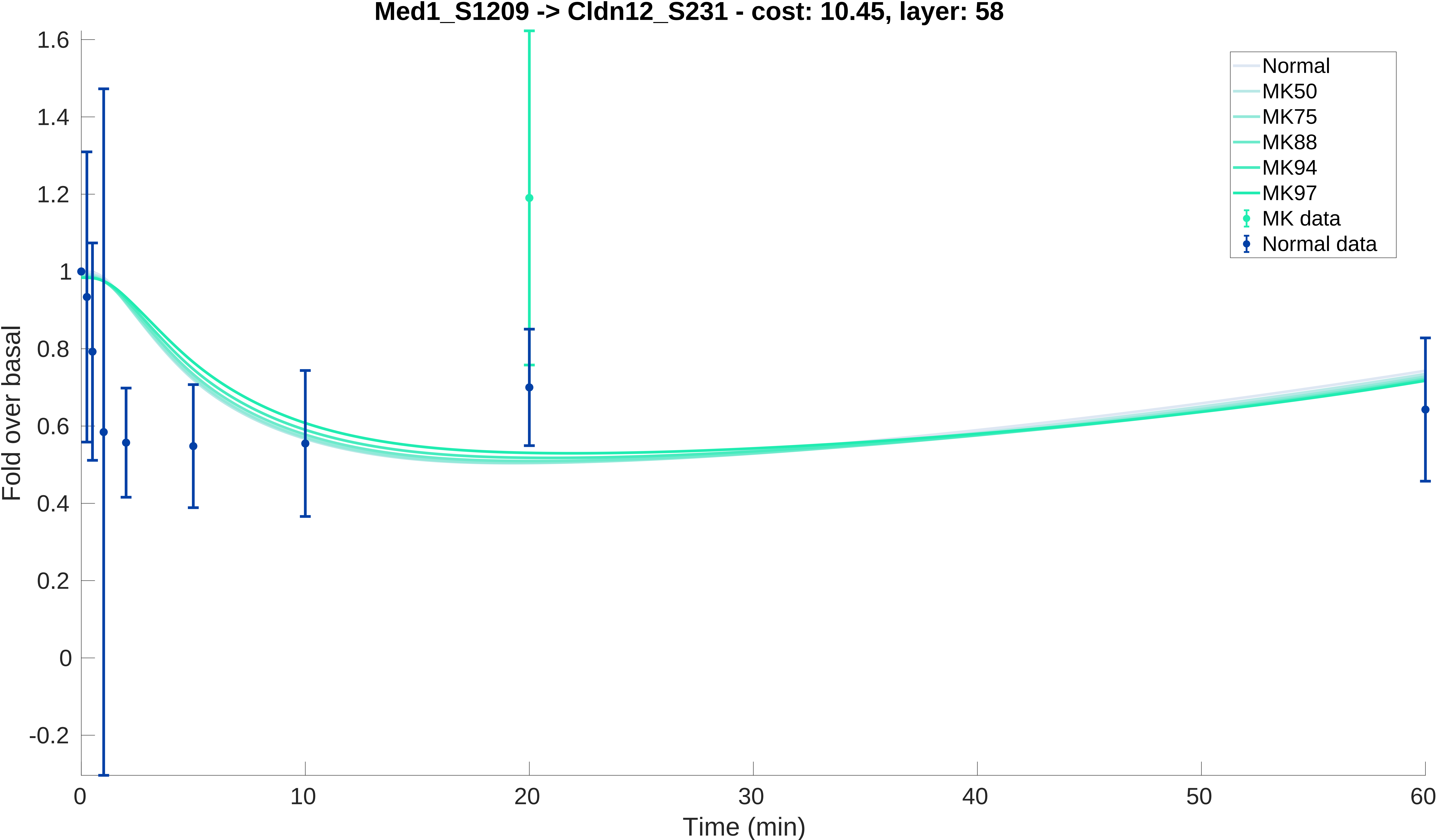
Med1_S1209 -> Cldn12_S231 - cost: 10.45, layer: 58
Fold over basal
Time (min)
1.6
1.4
1.2
1
0.8
0.6
0.4
0.2
0
-0.2
0
10
20
30
40
50
60
Normal
MK50
MK75
MK88
MK94
MK97
MK data
Normal data

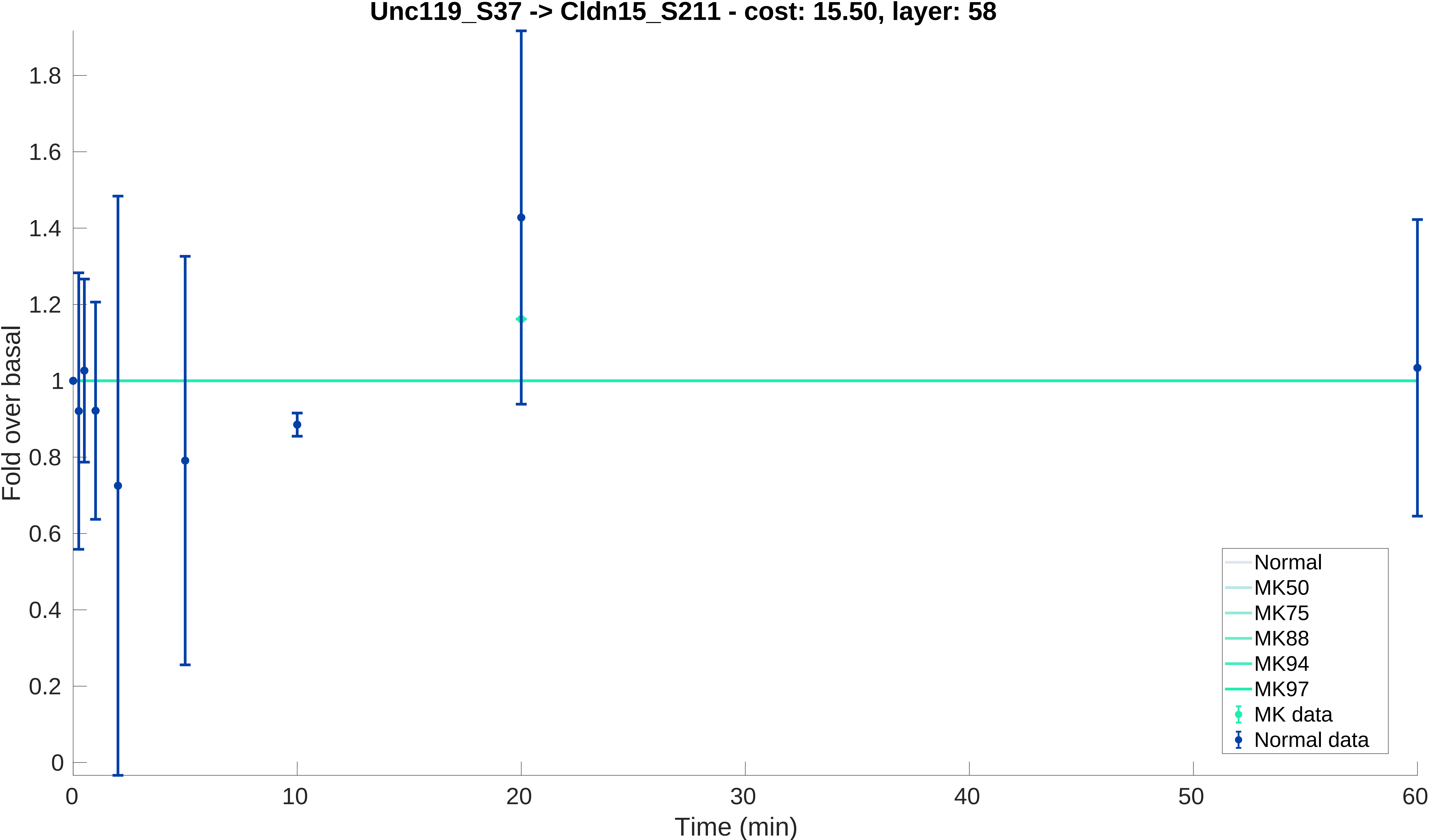

Unc119_S37 -> Cldn15_S211 - cost: 15.50, layer: 58
Fold over basal
Time (min)
0
0.2
0.4
0.6
0.8
1
1.2
1.4
1.6
1.8
0
10
20
30
40
50
60
Normal
MK50
MK75
MK88
MK94
MK97
MK data
Normal data

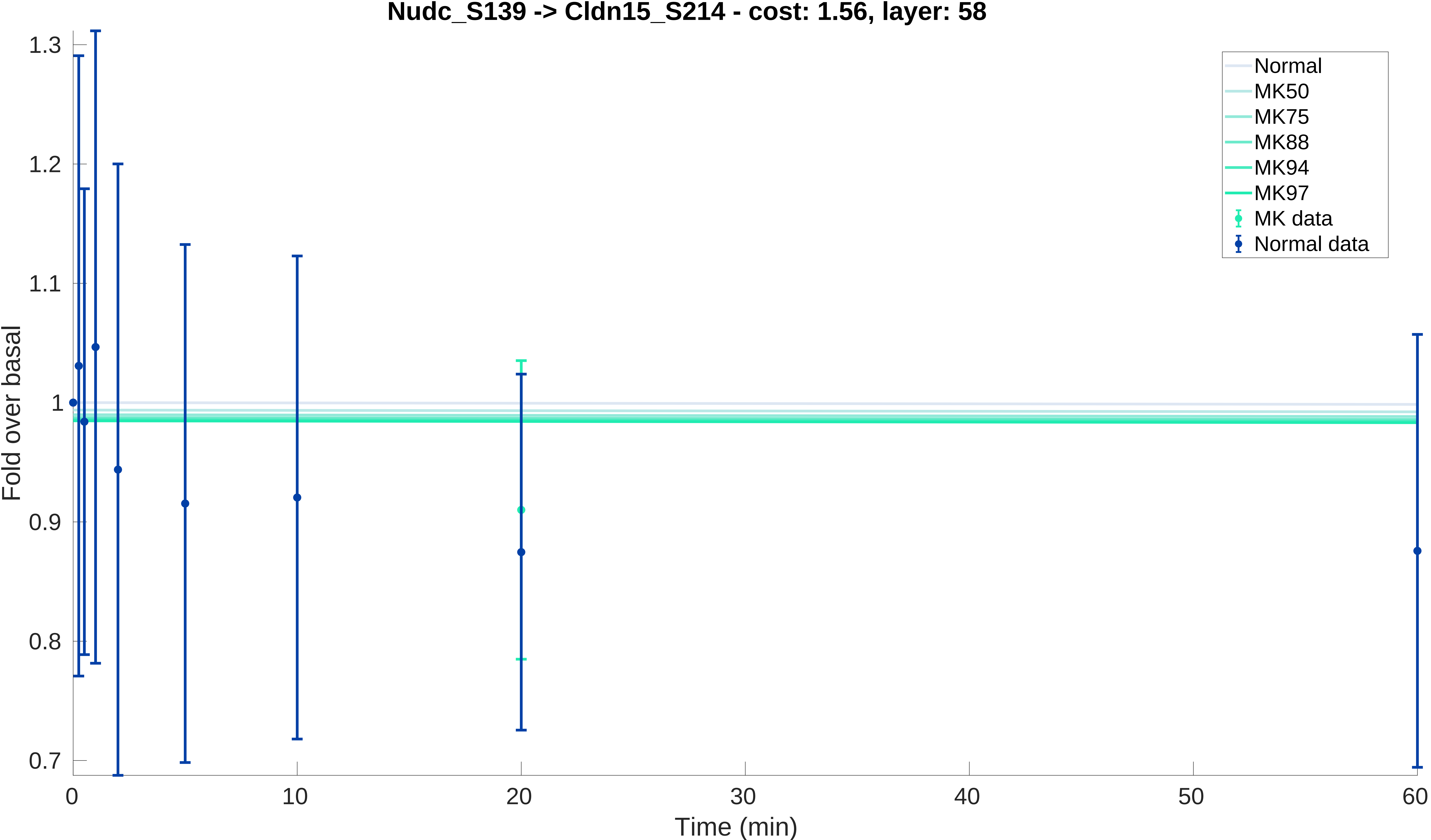
Nudc_S139 -> Cldn15_S214 - cost: 1.56, layer: 58
Fold over basal
Time (min)
1.3
1.2
1.1
1
0.9
0.8
0.7
0
10
20
30
40
50
60
Normal
MK50
MK75
MK88
MK94
MK97
MK data
Normal data

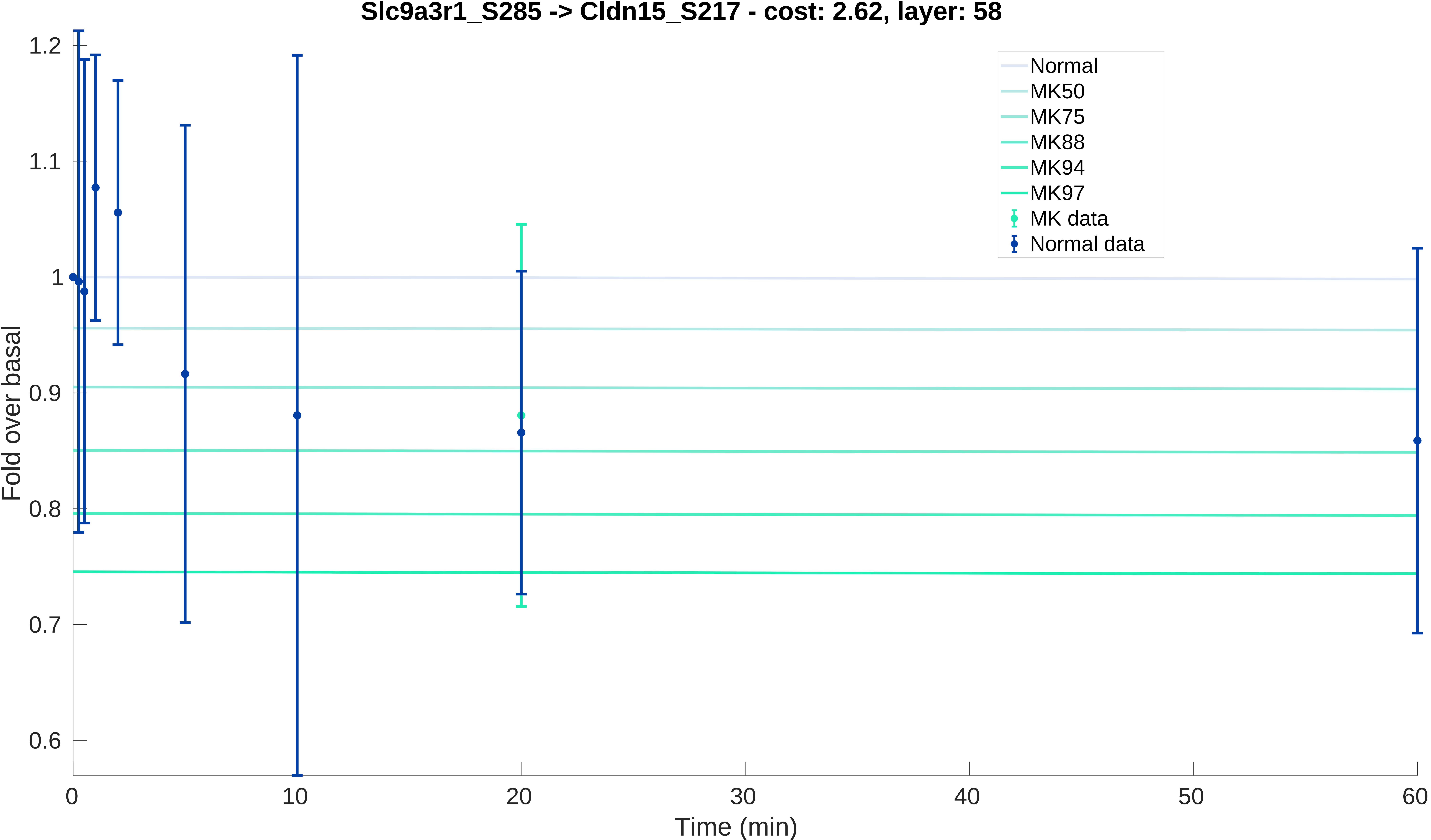
Slc9a3r1_S285 -> Cldn15_S217 - cost: 2.62, layer: 58
Fold over basal
Time (min)
1.2
1.1
1
0.9
0.8
0.7
0.6
0
10
20
30
40
50
60
Normal
MK50
MK75
MK88
MK94
MK97
MK data
Normal data

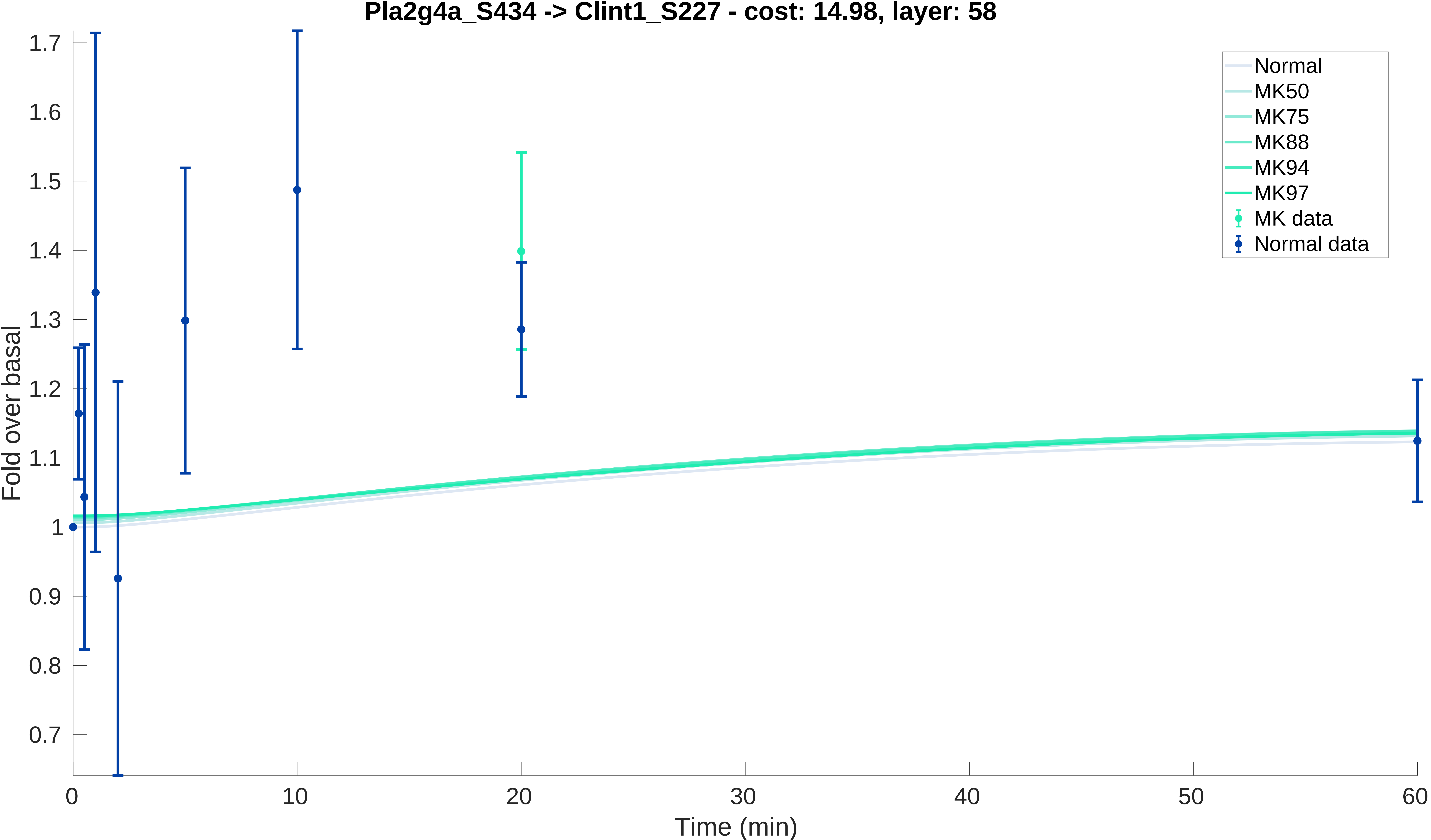

Pla2g4a_S434 -> Clint1_S227 - cost: 14.98, layer: 58
Normal
MK50
MK75
MK88
MK94
MK97
MK data
Normal data
Fold over basal
Time (min)
1.7
1.6
1.5
1.4
1.3
1.2
1.1
1
0.9
0.8
0.7
0
10
20
30
40
50
60

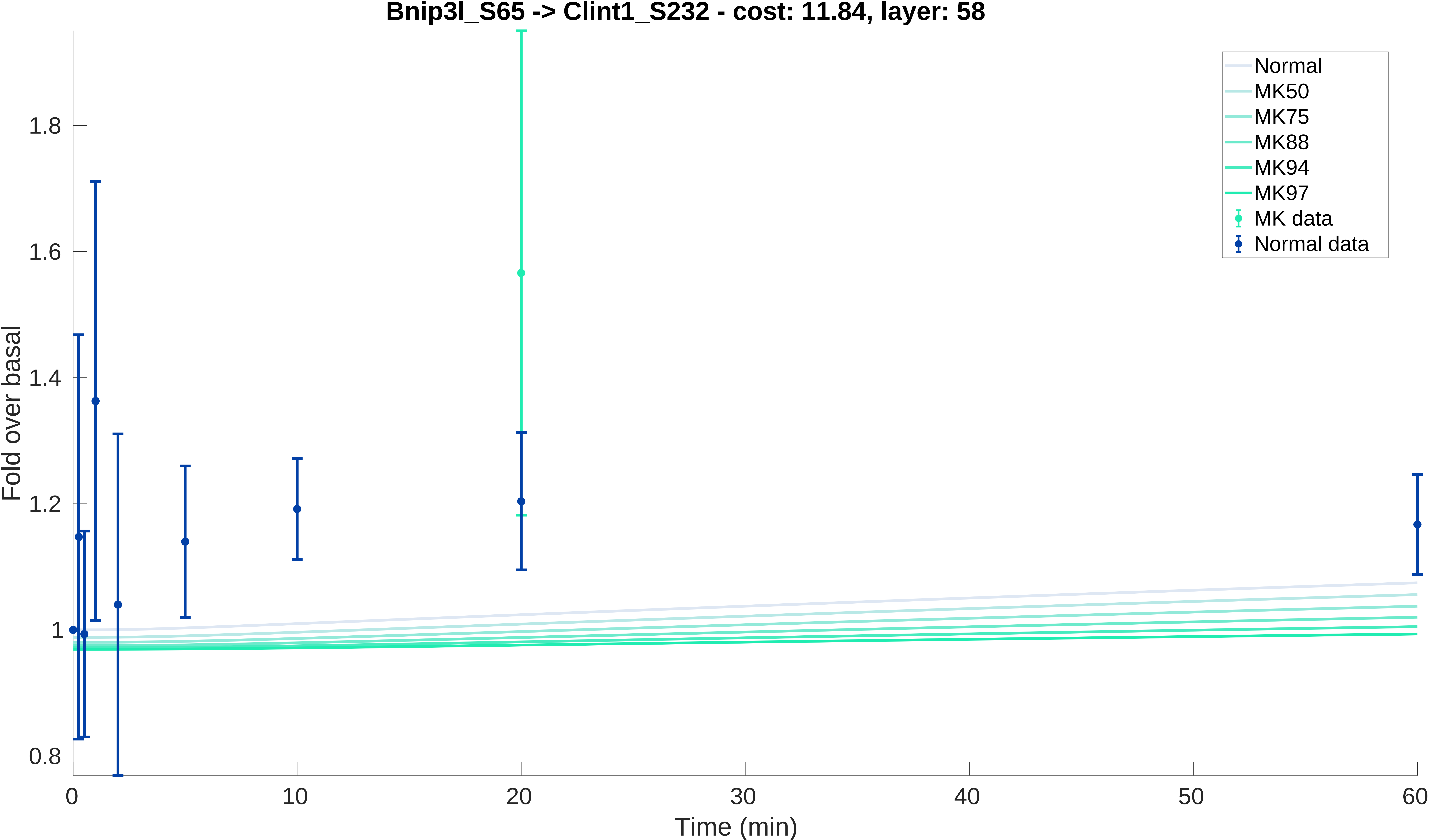

Bnip3l_S65 -> Clint1_S232 - cost: 11.84, layer: 58
Normal
MK50
MK75
MK88
MK94
MK97
MK data
Normal data
Fold over basal
0.8
1
1.2
1.4
1.6
1.8
0
10
20
30
40
50
60
Time (min)

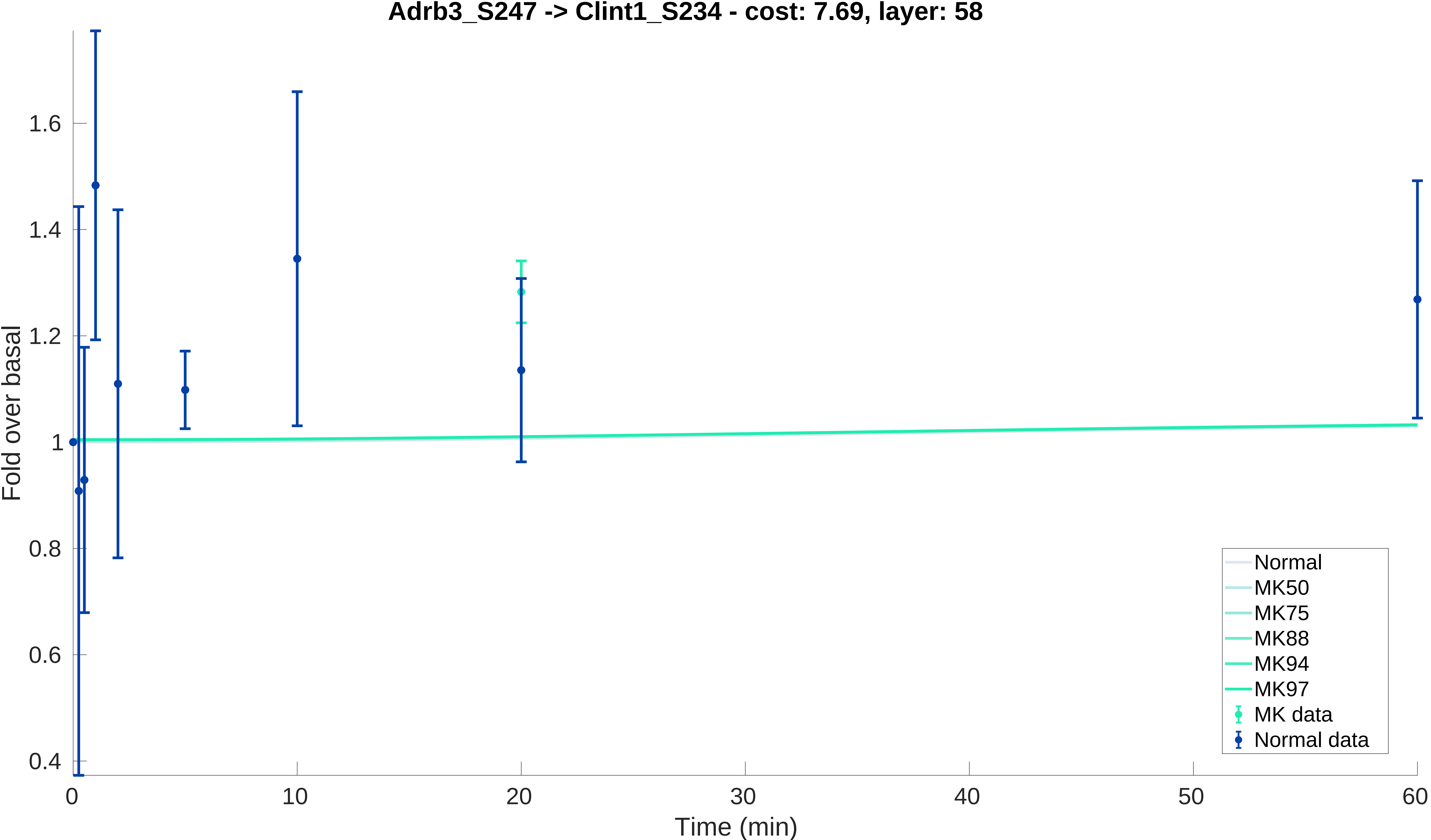

Adrb3_S247 -> Clint1_S234 - cost: 7.69, layer: 58
Fold over basal
Time (min)
0.4
0.6
0.8
1
1.2
1.4
1.6
0
10
20
30
40
50
60
Normal
MK50
MK75
MK88
MK94
MK97
MK data
Normal data

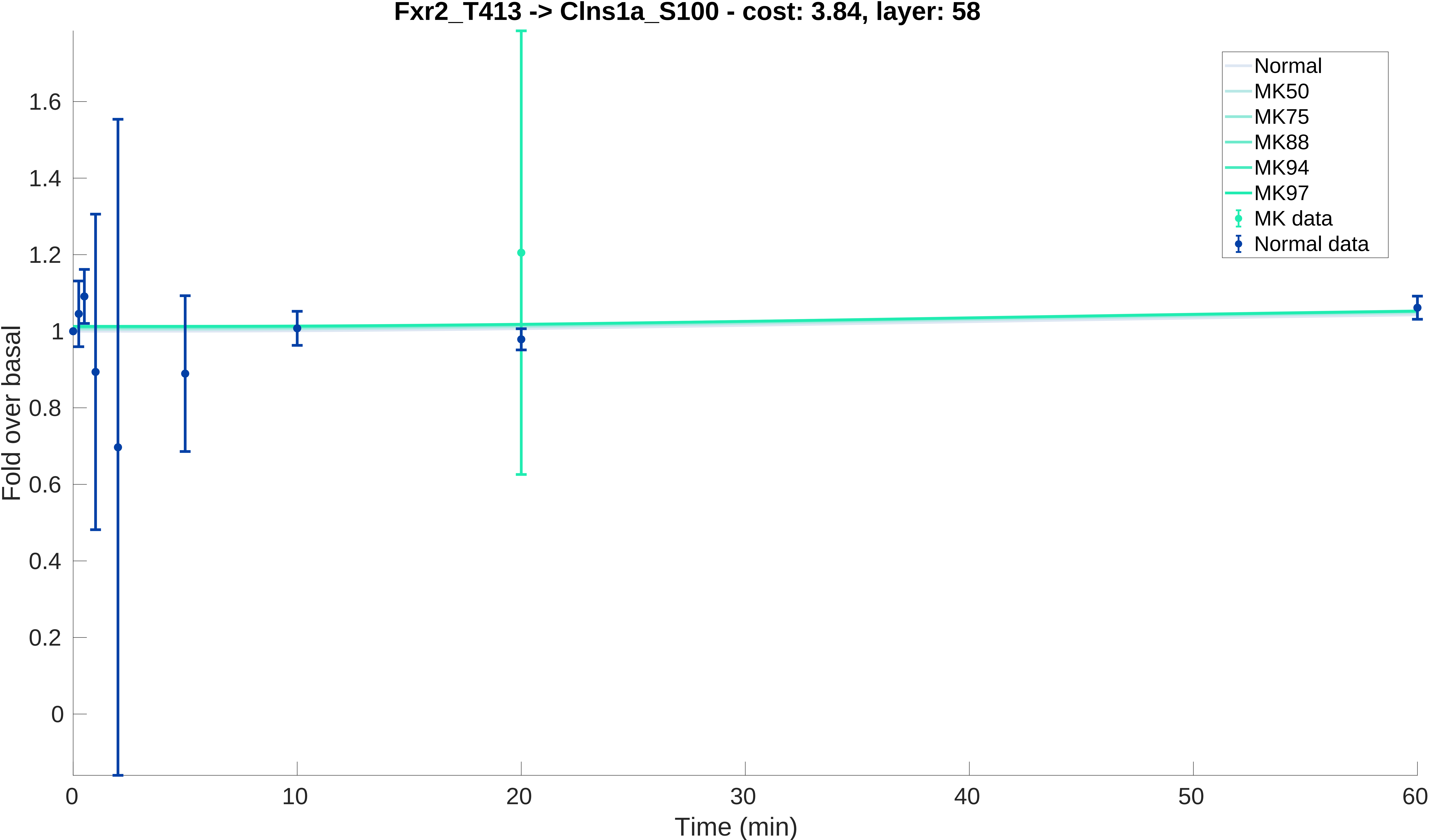

Fxr2_T413 -> Clns1a_S100 - cost: 3.84, layer: 58
Fold over basal
Time (min)
0
0.2
0.4
0.6
0.8
1
1.2
1.4
1.6
0
10
20
30
40
50
60
Normal
MK50
MK75
MK88
MK94
MK97
MK data
Normal data

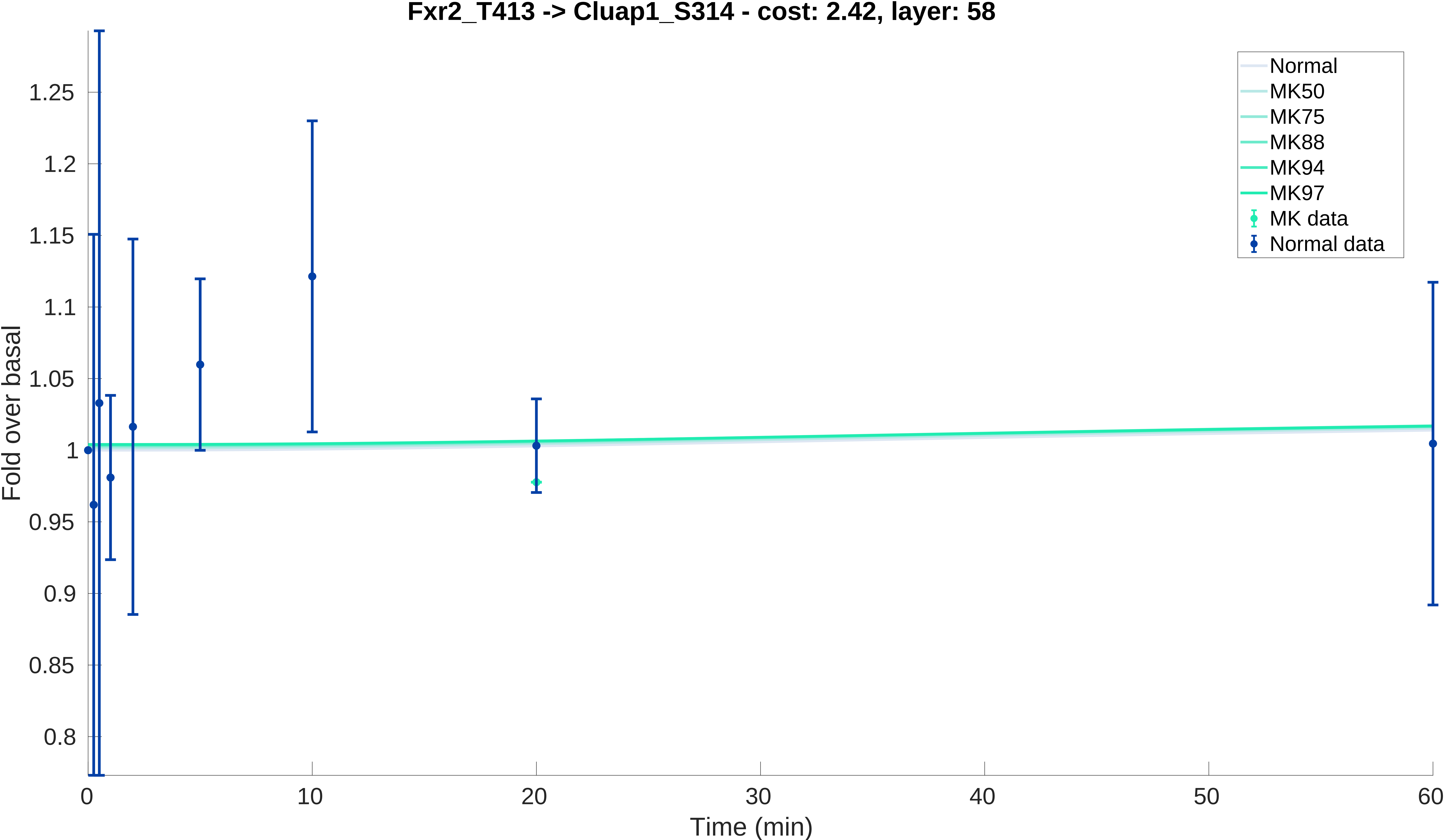
Fxr2_T413 -> Cluap1_S314 - cost: 2.42, layer: 58
Fold over basal
Time (min)
1.25
1.2
1.15
1.1
1.05
1
0.95
0.9
0.85
0.8
0
10
20
30
40
50
60
Normal
MK50
MK75
MK88
MK94
MK97
MK data
Normal data

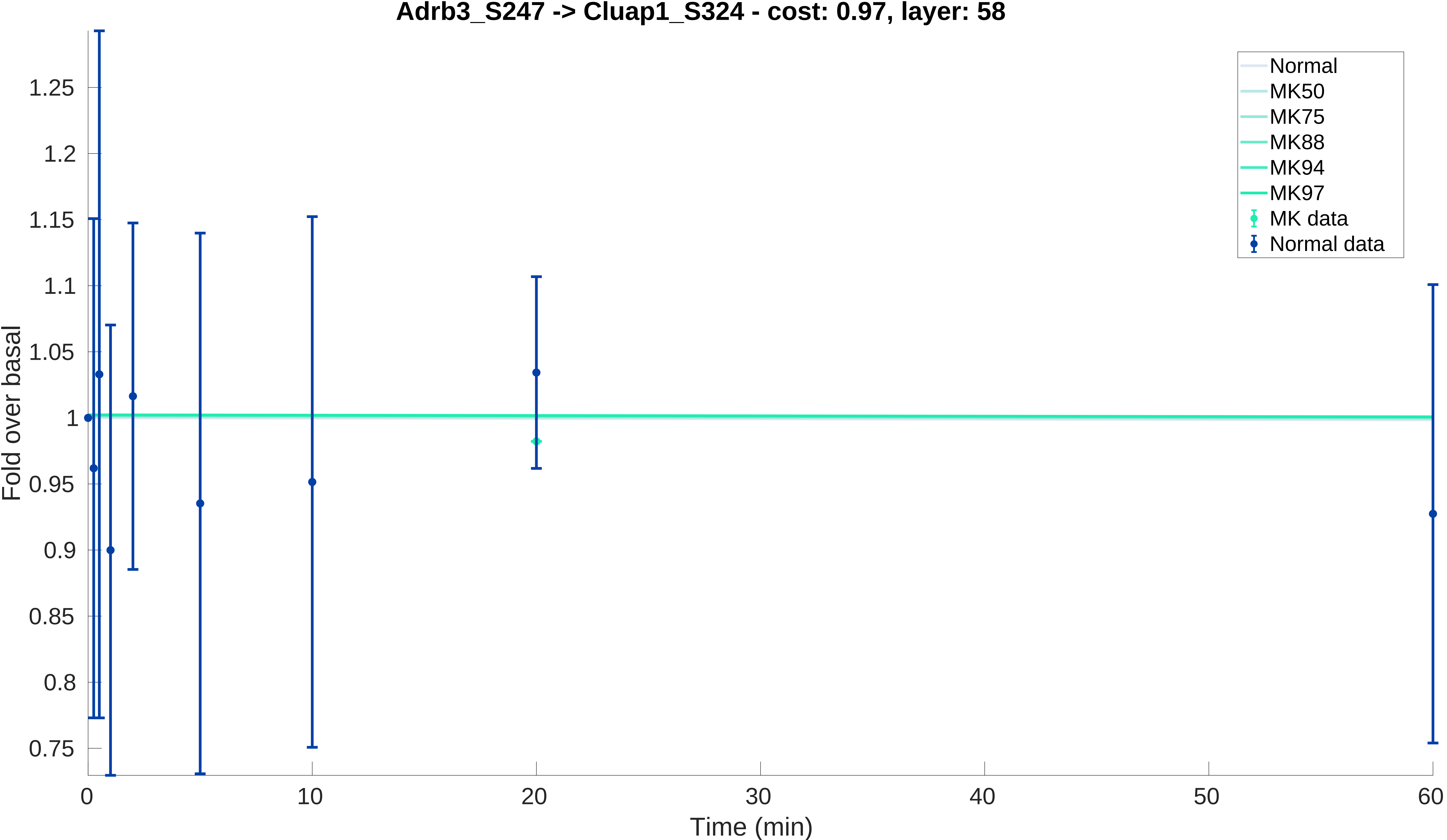

Adrb3_S247 -> Cluap1_S324 - cost: 0.97, layer: 58
Fold over basal
Time (min)
1.25
1.2
1.15
1.1
1.05
1
0.95
0.9
0.85
0.8
0.75
0
10
20
30
40
50
60
Normal
MK50
MK75
MK88
MK94
MK97
MK data
Normal data

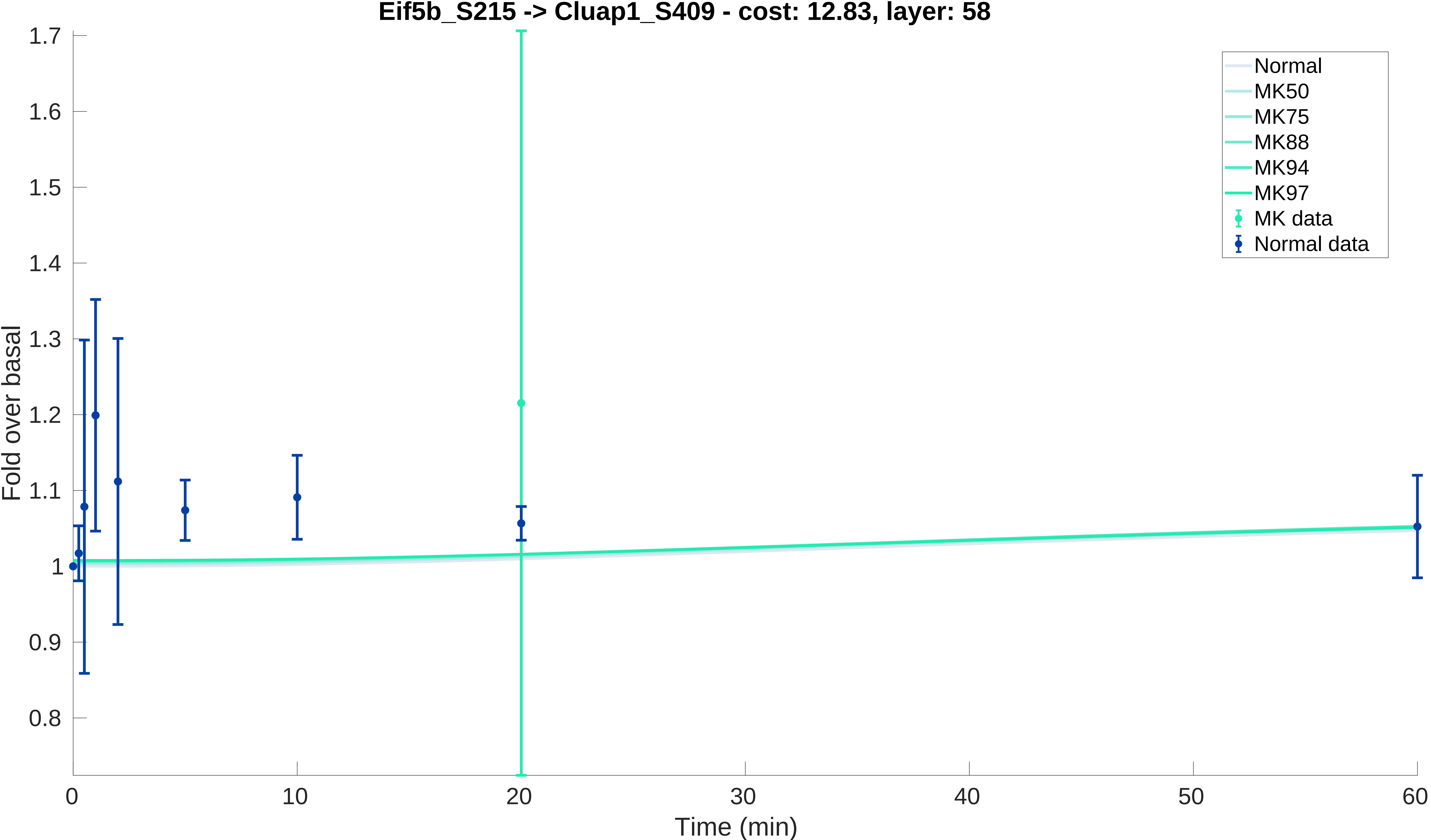

Eif5b_S215 -> Cluap1_S409 - cost: 12.83, layer: 58
Fold over basal
Time (min)
1.7
1.6
1.5
1.4
1.3
1.2
1.1
1
0.9
0.8
0
10
20
30
40
50
60
Normal
MK50
MK75
MK88
MK94
MK97
MK data
Normal data

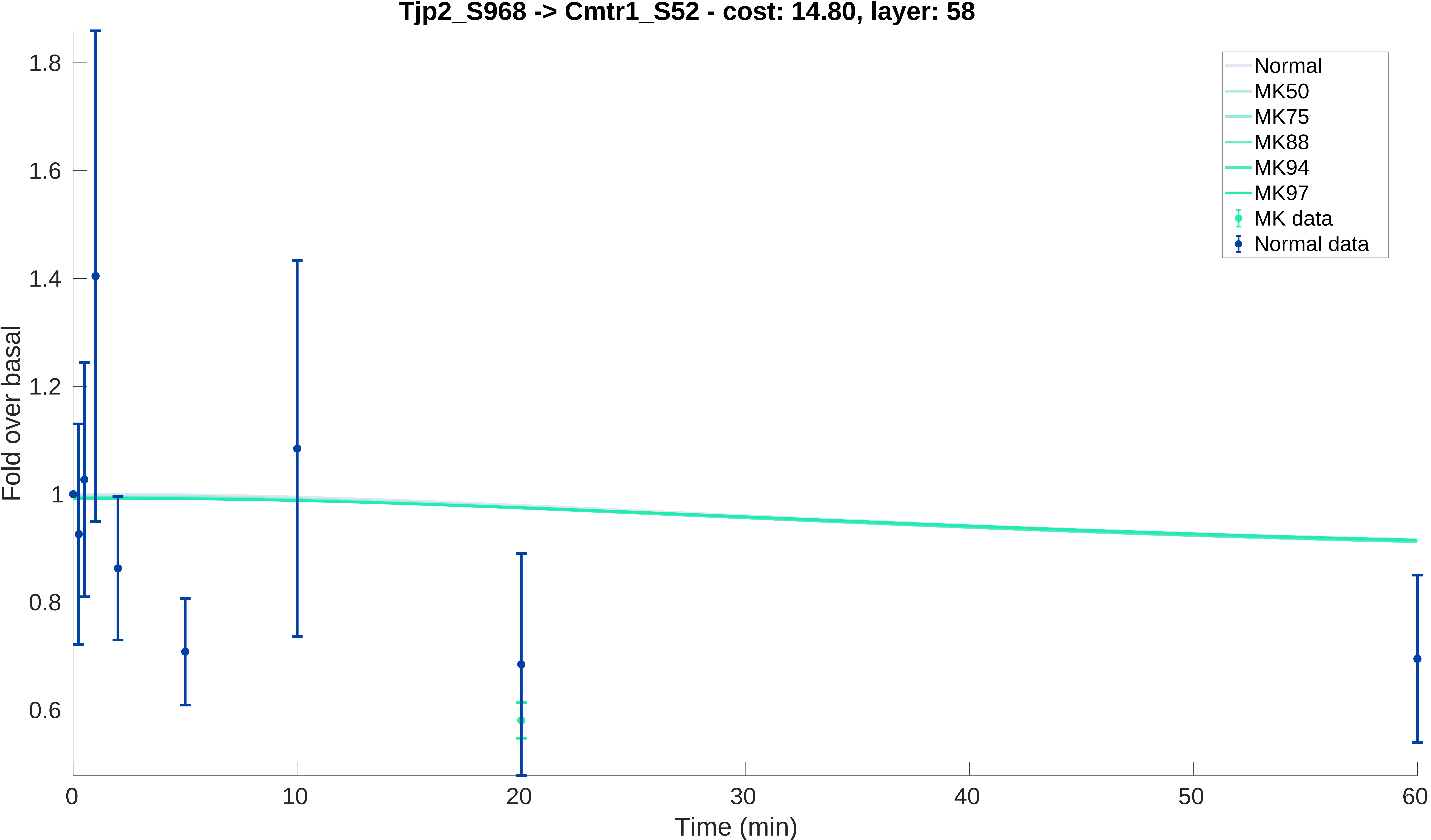
Tjp2_S968 -> Cmtr1_S52 - cost: 14.80, layer: 58
Fold over basal
Time (min)
1.8
1.6
1.4
1.2
1
0.8
0.6
0
10
20
30
40
50
60
Normal
MK50
MK75
MK88
MK94
MK97
MK data
Normal data

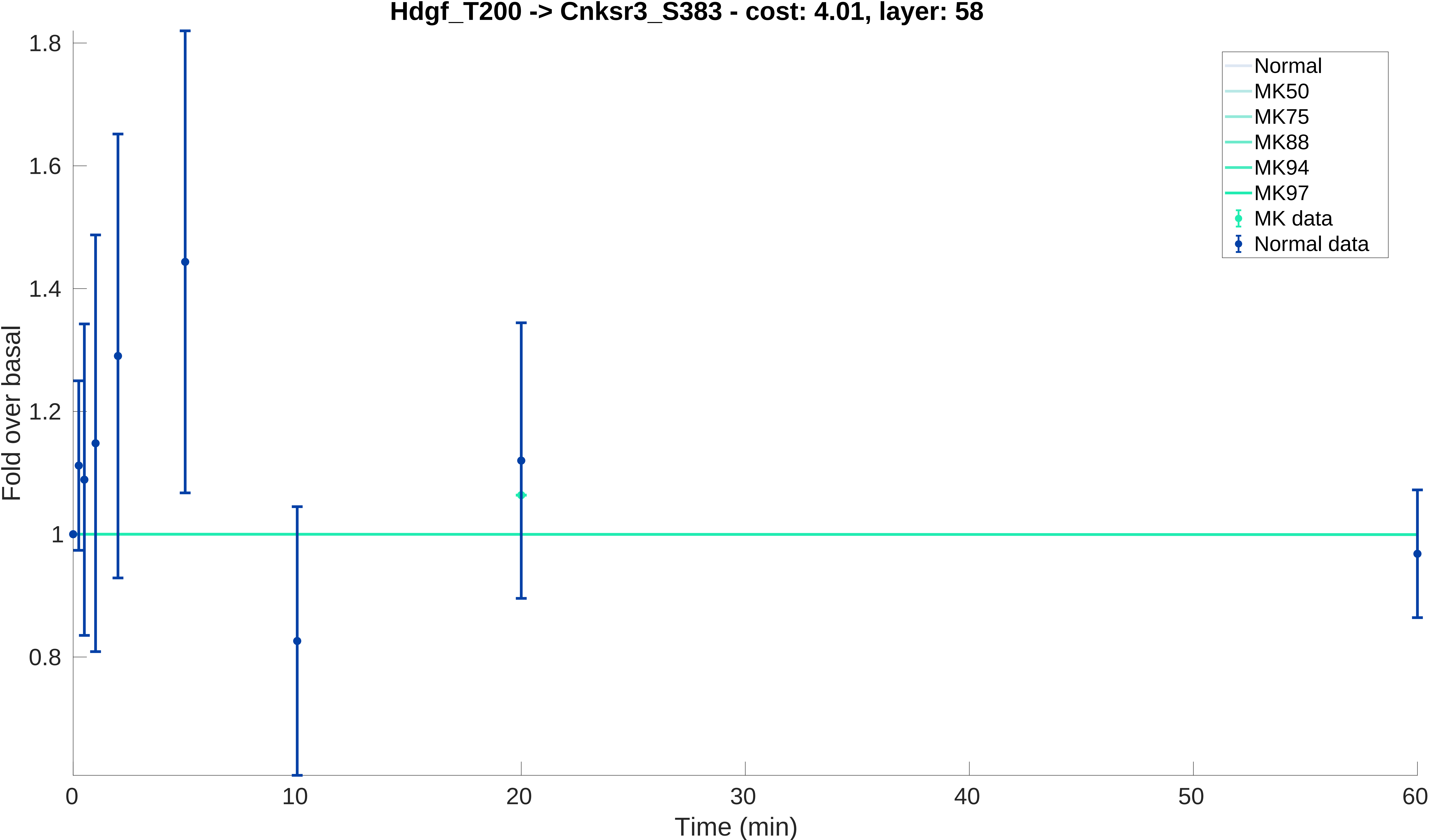

Hdgf_T200 -> Cnksr3_S383 - cost: 4.01, layer: 58
Fold over basal
1.8
1.6
1.4
1.2
1
0.8
0
10
20
30
40
50
60
Time (min)
Normal
MK50
MK75
MK88
MK94
MK97
MK data
Normal data

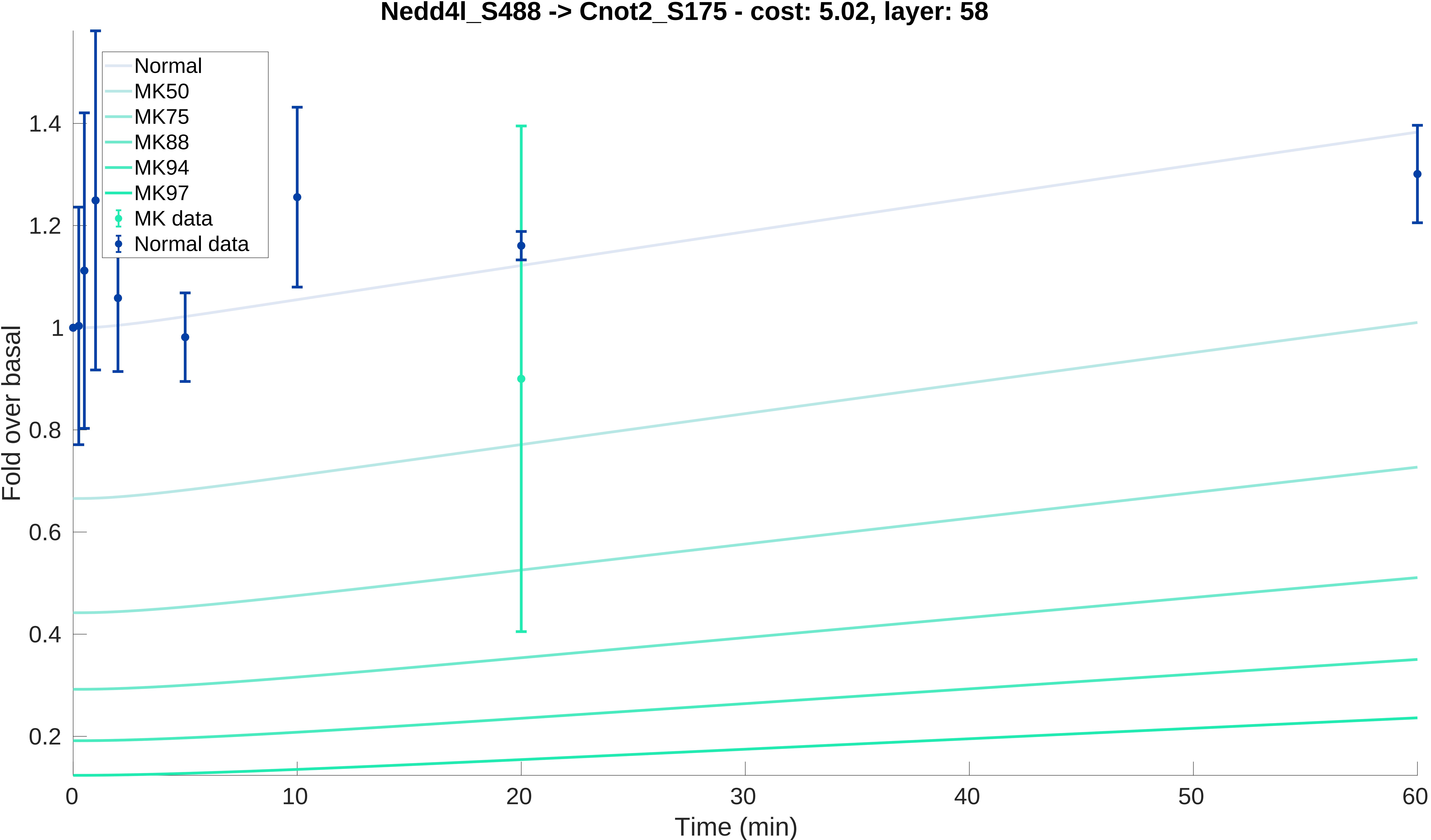
Nedd4l_S488 -> Cnot2_S175 - cost: 5.02, layer: 58
Normal
MK50
MK75
MK88
MK94
MK97
MK data
Normal data
Fold over basal
Time (min)
0.2
0.4
0.6
0.8
1
1.2
1.4
0
10
20
30
40
50
60

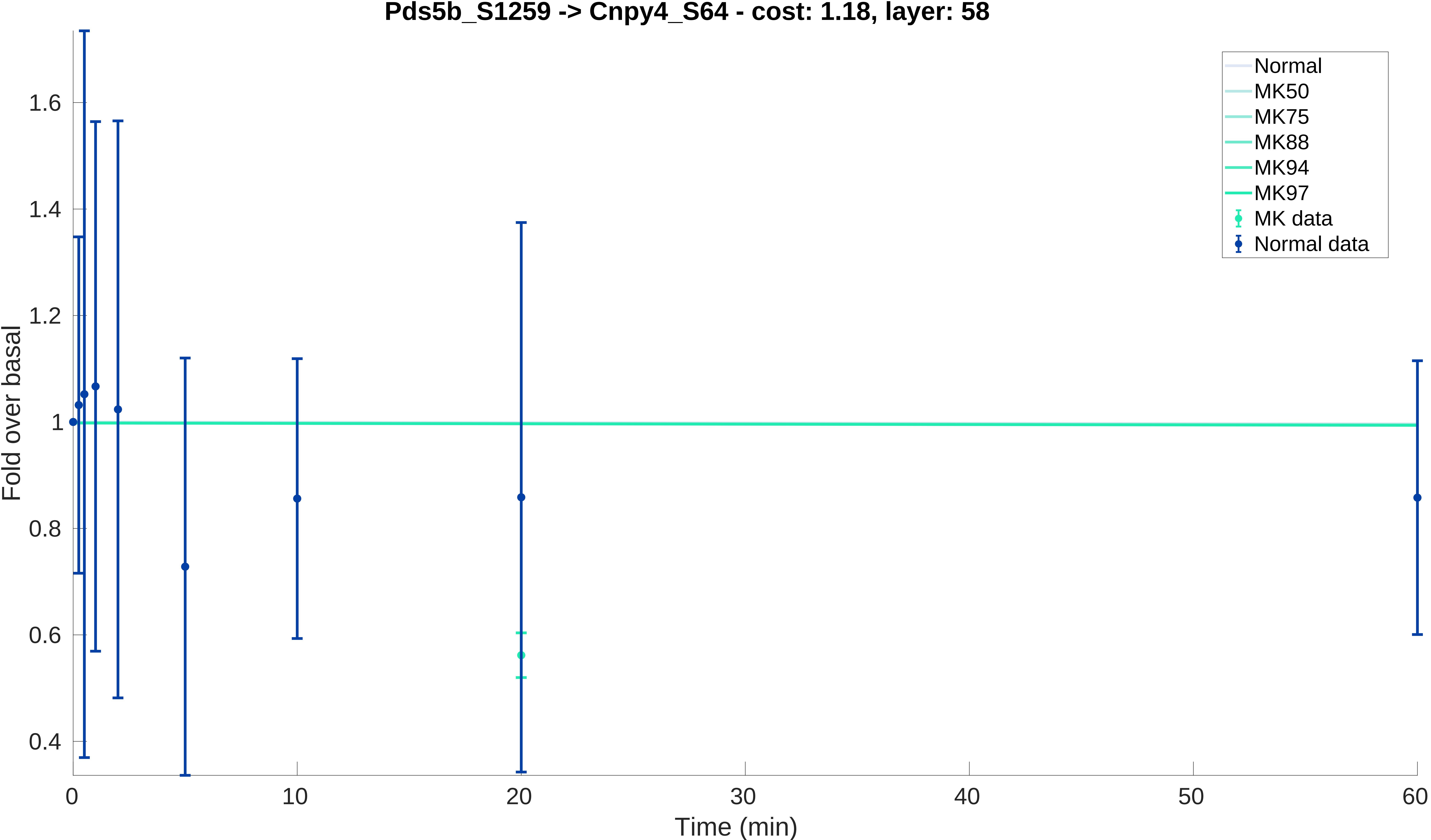

Pds5b_S1259 -> Cnpy4_S64 - cost: 1.18, layer: 58
Fold over basal
Time (min)
0
10
20
30
40
50
60
0.4
0.6
0.8
1
1.2
1.4
1.6
Normal
MK50
MK75
MK88
MK94
MK97
MK data
Normal data

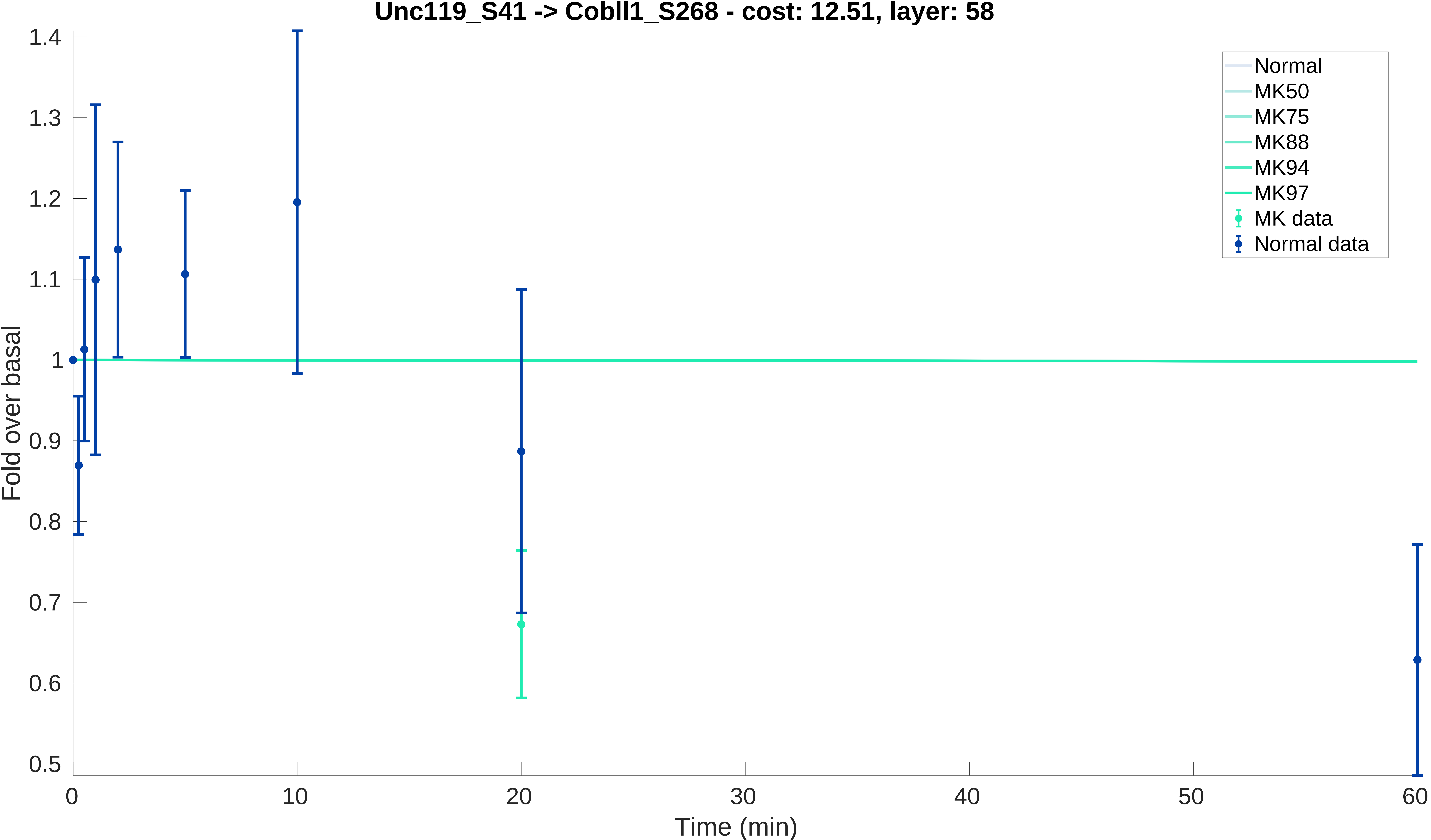

Unc119_S41 -> Cobll1_S268 - cost: 12.51, layer: 58
Fold over basal
Time (min)
Normal
MK50
MK75
MK88
MK94
MK97
MK data
Normal data
0
10
20
30
40
50
60
0.5
0.6
0.7
0.8
0.9
1
1.1
1.2
1.3
1.4

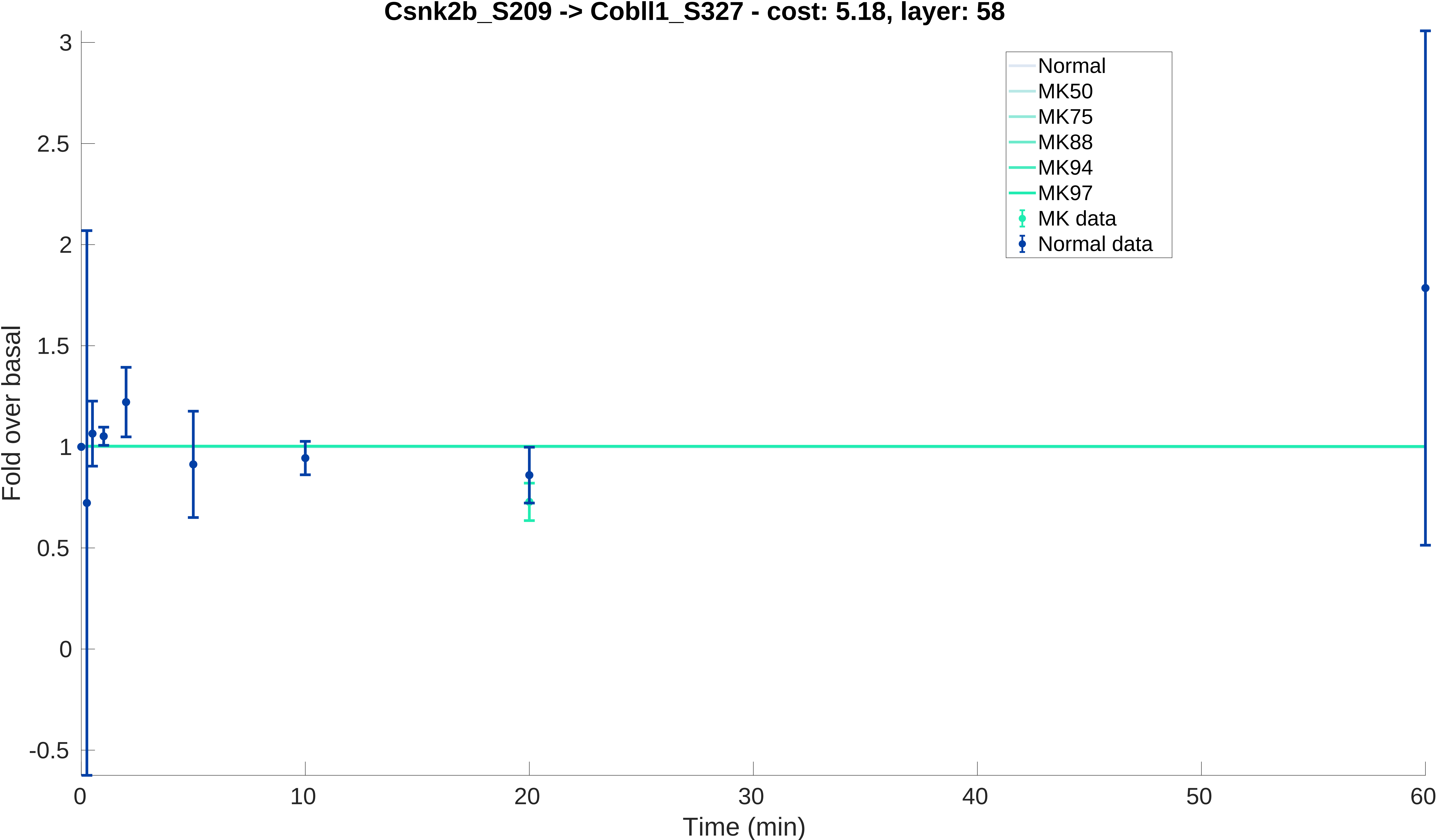

Csnk2b_S209 -> Cobll1_S327 - cost: 5.18, layer: 58
Normal
MK50
MK75
MK88
MK94
MK97
MK data
Normal data
Fold over basal
Time (min)
3
2.5
2
1.5
1
0.5
0
-0.5
0
10
20
30
40
50
60

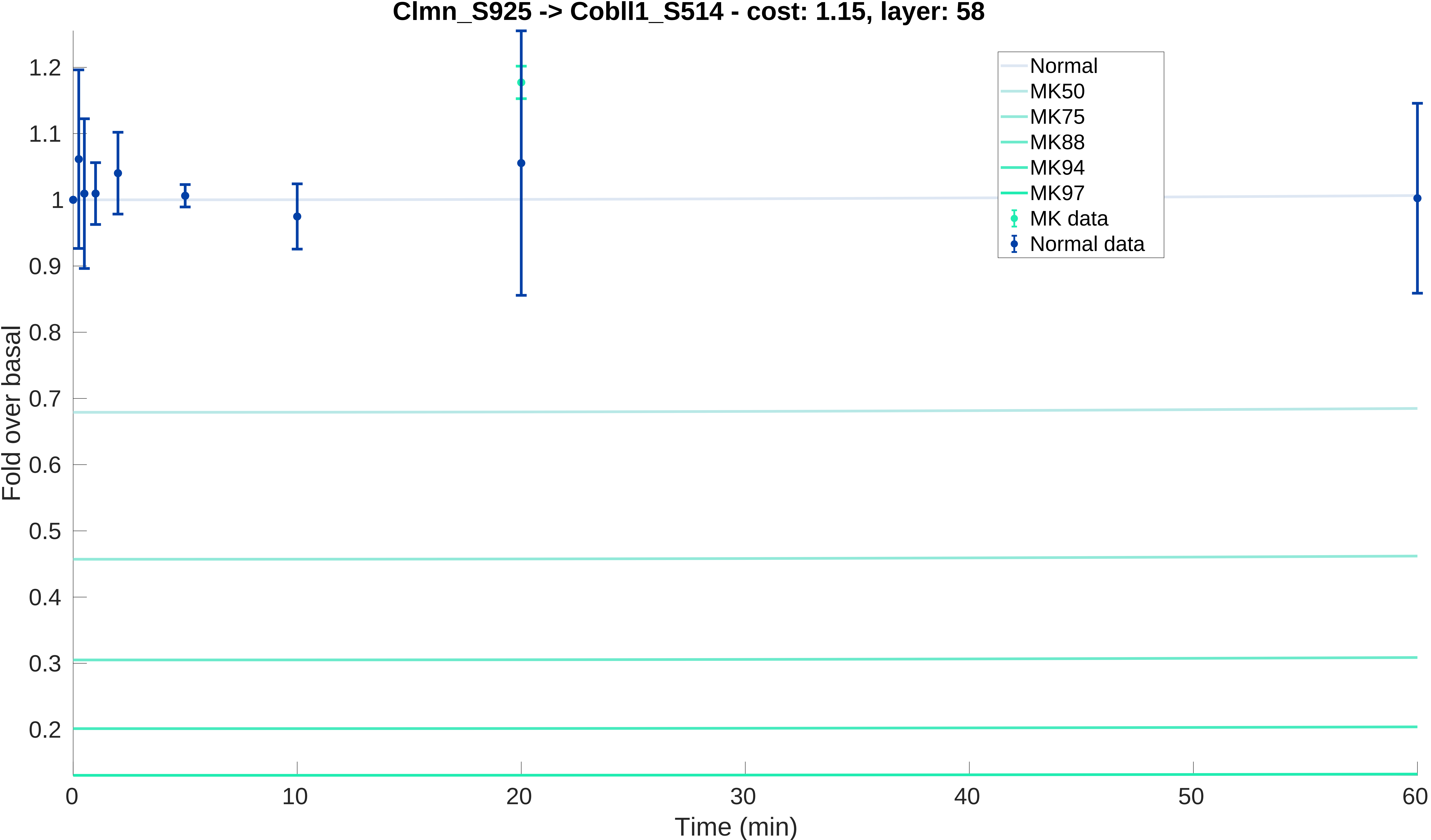

Clmn_S925 -> Cobll1_S514 - cost: 1.15, layer: 58
Fold over basal
Time (min)
0
10
20
30
40
50
60
0.2
0.3
0.4
0.5
0.6
0.7
0.8
0.9
1
1.1
1.2
Normal
MK50
MK75
MK88
MK94
MK97
MK data
Normal data

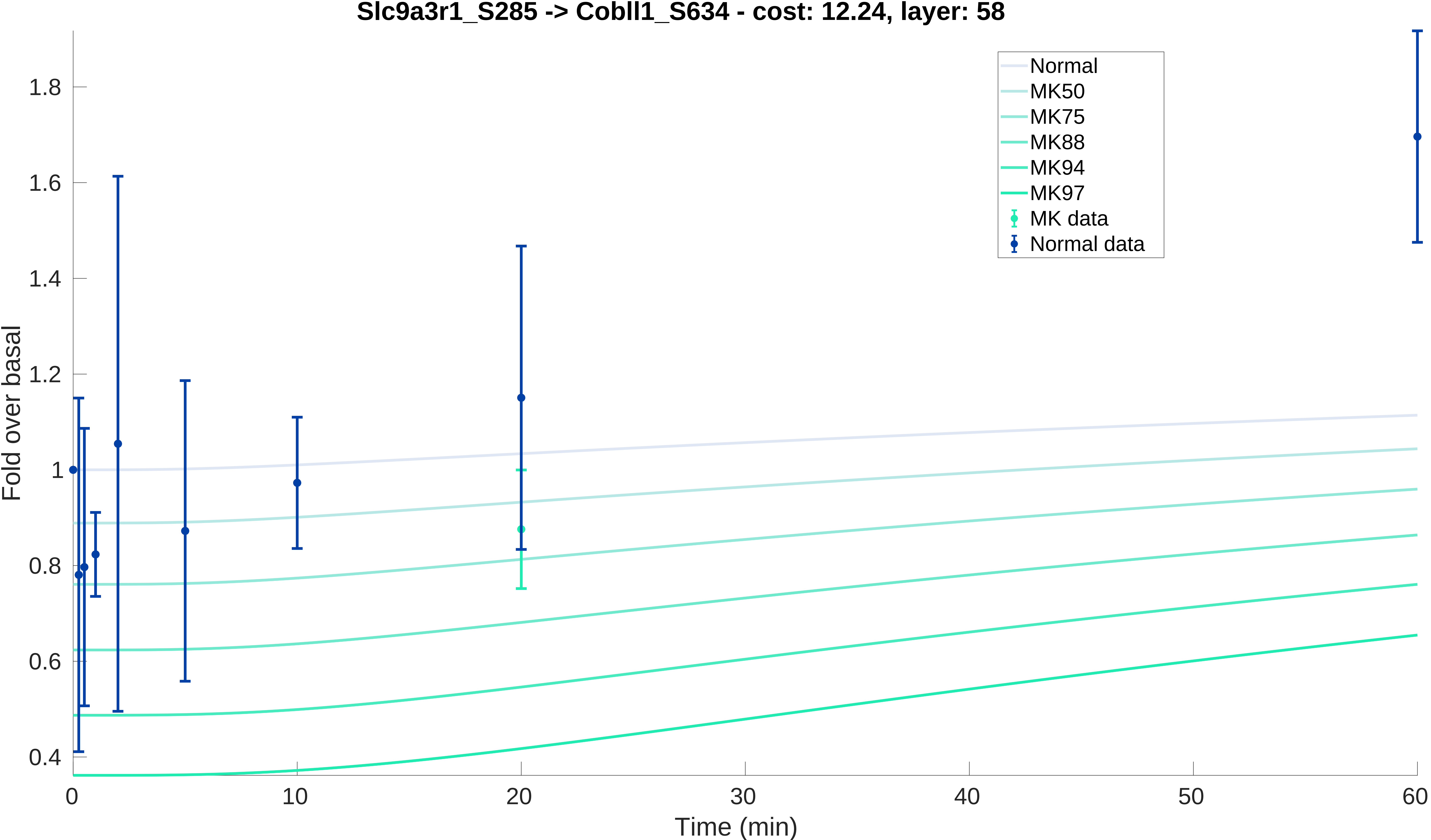

Slc9a3r1_S285 -> Cobll1_S634 - cost: 12.24, layer: 58
Fold over basal
Time (min)
0
10
20
30
40
50
60
0.4
0.6
0.8
1
1.2
1.4
1.6
1.8
Normal
MK50
MK75
MK88
MK94
MK97
MK data
Normal data

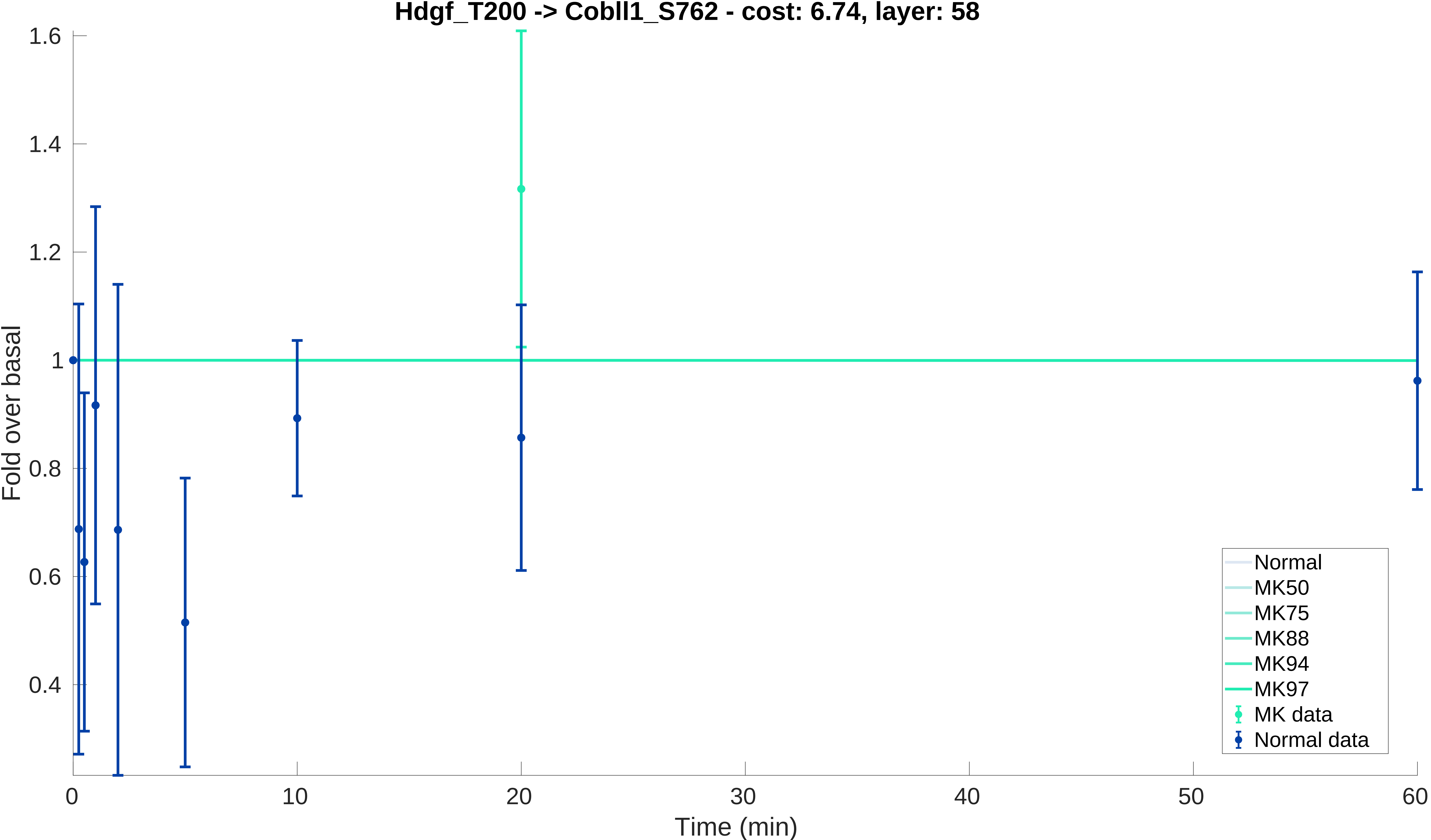

Hdgf_T200 -> Cobll1_S762 - cost: 6.74, layer: 58
Fold over basal
1.6
1.4
1.2
1
0.8
0.6
0.4
0
10
20
30
40
50
60
Time (min)
Normal
MK50
MK75
MK88
MK94
MK97
MK data
Normal data

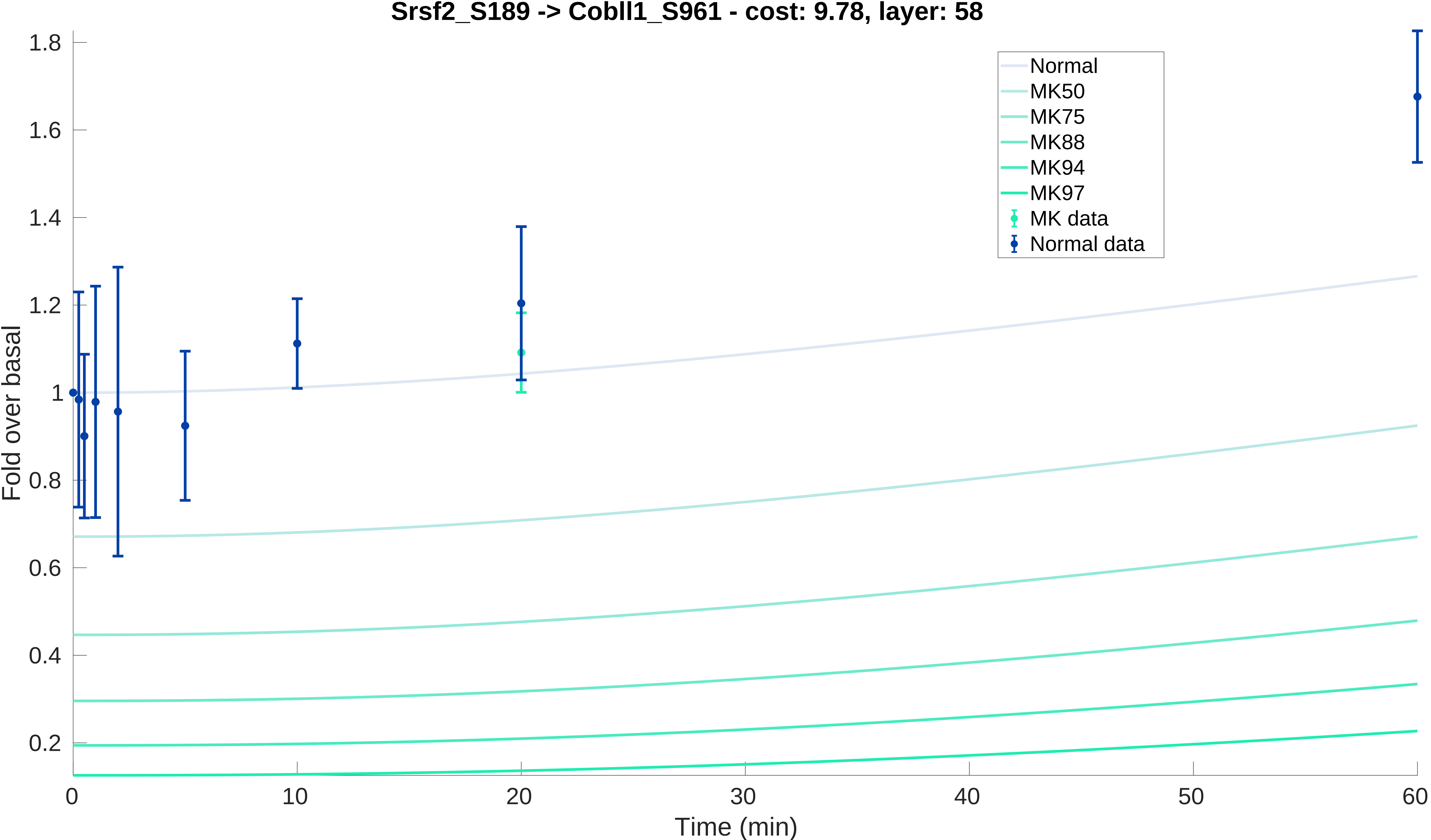

Srsf2_S189 -> Cobll1_S961 - cost: 9.78, layer: 58
Fold over basal
Time (min)
0.2
0.4
0.6
0.8
1
1.2
1.4
1.6
1.8
0
10
20
30
40
50
60
Normal
MK50
MK75
MK88
MK94
MK97
MK data
Normal data

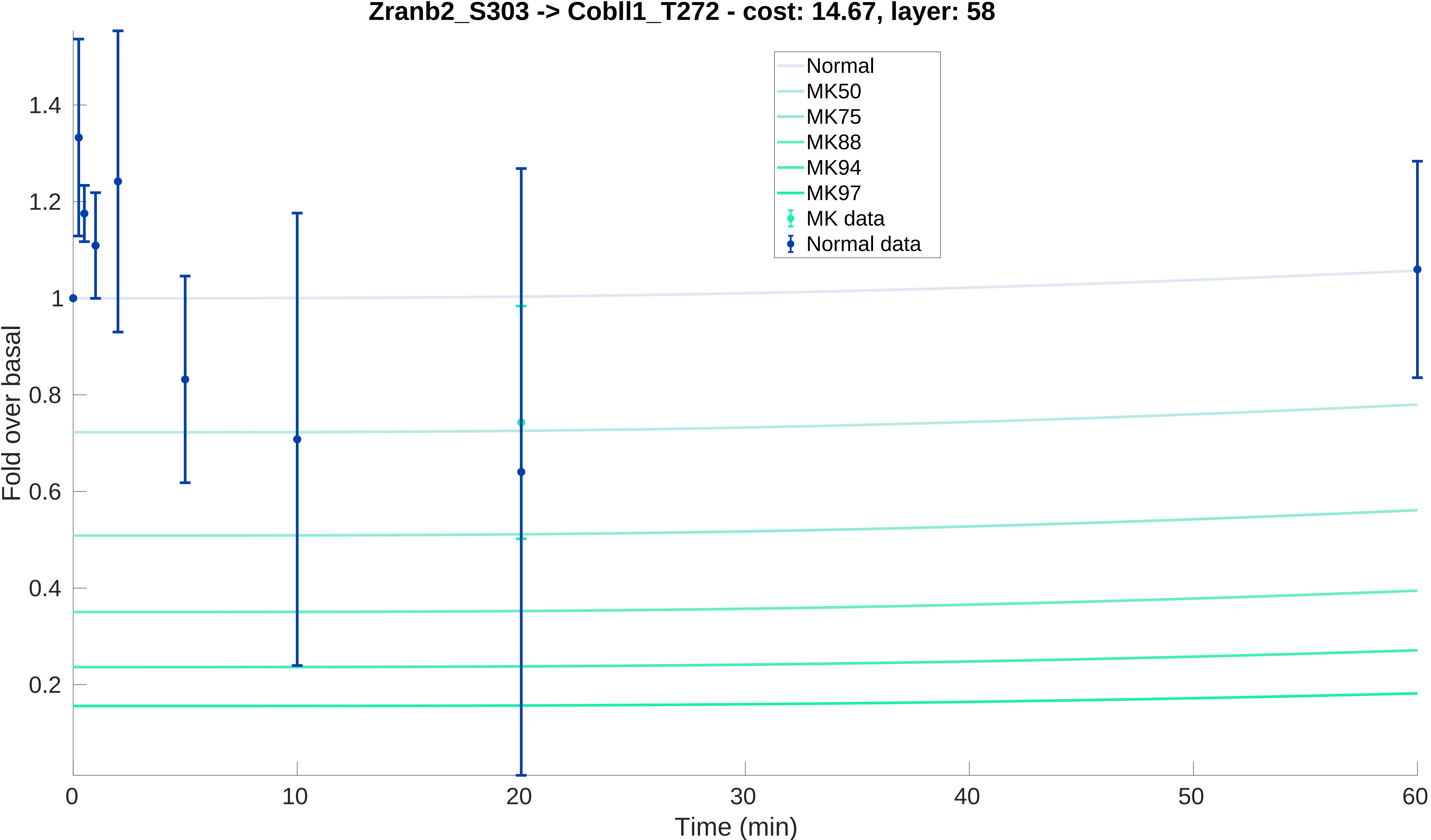

Zranb2_S303 -> Cobll1_T272 - cost: 14.67, layer: 58
Normal
MK50
MK75
MK88
MK94
MK97
MK data
Normal data
Fold over basal
Time (min)
0
10
20
30
40
50
60
0.2
0.4
0.6
0.8
1
1.2
1.4

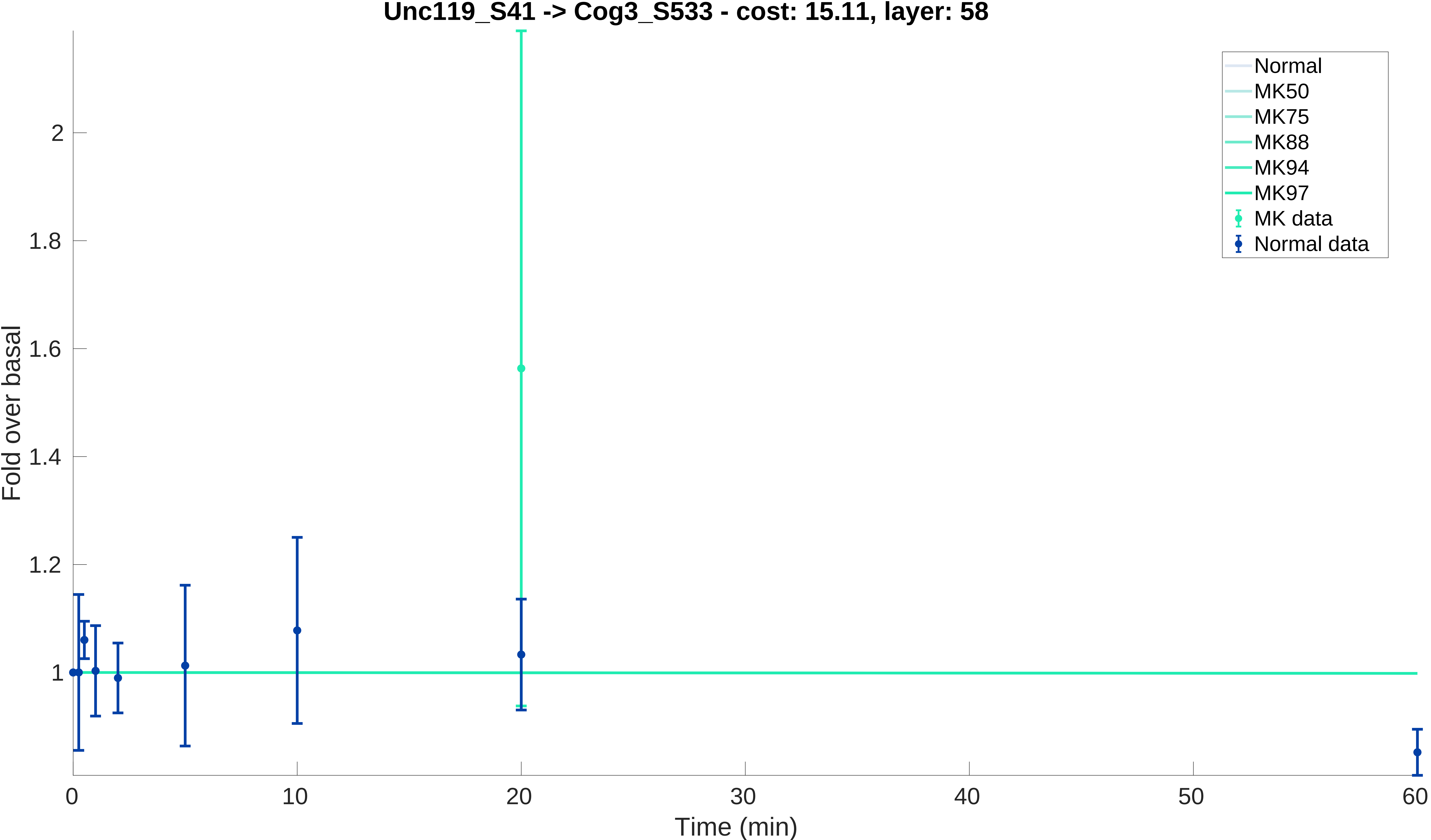
Unc119_S41 -> Cog3_S533 - cost: 15.11, layer: 58
Fold over basal
2
1.8
1.6
1.4
1.2
1
0
10
20
30
40
50
60
Time (min)
Normal
MK50
MK75
MK88
MK94
MK97
MK data
Normal data

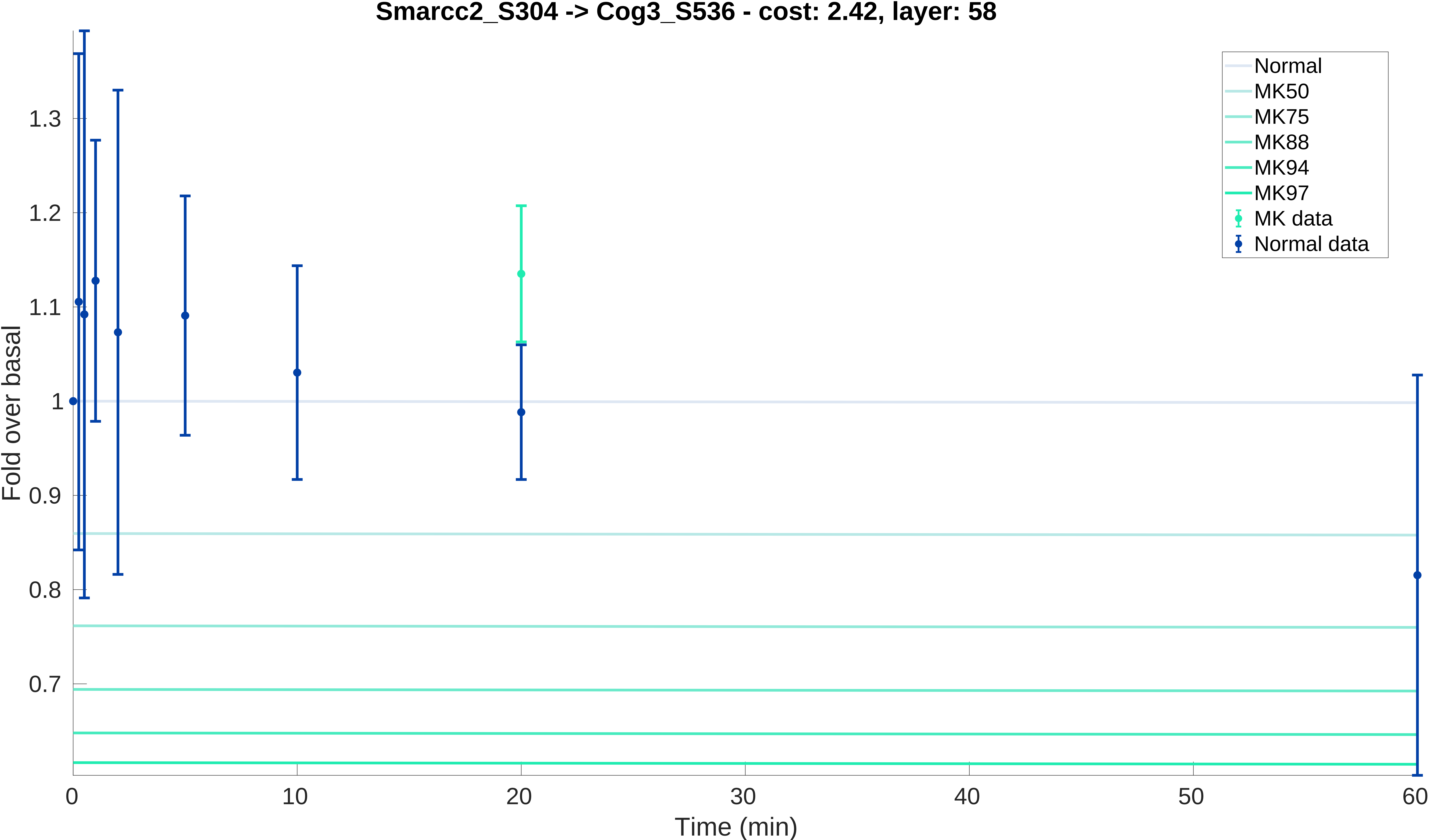
Smarcc2_S304 -> Cog3_S536 - cost: 2.42, layer: 58
Fold over basal
1.3
1.2
1.1
1
0.9
0.8
0.7
0
10
20
30
40
50
60
Time (min)
Normal
MK50
MK75
MK88
MK94
MK97
MK data
Normal data

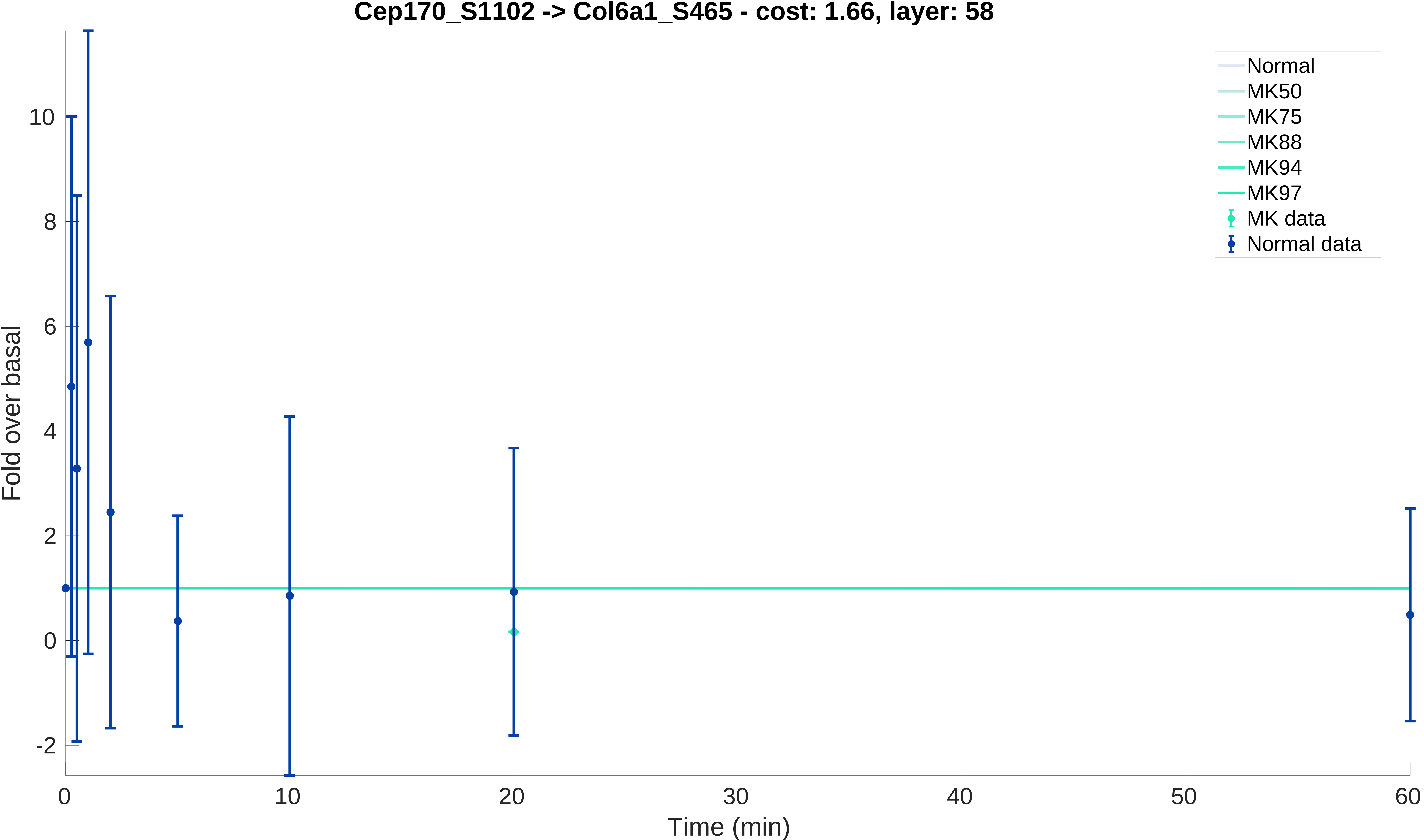

Cep170_S1102 -> Col6a1_S465 - cost: 1.66, layer: 58
Normal
MK50
MK75
MK88
MK94
MK97
MK data
Normal data
Fold over basal
Time (min)
10
8
6
4
2
0
-2
0
10
20
30
40
50
60

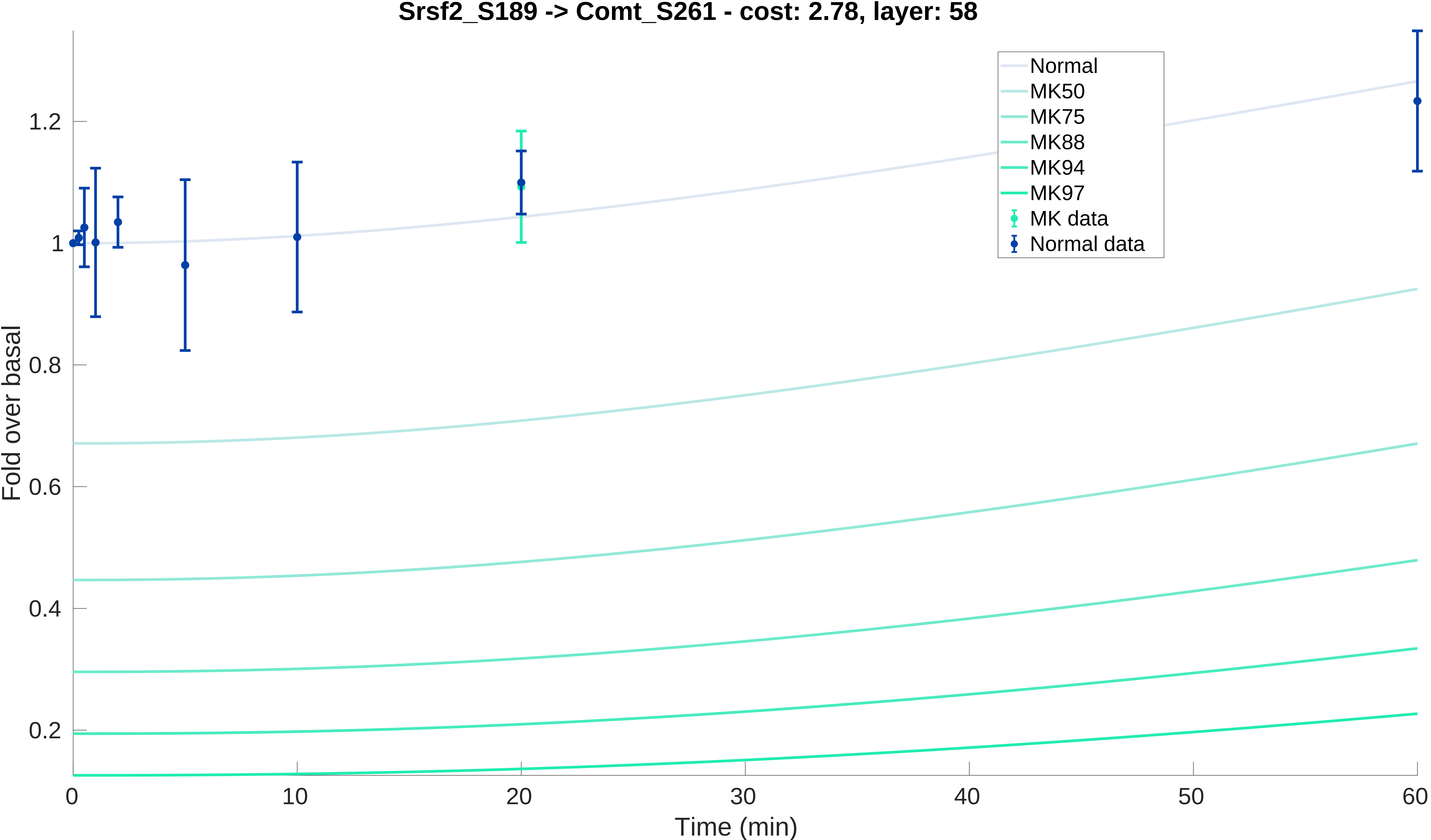
Srsf2_S189 -> Comt_S261 - cost: 2.78, layer: 58
Normal
MK50
MK75
MK88
MK94
MK97
MK data
Normal data
Fold over basal
1.2
1
0.8
0.6
0.4
0.2
0
10
20
30
40
50
60
Time (min)

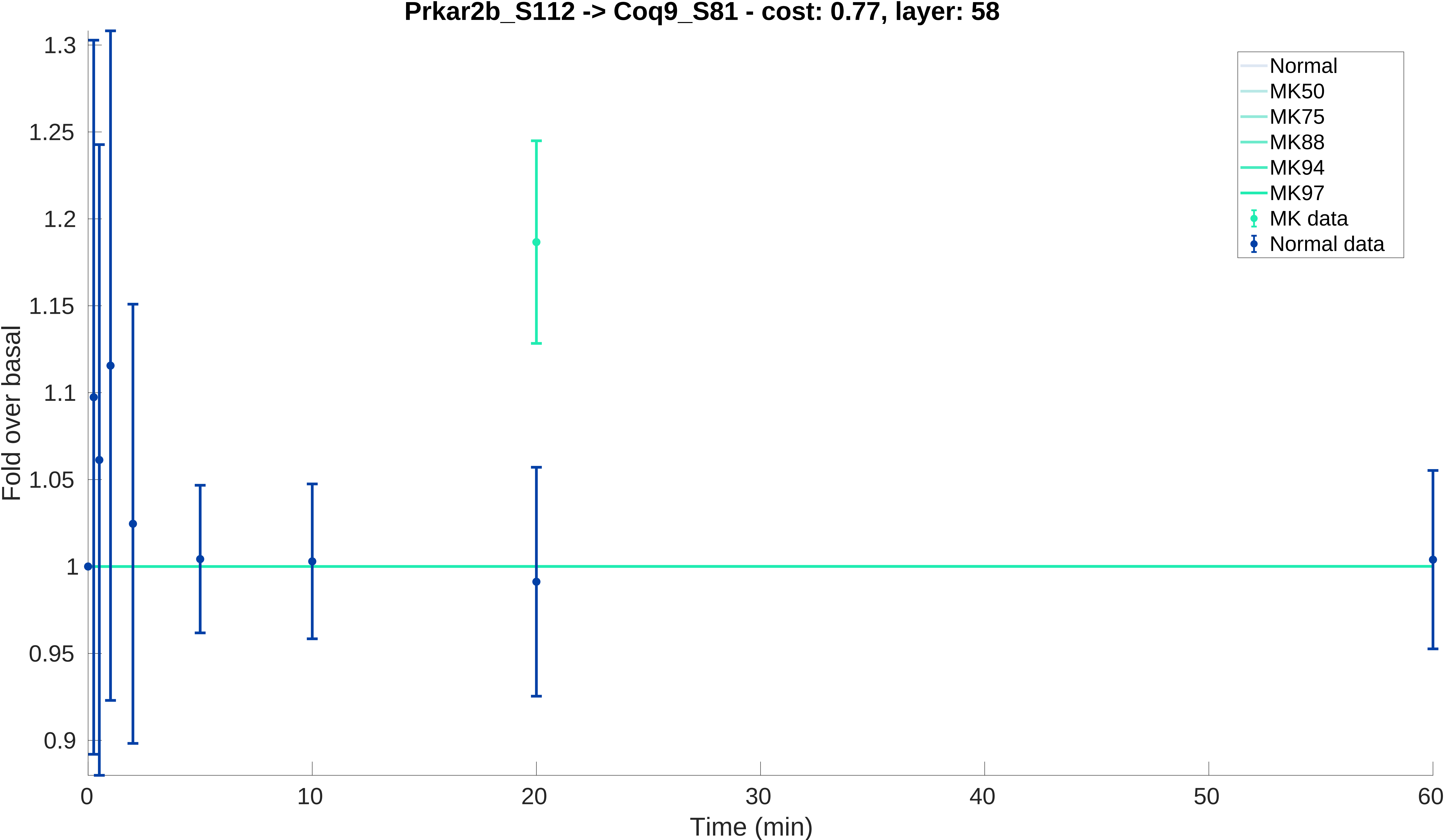

Prkar2b_S112 -> Coq9_S81 - cost: 0.77, layer: 58
Normal
MK50
MK75
MK88
MK94
MK97
MK data
Normal data
Fold over basal
Time (min)
1.3
1.25
1.2
1.15
1.1
1.05
1
0.95
0.9
0
10
20
30
40
50
60

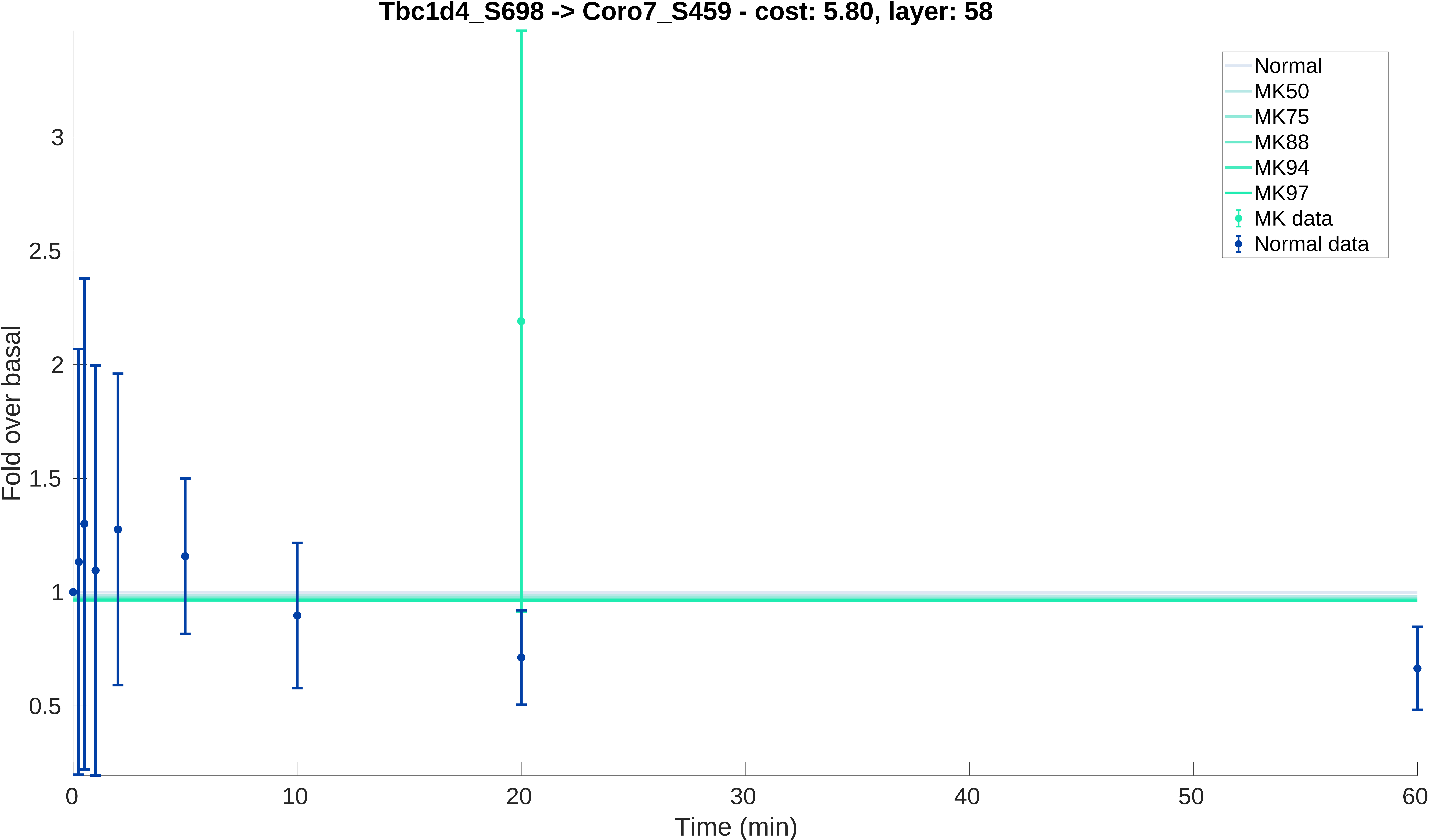

Tbc1d4_S698 -> Coro7_S459 - cost: 5.80, layer: 58
Fold over basal
Time (min)
0.5
1
1.5
2
2.5
3
0
10
20
30
40
50
60
Normal
MK50
MK75
MK88
MK94
MK97
MK data
Normal data

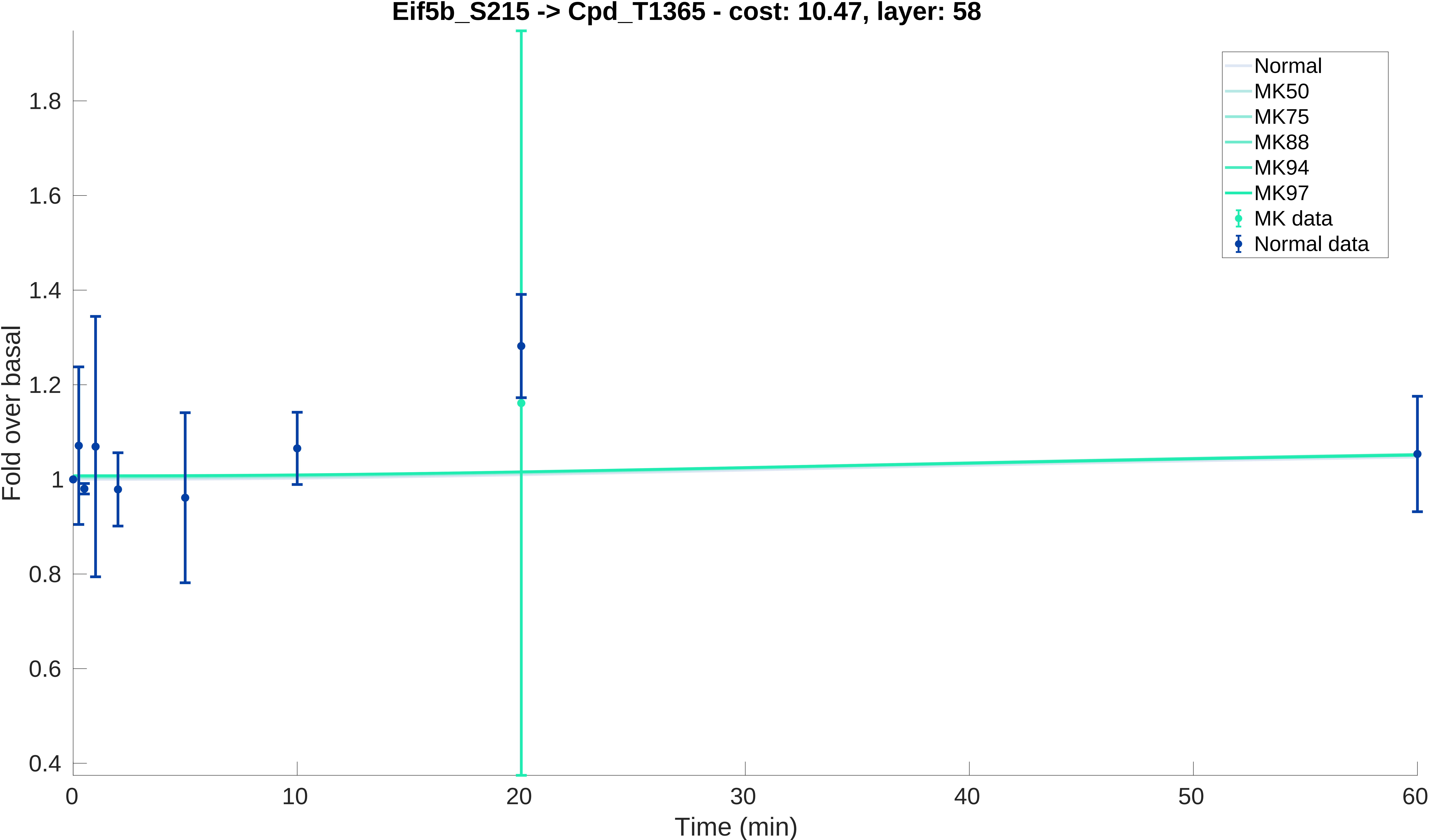
Eif5b_S215 -> Cpd_T1365 - cost: 10.47, layer: 58
Fold over basal
Time (min)
0.4
0.6
0.8
1
1.2
1.4
1.6
1.8
0
10
20
30
40
50
60
Normal
MK50
MK75
MK88
MK94
MK97
MK data
Normal data

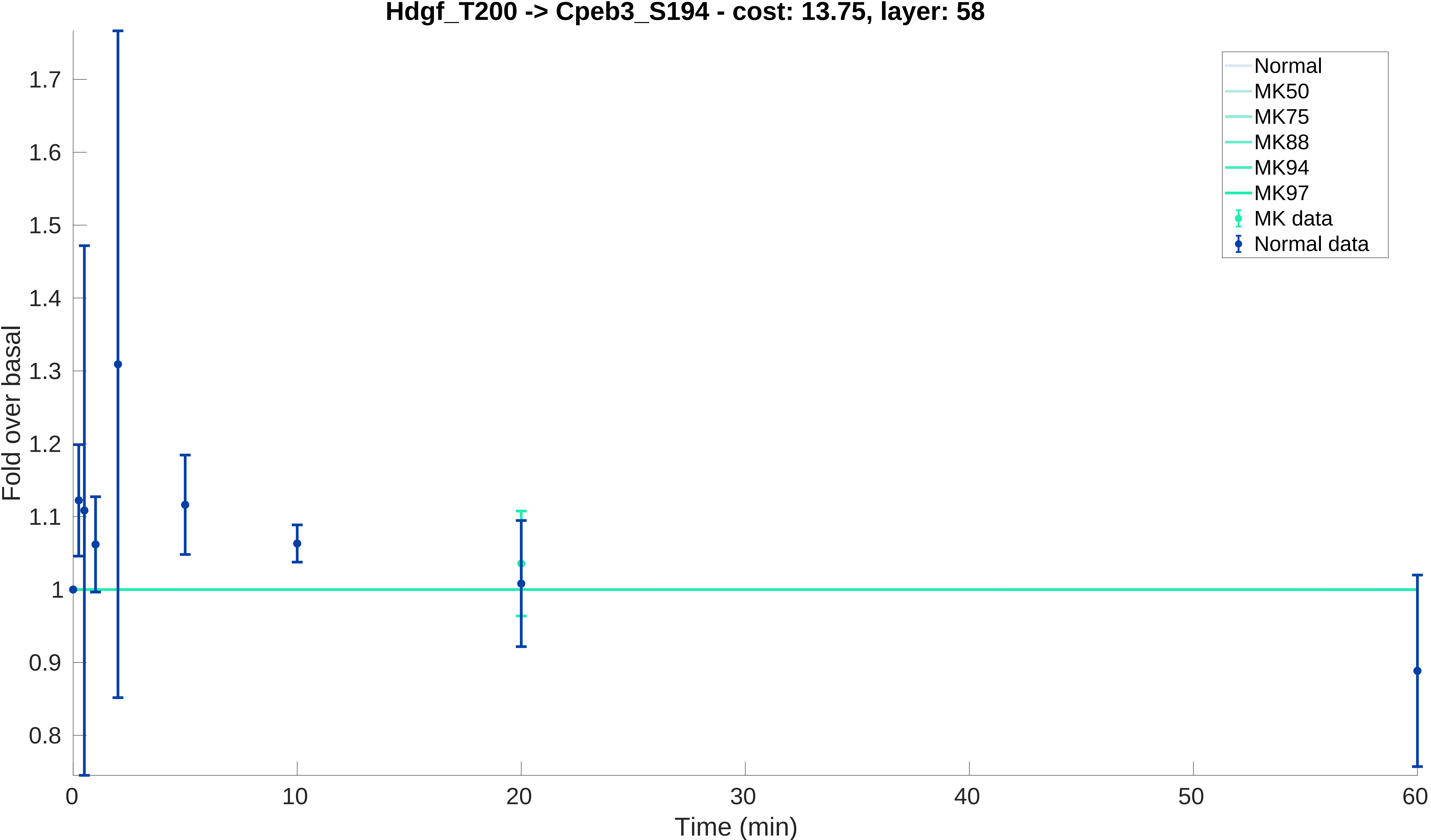

Hdgf_T200 -> Cpeb3_S194 - cost: 13.75, layer: 58
Fold over basal
Time (min)
1.7
1.6
1.5
1.4
1.3
1.2
1.1
1
0.9
0.8
0
10
20
30
40
50
60
Normal
MK50
MK75
MK88
MK94
MK97
MK data
Normal data

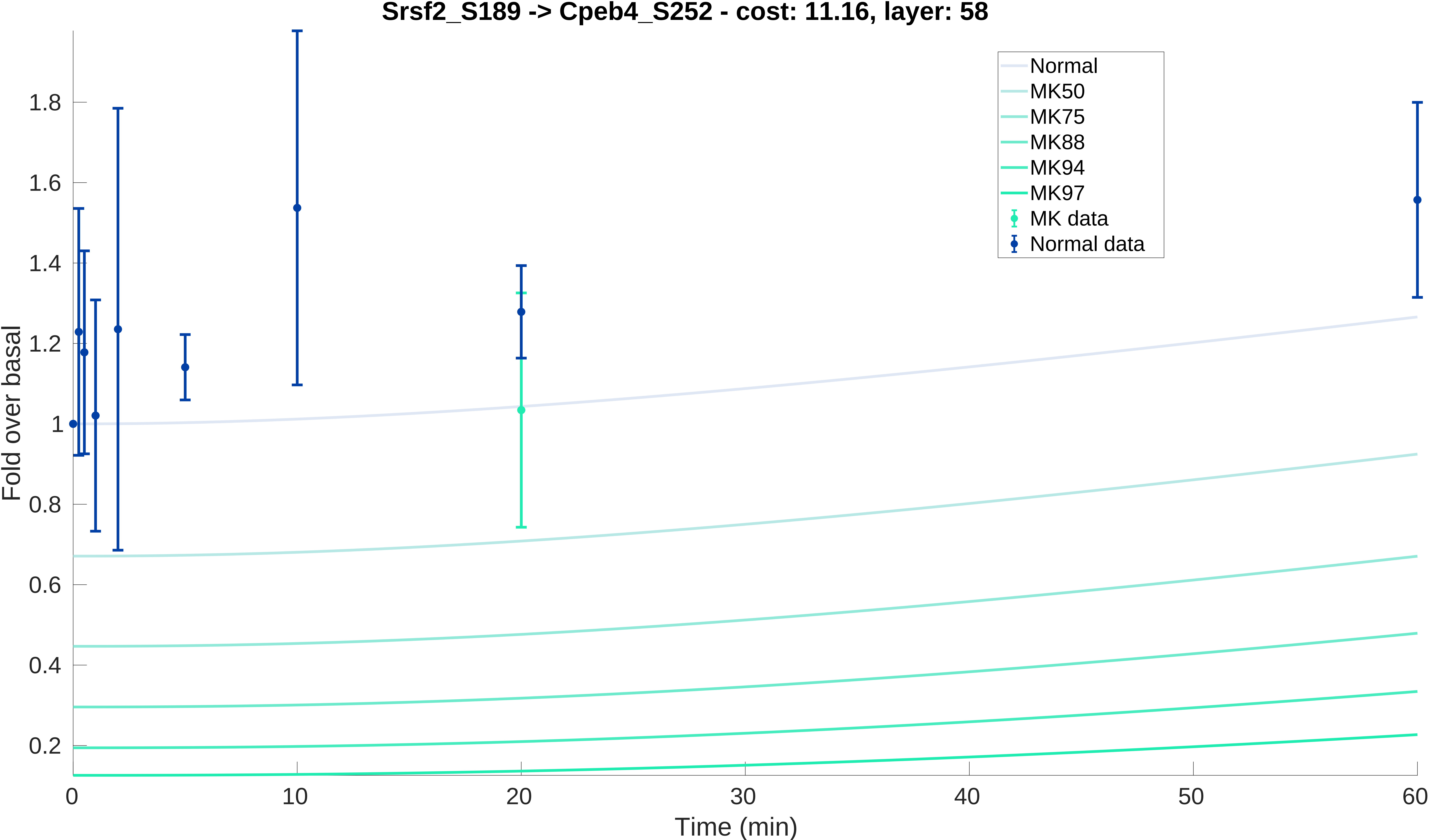

Srsf2_S189 -> Cpeb4_S252 - cost: 11.16, layer: 58
Normal
MK50
MK75
MK88
MK94
MK97
MK data
Normal data
Fold over basal
Time (min)
0
10
20
30
40
50
60
0.2
0.4
0.6
0.8
1
1.2
1.4
1.6
1.8

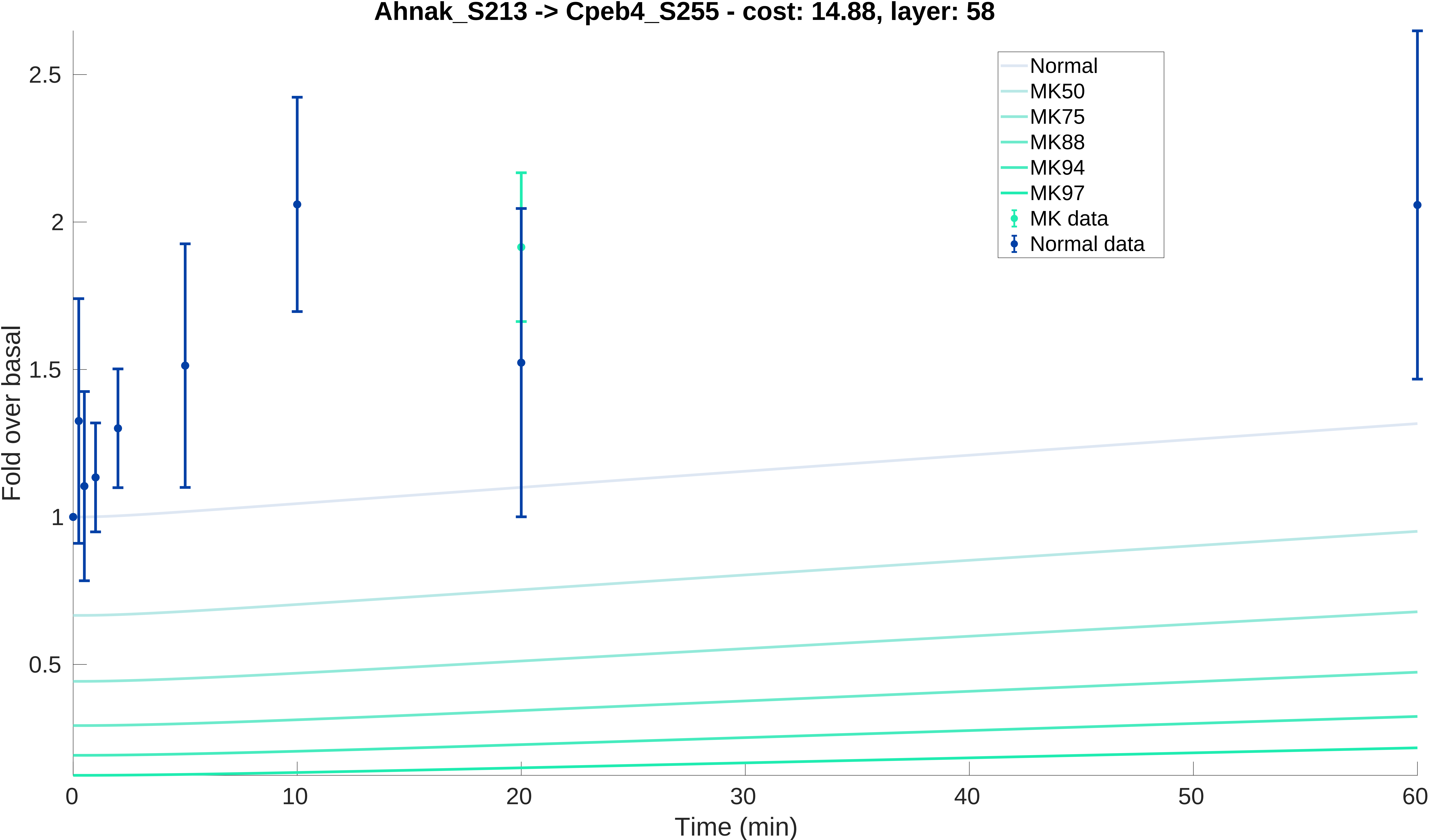

Ahnak_S213 -> Cpeb4_S255 - cost: 14.88, layer: 58
Fold over basal
Time (min)
Normal
MK50
MK75
MK88
MK94
MK97
MK data
Normal data
0.5
1
1.5
2
2.5
0
10
20
30
40
50
60

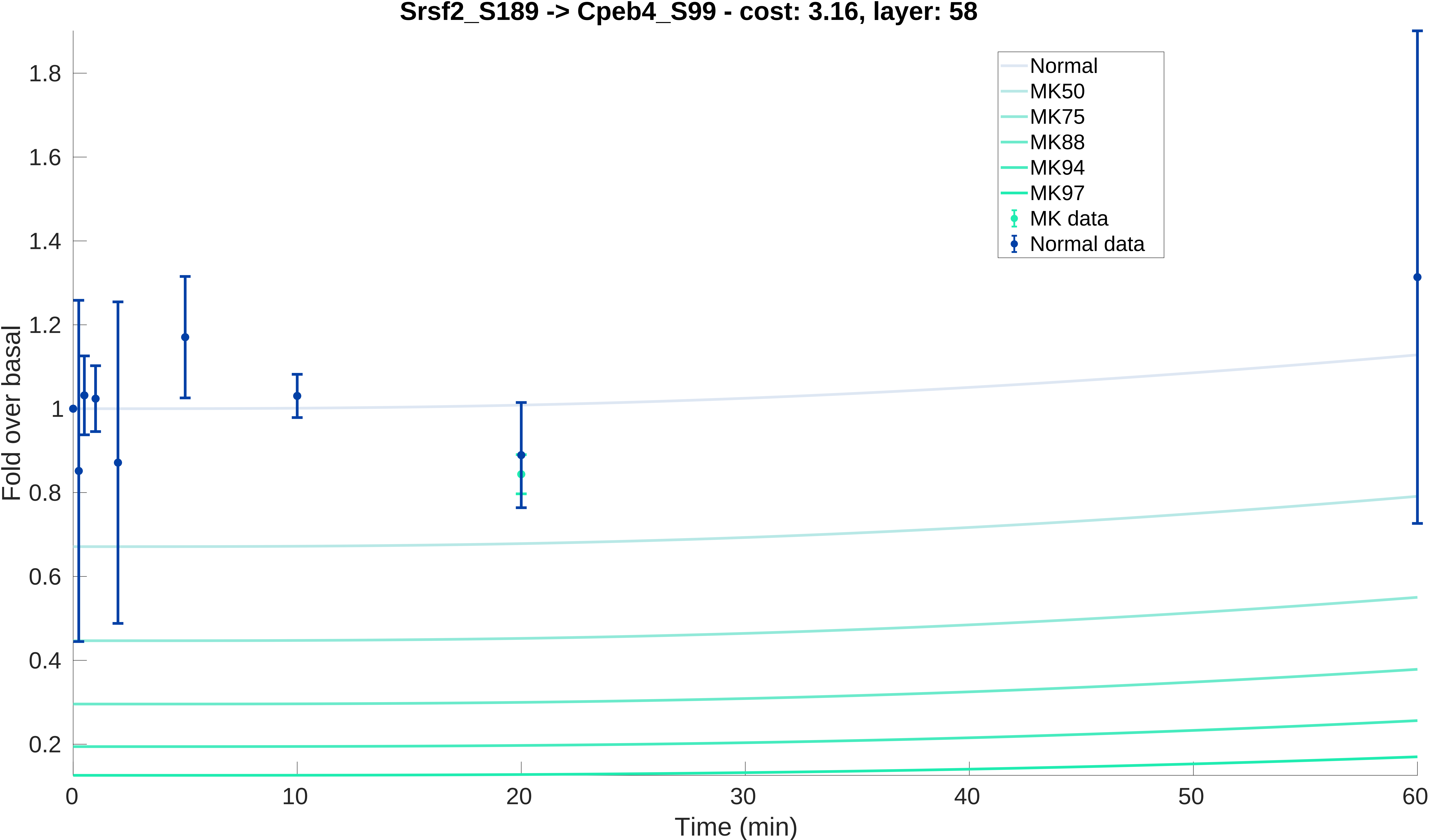

Srsf2_S189 -> Cpeb4_S99 - cost: 3.16, layer: 58
Normal
MK50
MK75
MK88
MK94
MK97
MK data
Normal data
Fold over basal
0.2
0.4
0.6
0.8
1
1.2
1.4
1.6
1.8
0
10
20
30
40
50
60
Time (min)

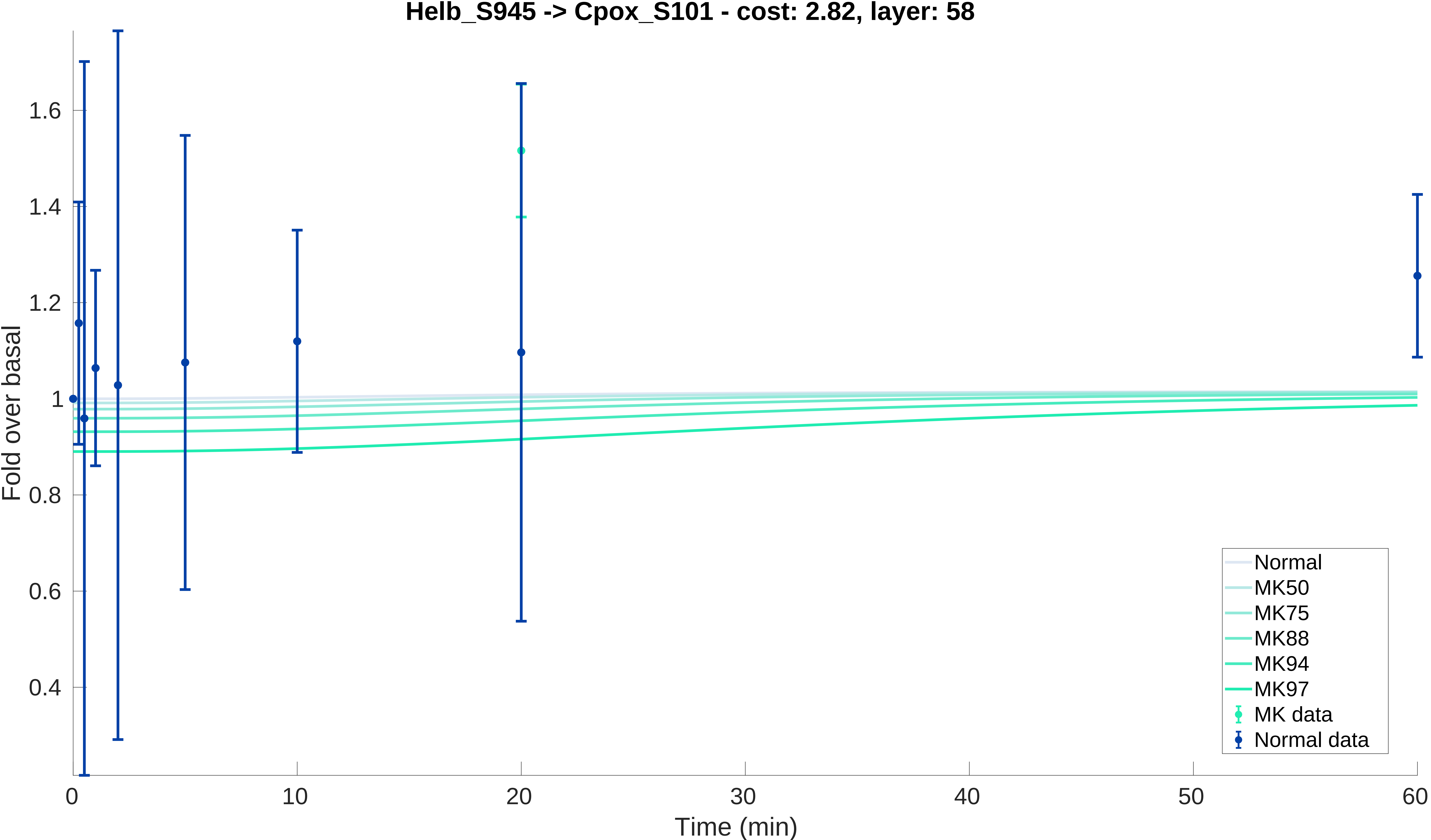
Helb_S945 -> Cpox_S101 - cost: 2.82, layer: 58
Fold over basal
Time (min)
0
10
20
30
40
50
60
0.4
0.6
0.8
1
1.2
1.4
1.6
Normal
MK50
MK75
MK88
MK94
MK97
MK data
Normal data

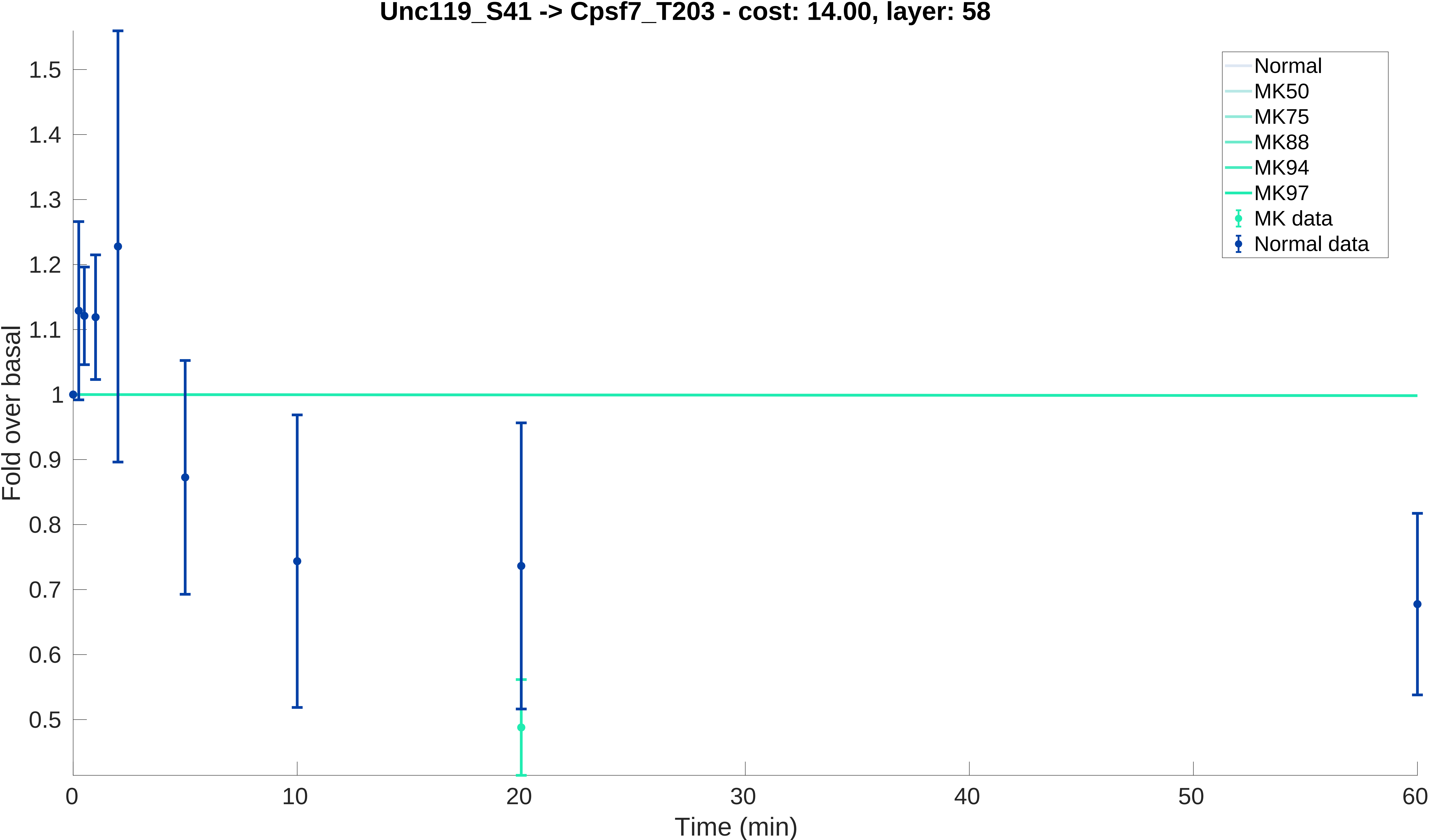

Unc119_S41 -> Cpsf7_T203 - cost: 14.00, layer: 58
Normal
MK50
MK75
MK88
MK94
MK97
MK data
Normal data
Fold over basal
1.5
1.4
1.3
1.2
1.1
1
0.9
0.8
0.7
0.6
0.5
0
10
20
30
40
50
60
Time (min)

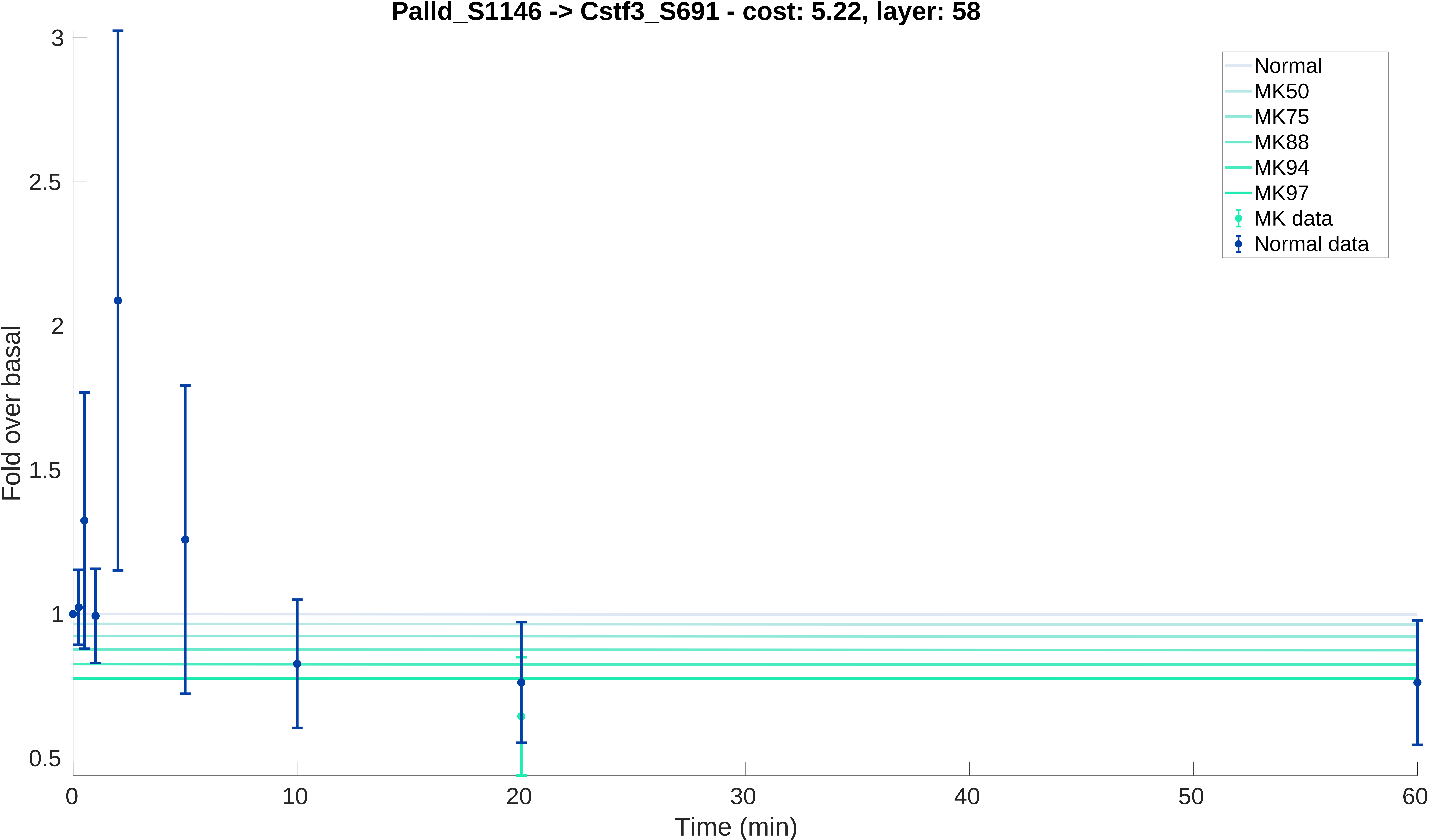

Palld_S1146 -> Cstf3_S691 - cost: 5.22, layer: 58
Fold over basal
Time (min)
0.5
1
1.5
2
2.5
3
0
10
20
30
40
50
60
Normal
MK50
MK75
MK88
MK94
MK97
MK data
Normal data

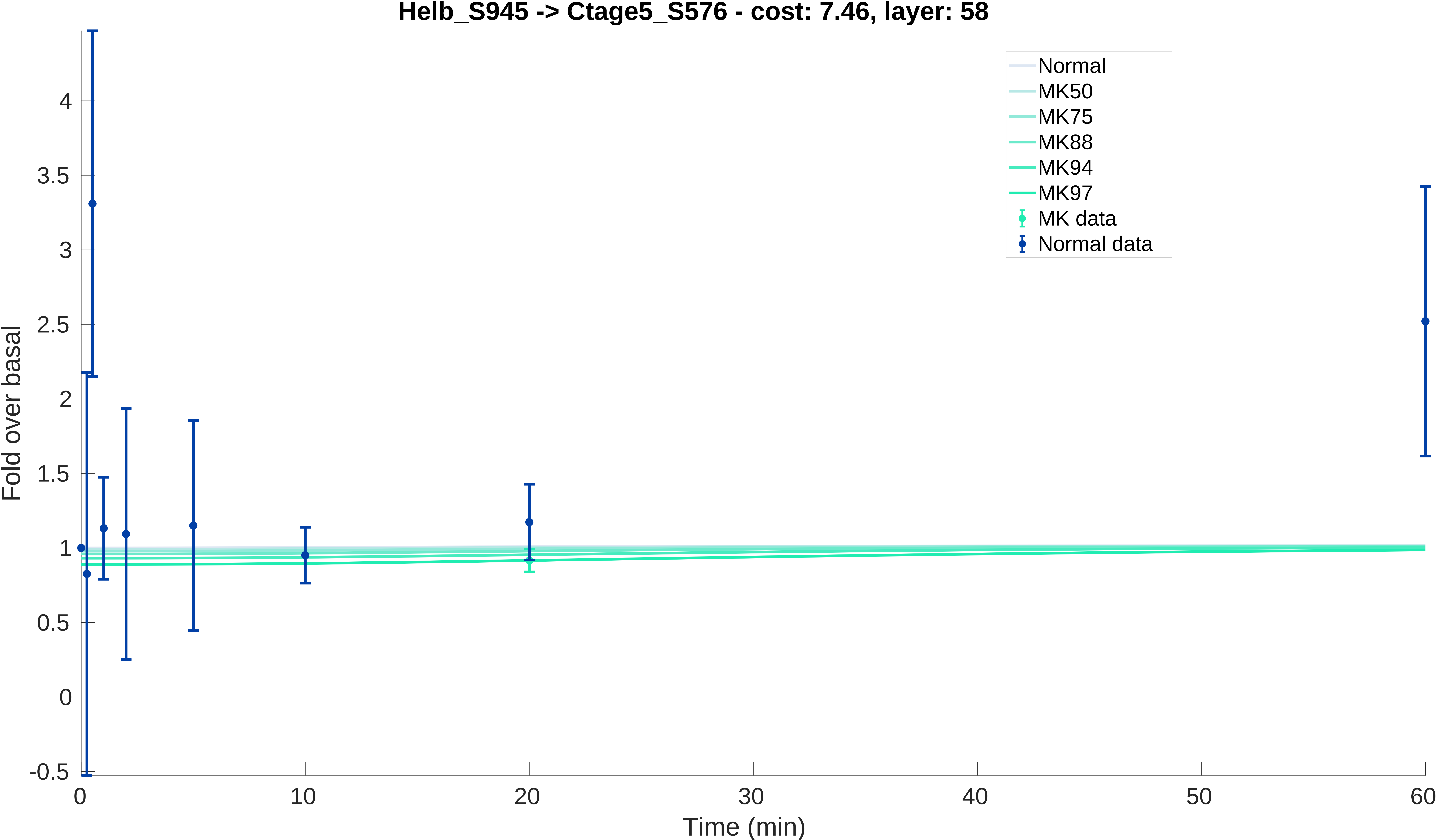

Helb_S945 -> Ctage5_S576 - cost: 7.46, layer: 58
Fold over basal
Time (min)
Normal
MK50
MK75
MK88
MK94
MK97
MK data
Normal data
-0.5
0
0.5
1
1.5
2
2.5
3
3.5
4
0
10
20
30
40
50
60

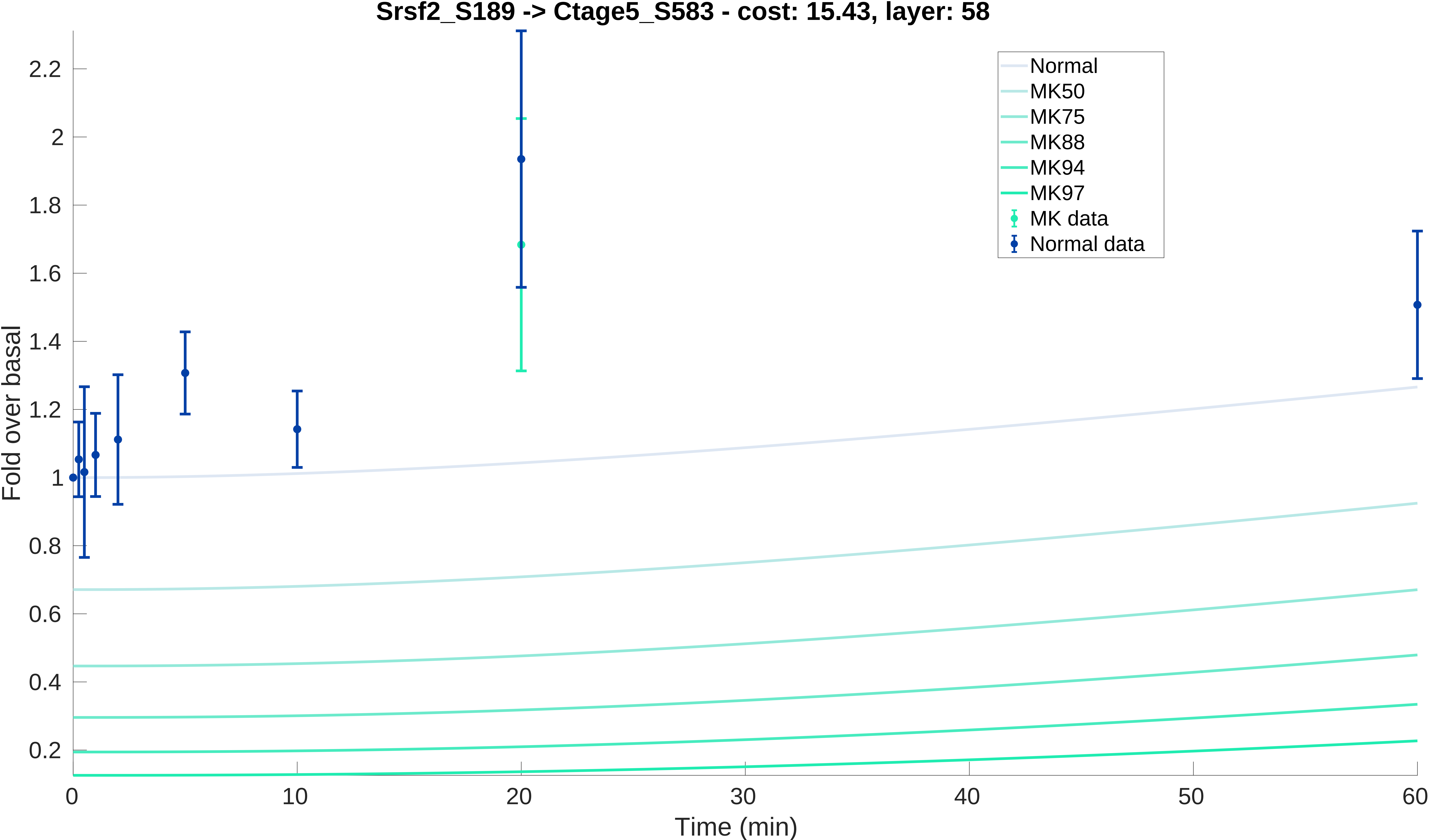

Srsf2_S189 -> Ctage5_S583 - cost: 15.43, layer: 58
Fold over basal
Time (min)
Normal
MK50
MK75
MK88
MK94
MK97
MK data
Normal data
2.2
2
1.8
1.6
1.4
1.2
1
0.8
0.6
0.4
0.2
0
10
20
30
40
50
60

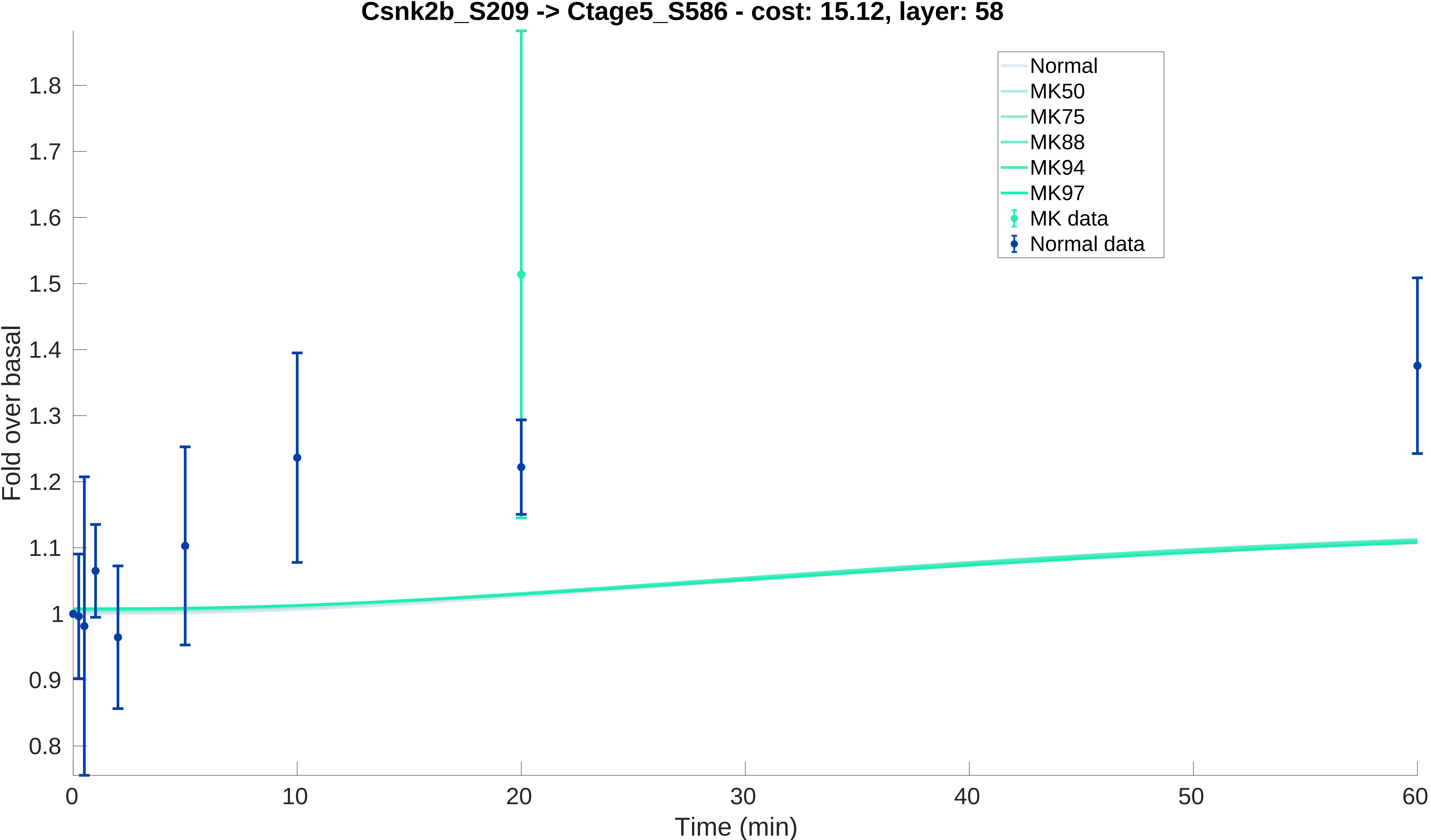

Csnk2b_S209 -> Ctage5_S586 - cost: 15.12, layer: 58
Fold over basal
Time (min)
0.8
0.9
1
1.1
1.2
1.3
1.4
1.5
1.6
1.7
1.8
0
10
20
30
40
50
60
Normal
MK50
MK75
MK88
MK94
MK97
MK data
Normal data

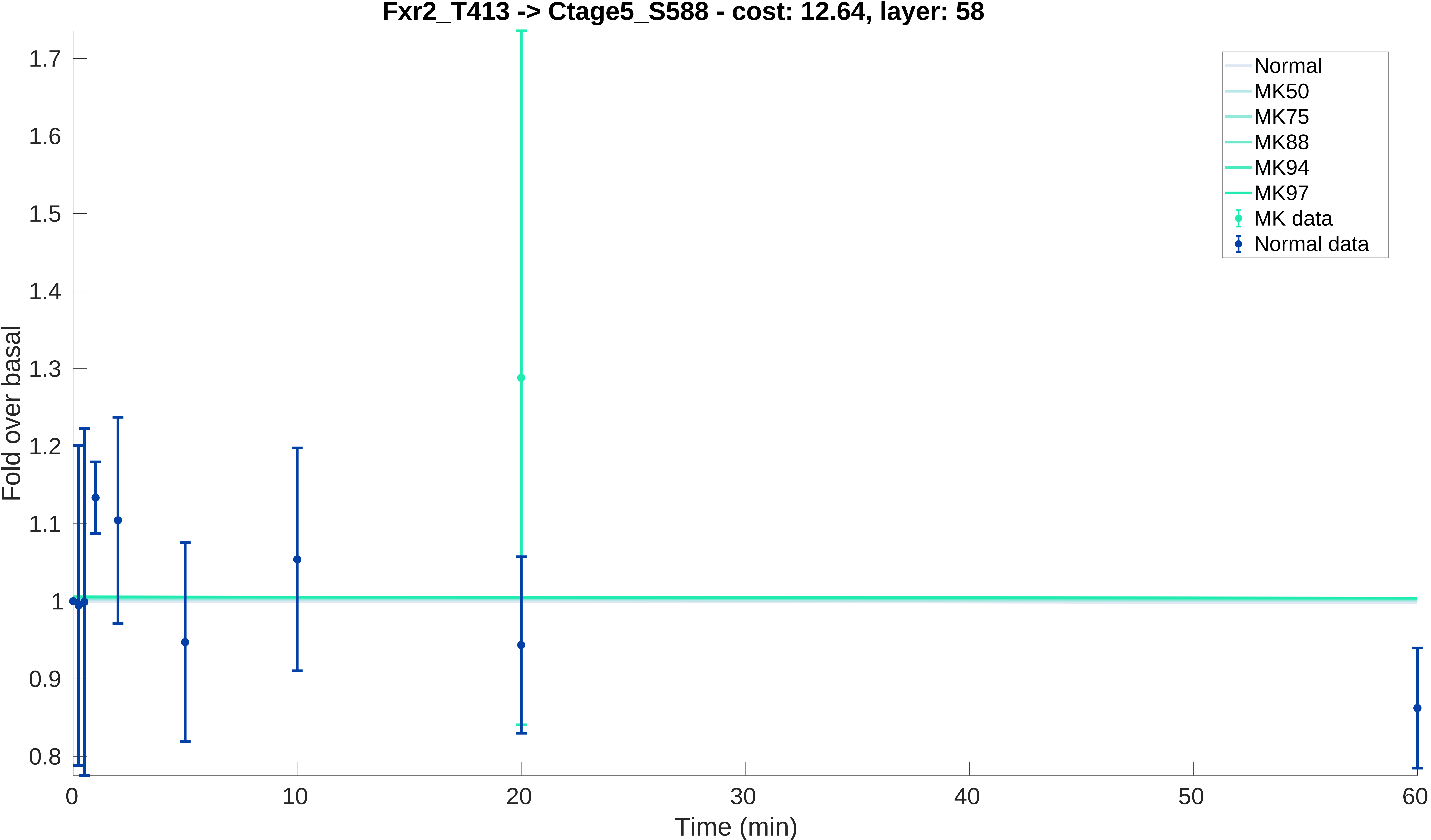

Fxr2_T413 -> Ctage5_S588 - cost: 12.64, layer: 58
Fold over basal
Time (min)
0.8
0.9
1
1.1
1.2
1.3
1.4
1.5
1.6
1.7
0
10
20
30
40
50
60
Normal
MK50
MK75
MK88
MK94
MK97
MK data
Normal data

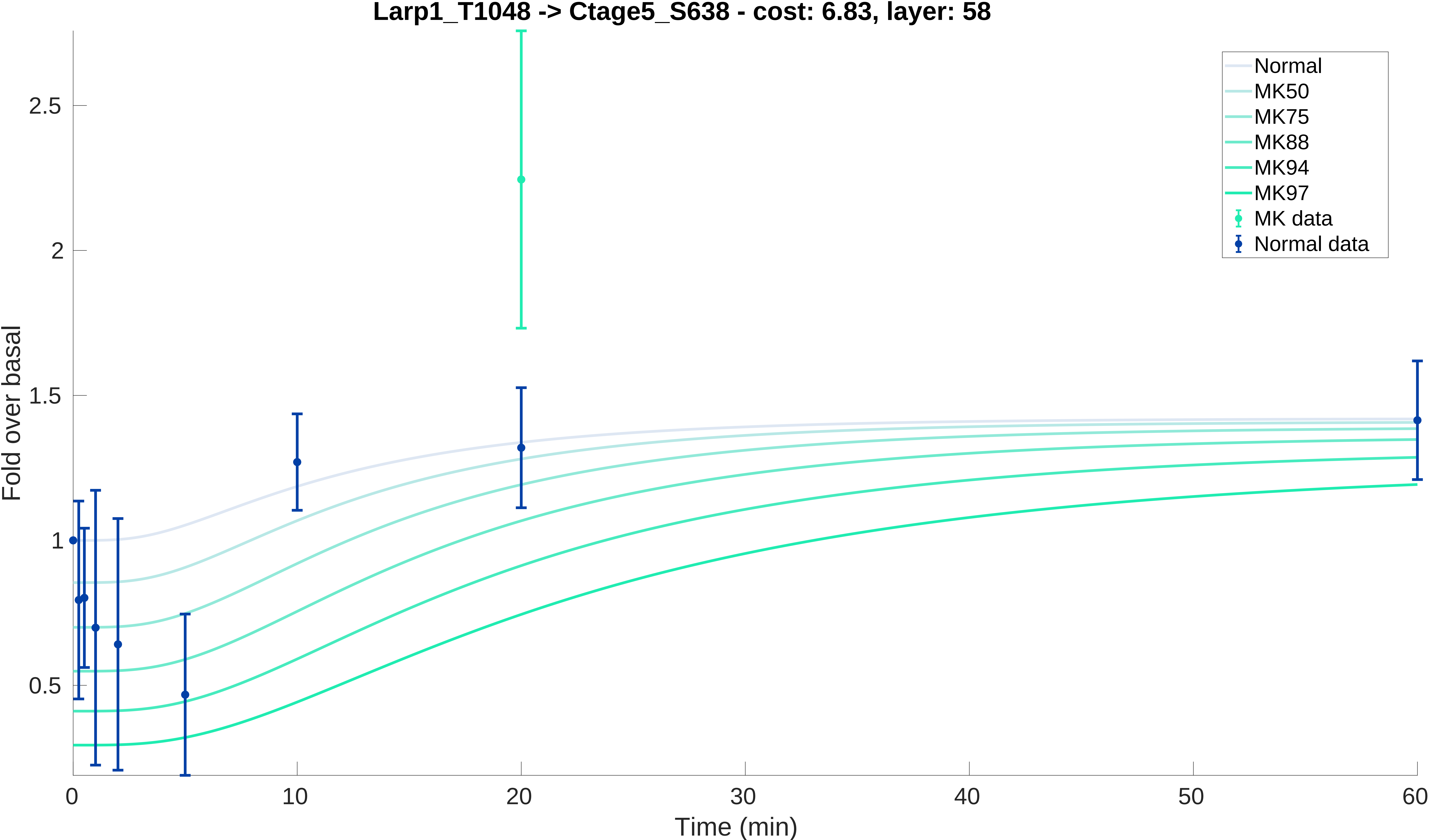
Larp1_T1048 -> Ctage5_S638 - cost: 6.83, layer: 58
Fold over basal
Time (min)
Normal
MK50
MK75
MK88
MK94
MK97
MK data
Normal data
0.5
1
1.5
2
2.5
0
10
20
30
40
50
60

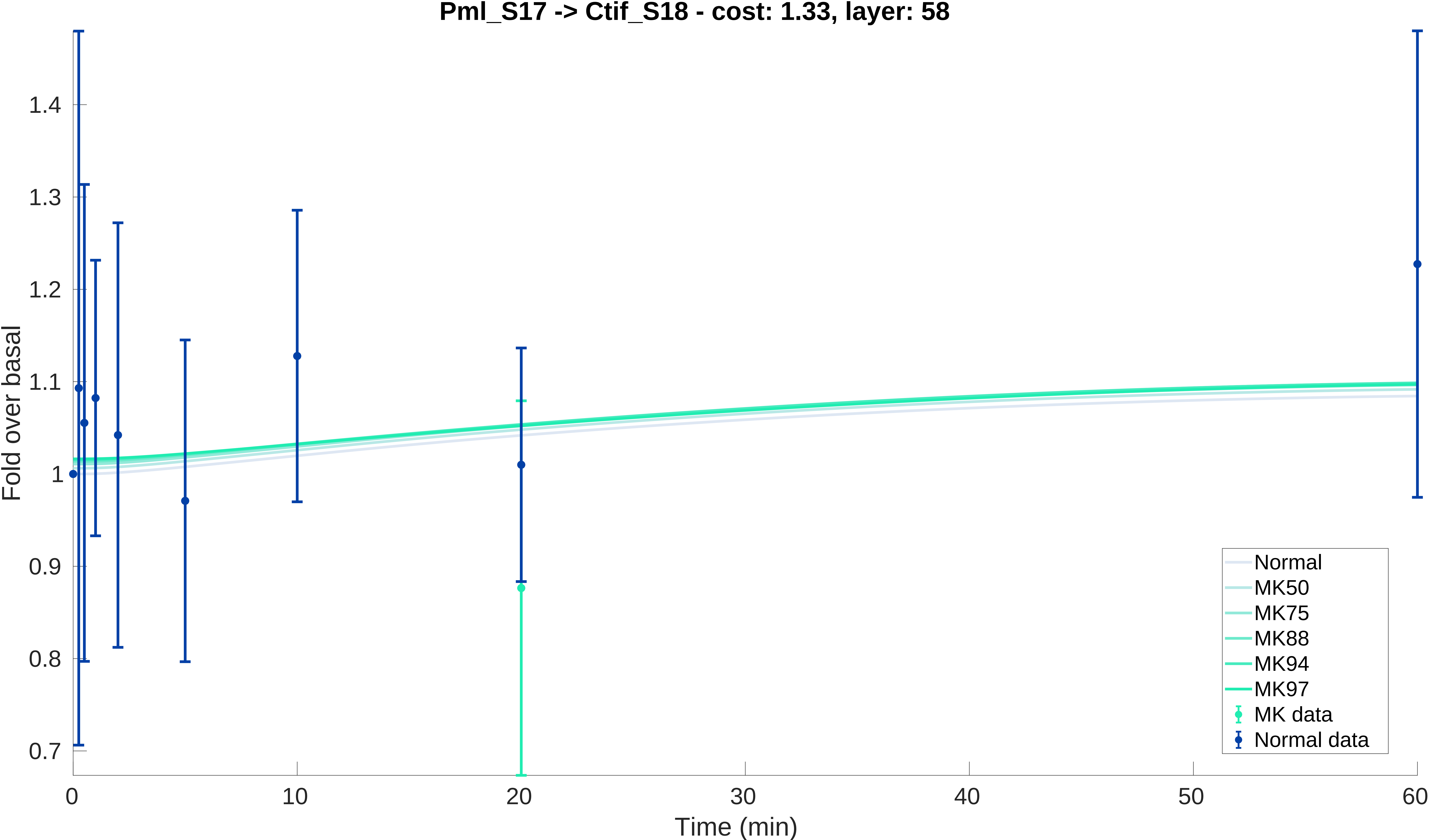

Pml_S17 -> Ctif_S18 - cost: 1.33, layer: 58
Fold over basal
1.4
1.3
1.2
1.1
1
0.9
0.8
0.7
0
10
20
30
40
50
60
Time (min)
Normal
MK50
MK75
MK88
MK94
MK97
MK data
Normal data

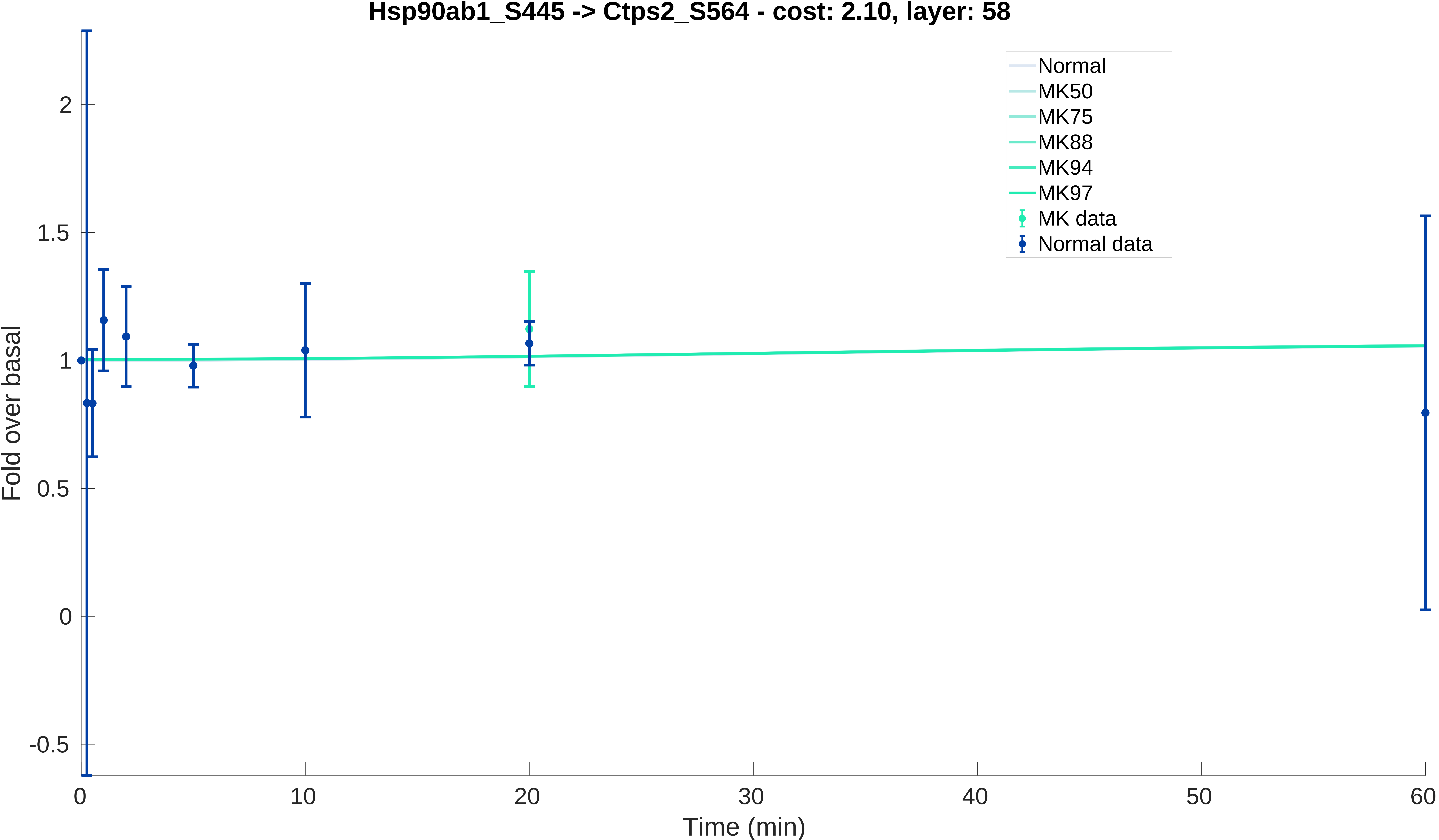

Hsp90ab1_S445 -> Ctps2_S564 - cost: 2.10, layer: 58
Normal
MK50
MK75
MK88
MK94
MK97
MK data
Normal data
Fold over basal
Time (min)
-0.5
0
0.5
1
1.5
2
0
10
20
30
40
50
60

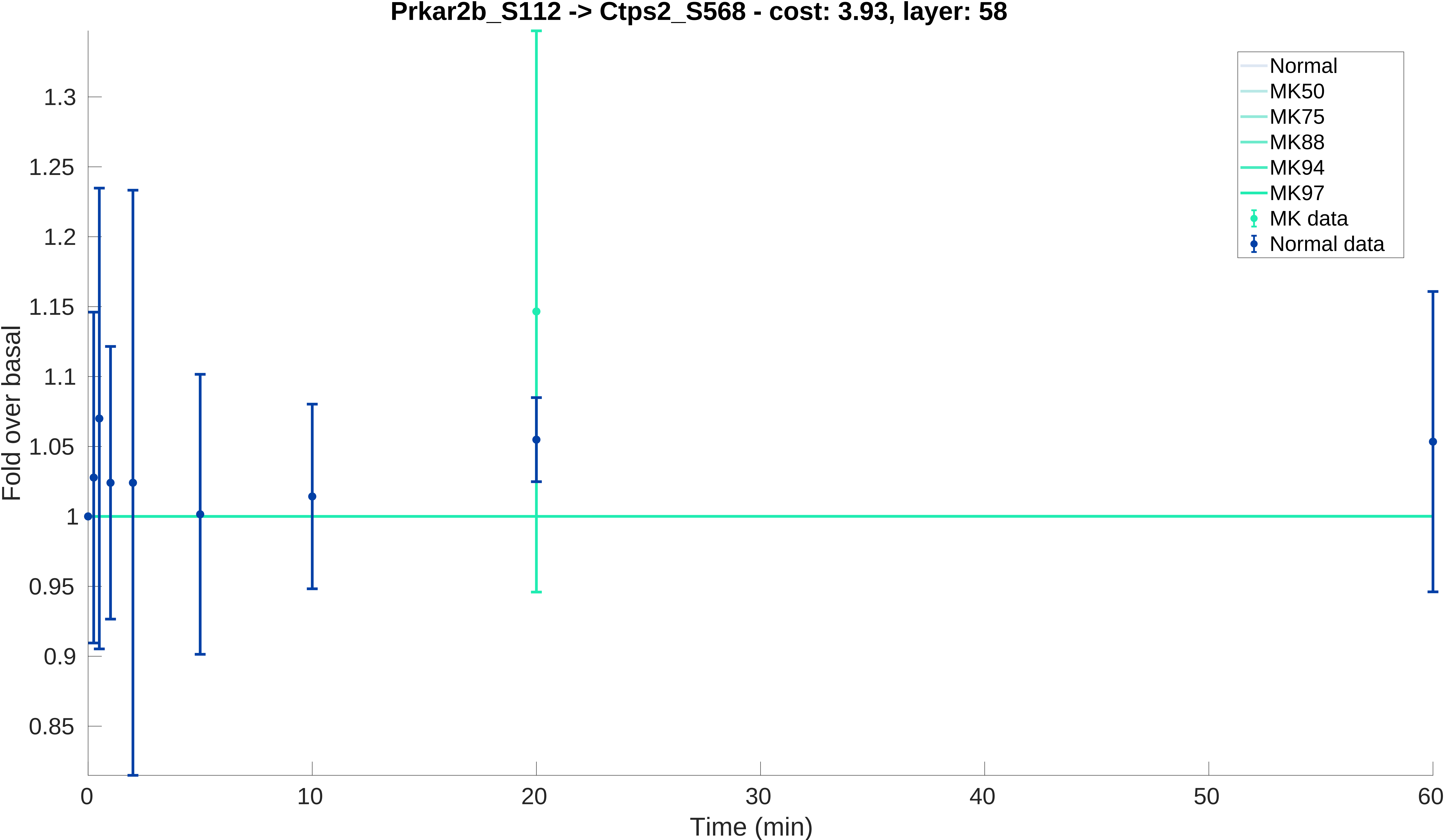
Prkar2b_S112 -> Ctps2_S568 - cost: 3.93, layer: 58
Fold over basal
Time (min)
1.3
1.25
1.2
1.15
1.1
1.05
1
0.95
0.9
0.85
0
10
20
30
40
50
60
Normal
MK50
MK75
MK88
MK94
MK97
MK data
Normal data

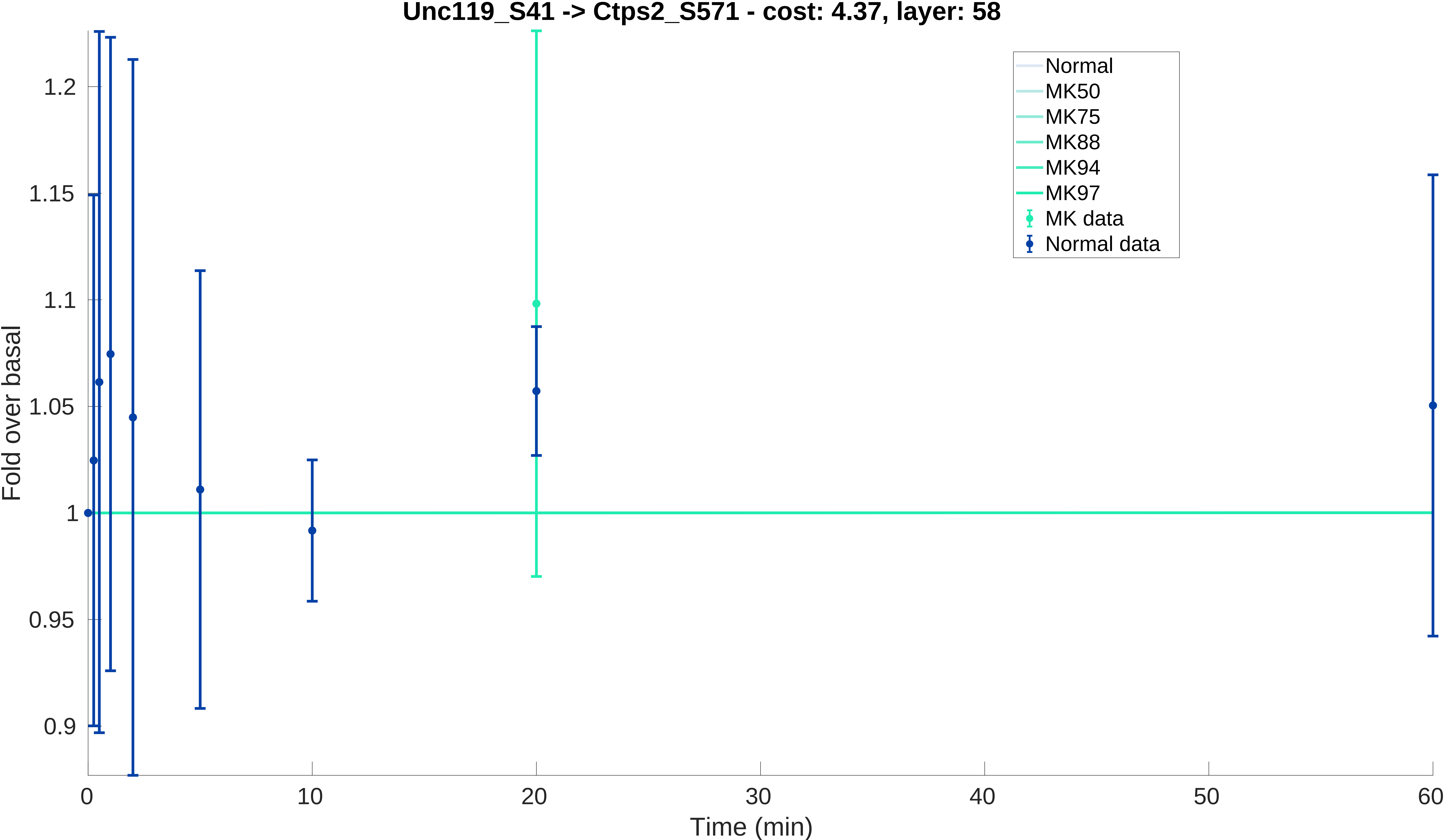

Unc119_S41 -> Ctps2_S571 - cost: 4.37, layer: 58
Fold over basal
Time (min)
1.2
1.15
1.1
1.05
1
0.95
0.9
0
10
20
30
40
50
60
Normal
MK50
MK75
MK88
MK94
MK97
MK data
Normal data

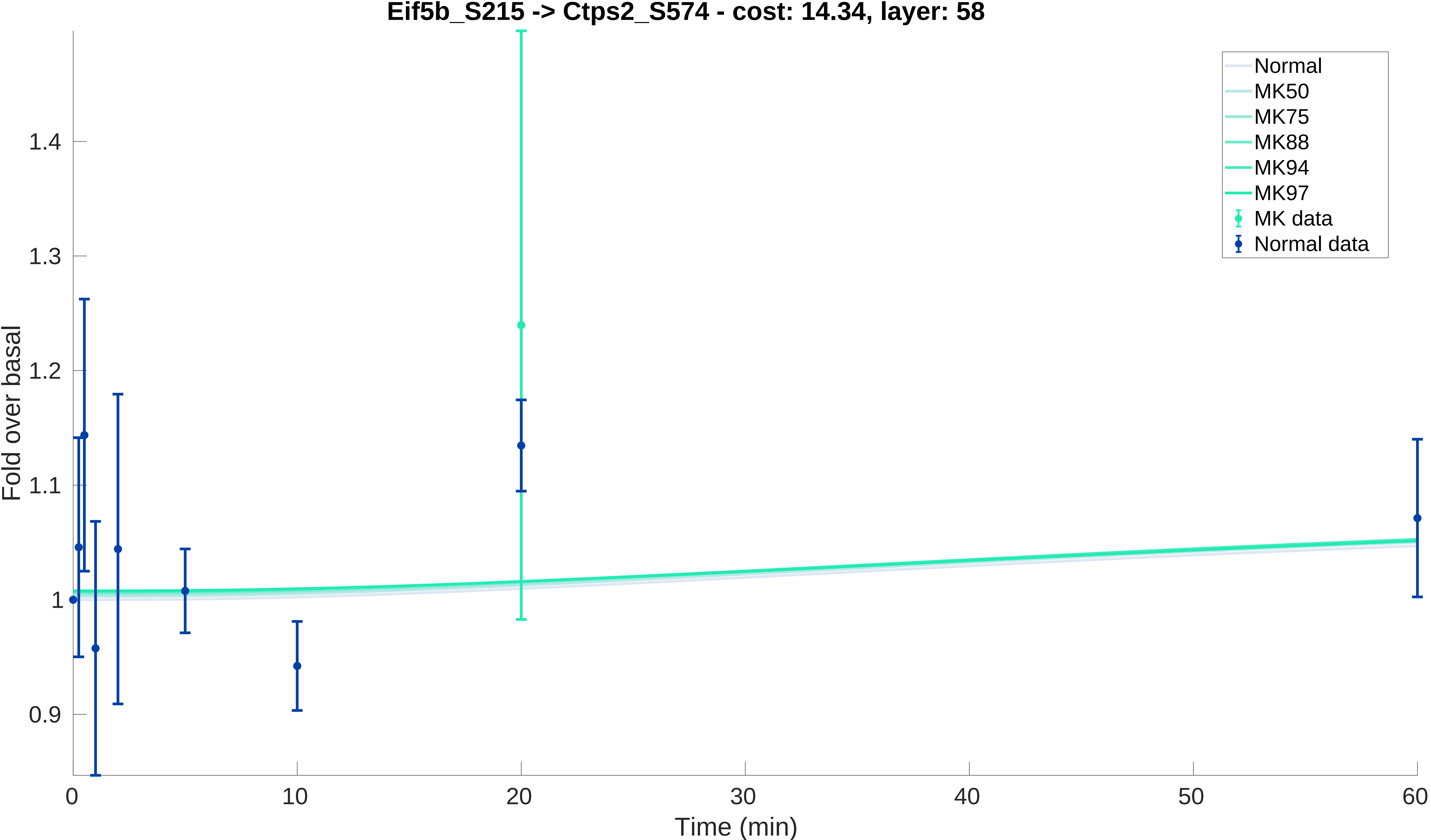

Eif5b_S215 -> Ctps2_S574 - cost: 14.34, layer: 58
Fold over basal
Time (min)
0.9
1
1.1
1.2
1.3
1.4
0
10
20
30
40
50
60
Normal
MK50
MK75
MK88
MK94
MK97
MK data
Normal data

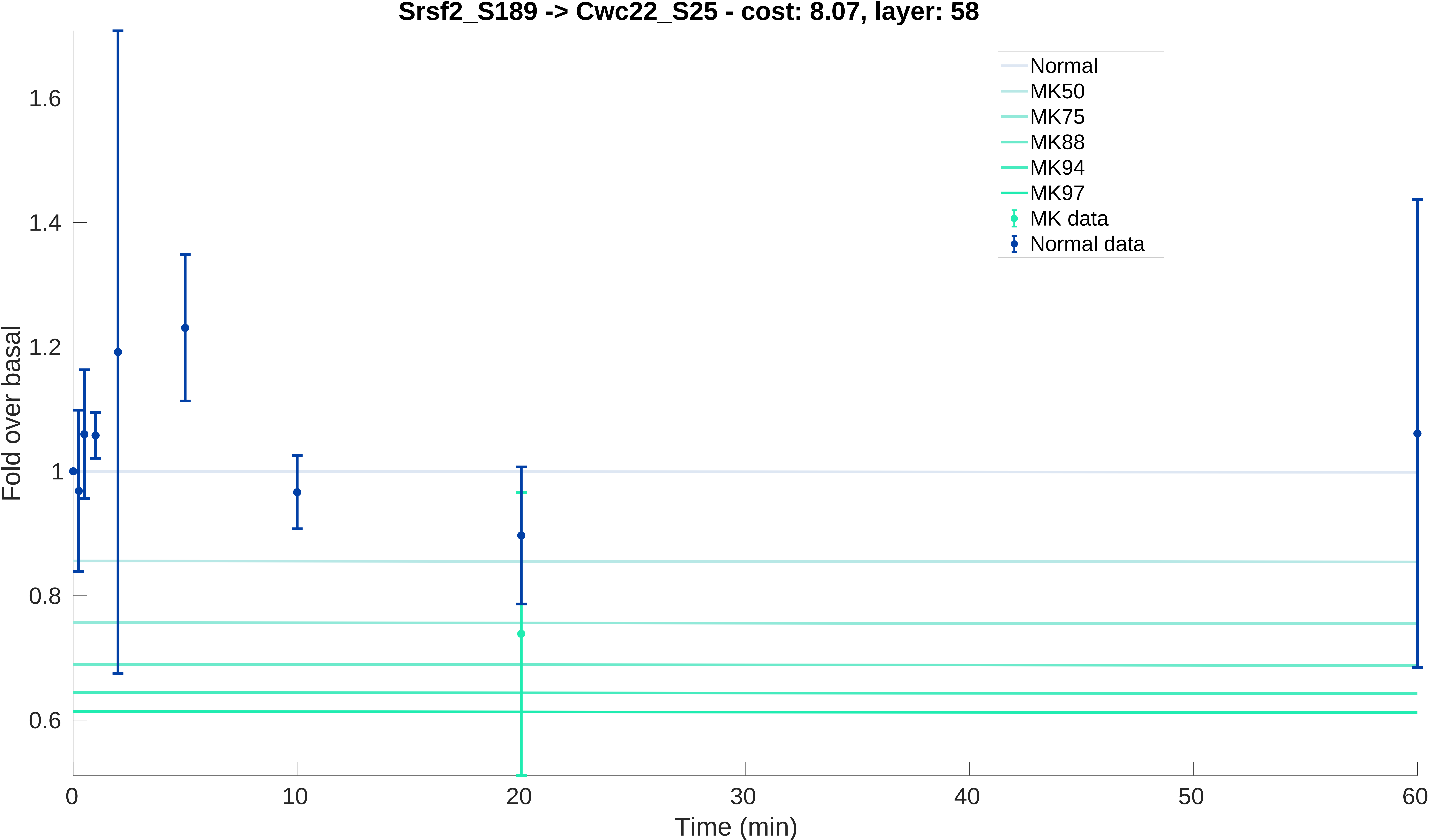

Srsf2_S189 -> Cwc22_S25 - cost: 8.07, layer: 58
Normal
MK50
MK75
MK88
MK94
MK97
MK data
Normal data
Fold over basal
1.6
1.4
1.2
1
0.8
0.6
0
10
20
30
40
50
60
Time (min)

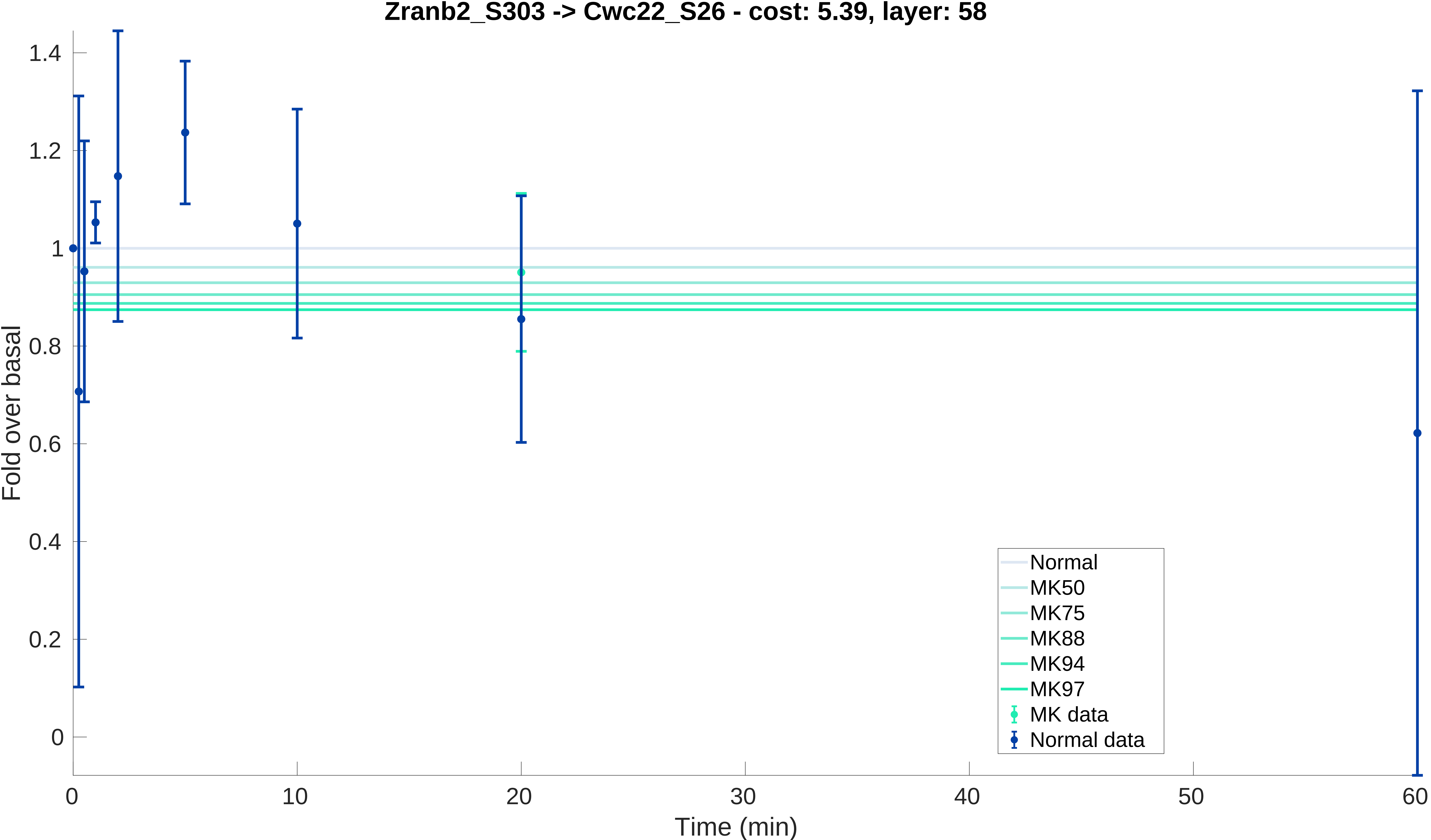

Zranb2_S303 -> Cwc22_S26 - cost: 5.39, layer: 58
Fold over basal
Time (min)
0
0.2
0.4
0.6
0.8
1
1.2
1.4
0
10
20
30
40
50
60
Normal
MK50
MK75
MK88
MK94
MK97
MK data
Normal data

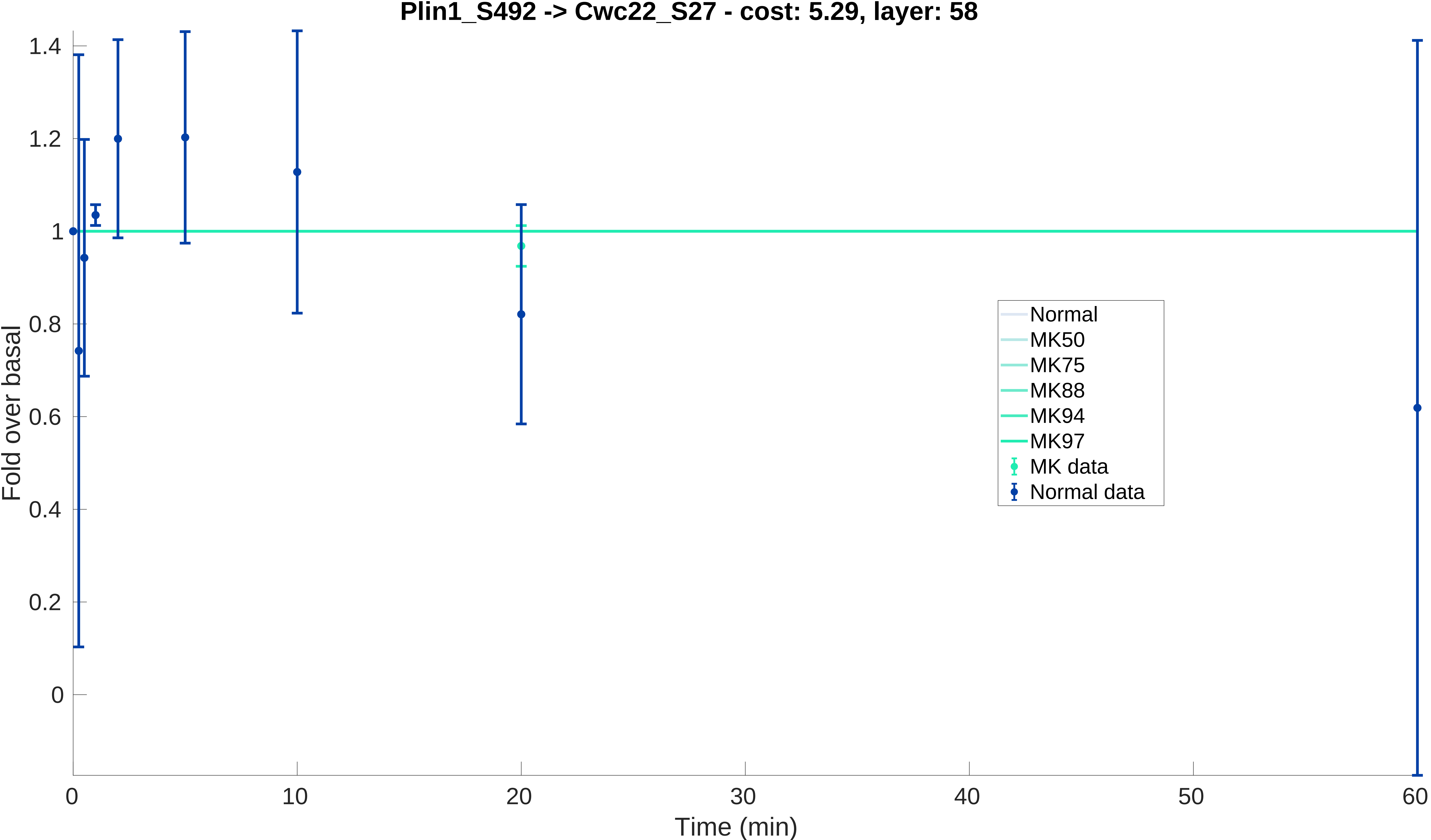

Plin1_S492 -> Cwc22_S27 - cost: 5.29, layer: 58
Fold over basal
1.4
1.2
1
0.8
0.6
0.4
0.2
0
0
10
20
30
40
50
60
Time (min)
Normal
MK50
MK75
MK88
MK94
MK97
MK data
Normal data

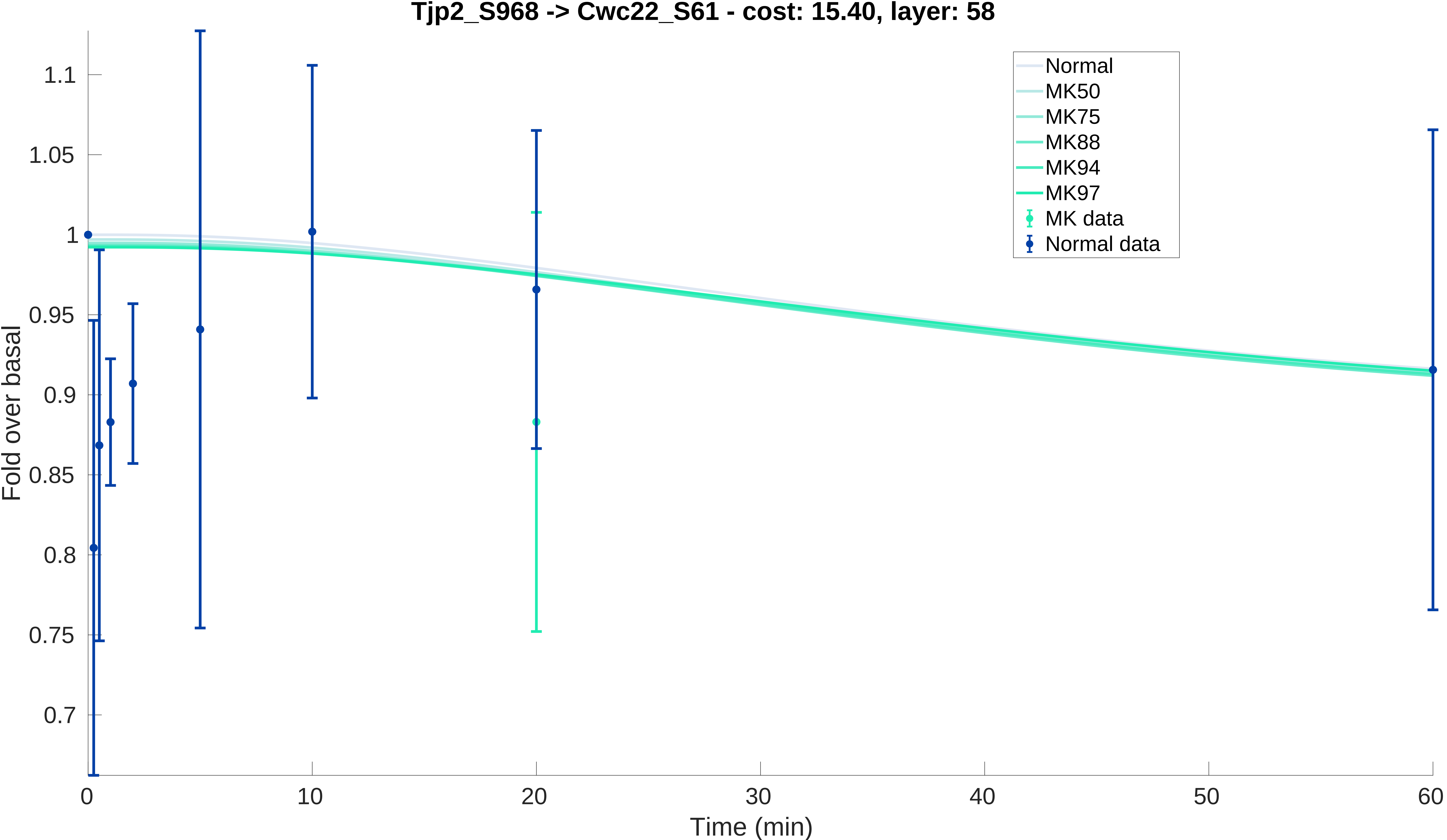

Tjp2_S968 -> Cwc22_S61 - cost: 15.40, layer: 58
Normal
MK50
MK75
MK88
MK94
MK97
MK data
Normal data
Fold over basal
Time (min)
1.1
1.05
1
0.95
0.9
0.85
0.8
0.75
0.7
0
10
20
30
40
50
60

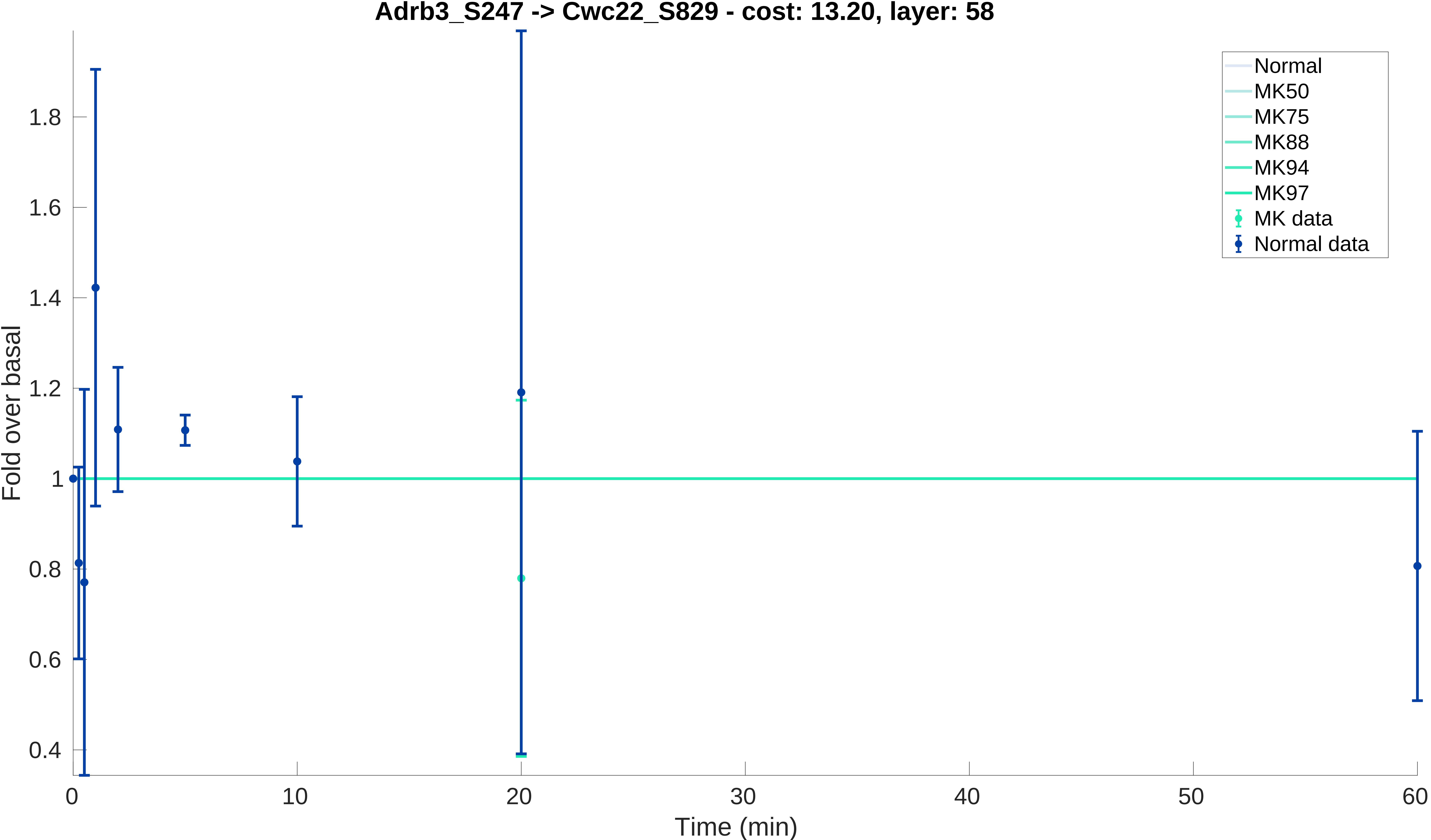

Adrb3_S247 -> Cwc22_S829 - cost: 13.20, layer: 58
Normal
MK50
MK75
MK88
MK94
MK97
MK data
Normal data
Fold over basal
1.8
1.6
1.4
1.2
1
0.8
0.6
0.4
0
10
20
30
40
50
60
Time (min)

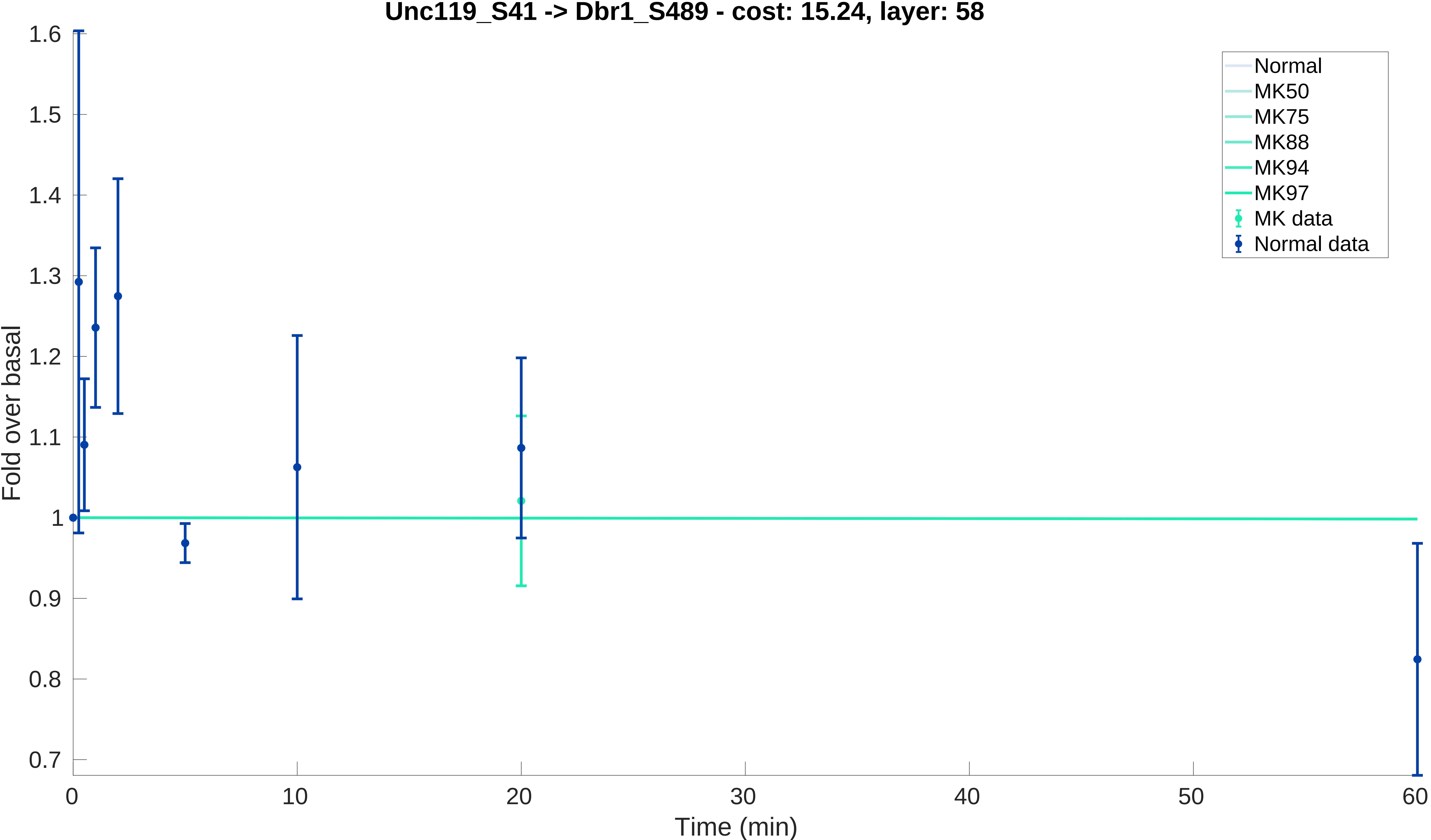
Unc119_S41 -> Dbr1_S489 - cost: 15.24, layer: 58
Fold over basal
Time (min)
0.7
0.8
0.9
1
1.1
1.2
1.3
1.4
1.5
1.6
0
10
20
30
40
50
60
Normal
MK50
MK75
MK88
MK94
MK97
MK data
Normal data

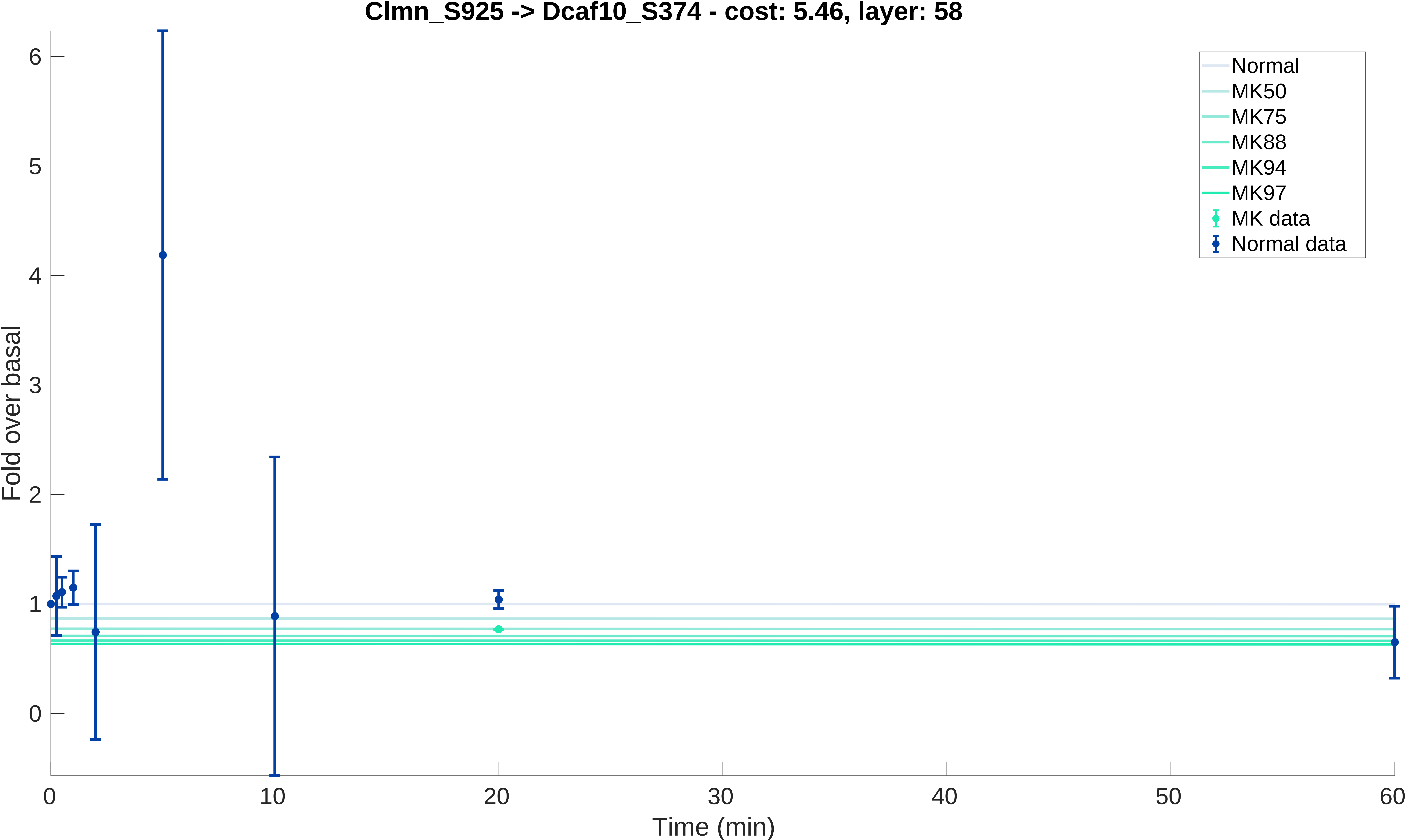
Clmn_S925 -> Dcaf10_S374 - cost: 5.46, layer: 58
Fold over basal
Time (min)
0
10
20
30
40
50
60
6
5
4
3
2
1
0
Normal
MK50
MK75
MK88
MK94
MK97
MK data
Normal data

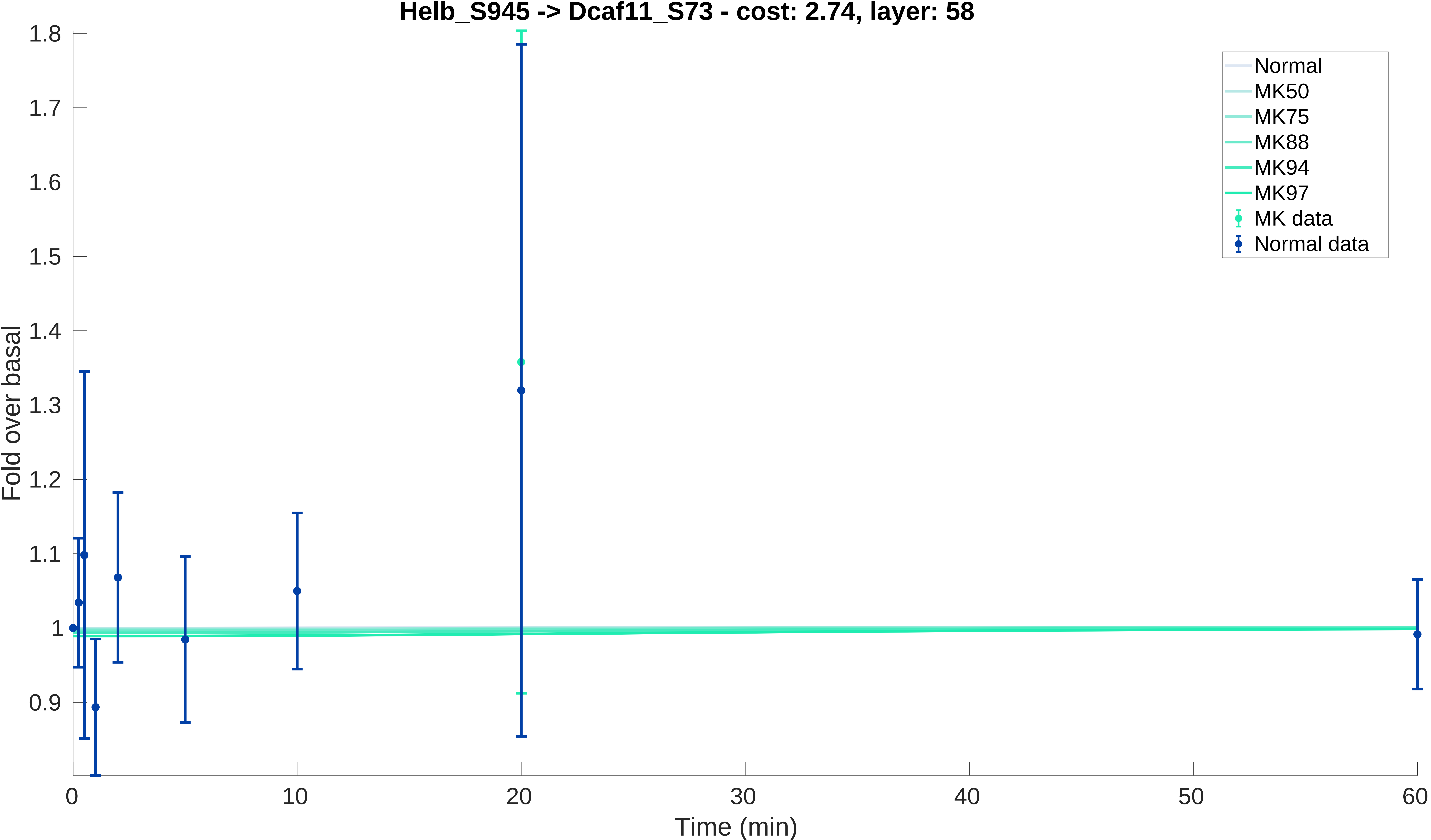

Helb_S945 -> Dcaf11_S73 - cost: 2.74, layer: 58
Fold over basal
Time (min)
Normal
MK50
MK75
MK88
MK94
MK97
MK data
Normal data
1.8
1.7
1.6
1.5
1.4
1.3
1.2
1.1
1
0.9
0
10
20
30
40
50
60

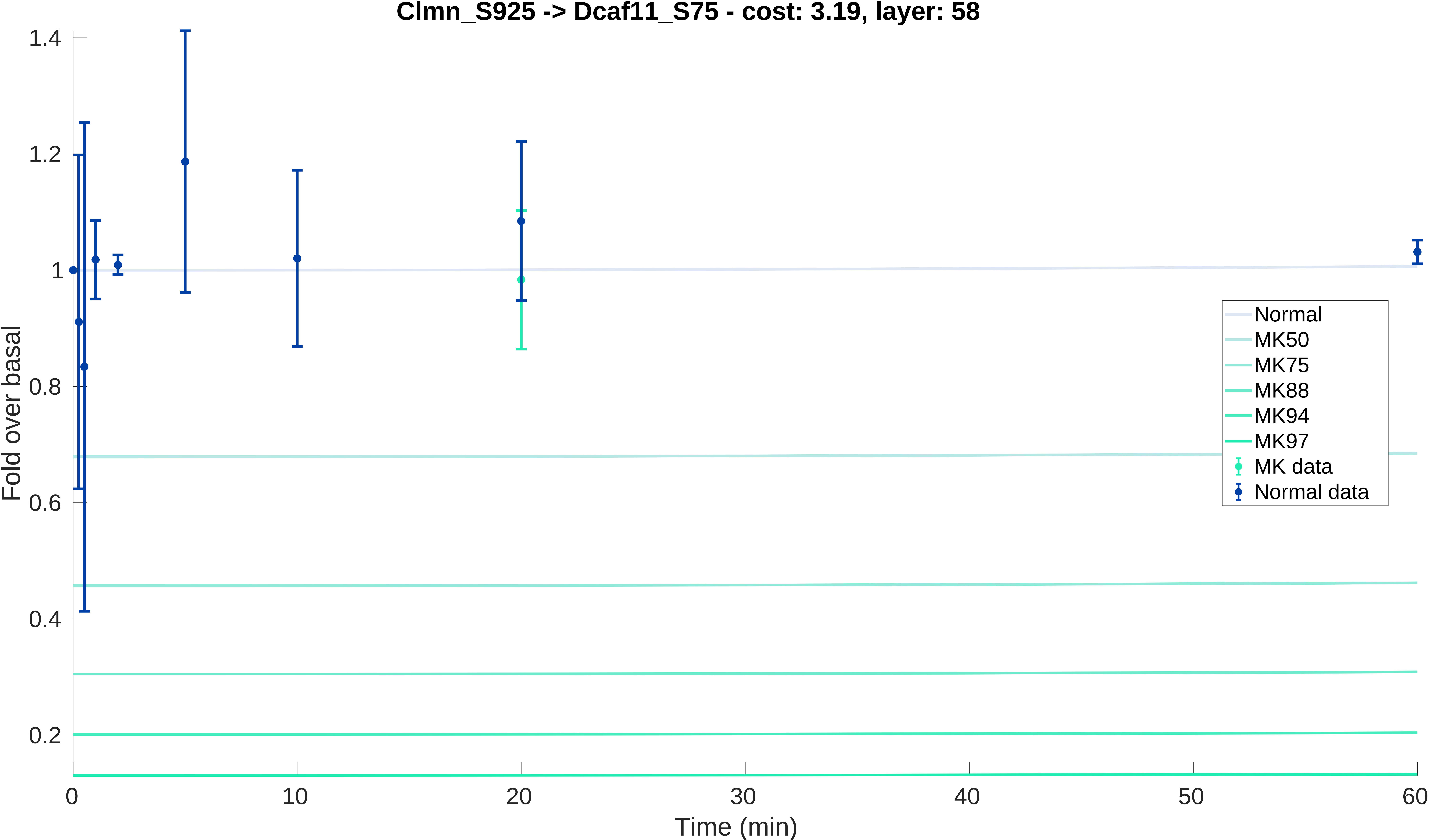
Clmn_S925 -> Dcaf11_S75 - cost: 3.19, layer: 58
Fold over basal
Time (min)
1.4
1.2
1
0.8
0.6
0.4
0.2
0
10
20
30
40
50
60
Normal
MK50
MK75
MK88
MK94
MK97
MK data
Normal data

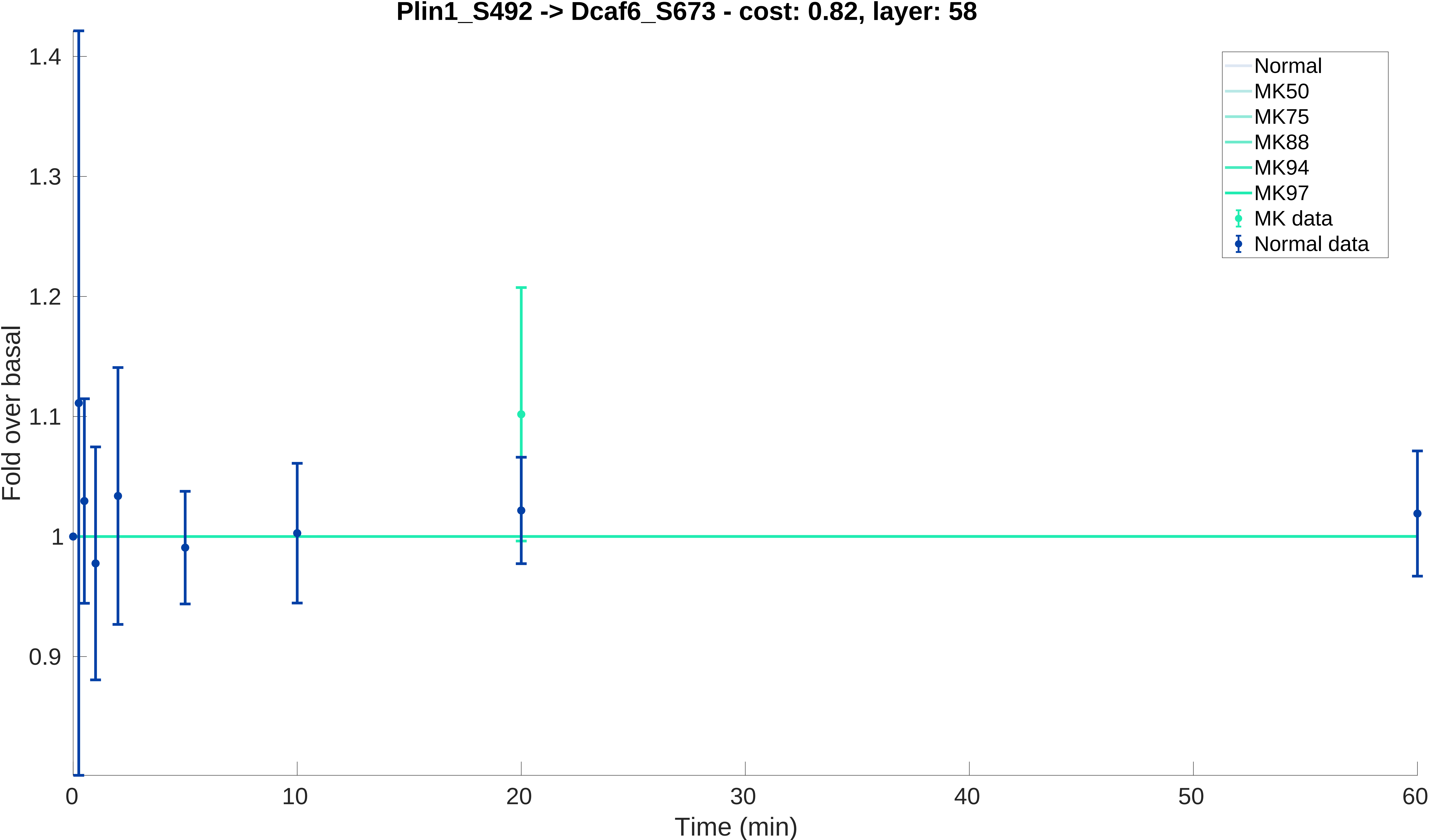

Plin1_S492 -> Dcaf6_S673 - cost: 0.82, layer: 58
Fold over basal
Time (min)
1.4
1.3
1.2
1.1
1
0.9
0
10
20
30
40
50
60
Normal
MK50
MK75
MK88
MK94
MK97
MK data
Normal data

Fxr2_T413 -> Dcaf6_T670 - cost: 0.59, layer: 58

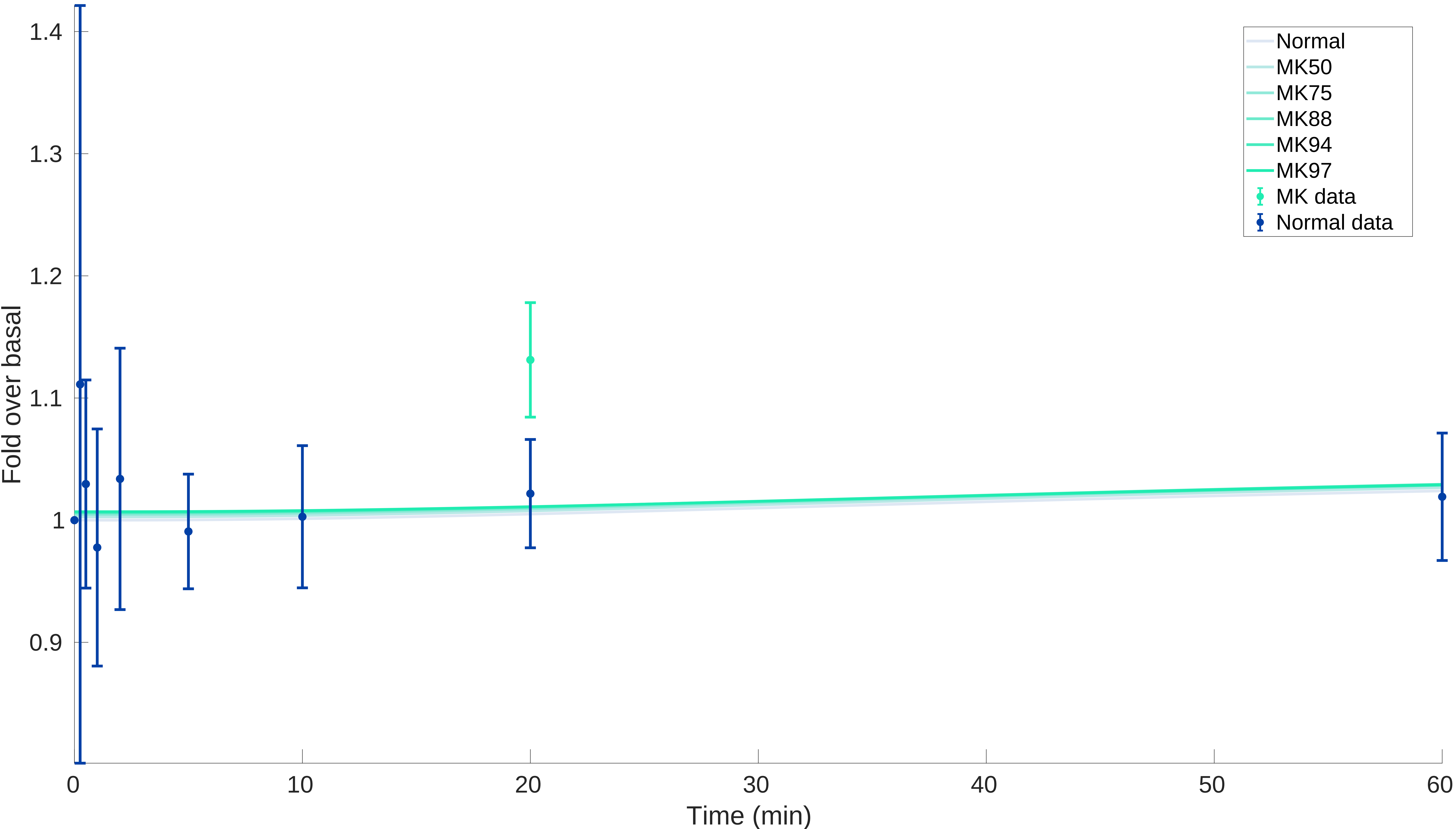

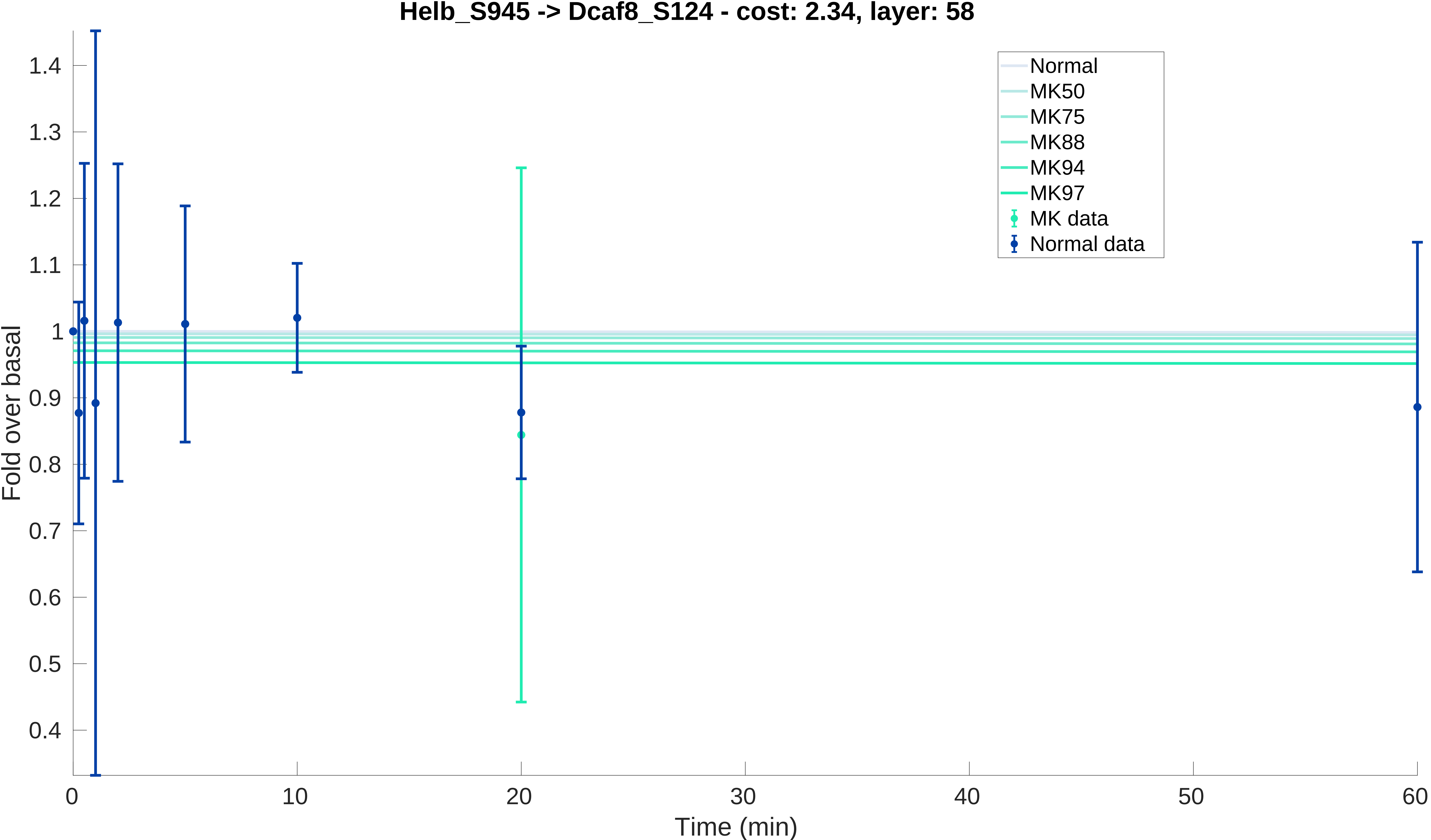

Helb_S945 -> Dcaf8_S124 - cost: 2.34, layer: 58
Normal
MK50
MK75
MK88
MK94
MK97
MK data
Normal data
Fold over basal
Time (min)
1.4
1.3
1.2
1.1
1
0.9
0.8
0.7
0.6
0.5
0.4
0
10
20
30
40
50
60

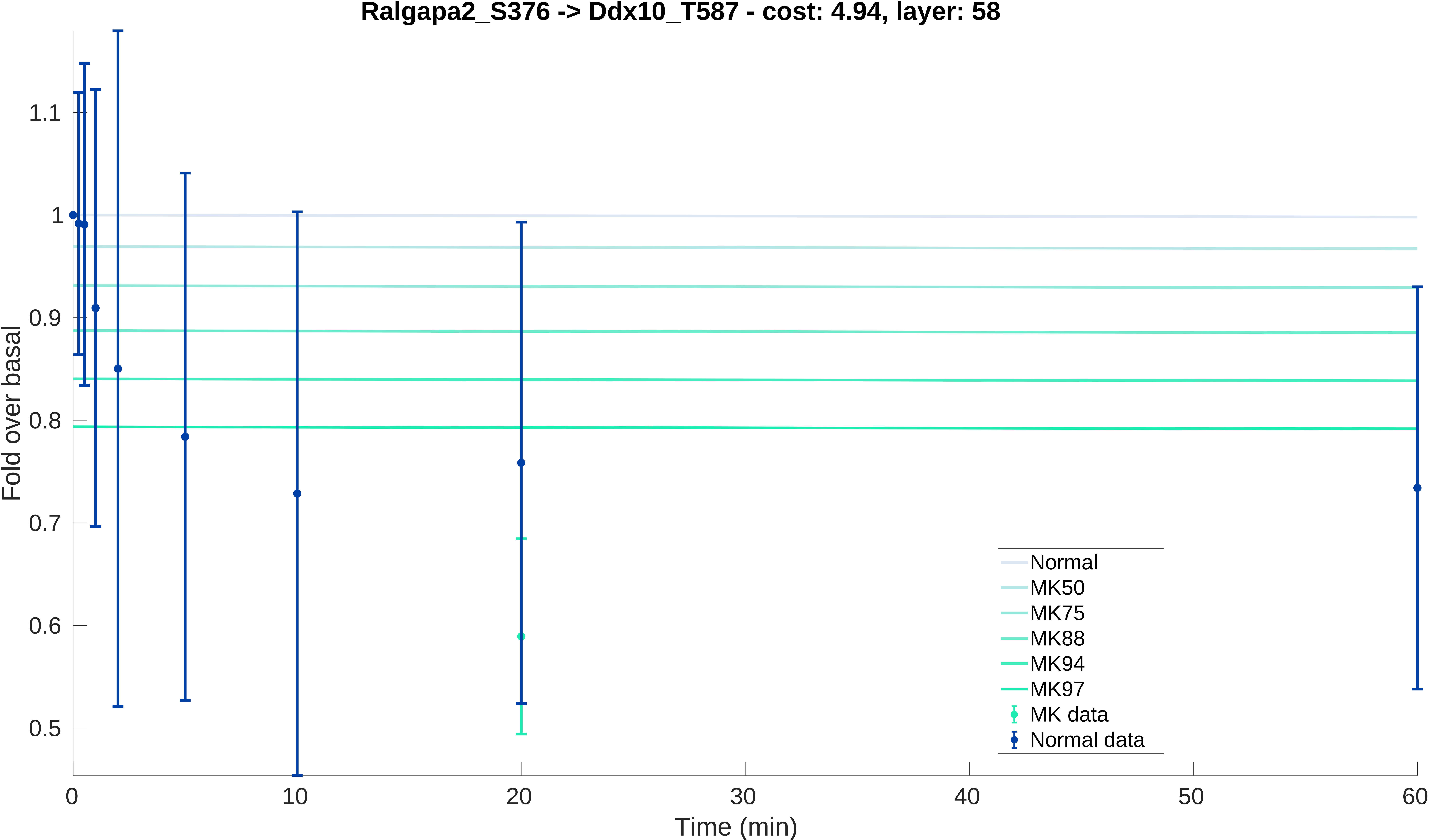
Ralgapa2_S376 -> Ddx10_T587 - cost: 4.94, layer: 58
Fold over basal
Time (min)
1.1
1
0.9
0.8
0.7
0.6
0.5
0
10
20
30
40
50
60
Normal
MK50
MK75
MK88
MK94
MK97
MK data
Normal data

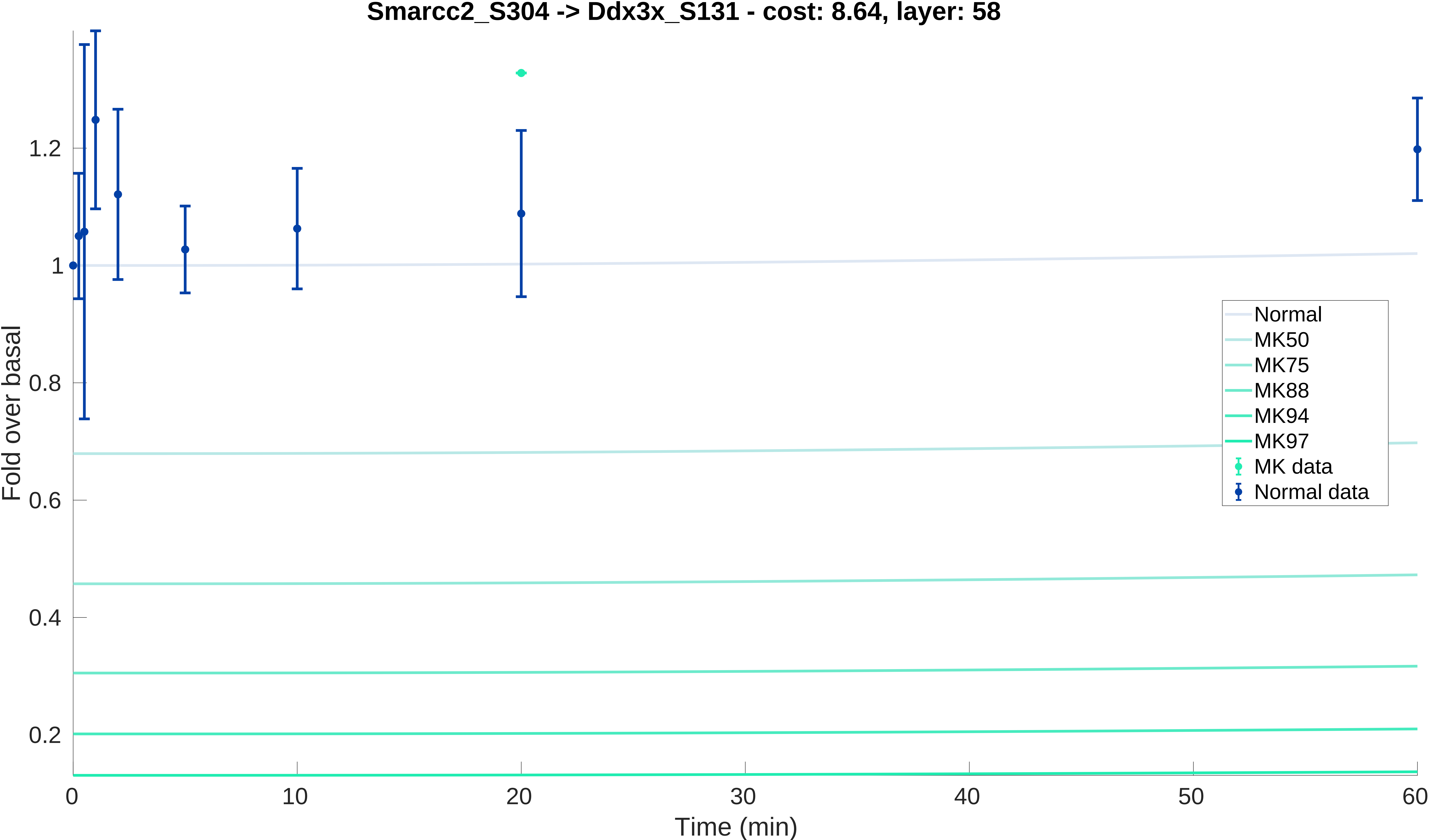

Smarcc2_S304 -> Ddx3x_S131 - cost: 8.64, layer: 58
Fold over basal
1.2
1
0.8
0.6
0.4
0.2
0
10
20
30
40
50
60
Time (min)
Normal
MK50
MK75
MK88
MK94
MK97
MK data
Normal data

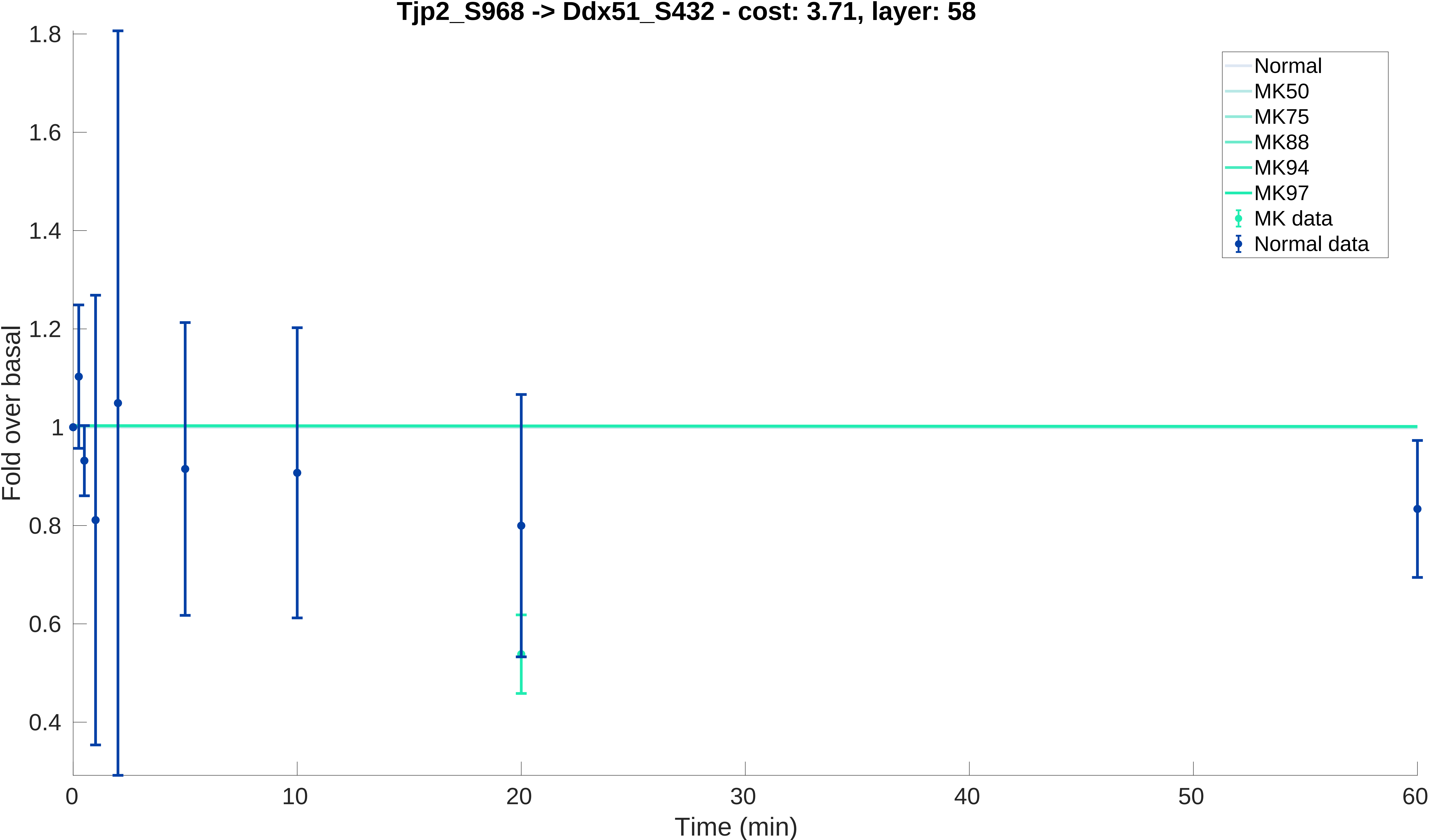

Tjp2_S968 -> Ddx51_S432 - cost: 3.71, layer: 58
Fold over basal
Time (min)
Normal
MK50
MK75
MK88
MK94
MK97
MK data
Normal data
1.8
1.6
1.4
1.2
1
0.8
0.6
0.4
0
10
20
30
40
50
60

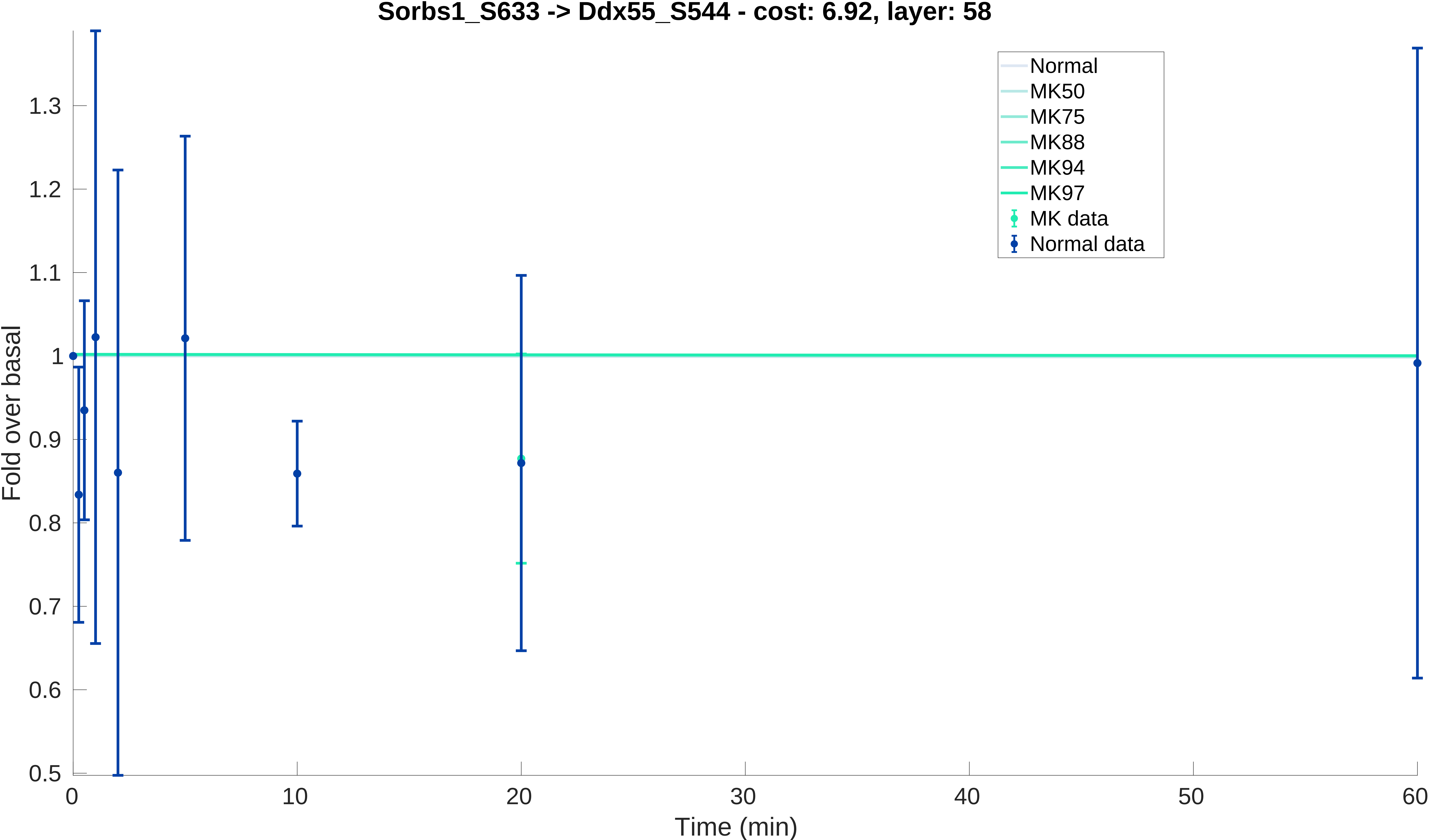

Sorbs1_S633 -> Ddx55_S544 - cost: 6.92, layer: 58
Normal
MK50
MK75
MK88
MK94
MK97
MK data
Normal data
Fold over basal
1.3
1.2
1.1
1
0.9
0.8
0.7
0.6
0.5
0
10
20
30
40
50
60
Time (min)

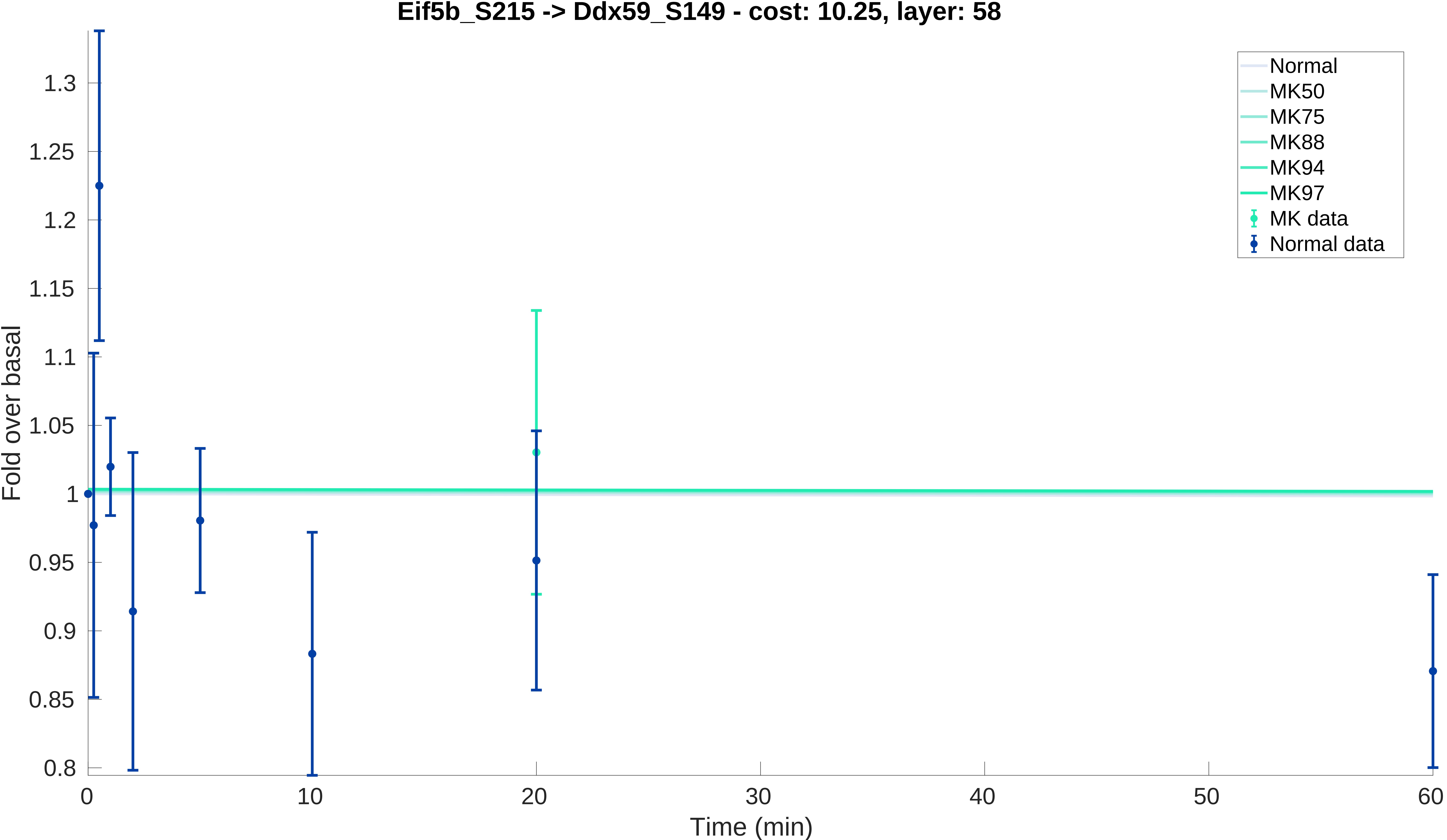

Eif5b_S215 -> Ddx59_S149 - cost: 10.25, layer: 58
Fold over basal
1.3
1.25
1.2
1.15
1.1
1.05
1
0.95
0.9
0.85
0.8
0
10
20
30
40
50
60
Time (min)
Normal
MK50
MK75
MK88
MK94
MK97
MK data
Normal data

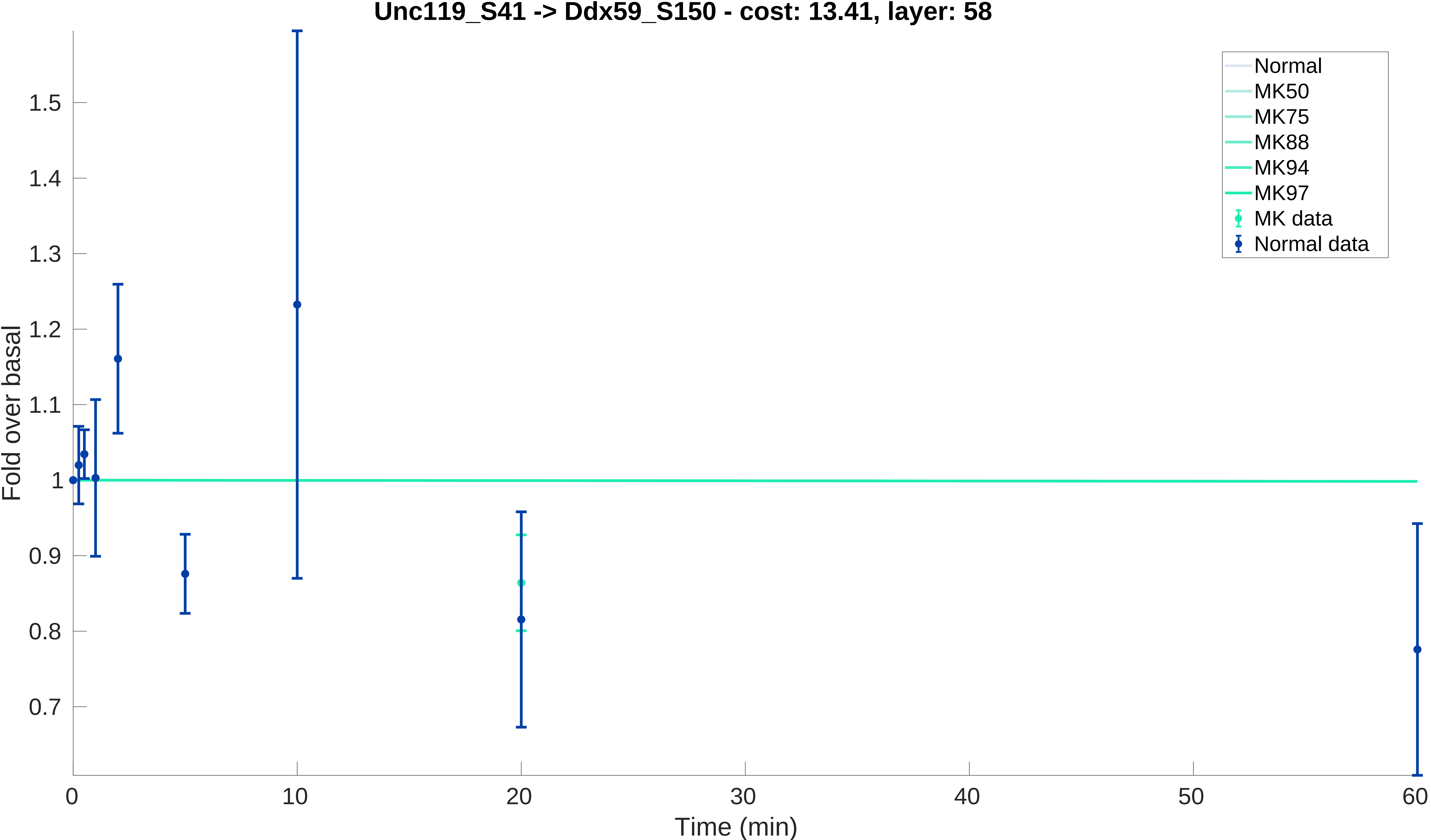

Unc119_S41 -> Ddx59_S150 - cost: 13.41, layer: 58
Fold over basal
Time (min)
1.5
1.4
1.3
1.2
1.1
1
0.9
0.8
0.7
0
10
20
30
40
50
60
Normal
MK50
MK75
MK88
MK94
MK97
MK data
Normal data

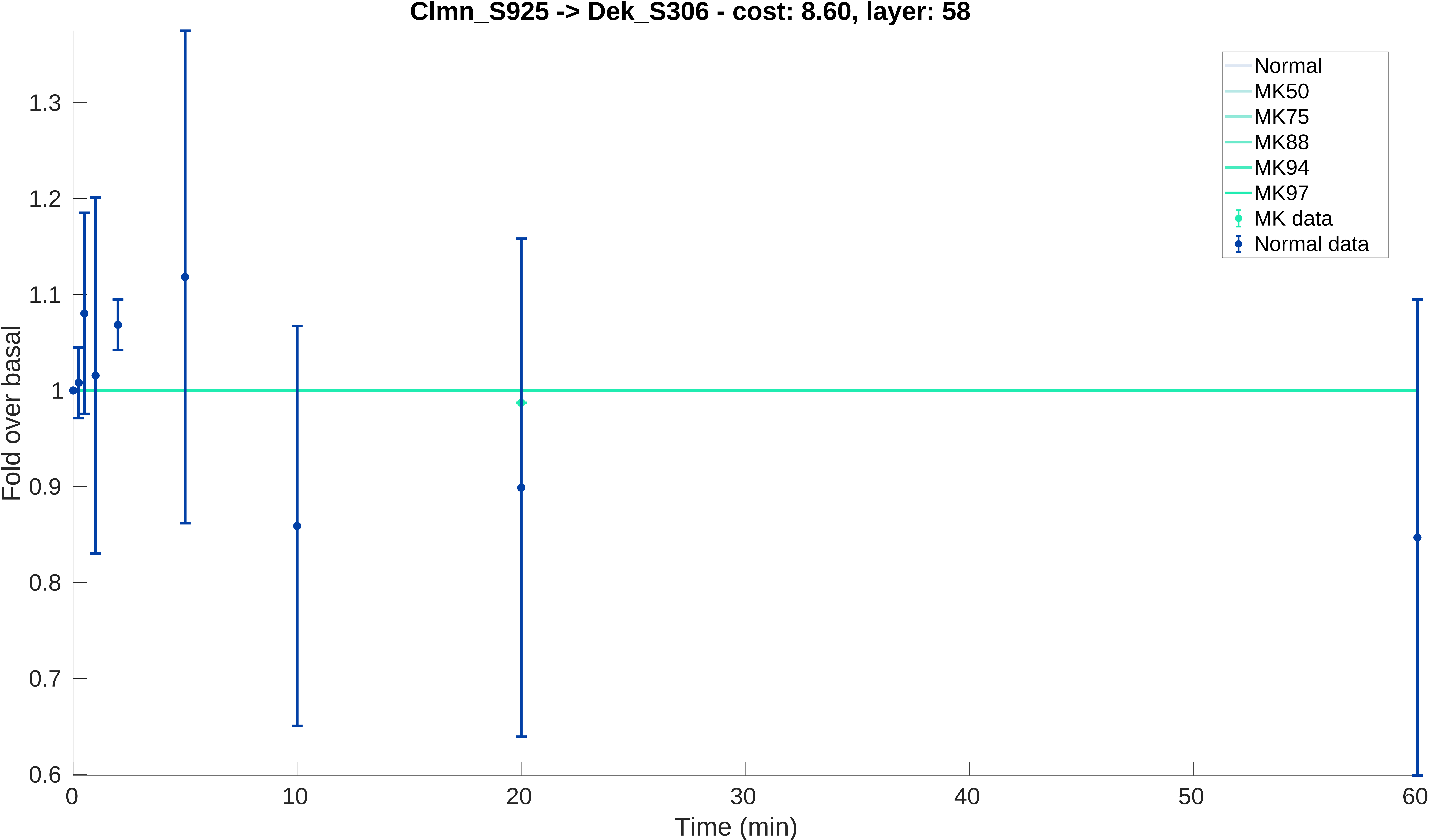
Clmn_S925 -> Dek_S306 - cost: 8.60, layer: 58
Fold over basal
1.3
1.2
1.1
1
0.9
0.8
0.7
0.6
0
10
20
30
40
50
60
Time (min)
Normal
MK50
MK75
MK88
MK94
MK97
MK data
Normal data

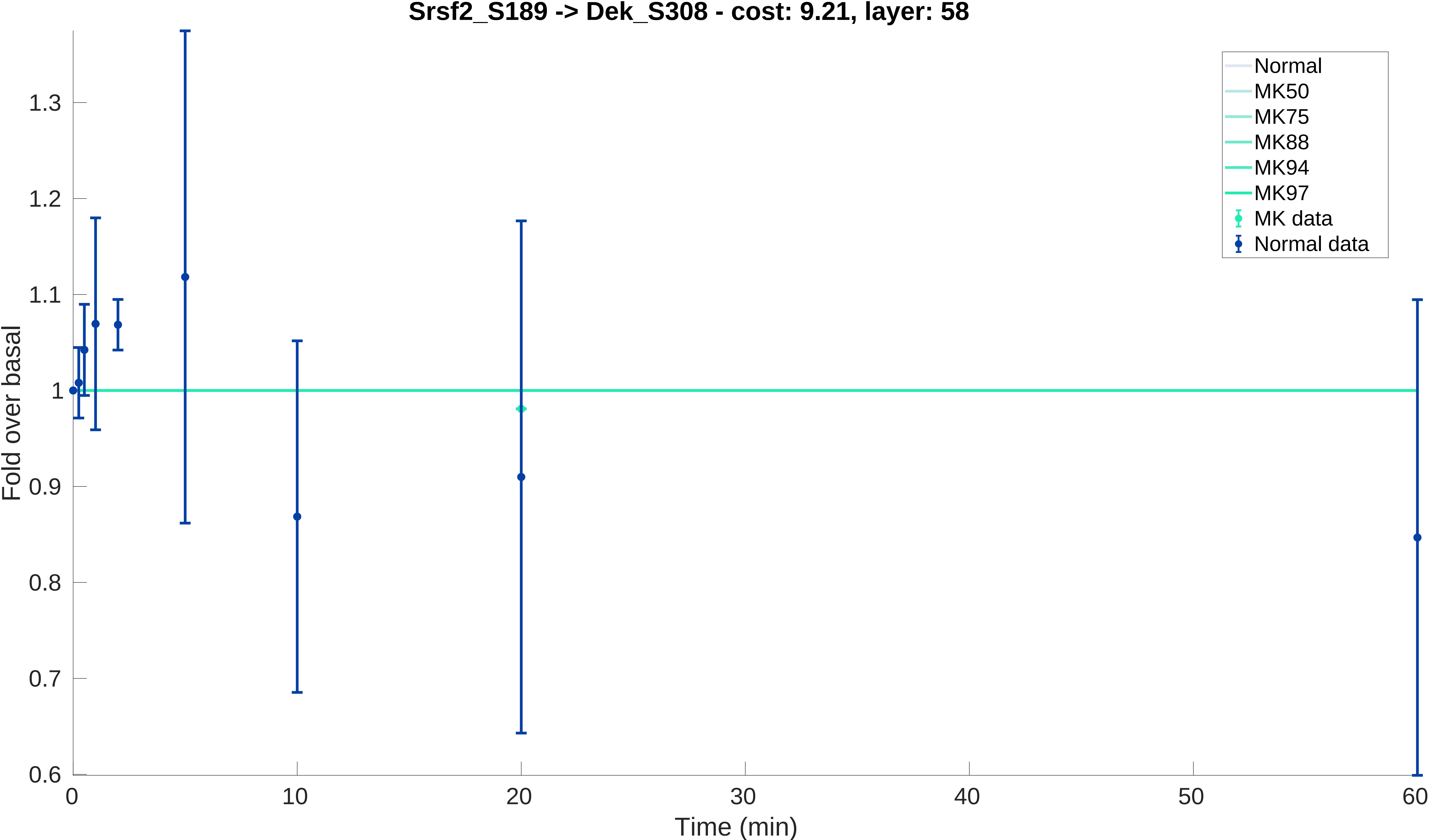

Srsf2_S189 -> Dek_S308 - cost: 9.21, layer: 58
Normal
MK50
MK75
MK88
MK94
MK97
MK data
Normal data
Fold over basal
Time (min)
0.6
0.7
0.8
0.9
1
1.1
1.2
1.3
0
10
20
30
40
50
60

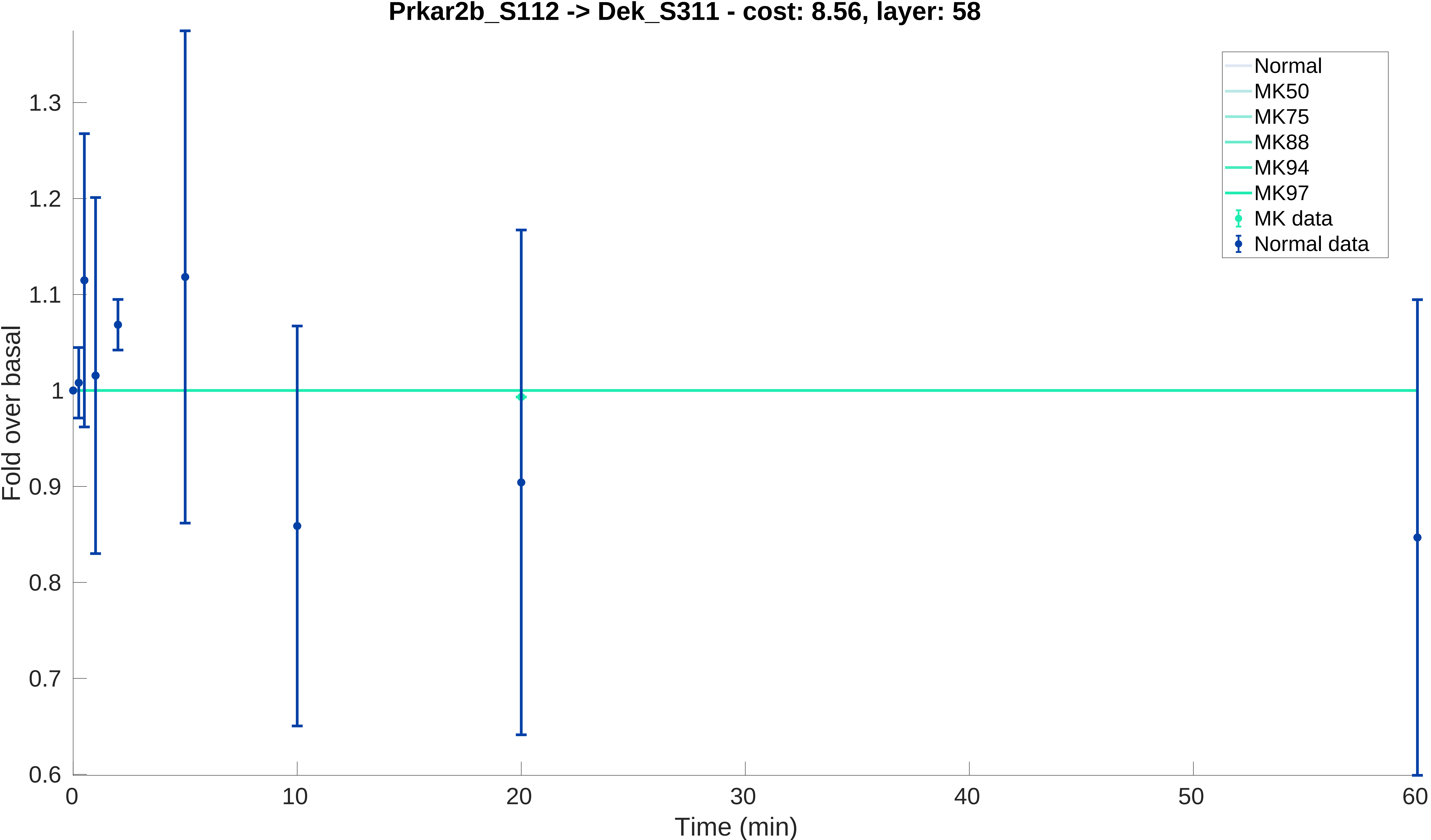

Prrkar2b_S112 -> Dek_S311 - cost: 8.56, layer: 58
Fold over basal
Time (min)
Normal
MK50
MK75
MK88
MK94
MK97
MK data
Normal data
0.6
0.7
0.8
0.9
1
1.1
1.2
1.3
0
10
20
30
40
50
60

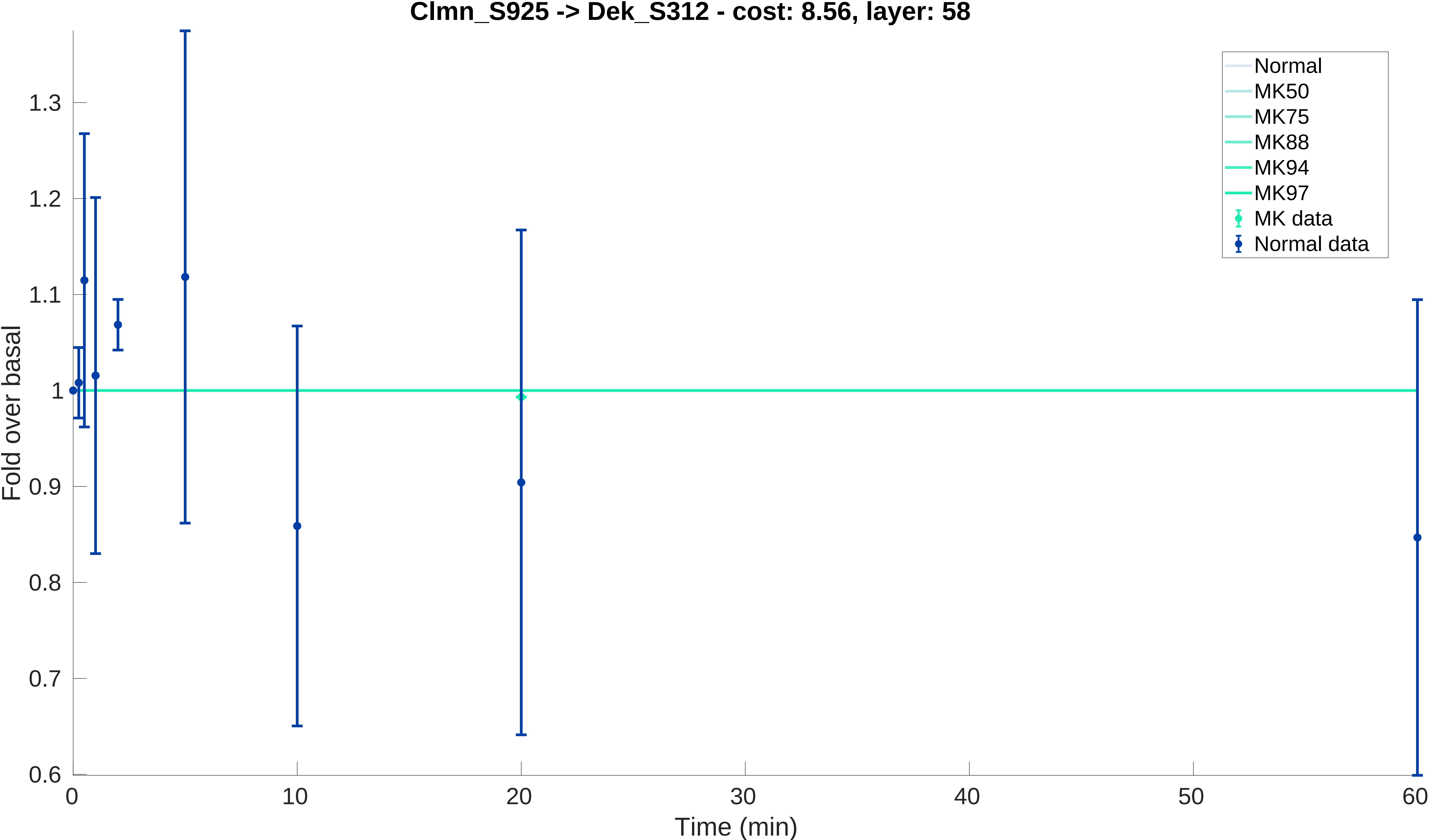

Clmn_S925 -> Dek_S312 - cost: 8.56, layer: 58
Fold over basal
Time (min)
1.3
1.2
1.1
1
0.9
0.8
0.7
0.6
0
10
20
30
40
50
60
Normal
MK50
MK75
MK88
MK94
MK97
MK data
Normal data

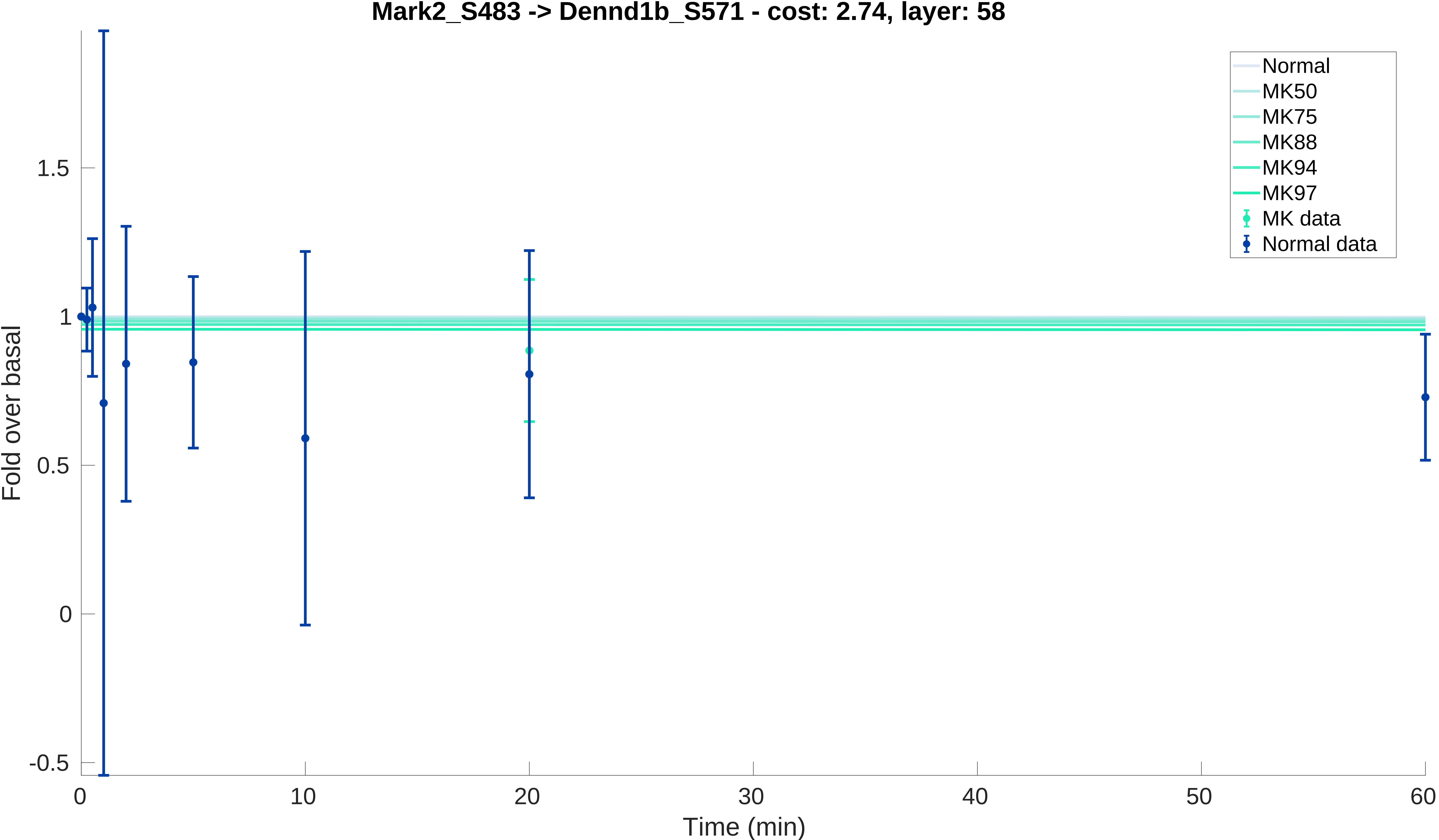

Mark2_S483 -> Dennd1b_S571 - cost: 2.74, layer: 58
Normal
MK50
MK75
MK88
MK94
MK97
MK data
Normal data
Fold over basal
1.5
1
0.5
0
-0.5
0
10
20
30
40
50
60
Time (min)

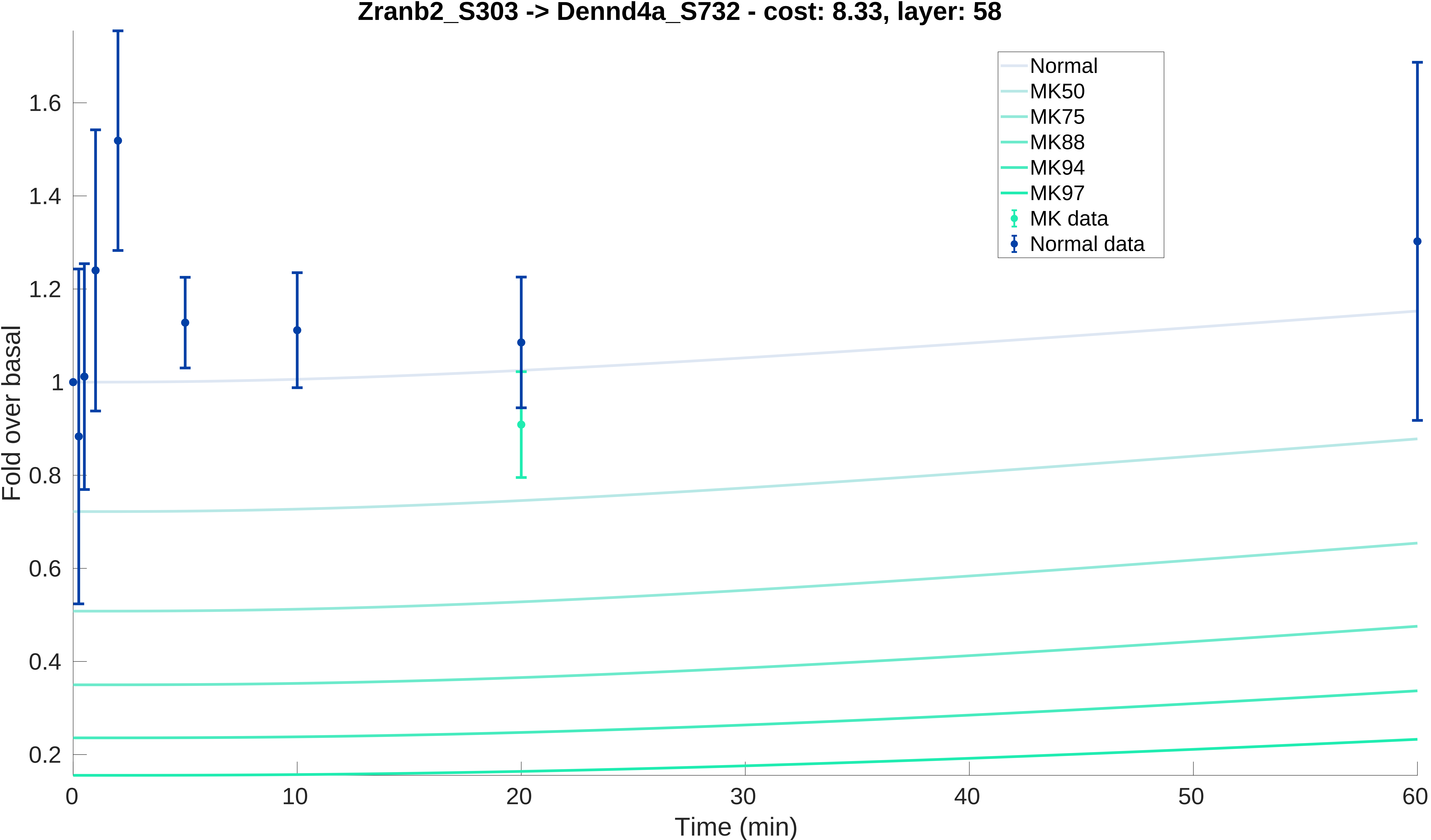

Zranb2_S303 -> Dennd4a_S732 - cost: 8.33, layer: 58
Normal
MK50
MK75
MK88
MK94
MK97
MK data
Normal data
Fold over basal
Time (min)
0.2
0.4
0.6
0.8
1
1.2
1.4
1.6
0
10
20
30
40
50
60

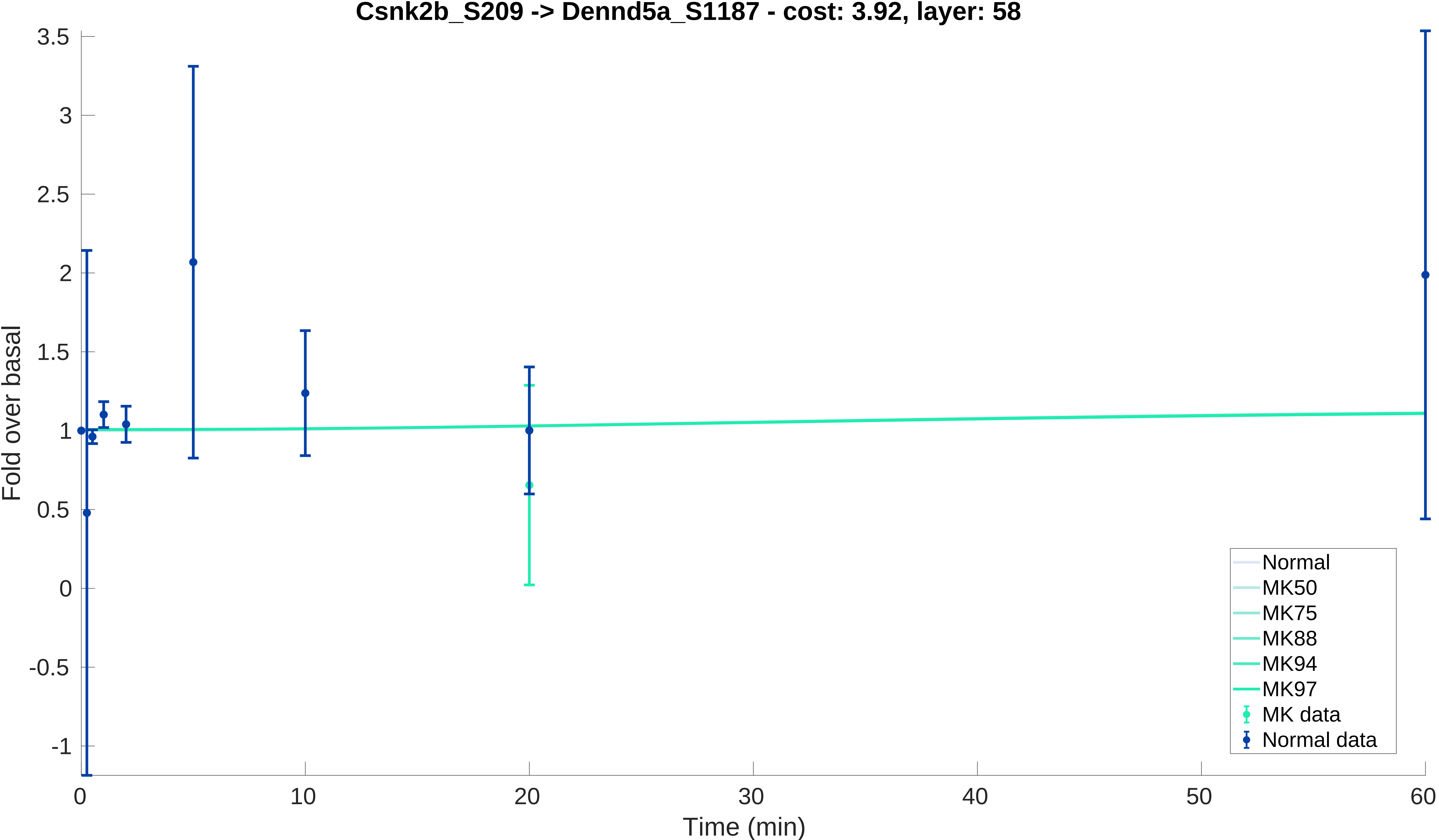

Csnk2b_S209 -> Dennd5a_S1187 - cost: 3.92, layer: 58
Fold over basal
3.5
3
2.5
2
1.5
1
0.5
0
-0.5
-1
0
10
20
30
40
50
60
Time (min)
Normal
MK50
MK75
MK88
MK94
MK97
MK data
Normal data

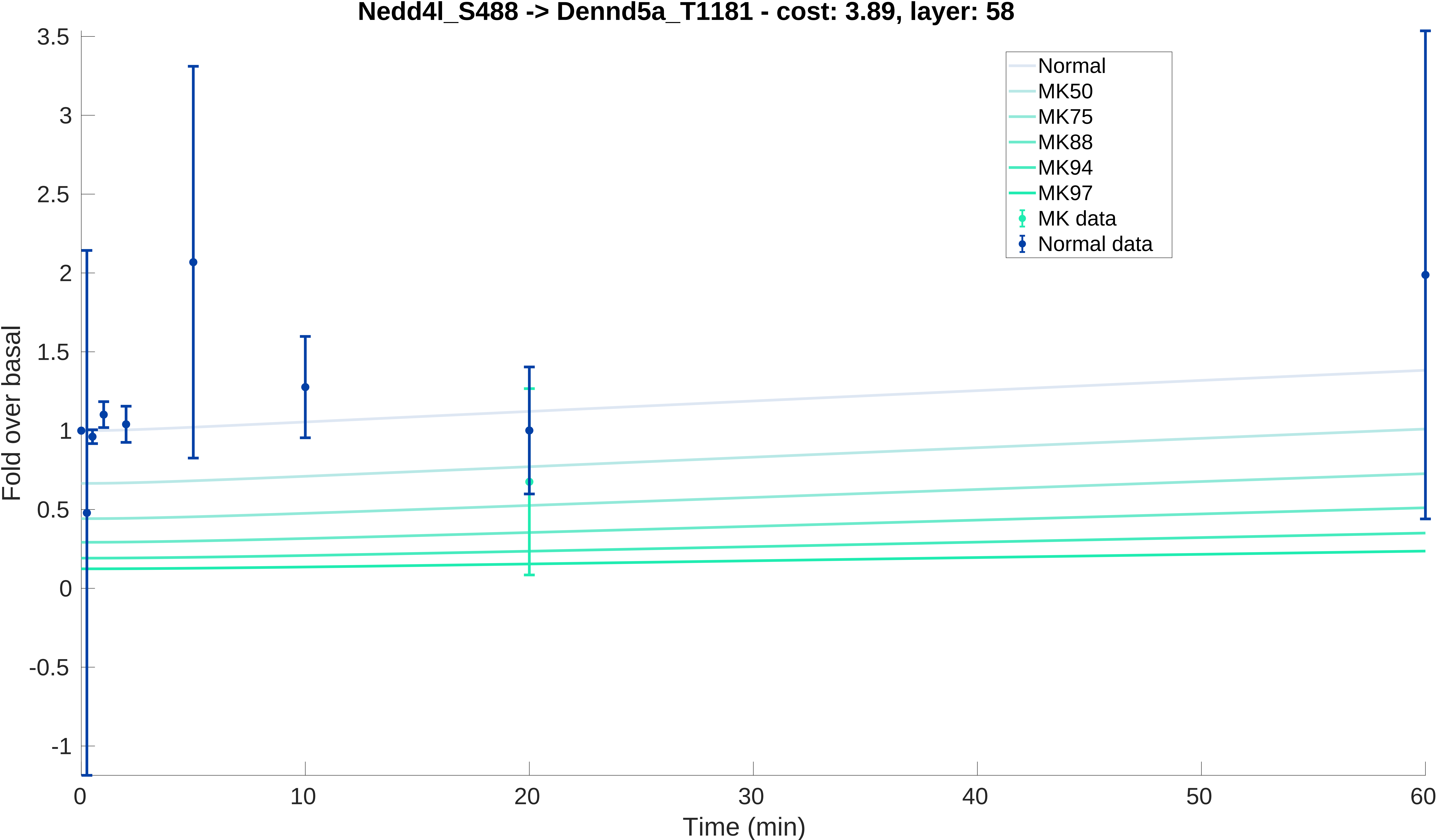

Nedd4l_S488 -> Dennd5a_T1181 - cost: 3.89, layer: 58
Fold over basal
Time (min)
3.5
3
2.5
2
1.5
1
0.5
0
-0.5
-1
0
10
20
30
40
50
60
Normal
MK50
MK75
MK88
MK94
MK97
MK data
Normal data

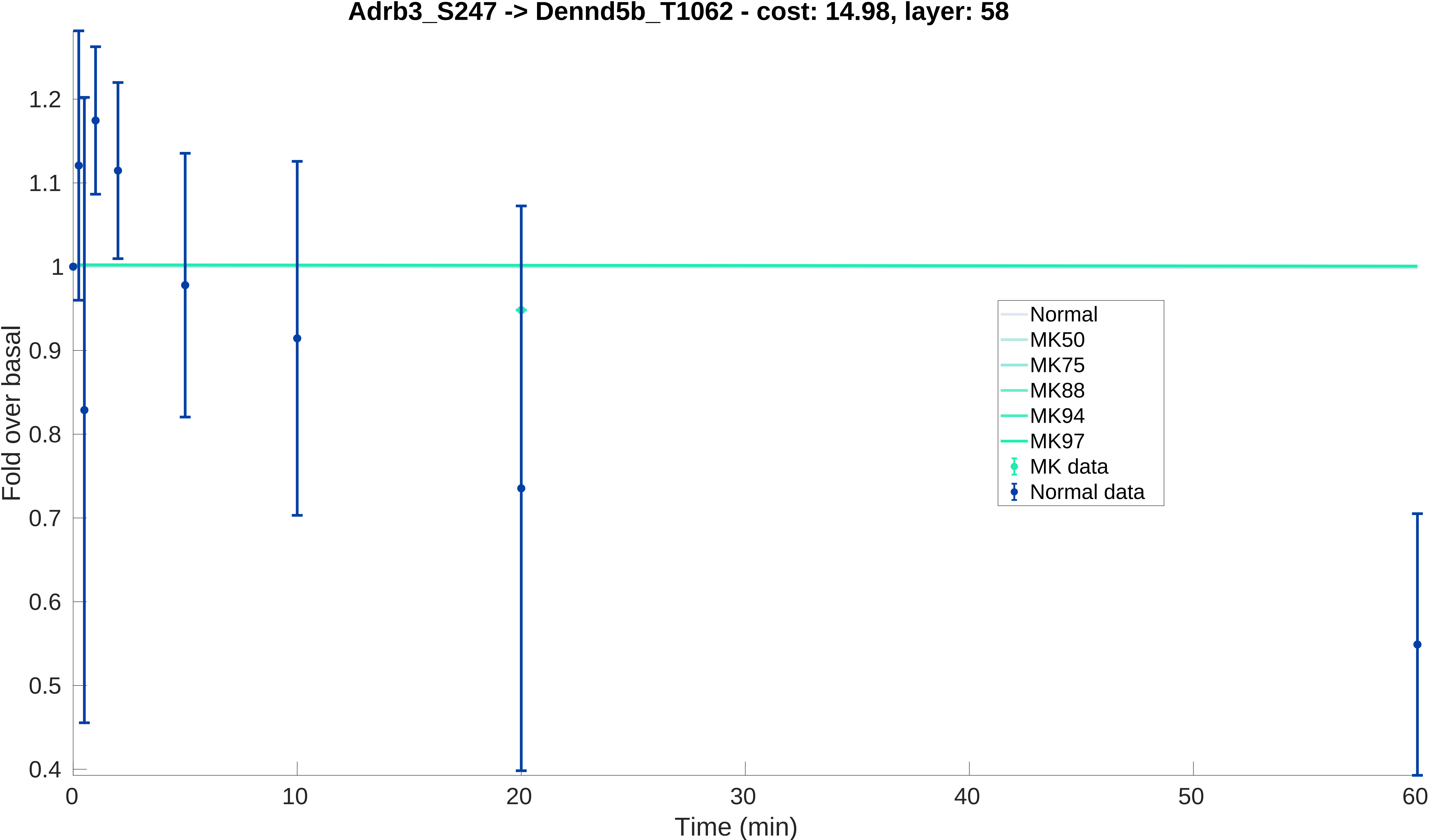

Adrb3_S247 -> Dennd5b_T1062 - cost: 14.98, layer: 58
Fold over basal
1.2
1.1
1
0.9
0.8
0.7
0.6
0.5
0.4
0
10
20
30
40
50
60
Time (min)
Normal
MK50
MK75
MK88
MK94
MK97
MK data
Normal data

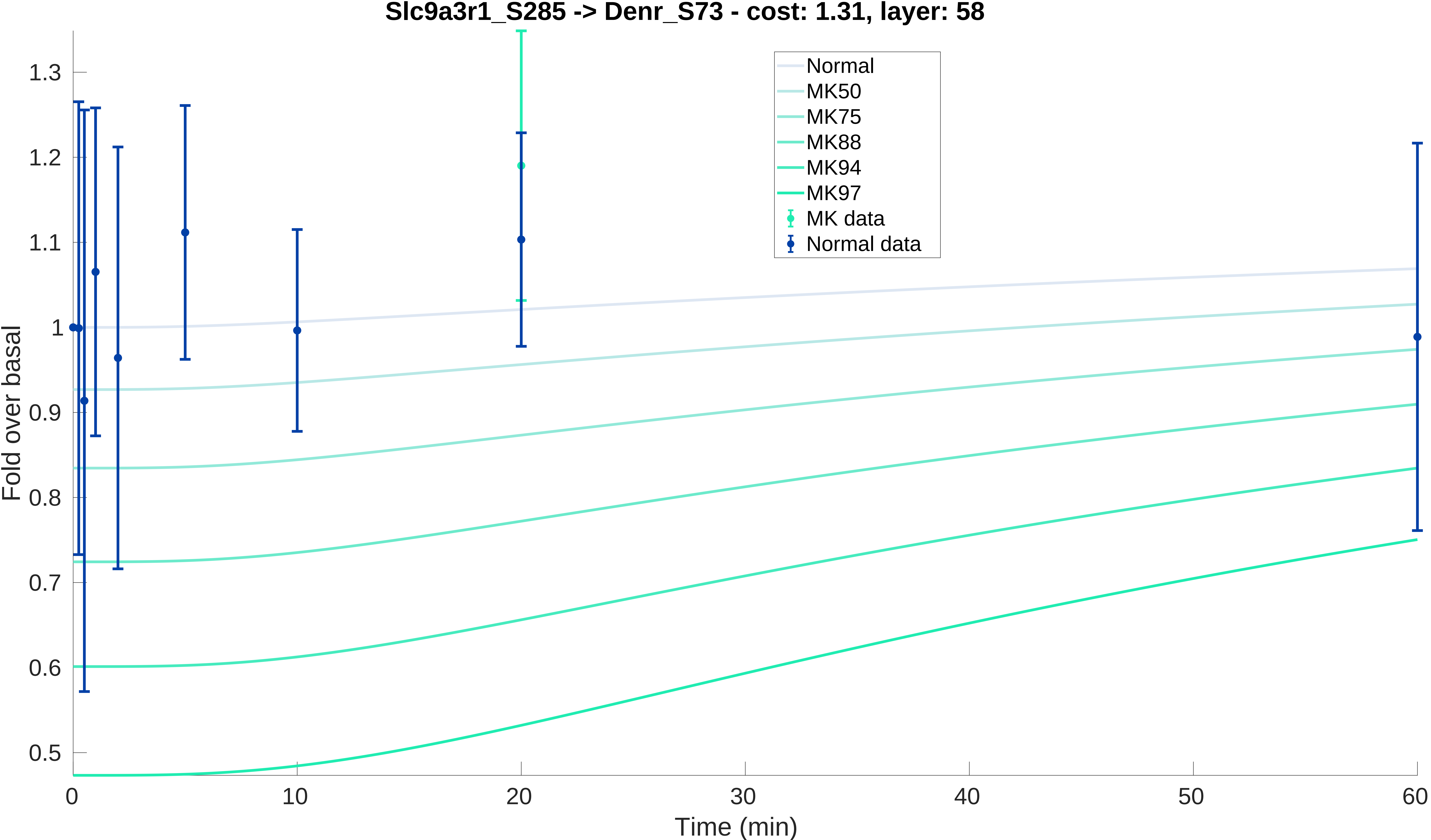

Slc9a3r1_S285 -> Denr_S73 - cost: 1.31, layer: 58
Normal
MK50
MK75
MK88
MK94
MK97
MK data
Normal data
Fold over basal
Time (min)
0.5
0.6
0.7
0.8
0.9
1
1.1
1.2
1.3
0
10
20
30
40
50
60

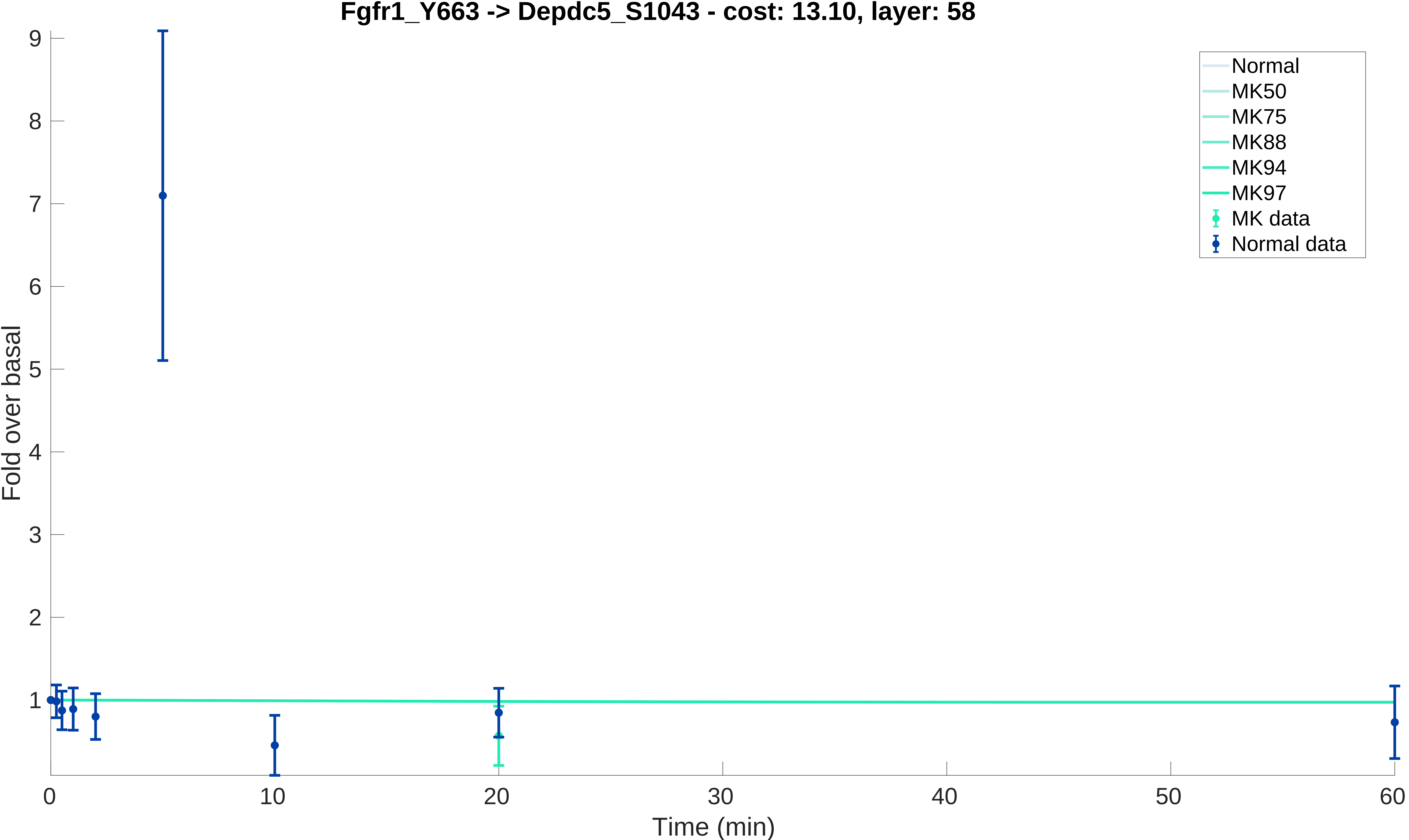
Fgfr1_Y663 -> Depdc5_S1043 - cost: 13.10, layer: 58
Fold over basal
Time (min)
0
10
20
30
40
50
60
1
2
3
4
5
6
7
8
9
Normal
MK50
MK75
MK88
MK94
MK97
MK data
Normal data

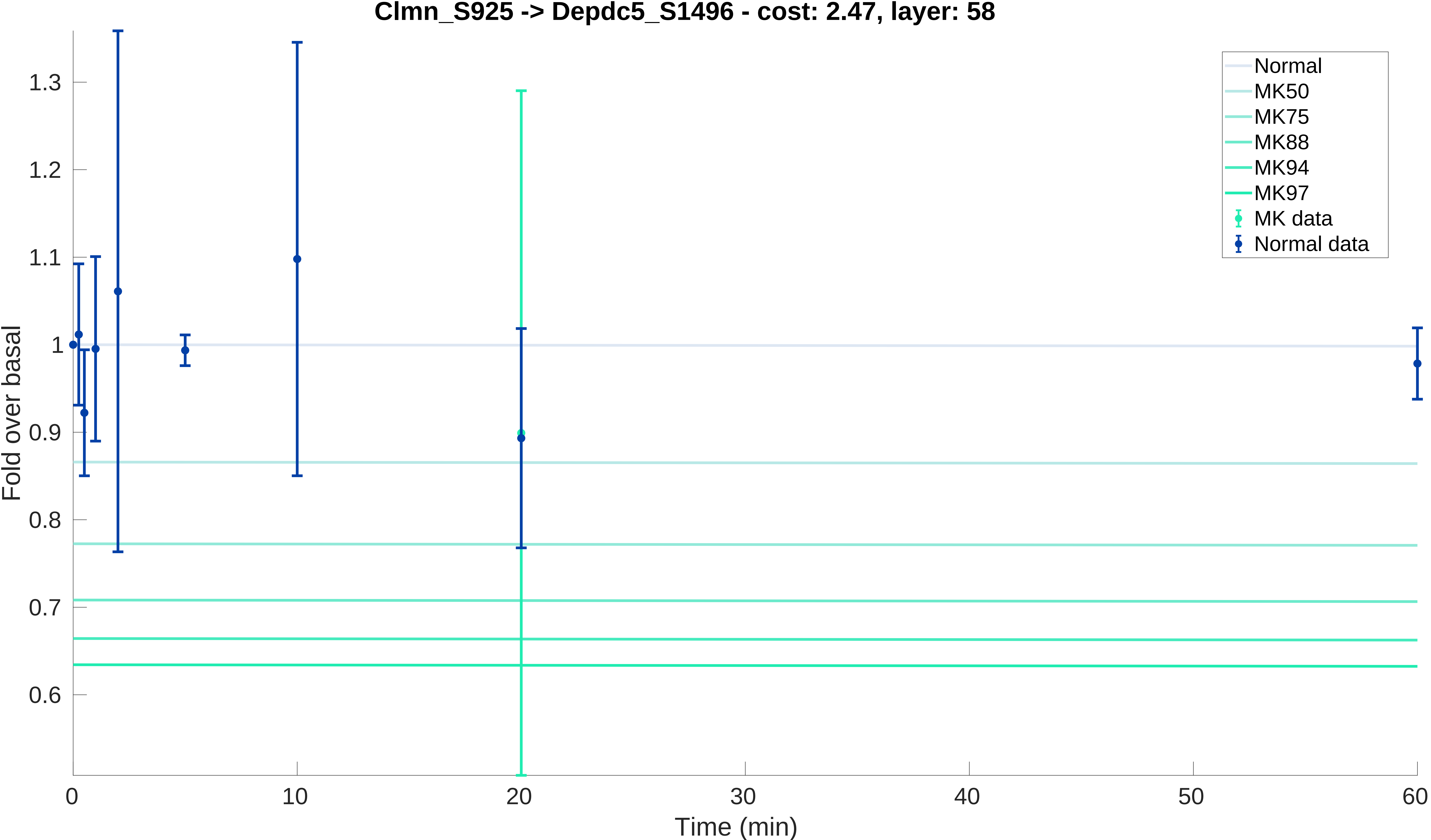

Clmn_S925 -> Depdc5_S1496 - cost: 2.47, layer: 58
Normal
MK50
MK75
MK88
MK94
MK97
MK data
Normal data
Fold over basal
1.3
1.2
1.1
1
0.9
0.8
0.7
0.6
0
10
20
30
40
50
60
Time (min)

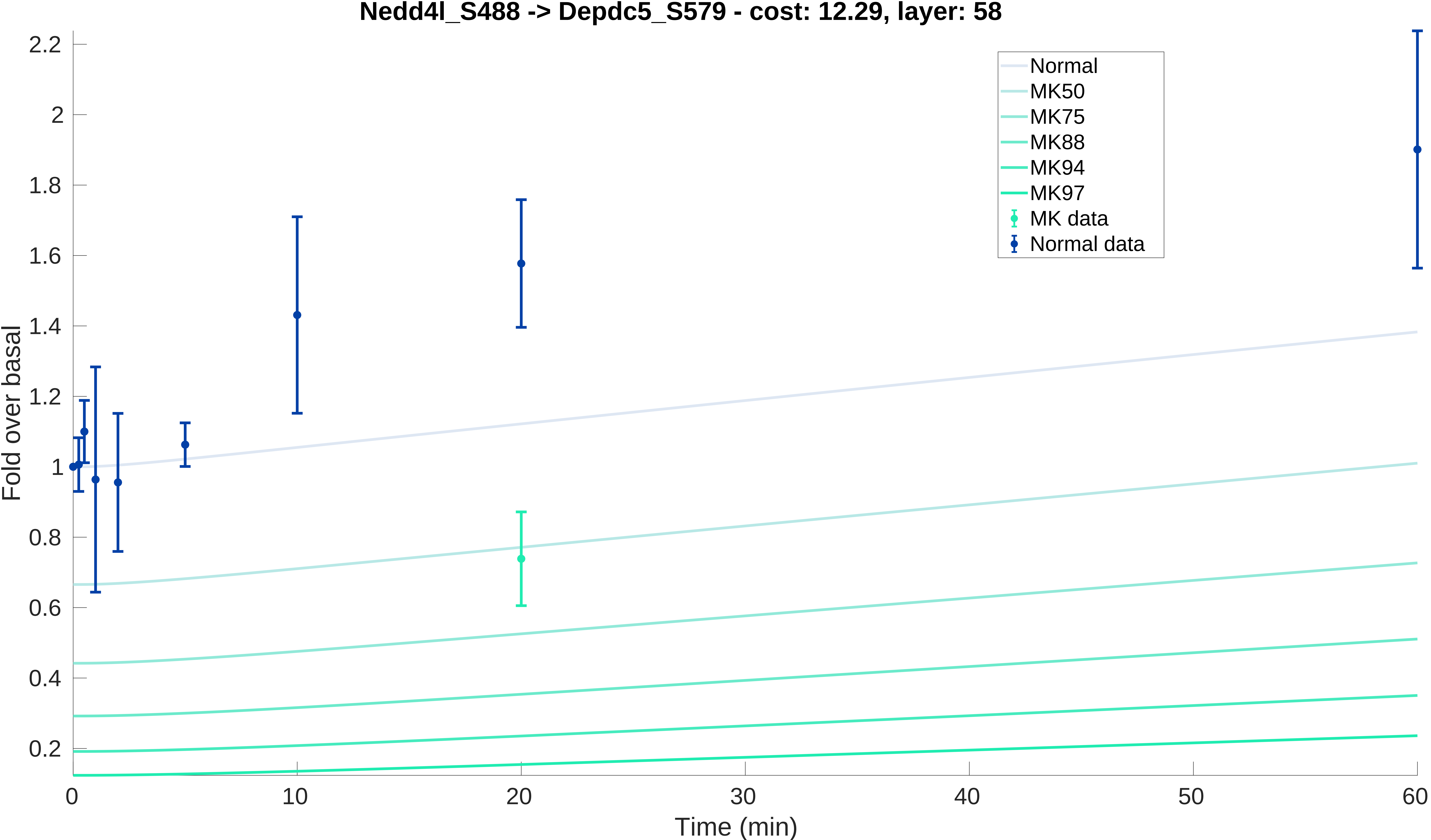

Nedd4l_S488 -> Depdc5_S579 - cost: 12.29, layer: 58
Fold over basal
Time (min)
Normal
MK50
MK75
MK88
MK94
MK97
MK data
Normal data
2.2
2
1.8
1.6
1.4
1.2
1
0.8
0.6
0.4
0.2
0
10
20
30
40
50
60

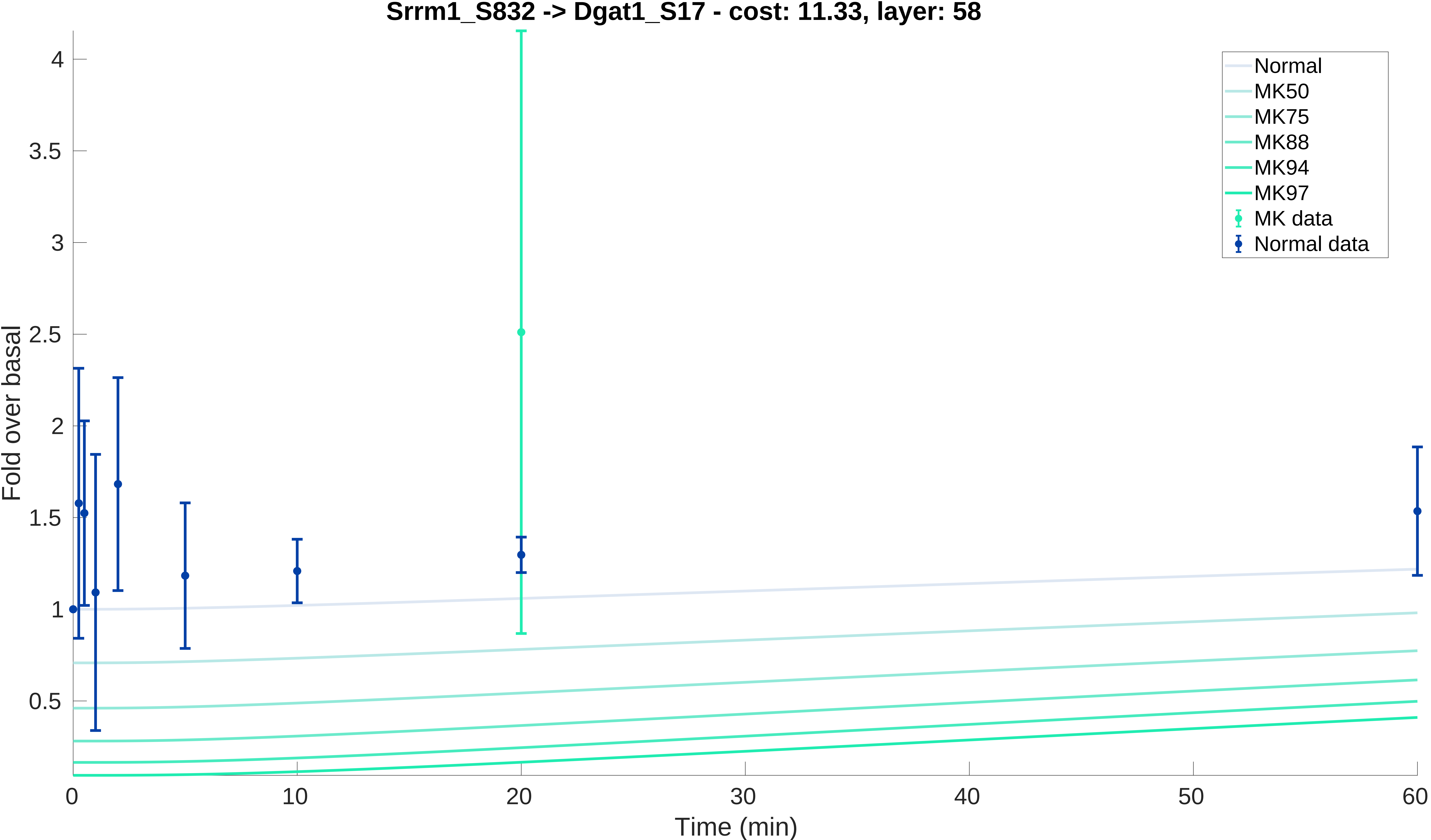

Srrm1_S832 -> Dgat1_S17 - cost: 11.33, layer: 58
Fold over basal
Time (min)
Normal
MK50
MK75
MK88
MK94
MK97
MK data
Normal data
0
10
20
30
40
50
60
0.5
1
1.5
2
2.5
3
3.5
4

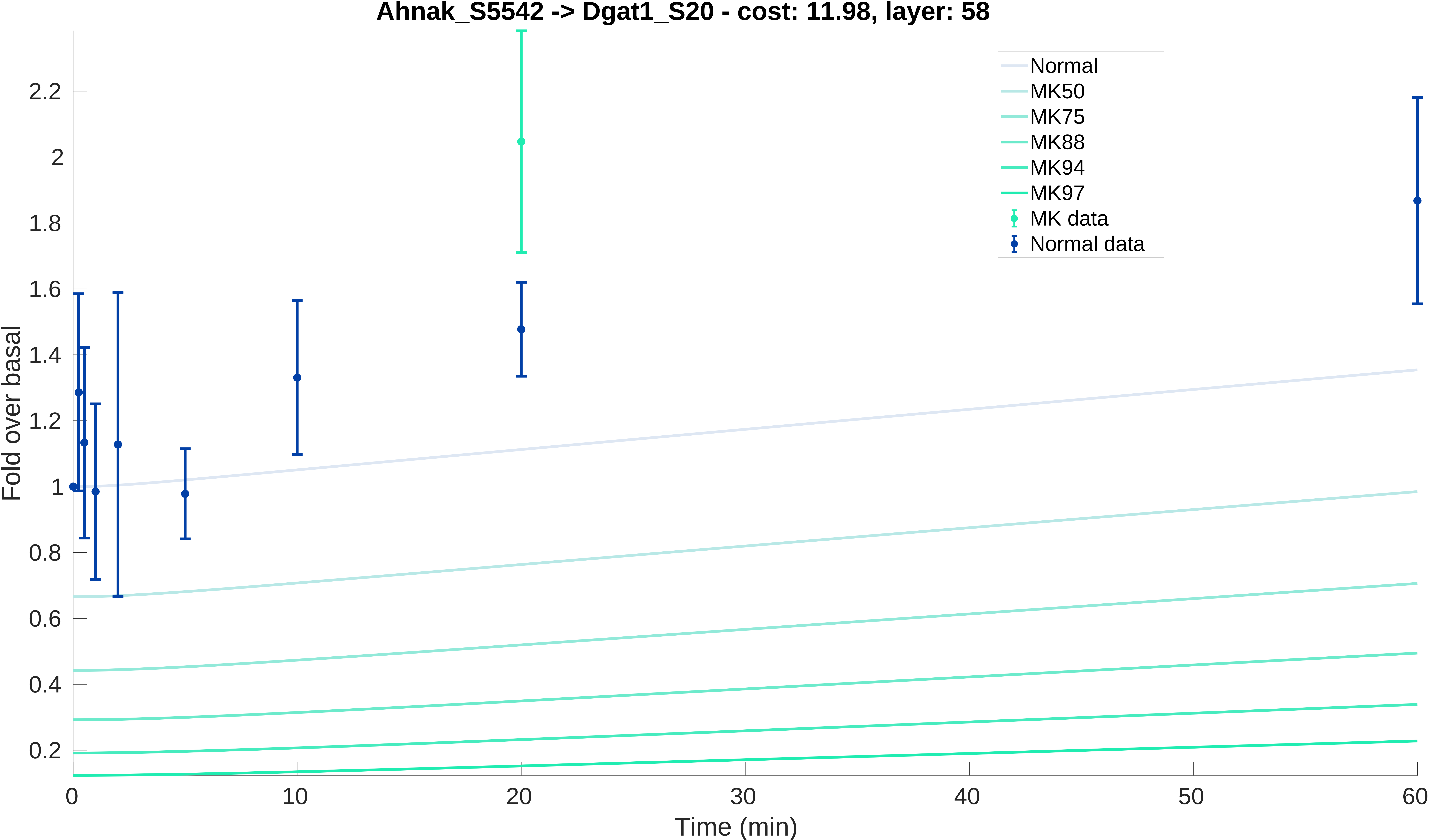

Ahnak_S5542 -> Dgat1_S20 - cost: 11.98, layer: 58
Fold over basal
Time (min)
0.2
0.4
0.6
0.8
1
1.2
1.4
1.6
1.8
2
2.2
0
10
20
30
40
50
60
Normal
MK50
MK75
MK88
MK94
MK97
MK data
Normal data

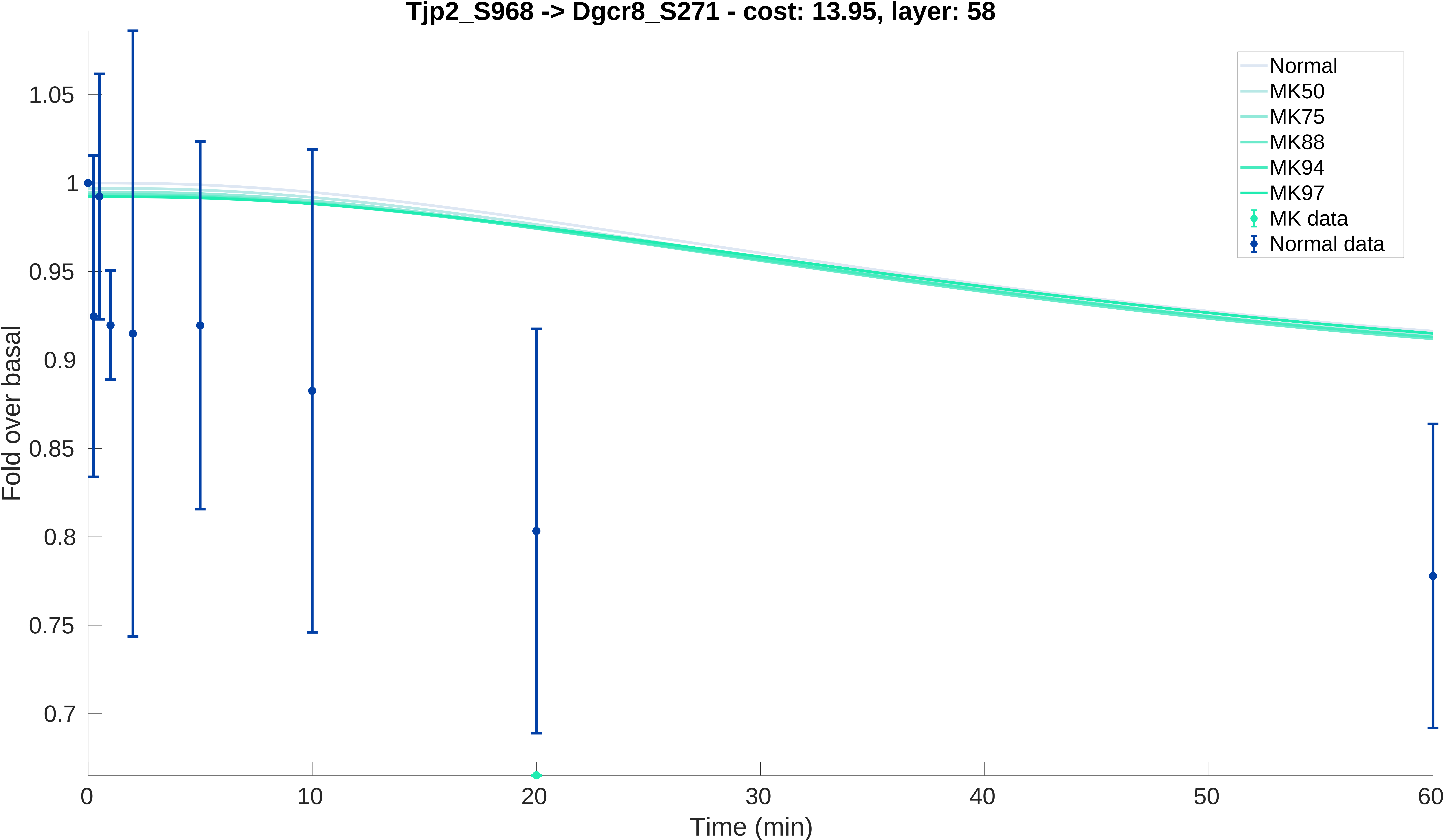

Tjp2_S968 -> Dgcr8_S271 - cost: 13.95, layer: 58
Normal
MK50
MK75
MK88
MK94
MK97
MK data
Normal data
Fold over basal
1.05
1
0.95
0.9
0.85
0.8
0.75
0.7
0
10
20
30
40
50
60
Time (min)

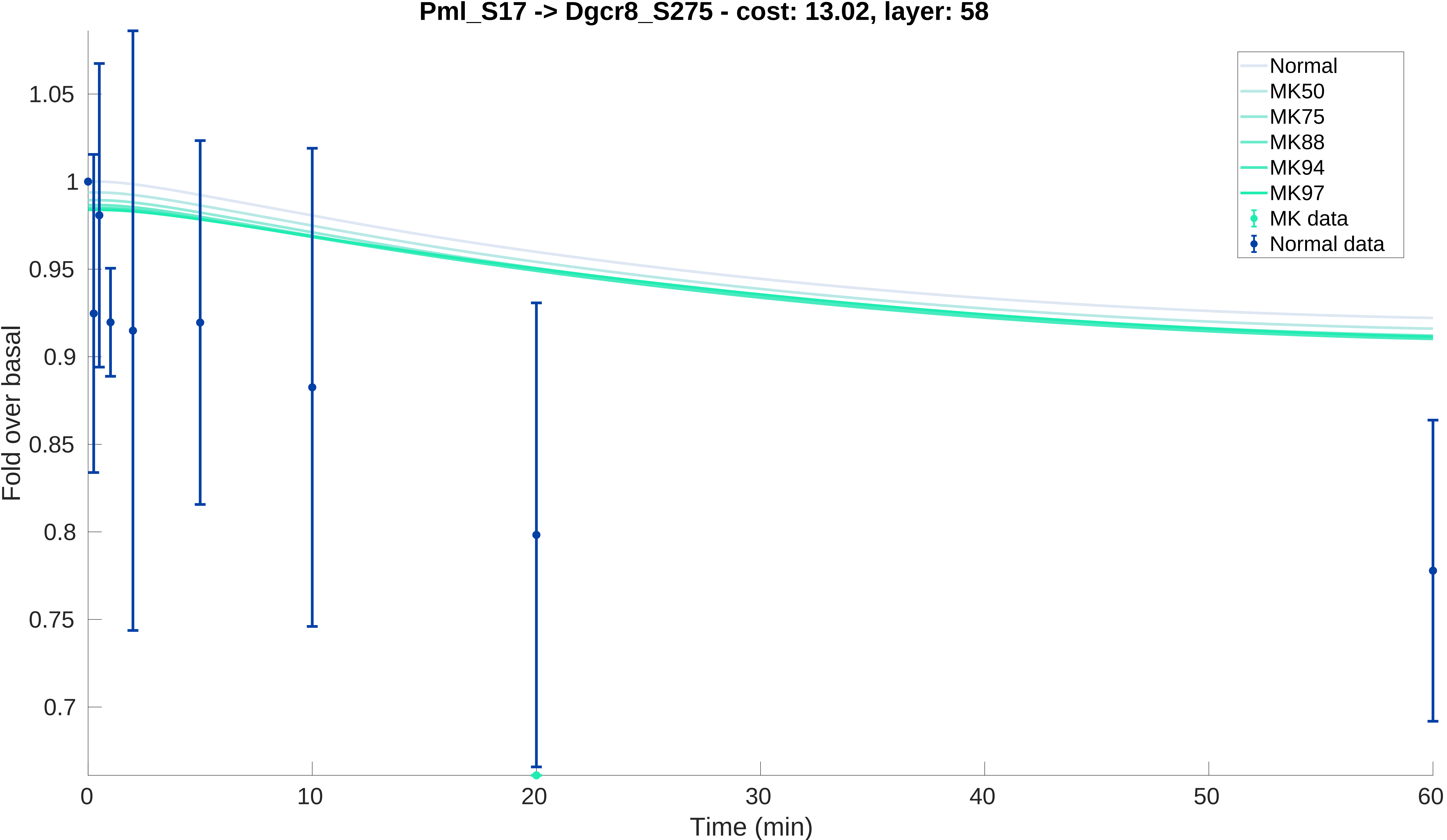

Pml_S17 -> Dgcr8_S275 - cost: 13.02, layer: 58
Fold over basal
Time (min)
1.05
1
0.95
0.9
0.85
0.8
0.75
0.7
0
10
20
30
40
50
60
Normal
MK50
MK75
MK88
MK94
MK97
MK data
Normal data

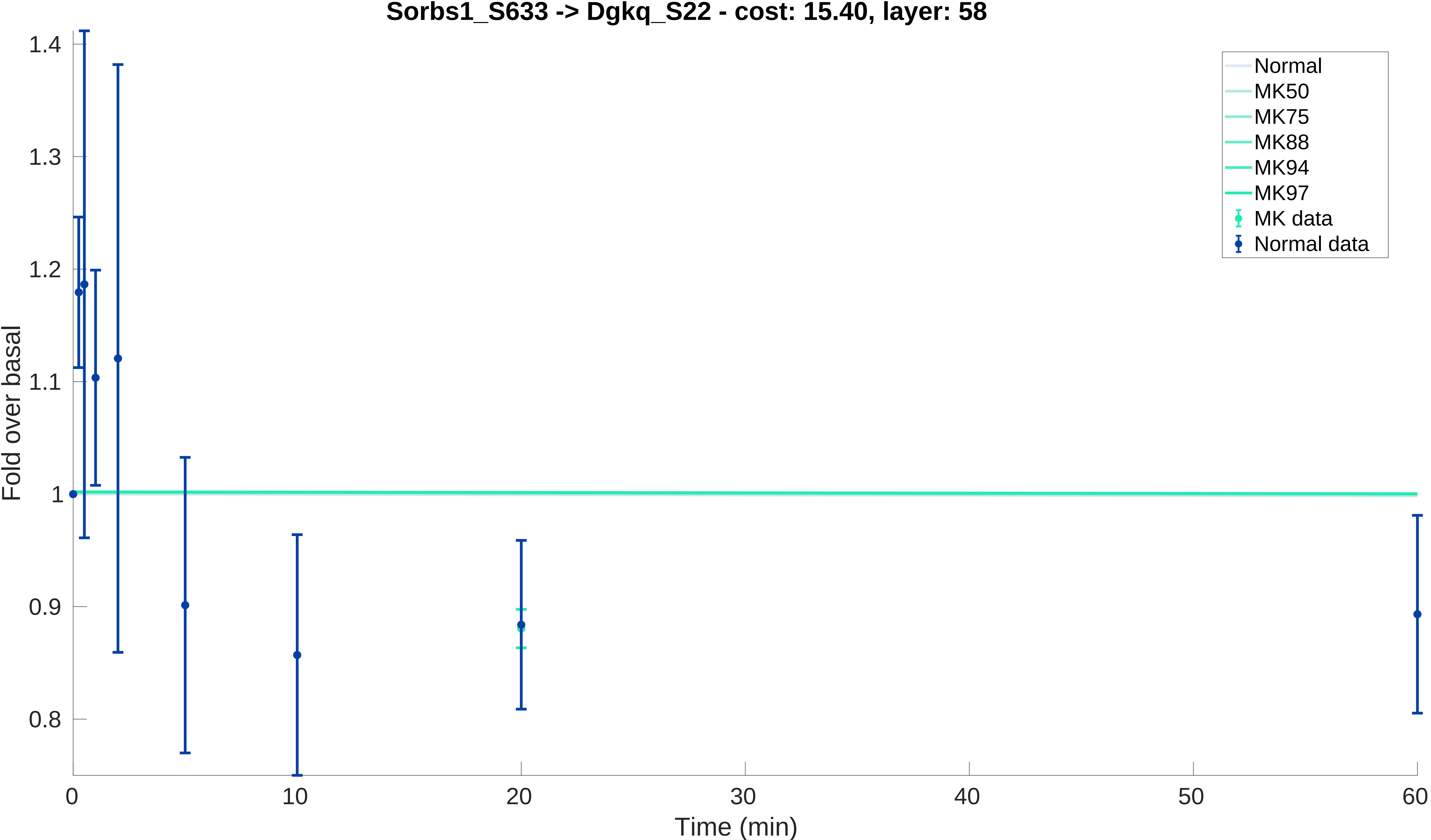

Sorbs1_S633 -> Dgkq_S22 - cost: 15.40, layer: 58
Fold over basal
Time (min)
1.4
1.3
1.2
1.1
1
0.9
0.8
0
10
20
30
40
50
60
Normal
MK50
MK75
MK88
MK94
MK97
MK data
Normal data

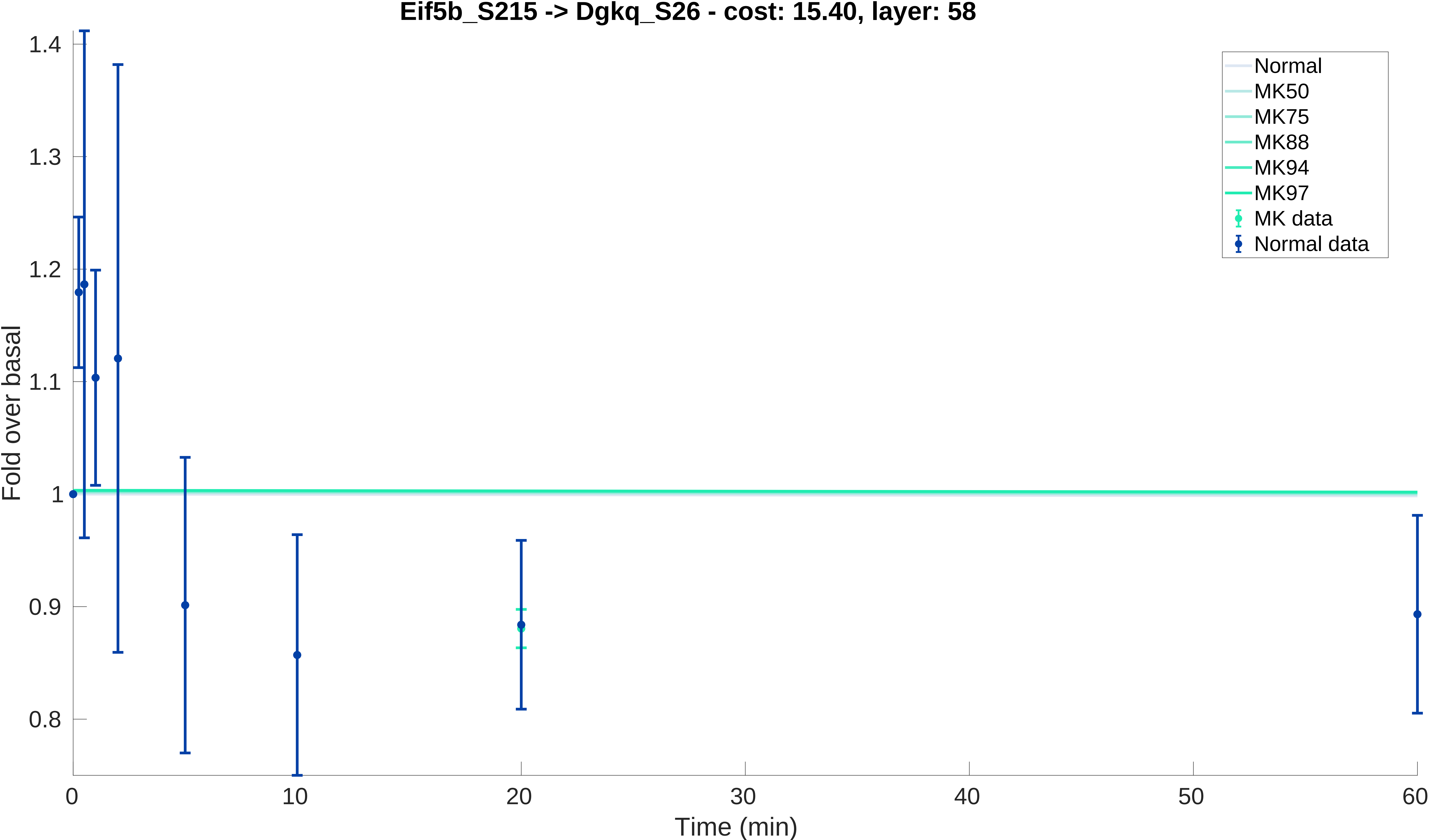

Eif5b_S215 -> Dgkq_S26 - cost: 15.40, layer: 58
Fold over basal
Time (min)
0.8
0.9
1
1.1
1.2
1.3
1.4
0
10
20
30
40
50
60
Normal
MK50
MK75
MK88
MK94
MK97
MK data
Normal data

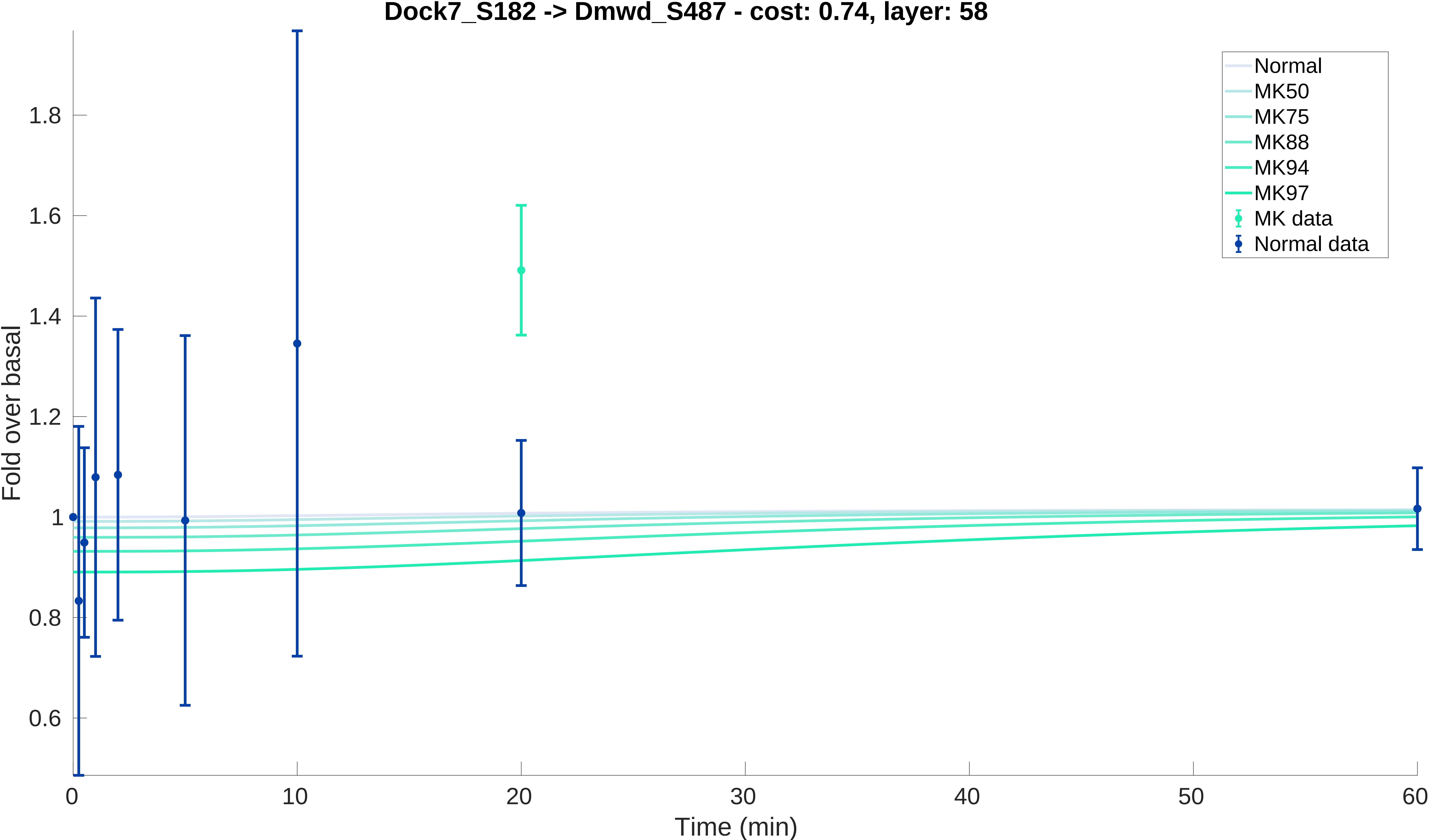

Dock7_S182 -> Dmwd_S487 - cost: 0.74, layer: 58
Fold over basal
Time (min)
0.6
0.8
1
1.2
1.4
1.6
1.8
0
10
20
30
40
50
60
Normal
MK50
MK75
MK88
MK94
MK97
MK data
Normal data

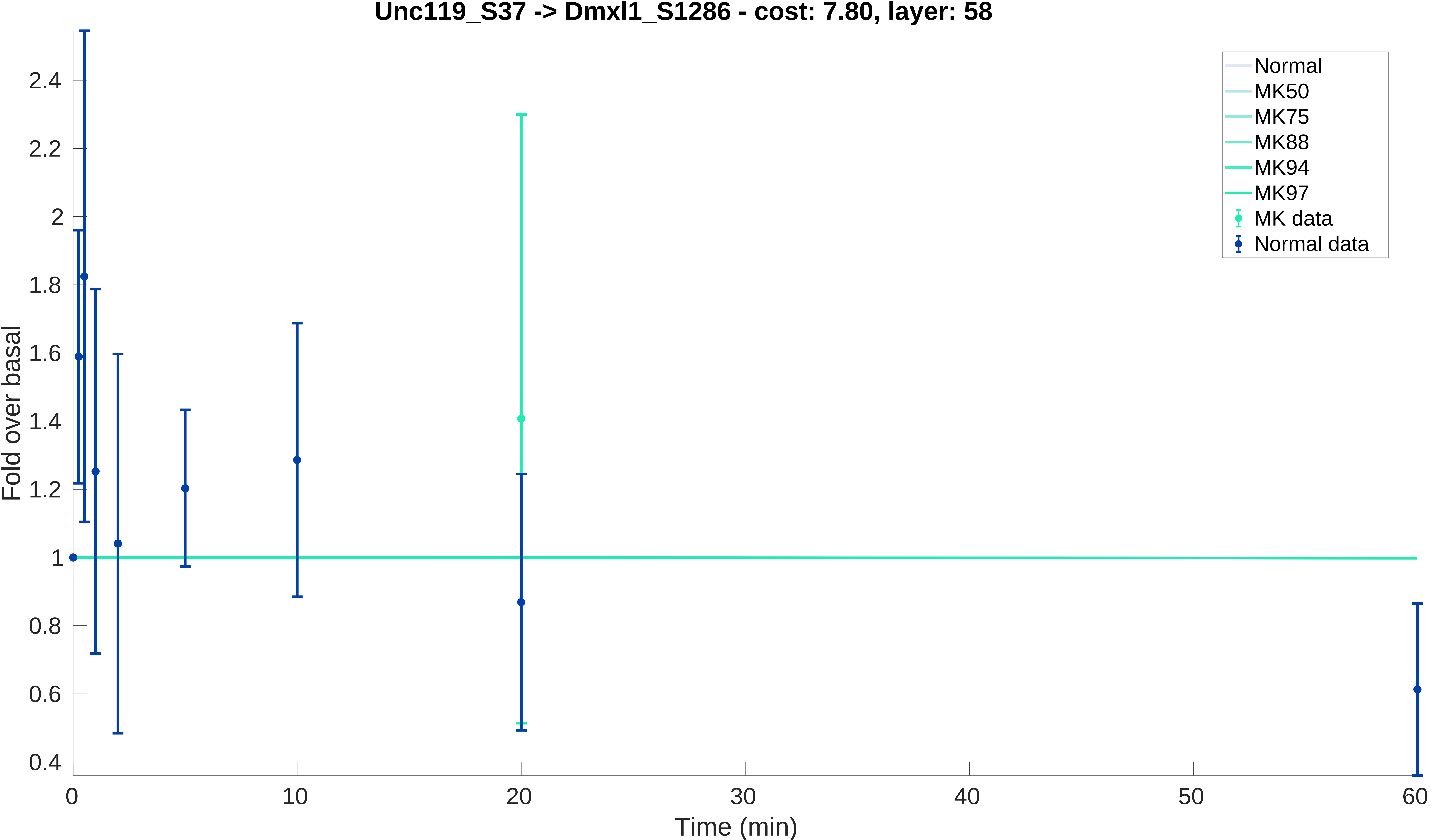

Unc119_S37 -> Dmxl1_S1286 - cost: 7.80, layer: 58
Fold over basal
Time (min)
0
10
20
30
40
50
60
0.4
0.6
0.8
1
1.2
1.4
1.6
1.8
2
2.2
2.4
Normal
MK50
MK75
MK88
MK94
MK97
MK data
Normal data

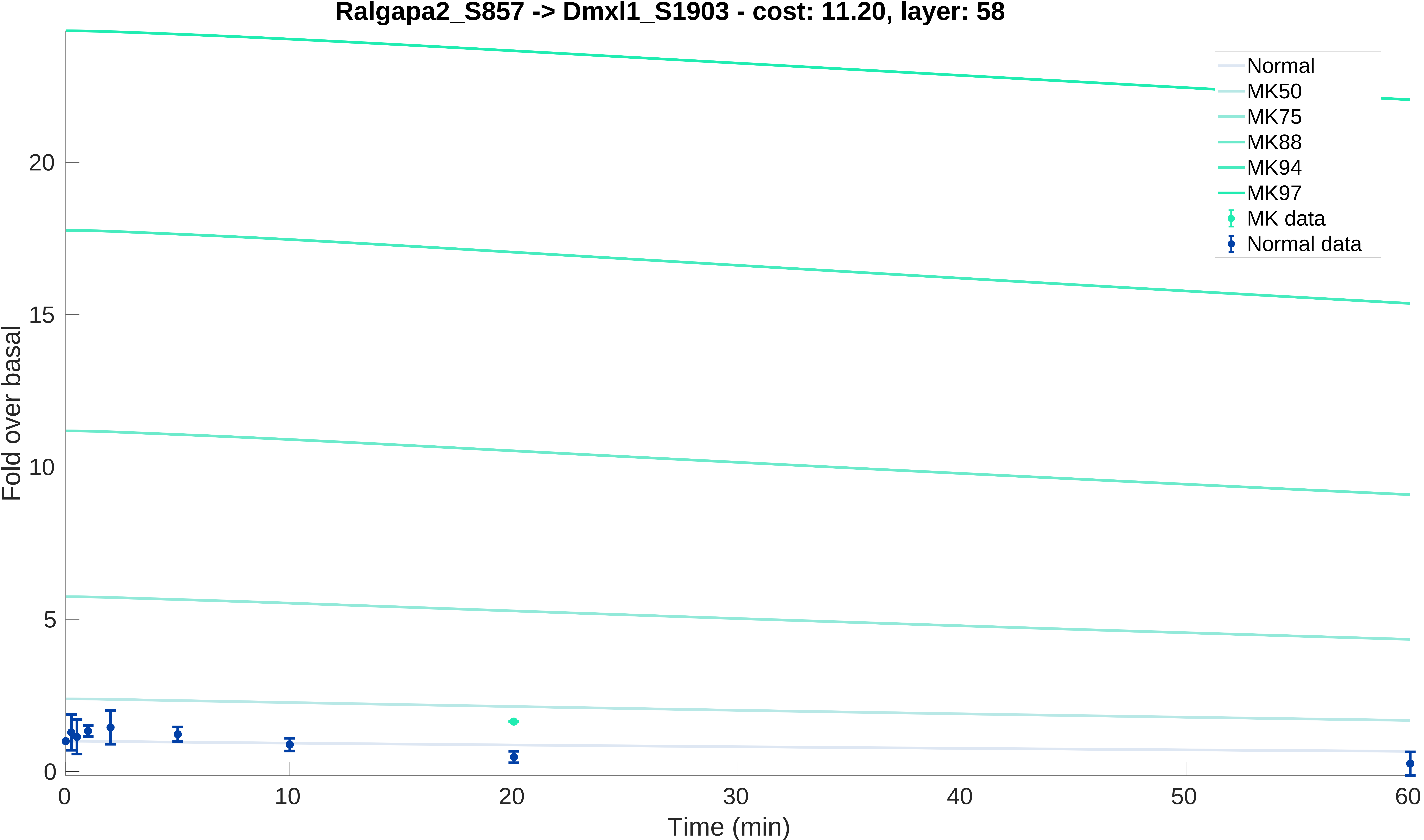

Ralgapa2_S857 -> Dmxl1_S1903 - cost: 11.20, layer: 58
Normal
MK50
MK75
MK88
MK94
MK97
MK data
Normal data
Fold over basal
0
5
10
15
20
Time (min)
0
10
20
30
40
50
60

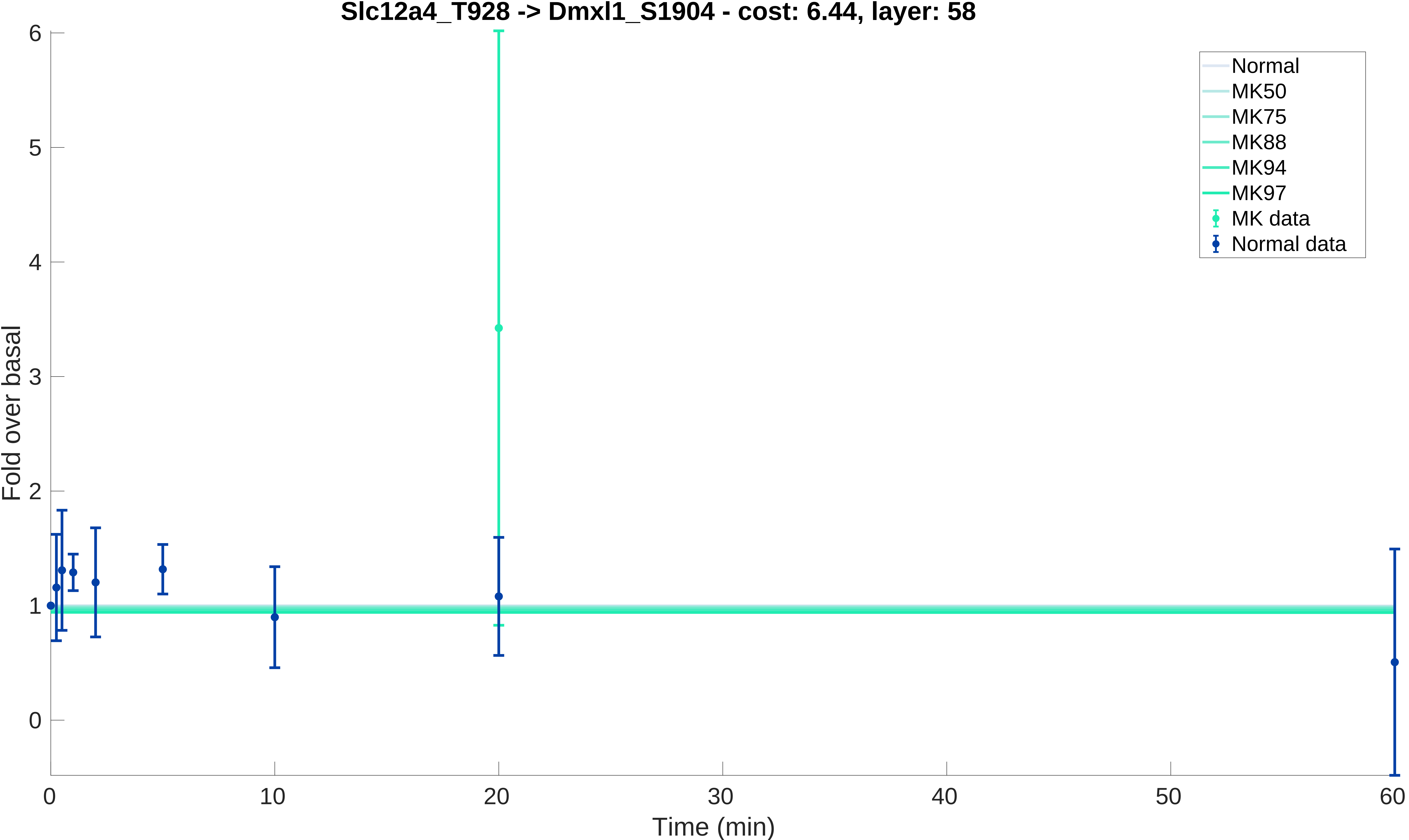

Slc12a4_T928 -> Dmxl1_S1904 - cost: 6.44, layer: 58
Fold over basal
Time (min)
0
1
2
3
4
5
6
0
10
20
30
40
50
60
Normal
MK50
MK75
MK88
MK94
MK97
MK data
Normal data

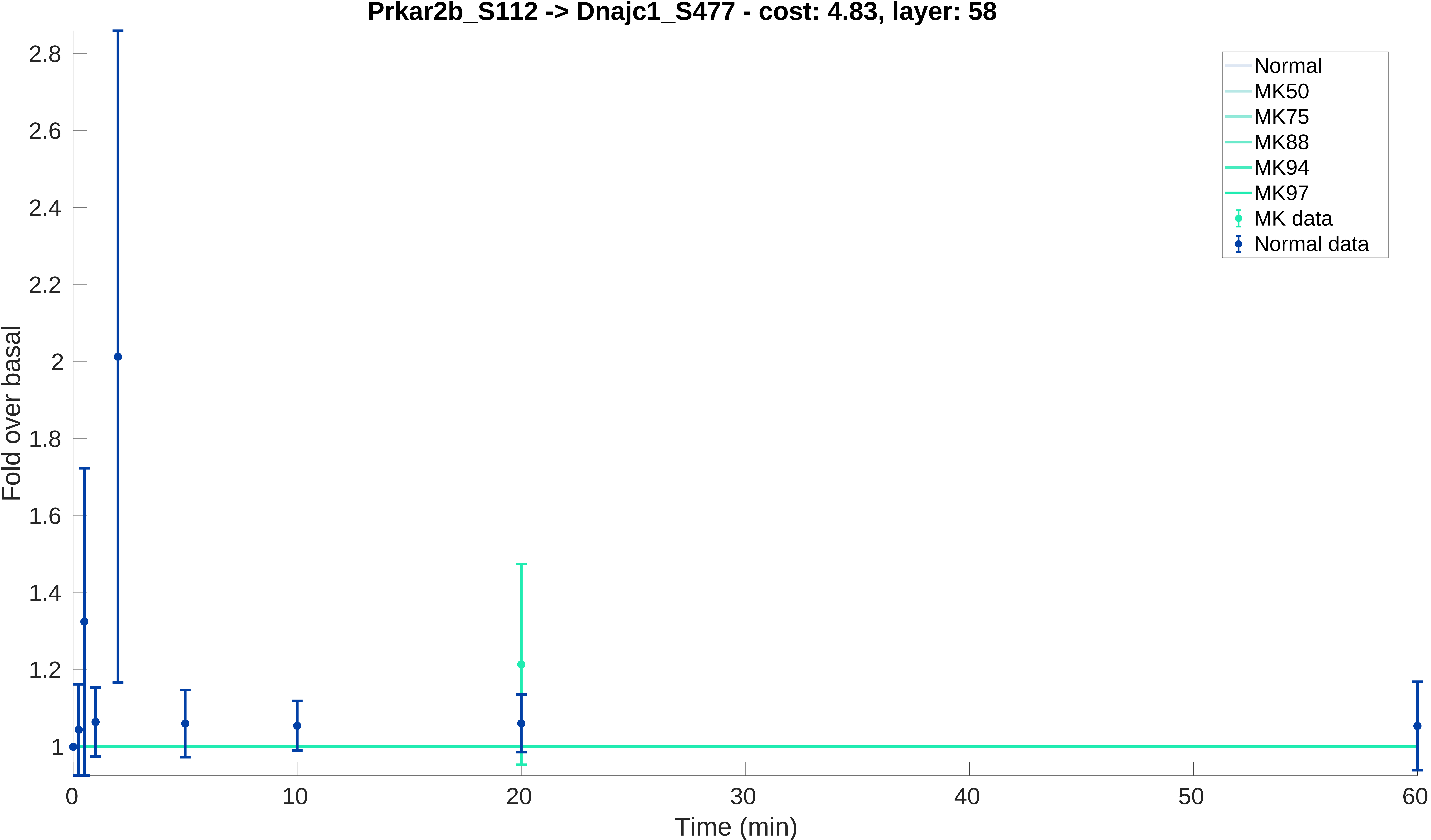

Prkar2b_S112 -> Dnajc1_S477 - cost: 4.83, layer: 58
Fold over basal
2.8
2.6
2.4
2.2
2
1.8
1.6
1.4
1.2
1
0
10
20
30
40
50
60
Time (min)
Normal
MK50
MK75
MK88
MK94
MK97
MK data
Normal data

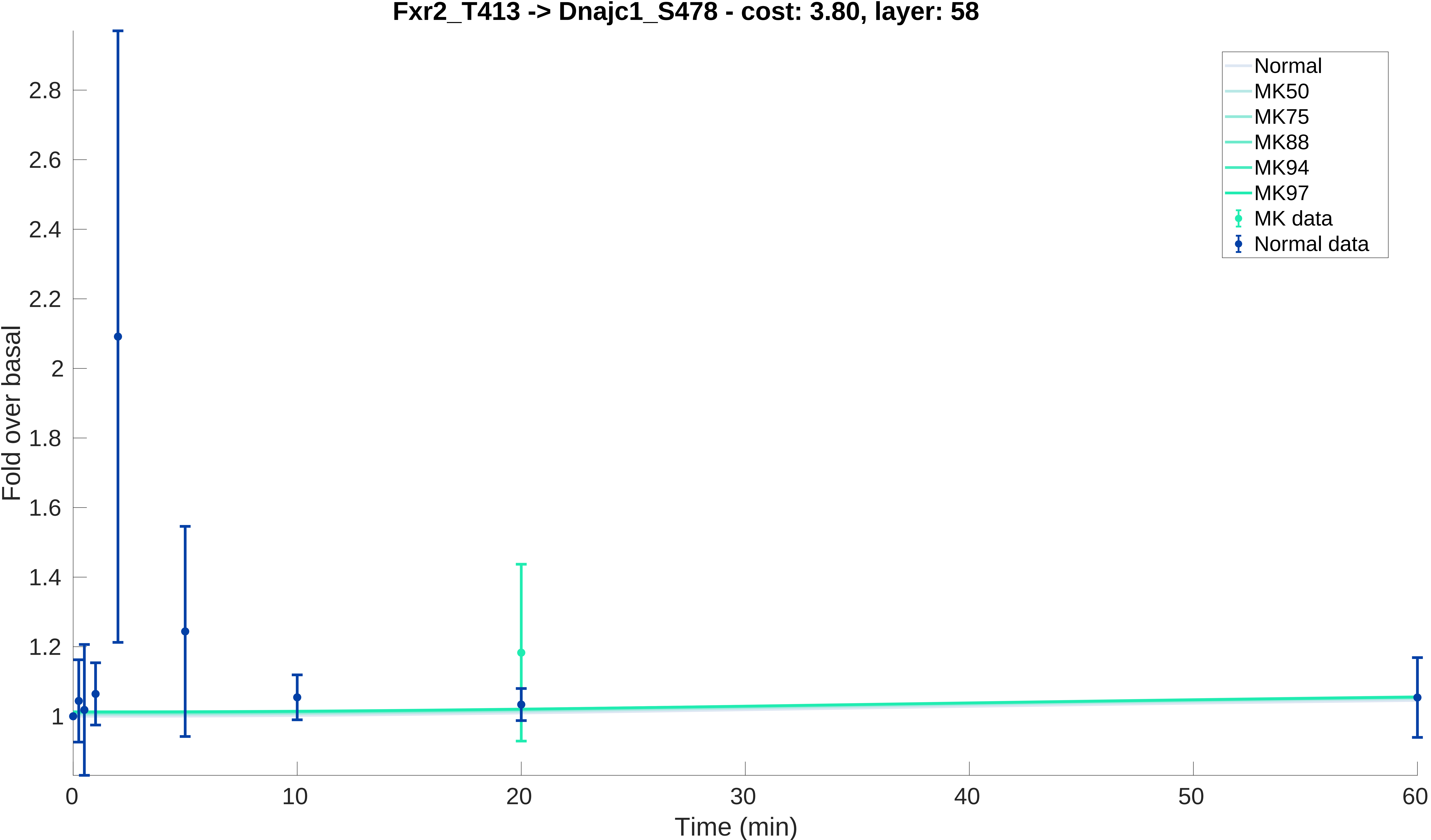

Fxr2_T413 -> Dnajc1_S478 - cost: 3.80, layer: 58
Fold over basal
Time (min)
2.8
2.6
2.4
2.2
2
1.8
1.6
1.4
1.2
1
0
10
20
30
40
50
60
Normal
MK50
MK75
MK88
MK94
MK97
MK data
Normal data

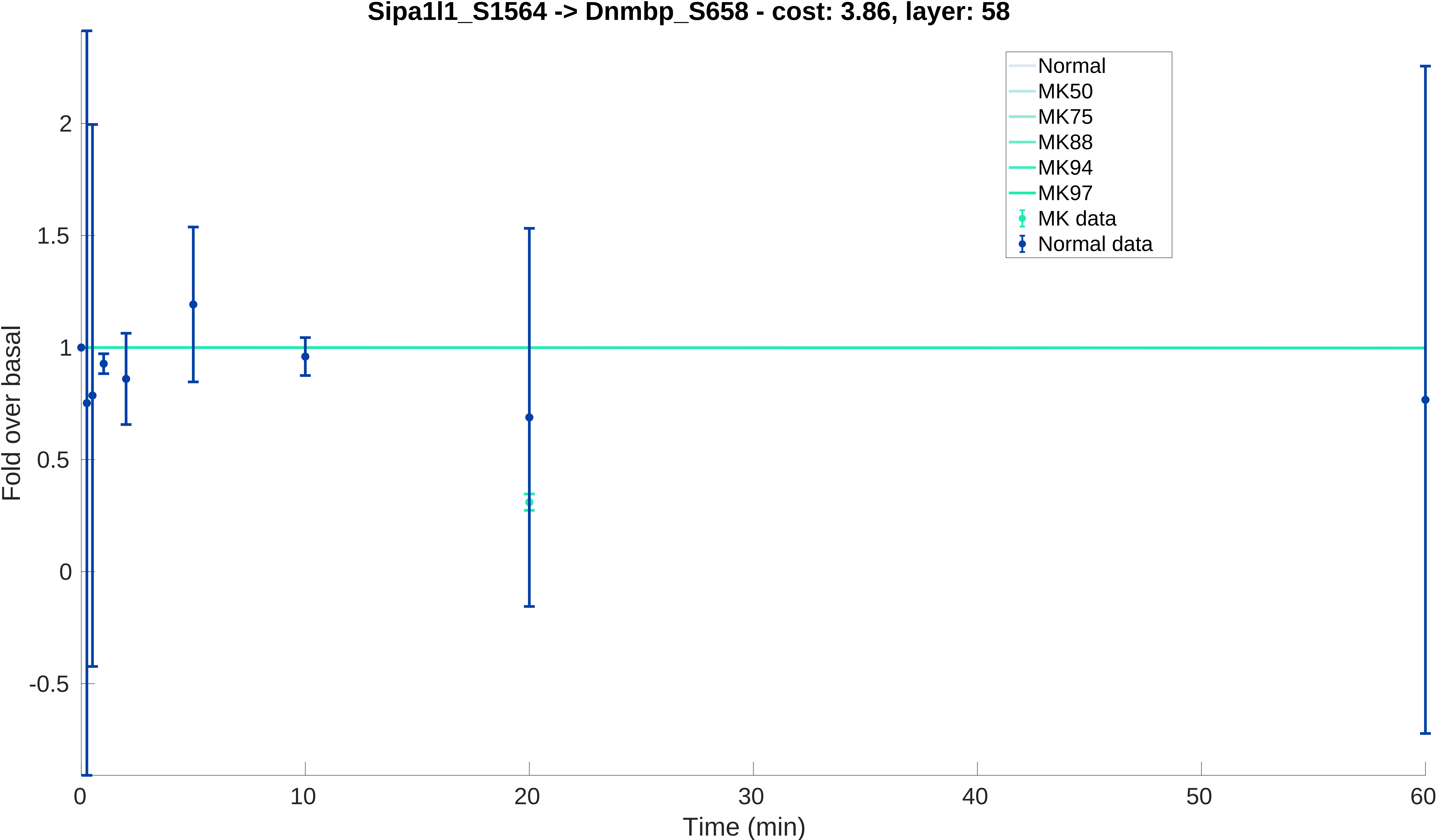

Sipa1l1_S1564 -> Dnmbp_S658 - cost: 3.86, layer: 58
Normal
MK50
MK75
MK88
MK94
MK97
MK data
Normal data
Fold over basal
Time (min)
2
1.5
1
0.5
0
-0.5
0
10
20
30
40
50
60

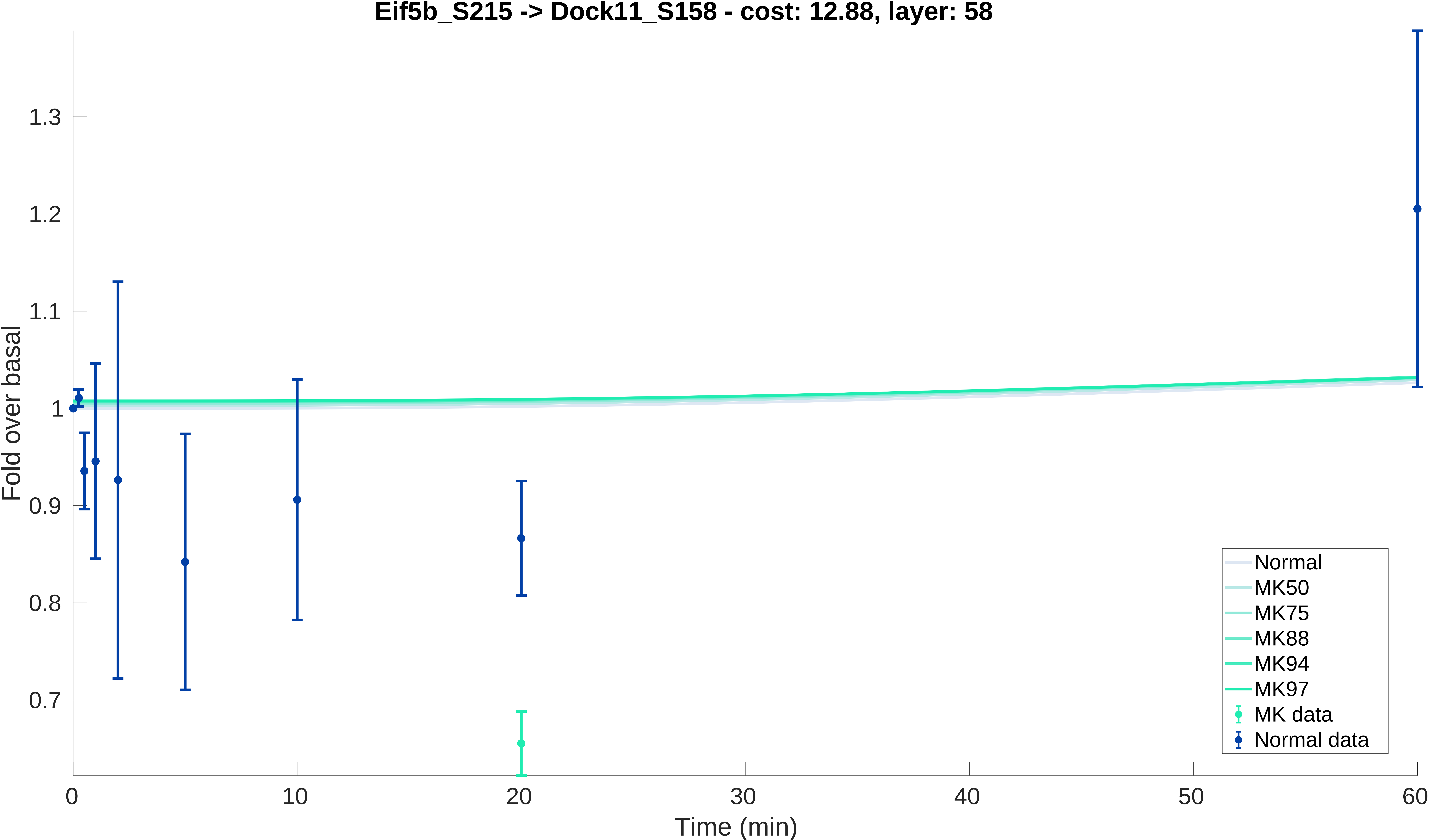

Eif5b_S215 -> Dock11_S158 - cost: 12.88, layer: 58
Fold over basal
Time (min)
0.7
0.8
0.9
1
1.1
1.2
1.3
0
10
20
30
40
50
60
Normal
MK50
MK75
MK88
MK94
MK97
MK data
Normal data

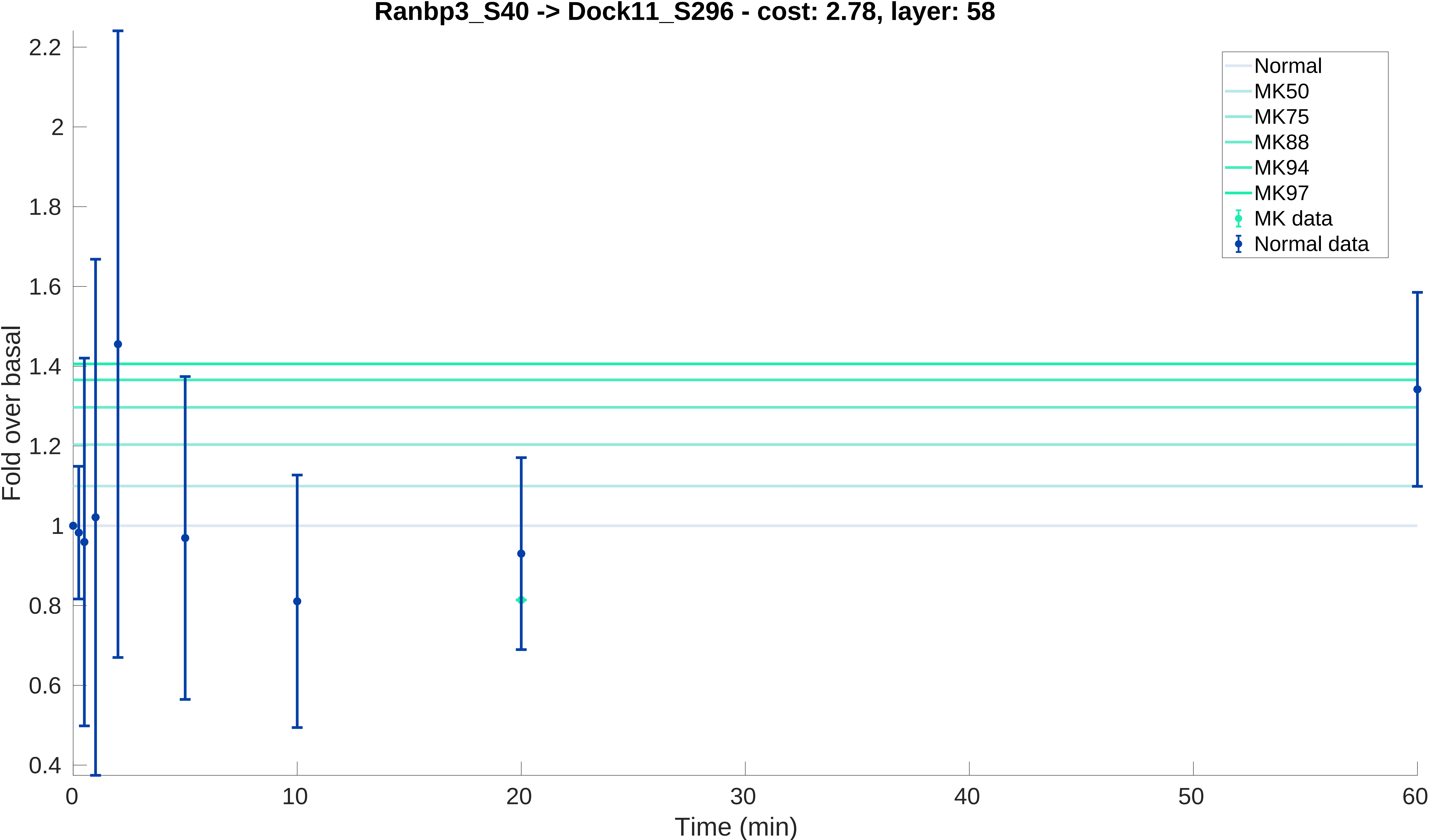

Ranbp3_S40 -> Dock11_S296 - cost: 2.78, layer: 58
Fold over basal
Time (min)
0.4
0.6
0.8
1
1.2
1.4
1.6
1.8
2
2.2
0
10
20
30
40
50
60
Normal
MK50
MK75
MK88
MK94
MK97
MK data
Normal data

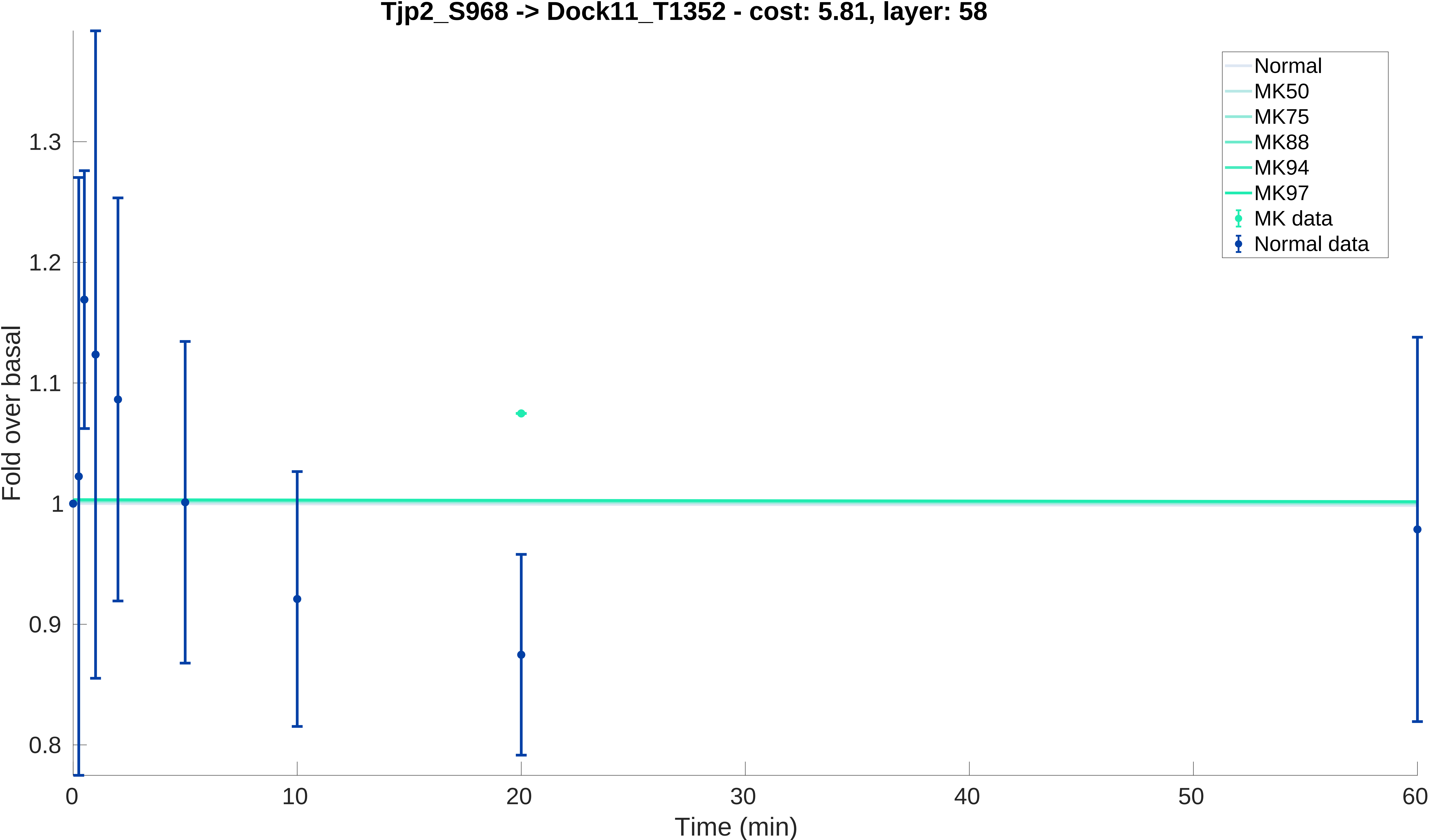
Tjp2_S968 -> Dock11_T1352 - cost: 5.81, layer: 58
Fold over basal
Time (min)
1.3
1.2
1.1
1
0.9
0.8
0
10
20
30
40
50
60
Normal
MK50
MK75
MK88
MK94
MK97
MK data
Normal data

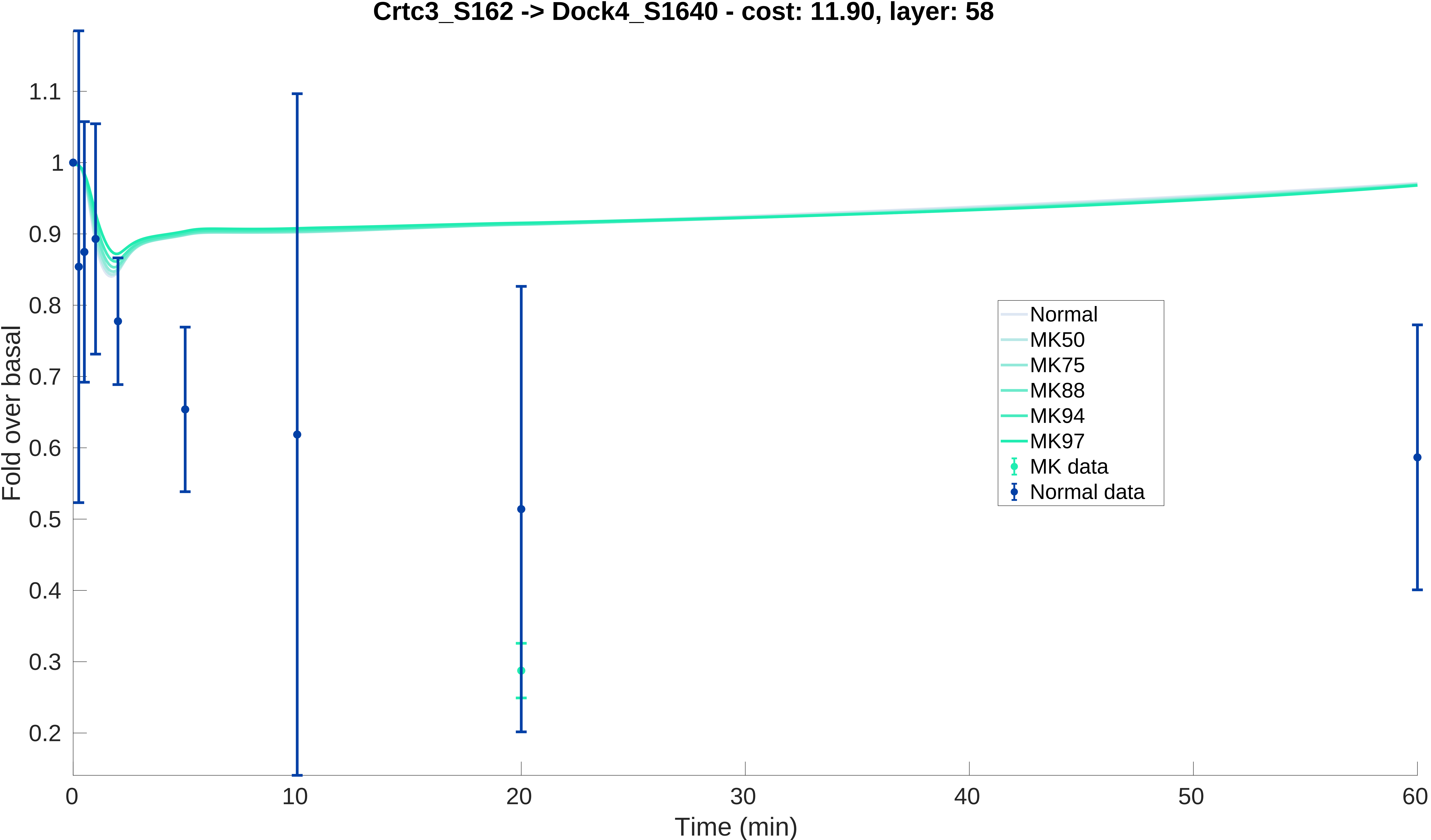

Crtc3_S162 -> Dock4_S1640 - cost: 11.90, layer: 58
Fold over basal
Time (min)
0
10
20
30
40
50
60
0.2
0.3
0.4
0.5
0.6
0.7
0.8
0.9
1
1.1
Normal
MK50
MK75
MK88
MK94
MK97
MK data
Normal data

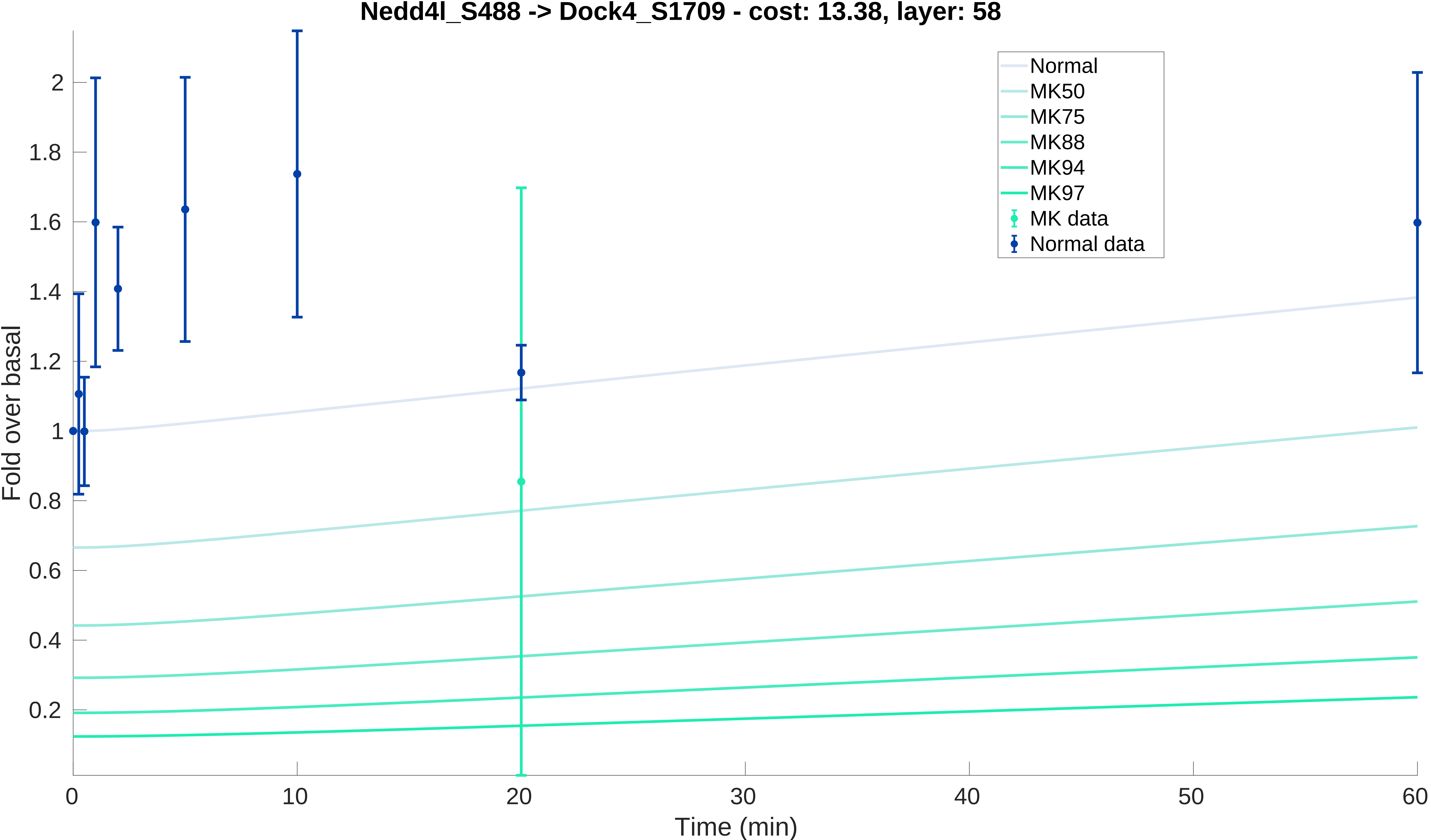
Nedd4l_S488 -> Dock4_S1709 - cost: 13.38, layer: 58
Normal
MK50
MK75
MK88
MK94
MK97
MK data
Normal data
Fold over basal
0.2
0.4
0.6
0.8
1
1.2
1.4
1.6
1.8
2
0
10
20
30
40
50
60
Time (min)

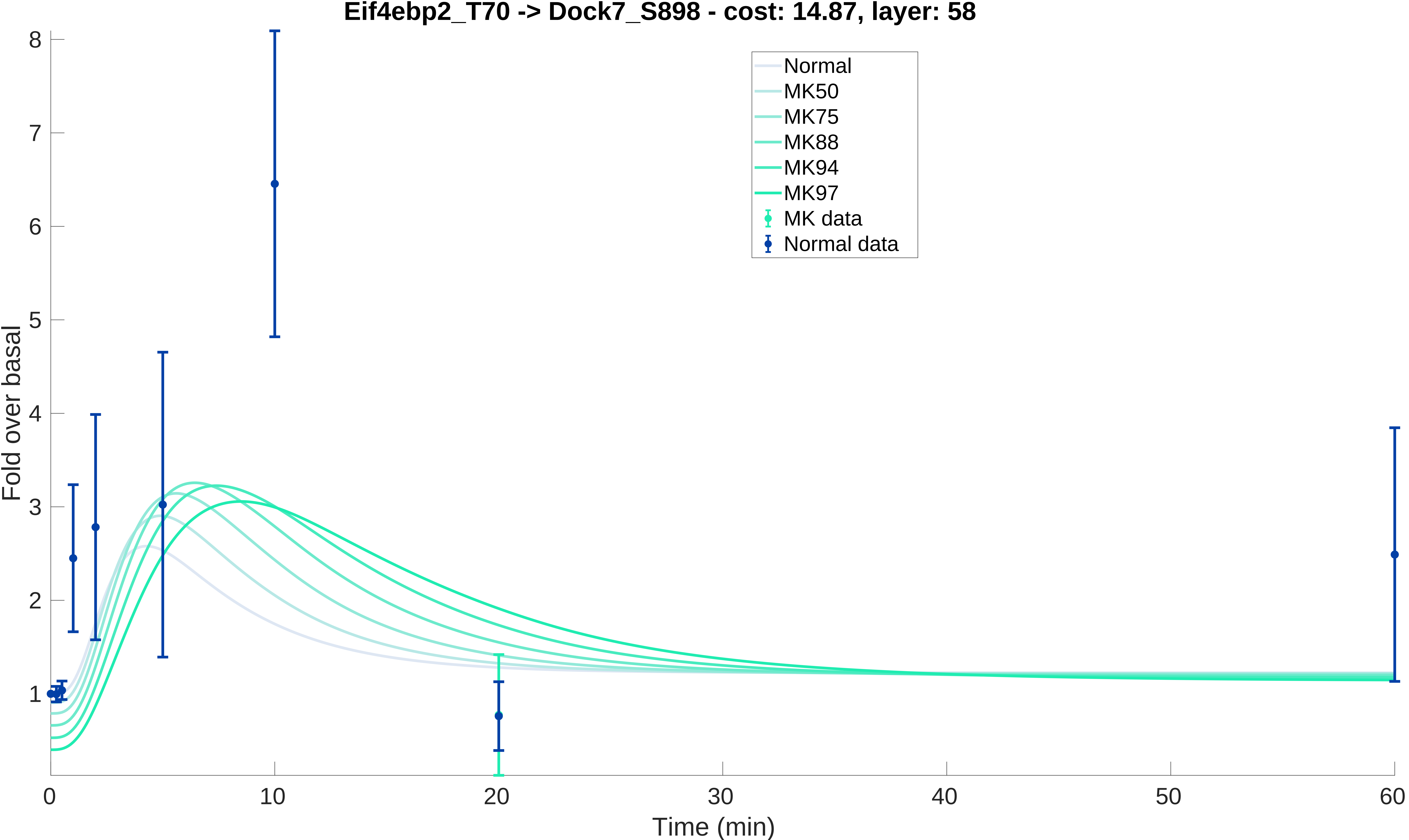

Eif4ebp2_T70 -> Dock7_S898 - cost: 14.87, layer: 58
Fold over basal
Time (min)
Normal
MK50
MK75
MK88
MK94
MK97
MK data
Normal data
0
10
20
30
40
50
60
1
2
3
4
5
6
7
8

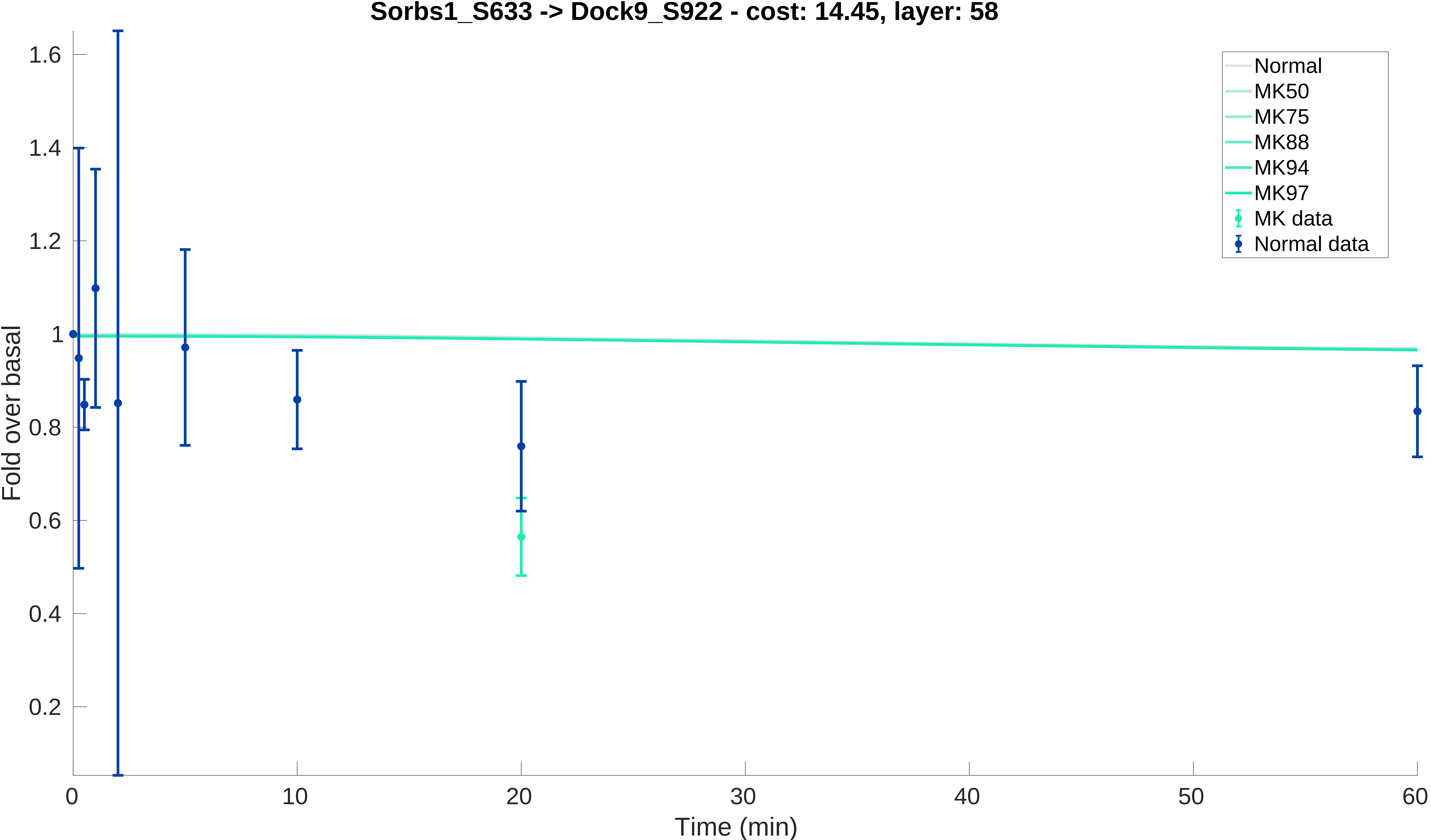

Sorbs1_S633 -> Dock9_S922 - cost: 14.45, layer: 58
Fold over basal
Time (min)
0
10
20
30
40
50
60
0.2
0.4
0.6
0.8
1
1.2
1.4
1.6
Normal
MK50
MK75
MK88
MK94
MK97
MK data
Normal data

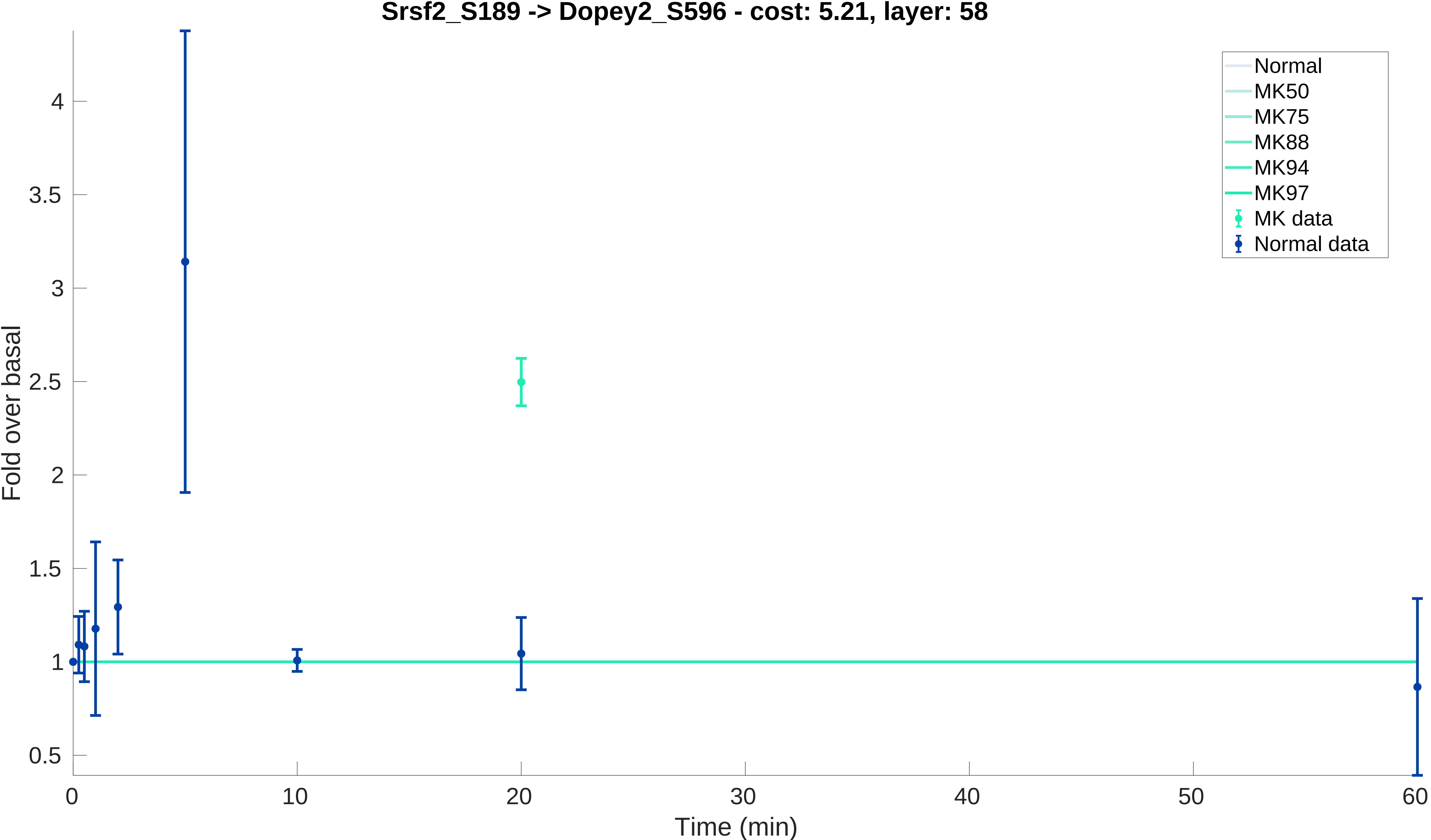
Srsf2_S189 -> Dopey2_S596 - cost: 5.21, layer: 58
Fold over basal
0.5
1
1.5
2
2.5
3
3.5
4
0
10
20
30
40
50
60
Time (min)
Normal
MK50
MK75
MK88
MK94
MK97
MK data
Normal data

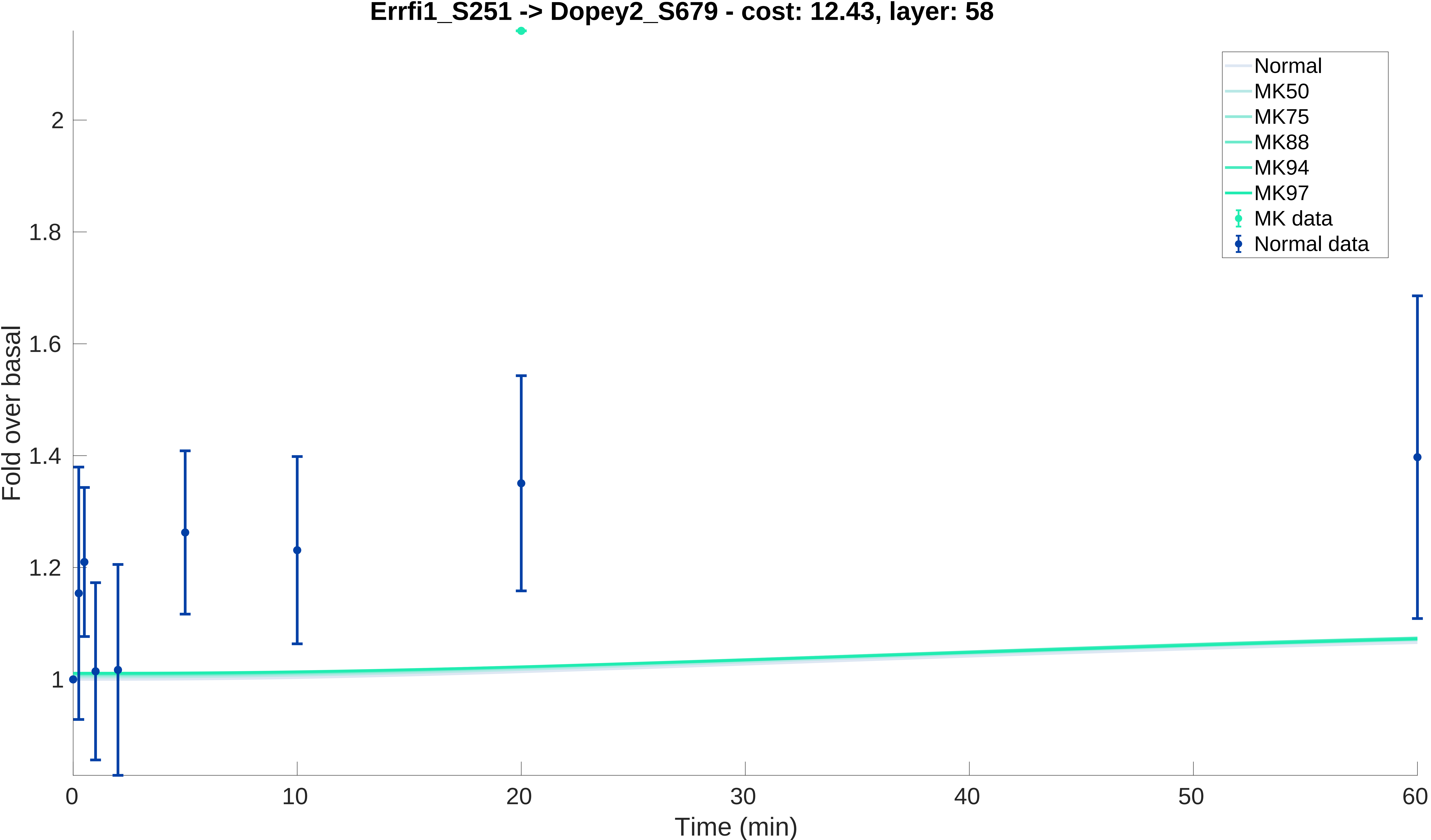

Errfi1_S251 -> Dopey2_S679 - cost: 12.43, layer: 58
Fold over basal
Time (min)
0
10
20
30
40
50
60
1
1.2
1.4
1.6
1.8
2
Normal
MK50
MK75
MK88
MK94
MK97
MK data
Normal data

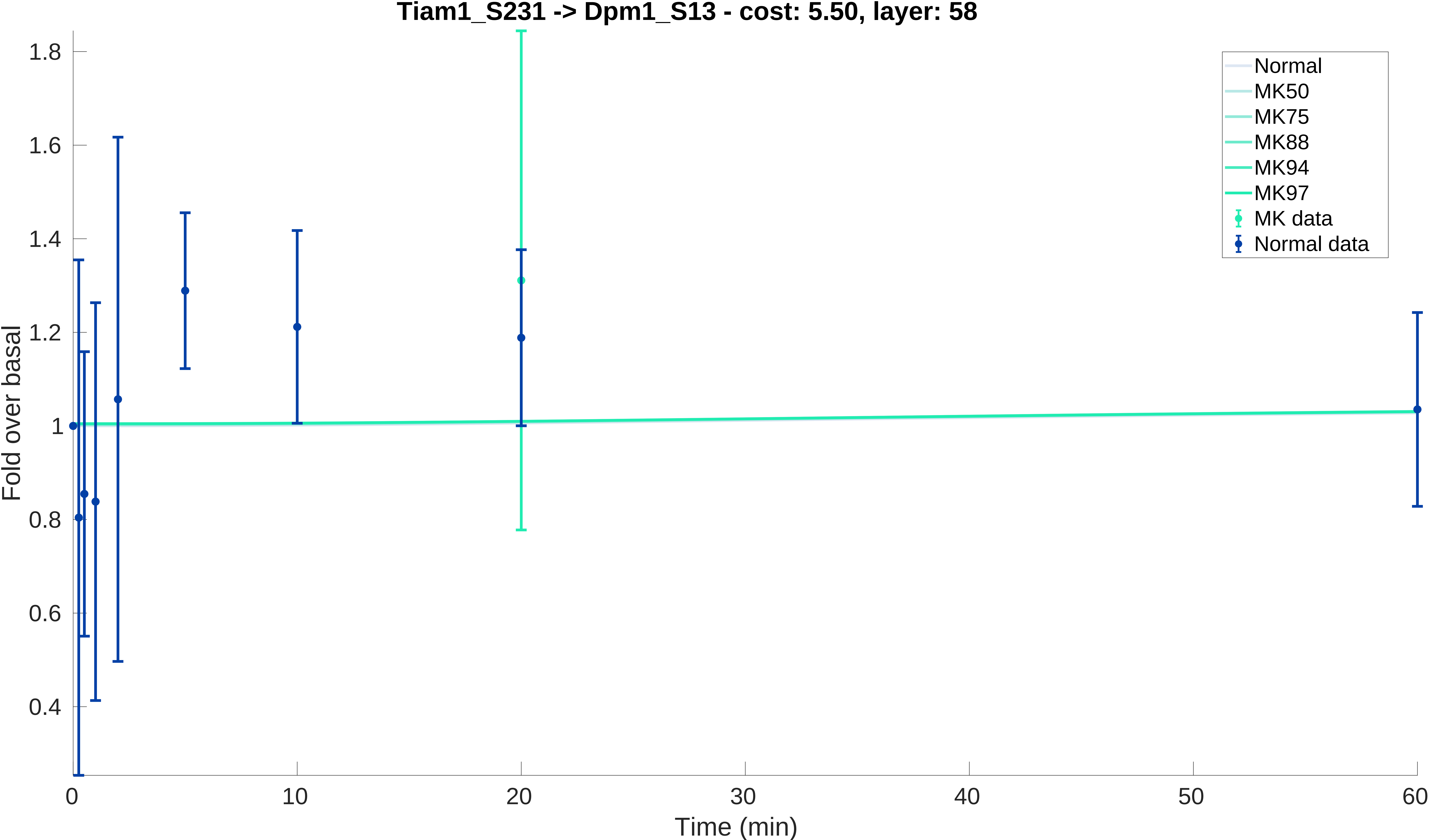

Tiam1_S231 -> Dpm1_S13 - cost: 5.50, layer: 58
Fold over basal
Time (min)
Normal
MK50
MK75
MK88
MK94
MK97
MK data
Normal data
1.8
1.6
1.4
1.2
1
0.8
0.6
0.4
0
10
20
30
40
50
60

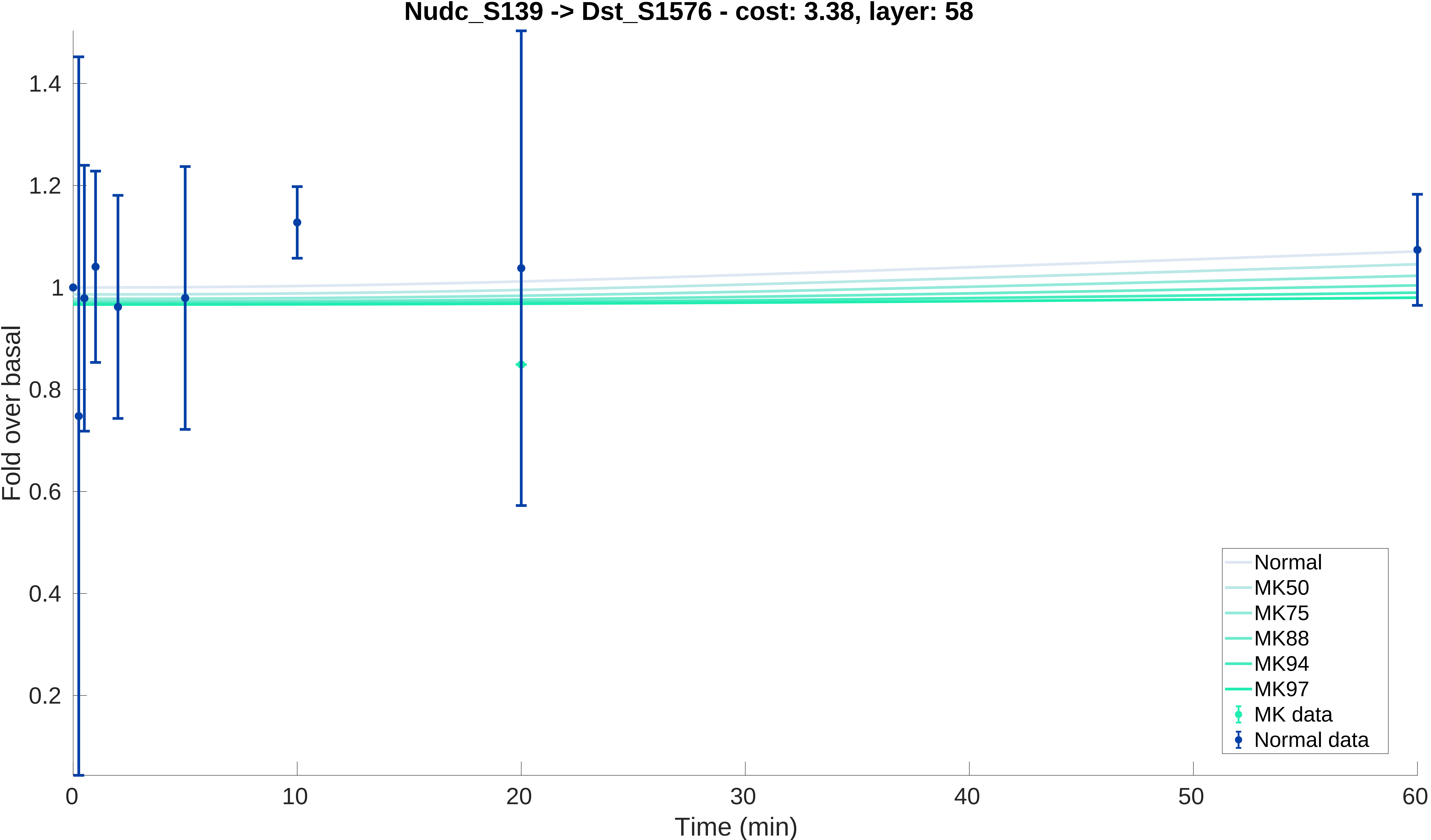

Nudc_S139 -> Dst_S1576 - cost: 3.38, layer: 58
Fold over basal
1.4
1.2
1
0.8
0.6
0.4
0.2
0
10
20
30
40
50
60
Time (min)
Normal
MK50
MK75
MK88
MK94
MK97
MK data
Normal data

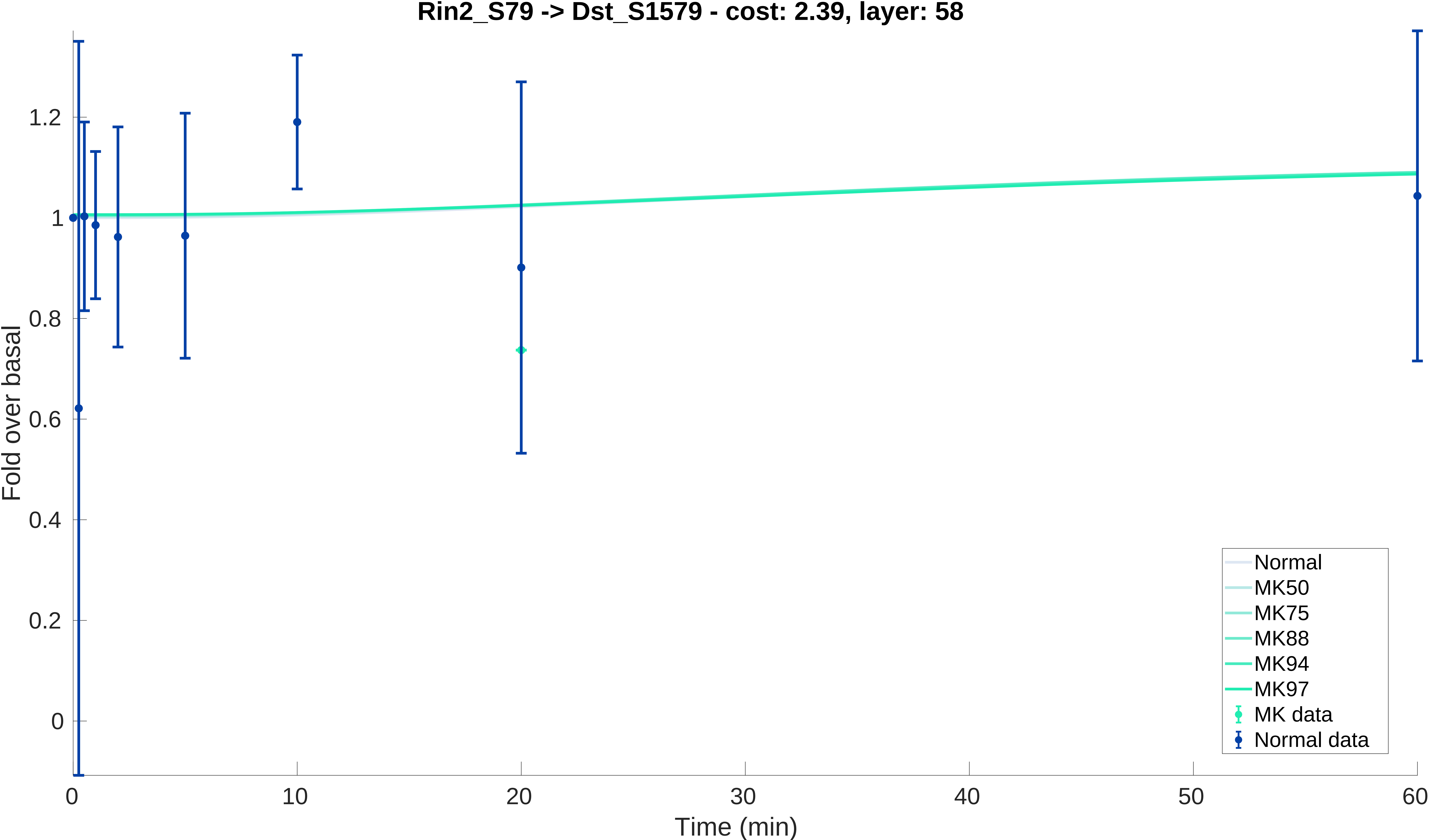

Rin2_S79 -> Dst_S1579 - cost: 2.39, layer: 58
Fold over basal
Time (min)
1.2
1
0.8
0.6
0.4
0.2
0
0
10
20
30
40
50
60
Normal
MK50
MK75
MK88
MK94
MK97
MK data
Normal data

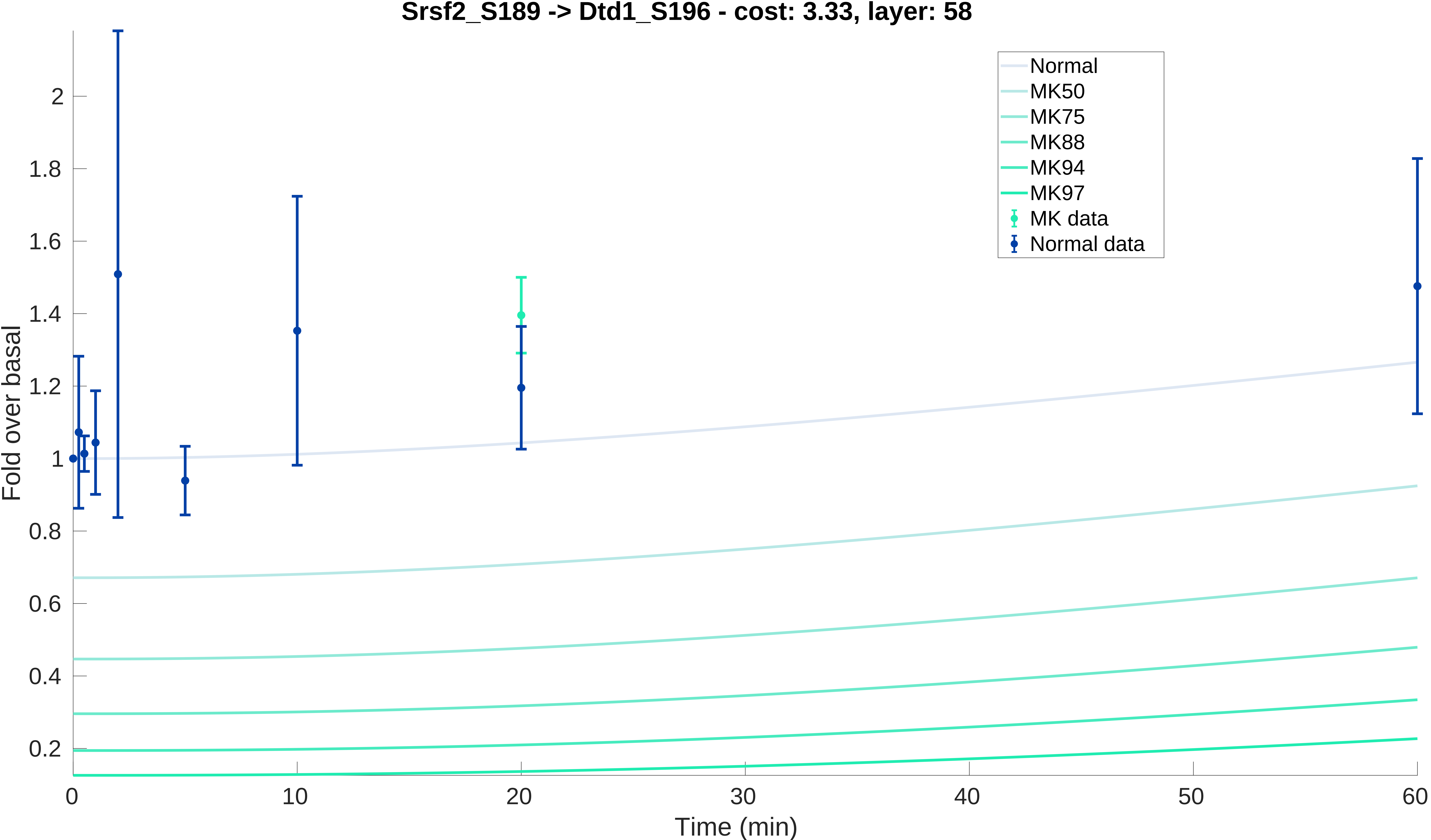
Srsf2_S189 -> Dtd1_S196 - cost: 3.33, layer: 58
Normal
MK50
MK75
MK88
MK94
MK97
MK data
Normal data
Fold over basal
Time (min)
0
10
20
30
40
50
60
0.2
0.4
0.6
0.8
1
1.2
1.4
1.6
1.8
2

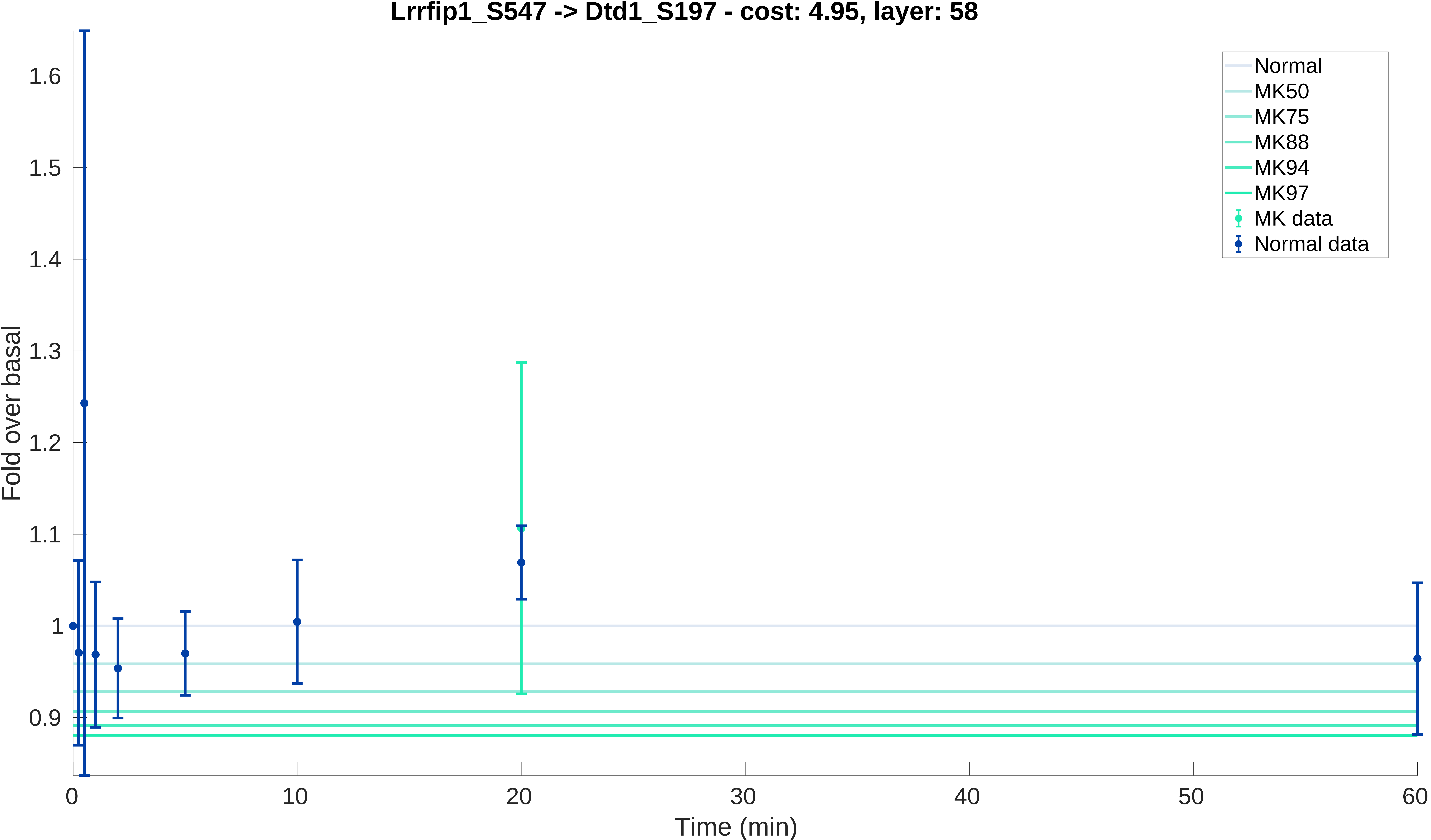

Lrrfip1_S547 -> Dtd1_S197 - cost: 4.95, layer: 58
Fold over basal
1.6
1.5
1.4
1.3
1.2
1.1
1
0.9
0
10
20
30
40
50
60
Time (min)
Normal
MK50
MK75
MK88
MK94
MK97
MK data
Normal data

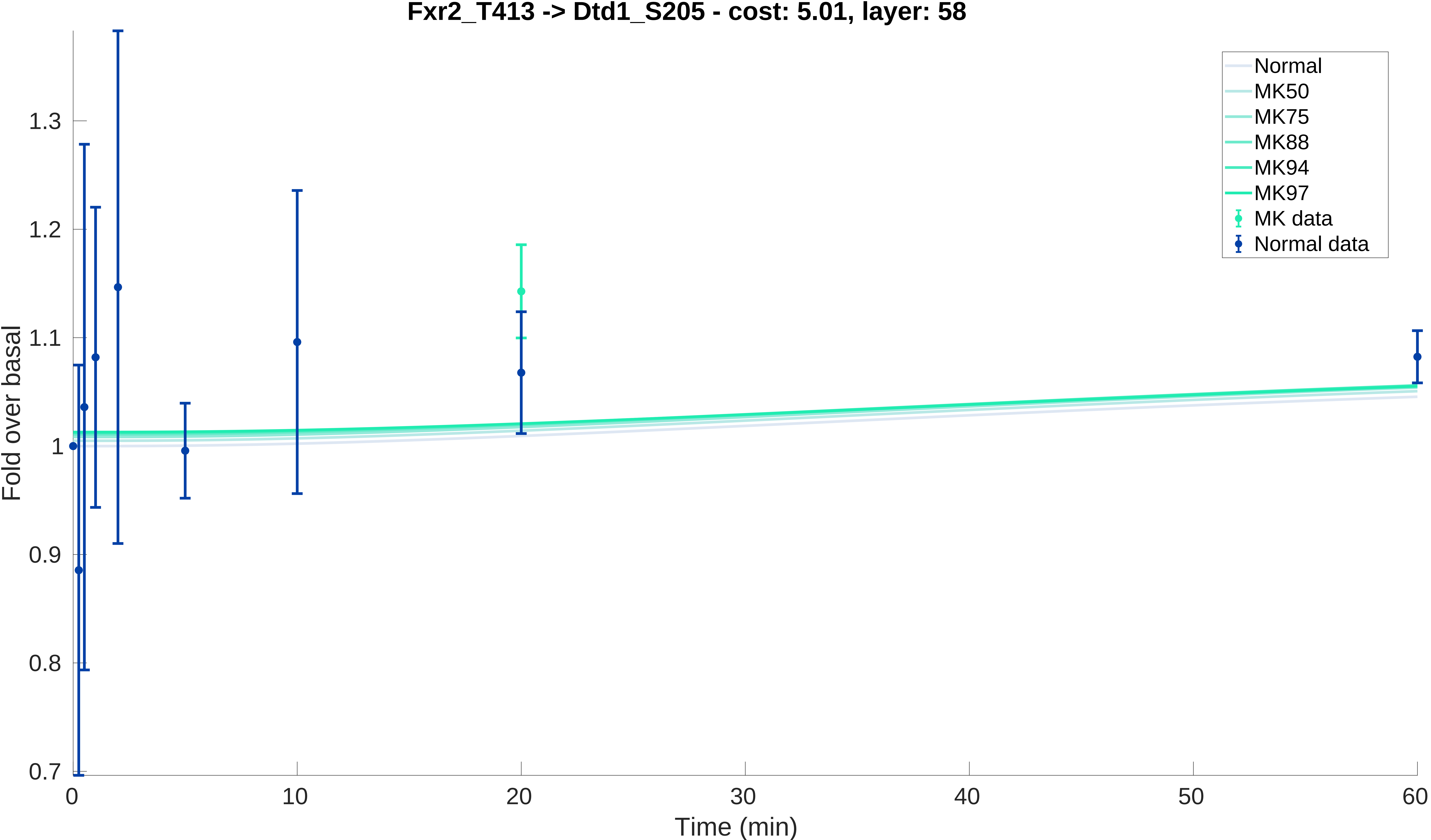

Fxr2_T413 -> Dtd1_S205 - cost: 5.01, layer: 58
Normal
MK50
MK75
MK88
MK94
MK97
MK data
Normal data
Fold over basal
1.3
1.2
1.1
1
0.9
0.8
0.7
0
10
20
30
40
50
60
Time (min)

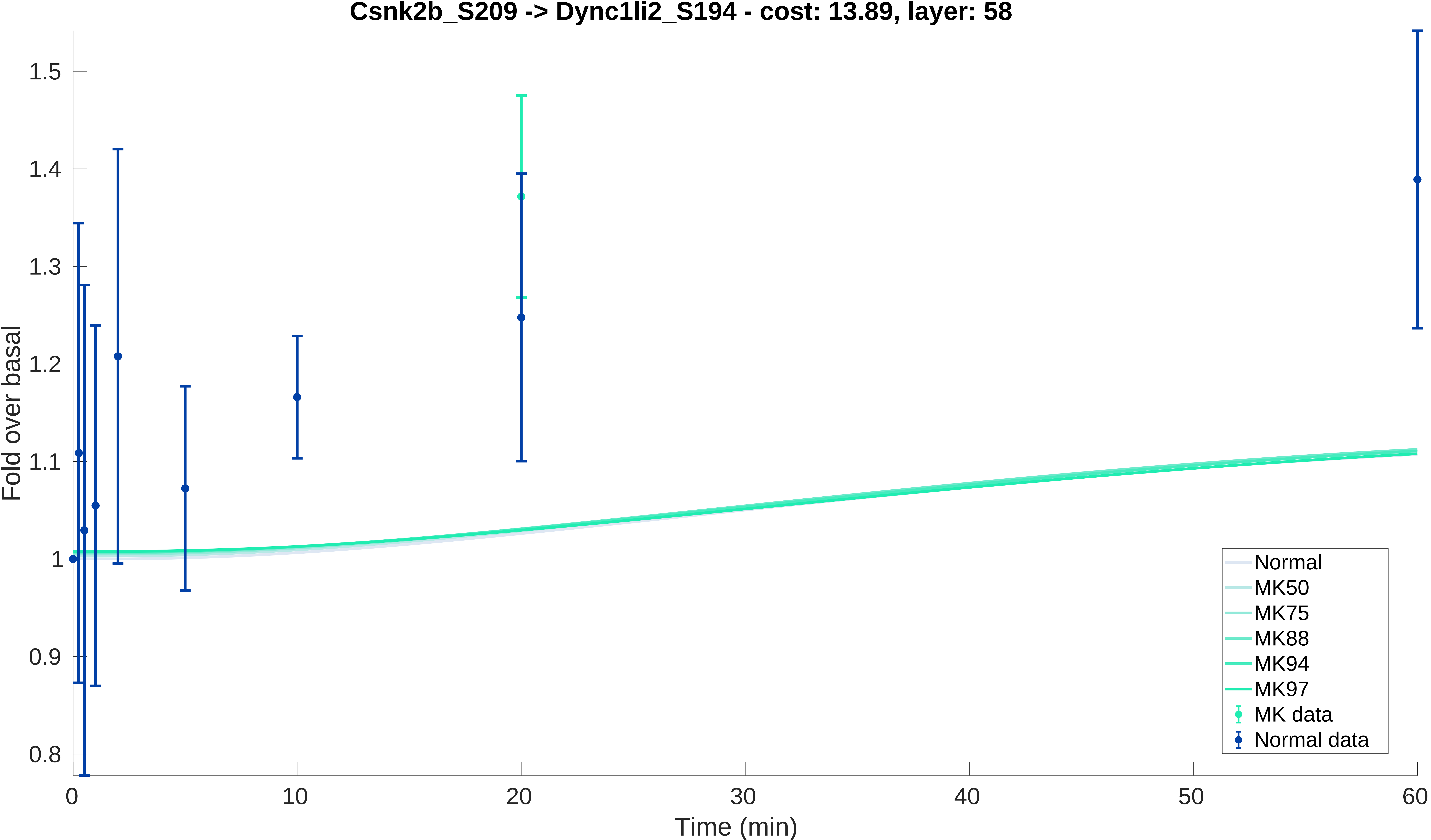

Csnk2b_S209 -> Dync1li2_S194 - cost: 13.89, layer: 58
Fold over basal
Time (min)
0.8
0.9
1
1.1
1.2
1.3
1.4
1.5
0
10
20
30
40
50
60
Normal
MK50
MK75
MK88
MK94
MK97
MK data
Normal data

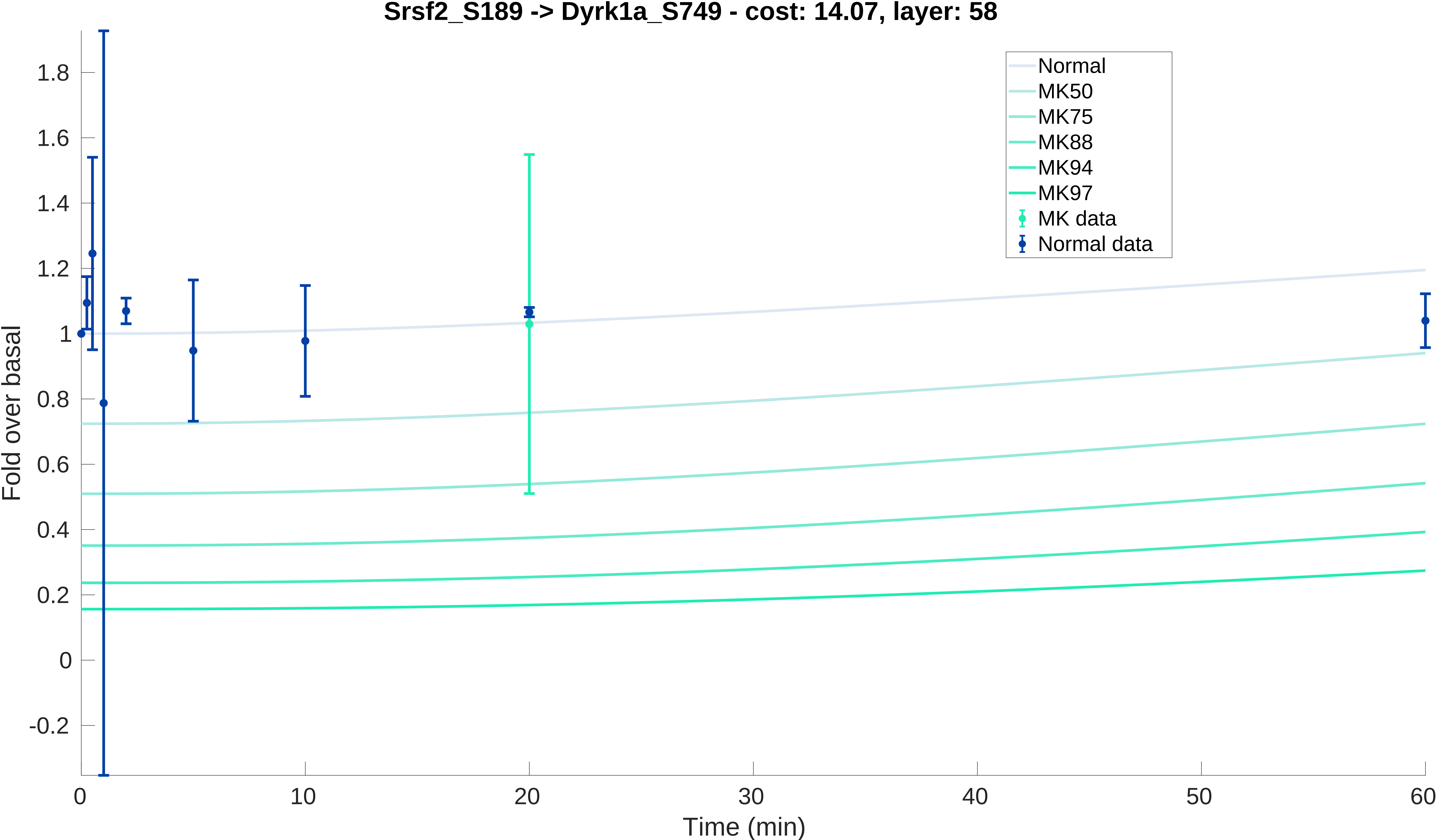

Srsf2_S189 -> Dyrk1a_S749 - cost: 14.07, layer: 58
Fold over basal
Time (min)
1.8
1.6
1.4
1.2
1
0.8
0.6
0.4
0.2
0
-0.2
0
10
20
30
40
50
60
Normal
MK50
MK75
MK88
MK94
MK97
MK data
Normal data

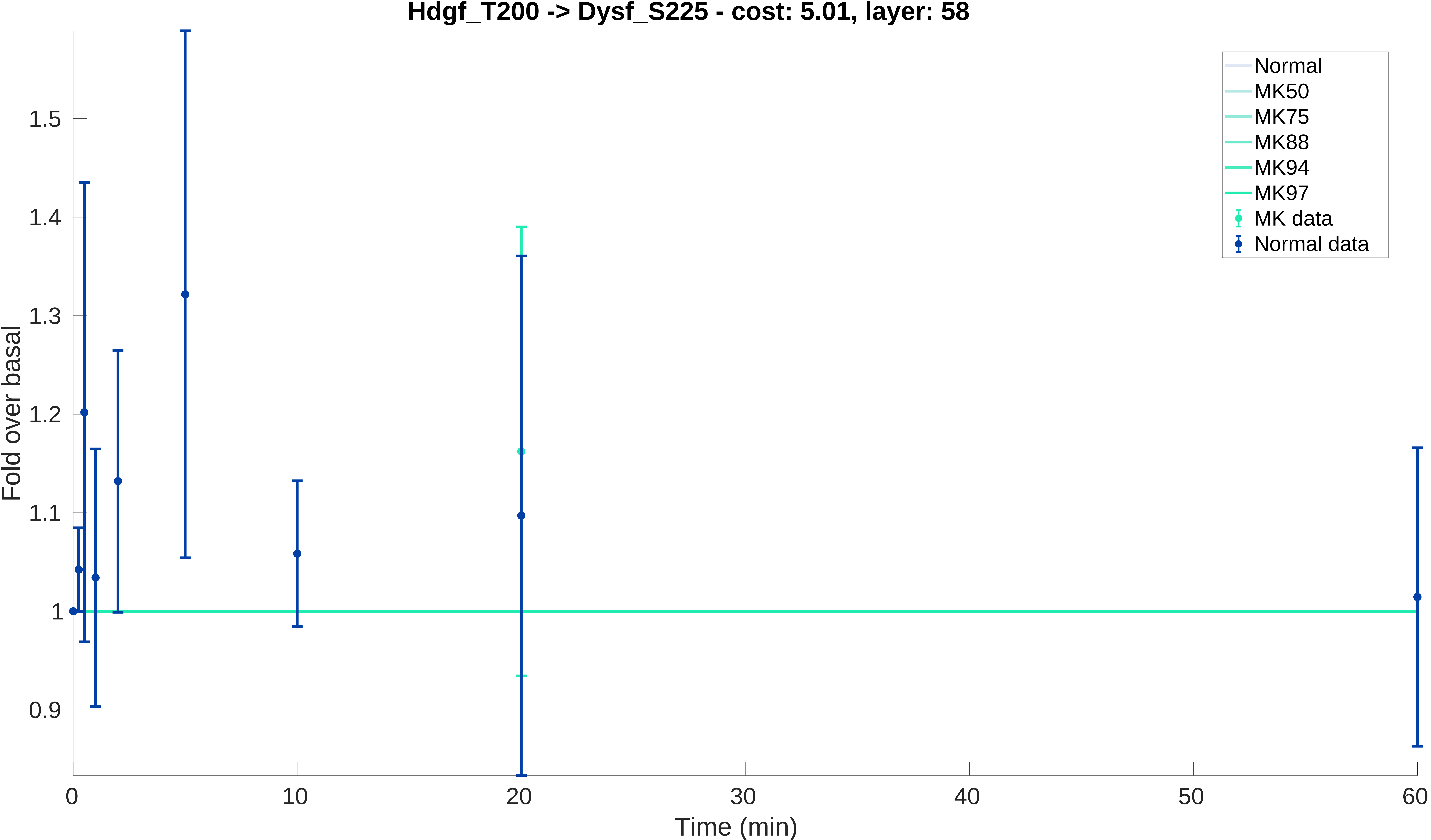

Hdgf_T200 -> Dysf_S225 - cost: 5.01, layer: 58
Fold over basal
Time (min)
0.9
1
1.1
1.2
1.3
1.4
1.5
0
10
20
30
40
50
60
Normal
MK50
MK75
MK88
MK94
MK97
MK data
Normal data

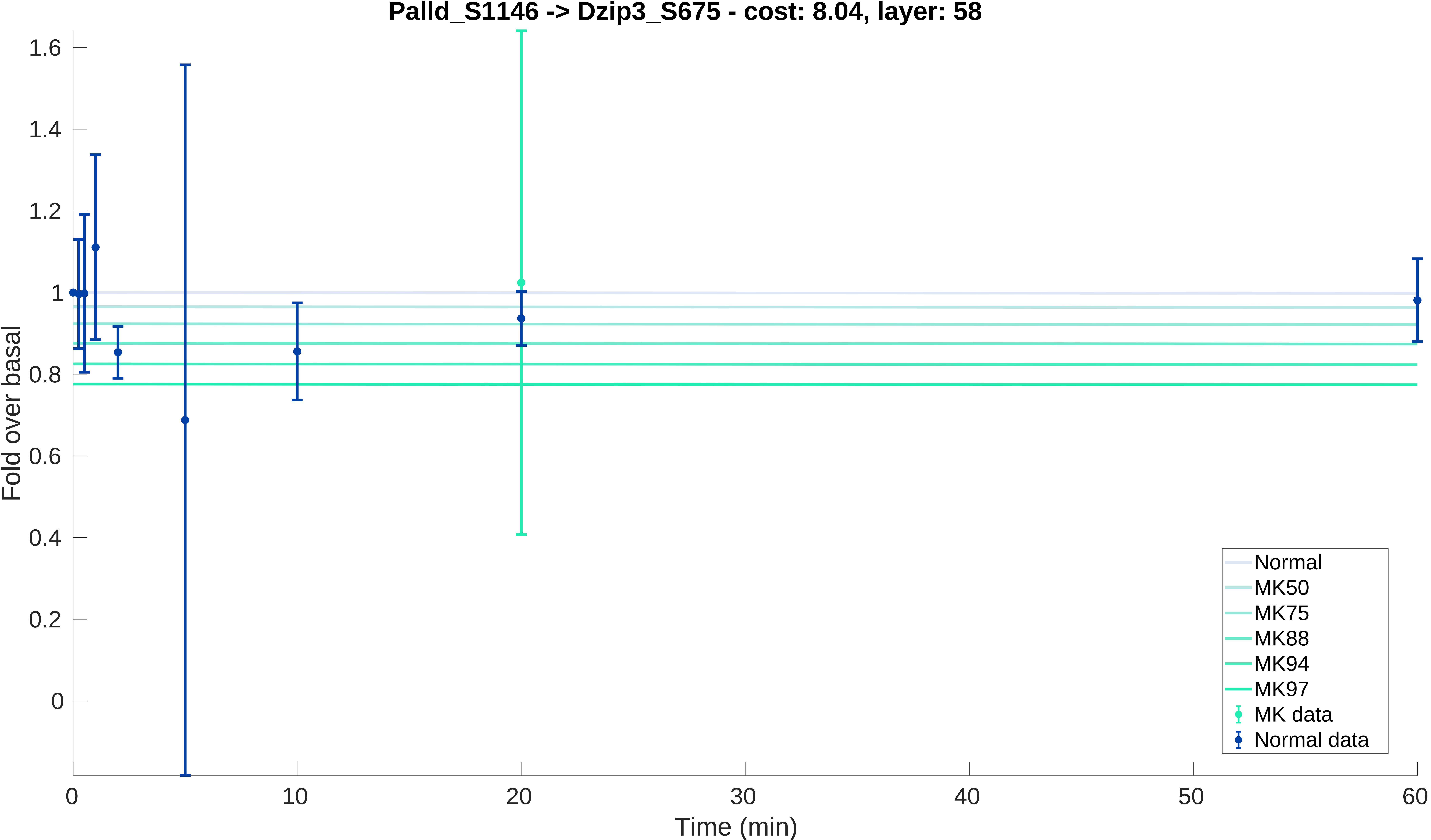

Palld_S1146 -> Dzip3_S675 - cost: 8.04, layer: 58
Fold over basal
Time (min)
1.6
1.4
1.2
1
0.8
0.6
0.4
0.2
0
0
10
20
30
40
50
60
Normal
MK50
MK75
MK88
MK94
MK97
MK data
Normal data

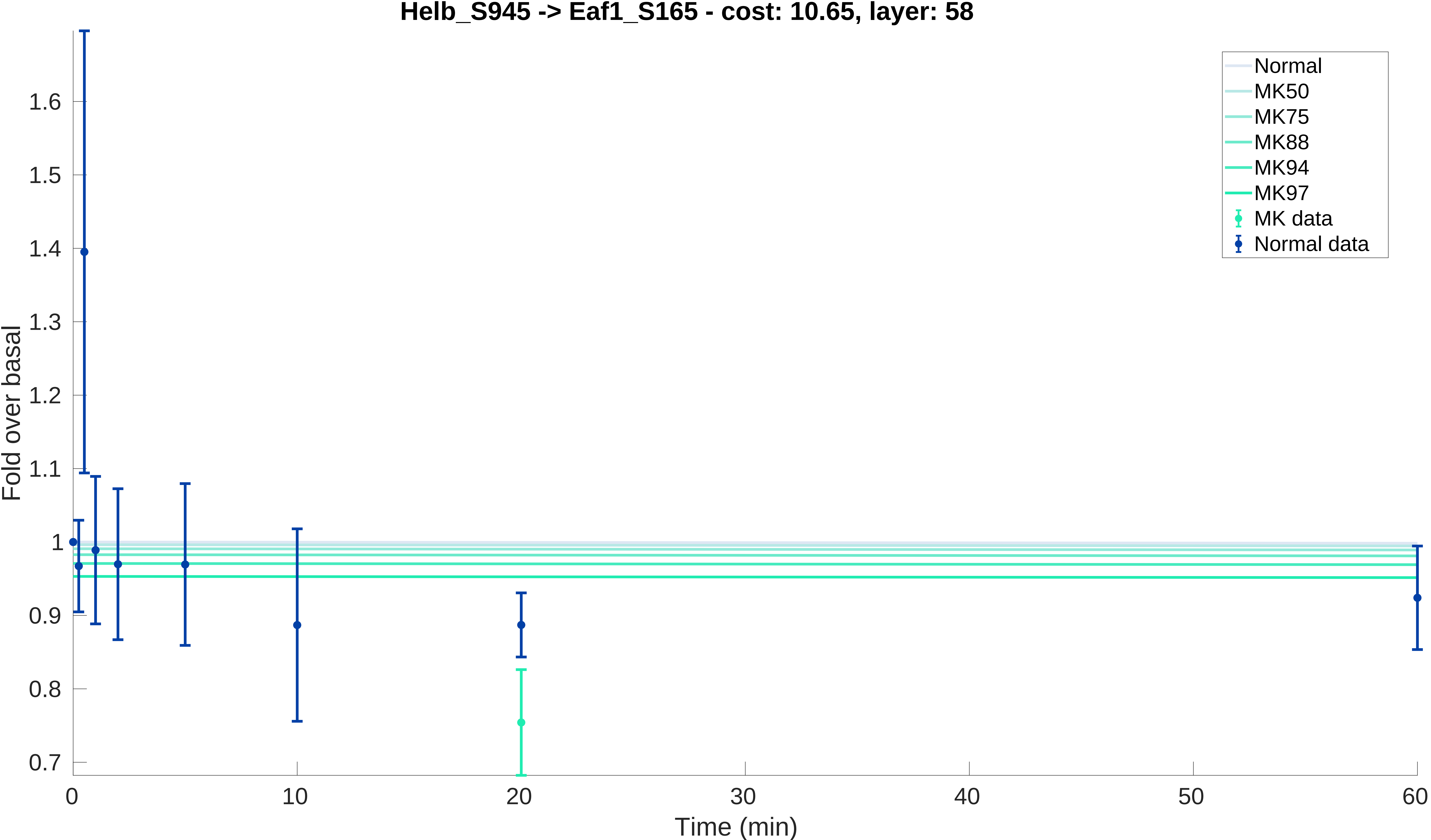

Helb_S945 -> Eaf1_S165 - cost: 10.65, layer: 58
Normal
MK50
MK75
MK88
MK94
MK97
MK data
Normal data
Fold over basal
Time (min)
1.6
1.5
1.4
1.3
1.2
1.1
1
0.9
0.8
0.7
0
10
20
30
40
50
60

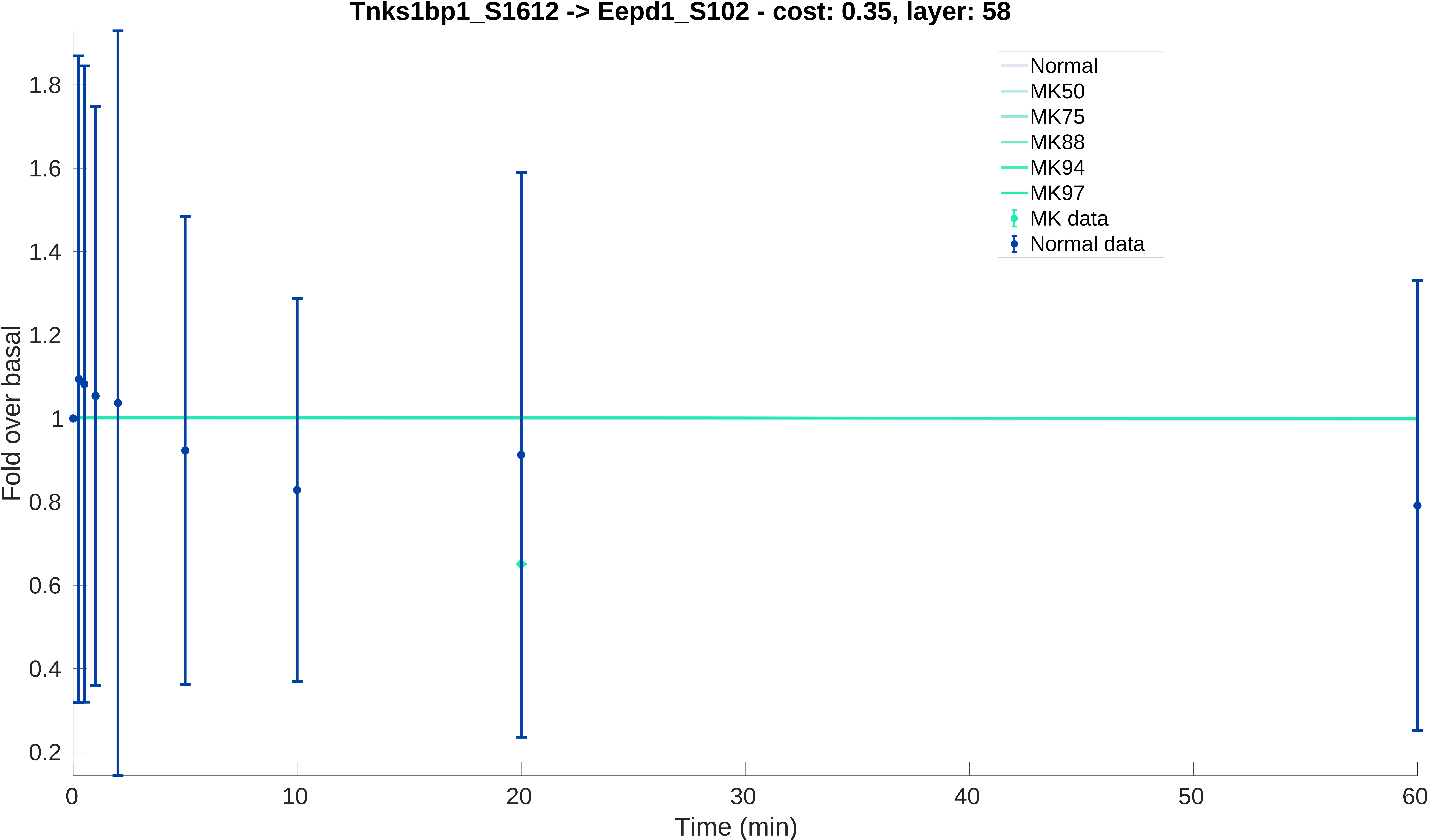

Tnks1bp1_S1612 -> Eepd1_S102 - cost: 0.35, layer: 58
Normal
MK50
MK75
MK88
MK94
MK97
MK data
Normal data
Fold over basal
1.8
1.6
1.4
1.2
1
0.8
0.6
0.4
0.2
0
10
20
30
40
50
60
Time (min)

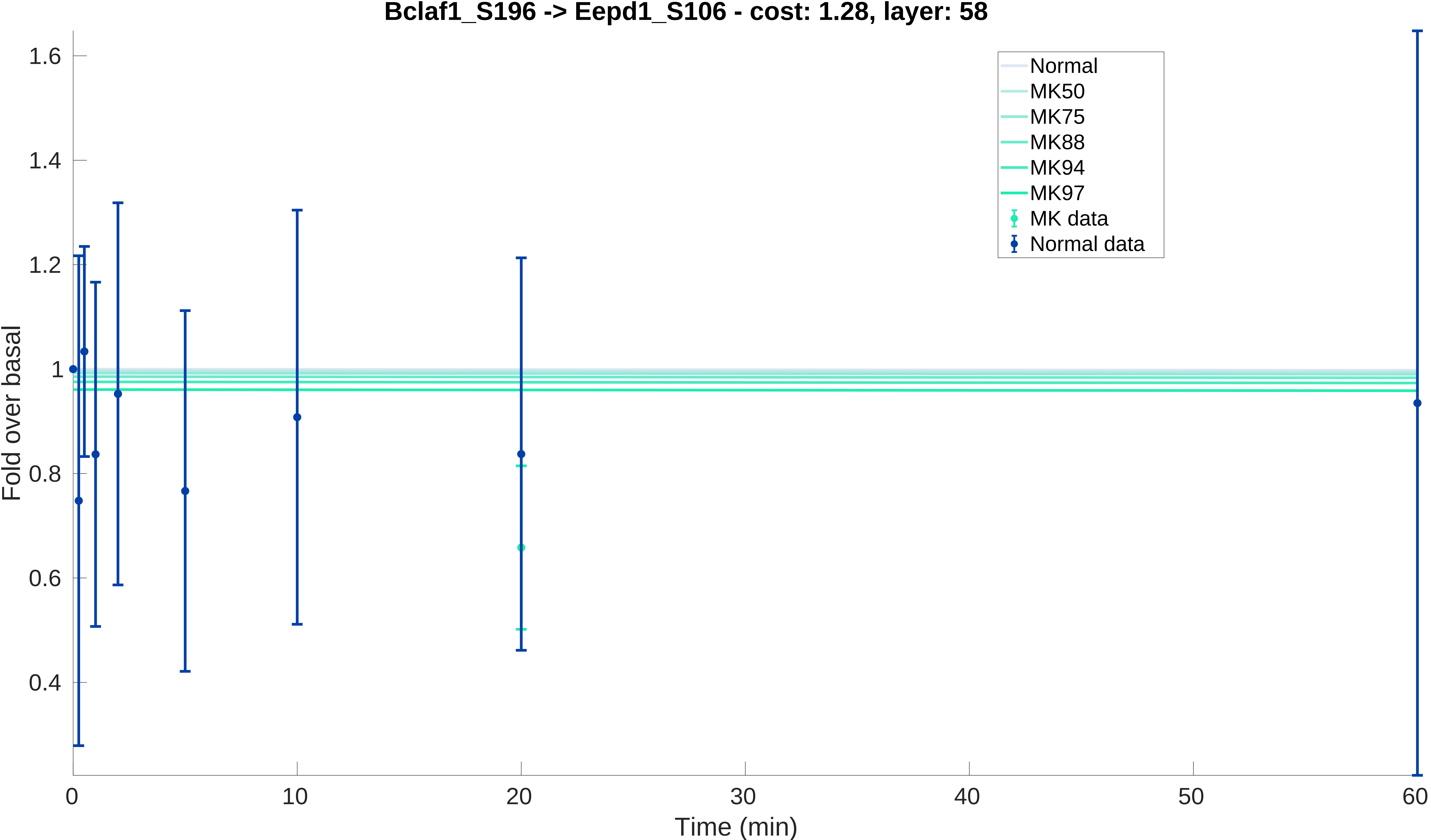

Bclaf1_S196 -> Eepd1_S106 - cost: 1.28, layer: 58
Fold over basal
Time (min)
1.6
1.4
1.2
1
0.8
0.6
0.4
0
10
20
30
40
50
60
Normal
MK50
MK75
MK88
MK94
MK97
MK data
Normal data

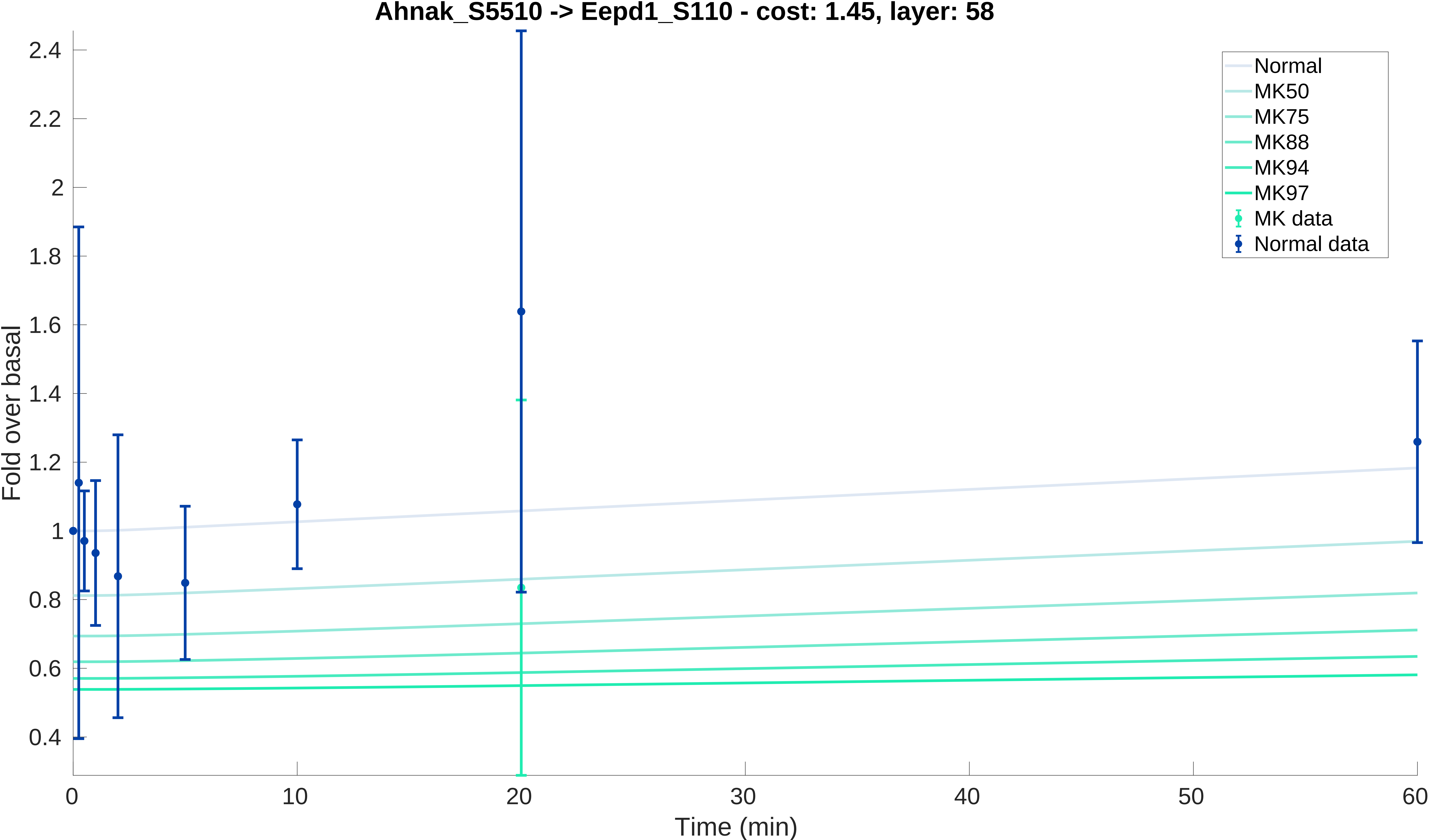

Ahnak_S5510 -> Eepd1_S110 - cost: 1.45, layer: 58
Fold over basal
Time (min)
Normal
MK50
MK75
MK88
MK94
MK97
MK data
Normal data
2.4
2.2
2
1.8
1.6
1.4
1.2
1
0.8
0.6
0.4
0
10
20
30
40
50
60

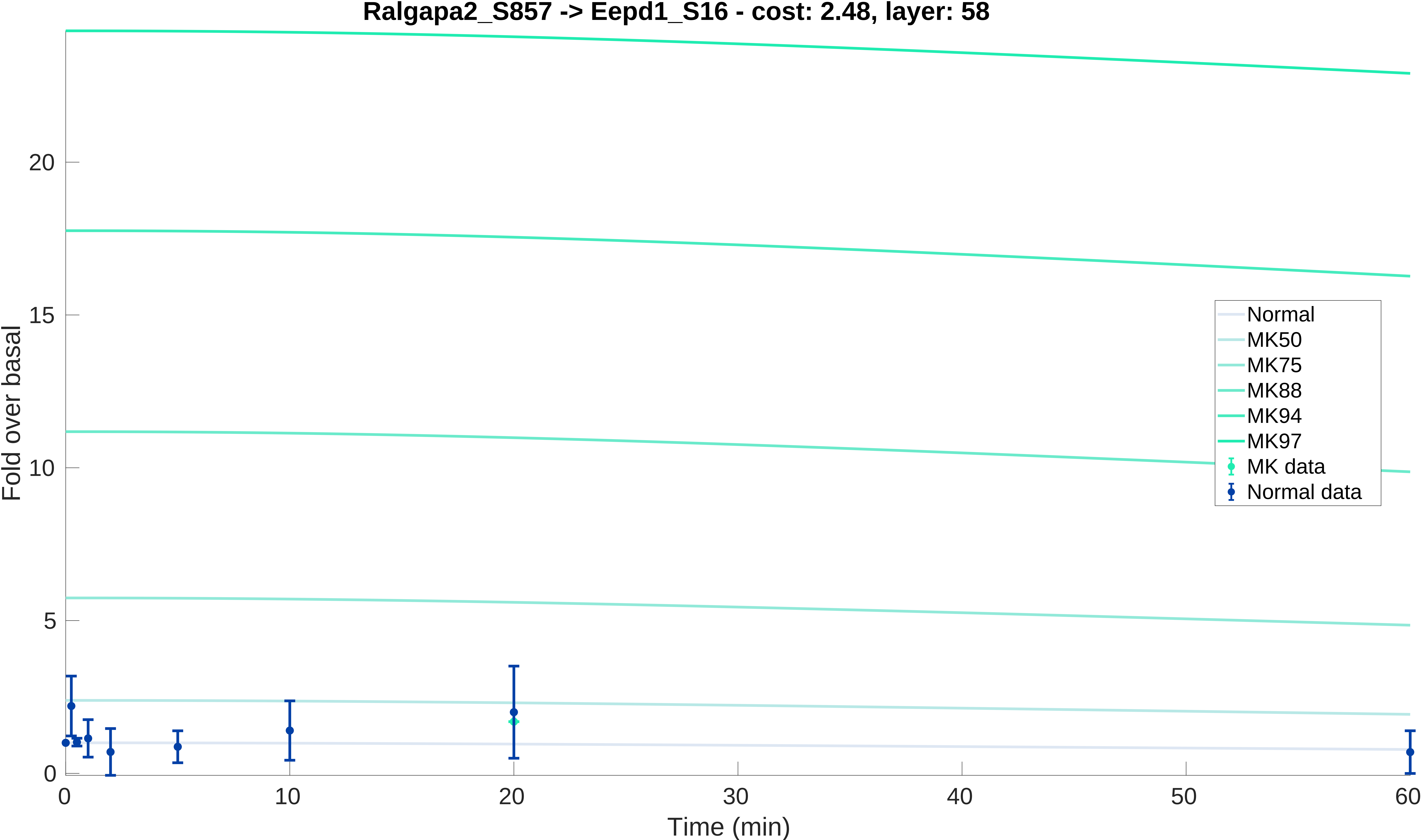

Ralgapa2_S857 -> Eepd1_S16 - cost: 2.48, layer: 58
Fold over basal
Time (min)
0
5
10
15
20
0
10
20
30
40
50
60
Normal
MK50
MK75
MK88
MK94
MK97
MK data
Normal data

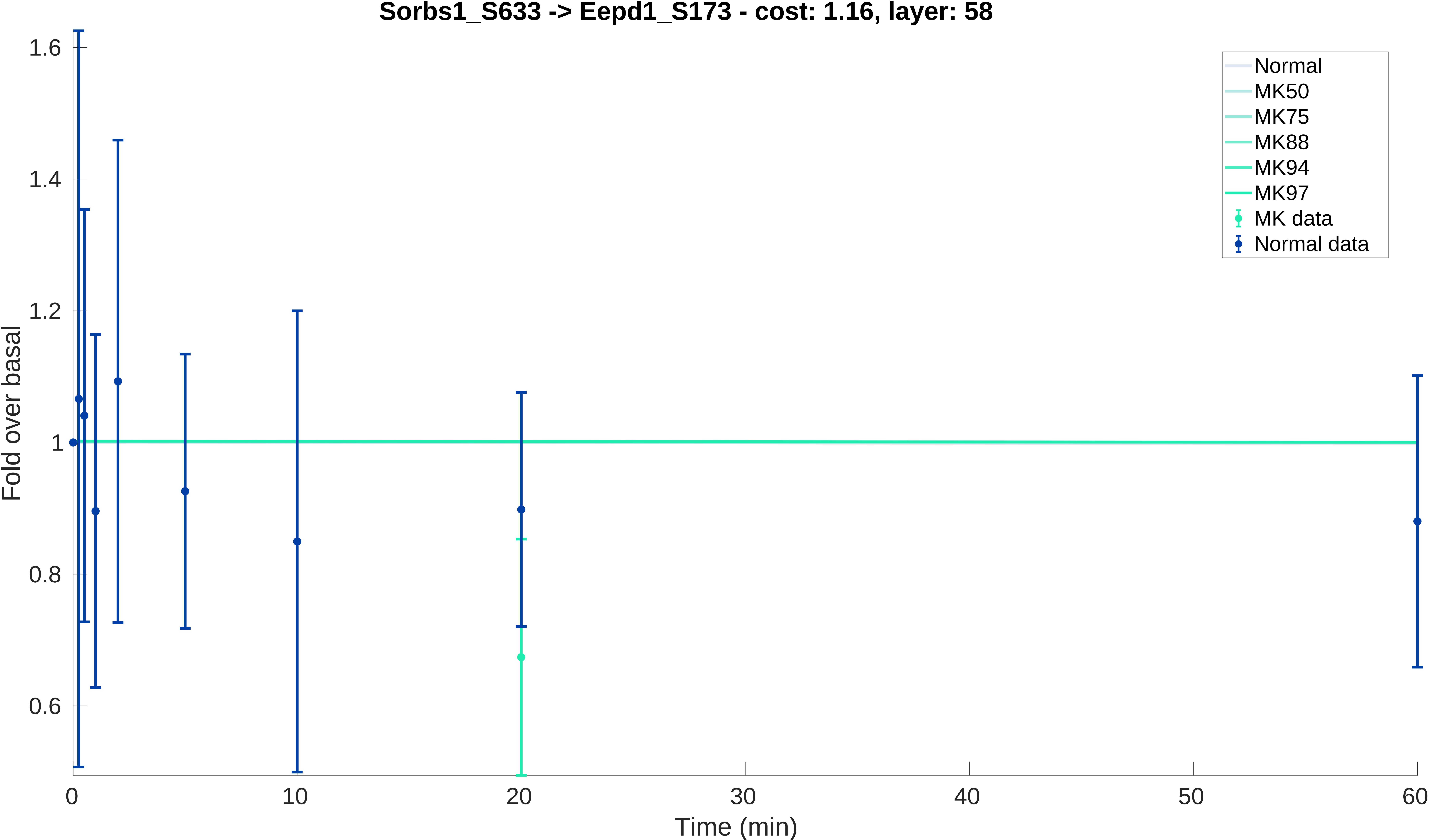

Sorbs1_S633 -> Eepd1_S173 - cost: 1.16, layer: 58
Fold over basal
Time (min)
0.6
0.8
1
1.2
1.4
1.6
0
10
20
30
40
50
60
Normal
MK50
MK75
MK88
MK94
MK97
MK data
Normal data

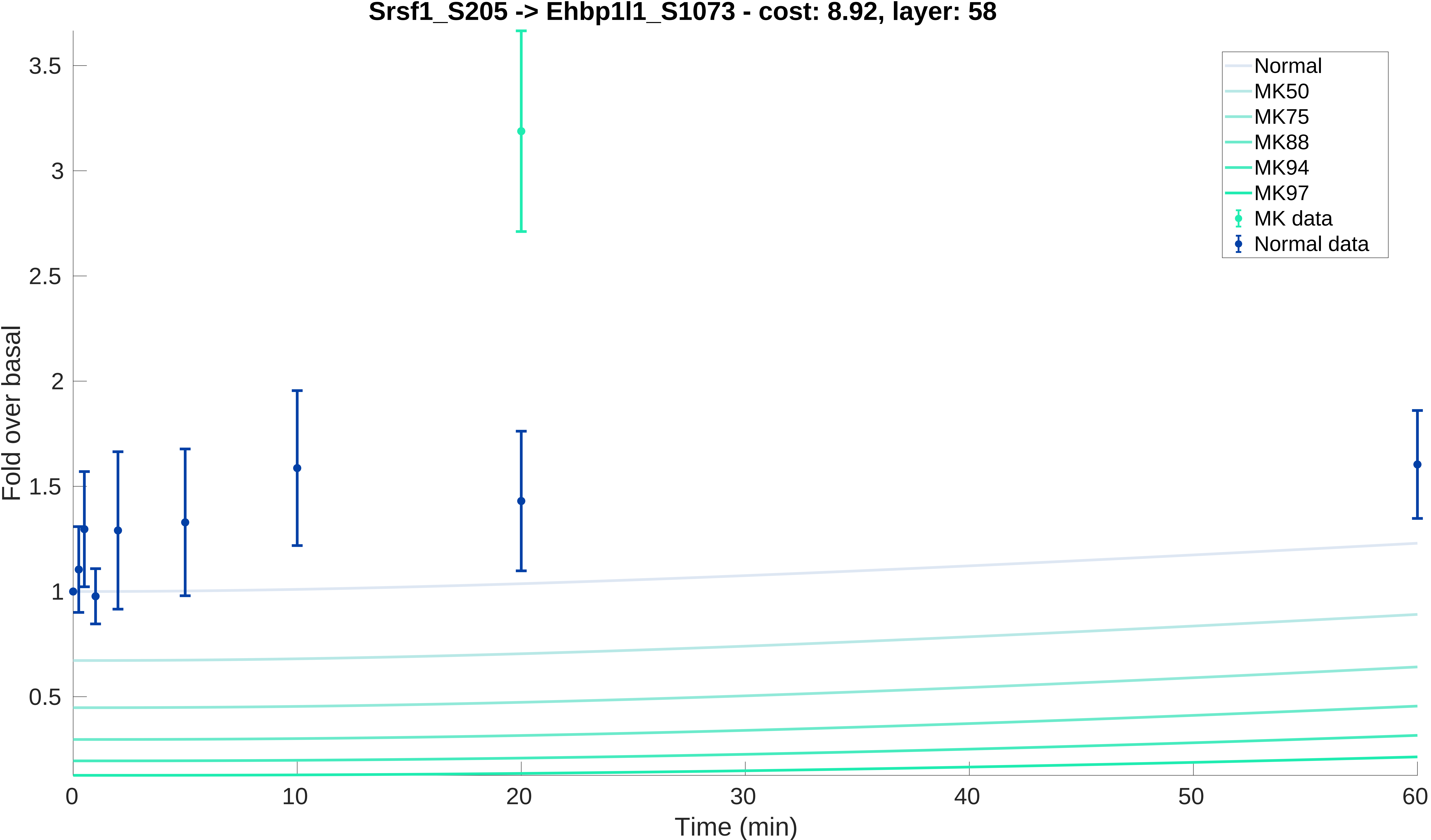

Srsf1_S205 -> Ehbp1l1_S1073 - cost: 8.92, layer: 58
Fold over basal
3.5
3
2.5
2
1.5
1
0.5
0
10
20
30
40
50
60
Time (min)
Normal
MK50
MK75
MK88
MK94
MK97
MK data
Normal data

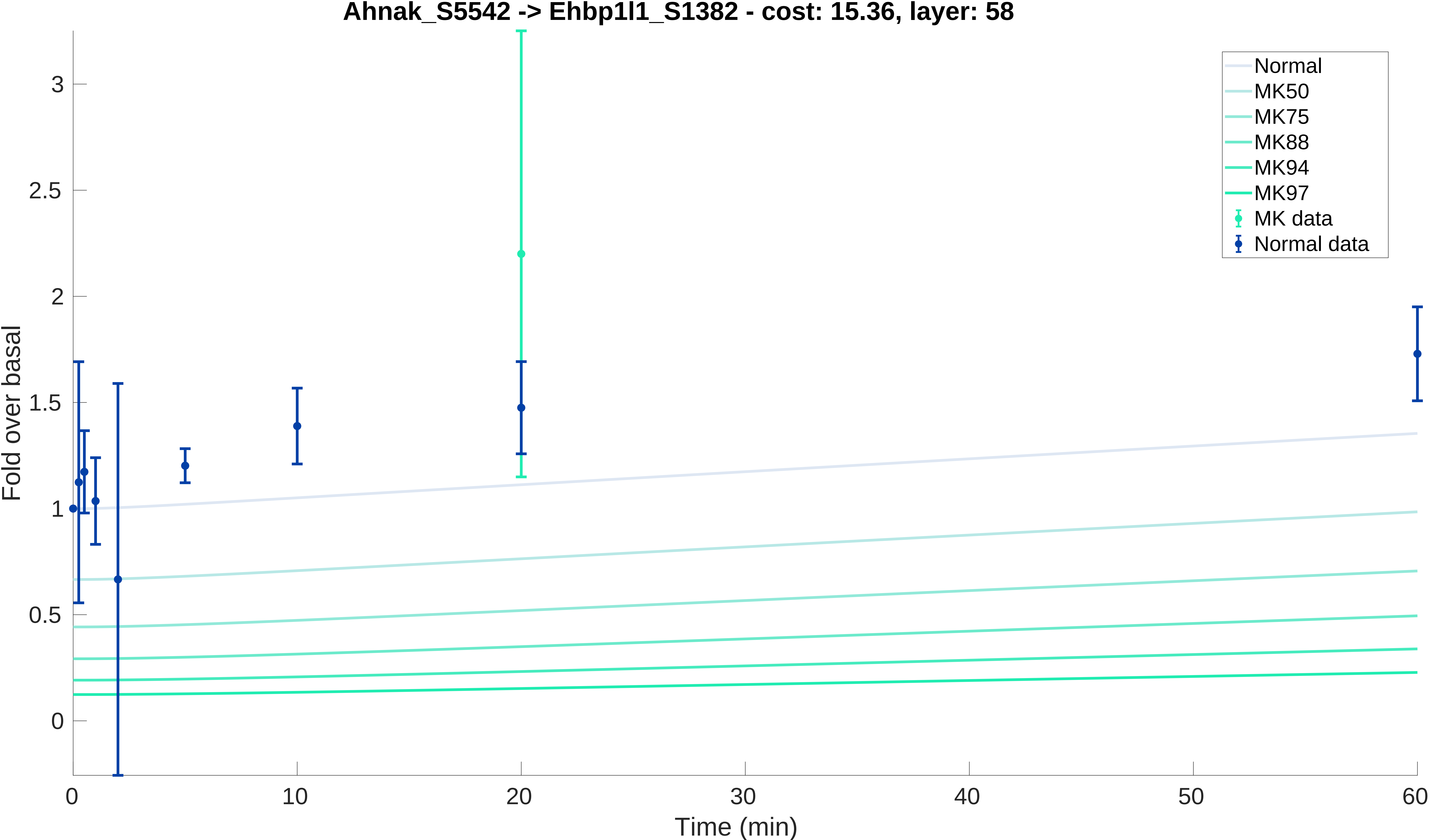

Ahnak_S5542 -> Ehbp1l1_S1382 - cost: 15.36, layer: 58
Normal
MK50
MK75
MK88
MK94
MK97
MK data
Normal data
Fold over basal
Time (min)
0
0.5
1
1.5
2
2.5
3
0
10
20
30
40
50
60

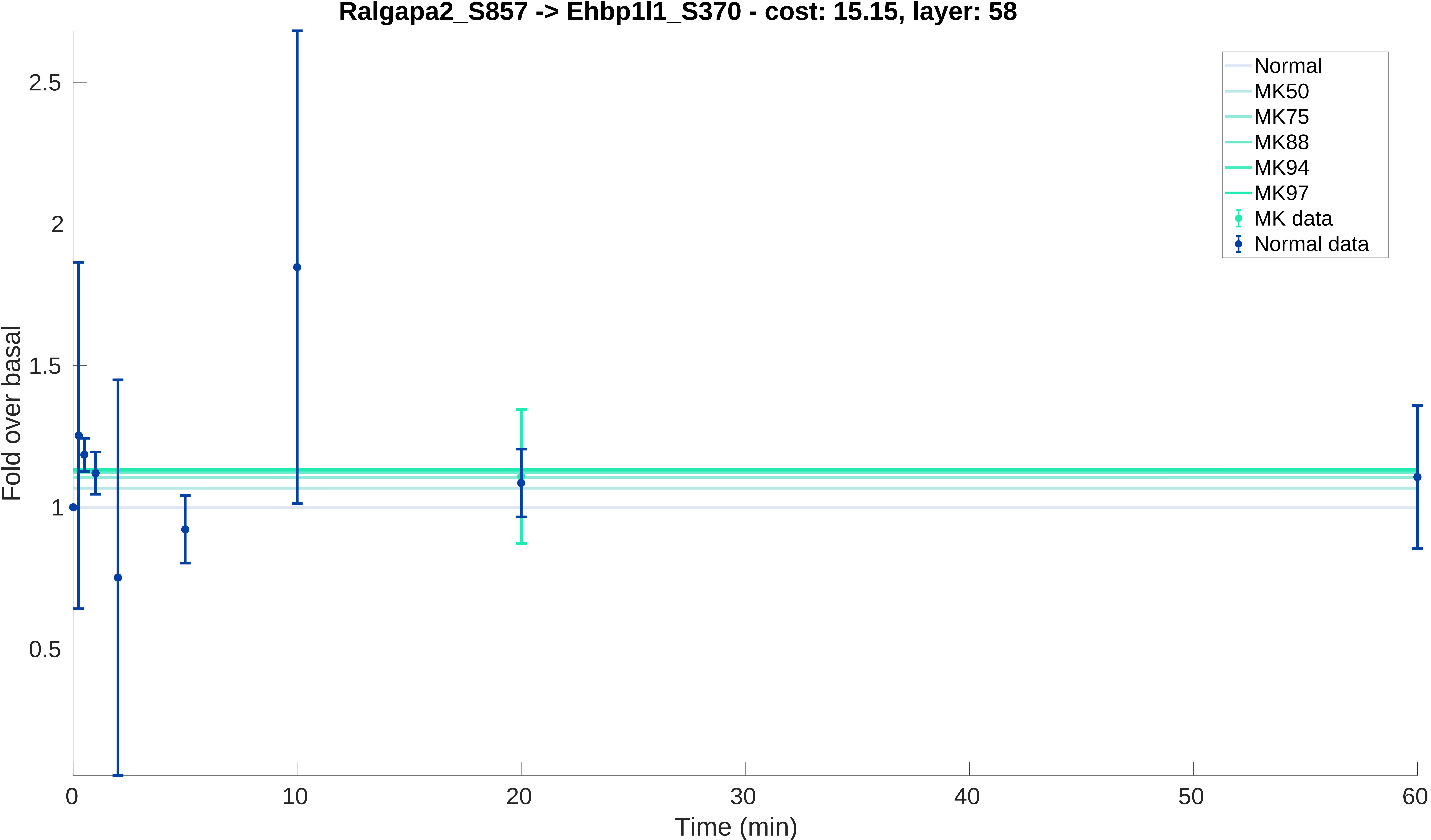

Ralgapa2_S857 -> Ehbp1l1_S370 - cost: 15.15, layer: 58
Fold over basal
Time (min)
0.5
1
1.5
2
2.5
0
10
20
30
40
50
60
Normal
MK50
MK75
MK88
MK94
MK97
MK data
Normal data

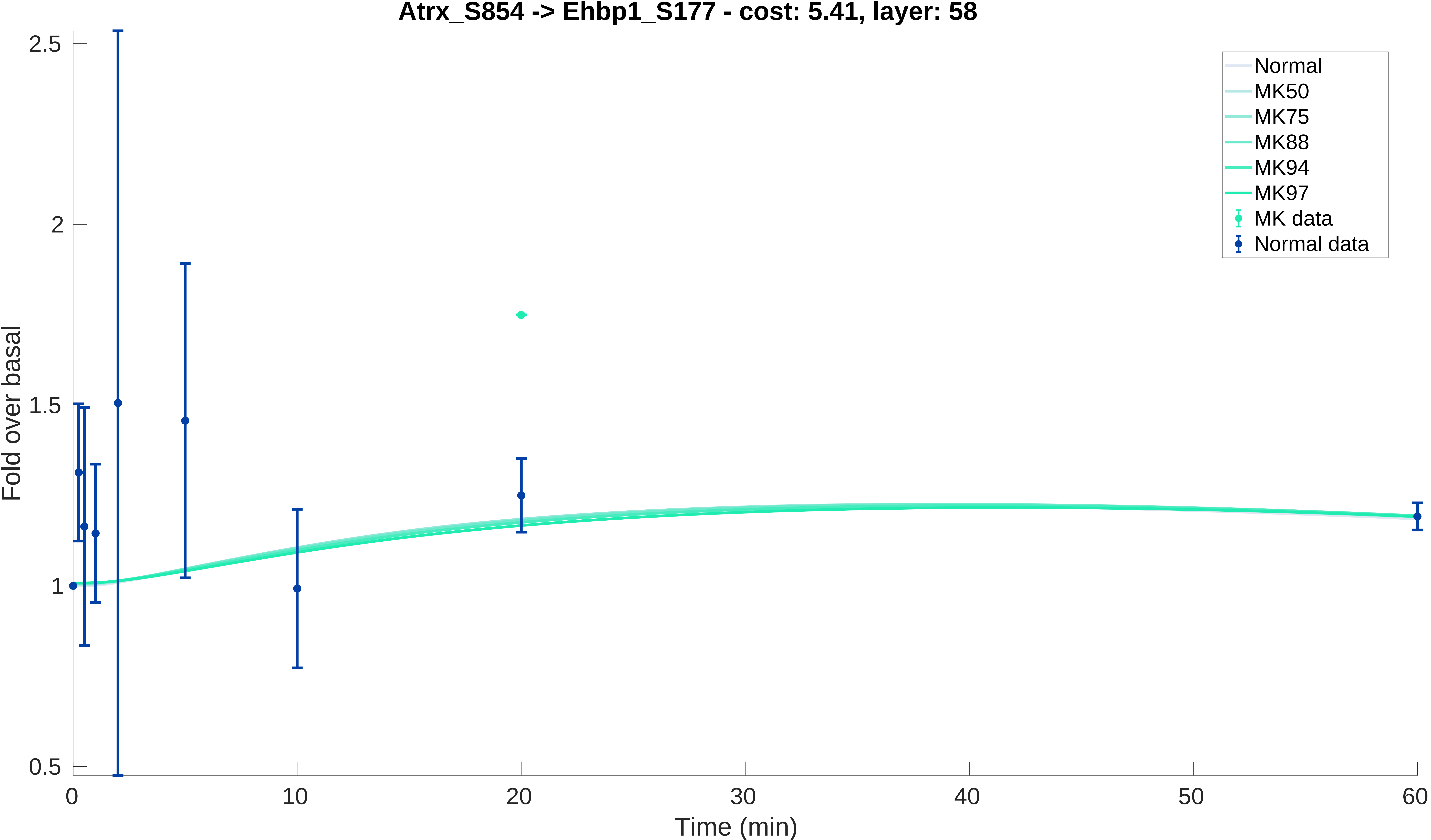

Atrx_S854 -> Ehbp1_S177 - cost: 5.41, layer: 58
Fold over basal
Time (min)
0.5
1
1.5
2
2.5
0
10
20
30
40
50
60
Normal
MK50
MK75
MK88
MK94
MK97
MK data
Normal data

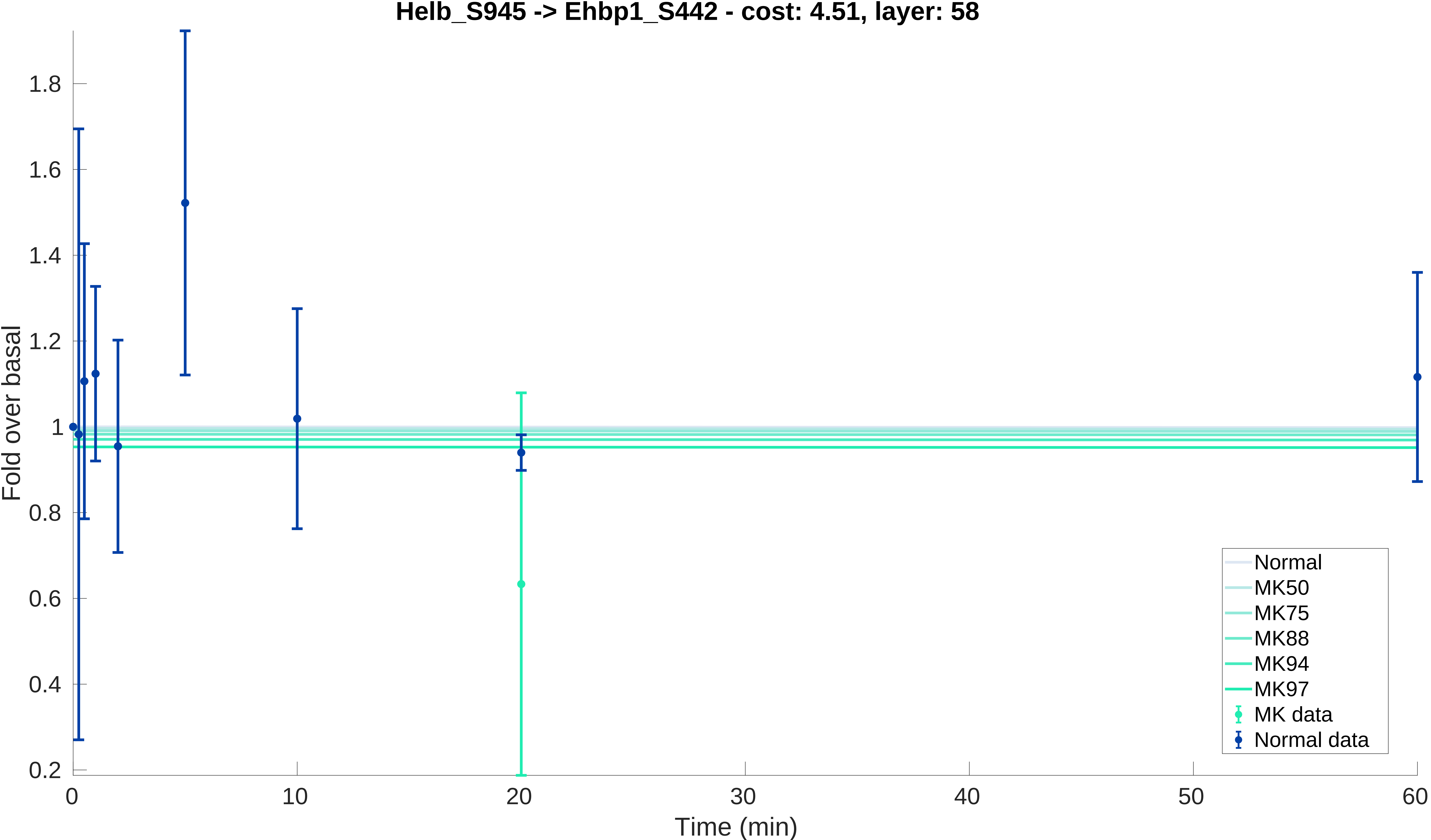

Helb_S945 -> Ehbp1_S442 - cost: 4.51, layer: 58
Fold over basal
1.8
1.6
1.4
1.2
1
0.8
0.6
0.4
0.2
0
10
20
30
40
50
60
Time (min)
Normal
MK50
MK75
MK88
MK94
MK97
MK data
Normal data

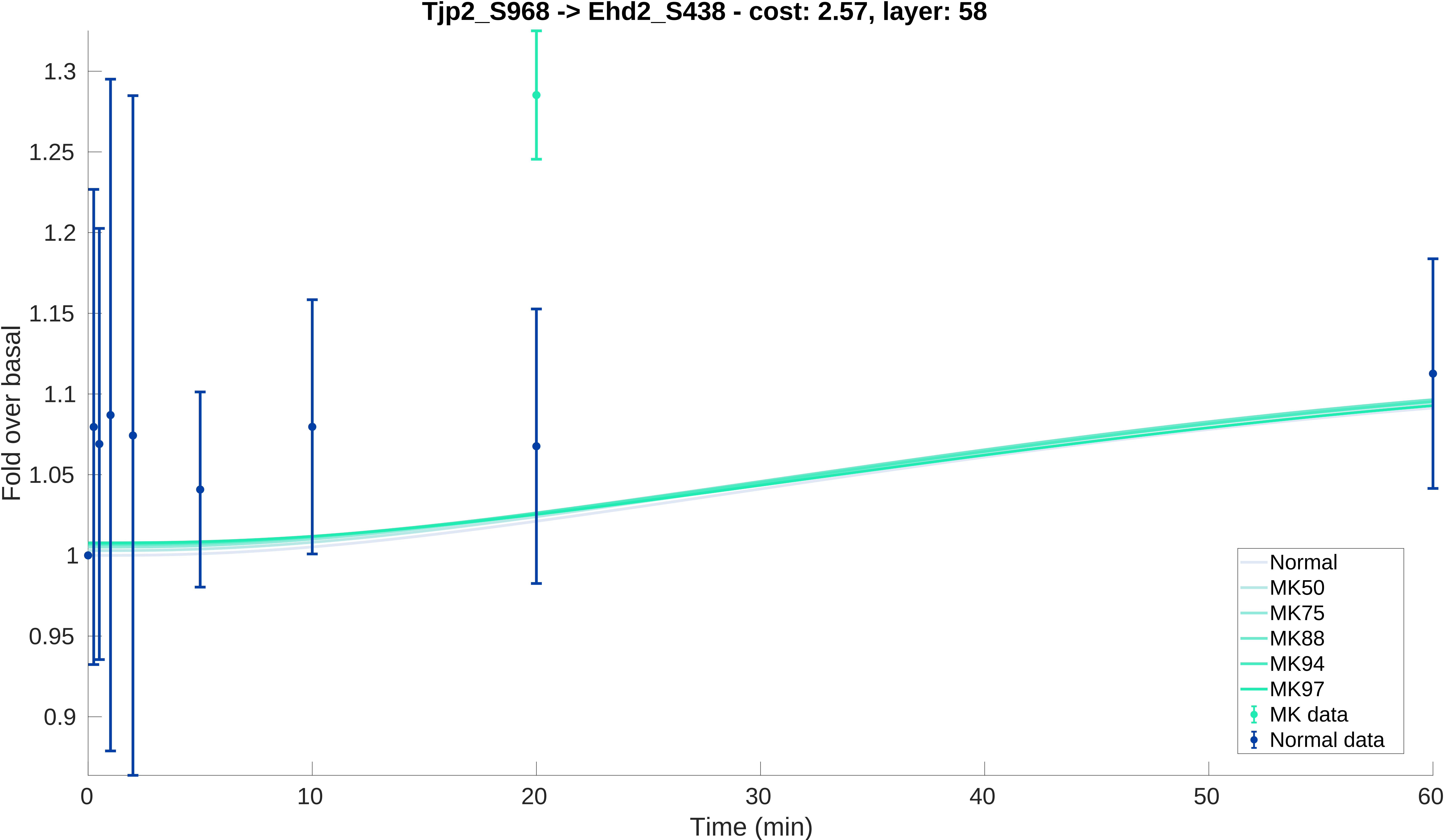
Tjp2_S968 -> Ehd2_S438 - cost: 2.57, layer: 58
Fold over basal
1.3
1.25
1.2
1.15
1.1
1.05
1
0.95
0.9
0
10
20
30
40
50
60
Time (min)
Normal
MK50
MK75
MK88
MK94
MK97
MK data
Normal data

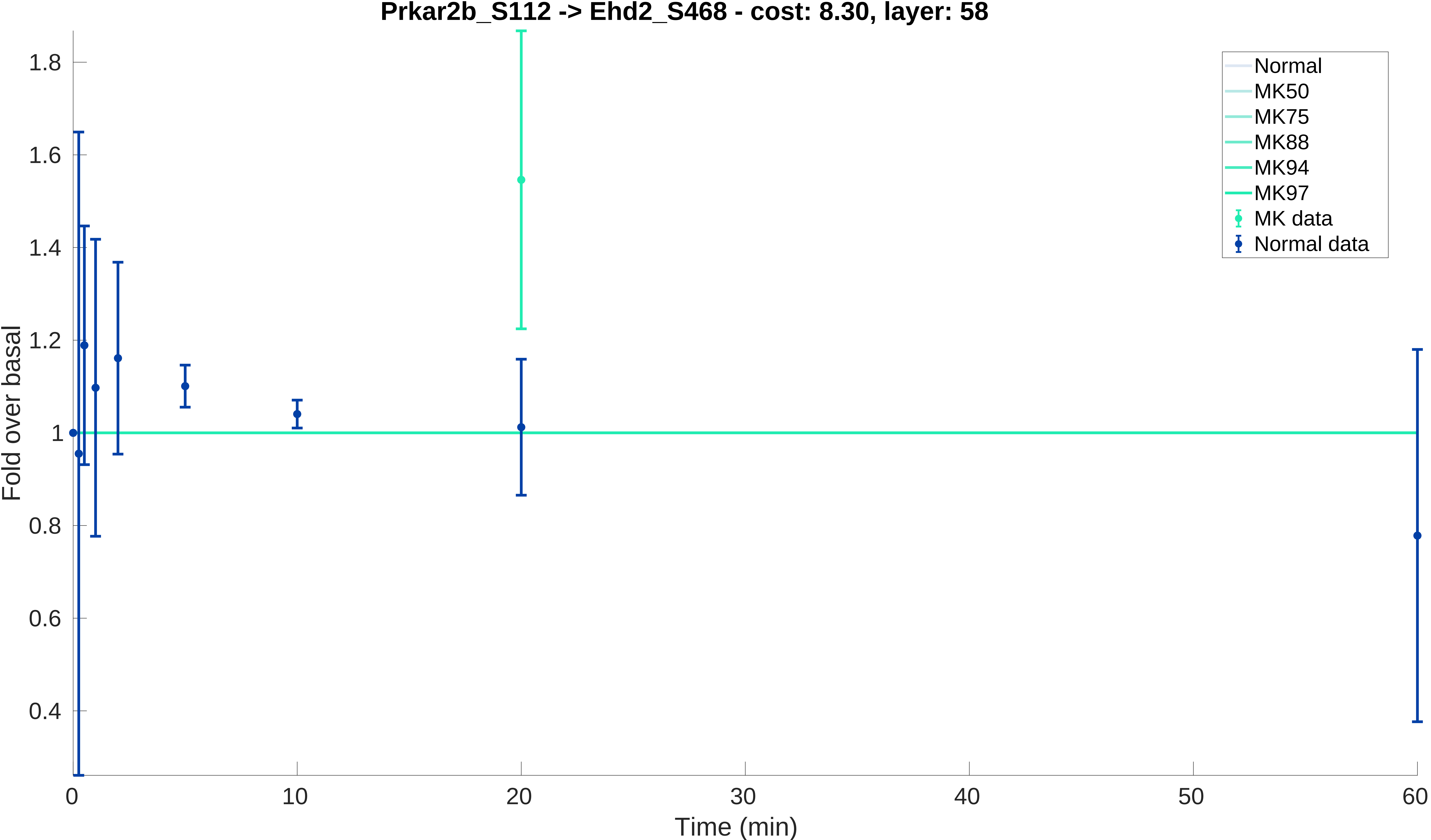
Prkar2b_S112 -> Ehd2_S468 - cost: 8.30, layer: 58
Fold over basal
Time (min)
1.8
1.6
1.4
1.2
1
0.8
0.6
0.4
0
10
20
30
40
50
60
Normal
MK50
MK75
MK88
MK94
MK97
MK data
Normal data

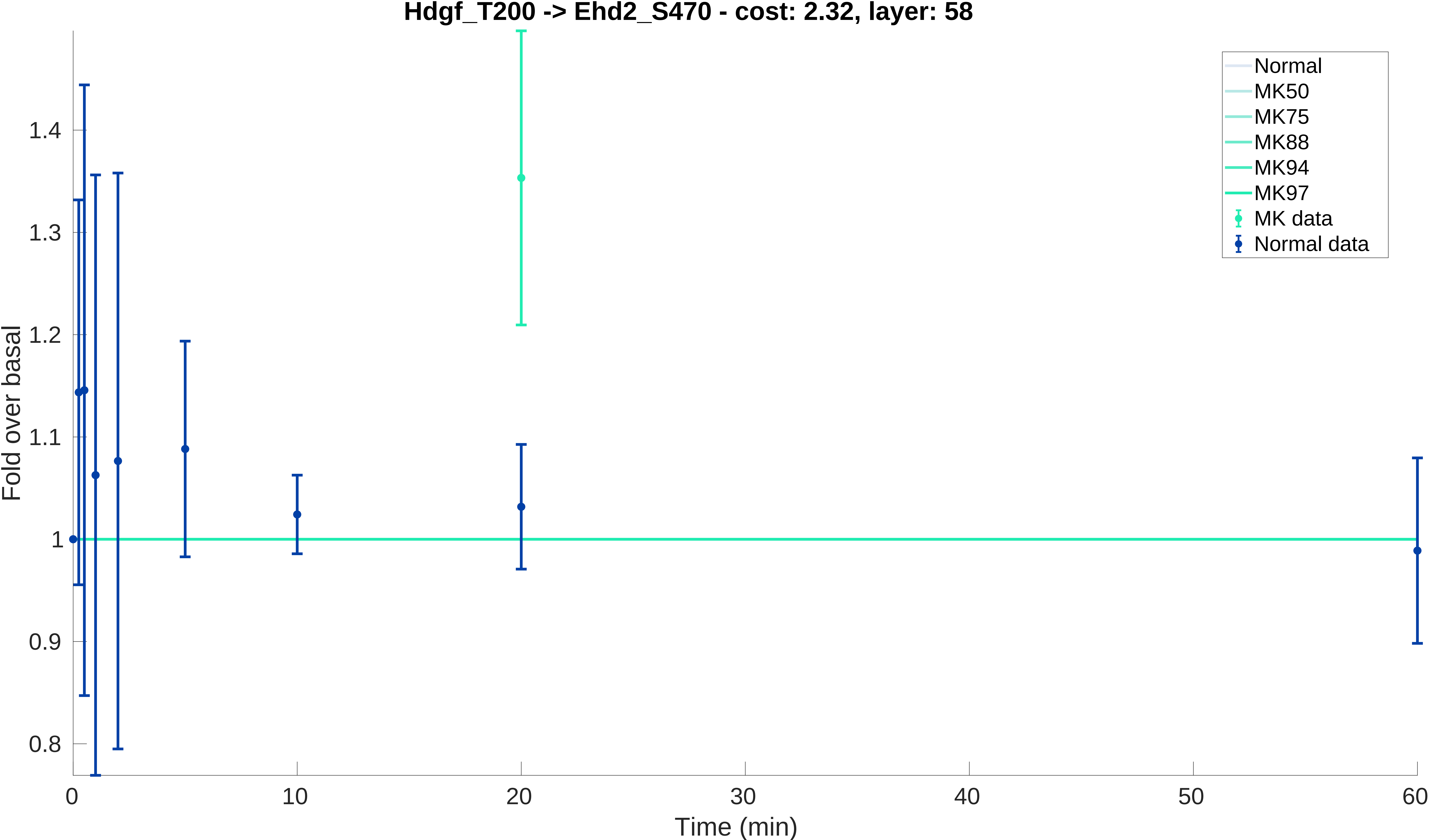

Hdgf_T200 -> Ehd2_S470 - cost: 2.32, layer: 58
Normal
MK50
MK75
MK88
MK94
MK97
MK data
Normal data
Fold over basal
1.4
1.3
1.2
1.1
1
0.9
0.8
0
10
20
30
40
50
60
Time (min)

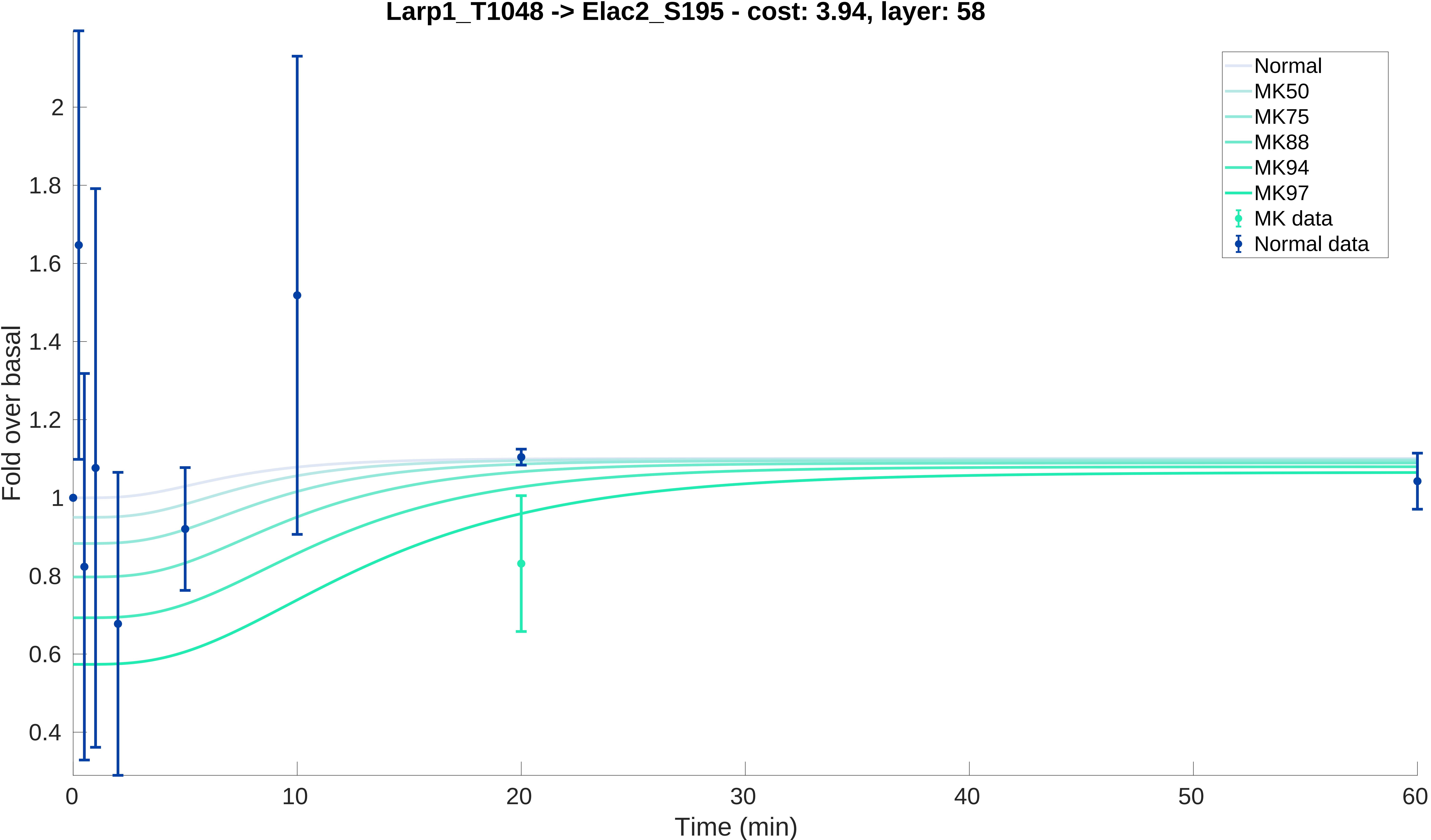

Larp1_T1048 -> Elac2_S195 - cost: 3.94, layer: 58
Fold over basal
Time (min)
0.4
0.6
0.8
1
1.2
1.4
1.6
1.8
2
0
10
20
30
40
50
60
Normal
MK50
MK75
MK88
MK94
MK97
MK data
Normal data

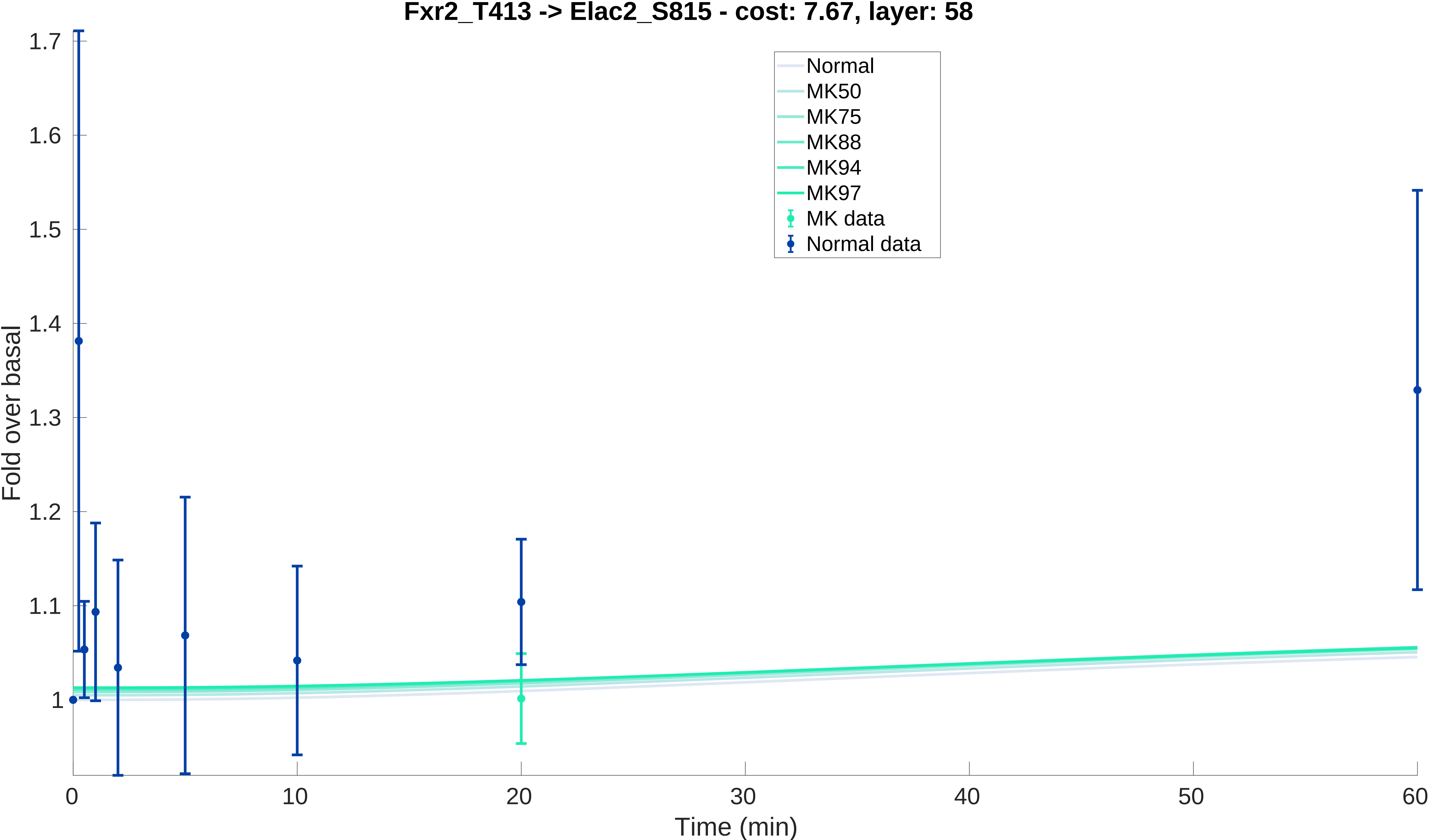

Fxr2_T413 -> Elac2_S815 - cost: 7.67, layer: 58
Normal
MK50
MK75
MK88
MK94
MK97
MK data
Normal data
Fold over basal
1.7
1.6
1.5
1.4
1.3
1.2
1.1
1
0
10
20
30
40
50
60
Time (min)

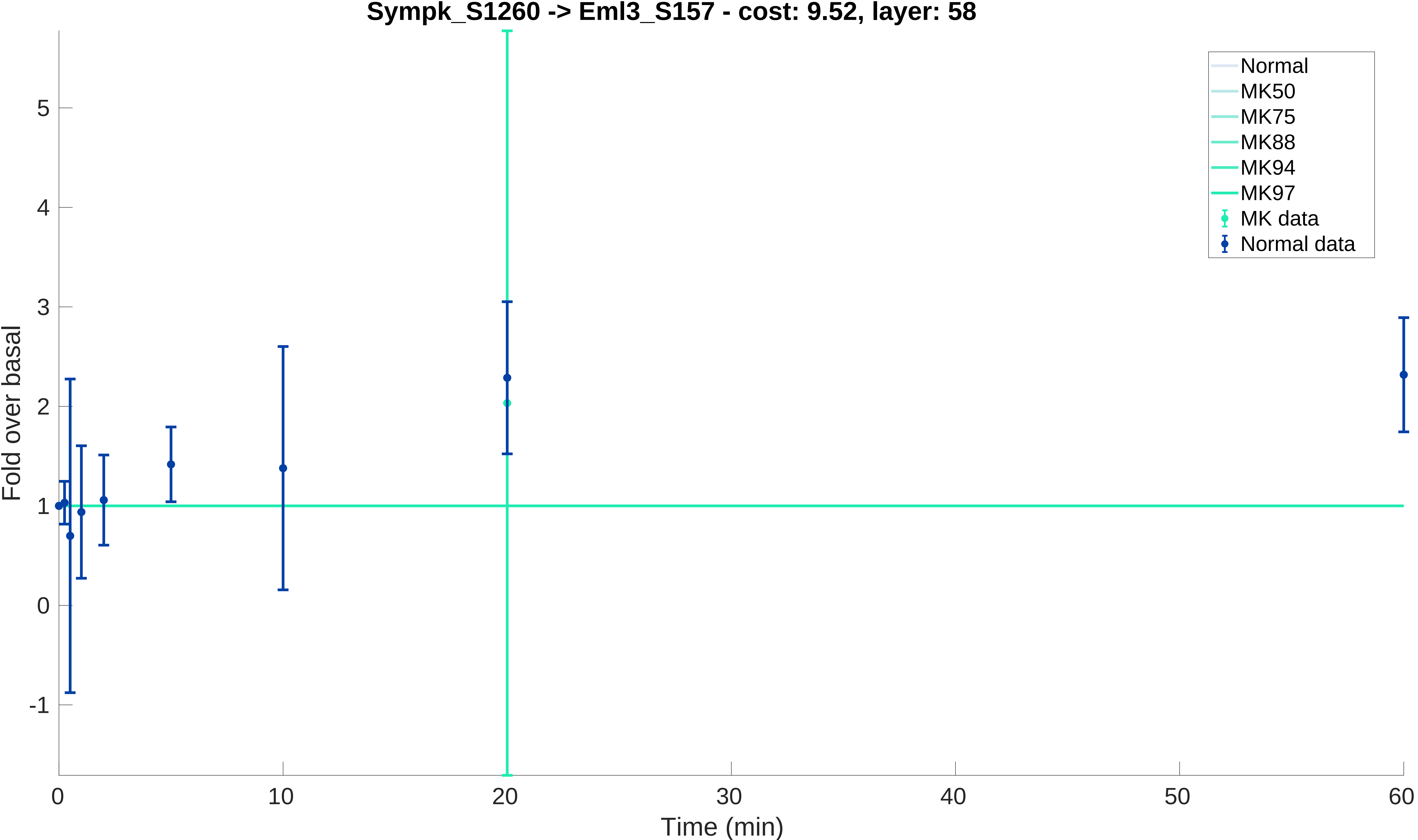
Sympk_S1260 -> Eml3_S157 - cost: 9.52, layer: 58
Fold over basal
Time (min)
5
4
3
2
1
0
-1
0
10
20
30
40
50
60
Normal
MK50
MK75
MK88
MK94
MK97
MK data
Normal data

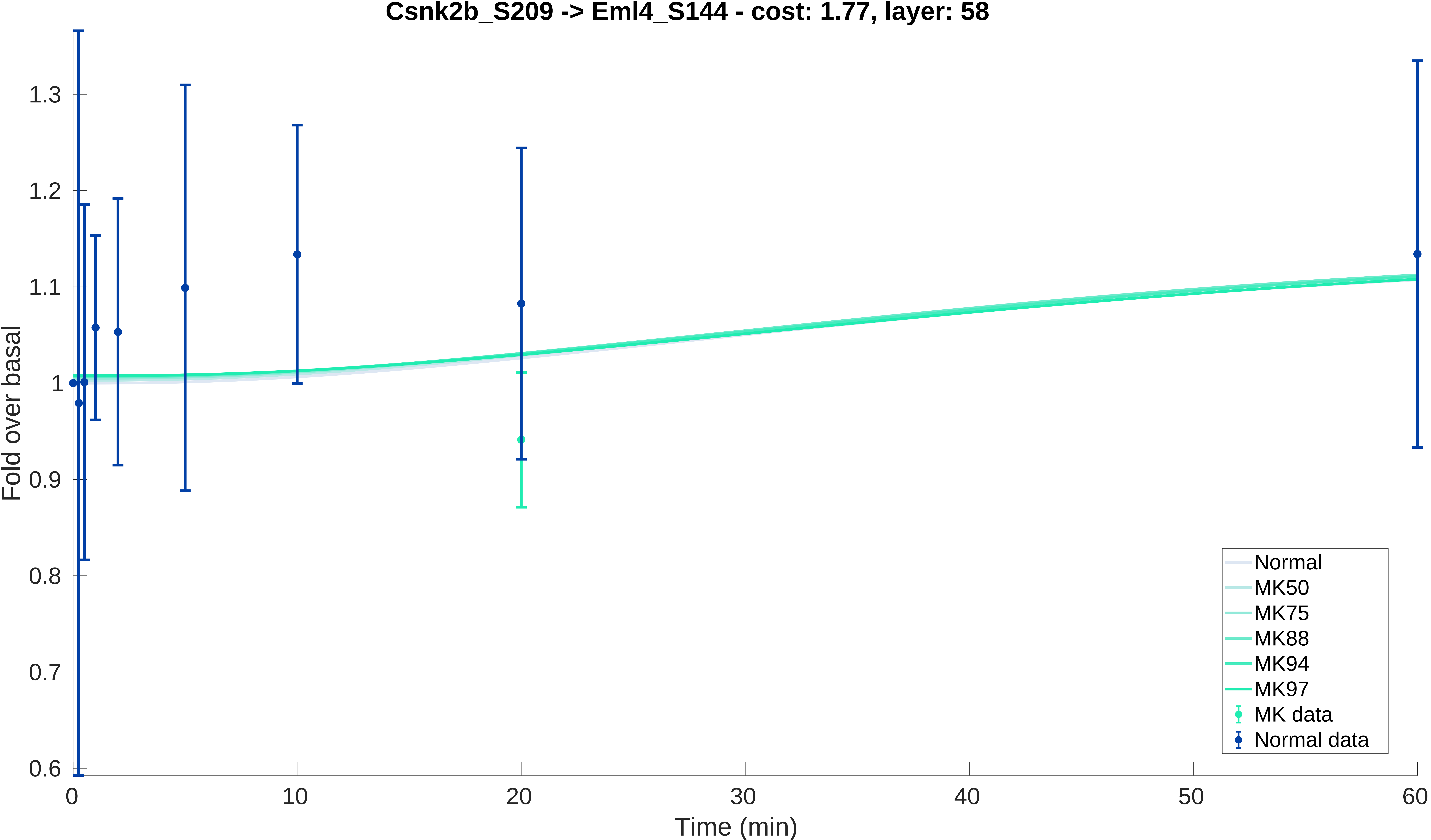

Csnk2b_S209 -> Eml4_S144 - cost: 1.77, layer: 58
Fold over basal
1.3
1.2
1.1
1
0.9
0.8
0.7
0.6
0
10
20
30
40
50
60
Time (min)
Normal
MK50
MK75
MK88
MK94
MK97
MK data
Normal data

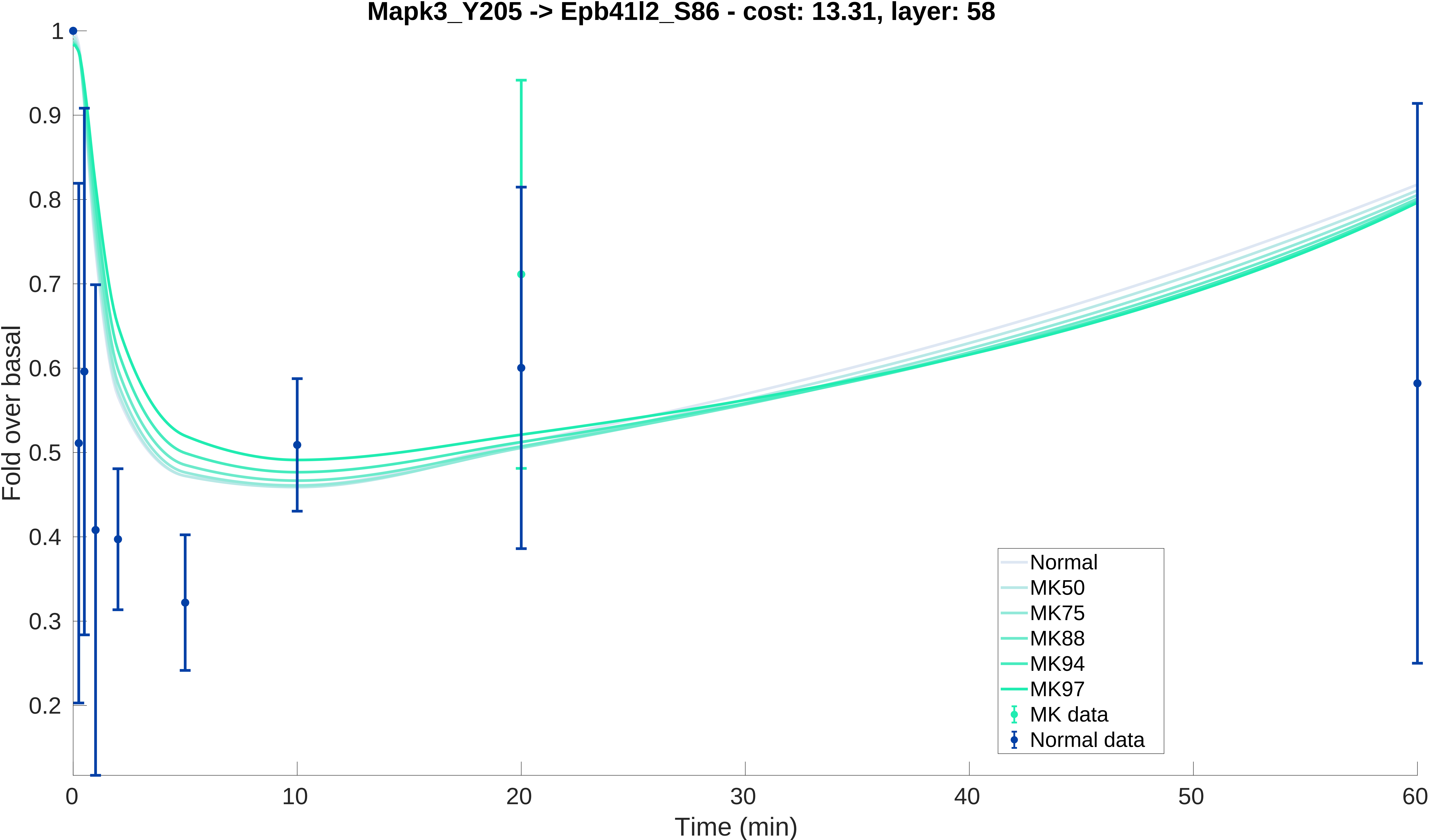

Mapk3_Y205 -> Epb41l2_S86 - cost: 13.31, layer: 58
Fold over basal
1
0.9
0.8
0.7
0.6
0.5
0.4
0.3
0.2
0
10
20
30
40
50
60
Time (min)
Normal
MK50
MK75
MK88
MK94
MK97
MK data
Normal data

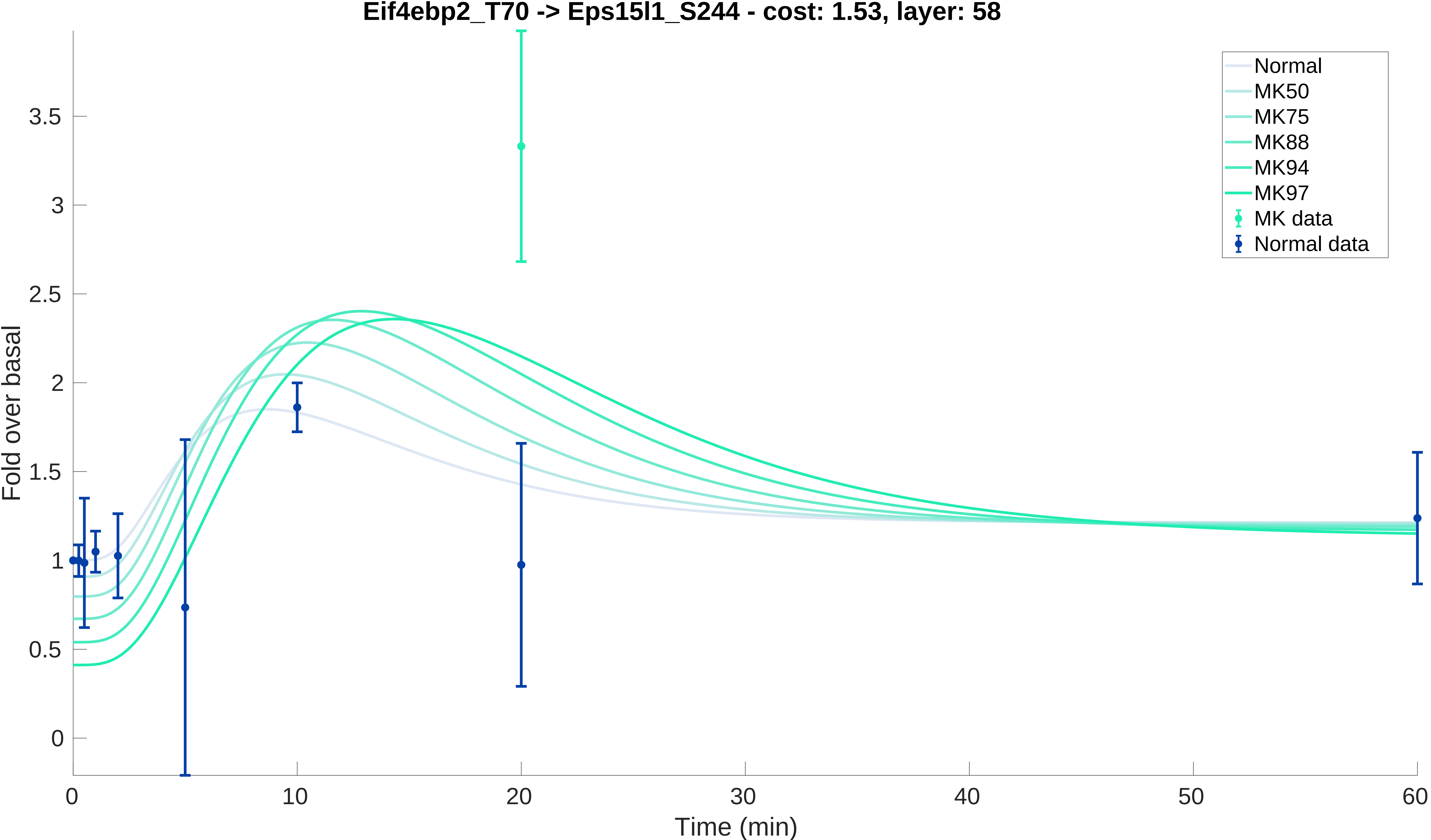

Eif4ebp2_T70 -> Eps15l1_S244 - cost: 1.53, layer: 58
Fold over basal
Time (min)
0
0.5
1
1.5
2
2.5
3
3.5
0
10
20
30
40
50
60
Normal
MK50
MK75
MK88
MK94
MK97
MK data
Normal data

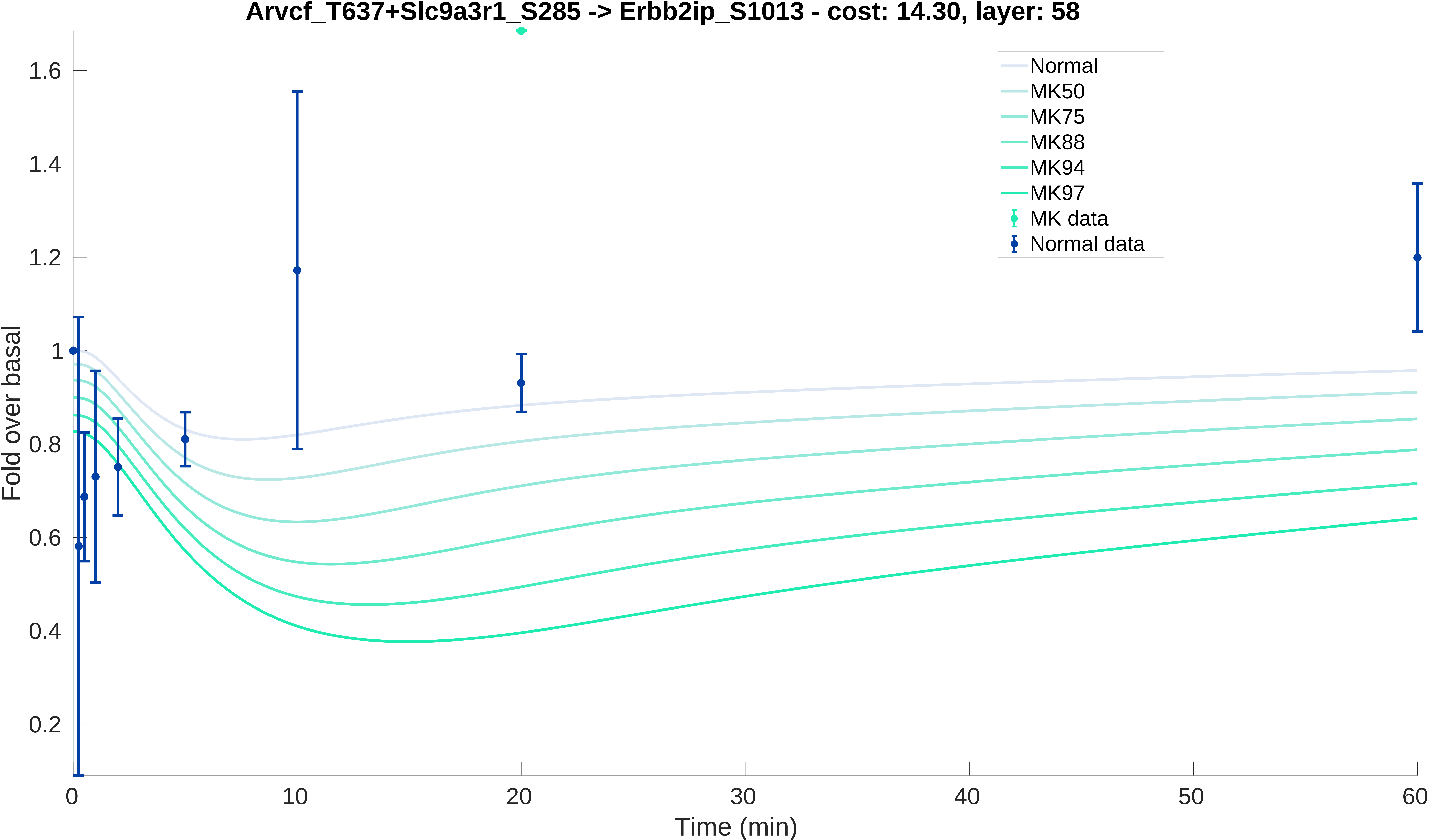

Arvcf_T637+Slc9a3r1_S285 -> Erbb2ip_S1013 - cost: 14.30, layer: 58
Normal
MK50
MK75
MK88
MK94
MK97
MK data
Normal data
Fold over basal
Time (min)
1.6
1.4
1.2
1
0.8
0.6
0.4
0.2
0
10
20
30
40
50
60

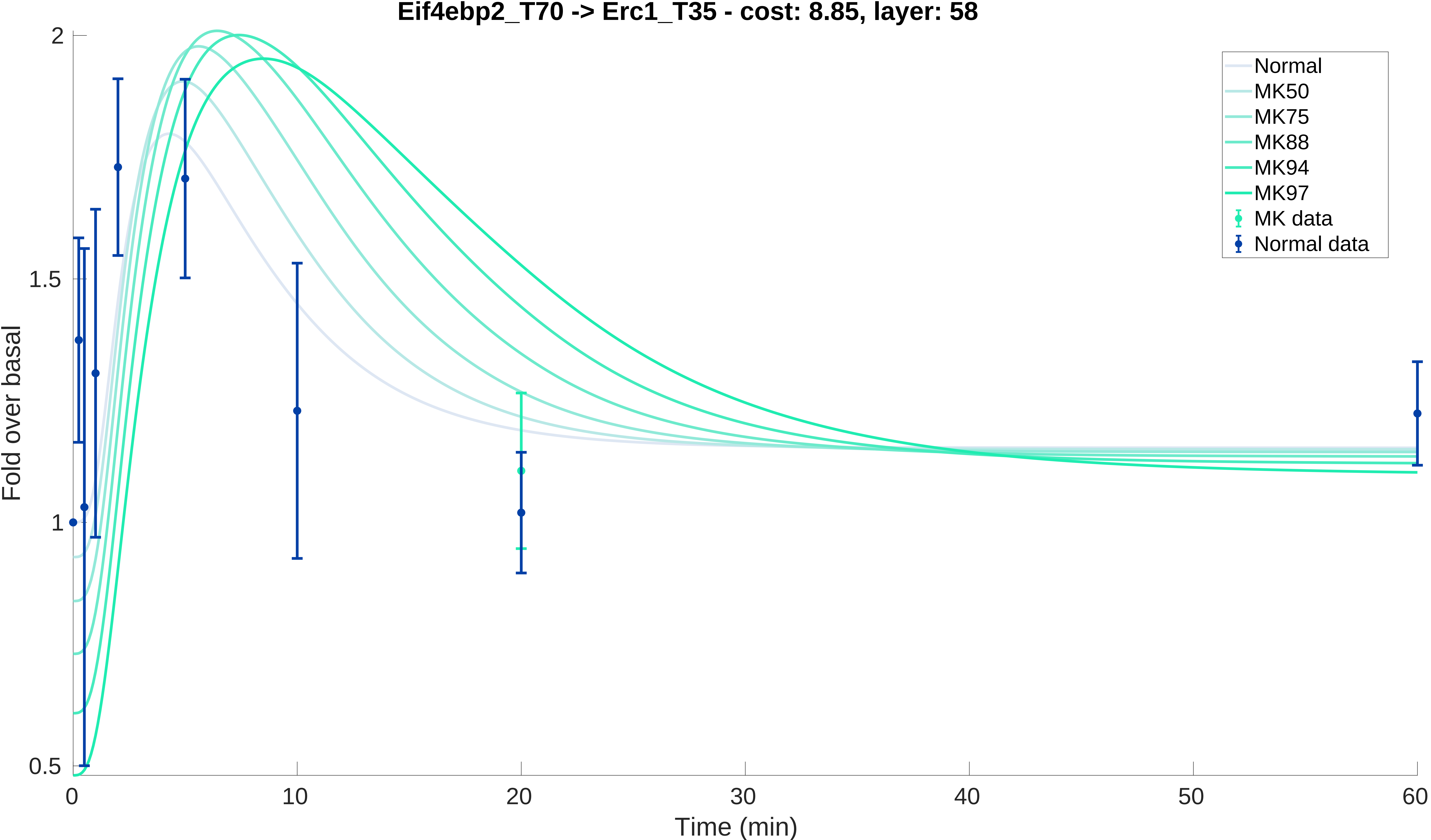
Eif4ebp2_T70 -> Erc1_T35 - cost: 8.85, layer: 58
Fold over basal
Time (min)
0
10
20
30
40
50
60
0.5
1
1.5
2
Normal
MK50
MK75
MK88
MK94
MK97
MK data
Normal data

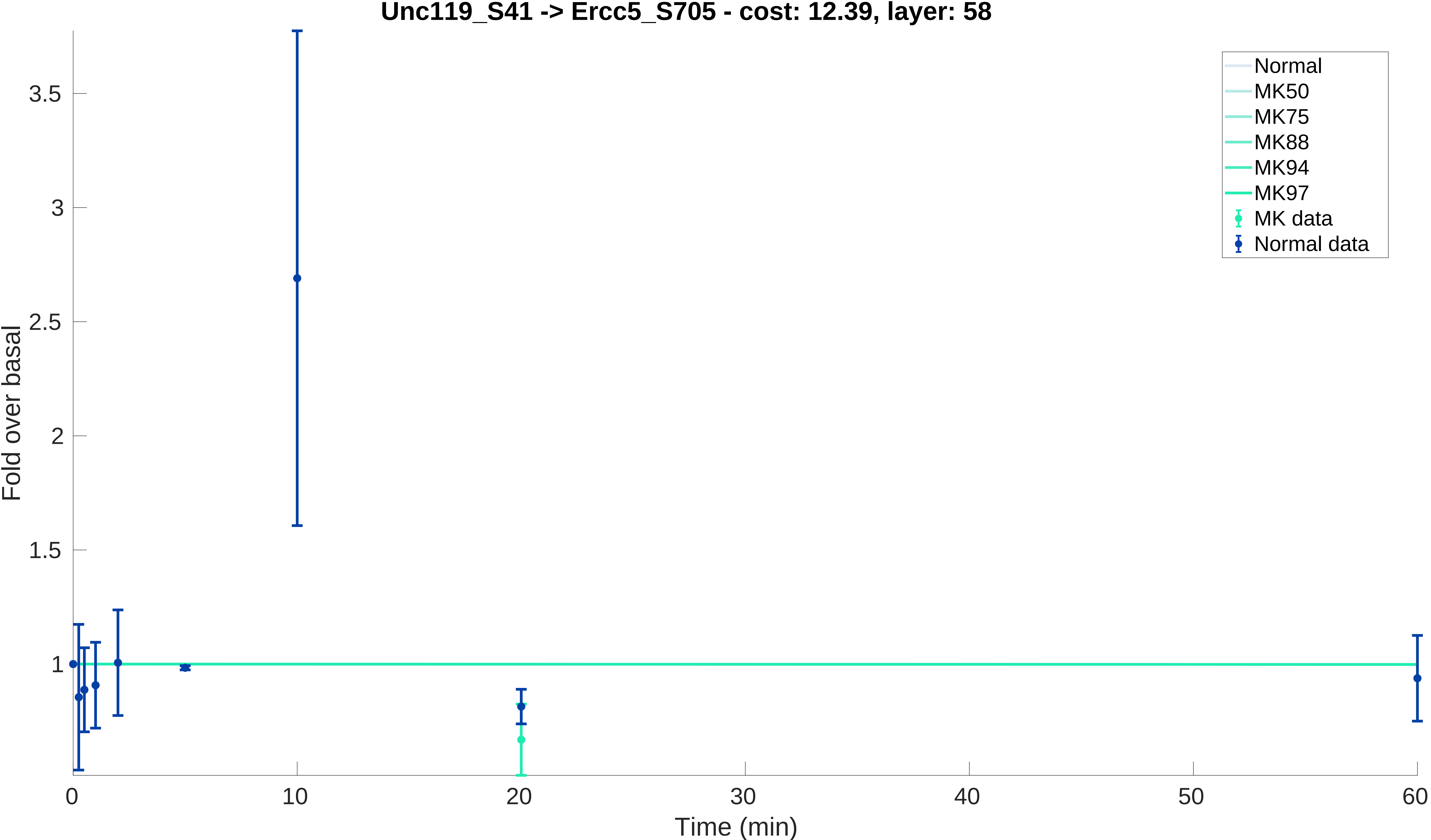
Unc119_S41 -> Ercc5_S705 - cost: 12.39, layer: 58
Fold over basal
Time (min)
3.5
3
2.5
2
1.5
1
0
10
20
30
40
50
60
Normal
MK50
MK75
MK88
MK94
MK97
MK data
Normal data

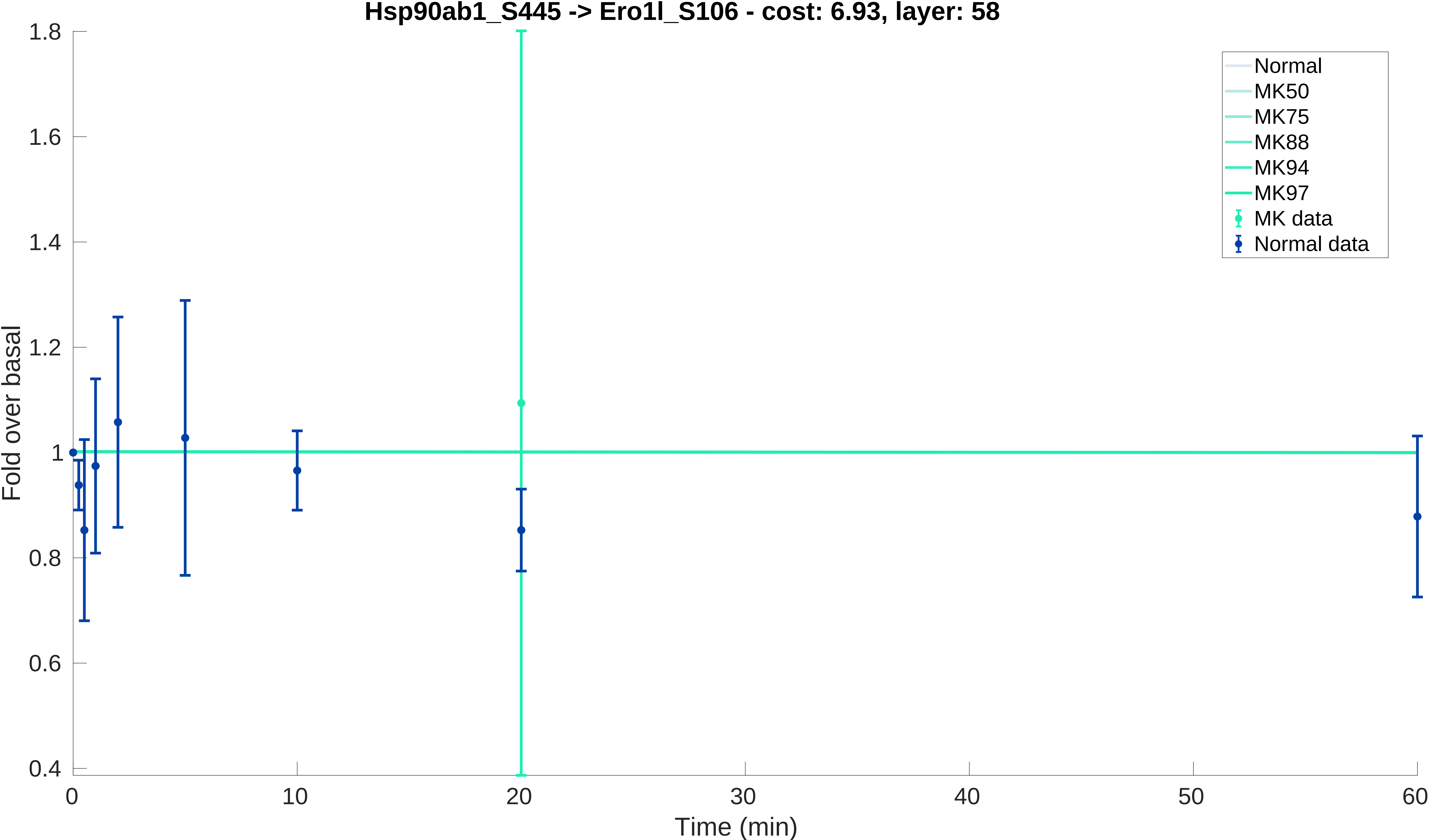

Hsp90ab1_S445 -> Ero1l_S106 - cost: 6.93, layer: 58
Fold over basal
Time (min)
1.8
1.6
1.4
1.2
1
0.8
0.6
0.4
0
10
20
30
40
50
60
Normal
MK50
MK75
MK88
MK94
MK97
MK data
Normal data

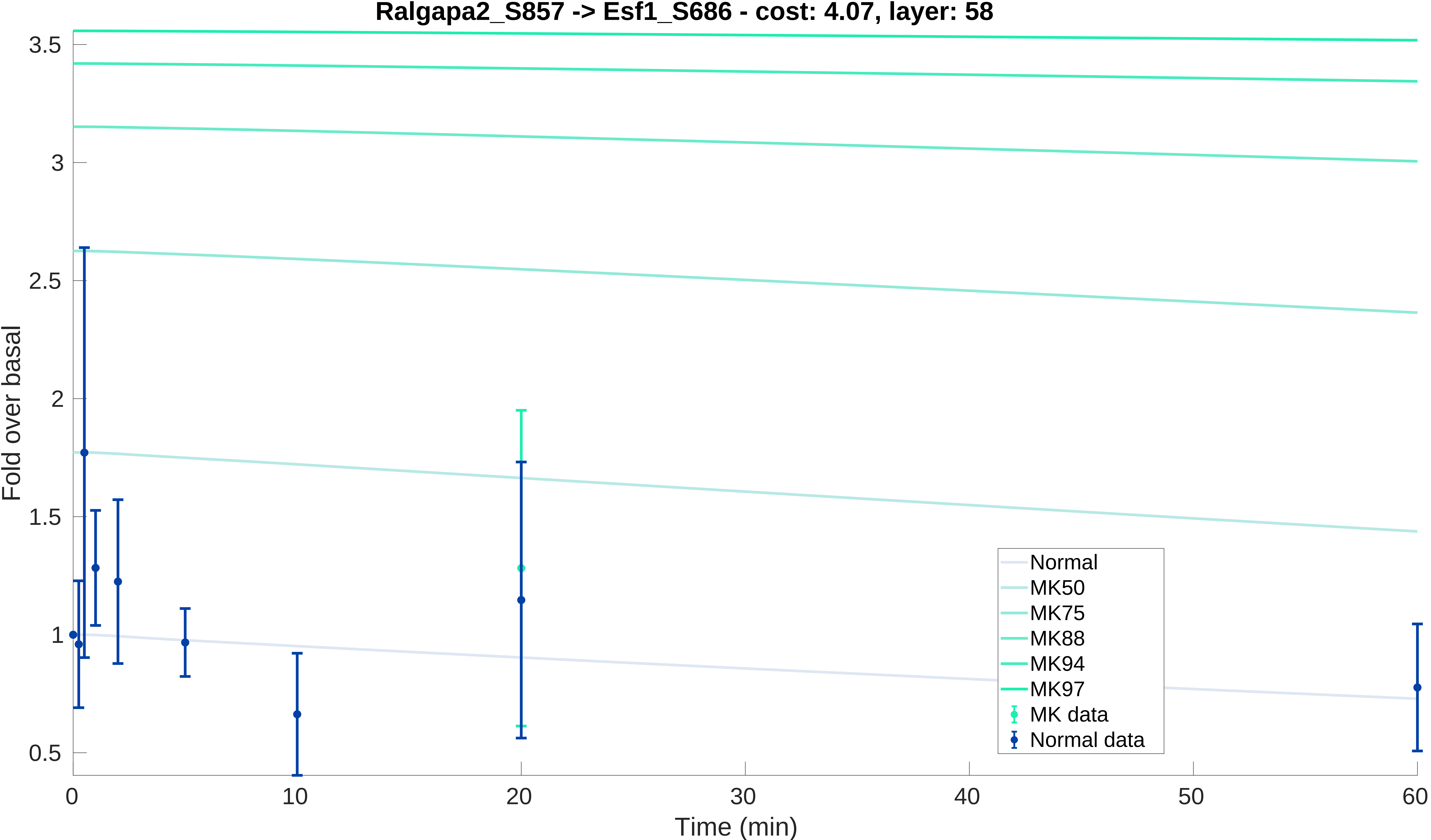
Ralgapa2_S857 -> Esf1_S686 - cost: 4.07, layer: 58
Fold over basal
Time (min)
0
10
20
30
40
50
60
0.5
1
1.5
2
2.5
3
3.5
Normal
MK50
MK75
MK88
MK94
MK97
MK data
Normal data

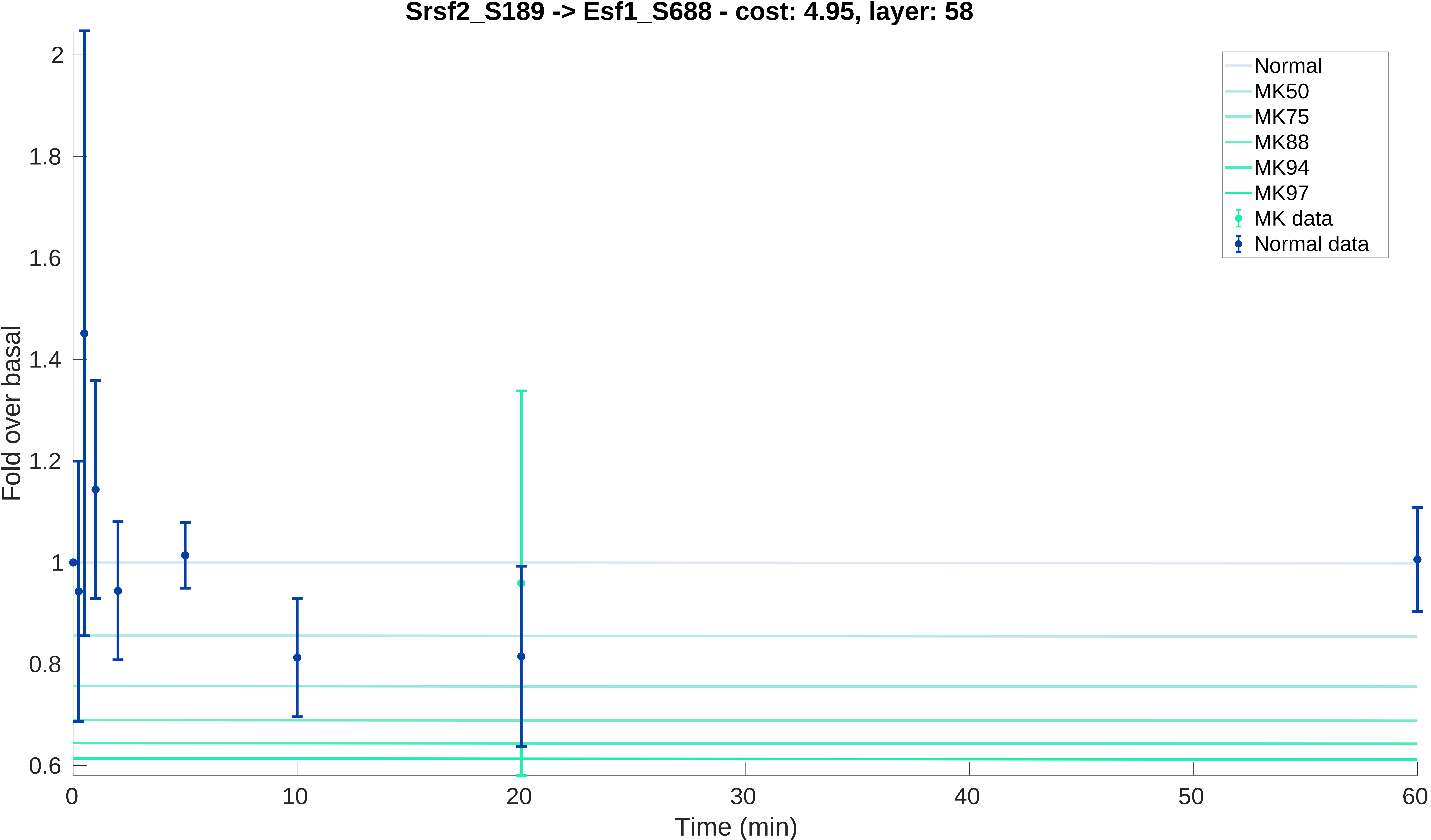

Srsf2_S189 -> Esf1_S688 - cost: 4.95, layer: 58
Fold over basal
Time (min)
0.6
0.8
1
1.2
1.4
1.6
1.8
2
0
10
20
30
40
50
60
Normal
MK50
MK75
MK88
MK94
MK97
MK data
Normal data

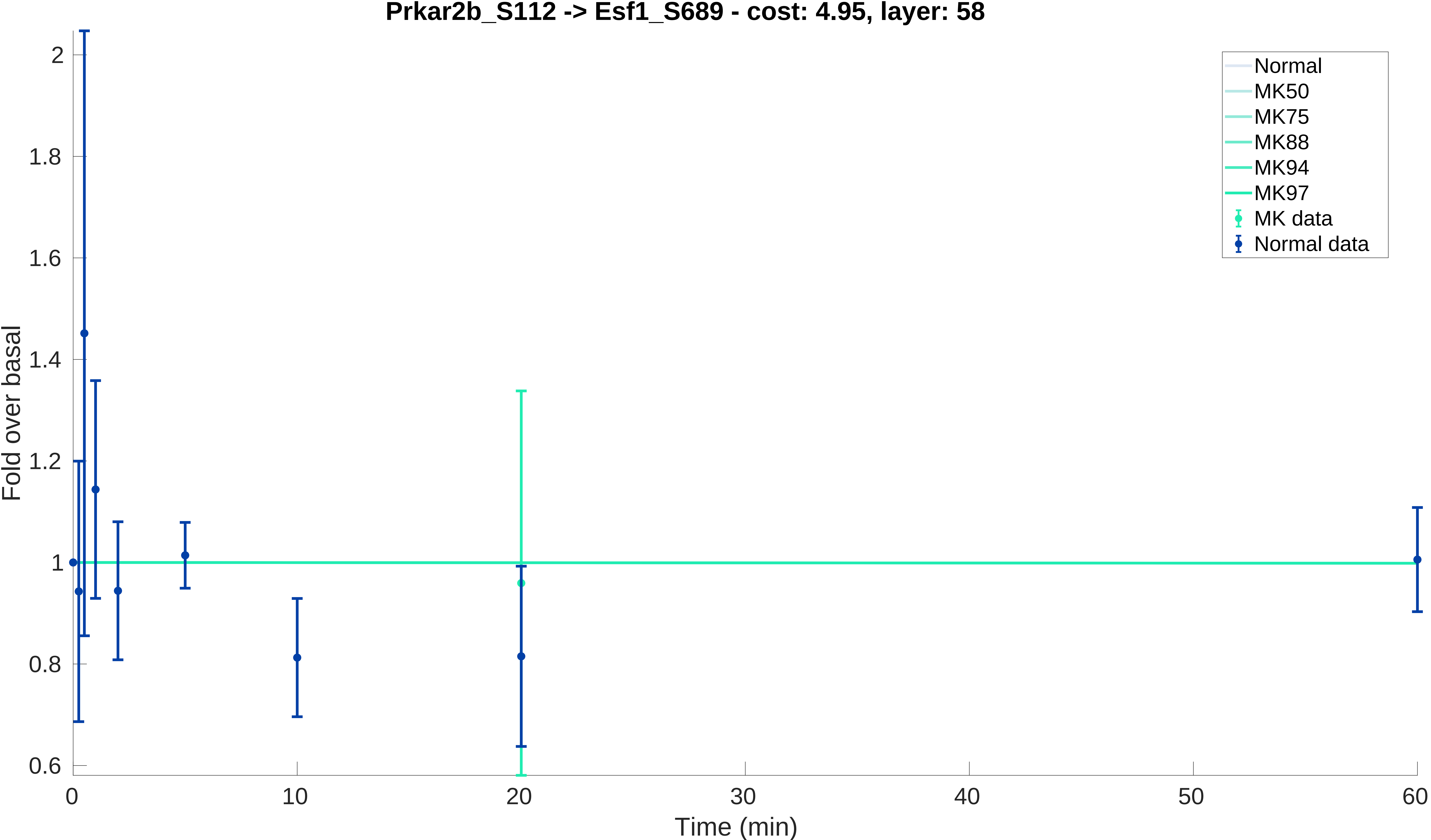

Prkar2b_S112 -> Esf1_S689 - cost: 4.95, layer: 58
Fold over basal
Time (min)
0.6
0.8
1
1.2
1.4
1.6
1.8
2
0
10
20
30
40
50
60
Normal
MK50
MK75
MK88
MK94
MK97
MK data
Normal data

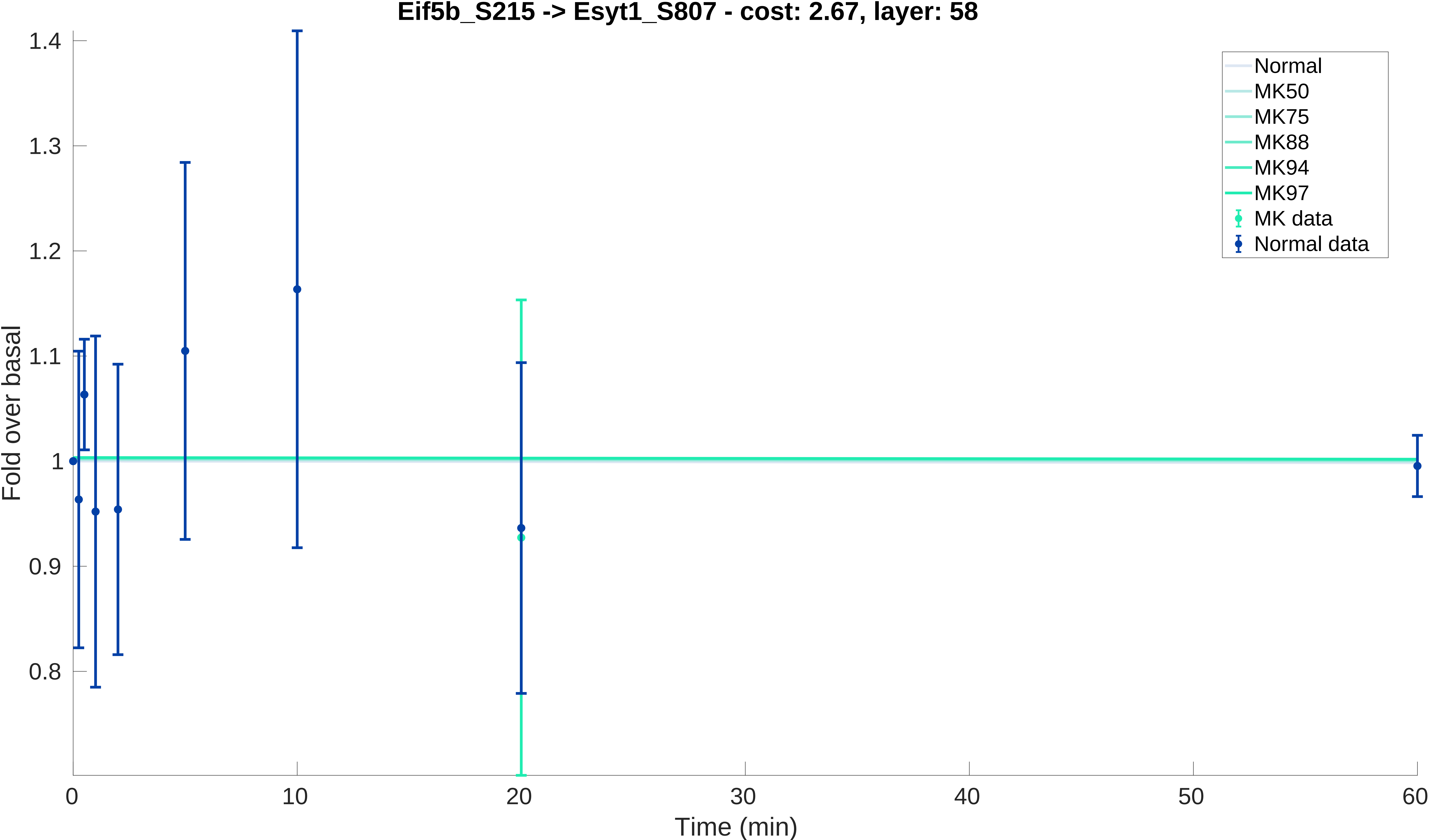

Eif5b_S215 -> Esyt1_S807 - cost: 2.67, layer: 58
Fold over basal
1.4
1.3
1.2
1.1
1
0.9
0.8
Time (min)
0
10
20
30
40
50
60
Normal
MK50
MK75
MK88
MK94
MK97
MK data
Normal data

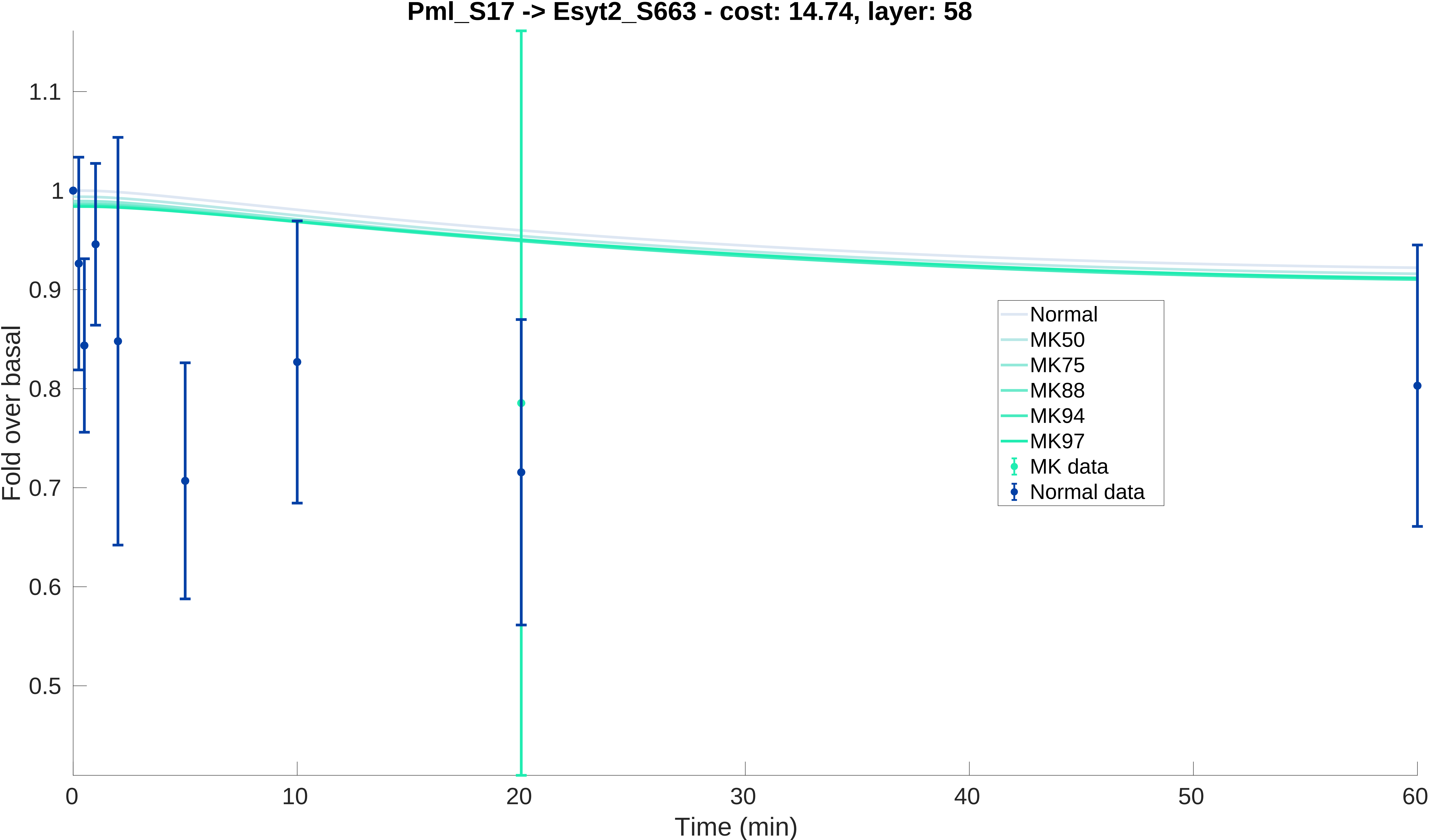

Pml_S17 -> Esyt2_S663 - cost: 14.74, layer: 58
Fold over basal
Time (min)
0.5
0.6
0.7
0.8
0.9
1
1.1
0
10
20
30
40
50
60
Normal
MK50
MK75
MK88
MK94
MK97
MK data
Normal data

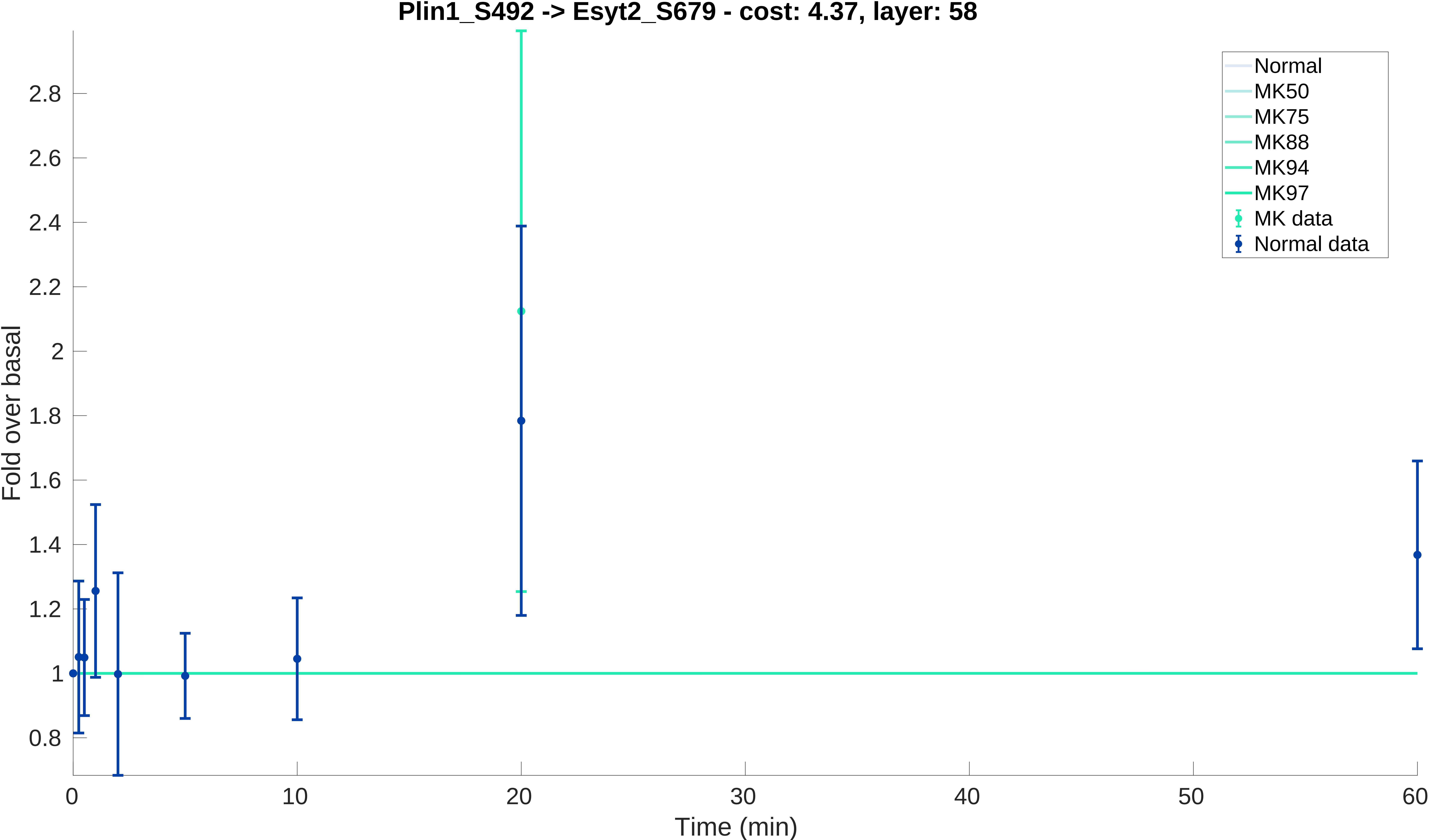

Plin1_S492 -> Esyt2_S679 - cost: 4.37, layer: 58
Normal
MK50
MK75
MK88
MK94
MK97
MK data
Normal data
Fold over basal
2.8
2.6
2.4
2.2
2
1.8
1.6
1.4
1.2
1
0.8
0
10
20
30
40
50
60
Time (min)

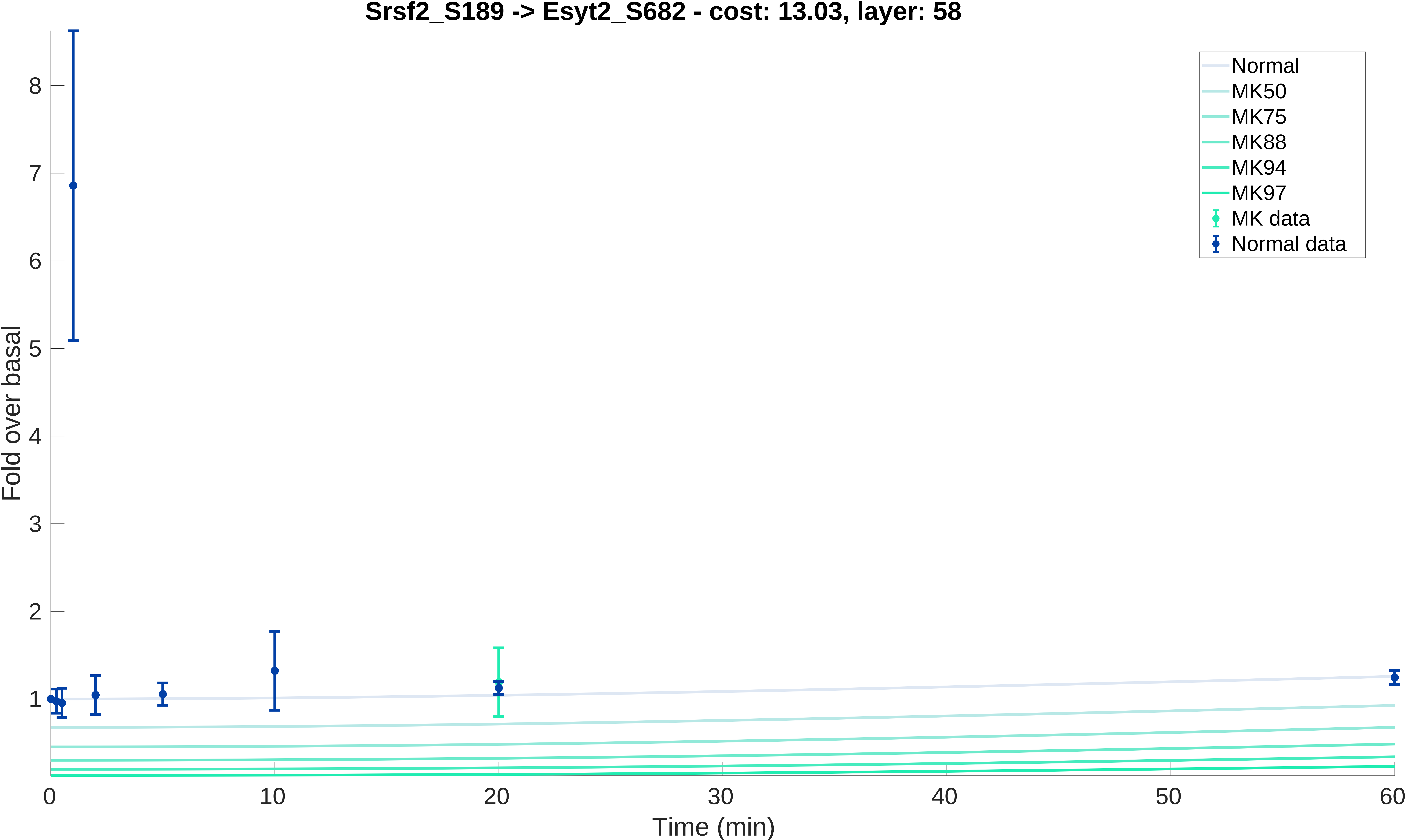
Srsf2_S189 -> Esyt2_S682 - cost: 13.03, layer: 58
Fold over basal
Time (min)
0
10
20
30
40
50
60
1
2
3
4
5
6
7
8
Normal
MK50
MK75
MK88
MK94
MK97
MK data
Normal data

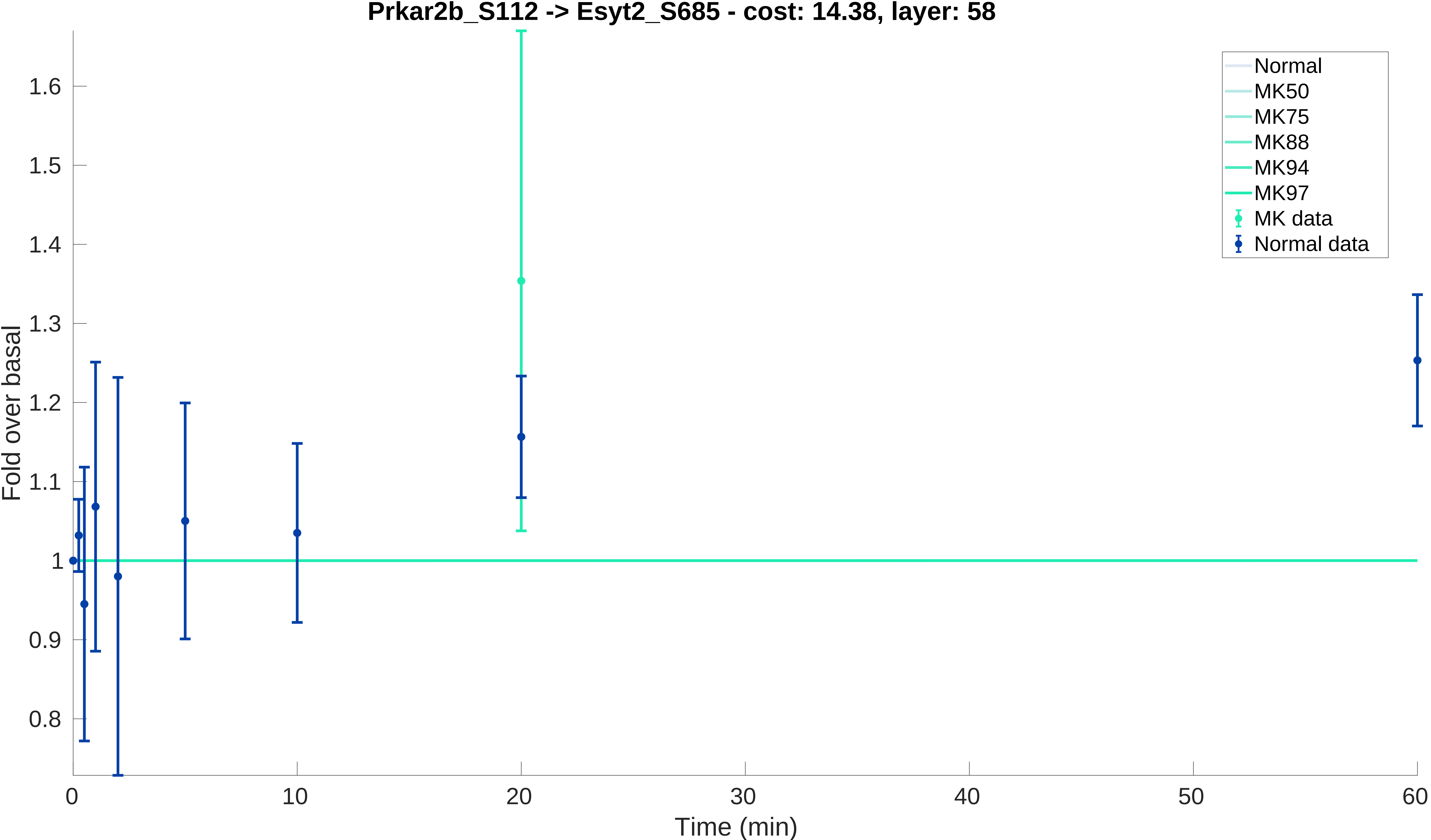

Prkar2b_S112 -> Esyt2_S685 - cost: 14.38, layer: 58
Fold over basal
Time (min)
1.6
1.5
1.4
1.3
1.2
1.1
1
0.9
0.8
0
10
20
30
40
50
60
Normal
MK50
MK75
MK88
MK94
MK97
MK data
Normal data

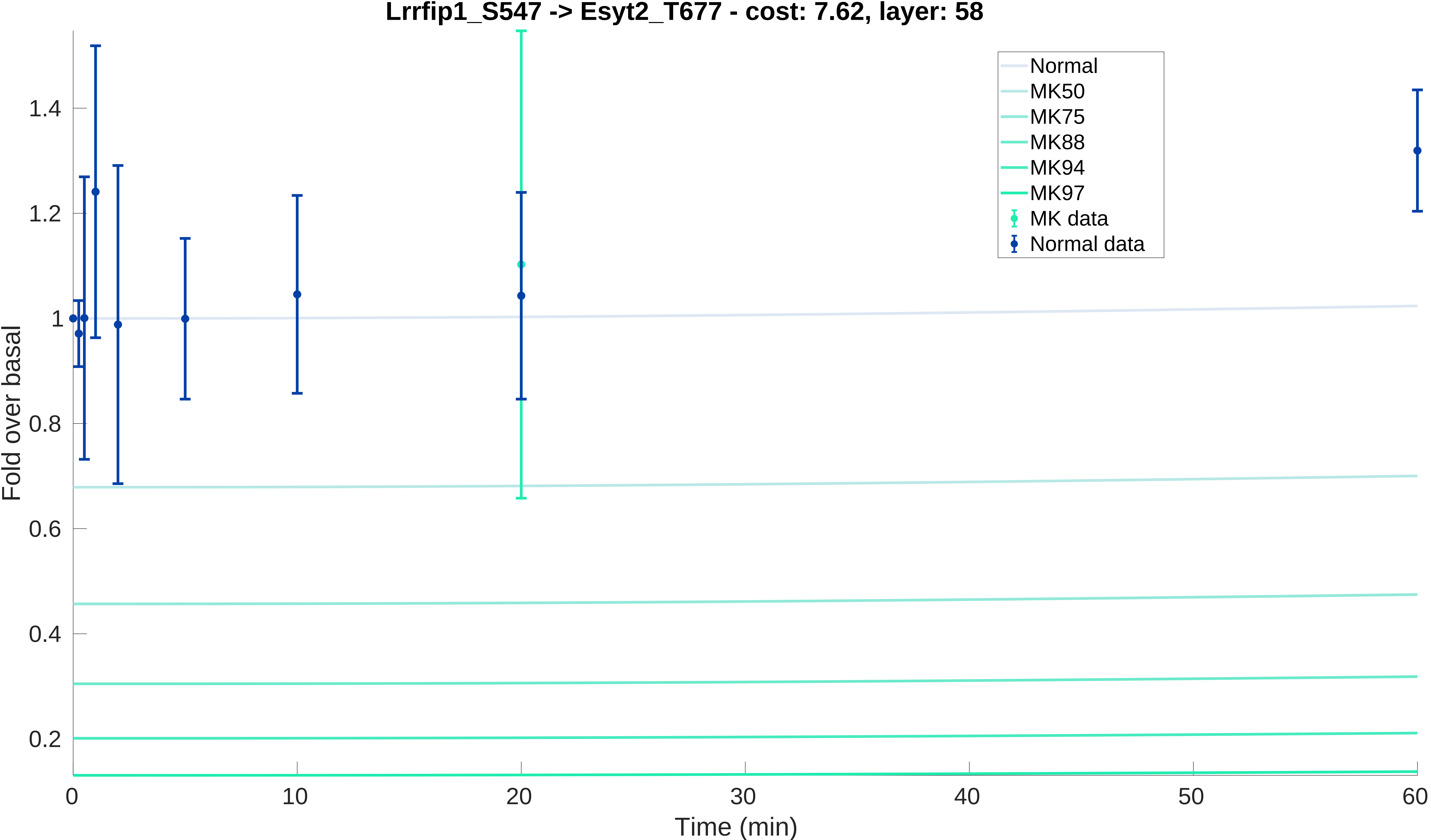

Lrrfip1_S547 -> Esyt2_T677 - cost: 7.62, layer: 58
Fold over basal
Time (min)
0.2
0.4
0.6
0.8
1
1.2
1.4
0
10
20
30
40
50
60
Normal
MK50
MK75
MK88
MK94
MK97
MK data
Normal data

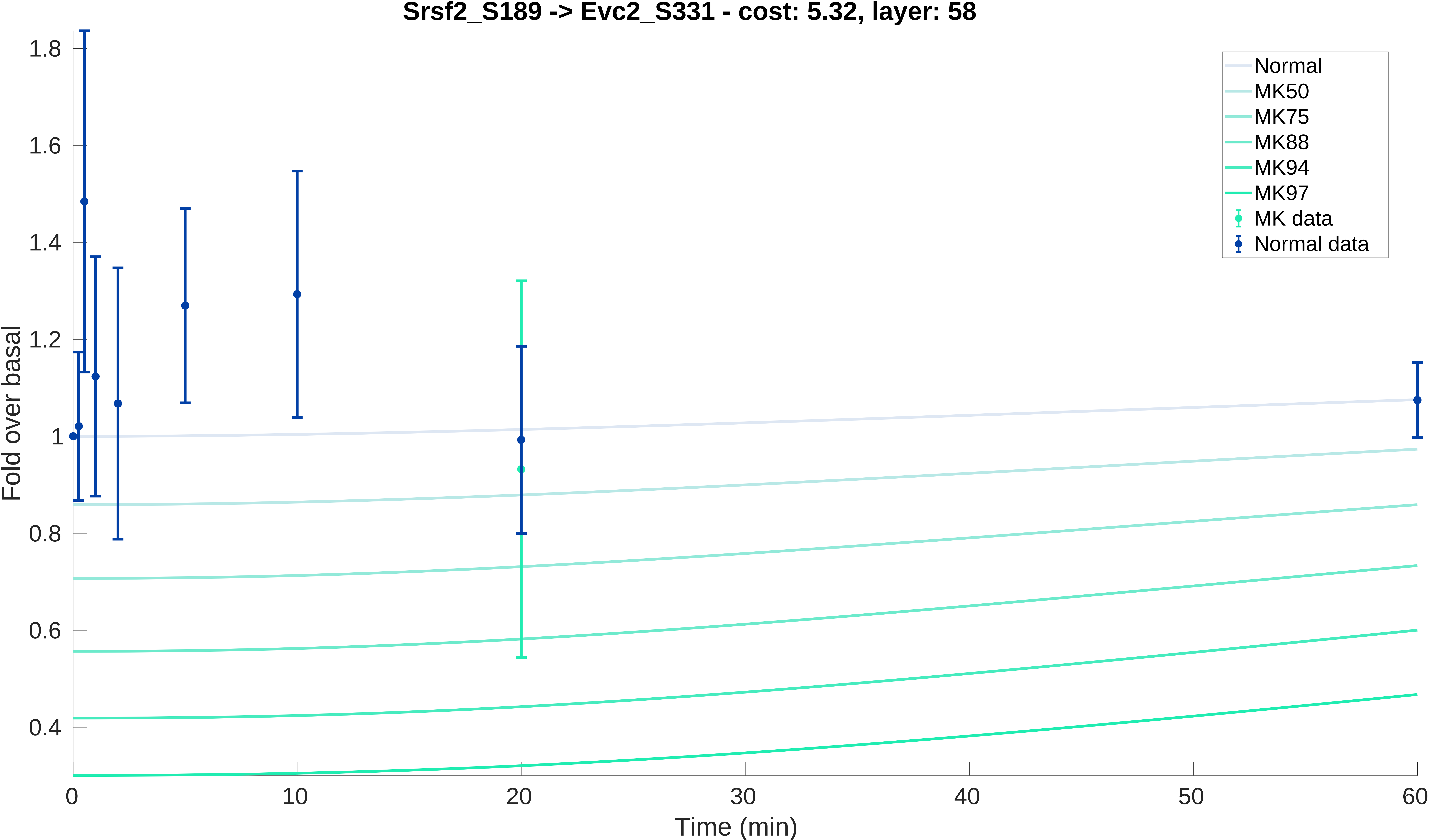
Srsf2_S189 -> Evc2_S331 - cost: 5.32, layer: 58
Fold over basal
Time (min)
0
10
20
30
40
50
60
0.4
0.6
0.8
1
1.2
1.4
1.6
1.8
Normal
MK50
MK75
MK88
MK94
MK97
MK data
Normal data

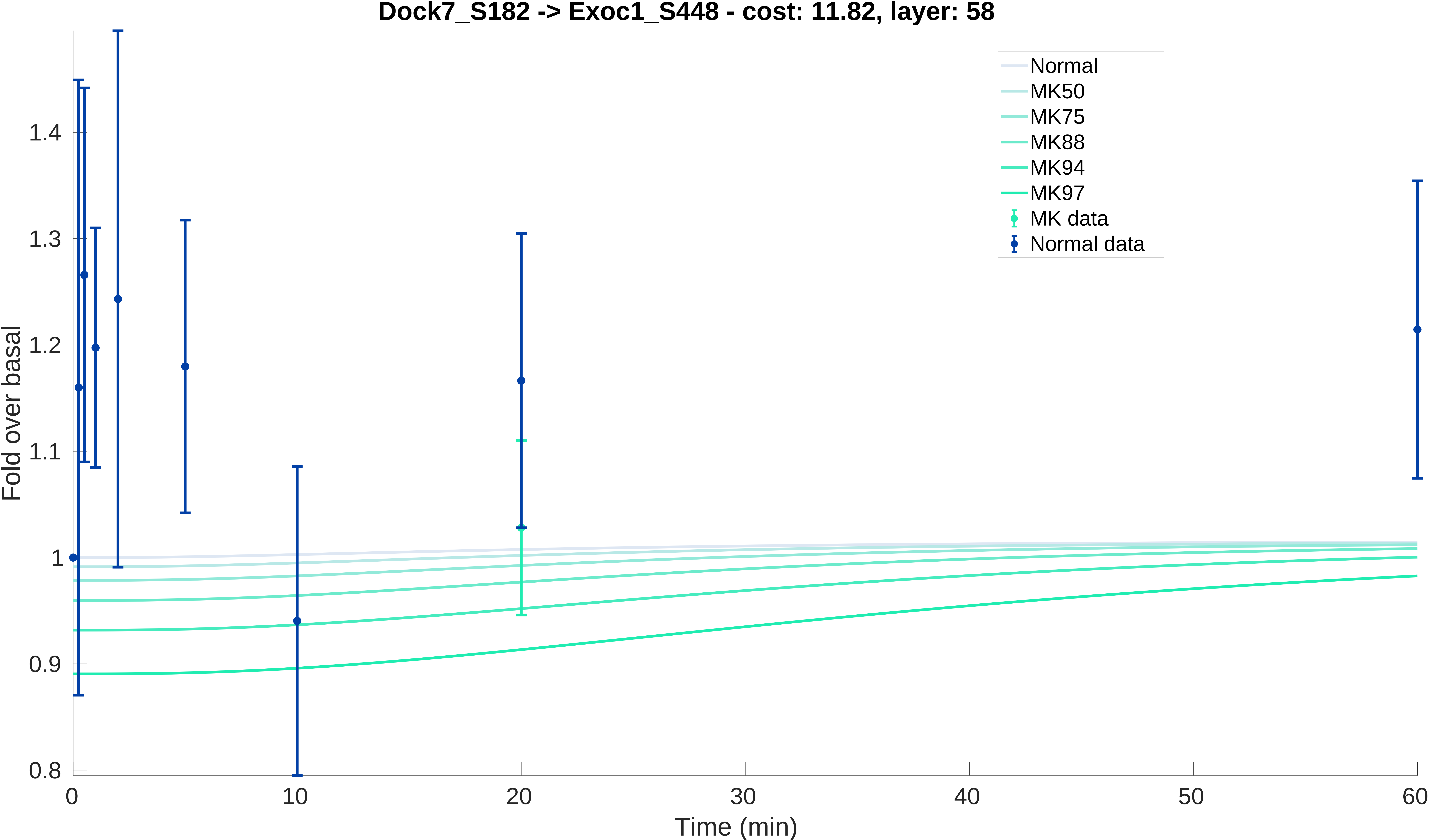

Dock7_S182 -> Exoc1_S448 - cost: 11.82, layer: 58
Normal
MK50
MK75
MK88
MK94
MK97
MK data
Normal data
Fold over basal
1.4
1.3
1.2
1.1
1
0.9
0.8
0
10
20
30
40
50
60
Time (min)

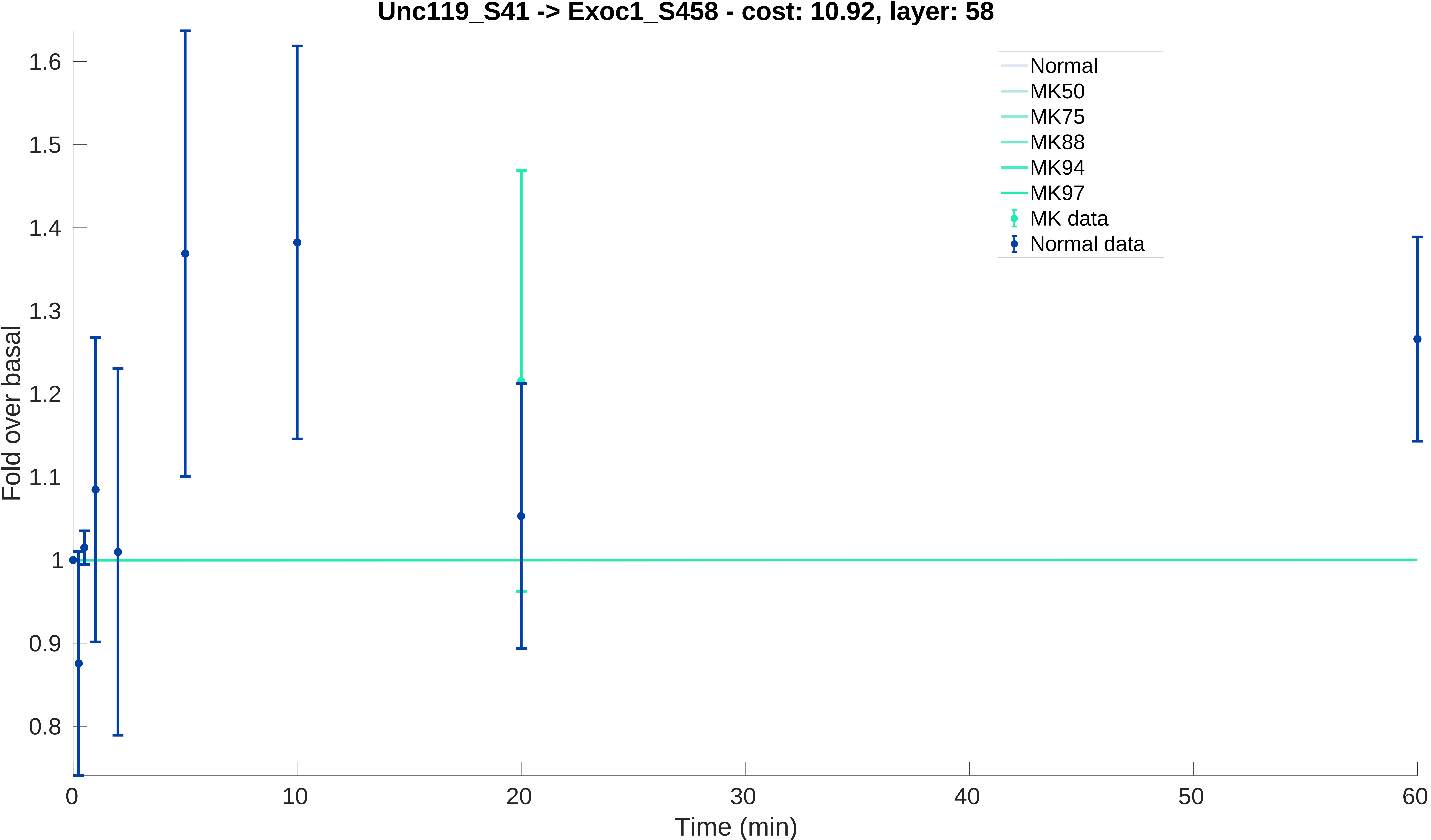

Unc119_S41 -> Exoc1_S458 - cost: 10.92, layer: 58
Fold over basal
1.6
1.5
1.4
1.3
1.2
1.1
1
0.9
0.8
0
10
20
30
40
50
60
Time (min)
Normal
MK50
MK75
MK88
MK94
MK97
MK data
Normal data

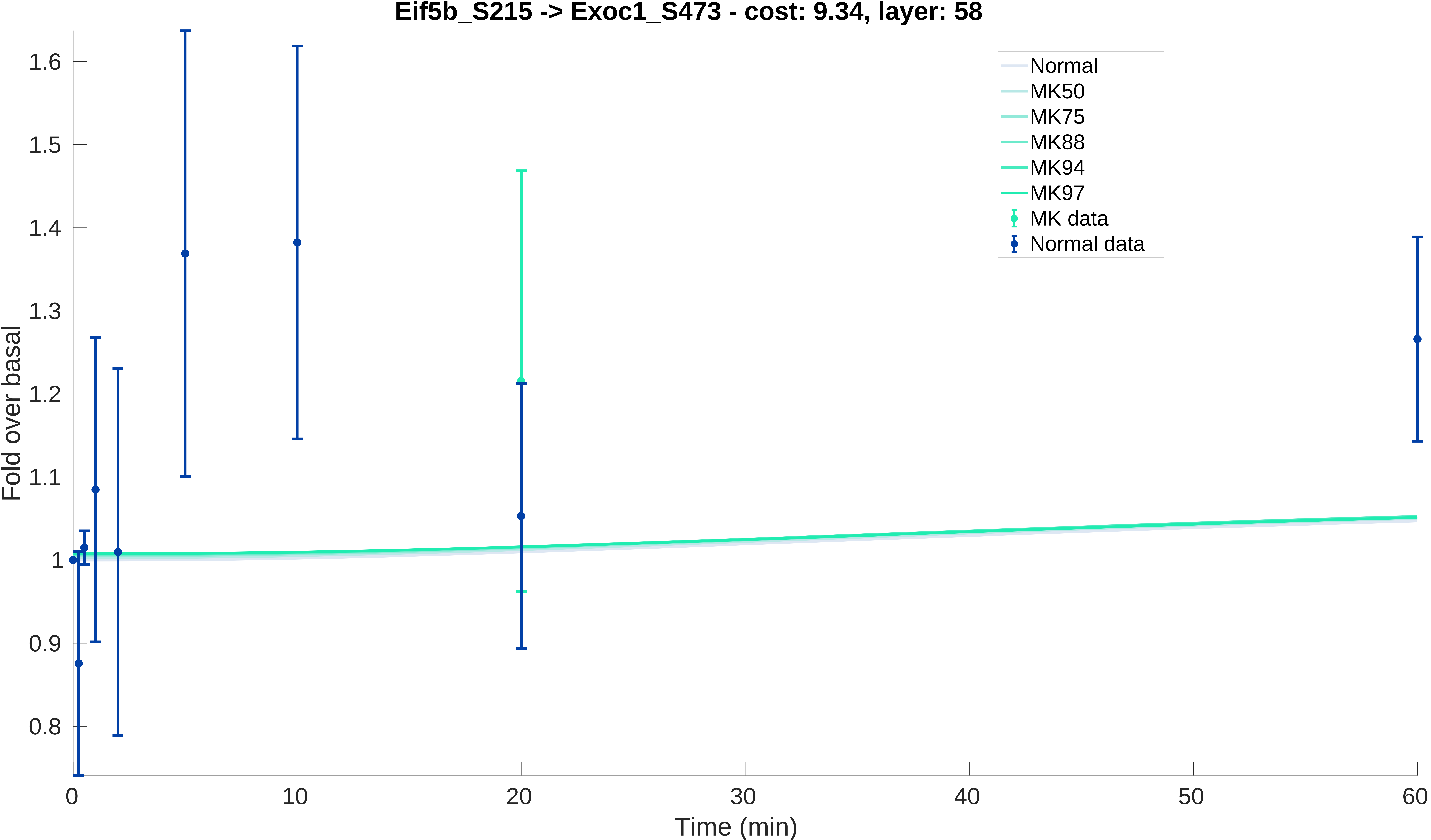

Eif5b_S215 -> Exoc1_S473 - cost: 9.34, layer: 58
Normal
MK50
MK75
MK88
MK94
MK97
MK data
Normal data
Fold over basal
Time (min)
1.6
1.5
1.4
1.3
1.2
1.1
1
0.9
0.8
0
10
20
30
40
50
60

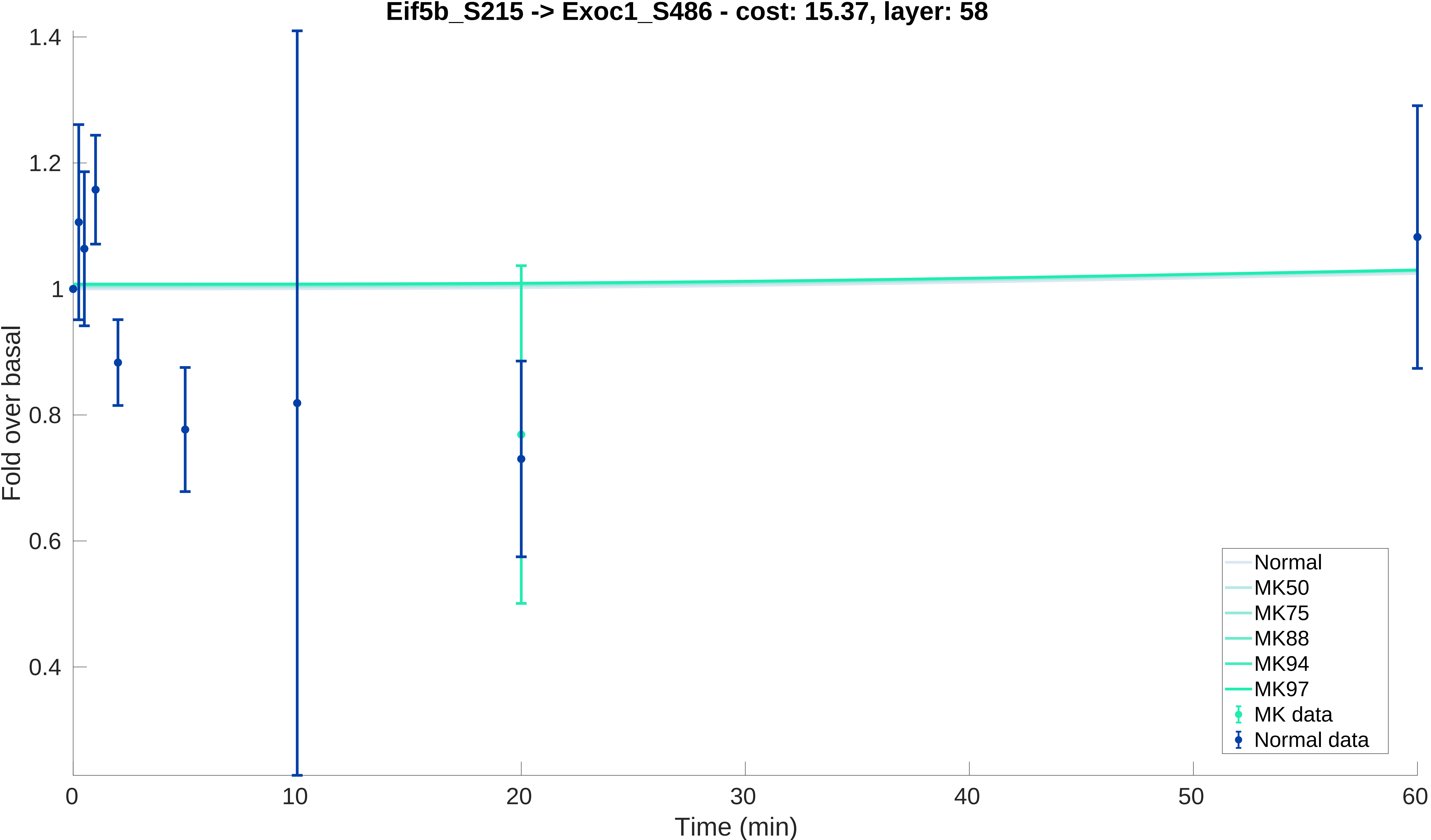
Eif5b_S215 -> Exoc1_S486 - cost: 15.37, layer: 58
Fold over basal
1.4
1.2
1
0.8
0.6
0.4
0
10
20
30
40
50
60
Time (min)
Normal
MK50
MK75
MK88
MK94
MK97
MK data
Normal data

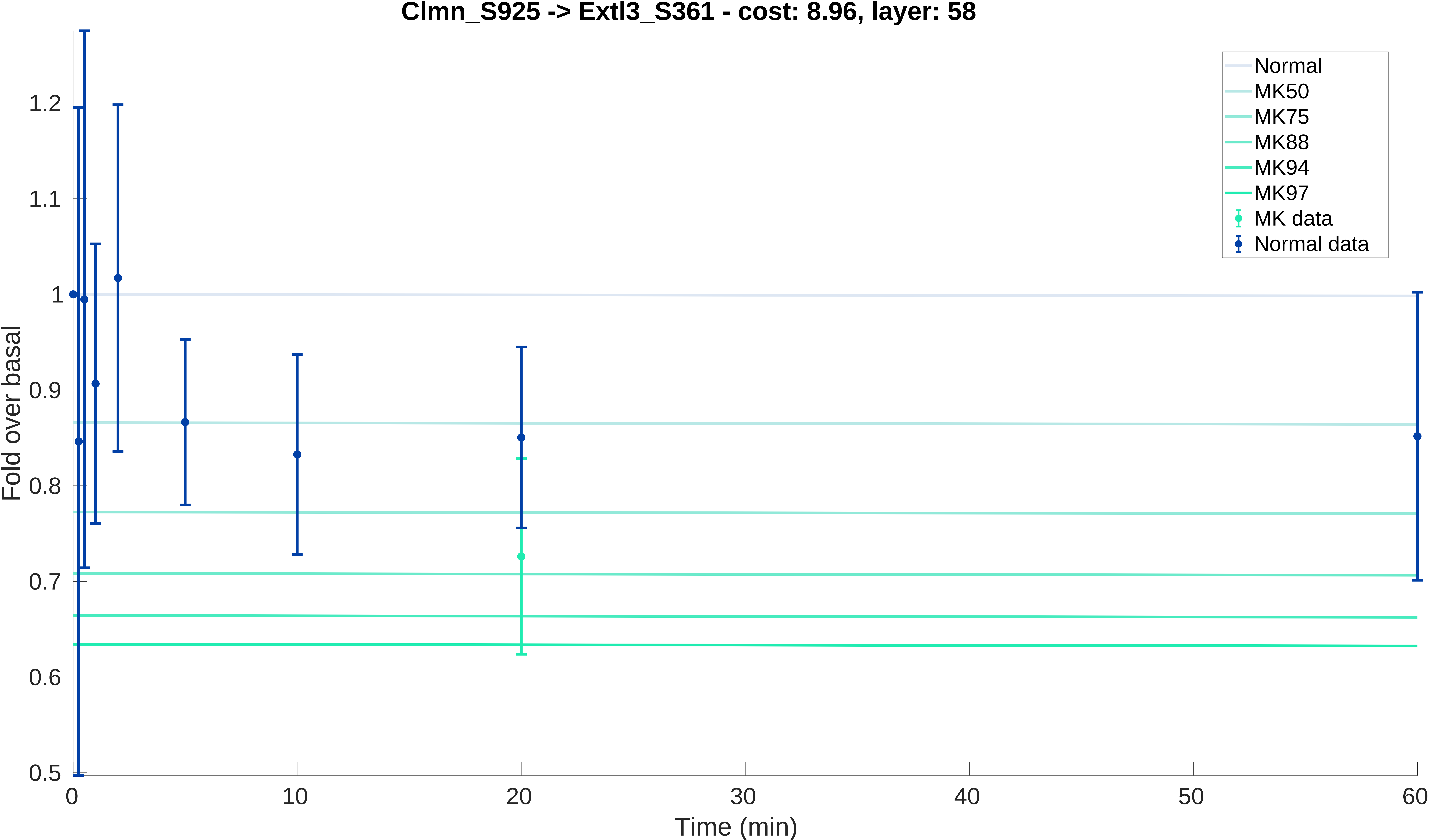

Clmn_S925 -> Extl3_S361 - cost: 8.96, layer: 58
Normal
MK50
MK75
MK88
MK94
MK97
MK data
Normal data
Fold over basal
Time (min)
0.5
0.6
0.7
0.8
0.9
1
1.1
1.2
0
10
20
30
40
50
60

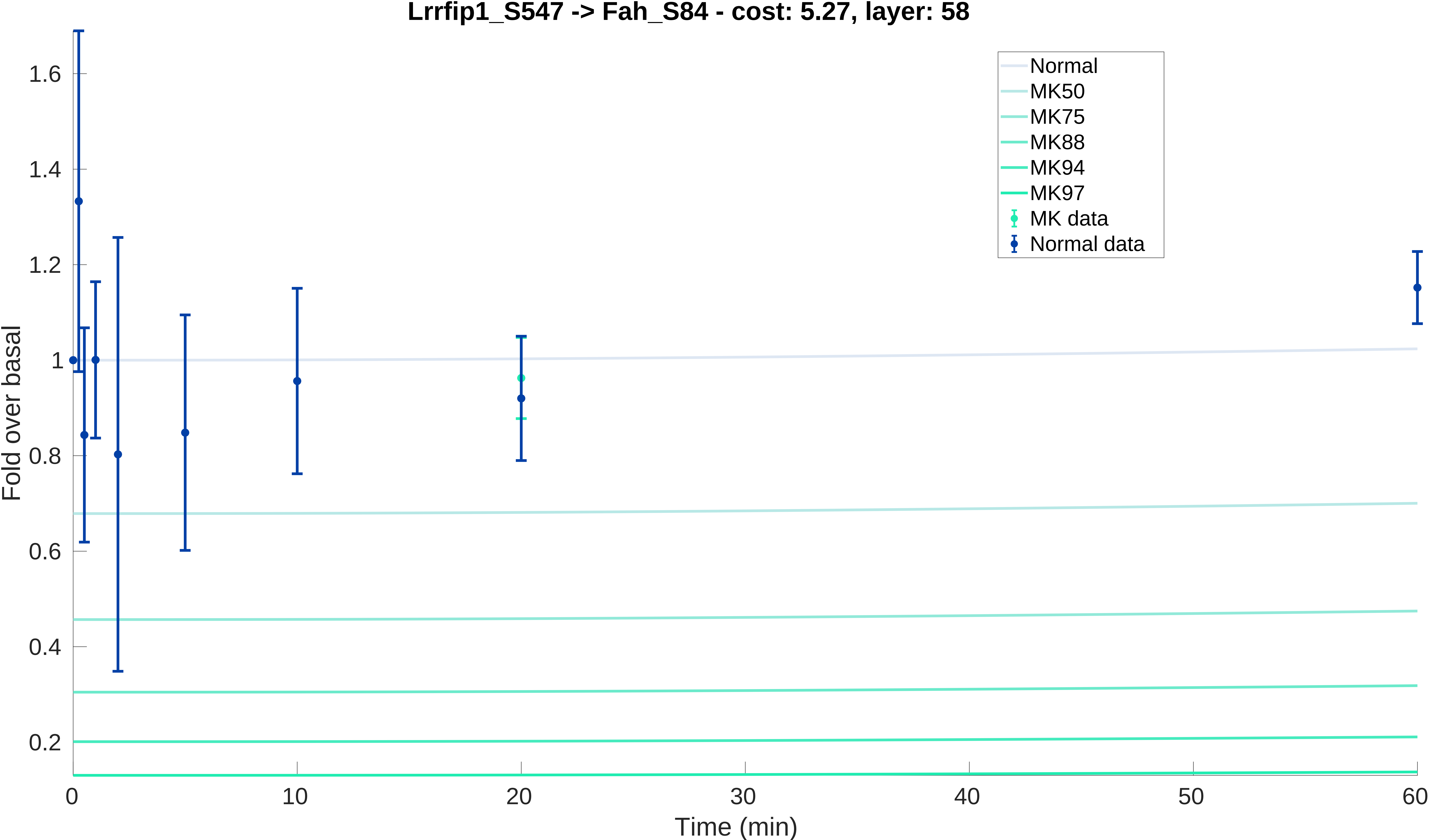

Lrrfip1_S547 -> Fah_S84 - cost: 5.27, layer: 58
Normal
MK50
MK75
MK88
MK94
MK97
MK data
Normal data
Fold over basal
Time (min)
0
10
20
30
40
50
60
0.2
0.4
0.6
0.8
1
1.2
1.4
1.6

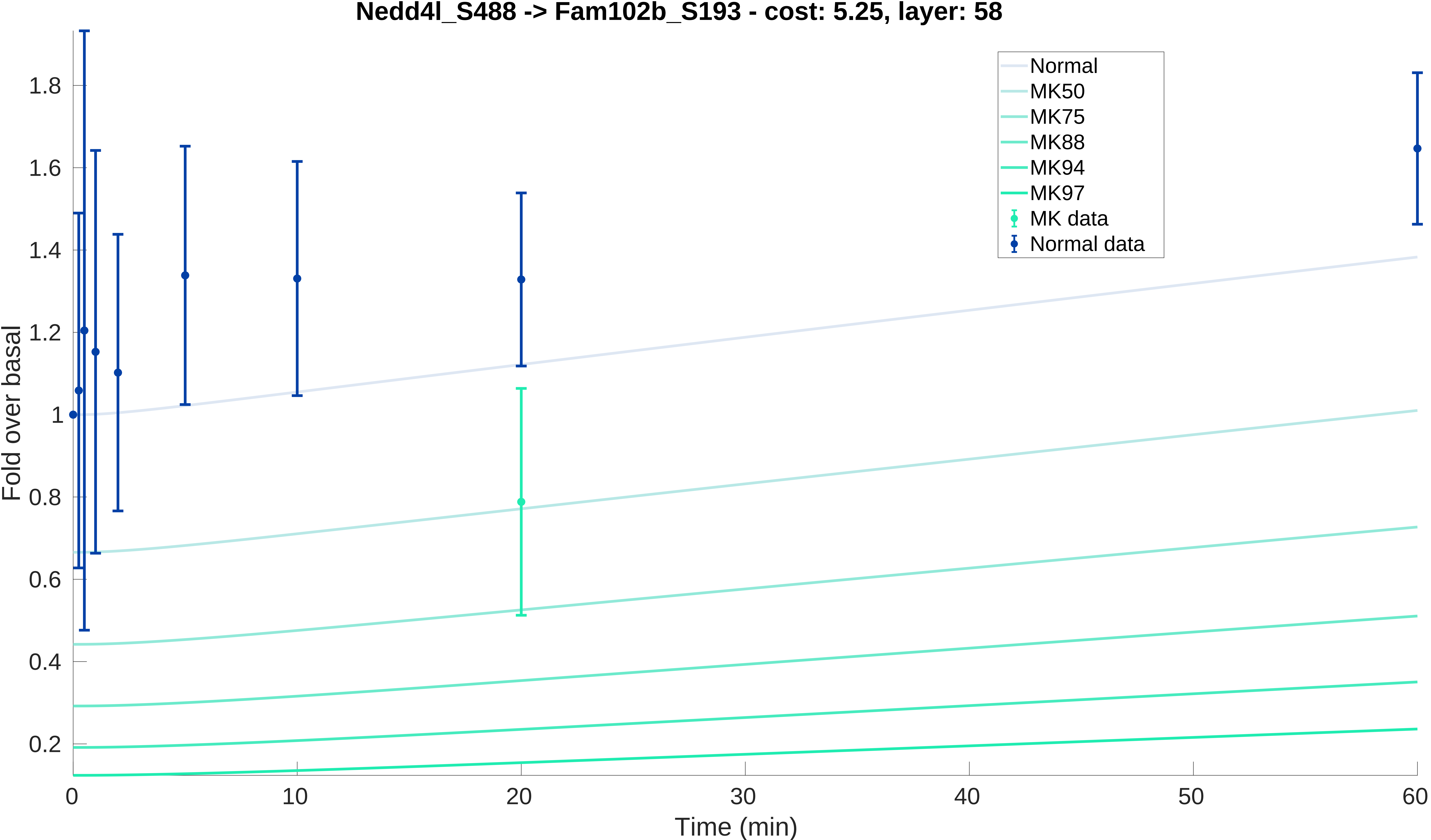

Nedd4l_S488 -> Fam102b_S193 - cost: 5.25, layer: 58
Fold over basal
1.8
1.6
1.4
1.2
1
0.8
0.6
0.4
0.2
0
10
20
30
40
50
60
Time (min)
Normal
MK50
MK75
MK88
MK94
MK97
MK data
Normal data

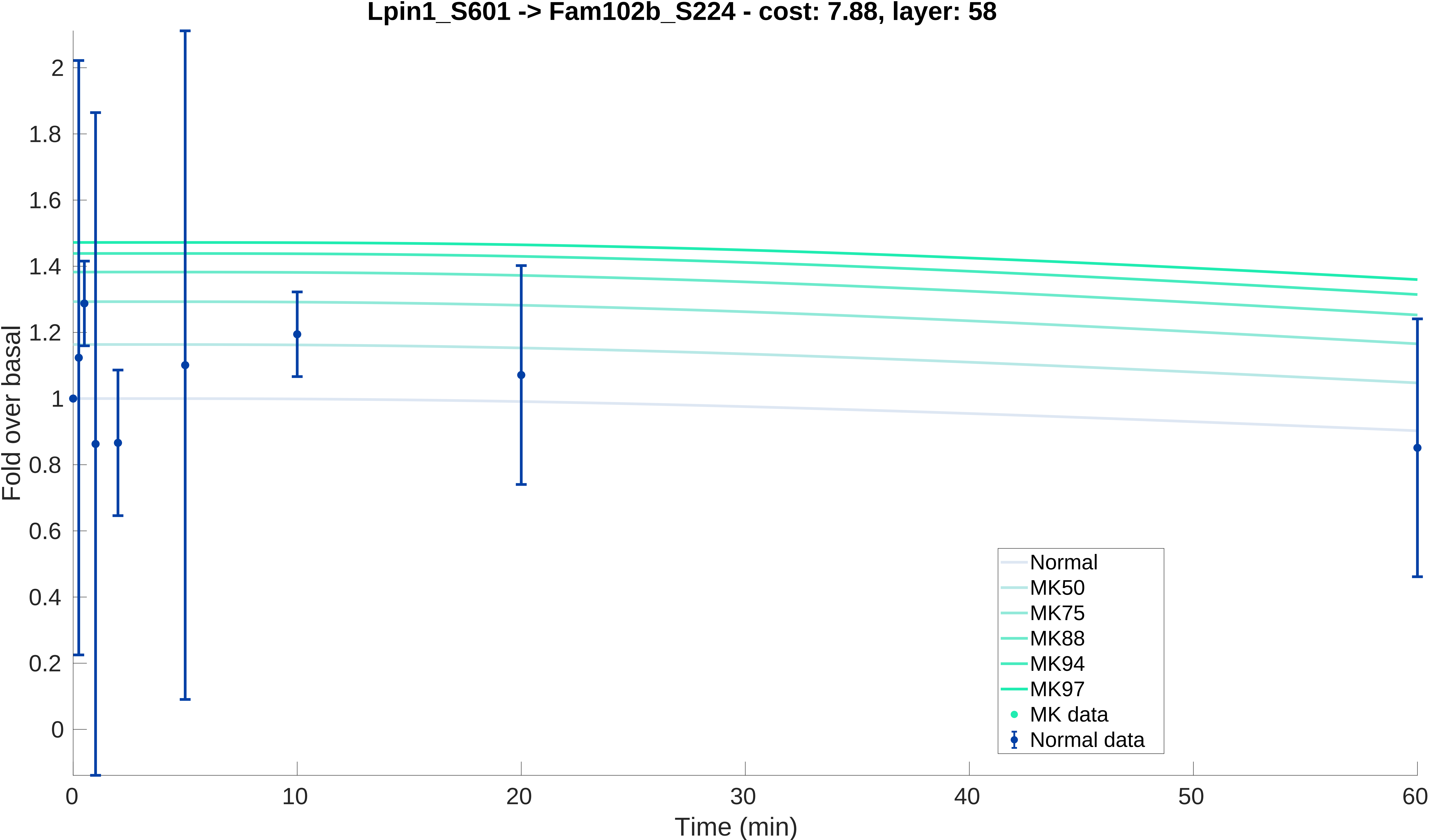

Lpin1_S601 -> Fam102b_S224 - cost: 7.88, layer: 58
Fold over basal
Time (min)
0
0.2
0.4
0.6
0.8
1
1.2
1.4
1.6
1.8
2
0
10
20
30
40
50
60
Normal
MK50
MK75
MK88
MK94
MK97
MK data
Normal data

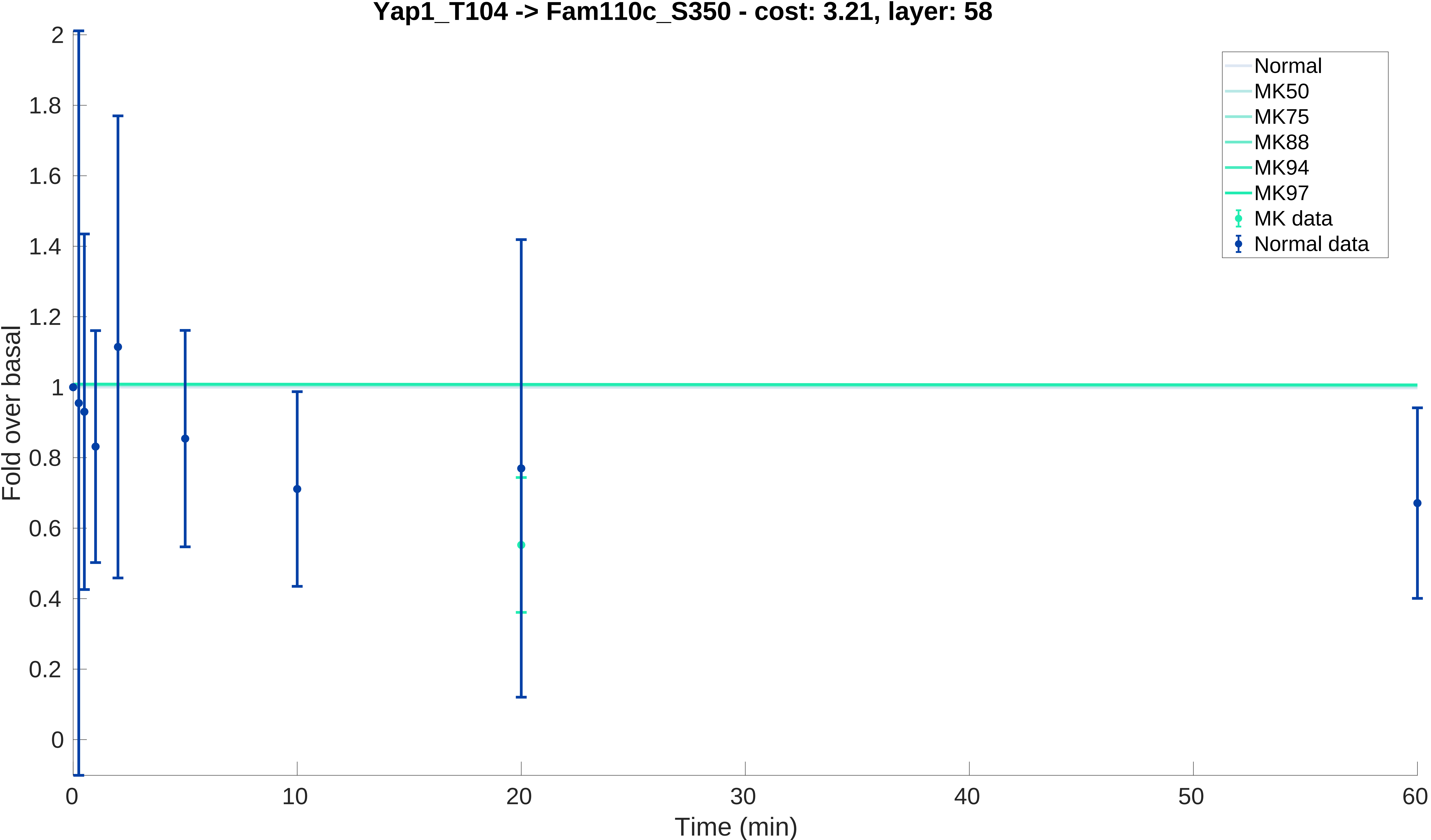
Yap1_T104 -> Fam110c_S350 - cost: 3.21, layer: 58
Fold over basal
Time (min)
Normal
MK50
MK75
MK88
MK94
MK97
MK data
Normal data
0
0.2
0.4
0.6
0.8
1
1.2
1.4
1.6
1.8
2
0
10
20
30
40
50
60

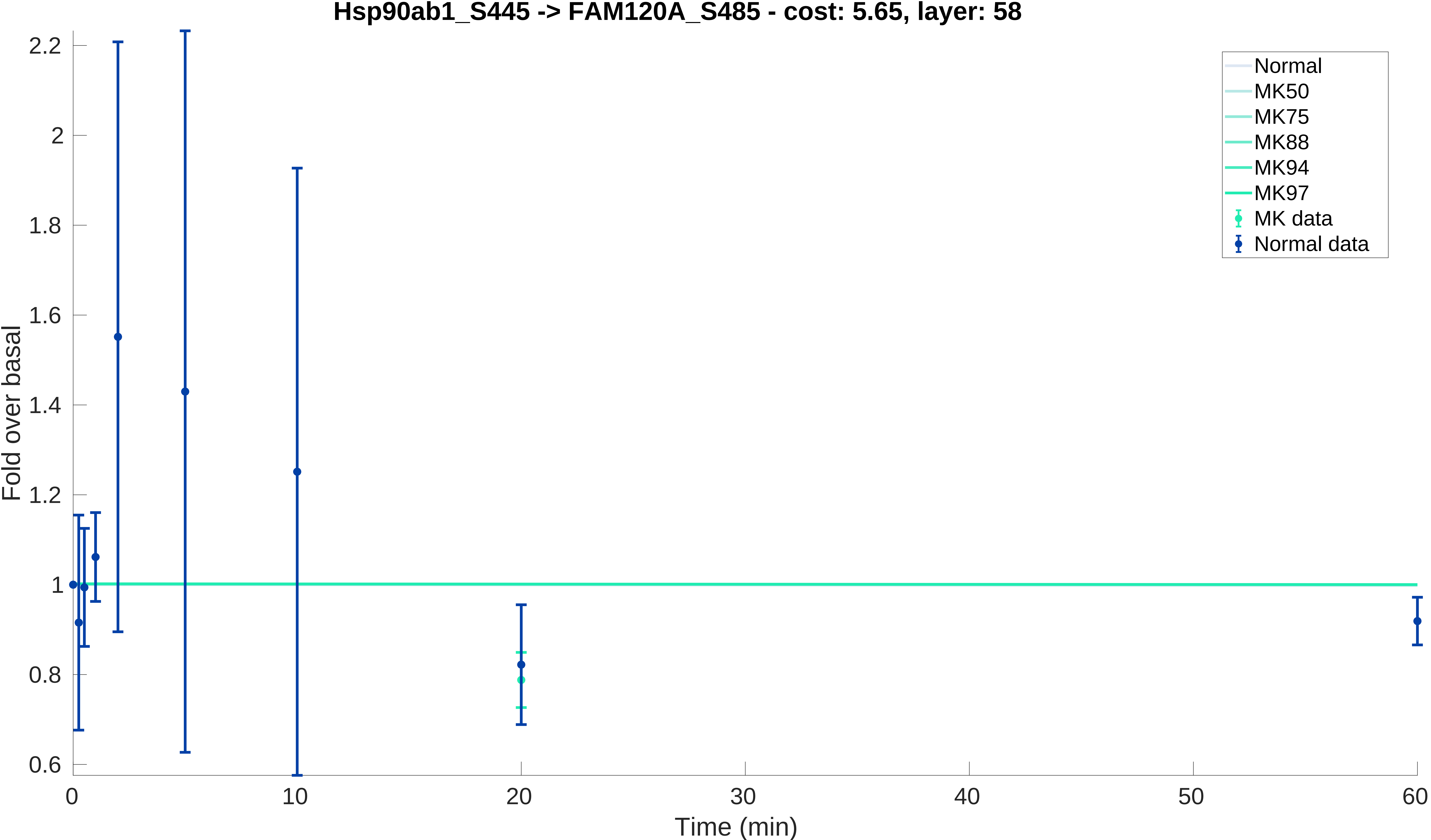
Hsp90ab1_S445 -> FAM120A_S485 - cost: 5.65, layer: 58
Fold over basal
Time (min)
2.2
2
1.8
1.6
1.4
1.2
1
0.8
0.6
0
10
20
30
40
50
60
Normal
MK50
MK75
MK88
MK94
MK97
MK data
Normal data

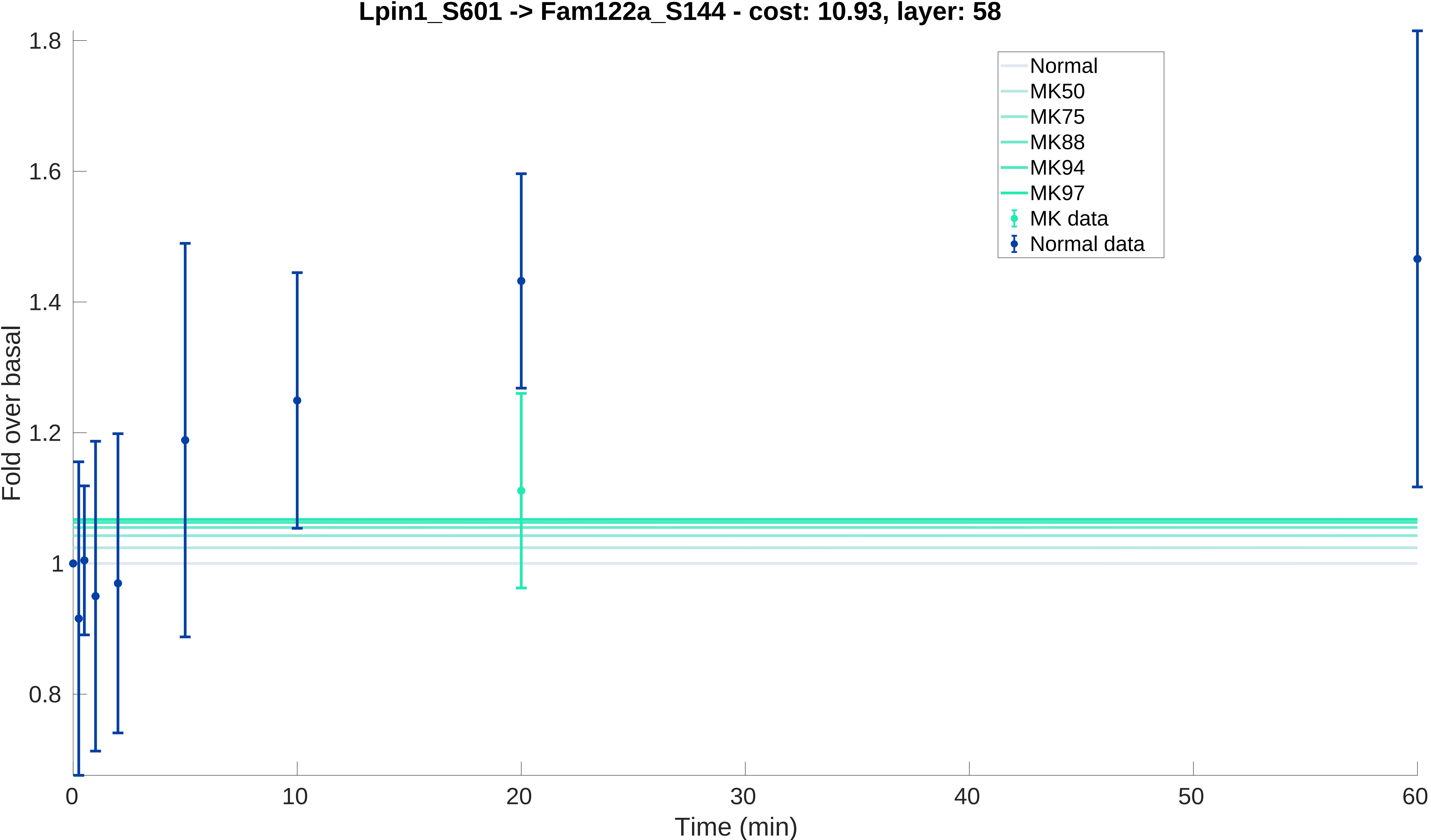

Lpin1_S601 -> Fam122a_S144 - cost: 10.93, layer: 58
Fold over basal
1.8
1.6
1.4
1.2
1
0.8
Time (min)
0
10
20
30
40
50
60
Normal
MK50
MK75
MK88
MK94
MK97
MK data
Normal data

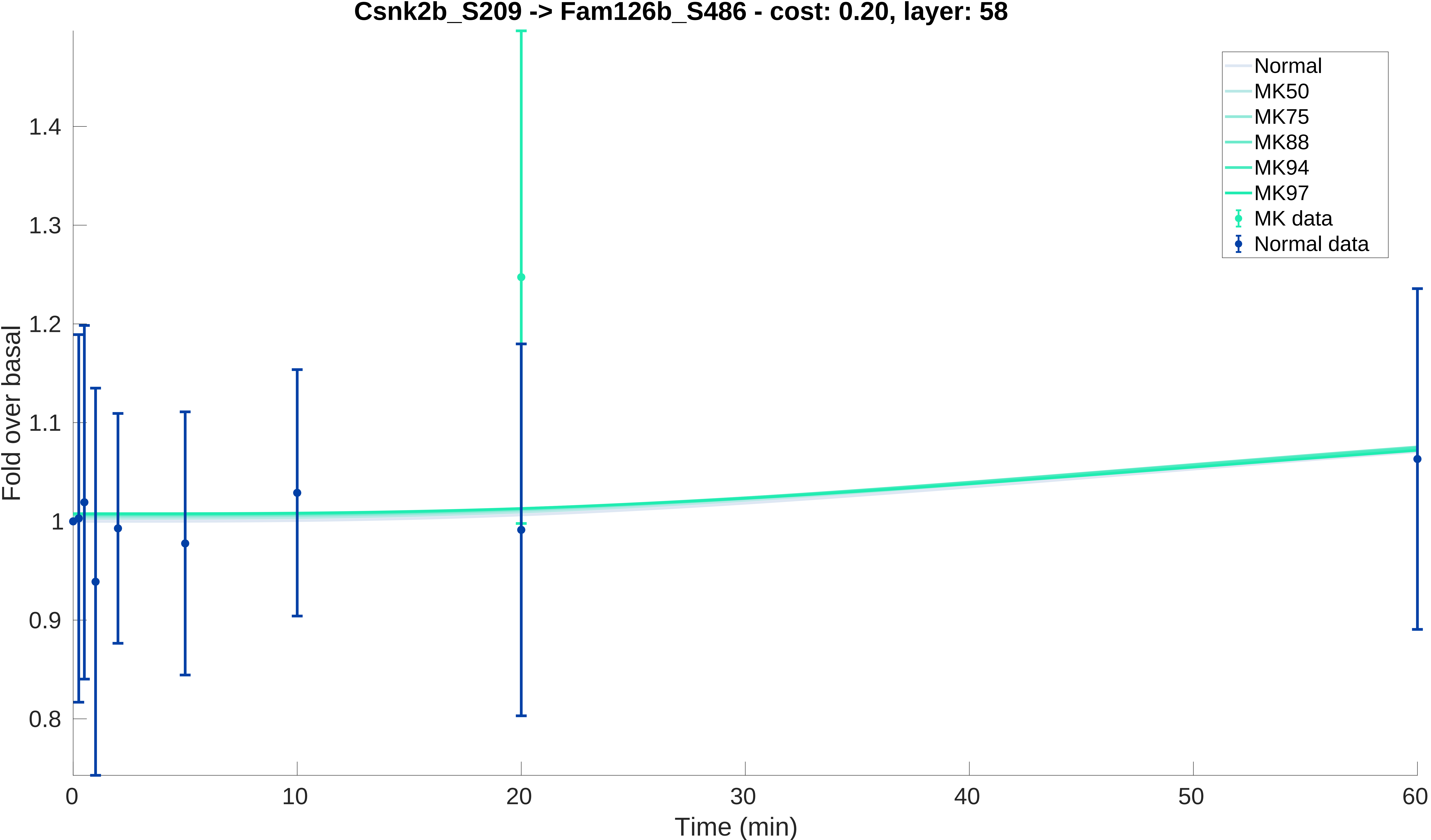

Csnk2b_S209 -> Fam126b_S486 - cost: 0.20, layer: 58
Fold over basal
Time (min)
1.4
1.3
1.2
1.1
1
0.9
0.8
0
10
20
30
40
50
60
Normal
MK50
MK75
MK88
MK94
MK97
MK data
Normal data

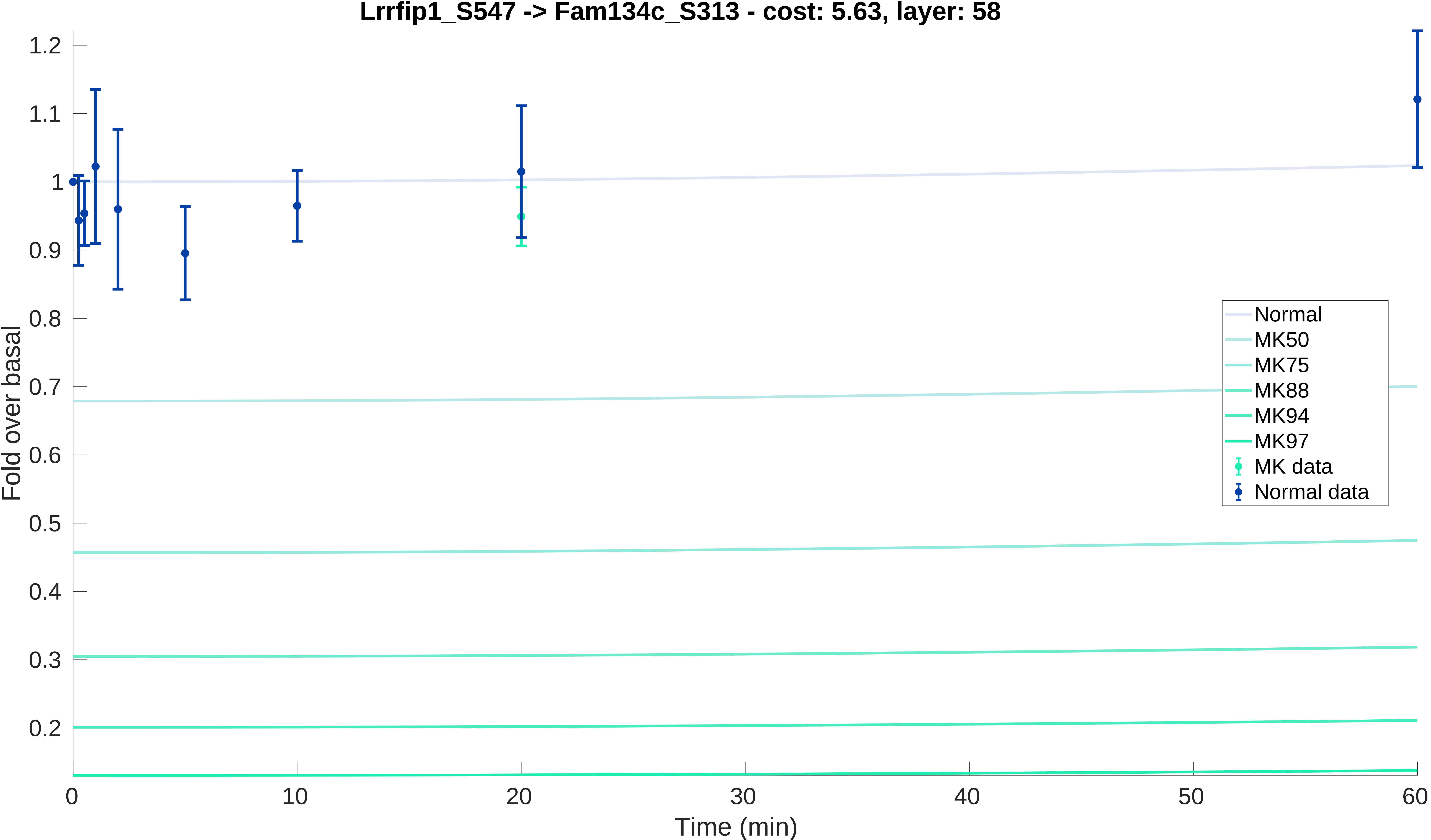

Lrrfip1_S547 -> Fam134c_S313 - cost: 5.63, layer: 58
Fold over basal
Time (min)
1.2
1.1
1
0.9
0.8
0.7
0.6
0.5
0.4
0.3
0.2
0
10
20
30
40
50
60
Normal
MK50
MK75
MK88
MK94
MK97
MK data
Normal data

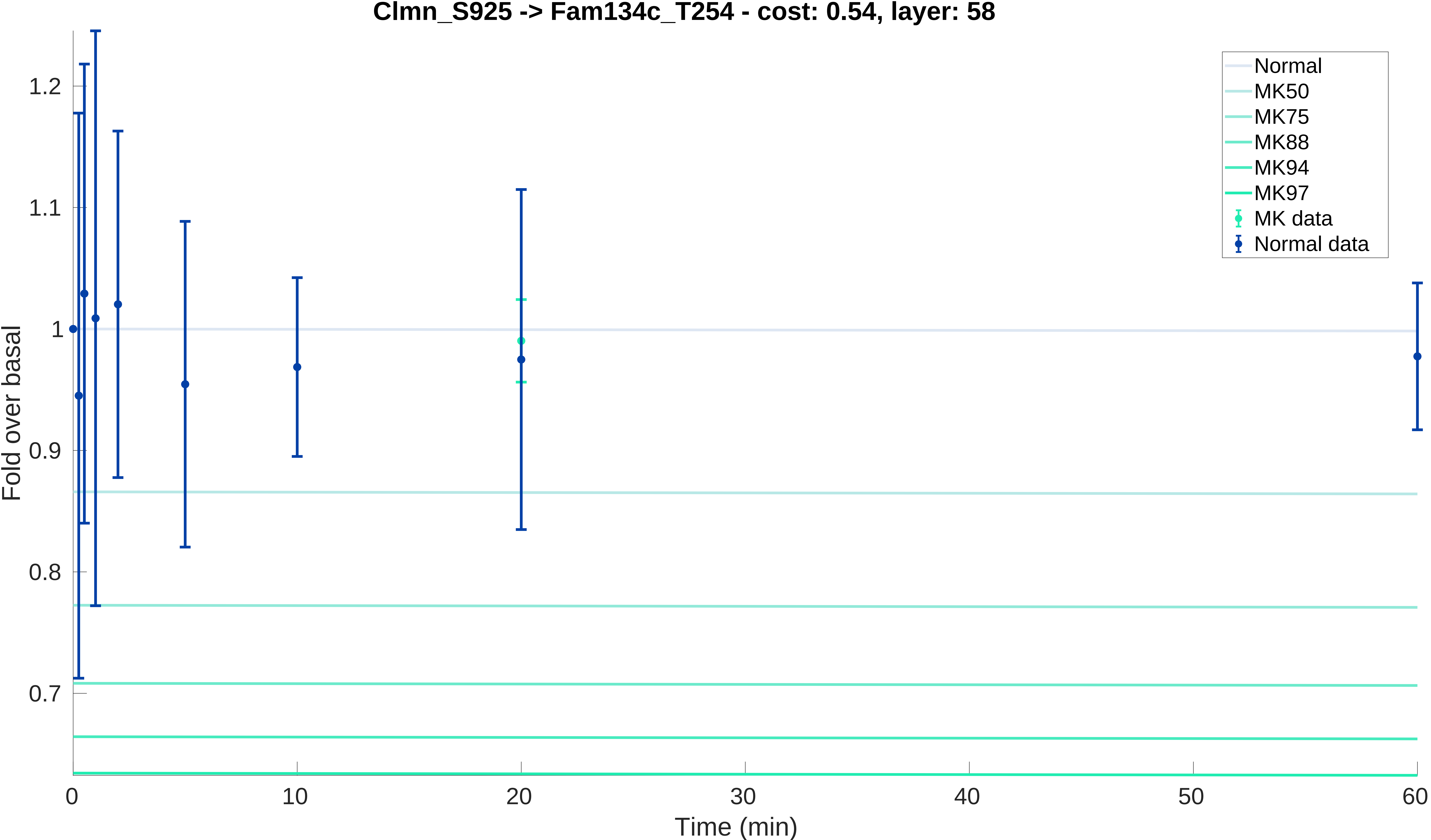

Clmn_S925 -> Fam134c_T254 - cost: 0.54, layer: 58
Fold over basal
Time (min)
0.7
0.8
0.9
1
1.1
1.2
0
10
20
30
40
50
60
Normal
MK50
MK75
MK88
MK94
MK97
MK data
Normal data

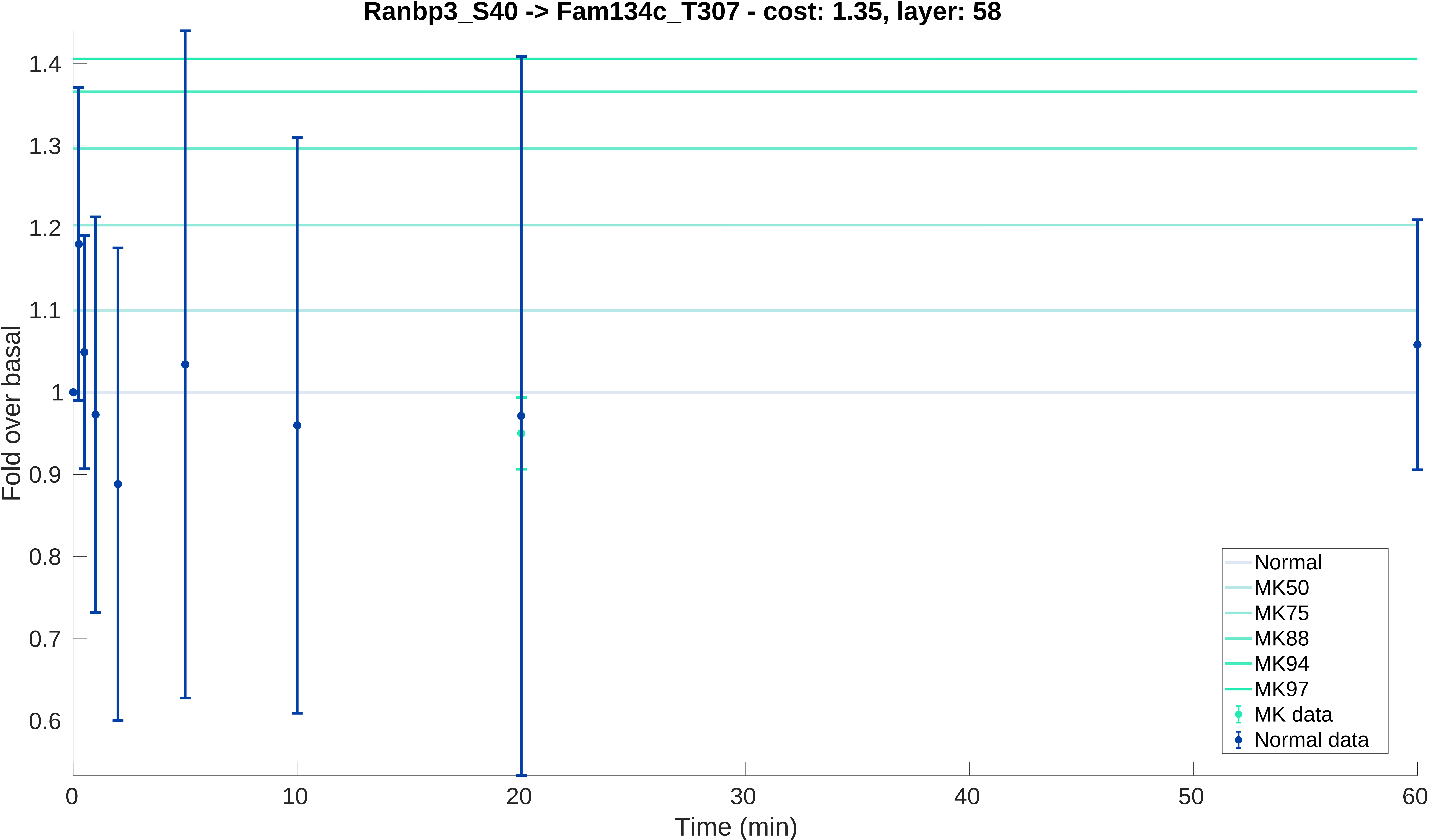

Ranbp3_S40 -> Fam134c_T307 - cost: 1.35, layer: 58
Fold over basal
Time (min)
0
10
20
30
40
50
60
0.6
0.7
0.8
0.9
1
1.1
1.2
1.3
1.4
Normal
MK50
MK75
MK88
MK94
MK97
MK data
Normal data

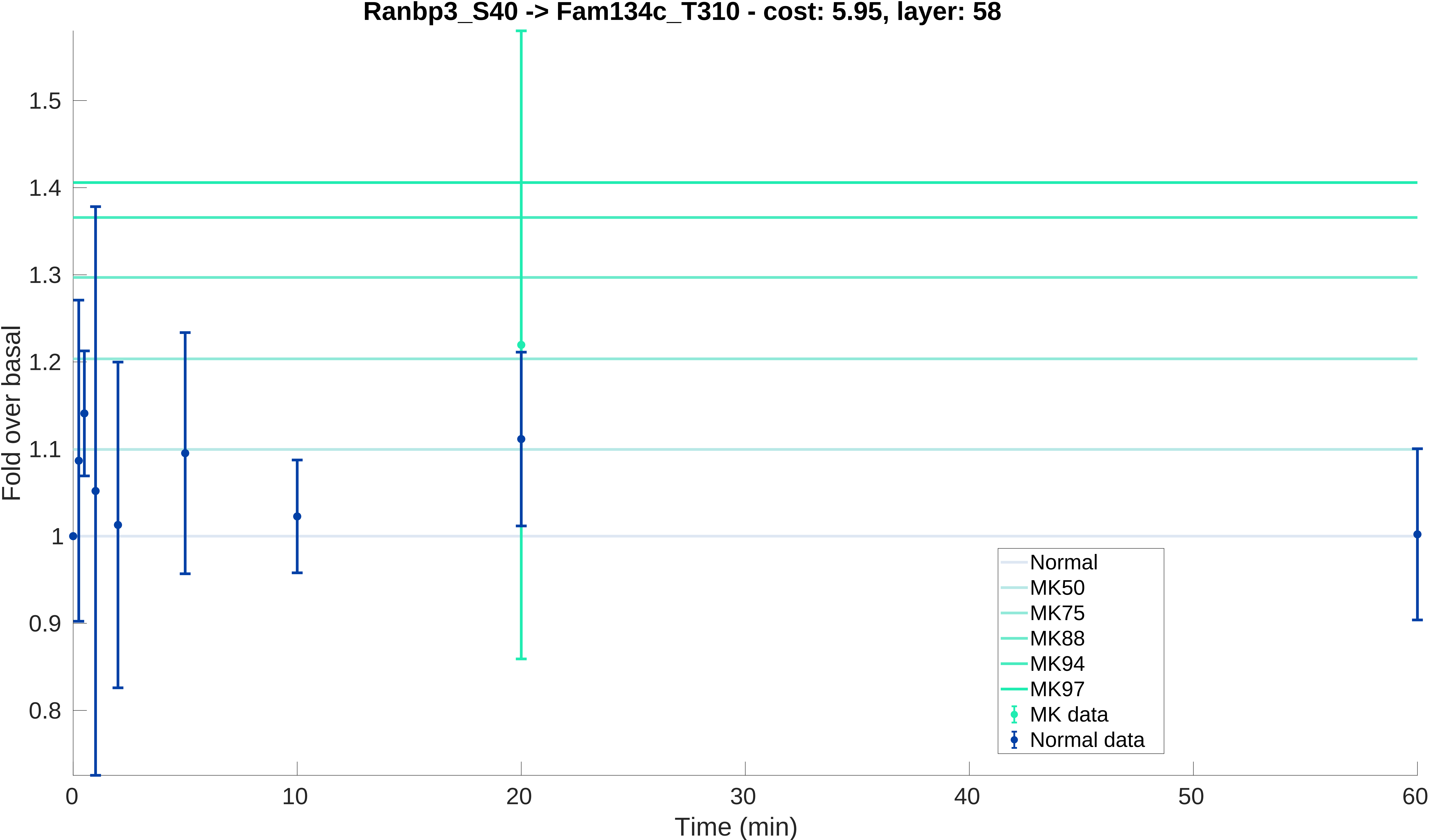

Ranbp3_S40 -> Fam134c_T310 - cost: 5.95, layer: 58
Fold over basal
Time (min)
1.5
1.4
1.3
1.2
1.1
1
0.9
0.8
0
10
20
30
40
50
60
Normal
MK50
MK75
MK88
MK94
MK97
MK data
Normal data

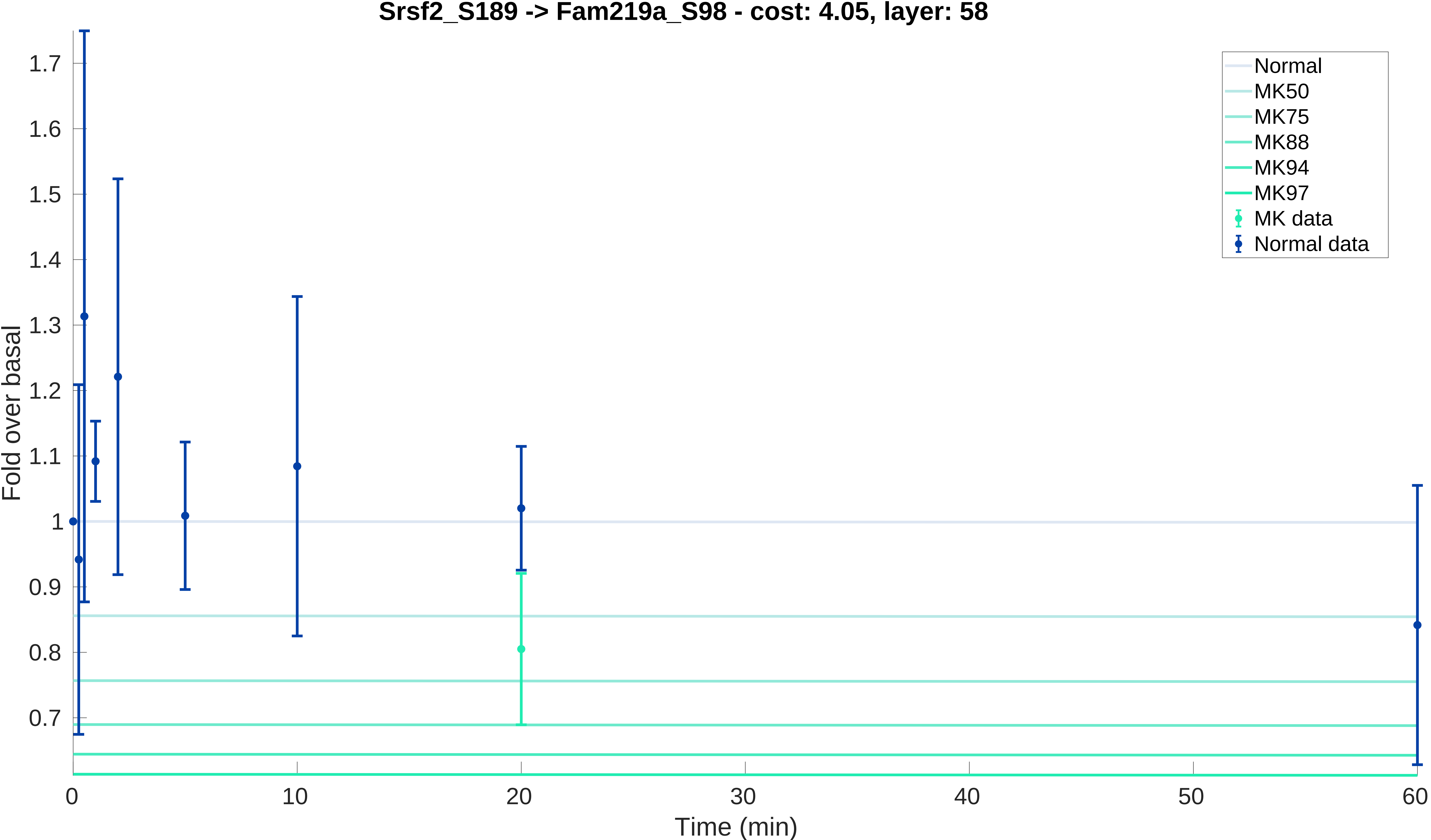

Srsf2_S189 -> Fam219a_S98 - cost: 4.05, layer: 58
Fold over basal
Time (min)
Normal
MK50
MK75
MK88
MK94
MK97
MK data
Normal data
0
10
20
30
40
50
60
0.7
0.8
0.9
1
1.1
1.2
1.3
1.4
1.5
1.6
1.7

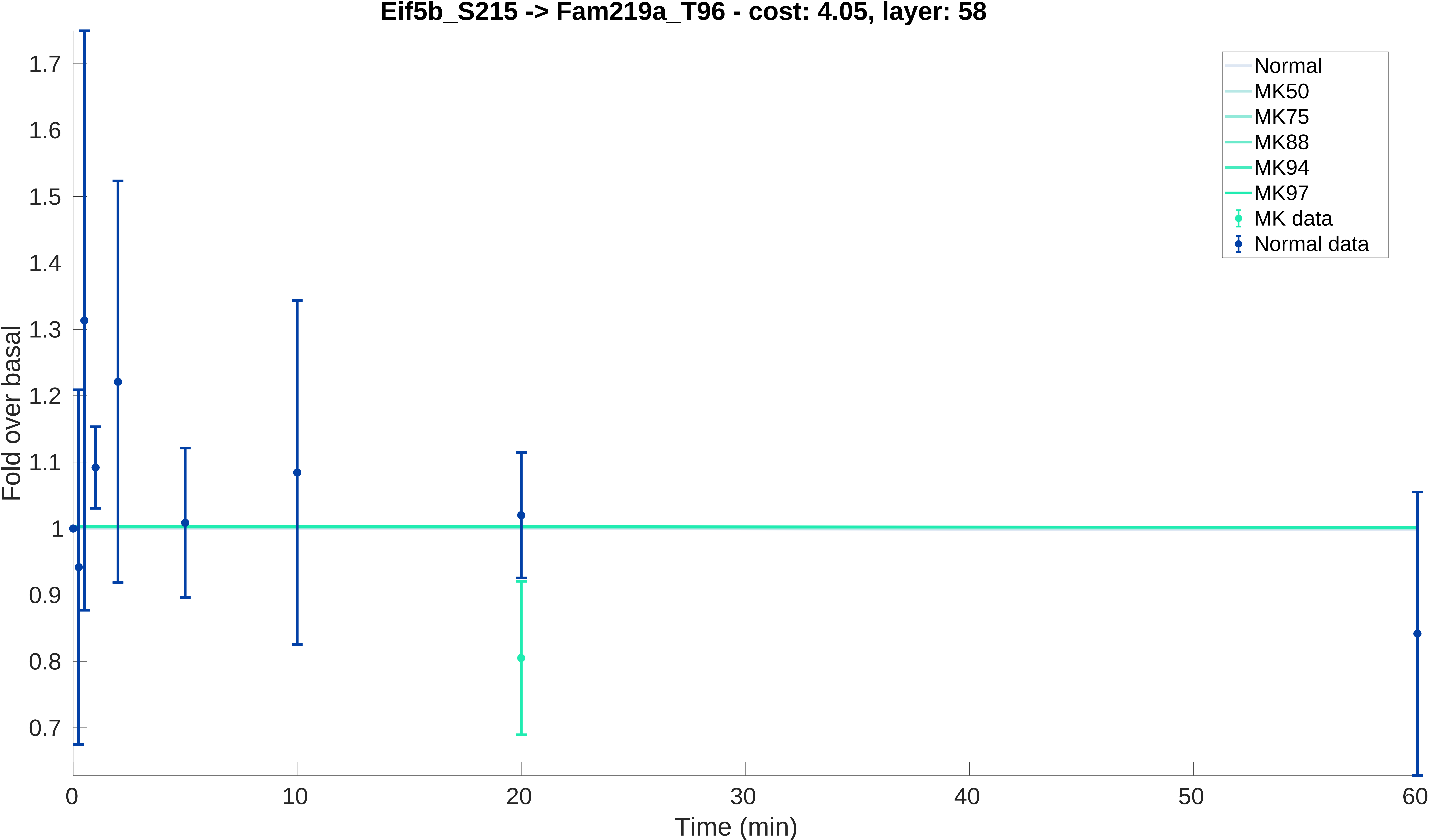

Eif5b_S215 -> Fam219a_T96 - cost: 4.05, layer: 58
Fold over basal
1.7
1.6
1.5
1.4
1.3
1.2
1.1
1
0.9
0.8
0.7
0
10
20
30
40
50
60
Time (min)
Normal
MK50
MK75
MK88
MK94
MK97
MK data
Normal data

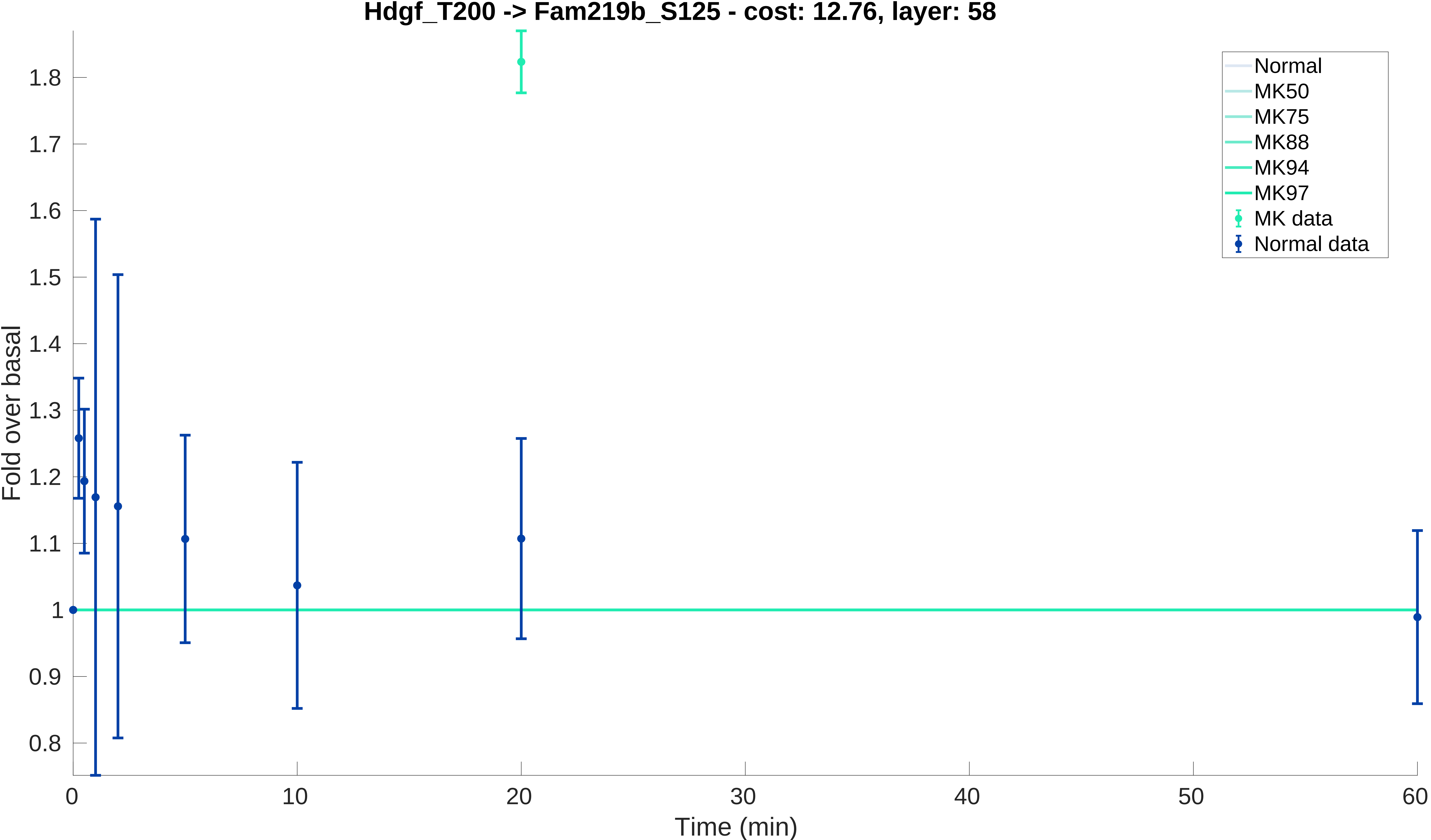
Hdgf_T200 -> Fam219b_S125 - cost: 12.76, layer: 58
Fold over basal
Time (min)
Normal
MK50
MK75
MK88
MK94
MK97
MK data
Normal data
0
10
20
30
40
50
60
0.8
0.9
1
1.1
1.2
1.3
1.4
1.5
1.6
1.7
1.8

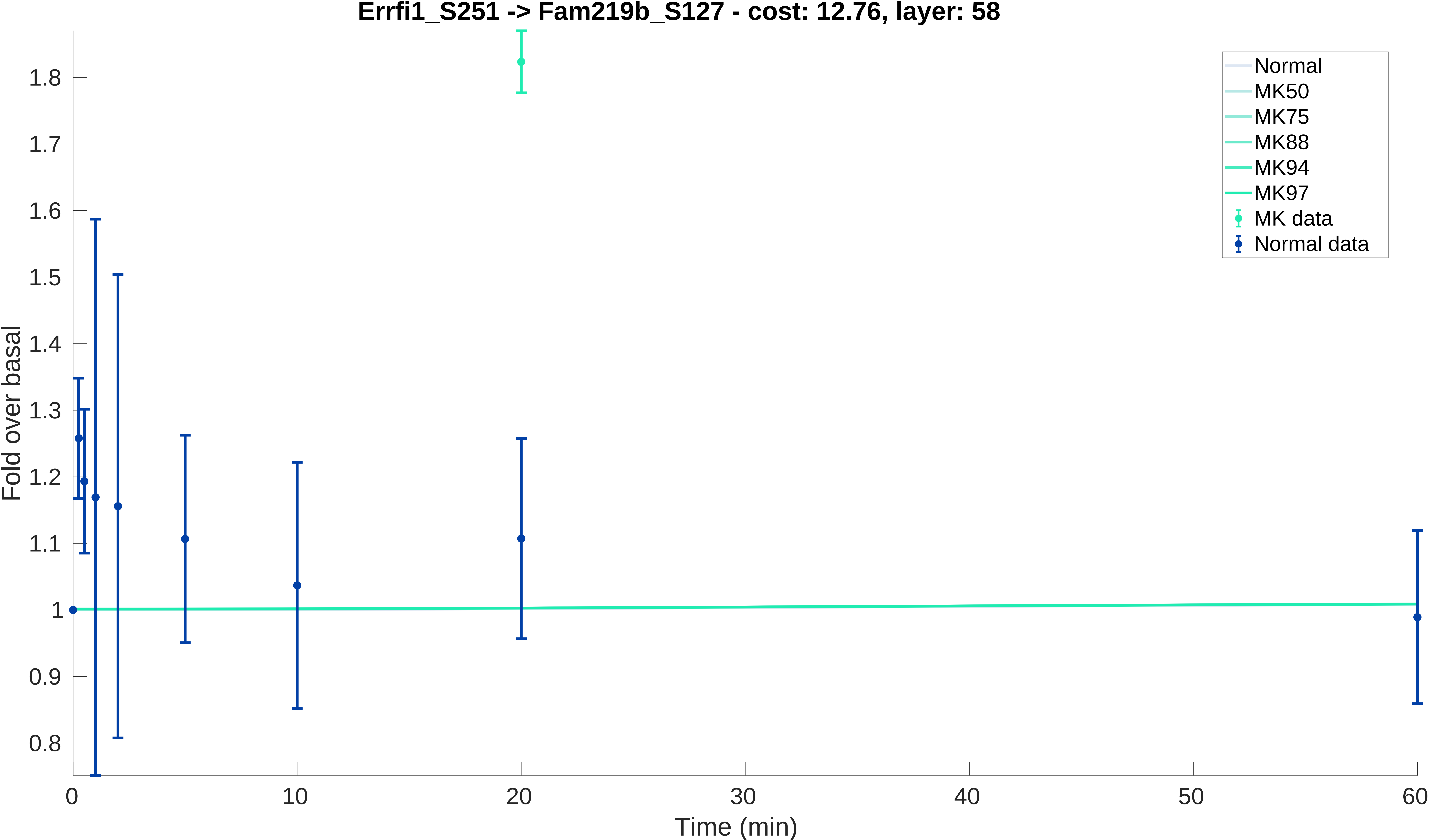
Errfi1_S251 -> Fam219b_S127 - cost: 12.76, layer: 58
Fold over basal
Time (min)
Normal
MK50
MK75
MK88
MK94
MK97
MK data
Normal data
1.8
1.7
1.6
1.5
1.4
1.3
1.2
1.1
1
0.9
0.8
0
10
20
30
40
50
60

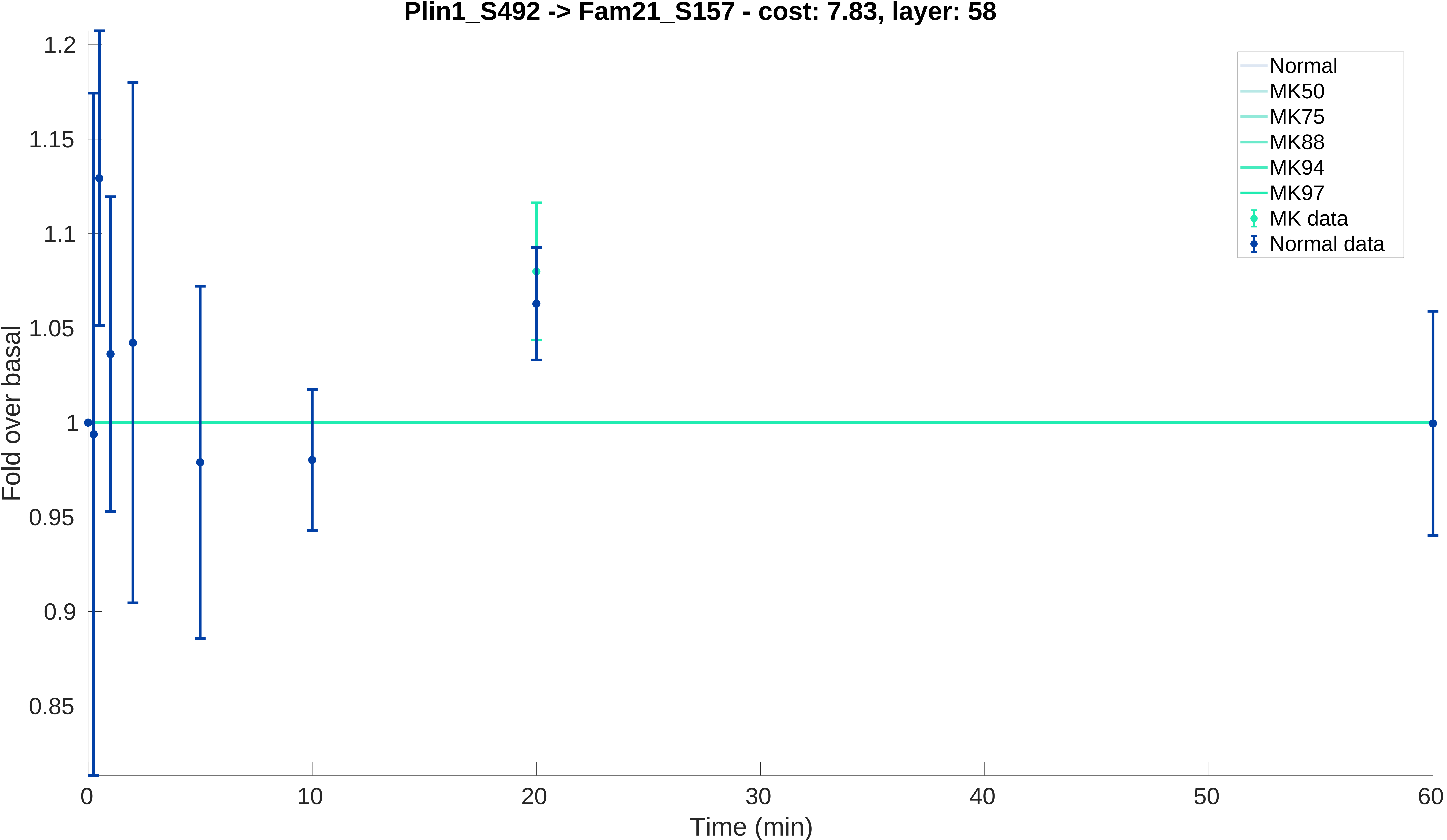

Plin1_S492 -> Fam21_S157 - cost: 7.83, layer: 58
Fold over basal
1.2
1.15
1.1
1.05
1
0.95
0.9
0.85
0
10
20
30
40
50
60
Time (min)
Normal
MK50
MK75
MK88
MK94
MK97
MK data
Normal data

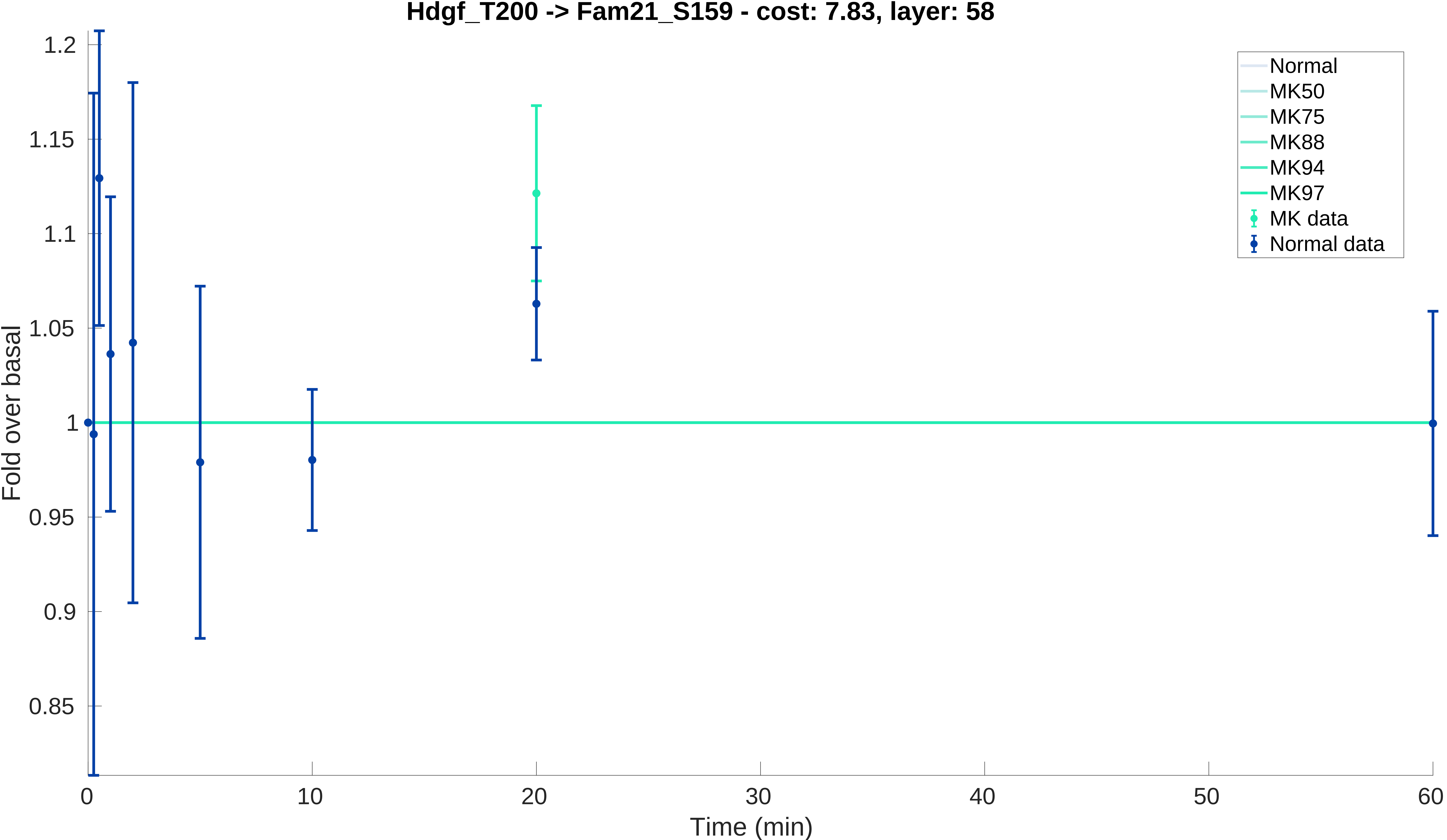
Hdgf_T200 -> Fam21_S159 - cost: 7.83, layer: 58
Fold over basal
1.2
1.15
1.1
1.05
1
0.95
0.9
0.85
0
10
20
30
40
50
60
Time (min)
Normal
MK50
MK75
MK88
MK94
MK97
MK data
Normal data

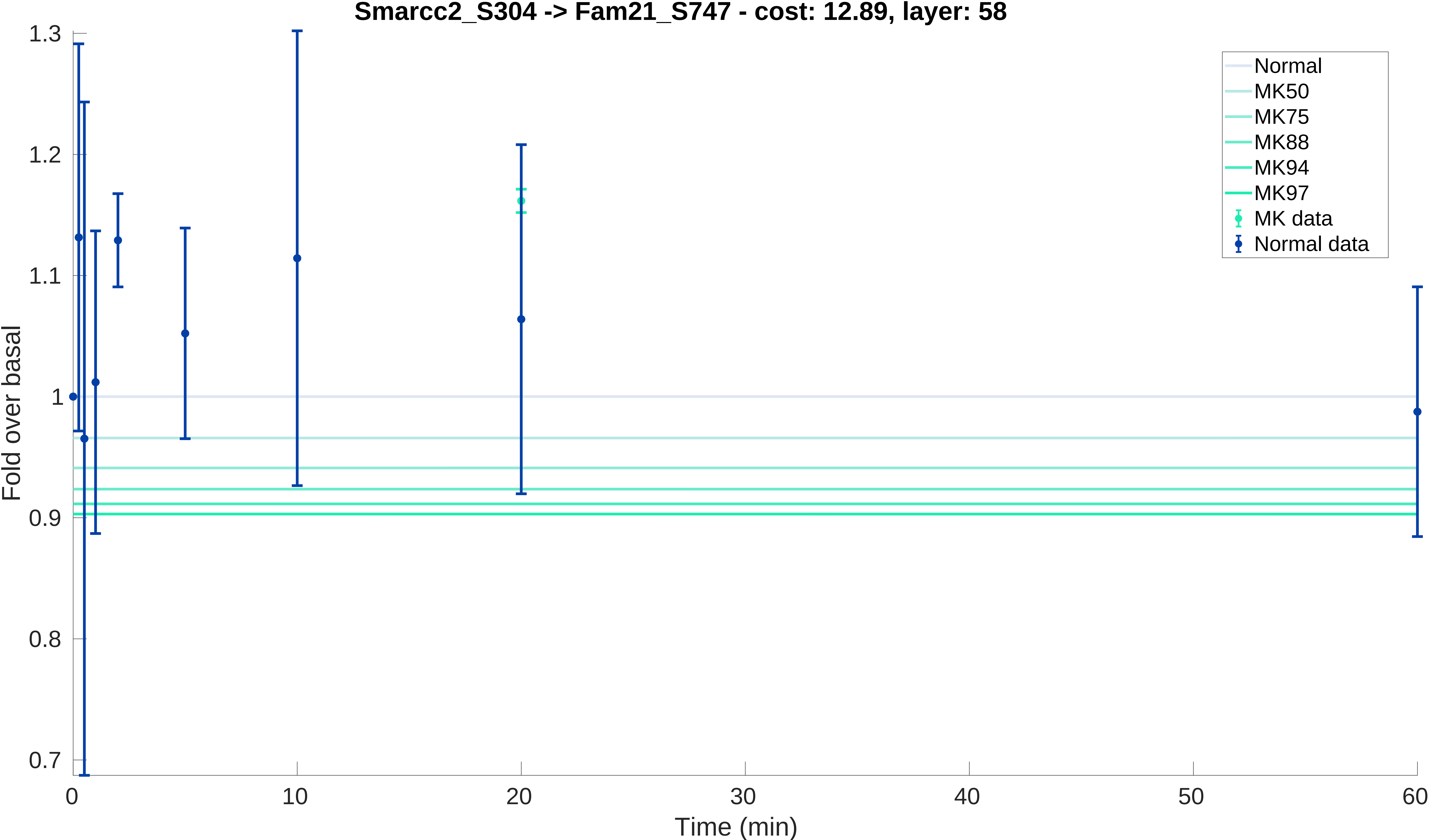

Smarcc2_S304 -> Fam21_S747 - cost: 12.89, layer: 58
Fold over basal
1.3
1.2
1.1
1
0.9
0.8
0.7
0
10
20
30
40
50
60
Time (min)
Normal
MK50
MK75
MK88
MK94
MK97
MK data
Normal data

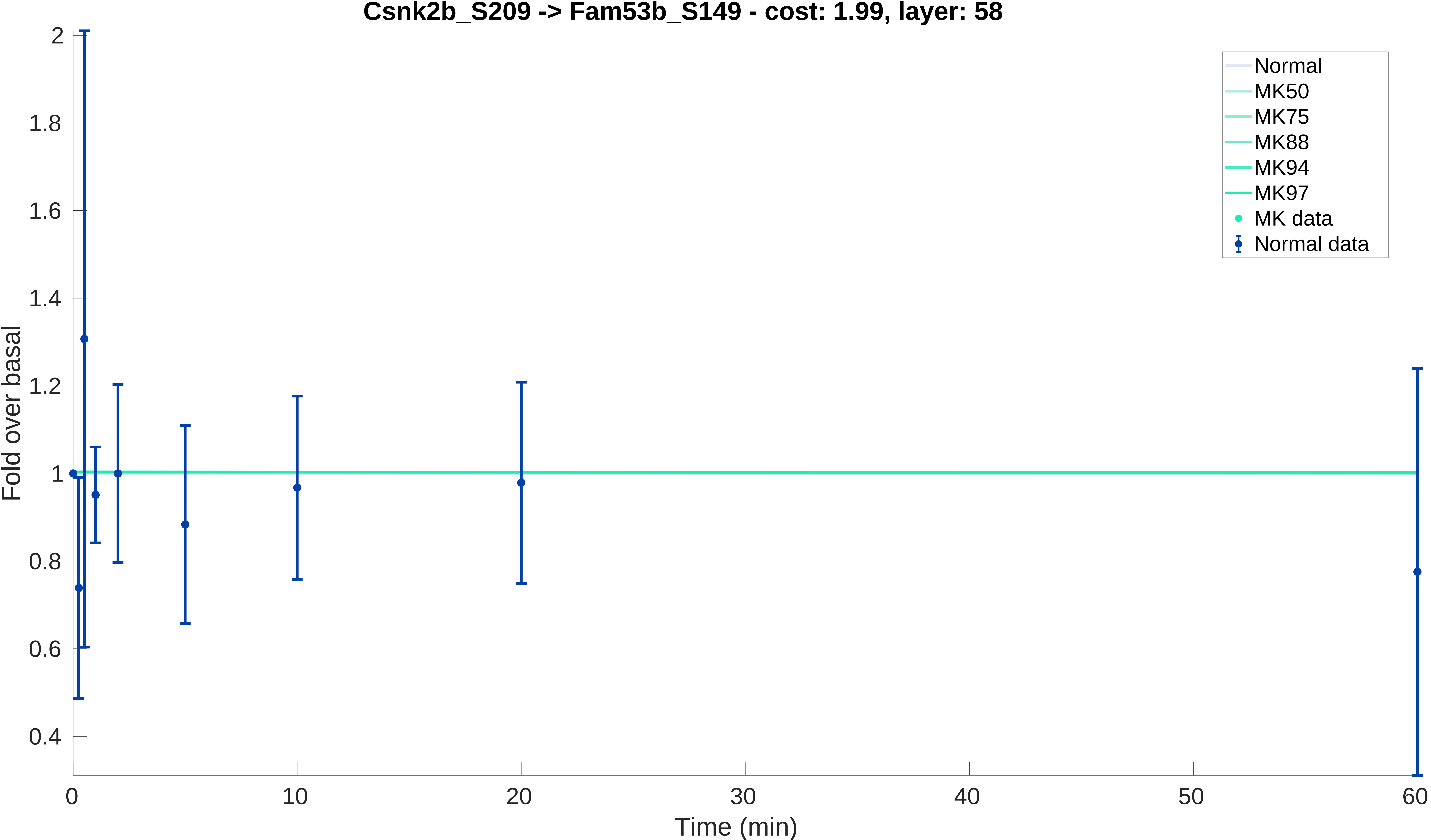

Csnk2b_S209 -> Fam53b_S149 - cost: 1.99, layer: 58
Fold over basal
Time (min)
2
1.8
1.6
1.4
1.2
1
0.8
0.6
0.4
0
10
20
30
40
50
60
Normal
MK50
MK75
MK88
MK94
MK97
MK data
Normal data

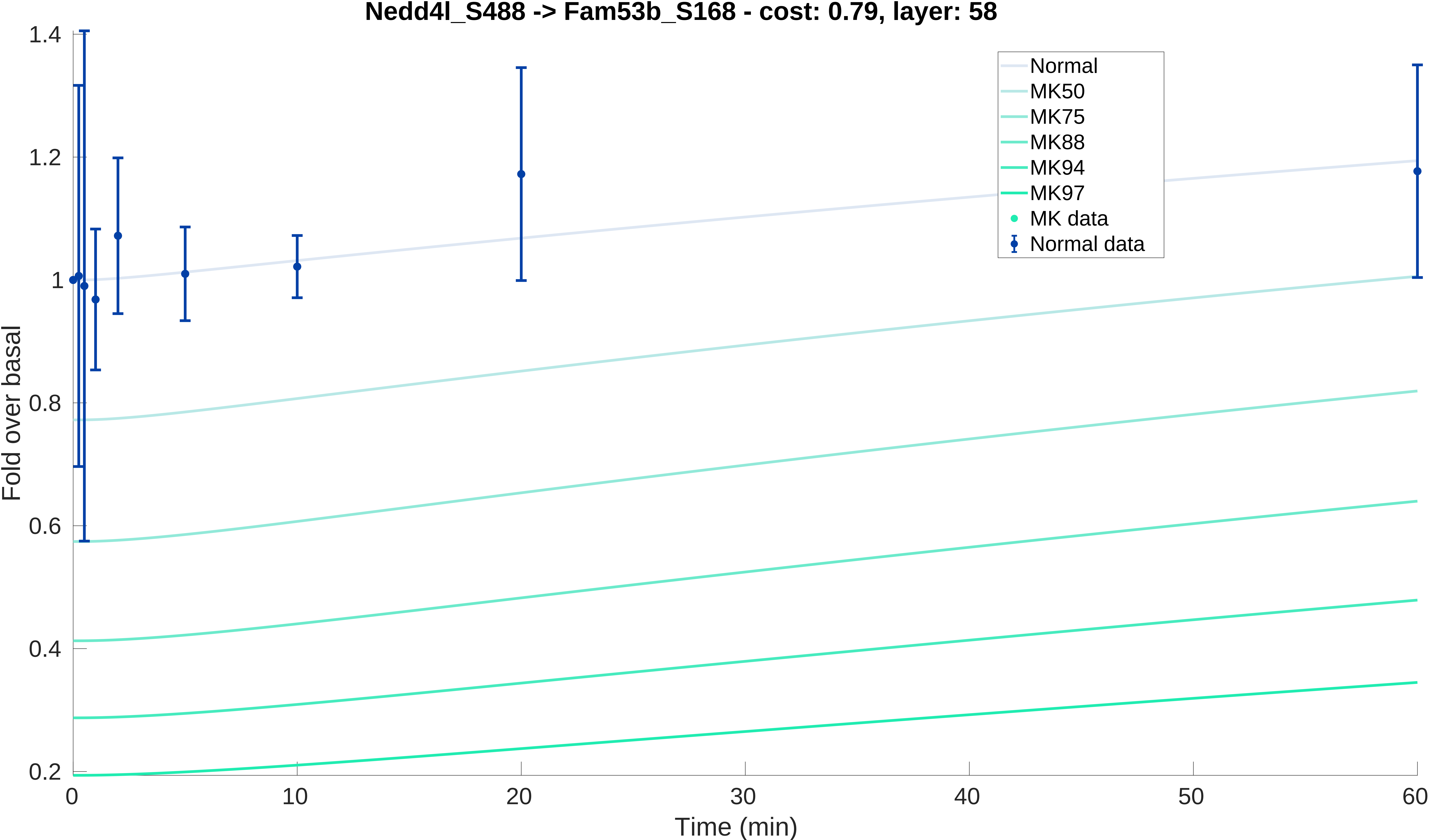

Nedd4l_S488 -> Fam53b_S168 - cost: 0.79, layer: 58
Fold over basal
Time (min)
0.2
0.4
0.6
0.8
1
1.2
1.4
0
10
20
30
40
50
60
Normal
MK50
MK75
MK88
MK94
MK97
MK data
Normal data

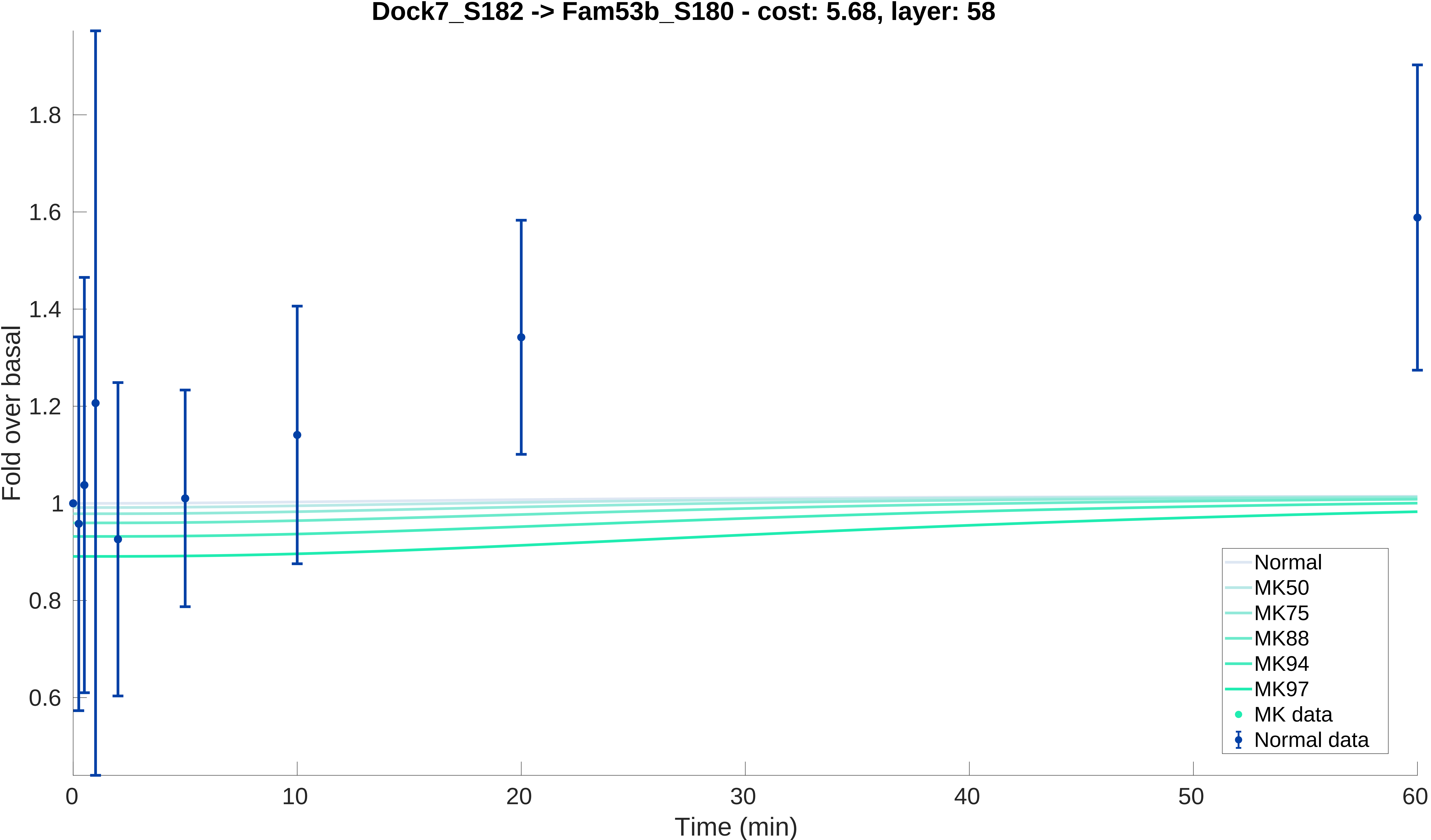
Dock7_S182 -> Fam53b_S180 - cost: 5.68, layer: 58
Fold over basal
1.8
1.6
1.4
1.2
1
0.8
0.6
0
10
20
30
40
50
60
Time (min)
Normal
MK50
MK75
MK88
MK94
MK97
MK data
Normal data

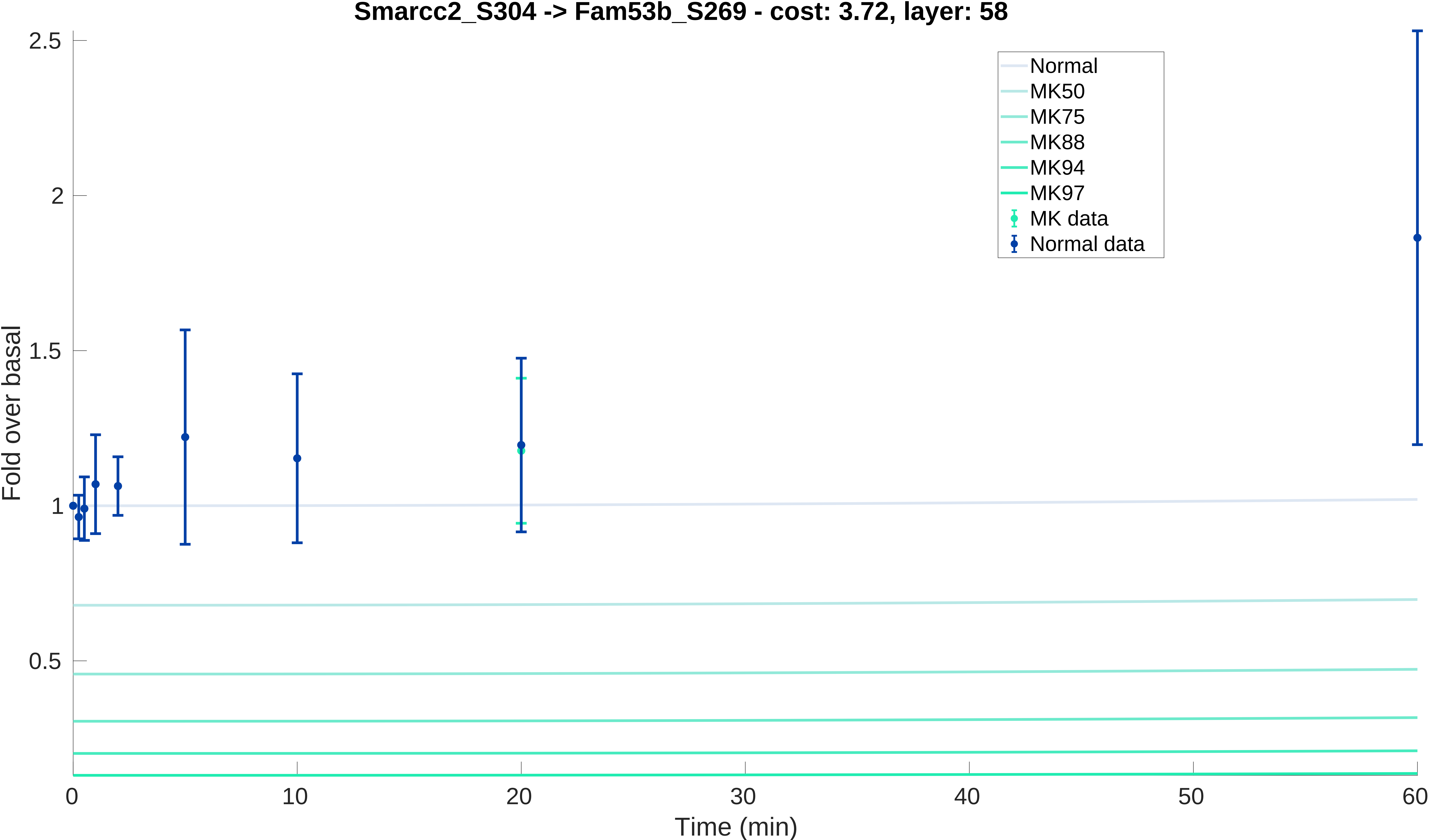

Smarcc2_S304 -> Fam53b_S269 - cost: 3.72, layer: 58
Fold over basal
2.5
2
1.5
1
0.5
0
10
20
30
40
50
60
Time (min)
Normal
MK50
MK75
MK88
MK94
MK97
MK data
Normal data

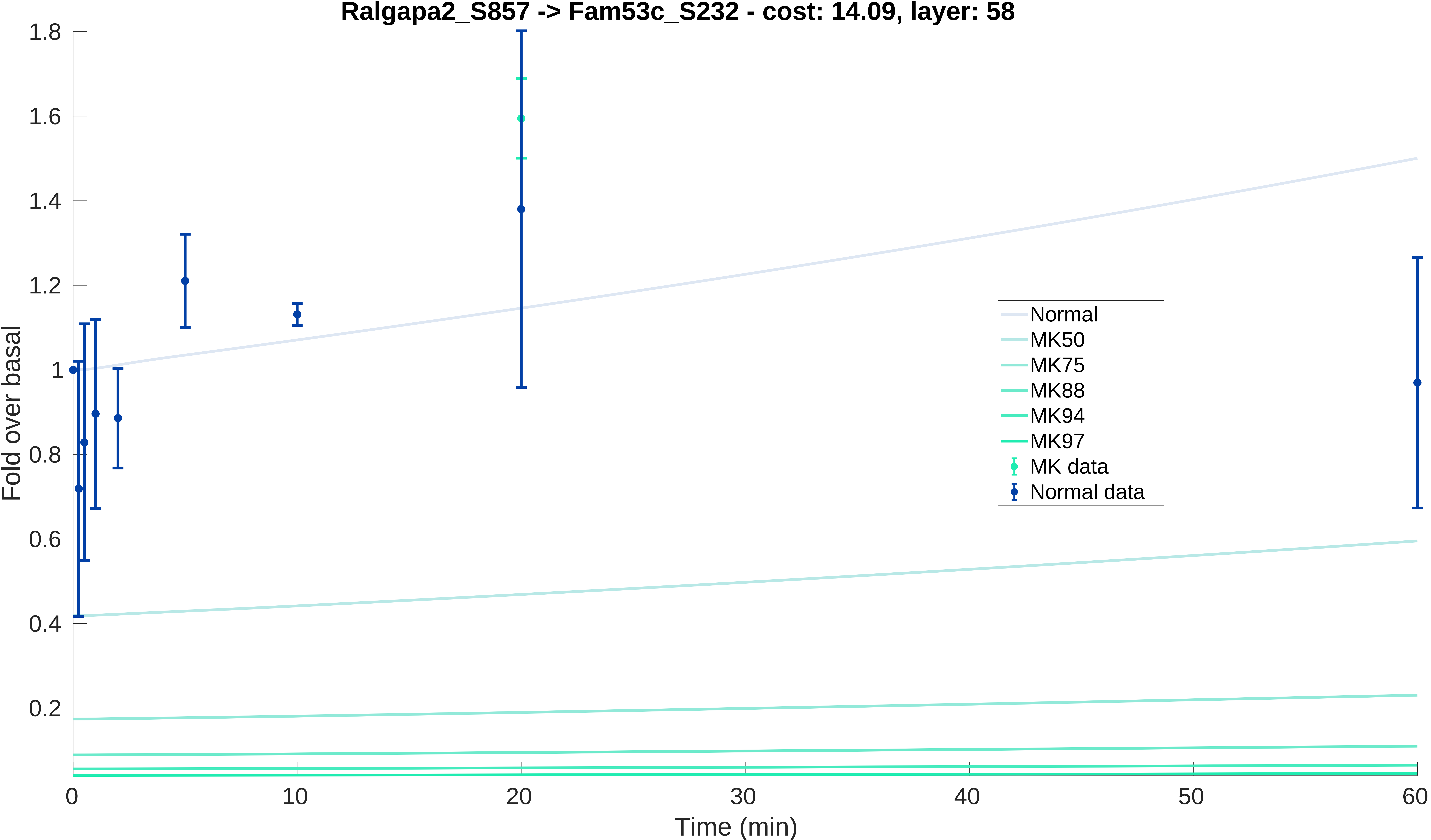
Ralgapa2_S857 -> Fam53c_S232 - cost: 14.09, layer: 58
1.8
1.6
1.4
1.2
1
0.8
0.6
0.4
0.2
Fold over basal
0
10
20
30
40
50
60
Time (min)
Normal
MK50
MK75
MK88
MK94
MK97
MK data
Normal data

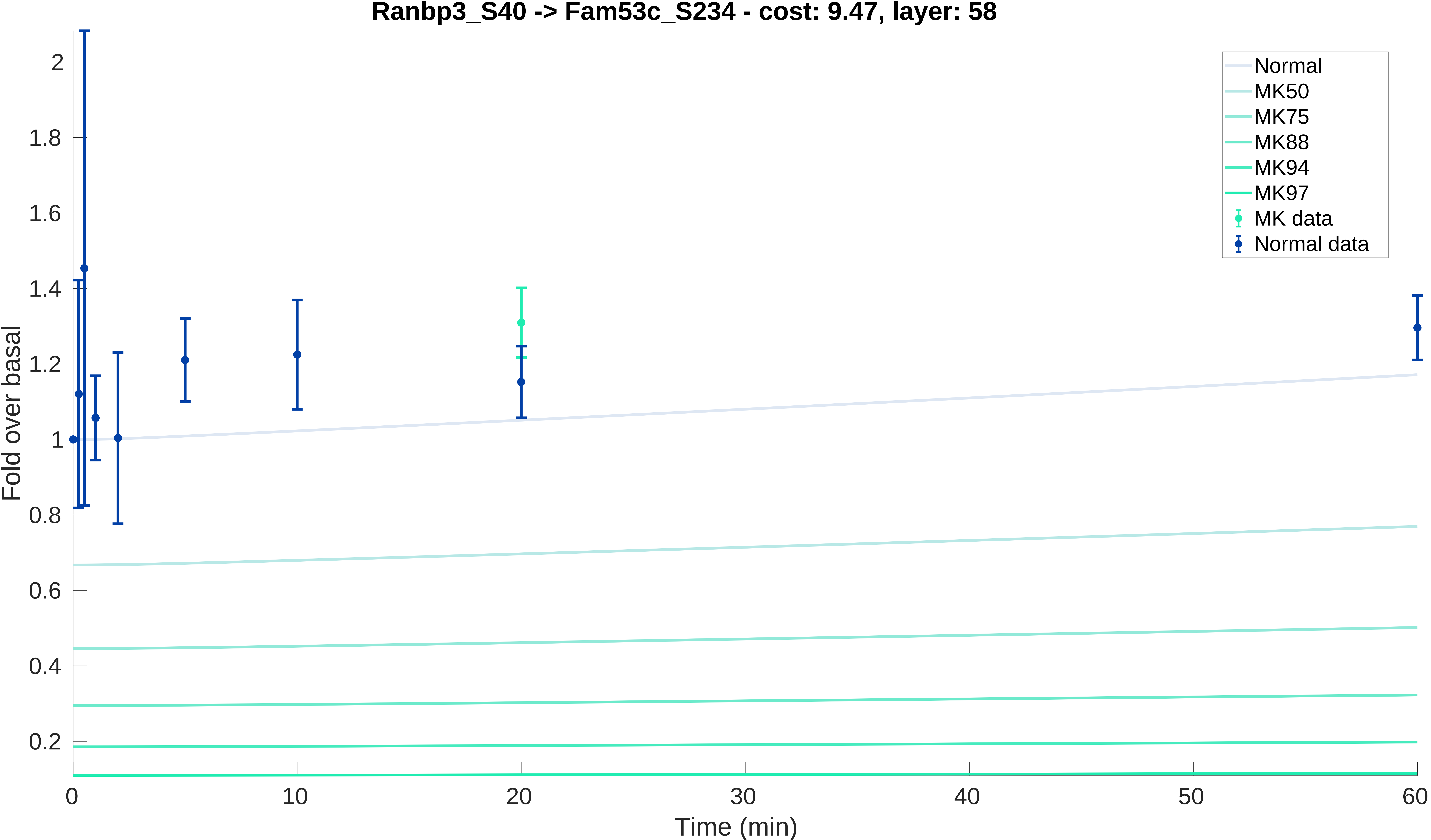
Ranbp3_S40 -> Fam53c_S234 - cost: 9.47, layer: 58
Fold over basal
Time (min)
0
10
20
30
40
50
60
0.2
0.4
0.6
0.8
1
1.2
1.4
1.6
1.8
2
Normal
MK50
MK75
MK88
MK94
MK97
MK data
Normal data

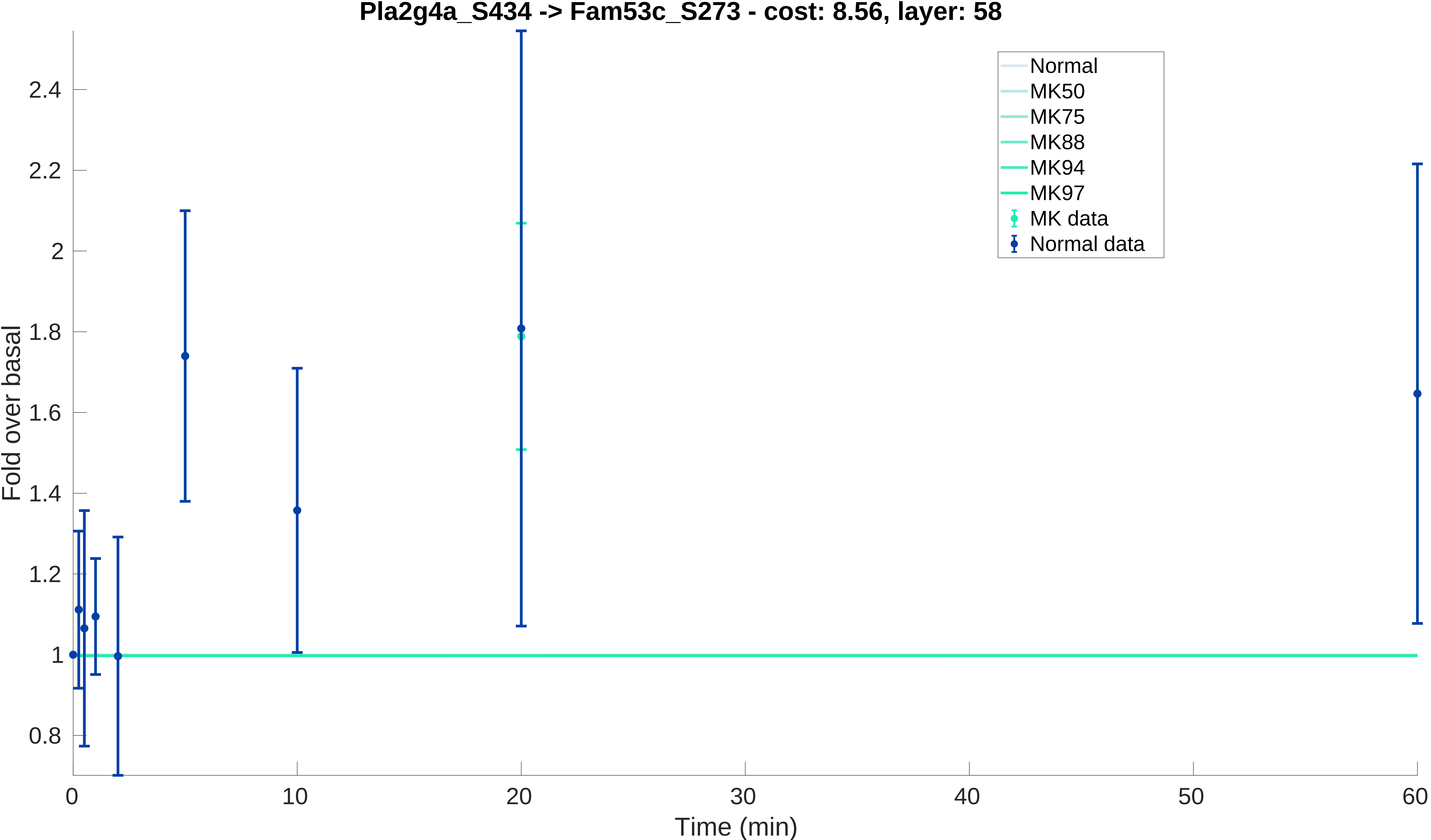

Pla2g4a_S434 -> Fam53c_S273 - cost: 8.56, layer: 58
Fold over basal
Time (min)
0.8
1
1.2
1.4
1.6
1.8
2
2.2
2.4
0
10
20
30
40
50
60
Normal
MK50
MK75
MK88
MK94
MK97
MK data
Normal data

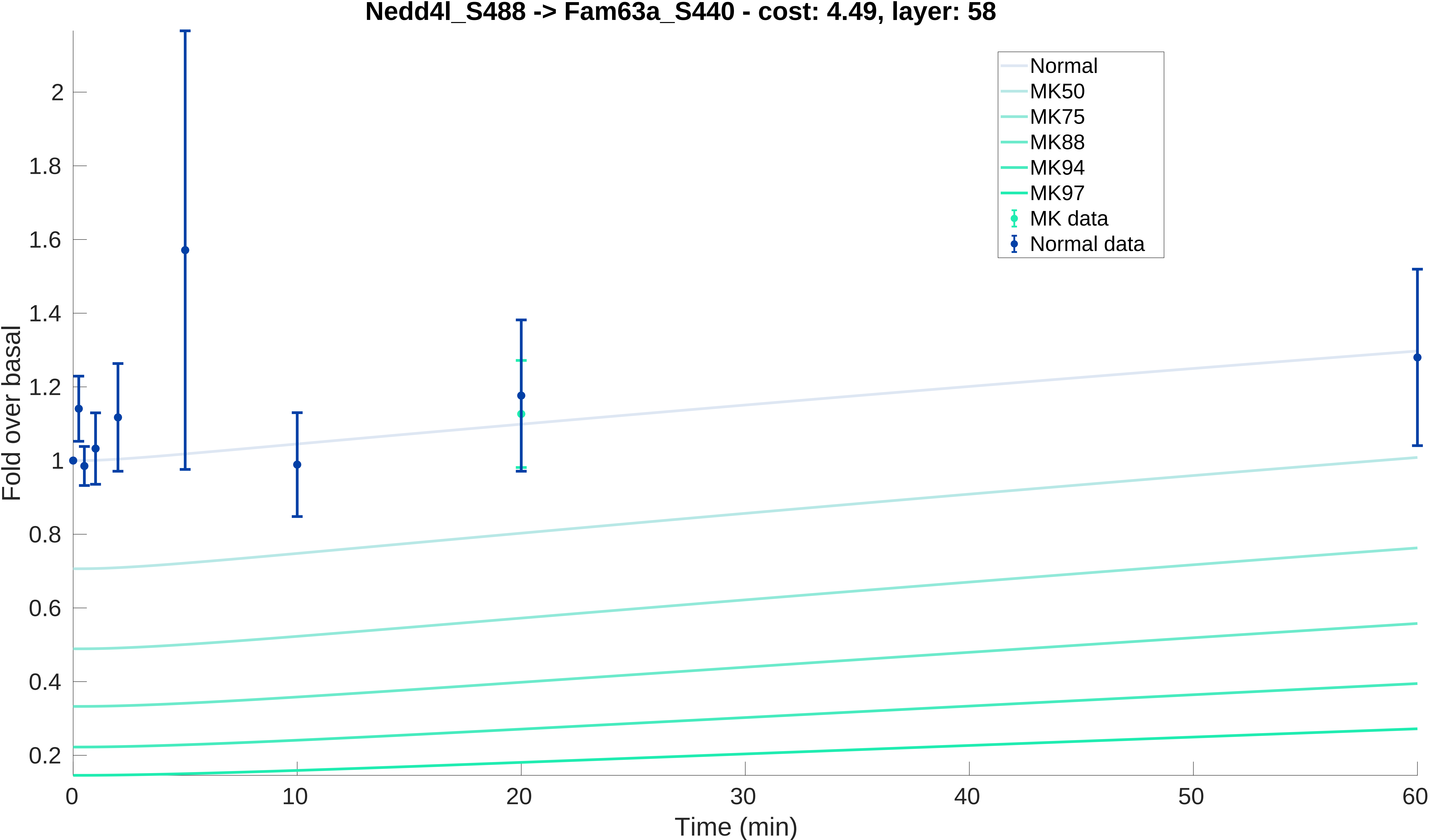

Nedd4l_S488 -> Fam63a_S440 - cost: 4.49, layer: 58
Fold over basal
Time (min)
Normal
MK50
MK75
MK88
MK94
MK97
MK data
Normal data
2
1.8
1.6
1.4
1.2
1
0.8
0.6
0.4
0.2
0
10
20
30
40
50
60

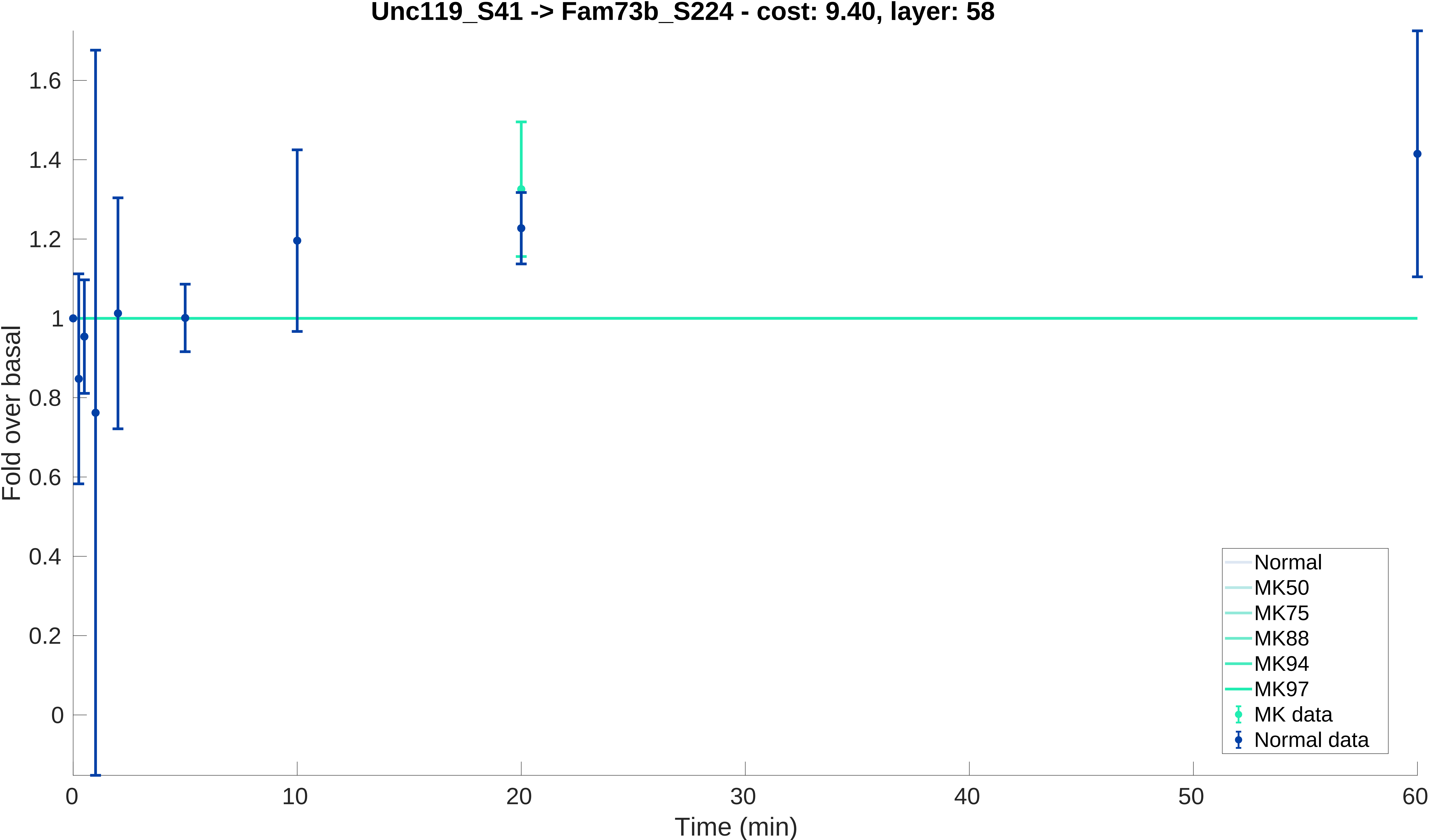

Unc119_S41 -> Fam73b_S224 - cost: 9.40, layer: 58
Fold over basal
1.6
1.4
1.2
1
0.8
0.6
0.4
0.2
0
Time (min)
0
10
20
30
40
50
60
Normal
MK50
MK75
MK88
MK94
MK97
MK data
Normal data

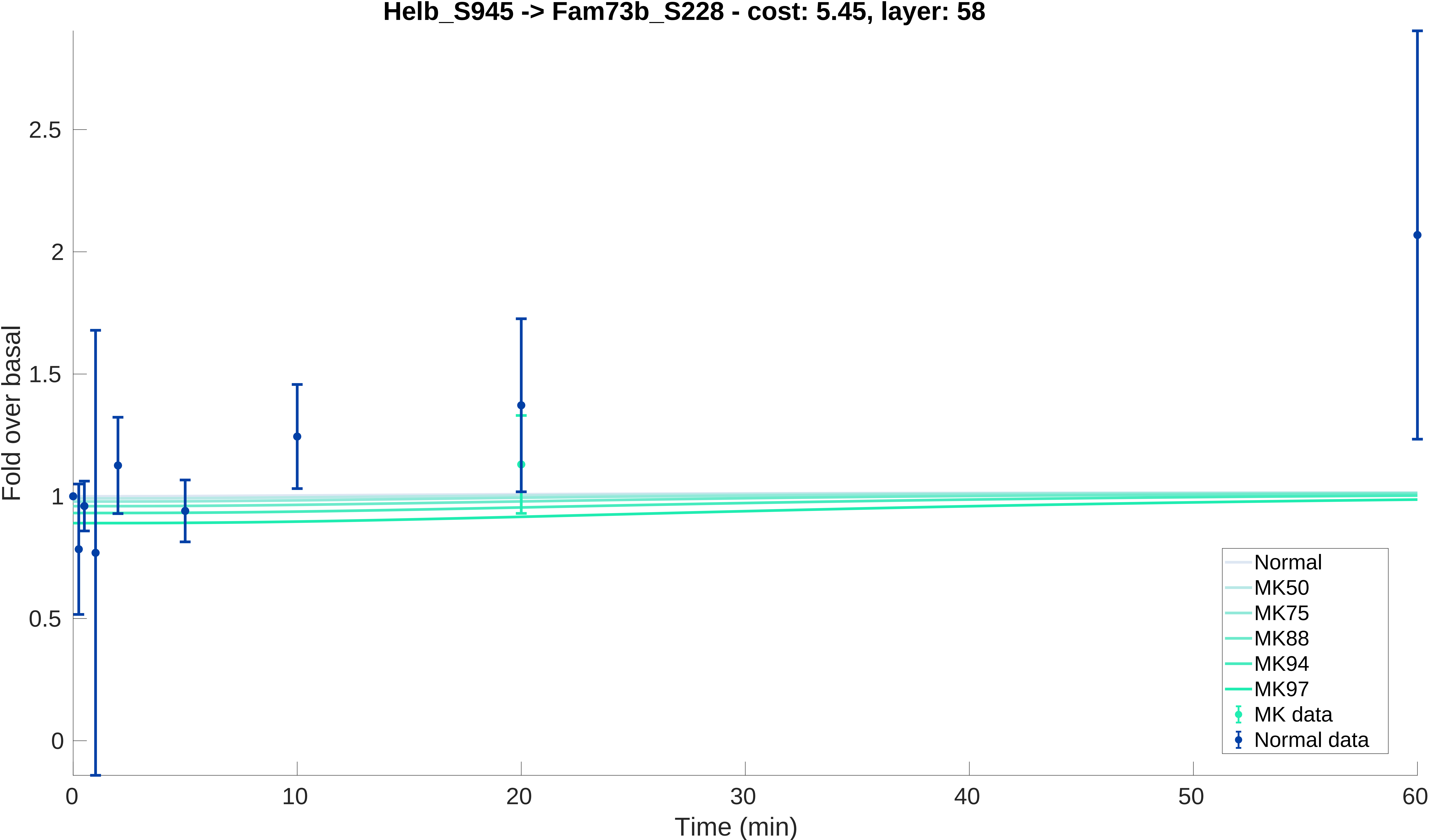

Helb_S945 -> Fam73b_S228 - cost: 5.45, layer: 58
Fold over basal
Time (min)
0
0.5
1
1.5
2
2.5
0
10
20
30
40
50
60
Normal
MK50
MK75
MK88
MK94
MK97
MK data
Normal data

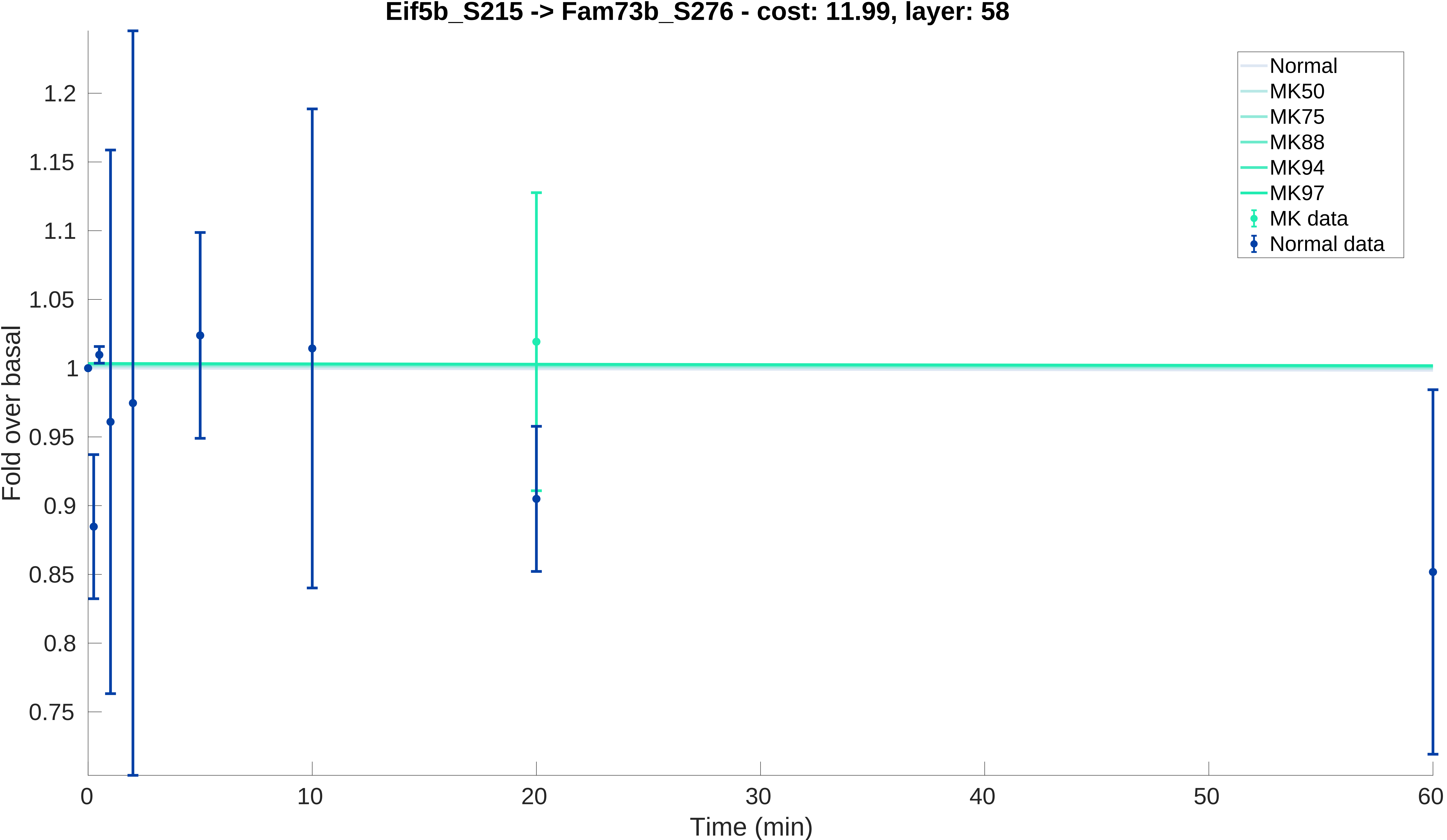

Eif5b_S215 -> Fam73b_S276 - cost: 11.99, layer: 58
Normal
MK50
MK75
MK88
MK94
MK97
MK data
Normal data
1.2
1.15
1.1
1.05
1
0.95
0.9
0.85
0.8
0.75
Fold over basal
0
10
20
30
40
50
60
Time (min)

#### Plin1_S492 -> Fam76b_S193 - cost: 2.85, layer: 58

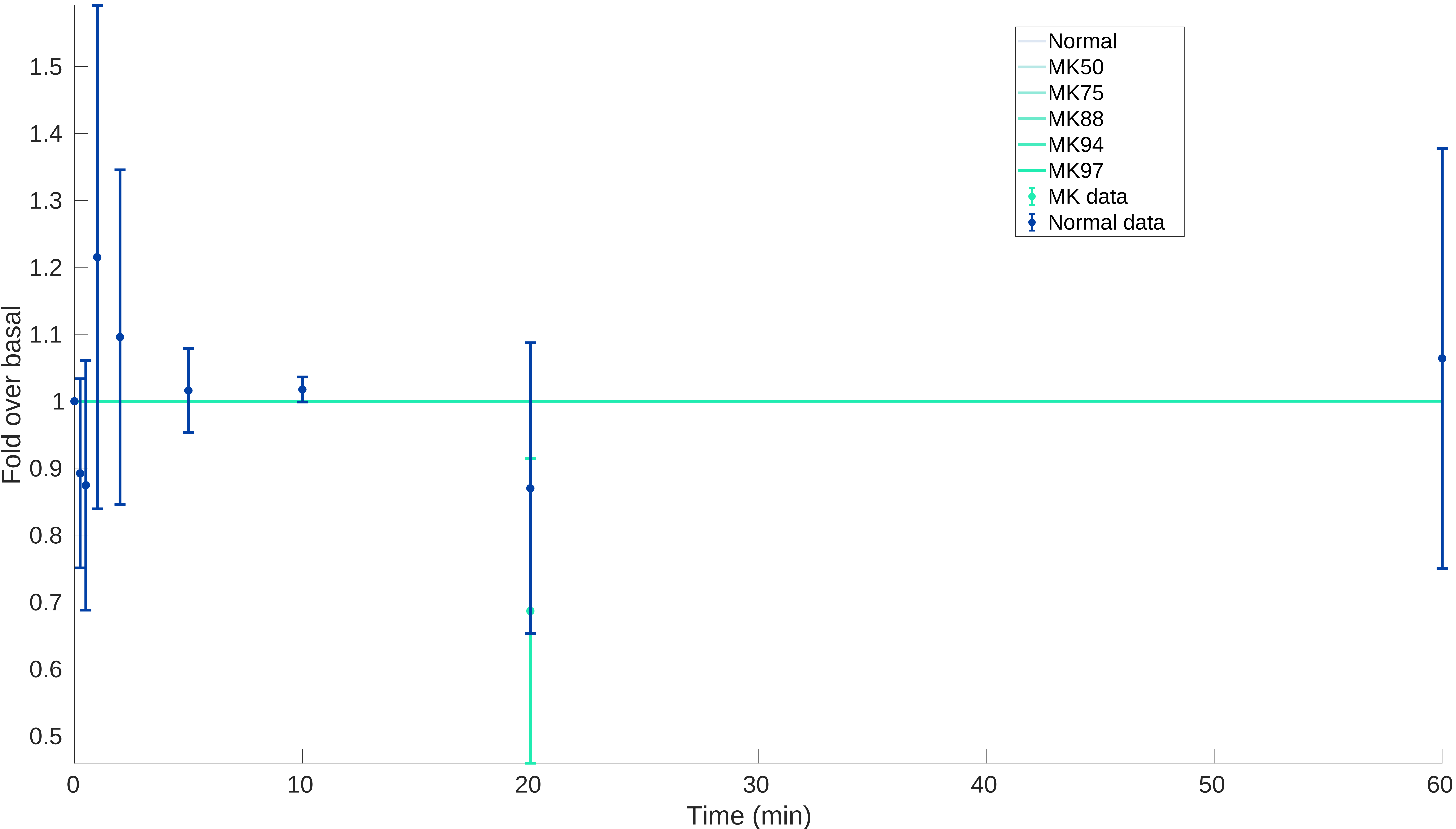

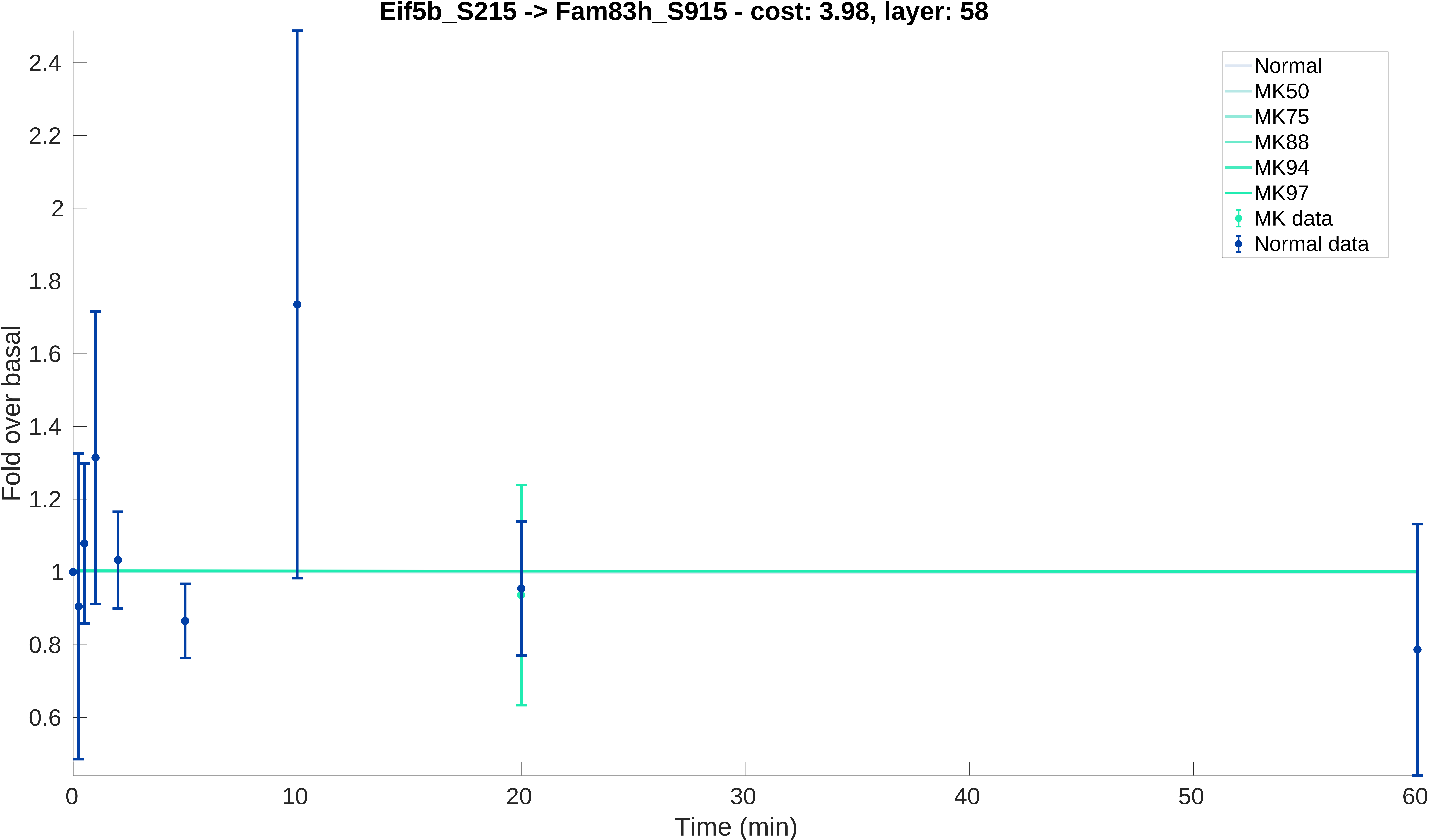

Eif5b_S215 -> Fam83h_S915 - cost: 3.98, layer: 58
Fold over basal
Time (min)
0.6
0.8
1
1.2
1.4
1.6
1.8
2
2.2
2.4
0
10
20
30
40
50
60
Normal
MK50
MK75
MK88
MK94
MK97
MK data
Normal data

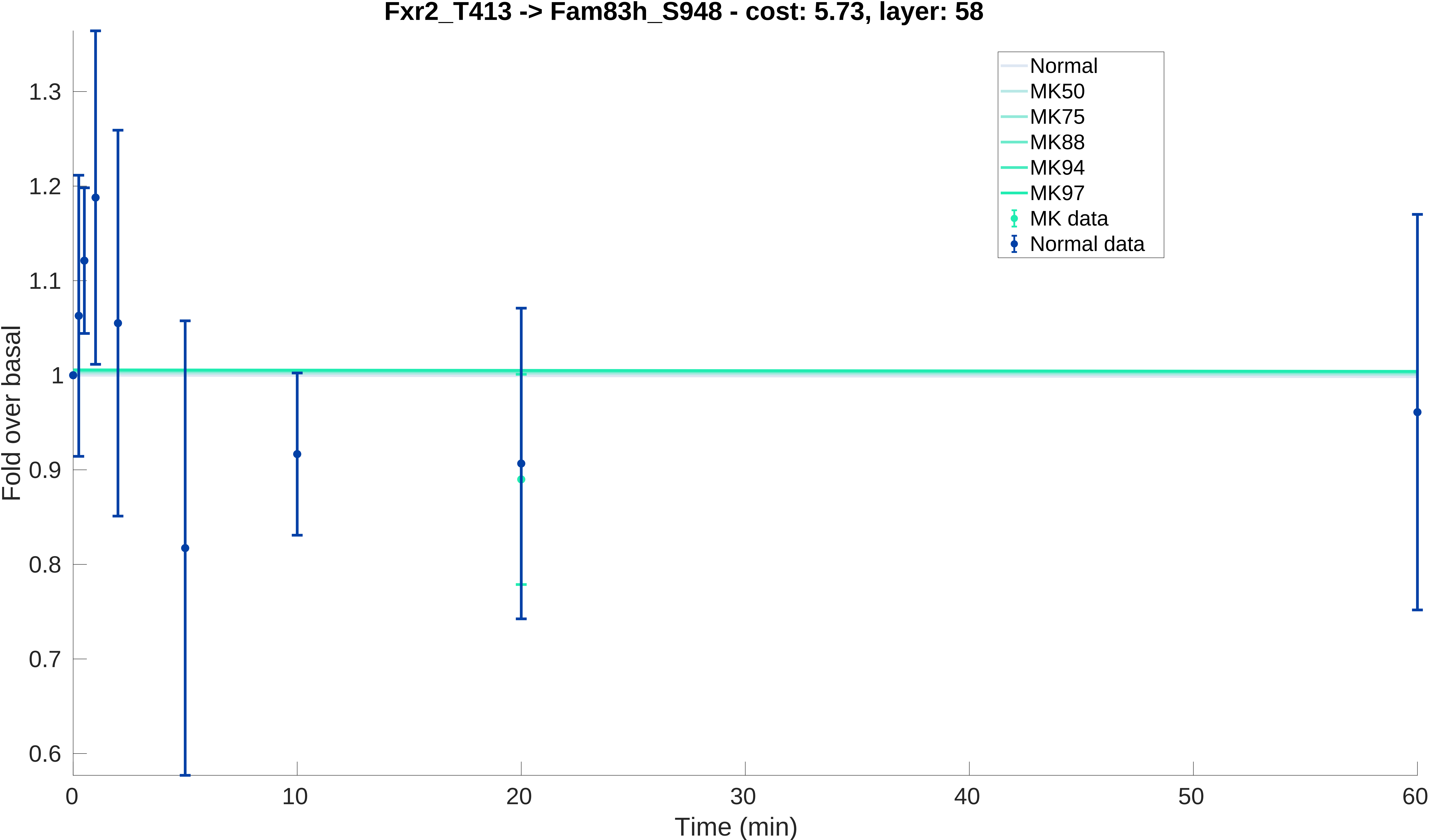

Fxr2_T413 -> Fam83h_S948 - cost: 5.73, layer: 58
Normal
MK50
MK75
MK88
MK94
MK97
MK data
Normal data
Fold over basal
1.3
1.2
1.1
1
0.9
0.8
0.7
0.6
0
10
20
30
40
50
60
Time (min)

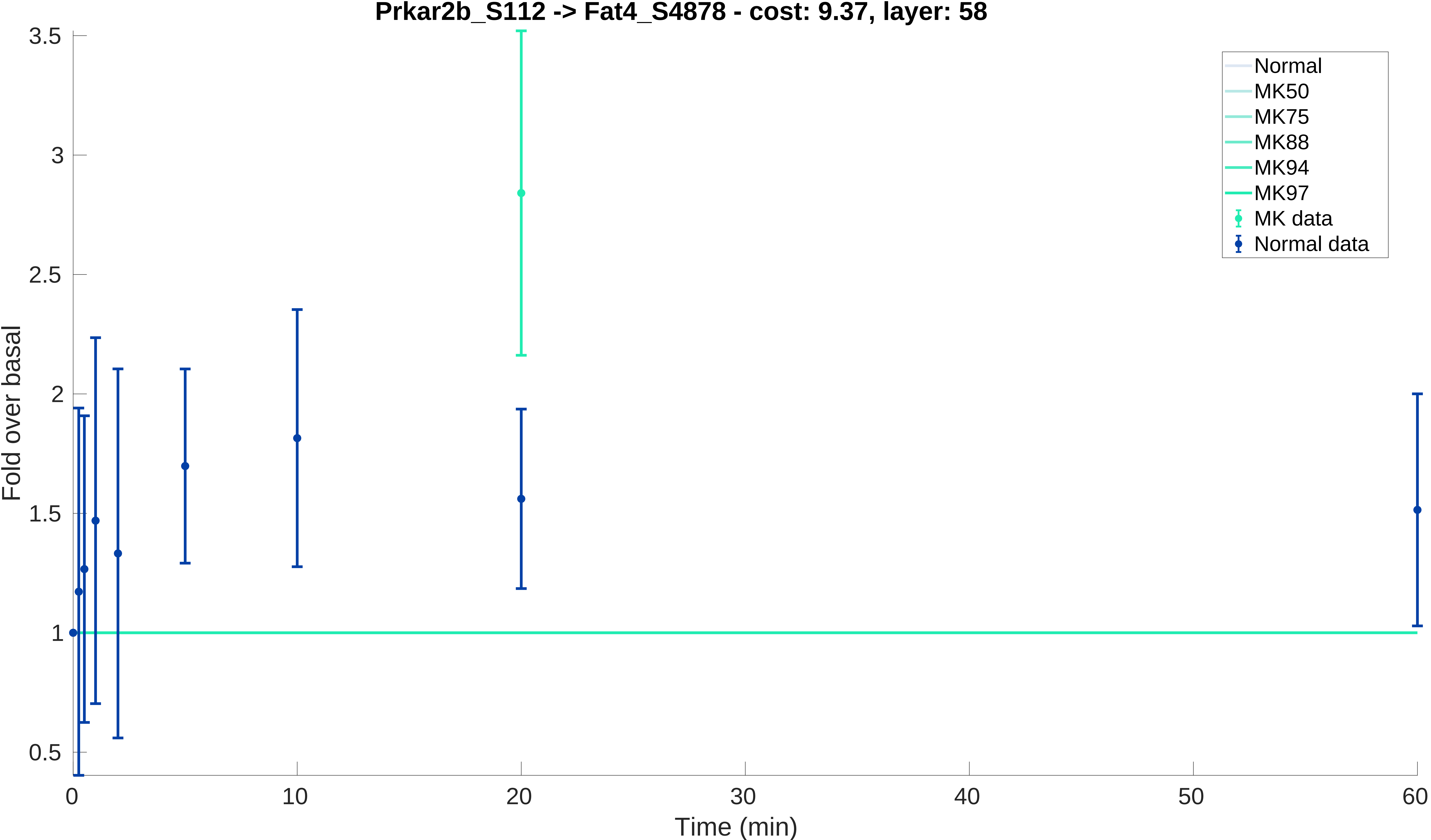

Prkar2b_S112 -> Fat4_S4878 - cost: 9.37, layer: 58
Fold over basal
Time (min)
3.5
3
2.5
2
1.5
1
0.5
0
10
20
30
40
50
60
Normal
MK50
MK75
MK88
MK94
MK97
MK data
Normal data

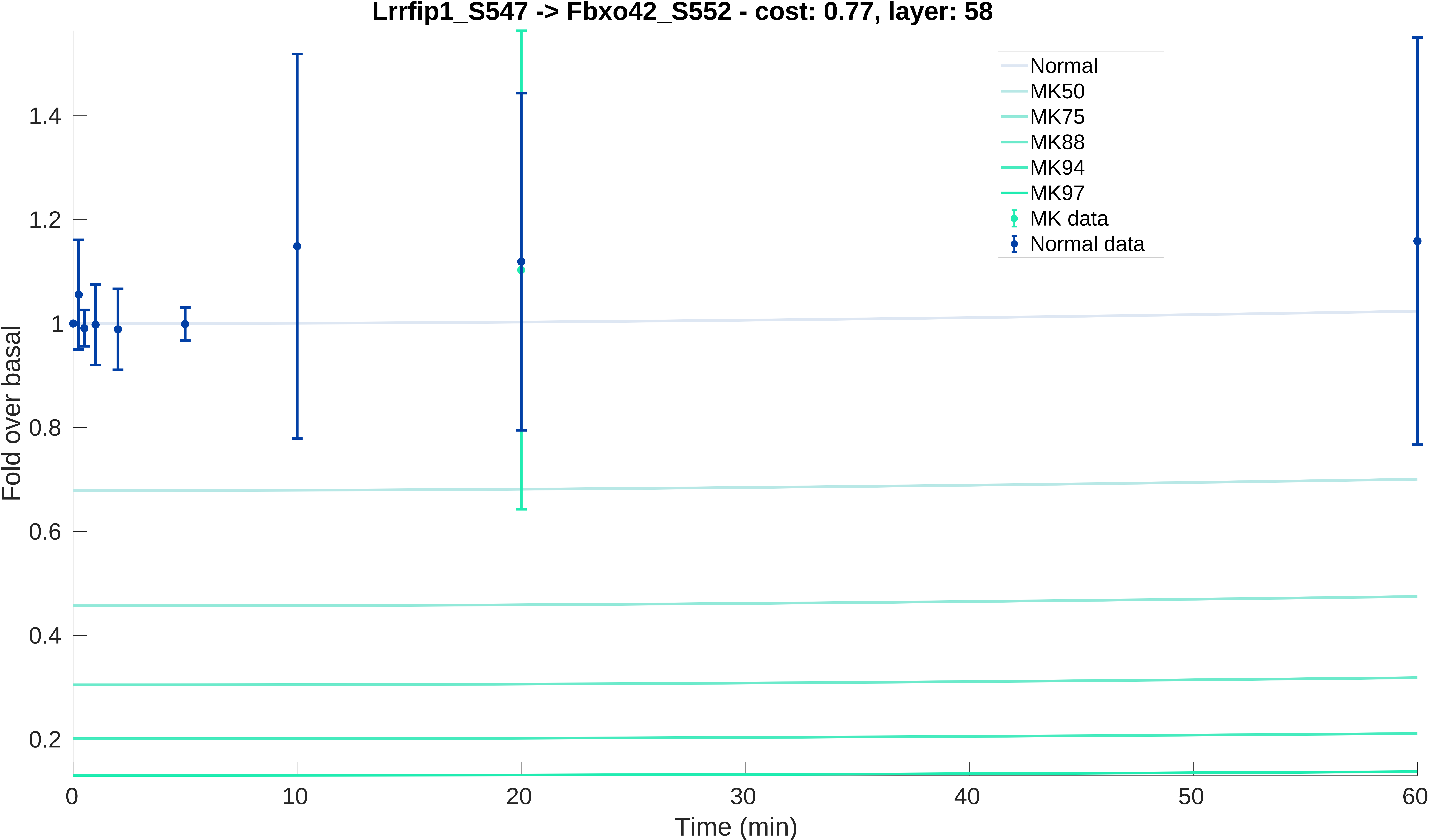

Lrrfip1_S547 -> Fbxo42_S552 - cost: 0.77, layer: 58
Normal
MK50
MK75
MK88
MK94
MK97
MK data
Normal data
Fold over basal
1.4
1.2
1
0.8
0.6
0.4
0.2
0
10
20
30
40
50
60
Time (min)

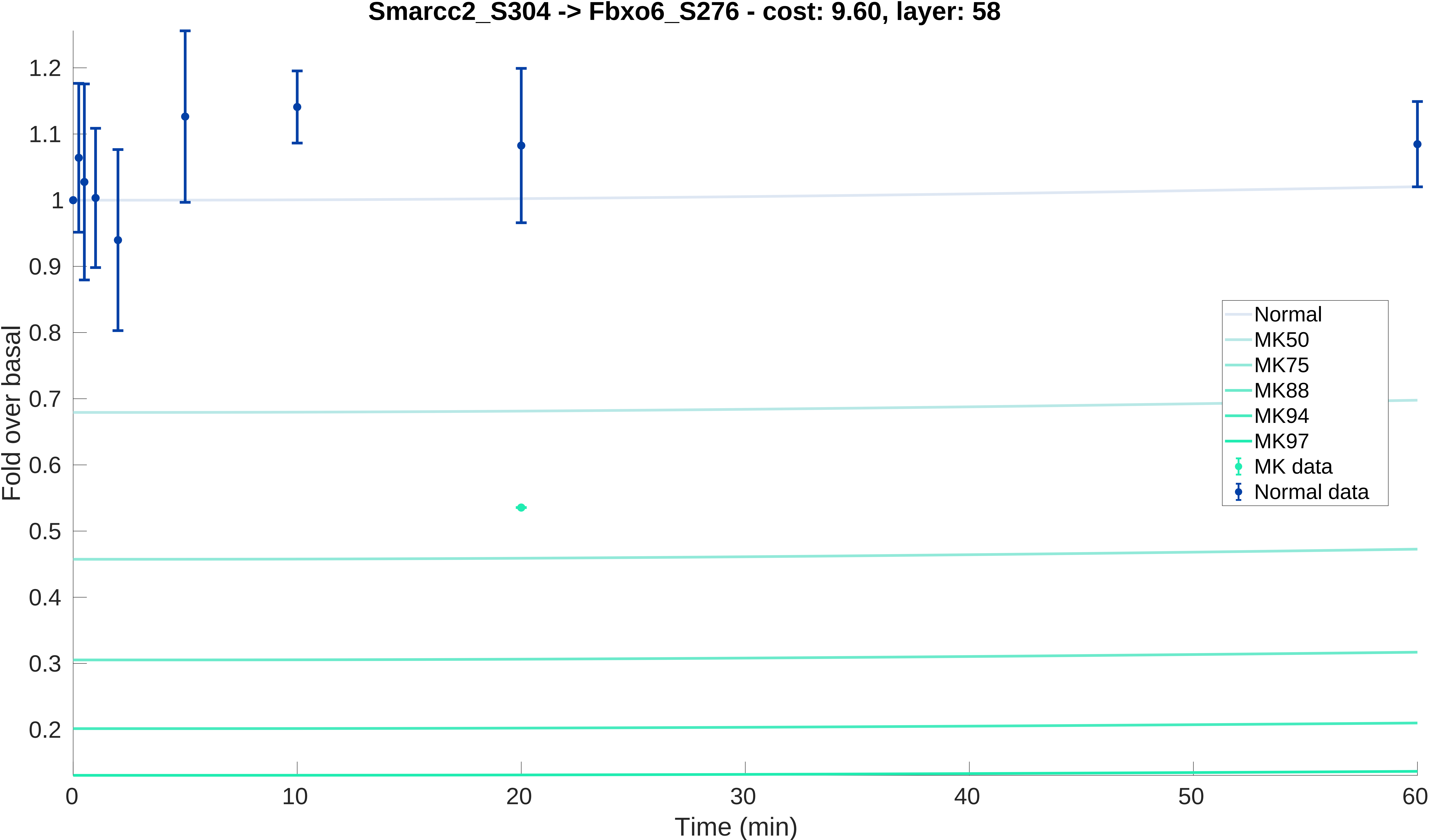

Smarcc2_S304 -> Fbxo6_S276 - cost: 9.60, layer: 58
Fold over basal
Time (min)
0
10
20
30
40
50
60
0.2
0.3
0.4
0.5
0.6
0.7
0.8
0.9
1
1.1
1.2
Normal
MK50
MK75
MK88
MK94
MK97
MK data
Normal data

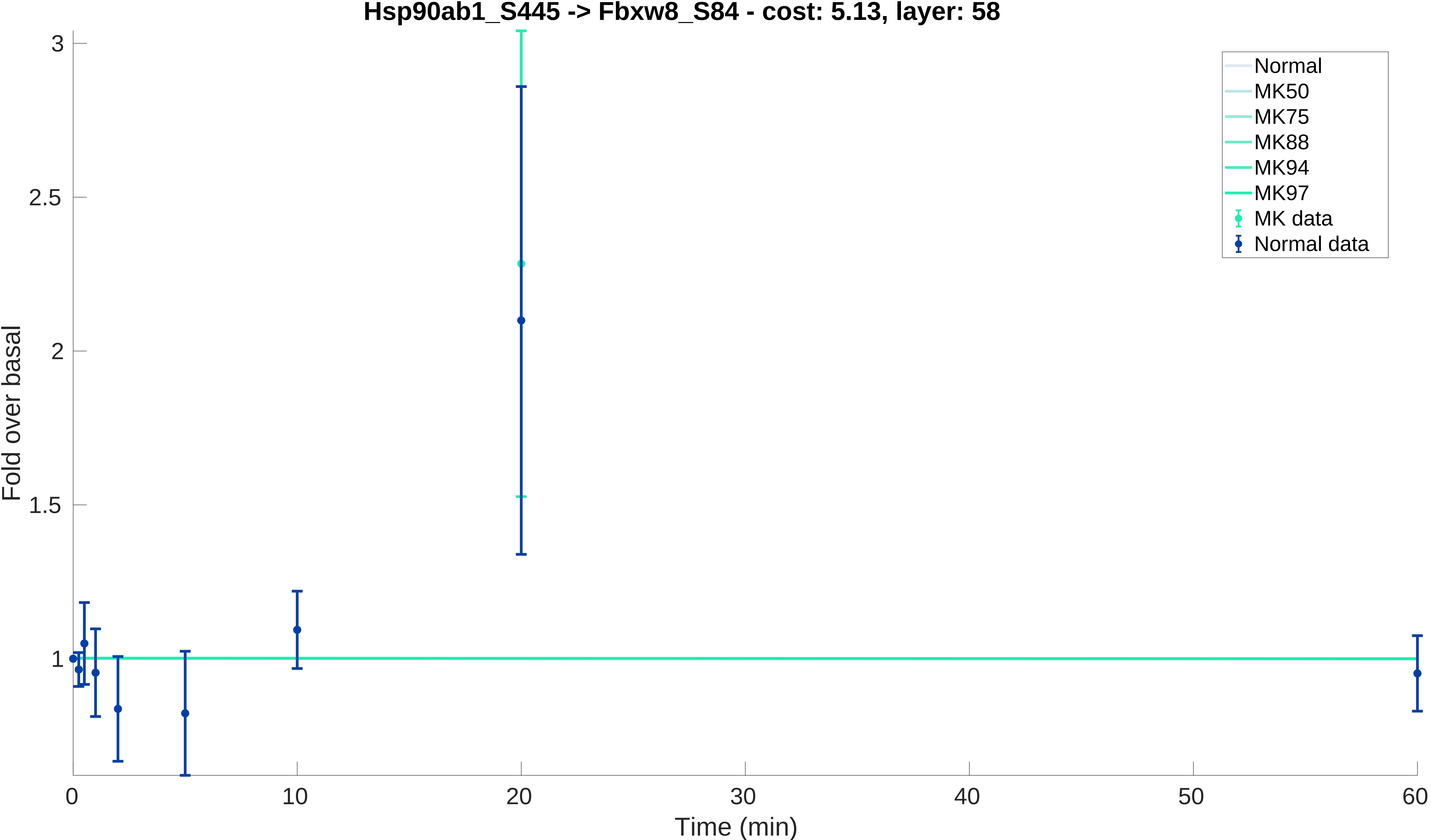

Hsp90ab1_S445 -> Fbxw8_S84 - cost: 5.13, layer: 58
Fold over basal
Time (min)
3
2.5
2
1.5
1
0
10
20
30
40
50
60
Normal
MK50
MK75
MK88
MK94
MK97
MK data
Normal data

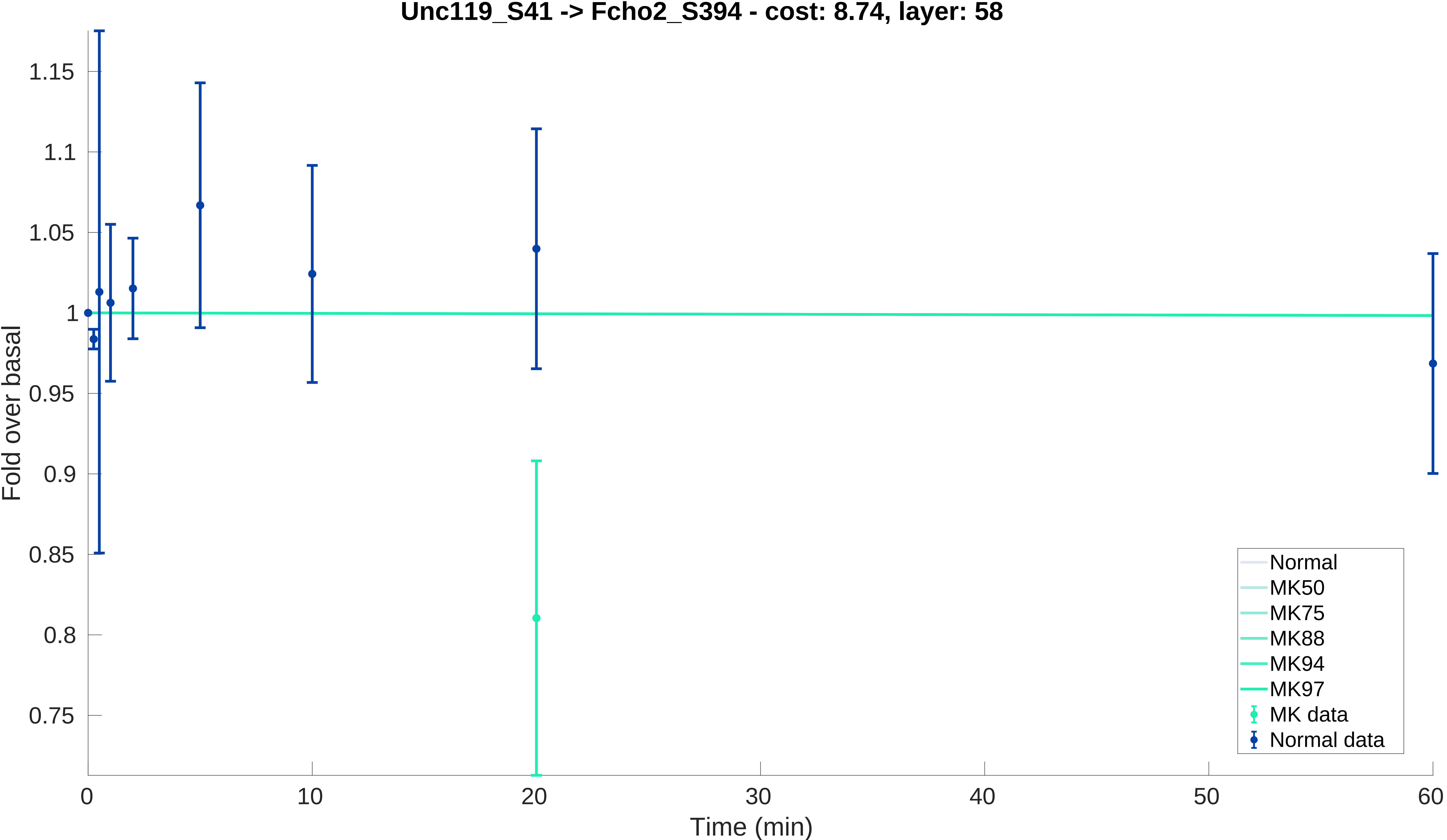

Unc119_S41 -> Fcho2_S394 - cost: 8.74, layer: 58
Fold over basal
1.15
1.1
1.05
1
0.95
0.9
0.85
0.8
0.75
0
10
20
30
40
50
60
Time (min)
Normal
MK50
MK75
MK88
MK94
MK97
MK data
Normal data

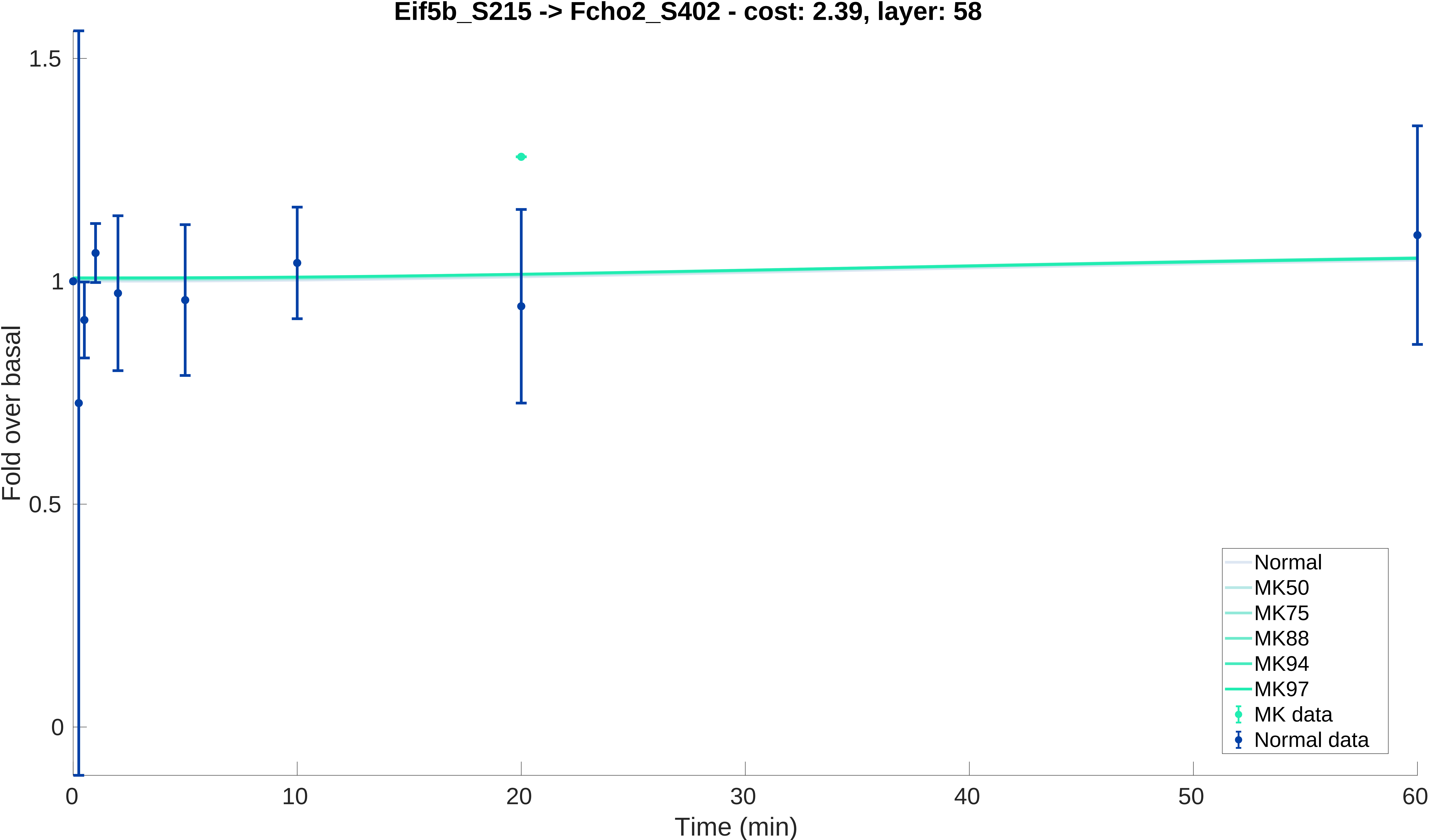

Eif5b_S215 -> Fcho2_S402 - cost: 2.39, layer: 58
Fold over basal
1.5
1
0.5
0
Time (min)
0
10
20
30
40
50
60
Normal
MK50
MK75
MK88
MK94
MK97
MK data
Normal data

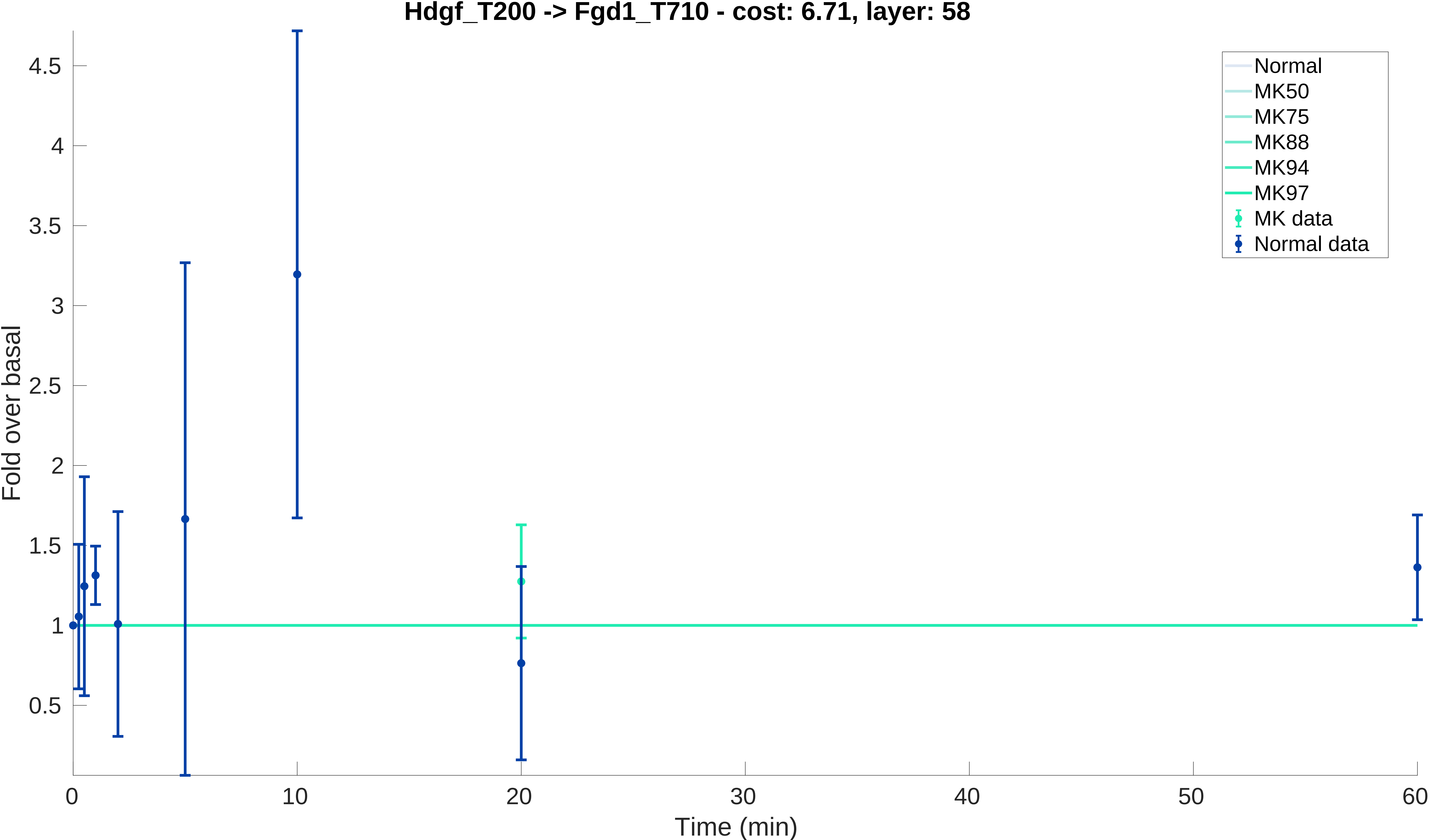

Hdgf_T200 -> Fgd1_T710 - cost: 6.71, layer: 58
Fold over basal
Time (min)
Normal
MK50
MK75
MK88
MK94
MK97
MK data
Normal data
0
10
20
30
40
50
60
0.5
1
1.5
2
2.5
3
3.5
4
4.5

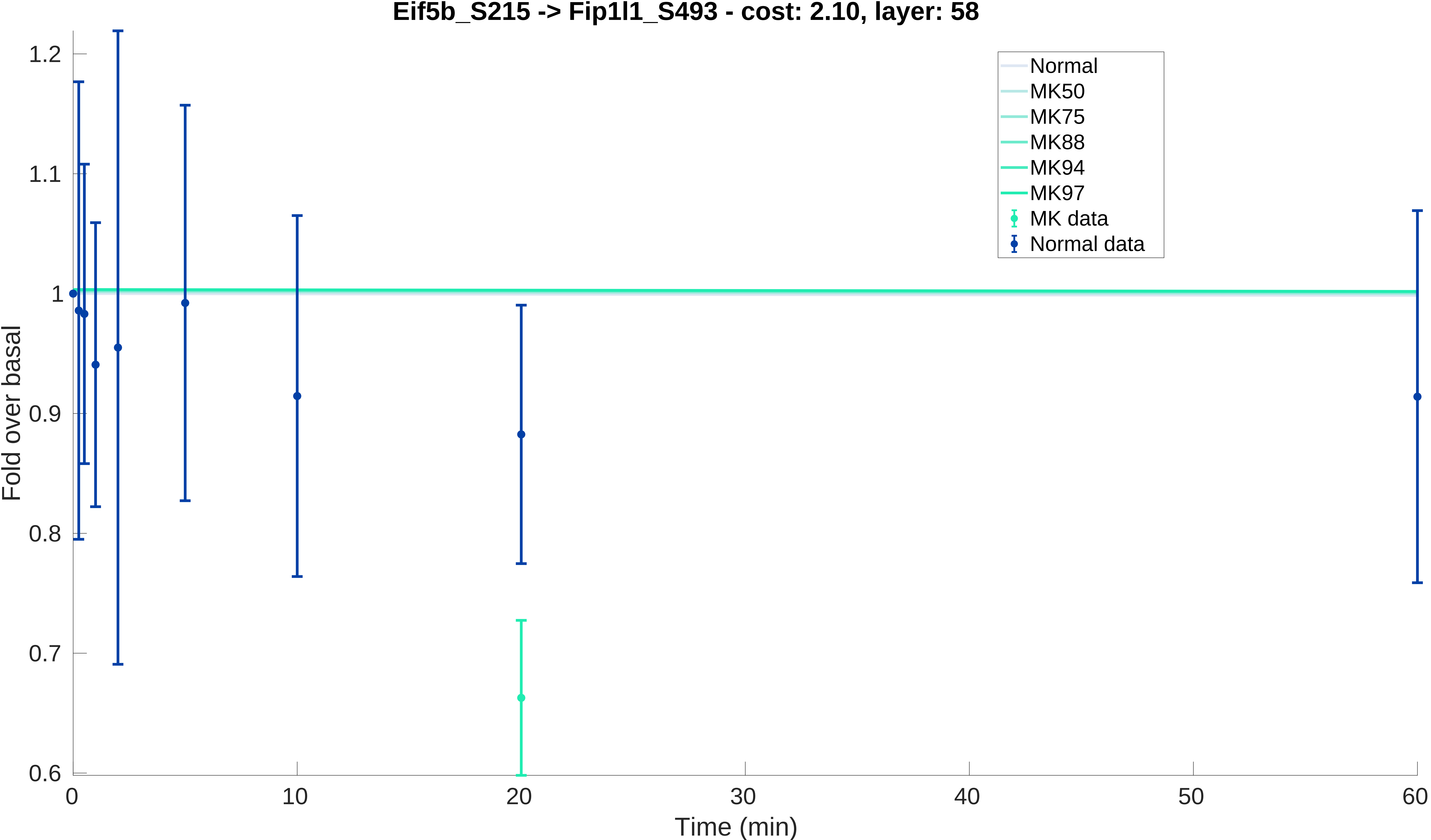

Eif5b_S215 -> Fip1l1_S493 - cost: 2.10, layer: 58
Fold over basal
Time (min)
Normal
MK50
MK75
MK88
MK94
MK97
MK data
Normal data
1.2
1.1
1
0.9
0.8
0.7
0.6
0
10
20
30
40
50
60

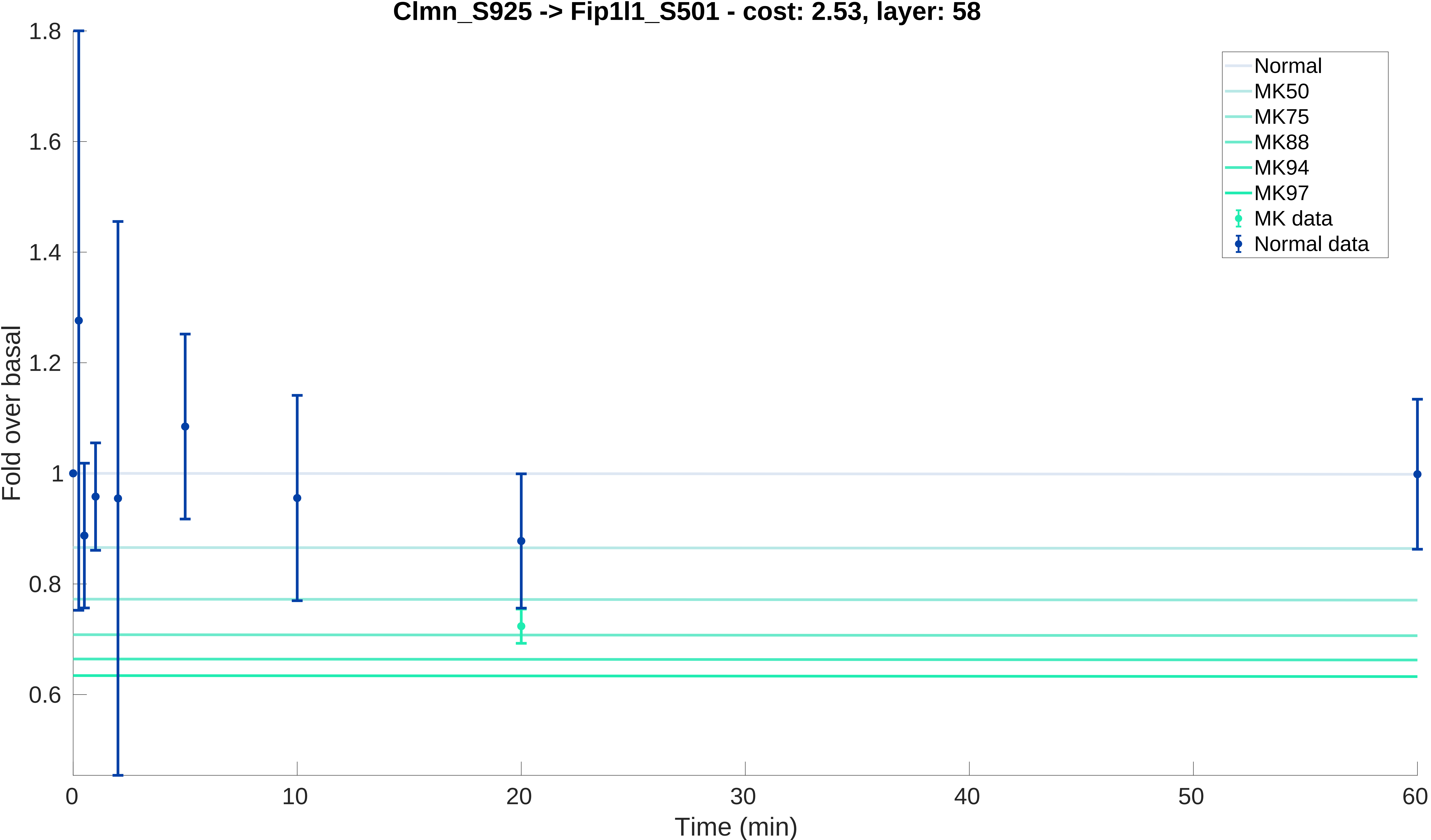

Clmn_S925 -> Fip1l1_S501 - cost: 2.53, layer: 58
Fold over basal
Time (min)
1.8
1.6
1.4
1.2
1
0.8
0.6
0
10
20
30
40
50
60
Normal
MK50
MK75
MK88
MK94
MK97
MK data
Normal data

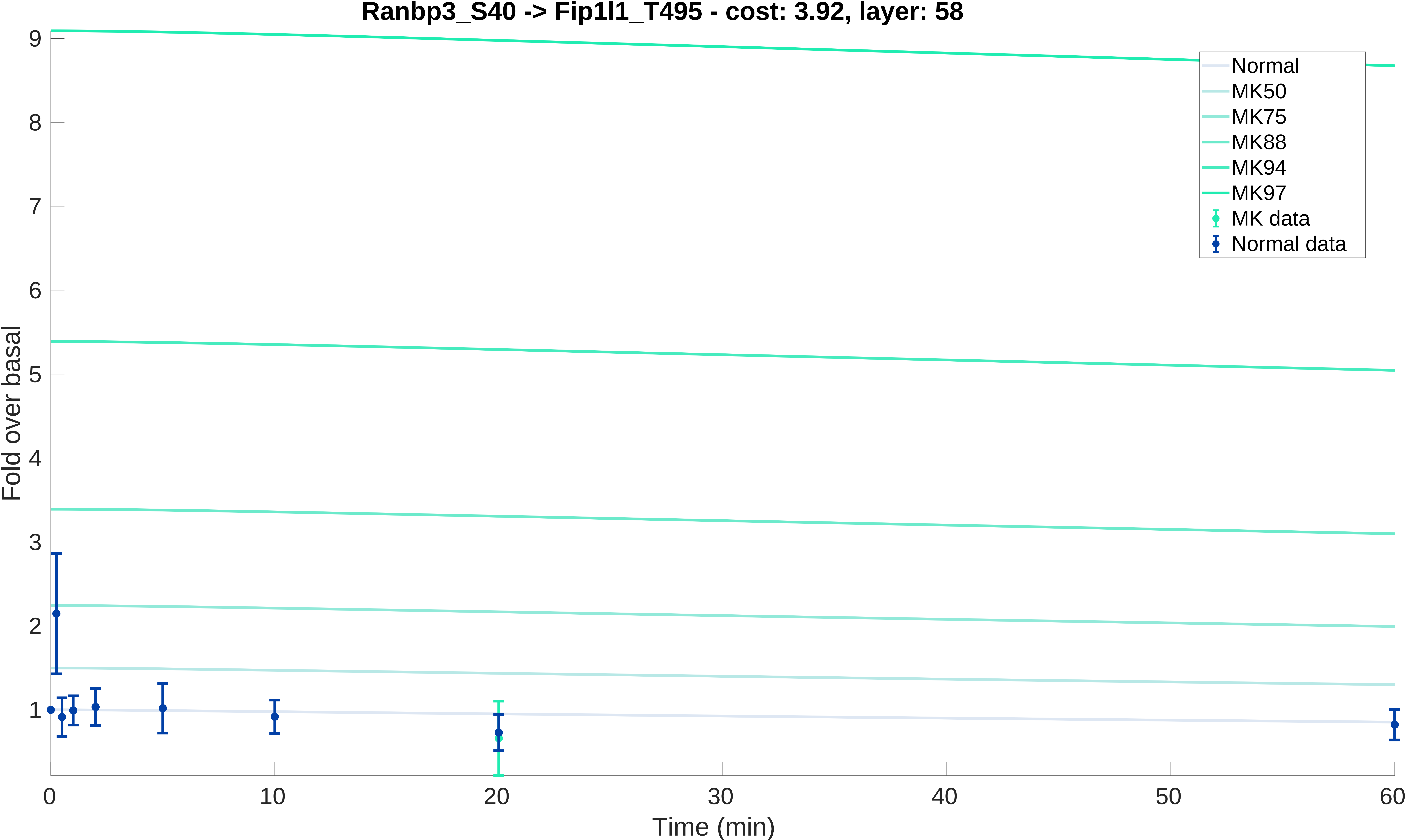

Ranbp3_S40 -> Fip1l1_T495 - cost: 3.92, layer: 58
Fold over basal
Time (min)
0
10
20
30
40
50
60
1
2
3
4
5
6
7
8
9
Normal
MK50
MK75
MK88
MK94
MK97
MK data
Normal data

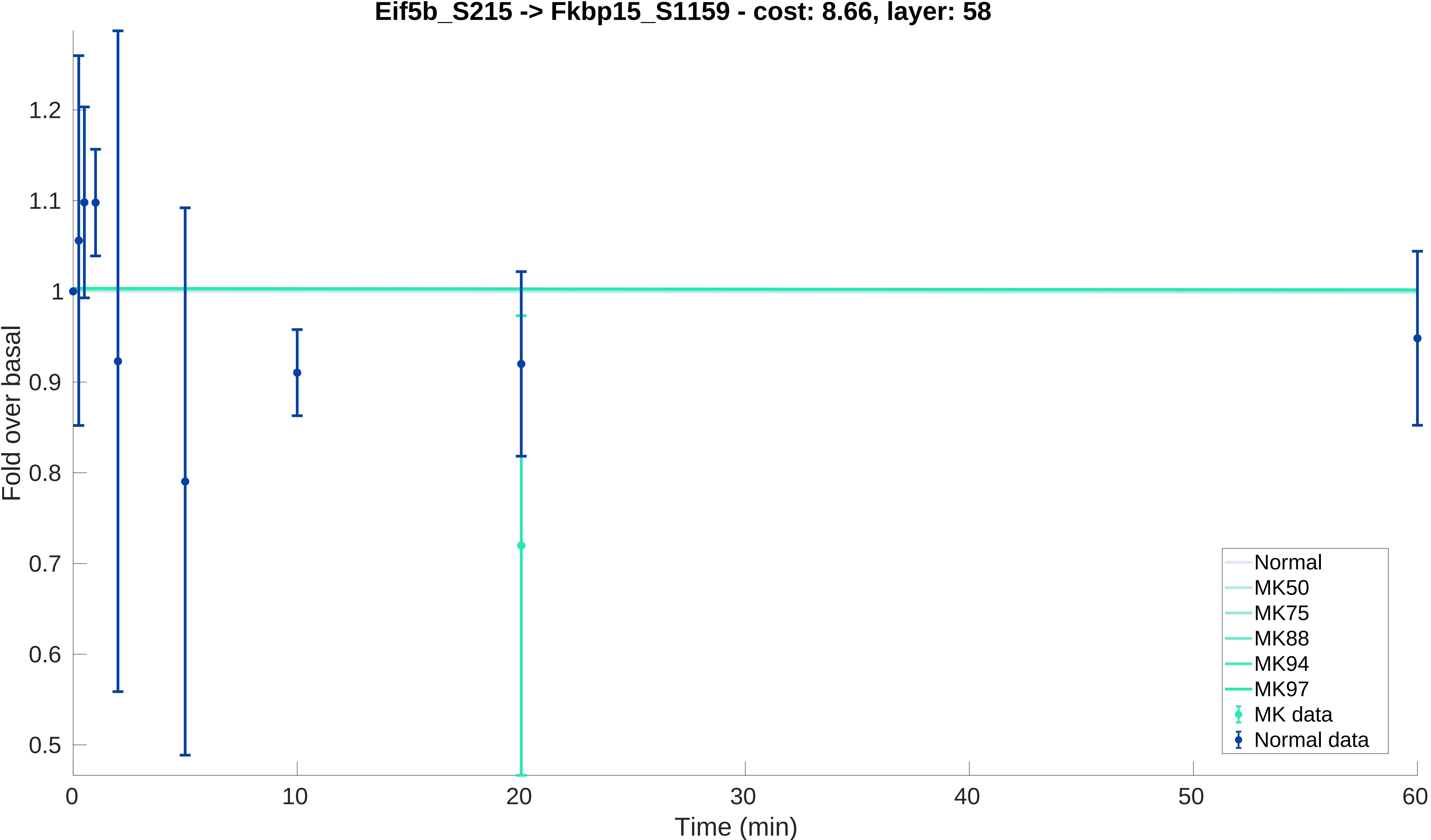

Eif5b_S215 -> Fkbp15_S1159 - cost: 8.66, layer: 58
Fold over basal
Time (min)
1.2
1.1
1
0.9
0.8
0.7
0.6
0.5
0
10
20
30
40
50
60
Normal
MK50
MK75
MK88
MK94
MK97
MK data
Normal data

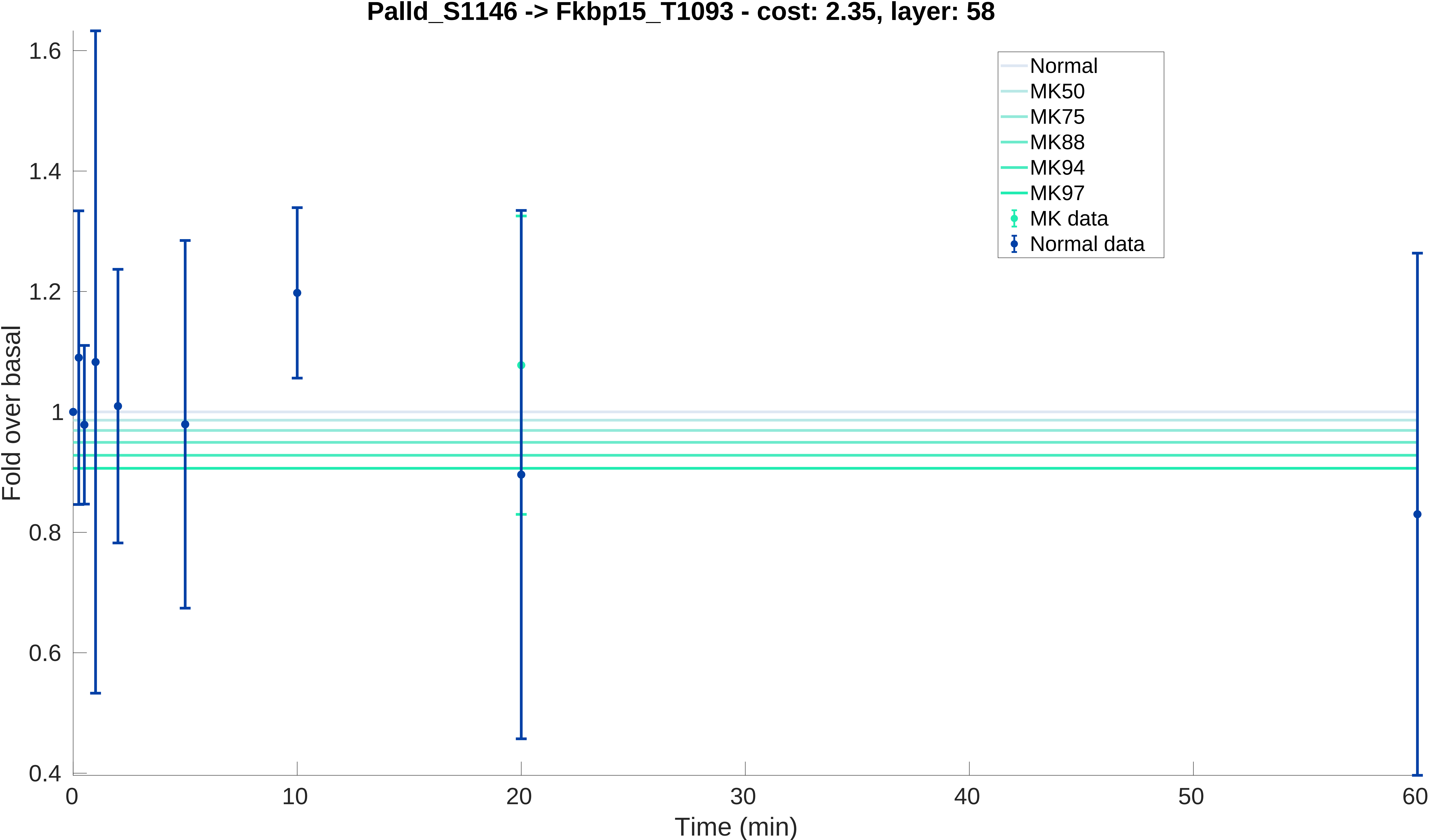

Palld_S1146 -> Fkbp15_T1093 - cost: 2.35, layer: 58
Fold over basal
Time (min)
Normal
MK50
MK75
MK88
MK94
MK97
MK data
Normal data
1.6
1.4
1.2
1
0.8
0.6
0.4
0
10
20
30
40
50
60

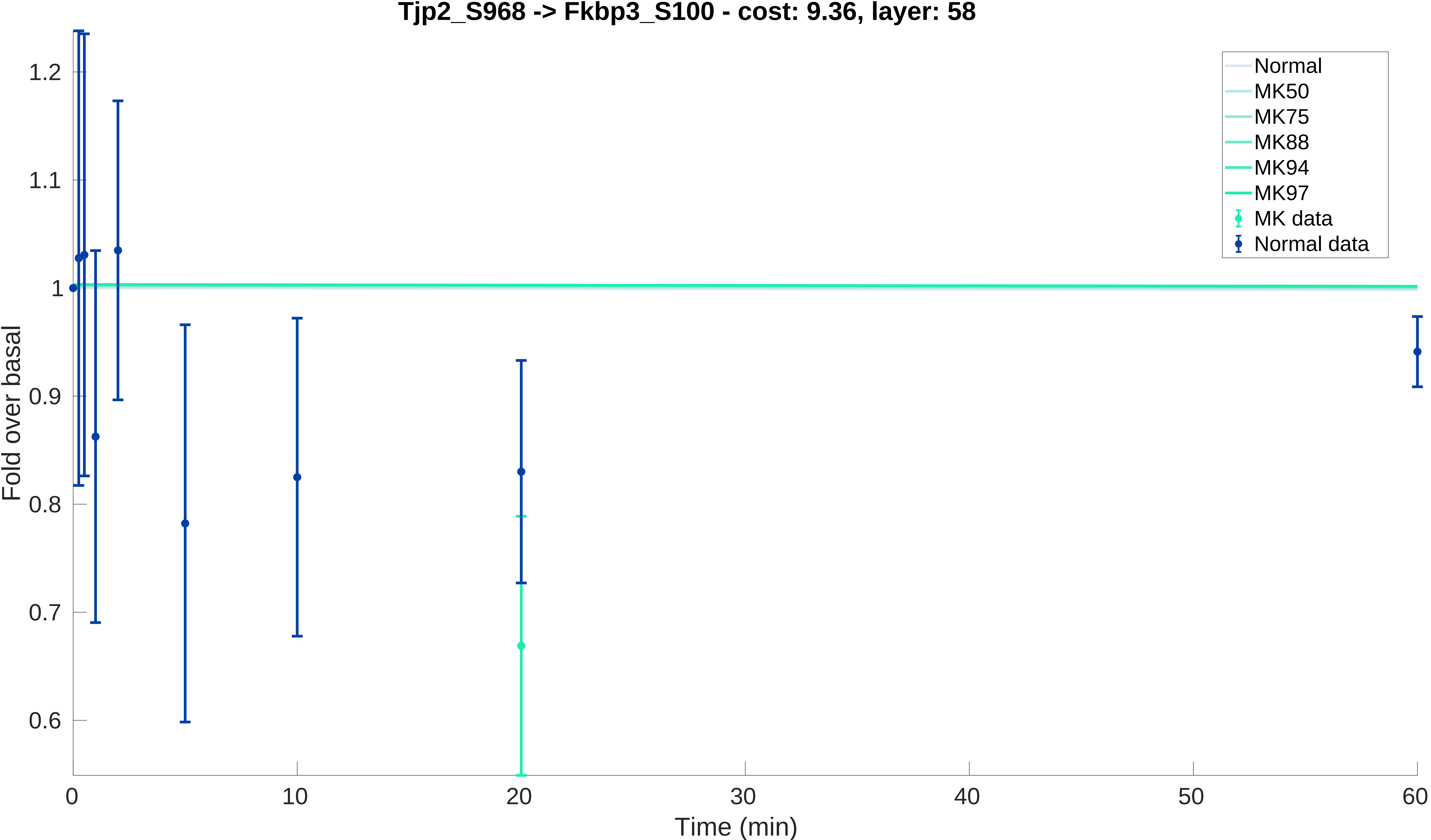

Tjp2_S968 -> Fkbp3_S100 - cost: 9.36, layer: 58
Normal
MK50
MK75
MK88
MK94
MK97
MK data
Normal data
Fold over basal
1.2
1.1
1
0.9
0.8
0.7
0.6
0
10
20
30
40
50
60
Time (min)

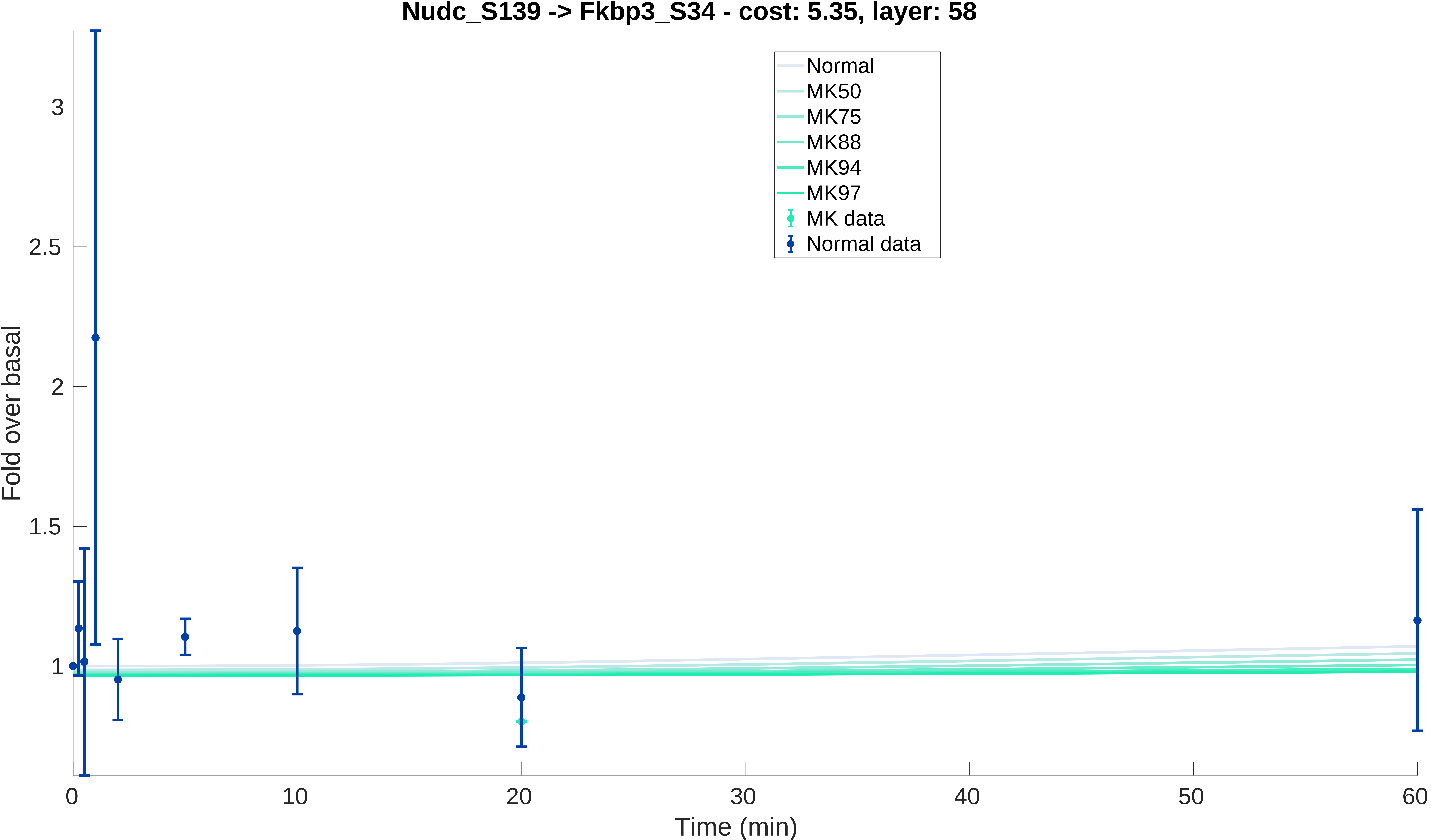
Nudc_S139 -> Fkbp3_S34 - cost: 5.35, layer: 58
Normal
MK50
MK75
MK88
MK94
MK97
MK data
Normal data
Fold over basal
3
2.5
2
1.5
1
0
10
20
30
40
50
60
Time (min)

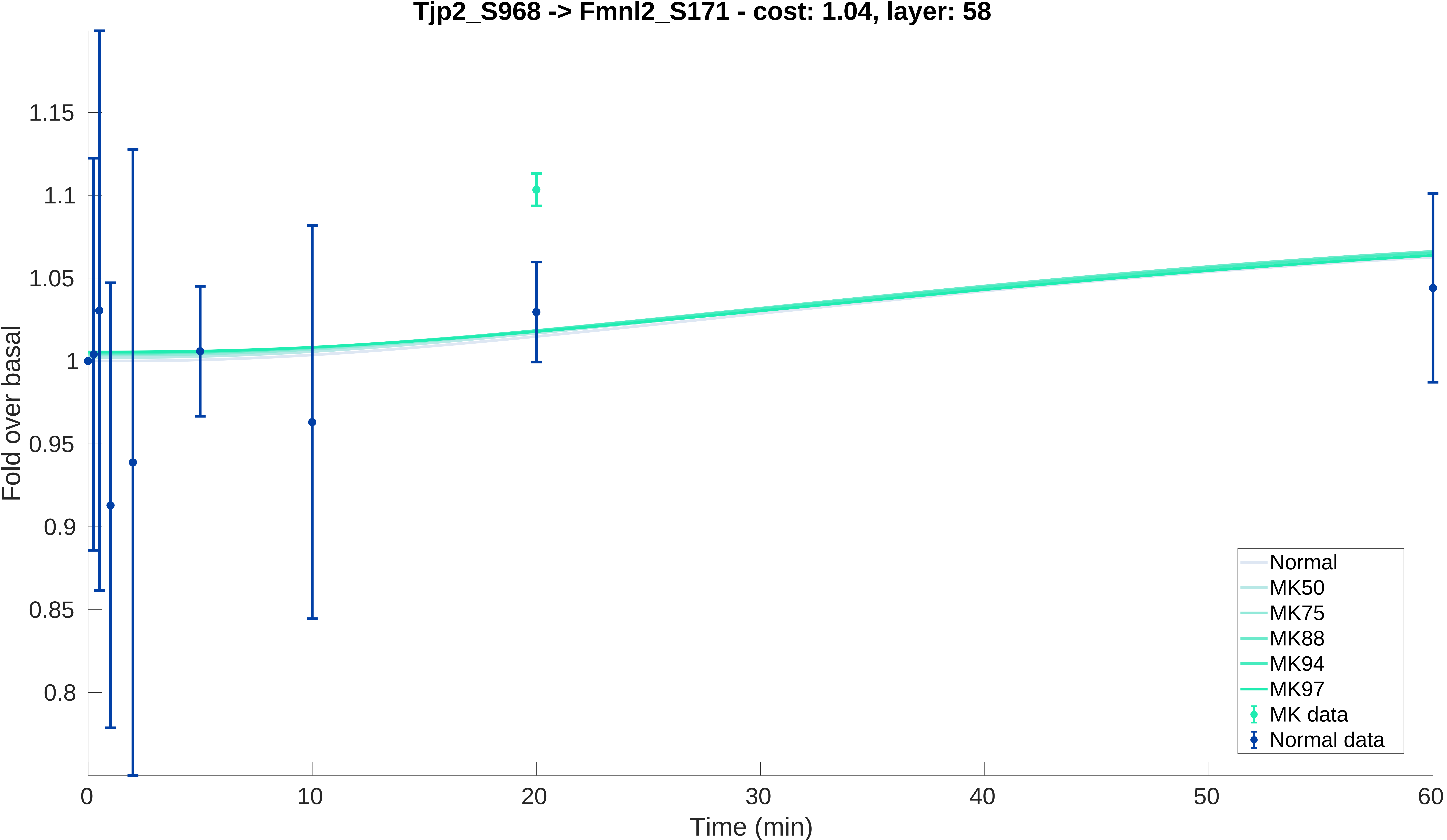

Tjp2_S968 -> Fmnl2_S171 - cost: 1.04, layer: 58
Fold over basal
1.15
1.1
1.05
1
0.95
0.9
0.85
0.8
0
10
20
30
40
50
60
Time (min)
Normal
MK50
MK75
MK88
MK94
MK97
MK data
Normal data

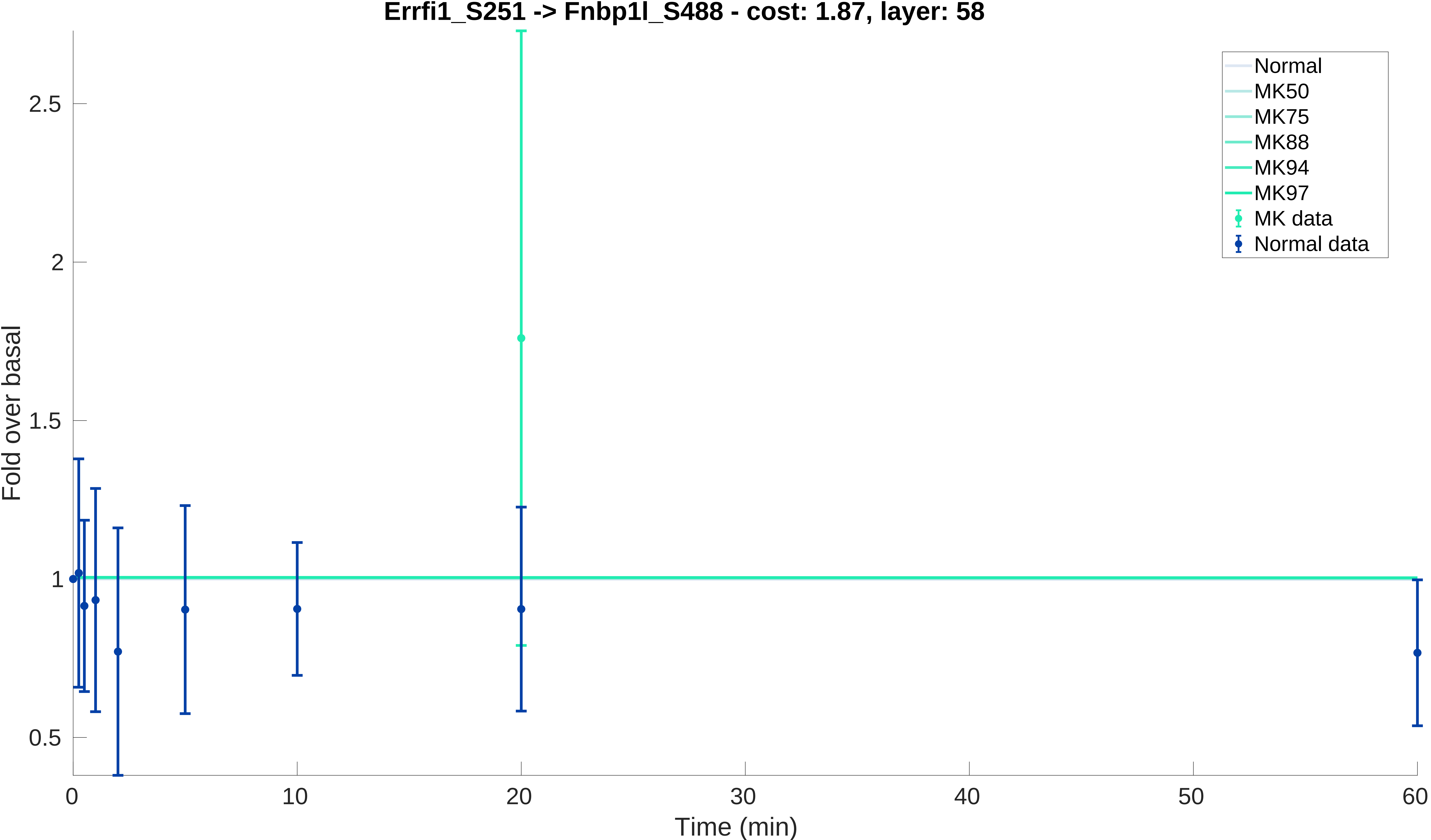

Errfi1_S251 -> Fnbp1l_S488 - cost: 1.87, layer: 58
Fold over basal
Time (min)
0.5
1
1.5
2
2.5
0
10
20
30
40
50
60
Normal
MK50
MK75
MK88
MK94
MK97
MK data
Normal data

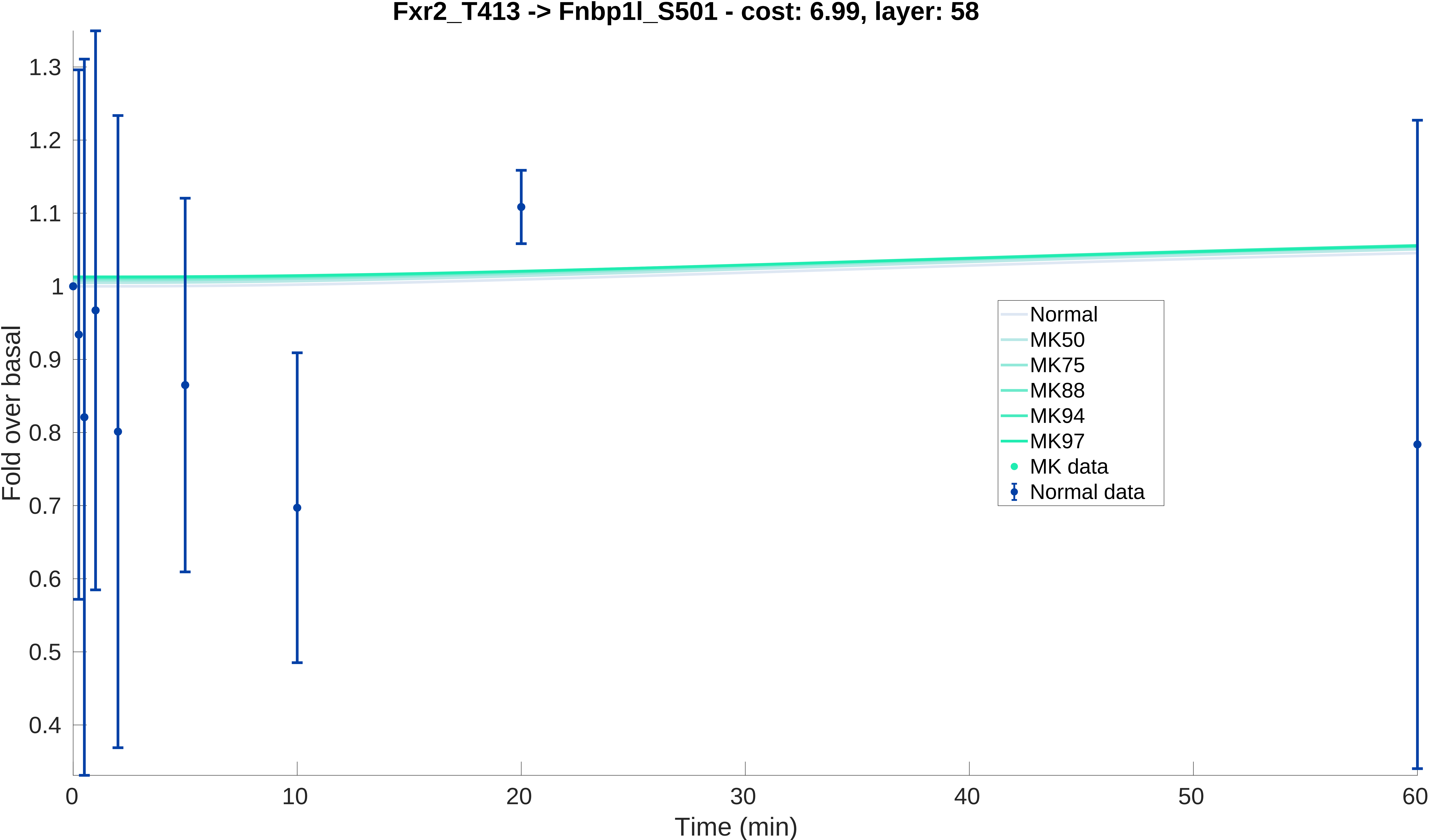

Fxr2_T413 -> Fnbp1l_S501 - cost: 6.99, layer: 58
Fold over basal
1.3
1.2
1.1
1
0.9
0.8
0.7
0.6
0.5
0.4
0
10
20
30
40
50
60
Time (min)
Normal
MK50
MK75
MK88
MK94
MK97
MK data
Normal data

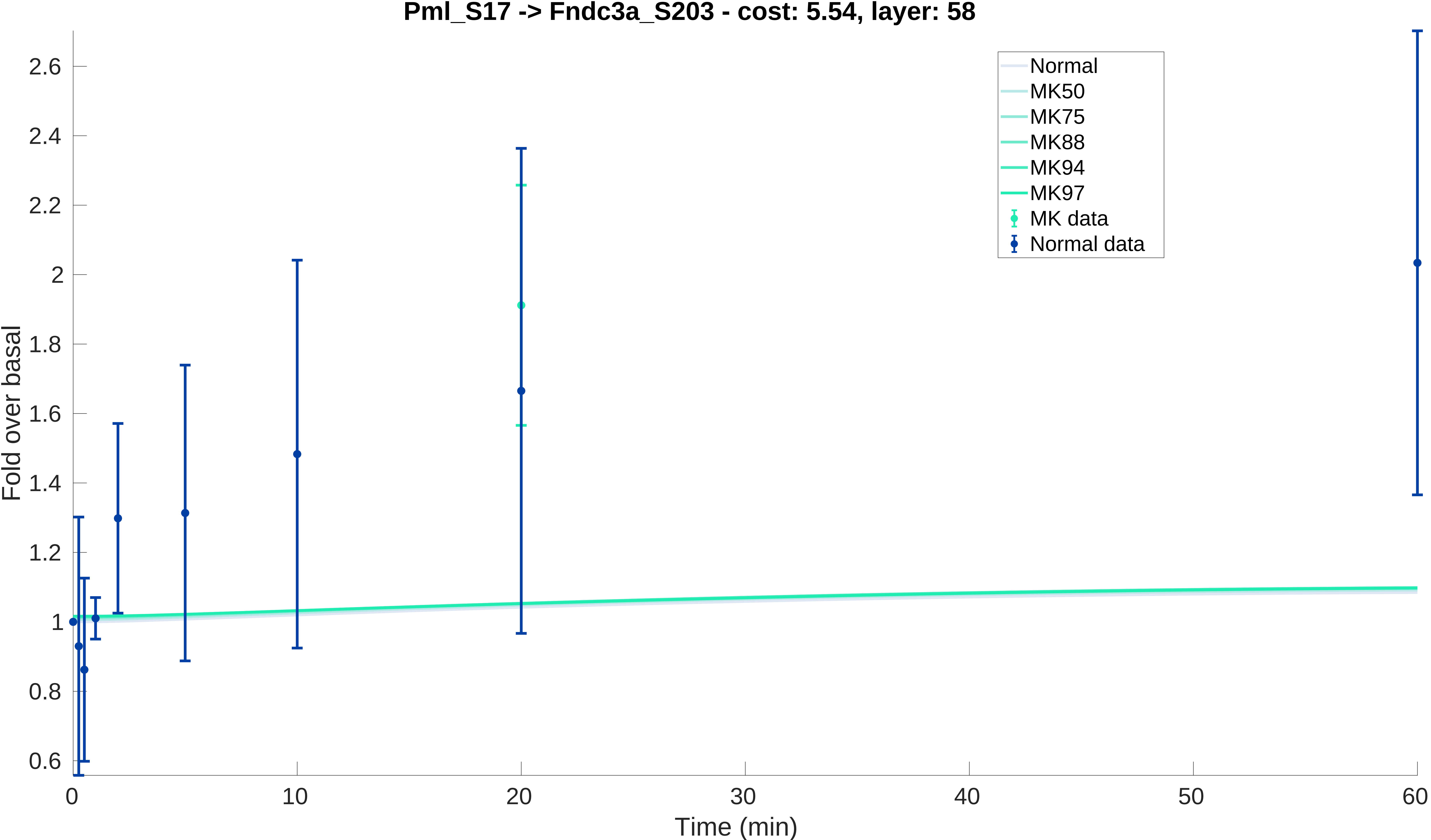

Pml_S17 -> Fndc3a_S203 - cost: 5.54, layer: 58
Normal
MK50
MK75
MK88
MK94
MK97
MK data
Normal data
Fold over basal
2.6
2.4
2.2
2
1.8
1.6
1.4
1.2
1
0.8
0.6
0
10
20
30
40
50
60
Time (min)

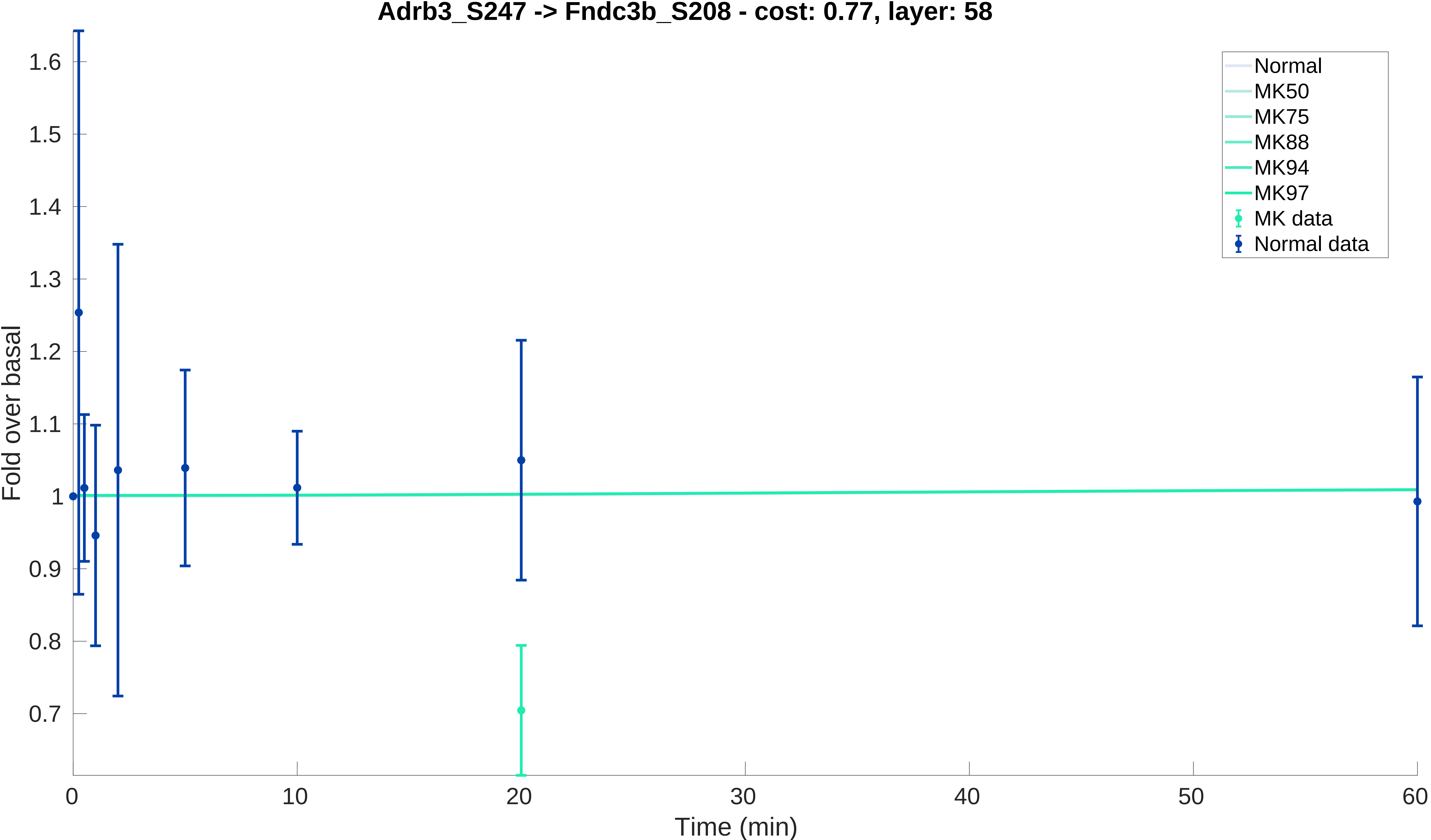

Adrb3_S247 -> Fndc3b_S208 - cost: 0.77, layer: 58
Fold over basal
1.6
1.5
1.4
1.3
1.2
1.1
1
0.9
0.8
0.7
0
10
20
30
40
50
60
Time (min)
Normal
MK50
MK75
MK88
MK94
MK97
MK data
Normal data

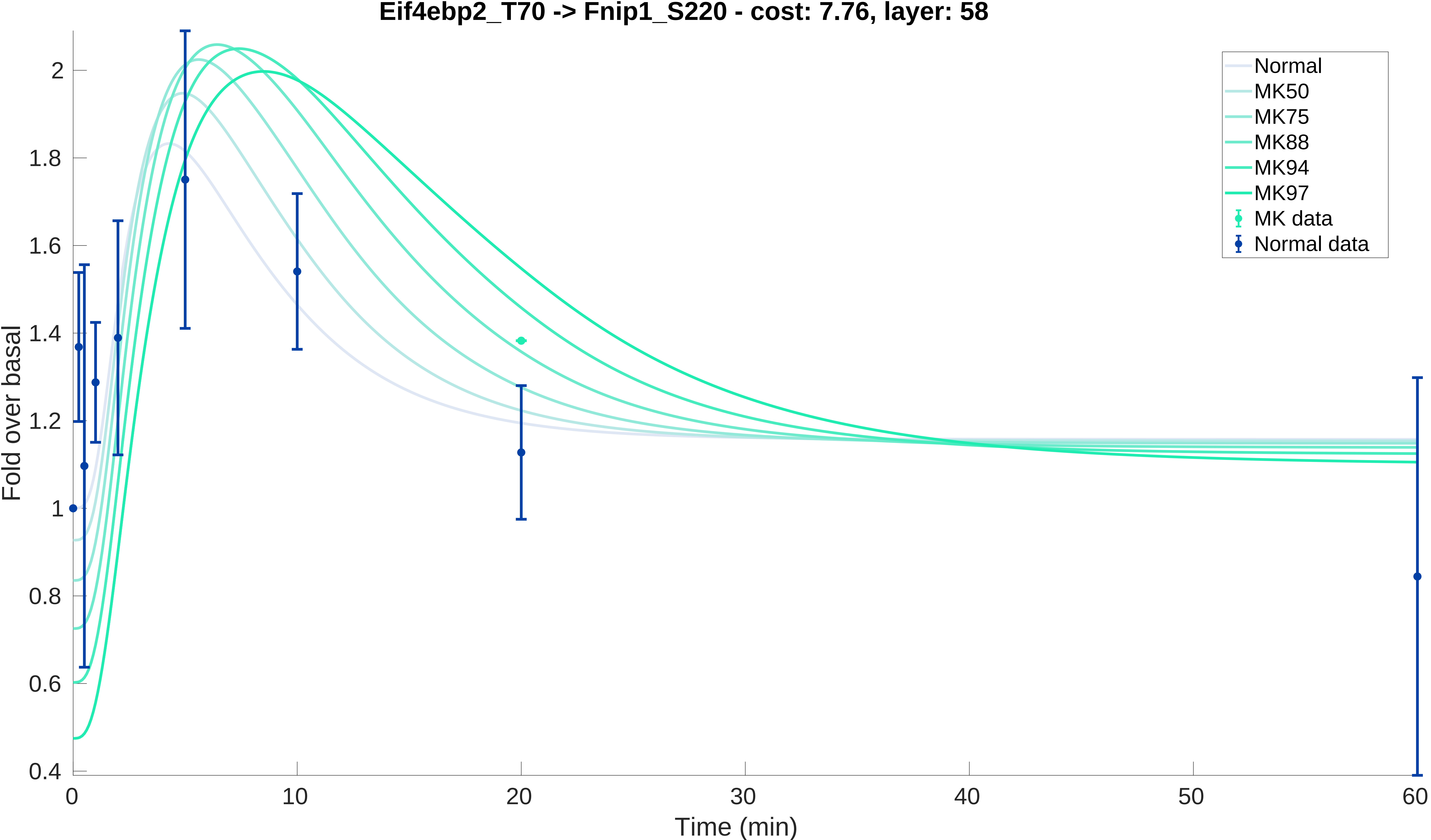
Eif4ebp2_T70 -> Fnip1_S220 - cost: 7.76, layer: 58
Fold over basal
Time (min)
0.4
0.6
0.8
1
1.2
1.4
1.6
1.8
2
0
10
20
30
40
50
60
Normal
MK50
MK75
MK88
MK94
MK97
MK data
Normal data

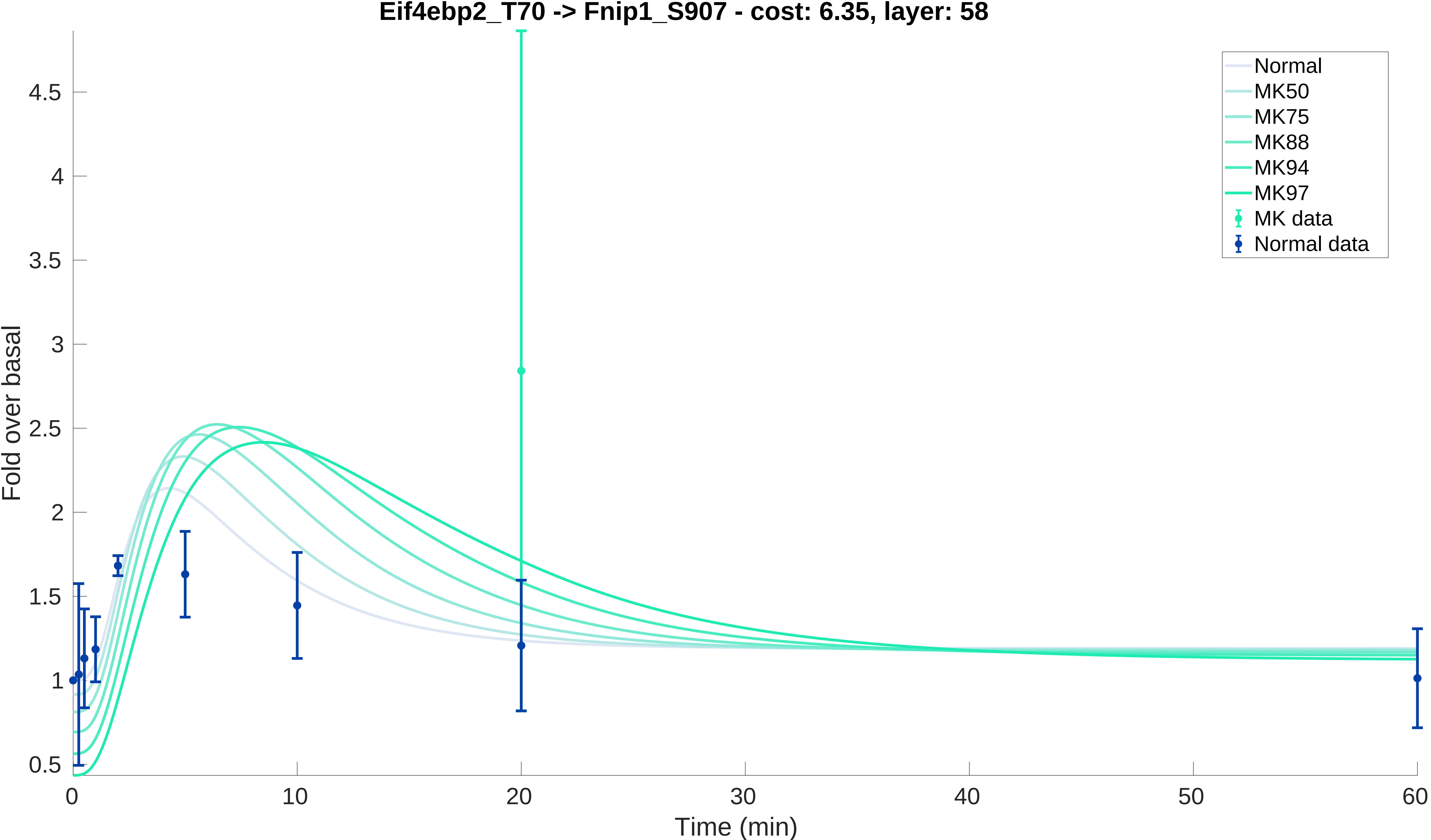

Eif4ebp2_T70 -> Fnip1_S907 - cost: 6.35, layer: 58
Fold over basal
Time (min)
0.5
1
1.5
2
2.5
3
3.5
4
4.5
0
10
20
30
40
50
60
Normal
MK50
MK75
MK88
MK94
MK97
MK data
Normal data

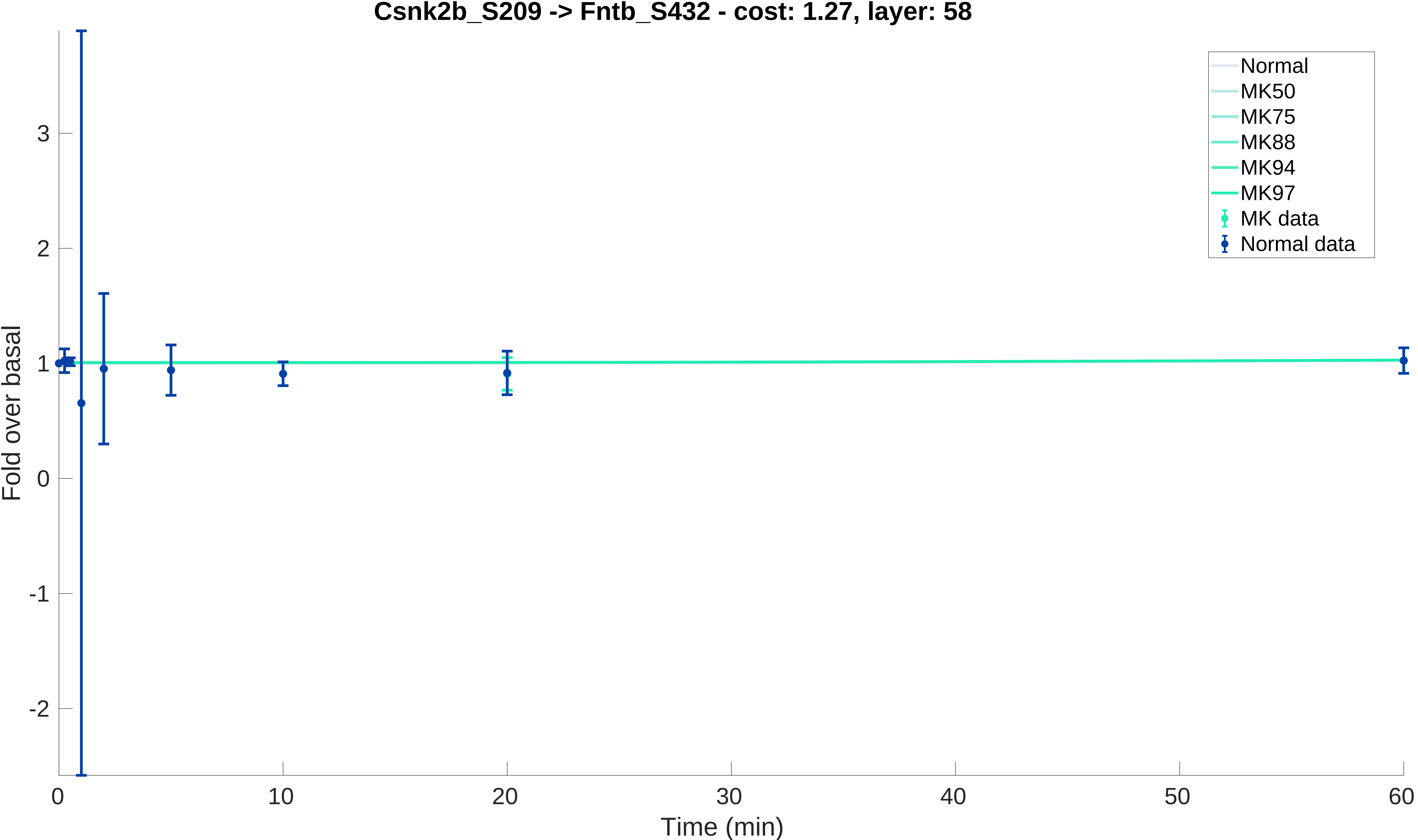

Csnk2b_S209 -> Fntb_S432 - cost: 1.27, layer: 58
Normal
MK50
MK75
MK88
MK94
MK97
MK data
Normal data
Fold over basal
3
2
1
0
-1
-2
0
10
20
30
40
50
60
Time (min)

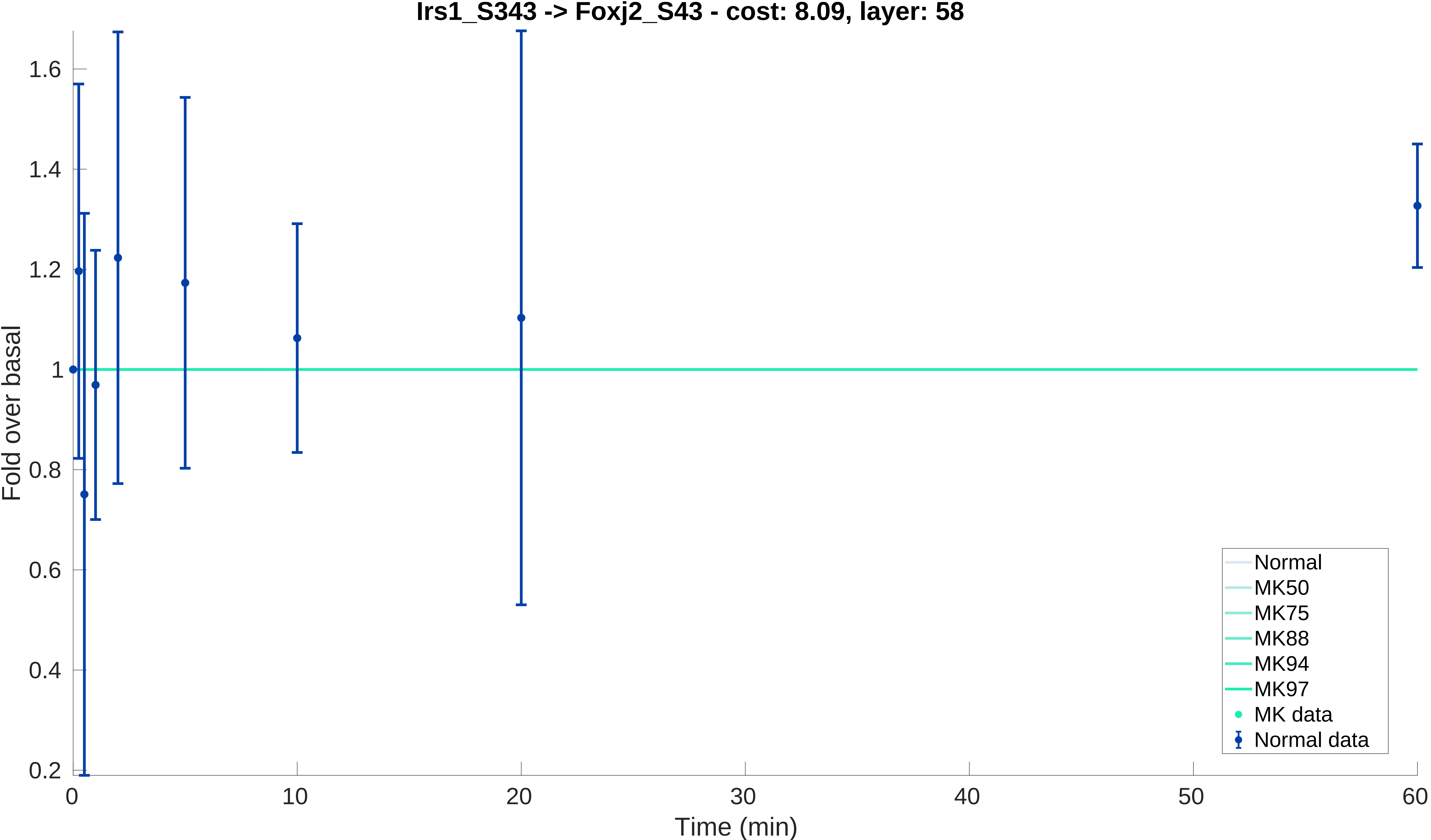

Irs1_S343 -> Foxj2_S43 - cost: 8.09, layer: 58
Fold over basal
Time (min)
0
10
20
30
40
50
60
0.2
0.4
0.6
0.8
1
1.2
1.4
1.6
Normal
MK50
MK75
MK88
MK94
MK97
MK data
Normal data

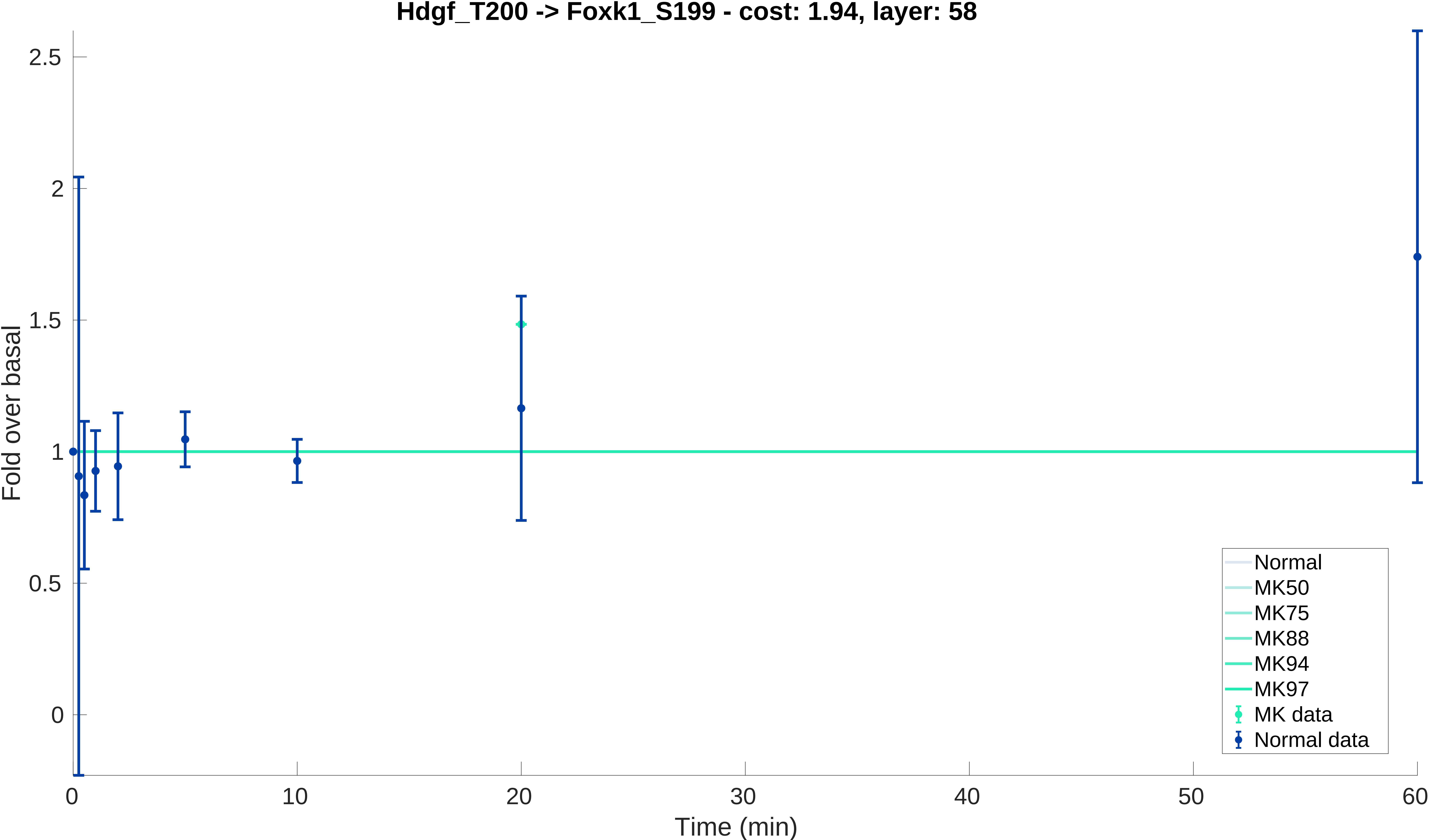

Hdgf_T200 -> Foxk1_S199 - cost: 1.94, layer: 58
Fold over basal
Time (min)
0
0.5
1
1.5
2
2.5
0
10
20
30
40
50
60
Normal
MK50
MK75
MK88
MK94
MK97
MK data
Normal data

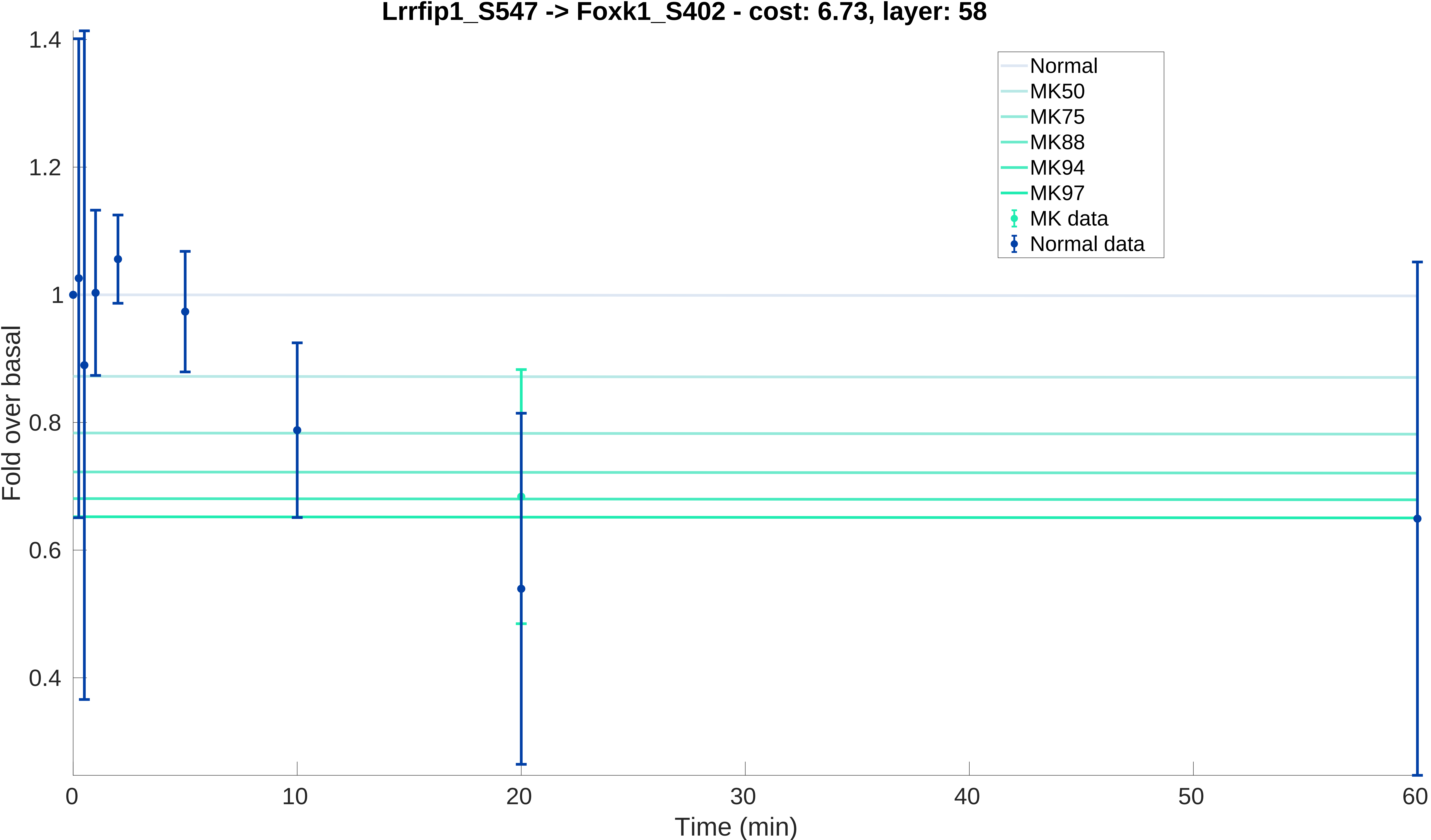

Lrrfip1_S547 -> Foxk1_S402 - cost: 6.73, layer: 58
Fold over basal
Time (min)
Normal
MK50
MK75
MK88
MK94
MK97
MK data
Normal data
1.4
1.2
1
0.8
0.6
0.4
0
10
20
30
40
50
60

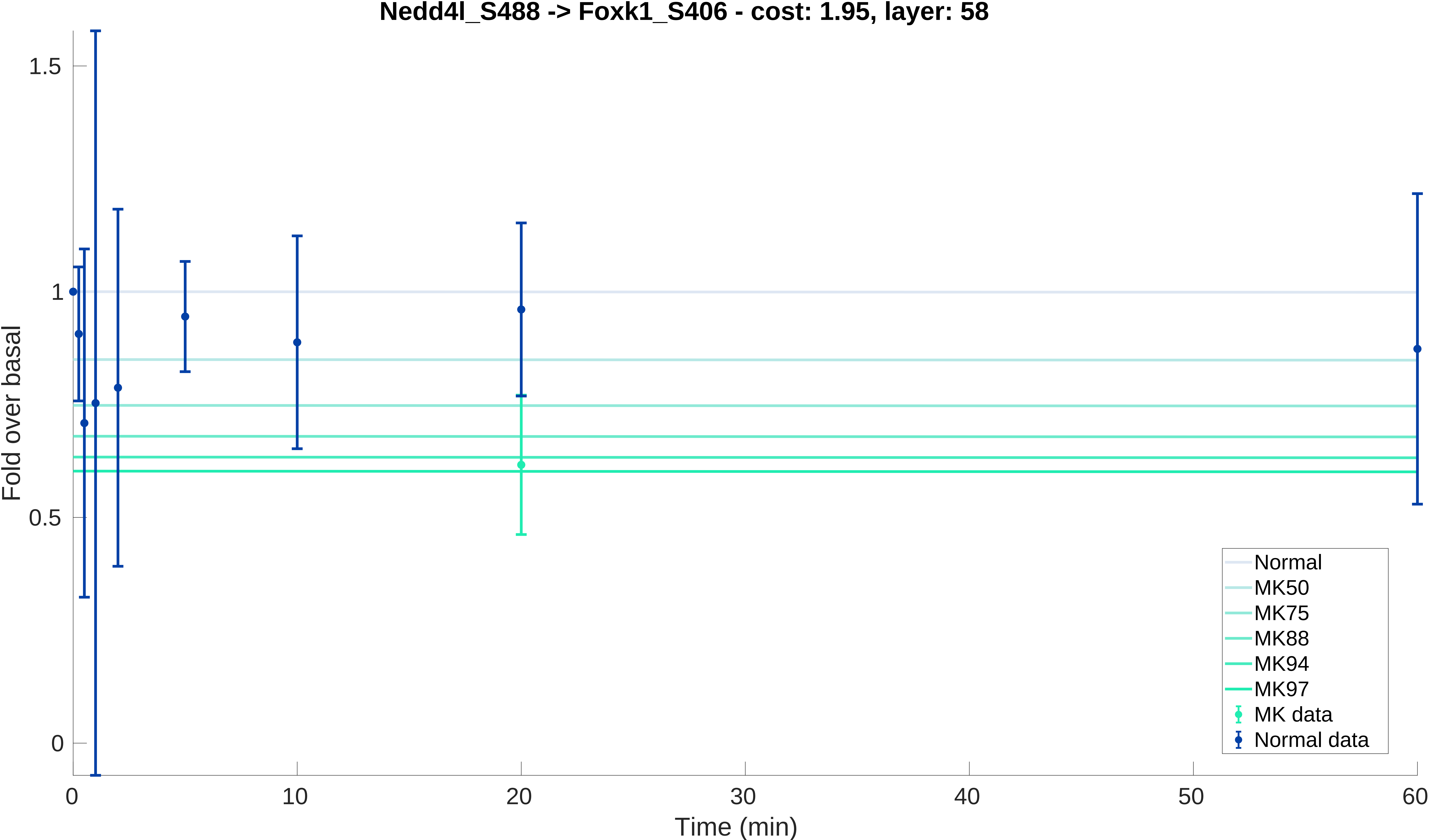

Nedd4l_S488 -> Foxk1_S406 - cost: 1.95, layer: 58
Fold over basal
Time (min)
0
0.5
1
1.5
0
10
20
30
40
50
60
Normal
MK50
MK75
MK88
MK94
MK97
MK data
Normal data

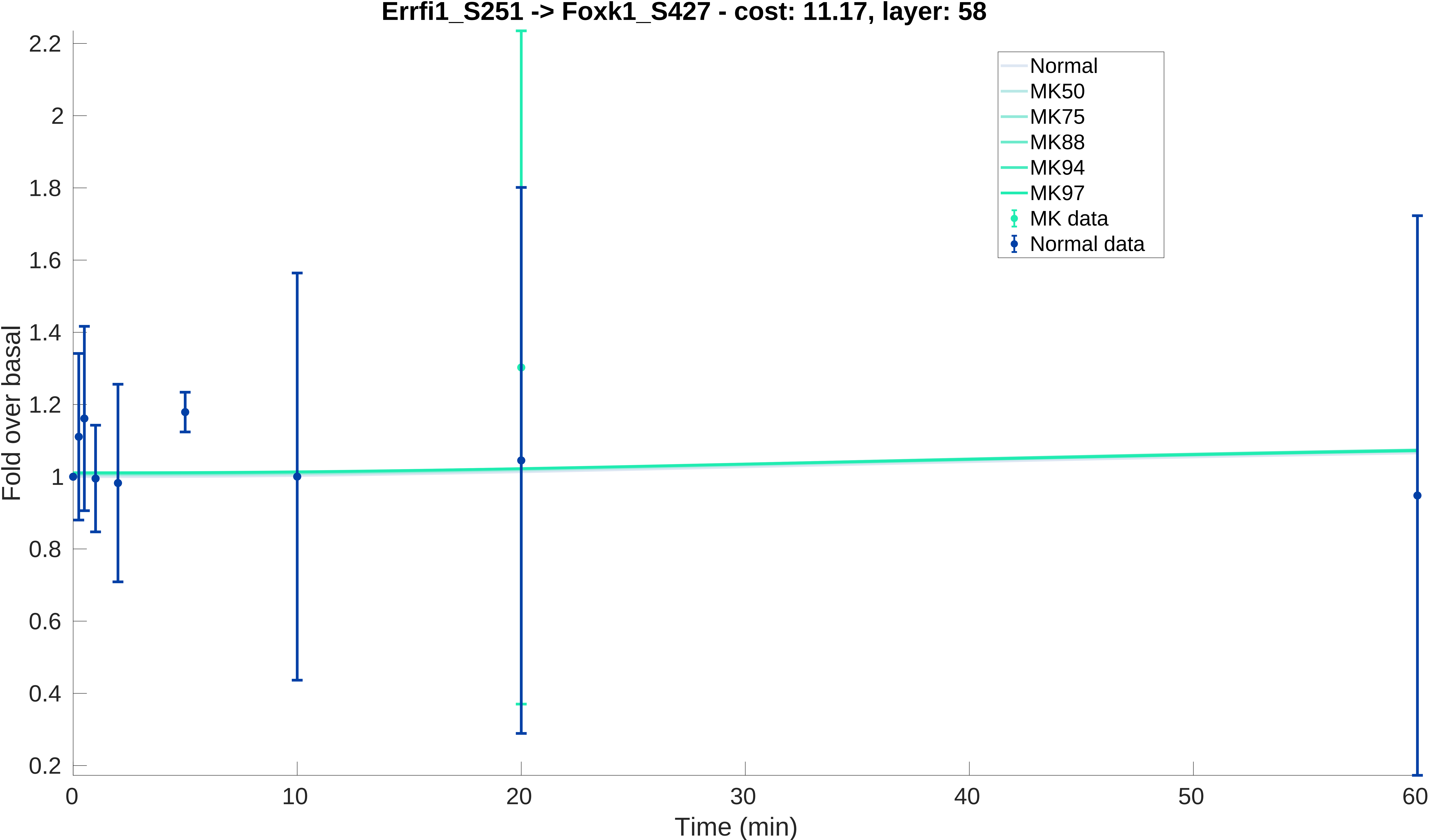

Errfi1_S251 -> Foxk1_S427 - cost: 11.17, layer: 58
Normal
MK50
MK75
MK88
MK94
MK97
MK data
Normal data
Fold over basal
Time (min)
0.2
0.4
0.6
0.8
1
1.2
1.4
1.6
1.8
2
2.2
0
10
20
30
40
50
60

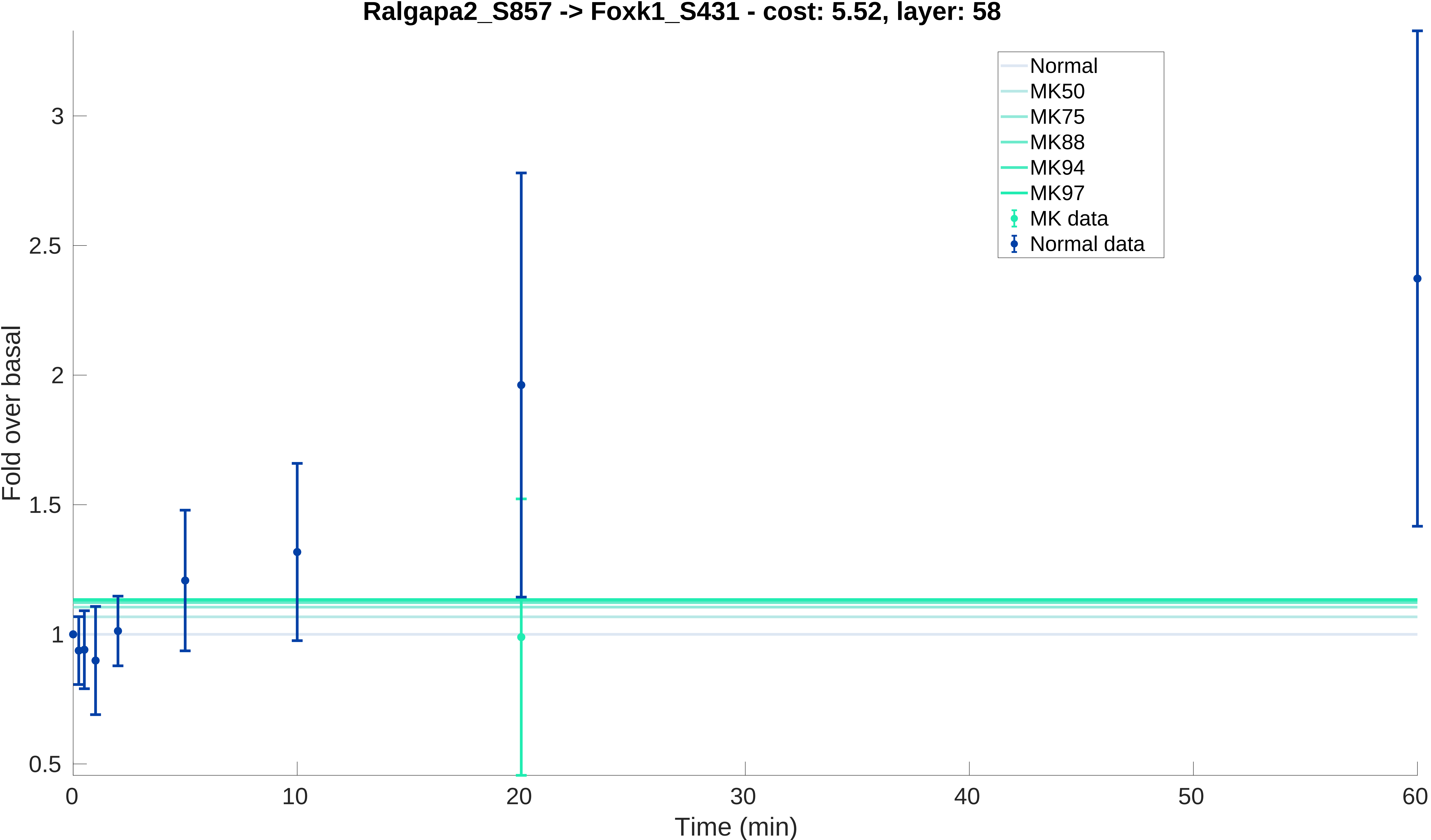

Ralgapa2_S857 -> Foxk1_S431 - cost: 5.52, layer: 58
Fold over basal
Time (min)
0.5
1
1.5
2
2.5
3
0
10
20
30
40
50
60
Normal
MK50
MK75
MK88
MK94
MK97
MK data
Normal data

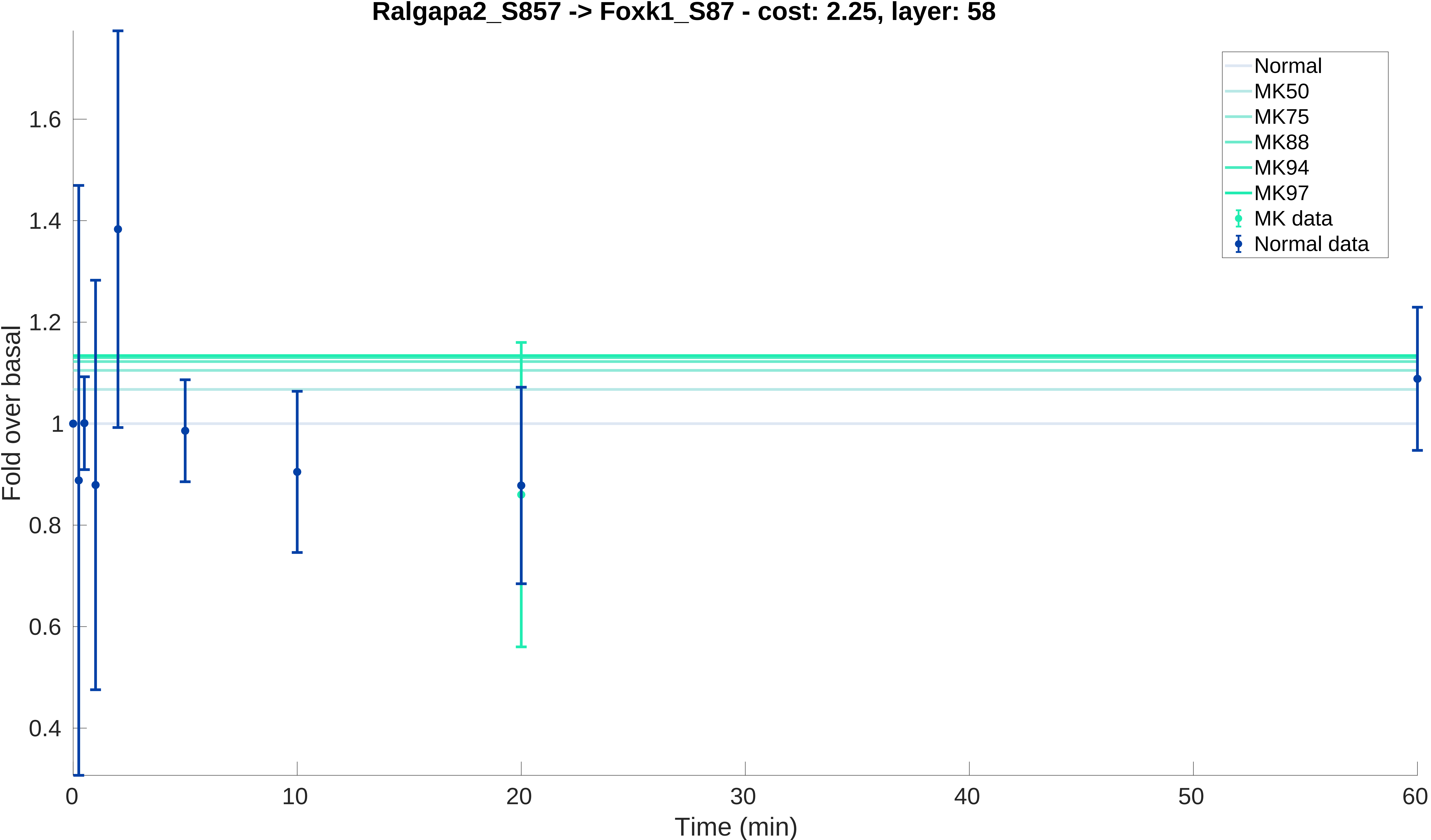

Ralgapa2_S857 -> Foxk1_S87 - cost: 2.25, layer: 58
Normal
MK50
MK75
MK88
MK94
MK97
MK data
Normal data
Fold over basal
1.6
1.4
1.2
1
0.8
0.6
0.4
0
10
20
30
40
50
60
Time (min)

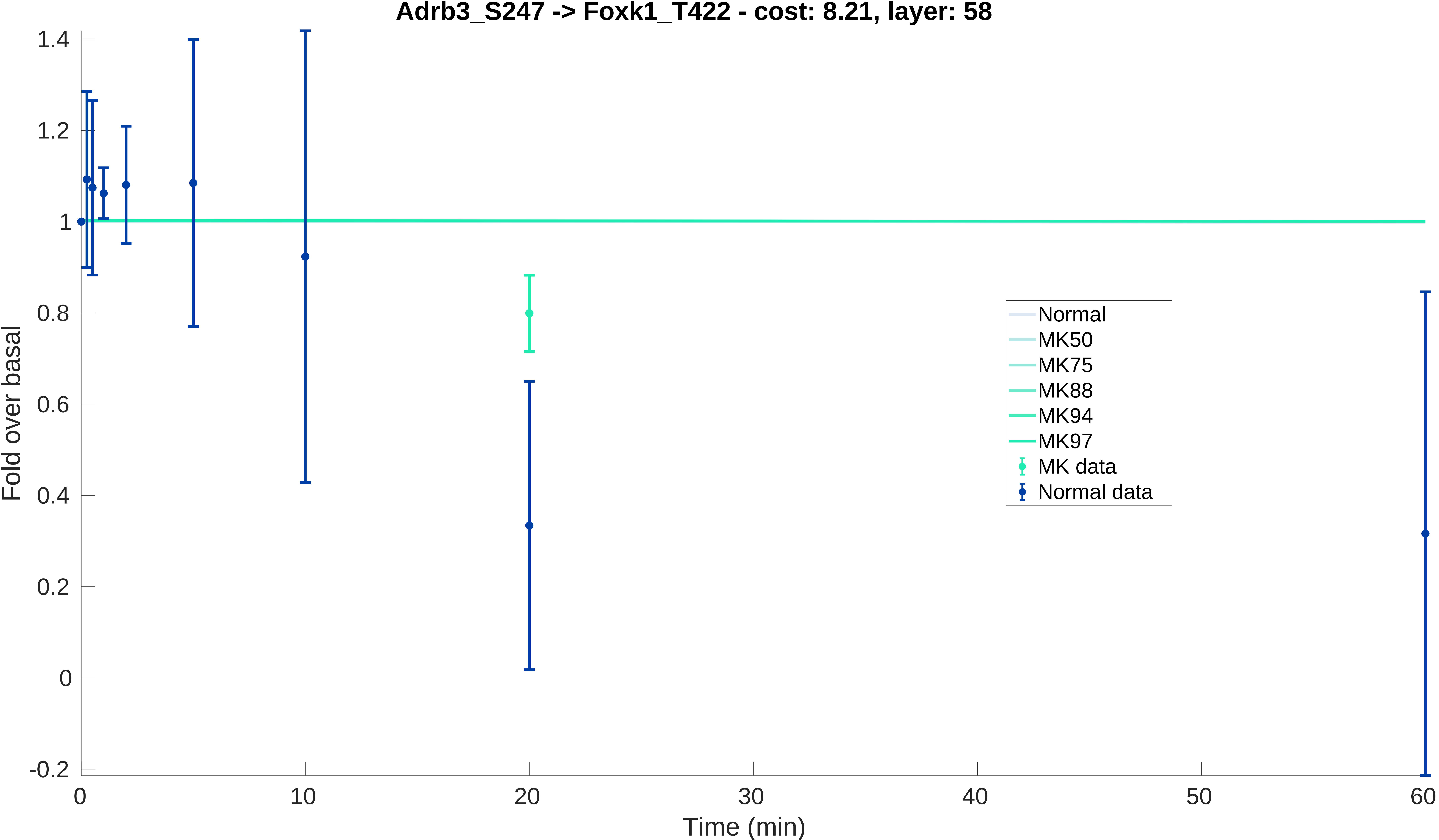

Adrb3_S247 -> Foxk1_T422 - cost: 8.21, layer: 58
Fold over basal
Time (min)
1.4
1.2
1
0.8
0.6
0.4
0.2
0
-0.2
0
10
20
30
40
50
60
Normal
MK50
MK75
MK88
MK94
MK97
MK data
Normal data

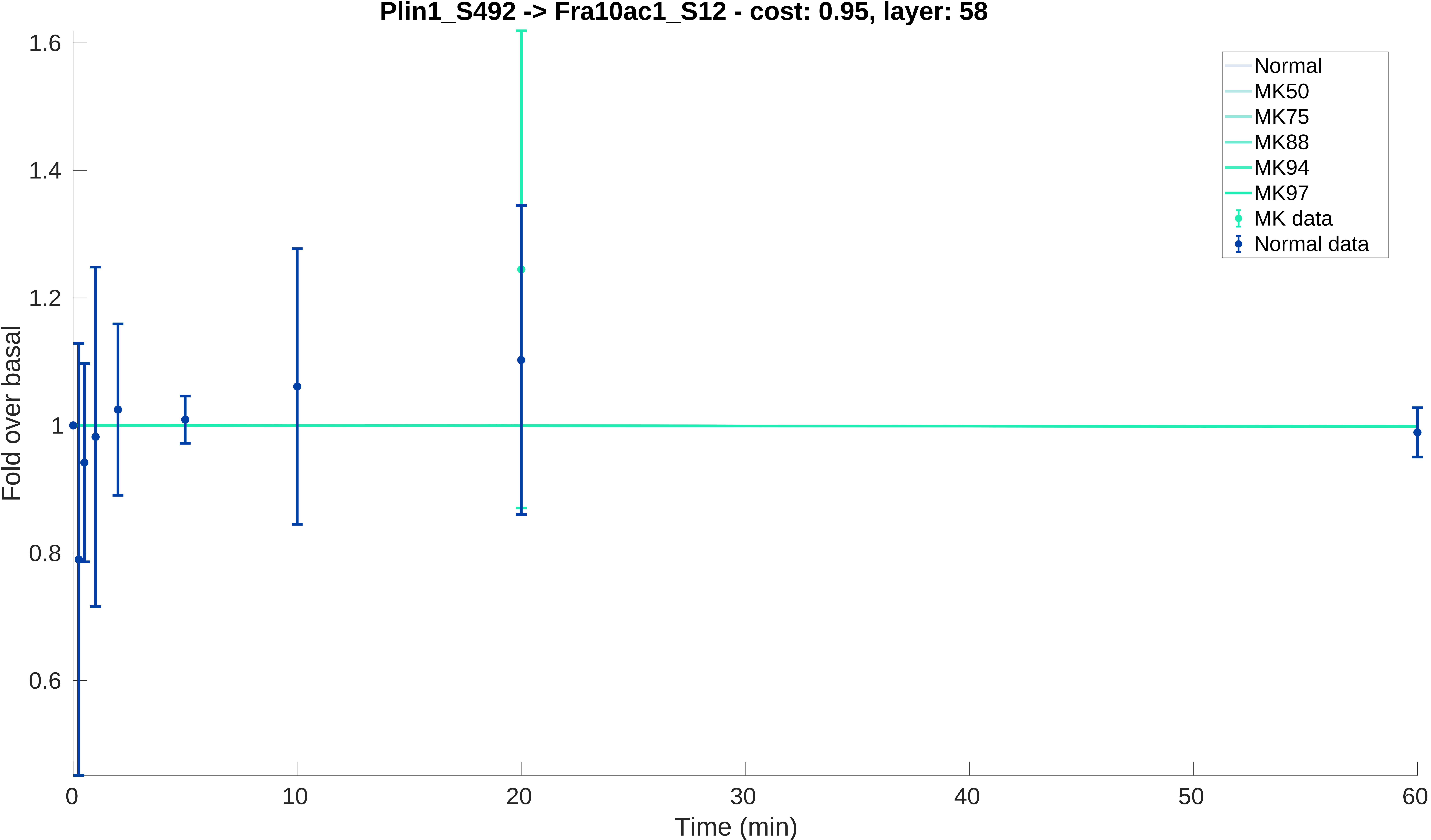

Plin1_S492 -> Fra10ac1_S12 - cost: 0.95, layer: 58
Fold over basal
Time (min)
1.6
1.4
1.2
1
0.8
0.6
0
10
20
30
40
50
60
Normal
MK50
MK75
MK88
MK94
MK97
MK data
Normal data

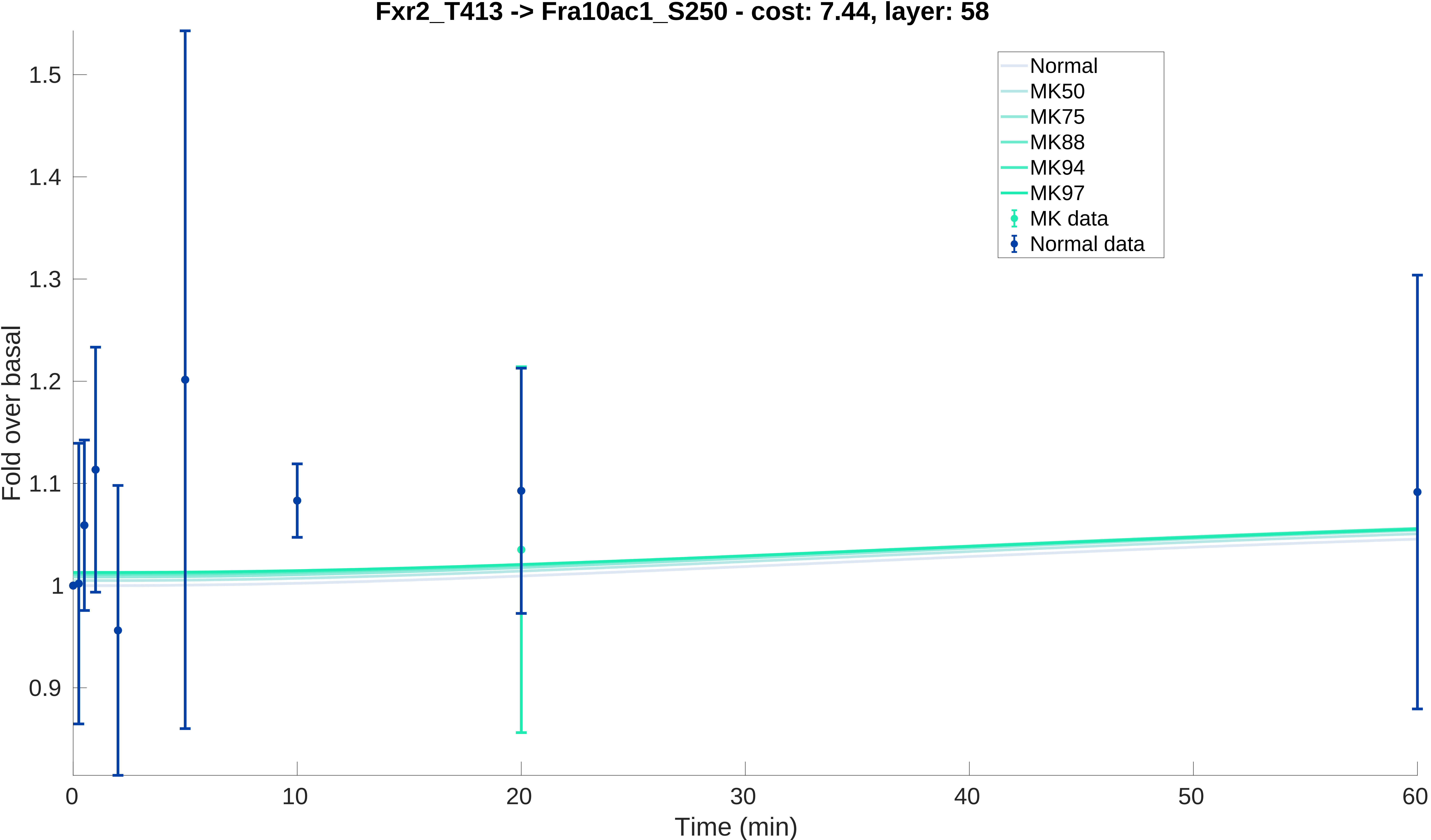

Fxr2_T413 -> Fra10ac1_S250 - cost: 7.44, layer: 58
Fold over basal
1.5
1.4
1.3
1.2
1.1
1
0.9
0
10
20
30
40
50
60
Time (min)
Normal
MK50
MK75
MK88
MK94
MK97
MK data
Normal data

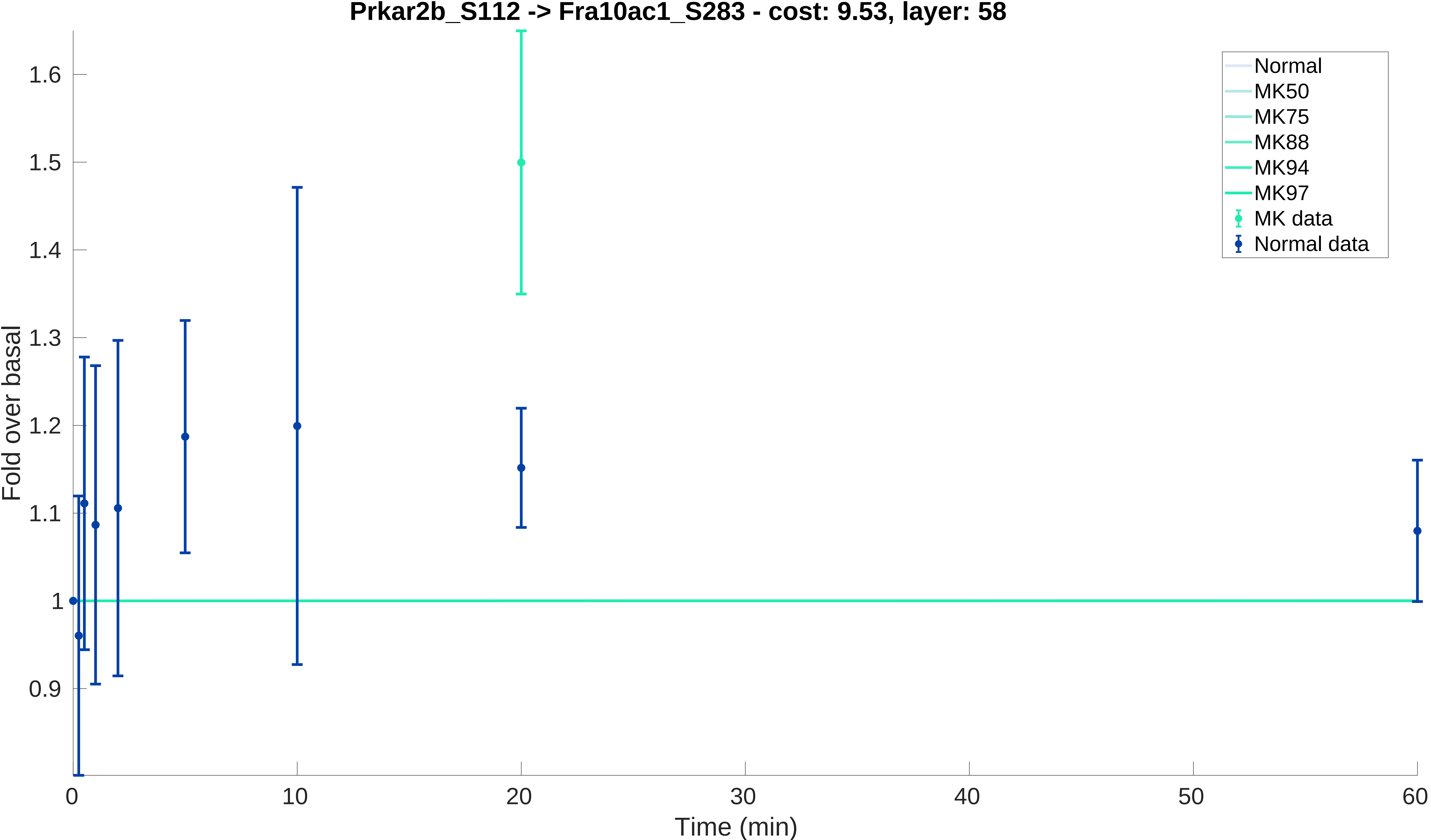

Prkar2b_S112 -> Fra10ac1_S283 - cost: 9.53, layer: 58
Normal
MK50
MK75
MK88
MK94
MK97
MK data
Normal data
Fold over basal
1.6
1.5
1.4
1.3
1.2
1.1
1
0.9
0
10
20
30
40
50
60
Time (min)

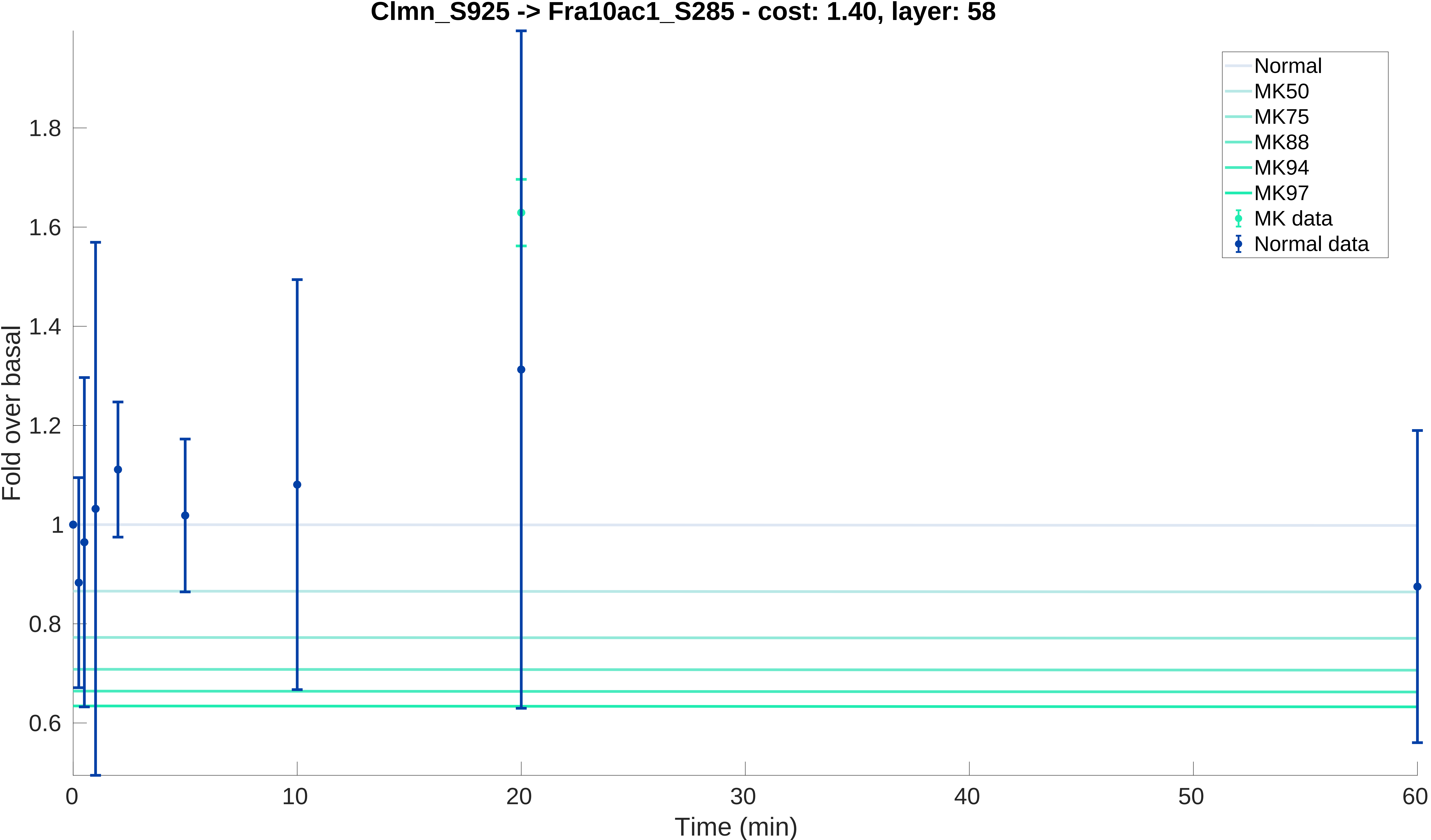

Clmn_S925 -> Fra10ac1_S285 - cost: 1.40, layer: 58
Normal
MK50
MK75
MK88
MK94
MK97
MK data
Normal data
Fold over basal
1.8
1.6
1.4
1.2
1
0.8
0.6
0
10
20
30
40
50
60
Time (min)

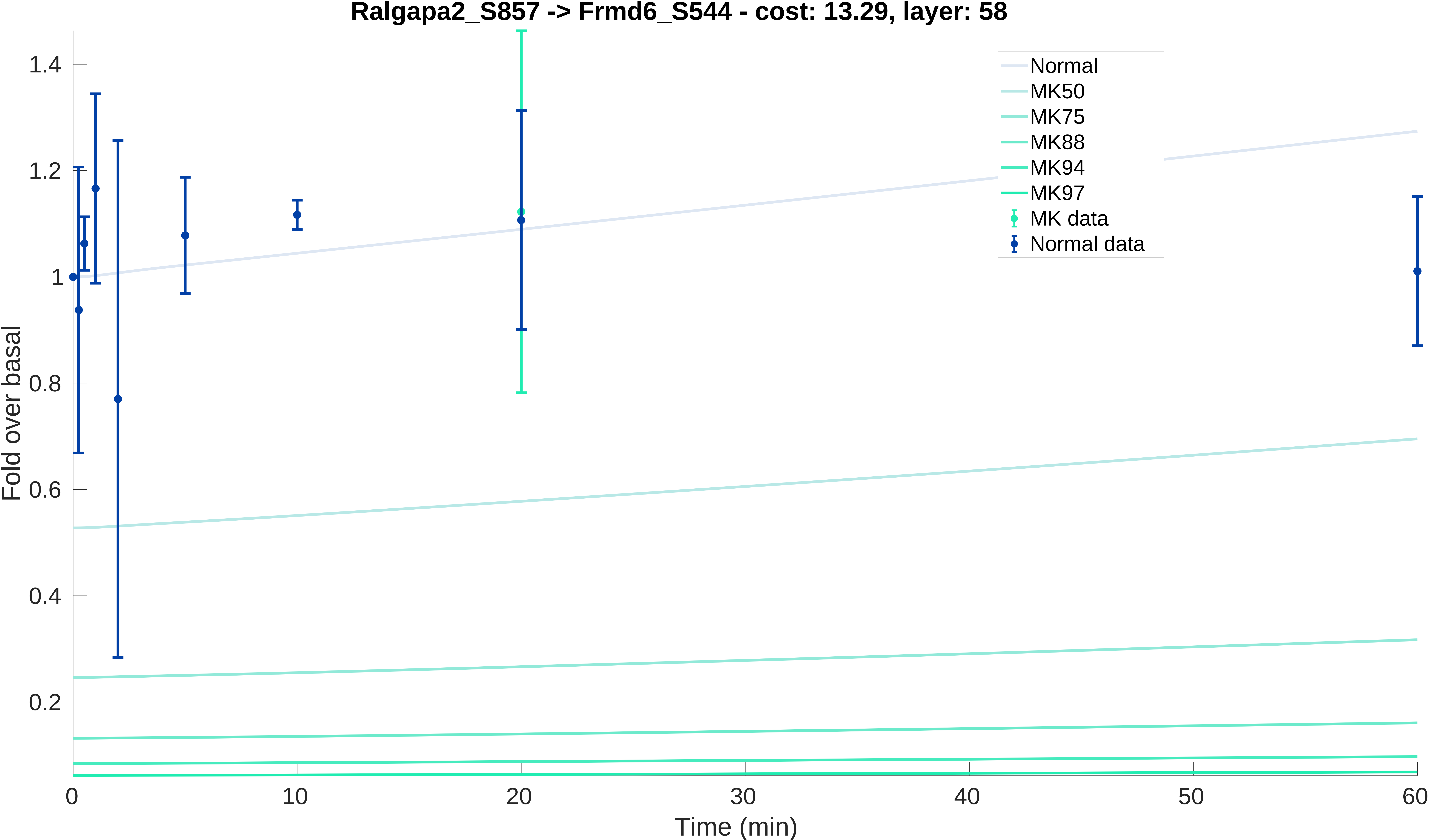

Ralgapa2_S857 -> Frmd6_S544 - cost: 13.29, layer: 58
Fold over basal
Time (min)
Normal
MK50
MK75
MK88
MK94
MK97
MK data
Normal data
0
0.2
0.4
0.6
0.8
1
1.2
1.4
0
10
20
30
40
50
60

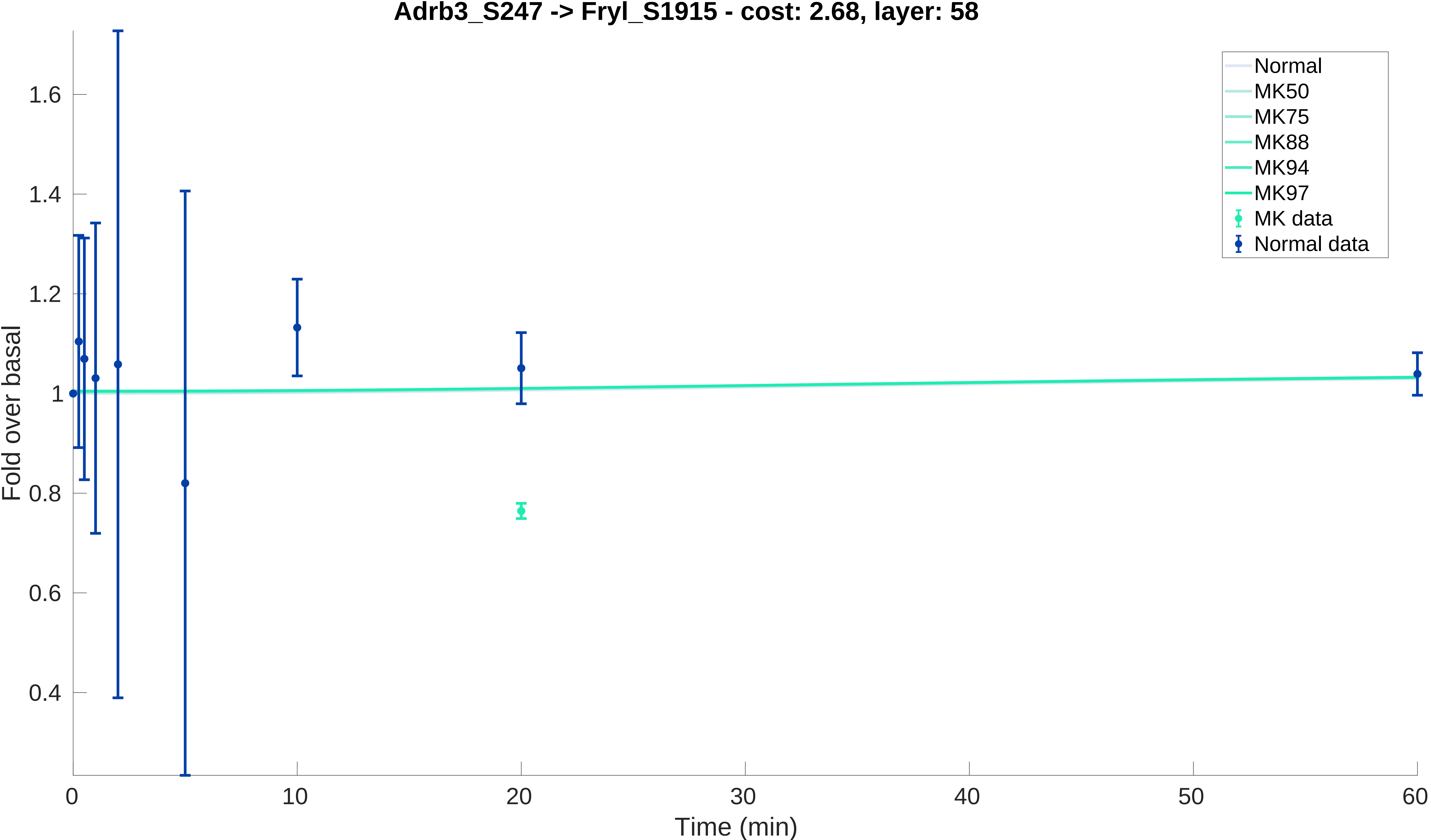

Adrb3_S247 -> Fryl_S1915 - cost: 2.68, layer: 58
Fold over basal
Time (min)
Normal
MK50
MK75
MK88
MK94
MK97
MK data
Normal data
0.4
0.6
0.8
1
1.2
1.4
1.6
0
10
20
30
40
50
60

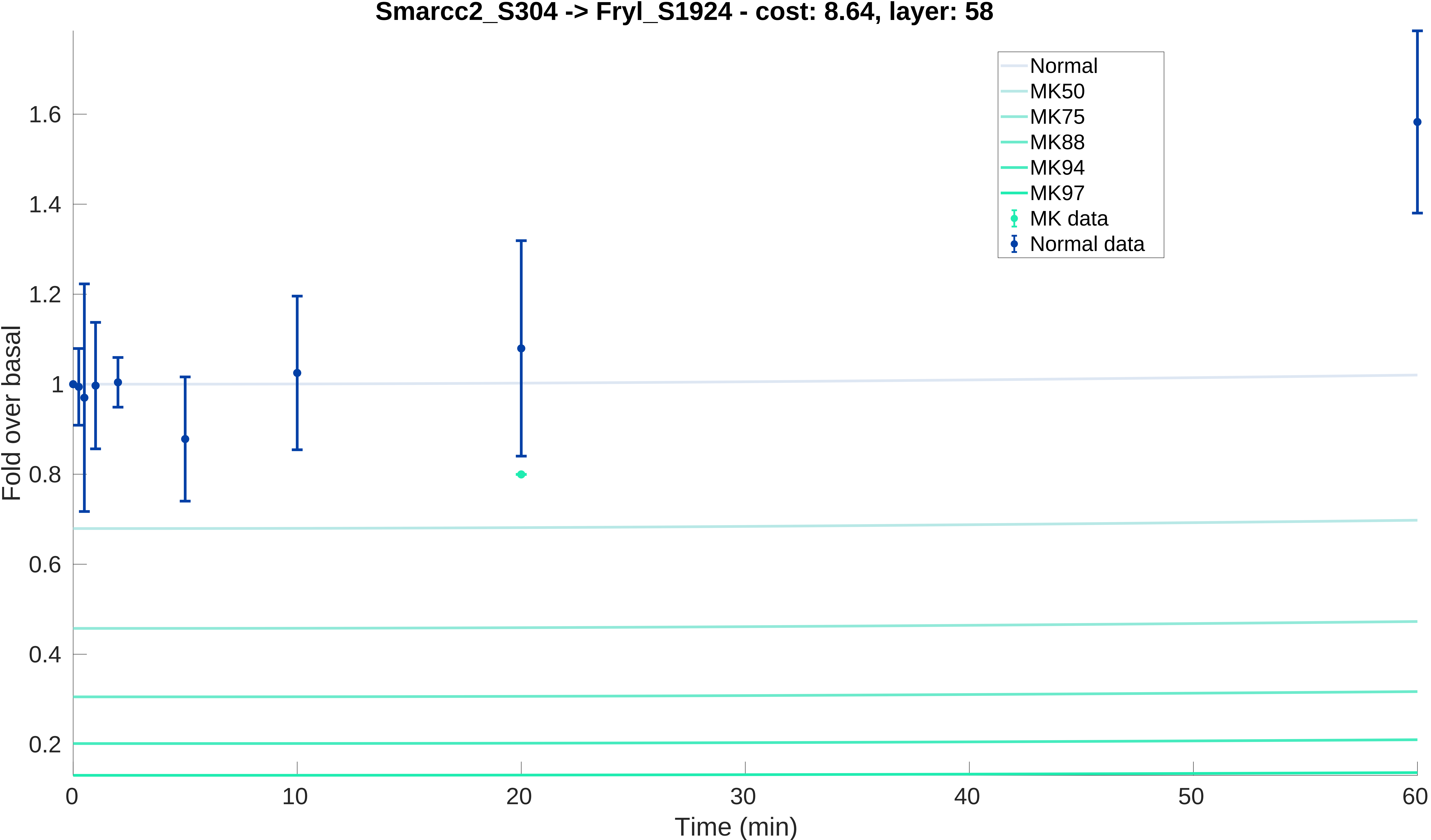
Smarcc2_S304 -> Fryl_S1924 - cost: 8.64, layer: 58
Normal
MK50
MK75
MK88
MK94
MK97
MK data
Normal data
1.6
1.4
1.2
1
0.8
0.6
0.4
0.2
Fold over basal
0
10
20
30
40
50
60
Time (min)

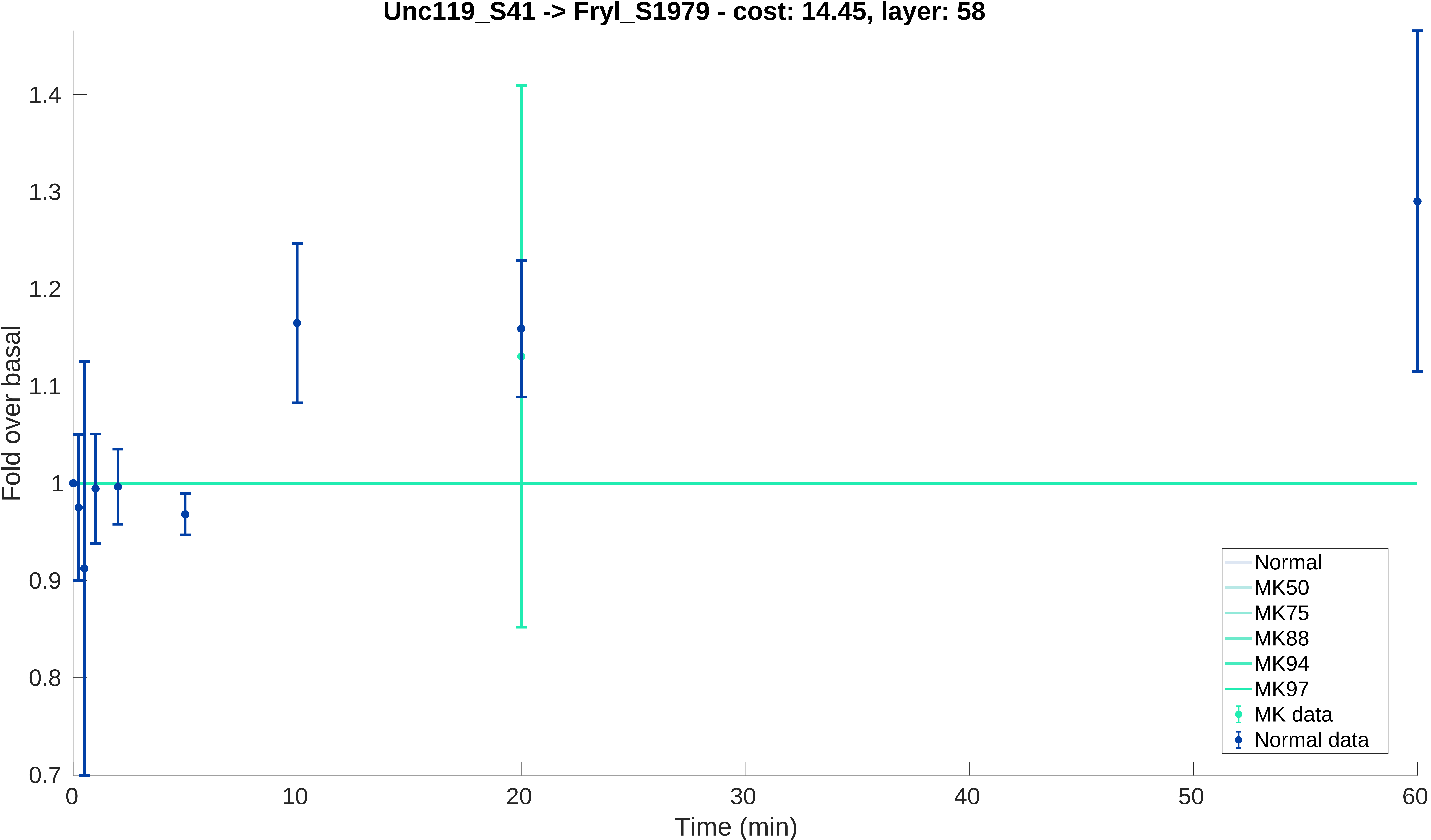

Unc119_S41 -> Fryl_S1979 - cost: 14.45, layer: 58
Fold over basal
1.4
1.3
1.2
1.1
1
0.9
0.8
0.7
0
10
20
30
40
50
60
Time (min)
Normal
MK50
MK75
MK88
MK94
MK97
MK data
Normal data

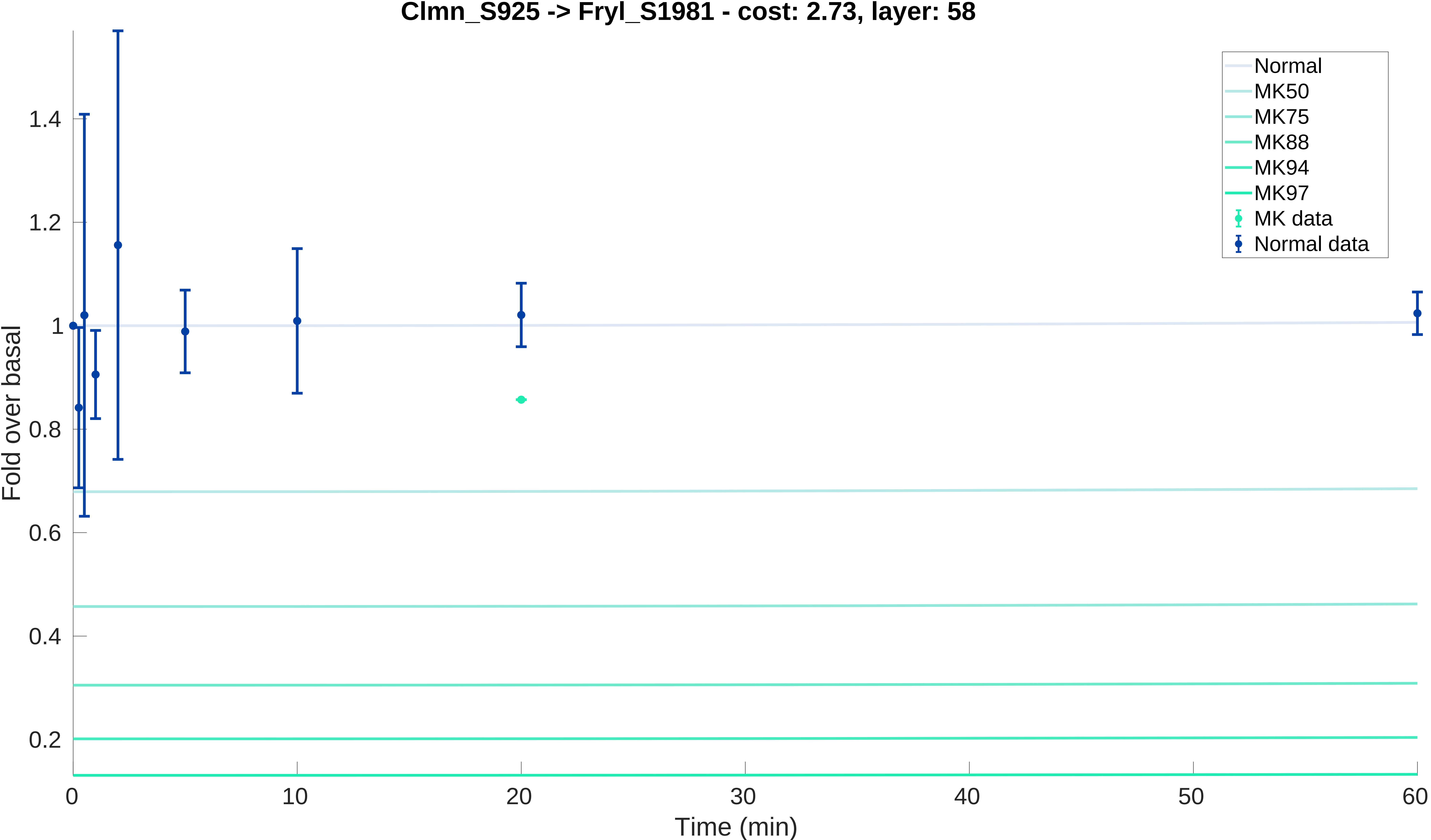
Clmn_S925 -> Fryl_S1981 - cost: 2.73, layer: 58
Fold over basal
1.4
1.2
1
0.8
0.6
0.4
0.2
0
10
20
30
40
50
60
Time (min)
Normal
MK50
MK75
MK88
MK94
MK97
MK data
Normal data

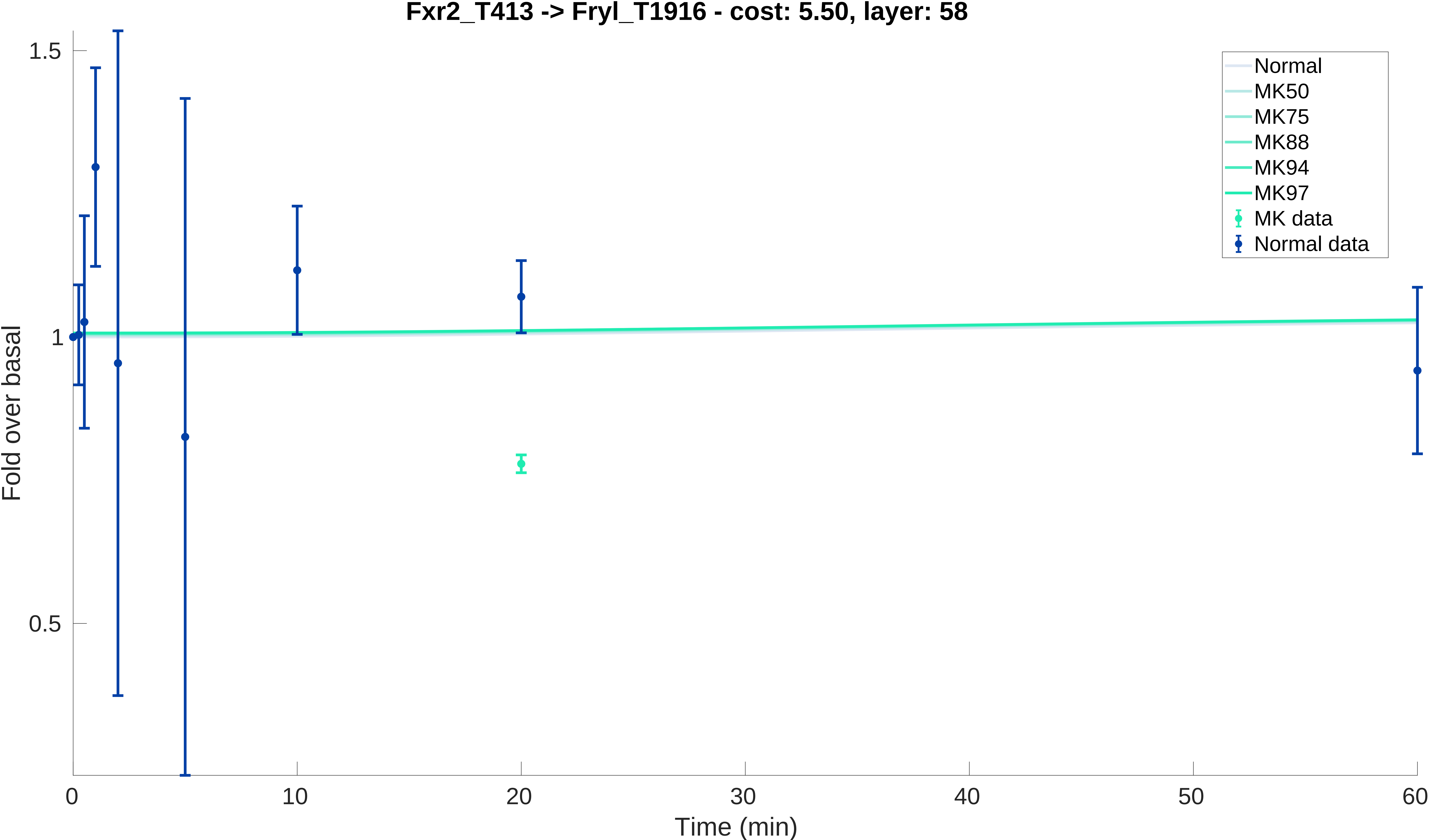

Fxr2_T413 -> Fryl_T1916 - cost: 5.50, layer: 58
Fold over basal
Time (min)
1.5
1
0.5
0
10
20
30
40
50
60
Normal
MK50
MK75
MK88
MK94
MK97
MK data
Normal data

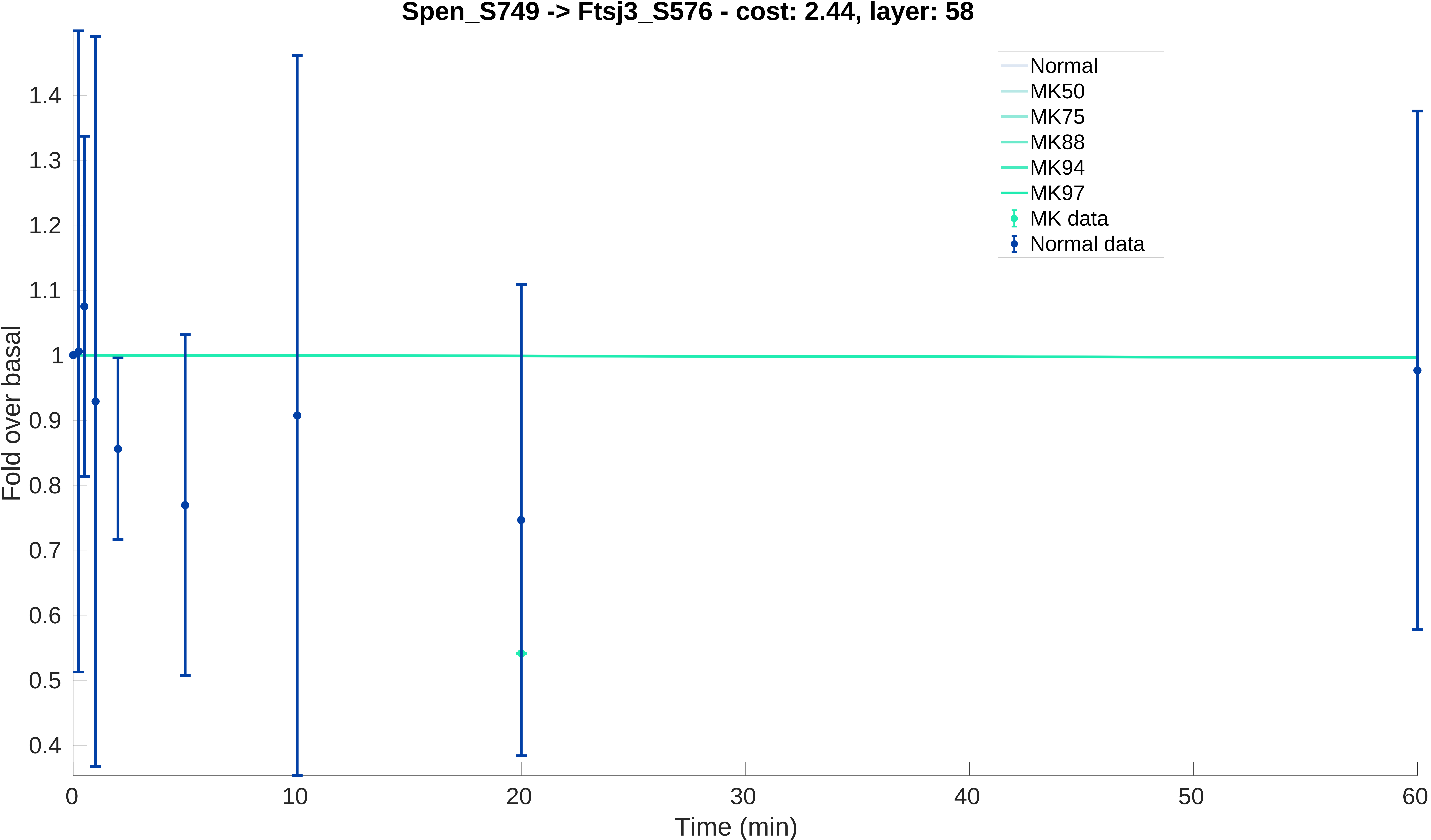
Spen_S749 -> Ftsj3_S576 - cost: 2.44, layer: 58
Normal
MK50
MK75
MK88
MK94
MK97
MK data
Normal data
Fold over basal
1.4
1.3
1.2
1.1
1
0.9
0.8
0.7
0.6
0.5
0.4
0
10
20
30
40
50
60
Time (min)

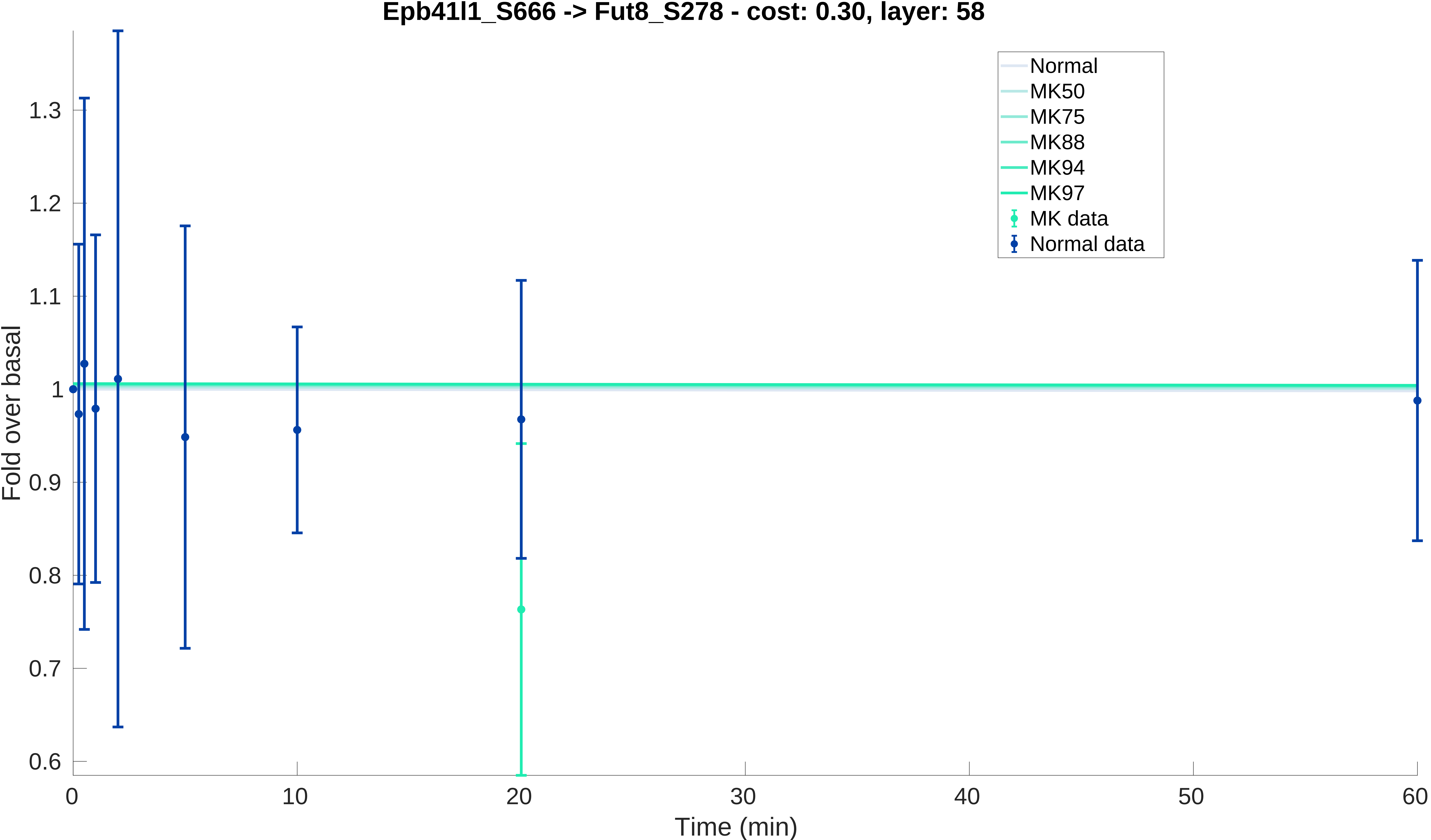

Epb41l1_S666 -> Fut8_S278 - cost: 0.30, layer: 58
Normal
MK50
MK75
MK88
MK94
MK97
MK data
Normal data
Fold over basal
1.3
1.2
1.1
1
0.9
0.8
0.7
0.6
0
10
20
30
40
50
60
Time (min)

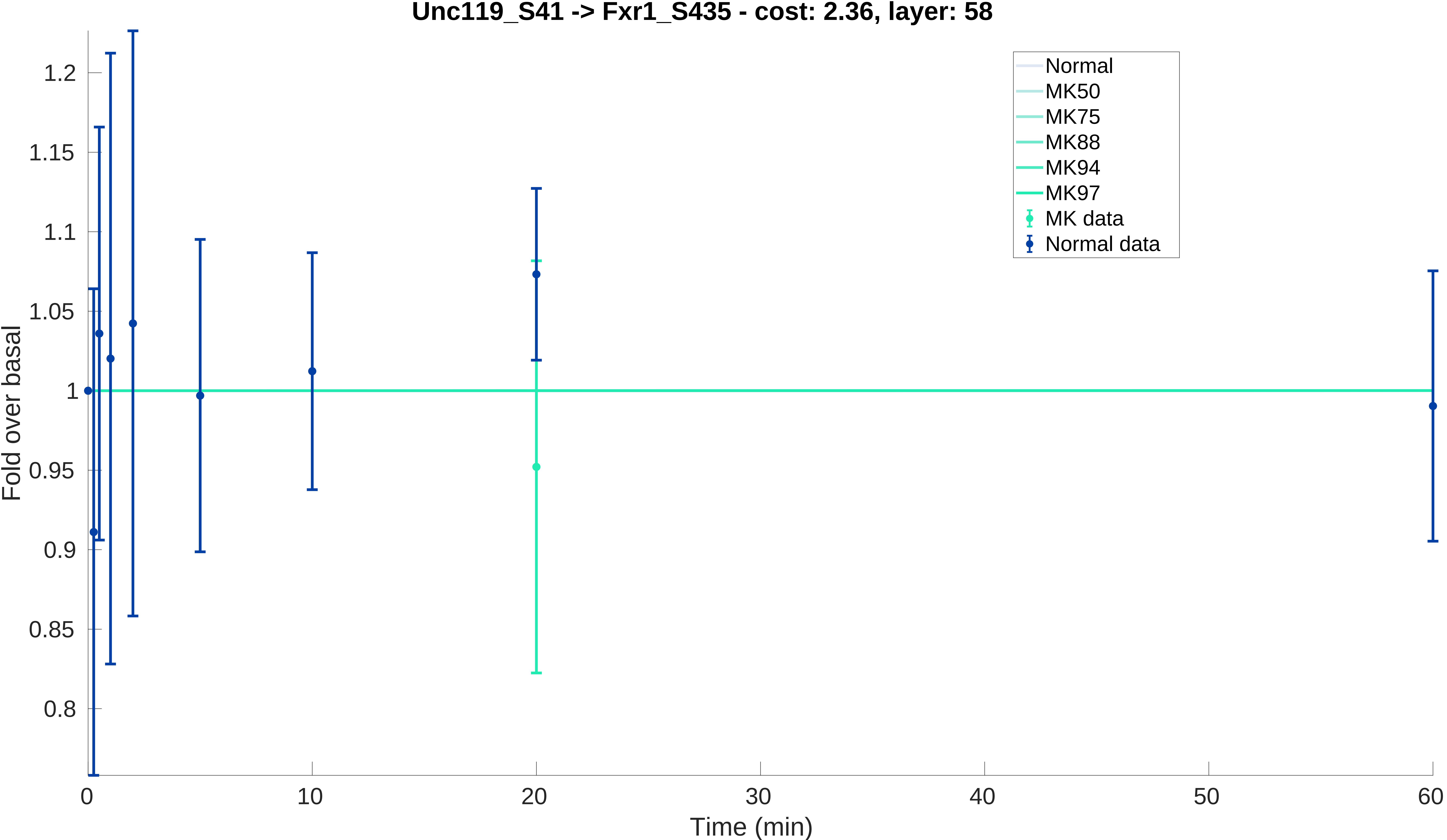

Unc119_S41 -> Fxr1_S435 - cost: 2.36, layer: 58
Normal
MK50
MK75
MK88
MK94
MK97
MK data
Normal data
Fold over basal
Time (min)
1.2
1.15
1.1
1.05
1
0.95
0.9
0.85
0.8
0
10
20
30
40
50
60

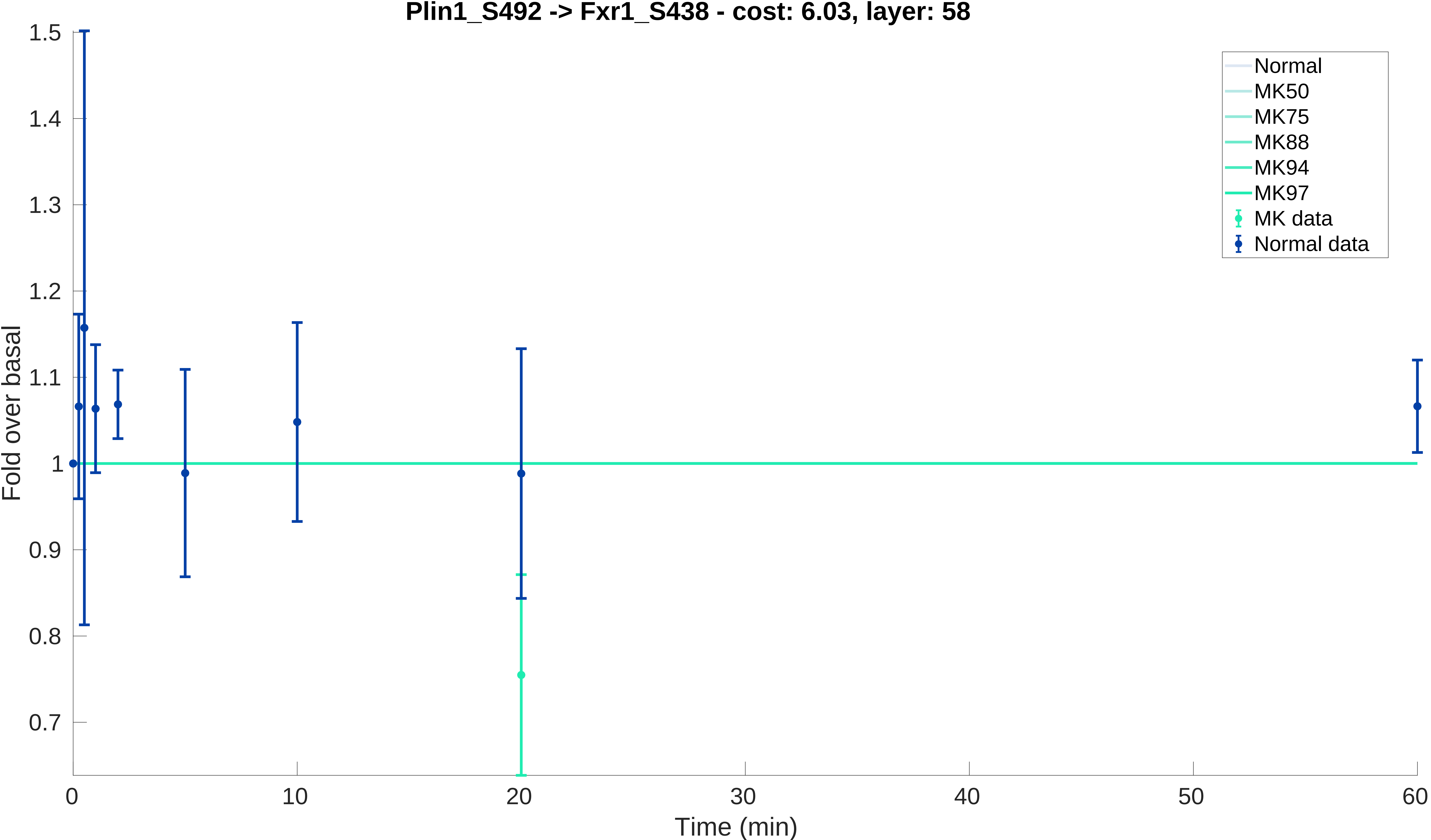

Plin1_S492 -> Fxr1_S438 - cost: 6.03, layer: 58
1.5
1.4
1.3
1.2
1.1
1
0.9
0.8
0.7
Fold over basal
0
10
20
30
40
50
60
Time (min)
Normal
MK50
MK75
MK88
MK94
MK97
MK data
Normal data

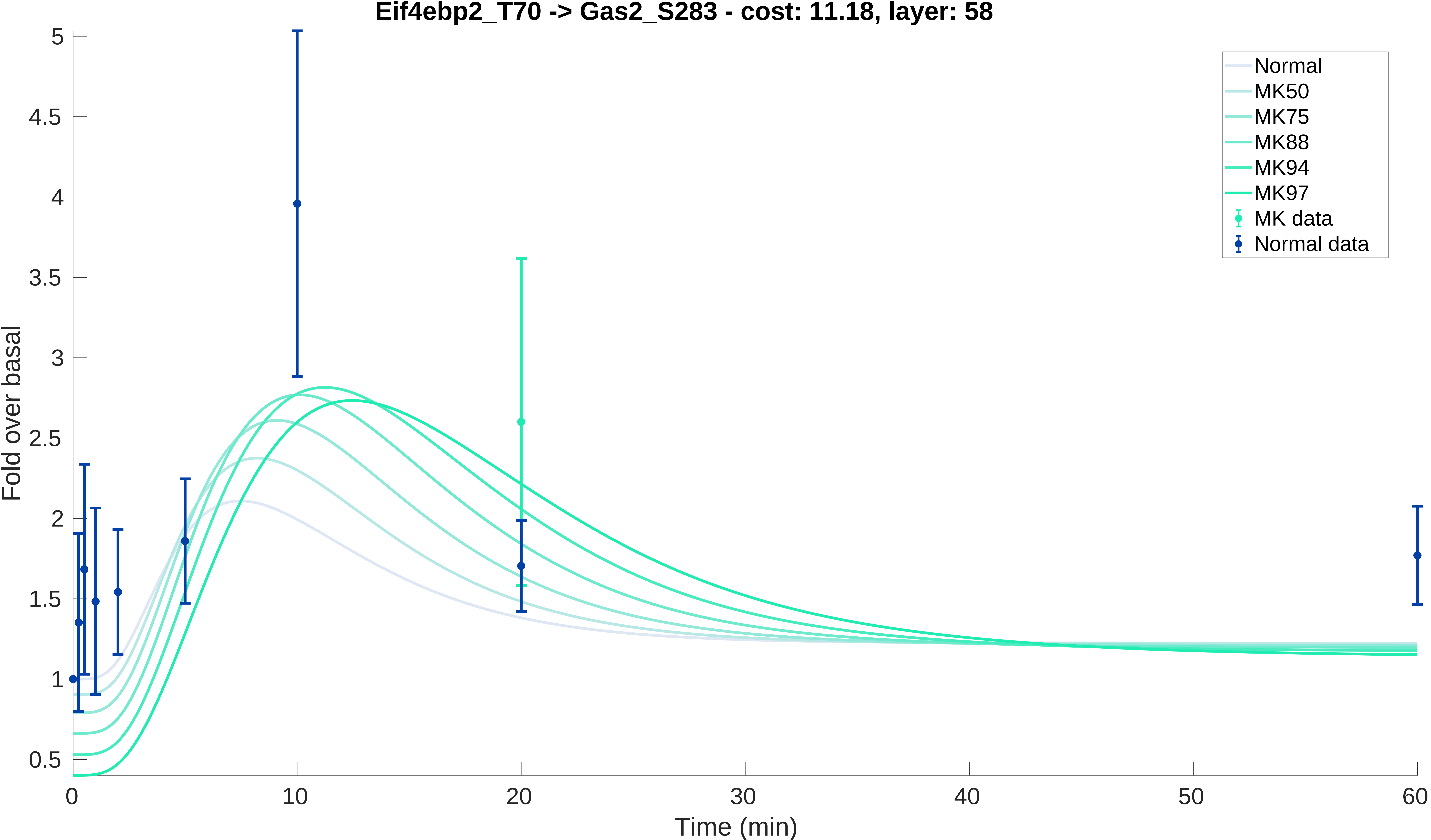

Eif4ebp2_T70 -> Gas2_S283 - cost: 11.18, layer: 58
Fold over basal
Time (min)
0.5
1
1.5
2
2.5
3
3.5
4
4.5
5
0
10
20
30
40
50
60
Normal
MK50
MK75
MK88
MK94
MK97
MK data
Normal data

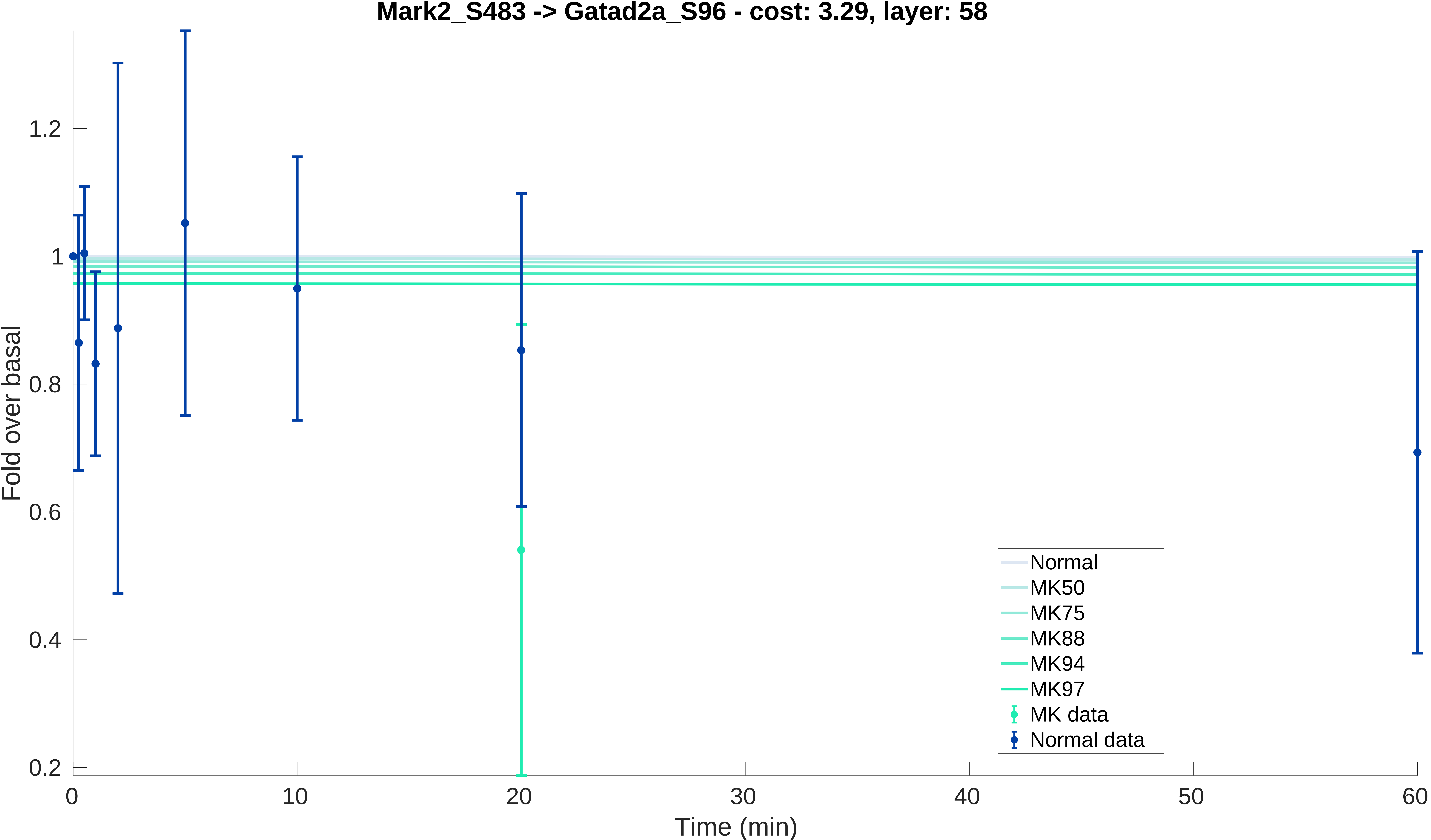

Mark2_S483 -> Gatad2a_S96 - cost: 3.29, layer: 58
Fold over basal
Time (min)
0.2
0.4
0.6
0.8
1
1.2
0
10
20
30
40
50
60
Normal
MK50
MK75
MK88
MK94
MK97
MK data
Normal data

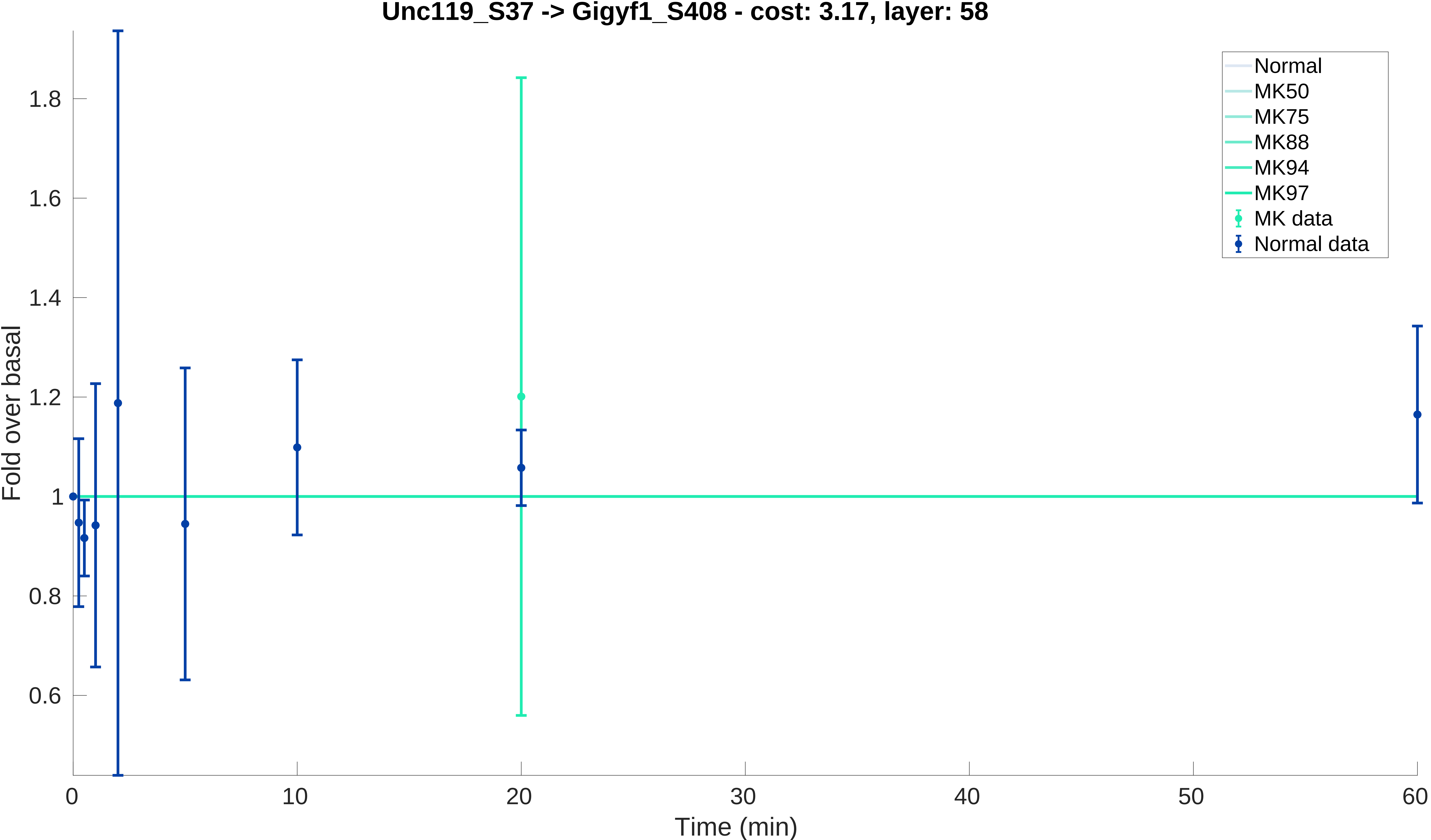

Unc119_S37 -> Gigyf1_S408 - cost: 3.17, layer: 58
Fold over basal
Time (min)
0.6
0.8
1
1.2
1.4
1.6
1.8
0
10
20
30
40
50
60
Normal
MK50
MK75
MK88
MK94
MK97
MK data
Normal data

### Eif4ebp2_T70 -> Glcci1_S397 - cost: 7.83, layer: 58

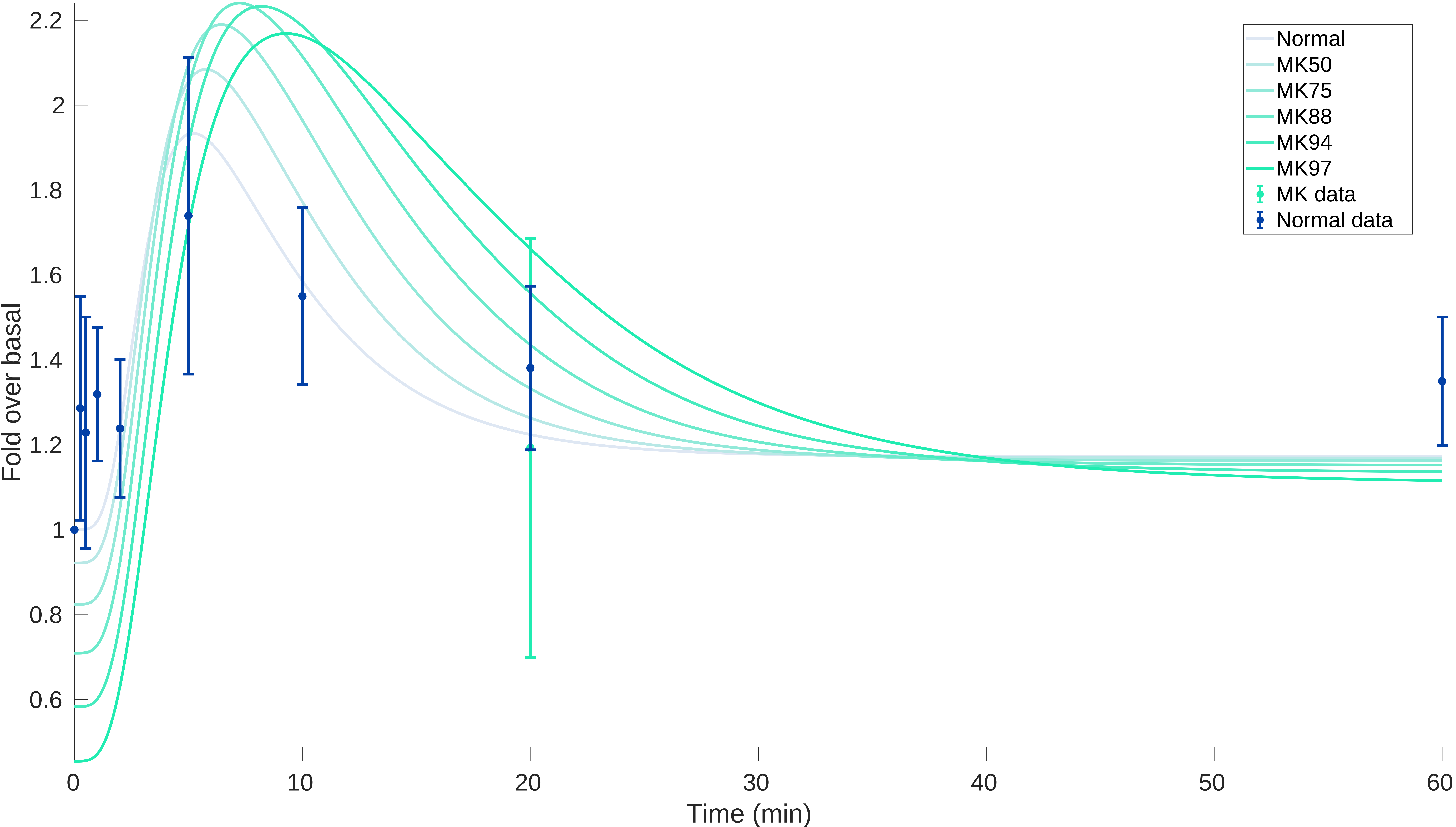

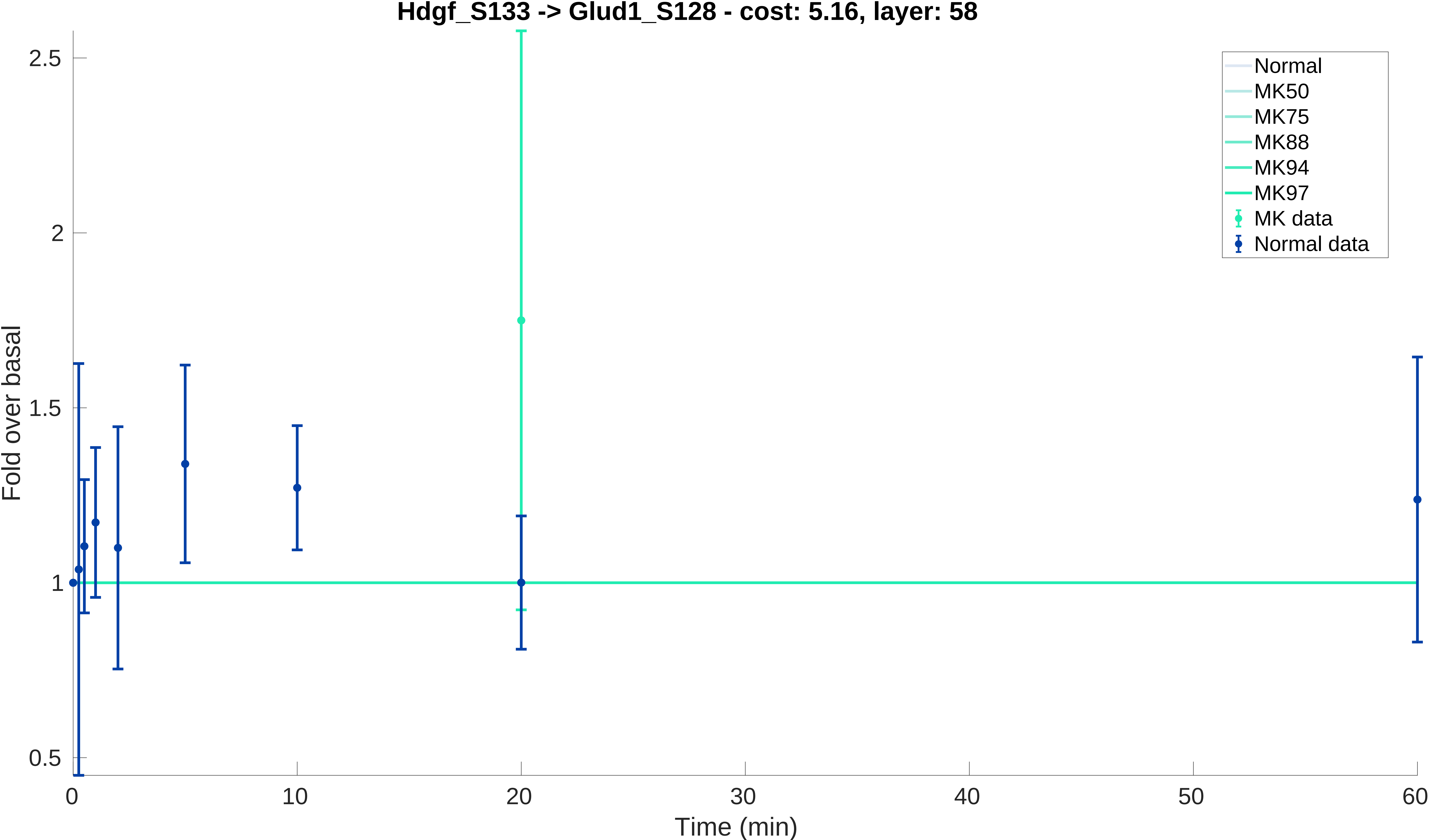

Hdgf_S133 -> Glud1_S128 - cost: 5.16, layer: 58
Normal
MK50
MK75
MK88
MK94
MK97
MK data
Normal data
Fold over basal
2.5
2
1.5
1
0.5
0
10
20
30
40
50
60
Time (min)

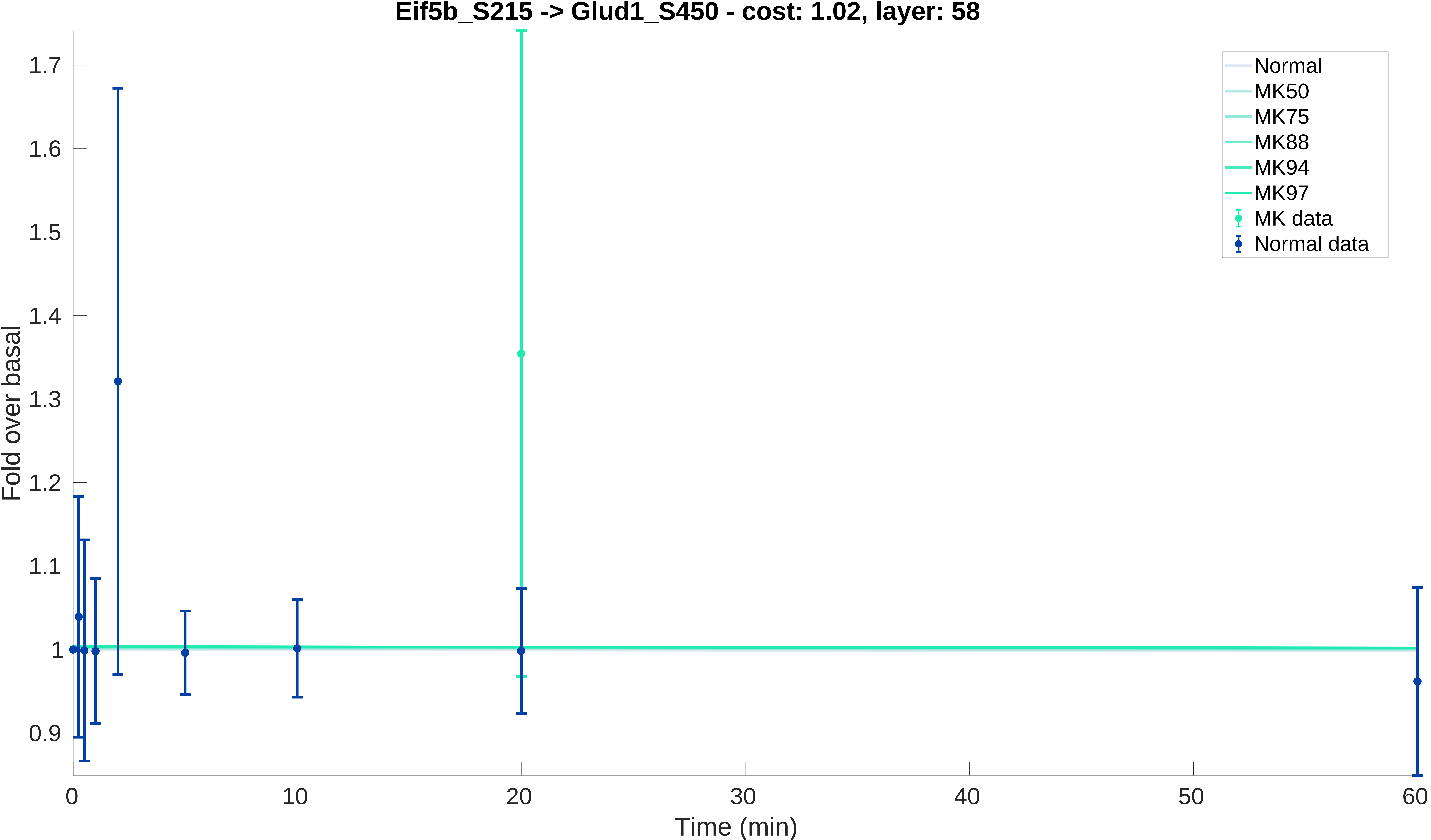

Eif5b_S215 -> Glud1_S450 - cost: 1.02, layer: 58
Fold over basal
Time (min)
1.7
1.6
1.5
1.4
1.3
1.2
1.1
1
0.9
0
10
20
30
40
50
60
Normal
MK50
MK75
MK88
MK94
MK97
MK data
Normal data

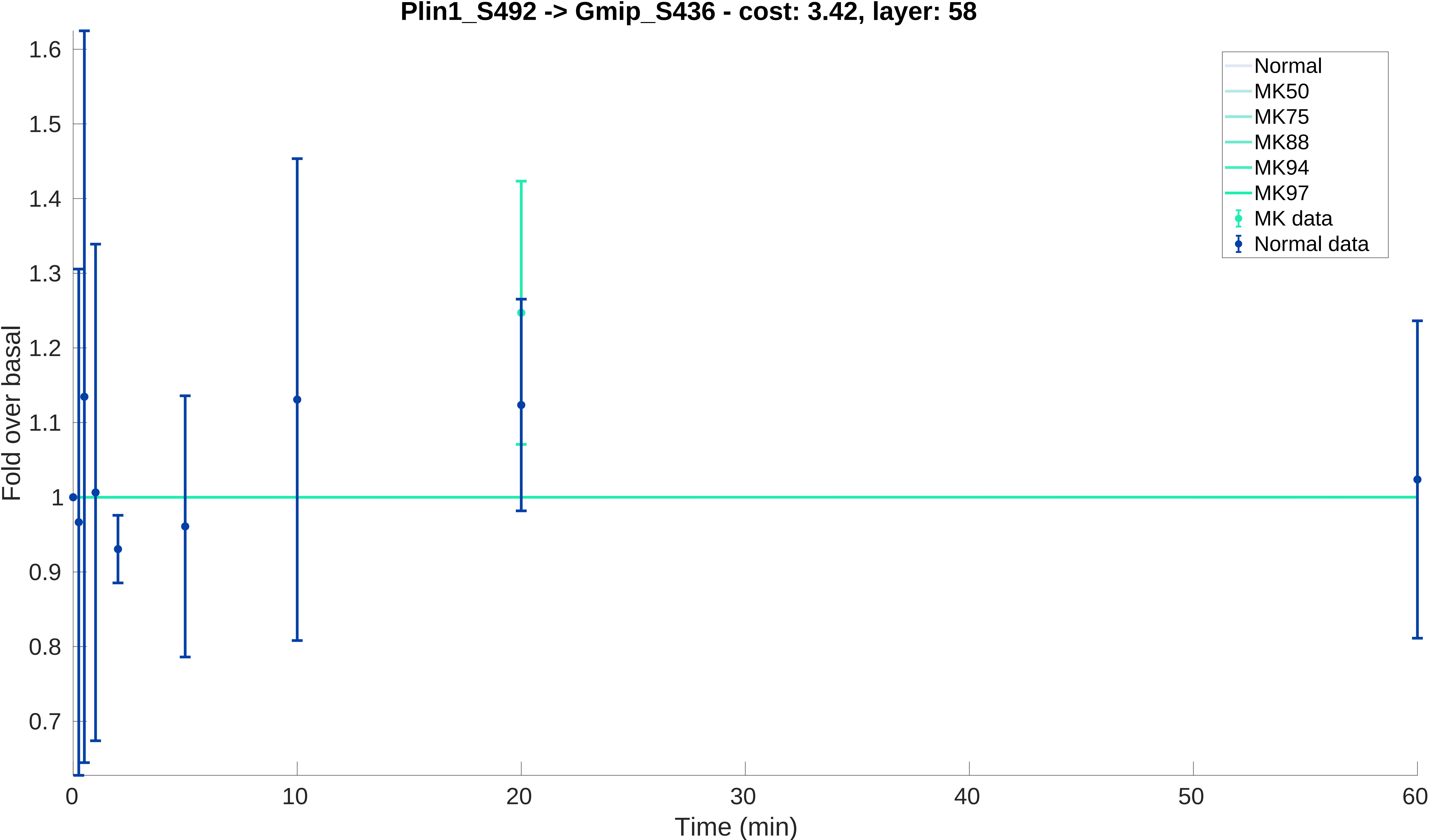

Plin1_S492 -> Gmip_S436 - cost: 3.42, layer: 58
Fold over basal
Time (min)
1.6
1.5
1.4
1.3
1.2
1.1
1
0.9
0.8
0.7
0
10
20
30
40
50
60
Normal
MK50
MK75
MK88
MK94
MK97
MK data
Normal data

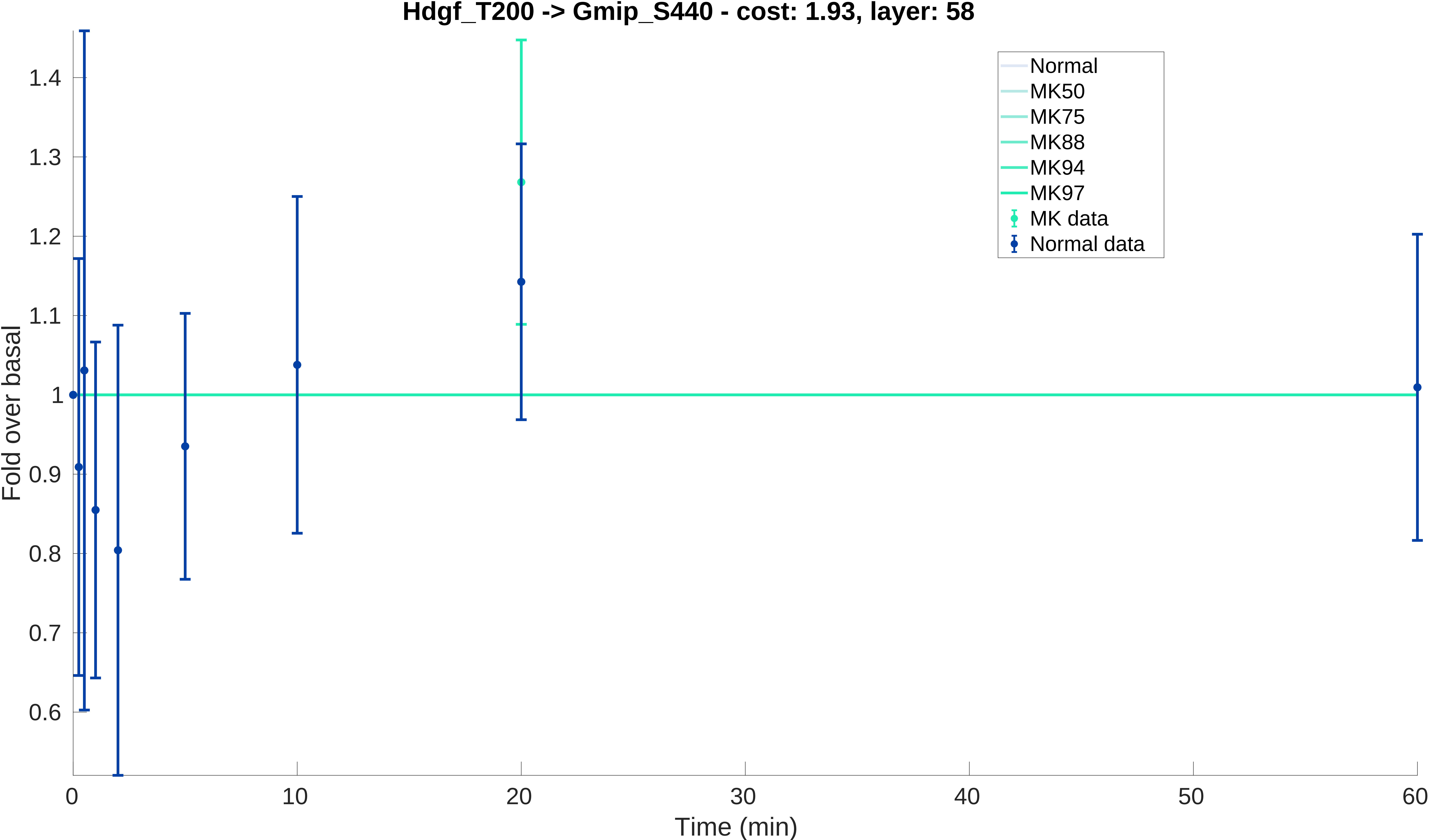

Hdgf_T200 -> Gmip_S440 - cost: 1.93, layer: 58
Normal
MK50
MK75
MK88
MK94
MK97
MK data
Normal data
Fold over basal
Time (min)
1.4
1.3
1.2
1.1
1
0.9
0.8
0.7
0.6
0
10
20
30
40
50
60

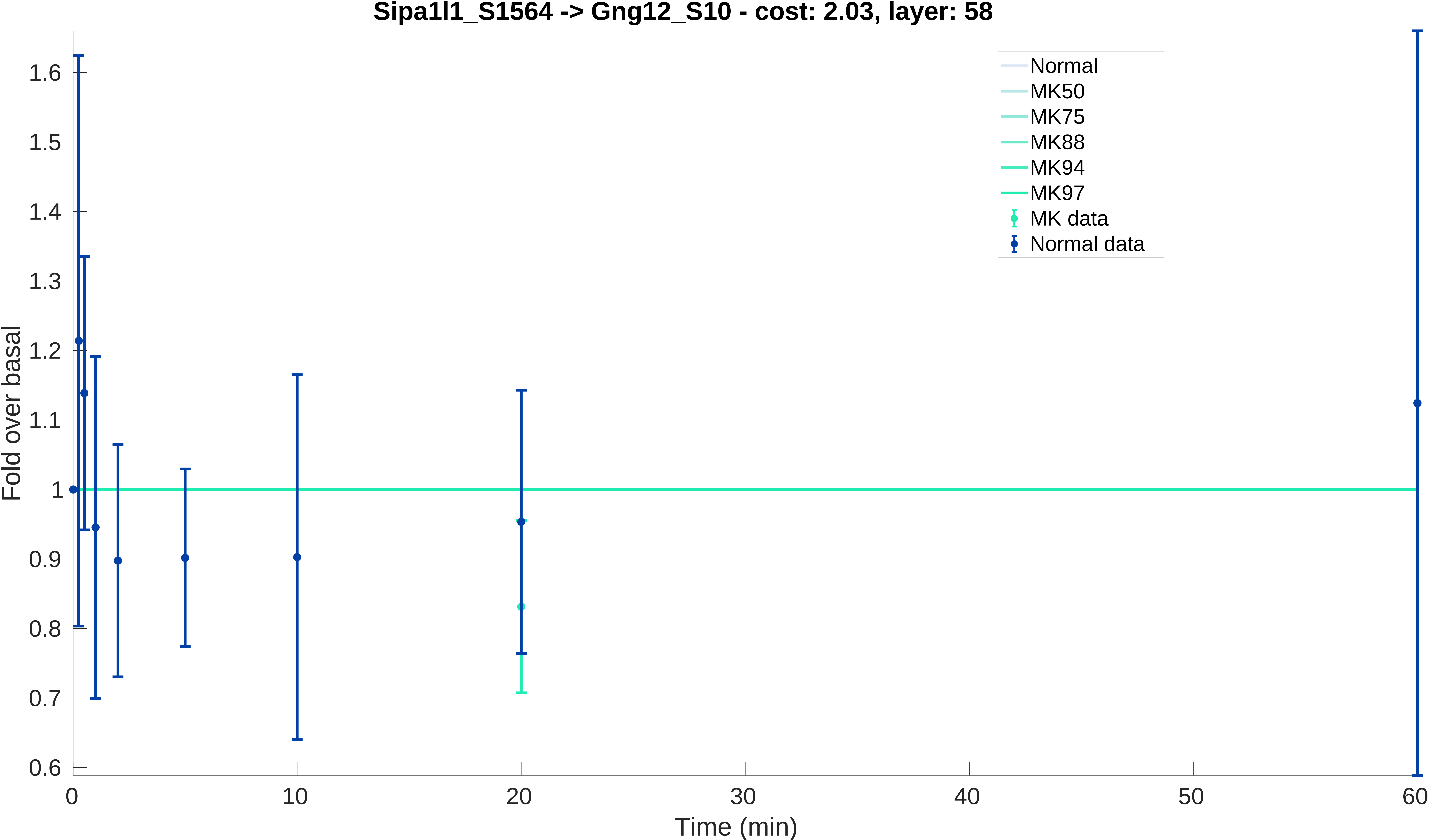

Sipa1l1_S1564 -> Gng12_S10 - cost: 2.03, layer: 58
Normal
MK50
MK75
MK88
MK94
MK97
MK data
Normal data
Fold over basal
0.6
0.7
0.8
0.9
1
1.1
1.2
1.3
1.4
1.5
1.6
0
10
20
30
40
50
60
Time (min)

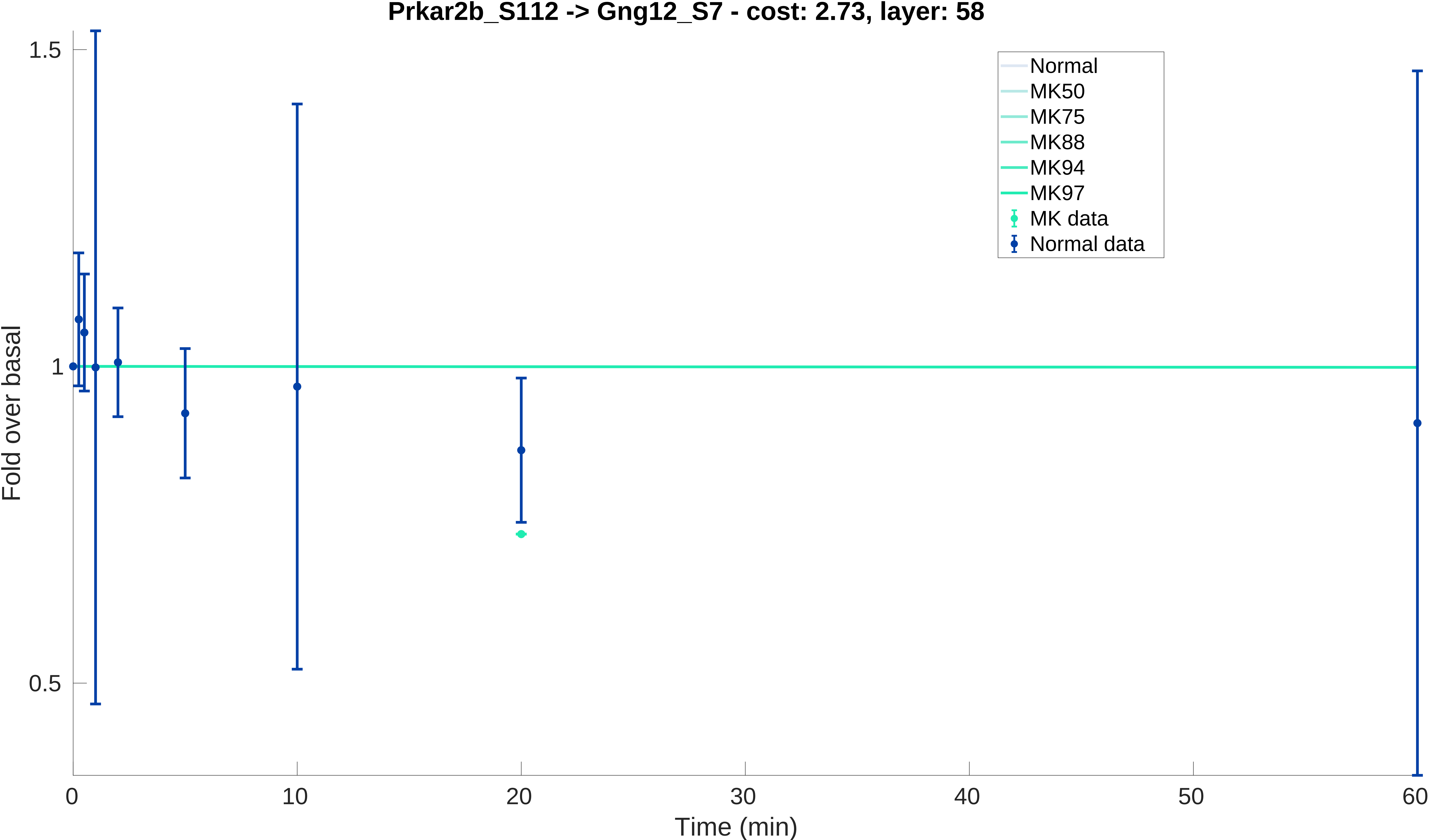

Prkar2b_S112 -> Gng12_S7 - cost: 2.73, layer: 58
Fold over basal
Time (min)
1.5
1
0.5
0
10
20
30
40
50
60
Normal
MK50
MK75
MK88
MK94
MK97
MK data
Normal data

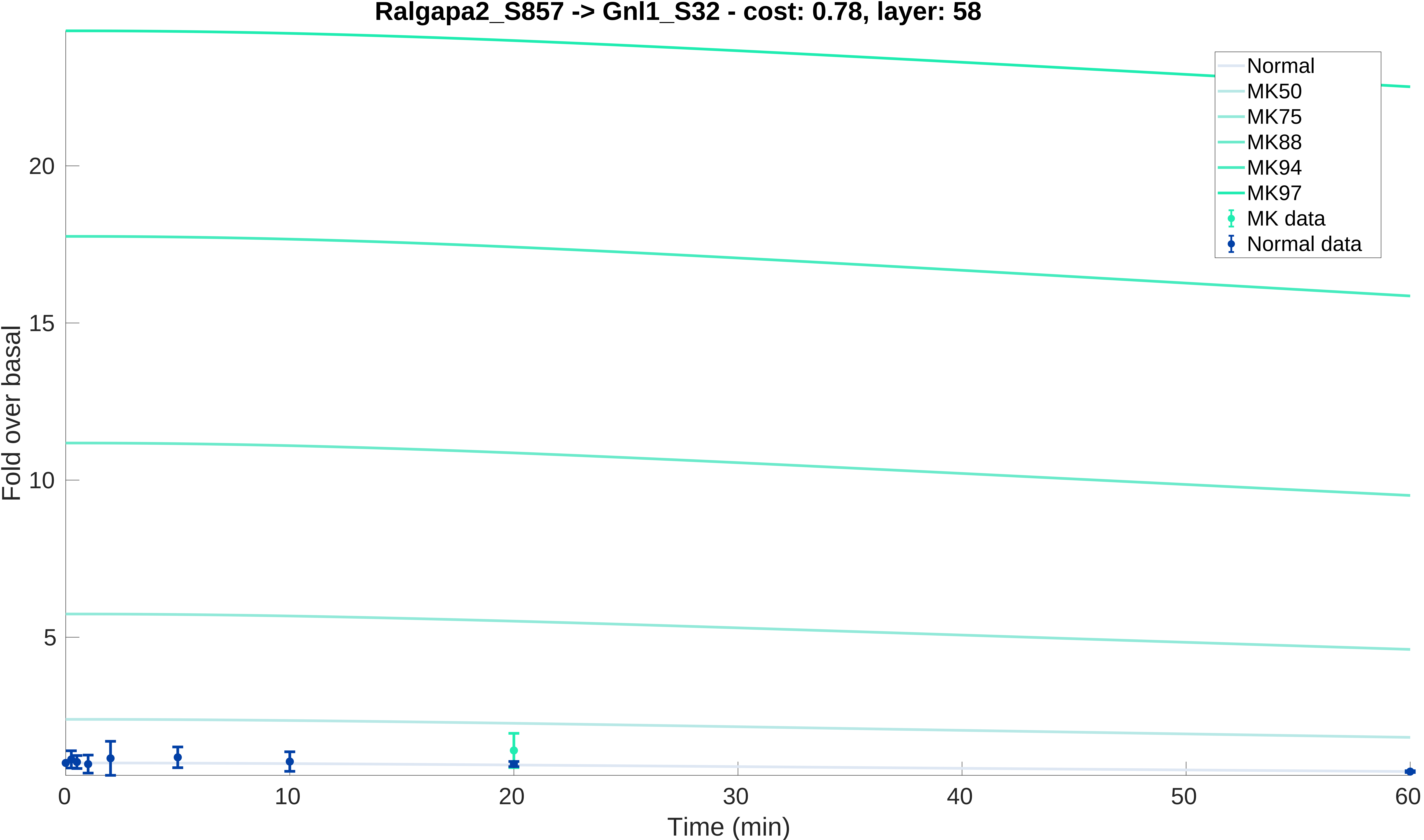
Ralgapa2_S857 -> Gnl1_S32 - cost: 0.78, layer: 58
Fold over basal
Time (min)
Normal
MK50
MK75
MK88
MK94
MK97
MK data
Normal data
0
10
20
30
40
50
60
5
10
15
20

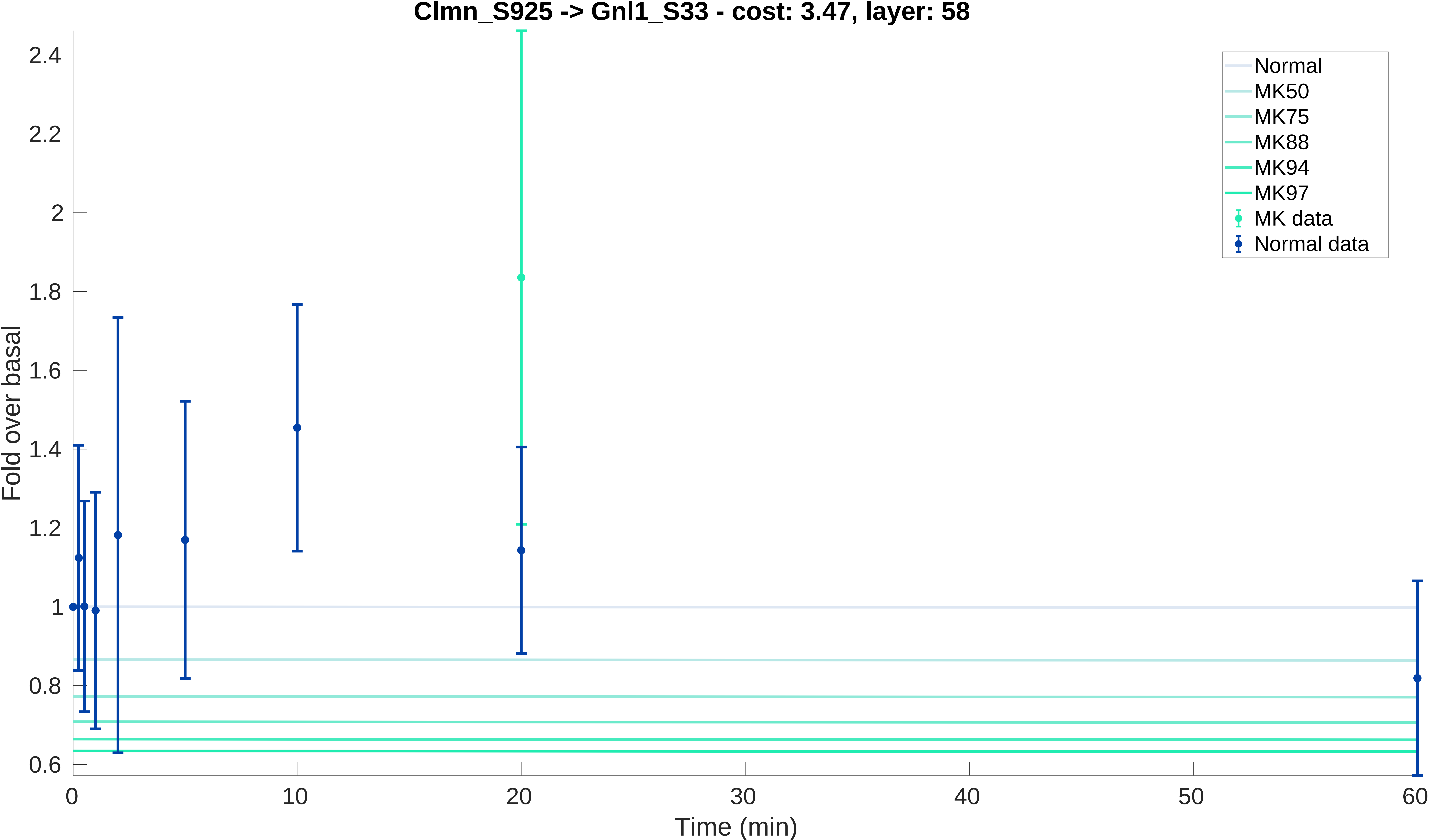

Clmn_S925 -> Gnl1_S33 - cost: 3.47, layer: 58
Fold over basal
2.4
2.2
2
1.8
1.6
1.4
1.2
1
0.8
0.6
0
10
20
30
40
50
60
Time (min)
Normal
MK50
MK75
MK88
MK94
MK97
MK data
Normal data

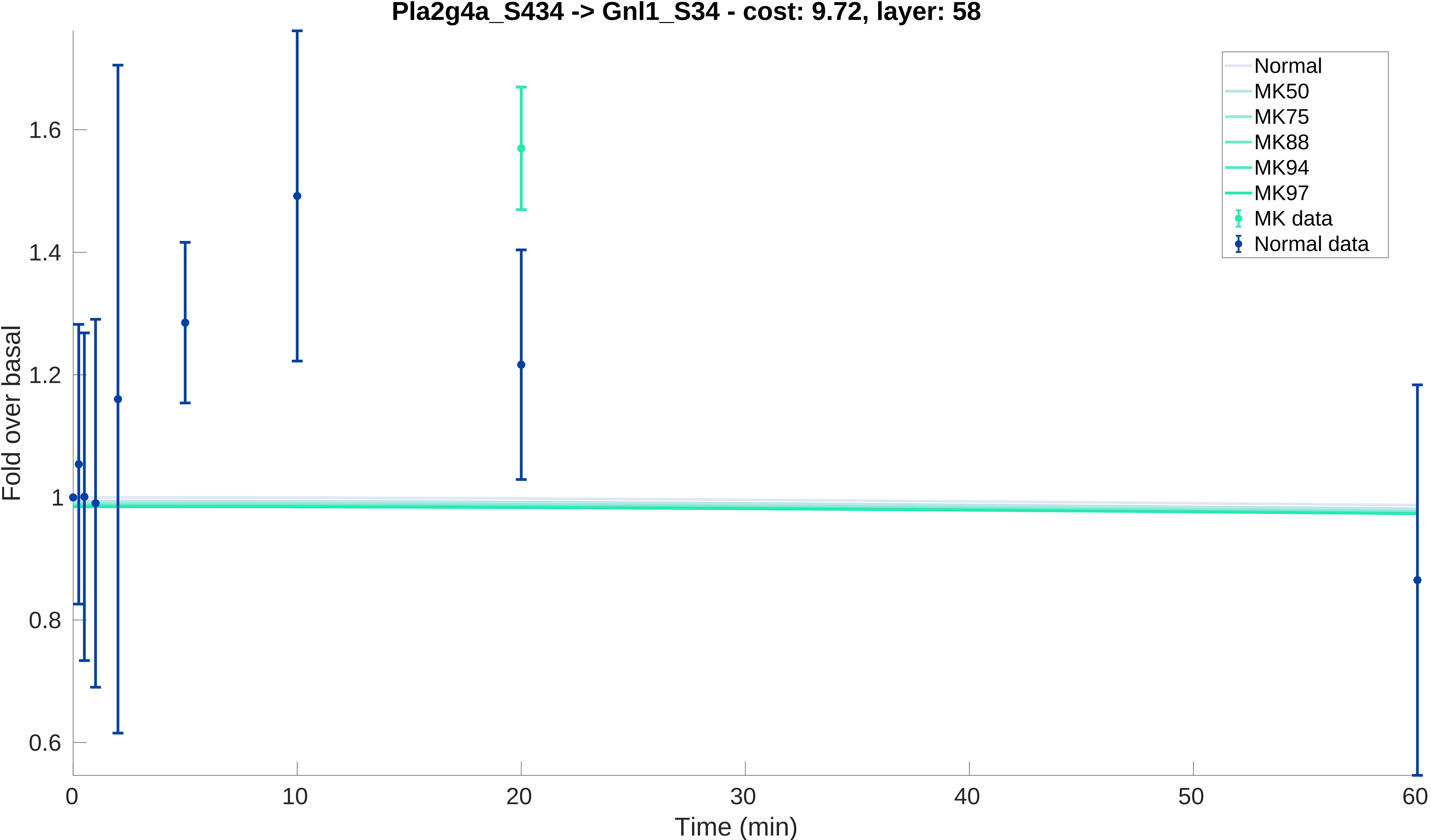

Pla2g4a_S434 -> Gnl1_S34 - cost: 9.72, layer: 58
Fold over basal
Time (min)
0.6
0.8
1
1.2
1.4
1.6
0
10
20
30
40
50
60
Normal
MK50
MK75
MK88
MK94
MK97
MK data
Normal data

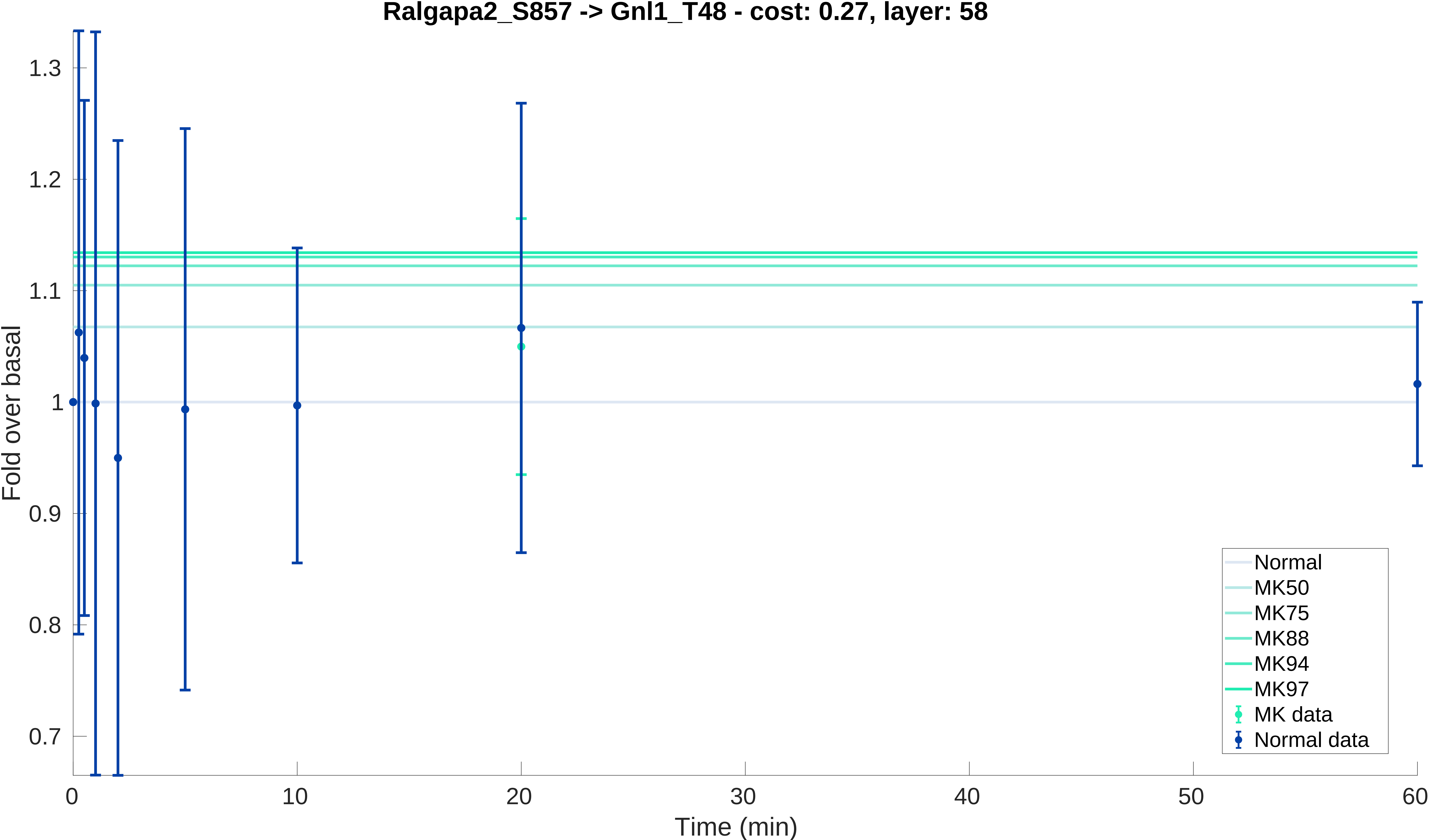
Ralgapa2_S857 -> Gnl1_T48 - cost: 0.27, layer: 58
Fold over basal
Time (min)
1.3
1.2
1.1
1
0.9
0.8
0.7
0
10
20
30
40
50
60
Normal
MK50
MK75
MK88
MK94
MK97
MK data
Normal data

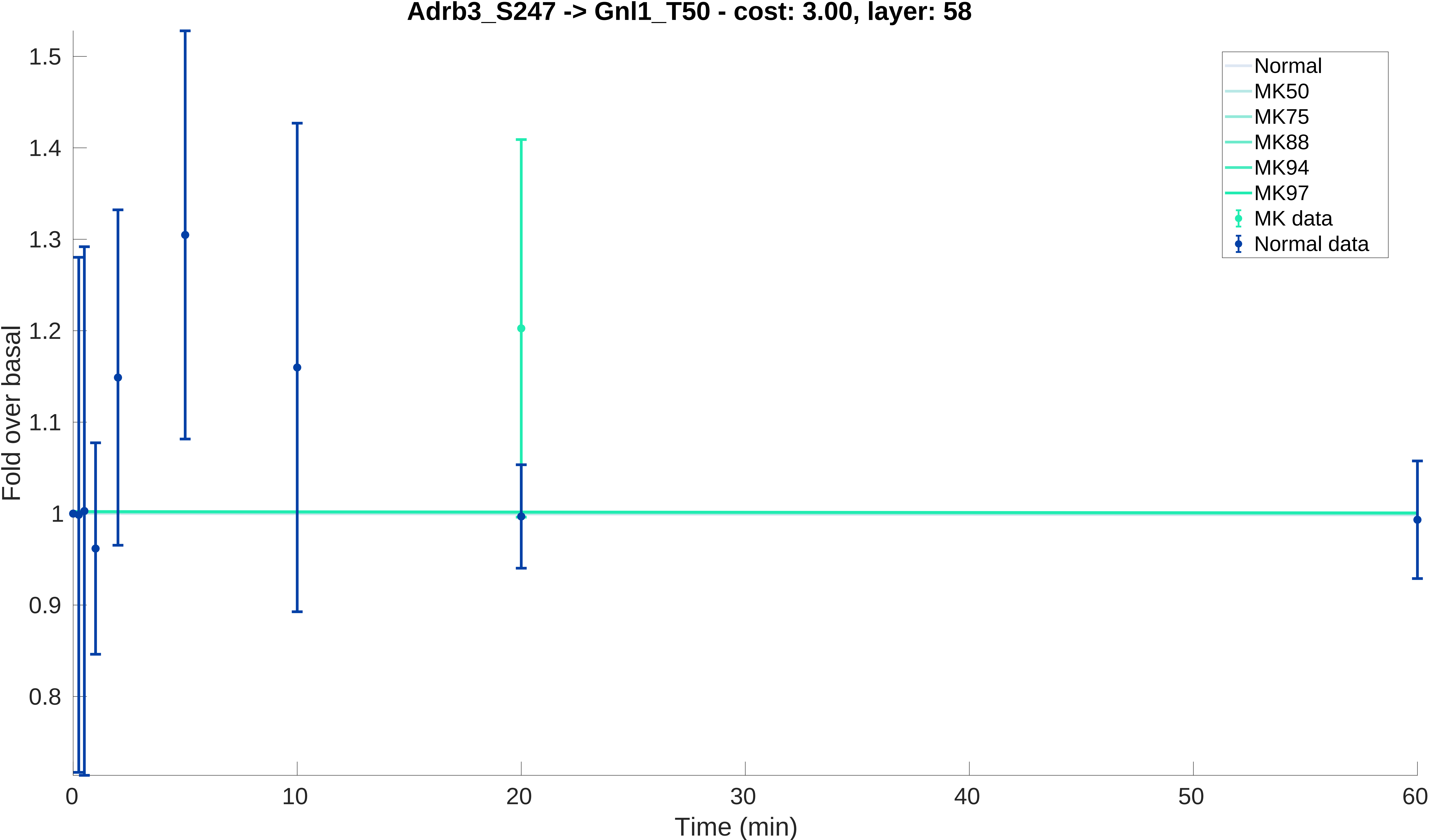

Adrb3_S247 -> Gnl1_T50 - cost: 3.00, layer: 58
Fold over basal
Time (min)
0
10
20
30
40
50
60
0.8
0.9
1
1.1
1.2
1.3
1.4
1.5
Normal
MK50
MK75
MK88
MK94
MK97
MK data
Normal data

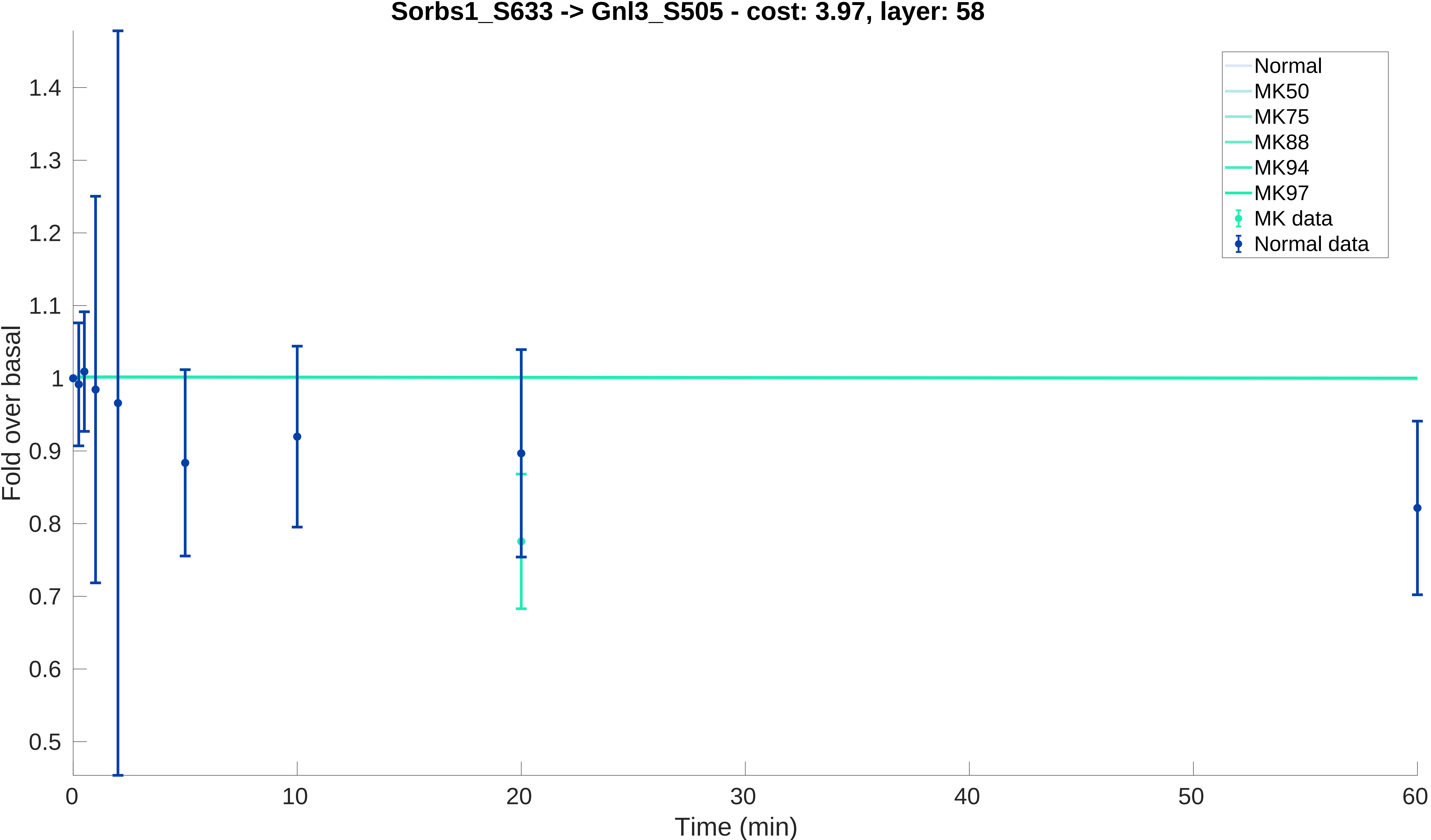

Sorbs1_S633 -> Gnl3_S505 - cost: 3.97, layer: 58
Fold over basal
Time (min)
1.4
1.3
1.2
1.1
1
0.9
0.8
0.7
0.6
0.5
0
10
20
30
40
50
60
Normal
MK50
MK75
MK88
MK94
MK97
MK data
Normal data

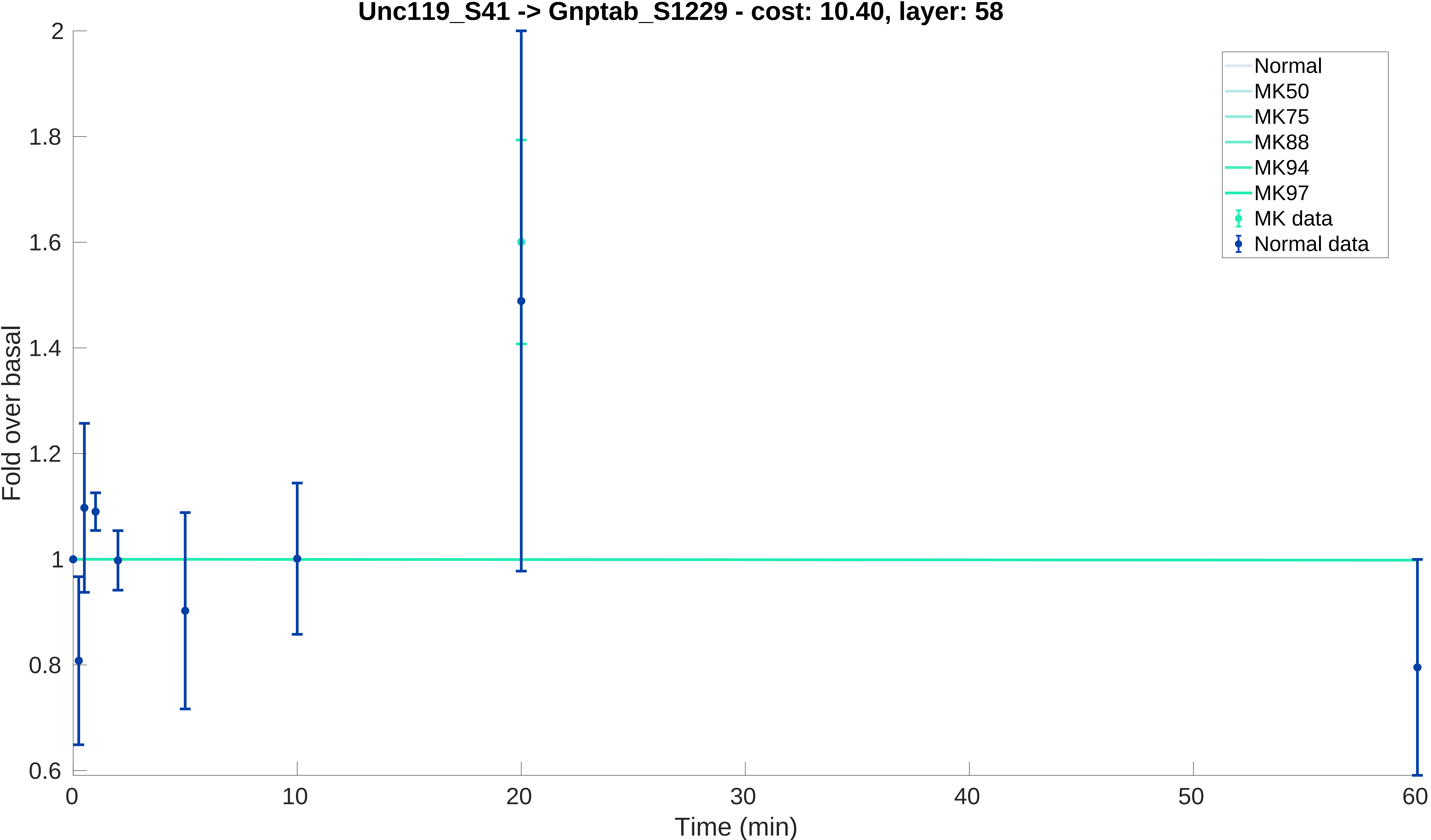

Unc119_S41 -> Gnptab_S1229 - cost: 10.40, layer: 58
Fold over basal
Time (min)
0.6
0.8
1
1.2
1.4
1.6
1.8
2
0
10
20
30
40
50
60
Normal
MK50
MK75
MK88
MK94
MK97
MK data
Normal data

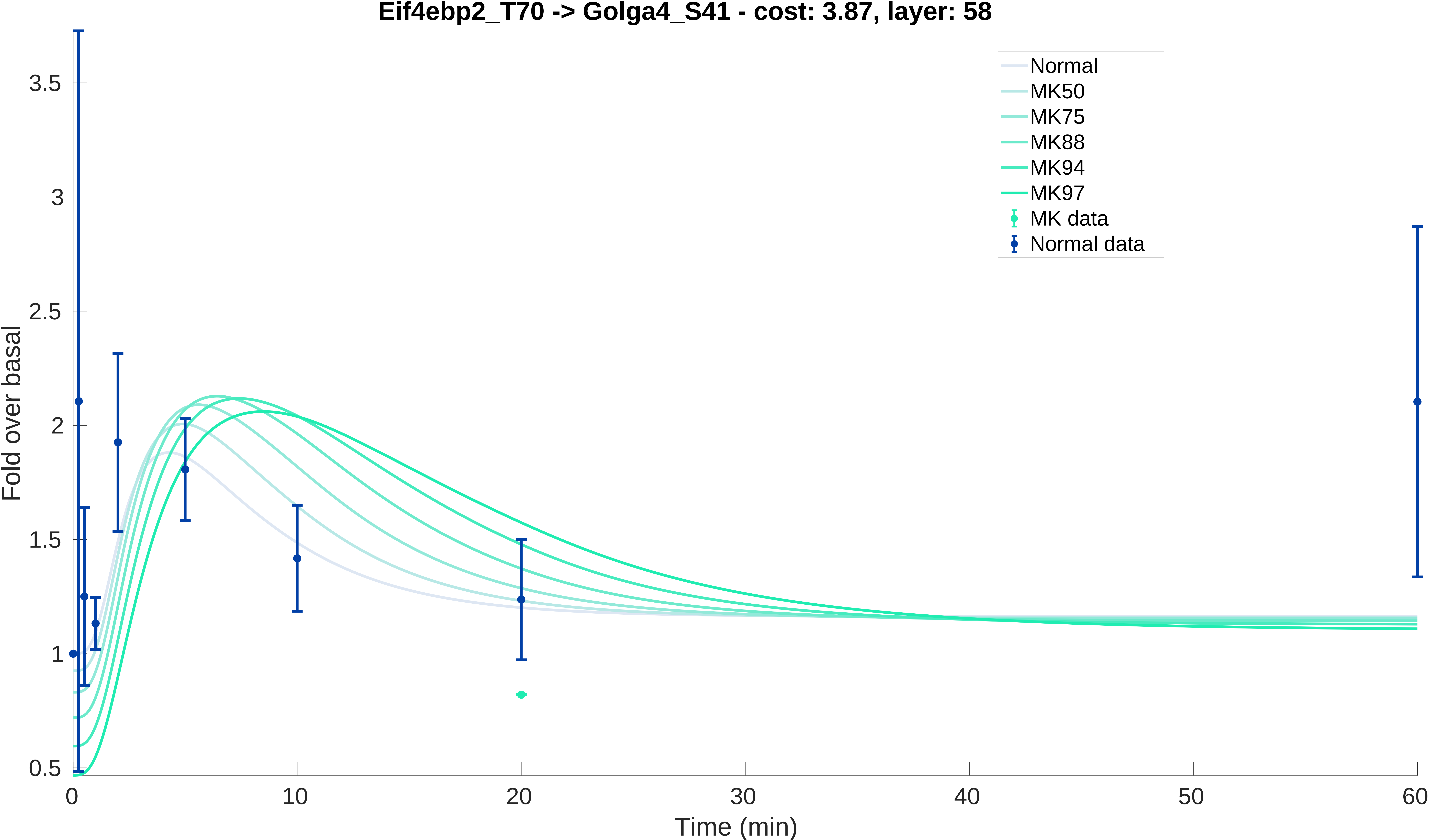

Eif4ebp2_T70 -> Golga4_S41 - cost: 3.87, layer: 58
Normal
MK50
MK75
MK88
MK94
MK97
MK data
Normal data
Fold over basal
Time (min)
0.5
1
1.5
2
2.5
3
3.5
0
10
20
30
40
50
60

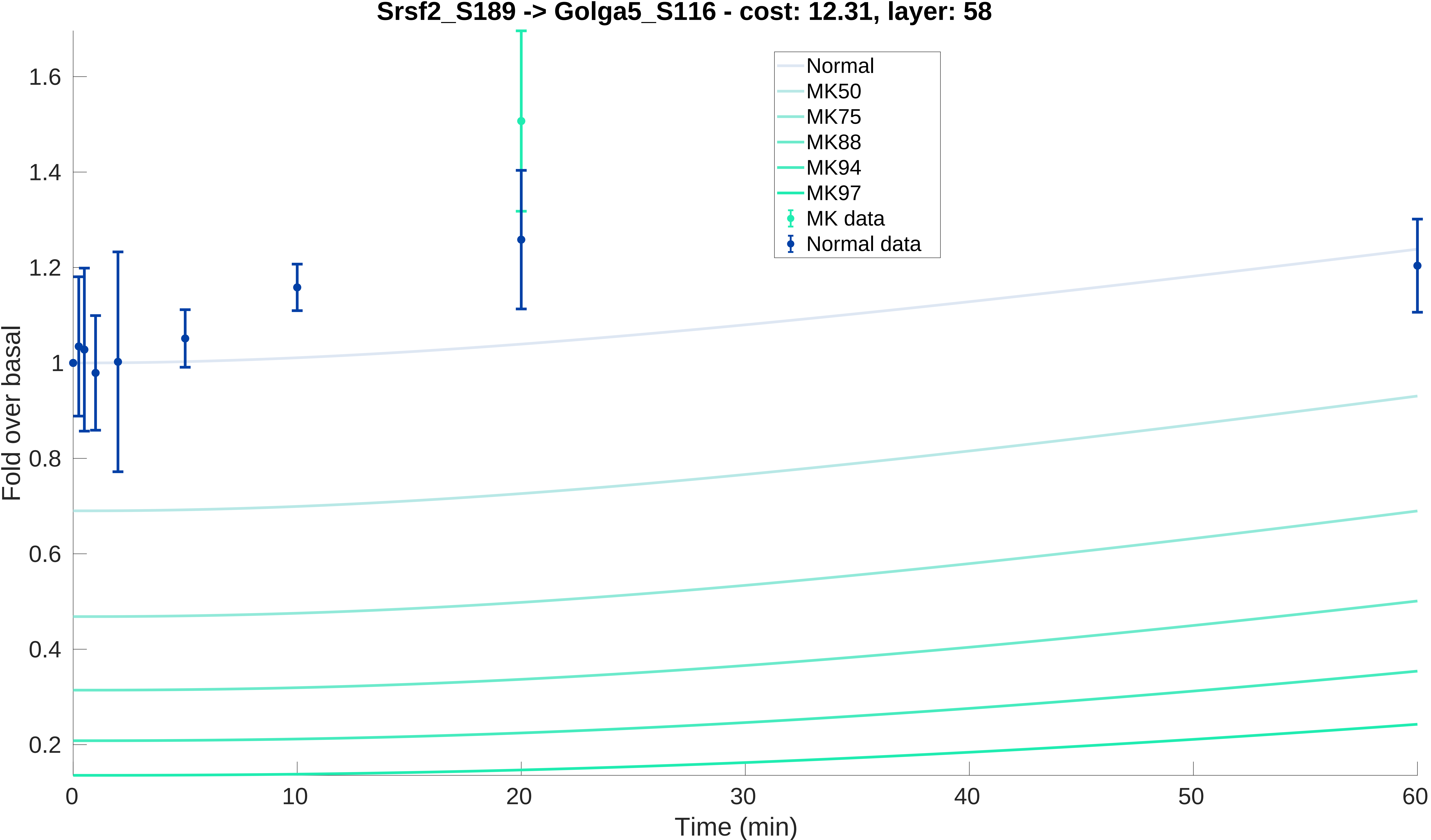

Srsf2_S189 -> Golga5_S116 - cost: 12.31, layer: 58
Normal
MK50
MK75
MK88
MK94
MK97
MK data
Normal data
Fold over basal
Time (min)
0
10
20
30
40
50
60
0.2
0.4
0.6
0.8
1
1.2
1.4
1.6

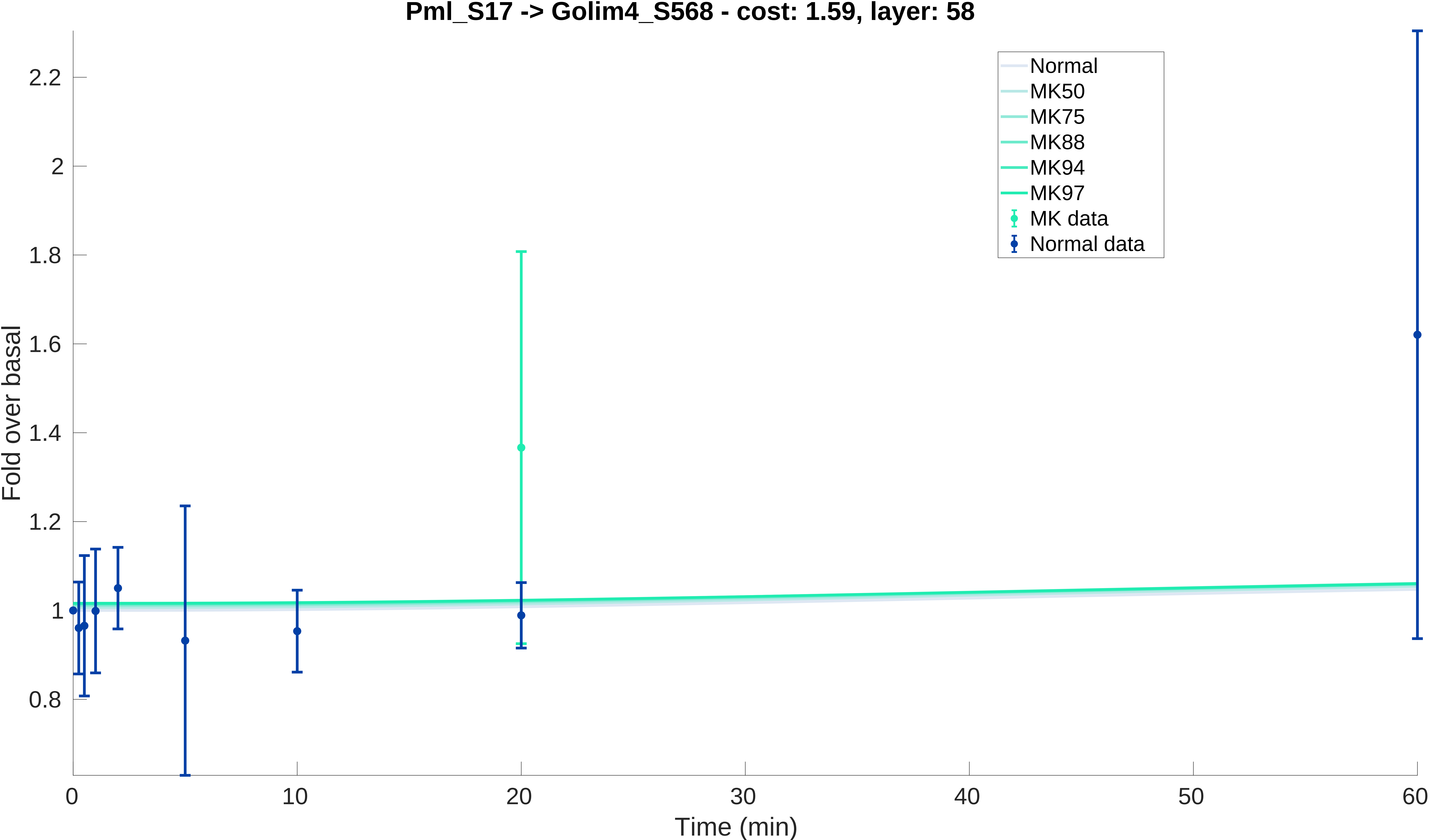

Pml_S17 -> Golim4_S568 - cost: 1.59, layer: 58
Normal
MK50
MK75
MK88
MK94
MK97
MK data
Normal data
Fold over basal
2.2
2
1.8
1.6
1.4
1.2
1
0.8
0
10
20
30
40
50
60
Time (min)

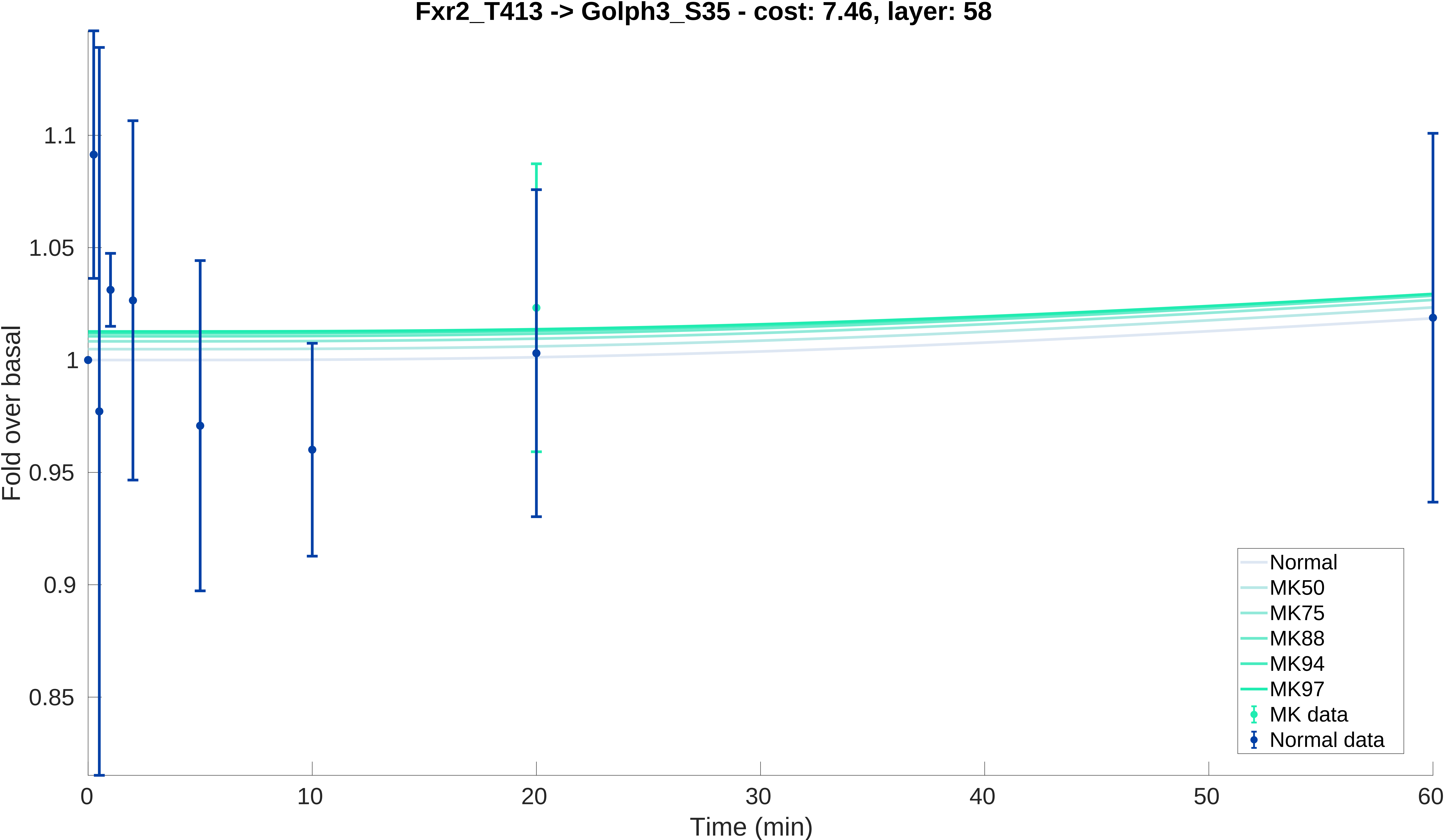

Fxr2_T413 -> Golph3_S35 - cost: 7.46, layer: 58
Fold over basal
Time (min)
1.1
1.05
1
0.95
0.9
0.85
0
10
20
30
40
50
60
Normal
MK50
MK75
MK88
MK94
MK97
MK data
Normal data

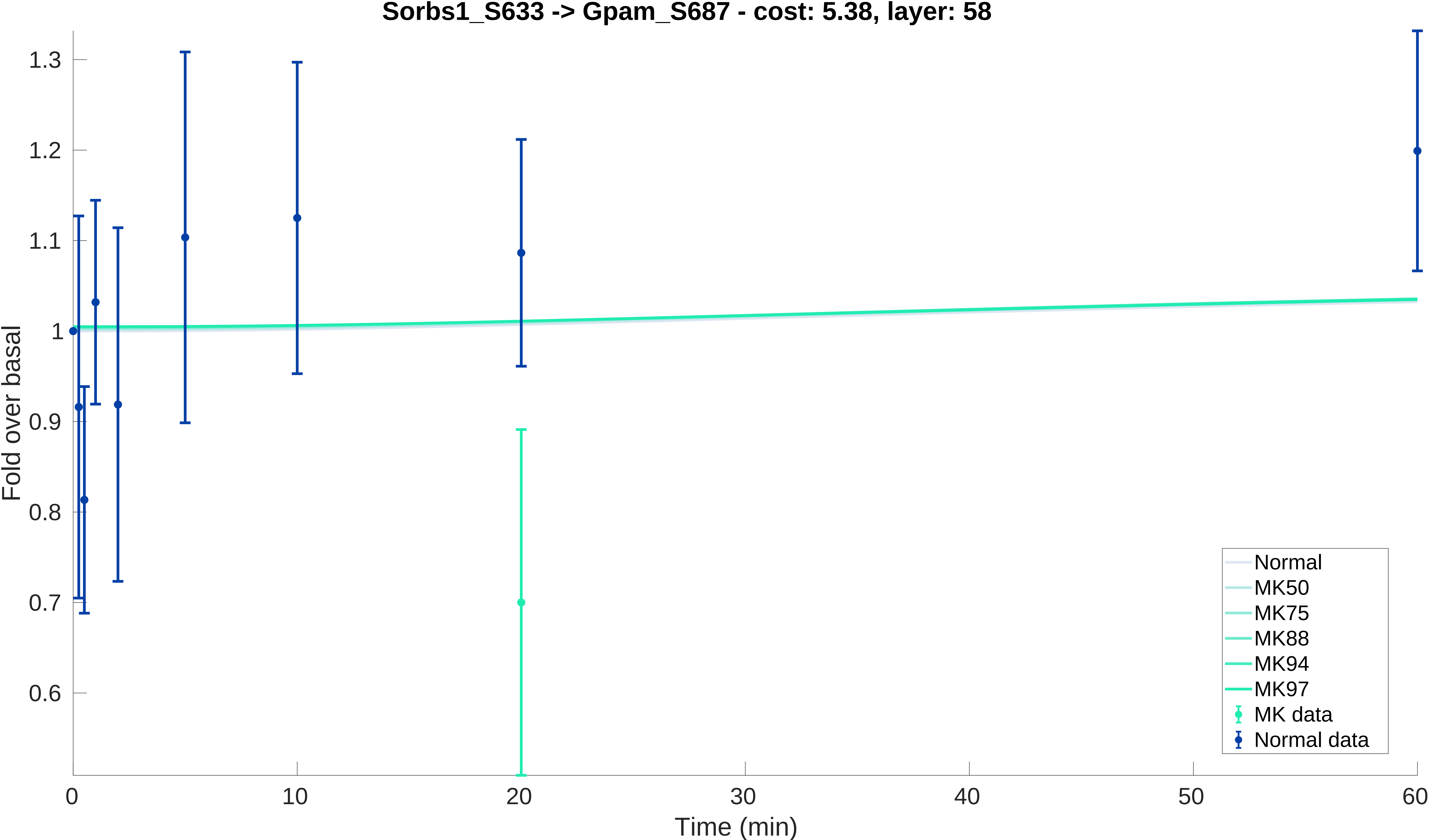
Sorbs1_S633 -> Gpam_S687 - cost: 5.38, layer: 58
Fold over basal
Time (min)
1.3
1.2
1.1
1
0.9
0.8
0.7
0.6
0
10
20
30
40
50
60
Normal
MK50
MK75
MK88
MK94
MK97
MK data
Normal data

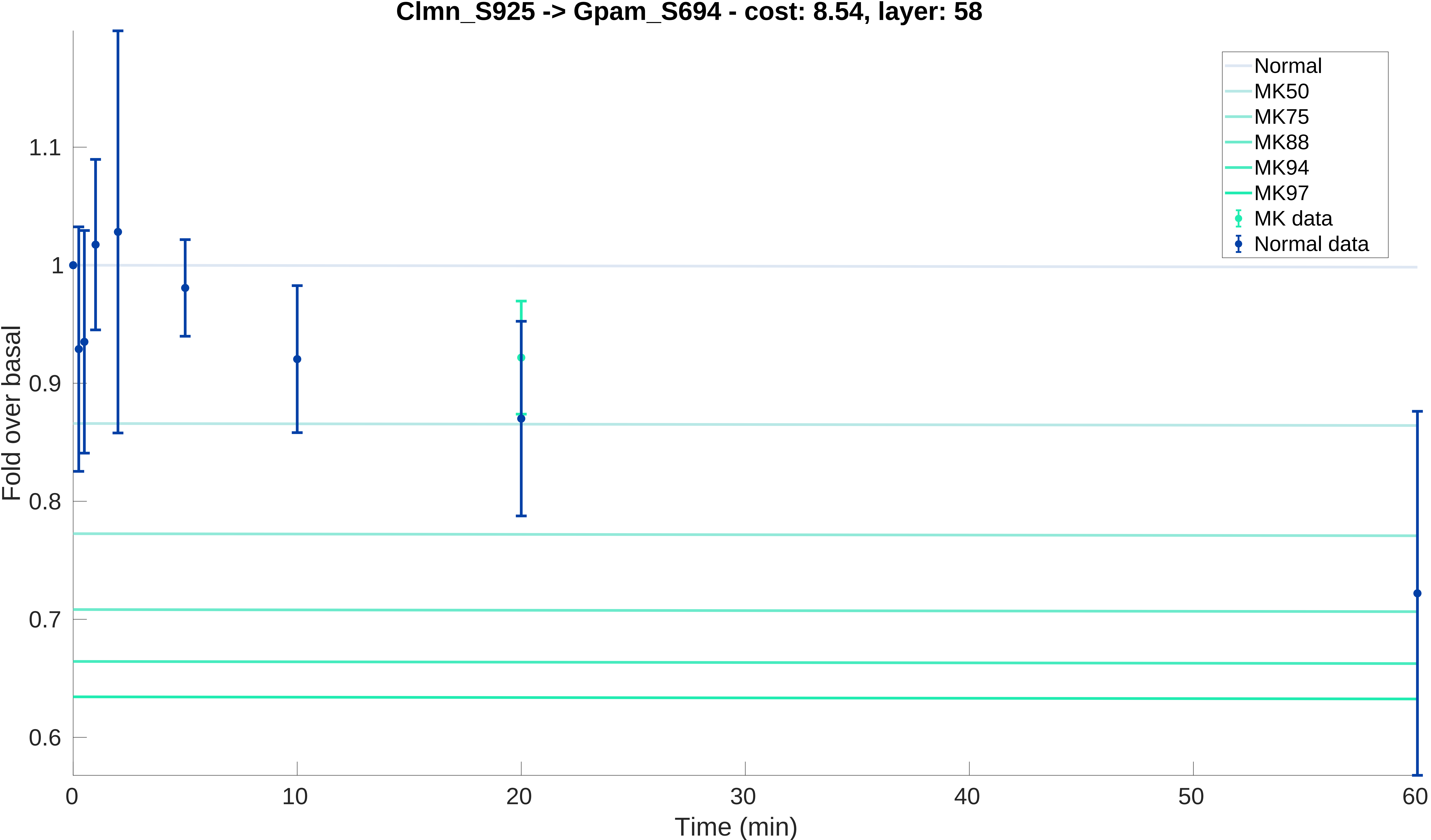
Clmn_S925 -> Gpam_S694 - cost: 8.54, layer: 58
Fold over basal
1.1
1
0.9
0.8
0.7
0.6
0
10
20
30
40
50
60
Time (min)
Normal
MK50
MK75
MK88
MK94
MK97
MK data
Normal data

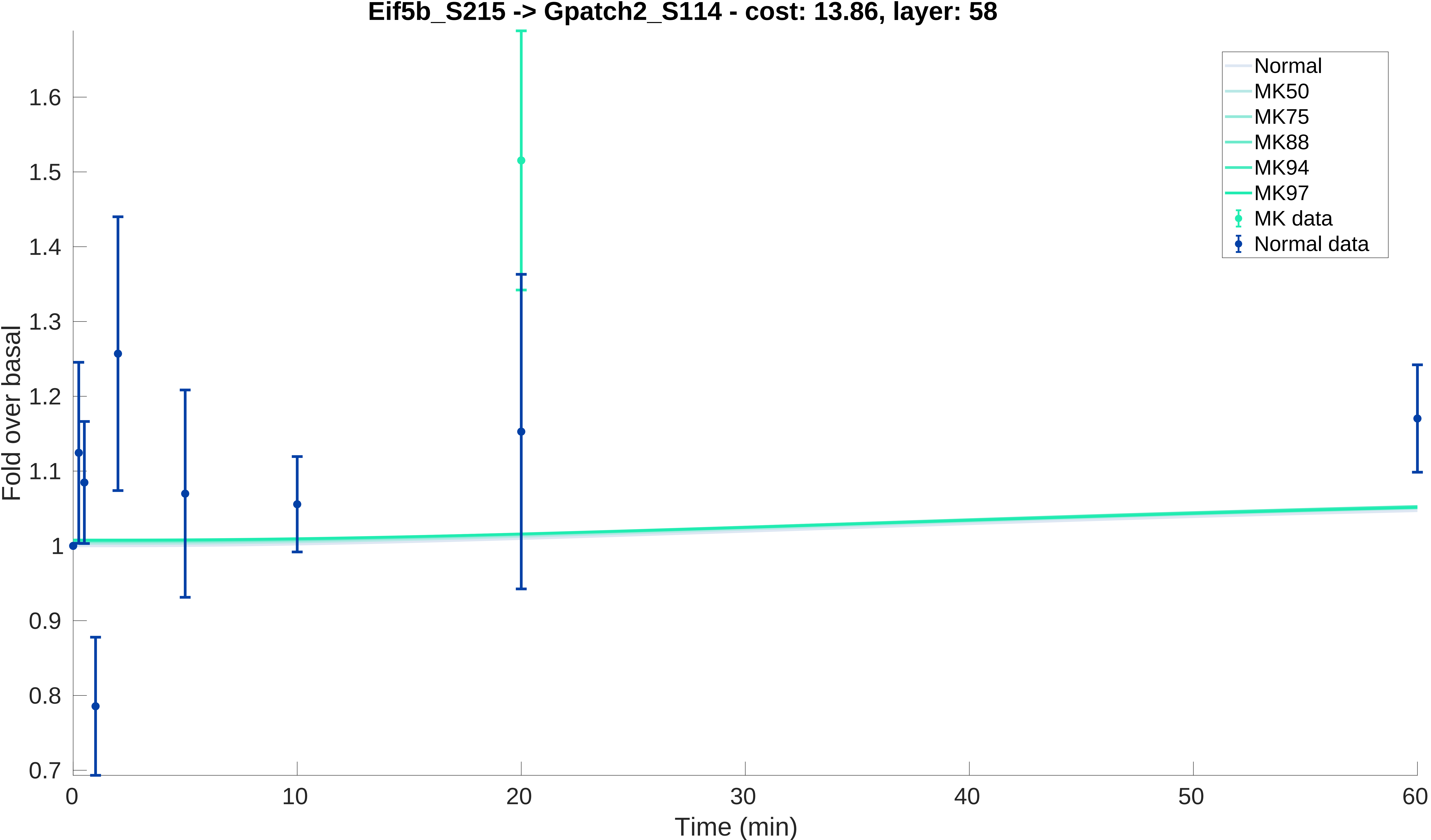

Eif5b_S215 -> Gpatch2_S114 - cost: 13.86, layer: 58
Fold over basal
Time (min)
0.7
0.8
0.9
1
1.1
1.2
1.3
1.4
1.5
1.6
0
10
20
30
40
50
60
Normal
MK50
MK75
MK88
MK94
MK97
MK data
Normal data

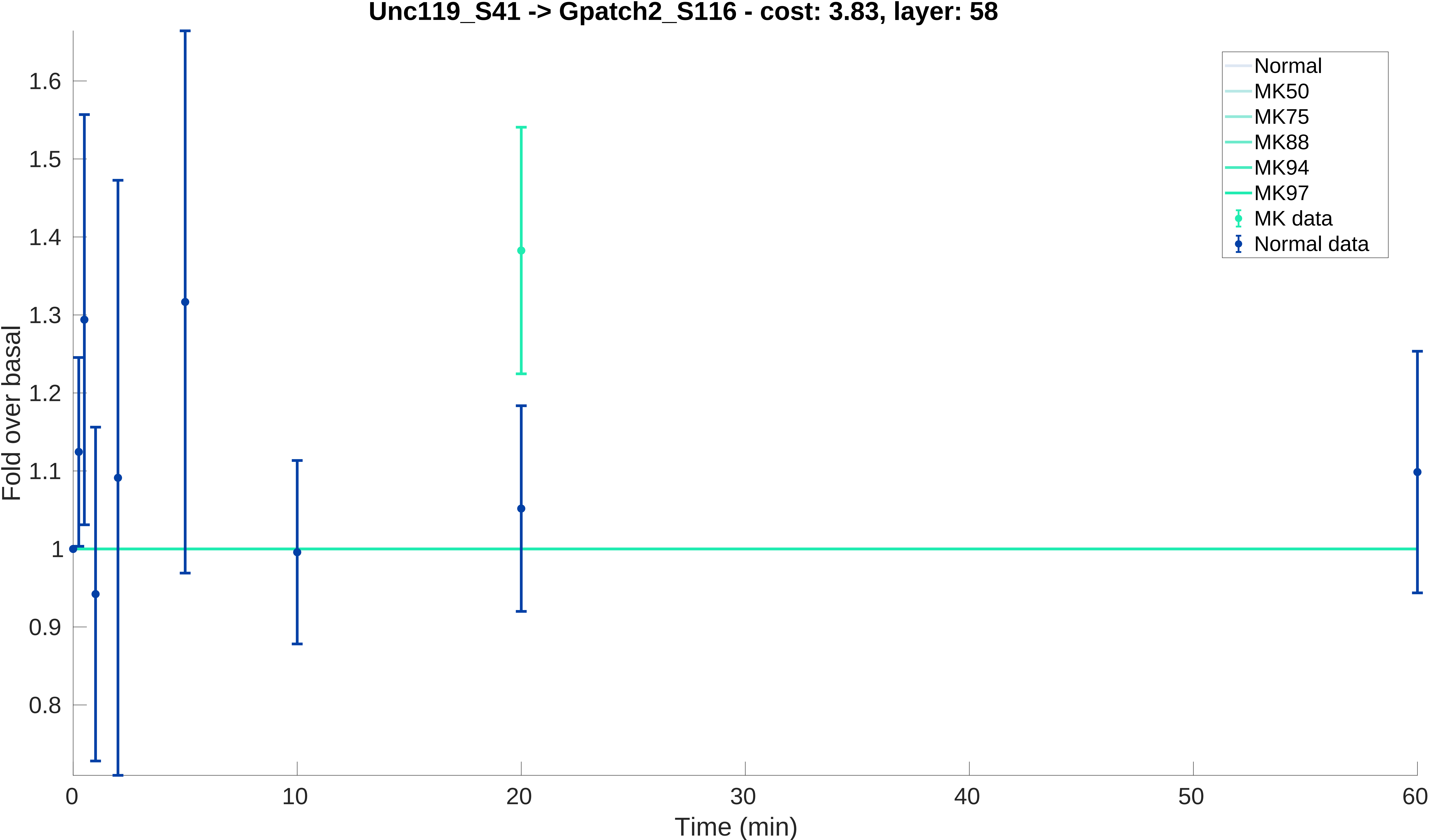

Unc119_S41 -> Gpatch2_S116 - cost: 3.83, layer: 58
Fold over basal
Time (min)
0.8
0.9
1
1.1
1.2
1.3
1.4
1.5
1.6
0
10
20
30
40
50
60
Normal
MK50
MK75
MK88
MK94
MK97
MK data
Normal data

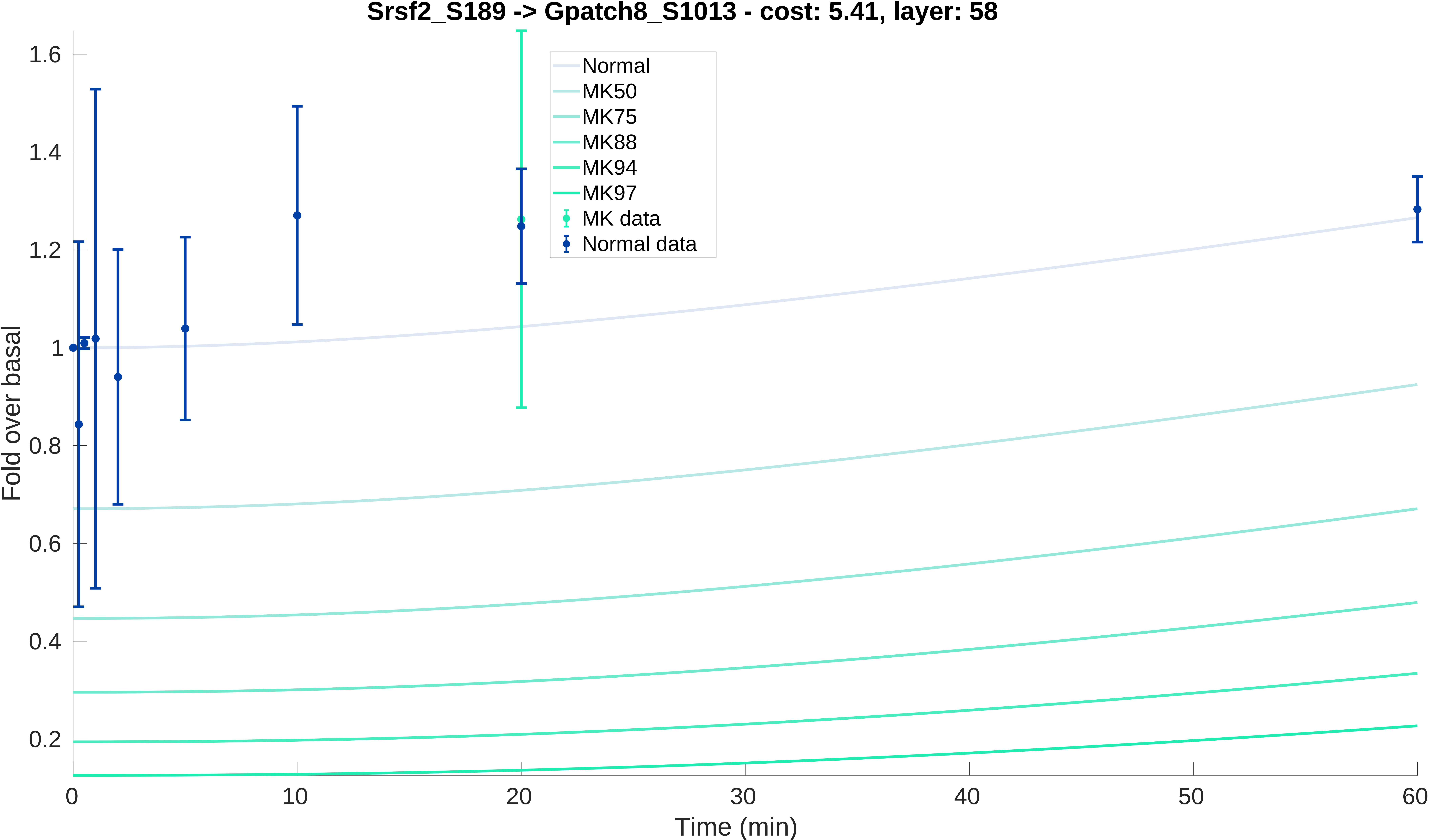

Srsf2_S189 -> Gpatch8_S1013 - cost: 5.41, layer: 58
Fold over basal
Time (min)
Normal
MK50
MK75
MK88
MK94
MK97
MK data
Normal data
0.2
0.4
0.6
0.8
1
1.2
1.4
1.6
0
10
20
30
40
50
60

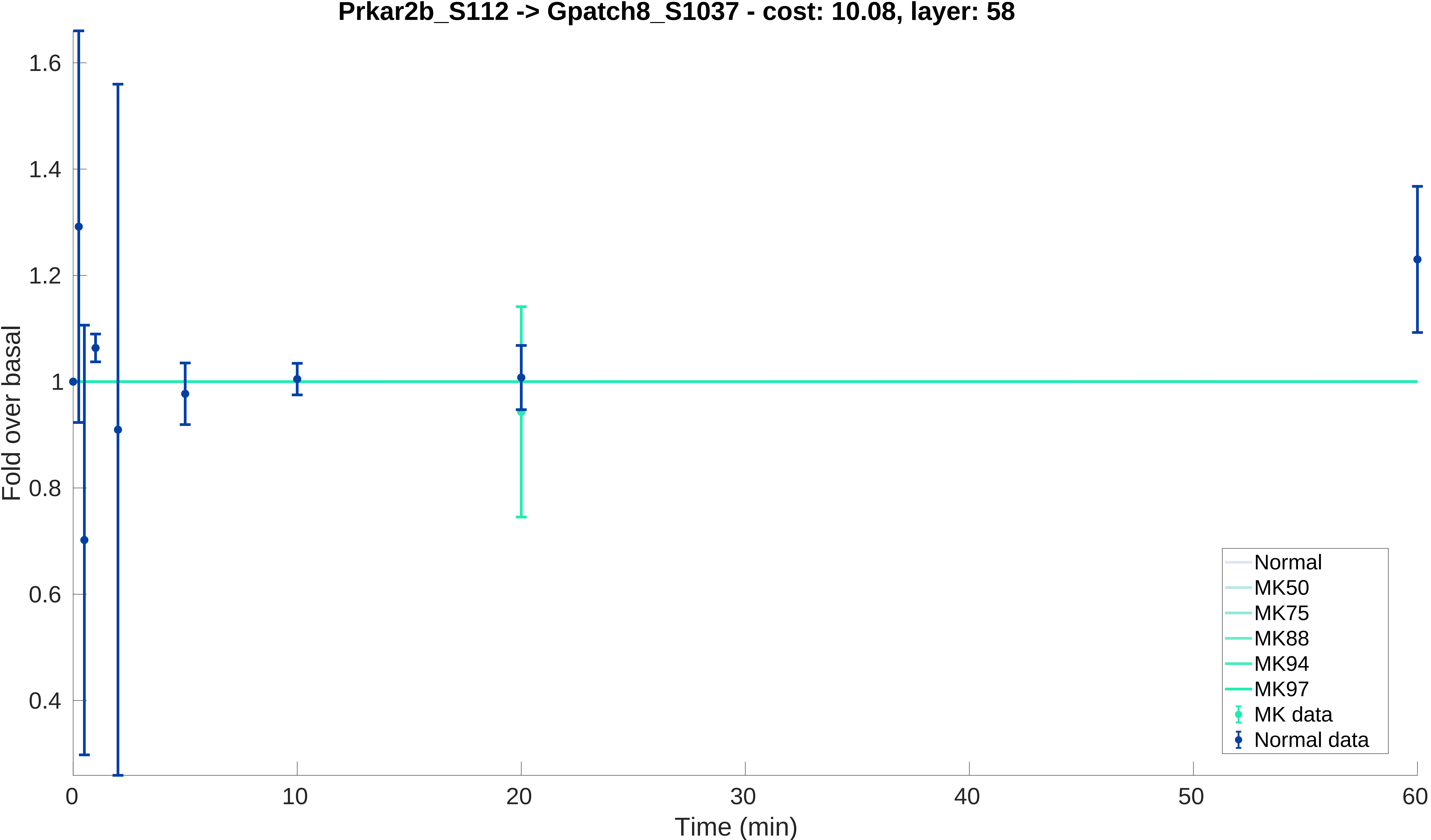

Prkar2b_S112 -> Gpatch8_S1037 - cost: 10.08, layer: 58
Fold over basal
Time (min)
1.6
1.4
1.2
1
0.8
0.6
0.4
0
10
20
30
40
50
60
Normal
MK50
MK75
MK88
MK94
MK97
MK data
Normal data

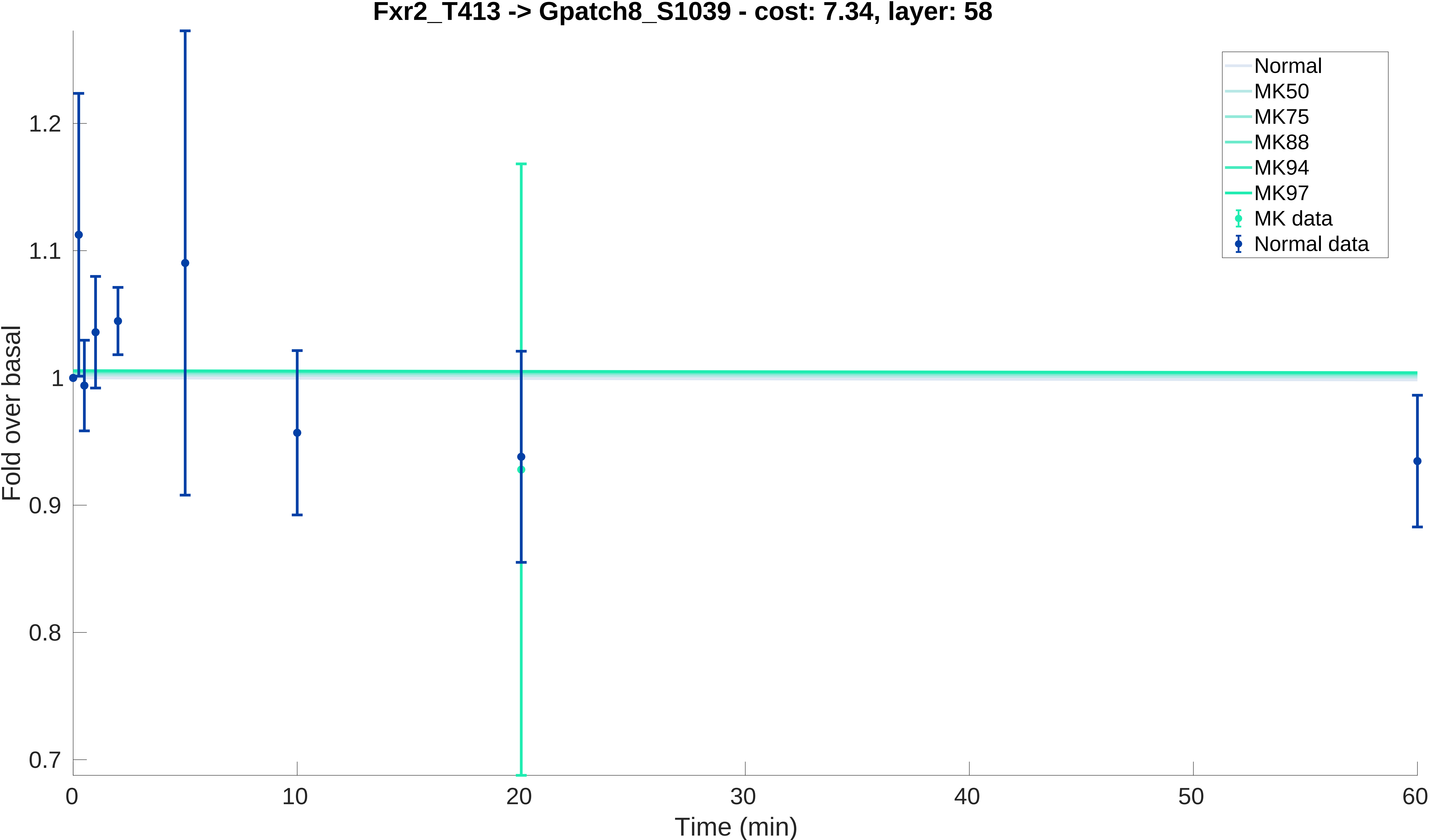

Fxr2_T413 -> Gpatch8_S1039 - cost: 7.34, layer: 58
Normal
MK50
MK75
MK88
MK94
MK97
MK data
Normal data
Fold over basal
1.2
1.1
1
0.9
0.8
0.7
0
10
20
30
40
50
60
Time (min)

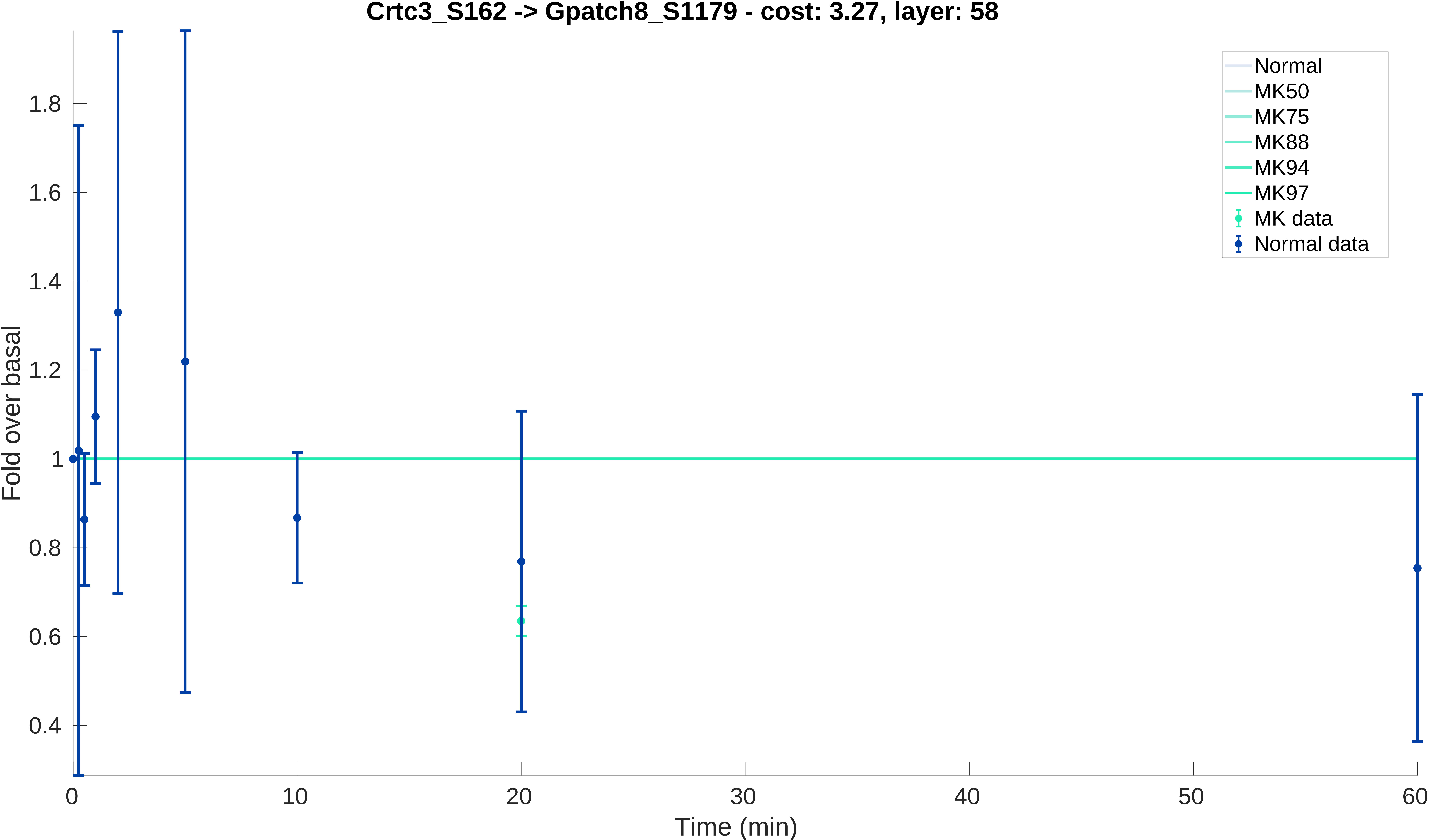

Crtc3_S162 -> Gpatch8_S1179 - cost: 3.27, layer: 58
Fold over basal
Time (min)
0.4
0.6
0.8
1
1.2
1.4
1.6
1.8
0
10
20
30
40
50
60
Normal
MK50
MK75
MK88
MK94
MK97
MK data
Normal data

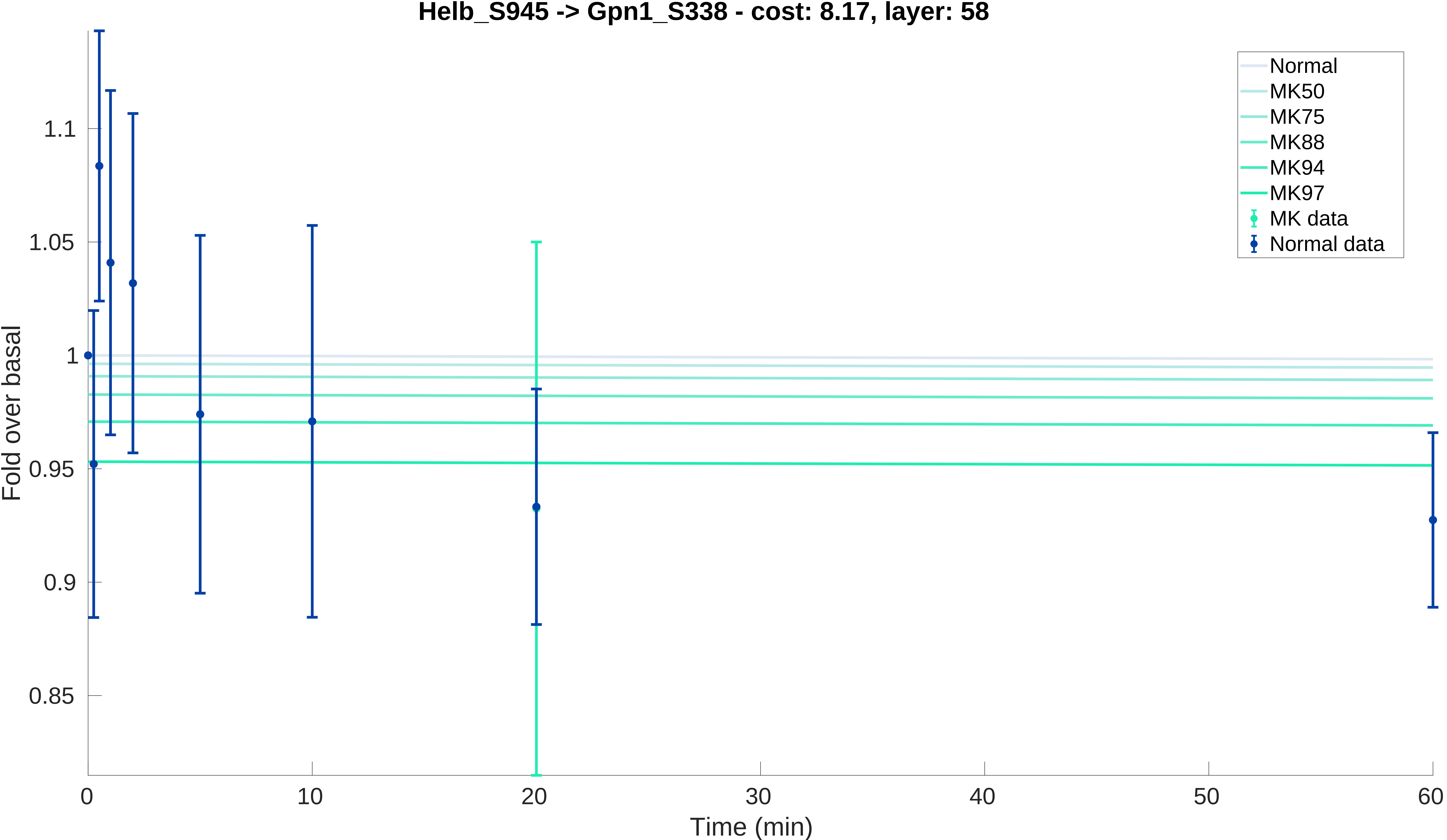
Helb_S945 -> Gpn1_S338 - cost: 8.17, layer: 58
Normal
MK50
MK75
MK88
MK94
MK97
MK data
Normal data
Fold over basal
1.1
1.05
1
0.95
0.9
0.85
0
10
20
30
40
50
60
Time (min)

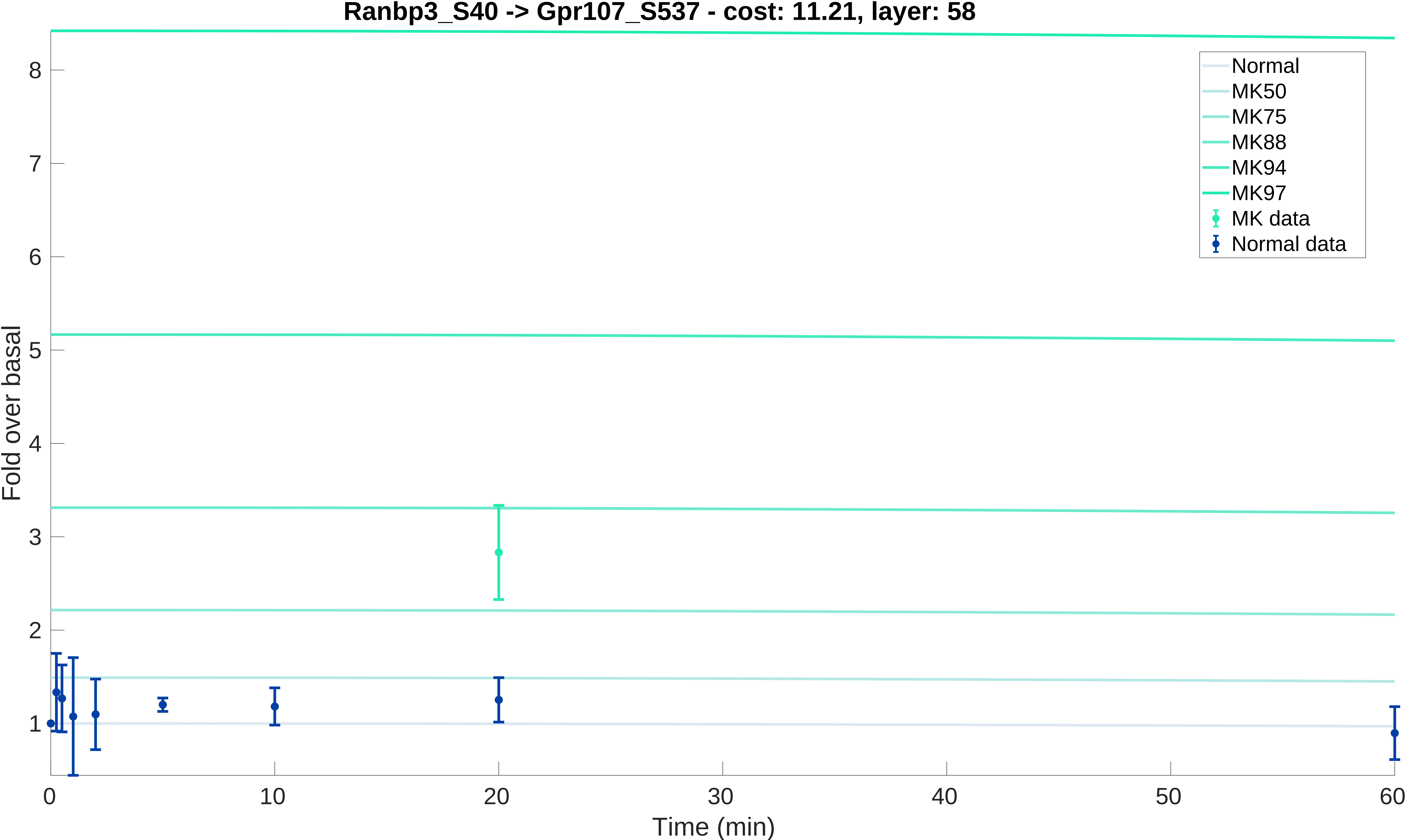

Ranbp3_S40 -> Gpr107_S537 - cost: 11.21, layer: 58
Fold over basal
Time (min)
0
10
20
30
40
50
60
1
2
3
4
5
6
7
8
Normal
MK50
MK75
MK88
MK94
MK97
MK data
Normal data

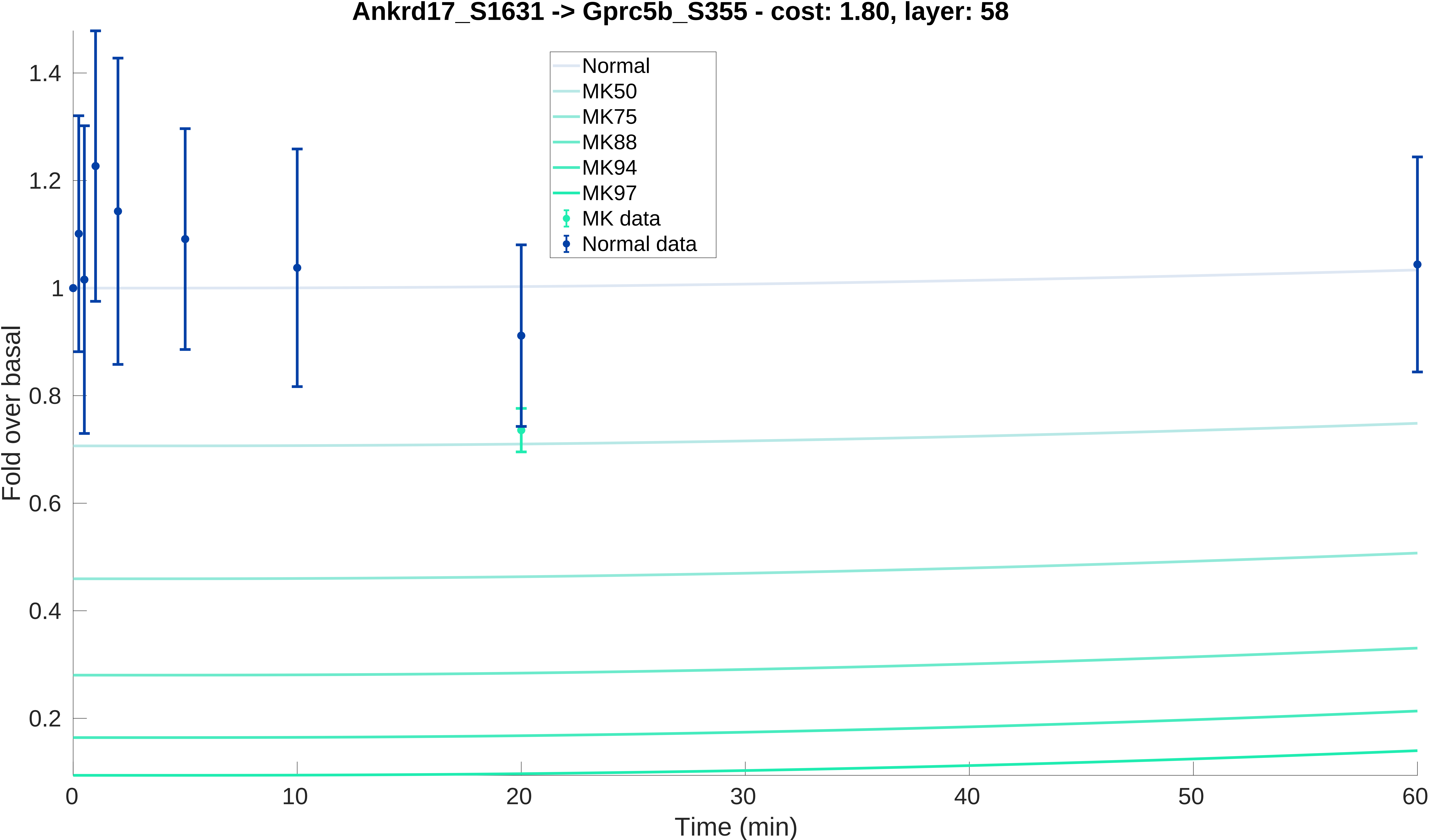

Ankrd17_S1631 -> Gprc5b_S355 - cost: 1.80, layer: 58
Normal
MK50
MK75
MK88
MK94
MK97
MK data
Normal data
Fold over basal
Time (min)
0
10
20
30
40
50
60
0.2
0.4
0.6
0.8
1
1.2
1.4

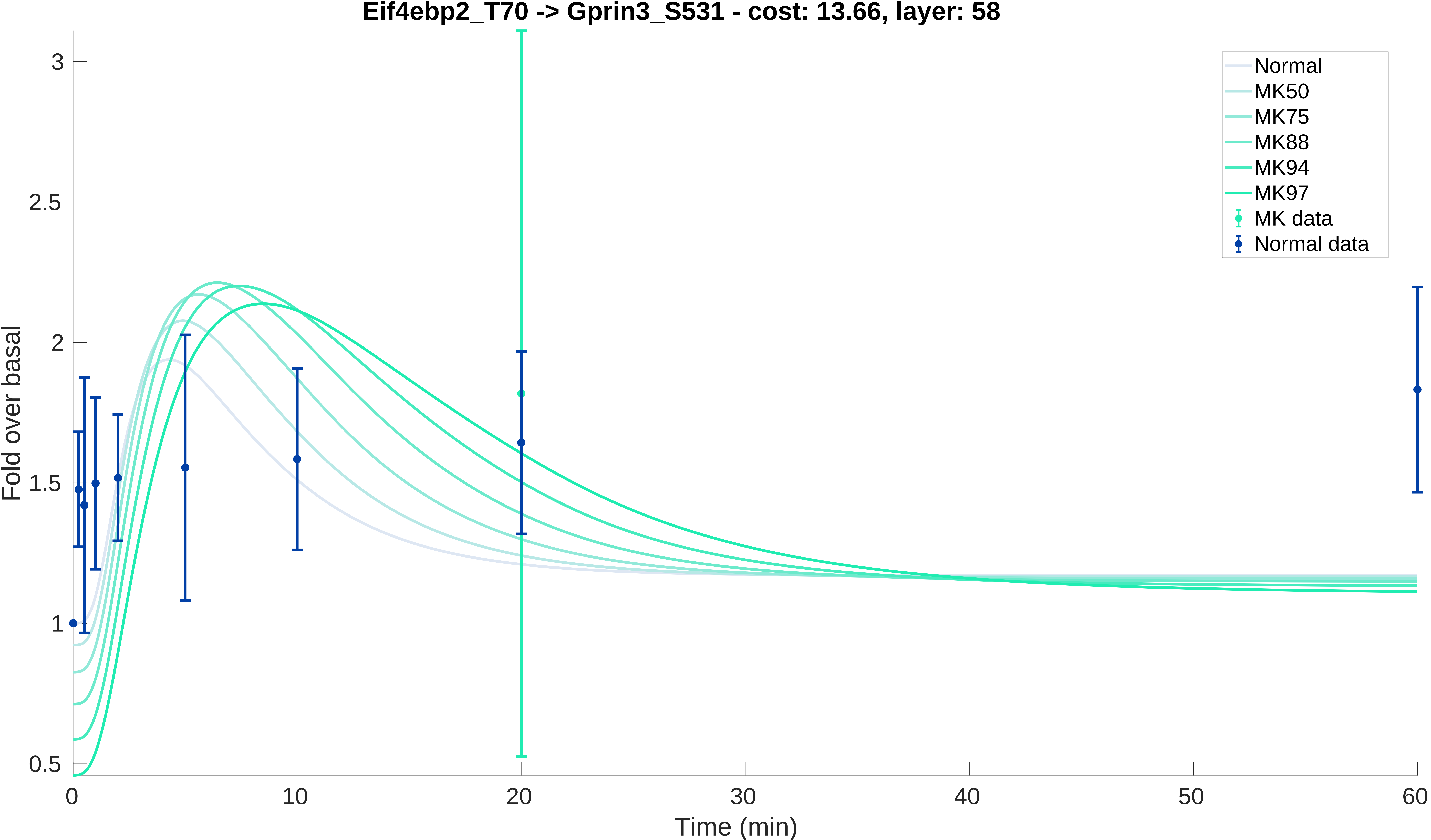

Eif4ebp2_T70 -> Gprin3_S531 - cost: 13.66, layer: 58
Normal
MK50
MK75
MK88
MK94
MK97
MK data
Normal data
Fold over basal
Time (min)
0.5
1
1.5
2
2.5
3
0
10
20
30
40
50
60

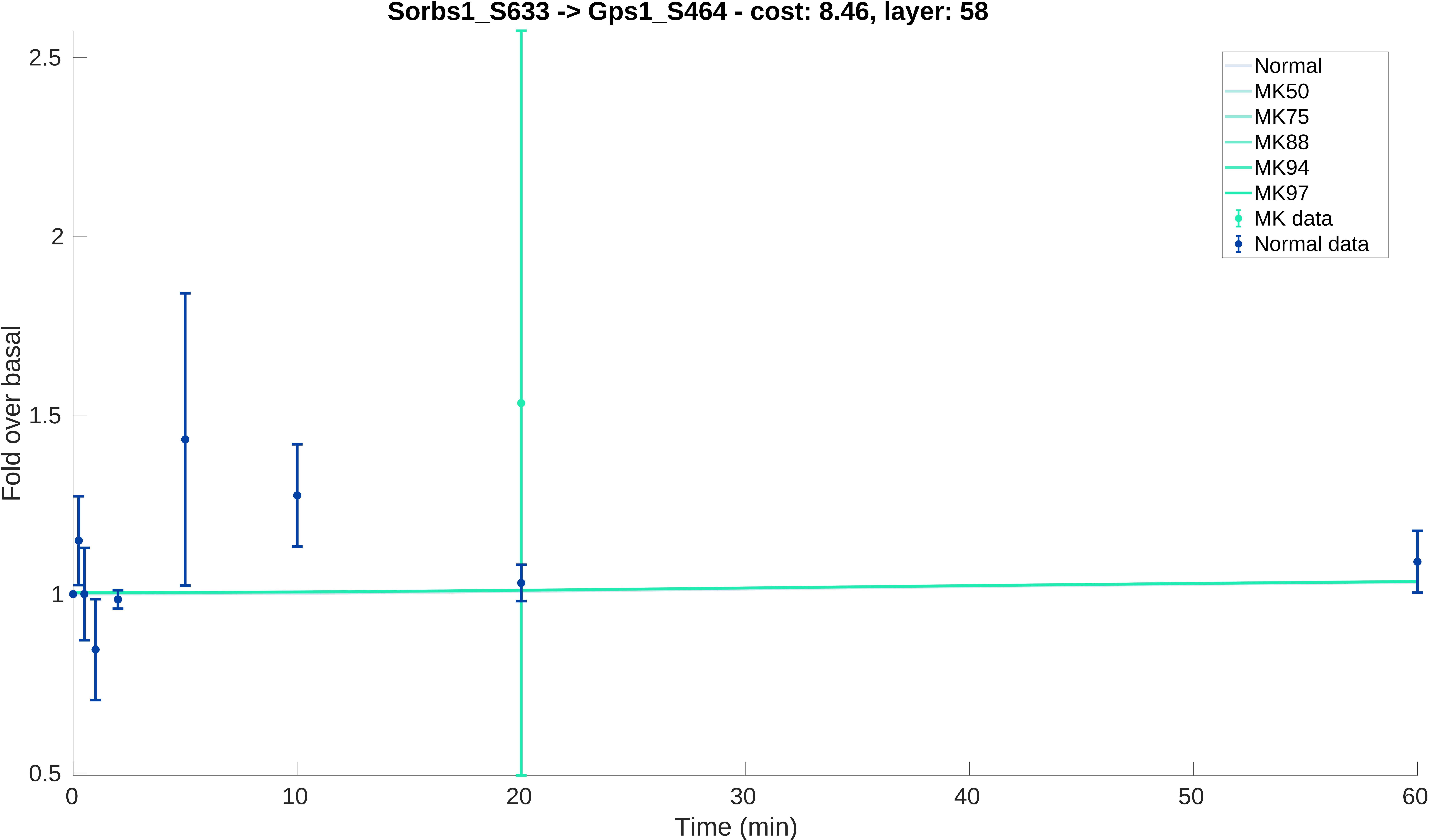

Sorbs1_S633 -> Gps1_S464 - cost: 8.46, layer: 58
Normal
MK50
MK75
MK88
MK94
MK97
MK data
Normal data
Fold over basal
Time (min)
2.5
2
1.5
1
0.5
0
10
20
30
40
50
60

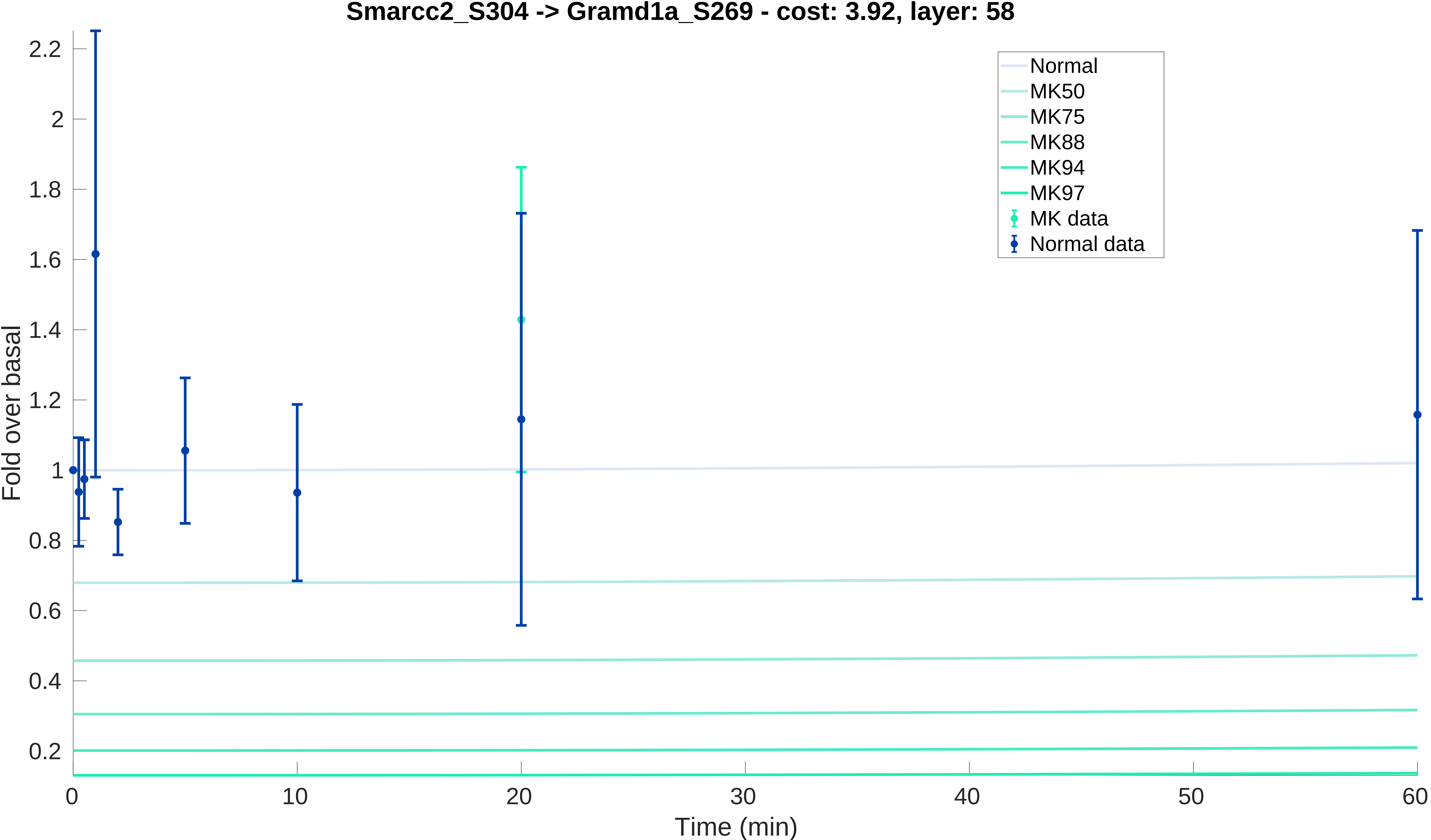
Smarcc2_S304 -> Gramd1a_S269 - cost: 3.92, layer: 58
Fold over basal
2.2
2
1.8
1.6
1.4
1.2
1
0.8
0.6
0.4
0.2
0
10
20
30
40
50
60
Time (min)
Normal
MK50
MK75
MK88
MK94
MK97
MK data
Normal data

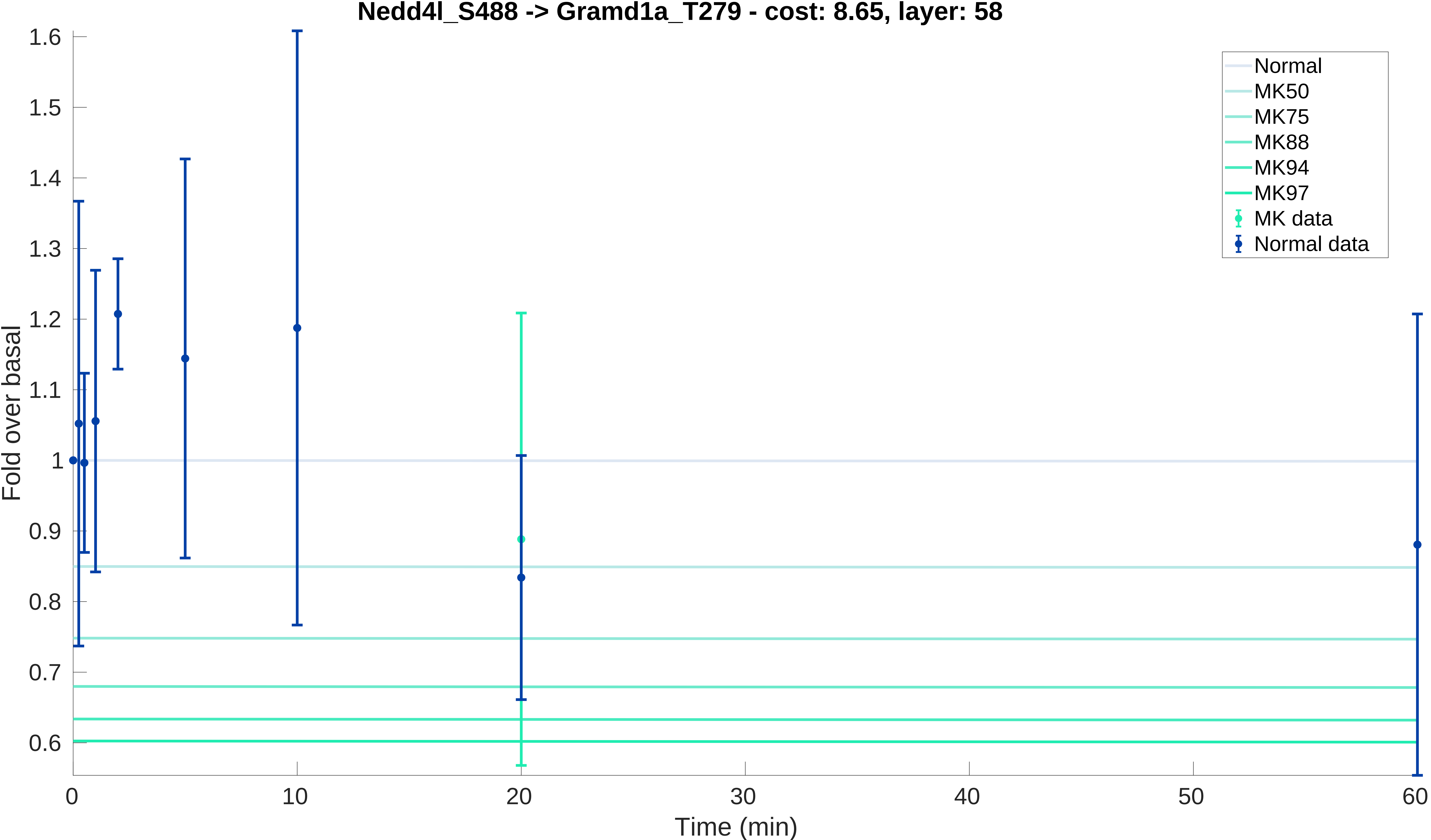
Nedd4l_S488 -> Gramd1a_T279 - cost: 8.65, layer: 58
Fold over basal
1.6
1.5
1.4
1.3
1.2
1.1
1
0.9
0.8
0.7
0.6
0
10
20
30
40
50
60
Time (min)
Normal
MK50
MK75
MK88
MK94
MK97
MK data
Normal data

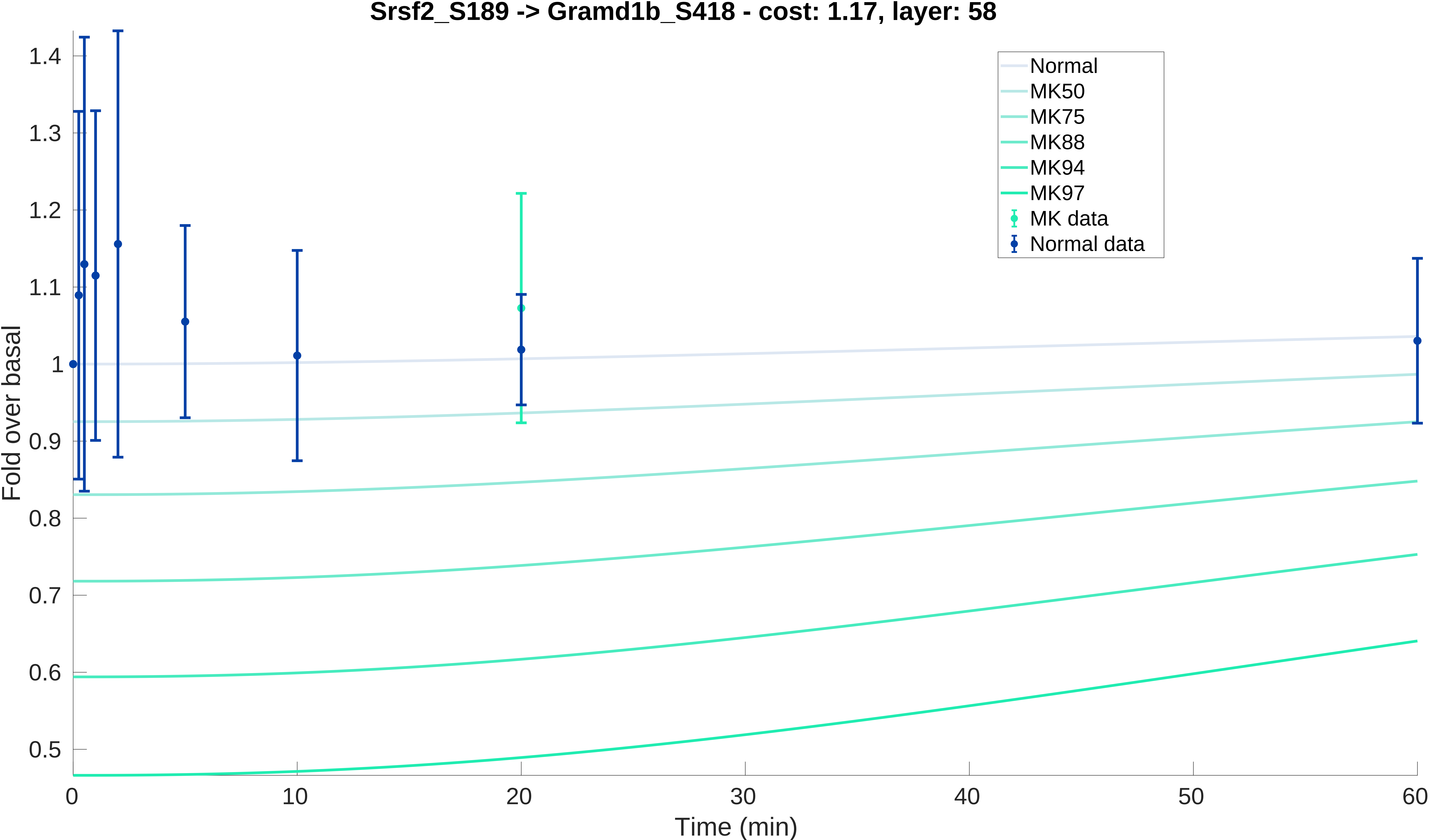

Srsf2_S189 -> Gramd1b_S418 - cost: 1.17, layer: 58
Fold over basal
Time (min)
Normal
MK50
MK75
MK88
MK94
MK97
MK data
Normal data
0.5
0.6
0.7
0.8
0.9
1
1.1
1.2
1.3
1.4
0
10
20
30
40
50
60

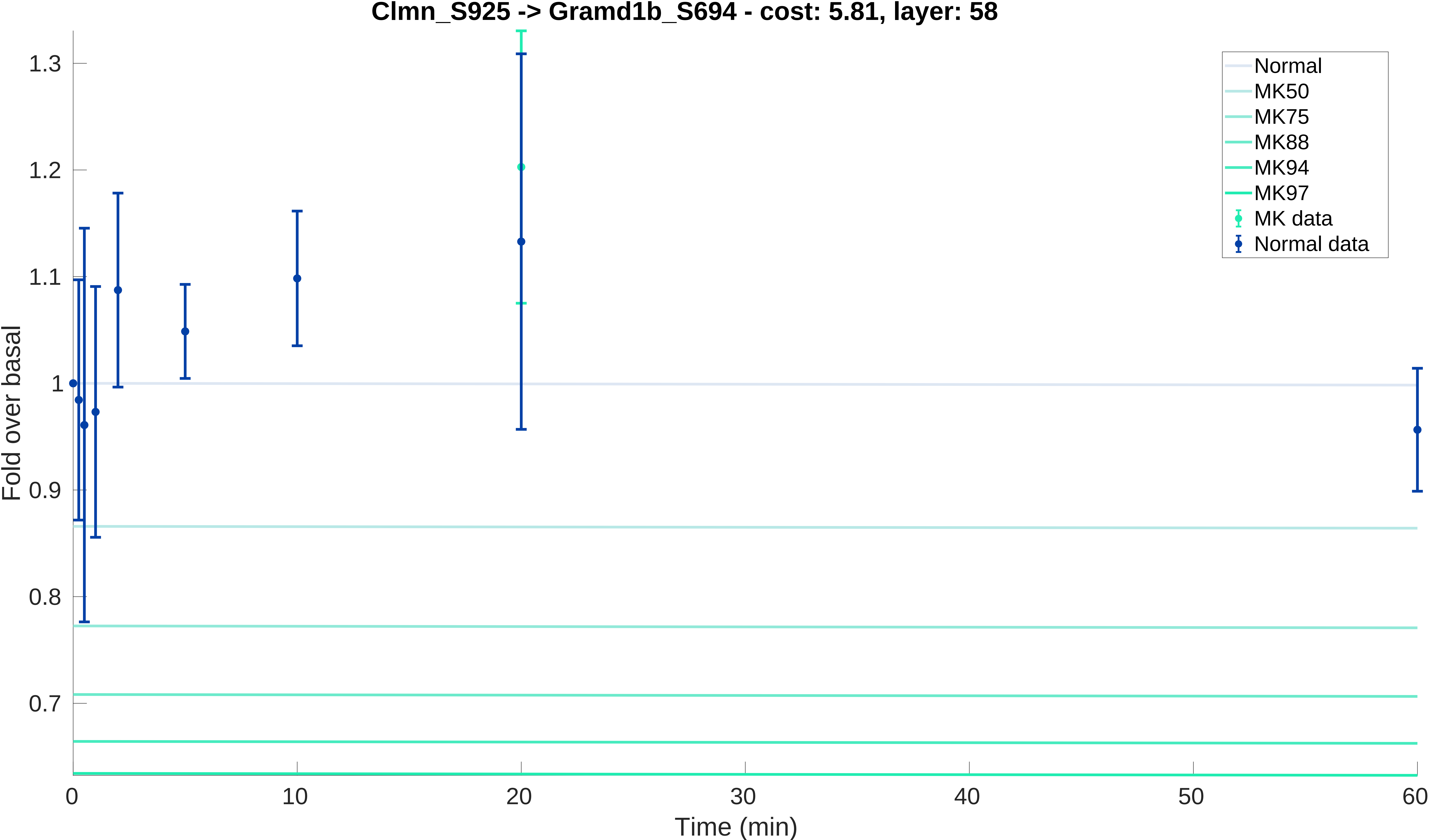

Clmn_S925 -> Gramd1b_S694 - cost: 5.81, layer: 58
Fold over basal
Time (min)
1.3
1.2
1.1
1
0.9
0.8
0.7
0
10
20
30
40
50
60
Normal
MK50
MK75
MK88
MK94
MK97
MK data
Normal data

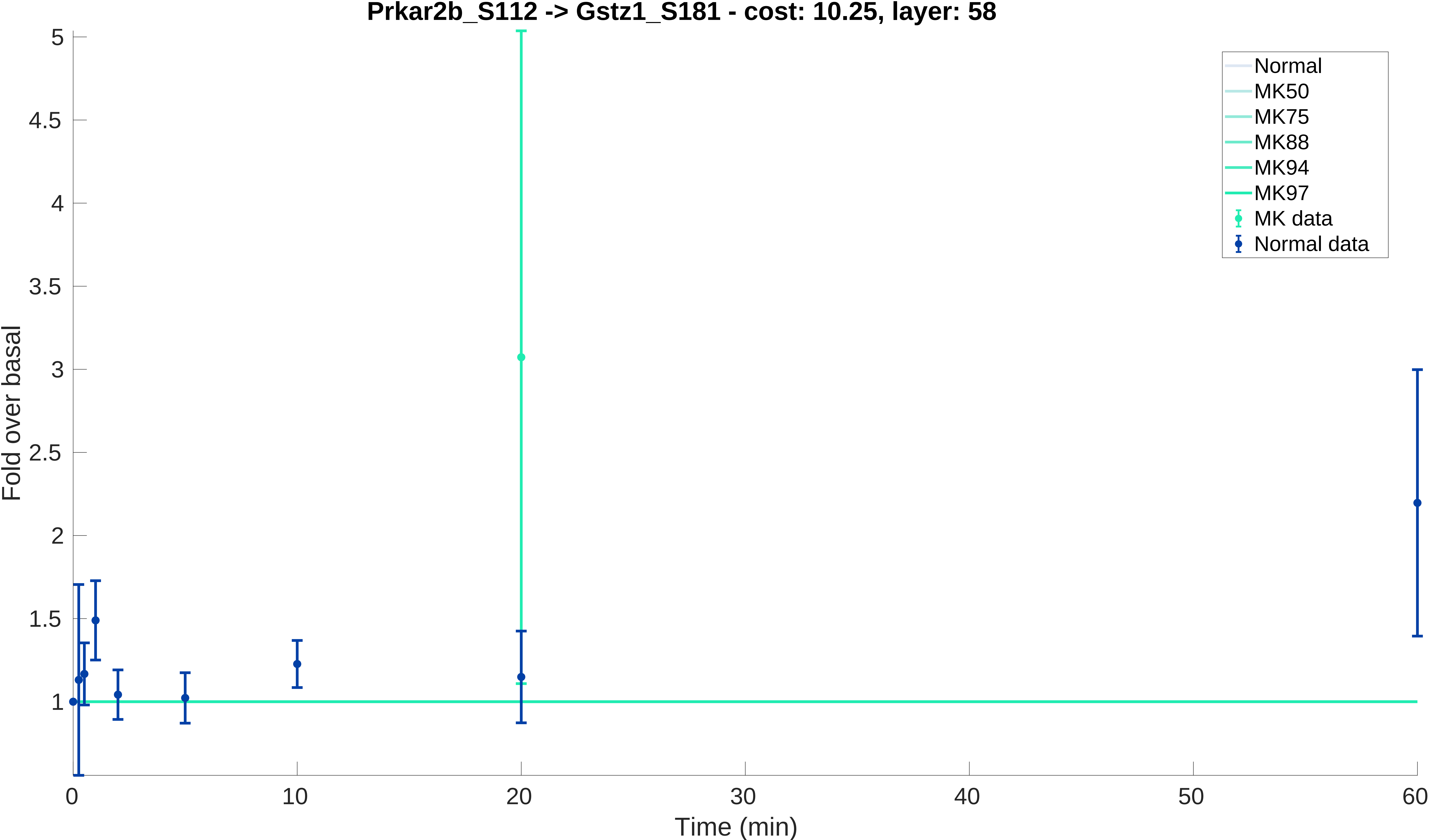

Prkar2b_S112 -> Gstz1_S181 - cost: 10.25, layer: 58
Fold over basal
Time (min)
0
10
20
30
40
50
60
1
1.5
2
2.5
3
3.5
4
4.5
5
Normal
MK50
MK75
MK88
MK94
MK97
MK data
Normal data

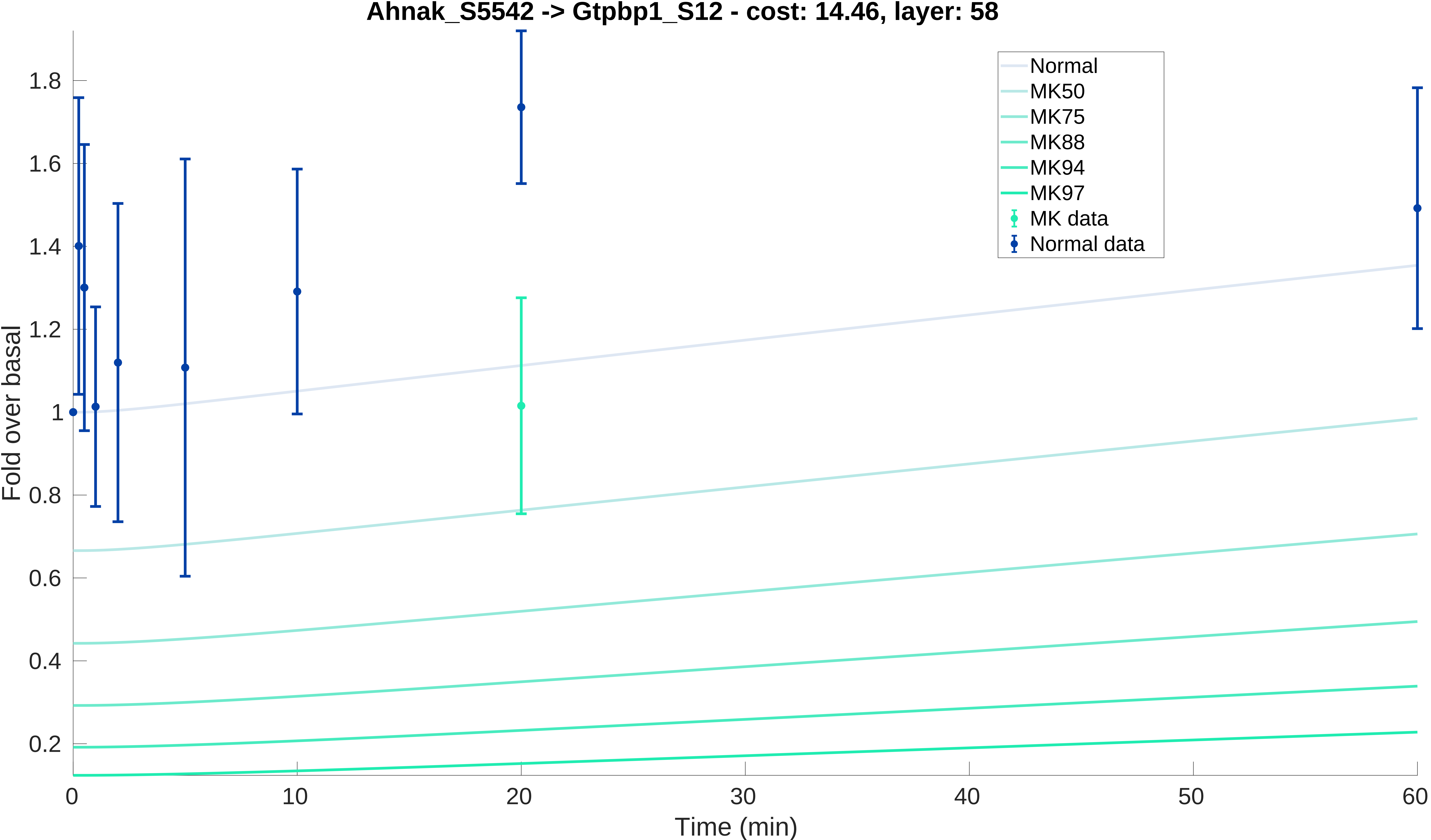

Ahnak_S5542 -> Gtpbp1_S12 - cost: 14.46, layer: 58
Normal
MK50
MK75
MK88
MK94
MK97
MK data
Normal data
Fold over basal
Time (min)
0
10
20
30
40
50
60
0.2
0.4
0.6
0.8
1
1.2
1.4
1.6
1.8

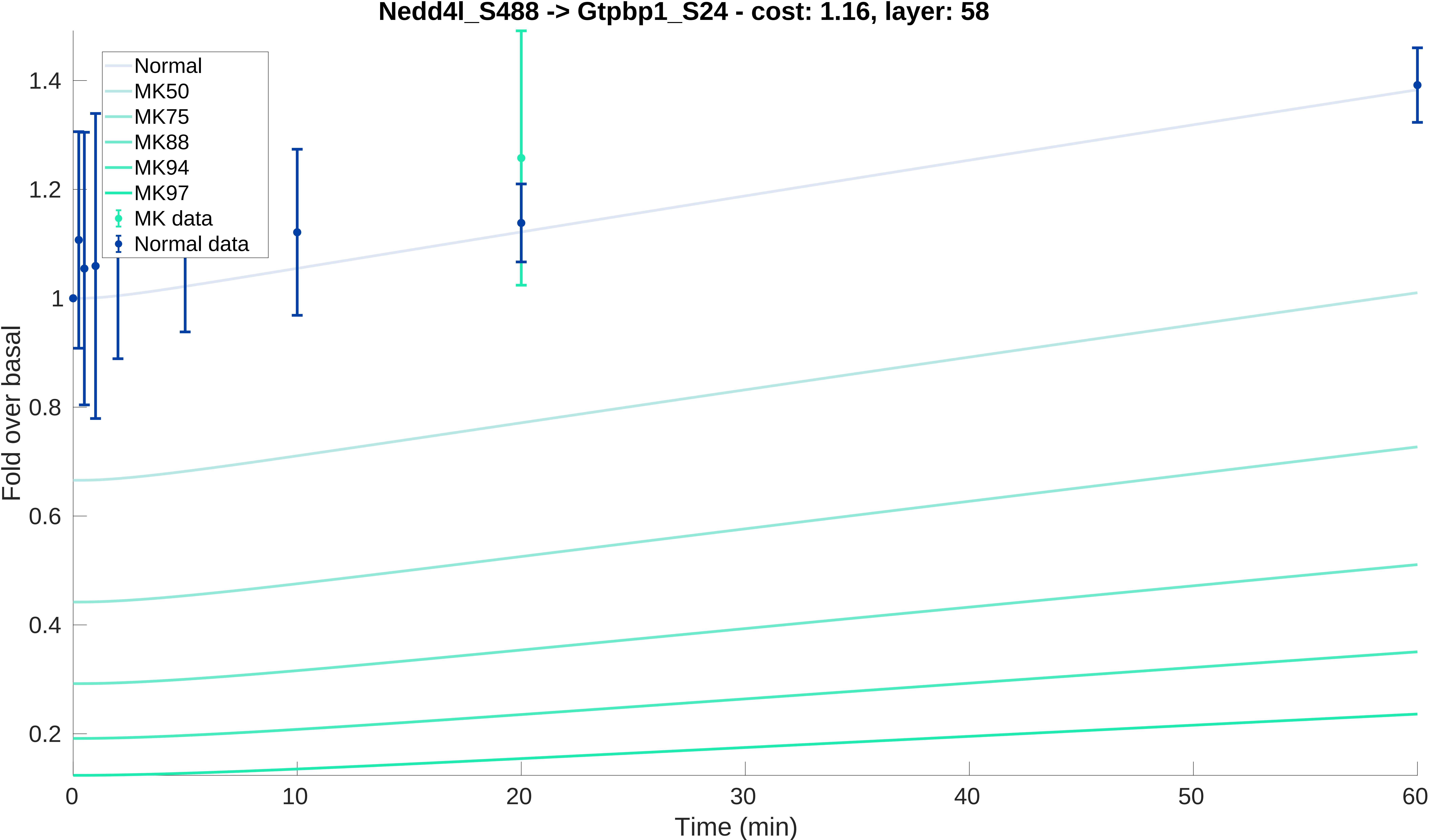

Nedd4l_S488 -> Gtpbp1_S24 - cost: 1.16, layer: 58
Normal
MK50
MK75
MK88
MK94
MK97
MK data
Normal data
Fold over basal
Time (min)
0.2
0.4
0.6
0.8
1
1.2
1.4
0
10
20
30
40
50
60

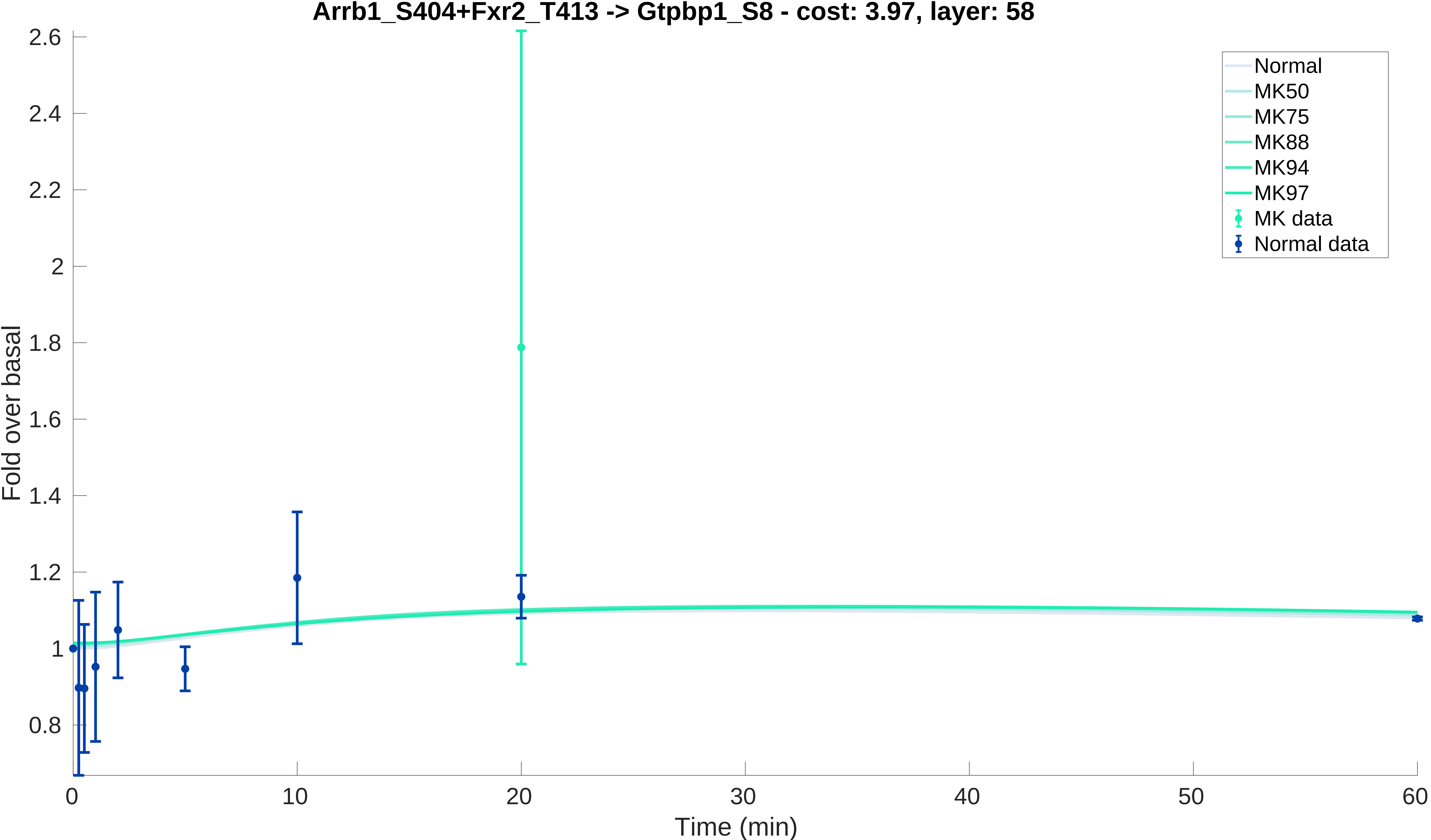

Arrb1_S404+Fxr2_T413 -> Gtpbp1_S8 - cost: 3.97, layer: 58
Fold over basal
2.6
2.4
2.2
2
1.8
1.6
1.4
1.2
1
0.8
0
10
20
30
40
50
60
Time (min)
Normal
MK50
MK75
MK88
MK94
MK97
MK data
Normal data

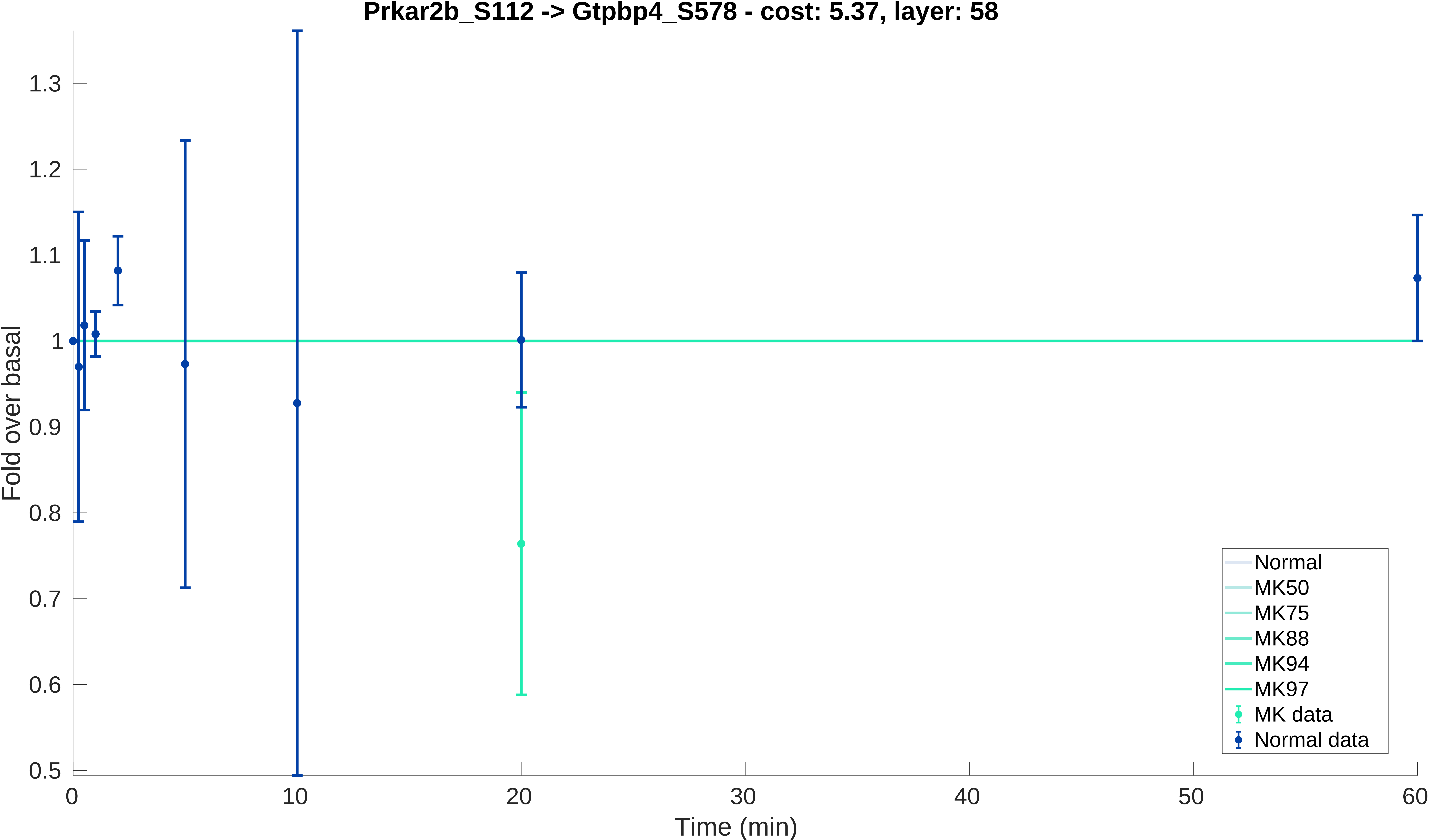

Prkar2b_S112 -> Gtpbp4_S578 - cost: 5.37, layer: 58
Fold over basal
1.3
1.2
1.1
1
0.9
0.8
0.7
0.6
0.5
0
10
20
30
40
50
60
Time (min)
Normal
MK50
MK75
MK88
MK94
MK97
MK data
Normal data

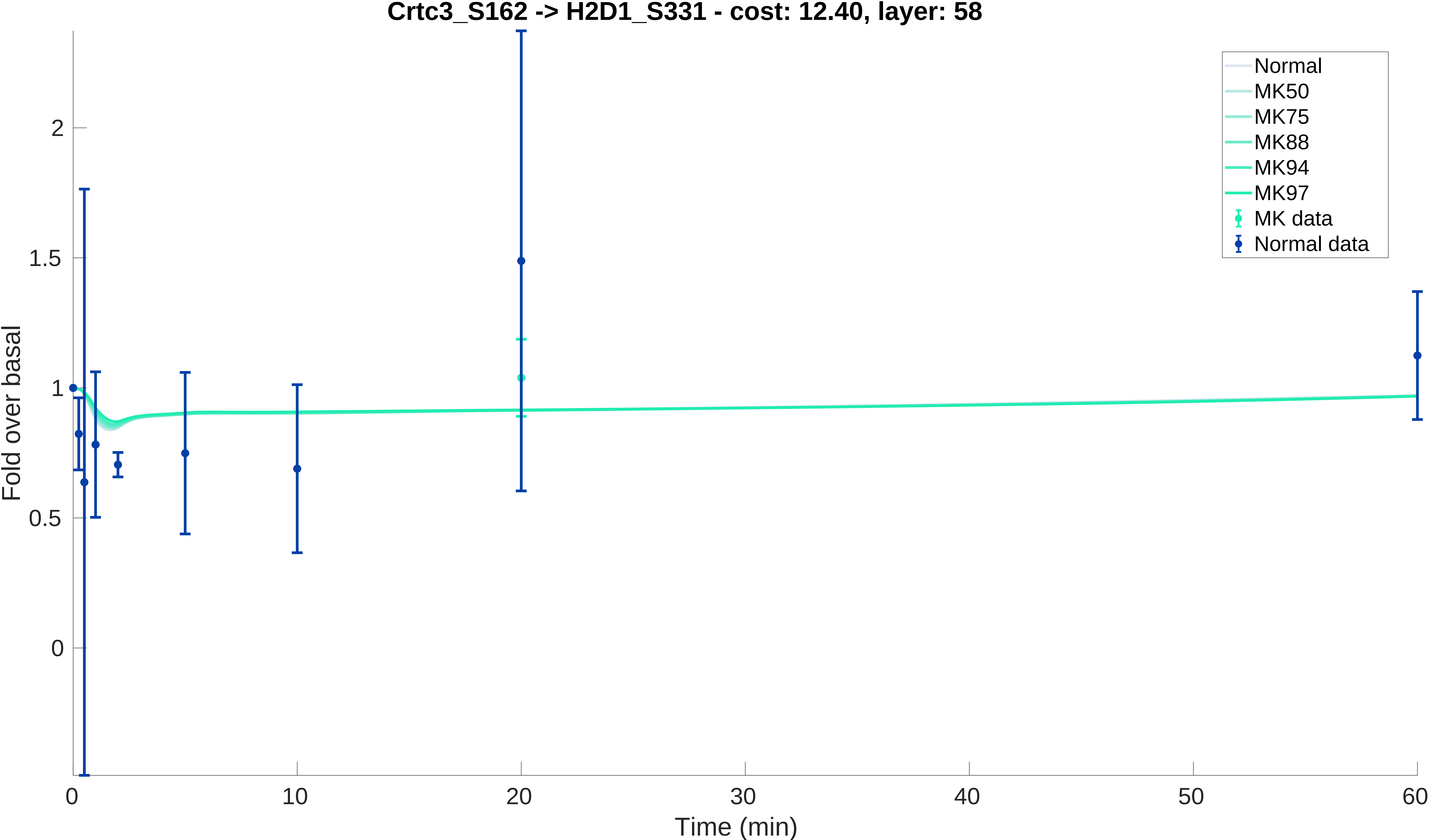

Crtc3_S162 -> H2D1_S331 - cost: 12.40, layer: 58
Normal
MK50
MK75
MK88
MK94
MK97
MK data
Normal data
Fold over basal
Time (min)
0
10
20
30
40
50
60
0
0.5
1
1.5
2

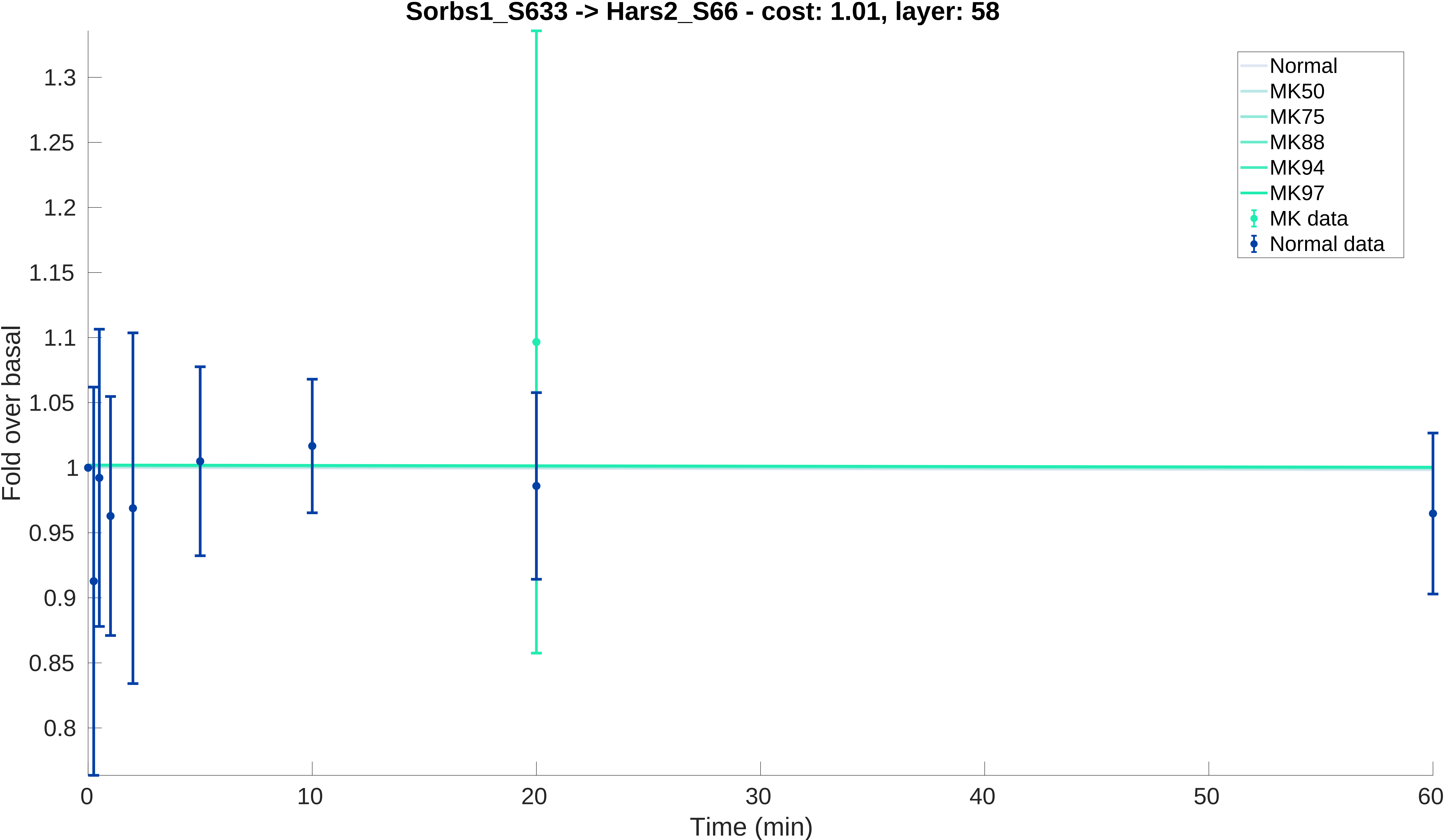

Sorbs1_S633 -> Hars2_S66 - cost: 1.01, layer: 58
Fold over basal
Time (min)
Normal
MK50
MK75
MK88
MK94
MK97
MK data
Normal data
1.3
1.25
1.2
1.15
1.1
1.05
1
0.95
0.9
0.85
0.8
0
10
20
30
40
50
60

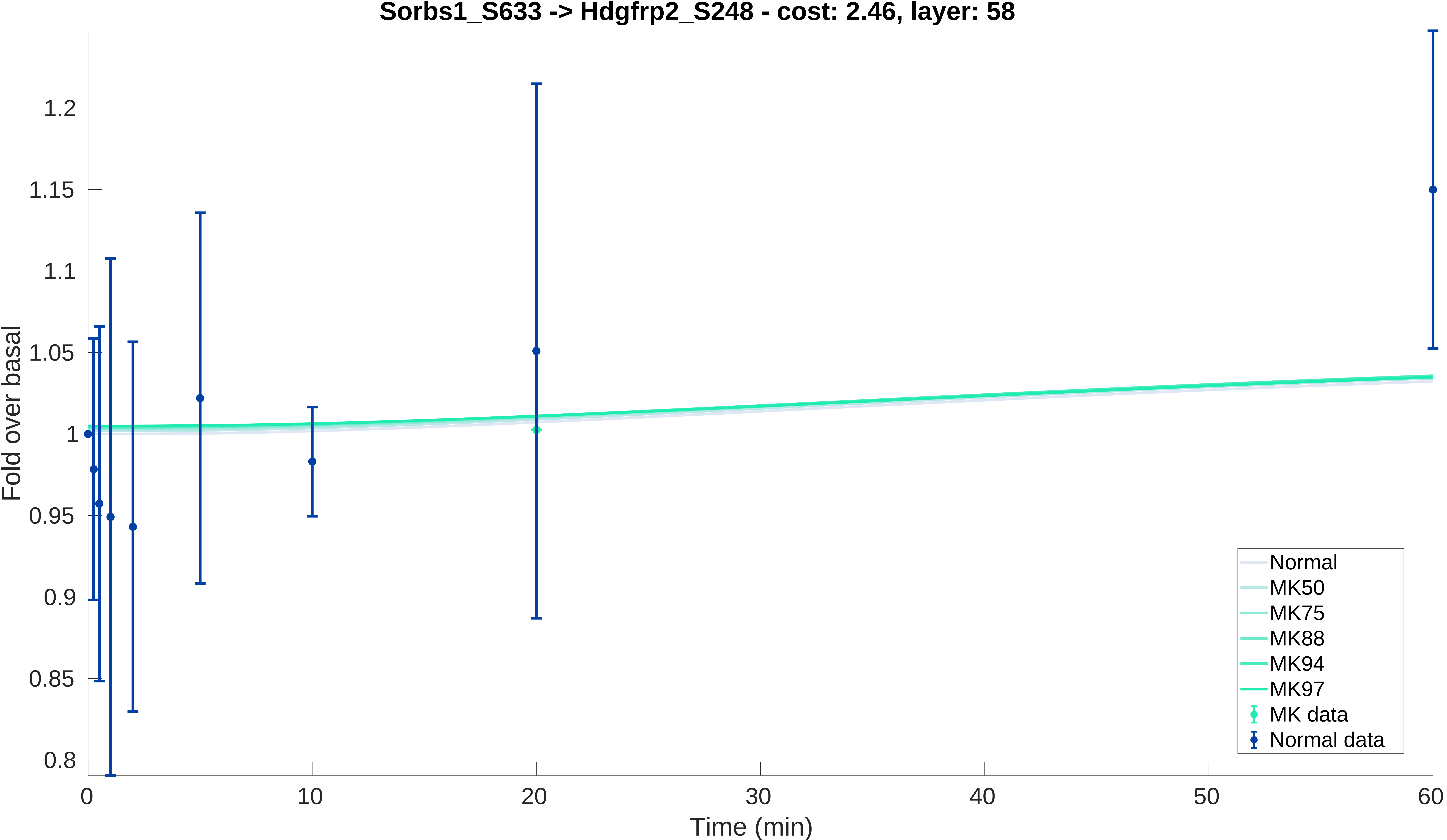

Sorbs1_S633 -> Hdgfrp2_S248 - cost: 2.46, layer: 58
Fold over basal
1.2
1.15
1.1
1.05
1
0.95
0.9
0.85
0.8
0
10
20
30
40
50
60
Time (min)
Normal
MK50
MK75
MK88
MK94
MK97
MK data
Normal data

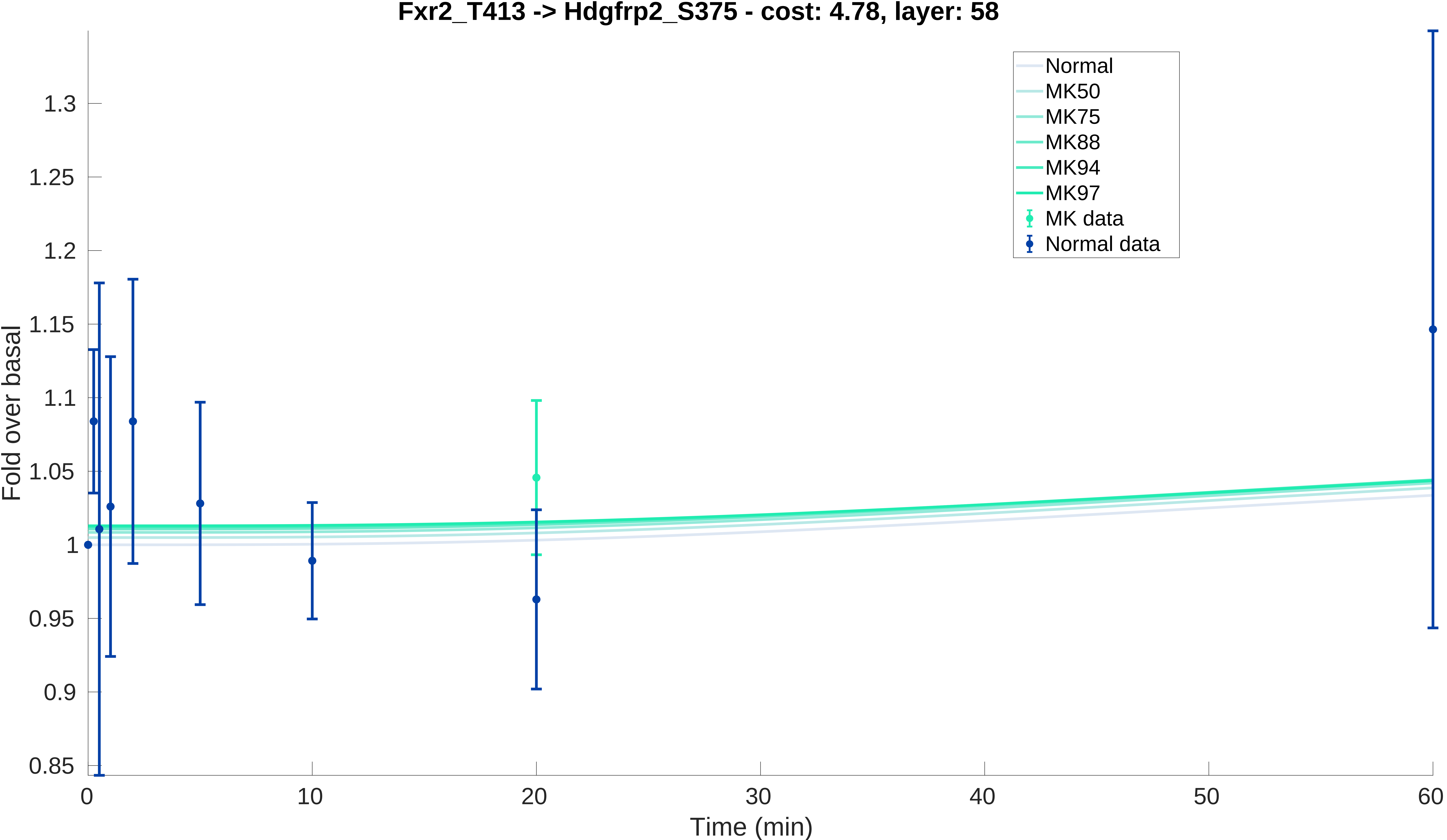

Fxr2_T413 -> Hdgfrp2_S375 - cost: 4.78, layer: 58
Fold over basal
Time (min)
1.3
1.25
1.2
1.15
1.1
1.05
1
0.95
0.9
0.85
0
10
20
30
40
50
60
Normal
MK50
MK75
MK88
MK94
MK97
MK data
Normal data

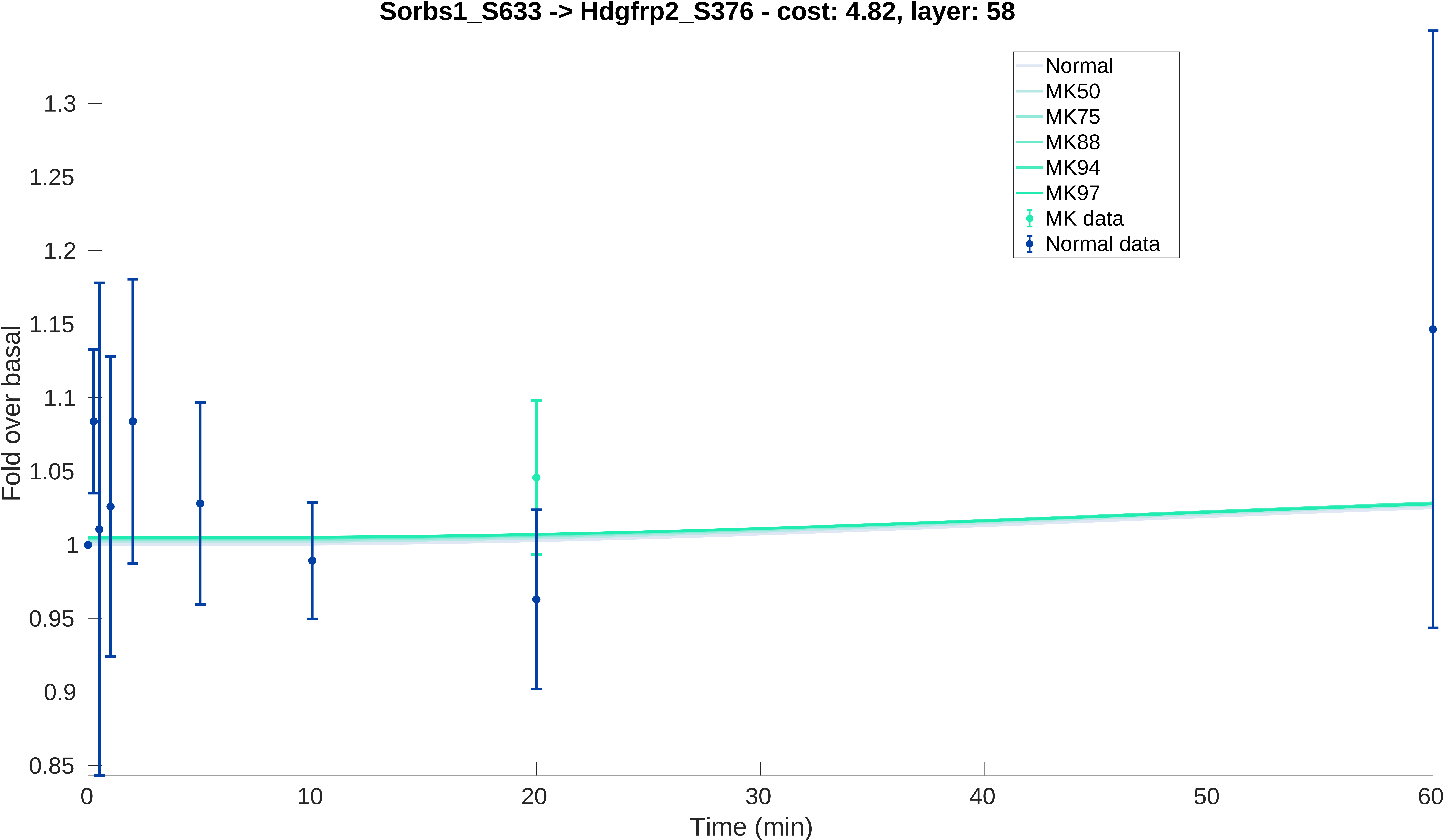
Sorbs1_S633 -> Hdgfrp2_S376 - cost: 4.82, layer: 58
Fold over basal
Time (min)
1.3
1.25
1.2
1.15
1.1
1.05
1
0.95
0.9
0.85
0
10
20
30
40
50
60
Normal
MK50
MK75
MK88
MK94
MK97
MK data
Normal data

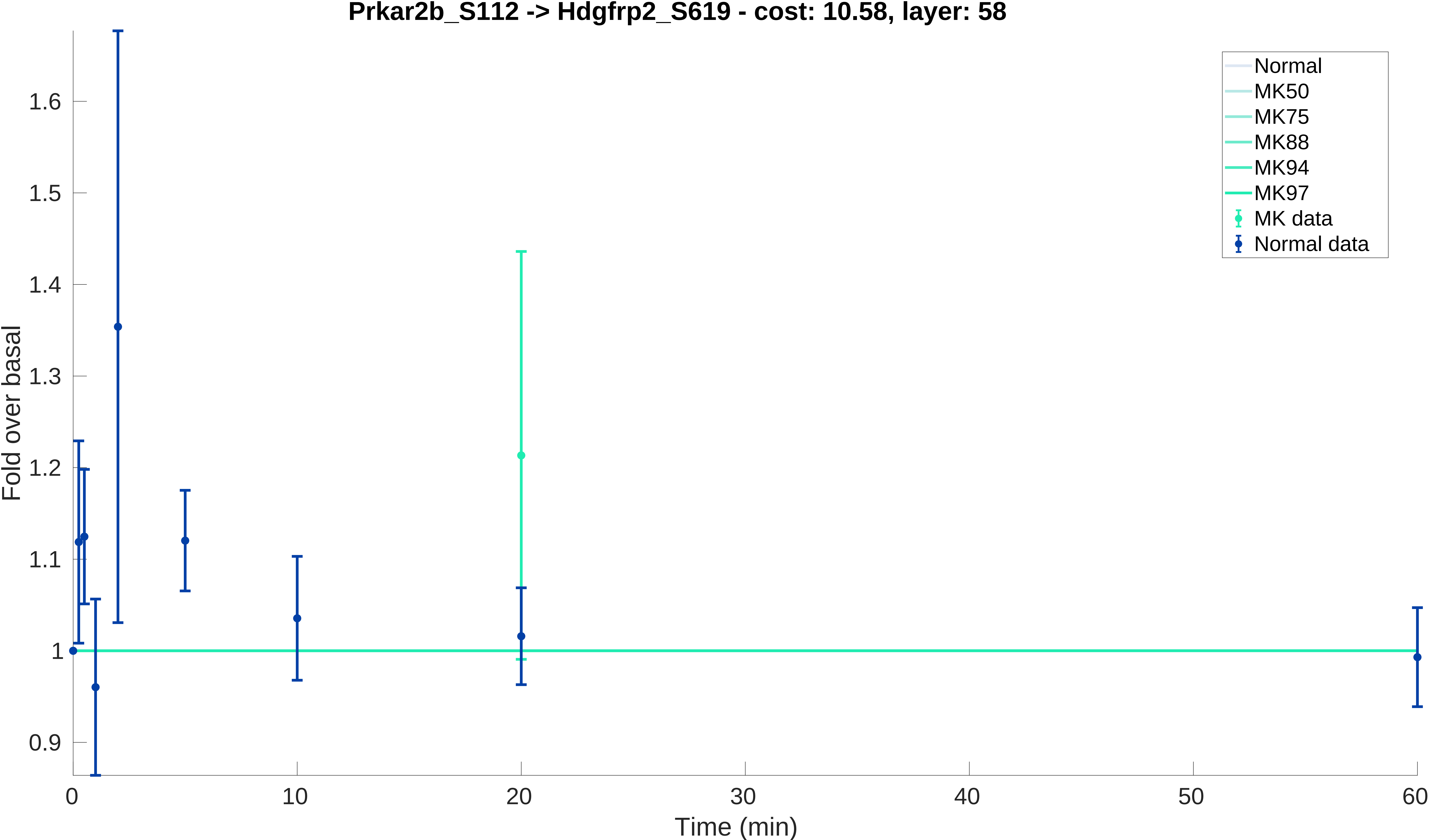

Prkar2b_S112 -> Hdgfrp2_S619 - cost: 10.58, layer: 58
Fold over basal
1.6
1.5
1.4
1.3
1.2
1.1
1
0.9
0
10
20
30
40
50
60
Time (min)
Normal
MK50
MK75
MK88
MK94
MK97
MK data
Normal data

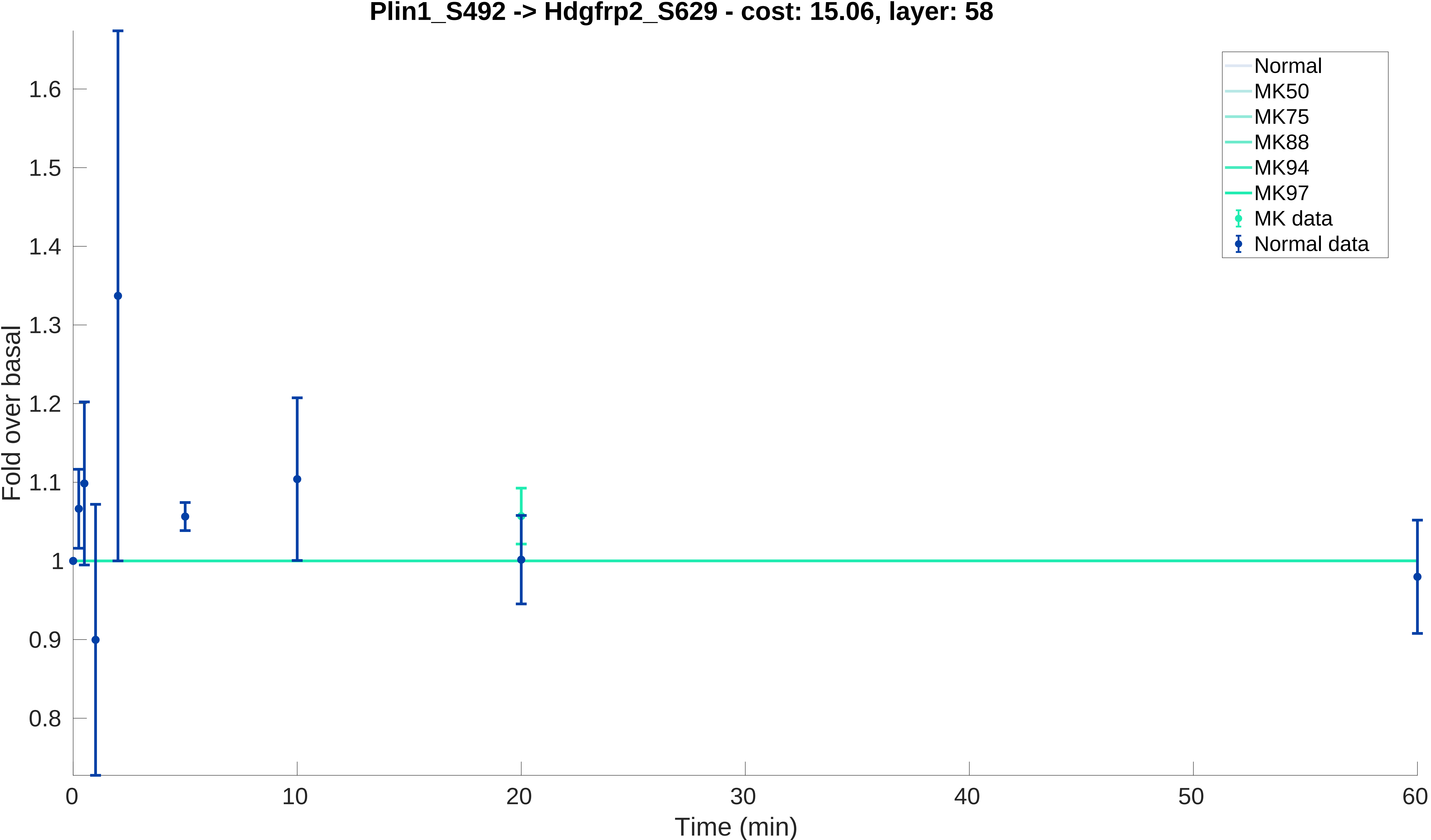
Plin1_S492 -> Hdgfrp2_S629 - cost: 15.06, layer: 58
Fold over basal
Time (min)
0.8
0.9
1
1.1
1.2
1.3
1.4
1.5
1.6
0
10
20
30
40
50
60
Normal
MK50
MK75
MK88
MK94
MK97
MK data
Normal data

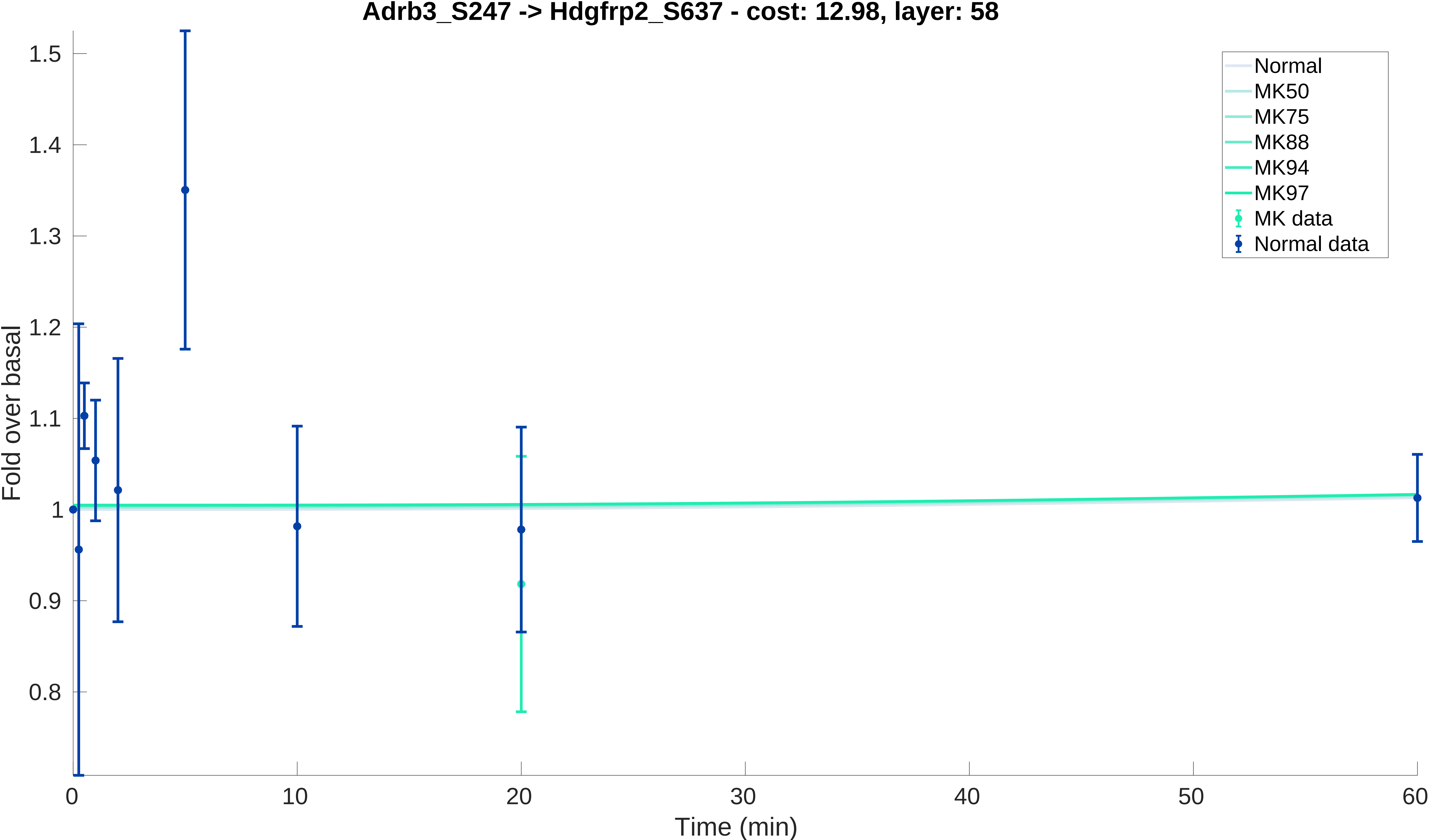

Adrb3_S247 -> Hdgfrp2_S637 - cost: 12.98, layer: 58
Fold over basal
Time (min)
1.5
1.4
1.3
1.2
1.1
1
0.9
0.8
0
10
20
30
40
50
60
Normal
MK50
MK75
MK88
MK94
MK97
MK data
Normal data

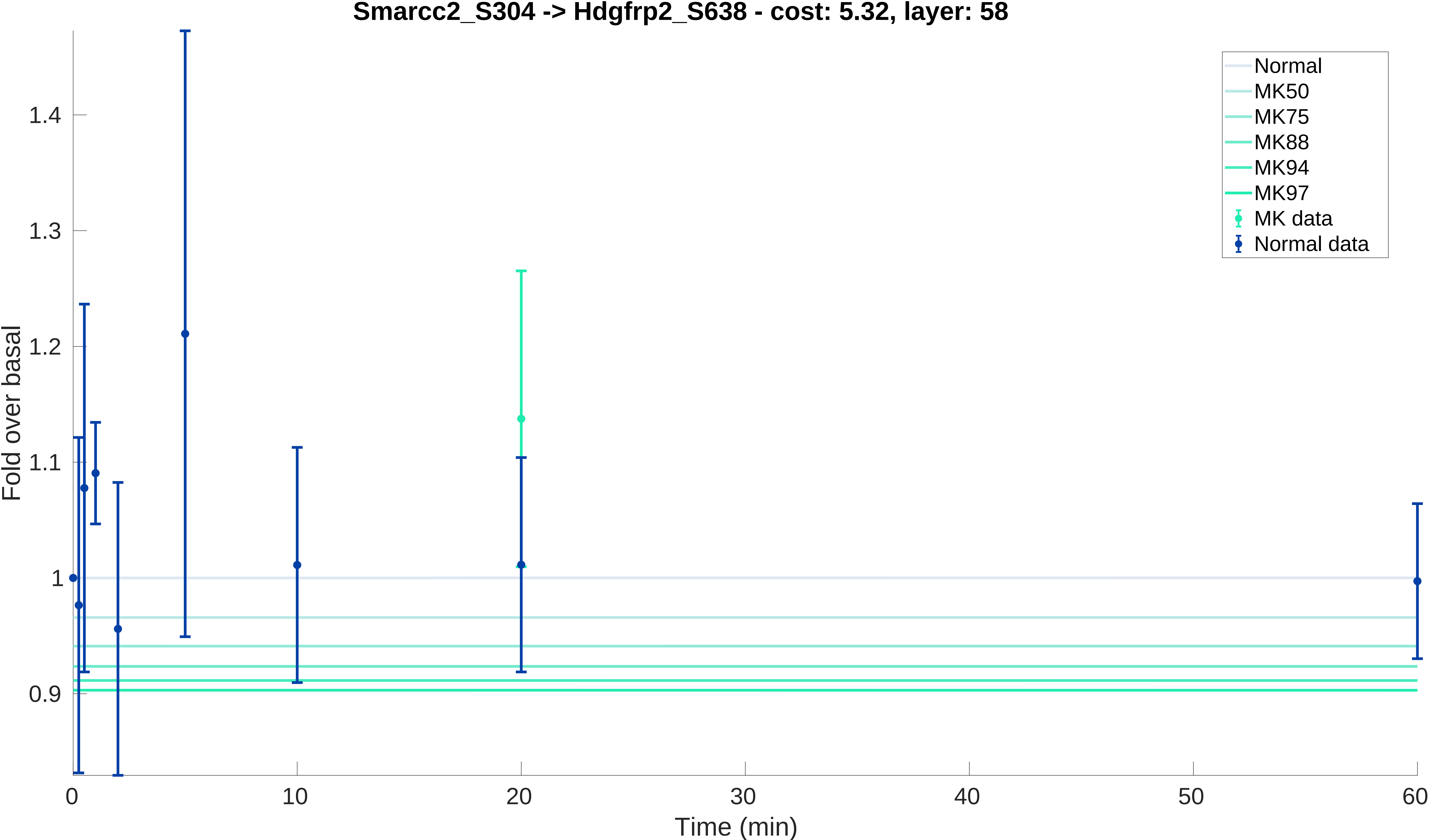

Smarcc2_S304 -> Hdgfrp2_S638 - cost: 5.32, layer: 58
Fold over basal
Time (min)
1.4
1.3
1.2
1.1
1
0.9
0
10
20
30
40
50
60
Normal
MK50
MK75
MK88
MK94
MK97
MK data
Normal data

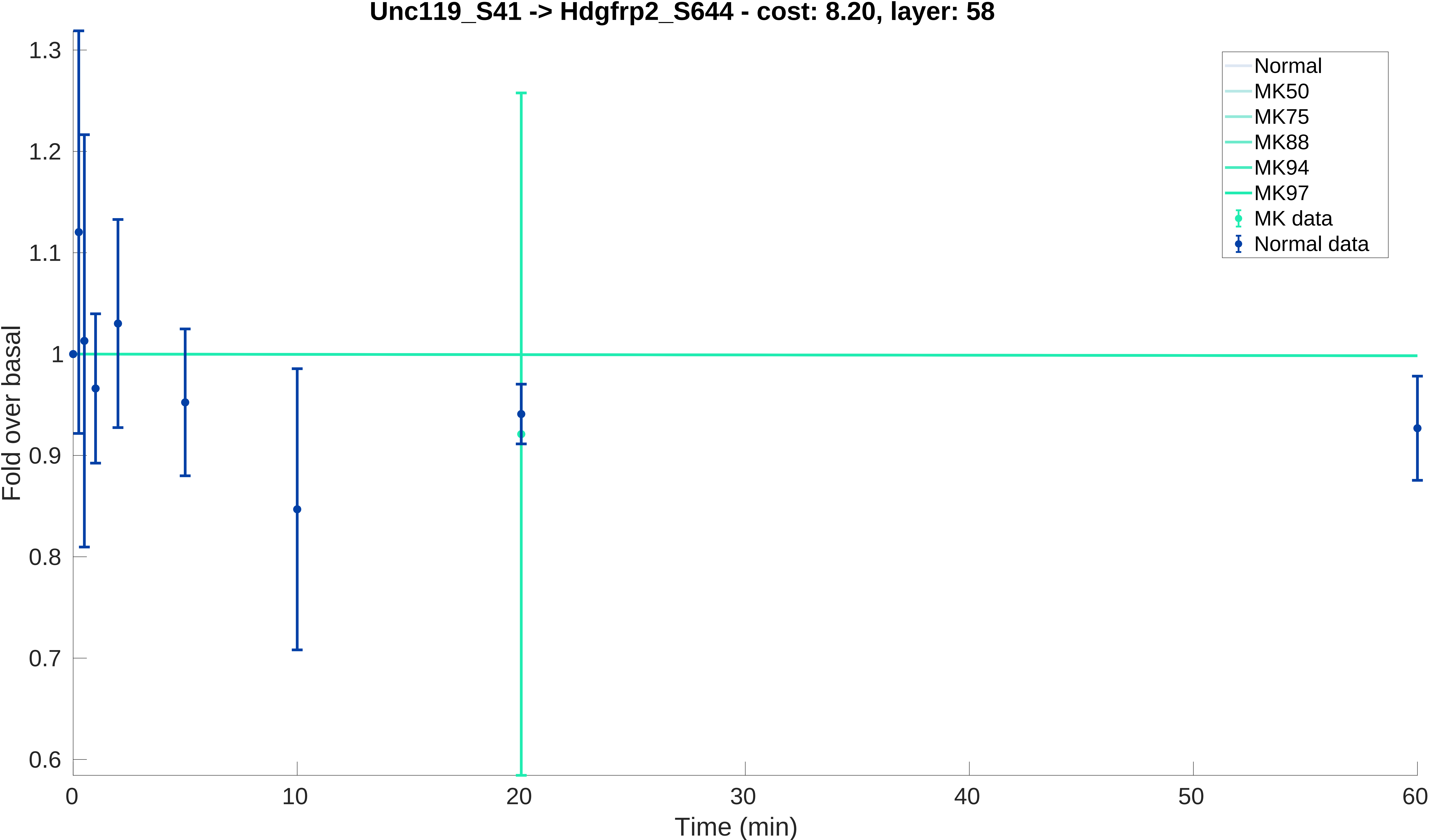

Unc119_S41 -> Hdgfrp2_S644 - cost: 8.20, layer: 58
Fold over basal
Time (min)
1.3
1.2
1.1
1
0.9
0.8
0.7
0.6
0
10
20
30
40
50
60
Normal
MK50
MK75
MK88
MK94
MK97
MK data
Normal data

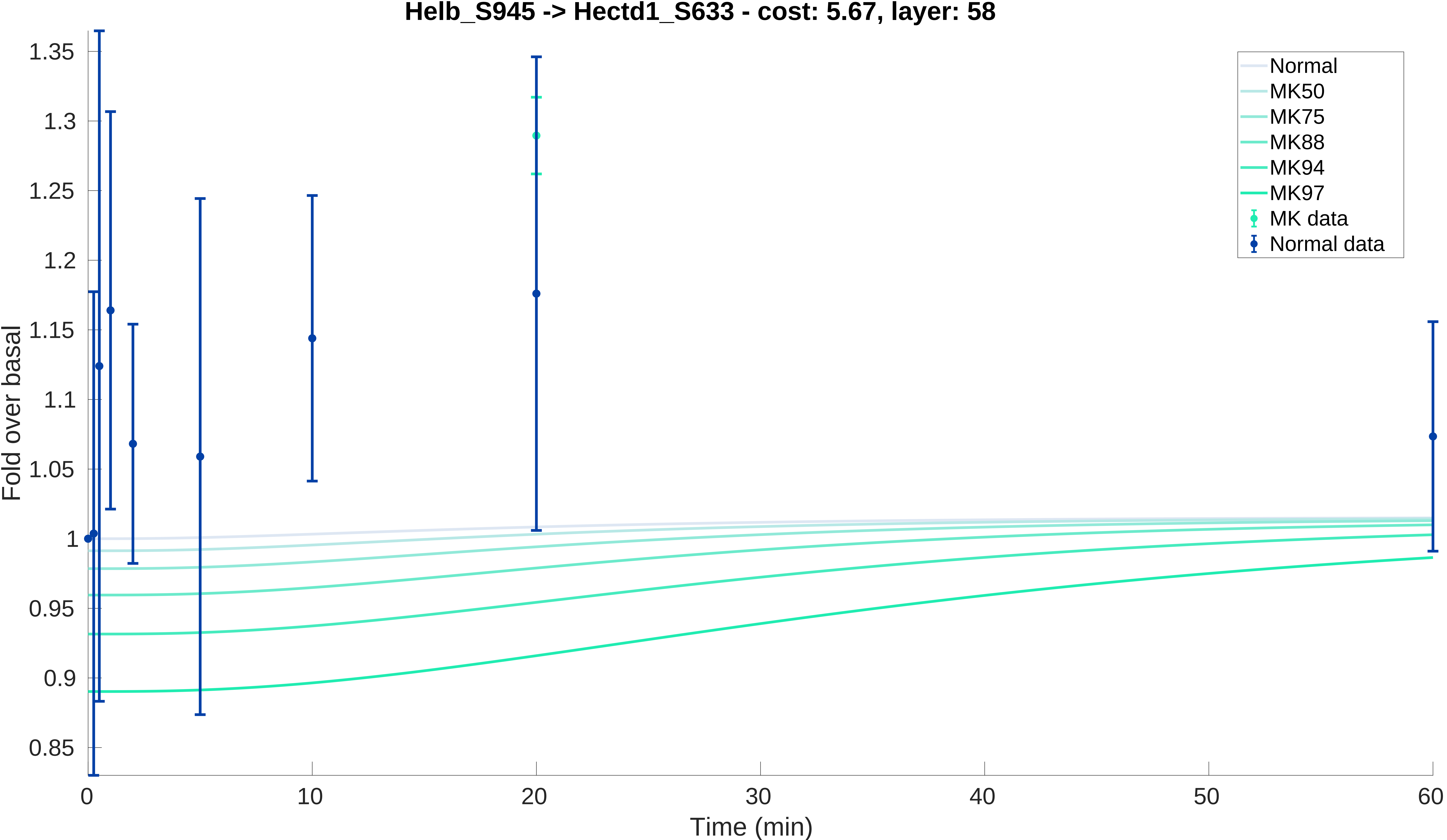

Helb_S945 -> Hectd1_S633 - cost: 5.67, layer: 58
Fold over basal
Time (min)
1.35
1.3
1.25
1.2
1.15
1.1
1.05
1
0.95
0.9
0.85
0
10
20
30
40
50
60
Normal
MK50
MK75
MK88
MK94
MK97
MK data
Normal data

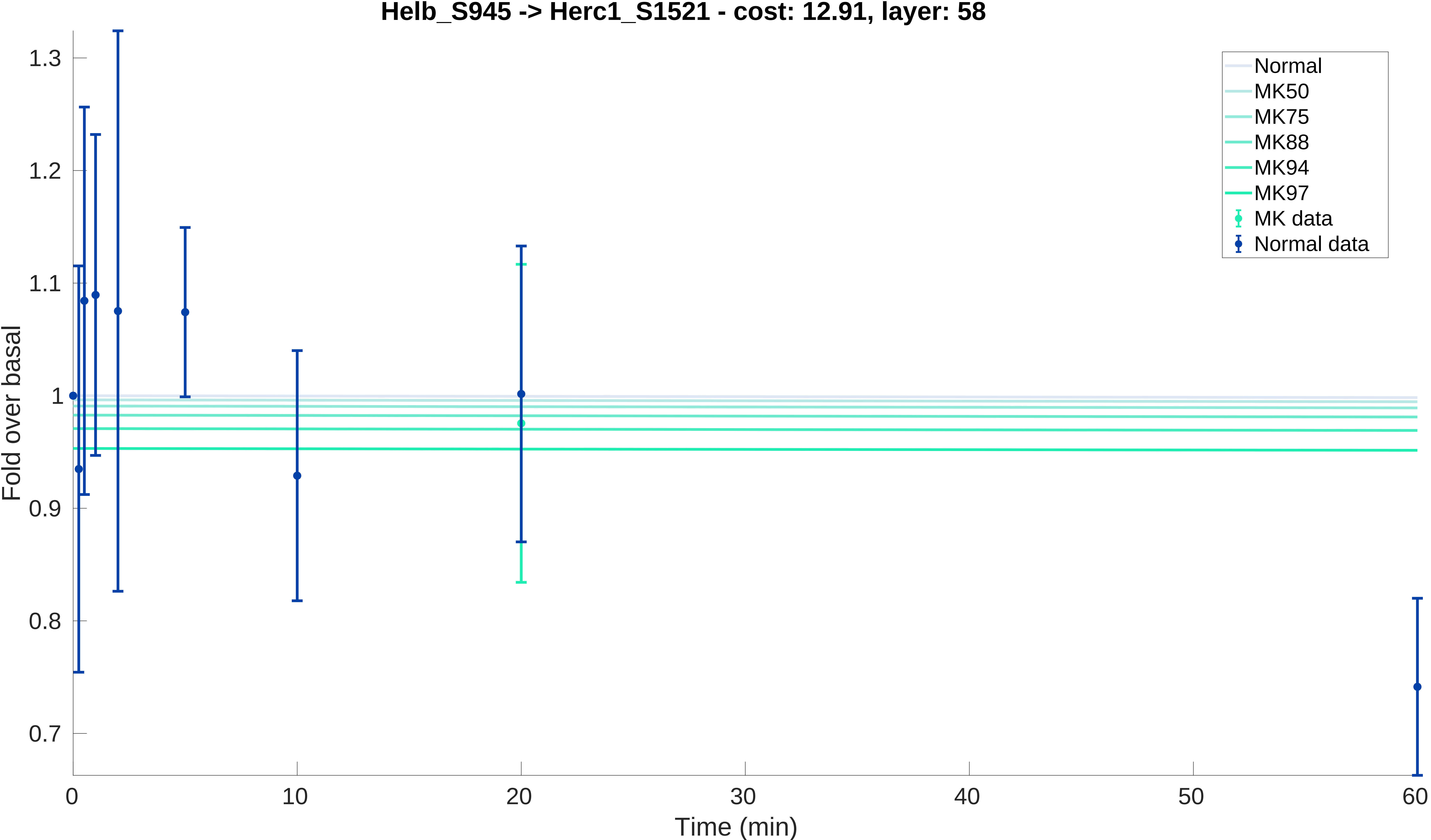
Helb_S945 -> Herc1_S1521 - cost: 12.91, layer: 58
Fold over basal
Time (min)
Normal
MK50
MK75
MK88
MK94
MK97
MK data
Normal data
1.3
1.2
1.1
1
0.9
0.8
0.7
0
10
20
30
40
50
60

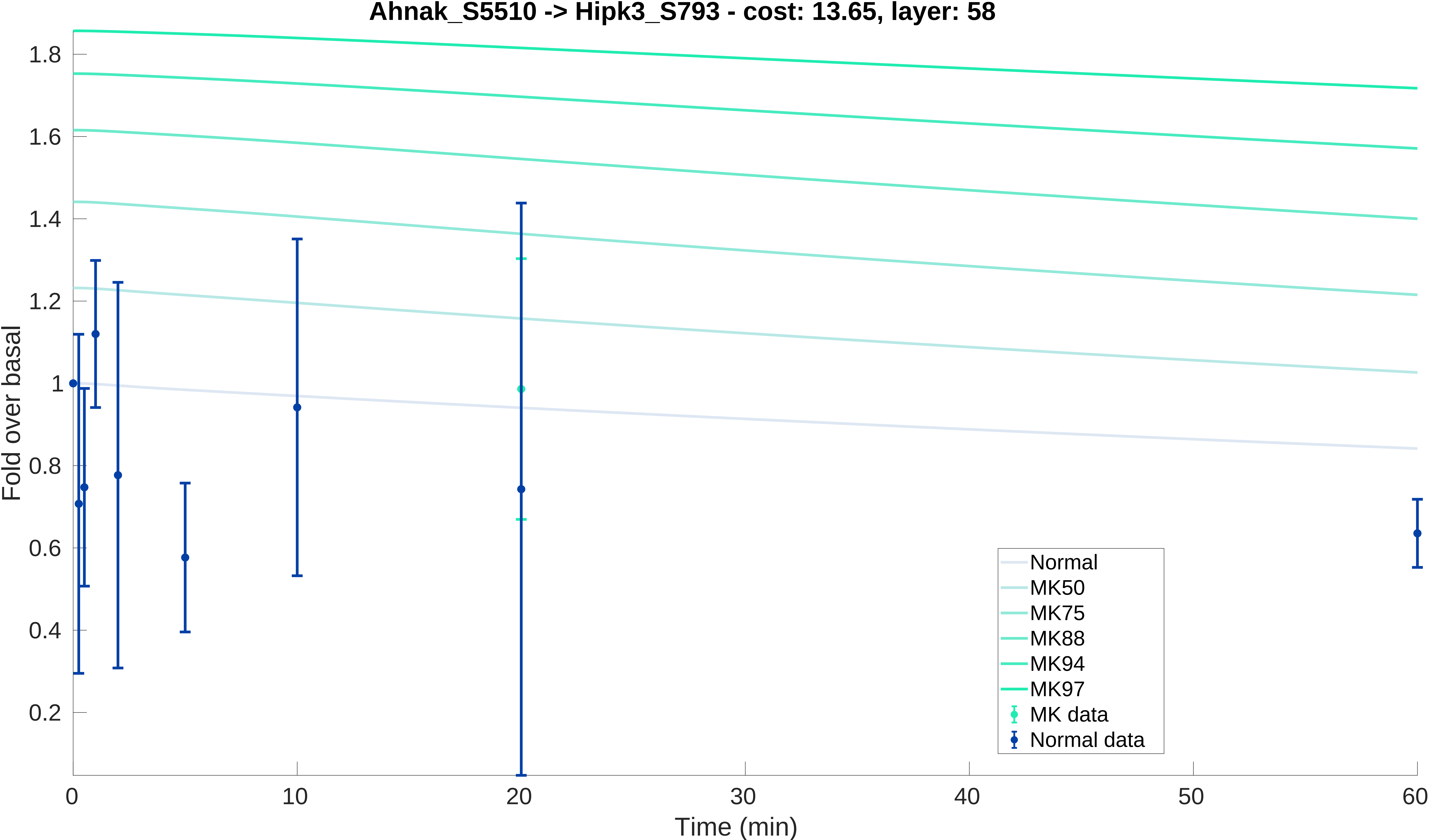

Ahnak_S5510 -> Hipk3_S793 - cost: 13.65, layer: 58
Fold over basal
Time (min)
0
10
20
30
40
50
60
0.2
0.4
0.6
0.8
1
1.2
1.4
1.6
1.8
Normal
MK50
MK75
MK88
MK94
MK97
MK data
Normal data

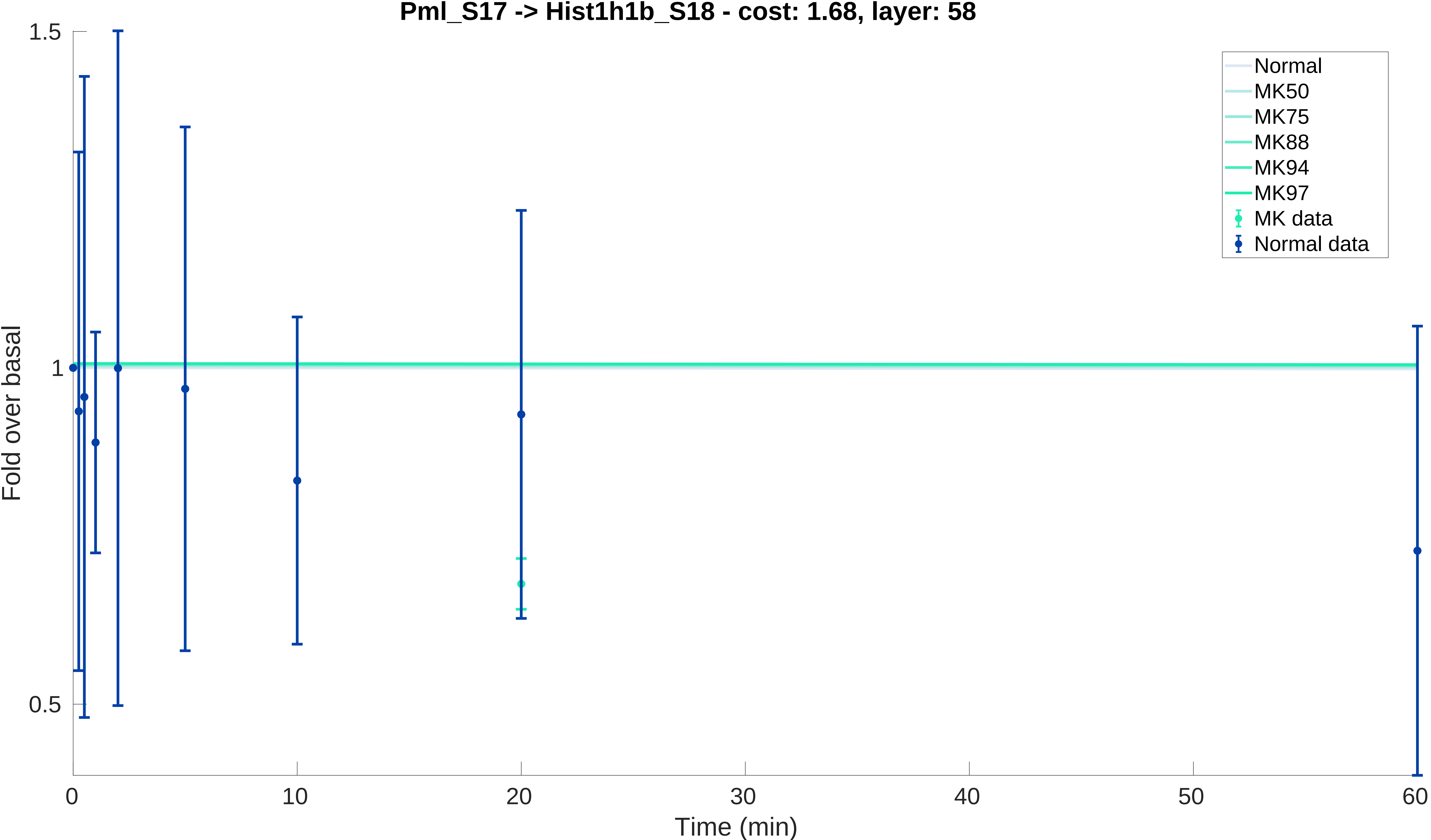

Pml_S17 -> Hist1h1b_S18 - cost: 1.68, layer: 58
1.5
1
0.5
Fold over basal
0
10
20
30
40
50
60
Time (min)
Normal
MK50
MK75
MK88
MK94
MK97
MK data
Normal data

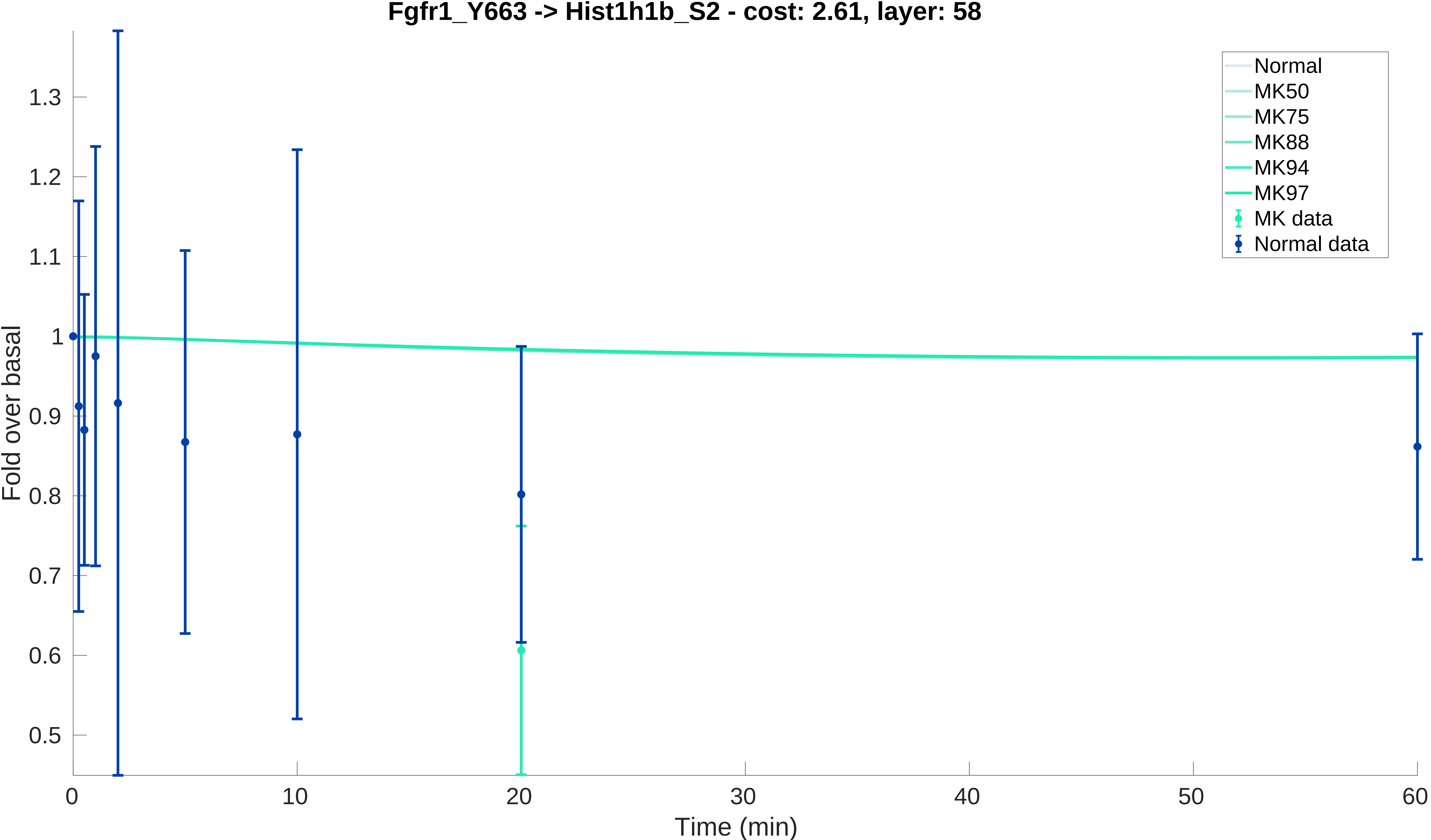

Fgfr1_Y663 -> Hist1h1b_S2 - cost: 2.61, layer: 58
Fold over basal
Time (min)
Normal
MK50
MK75
MK88
MK94
MK97
MK data
Normal data
1.3
1.2
1.1
1
0.9
0.8
0.7
0.6
0.5
0
10
20
30
40
50
60

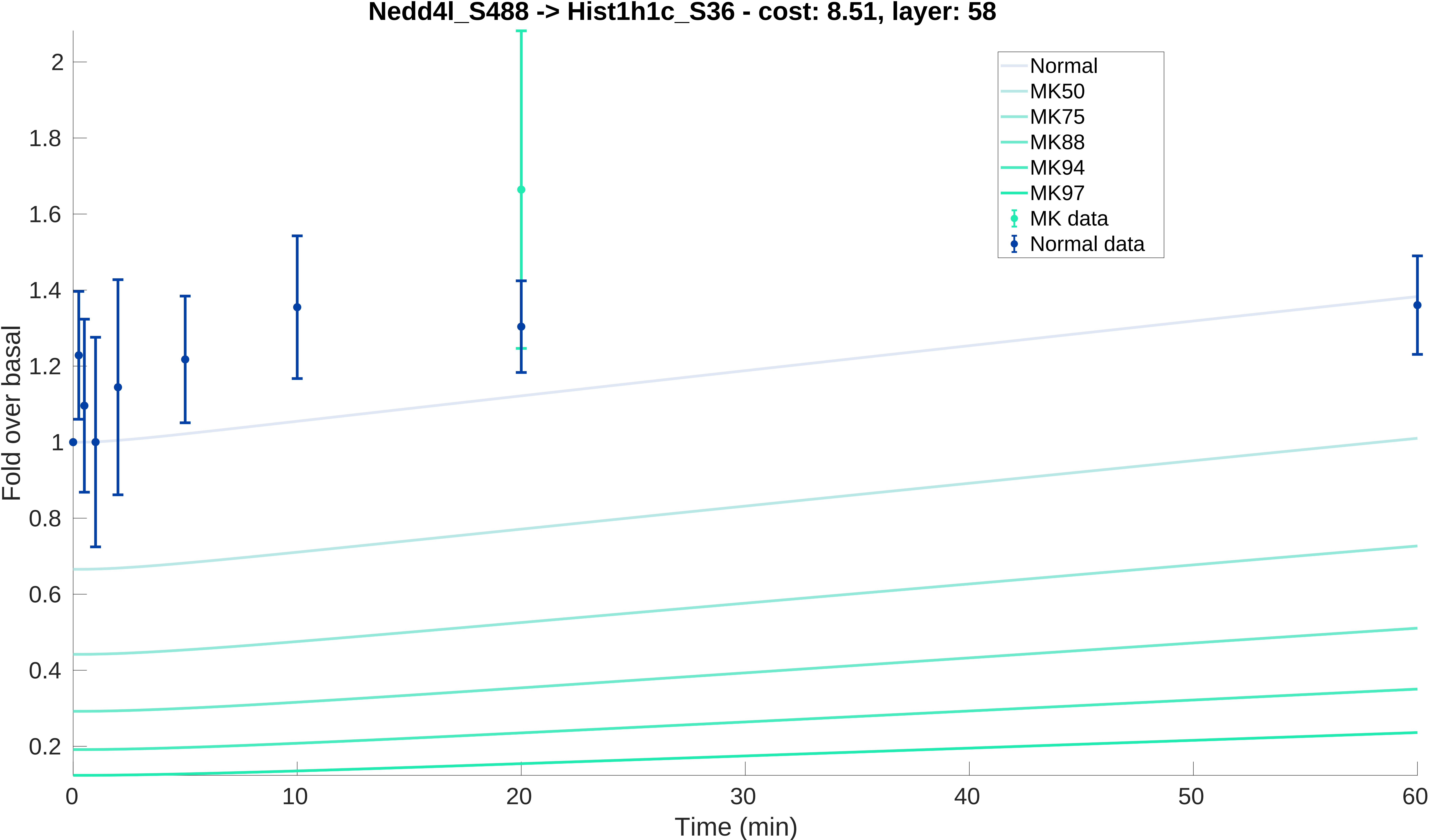

Nedd4l_S488 -> Hist1h1c_S36 - cost: 8.51, layer: 58
Fold over basal
Time (min)
0
10
20
30
40
50
60
0.2
0.4
0.6
0.8
1
1.2
1.4
1.6
1.8
2
Normal
MK50
MK75
MK88
MK94
MK97
MK data
Normal data

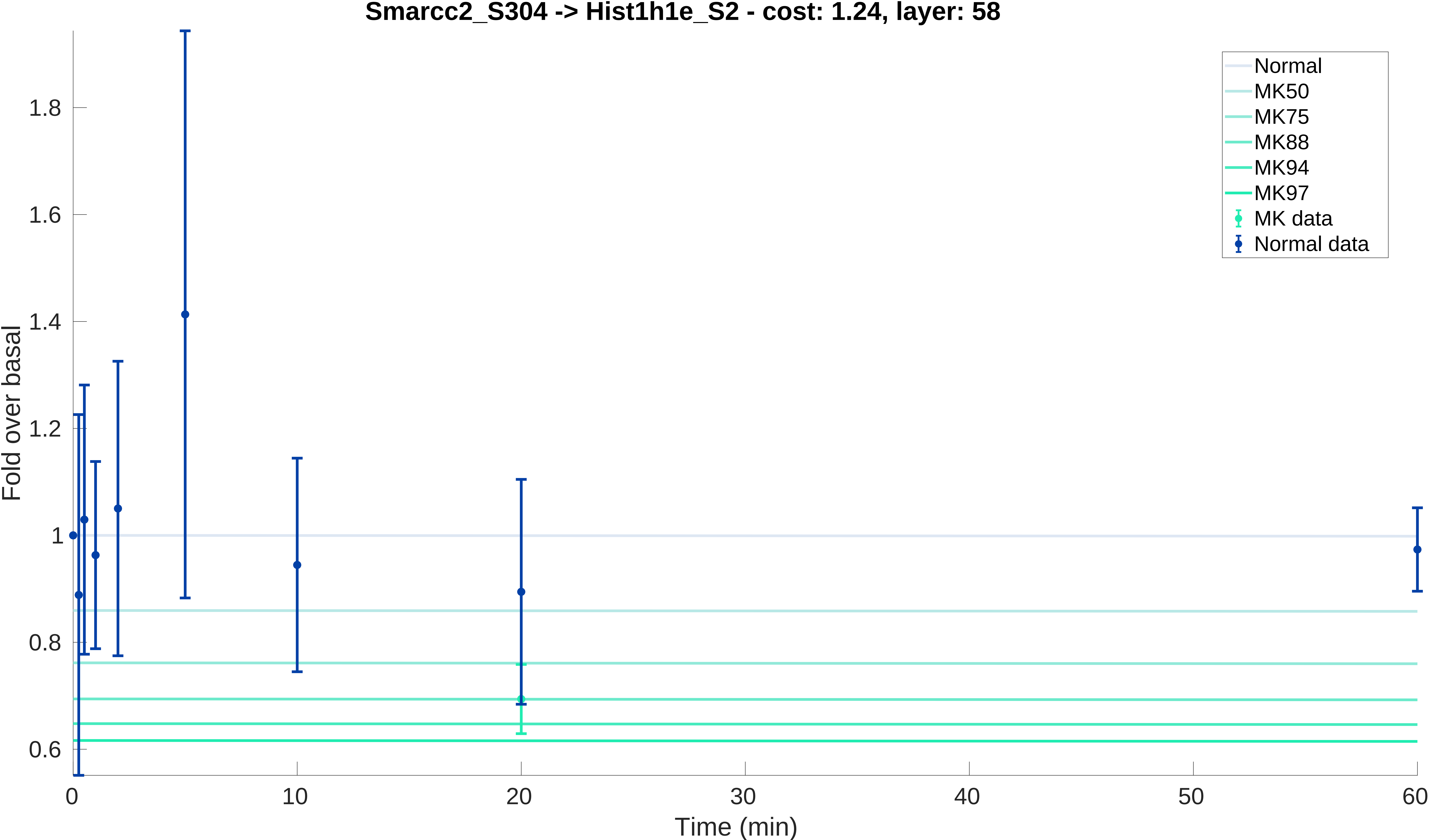

Smarcc2_S304 -> Hist1h1e_S2 - cost: 1.24, layer: 58
Fold over basal
Time (min)
1.8
1.6
1.4
1.2
1
0.8
0.6
0
10
20
30
40
50
60
Normal
MK50
MK75
MK88
MK94
MK97
MK data
Normal data

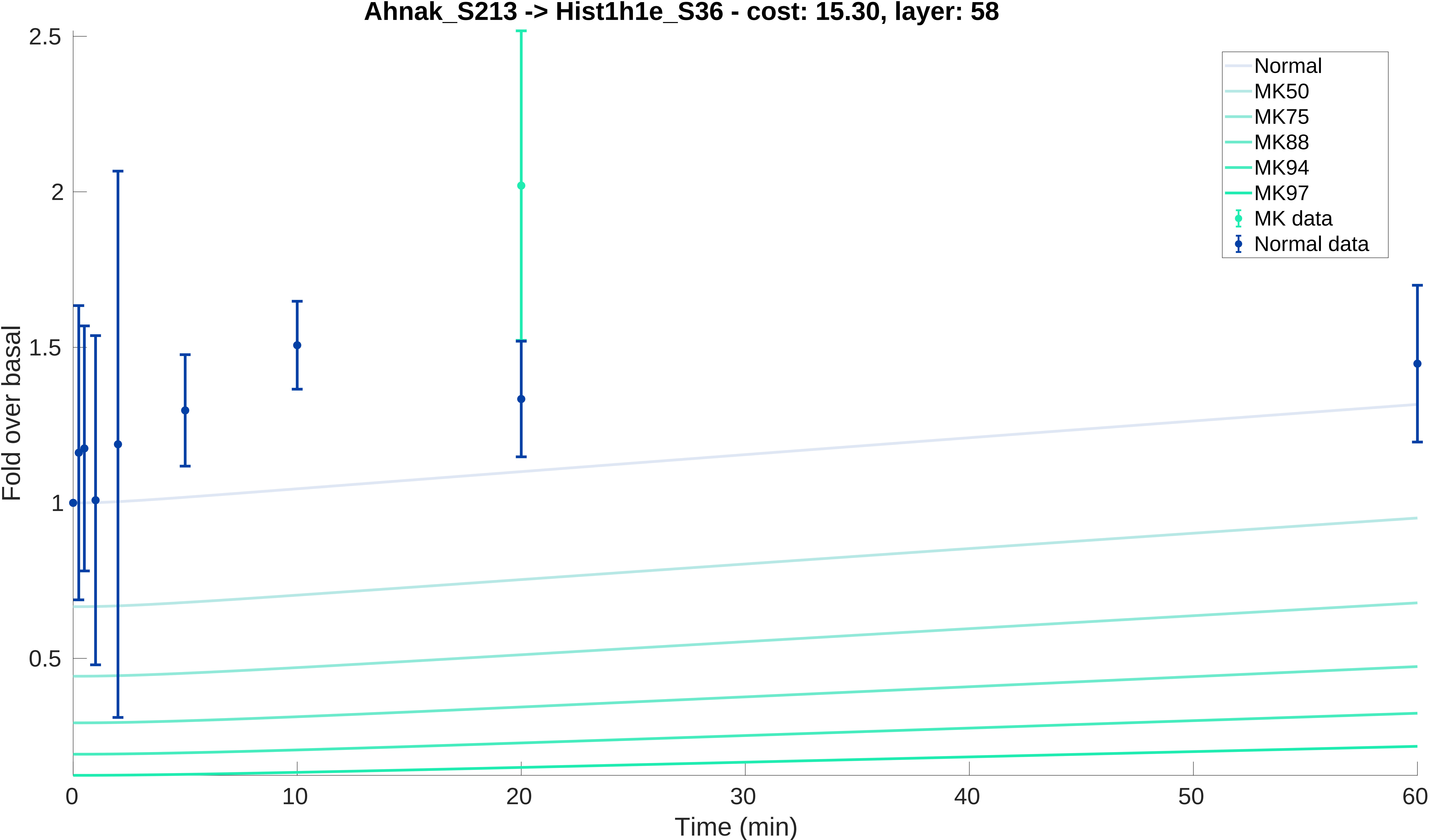

Ahnak_S213 -> Hist1h1e_S36 - cost: 15.30, layer: 58
Fold over basal
Time (min)
0
10
20
30
40
50
60
0.5
1
1.5
2
2.5
Normal
MK50
MK75
MK88
MK94
MK97
MK data
Normal data

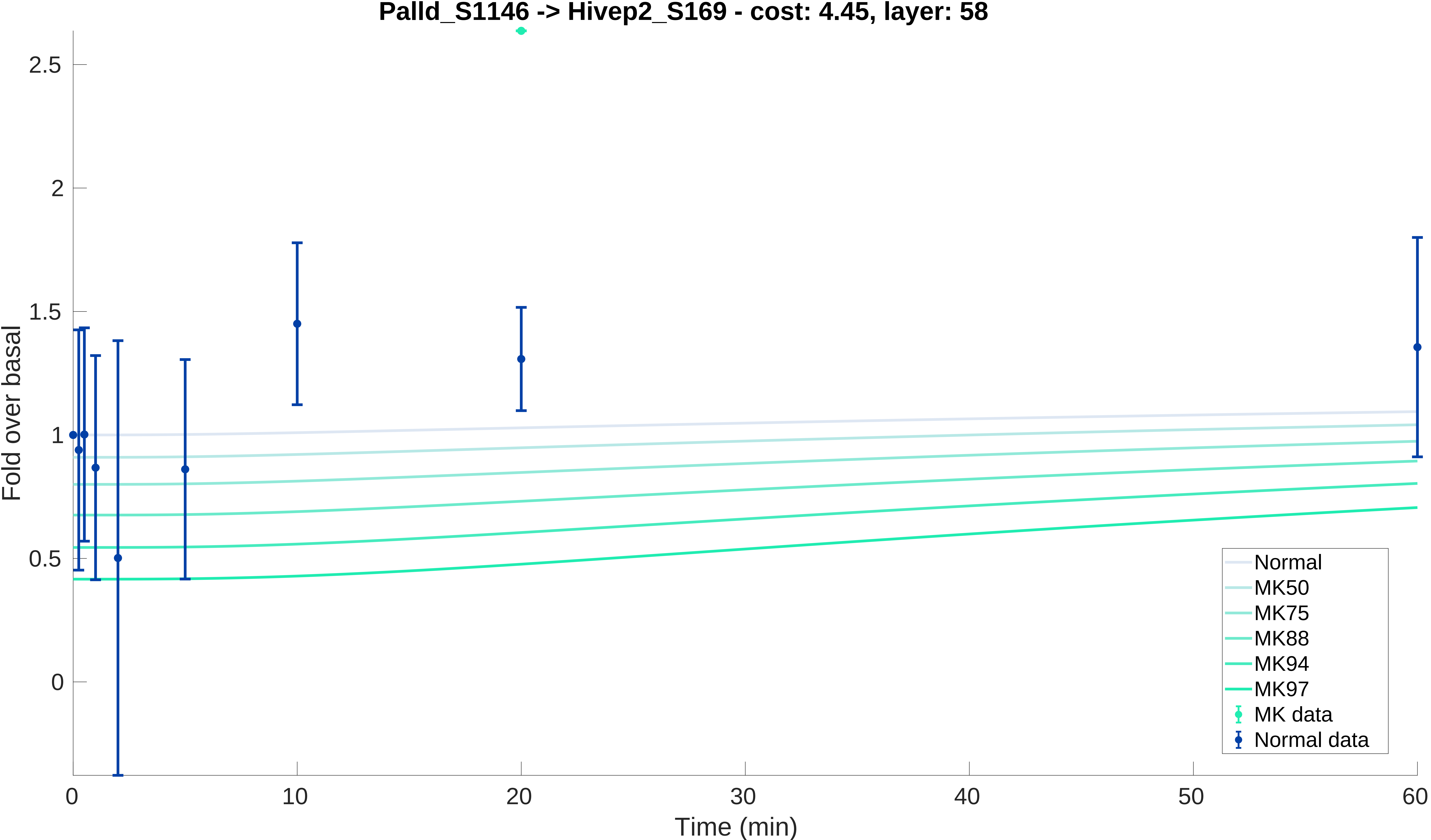

Palld_S1146 -> Hivep2_S169 - cost: 4.45, layer: 58
Fold over basal
Time (min)
0
0.5
1
1.5
2
2.5
0
10
20
30
40
50
60
Normal
MK50
MK75
MK88
MK94
MK97
MK data
Normal data

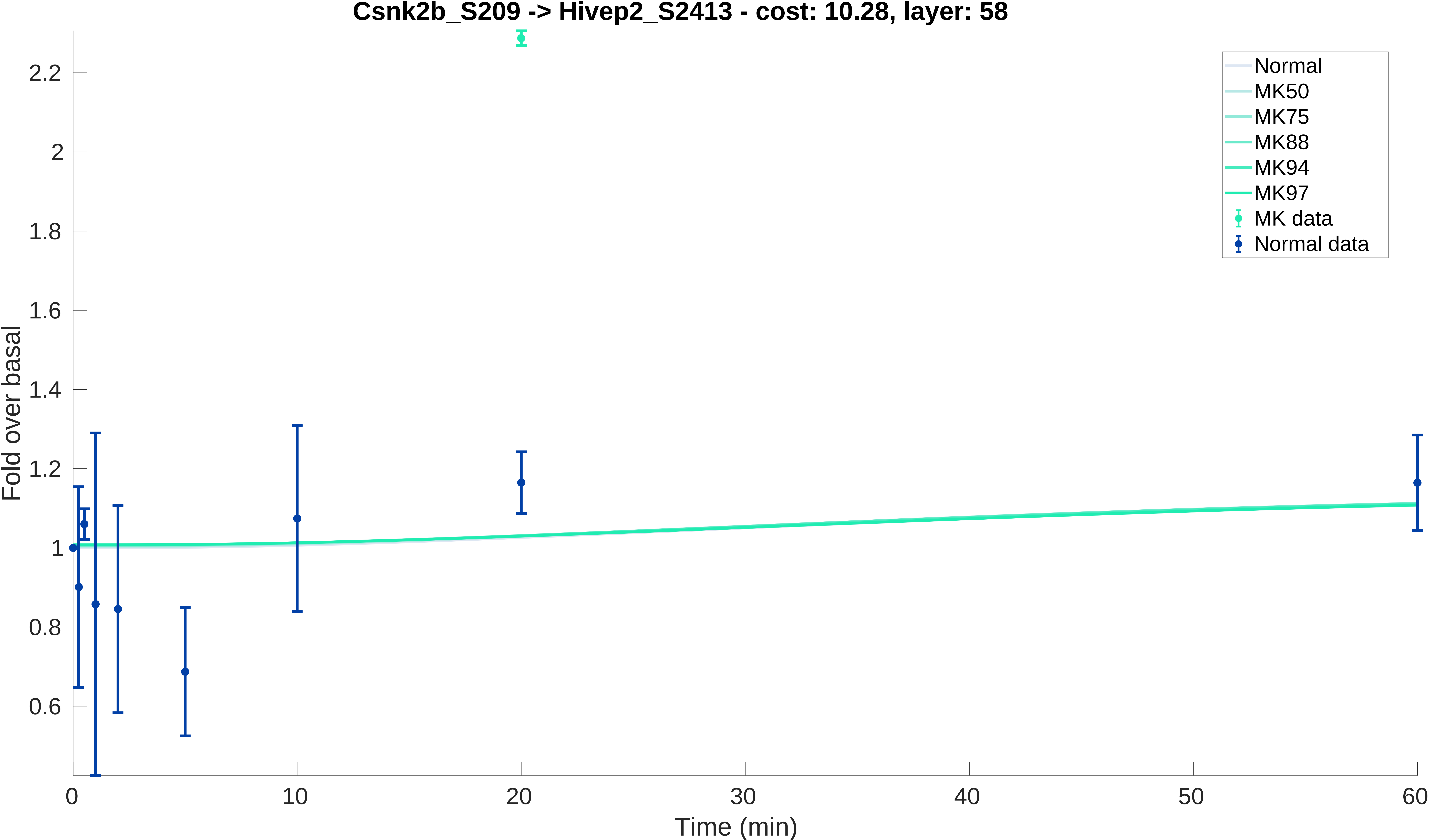

Csnk2b_S209 -> Hivep2_S2413 - cost: 10.28, layer: 58
Fold over basal
Time (min)
0
10
20
30
40
50
60
0.6
0.8
1
1.2
1.4
1.6
1.8
2
2.2
Normal
MK50
MK75
MK88
MK94
MK97
MK data
Normal data

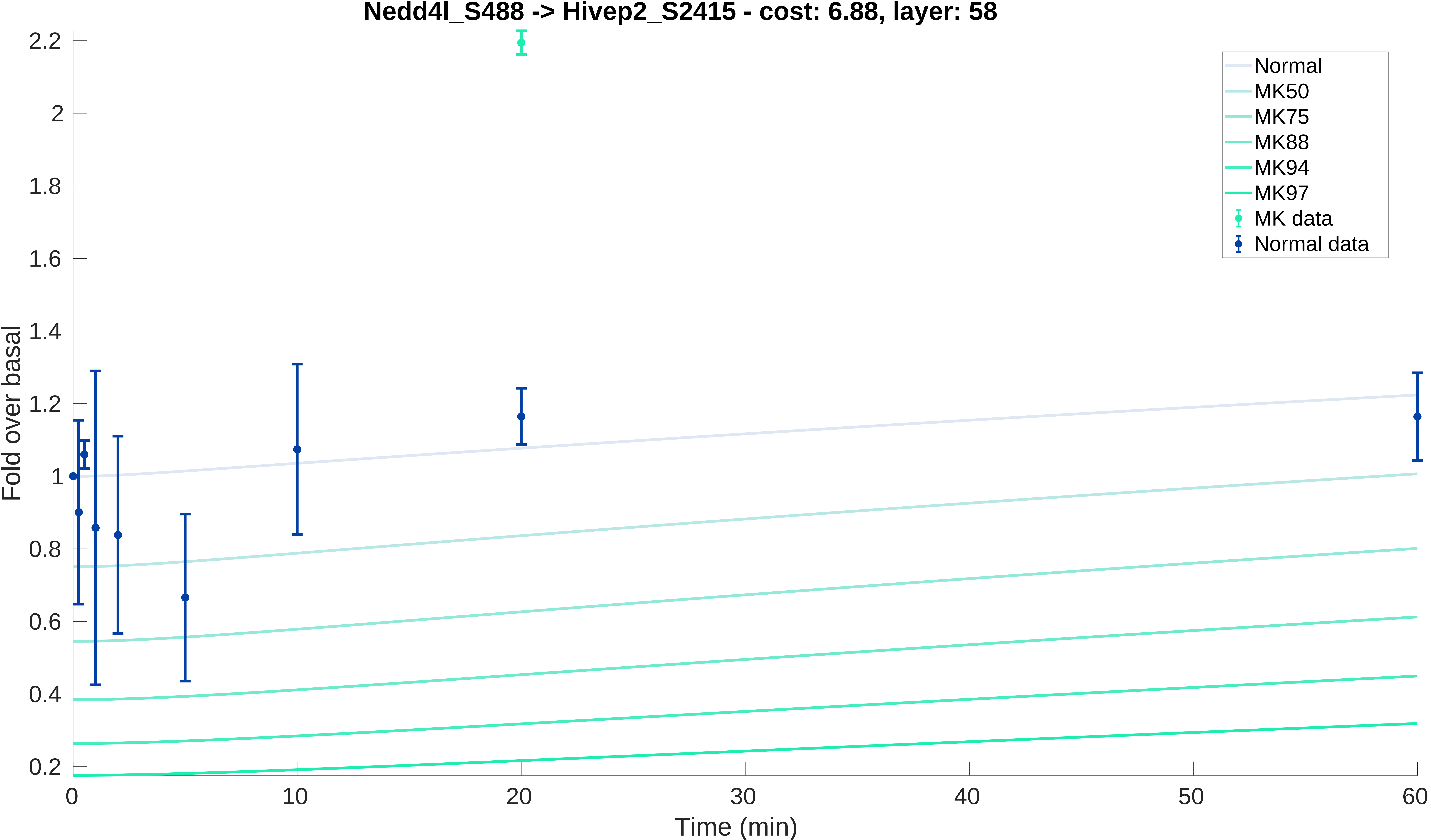

Nedd4l_S488 -> Hivep2_S2415 - cost: 6.88, layer: 58
Fold over basal
Time (min)
2.2
2
1.8
1.6
1.4
1.2
1
0.8
0.6
0.4
0.2
0
10
20
30
40
50
60
Normal
MK50
MK75
MK88
MK94
MK97
MK data
Normal data

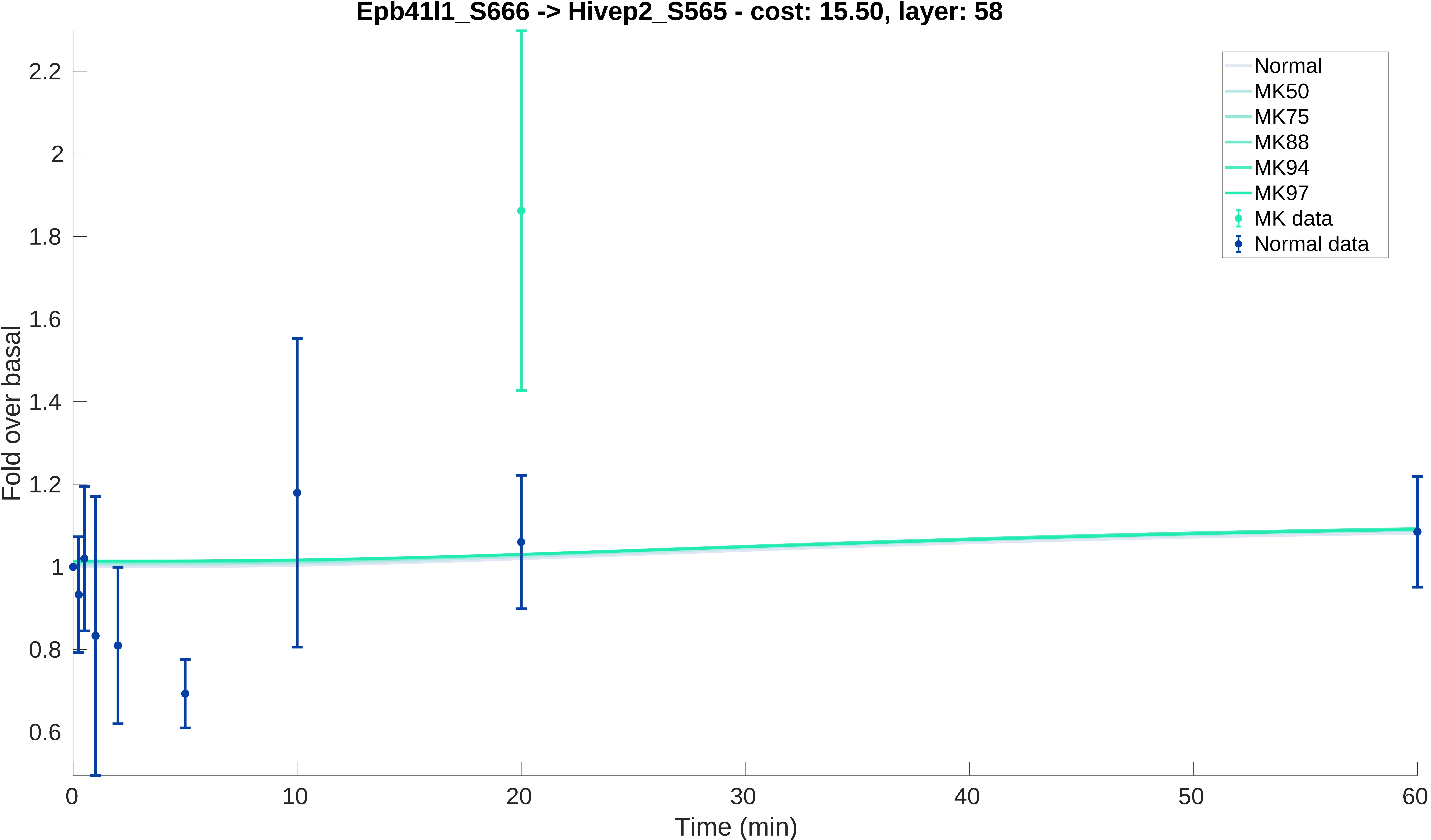
Epb41l1_S666 -> Hivep2_S565 - cost: 15.50, layer: 58
Fold over basal
Time (min)
0.6
0.8
1
1.2
1.4
1.6
1.8
2
2.2
0
10
20
30
40
50
60
Normal
MK50
MK75
MK88
MK94
MK97
MK data
Normal data

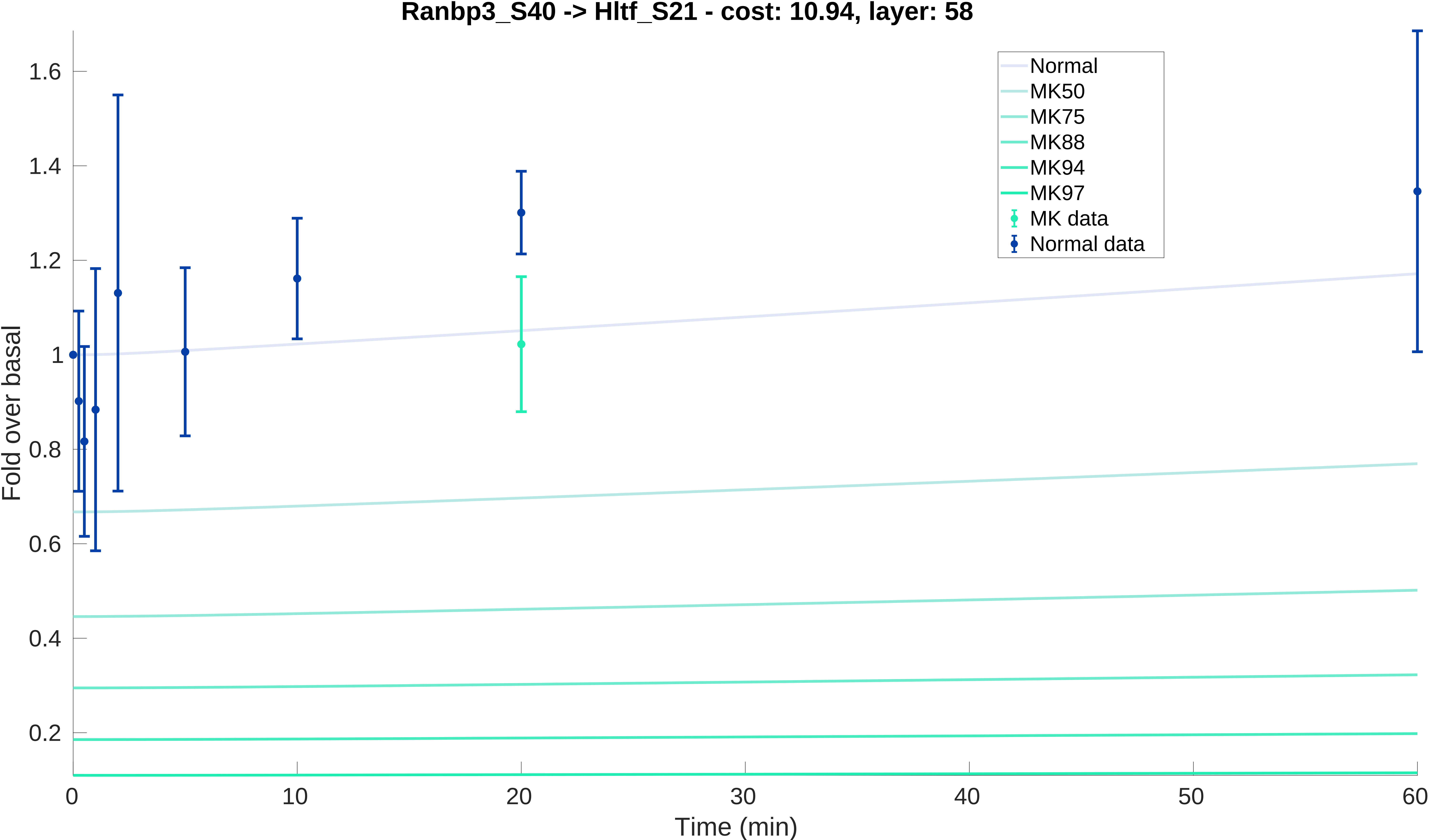

Ranbp3_S40 -> Hltf_S21 - cost: 10.94, layer: 58
Normal
MK50
MK75
MK88
MK94
MK97
MK data
Normal data
Fold over basal
1.6
1.4
1.2
1
0.8
0.6
0.4
0.2
0
10
20
30
40
50
60
Time (min)

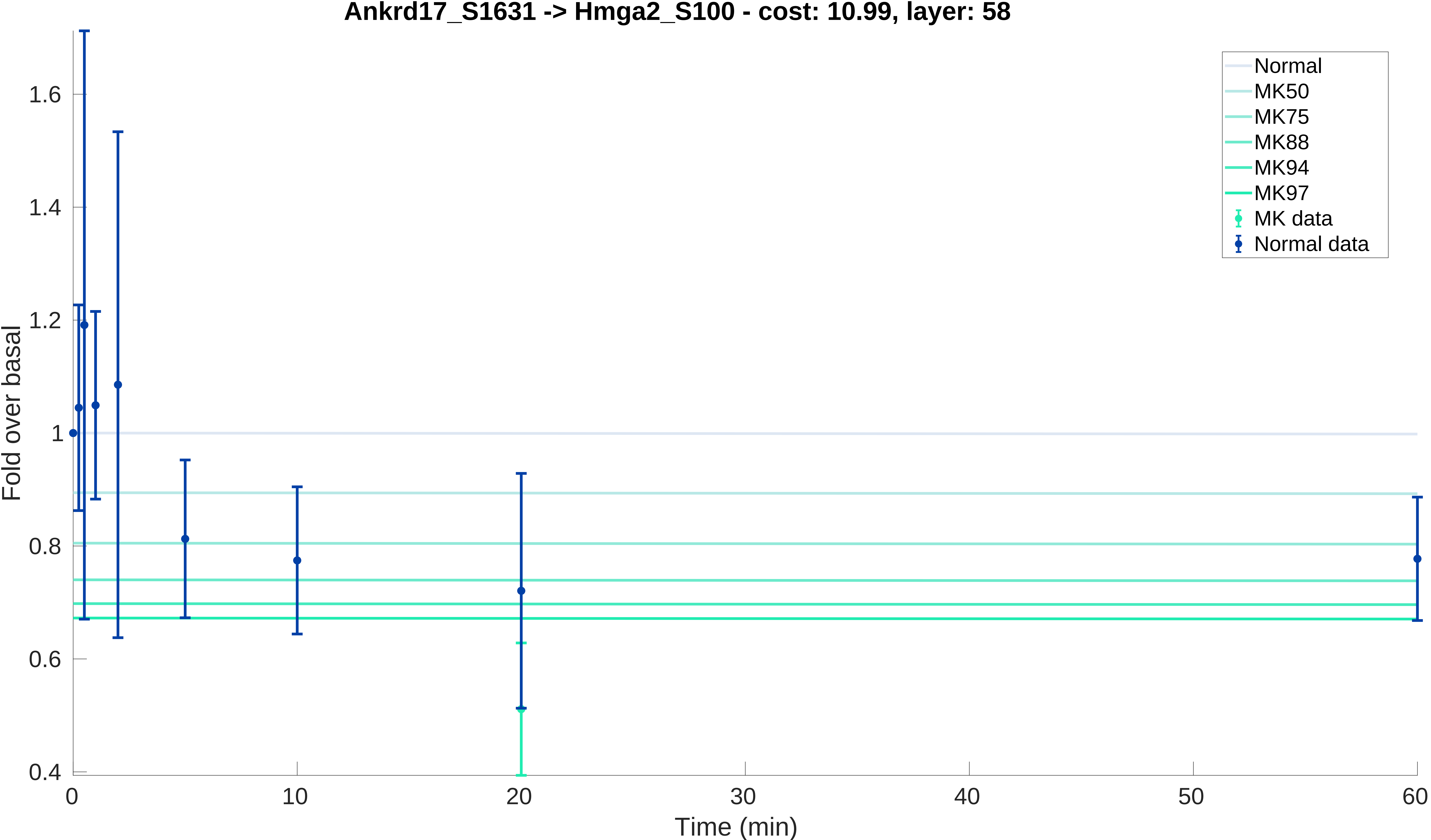

Ankrd17_S1631 -> Hmga2_S100 - cost: 10.99, layer: 58
Fold over basal
Time (min)
0.4
0.6
0.8
1
1.2
1.4
1.6
0
10
20
30
40
50
60
Normal
MK50
MK75
MK88
MK94
MK97
MK data
Normal data

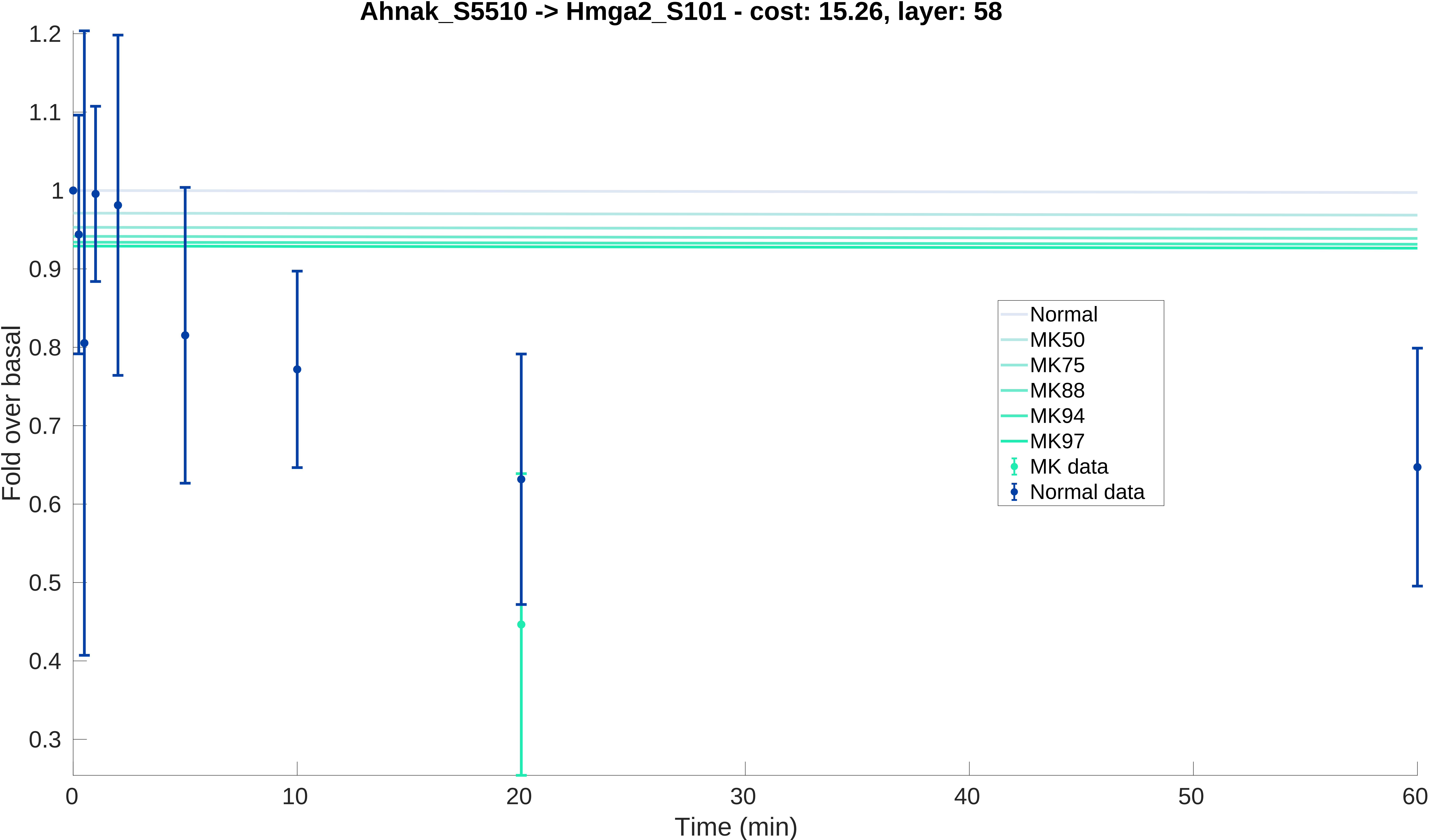

Ahnak_S5510 -> Hmga2_S101 - cost: 15.26, layer: 58
Fold over basal
Time (min)
0
10
20
30
40
50
60
0.3
0.4
0.5
0.6
0.7
0.8
0.9
1
1.1
1.2
Normal
MK50
MK75
MK88
MK94
MK97
MK data
Normal data

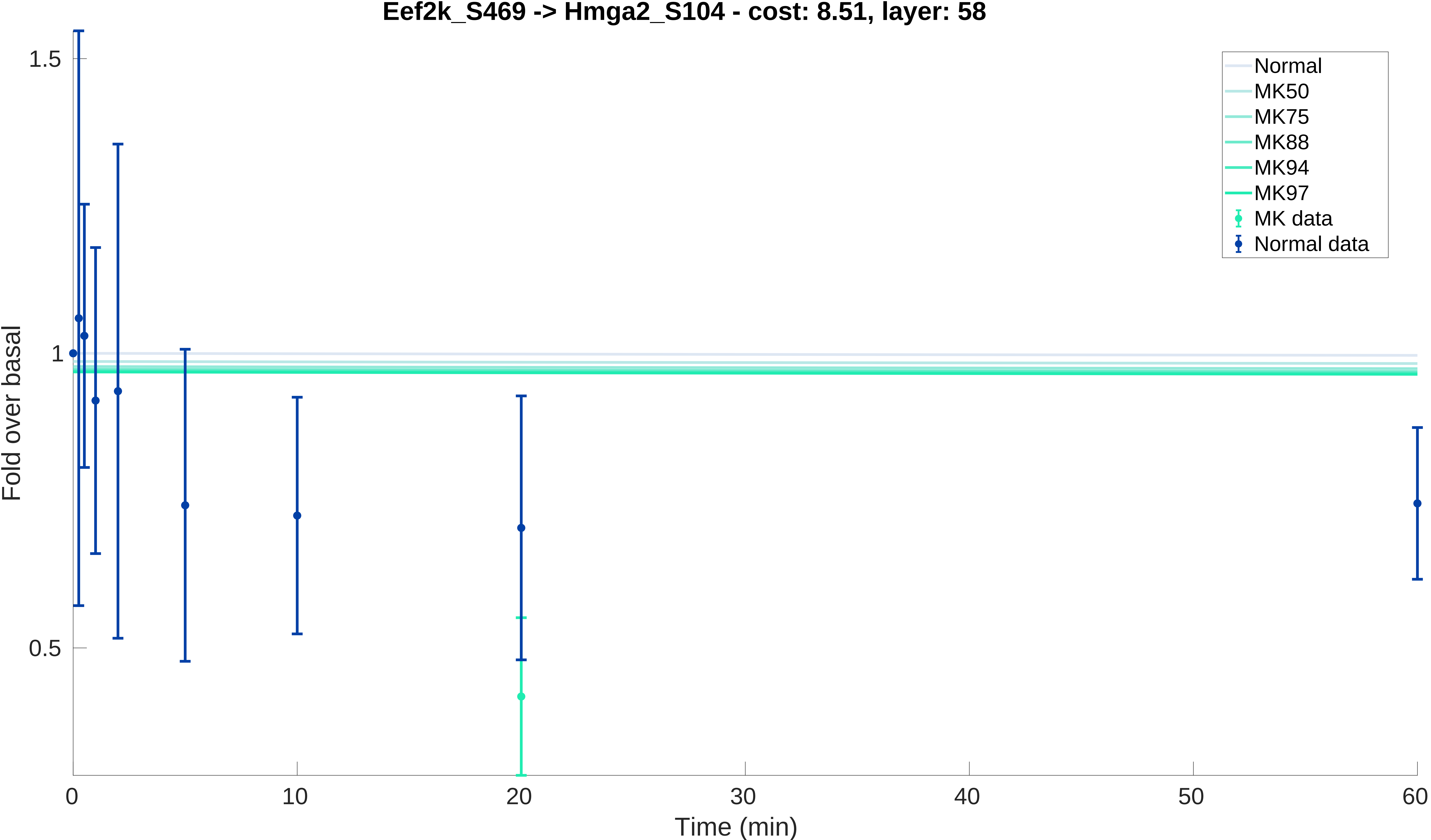

Eef2k_S469 -> Hmga2_S104 - cost: 8.51, layer: 58
Fold over basal
1.5
1
0.5
0
10
20
30
40
50
60
Time (min)
Normal
MK50
MK75
MK88
MK94
MK97
MK data
Normal data

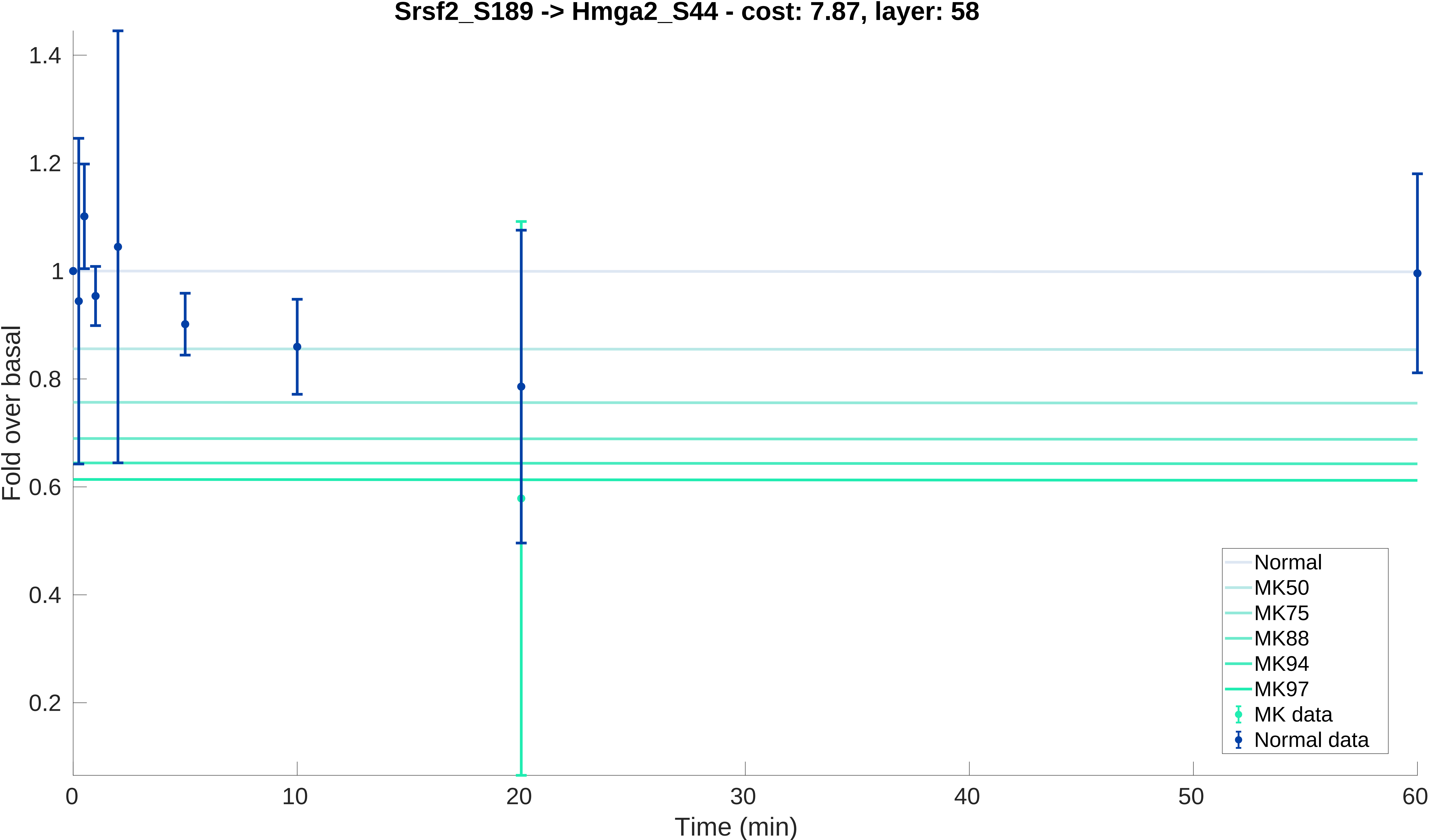

Srsf2_S189 -> Hmga2_S44 - cost: 7.87, layer: 58
1.4
1.2
1
0.8
0.6
0.4
0.2
Fold over basal
0
10
20
30
40
50
60
Time (min)
Normal
MK50
MK75
MK88
MK94
MK97
MK data
Normal data

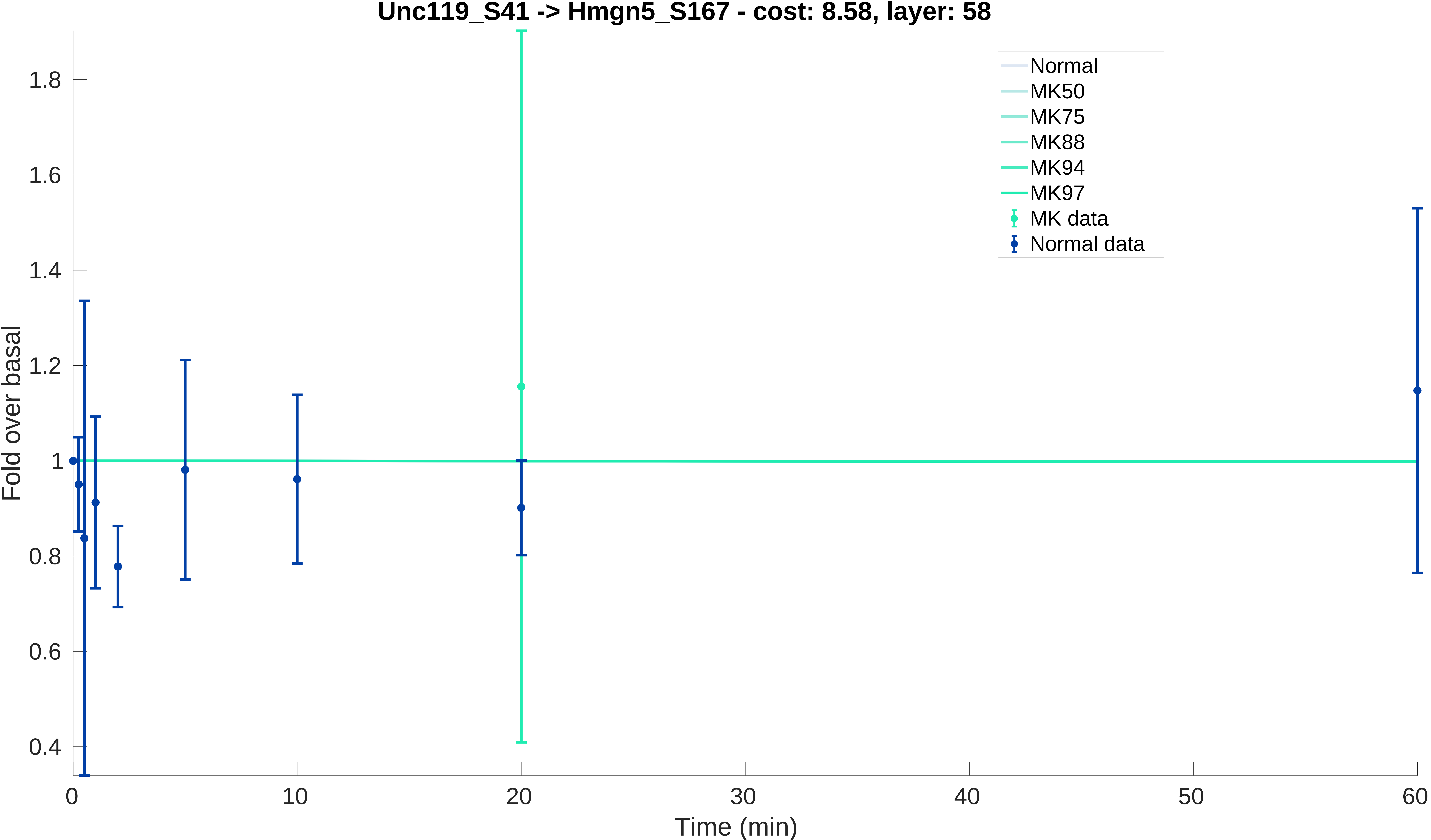

Unc119_S41 -> Hmgn5_S167 - cost: 8.58, layer: 58
Normal
MK50
MK75
MK88
MK94
MK97
MK data
Normal data
Fold over basal
Time (min)
1.8
1.6
1.4
1.2
1
0.8
0.6
0.4
0
10
20
30
40
50
60

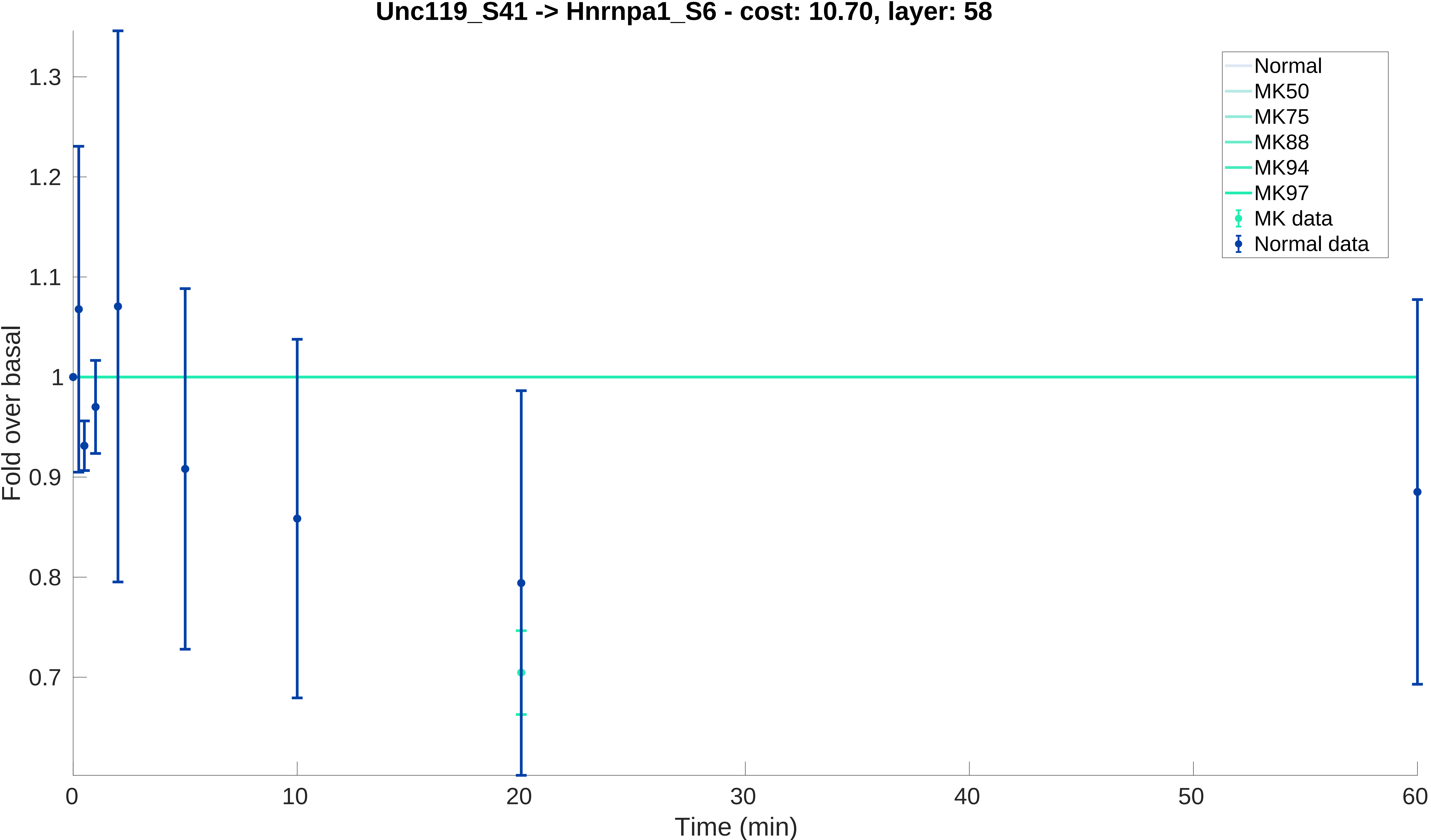

Unc119_S41 -> Hnrnpa1_S6 - cost: 10.70, layer: 58
Fold over basal
Time (min)
1.3
1.2
1.1
1
0.9
0.8
0.7
0
10
20
30
40
50
60
Normal
MK50
MK75
MK88
MK94
MK97
MK data
Normal data

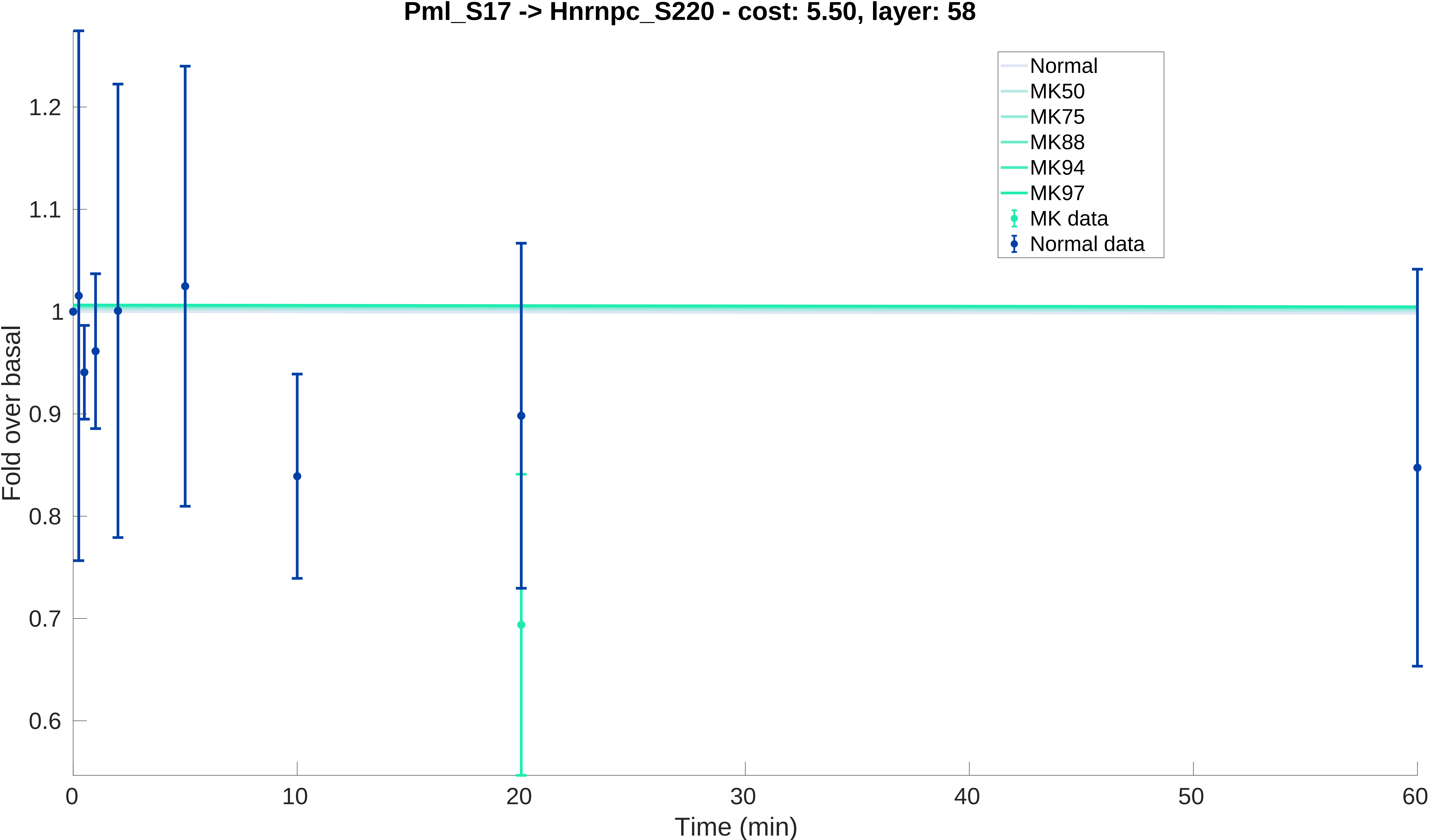

Pml_S17 -> Hnrnpc_S220 - cost: 5.50, layer: 58
Normal
MK50
MK75
MK88
MK94
MK97
MK data
Normal data
Fold over basal
Time (min)
1.2
1.1
1
0.9
0.8
0.7
0.6
0
10
20
30
40
50
60

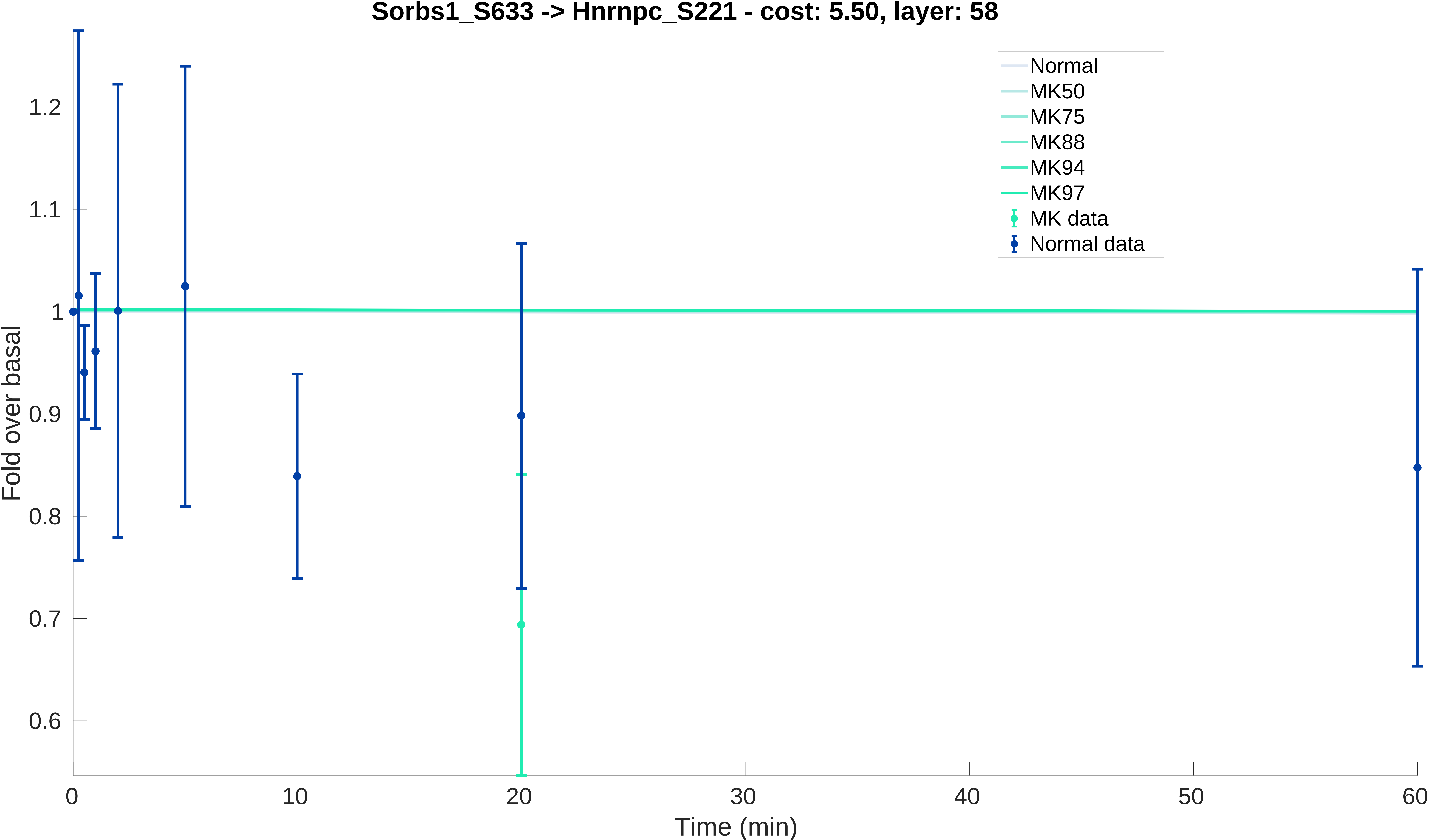
Sorbs1_S633 -> Hnrnpc_S221 - cost: 5.50, layer: 58
Normal
MK50
MK75
MK88
MK94
MK97
MK data
Normal data
Fold over basal
Time (min)
1.2
1.1
1
0.9
0.8
0.7
0.6
0
10
20
30
40
50
60

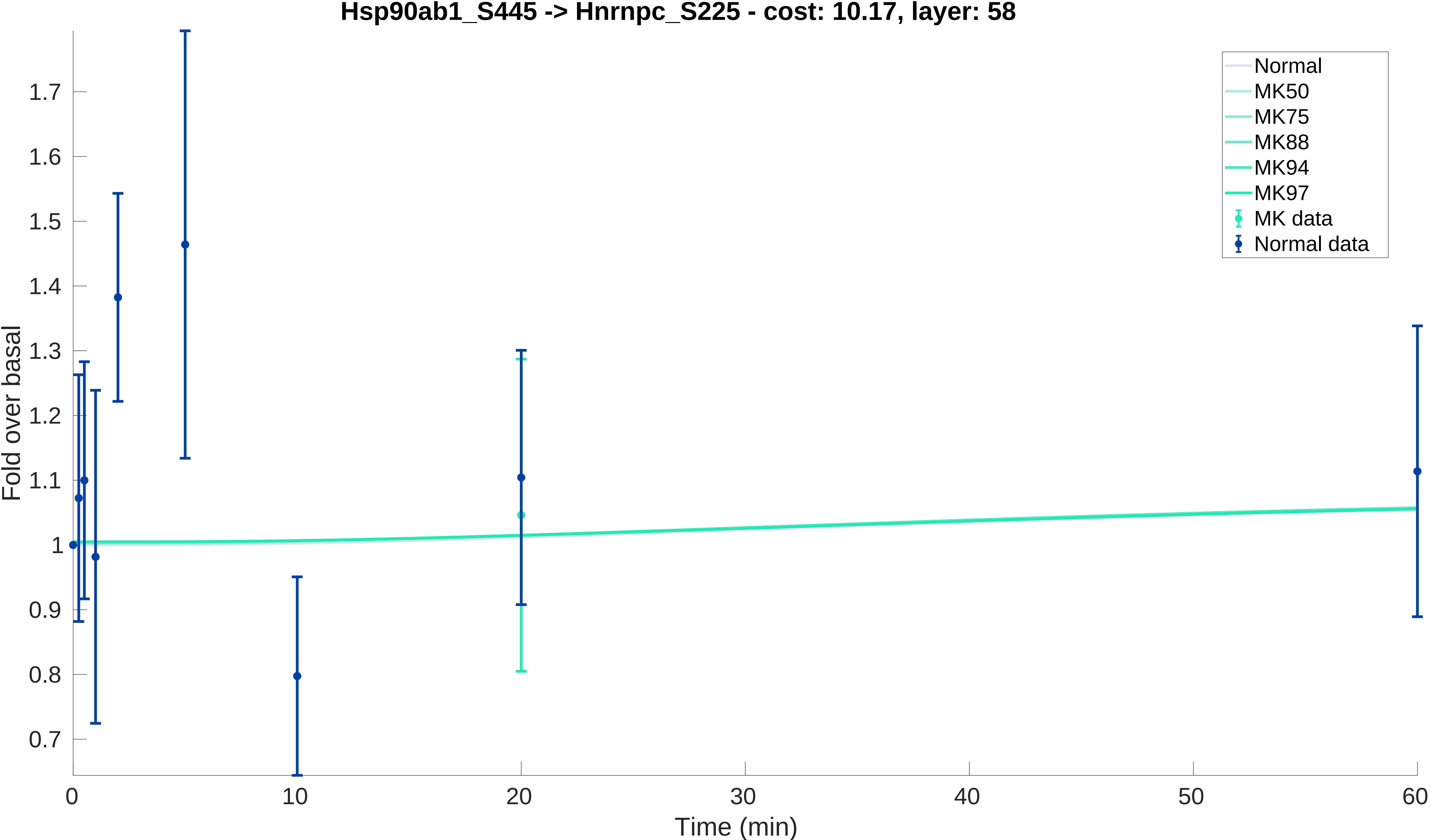
Hsp90ab1_S445 -> Hnrnpc_S225 - cost: 10.17, layer: 58
Fold over basal
Time (min)
0.7
0.8
0.9
1
1.1
1.2
1.3
1.4
1.5
1.6
1.7
0
10
20
30
40
50
60
Normal
MK50
MK75
MK88
MK94
MK97
MK data
Normal data

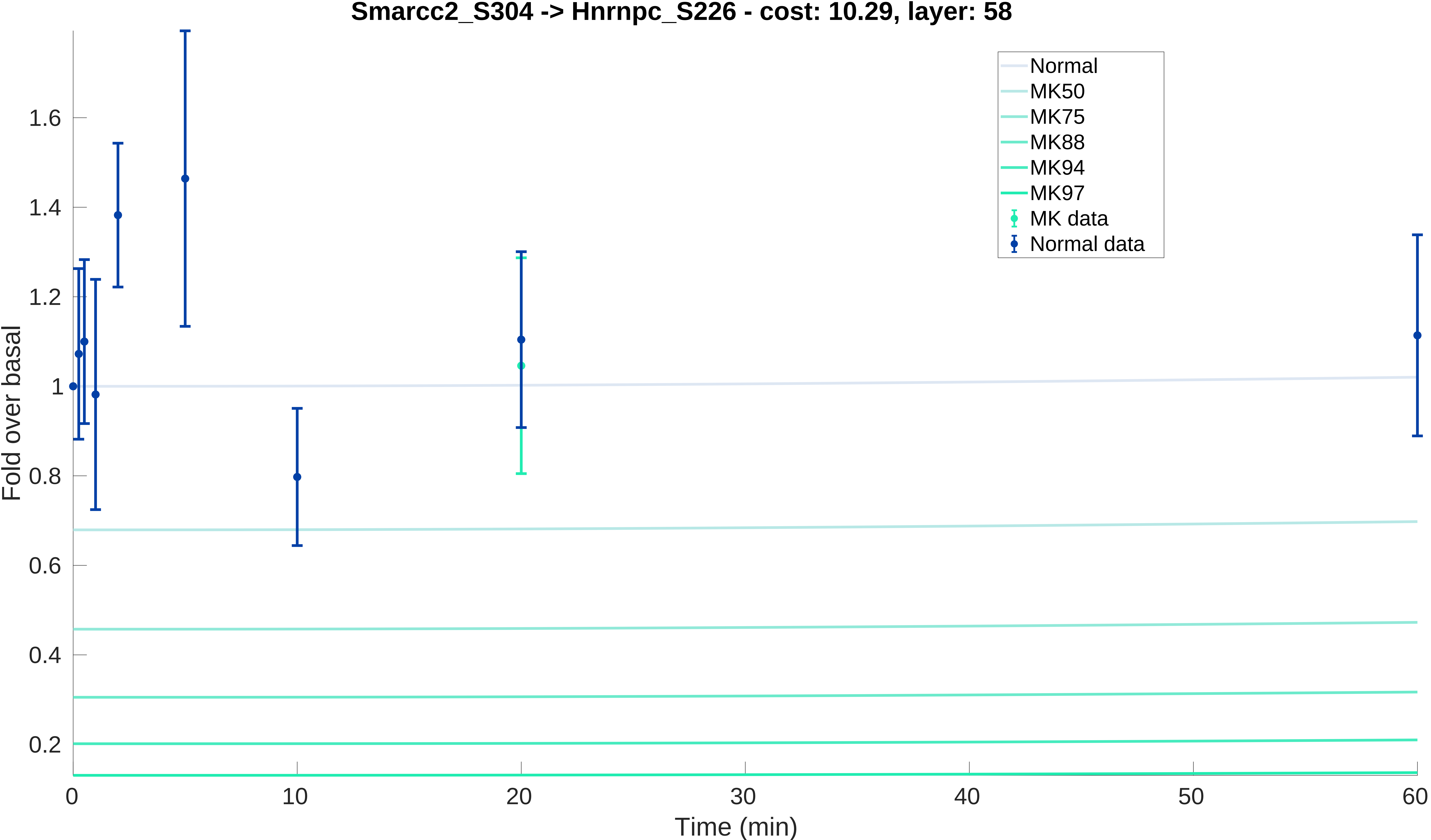
Smarcc2_S304 -> Hnrnpc_S226 - cost: 10.29, layer: 58
Fold over basal
Time (min)
0.2
0.4
0.6
0.8
1
1.2
1.4
1.6
0
10
20
30
40
50
60
Normal
MK50
MK75
MK88
MK94
MK97
MK data
Normal data

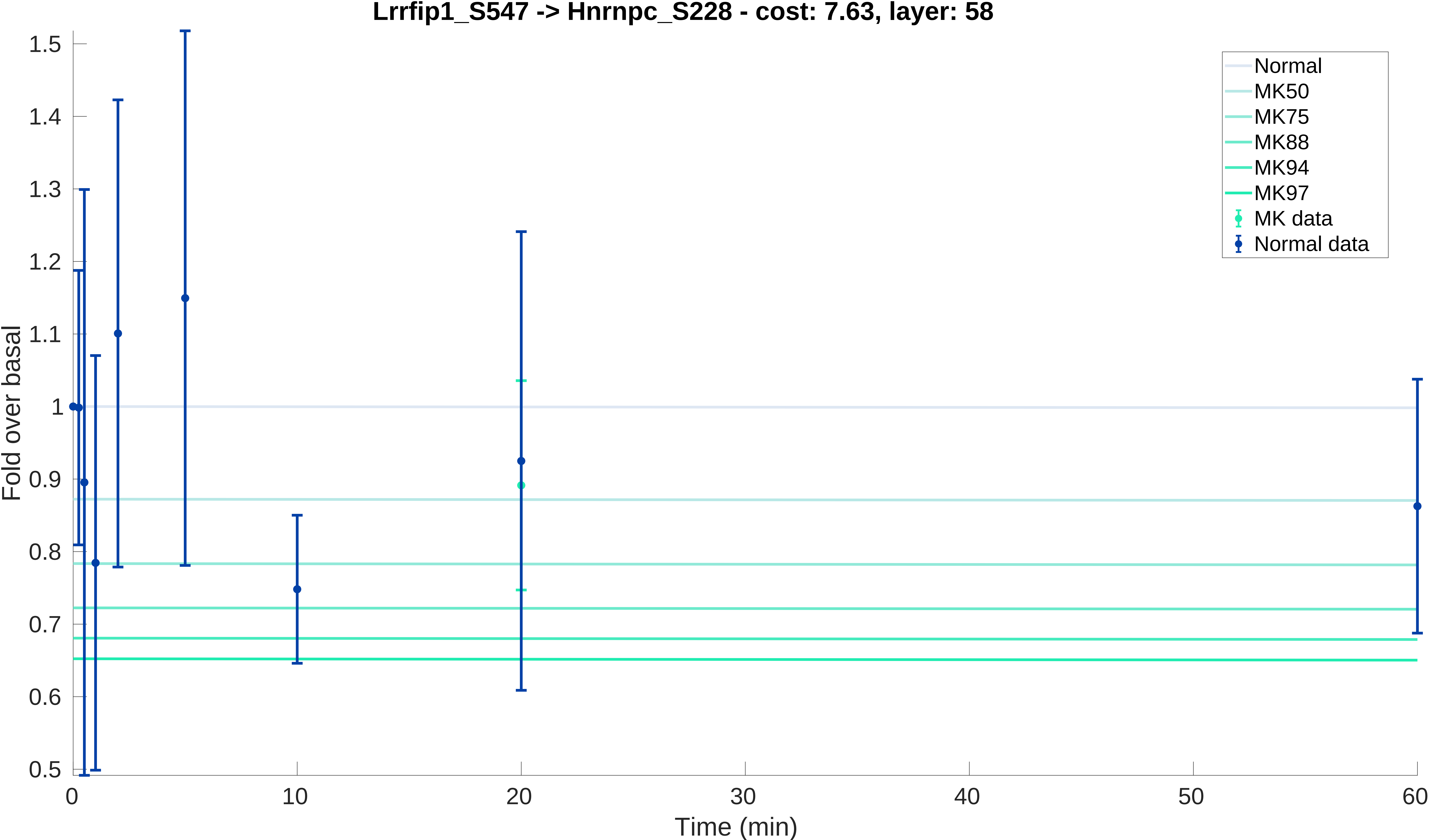

Lrrfip1_S547 -> Hnrnpc_S228 - cost: 7.63, layer: 58
Fold over basal
Time (min)
1.5
1.4
1.3
1.2
1.1
1
0.9
0.8
0.7
0.6
0.5
0
10
20
30
40
50
60
Normal
MK50
MK75
MK88
MK94
MK97
MK data
Normal data

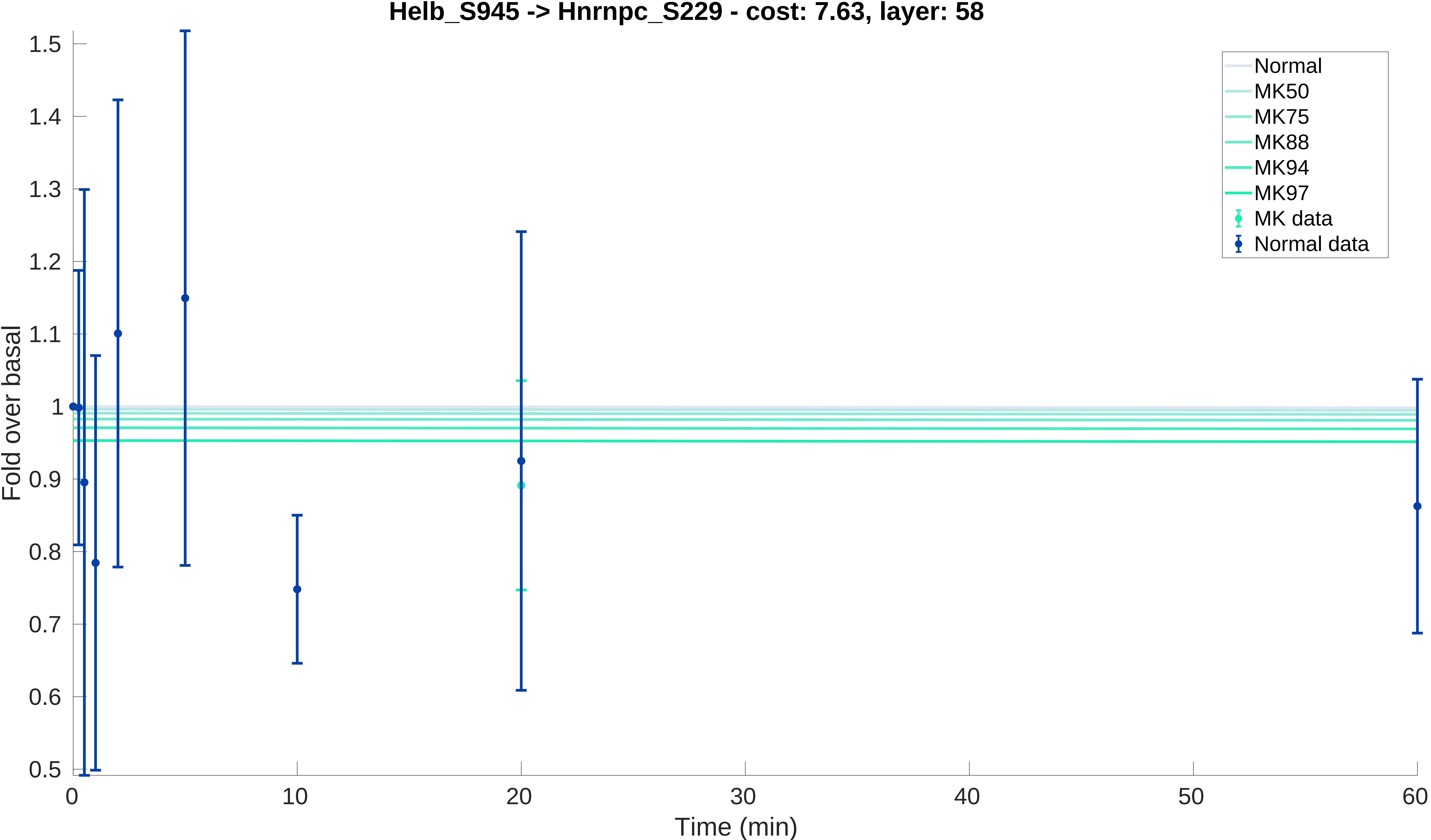

Helb_S945 -> Hnrnpc_S229 - cost: 7.63, layer: 58
Fold over basal
Time (min)
1.5
1.4
1.3
1.2
1.1
1
0.9
0.8
0.7
0.6
0.5
0
10
20
30
40
50
60
Normal
MK50
MK75
MK88
MK94
MK97
MK data
Normal data

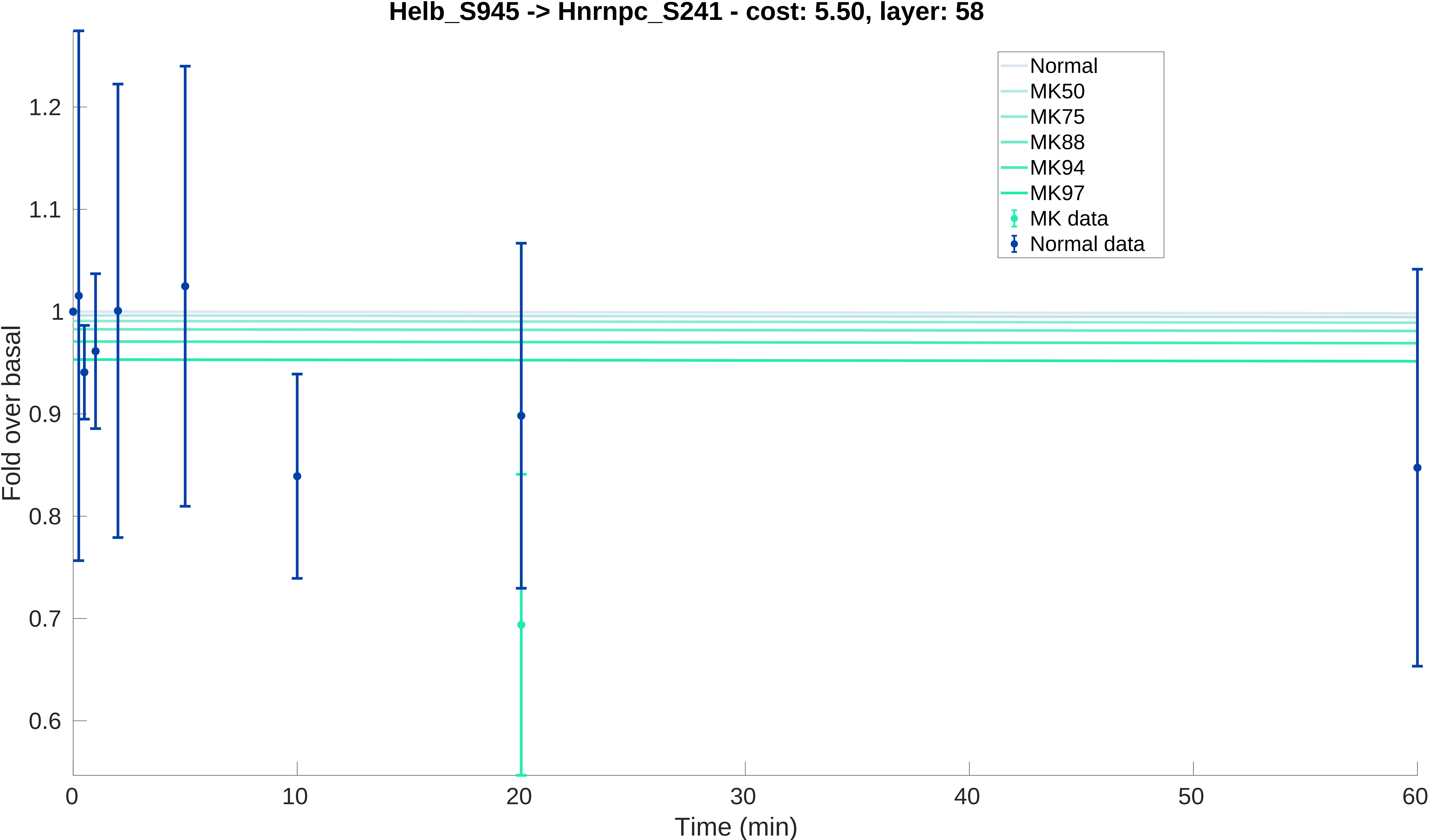

Helb_S945 -> Hnrnpc_S241 - cost: 5.50, layer: 58
Normal
MK50
MK75
MK88
MK94
MK97
MK data
Normal data
Fold over basal
1.2
1.1
1
0.9
0.8
0.7
0.6
0
10
20
30
40
50
60
Time (min)

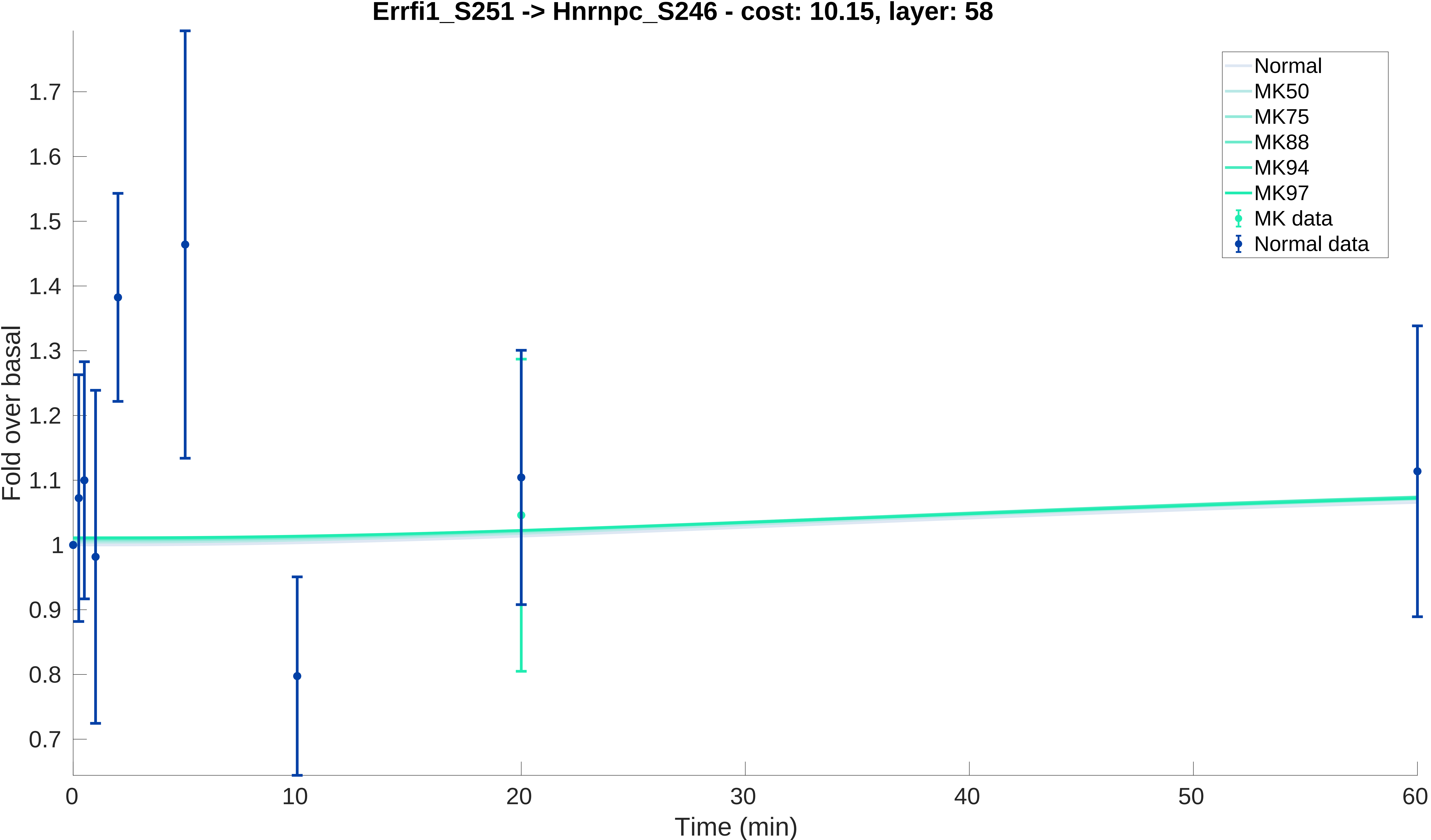

Errfi1_S251 -> Hnrnpc_S246 - cost: 10.15, layer: 58
Normal
MK50
MK75
MK88
MK94
MK97
MK data
Normal data
Fold over basal
1.7
1.6
1.5
1.4
1.3
1.2
1.1
1
0.9
0.8
0.7
0
10
20
30
40
50
60
Time (min)

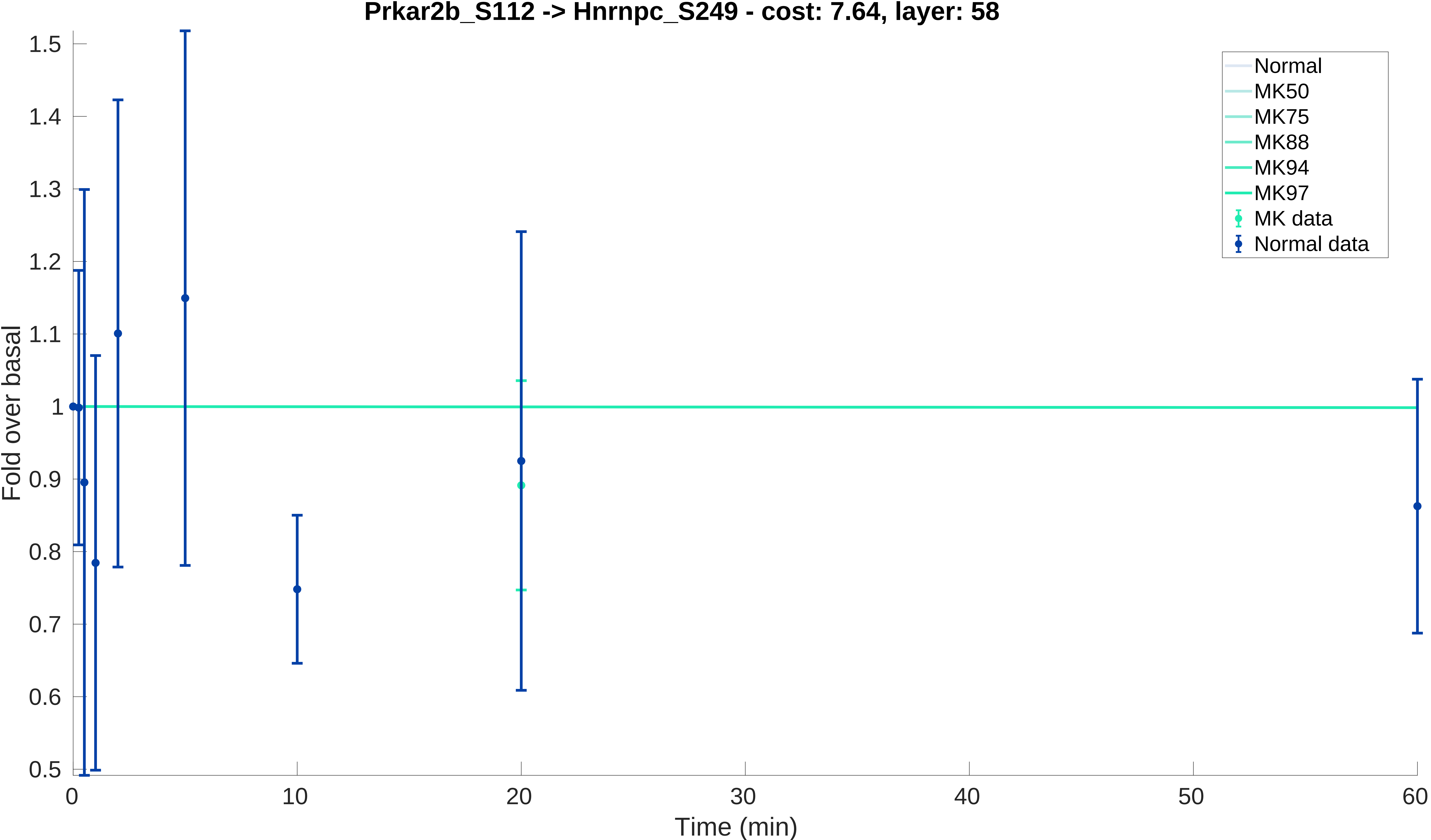

Prkar2b_S112 -> Hnrnpc_S249 - cost: 7.64, layer: 58
Fold over basal
Time (min)
0.5
0.6
0.7
0.8
0.9
1
1.1
1.2
1.3
1.4
1.5
0
10
20
30
40
50
60
Normal
MK50
MK75
MK88
MK94
MK97
MK data
Normal data

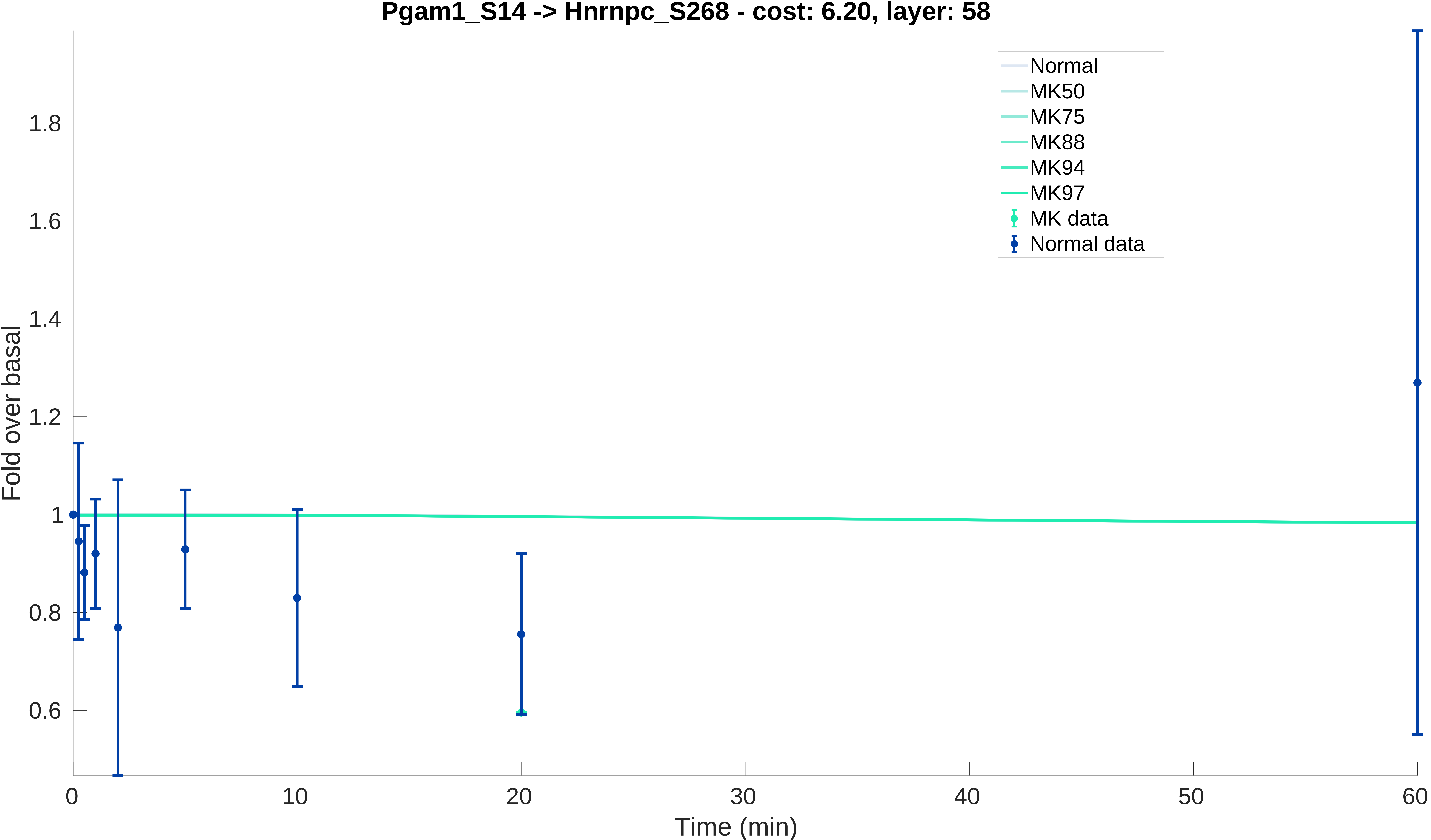

Pgam1_S14 -> Hnrnpc_S268 - cost: 6.20, layer: 58
Normal
MK50
MK75
MK88
MK94
MK97
MK data
Normal data
Fold over basal
1.8
1.6
1.4
1.2
1
0.8
0.6
0
10
20
30
40
50
60
Time (min)

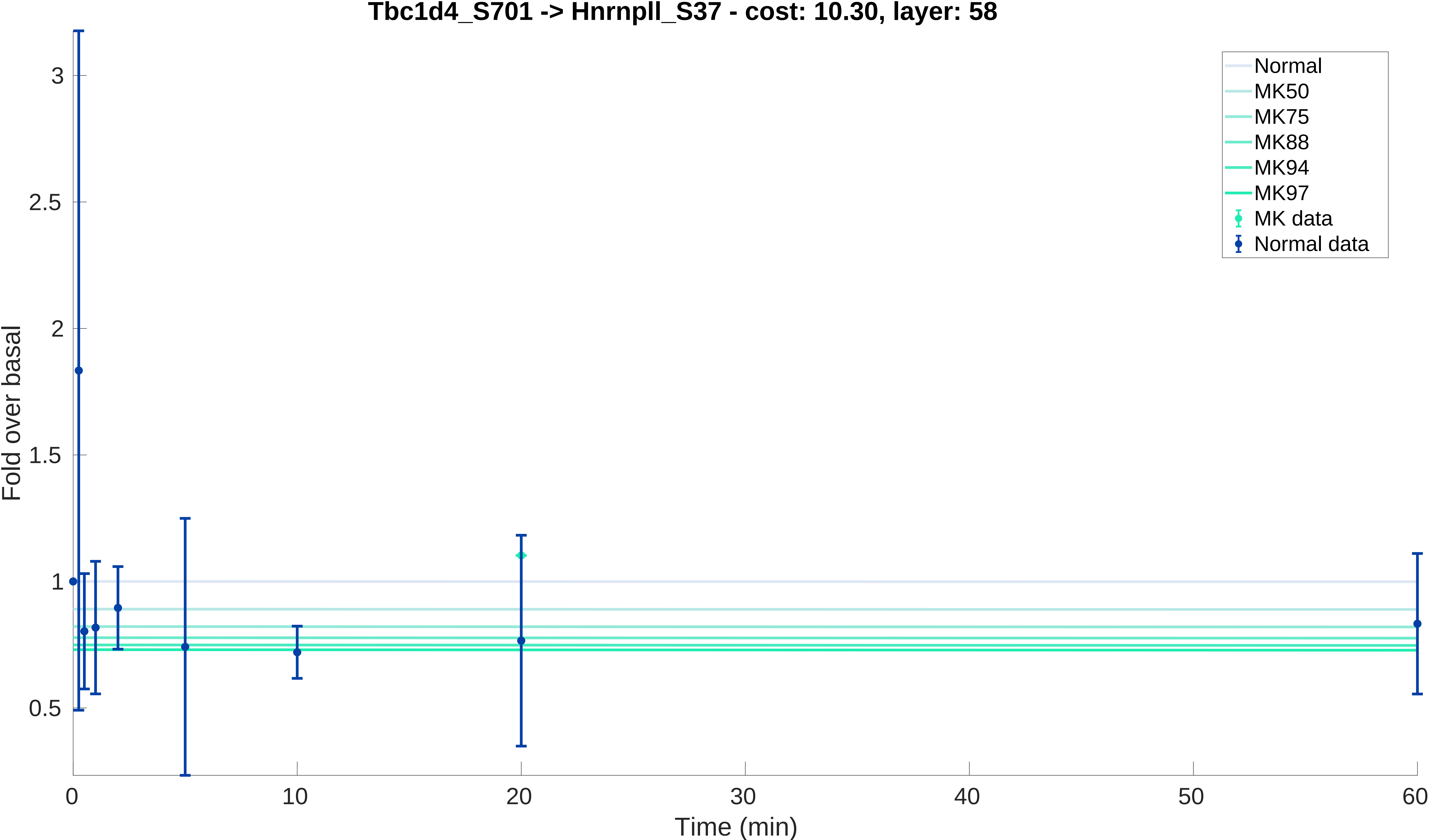

Tbc1d4_S701 -> Hnrnpll_S37 - cost: 10.30, layer: 58
Fold over basal
Time (min)
0.5
1
1.5
2
2.5
3
0
10
20
30
40
50
60
Normal
MK50
MK75
MK88
MK94
MK97
MK data
Normal data

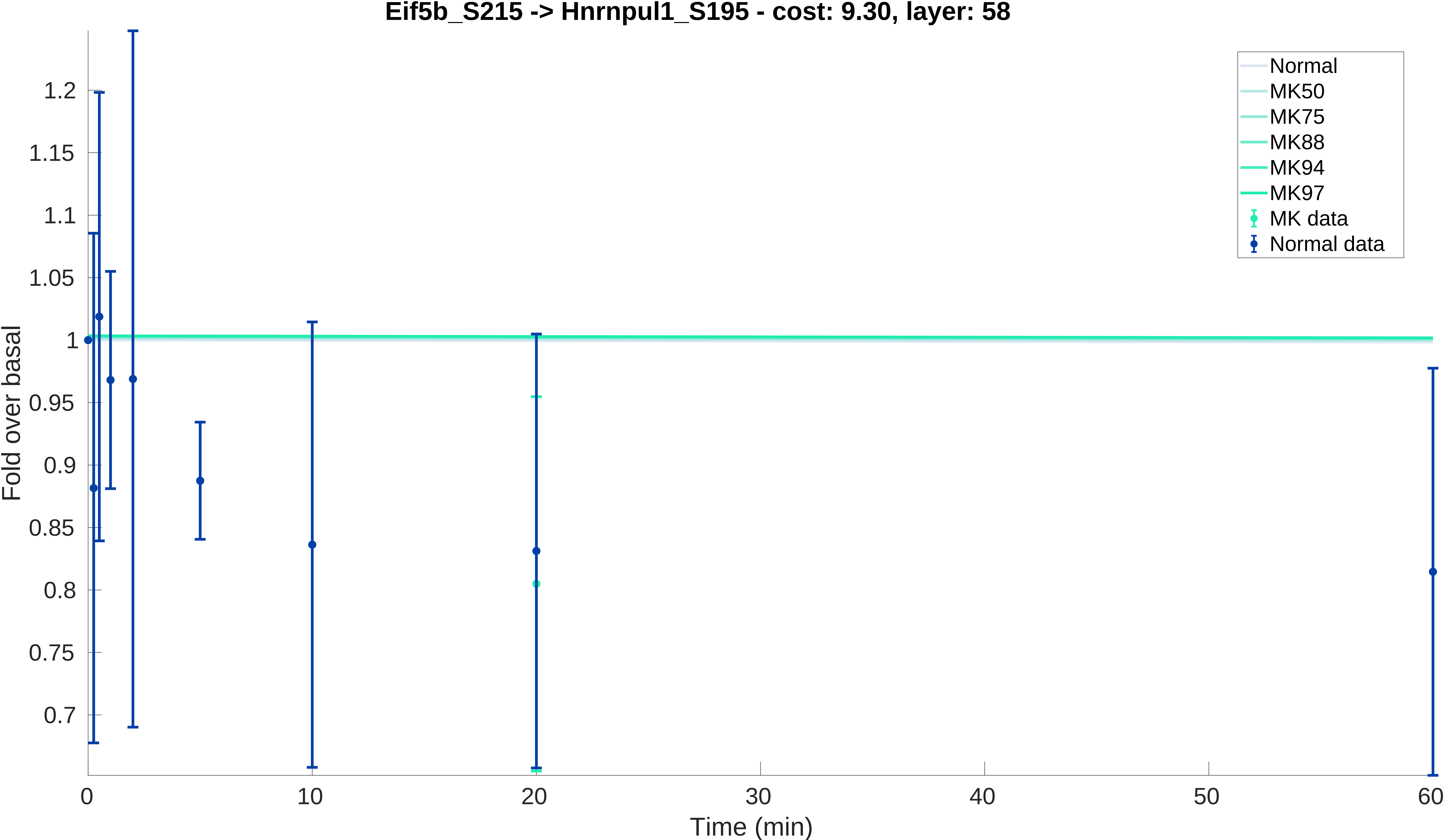

Eif5b_S215 -> Hnrnpul1_S195 - cost: 9.30, layer: 58
Normal
MK50
MK75
MK88
MK94
MK97
MK data
Normal data
Fold over basal
1.2
1.15
1.1
1.05
1
0.95
0.9
0.85
0.8
0.75
0.7
0
10
20
30
40
50
60
Time (min)

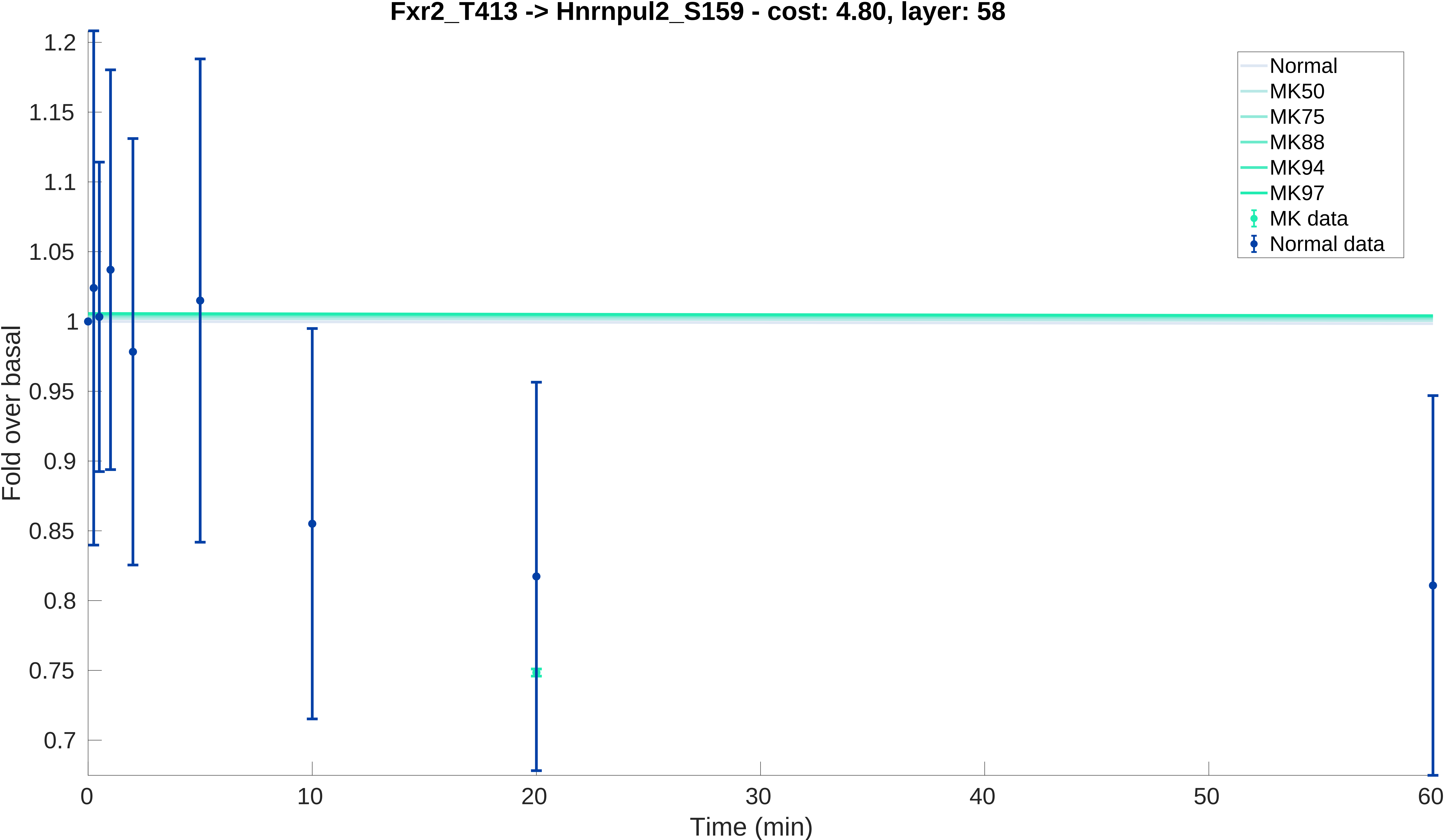

Fxr2_T413 -> Hnrnpul2_S159 - cost: 4.80, layer: 58
Fold over basal
Time (min)
Normal
MK50
MK75
MK88
MK94
MK97
MK data
Normal data
1.2
1.15
1.1
1.05
1
0.95
0.9
0.85
0.8
0.75
0.7
0
10
20
30
40
50
60

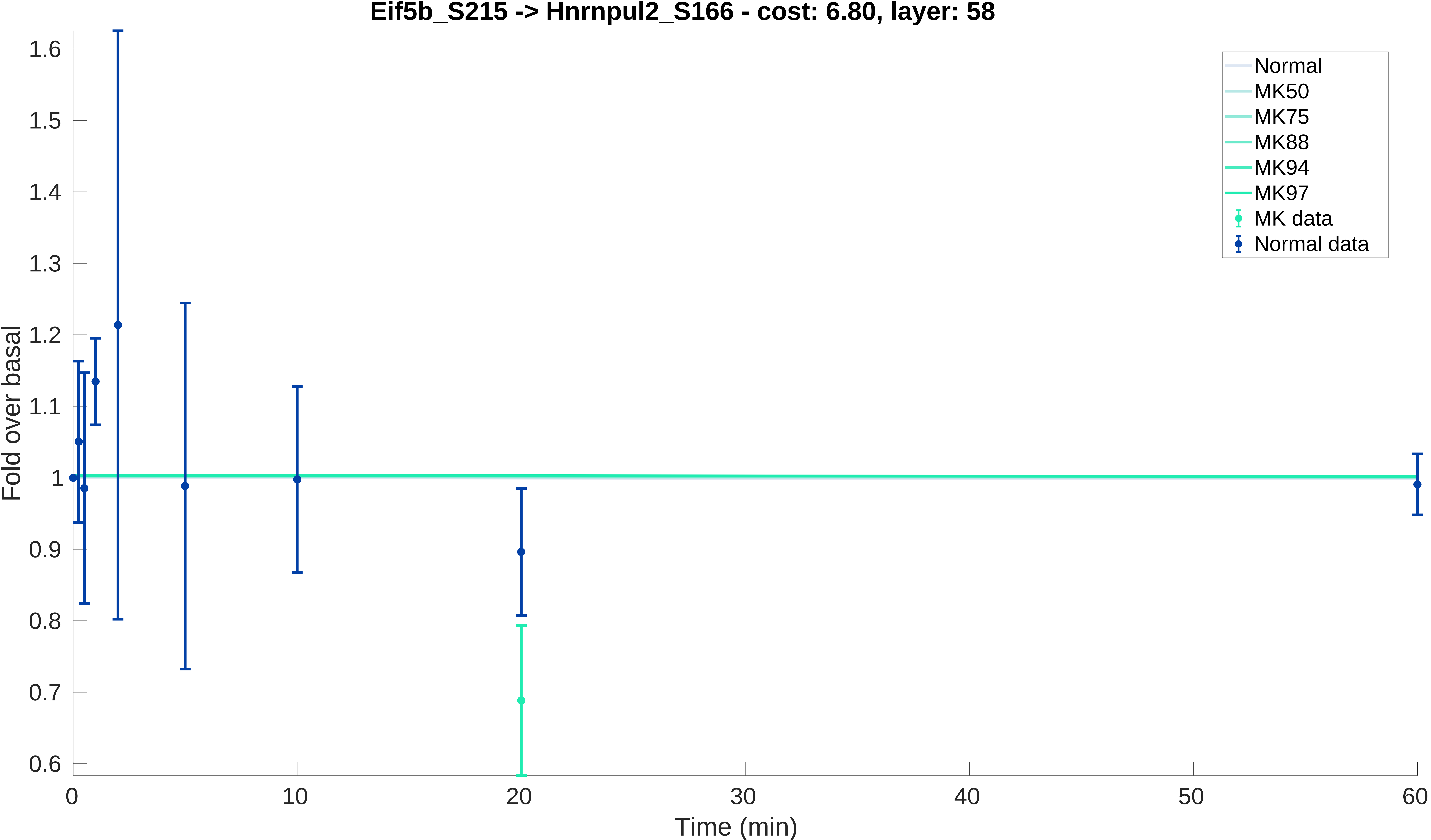
Eif5b_S215 -> Hnrnpul2_S166 - cost: 6.80, layer: 58
Fold over basal
Time (min)
1.6
1.5
1.4
1.3
1.2
1.1
1
0.9
0.8
0.7
0.6
0
10
20
30
40
50
60
Normal
MK50
MK75
MK88
MK94
MK97
MK data
Normal data

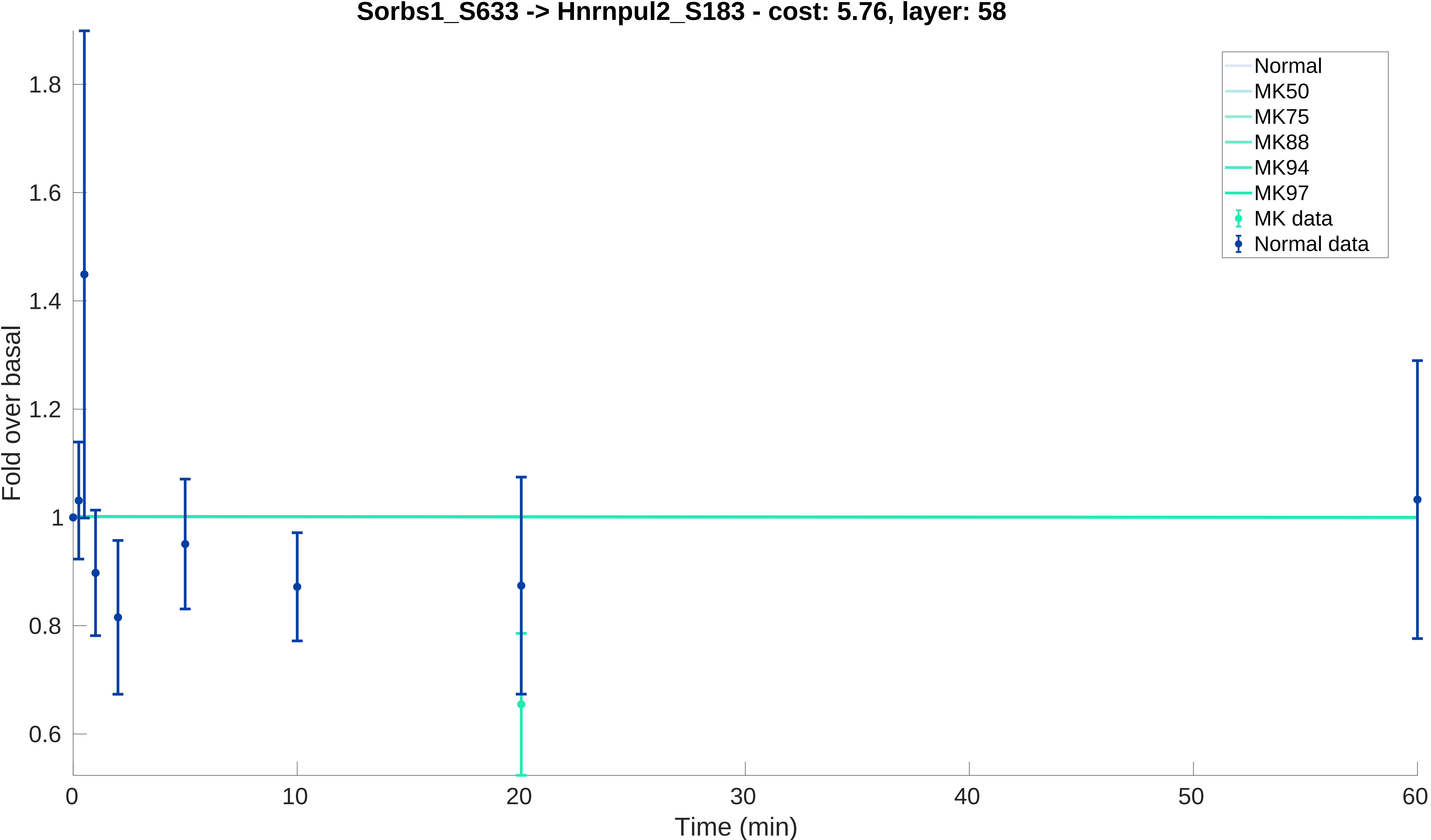

Sorbs1_S633 -> Hnrnpul2_S183 - cost: 5.76, layer: 58
Fold over basal
Time (min)
0.6
0.8
1
1.2
1.4
1.6
1.8
0
10
20
30
40
50
60
Normal
MK50
MK75
MK88
MK94
MK97
MK data
Normal data

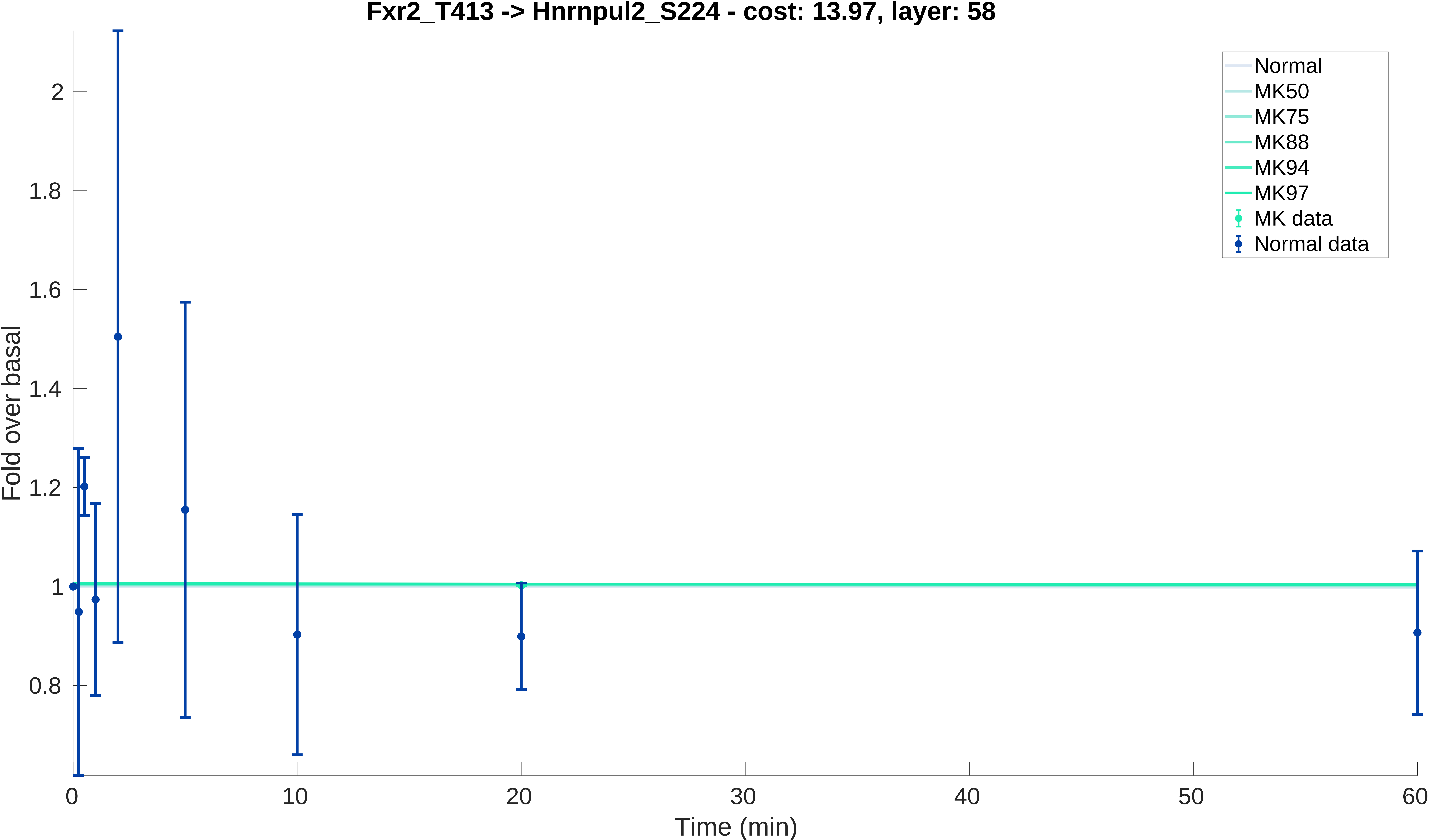
Fxr2_T413 -> Hnrnpul2_S224 - cost: 13.97, layer: 58
Fold over basal
Time (min)
0.8
1
1.2
1.4
1.6
1.8
2
0
10
20
30
40
50
60
Normal
MK50
MK75
MK88
MK94
MK97
MK data
Normal data

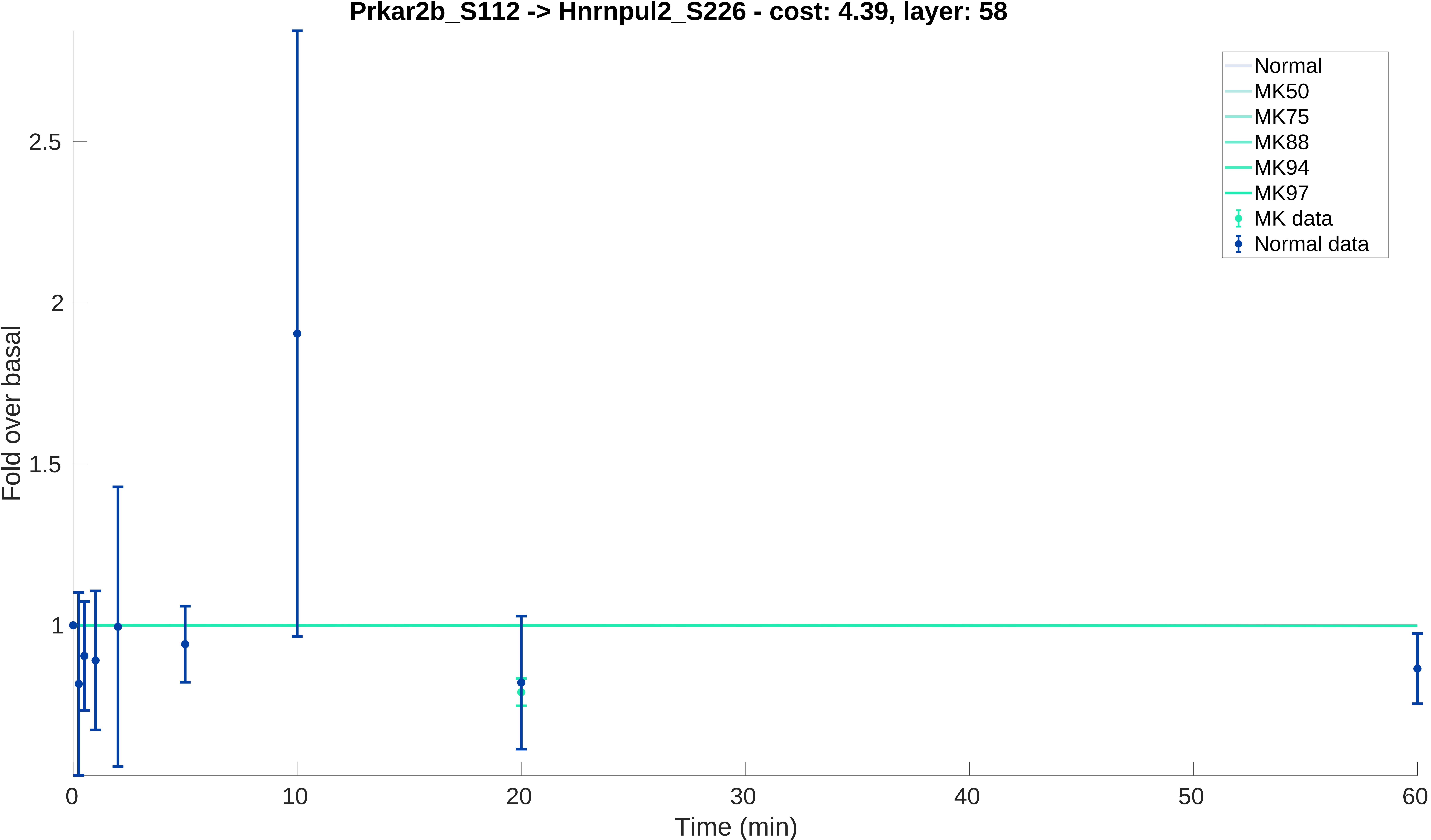

Prkar2b_S112 -> Hnrnpul2_S226 - cost: 4.39, layer: 58
Fold over basal
Time (min)
2.5
2
1.5
1
0
10
20
30
40
50
60
Normal
MK50
MK75
MK88
MK94
MK97
MK data
Normal data

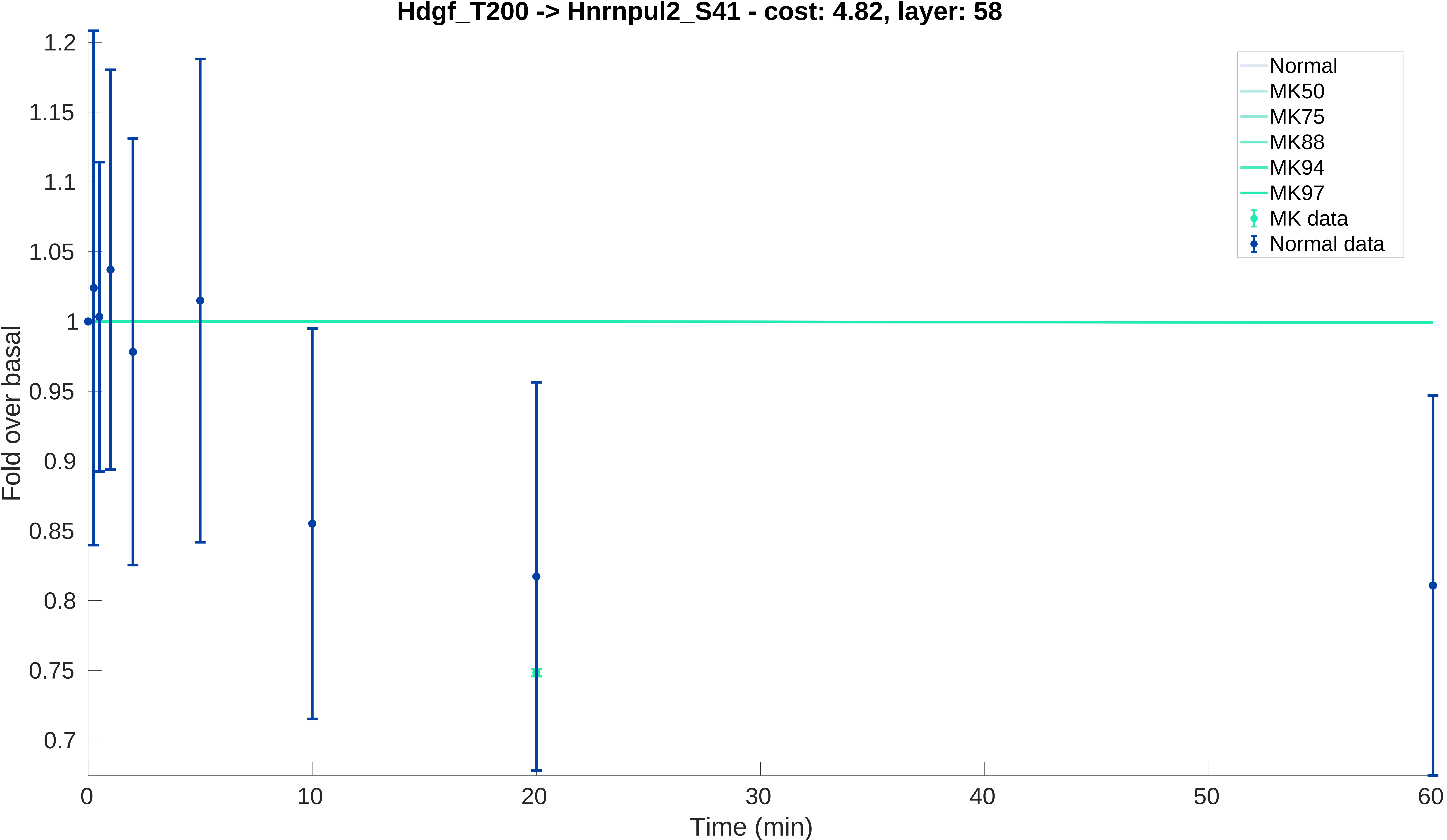
Hdgf_T200 -> Hnrnpul2_S41 - cost: 4.82, layer: 58
Fold over basal
Time (min)
1.2
1.15
1.1
1.05
1
0.95
0.9
0.85
0.8
0.75
0.7
0
10
20
30
40
50
60
Normal
MK50
MK75
MK88
MK94
MK97
MK data
Normal data

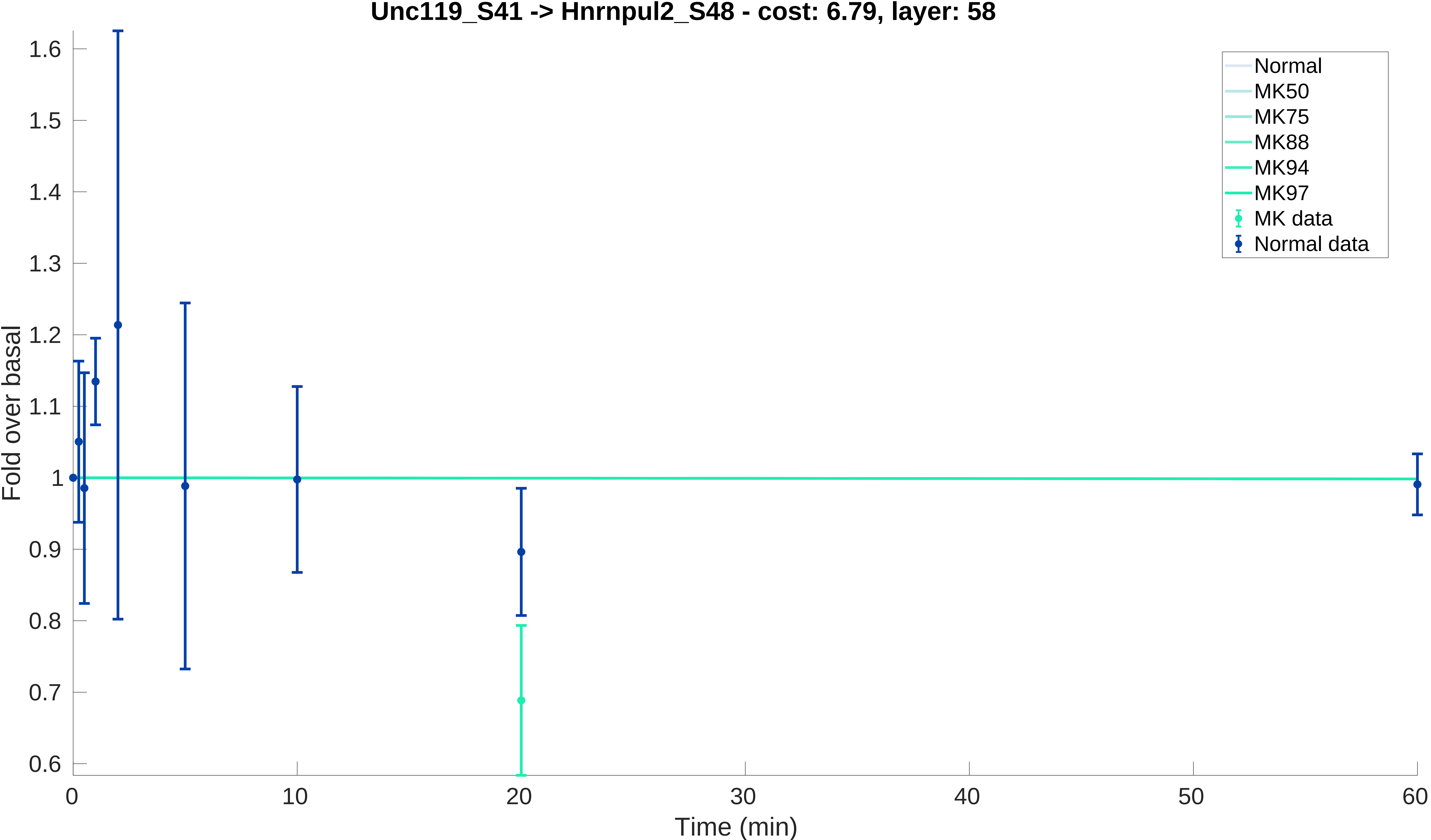
Unc119_S41 -> Hnrnpul2_S48 - cost: 6.79, layer: 58
Fold over basal
1.6
1.5
1.4
1.3
1.2
1.1
1
0.9
0.8
0.7
0.6
0
10
20
30
40
50
60
Time (min)
Normal
MK50
MK75
MK88
MK94
MK97
MK data
Normal data

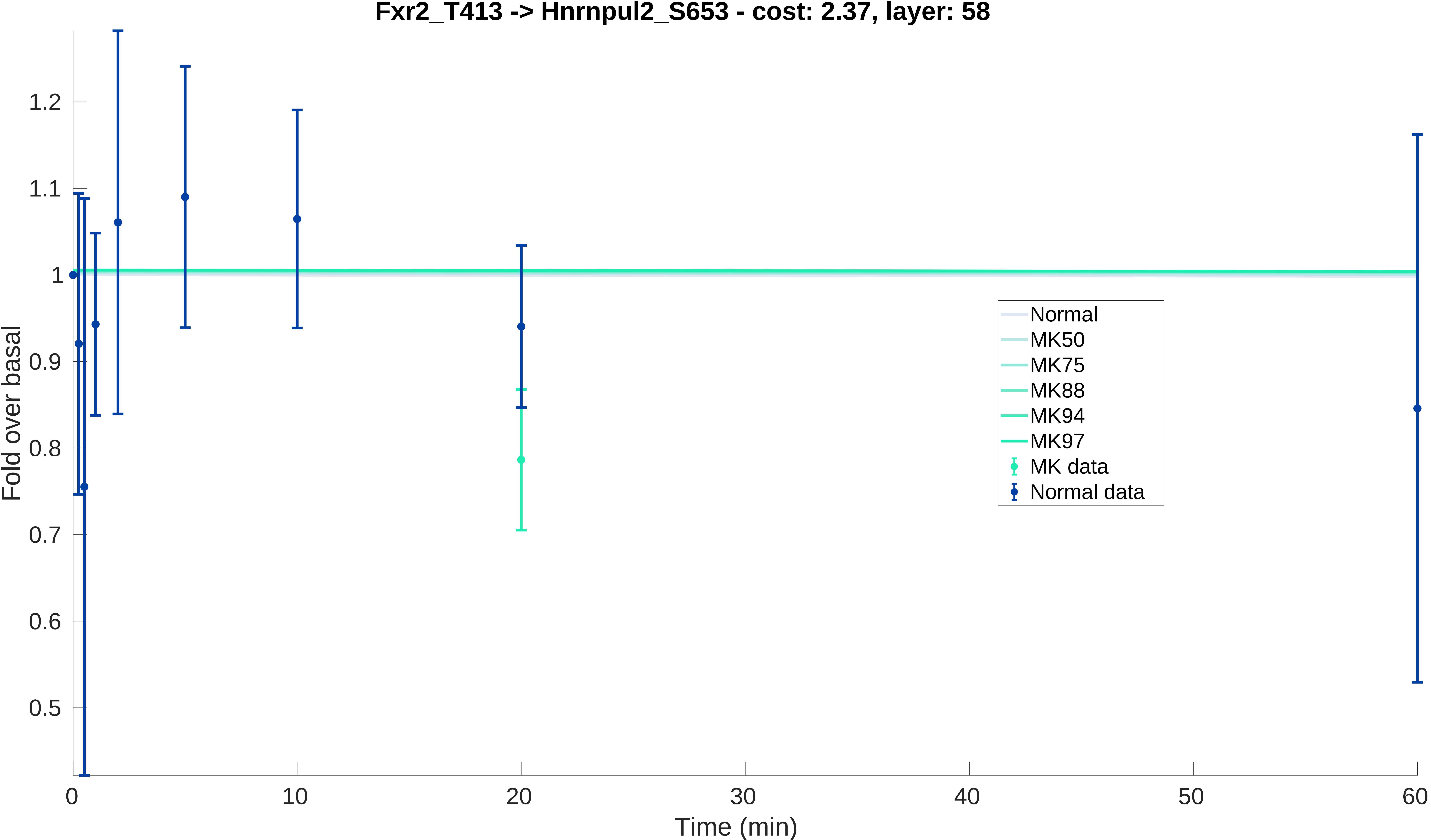

Fxr2_T413 -> Hnrnpul2_S653 - cost: 2.37, layer: 58
Fold over basal
1.2
1.1
1
0.9
0.8
0.7
0.6
0.5
0
10
20
30
40
50
60
Time (min)
Normal
MK50
MK75
MK88
MK94
MK97
MK data
Normal data

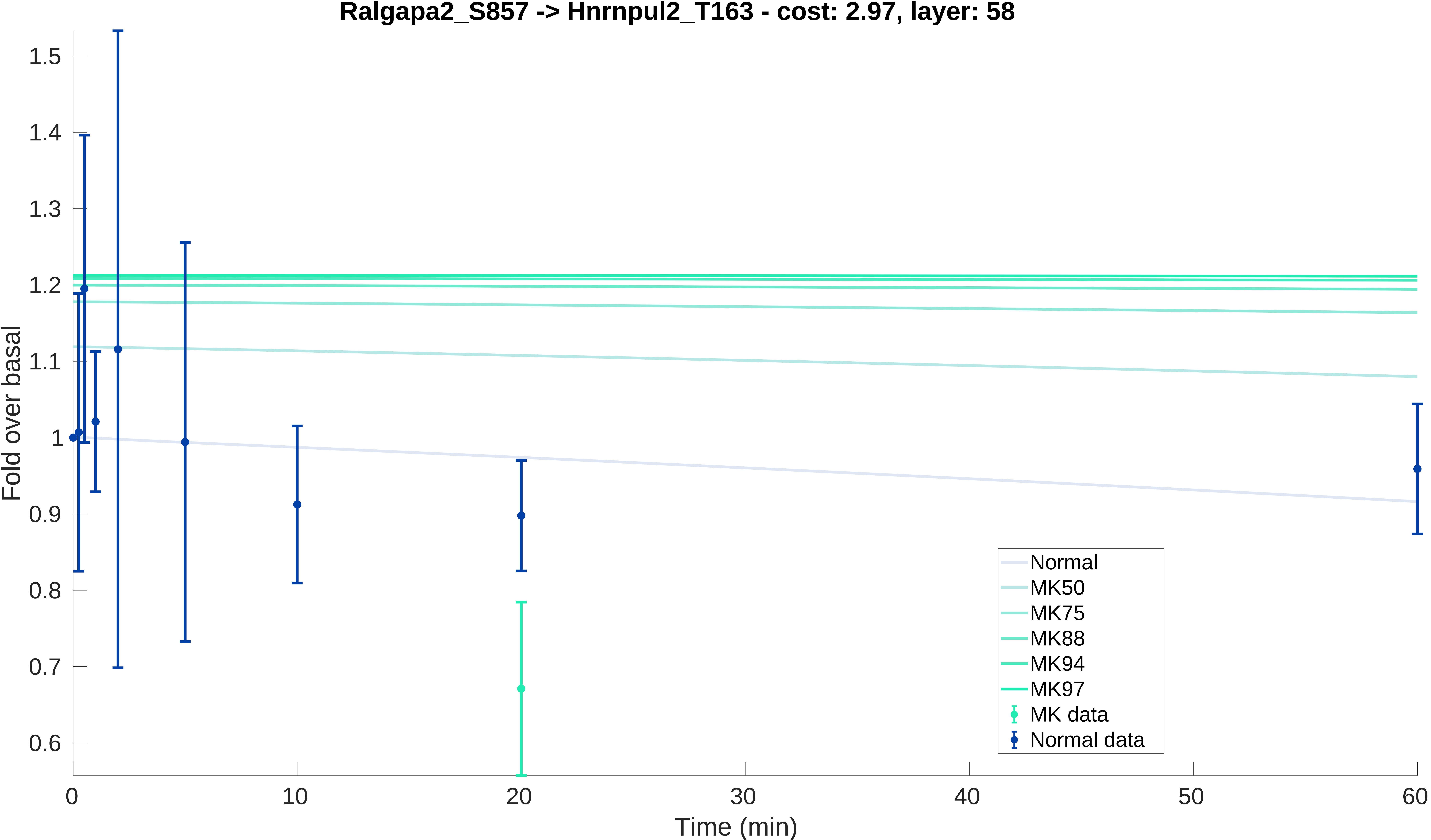

Ralgapa2_S857 -> Hnrnpul2_T163 - cost: 2.97, layer: 58
Fold over basal
Time (min)
1.5
1.4
1.3
1.2
1.1
1
0.9
0.8
0.7
0.6
0
10
20
30
40
50
60
Normal
MK50
MK75
MK88
MK94
MK97
MK data
Normal data

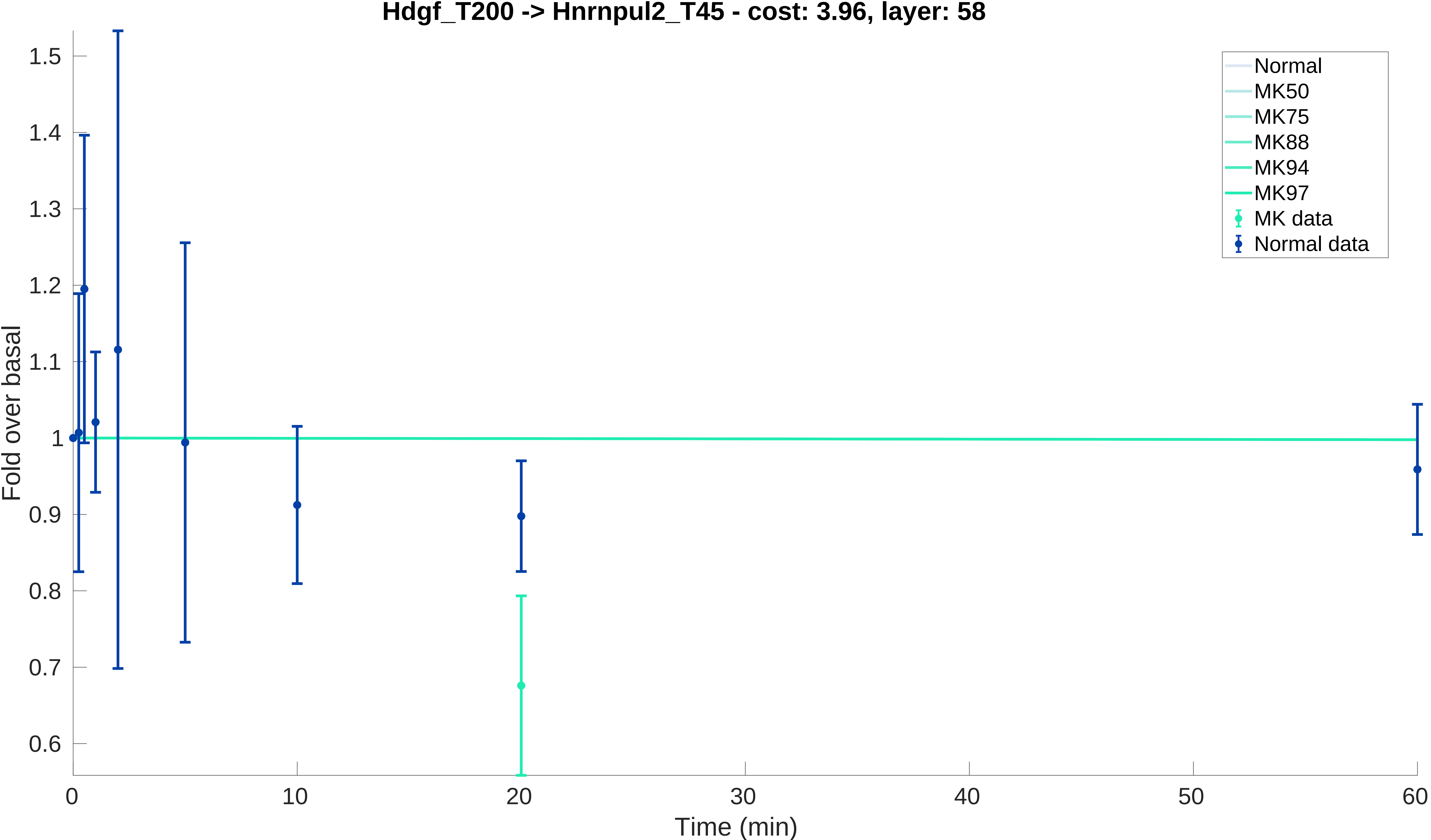

Hdgf_T200 -> Hnrnpul2_T45 - cost: 3.96, layer: 58
Fold over basal
1.5
1.4
1.3
1.2
1.1
1
0.9
0.8
0.7
0.6
0
10
20
30
40
50
60
Time (min)
Normal
MK50
MK75
MK88
MK94
MK97
MK data
Normal data

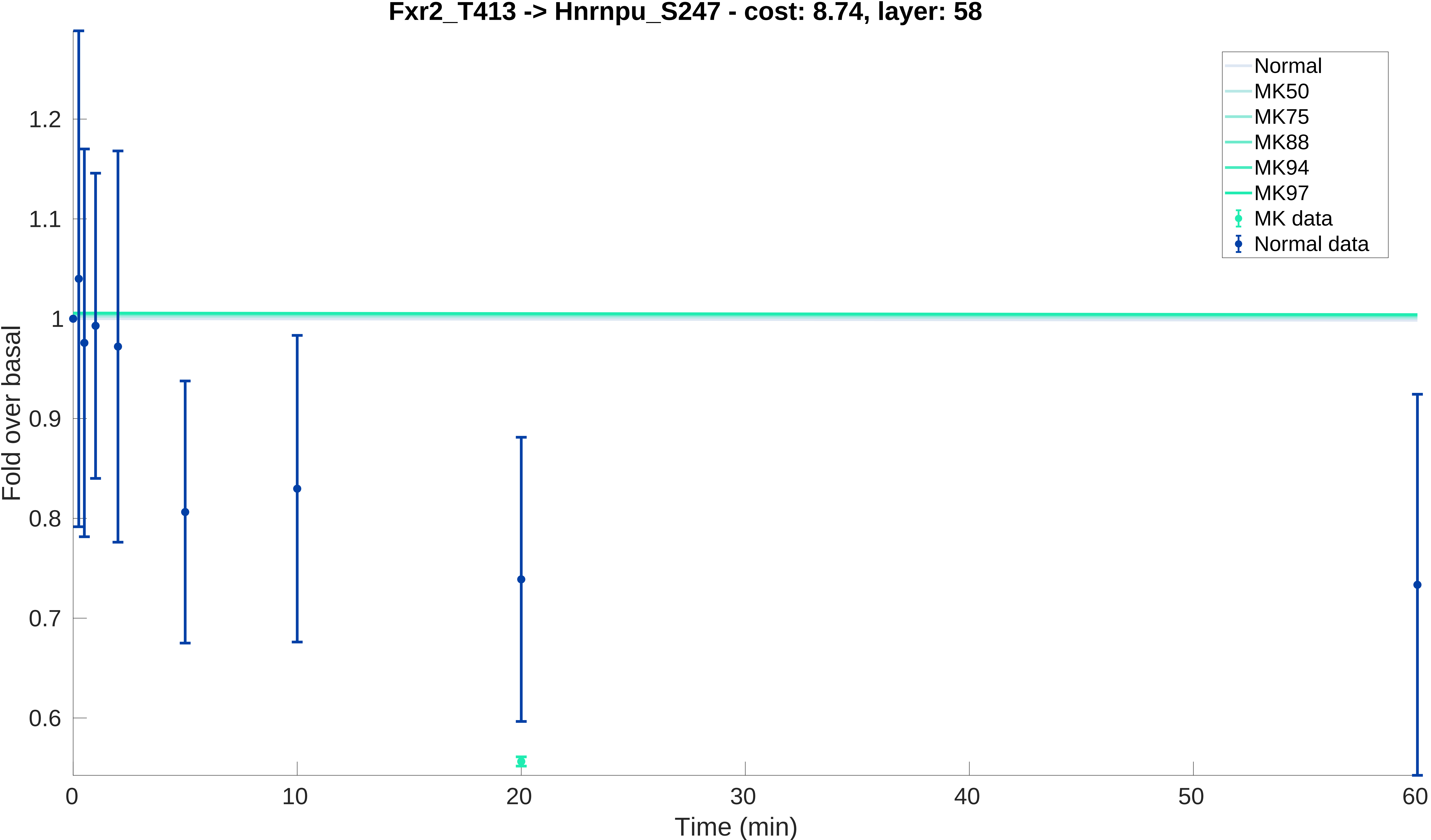

Fxr2_T413 -> Hnrnpu_S247 - cost: 8.74, layer: 58
Normal
MK50
MK75
MK88
MK94
MK97
MK data
Normal data
Fold over basal
1.2
1.1
1
0.9
0.8
0.7
0.6
0
10
20
30
40
50
60
Time (min)

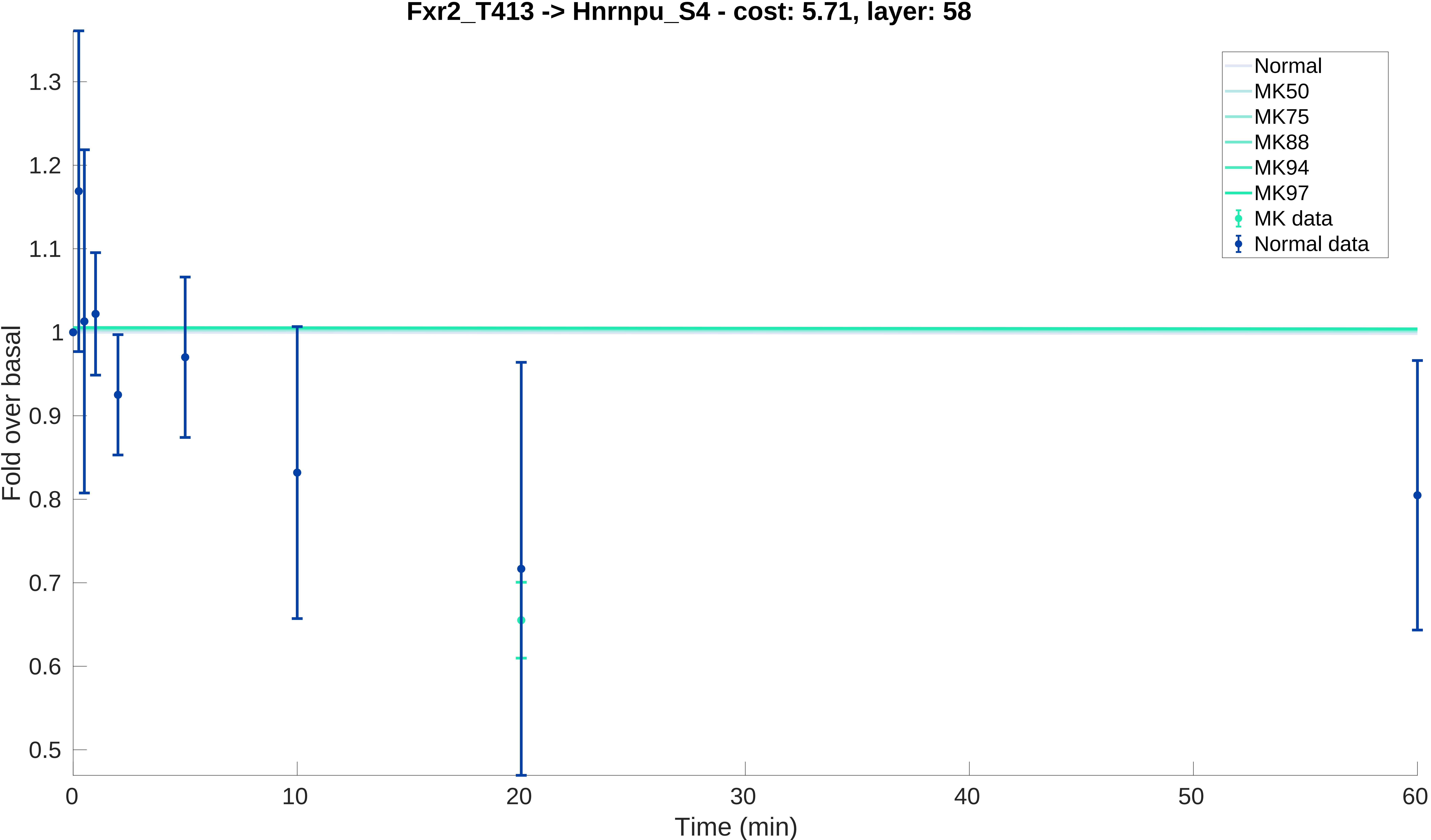

Fxr2_T413 -> Hnrnpu_S4 - cost: 5.71, layer: 58
Fold over basal
Time (min)
1.3
1.2
1.1
1
0.9
0.8
0.7
0.6
0.5
0
10
20
30
40
50
60
Normal
MK50
MK75
MK88
MK94
MK97
MK data
Normal data

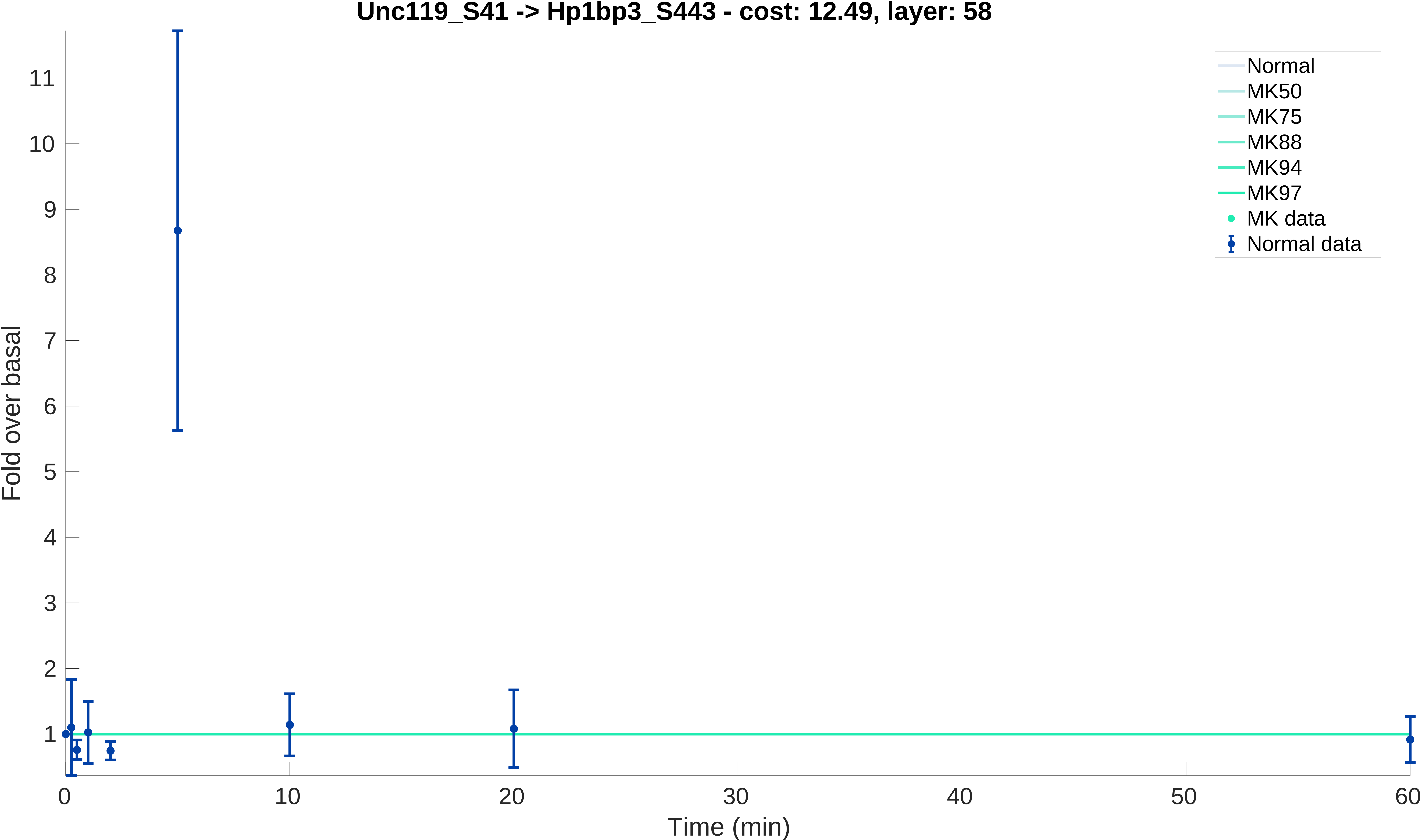
Unc119_S41 -> Hp1bp3_S443 - cost: 12.49, layer: 58
Fold over basal
Time (min)
0
10
20
30
40
50
60
1
2
3
4
5
6
7
8
9
10
11
Normal
MK50
MK75
MK88
MK94
MK97
MK data
Normal data

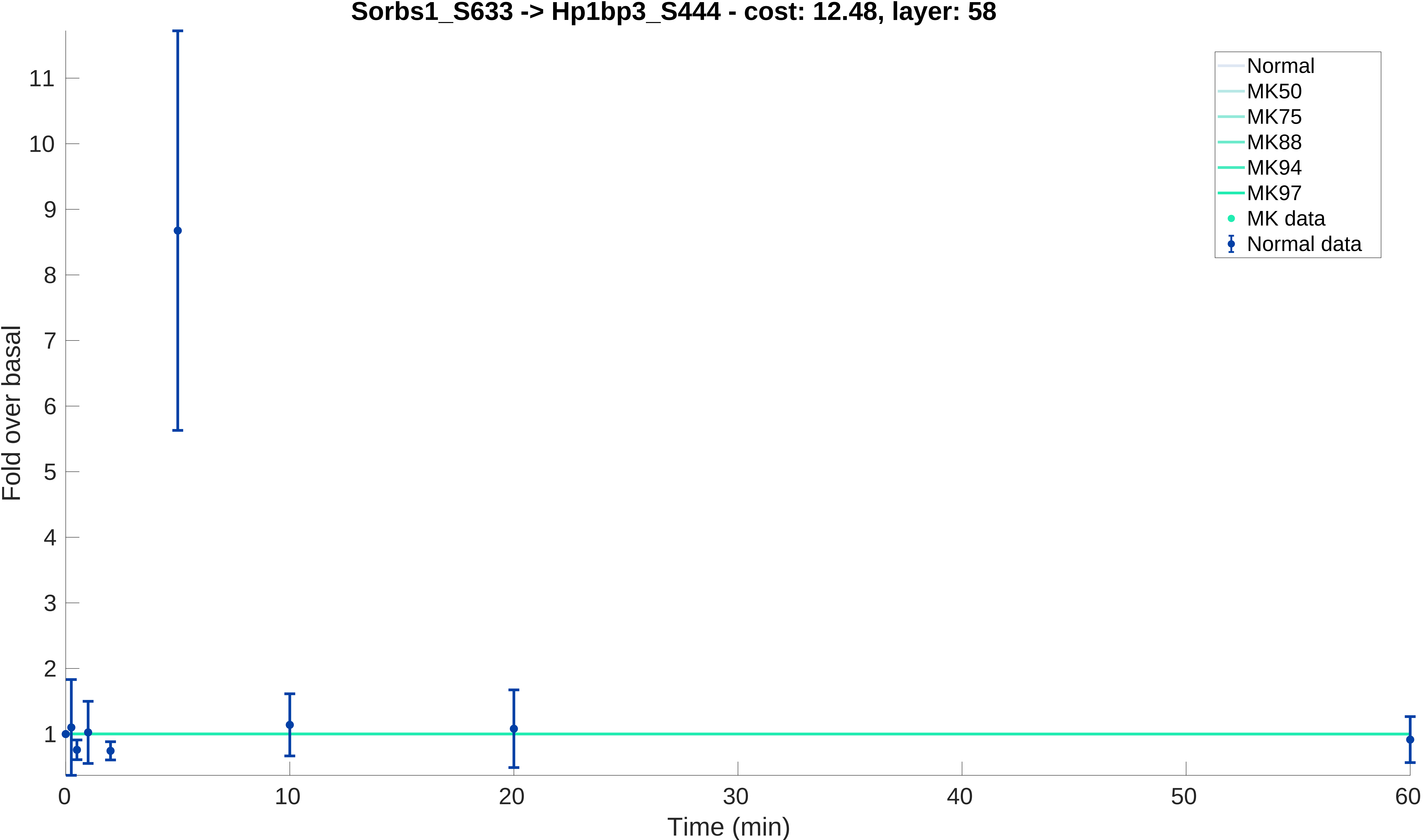

Sorbs1_S633 -> Hp1bp3_S444 - cost: 12.48, layer: 58
Fold over basal
Time (min)
0
10
20
30
40
50
60
1
2
3
4
5
6
7
8
9
10
11
Normal
MK50
MK75
MK88
MK94
MK97
MK data
Normal data

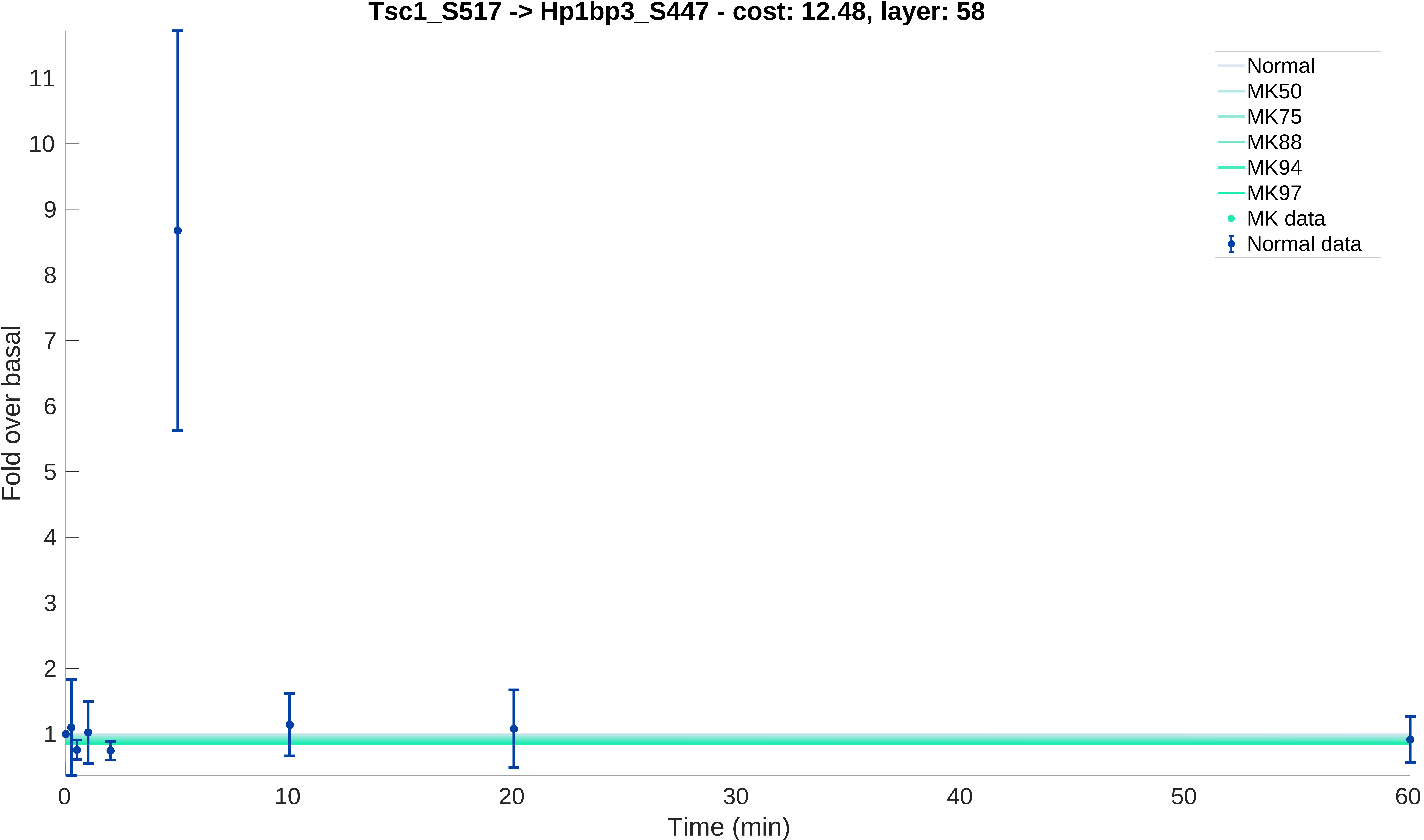
Tsc1_S517 -> Hp1bp3_S447 - cost: 12.48, layer: 58
Fold over basal
Time (min)
Normal
MK50
MK75
MK88
MK94
MK97
MK data
Normal data
0
10
20
30
40
50
60
1
2
3
4
5
6
7
8
9
10
11

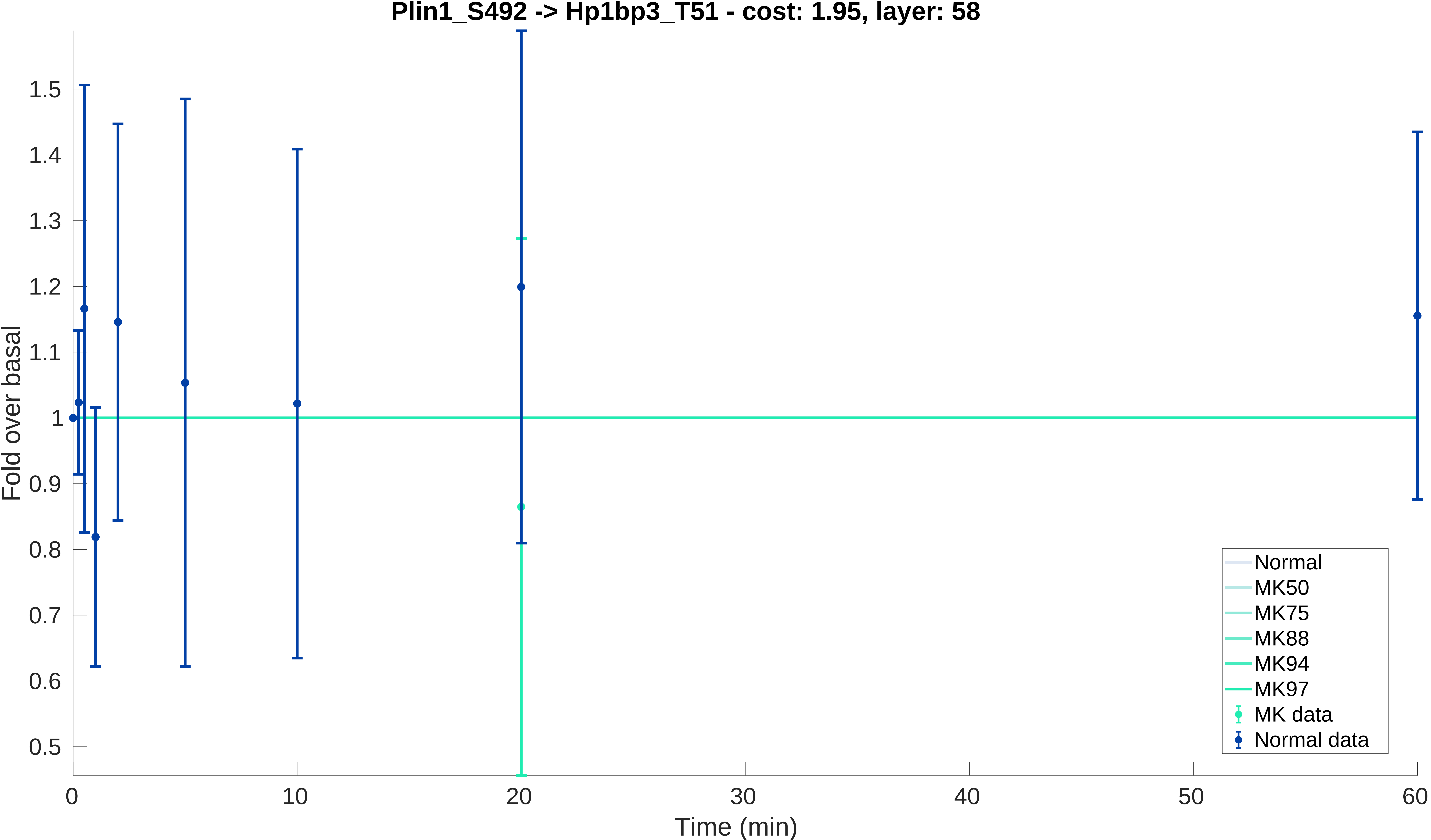

Plin1_S492 -> Hp1bp3_T51 - cost: 1.95, layer: 58
Fold over basal
Time (min)
0.5
0.6
0.7
0.8
0.9
1
1.1
1.2
1.3
1.4
1.5
0
10
20
30
40
50
60
Normal
MK50
MK75
MK88
MK94
MK97
MK data
Normal data

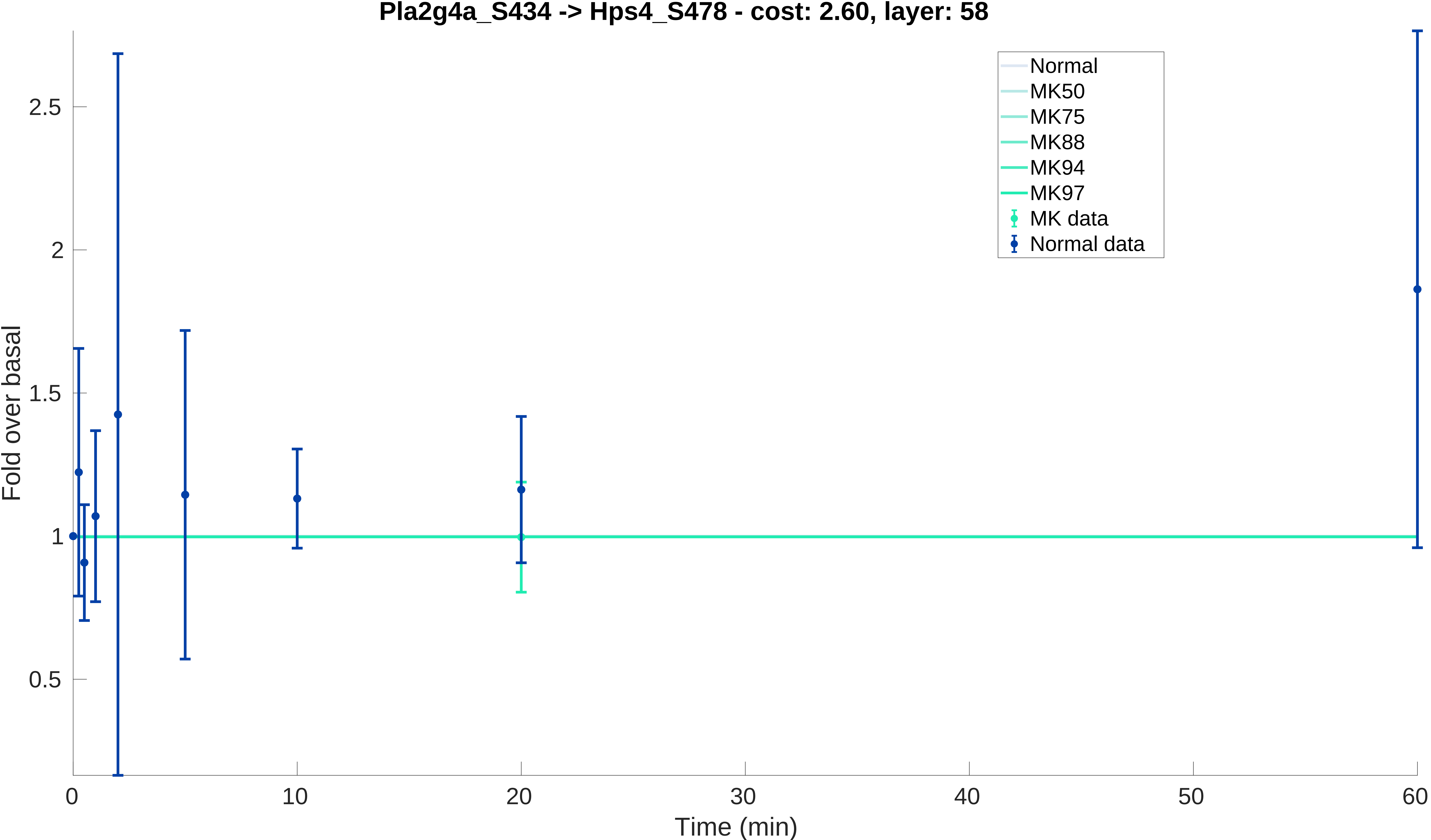

Pla2g4a_S434 -> Hps4_S478 - cost: 2.60, layer: 58
Normal
MK50
MK75
MK88
MK94
MK97
MK data
Normal data
Fold over basal
2.5
2
1.5
1
0.5
0
10
20
30
40
50
60
Time (min)

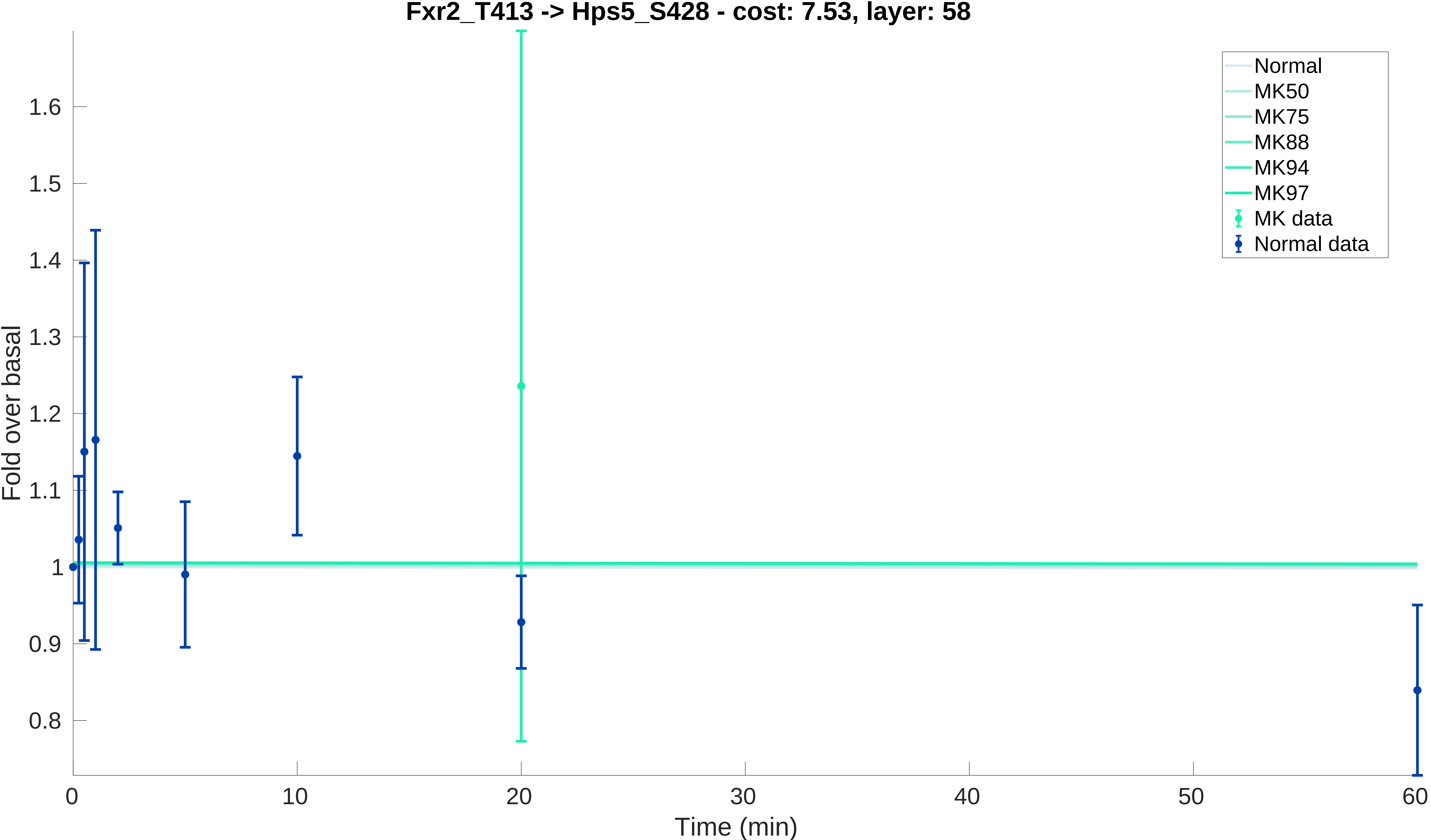

Fxr2_T413 -> Hps5_S428 - cost: 7.53, layer: 58
Fold over basal
Time (min)
Normal
MK50
MK75
MK88
MK94
MK97
MK data
Normal data
1.6
1.5
1.4
1.3
1.2
1.1
1
0.9
0.8
0
10
20
30
40
50
60

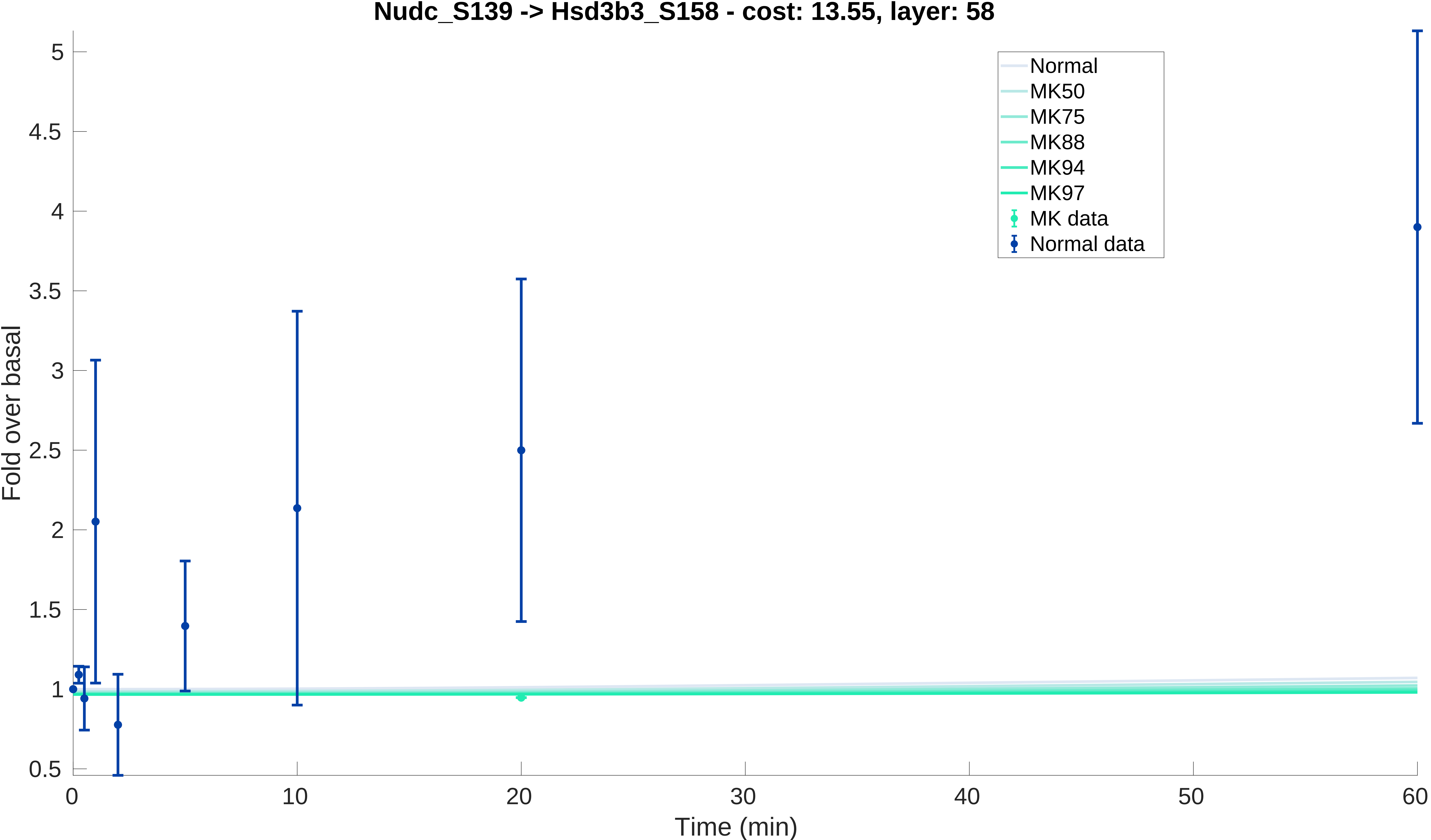

Nudc_S139 -> Hsd3b3_S158 - cost: 13.55, layer: 58
Normal
MK50
MK75
MK88
MK94
MK97
MK data
Normal data
Fold over basal
Time (min)
0.5
1
1.5
2
2.5
3
3.5
4
4.5
5
0
10
20
30
40
50
60

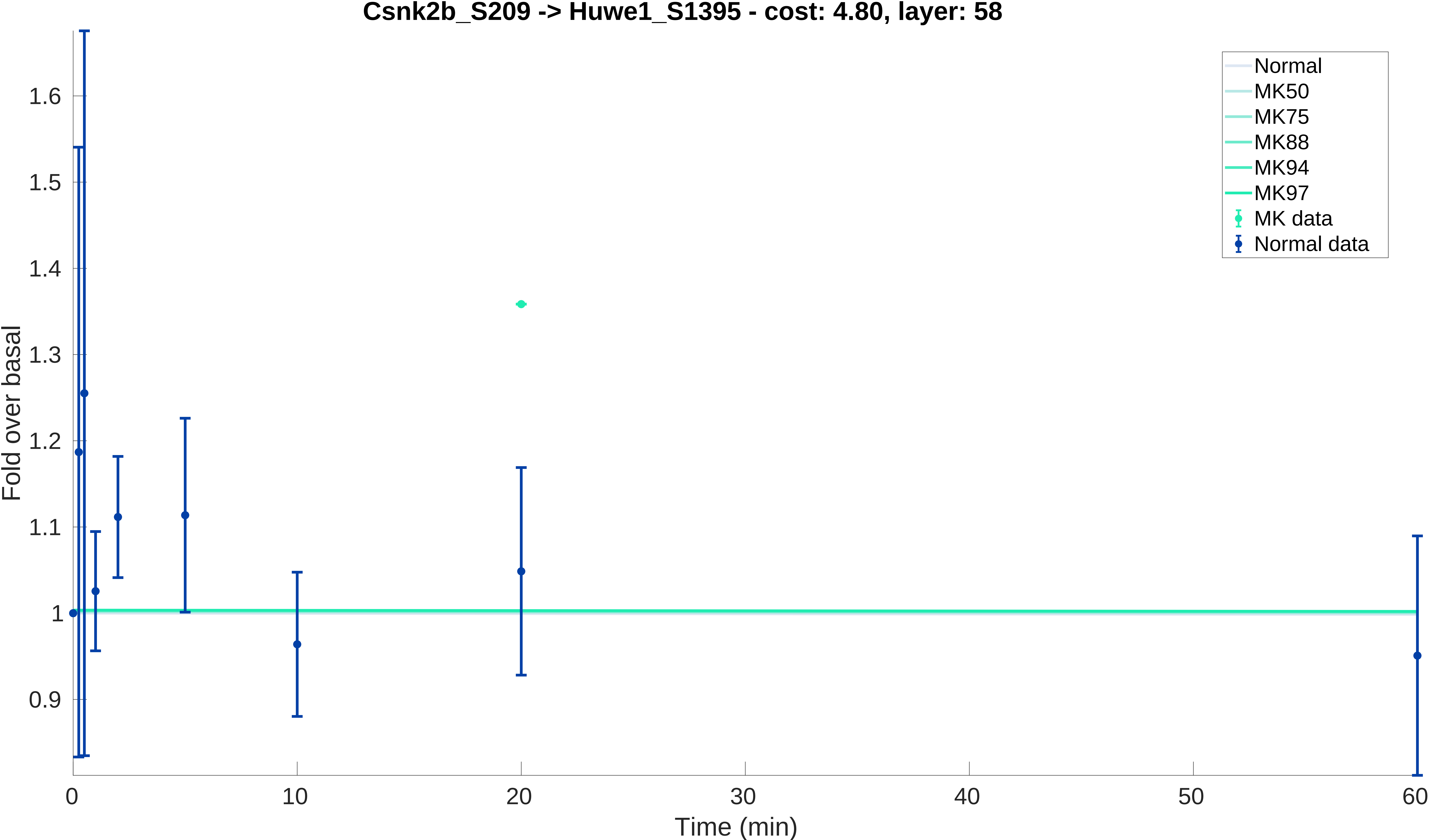

Csnk2b_S209 -> Huwe1_S1395 - cost: 4.80, layer: 58
Fold over basal
Time (min)
Normal
MK50
MK75
MK88
MK94
MK97
MK data
Normal data
1.6
1.5
1.4
1.3
1.2
1.1
1
0.9
0
10
20
30
40
50
60

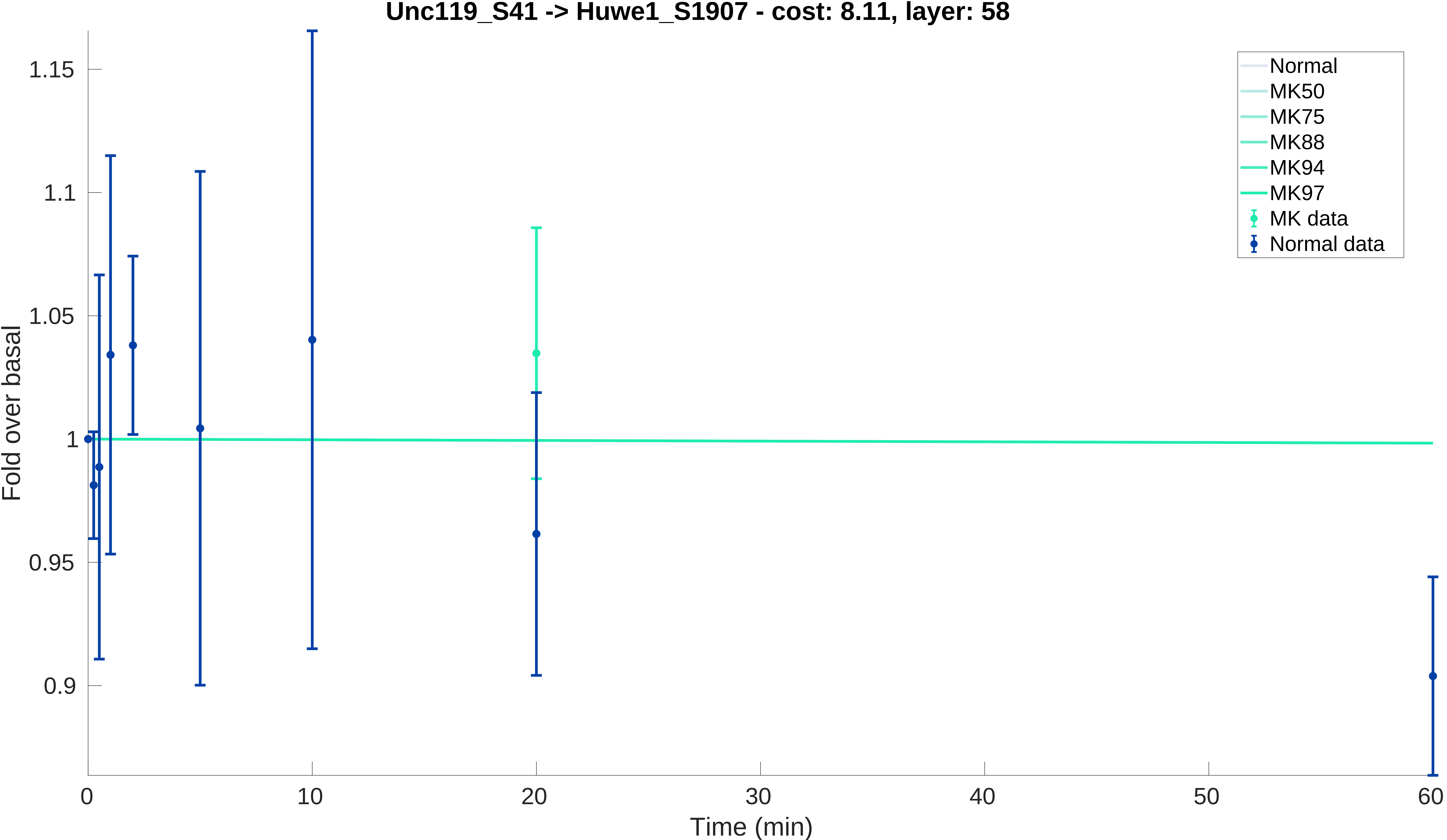

Unc119_S41 -> Huwe1_S1907 - cost: 8.11, layer: 58
Fold over basal
1.15
1.1
1.05
1
0.95
0.9
0
10
20
30
40
50
60
Time (min)
Normal
MK50
MK75
MK88
MK94
MK97
MK data
Normal data

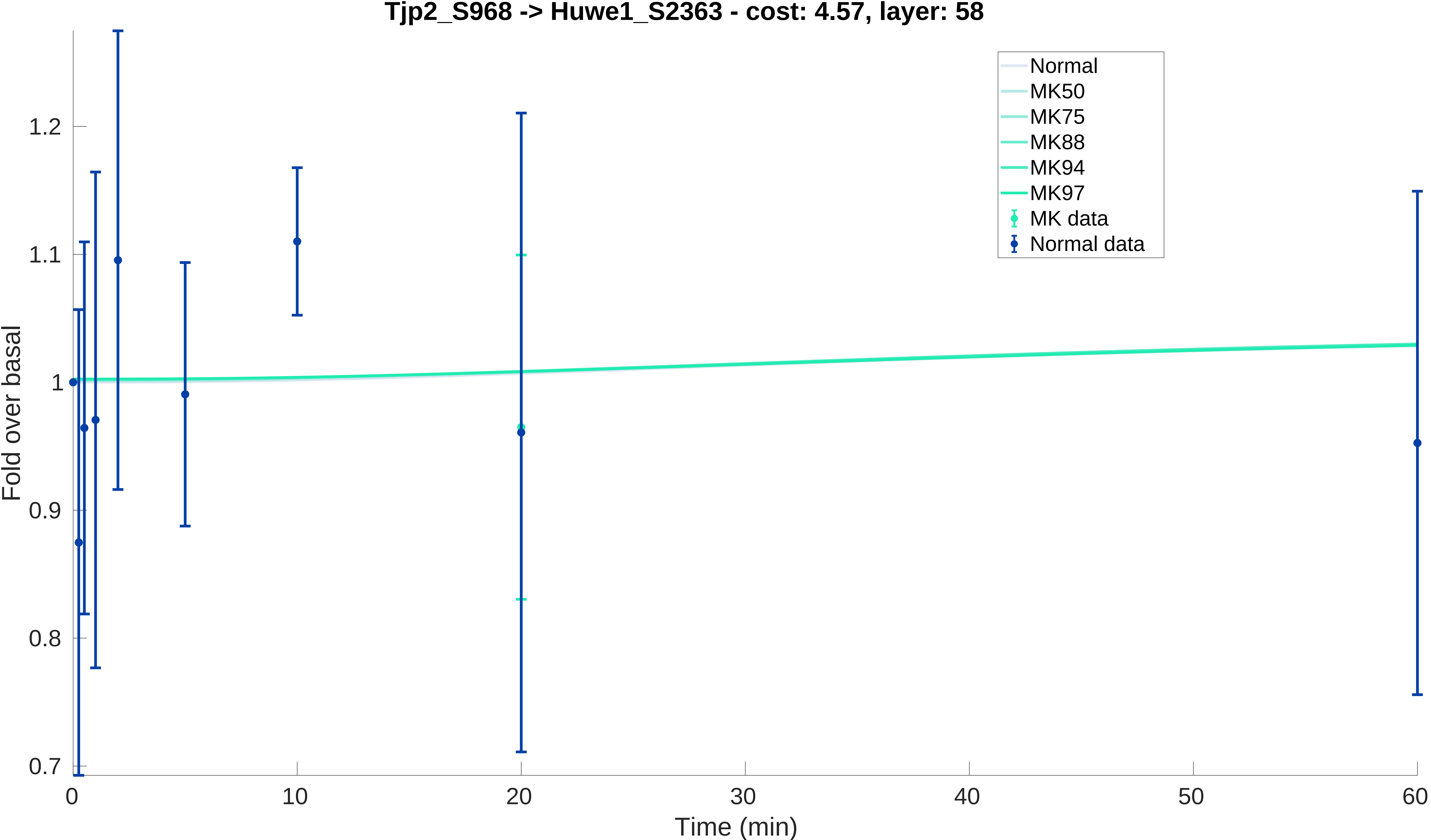
Tjp2_S968 -> Huwe1_S2363 - cost: 4.57, layer: 58
Fold over basal
Time (min)
0
10
20
30
40
50
60
0.7
0.8
0.9
1
1.1
1.2
Normal
MK50
MK75
MK88
MK94
MK97
MK data
Normal data

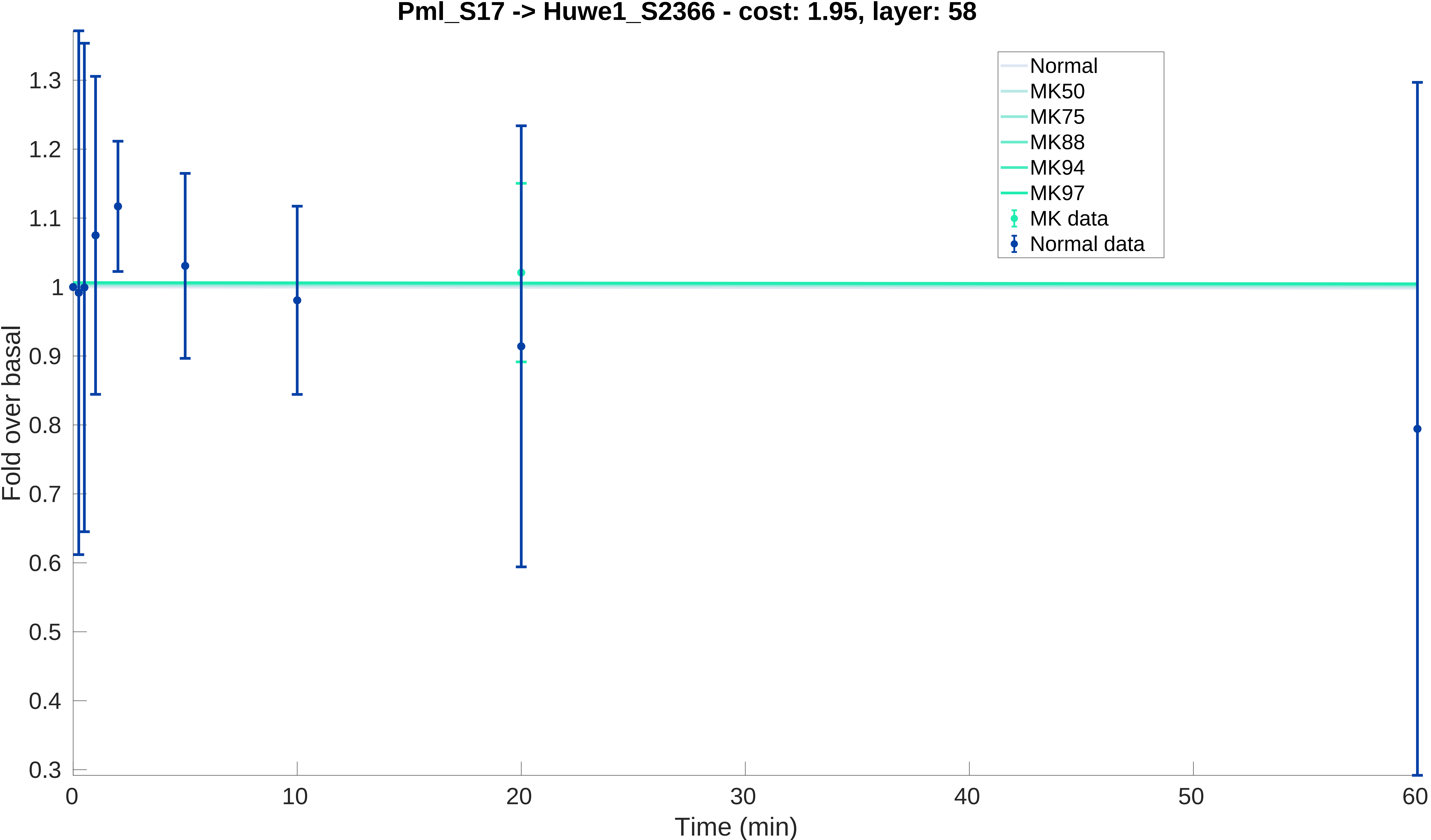
Pml_S17 -> Huwe1_S2366 - cost: 1.95, layer: 58
Normal
MK50
MK75
MK88
MK94
MK97
MK data
Normal data
Fold over basal
Time (min)
0.3
0.4
0.5
0.6
0.7
0.8
0.9
1
1.1
1.2
1.3
0
10
20
30
40
50
60

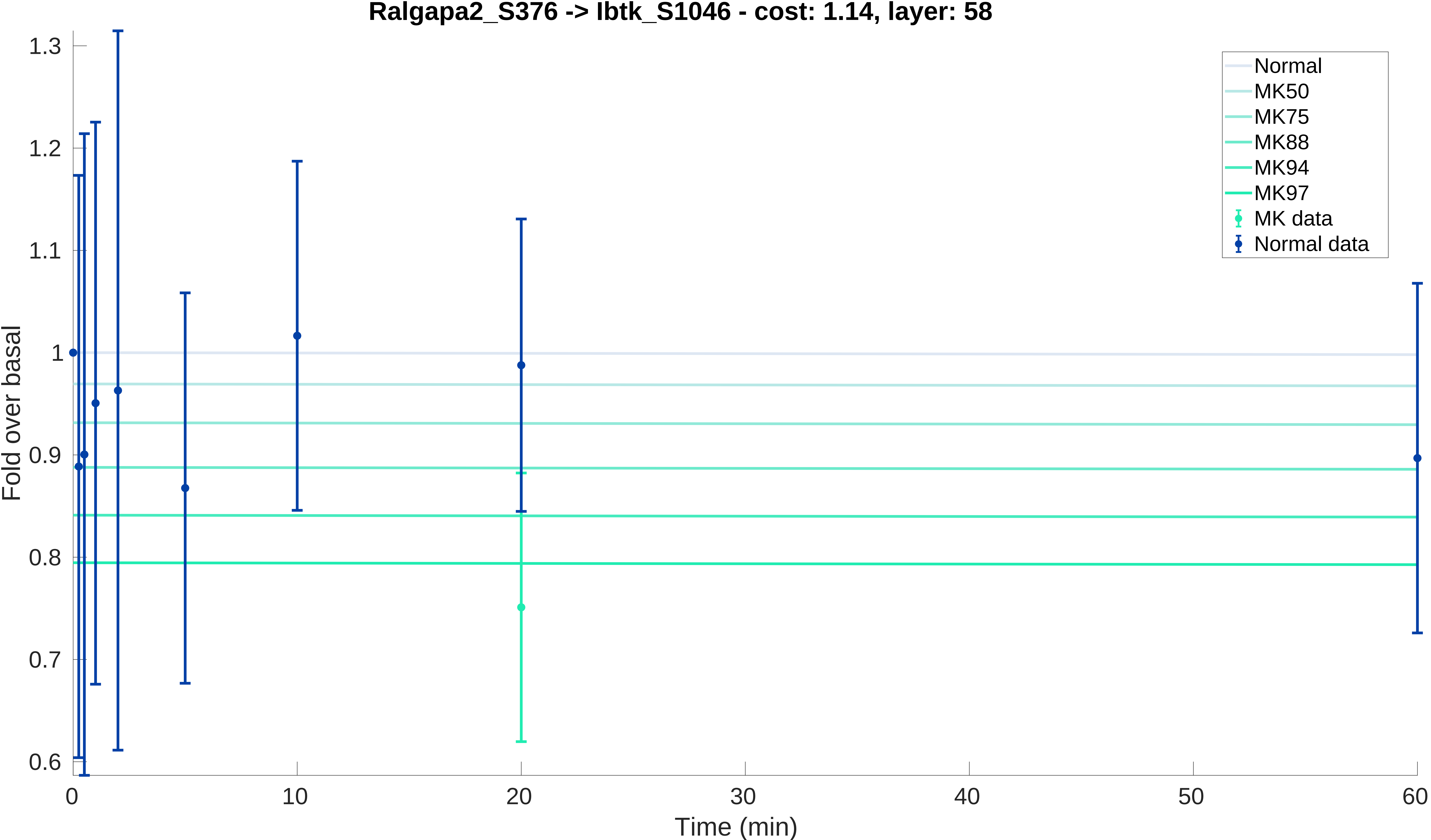
Ralgapa2_S376 -> Ibtk_S1046 - cost: 1.14, layer: 58
Fold over basal
Time (min)
0
10
20
30
40
50
60
0.6
0.7
0.8
0.9
1
1.1
1.2
1.3
Normal
MK50
MK75
MK88
MK94
MK97
MK data
Normal data

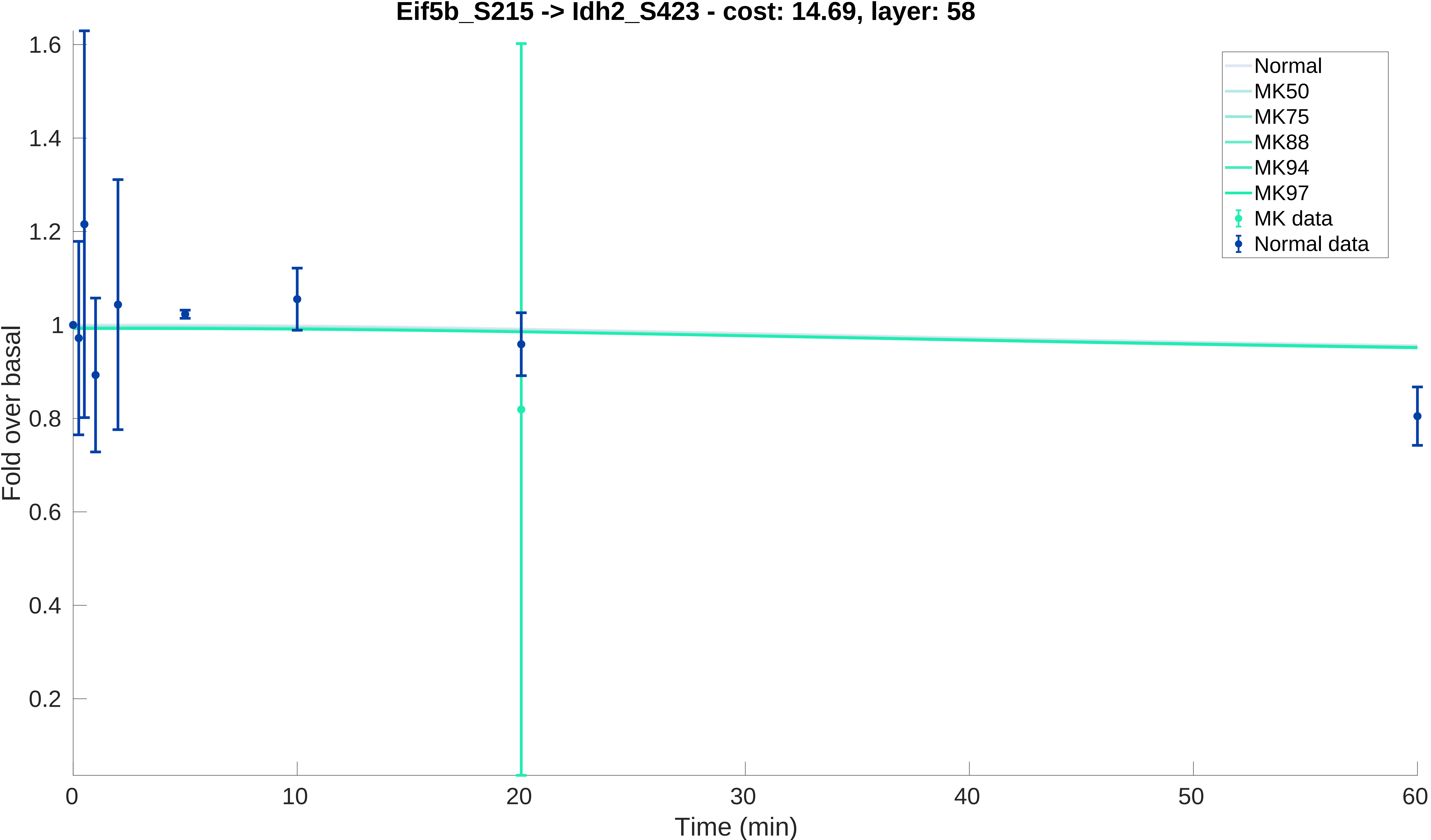

Eif5b_S215 -> Idh2_S423 - cost: 14.69, layer: 58
Fold over basal
Time (min)
1.6
1.4
1.2
1
0.8
0.6
0.4
0.2
0
10
20
30
40
50
60
Normal
MK50
MK75
MK88
MK94
MK97
MK data
Normal data

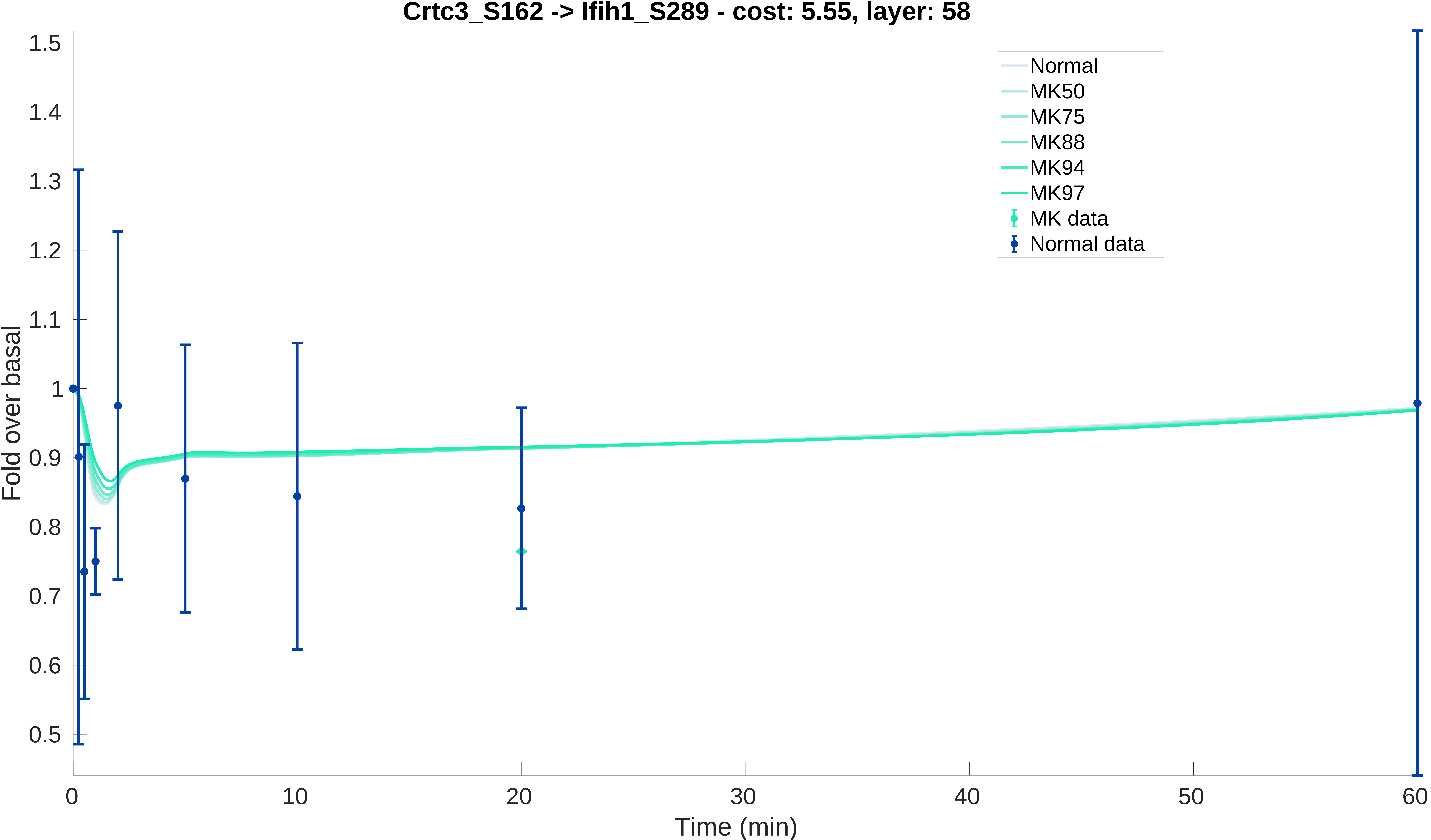

Crtc3_S162 -> Ifih1_S289 - cost: 5.55, layer: 58
Fold over basal
Time (min)
Normal
MK50
MK75
MK88
MK94
MK97
MK data
Normal data
1.5
1.4
1.3
1.2
1.1
1
0.9
0.8
0.7
0.6
0.5
0
10
20
30
40
50
60

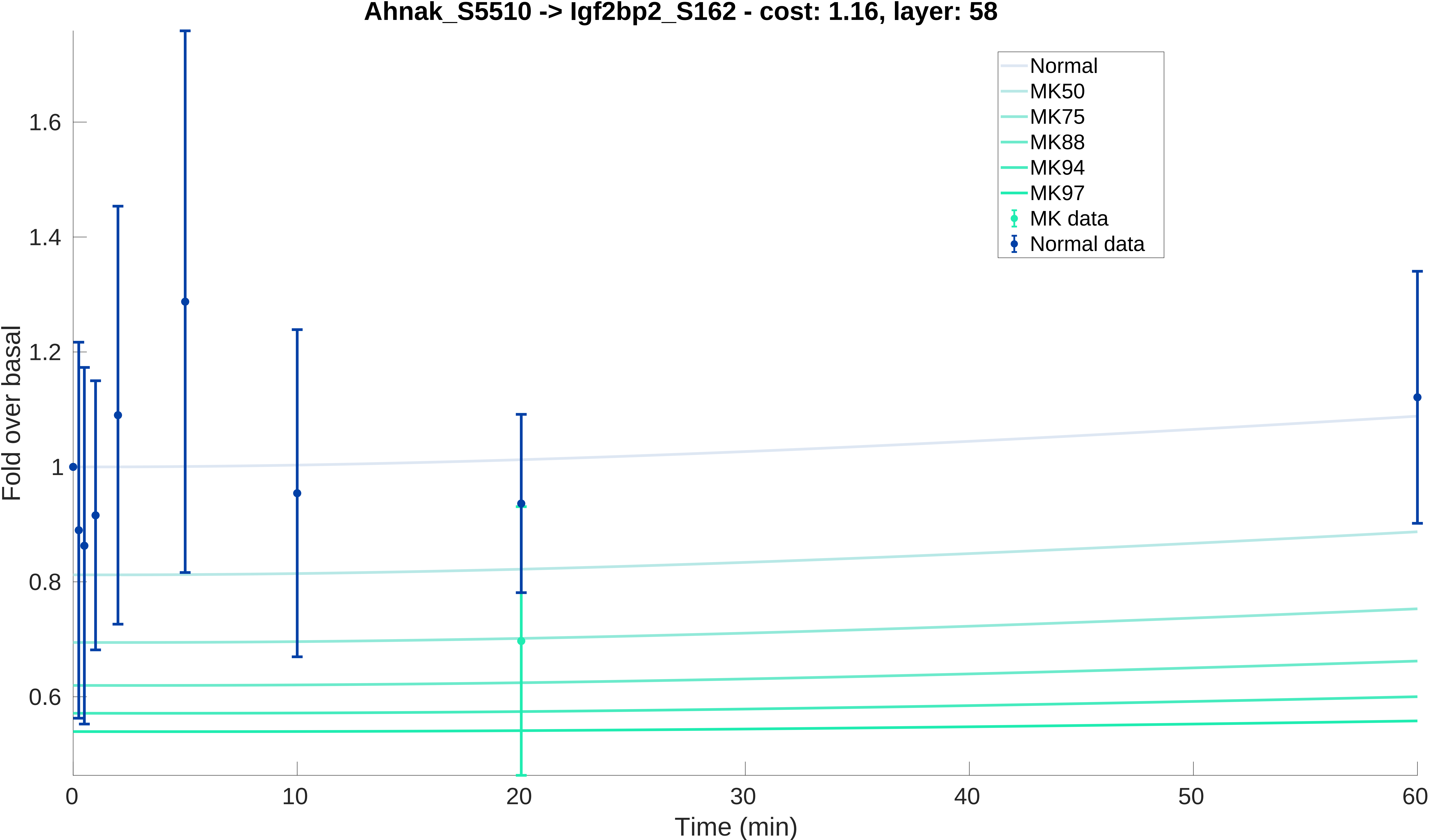

Ahnak_S5510 -> Igf2bp2_S162 - cost: 1.16, layer: 58
Fold over basal
1.6
1.4
1.2
1
0.8
0.6
0
10
20
30
40
50
60
Time (min)
Normal
MK50
MK75
MK88
MK94
MK97
MK data
Normal data

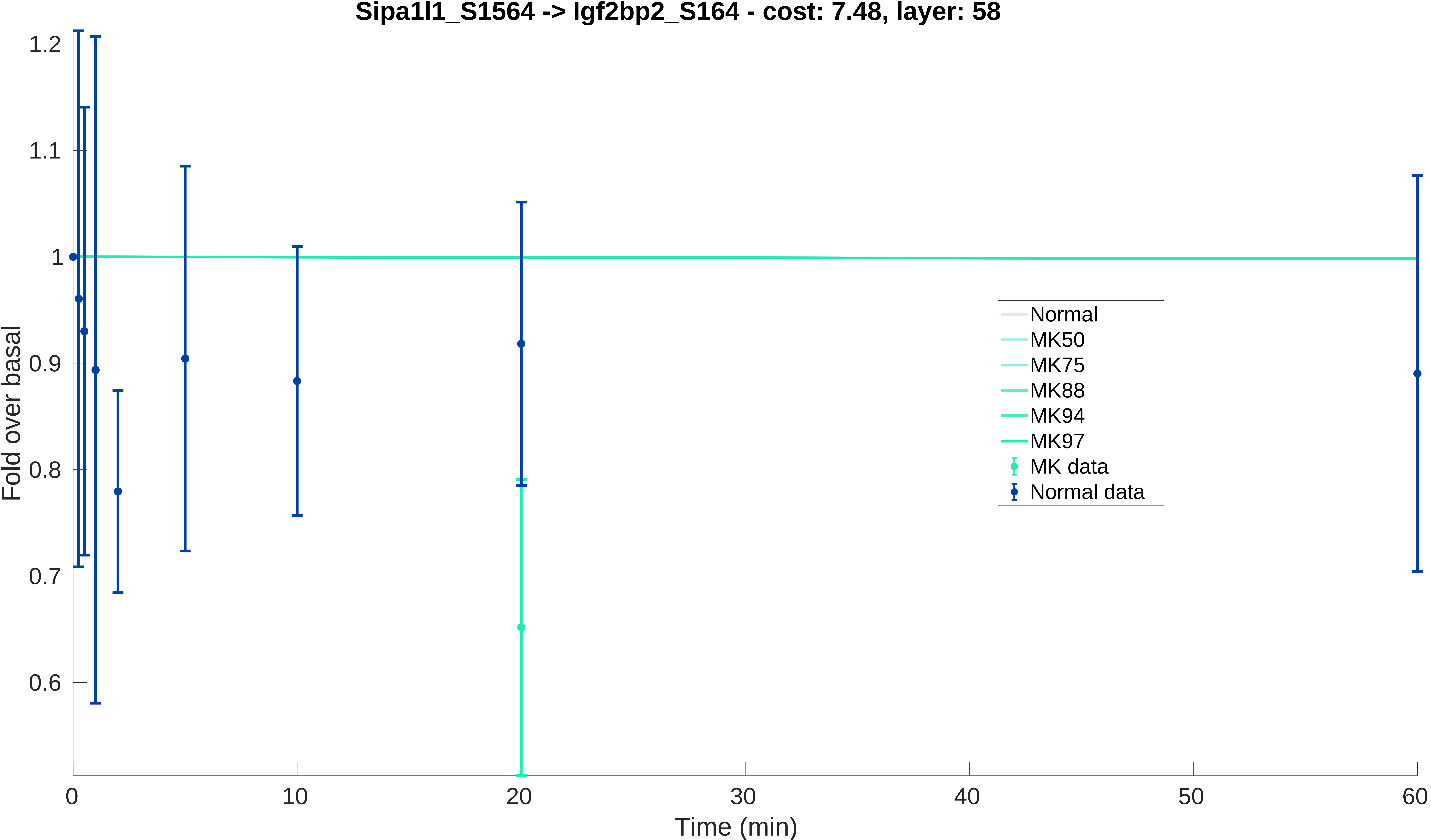

Sipa1l1_S1564 -> Igf2bp2_S164 - cost: 7.48, layer: 58
Fold over basal
Time (min)
1.2
1.1
1
0.9
0.8
0.7
0.6
0
10
20
30
40
50
60
Normal
MK50
MK75
MK88
MK94
MK97
MK data
Normal data

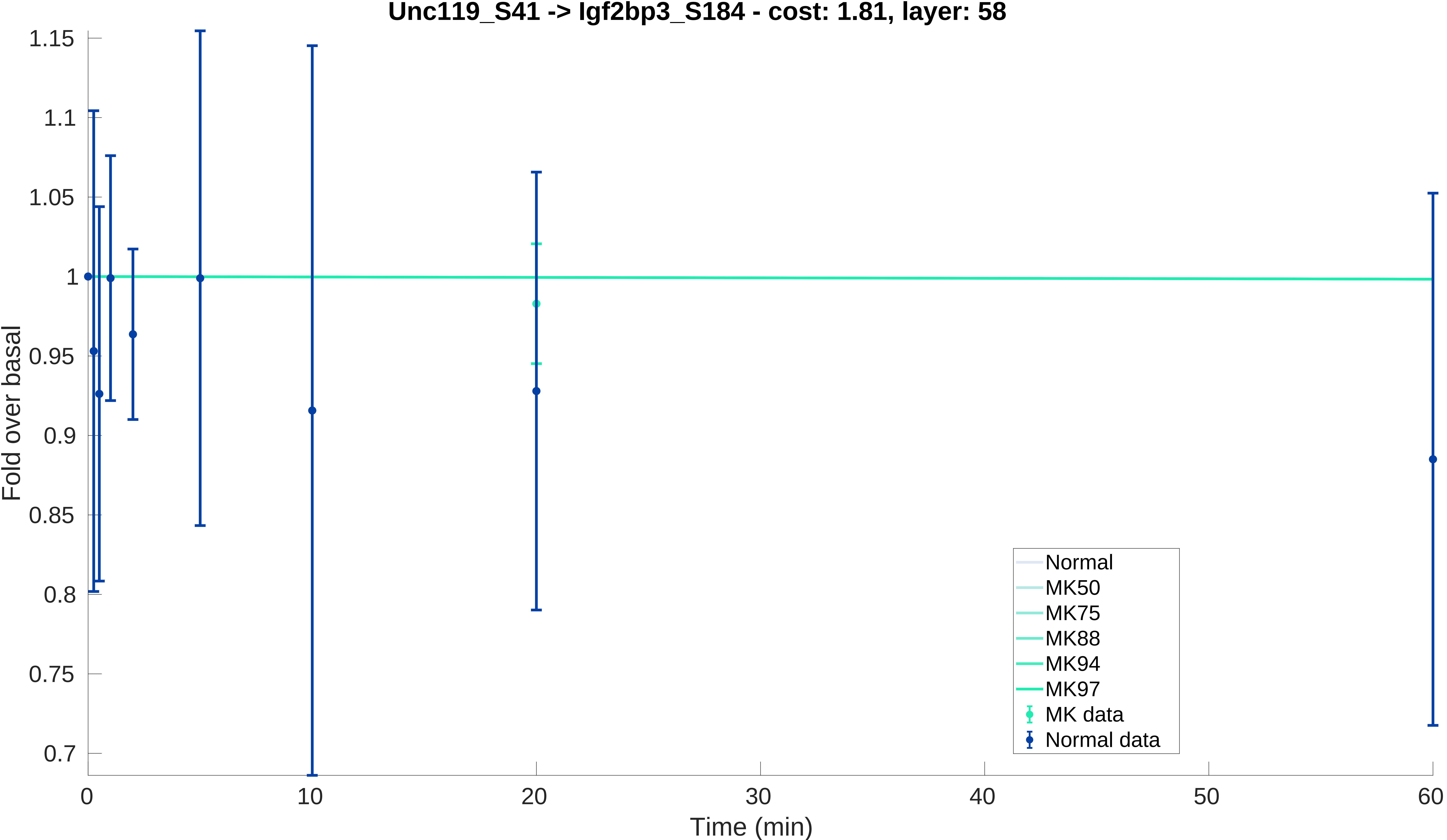

Unc119_S41 -> Igf2bp3_S184 - cost: 1.81, layer: 58
Fold over basal
Time (min)
1.15
1.1
1.05
1
0.95
0.9
0.85
0.8
0.75
0.7
0
10
20
30
40
50
60
Normal
MK50
MK75
MK88
MK94
MK97
MK data
Normal data

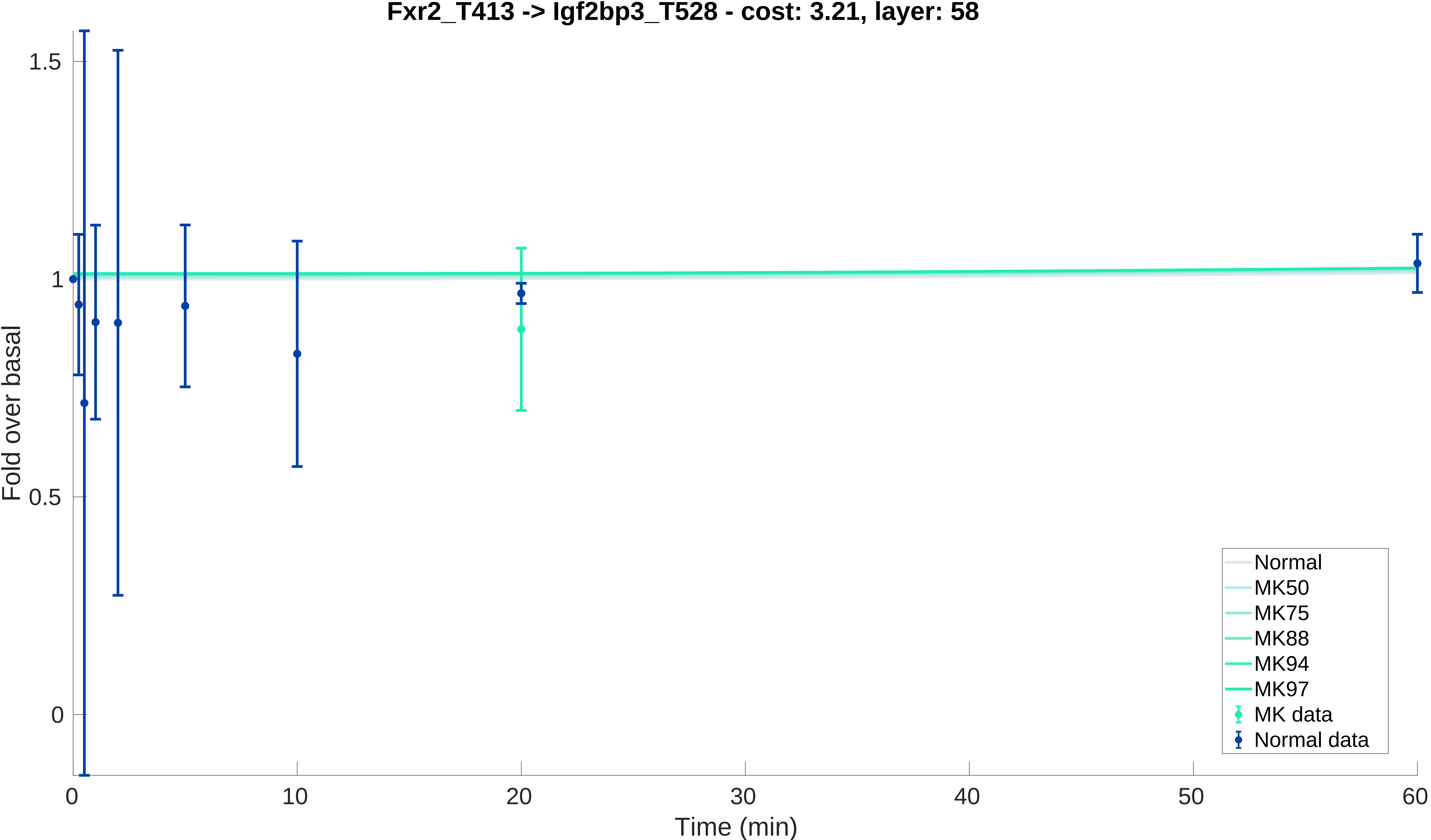

Fxr2_T413 -> Igf2bp3_T528 - cost: 3.21, layer: 58
Fold over basal
1.5
1
0.5
0
0
10
20
30
40
50
60
Time (min)
Normal
MK50
MK75
MK88
MK94
MK97
MK data
Normal data

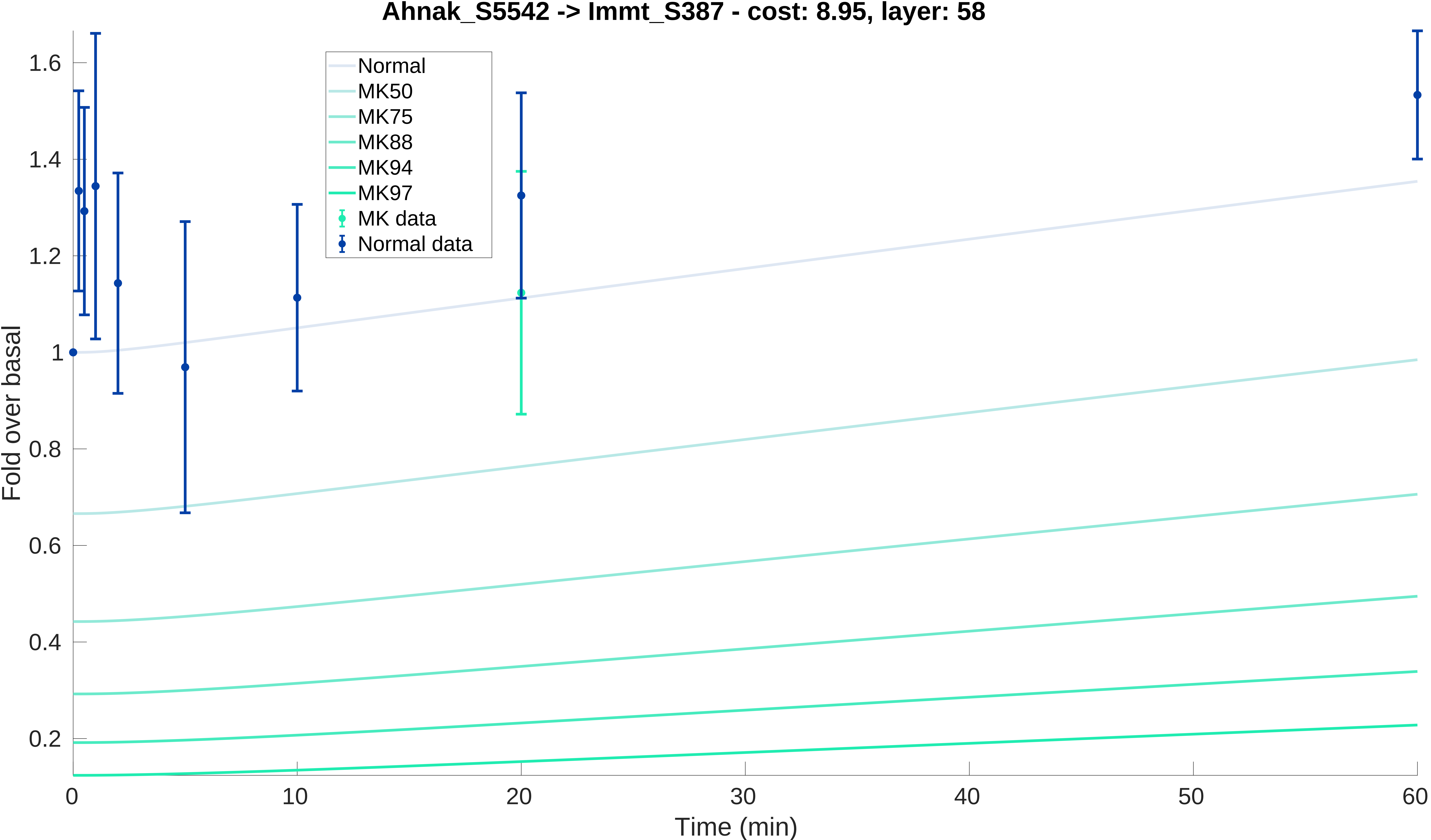
Ahnak_S5542 -> Immt_S387 - cost: 8.95, layer: 58
Normal
MK50
MK75
MK88
MK94
MK97
MK data
Normal data
Fold over basal
1.6
1.4
1.2
1
0.8
0.6
0.4
0.2
0
10
20
30
40
50
60
Time (min)

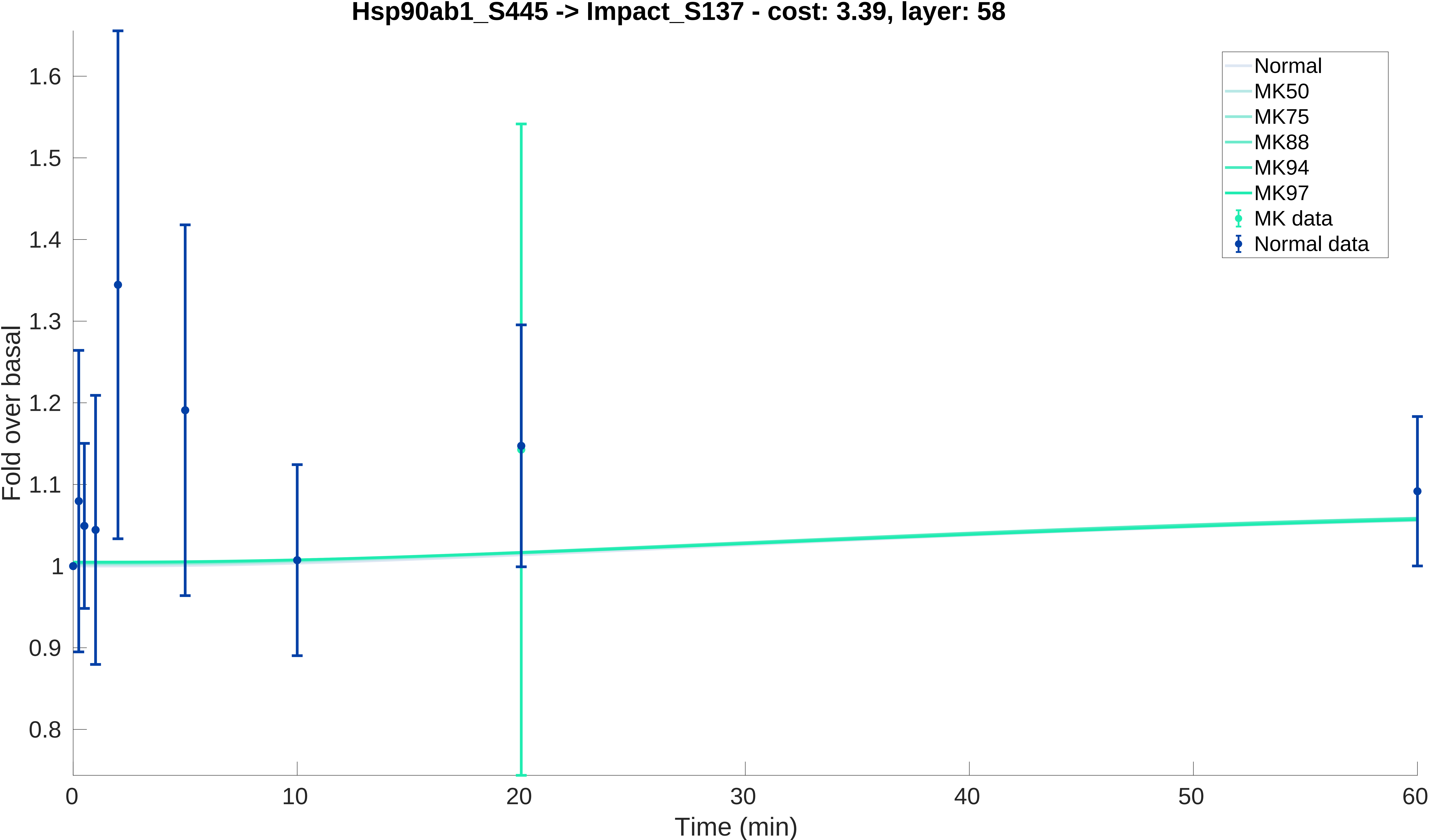

Hsp90ab1_S445 -> Impact_S137 - cost: 3.39, layer: 58
Fold over basal
Time (min)
0.8
0.9
1
1.1
1.2
1.3
1.4
1.5
1.6
0
10
20
30
40
50
60
Normal
MK50
MK75
MK88
MK94
MK97
MK data
Normal data

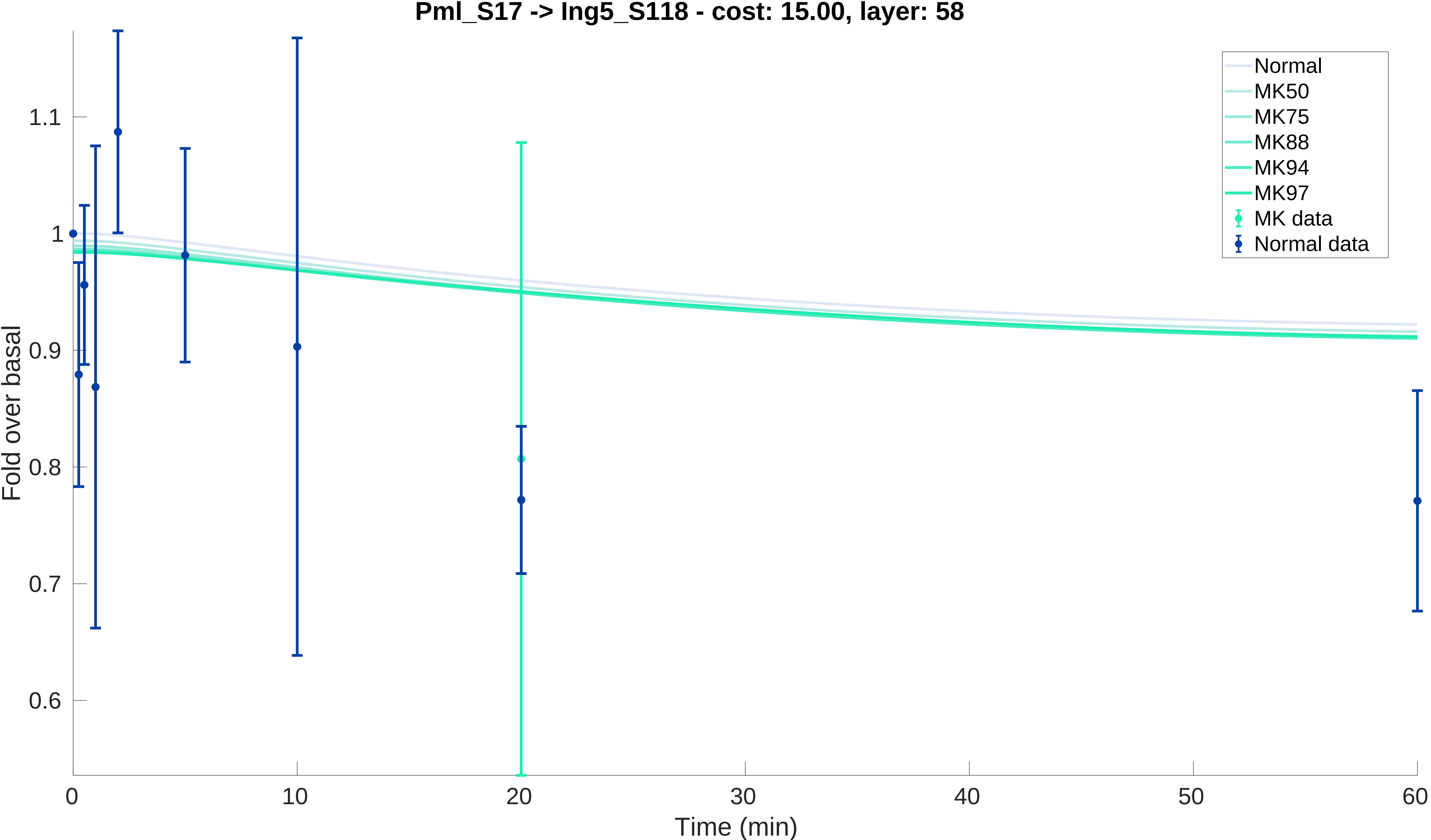

Pml_S17 -> Ing5_S118 - cost: 15.00, layer: 58
Fold over basal
Time (min)
0.6
0.7
0.8
0.9
1
1.1
0
10
20
30
40
50
60
Normal
MK50
MK75
MK88
MK94
MK97
MK data
Normal data

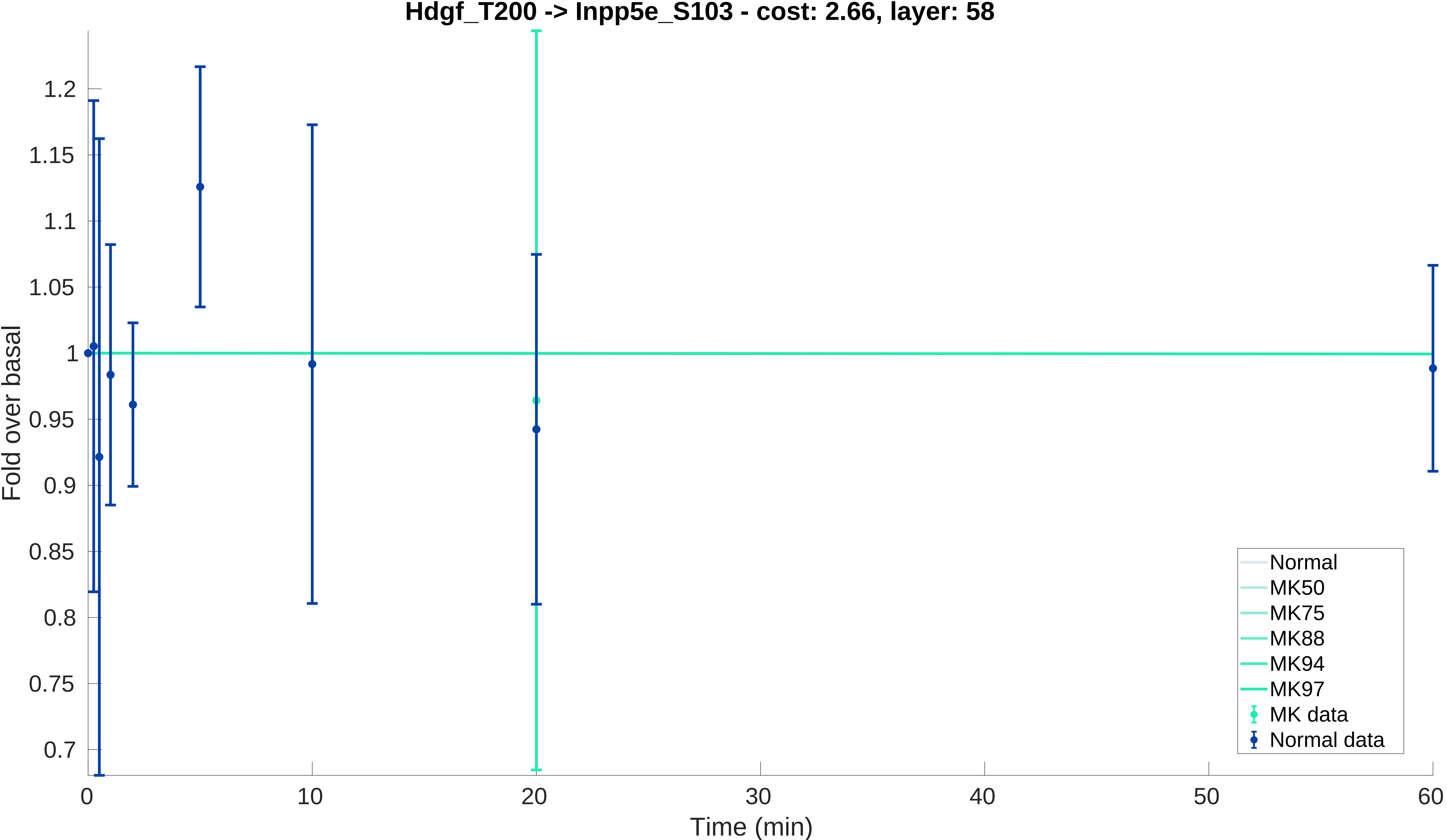

Hdgf_T200 -> Inpp5e_S103 - cost: 2.66, layer: 58
Fold over basal
1.2
1.15
1.1
1.05
1
0.95
0.9
0.85
0.8
0.75
0.7
0
10
20
30
40
50
60
Time (min)
Normal
MK50
MK75
MK88
MK94
MK97
MK data
Normal data

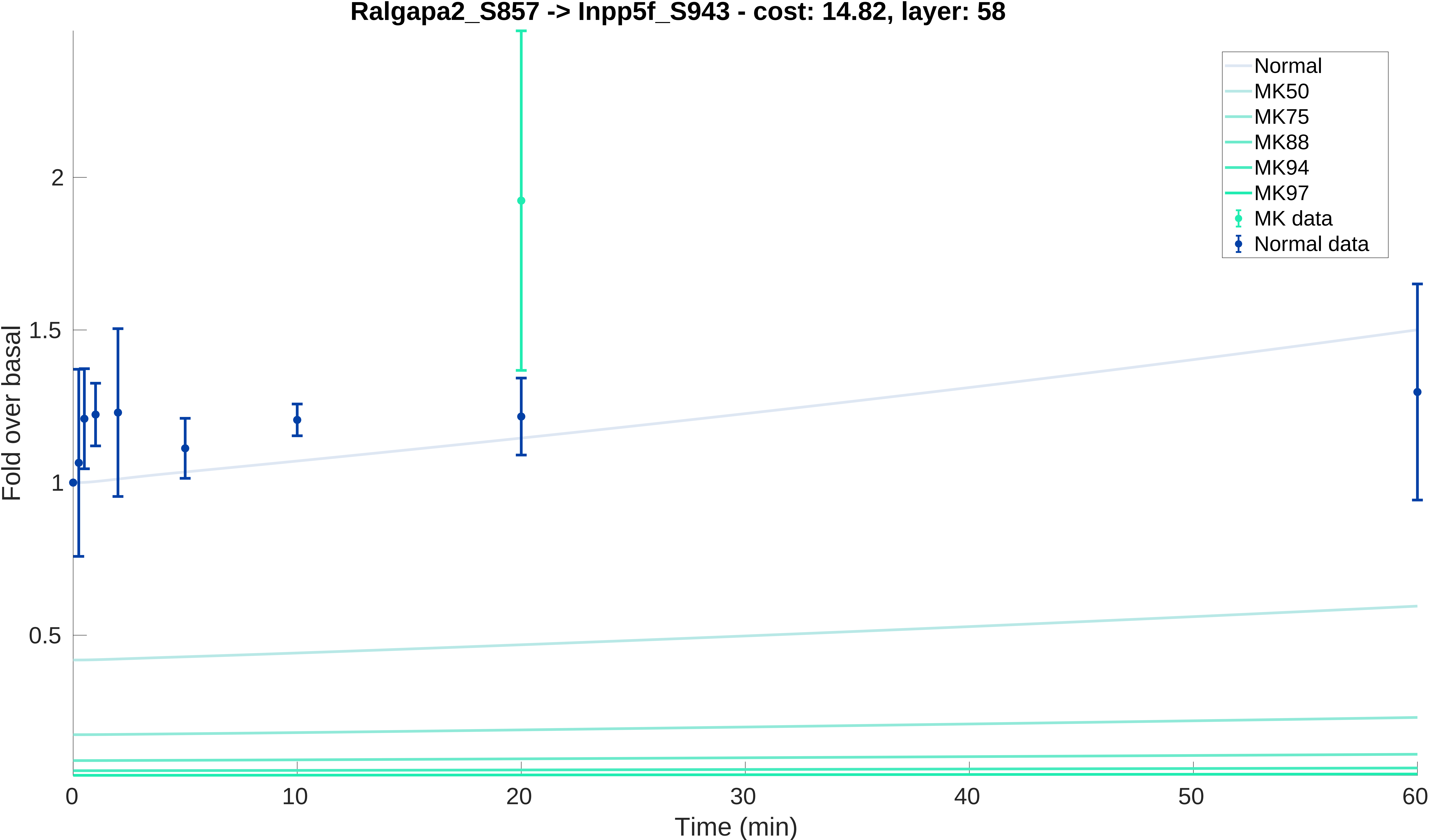

Ralgapa2_S857 -> Inpp5f_S943 - cost: 14.82, layer: 58
Fold over basal
Time (min)
0.5
1
1.5
2
0
10
20
30
40
50
60
Normal
MK50
MK75
MK88
MK94
MK97
MK data
Normal data

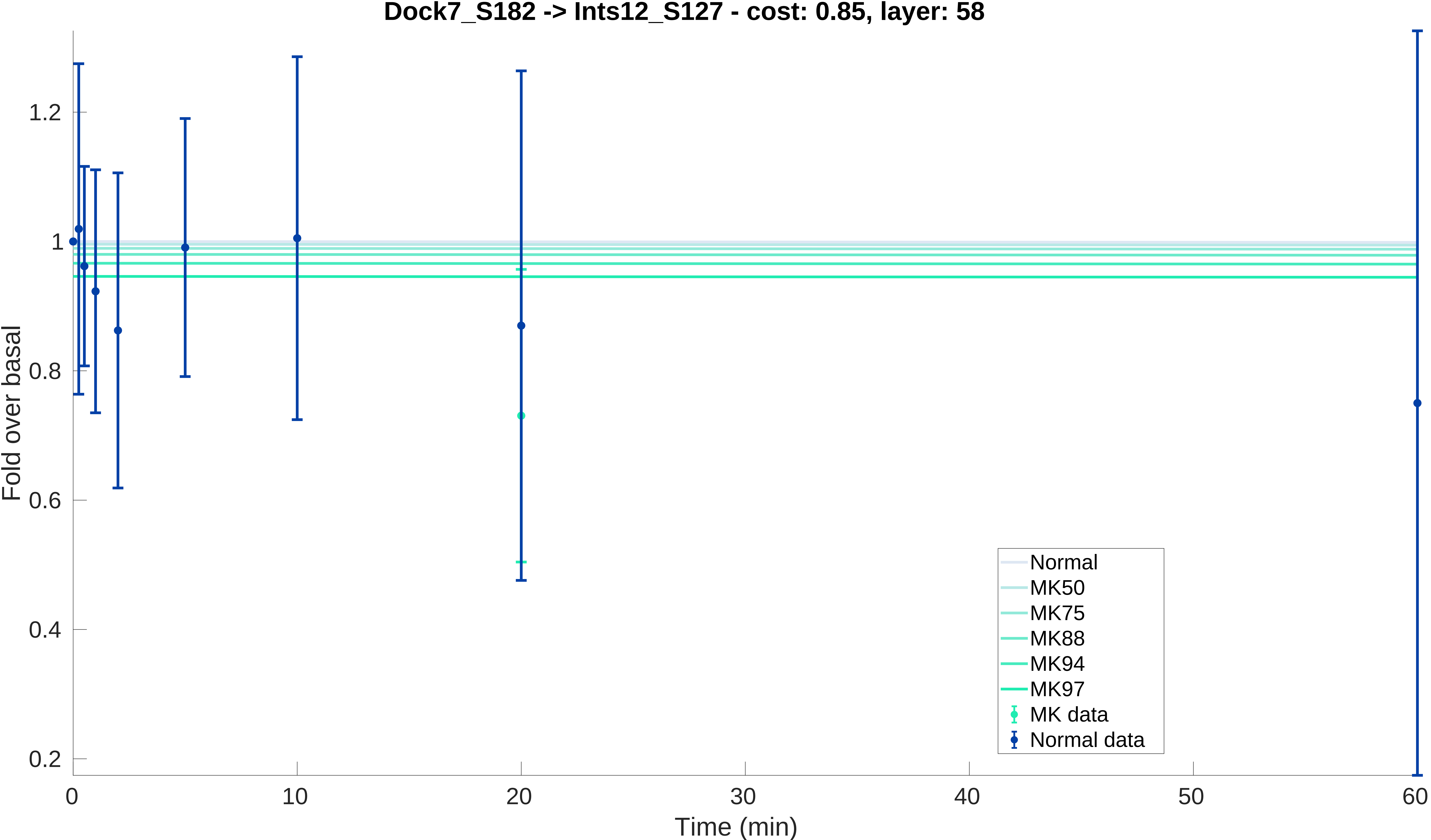

Dock7_S182 -> Ints12_S127 - cost: 0.85, layer: 58
Fold over basal
Time (min)
0.2
0.4
0.6
0.8
1
1.2
0
10
20
30
40
50
60
Normal
MK50
MK75
MK88
MK94
MK97
MK data
Normal data

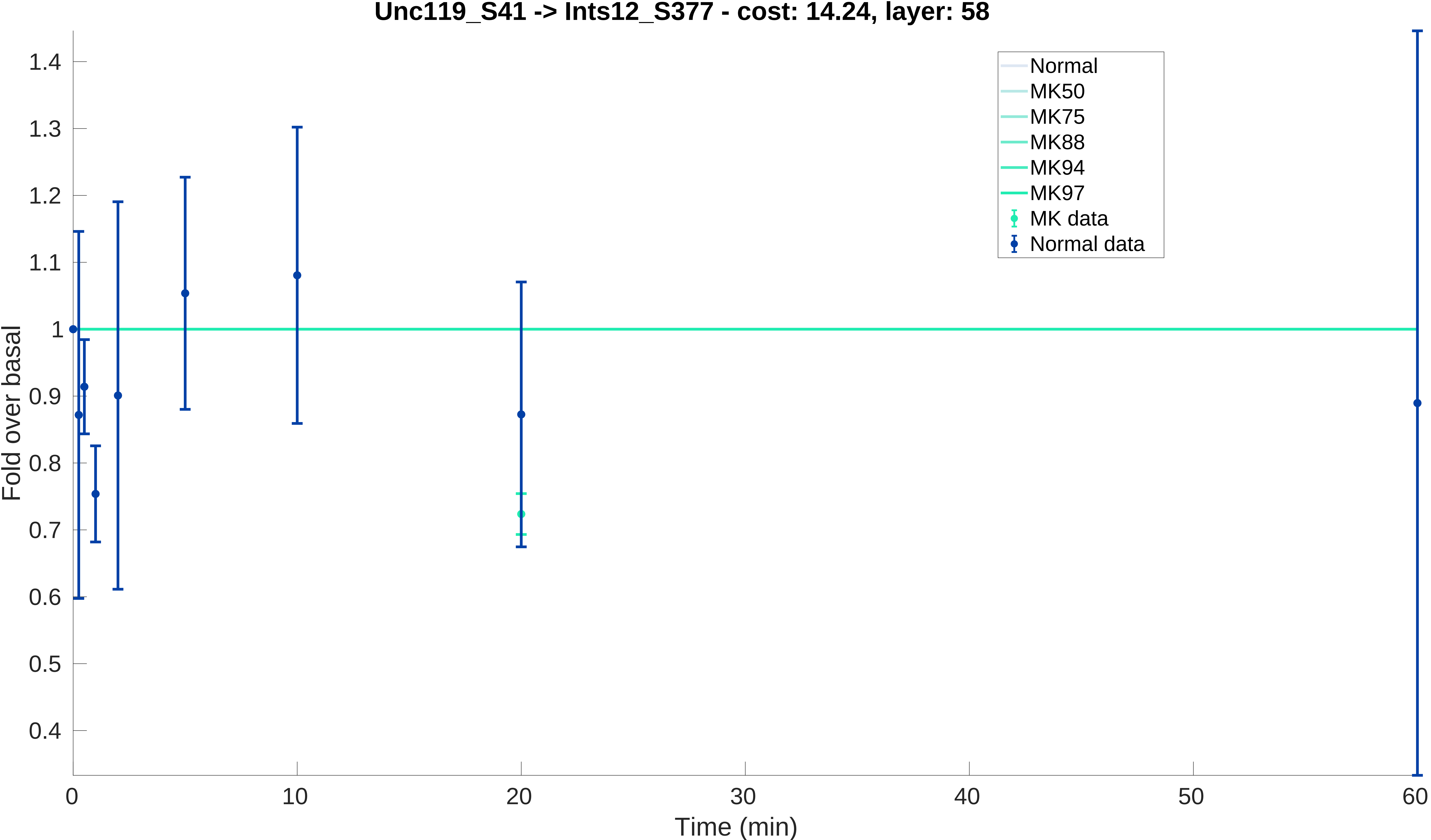

Unc119_S41 -> Ints12_S377 - cost: 14.24, layer: 58
Fold over basal
1.4
1.3
1.2
1.1
1
0.9
0.8
0.7
0.6
0.5
0.4
0
10
20
30
40
50
60
Time (min)
Normal
MK50
MK75
MK88
MK94
MK97
MK data
Normal data

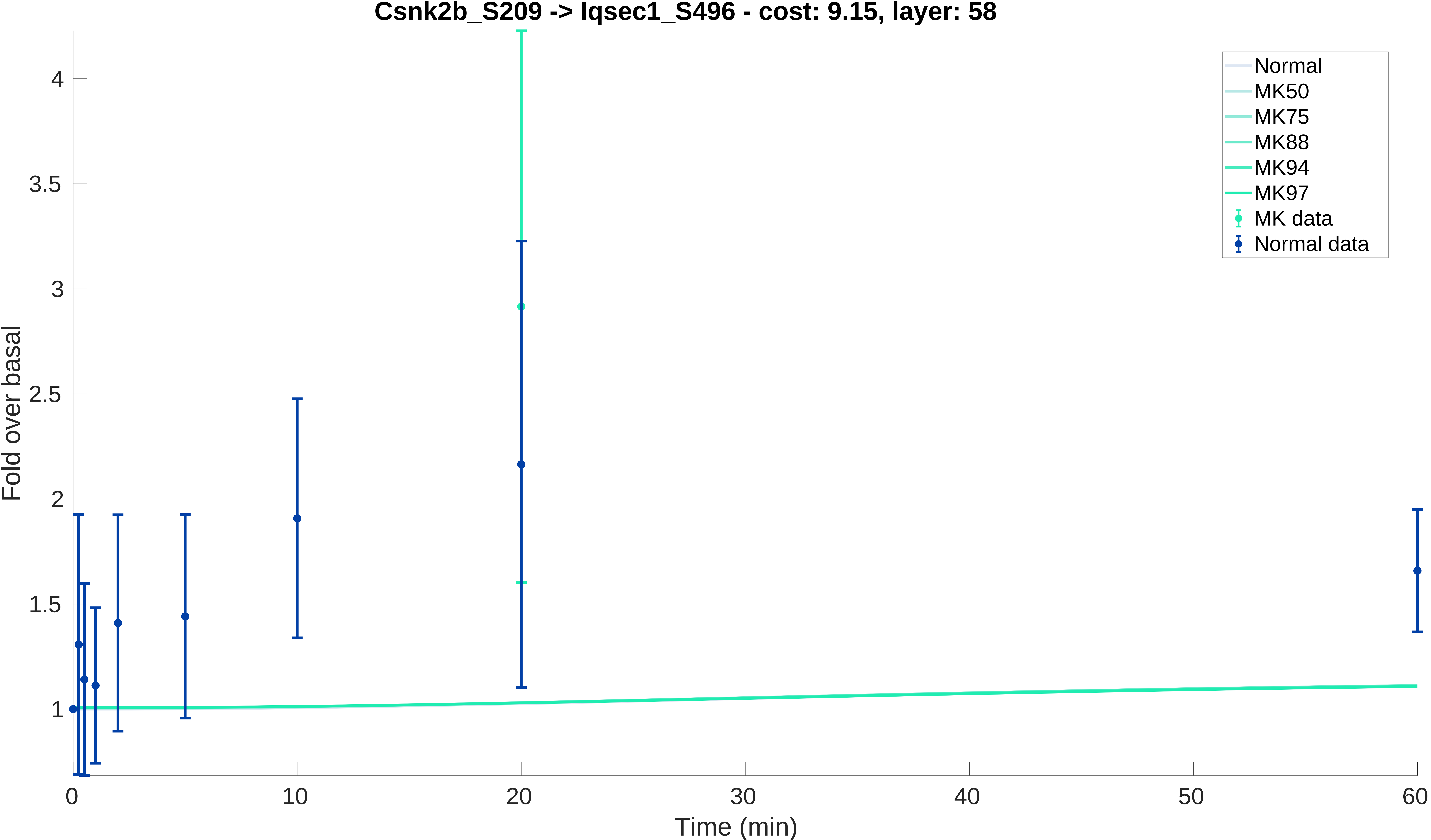
Csnk2b_S209 -> Iqsec1_S496 - cost: 9.15, layer: 58
Fold over basal
Time (min)
0
10
20
30
40
50
60
1
1.5
2
2.5
3
3.5
4
Normal
MK50
MK75
MK88
MK94
MK97
MK data
Normal data

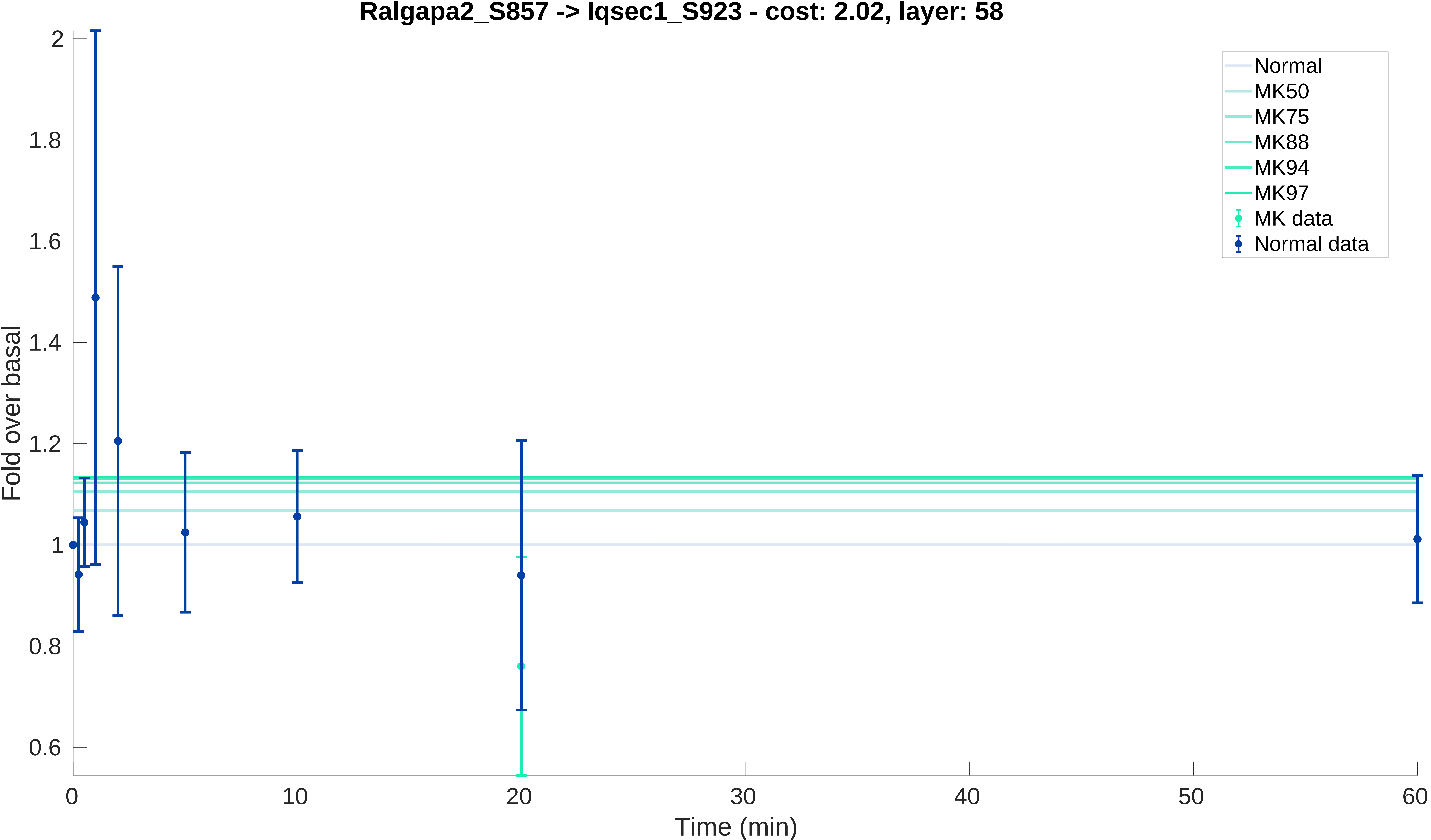
Ralgapa2_S857 -> Iqsec1_S923 - cost: 2.02, layer: 58
Fold over basal
Time (min)
0
10
20
30
40
50
60
0.6
0.8
1
1.2
1.4
1.6
1.8
2
Normal
MK50
MK75
MK88
MK94
MK97
MK data
Normal data

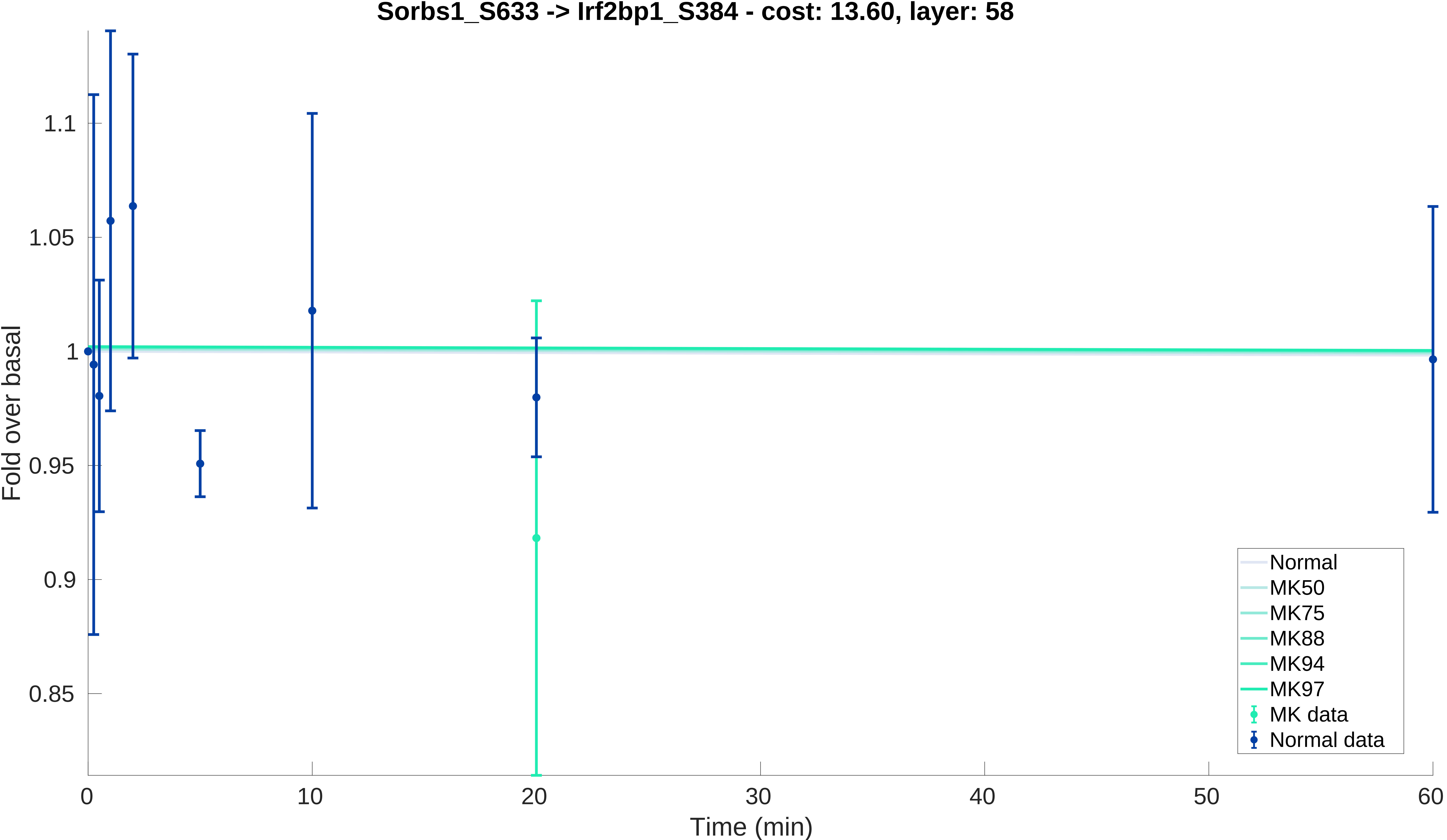

Sorbs1_S633 -> Irf2bp1_S384 - cost: 13.60, layer: 58
Fold over basal
Time (min)
1.1
1.05
1
0.95
0.9
0.85
0
10
20
30
40
50
60
Normal
MK50
MK75
MK88
MK94
MK97
MK data
Normal data

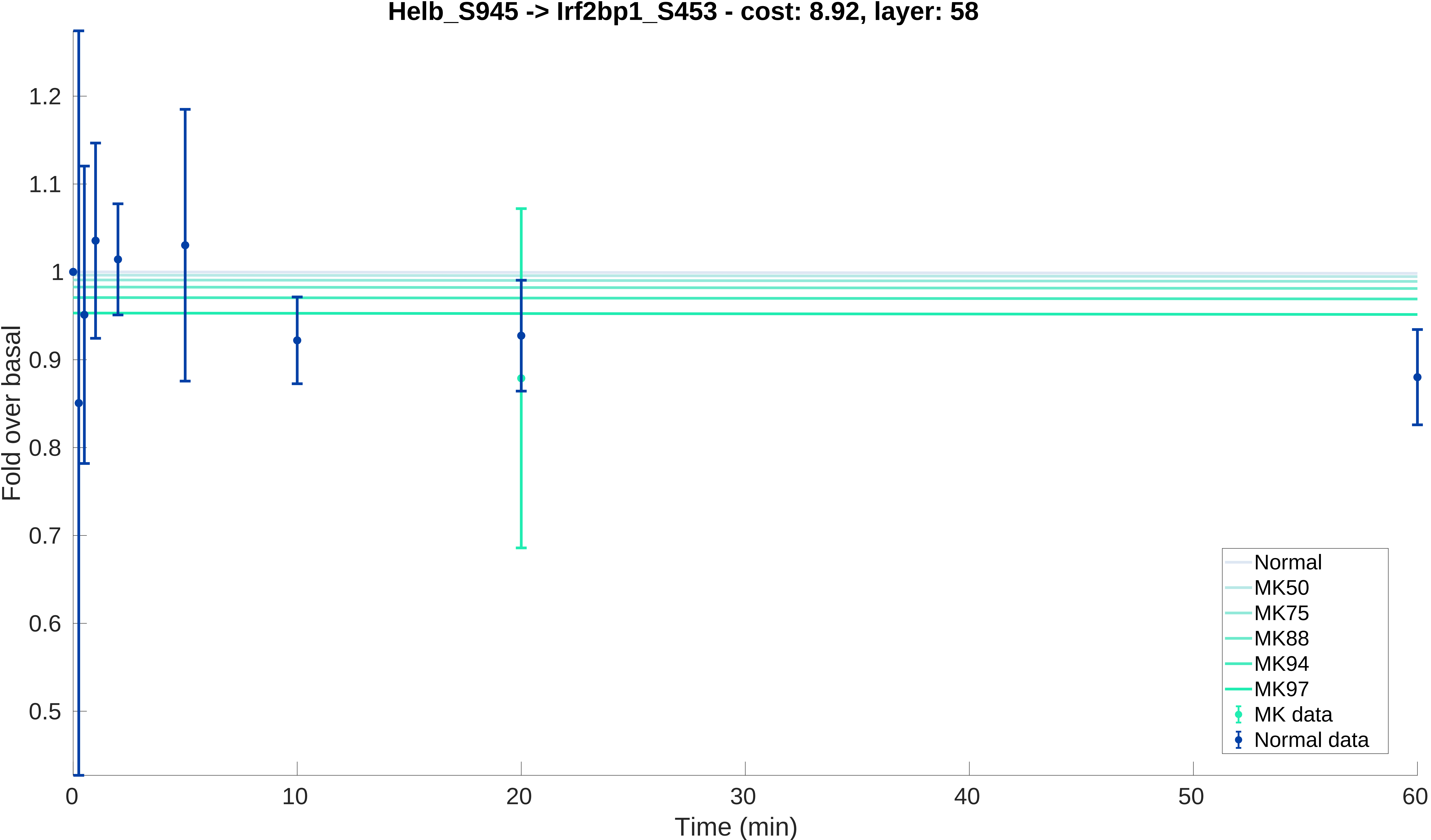

Helb_S945 -> Irf2bp1_S453 - cost: 8.92, layer: 58
Fold over basal
1.2
1.1
1
0.9
0.8
0.7
0.6
0.5
0
10
20
30
40
50
60
Time (min)
Normal
MK50
MK75
MK88
MK94
MK97
MK data
Normal data

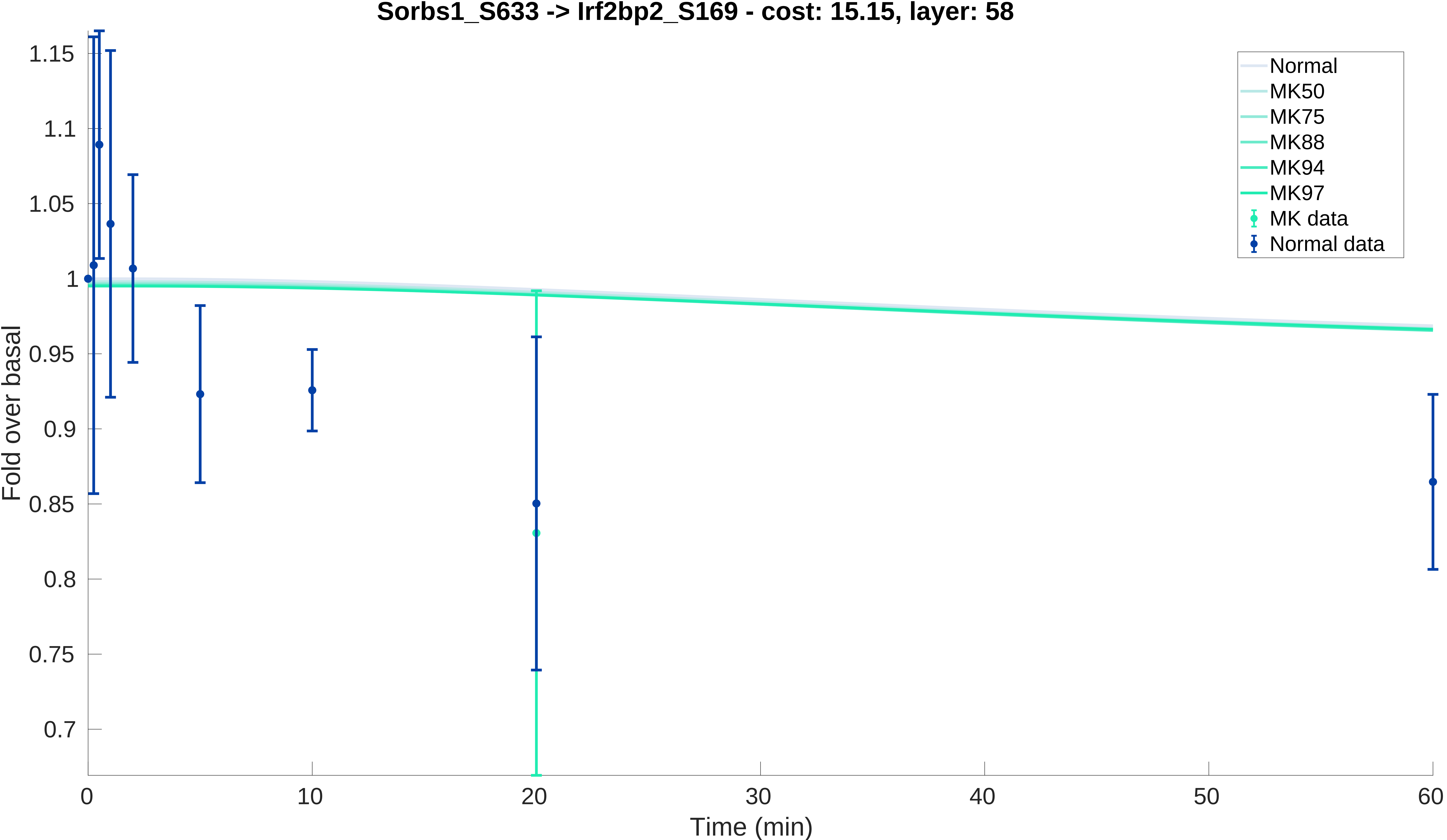

Sorbs1_S633 -> Irf2bp2_S169 - cost: 15.15, layer: 58
Fold over basal
Time (min)
1.15
1.1
1.05
1
0.95
0.9
0.85
0.8
0.75
0.7
0
10
20
30
40
50
60
Normal
MK50
MK75
MK88
MK94
MK97
MK data
Normal data

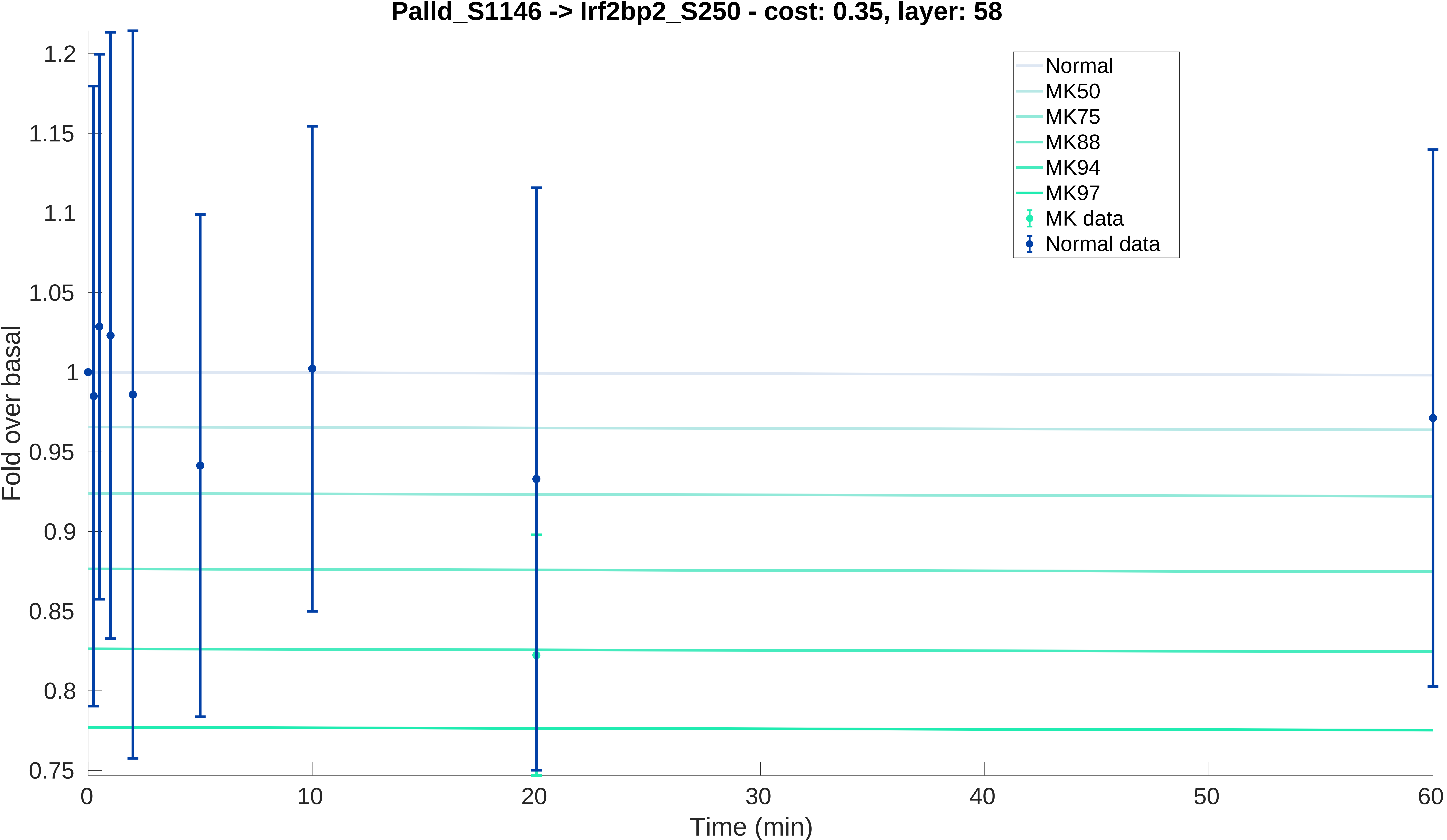

Palld_S1146 -> Irf2bp2_S250 - cost: 0.35, layer: 58
Normal
MK50
MK75
MK88
MK94
MK97
MK data
Normal data
Fold over basal
Time (min)
1.2
1.15
1.1
1.05
1
0.95
0.9
0.85
0.8
0.75
0
10
20
30
40
50
60

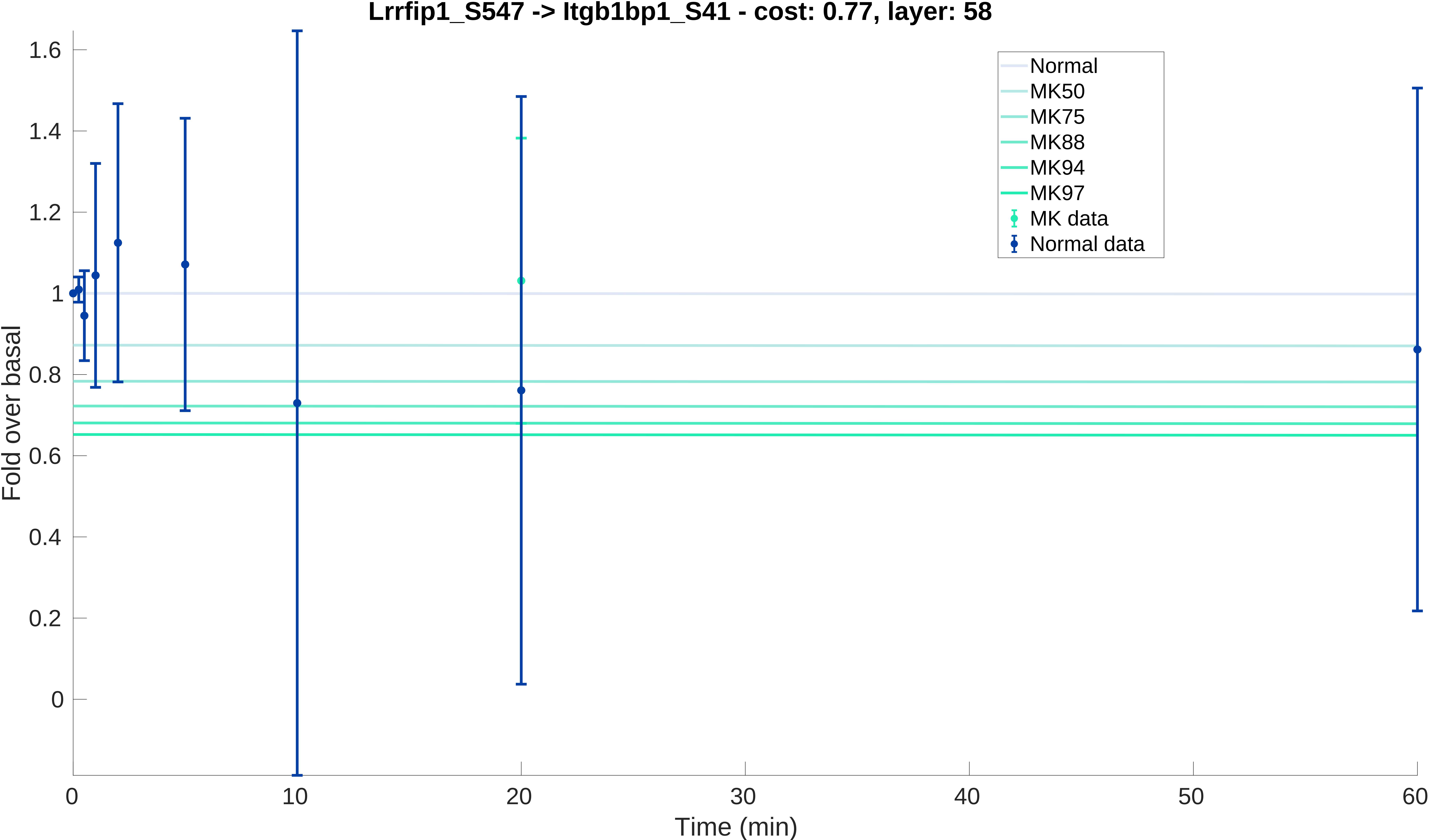

Lrrfip1_S547 -> Itgb1bp1_S41 - cost: 0.77, layer: 58
Fold over basal
Time (min)
1.6
1.4
1.2
1
0.8
0.6
0.4
0.2
0
0
10
20
30
40
50
60
Normal
MK50
MK75
MK88
MK94
MK97
MK data
Normal data

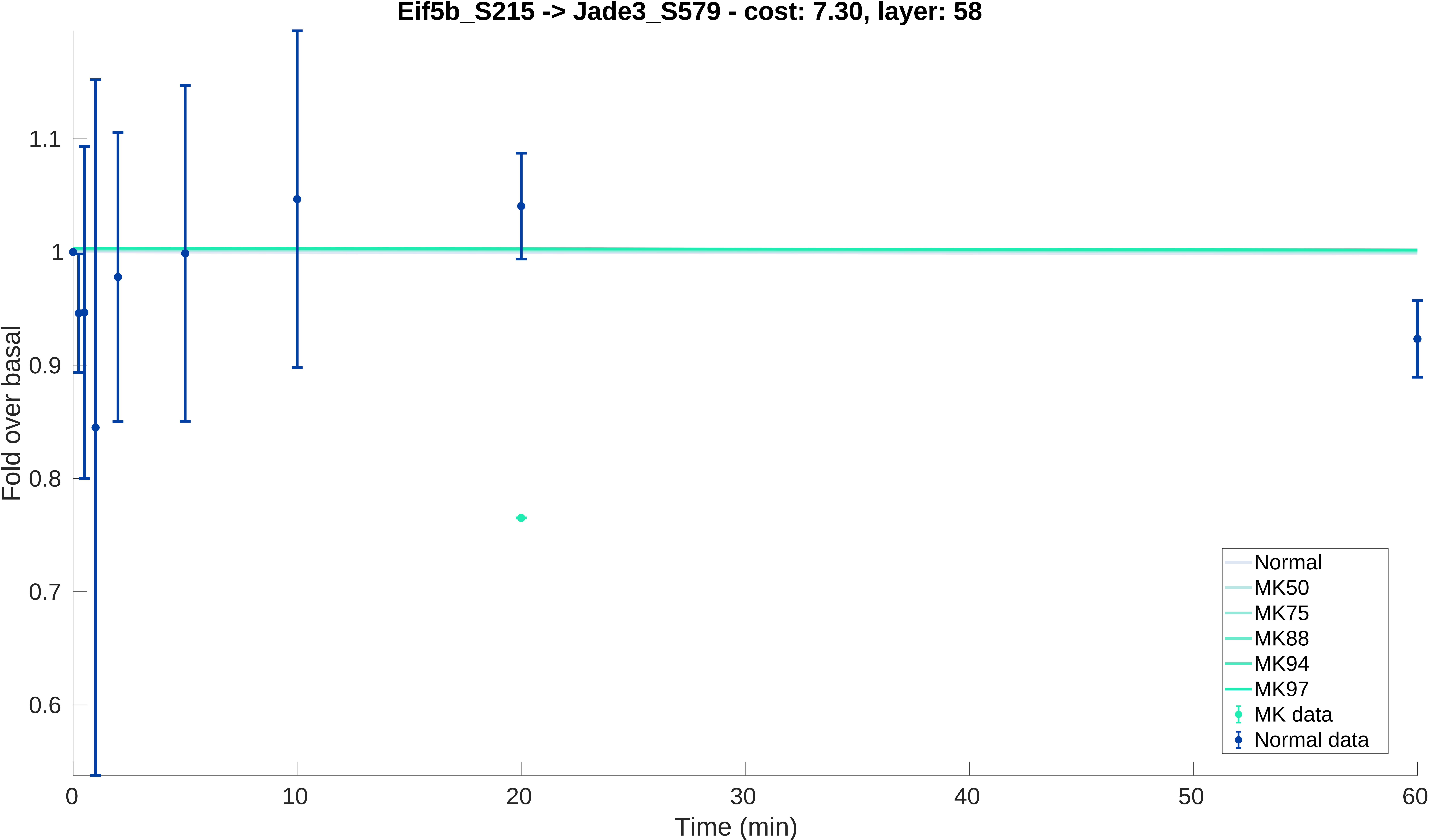
Eif5b_S215 -> Jade3_S579 - cost: 7.30, layer: 58
Fold over basal
Time (min)
1.1
1
0.9
0.8
0.7
0.6
0
10
20
30
40
50
60
Normal
MK50
MK75
MK88
MK94
MK97
MK data
Normal data

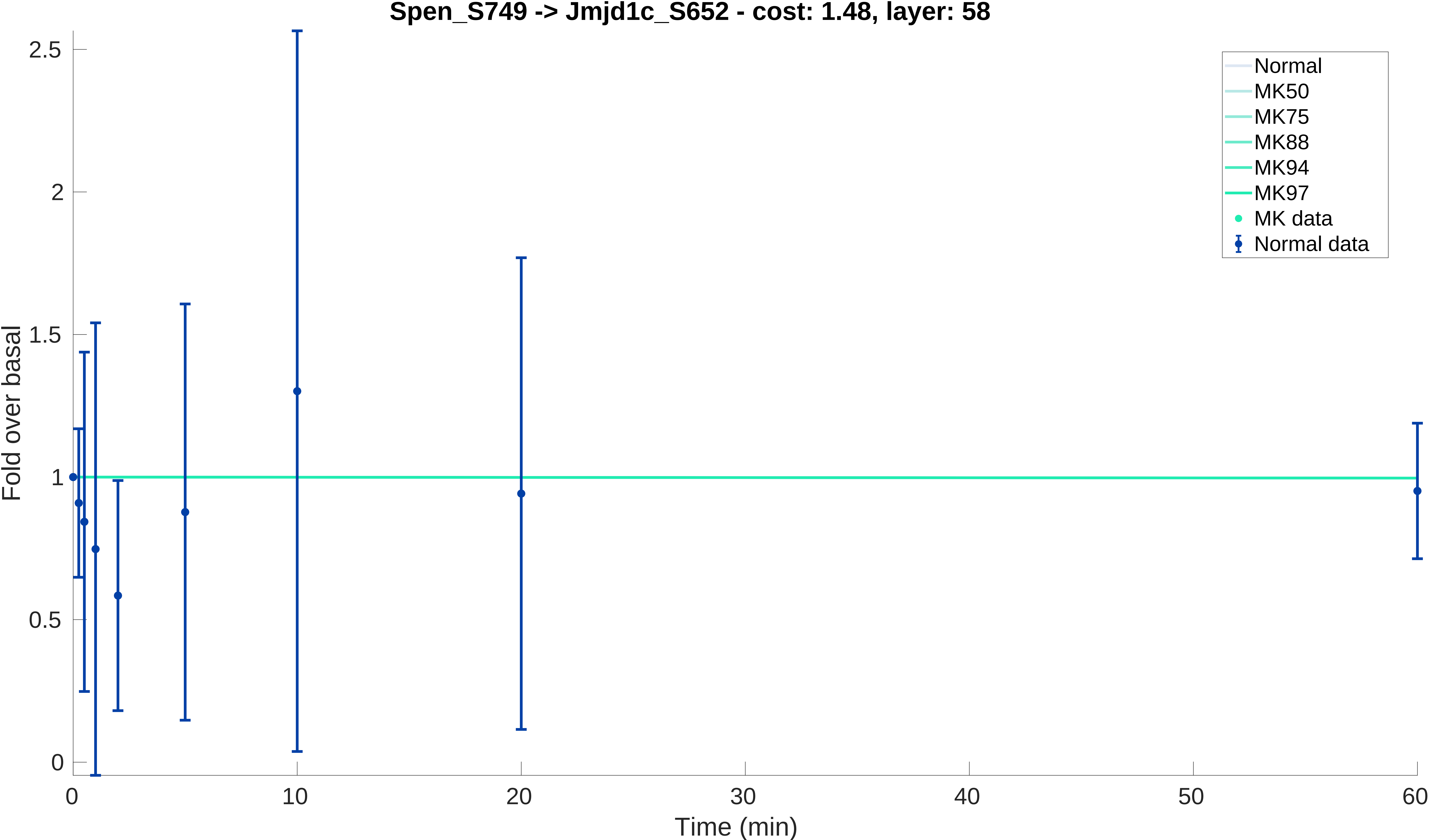
Spen_S749 -> Jmjd1c_S652 - cost: 1.48, layer: 58
Normal
MK50
MK75
MK88
MK94
MK97
MK data
Normal data
Fold over basal
Time (min)
0
0.5
1
1.5
2
2.5
0
10
20
30
40
50
60

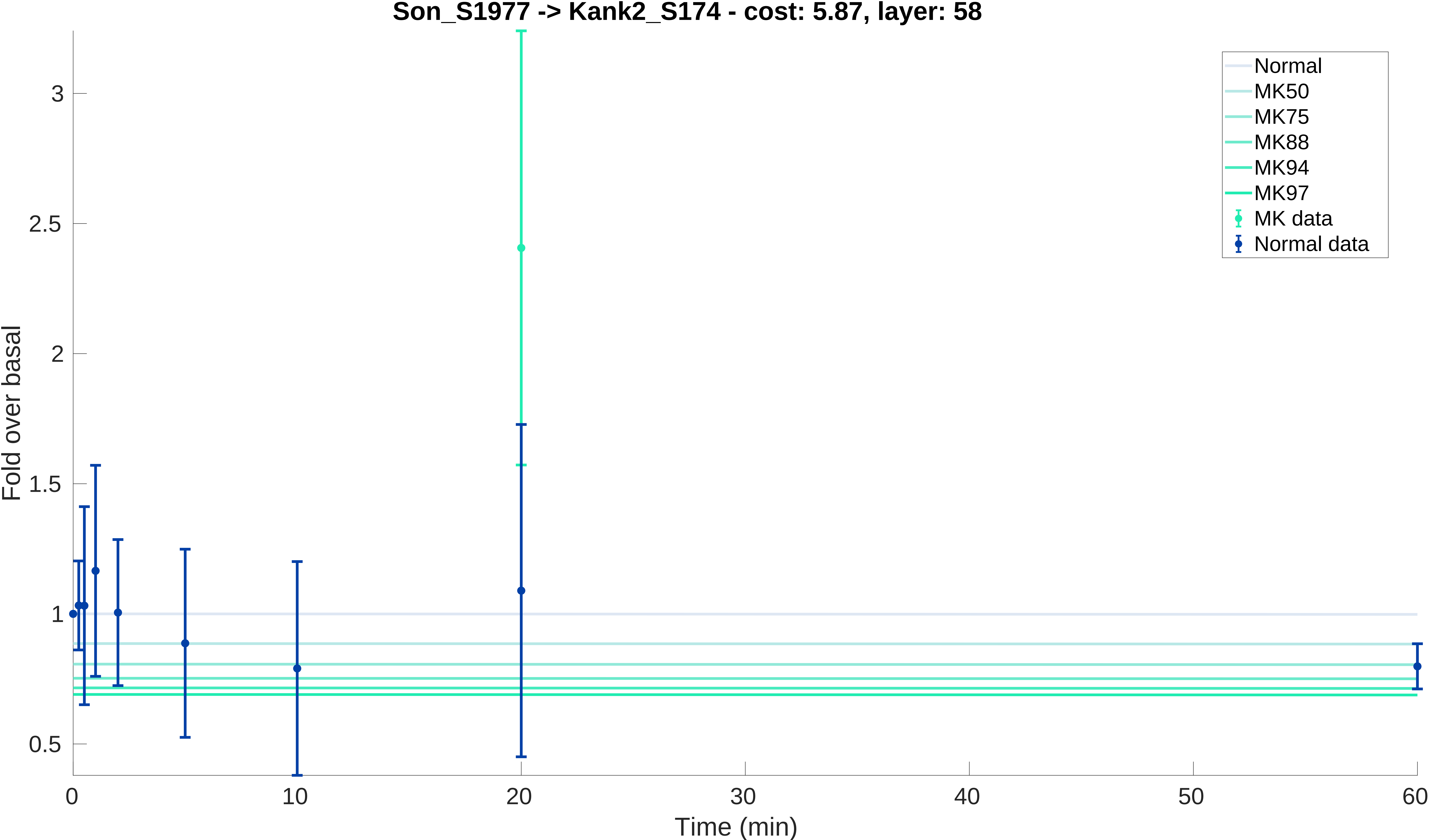
Son_S1977 -> Kank2_S174 - cost: 5.87, layer: 58
Fold over basal
3
2.5
2
1.5
1
0.5
0
10
20
30
40
50
60
Time (min)
Normal
MK50
MK75
MK88
MK94
MK97
MK data
Normal data

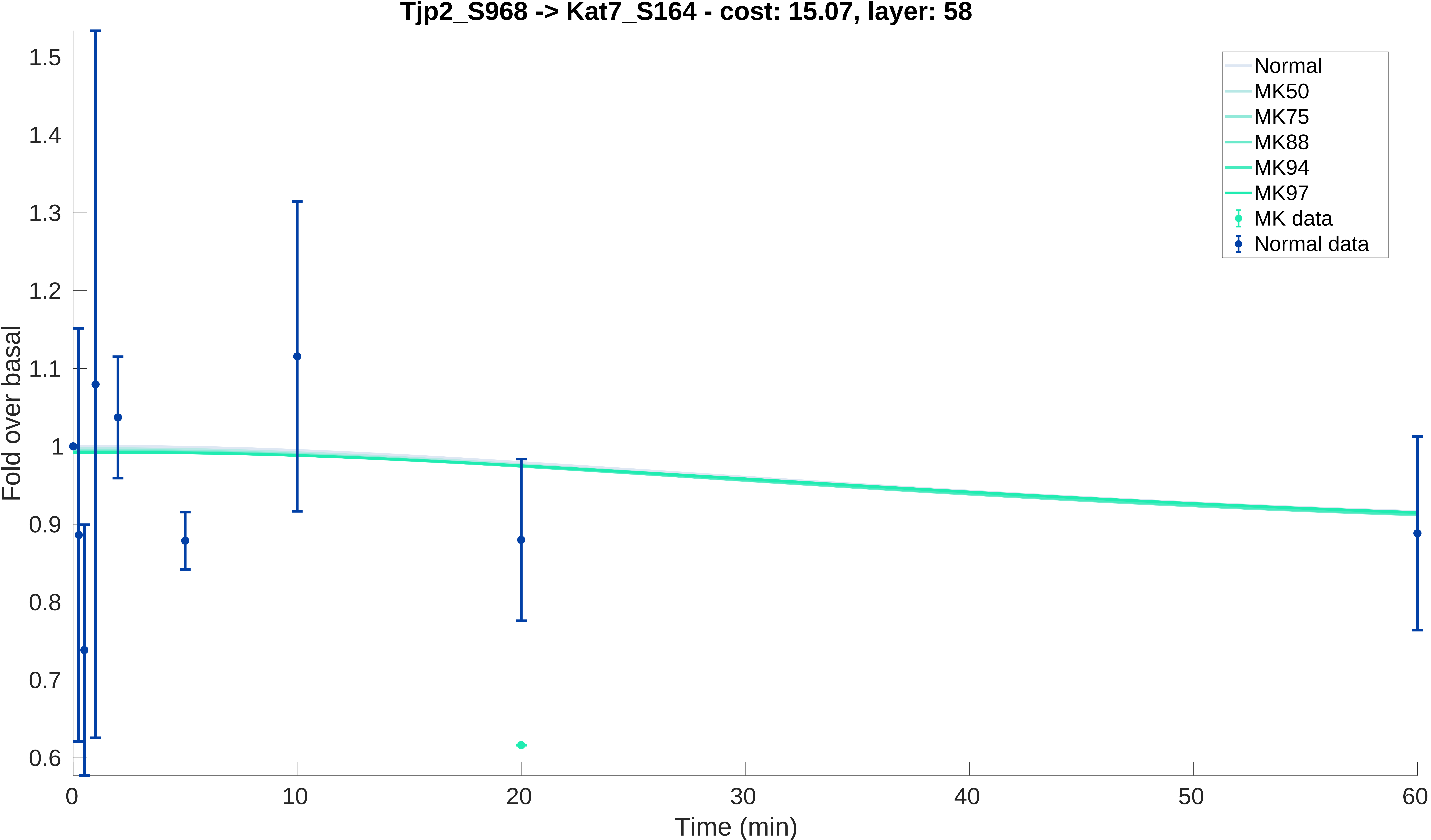

Tjp2_S968 -> Kat7_S164 - cost: 15.07, layer: 58
Fold over basal
Time (min)
0.6
0.7
0.8
0.9
1
1.1
1.2
1.3
1.4
1.5
0
10
20
30
40
50
60
Normal
MK50
MK75
MK88
MK94
MK97
MK data
Normal data

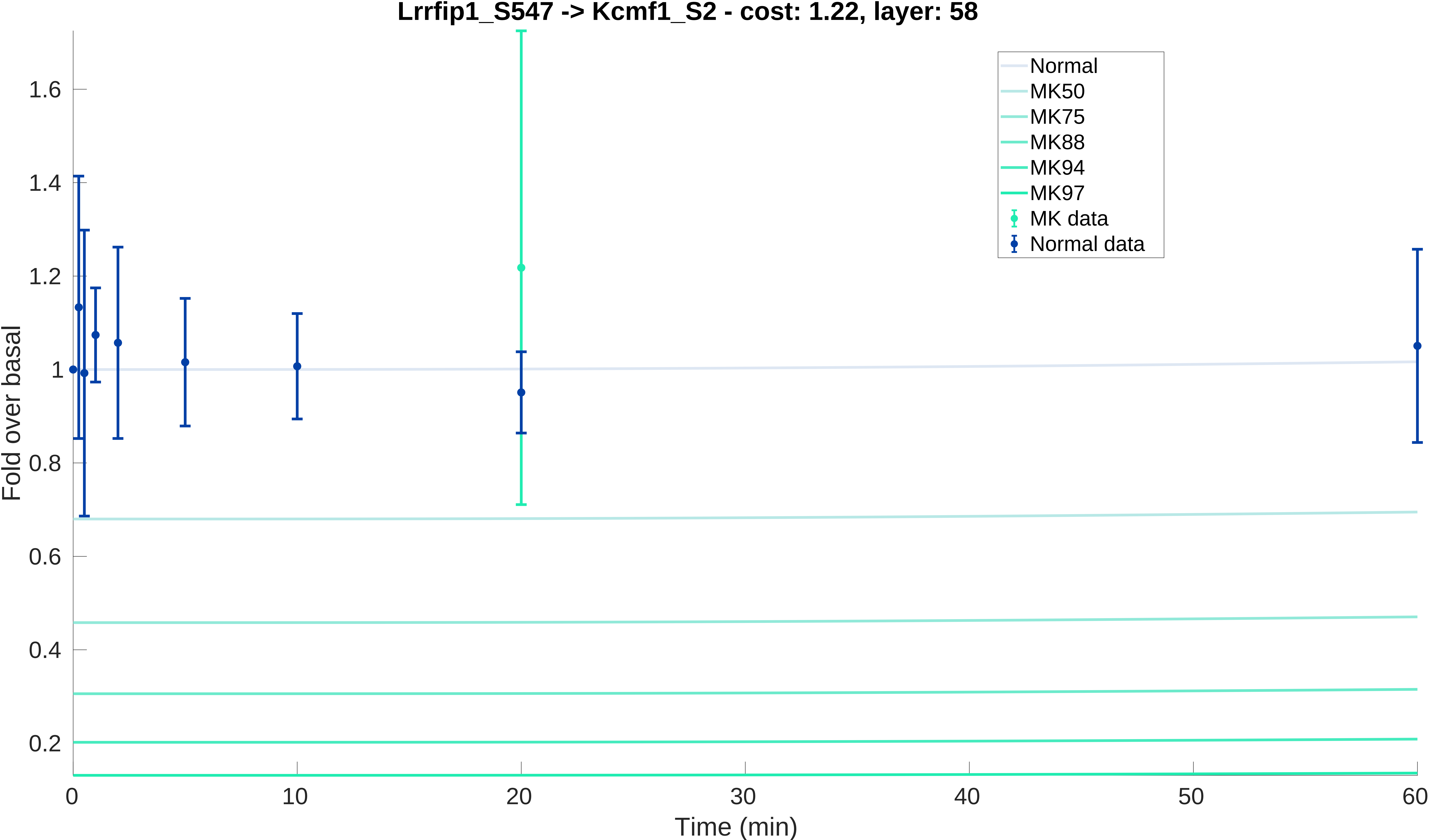

Lrrfip1_S547 -> Kcmf1_S2 - cost: 1.22, layer: 58
Normal
MK50
MK75
MK88
MK94
MK97
MK data
Normal data
Fold over basal
Time (min)
0.2
0.4
0.6
0.8
1
1.2
1.4
1.6
0
10
20
30
40
50
60

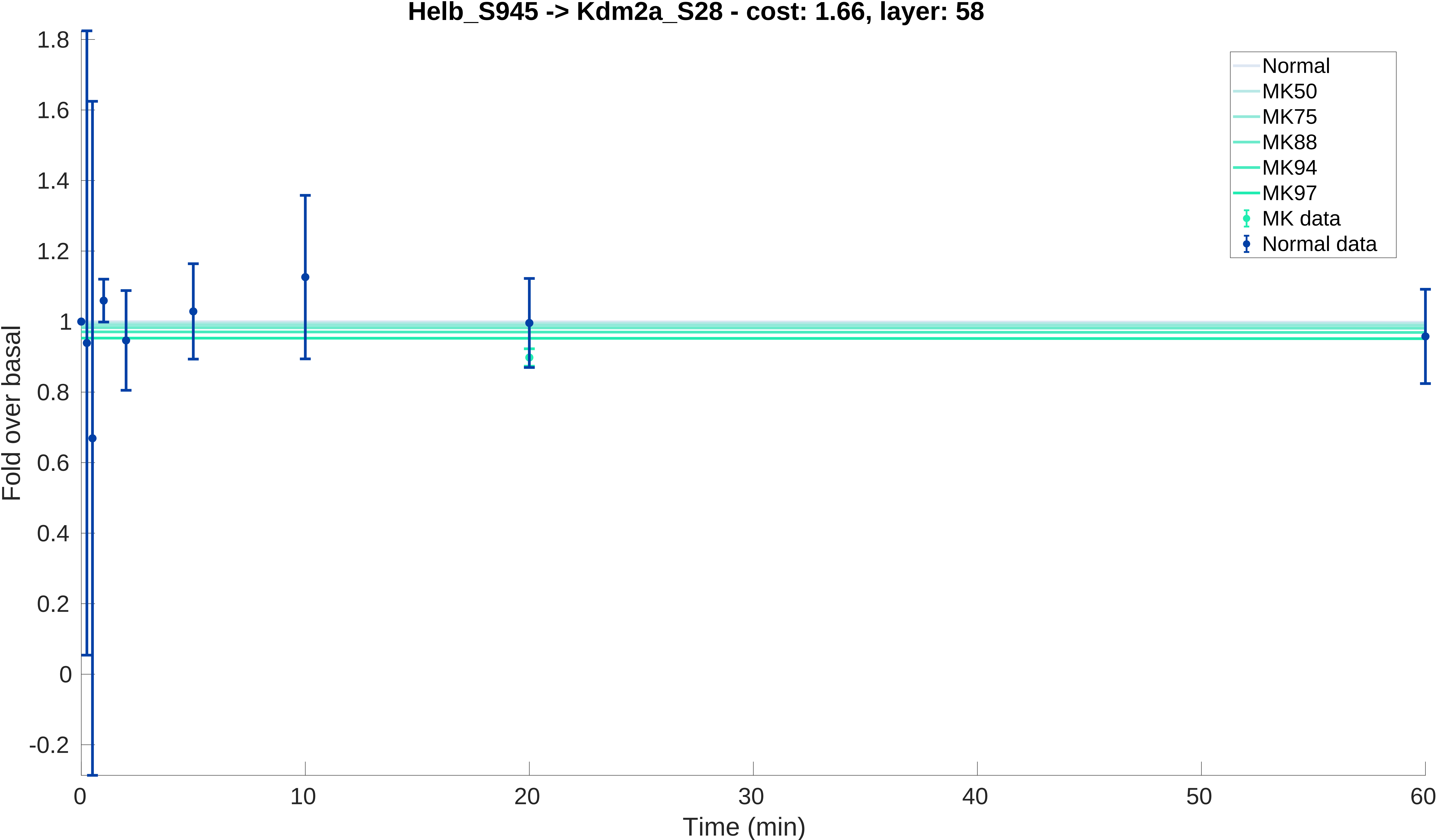

Helb_S945 -> Kdm2a_S28 - cost: 1.66, layer: 58
Fold over basal
1.8
1.6
1.4
1.2
1
0.8
0.6
0.4
0.2
0
-0.2
Time (min)
0
10
20
30
40
50
60
Normal
MK50
MK75
MK88
MK94
MK97
MK data
Normal data

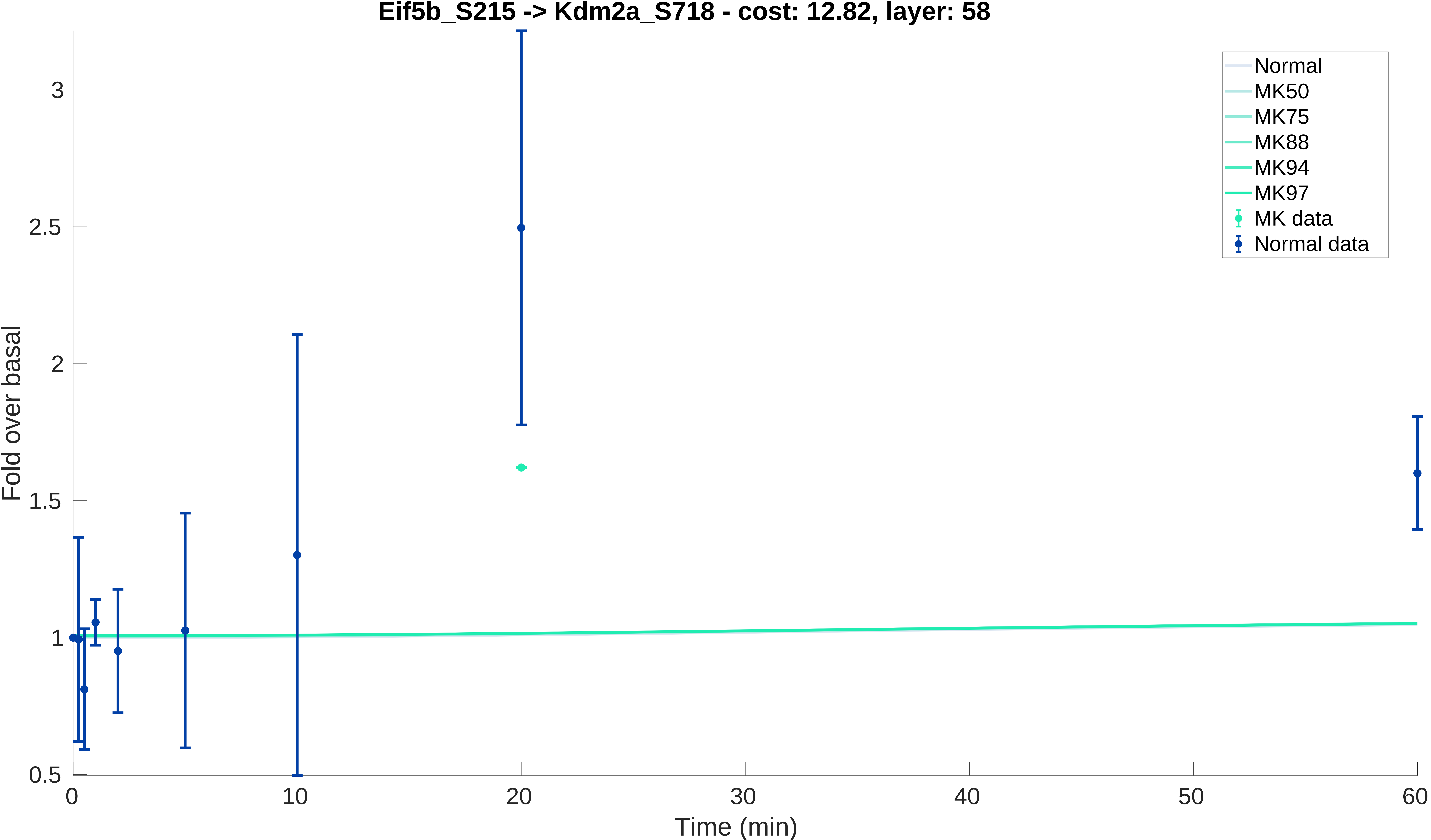

Eif5b_S215 -> Kdm2a_S718 - cost: 12.82, layer: 58
Fold over basal
Time (min)
0.5
1
1.5
2
2.5
3
0
10
20
30
40
50
60
Normal
MK50
MK75
MK88
MK94
MK97
MK data
Normal data

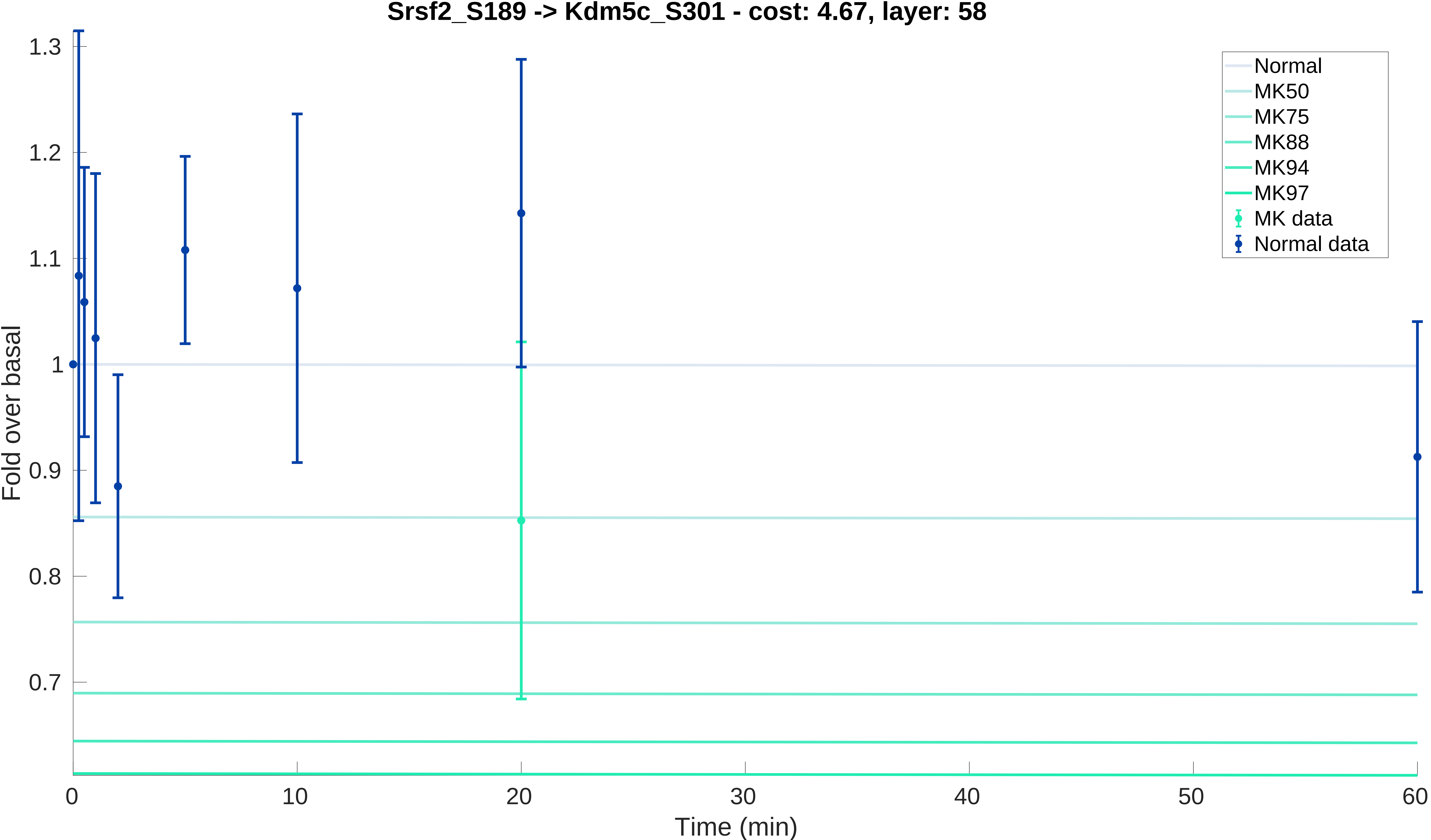
Srsf2_S189 -> Kdm5c_S301 - cost: 4.67, layer: 58
Fold over basal
1.3
1.2
1.1
1
0.9
0.8
0.7
0
10
20
30
40
50
60
Time (min)
Normal
MK50
MK75
MK88
MK94
MK97
MK data
Normal data

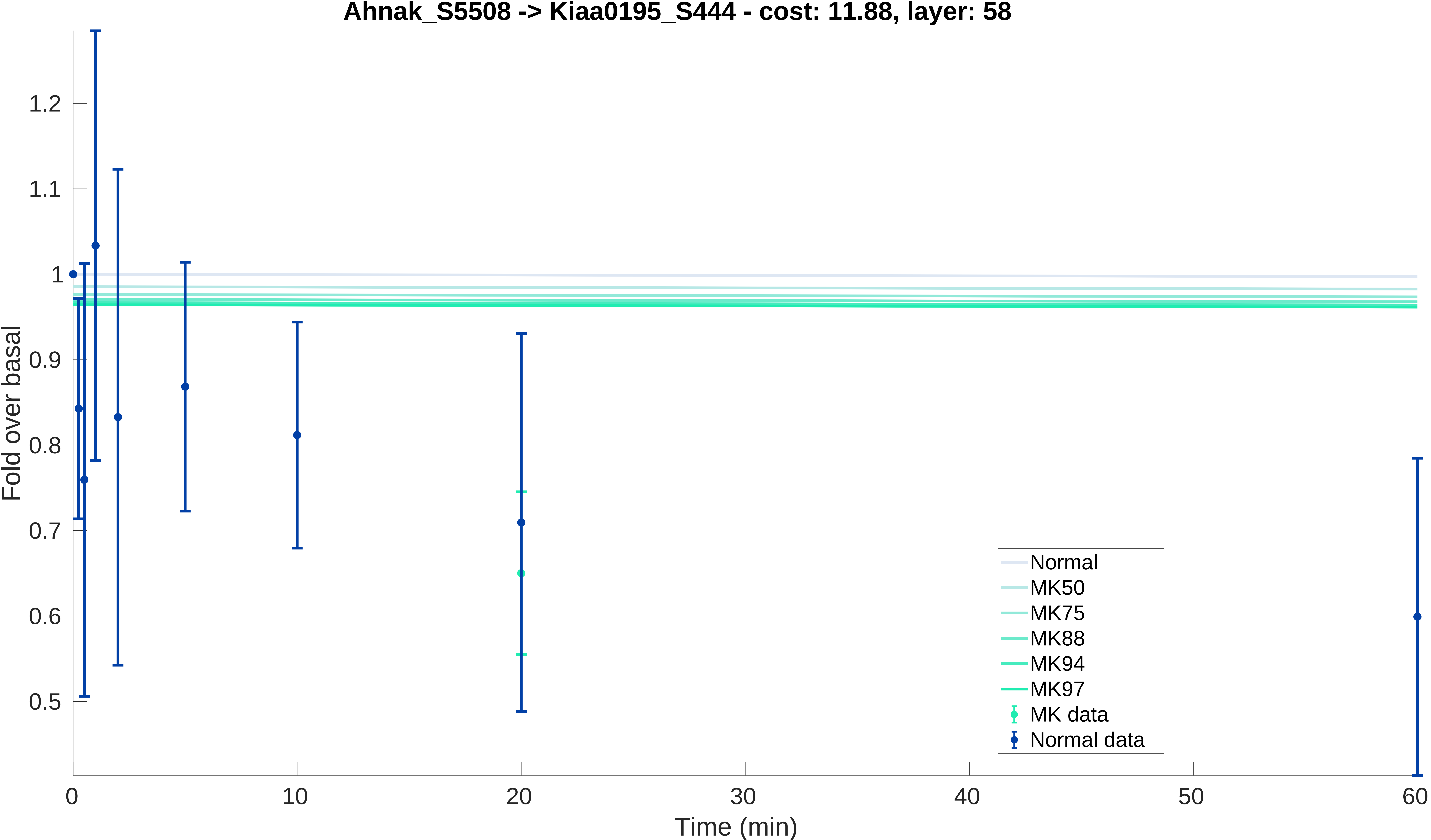

Ahnak_S5508 -> Kiaa0195_S444 - cost: 11.88, layer: 58
Fold over basal
1.2
1.1
1
0.9
0.8
0.7
0.6
0.5
0
10
20
30
40
50
60
Time (min)
Normal
MK50
MK75
MK88
MK94
MK97
MK data
Normal data

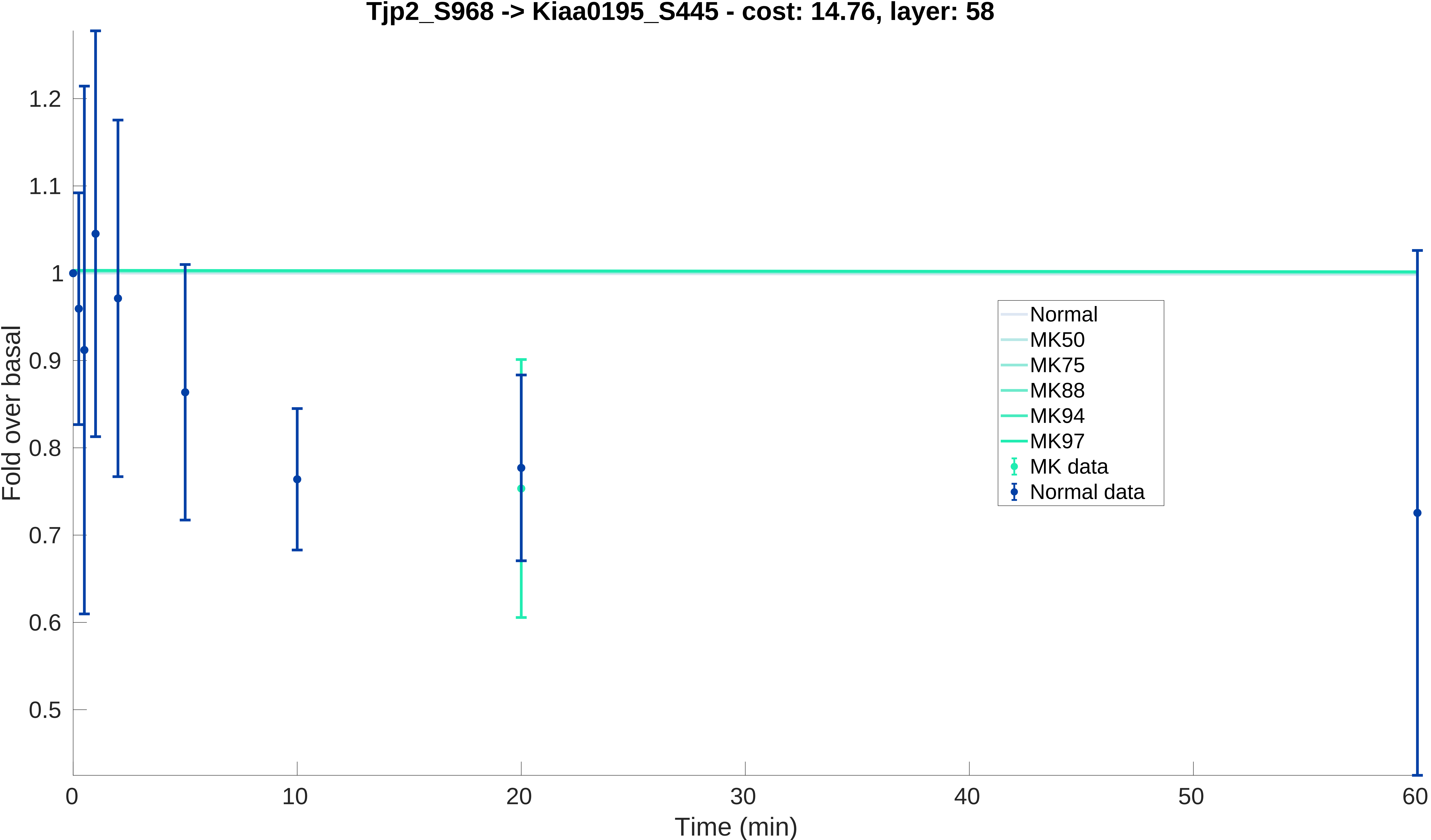

Tjp2_S968 -> Kiaa0195_S445 - cost: 14.76, layer: 58
Fold over basal
Time (min)
1.2
1.1
1
0.9
0.8
0.7
0.6
0.5
0
10
20
30
40
50
60
Normal
MK50
MK75
MK88
MK94
MK97
MK data
Normal data

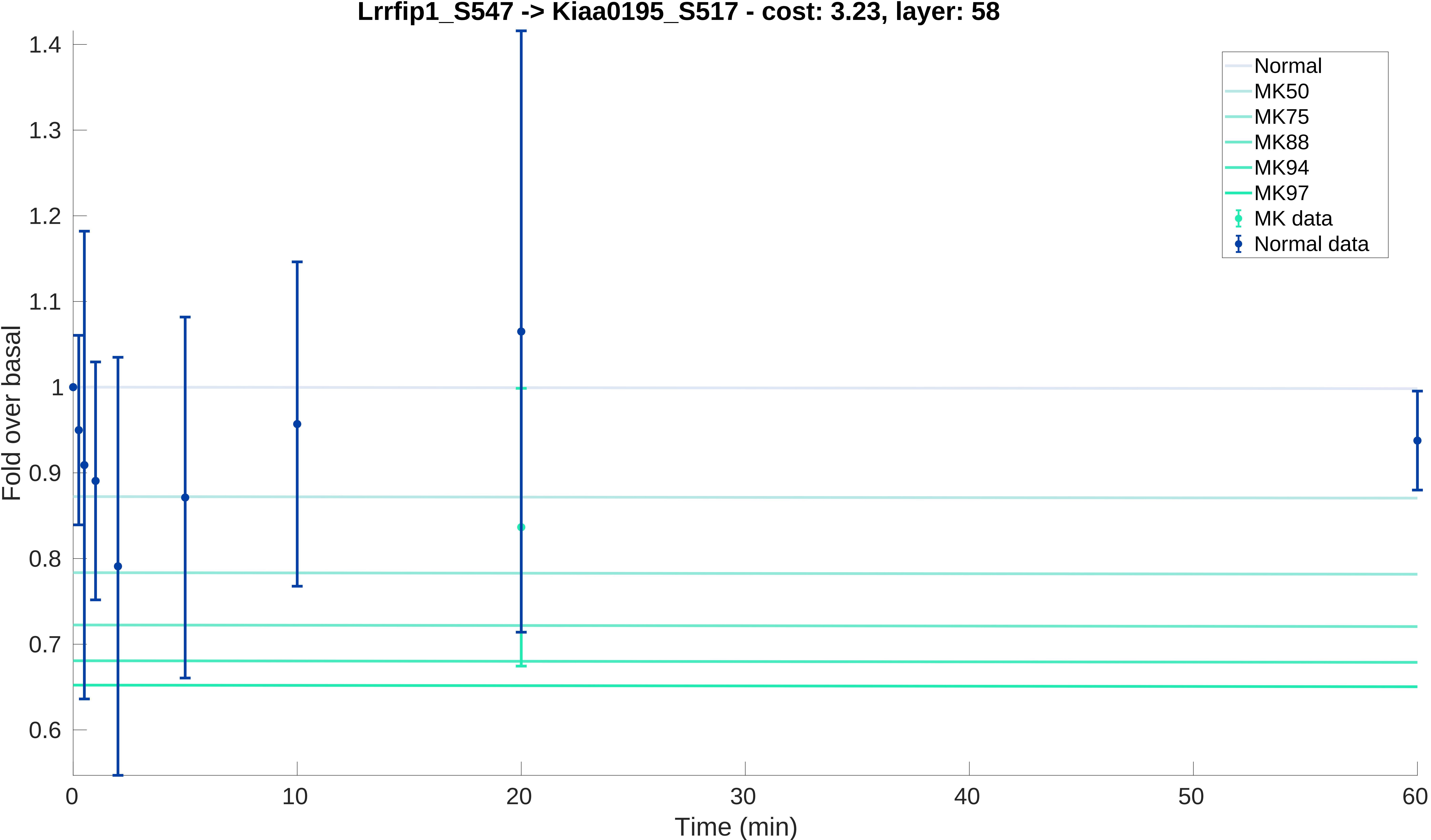

Lrrfip1_S547 -> Kiaa0195_S517 - cost: 3.23, layer: 58
Fold over basal
Time (min)
1.4
1.3
1.2
1.1
1
0.9
0.8
0.7
0.6
0
10
20
30
40
50
60
Normal
MK50
MK75
MK88
MK94
MK97
MK data
Normal data

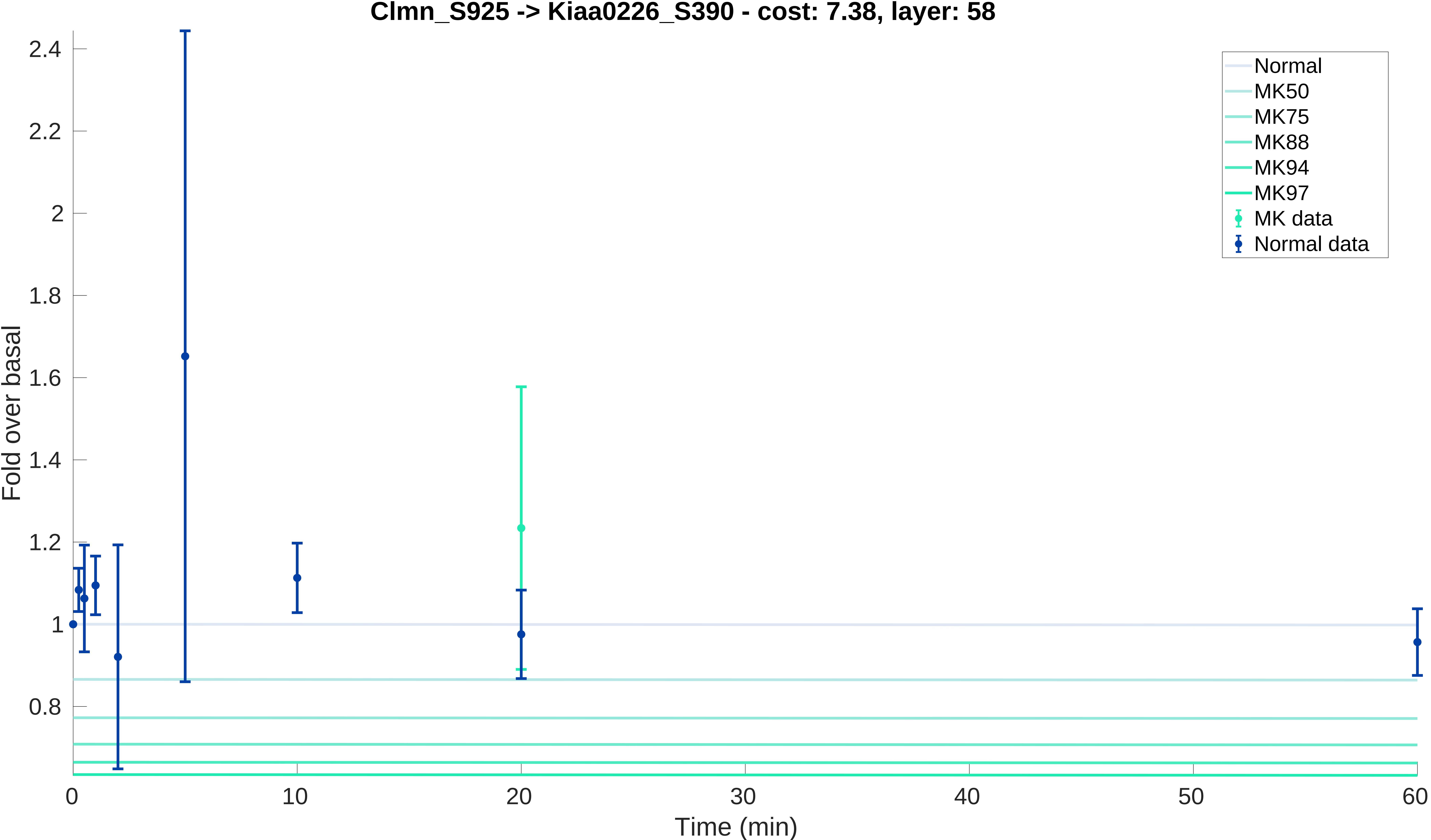

Clmn_S925 -> Kiaa0226_S390 - cost: 7.38, layer: 58
Fold over basal
2.4
2.2
2
1.8
1.6
1.4
1.2
1
0.8
0
10
20
30
40
50
60
Time (min)
Normal
MK50
MK75
MK88
MK94
MK97
MK data
Normal data

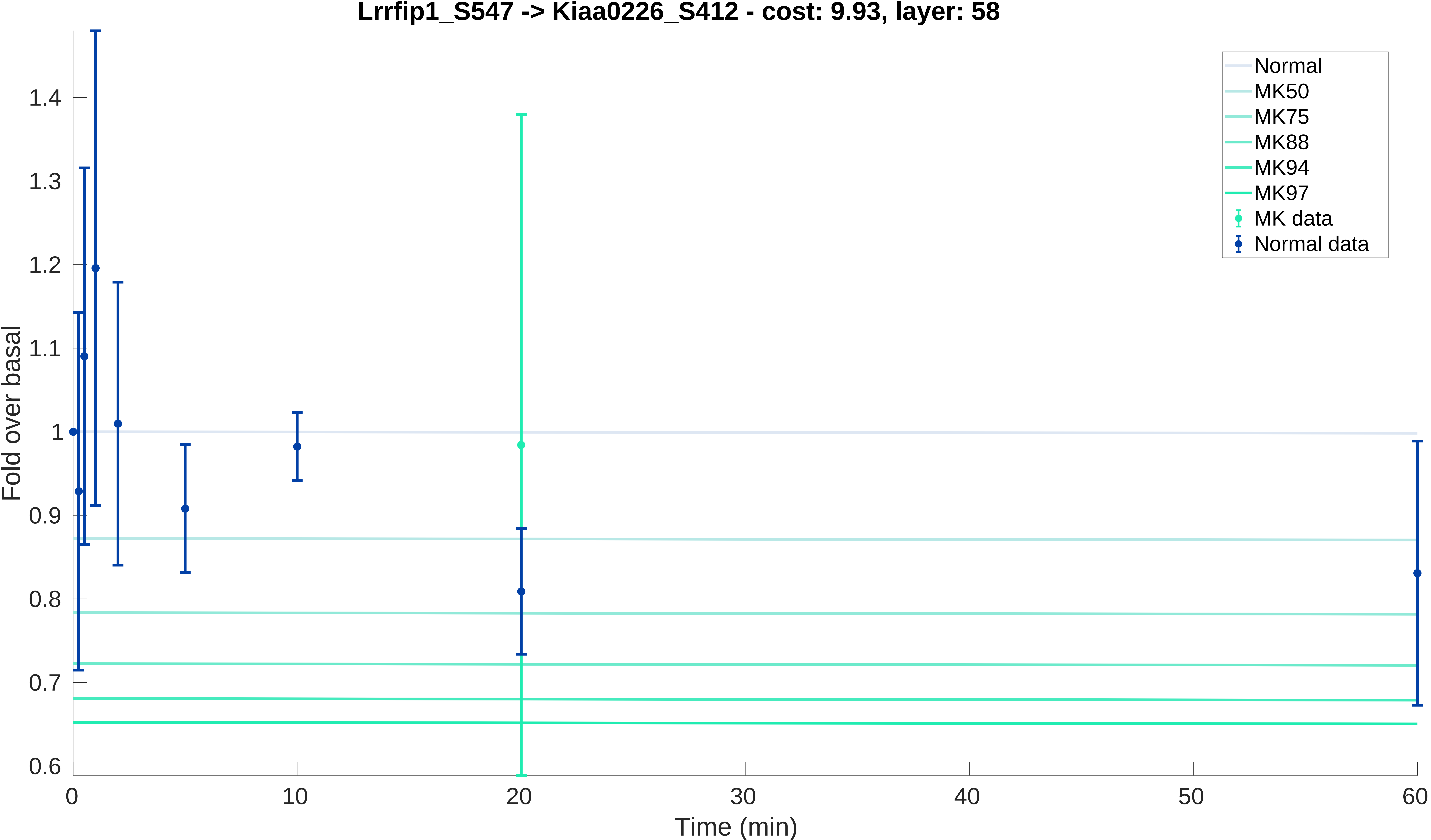
Lrrfip1_S547 -> Kiaa0226_S412 - cost: 9.93, layer: 58
Fold over basal
Time (min)
1.4
1.3
1.2
1.1
1
0.9
0.8
0.7
0.6
0
10
20
30
40
50
60
Normal
MK50
MK75
MK88
MK94
MK97
MK data
Normal data

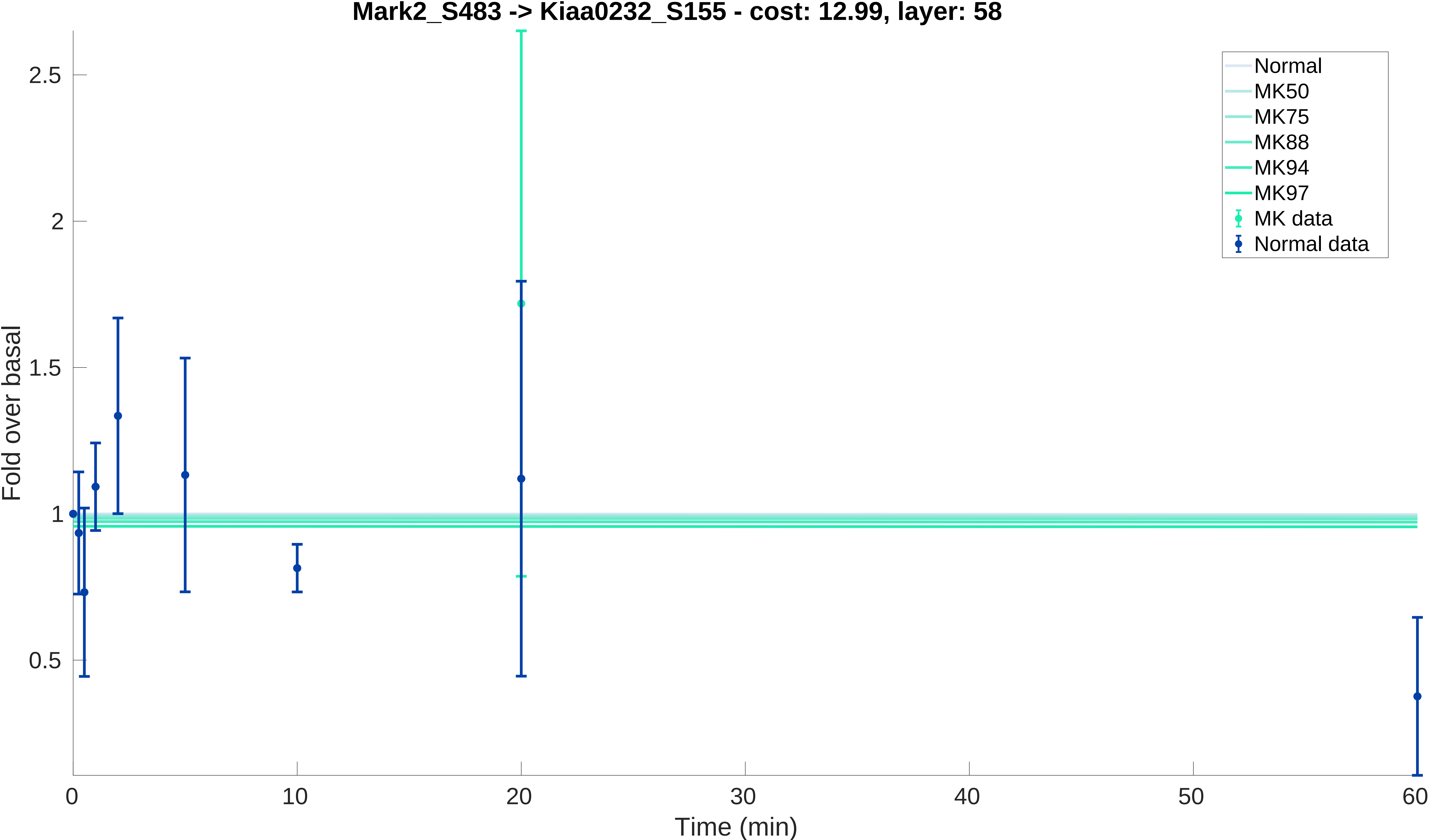

Mark2_S483 -> Kiaa0232_S155 - cost: 12.99, layer: 58
Fold over basal
Time (min)
0.5
1
1.5
2
2.5
0
10
20
30
40
50
60
Normal
MK50
MK75
MK88
MK94
MK97
MK data
Normal data

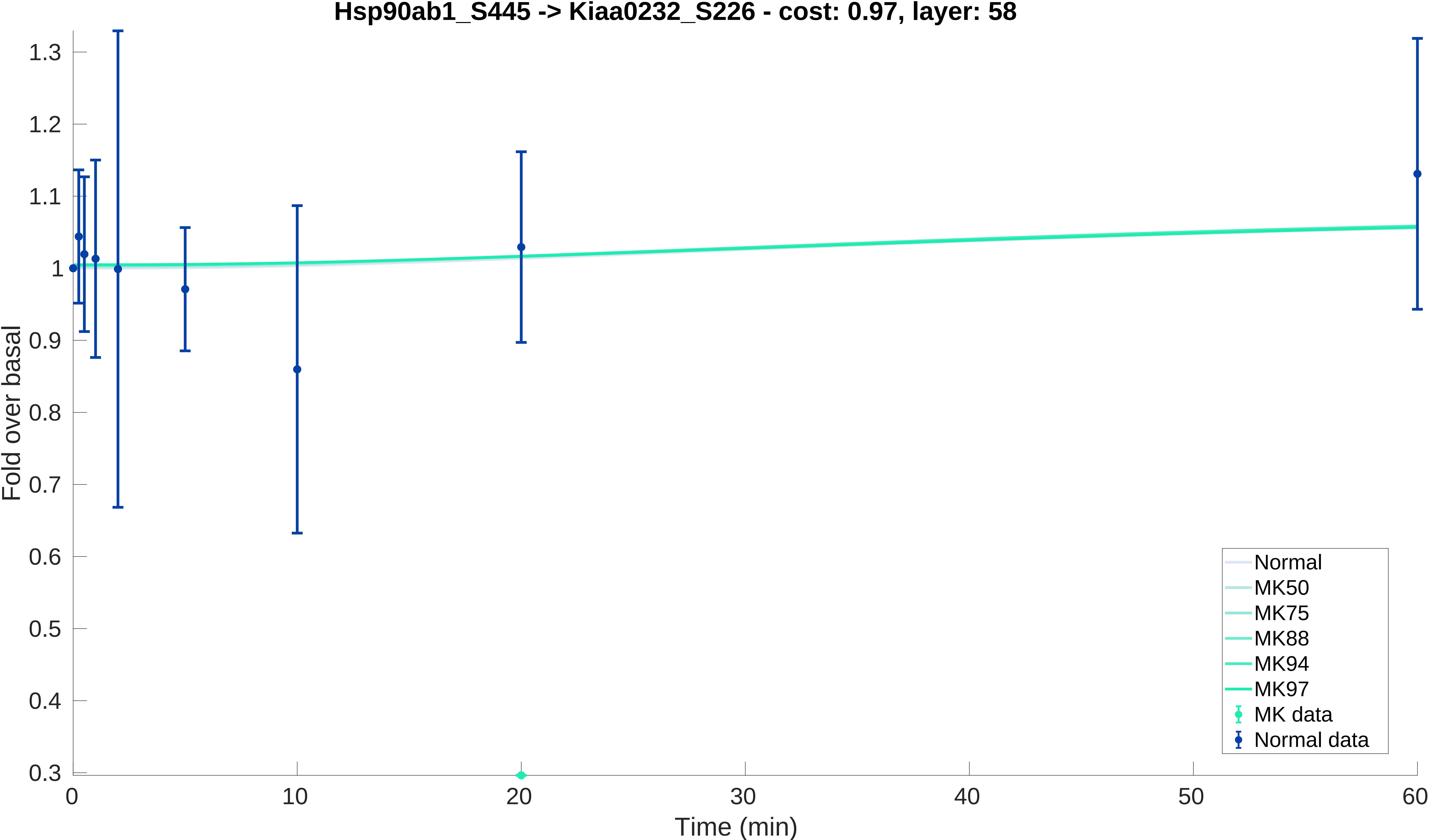

Hsp90ab1_S445 -> Kiaa0232_S226 - cost: 0.97, layer: 58
Fold over basal
1.3
1.2
1.1
1
0.9
0.8
0.7
0.6
0.5
0.4
0.3
0
10
20
30
40
50
60
Time (min)
Normal
MK50
MK75
MK88
MK94
MK97
MK data
Normal data

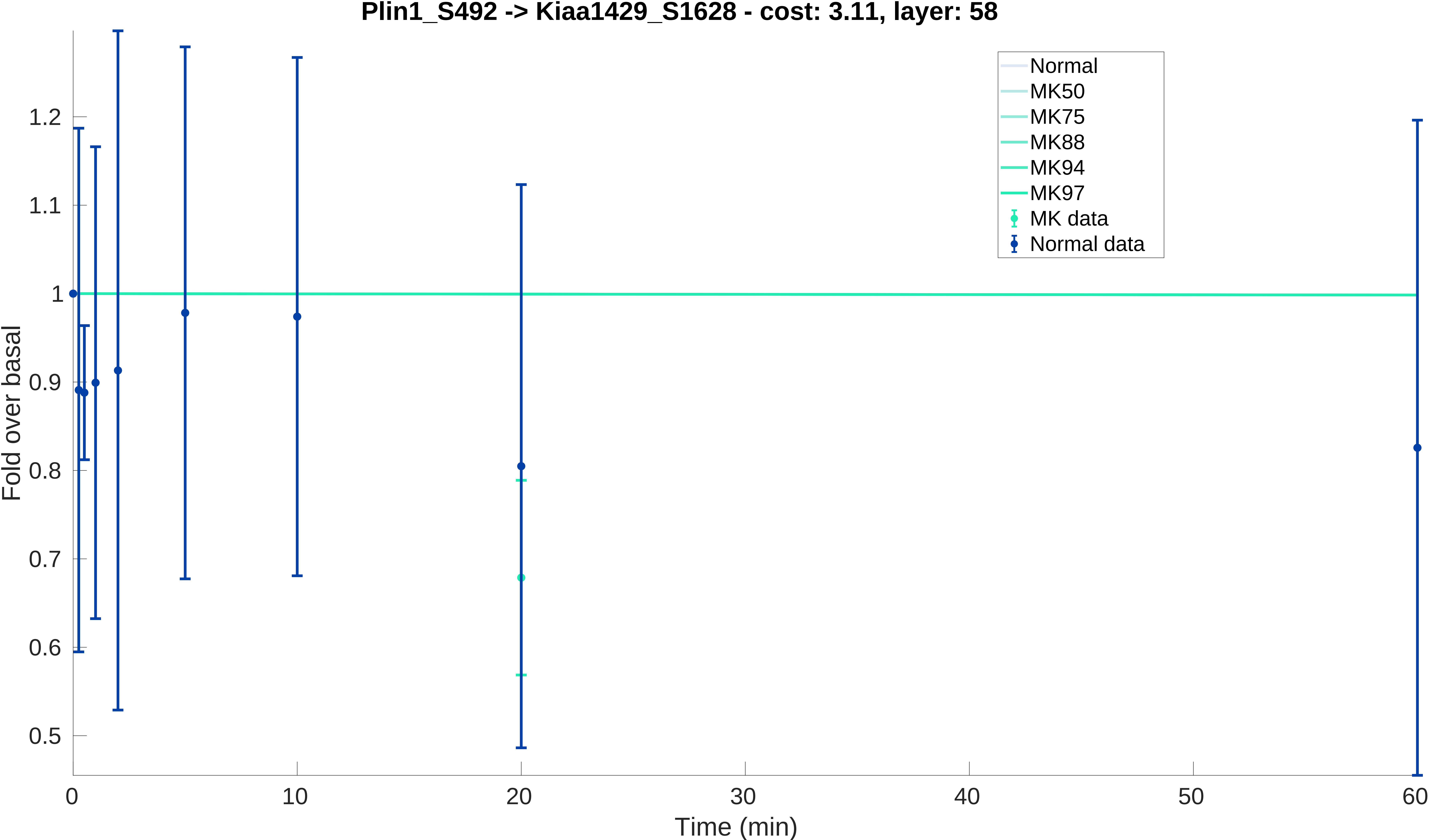
Plin1_S492 -> Kiaa1429_S1628 - cost: 3.11, layer: 58
Normal
MK50
MK75
MK88
MK94
MK97
MK data
Normal data
Fold over basal
1.2
1.1
1
0.9
0.8
0.7
0.6
0.5
0
10
20
30
40
50
60
Time (min)

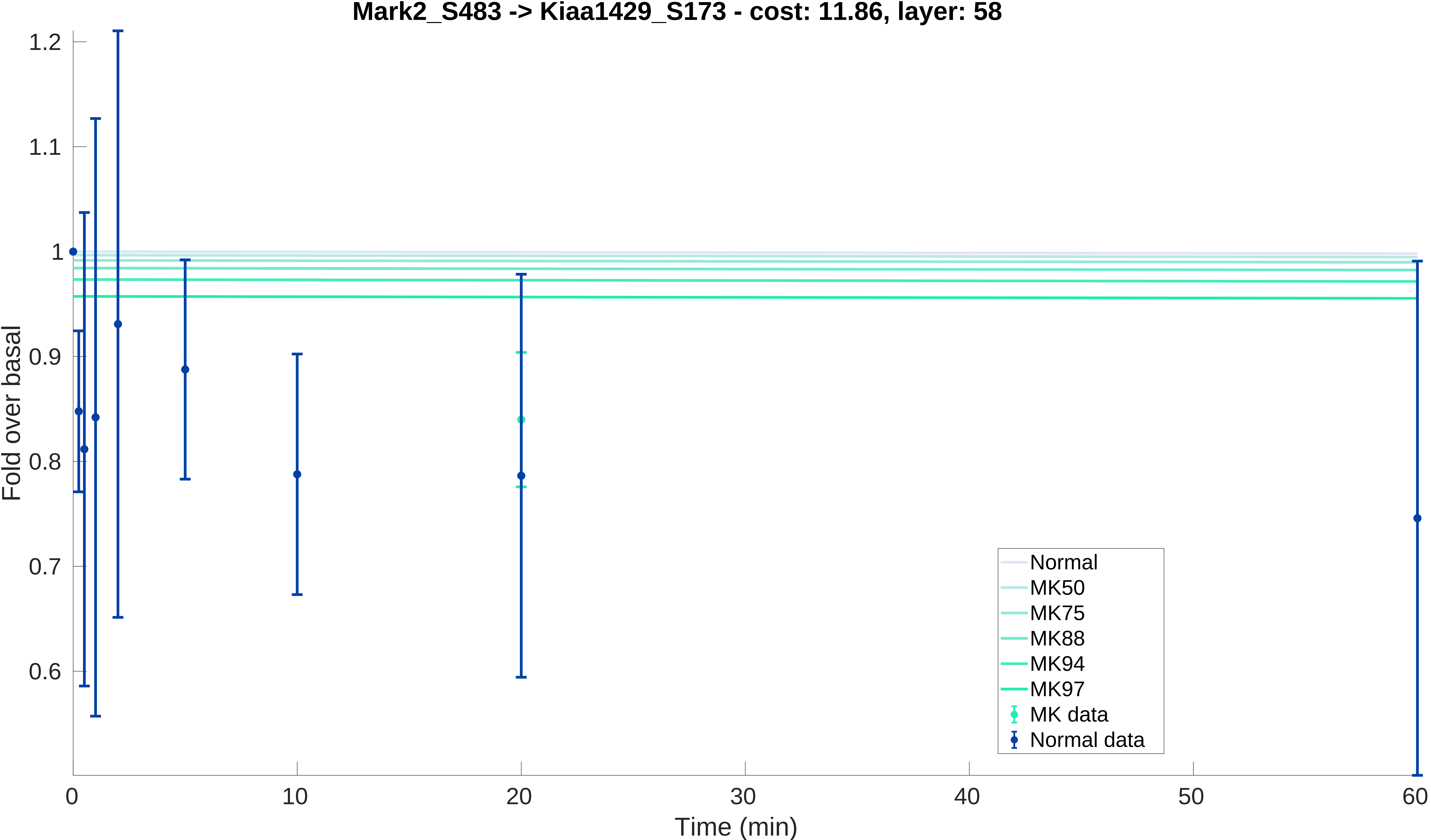

Mark2_S483 -> Kiaa1429_S173 - cost: 11.86, layer: 58
Fold over basal
Time (min)
1.2
1.1
1
0.9
0.8
0.7
0.6
0
10
20
30
40
50
60
Normal
MK50
MK75
MK88
MK94
MK97
MK data
Normal data

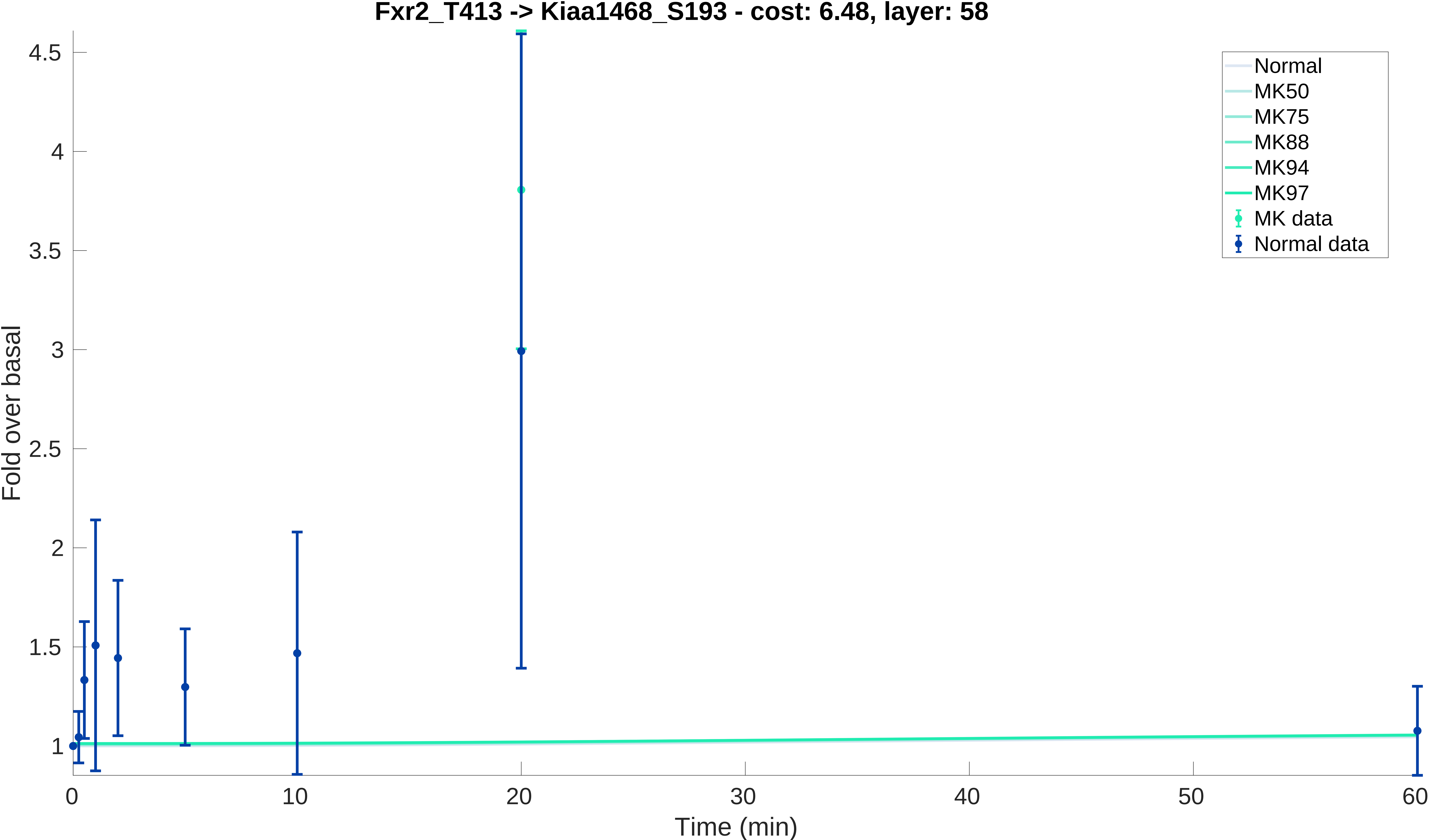

Fxr2_T413 -> Kiaa1468_S193 - cost: 6.48, layer: 58
Fold over basal
Time (min)
4.5
4
3.5
3
2.5
2
1.5
1
0
10
20
30
40
50
60
Normal
MK50
MK75
MK88
MK94
MK97
MK data
Normal data

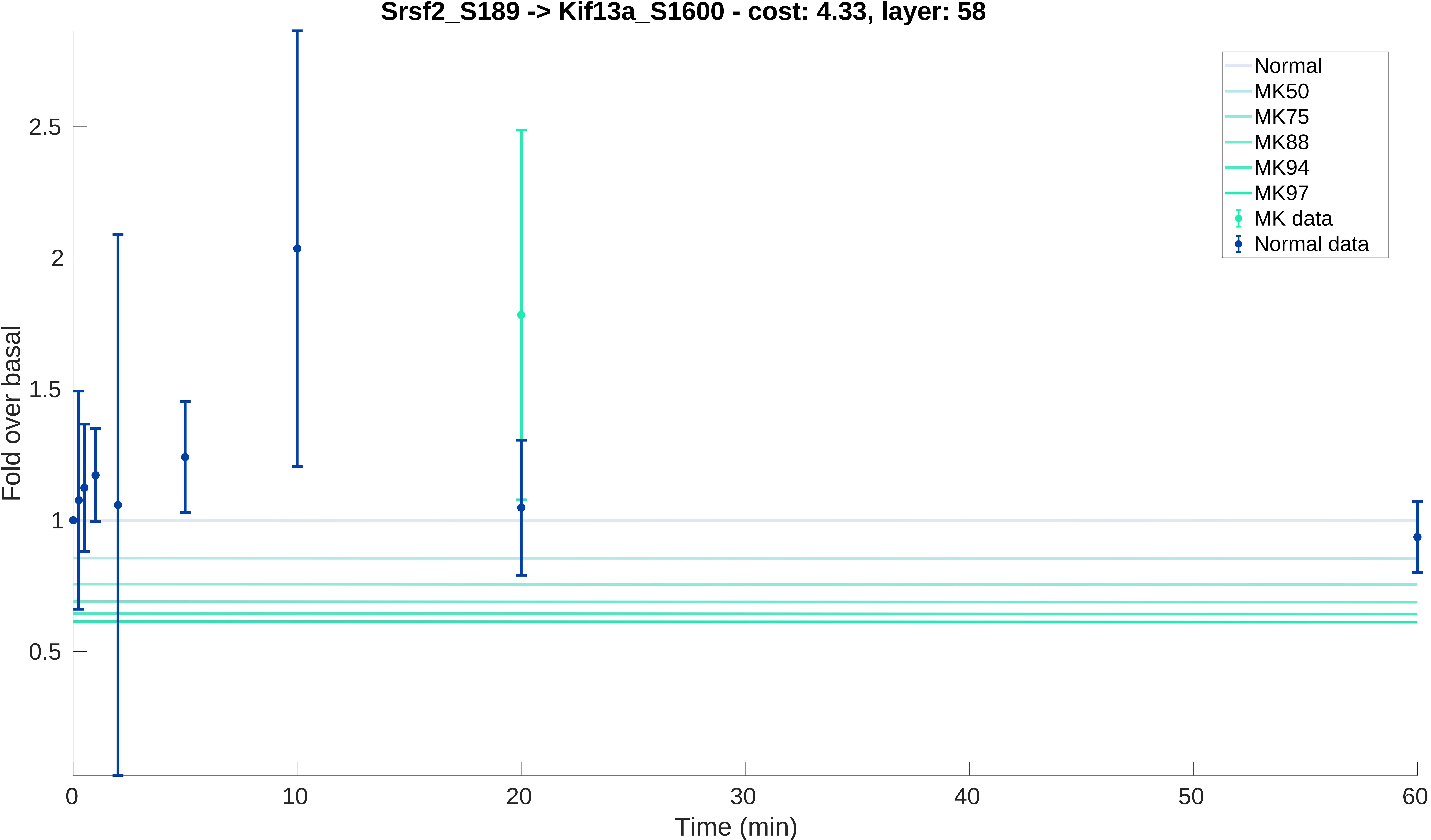
Srsf2_S189 -> Kif13a_S1600 - cost: 4.33, layer: 58
Fold over basal
Time (min)
0.5
1
1.5
2
2.5
0
10
20
30
40
50
60
Normal
MK50
MK75
MK88
MK94
MK97
MK data
Normal data

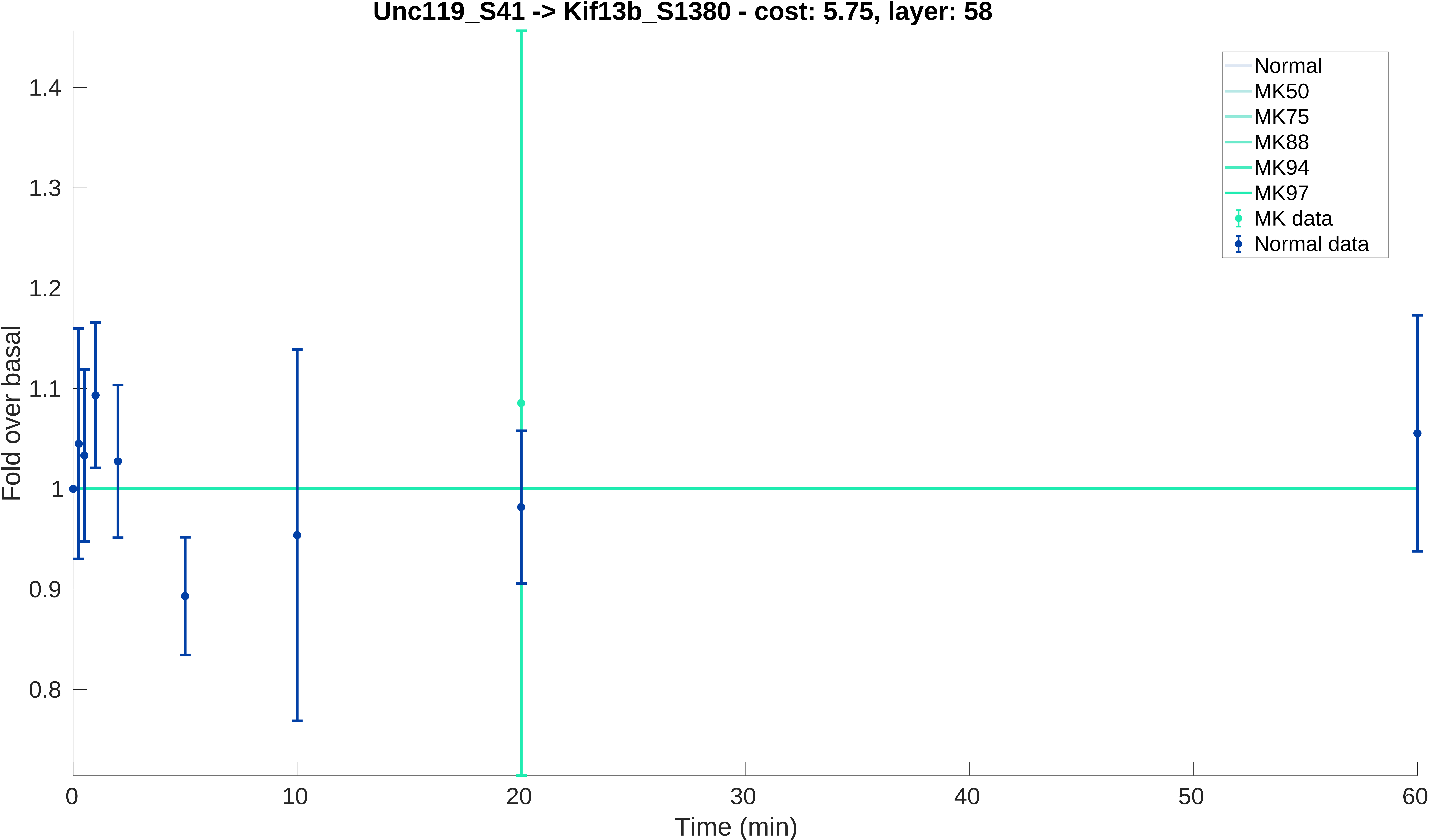

Unc119_S41 -> Kif13b_S1380 - cost: 5.75, layer: 58
Fold over basal
Time (min)
1.4
1.3
1.2
1.1
1
0.9
0.8
0
10
20
30
40
50
60
Normal
MK50
MK75
MK88
MK94
MK97
MK data
Normal data

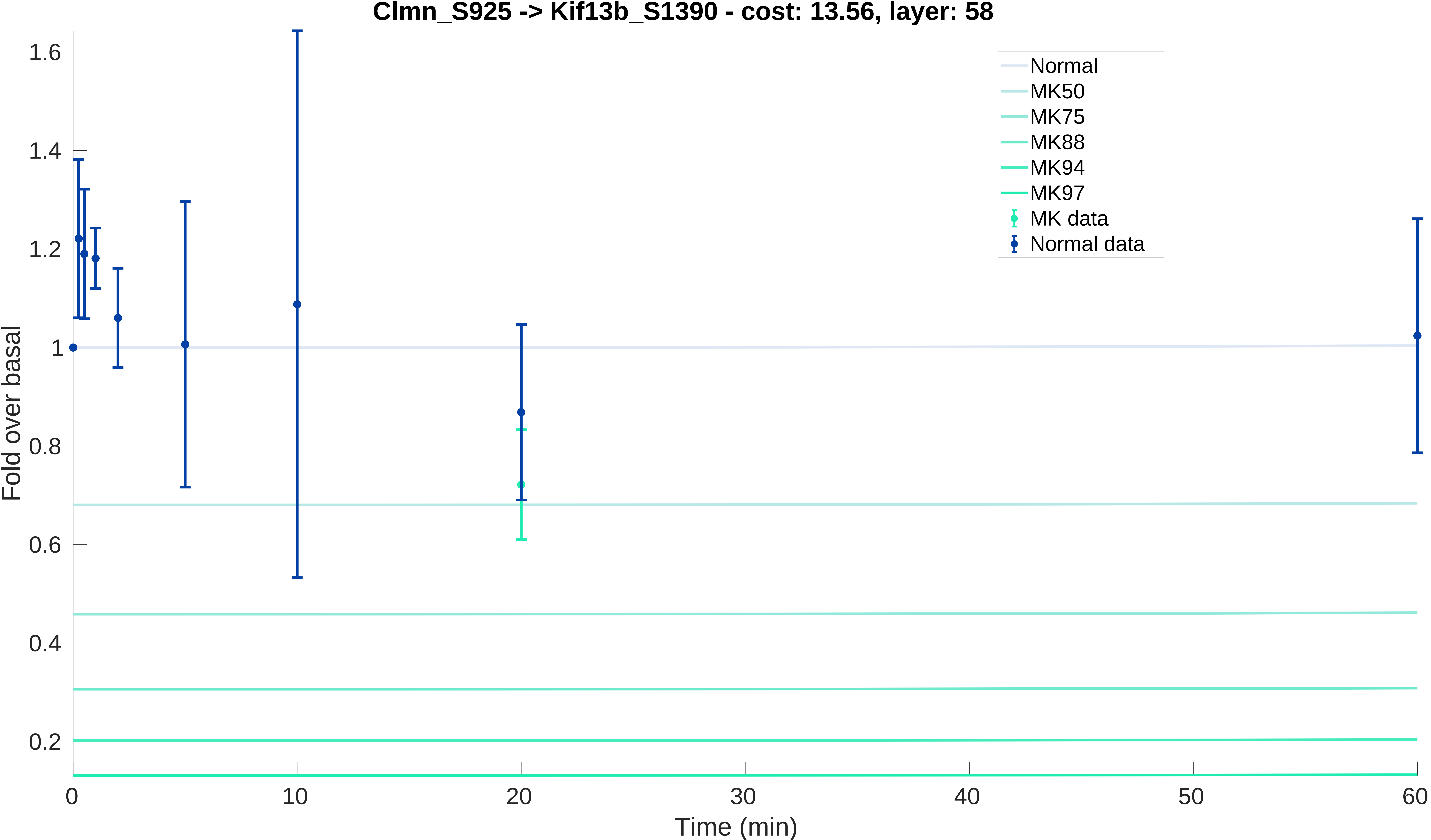

Clmn_S925 -> Kif13b_S1390 - cost: 13.56, layer: 58
Fold over basal
Time (min)
0
10
20
30
40
50
60
0.2
0.4
0.6
0.8
1
1.2
1.4
1.6
Normal
MK50
MK75
MK88
MK94
MK97
MK data
Normal data

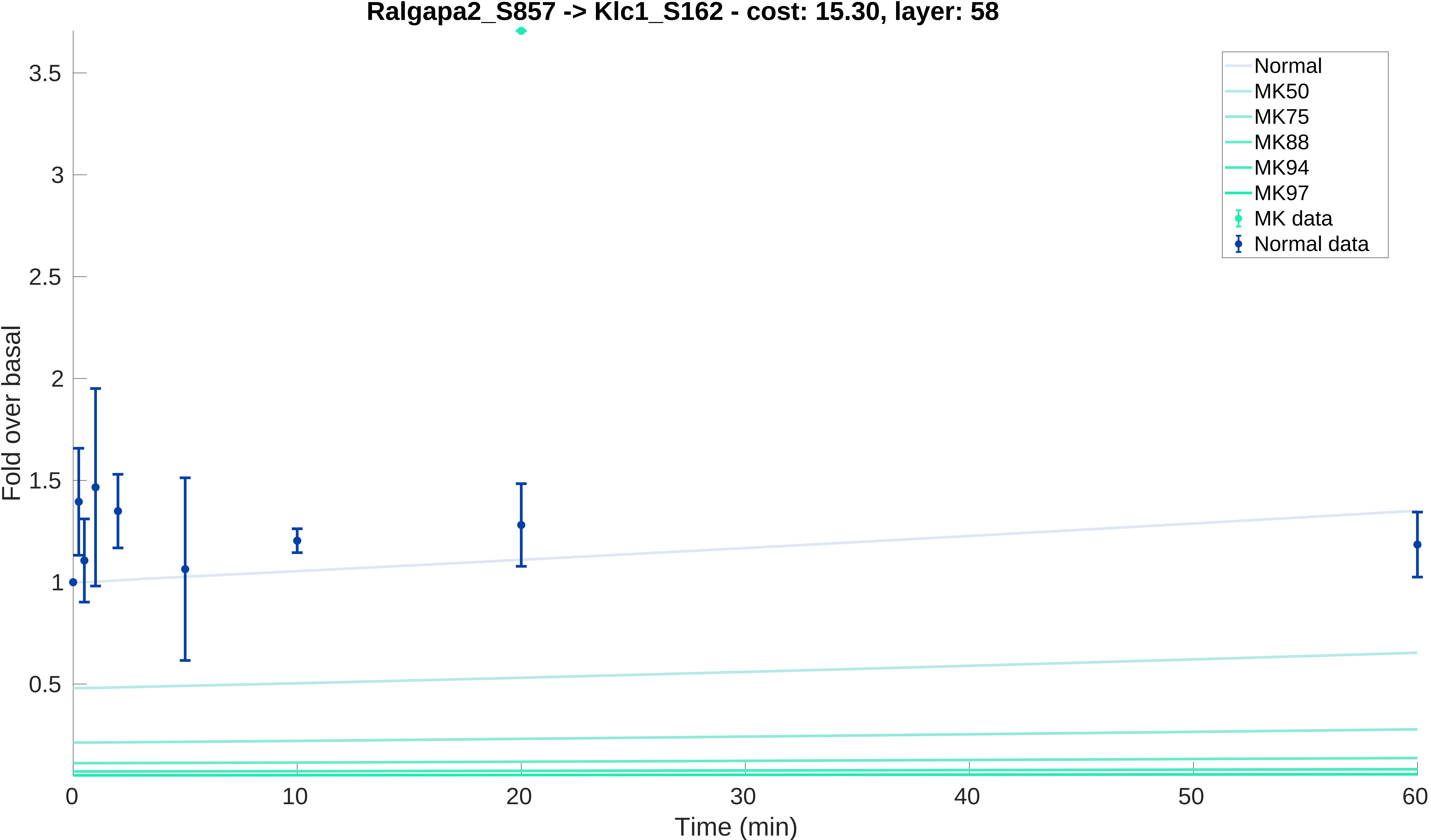
Ralgapa2_S857 -> Klc1_S162 - cost: 15.30, layer: 58
Fold over basal
Time (min)
0
10
20
30
40
50
60
0.5
1
1.5
2
2.5
3
3.5
Normal
MK50
MK75
MK88
MK94
MK97
MK data
Normal data

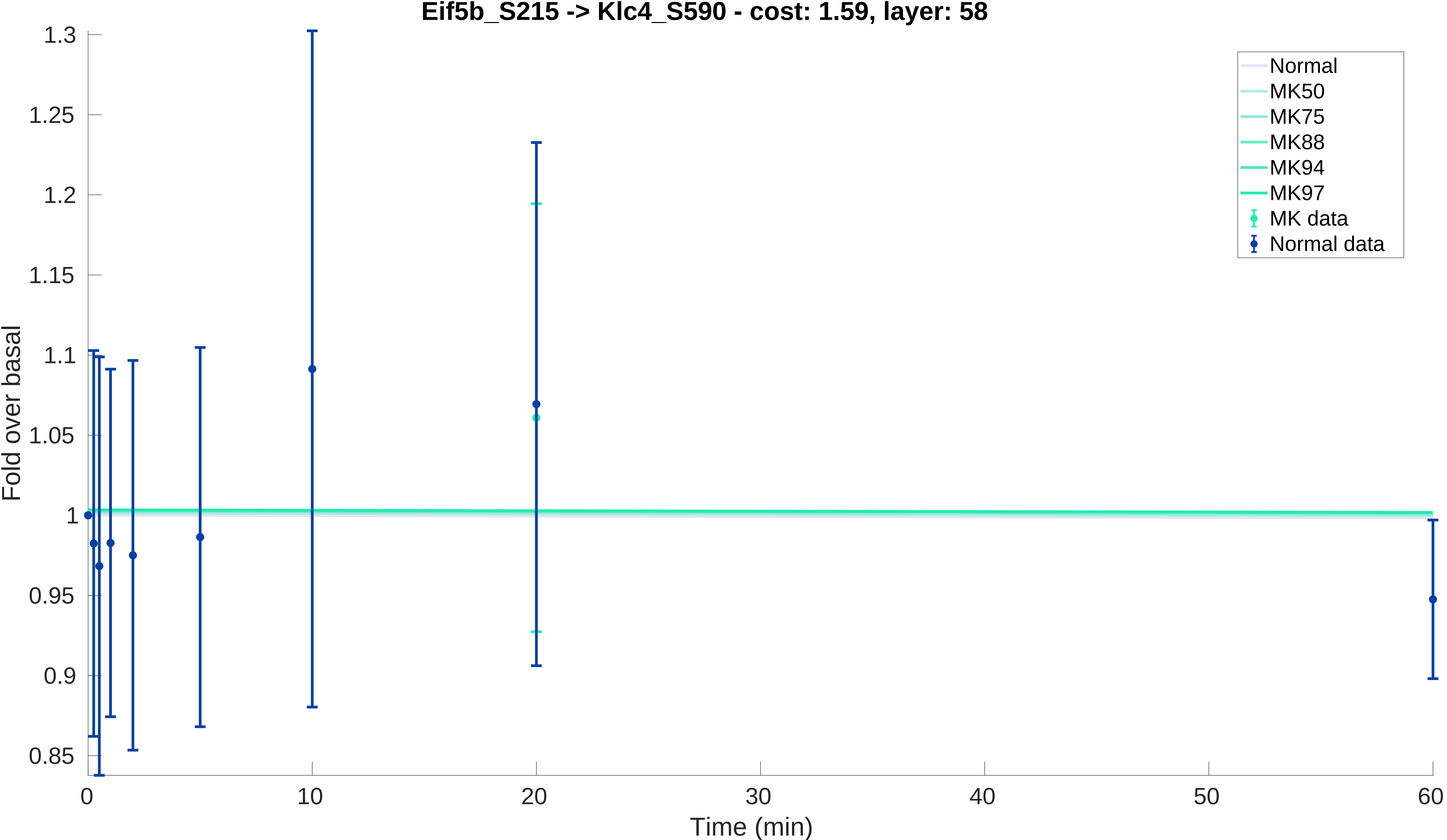

Eif5b_S215 -> Klc4_S590 - cost: 1.59, layer: 58
Fold over basal
1.3
1.25
1.2
1.15
1.1
1.05
1
0.95
0.9
0.85
0
10
20
30
40
50
60
Time (min)
Normal
MK50
MK75
MK88
MK94
MK97
MK data
Normal data

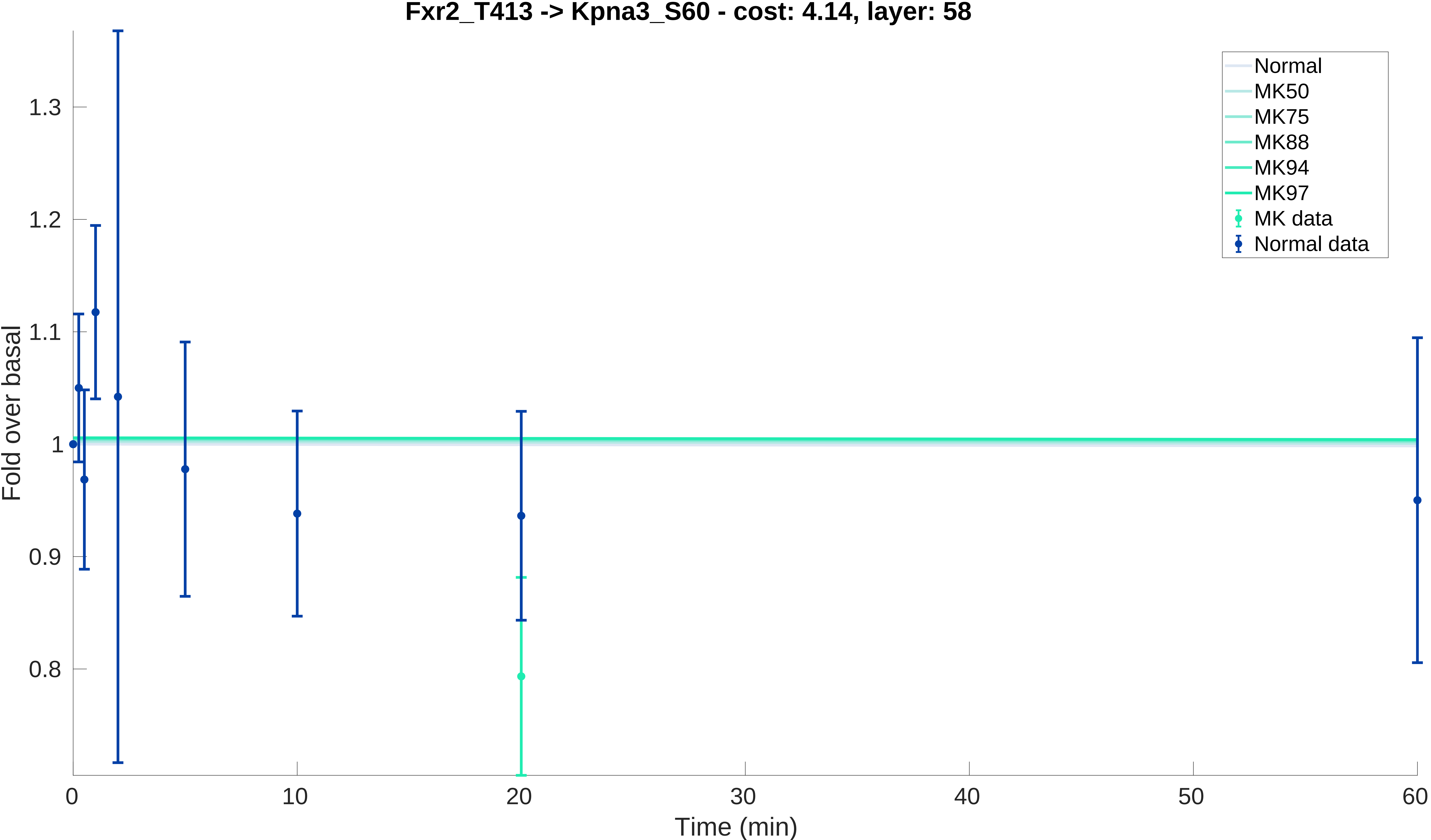

Fxr2_T413 -> Kpna3_S60 - cost: 4.14, layer: 58
Normal
MK50
MK75
MK88
MK94
MK97
MK data
Normal data
Fold over basal
1.3
1.2
1.1
1
0.9
0.8
0
10
20
30
40
50
60
Time (min)

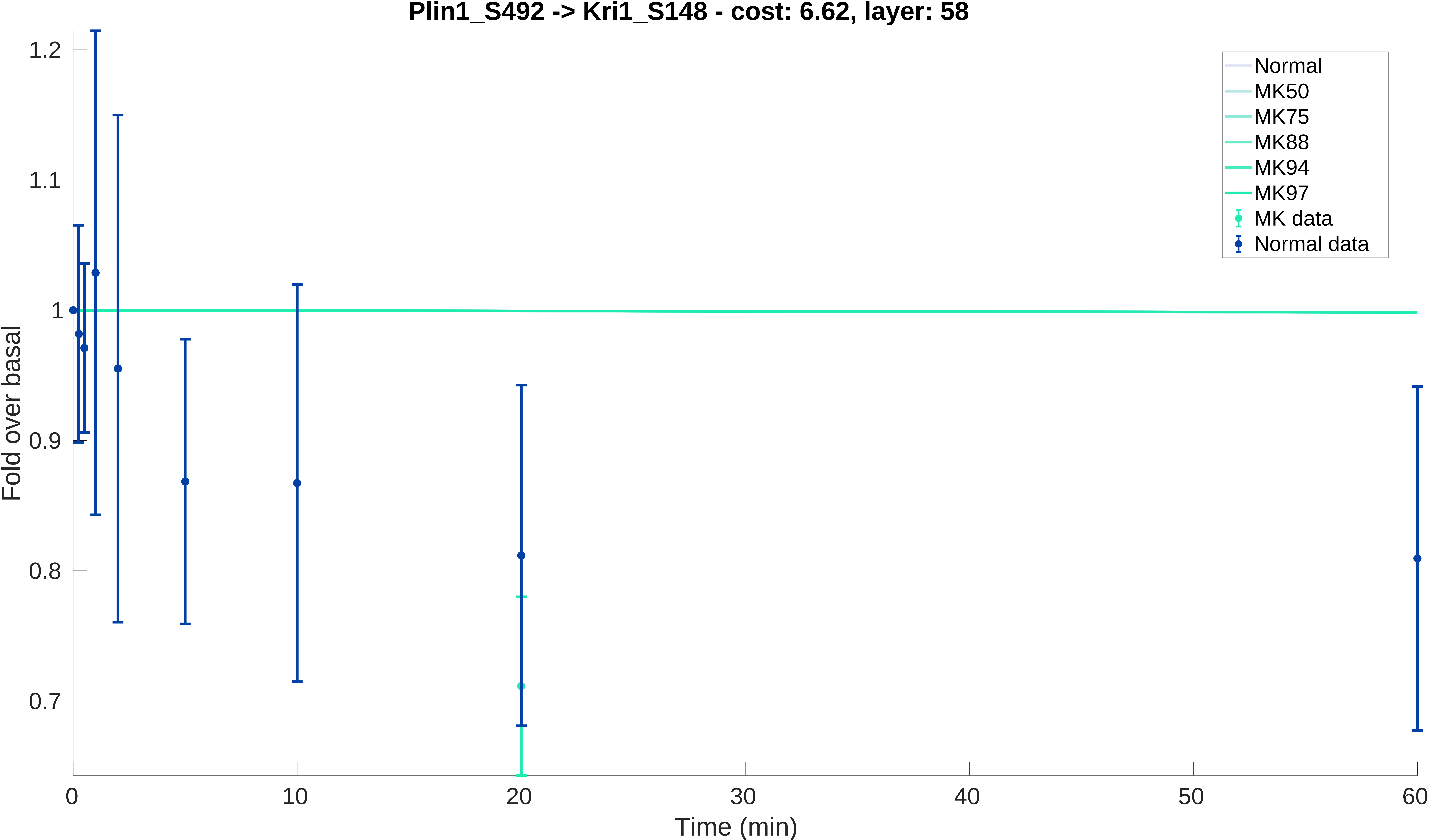

Plin1_S492 -> Kri1_S148 - cost: 6.62, layer: 58
Fold over basal
Time (min)
1.2
1.1
1
0.9
0.8
0.7
0
10
20
30
40
50
60
Normal
MK50
MK75
MK88
MK94
MK97
MK data
Normal data

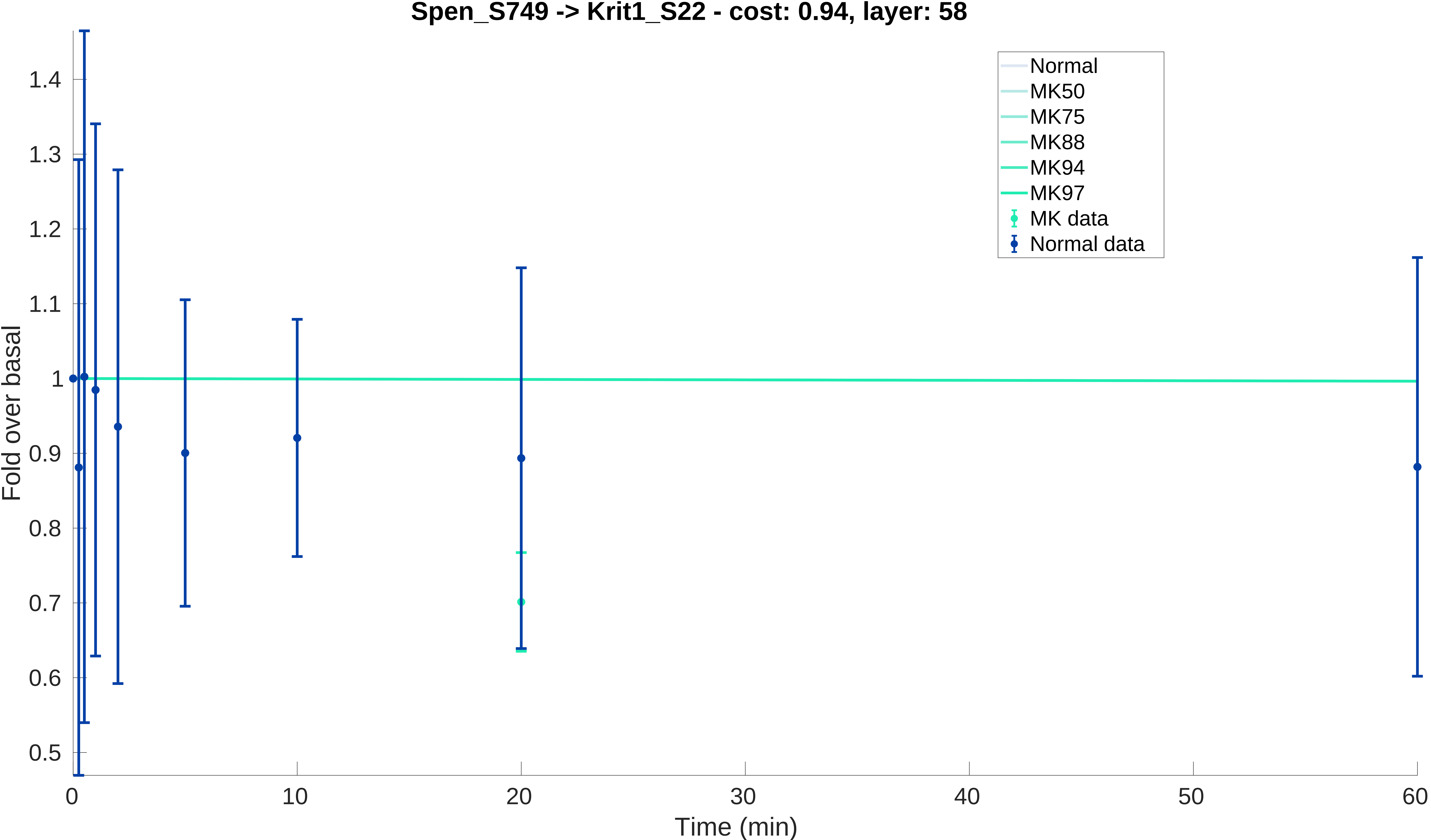

Spen_S749 -> Krit1_S22 - cost: 0.94, layer: 58
Normal
MK50
MK75
MK88
MK94
MK97
MK data
Normal data
Fold over basal
1.4
1.3
1.2
1.1
1
0.9
0.8
0.7
0.6
0.5
0
10
20
30
40
50
60
Time (min)

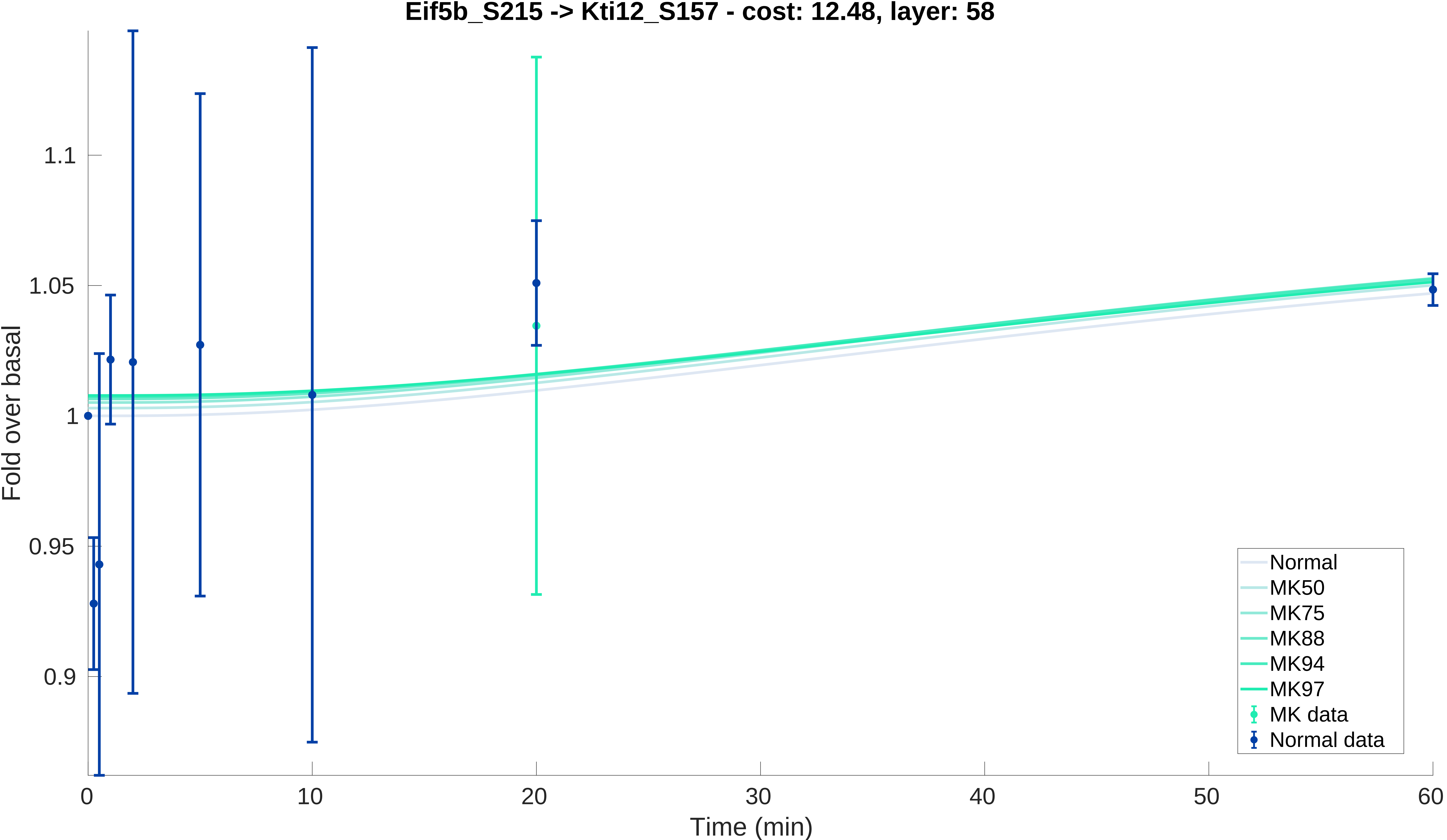

Eif5b_S215 -> Kti12_S157 - cost: 12.48, layer: 58
Fold over basal
1.1
1.05
1
0.95
0.9
0
10
20
30
40
50
60
Time (min)
Normal
MK50
MK75
MK88
MK94
MK97
MK data
Normal data

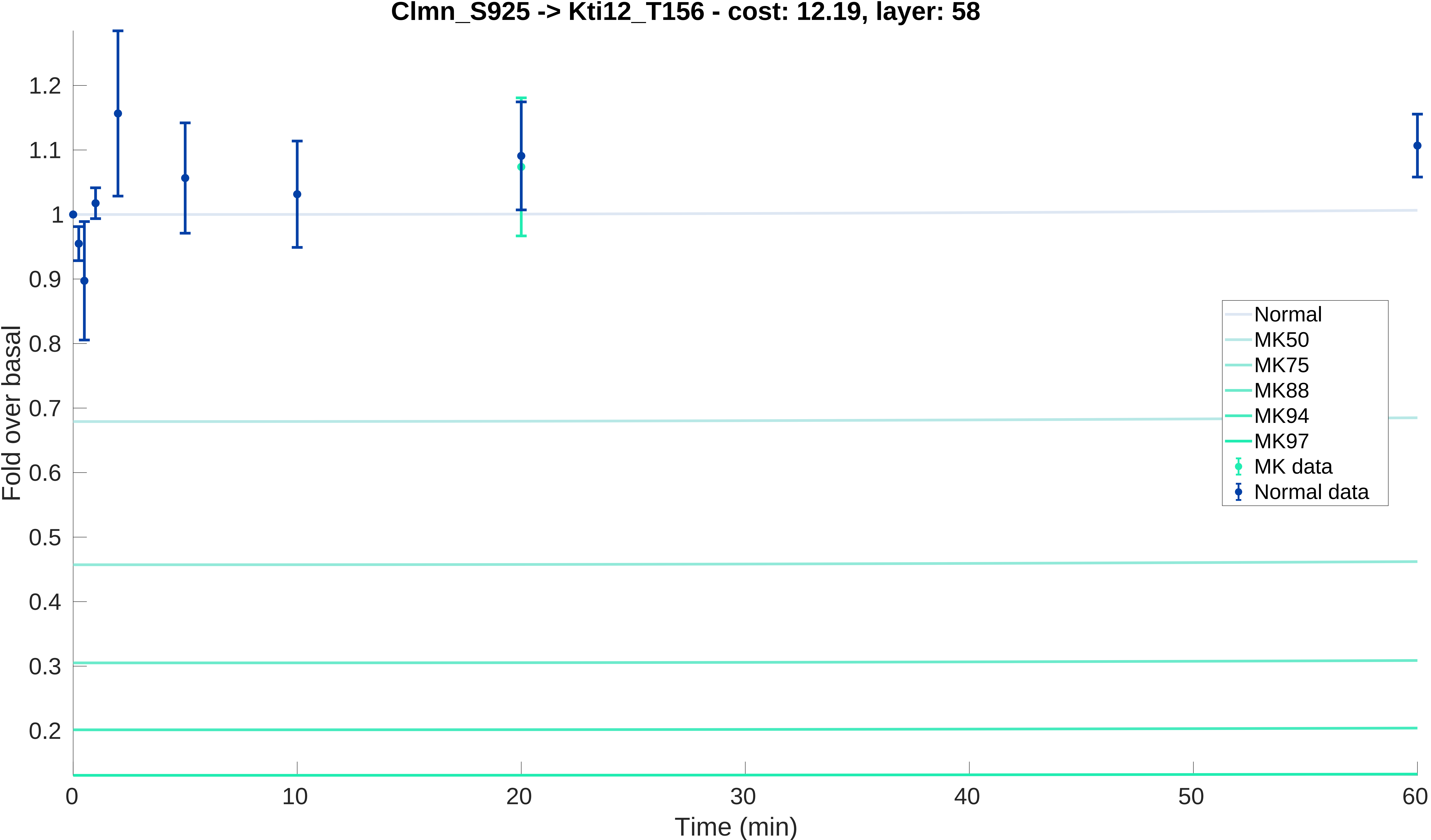

Clmn_S925 -> Kti12_T156 - cost: 12.19, layer: 58
Fold over basal
Time (min)
0
10
20
30
40
50
60
0.2
0.3
0.4
0.5
0.6
0.7
0.8
0.9
1
1.1
1.2
Normal
MK50
MK75
MK88
MK94
MK97
MK data
Normal data

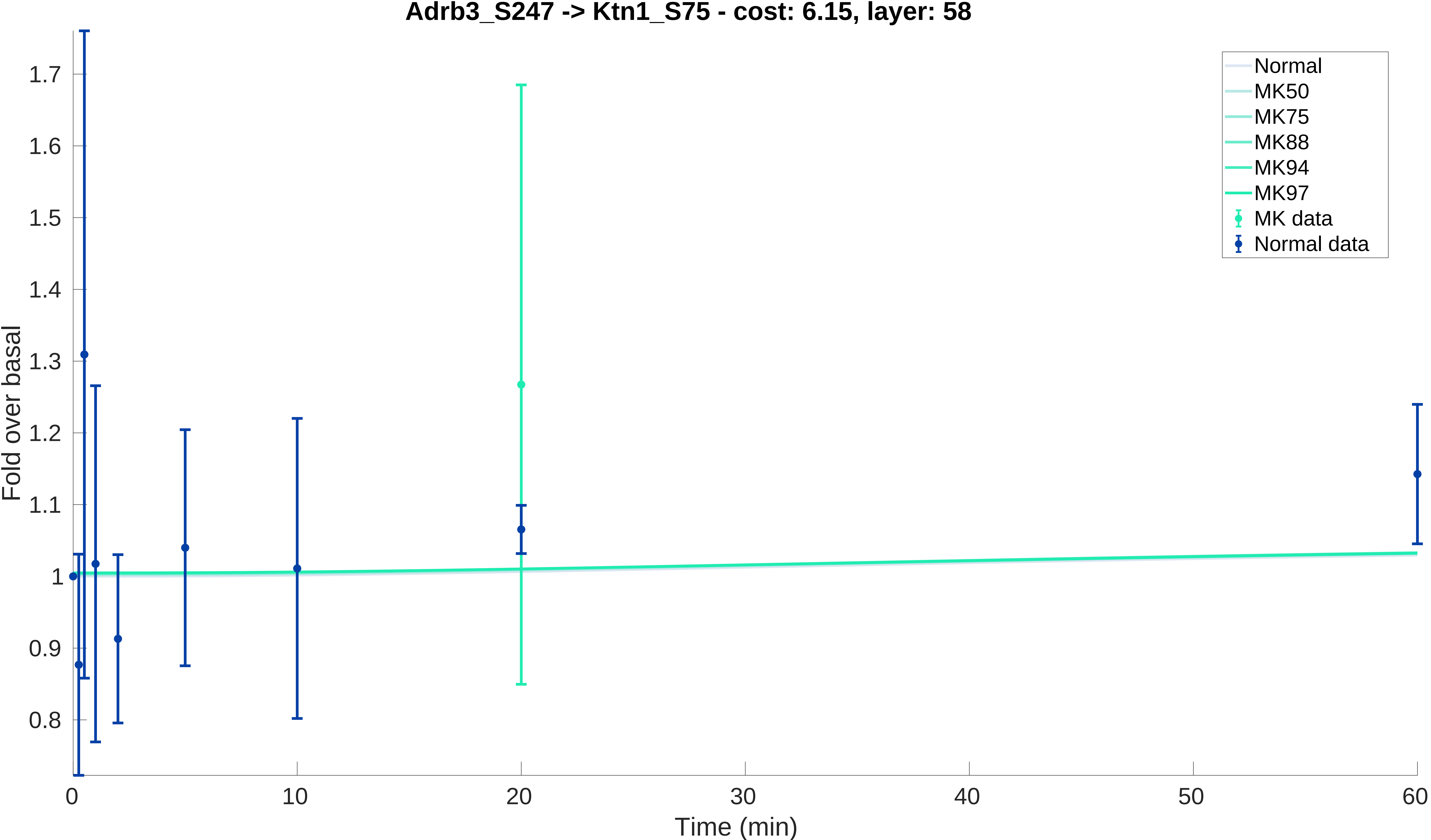

Adrb3_S247 -> Ktn1_S75 - cost: 6.15, layer: 58
Fold over basal
1.7
1.6
1.5
1.4
1.3
1.2
1.1
1
0.9
0.8
0
10
20
30
40
50
60
Time (min)
Normal
MK50
MK75
MK88
MK94
MK97
MK data
Normal data

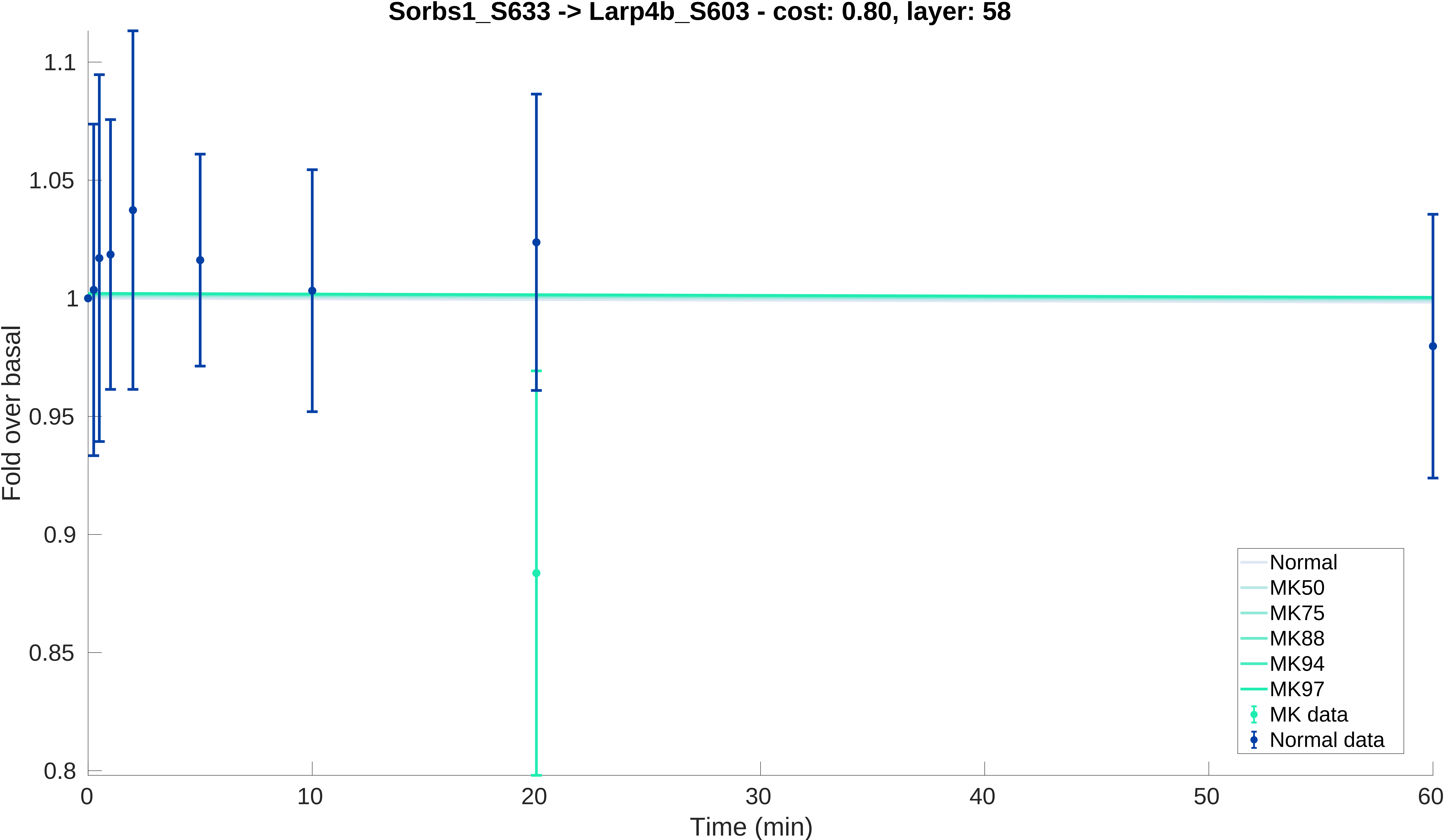
Sorbs1_S633 -> Larp4b_S603 - cost: 0.80, layer: 58
Fold over basal
1.1
1.05
1
0.95
0.9
0.85
0.8
0
10
20
30
40
50
60
Time (min)
Normal
MK50
MK75
MK88
MK94
MK97
MK data
Normal data

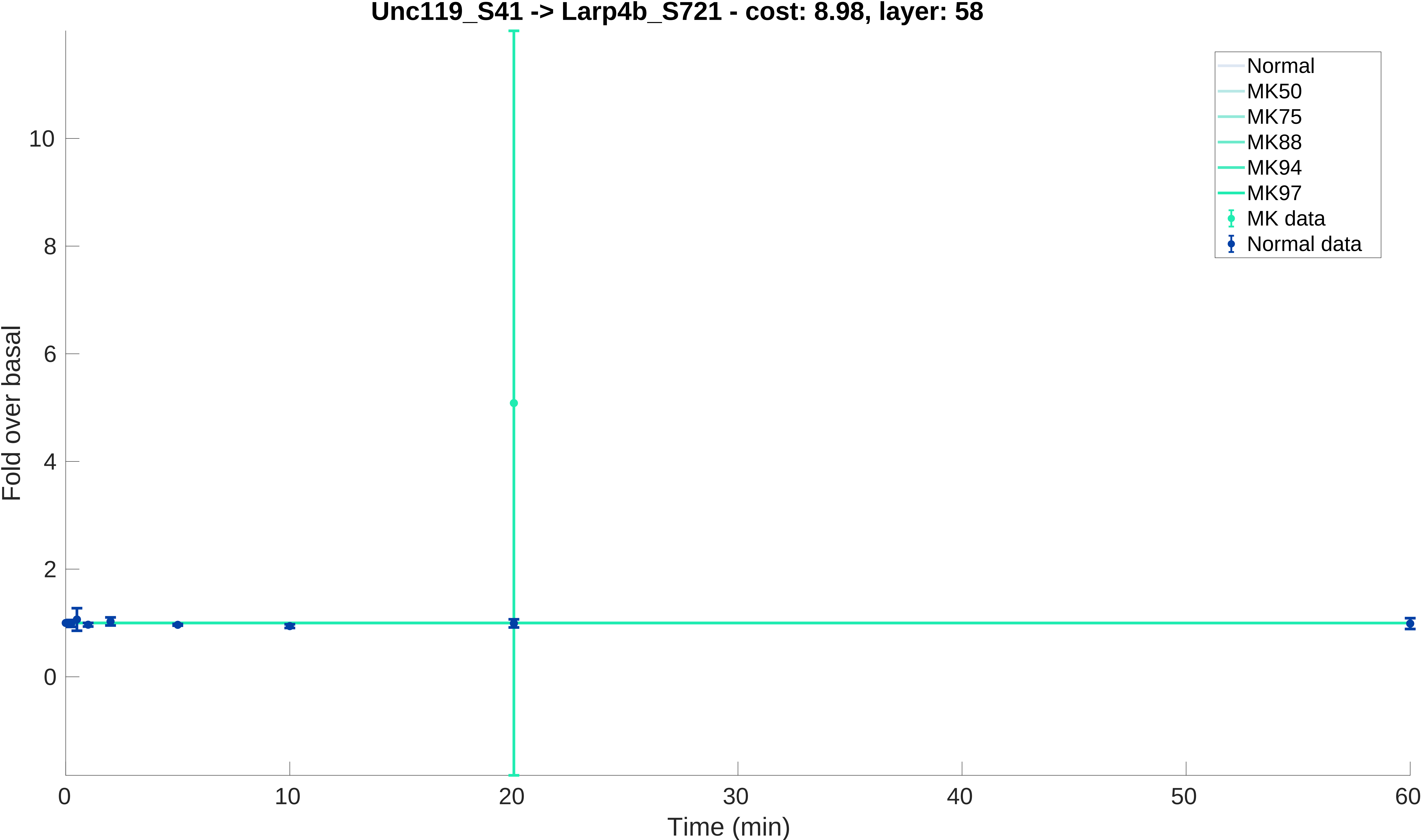

Unc119_S41 -> Larp4b_S721 - cost: 8.98, layer: 58
Fold over basal
Time (min)
Normal
MK50
MK75
MK88
MK94
MK97
MK data
Normal data
0
2
4
6
8
10
0
10
20
30
40
50
60

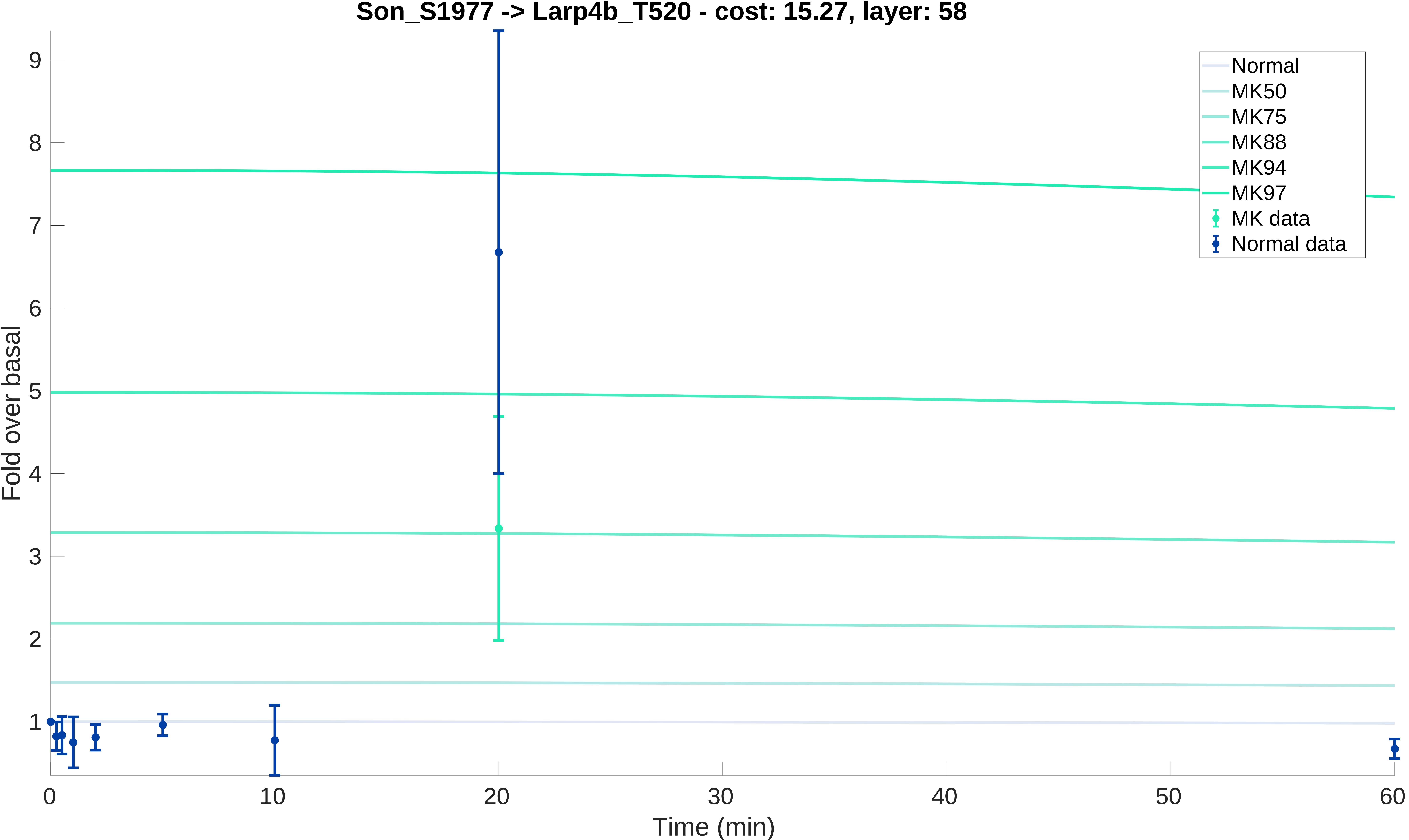
Son_S1977 -> Larp4b_T520 - cost: 15.27, layer: 58
Fold over basal
Time (min)
0
10
20
30
40
50
60
1
2
3
4
5
6
7
8
9
Normal
MK50
MK75
MK88
MK94
MK97
MK data
Normal data

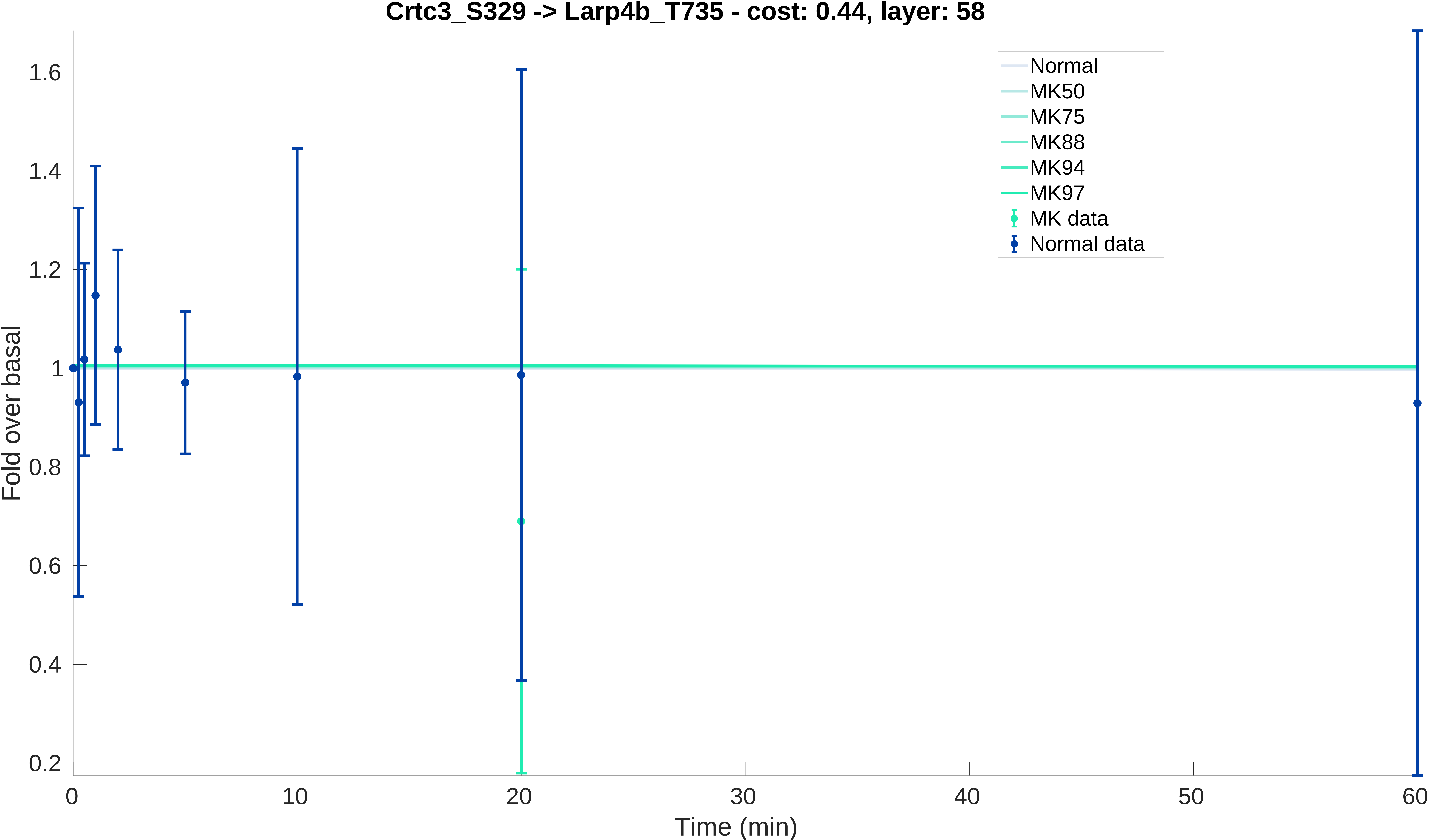

Crtc3_S329 -> Larp4b_T735 - cost: 0.44, layer: 58
Fold over basal
Time (min)
1.6
1.4
1.2
1
0.8
0.6
0.4
0.2
0
10
20
30
40
50
60
Normal
MK50
MK75
MK88
MK94
MK97
MK data
Normal data

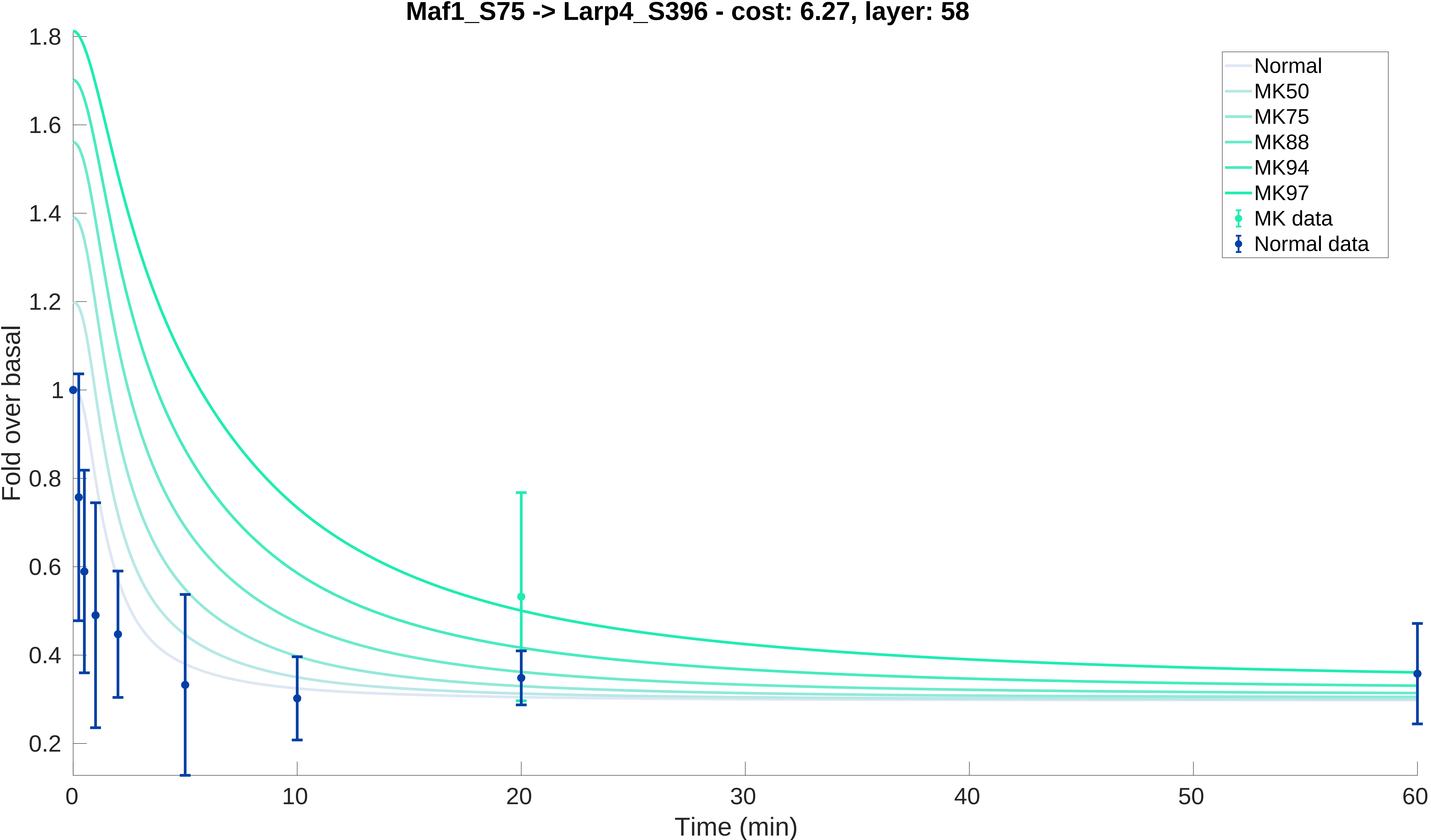

Maf1_S75 -> Larp4_S396 - cost: 6.27, layer: 58
Fold over basal
Time (min)
0
10
20
30
40
50
60
0.2
0.4
0.6
0.8
1
1.2
1.4
1.6
1.8
Normal
MK50
MK75
MK88
MK94
MK97
MK data
Normal data

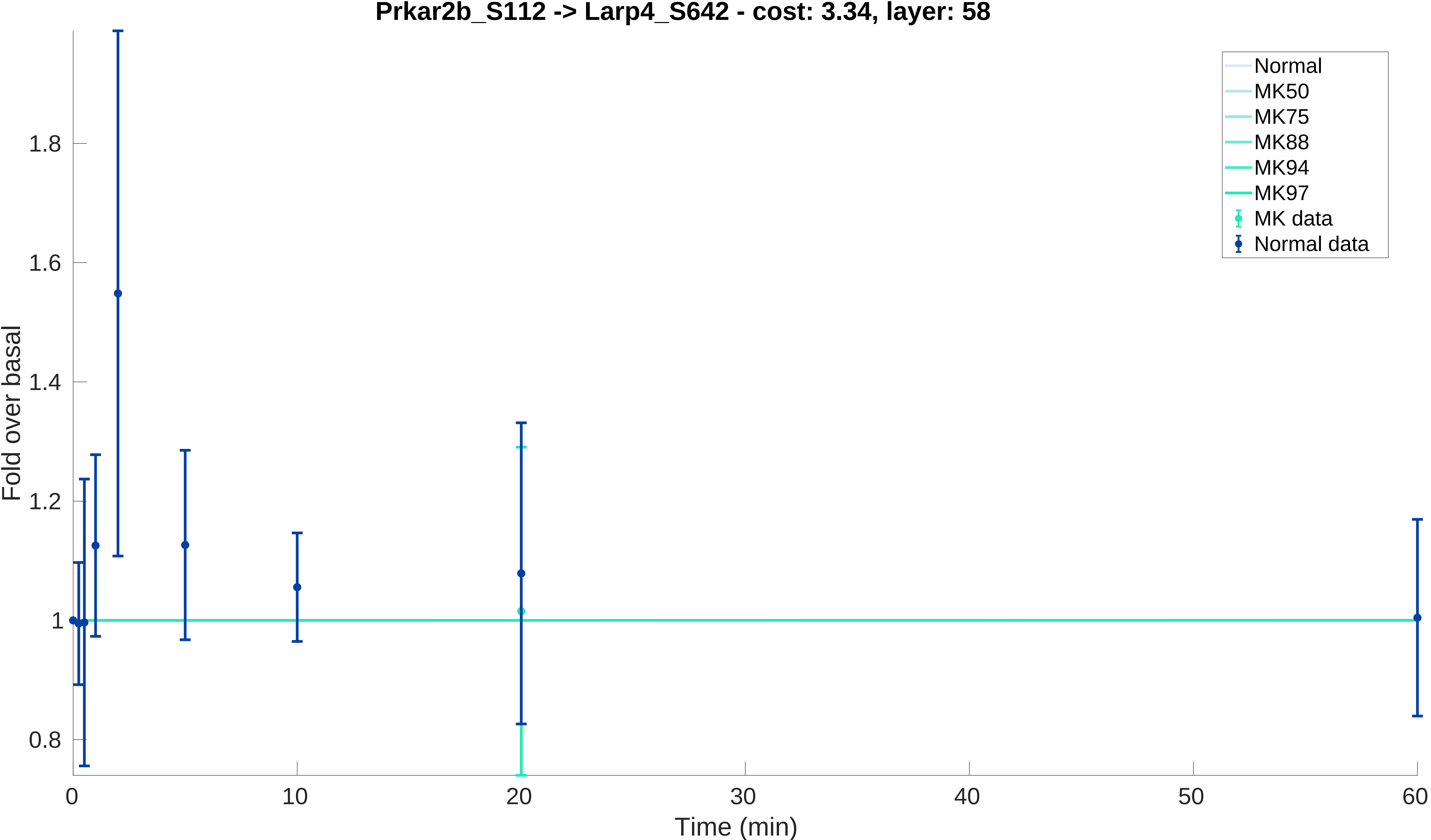

Prkar2b_S112 -> Larp4_S642 - cost: 3.34, layer: 58
Fold over basal
Time (min)
0.8
1
1.2
1.4
1.6
1.8
0
10
20
30
40
50
60
Normal
MK50
MK75
MK88
MK94
MK97
MK data
Normal data

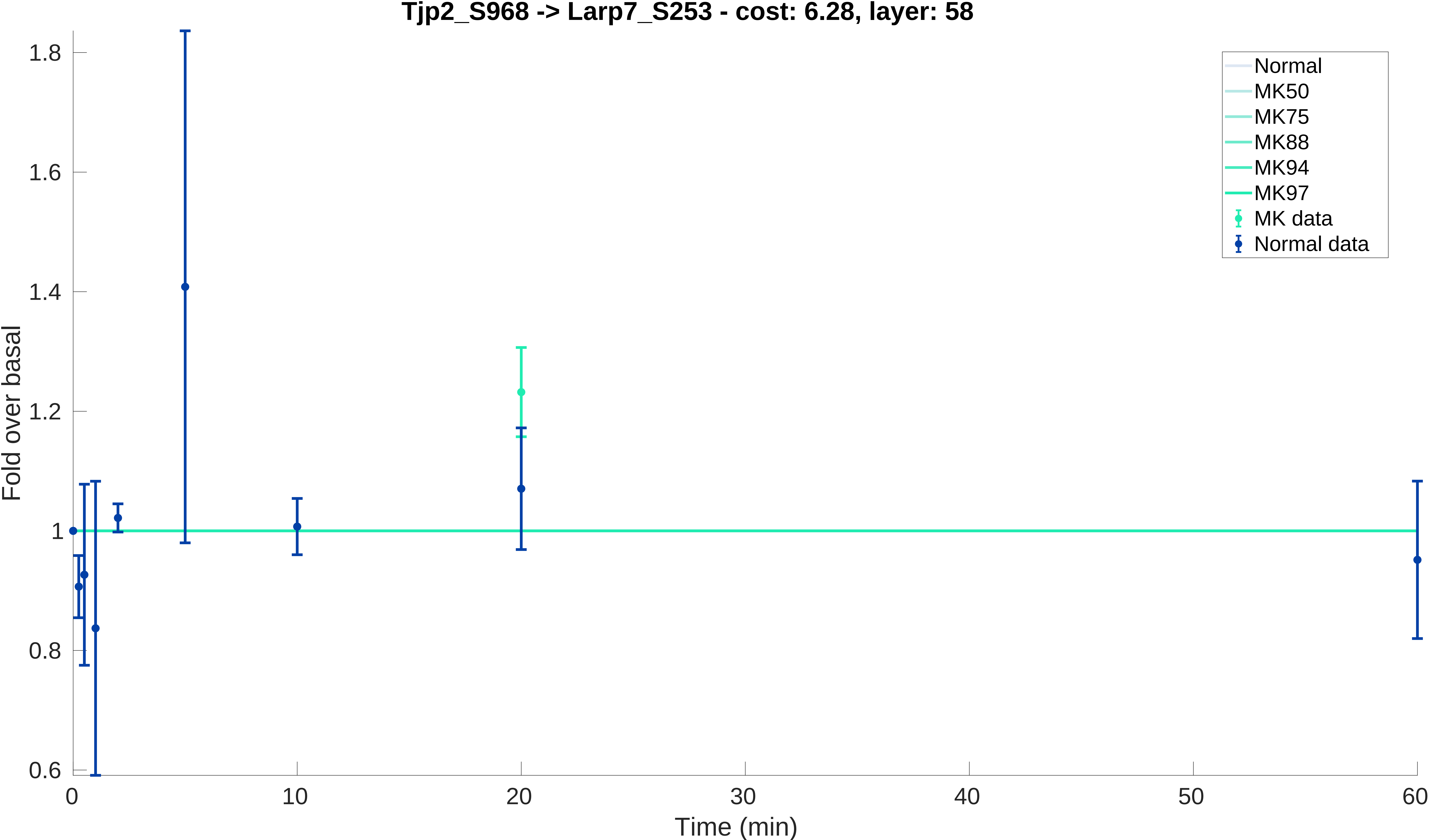

Tjp2_S968 -> Larp7_S253 - cost: 6.28, layer: 58
Fold over basal
Time (min)
1.8
1.6
1.4
1.2
1
0.8
0.6
0
10
20
30
40
50
60
Normal
MK50
MK75
MK88
MK94
MK97
MK data
Normal data

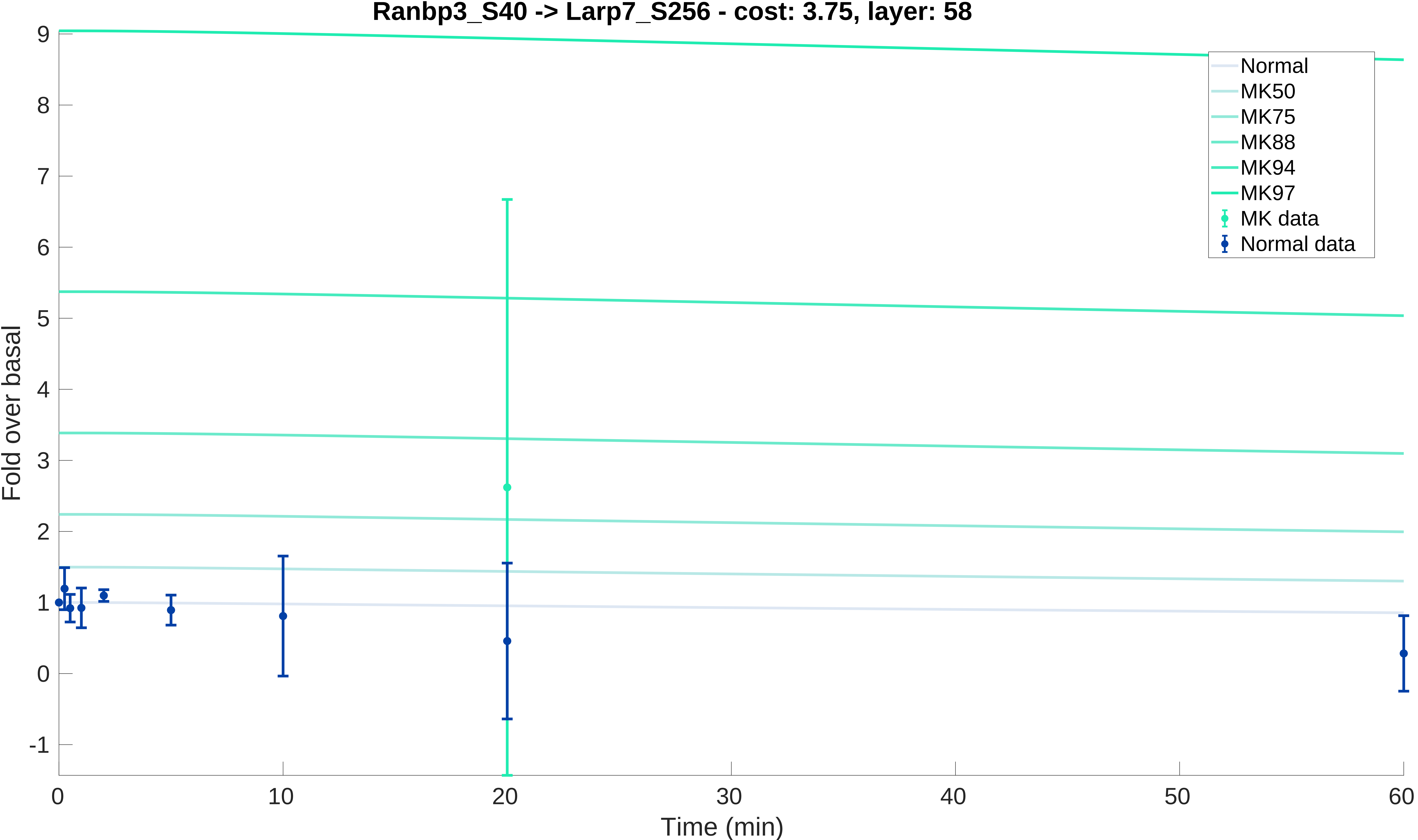

Ranbp3_S40 -> Larp7_S256 - cost: 3.75, layer: 58
Fold over basal
Time (min)
9
8
7
6
5
4
3
2
1
0
-1
0
10
20
30
40
50
60
Normal
MK50
MK75
MK88
MK94
MK97
MK data
Normal data

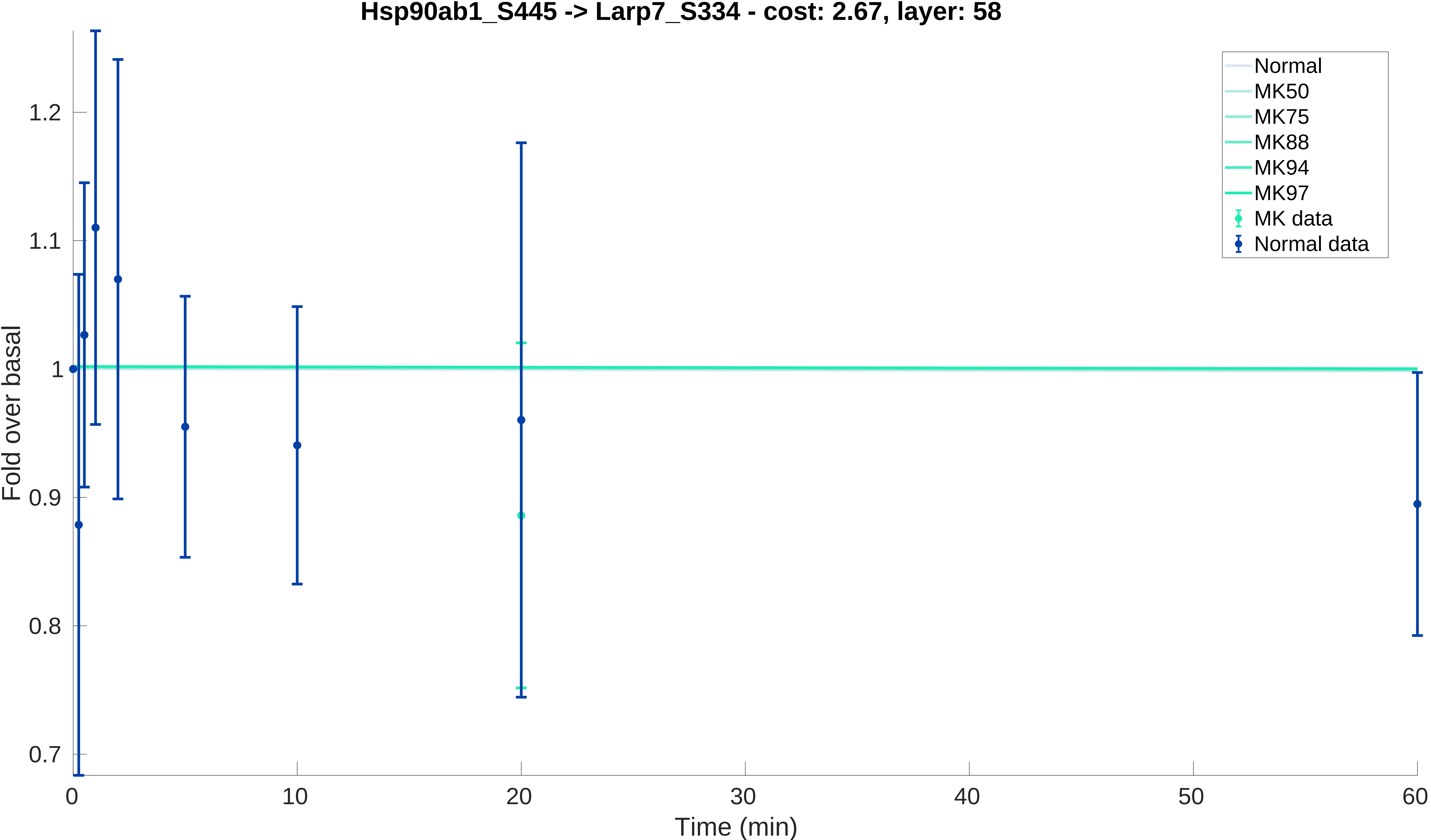
Hsp90ab1_S445 -> Larp7_S334 - cost: 2.67, layer: 58
Fold over basal
1.2
1.1
1
0.9
0.8
0.7
0
10
20
30
40
50
60
Time (min)
Normal
MK50
MK75
MK88
MK94
MK97
MK data
Normal data

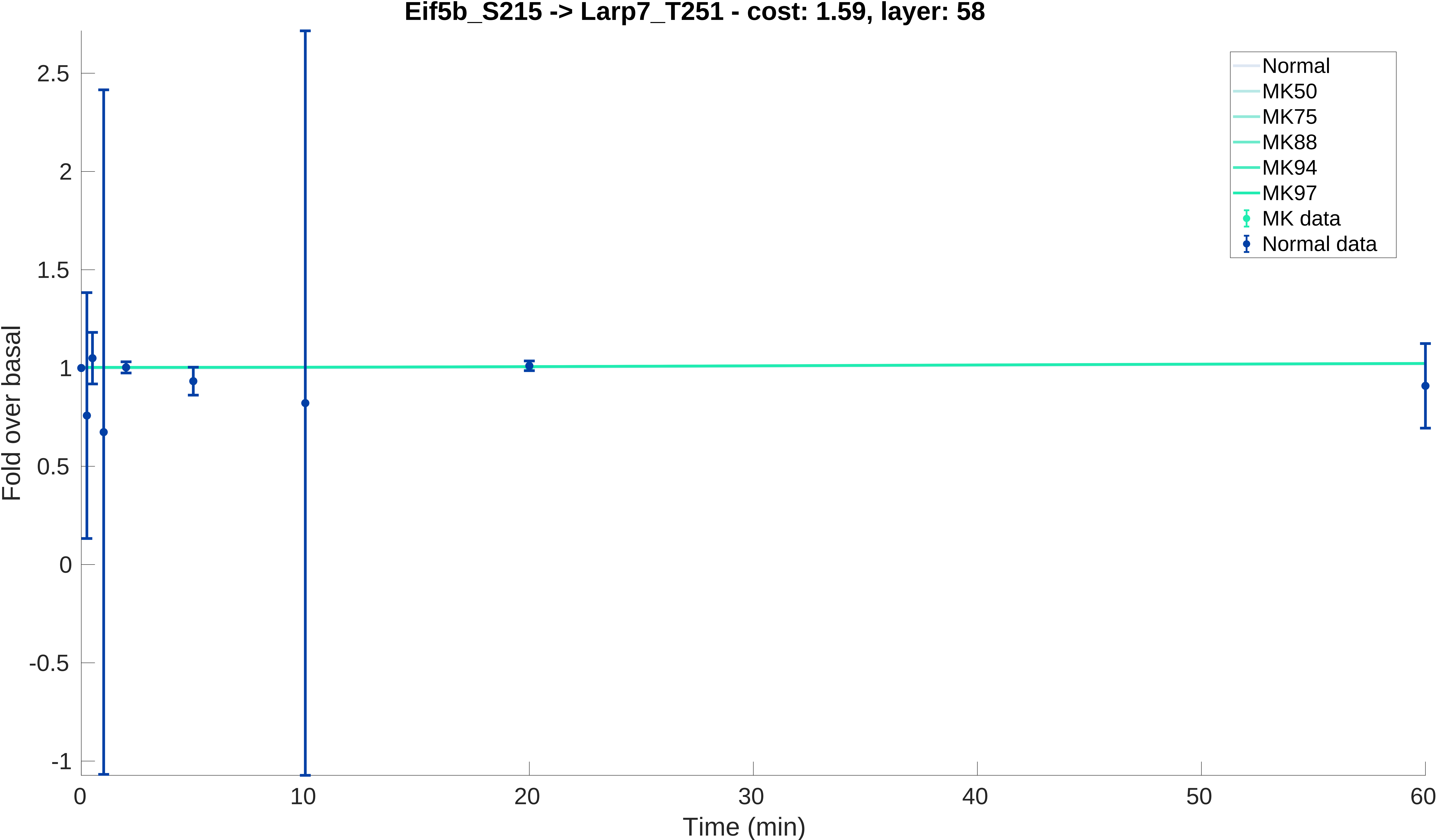

Eif5b_S215 -> Larp7_T251 - cost: 1.59, layer: 58
Normal
MK50
MK75
MK88
MK94
MK97
MK data
Normal data
Fold over basal
Time (min)
2.5
2
1.5
1
0.5
0
-0.5
-1
0
10
20
30
40
50
60

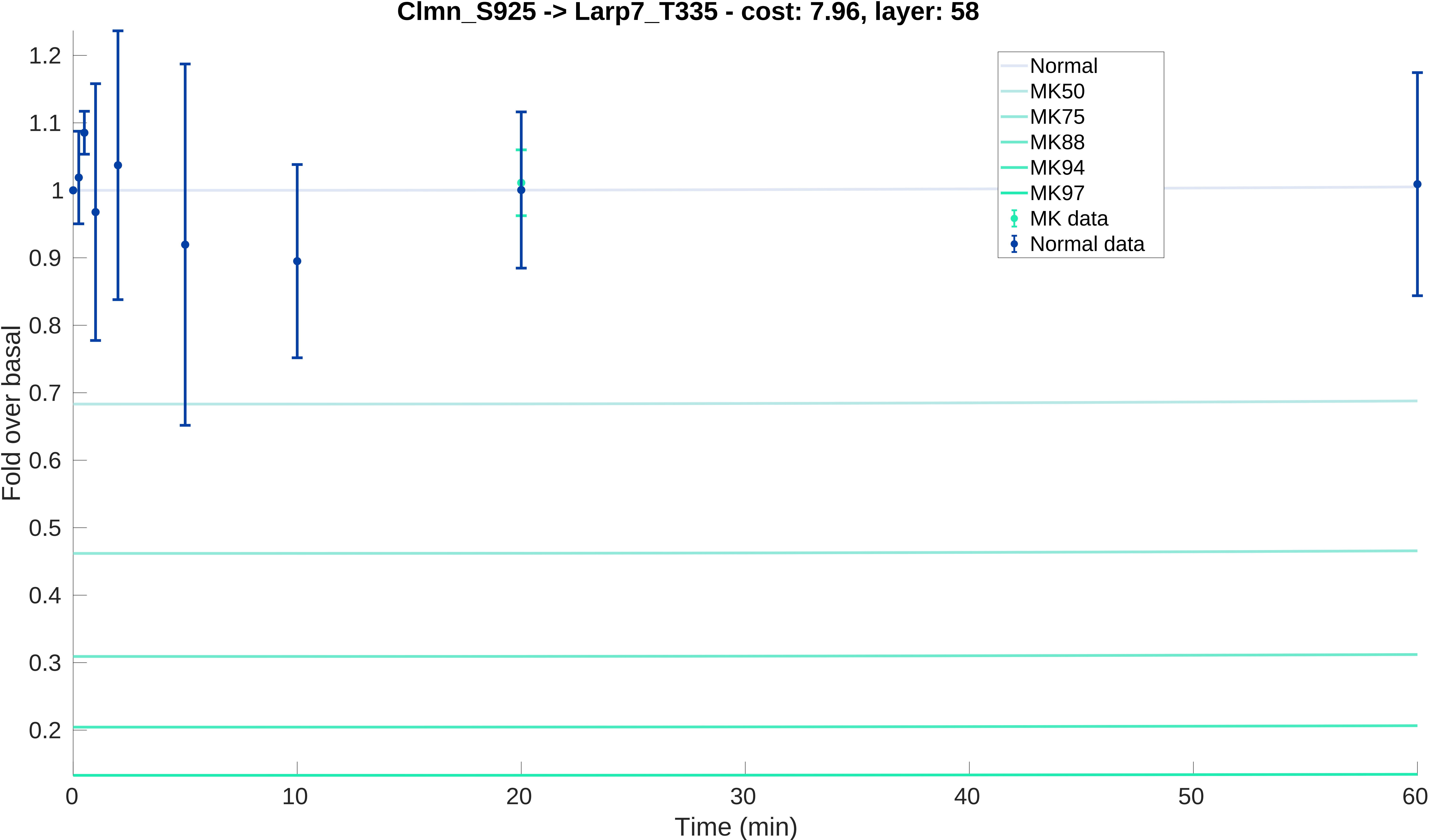

Clmn_S925 -> Larp7_T335 - cost: 7.96, layer: 58
Fold over basal
Time (min)
Normal
MK50
MK75
MK88
MK94
MK97
MK data
Normal data
1.2
1.1
1
0.9
0.8
0.7
0.6
0.5
0.4
0.3
0.2
0
10
20
30
40
50
60

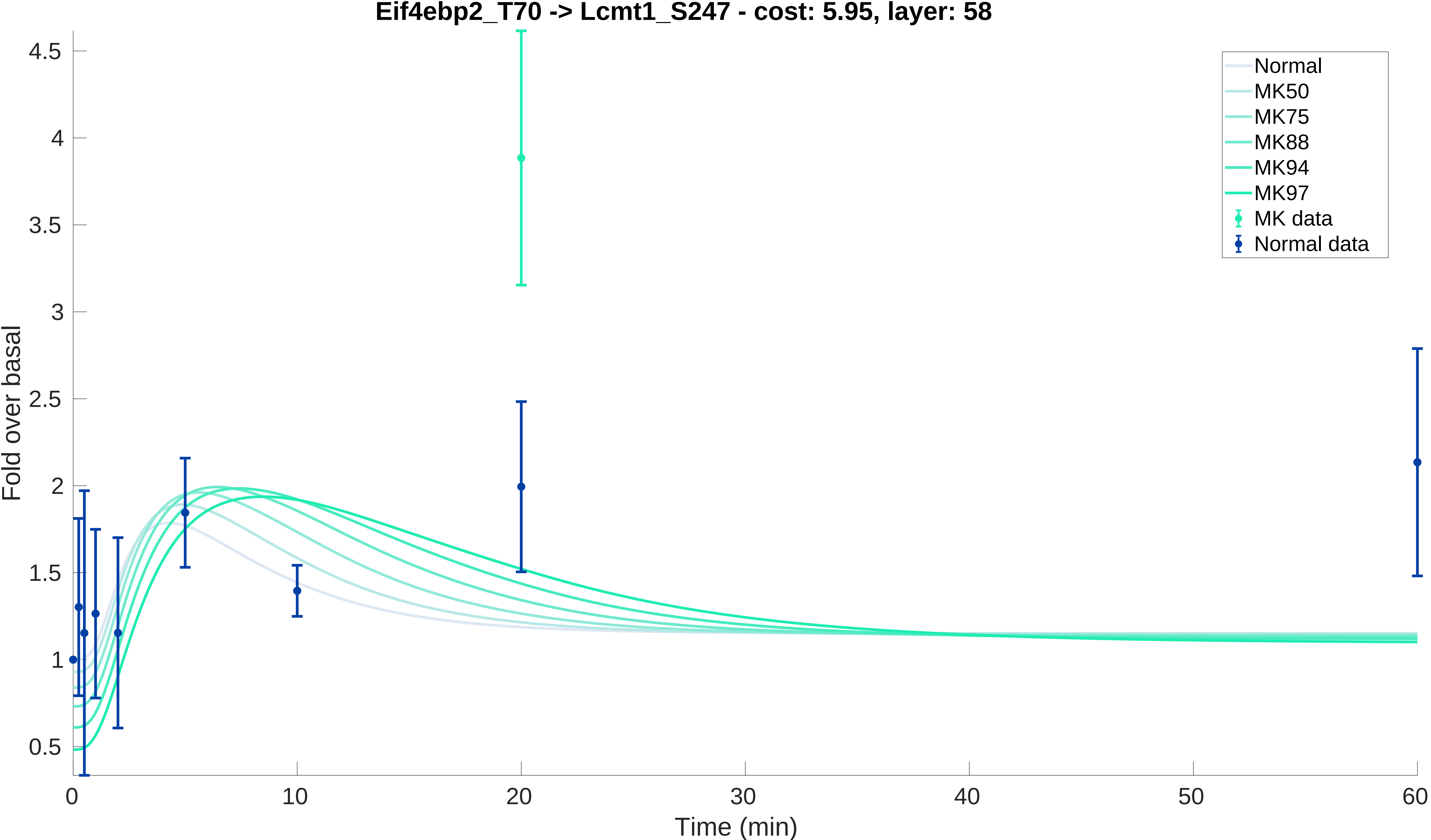

Eif4ebp2_T70 -> Lcmt1_S247 - cost: 5.95, layer: 58
Normal
MK50
MK75
MK88
MK94
MK97
MK data
Normal data
Fold over basal
Time (min)
0.5
1
1.5
2
2.5
3
3.5
4
4.5
0
10
20
30
40
50
60

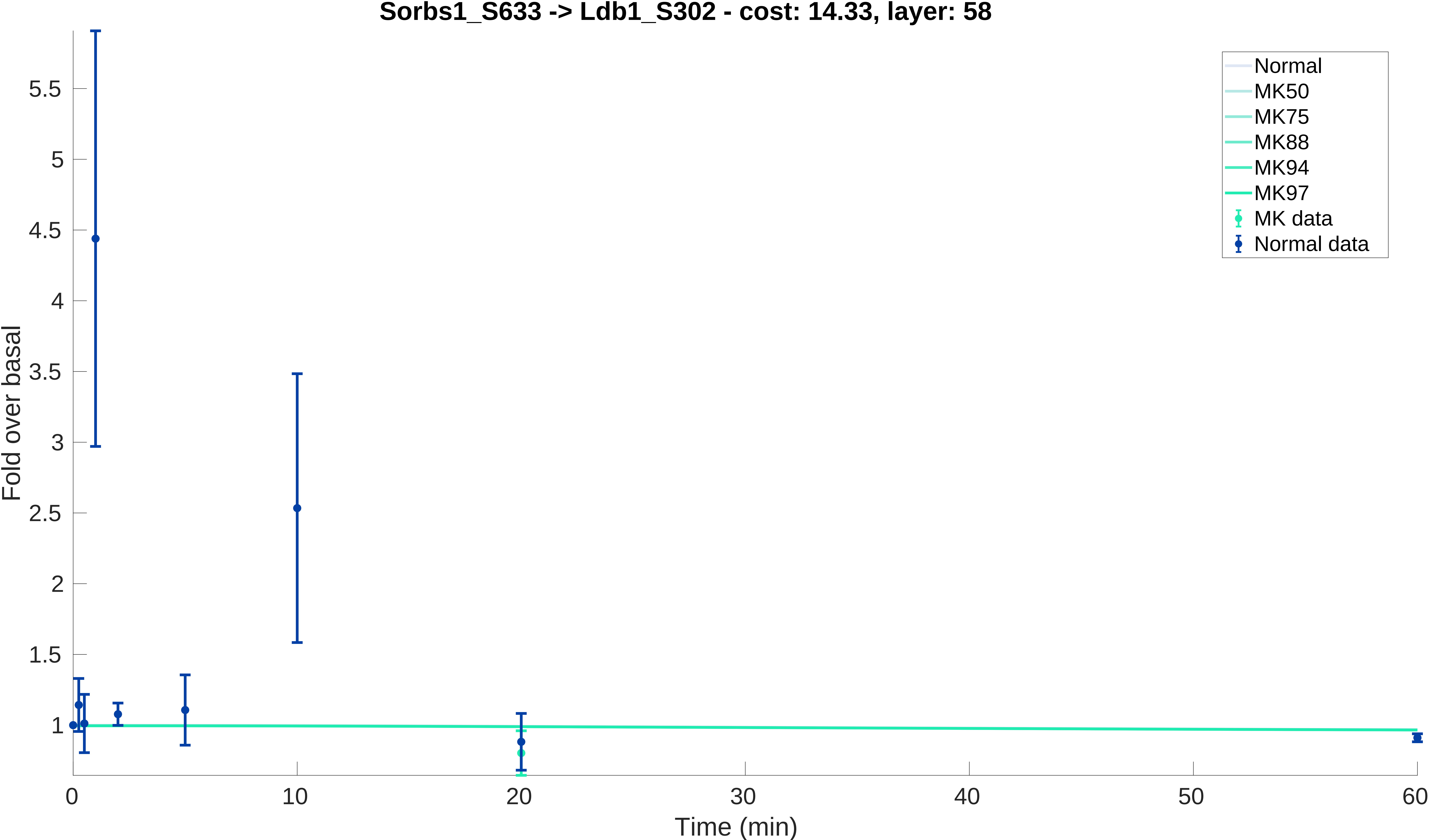
Sorbs1_S633 -> Ldb1_S302 - cost: 14.33, layer: 58
Fold over basal
Time (min)
Normal
MK50
MK75
MK88
MK94
MK97
MK data
Normal data
5.5
5
4.5
4
3.5
3
2.5
2
1.5
1
0
10
20
30
40
50
60

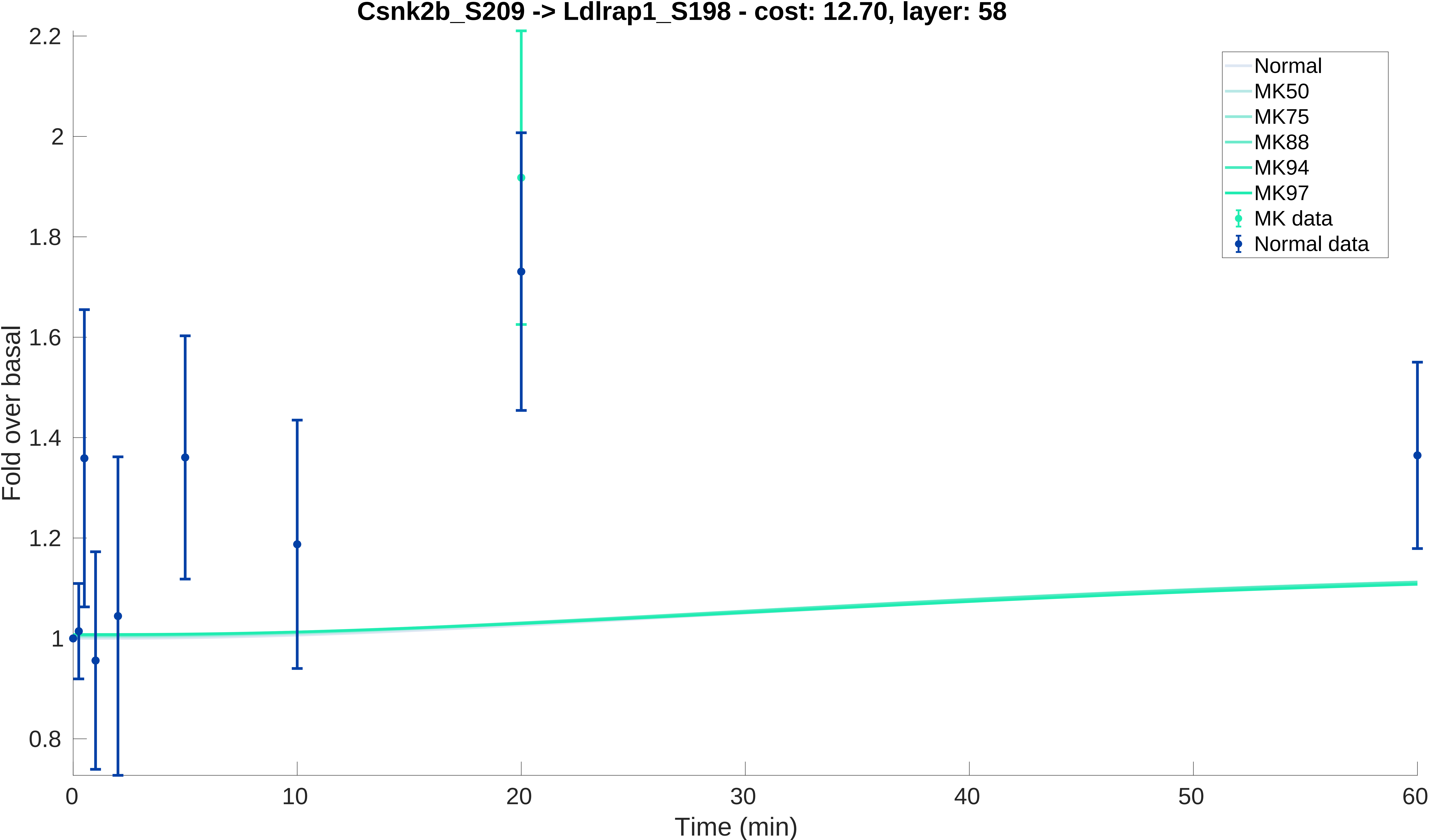

Csnk2b_S209 -> Ldlrap1_S198 - cost: 12.70, layer: 58
Fold over basal
Time (min)
2.2
2
1.8
1.6
1.4
1.2
1
0.8
0
10
20
30
40
50
60
Normal
MK50
MK75
MK88
MK94
MK97
MK data
Normal data

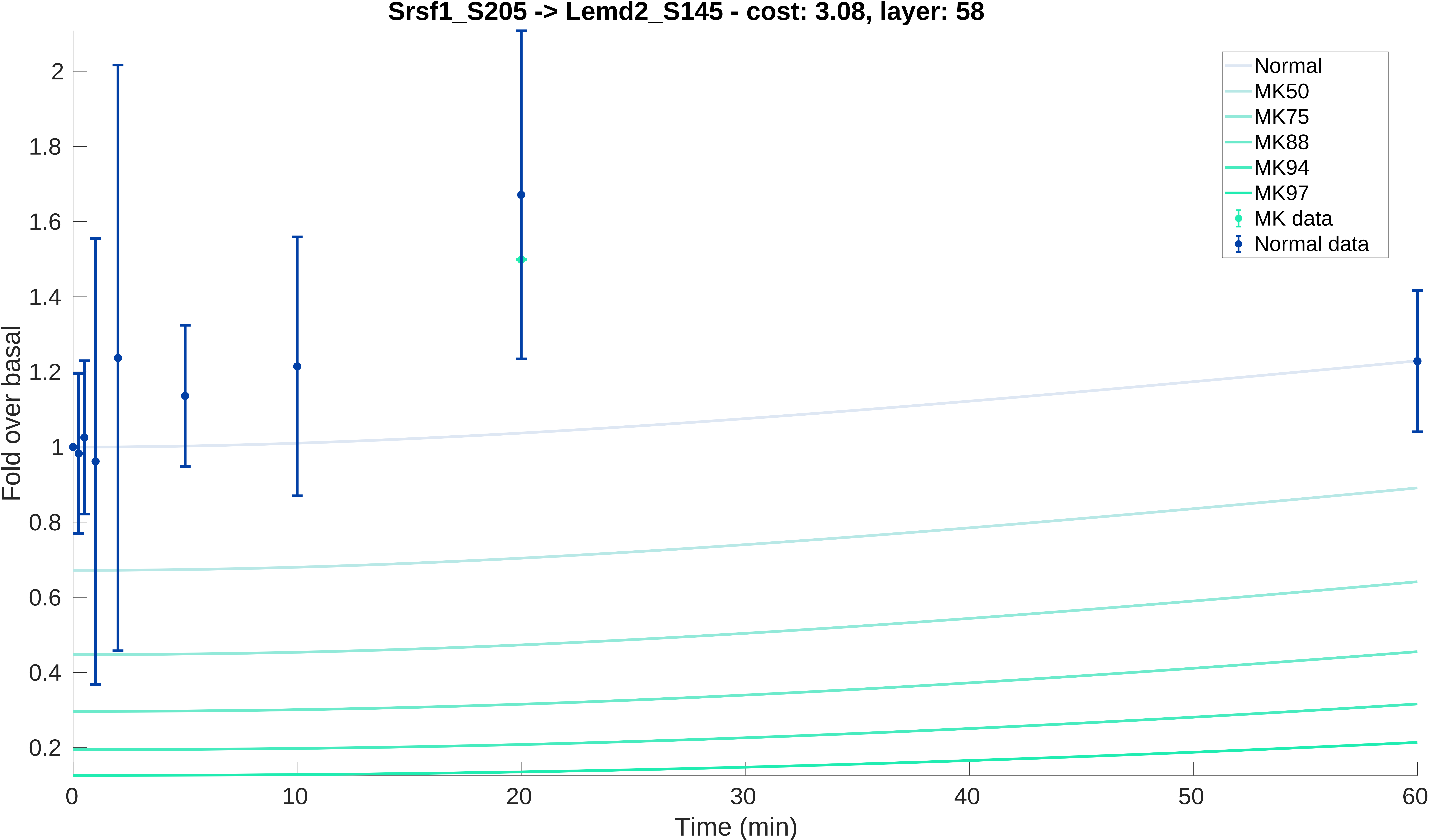

Srsf1_S205 -> Lemd2_S145 - cost: 3.08, layer: 58
Fold over basal
Time (min)
0
10
20
30
40
50
60
0.2
0.4
0.6
0.8
1
1.2
1.4
1.6
1.8
2
Normal
MK50
MK75
MK88
MK94
MK97
MK data
Normal data

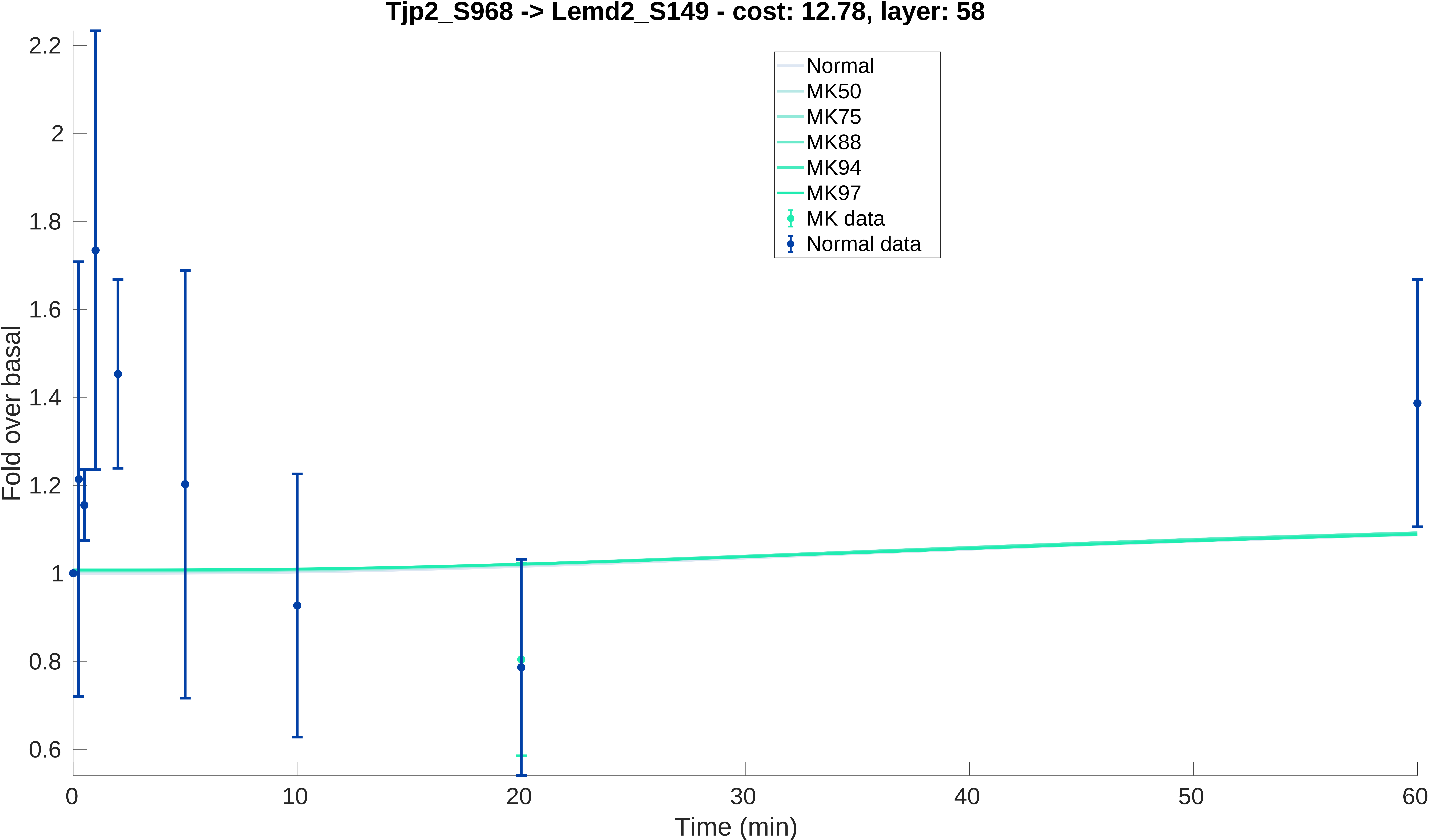

Tjp2_S968 -> Lemd2_S149 - cost: 12.78, layer: 58
Normal
MK50
MK75
MK88
MK94
MK97
MK data
Normal data
Fold over basal
Time (min)
0.6
0.8
1
1.2
1.4
1.6
1.8
2
2.2
0
10
20
30
40
50
60

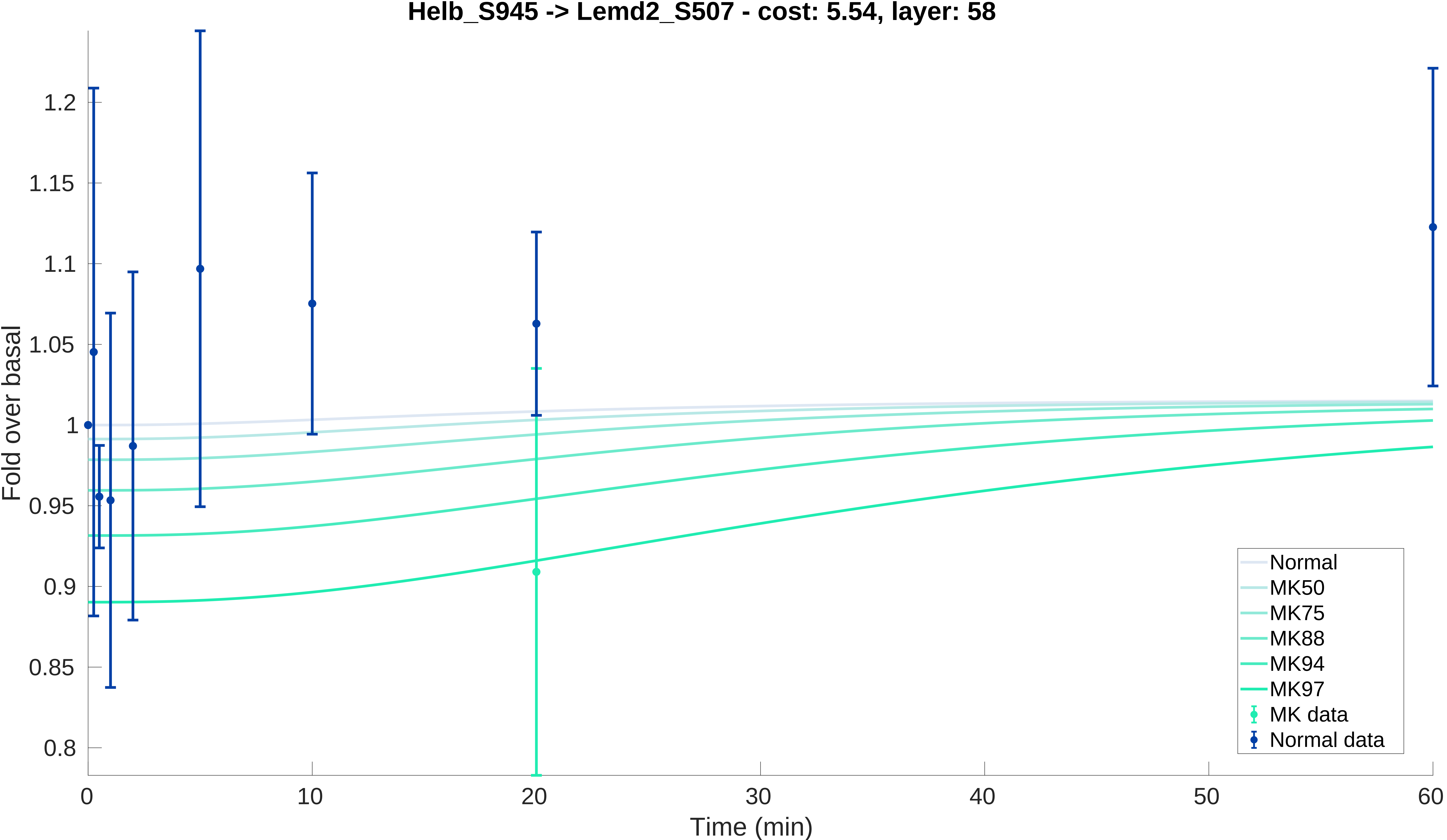
Helb_S945 -> Lemd2_S507 - cost: 5.54, layer: 58
Fold over basal
1.2
1.15
1.1
1.05
1
0.95
0.9
0.85
0.8
0
10
20
30
40
50
60
Time (min)
Normal
MK50
MK75
MK88
MK94
MK97
MK data
Normal data

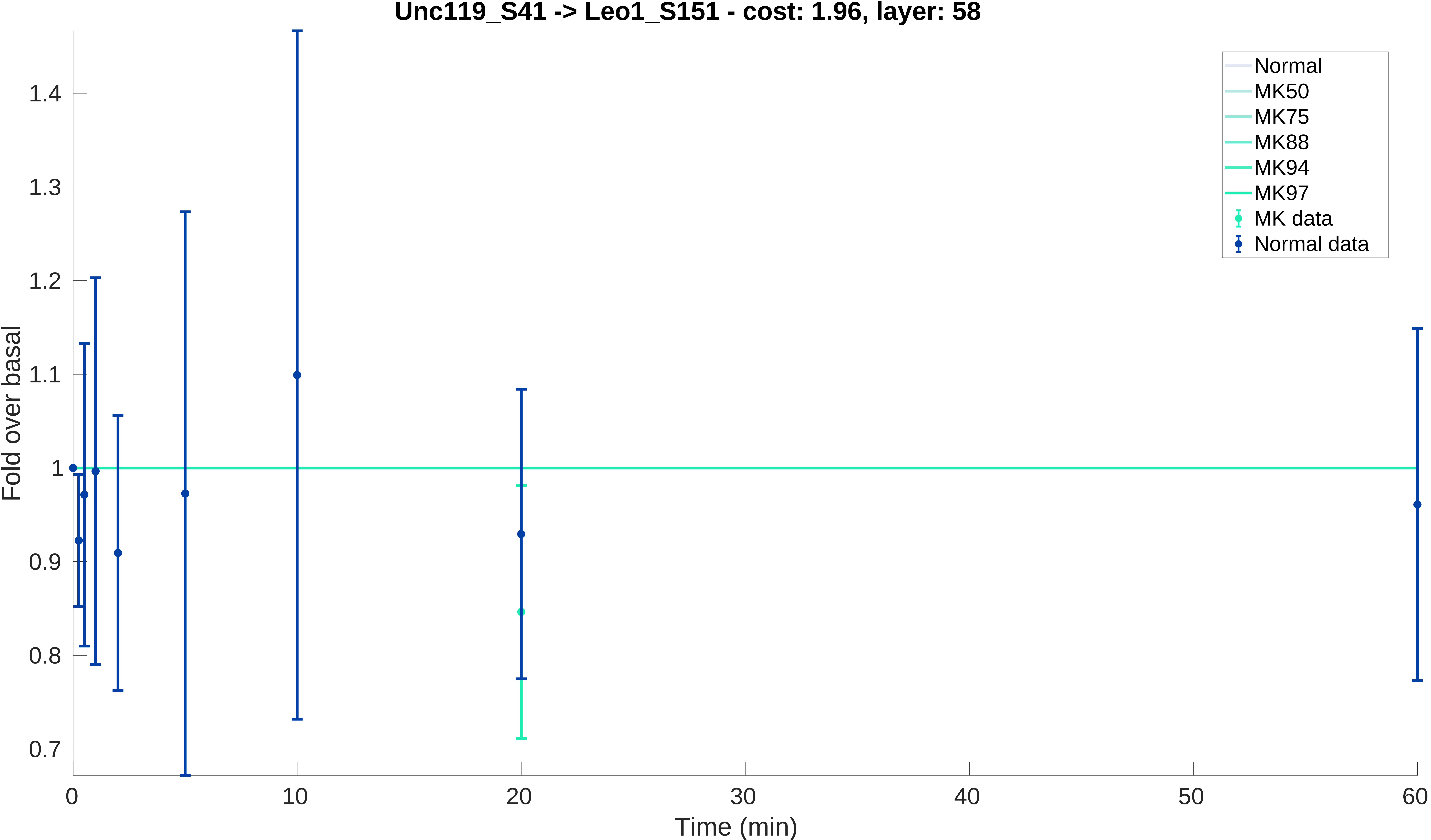

Unc119_S41 -> Leo1_S151 - cost: 1.96, layer: 58
Fold over basal
Time (min)
0.7
0.8
0.9
1
1.1
1.2
1.3
1.4
0
10
20
30
40
50
60
Normal
MK50
MK75
MK88
MK94
MK97
MK data
Normal data

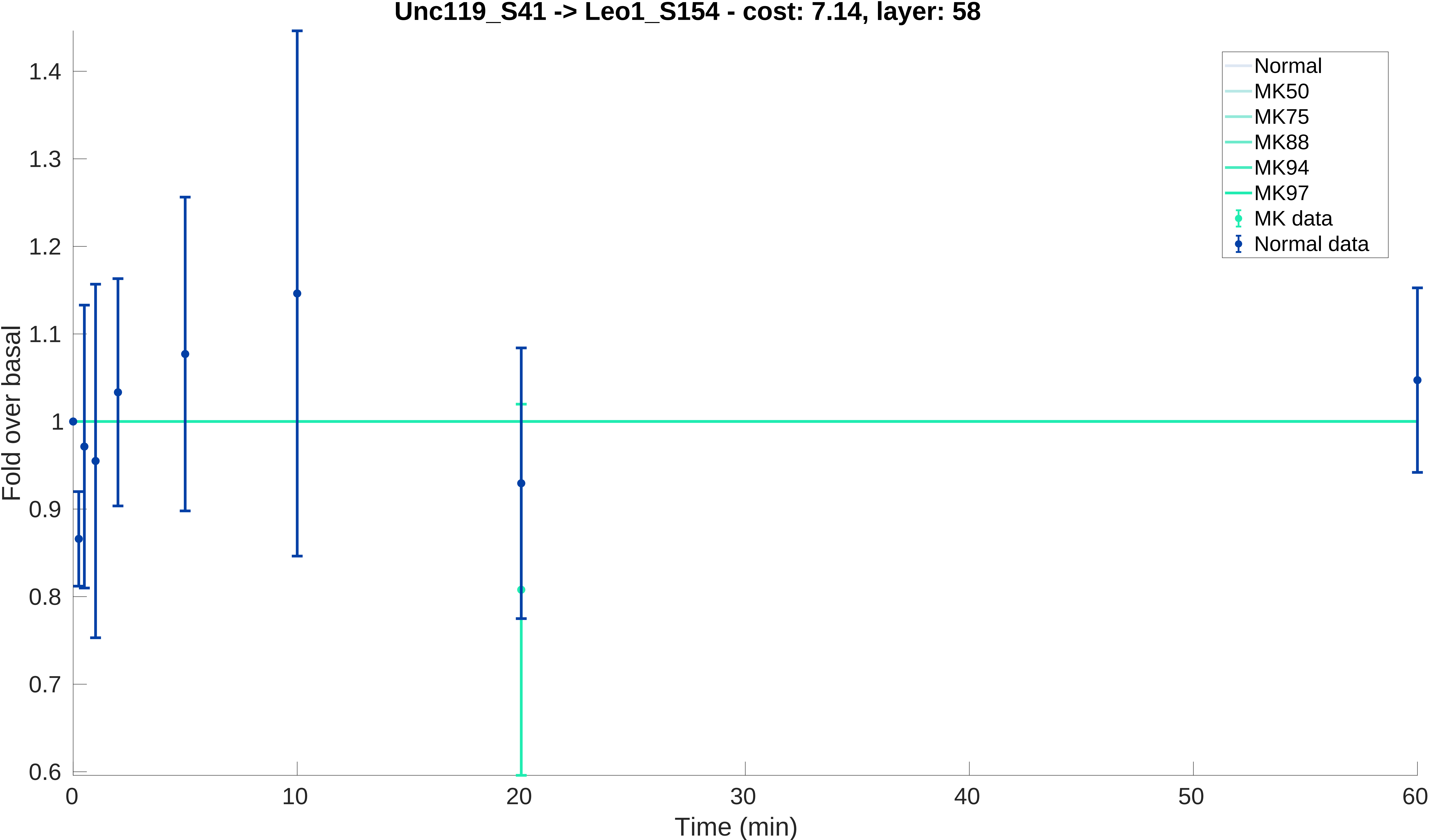

Unc119_S41 -> Leo1_S154 - cost: 7.14, layer: 58
Fold over basal
Time (min)
Normal
MK50
MK75
MK88
MK94
MK97
MK data
Normal data
1.4
1.3
1.2
1.1
1
0.9
0.8
0.7
0.6
0
10
20
30
40
50
60

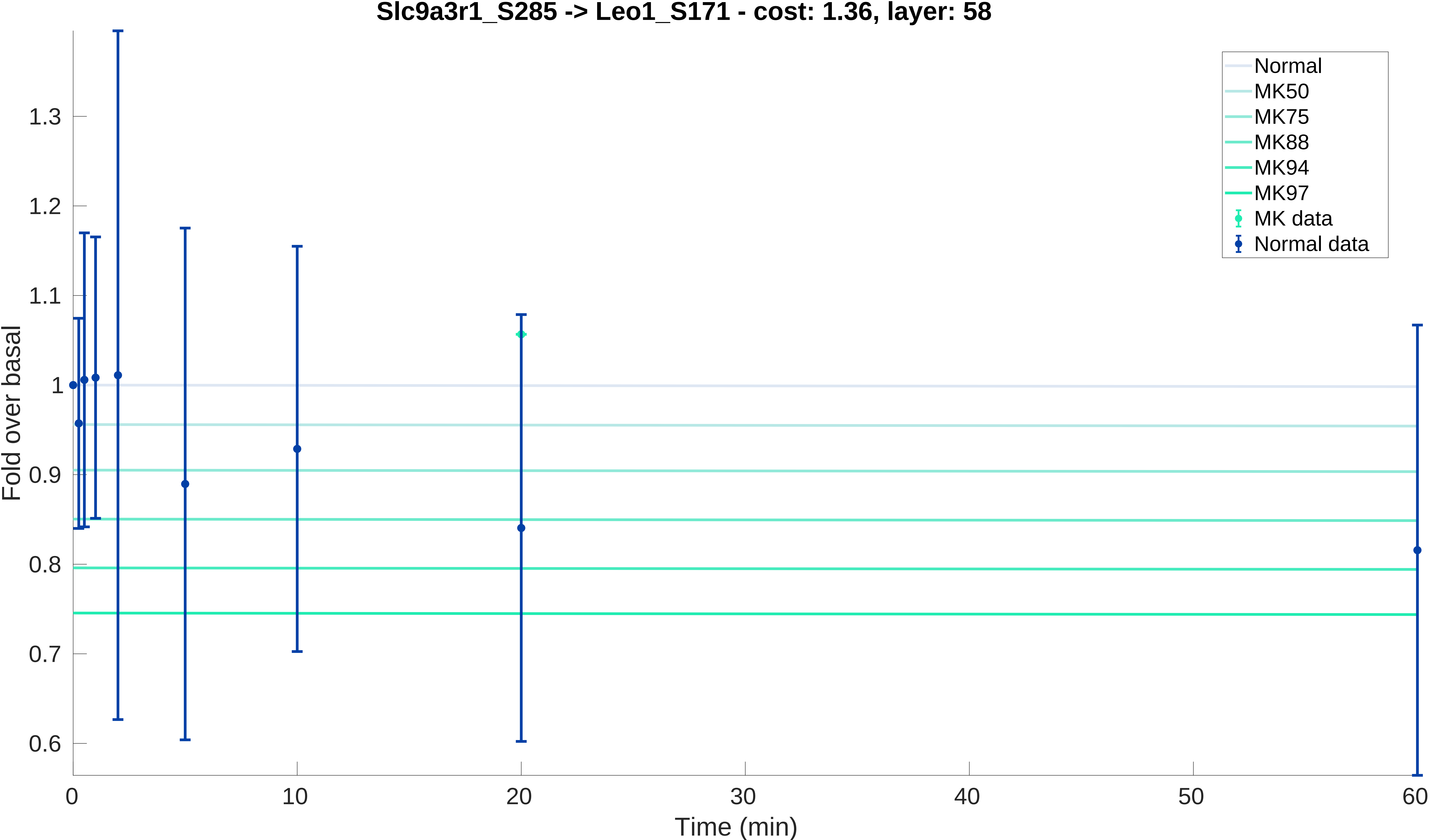

Slc9a3r1_S285 -> Leo1_S171 - cost: 1.36, layer: 58
Normal
MK50
MK75
MK88
MK94
MK97
MK data
Normal data
Fold over basal
1.3
1.2
1.1
1
0.9
0.8
0.7
0.6
0
10
20
30
40
50
60
Time (min)

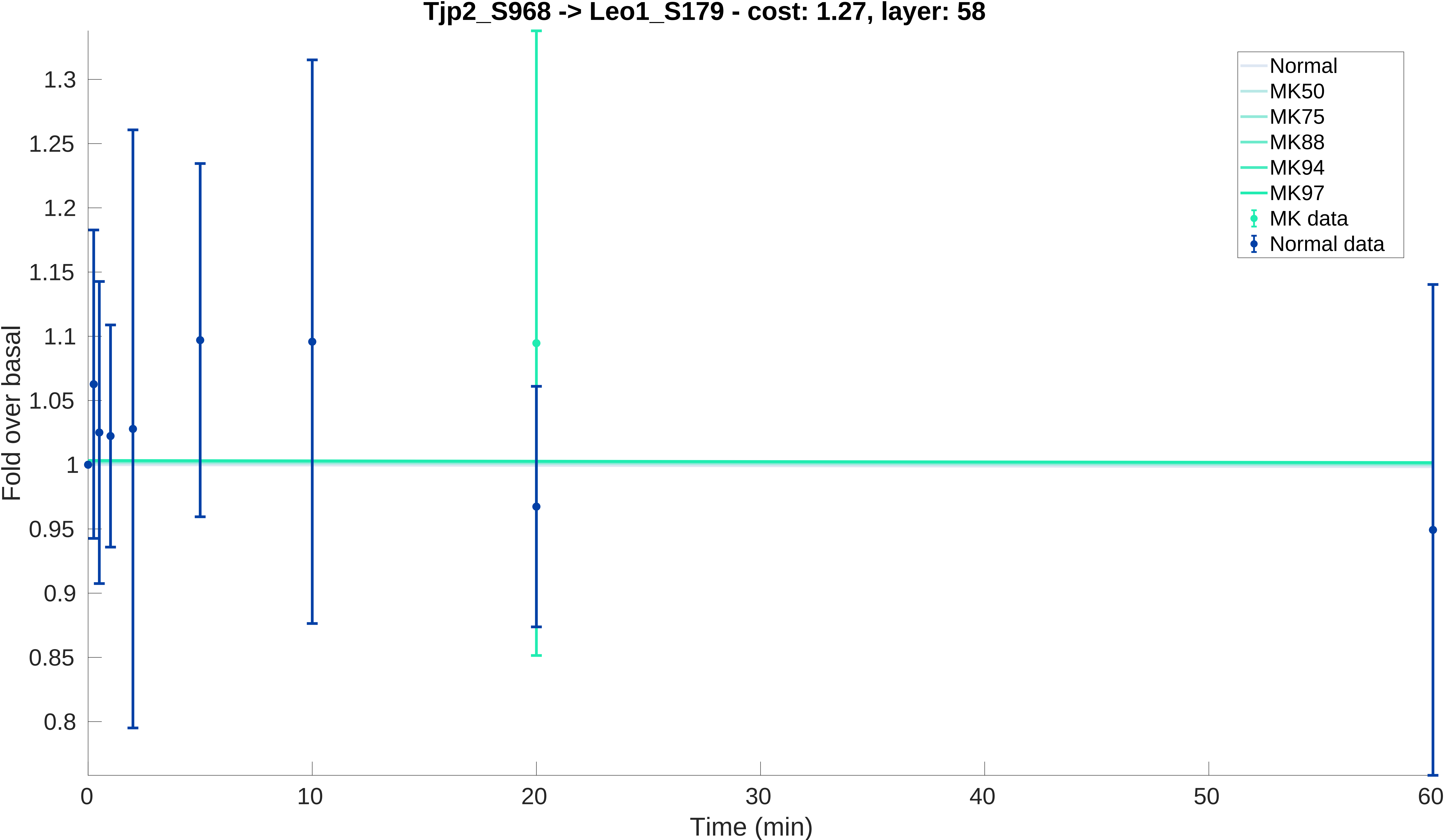

Tjp2_S968 -> Leo1_S179 - cost: 1.27, layer: 58
Normal
MK50
MK75
MK88
MK94
MK97
MK data
Normal data
Fold over basal
1.3
1.25
1.2
1.15
1.1
1.05
1
0.95
0.9
0.85
0.8
0
10
20
30
40
50
60
Time (min)

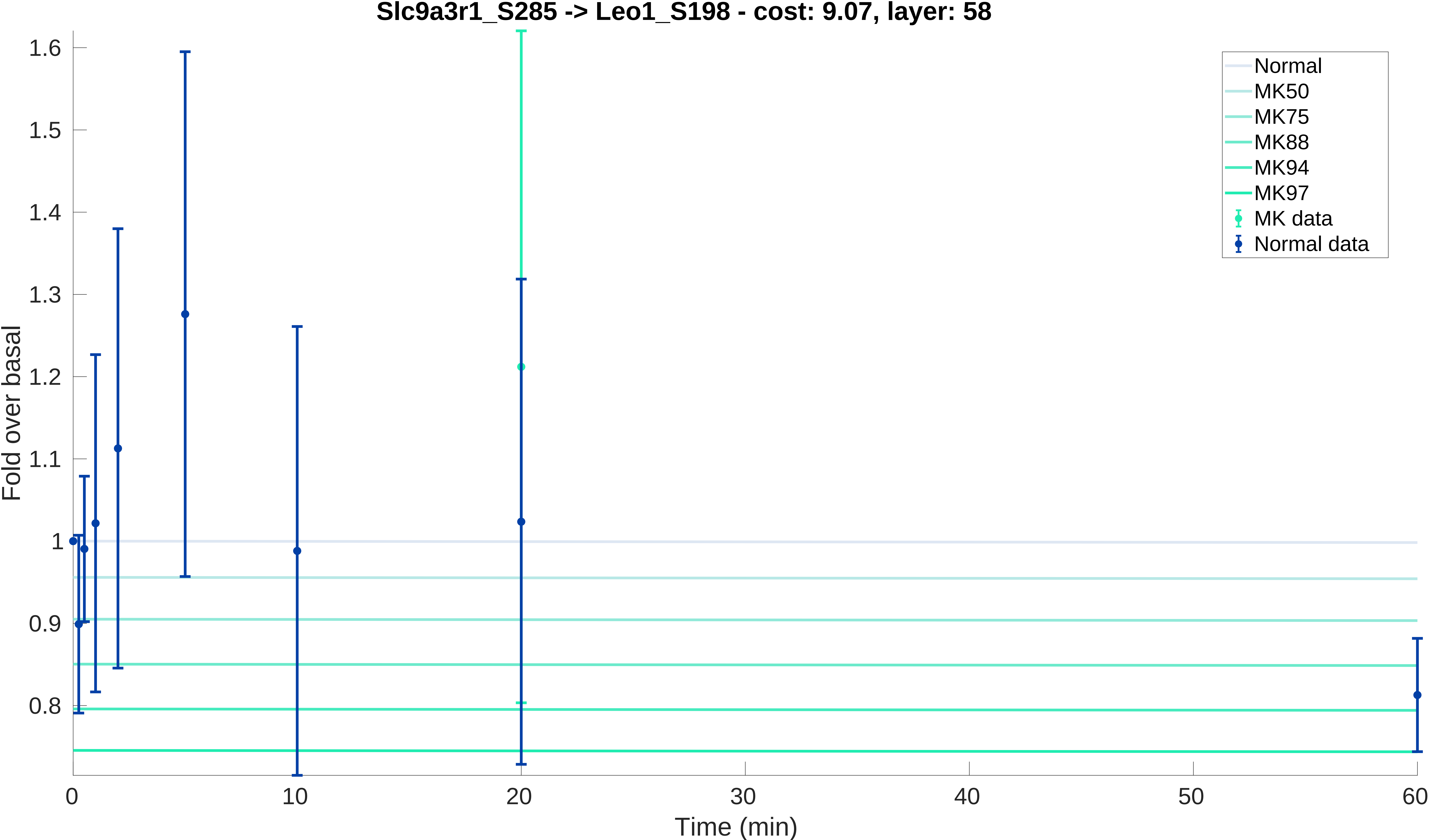

Slc9a3r1_S285 -> Leo1_S198 - cost: 9.07, layer: 58
Fold over basal
Time (min)
1.6
1.5
1.4
1.3
1.2
1.1
1
0.9
0.8
0
10
20
30
40
50
60
Normal
MK50
MK75
MK88
MK94
MK97
MK data
Normal data

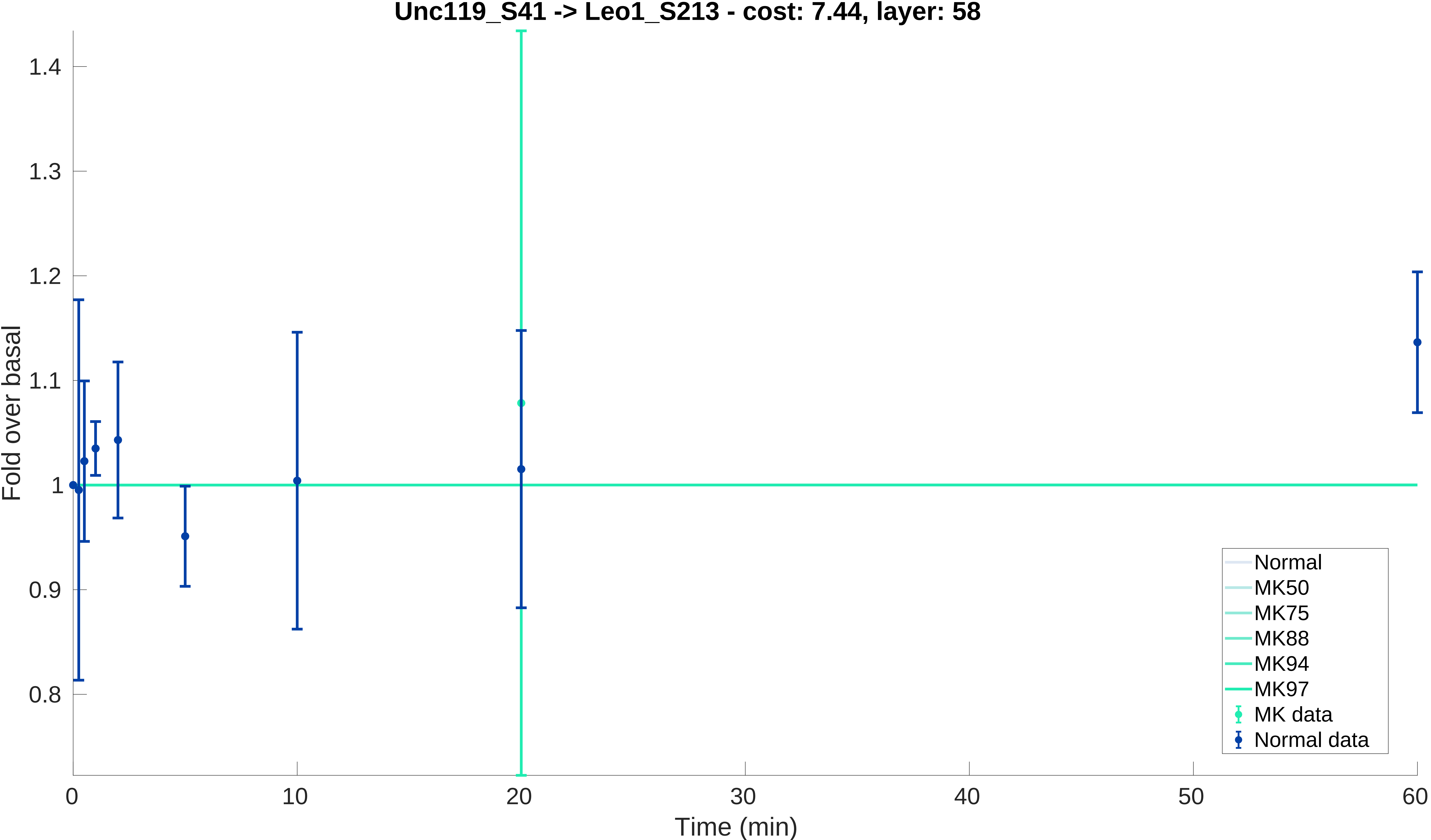

Unc119_S41 -> Leo1_S213 - cost: 7.44, layer: 58
Fold over basal
1.4
1.3
1.2
1.1
1
0.9
0.8
0
10
20
30
40
50
60
Time (min)
Normal
MK50
MK75
MK88
MK94
MK97
MK data
Normal data

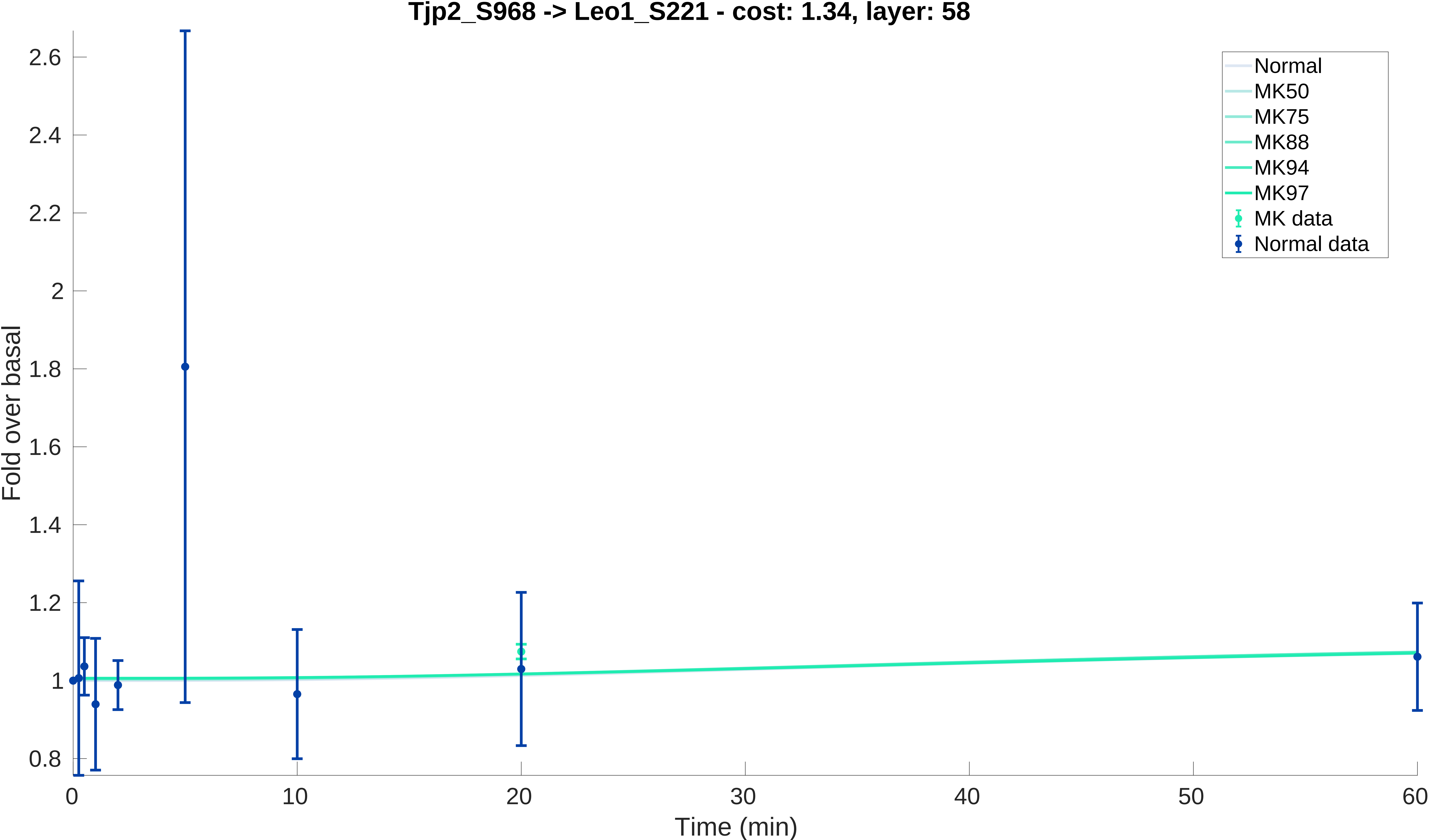

Tjp2_S968 -> Leo1_S221 - cost: 1.34, layer: 58
Fold over basal
Time (min)
0.8
1
1.2
1.4
1.6
1.8
2
2.2
2.4
2.6
0
10
20
30
40
50
60
Normal
MK50
MK75
MK88
MK94
MK97
MK data
Normal data

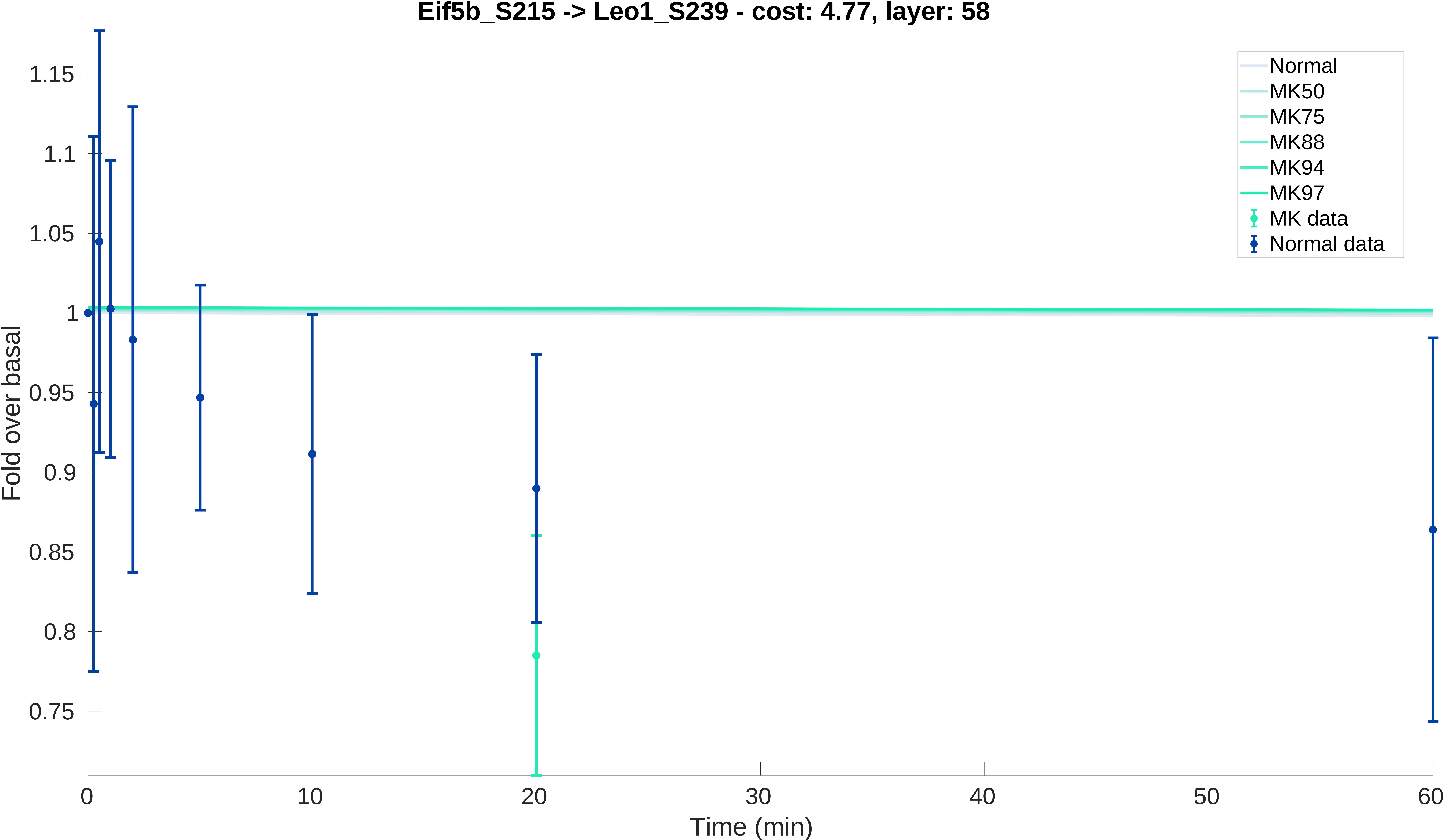

Eif5b_S215 -> Leo1_S239 - cost: 4.77, layer: 58
Normal
MK50
MK75
MK88
MK94
MK97
MK data
Normal data
Fold over basal
1.15
1.1
1.05
1
0.95
0.9
0.85
0.8
0.75
0
10
20
30
40
50
60
Time (min)

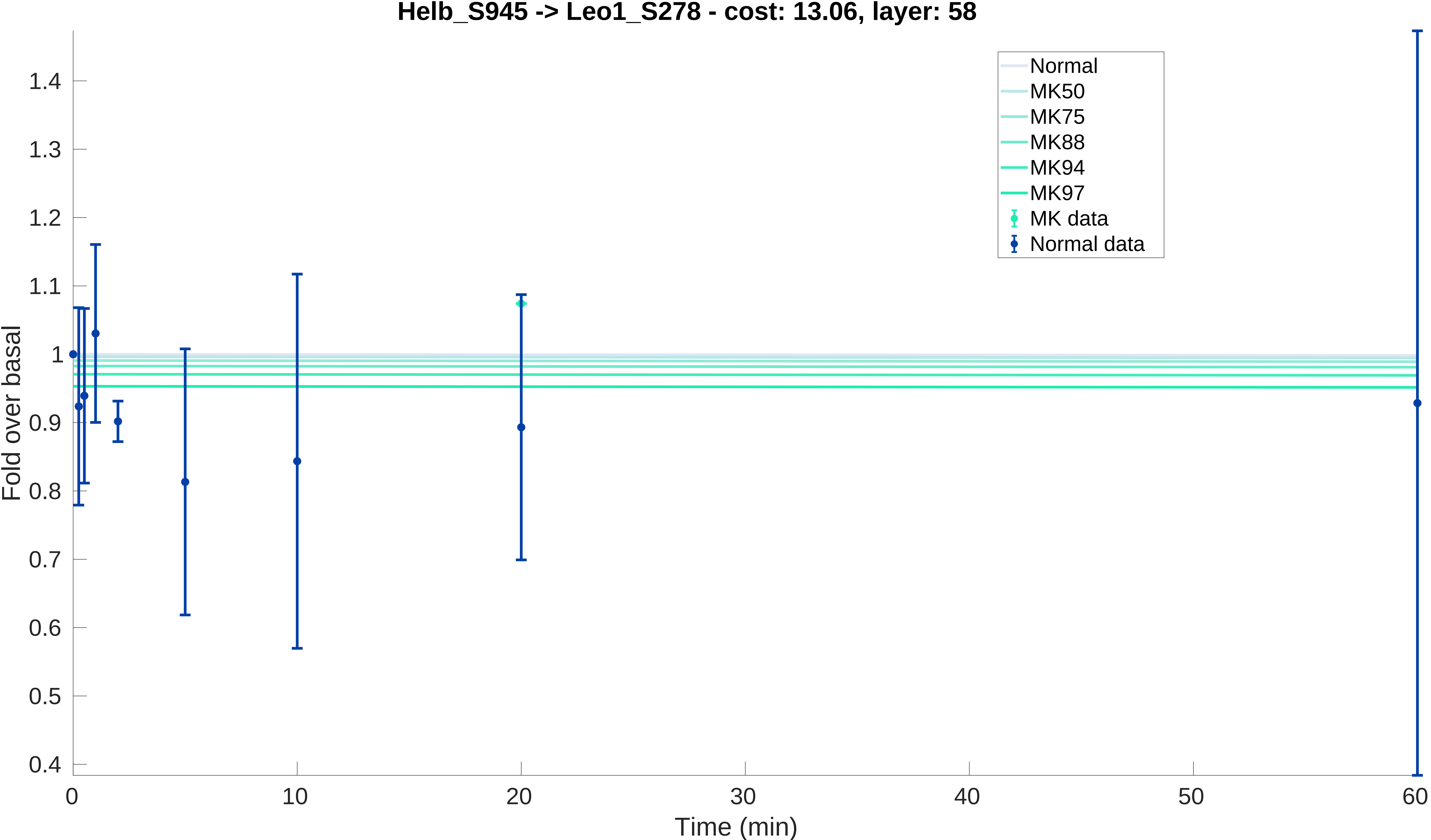

Helb_S945 -> Leo1_S278 - cost: 13.06, layer: 58
Normal
MK50
MK75
MK88
MK94
MK97
MK data
Normal data
Fold over basal
Time (min)
0
10
20
30
40
50
60
0.4
0.5
0.6
0.7
0.8
0.9
1
1.1
1.2
1.3
1.4

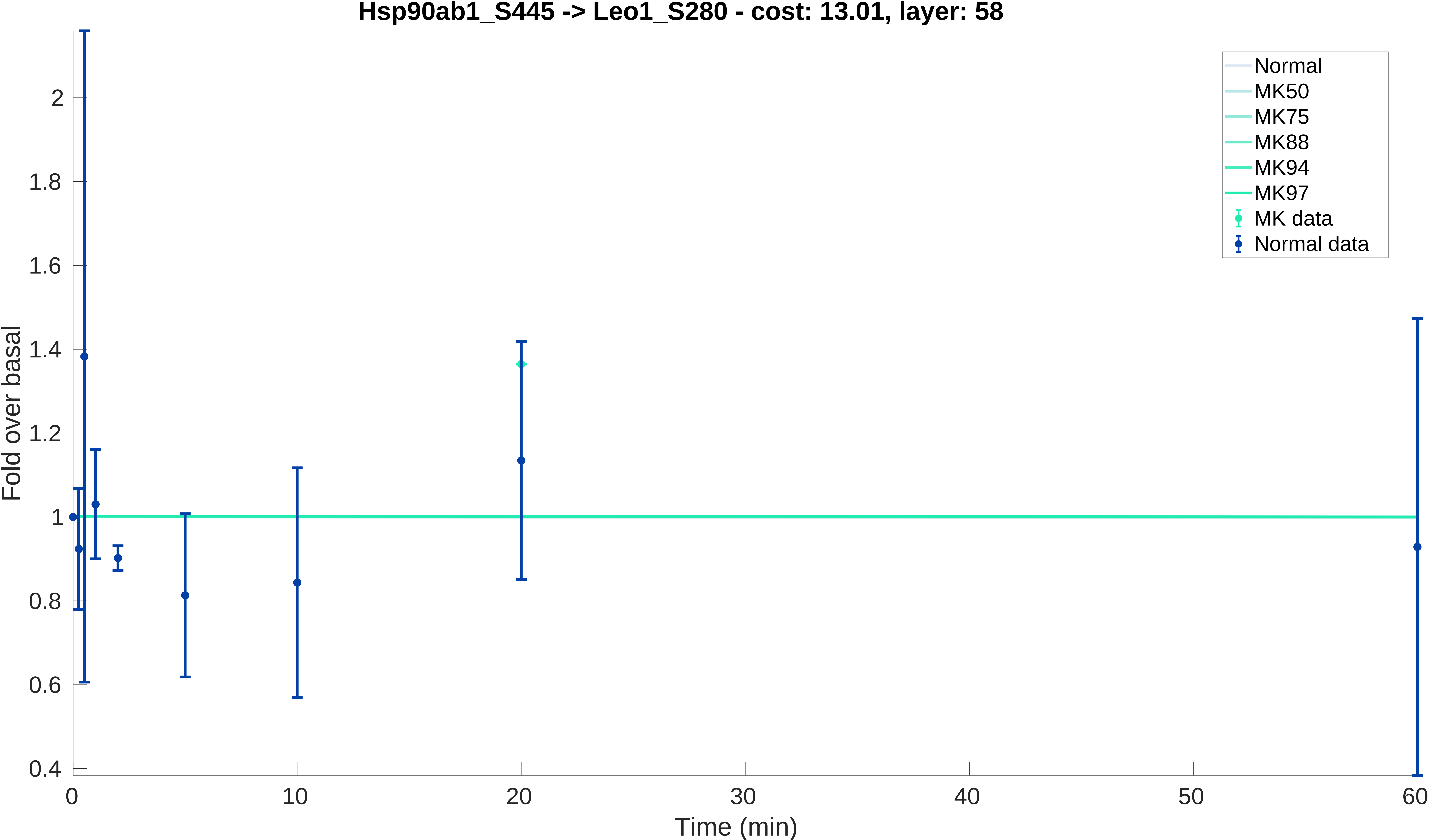

Hsp90ab1_S445 -> Leo1_S280 - cost: 13.01, layer: 58
Fold over basal
Time (min)
0.4
0.6
0.8
1
1.2
1.4
1.6
1.8
2
0
10
20
30
40
50
60
Normal
MK50
MK75
MK88
MK94
MK97
MK data
Normal data

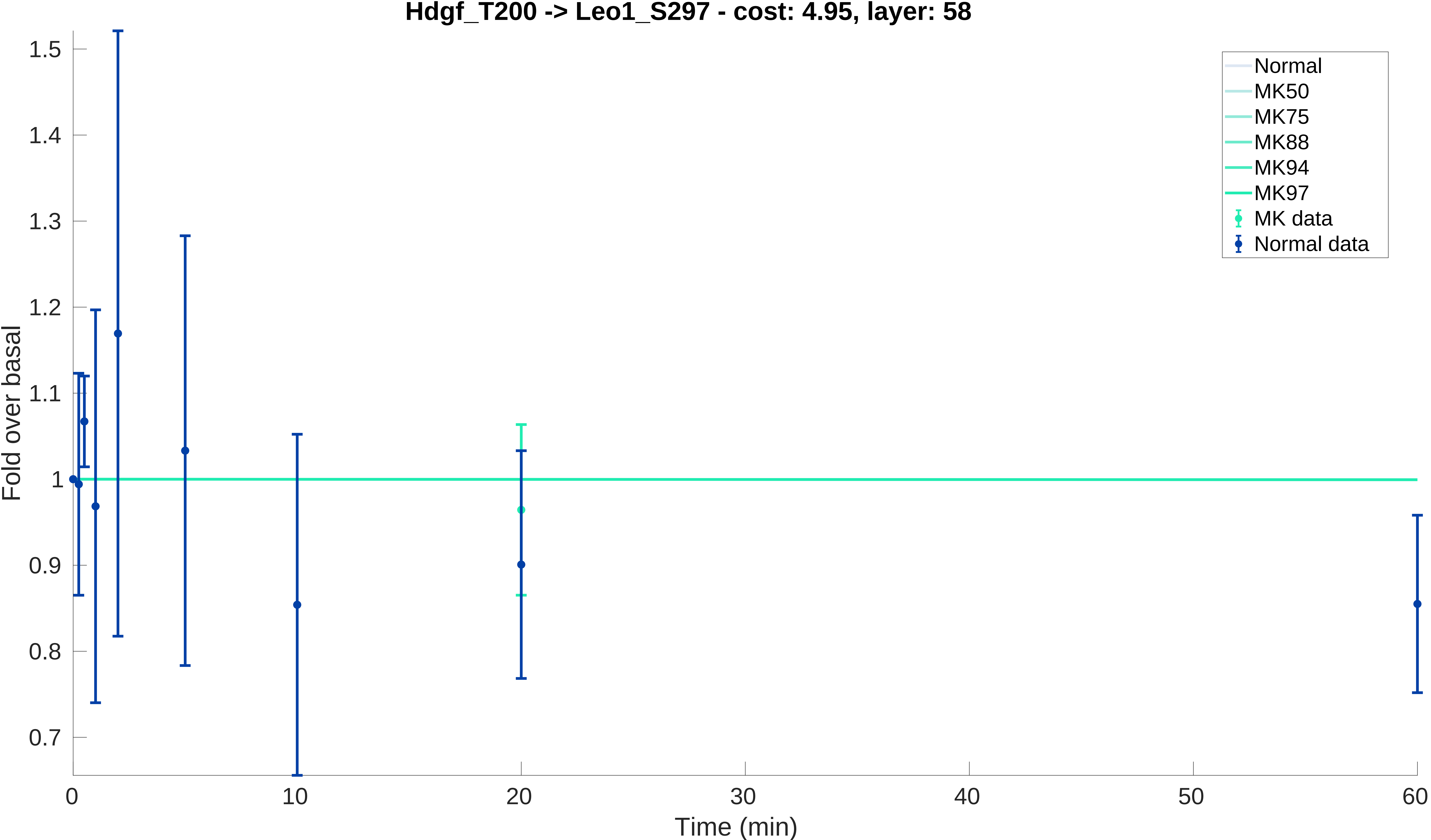

Hdgf_T200 -> Leo1_S297 - cost: 4.95, layer: 58
Normal
MK50
MK75
MK88
MK94
MK97
MK data
Normal data
Fold over basal
Time (min)
0
10
20
30
40
50
60
0.7
0.8
0.9
1
1.1
1.2
1.3
1.4
1.5

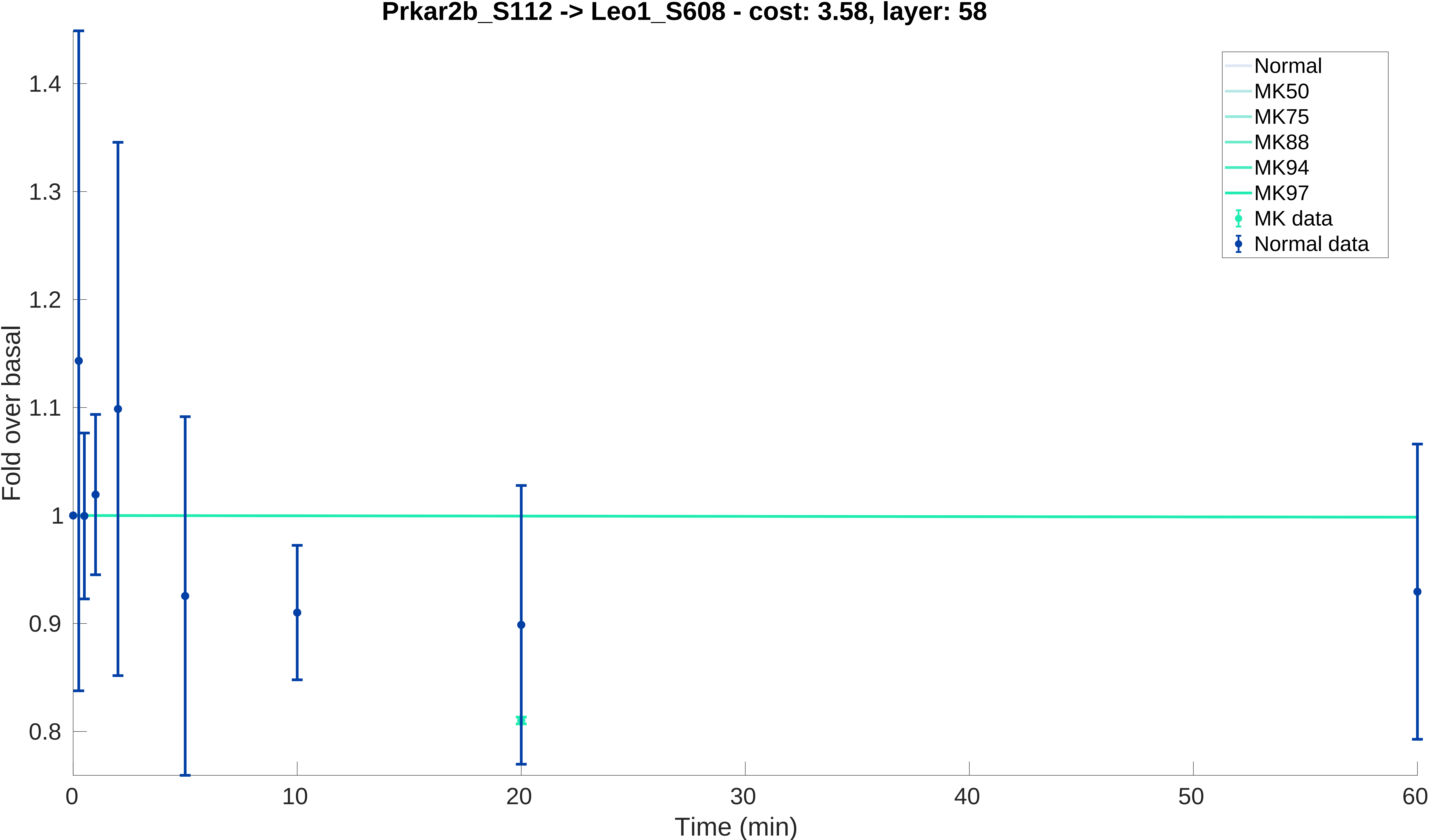

Prkar2b_S112 -> Leo1_S608 - cost: 3.58, layer: 58
Fold over basal
Time (min)
1.4
1.3
1.2
1.1
1
0.9
0.8
0
10
20
30
40
50
60
Normal
MK50
MK75
MK88
MK94
MK97
MK data
Normal data

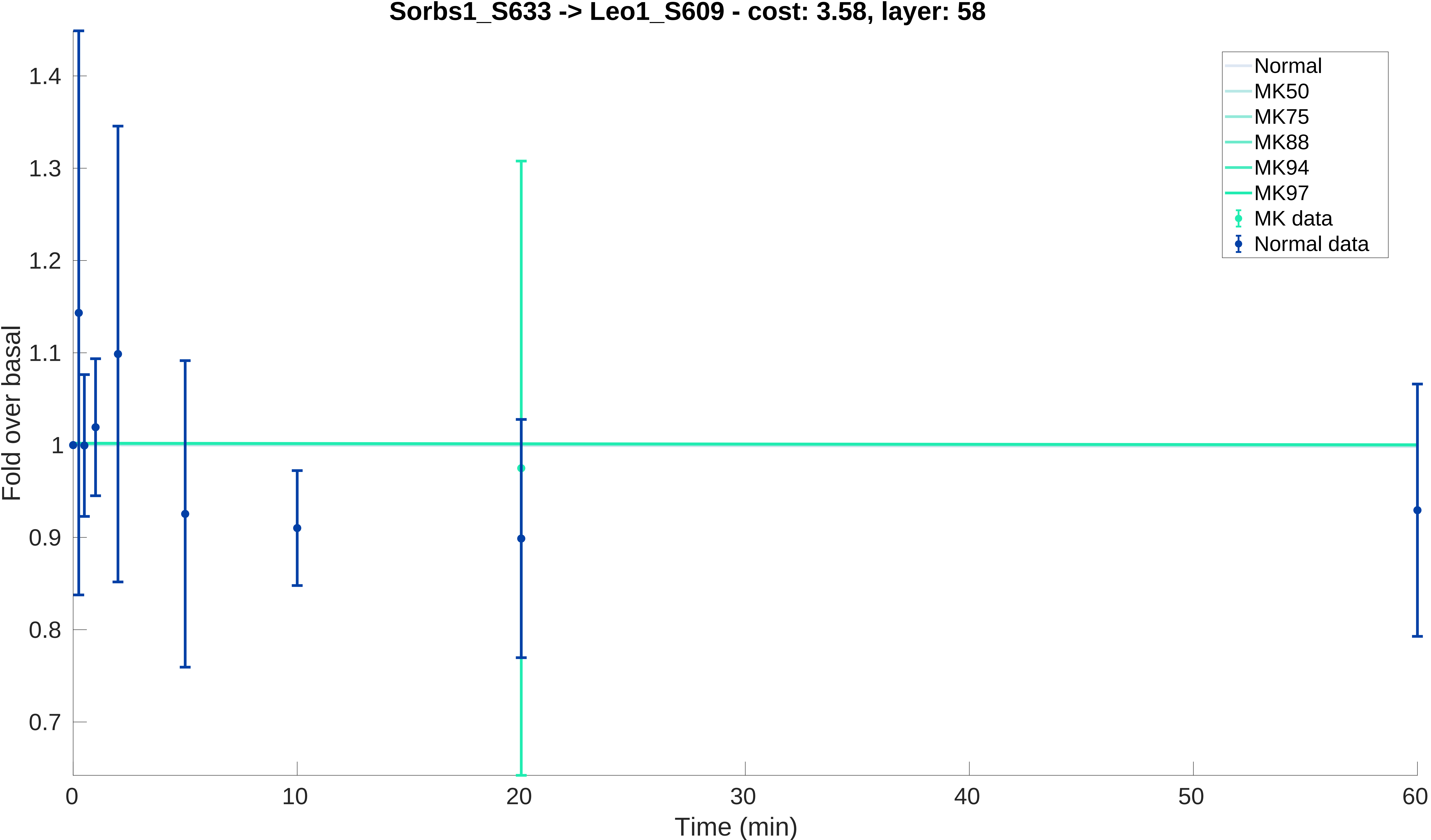

Sorbs1_S633 -> Leo1_S609 - cost: 3.58, layer: 58
Fold over basal
Time (min)
1.4
1.3
1.2
1.1
1
0.9
0.8
0.7
0
10
20
30
40
50
60
Normal
MK50
MK75
MK88
MK94
MK97
MK data
Normal data

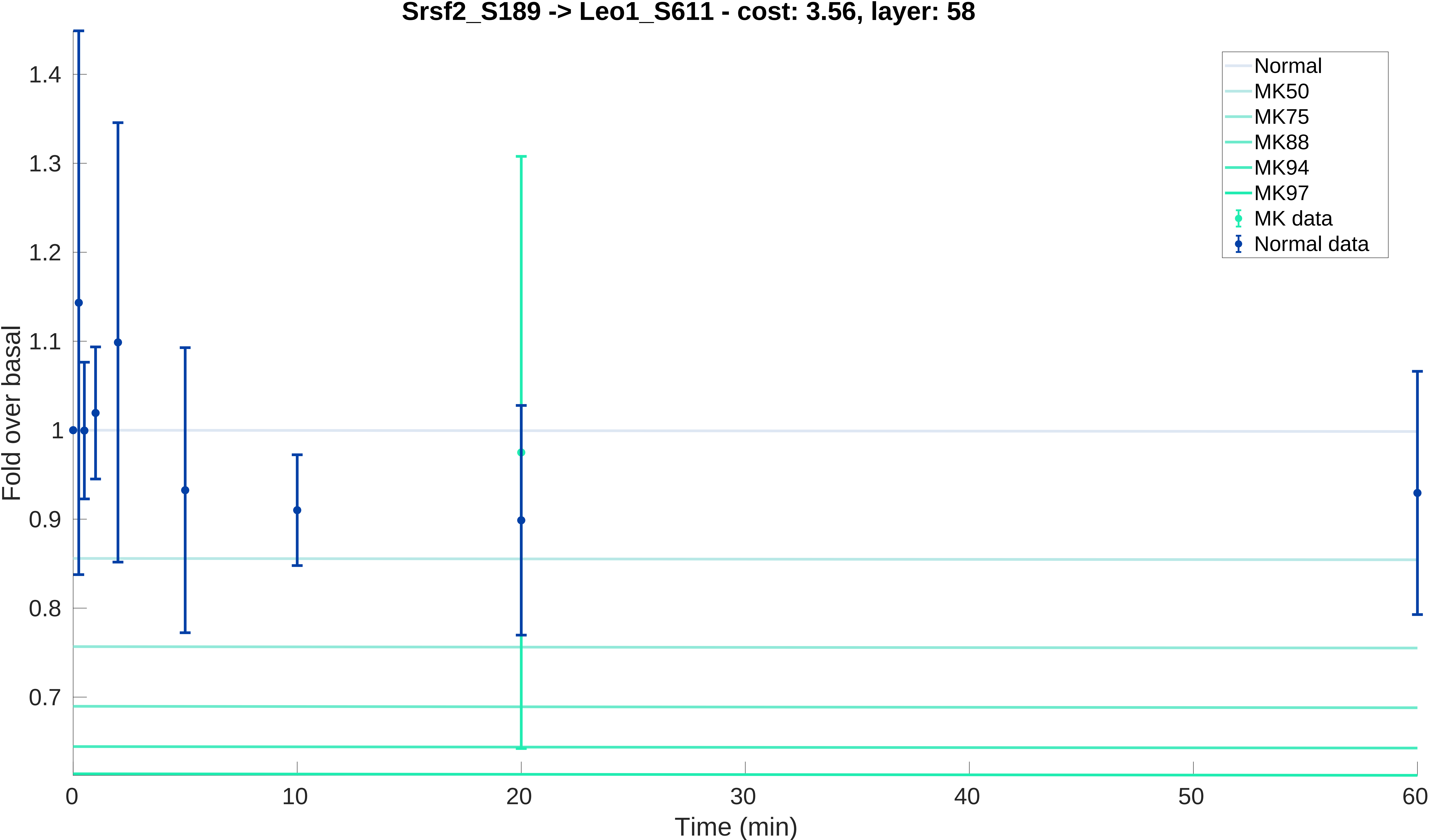

Srsf2_S189 -> Leo1_S611 - cost: 3.56, layer: 58
Fold over basal
Time (min)
1.4
1.3
1.2
1.1
1
0.9
0.8
0.7
0
10
20
30
40
50
60
Normal
MK50
MK75
MK88
MK94
MK97
MK data
Normal data

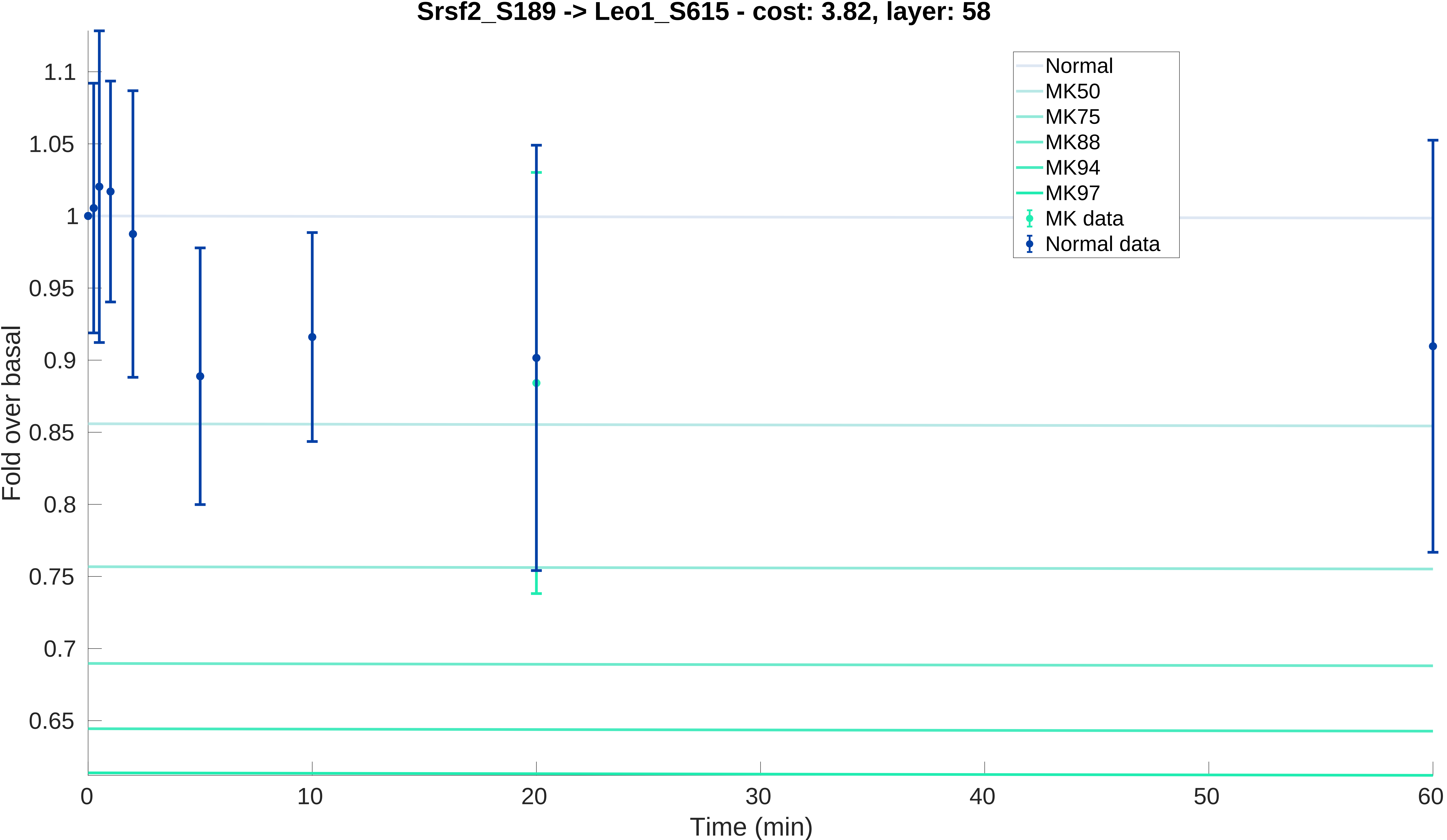

Srsf2_S189 -> Leo1_S615 - cost: 3.82, layer: 58
Normal
MK50
MK75
MK88
MK94
MK97
MK data
Normal data
Fold over basal
Time (min)
1.1
1.05
1
0.95
0.9
0.85
0.8
0.75
0.7
0.65
0
10
20
30
40
50
60

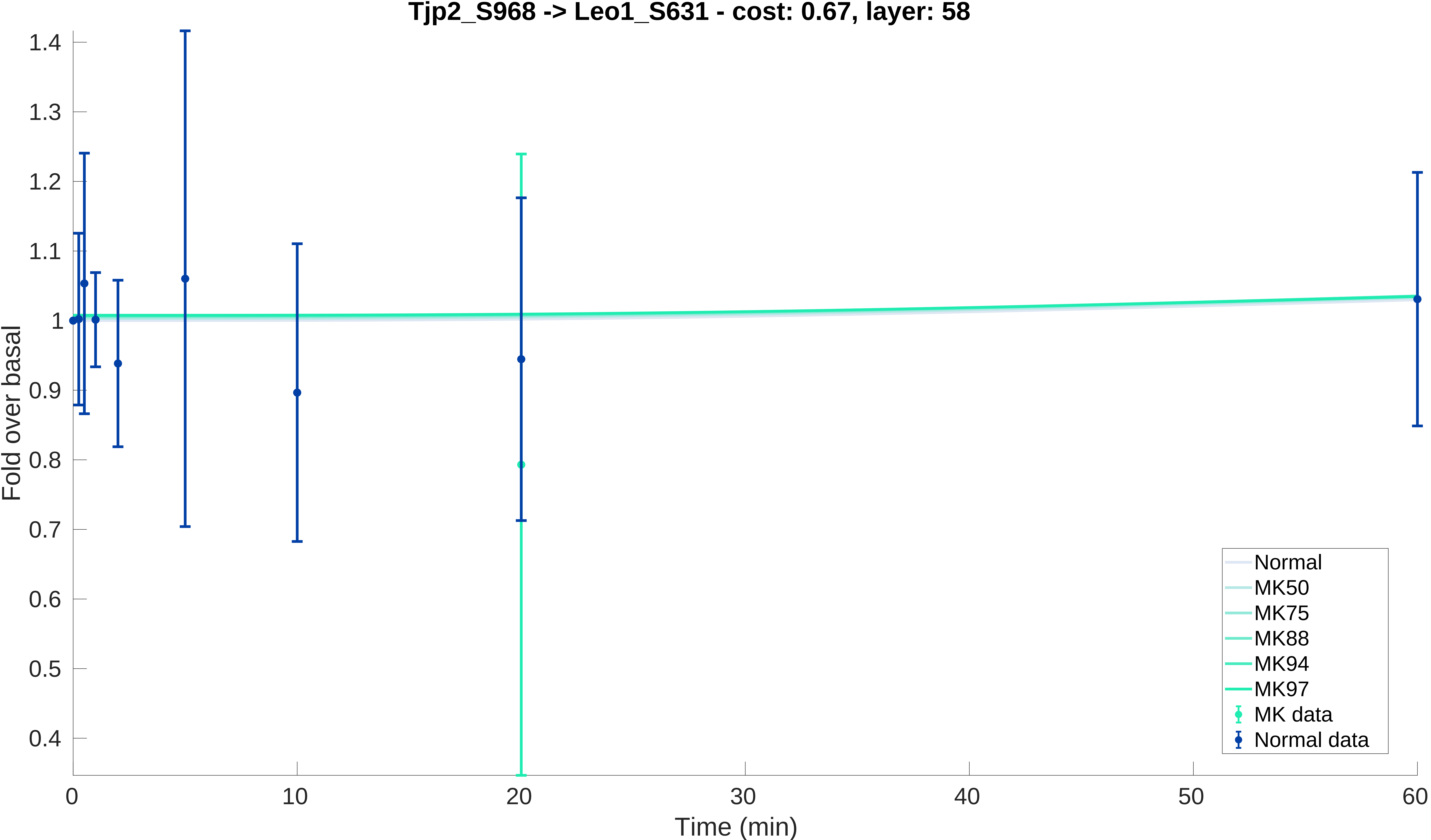

Tjp2_S968 -> Leo1_S631 - cost: 0.67, layer: 58
Fold over basal
Time (min)
1.4
1.3
1.2
1.1
1
0.9
0.8
0.7
0.6
0.5
0.4
0
10
20
30
40
50
60
Normal
MK50
MK75
MK88
MK94
MK97
MK data
Normal data

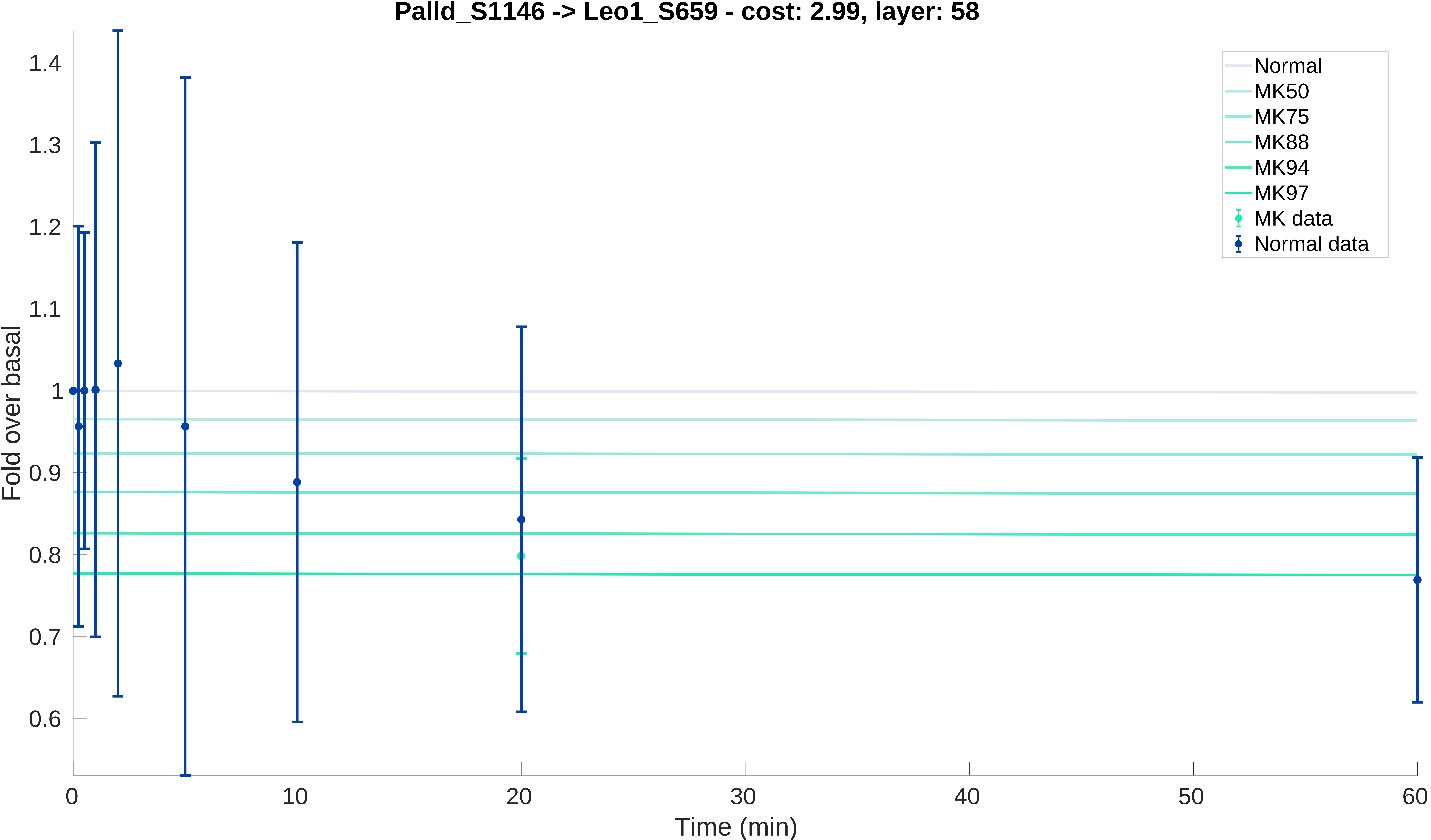

Palld_S1146 -> Leo1_S659 - cost: 2.99, layer: 58
Normal
MK50
MK75
MK88
MK94
MK97
MK data
Normal data
Fold over basal
Time (min)
1.4
1.3
1.2
1.1
1
0.9
0.8
0.7
0.6
0
10
20
30
40
50
60

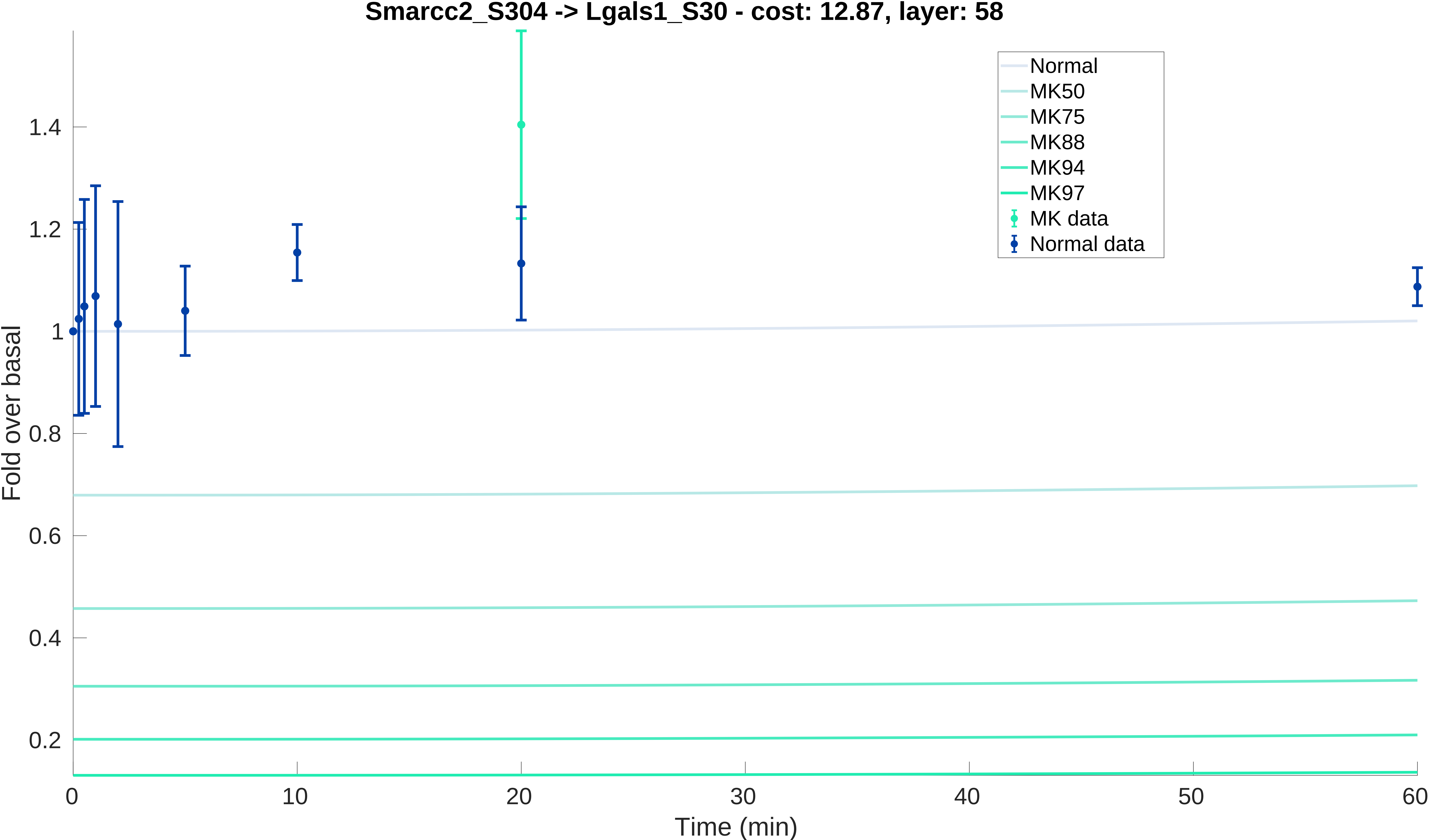

Smarcc2_S304 -> Lgals1_S30 - cost: 12.87, layer: 58
Normal
MK50
MK75
MK88
MK94
MK97
MK data
Normal data
Fold over basal
0.2
0.4
0.6
0.8
1
1.2
1.4
Time (min)
0
10
20
30
40
50
60

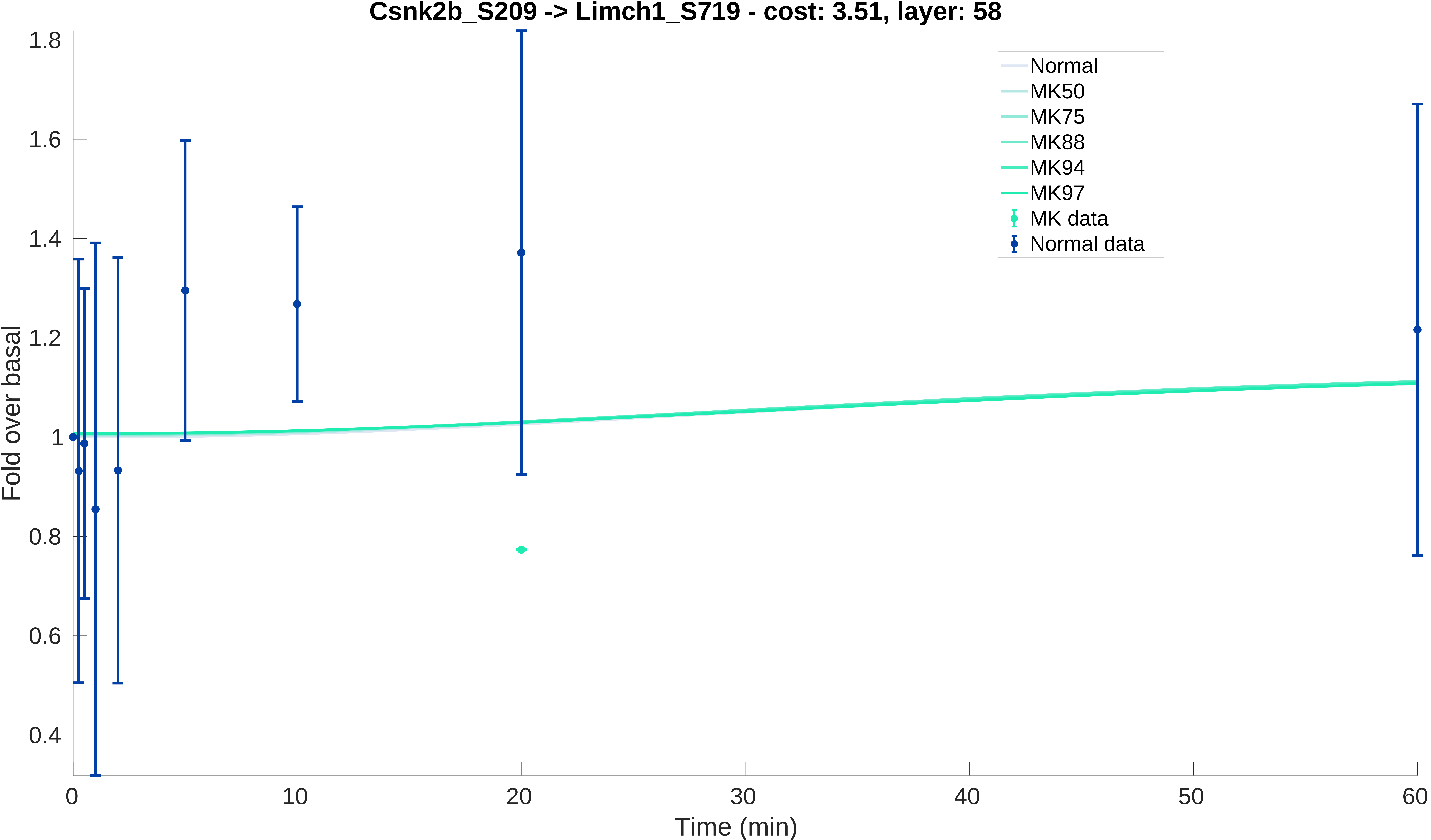

Csnk2b_S209 -> Limch1_S719 - cost: 3.51, layer: 58
Fold over basal
Time (min)
1.8
1.6
1.4
1.2
1
0.8
0.6
0.4
0
10
20
30
40
50
60
Normal
MK50
MK75
MK88
MK94
MK97
MK data
Normal data

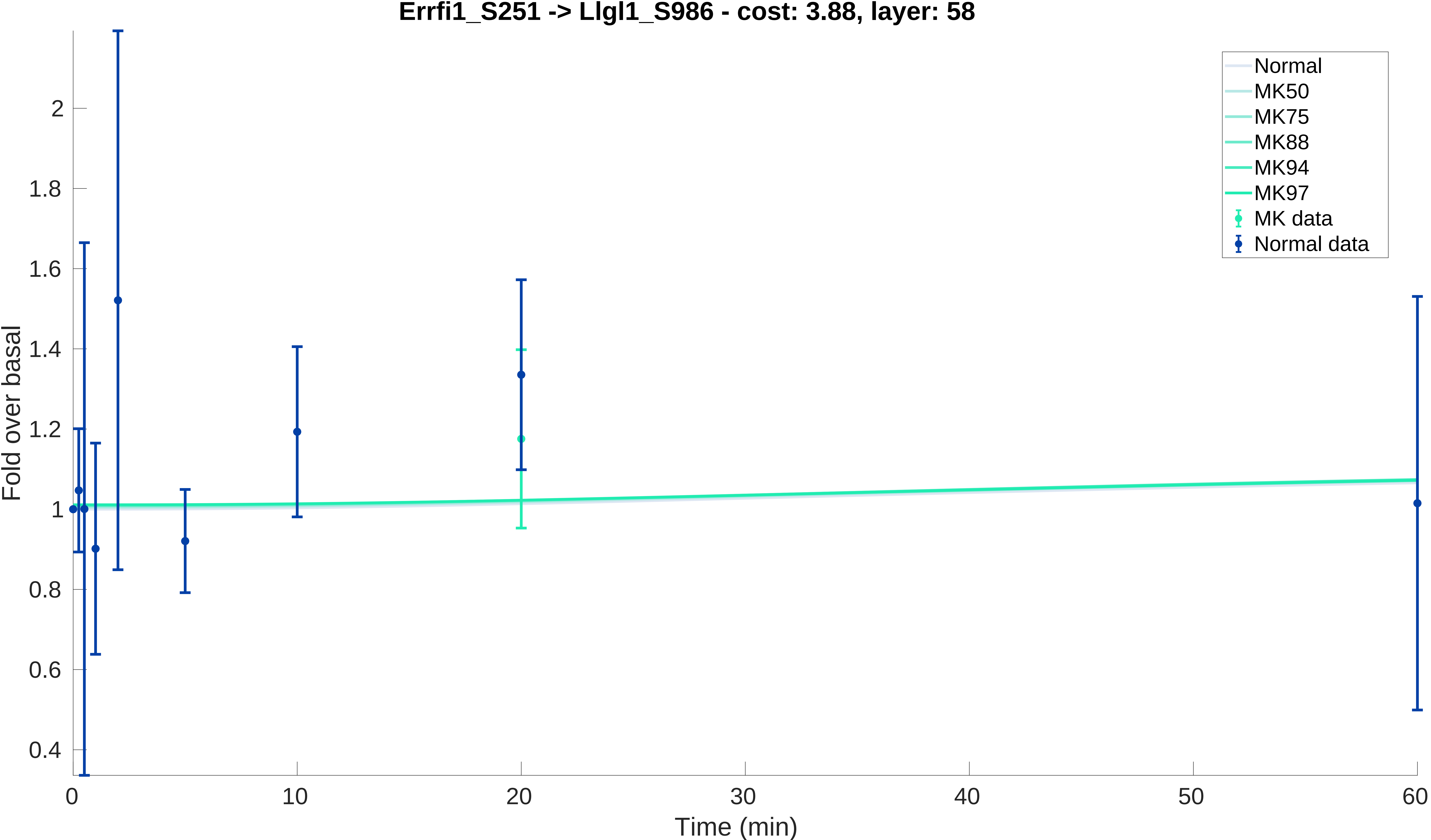

Errfi1_S251 -> Llgl1_S986 - cost: 3.88, layer: 58
Normal
MK50
MK75
MK88
MK94
MK97
MK data
Normal data
Fold over basal
0.4
0.6
0.8
1
1.2
1.4
1.6
1.8
2
0
10
20
30
40
50
60
Time (min)

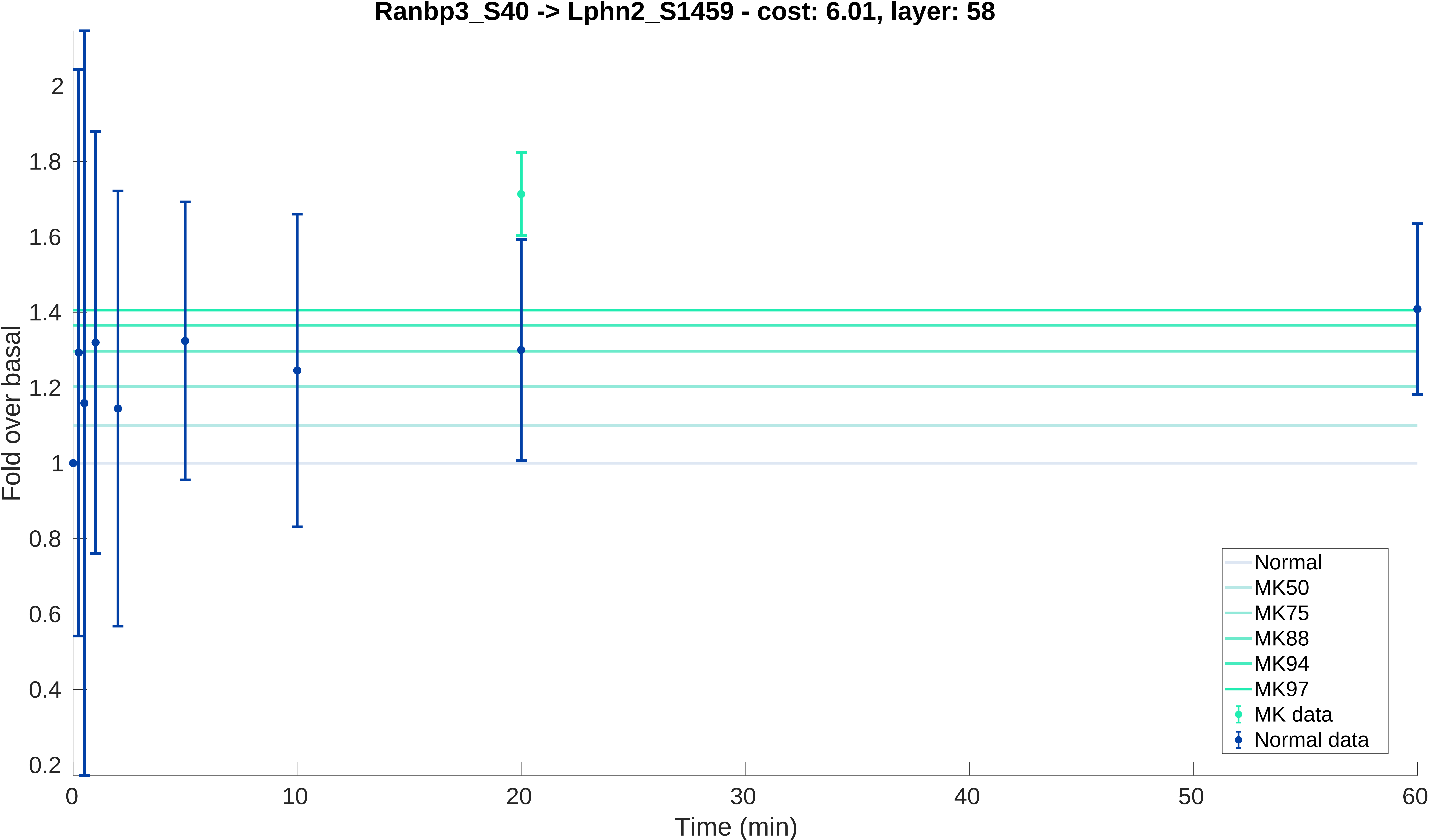

Ranbp3_S40 -> Lphn2_S1459 - cost: 6.01, layer: 58
Fold over basal
Time (min)
0
10
20
30
40
50
60
0.2
0.4
0.6
0.8
1
1.2
1.4
1.6
1.8
2
Normal
MK50
MK75
MK88
MK94
MK97
MK data
Normal data

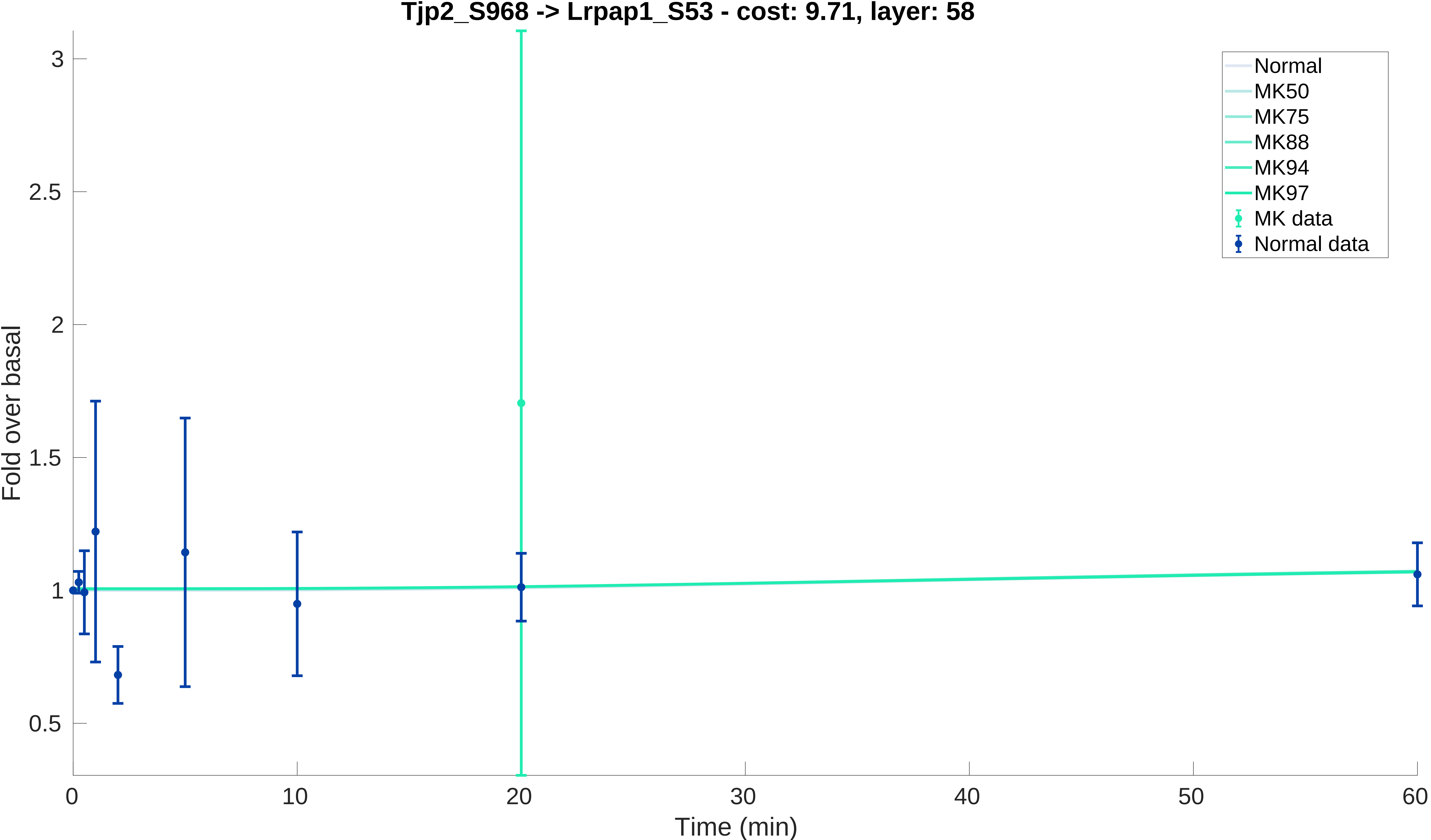

Tjp2_S968 -> Lrpap1_S53 - cost: 9.71, layer: 58
Fold over basal
Time (min)
0.5
1
1.5
2
2.5
3
0
10
20
30
40
50
60
Normal
MK50
MK75
MK88
MK94
MK97
MK data
Normal data

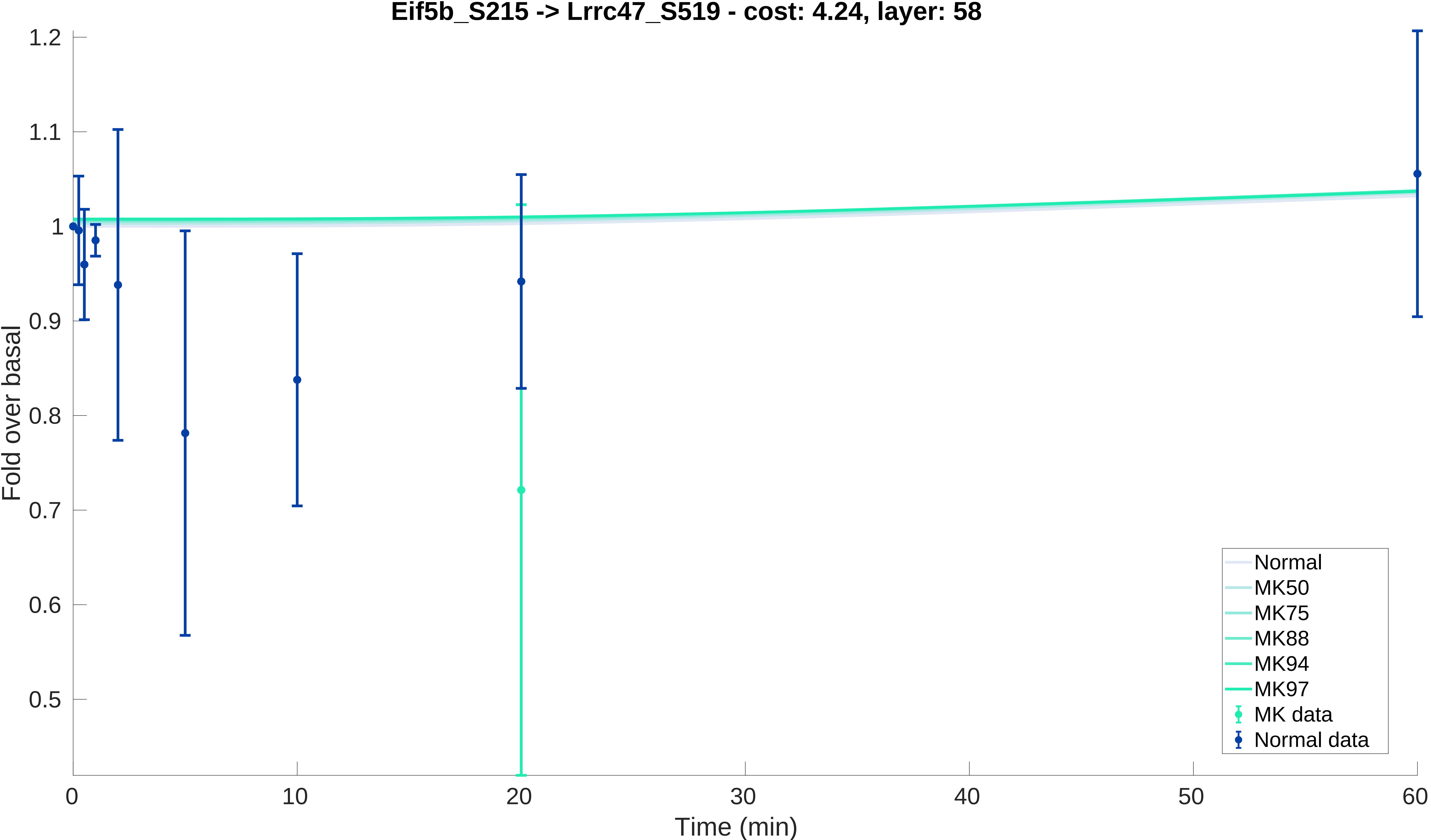
Eif5b_S215 -> Lrrc47_S519 - cost: 4.24, layer: 58
Fold over basal
Time (min)
1.2
1.1
1
0.9
0.8
0.7
0.6
0.5
0
10
20
30
40
50
60
Normal
MK50
MK75
MK88
MK94
MK97
MK data
Normal data

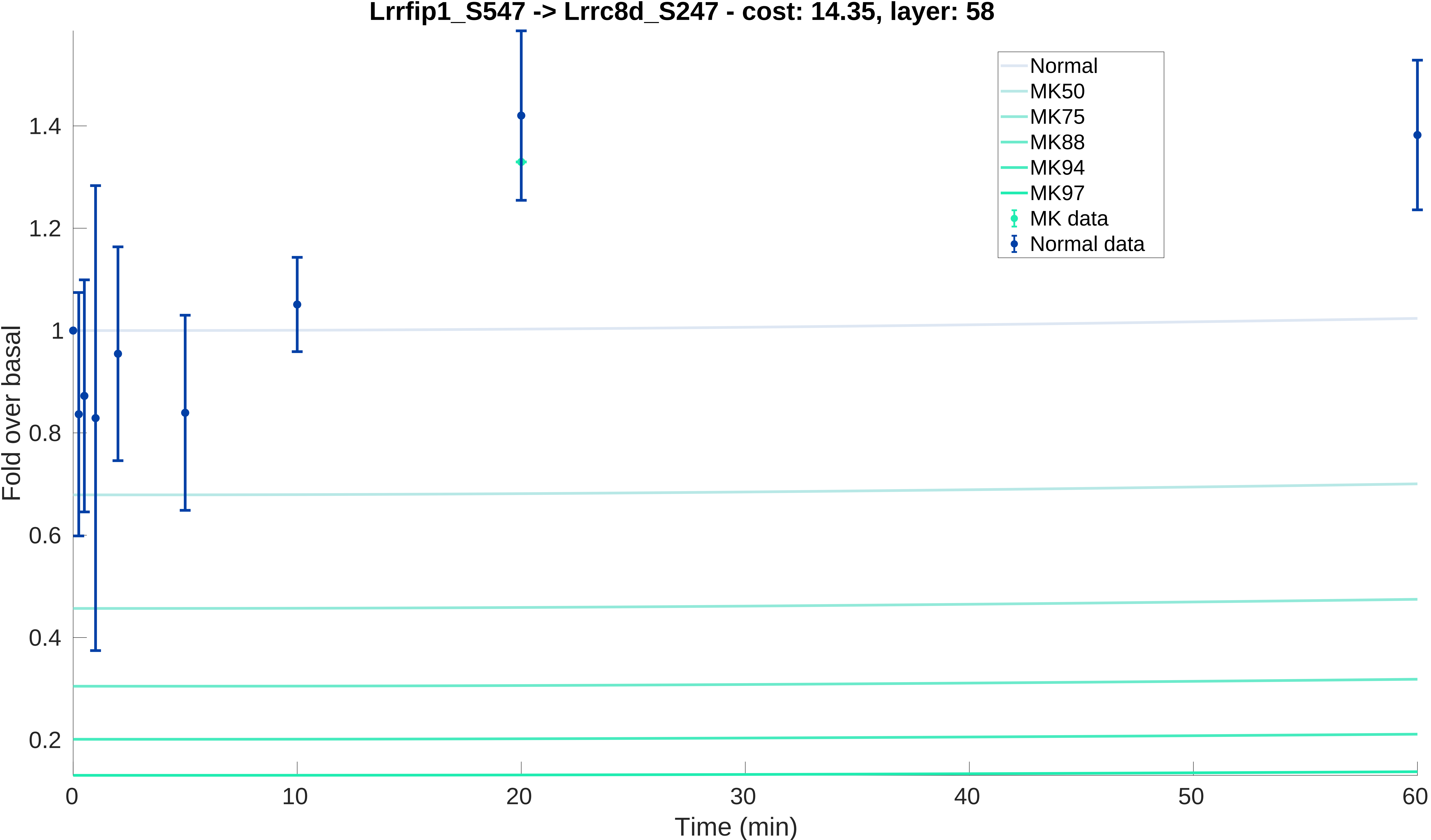

Lrrfip1_S547 -> Lrrc8d_S247 - cost: 14.35, layer: 58
Normal
MK50
MK75
MK88
MK94
MK97
MK data
Normal data
Fold over basal
1.4
1.2
1
0.8
0.6
0.4
0.2
0
10
20
30
40
50
60
Time (min)

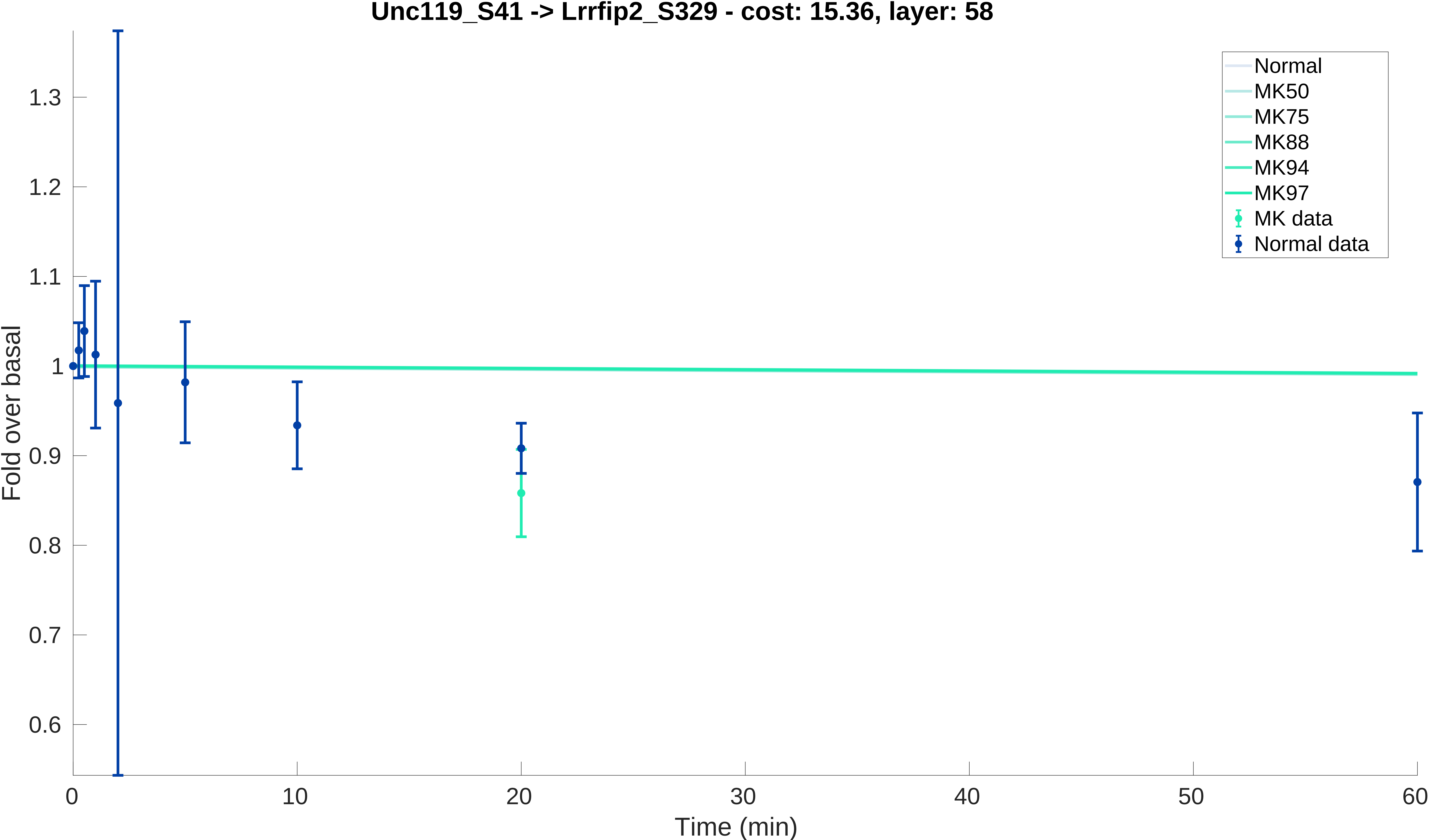

Unc119_S41 -> Lrrfip2_S329 - cost: 15.36, layer: 58
Fold over basal
1.3
1.2
1.1
1
0.9
0.8
0.7
0.6
0
10
20
30
40
50
60
Time (min)
Normal
MK50
MK75
MK88
MK94
MK97
MK data
Normal data

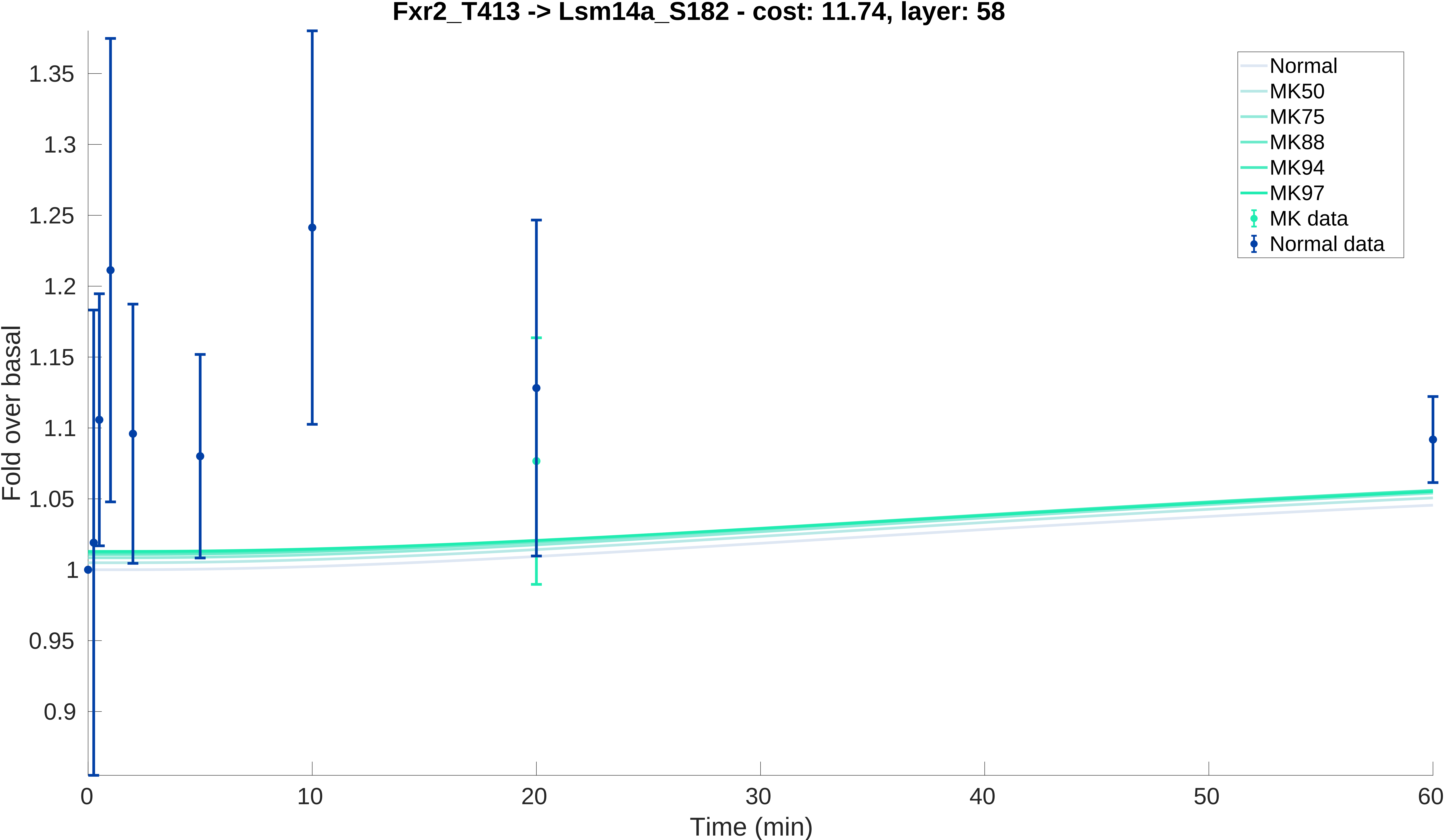

Fxr2_T413 -> Lsm14a_S182 - cost: 11.74, layer: 58
Fold over basal
1.35
1.3
1.25
1.2
1.15
1.1
1.05
1
0.95
0.9
0
10
20
30
40
50
60
Time (min)
Normal
MK50
MK75
MK88
MK94
MK97
MK data
Normal data

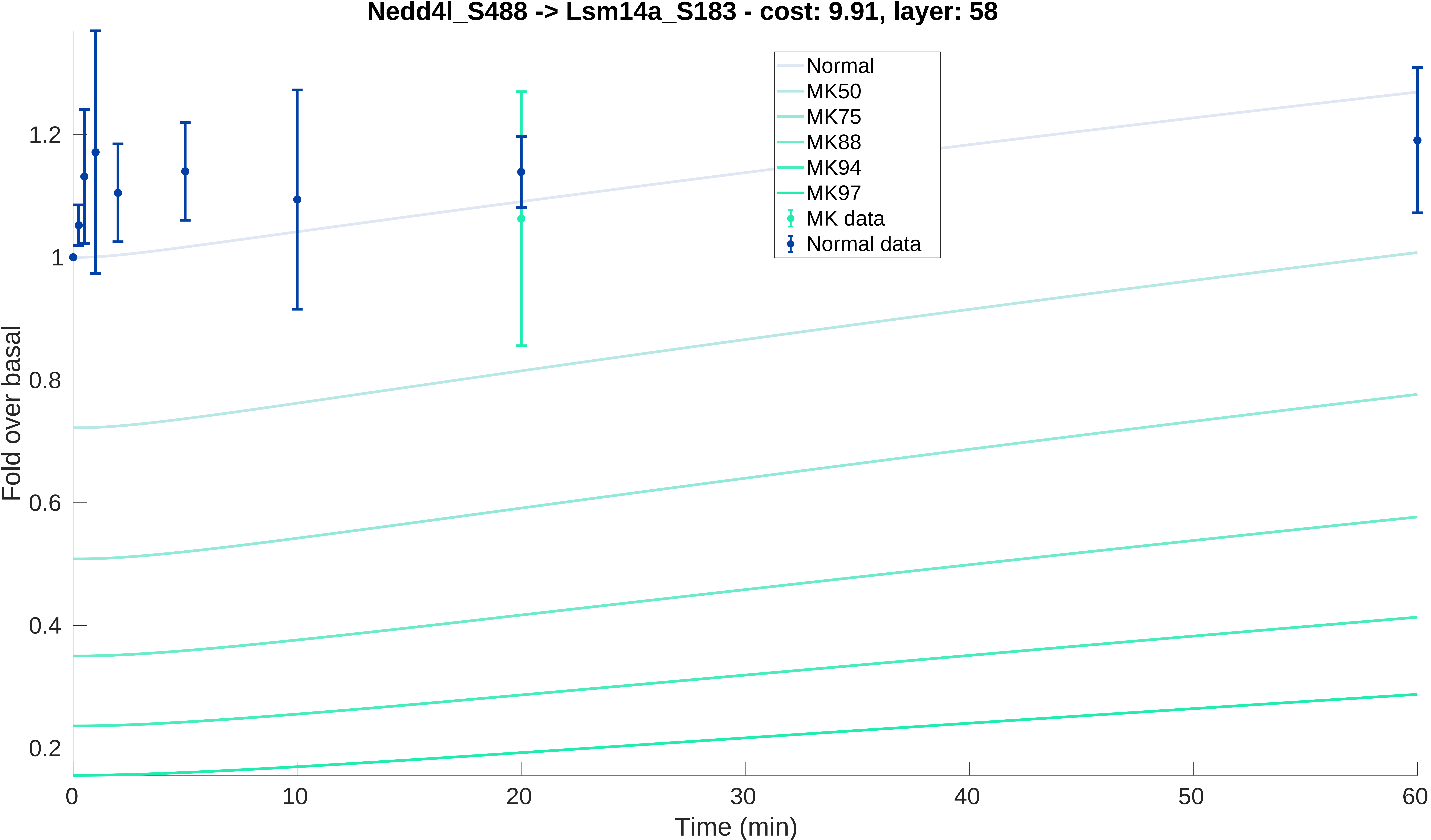

Nedd4l_S488 -> Lsm14a_S183 - cost: 9.91, layer: 58
Normal
MK50
MK75
MK88
MK94
MK97
MK data
Normal data
Fold over basal
1.2
1
0.8
0.6
0.4
0.2
0
10
20
30
40
50
60
Time (min)

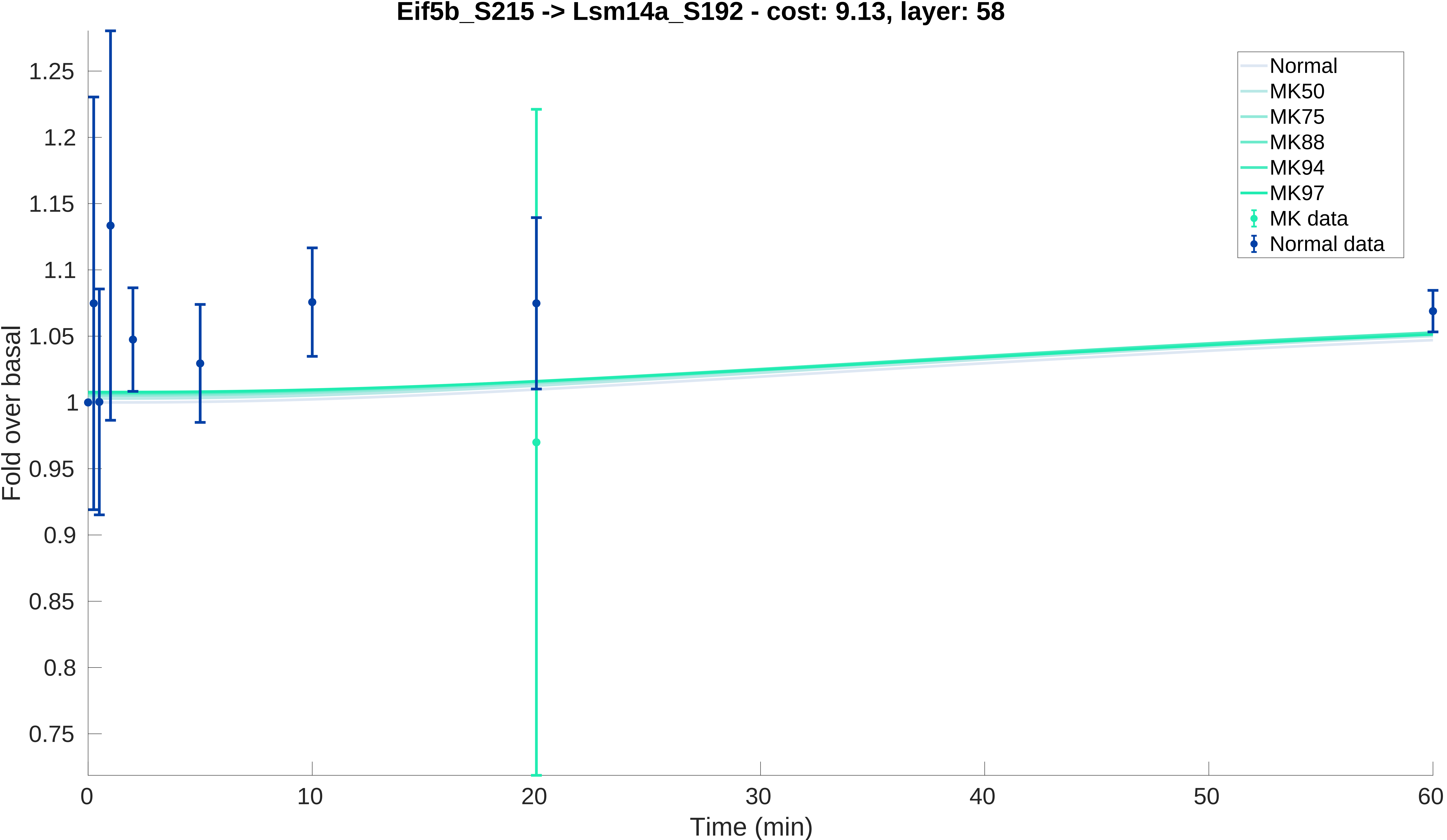

Eif5b_S215 -> Lsm14a_S192 - cost: 9.13, layer: 58
Fold over basal
Time (min)
1.25
1.2
1.15
1.1
1.05
1
0.95
0.9
0.85
0.8
0.75
0
10
20
30
40
50
60
Normal
MK50
MK75
MK88
MK94
MK97
MK data
Normal data

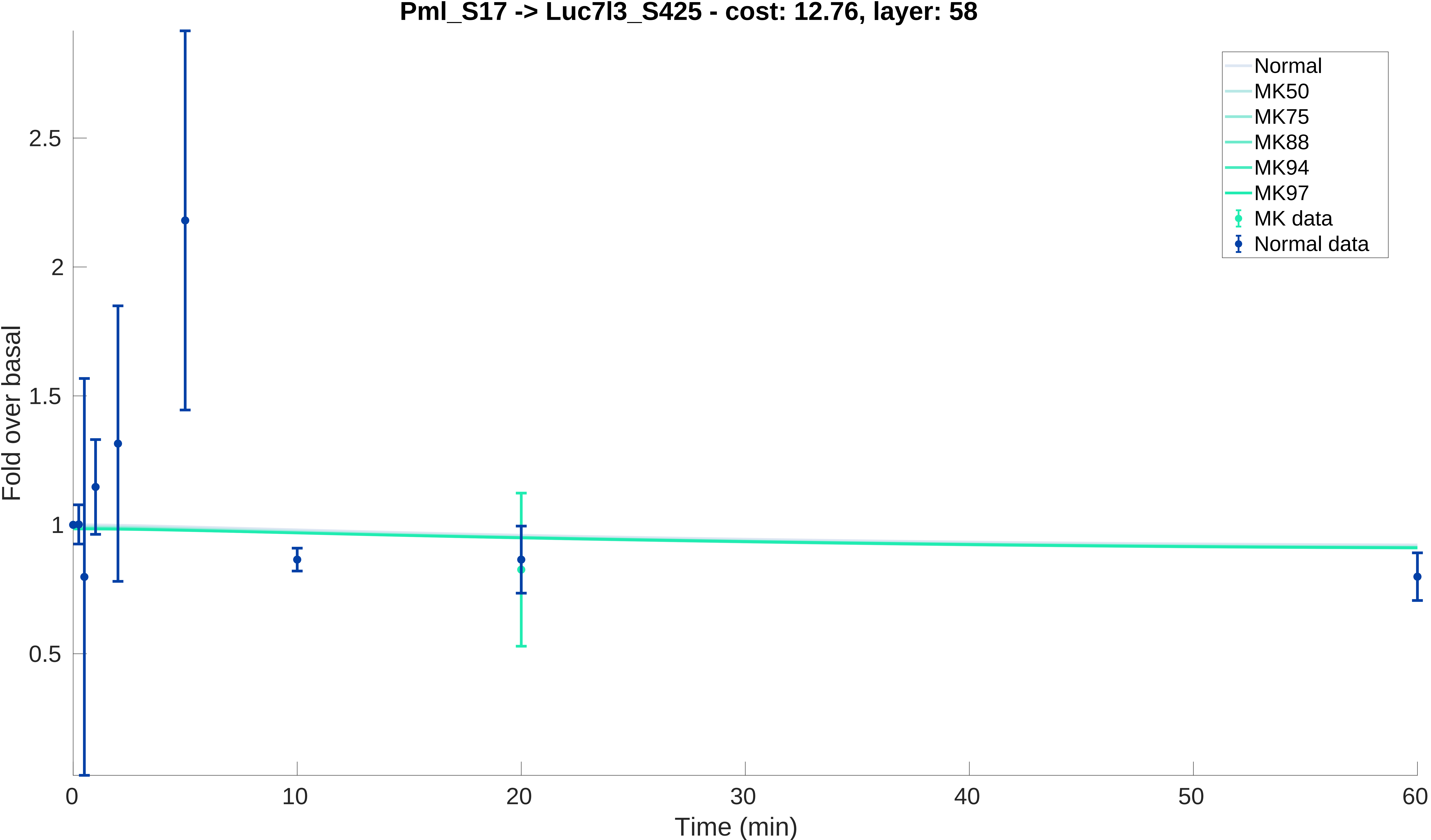

Pml_S17 -> Luc7l3_S425 - cost: 12.76, layer: 58
Normal
MK50
MK75
MK88
MK94
MK97
MK data
Normal data
Fold over basal
2.5
2
1.5
1
0.5
0
10
20
30
40
50
60
Time (min)

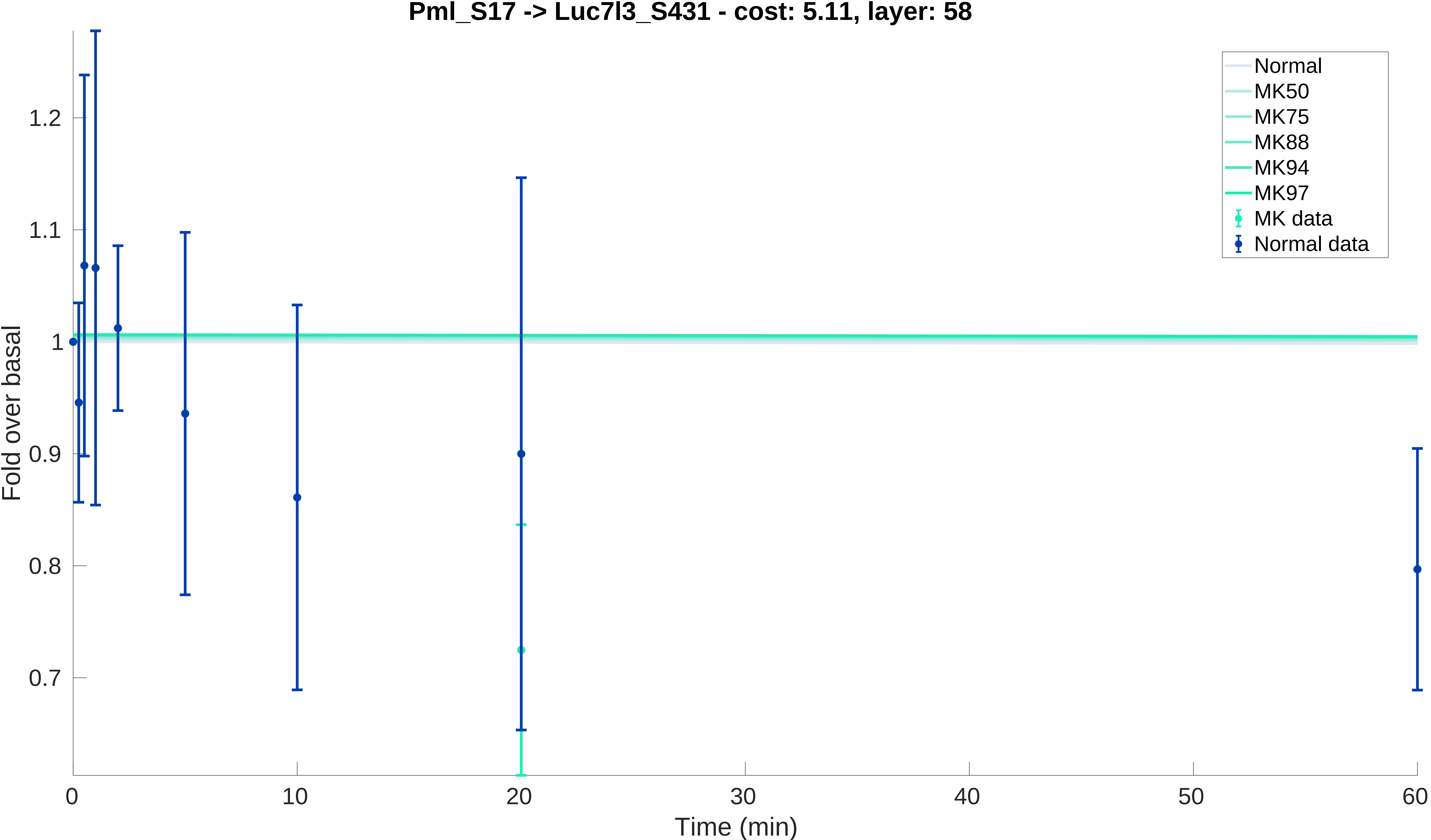

Pml_S17 -> Luc7l3_S431 - cost: 5.11, layer: 58
Fold over basal
Time (min)
0
10
20
30
40
50
60
0.7
0.8
0.9
1
1.1
1.2
Normal
MK50
MK75
MK88
MK94
MK97
MK data
Normal data

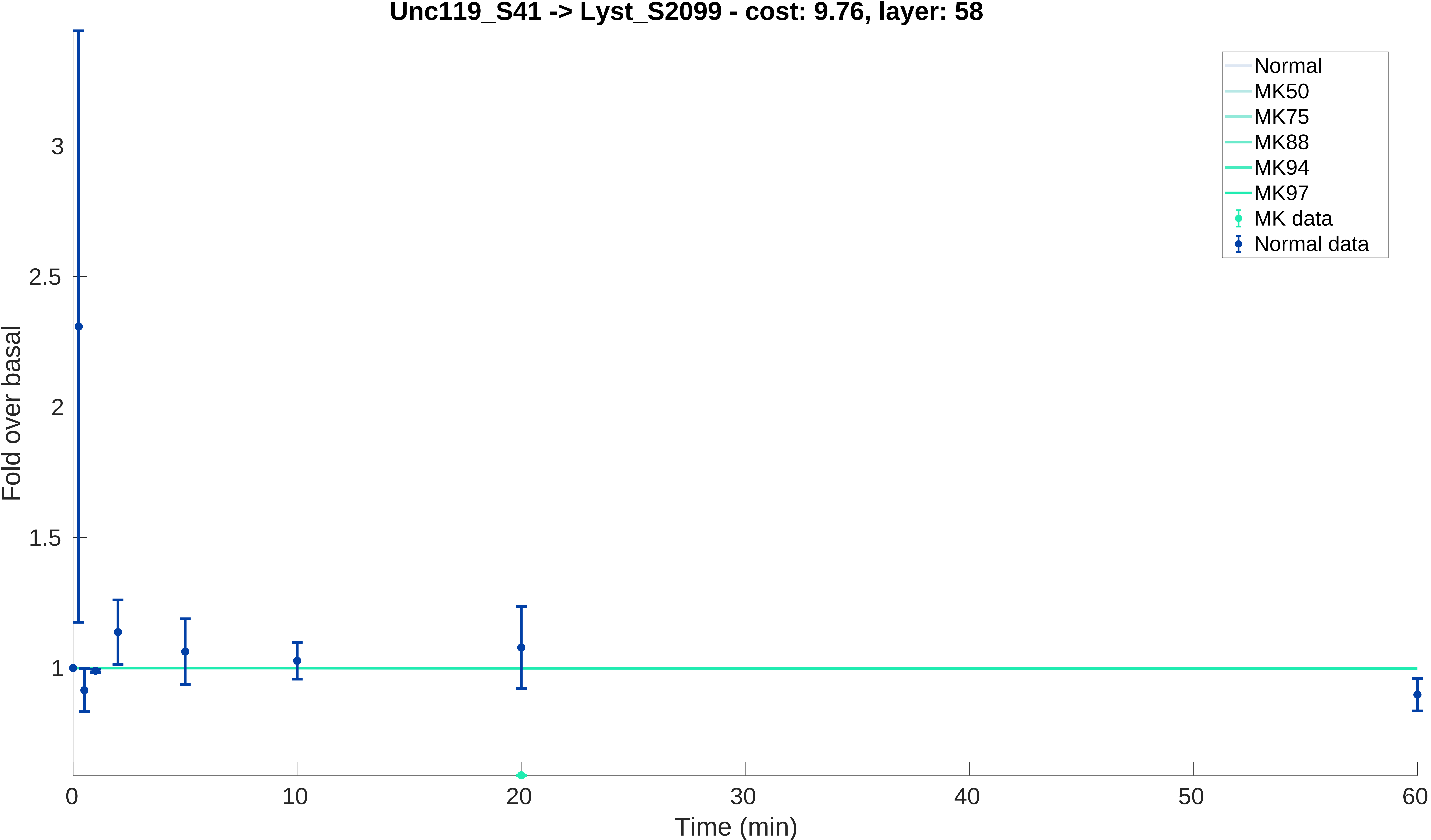

Unc119_S41 -> Lyst_S2099 - cost: 9.76, layer: 58
Normal
MK50
MK75
MK88
MK94
MK97
MK data
Normal data
Fold over basal
Time (min)
3
2.5
2
1.5
1
0
10
20
30
40
50
60

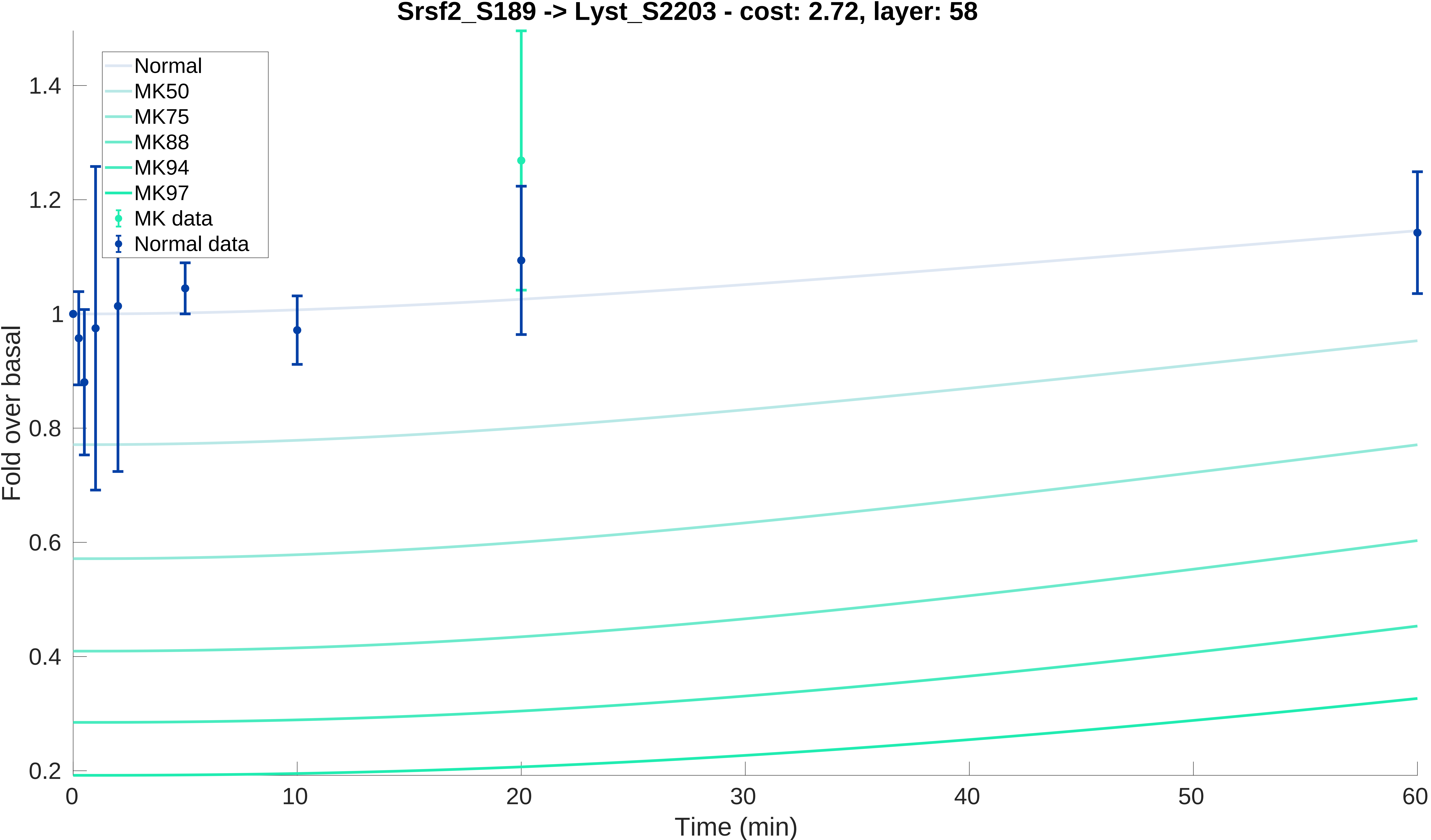
Srsf2_S189 -> Lyst_S2203 - cost: 2.72, layer: 58
Normal
MK50
MK75
MK88
MK94
MK97
MK data
Normal data
Fold over basal
1.4
1.2
1
0.8
0.6
0.4
0.2
0
10
20
30
40
50
60
Time (min)

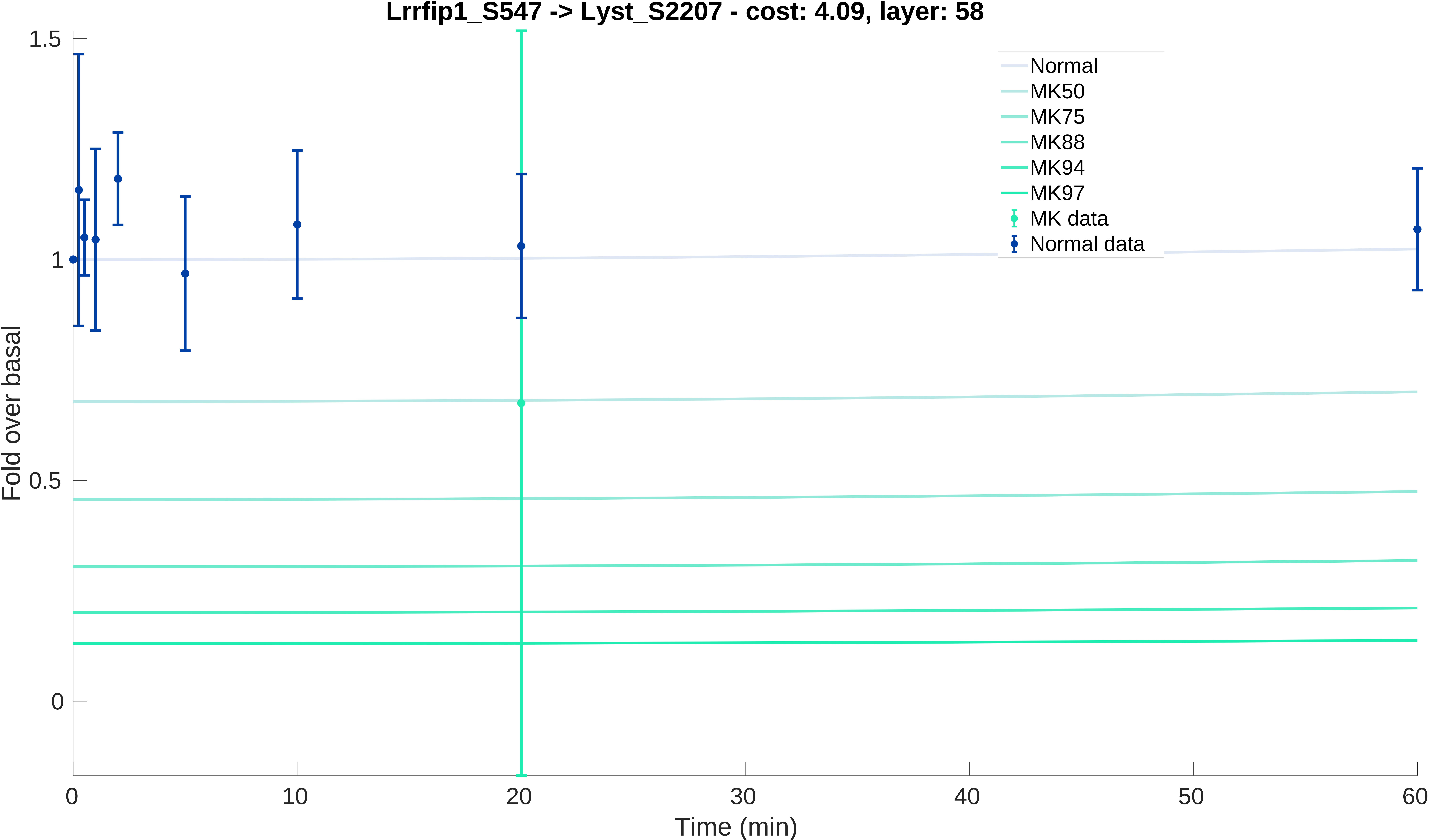

Lrrfip1_S547 -> Lyst_S2207 - cost: 4.09, layer: 58
Fold over basal
Time (min)
1.5
1
0.5
0
0
10
20
30
40
50
60
Normal
MK50
MK75
MK88
MK94
MK97
MK data
Normal data

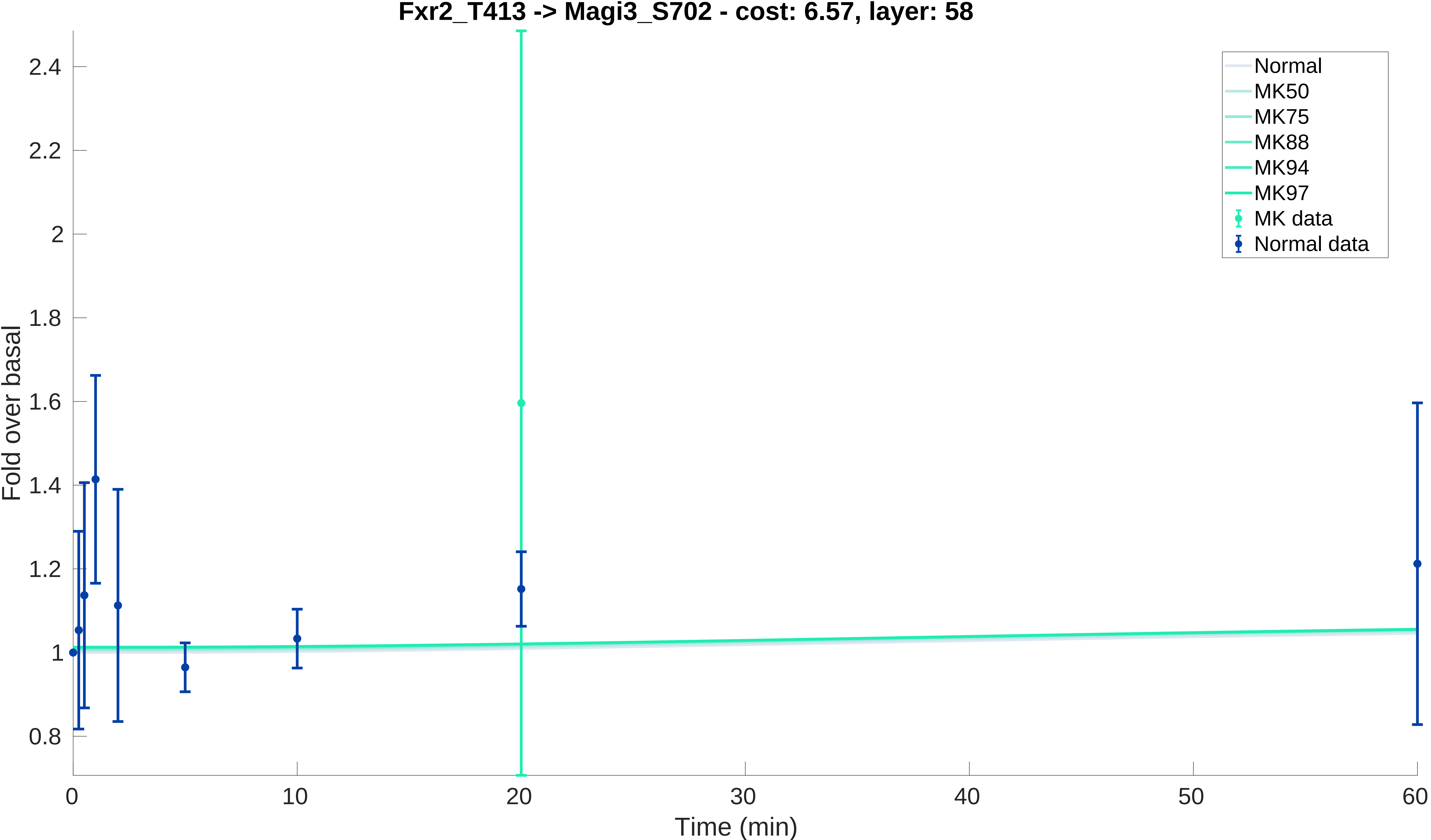

Fxr2_T413 -> Magi3_S702 - cost: 6.57, layer: 58
Normal
MK50
MK75
MK88
MK94
MK97
MK data
Normal data
Fold over basal
2.4
2.2
2
1.8
1.6
1.4
1.2
1
0.8
0
10
20
30
40
50
60
Time (min)

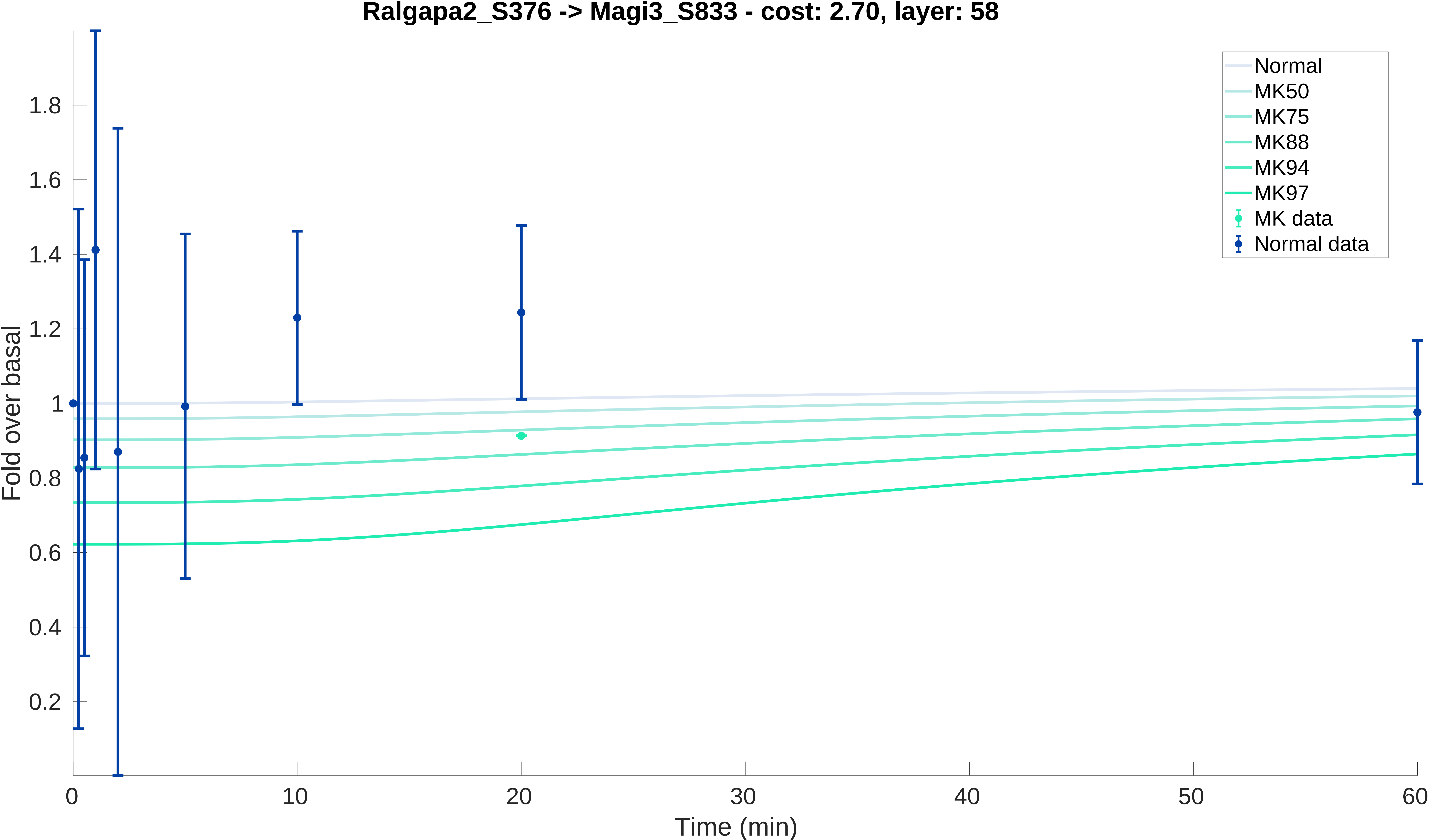

Ralgapa2_S376 -> Magi3_S833 - cost: 2.70, layer: 58
Normal
MK50
MK75
MK88
MK94
MK97
MK data
Normal data
Fold over basal
Time (min)
0
10
20
30
40
50
60
0.2
0.4
0.6
0.8
1
1.2
1.4
1.6
1.8

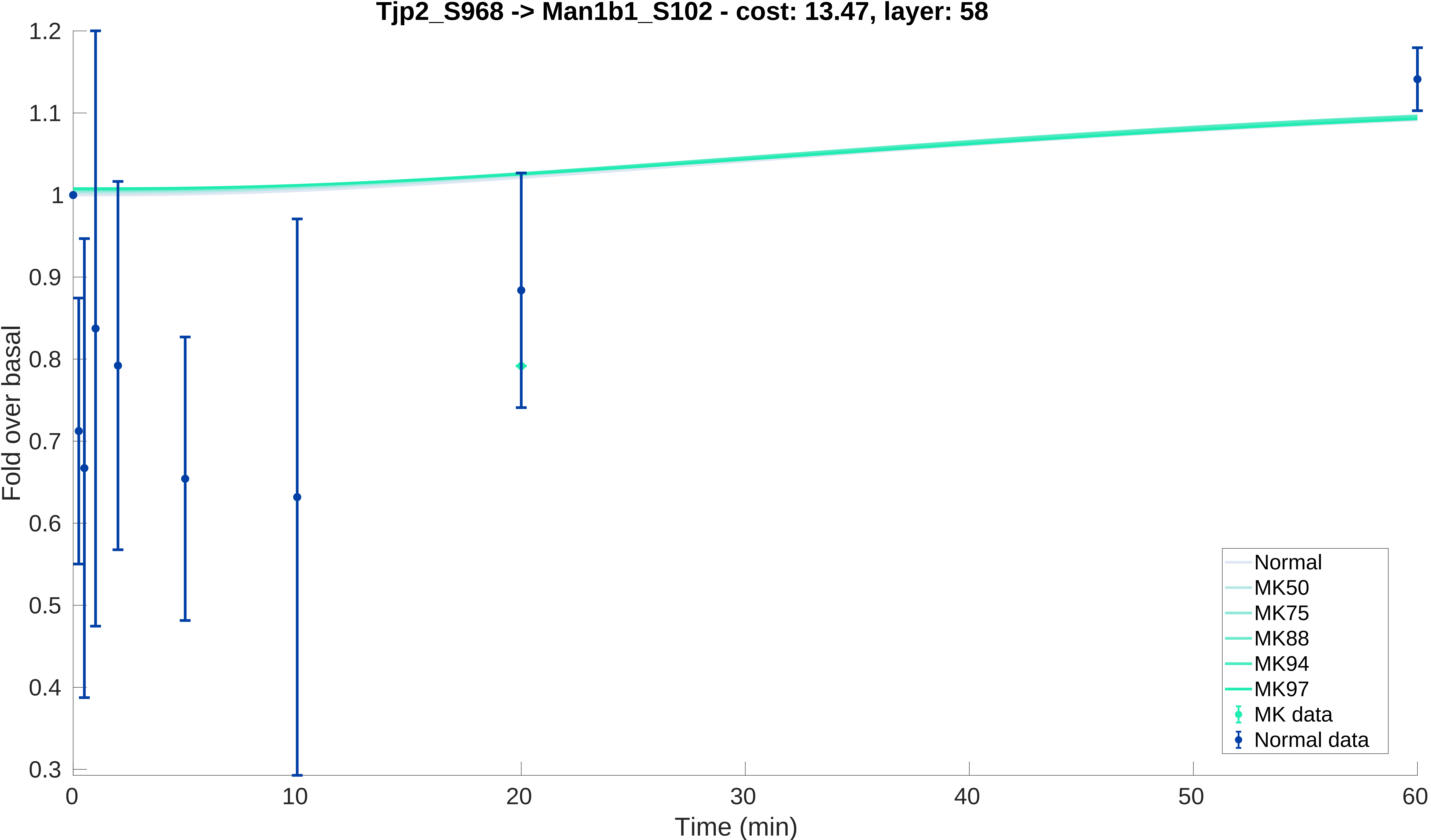

Tjp2_S968 -> Man1b1_S102 - cost: 13.47, layer: 58
Fold over basal
Time (min)
1.2
1.1
1
0.9
0.8
0.7
0.6
0.5
0.4
0.3
0
10
20
30
40
50
60
Normal
MK50
MK75
MK88
MK94
MK97
MK data
Normal data

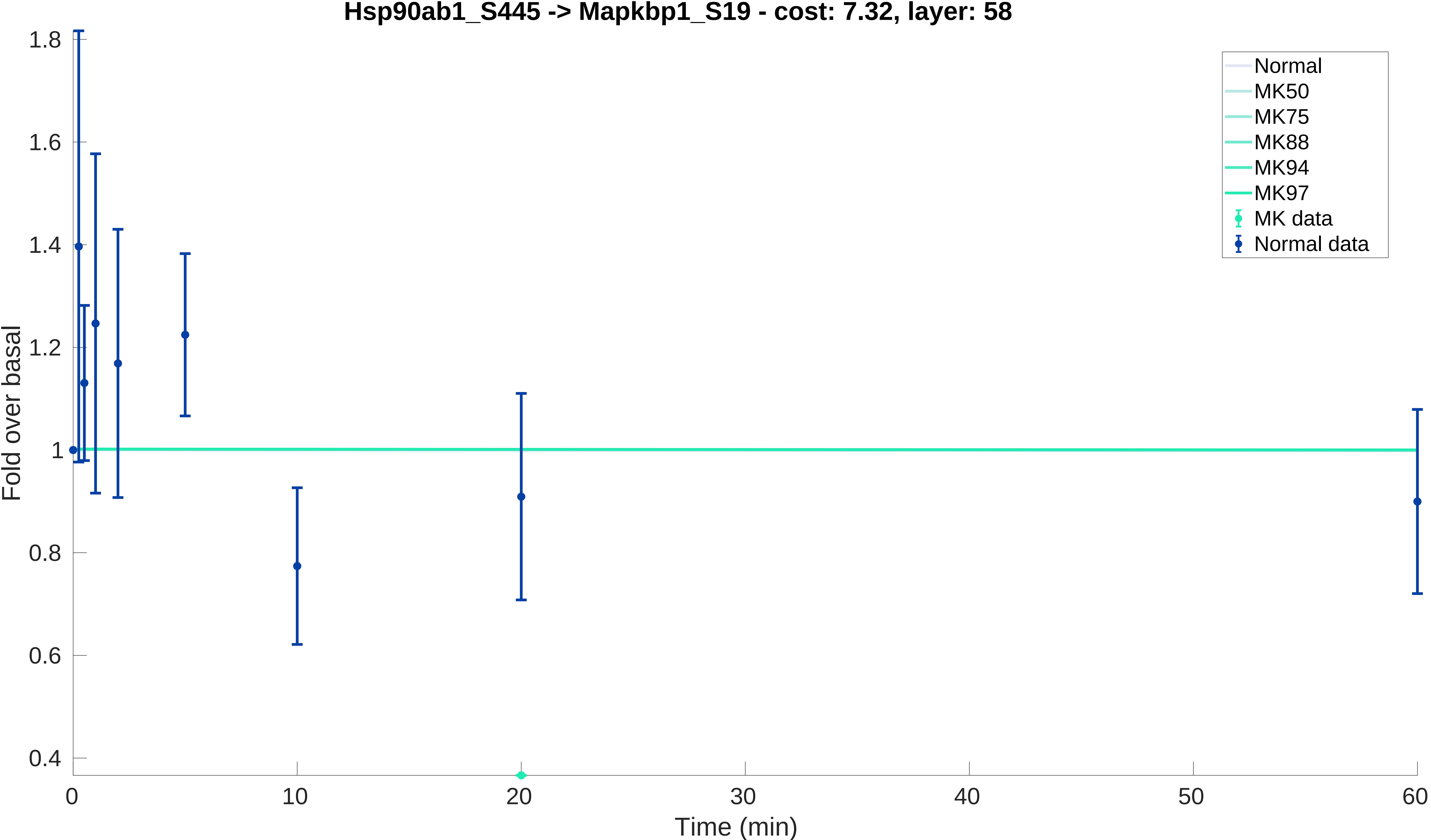

Hsp90ab1_S445 -> Mapkbp1_S19 - cost: 7.32, layer: 58
Fold over basal
Time (min)
0.4
0.6
0.8
1
1.2
1.4
1.6
1.8
0
10
20
30
40
50
60
Normal
MK50
MK75
MK88
MK94
MK97
MK data
Normal data

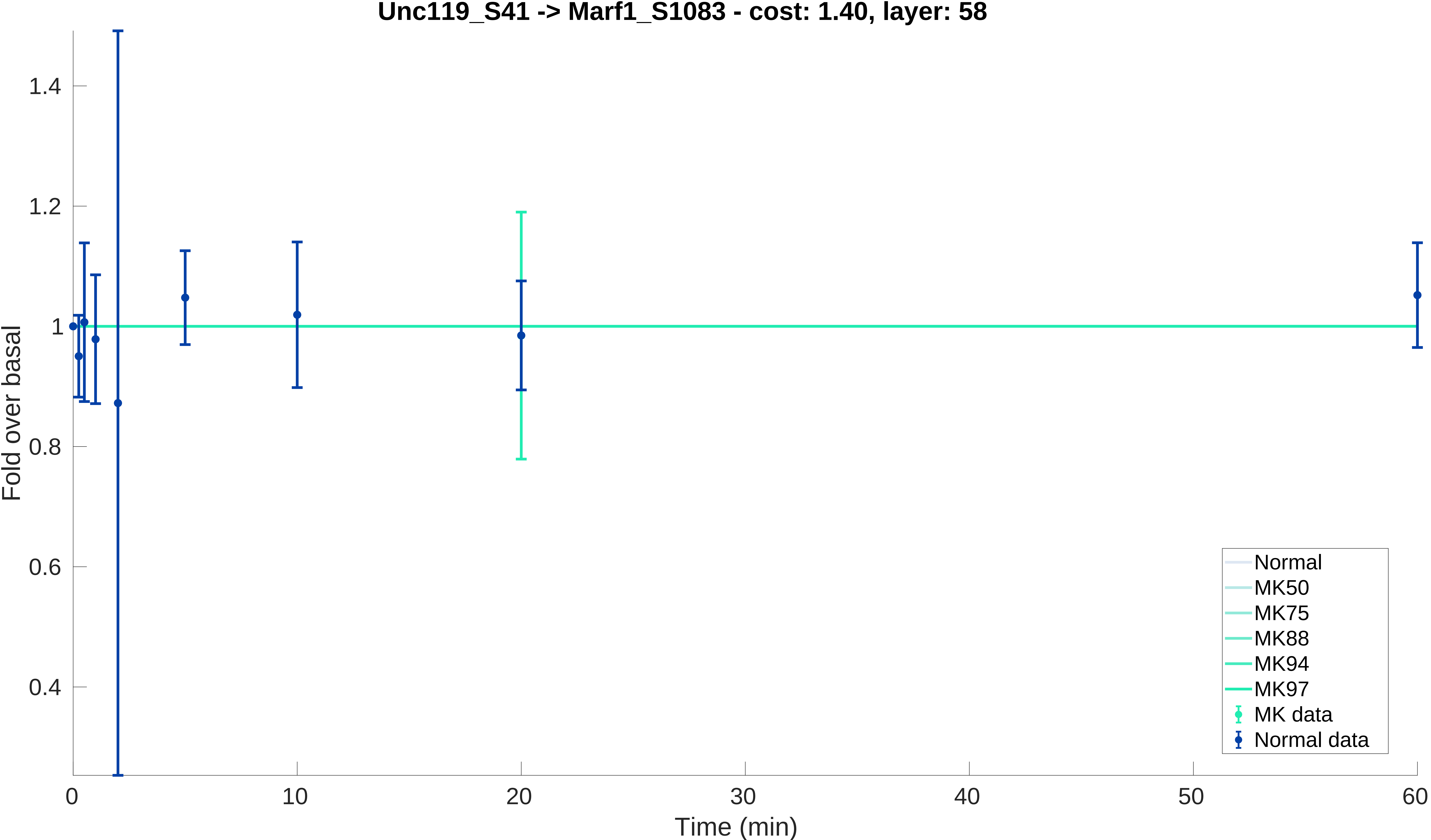

Unc119_S41 -> Marf1_S1083 - cost: 1.40, layer: 58
Fold over basal
1.4
1.2
1
0.8
0.6
0.4
0
10
20
30
40
50
60
Time (min)
Normal
MK50
MK75
MK88
MK94
MK97
MK data
Normal data

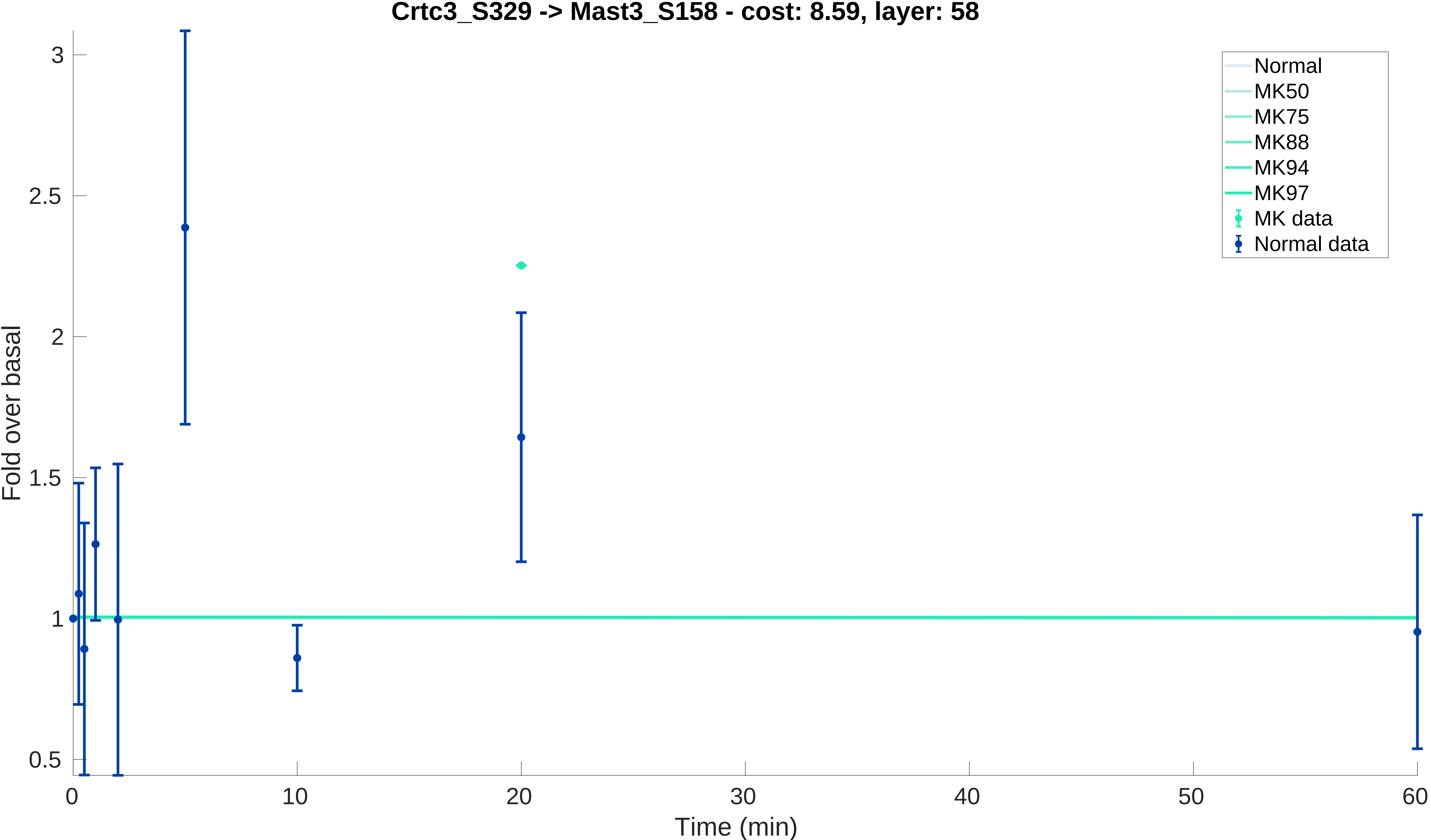

Crtc3_S329 -> Mast3_S158 - cost: 8.59, layer: 58
Fold over basal
Time (min)
0.5
1
1.5
2
2.5
3
0
10
20
30
40
50
60
Normal
MK50
MK75
MK88
MK94
MK97
MK data
Normal data

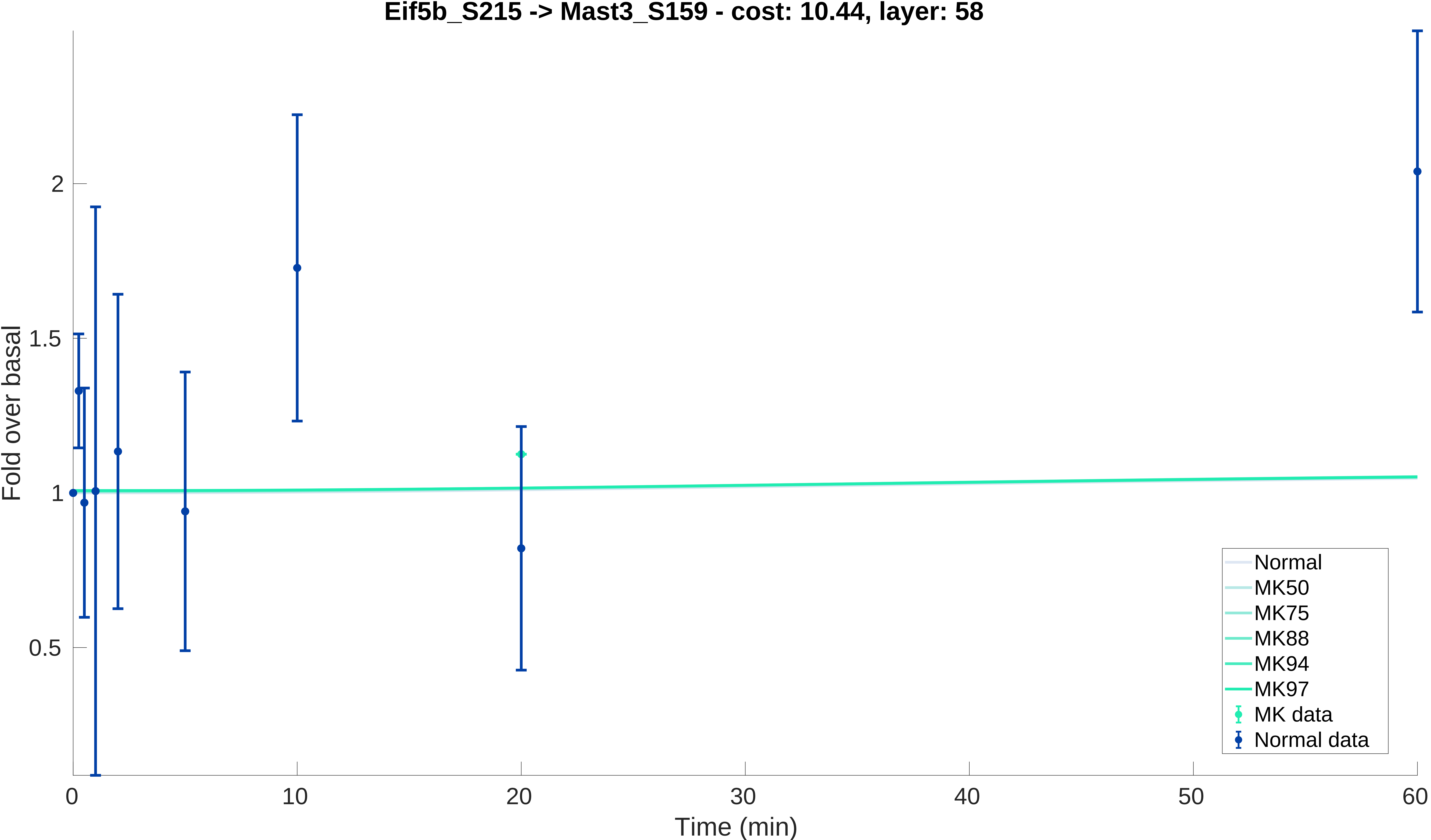
Eif5b_S215 -> Mast3_S159 - cost: 10.44, layer: 58
Fold over basal
Time (min)
0
10
20
30
40
50
60
0.5
1
1.5
2
Normal
MK50
MK75
MK88
MK94
MK97
MK data
Normal data

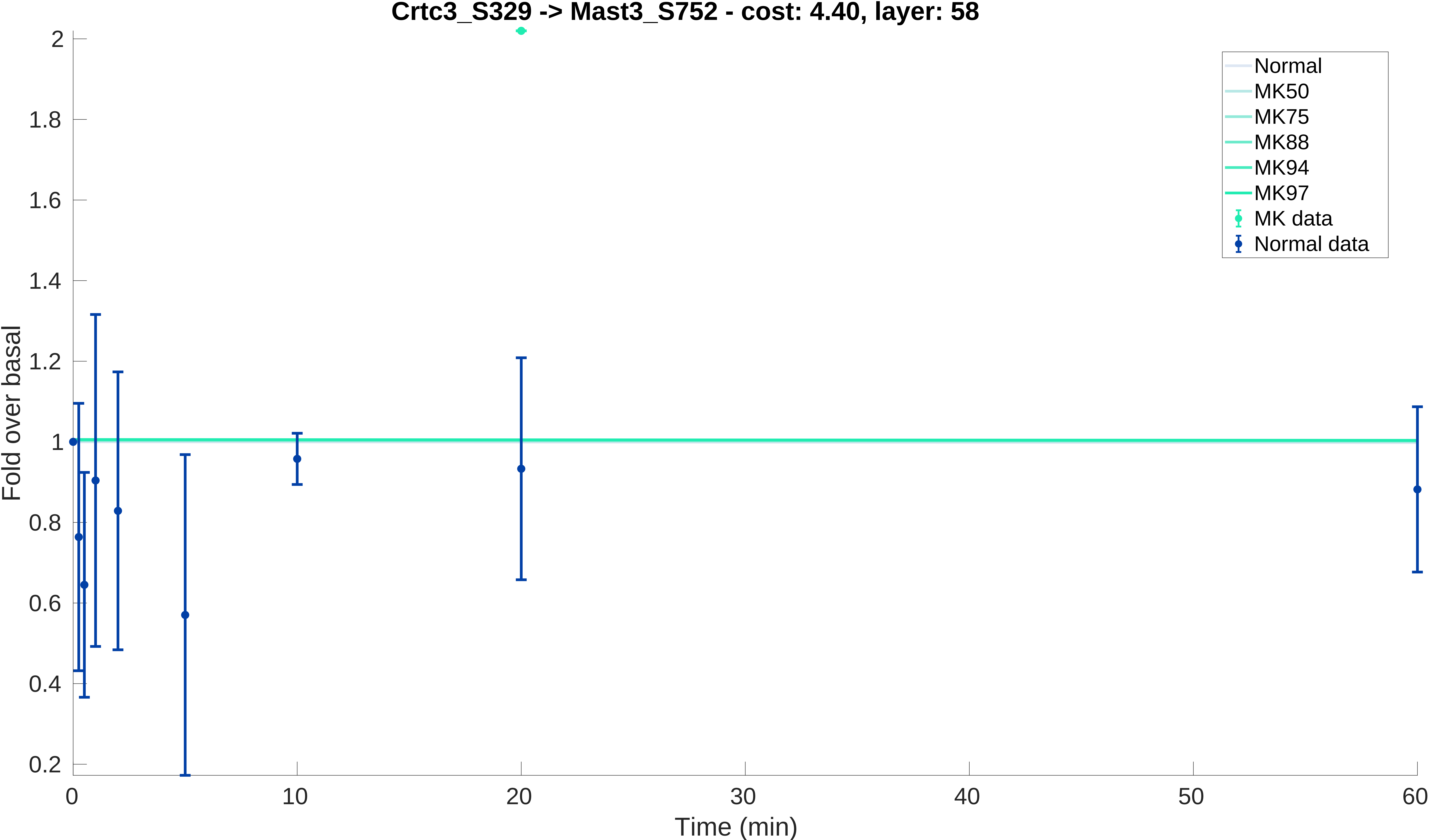

Crtc3_S329 -> Mast3_S752 - cost: 4.40, layer: 58
Fold over basal
Time (min)
Normal
MK50
MK75
MK88
MK94
MK97
MK data
Normal data
0.2
0.4
0.6
0.8
1
1.2
1.4
1.6
1.8
2
0
10
20
30
40
50
60

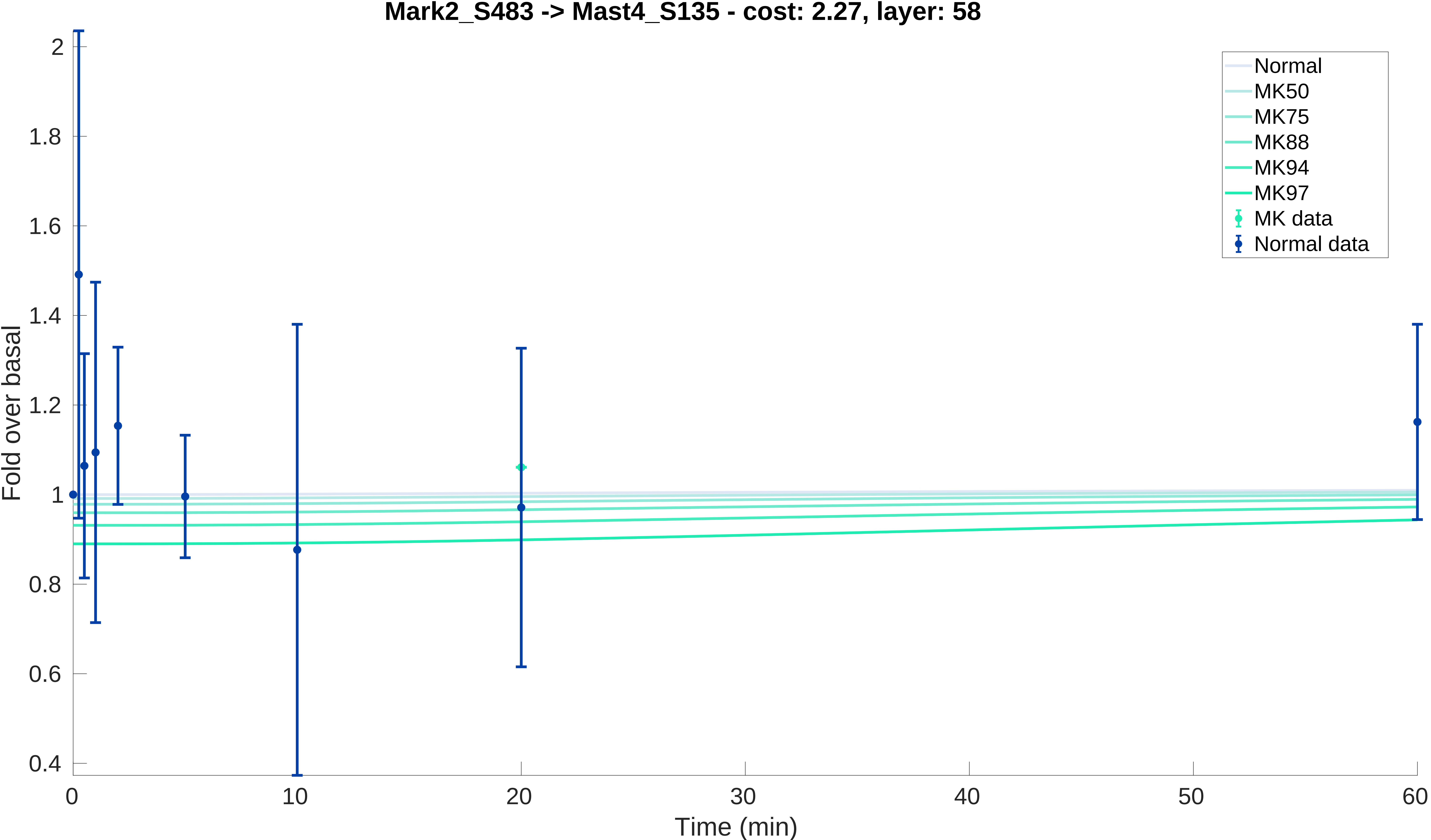

Mark2_S483 -> Mast4_S135 - cost: 2.27, layer: 58
Normal
MK50
MK75
MK88
MK94
MK97
MK data
Normal data
Fold over basal
Time (min)
0.4
0.6
0.8
1
1.2
1.4
1.6
1.8
2
0
10
20
30
40
50
60

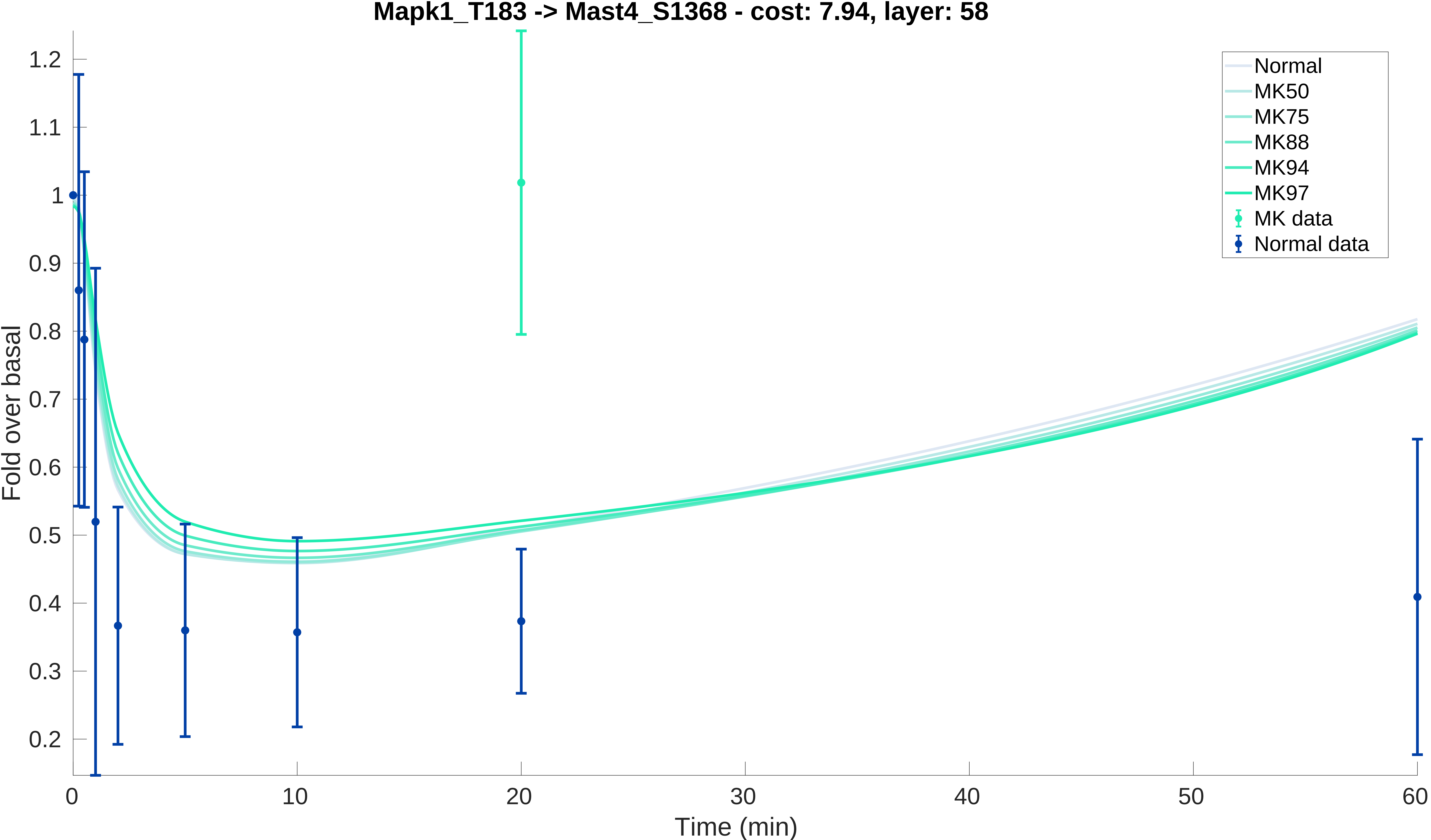

Mapk1_T183 -> Mast4_S1368 - cost: 7.94, layer: 58
Fold over basal
Time (min)
0
10
20
30
40
50
60
0.2
0.3
0.4
0.5
0.6
0.7
0.8
0.9
1
1.1
1.2
Normal
MK50
MK75
MK88
MK94
MK97
MK data
Normal data

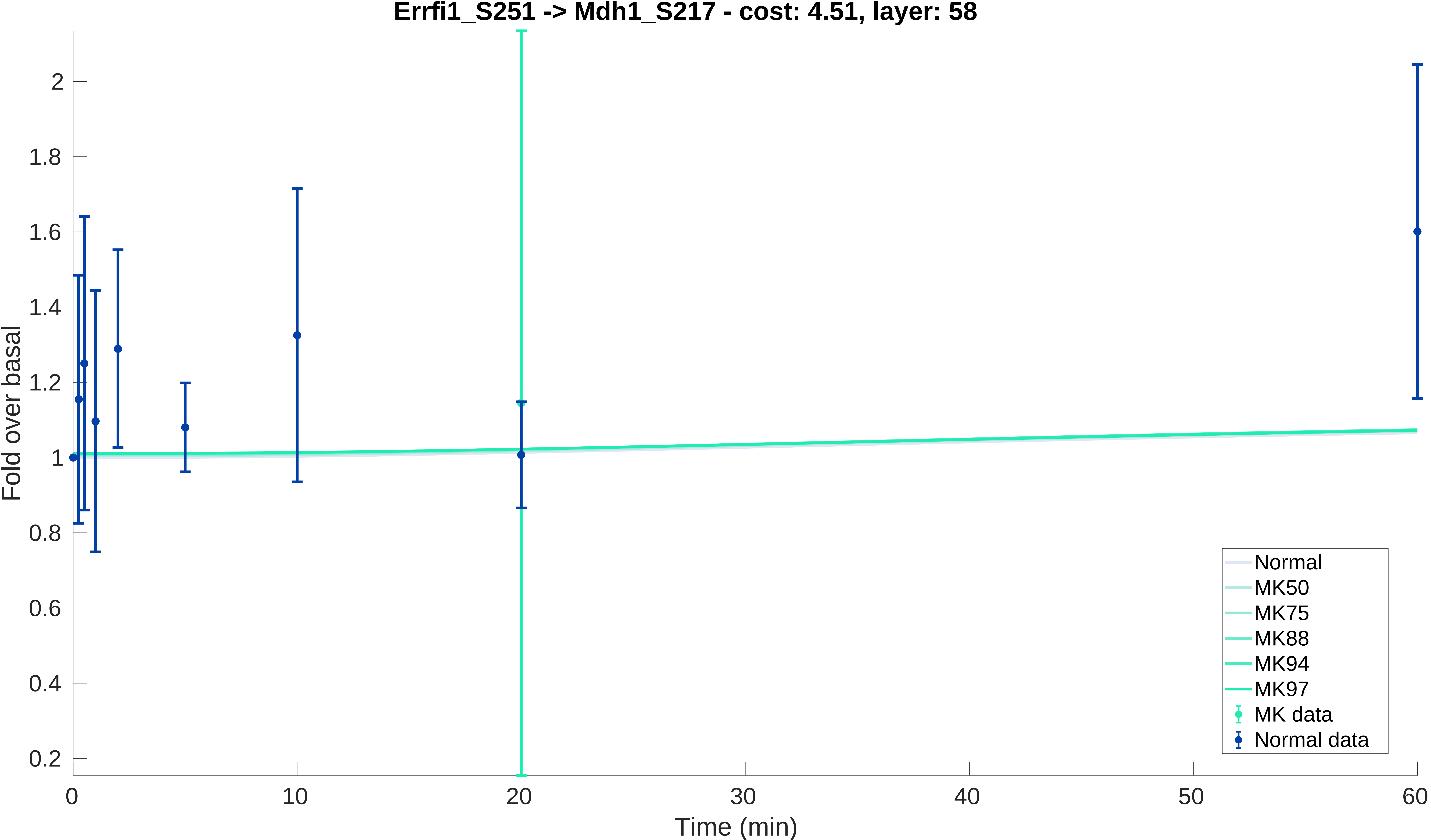
Errfi1_S251 -> Mdh1_S217 - cost: 4.51, layer: 58
Fold over basal
2
1.8
1.6
1.4
1.2
1
0.8
0.6
0.4
0.2
0
10
20
30
40
50
60
Time (min)
Normal
MK50
MK75
MK88
MK94
MK97
MK data
Normal data

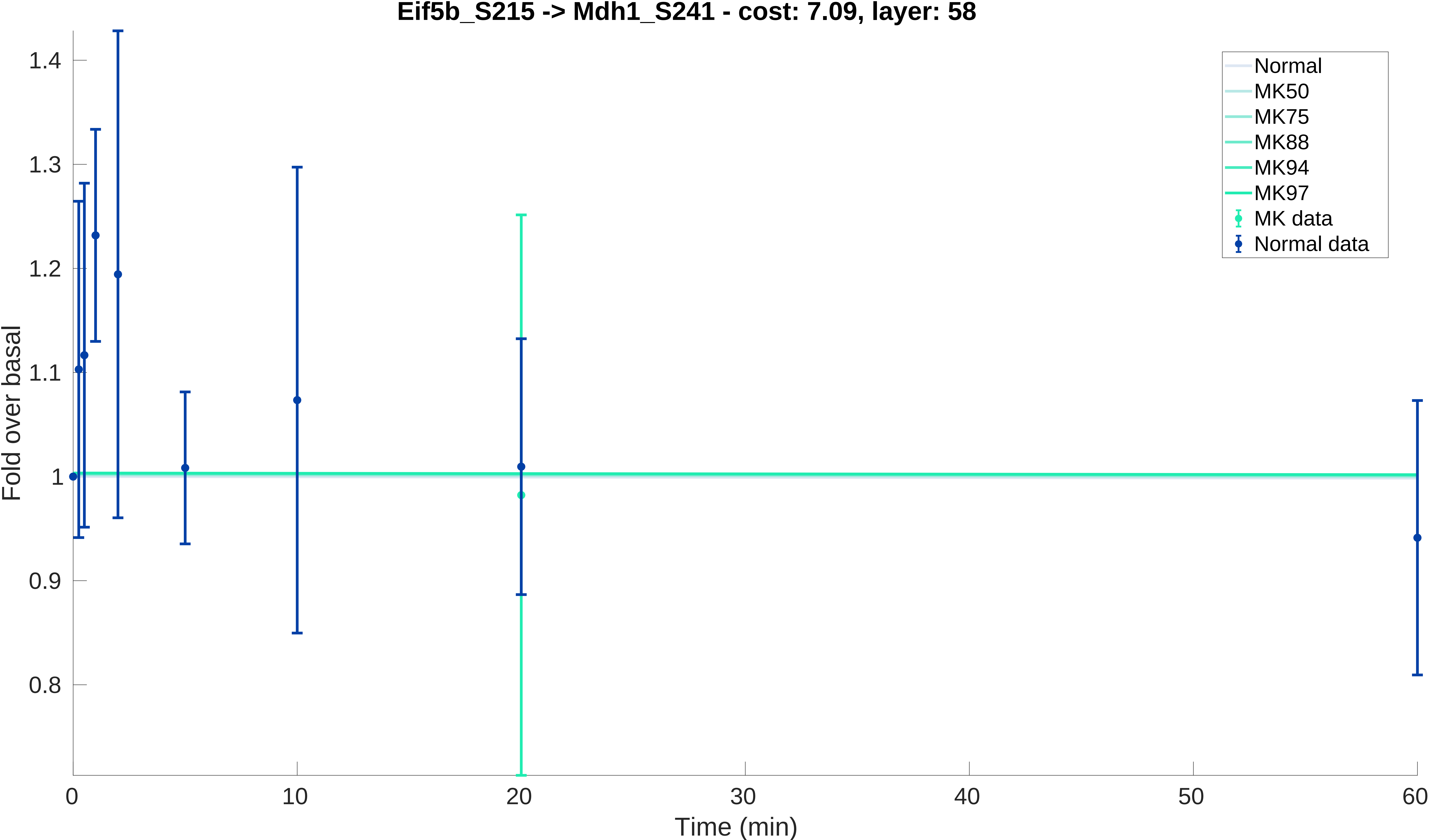

Eif5b_S215 -> Mdh1_S241 - cost: 7.09, layer: 58
1.4
1.3
1.2
1.1
1
0.9
0.8
Fold over basal
0
10
20
30
40
50
60
Time (min)
Normal
MK50
MK75
MK88
MK94
MK97
MK data
Normal data

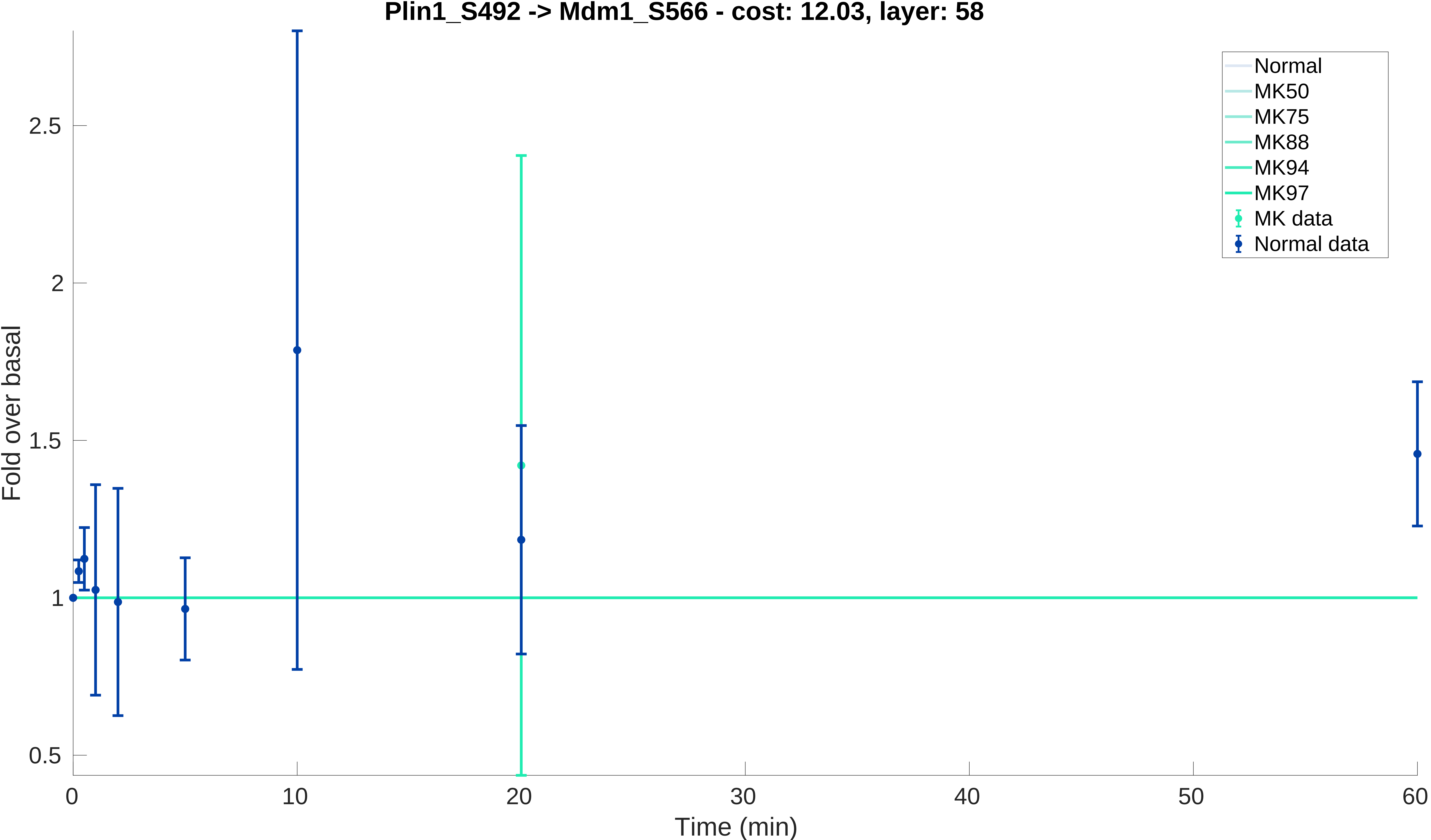

Plin1_S492 -> Mdm1_S566 - cost: 12.03, layer: 58
Normal
MK50
MK75
MK88
MK94
MK97
MK data
Normal data
Fold over basal
2.5
2
1.5
1
0.5
0
10
20
30
40
50
60
Time (min)

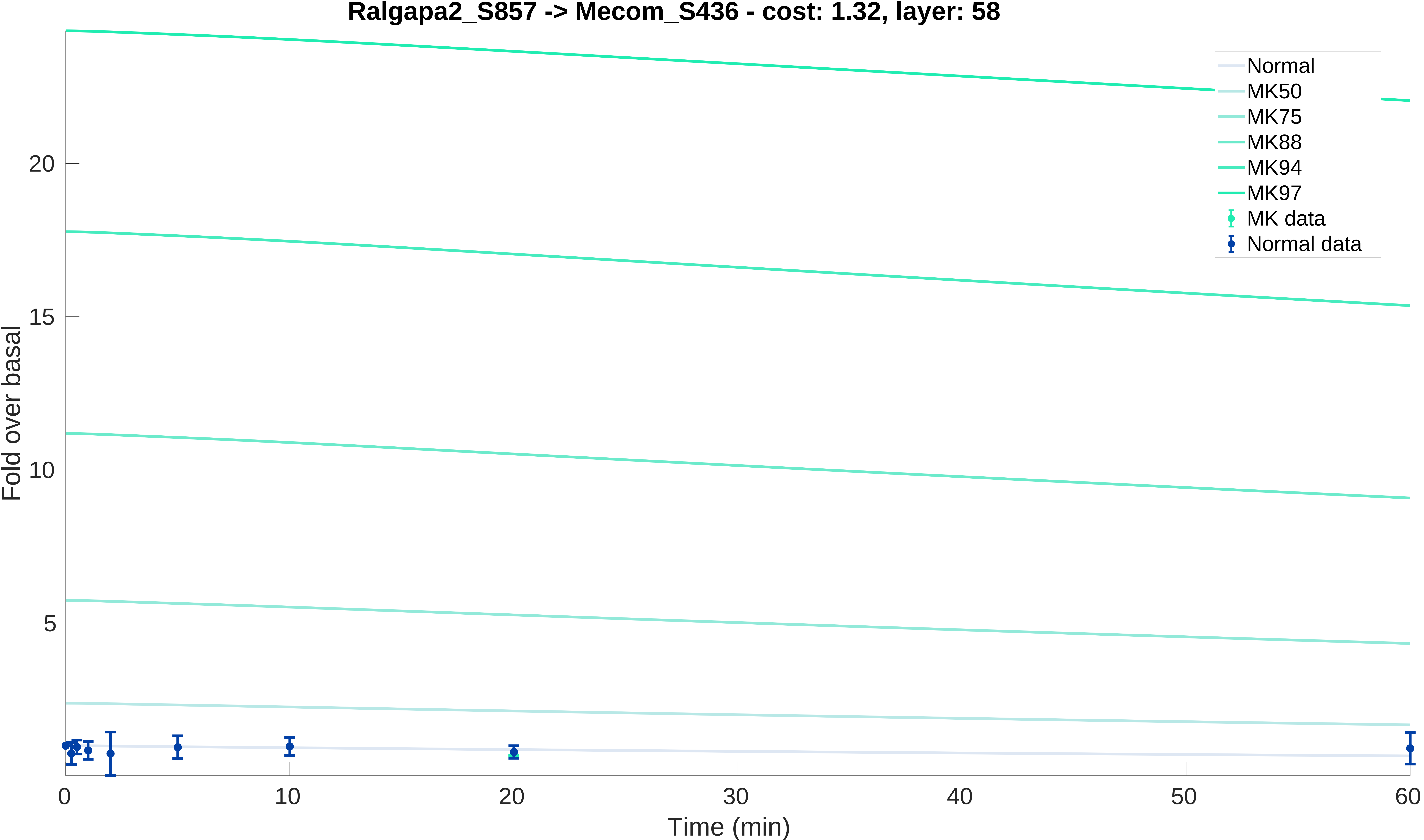

Ralgapa2_S857 -> Mecom_S436 - cost: 1.32, layer: 58
Normal
MK50
MK75
MK88
MK94
MK97
MK data
Normal data
Fold over basal
20
15
10
5
0
10
20
30
40
50
60
Time (min)

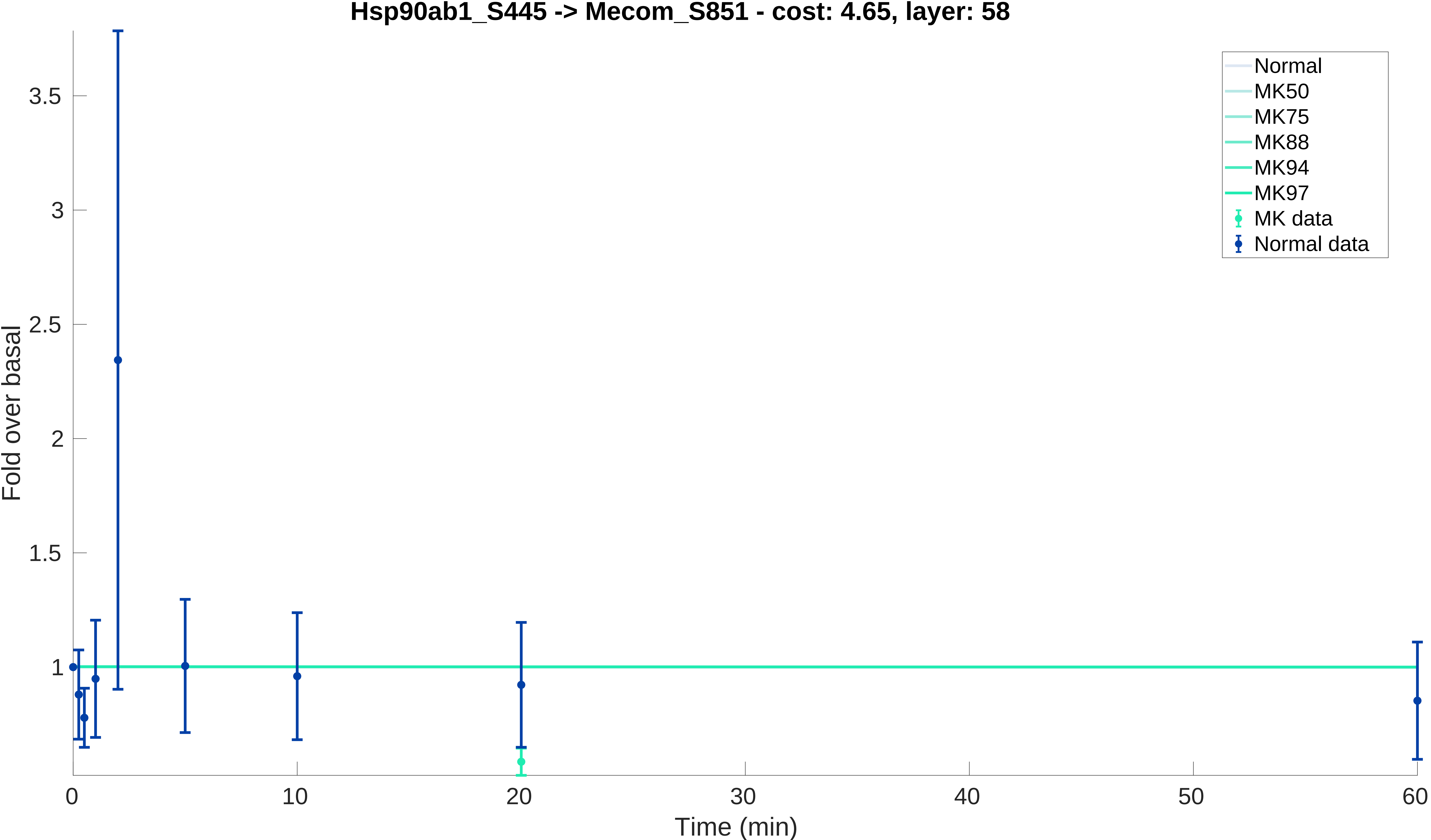

Hsp90ab1_S445 -> Mecom_S851 - cost: 4.65, layer: 58
Fold over basal
Time (min)
3.5
3
2.5
2
1.5
1
0
10
20
30
40
50
60
Normal
MK50
MK75
MK88
MK94
MK97
MK data
Normal data

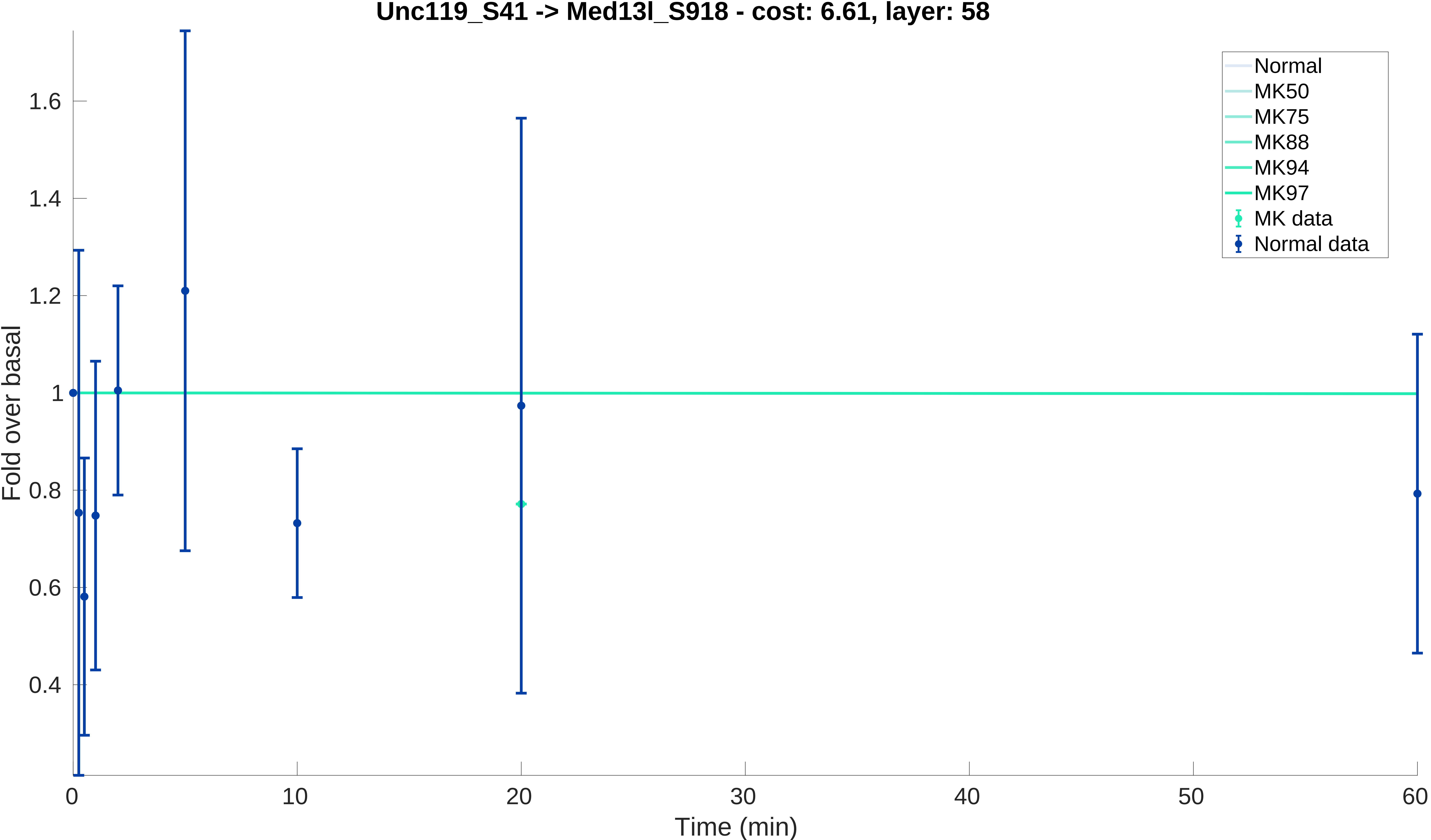

Unc119_S41 -> Med13l_S918 - cost: 6.61, layer: 58
Fold over basal
1.6
1.4
1.2
1
0.8
0.6
0.4
0
10
20
30
40
50
60
Time (min)
Normal
MK50
MK75
MK88
MK94
MK97
MK data
Normal data

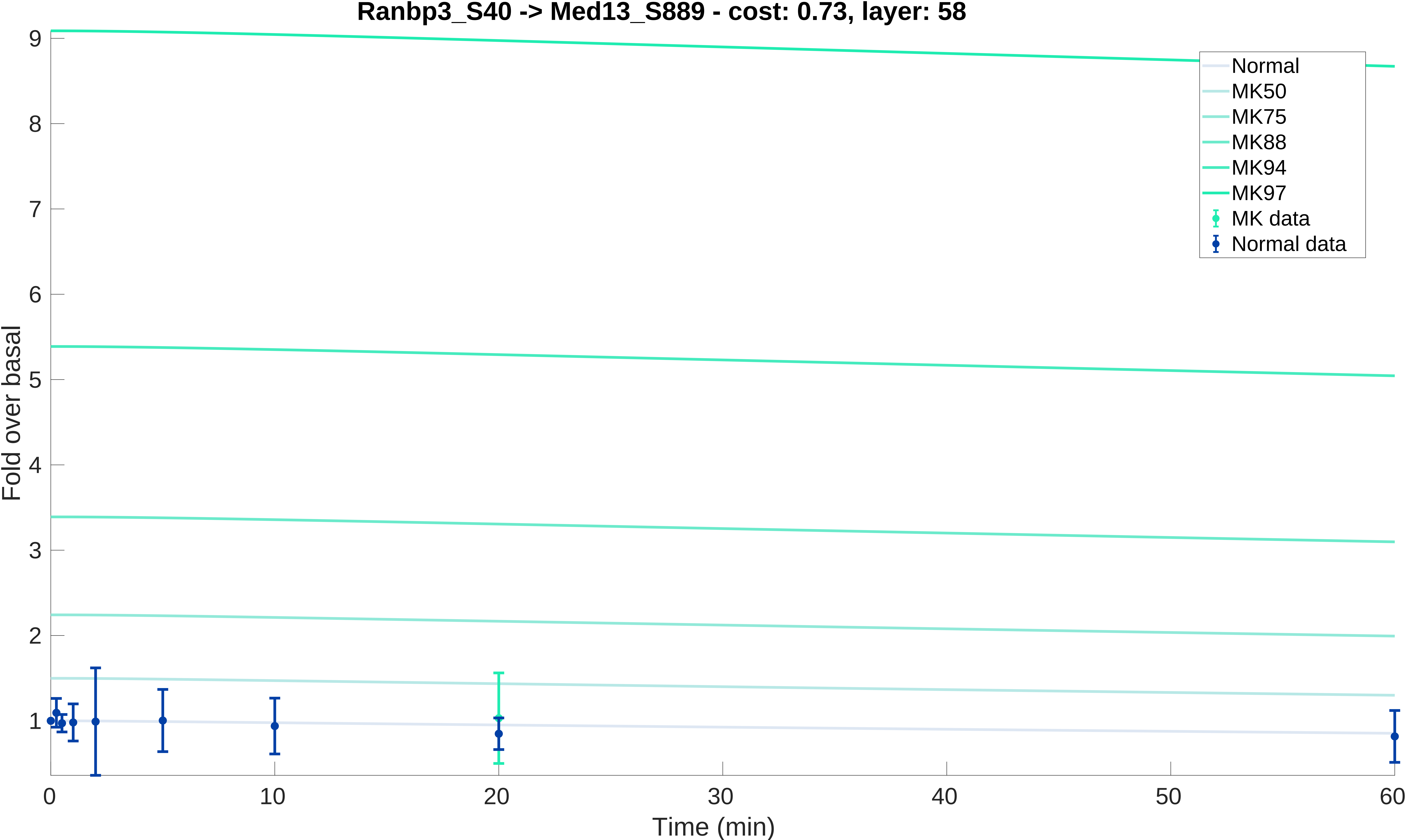

Ranbp3_S40 -> Med13_S889 - cost: 0.73, layer: 58
Fold over basal
Time (min)
0
10
20
30
40
50
60
1
2
3
4
5
6
7
8
9
Normal
MK50
MK75
MK88
MK94
MK97
MK data
Normal data

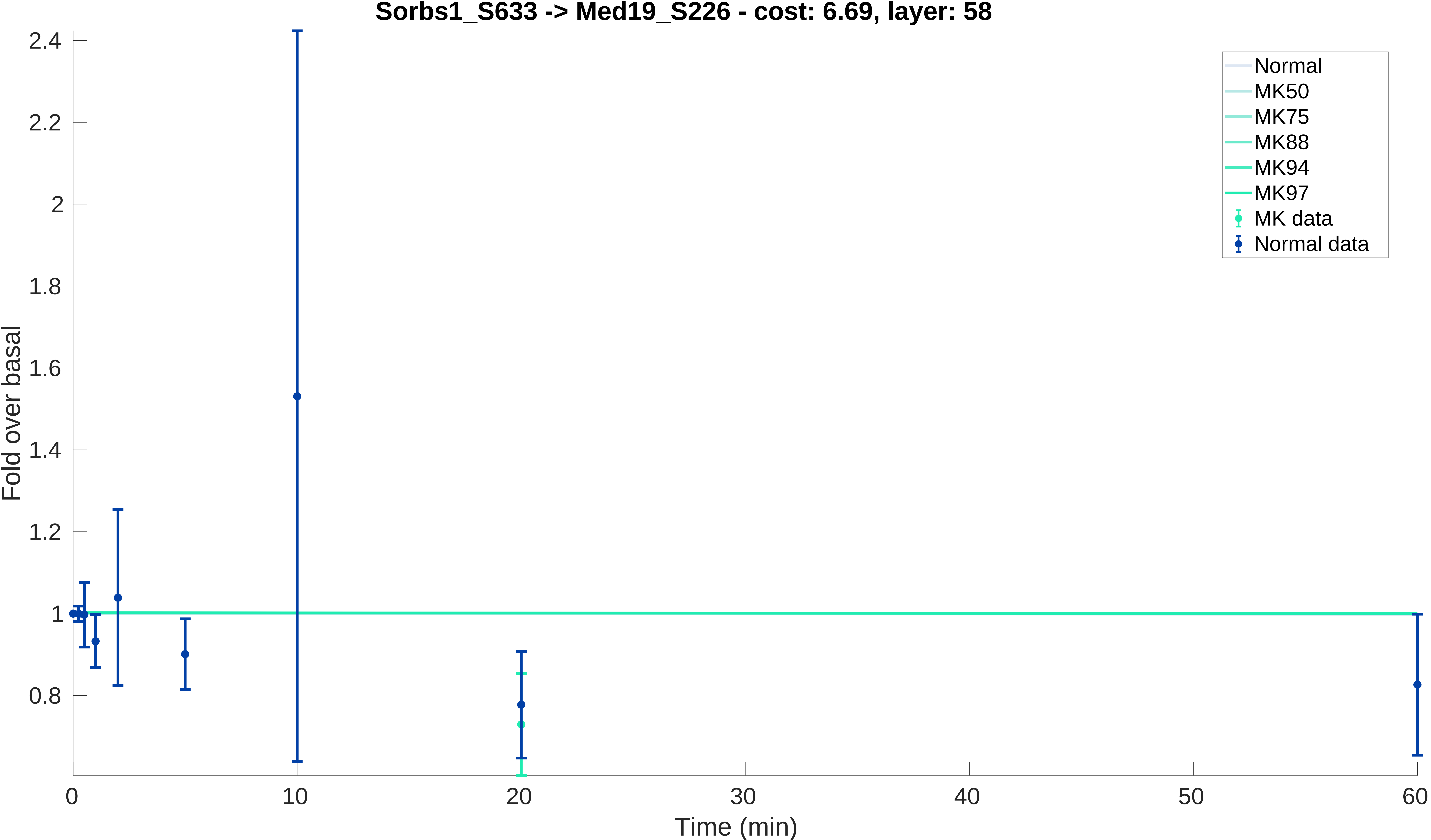

Sorbs1_S633 -> Med19_S226 - cost: 6.69, layer: 58
Fold over basal
Time (min)
2.4
2.2
2
1.8
1.6
1.4
1.2
1
0.8
0
10
20
30
40
50
60
Normal
MK50
MK75
MK88
MK94
MK97
MK data
Normal data

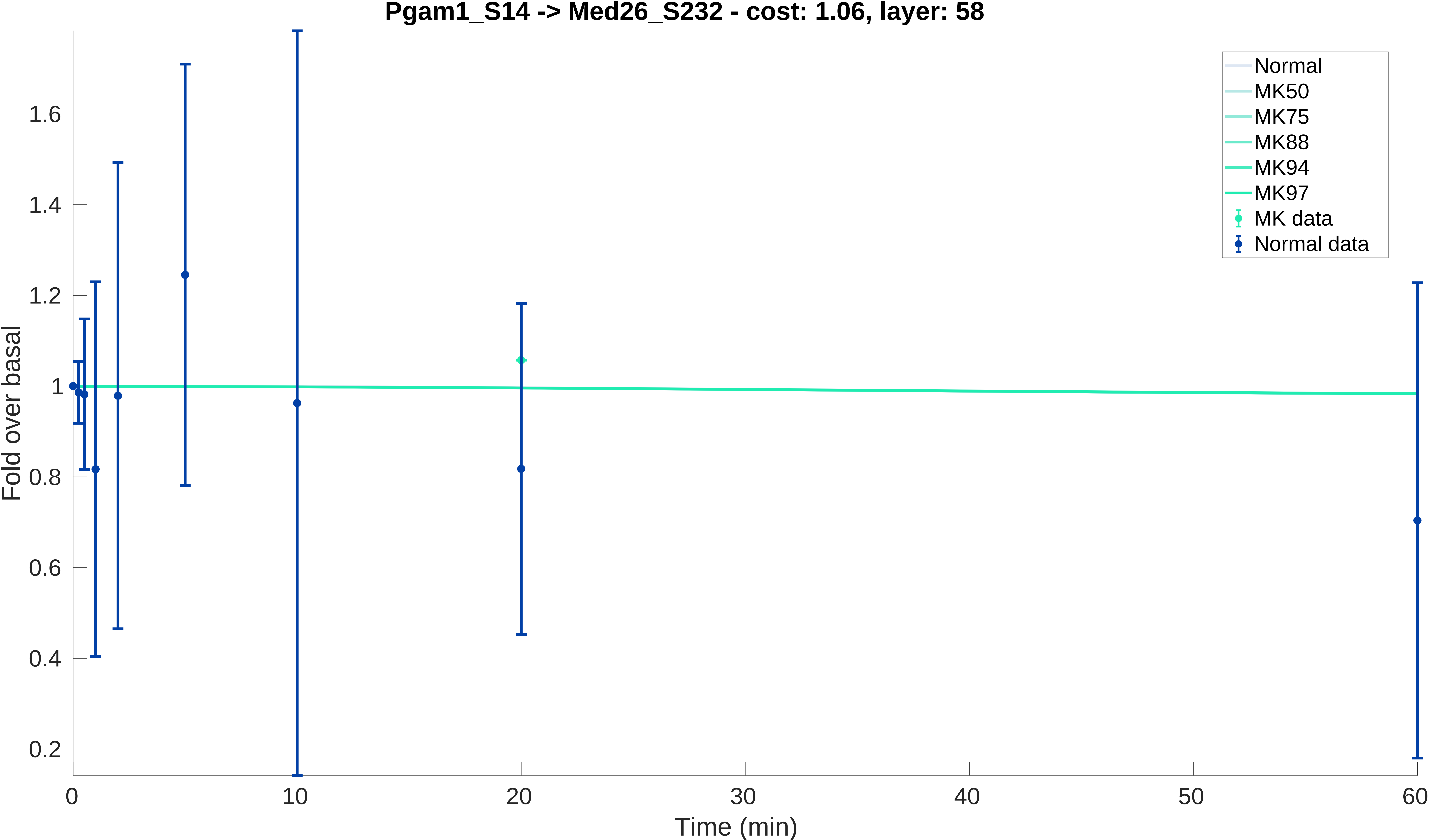

Pgam1_S14 -> Med26_S232 - cost: 1.06, layer: 58
Fold over basal
Time (min)
0
10
20
30
40
50
60
0.2
0.4
0.6
0.8
1
1.2
1.4
1.6
Normal
MK50
MK75
MK88
MK94
MK97
MK data
Normal data

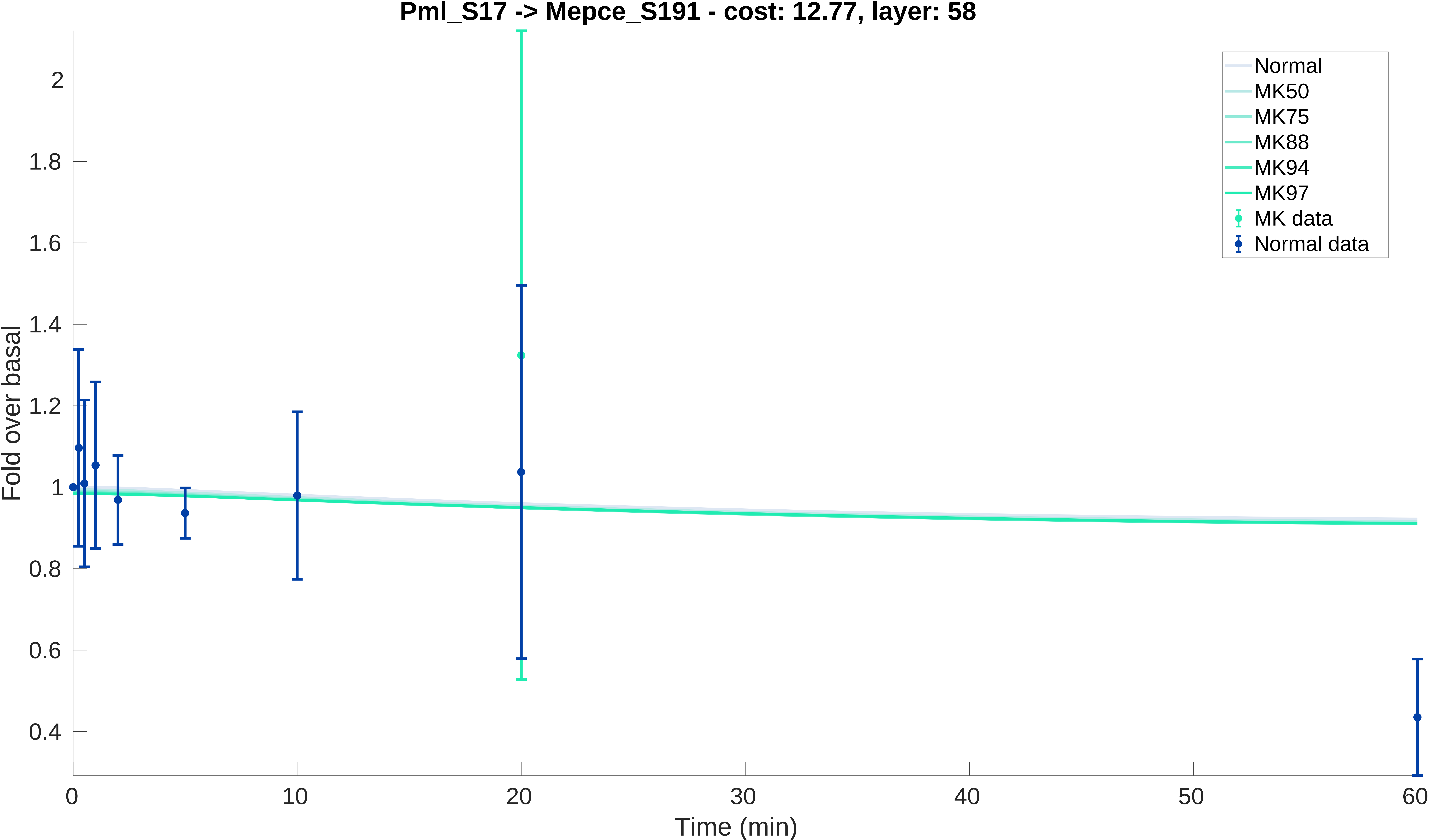

Pml_S17 -> Mepce_S191 - cost: 12.77, layer: 58
Fold over basal
Time (min)
0
10
20
30
40
50
60
0.4
0.6
0.8
1
1.2
1.4
1.6
1.8
2
Normal
MK50
MK75
MK88
MK94
MK97
MK data
Normal data

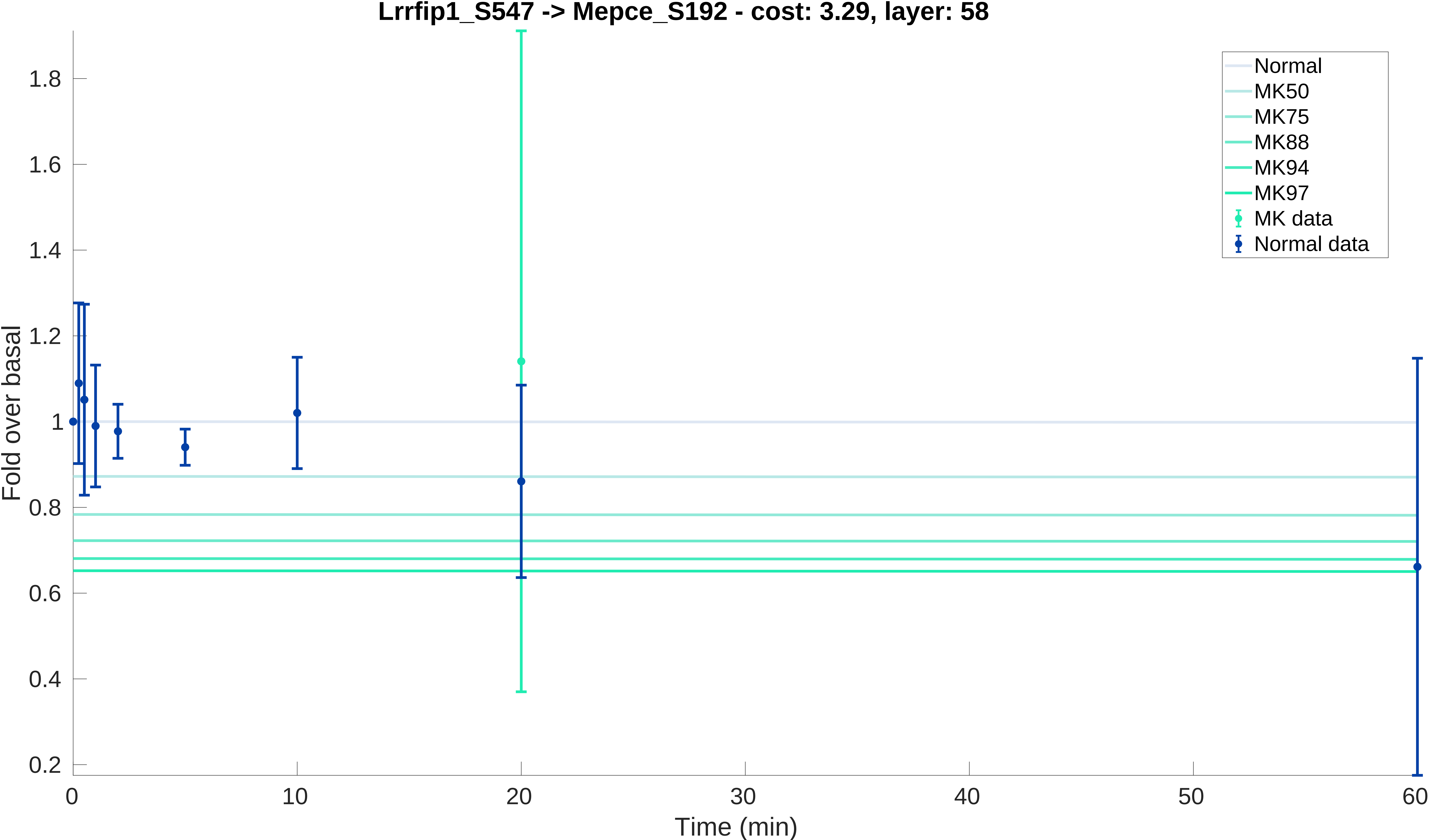
Lrrfip1_S547 -> Mepce_S192 - cost: 3.29, layer: 58
Fold over basal
Time (min)
0.2
0.4
0.6
0.8
1
1.2
1.4
1.6
1.8
0
10
20
30
40
50
60
Normal
MK50
MK75
MK88
MK94
MK97
MK data
Normal data

Tjp2_S968 -> Mepce_T188 - cost: 14.70, layer: 58

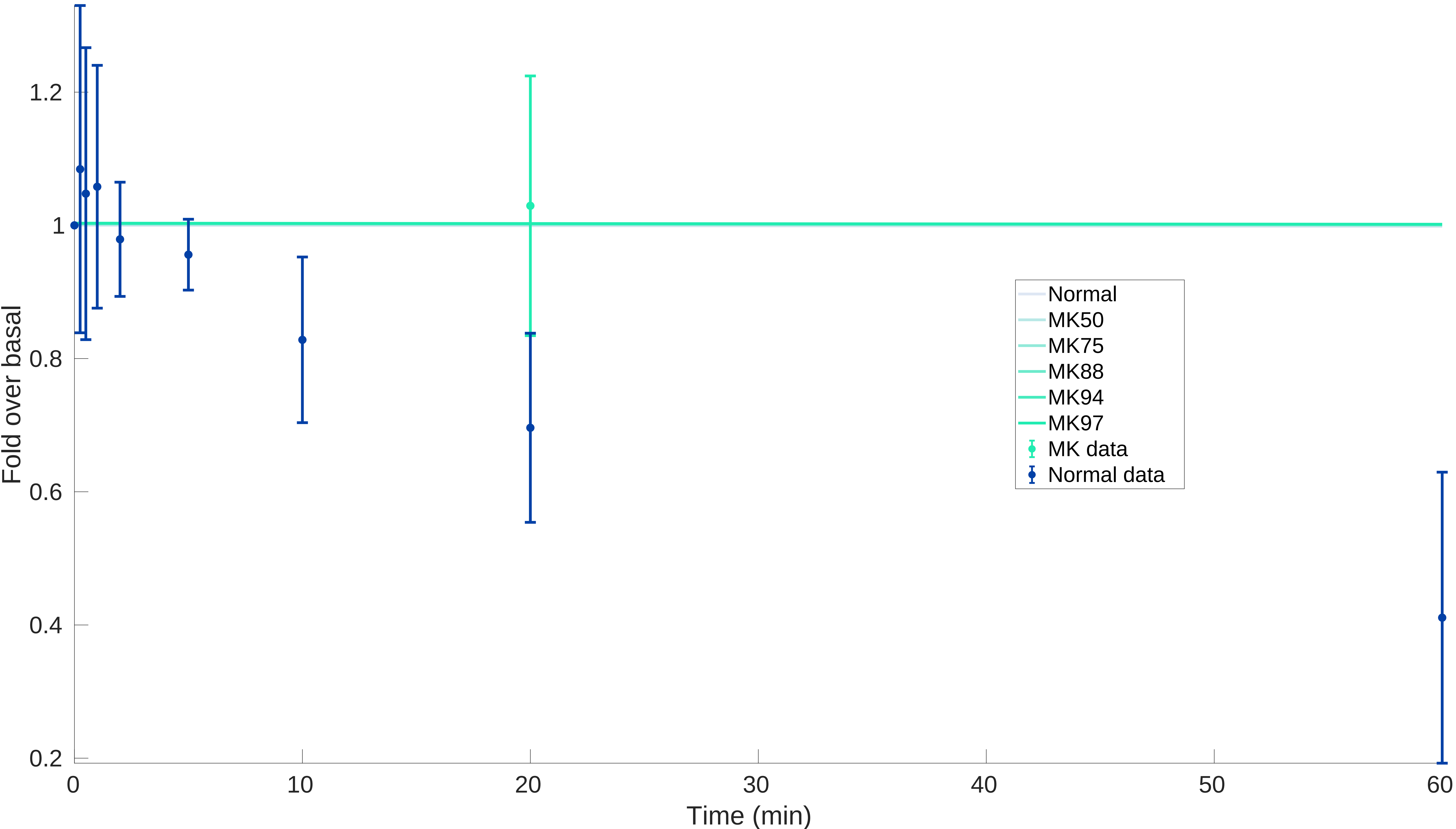

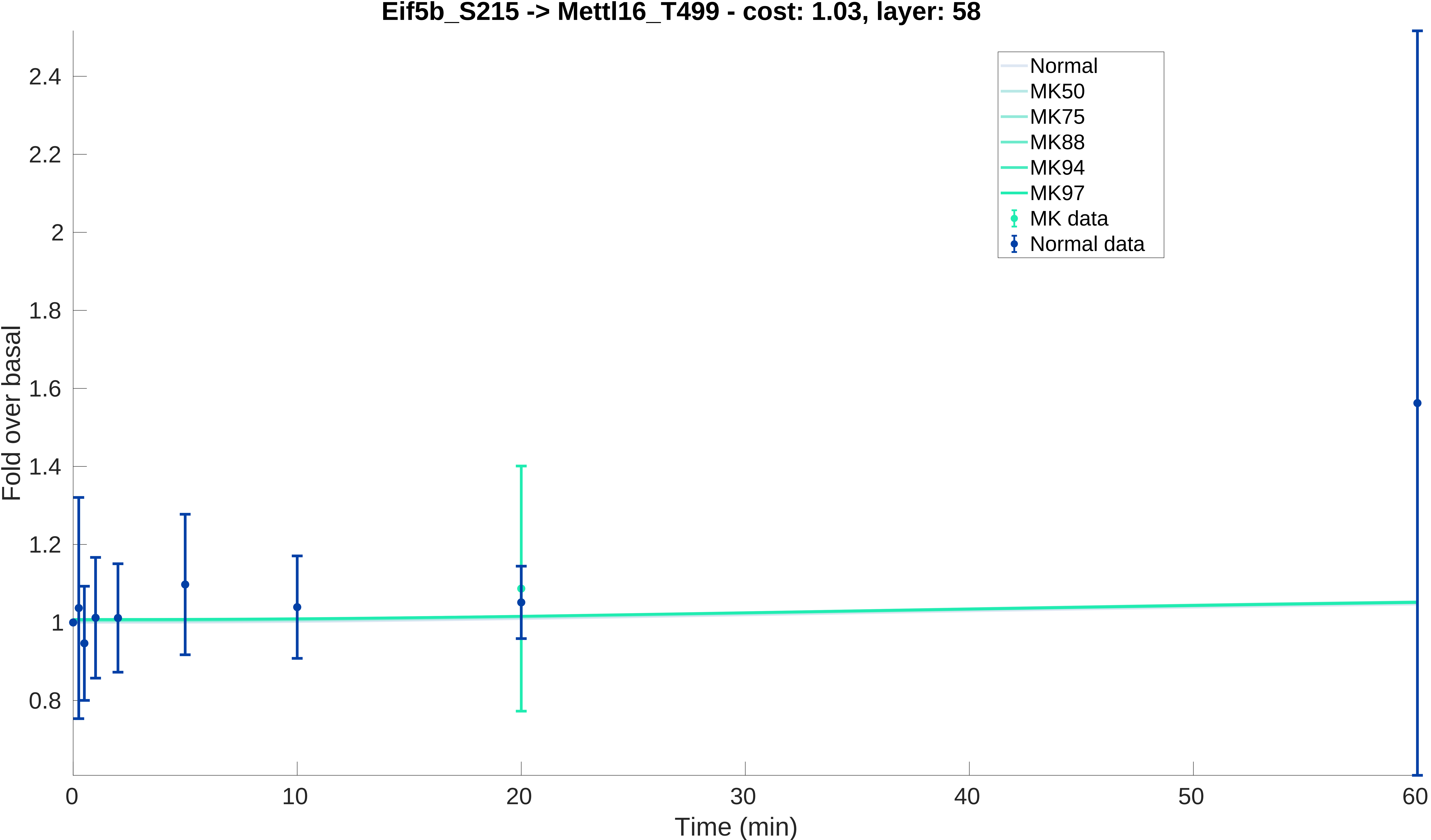
Eif5b_S215 -> Mettl16_T499 - cost: 1.03, layer: 58
Normal
MK50
MK75
MK88
MK94
MK97
MK data
Normal data
Fold over basal
Time (min)
0.8
1
1.2
1.4
1.6
1.8
2
2.2
2.4
0
10
20
30
40
50
60

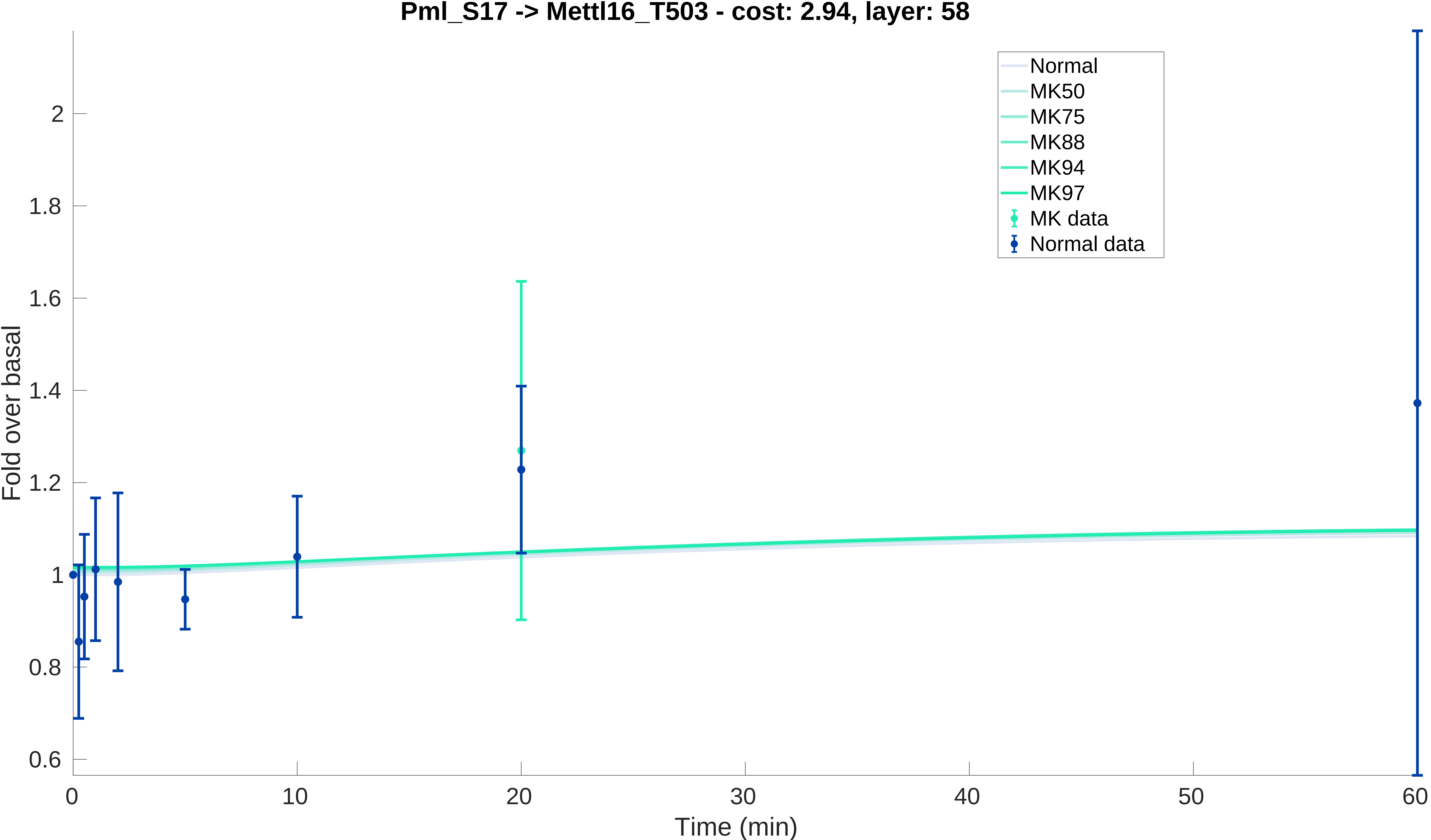

Pml_S17 -> Mettl16_T503 - cost: 2.94, layer: 58
Normal
MK50
MK75
MK88
MK94
MK97
MK data
Normal data
Fold over basal
Time (min)
0.6
0.8
1
1.2
1.4
1.6
1.8
2
0
10
20
30
40
50
60

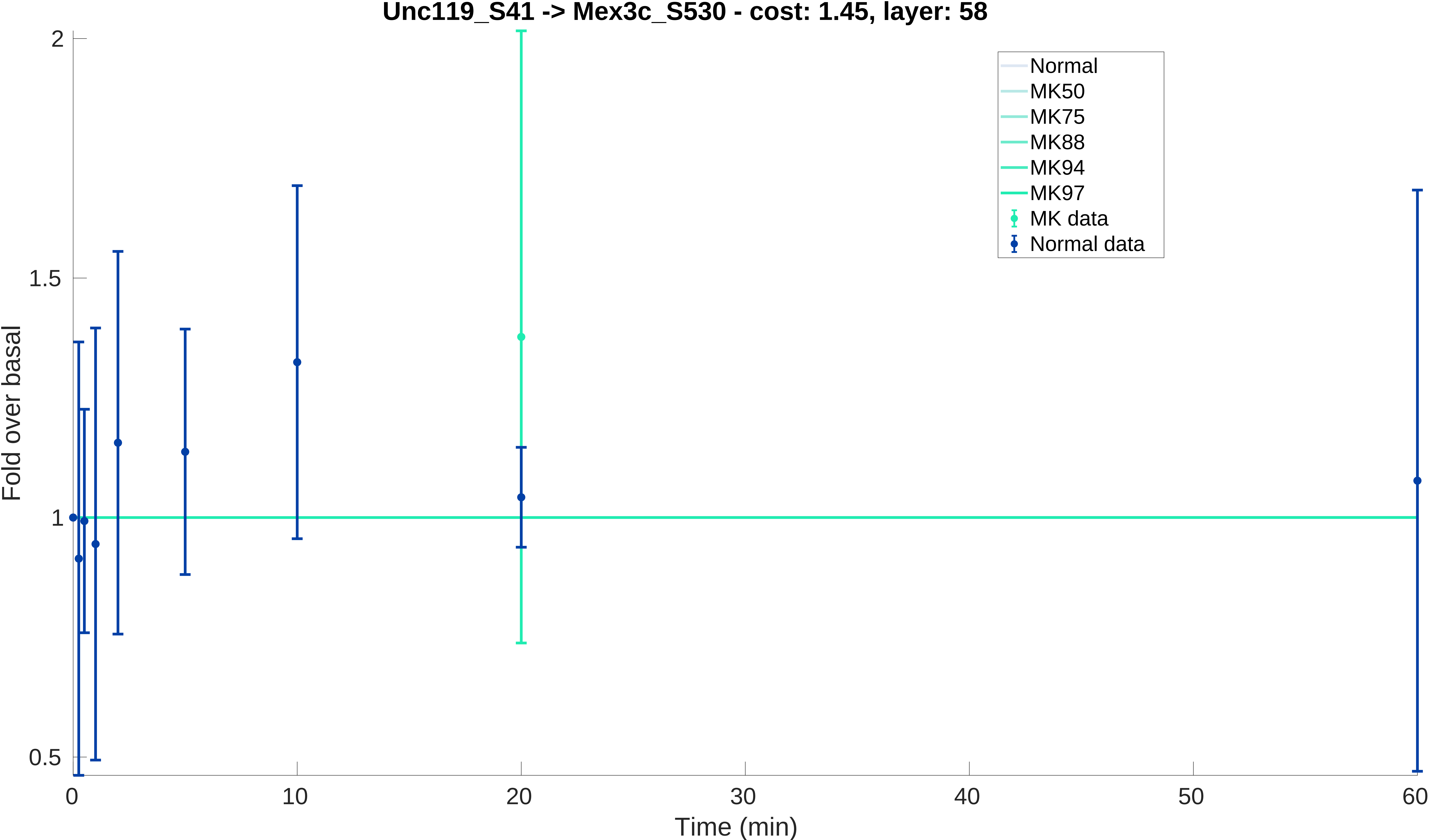

Unc119_S41 -> Mex3c_S530 - cost: 1.45, layer: 58
Fold over basal
Time (min)
0.5
1
1.5
2
0
10
20
30
40
50
60
Normal
MK50
MK75
MK88
MK94
MK97
MK data
Normal data

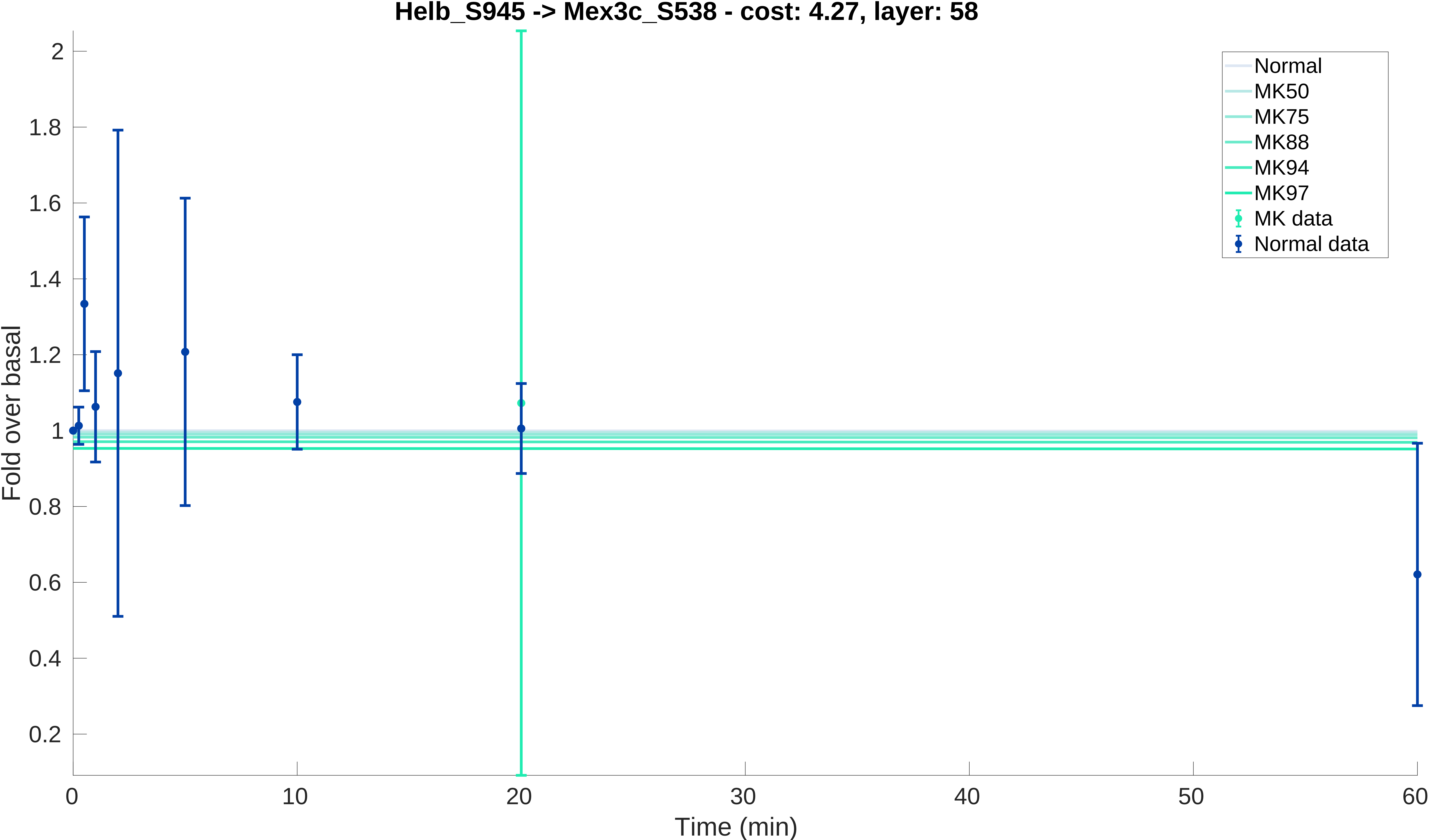
Helb_S945 -> Mex3c_S538 - cost: 4.27, layer: 58
Fold over basal
Time (min)
0.2
0.4
0.6
0.8
1
1.2
1.4
1.6
1.8
2
0
10
20
30
40
50
60
Normal
MK50
MK75
MK88
MK94
MK97
MK data
Normal data

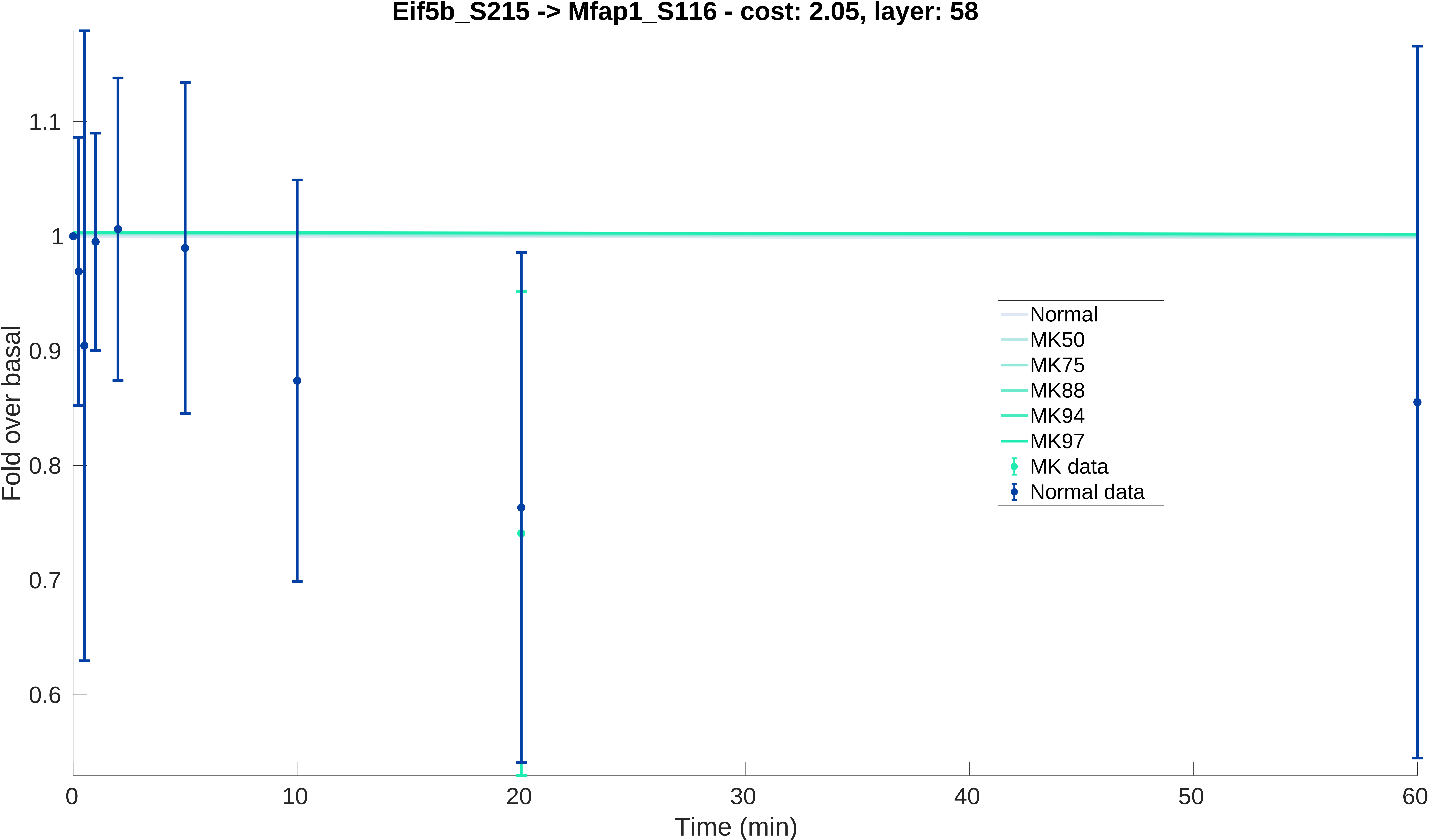
Eif5b_S215 -> Mfap1_S116 - cost: 2.05, layer: 58
Fold over basal
1.1
1
0.9
0.8
0.7
0.6
0
10
20
30
40
50
60
Time (min)
Normal
MK50
MK75
MK88
MK94
MK97
MK data
Normal data

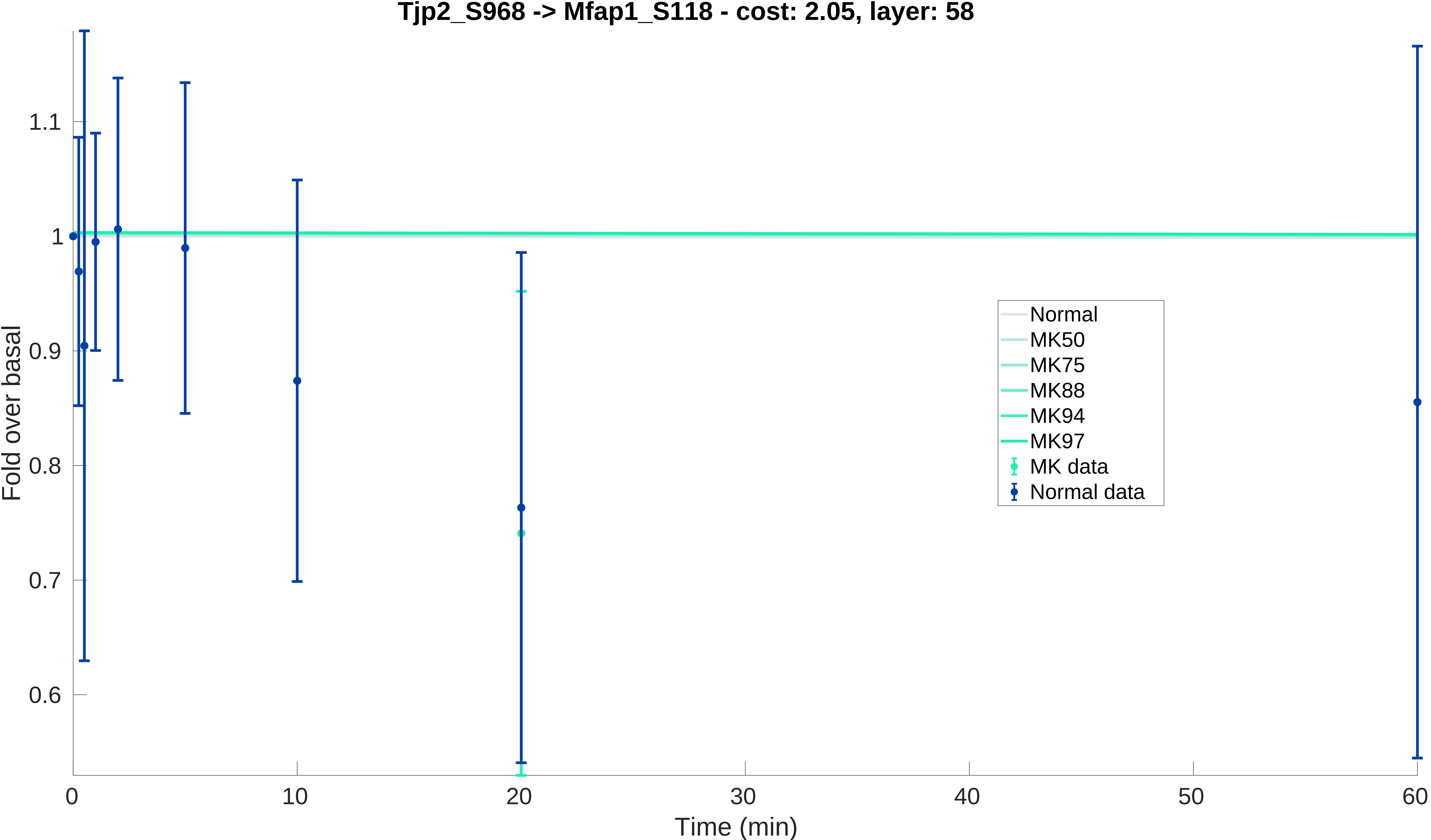

Tjp2_S968 -> Mfap1_S118 - cost: 2.05, layer: 58
Fold over basal
Time (min)
0.6
0.7
0.8
0.9
1
1.1
0
10
20
30
40
50
60
Normal
MK50
MK75
MK88
MK94
MK97
MK data
Normal data

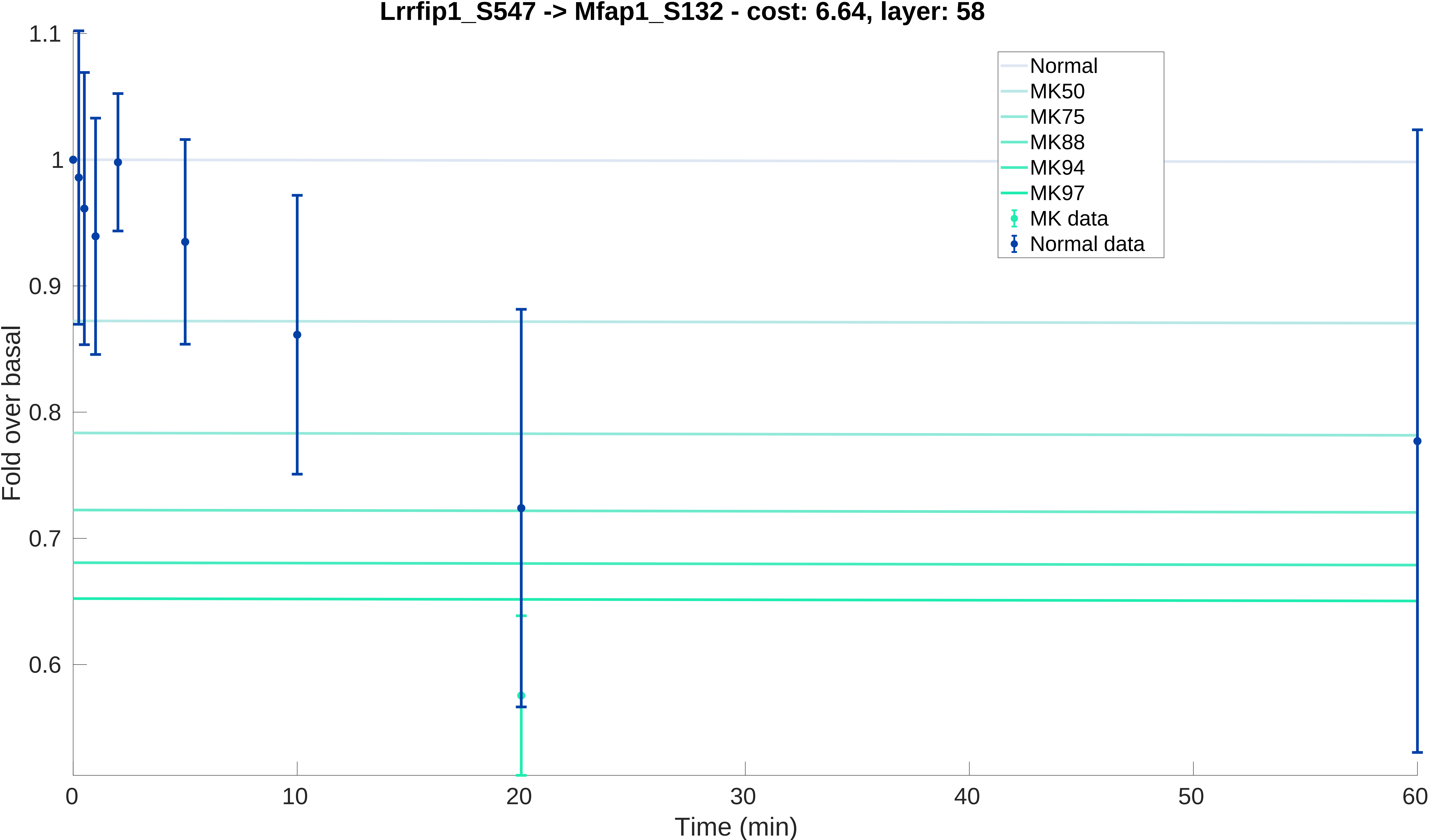

Lrrfip1_S547 -> Mfap1_S132 - cost: 6.64, layer: 58
Fold over basal
Time (min)
1.1
1
0.9
0.8
0.7
0.6
0
10
20
30
40
50
60
Normal
MK50
MK75
MK88
MK94
MK97
MK data
Normal data

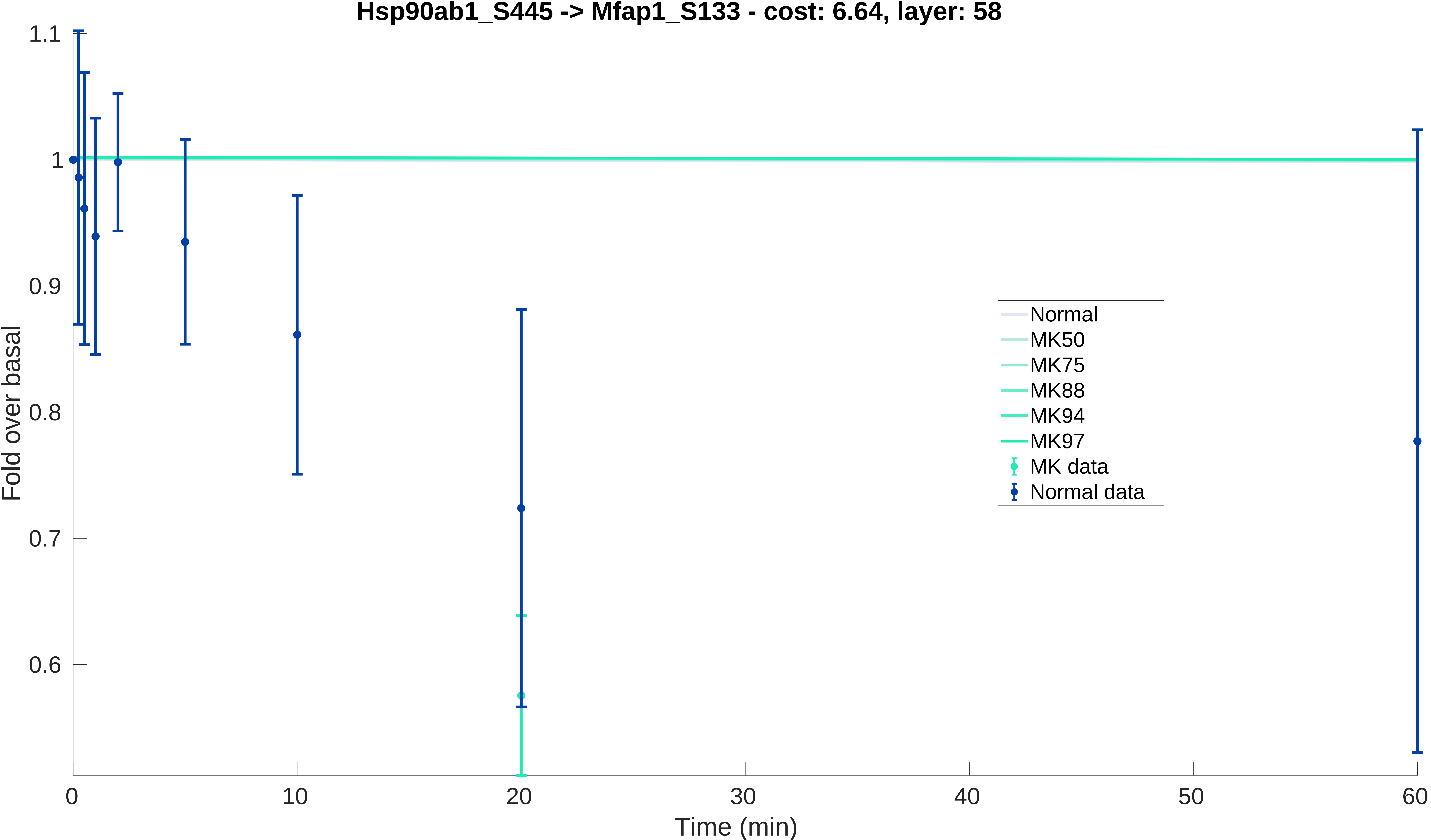

Hsp90ab1_S445 -> Mfap1_S133 - cost: 6.64, layer: 58
Fold over basal
Time (min)
1.1
1
0.9
0.8
0.7
0.6
0
10
20
30
40
50
60
Normal
MK50
MK75
MK88
MK94
MK97
MK data
Normal data

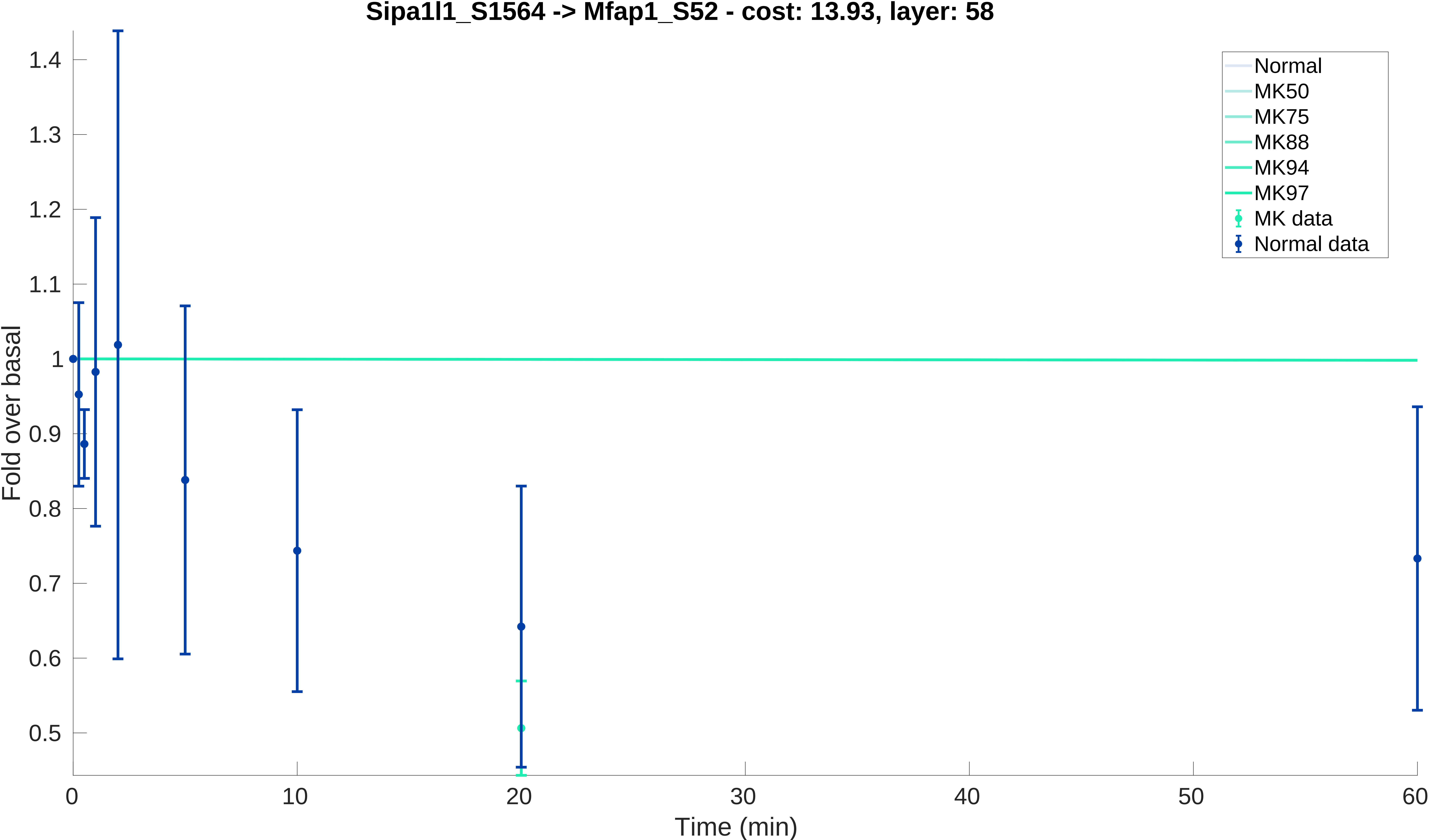

Sipa1l1_S1564 -> Mfap1_S52 - cost: 13.93, layer: 58
Fold over basal
1.4
1.3
1.2
1.1
1
0.9
0.8
0.7
0.6
0.5
0
10
20
30
40
50
60
Time (min)
Normal
MK50
MK75
MK88
MK94
MK97
MK data
Normal data

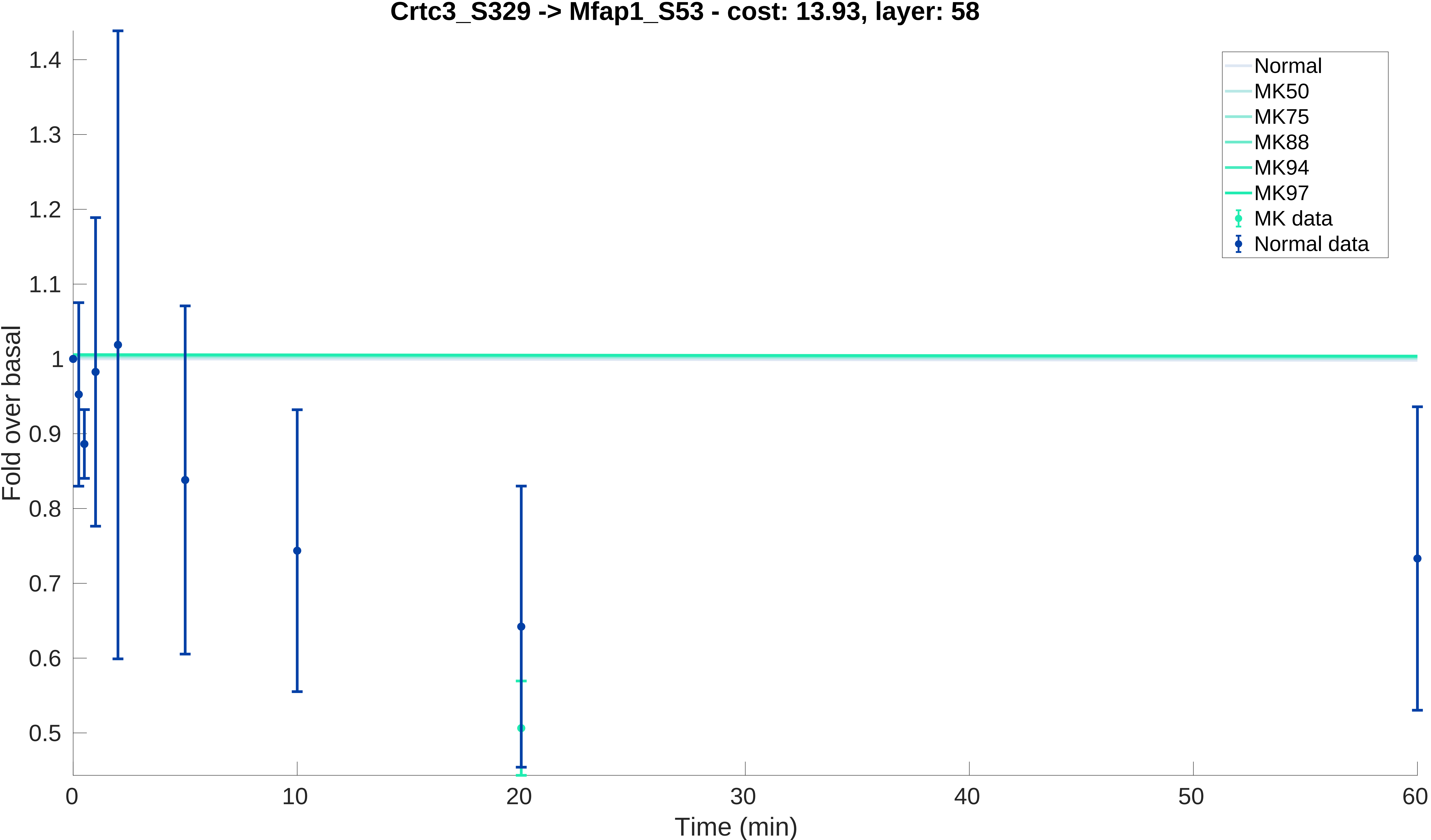

Crtc3_S329 -> Mfap1_S53 - cost: 13.93, layer: 58
Fold over basal
Time (min)
1.4
1.3
1.2
1.1
1
0.9
0.8
0.7
0.6
0.5
0
10
20
30
40
50
60
Normal
MK50
MK75
MK88
MK94
MK97
MK data
Normal data

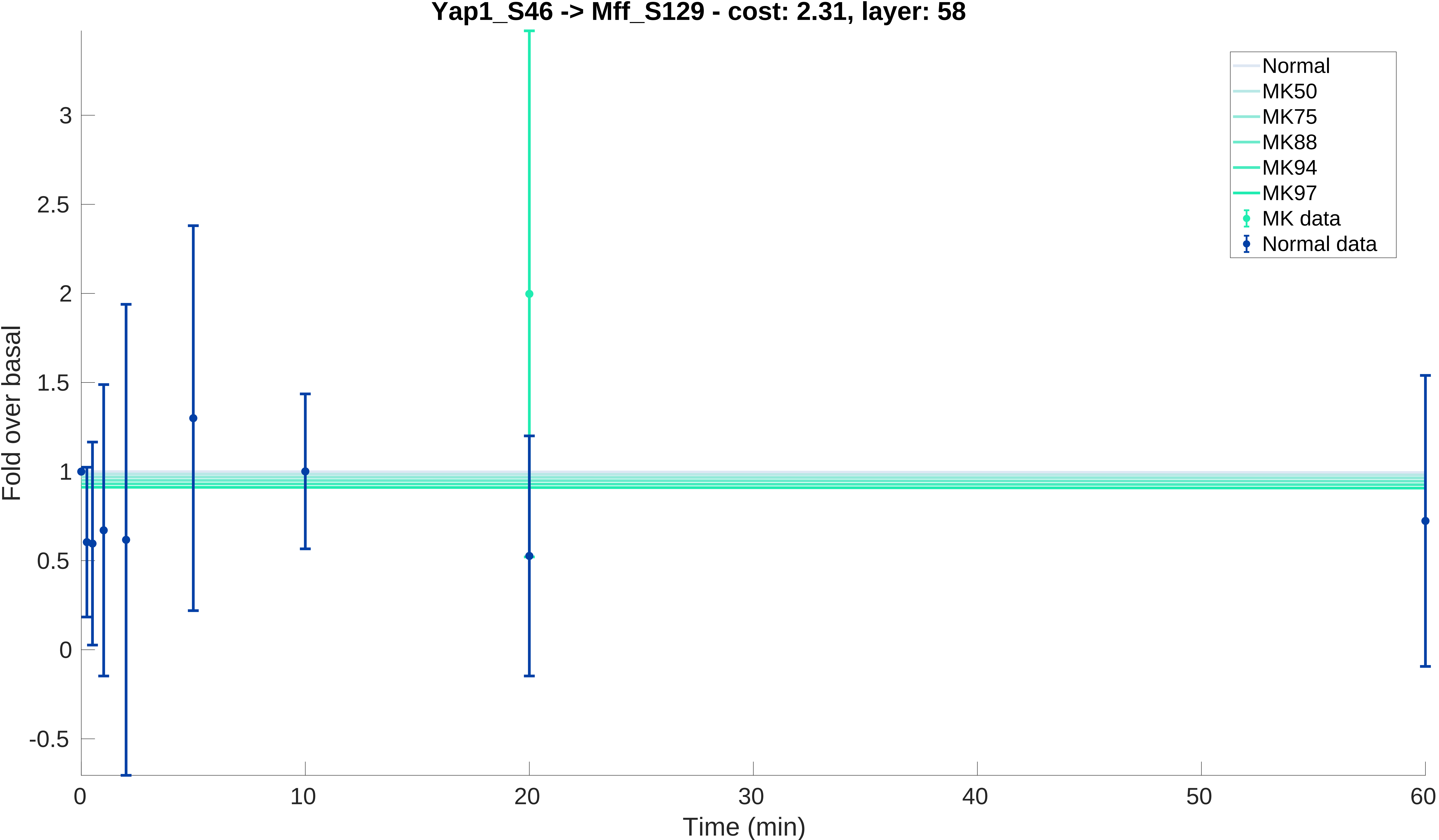

Yap1_S46 -> Mff_S129 - cost: 2.31, layer: 58
Normal
MK50
MK75
MK88
MK94
MK97
MK data
Normal data
Fold over basal
Time (min)
3
2.5
2
1.5
1
0.5
0
-0.5
0
10
20
30
40
50
60

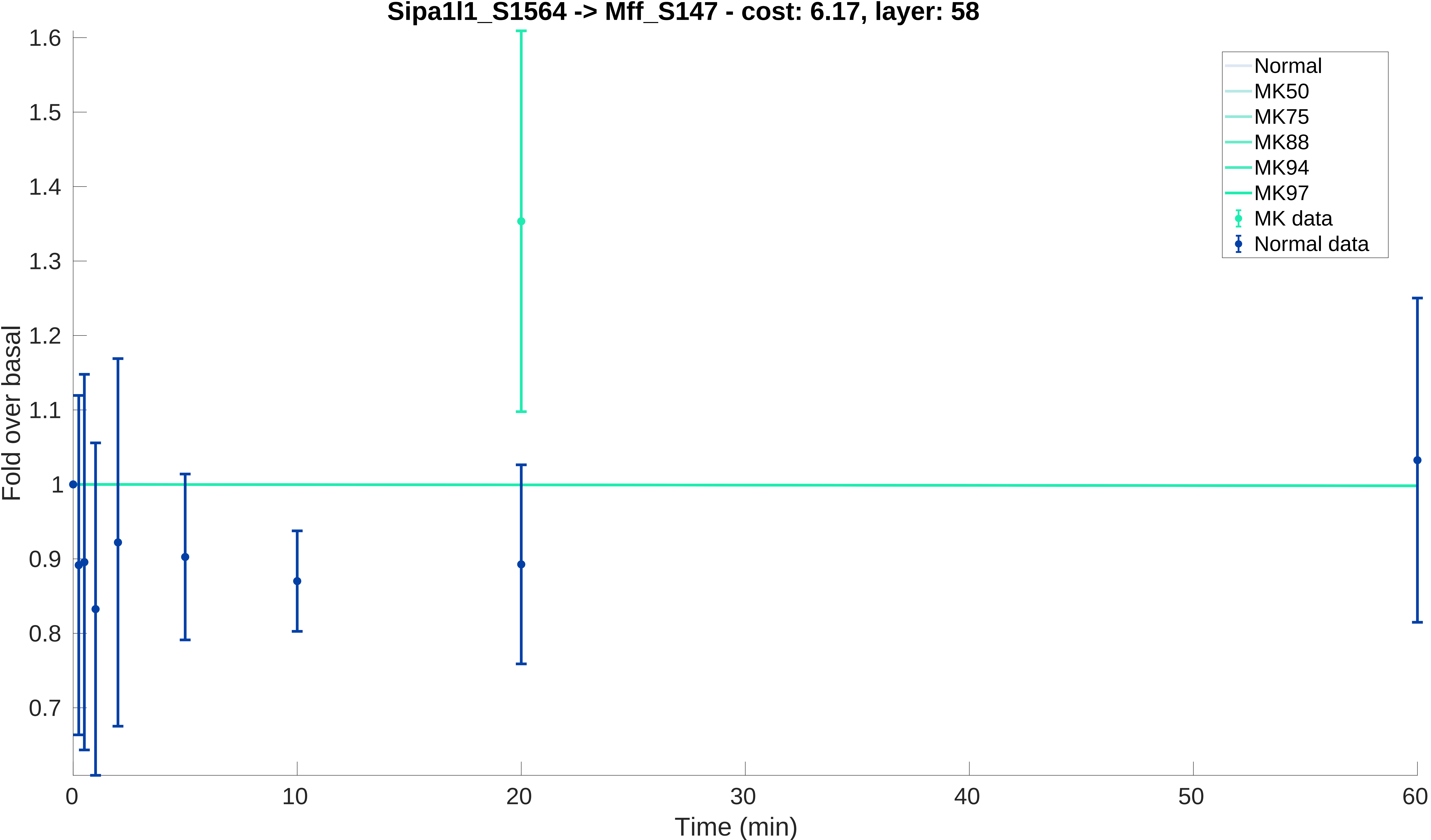

Sipa1l1_S1564 -> Mff_S147 - cost: 6.17, layer: 58
Fold over basal
Time (min)
0.7
0.8
0.9
1
1.1
1.2
1.3
1.4
1.5
1.6
0
10
20
30
40
50
60
Normal
MK50
MK75
MK88
MK94
MK97
MK data
Normal data

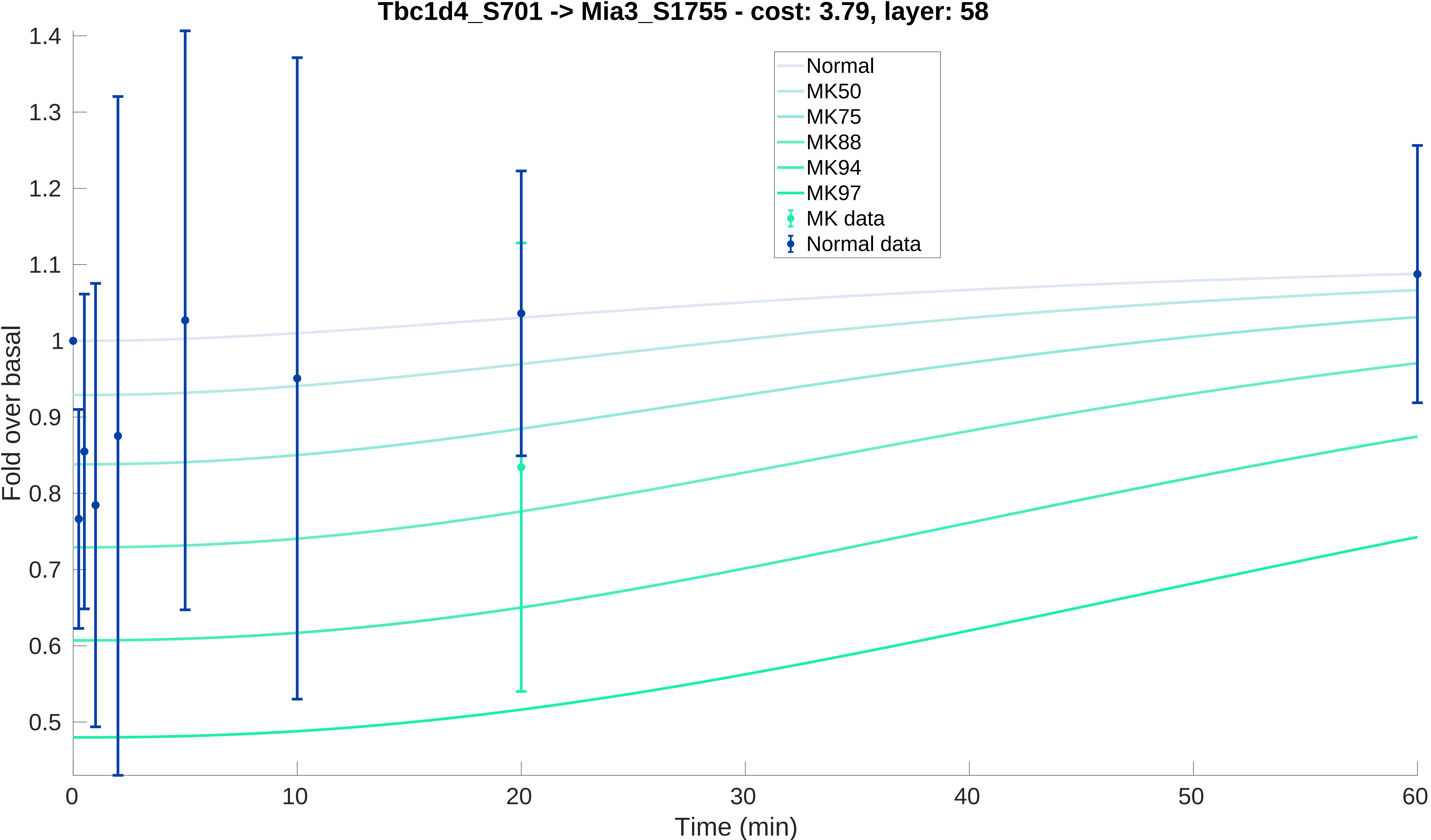
Tbc1d4_S701 -> Mia3_S1755 - cost: 3.79, layer: 58
Normal
MK50
MK75
MK88
MK94
MK97
MK data
Normal data
Fold over basal
Time (min)
1.4
1.3
1.2
1.1
1
0.9
0.8
0.7
0.6
0.5
0
10
20
30
40
50
60

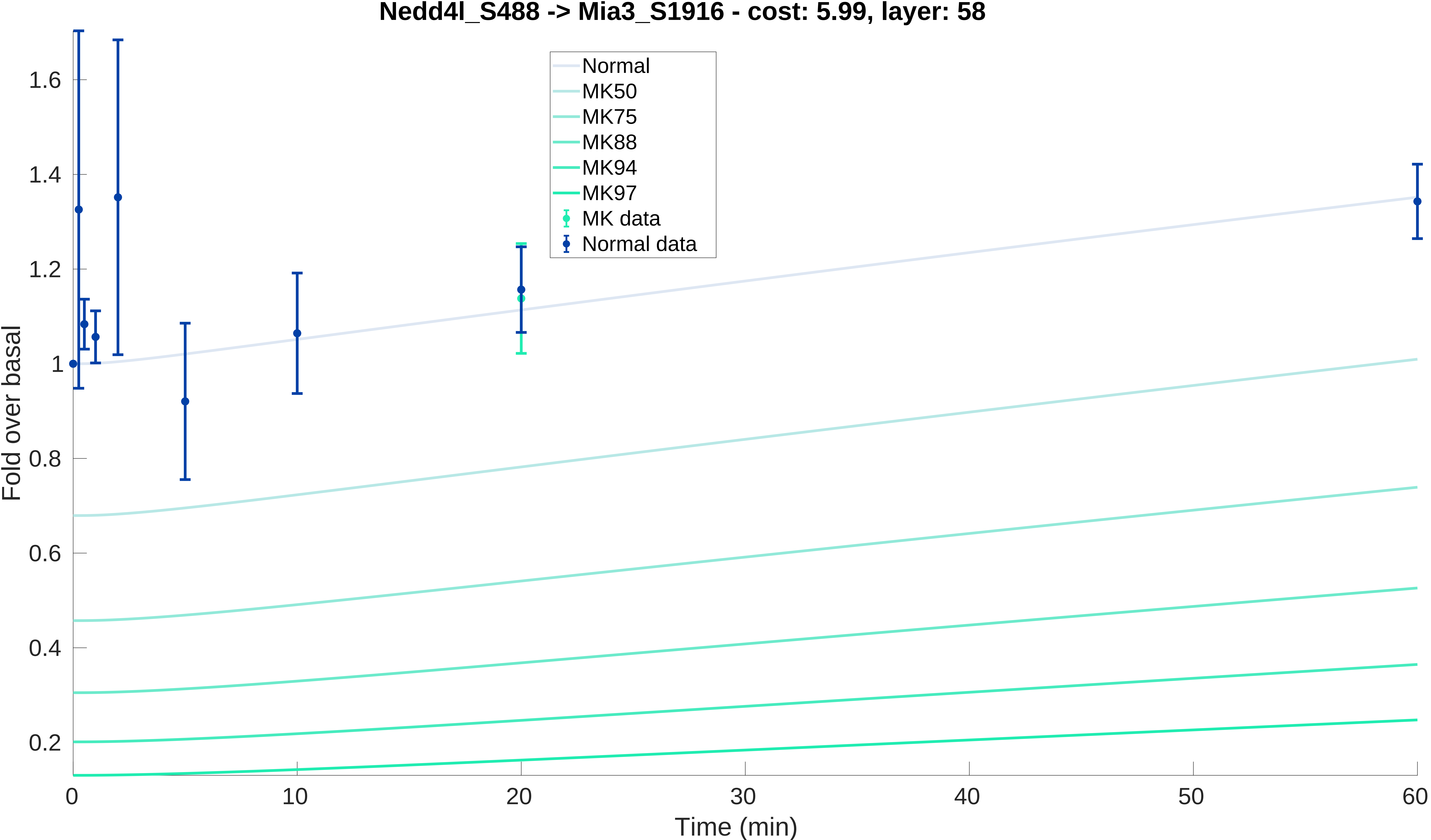

Nedd4l_S488 -> Mia3_S1916 - cost: 5.99, layer: 58
Normal
MK50
MK75
MK88
MK94
MK97
MK data
Normal data
Fold over basal
0.2
0.4
0.6
0.8
1
1.2
1.4
1.6
0
10
20
30
40
50
60
Time (min)

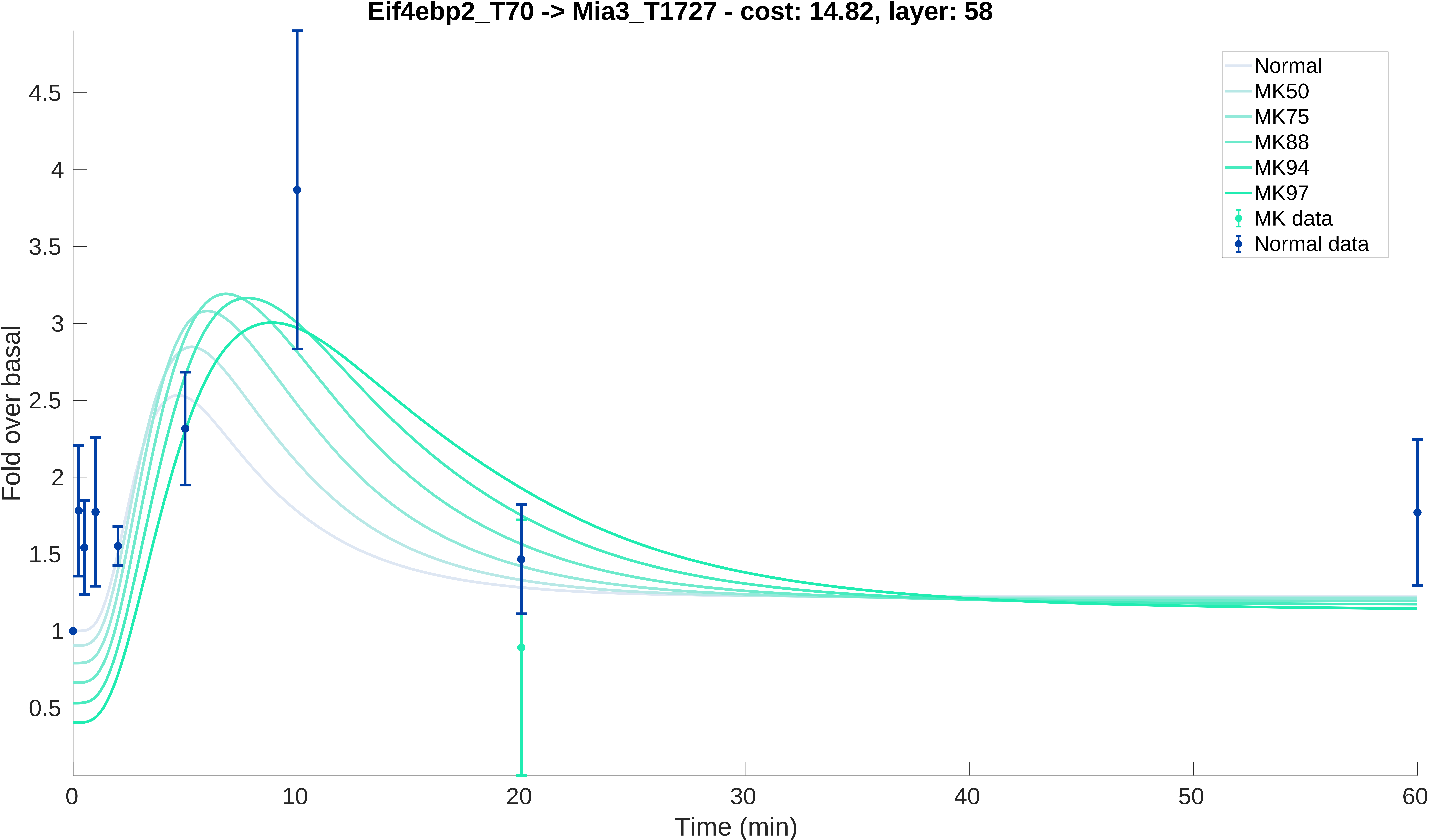
Eif4ebp2_T70 -> Mia3_T1727 - cost: 14.82, layer: 58
Fold over basal
Time (min)
0.5
1
1.5
2
2.5
3
3.5
4
4.5
0
10
20
30
40
50
60
Normal
MK50
MK75
MK88
MK94
MK97
MK data
Normal data

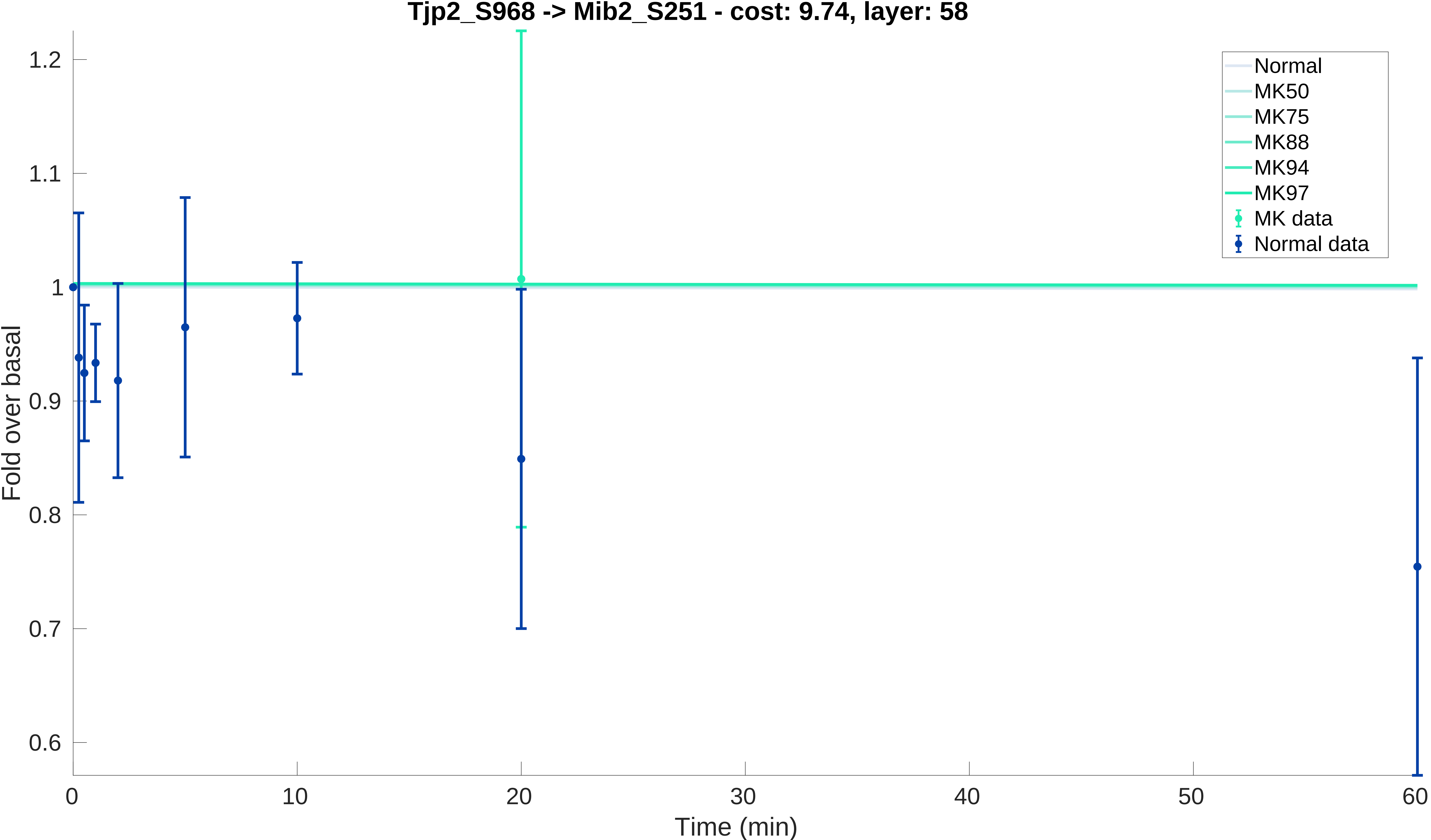

Tjp2_S968 -> Mib2_S251 - cost: 9.74, layer: 58
Fold over basal
Time (min)
1.2
1.1
1
0.9
0.8
0.7
0.6
0
10
20
30
40
50
60
Normal
MK50
MK75
MK88
MK94
MK97
MK data
Normal data

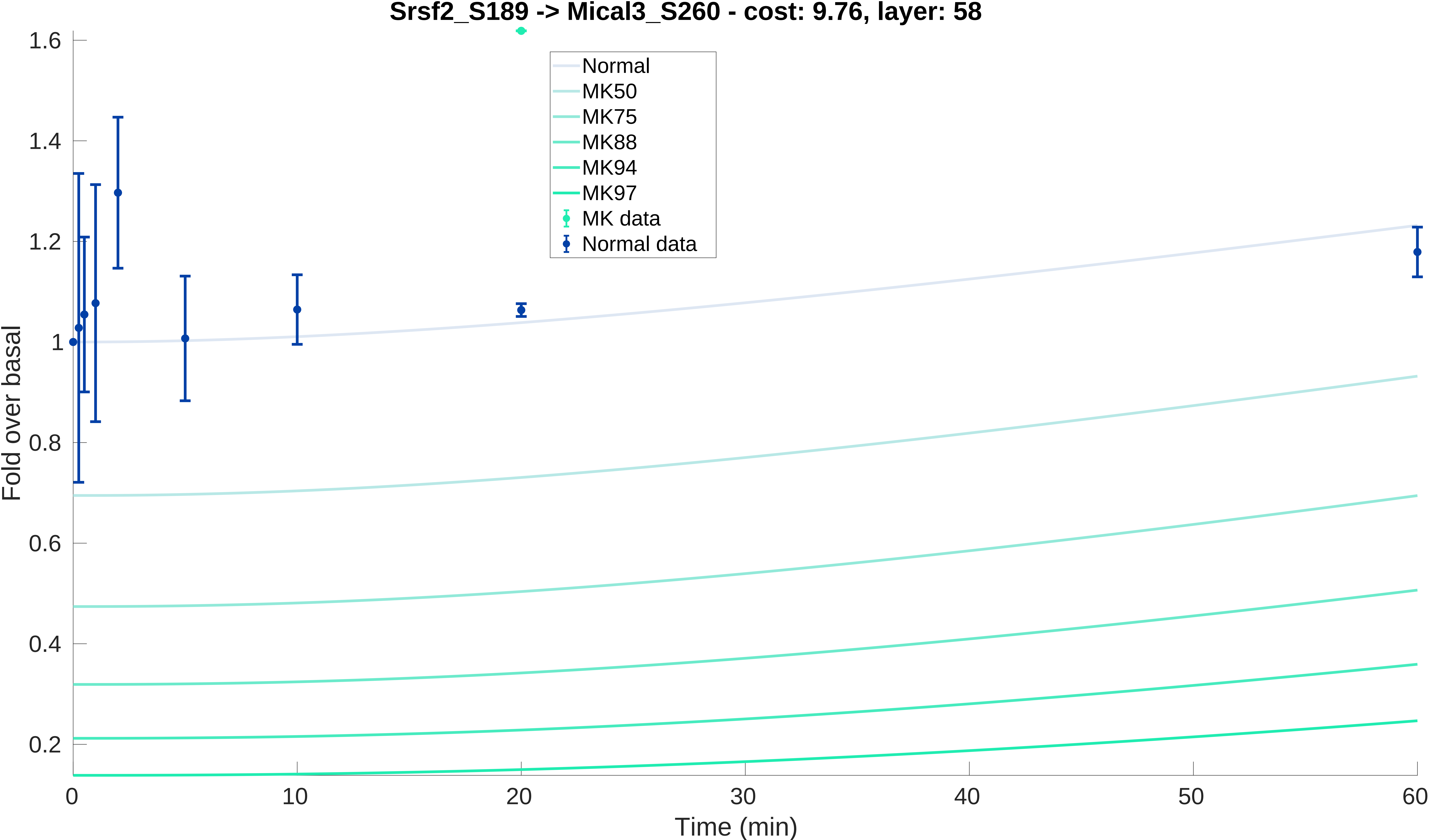

Srsf2_S189 -> Mical3_S260 - cost: 9.76, layer: 58
Fold over basal
Time (min)
Normal
MK50
MK75
MK88
MK94
MK97
MK data
Normal data
1.6
1.4
1.2
1
0.8
0.6
0.4
0.2
0
10
20
30
40
50
60

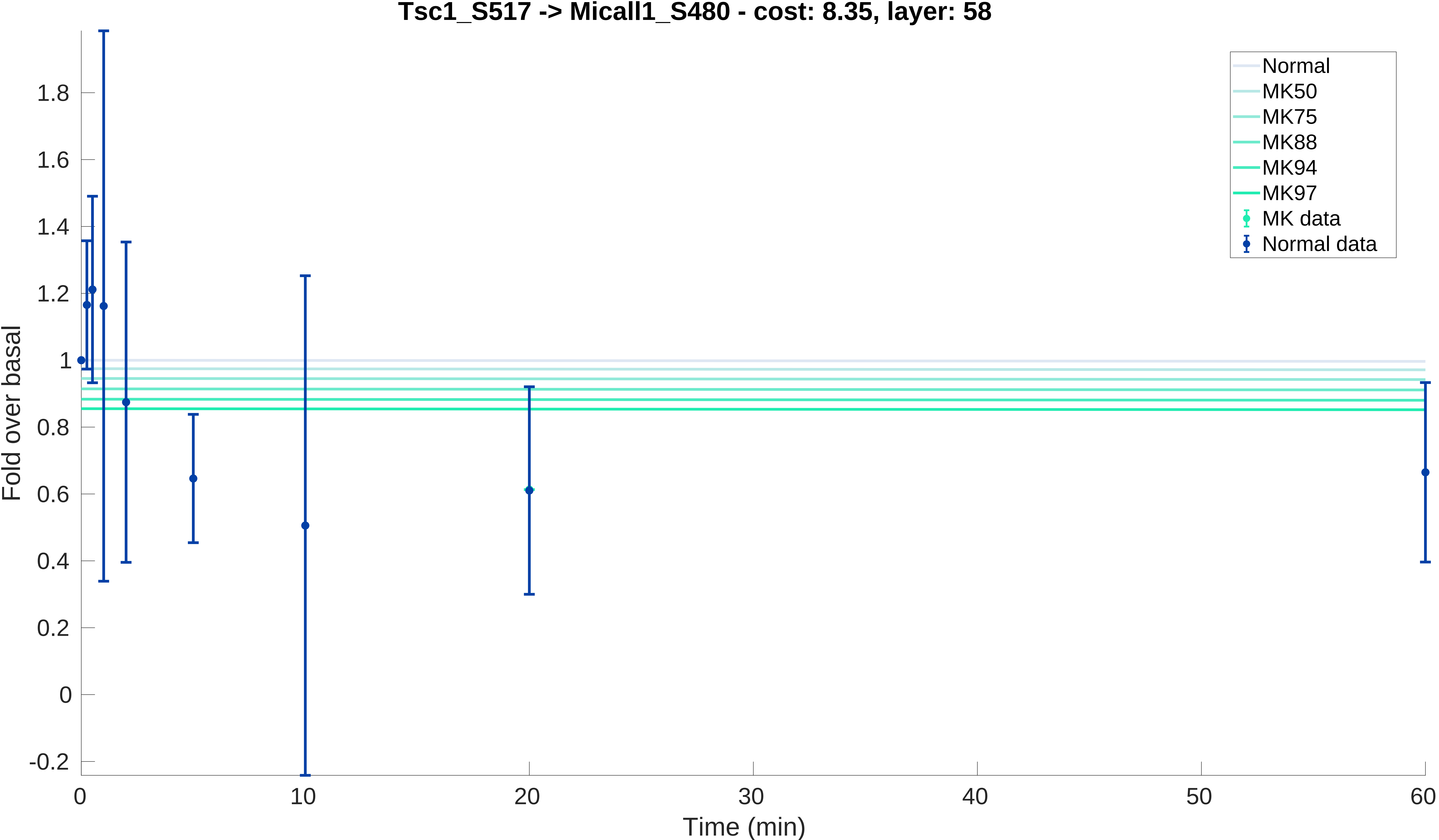

Tsc1_S517 -> Micall1_S480 - cost: 8.35, layer: 58
Normal
MK50
MK75
MK88
MK94
MK97
MK data
Normal data
Fold over basal
Time (min)
1.8
1.6
1.4
1.2
1
0.8
0.6
0.4
0.2
0
-0.2
0
10
20
30
40
50
60

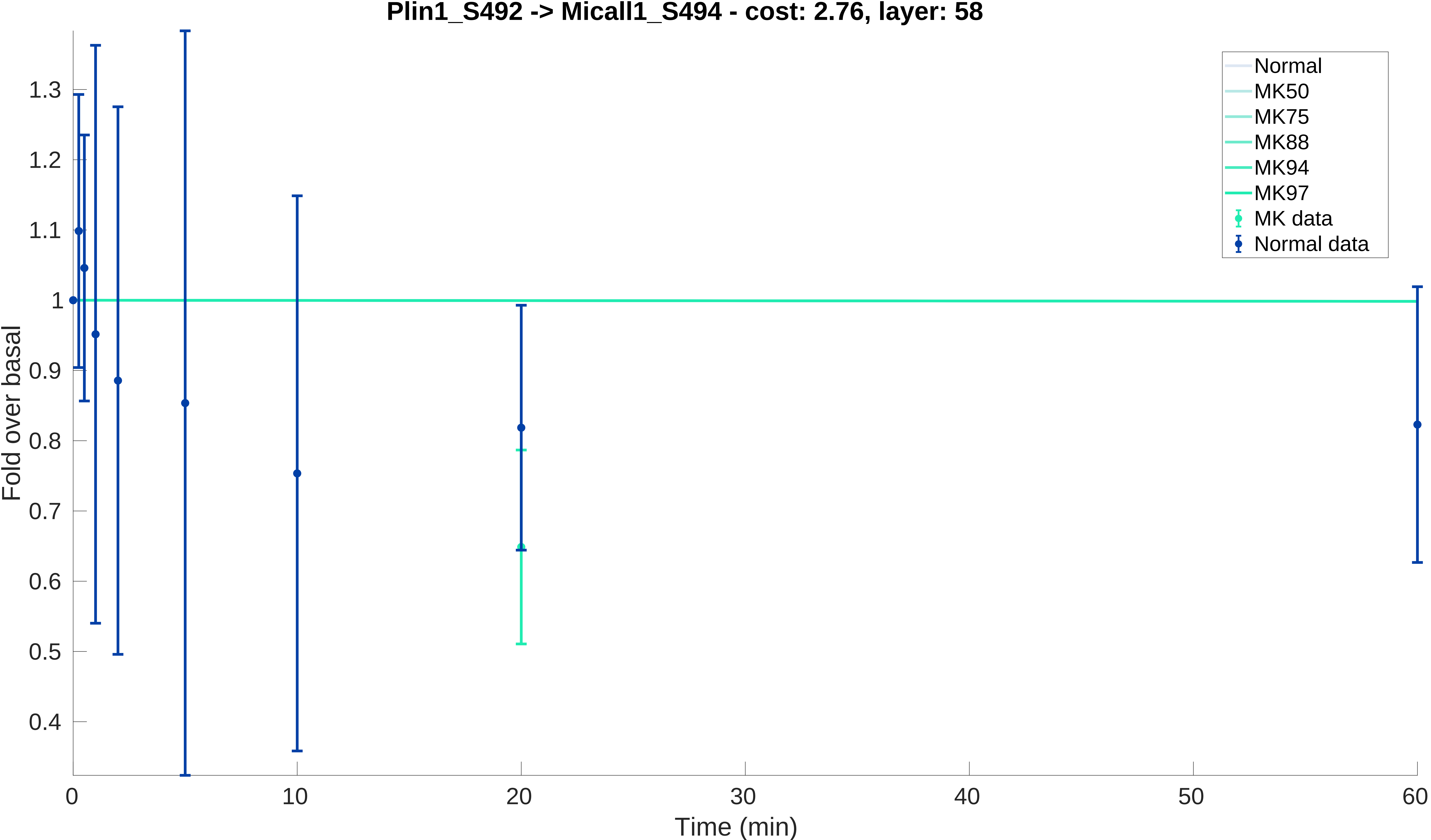

Plin1_S492 -> Micall1_S494 - cost: 2.76, layer: 58
Fold over basal
1.3
1.2
1.1
1
0.9
0.8
0.7
0.6
0.5
0.4
0
10
20
30
40
50
60
Time (min)
Normal
MK50
MK75
MK88
MK94
MK97
MK data
Normal data

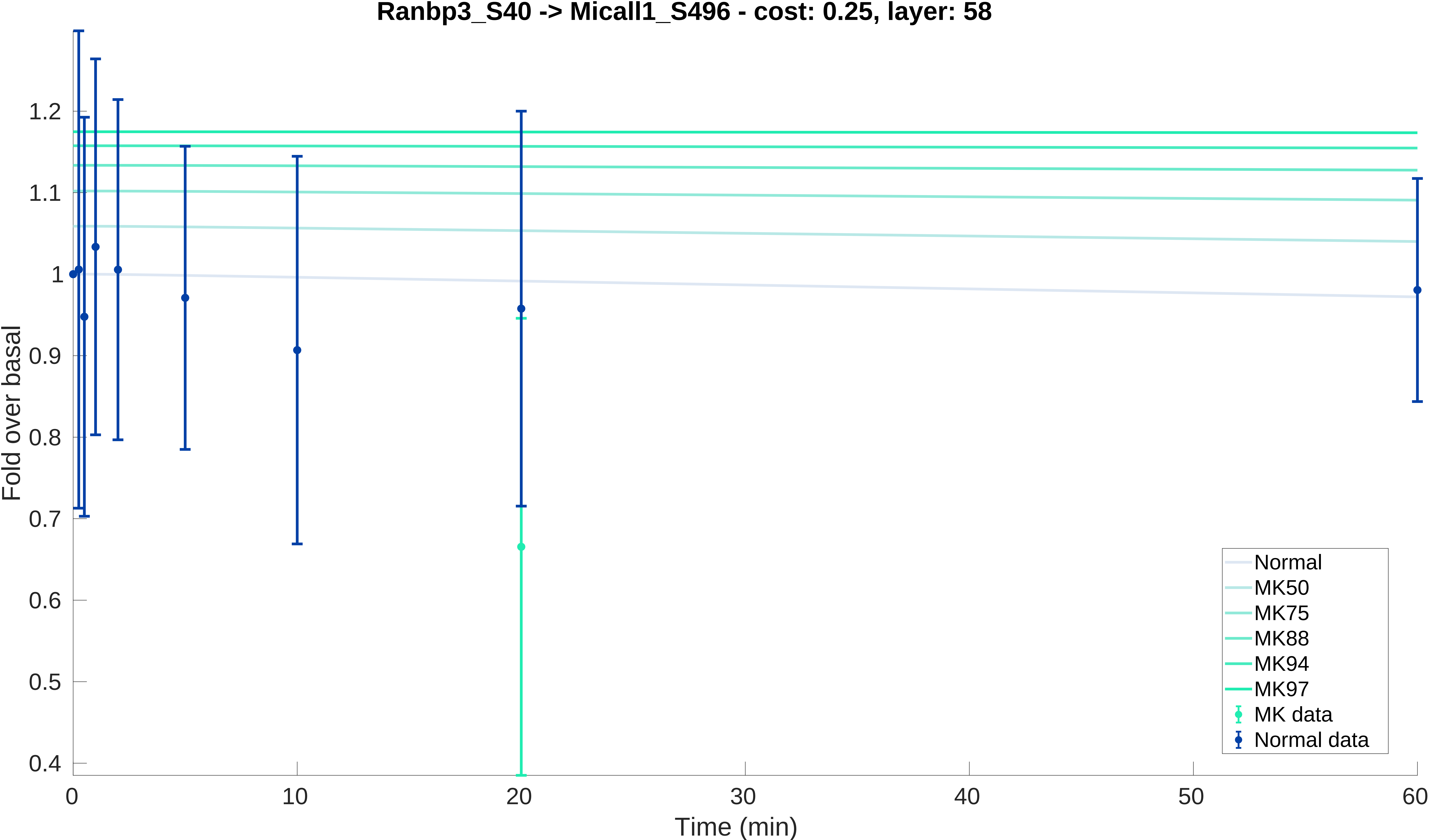

Ranbp3_S40 -> Micall1_S496 - cost: 0.25, layer: 58
Fold over basal
Time (min)
0.4
0.5
0.6
0.7
0.8
0.9
1
1.1
1.2
0
10
20
30
40
50
60
Normal
MK50
MK75
MK88
MK94
MK97
MK data
Normal data

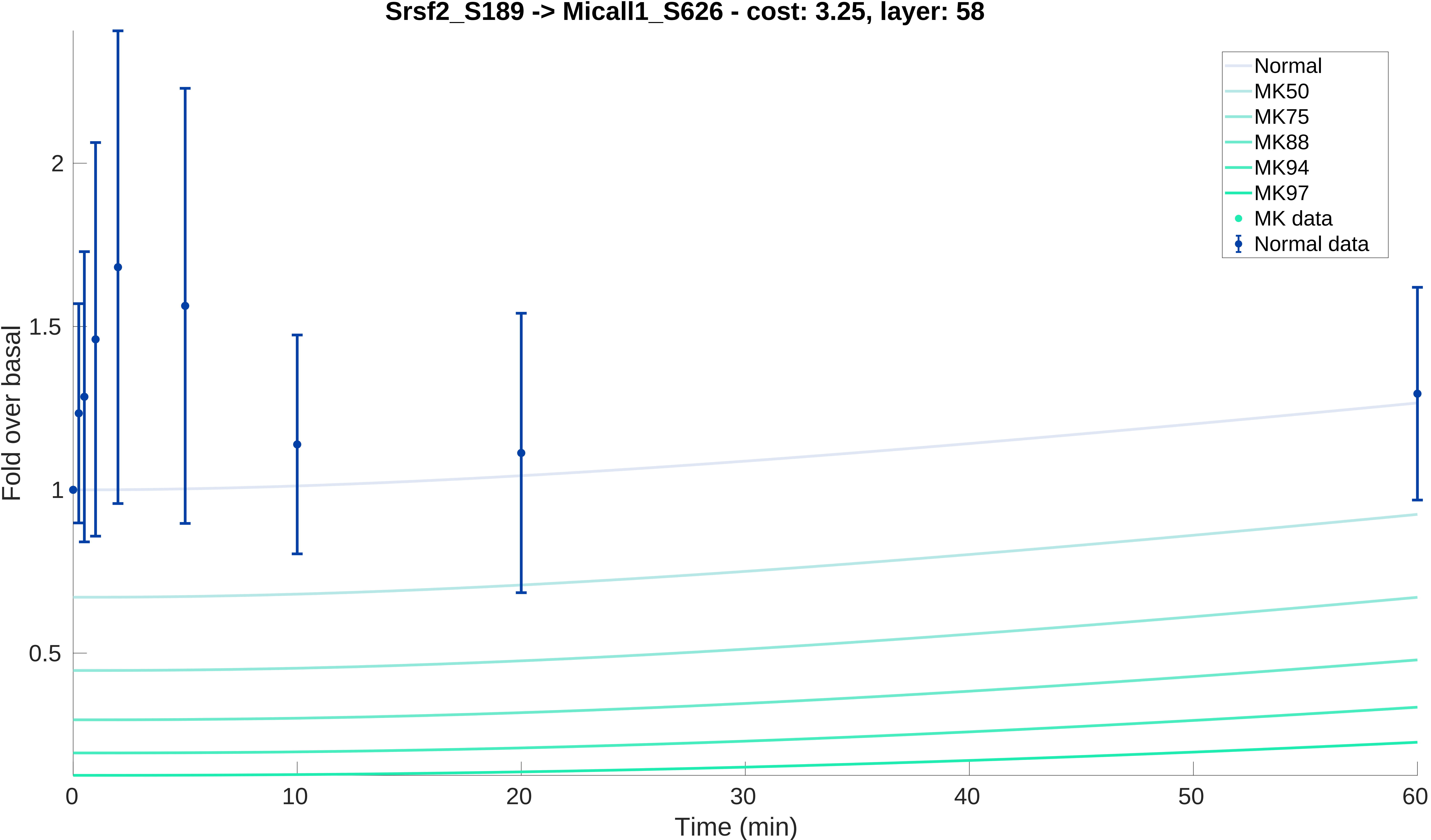

Srsf2_S189 -> Micall1_S626 - cost: 3.25, layer: 58
Fold over basal
Time (min)
0
10
20
30
40
50
60
0.5
1
1.5
2
Normal
MK50
MK75
MK88
MK94
MK97
MK data
Normal data

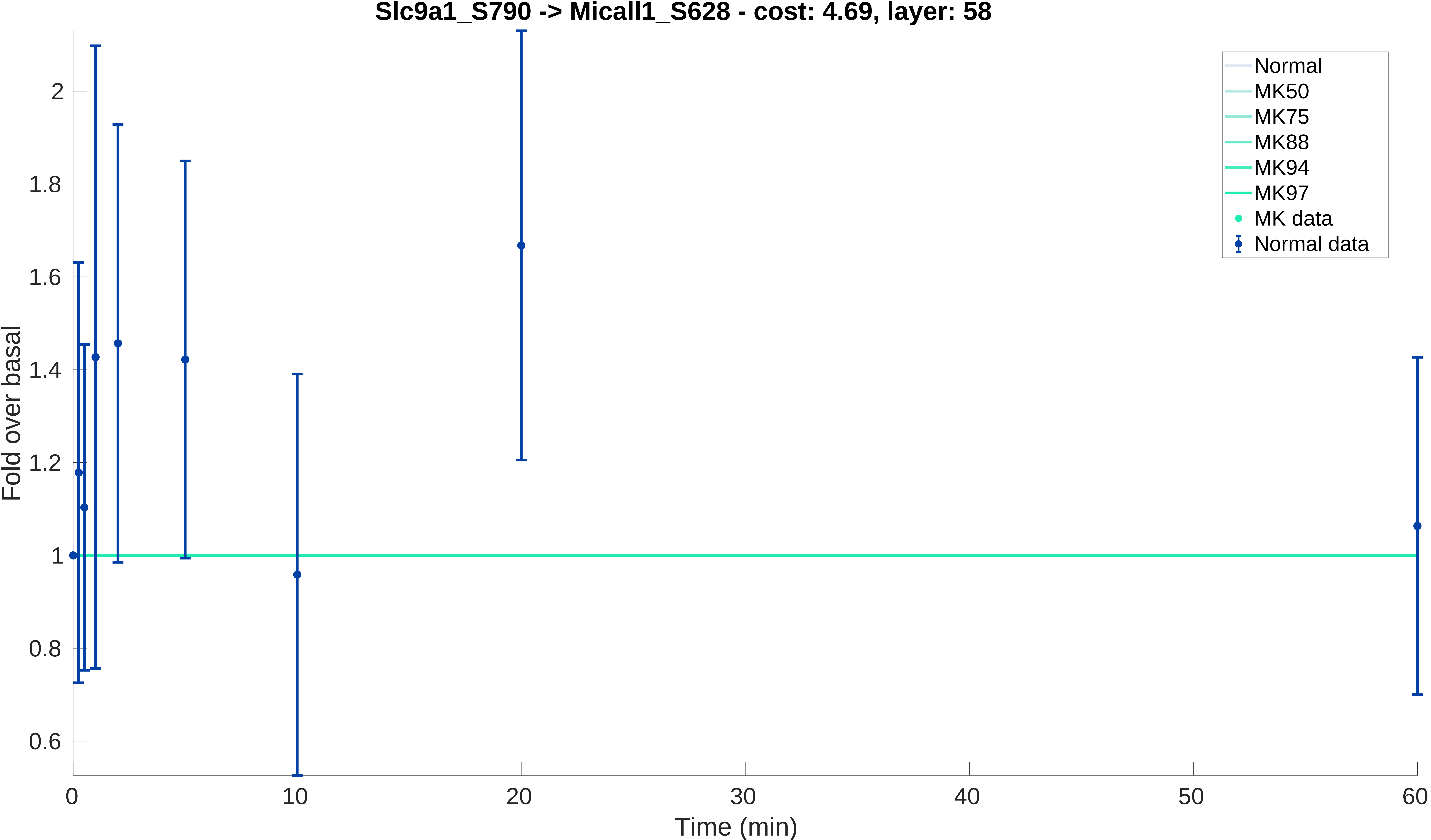

Slc9a1_S790 -> Micall1_S628 - cost: 4.69, layer: 58
Fold over basal
Time (min)
0.6
0.8
1
1.2
1.4
1.6
1.8
2
0
10
20
30
40
50
60
Normal
MK50
MK75
MK88
MK94
MK97
MK data
Normal data

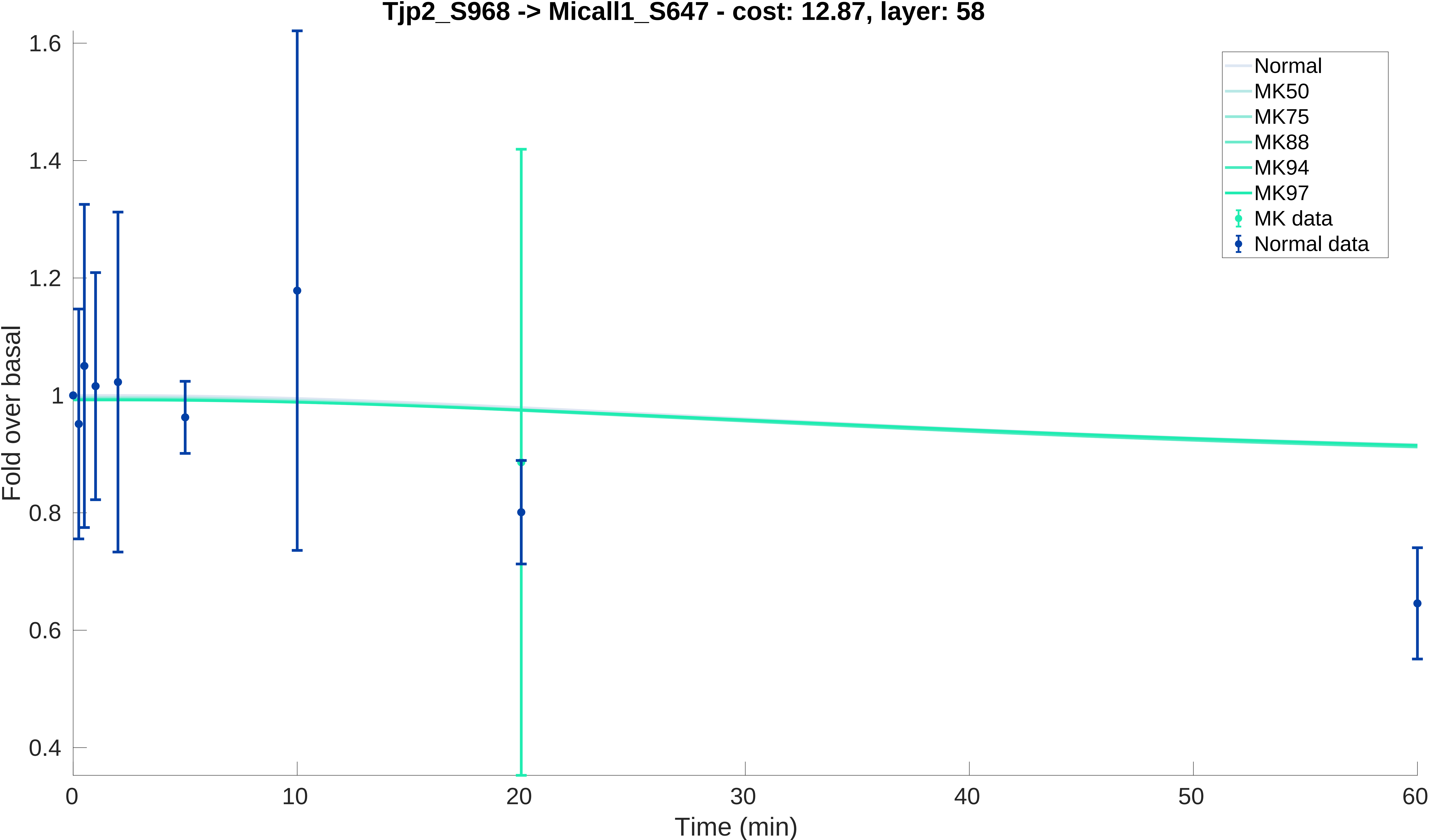
Tjp2_S968 -> Micall1_S647 - cost: 12.87, layer: 58
Fold over basal
Time (min)
1.6
1.4
1.2
1
0.8
0.6
0.4
0
10
20
30
40
50
60
Normal
MK50
MK75
MK88
MK94
MK97
MK data
Normal data

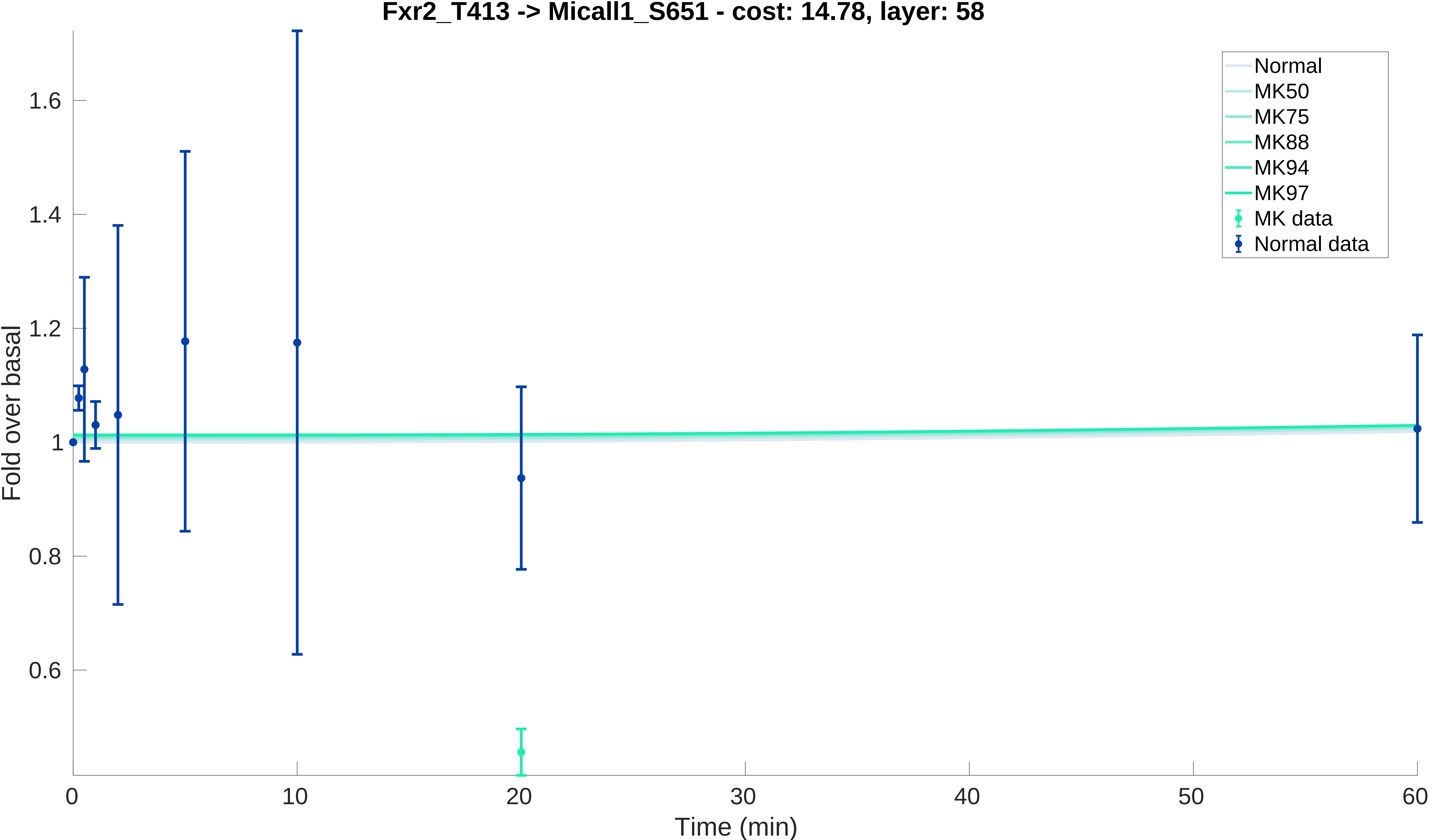

Fxr2_T413 -> Micall1_S651 - cost: 14.78, layer: 58
Fold over basal
Time (min)
0.6
0.8
1
1.2
1.4
1.6
0
10
20
30
40
50
60
Normal
MK50
MK75
MK88
MK94
MK97
MK data
Normal data

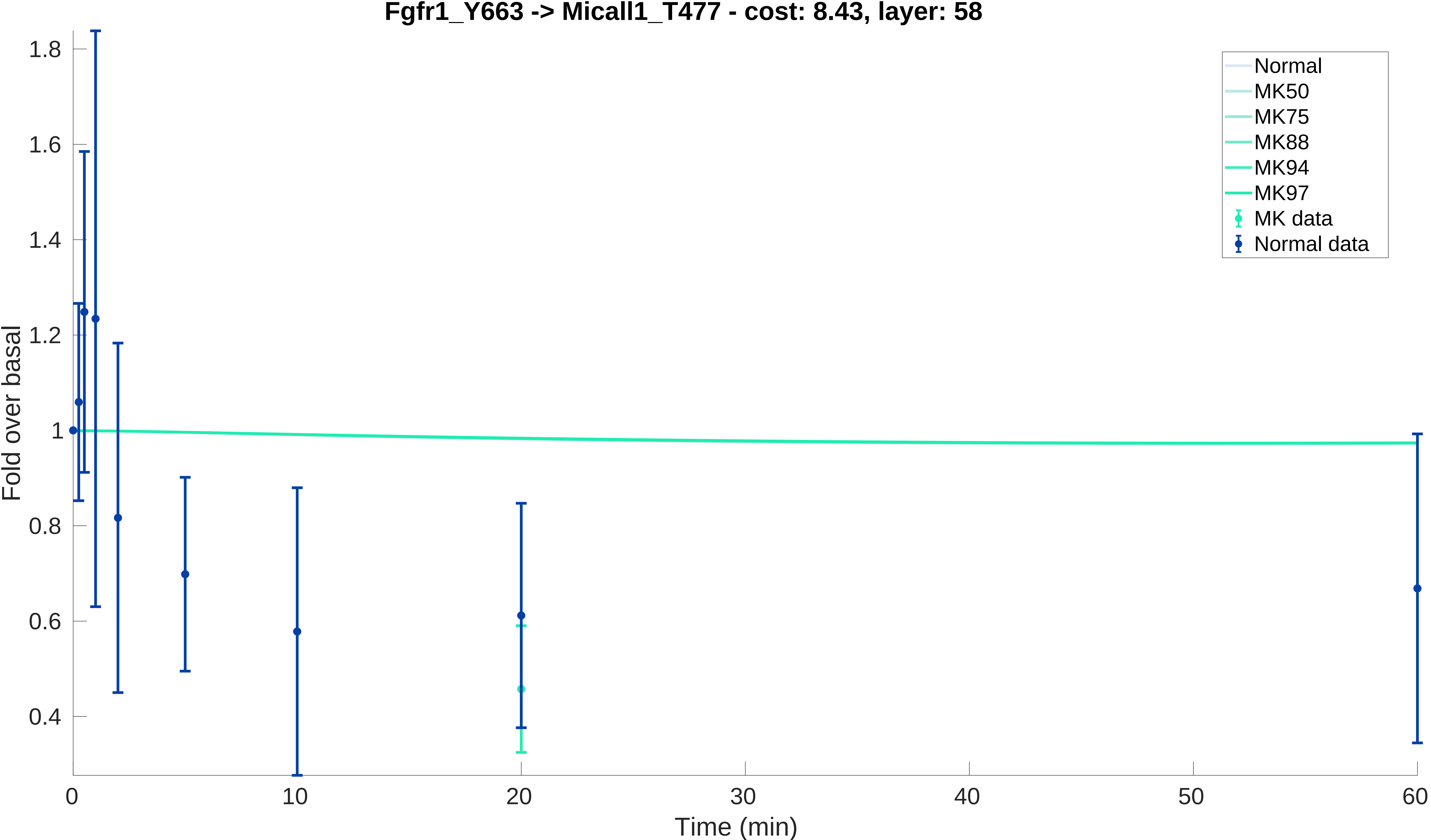
Fgfr1_Y663 -> Micall1_T477 - cost: 8.43, layer: 58
Fold over basal
Time (min)
1.8
1.6
1.4
1.2
1
0.8
0.6
0.4
0
10
20
30
40
50
60
Normal
MK50
MK75
MK88
MK94
MK97
MK data
Normal data

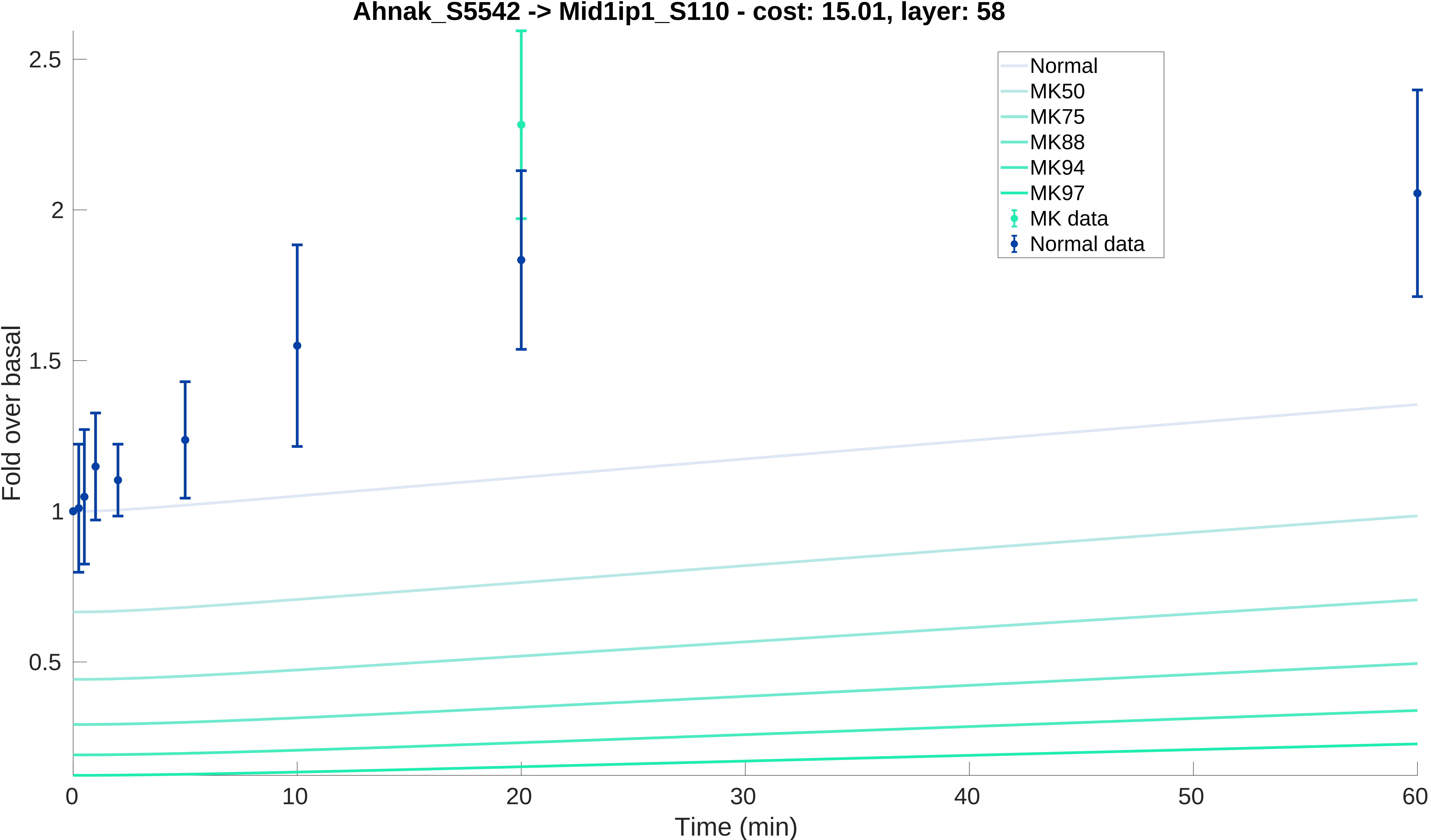

Ahnak_S5542 -> Mid1ip1_S110 - cost: 15.01, layer: 58
Fold over basal
Time (min)
0.5
1
1.5
2
2.5
0
10
20
30
40
50
60
Normal
MK50
MK75
MK88
MK94
MK97
MK data
Normal data

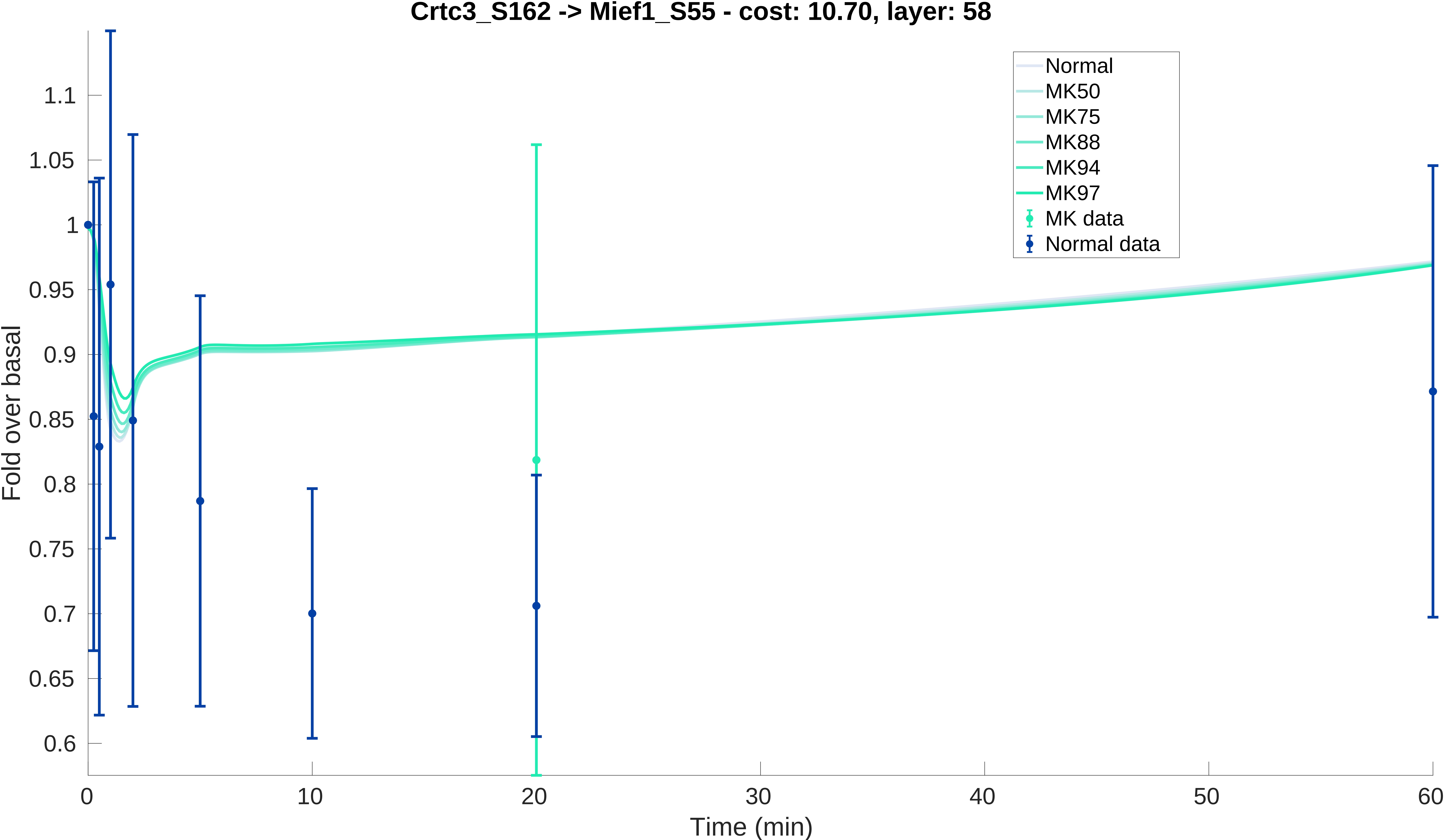

Crtc3_S162 -> Mief1_S55 - cost: 10.70, layer: 58
Normal
MK50
MK75
MK88
MK94
MK97
MK data
Normal data
Fold over basal
Time (min)
0
10
20
30
40
50
60
0.6
0.65
0.7
0.75
0.8
0.85
0.9
0.95
1
1.05
1.1

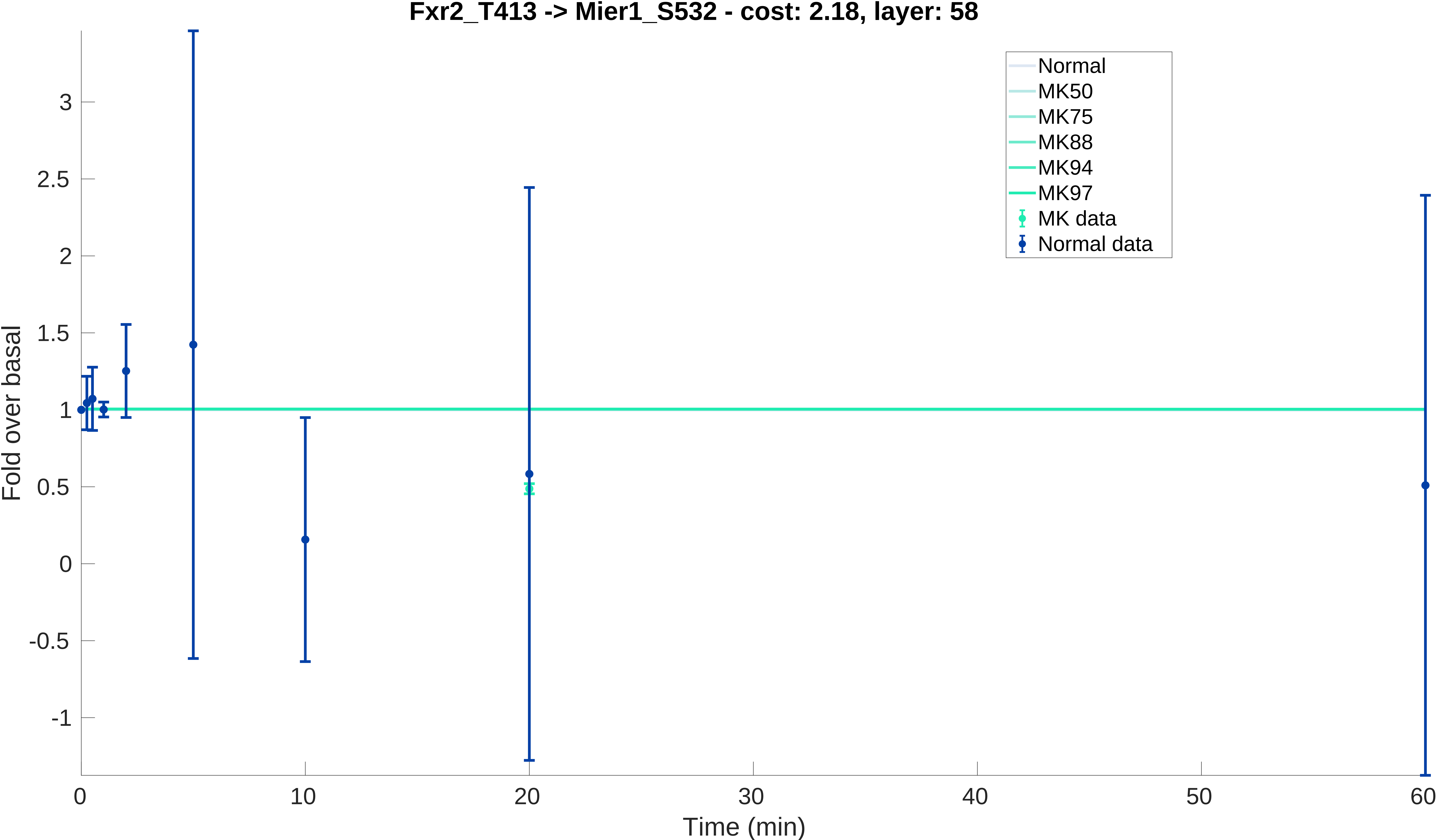

Fxr2_T413 -> Mier1_S532 - cost: 2.18, layer: 58
Fold over basal
3
2.5
2
1.5
1
0.5
0
-0.5
-1
0
10
20
30
40
50
60
Time (min)
Normal
MK50
MK75
MK88
MK94
MK97
MK data
Normal data

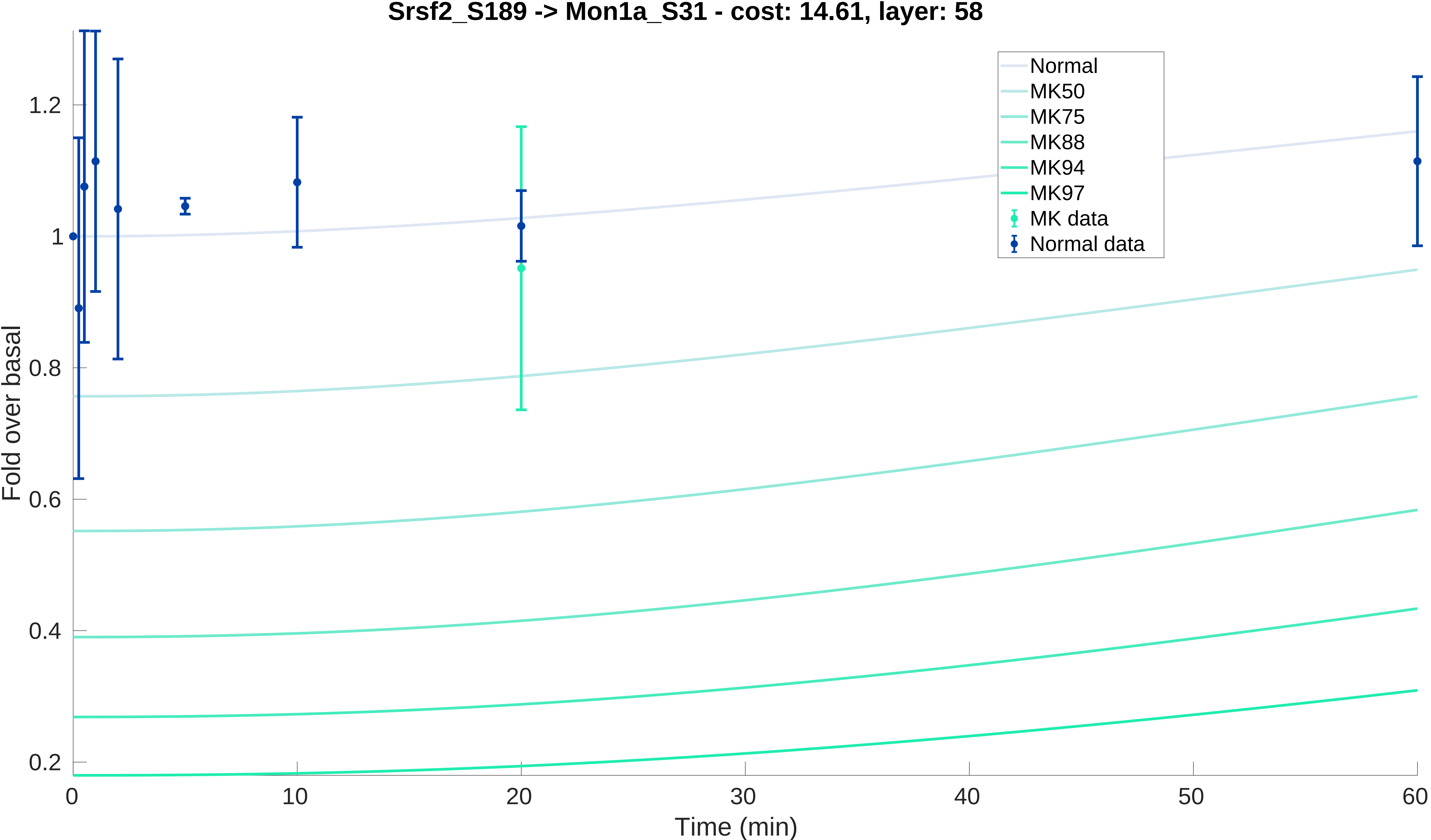

Srsf2_S189 -> Mon1a_S31 - cost: 14.61, layer: 58
Fold over basal
Time (min)
0.2
0.4
0.6
0.8
1
1.2
0
10
20
30
40
50
60
Normal
MK50
MK75
MK88
MK94
MK97
MK data
Normal data

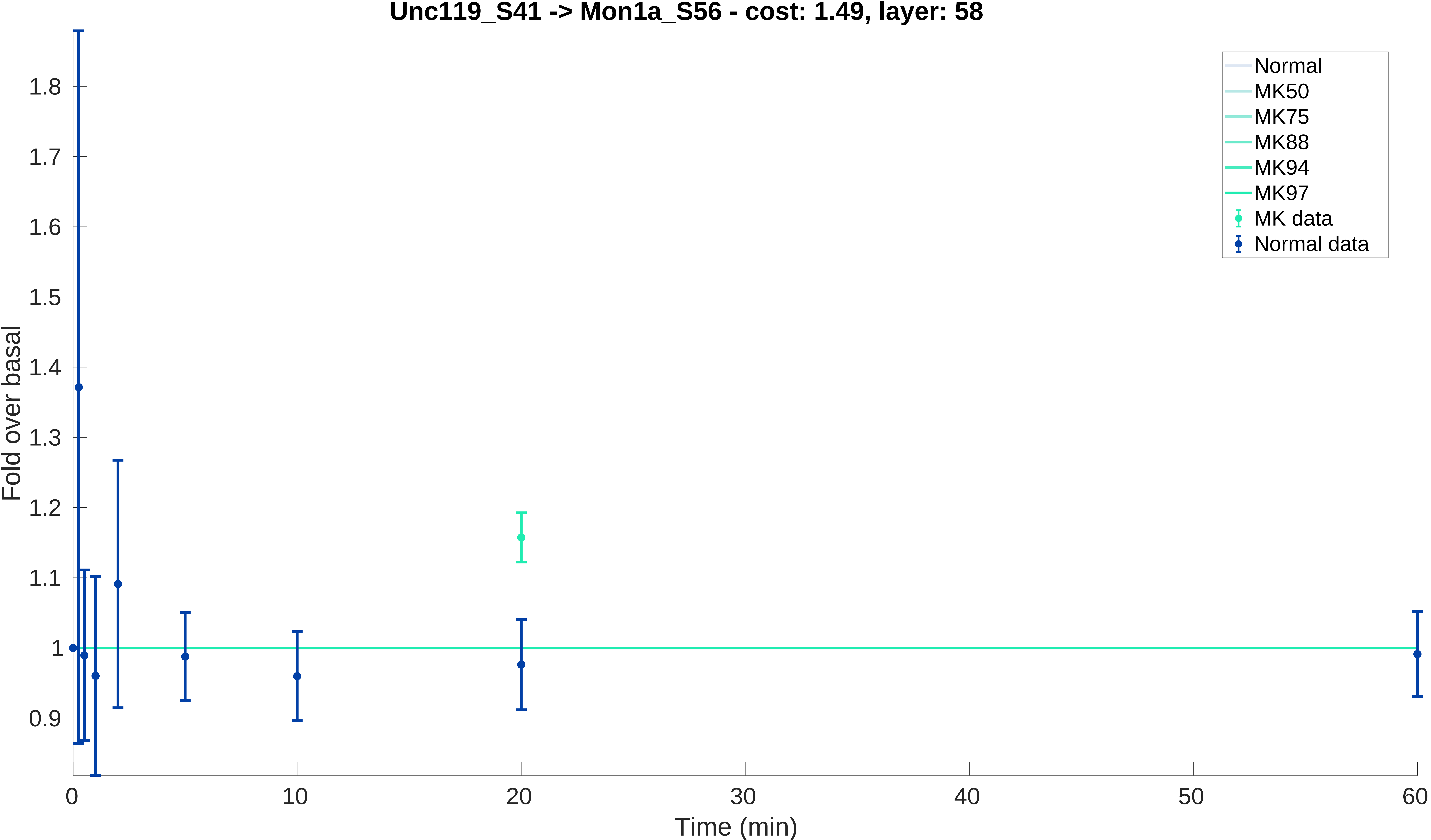

Unc119_S41 -> Mon1a_S56 - cost: 1.49, layer: 58
Normal
MK50
MK75
MK88
MK94
MK97
MK data
Normal data
Fold over basal
Time (min)
0.9
1
1.1
1.2
1.3
1.4
1.5
1.6
1.7
1.8
0
10
20
30
40
50
60

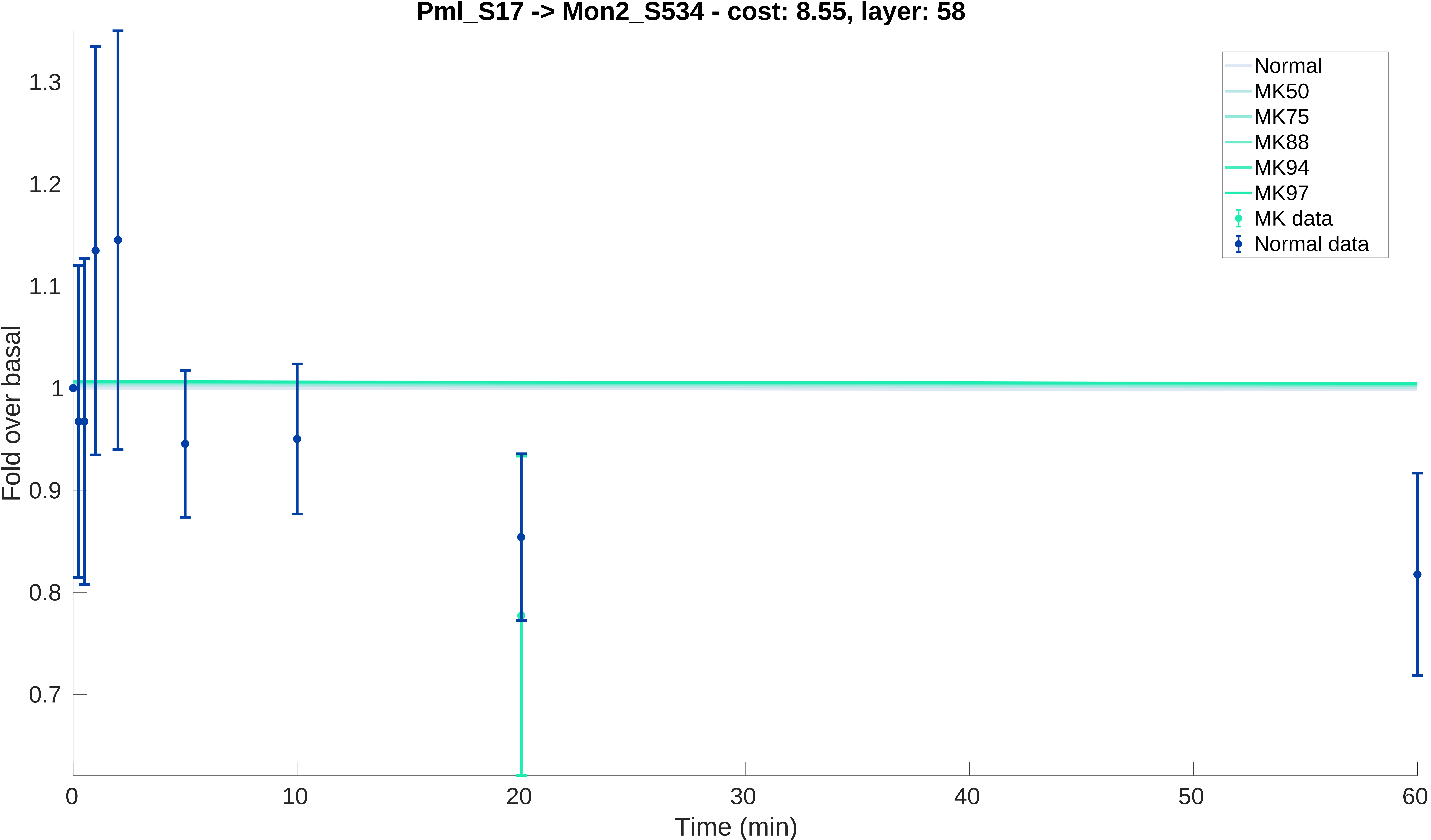

Pml_S17 -> Mon2_S534 - cost: 8.55, layer: 58
Fold over basal
1.3
1.2
1.1
1
0.9
0.8
0.7
0
10
20
30
40
50
60
Time (min)
Normal
MK50
MK75
MK88
MK94
MK97
MK data
Normal data

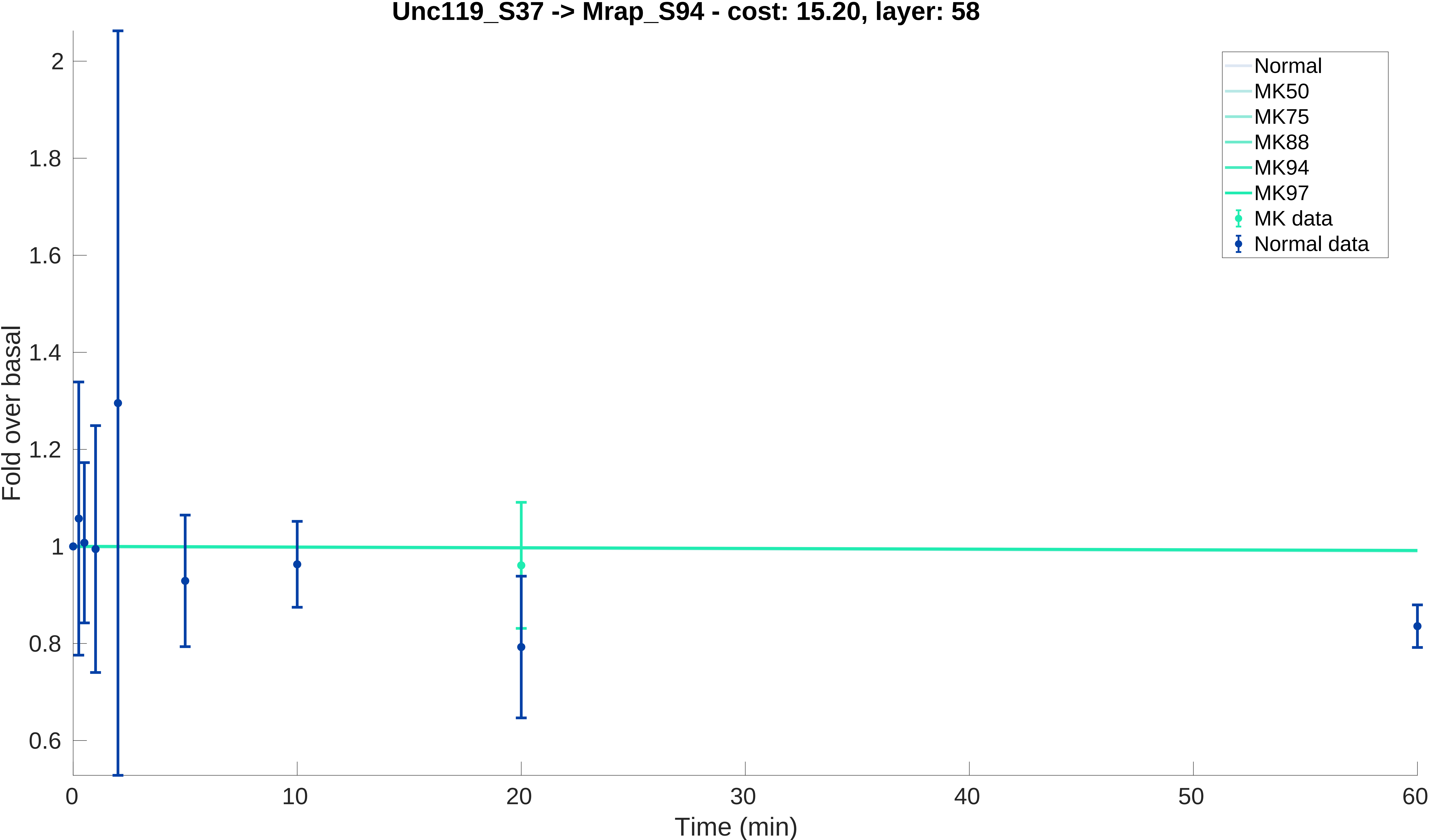
Unc119_S37 -> Mrap_S94 - cost: 15.20, layer: 58
Fold over basal
Time (min)
2
1.8
1.6
1.4
1.2
1
0.8
0.6
0
10
20
30
40
50
60
Normal
MK50
MK75
MK88
MK94
MK97
MK data
Normal data

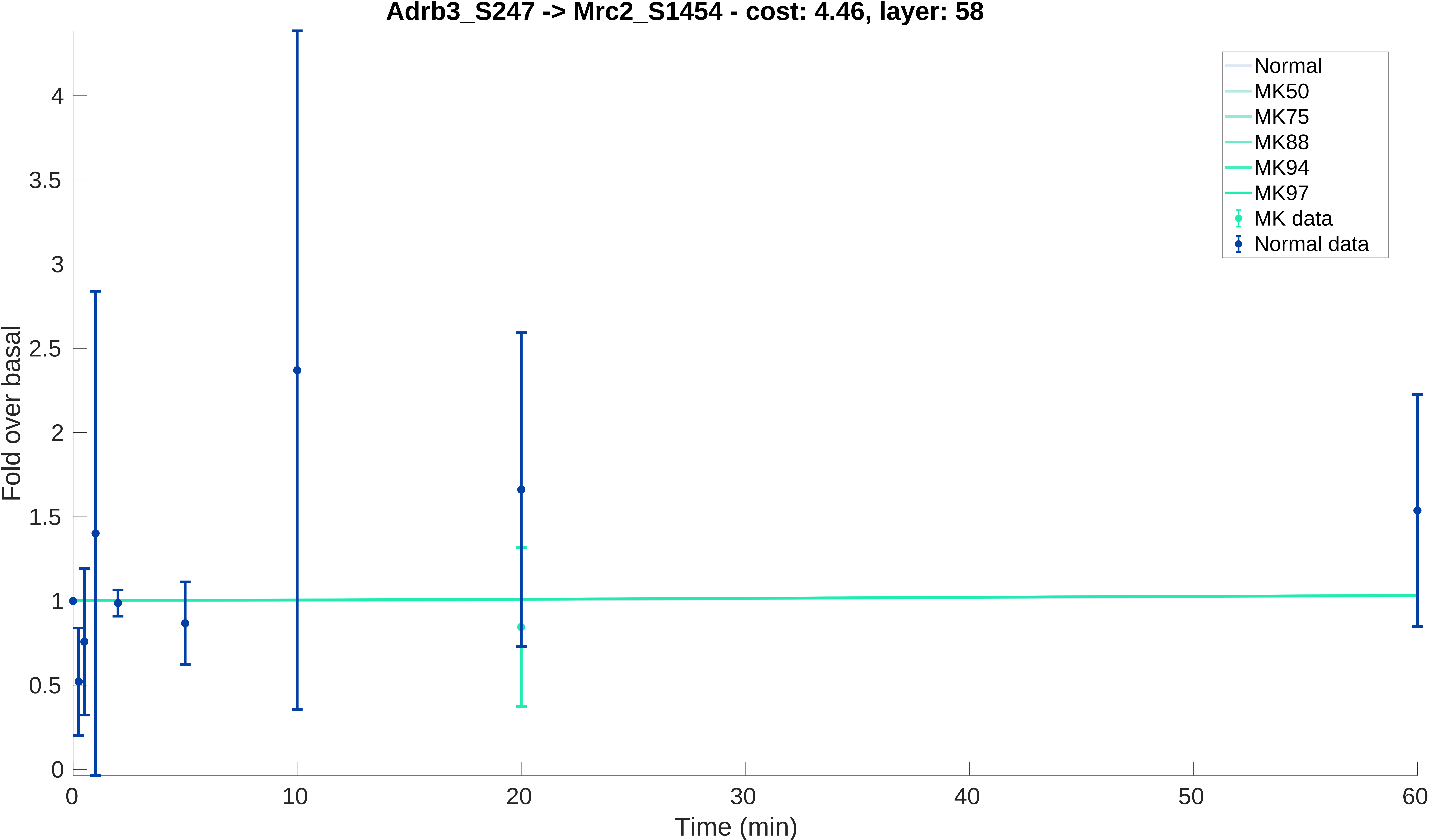
Adrb3_S247 -> Mrc2_S1454 - cost: 4.46, layer: 58
Fold over basal
Time (min)
0
0.5
1
1.5
2
2.5
3
3.5
4
0
10
20
30
40
50
60
Normal
MK50
MK75
MK88
MK94
MK97
MK data
Normal data

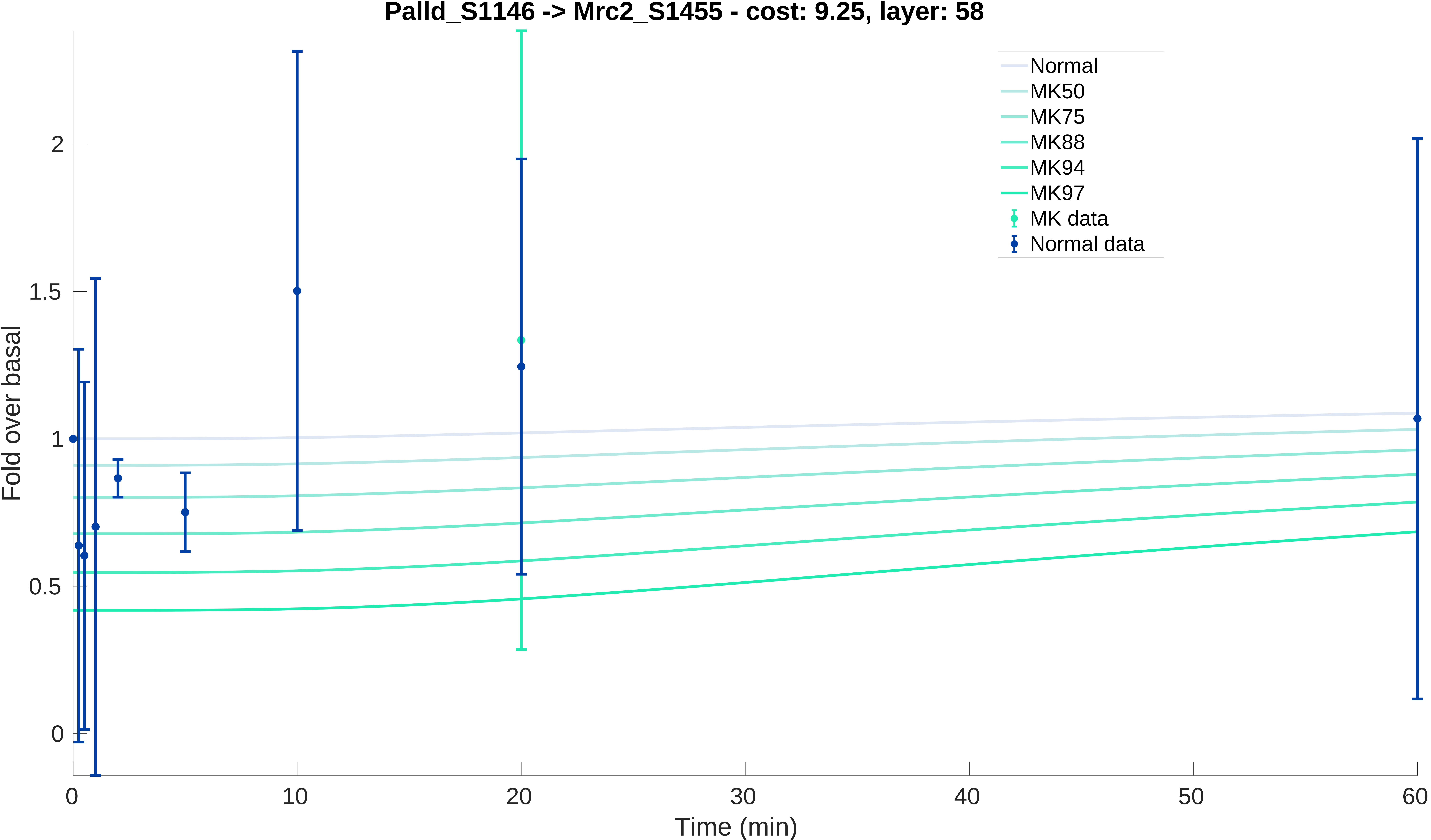

Palld_S1146 -> Mrc2_S1455 - cost: 9.25, layer: 58
Normal
MK50
MK75
MK88
MK94
MK97
MK data
Normal data
Fold over basal
Time (min)
0
0.5
1
1.5
2
0
10
20
30
40
50
60

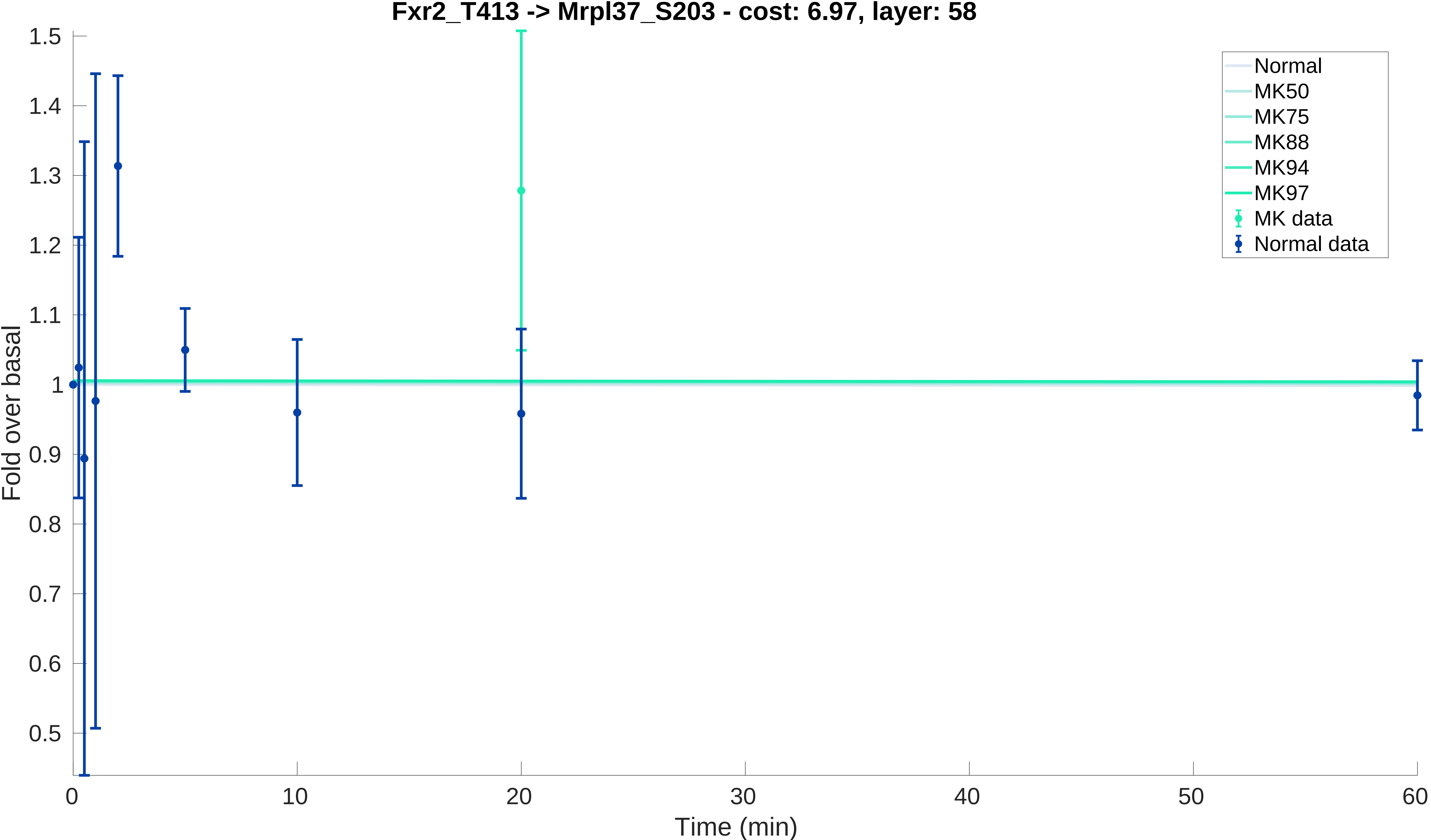

Fxr2_T413 -> Mrpl37_S203 - cost: 6.97, layer: 58
Fold over basal
Time (min)
1.5
1.4
1.3
1.2
1.1
1
0.9
0.8
0.7
0.6
0.5
0
10
20
30
40
50
60
Normal
MK50
MK75
MK88
MK94
MK97
MK data
Normal data

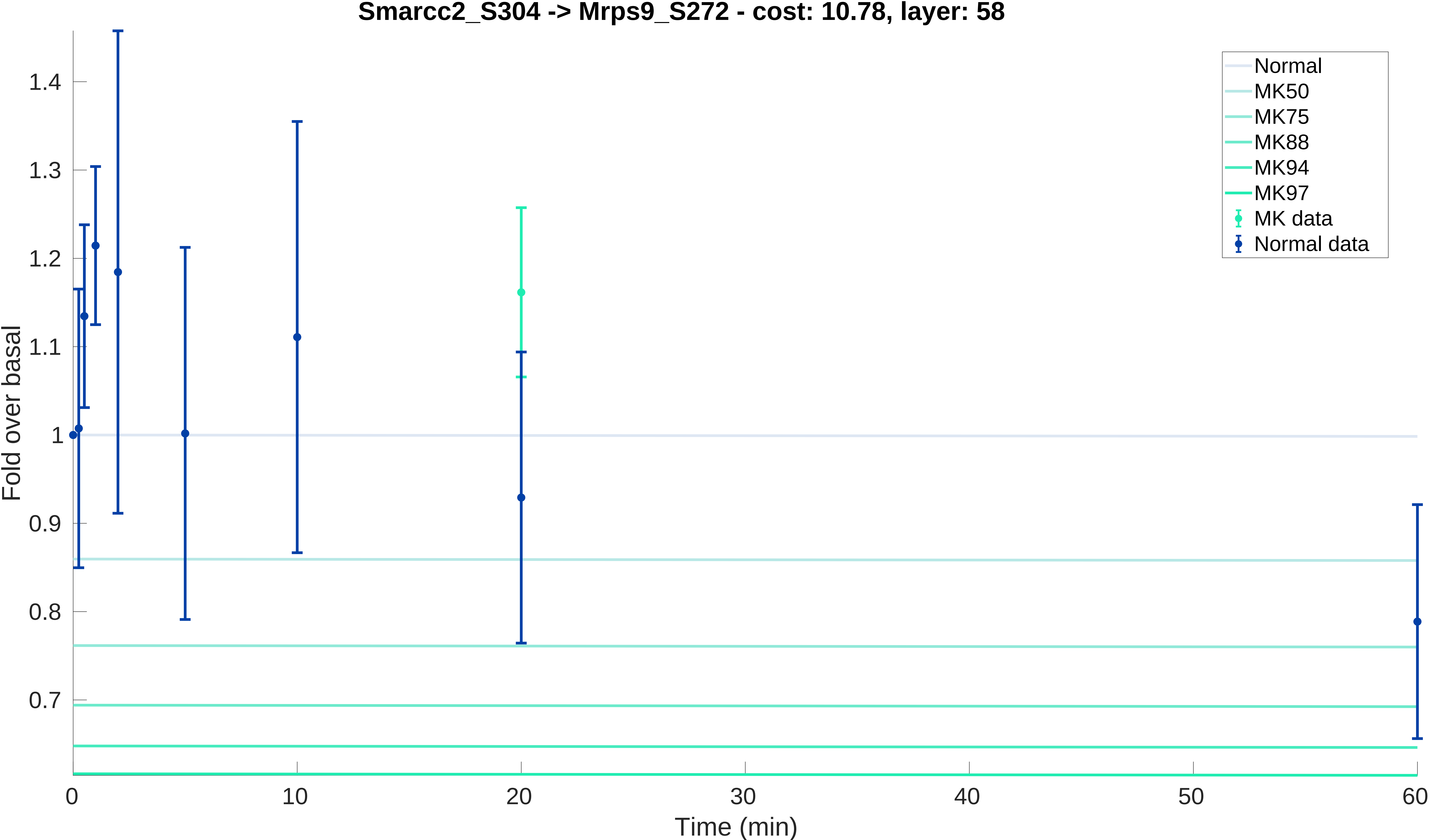
Smarcc2_S304 -> Mrps9_S272 - cost: 10.78, layer: 58
Fold over basal
1.4
1.3
1.2
1.1
1
0.9
0.8
0.7
0
10
20
30
40
50
60
Time (min)
Normal
MK50
MK75
MK88
MK94
MK97
MK data
Normal data

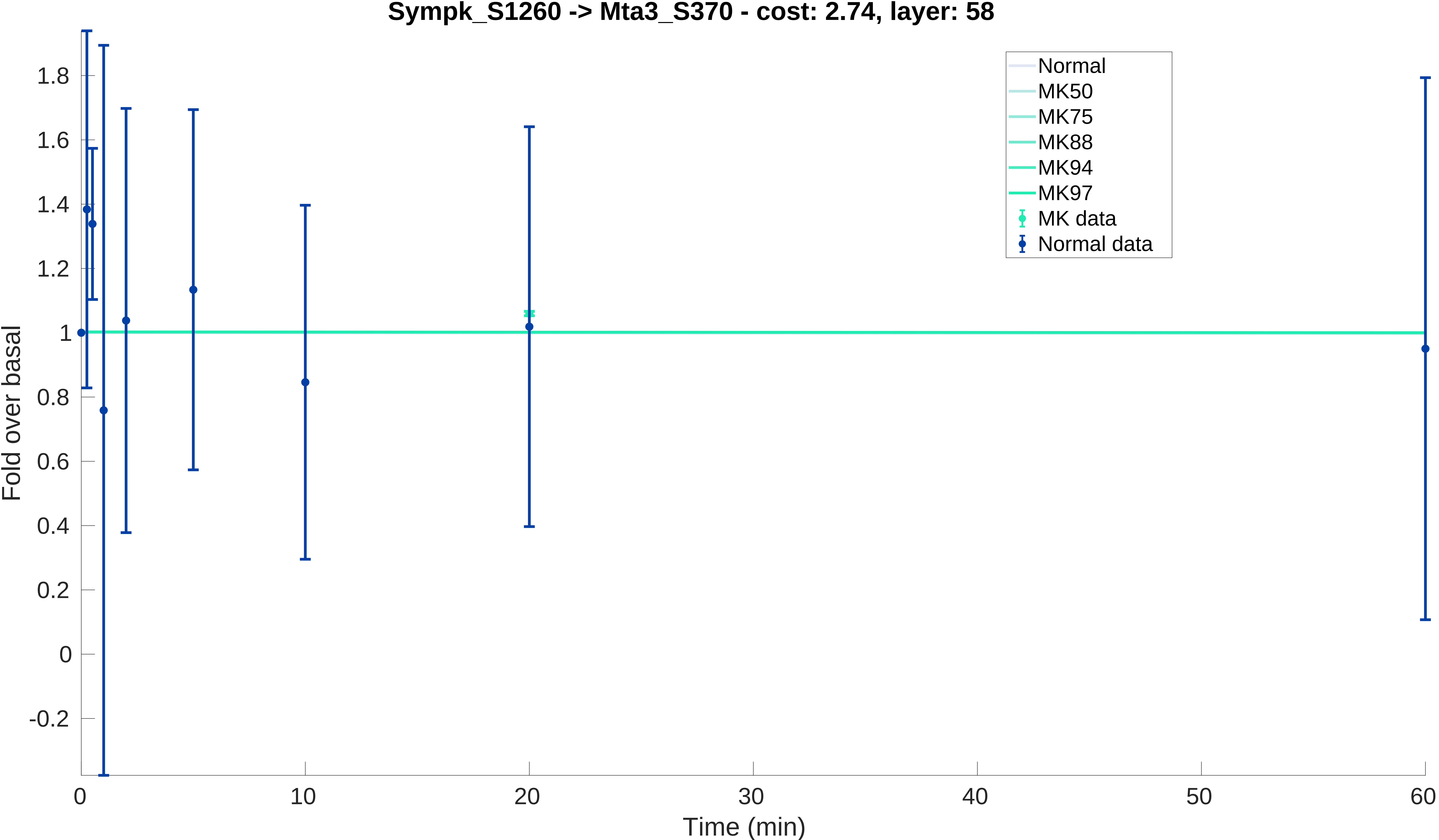
Sympk_S1260 -> Mta3_S370 - cost: 2.74, layer: 58
Normal
MK50
MK75
MK88
MK94
MK97
MK data
Normal data
Fold over basal
Time (min)
1.8
1.6
1.4
1.2
1
0.8
0.6
0.4
0.2
0
-0.2
0
10
20
30
40
50
60

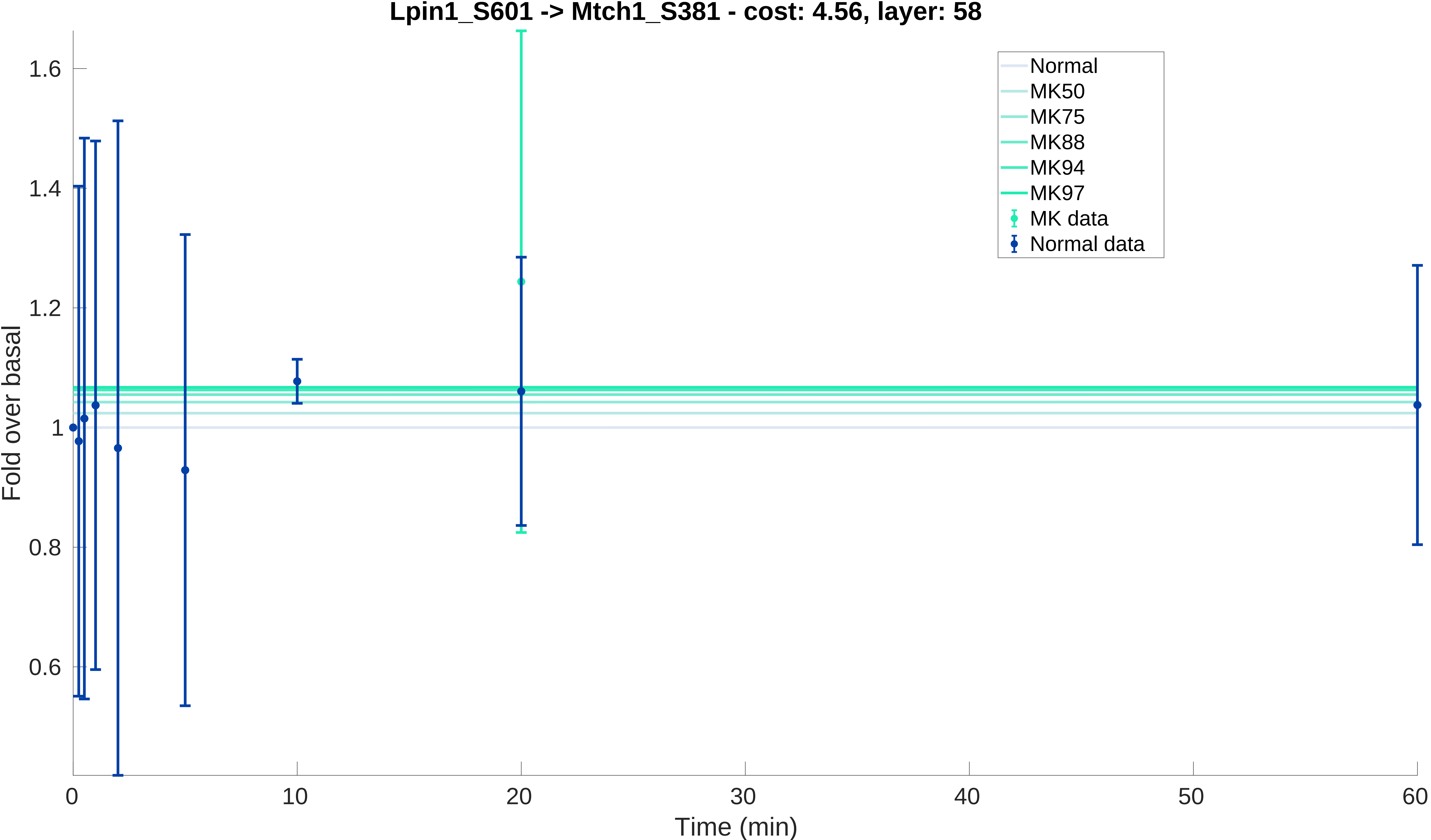

Lpin1_S601 -> Mtch1_S381 - cost: 4.56, layer: 58
Fold over basal
Time (min)
1.6
1.4
1.2
1
0.8
0.6
0
10
20
30
40
50
60
Normal
MK50
MK75
MK88
MK94
MK97
MK data
Normal data

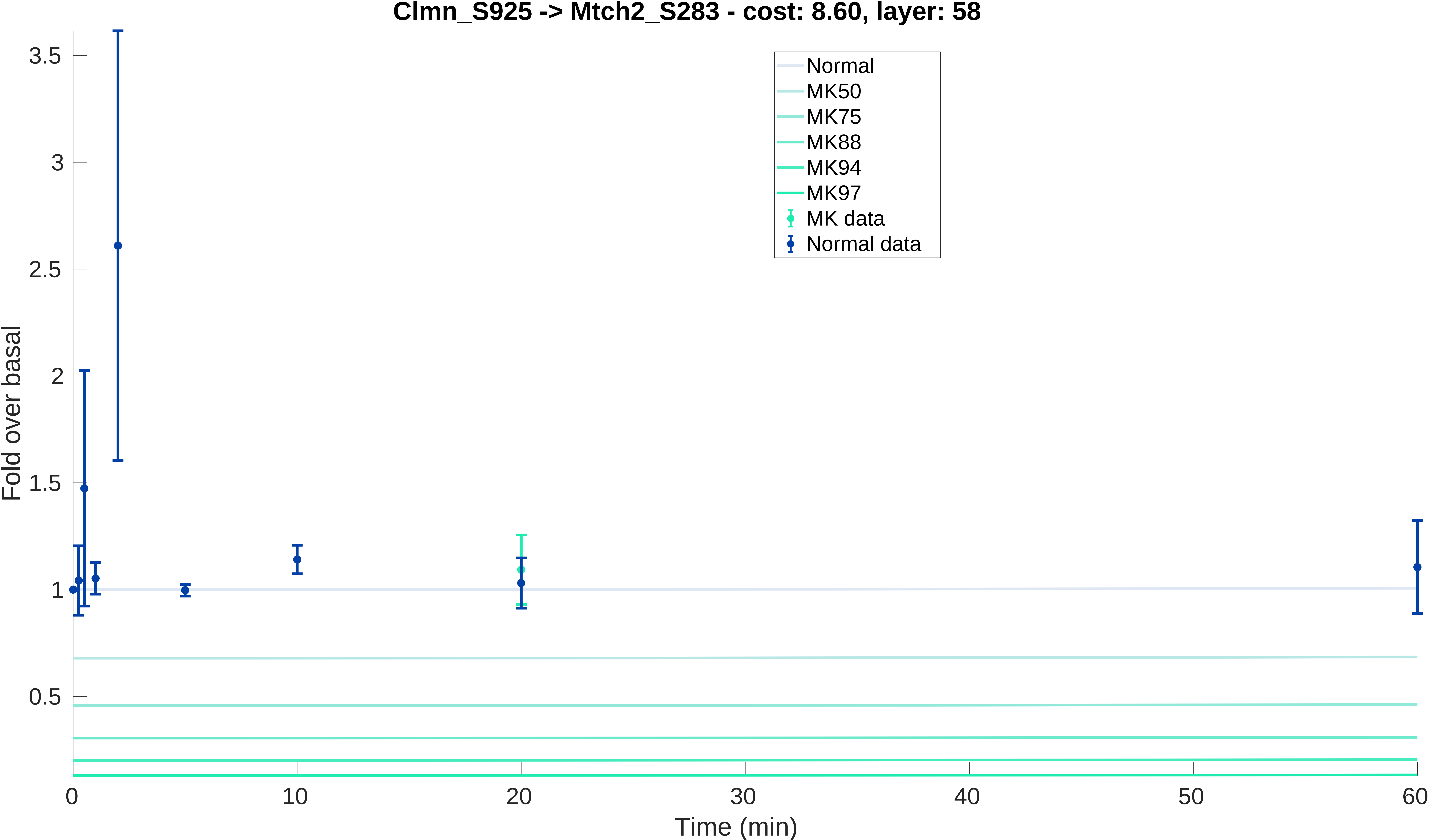
Clmn_S925 -> Mtch2_S283 - cost: 8.60, layer: 58
Fold over basal
3.5
3
2.5
2
1.5
1
0.5
0
10
20
30
40
50
60
Time (min)
Normal
MK50
MK75
MK88
MK94
MK97
MK data
Normal data

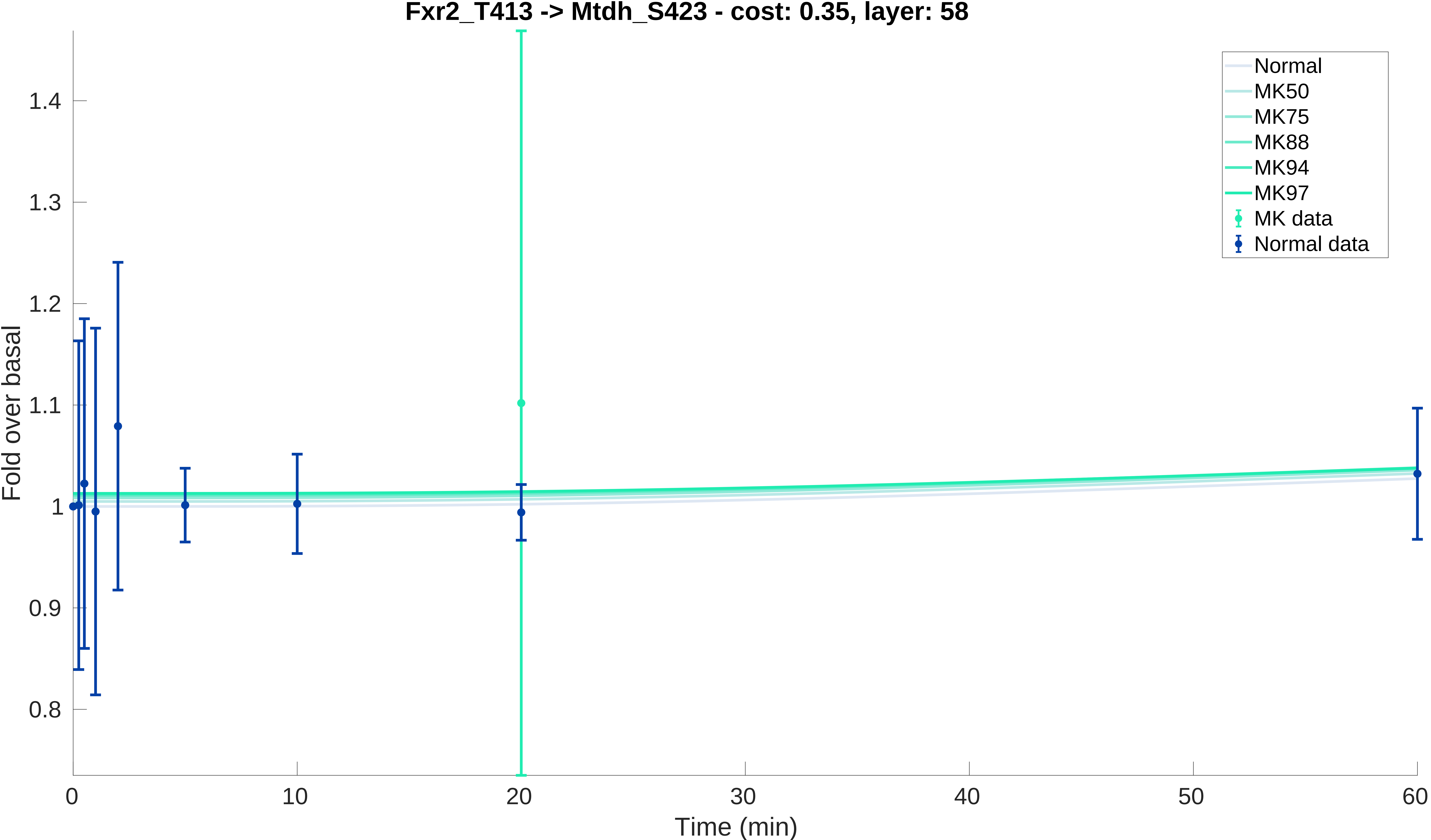
Fxr2_T413 -> Mtdh_S423 - cost: 0.35, layer: 58
Normal
MK50
MK75
MK88
MK94
MK97
MK data
Normal data
Fold over basal
1.4
1.3
1.2
1.1
1
0.9
0.8
0
10
20
30
40
50
60
Time (min)

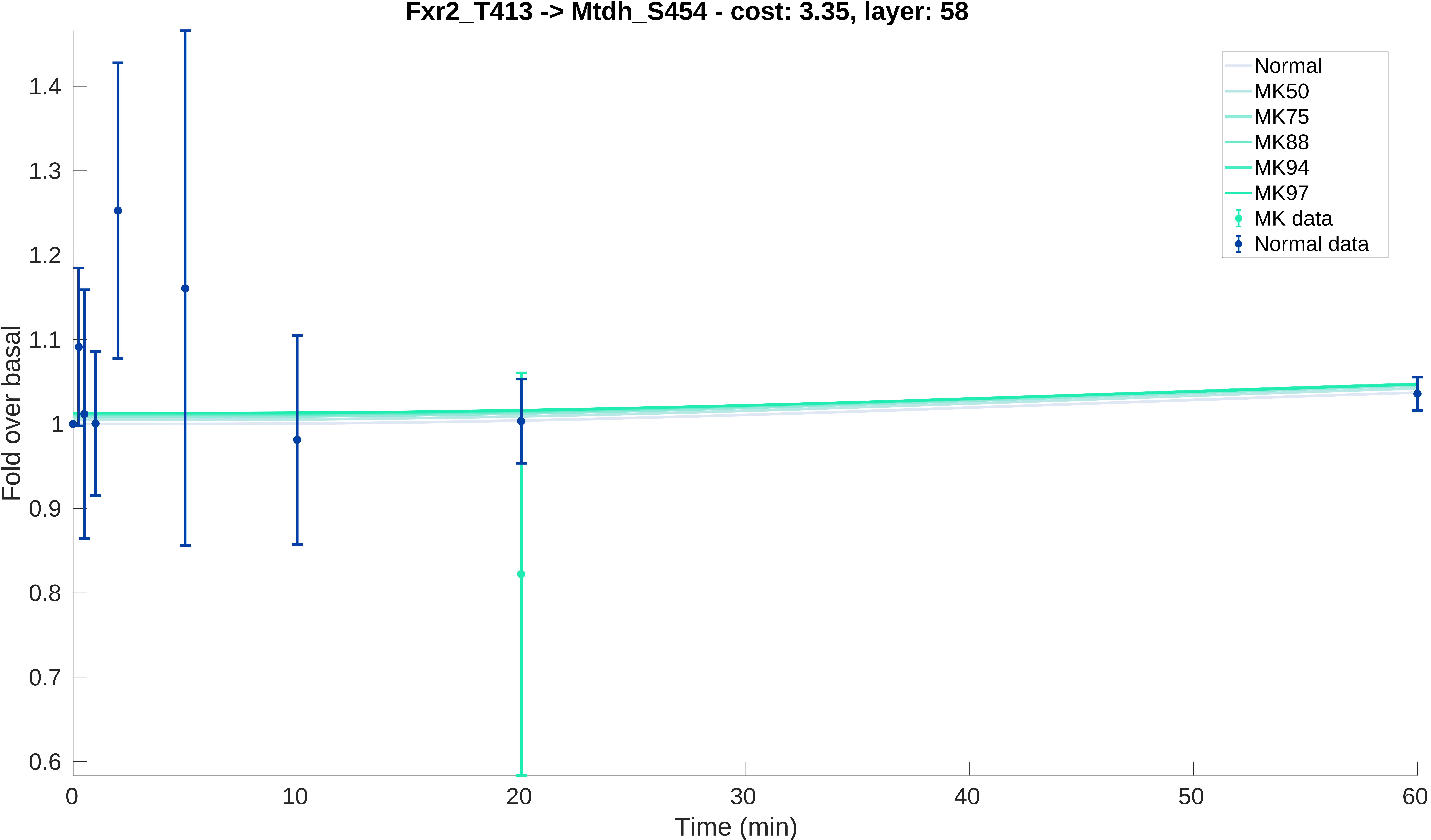

Fxr2_T413 -> Mtdh_S454 - cost: 3.35, layer: 58
Fold over basal
Time (min)
0.6
0.7
0.8
0.9
1
1.1
1.2
1.3
1.4
0
10
20
30
40
50
60
Normal
MK50
MK75
MK88
MK94
MK97
MK data
Normal data

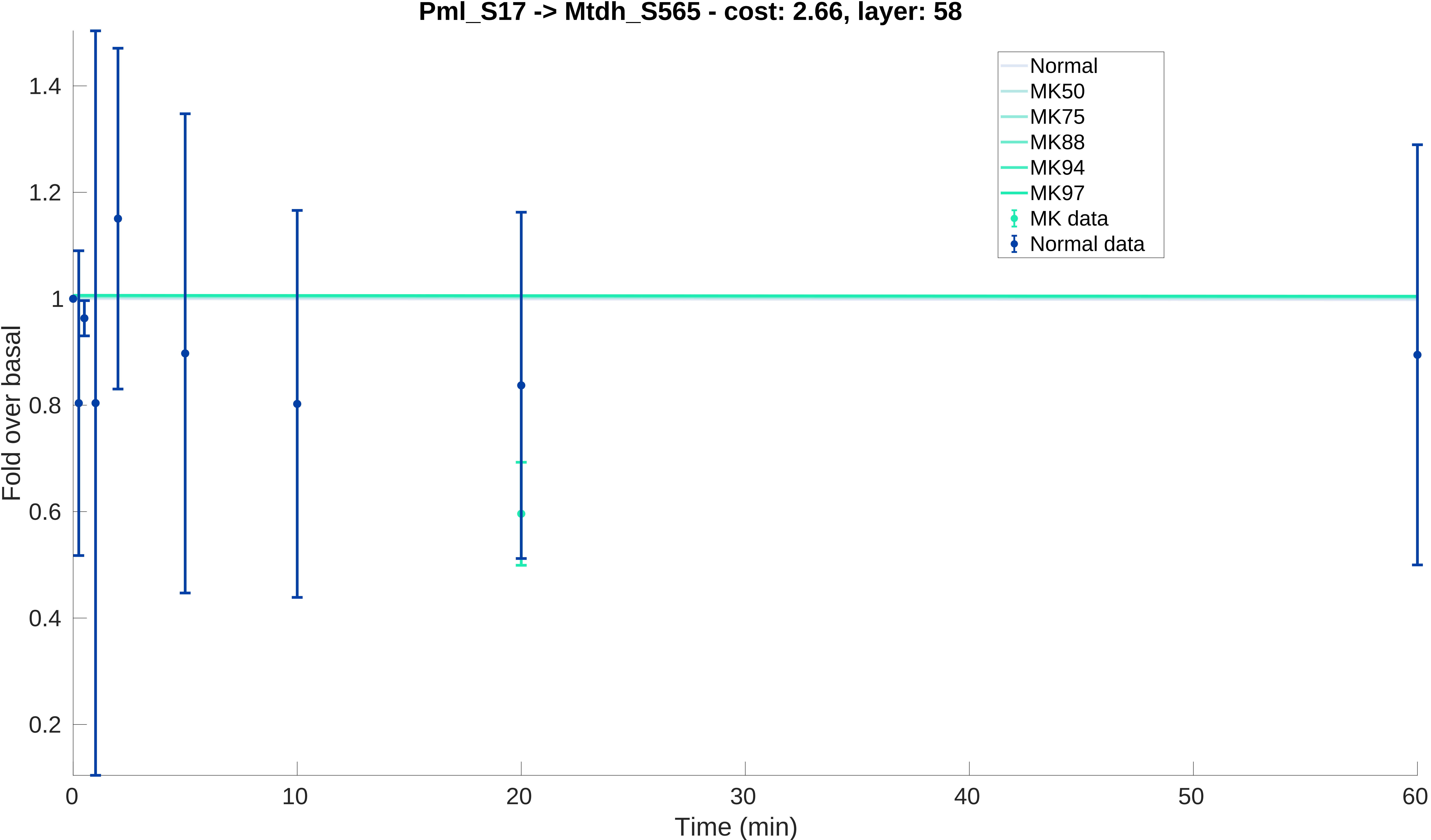

Pml_S17 -> Mtdh_S565 - cost: 2.66, layer: 58
Normal
MK50
MK75
MK88
MK94
MK97
MK data
Normal data
Fold over basal
1.4
1.2
1
0.8
0.6
0.4
0.2
0
10
20
30
40
50
60
Time (min)

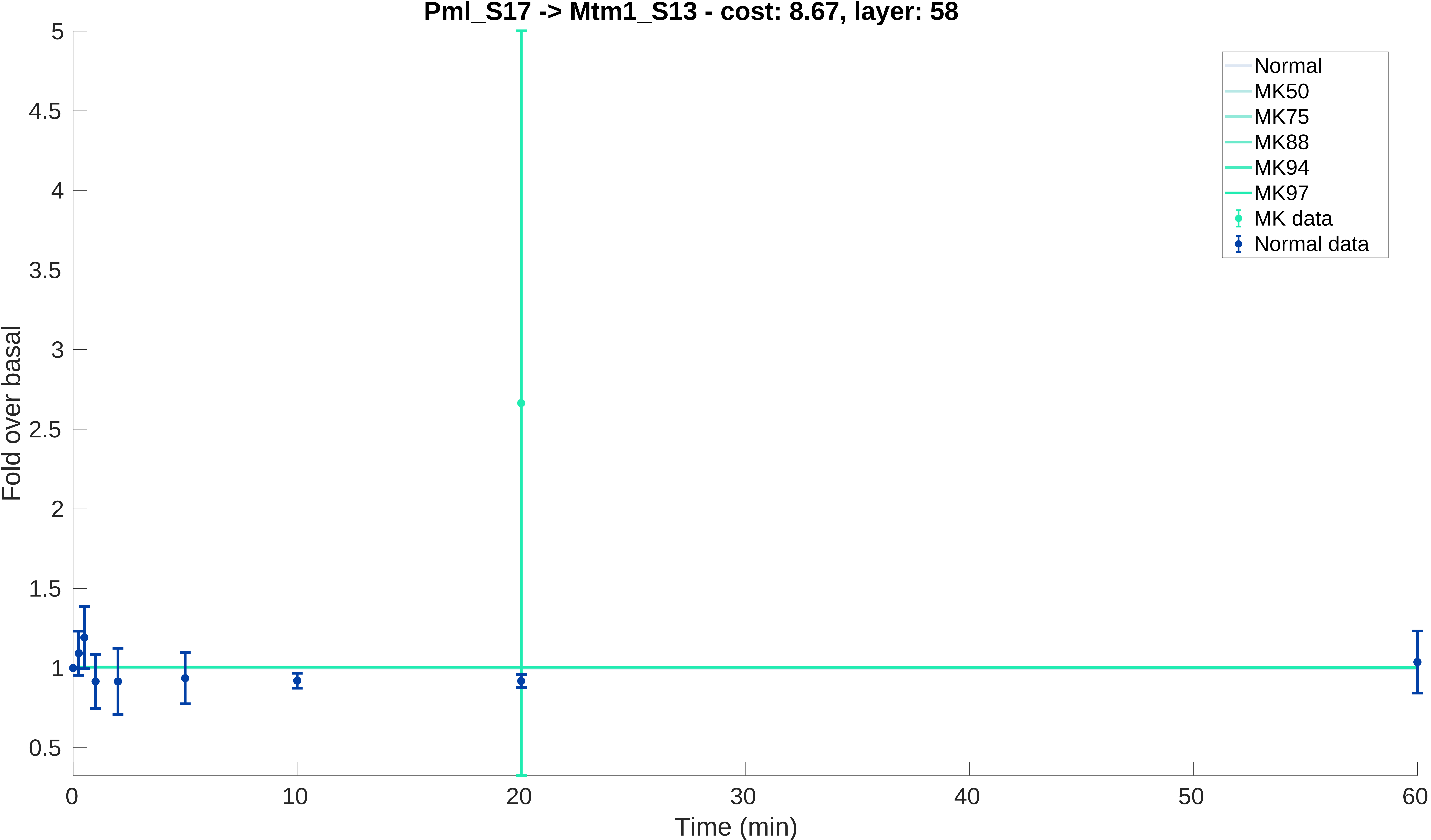

Pml_S17 -> Mtm1_S13 - cost: 8.67, layer: 58
Fold over basal
Time (min)
0.5
1
1.5
2
2.5
3
3.5
4
4.5
5
0
10
20
30
40
50
60
Normal
MK50
MK75
MK88
MK94
MK97
MK data
Normal data

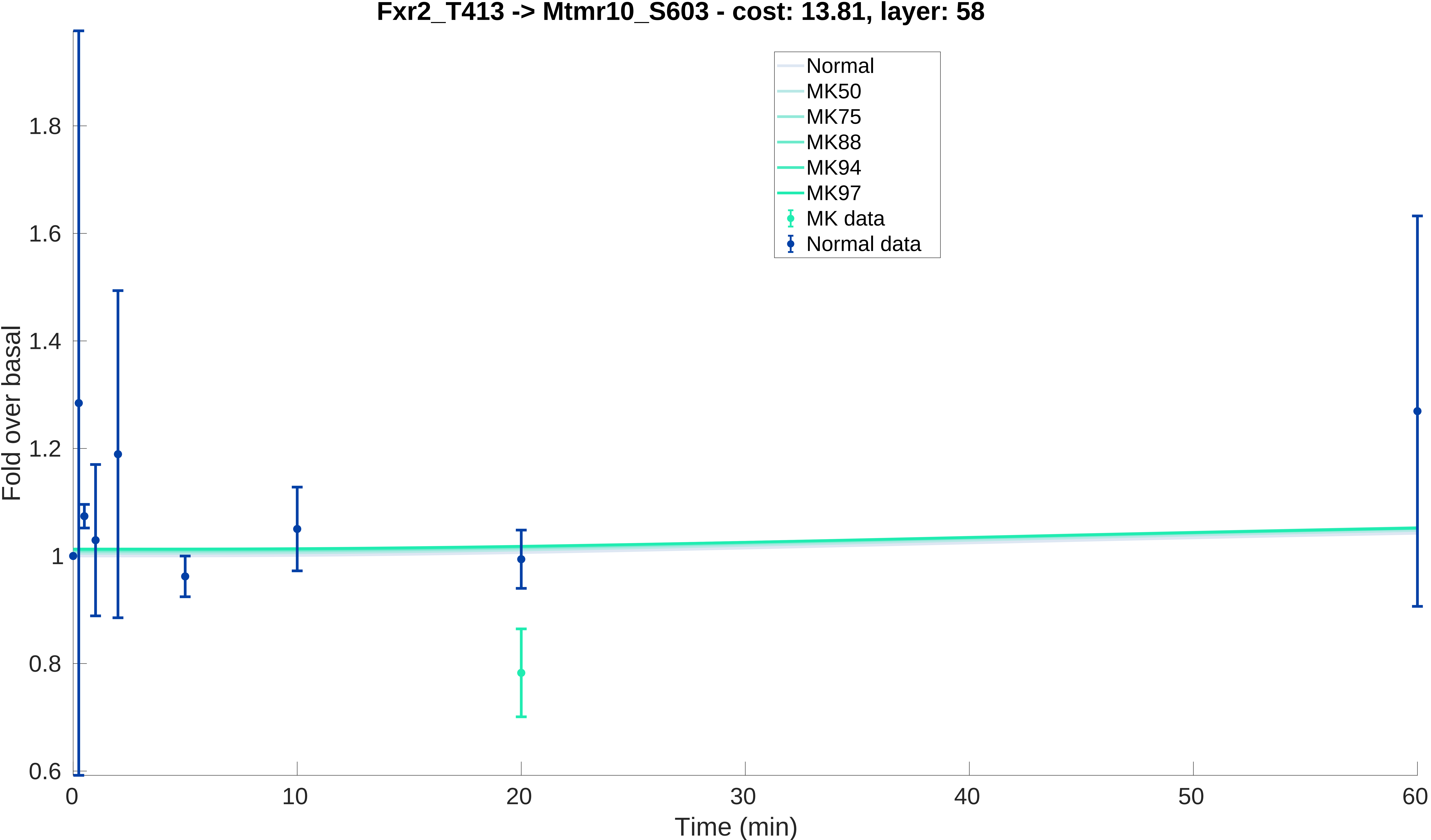
Fxr2_T413 -> Mtmr10_S603 - cost: 13.81, layer: 58
Normal
MK50
MK75
MK88
MK94
MK97
MK data
Normal data
Fold over basal
1.8
1.6
1.4
1.2
1
0.8
0.6
0
10
20
30
40
50
60
Time (min)

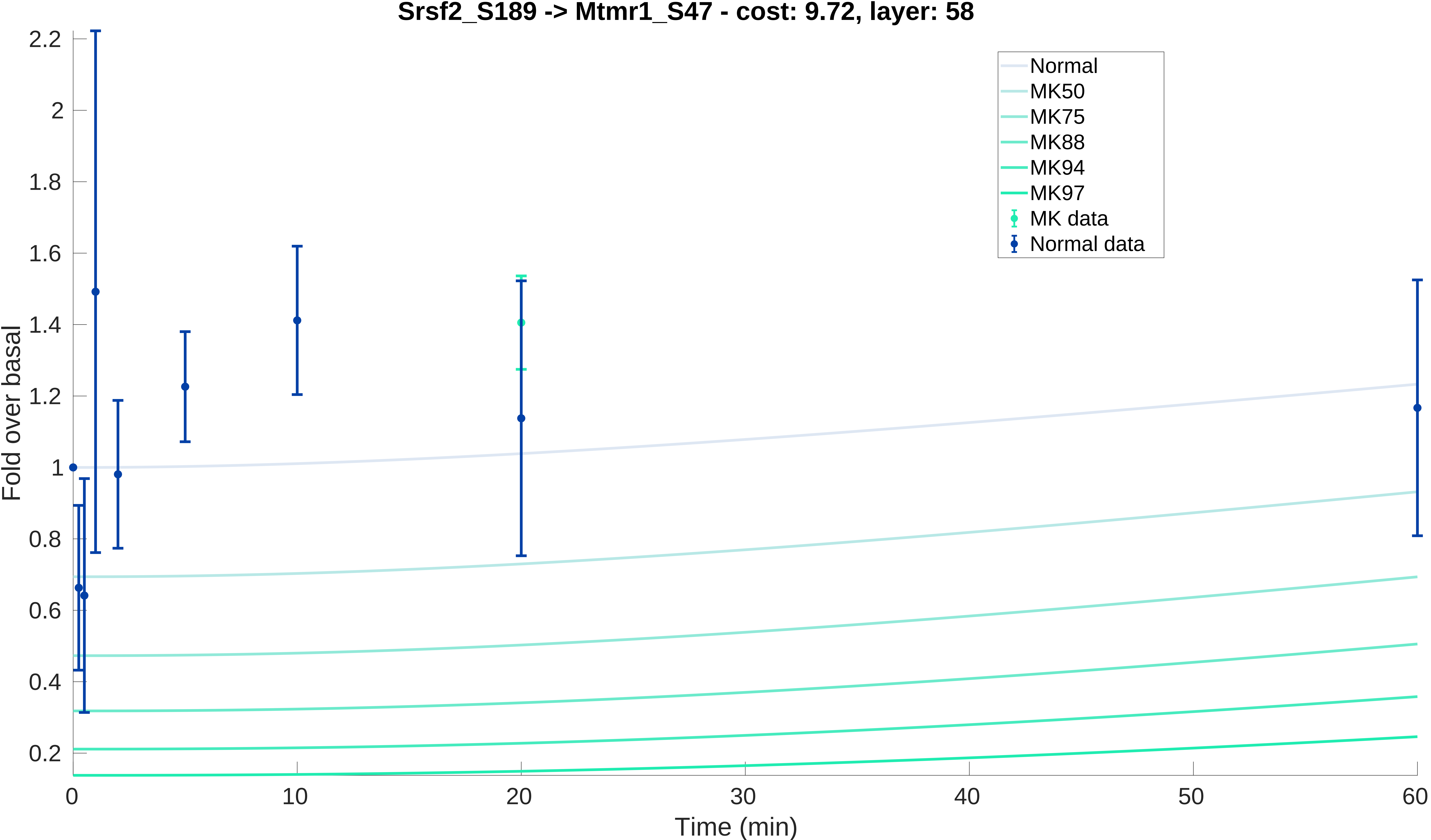

Srsf2_S189 -> Mtmr1_S47 - cost: 9.72, layer: 58
Normal
MK50
MK75
MK88
MK94
MK97
MK data
Normal data
Fold over basal
2.2
2
1.8
1.6
1.4
1.2
1
0.8
0.6
0.4
0.2
0
10
20
30
40
50
60
Time (min)

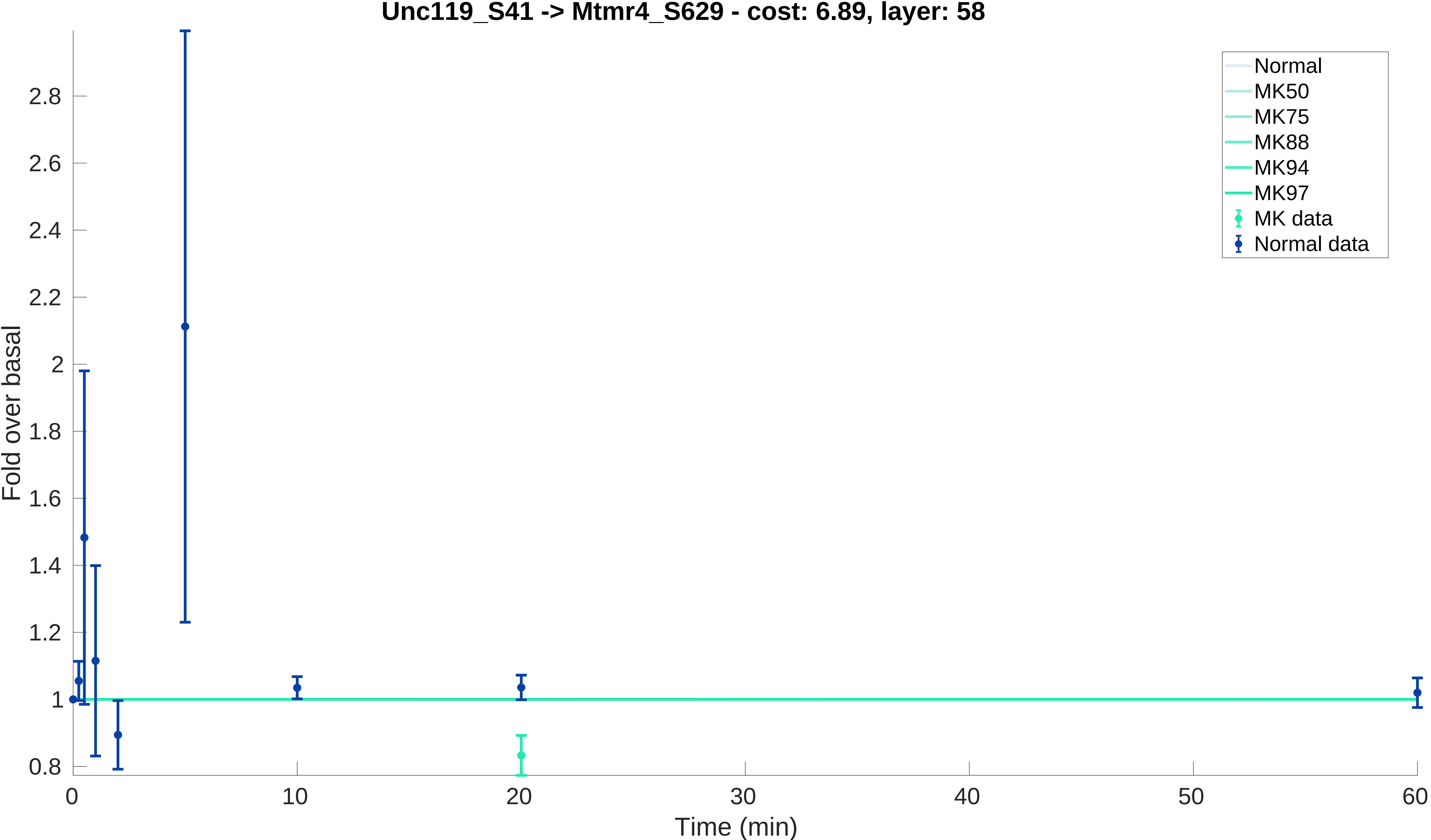
Unc119_S41 -> Mtmr4_S629 - cost: 6.89, layer: 58
Fold over basal
2.8
2.6
2.4
2.2
2
1.8
1.6
1.4
1.2
1
0.8
0
10
20
30
40
50
60
Time (min)
Normal
MK50
MK75
MK88
MK94
MK97
MK data
Normal data

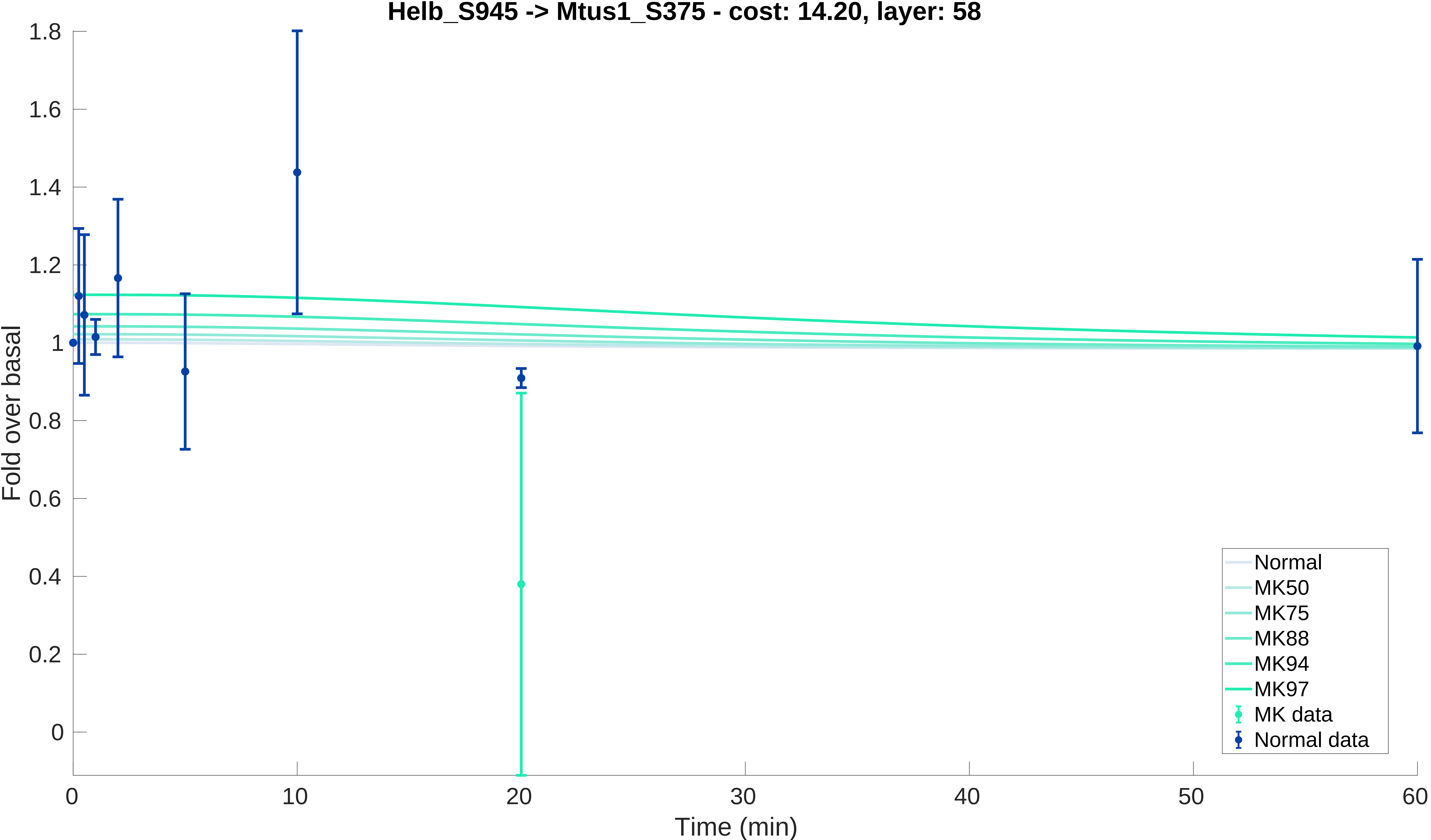

Helb_S945 -> Mtus1_S375 - cost: 14.20, layer: 58
Fold over basal
Time (min)
1.8
1.6
1.4
1.2
1
0.8
0.6
0.4
0.2
0
0
10
20
30
40
50
60
Normal
MK50
MK75
MK88
MK94
MK97
MK data
Normal data

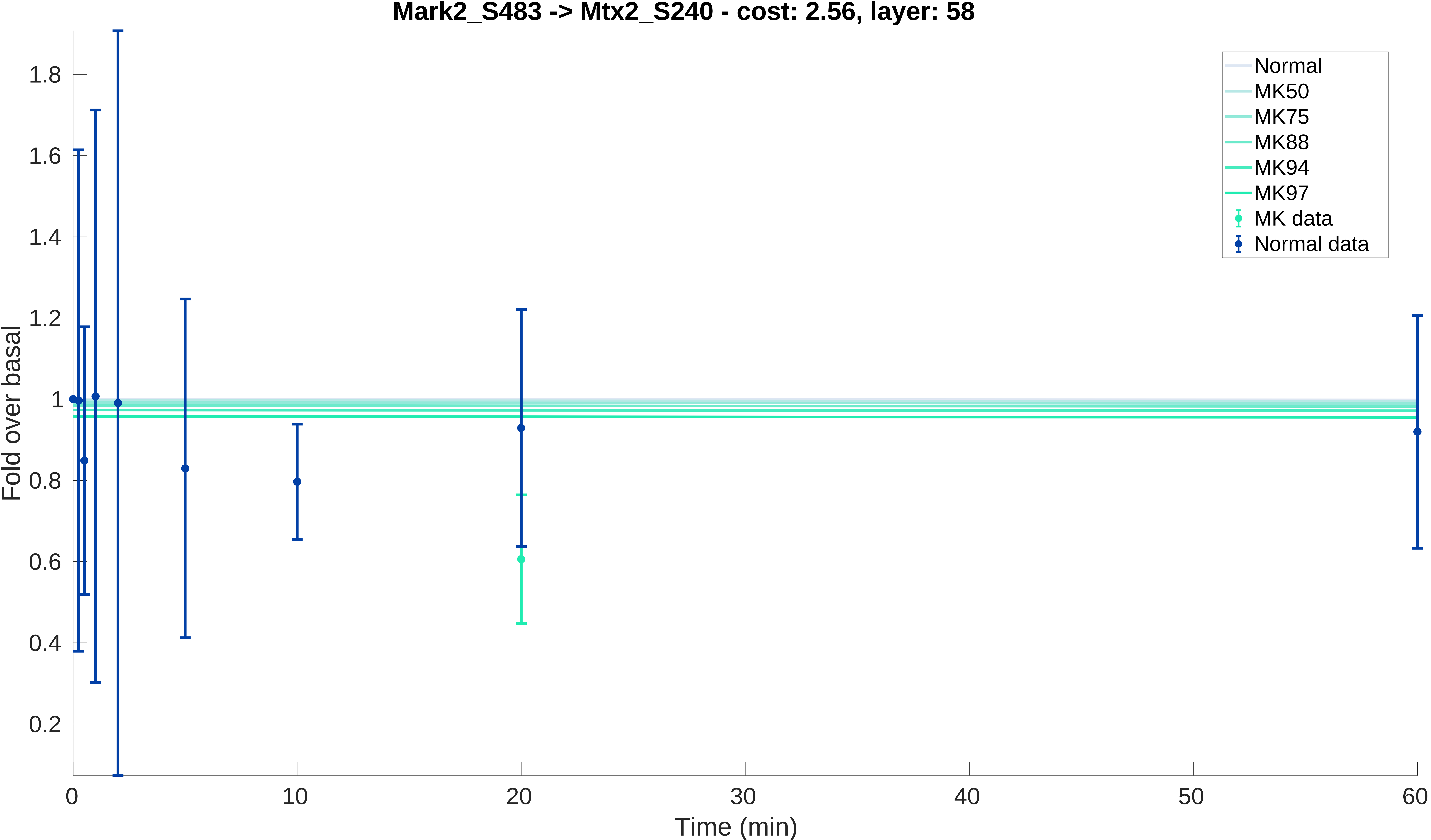
Mark2_S483 -> Mtx2_S240 - cost: 2.56, layer: 58
Normal
MK50
MK75
MK88
MK94
MK97
MK data
Normal data
Fold over basal
1.8
1.6
1.4
1.2
1
0.8
0.6
0.4
0.2
0
10
20
30
40
50
60
Time (min)

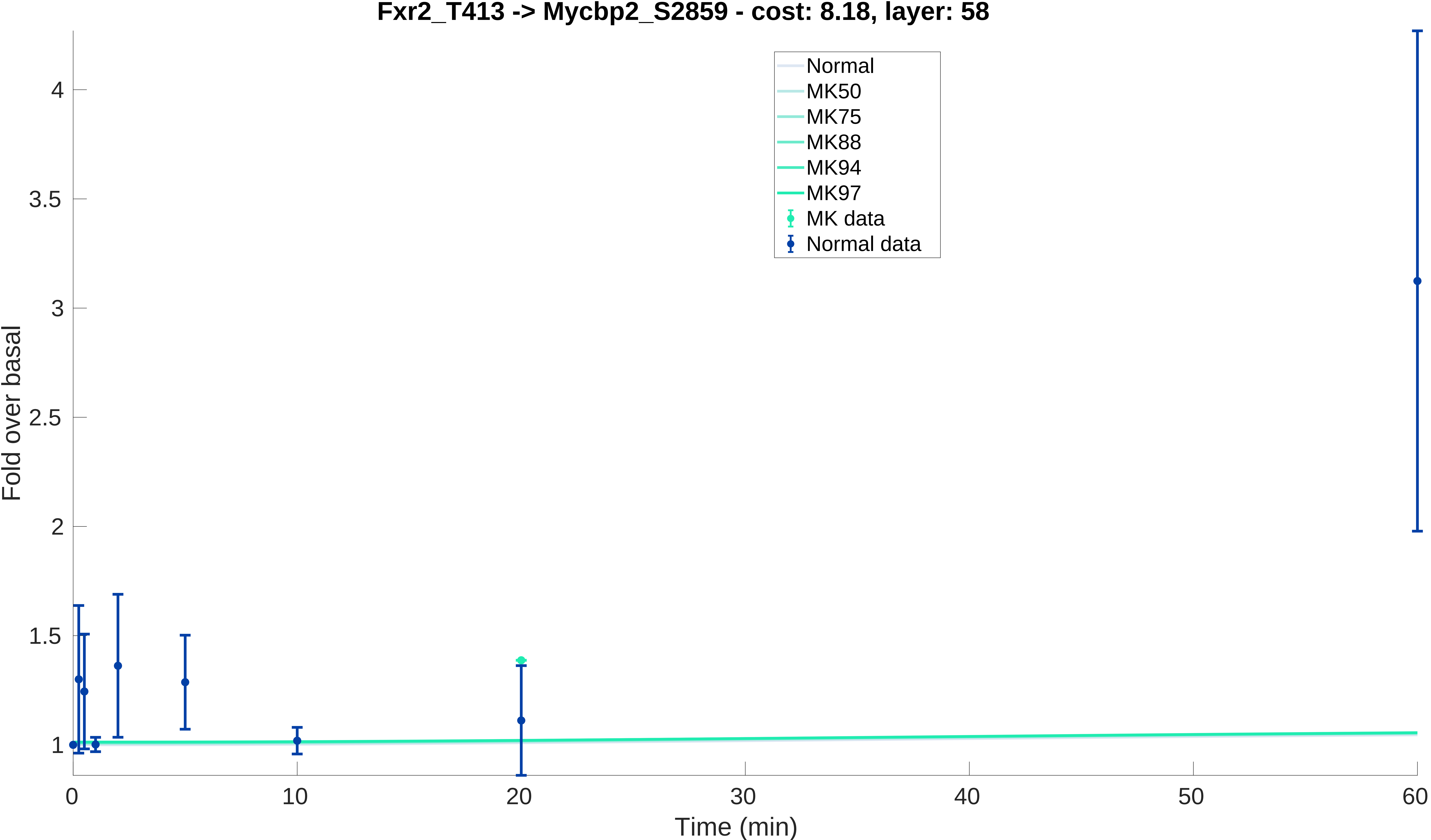

Fxr2_T413 -> Mycbp2_S2859 - cost: 8.18, layer: 58
Fold over basal
Time (min)
0
10
20
30
40
50
60
1
1.5
2
2.5
3
3.5
4
Normal
MK50
MK75
MK88
MK94
MK97
MK data
Normal data

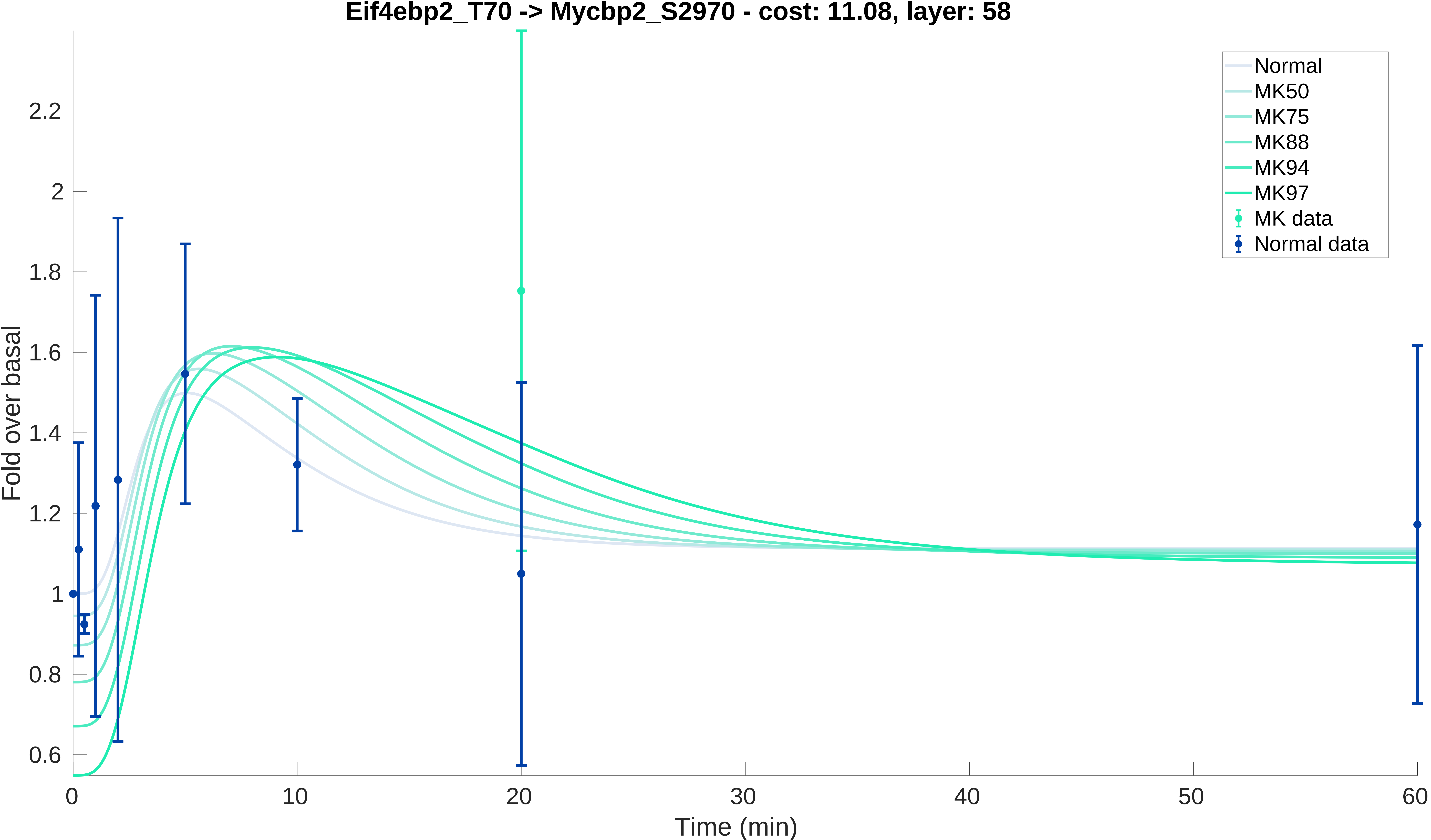

Eif4ebp2_T70 -> Mycbp2_S2970 - cost: 11.08, layer: 58
Fold over basal
Time (min)
Normal
MK50
MK75
MK88
MK94
MK97
MK data
Normal data
0.6
0.8
1
1.2
1.4
1.6
1.8
2
2.2
0
10
20
30
40
50
60

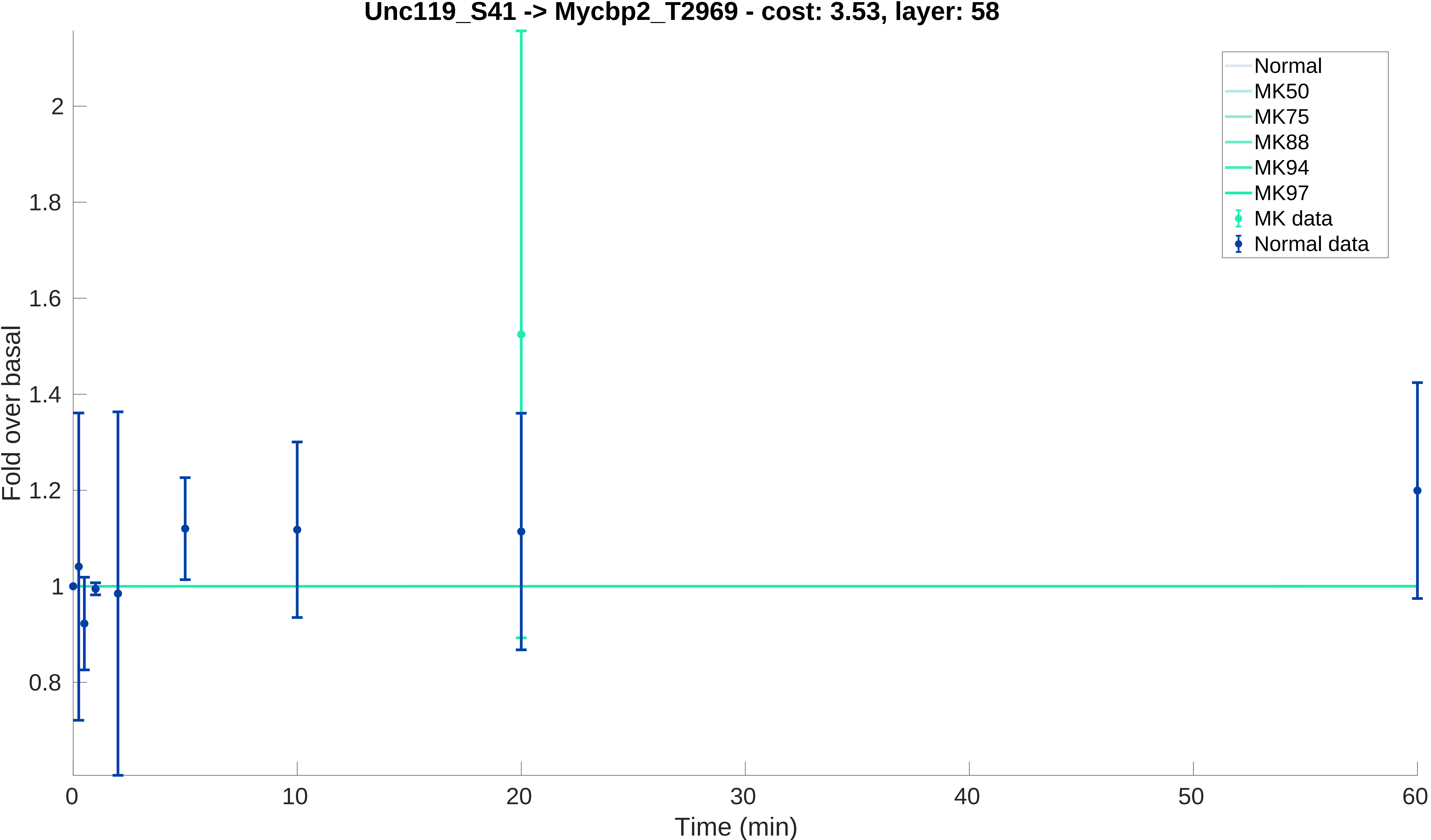

Unc119_S41 -> Mycbp2_T2969 - cost: 3.53, layer: 58
Fold over basal
Time (min)
0.8
1
1.2
1.4
1.6
1.8
2
0
10
20
30
40
50
60
Normal
MK50
MK75
MK88
MK94
MK97
MK data
Normal data

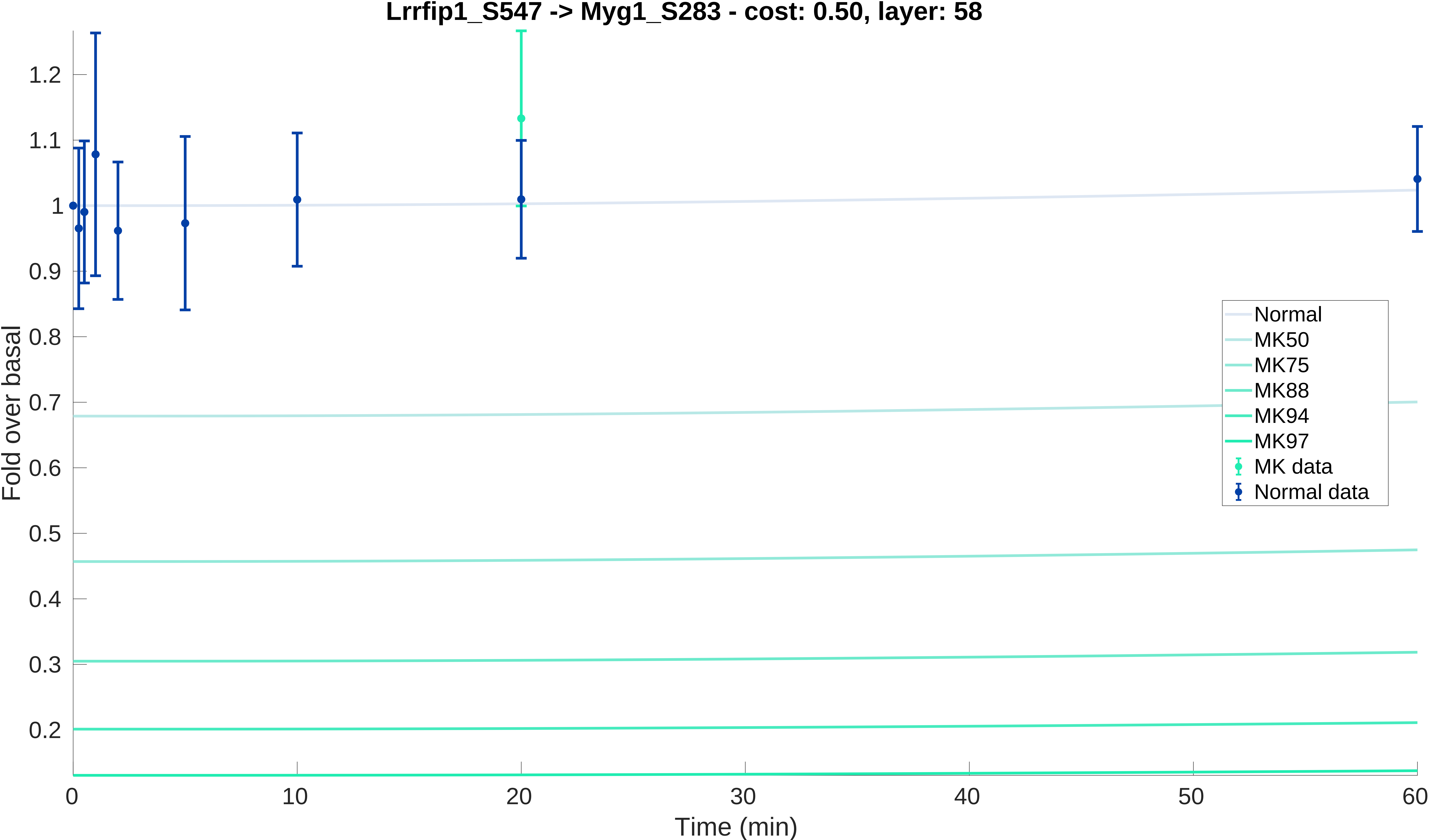
Lrrfip1_S547 -> Myg1_S283 - cost: 0.50, layer: 58
Fold over basal
Time (min)
0
10
20
30
40
50
60
0.2
0.3
0.4
0.5
0.6
0.7
0.8
0.9
1
1.1
1.2
Normal
MK50
MK75
MK88
MK94
MK97
MK data
Normal data

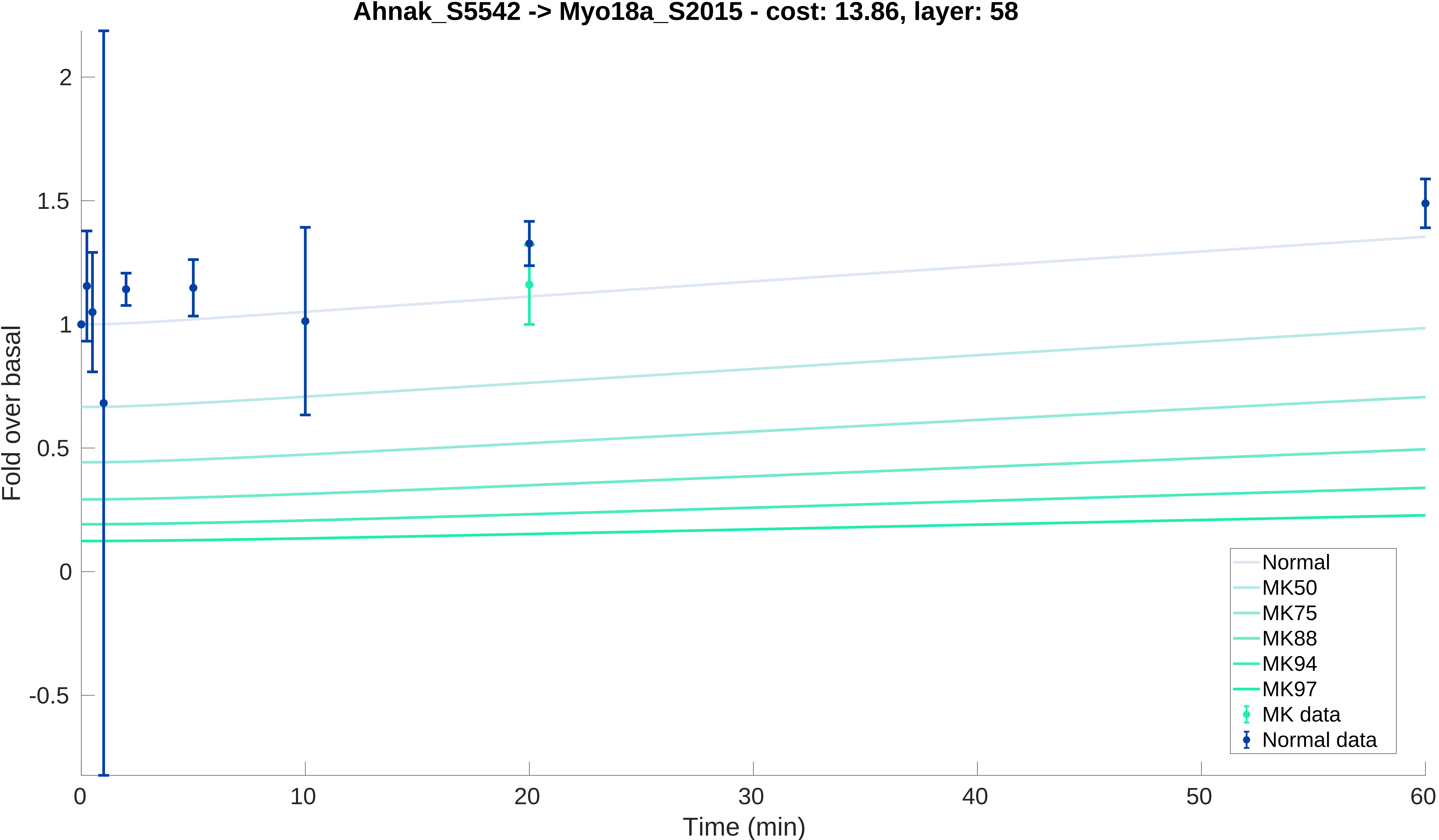

Ahnak_S5542 -> Myo18a_S2015 - cost: 13.86, layer: 58
Fold over basal
Time (min)
2
1.5
1
0.5
0
-0.5
0
10
20
30
40
50
60
Normal
MK50
MK75
MK88
MK94
MK97
MK data
Normal data

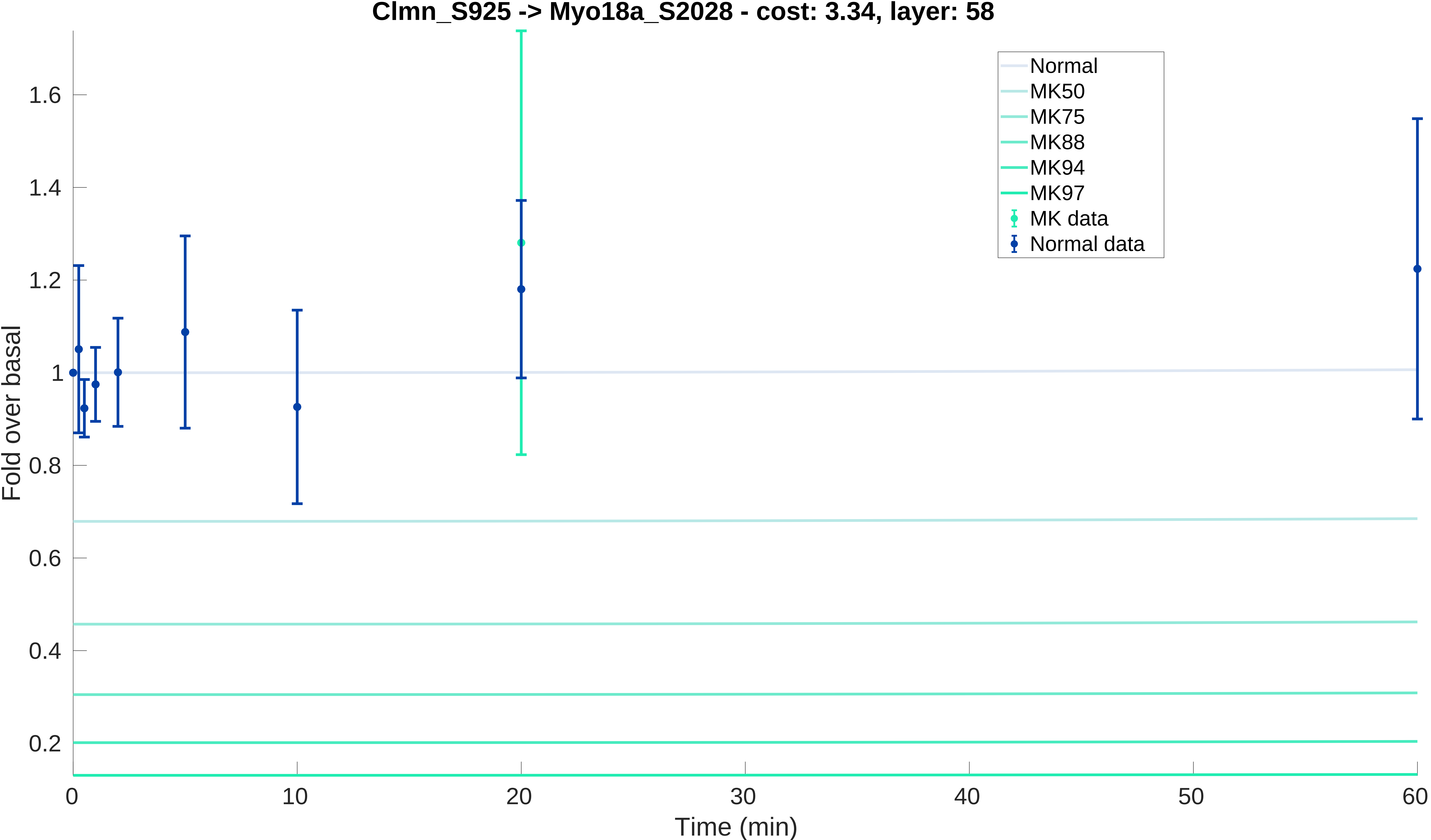

Clmn_S925 -> Myo18a_S2028 - cost: 3.34, layer: 58
Normal
MK50
MK75
MK88
MK94
MK97
MK data
Normal data
Fold over basal
Time (min)
0.2
0.4
0.6
0.8
1
1.2
1.4
1.6
0
10
20
30
40
50
60

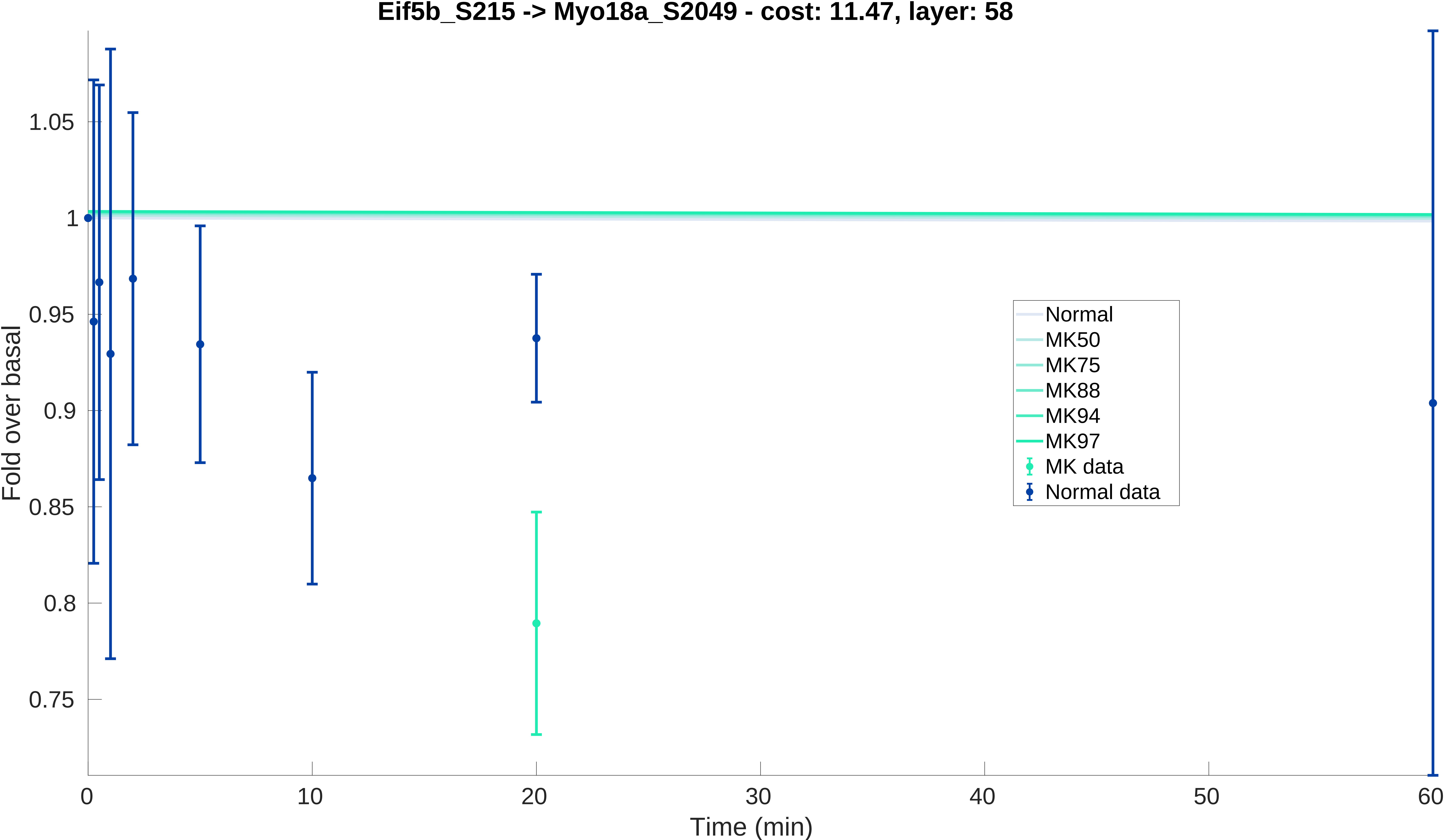

Eif5b_S215 -> Myo18a_S2049 - cost: 11.47, layer: 58
Fold over basal
1.05
1
0.95
0.9
0.85
0.8
0.75
0
10
20
30
40
50
60
Time (min)
Normal
MK50
MK75
MK88
MK94
MK97
MK data
Normal data

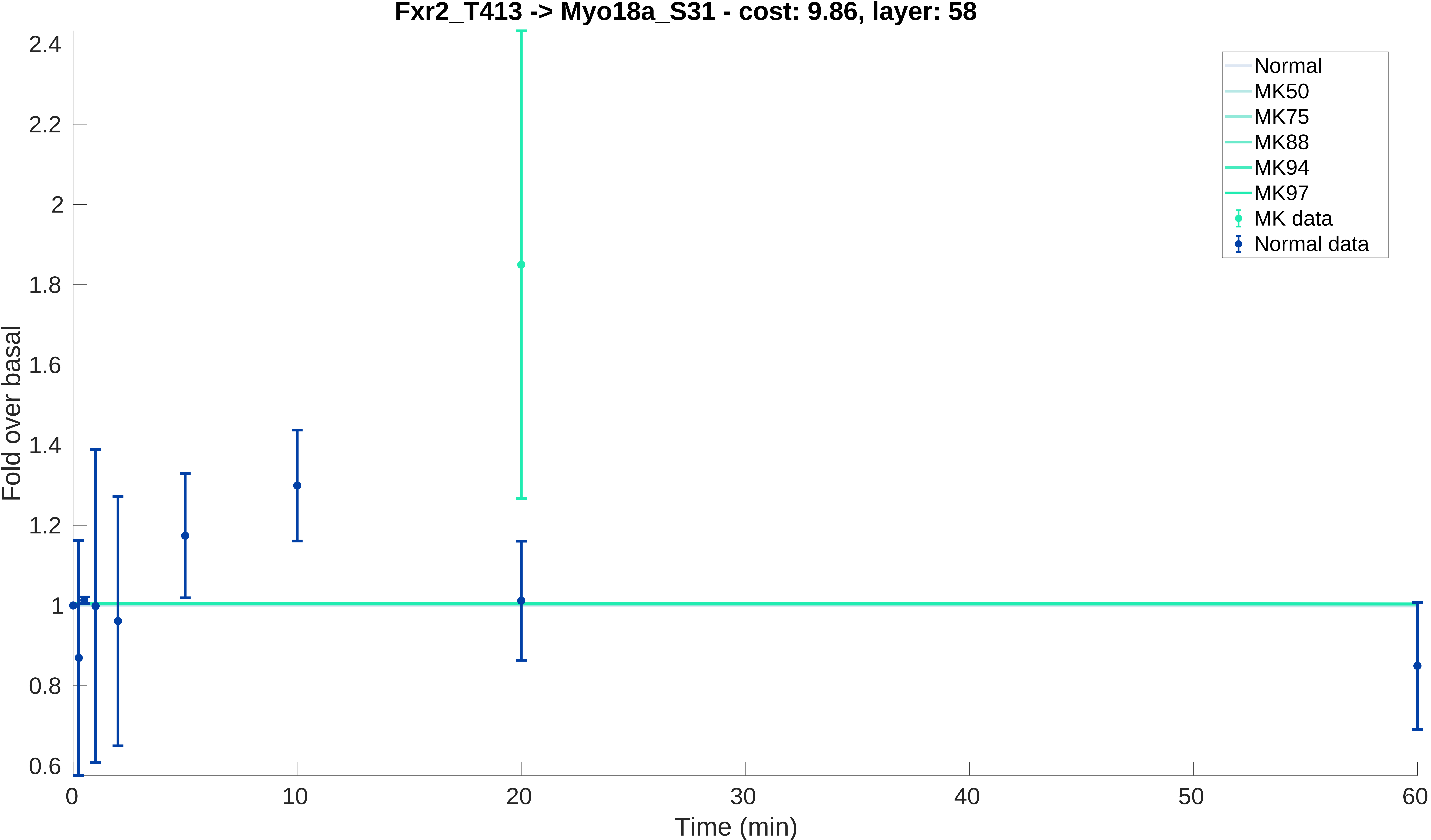

Fxr2_T413 -> Myo18a_S31 - cost: 9.86, layer: 58
Fold over basal
Time (min)
2.4
2.2
2
1.8
1.6
1.4
1.2
1
0.8
0.6
0
10
20
30
40
50
60
Normal
MK50
MK75
MK88
MK94
MK97
MK data
Normal data

#### Helb_S945 -> Myo18a_S59 - cost: 2.97, layer: 58

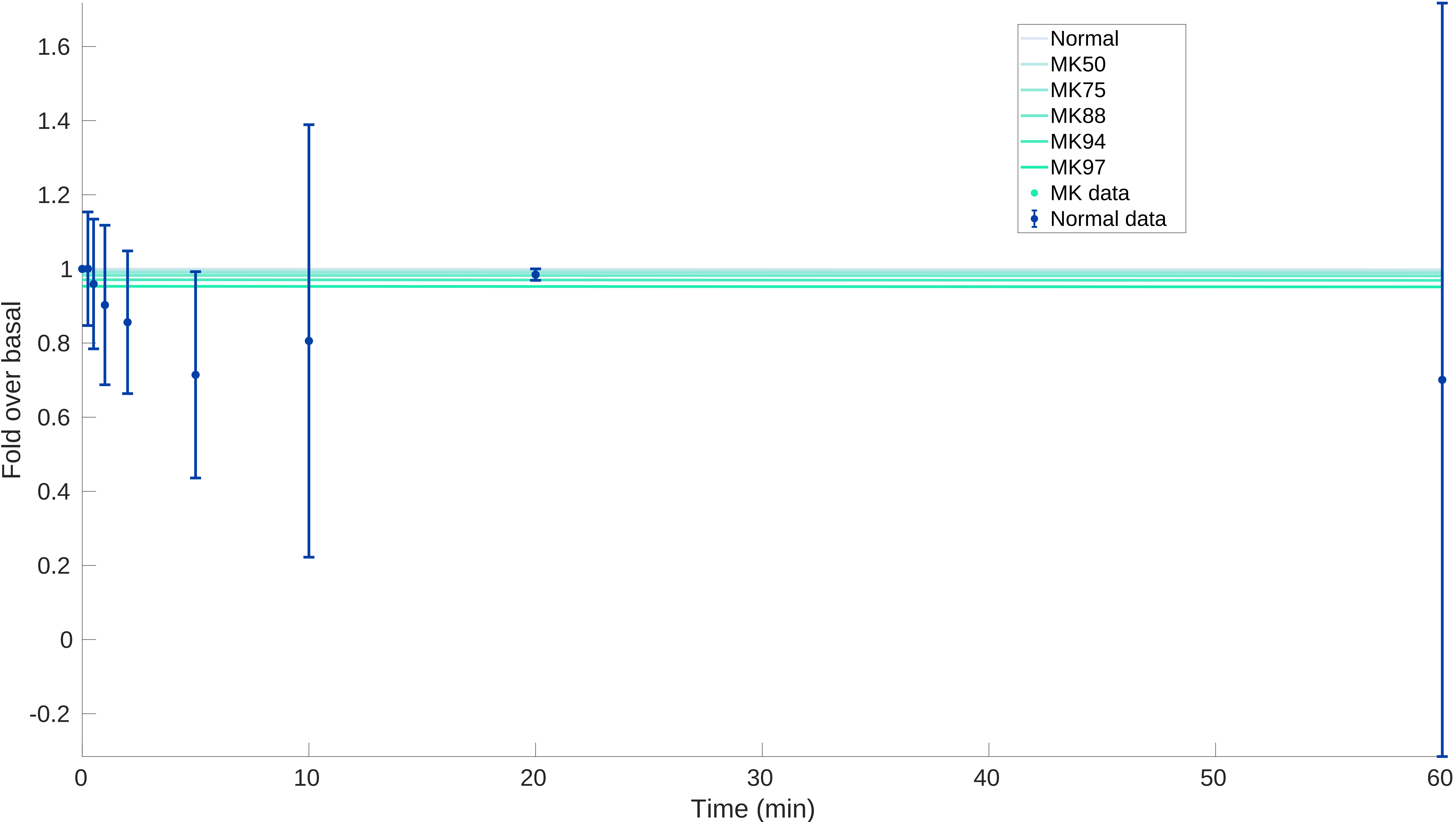

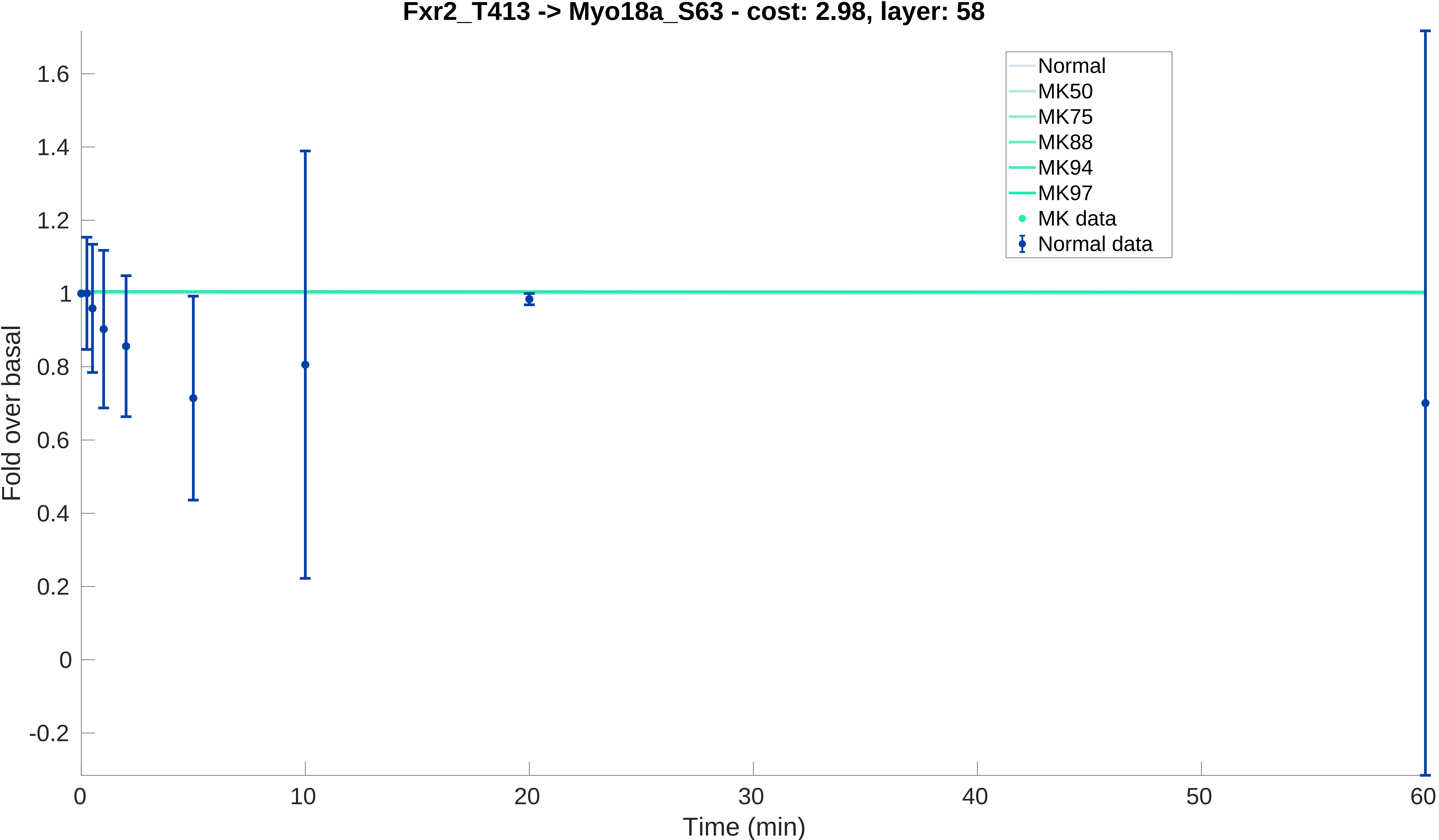
Fxr2_T413 -> Myo18a_S63 - cost: 2.98, layer: 58
Fold over basal
Time (min)
1.6
1.4
1.2
1
0.8
0.6
0.4
0.2
0
-0.2
0
10
20
30
40
50
60
Normal
MK50
MK75
MK88
MK94
MK97
MK data
Normal data

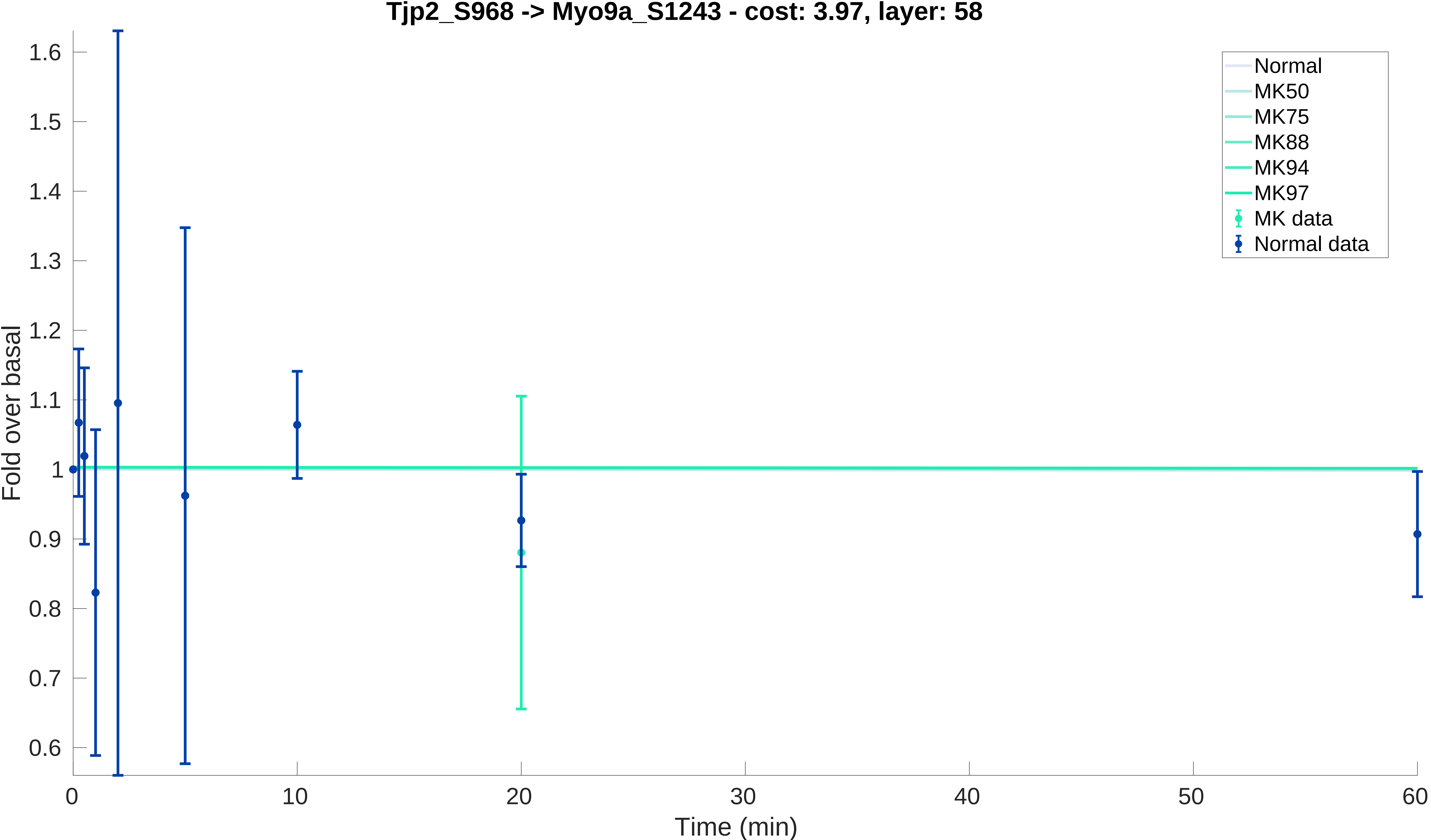

Tjp2_S968 -> Myo9a_S1243 - cost: 3.97, layer: 58
Fold over basal
Time (min)
1.6
1.5
1.4
1.3
1.2
1.1
1
0.9
0.8
0.7
0.6
0
10
20
30
40
50
60
Normal
MK50
MK75
MK88
MK94
MK97
MK data
Normal data

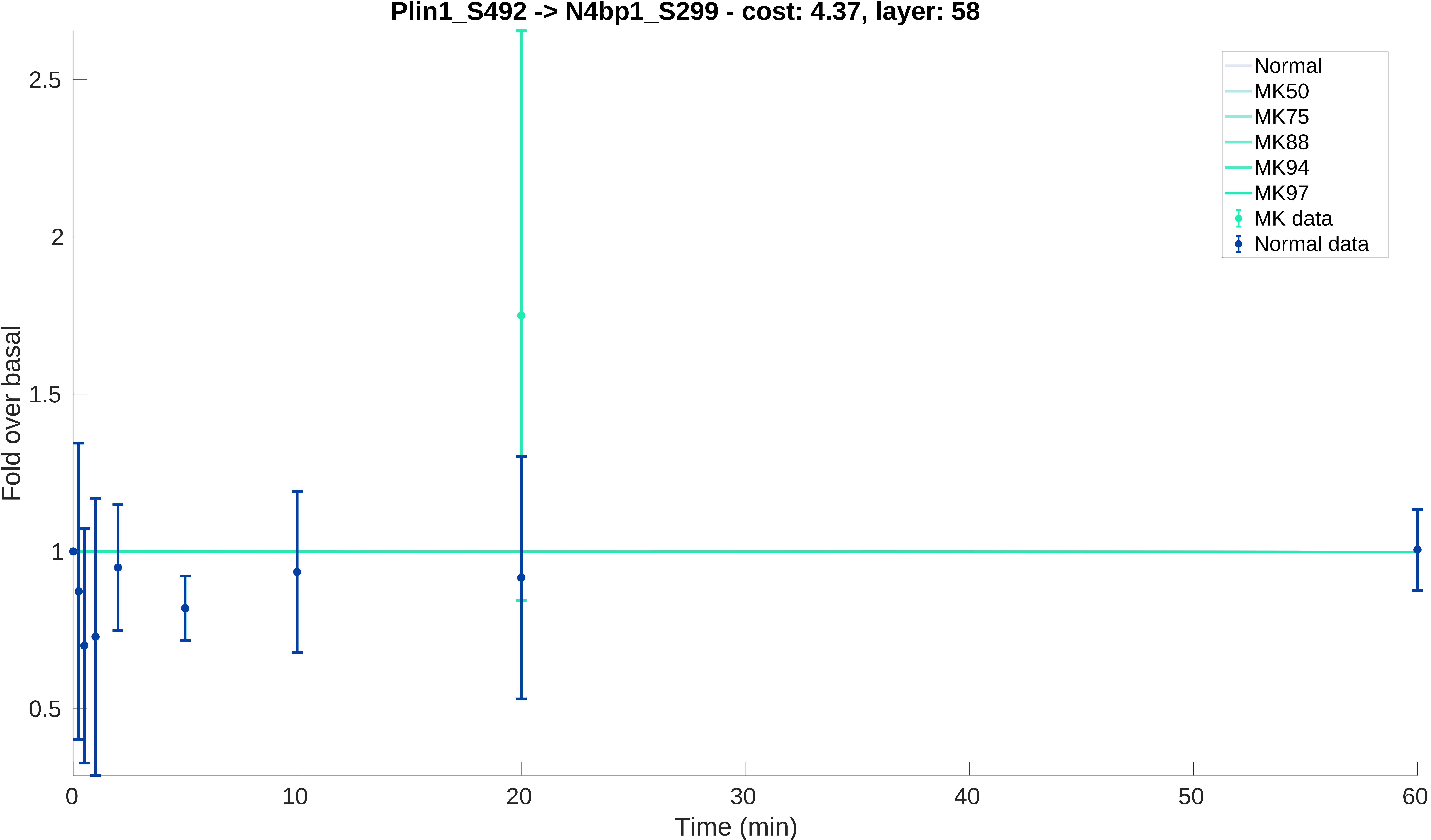
Plin1_S492 -> N4bp1_S299 - cost: 4.37, layer: 58
Normal
MK50
MK75
MK88
MK94
MK97
MK data
Normal data
Fold over basal
2.5
2
1.5
1
0.5
0
10
20
30
40
50
60
Time (min)

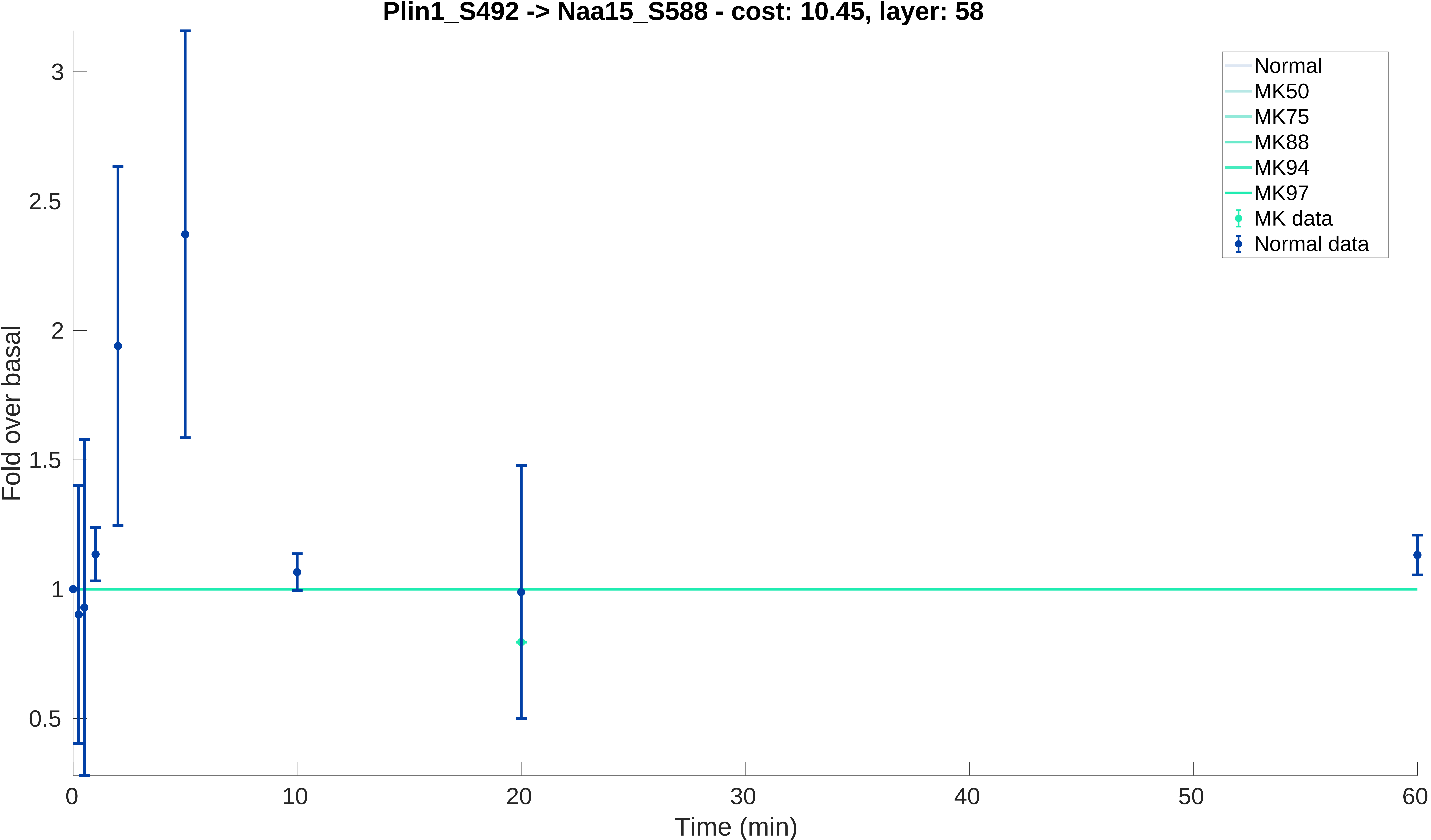
Plin1_S492 -> Naa15_S588 - cost: 10.45, layer: 58
Fold over basal
Time (min)
0.5
1
1.5
2
2.5
3
0
10
20
30
40
50
60
Normal
MK50
MK75
MK88
MK94
MK97
MK data
Normal data

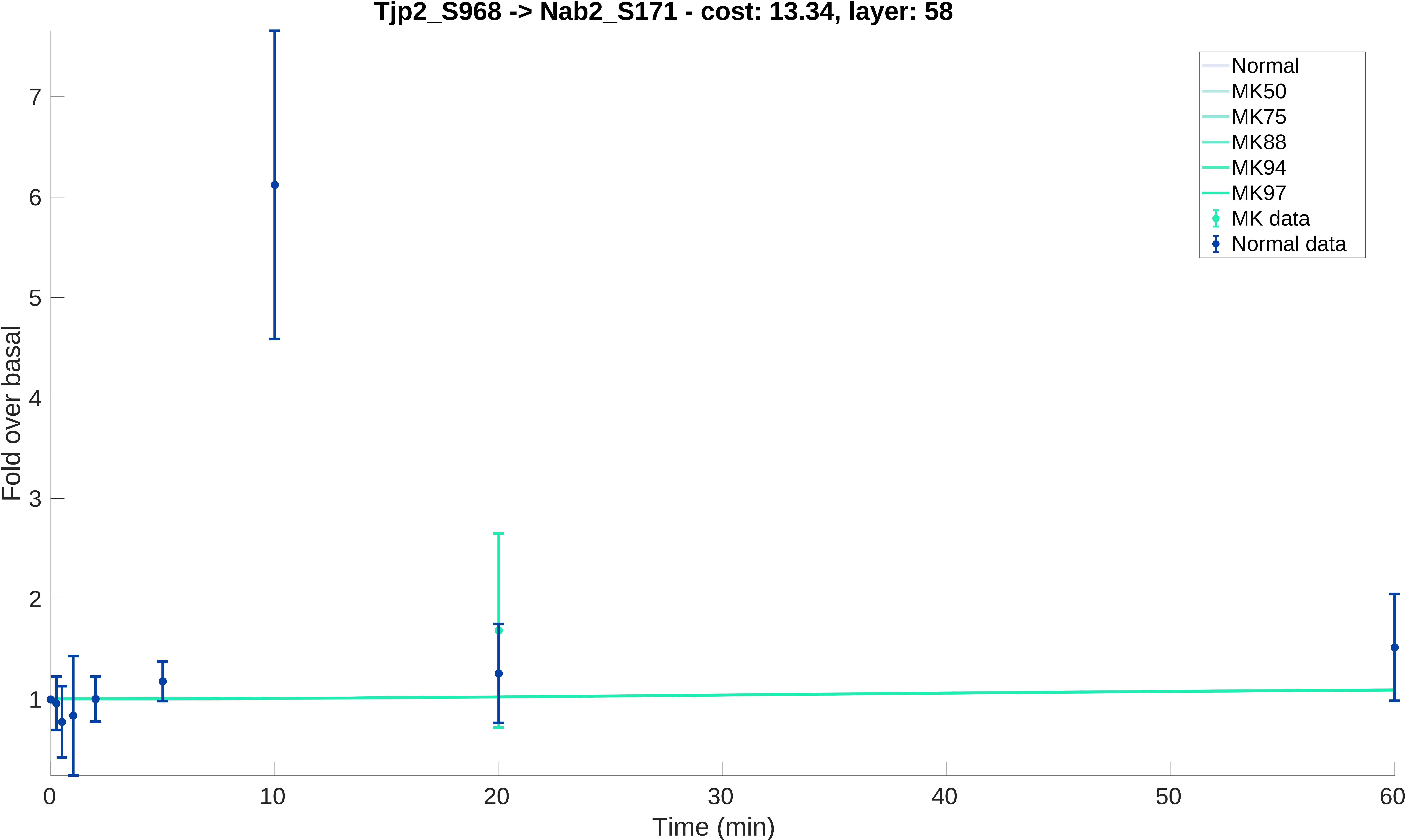

Tjp2_S968 -> Nab2_S171 - cost: 13.34, layer: 58
Fold over basal
Time (min)
0
10
20
30
40
50
60
1
2
3
4
5
6
7
Normal
MK50
MK75
MK88
MK94
MK97
MK data
Normal data

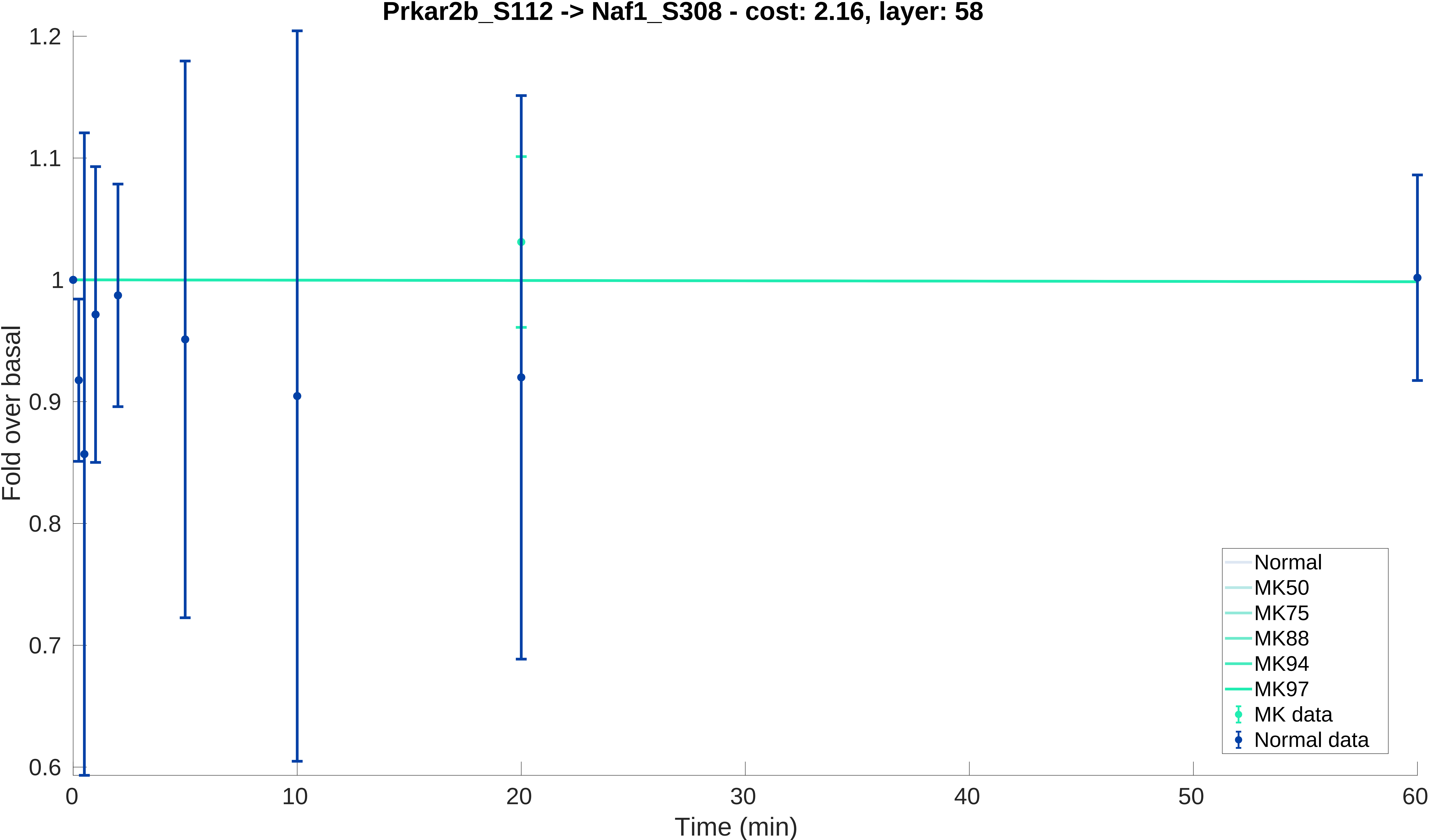

Prkar2b_S112 -> Naf1_S308 - cost: 2.16, layer: 58
Fold over basal
1.2
1.1
1
0.9
0.8
0.7
0.6
0
10
20
30
40
50
60
Time (min)
Normal
MK50
MK75
MK88
MK94
MK97
MK data
Normal data

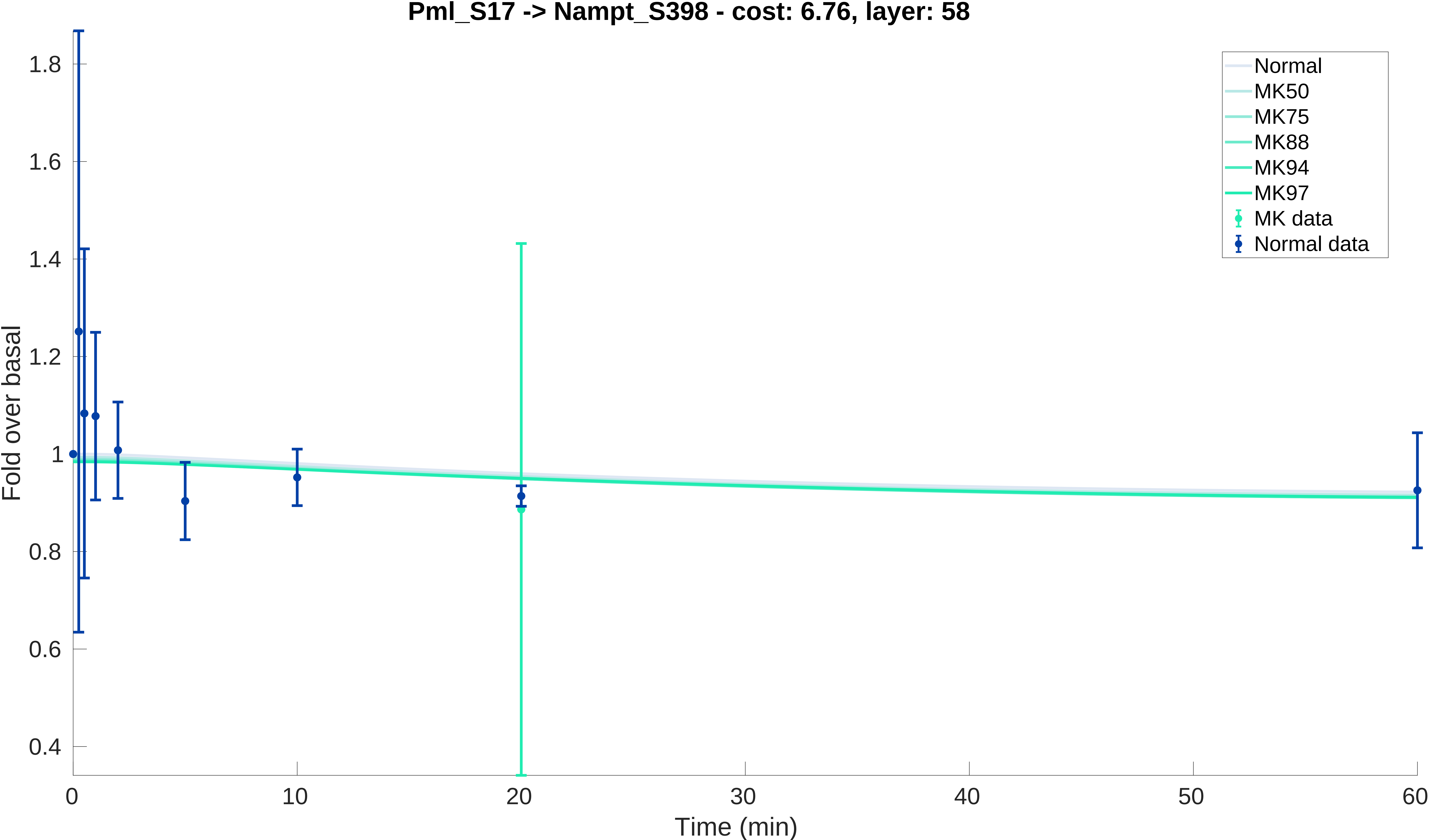

Pml_S17 -> Nampt_S398 - cost: 6.76, layer: 58
Normal
MK50
MK75
MK88
MK94
MK97
MK data
Normal data
Fold over basal
1.8
1.6
1.4
1.2
1
0.8
0.6
0.4
0
10
20
30
40
50
60
Time (min)

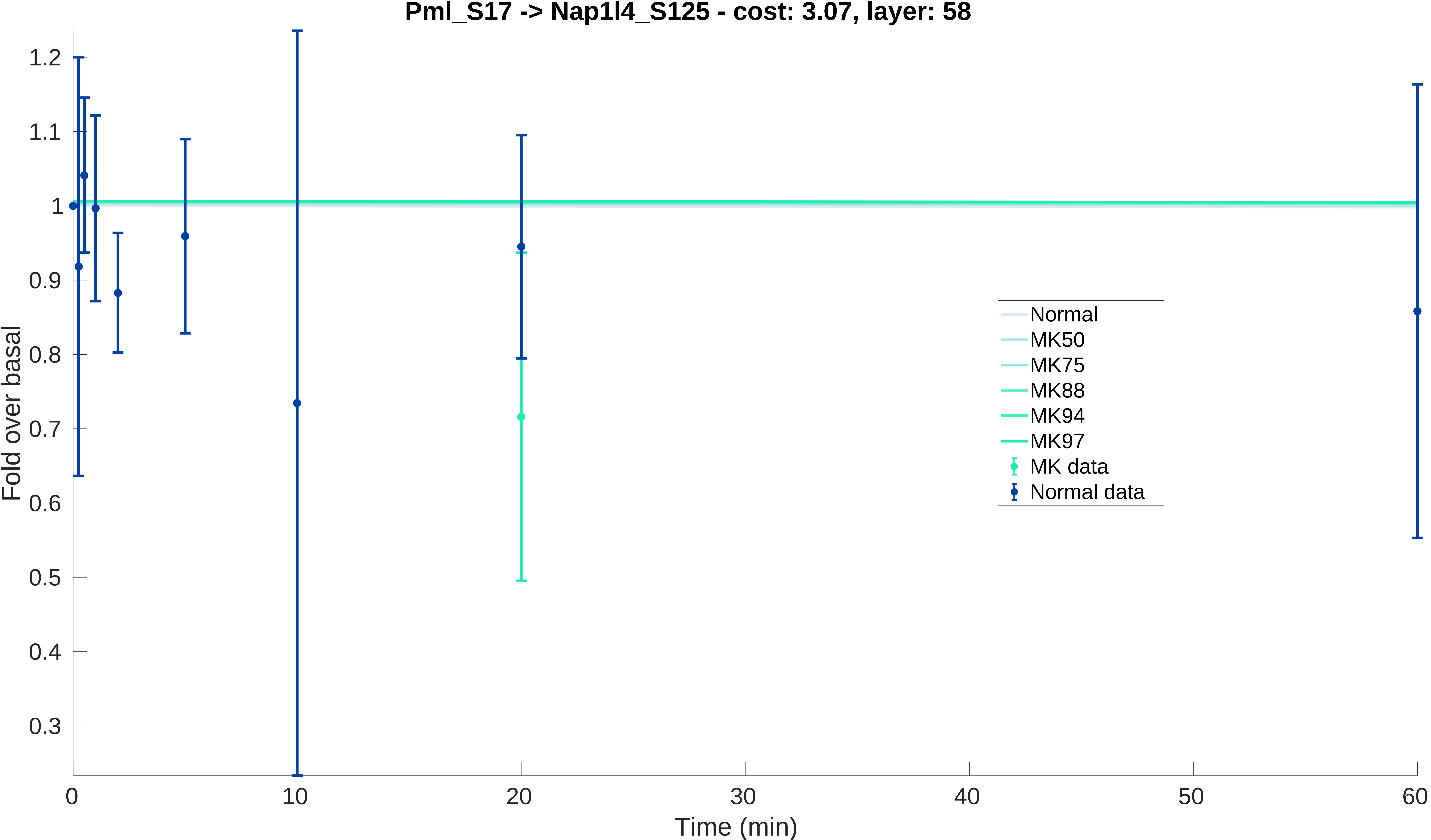

Pml_S17 -> Nap1l4_S125 - cost: 3.07, layer: 58
Fold over basal
Time (min)
1.2
1.1
1
0.9
0.8
0.7
0.6
0.5
0.4
0.3
0
10
20
30
40
50
60
Normal
MK50
MK75
MK88
MK94
MK97
MK data
Normal data

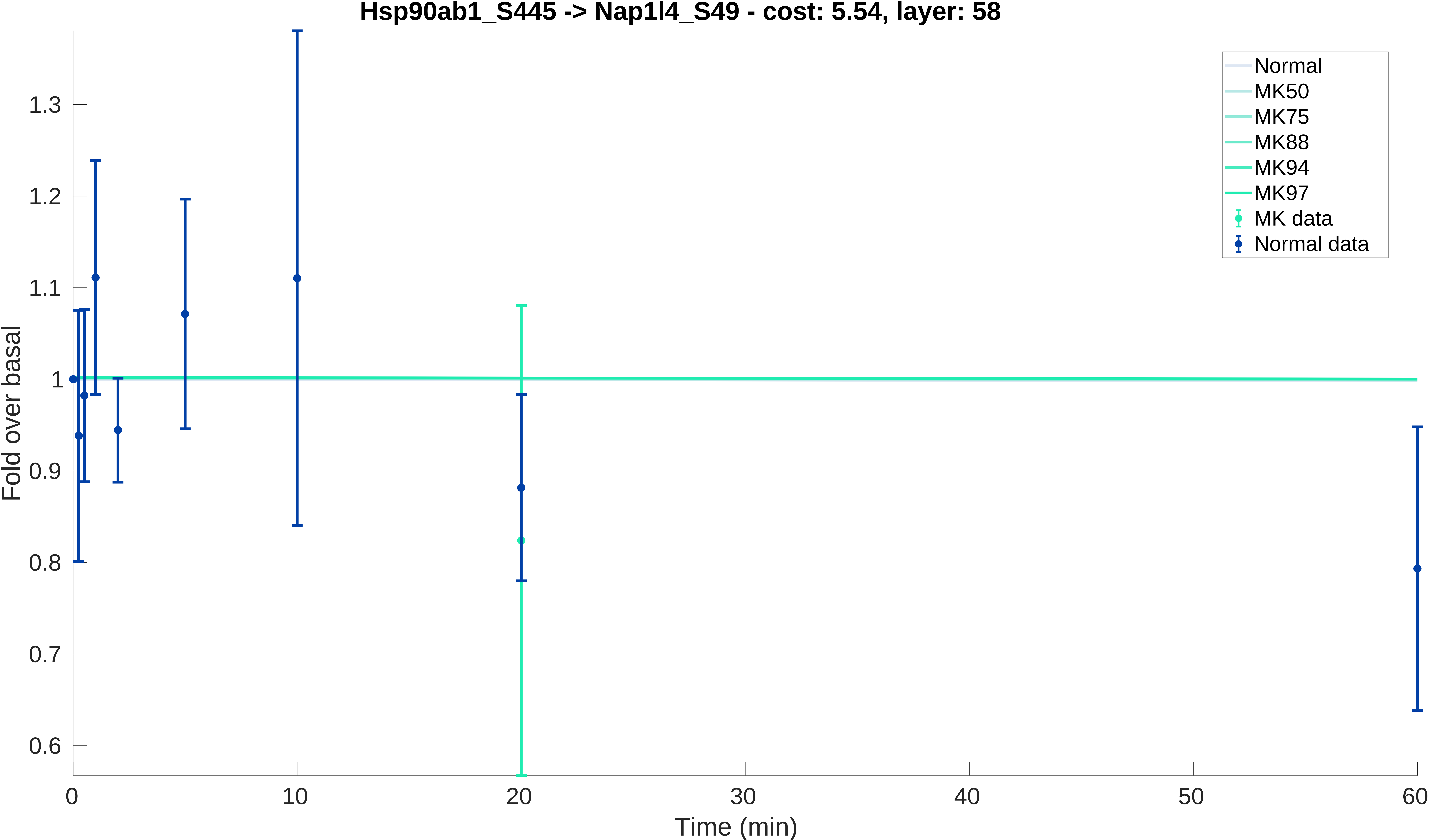

Hsp90ab1_S445 -> Nap1l4_S49 - cost: 5.54, layer: 58
Normal
MK50
MK75
MK88
MK94
MK97
MK data
Normal data
Fold over basal
1.3
1.2
1.1
1
0.9
0.8
0.7
0.6
0
10
20
30
40
50
60
Time (min)

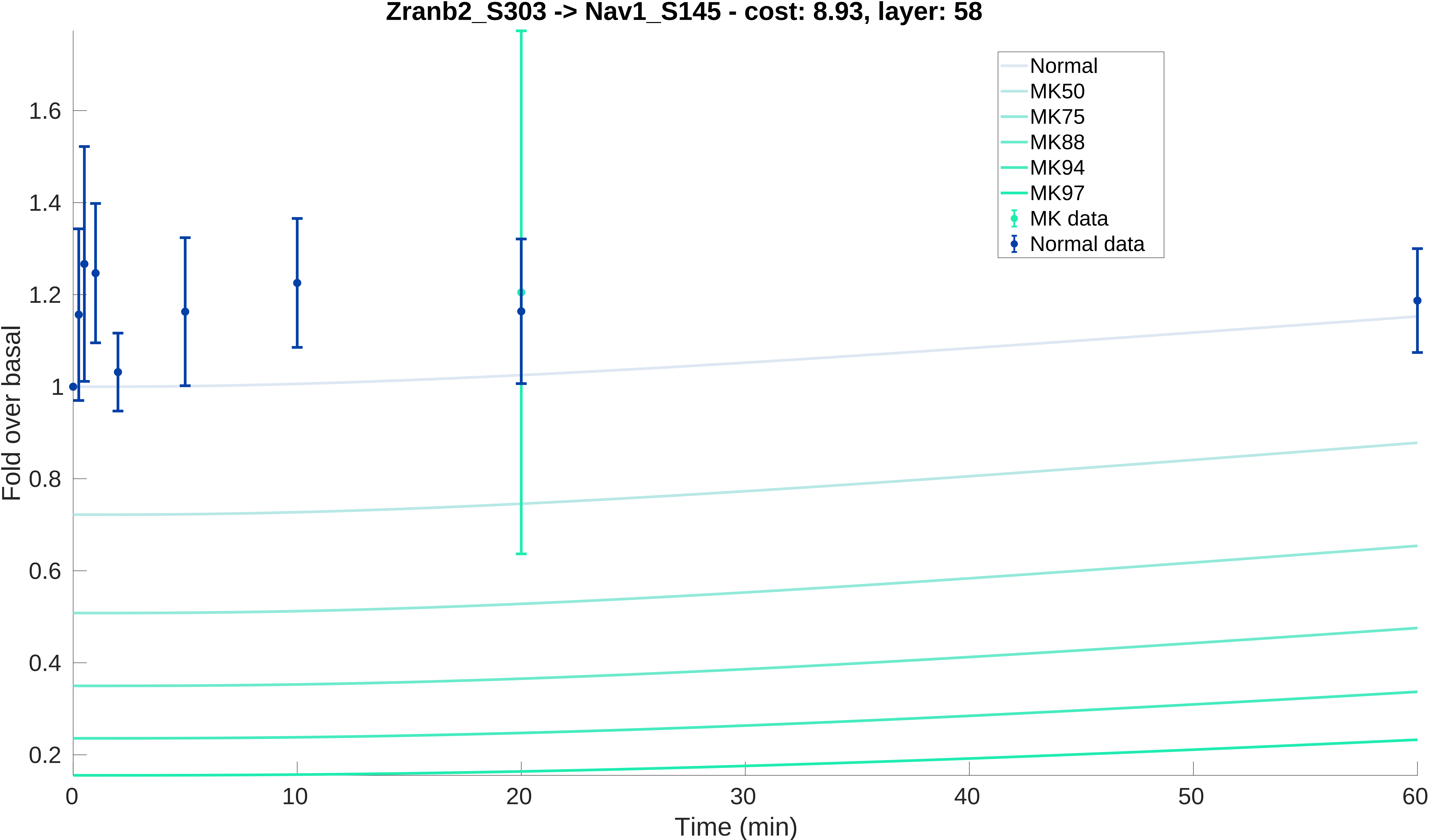

Zranb2_S303 -> Nav1_S145 - cost: 8.93, layer: 58
Fold over basal
Time (min)
1.6
1.4
1.2
1
0.8
0.6
0.4
0.2
0
10
20
30
40
50
60
Normal
MK50
MK75
MK88
MK94
MK97
MK data
Normal data

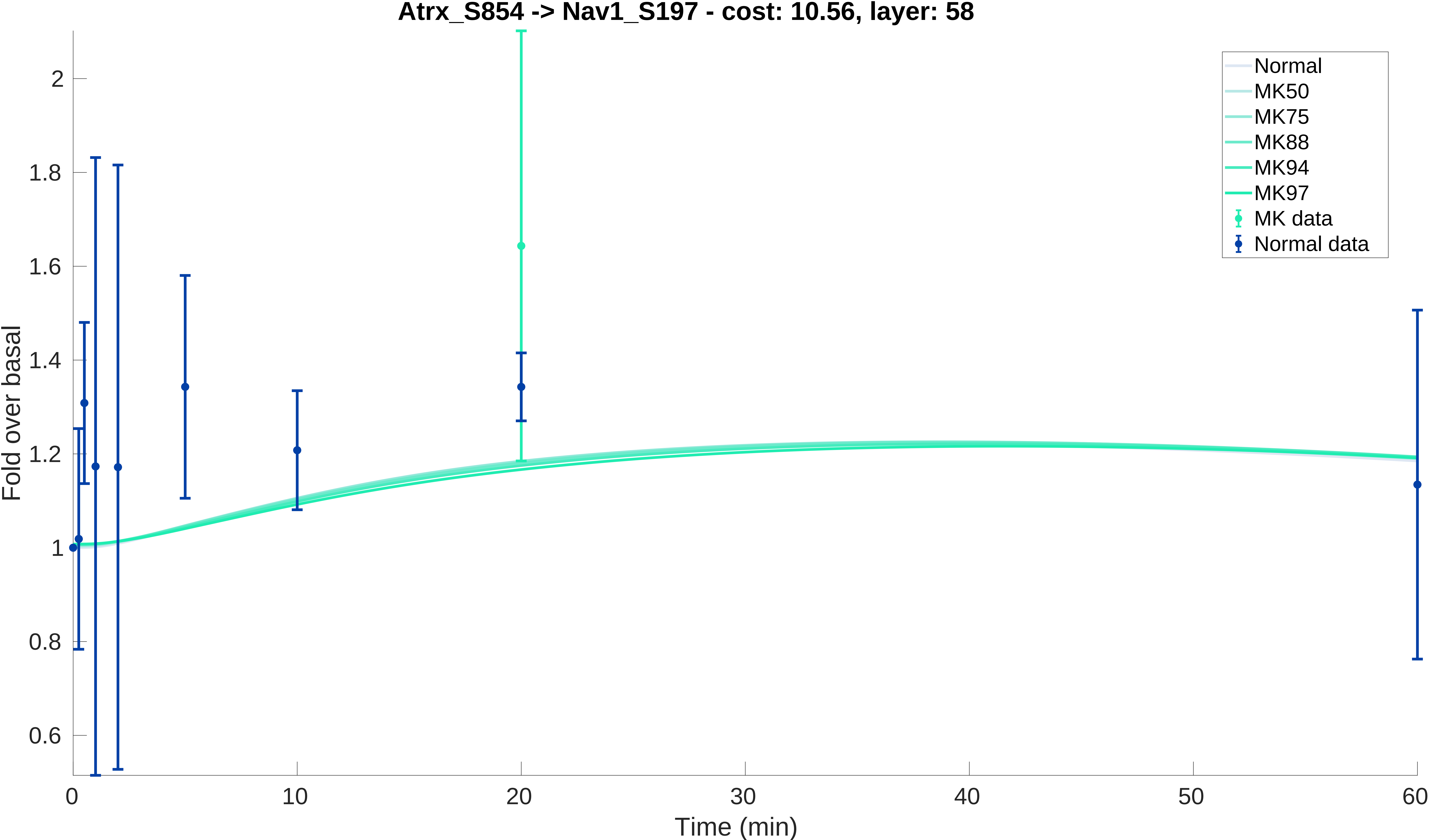

Atrx_S854 -> Nav1_S197 - cost: 10.56, layer: 58
Fold over basal
Time (min)
0.6
0.8
1
1.2
1.4
1.6
1.8
2
0
10
20
30
40
50
60
Normal
MK50
MK75
MK88
MK94
MK97
MK data
Normal data

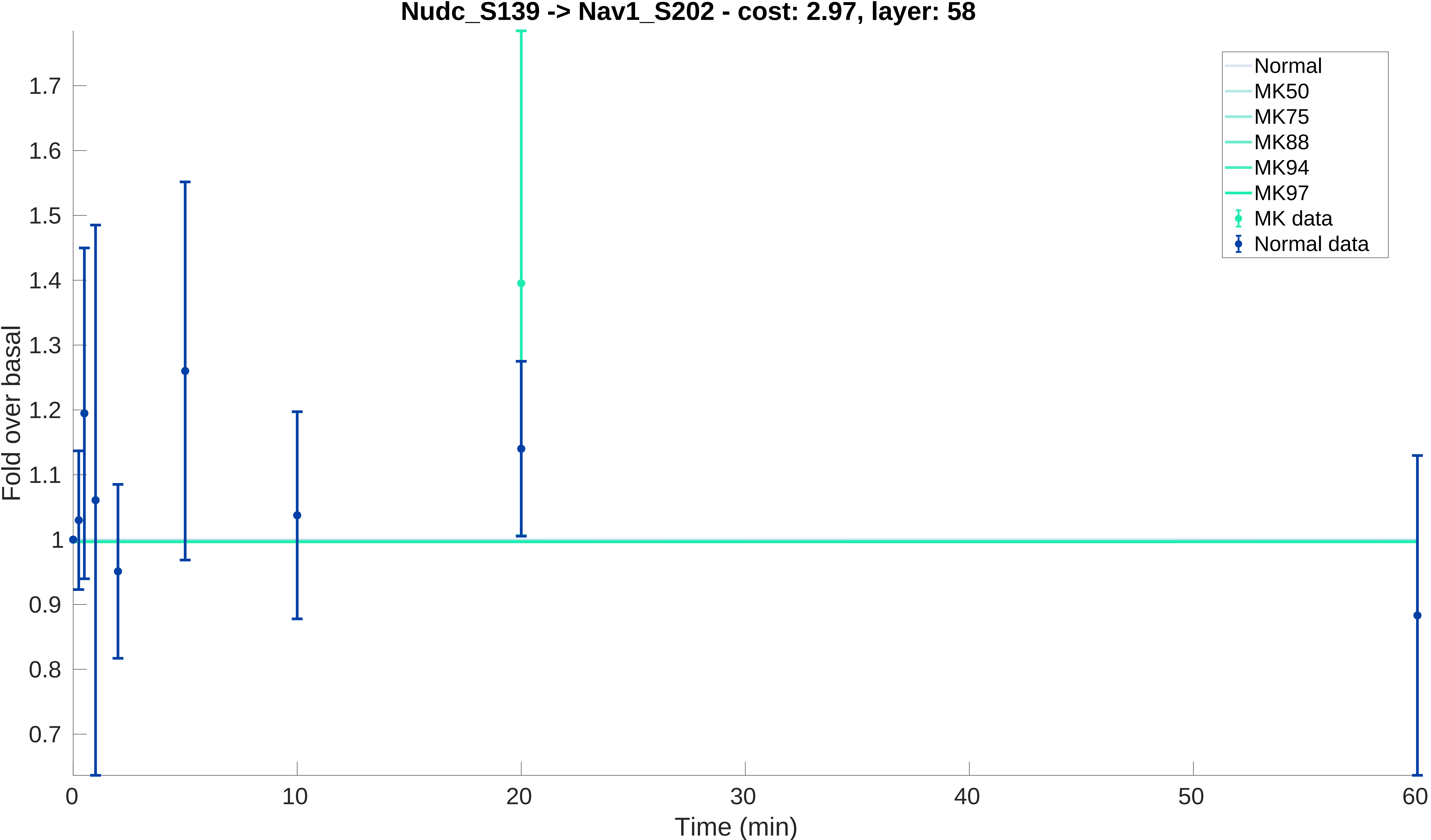
Nudc_S139 -> Nav1_S202 - cost: 2.97, layer: 58
Fold over basal
Time (min)
0.7
0.8
0.9
1
1.1
1.2
1.3
1.4
1.5
1.6
1.7
0
10
20
30
40
50
60
Normal
MK50
MK75
MK88
MK94
MK97
MK data
Normal data

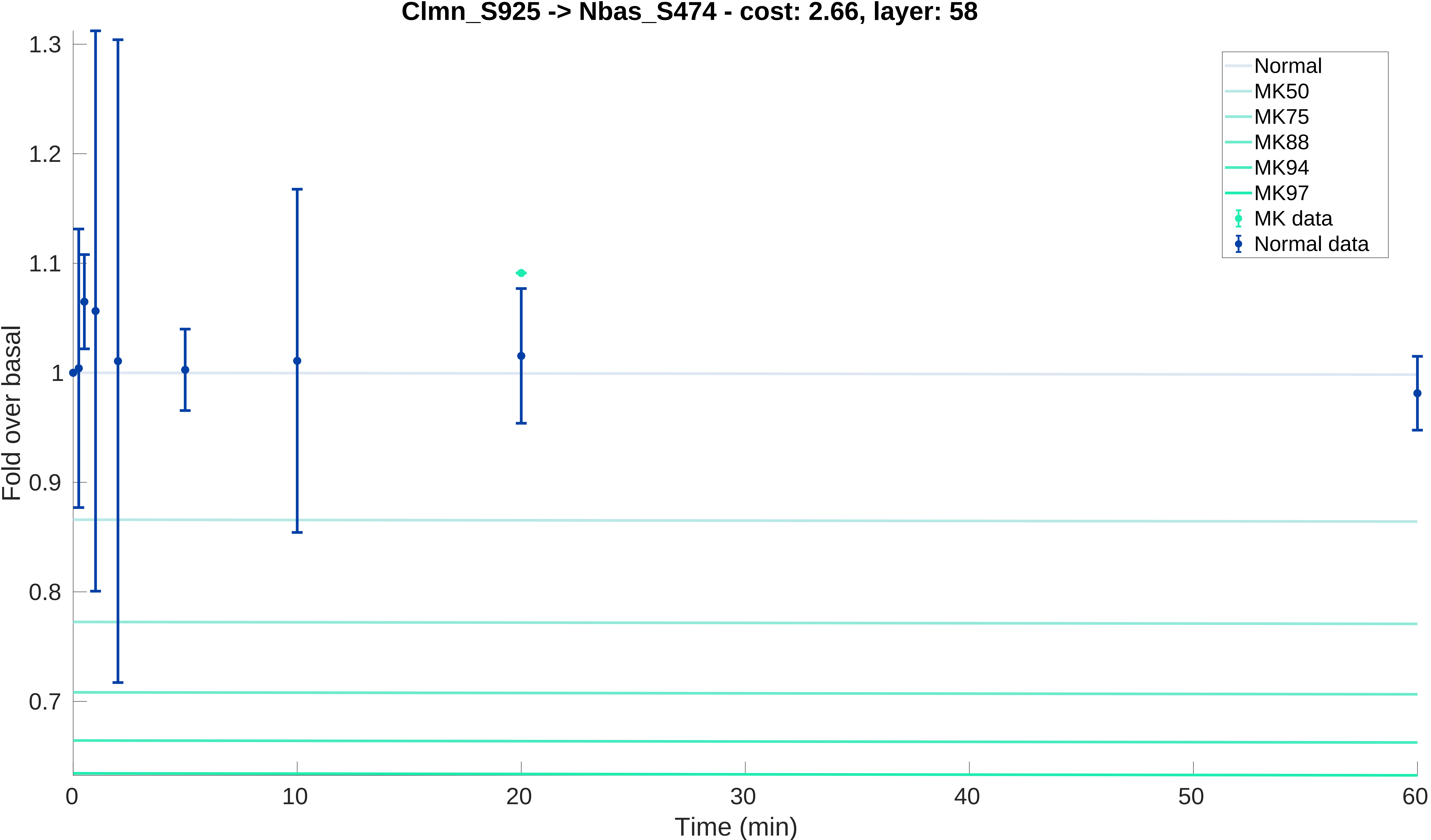

Clmn_S925 -> Nbas_S474 - cost: 2.66, layer: 58
Fold over basal
Time (min)
1.3
1.2
1.1
1
0.9
0.8
0.7
0
10
20
30
40
50
60
Normal
MK50
MK75
MK88
MK94
MK97
MK data
Normal data

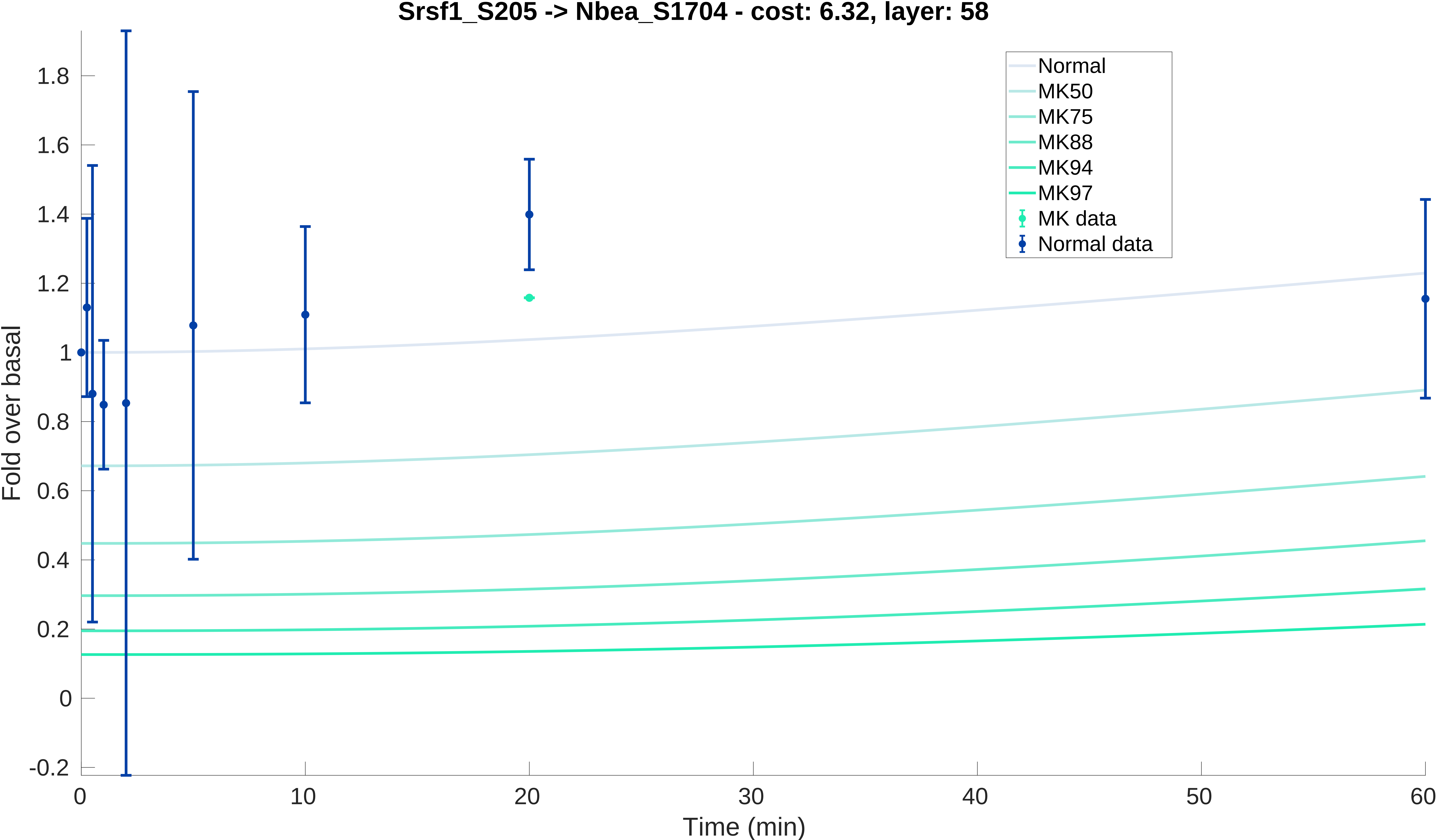

Srsf1_S205 -> Nbea_S1704 - cost: 6.32, layer: 58
Normal
MK50
MK75
MK88
MK94
MK97
MK data
Normal data
Fold over basal
Time (min)
1.8
1.6
1.4
1.2
1
0.8
0.6
0.4
0.2
0
-0.2
0
10
20
30
40
50
60

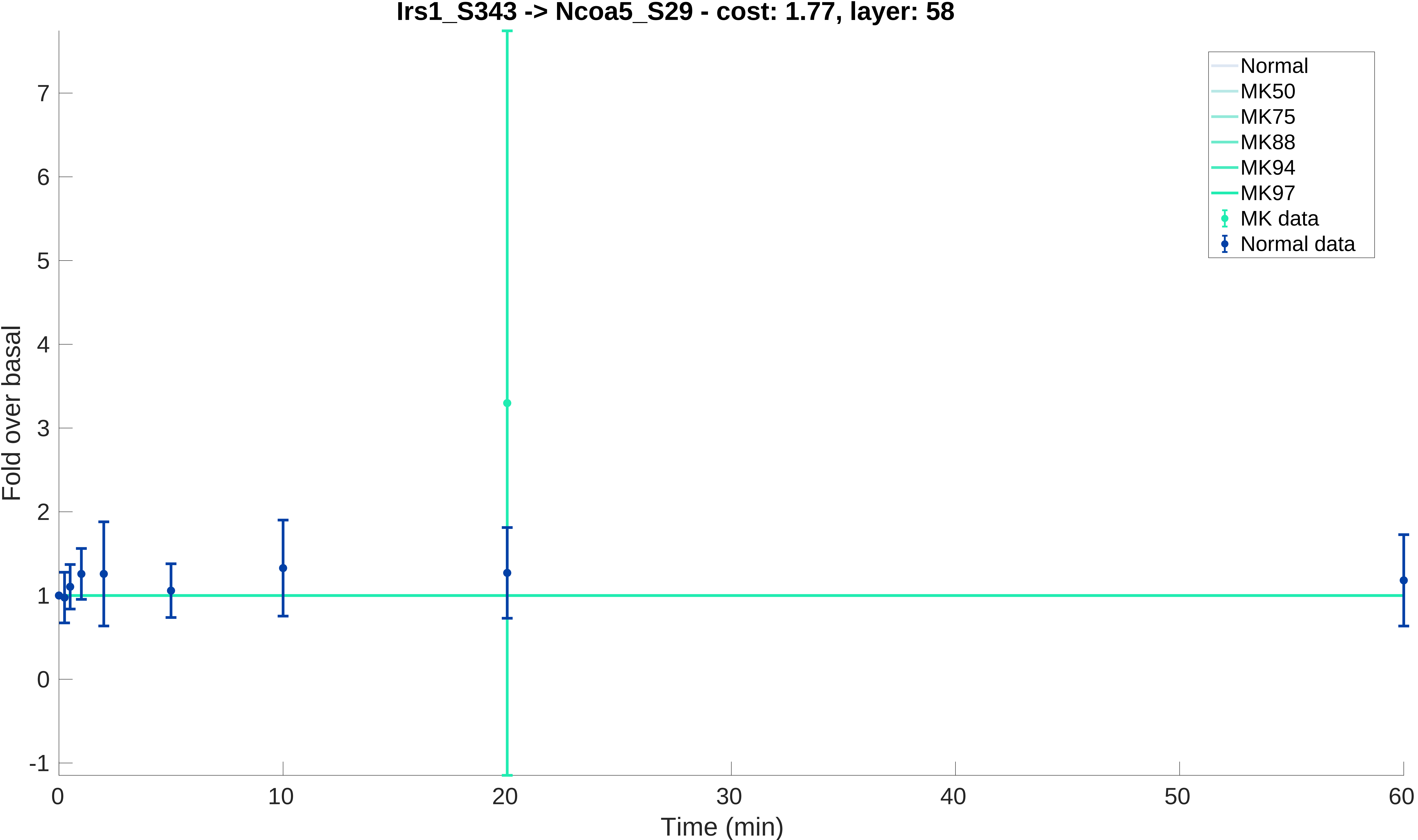

Irs1_S343 -> Ncoa5_S29 - cost: 1.77, layer: 58
Normal
MK50
MK75
MK88
MK94
MK97
MK data
Normal data
Fold over basal
Time (min)
-1
0
1
2
3
4
5
6
7
0
10
20
30
40
50
60

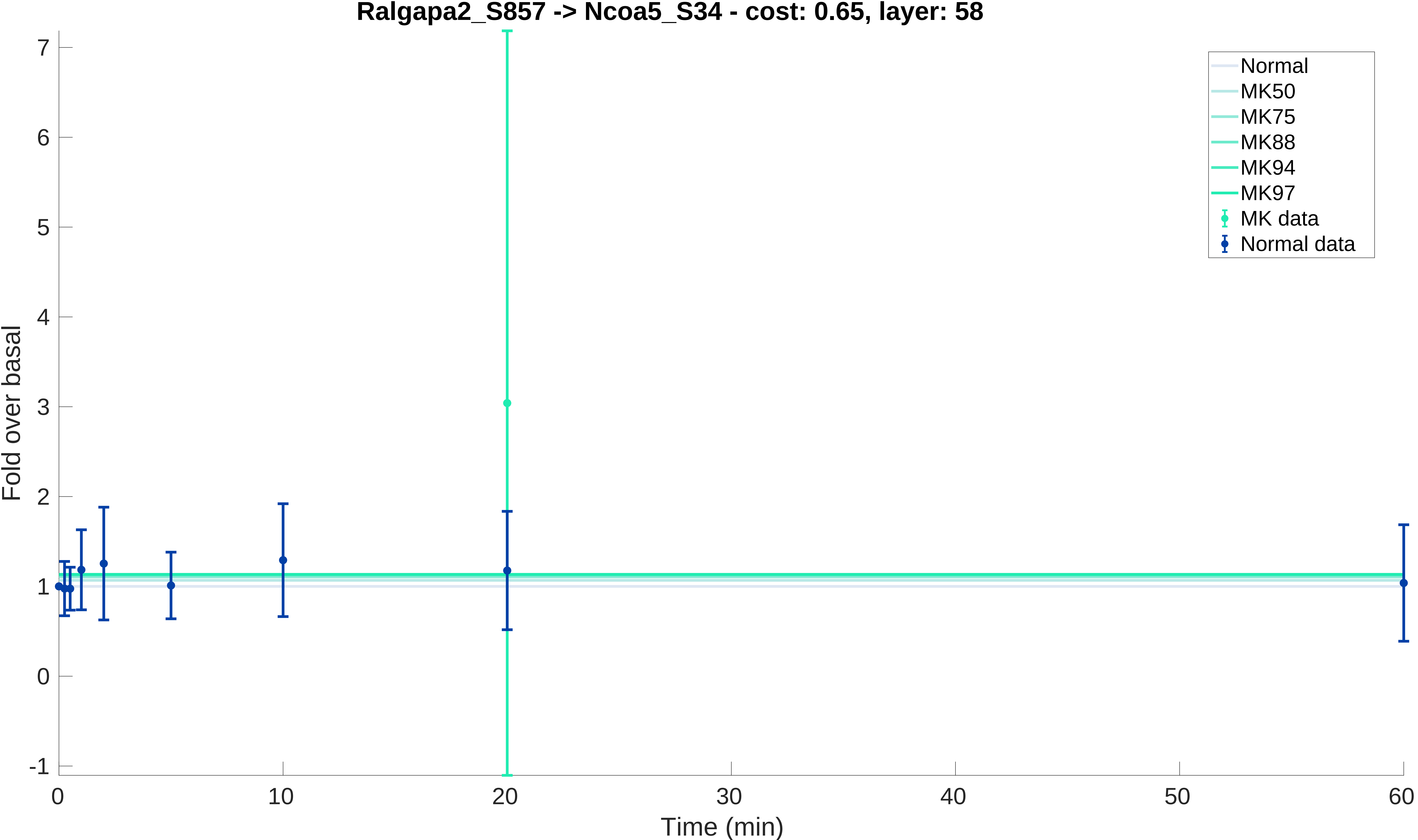
Ralgapa2_S857 -> Ncoa5_S34 - cost: 0.65, layer: 58
Fold over basal
Time (min)
7
6
5
4
3
2
1
0
-1
0
10
20
30
40
50
60
Normal
MK50
MK75
MK88
MK94
MK97
MK data
Normal data

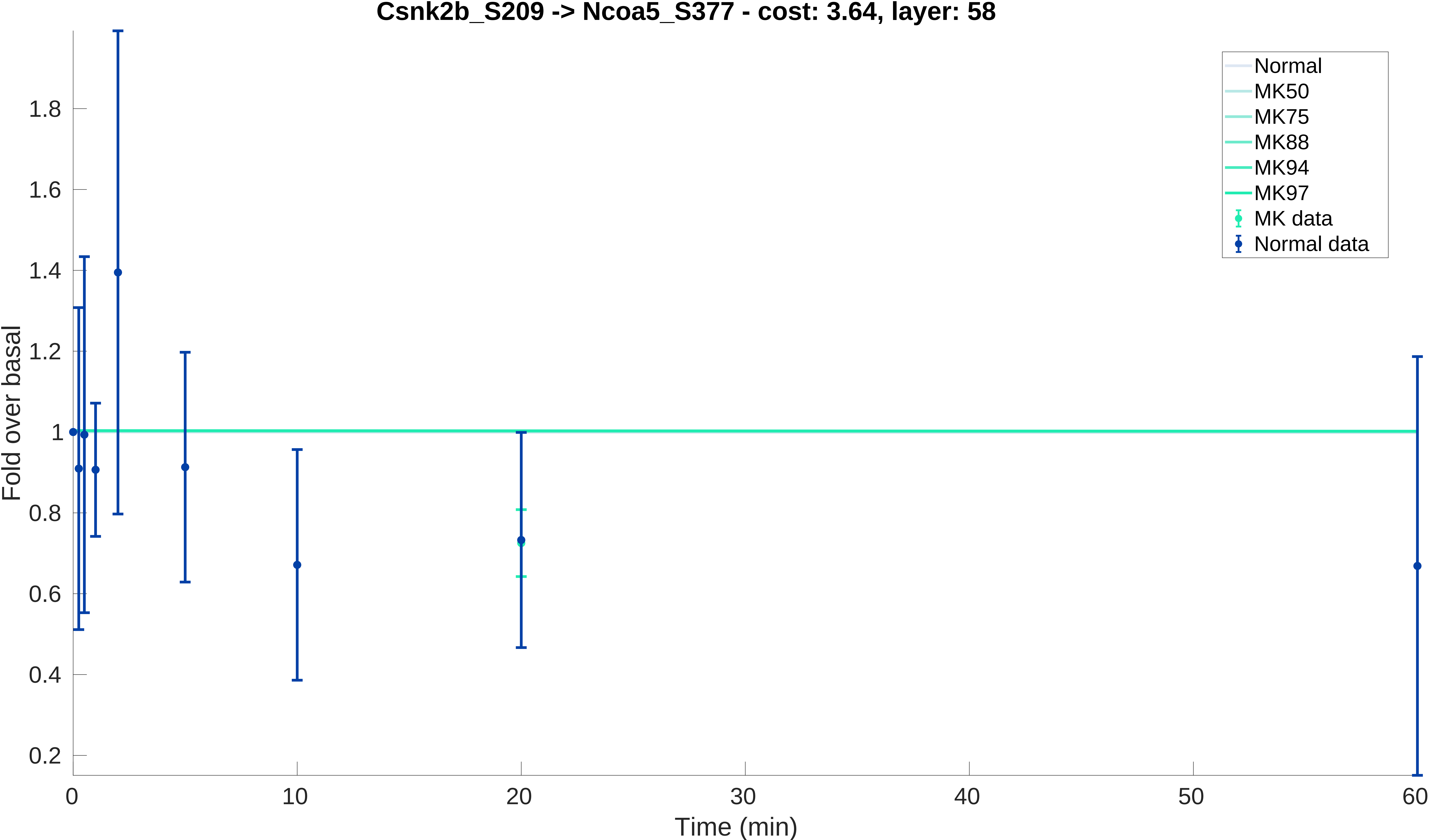

Csnk2b_S209 -> Ncoa5_S377 - cost: 3.64, layer: 58
Fold over basal
Time (min)
1.8
1.6
1.4
1.2
1
0.8
0.6
0.4
0.2
0
10
20
30
40
50
60
Normal
MK50
MK75
MK88
MK94
MK97
MK data
Normal data

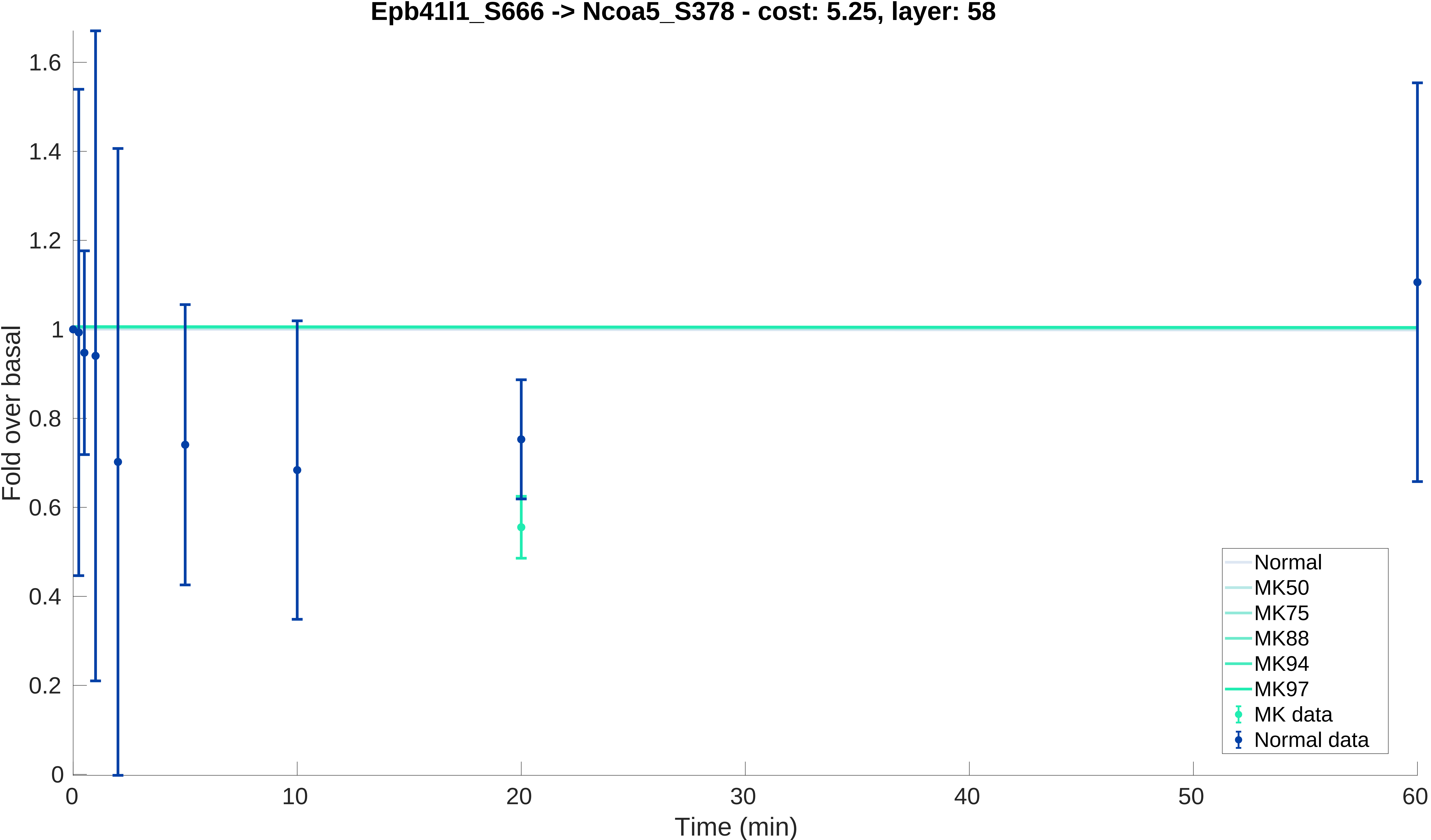
Epb41l1_S666 -> Ncoa5_S378 - cost: 5.25, layer: 58
Fold over basal
Time (min)
0
0.2
0.4
0.6
0.8
1
1.2
1.4
1.6
0
10
20
30
40
50
60
Normal
MK50
MK75
MK88
MK94
MK97
MK data
Normal data

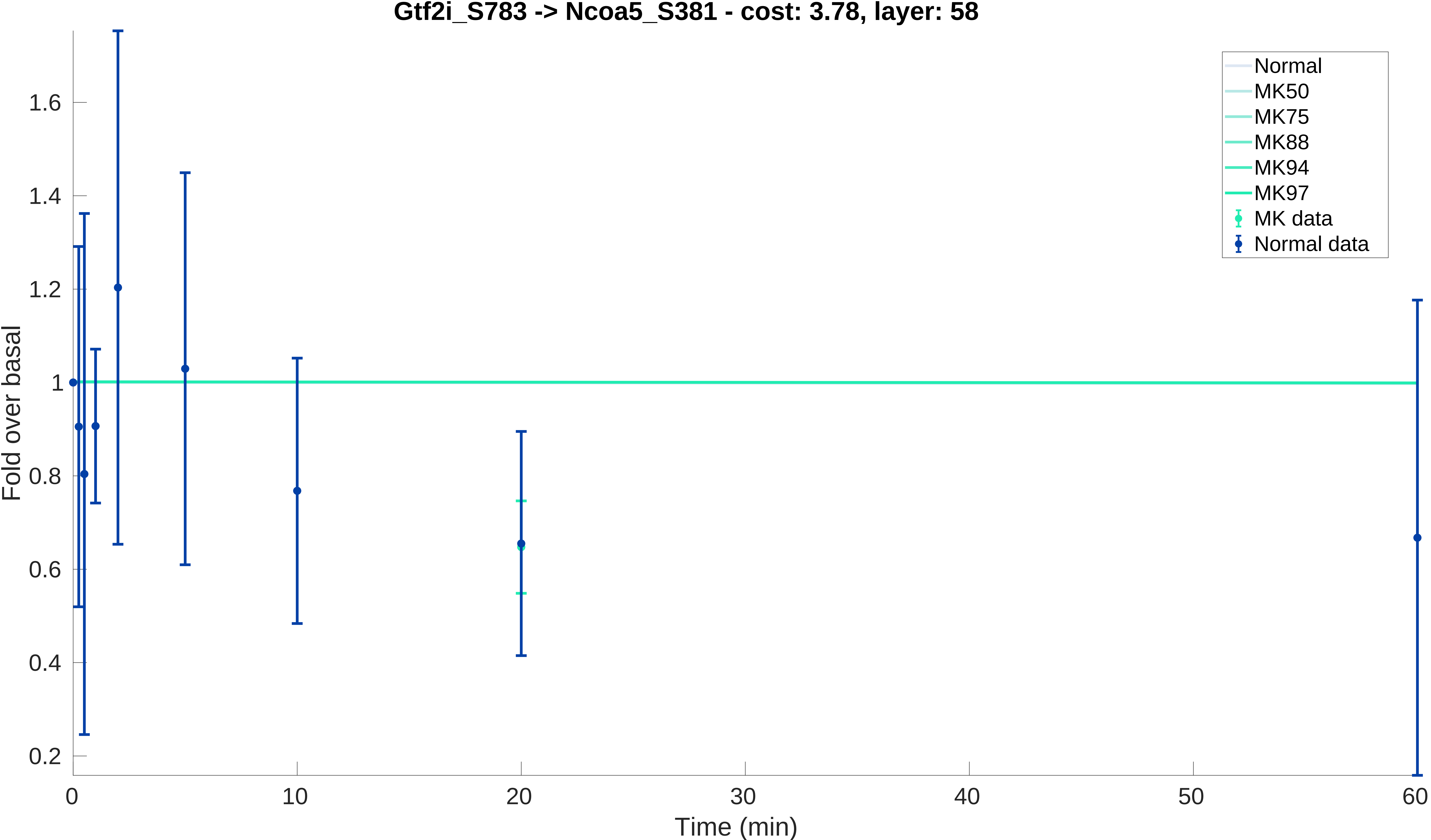

Gtf2i_S783 -> Ncoa5_S381 - cost: 3.78, layer: 58
Fold over basal
Time (min)
0.2
0.4
0.6
0.8
1
1.2
1.4
1.6
0
10
20
30
40
50
60
Normal
MK50
MK75
MK88
MK94
MK97
MK data
Normal data

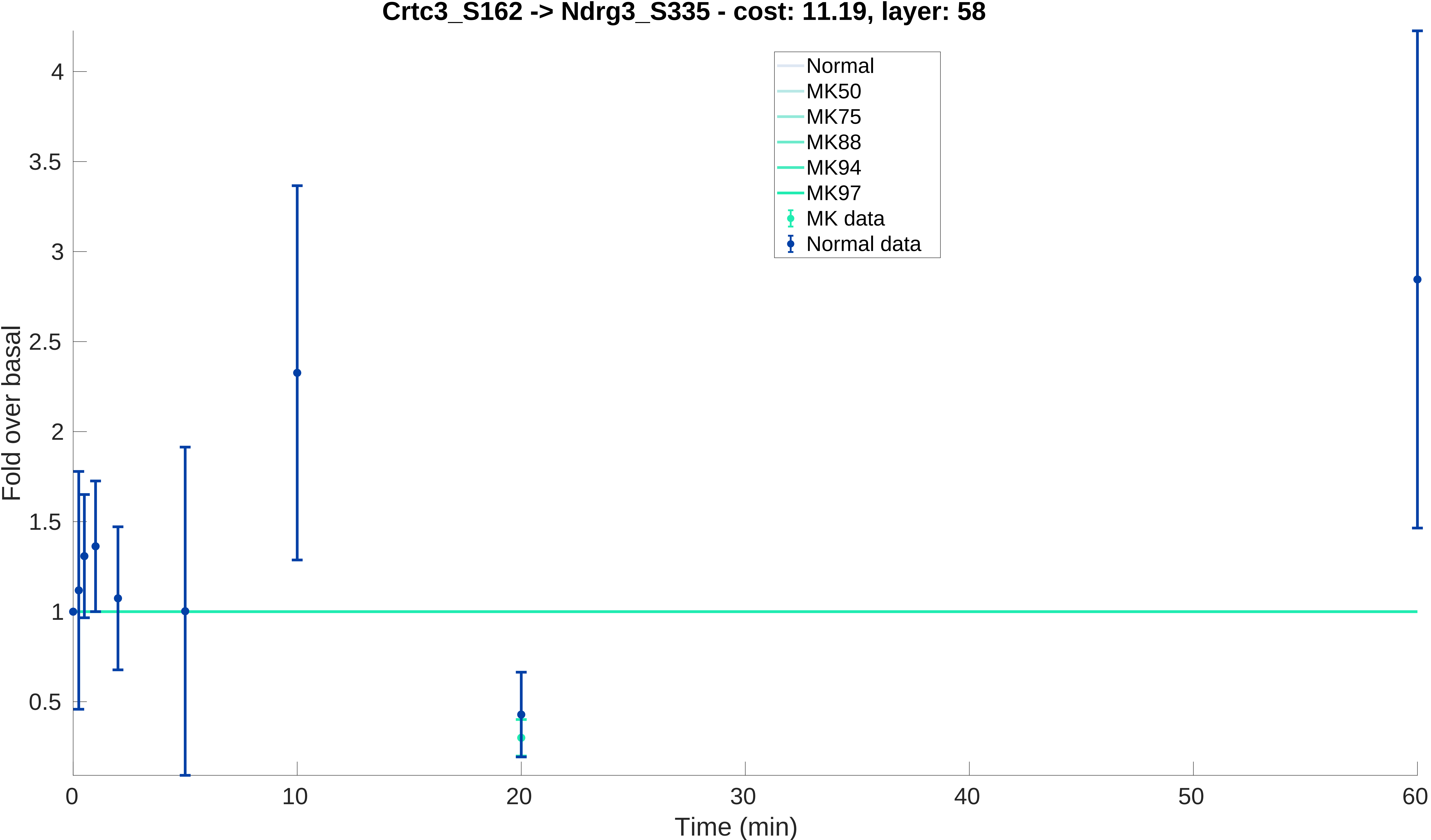
Crtc3_S162 -> Ndrg3_S335 - cost: 11.19, layer: 58
Normal
MK50
MK75
MK88
MK94
MK97
MK data
Normal data
Fold over basal
4
3.5
3
2.5
2
1.5
1
0.5
0
10
20
30
40
50
60
Time (min)

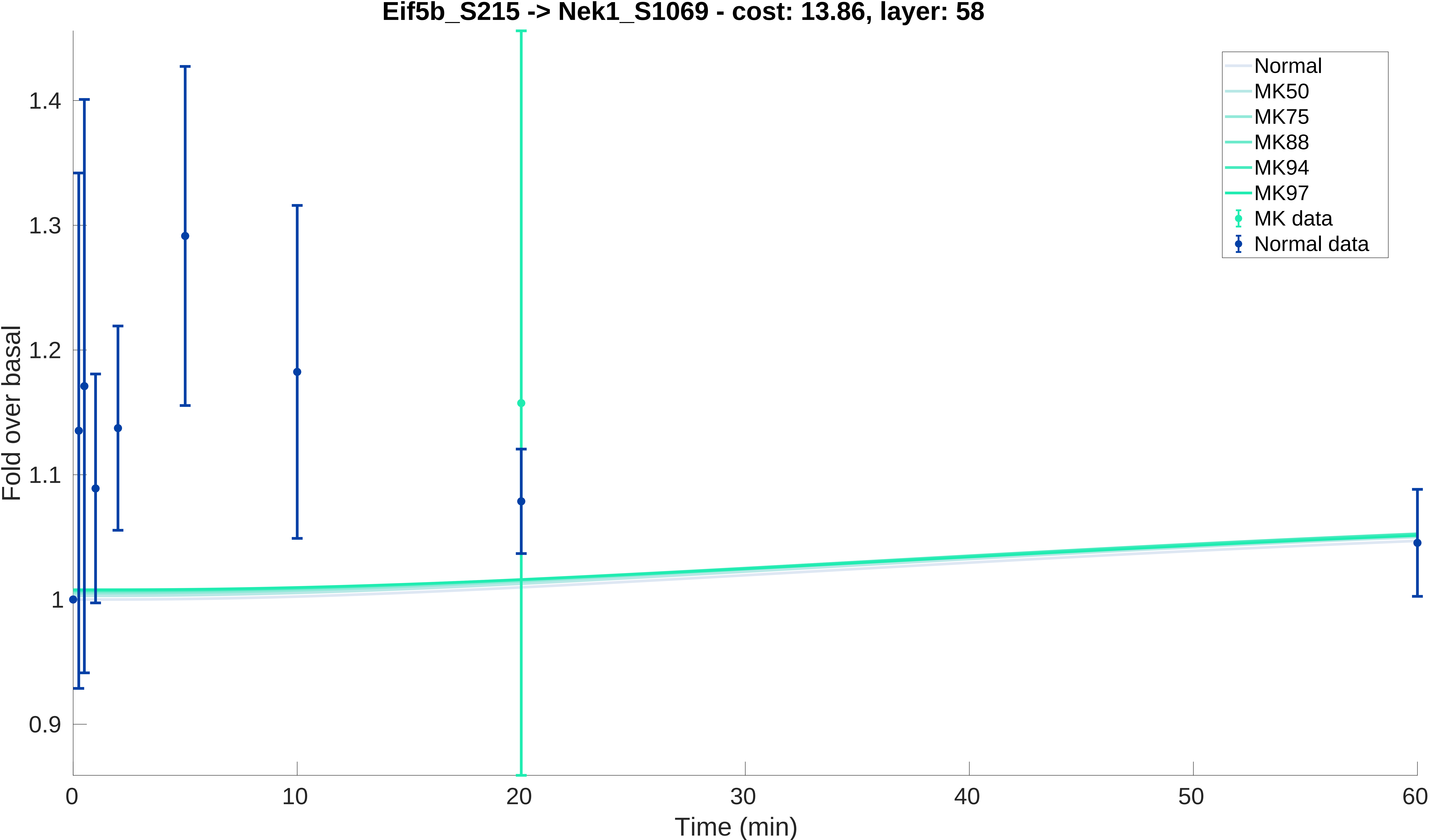
Eif5b_S215 -> Nek1_S1069 - cost: 13.86, layer: 58
Fold over basal
1.4
1.3
1.2
1.1
1
0.9
0
10
20
30
40
50
60
Time (min)
Normal
MK50
MK75
MK88
MK94
MK97
MK data
Normal data

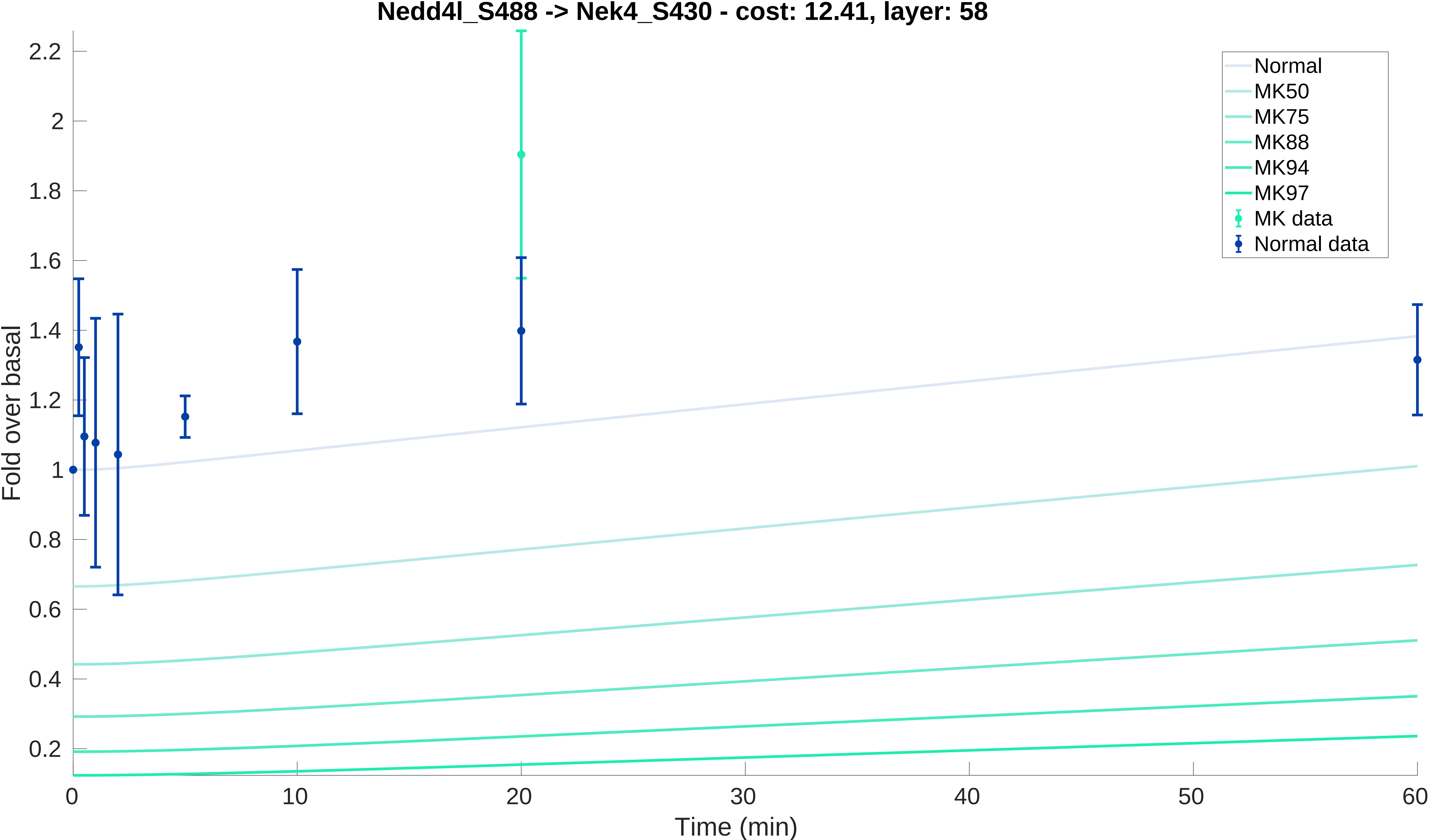

Nedd4l_S488 -> Nek4_S430 - cost: 12.41, layer: 58
Fold over basal
Time (min)
0
10
20
30
40
50
60
0.2
0.4
0.6
0.8
1
1.2
1.4
1.6
1.8
2
2.2
Normal
MK50
MK75
MK88
MK94
MK97
MK data
Normal data

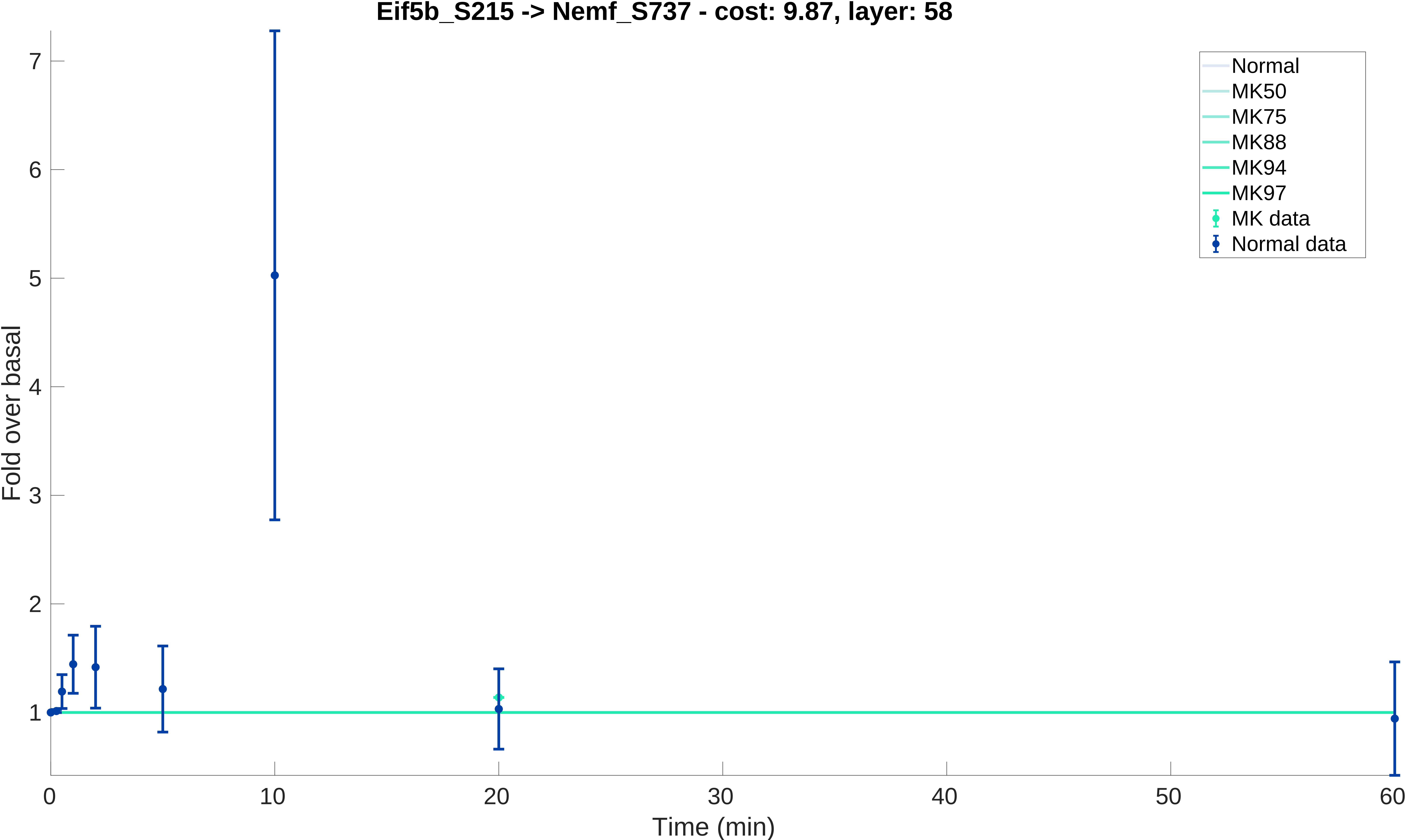
Eif5b_S215 -> Nemf_S737 - cost: 9.87, layer: 58
Fold over basal
Time (min)
0
10
20
30
40
50
60
1
2
3
4
5
6
7
Normal
MK50
MK75
MK88
MK94
MK97
MK data
Normal data

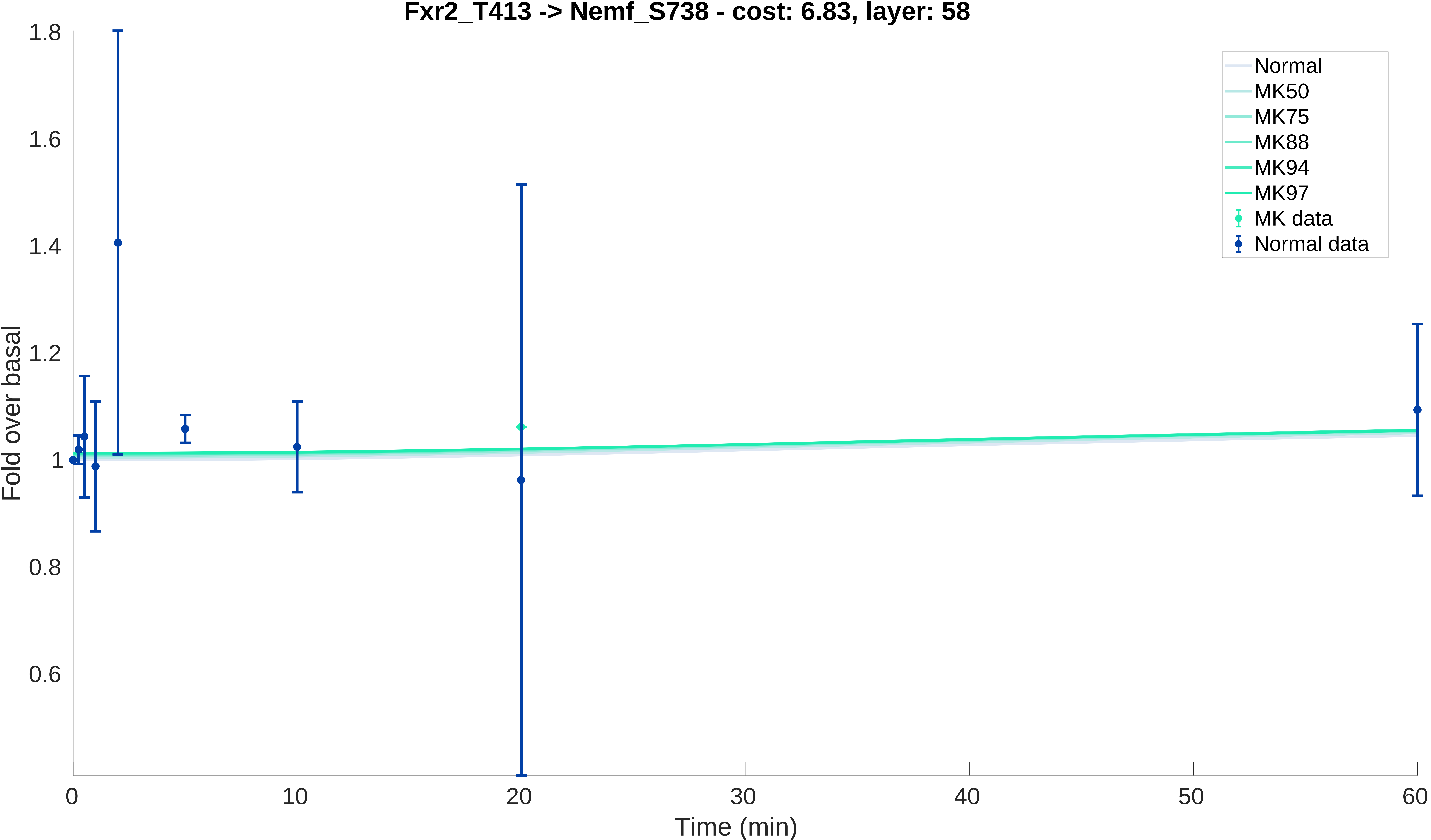

Fxr2_T413 -> Nemf_S738 - cost: 6.83, layer: 58
Fold over basal
1.8
1.6
1.4
1.2
1
0.8
0.6
0
10
20
30
40
50
60
Time (min)
Normal
MK50
MK75
MK88
MK94
MK97
MK data
Normal data

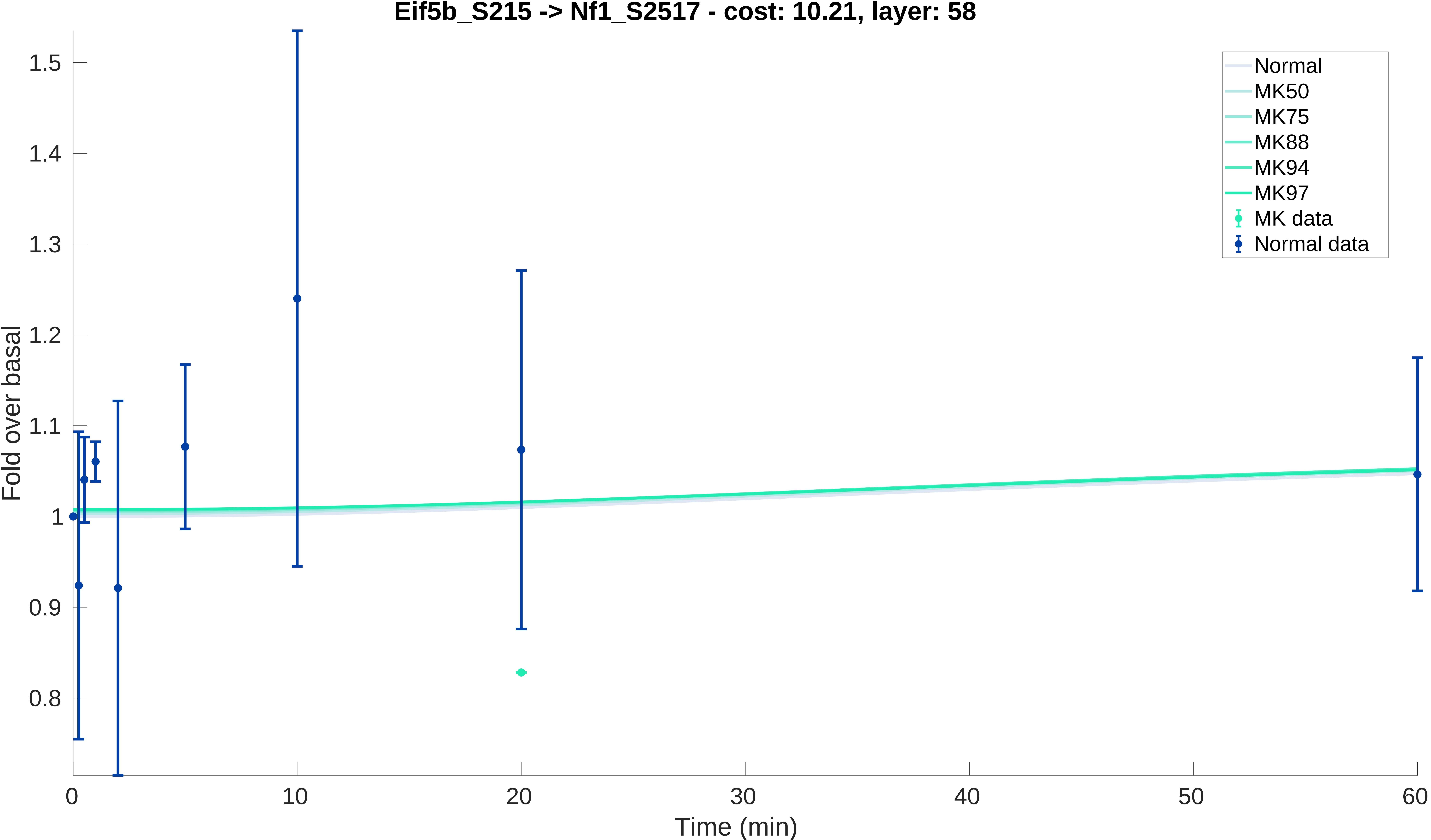
Eif5b_S215 -> Nf1_S2517 - cost: 10.21, layer: 58
Fold over basal
1.5
1.4
1.3
1.2
1.1
1
0.9
0.8
0
10
20
30
40
50
60
Time (min)
Normal
MK50
MK75
MK88
MK94
MK97
MK data
Normal data

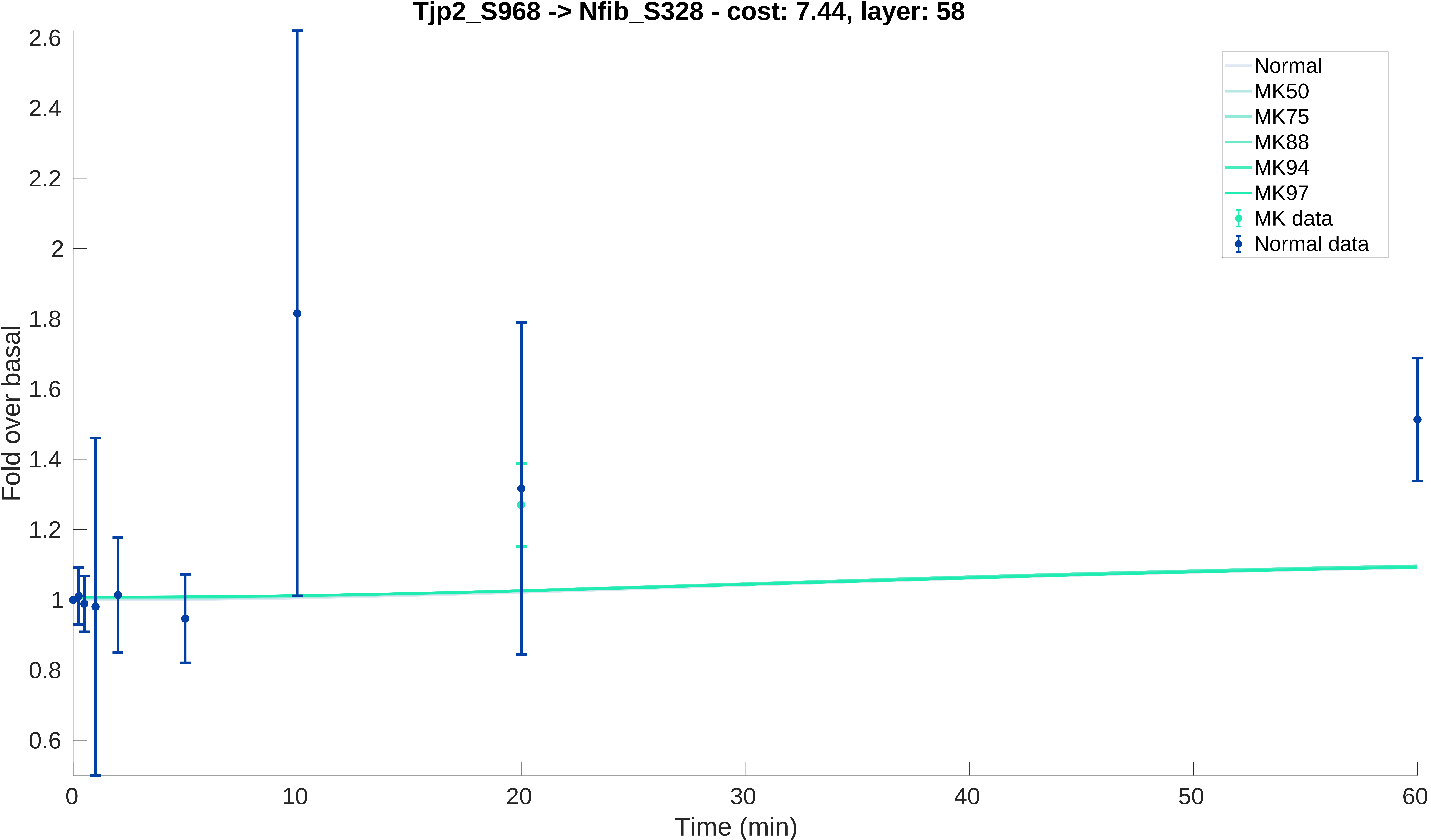

Tjp2_S968 -> Nfib_S328 - cost: 7.44, layer: 58
Fold over basal
Time (min)
Normal
MK50
MK75
MK88
MK94
MK97
MK data
Normal data
2.6
2.4
2.2
2
1.8
1.6
1.4
1.2
1
0.8
0.6
0
10
20
30
40
50
60

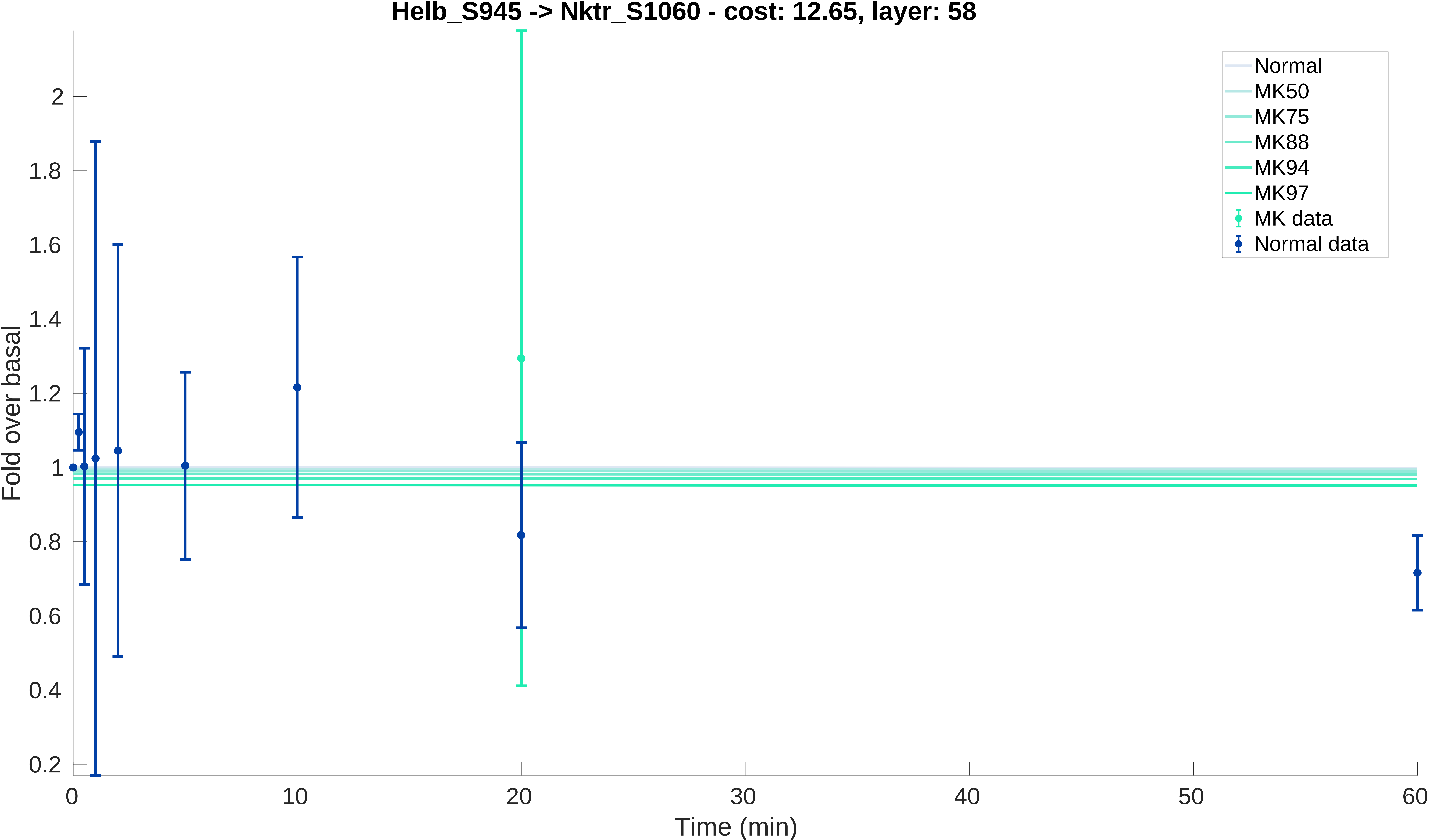

Helb_S945 -> Nktr_S1060 - cost: 12.65, layer: 58
Fold over basal
Time (min)
0.2
0.4
0.6
0.8
1
1.2
1.4
1.6
1.8
2
0
10
20
30
40
50
60
Normal
MK50
MK75
MK88
MK94
MK97
MK data
Normal data

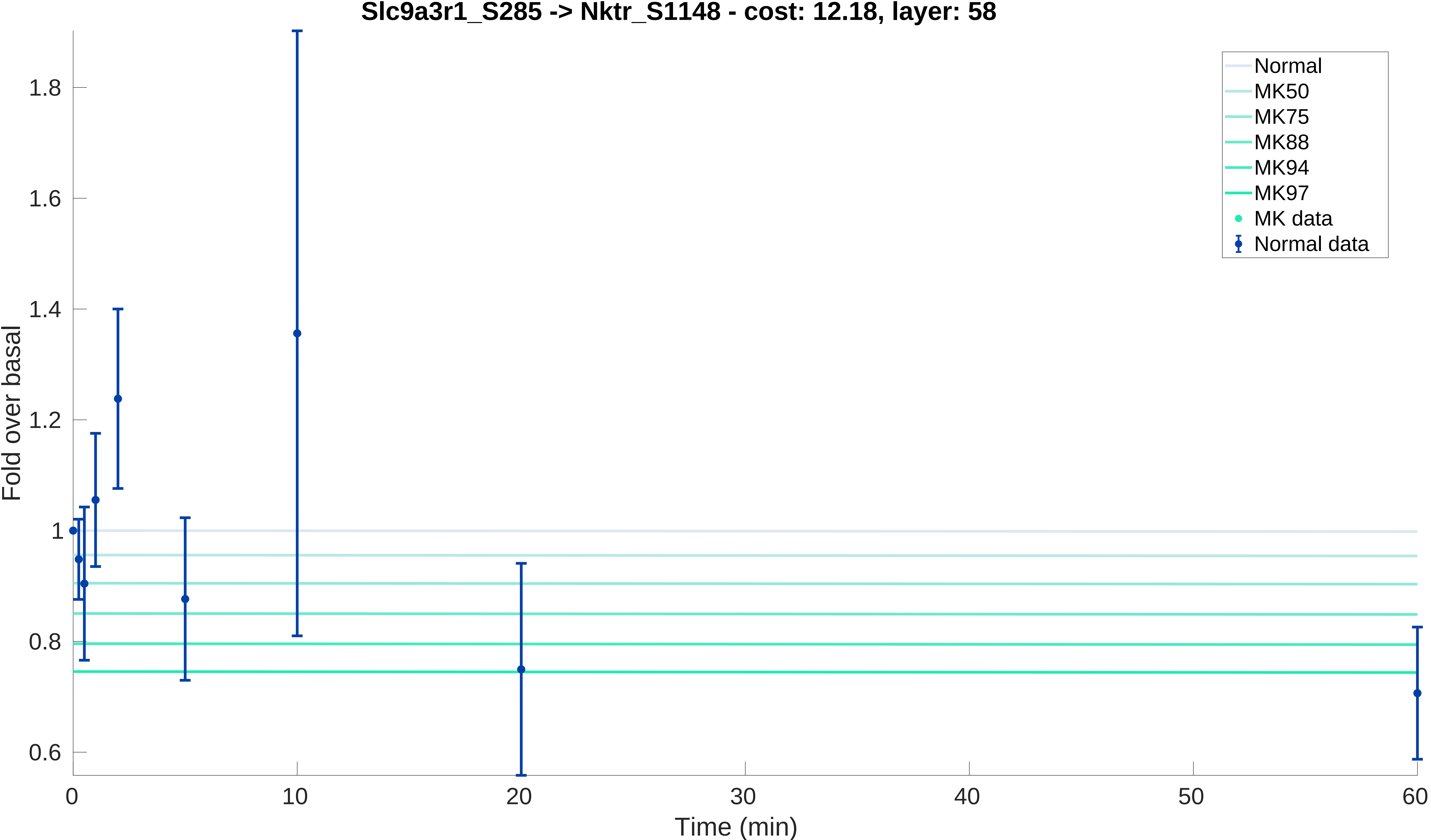

Slc9a3r1_S285 -> Nktr_S1148 - cost: 12.18, layer: 58
Fold over basal
Time (min)
0.6
0.8
1
1.2
1.4
1.6
1.8
0
10
20
30
40
50
60
Normal
MK50
MK75
MK88
MK94
MK97
MK data
Normal data

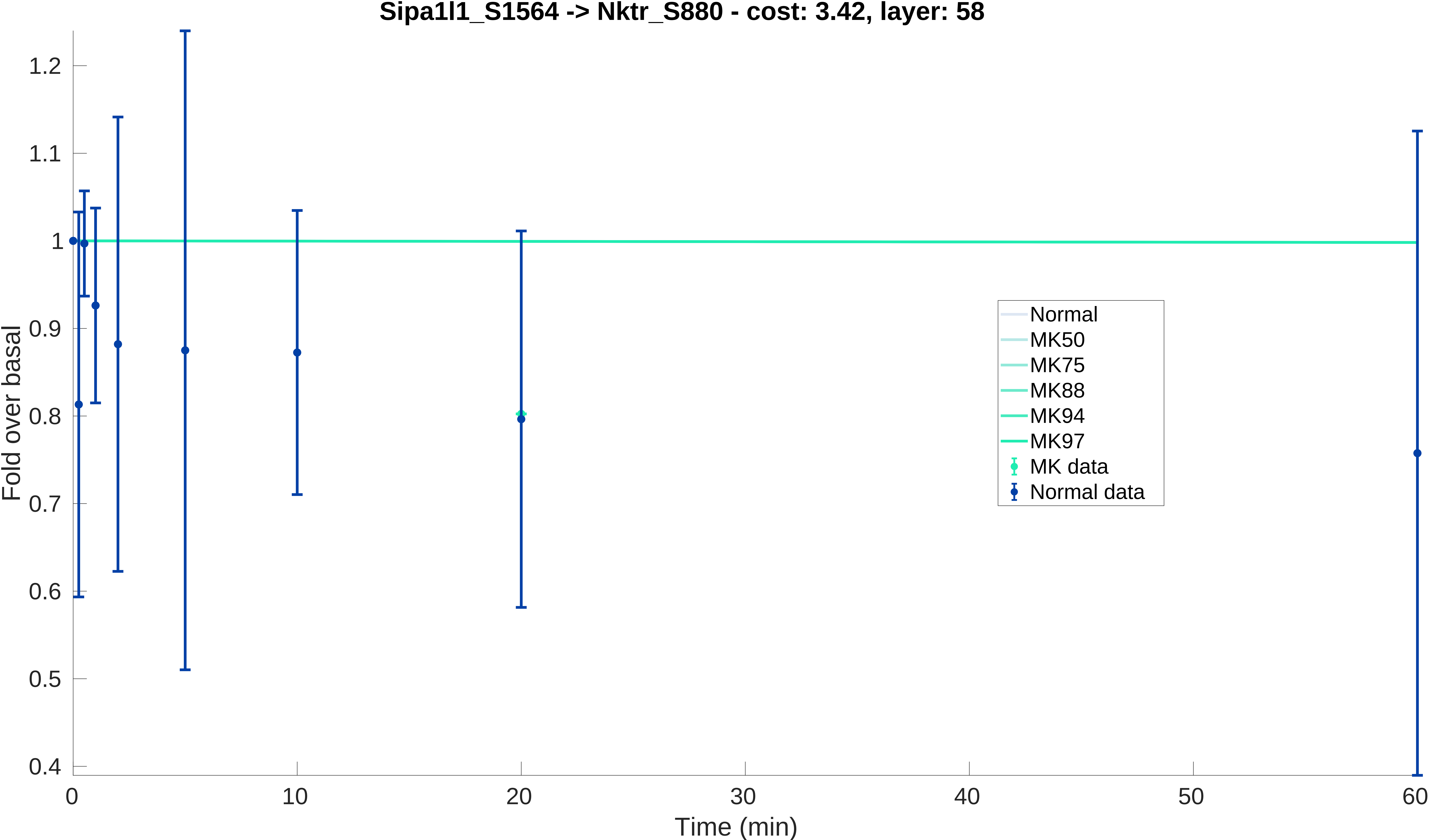
Sipa1l1_S1564 -> Nktr_S880 - cost: 3.42, layer: 58
Fold over basal
1.2
1.1
1
0.9
0.8
0.7
0.6
0.5
0.4
0
10
20
30
40
50
60
Time (min)
Normal
MK50
MK75
MK88
MK94
MK97
MK data
Normal data

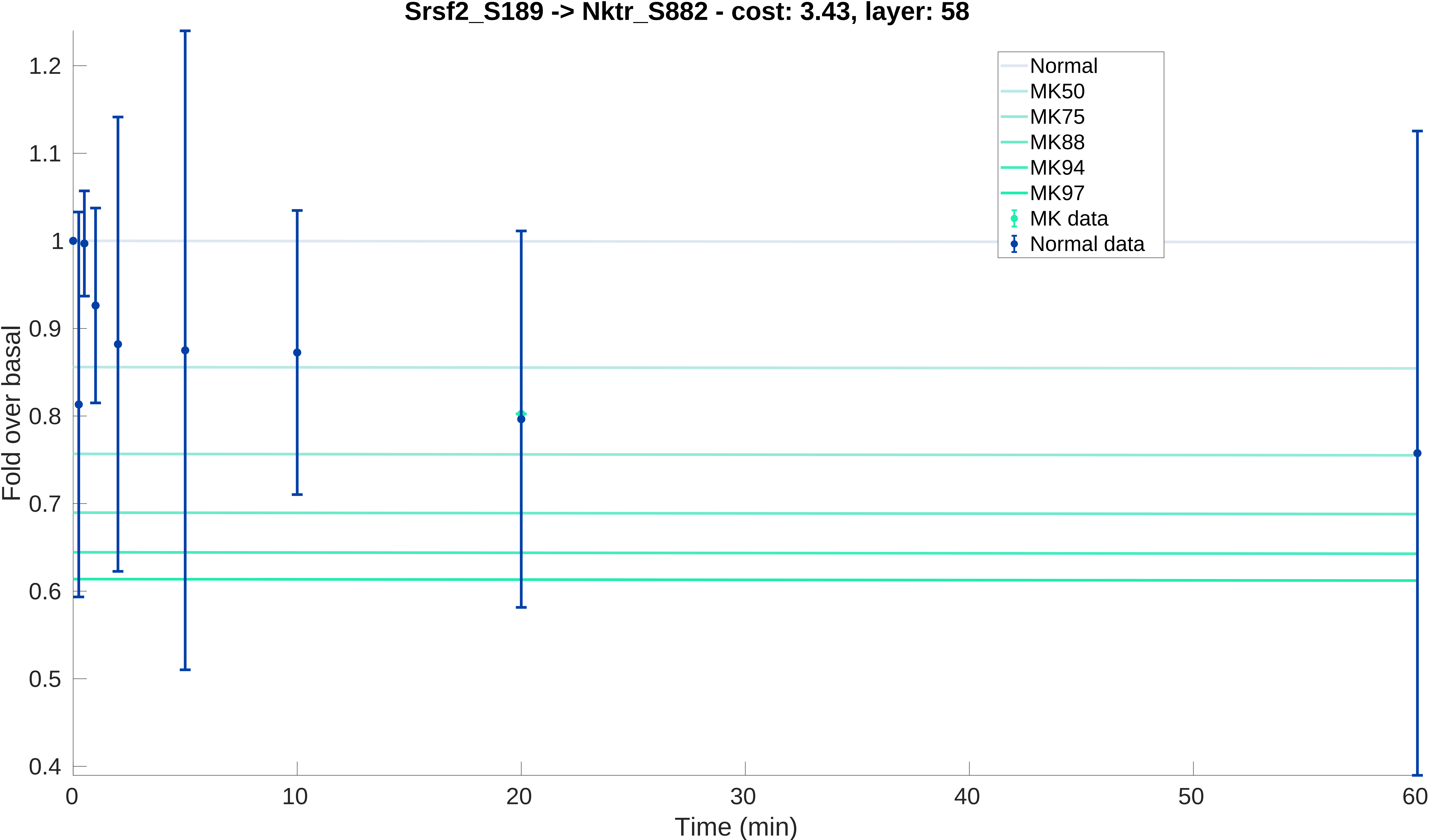

Srsf2_S189 -> Nktr_S882 - cost: 3.43, layer: 58
Normal
MK50
MK75
MK88
MK94
MK97
MK data
Normal data
Fold over basal
Time (min)
1.2
1.1
1
0.9
0.8
0.7
0.6
0.5
0.4
0
10
20
30
40
50
60

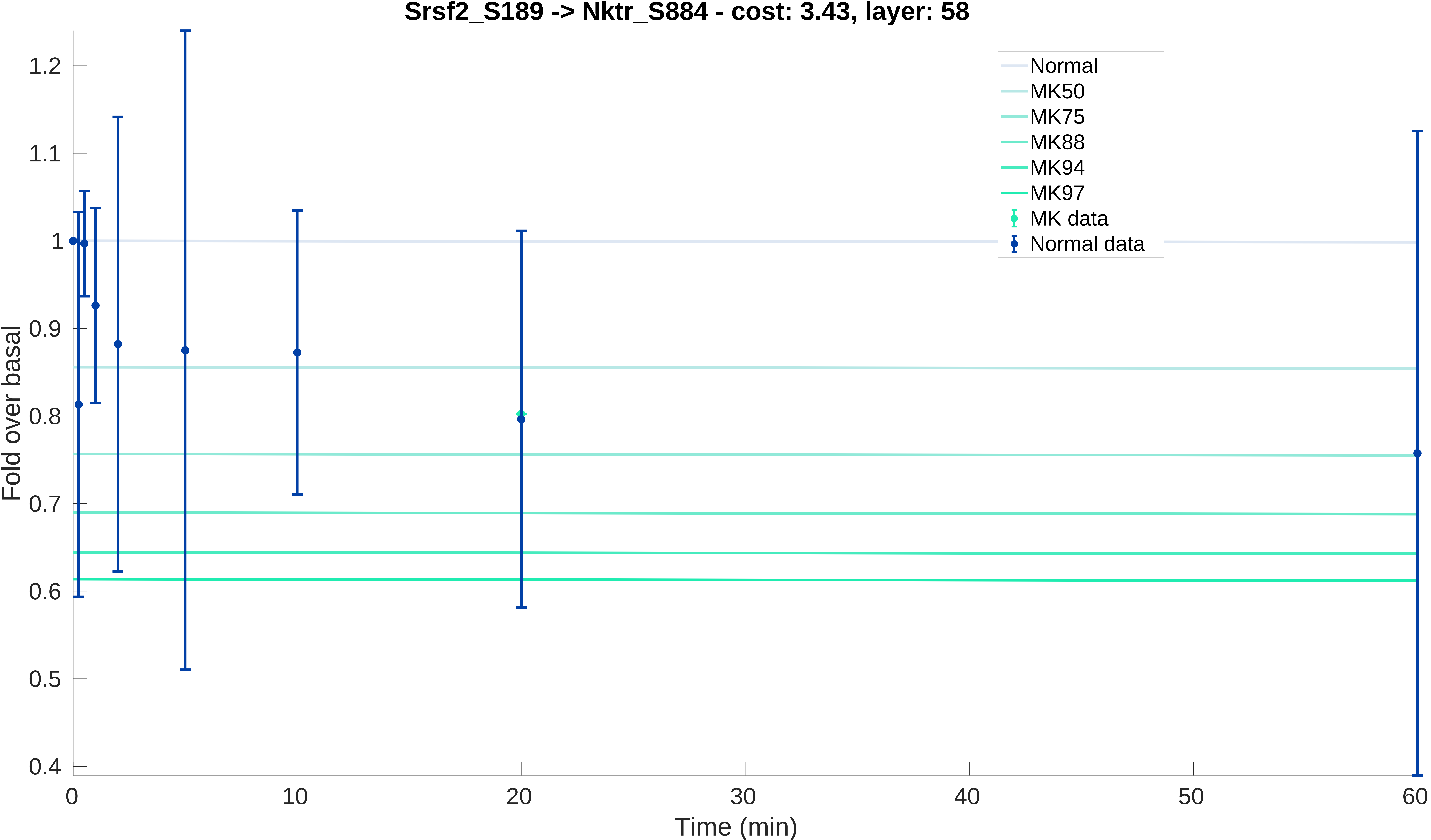

Srsf2_S189 -> Nktr_S884 - cost: 3.43, layer: 58
Normal
MK50
MK75
MK88
MK94
MK97
MK data
Normal data
Fold over basal
Time (min)
1.2
1.1
1
0.9
0.8
0.7
0.6
0.5
0.4
0
10
20
30
40
50
60

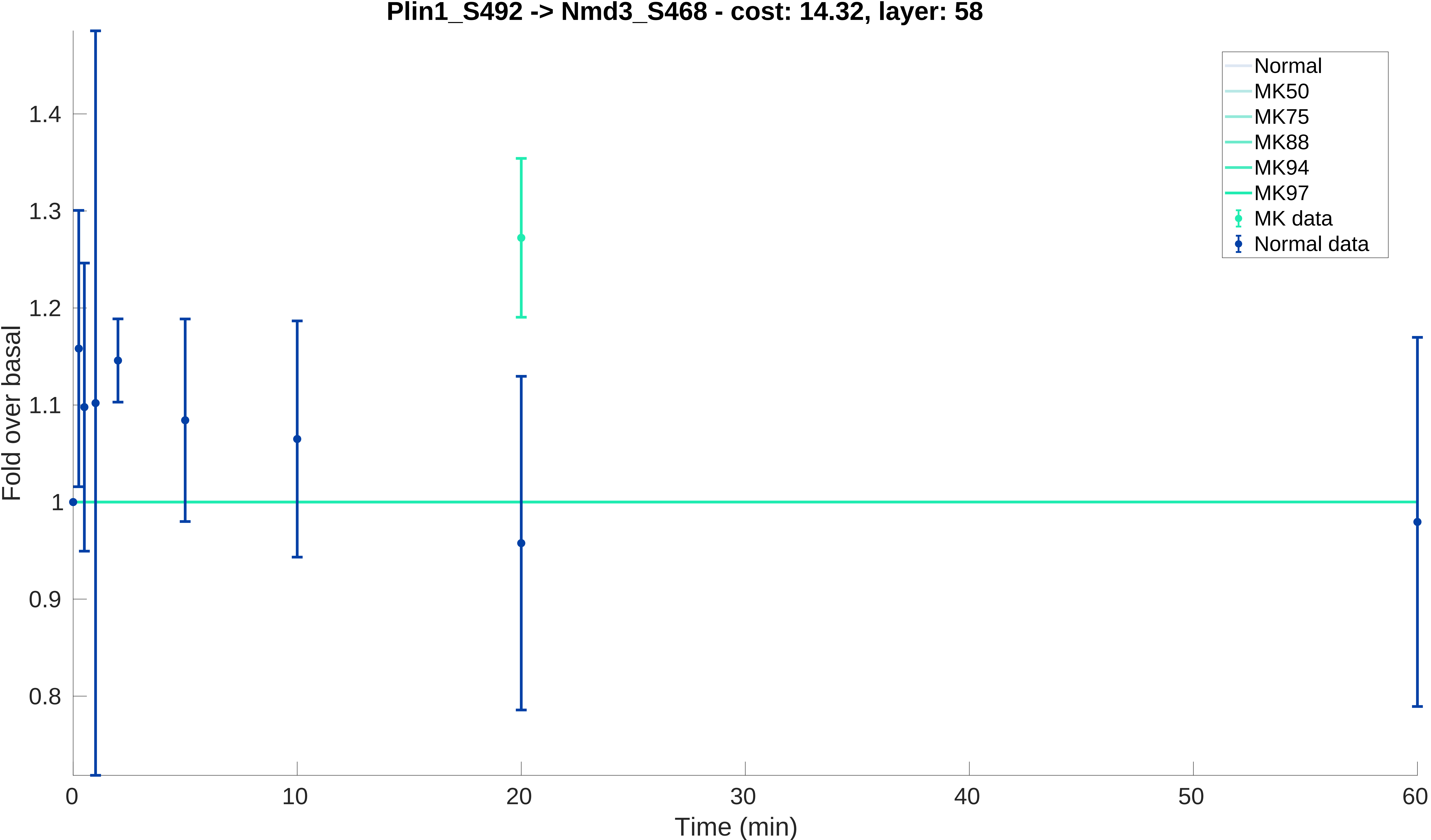

Plin1_S492 -> Nmd3_S468 - cost: 14.32, layer: 58
Normal
MK50
MK75
MK88
MK94
MK97
MK data
Normal data
Fold over basal
1.4
1.3
1.2
1.1
1
0.9
0.8
0
10
20
30
40
50
60
Time (min)

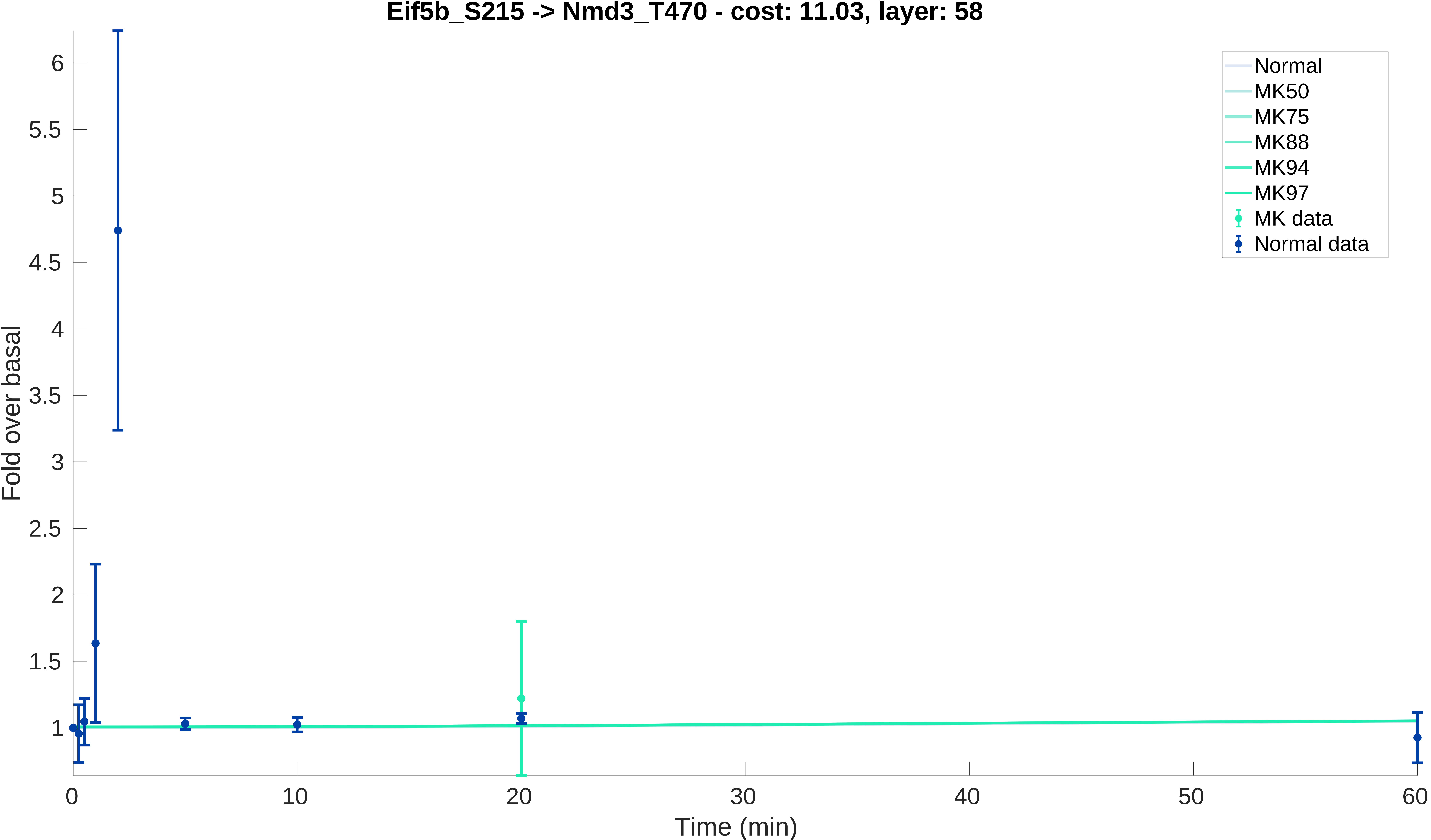

Eif5b_S215 -> Nmd3_T470 - cost: 11.03, layer: 58
Fold over basal
Time (min)
6
5.5
5
4.5
4
3.5
3
2.5
2
1.5
1
0
10
20
30
40
50
60
Normal
MK50
MK75
MK88
MK94
MK97
MK data
Normal data

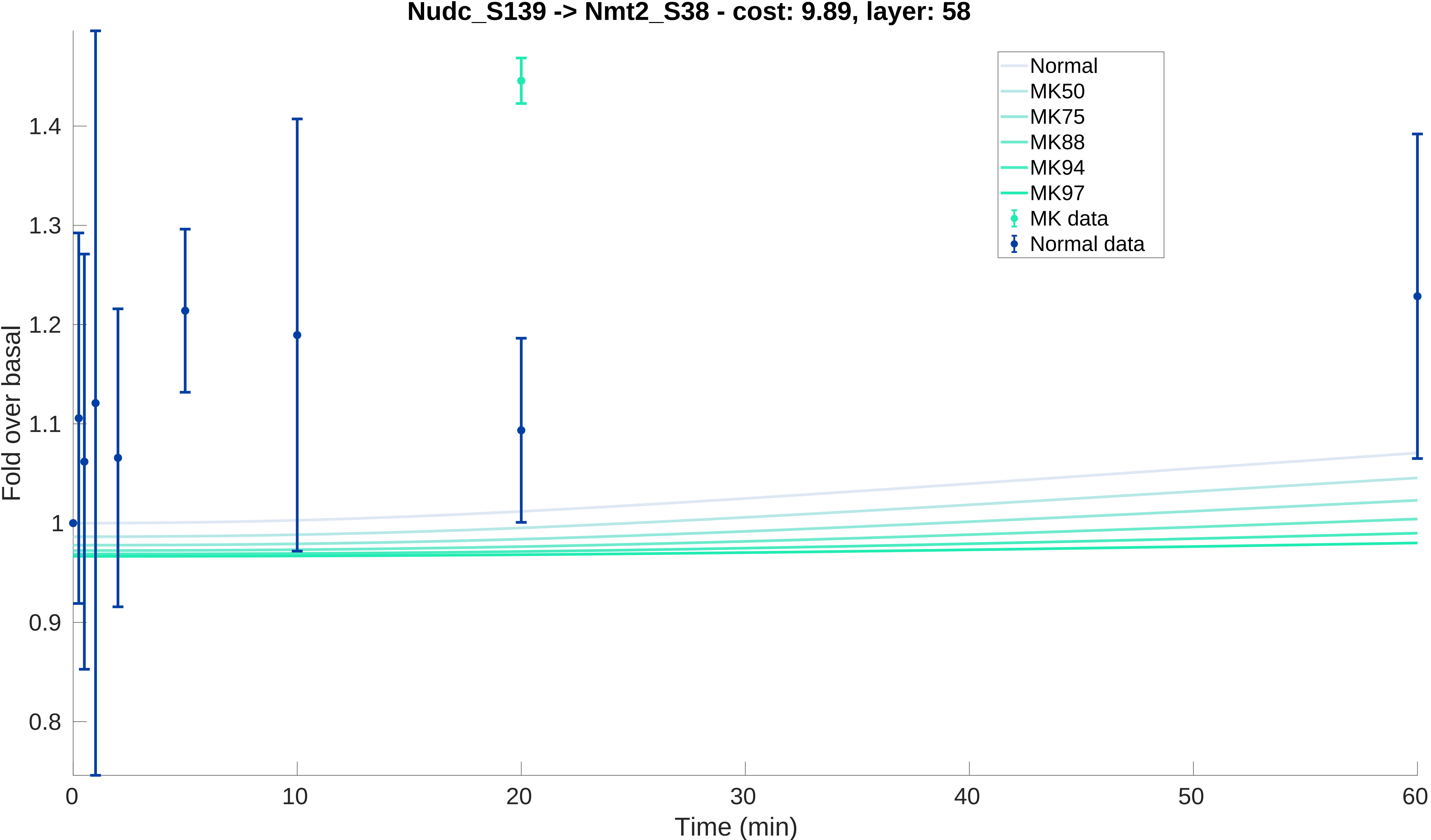
Nudc_S139 -> Nmt2_S38 - cost: 9.89, layer: 58
Normal
MK50
MK75
MK88
MK94
MK97
MK data
Normal data
Fold over basal
Time (min)
0.8
0.9
1
1.1
1.2
1.3
1.4
0
10
20
30
40
50
60

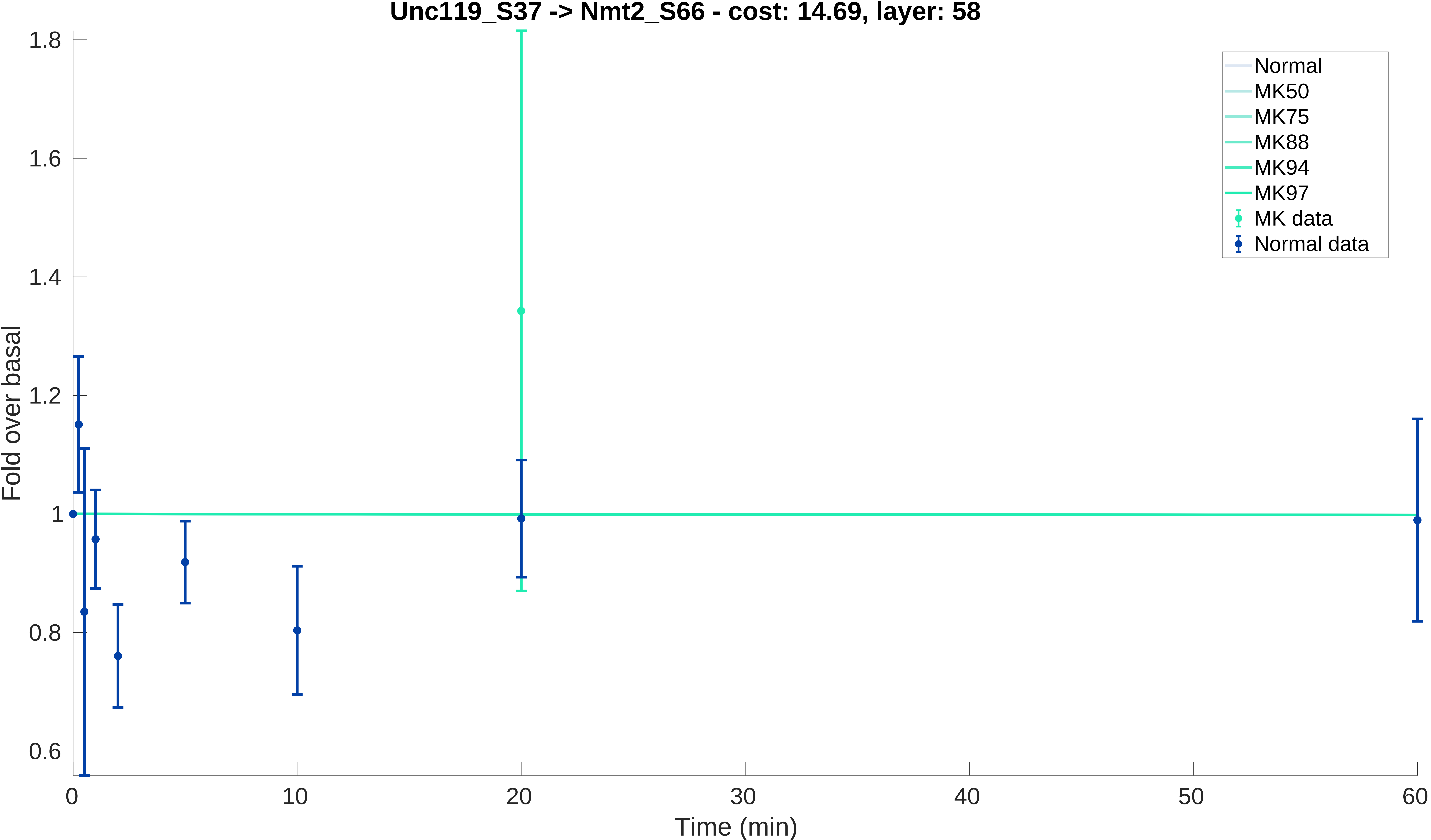

Unc119_S37 -> Nmt2_S66 - cost: 14.69, layer: 58
Fold over basal
Time (min)
0.6
0.8
1
1.2
1.4
1.6
1.8
0
10
20
30
40
50
60
Normal
MK50
MK75
MK88
MK94
MK97
MK data
Normal data

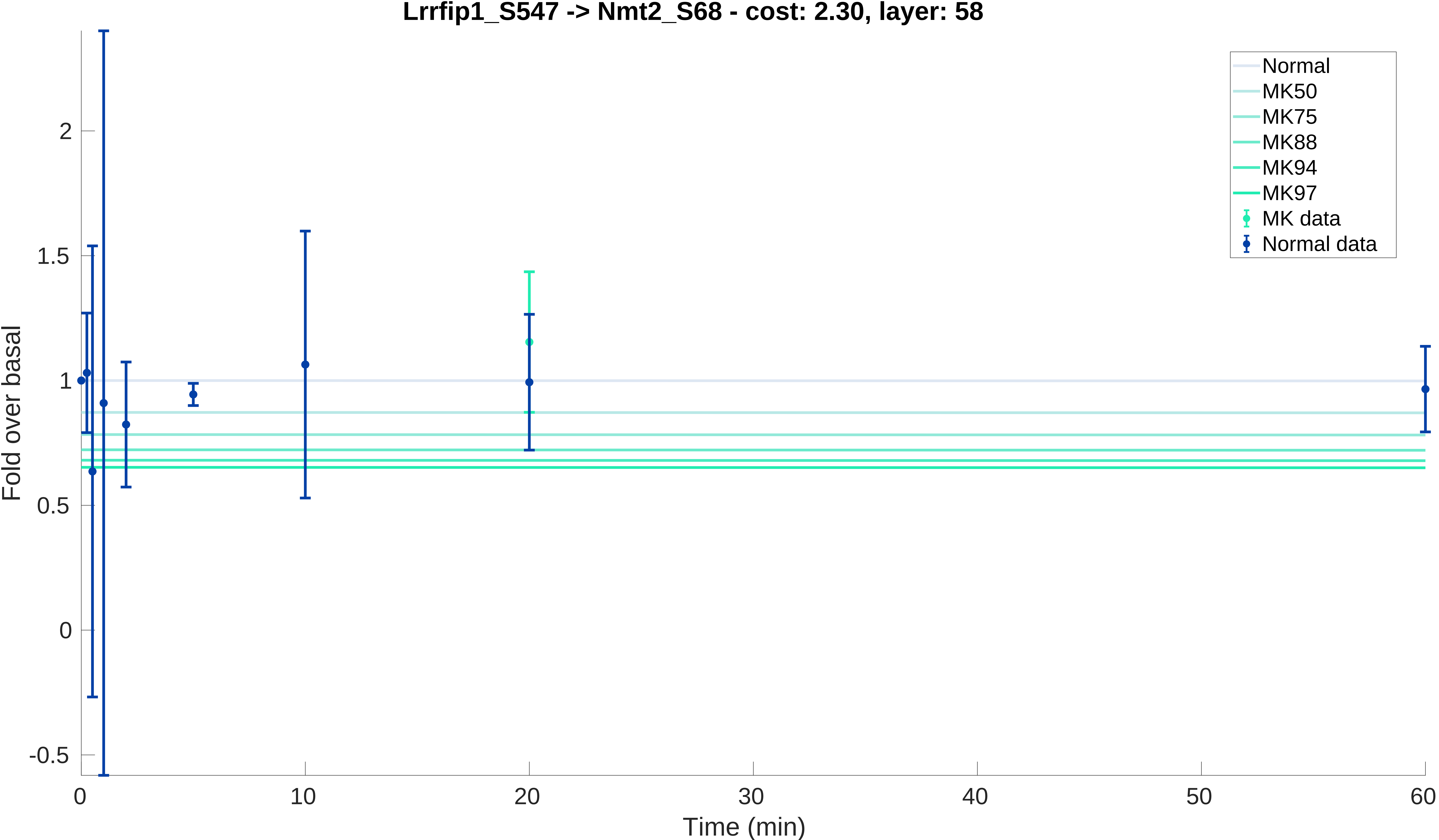

Lrrfip1_S547 -> Nmt2_S68 - cost: 2.30, layer: 58
Fold over basal
Time (min)
-0.5
0
0.5
1
1.5
2
0
10
20
30
40
50
60
Normal
MK50
MK75
MK88
MK94
MK97
MK data
Normal data

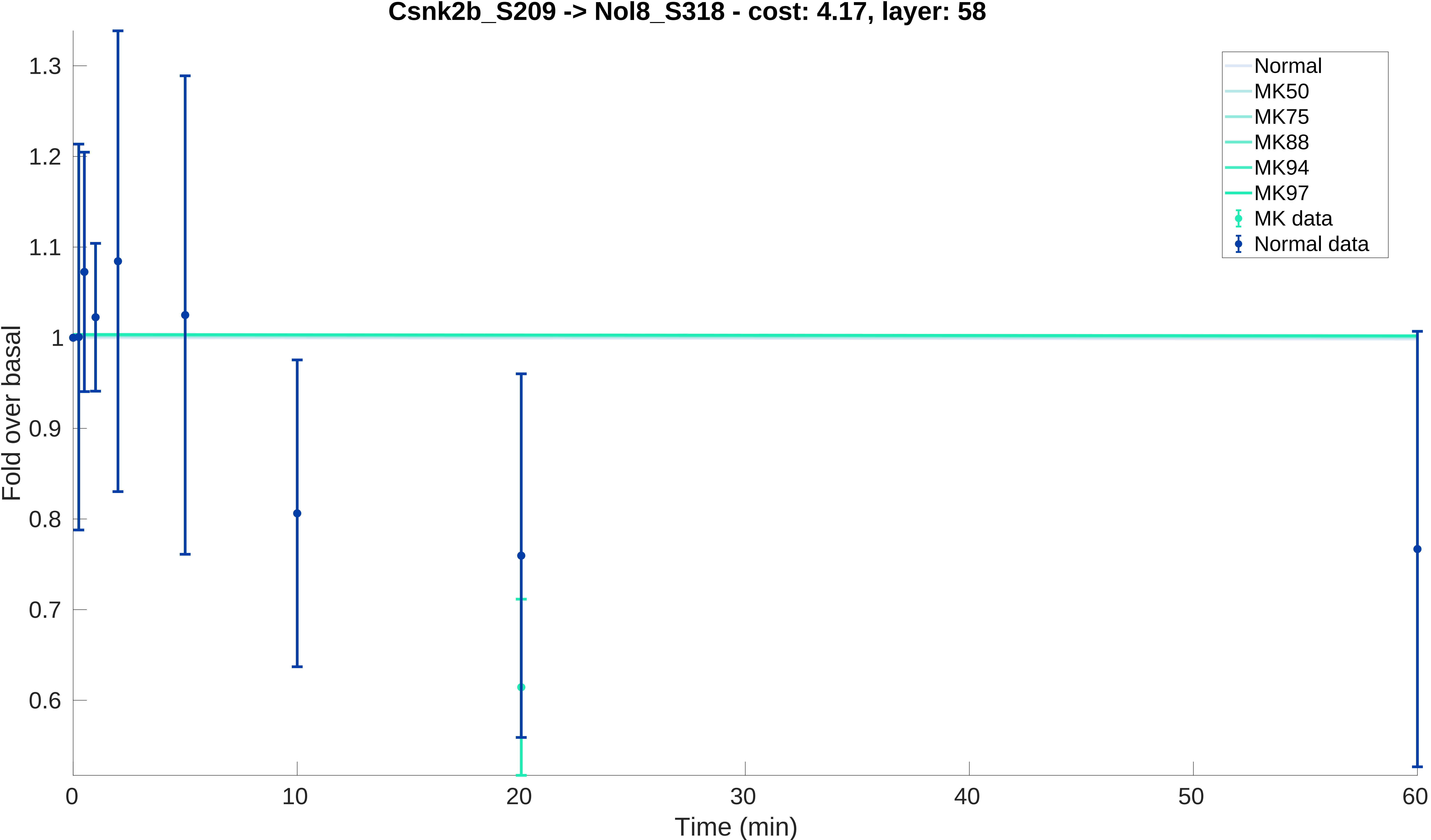
Csnk2b_S209 -> Nol8_S318 - cost: 4.17, layer: 58
Normal
MK50
MK75
MK88
MK94
MK97
MK data
Normal data
Fold over basal
Time (min)
0
10
20
30
40
50
60
0.6
0.7
0.8
0.9
1
1.1
1.2
1.3

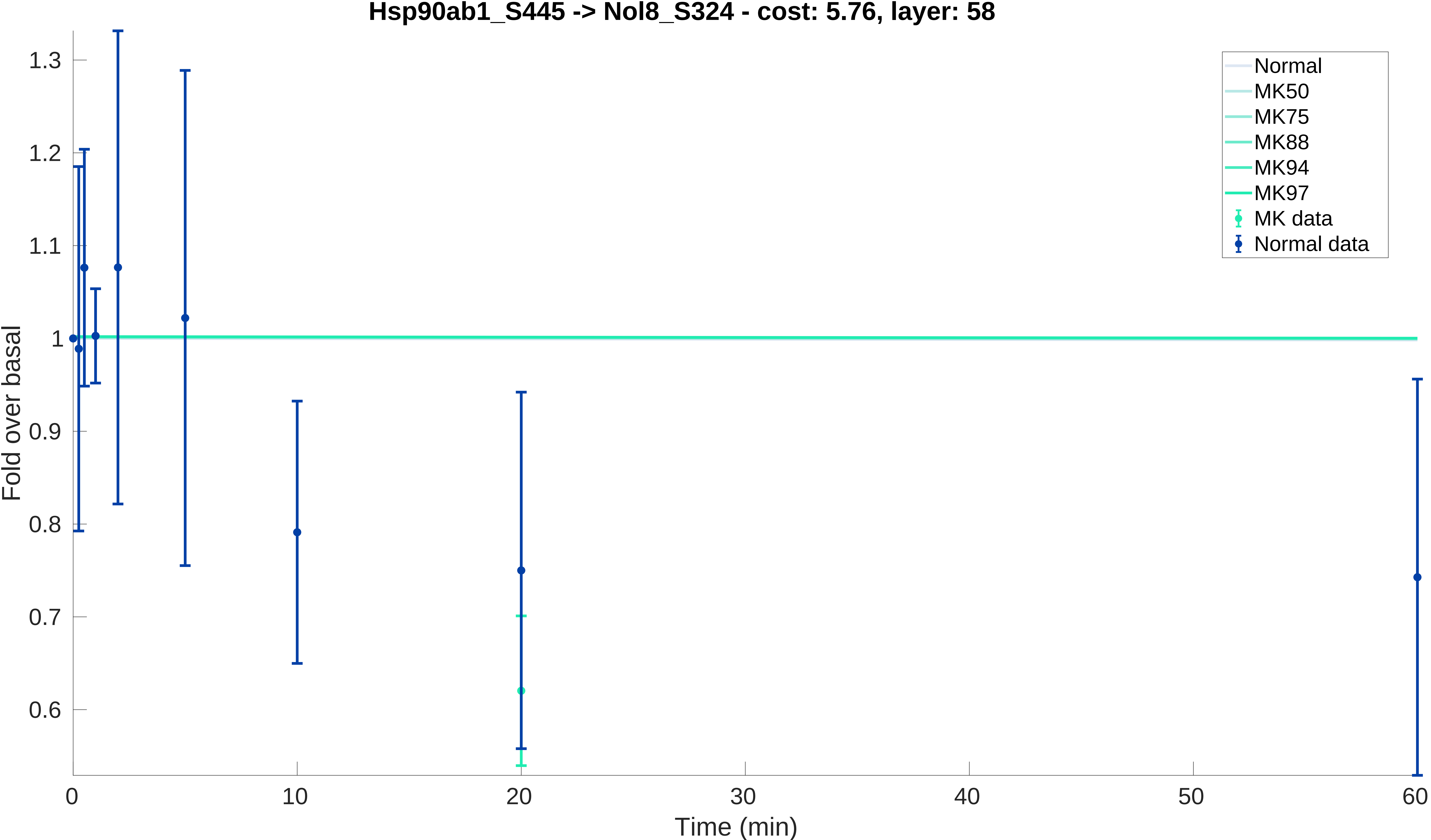
Hsp90ab1_S445 -> Nol8_S324 - cost: 5.76, layer: 58
Fold over basal
Time (min)
0.6
0.7
0.8
0.9
1
1.1
1.2
1.3
0
10
20
30
40
50
60
Normal
MK50
MK75
MK88
MK94
MK97
MK data
Normal data

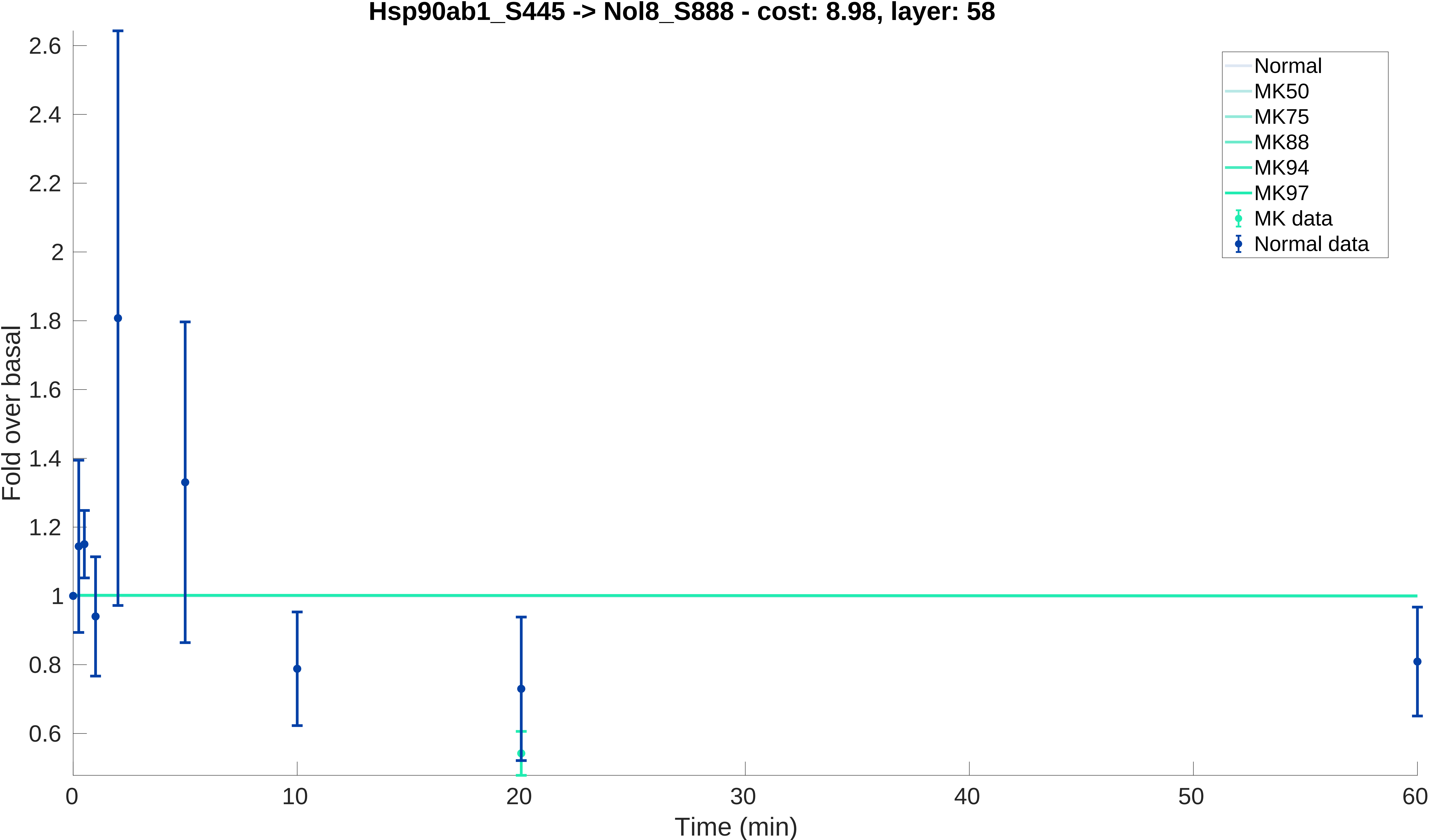

Hsp90ab1_S445 -> Nol8_S888 - cost: 8.98, layer: 58
Fold over basal
Time (min)
Normal
MK50
MK75
MK88
MK94
MK97
MK data
Normal data
2.6
2.4
2.2
2
1.8
1.6
1.4
1.2
1
0.8
0.6
0
10
20
30
40
50
60

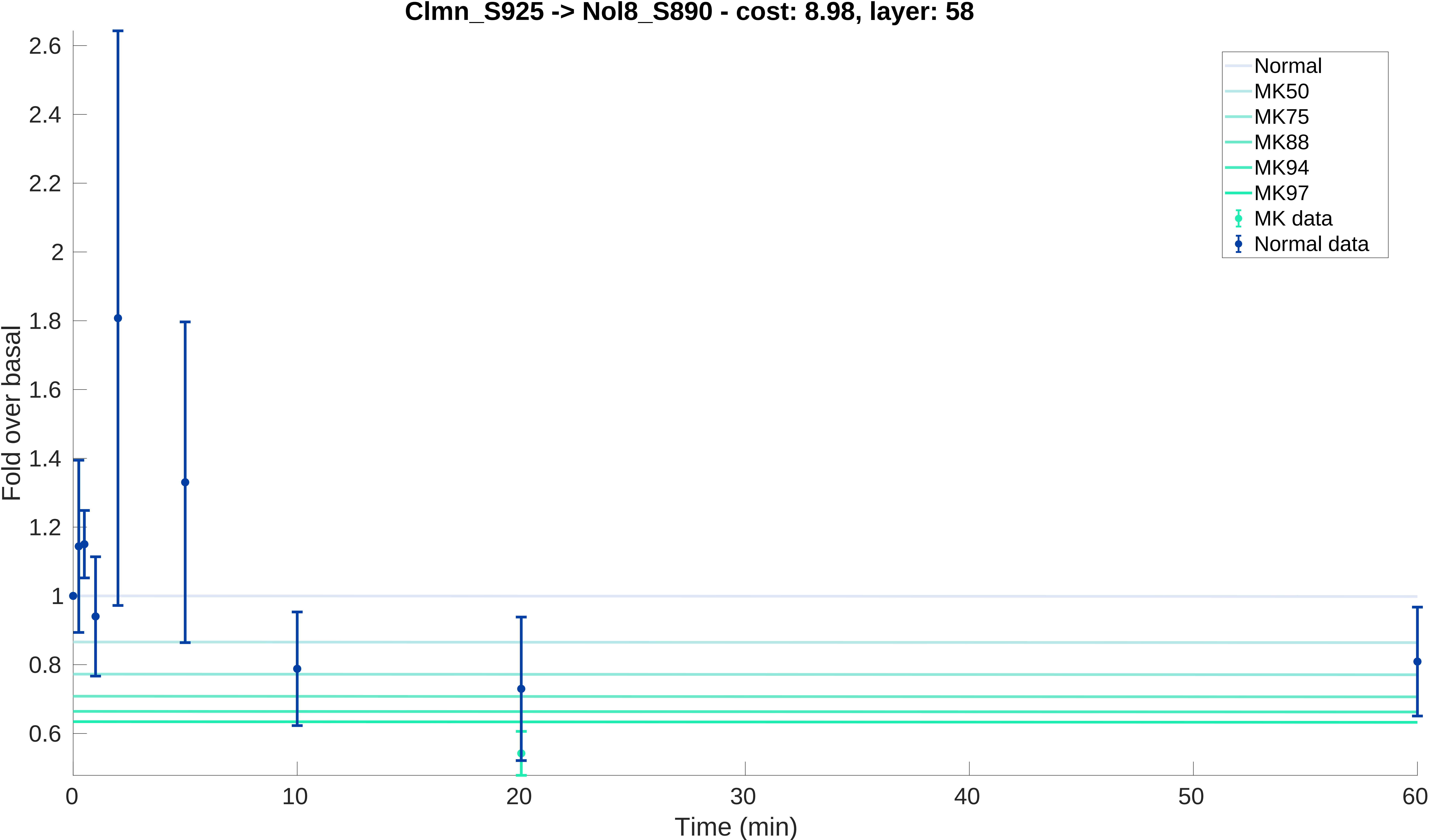

Clmn_S925 -> Nol8_S890 - cost: 8.98, layer: 58
Fold over basal
Time (min)
2.6
2.4
2.2
2
1.8
1.6
1.4
1.2
1
0.8
0.6
0
10
20
30
40
50
60
Normal
MK50
MK75
MK88
MK94
MK97
MK data
Normal data

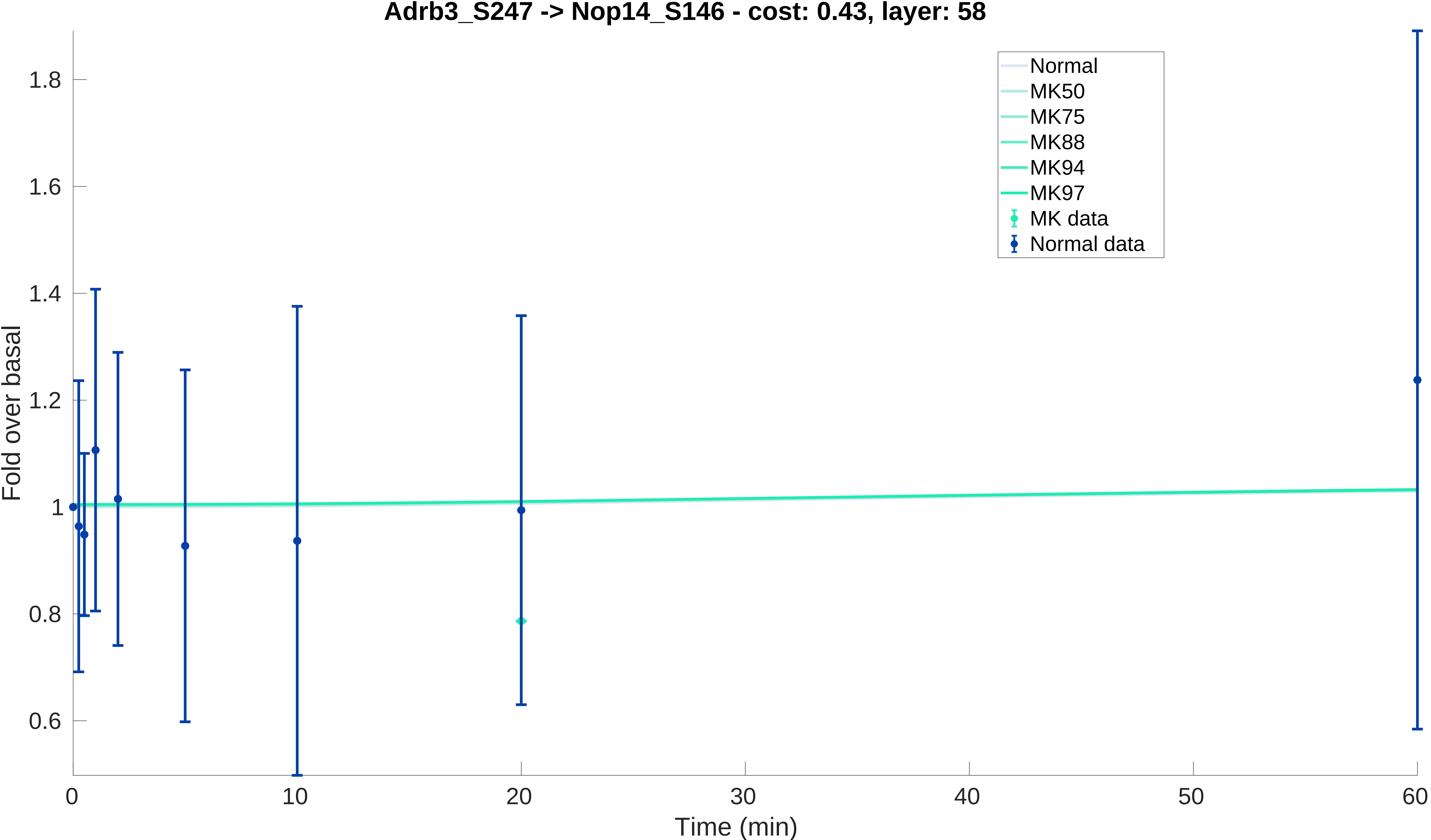

Adrb3_S247 -> Nop14_S146 - cost: 0.43, layer: 58
Normal
MK50
MK75
MK88
MK94
MK97
MK data
Normal data
Fold over basal
Time (min)
1.8
1.6
1.4
1.2
1
0.8
0.6
0
10
20
30
40
50
60

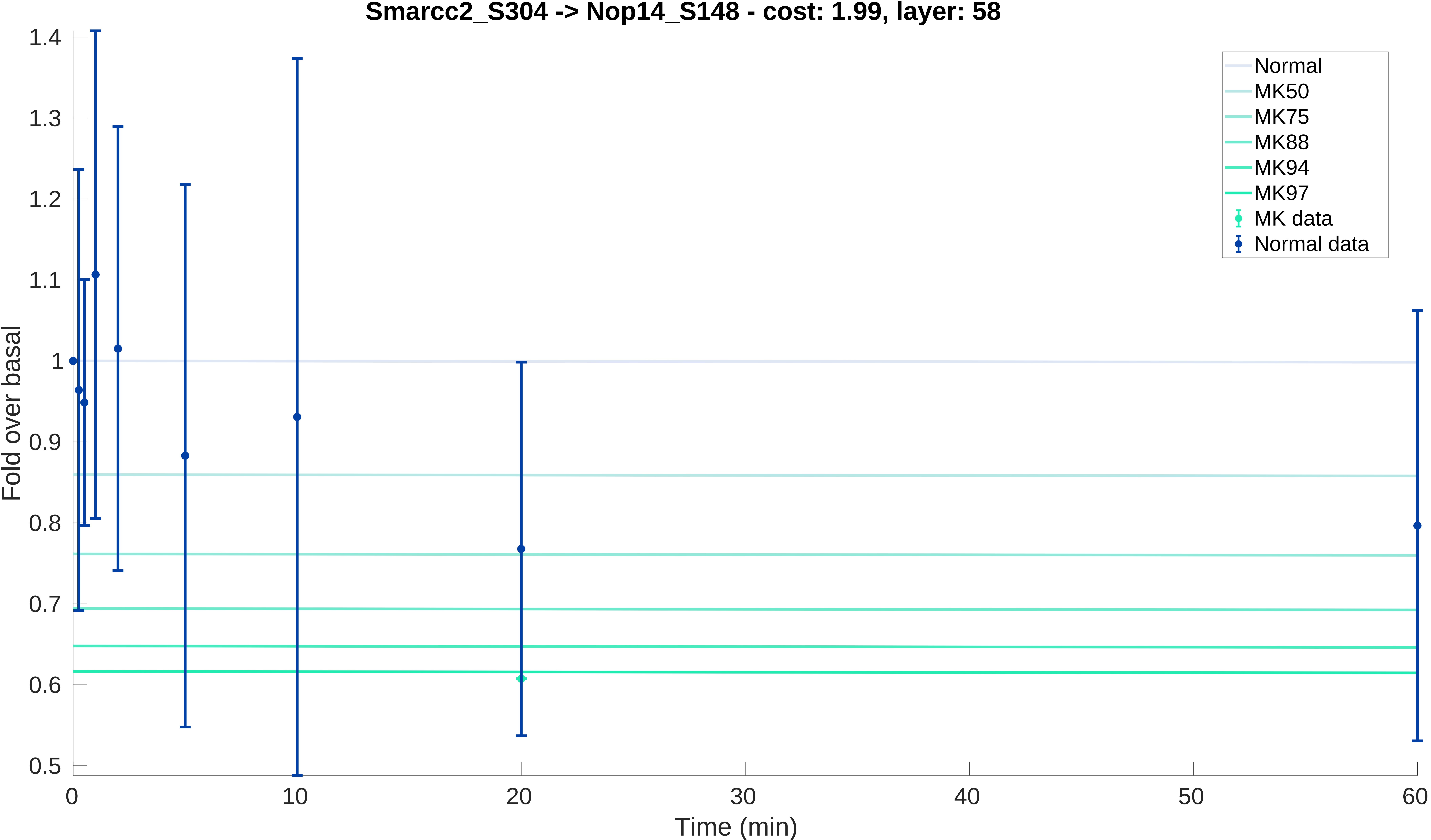

Smarcc2_S304 -> Nop14_S148 - cost: 1.99, layer: 58
Fold over basal
Time (min)
1.4
1.3
1.2
1.1
1
0.9
0.8
0.7
0.6
0.5
0
10
20
30
40
50
60
Normal
MK50
MK75
MK88
MK94
MK97
MK data
Normal data

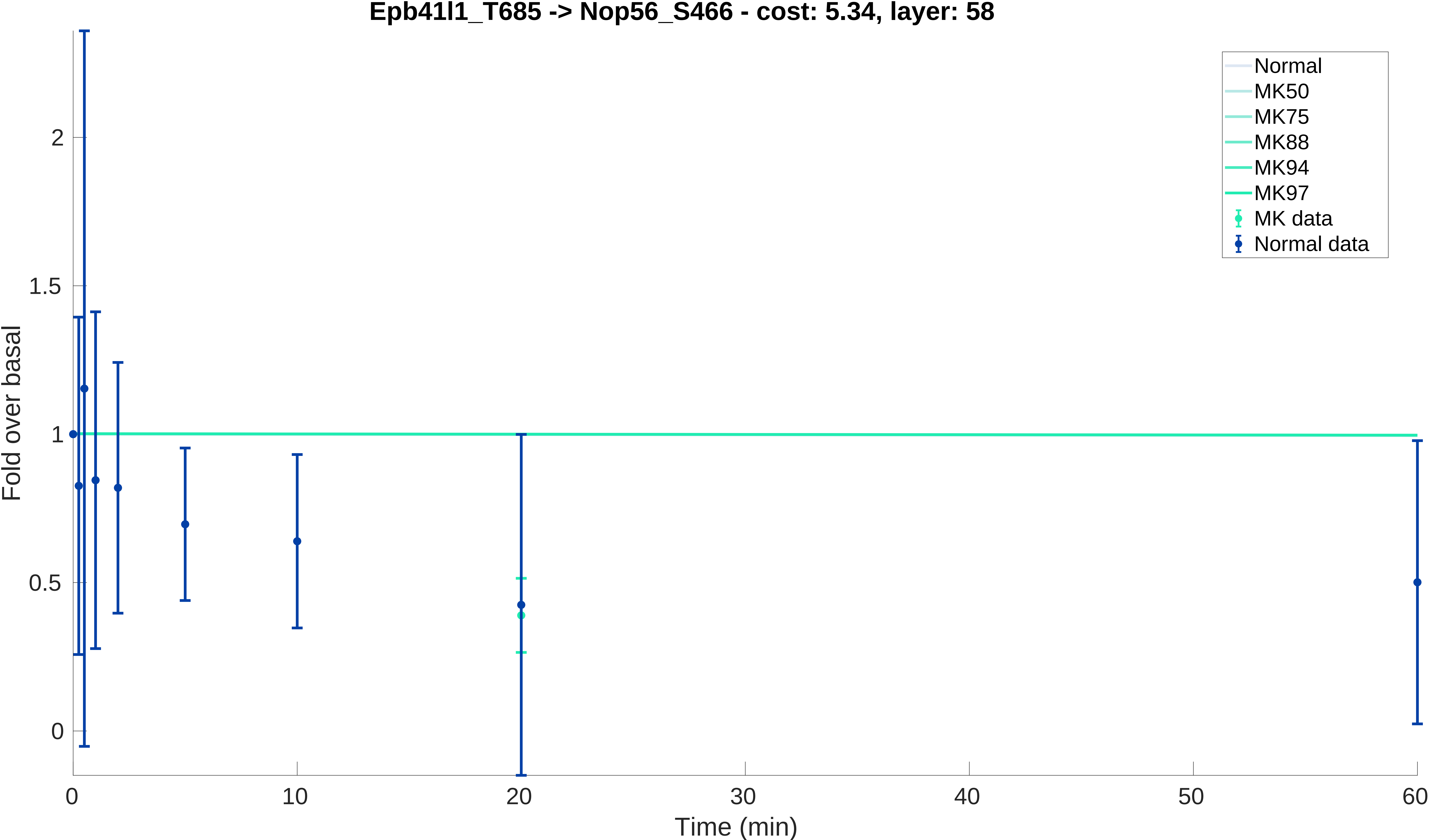

Epb41l1_T685 -> Nop56_S466 - cost: 5.34, layer: 58
Normal
MK50
MK75
MK88
MK94
MK97
MK data
Normal data
Fold over basal
Time (min)
0
0.5
1
1.5
2
0
10
20
30
40
50
60

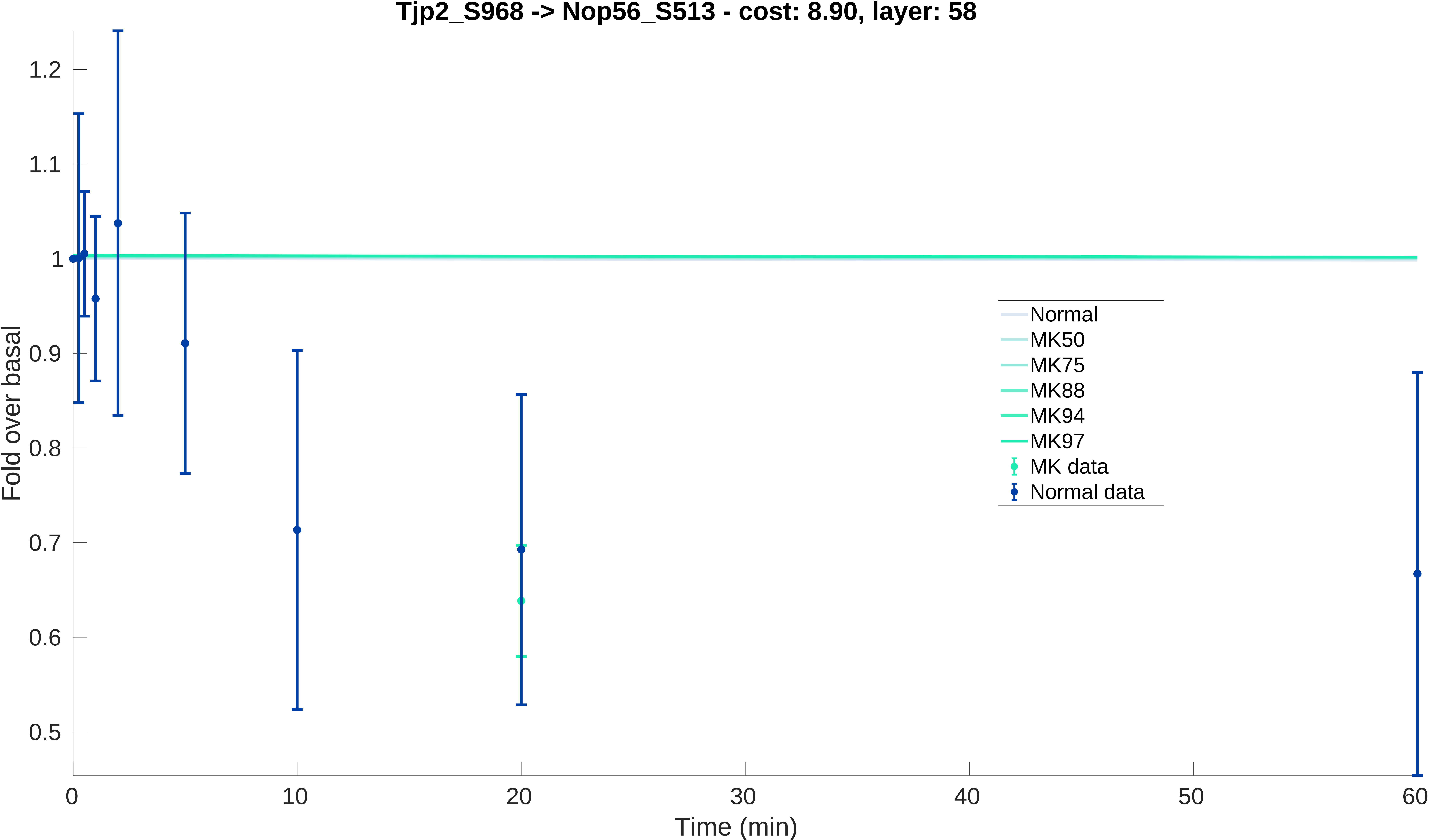

Tjp2_S968 -> Nop56_S513 - cost: 8.90, layer: 58
Fold over basal
Time (min)
1.2
1.1
1
0.9
0.8
0.7
0.6
0.5
0
10
20
30
40
50
60
Normal
MK50
MK75
MK88
MK94
MK97
MK data
Normal data

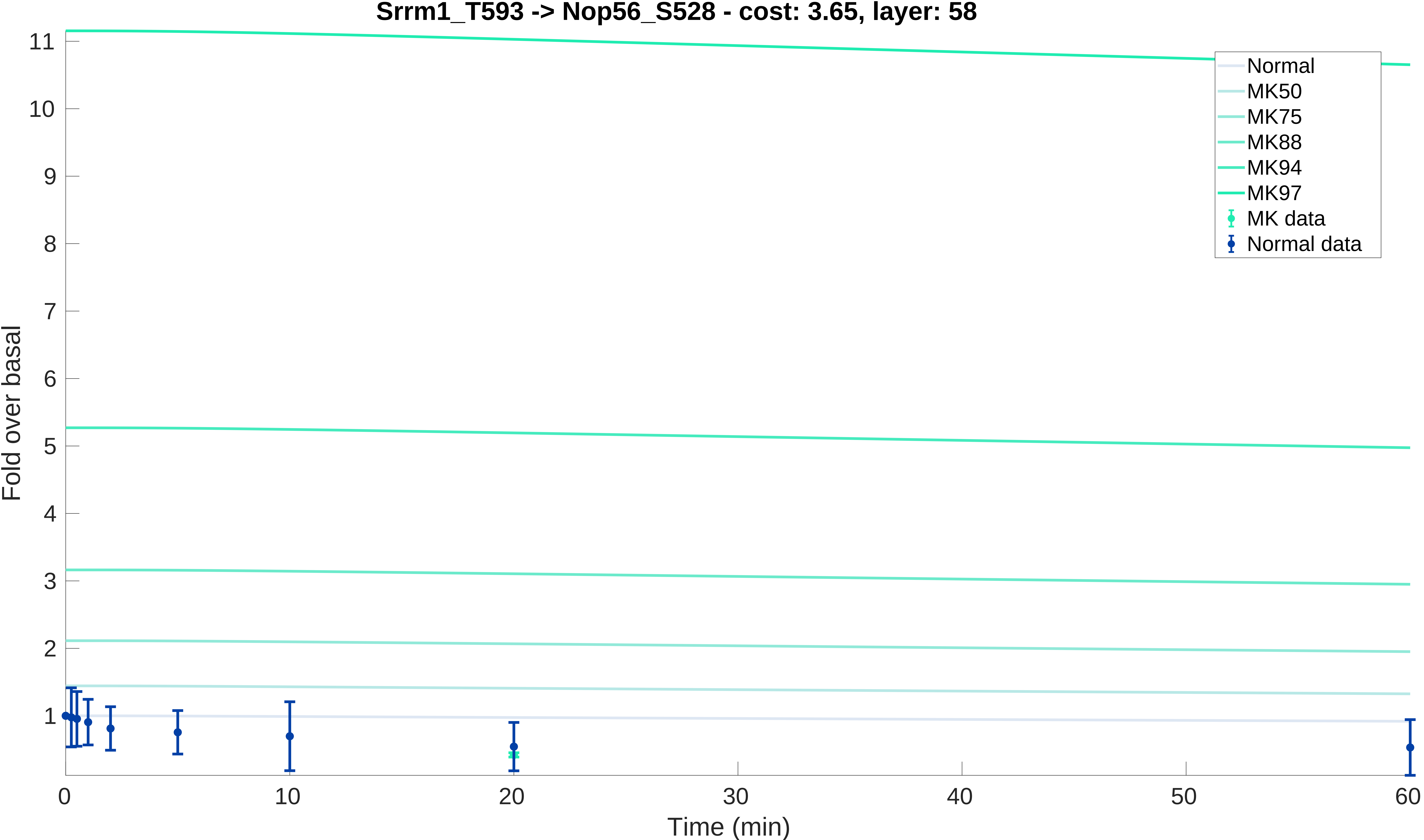

Srrm1_T593 -> Nop56_S528 - cost: 3.65, layer: 58
Fold over basal
Time (min)
Normal
MK50
MK75
MK88
MK94
MK97
MK data
Normal data
11
10
9
8
7
6
5
4
3
2
1
0
10
20
30
40
50
60

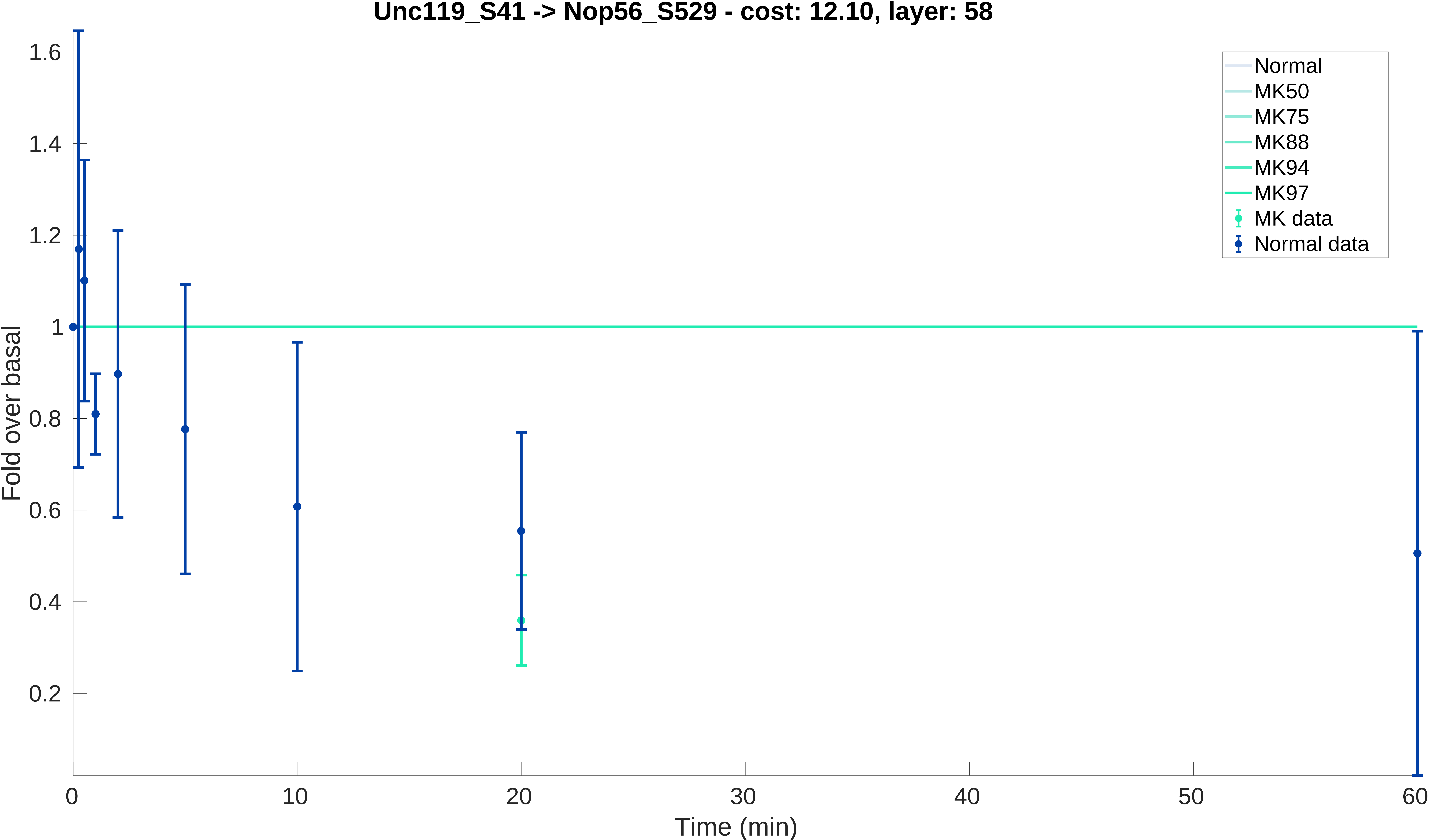

Unc119_S41 -> Nop56_S529 - cost: 12.10, layer: 58
Fold over basal
1.6
1.4
1.2
1
0.8
0.6
0.4
0.2
0
10
20
30
40
50
60
Time (min)
Normal
MK50
MK75
MK88
MK94
MK97
MK data
Normal data

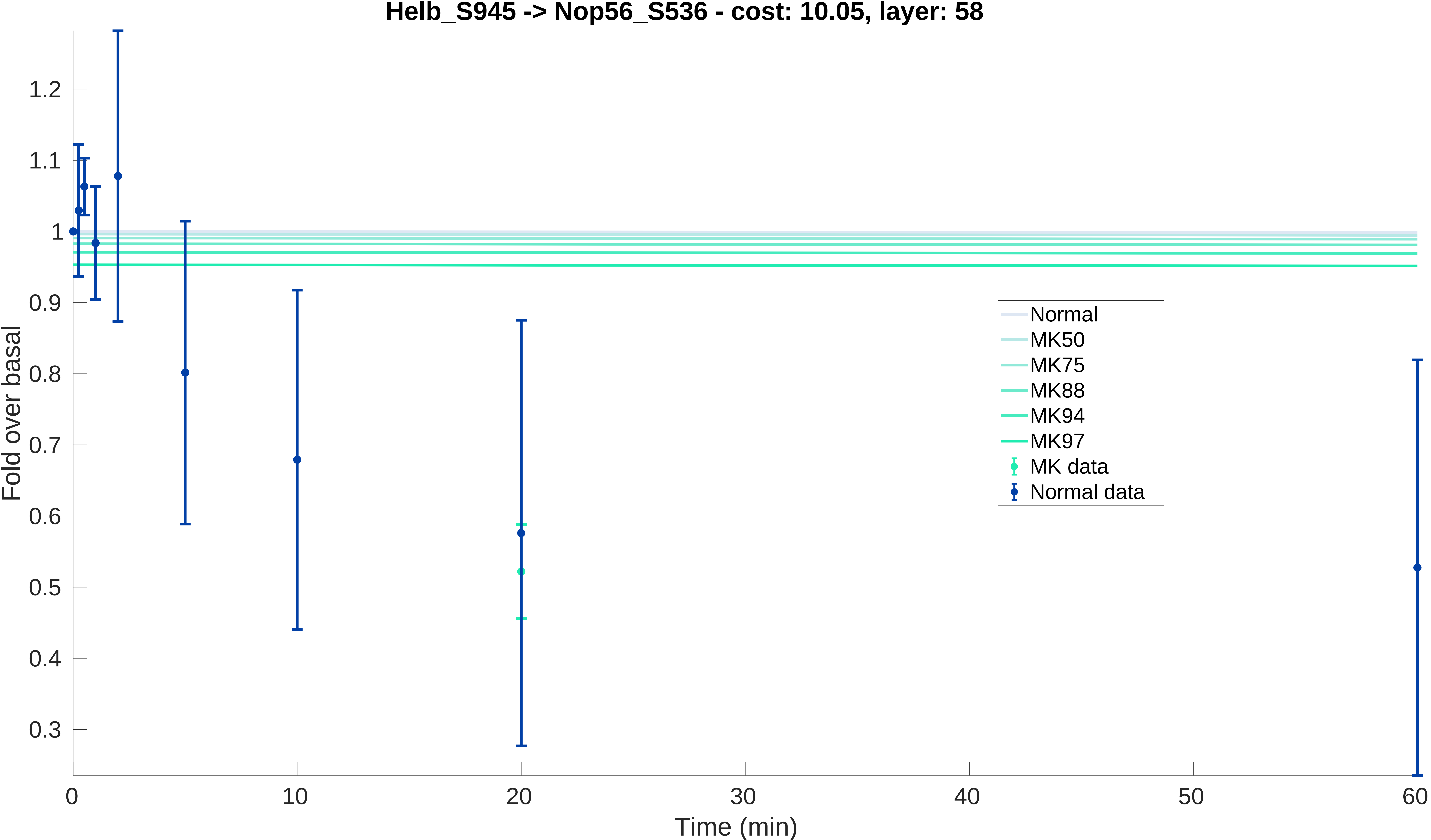

Helb_S945 -> Nop56_S536 - cost: 10.05, layer: 58
Fold over basal
1.2
1.1
1
0.9
0.8
0.7
0.6
0.5
0.4
0.3
0
10
20
30
40
50
60
Time (min)
Normal
MK50
MK75
MK88
MK94
MK97
MK data
Normal data

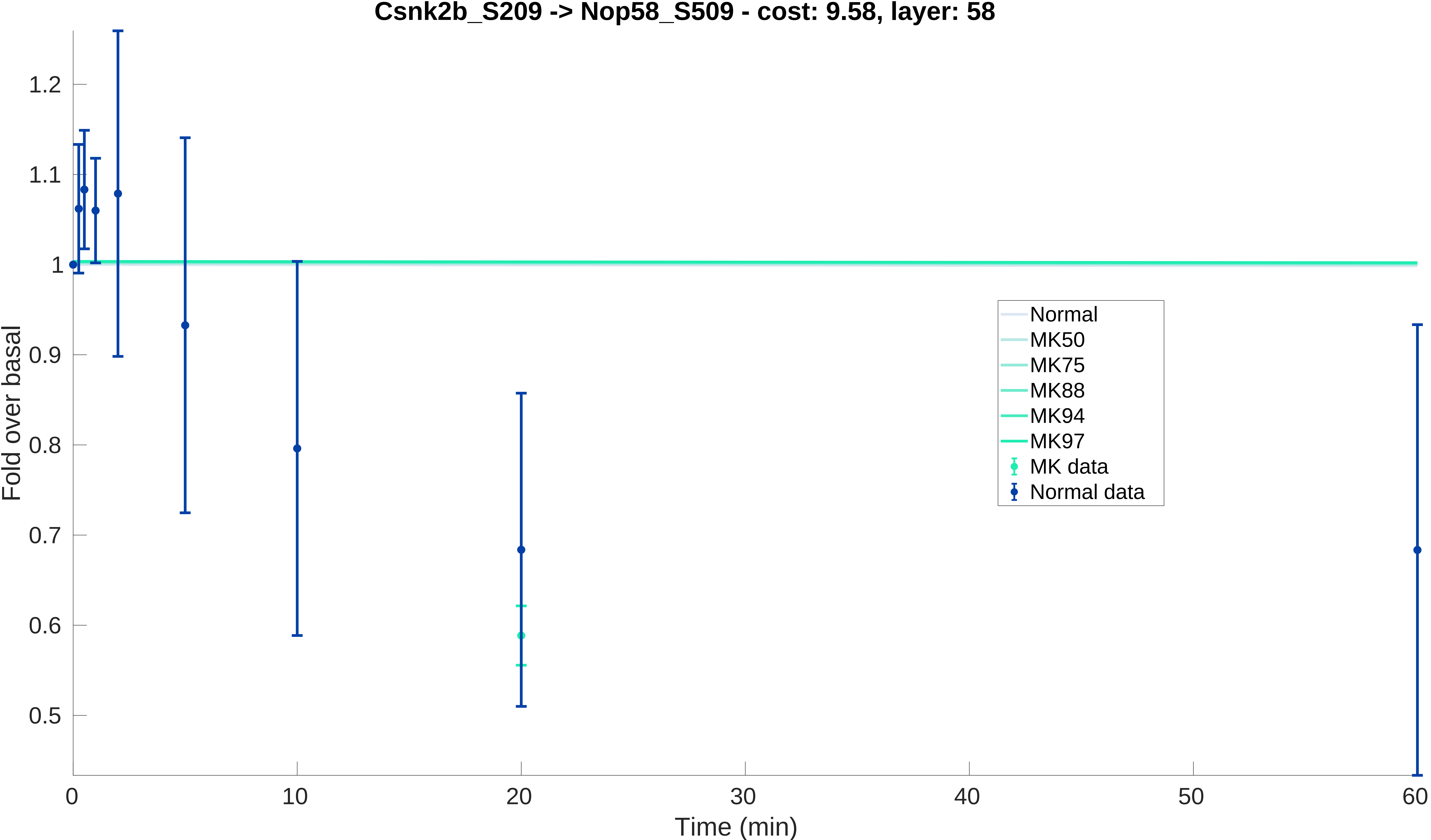

Csnk2b_S209 -> Nop58_S509 - cost: 9.58, layer: 58
Fold over basal
1.2
1.1
1
0.9
0.8
0.7
0.6
0.5
0
10
20
30
40
50
60
Time (min)
Normal
MK50
MK75
MK88
MK94
MK97
MK data
Normal data

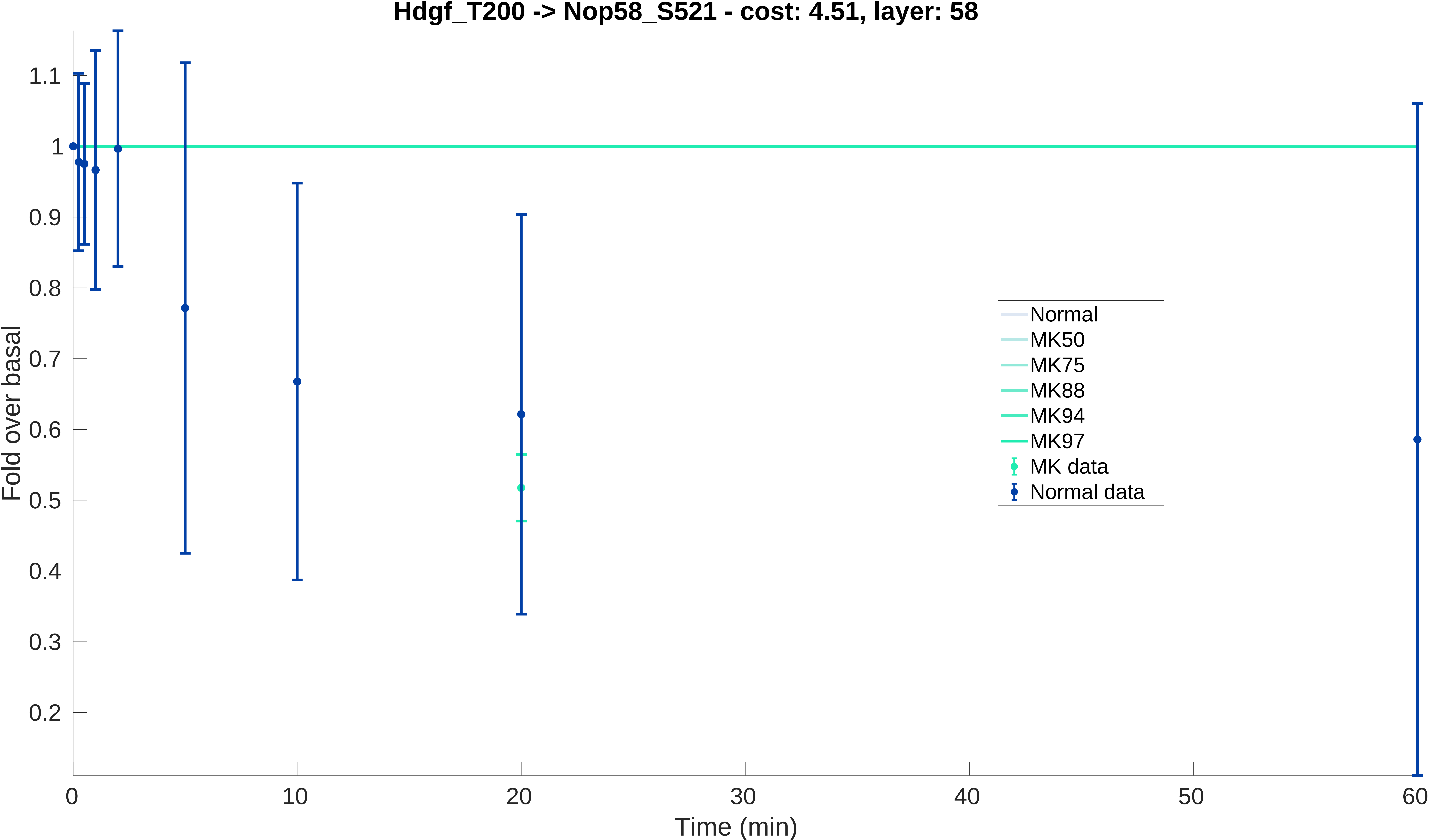
Hdgf_T200 -> Nop58_S521 - cost: 4.51, layer: 58
Fold over basal
Time (min)
0
10
20
30
40
50
60
0.2
0.3
0.4
0.5
0.6
0.7
0.8
0.9
1
1.1
Normal
MK50
MK75
MK88
MK94
MK97
MK data
Normal data

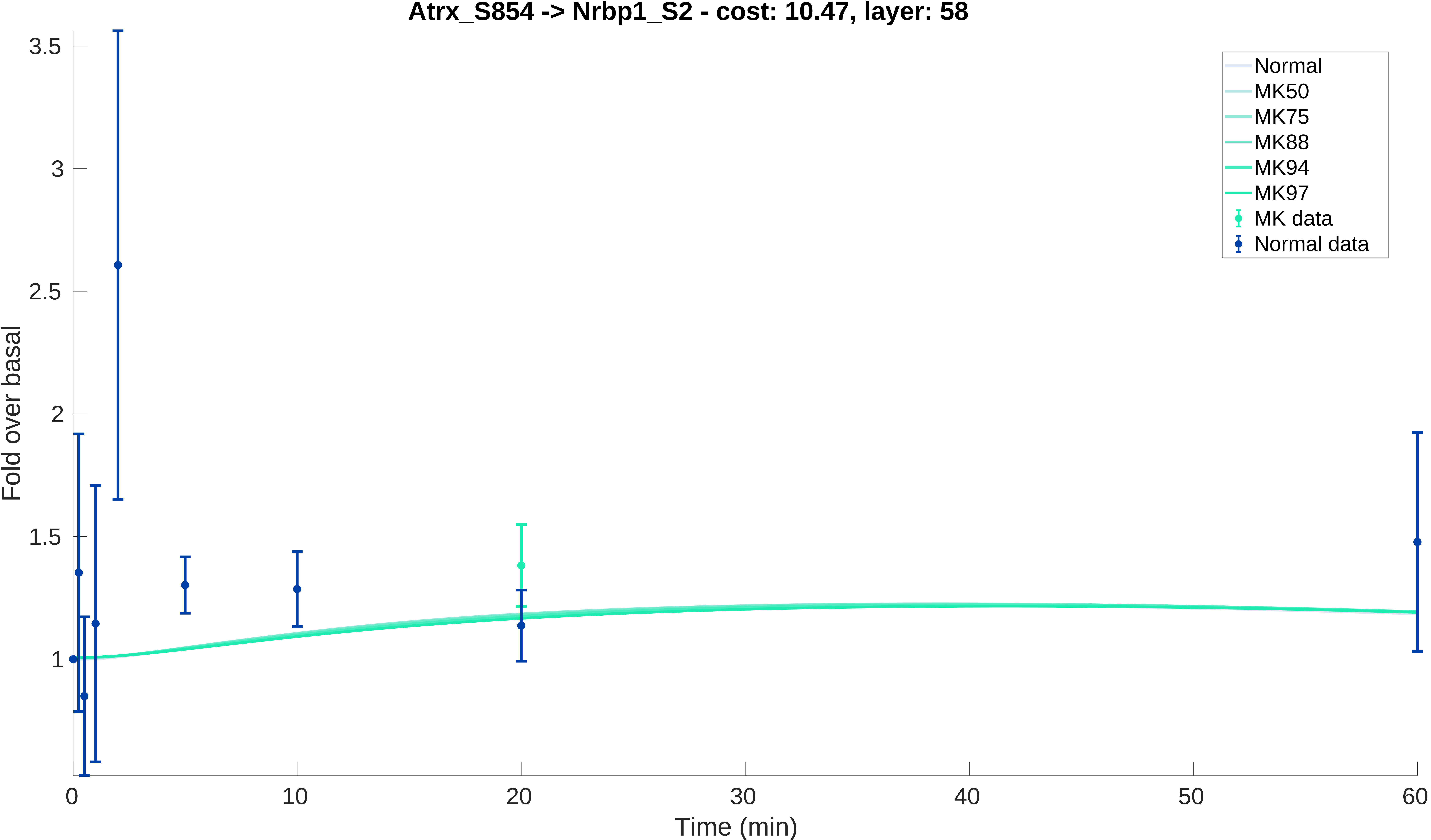

Atrx_S854 -> Nrbp1_S2 - cost: 10.47, layer: 58
Fold over basal
Time (min)
0
10
20
30
40
50
60
1
1.5
2
2.5
3
3.5
Normal
MK50
MK75
MK88
MK94
MK97
MK data
Normal data

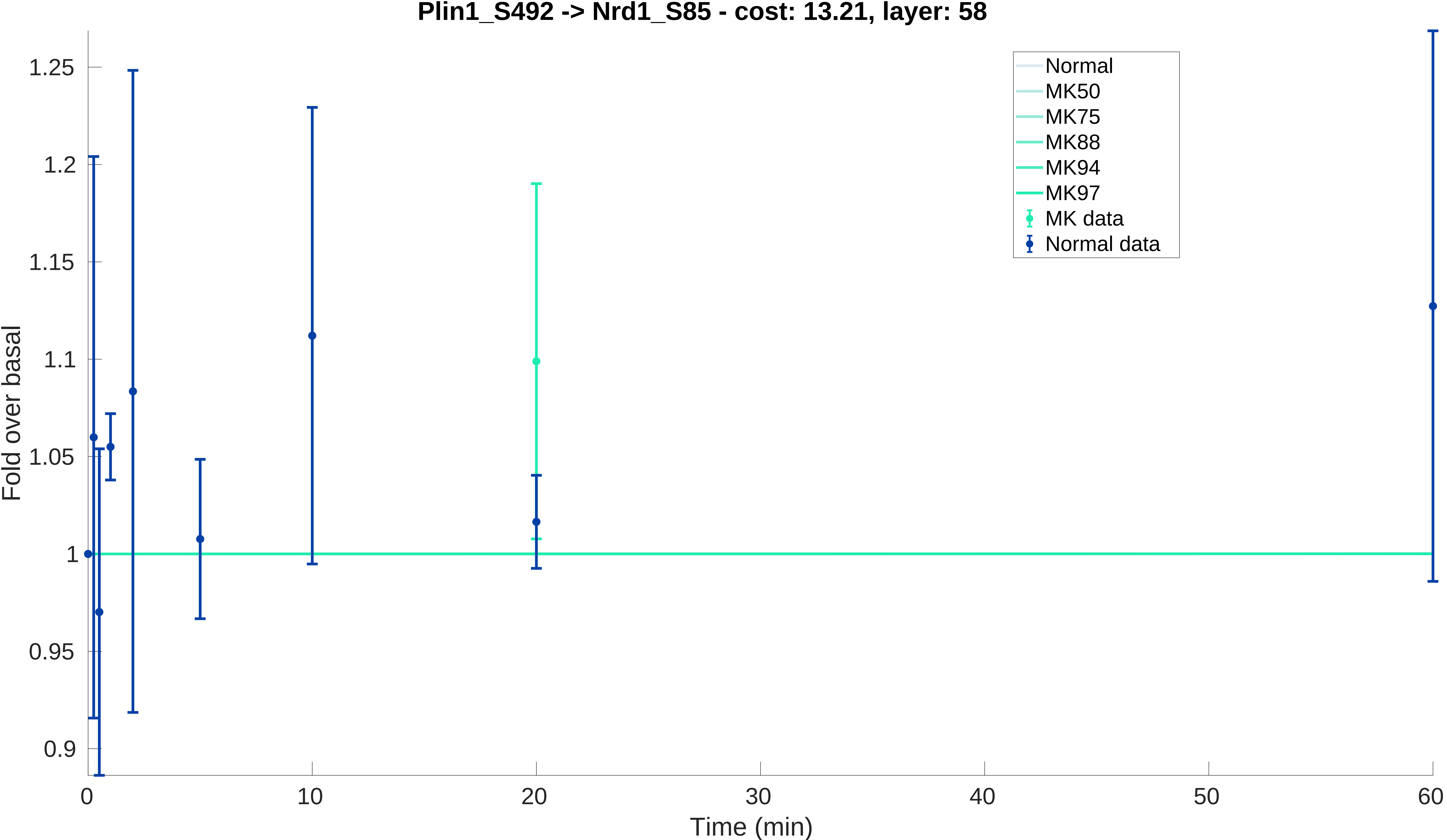

Plin1_S492 -> Nrd1_S85 - cost: 13.21, layer: 58
Normal
MK50
MK75
MK88
MK94
MK97
MK data
Normal data
Fold over basal
1.25
1.2
1.15
1.1
1.05
1
0.95
0.9
0
10
20
30
40
50
60
Time (min)

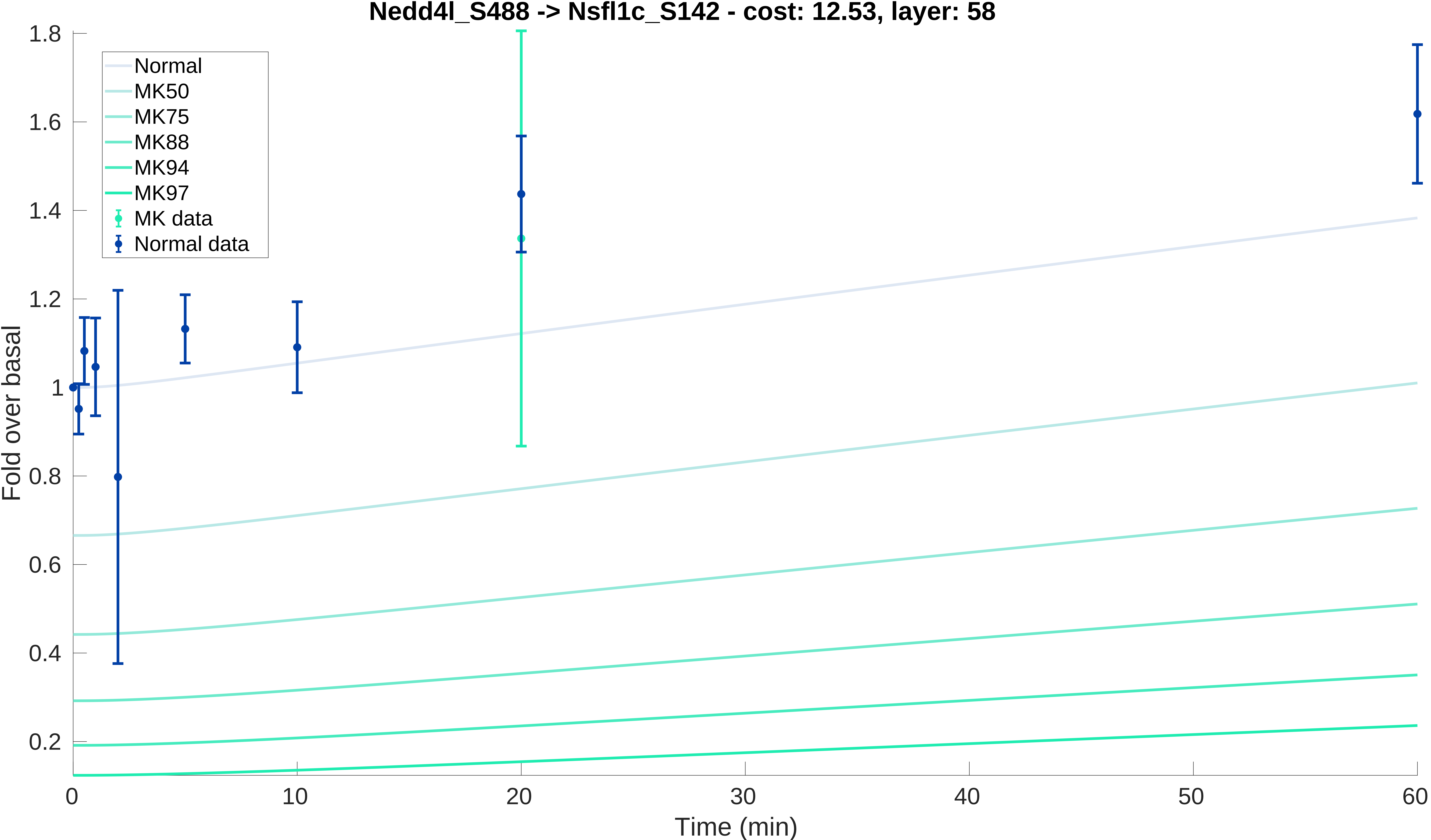

Nedd4l_S488 -> Nsfl1c_S142 - cost: 12.53, layer: 58
Fold over basal
Time (min)
Normal
MK50
MK75
MK88
MK94
MK97
MK data
Normal data
1.8
1.6
1.4
1.2
1
0.8
0.6
0.4
0.2
0
10
20
30
40
50
60

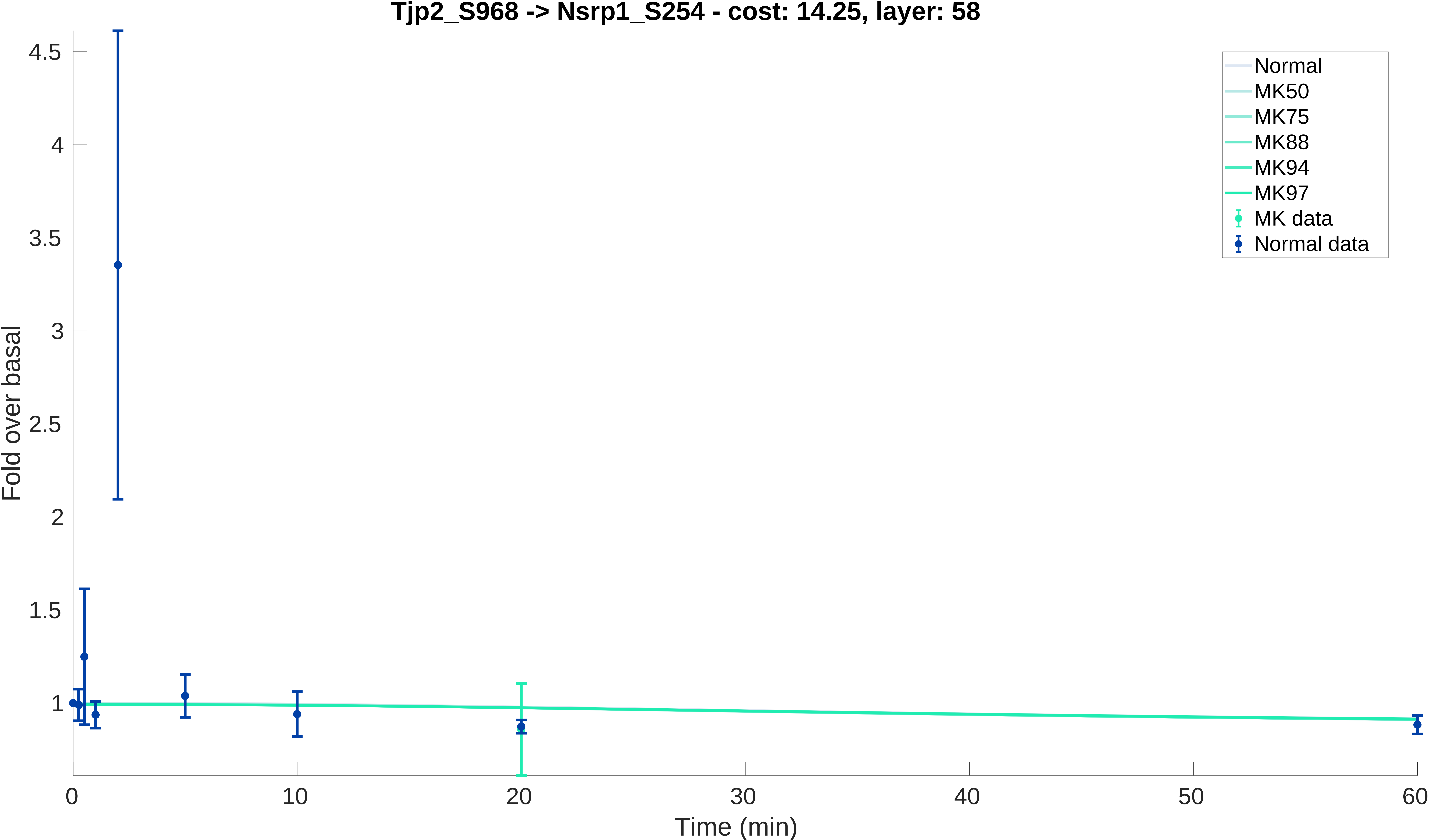

Tjp2_S968 -> Nsrp1_S254 - cost: 14.25, layer: 58
Fold over basal
Time (min)
4.5
4
3.5
3
2.5
2
1.5
1
0
10
20
30
40
50
60
Normal
MK50
MK75
MK88
MK94
MK97
MK data
Normal data

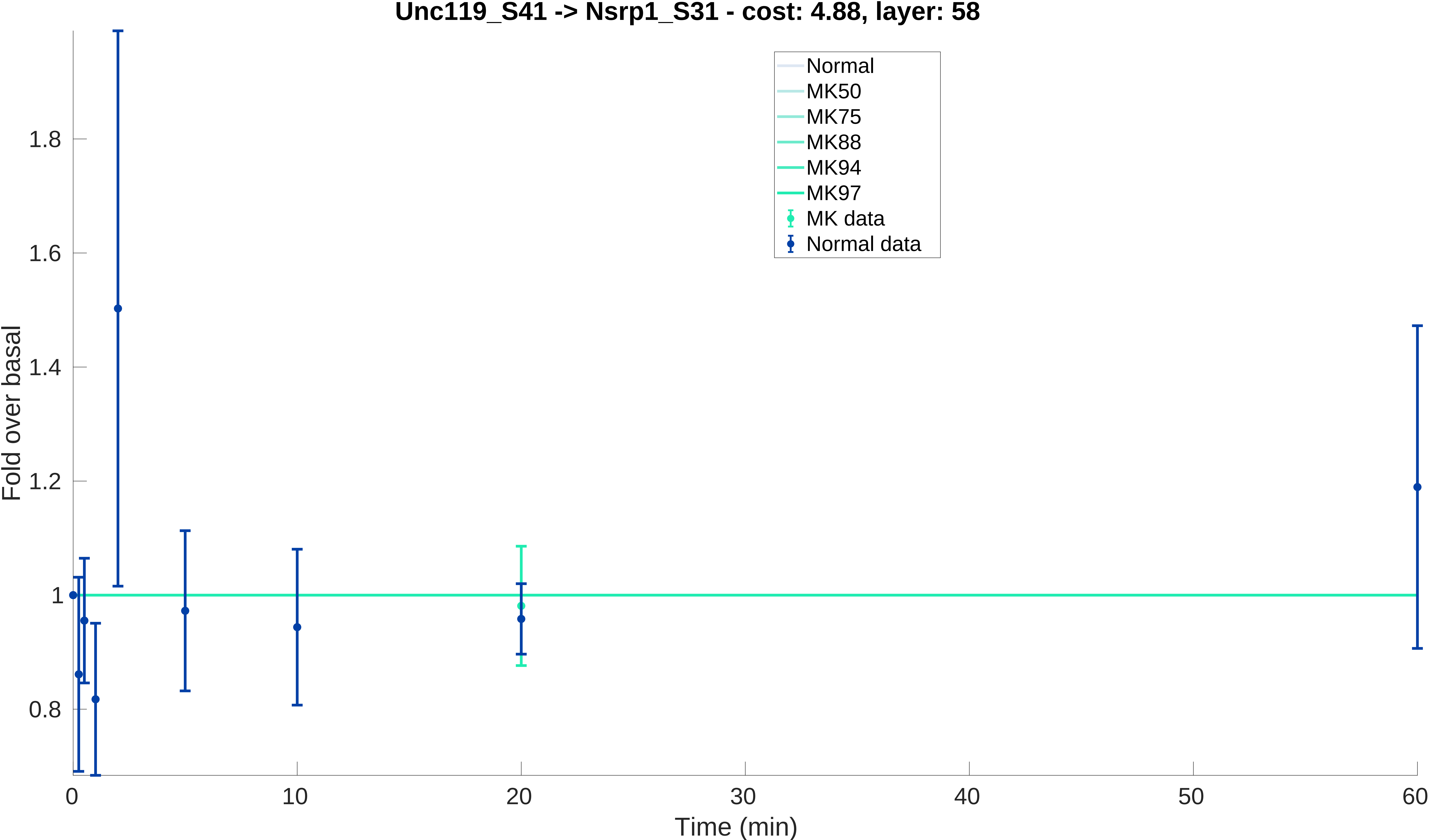

Unc119_S41 -> Nsrp1_S31 - cost: 4.88, layer: 58
Normal
MK50
MK75
MK88
MK94
MK97
MK data
Normal data
Fold over basal
0.8
1
1.2
1.4
1.6
1.8
0
10
20
30
40
50
60
Time (min)

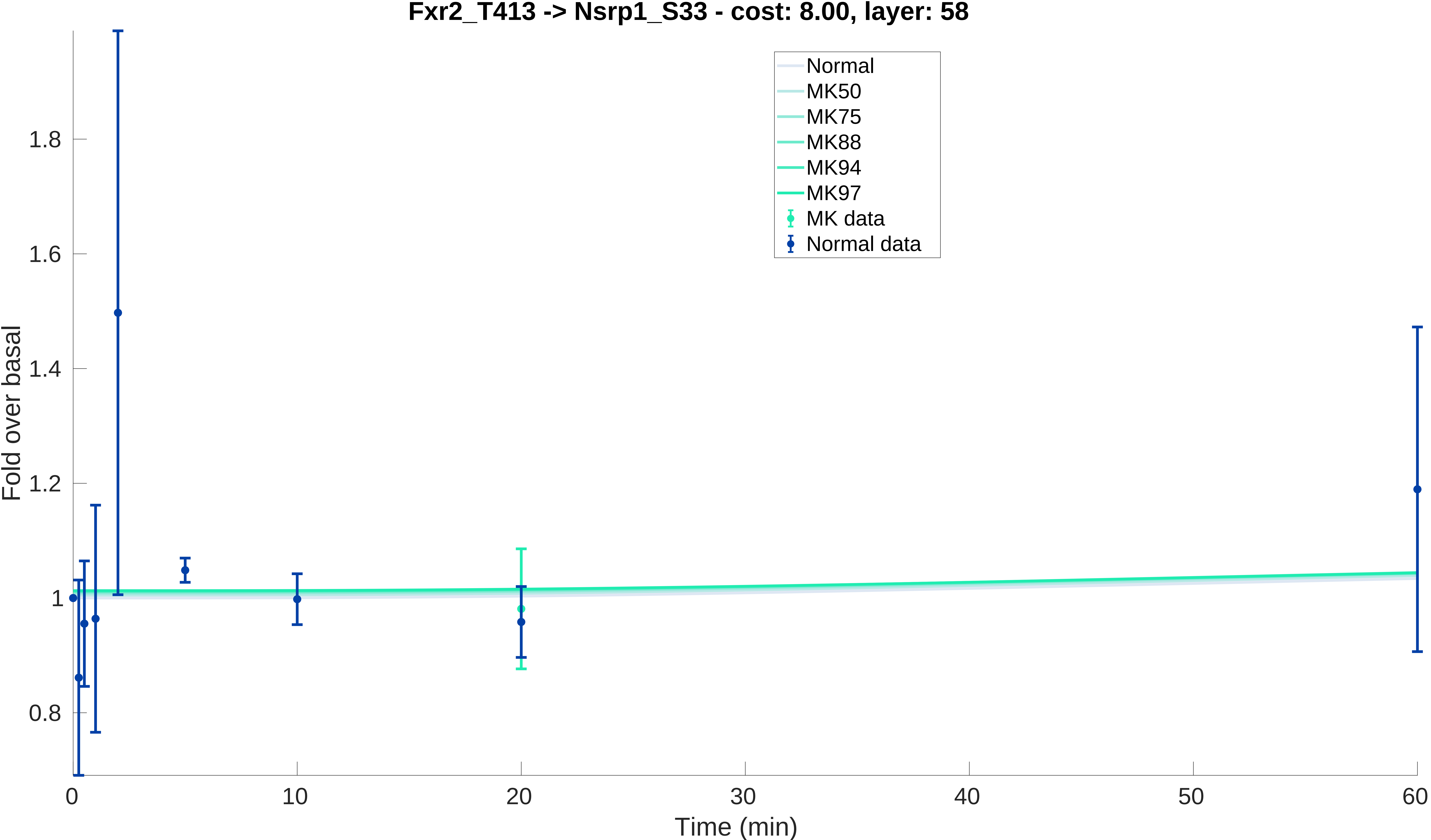

Fxr2_T413 -> Nsrp1_S33 - cost: 8.00, layer: 58
Normal
MK50
MK75
MK88
MK94
MK97
MK data
Normal data
Fold over basal
1.8
1.6
1.4
1.2
1
0.8
0
10
20
30
40
50
60
Time (min)

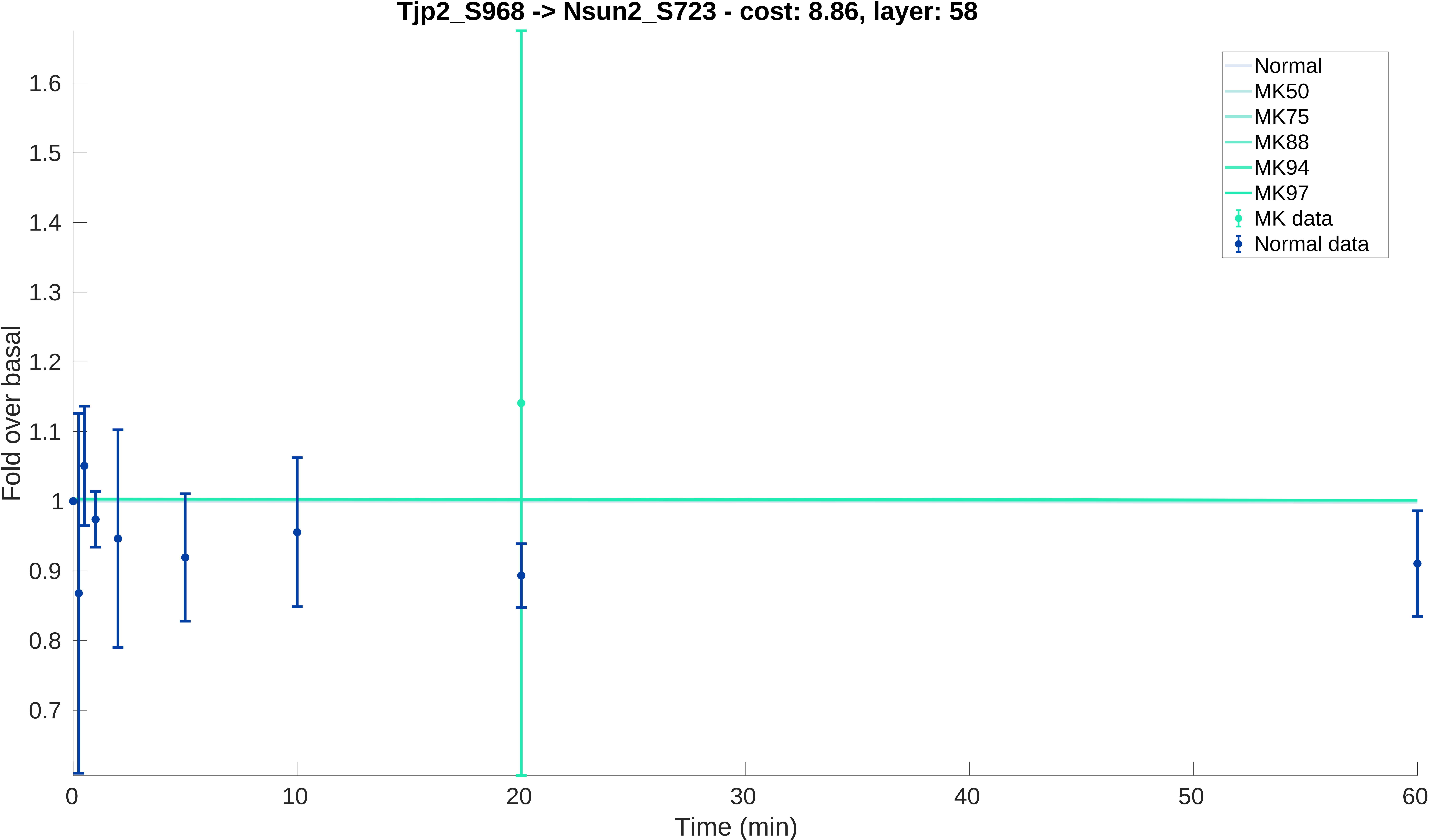

Tjp2_S968 -> Nsun2_S723 - cost: 8.86, layer: 58
Fold over basal
Time (min)
1.6
1.5
1.4
1.3
1.2
1.1
1
0.9
0.8
0.7
0
10
20
30
40
50
60
Normal
MK50
MK75
MK88
MK94
MK97
MK data
Normal data

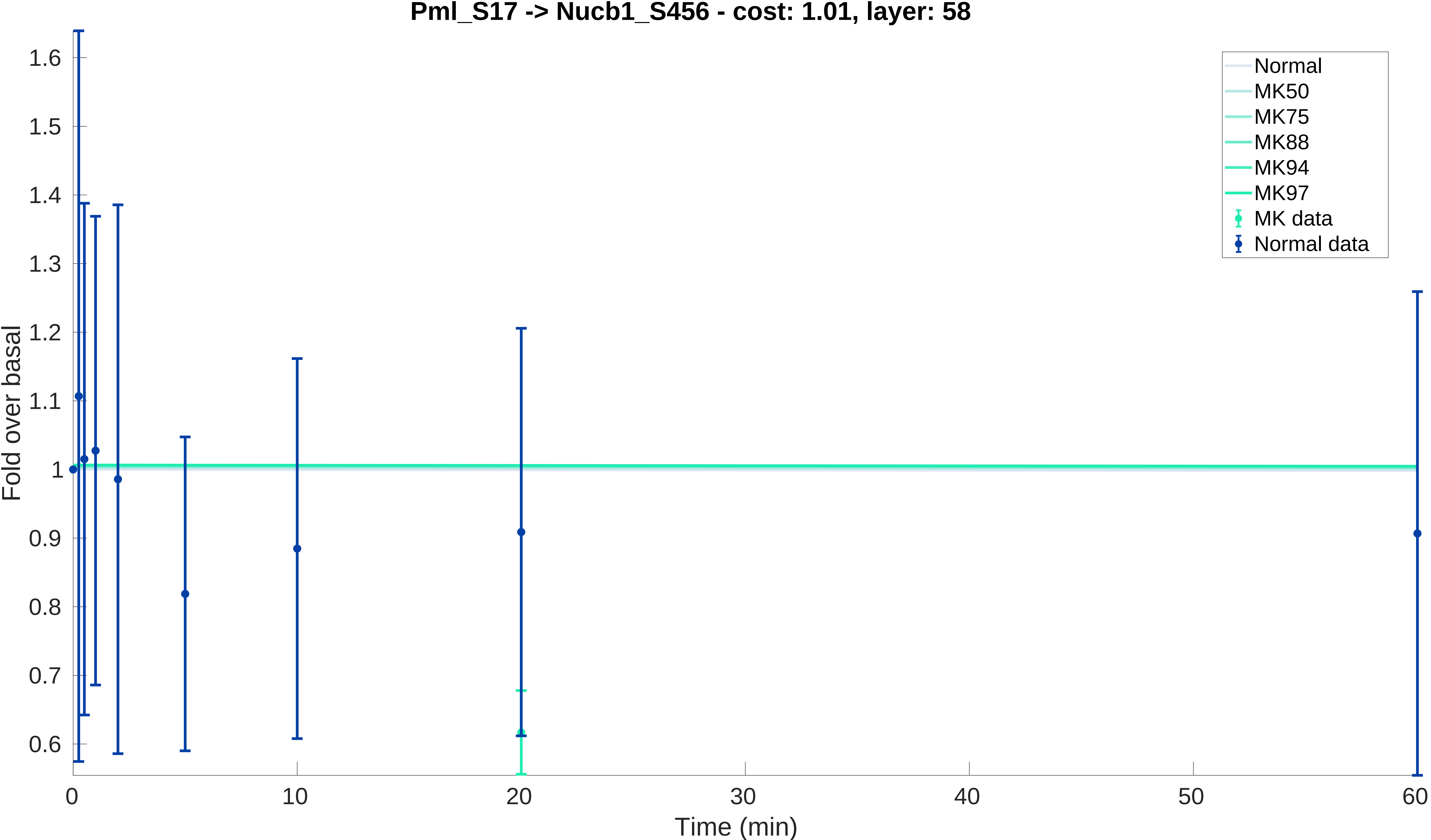

Pml_S17 -> Nucb1_S456 - cost: 1.01, layer: 58
Fold over basal
1.6
1.5
1.4
1.3
1.2
1.1
1
0.9
0.8
0.7
0.6
0
10
20
30
40
50
60
Time (min)
Normal
MK50
MK75
MK88
MK94
MK97
MK data
Normal data

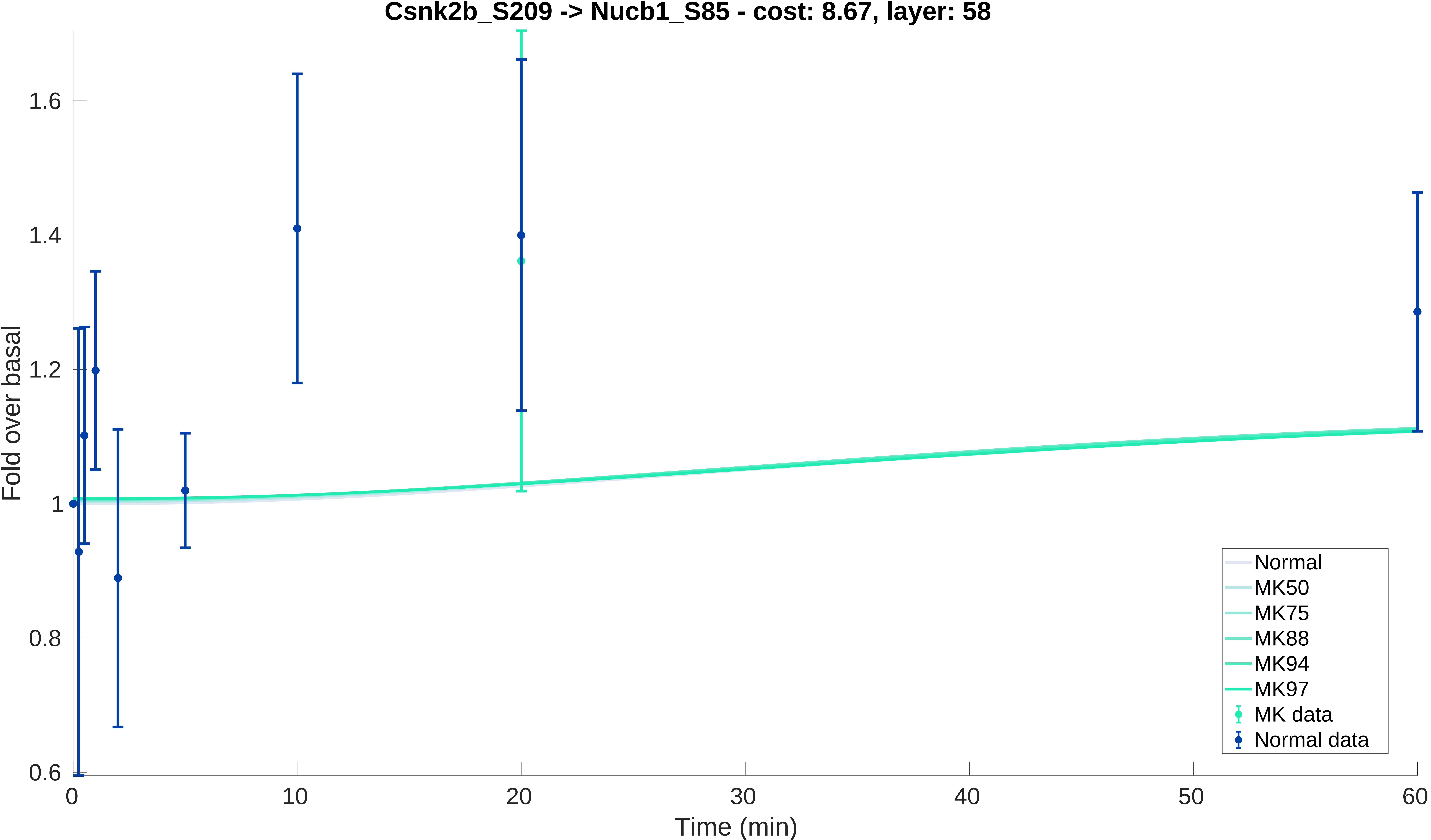

Csnk2b_S209 -> Nucb1_S85 - cost: 8.67, layer: 58
Fold over basal
1.6
1.4
1.2
1
0.8
0.6
0
10
20
30
40
50
60
Time (min)
Normal
MK50
MK75
MK88
MK94
MK97
MK data
Normal data

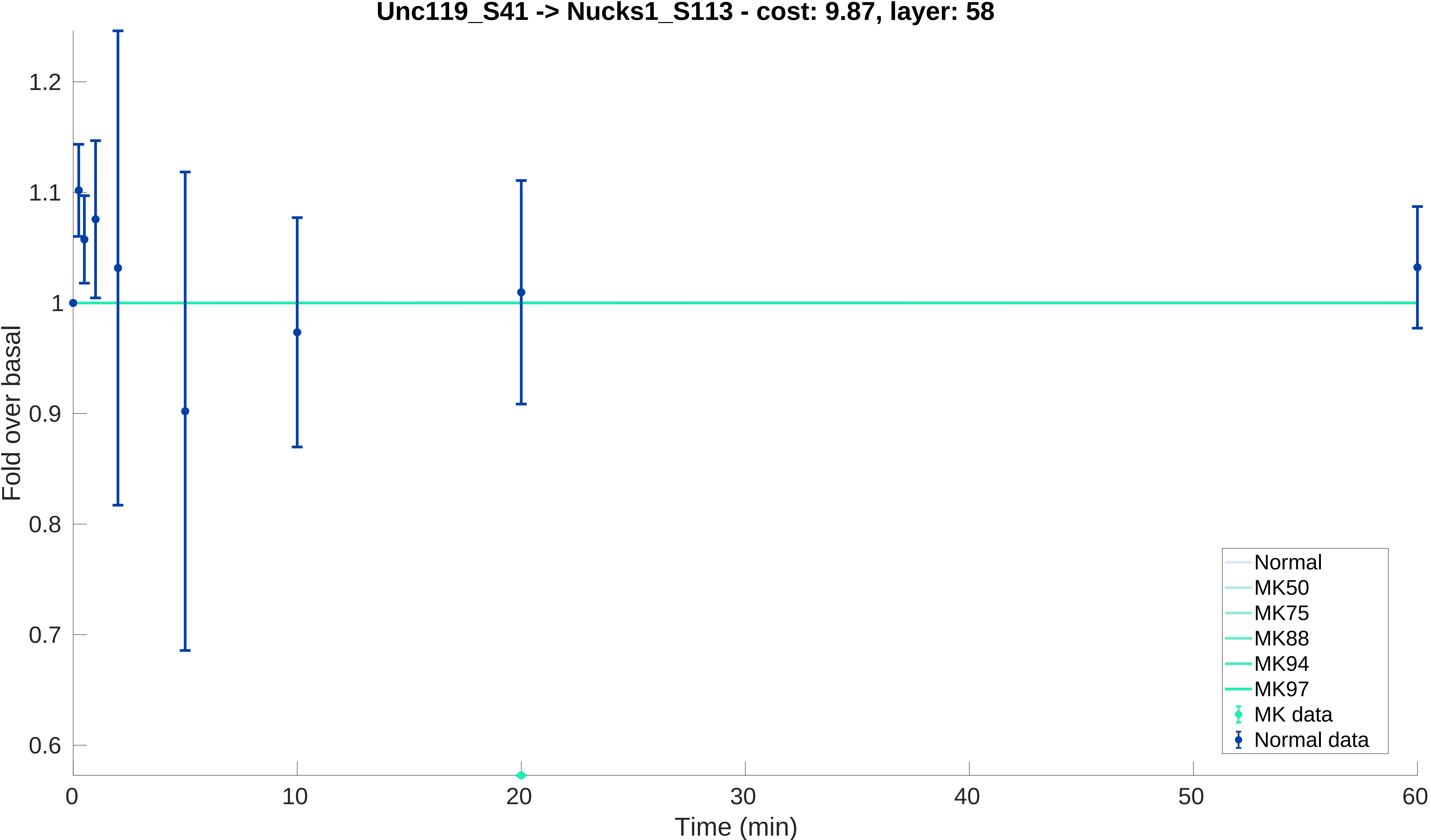

Unc119_S41 -> Nucks1_S113 - cost: 9.87, layer: 58
Fold over basal
1.2
1.1
1
0.9
0.8
0.7
0.6
0
10
20
30
40
50
60
Time (min)
Normal
MK50
MK75
MK88
MK94
MK97
MK data
Normal data

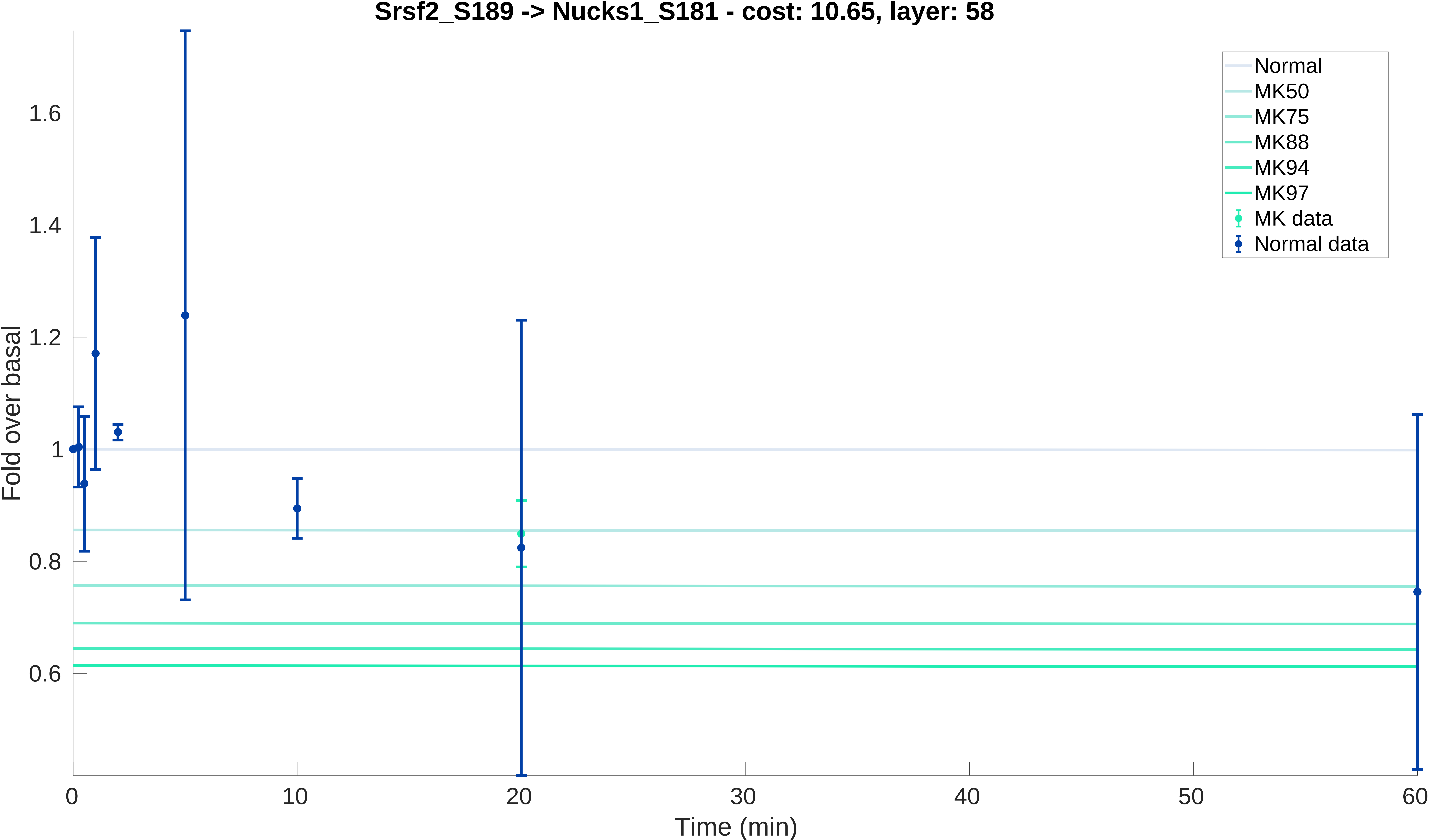

Srsf2_S189 -> Nucks1_S181 - cost: 10.65, layer: 58
Fold over basal
1.6
1.4
1.2
1
0.8
0.6
0
10
20
30
40
50
60
Time (min)
Normal
MK50
MK75
MK88
MK94
MK97
MK data
Normal data

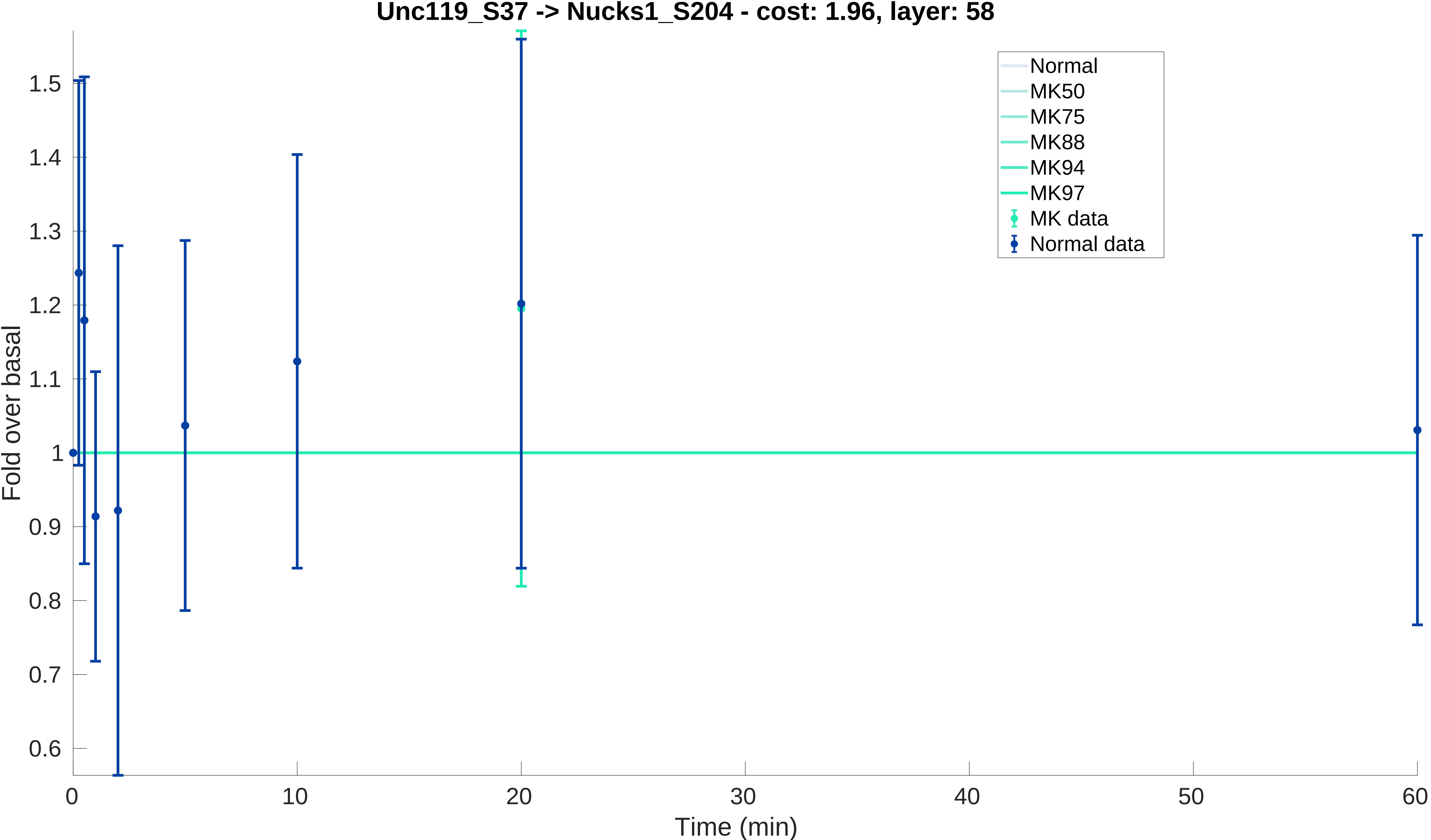

Unc119_S37 -> Nucks1_S204 - cost: 1.96, layer: 58
Fold over basal
Time (min)
0.6
0.7
0.8
0.9
1
1.1
1.2
1.3
1.4
1.5
0
10
20
30
40
50
60
Normal
MK50
MK75
MK88
MK94
MK97
MK data
Normal data

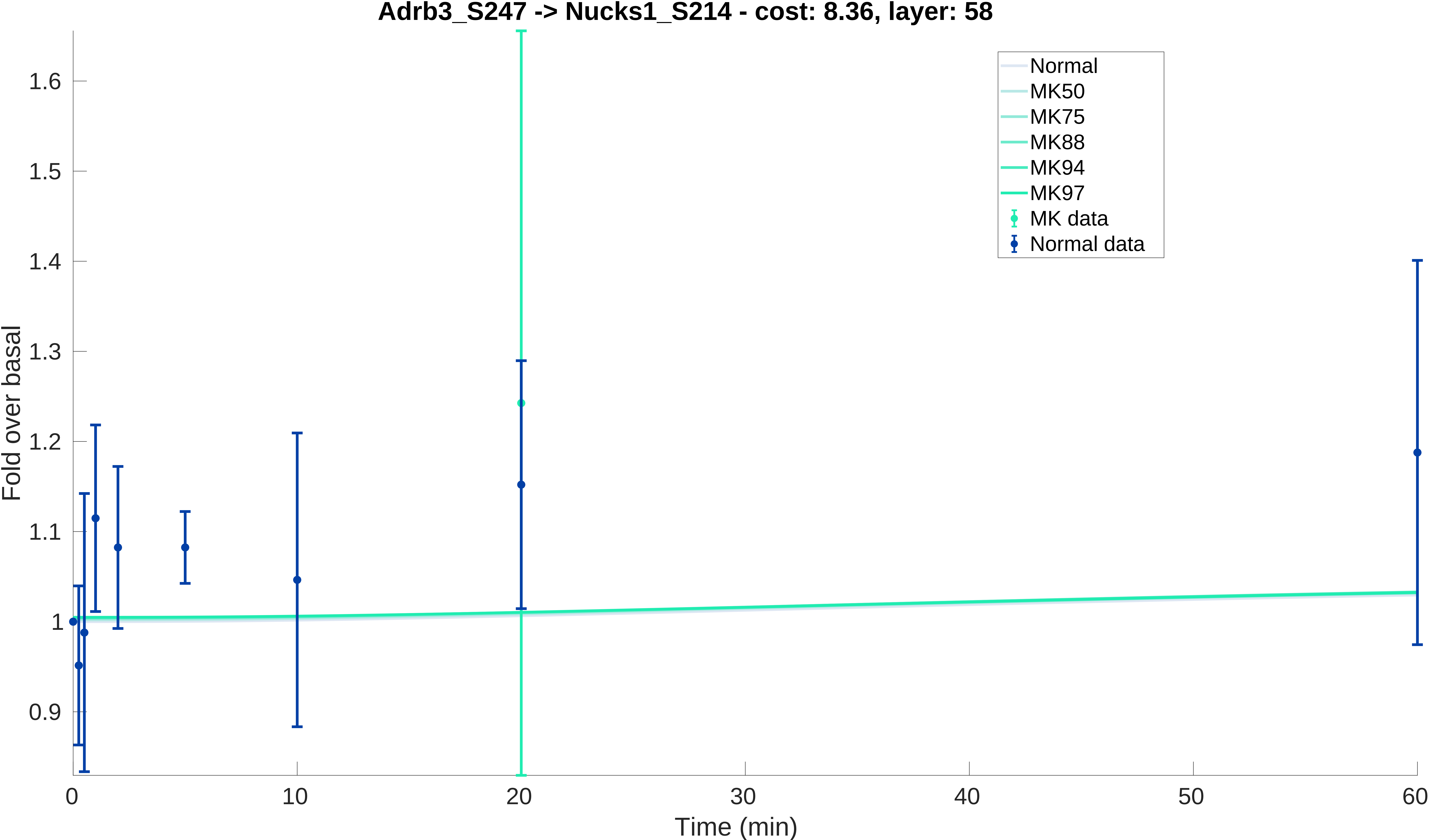

Adrb3_S247 -> Nucks1_S214 - cost: 8.36, layer: 58
Normal
MK50
MK75
MK88
MK94
MK97
MK data
Normal data
Fold over basal
Time (min)
1.6
1.5
1.4
1.3
1.2
1.1
1
0.9
0
10
20
30
40
50
60

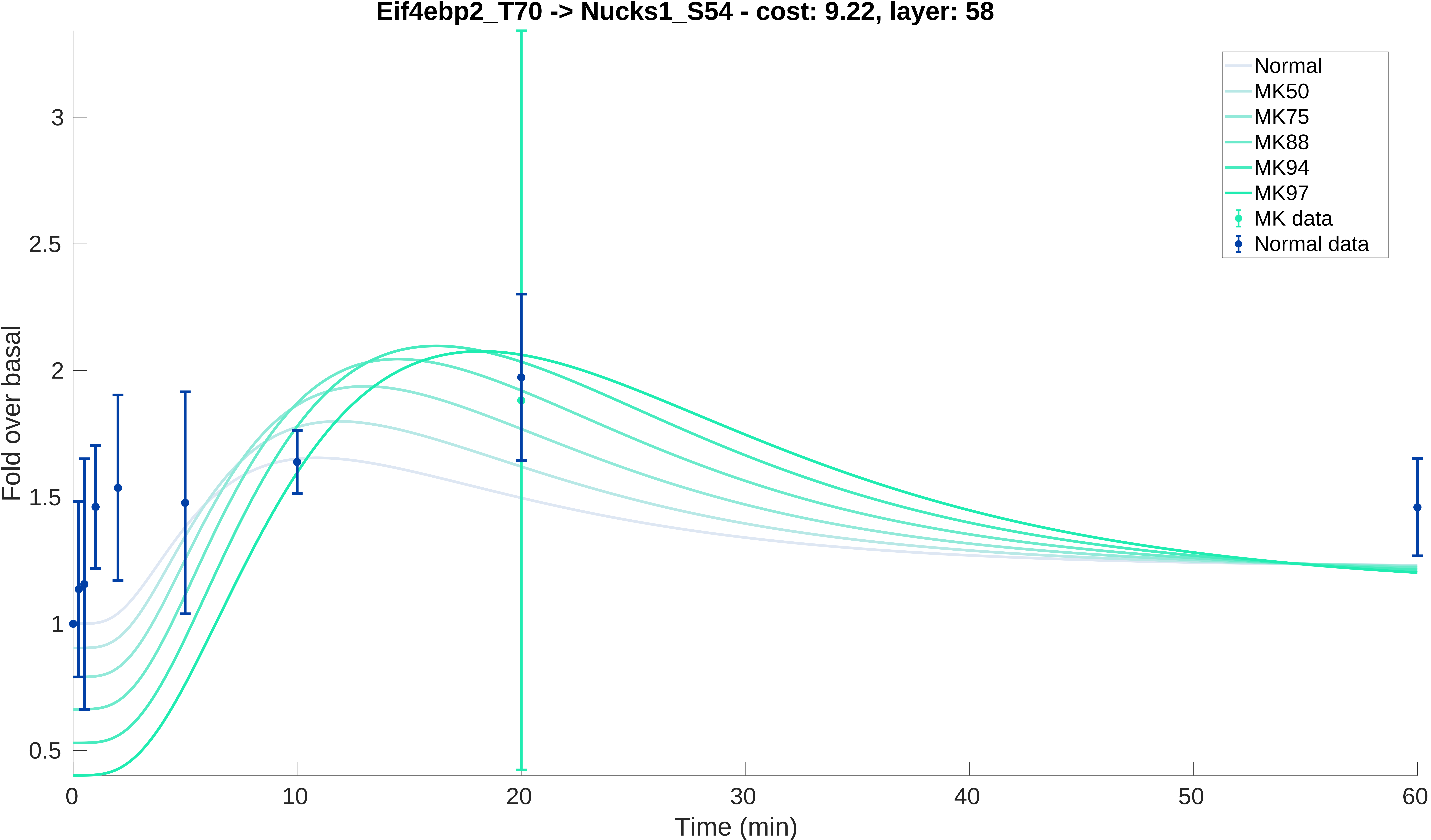

Eif4ebp2_T70 -> Nucks1_S54 - cost: 9.22, layer: 58
Normal
MK50
MK75
MK88
MK94
MK97
MK data
Normal data
Fold over basal
Time (min)
0.5
1
1.5
2
2.5
3
0
10
20
30
40
50
60

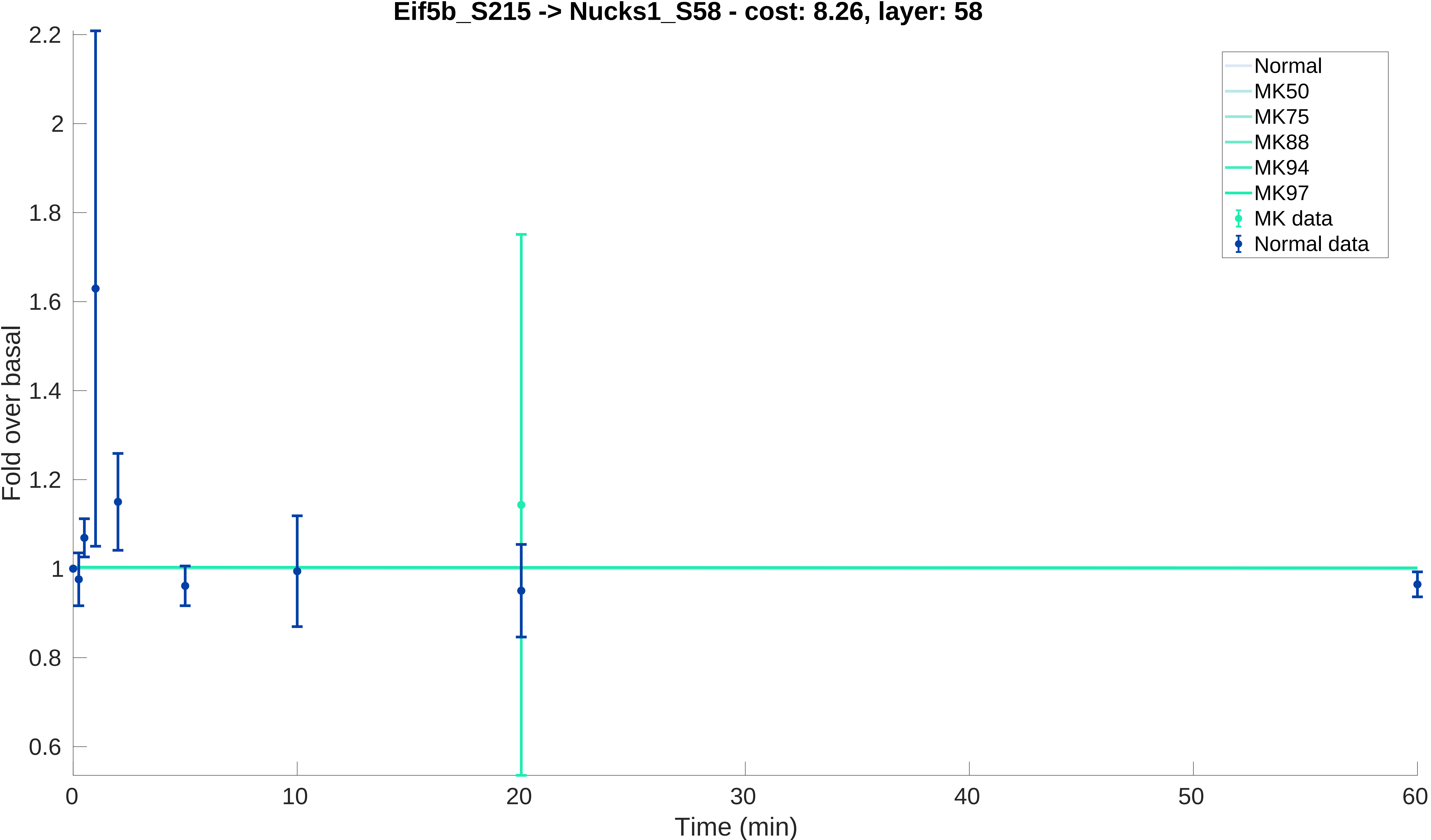
Eif5b_S215 -> Nucks1_S58 - cost: 8.26, layer: 58
Fold over basal
Time (min)
0.6
0.8
1
1.2
1.4
1.6
1.8
2
2.2
0
10
20
30
40
50
60
Normal
MK50
MK75
MK88
MK94
MK97
MK data
Normal data

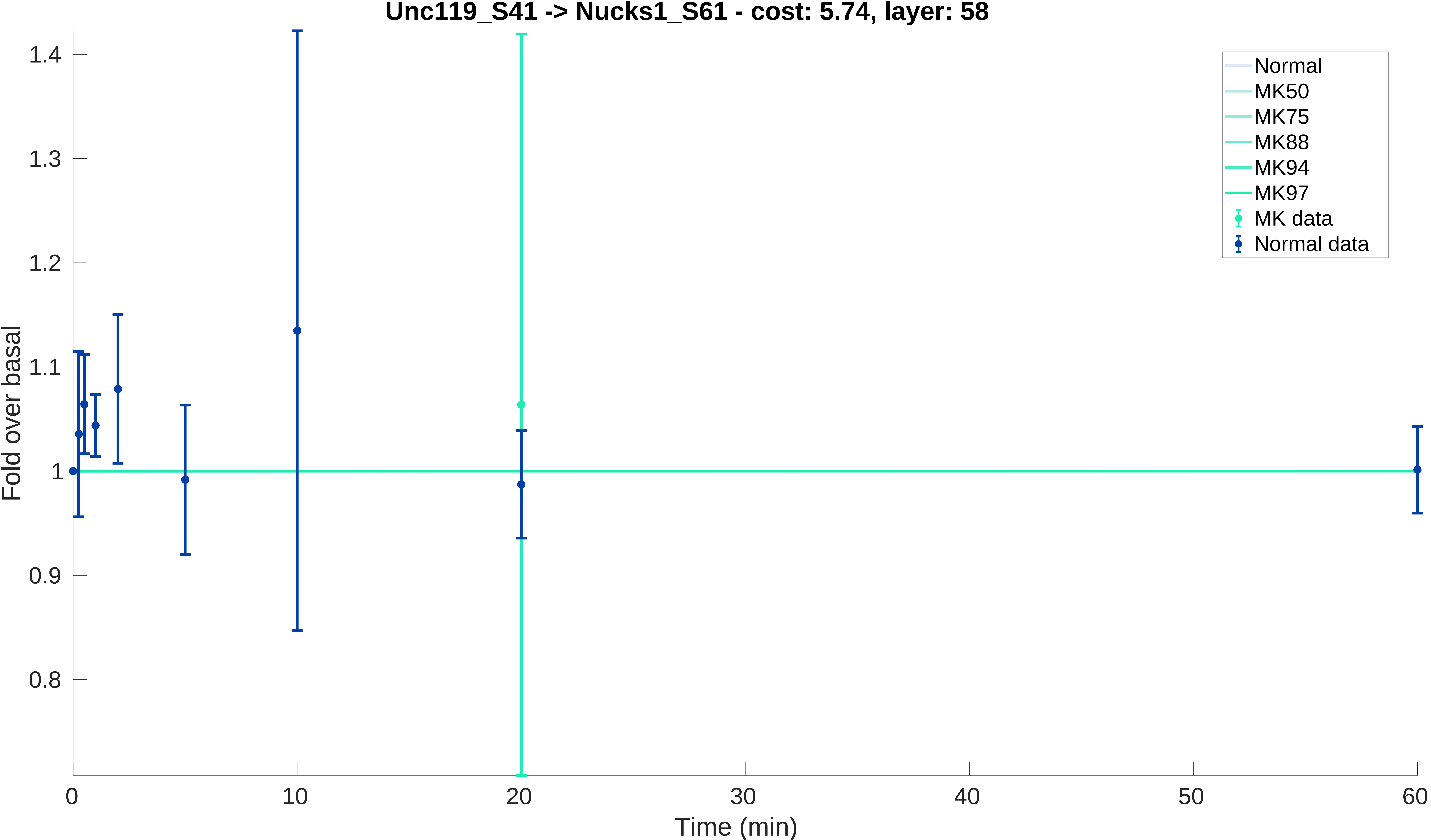

Unc119_S41 -> Nucks1_S61 - cost: 5.74, layer: 58
Fold over basal
Time (min)
1.4
1.3
1.2
1.1
1
0.9
0.8
0
10
20
30
40
50
60
Normal
MK50
MK75
MK88
MK94
MK97
MK data
Normal data

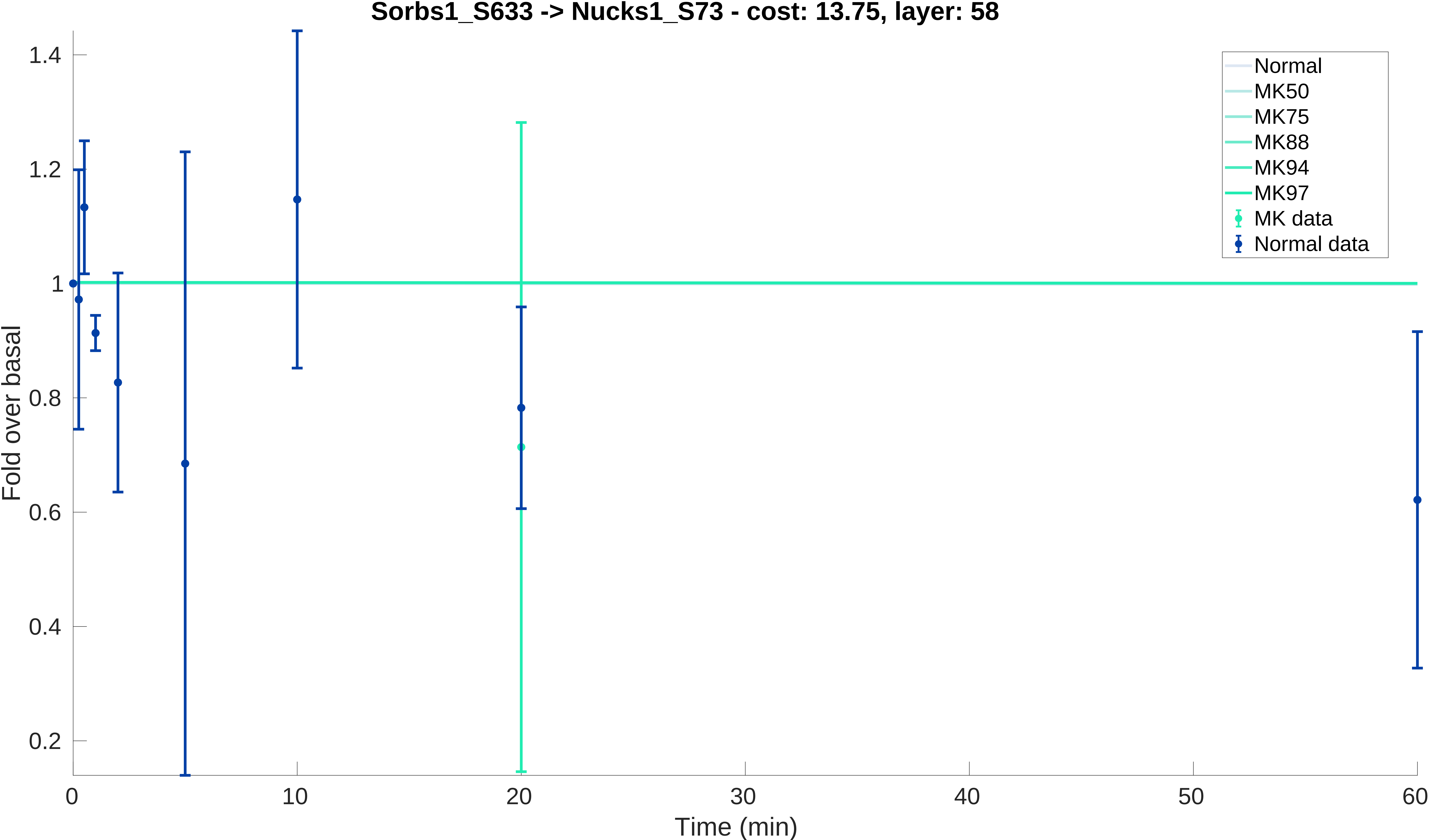
Sorbs1_S633 -> Nucks1_S73 - cost: 13.75, layer: 58
Fold over basal
Time (min)
0.2
0.4
0.6
0.8
1
1.2
1.4
0
10
20
30
40
50
60
Normal
MK50
MK75
MK88
MK94
MK97
MK data
Normal data

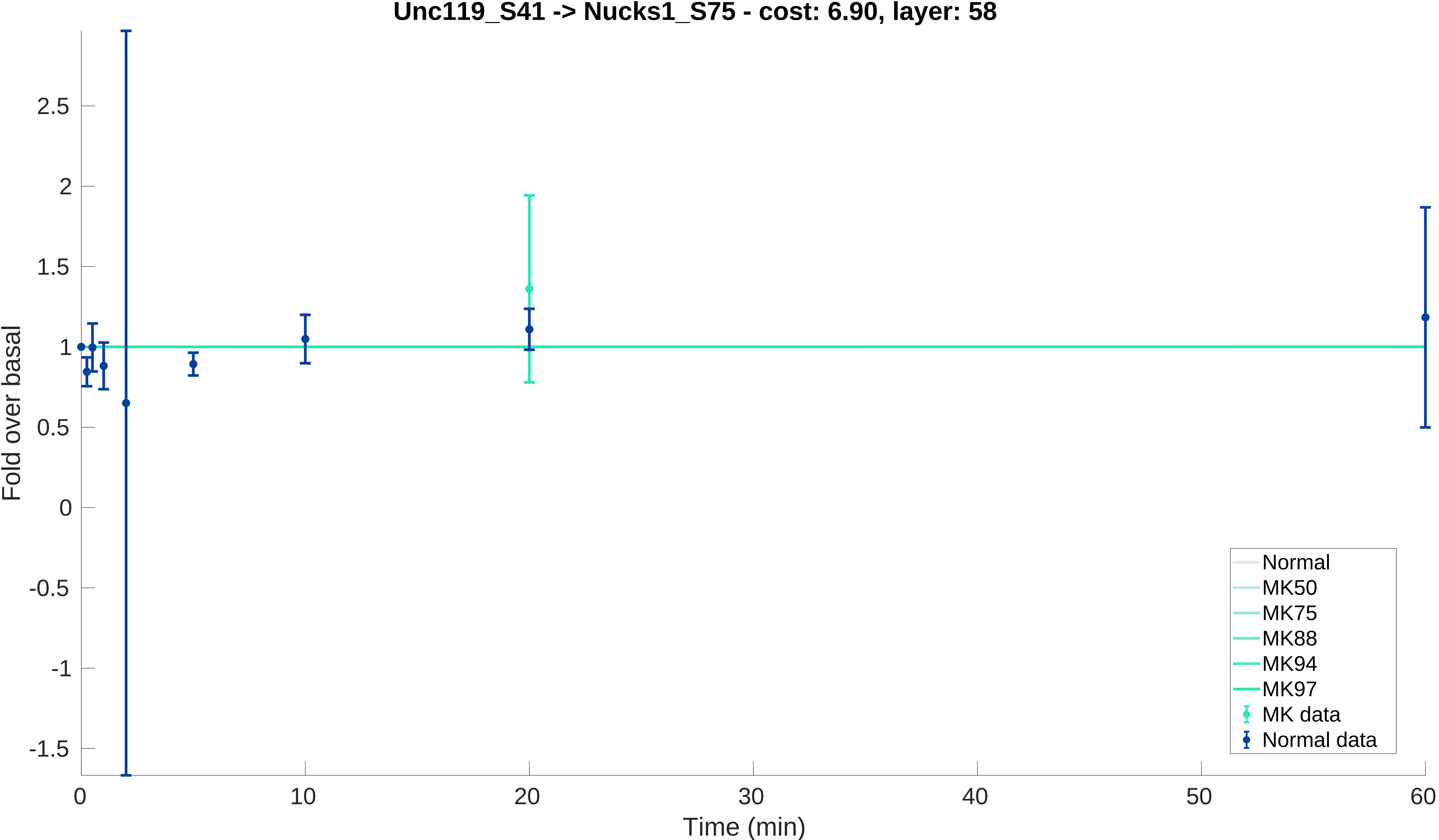

Unc119_S41 -> Nucks1_S75 - cost: 6.90, layer: 58
Fold over basal
Time (min)
2.5
2
1.5
1
0.5
0
-0.5
-1
-1.5
0
10
20
30
40
50
60
Normal
MK50
MK75
MK88
MK94
MK97
MK data
Normal data

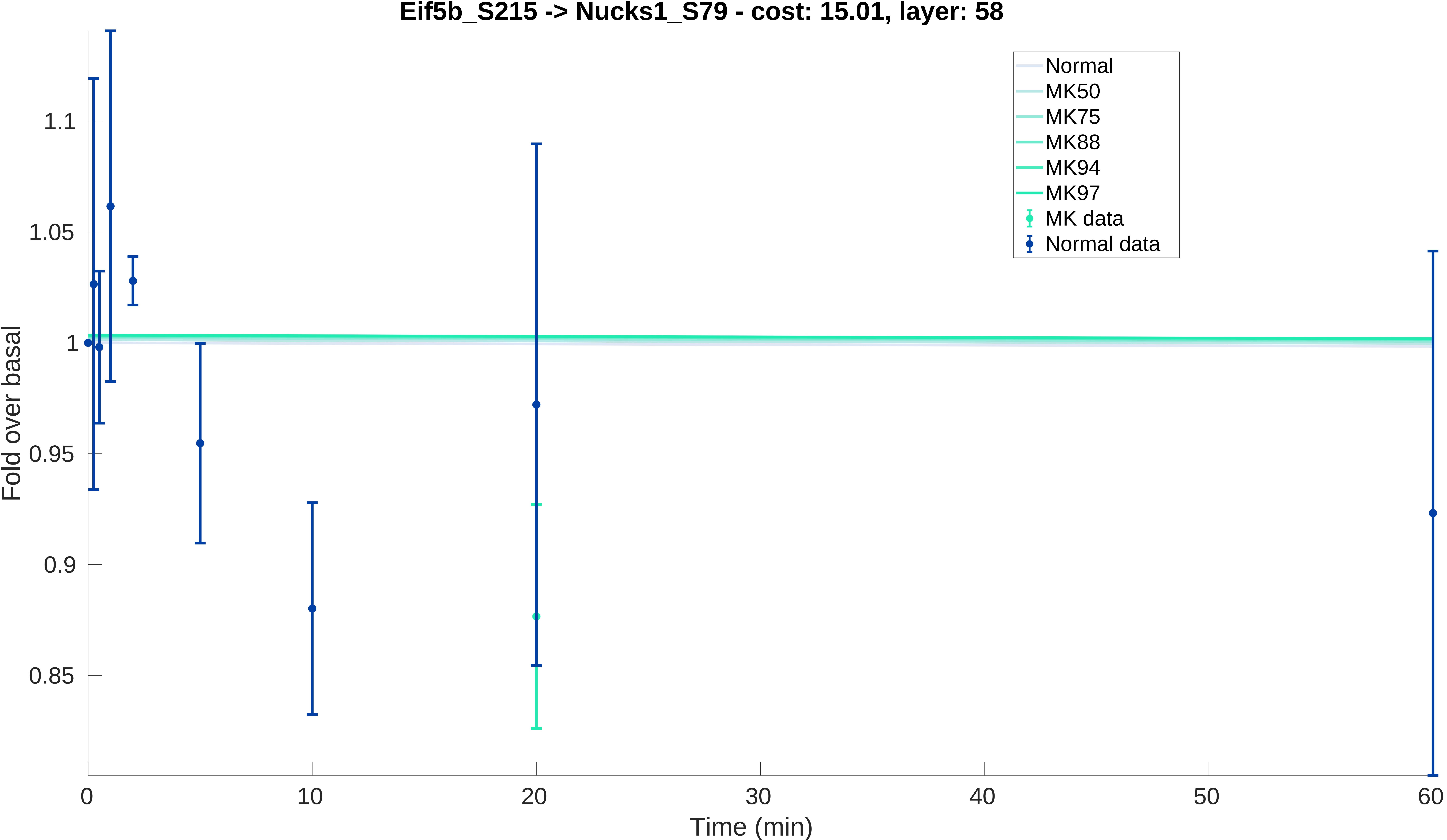

Eif5b_S215 -> Nucks1_S79 - cost: 15.01, layer: 58
Normal
MK50
MK75
MK88
MK94
MK97
MK data
Normal data
Fold over basal
1.1
1.05
1
0.95
0.9
0.85
0
10
20
30
40
50
60
Time (min)

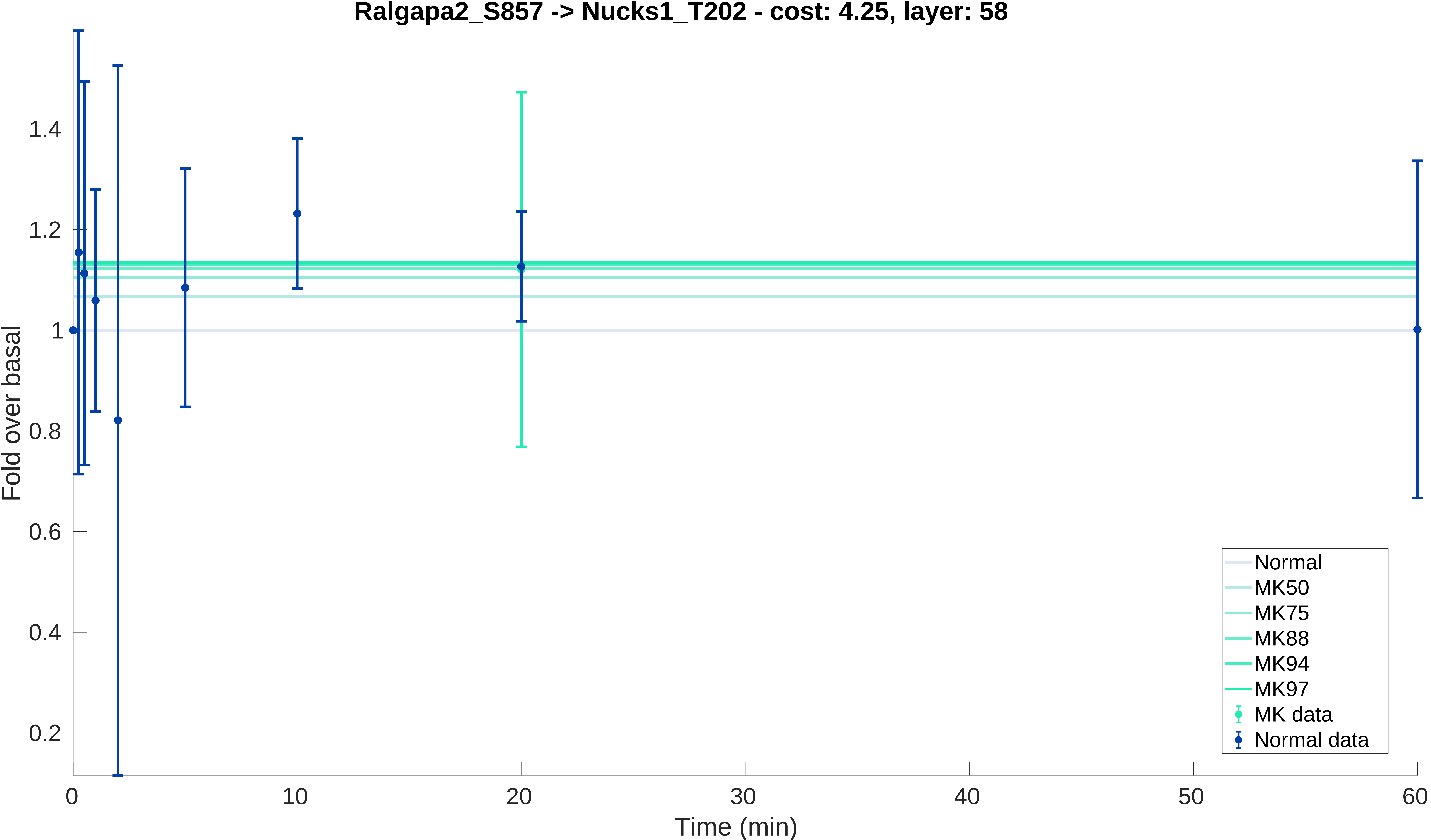

Ralgapa2_S857 -> Nucks1_T202 - cost: 4.25, layer: 58
Fold over basal
Time (min)
1.4
1.2
1
0.8
0.6
0.4
0.2
0
10
20
30
40
50
60
Normal
MK50
MK75
MK88
MK94
MK97
MK data
Normal data

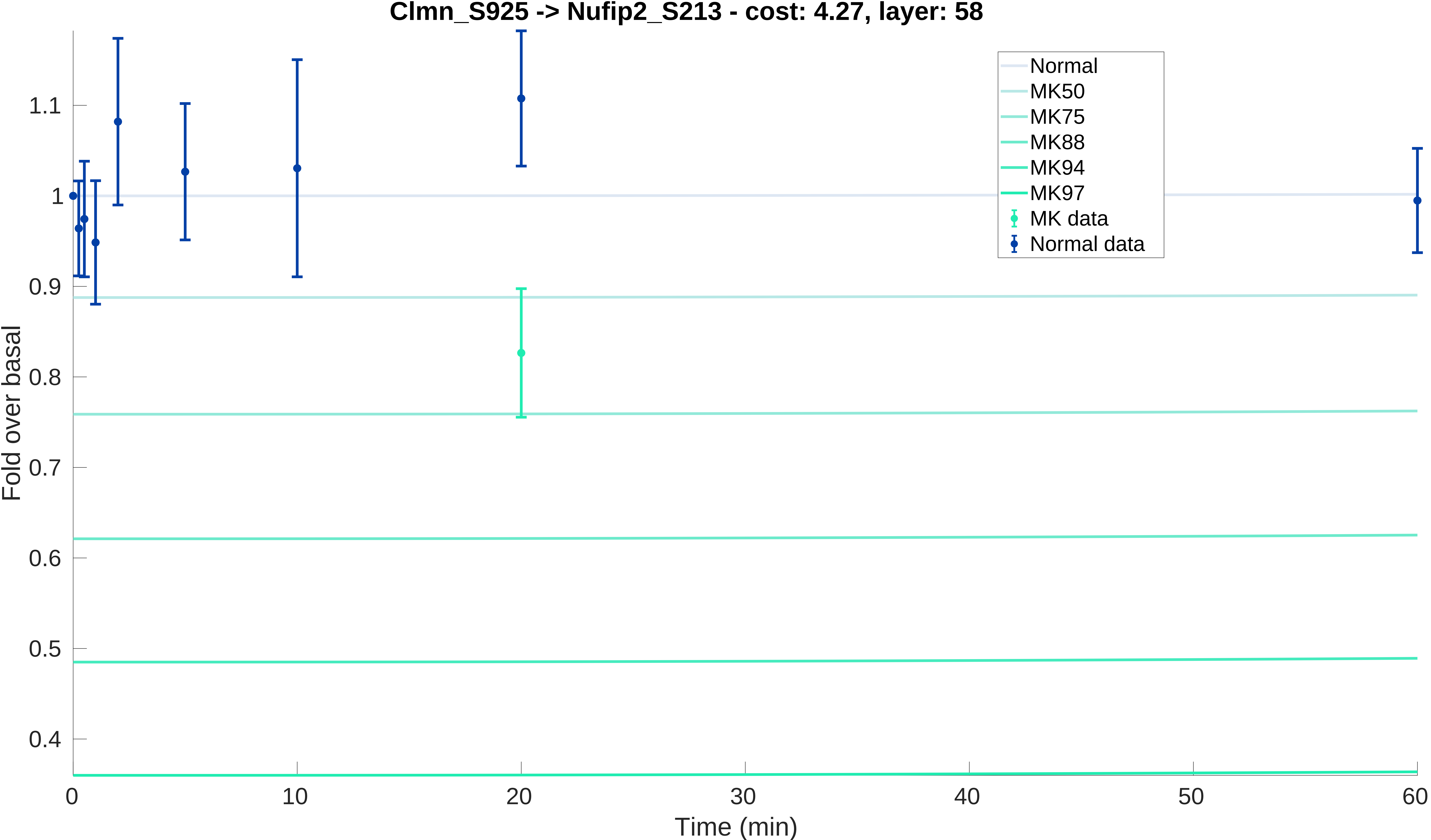
Clmn_S925 -> Nufip2_S213 - cost: 4.27, layer: 58
Fold over basal
1.1
1
0.9
0.8
0.7
0.6
0.5
0.4
0
10
20
30
40
50
60
Time (min)
Normal
MK50
MK75
MK88
MK94
MK97
MK data
Normal data

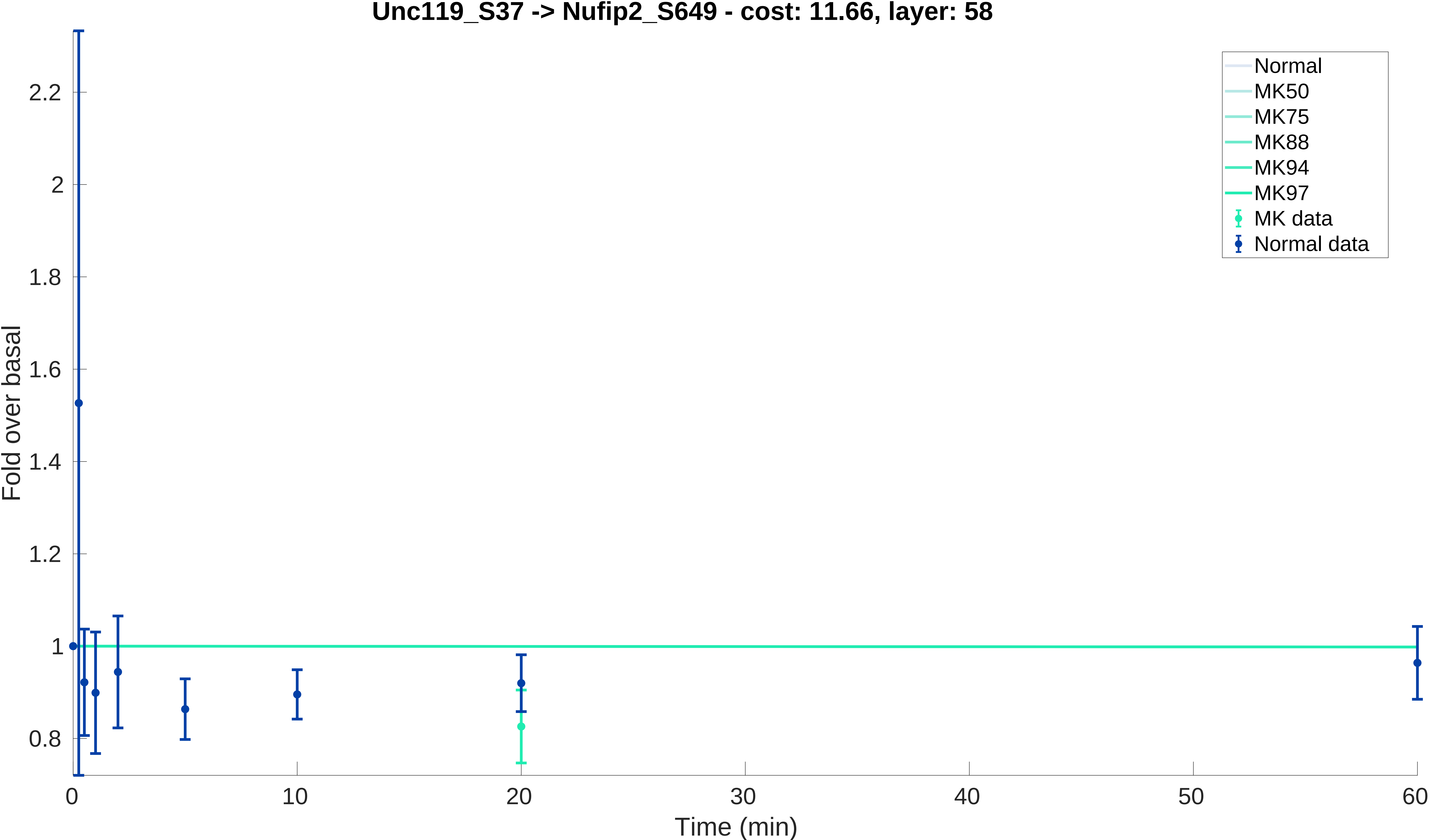

Unc119_S37 -> Nufip2_S649 - cost: 11.66, layer: 58
Normal
MK50
MK75
MK88
MK94
MK97
MK data
Normal data
Fold over basal
Time (min)
0.8
1
1.2
1.4
1.6
1.8
2
2.2
0
10
20
30
40
50
60

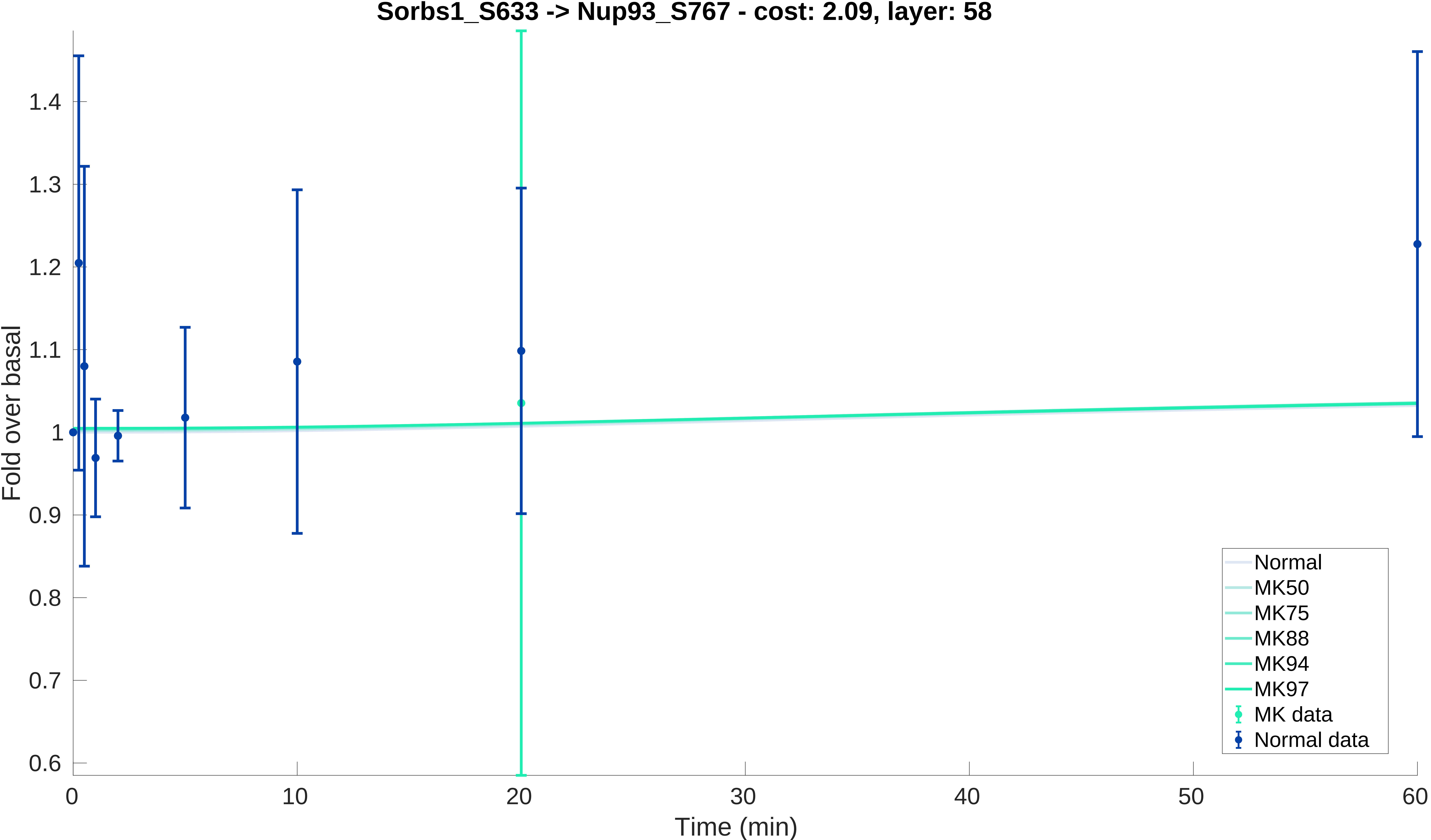

Sorbs1_S633 -> Nup93_S767 - cost: 2.09, layer: 58
Fold over basal
1.4
1.3
1.2
1.1
1
0.9
0.8
0.7
0.6
0
10
20
30
40
50
60
Time (min)
Normal
MK50
MK75
MK88
MK94
MK97
MK data
Normal data

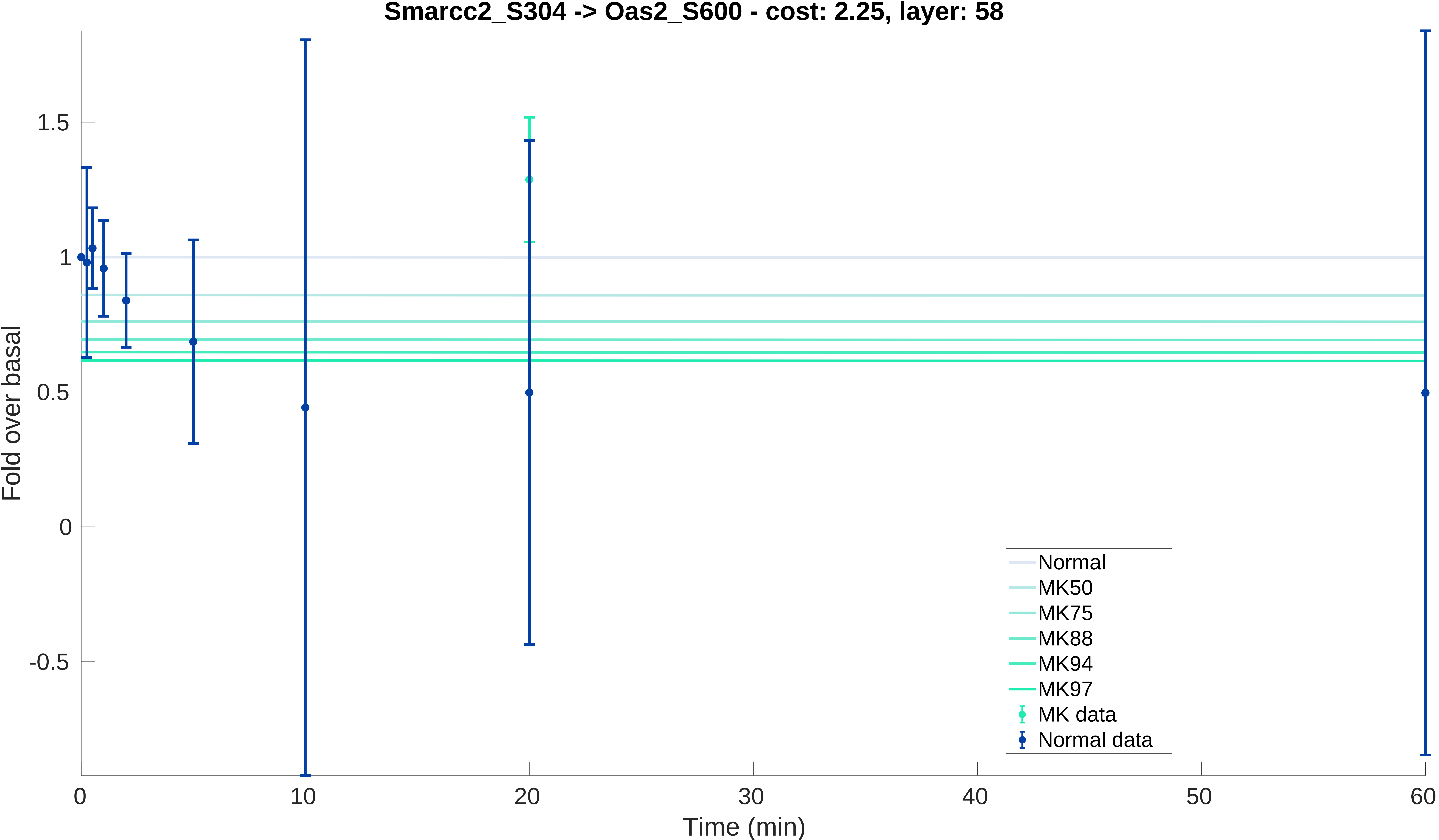

Smarcc2_S304 -> Oas2_S600 - cost: 2.25, layer: 58
1.5
1
0.5
0
-0.5
Fold over basal
0
10
20
30
40
50
60
Time (min)
Normal
MK50
MK75
MK88
MK94
MK97
MK data
Normal data

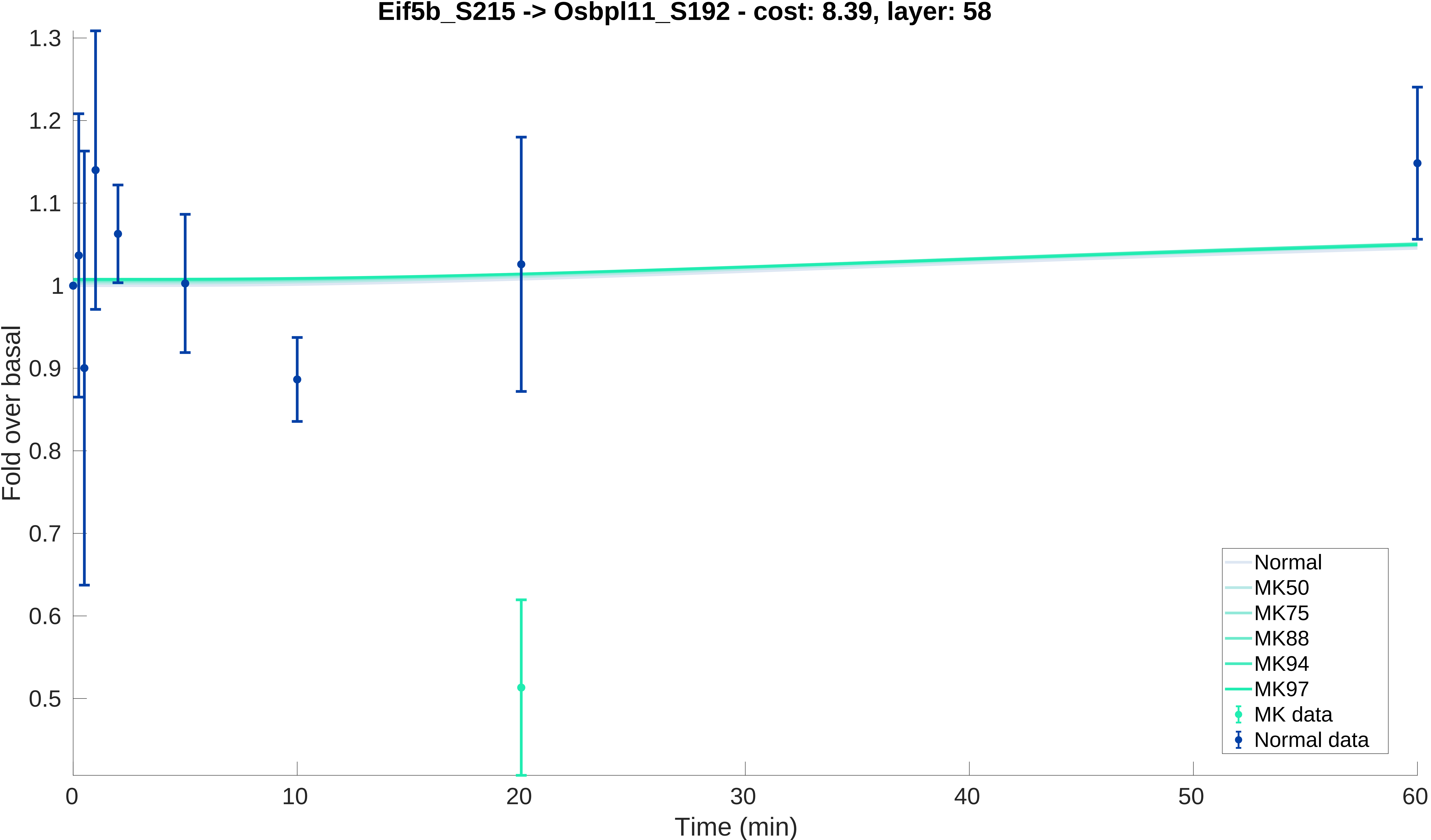

Eif5b_S215 -> Osbpl11_S192 - cost: 8.39, layer: 58
Fold over basal
Time (min)
1.3
1.2
1.1
1
0.9
0.8
0.7
0.6
0.5
0
10
20
30
40
50
60
Normal
MK50
MK75
MK88
MK94
MK97
MK data
Normal data

Smarcc2_S304 -> Osbpl3_S33 - cost: 9.63, layer: 58

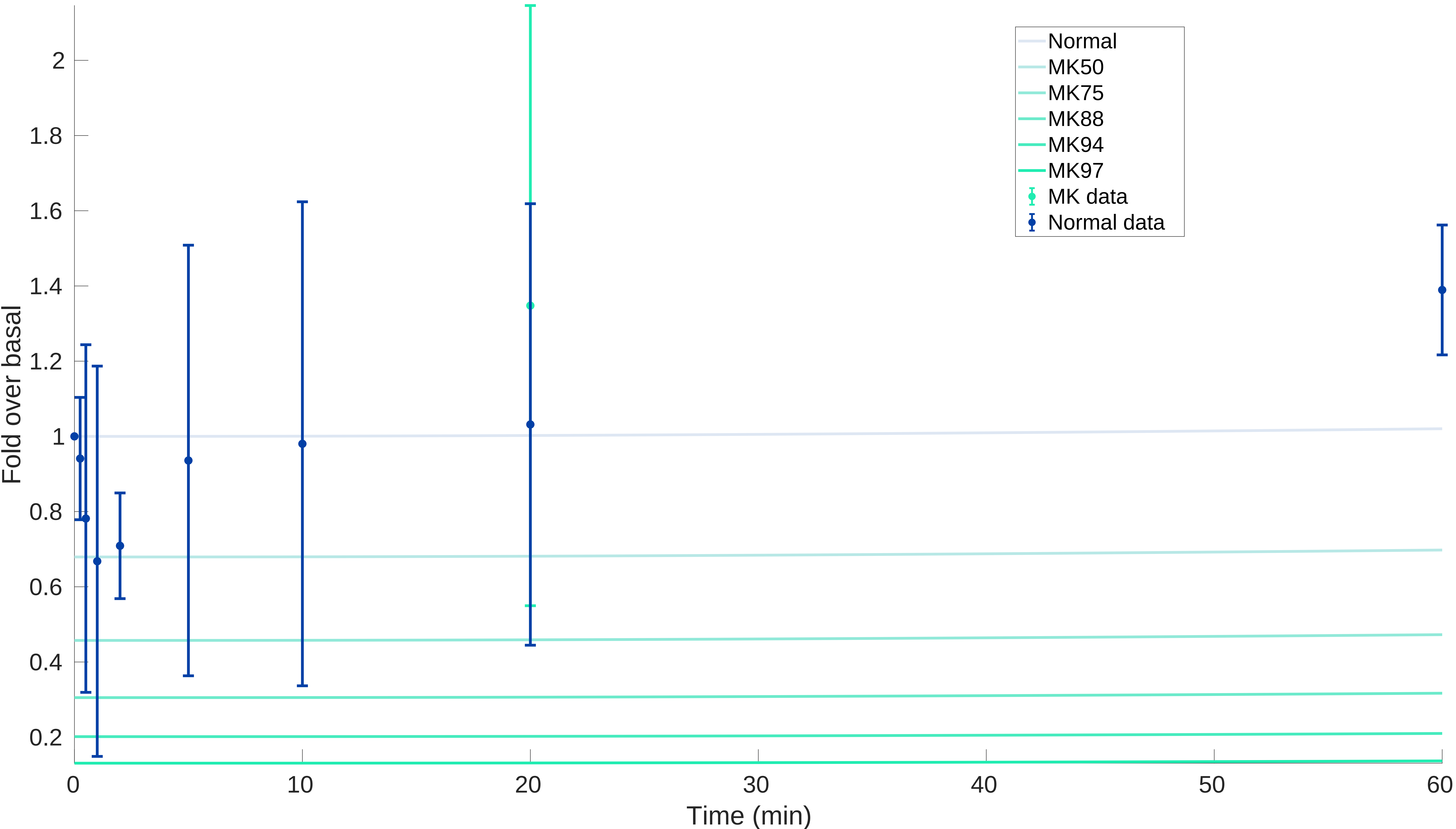

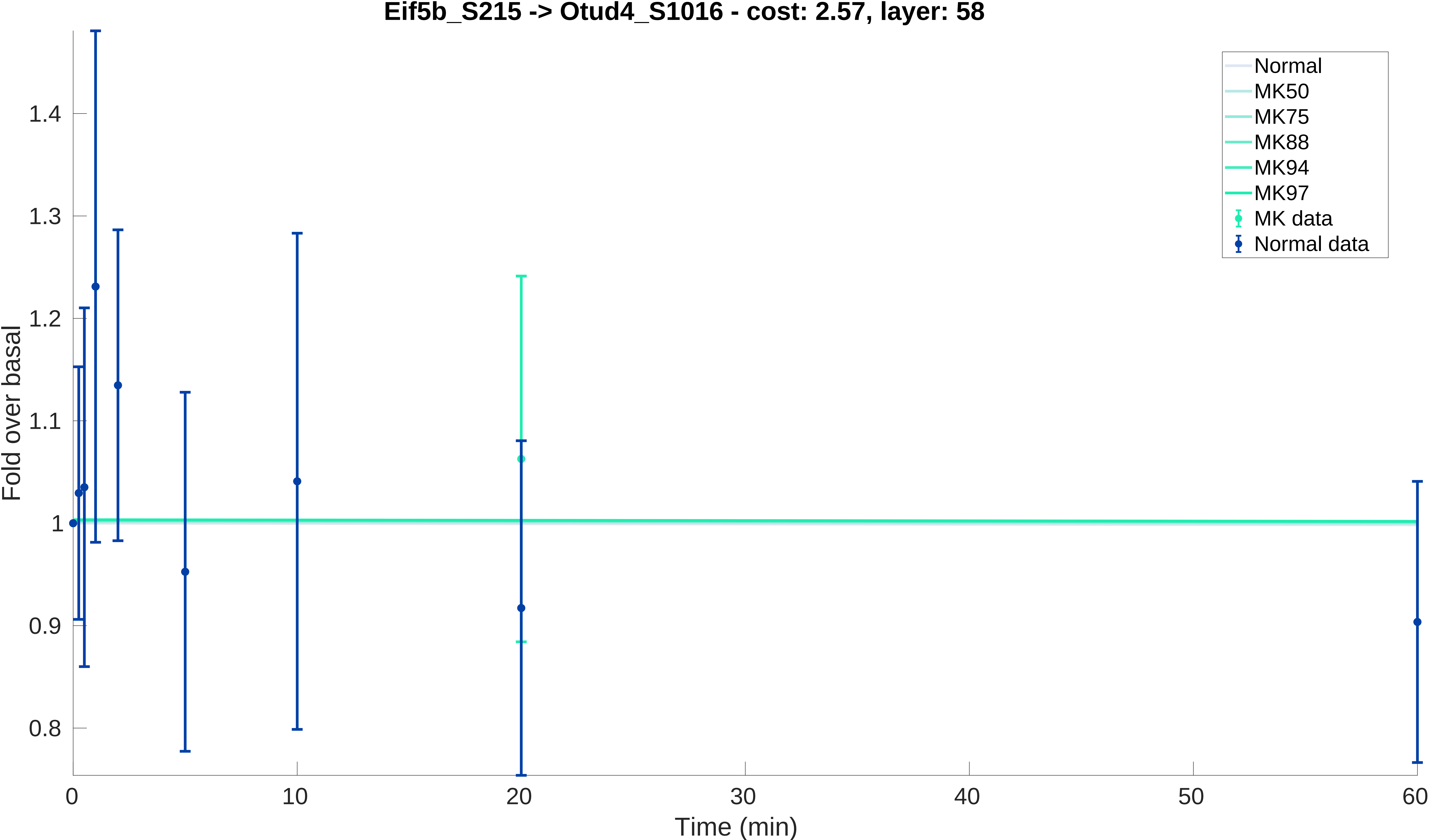

Eif5b_S215 -> Otud4_S1016 - cost: 2.57, layer: 58
Normal
MK50
MK75
MK88
MK94
MK97
MK data
Normal data
Fold over basal
1.4
1.3
1.2
1.1
1
0.9
0.8
0
10
20
30
40
50
60
Time (min)

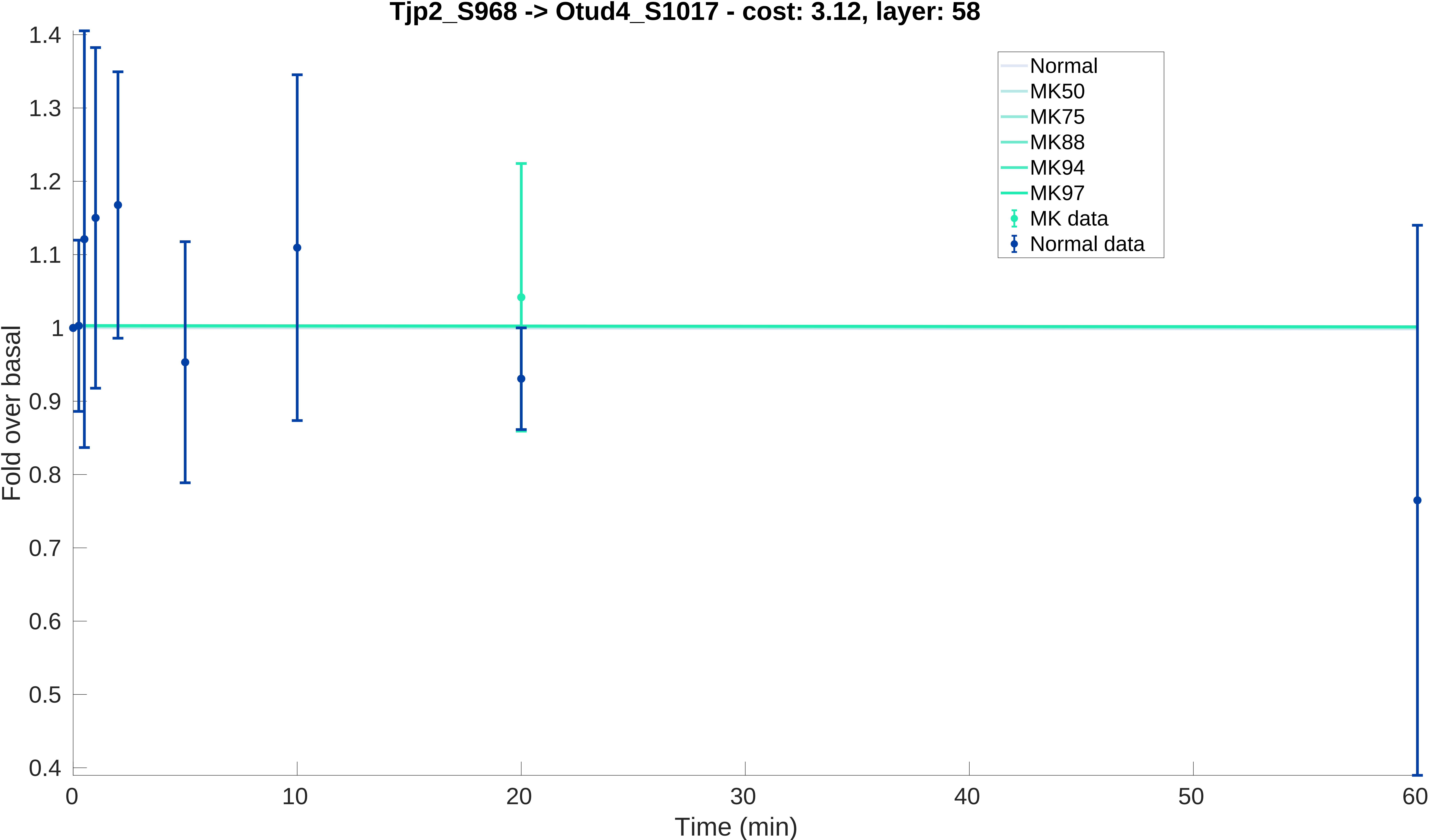

Tjp2_S968 -> Otud4_S1017 - cost: 3.12, layer: 58
Fold over basal
Time (min)
1.4
1.3
1.2
1.1
1
0.9
0.8
0.7
0.6
0.5
0.4
0
10
20
30
40
50
60
Normal
MK50
MK75
MK88
MK94
MK97
MK data
Normal data

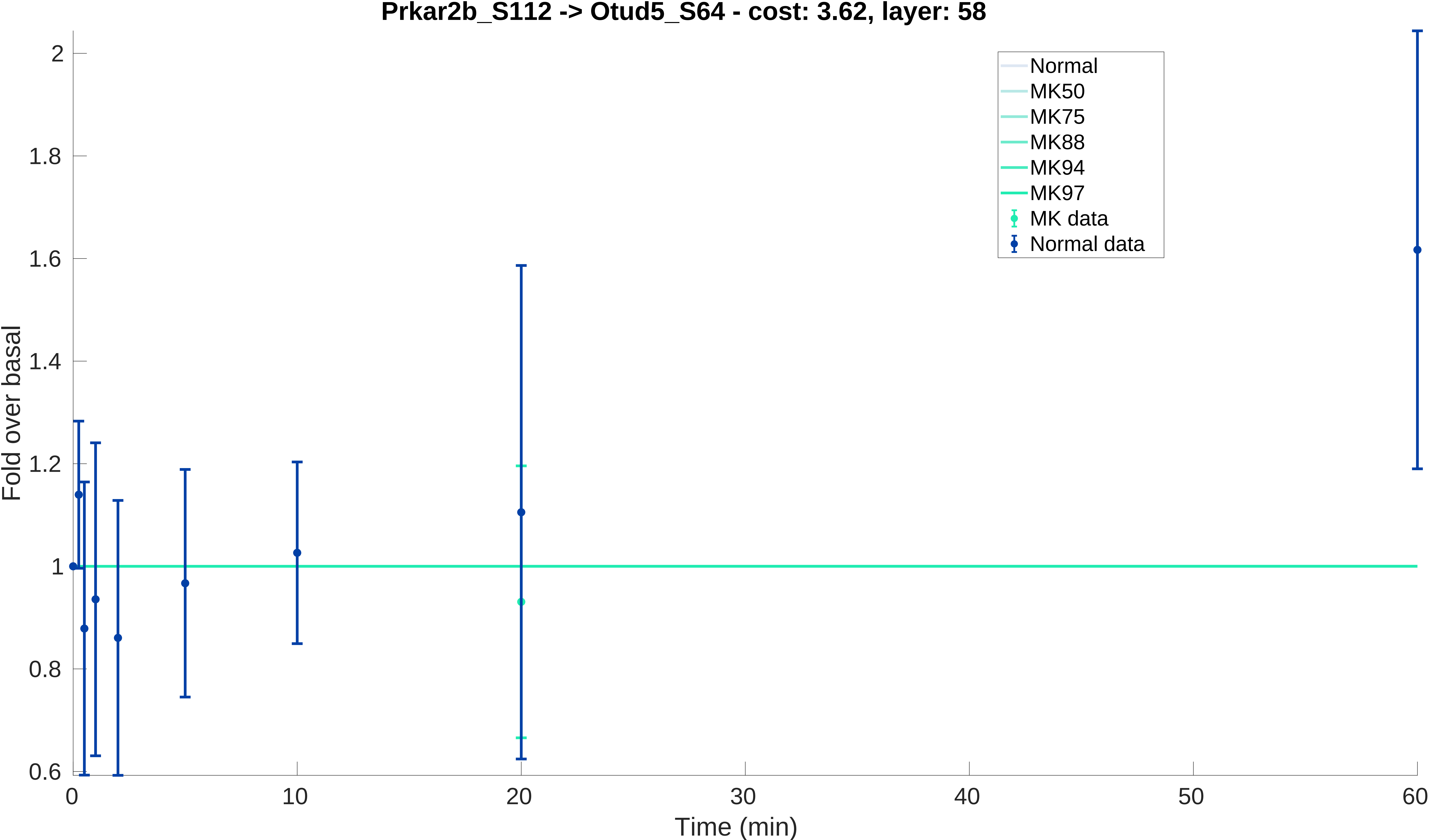
Prkar2b_S112 -> Otud5_S64 - cost: 3.62, layer: 58
Normal
MK50
MK75
MK88
MK94
MK97
MK data
Normal data
Fold over basal
Time (min)
0.6
0.8
1
1.2
1.4
1.6
1.8
2
0
10
20
30
40
50
60

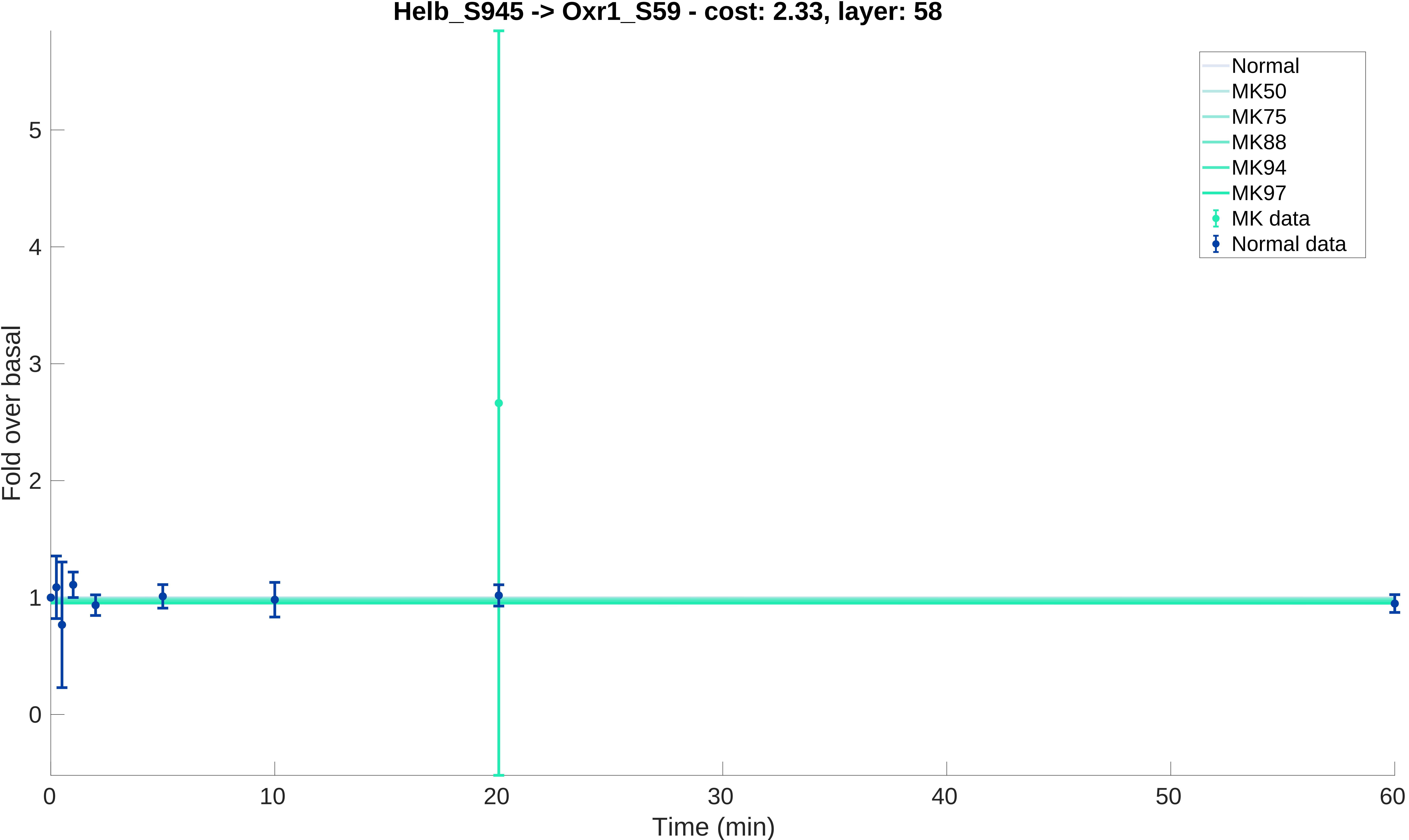

Helb_S945 -> Oxr1_S59 - cost: 2.33, layer: 58
Fold over basal
Time (min)
0
10
20
30
40
50
60
0
1
2
3
4
5
Normal
MK50
MK75
MK88
MK94
MK97
MK data
Normal data

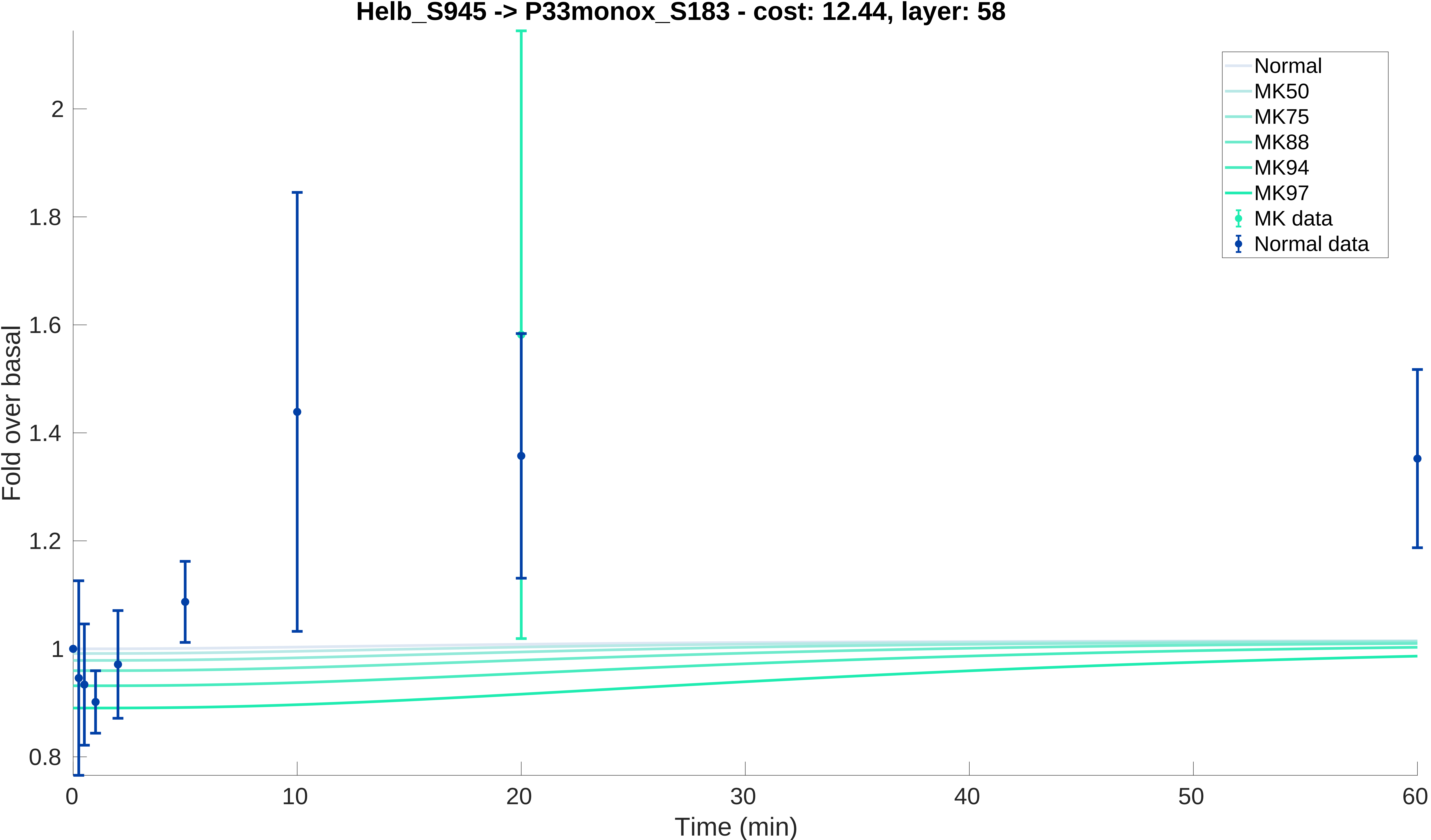

Helb_S945 -> P33monox_S183 - cost: 12.44, layer: 58
Fold over basal
Time (min)
0
10
20
30
40
50
60
0.8
1
1.2
1.4
1.6
1.8
2
Normal
MK50
MK75
MK88
MK94
MK97
MK data
Normal data

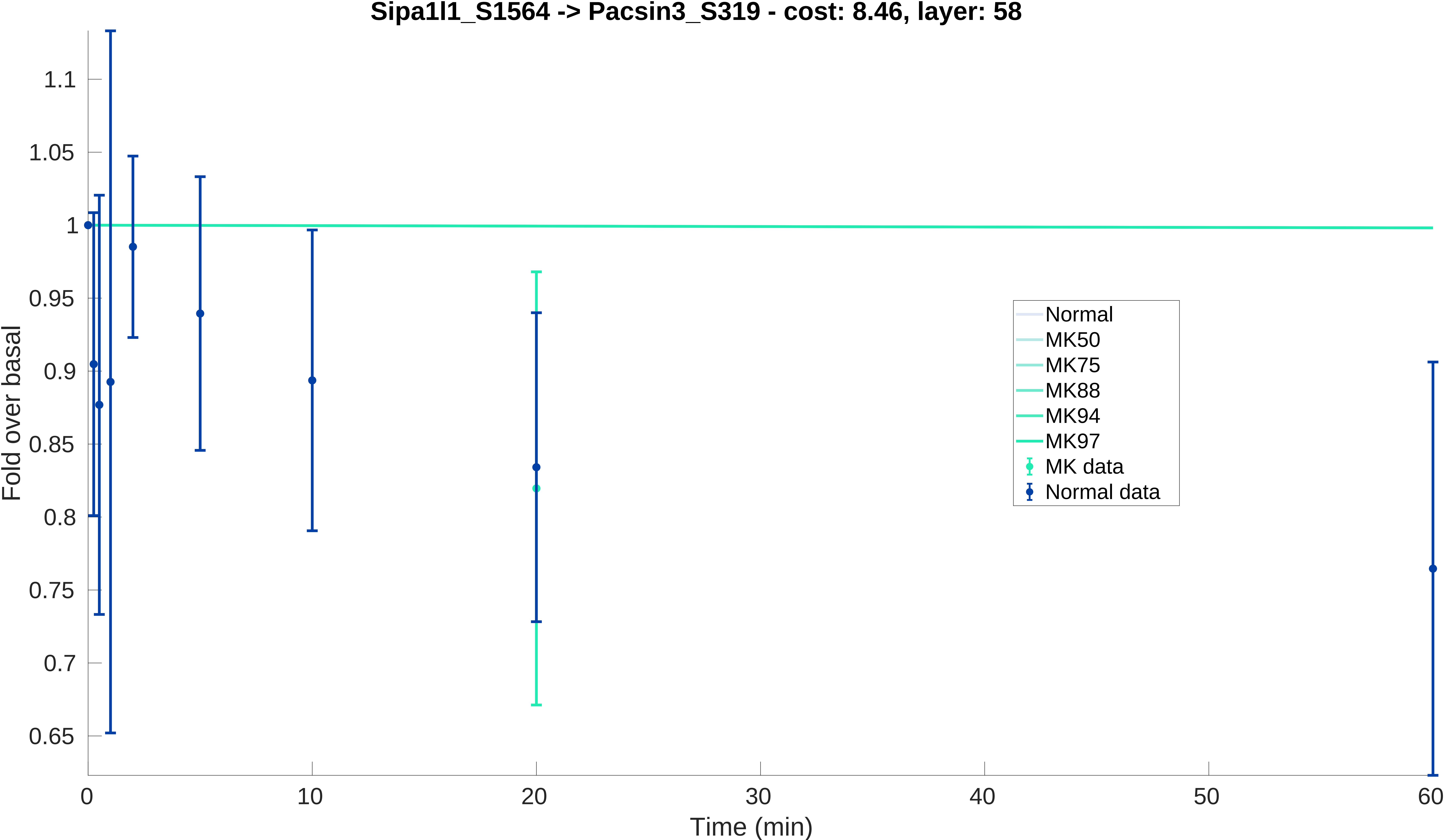
Sipa1l1_S1564 -> Pacsin3_S319 - cost: 8.46, layer: 58
Fold over basal
1.1
1.05
1
0.95
0.9
0.85
0.8
0.75
0.7
0.65
0
10
20
30
40
50
60
Time (min)
Normal
MK50
MK75
MK88
MK94
MK97
MK data
Normal data

Plin1_S492 -> Pacsin3_S354 - cost: 9.89, layer: 58

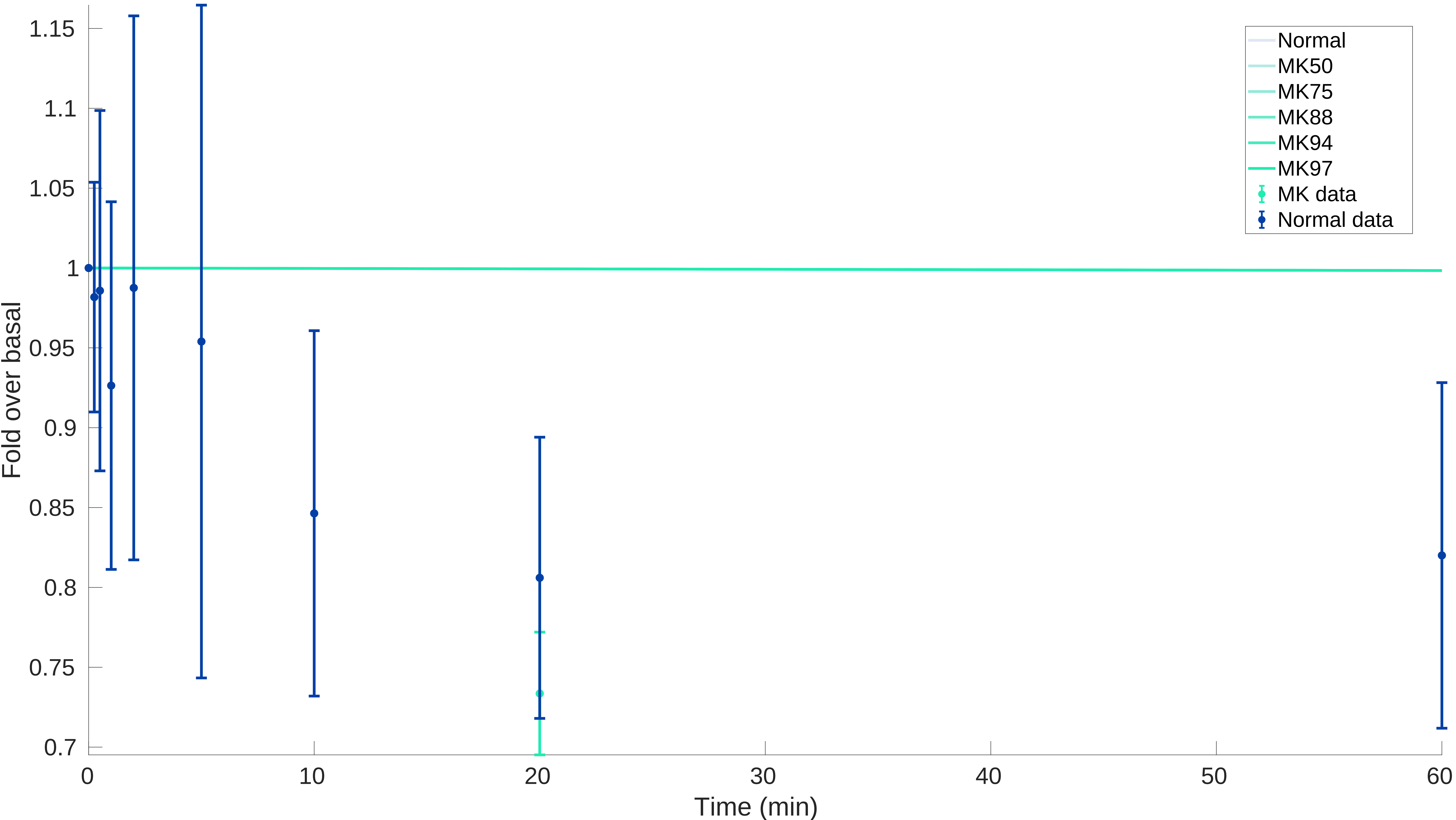

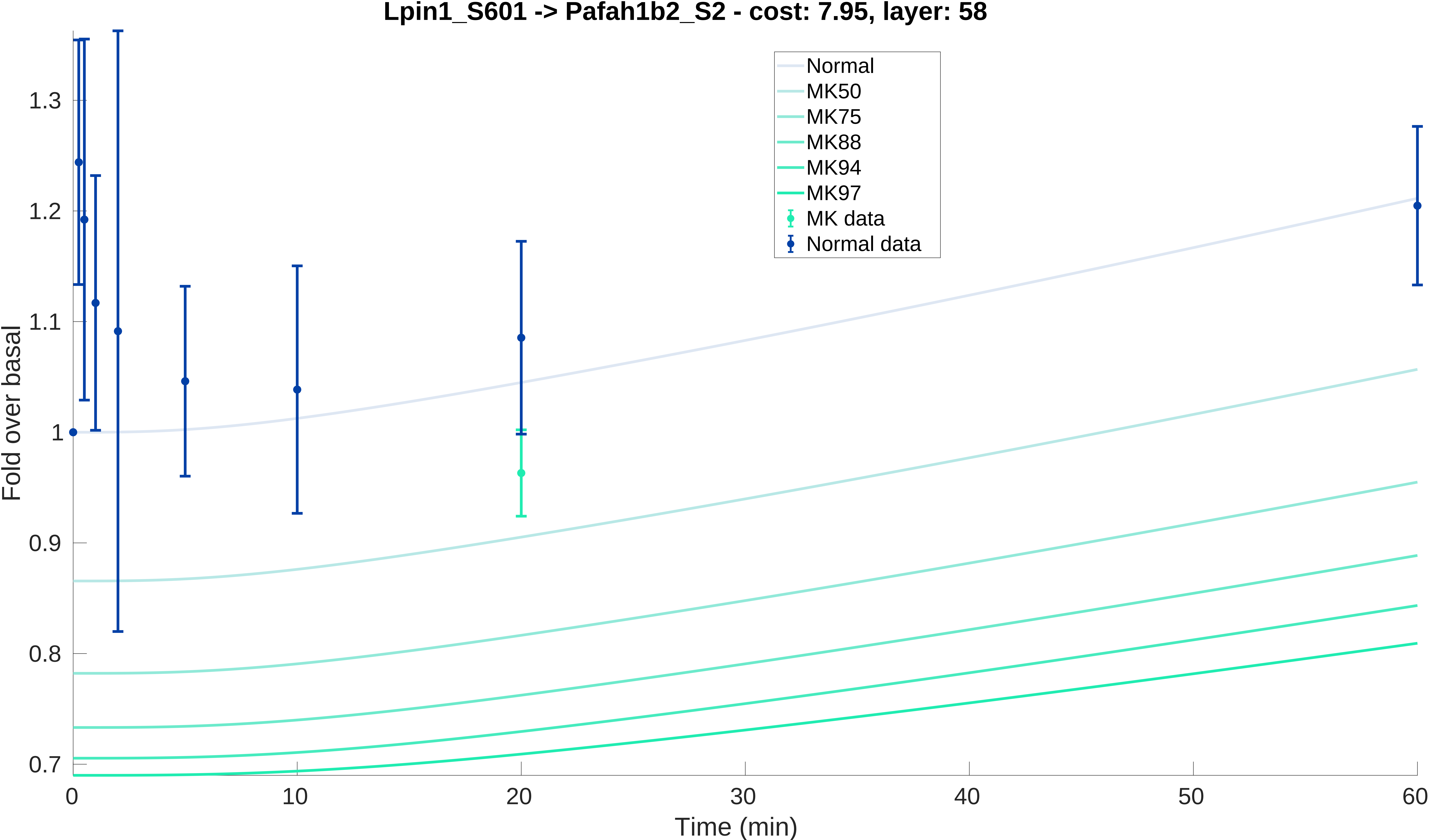

Lpin1_S601 -> Pafah1b2_S2 - cost: 7.95, layer: 58
Normal
MK50
MK75
MK88
MK94
MK97
MK data
Normal data
1.3
1.2
1.1
1
0.9
0.8
0.7
Fold over basal
0
10
20
30
40
50
60
Time (min)

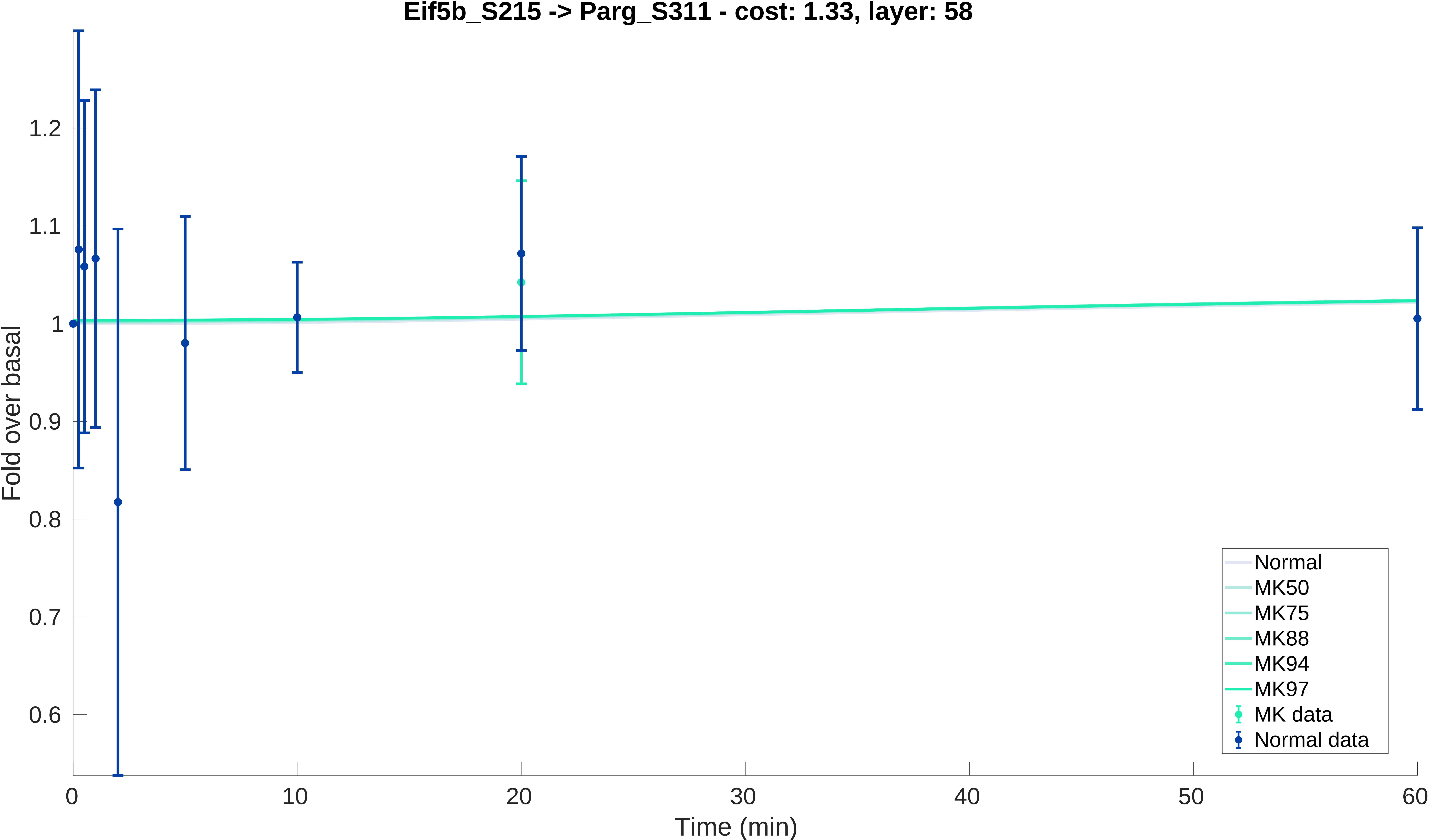

Eif5b_S215 -> Parg_S311 - cost: 1.33, layer: 58
Fold over basal
1.2
1.1
1
0.9
0.8
0.7
0.6
0
10
20
30
40
50
60
Time (min)
Normal
MK50
MK75
MK88
MK94
MK97
MK data
Normal data

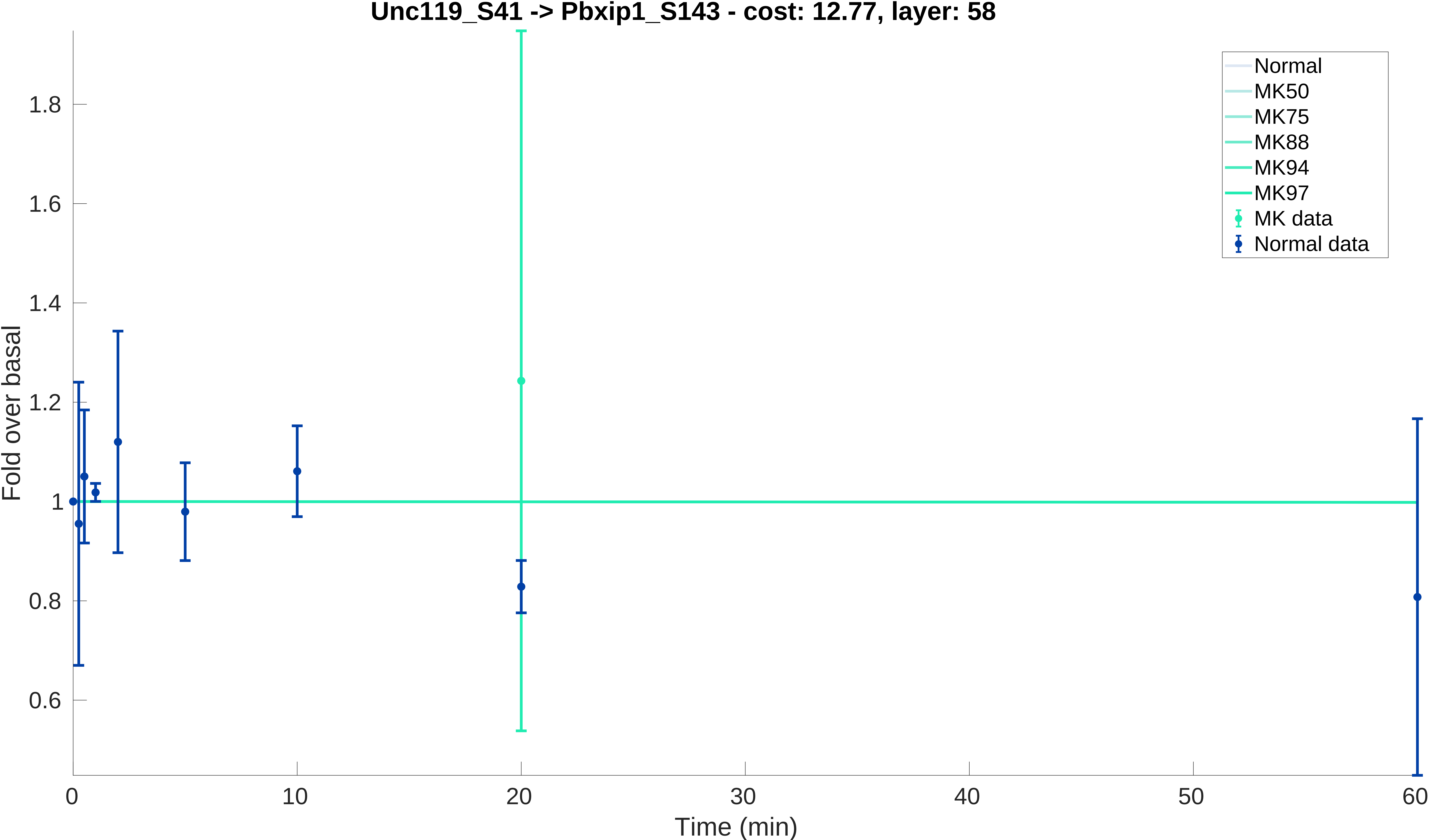

Unc119_S41 -> Pbxip1_S143 - cost: 12.77, layer: 58
Fold over basal
Time (min)
1.8
1.6
1.4
1.2
1
0.8
0.6
0
10
20
30
40
50
60
Normal
MK50
MK75
MK88
MK94
MK97
MK data
Normal data

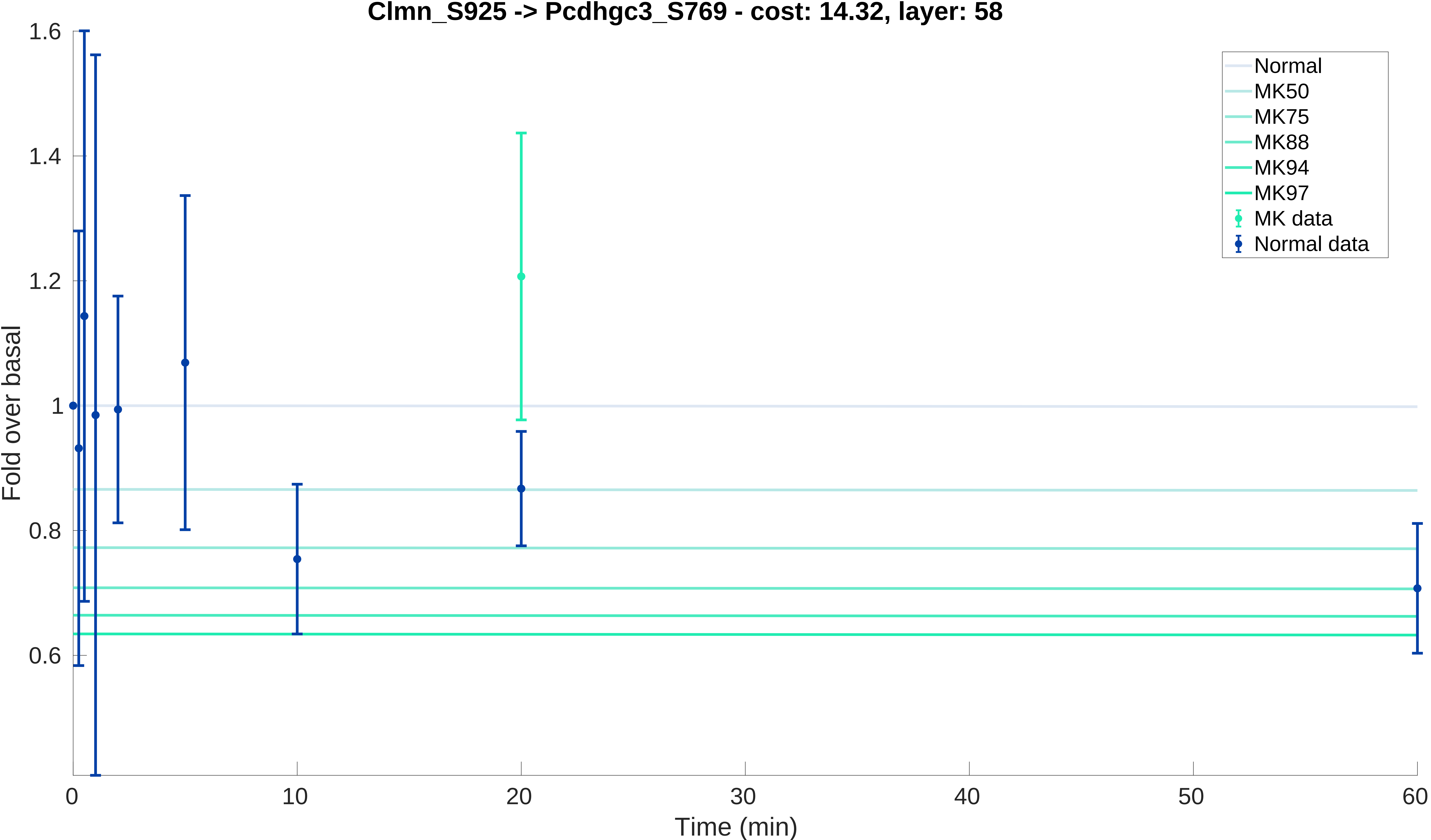

Clmn_S925 -> Pcdhgc3_S769 - cost: 14.32, layer: 58
Fold over basal
Time (min)
1.6
1.4
1.2
1
0.8
0.6
0
10
20
30
40
50
60
Normal
MK50
MK75
MK88
MK94
MK97
MK data
Normal data

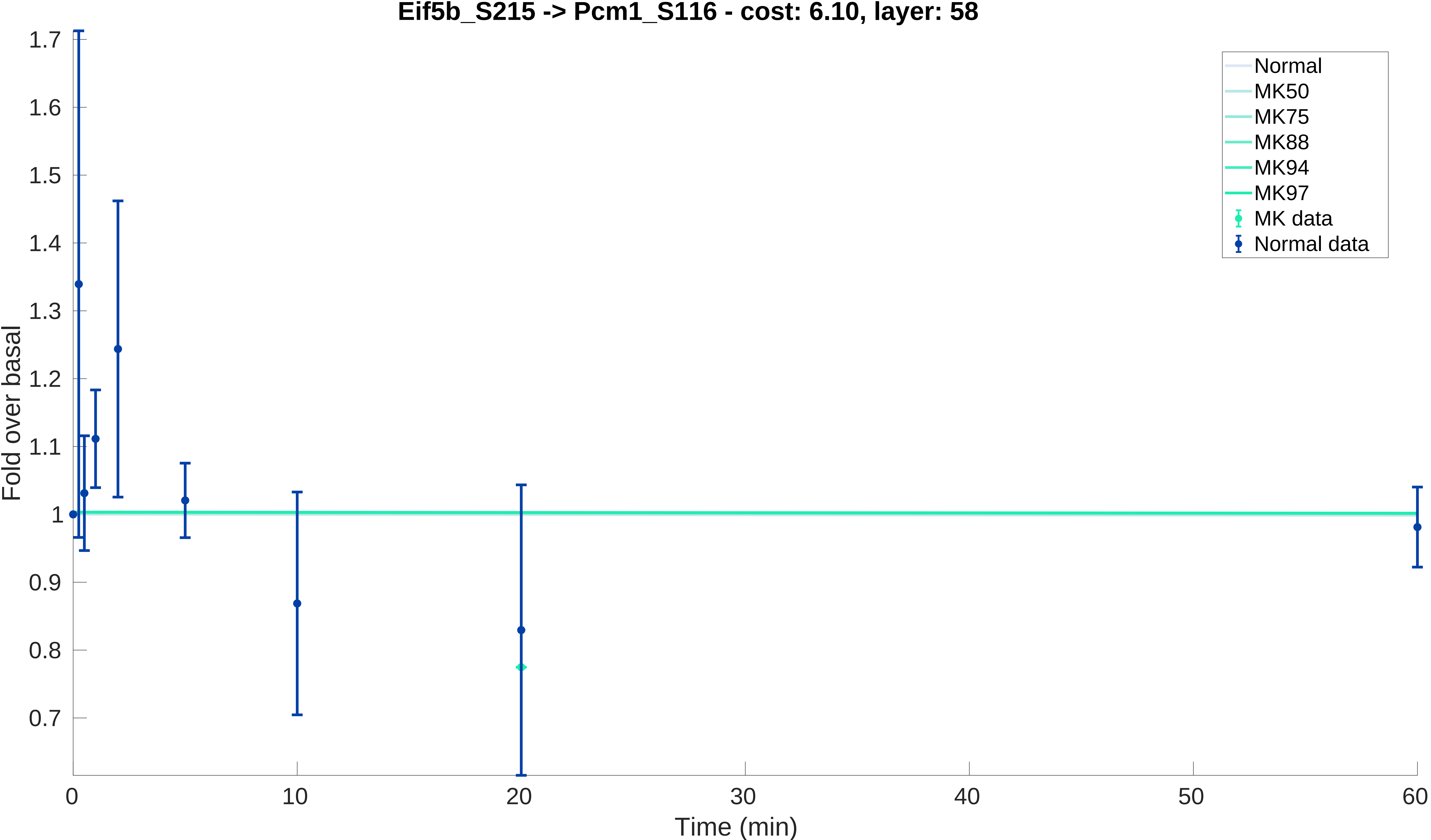
Eif5b_S215 -> Pcm1_S116 - cost: 6.10, layer: 58
Fold over basal
Time (min)
0.7
0.8
0.9
1
1.1
1.2
1.3
1.4
1.5
1.6
1.7
0
10
20
30
40
50
60
Normal
MK50
MK75
MK88
MK94
MK97
MK data
Normal data

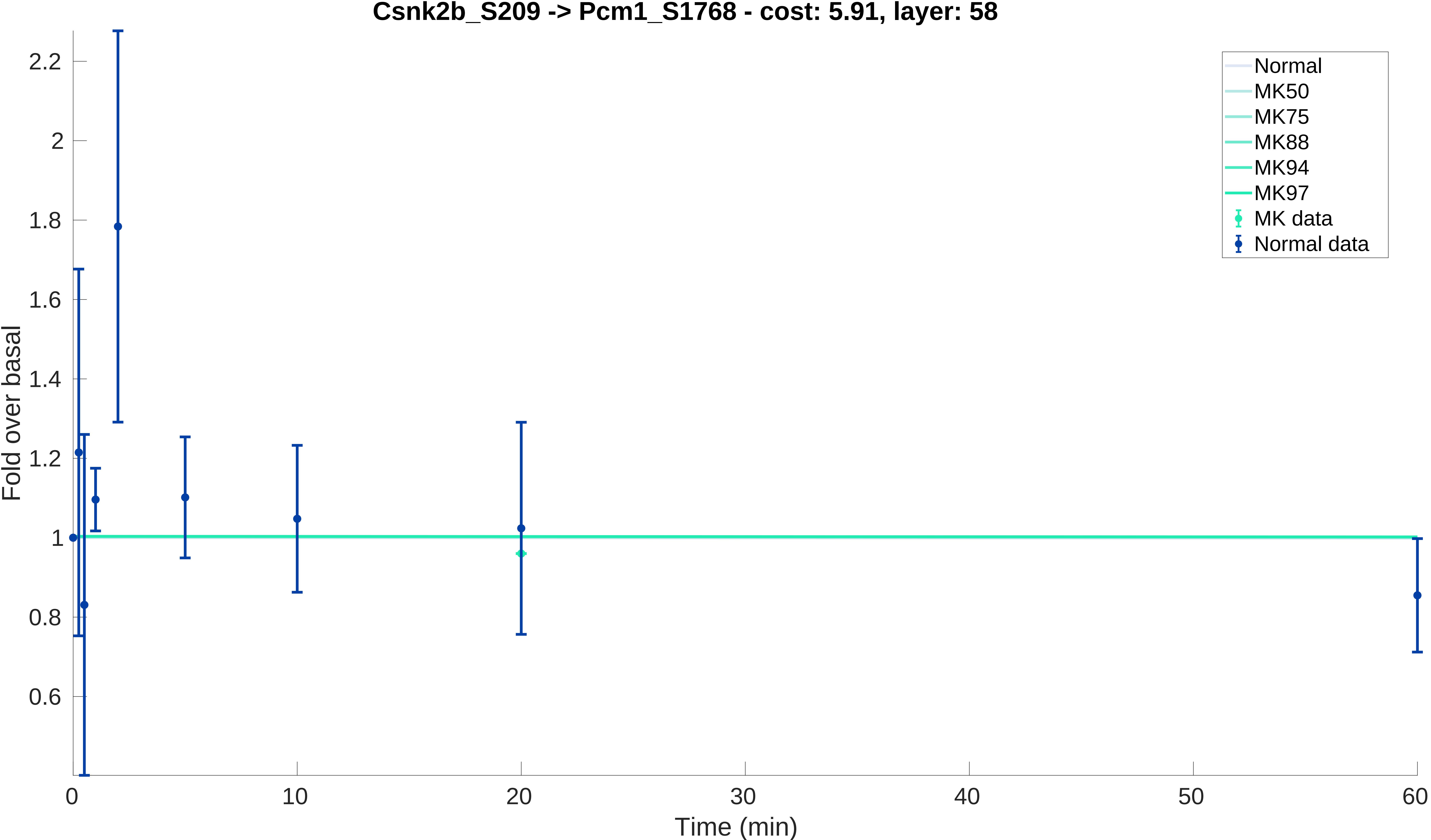

Csnk2b_S209 -> Pcm1_S1768 - cost: 5.91, layer: 58
Normal
MK50
MK75
MK88
MK94
MK97
MK data
Normal data
Fold over basal
Time (min)
0
10
20
30
40
50
60
0.6
0.8
1
1.2
1.4
1.6
1.8
2
2.2

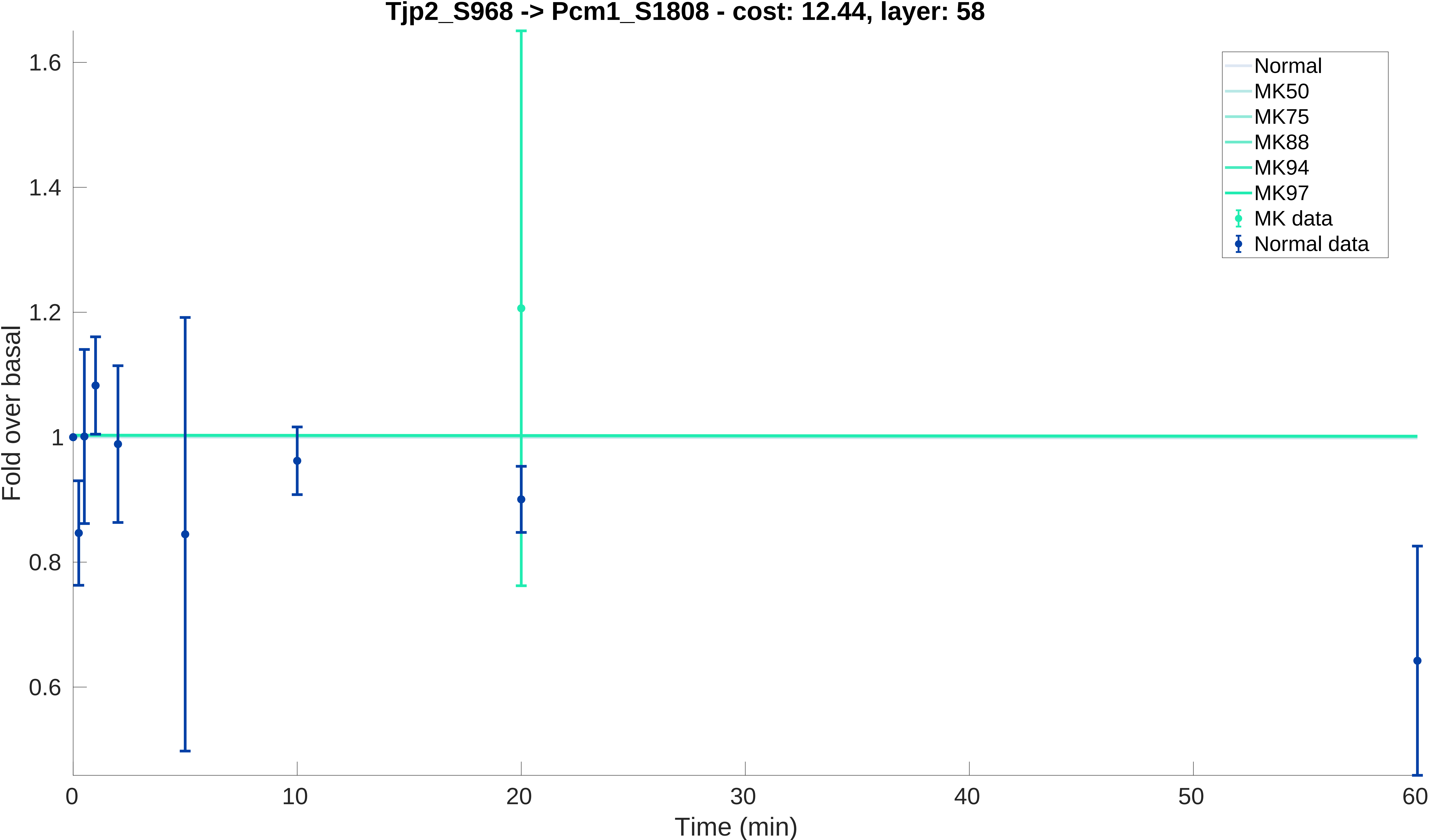
Tjp2_S968 -> Pcm1_S1808 - cost: 12.44, layer: 58
Fold over basal
Time (min)
1.6
1.4
1.2
1
0.8
0.6
0
10
20
30
40
50
60
Normal
MK50
MK75
MK88
MK94
MK97
MK data
Normal data

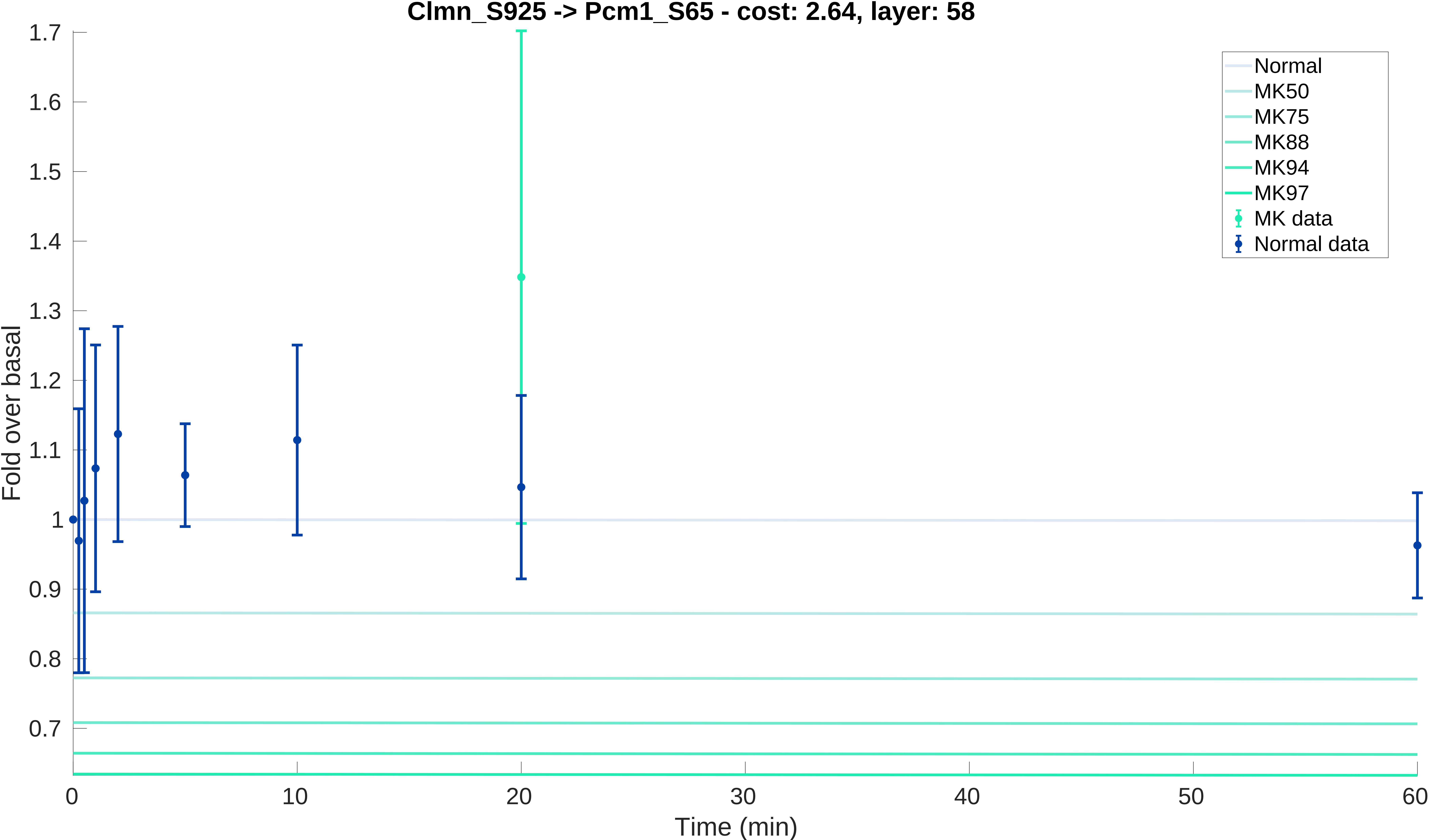

Clmn_S925 -> Pcm1_S65 - cost: 2.64, layer: 58
Fold over basal
Time (min)
1.7
1.6
1.5
1.4
1.3
1.2
1.1
1
0.9
0.8
0.7
0
10
20
30
40
50
60
Normal
MK50
MK75
MK88
MK94
MK97
MK data
Normal data

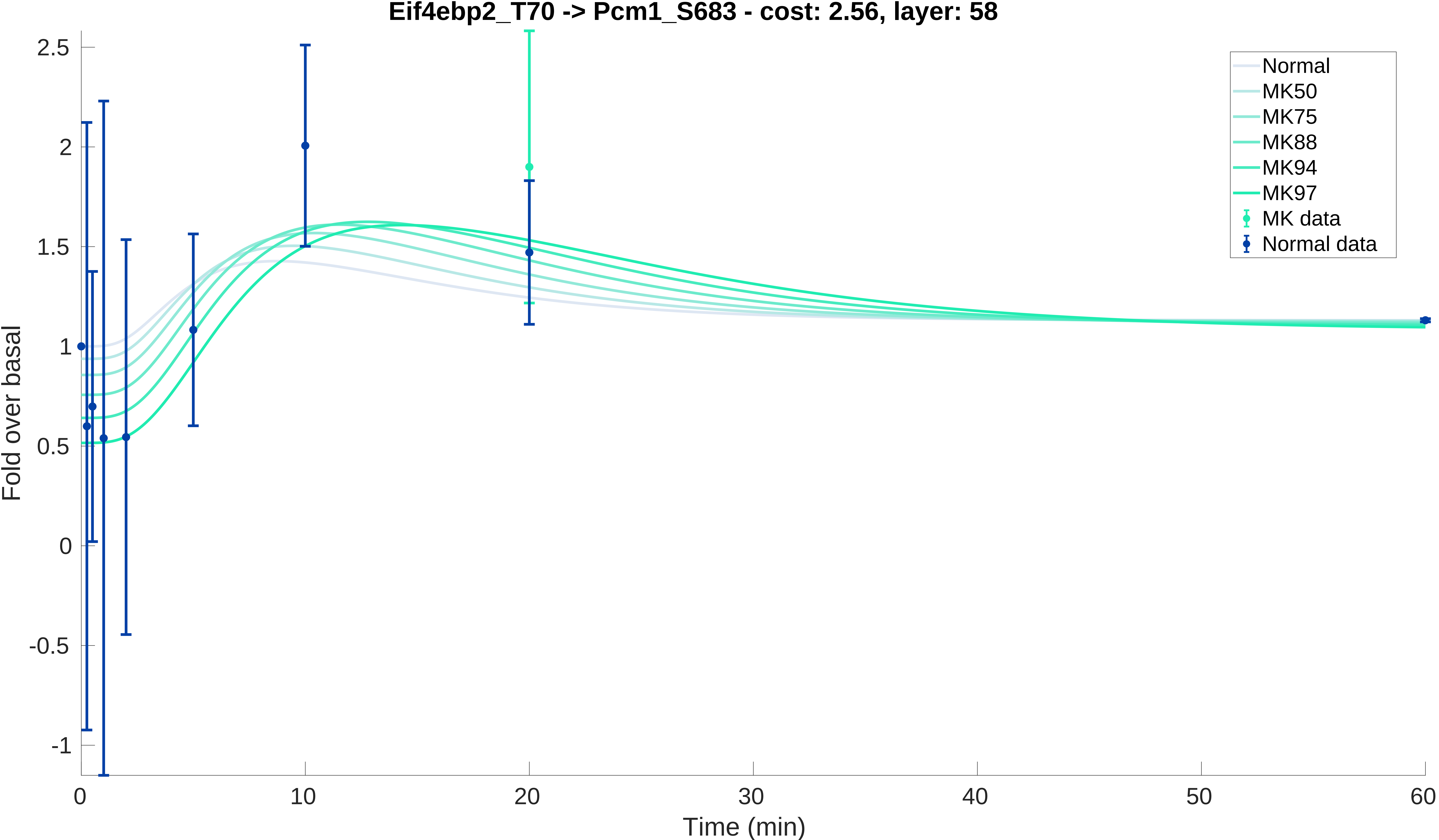

Eif4ebp2_T70 -> Pcm1_S683 - cost: 2.56, layer: 58
Fold over basal
Time (min)
2.5
2
1.5
1
0.5
0
-0.5
-1
0
10
20
30
40
50
60
Normal
MK50
MK75
MK88
MK94
MK97
MK data
Normal data

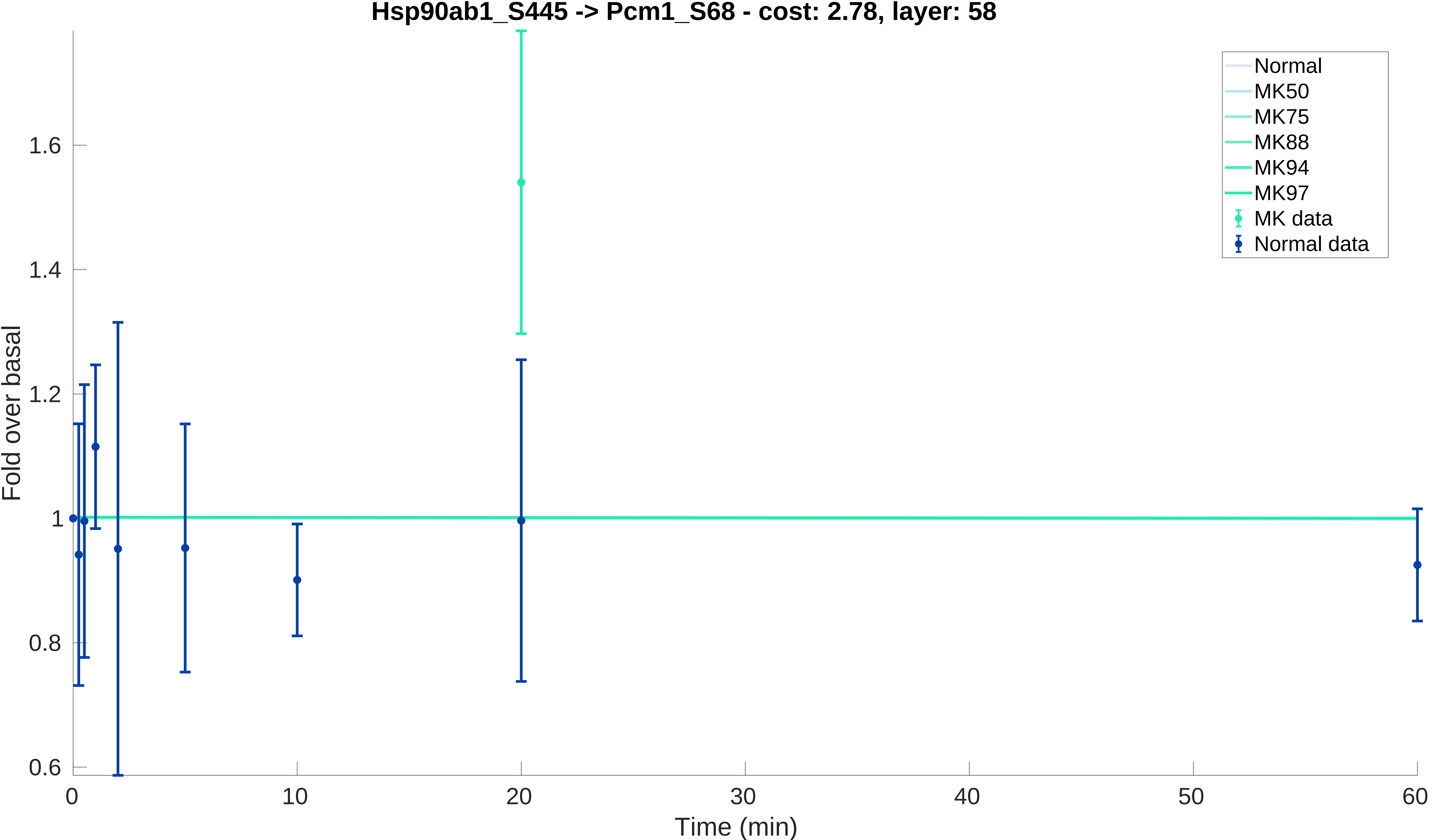

Hsp90ab1_S445 -> Pcm1_S68 - cost: 2.78, layer: 58
Fold over basal
1.6
1.4
1.2
1
0.8
0.6
0
10
20
30
40
50
60
Time (min)
Normal
MK50
MK75
MK88
MK94
MK97
MK data
Normal data

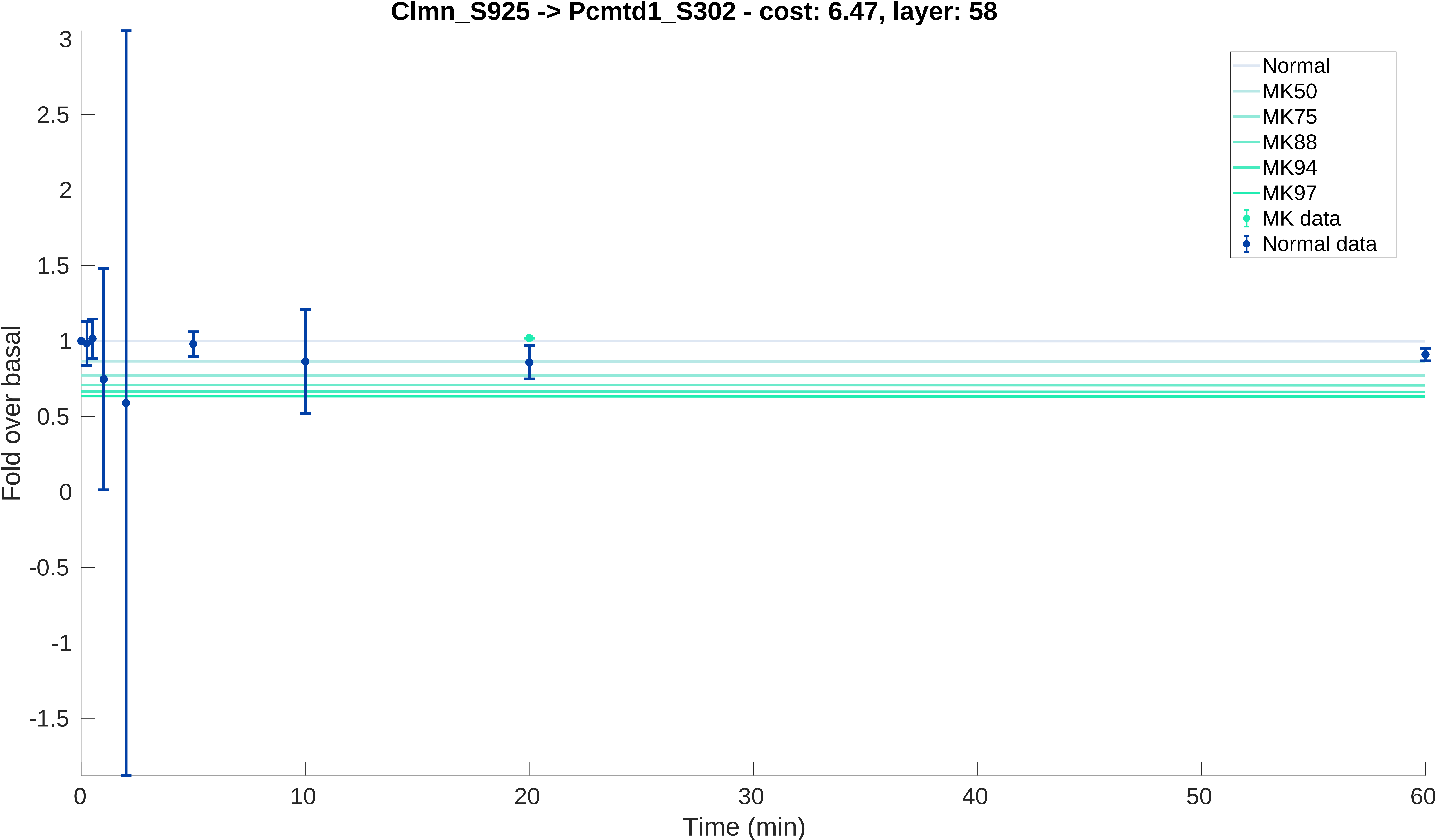
Clmn_S925 -> Pcmtd1_S302 - cost: 6.47, layer: 58
Fold over basal
Time (min)
Normal
MK50
MK75
MK88
MK94
MK97
MK data
Normal data
3
2.5
2
1.5
1
0.5
0
-0.5
-1
-1.5
0
10
20
30
40
50
60

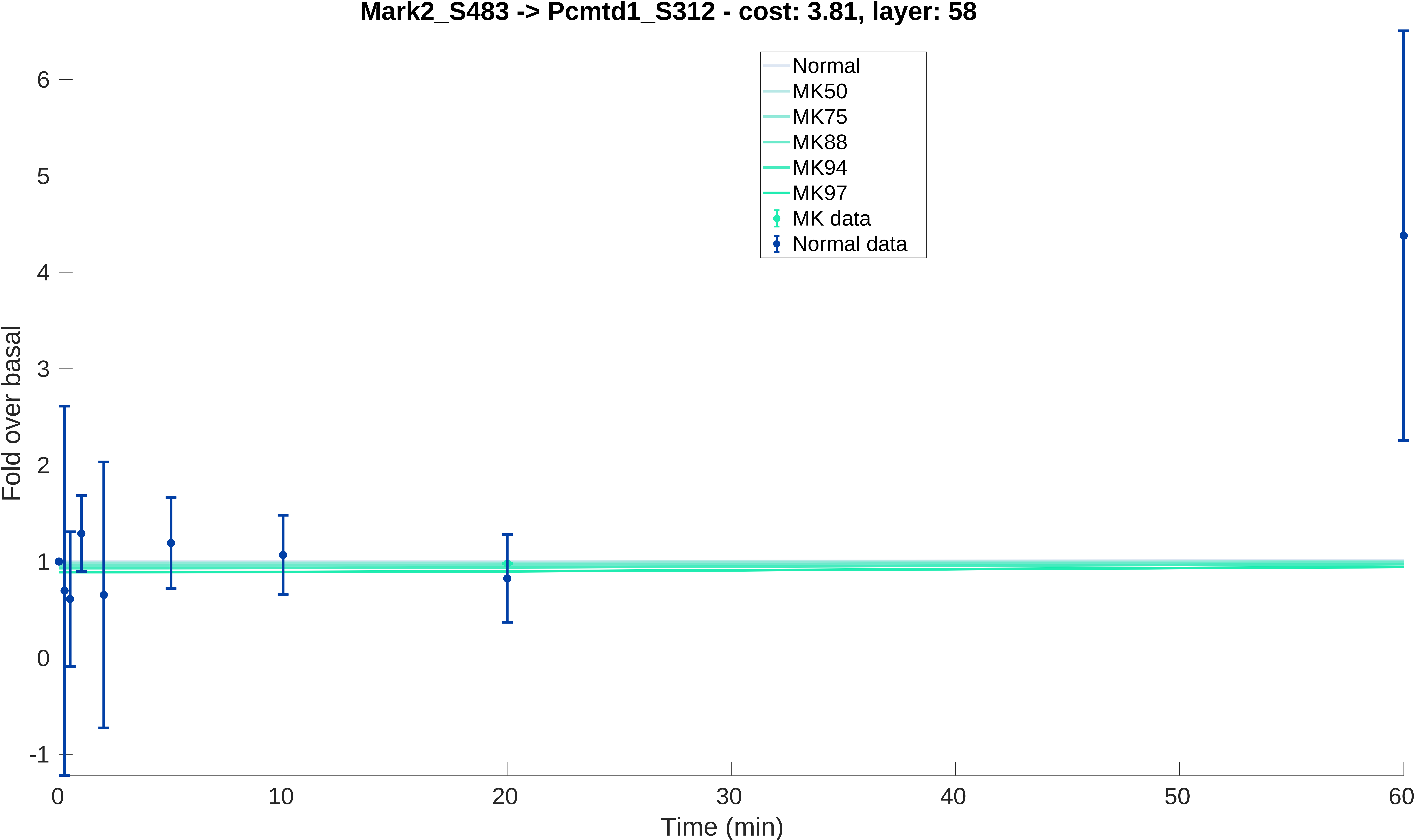

Mark2_S483 -> Pcmtd1_S312 - cost: 3.81, layer: 58
Normal
MK50
MK75
MK88
MK94
MK97
MK data
Normal data
Fold over basal
Time (min)
-1
0
1
2
3
4
5
6
0
10
20
30
40
50
60

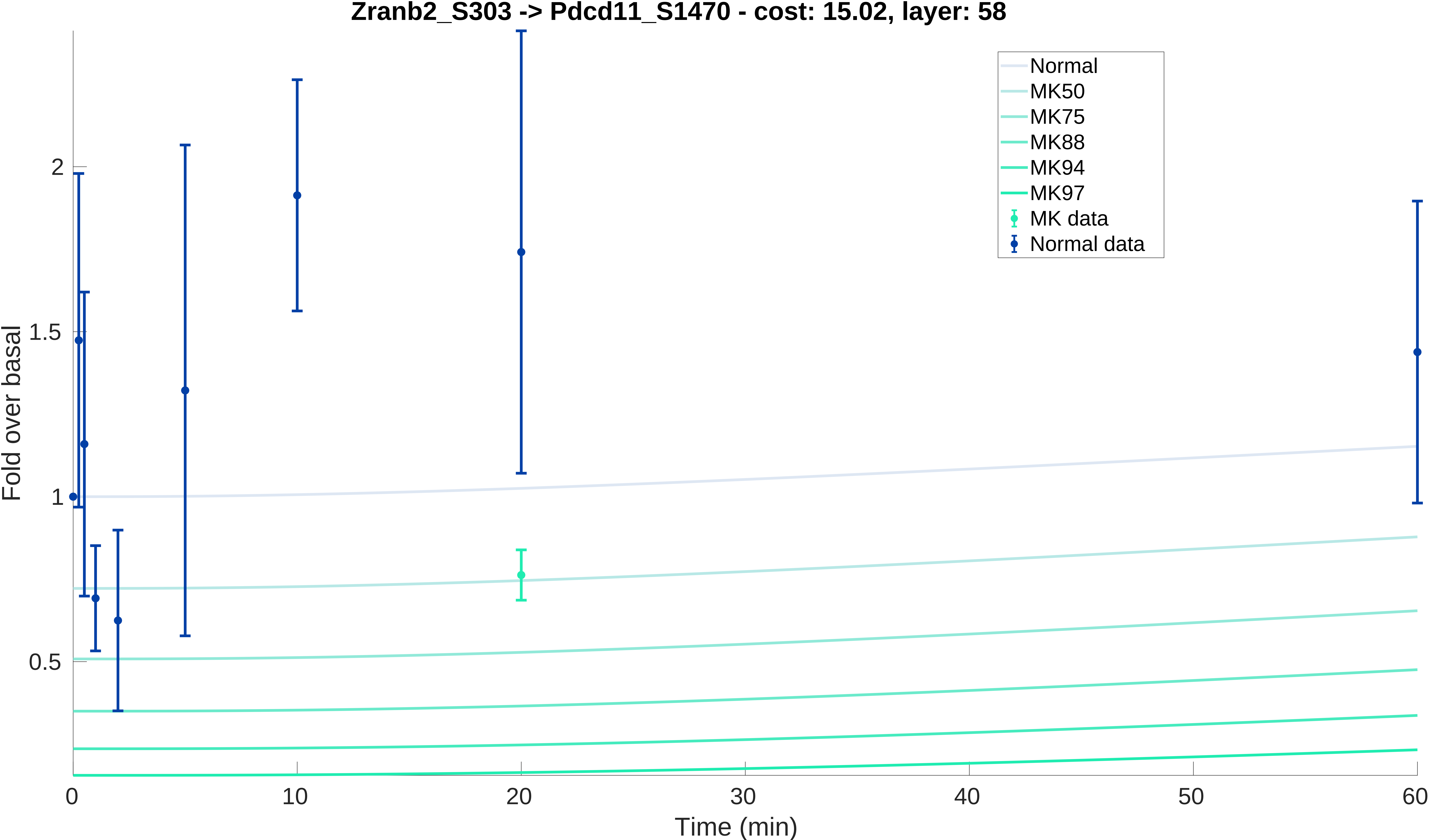

Zranb2_S303 -> Pdcd11_S1470 - cost: 15.02, layer: 58
Fold over basal
Time (min)
0
10
20
30
40
50
60
0.5
1
1.5
2
Normal
MK50
MK75
MK88
MK94
MK97
MK data
Normal data

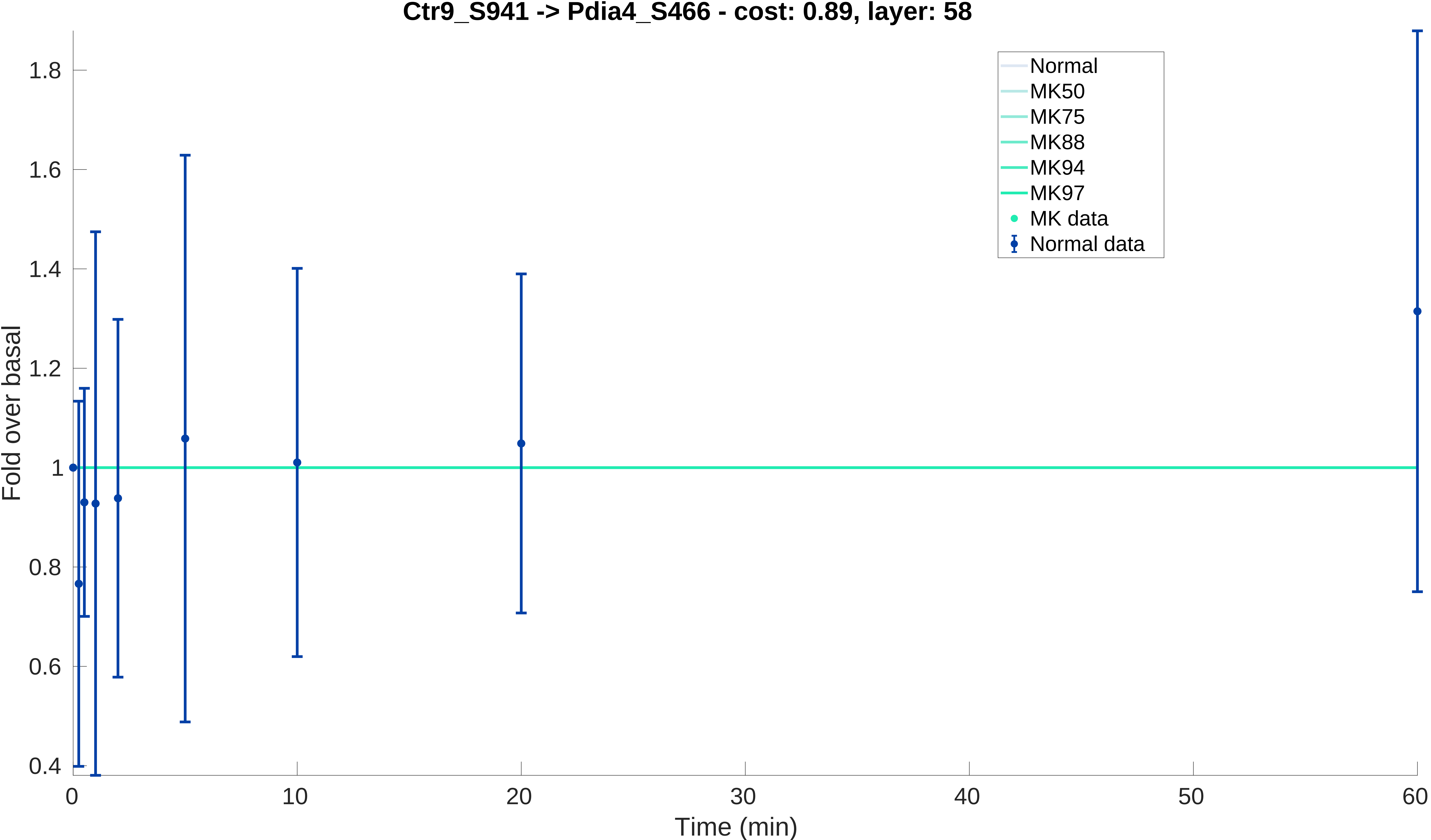

Ctr9_S941 -> Pdia4_S466 - cost: 0.89, layer: 58
Normal
MK50
MK75
MK88
MK94
MK97
MK data
Normal data
Fold over basal
Time (min)
0.4
0.6
0.8
1
1.2
1.4
1.6
1.8
0
10
20
30
40
50
60

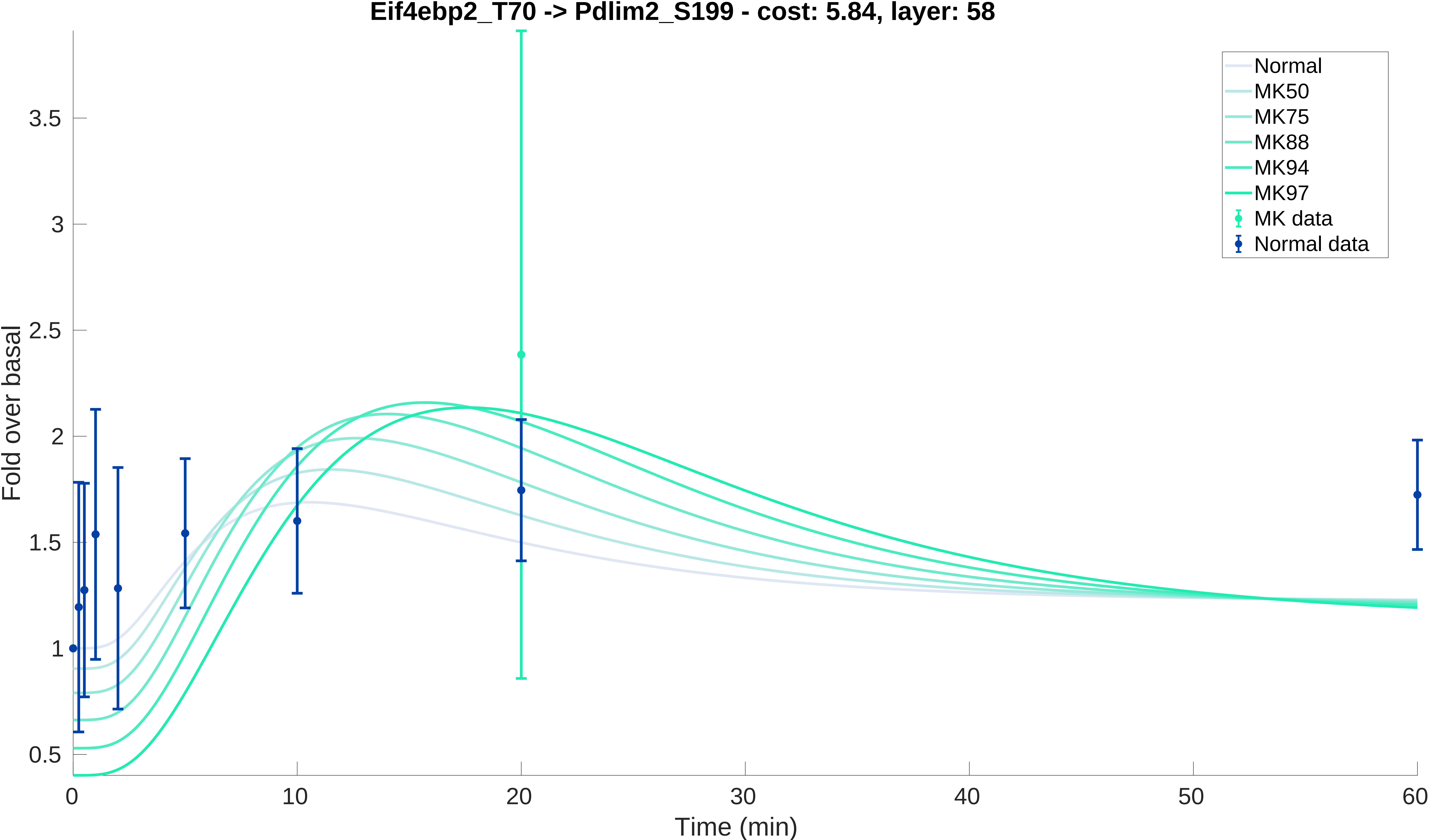

Eif4ebp2_T70 -> Pdlim2_S199 - cost: 5.84, layer: 58
Fold over basal
3.5
3
2.5
2
1.5
1
0.5
0
10
20
30
40
50
60
Time (min)
Normal
MK50
MK75
MK88
MK94
MK97
MK data
Normal data

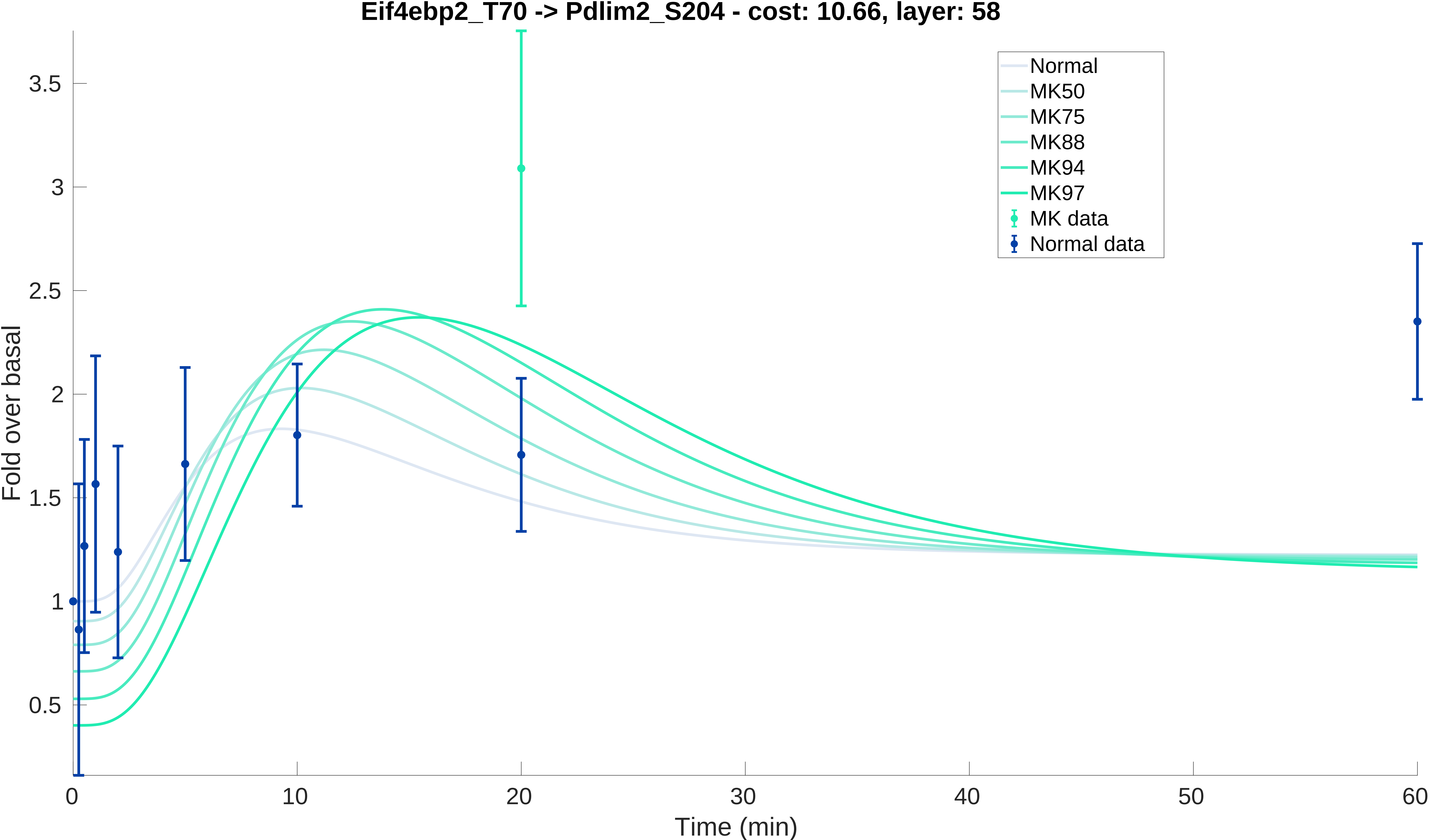

Eif4ebp2_T70 -> Pdlim2_S204 - cost: 10.66, layer: 58
Normal
MK50
MK75
MK88
MK94
MK97
MK data
Normal data
Fold over basal
0.5
1
1.5
2
2.5
3
3.5
0
10
20
30
40
50
60
Time (min)

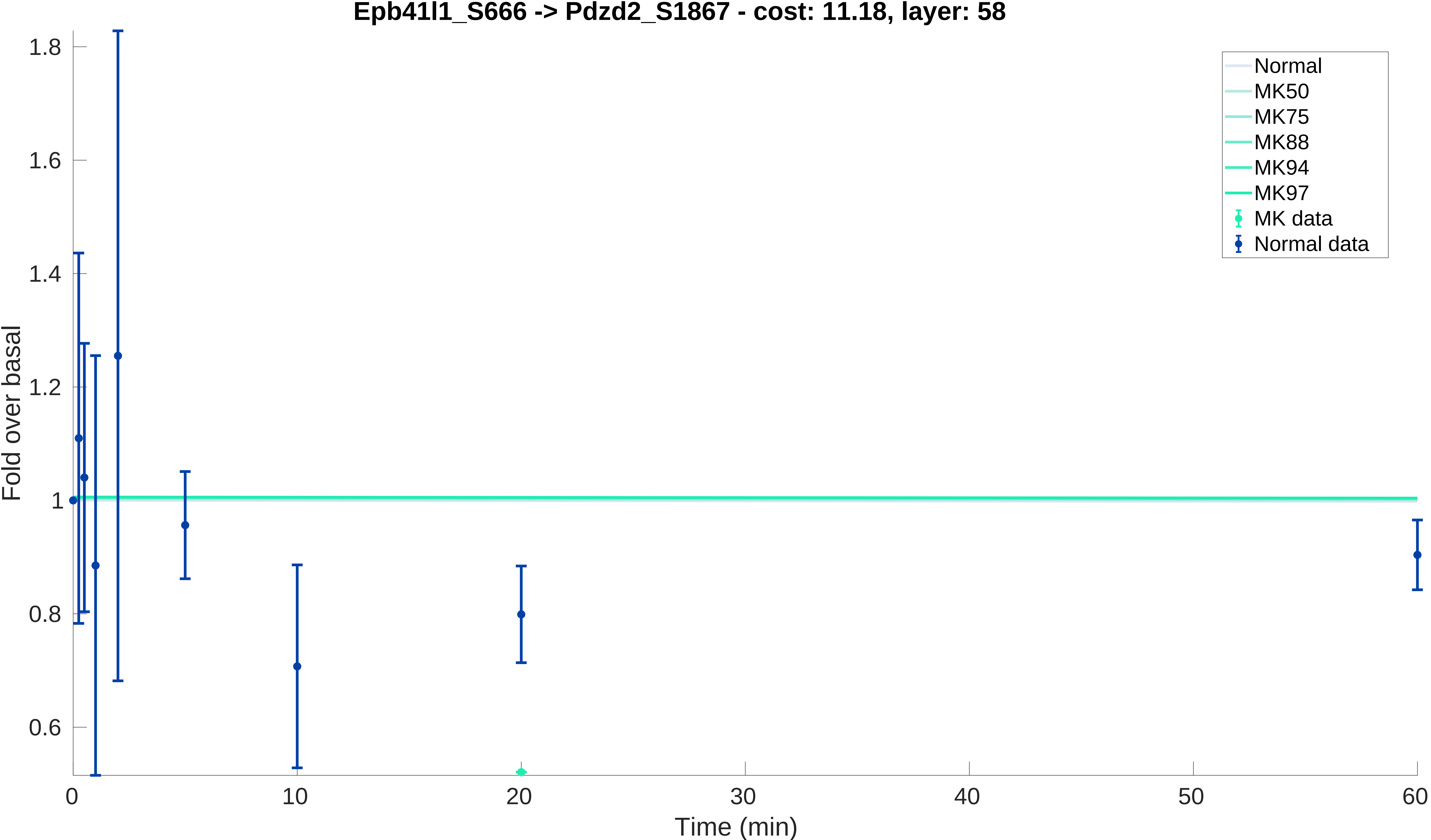

Epb41l1_S666 -> Pdzd2_S1867 - cost: 11.18, layer: 58
Fold over basal
Time (min)
1.8
1.6
1.4
1.2
1
0.8
0.6
0
10
20
30
40
50
60
Normal
MK50
MK75
MK88
MK94
MK97
MK data
Normal data

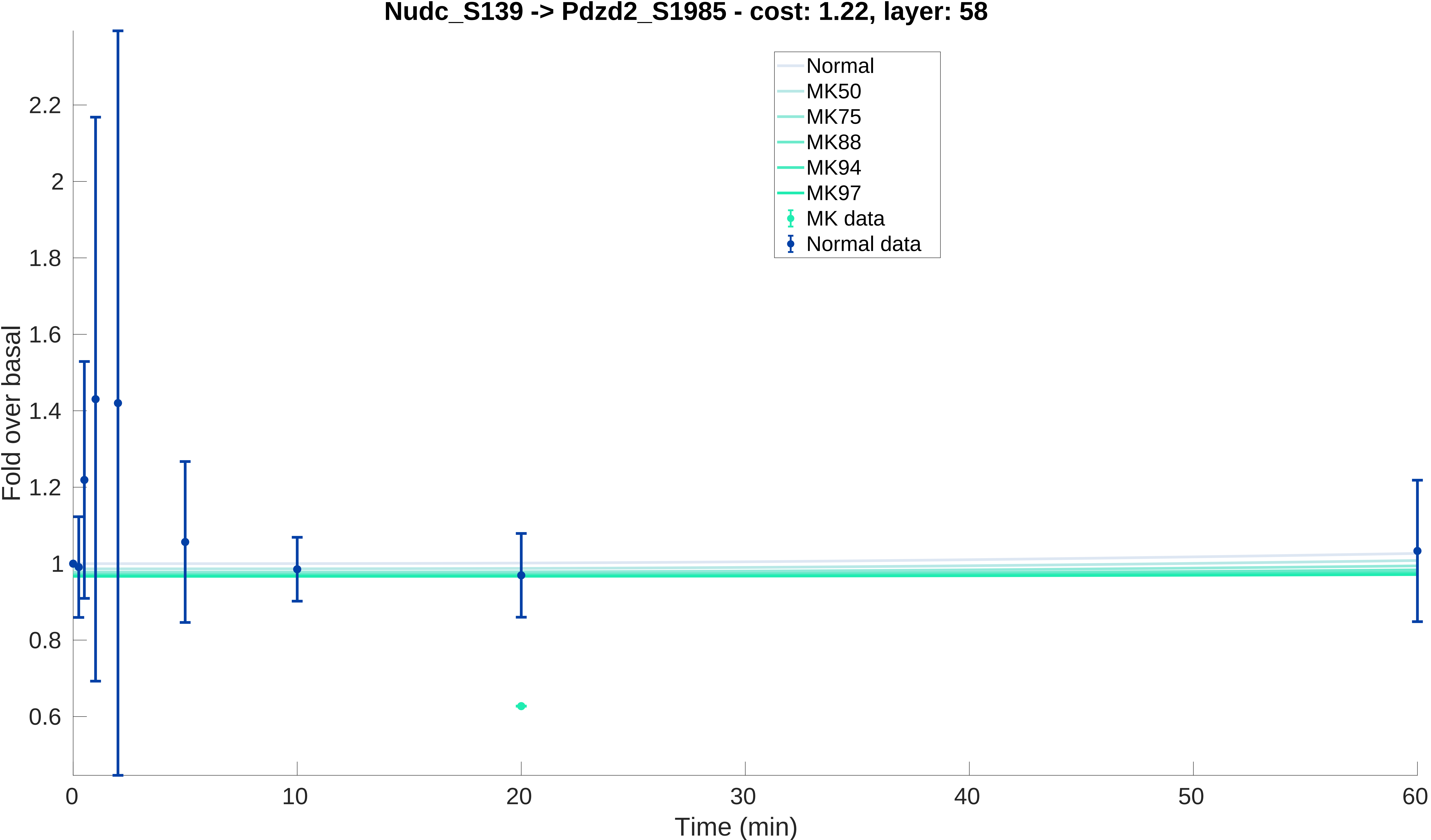
Nudc_S139 -> Pdzd2_S1985 - cost: 1.22, layer: 58
Normal
MK50
MK75
MK88
MK94
MK97
MK data
Normal data
Fold over basal
2.2
2
1.8
1.6
1.4
1.2
1
0.8
0.6
0
10
20
30
40
50
60
Time (min)

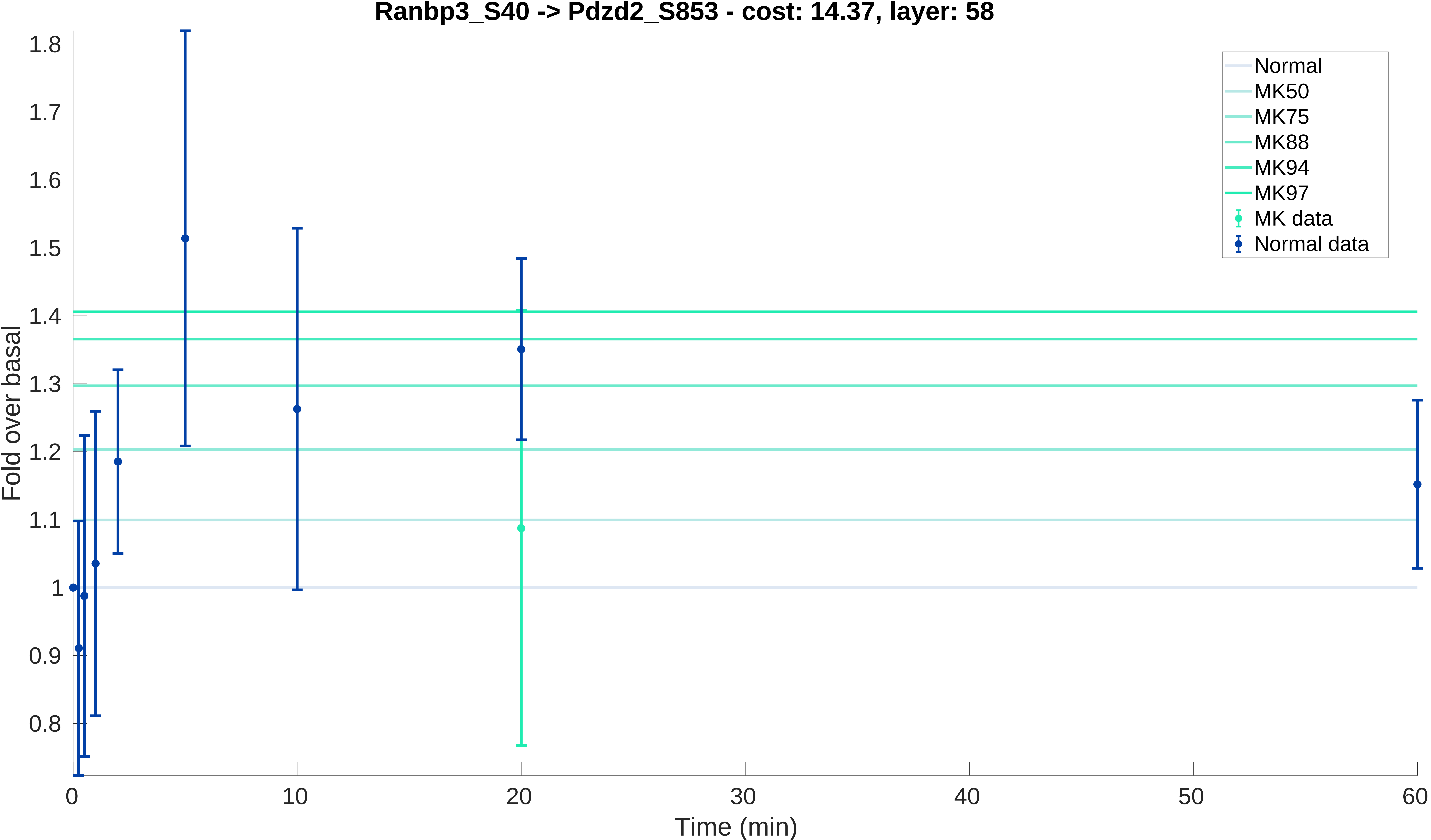

Ranbp3_S40 -> Pdzd2_S853 - cost: 14.37, layer: 58
Fold over basal
Time (min)
0
10
20
30
40
50
60
0.8
0.9
1
1.1
1.2
1.3
1.4
1.5
1.6
1.7
1.8
Normal
MK50
MK75
MK88
MK94
MK97
MK data
Normal data

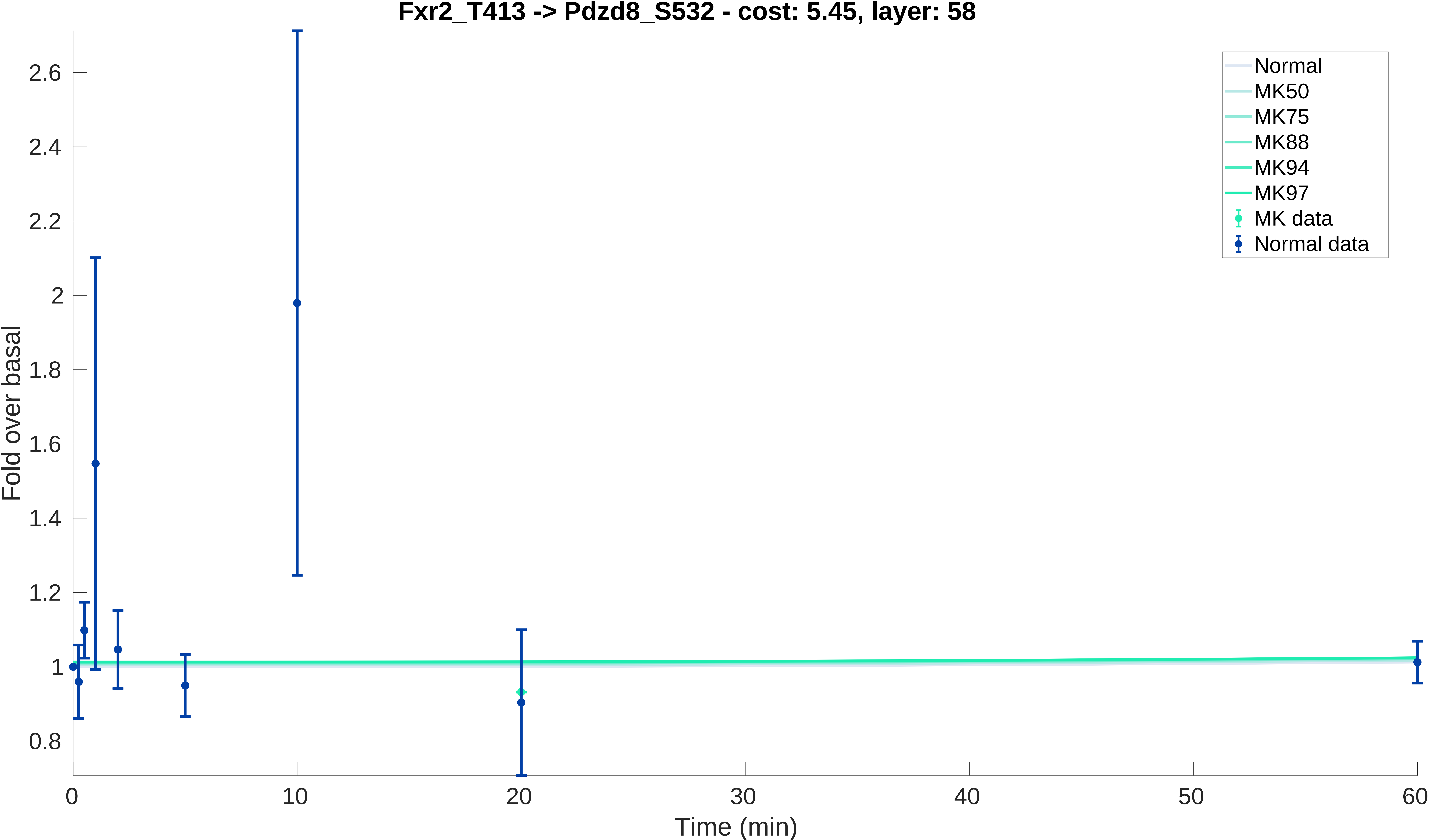

Fxr2_T413 -> Pdzd8_S532 - cost: 5.45, layer: 58
Fold over basal
2.6
2.4
2.2
2
1.8
1.6
1.4
1.2
1
0.8
0
10
20
30
40
50
60
Time (min)
Normal
MK50
MK75
MK88
MK94
MK97
MK data
Normal data

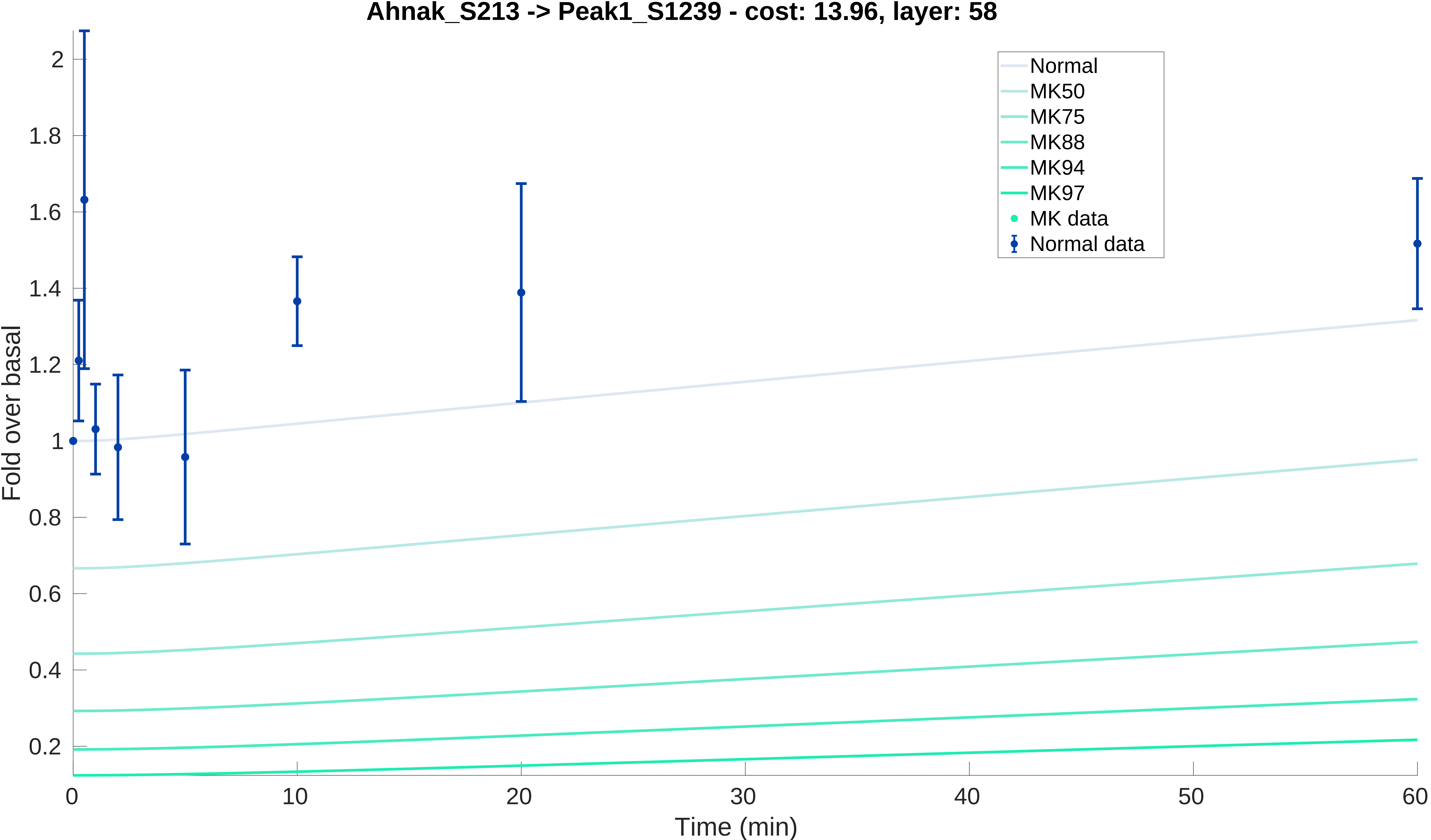

Ahnak_S213 -> Peak1_S1239 - cost: 13.96, layer: 58
Fold over basal
Time (min)
0.2
0.4
0.6
0.8
1
1.2
1.4
1.6
1.8
2
0
10
20
30
40
50
60
Normal
MK50
MK75
MK88
MK94
MK97
MK data
Normal data

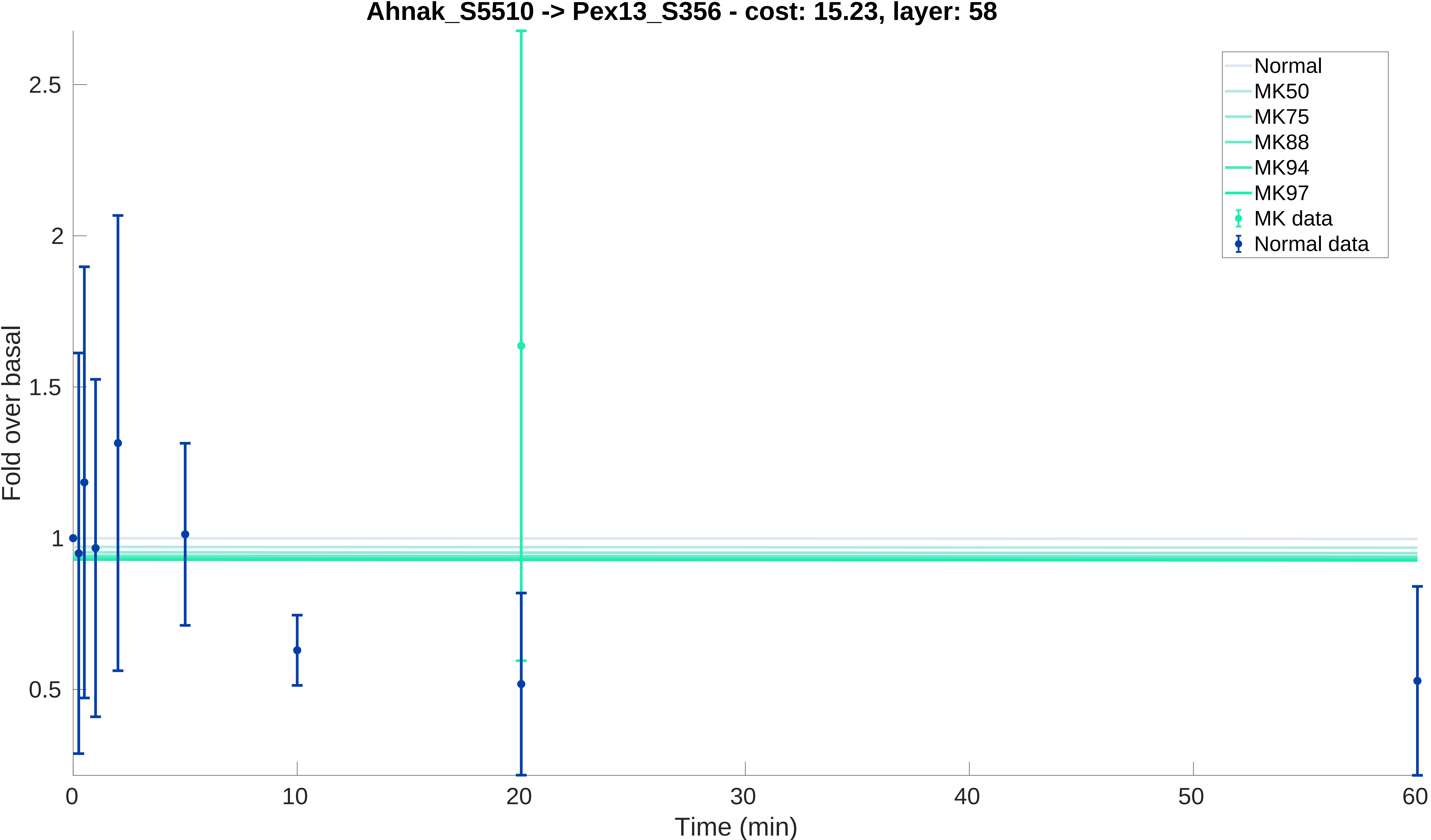

Ahnak_S5510 -> Pex13_S356 - cost: 15.23, layer: 58
Fold over basal
Time (min)
0.5
1
1.5
2
2.5
0
10
20
30
40
50
60
Normal
MK50
MK75
MK88
MK94
MK97
MK data
Normal data

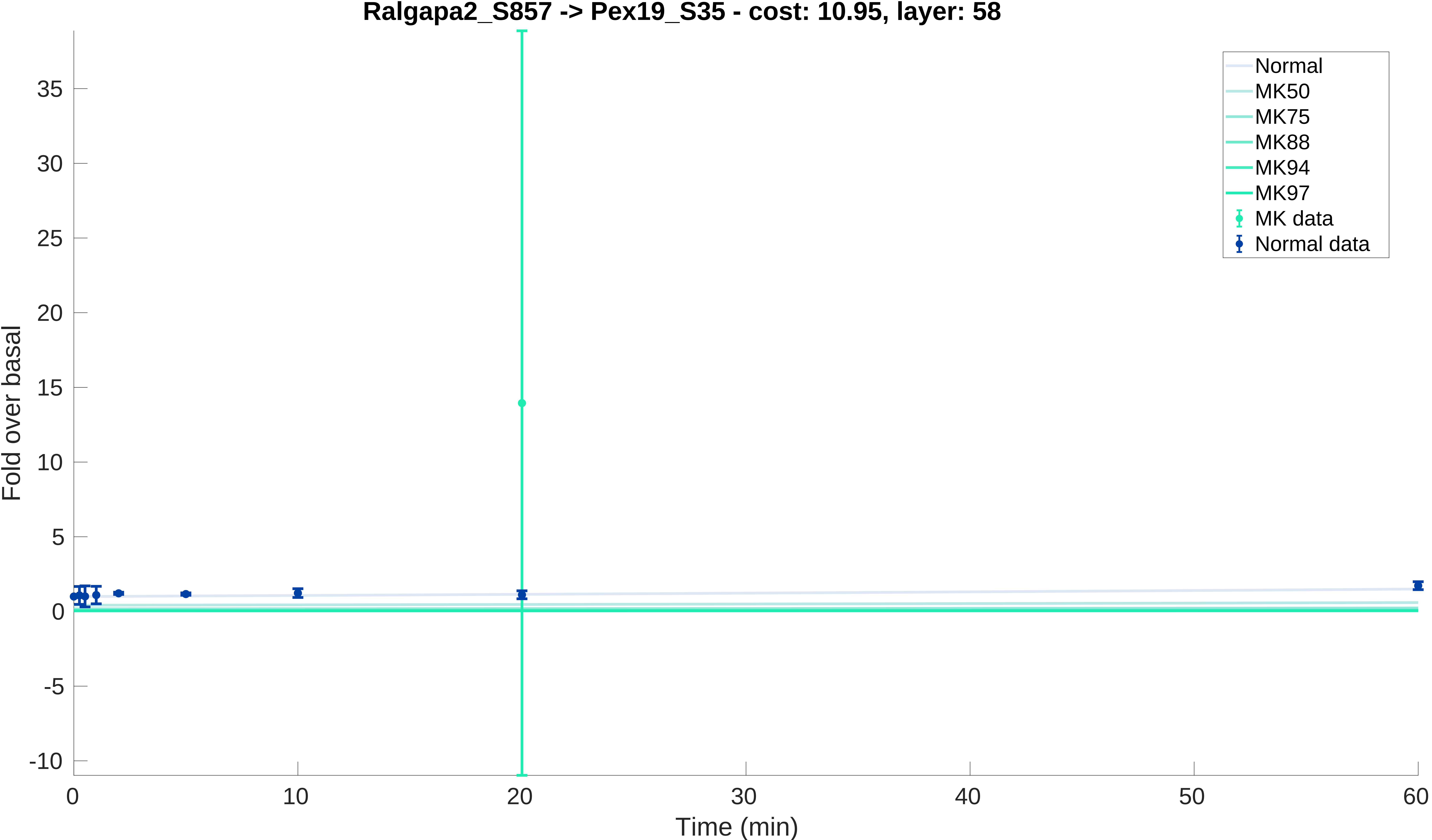

Ralgapa2_S857 -> Pex19_S35 - cost: 10.95, layer: 58
Fold over basal
Time (min)
35
30
25
20
15
10
5
0
-5
-10
0
10
20
30
40
50
60
Normal
MK50
MK75
MK88
MK94
MK97
MK data
Normal data

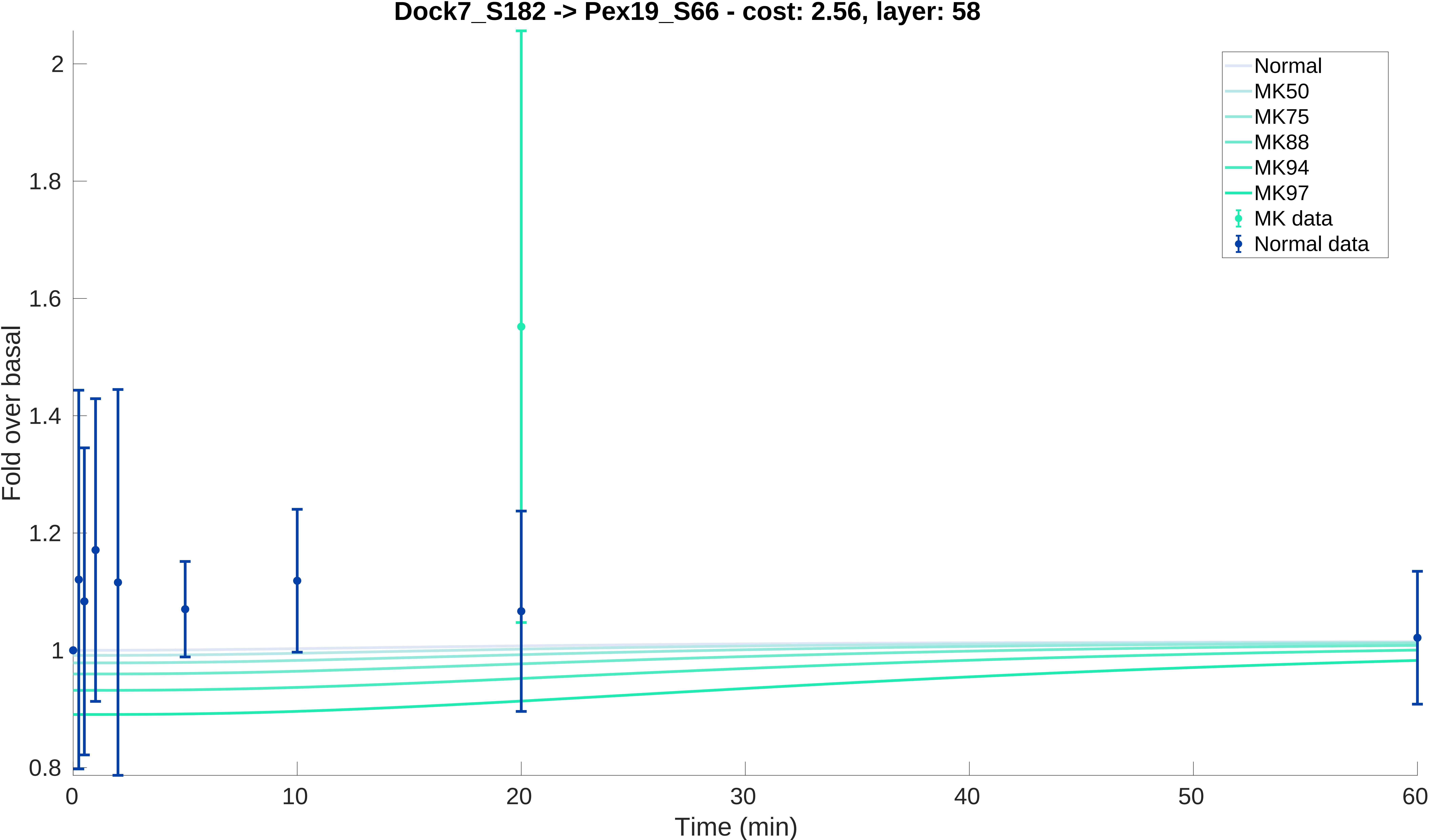

Dock7_S182 -> Pex19_S66 - cost: 2.56, layer: 58
Fold over basal
Time (min)
0.8
1
1.2
1.4
1.6
1.8
2
0
10
20
30
40
50
60
Normal
MK50
MK75
MK88
MK94
MK97
MK data
Normal data

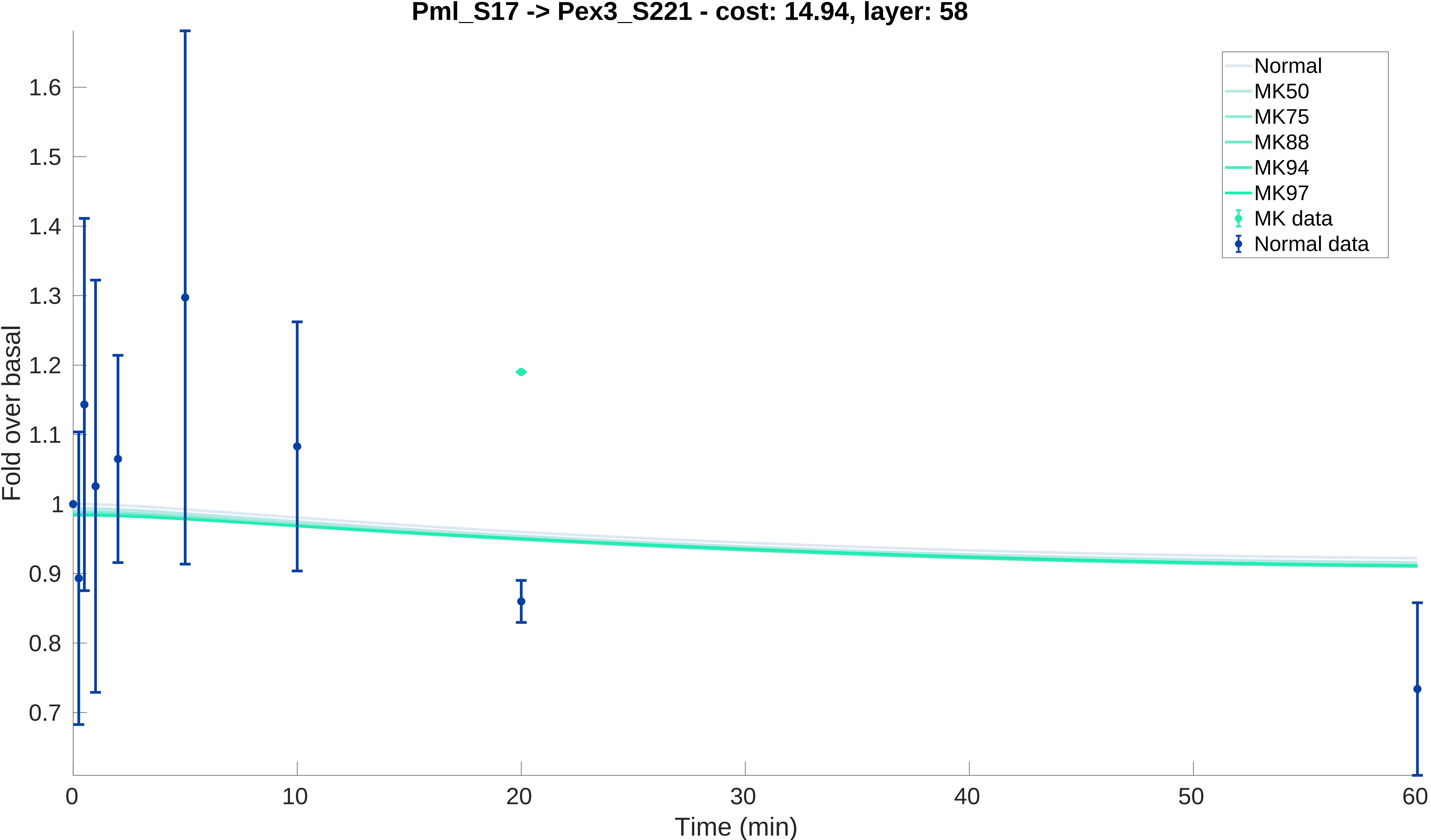
Pml_S17 -> Pex3_S221 - cost: 14.94, layer: 58
Fold over basal
Time (min)
0.7
0.8
0.9
1
1.1
1.2
1.3
1.4
1.5
1.6
0
10
20
30
40
50
60
Normal
MK50
MK75
MK88
MK94
MK97
MK data
Normal data

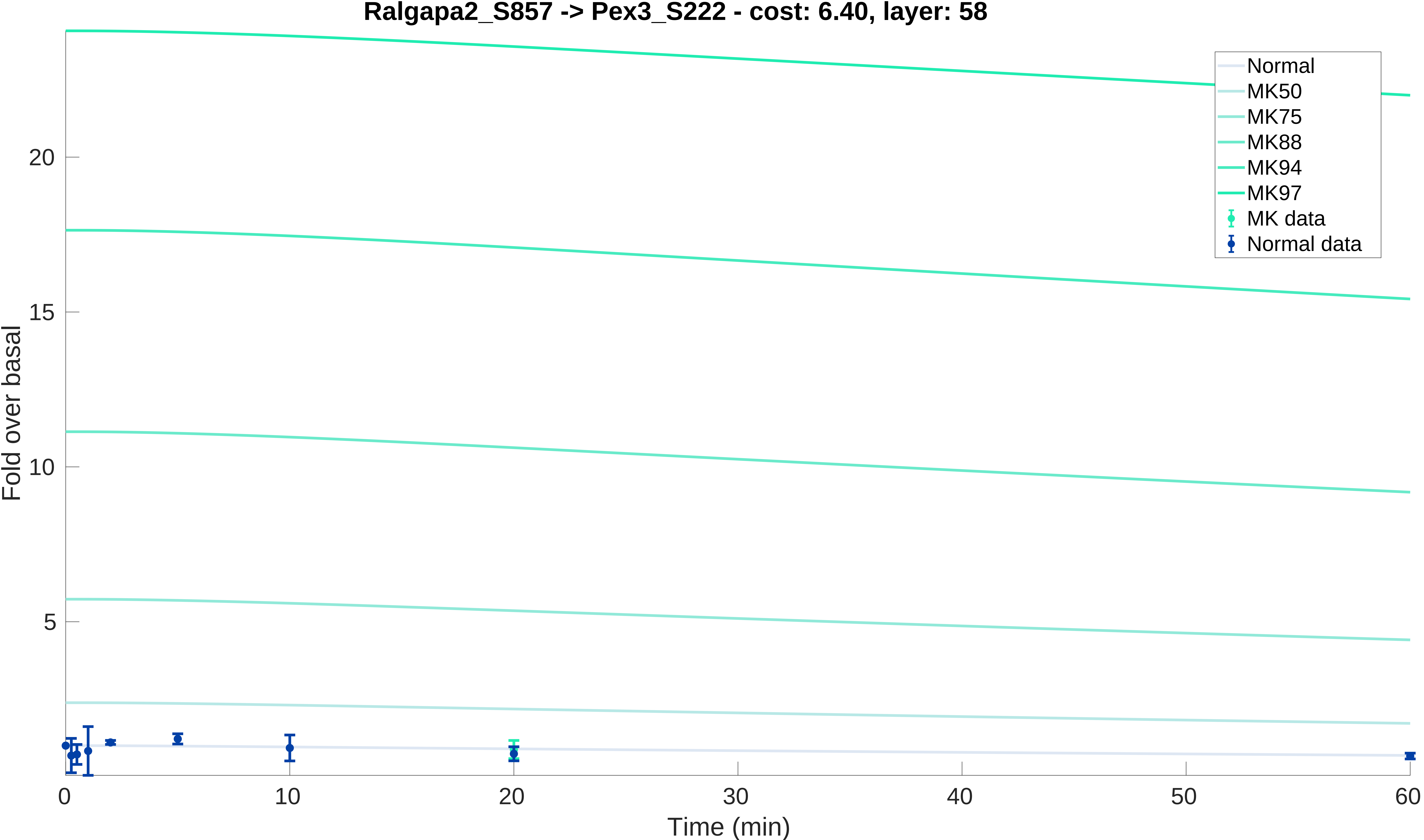

Ralgapa2_S857 -> Pex3_S222 - cost: 6.40, layer: 58
Fold over basal
Time (min)
0
10
20
30
40
50
60
5
10
15
20
Normal
MK50
MK75
MK88
MK94
MK97
MK data
Normal data

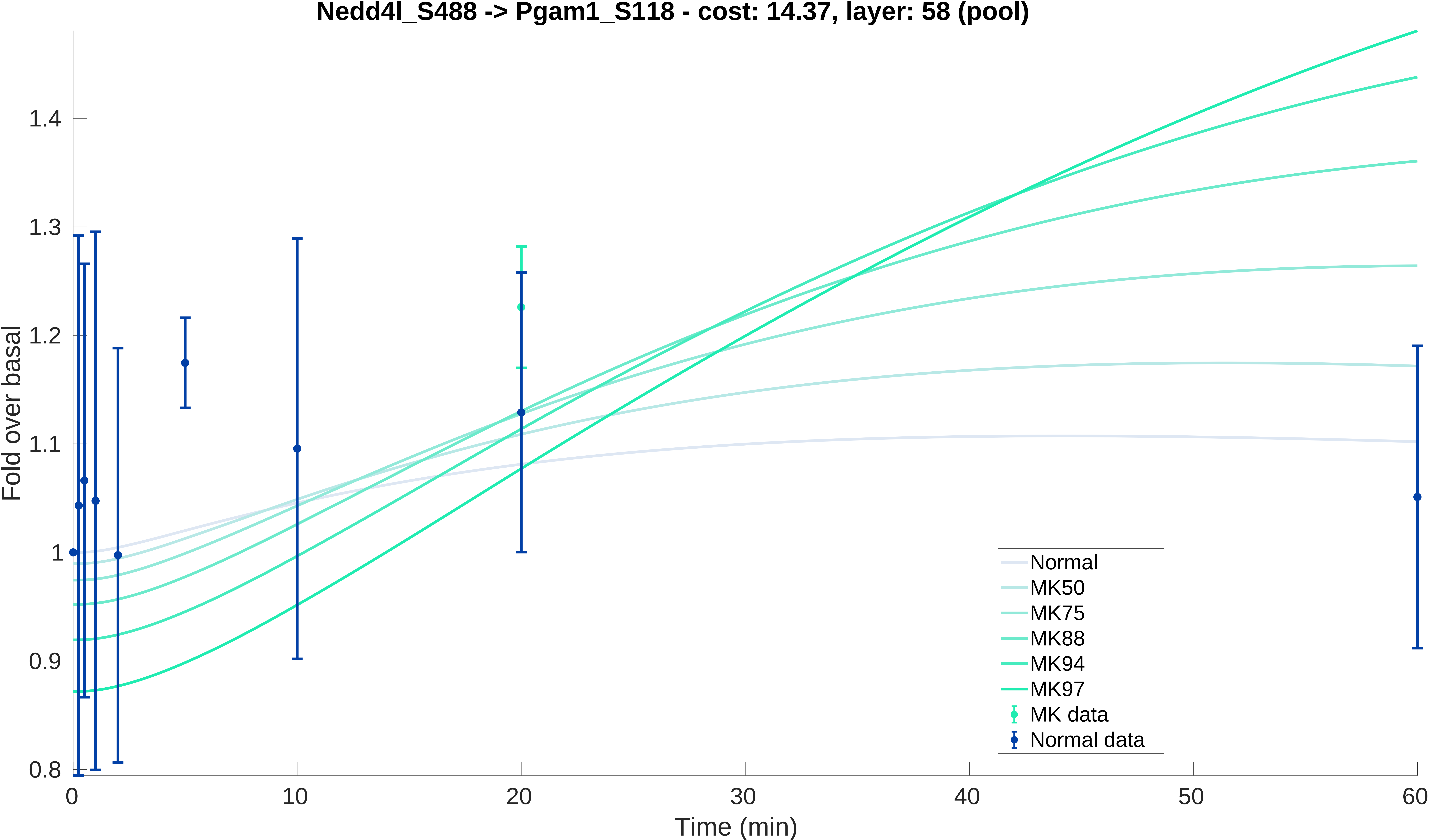
Nedd4l_S488 -> Pgam1_S118 - cost: 14.37, layer: 58 (pool)
Fold over basal
Time (min)
0.8
0.9
1
1.1
1.2
1.3
1.4
0
10
20
30
40
50
60
Normal
MK50
MK75
MK88
MK94
MK97
MK data
Normal data

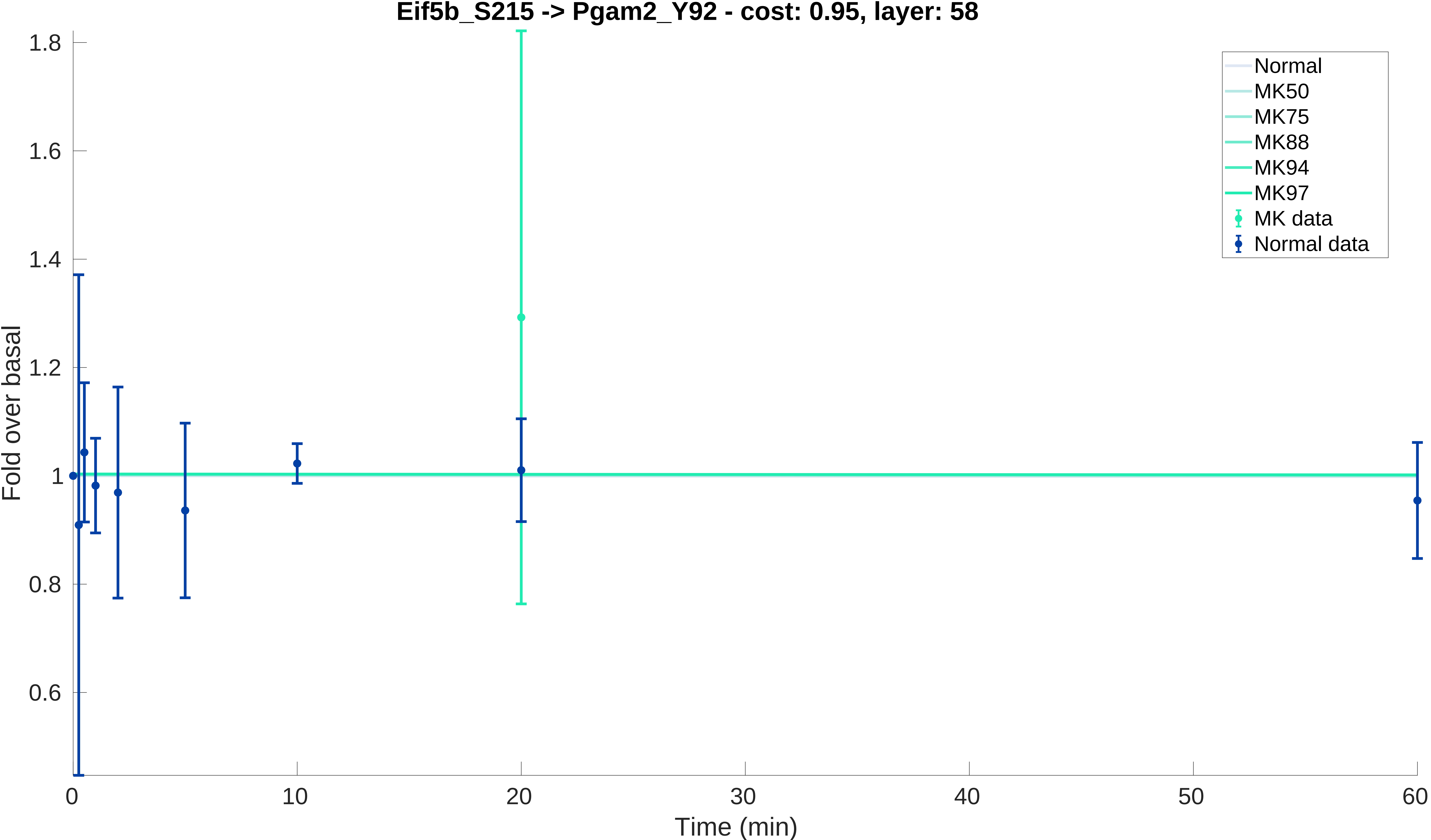
Eif5b_S215 -> Pgam2_Y92 - cost: 0.95, layer: 58
Fold over basal
Time (min)
0.6
0.8
1
1.2
1.4
1.6
1.8
0
10
20
30
40
50
60
Normal
MK50
MK75
MK88
MK94
MK97
MK data
Normal data

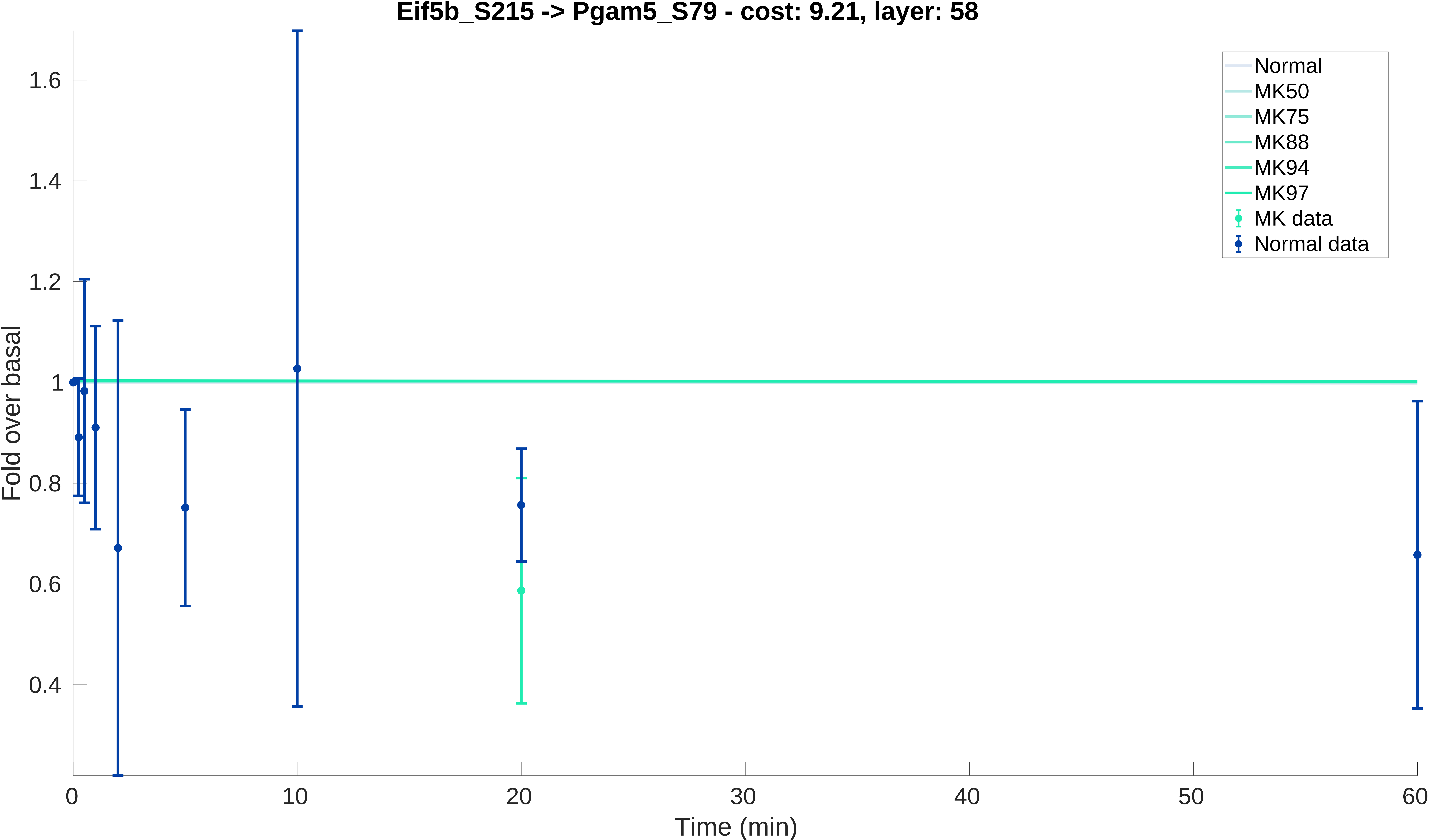
Eif5b_S215 -> Pgam5_S79 - cost: 9.21, layer: 58
Fold over basal
Time (min)
Normal
MK50
MK75
MK88
MK94
MK97
MK data
Normal data
0
10
20
30
40
50
60
0.4
0.6
0.8
1
1.2
1.4
1.6

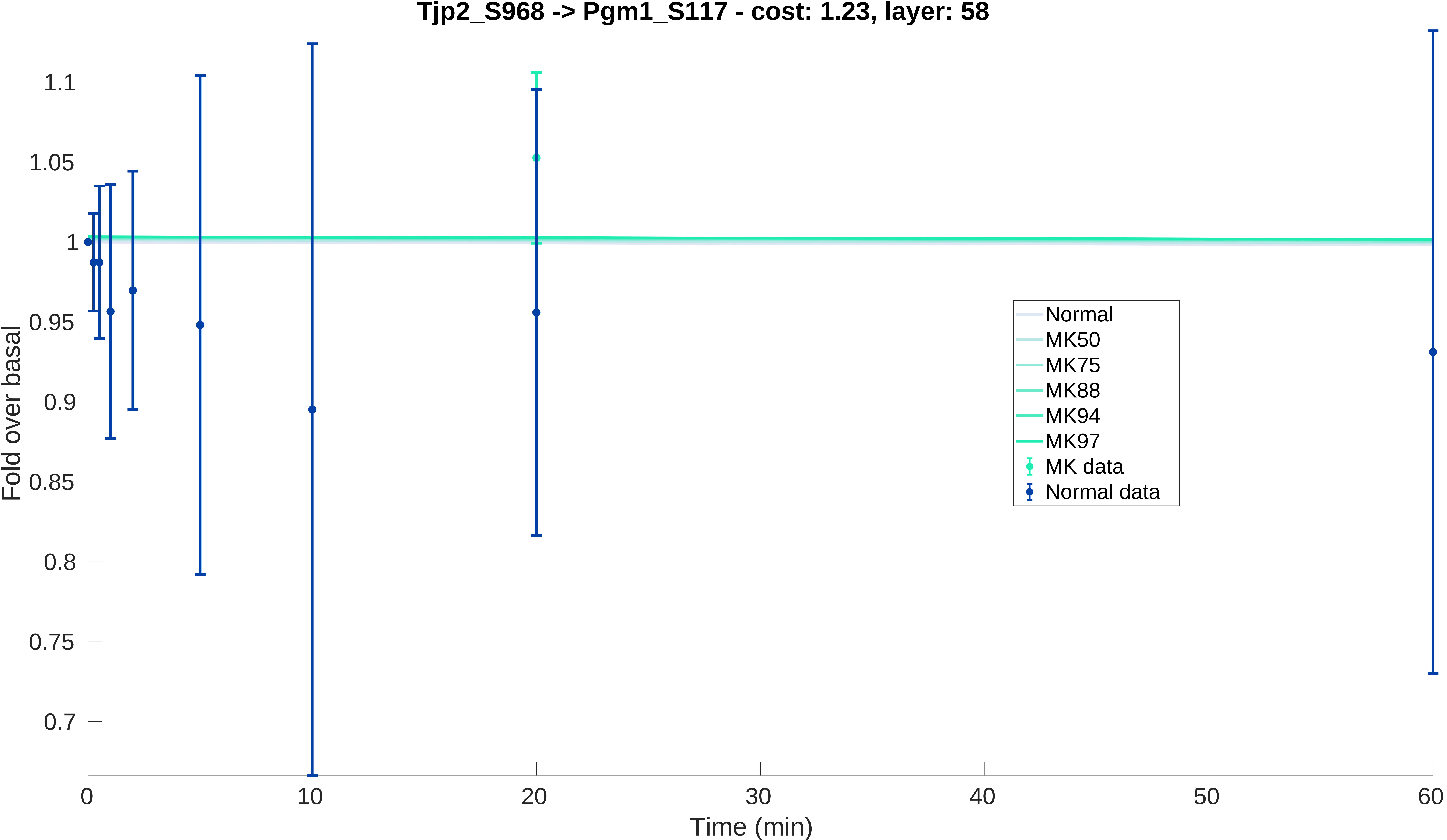

Tjp2_S968 -> Pgm1_S117 - cost: 1.23, layer: 58
Fold over basal
1.1
1.05
1
0.95
0.9
0.85
0.8
0.75
0.7
0
10
20
30
40
50
60
Time (min)
Normal
MK50
MK75
MK88
MK94
MK97
MK data
Normal data

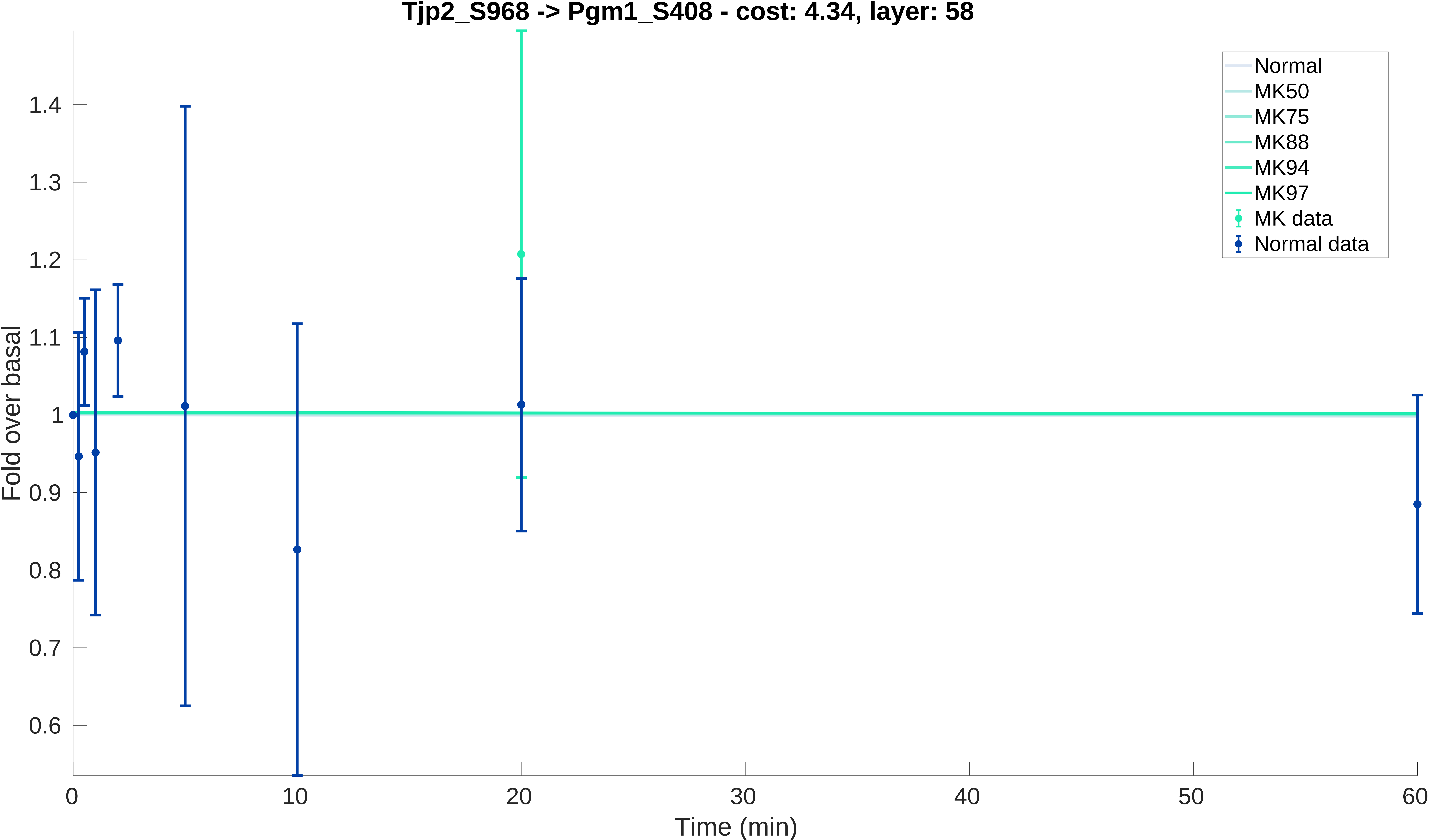
Tjp2_S968 -> Pgm1_S408 - cost: 4.34, layer: 58
Normal
MK50
MK75
MK88
MK94
MK97
MK data
Normal data
Fold over basal
1.4
1.3
1.2
1.1
1
0.9
0.8
0.7
0.6
0
10
20
30
40
50
60
Time (min)

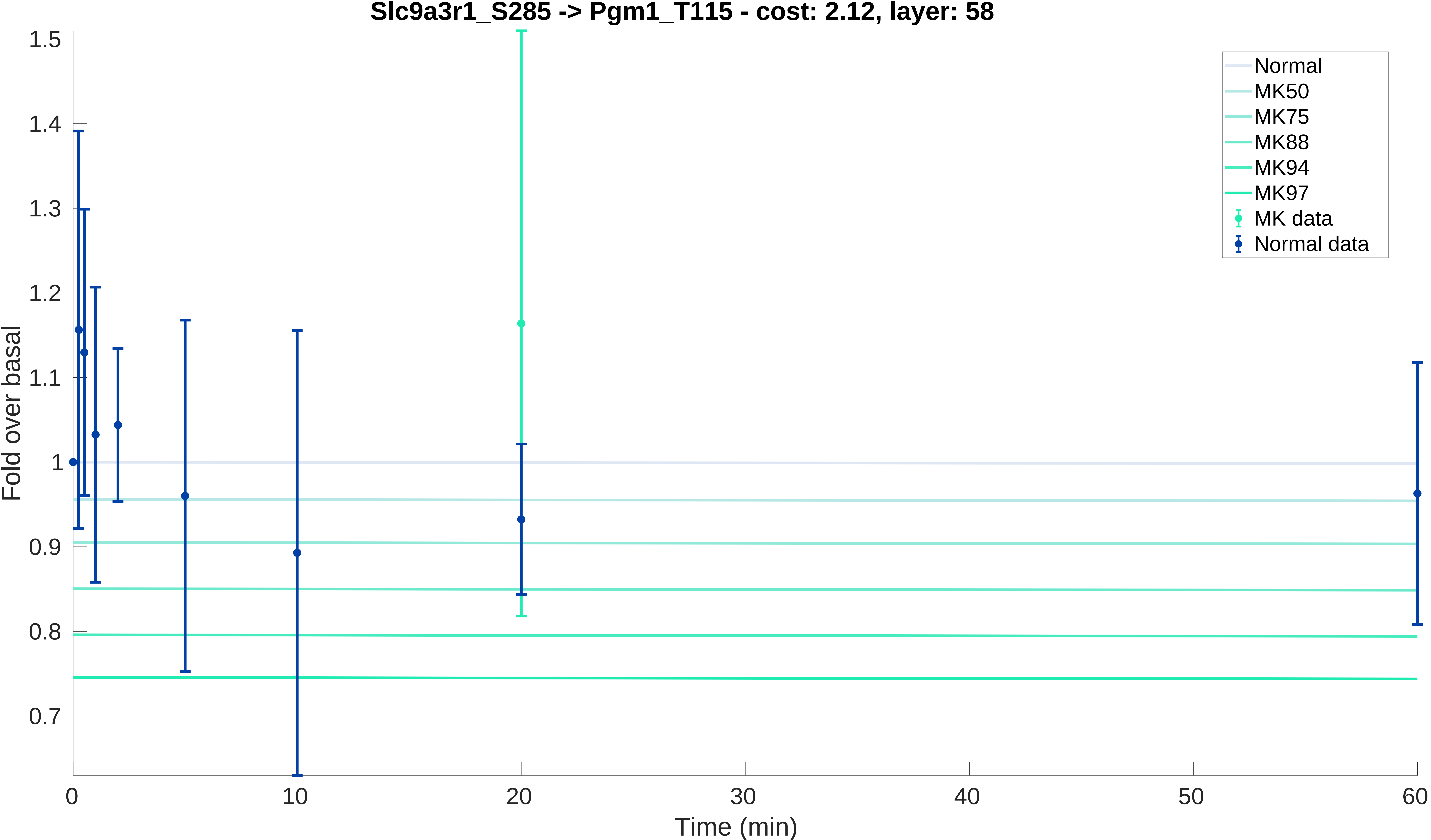
Slc9a3r1_S285 -> Pgm1_T115 - cost: 2.12, layer: 58
Fold over basal
Time (min)
1.5
1.4
1.3
1.2
1.1
1
0.9
0.8
0.7
0
10
20
30
40
50
60
Normal
MK50
MK75
MK88
MK94
MK97
MK data
Normal data

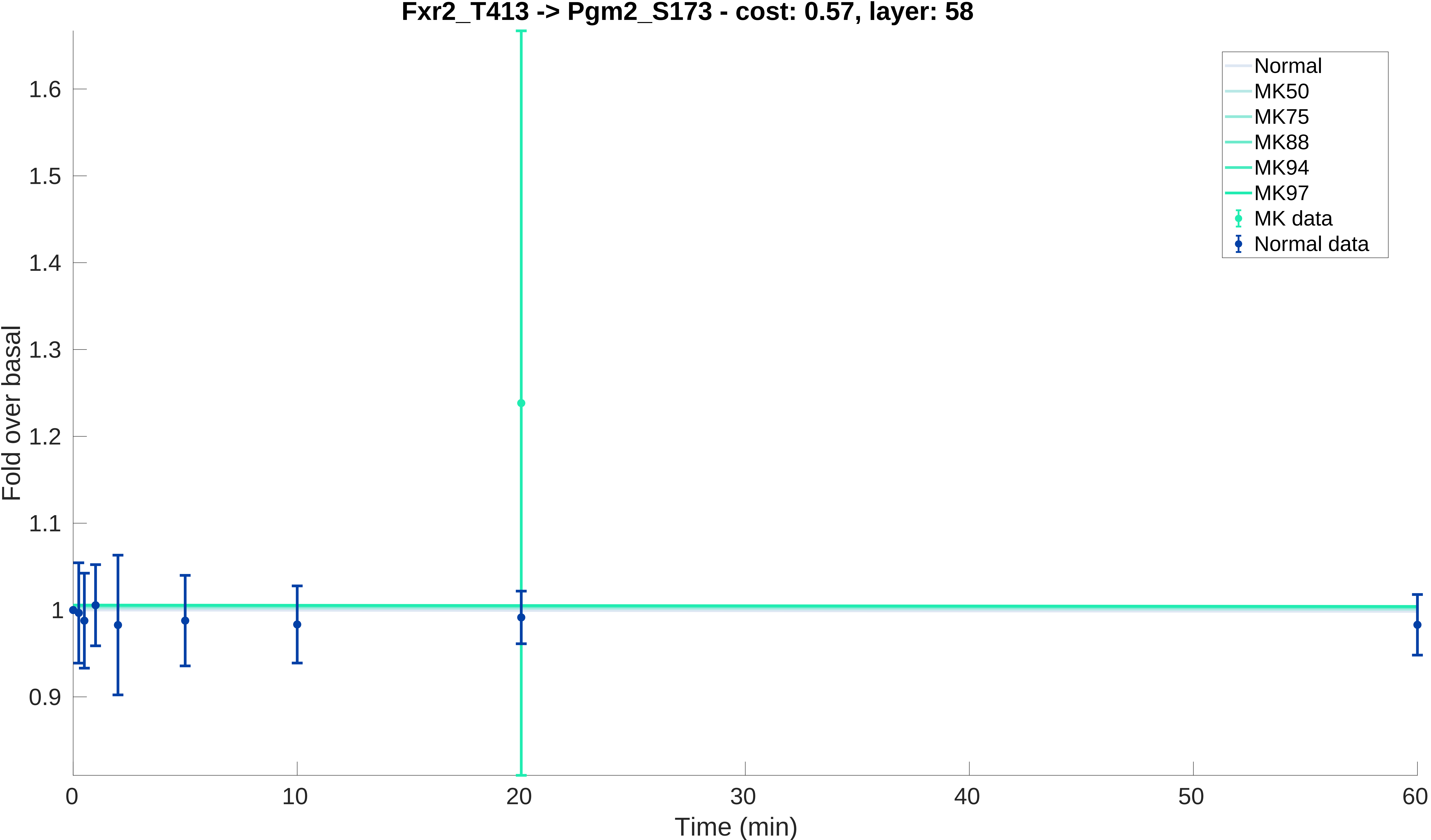

Fxr2_T413 -> Pgm2_S173 - cost: 0.57, layer: 58
Fold over basal
Time (min)
Normal
MK50
MK75
MK88
MK94
MK97
MK data
Normal data
1.6
1.5
1.4
1.3
1.2
1.1
1
0.9
0
10
20
30
40
50
60

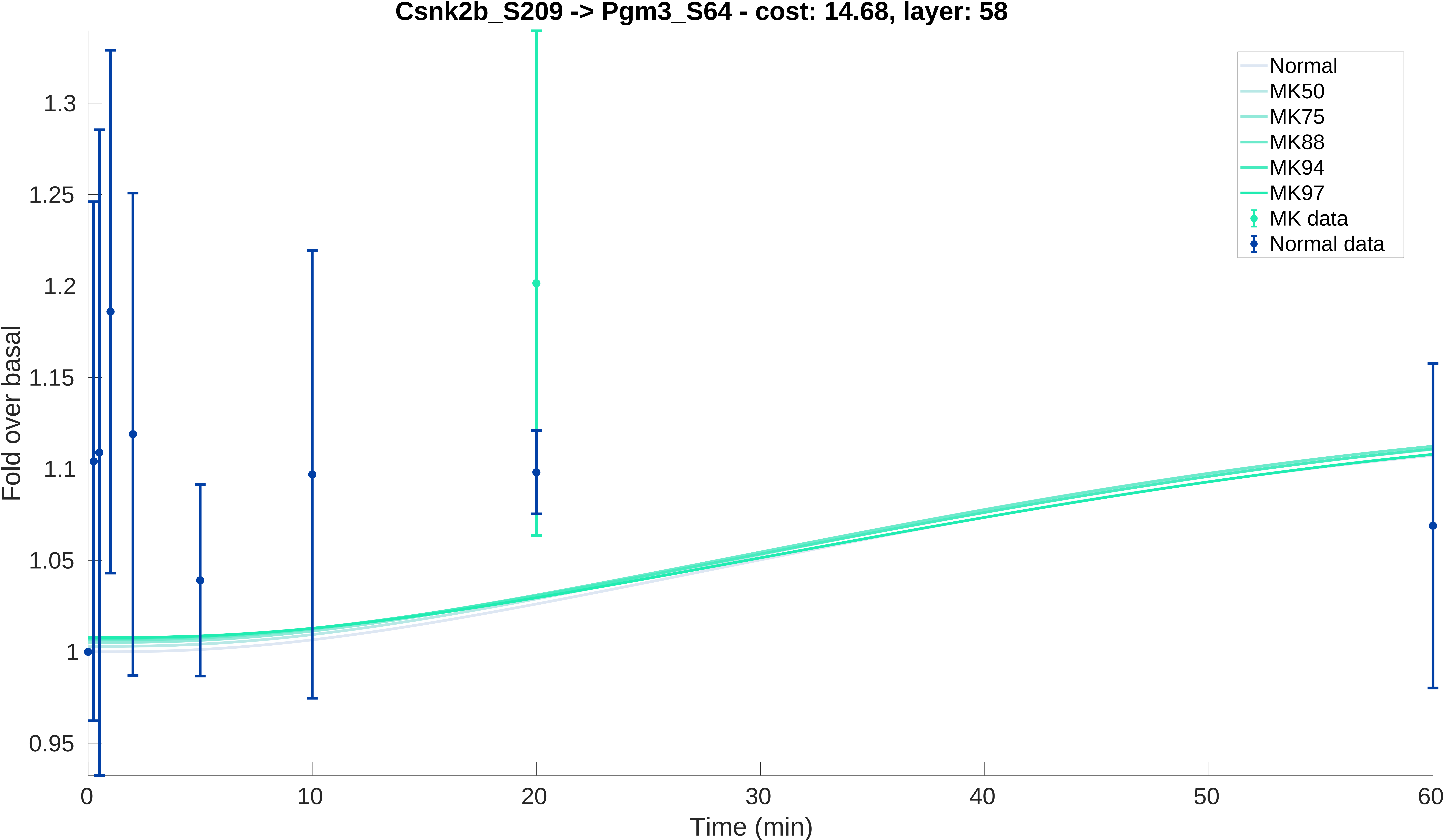

Csnk2b_S209 -> Pgm3_S64 - cost: 14.68, layer: 58
Fold over basal
1.3
1.25
1.2
1.15
1.1
1.05
1
0.95
Time (min)
0
10
20
30
40
50
60
Normal
MK50
MK75
MK88
MK94
MK97
MK data
Normal data

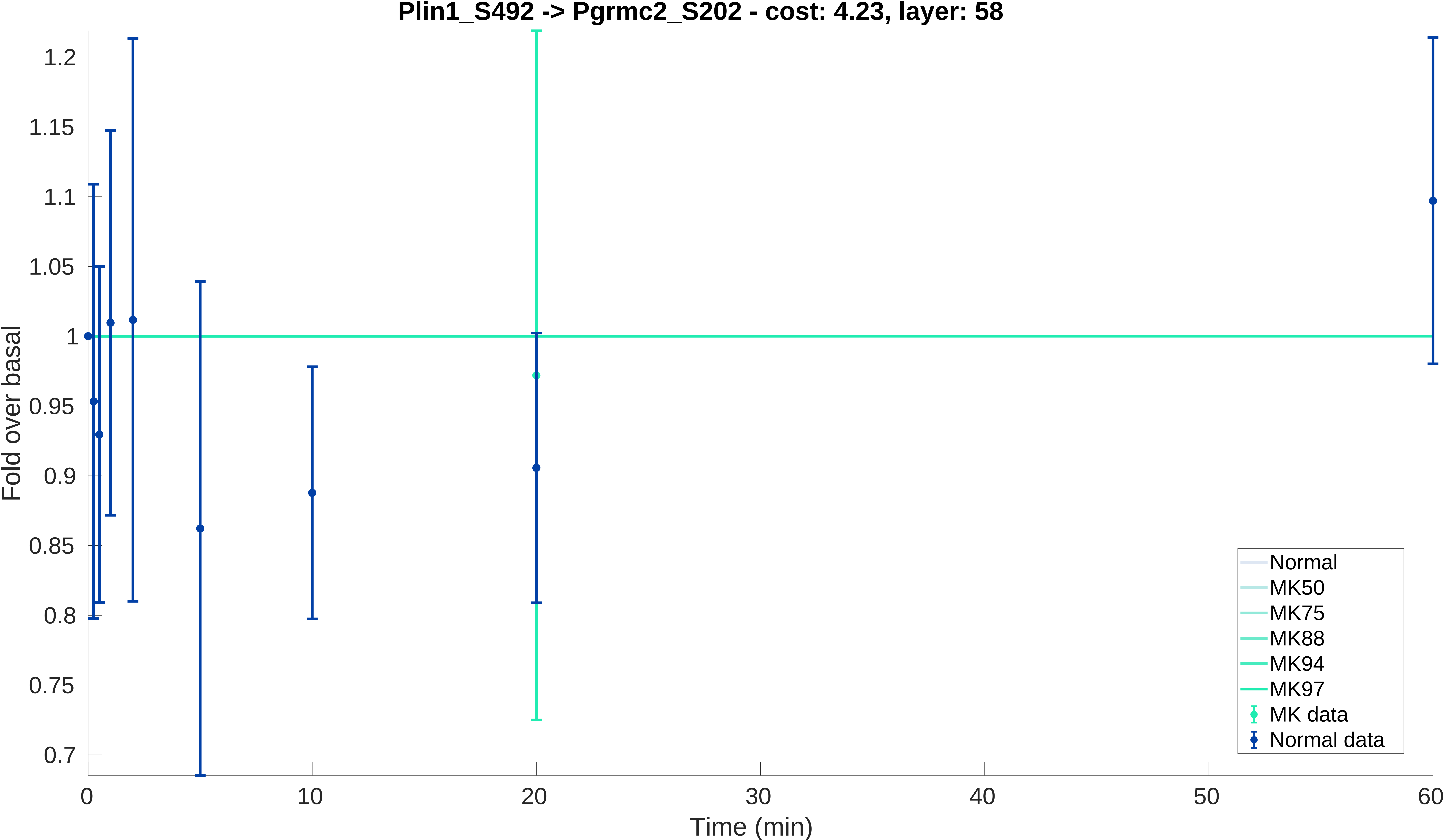
Plin1_S492 -> Pgrmc2_S202 - cost: 4.23, layer: 58
Fold over basal
1.2
1.15
1.1
1.05
1
0.95
0.9
0.85
0.8
0.75
0.7
0
10
20
30
40
50
60
Time (min)
Normal
MK50
MK75
MK88
MK94
MK97
MK data
Normal data

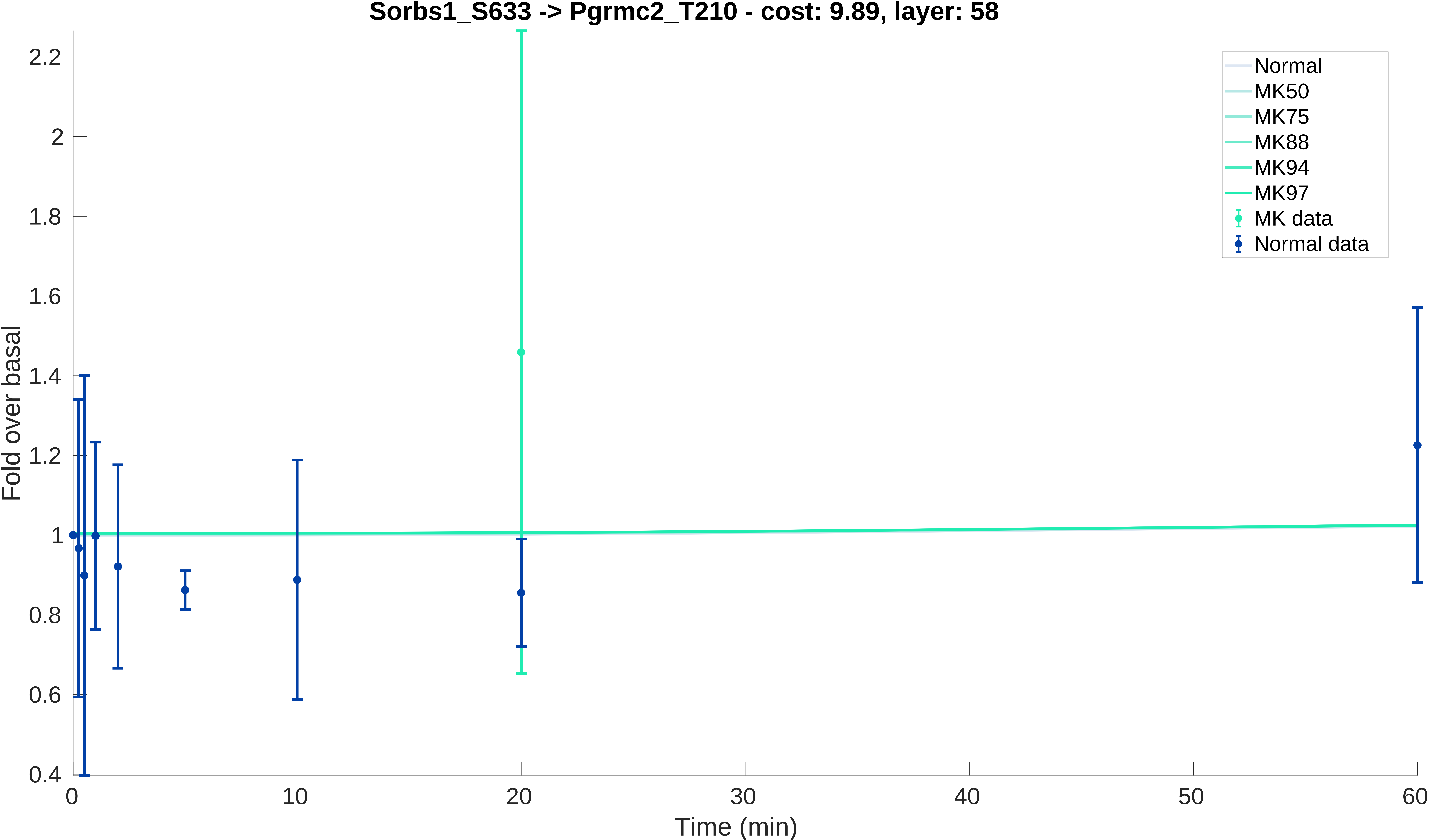

Sorbs1_S633 -> Pgrmc2_T210 - cost: 9.89, layer: 58
Fold over basal
Time (min)
0.4
0.6
0.8
1
1.2
1.4
1.6
1.8
2
2.2
0
10
20
30
40
50
60
Normal
MK50
MK75
MK88
MK94
MK97
MK data
Normal data

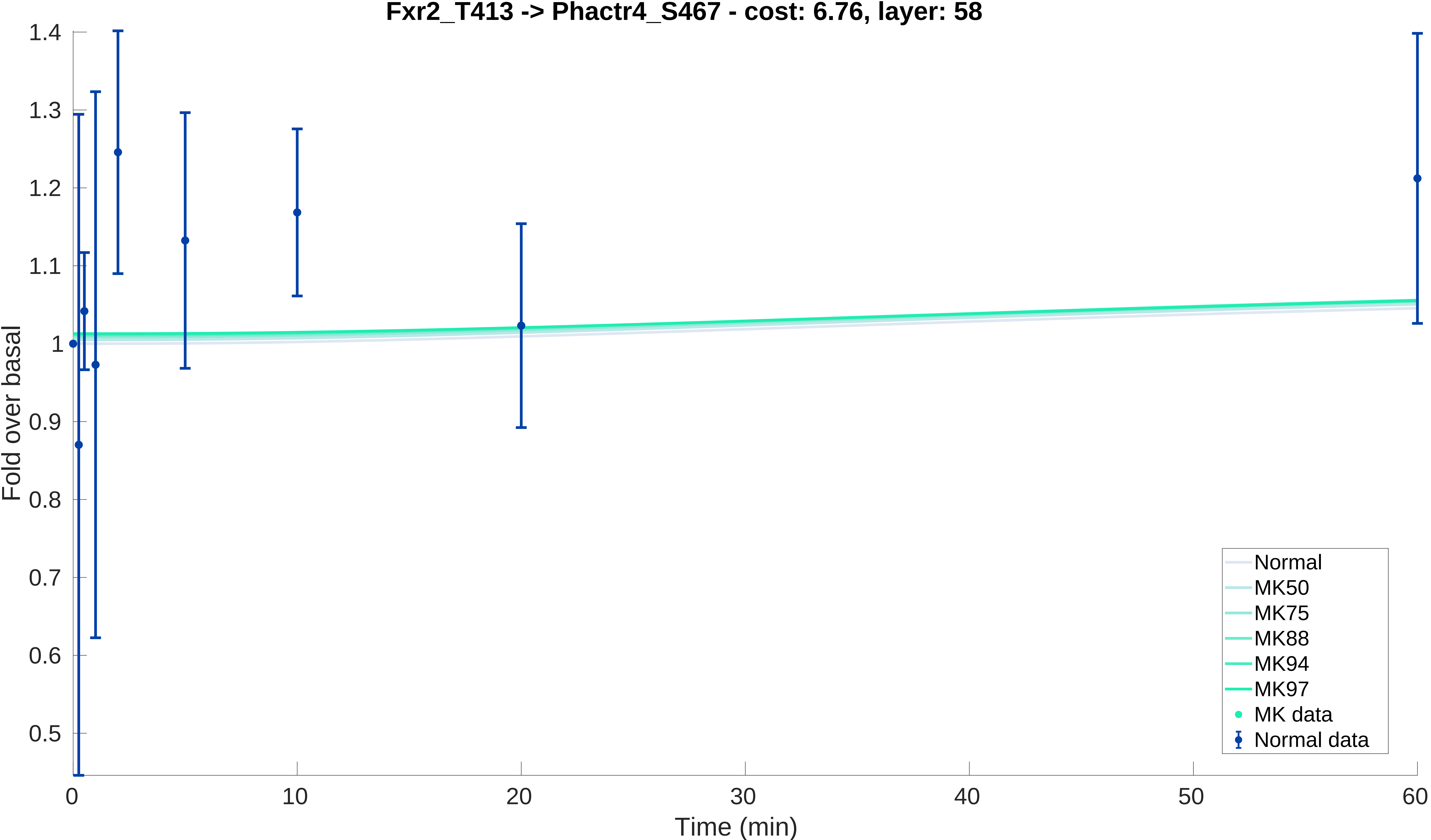

Fxr2_T413 -> Phactr4_S467 - cost: 6.76, layer: 58
Fold over basal
Time (min)
1.4
1.3
1.2
1.1
1
0.9
0.8
0.7
0.6
0.5
0
10
20
30
40
50
60
Normal
MK50
MK75
MK88
MK94
MK97
MK data
Normal data

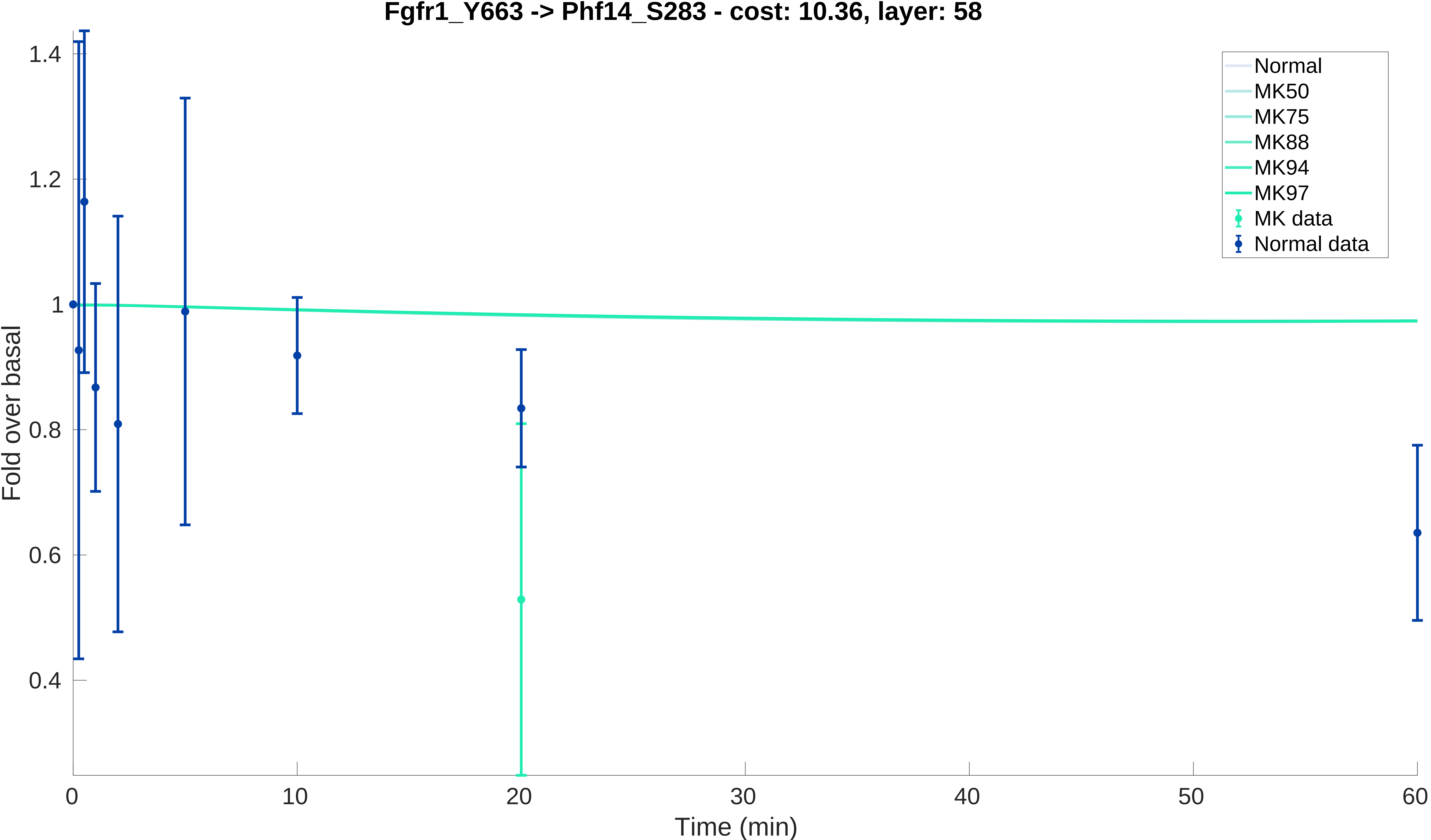
Fgfr1_Y663 -> Phf14_S283 - cost: 10.36, layer: 58
Fold over basal
Time (min)
0
10
20
30
40
50
60
0.4
0.6
0.8
1
1.2
1.4
Normal
MK50
MK75
MK88
MK94
MK97
MK data
Normal data

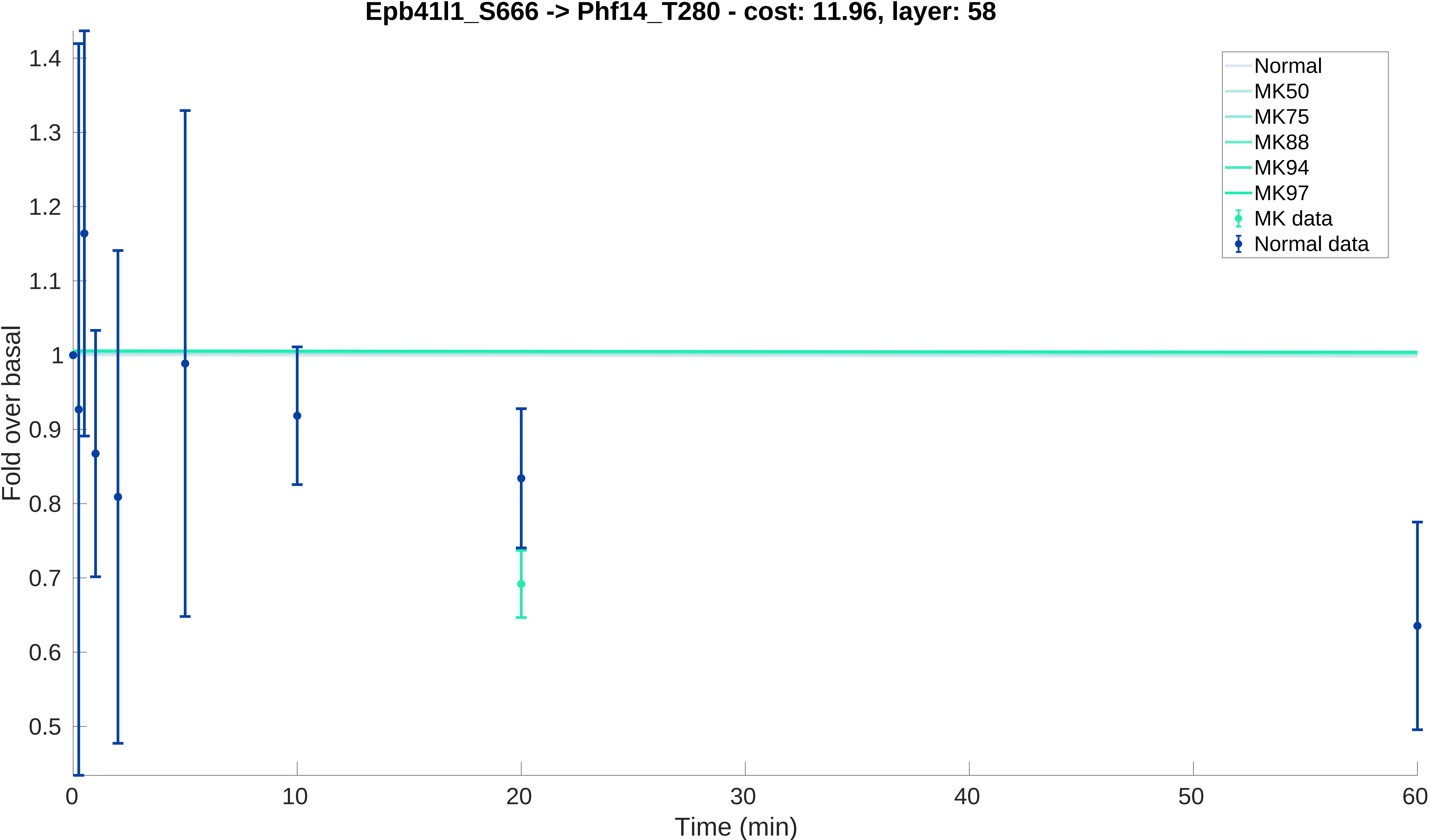

Epb41l1_S666 -> Phf14_T280 - cost: 11.96, layer: 58
Fold over basal
Time (min)
0.5
0.6
0.7
0.8
0.9
1
1.1
1.2
1.3
1.4
0
10
20
30
40
50
60
Normal
MK50
MK75
MK88
MK94
MK97
MK data
Normal data

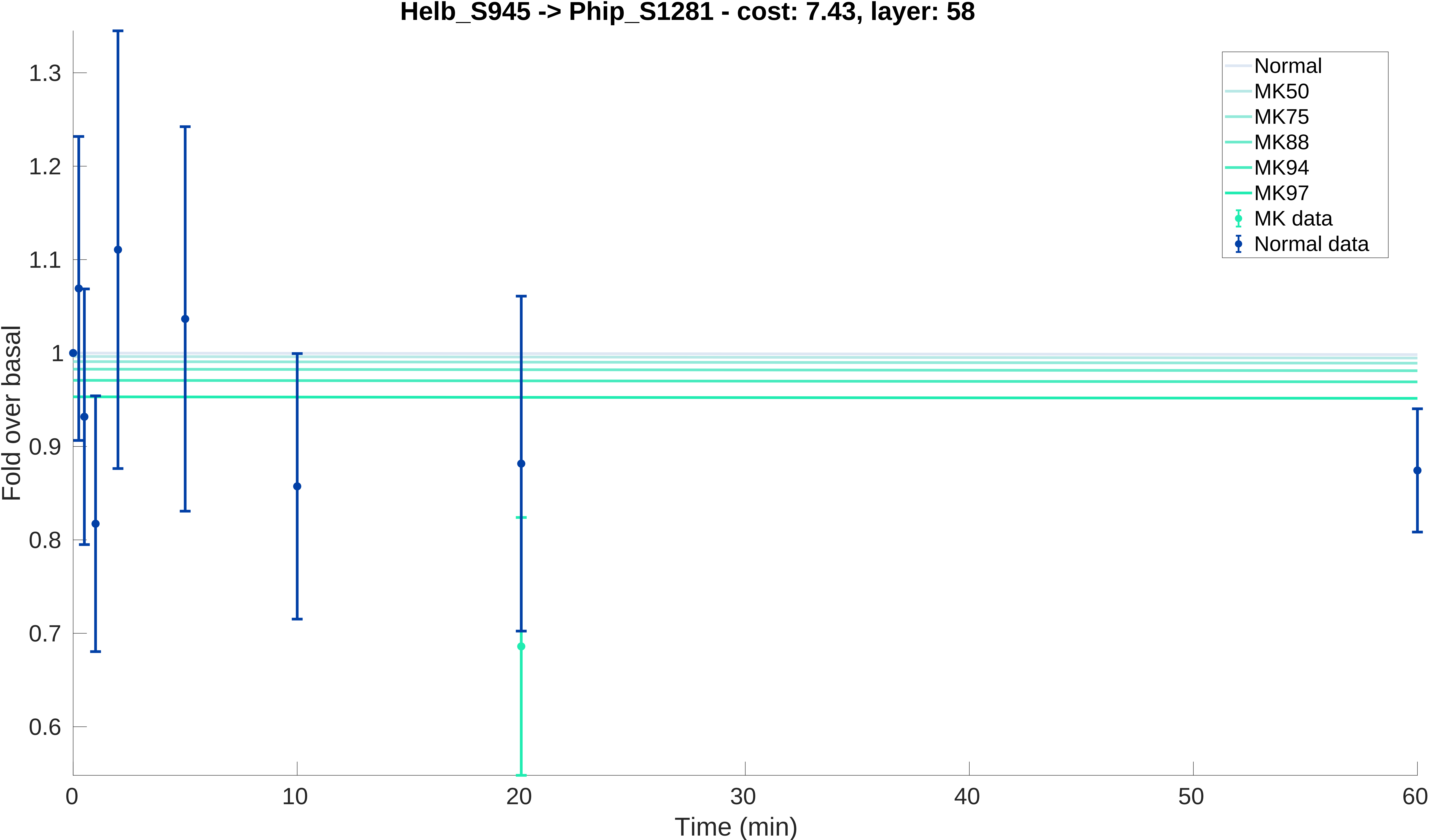

Helb_S945 -> Phip_S1281 - cost: 7.43, layer: 58
Normal
MK50
MK75
MK88
MK94
MK97
MK data
Normal data
Fold over basal
1.3
1.2
1.1
1
0.9
0.8
0.7
0.6
0
10
20
30
40
50
60
Time (min)

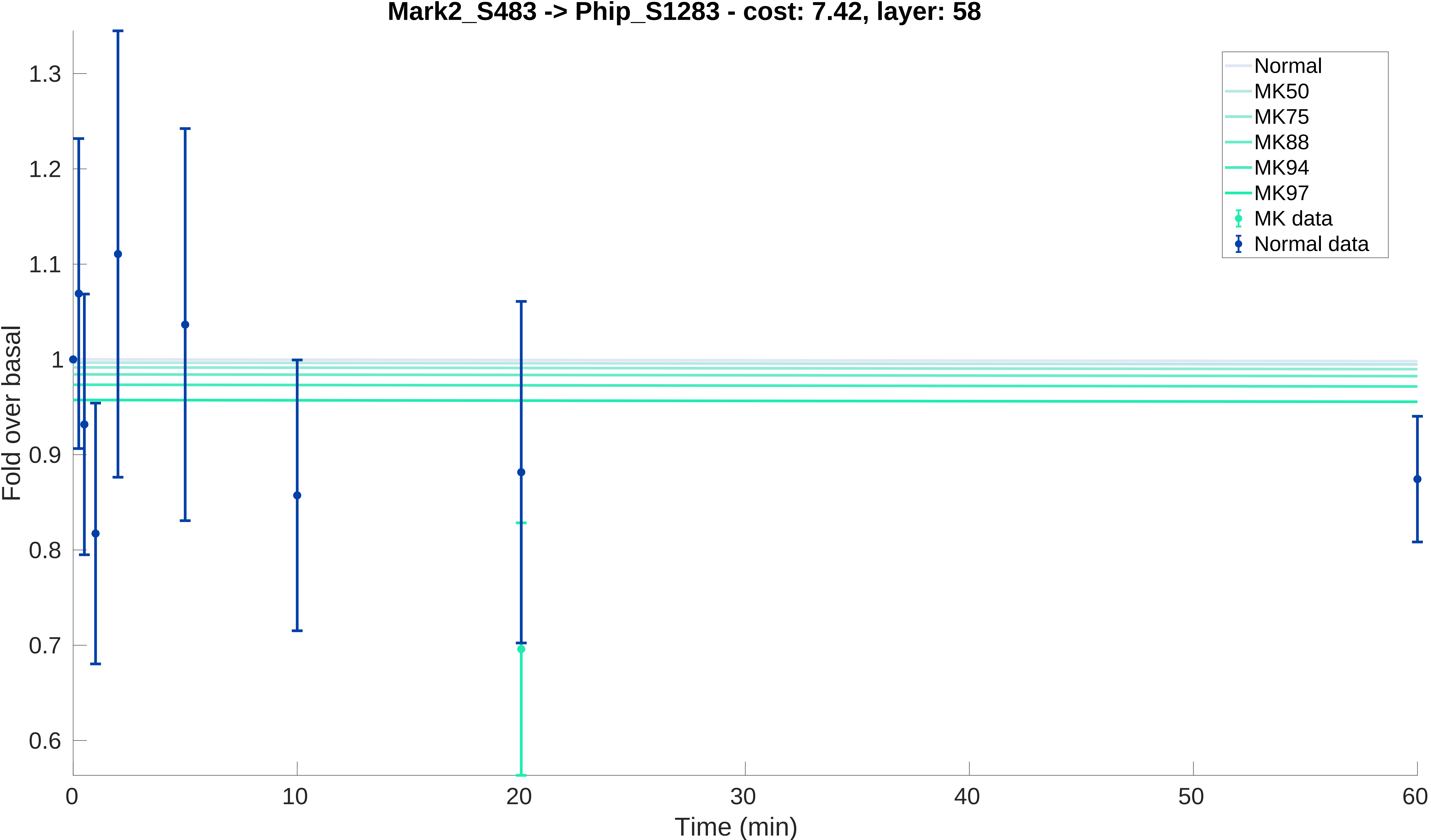

Mark2_S483 -> Phip_S1283 - cost: 7.42, layer: 58
Fold over basal
1.3
1.2
1.1
1
0.9
0.8
0.7
0.6
0
10
20
30
40
50
60
Time (min)
Normal
MK50
MK75
MK88
MK94
MK97
MK data
Normal data

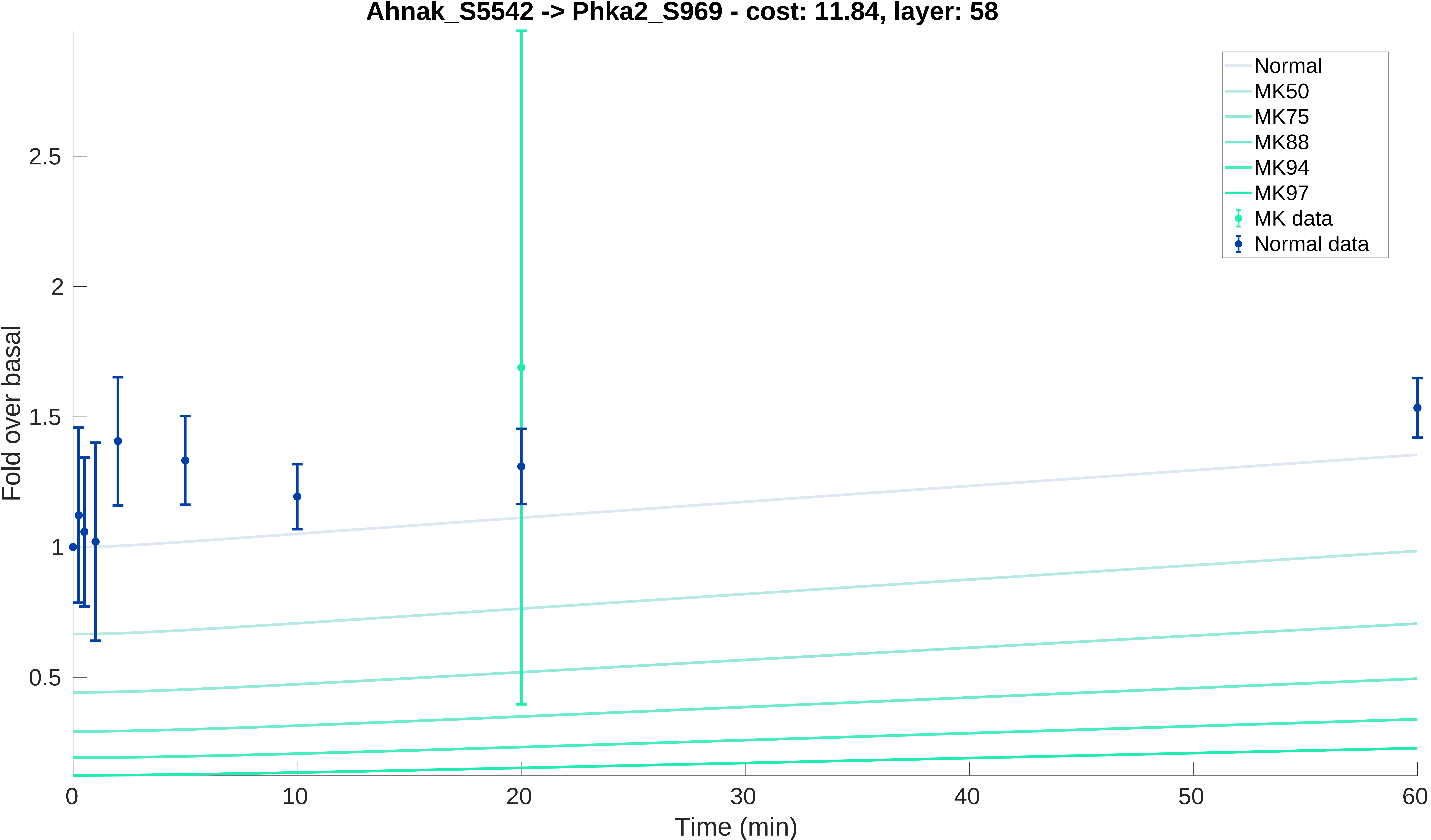
Ahnak_S5542 -> Phka2_S969 - cost: 11.84, layer: 58
Normal
MK50
MK75
MK88
MK94
MK97
MK data
Normal data
Fold over basal
Time (min)
0.5
1
1.5
2
2.5
0
10
20
30
40
50
60

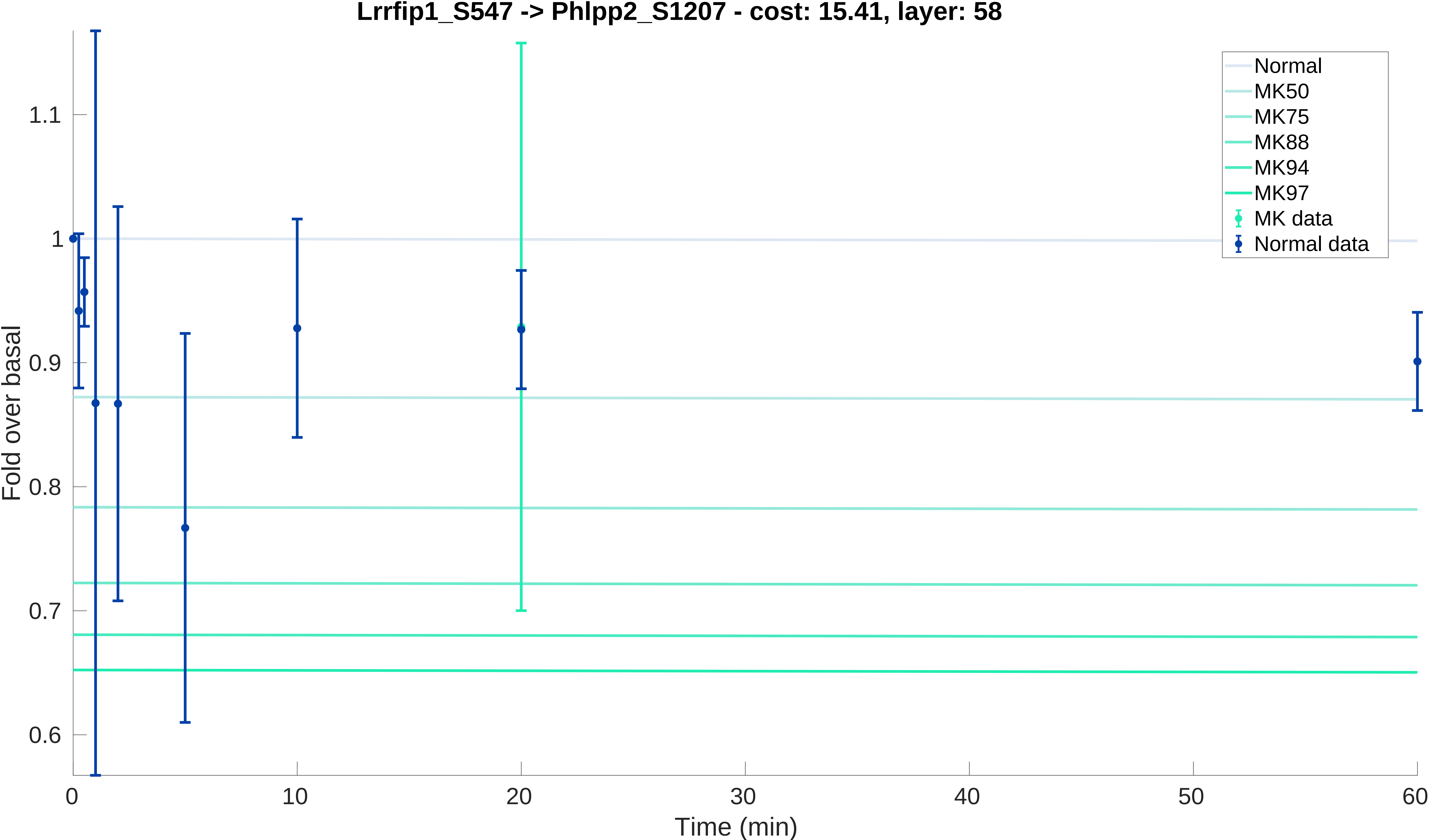

Lrrfip1_S547 -> Phlpp2_S1207 - cost: 15.41, layer: 58
Normal
MK50
MK75
MK88
MK94
MK97
MK data
Normal data
Fold over basal
1.1
1
0.9
0.8
0.7
0.6
0
10
20
30
40
50
60
Time (min)

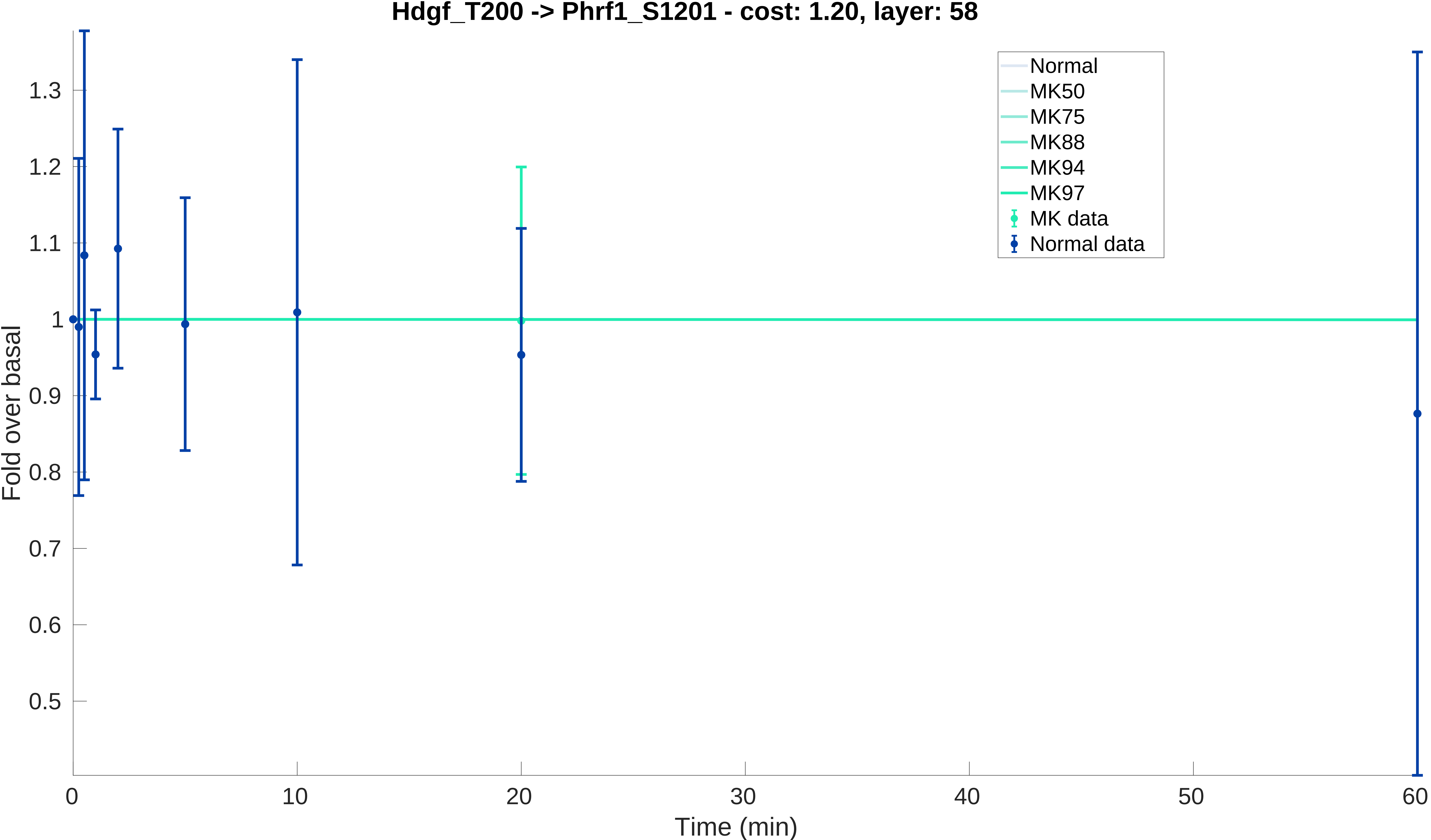

Hdgf_T200 -> Phrf1_S1201 - cost: 1.20, layer: 58
Normal
MK50
MK75
MK88
MK94
MK97
MK data
Normal data
Fold over basal
1.3
1.2
1.1
1
0.9
0.8
0.7
0.6
0.5
0
10
20
30
40
50
60
Time (min)

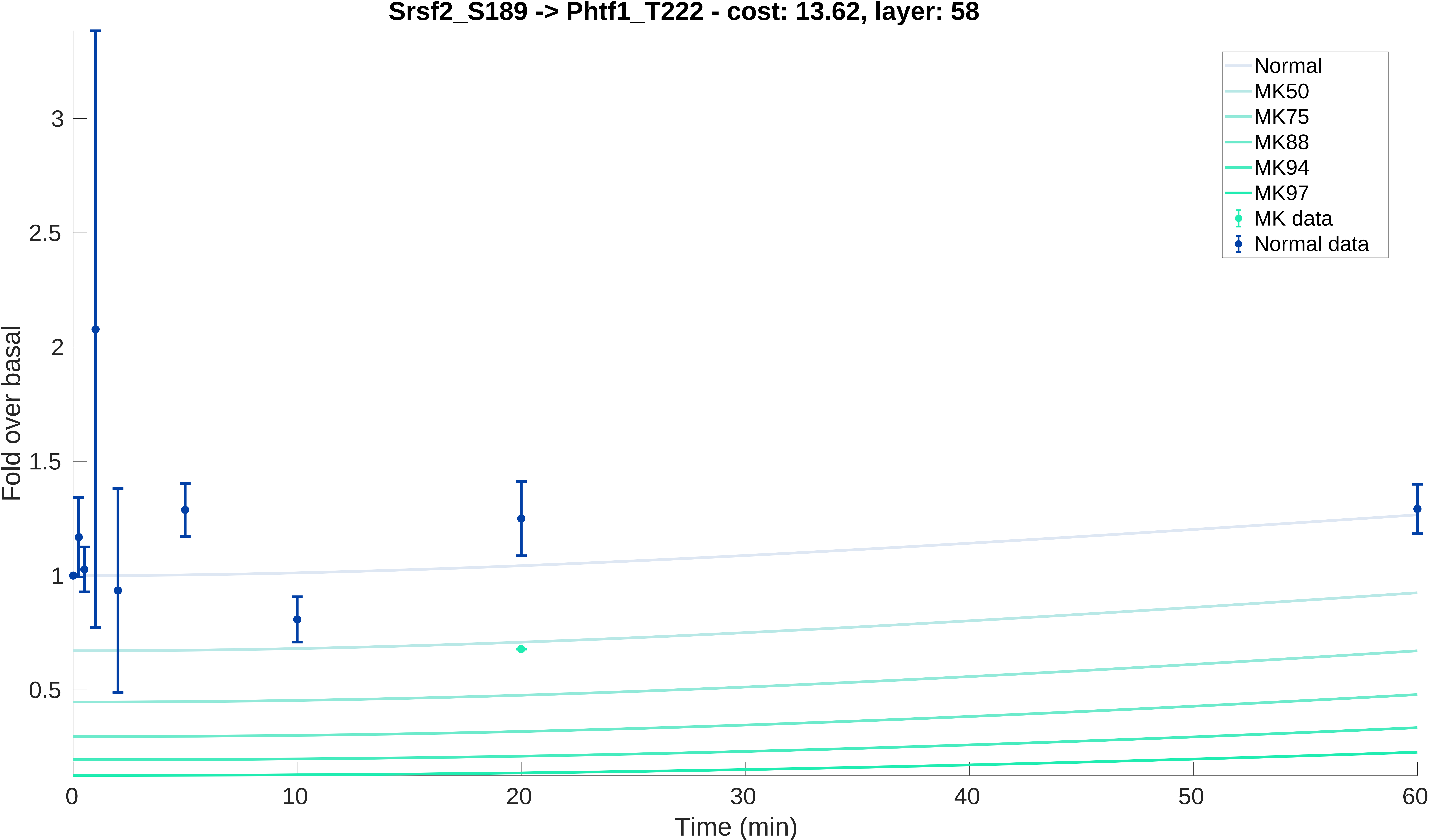
Srsf2_S189 -> Phtf1_T222 - cost: 13.62, layer: 58
Fold over basal
Time (min)
0
10
20
30
40
50
60
0.5
1
1.5
2
2.5
3
Normal
MK50
MK75
MK88
MK94
MK97
MK data
Normal data

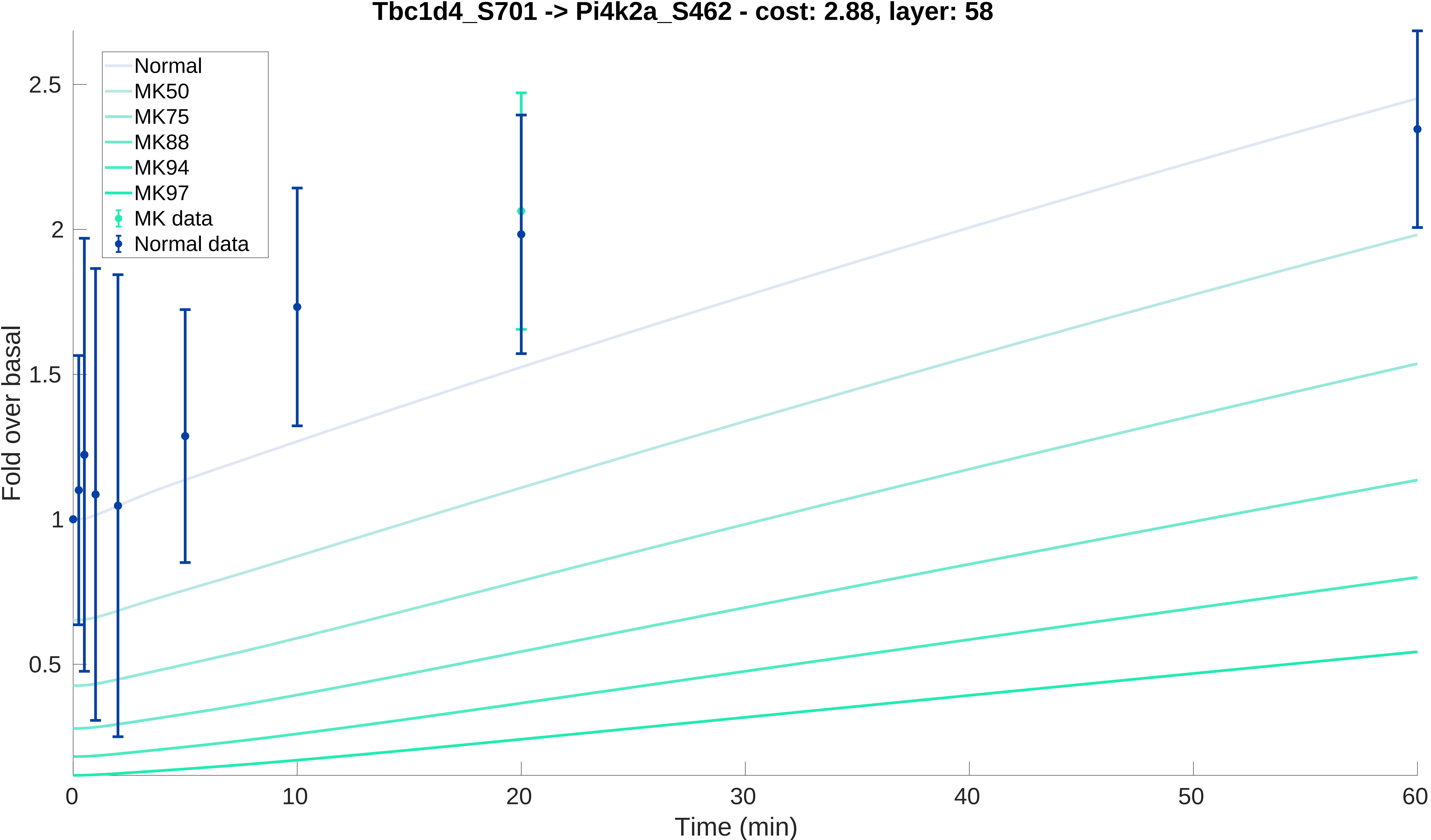

Tbc1d4_S701 -> Pi4k2a_S462 - cost: 2.88, layer: 58
Normal
MK50
MK75
MK88
MK94
MK97
MK data
Normal data
Fold over basal
0.5
1
1.5
2
2.5
0
10
20
30
40
50
60
Time (min)

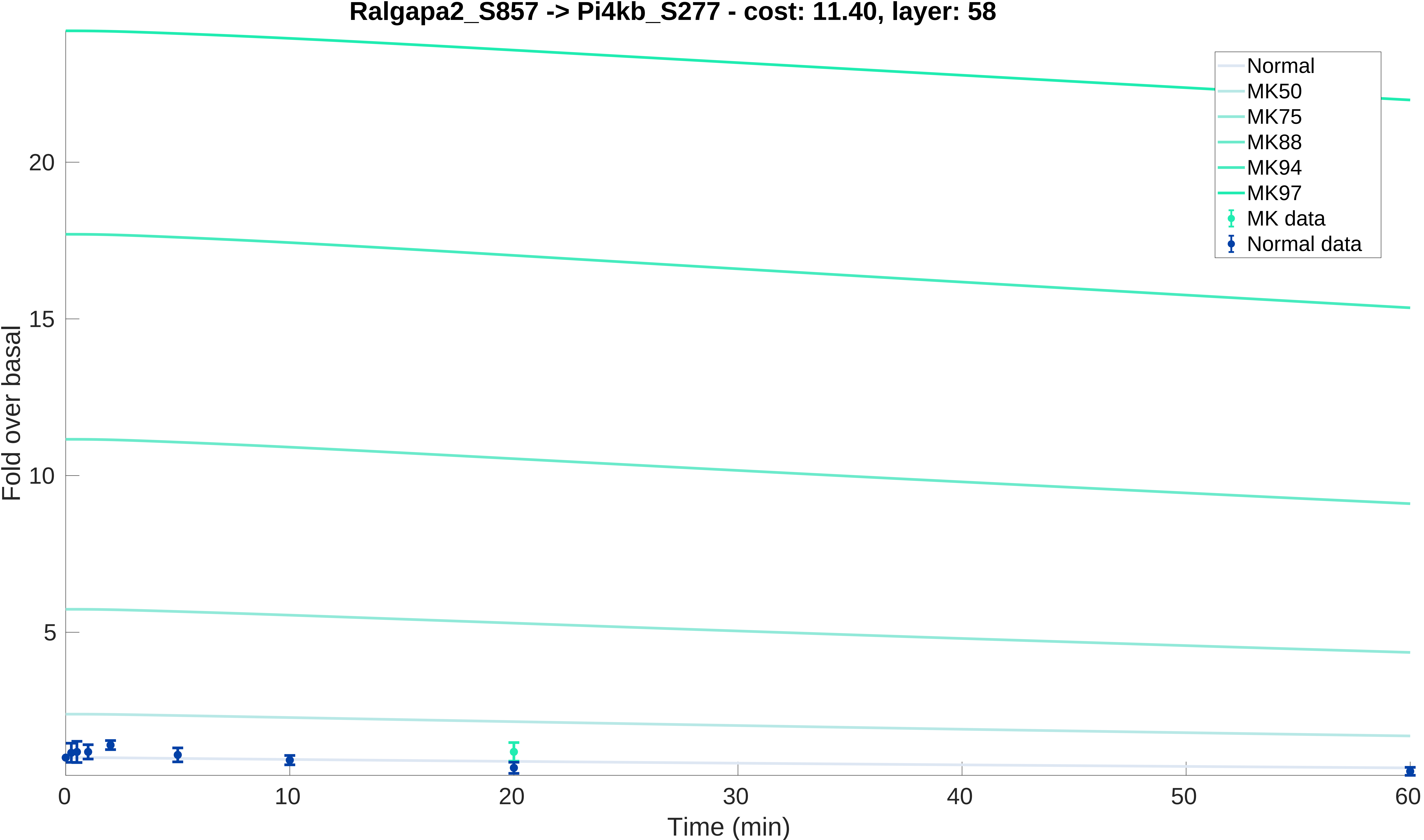
Ralgapa2_S857 -> Pi4kb_S277 - cost: 11.40, layer: 58
Normal
MK50
MK75
MK88
MK94
MK97
MK data
Normal data
Fold over basal
Time (min)
0
10
20
30
40
50
60
5
10
15
20

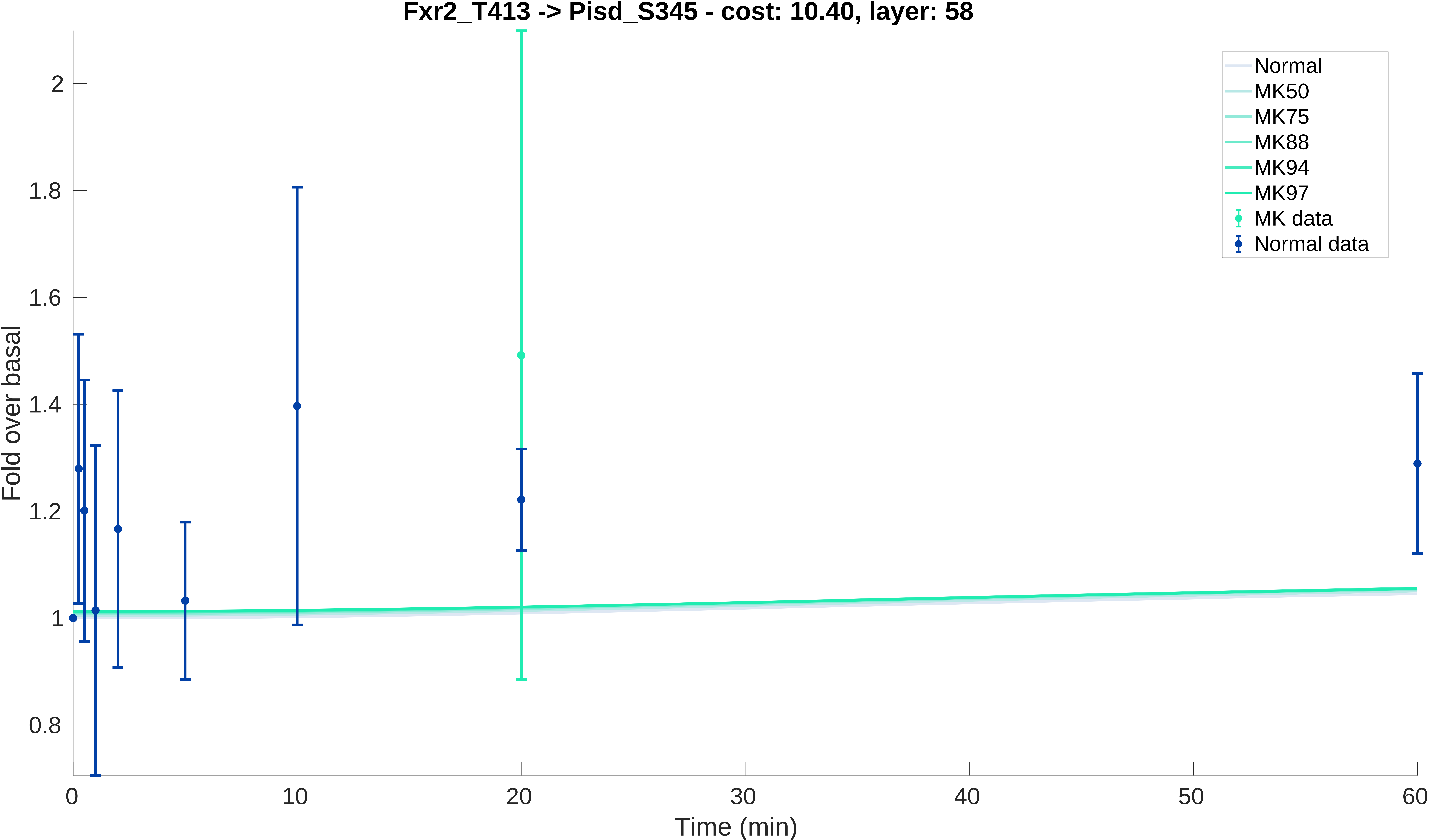
Fxr2_T413 -> Pisd_S345 - cost: 10.40, layer: 58
Fold over basal
Time (min)
0.8
1
1.2
1.4
1.6
1.8
2
0
10
20
30
40
50
60
Normal
MK50
MK75
MK88
MK94
MK97
MK data
Normal data

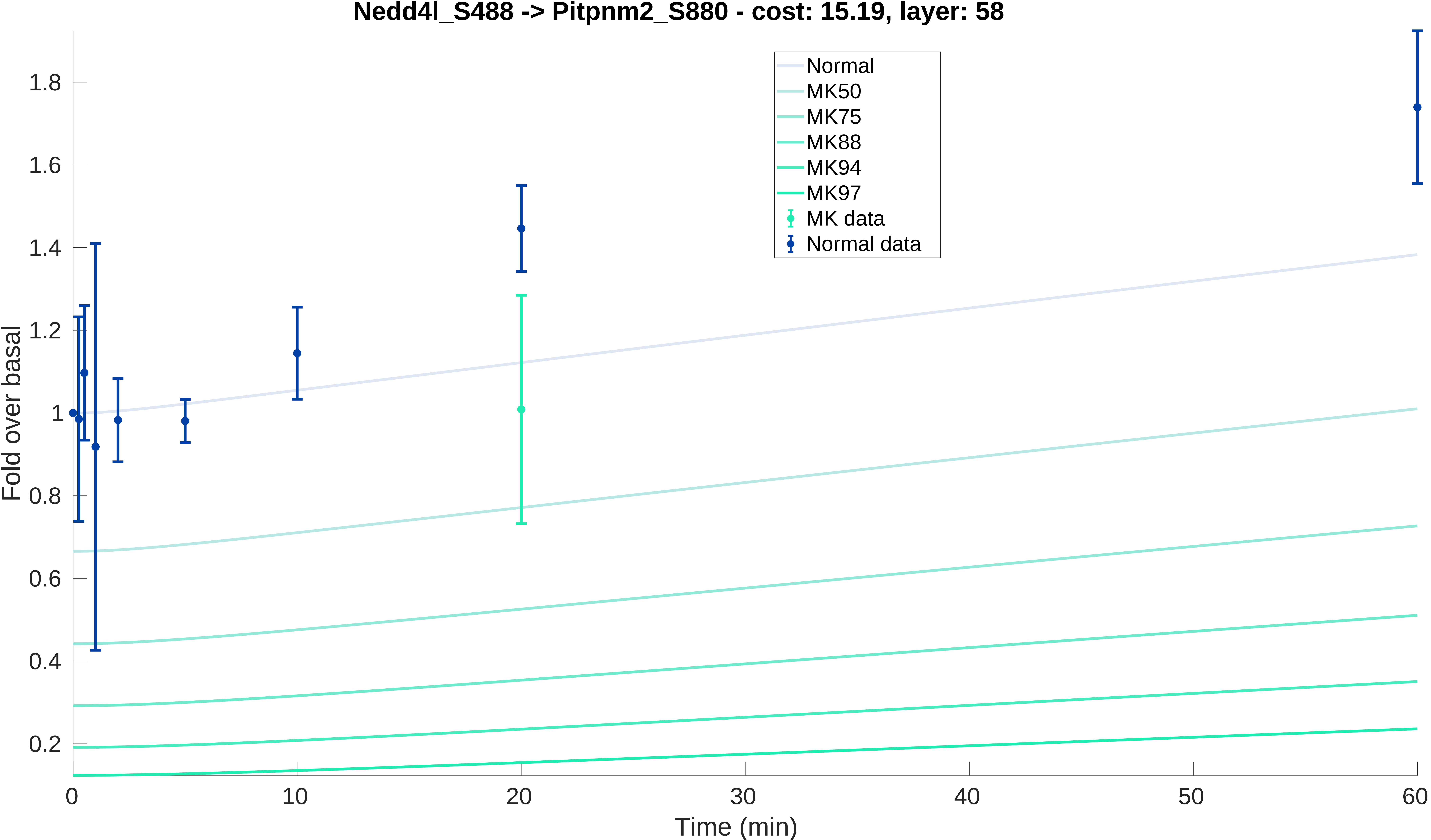

Nedd4l_S488 -> Pitpnm2_S880 - cost: 15.19, layer: 58
Normal
MK50
MK75
MK88
MK94
MK97
MK data
Normal data
1.8
1.6
1.4
1.2
1
0.8
0.6
0.4
0.2
Fold over basal
0
10
20
30
40
50
60
Time (min)

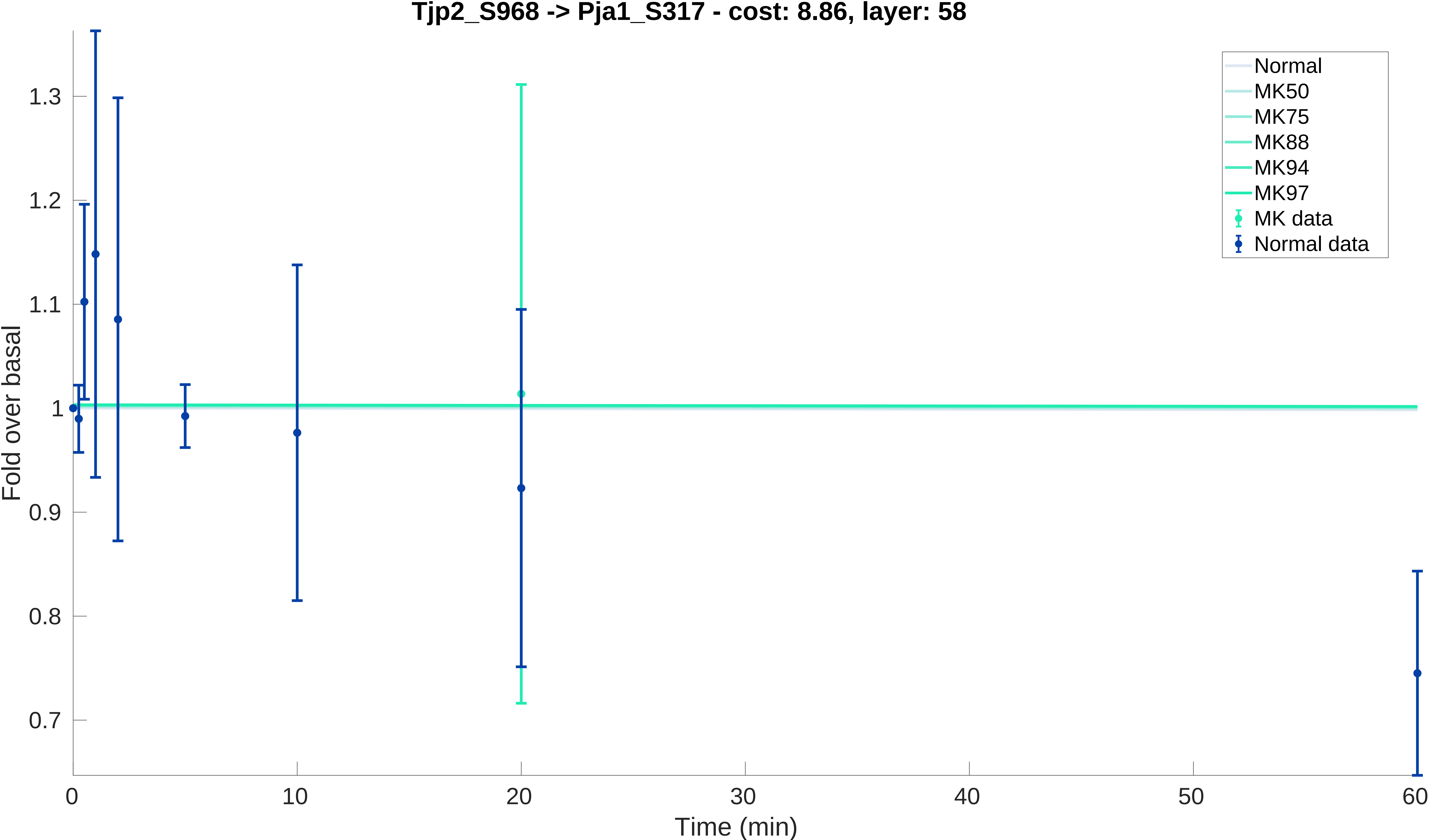

Tjp2_S968 -> Pja1_S317 - cost: 8.86, layer: 58
Normal
MK50
MK75
MK88
MK94
MK97
MK data
Normal data
Fold over basal
Time (min)
1.3
1.2
1.1
1
0.9
0.8
0.7
0
10
20
30
40
50
60

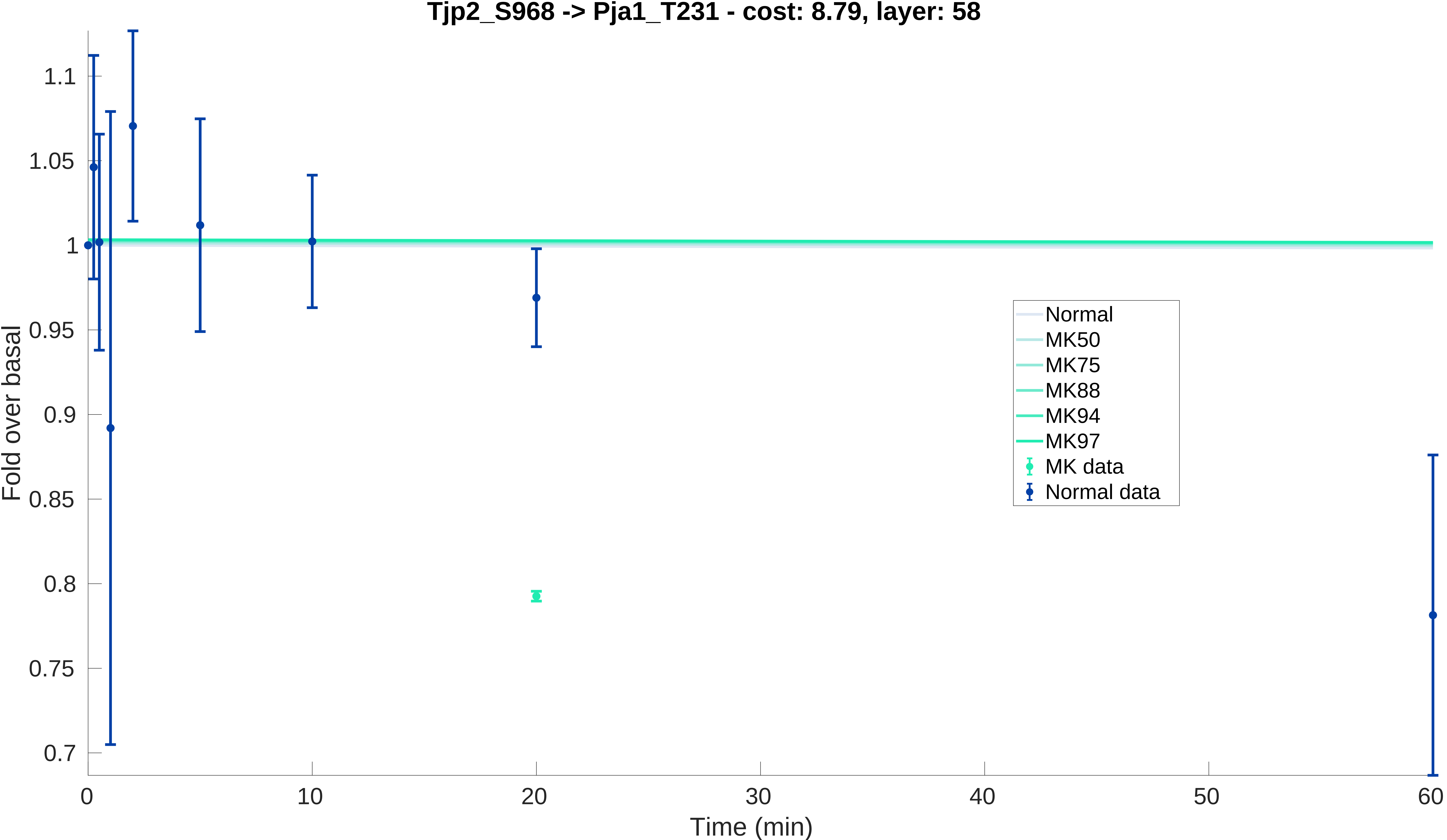

Tjp2_S968 -> Pja1_T231 - cost: 8.79, layer: 58
Fold over basal
Time (min)
1.1
1.05
1
0.95
0.9
0.85
0.8
0.75
0.7
0
10
20
30
40
50
60
Normal
MK50
MK75
MK88
MK94
MK97
MK data
Normal data

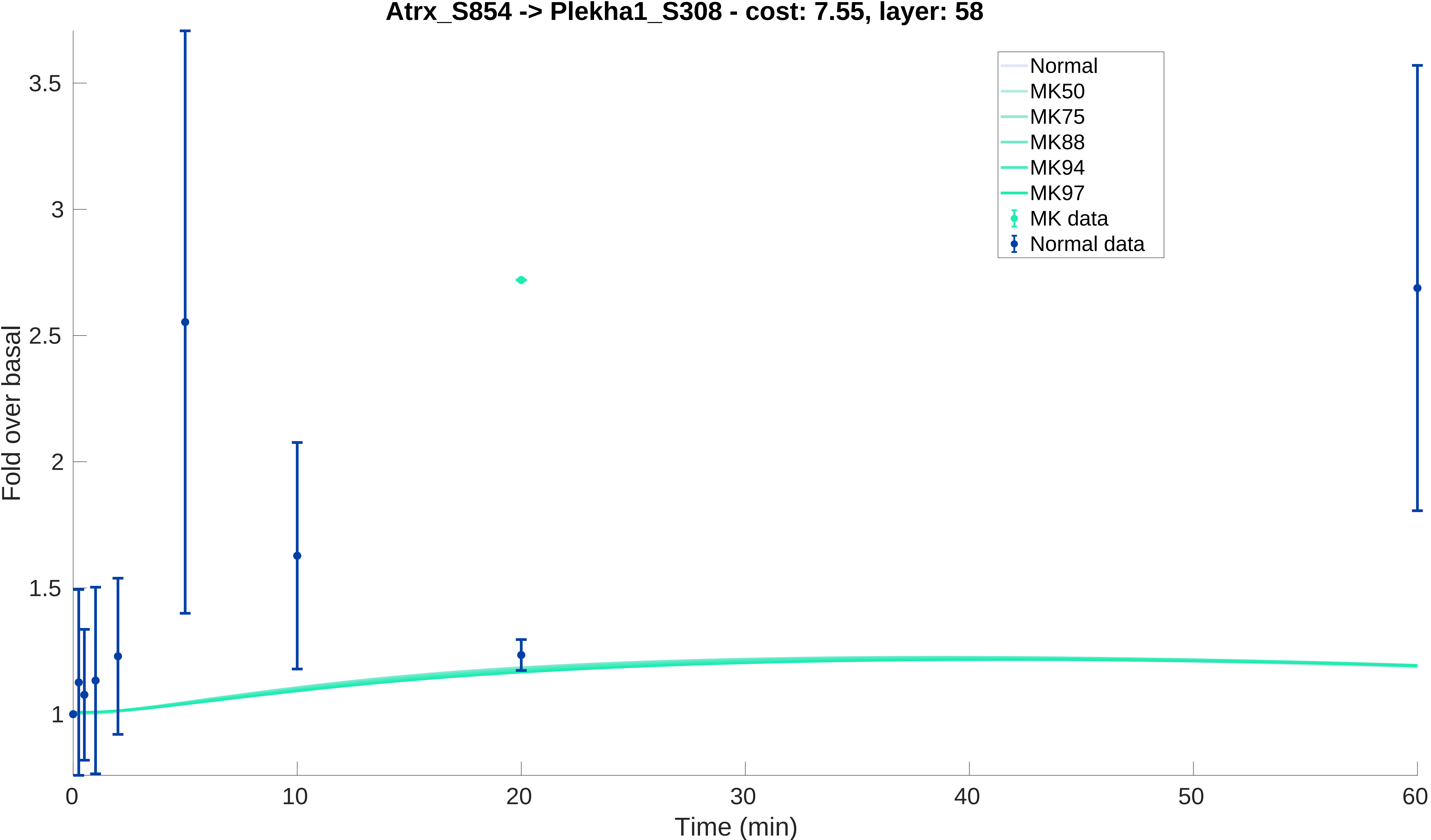

Atrx_S854 -> Plekha1_S308 - cost: 7.55, layer: 58
Normal
MK50
MK75
MK88
MK94
MK97
MK data
Normal data
Fold over basal
Time (min)
0
10
20
30
40
50
60
1
1.5
2
2.5
3
3.5

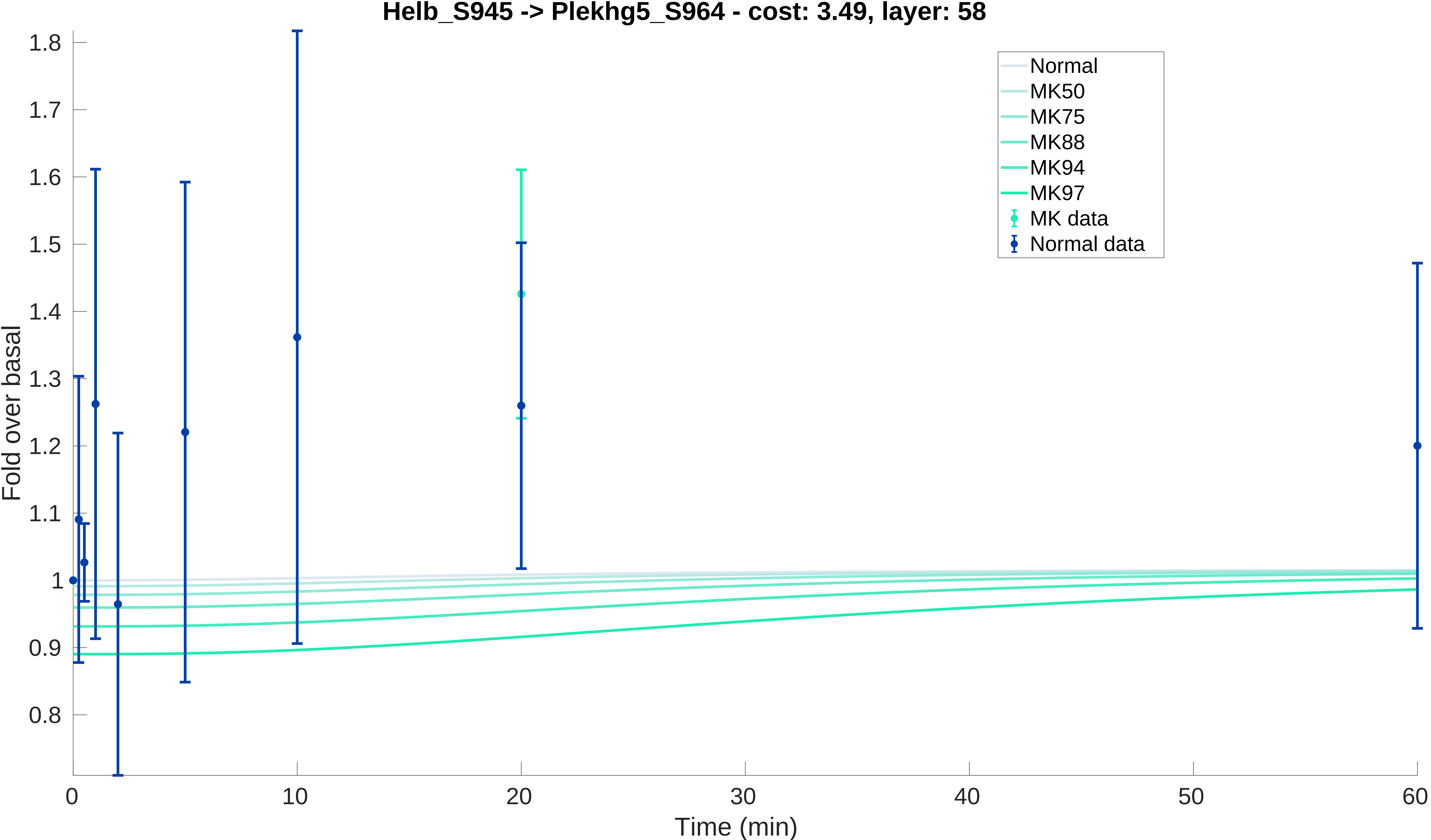

Helb_S945 -> Plekhg5_S964 - cost: 3.49, layer: 58
Fold over basal
Time (min)
0.8
0.9
1
1.1
1.2
1.3
1.4
1.5
1.6
1.7
1.8
0
10
20
30
40
50
60
Normal
MK50
MK75
MK88
MK94
MK97
MK data
Normal data

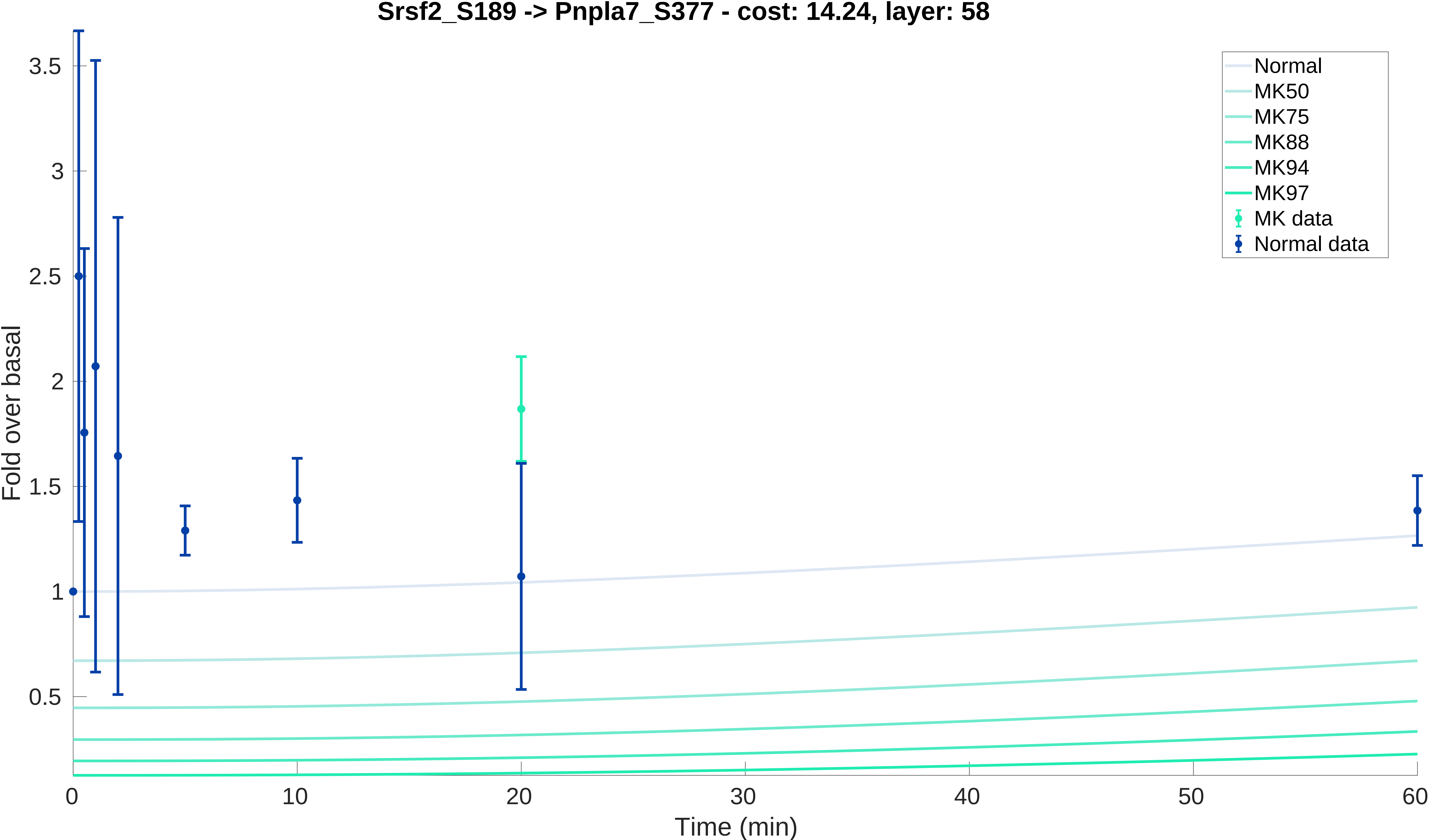

Srsf2_S189 -> Pnpla7_S377 - cost: 14.24, layer: 58
Fold over basal
3.5
3
2.5
2
1.5
1
0.5
0
10
20
30
40
50
60
Time (min)
Normal
MK50
MK75
MK88
MK94
MK97
MK data
Normal data

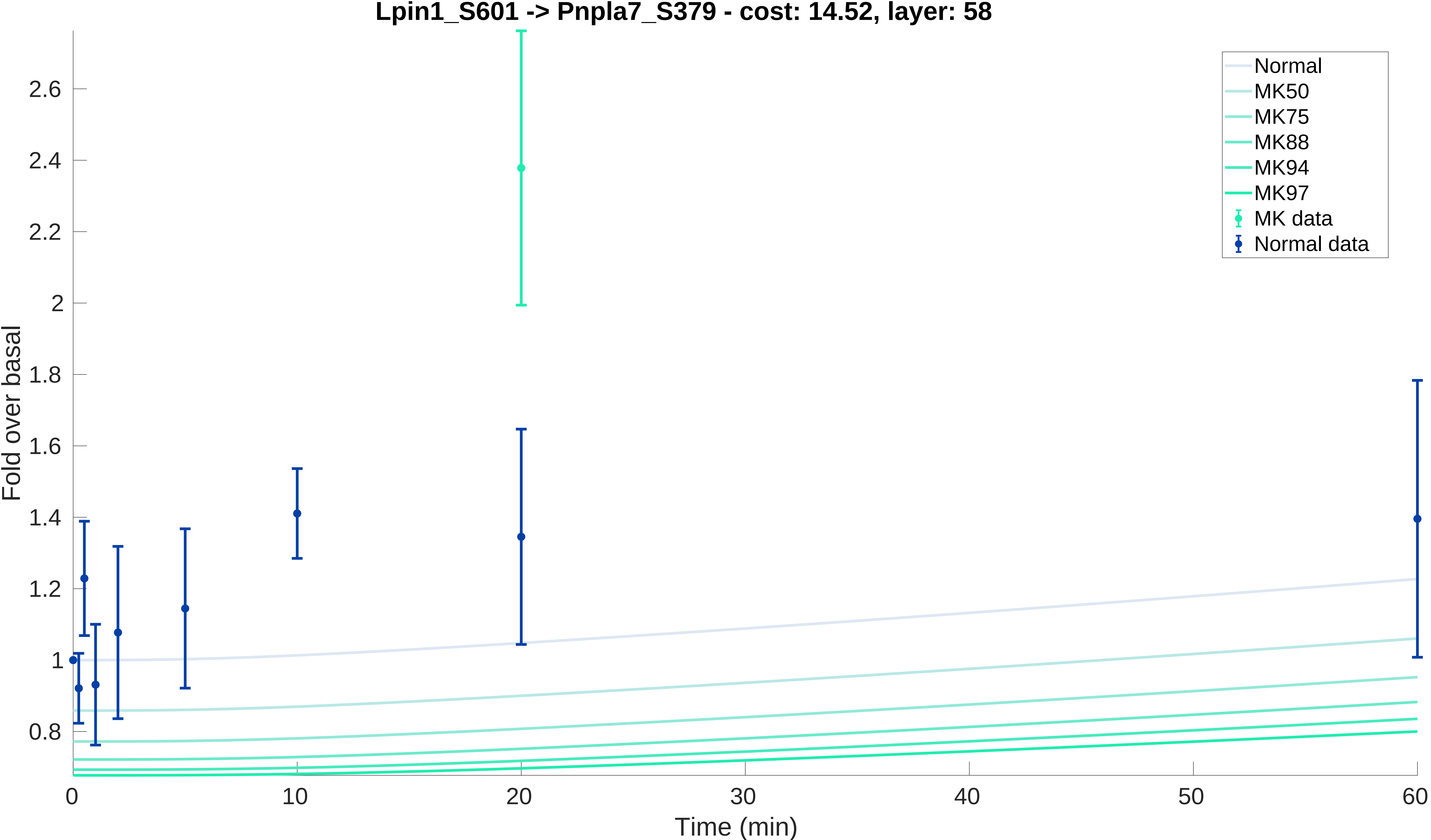

Lpin1_S601 -> Pnpla7_S379 - cost: 14.52, layer: 58
Fold over basal
Time (min)
0
10
20
30
40
50
60
0.8
1
1.2
1.4
1.6
1.8
2
2.2
2.4
2.6
Normal
MK50
MK75
MK88
MK94
MK97
MK data
Normal data

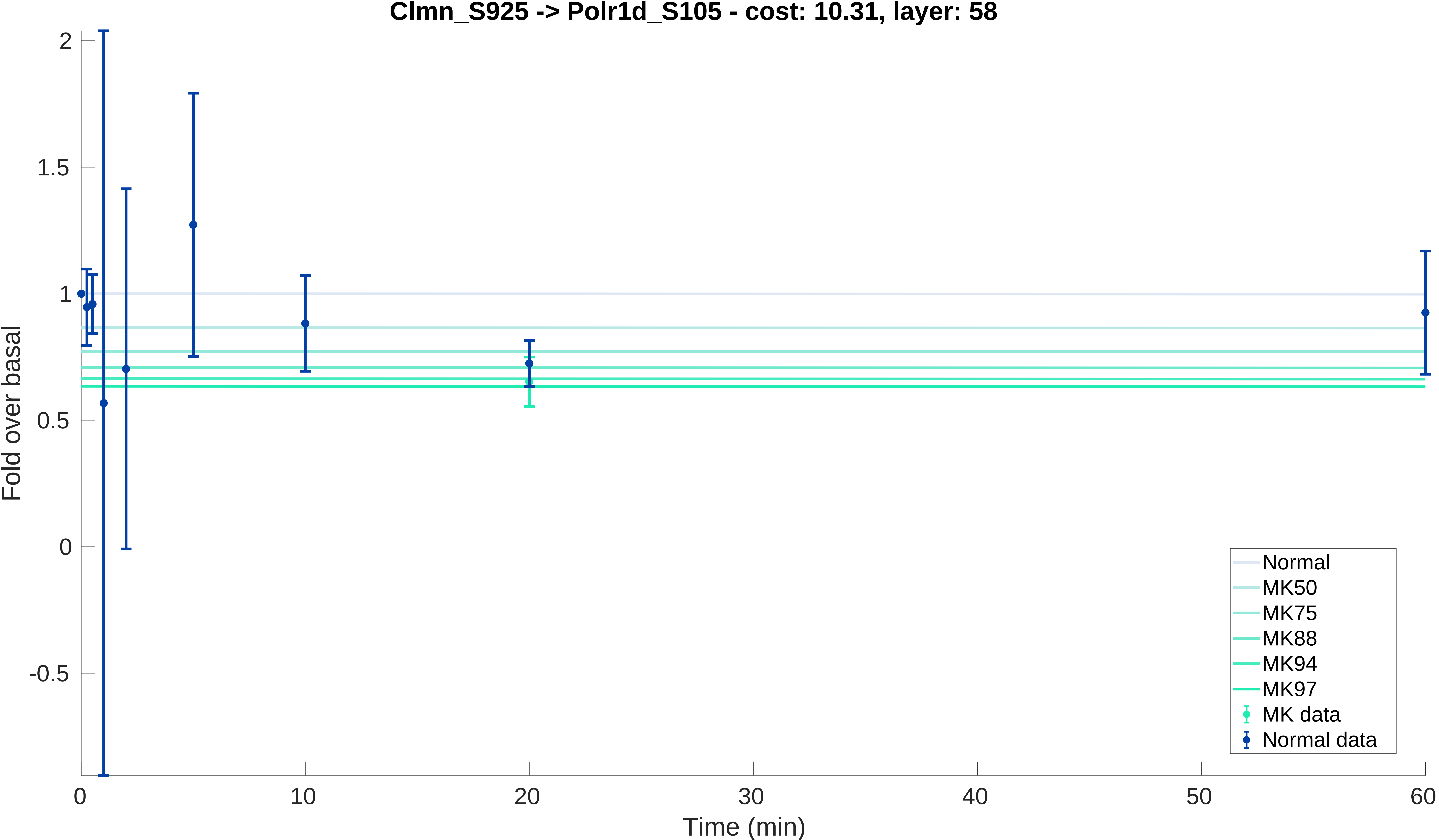
Clmn_S925 -> Polr1d_S105 - cost: 10.31, layer: 58
Fold over basal
2
1.5
1
0.5
0
-0.5
0
10
20
30
40
50
60
Time (min)
Normal
MK50
MK75
MK88
MK94
MK97
MK data
Normal data

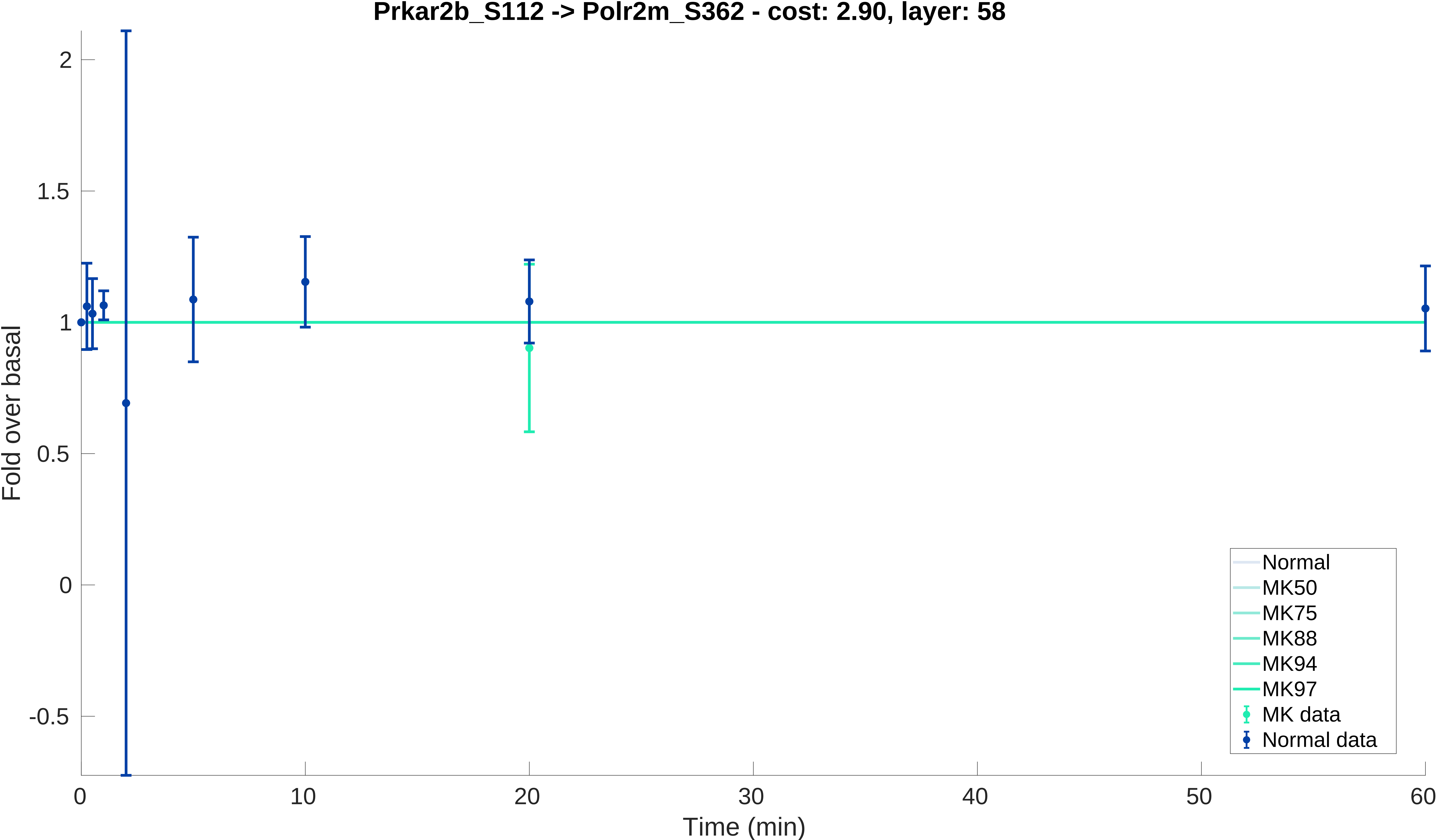

Prkar2b_S112 -> Polr2m_S362 - cost: 2.90, layer: 58
Fold over basal
Time (min)
2
1.5
1
0.5
0
-0.5
0
10
20
30
40
50
60
Normal
MK50
MK75
MK88
MK94
MK97
MK data
Normal data

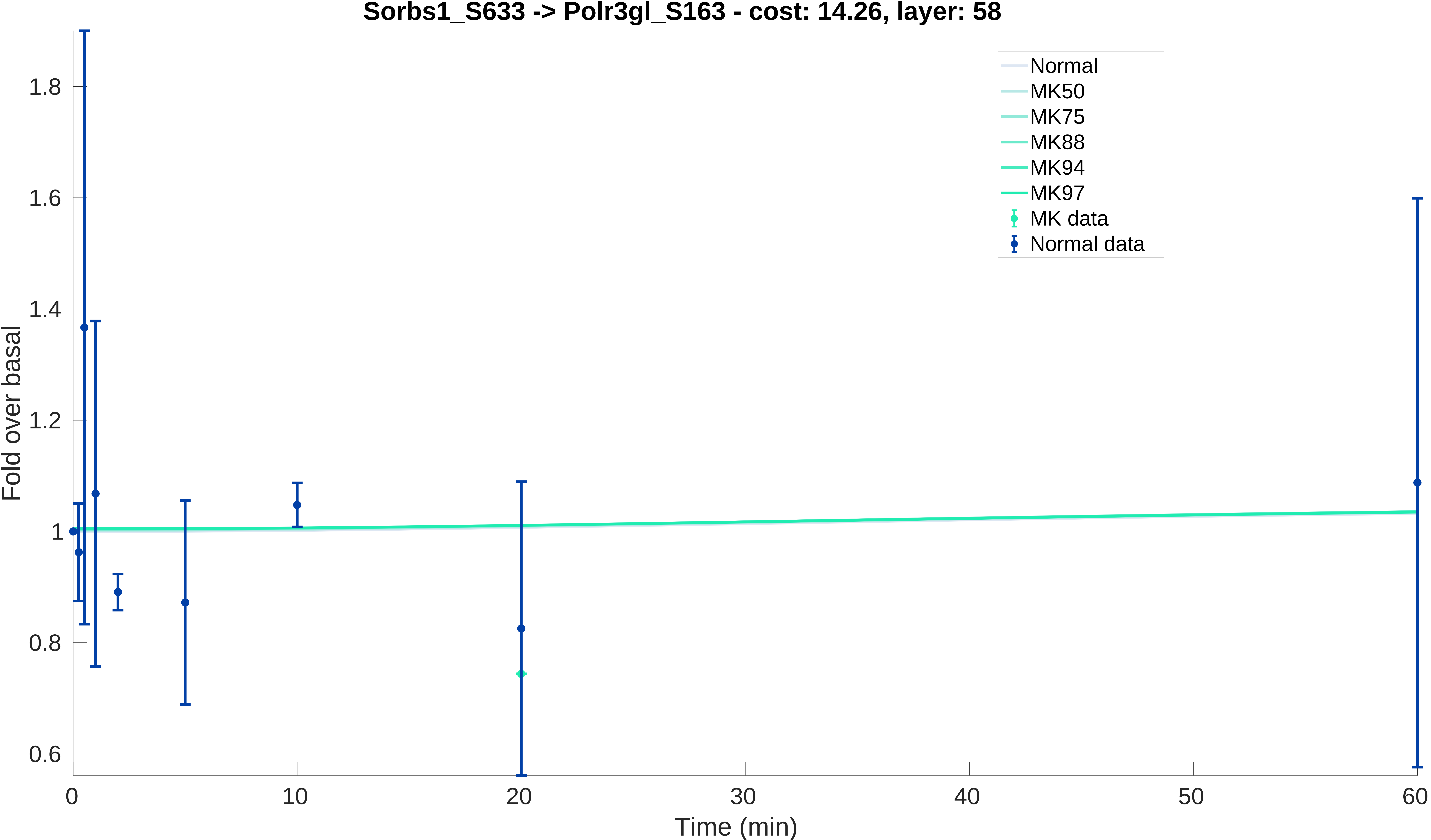

Sorbs1_S633 -> Polr3gl_S163 - cost: 14.26, layer: 58
Normal
MK50
MK75
MK88
MK94
MK97
MK data
Normal data
Fold over basal
1.8
1.6
1.4
1.2
1
0.8
0.6
0
10
20
30
40
50
60
Time (min)

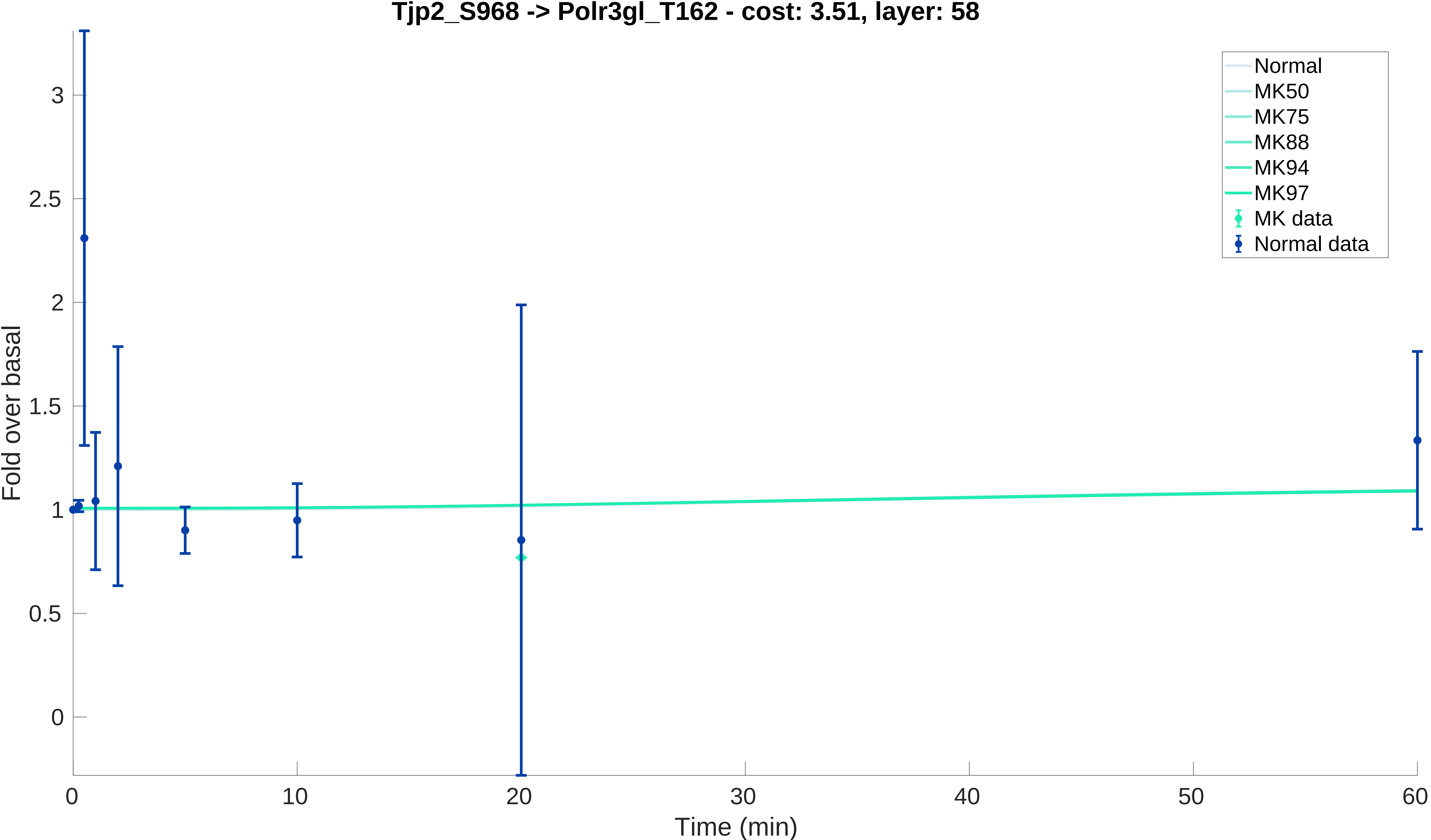

Tjp2_S968 -> Polr3gl_T162 - cost: 3.51, layer: 58
Fold over basal
Time (min)
0
0.5
1
1.5
2
2.5
3
0
10
20
30
40
50
60
Normal
MK50
MK75
MK88
MK94
MK97
MK data
Normal data

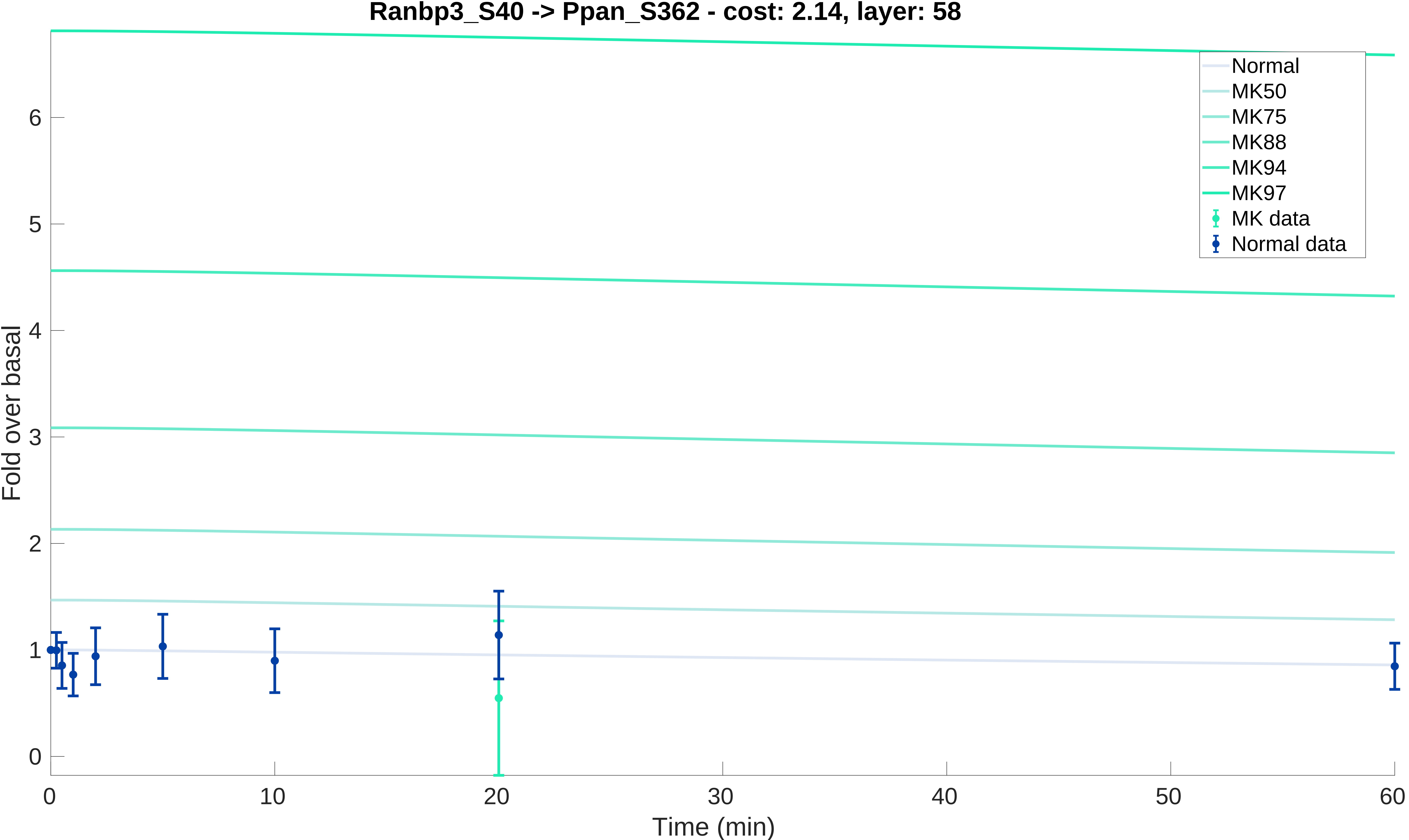

Ranbp3_S40 -> Ppan_S362 - cost: 2.14, layer: 58
Fold over basal
Time (min)
0
1
2
3
4
5
6
0
10
20
30
40
50
60
Normal
MK50
MK75
MK88
MK94
MK97
MK data
Normal data

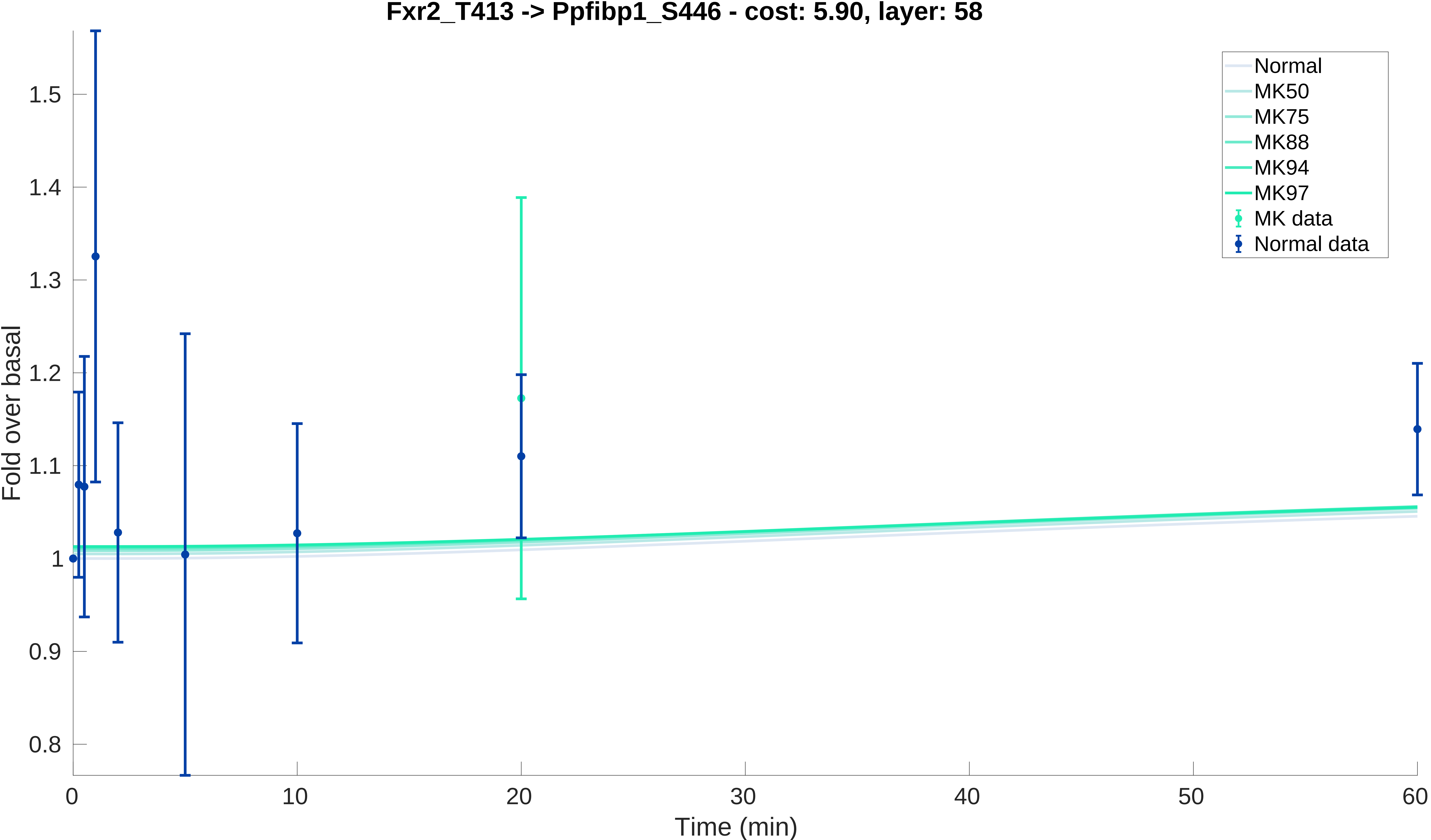

Fxr2_T413 -> Ppfibp1_S446 - cost: 5.90, layer: 58
Normal
MK50
MK75
MK88
MK94
MK97
MK data
Normal data
Fold over basal
1.5
1.4
1.3
1.2
1.1
1
0.9
0.8
0
10
20
30
40
50
60
Time (min)

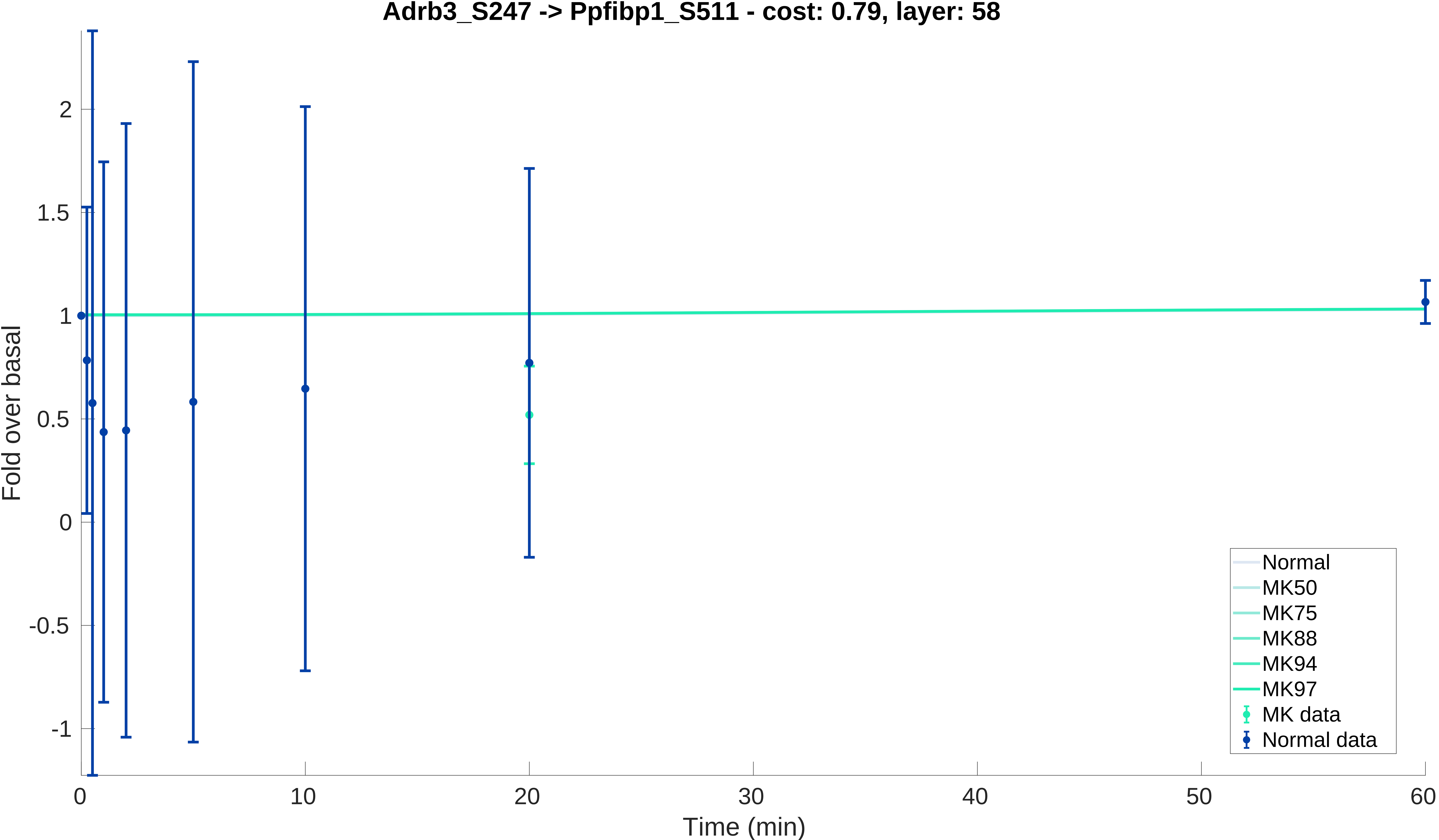

Adrb3_S247 -> Ppfibp1_S511 - cost: 0.79, layer: 58
Fold over basal
Time (min)
2
1.5
1
0.5
0
-0.5
-1
0
10
20
30
40
50
60
Normal
MK50
MK75
MK88
MK94
MK97
MK data
Normal data

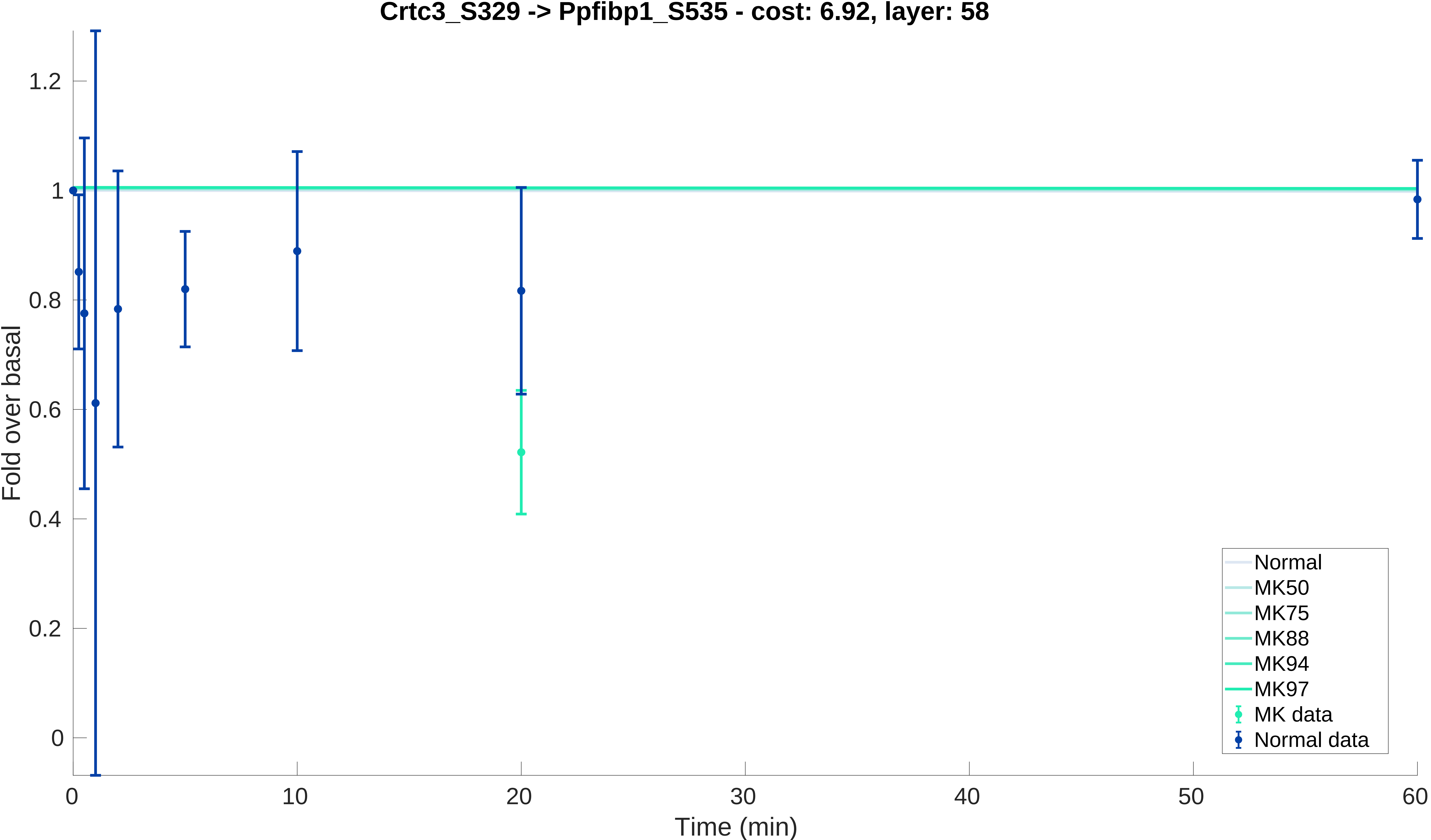

Crtc3_S329 -> Ppfibp1_S535 - cost: 6.92, layer: 58
Fold over basal
1.2
1
0.8
0.6
0.4
0.2
0
Time (min)
0
10
20
30
40
50
60
Normal
MK50
MK75
MK88
MK94
MK97
MK data
Normal data

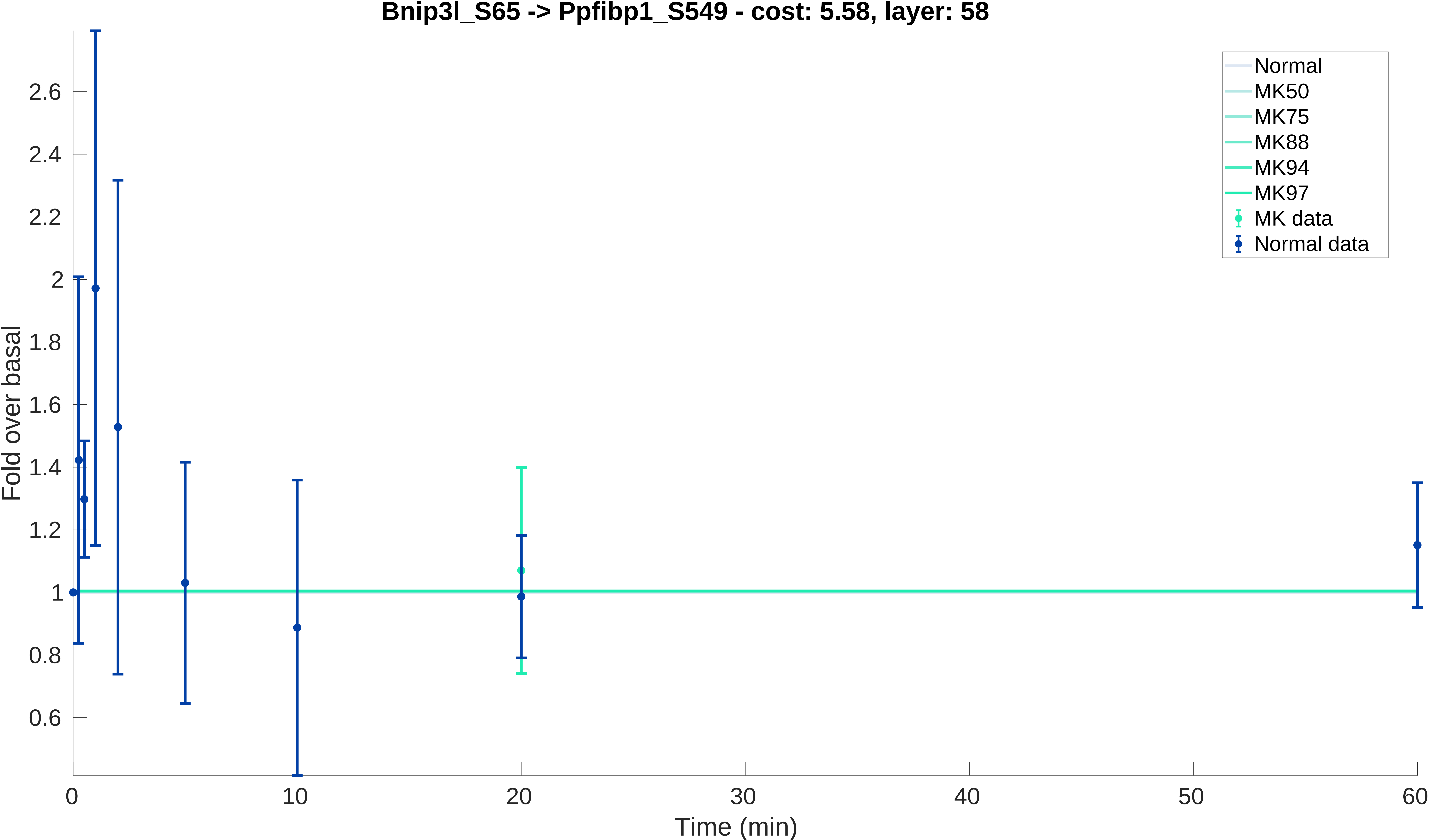

Bnip3l_S65 -> Ppfibp1_S549 - cost: 5.58, layer: 58
Fold over basal
Time (min)
0.6
0.8
1
1.2
1.4
1.6
1.8
2
2.2
2.4
2.6
0
10
20
30
40
50
60
Normal
MK50
MK75
MK88
MK94
MK97
MK data
Normal data

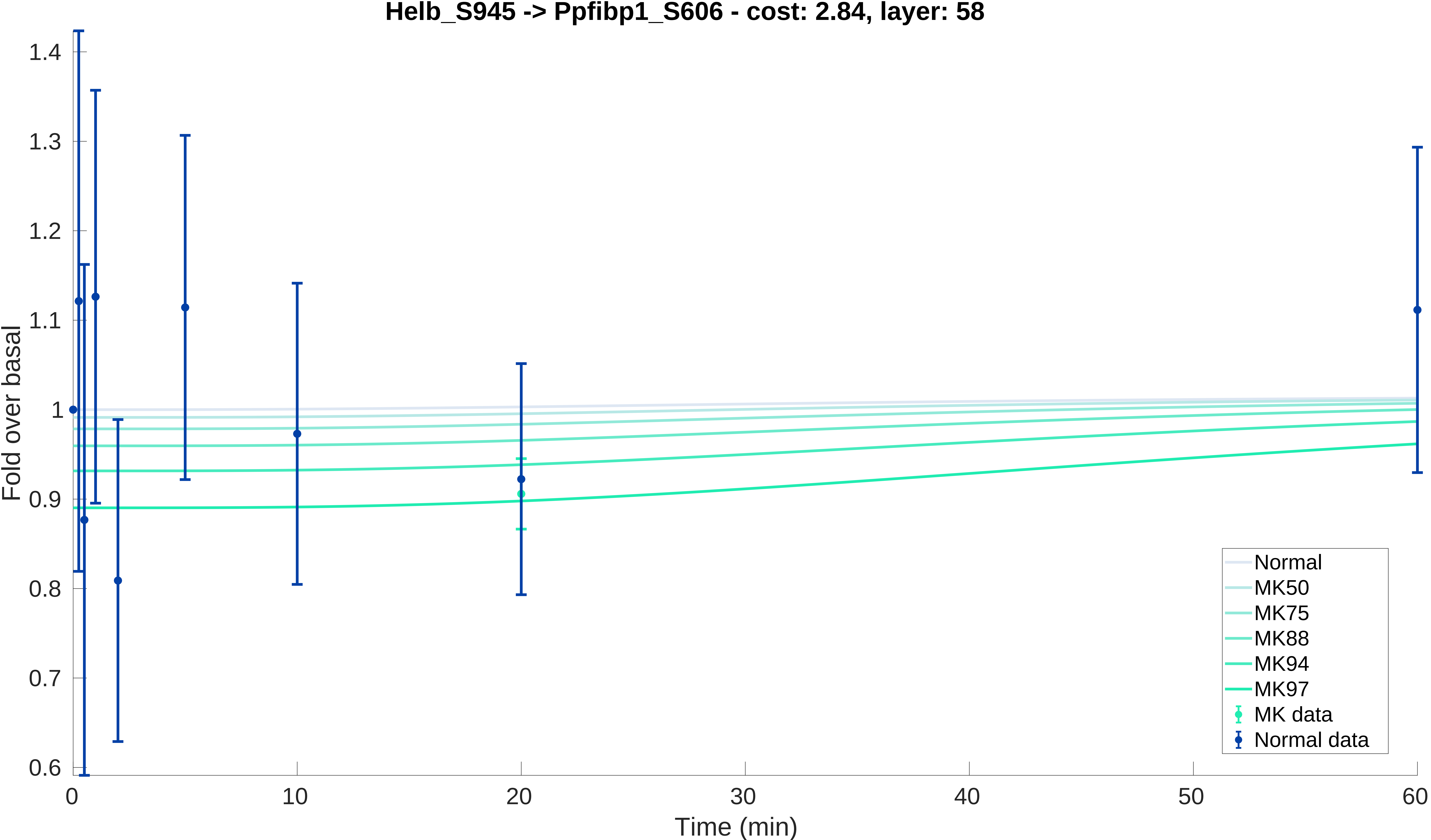

Helb_S945 -> Ppfibp1_S606 - cost: 2.84, layer: 58
Fold over basal
Time (min)
1.4
1.3
1.2
1.1
1
0.9
0.8
0.7
0.6
0
10
20
30
40
50
60
Normal
MK50
MK75
MK88
MK94
MK97
MK data
Normal data

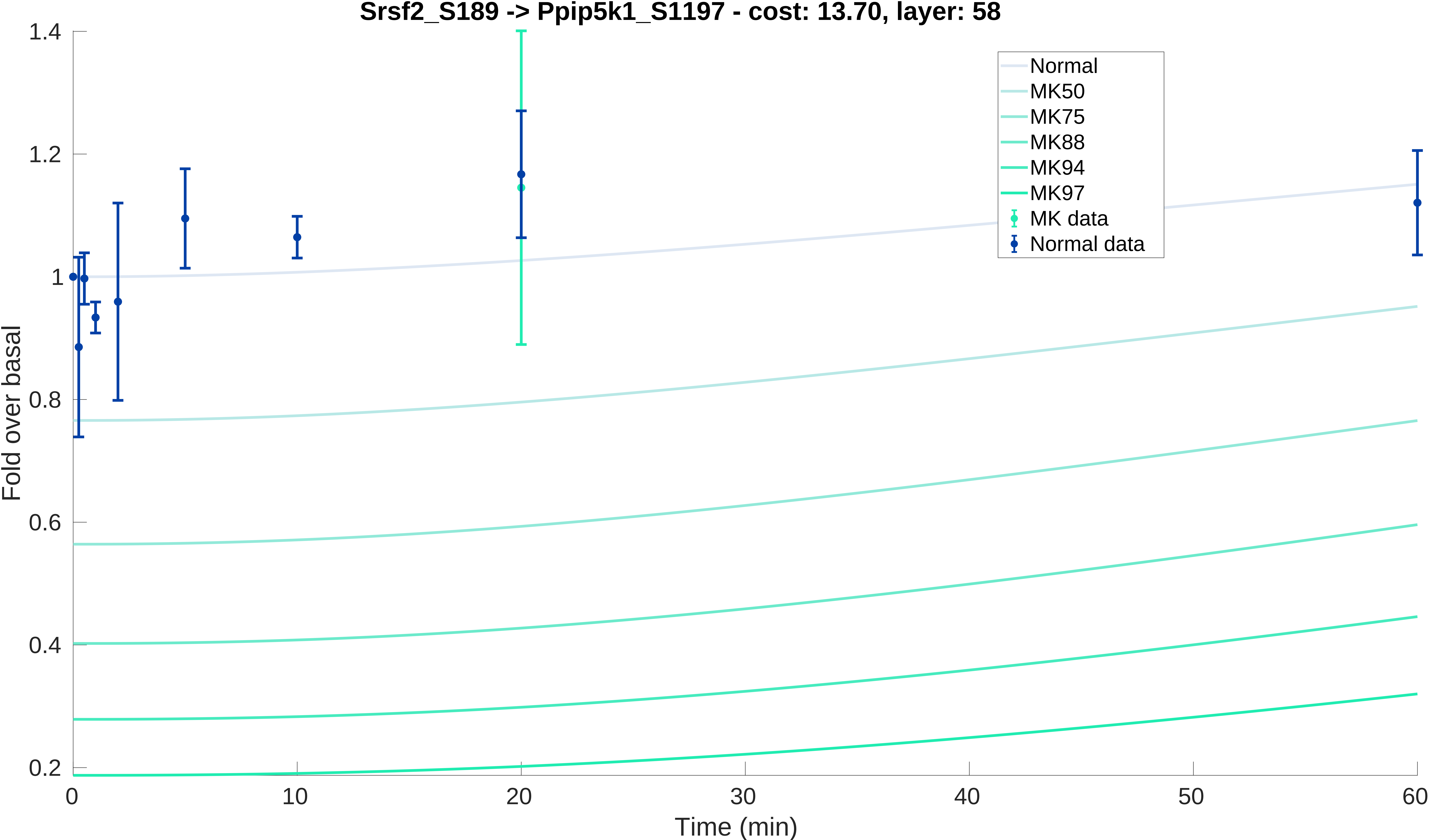

Srsf2_S189 -> Ppip5k1_S1197 - cost: 13.70, layer: 58
Fold over basal
Time (min)
Normal
MK50
MK75
MK88
MK94
MK97
MK data
Normal data
1.4
1.2
1
0.8
0.6
0.4
0.2
0
10
20
30
40
50
60

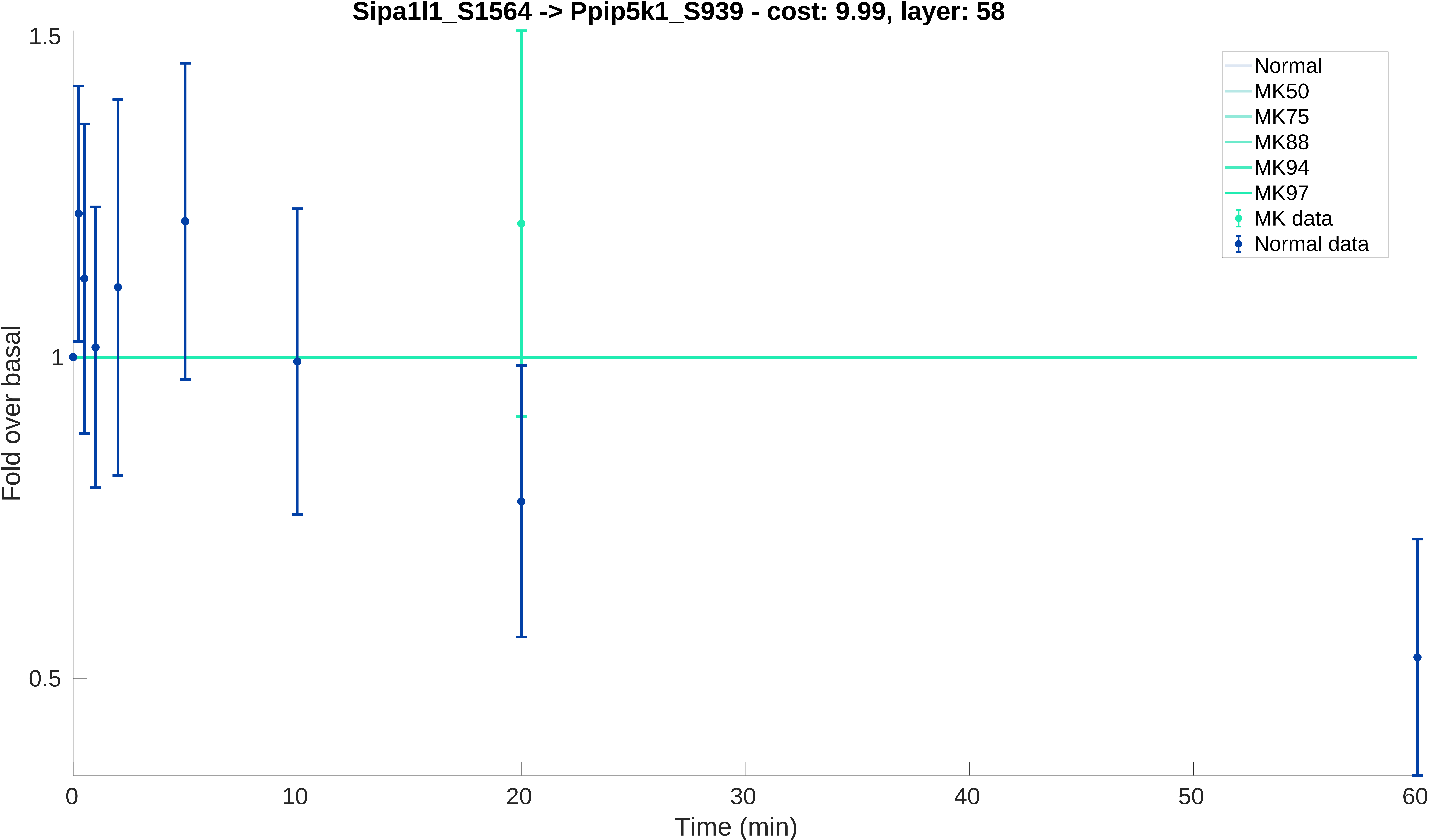

Sipa1l1_S1564 -> Ppip5k1_S939 - cost: 9.99, layer: 58
Fold over basal
Time (min)
1.5
1
0.5
0
10
20
30
40
50
60
Normal
MK50
MK75
MK88
MK94
MK97
MK data
Normal data

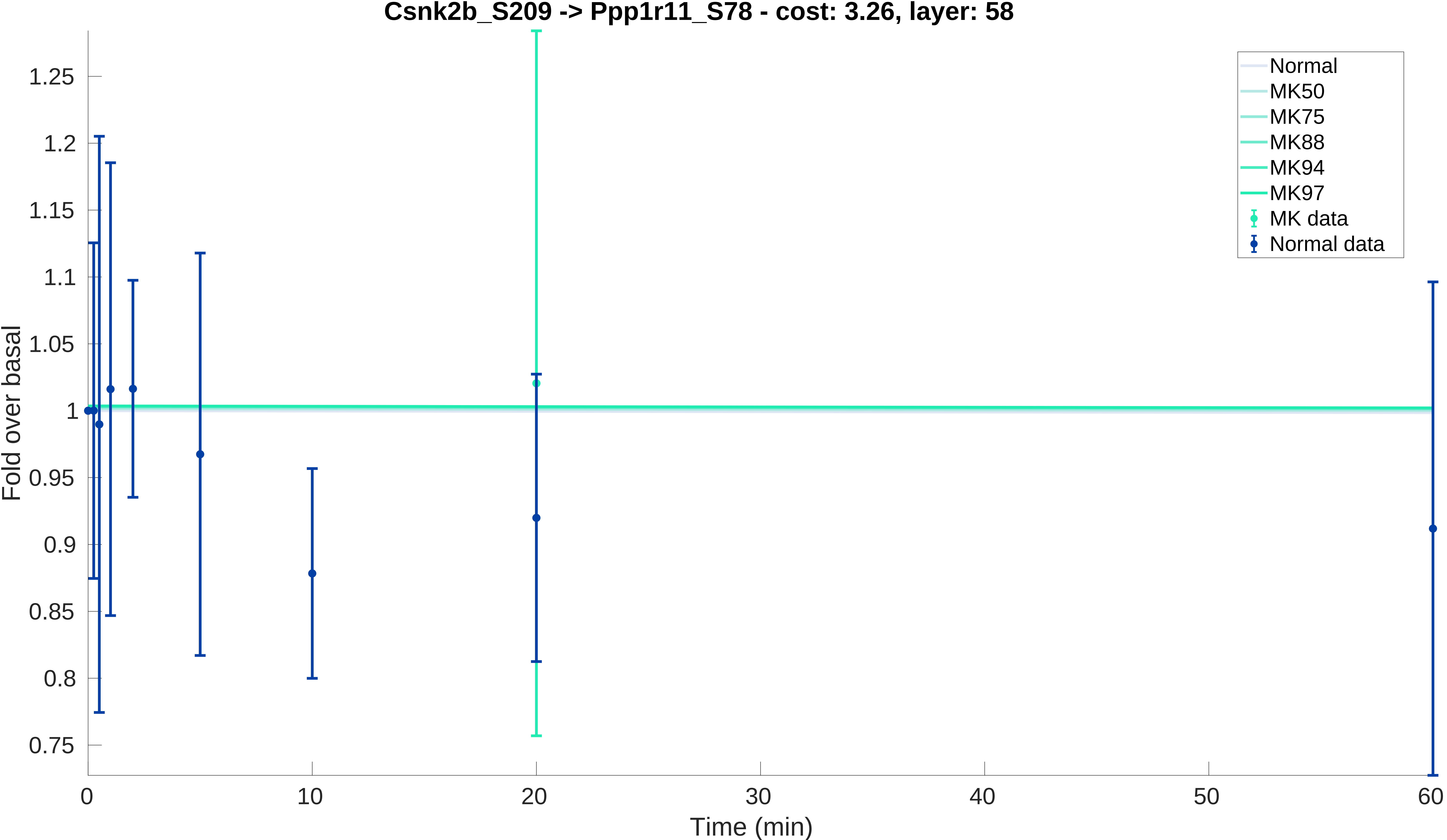
Csnk2b_S209 -> Ppp1r11_S78 - cost: 3.26, layer: 58
Fold over basal
1.25
1.2
1.15
1.1
1.05
1
0.95
0.9
0.85
0.8
0.75
0
10
20
30
40
50
60
Time (min)
Normal
MK50
MK75
MK88
MK94
MK97
MK data
Normal data

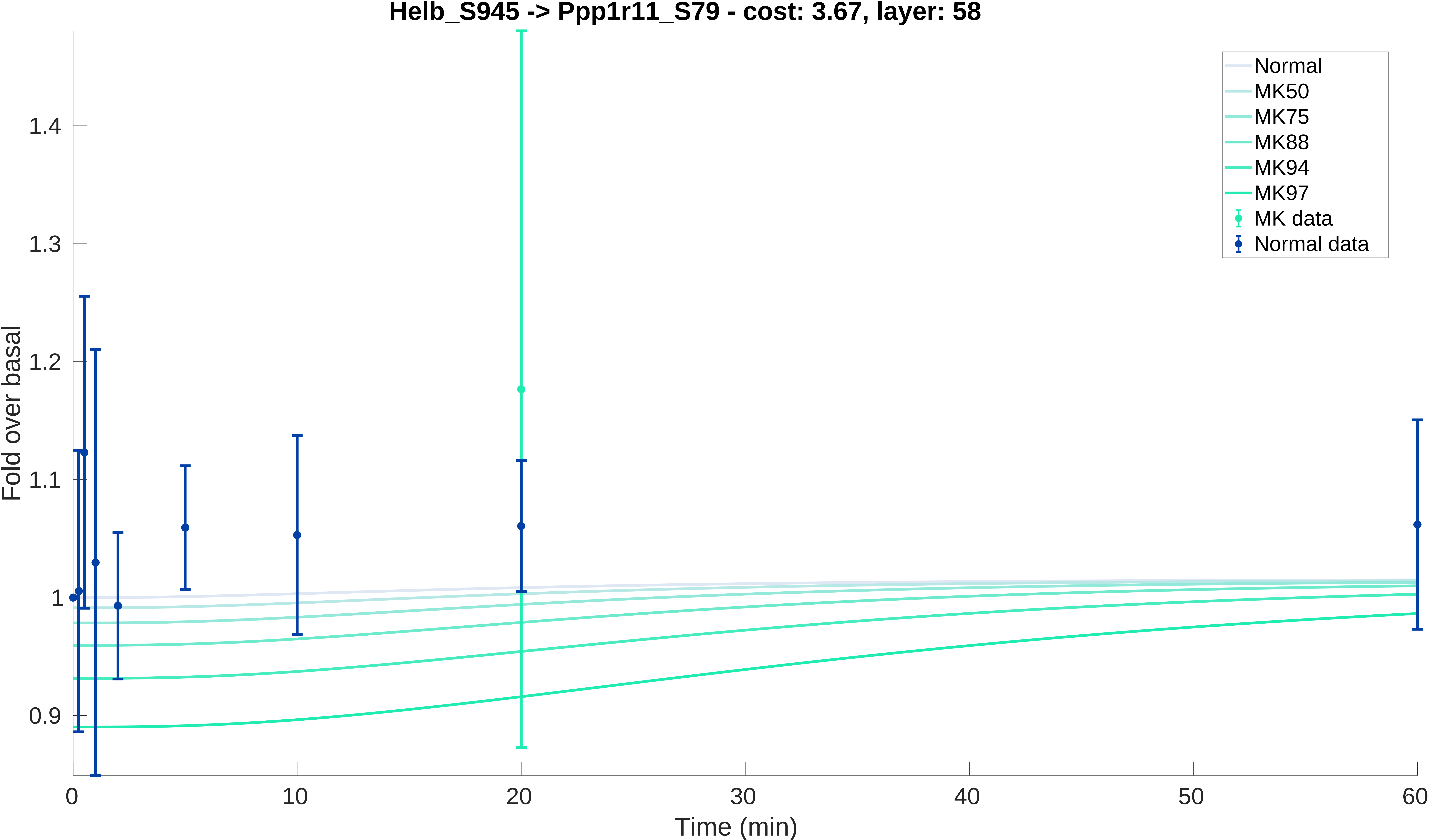

Helb_S945 -> Ppp1r11_S79 - cost: 3.67, layer: 58
Fold over basal
Time (min)
1.4
1.3
1.2
1.1
1
0.9
0
10
20
30
40
50
60
Normal
MK50
MK75
MK88
MK94
MK97
MK data
Normal data

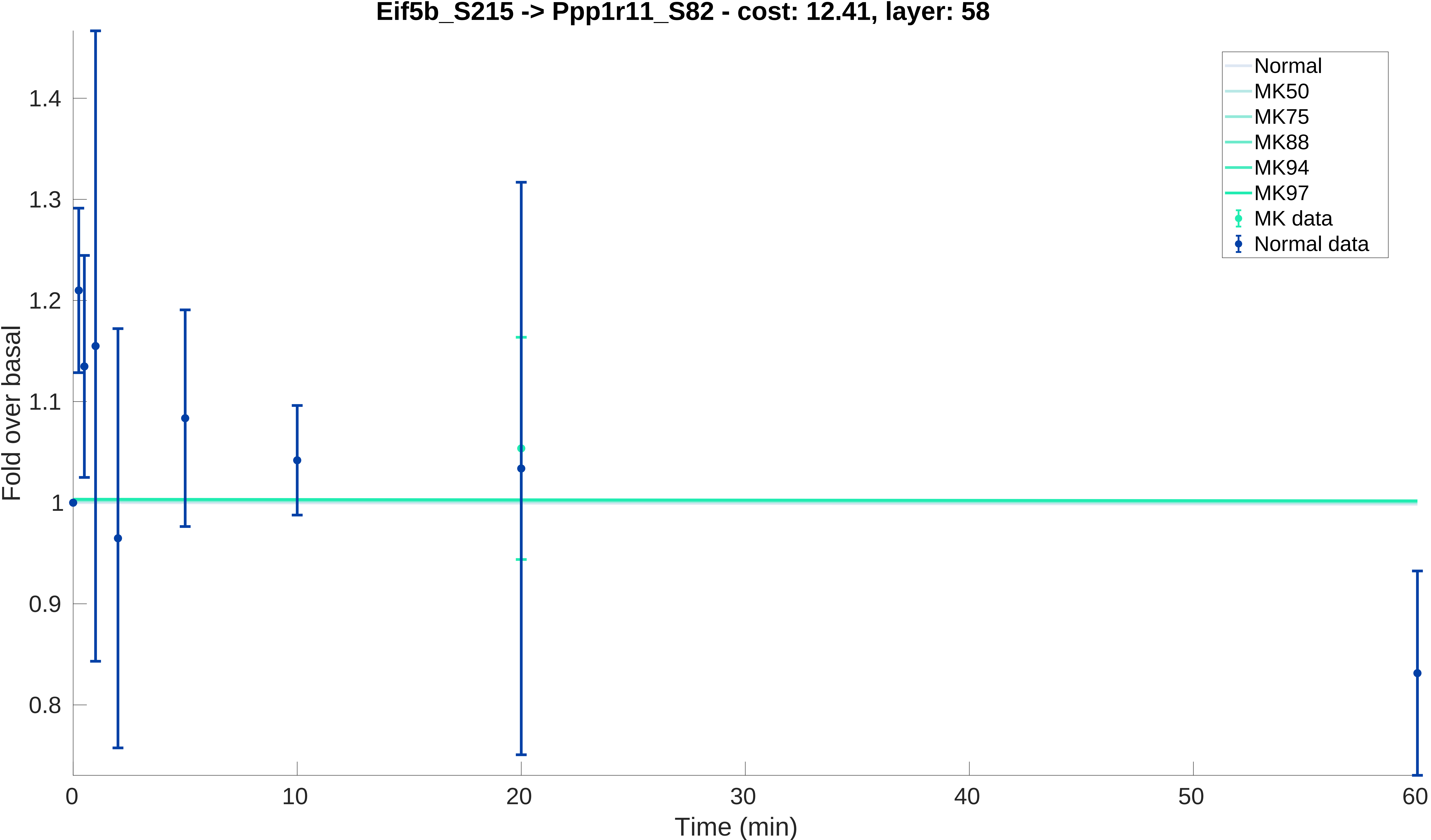

Eif5b_S215 -> Ppp1r11_S82 - cost: 12.41, layer: 58
Fold over basal
Time (min)
0
10
20
30
40
50
60
0.8
0.9
1
1.1
1.2
1.3
1.4
Normal
MK50
MK75
MK88
MK94
MK97
MK data
Normal data

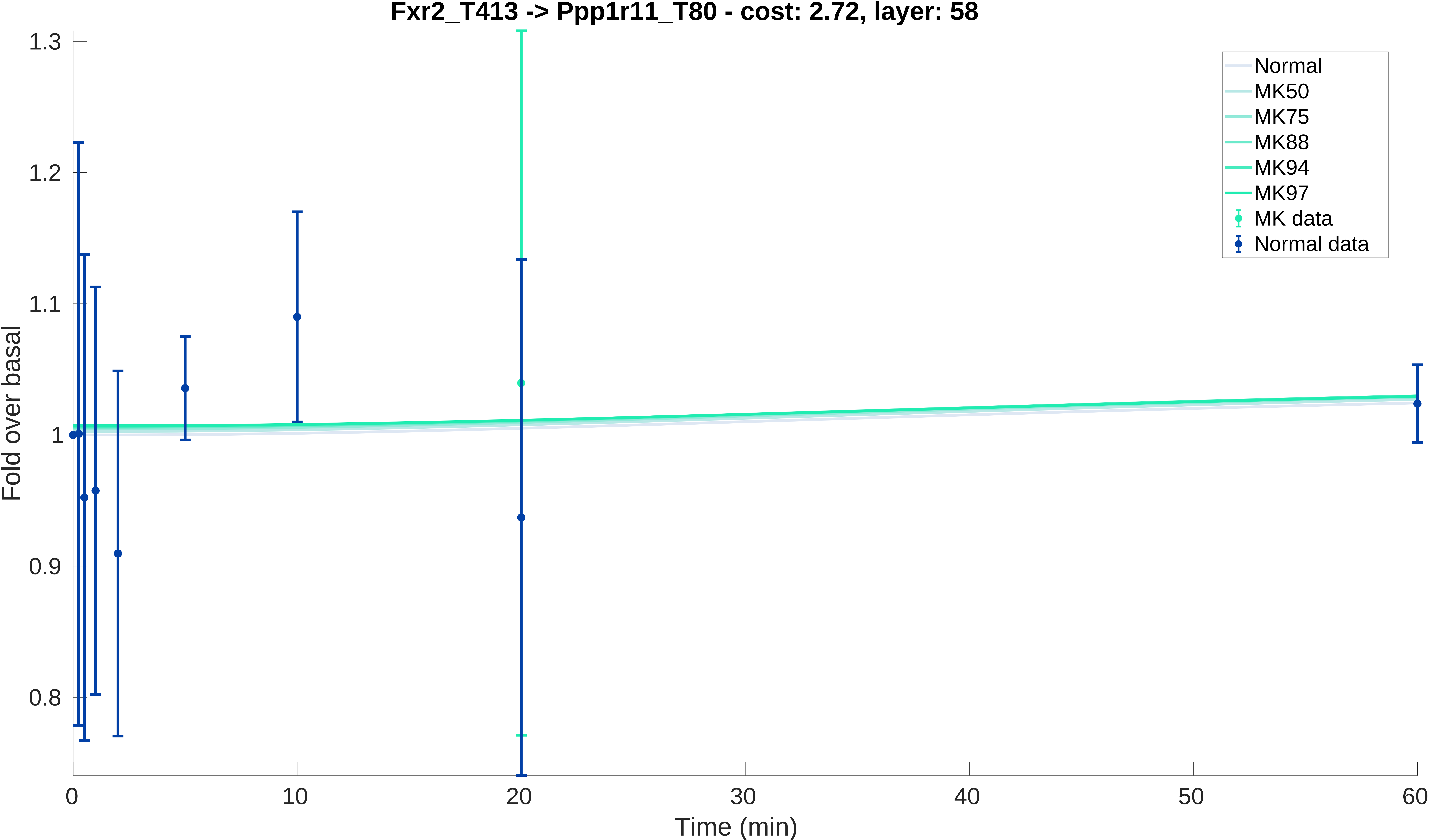
Fxr2_T413 -> Ppp1r11_T80 - cost: 2.72, layer: 58
Fold over basal
Time (min)
1.3
1.2
1.1
1
0.9
0.8
0
10
20
30
40
50
60
Normal
MK50
MK75
MK88
MK94
MK97
MK data
Normal data

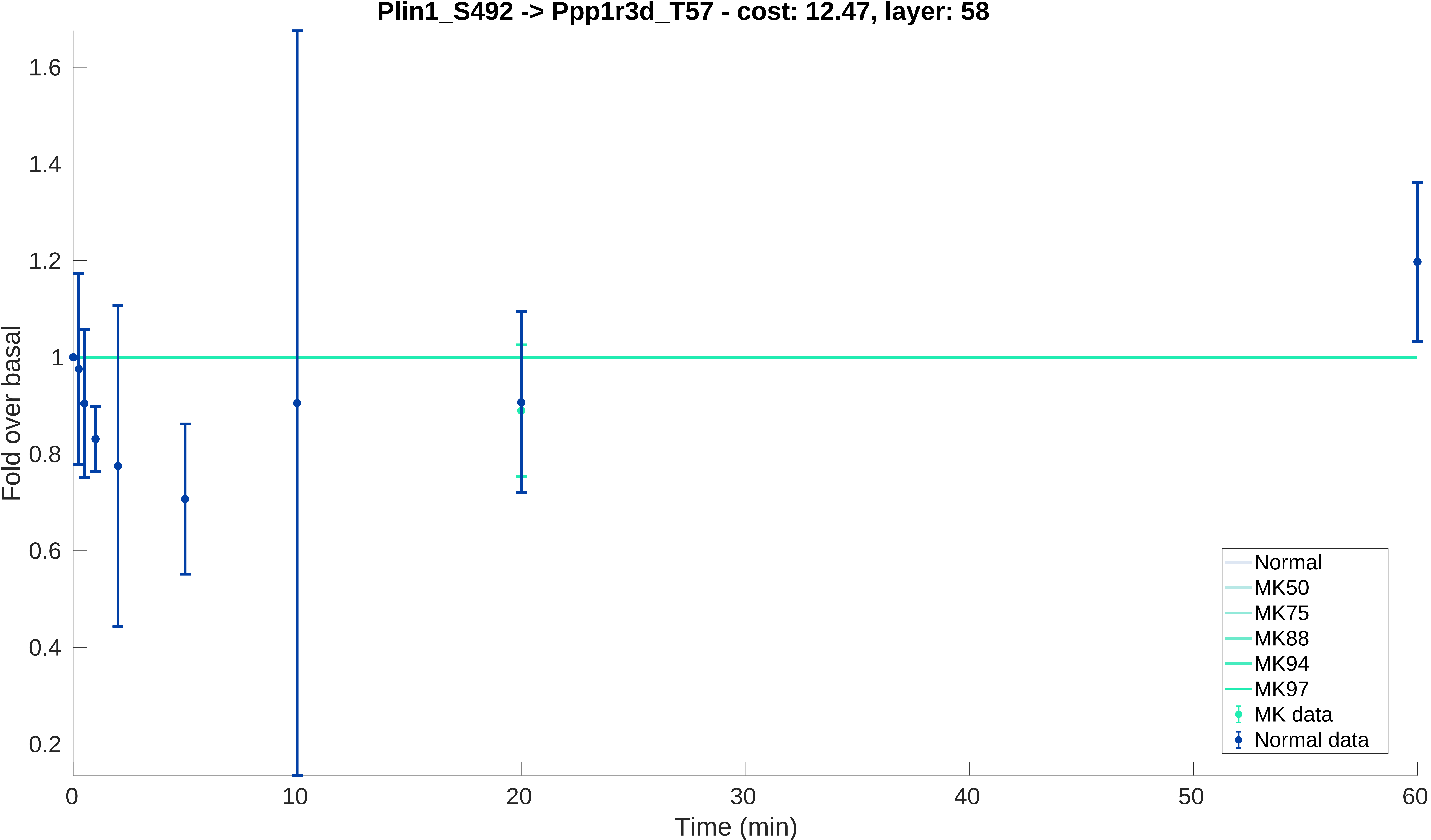

Plin1_S492 -> Ppp1r3d_T57 - cost: 12.47, layer: 58
Fold over basal
Time (min)
0.2
0.4
0.6
0.8
1
1.2
1.4
1.6
0
10
20
30
40
50
60
Normal
MK50
MK75
MK88
MK94
MK97
MK data
Normal data

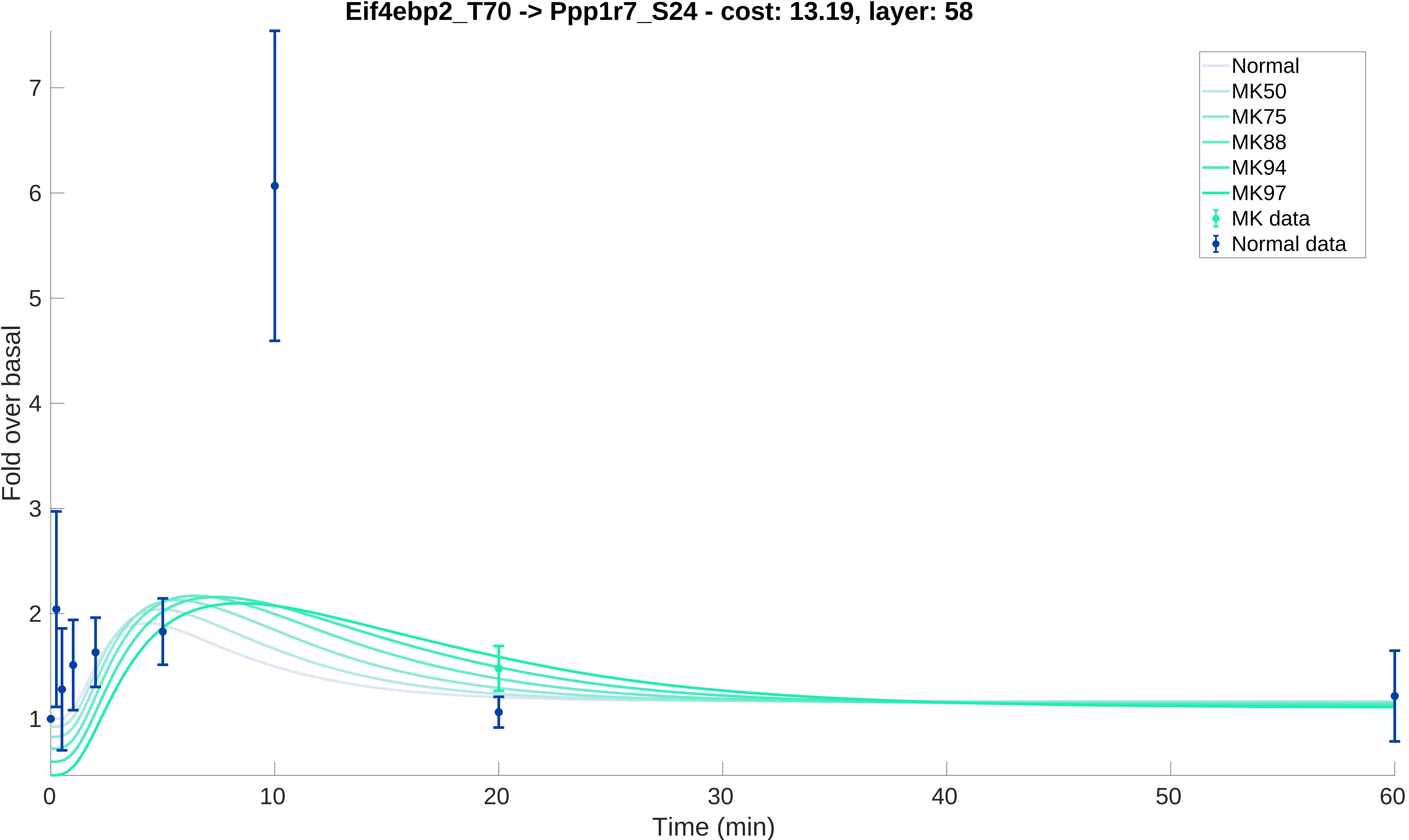
Eif4ebp2_T70 -> Ppp1r7_S24 - cost: 13.19, layer: 58
Fold over basal
Time (min)
0
10
20
30
40
50
60
1
2
3
4
5
6
7
Normal
MK50
MK75
MK88
MK94
MK97
MK data
Normal data

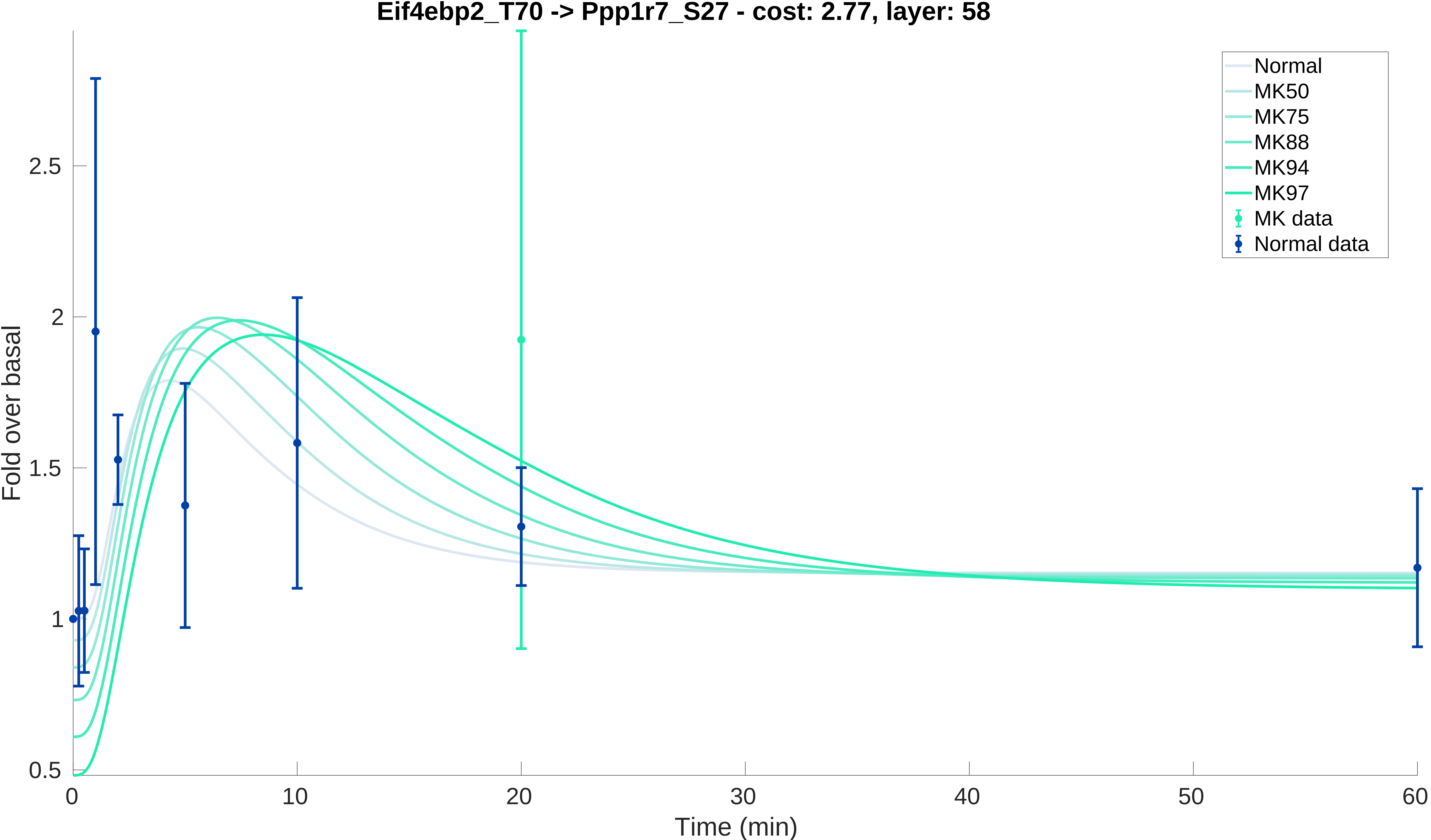

Eif4ebp2_T70 -> Ppp1r7_S27 - cost: 2.77, layer: 58
Normal
MK50
MK75
MK88
MK94
MK97
MK data
Normal data
Fold over basal
Time (min)
0.5
1
1.5
2
2.5
0
10
20
30
40
50
60

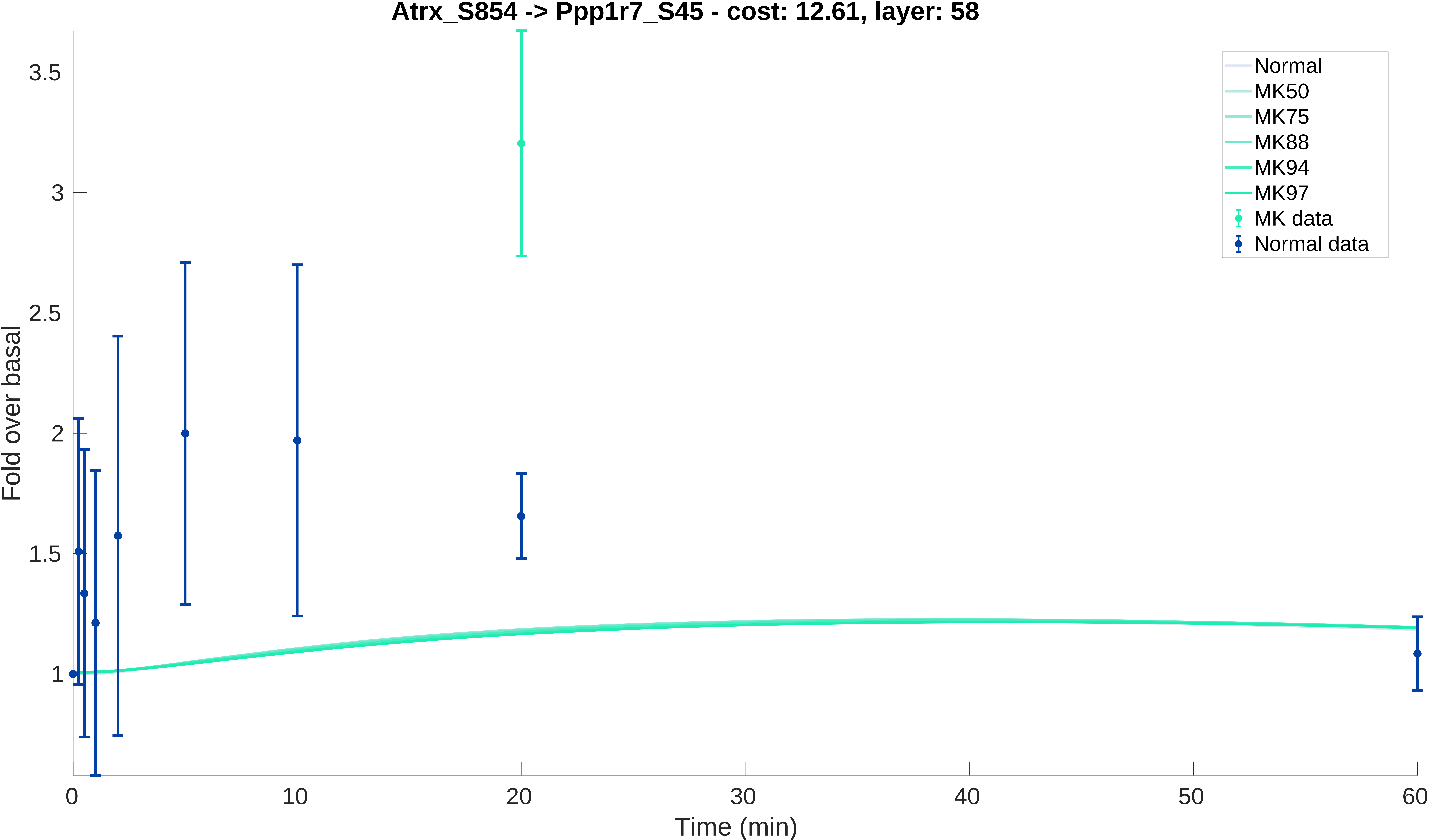

Atrx_S854 -> Ppp1r7_S45 - cost: 12.61, layer: 58
Fold over basal
Time (min)
3.5
3
2.5
2
1.5
1
0
10
20
30
40
50
60
Normal
MK50
MK75
MK88
MK94
MK97
MK data
Normal data

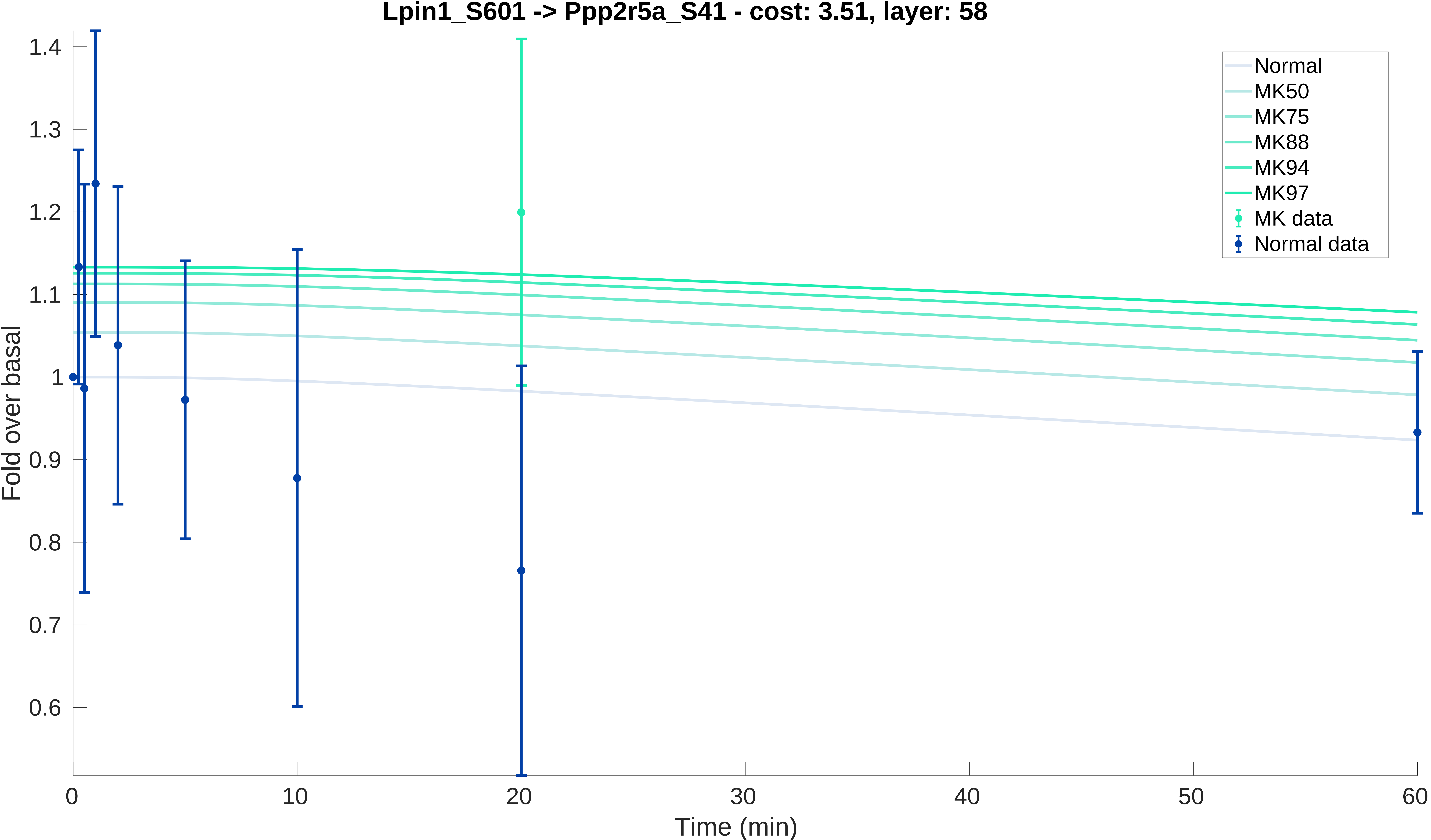

Lpin1_S601 -> Ppp2r5a_S41 - cost: 3.51, layer: 58
Fold over basal
1.4
1.3
1.2
1.1
1
0.9
0.8
0.7
0.6
0
10
20
30
40
50
60
Time (min)
Normal
MK50
MK75
MK88
MK94
MK97
MK data
Normal data

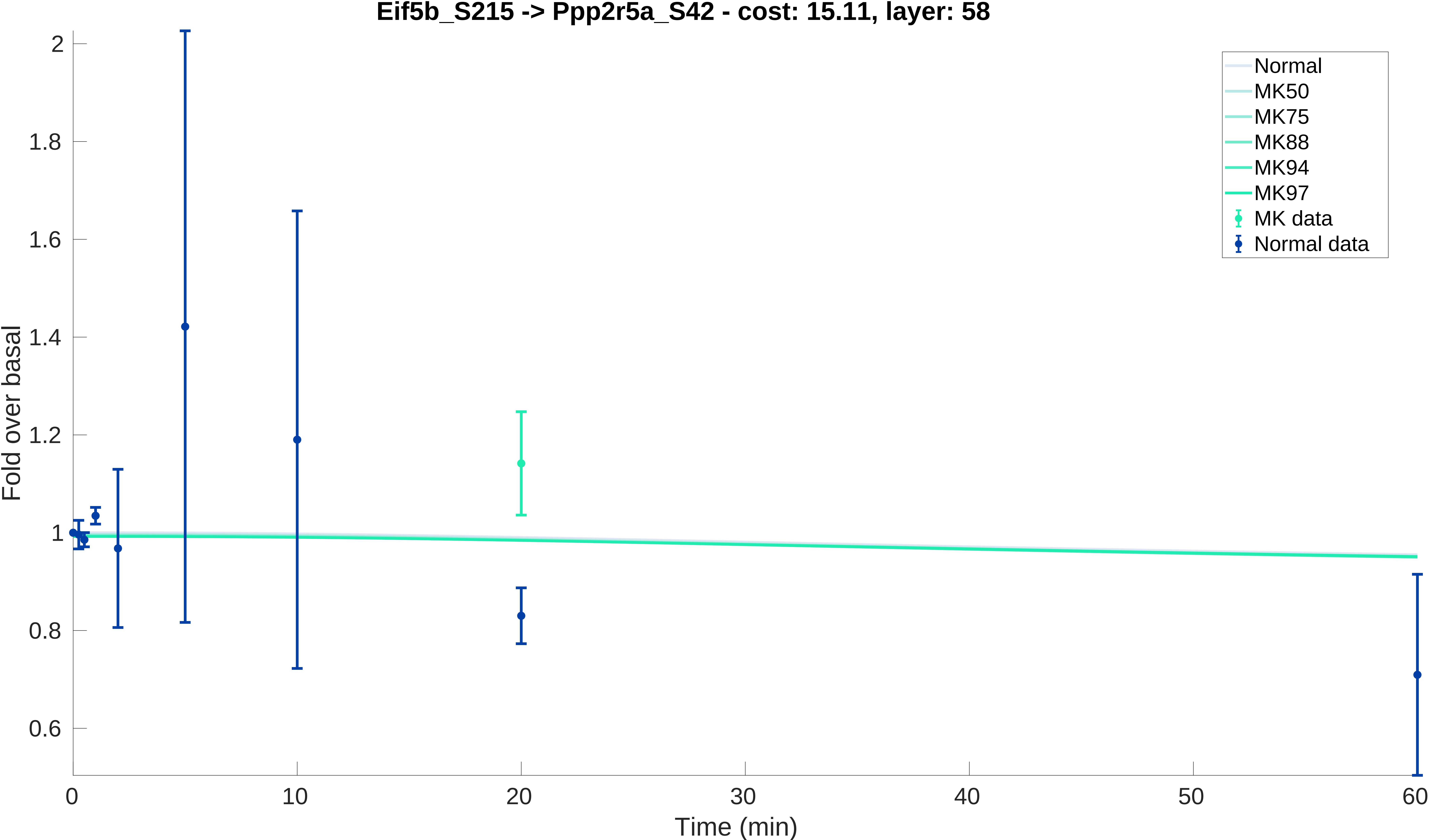

Eif5b_S215 -> Ppp2r5a_S42 - cost: 15.11, layer: 58
Fold over basal
Time (min)
0
10
20
30
40
50
60
0.6
0.8
1
1.2
1.4
1.6
1.8
2
Normal
MK50
MK75
MK88
MK94
MK97
MK data
Normal data

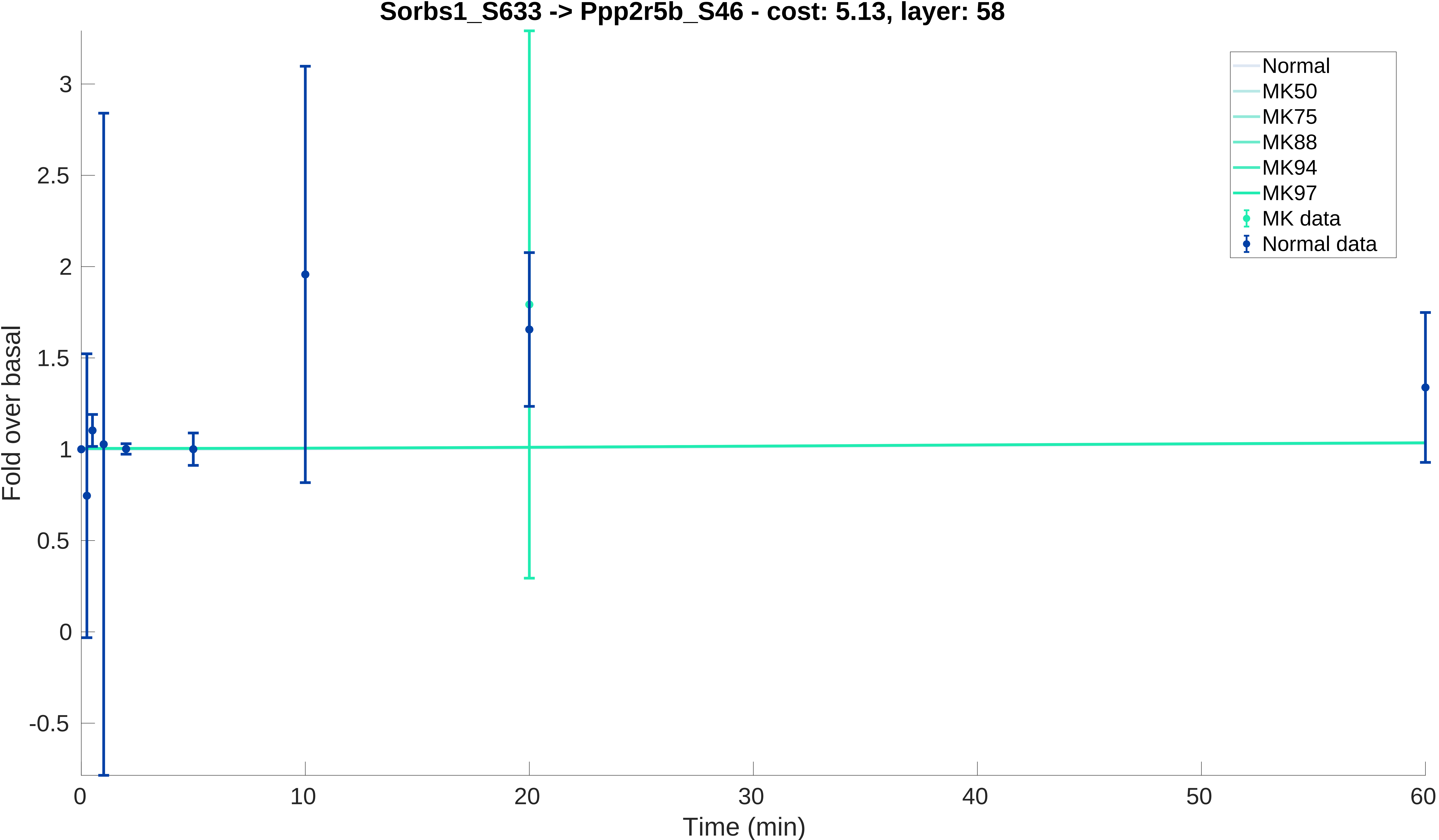

Sorbs1_S633 -> Ppp2r5b_S46 - cost: 5.13, layer: 58
Fold over basal
Time (min)
3
2.5
2
1.5
1
0.5
0
-0.5
0
10
20
30
40
50
60
Normal
MK50
MK75
MK88
MK94
MK97
MK data
Normal data

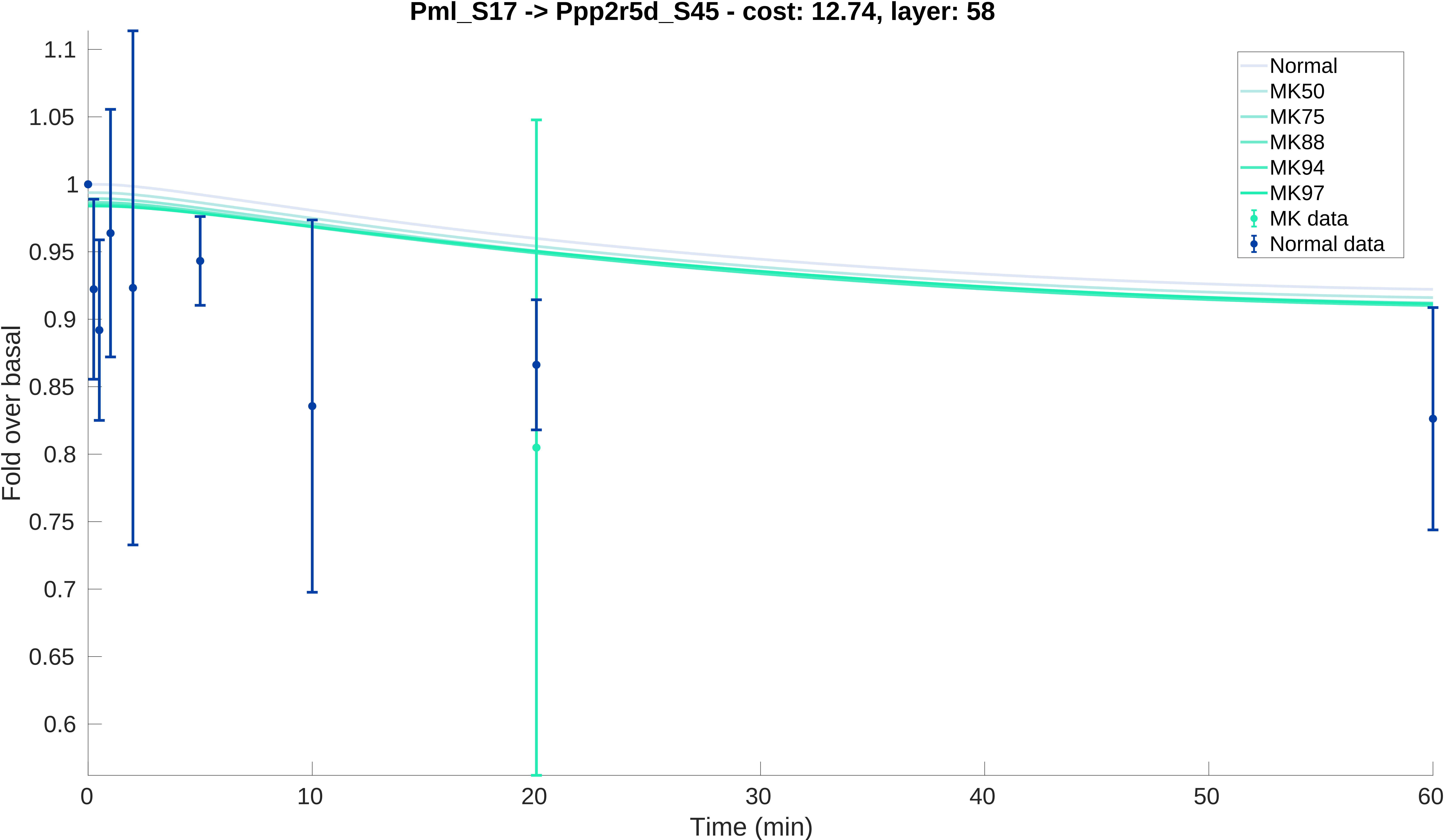

Pml_S17 -> Ppp2r5d_S45 - cost: 12.74, layer: 58
Fold over basal
Time (min)
1.1
1.05
1
0.95
0.9
0.85
0.8
0.75
0.7
0.65
0.6
0
10
20
30
40
50
60
Normal
MK50
MK75
MK88
MK94
MK97
MK data
Normal data

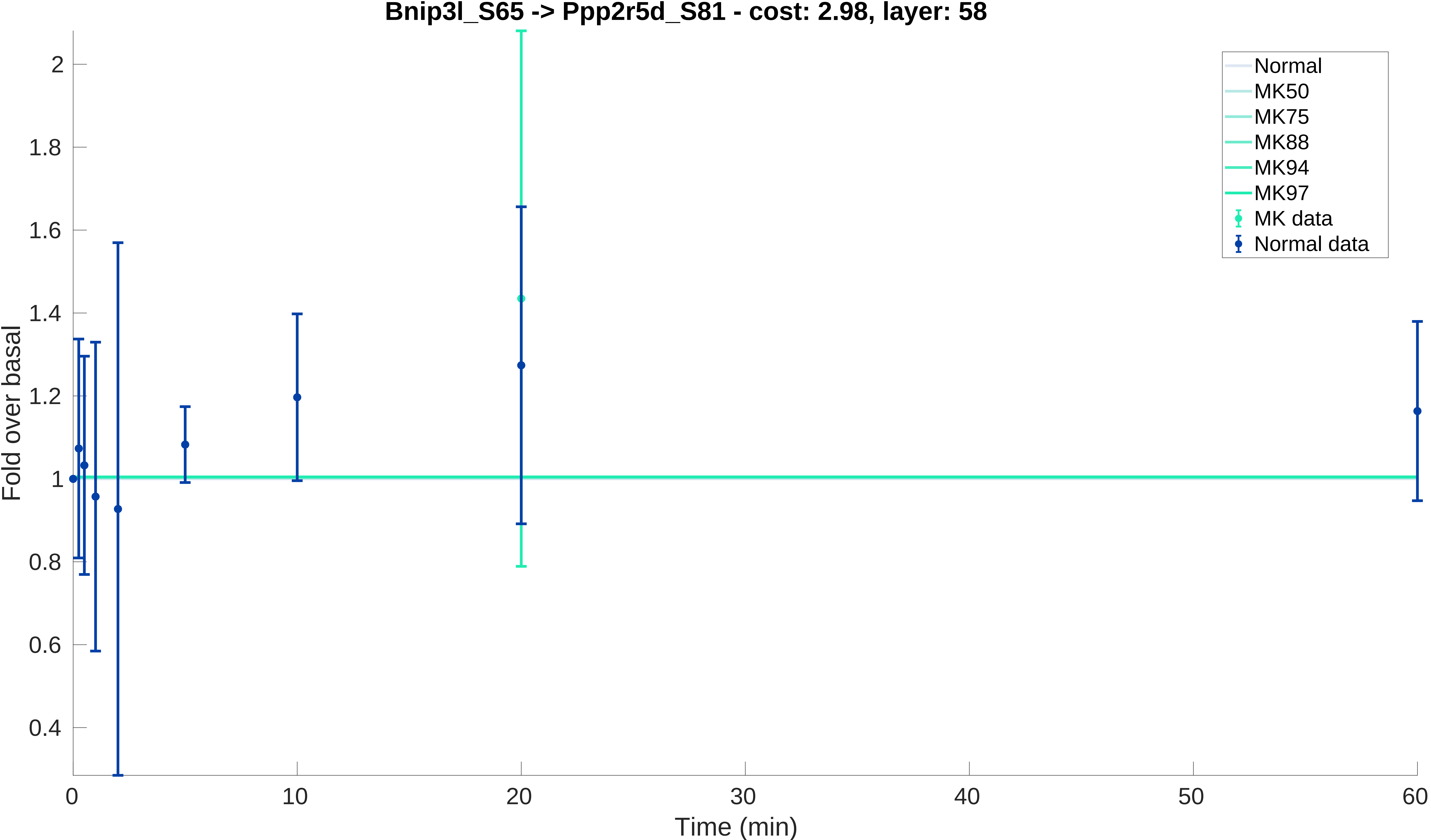
Bnip3l_S65 -> Ppp2r5d_S81 - cost: 2.98, layer: 58
Fold over basal
Time (min)
0
10
20
30
40
50
60
0.4
0.6
0.8
1
1.2
1.4
1.6
1.8
2
Normal
MK50
MK75
MK88
MK94
MK97
MK data
Normal data

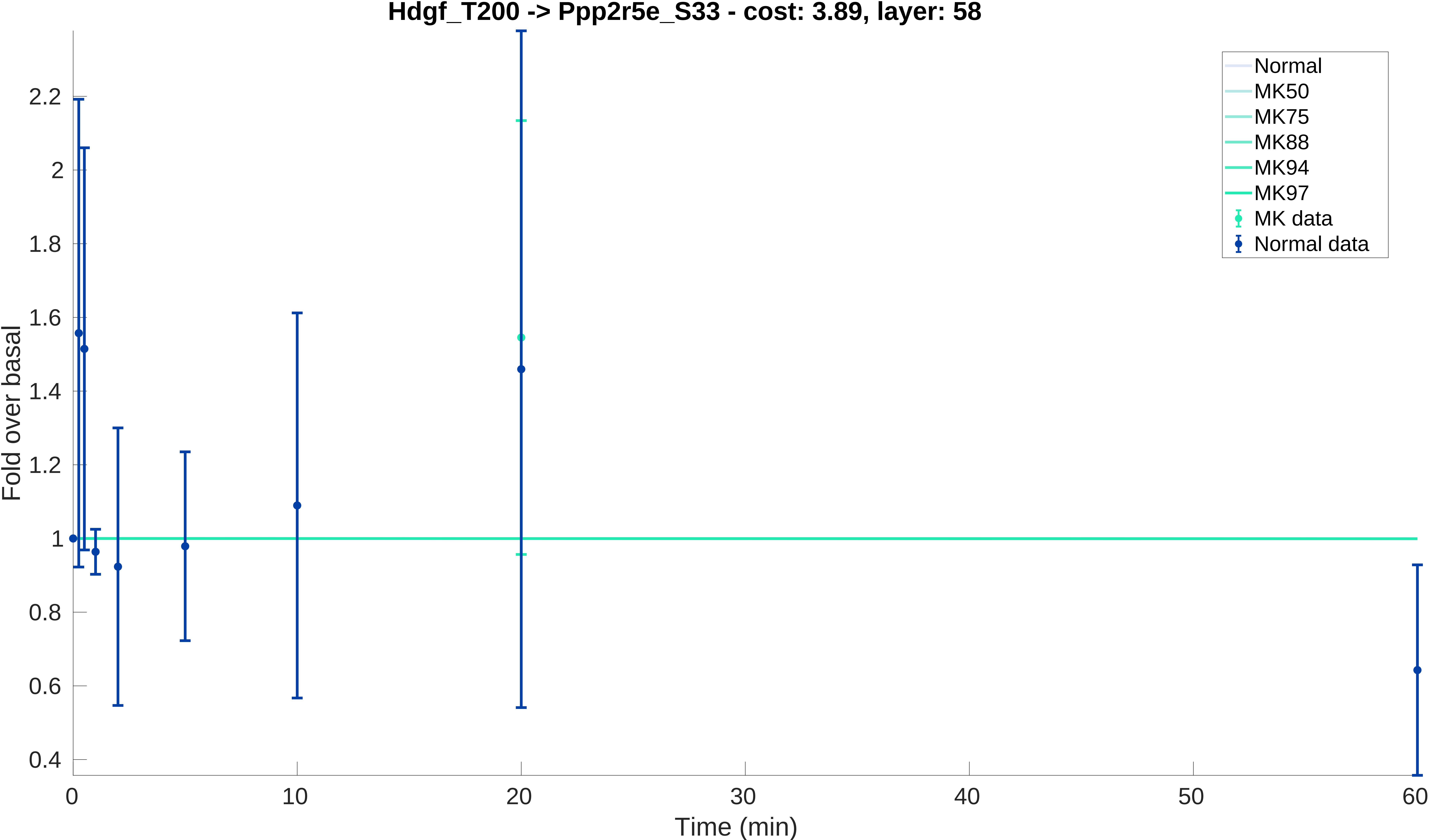

Hdgf_T200 -> Ppp2r5e_S33 - cost: 3.89, layer: 58
Fold over basal
Time (min)
0.4
0.6
0.8
1
1.2
1.4
1.6
1.8
2
2.2
0
10
20
30
40
50
60
Normal
MK50
MK75
MK88
MK94
MK97
MK data
Normal data

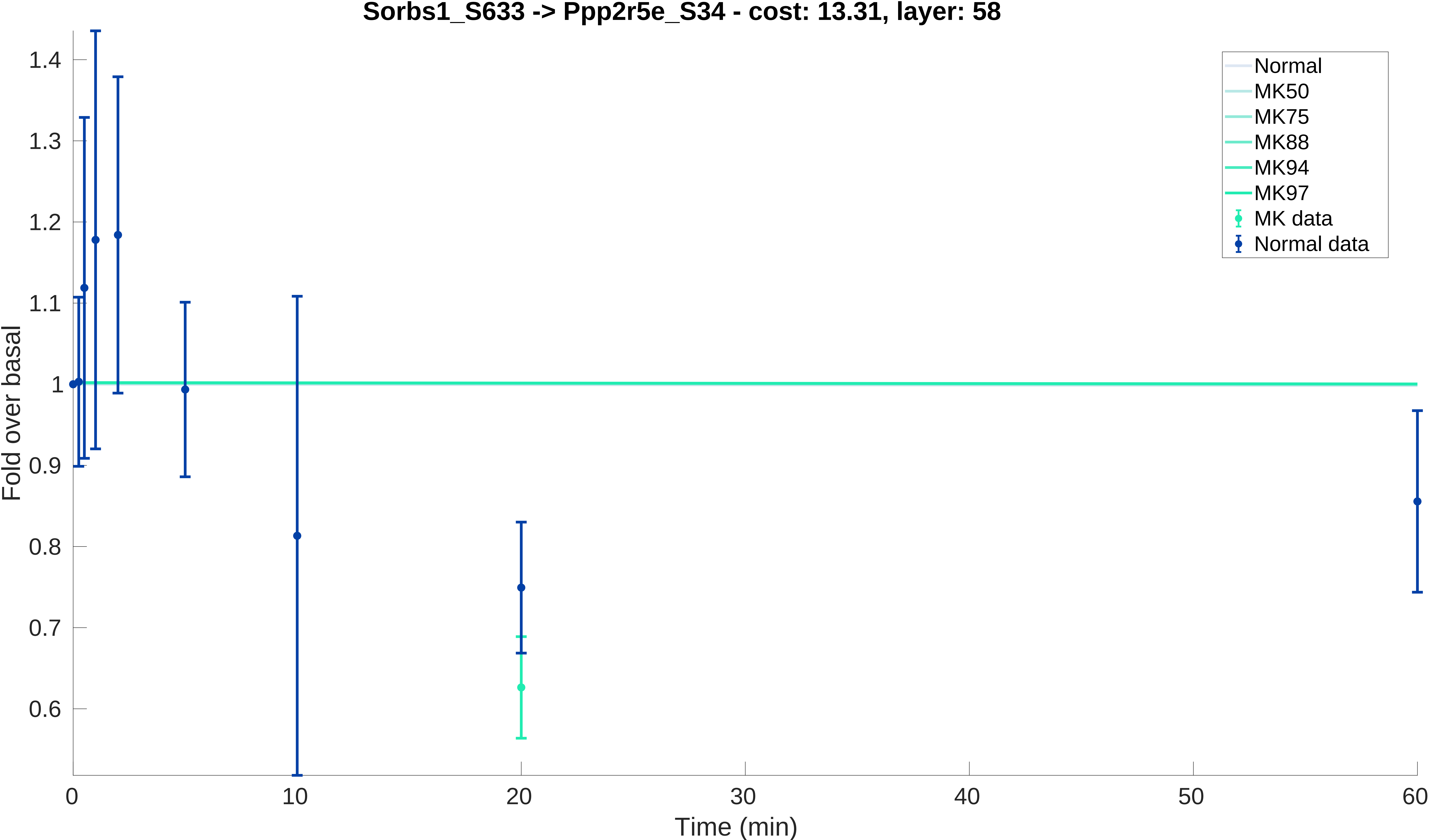

Sorbs1_S633 -> Ppp2r5e_S34 - cost: 13.31, layer: 58
Fold over basal
Time (min)
0
10
20
30
40
50
60
0.6
0.7
0.8
0.9
1
1.1
1.2
1.3
1.4
Normal
MK50
MK75
MK88
MK94
MK97
MK data
Normal data

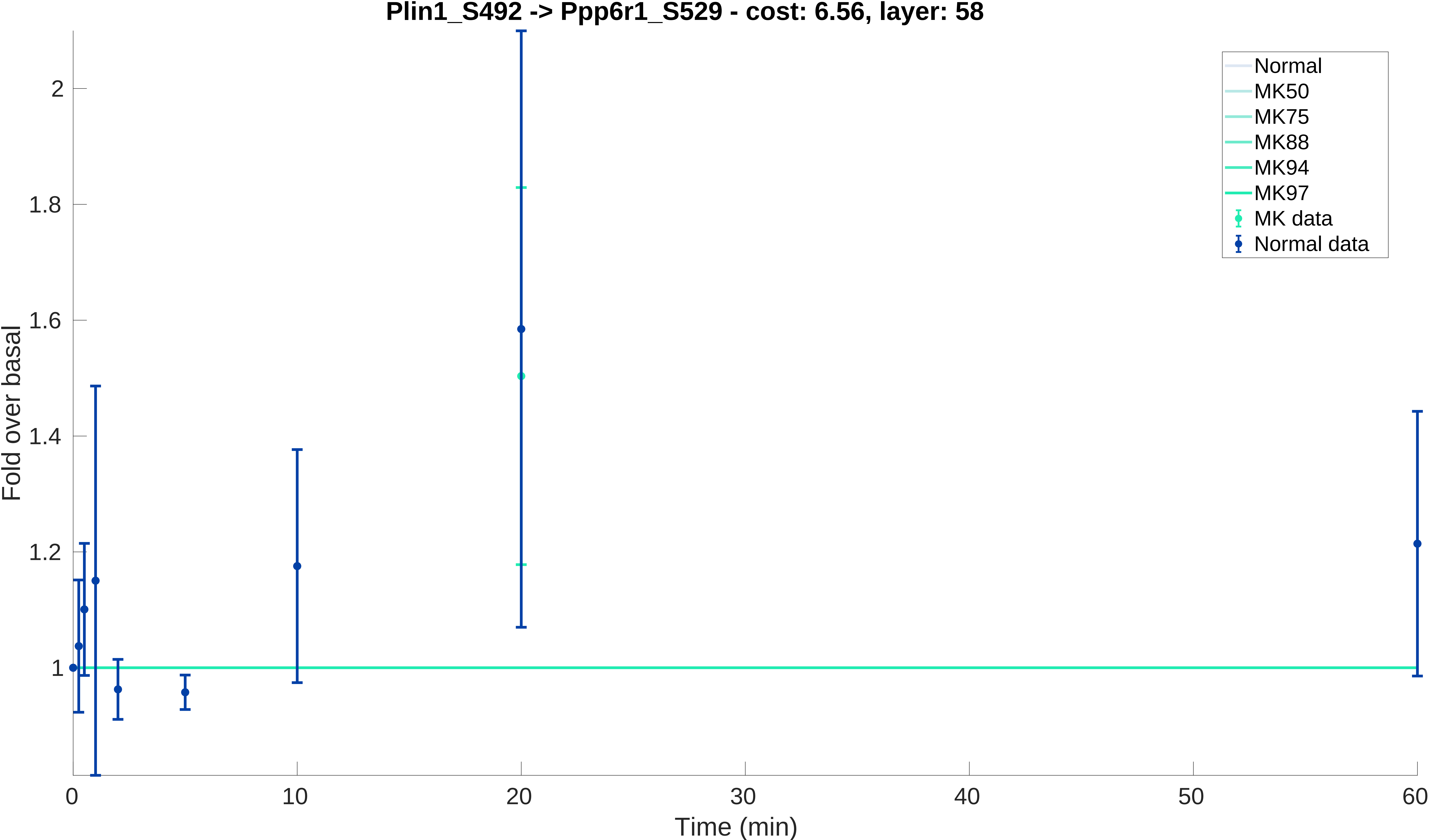

Plin1_S492 -> Ppp6r1_S529 - cost: 6.56, layer: 58
Normal
MK50
MK75
MK88
MK94
MK97
MK data
Normal data
Fold over basal
2
1.8
1.6
1.4
1.2
1
0
10
20
30
40
50
60
Time (min)

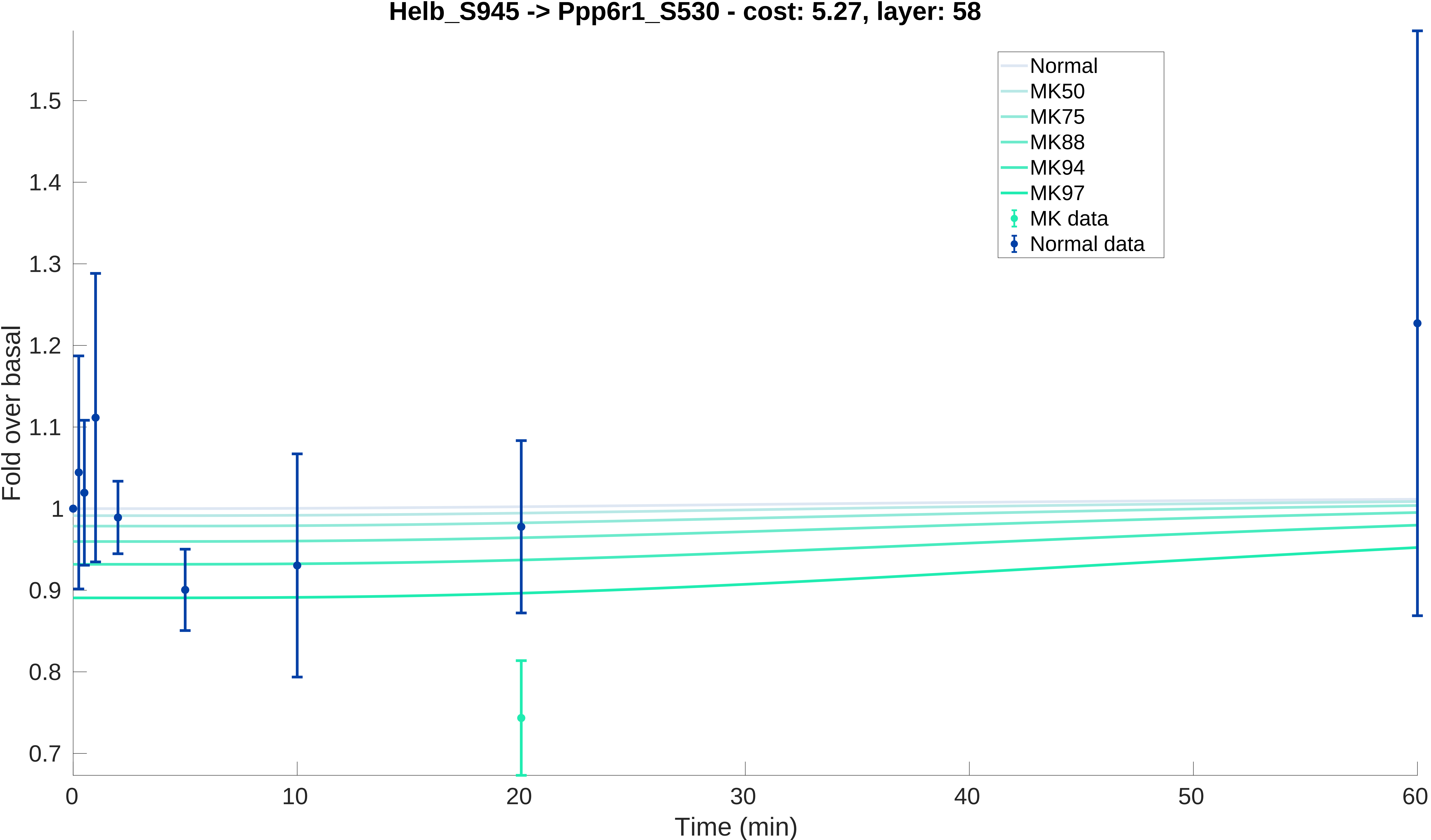

Helb_S945 -> Ppp6r1_S530 - cost: 5.27, layer: 58
Fold over basal
Time (min)
0.7
0.8
0.9
1
1.1
1.2
1.3
1.4
1.5
0
10
20
30
40
50
60
Normal
MK50
MK75
MK88
MK94
MK97
MK data
Normal data

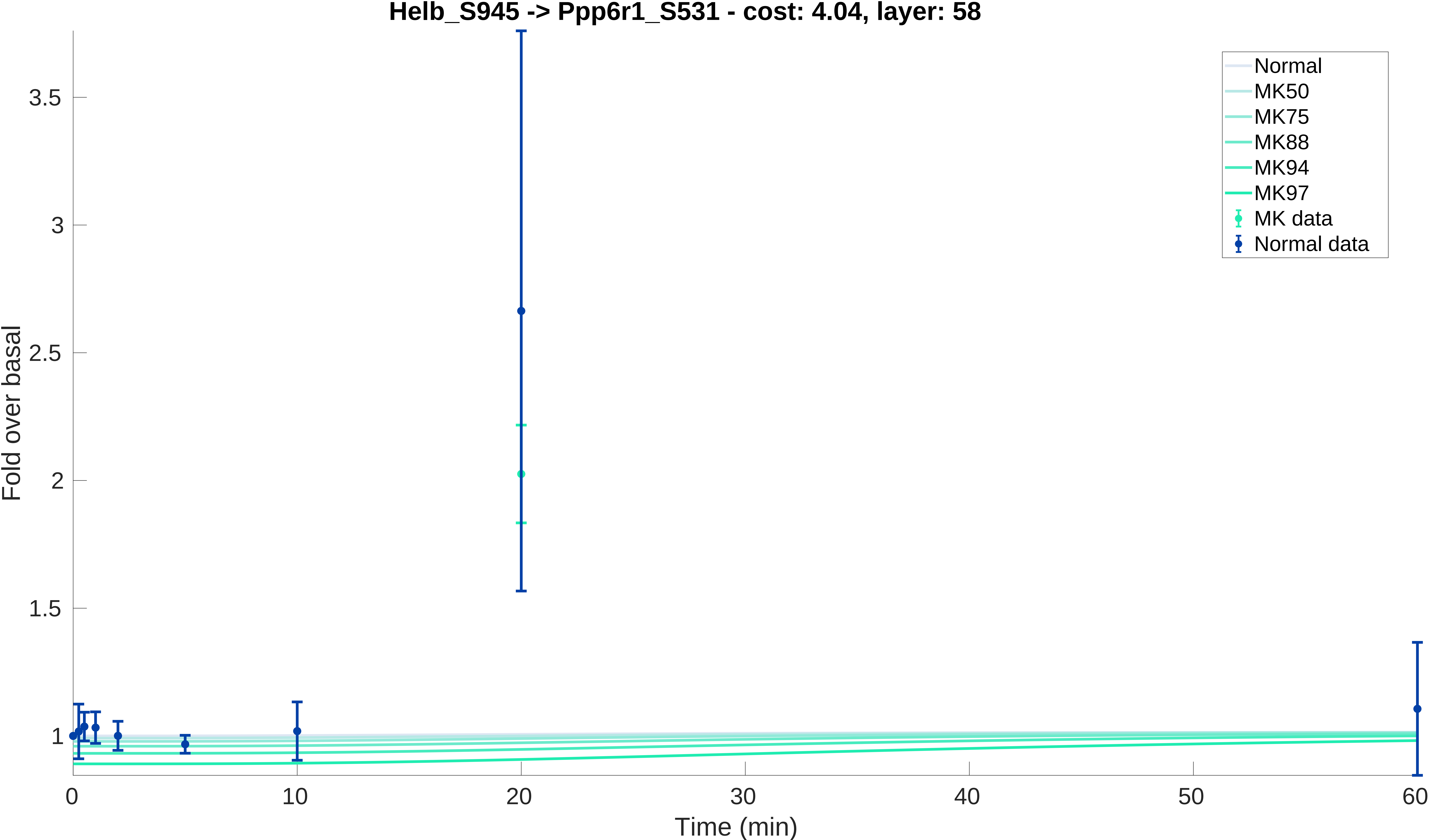
Helb_S945 -> Ppp6r1_S531 - cost: 4.04, layer: 58
Fold over basal
3.5
3
2.5
2
1.5
1
0
10
20
30
40
50
60
Time (min)
Normal
MK50
MK75
MK88
MK94
MK97
MK data
Normal data

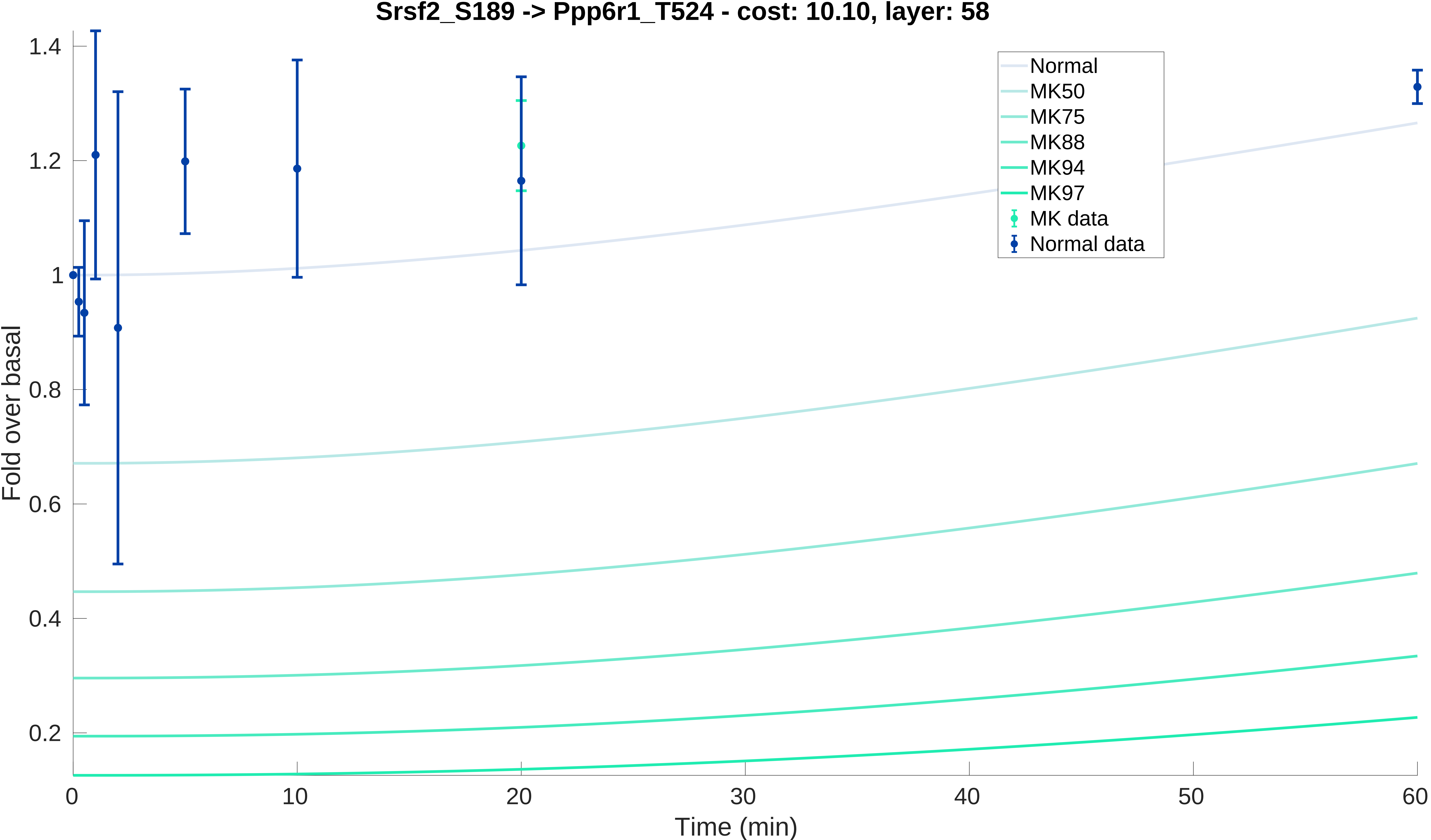

Srsf2_S189 -> Ppp6r1_T524 - cost: 10.10, layer: 58
Normal
MK50
MK75
MK88
MK94
MK97
MK data
Normal data
Fold over basal
Time (min)
0.2
0.4
0.6
0.8
1
1.2
1.4
0
10
20
30
40
50
60

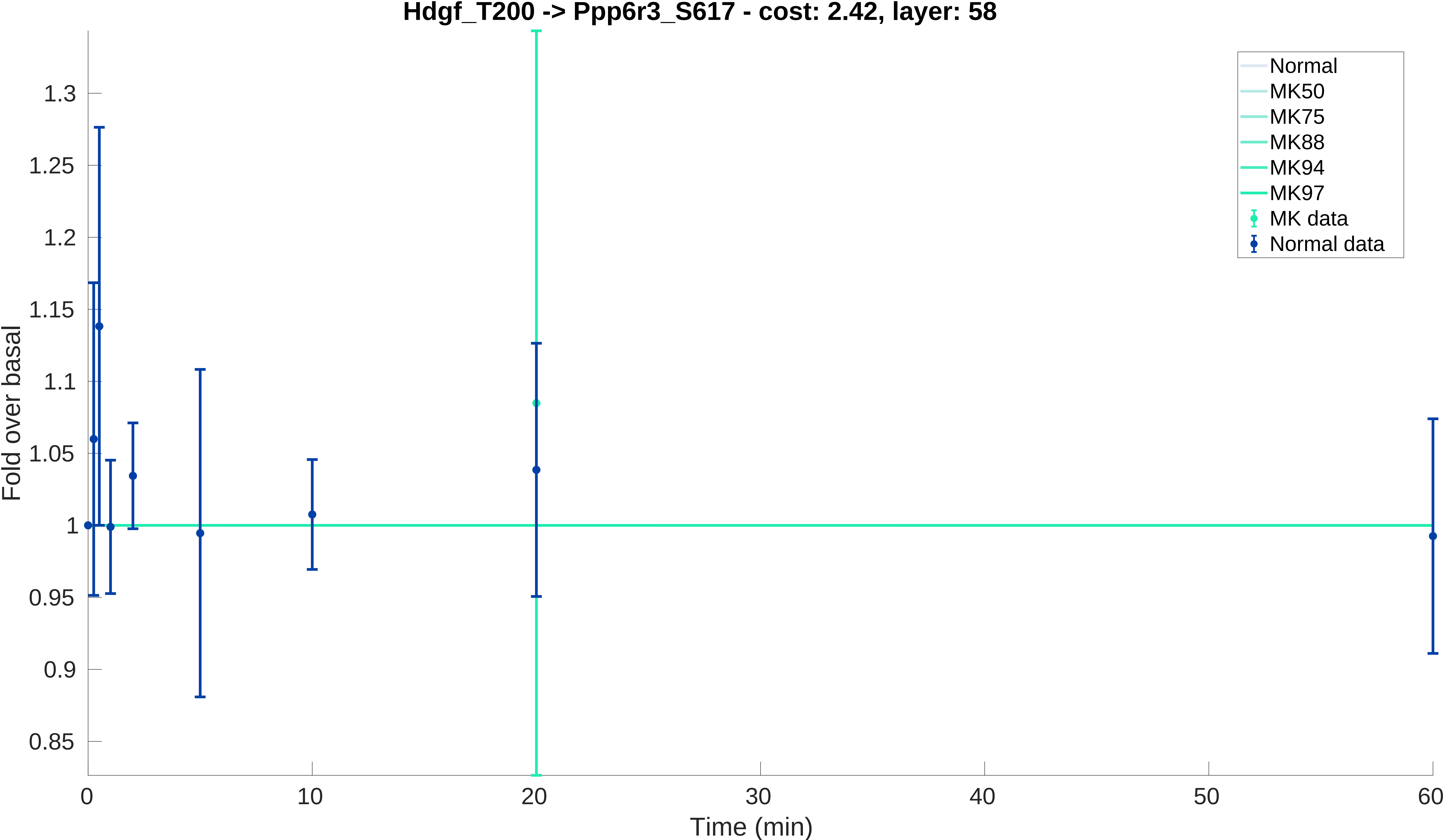

Hdgf_T200 -> Ppp6r3_S617 - cost: 2.42, layer: 58
Normal
MK50
MK75
MK88
MK94
MK97
MK data
Normal data
Fold over basal
1.3
1.25
1.2
1.15
1.1
1.05
1
0.95
0.9
0.85
0
10
20
30
40
50
60
Time (min)

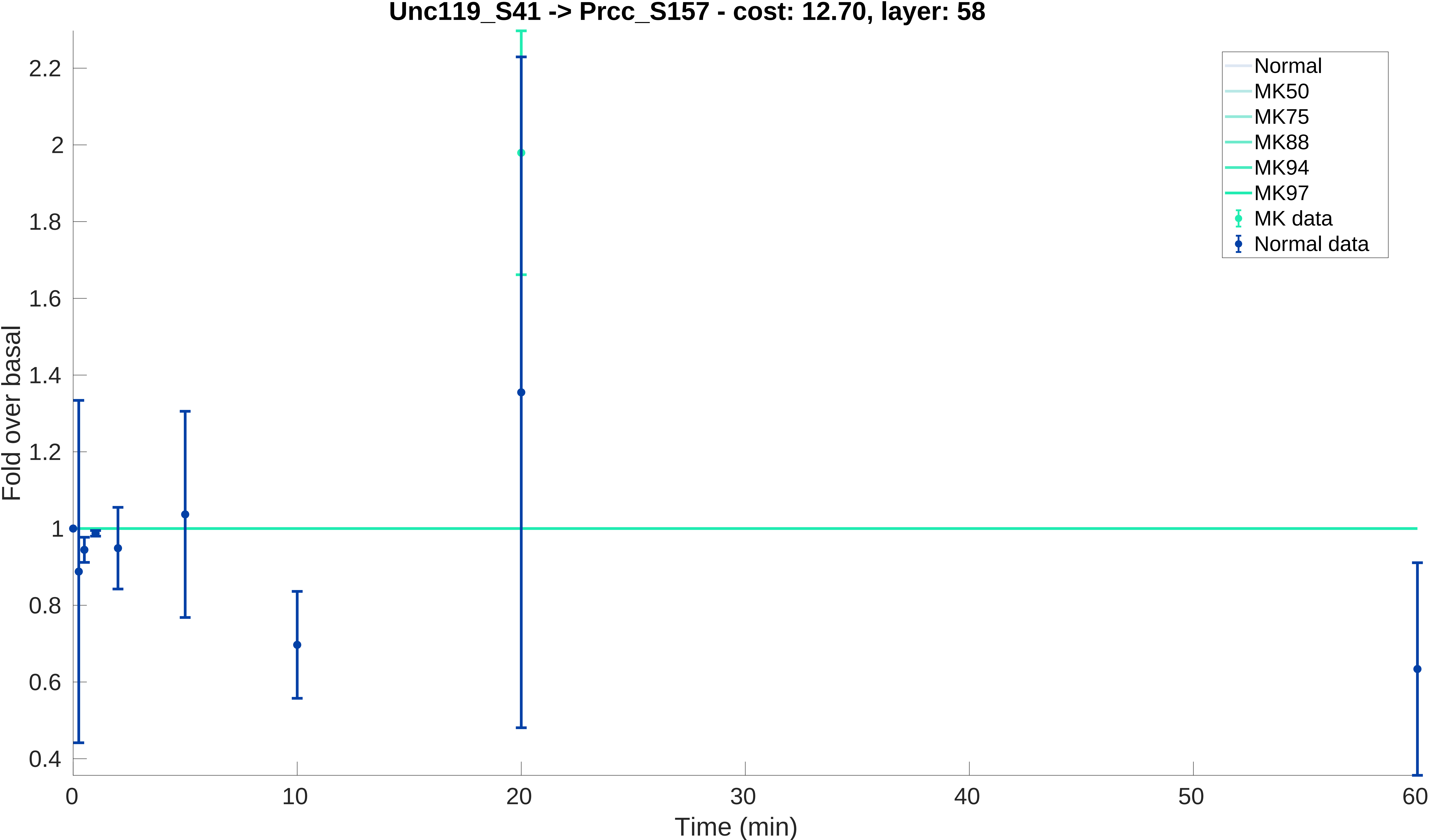

Unc119_S41 -> Prcc_S157 - cost: 12.70, layer: 58
Fold over basal
Time (min)
0.4
0.6
0.8
1
1.2
1.4
1.6
1.8
2
2.2
0
10
20
30
40
50
60
Normal
MK50
MK75
MK88
MK94
MK97
MK data
Normal data

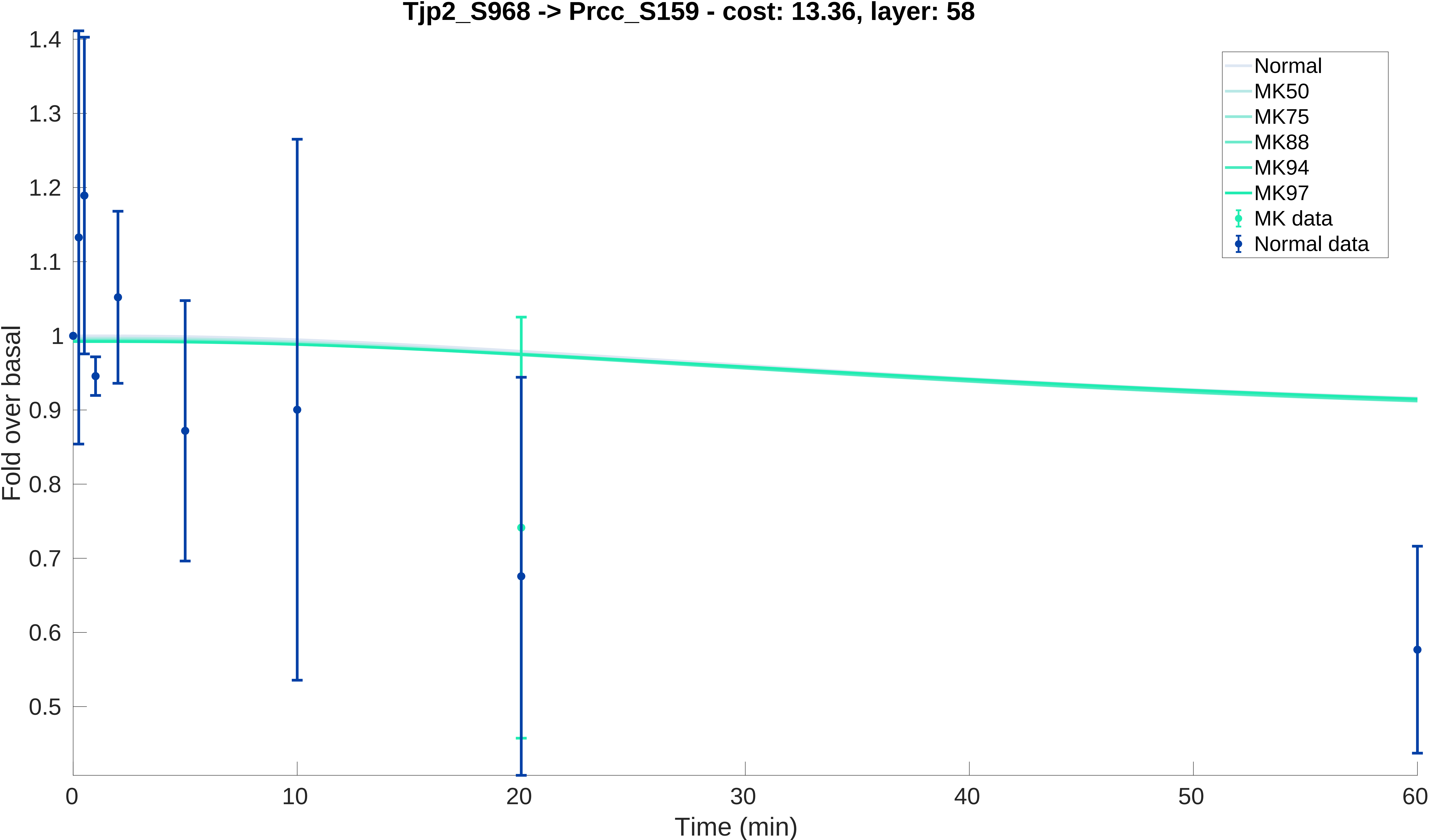
Tjp2_S968 -> Prcc_S159 - cost: 13.36, layer: 58
Fold over basal
Time (min)
1.4
1.3
1.2
1.1
1
0.9
0.8
0.7
0.6
0.5
0
10
20
30
40
50
60
Normal
MK50
MK75
MK88
MK94
MK97
MK data
Normal data

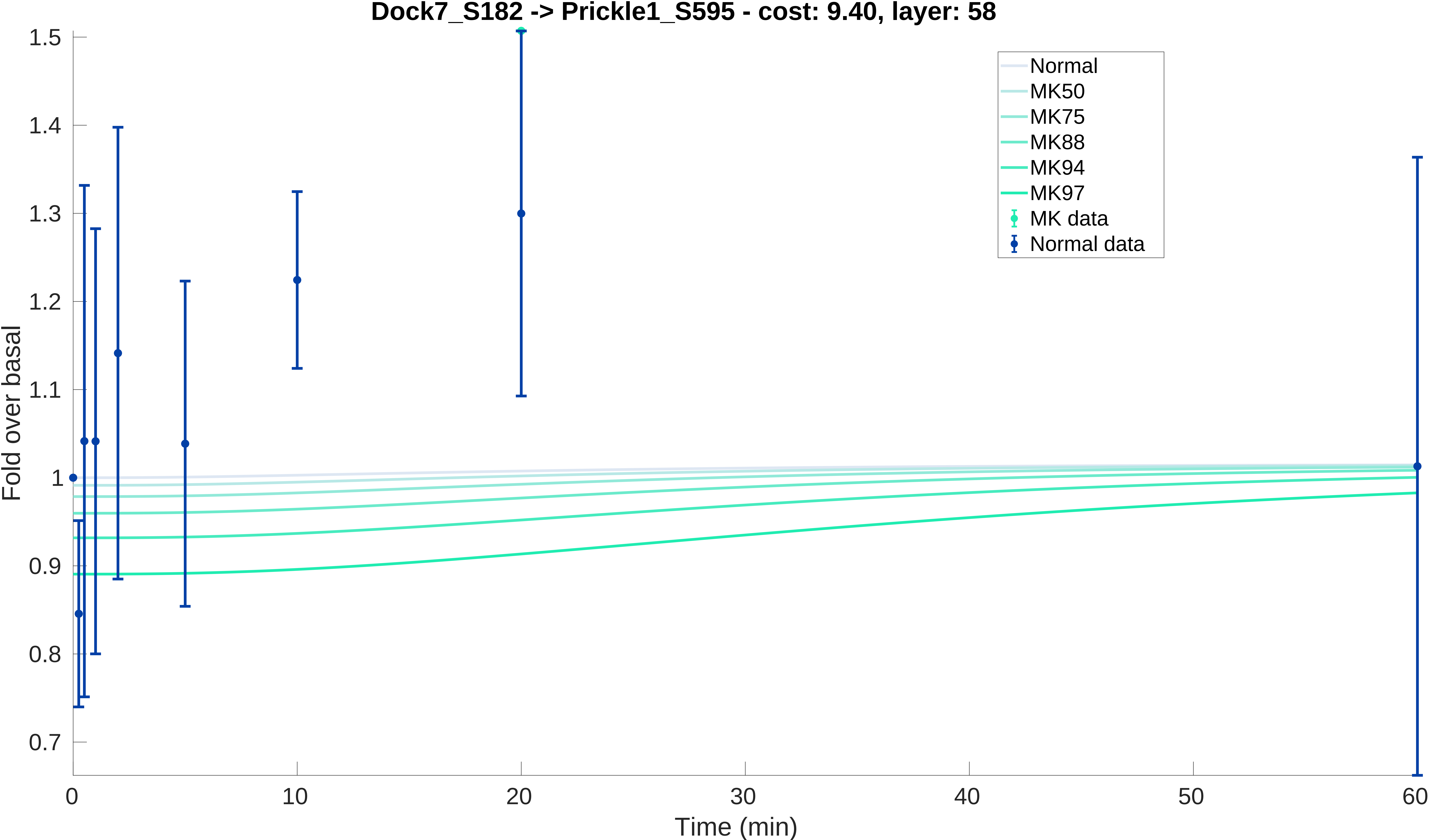

Dock7_S182 -> Prickle1_S595 - cost: 9.40, layer: 58
Fold over basal
Time (min)
Normal
MK50
MK75
MK88
MK94
MK97
MK data
Normal data
1.5
1.4
1.3
1.2
1.1
1
0.9
0.8
0.7
0
10
20
30
40
50
60

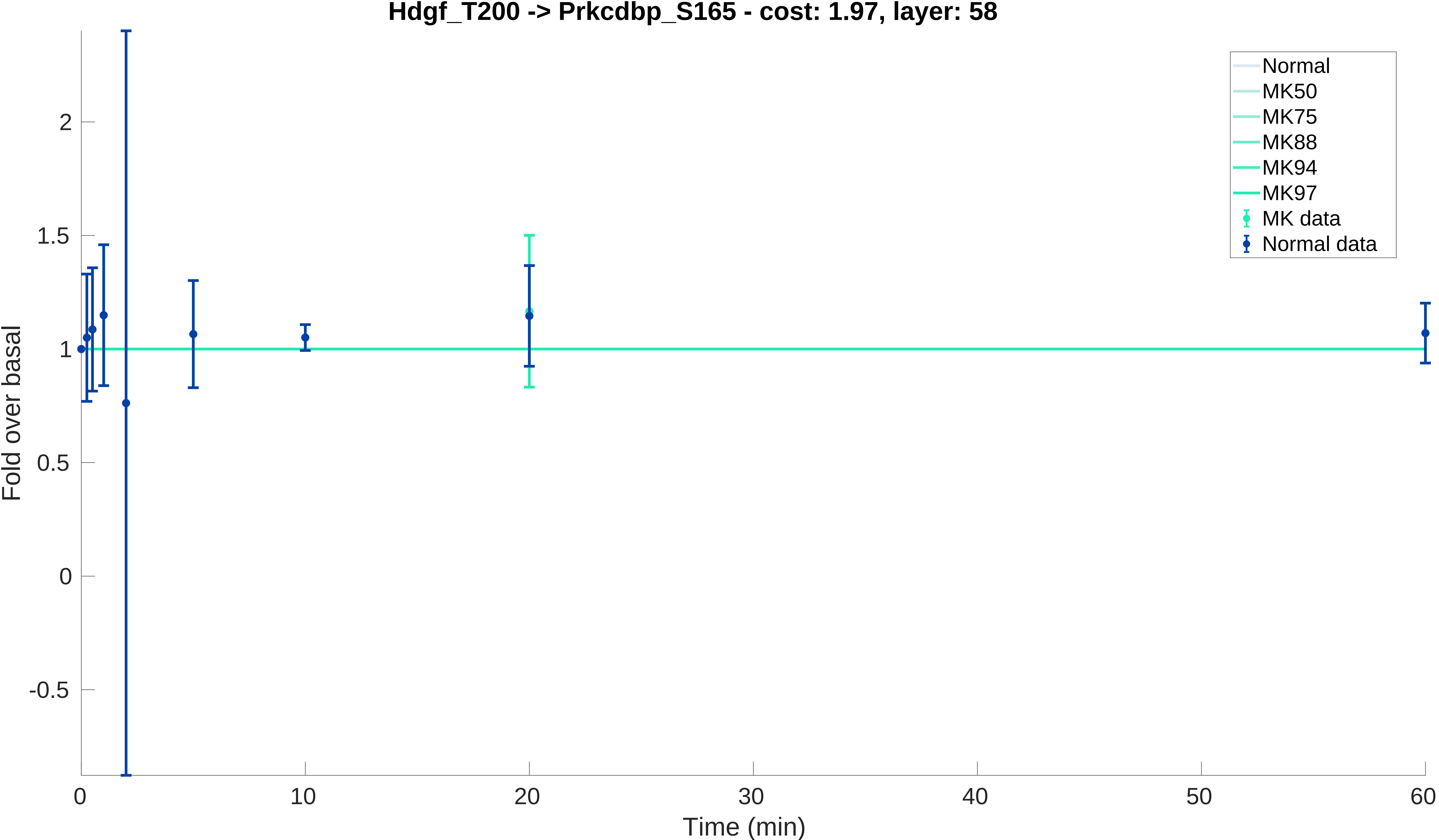

Hdgf_T200 -> Prkcdbp_S165 - cost: 1.97, layer: 58
Normal
MK50
MK75
MK88
MK94
MK97
MK data
Normal data
Fold over basal
Time (min)
2
1.5
1
0.5
0
-0.5
0
10
20
30
40
50
60

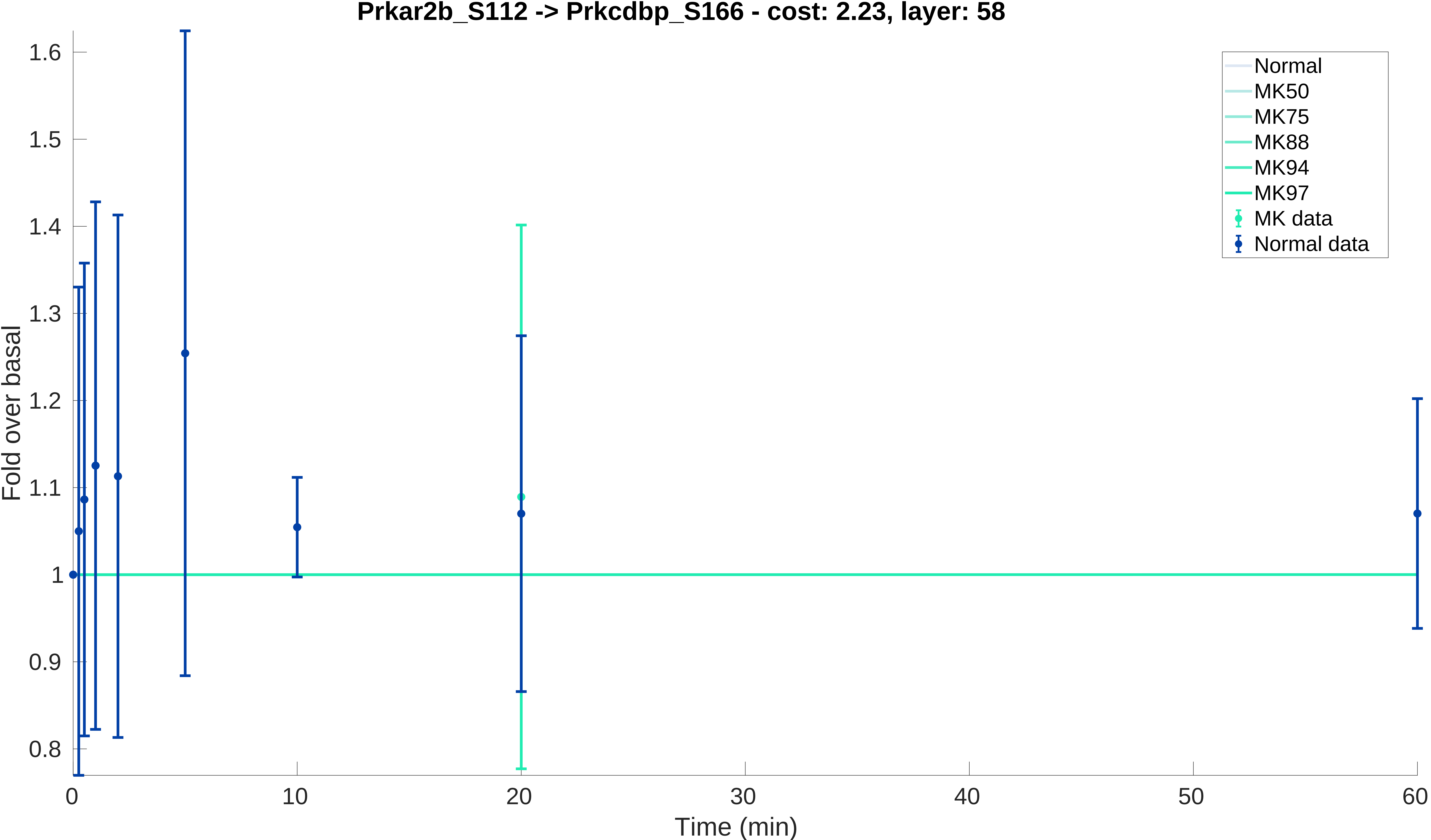

Prkar2b_S112 -> Prkcdbp_S166 - cost: 2.23, layer: 58
Fold over basal
Time (min)
1.6
1.5
1.4
1.3
1.2
1.1
1
0.9
0.8
0
10
20
30
40
50
60
Normal
MK50
MK75
MK88
MK94
MK97
MK data
Normal data

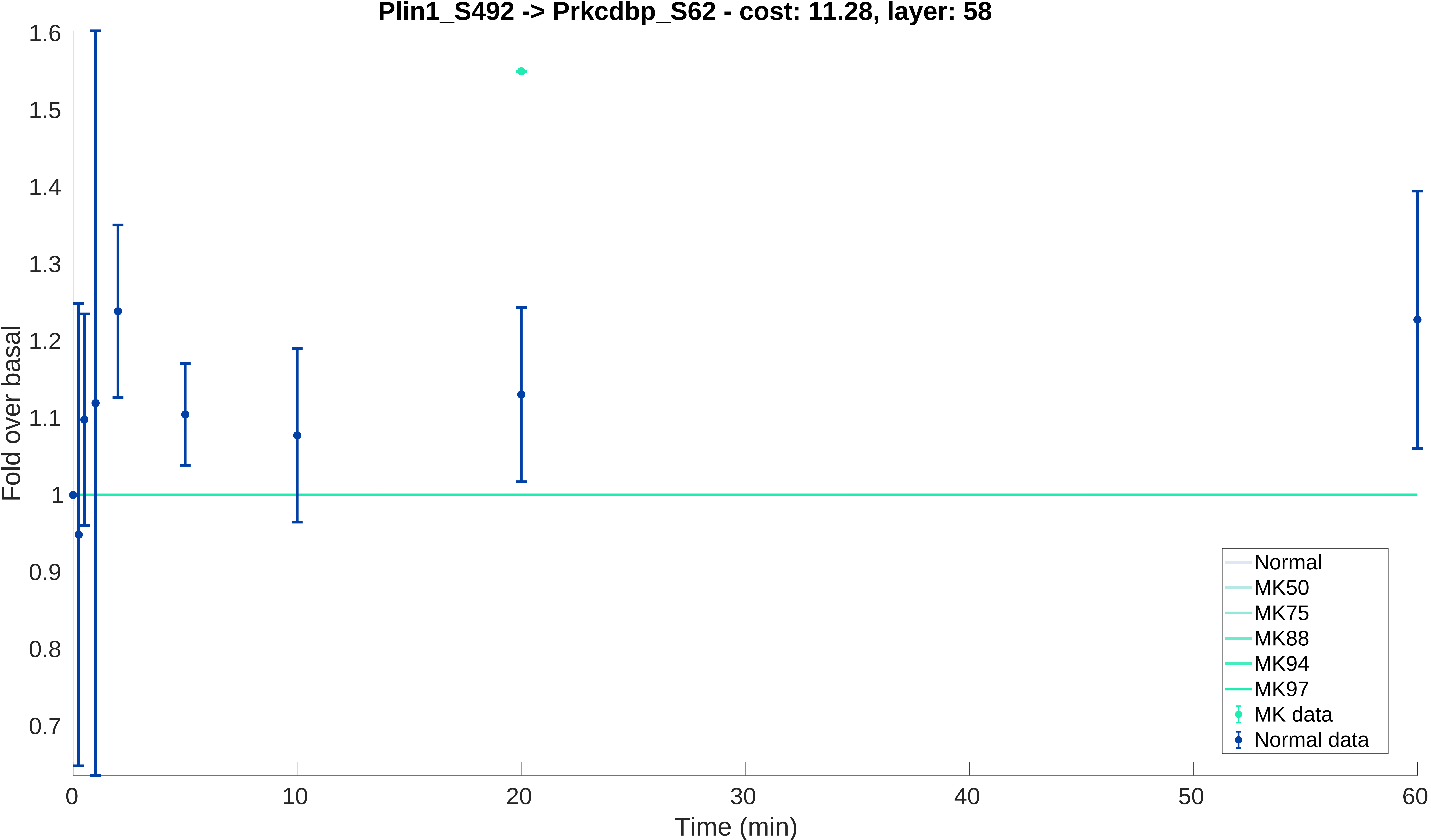

Plin1_S492 -> Prkcdbp_S62 - cost: 11.28, layer: 58
Fold over basal
1.6
1.5
1.4
1.3
1.2
1.1
1
0.9
0.8
0.7
0
10
20
30
40
50
60
Time (min)
Normal
MK50
MK75
MK88
MK94
MK97
MK data
Normal data

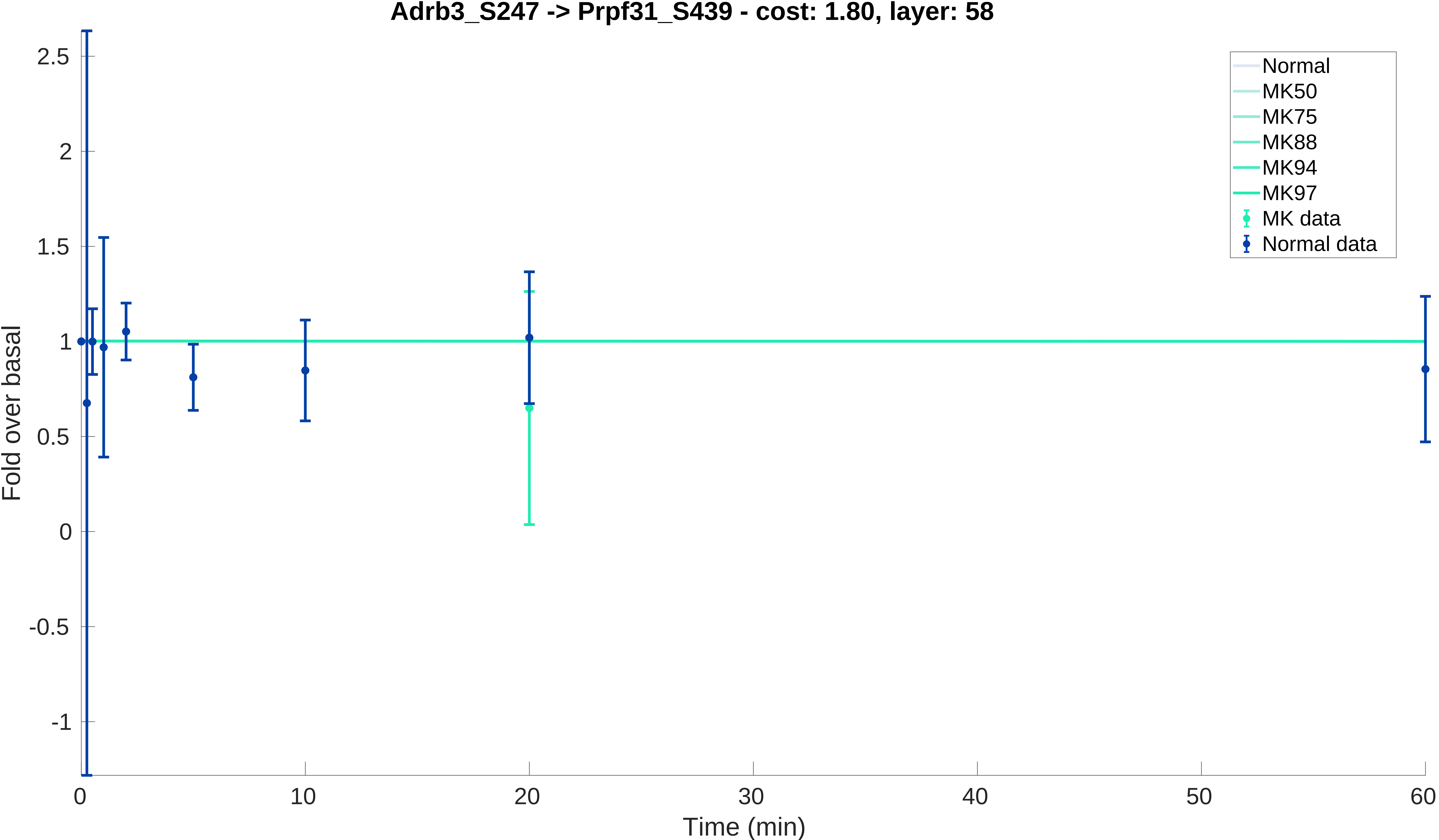
Adrb3_S247 -> Prpf31_S439 - cost: 1.80, layer: 58
Fold over basal
Time (min)
2.5
2
1.5
1
0.5
0
-0.5
-1
0
10
20
30
40
50
60
Normal
MK50
MK75
MK88
MK94
MK97
MK data
Normal data

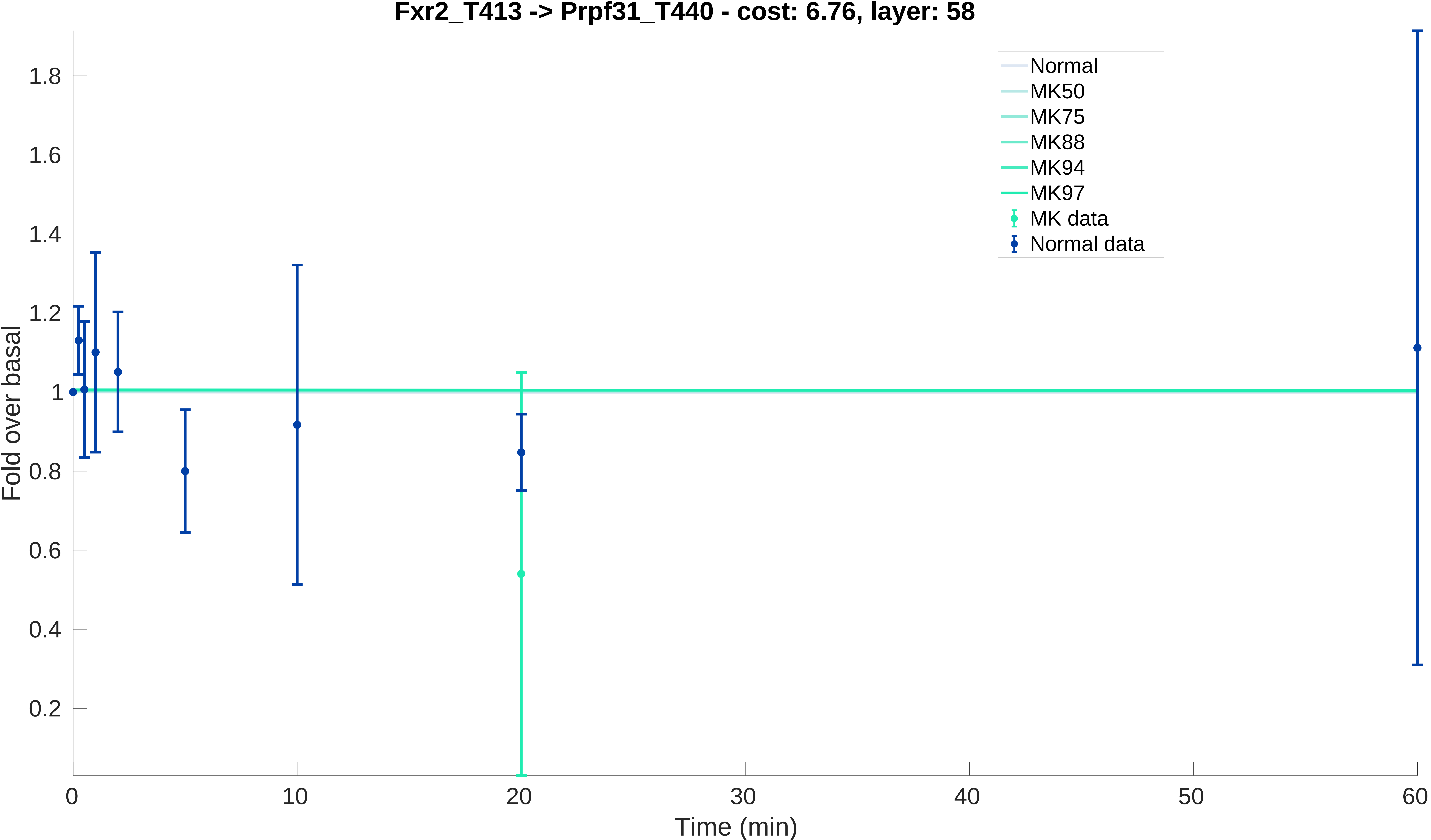

Fxr2_T413 -> Prpf31_T440 - cost: 6.76, layer: 58
Normal
MK50
MK75
MK88
MK94
MK97
MK data
Normal data
Fold over basal
1.8
1.6
1.4
1.2
1
0.8
0.6
0.4
0.2
0
10
20
30
40
50
60
Time (min)

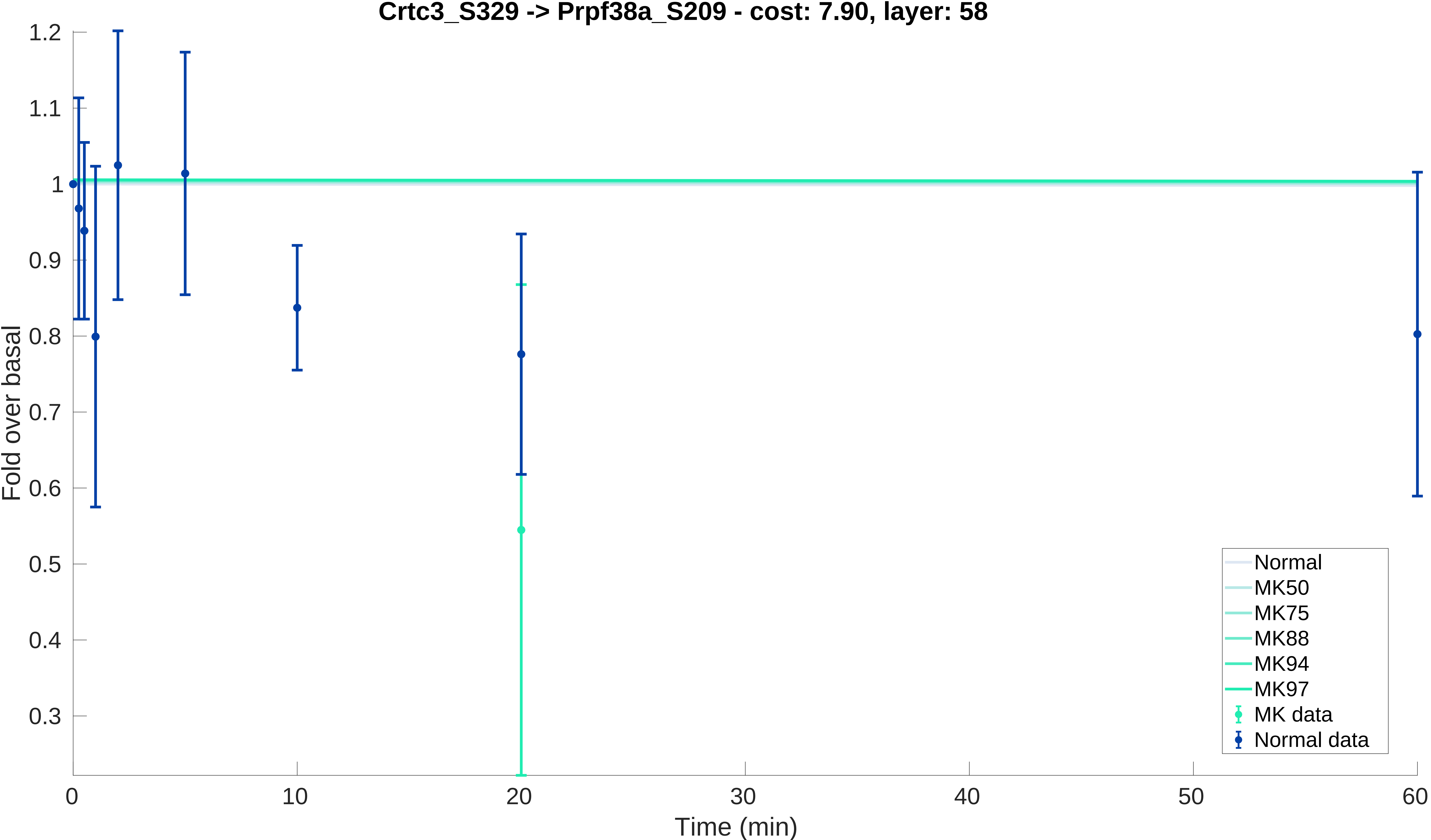

Crtc3_S329 -> Prpf38a_S209 - cost: 7.90, layer: 58
Fold over basal
1.2
1.1
1
0.9
0.8
0.7
0.6
0.5
0.4
0.3
0
10
20
30
40
50
60
Time (min)
Normal
MK50
MK75
MK88
MK94
MK97
MK data
Normal data

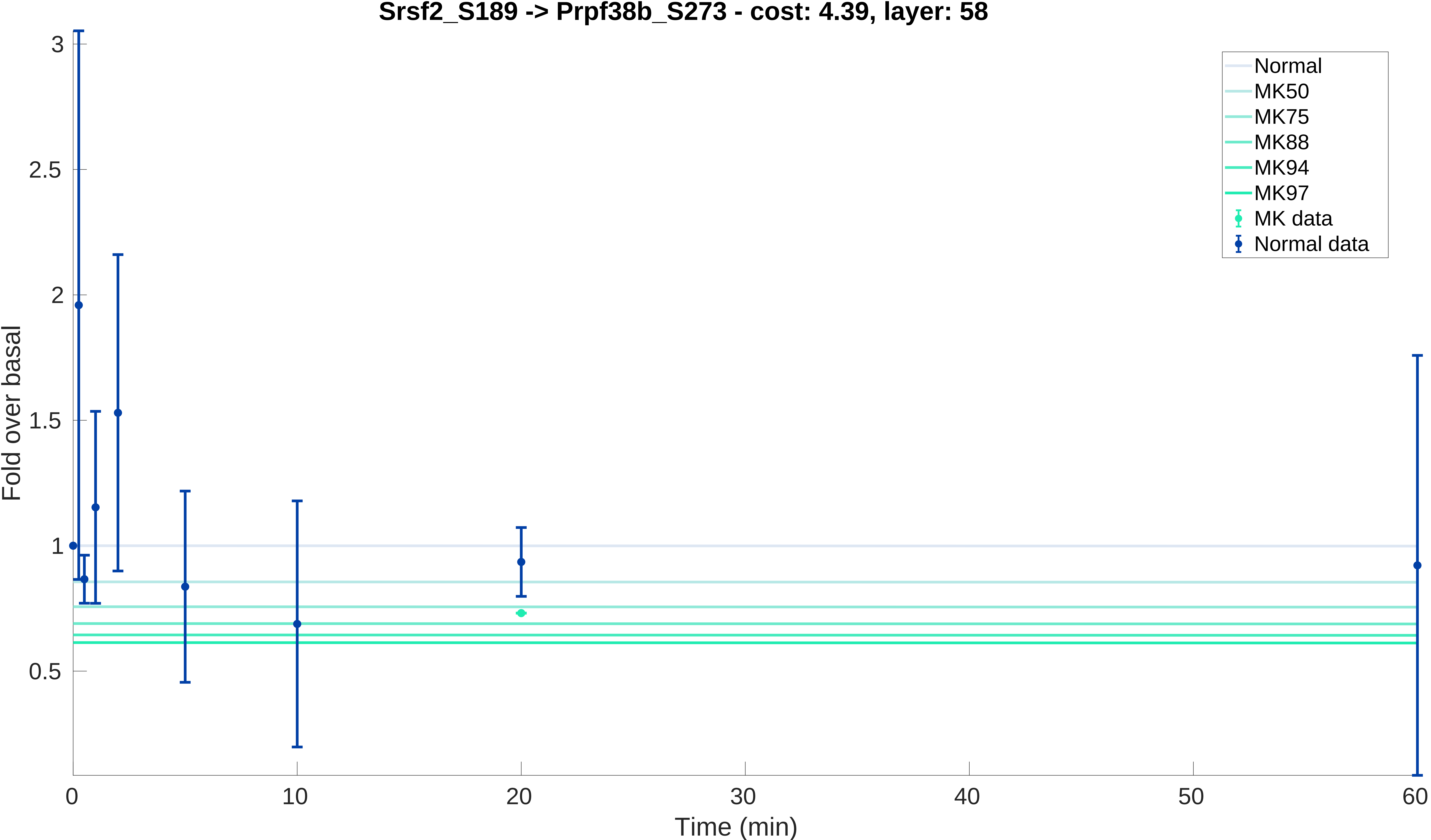

Srsf2_S189 -> Prpf38b_S273 - cost: 4.39, layer: 58
Fold over basal
Time (min)
0.5
1
1.5
2
2.5
3
0
10
20
30
40
50
60
Normal
MK50
MK75
MK88
MK94
MK97
MK data
Normal data

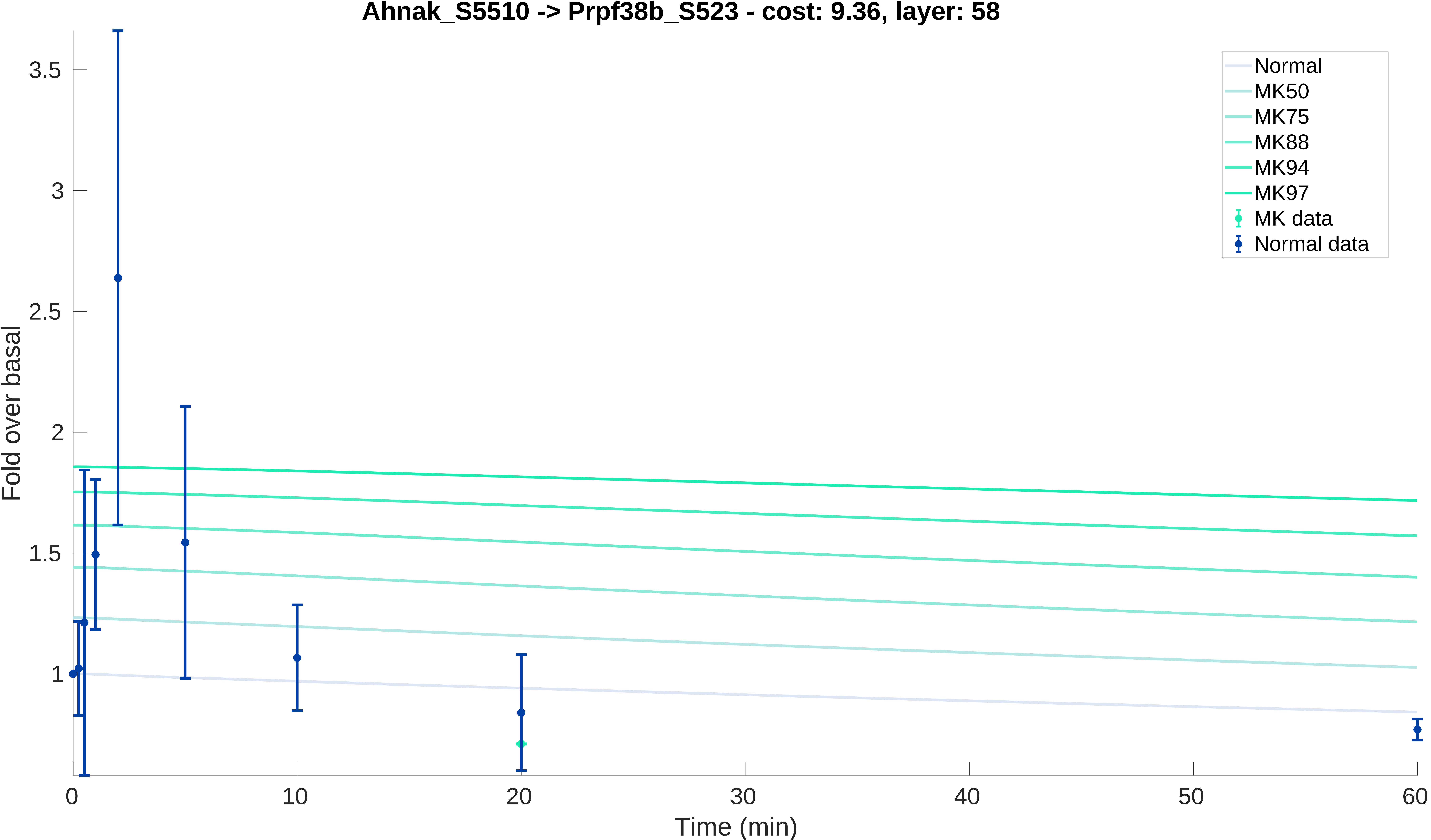

Ahnak_S5510 -> Prpf38b_S523 - cost: 9.36, layer: 58
Fold over basal
Time (min)
0
10
20
30
40
50
60
1
1.5
2
2.5
3
3.5
Normal
MK50
MK75
MK88
MK94
MK97
MK data
Normal data

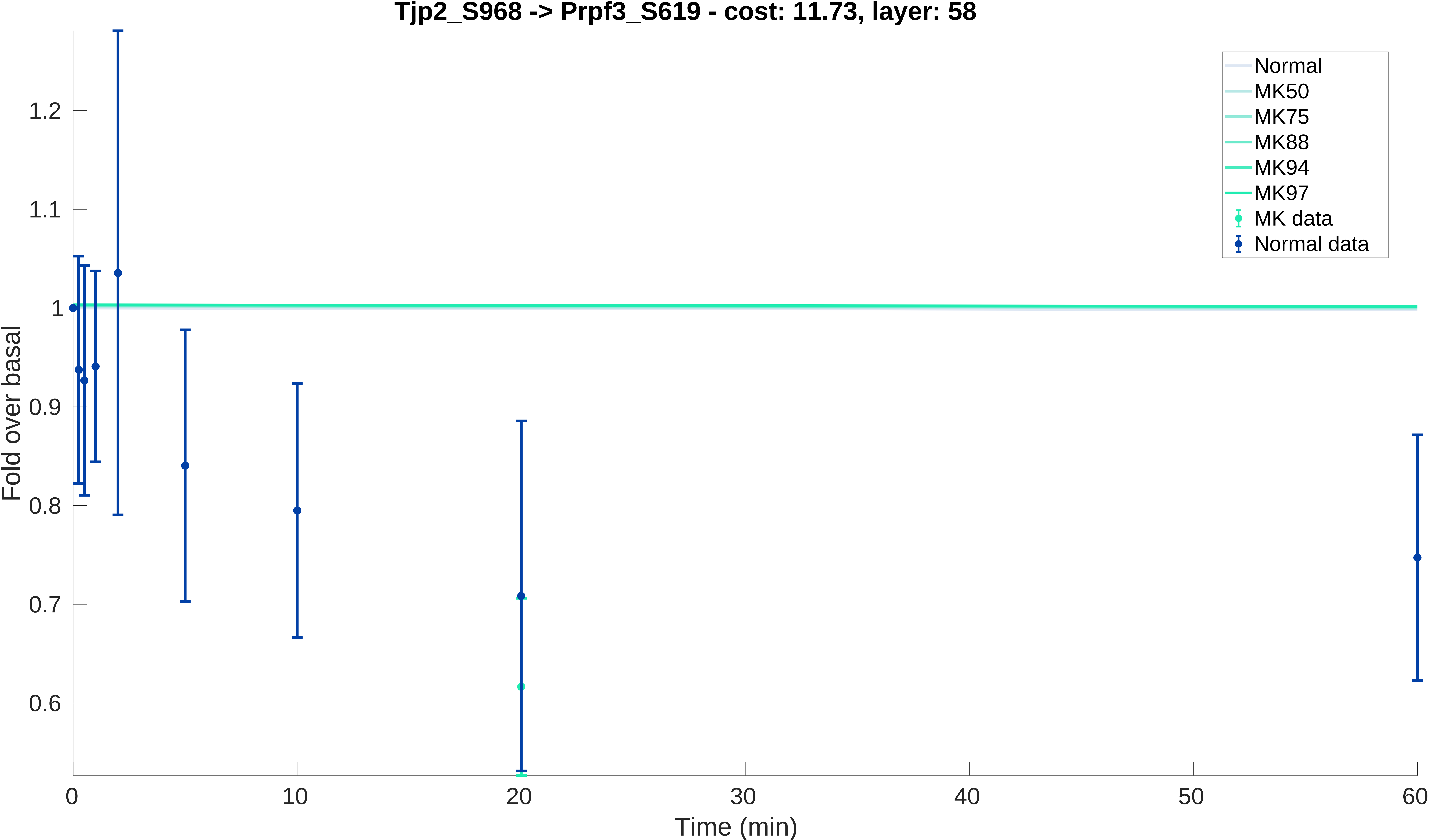
Tjp2_S968 -> Prpf3_S619 - cost: 11.73, layer: 58
Normal
MK50
MK75
MK88
MK94
MK97
MK data
Normal data
Fold over basal
1.2
1.1
1
0.9
0.8
0.7
0.6
0
10
20
30
40
50
60
Time (min)

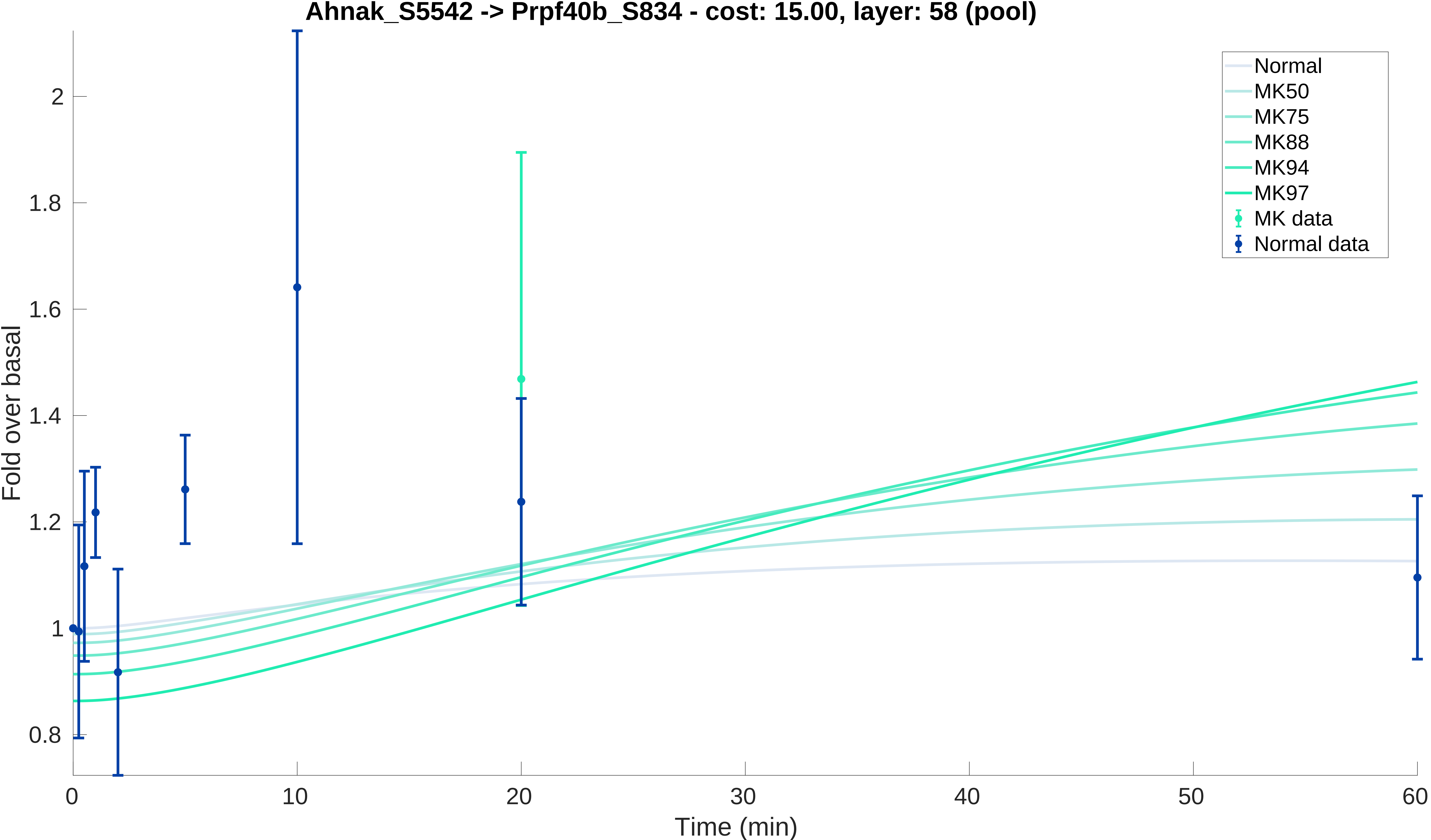
Ahnak_S5542 -> Prpf40b_S834 - cost: 15.00, layer: 58 (pool)
Fold over basal
Time (min)
0.8
1
1.2
1.4
1.6
1.8
2
0
10
20
30
40
50
60
Normal
MK50
MK75
MK88
MK94
MK97
MK data
Normal data

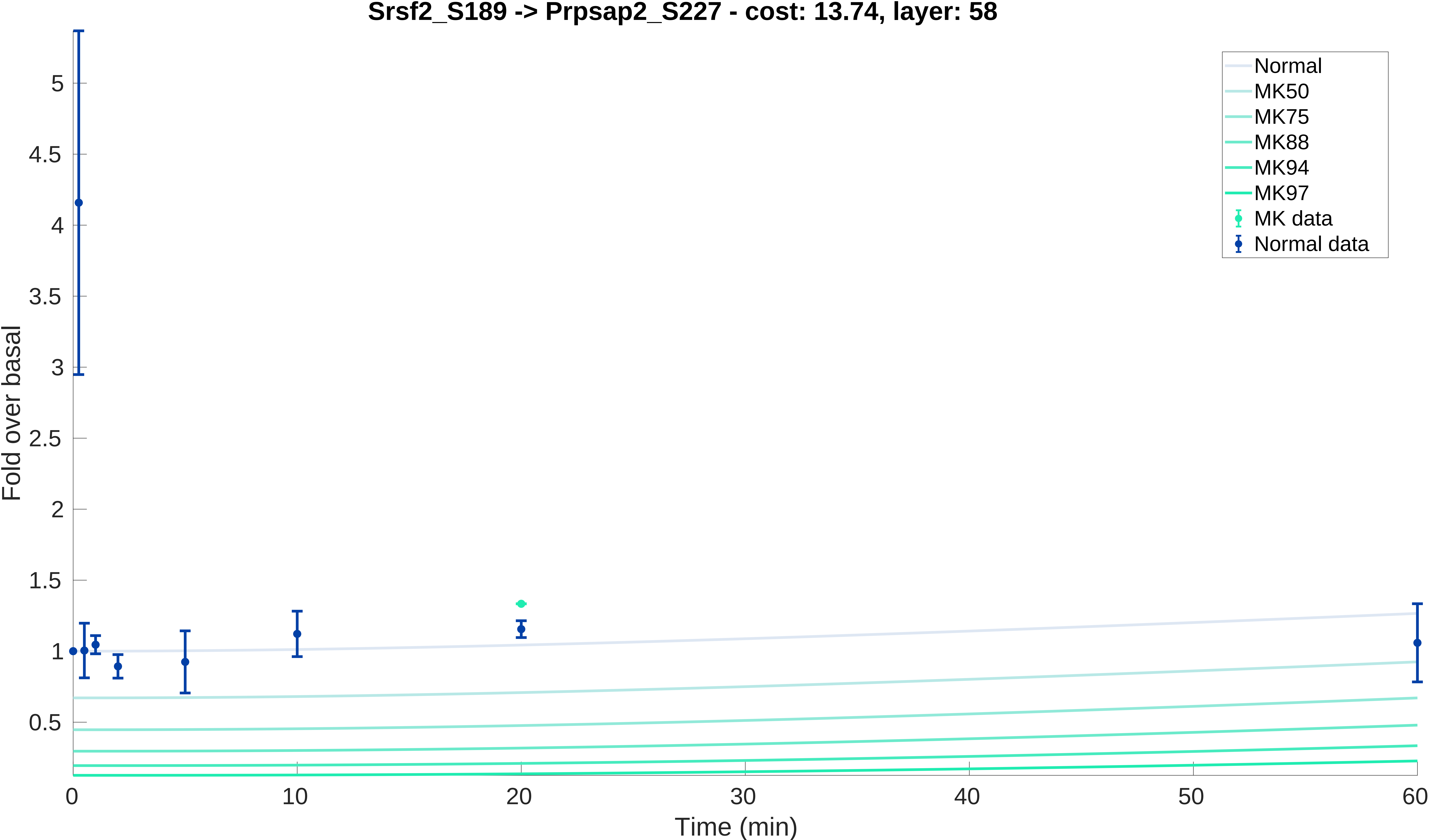

Srsf2_S189 -> Prpsap2_S227 - cost: 13.74, layer: 58
Fold over basal
Time (min)
0
10
20
30
40
50
60
0.5
1
1.5
2
2.5
3
3.5
4
4.5
5
Normal
MK50
MK75
MK88
MK94
MK97
MK data
Normal data

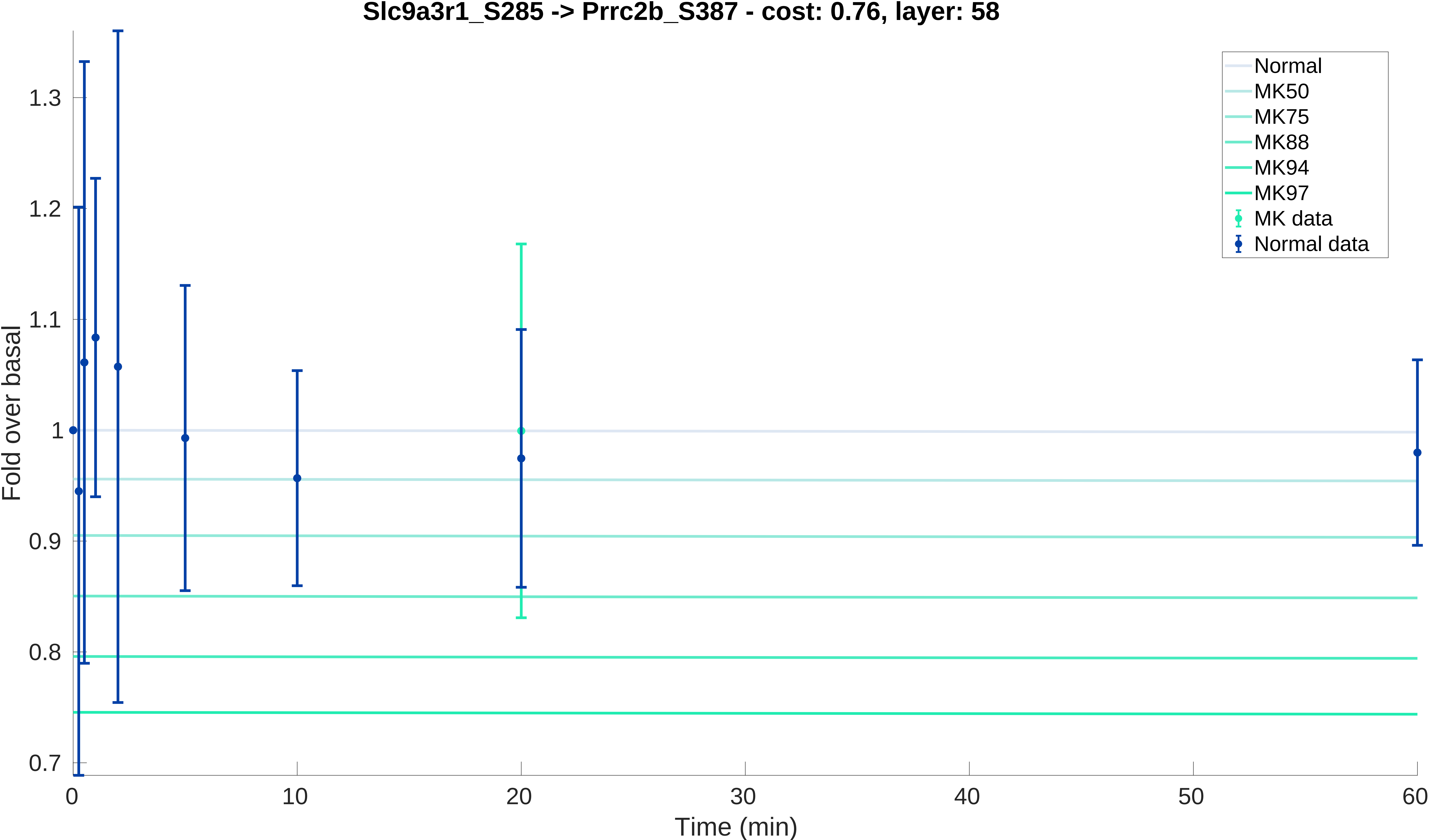

Slc9a3r1_S285 -> Prrc2b_S387 - cost: 0.76, layer: 58
Fold over basal
Time (min)
1.3
1.2
1.1
1
0.9
0.8
0.7
0
10
20
30
40
50
60
Normal
MK50
MK75
MK88
MK94
MK97
MK data
Normal data

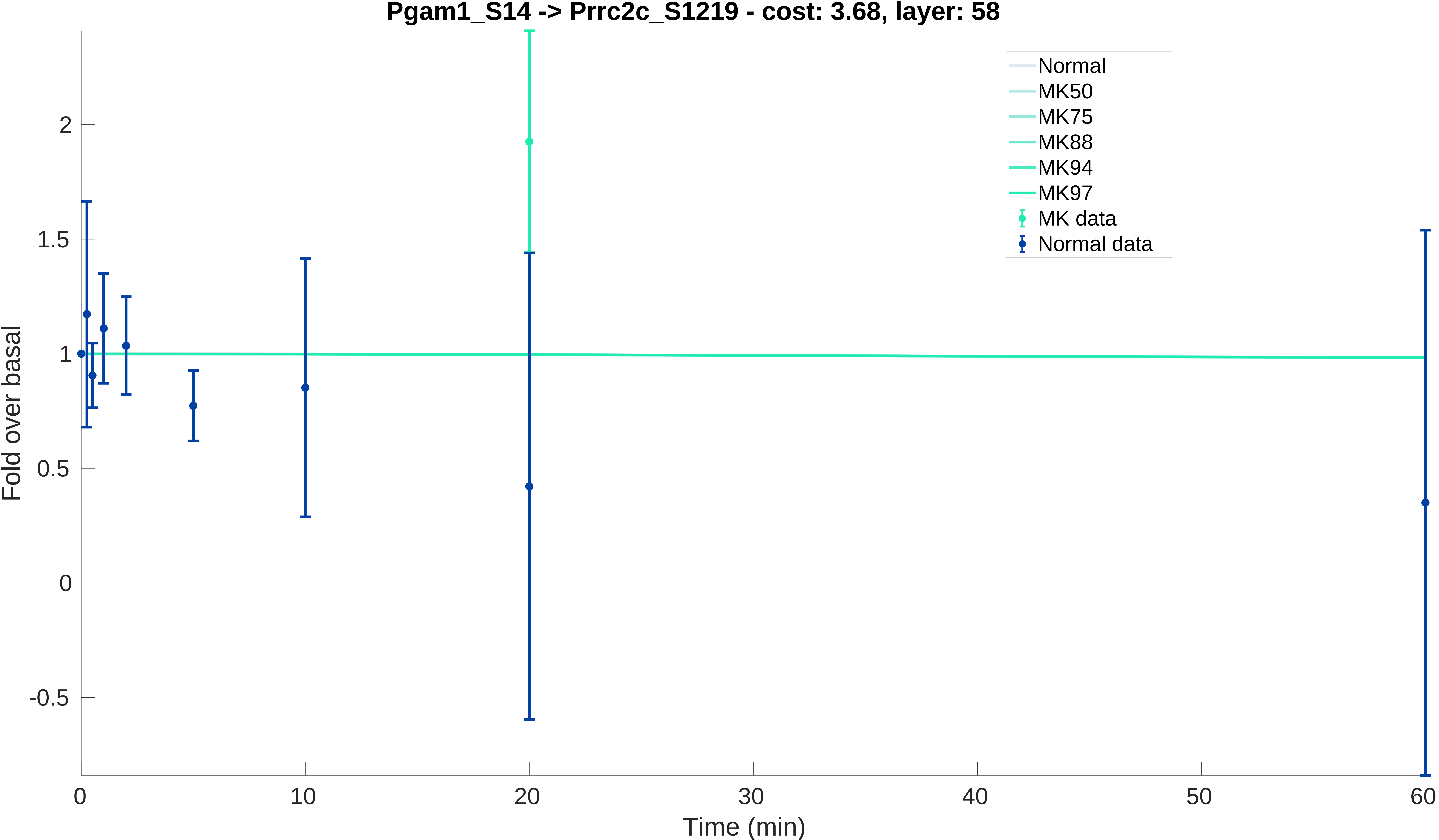

Pgam1_S14 -> Prrc2c_S1219 - cost: 3.68, layer: 58
Fold over basal
Time (min)
2
1.5
1
0.5
0
-0.5
0
10
20
30
40
50
60
Normal
MK50
MK75
MK88
MK94
MK97
MK data
Normal data

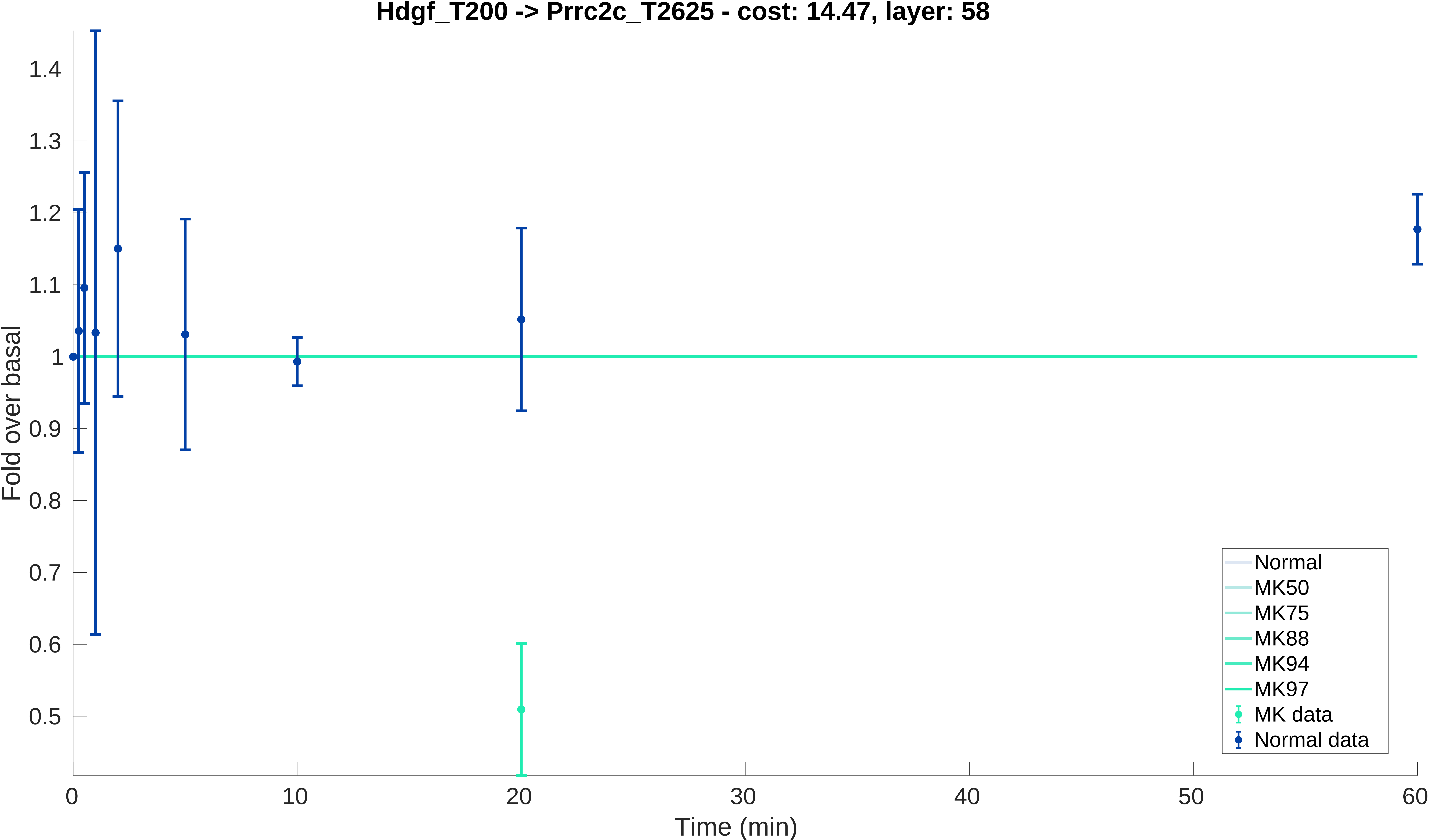
Hdgf_T200 -> Prrc2c_T2625 - cost: 14.47, layer: 58
Fold over basal
Time (min)
0
10
20
30
40
50
60
0.5
0.6
0.7
0.8
0.9
1
1.1
1.2
1.3
1.4
Normal
MK50
MK75
MK88
MK94
MK97
MK data
Normal data

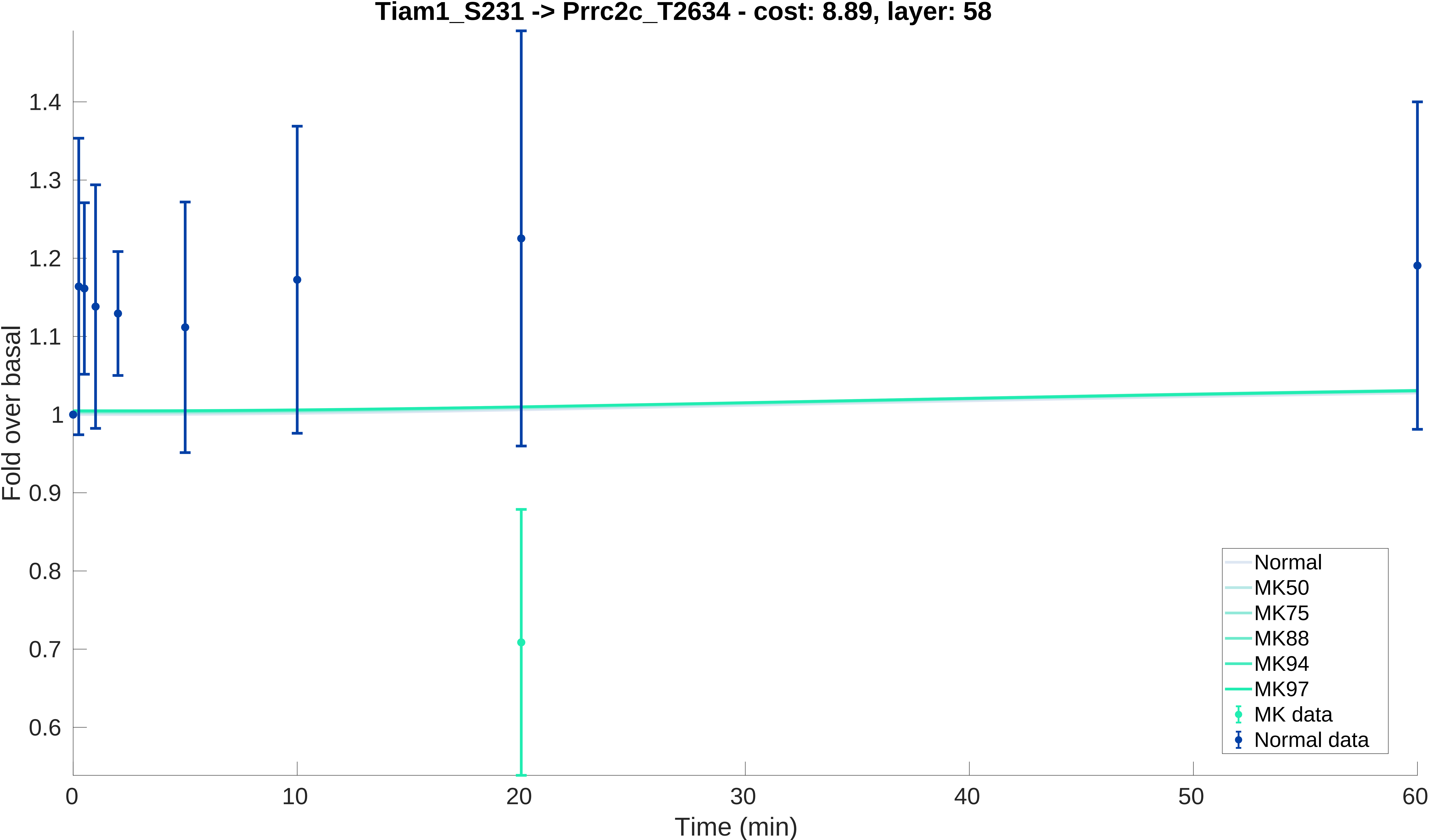

Tiam1_S231 -> Prrc2c_T2634 - cost: 8.89, layer: 58
Fold over basal
Time (min)
1.4
1.3
1.2
1.1
1
0.9
0.8
0.7
0.6
0
10
20
30
40
50
60
Normal
MK50
MK75
MK88
MK94
MK97
MK data
Normal data

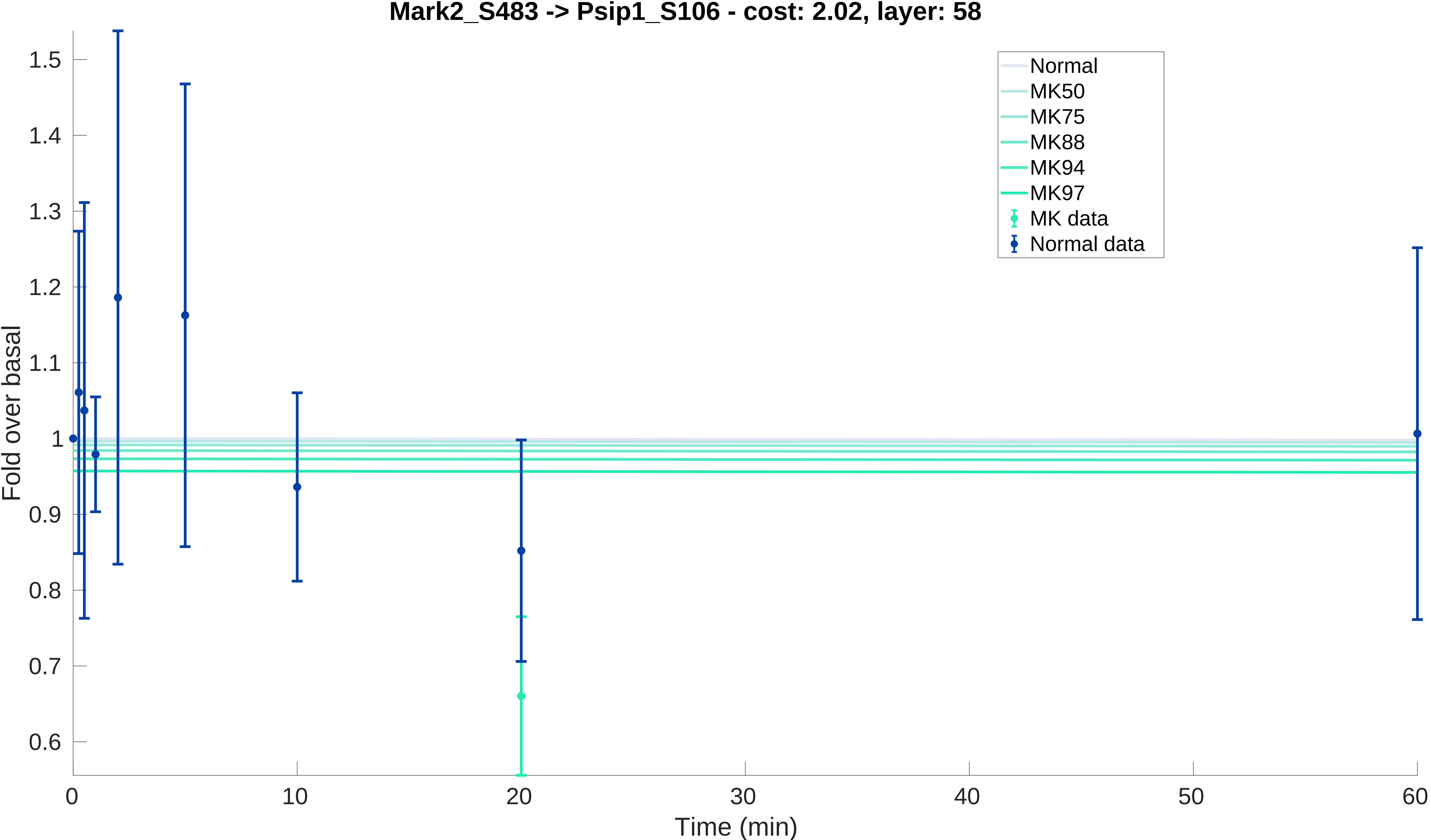

Mark2_S483 -> Psip1_S106 - cost: 2.02, layer: 58
Fold over basal
Time (min)
1.5
1.4
1.3
1.2
1.1
1
0.9
0.8
0.7
0.6
0
10
20
30
40
50
60
Normal
MK50
MK75
MK88
MK94
MK97
MK data
Normal data

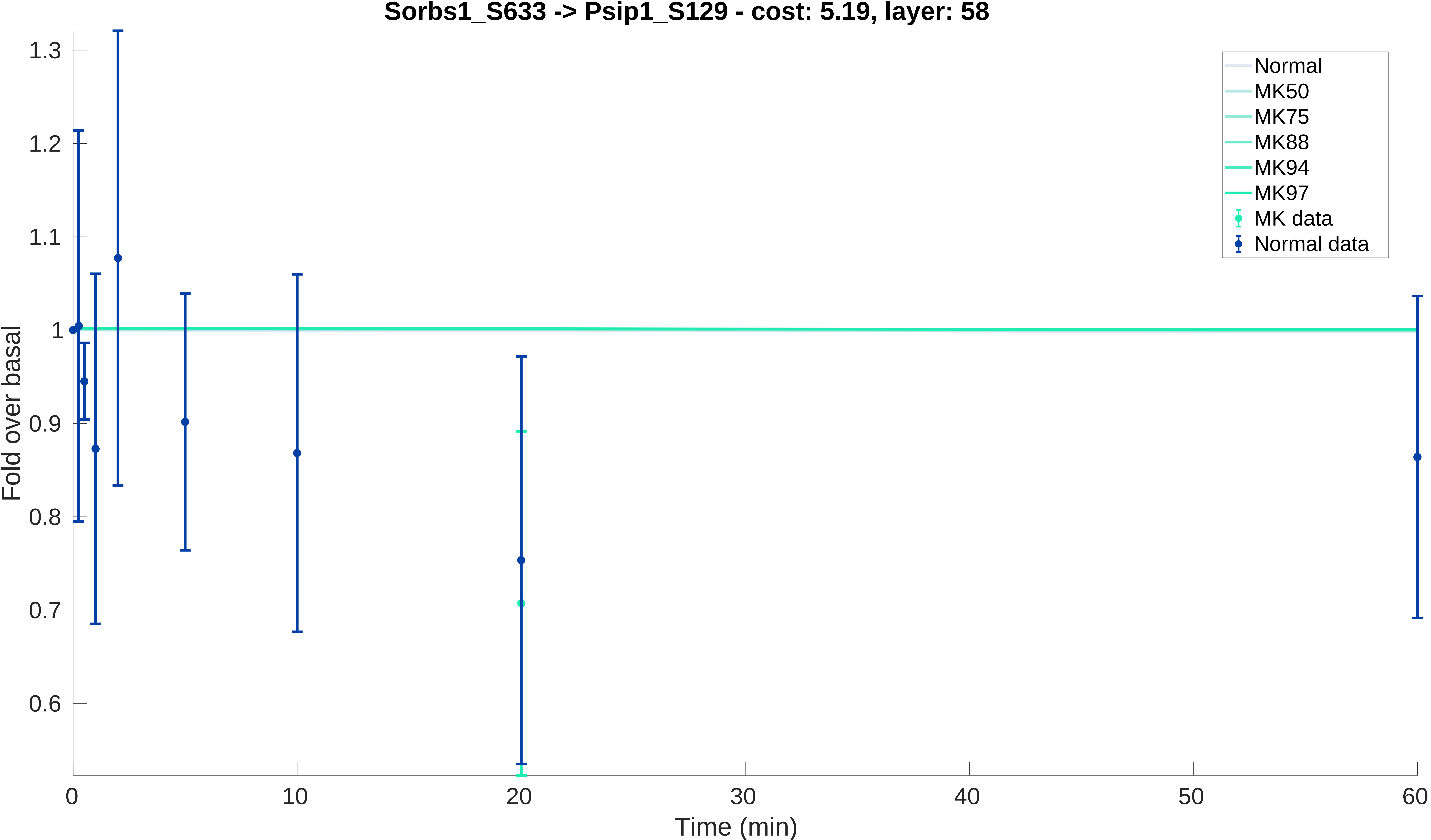

Sorbs1_S633 -> Psip1_S129 - cost: 5.19, layer: 58
Fold over basal
Time (min)
0.6
0.7
0.8
0.9
1
1.1
1.2
1.3
0
10
20
30
40
50
60
Normal
MK50
MK75
MK88
MK94
MK97
MK data
Normal data

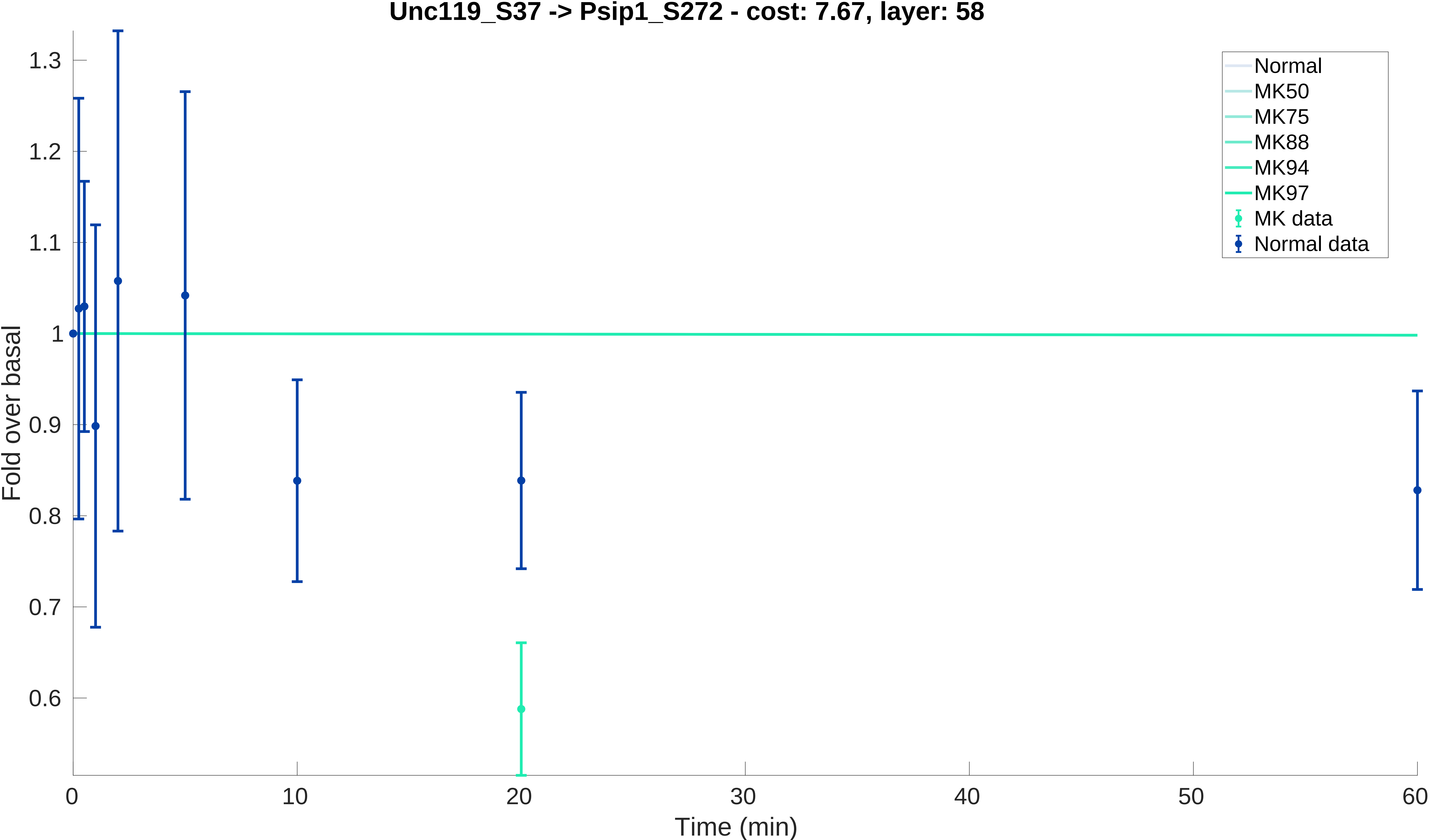
Unc119_S37 -> Psip1_S272 - cost: 7.67, layer: 58
Fold over basal
Time (min)
0
10
20
30
40
50
60
0.6
0.7
0.8
0.9
1
1.1
1.2
1.3
Normal
MK50
MK75
MK88
MK94
MK97
MK data
Normal data

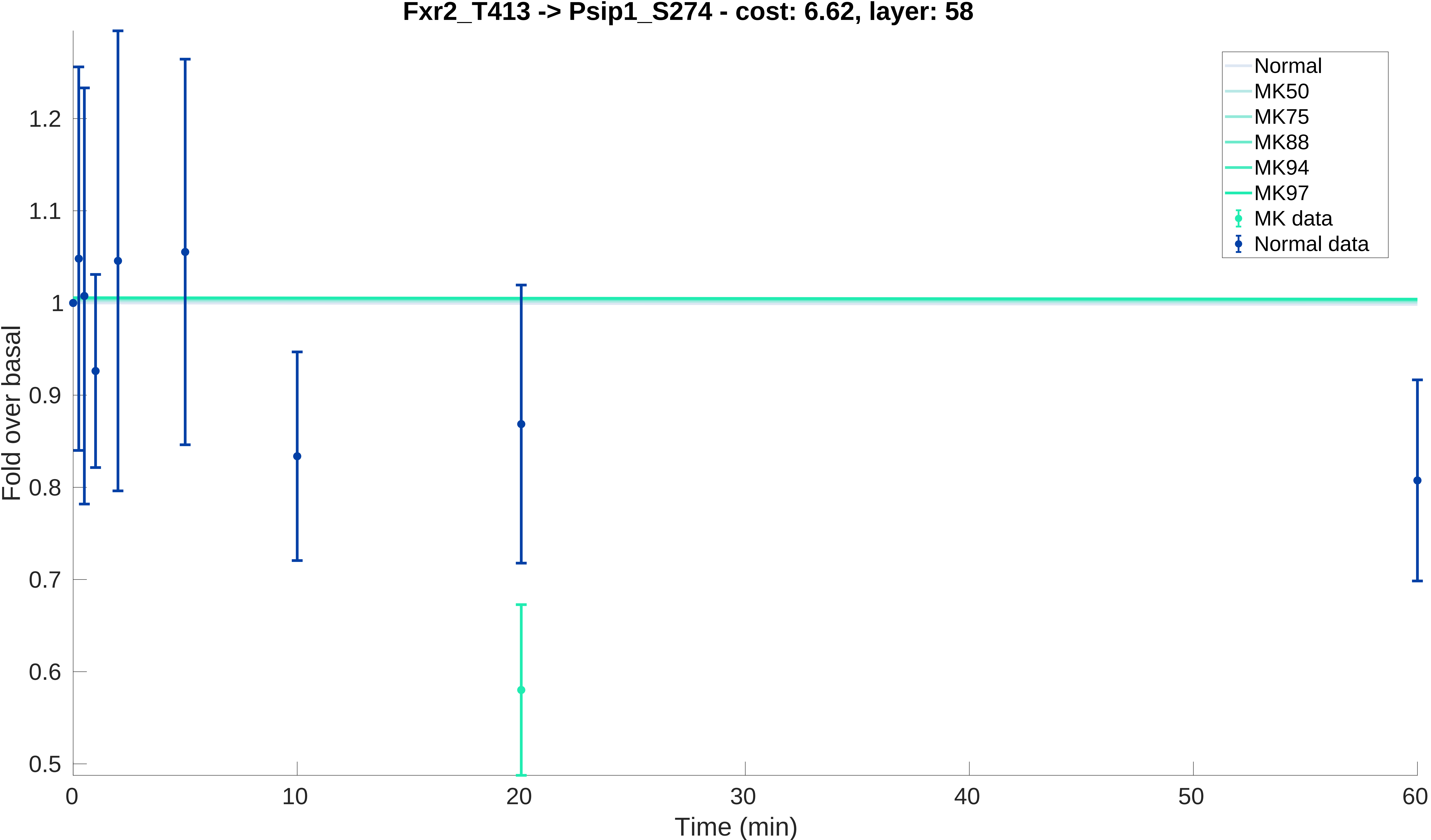

Fxr2_T413 -> Psip1_S274 - cost: 6.62, layer: 58
Fold over basal
Time (min)
0.5
0.6
0.7
0.8
0.9
1
1.1
1.2
0
10
20
30
40
50
60
Normal
MK50
MK75
MK88
MK94
MK97
MK data
Normal data

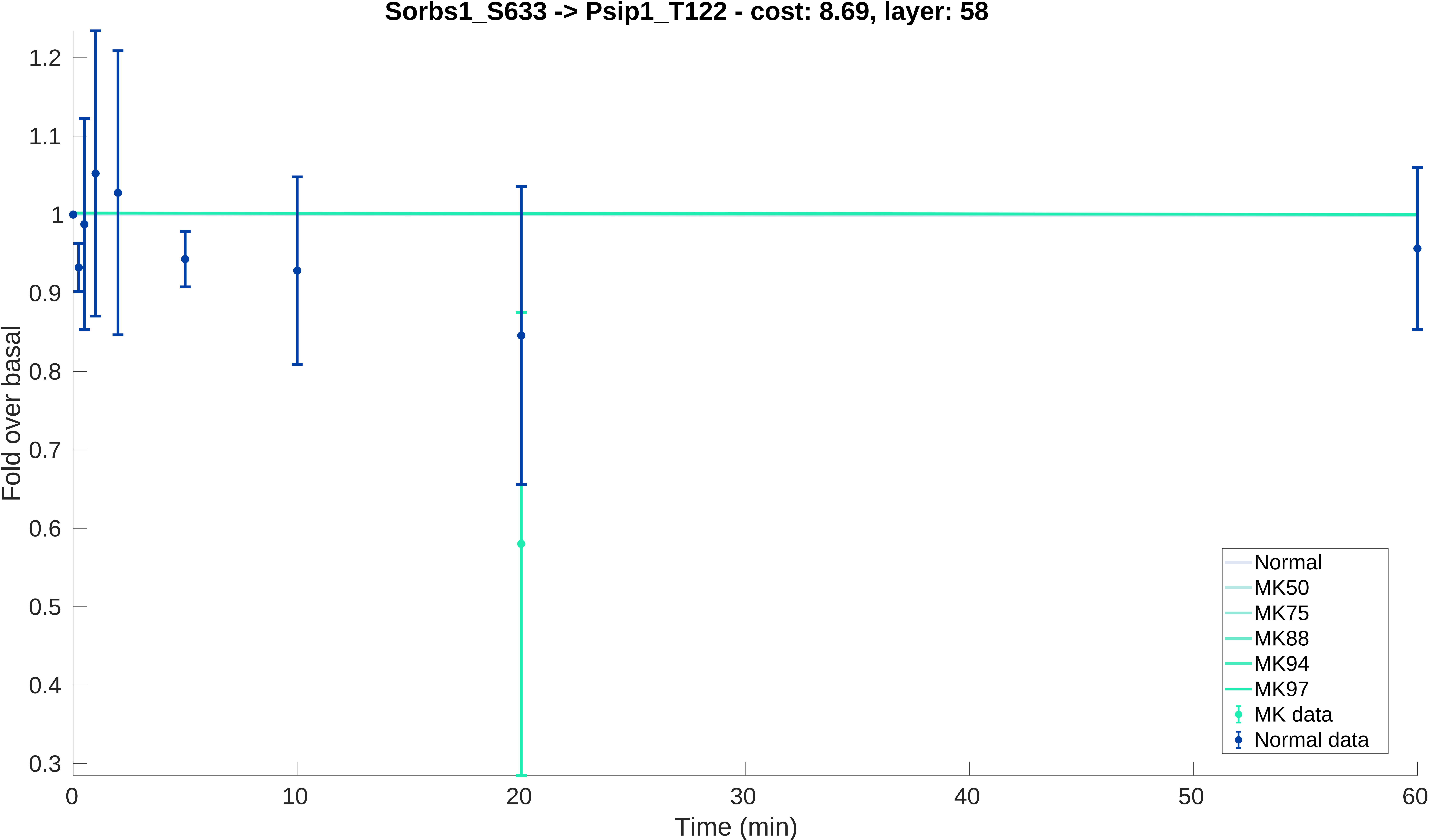

Sorbs1_S633 -> Psip1_T122 - cost: 8.69, layer: 58
Fold over basal
Time (min)
1.2
1.1
1
0.9
0.8
0.7
0.6
0.5
0.4
0.3
0
10
20
30
40
50
60
Normal
MK50
MK75
MK88
MK94
MK97
MK data
Normal data

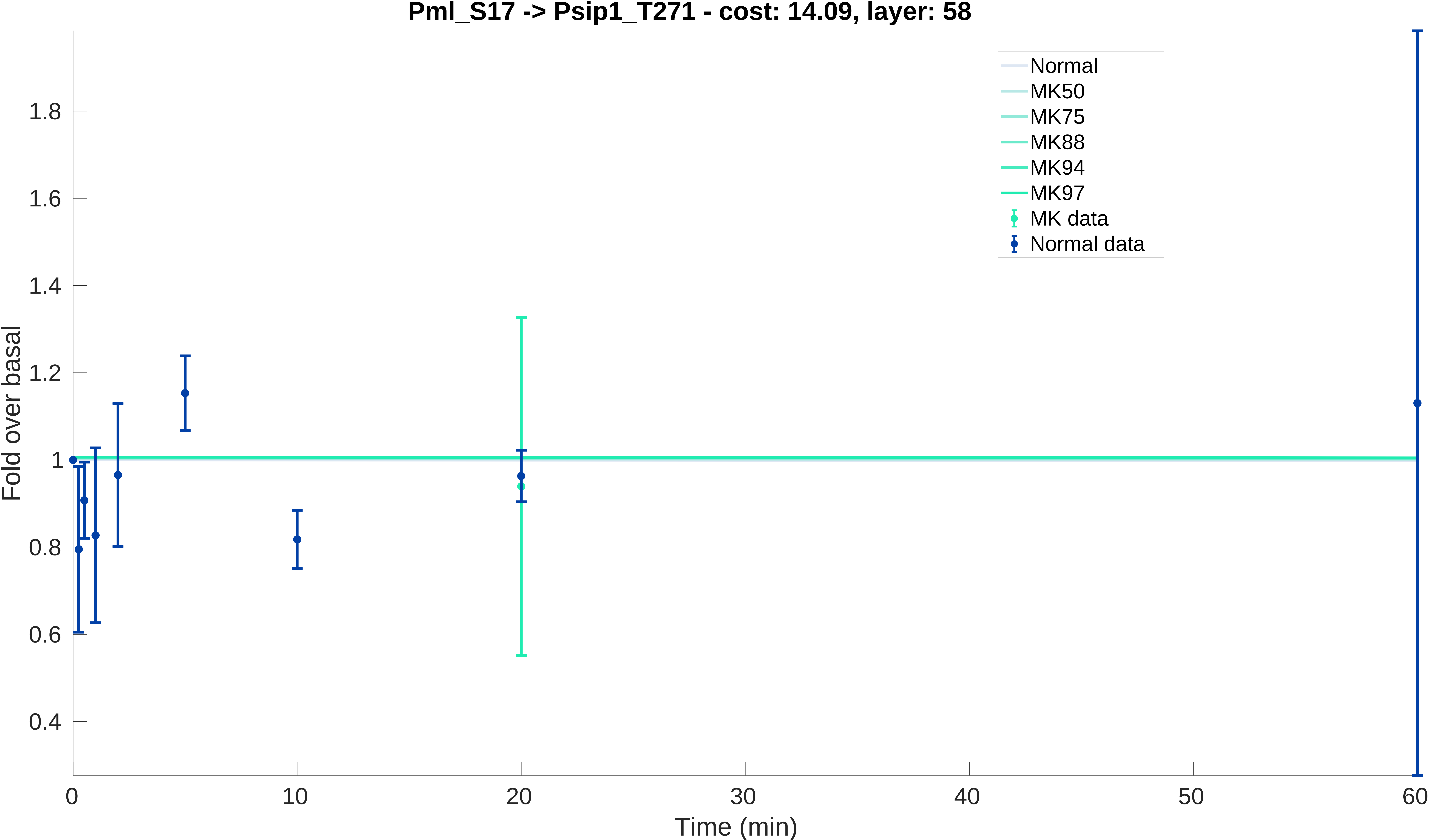

Pml_S17 -> Psip1_T271 - cost: 14.09, layer: 58
Normal
MK50
MK75
MK88
MK94
MK97
MK data
Normal data
Fold over basal
Time (min)
0.4
0.6
0.8
1
1.2
1.4
1.6
1.8
0
10
20
30
40
50
60

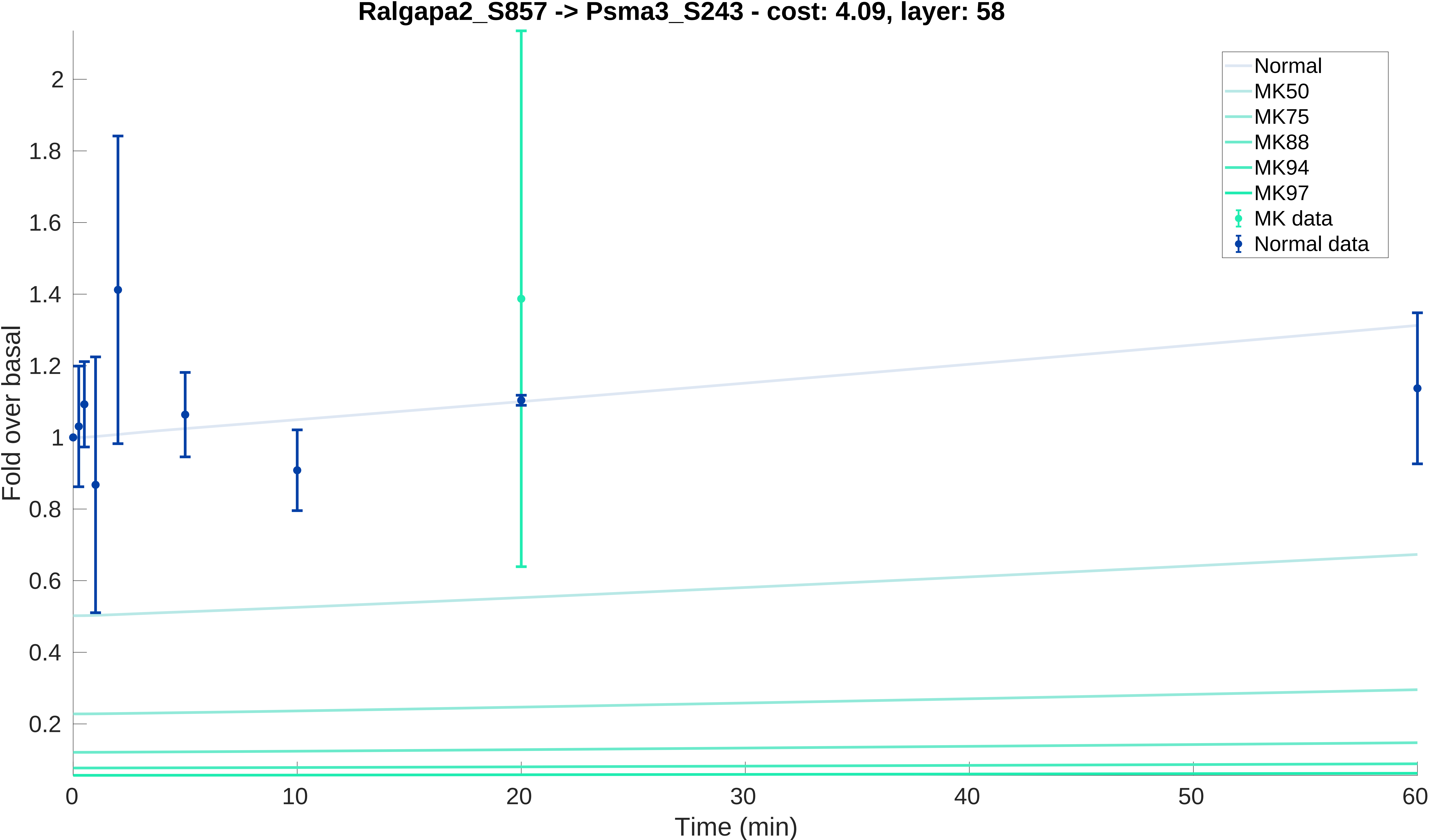

Ralgapa2_S857 -> Psma3_S243 - cost: 4.09, layer: 58
Normal
MK50
MK75
MK88
MK94
MK97
MK data
Normal data
Fold over basal
Time (min)
0
10
20
30
40
50
60
0.2
0.4
0.6
0.8
1
1.2
1.4
1.6
1.8
2

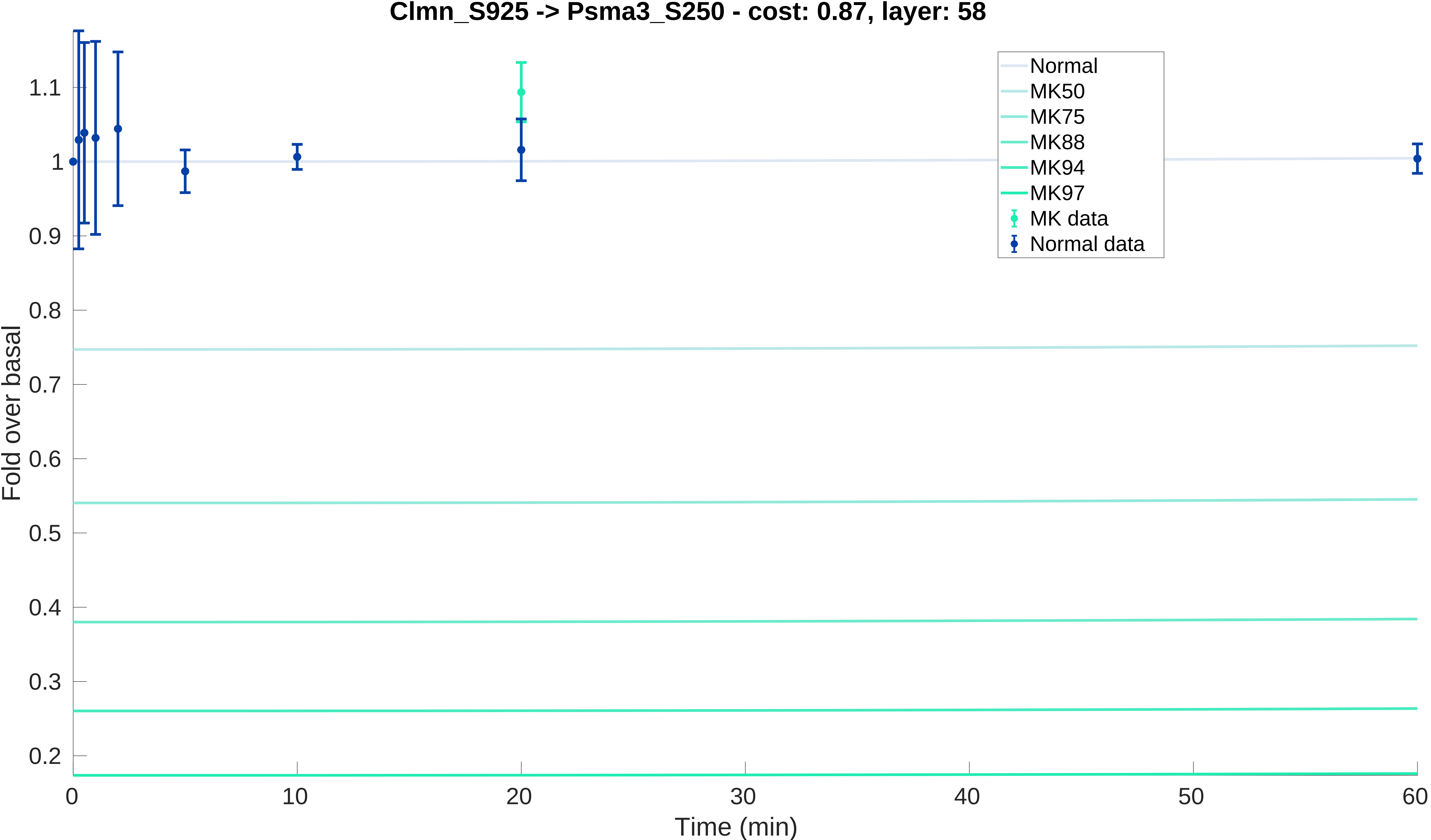

Clmn_S925 -> Psma3_S250 - cost: 0.87, layer: 58
Normal
MK50
MK75
MK88
MK94
MK97
MK data
Normal data
Fold over basal
Time (min)
0.2
0.3
0.4
0.5
0.6
0.7
0.8
0.9
1
1.1
0
10
20
30
40
50
60

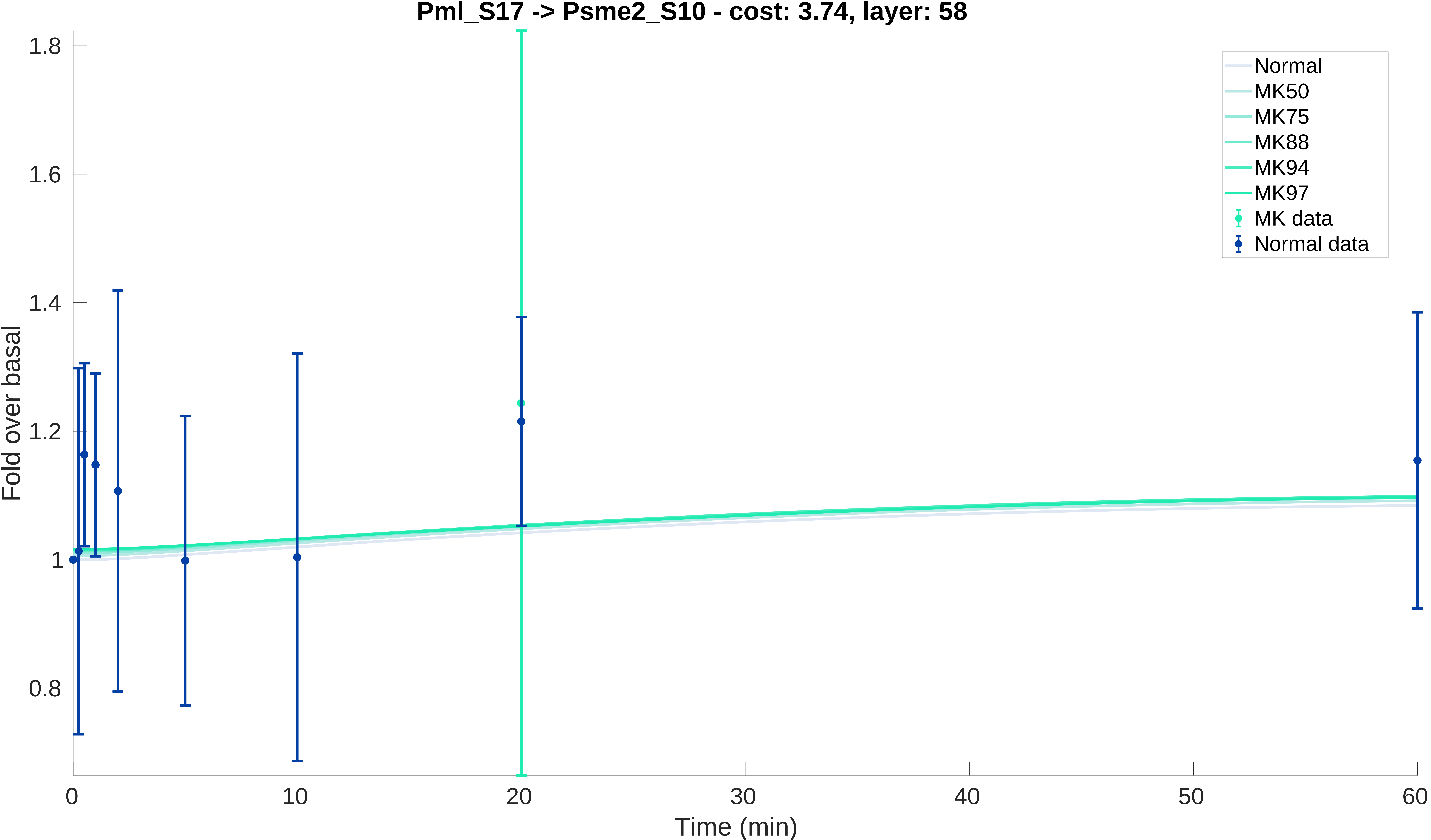
Pml_S17 -> Psme2_S10 - cost: 3.74, layer: 58
Fold over basal
Time (min)
1.8
1.6
1.4
1.2
1
0.8
0
10
20
30
40
50
60
Normal
MK50
MK75
MK88
MK94
MK97
MK data
Normal data

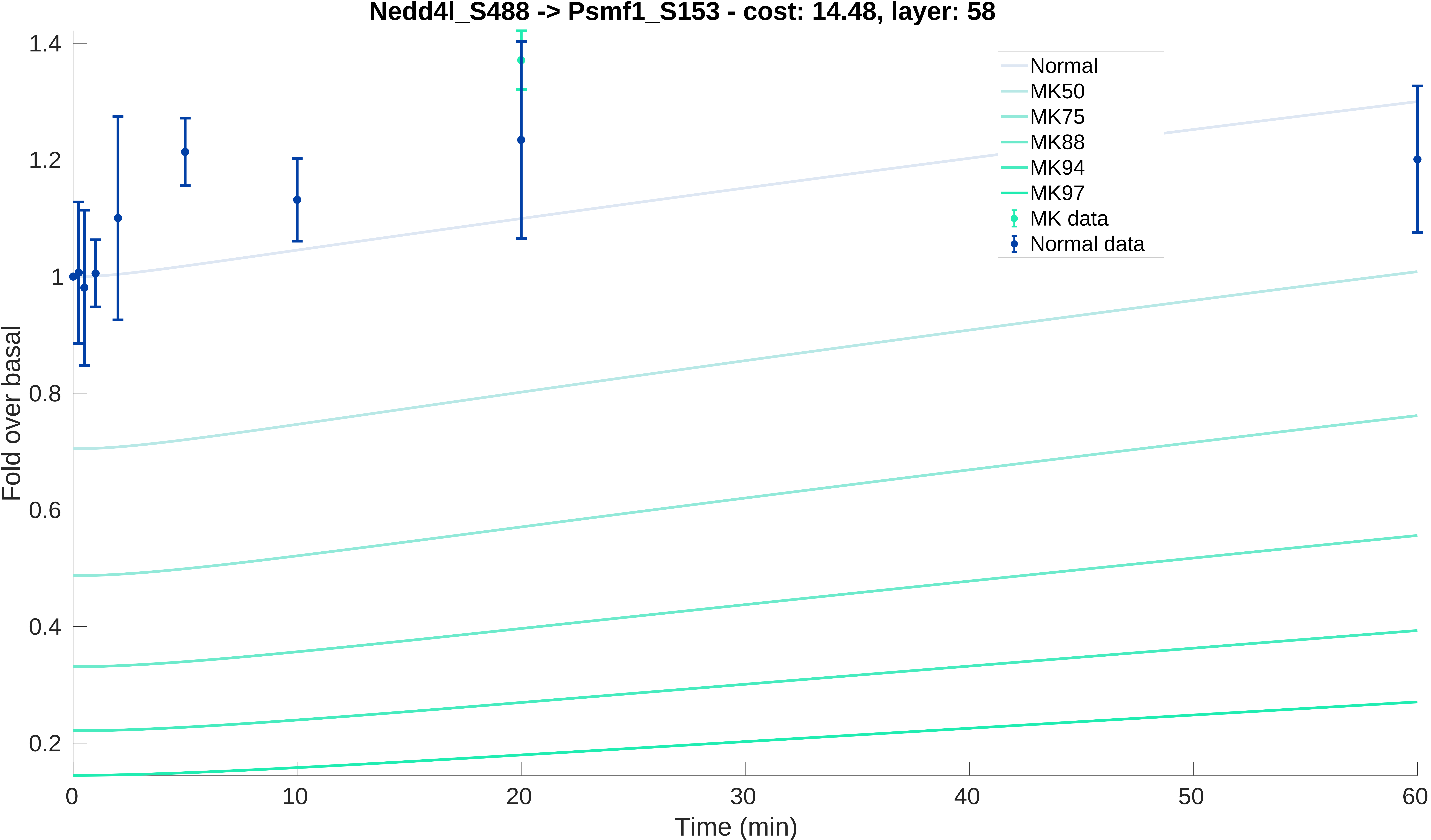
Nedd4l_S488 -> Psmf1_S153 - cost: 14.48, layer: 58
Fold over basal
Time (min)
1.4
1.2
1
0.8
0.6
0.4
0.2
0
10
20
30
40
50
60
Normal
MK50
MK75
MK88
MK94
MK97
MK data
Normal data

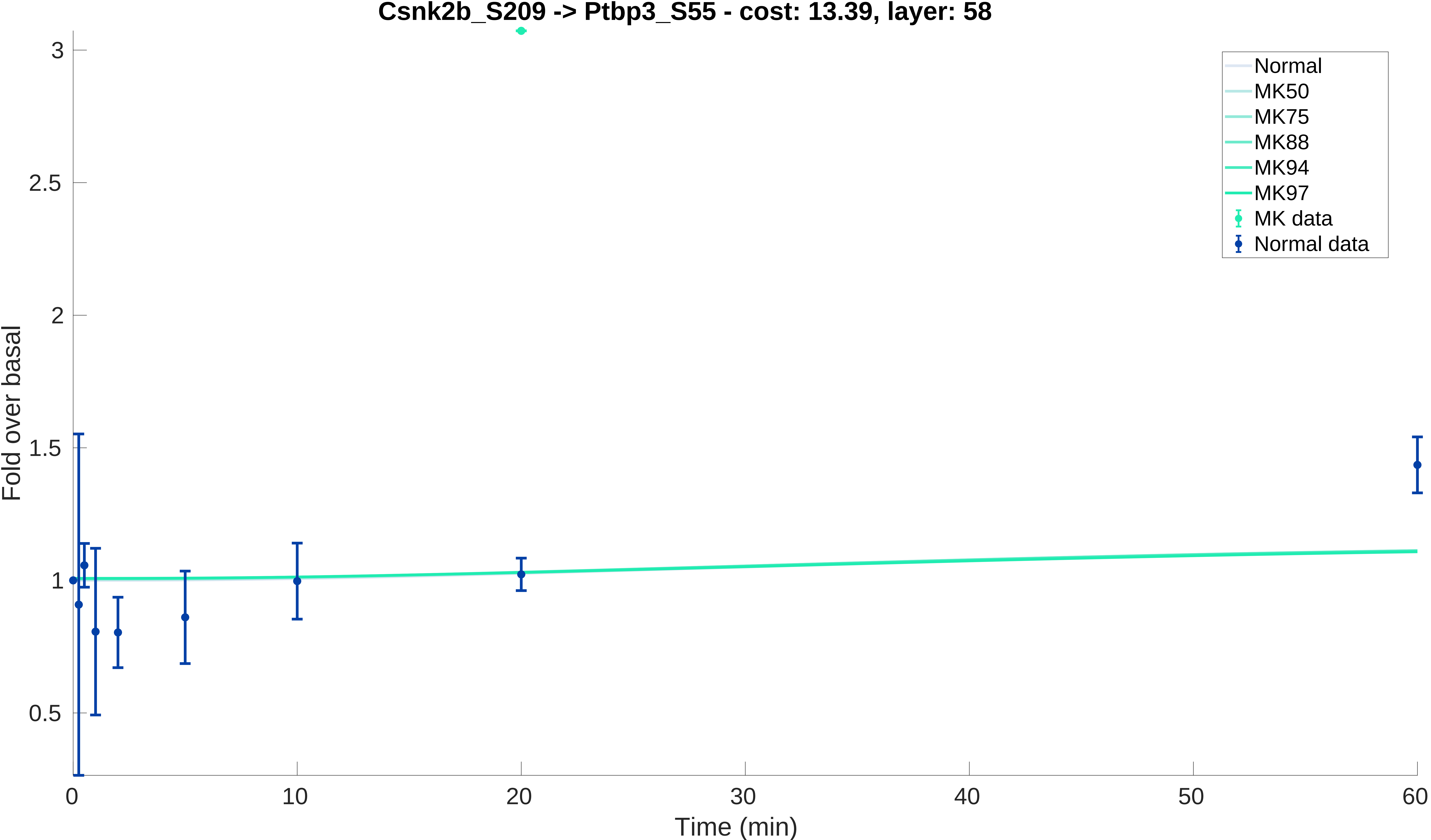

Csnk2b_S209 -> Ptbp3_S55 - cost: 13.39, layer: 58
Fold over basal
Time (min)
0.5
1
1.5
2
2.5
3
0
10
20
30
40
50
60
Normal
MK50
MK75
MK88
MK94
MK97
MK data
Normal data

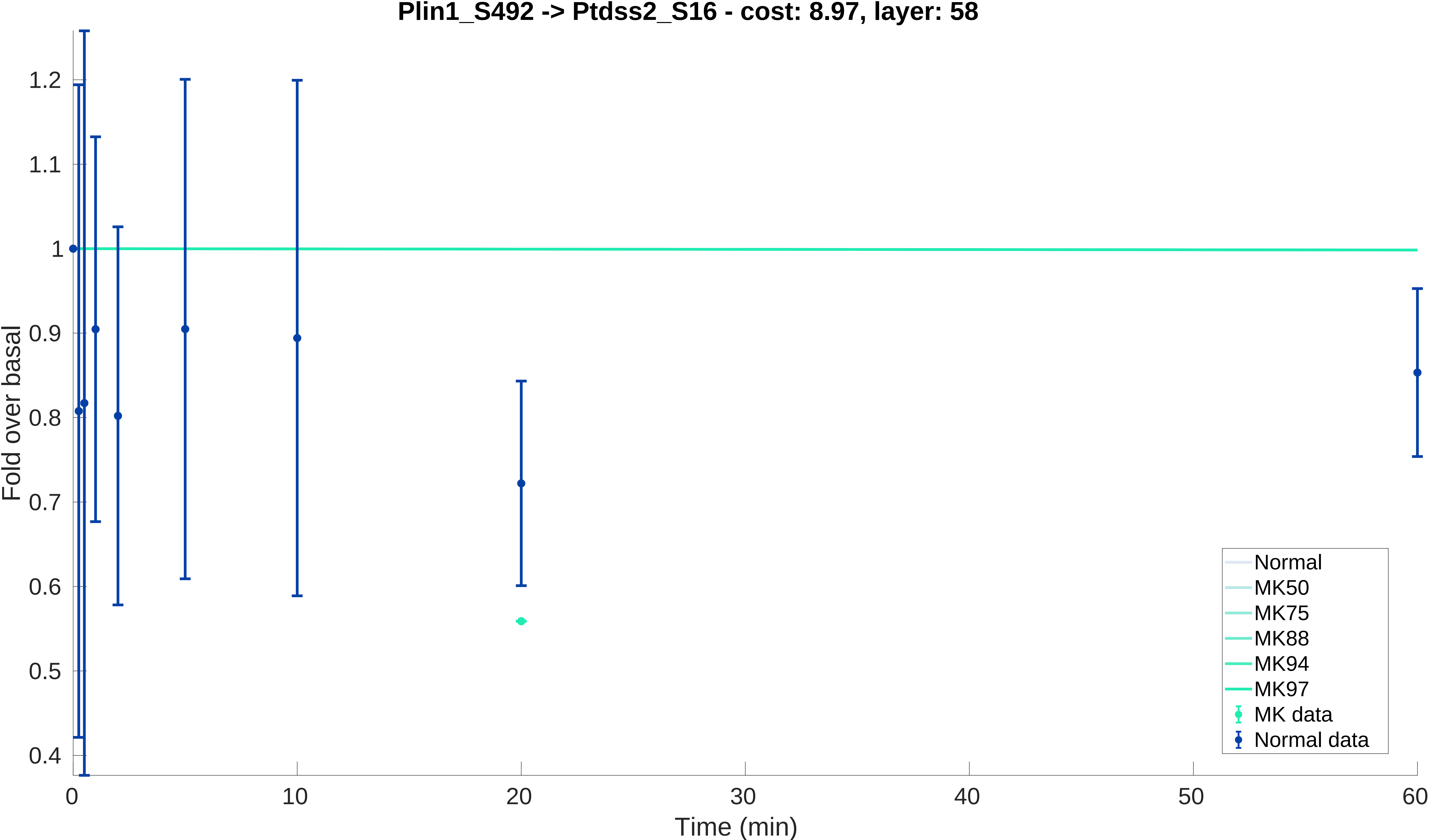

Plin1_S492 -> Ptdss2_S16 - cost: 8.97, layer: 58
Fold over basal
1.2
1.1
1
0.9
0.8
0.7
0.6
0.5
0.4
0
10
20
30
40
50
60
Time (min)
Normal
MK50
MK75
MK88
MK94
MK97
MK data
Normal data

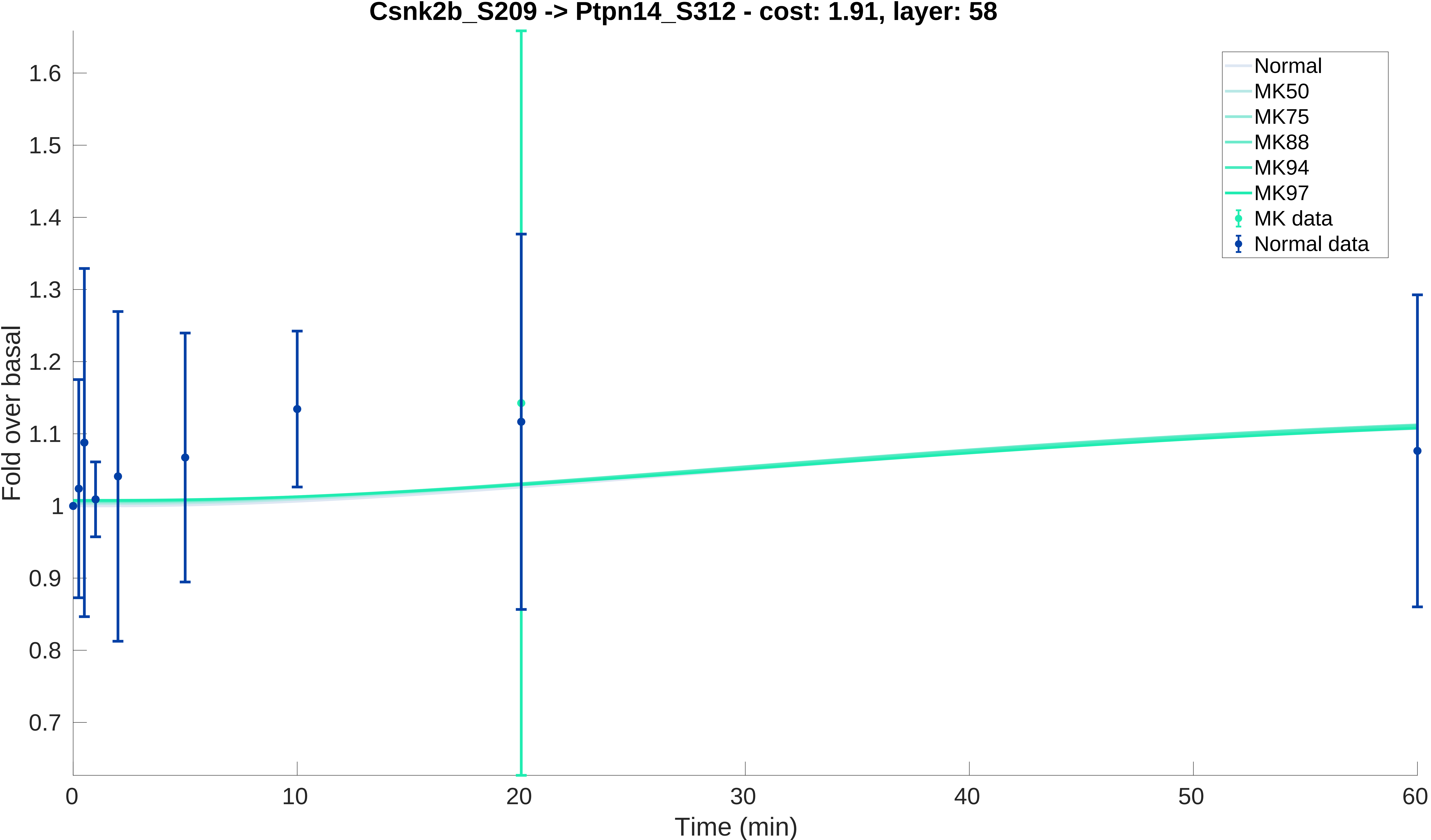

Csnk2b_S209 -> Ptpn14_S312 - cost: 1.91, layer: 58
Fold over basal
Time (min)
0
10
20
30
40
50
60
0.7
0.8
0.9
1
1.1
1.2
1.3
1.4
1.5
1.6
Normal
MK50
MK75
MK88
MK94
MK97
MK data
Normal data

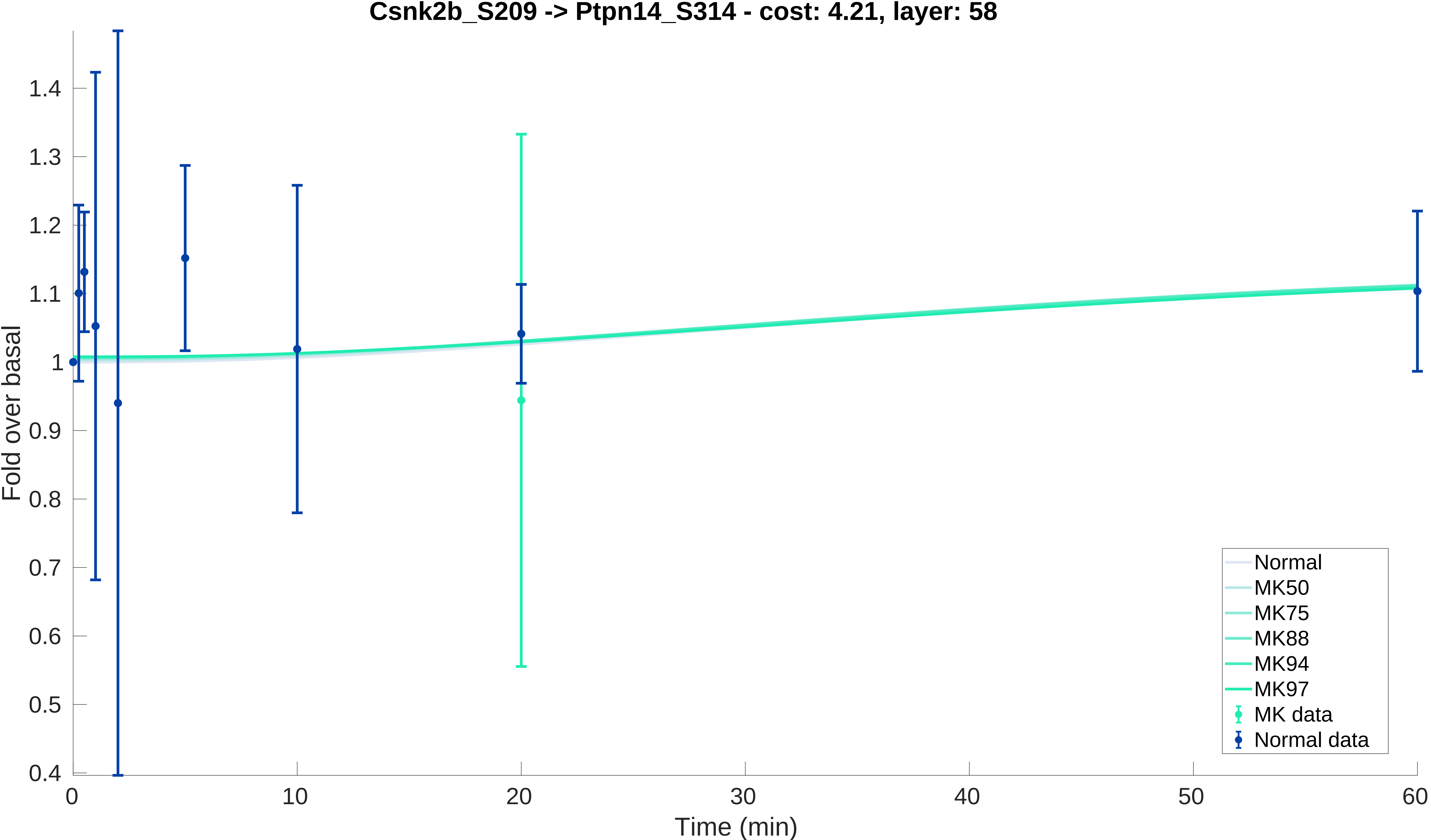
Csnk2b_S209 -> Ptpn14_S314 - cost: 4.21, layer: 58
Fold over basal
Time (min)
1.4
1.3
1.2
1.1
1
0.9
0.8
0.7
0.6
0.5
0.4
0
10
20
30
40
50
60
Normal
MK50
MK75
MK88
MK94
MK97
MK data
Normal data

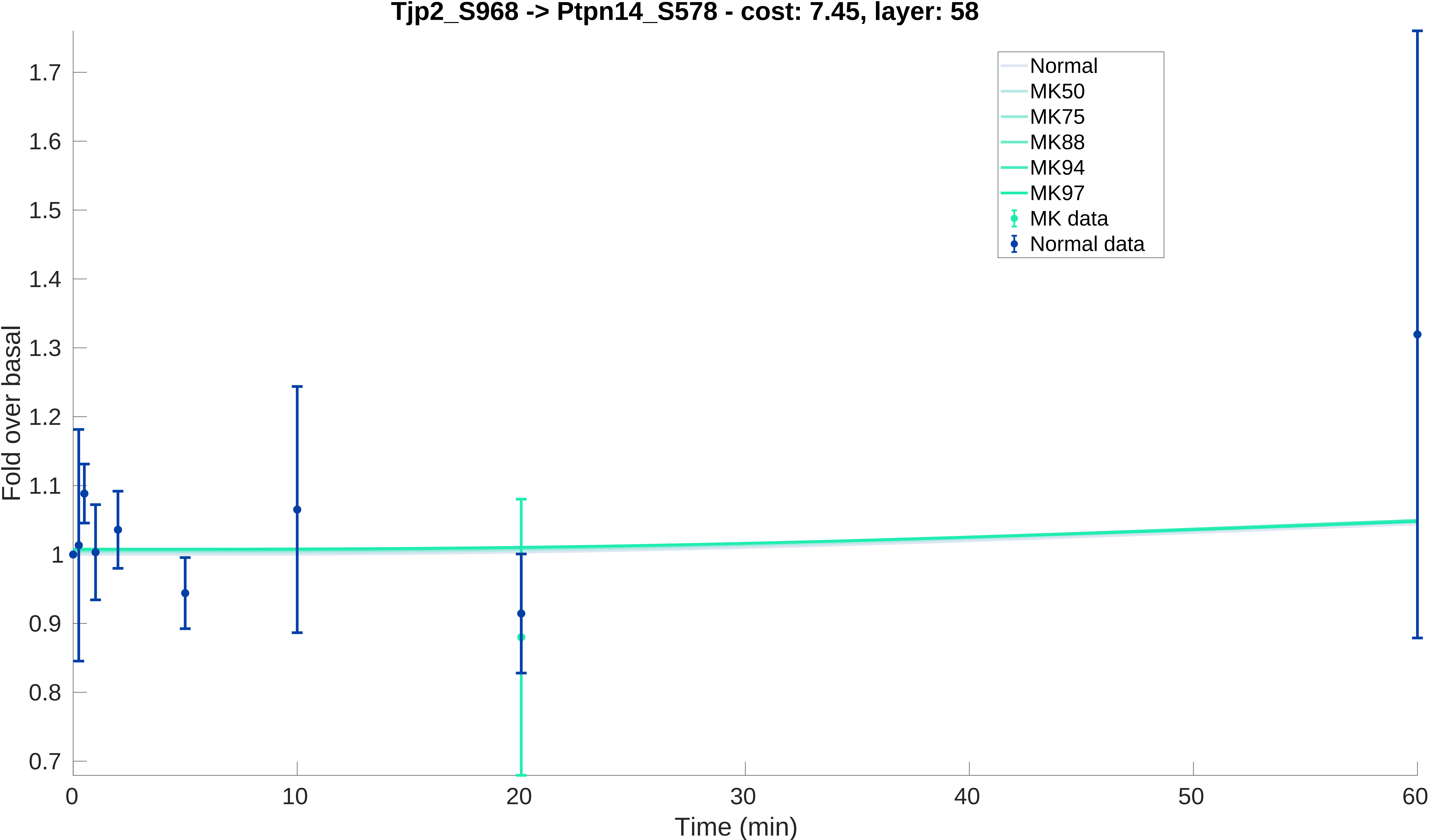

Tjp2_S968 -> Ptpn14_S578 - cost: 7.45, layer: 58
Normal
MK50
MK75
MK88
MK94
MK97
MK data
Normal data
Fold over basal
1.7
1.6
1.5
1.4
1.3
1.2
1.1
1
0.9
0.8
0.7
0
10
20
30
40
50
60
Time (min)

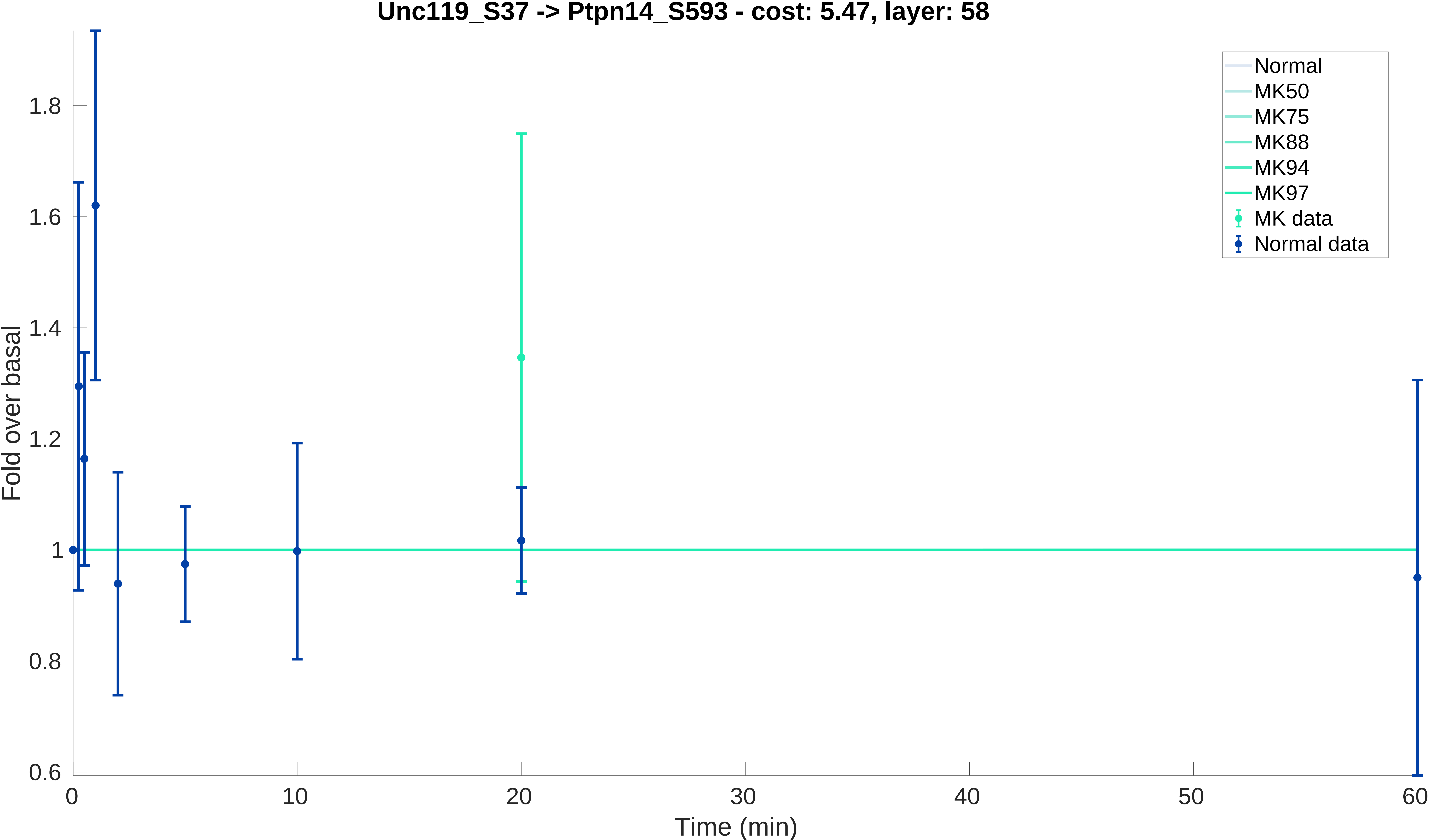

Unc119_S37 -> Ptpn14_S593 - cost: 5.47, layer: 58
Fold over basal
Time (min)
0.6
0.8
1
1.2
1.4
1.6
1.8
0
10
20
30
40
50
60
Normal
MK50
MK75
MK88
MK94
MK97
MK data
Normal data

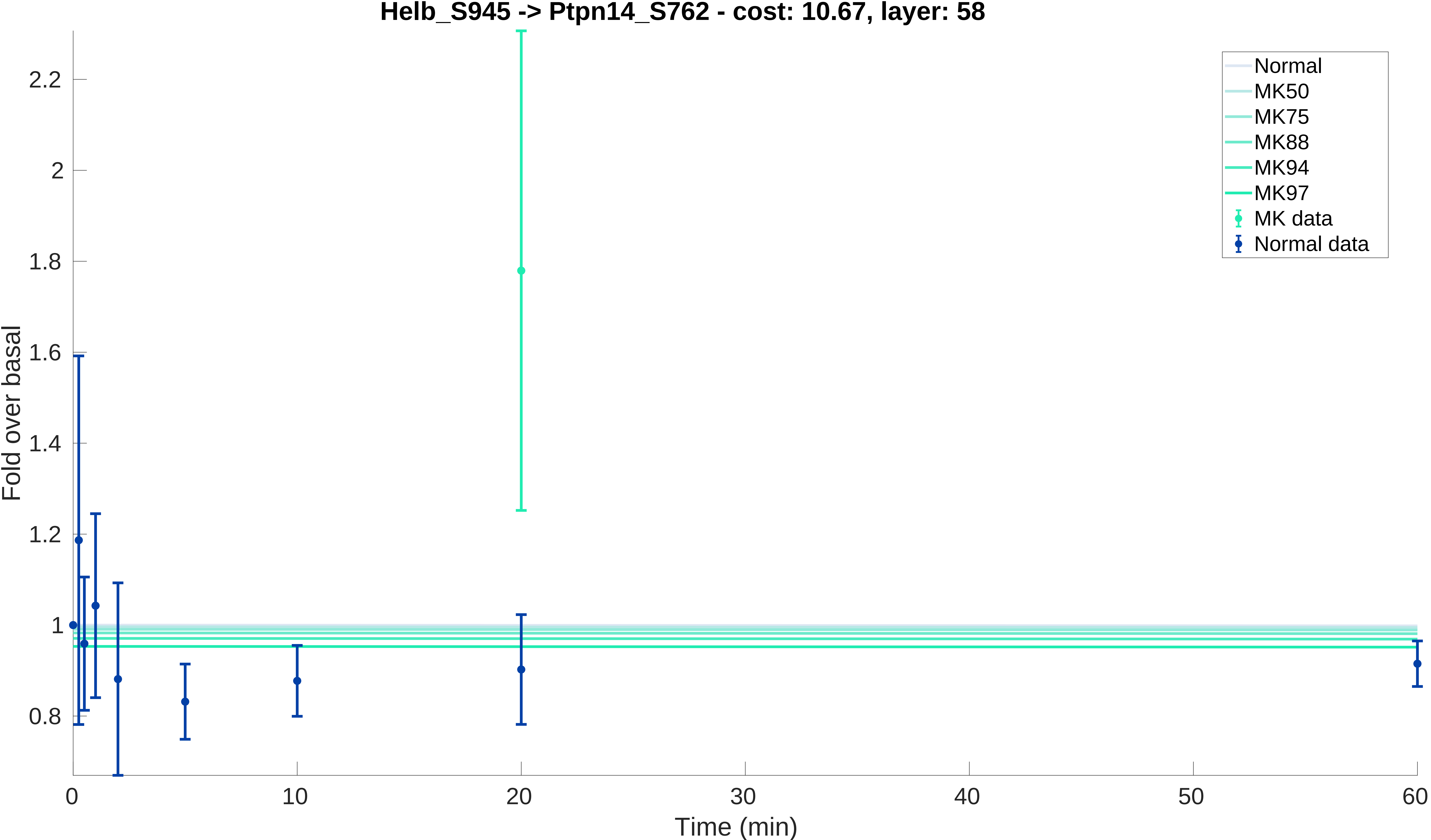

Helb_S945 -> Ptpn14_S762 - cost: 10.67, layer: 58
Fold over basal
2.2
2
1.8
1.6
1.4
1.2
1
0.8
0
10
20
30
40
50
60
Time (min)
Normal
MK50
MK75
MK88
MK94
MK97
MK data
Normal data

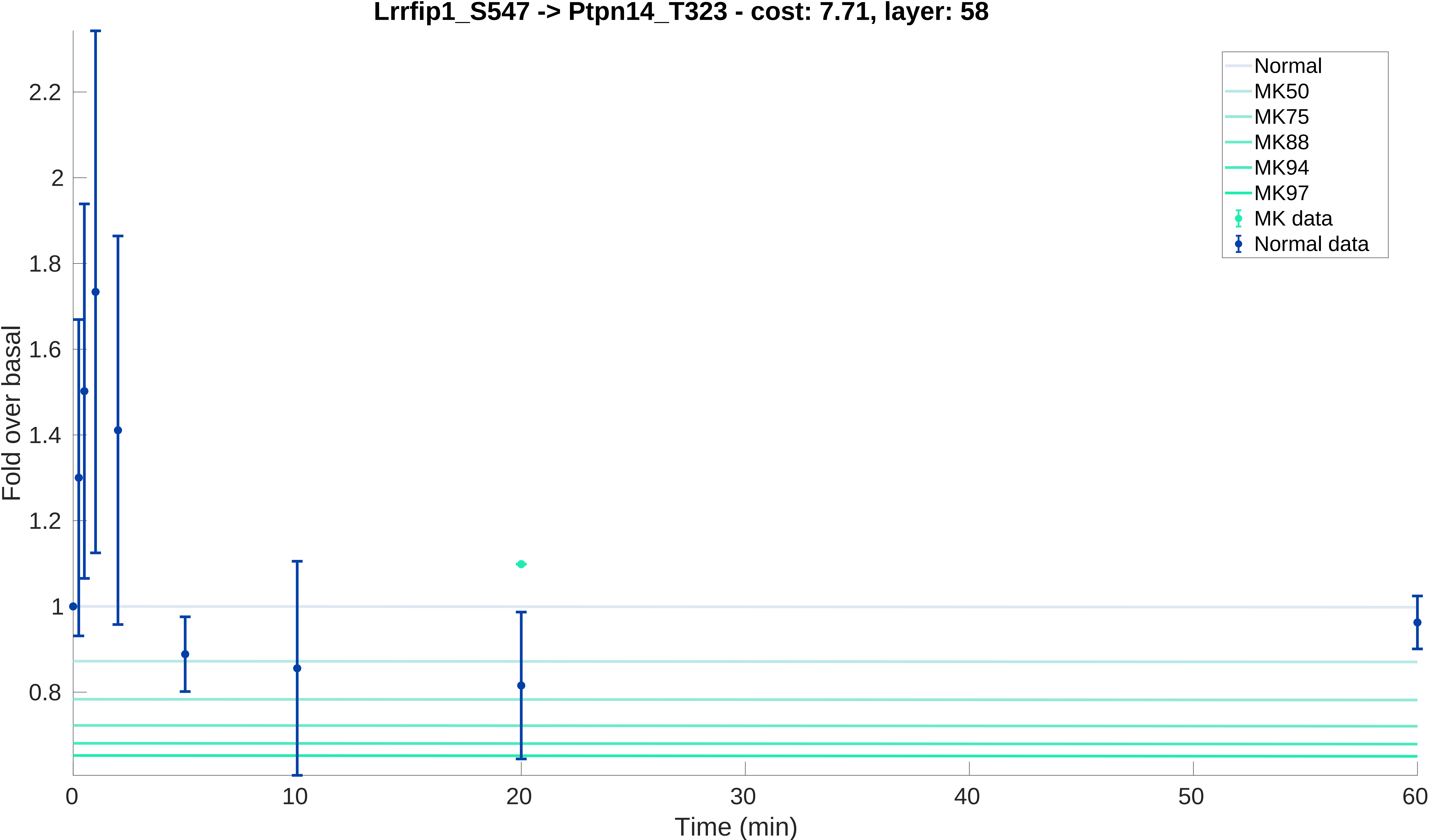

Lrrfip1_S547 -> Ptpn14_T323 - cost: 7.71, layer: 58
Fold over basal
Time (min)
0
10
20
30
40
50
60
0.8
1
1.2
1.4
1.6
1.8
2
2.2
Normal
MK50
MK75
MK88
MK94
MK97
MK data
Normal data

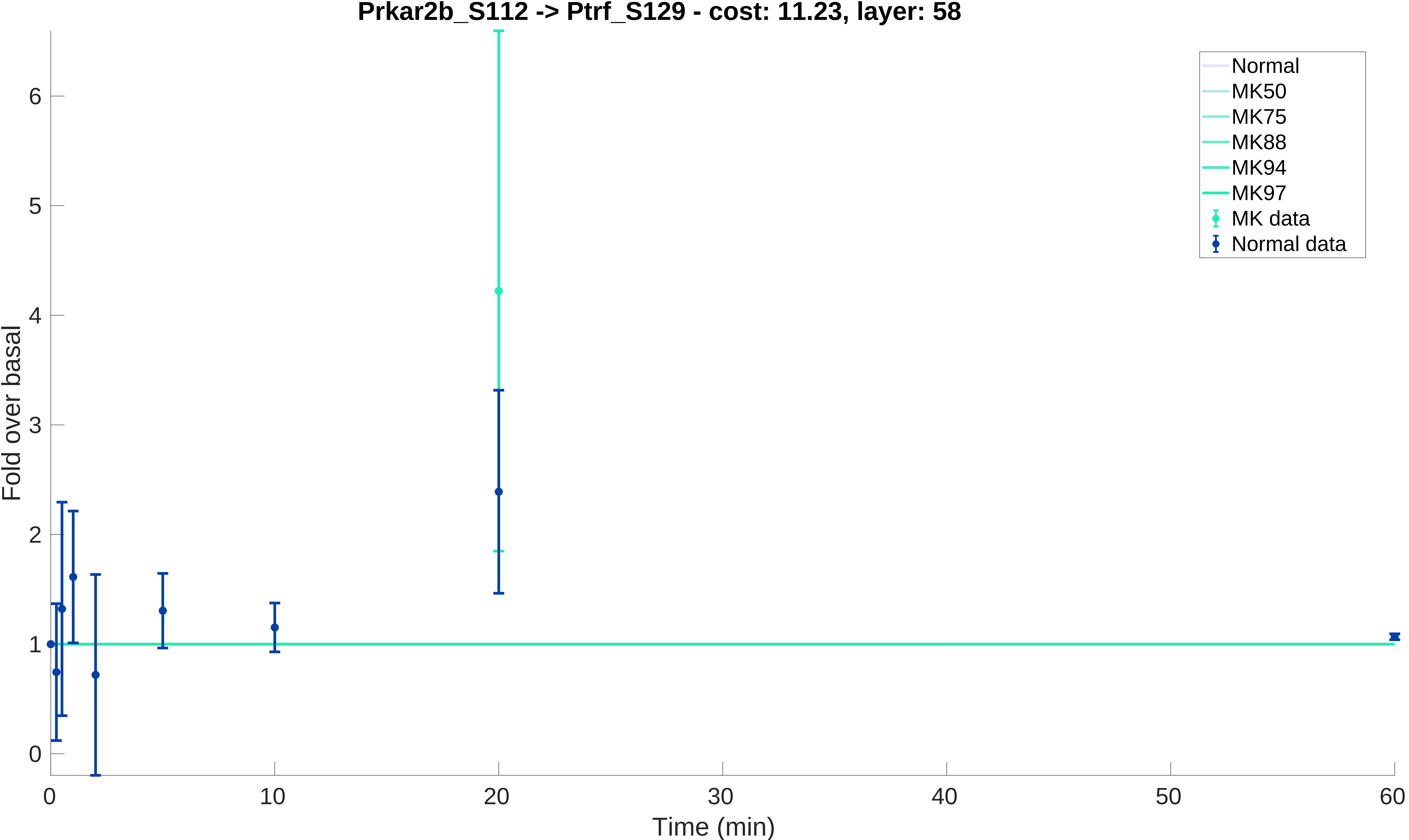

Prkar2b_S112 -> Ptrf_S129 - cost: 11.23, layer: 58
Normal
MK50
MK75
MK88
MK94
MK97
MK data
Normal data
Fold over basal
Time (min)
0
1
2
3
4
5
6
0
10
20
30
40
50
60

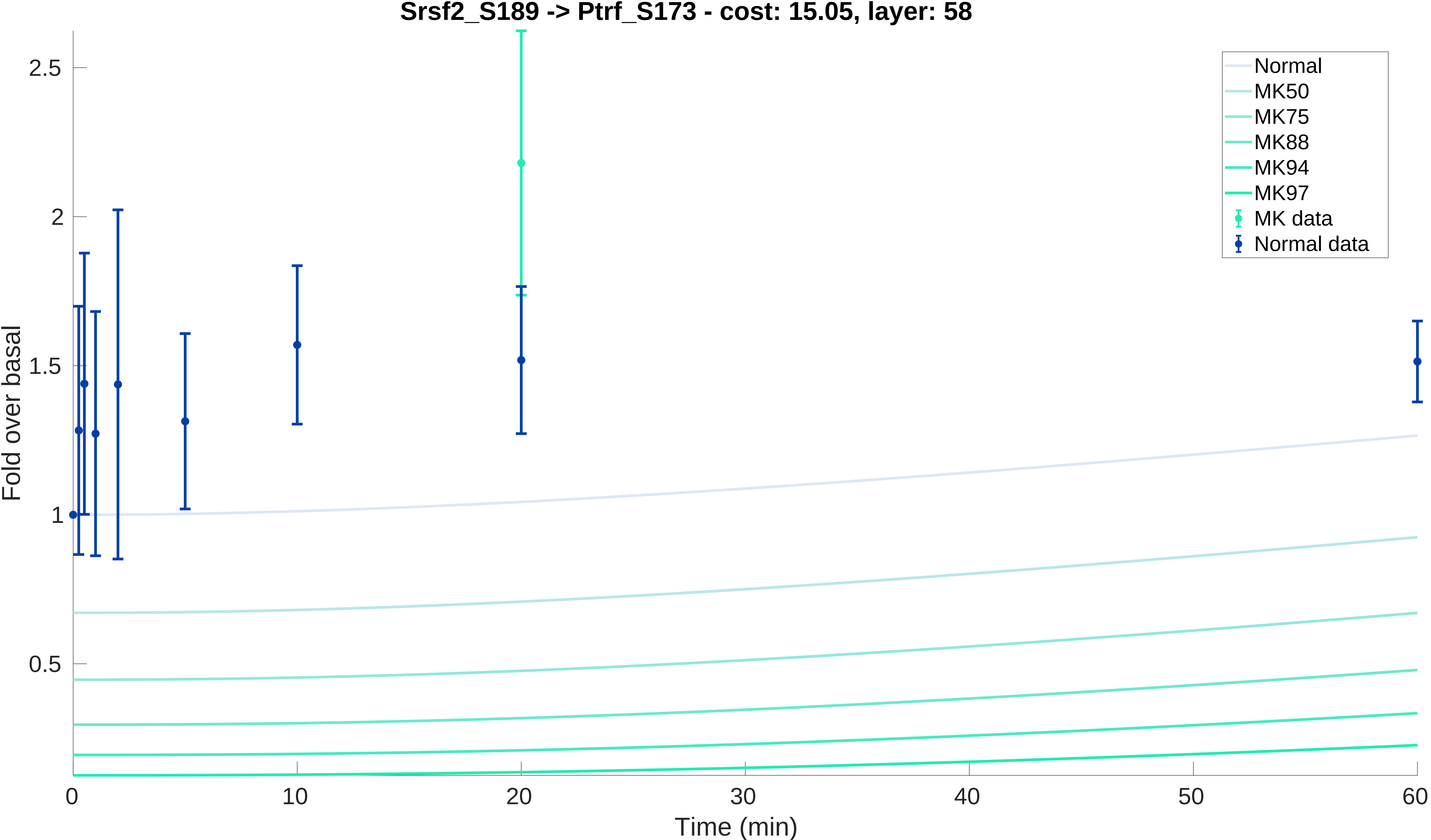

Srsf2_S189 -> Ptrf_S173 - cost: 15.05, layer: 58
Fold over basal
Time (min)
0
10
20
30
40
50
60
0.5
1
1.5
2
2.5
Normal
MK50
MK75
MK88
MK94
MK97
MK data
Normal data

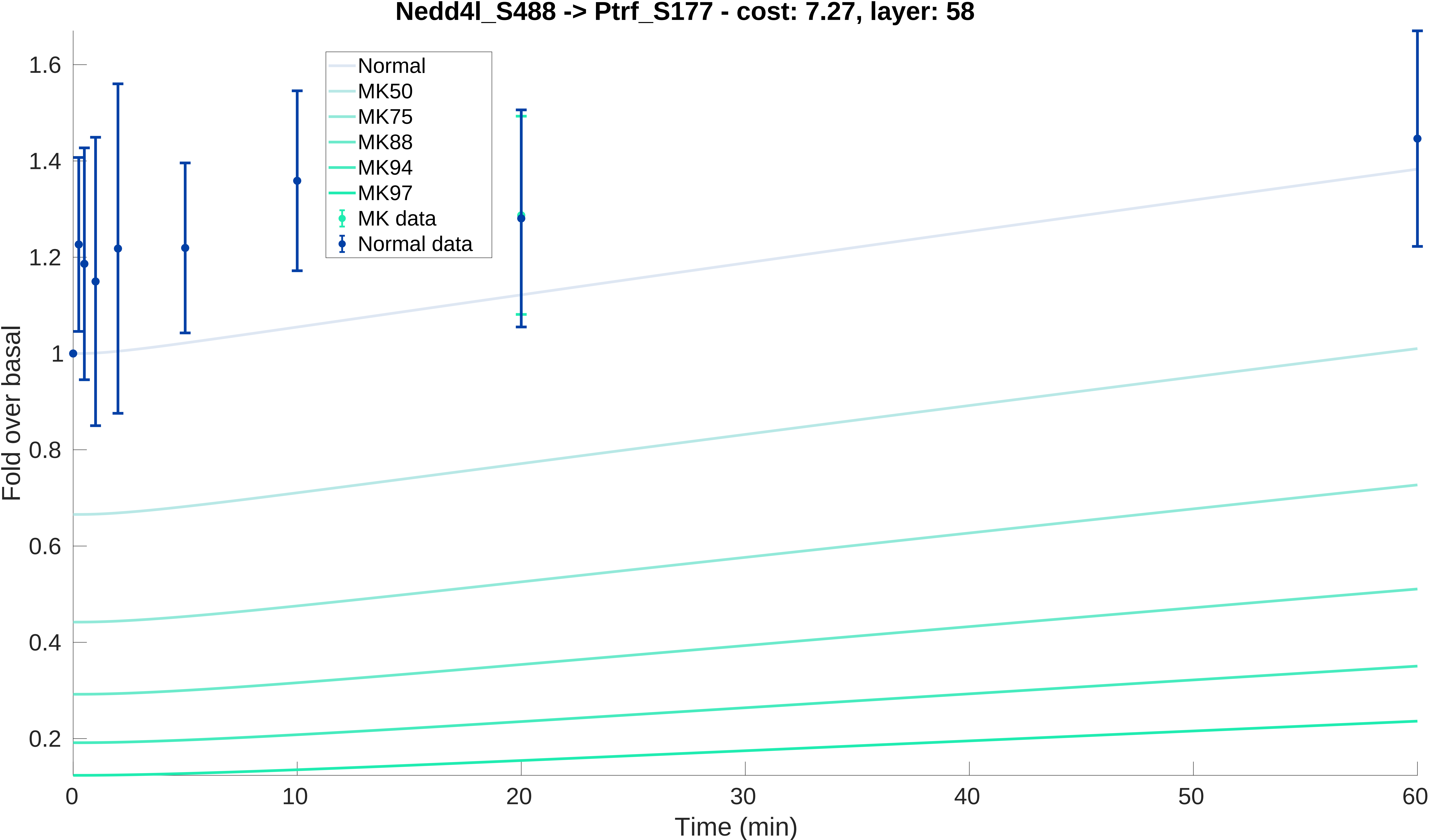

Nedd4l_S488 -> Ptrf_S177 - cost: 7.27, layer: 58
Normal
MK50
MK75
MK88
MK94
MK97
MK data
Normal data
Fold over basal
1.6
1.4
1.2
1
0.8
0.6
0.4
0.2
0
10
20
30
40
50
60
Time (min)

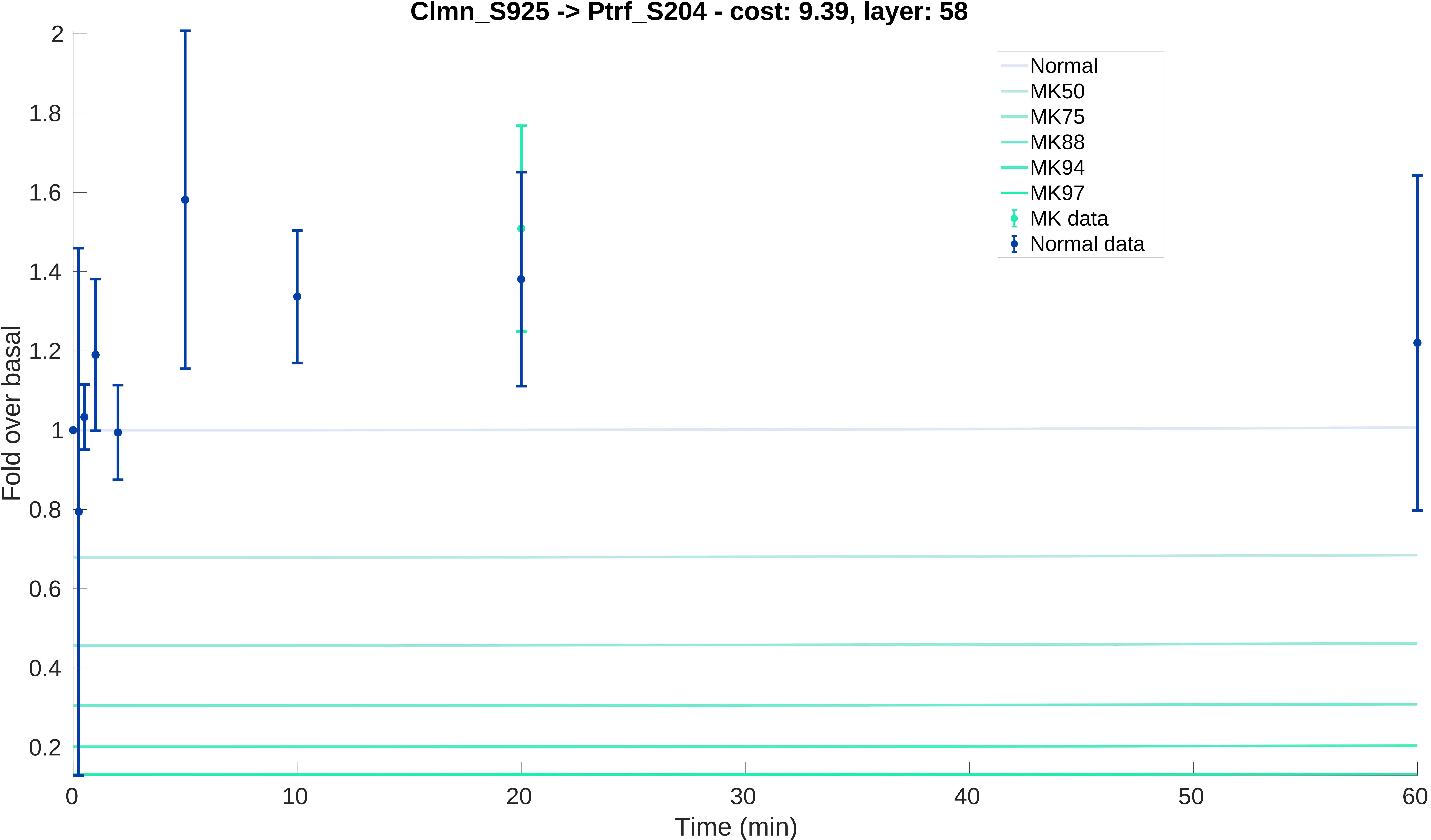

Clmn_S925 -> Ptrf_S204 - cost: 9.39, layer: 58
Fold over basal
Time (min)
2
1.8
1.6
1.4
1.2
1
0.8
0.6
0.4
0.2
0
10
20
30
40
50
60
Normal
MK50
MK75
MK88
MK94
MK97
MK data
Normal data

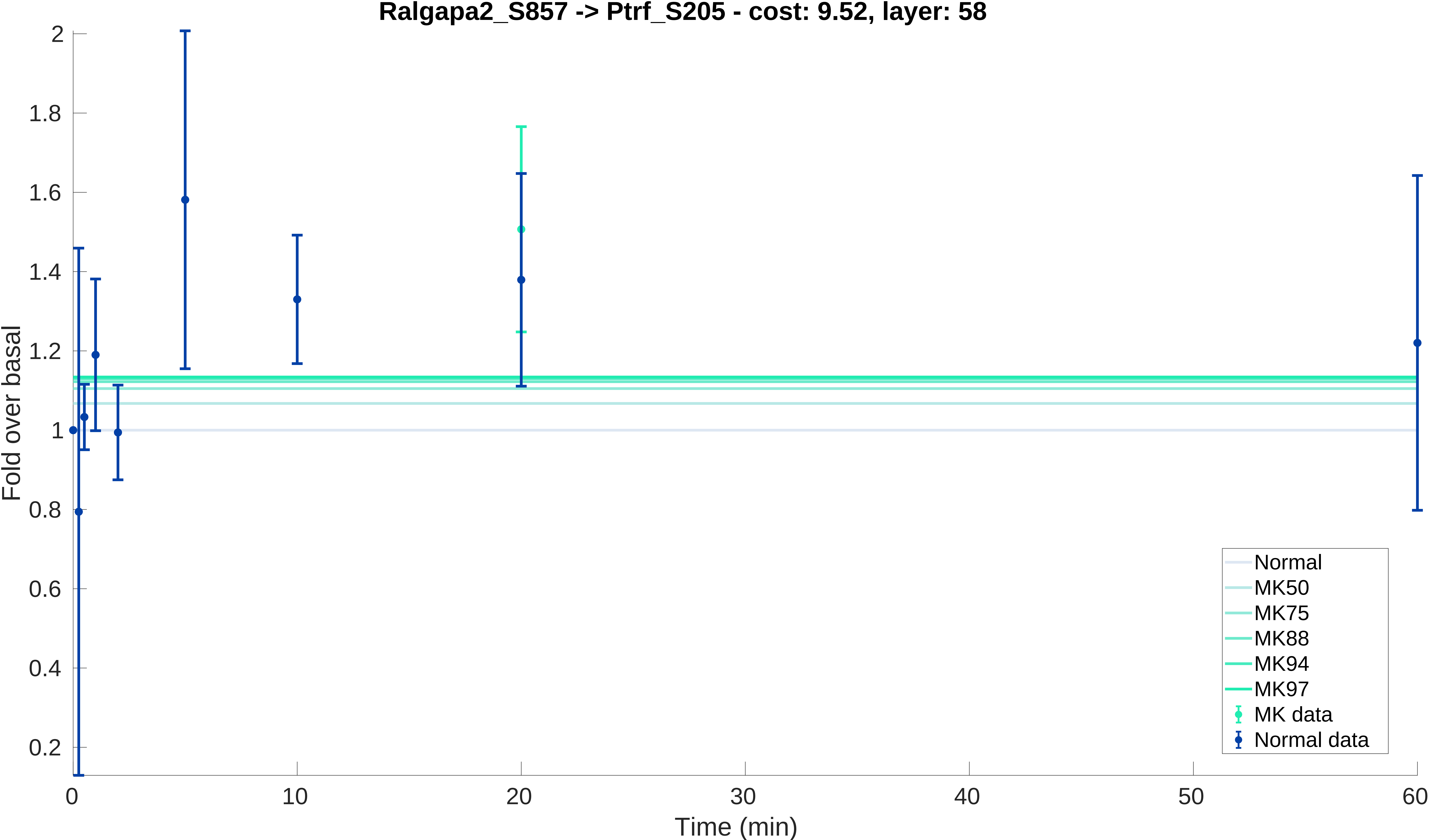

Ralgapa2_S857 -> Ptrf_S205 - cost: 9.52, layer: 58
Fold over basal
Time (min)
0
10
20
30
40
50
60
0.2
0.4
0.6
0.8
1
1.2
1.4
1.6
1.8
2
Normal
MK50
MK75
MK88
MK94
MK97
MK data
Normal data

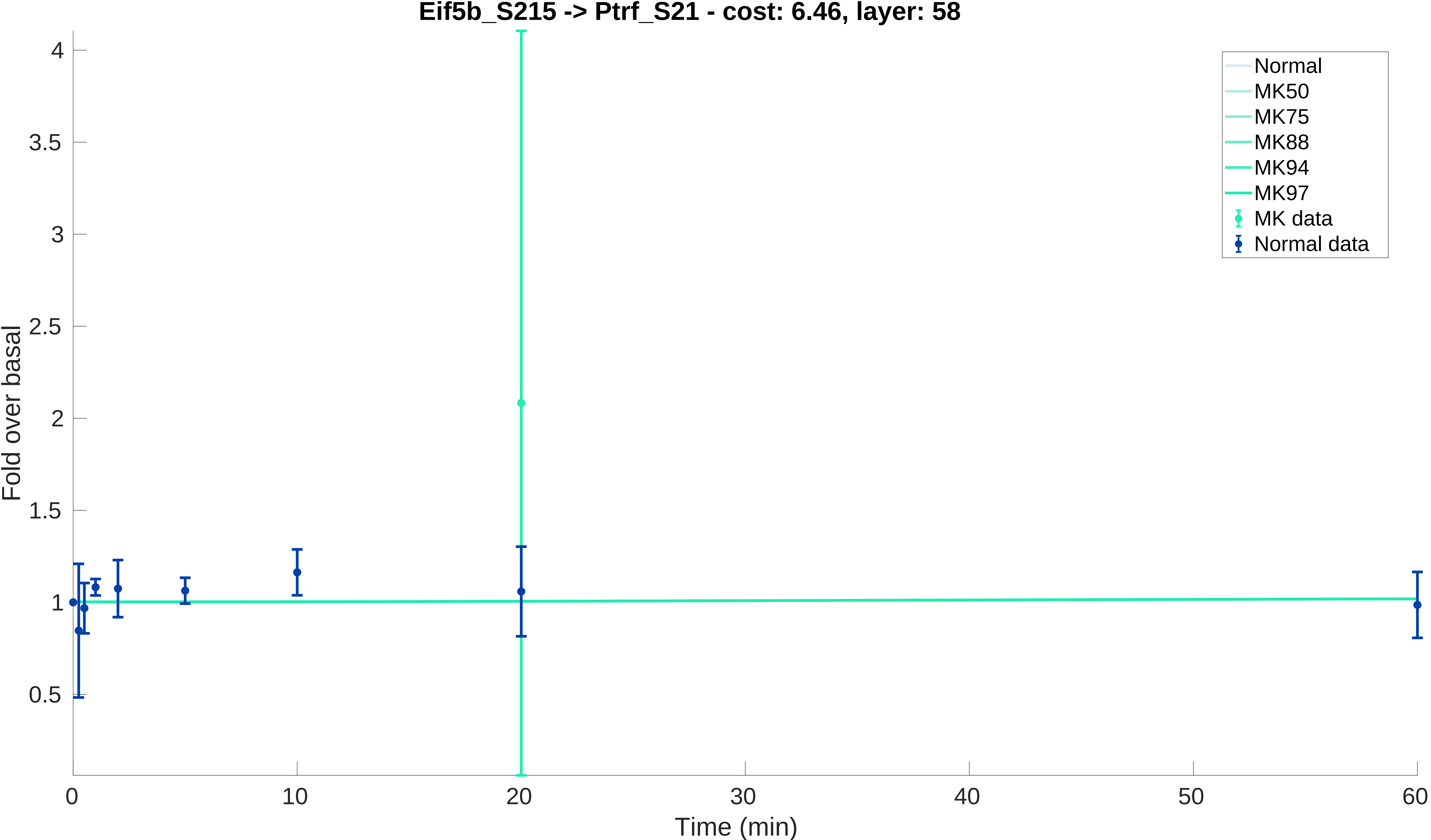

Eif5b_S215 -> Ptrf_S21 - cost: 6.46, layer: 58
Fold over basal
Time (min)
0.5
1
1.5
2
2.5
3
3.5
4
0
10
20
30
40
50
60
Normal
MK50
MK75
MK88
MK94
MK97
MK data
Normal data

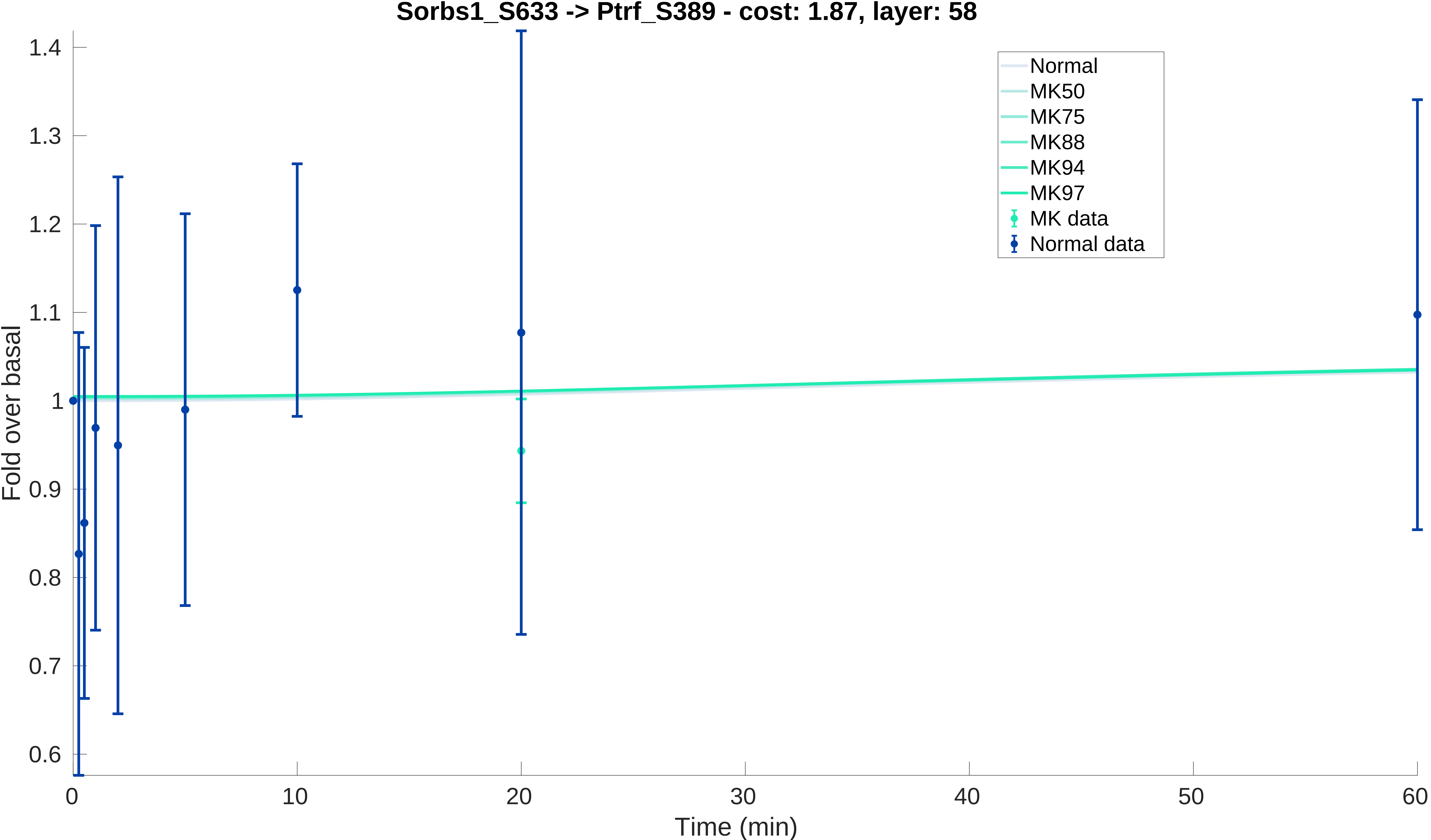

Sorbs1_S633 -> Ptrf_S389 - cost: 1.87, layer: 58
Fold over basal
Time (min)
Normal
MK50
MK75
MK88
MK94
MK97
MK data
Normal data
1.4
1.3
1.2
1.1
1
0.9
0.8
0.7
0.6
0
10
20
30
40
50
60

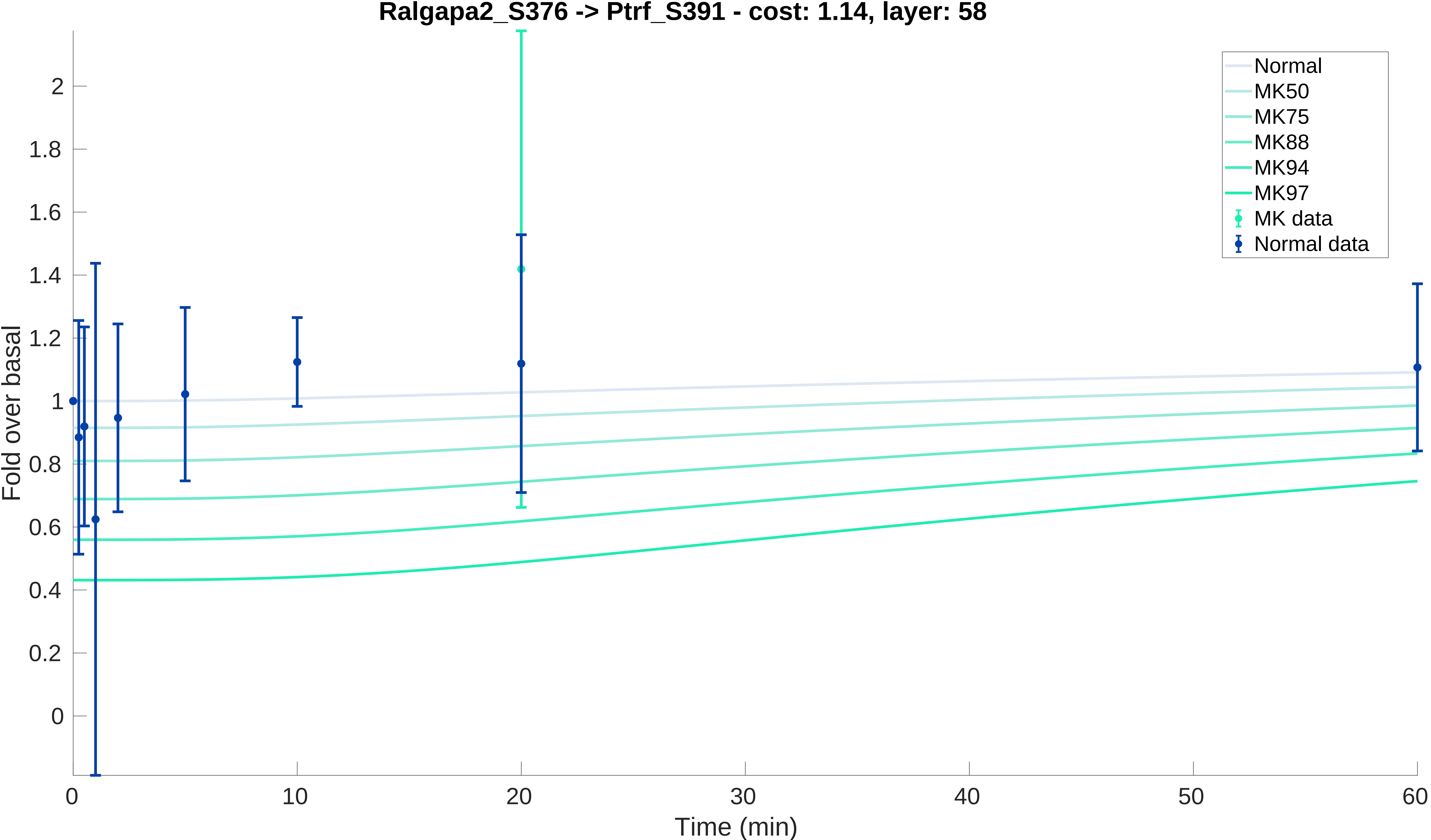

Ralgapa2_S376 -> Ptrf_S391 - cost: 1.14, layer: 58
Normal
MK50
MK75
MK88
MK94
MK97
MK data
Normal data
Fold over basal
2
1.8
1.6
1.4
1.2
1
0.8
0.6
0.4
0.2
0
0
10
20
30
40
50
60
Time (min)

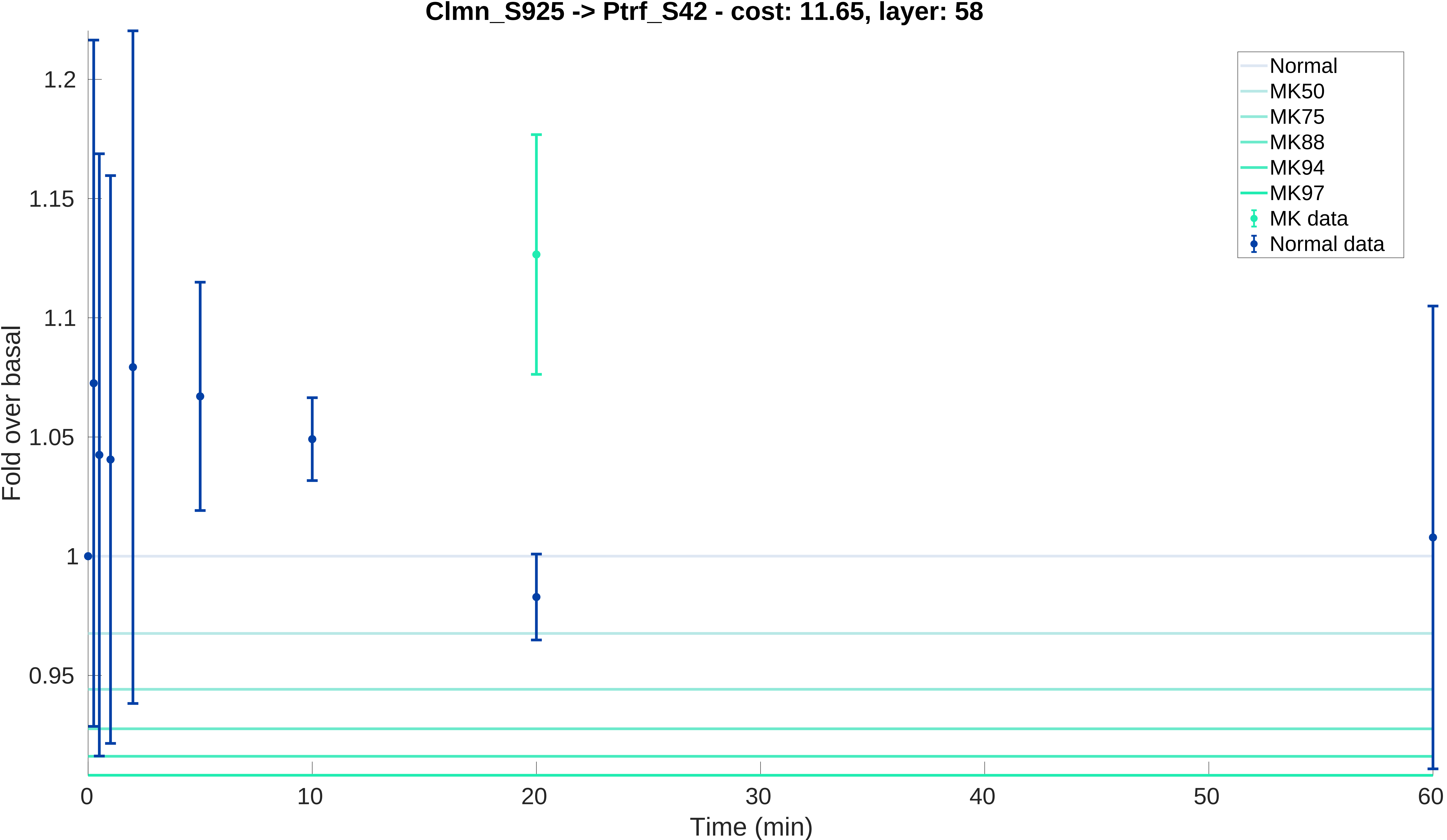
Clmn_S925 -> Ptrf_S42 - cost: 11.65, layer: 58
Fold over basal
1.2
1.15
1.1
1.05
1
0.95
Time (min)
0
10
20
30
40
50
60
Normal
MK50
MK75
MK88
MK94
MK97
MK data
Normal data

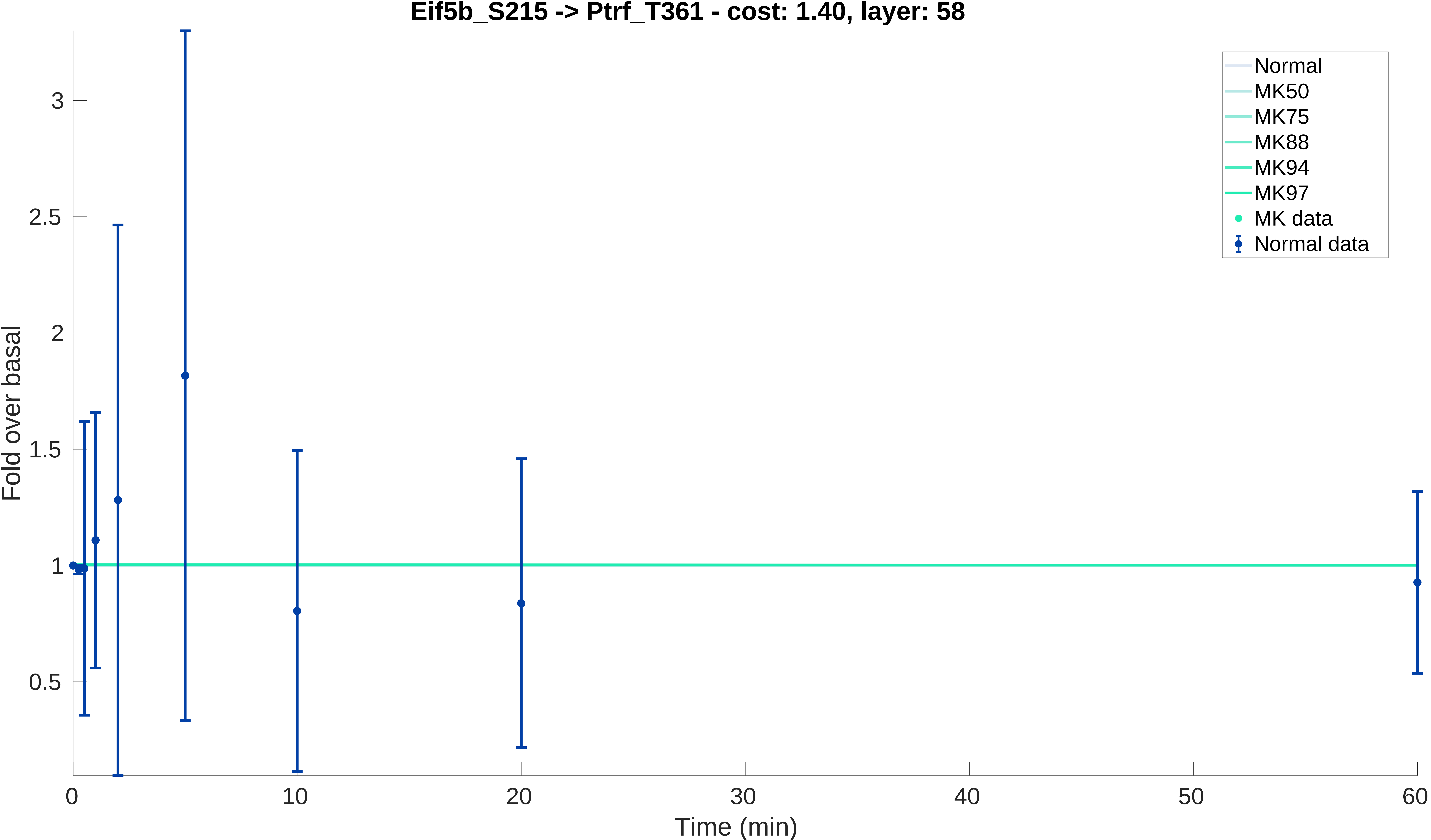
Eif5b_S215 -> Ptrf_T361 - cost: 1.40, layer: 58
Fold over basal
3
2.5
2
1.5
1
0.5
0
10
20
30
40
50
60
Time (min)
Normal
MK50
MK75
MK88
MK94
MK97
MK data
Normal data

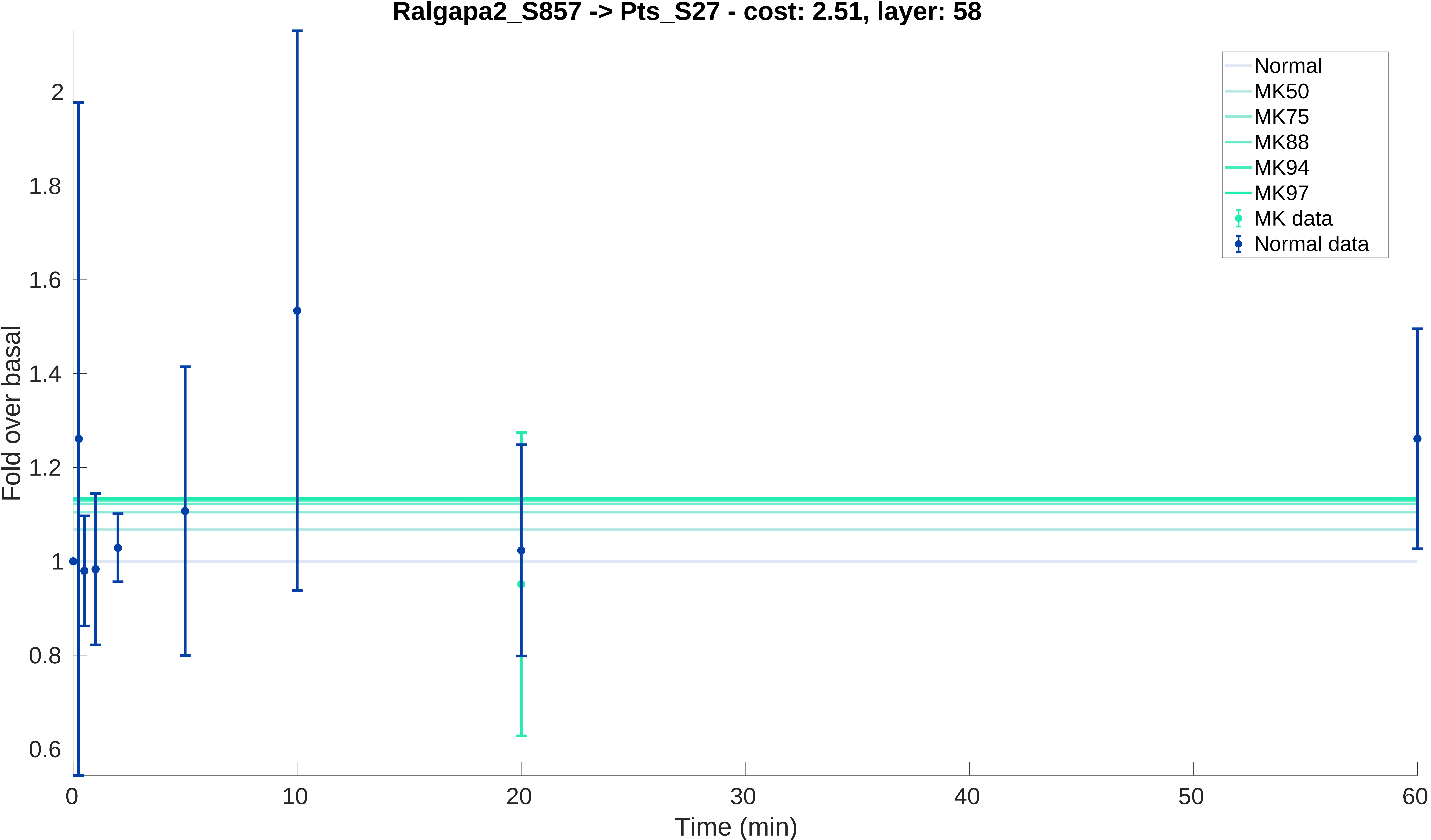

Ralgapa2_S857 -> Pts_S27 - cost: 2.51, layer: 58
Normal
MK50
MK75
MK88
MK94
MK97
MK data
Normal data
Fold over basal
2
1.8
1.6
1.4
1.2
1
0.8
0.6
0
10
20
30
40
50
60
Time (min)

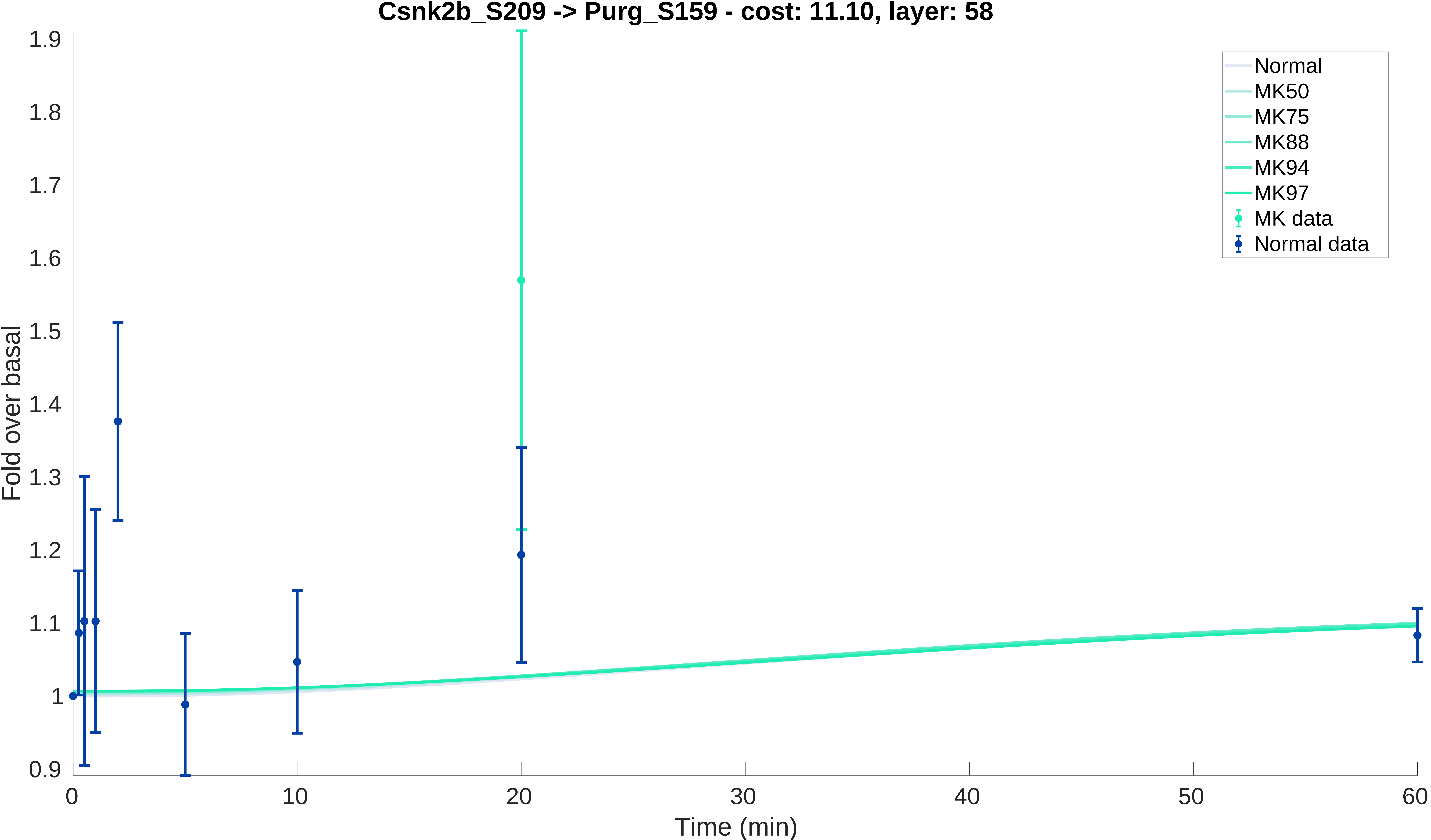

Csnk2b_S209 -> Purg_S159 - cost: 11.10, layer: 58
Fold over basal
Time (min)
0.9
1
1.1
1.2
1.3
1.4
1.5
1.6
1.7
1.8
1.9
0
10
20
30
40
50
60
Normal
MK50
MK75
MK88
MK94
MK97
MK data
Normal data

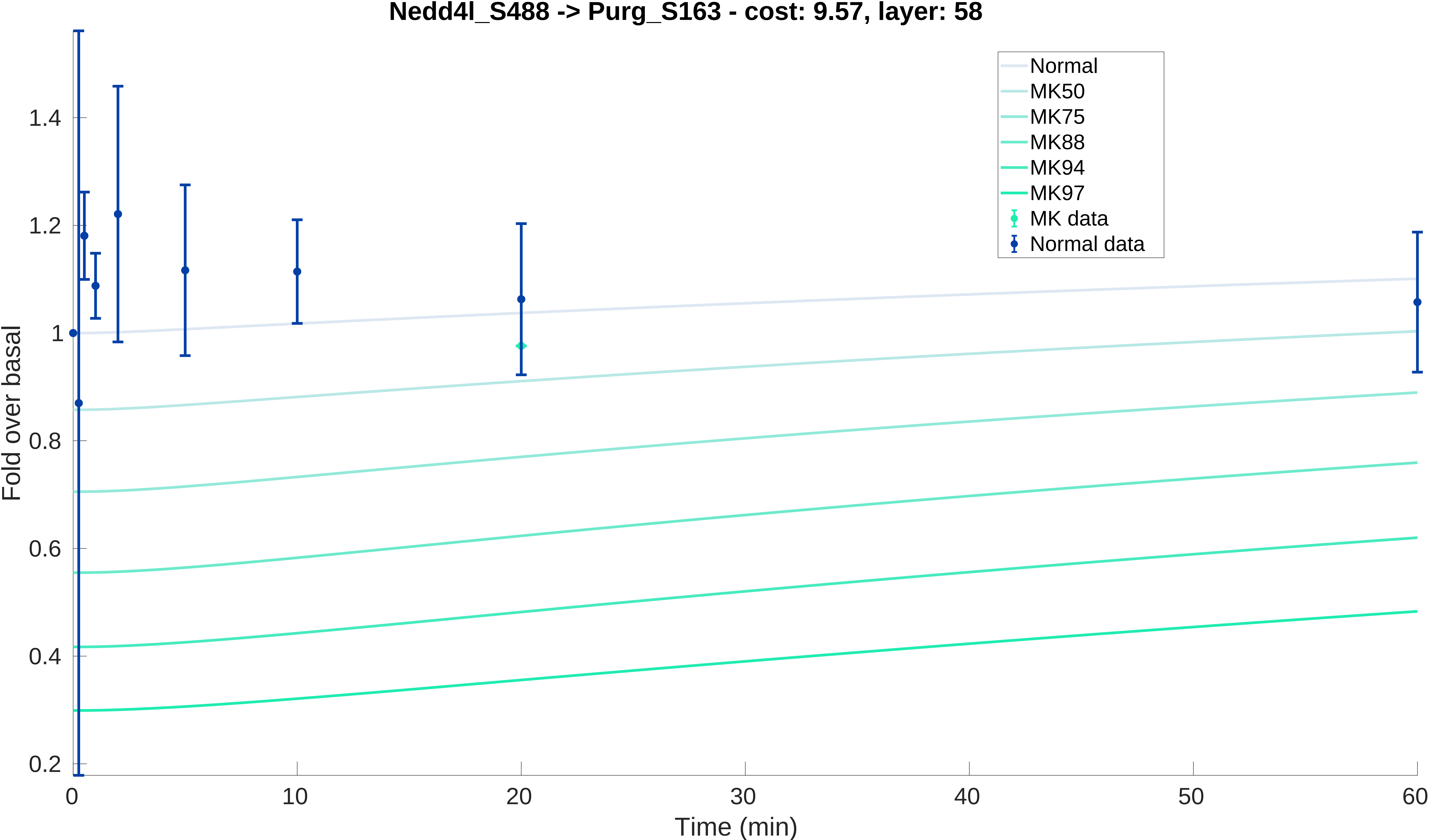
Nedd4l_S488 -> Purg_S163 - cost: 9.57, layer: 58
Normal
MK50
MK75
MK88
MK94
MK97
MK data
Normal data
Fold over basal
Time (min)
0.2
0.4
0.6
0.8
1
1.2
1.4
0
10
20
30
40
50
60

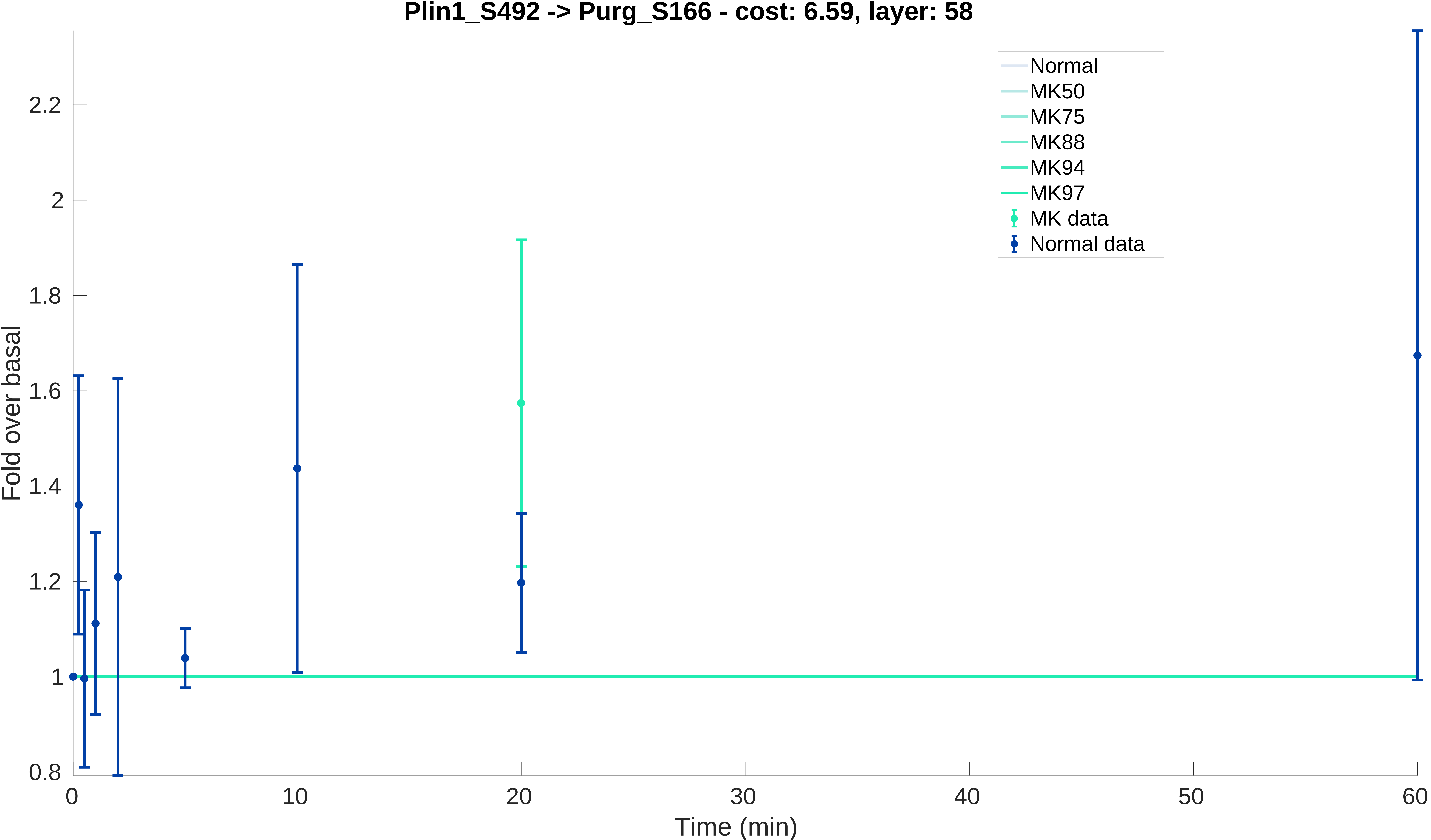

Plin1_S492 -> Purg_S166 - cost: 6.59, layer: 58
Normal
MK50
MK75
MK88
MK94
MK97
MK data
Normal data
Fold over basal
0.8
1
1.2
1.4
1.6
1.8
2
2.2
0
10
20
30
40
50
60
Time (min)

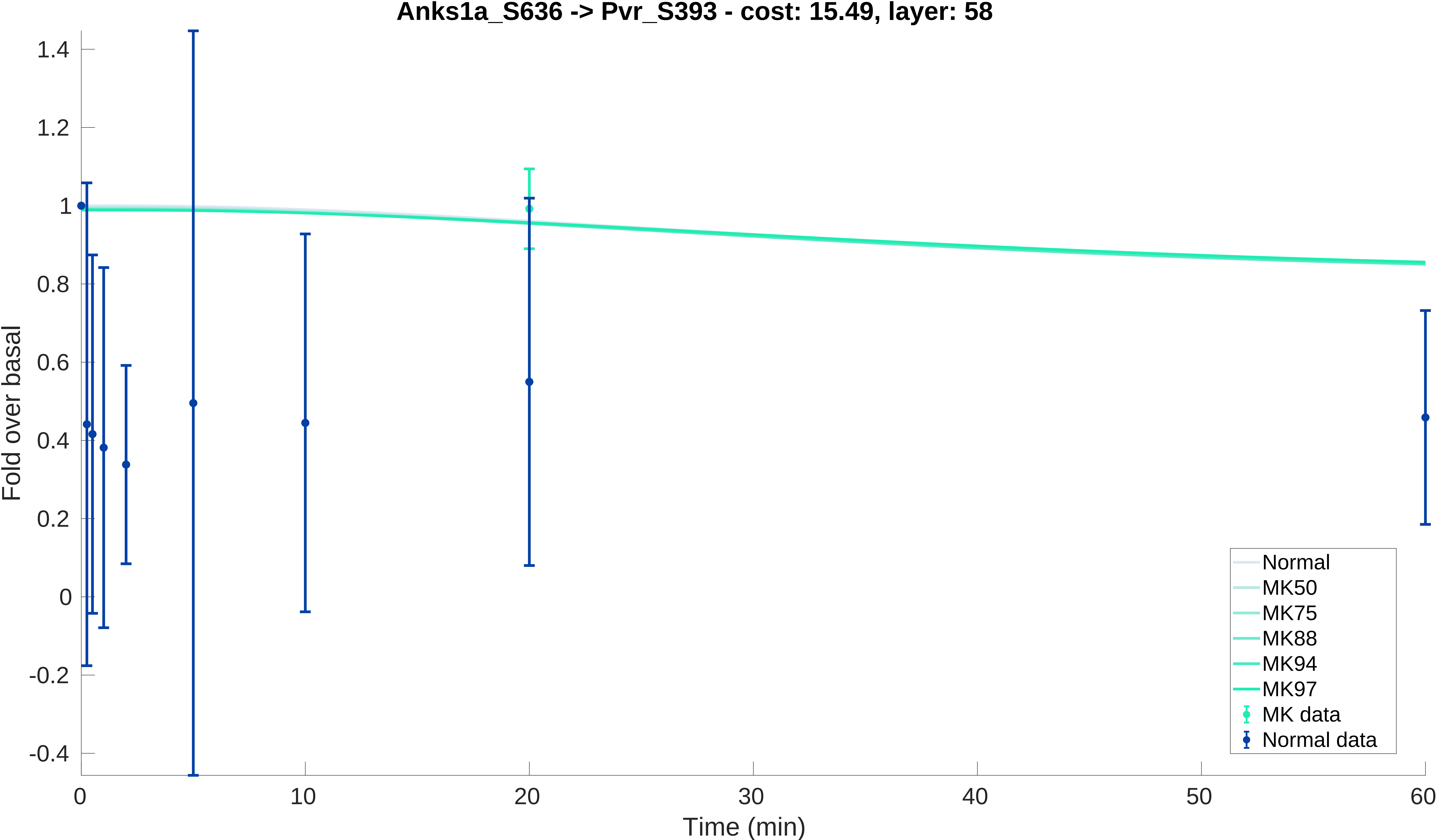

Anks1a_S636 -> Pvr_S393 - cost: 15.49, layer: 58
Fold over basal
Time (min)
1.4
1.2
1
0.8
0.6
0.4
0.2
0
-0.2
-0.4
0
10
20
30
40
50
60
Normal
MK50
MK75
MK88
MK94
MK97
MK data
Normal data

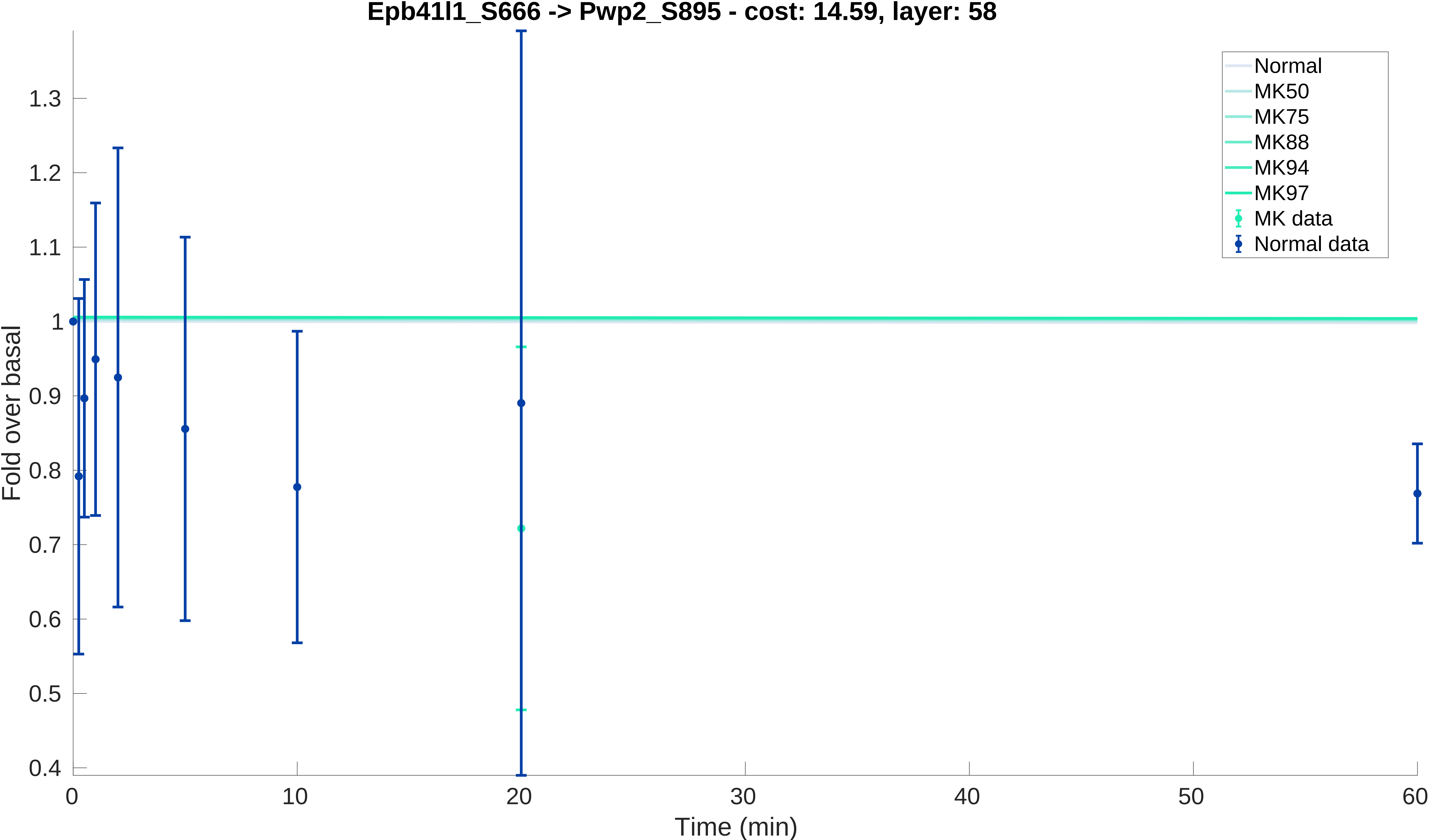
Epb41l1_S666 -> Pwp2_S895 - cost: 14.59, layer: 58
Fold over basal
Time (min)
1.3
1.2
1.1
1
0.9
0.8
0.7
0.6
0.5
0.4
0
10
20
30
40
50
60
Normal
MK50
MK75
MK88
MK94
MK97
MK data
Normal data

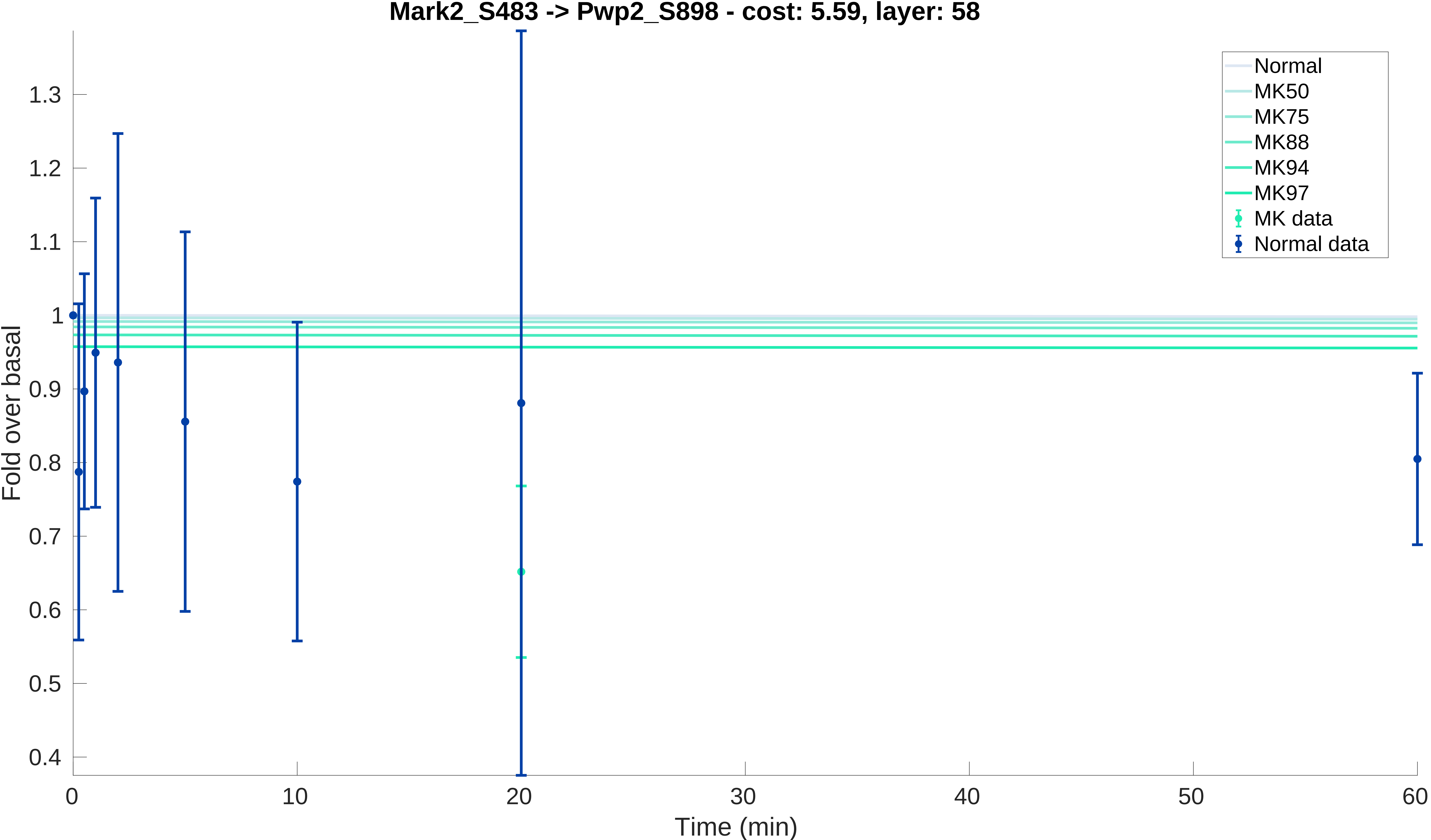
Mark2_S483 -> Pwp2_S898 - cost: 5.59, layer: 58
Normal
MK50
MK75
MK88
MK94
MK97
MK data
Normal data
Fold over basal
1.3
1.2
1.1
1
0.9
0.8
0.7
0.6
0.5
0.4
0
10
20
30
40
50
60
Time (min)

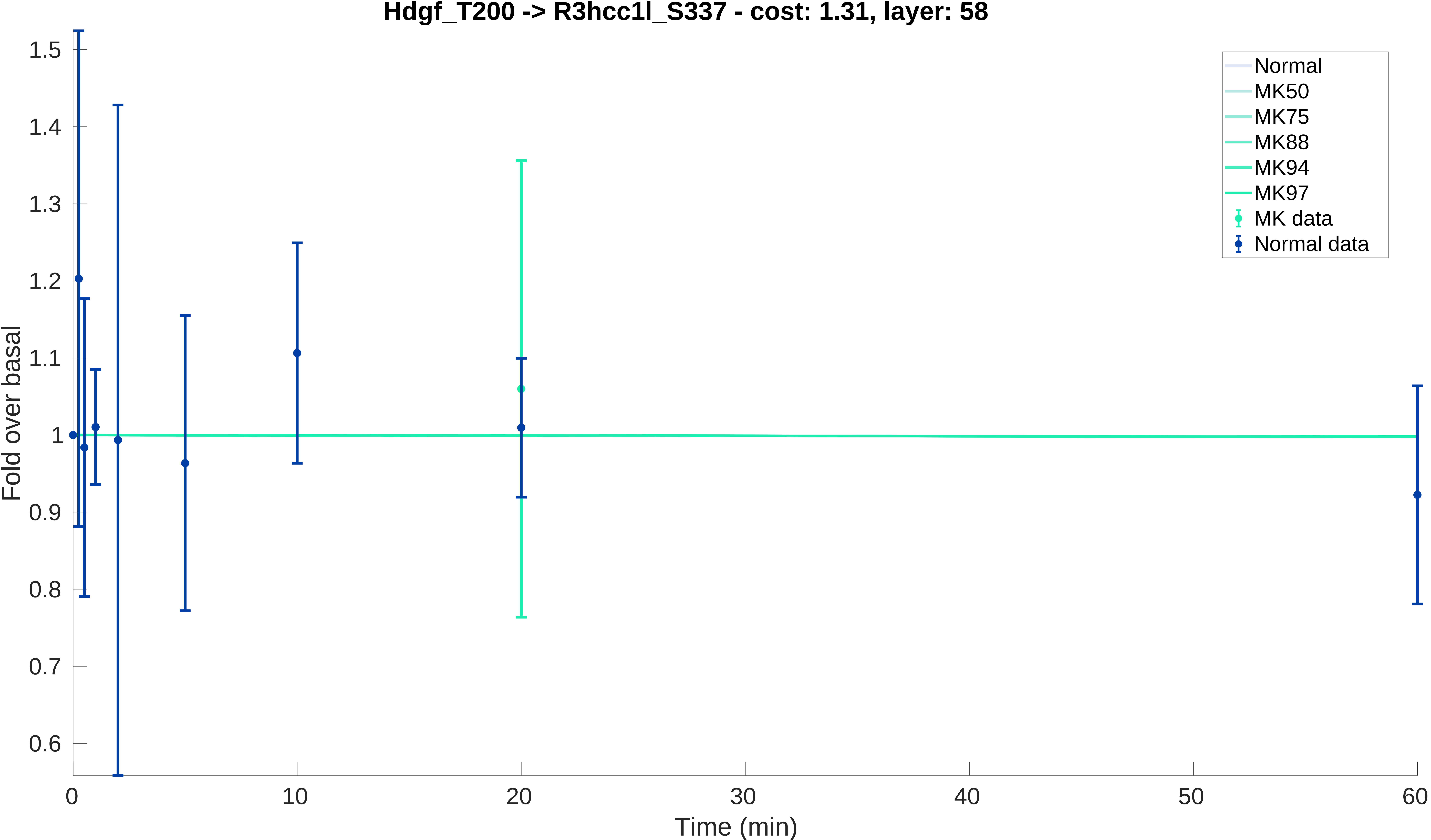

Hdgf_T200 -> R3hcc1l_S337 - cost: 1.31, layer: 58
Normal
MK50
MK75
MK88
MK94
MK97
MK data
Normal data
Fold over basal
Time (min)
1.5
1.4
1.3
1.2
1.1
1
0.9
0.8
0.7
0.6
0
10
20
30
40
50
60

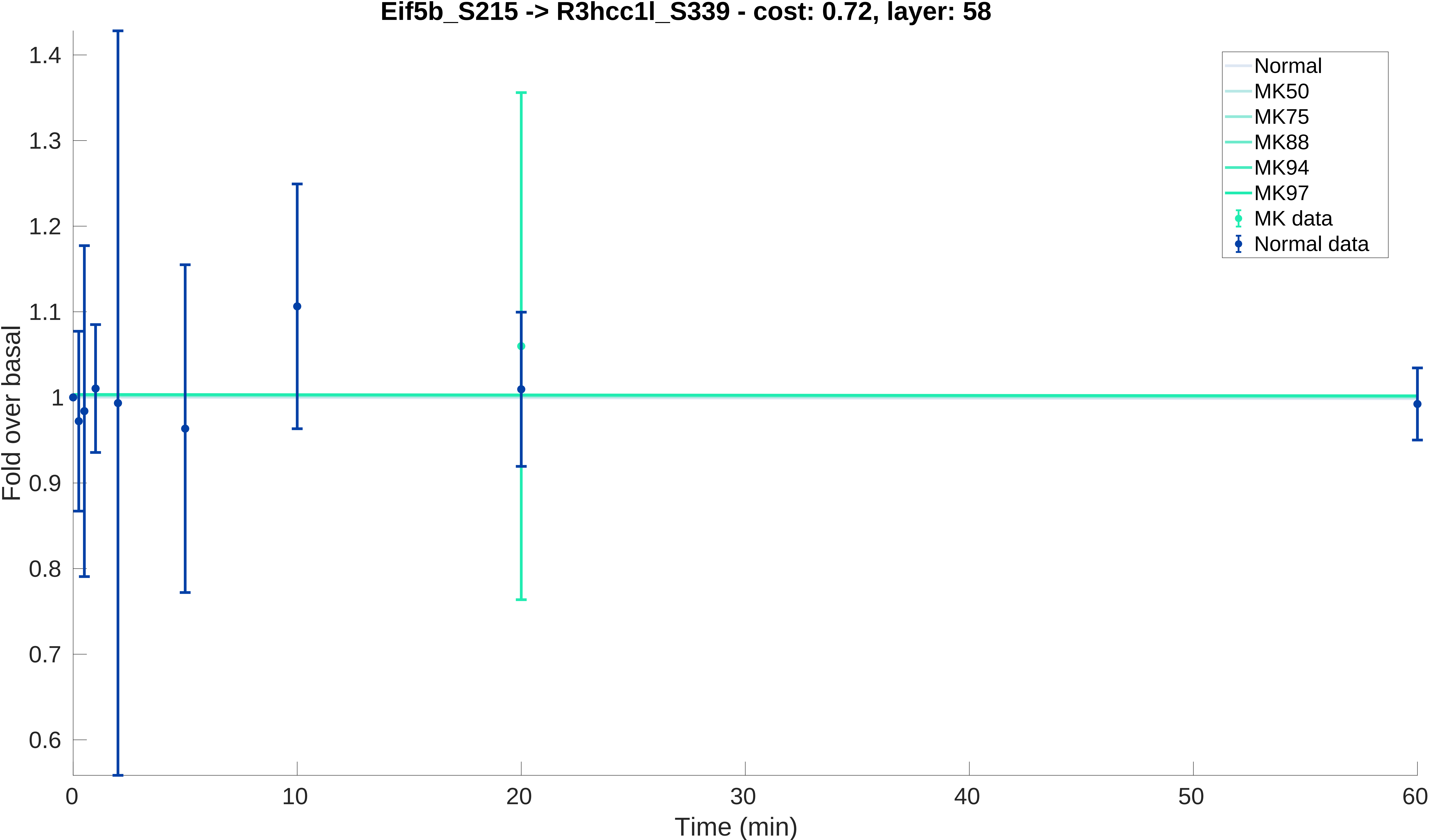

Eif5b_S215 -> R3hcc1l_S339 - cost: 0.72, layer: 58
Fold over basal
Time (min)
Normal
MK50
MK75
MK88
MK94
MK97
MK data
Normal data
1.4
1.3
1.2
1.1
1
0.9
0.8
0.7
0.6
0
10
20
30
40
50
60

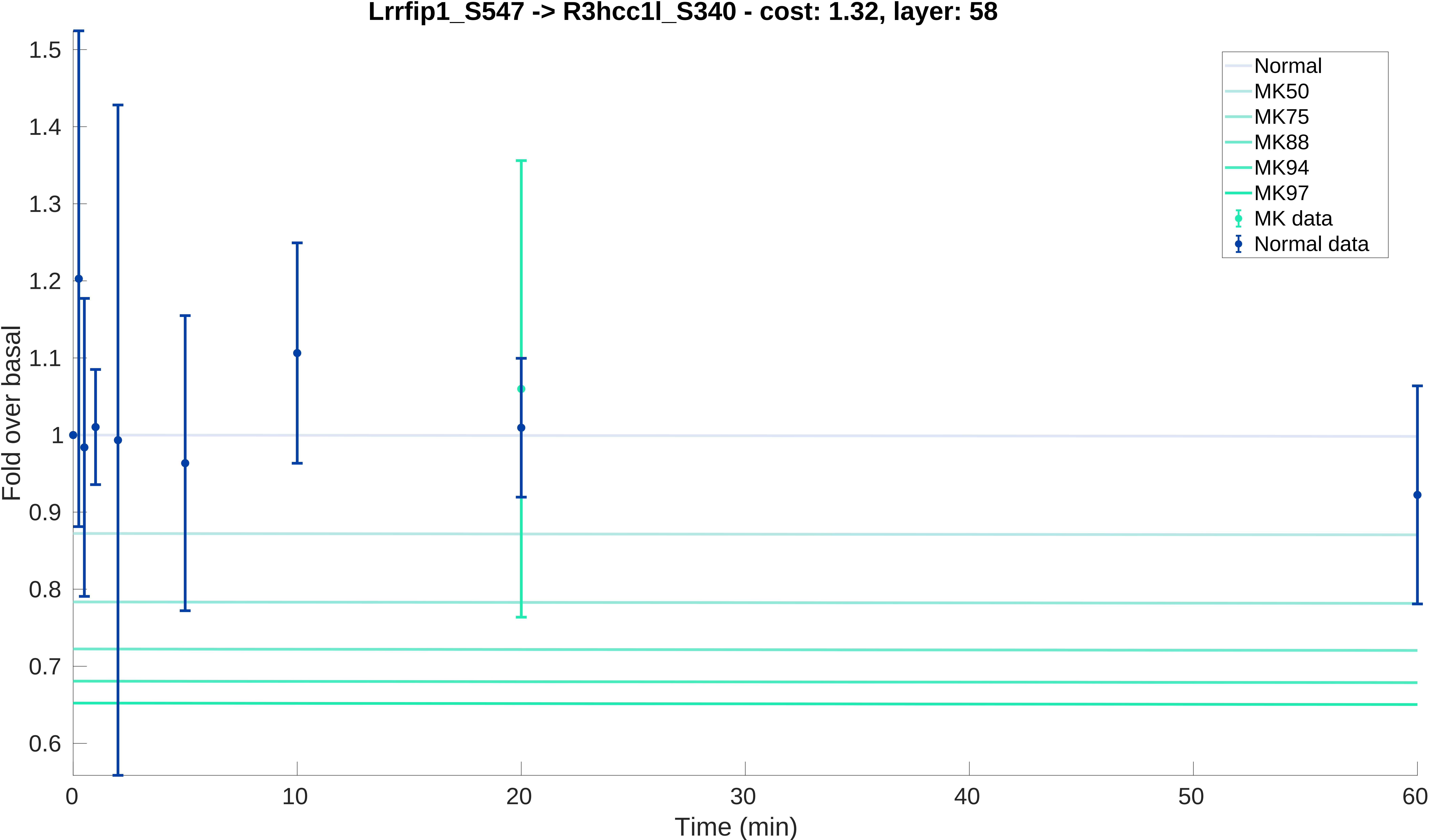
Lrrfip1_S547 -> R3hcc1l_S340 - cost: 1.32, layer: 58
Fold over basal
1.5
1.4
1.3
1.2
1.1
1
0.9
0.8
0.7
0.6
Time (min)
0
10
20
30
40
50
60
Normal
MK50
MK75
MK88
MK94
MK97
MK data
Normal data

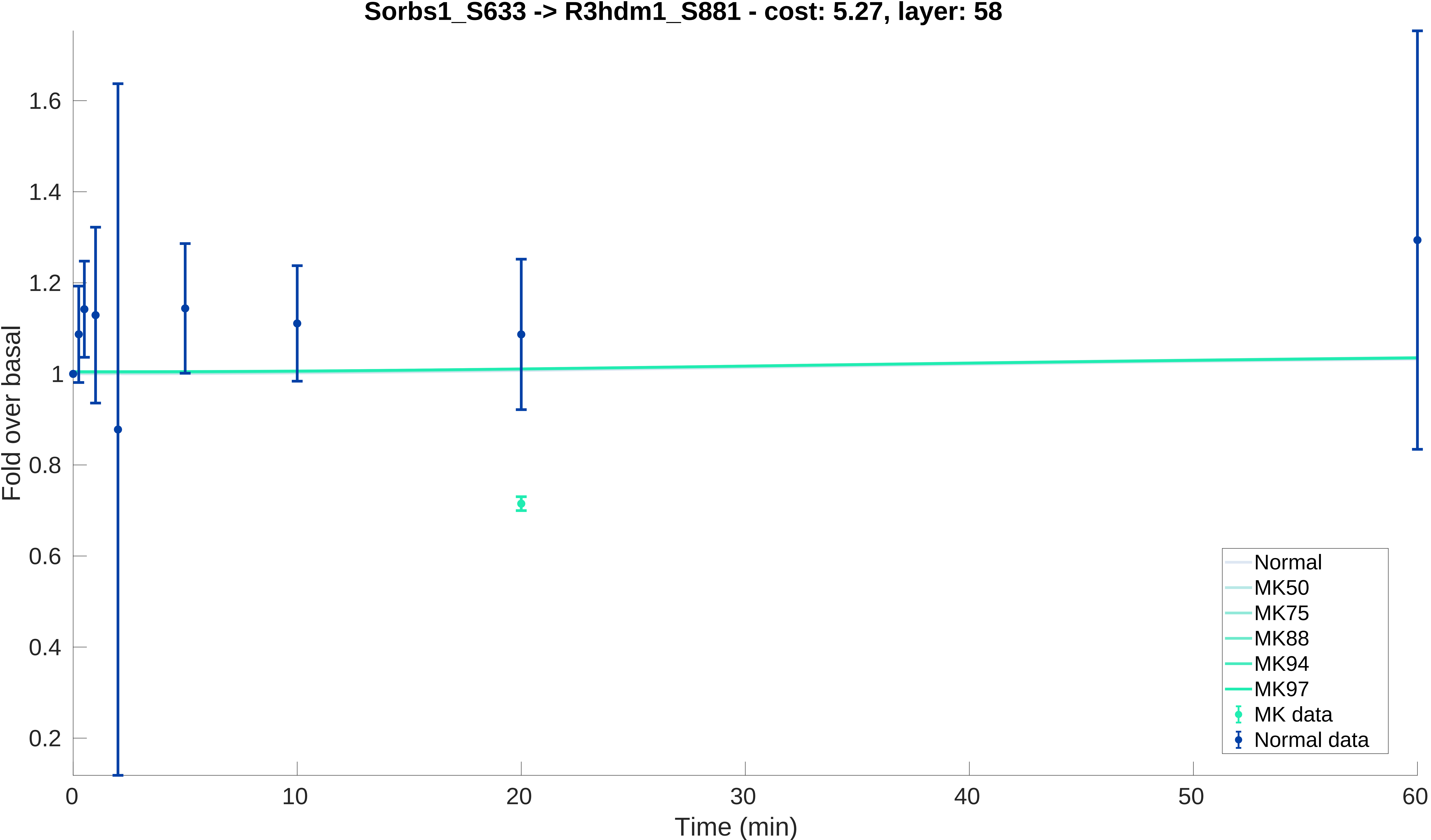

Sorbs1_S633 -> R3hdm1_S881 - cost: 5.27, layer: 58
Fold over basal
1.6
1.4
1.2
1
0.8
0.6
0.4
0.2
0
10
20
30
40
50
60
Time (min)
Normal
MK50
MK75
MK88
MK94
MK97
MK data
Normal data

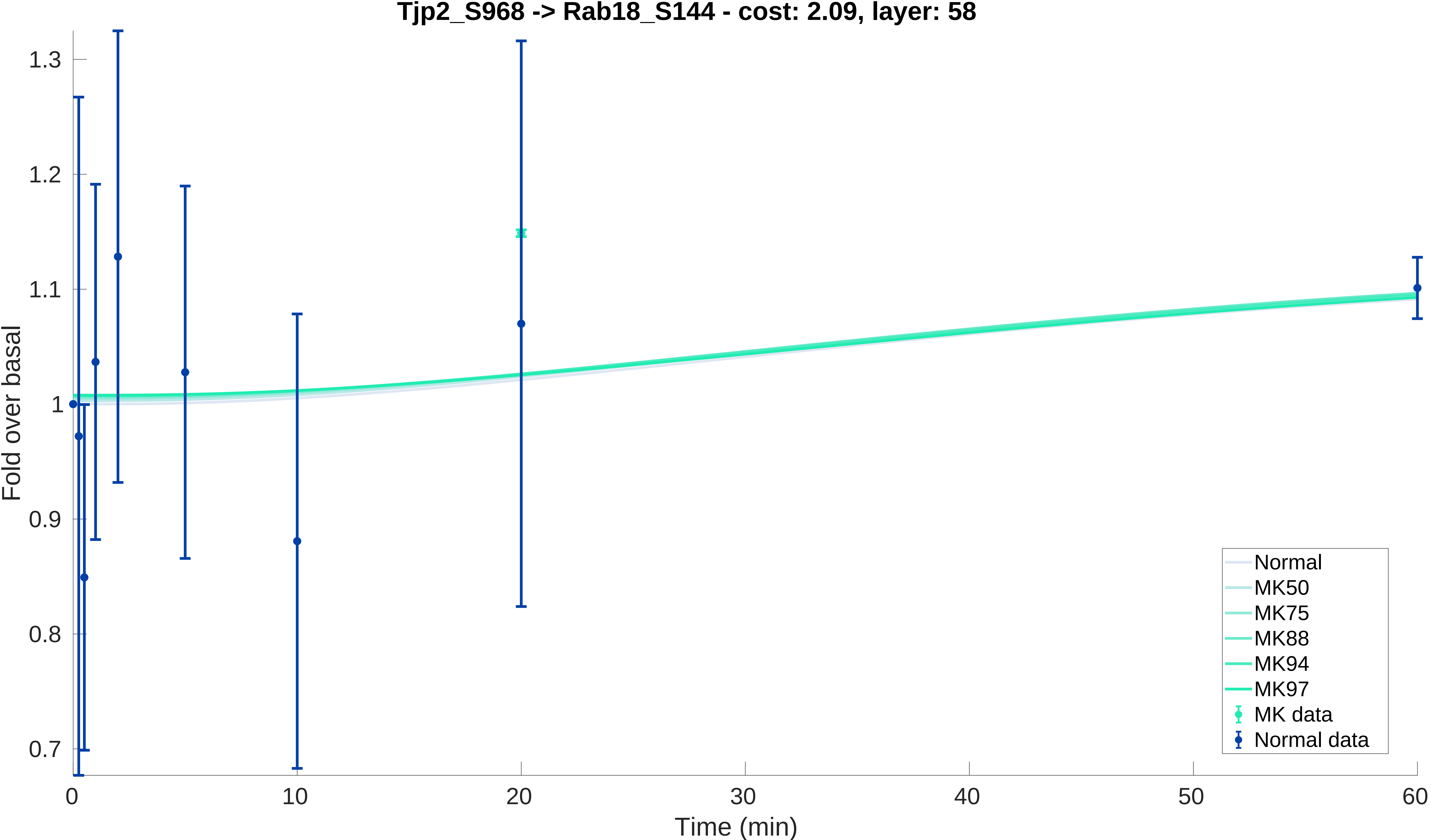

Tjp2_S968 -> Rab18_S144 - cost: 2.09, layer: 58
Fold over basal
Time (min)
1.3
1.2
1.1
1
0.9
0.8
0.7
0
10
20
30
40
50
60
Normal
MK50
MK75
MK88
MK94
MK97
MK data
Normal data

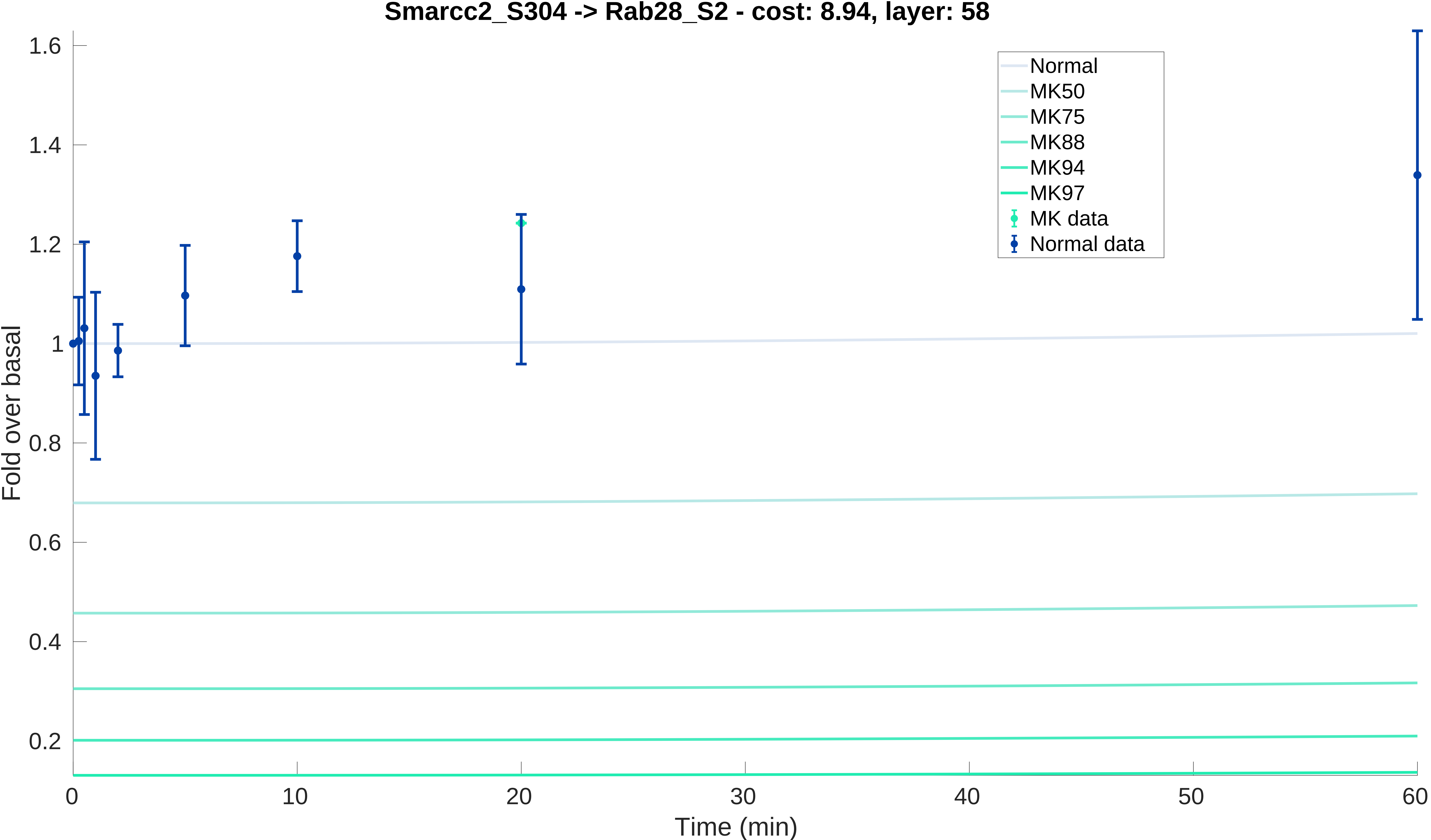

Smarcc2_S304 -> Rab28_S2 - cost: 8.94, layer: 58
Normal
MK50
MK75
MK88
MK94
MK97
MK data
Normal data
Fold over basal
Time (min)
1.6
1.4
1.2
1
0.8
0.6
0.4
0.2
0
10
20
30
40
50
60

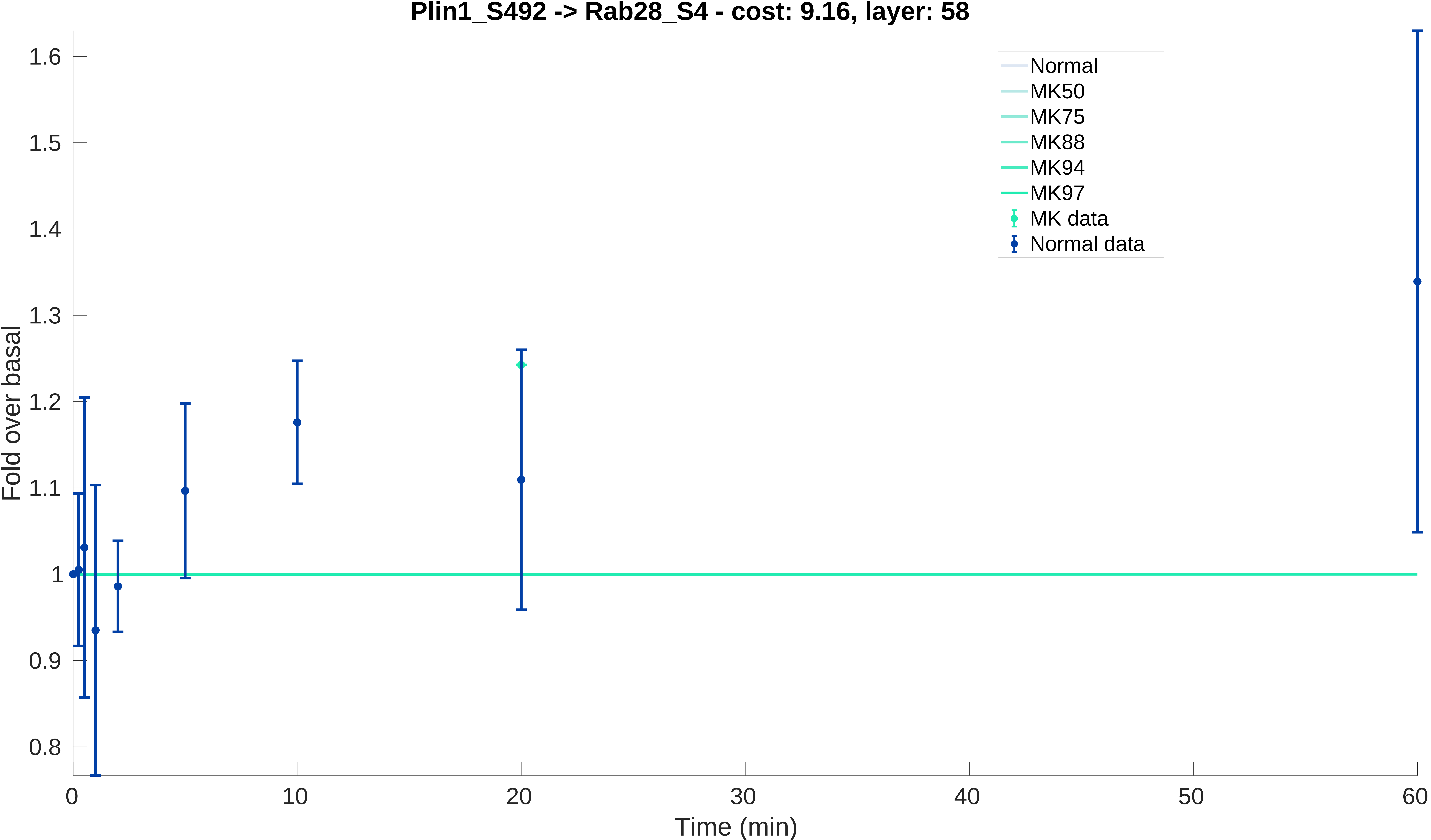
Plin1_S492 -> Rab28_S4 - cost: 9.16, layer: 58
Fold over basal
Time (min)
0.8
0.9
1
1.1
1.2
1.3
1.4
1.5
1.6
0
10
20
30
40
50
60
Normal
MK50
MK75
MK88
MK94
MK97
MK data
Normal data

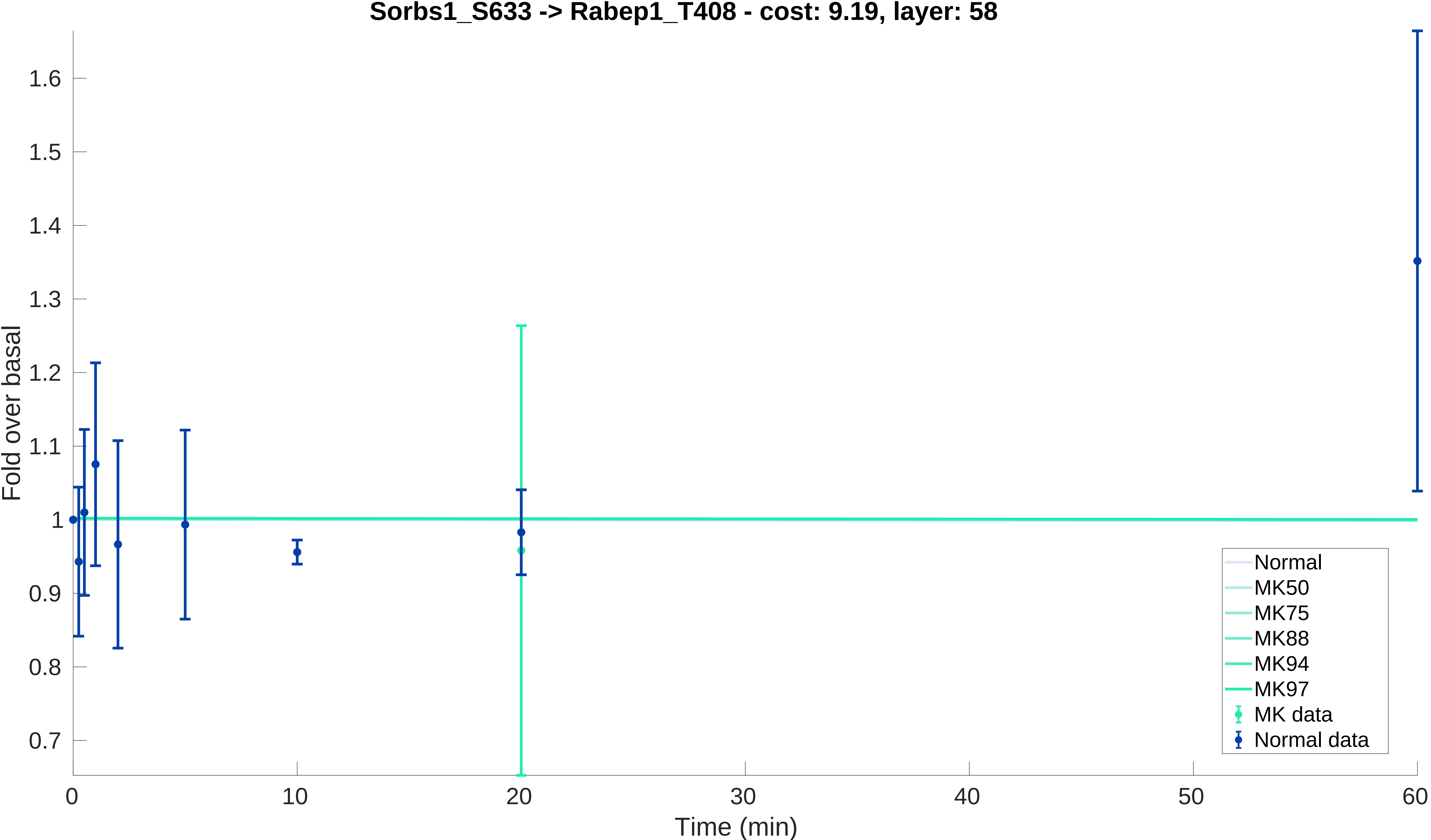
Sorbs1_S633 -> Rabep1_T408 - cost: 9.19, layer: 58
Fold over basal
1.6
1.5
1.4
1.3
1.2
1.1
1
0.9
0.8
0.7
0
10
20
30
40
50
60
Time (min)
Normal
MK50
MK75
MK88
MK94
MK97
MK data
Normal data

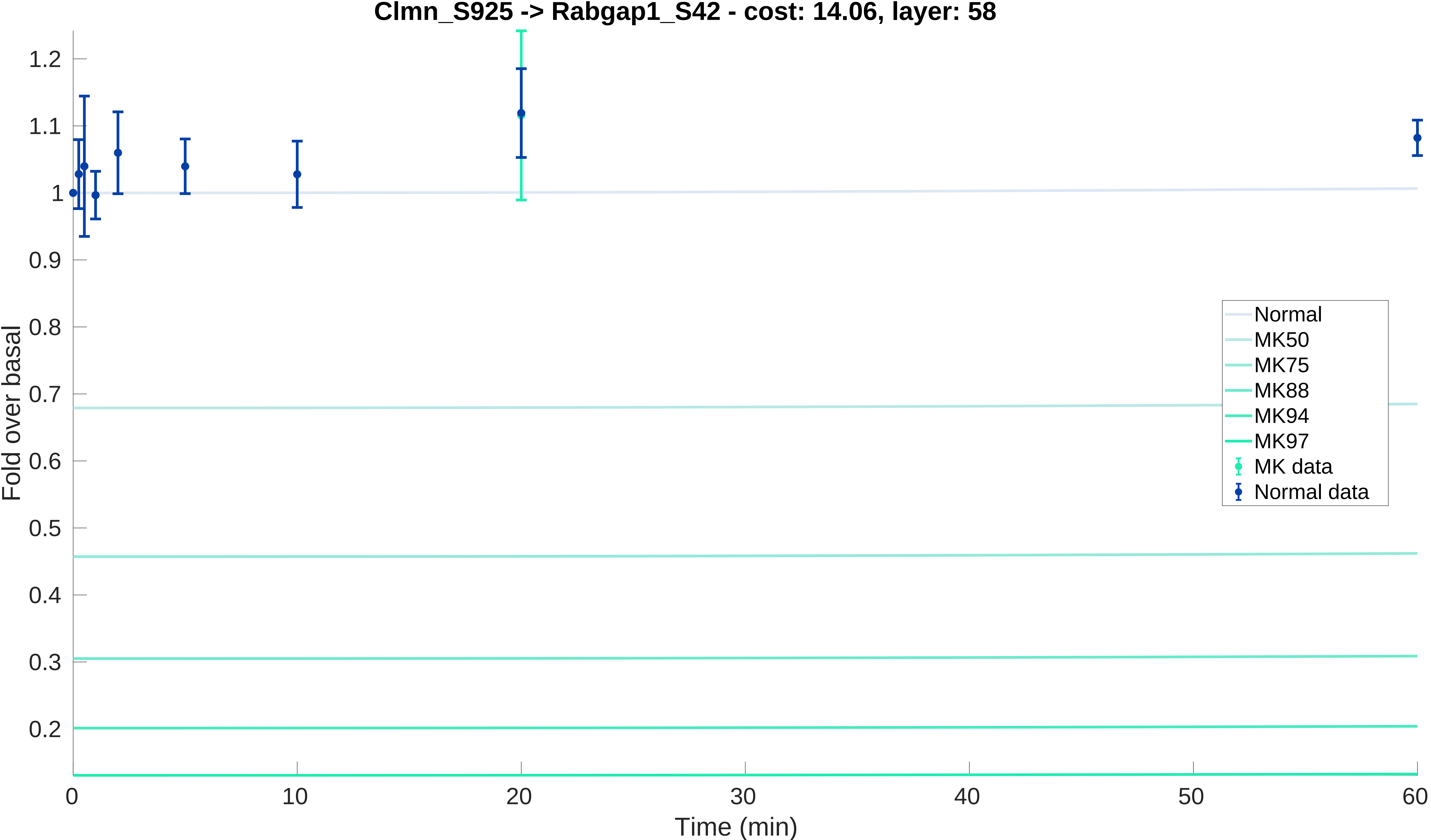

Clmn_S925 -> Rabgap1_S42 - cost: 14.06, layer: 58
Fold over basal
1.2
1.1
1
0.9
0.8
0.7
0.6
0.5
0.4
0.3
0.2
0
10
20
30
40
50
60
Time (min)
Normal
MK50
MK75
MK88
MK94
MK97
MK data
Normal data

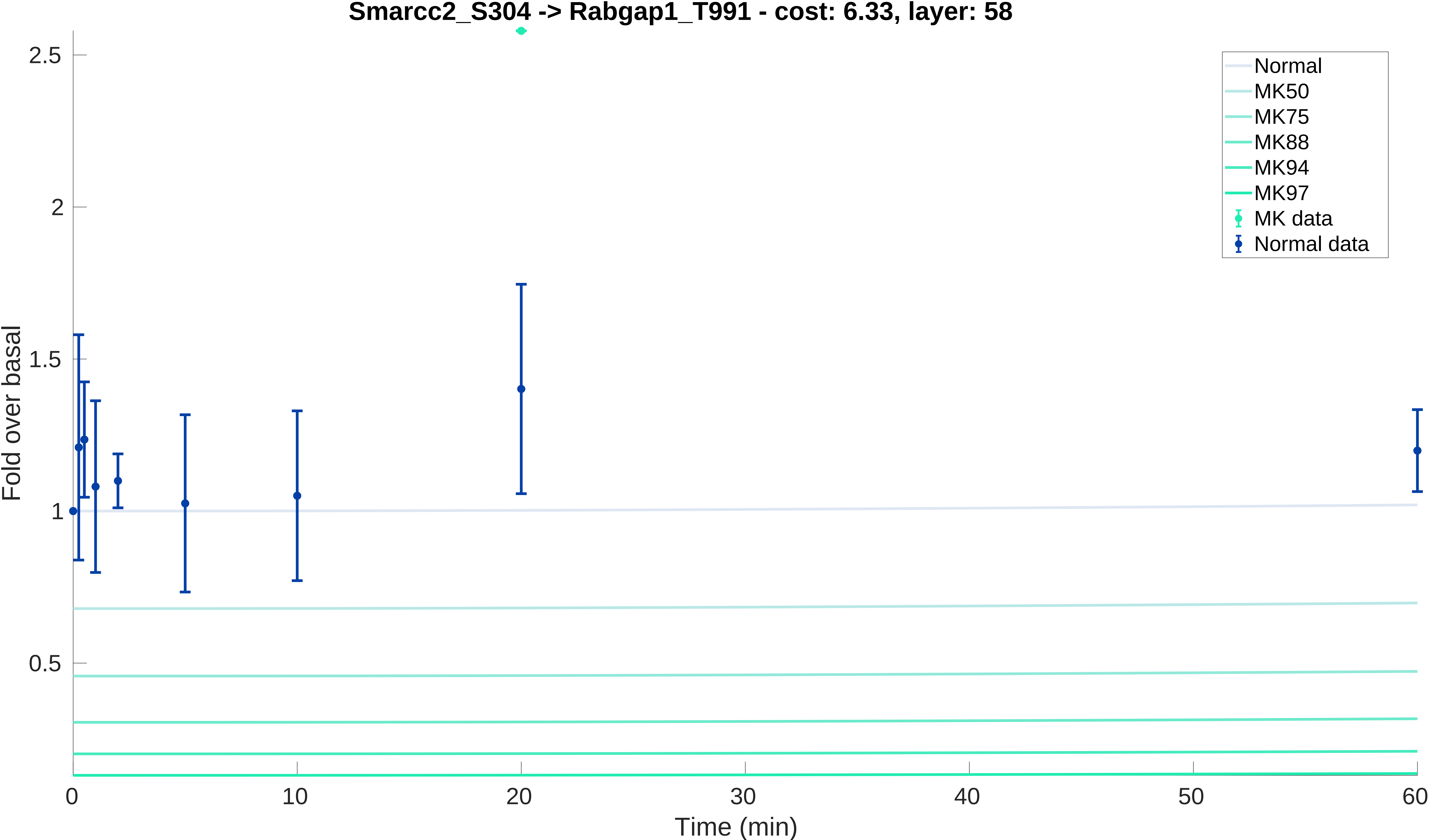

Smarcc2_S304 -> Rabgap1_T991 - cost: 6.33, layer: 58
Fold over basal
2.5
2
1.5
1
0.5
0
10
20
30
40
50
60
Time (min)
Normal
MK50
MK75
MK88
MK94
MK97
MK data
Normal data

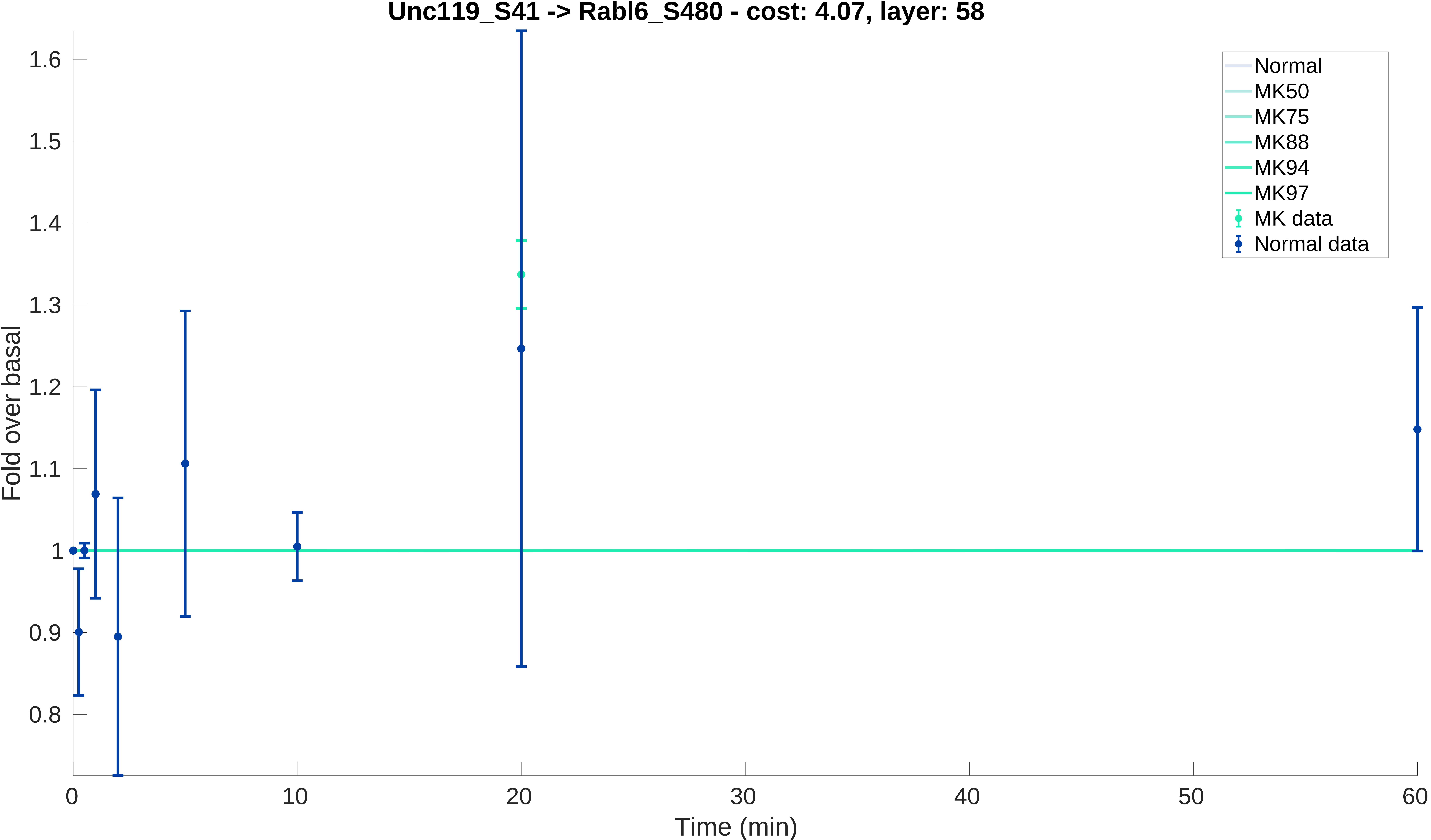

Unc119_S41 -> Rabl6_S480 - cost: 4.07, layer: 58
Fold over basal
Time (min)
1.6
1.5
1.4
1.3
1.2
1.1
1
0.9
0.8
0
10
20
30
40
50
60
Normal
MK50
MK75
MK88
MK94
MK97
MK data
Normal data

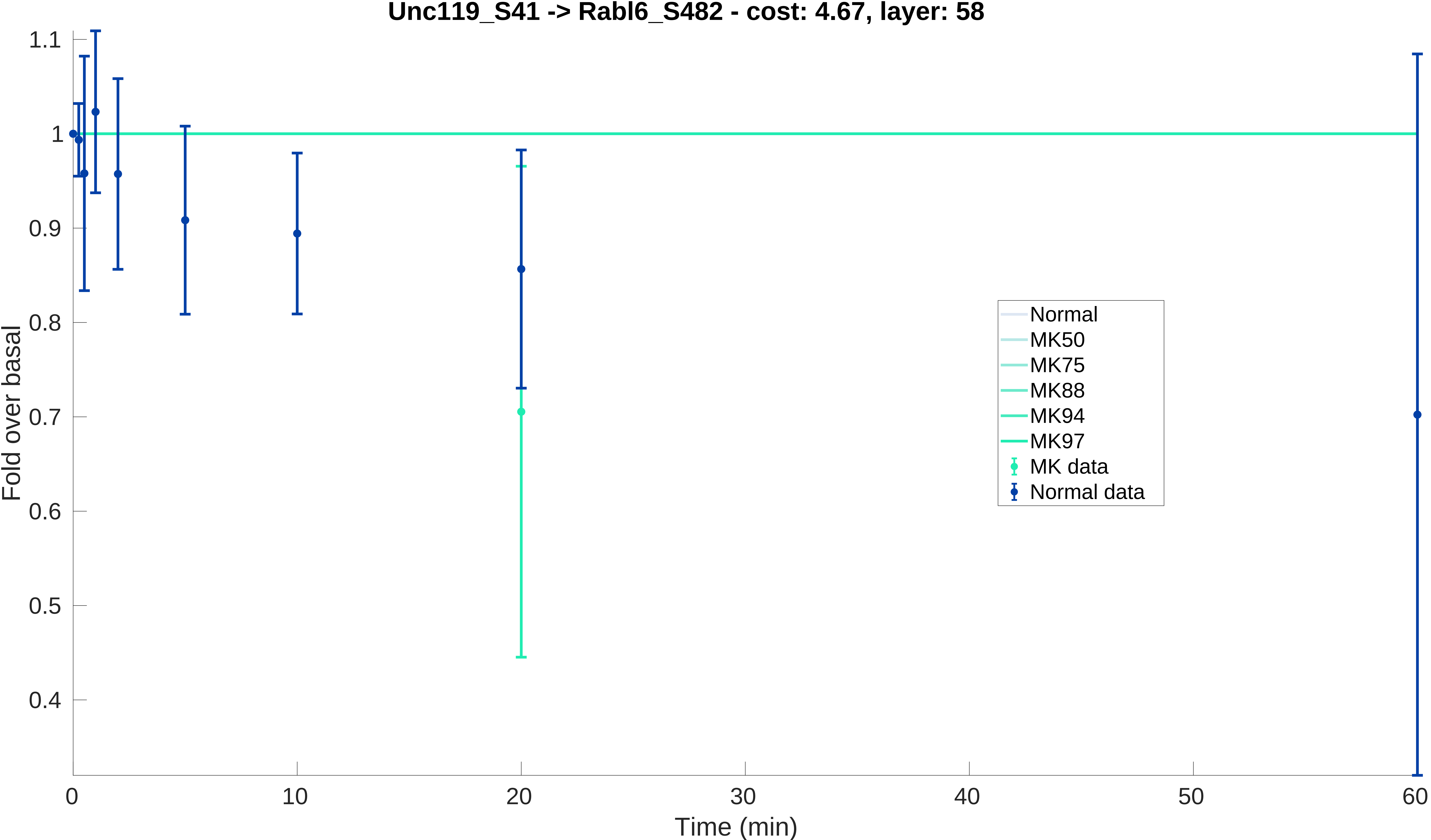
Unc119_S41 -> Rabl6_S482 - cost: 4.67, layer: 58
Fold over basal
1.1
1
0.9
0.8
0.7
0.6
0.5
0.4
0
10
20
30
40
50
60
Time (min)
Normal
MK50
MK75
MK88
MK94
MK97
MK data
Normal data

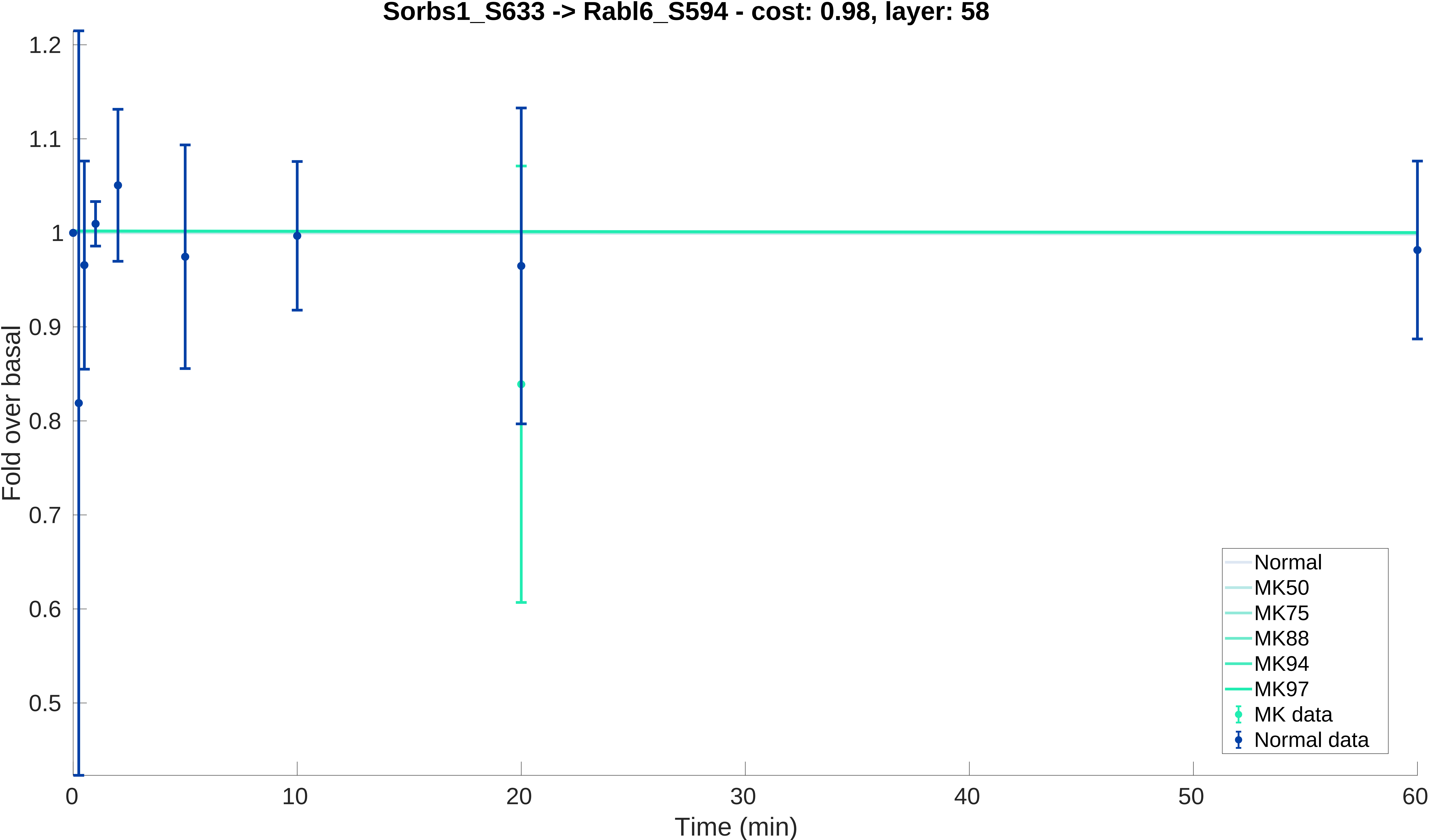
Sorbs1_S633 -> Rabl6_S594 - cost: 0.98, layer: 58
Fold over basal
1.2
1.1
1
0.9
0.8
0.7
0.6
0.5
0
10
20
30
40
50
60
Time (min)
Normal
MK50
MK75
MK88
MK94
MK97
MK data
Normal data

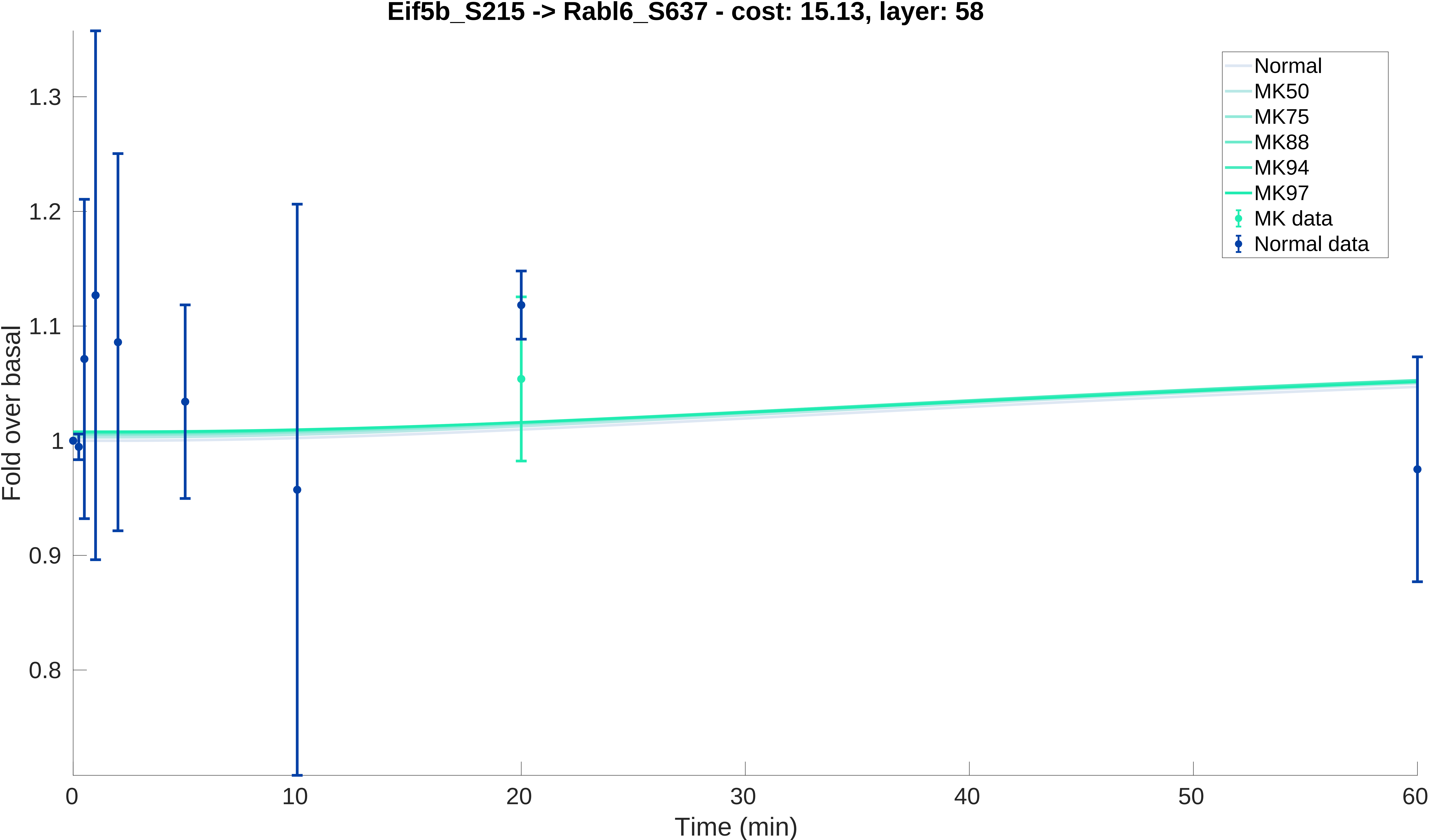
Eif5b_S215 -> Rabl6_S637 - cost: 15.13, layer: 58
Fold over basal
1.3
1.2
1.1
1
0.9
0.8
0
10
20
30
40
50
60
Time (min)
Normal
MK50
MK75
MK88
MK94
MK97
MK data
Normal data

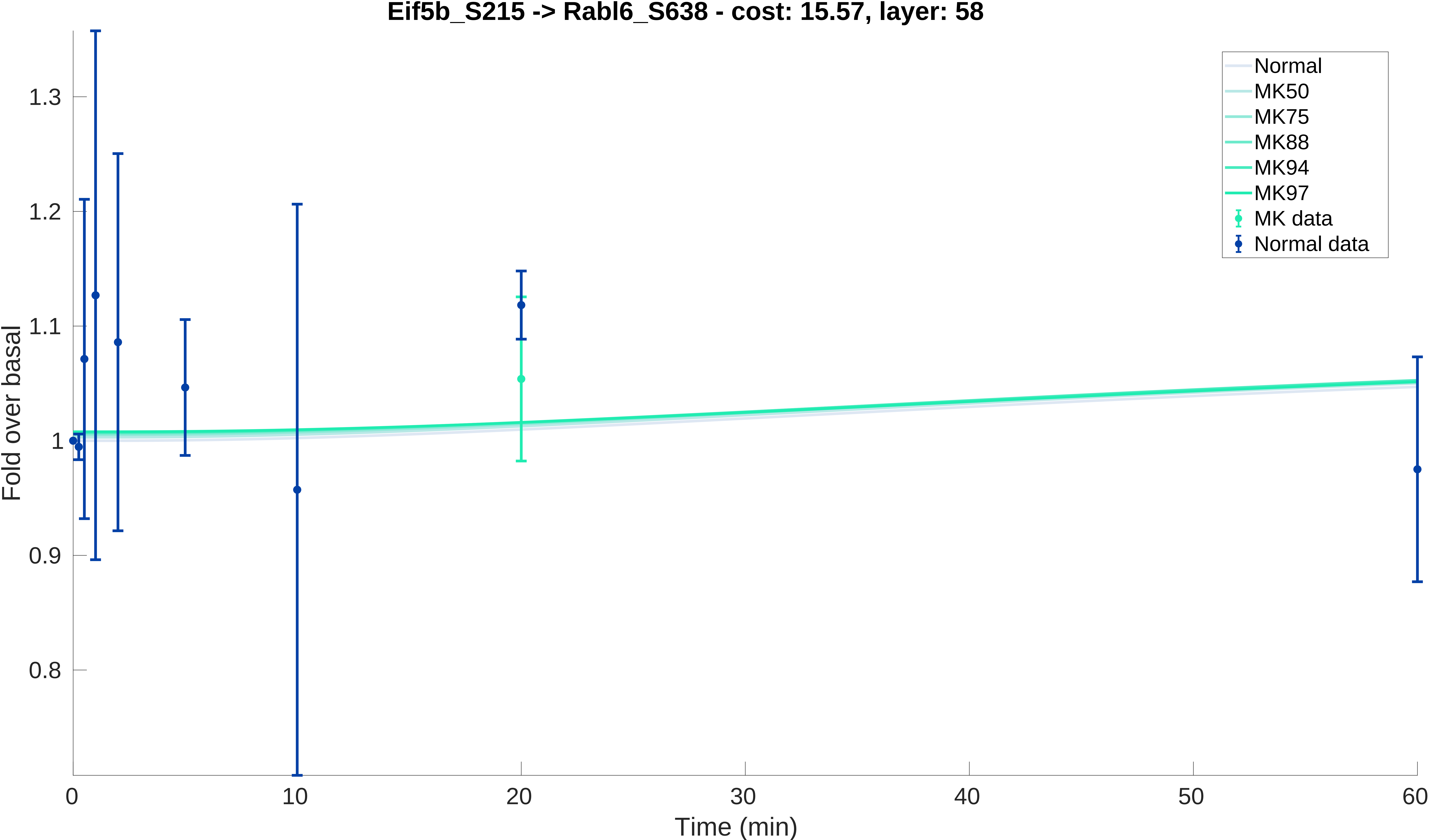

Eif5b_S215 -> Rabl6_S638 - cost: 15.57, layer: 58
Normal
MK50
MK75
MK88
MK94
MK97
MK data
Normal data
Fold over basal
Time (min)
1.3
1.2
1.1
1
0.9
0.8
0
10
20
30
40
50
60

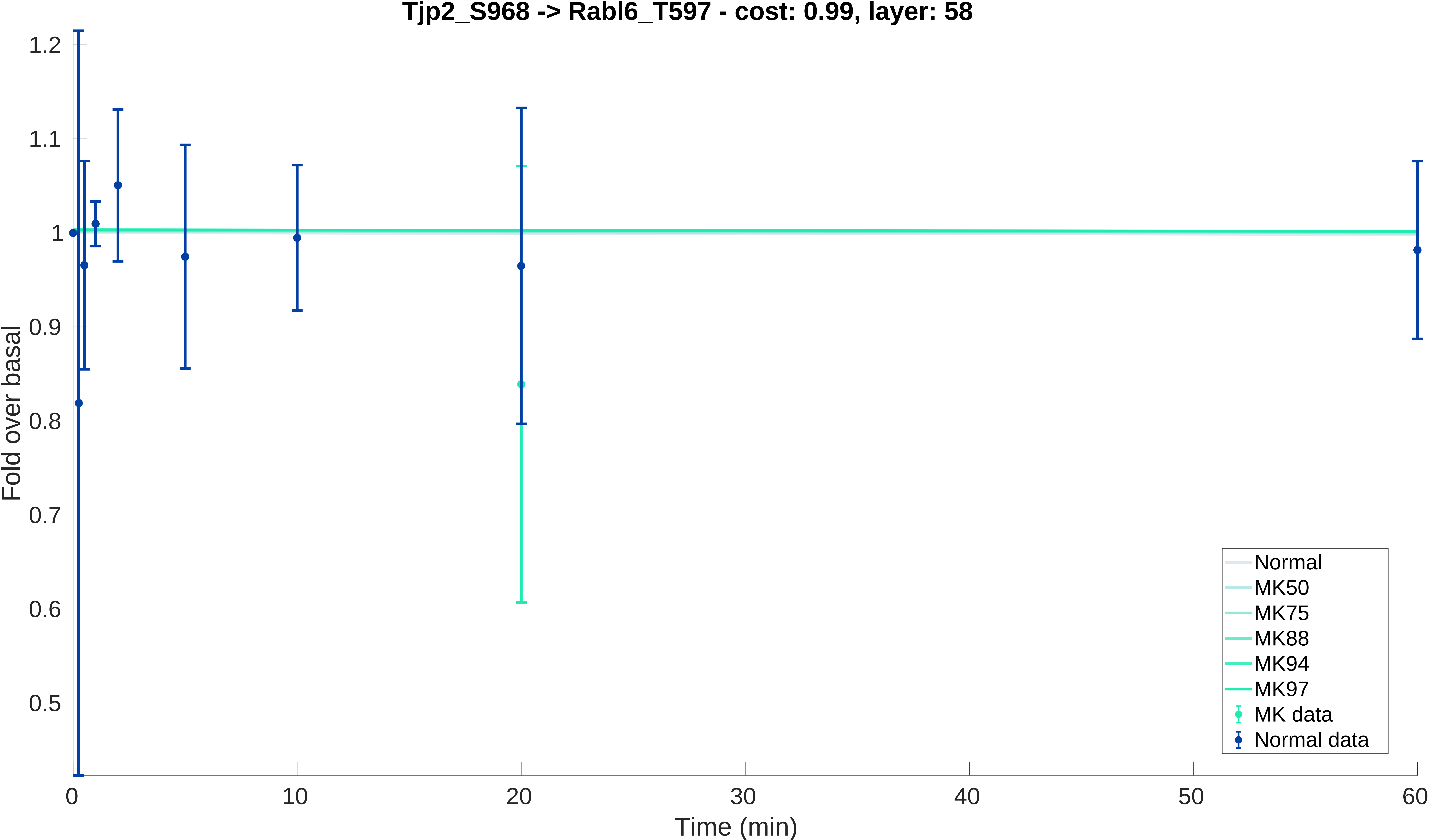
Tjp2_S968 -> Rabl6_T597 - cost: 0.99, layer: 58
Fold over basal
Time (min)
1.2
1.1
1
0.9
0.8
0.7
0.6
0.5
0
10
20
30
40
50
60
Normal
MK50
MK75
MK88
MK94
MK97
MK data
Normal data

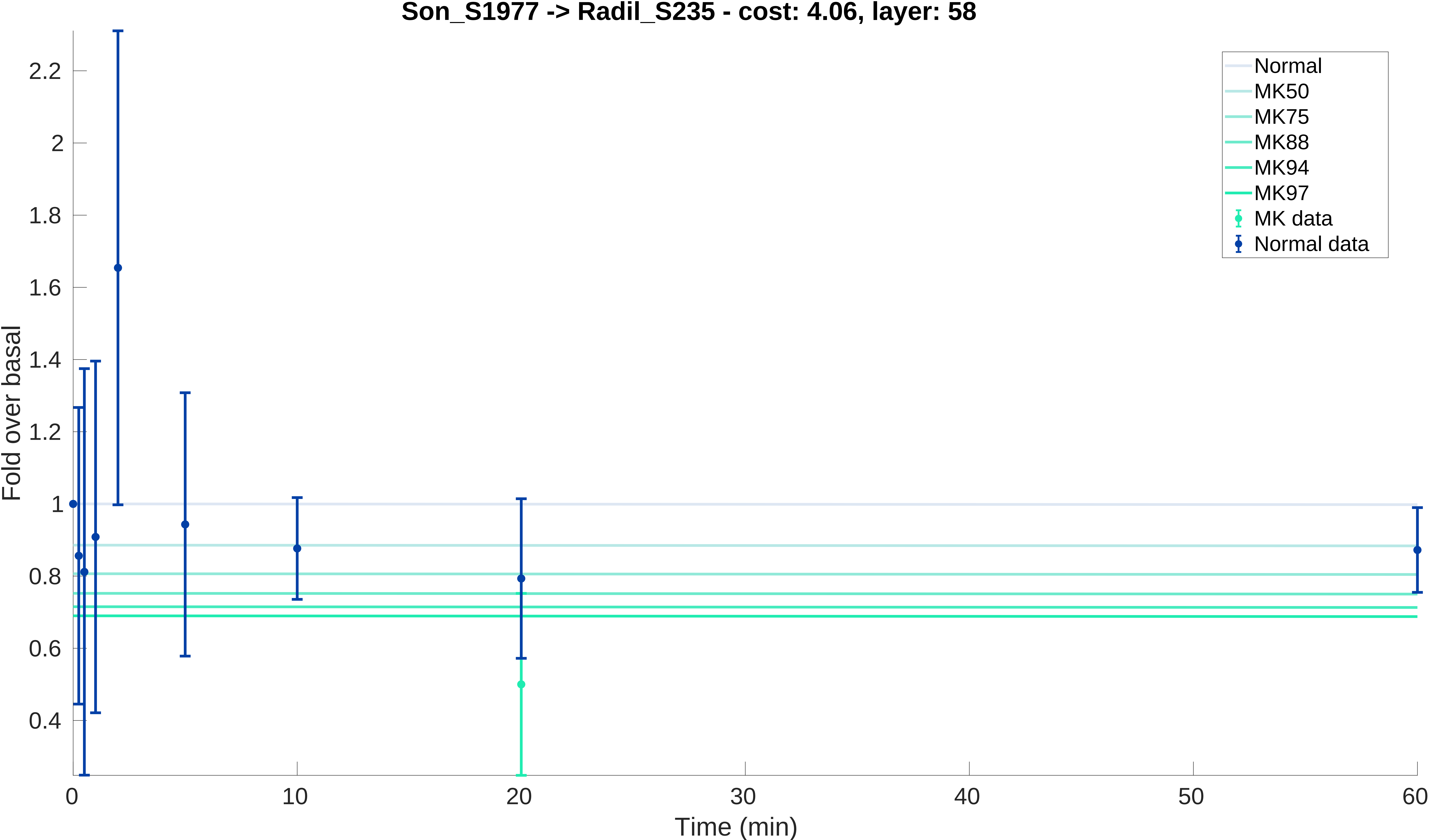
Son_S1977 -> Radil_S235 - cost: 4.06, layer: 58
Fold over basal
Time (min)
0.4
0.6
0.8
1
1.2
1.4
1.6
1.8
2
2.2
0
10
20
30
40
50
60
Normal
MK50
MK75
MK88
MK94
MK97
MK data
Normal data

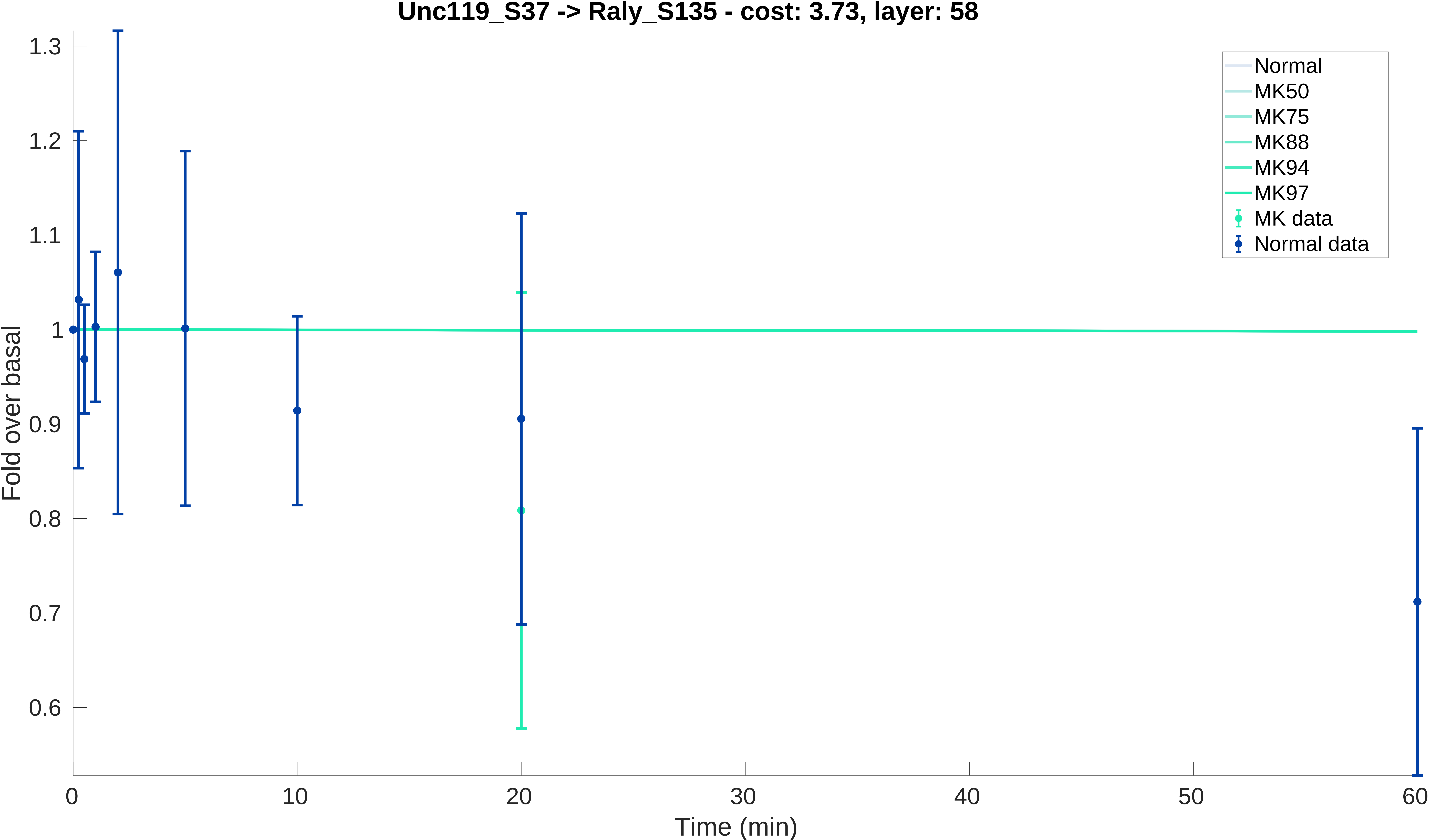

Unc119_S37 -> Raly_S135 - cost: 3.73, layer: 58
Fold over basal
Time (min)
1.3
1.2
1.1
1
0.9
0.8
0.7
0.6
0
10
20
30
40
50
60
Normal
MK50
MK75
MK88
MK94
MK97
MK data
Normal data

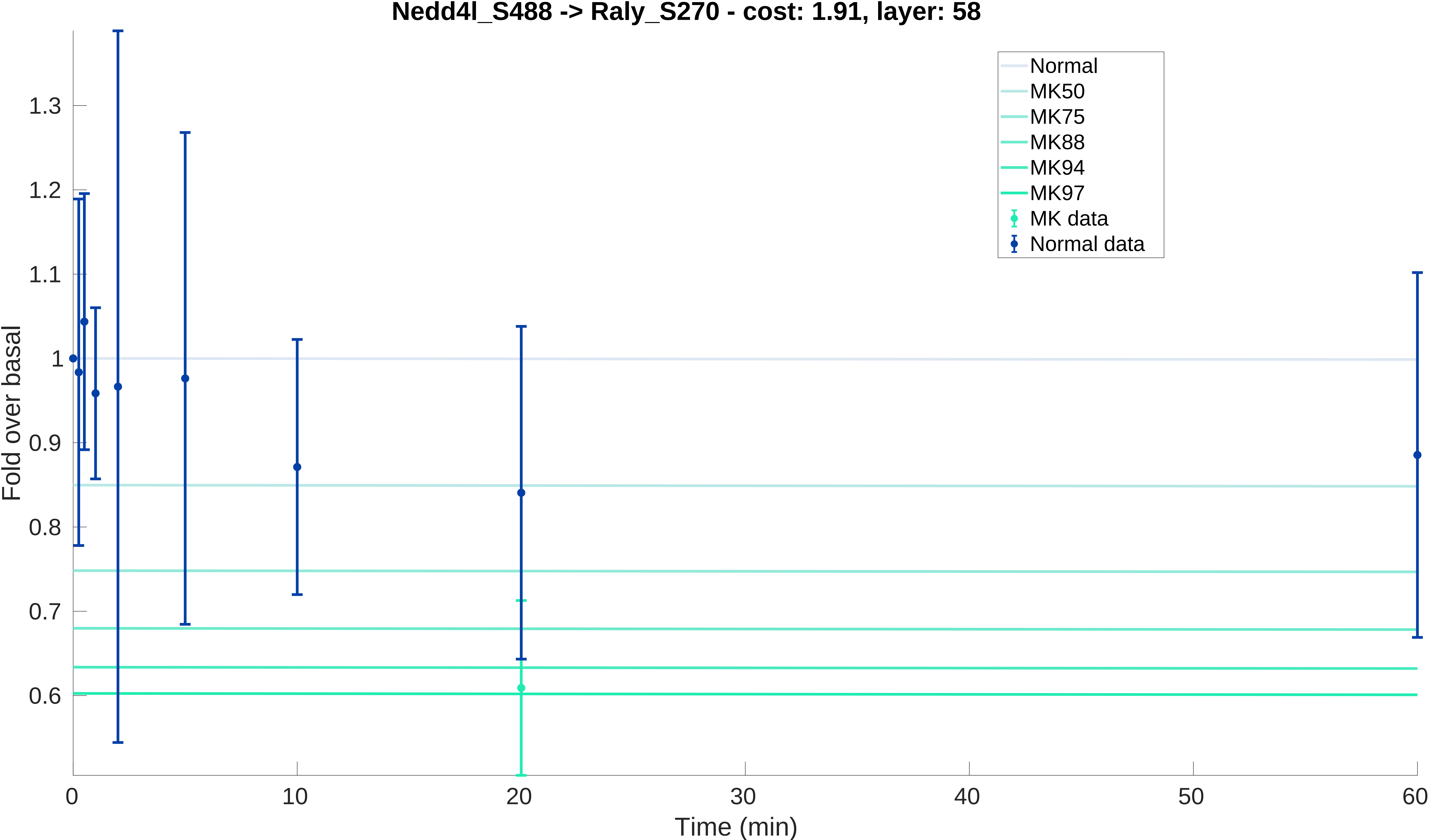
Nedd4l_S488 -> Raly_S270 - cost: 1.91, layer: 58
Normal
MK50
MK75
MK88
MK94
MK97
MK data
Normal data
Fold over basal
1.3
1.2
1.1
1
0.9
0.8
0.7
0.6
0
10
20
30
40
50
60
Time (min)

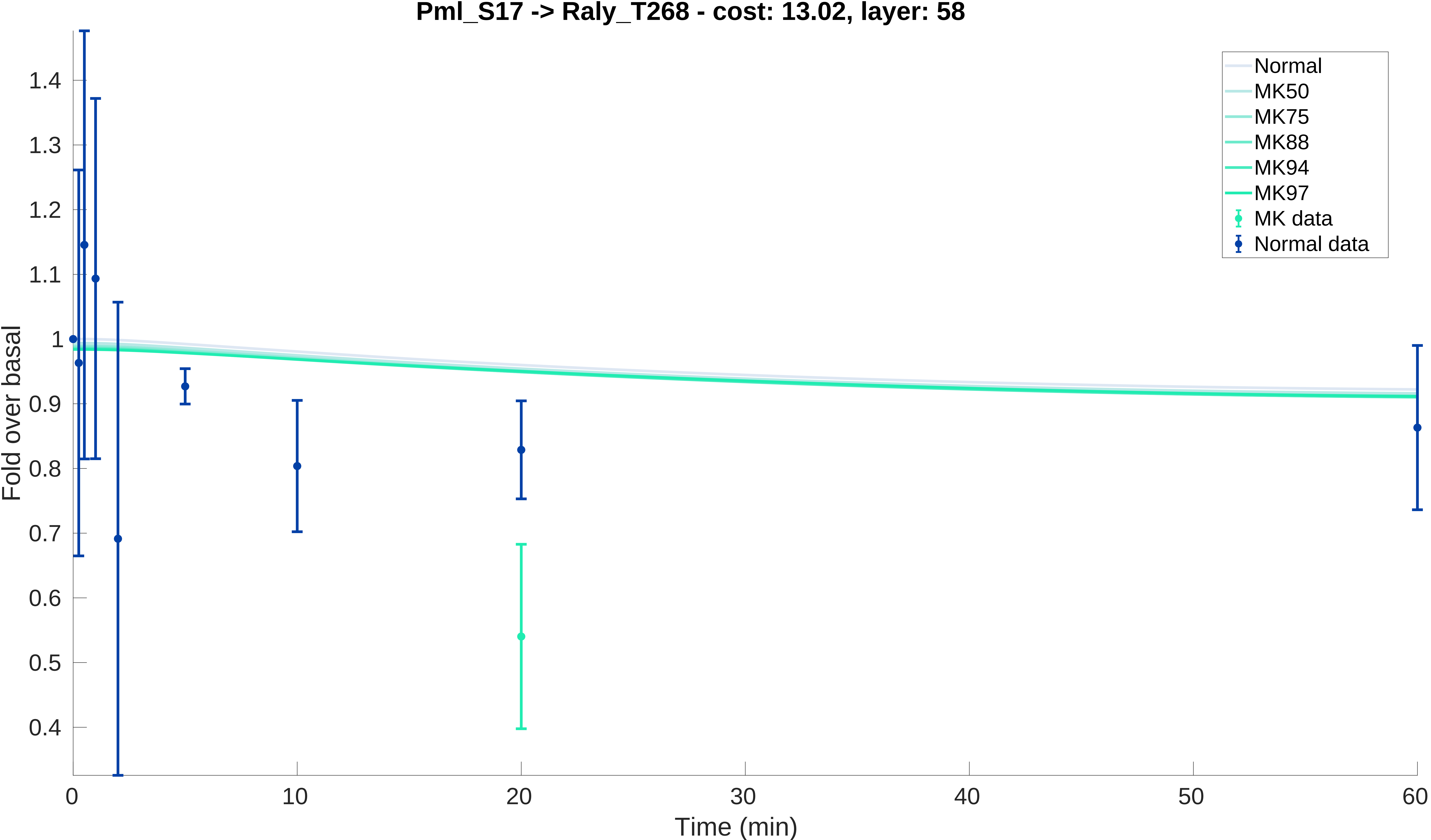

Pml_S17 -> Raly_T268 - cost: 13.02, layer: 58
Fold over basal
Time (min)
0
10
20
30
40
50
60
0.4
0.5
0.6
0.7
0.8
0.9
1
1.1
1.2
1.3
1.4
Normal
MK50
MK75
MK88
MK94
MK97
MK data
Normal data

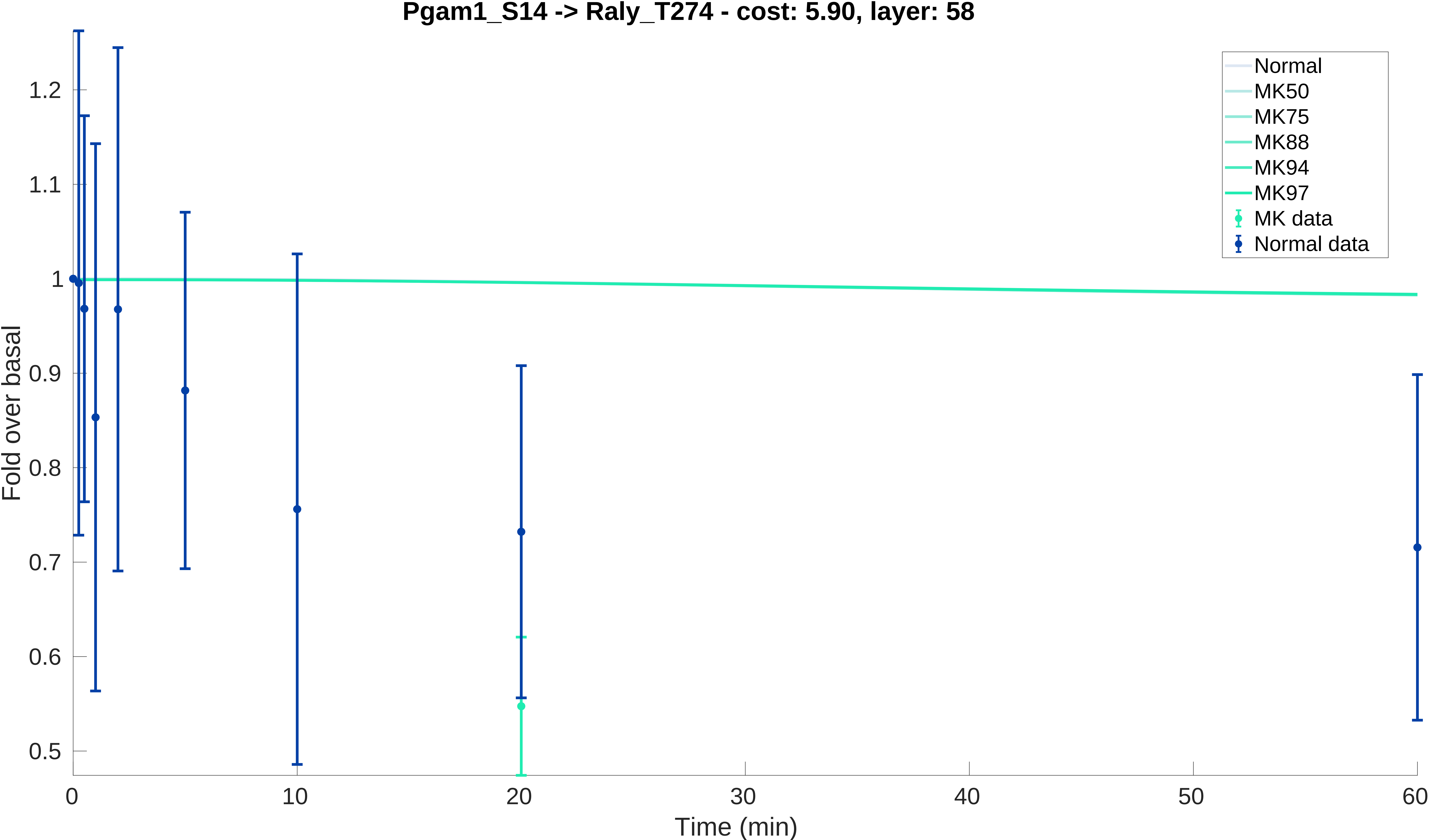

Pgam1_S14 -> Raly_T274 - cost: 5.90, layer: 58
Normal
MK50
MK75
MK88
MK94
MK97
MK data
Normal data
Fold over basal
0.5
0.6
0.7
0.8
0.9
1
1.1
1.2
0
10
20
30
40
50
60
Time (min)

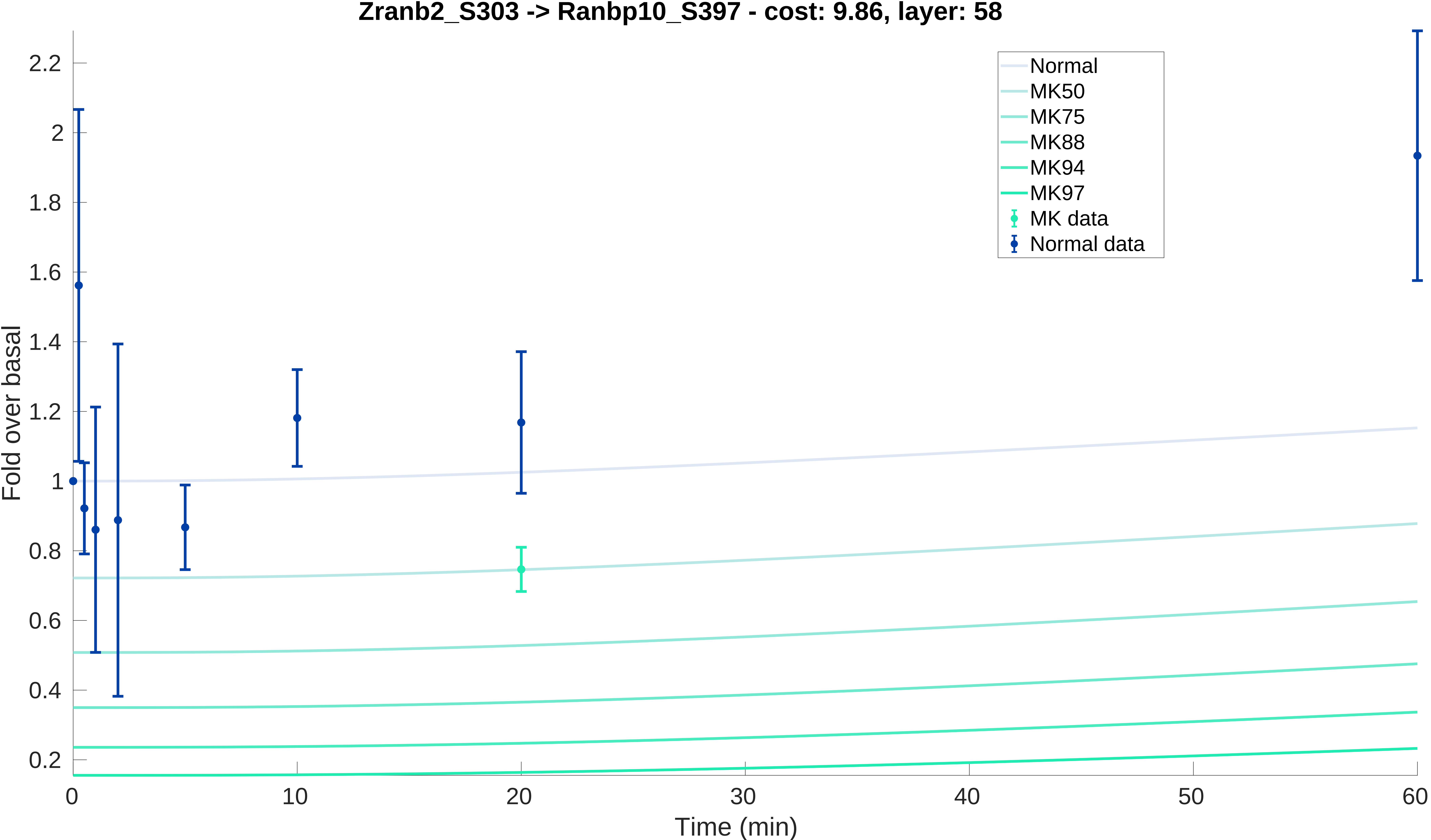

Zranb2_S303 -> Ranbp10_S397 - cost: 9.86, layer: 58
Normal
MK50
MK75
MK88
MK94
MK97
MK data
Normal data
Fold over basal
Time (min)
0.2
0.4
0.6
0.8
1
1.2
1.4
1.6
1.8
2
2.2
0
10
20
30
40
50
60

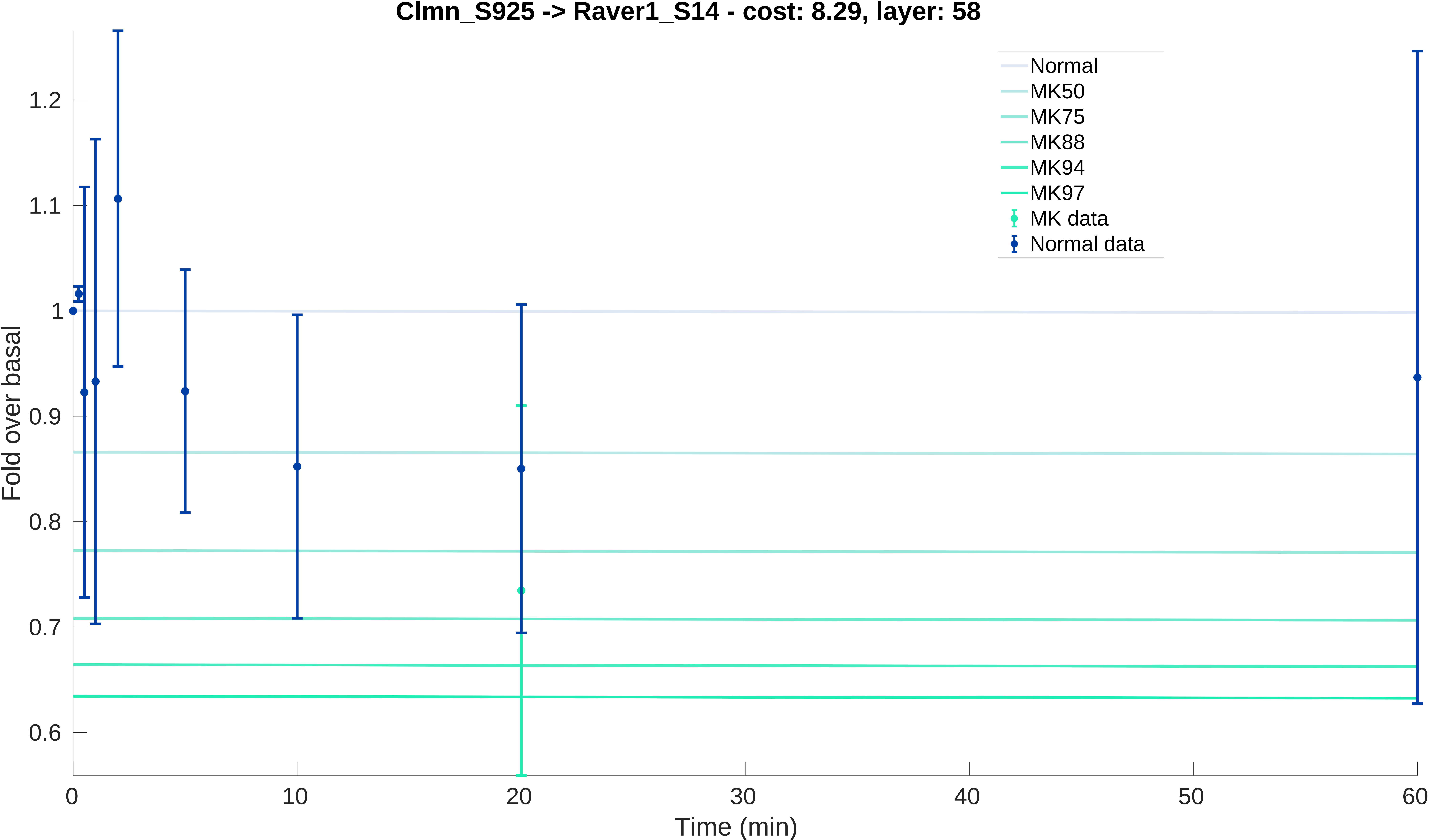

Clmn_S925 -> Raver1_S14 - cost: 8.29, layer: 58
Normal
MK50
MK75
MK88
MK94
MK97
MK data
Normal data
Fold over basal
Time (min)
1.2
1.1
1
0.9
0.8
0.7
0.6
0
10
20
30
40
50
60

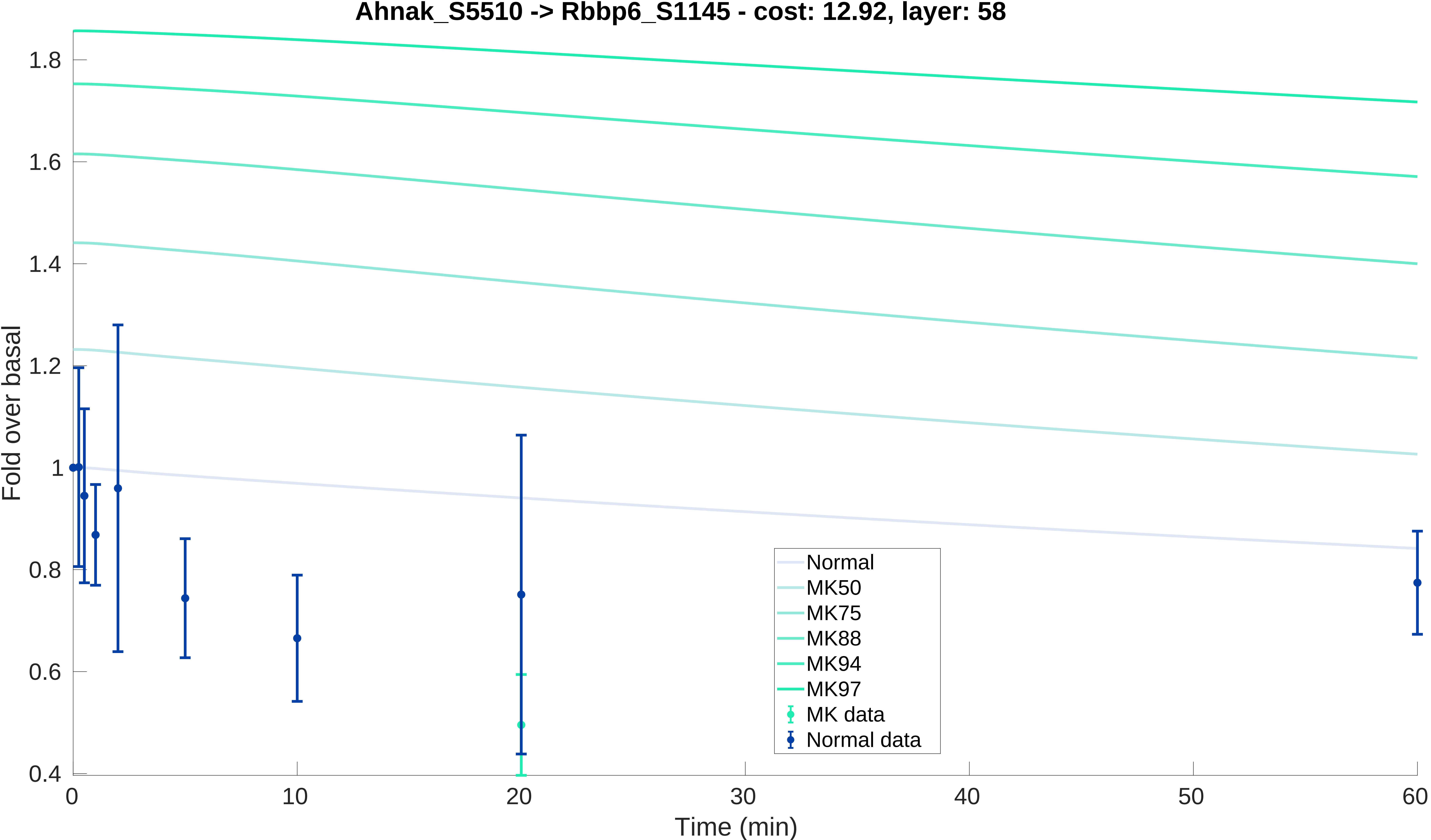

Ahnak_S5510 -> Rbbp6_S1145 - cost: 12.92, layer: 58
Fold over basal
Time (min)
0.4
0.6
0.8
1
1.2
1.4
1.6
1.8
0
10
20
30
40
50
60
Normal
MK50
MK75
MK88
MK94
MK97
MK data
Normal data

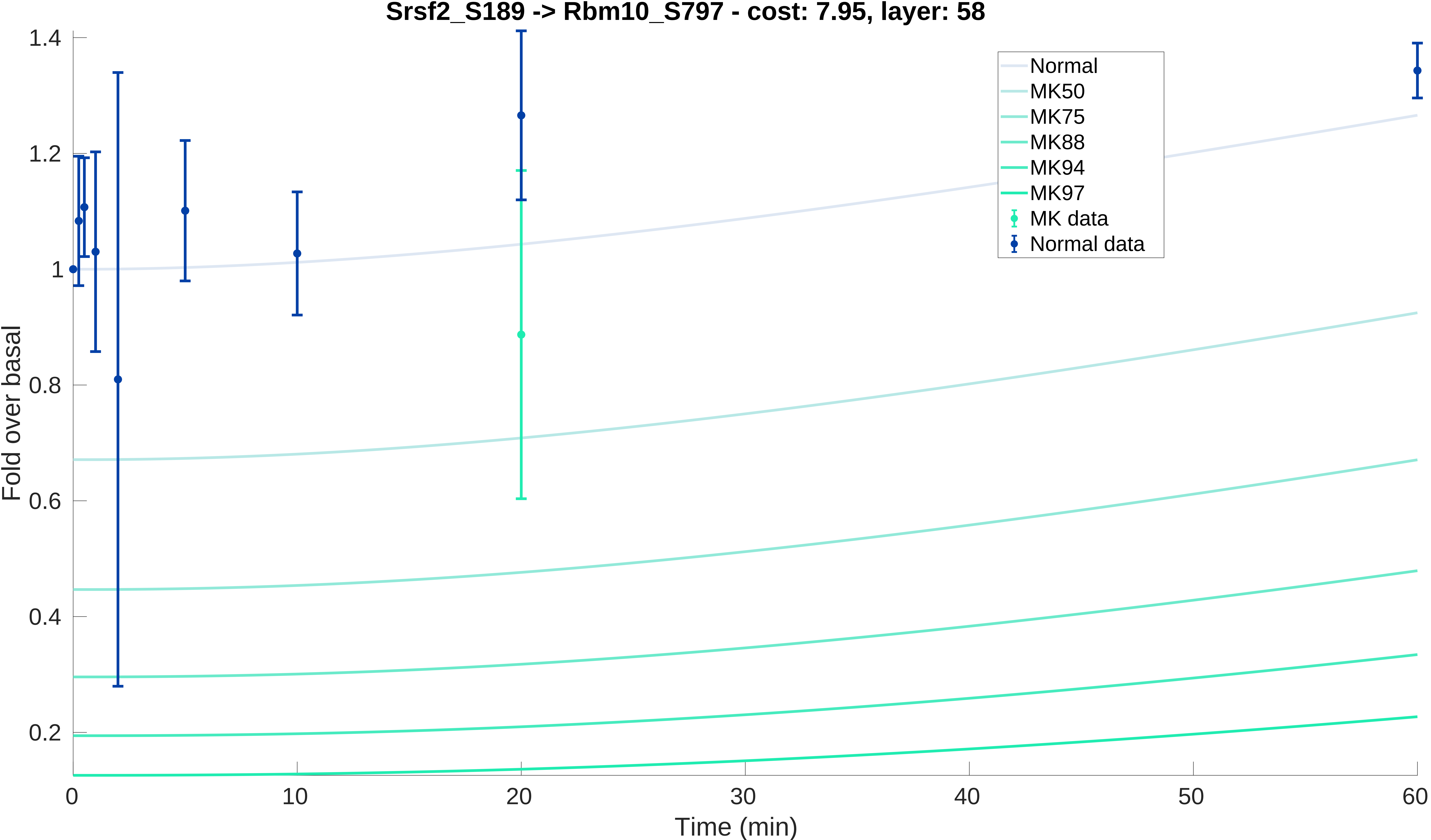

Srsf2_S189 -> Rbm10_S797 - cost: 7.95, layer: 58
Fold over basal
Time (min)
0
10
20
30
40
50
60
0.2
0.4
0.6
0.8
1
1.2
1.4
Normal
MK50
MK75
MK88
MK94
MK97
MK data
Normal data

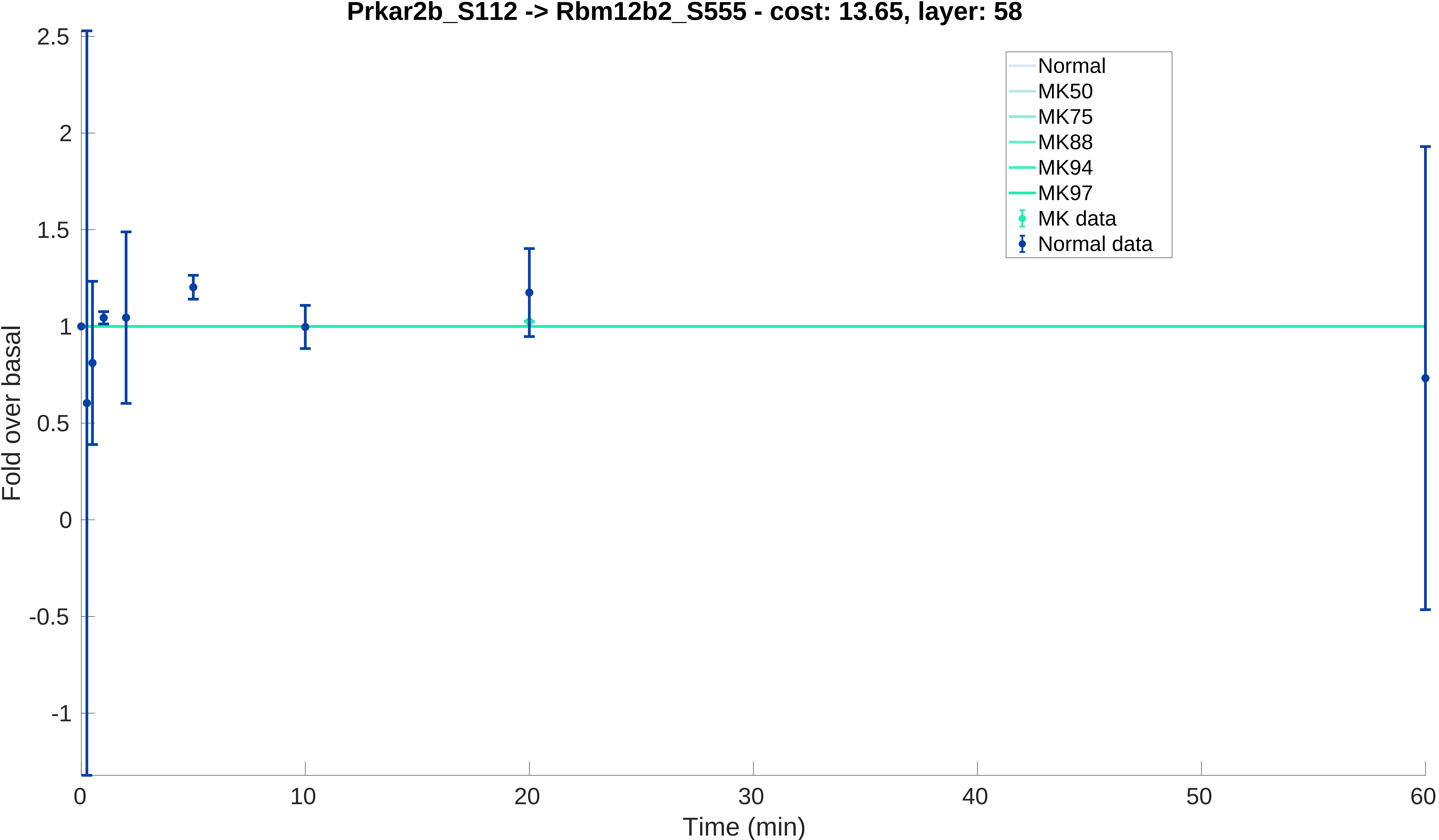

Prkar2b_S112 -> Rbm12b2_S555 - cost: 13.65, layer: 58
Fold over basal
Time (min)
2.5
2
1.5
1
0.5
0
-0.5
-1
0
10
20
30
40
50
60
Normal
MK50
MK75
MK88
MK94
MK97
MK data
Normal data

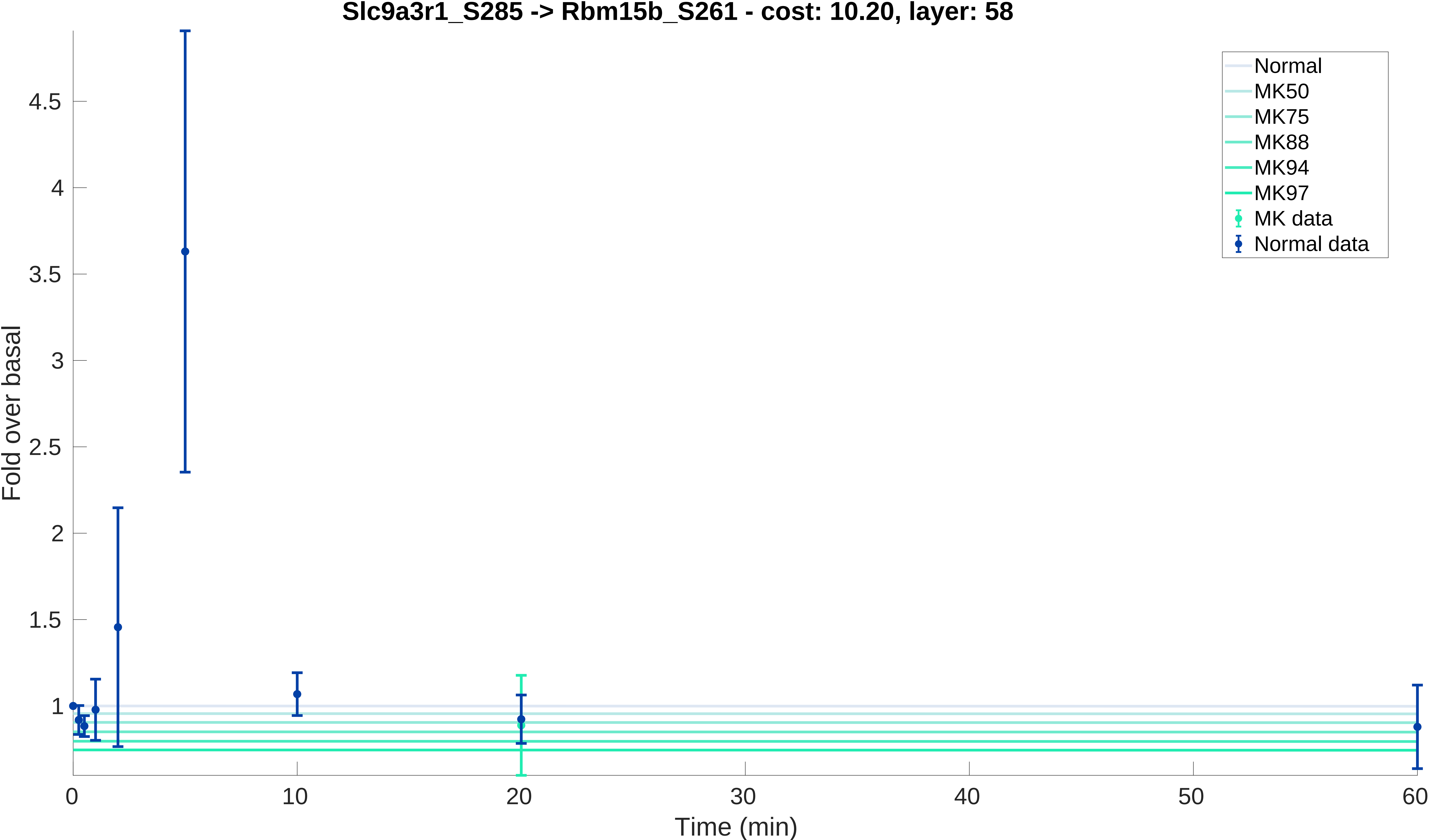

Slc9a3r1_S285 -> Rbm15b_S261 - cost: 10.20, layer: 58
Fold over basal
Time (min)
0
10
20
30
40
50
60
1
1.5
2
2.5
3
3.5
4
4.5
Normal
MK50
MK75
MK88
MK94
MK97
MK data
Normal data

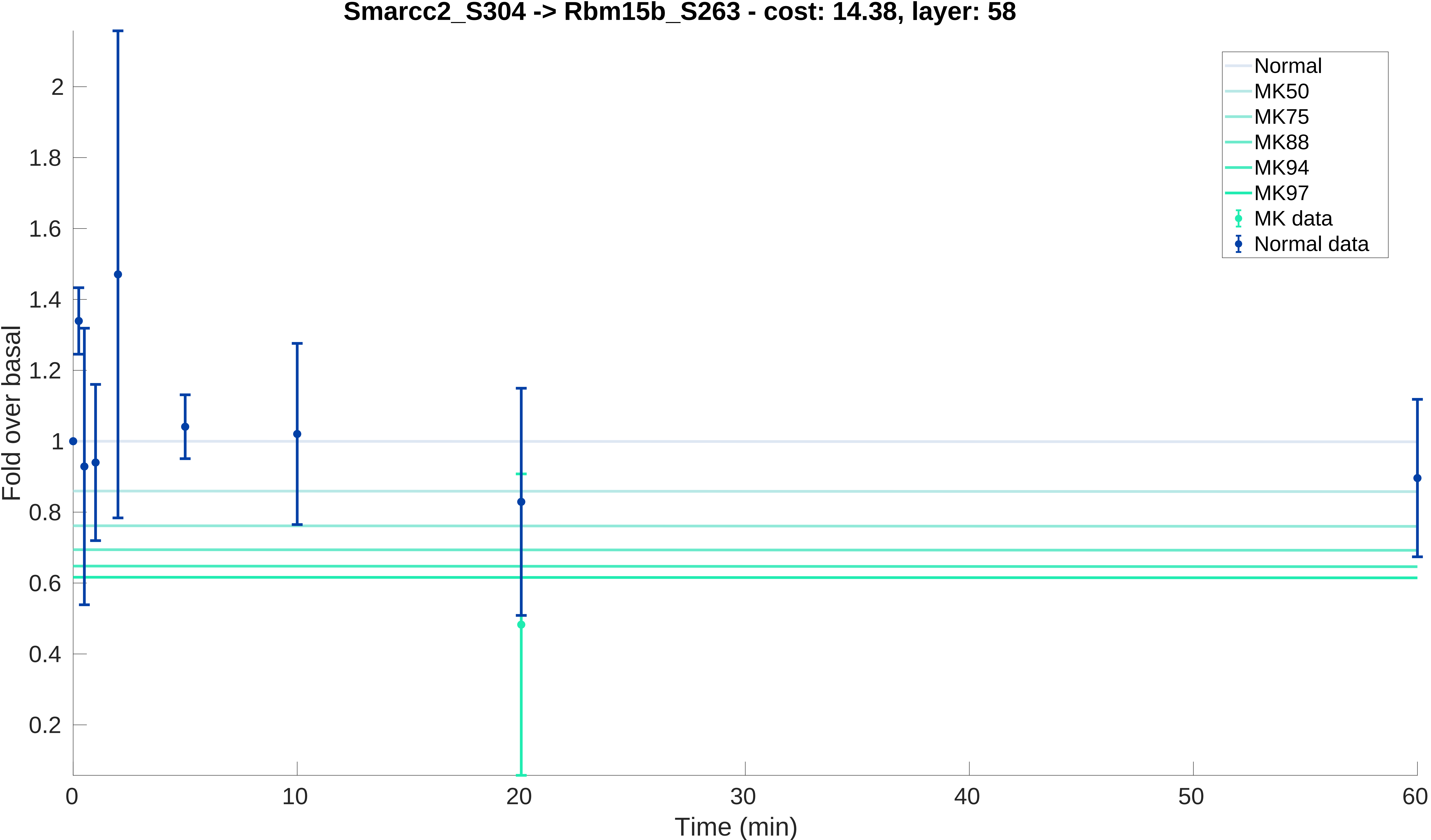

Smarcc2_S304 -> Rbm15b_S263 - cost: 14.38, layer: 58
Fold over basal
Time (min)
0
10
20
30
40
50
60
0.2
0.4
0.6
0.8
1
1.2
1.4
1.6
1.8
2
Normal
MK50
MK75
MK88
MK94
MK97
MK data
Normal data

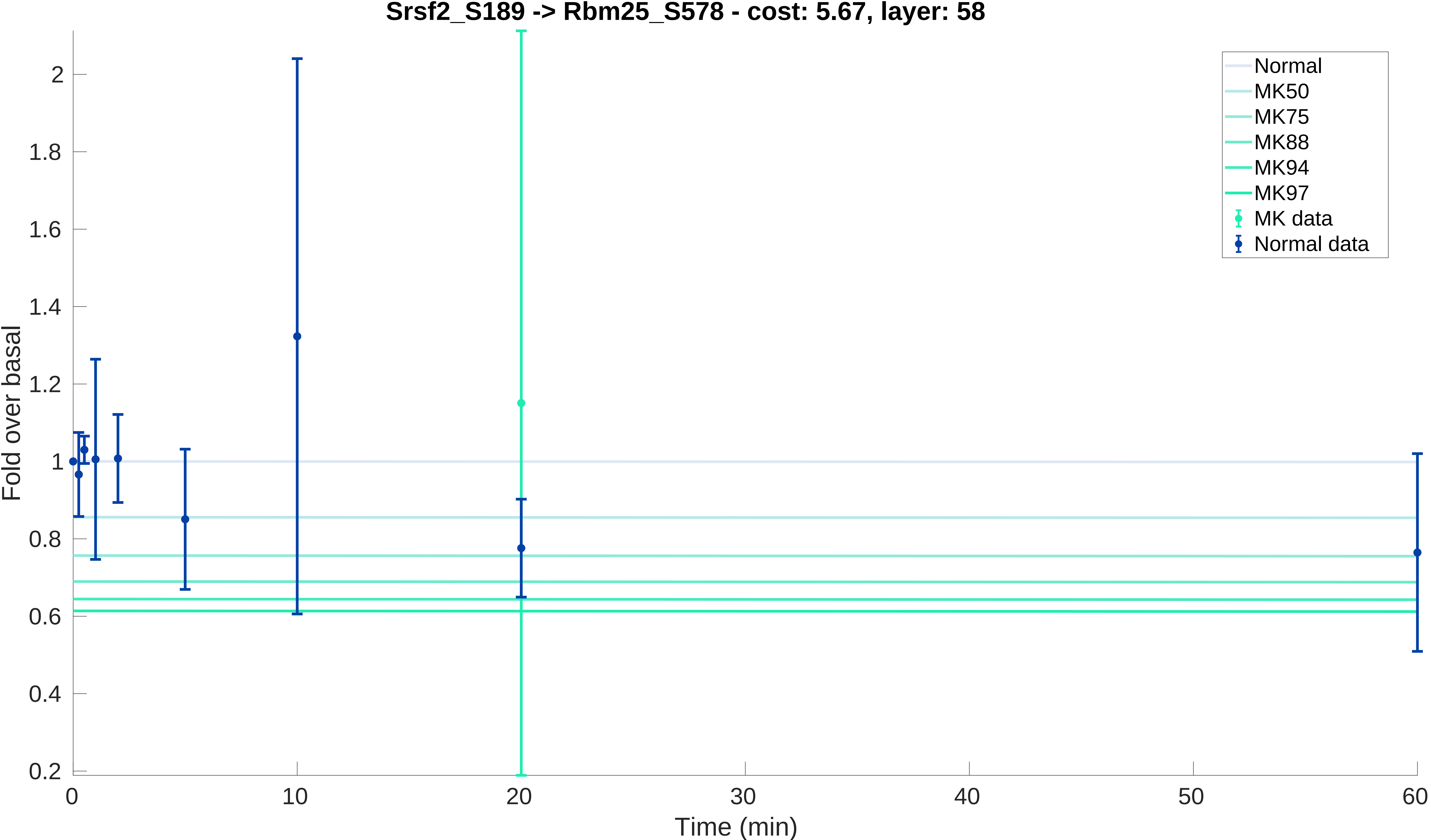

Srsf2_S189 -> Rbm25_S578 - cost: 5.67, layer: 58
Fold over basal
Time (min)
0.2
0.4
0.6
0.8
1
1.2
1.4
1.6
1.8
2
0
10
20
30
40
50
60
Normal
MK50
MK75
MK88
MK94
MK97
MK data
Normal data

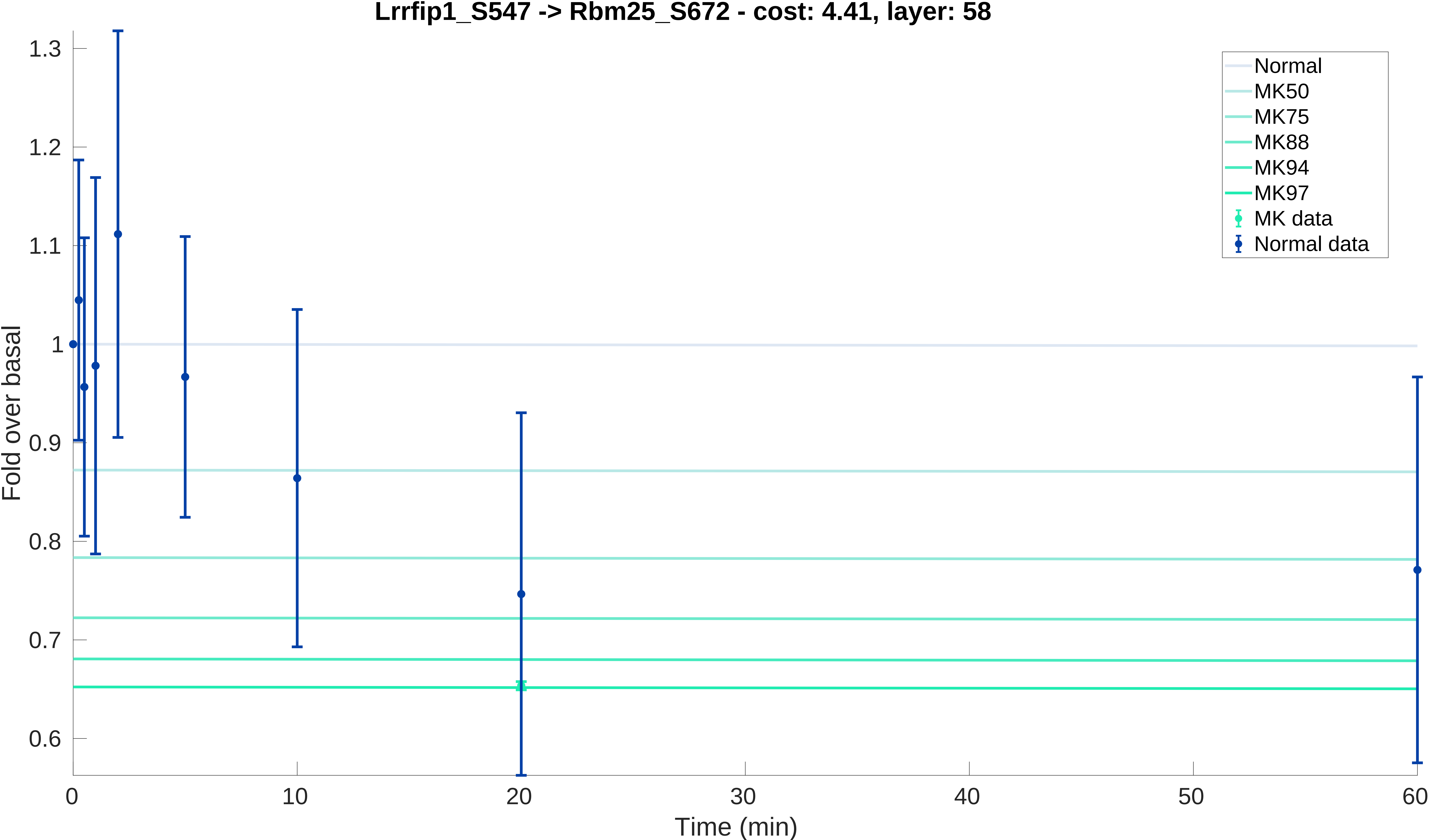

Lrrfip1_S547 -> Rbm25_S672 - cost: 4.41, layer: 58
Normal
MK50
MK75
MK88
MK94
MK97
MK data
Normal data
Fold over basal
Time (min)
1.3
1.2
1.1
1
0.9
0.8
0.7
0.6
0
10
20
30
40
50
60

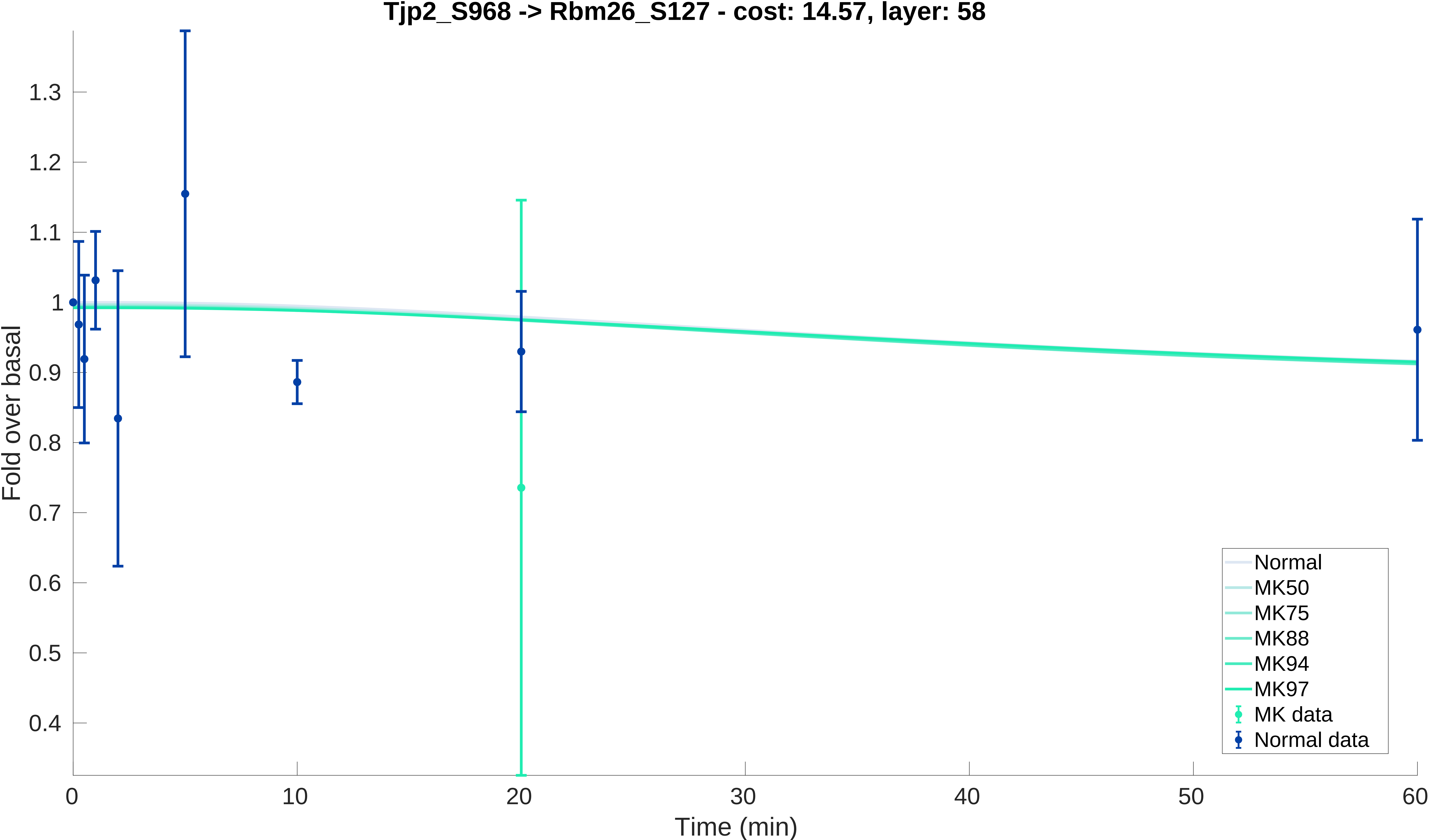

Tjp2_S968 -> Rbm26_S127 - cost: 14.57, layer: 58
Fold over basal
Time (min)
0.4
0.5
0.6
0.7
0.8
0.9
1
1.1
1.2
1.3
0
10
20
30
40
50
60
Normal
MK50
MK75
MK88
MK94
MK97
MK data
Normal data

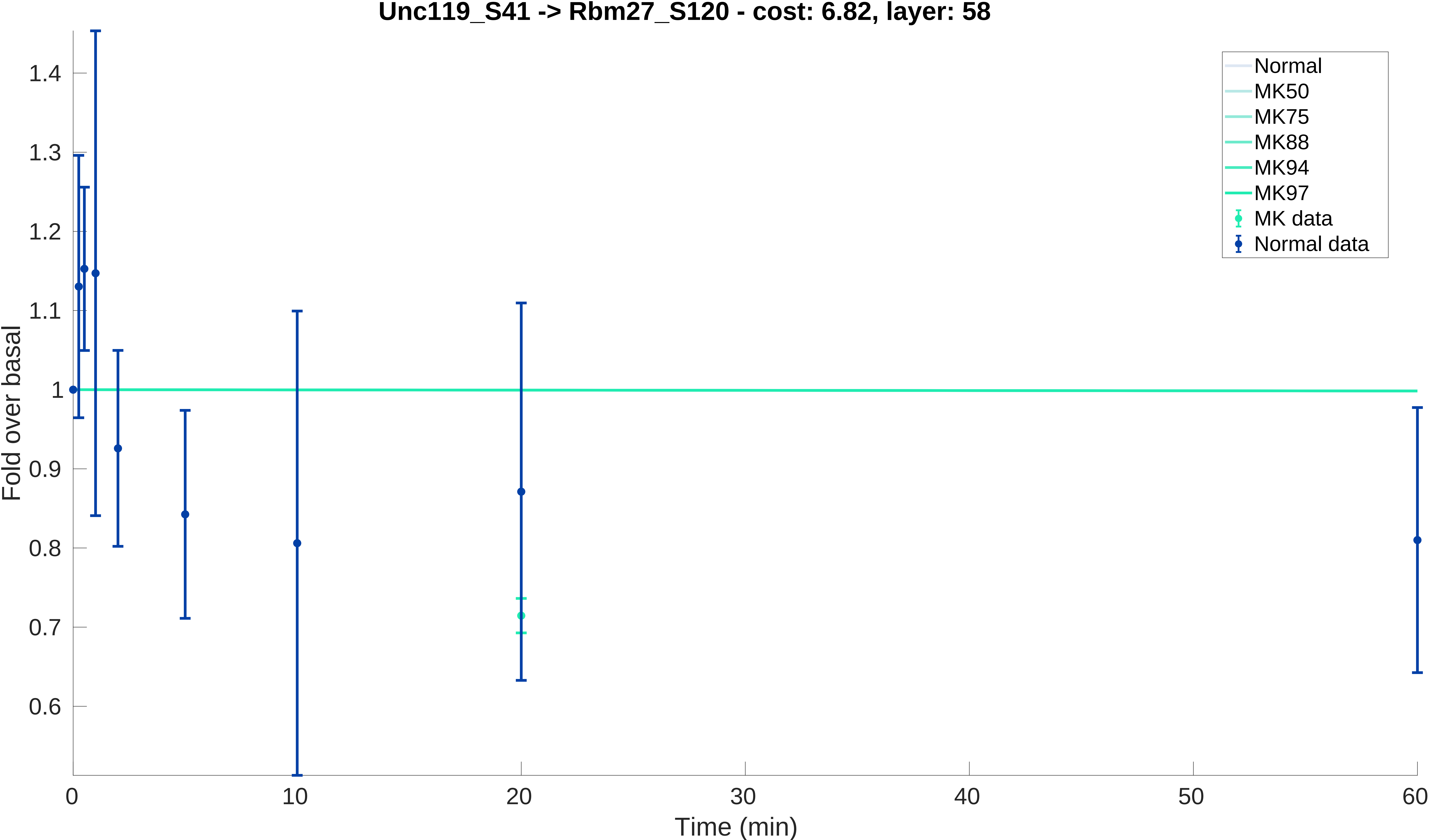

Unc119_S41 -> Rbm27_S120 - cost: 6.82, layer: 58
Fold over basal
1.4
1.3
1.2
1.1
1
0.9
0.8
0.7
0.6
0
10
20
30
40
50
60
Time (min)
Normal
MK50
MK75
MK88
MK94
MK97
MK data
Normal data

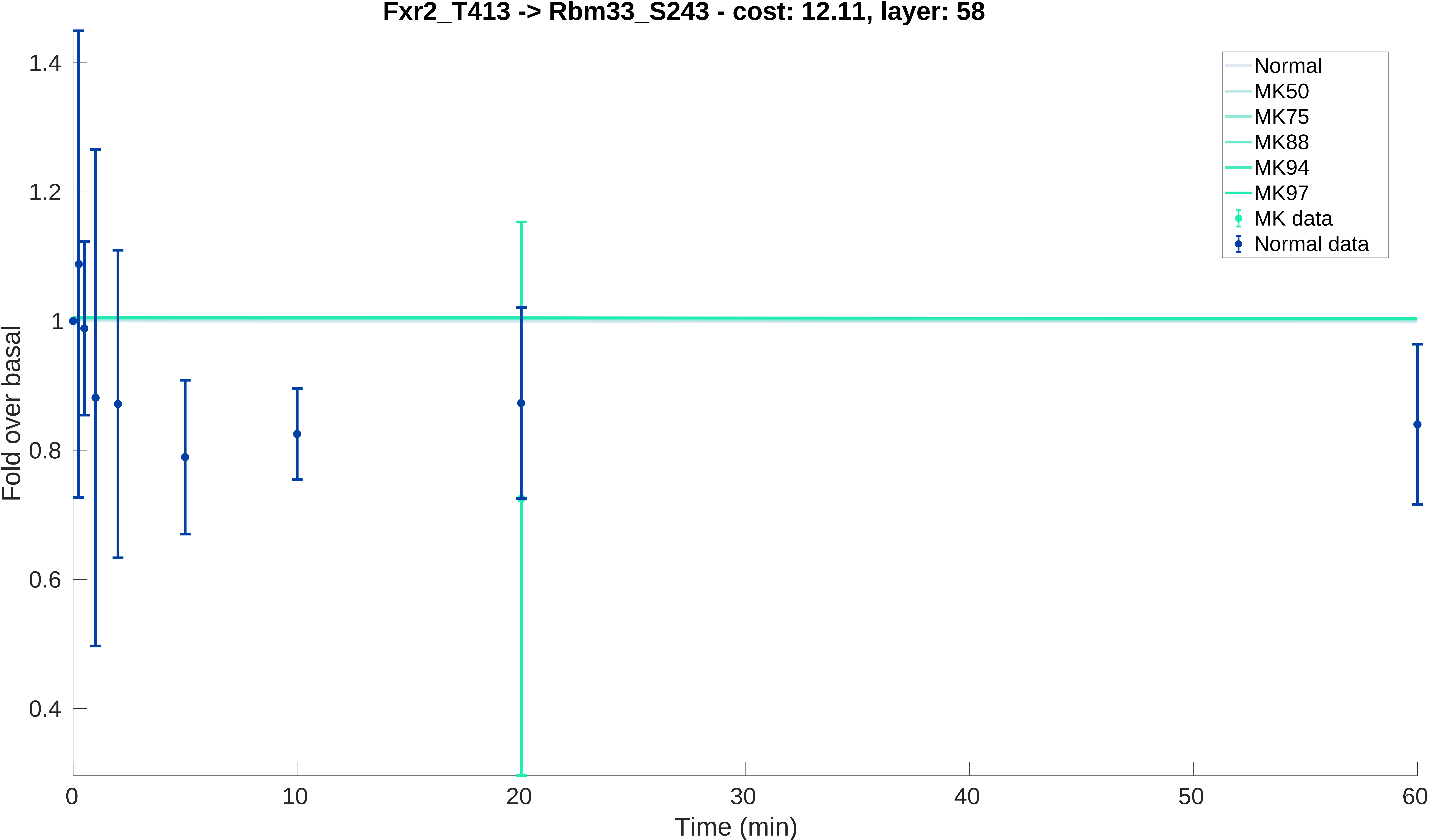

Fxr2_T413 -> Rbm33_S243 - cost: 12.11, layer: 58
Normal
MK50
MK75
MK88
MK94
MK97
MK data
Normal data
Fold over basal
1.4
1.2
1
0.8
0.6
0.4
0
10
20
30
40
50
60
Time (min)

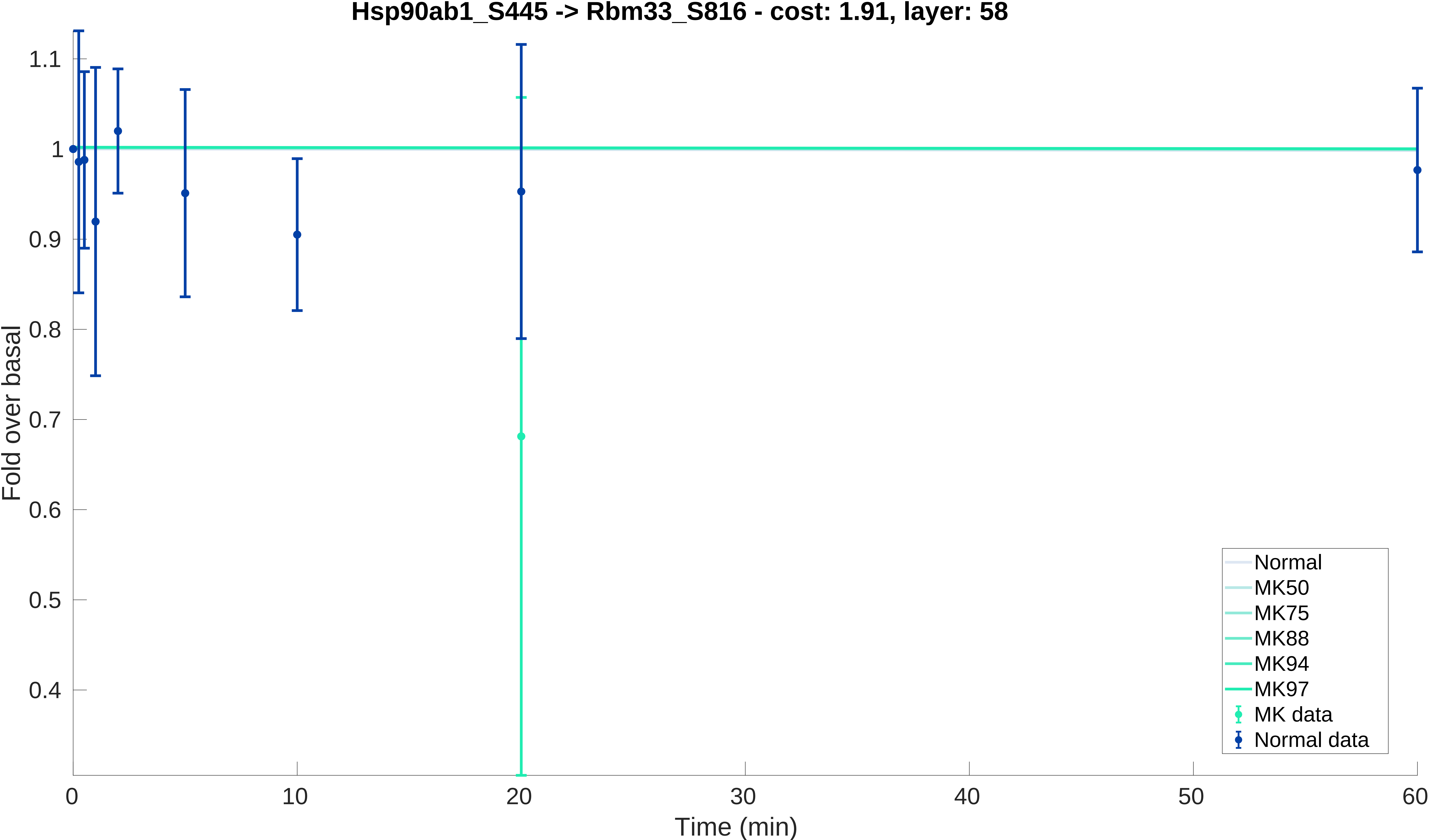

Hsp90ab1_S445 -> Rbm33_S816 - cost: 1.91, layer: 58
Fold over basal
Time (min)
1.1
1
0.9
0.8
0.7
0.6
0.5
0.4
0
10
20
30
40
50
60
Normal
MK50
MK75
MK88
MK94
MK97
MK data
Normal data

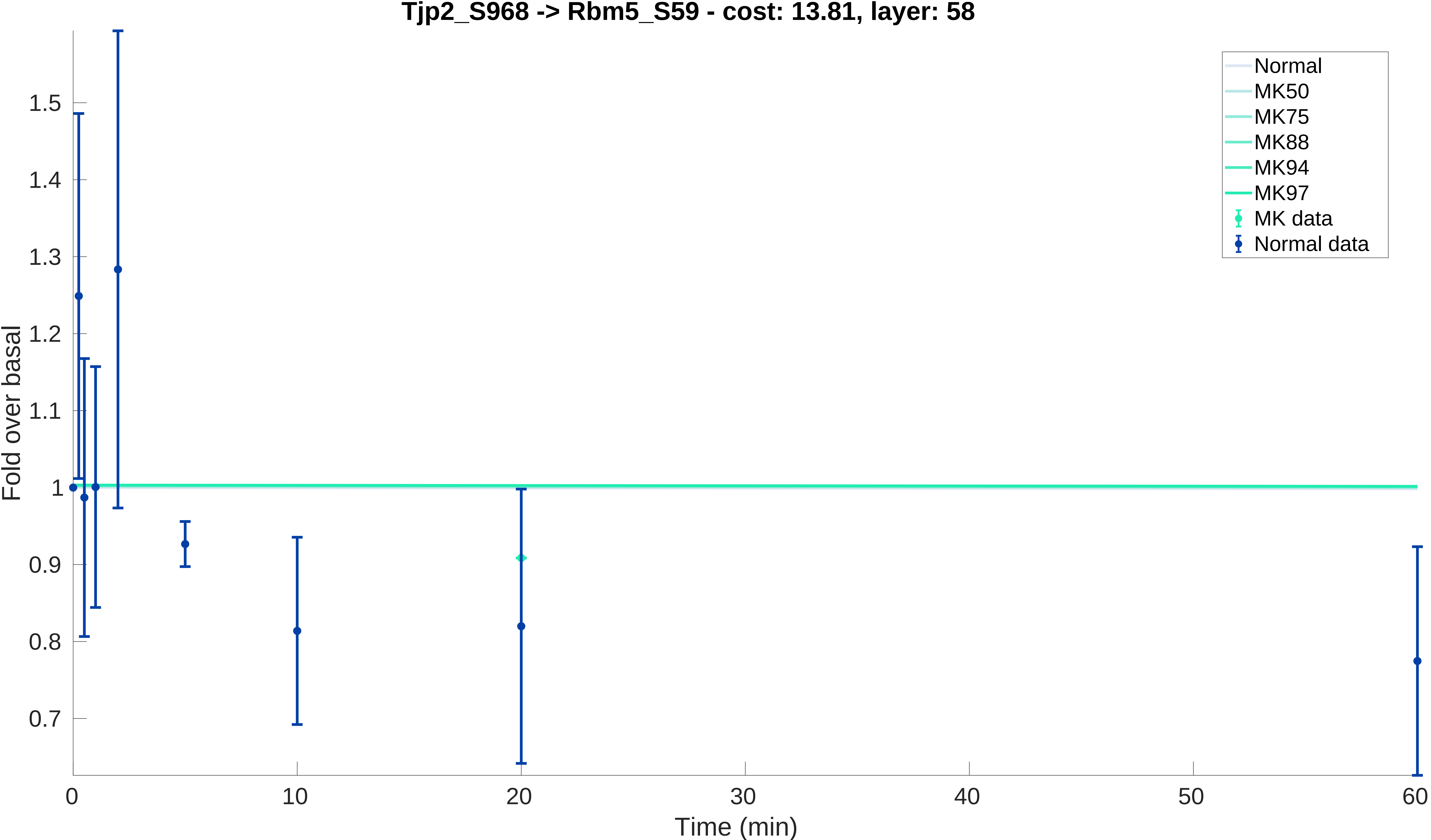
Tjp2_S968 -> Rbm5_S59 - cost: 13.81, layer: 58
Fold over basal
Time (min)
Normal
MK50
MK75
MK88
MK94
MK97
MK data
Normal data
0.7
0.8
0.9
1
1.1
1.2
1.3
1.4
1.5
0
10
20
30
40
50
60

Eif5b_S215 -> Rbm5_S69 - cost: 2.58, layer: 58

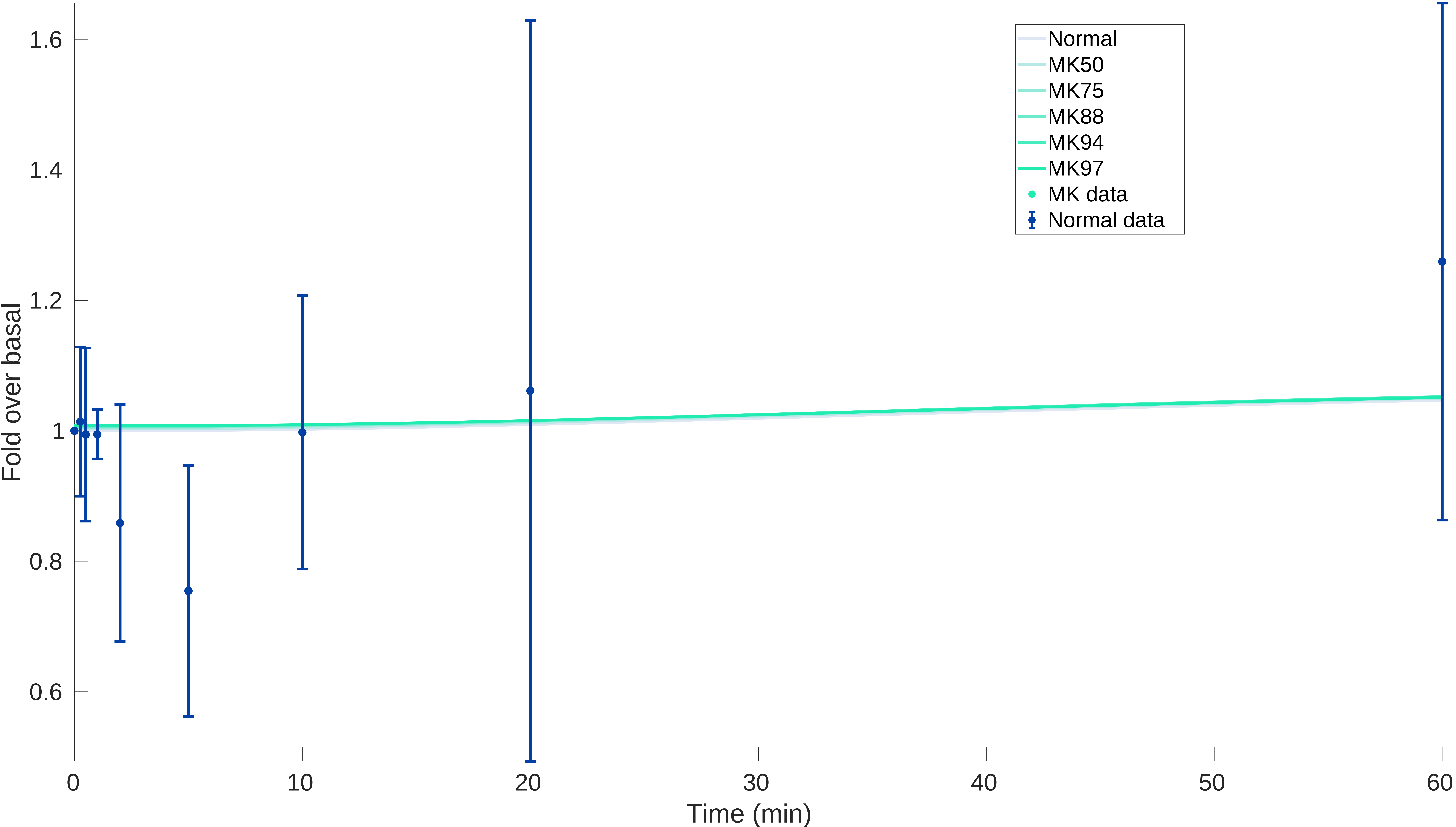

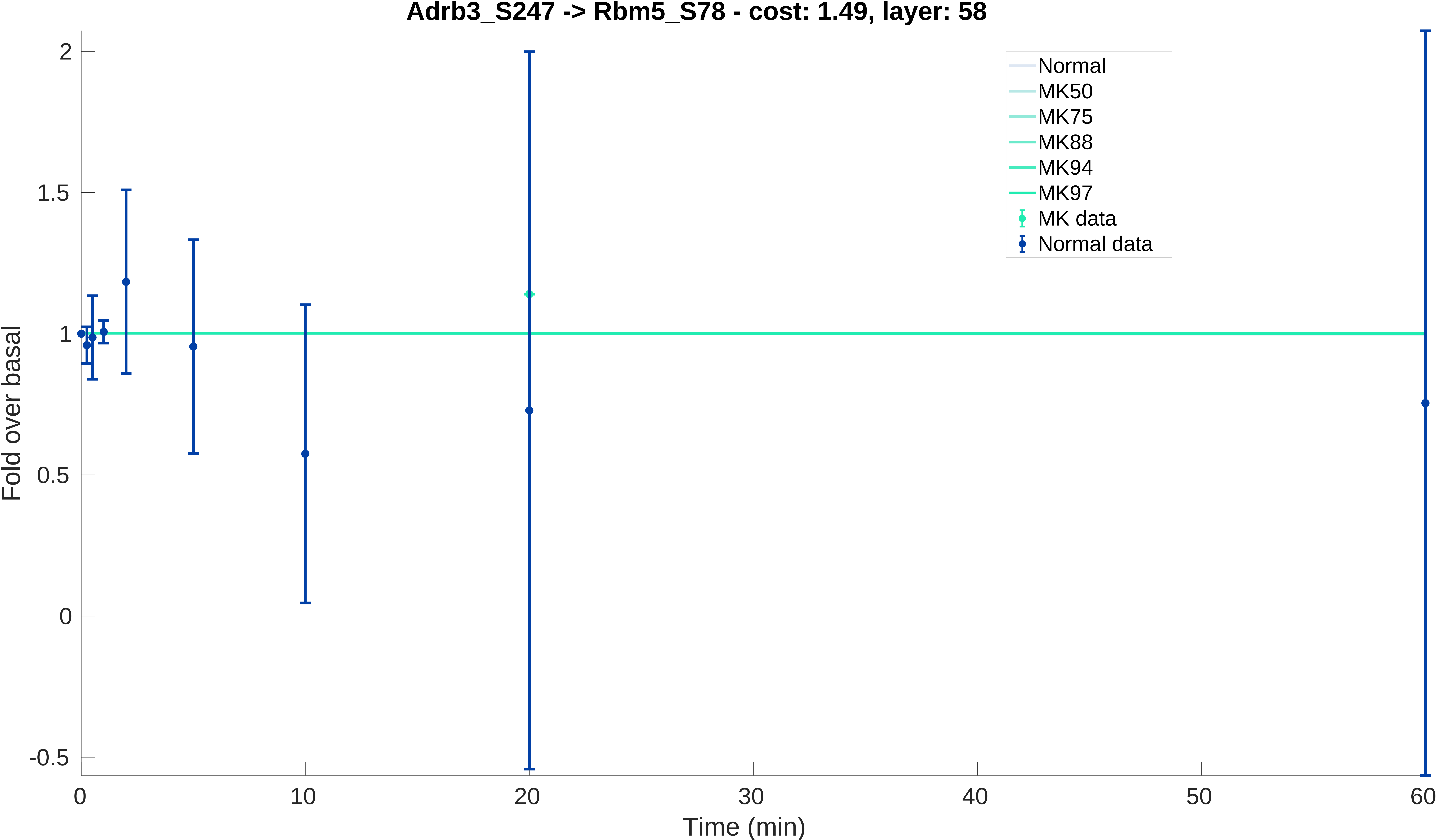

Adrb3_S247 -> Rbm5_S78 - cost: 1.49, layer: 58
Normal
MK50
MK75
MK88
MK94
MK97
MK data
Normal data
Fold over basal
Time (min)
2
1.5
1
0.5
0
-0.5
0
10
20
30
40
50
60

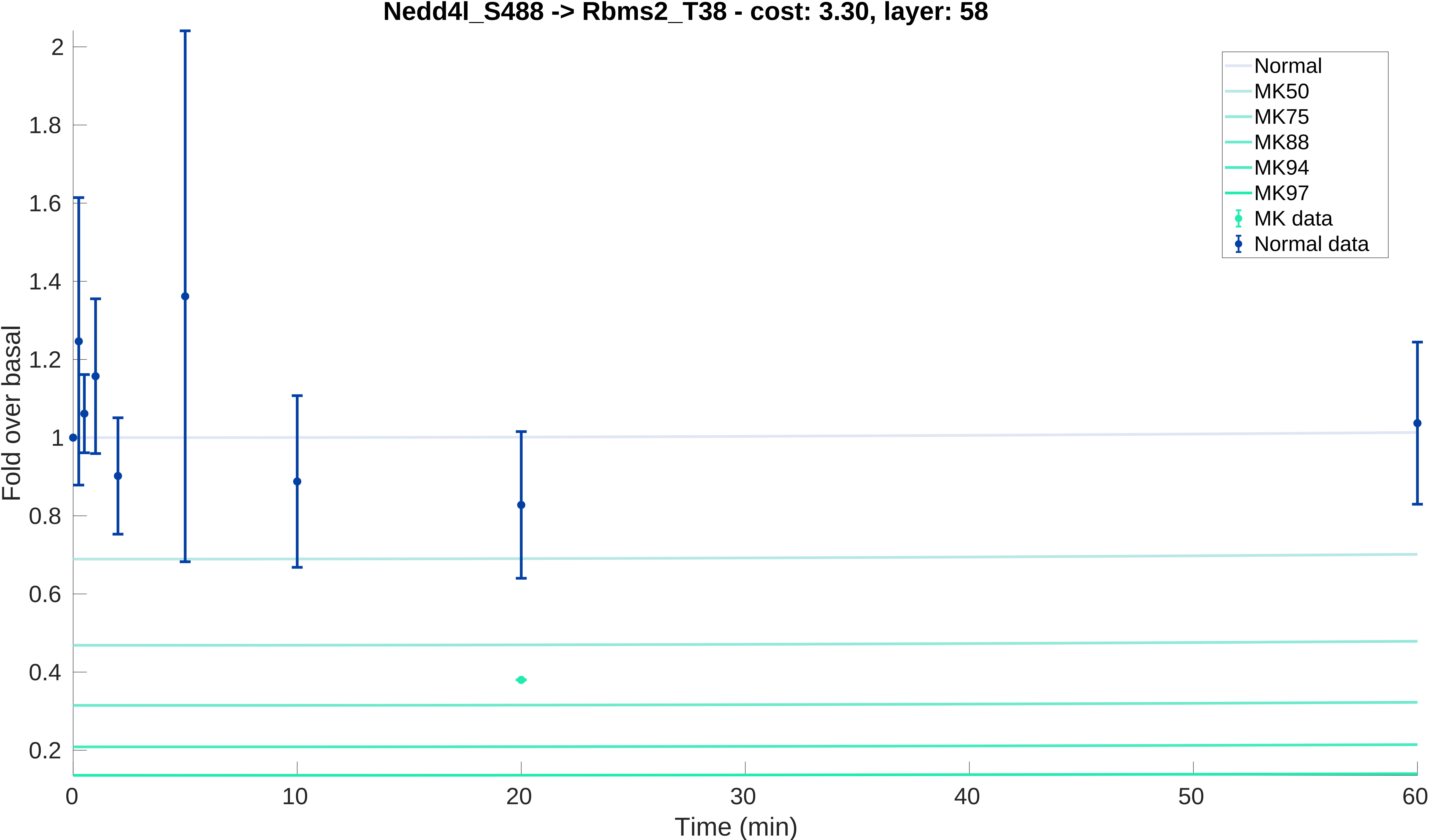

Nedd4l_S488 -> Rbms2_T38 - cost: 3.30, layer: 58
Fold over basal
Time (min)
Normal
MK50
MK75
MK88
MK94
MK97
MK data
Normal data
0.2
0.4
0.6
0.8
1
1.2
1.4
1.6
1.8
2
0
10
20
30
40
50
60

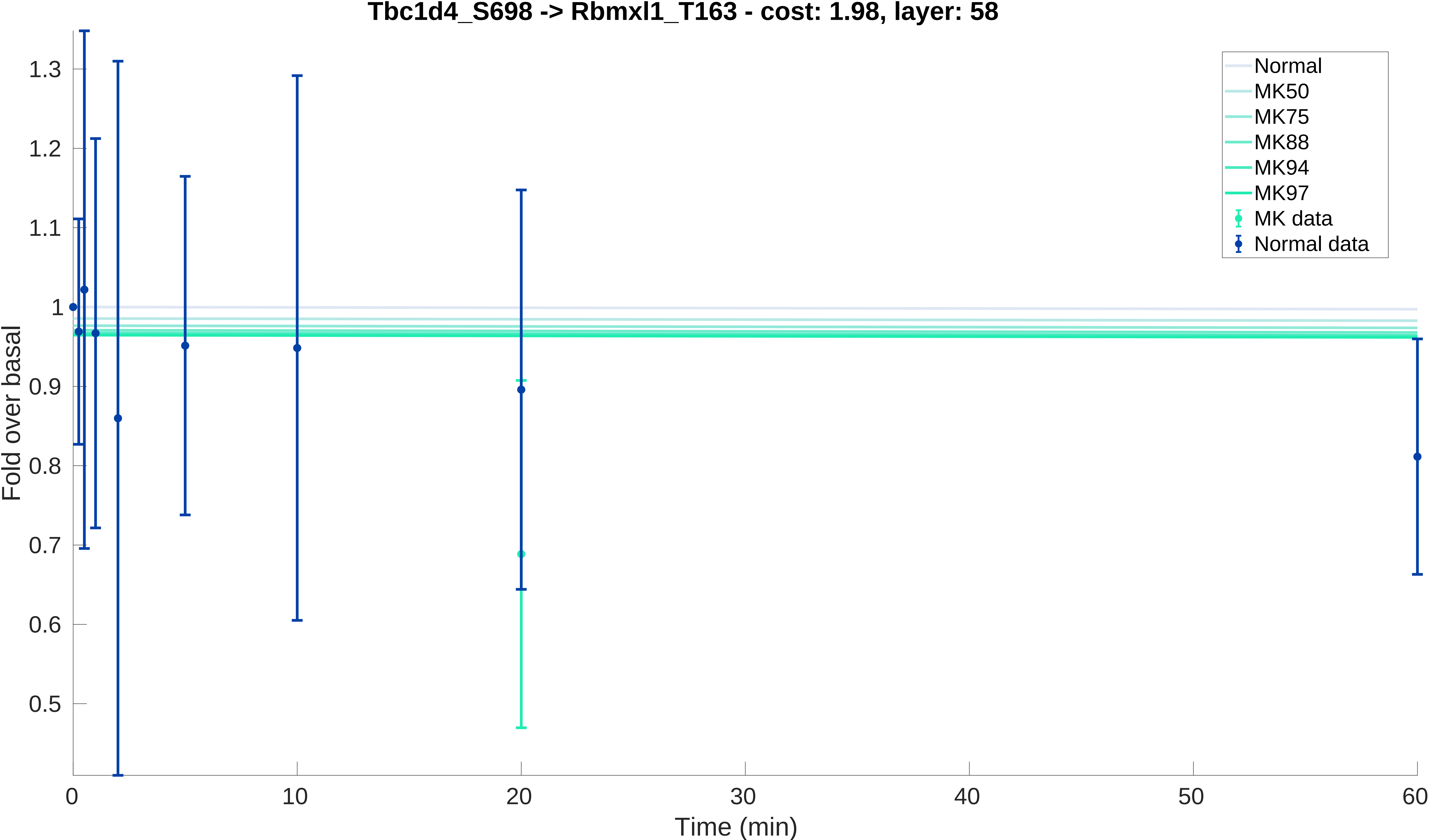

Tbc1d4_S698 -> Rbmxl1_T163 - cost: 1.98, layer: 58
Fold over basal
Time (min)
Normal
MK50
MK75
MK88
MK94
MK97
MK data
Normal data
1.3
1.2
1.1
1
0.9
0.8
0.7
0.6
0.5
0
10
20
30
40
50
60

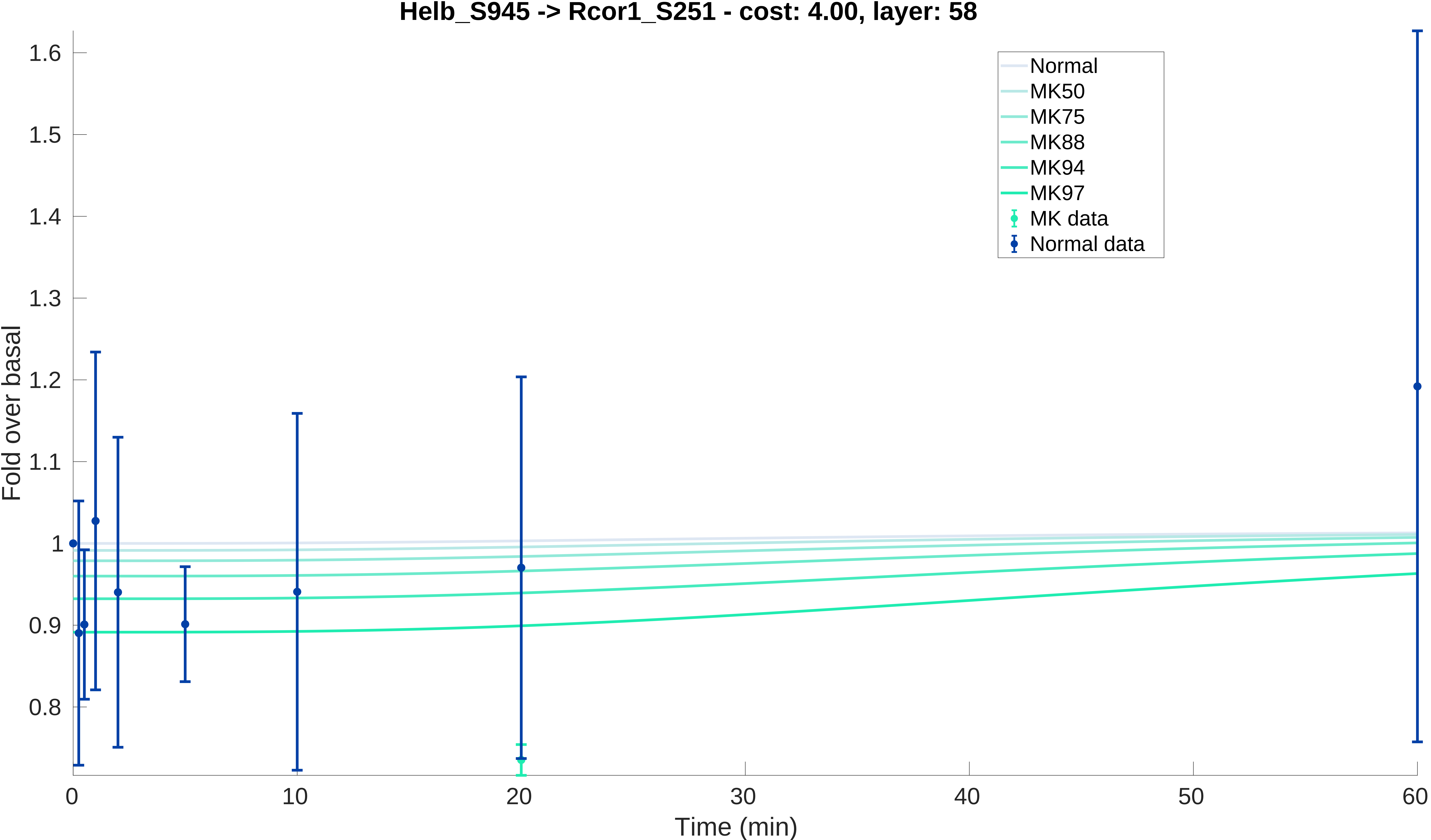

Helb_S945 -> Rcor1_S251 - cost: 4.00, layer: 58
Fold over basal
1.6
1.5
1.4
1.3
1.2
1.1
1
0.9
0.8
0
10
20
30
40
50
60
Time (min)
Normal
MK50
MK75
MK88
MK94
MK97
MK data
Normal data

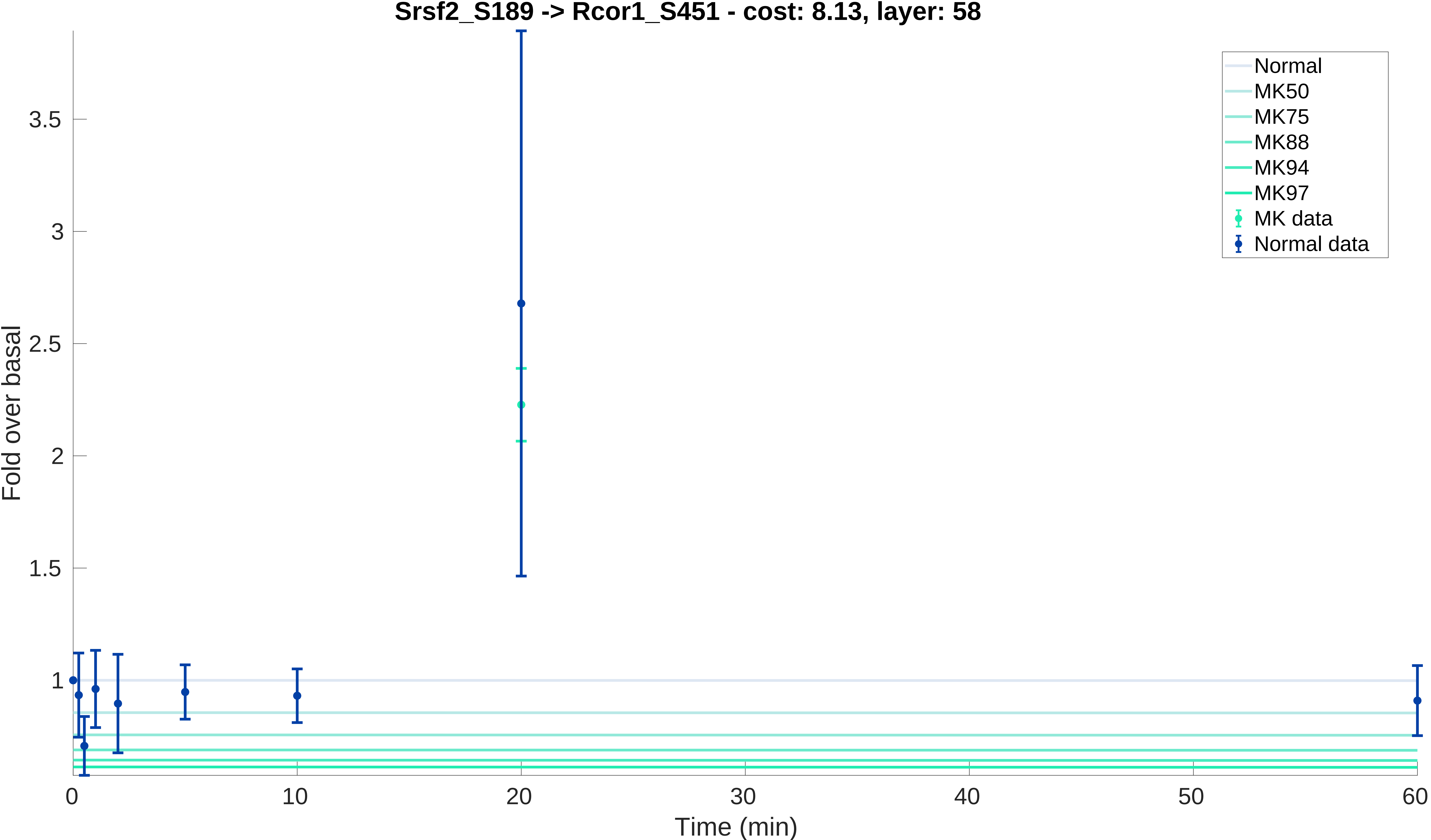

Srsf2_S189 -> Rcor1_S451 - cost: 8.13, layer: 58
Fold over basal
Time (min)
Normal
MK50
MK75
MK88
MK94
MK97
MK data
Normal data
3.5
3
2.5
2
1.5
1
0
10
20
30
40
50
60

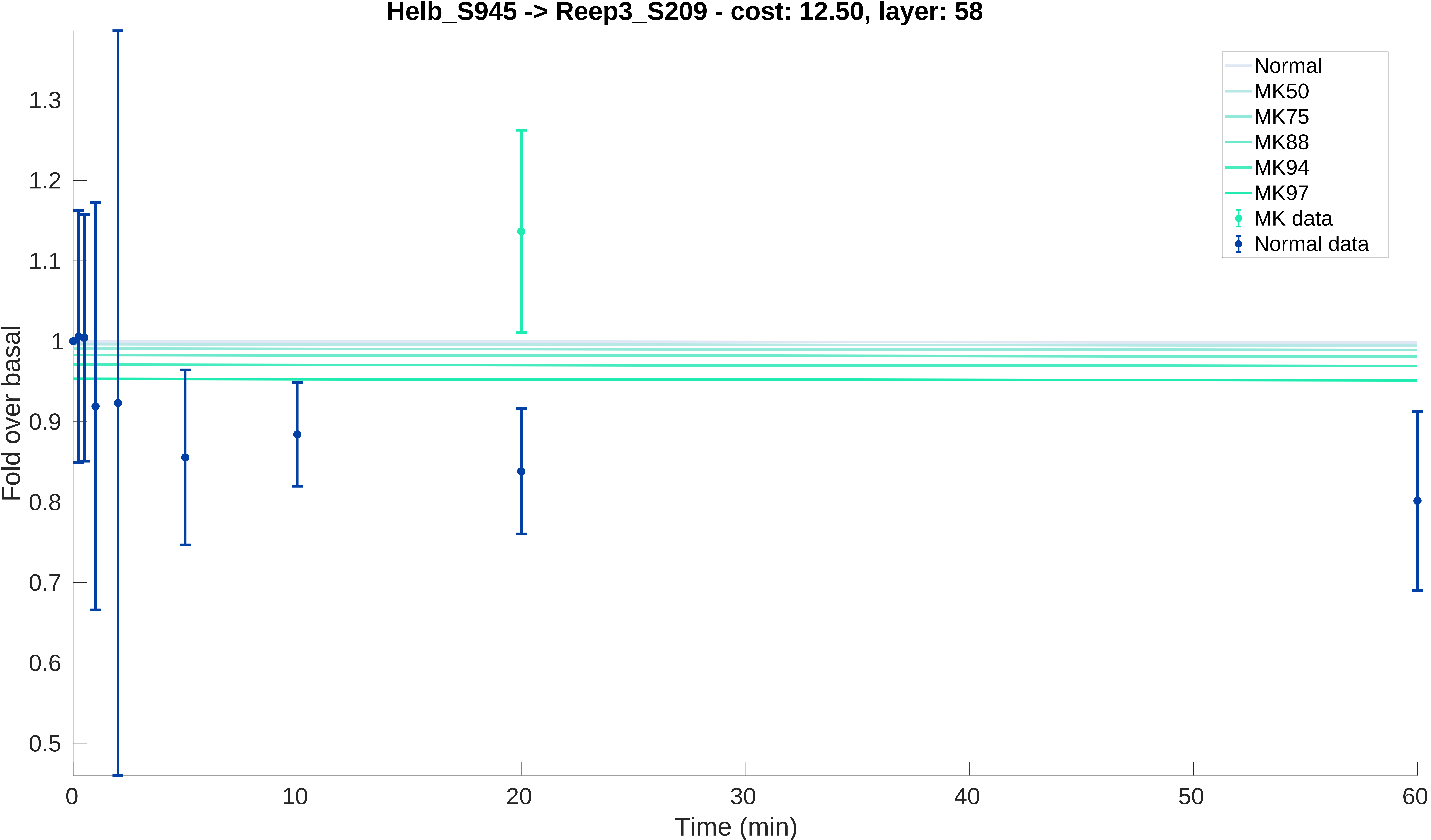
Helb_S945 -> Reep3_S209 - cost: 12.50, layer: 58
Normal
MK50
MK75
MK88
MK94
MK97
MK data
Normal data
Fold over basal
1.3
1.2
1.1
1
0.9
0.8
0.7
0.6
0.5
0
10
20
30
40
50
60
Time (min)

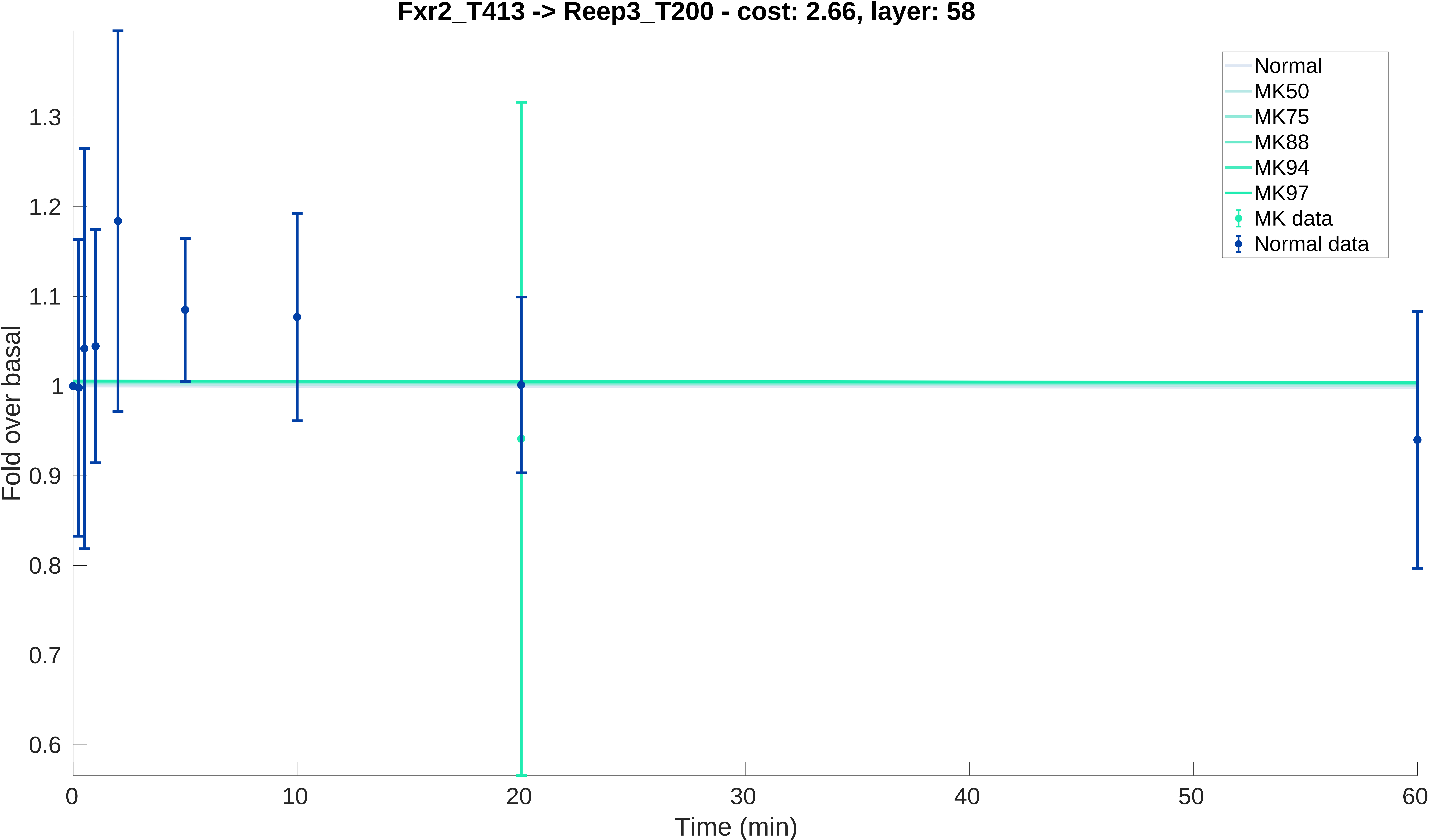

Fxr2_T413 -> Reep3_T200 - cost: 2.66, layer: 58
Normal
MK50
MK75
MK88
MK94
MK97
MK data
Normal data
1.3
1.2
1.1
1
0.9
0.8
0.7
0.6
Fold over basal
0
10
20
30
40
50
60
Time (min)

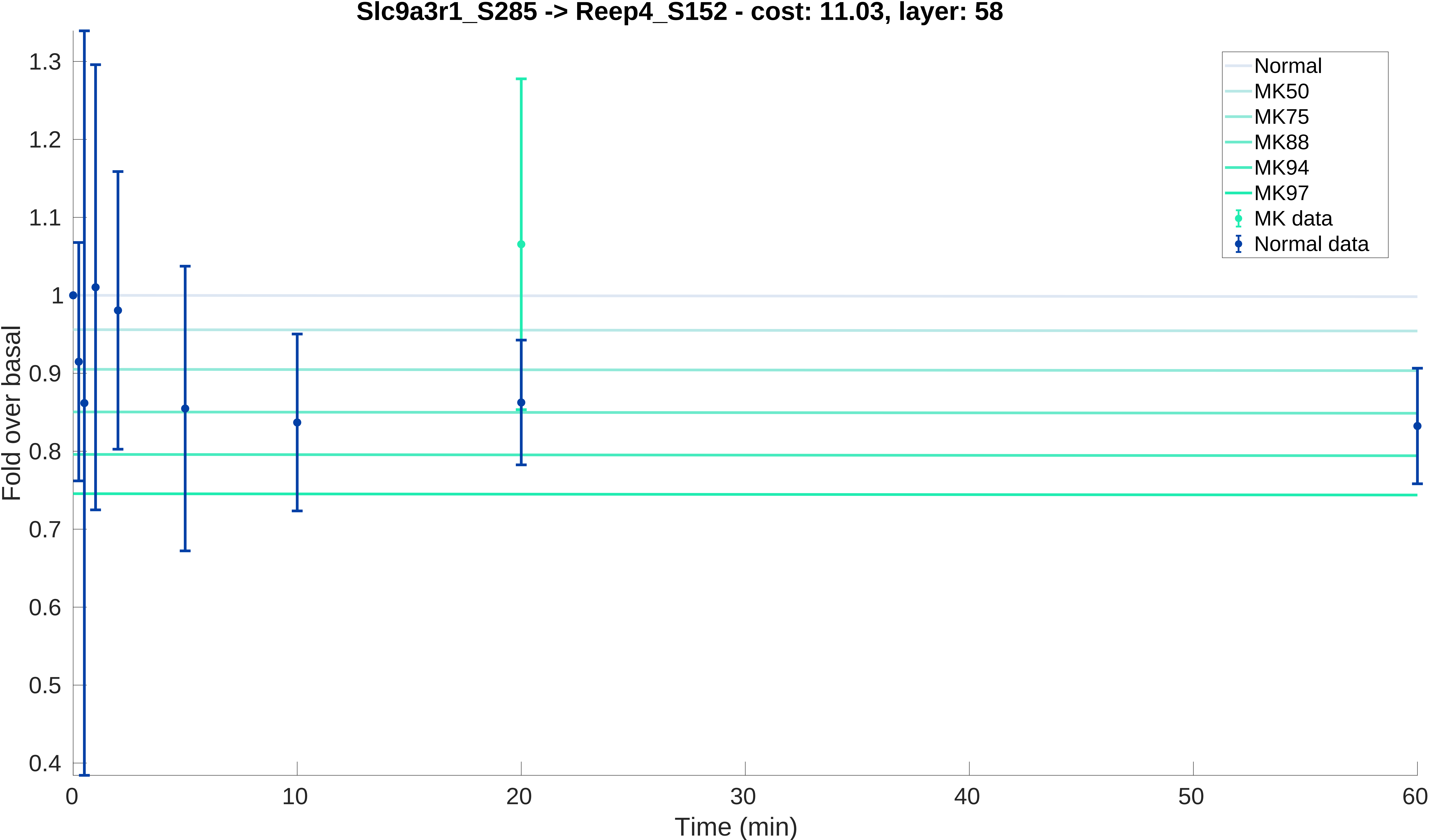

Slc9a3r1_S285 -> Reep4_S152 - cost: 11.03, layer: 58
Fold over basal
Time (min)
1.3
1.2
1.1
1
0.9
0.8
0.7
0.6
0.5
0.4
0
10
20
30
40
50
60
Normal
MK50
MK75
MK88
MK94
MK97
MK data
Normal data

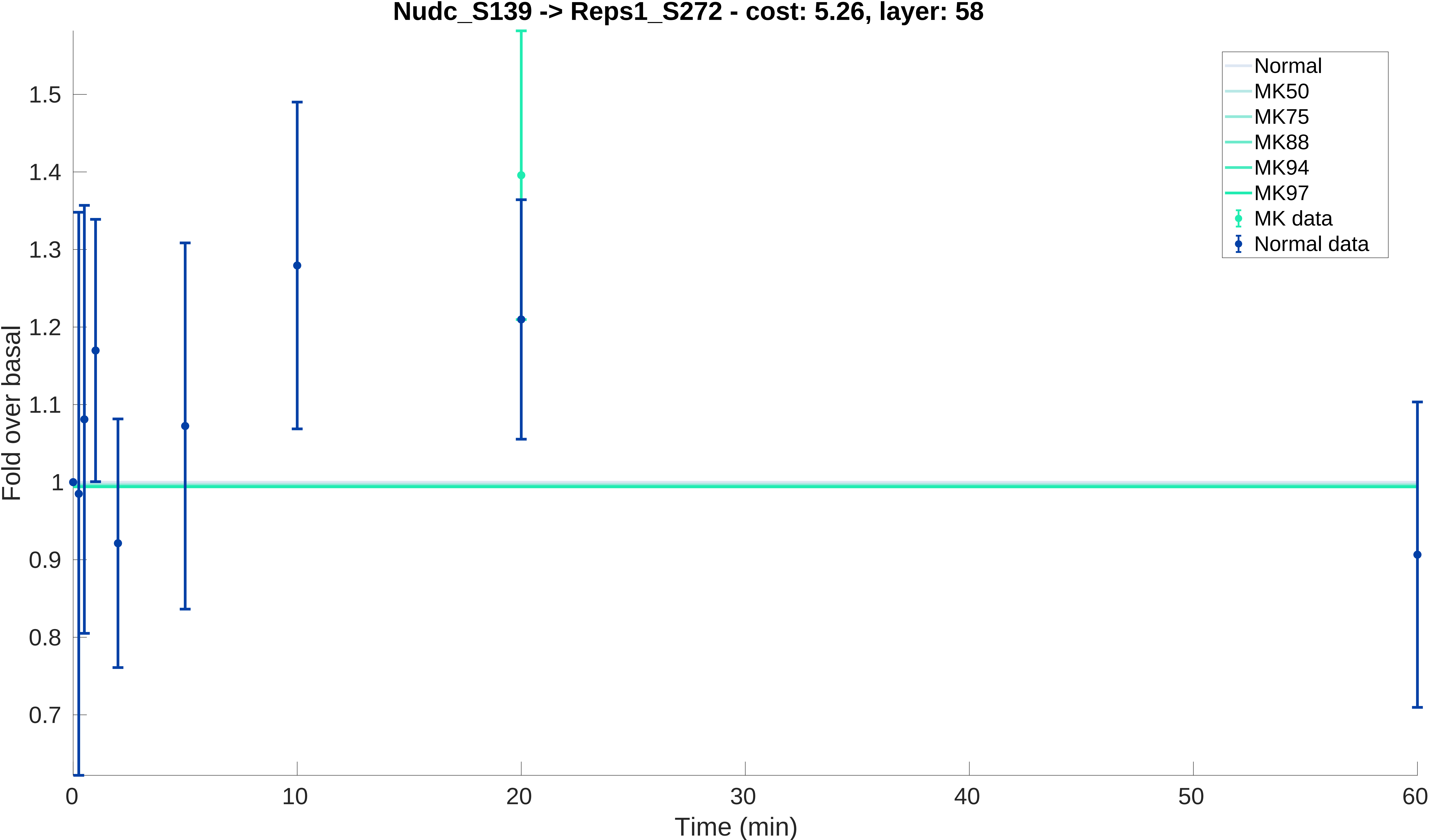

Nudc_S139 -> Reps1_S272 - cost: 5.26, layer: 58
Normal
MK50
MK75
MK88
MK94
MK97
MK data
Normal data
Fold over basal
1.5
1.4
1.3
1.2
1.1
1
0.9
0.8
0.7
0
10
20
30
40
50
60
Time (min)

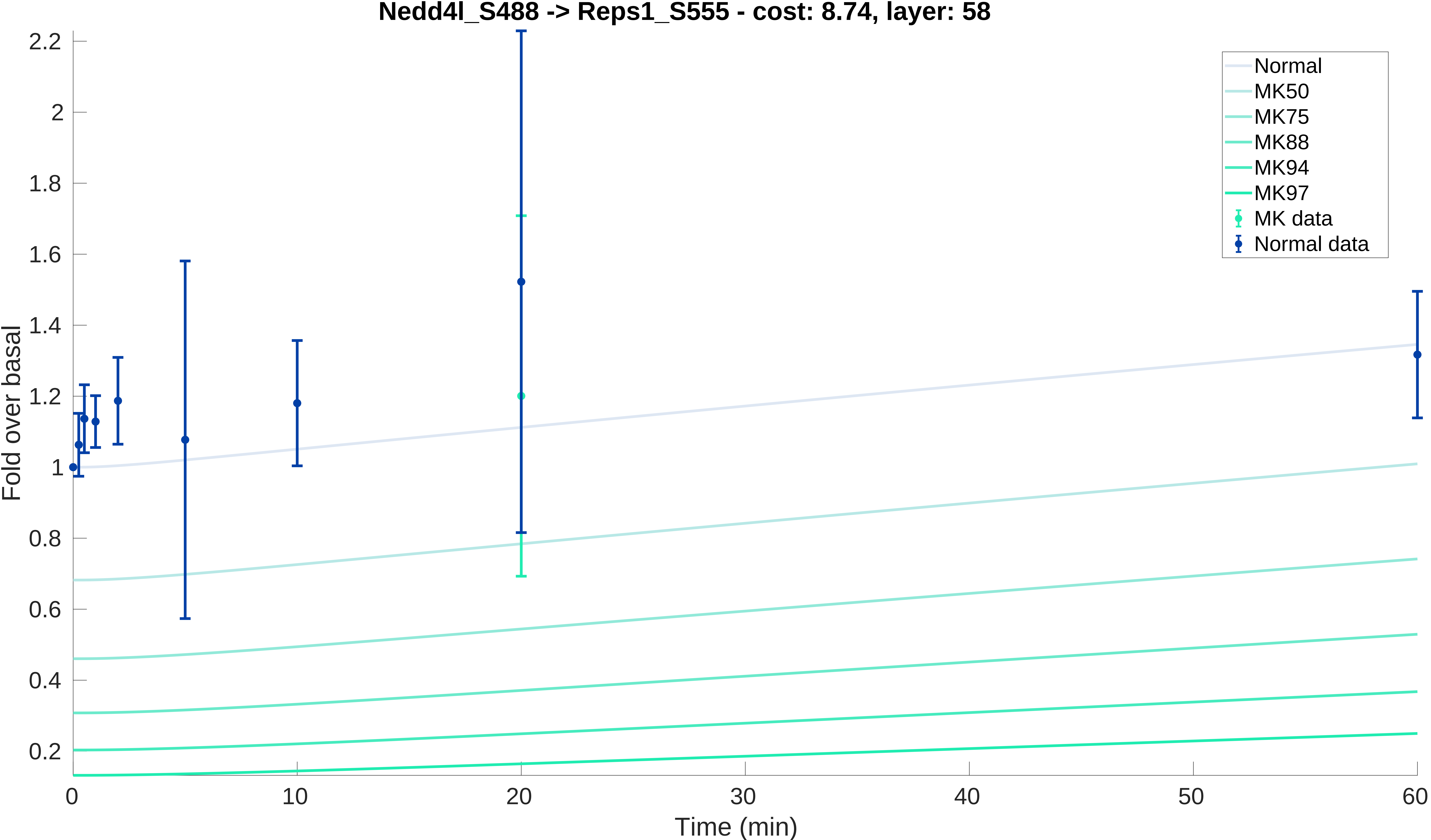

Nedd4l_S488 -> Reps1_S555 - cost: 8.74, layer: 58
Fold over basal
Time (min)
0
10
20
30
40
50
60
0.2
0.4
0.6
0.8
1
1.2
1.4
1.6
1.8
2
2.2
Normal
MK50
MK75
MK88
MK94
MK97
MK data
Normal data

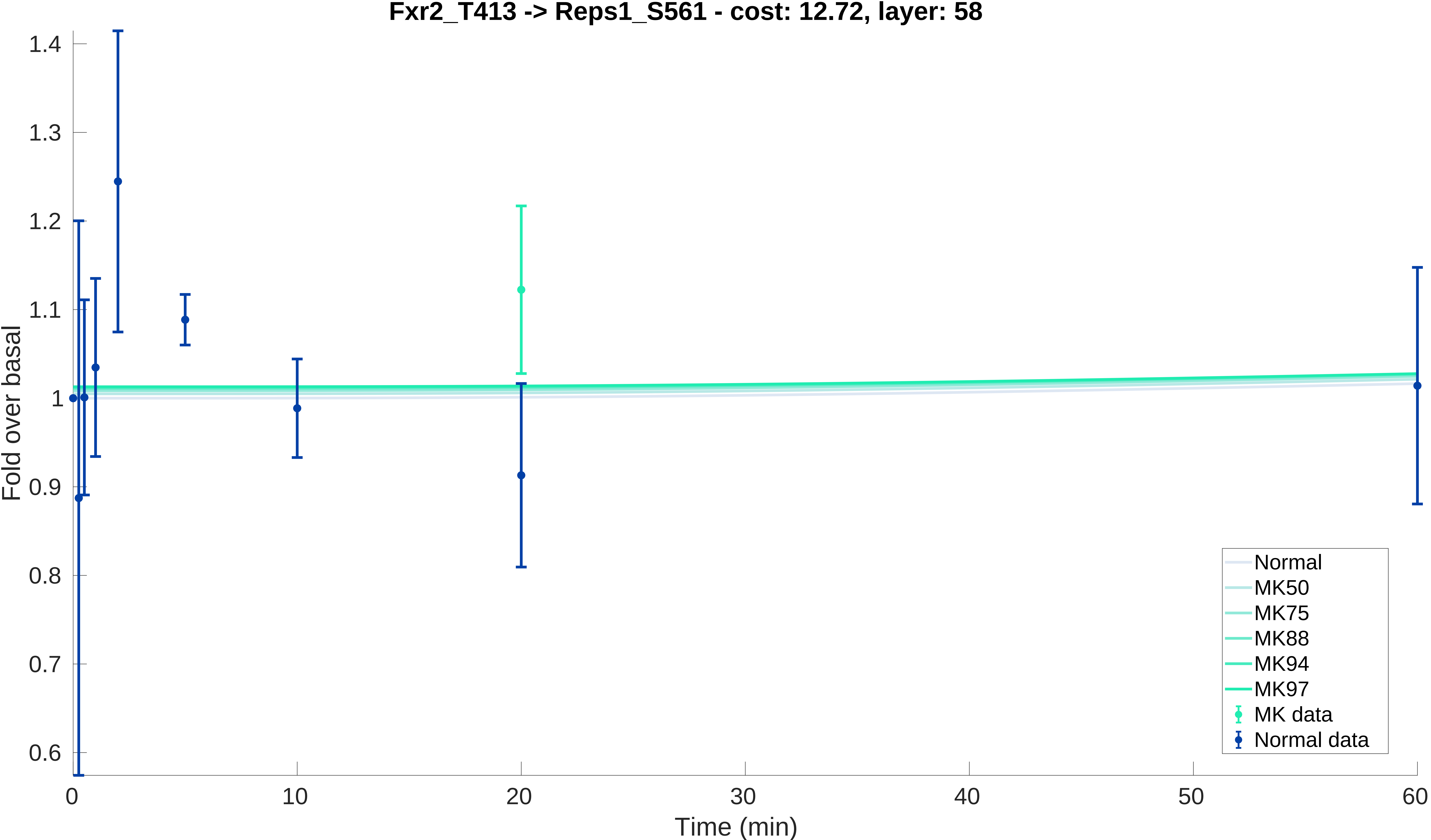

Fxr2_T413 -> Reps1_S561 - cost: 12.72, layer: 58
Fold over basal
Time (min)
1.4
1.3
1.2
1.1
1
0.9
0.8
0.7
0.6
0
10
20
30
40
50
60
Normal
MK50
MK75
MK88
MK94
MK97
MK data
Normal data

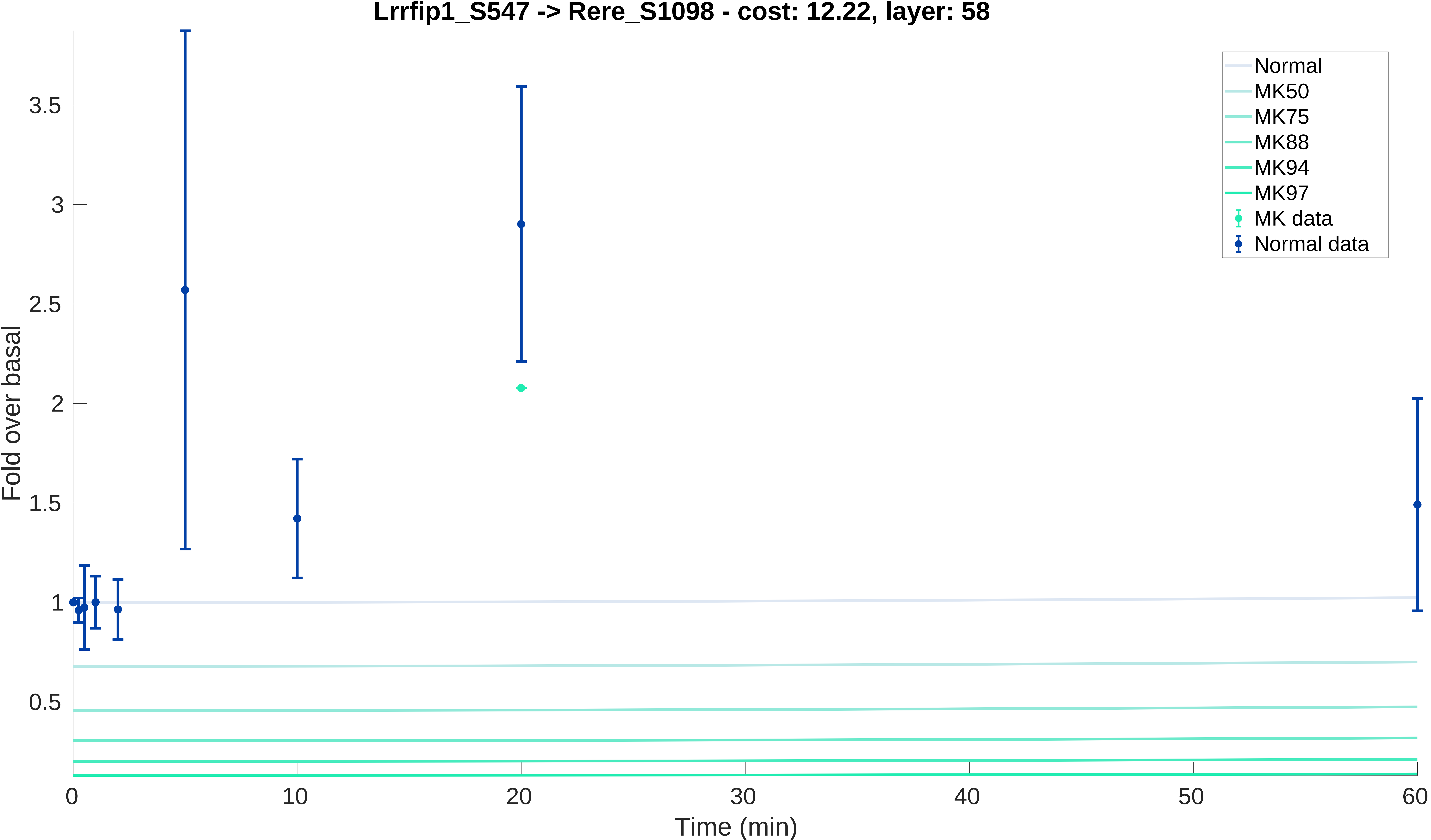

Lrrfip1_S547 -> Rere_S1098 - cost: 12.22, layer: 58
Fold over basal
Time (min)
0.5
1
1.5
2
2.5
3
3.5
0
10
20
30
40
50
60
Normal
MK50
MK75
MK88
MK94
MK97
MK data
Normal data

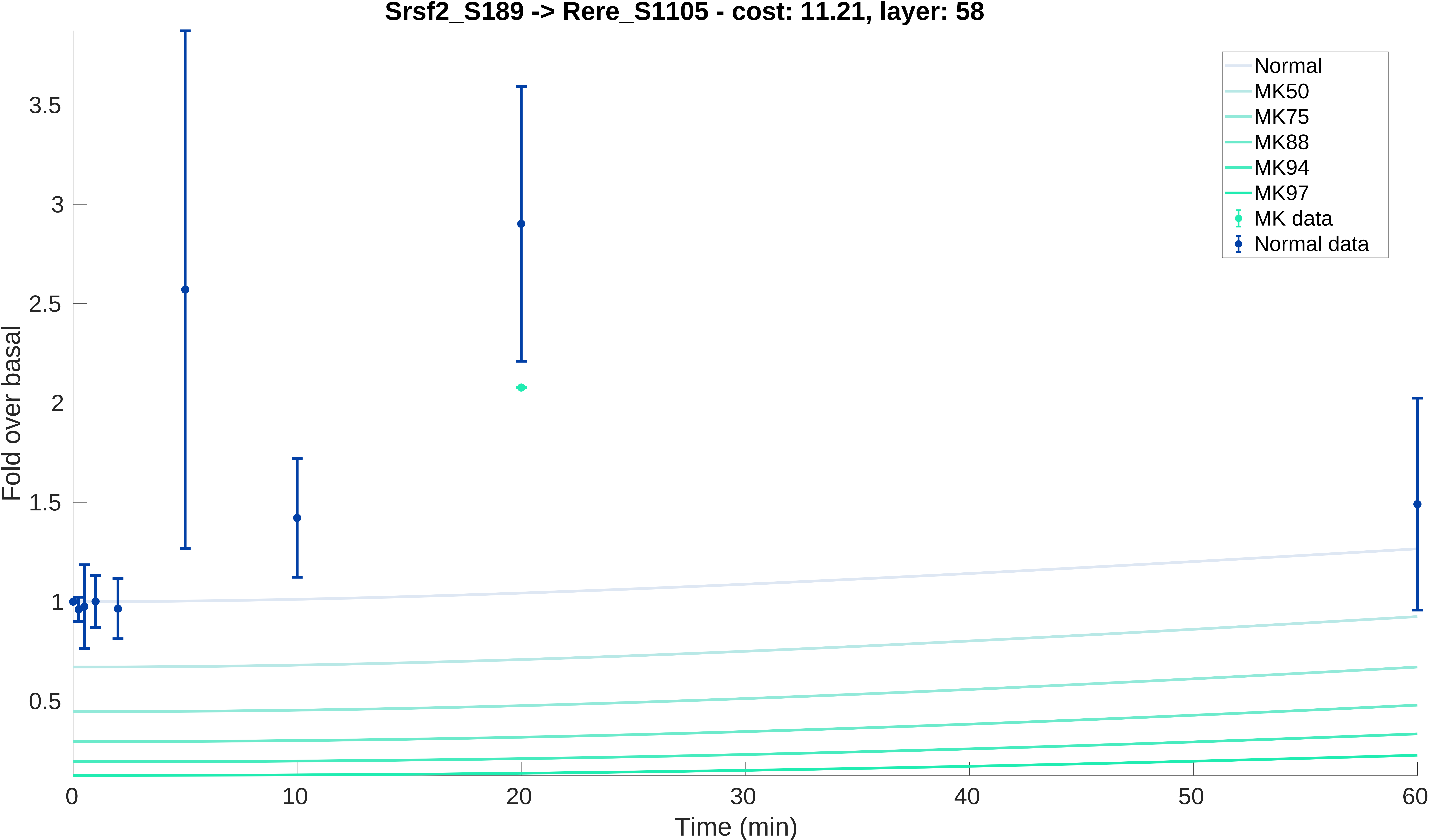

Srsf2_S189 -> Rere_S1105 - cost: 11.21, layer: 58
Fold over basal
Time (min)
0
10
20
30
40
50
60
0.5
1
1.5
2
2.5
3
3.5
Normal
MK50
MK75
MK88
MK94
MK97
MK data
Normal data

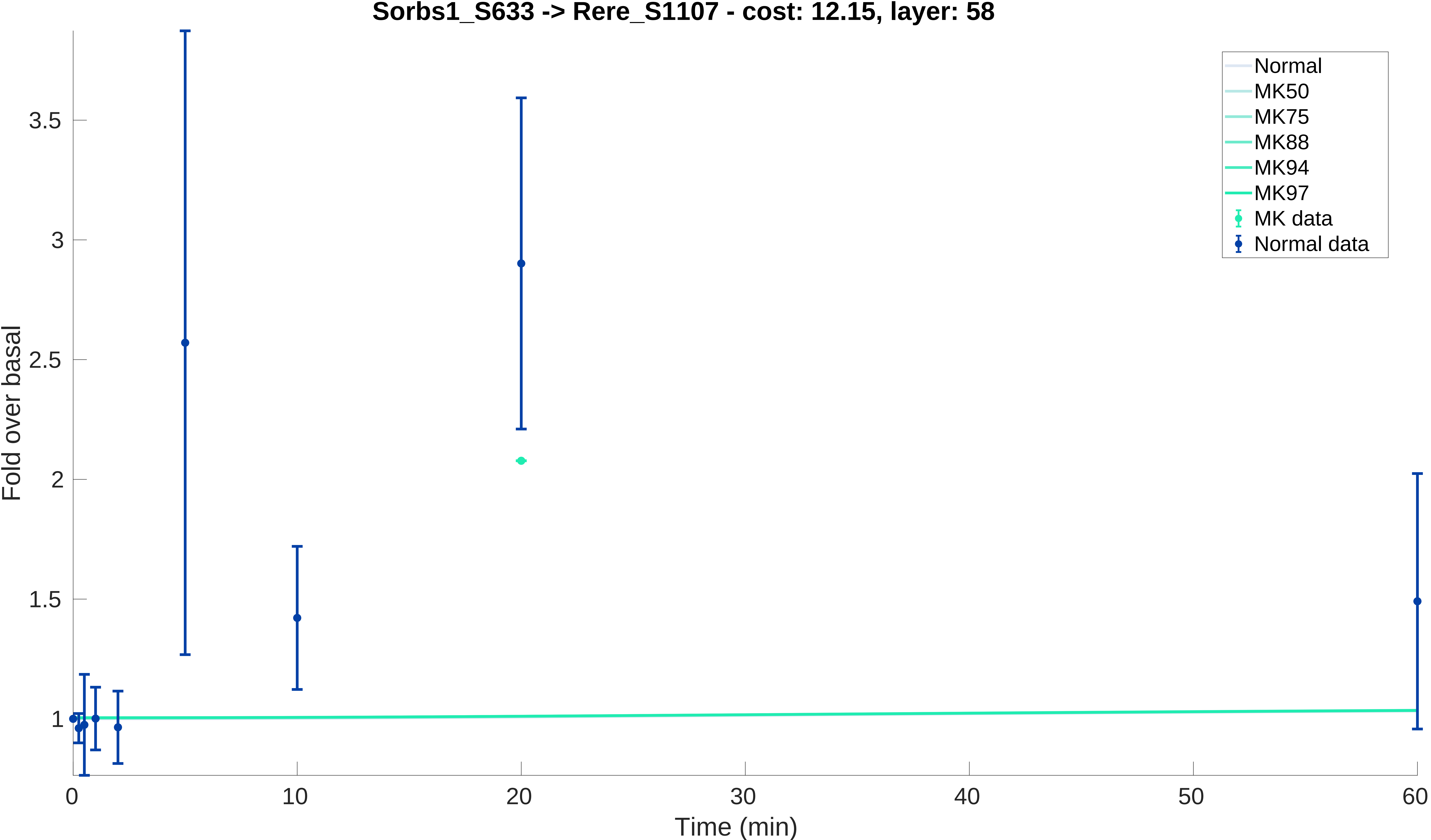

Sorbs1_S633 -> Rere_S1107 - cost: 12.15, layer: 58
Fold over basal
Time (min)
Normal
MK50
MK75
MK88
MK94
MK97
MK data
Normal data
3.5
3
2.5
2
1.5
1
0
10
20
30
40
50
60

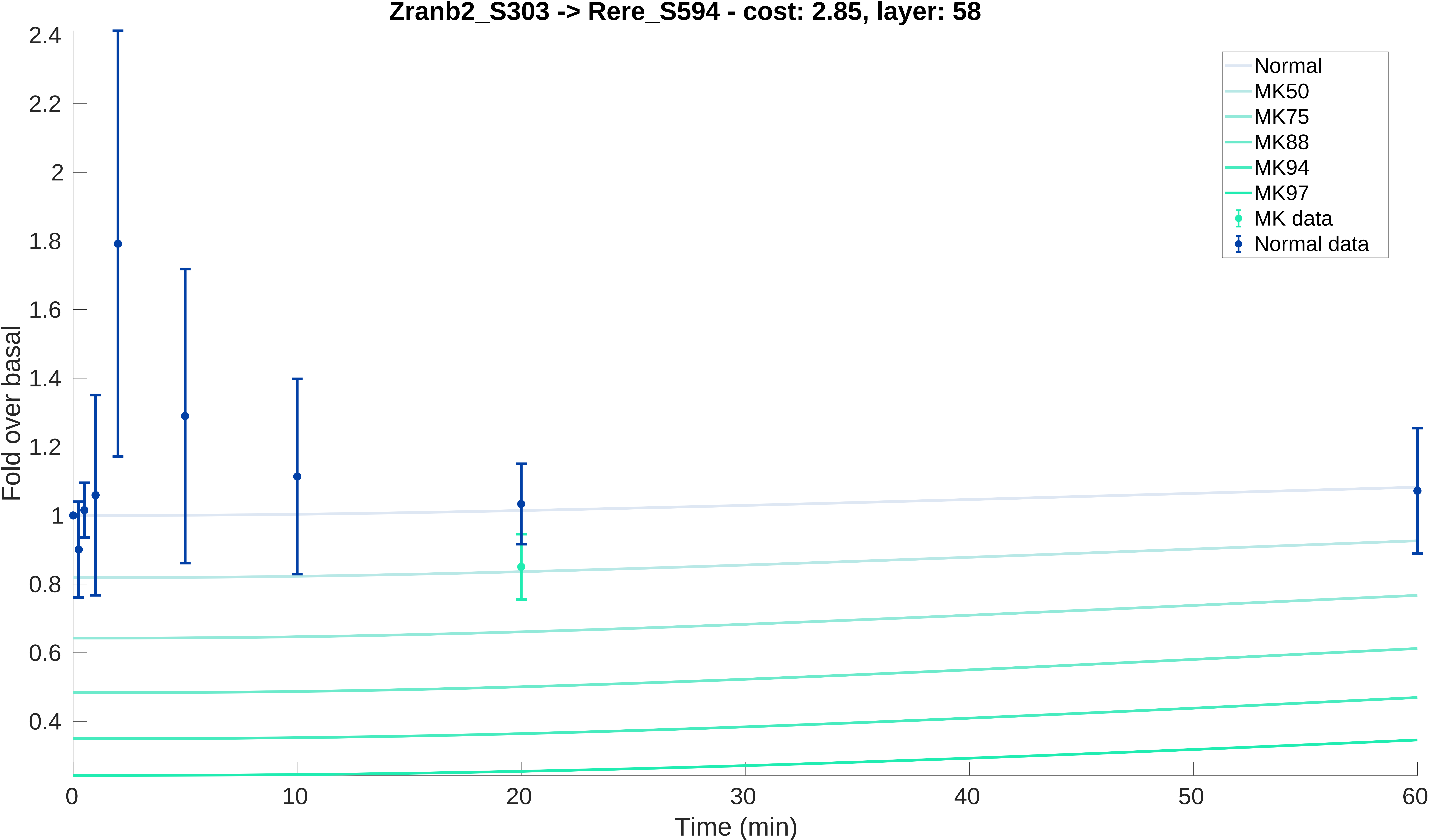

Zranb2_S303 -> Rere_S594 - cost: 2.85, layer: 58
Fold over basal
2.4
2.2
2
1.8
1.6
1.4
1.2
1
0.8
0.6
0.4
0
10
20
30
40
50
60
Time (min)
Normal
MK50
MK75
MK88
MK94
MK97
MK data
Normal data

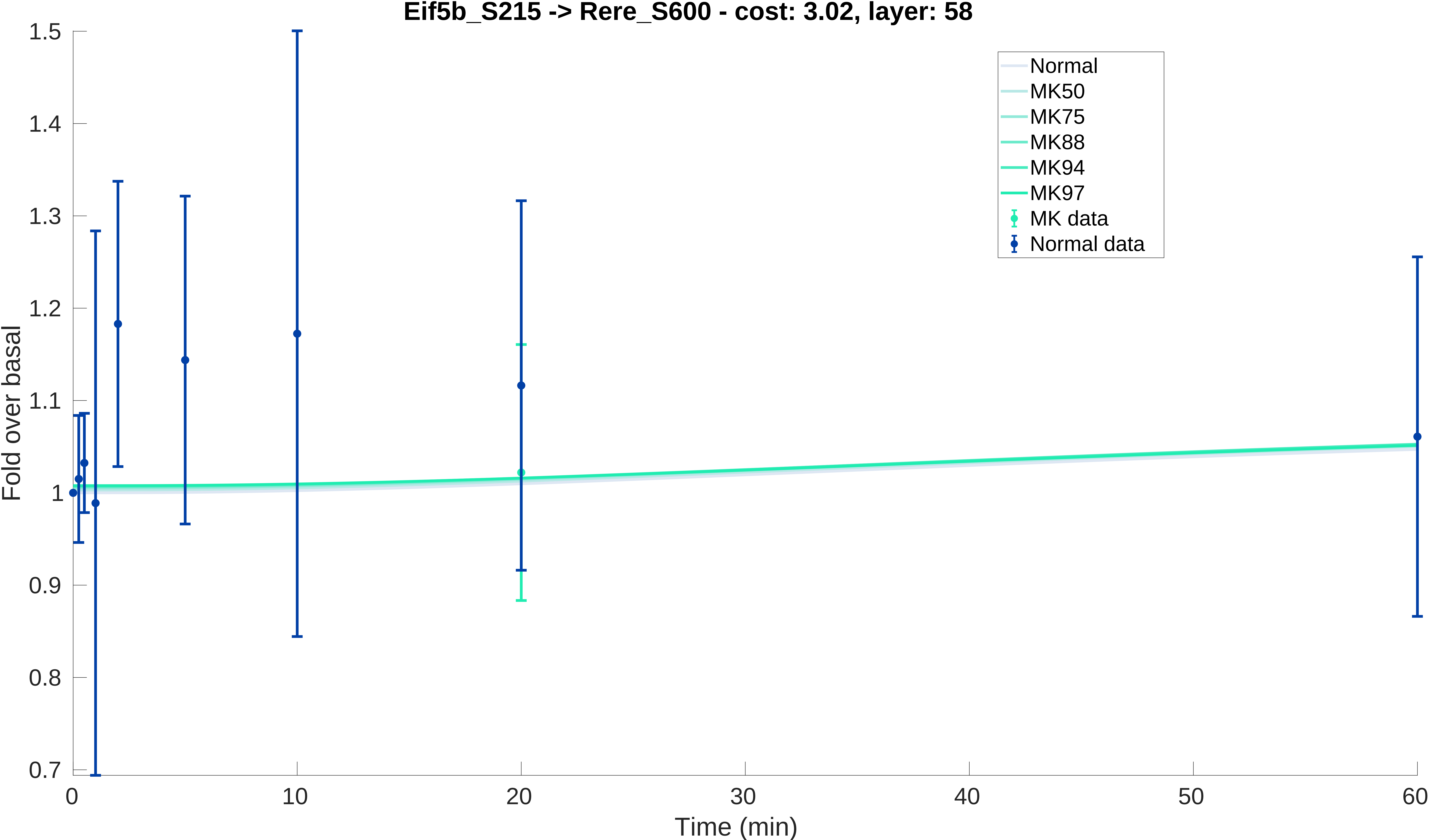

Eif5b_S215 -> Rere_S600 - cost: 3.02, layer: 58
Fold over basal
Time (min)
Normal
MK50
MK75
MK88
MK94
MK97
MK data
Normal data
1.5
1.4
1.3
1.2
1.1
1
0.9
0.8
0.7
0
10
20
30
40
50
60

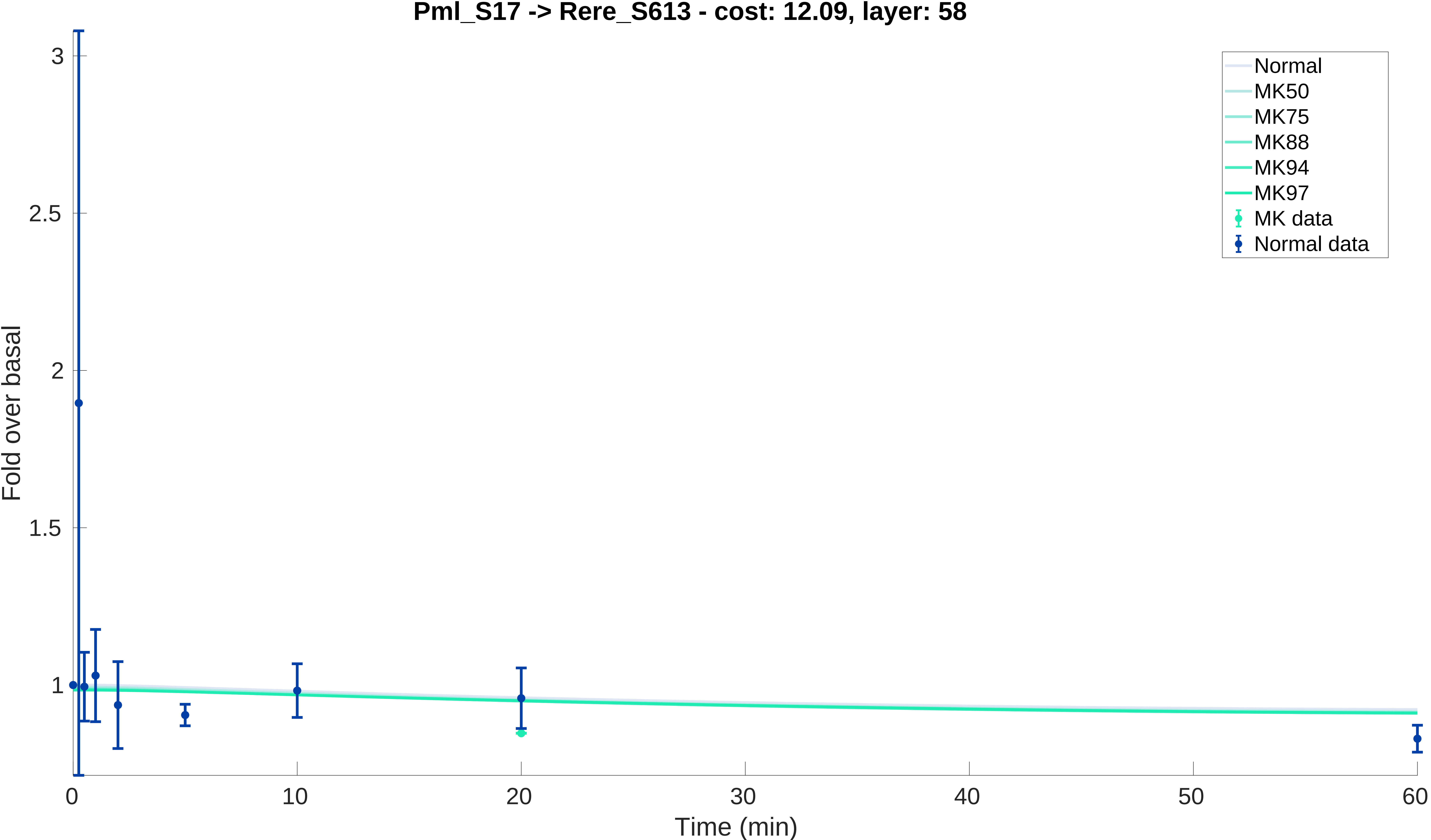

Pml_S17 -> Rere_S613 - cost: 12.09, layer: 58
Fold over basal
Time (min)
3
2.5
2
1.5
1
0
10
20
30
40
50
60
Normal
MK50
MK75
MK88
MK94
MK97
MK data
Normal data

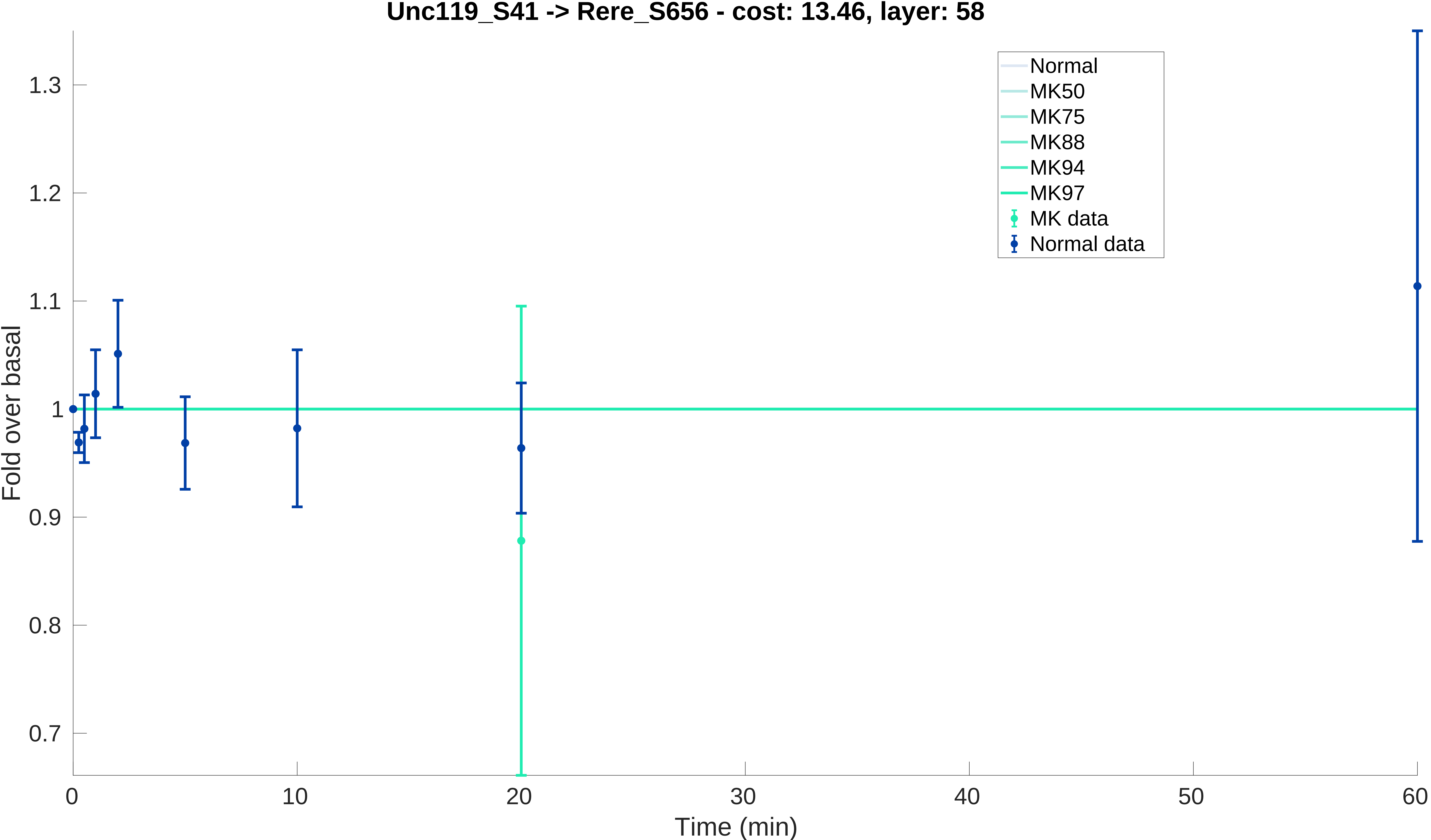
Unc119_S41 -> Rere_S656 - cost: 13.46, layer: 58
Normal
MK50
MK75
MK88
MK94
MK97
MK data
Normal data
Fold over basal
1.3
1.2
1.1
1
0.9
0.8
0.7
0
10
20
30
40
50
60
Time (min)

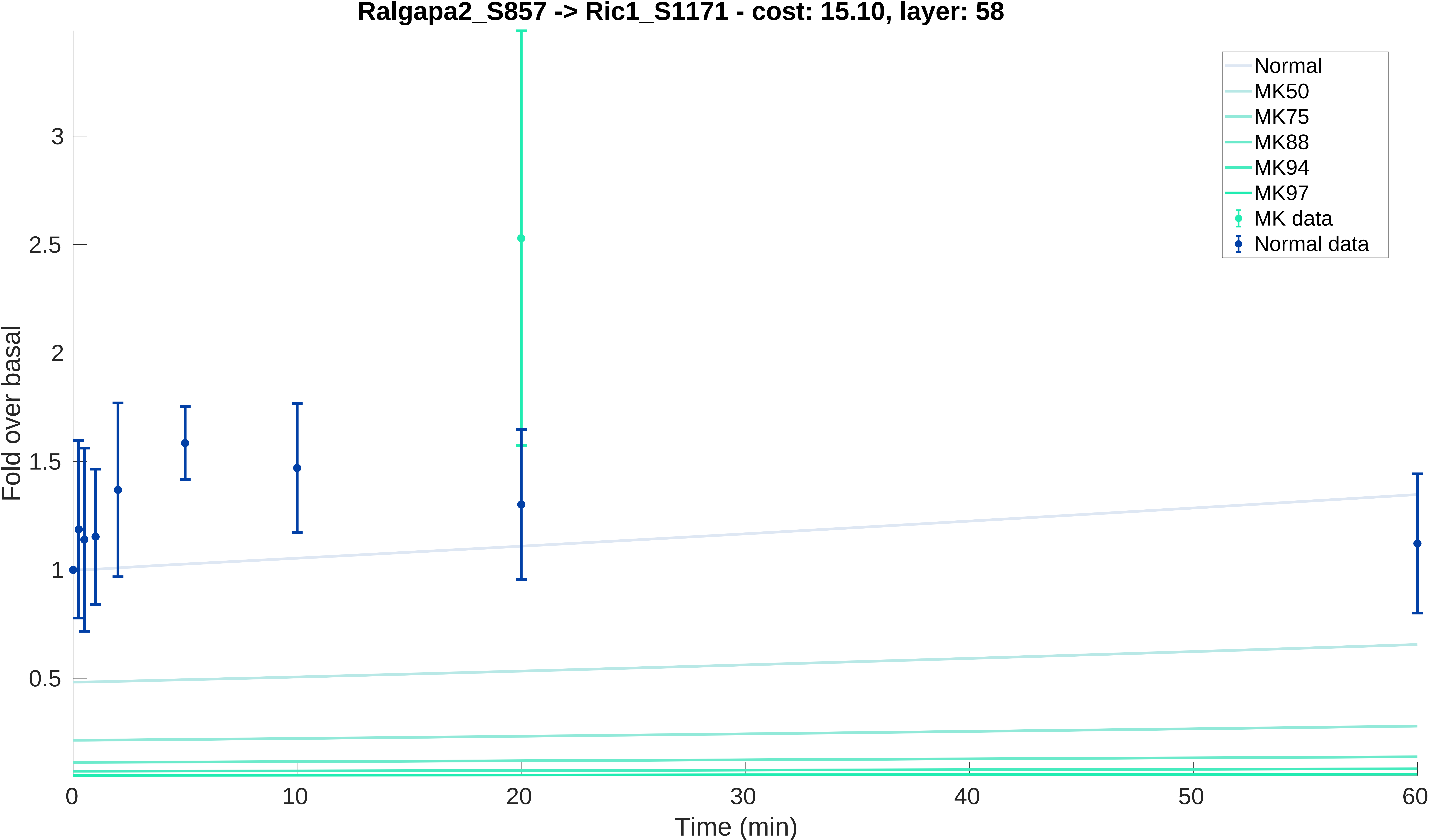

Ralgapa2_S857 -> Ric1_S1171 - cost: 15.10, layer: 58
Normal
MK50
MK75
MK88
MK94
MK97
MK data
Normal data
Fold over basal
0.5
1
1.5
2
2.5
3
Time (min)
0
10
20
30
40
50
60

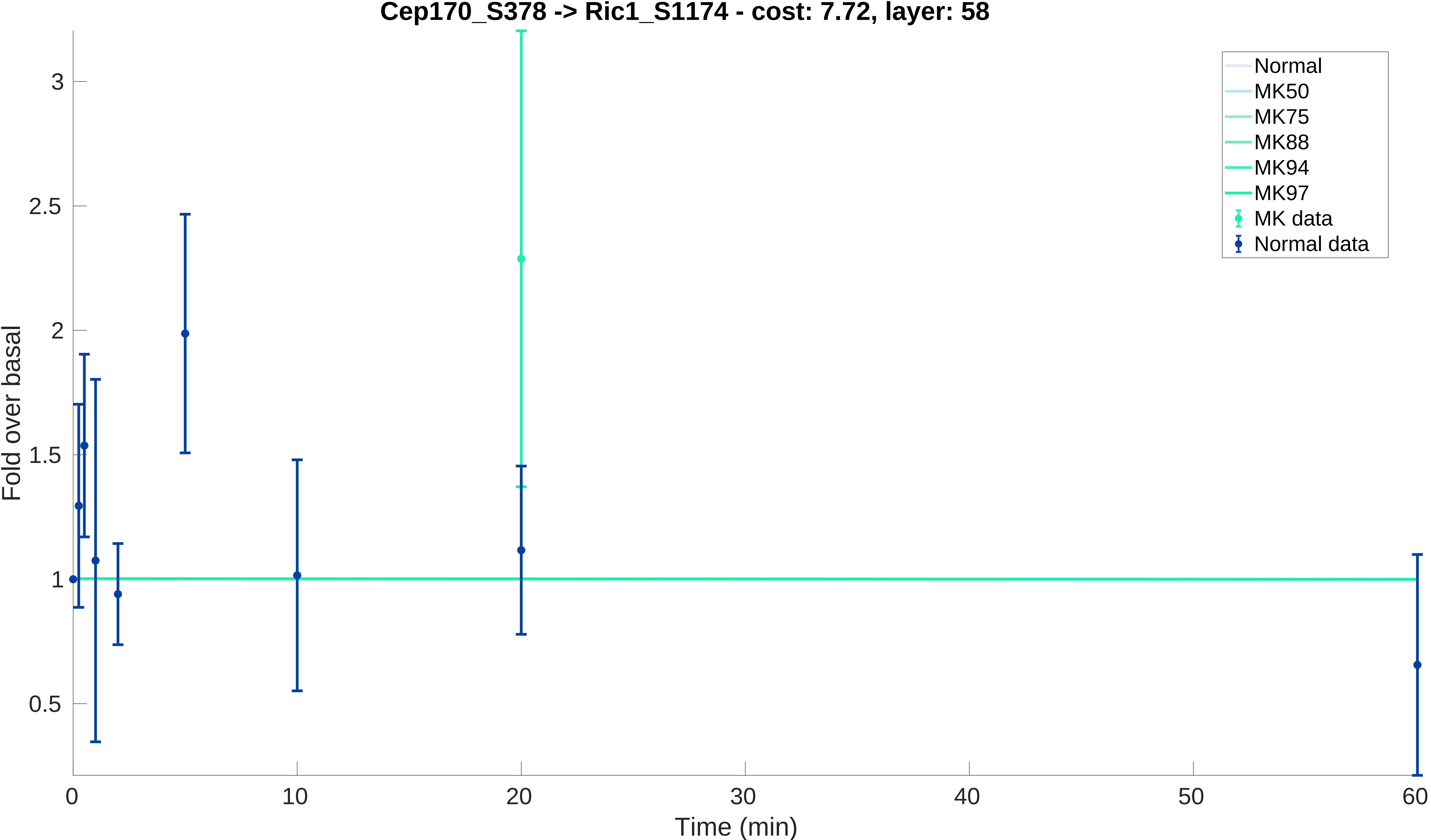

Cep170_S378 -> Ric1_S1174 - cost: 7.72, layer: 58
Fold over basal
Time (min)
0.5
1
1.5
2
2.5
3
0
10
20
30
40
50
60
Normal
MK50
MK75
MK88
MK94
MK97
MK data
Normal data

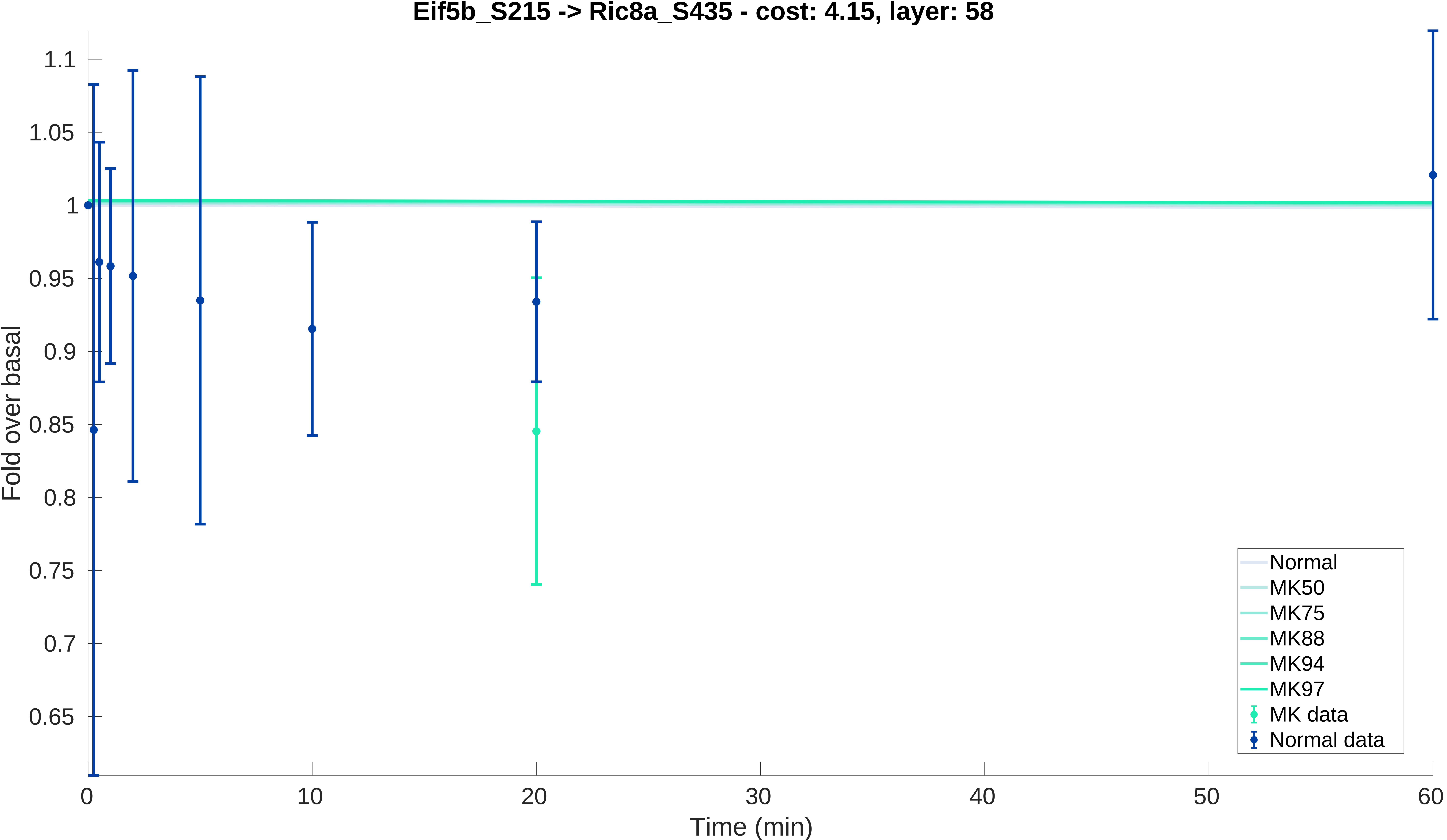

Eif5b_S215 -> Ric8a_S435 - cost: 4.15, layer: 58
Fold over basal
Time (min)
1.1
1.05
1
0.95
0.9
0.85
0.8
0.75
0.7
0.65
0
10
20
30
40
50
60
Normal
MK50
MK75
MK88
MK94
MK97
MK data
Normal data

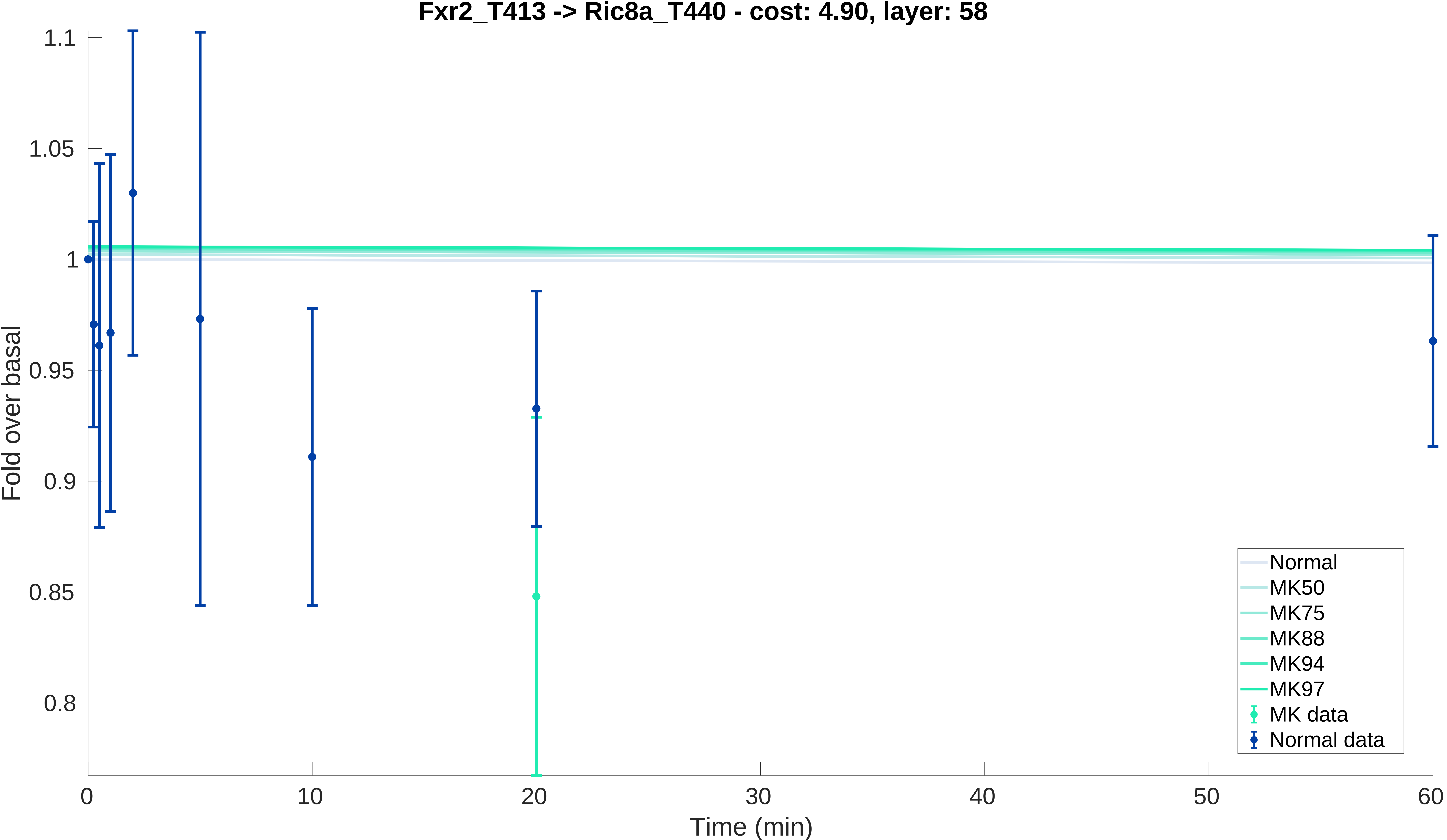

Fxr2_T413 -> Ric8a_T440 - cost: 4.90, layer: 58
Fold over basal
Time (min)
1.1
1.05
1
0.95
0.9
0.85
0.8
0
10
20
30
40
50
60
Normal
MK50
MK75
MK88
MK94
MK97
MK data
Normal data

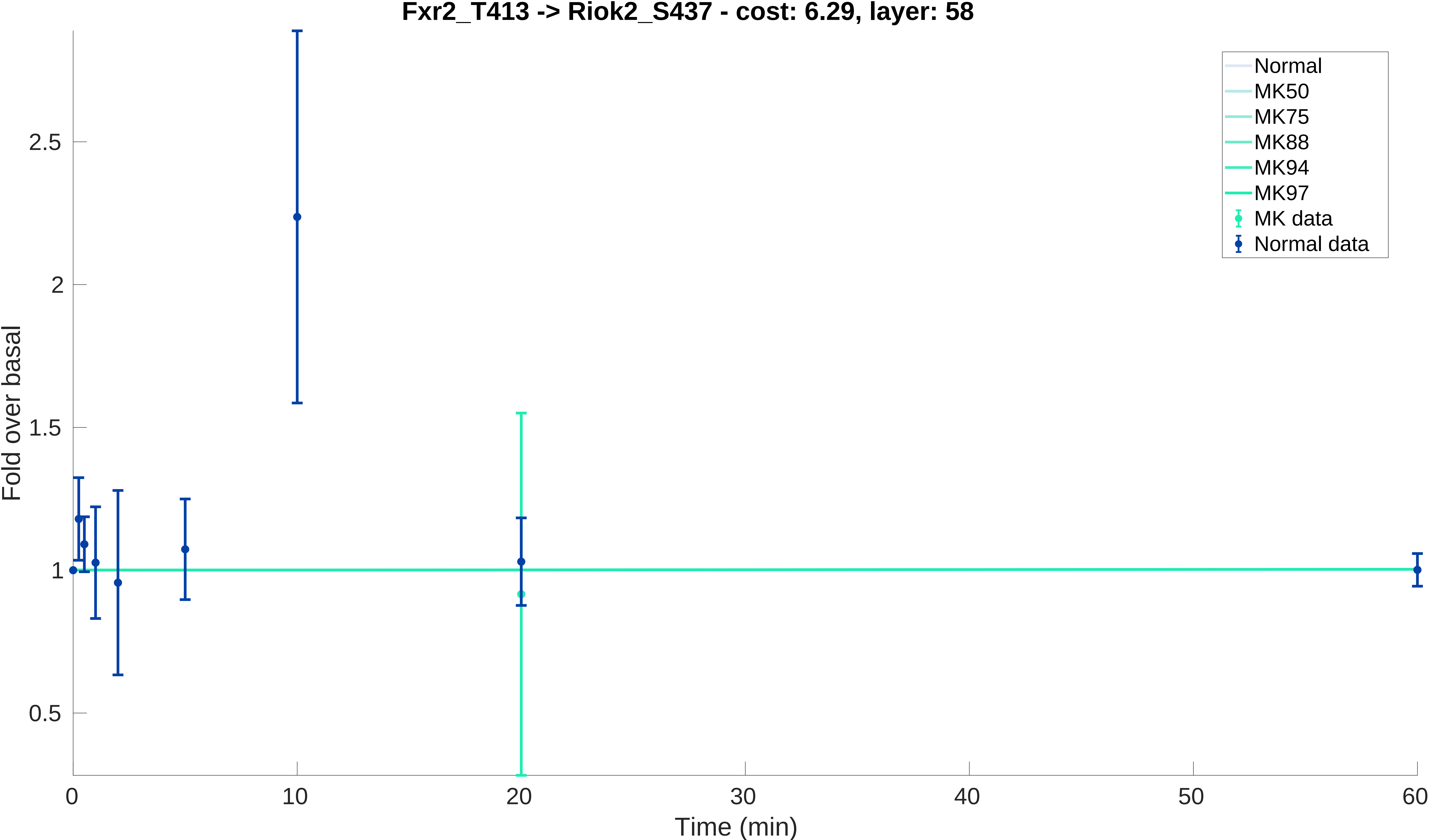

Fxr2_T413 -> Riok2_S437 - cost: 6.29, layer: 58
Normal
MK50
MK75
MK88
MK94
MK97
MK data
Normal data
Fold over basal
0.5
1
1.5
2
2.5
0
10
20
30
40
50
60
Time (min)

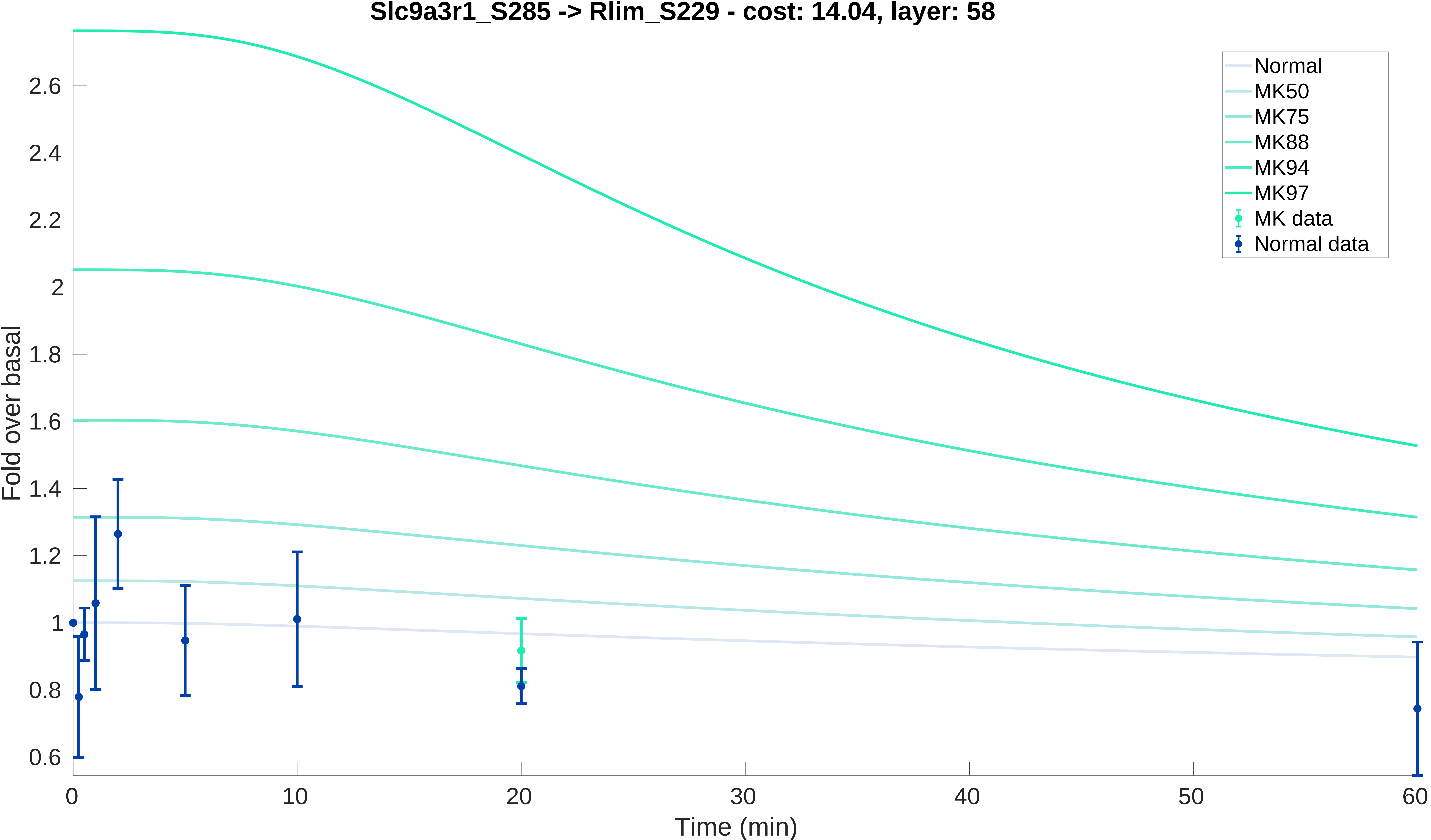

Slc9a3r1_S285 -> Rlim_S229 - cost: 14.04, layer: 58
Fold over basal
Time (min)
0.6
0.8
1
1.2
1.4
1.6
1.8
2
2.2
2.4
2.6
0
10
20
30
40
50
60
Normal
MK50
MK75
MK88
MK94
MK97
MK data
Normal data

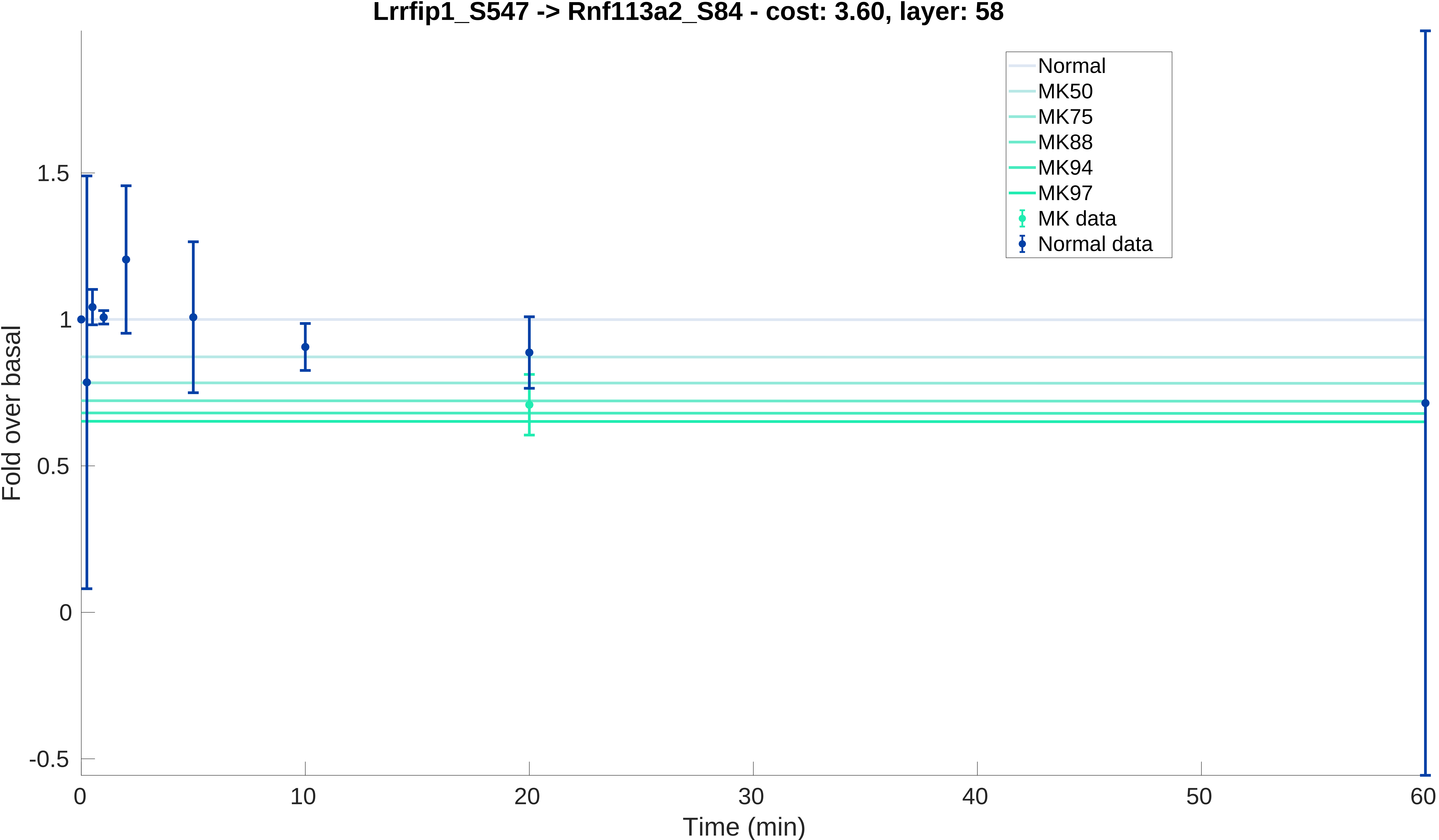
Lrrfip1_S547 -> Rnf113a2_S84 - cost: 3.60, layer: 58
Fold over basal
Time (min)
1.5
1
0.5
0
-0.5
0
10
20
30
40
50
60
Normal
MK50
MK75
MK88
MK94
MK97
MK data
Normal data

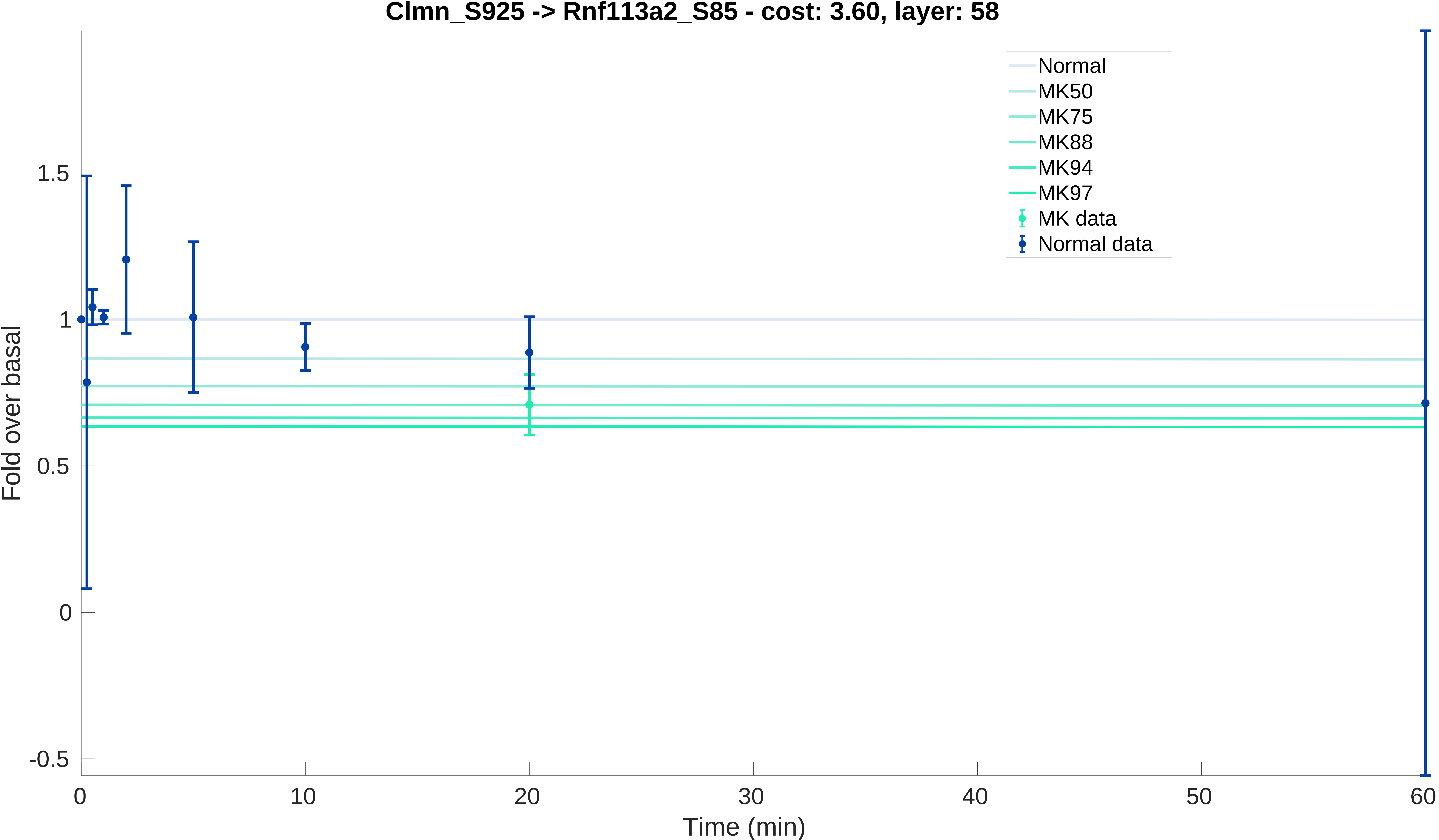

Clmn_S925 -> Rnf113a2_S85 - cost: 3.60, layer: 58
Normal
MK50
MK75
MK88
MK94
MK97
MK data
Normal data
1.5
1
0.5
0
-0.5
Fold over basal
0
10
20
30
40
50
60
Time (min)

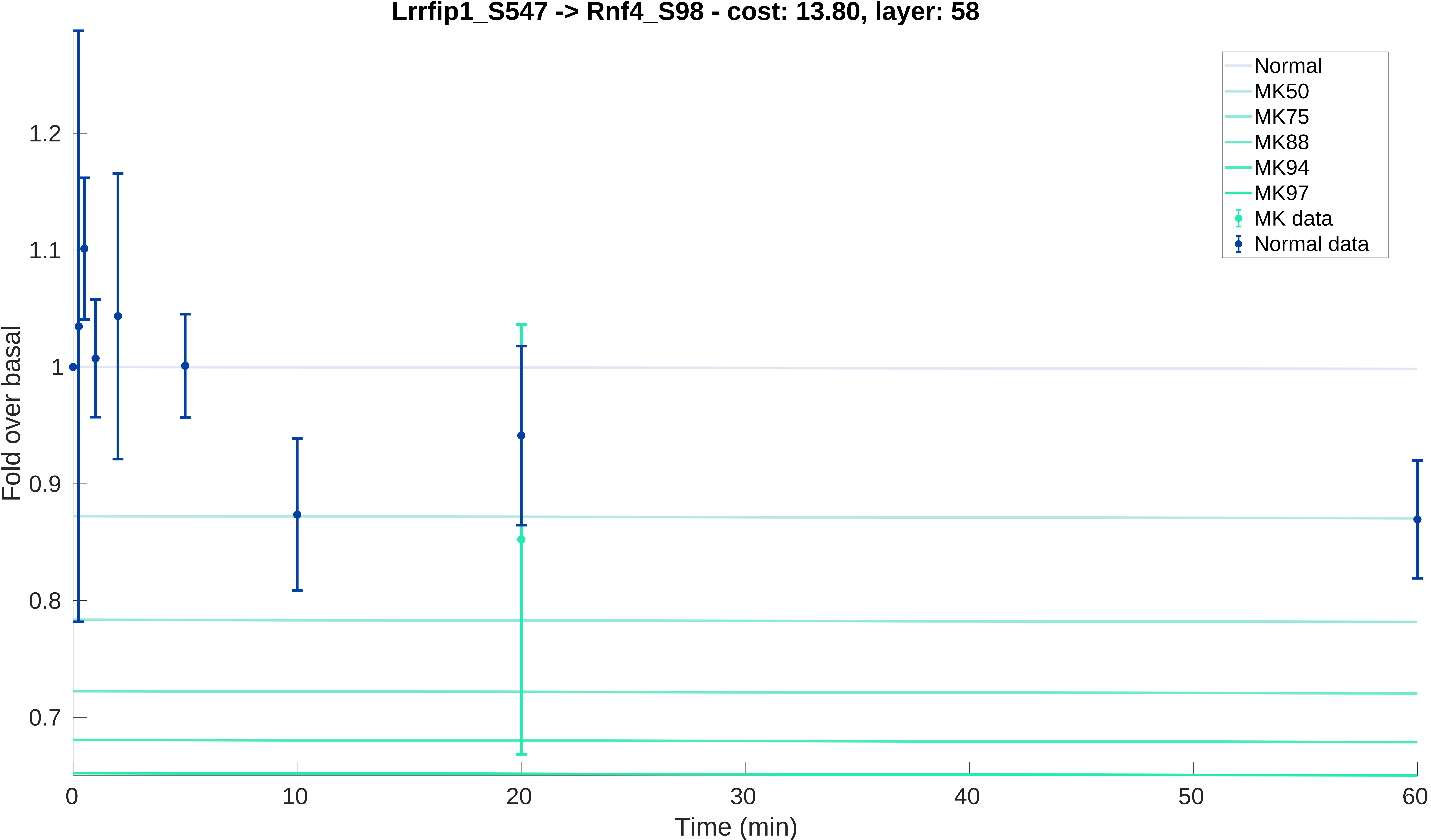

Lrrfip1_S547 -> Rnf4_S98 - cost: 13.80, layer: 58
Normal
MK50
MK75
MK88
MK94
MK97
MK data
Normal data
Fold over basal
1.2
1.1
1
0.9
0.8
0.7
0
10
20
30
40
50
60
Time (min)

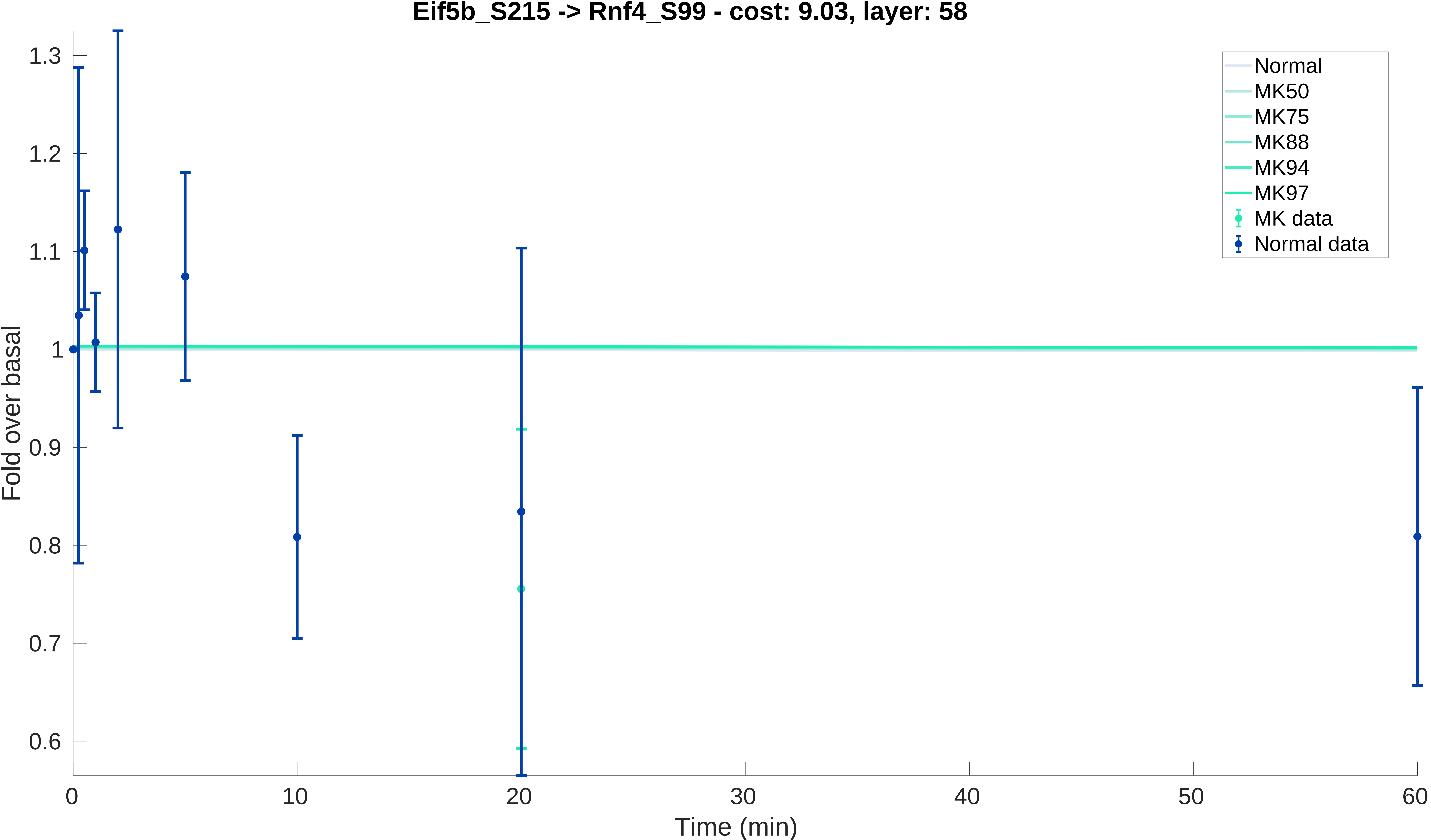

Eif5b_S215 -> Rnf4_S99 - cost: 9.03, layer: 58
Fold over basal
Time (min)
1.3
1.2
1.1
1
0.9
0.8
0.7
0.6
0
10
20
30
40
50
60
Normal
MK50
MK75
MK88
MK94
MK97
MK data
Normal data

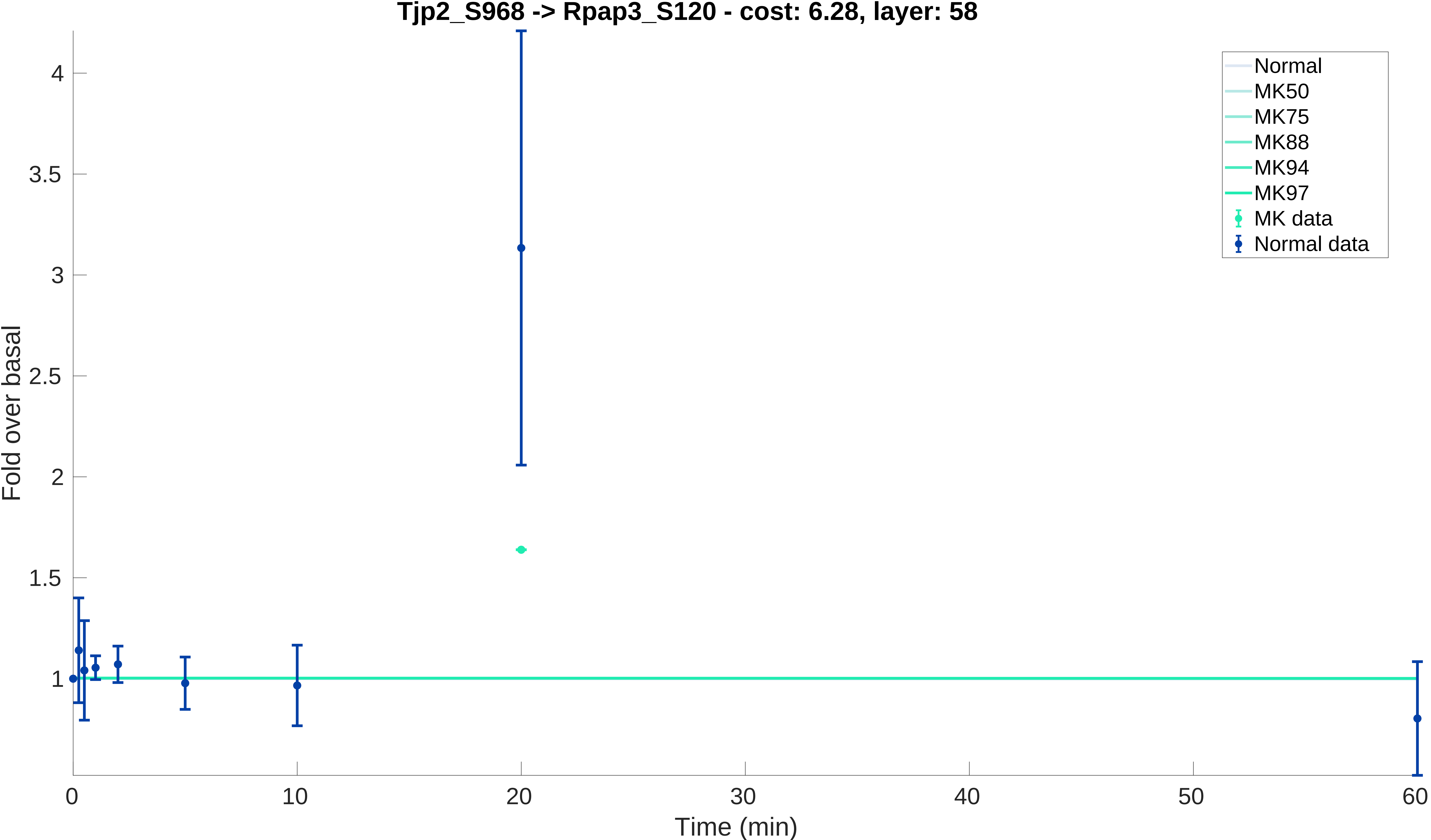
Tjp2_S968 -> Rpap3_S120 - cost: 6.28, layer: 58
Fold over basal
Time (min)
0
10
20
30
40
50
60
1
1.5
2
2.5
3
3.5
4
Normal
MK50
MK75
MK88
MK94
MK97
MK data
Normal data

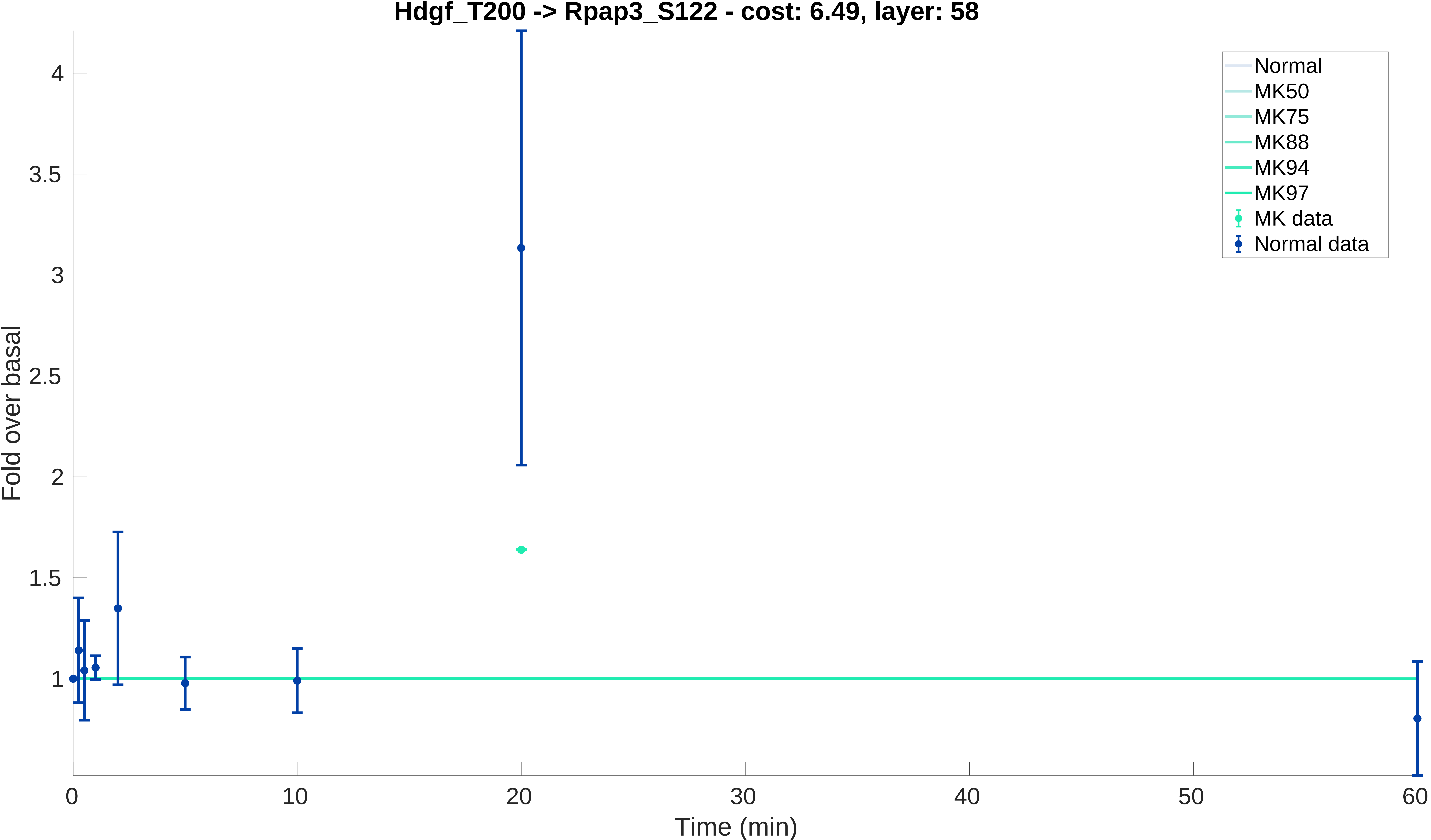
Hdgf_T200 -> Rpap3_S122 - cost: 6.49, layer: 58
Fold over basal
Time (min)
4
3.5
3
2.5
2
1.5
1
0
10
20
30
40
50
60
Normal
MK50
MK75
MK88
MK94
MK97
MK data
Normal data

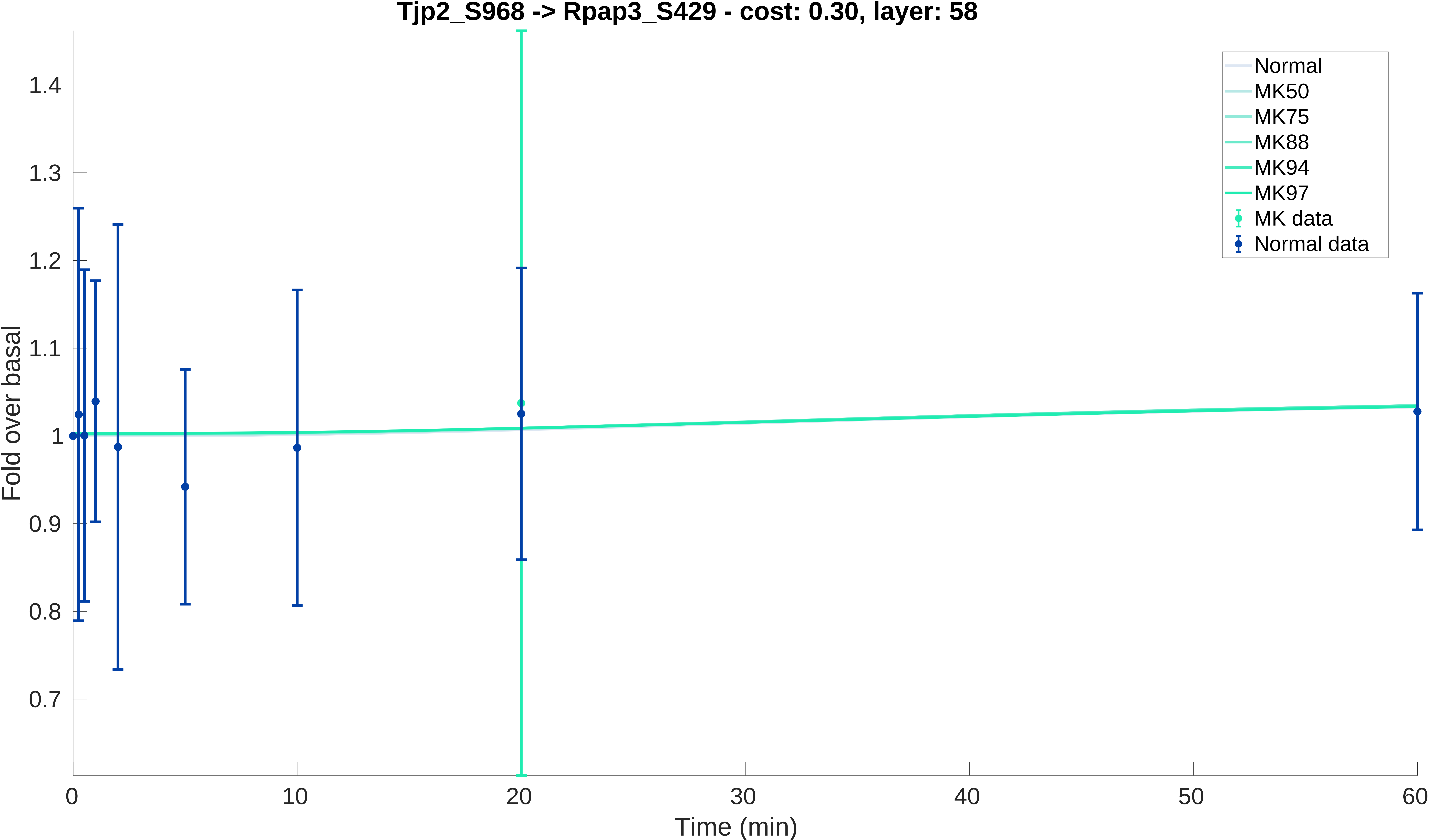
Tjp2_S968 -> Rpap3_S429 - cost: 0.30, layer: 58
Fold over basal
Time (min)
1.4
1.3
1.2
1.1
1
0.9
0.8
0.7
0
10
20
30
40
50
60
Normal
MK50
MK75
MK88
MK94
MK97
MK data
Normal data

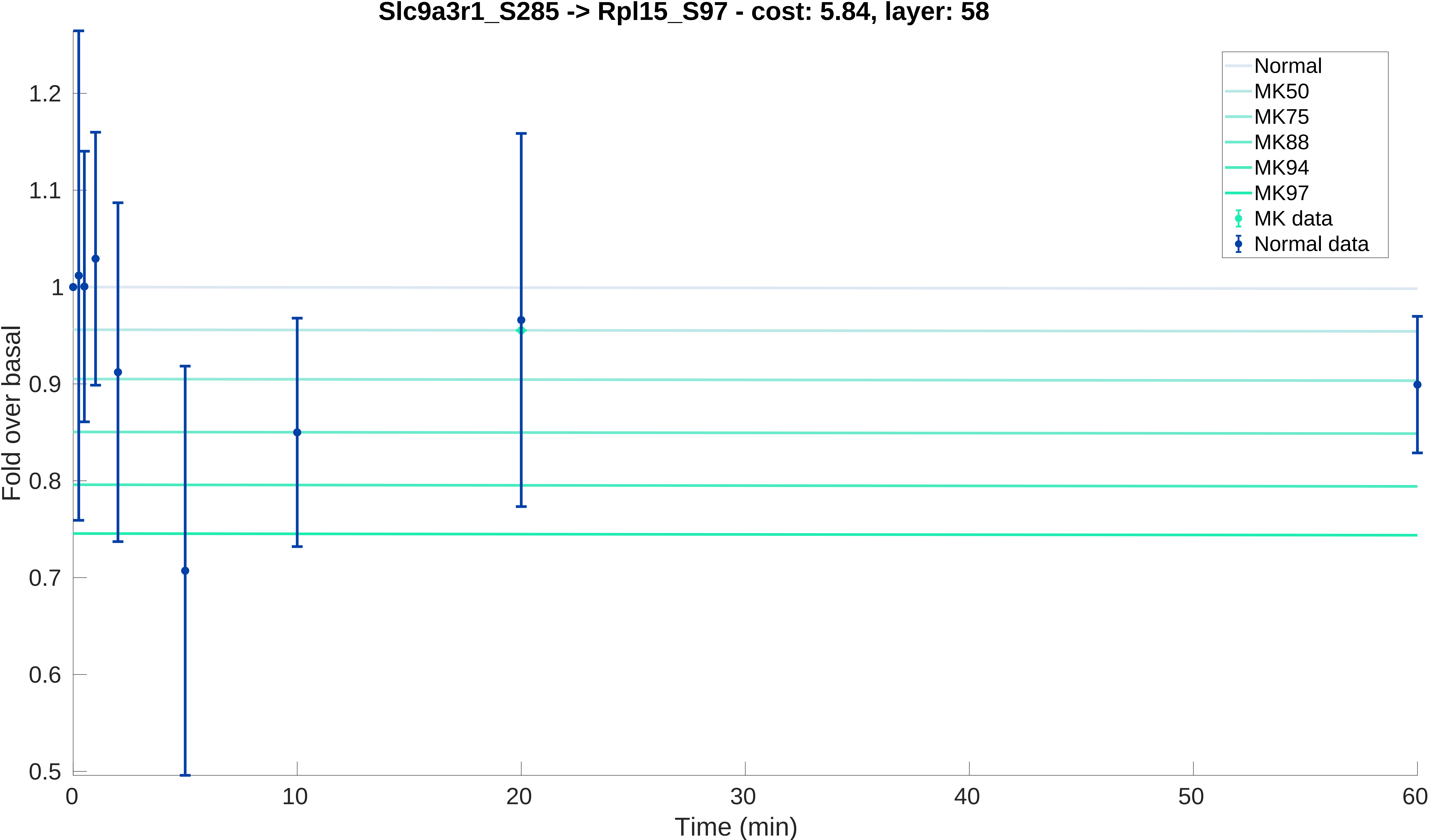

Slc9a3r1_S285 -> Rpl15_S97 - cost: 5.84, layer: 58
Normal
MK50
MK75
MK88
MK94
MK97
MK data
Normal data
Fold over basal
Time (min)
0.5
0.6
0.7
0.8
0.9
1
1.1
1.2
0
10
20
30
40
50
60

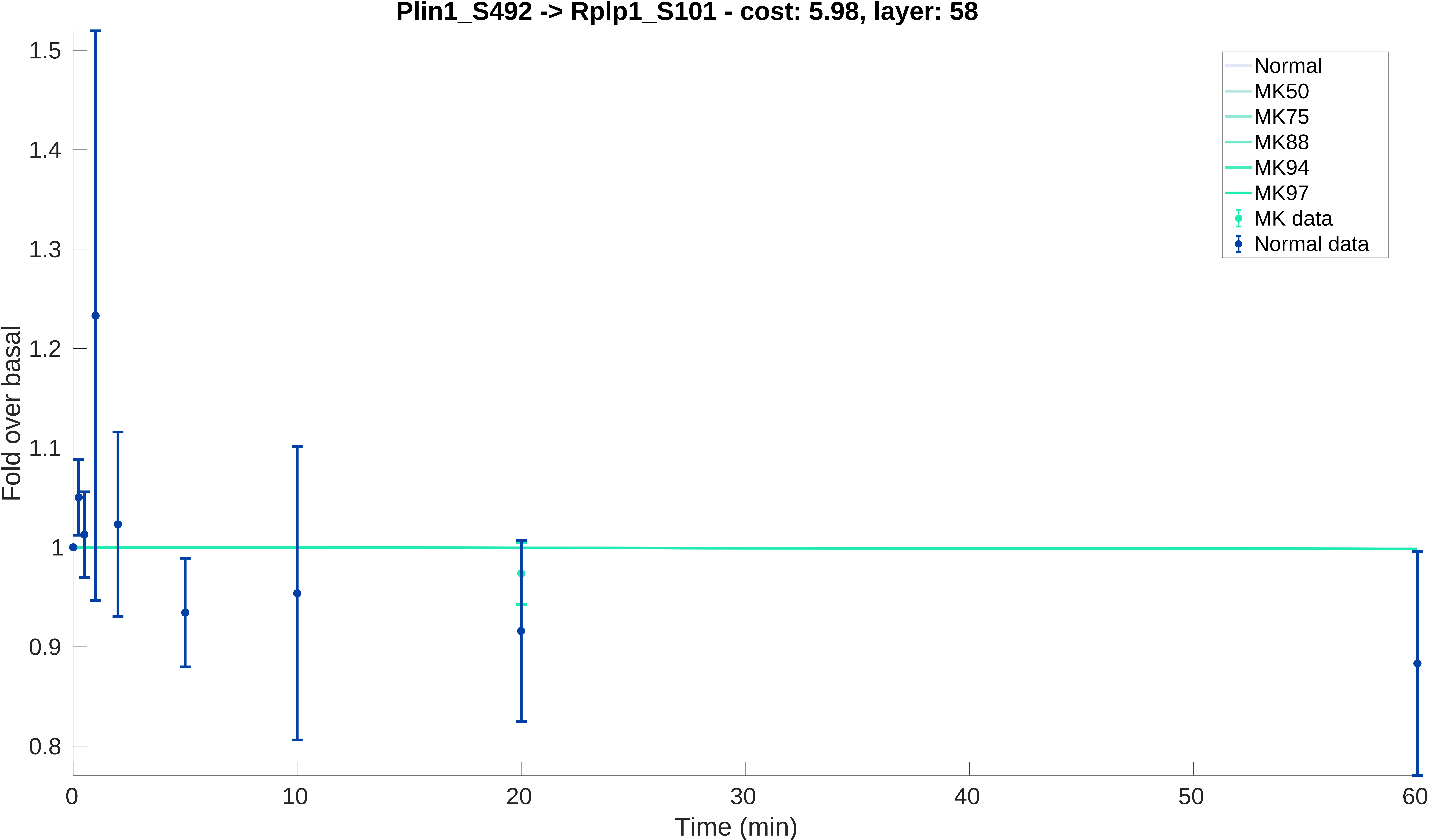
Plin1_S492 -> Rplp1_S101 - cost: 5.98, layer: 58
Fold over basal
Time (min)
1.5
1.4
1.3
1.2
1.1
1
0.9
0.8
0
10
20
30
40
50
60
Normal
MK50
MK75
MK88
MK94
MK97
MK data
Normal data

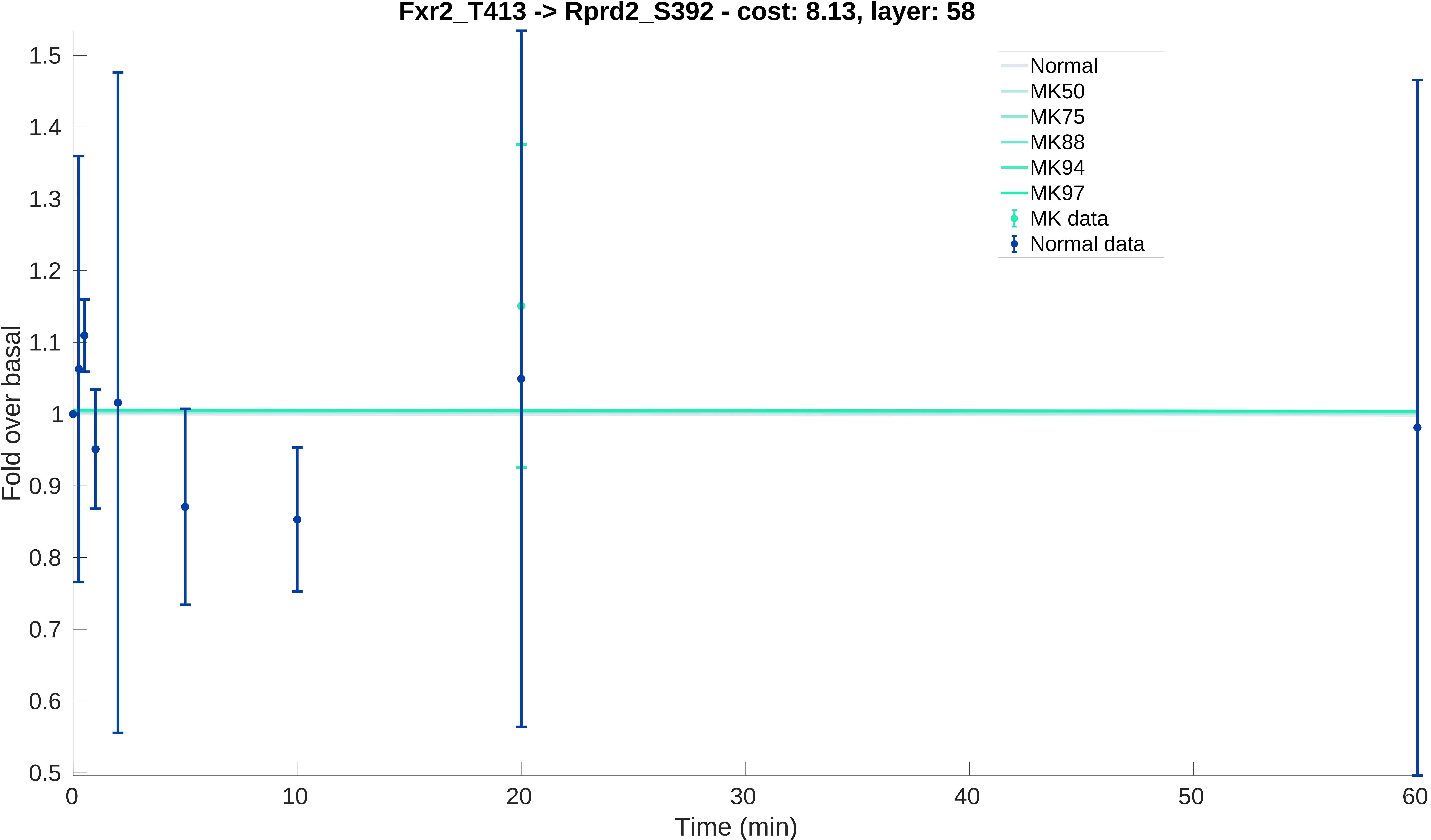

Fxr2_T413 -> Rprd2_S392 - cost: 8.13, layer: 58
Normal
MK50
MK75
MK88
MK94
MK97
MK data
Normal data
Fold over basal
Time (min)
1.5
1.4
1.3
1.2
1.1
1
0.9
0.8
0.7
0.6
0.5
0
10
20
30
40
50
60

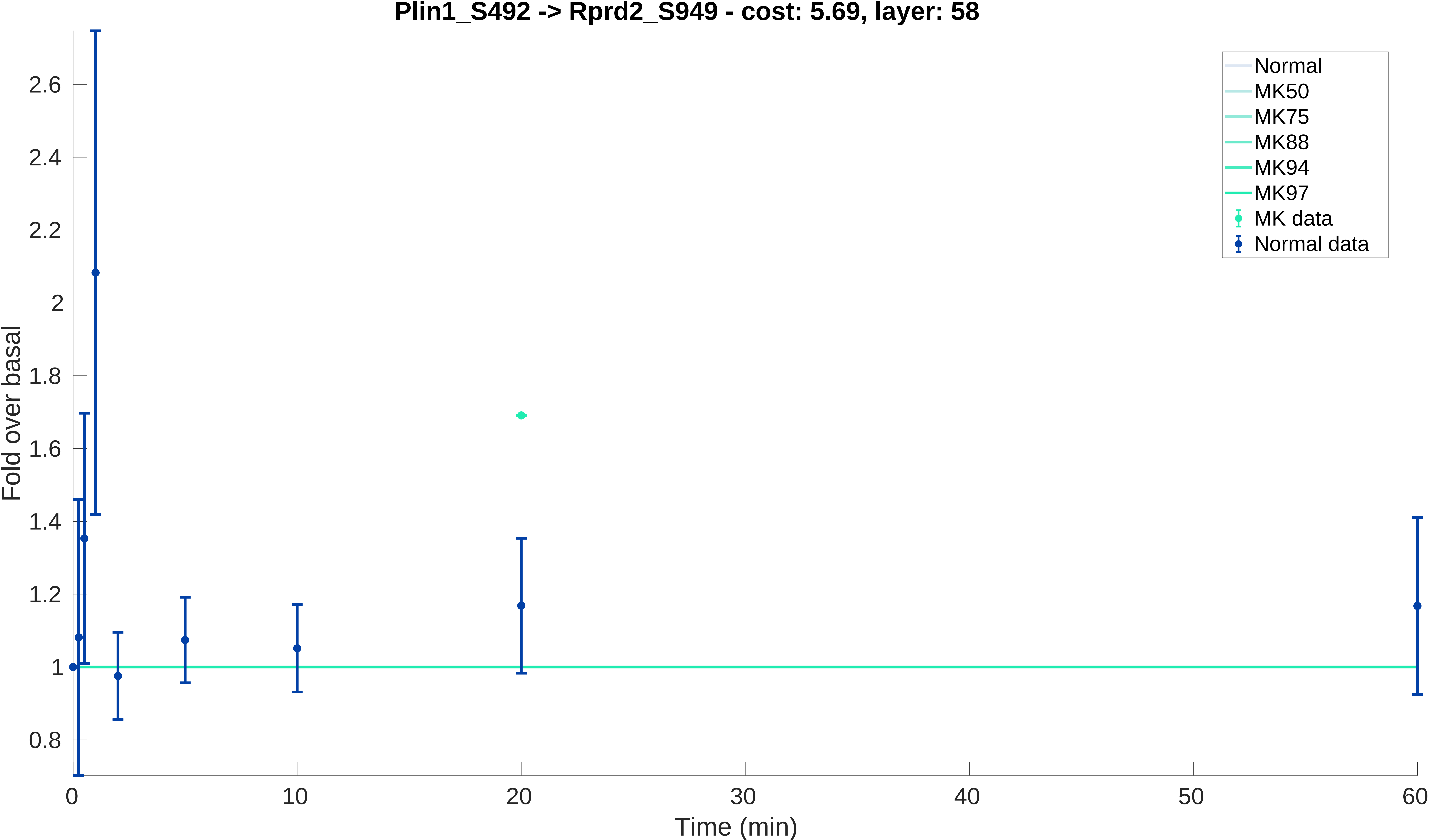
Plin1_S492 -> Rprd2_S949 - cost: 5.69, layer: 58
Fold over basal
Time (min)
Normal
MK50
MK75
MK88
MK94
MK97
MK data
Normal data
2.6
2.4
2.2
2
1.8
1.6
1.4
1.2
1
0.8
0
10
20
30
40
50
60

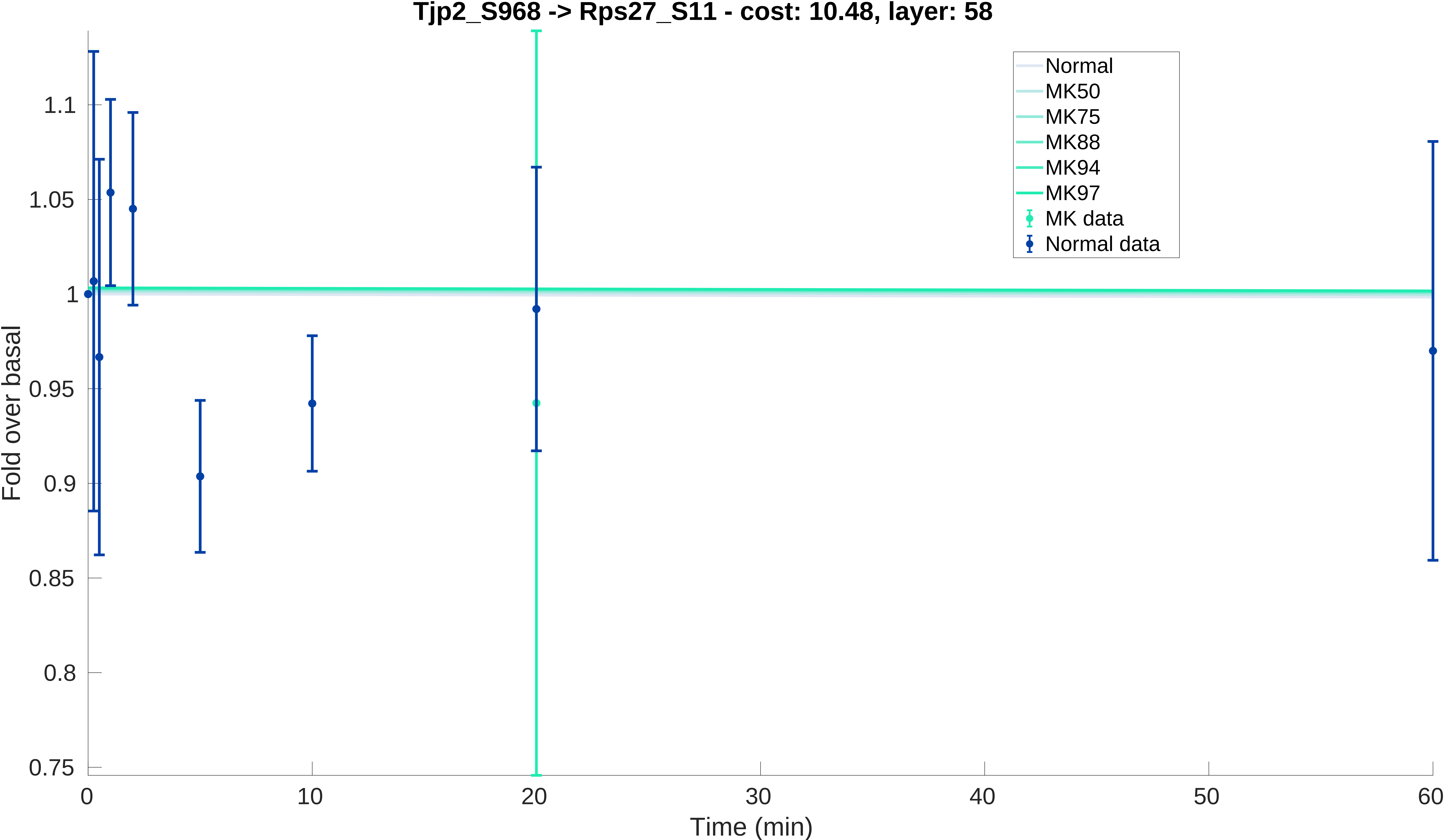

Tjp2_S968 -> Rps27_S11 - cost: 10.48, layer: 58
Normal
MK50
MK75
MK88
MK94
MK97
MK data
Normal data
Fold over basal
1.1
1.05
1
0.95
0.9
0.85
0.8
0.75
0
10
20
30
40
50
60
Time (min)

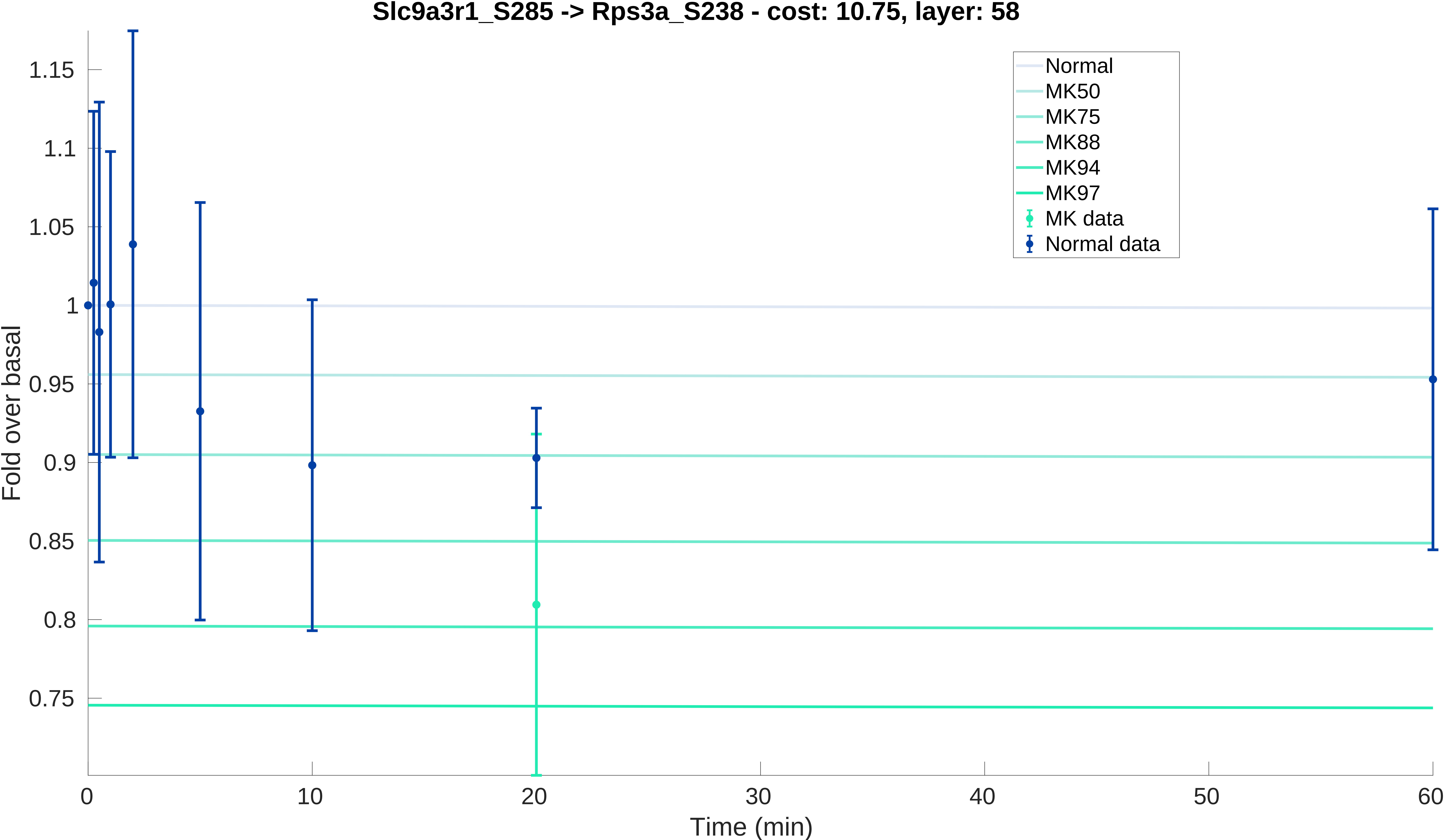

Slc9a3r1_S285 -> Rps3a_S238 - cost: 10.75, layer: 58
Normal
MK50
MK75
MK88
MK94
MK97
MK data
Normal data
Fold over basal
Time (min)
1.15
1.1
1.05
1
0.95
0.9
0.85
0.8
0.75
0
10
20
30
40
50
60

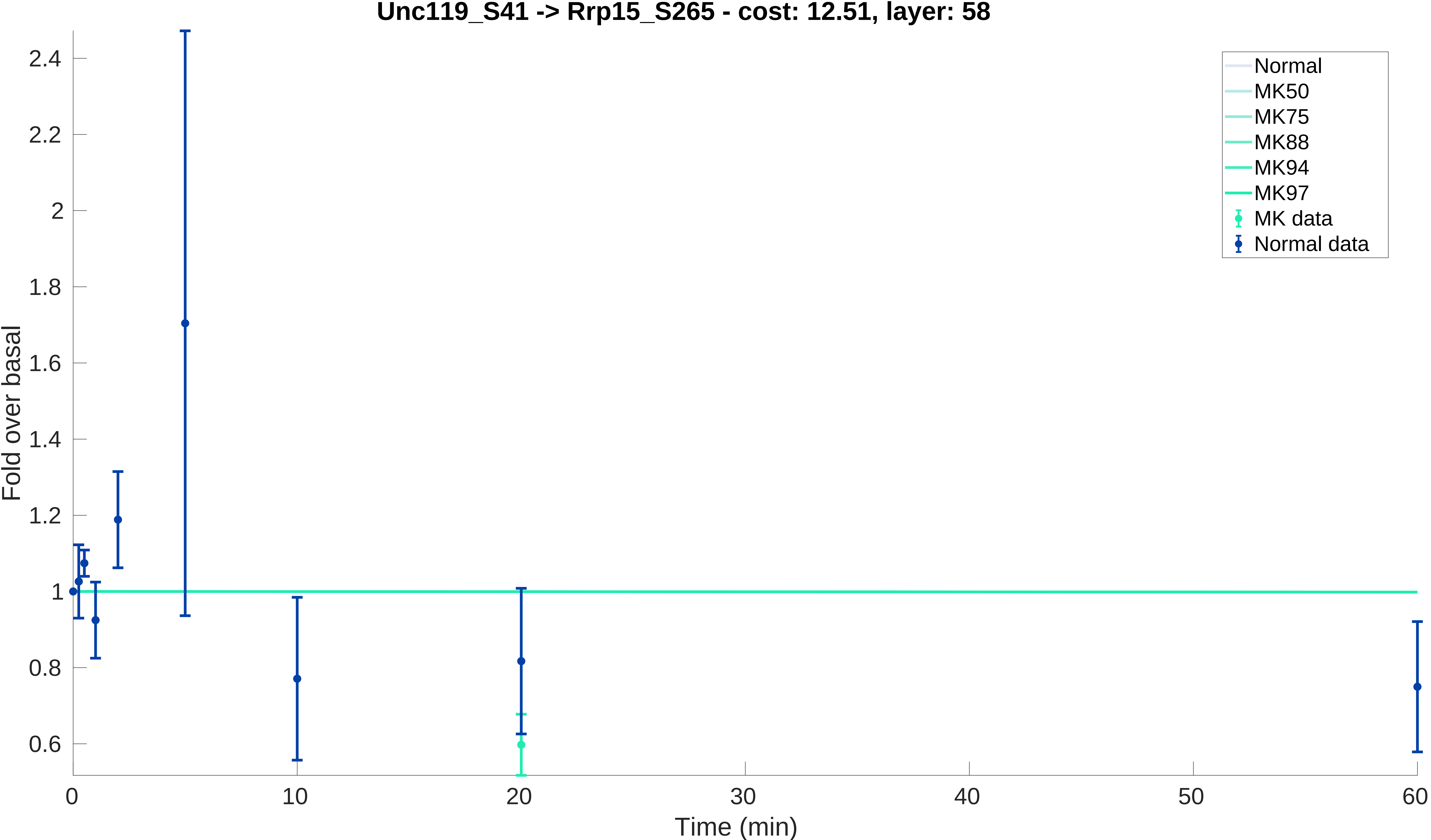
Unc119_S41 -> Rrp15_S265 - cost: 12.51, layer: 58
Fold over basal
Time (min)
2.4
2.2
2
1.8
1.6
1.4
1.2
1
0.8
0.6
0
10
20
30
40
50
60
Normal
MK50
MK75
MK88
MK94
MK97
MK data
Normal data

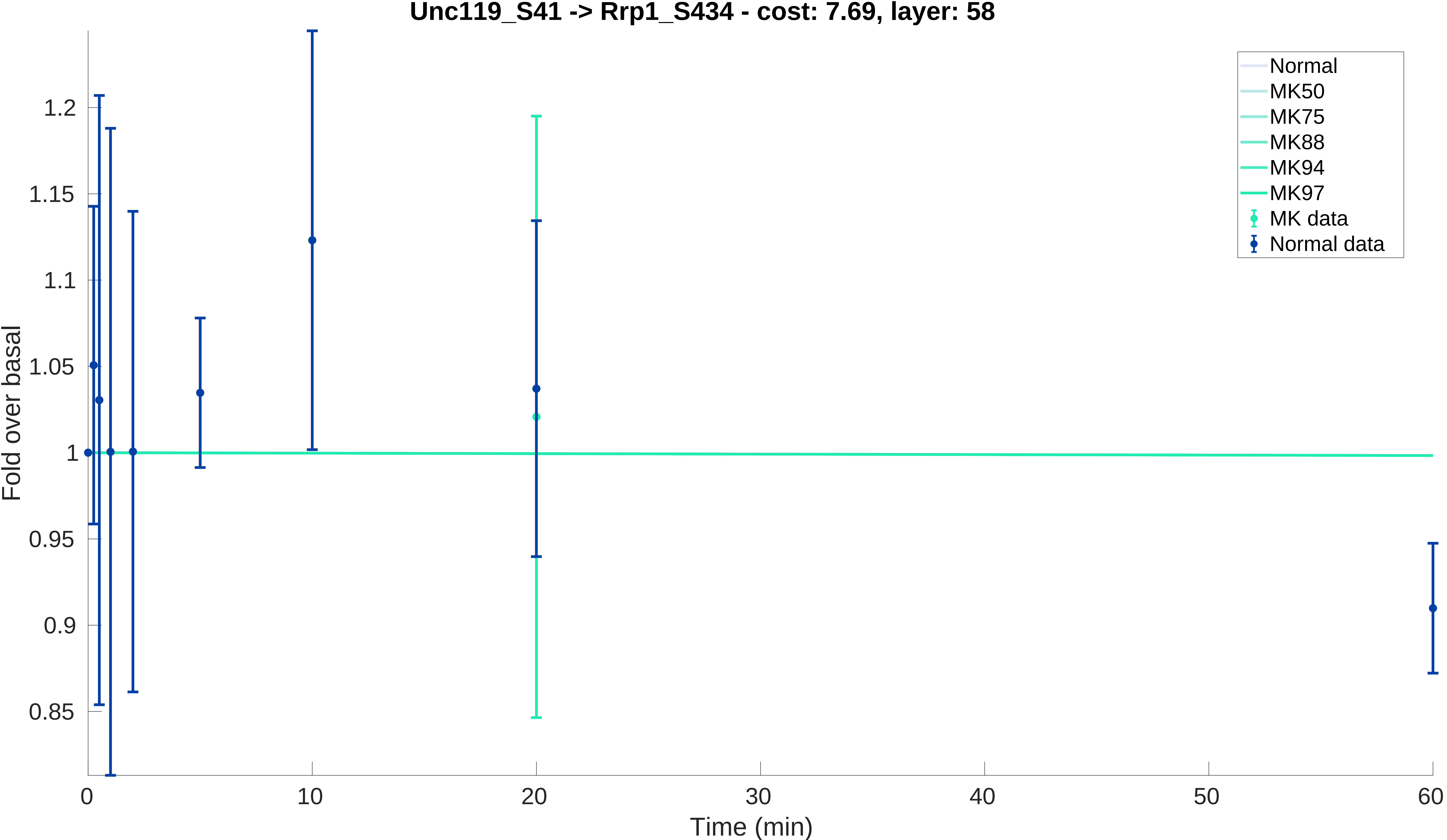

Unc119_S41 -> Rrp1_S434 - cost: 7.69, layer: 58
Fold over basal
Time (min)
1.2
1.15
1.1
1.05
1
0.95
0.9
0.85
0
10
20
30
40
50
60
Normal
MK50
MK75
MK88
MK94
MK97
MK data
Normal data

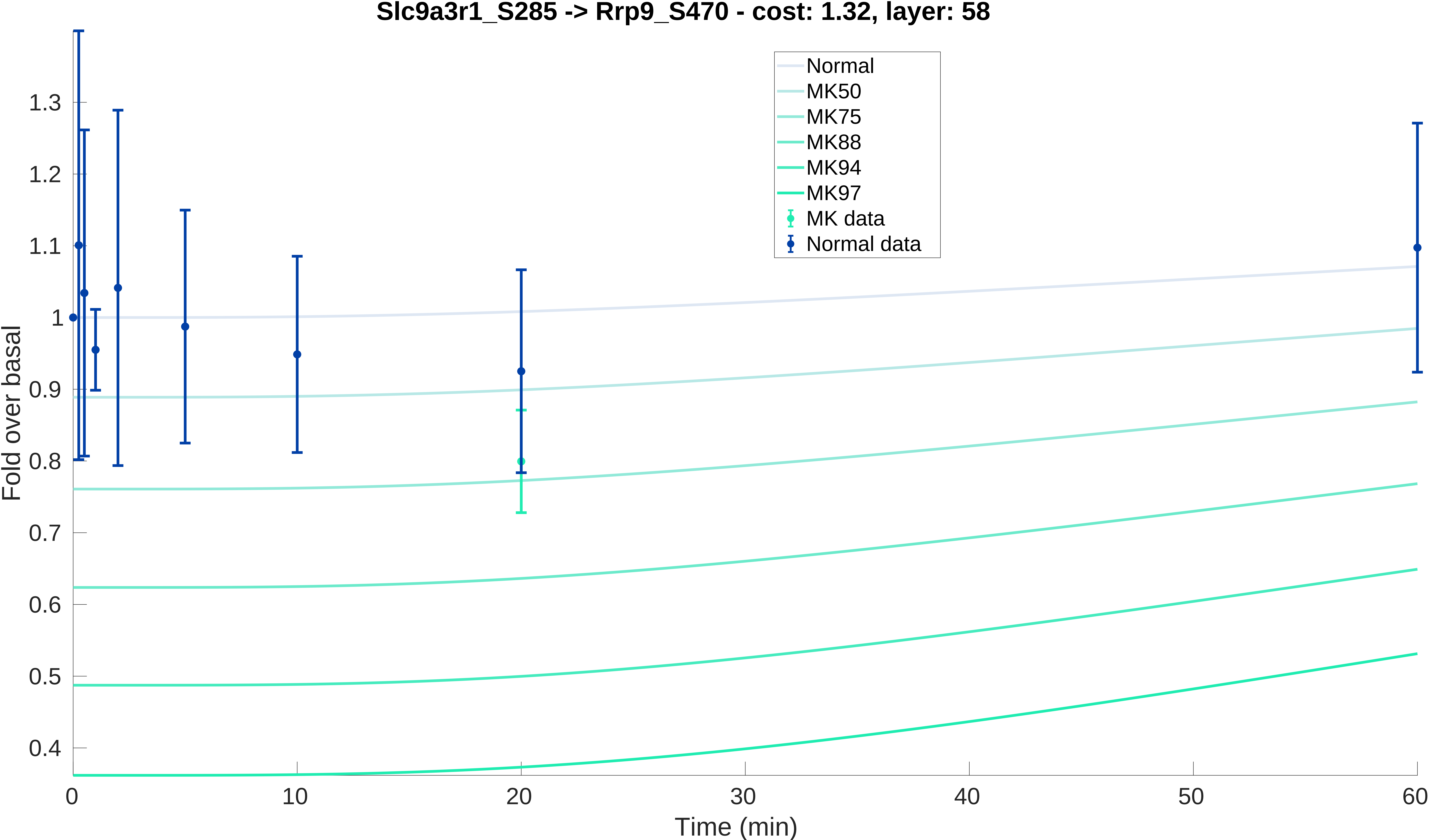

Slc9a3r1_S285 -> Rrp9_S470 - cost: 1.32, layer: 58
Normal
MK50
MK75
MK88
MK94
MK97
MK data
Normal data
Fold over basal
1.3
1.2
1.1
1
0.9
0.8
0.7
0.6
0.5
0.4
0
10
20
30
40
50
60
Time (min)

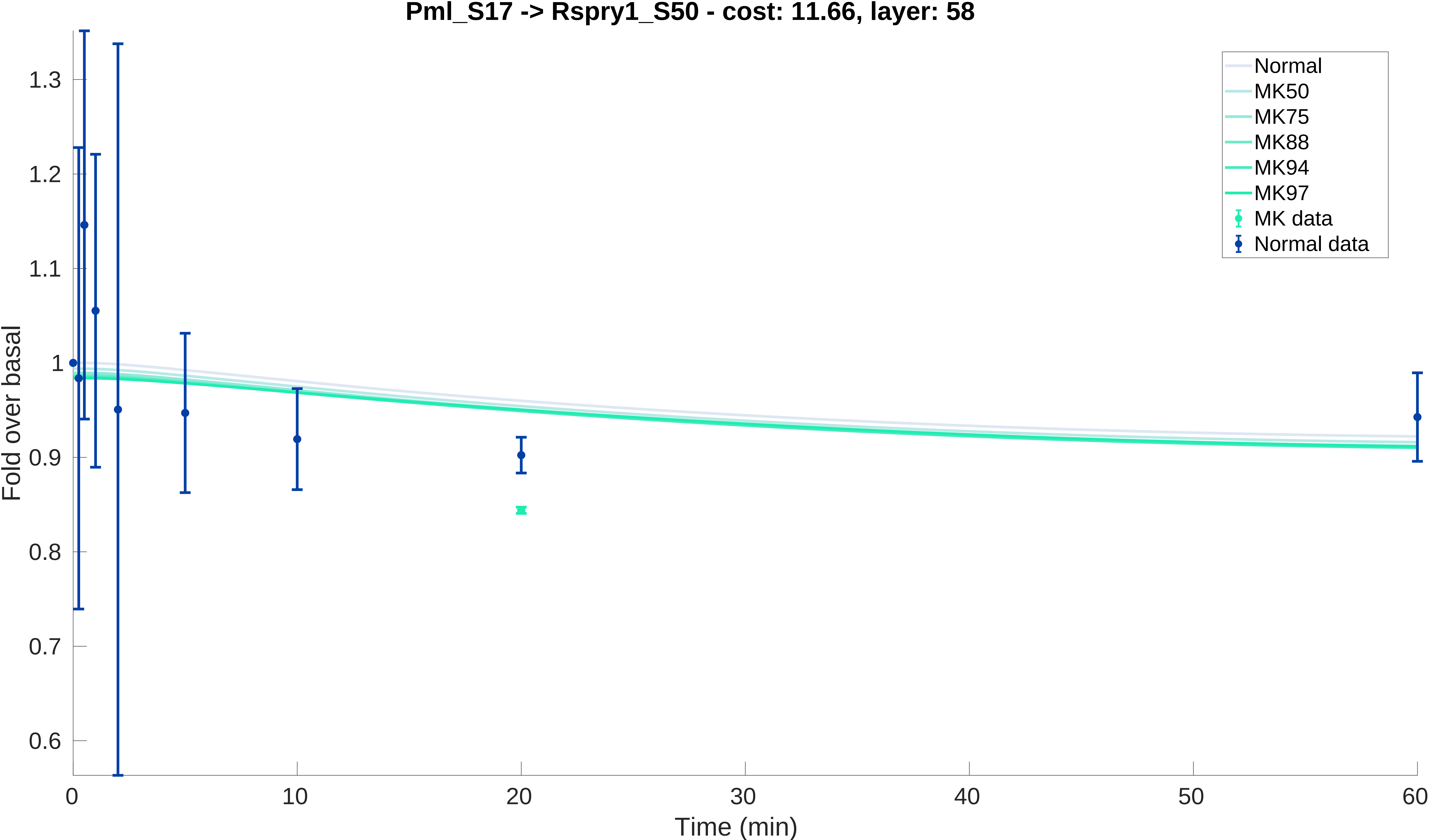
Pml_S17 -> Rspry1_S50 - cost: 11.66, layer: 58
Normal
MK50
MK75
MK88
MK94
MK97
MK data
Normal data
Fold over basal
1.3
1.2
1.1
1
0.9
0.8
0.7
0.6
0
10
20
30
40
50
60
Time (min)

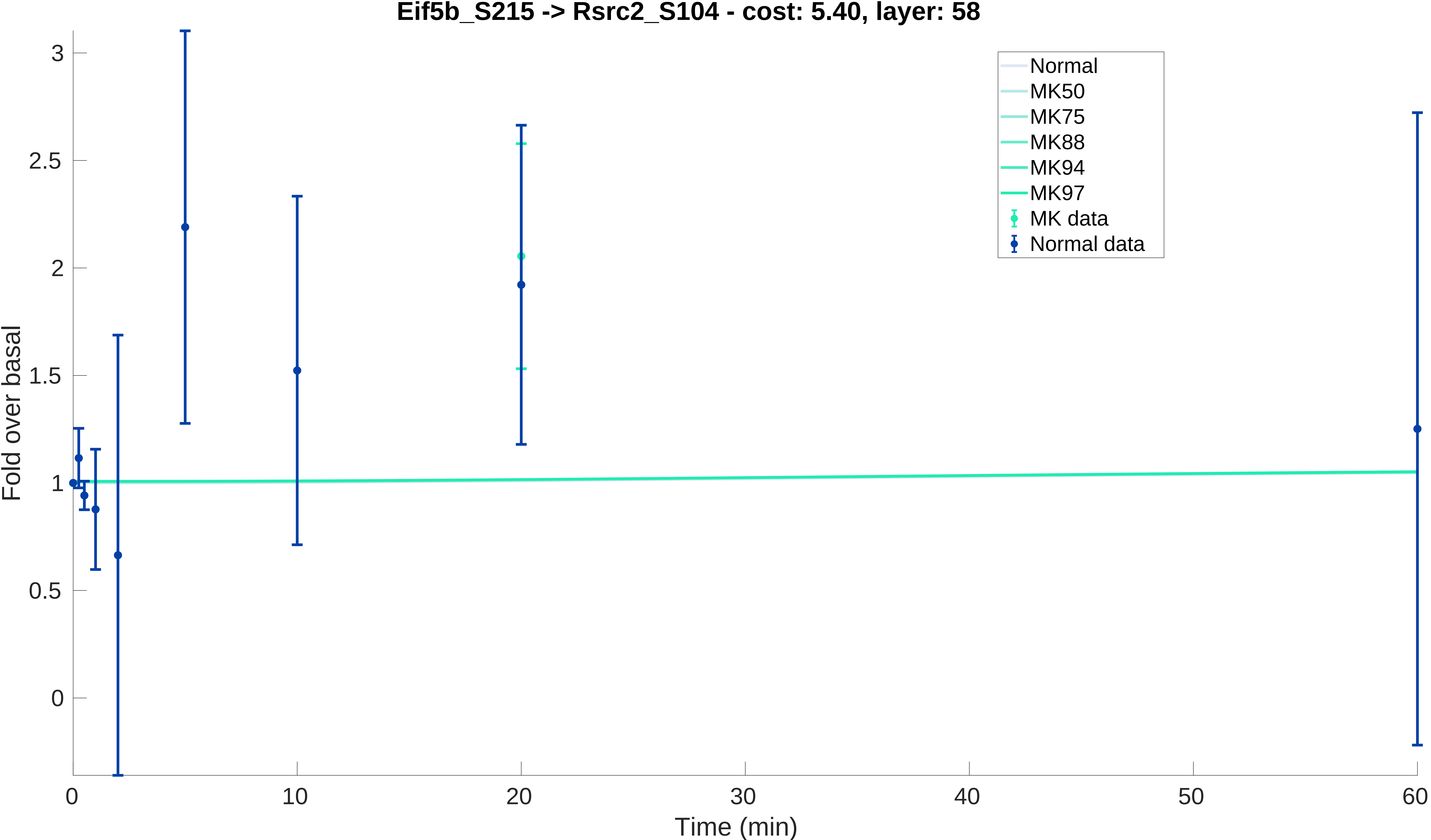

Eif5b_S215 -> Rsrc2_S104 - cost: 5.40, layer: 58
Normal
MK50
MK75
MK88
MK94
MK97
MK data
Normal data
Fold over basal
Time (min)
3
2.5
2
1.5
1
0.5
0
0
10
20
30
40
50
60

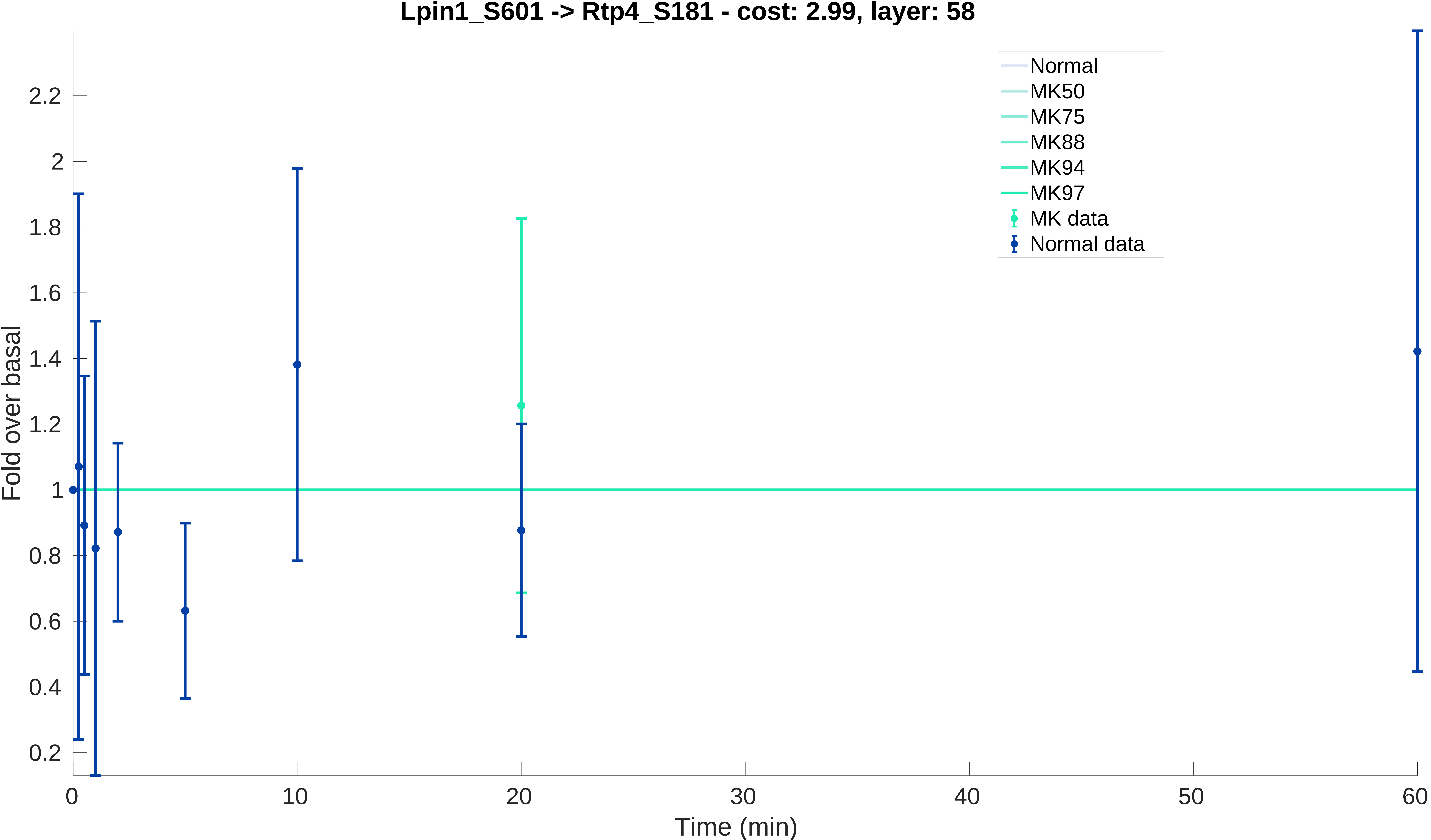

Lpin1_S601 -> Rtp4_S181 - cost: 2.99, layer: 58
Normal
MK50
MK75
MK88
MK94
MK97
MK data
Normal data
Fold over basal
Time (min)
0.2
0.4
0.6
0.8
1
1.2
1.4
1.6
1.8
2
2.2
0
10
20
30
40
50
60

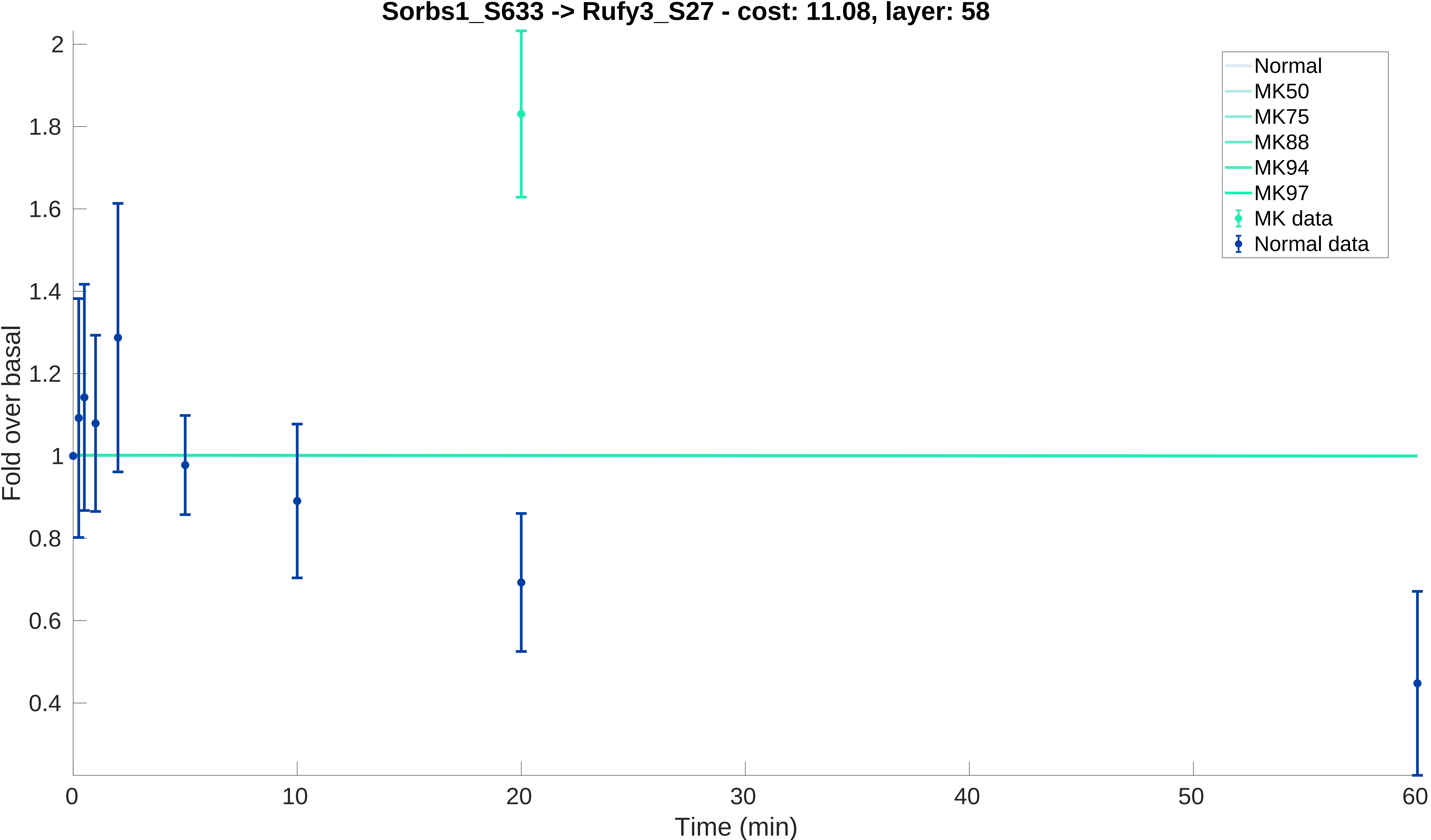

Sorbs1_S633 -> Rufy3_S27 - cost: 11.08, layer: 58
Normal
MK50
MK75
MK88
MK94
MK97
MK data
Normal data
Fold over basal
Time (min)
0
10
20
30
40
50
60
0.4
0.6
0.8
1
1.2
1.4
1.6
1.8
2

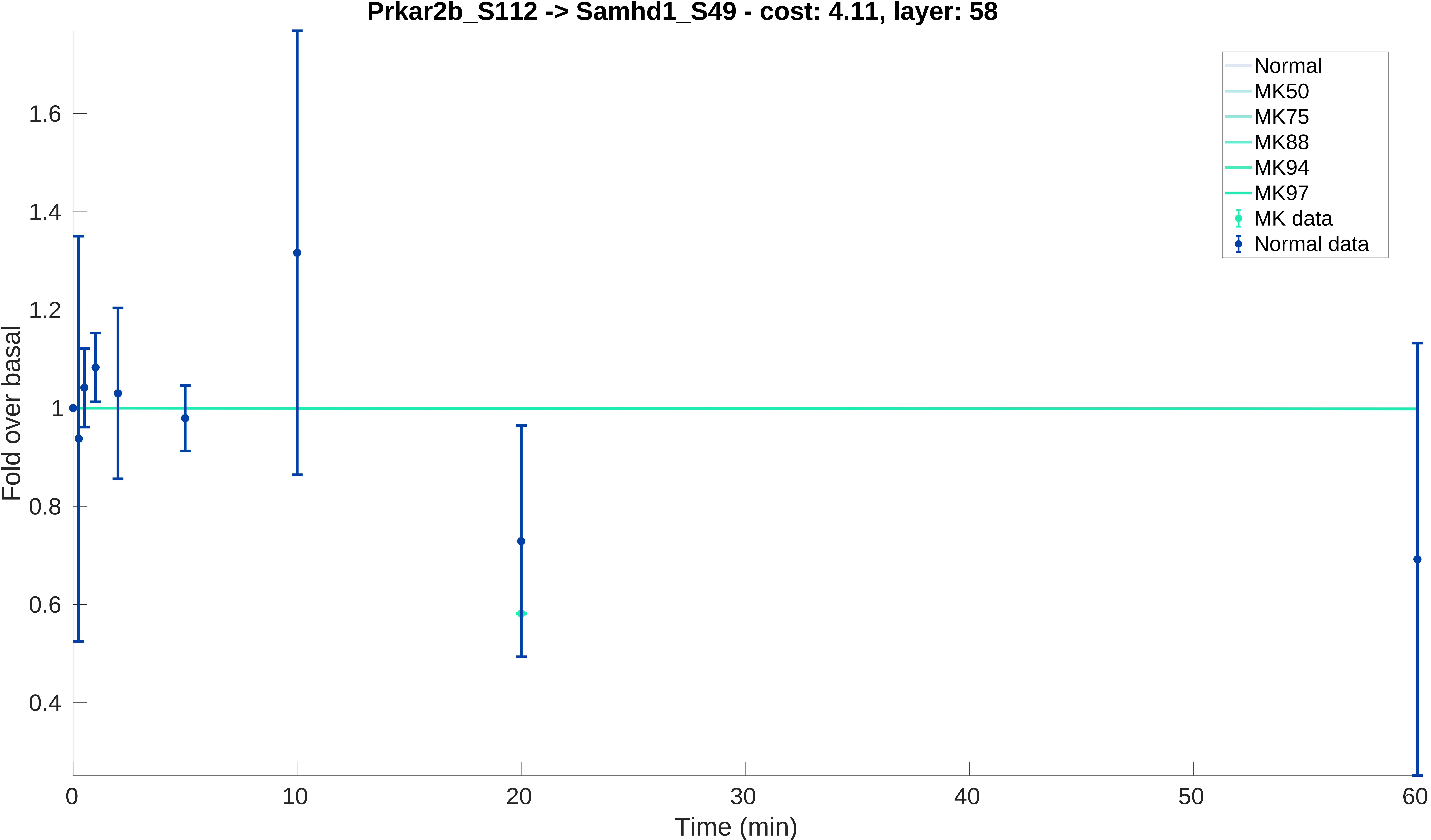

Prkar2b_S112 -> Samhd1_S49 - cost: 4.11, layer: 58
Fold over basal
1.6
1.4
1.2
1
0.8
0.6
0.4
0
10
20
30
40
50
60
Time (min)
Normal
MK50
MK75
MK88
MK94
MK97
MK data
Normal data

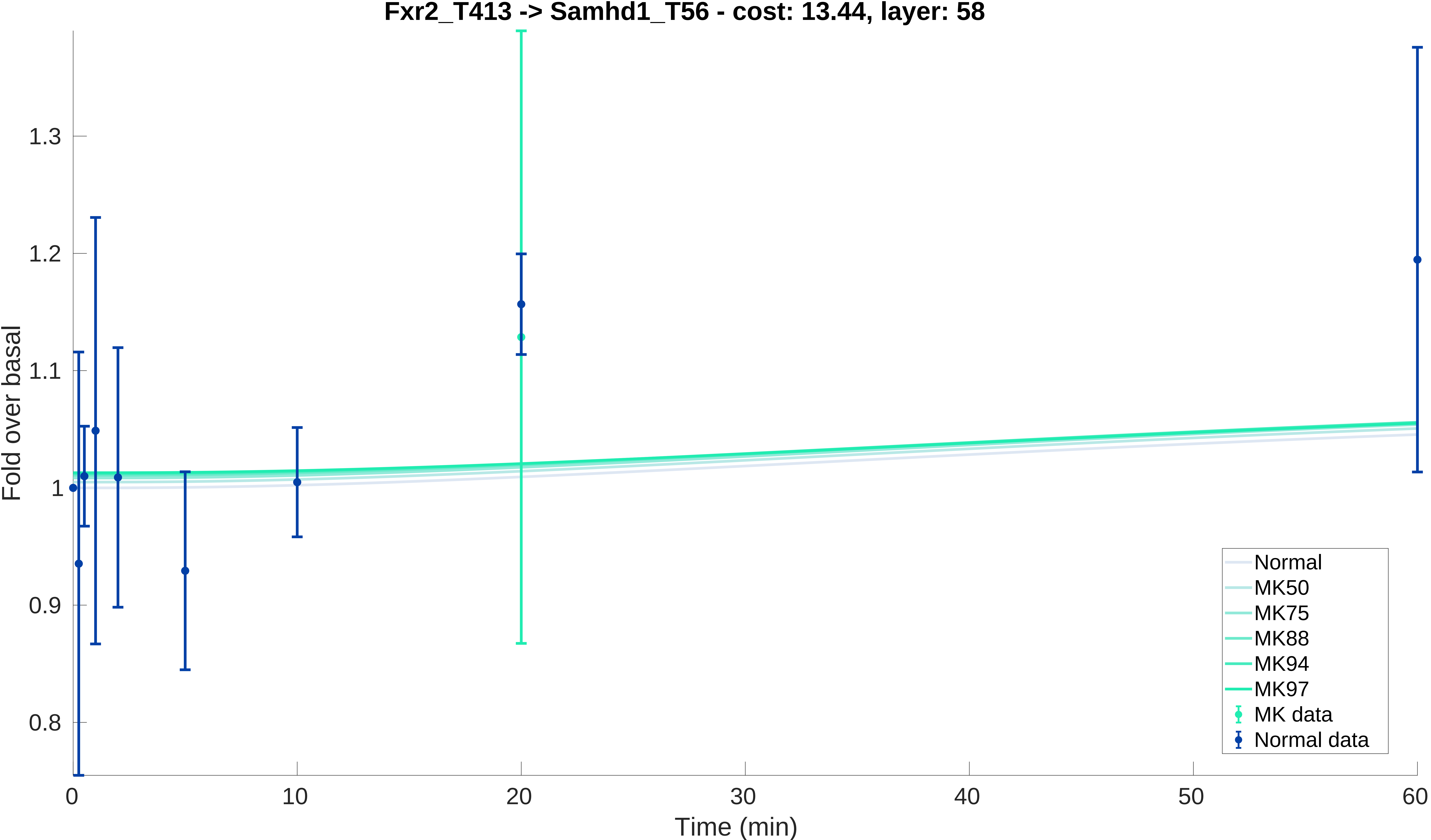

Fxr2_T413 -> Samhd1_T56 - cost: 13.44, layer: 58
Fold over basal
1.3
1.2
1.1
1
0.9
0.8
0
10
20
30
40
50
60
Time (min)
Normal
MK50
MK75
MK88
MK94
MK97
MK data
Normal data

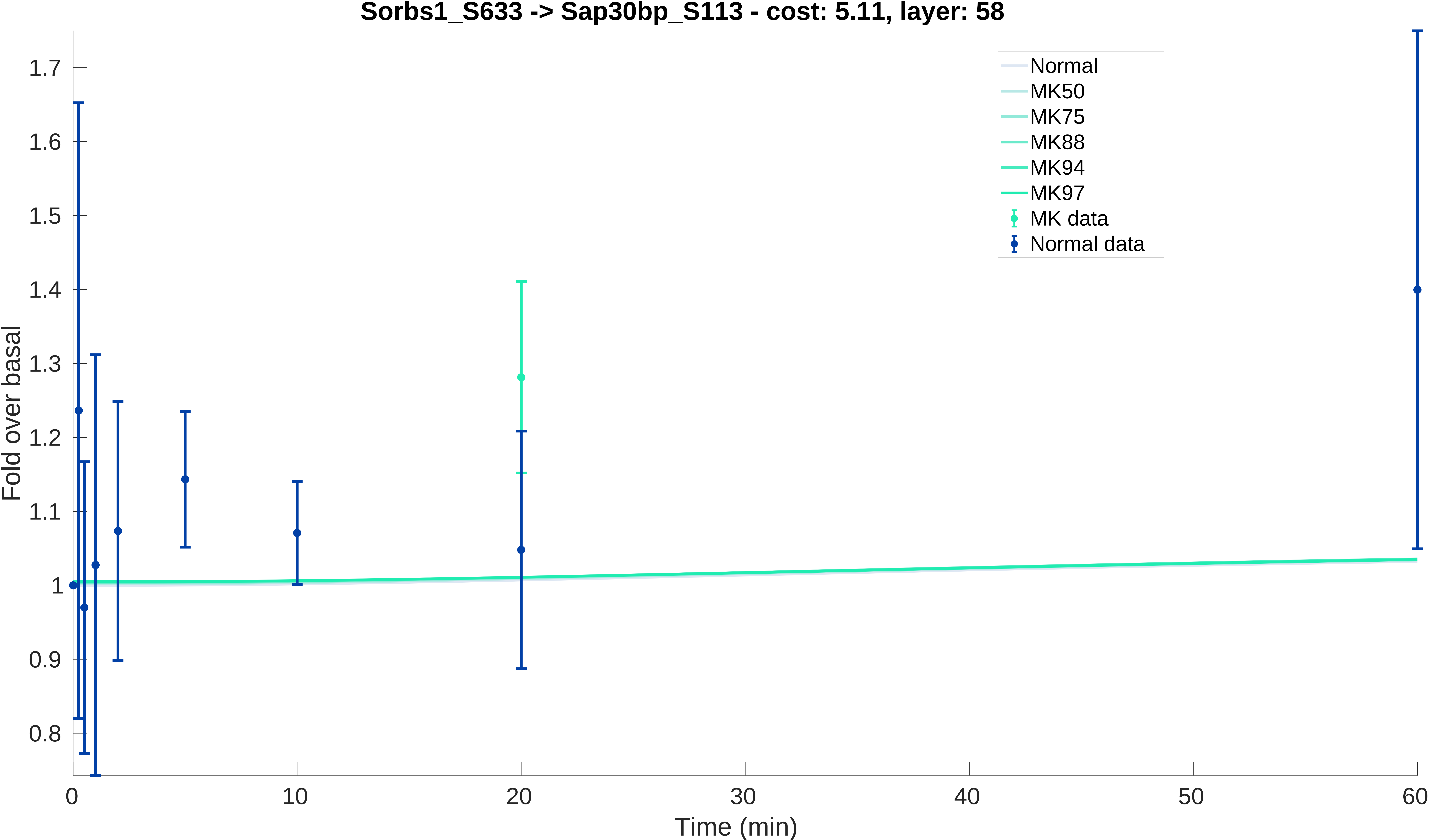
Sorbs1_S633 -> Sap30bp_S113 - cost: 5.11, layer: 58
Fold over basal
Time (min)
0.8
0.9
1
1.1
1.2
1.3
1.4
1.5
1.6
1.7
0
10
20
30
40
50
60
Normal
MK50
MK75
MK88
MK94
MK97
MK data
Normal data

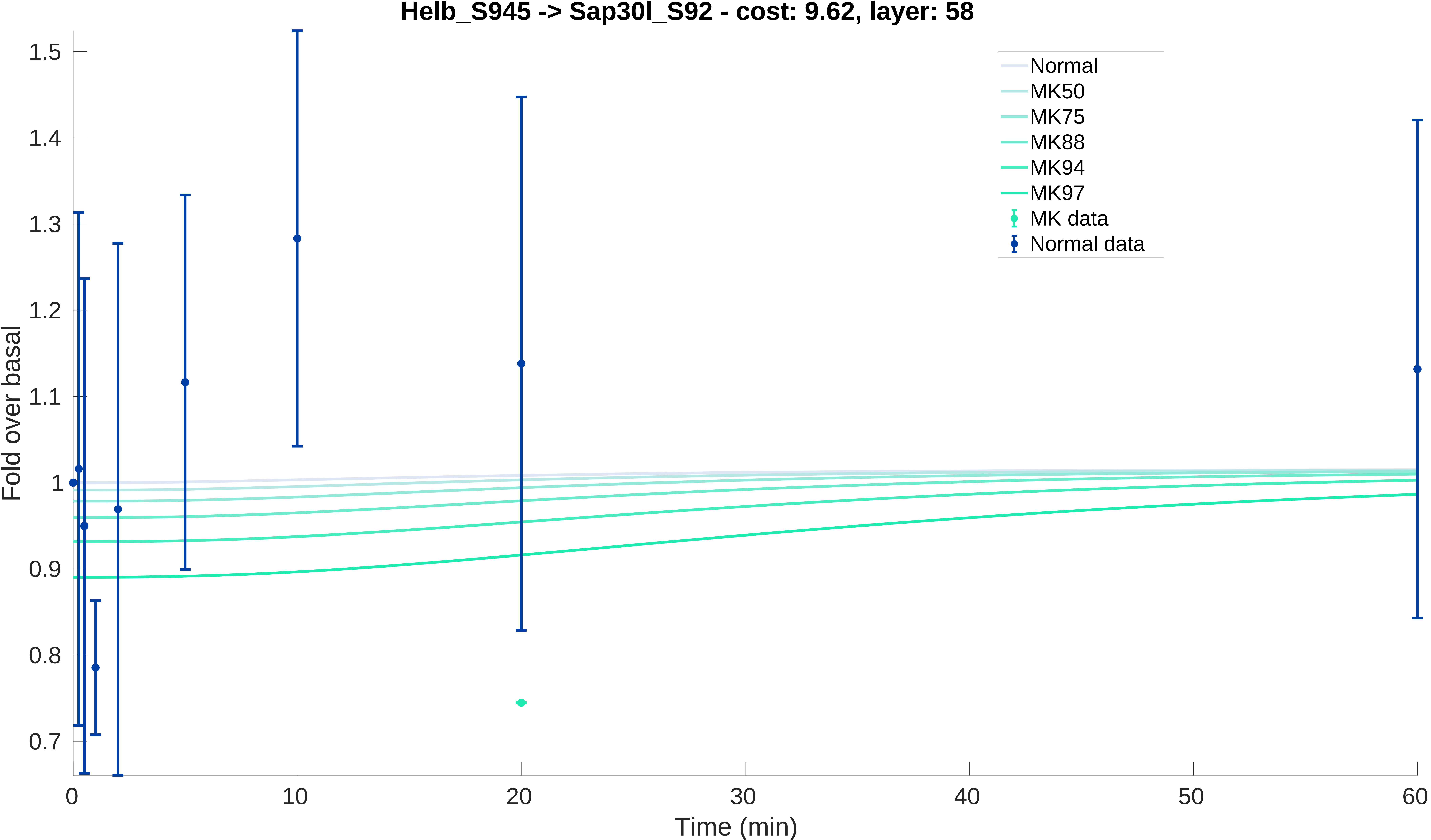
Helb_S945 -> Sap30l_S92 - cost: 9.62, layer: 58
Normal
MK50
MK75
MK88
MK94
MK97
MK data
Normal data
Fold over basal
Time (min)
1.5
1.4
1.3
1.2
1.1
1
0.9
0.8
0.7
0
10
20
30
40
50
60

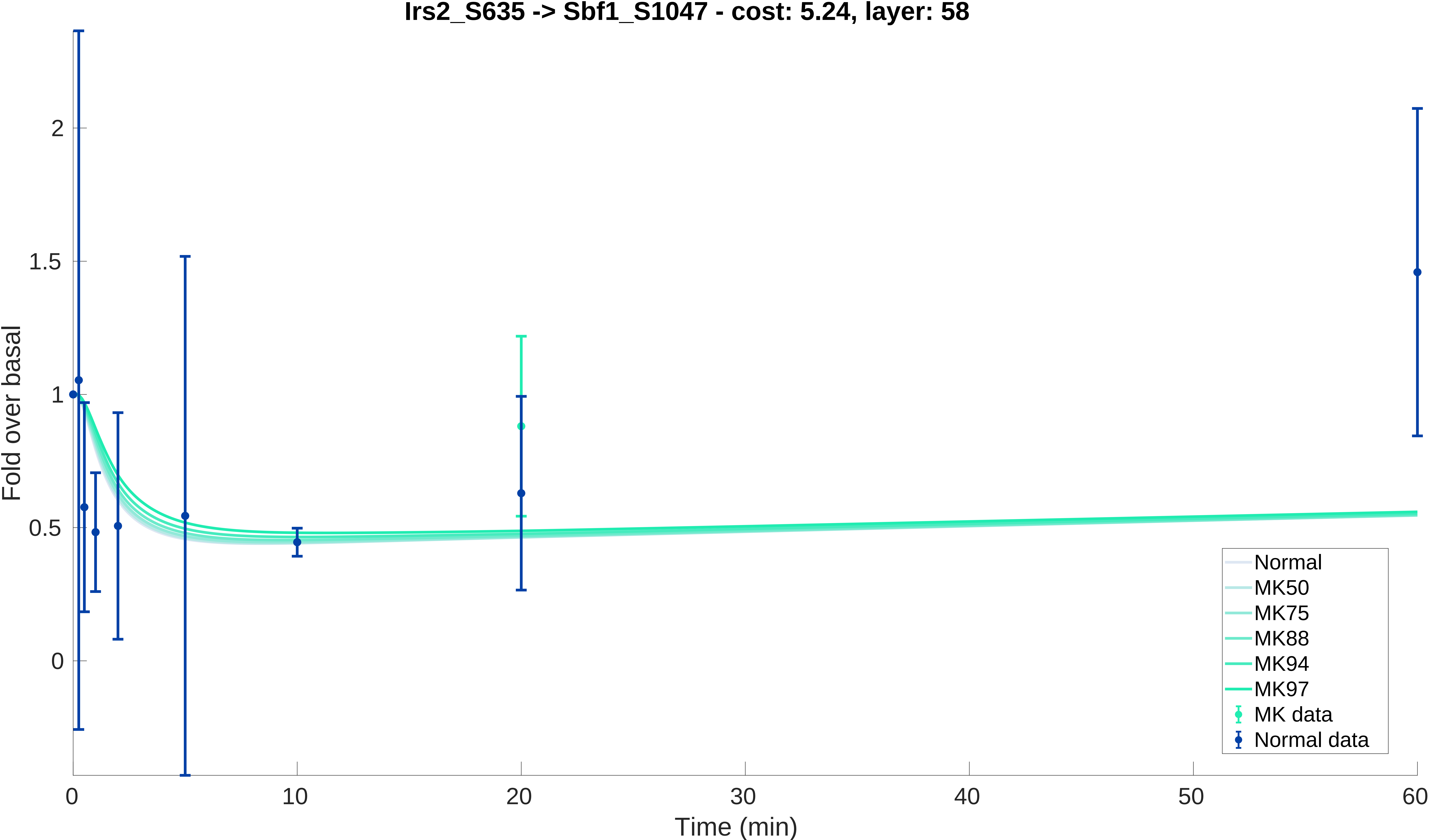

Irs2_S635 -> Sbf1_S1047 - cost: 5.24, layer: 58
Fold over basal
Time (min)
0
0.5
1
1.5
2
0
10
20
30
40
50
60
Normal
MK50
MK75
MK88
MK94
MK97
MK data
Normal data

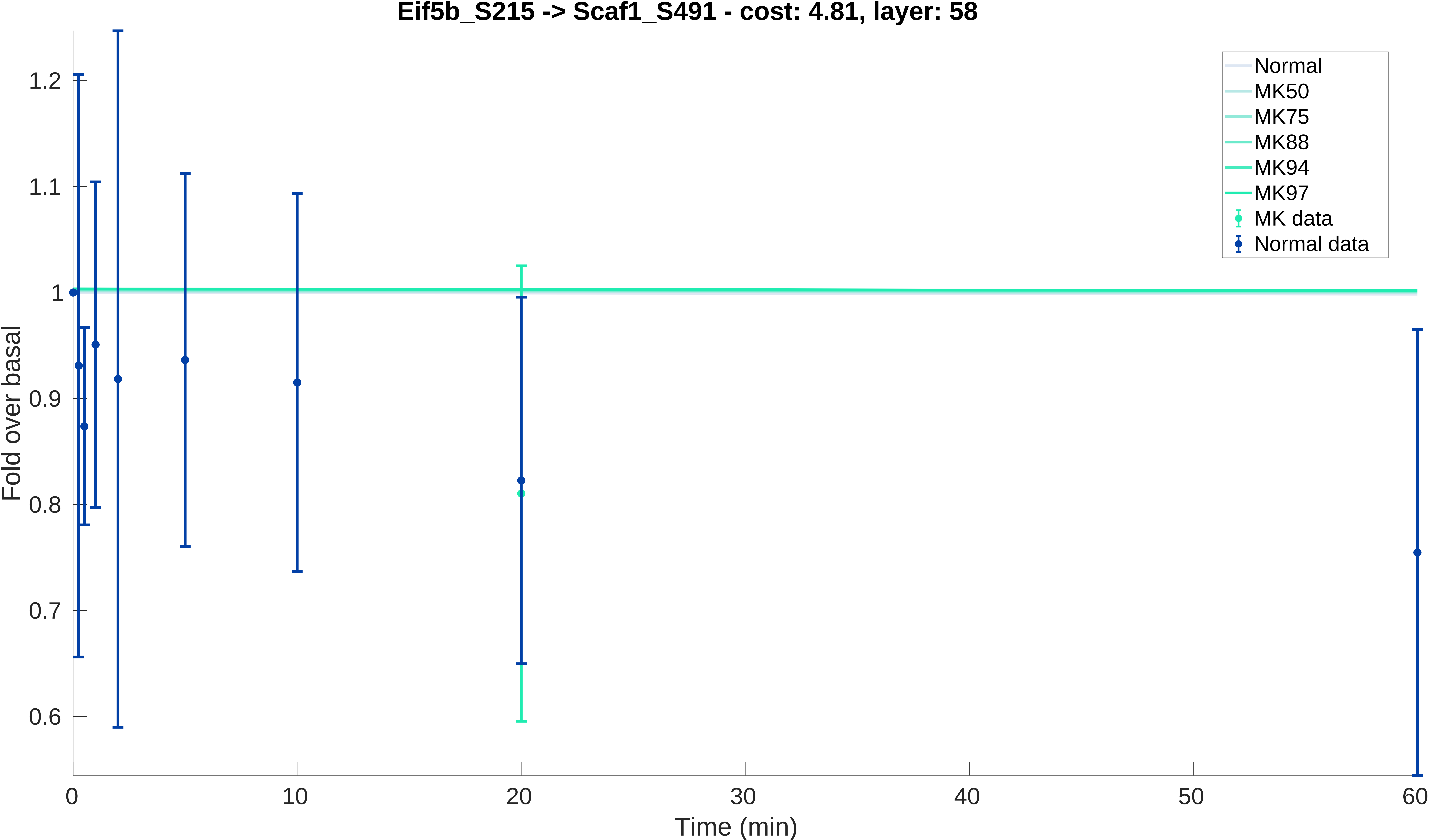

Eif5b_S215 -> Scaf1_S491 - cost: 4.81, layer: 58
Fold over basal
1.2
1.1
1
0.9
0.8
0.7
0.6
0
10
20
30
40
50
60
Time (min)
Normal
MK50
MK75
MK88
MK94
MK97
MK data
Normal data

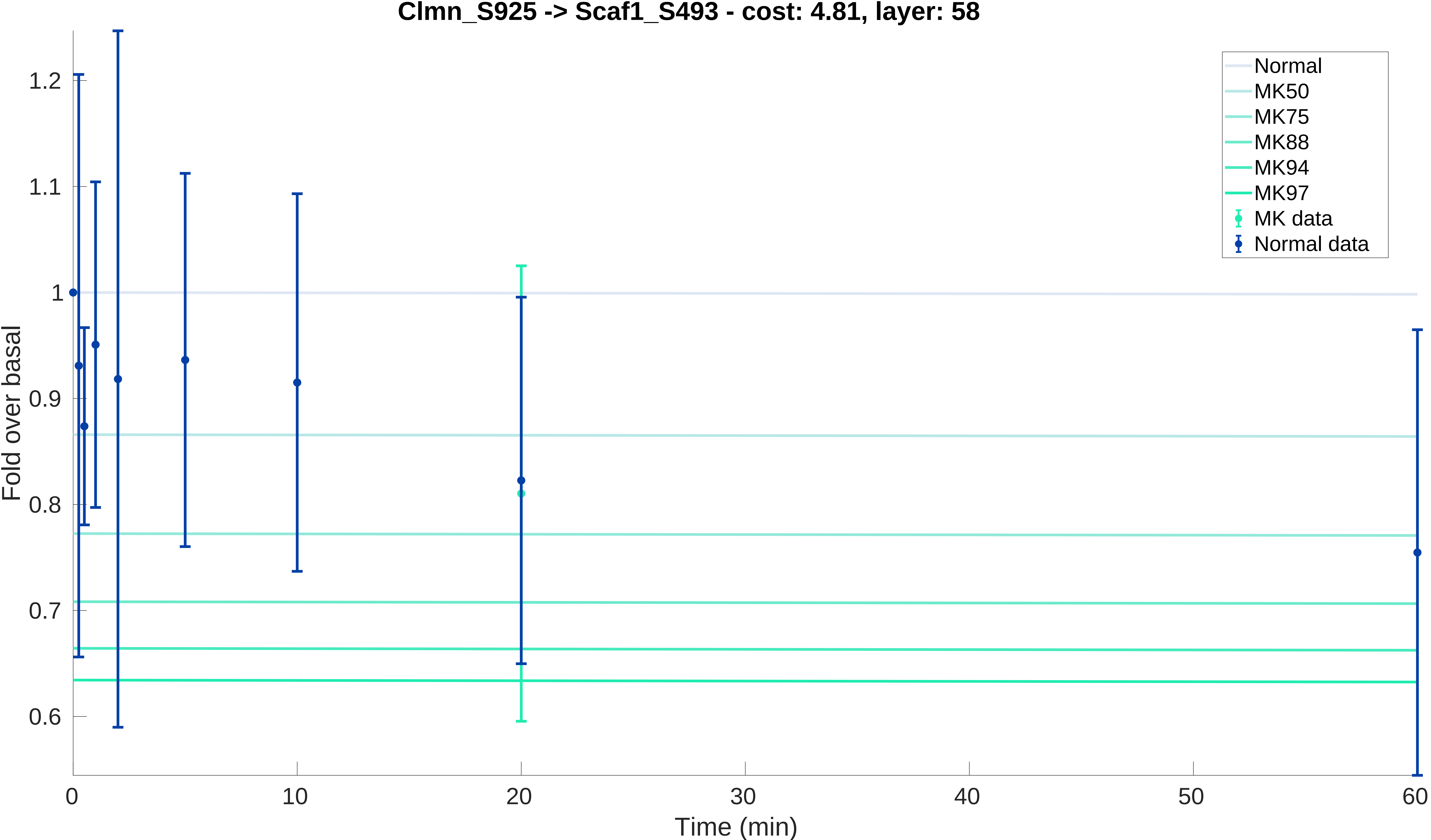

Clmn_S925 -> Scaf1_S493 - cost: 4.81, layer: 58
Fold over basal
Time (min)
0.6
0.7
0.8
0.9
1
1.1
1.2
0
10
20
30
40
50
60
Normal
MK50
MK75
MK88
MK94
MK97
MK data
Normal data

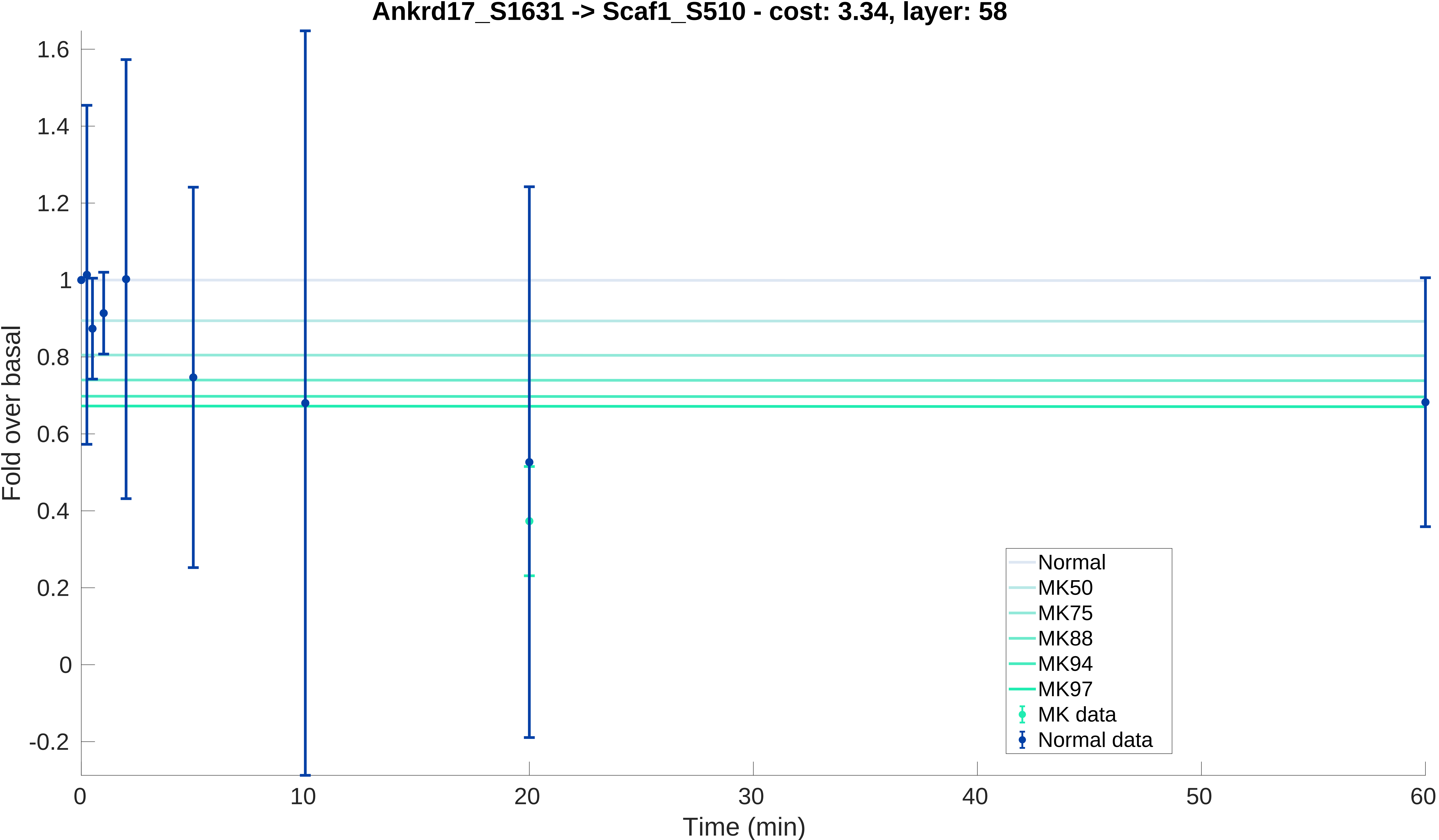
Ankrd17_S1631 -> Scaf1_S510 - cost: 3.34, layer: 58
Fold over basal
Time (min)
1.6
1.4
1.2
1
0.8
0.6
0.4
0.2
0
-0.2
0
10
20
30
40
50
60
Normal
MK50
MK75
MK88
MK94
MK97
MK data
Normal data

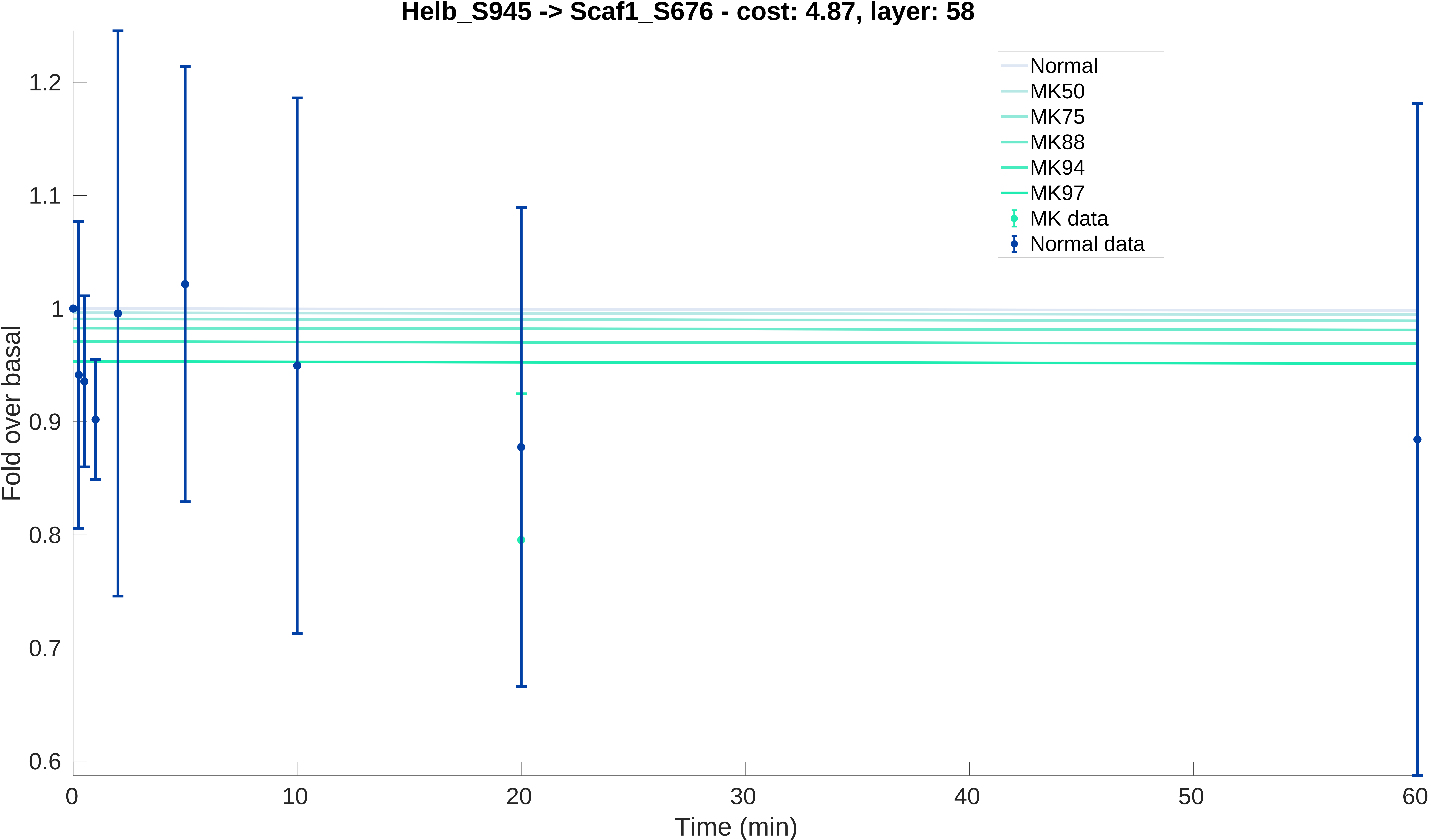

Helb_S945 -> Scaf1_S676 - cost: 4.87, layer: 58
Normal
MK50
MK75
MK88
MK94
MK97
MK data
Normal data
Fold over basal
1.2
1.1
1
0.9
0.8
0.7
0.6
0
10
20
30
40
50
60
Time (min)

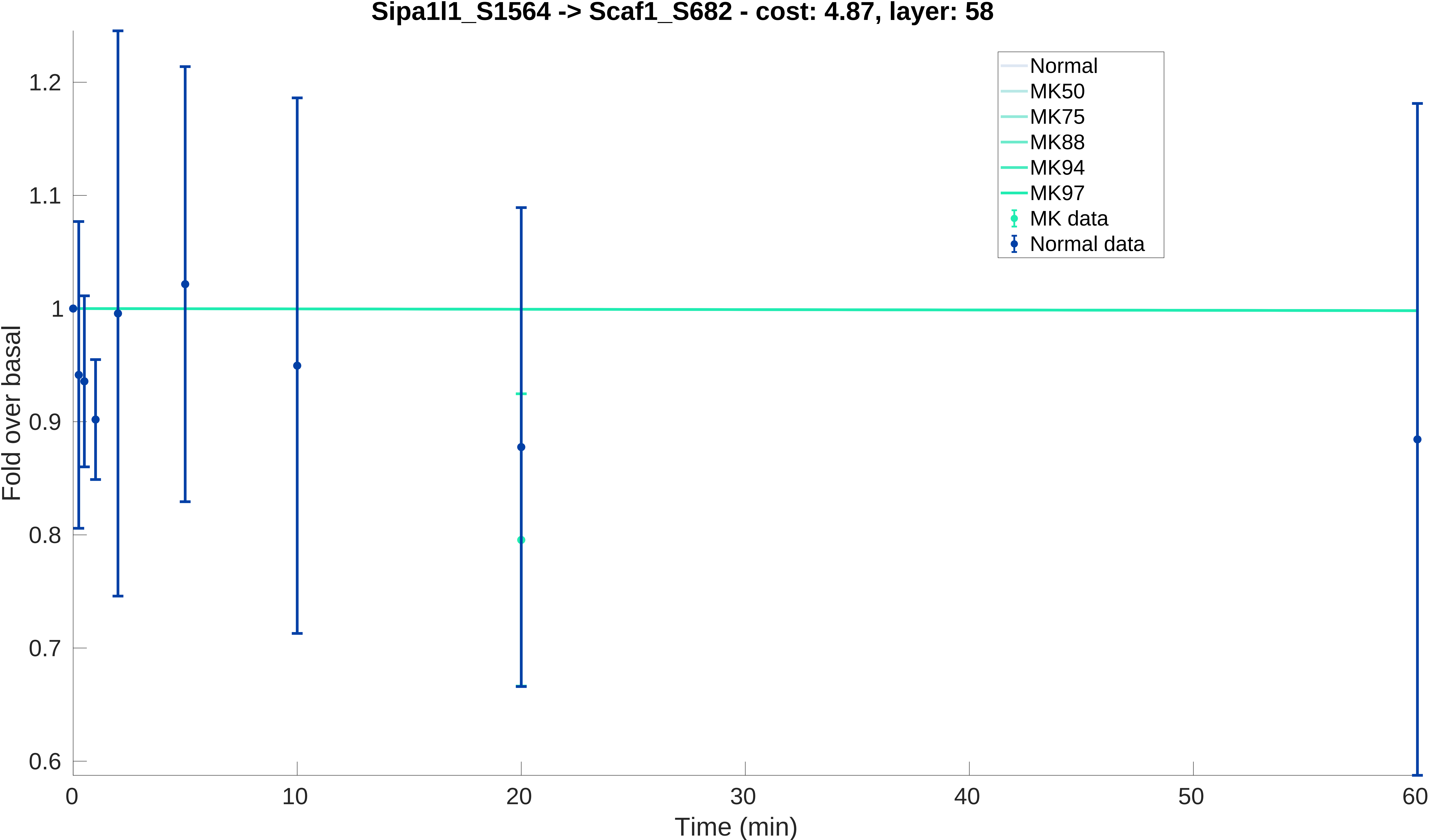

Sipa1l1_S1564 -> Scaf1_S682 - cost: 4.87, layer: 58
Normal
MK50
MK75
MK88
MK94
MK97
MK data
Normal data
Fold over basal
Time (min)
1.2
1.1
1
0.9
0.8
0.7
0.6
0
10
20
30
40
50
60

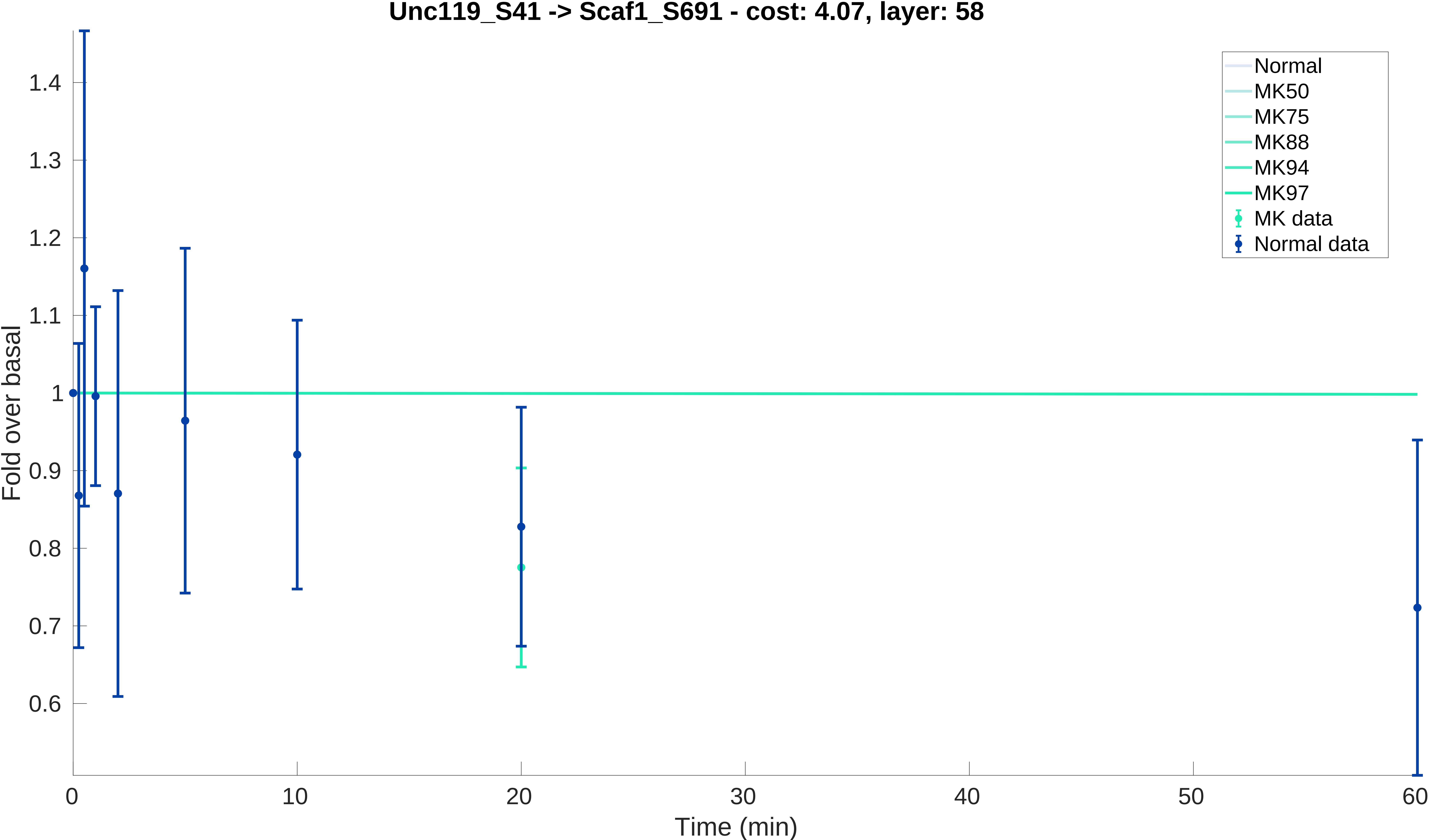

Unc119_S41 -> Scaf1_S691 - cost: 4.07, layer: 58
Fold over basal
Time (min)
Normal
MK50
MK75
MK88
MK94
MK97
MK data
Normal data
1.4
1.3
1.2
1.1
1
0.9
0.8
0.7
0.6
0
10
20
30
40
50
60

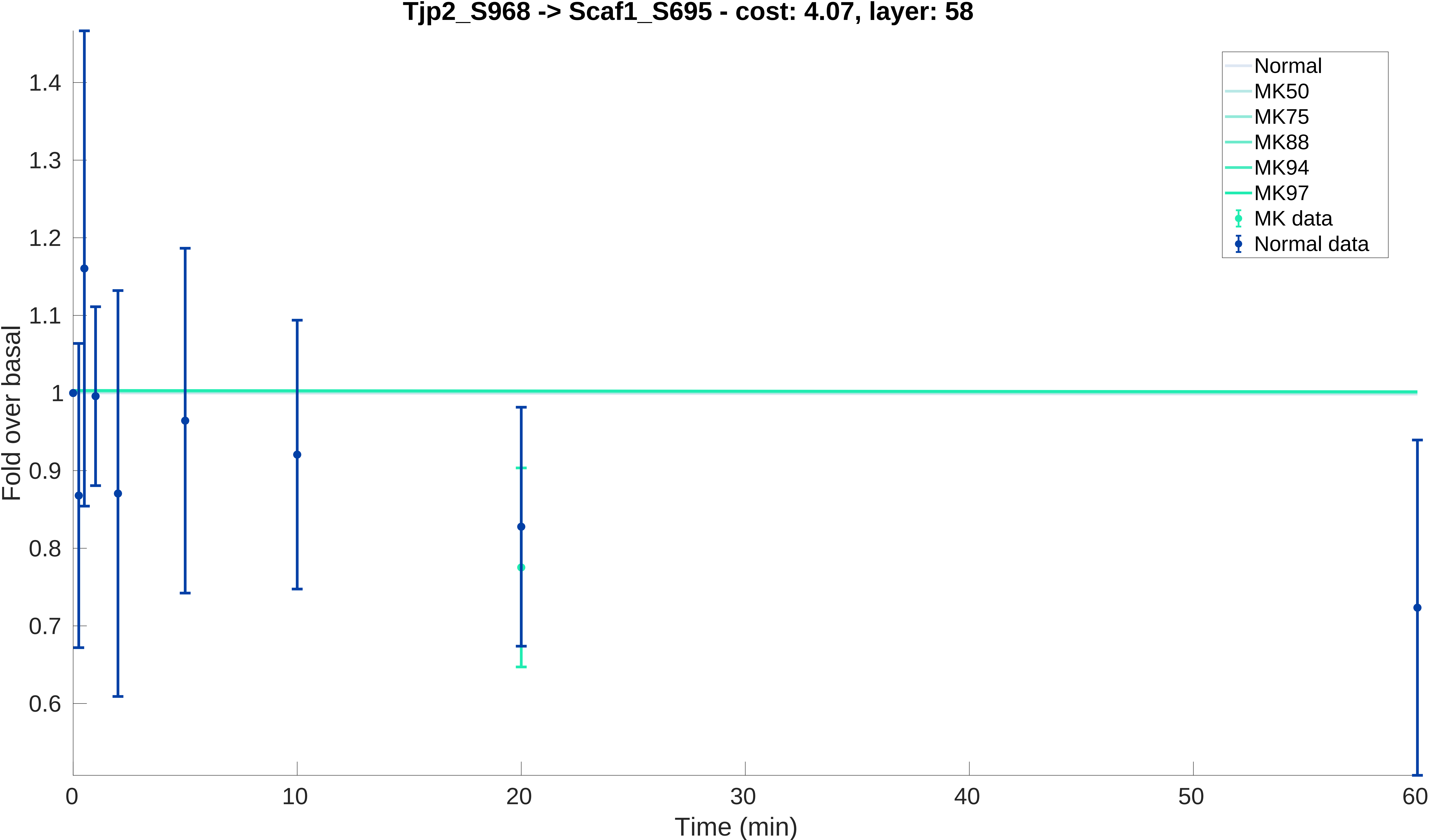

Tjp2_S968 -> Scaf1_S695 - cost: 4.07, layer: 58
Fold over basal
1.4
1.3
1.2
1.1
1
0.9
0.8
0.7
0.6
0
10
20
30
40
50
60
Time (min)
Normal
MK50
MK75
MK88
MK94
MK97
MK data
Normal data

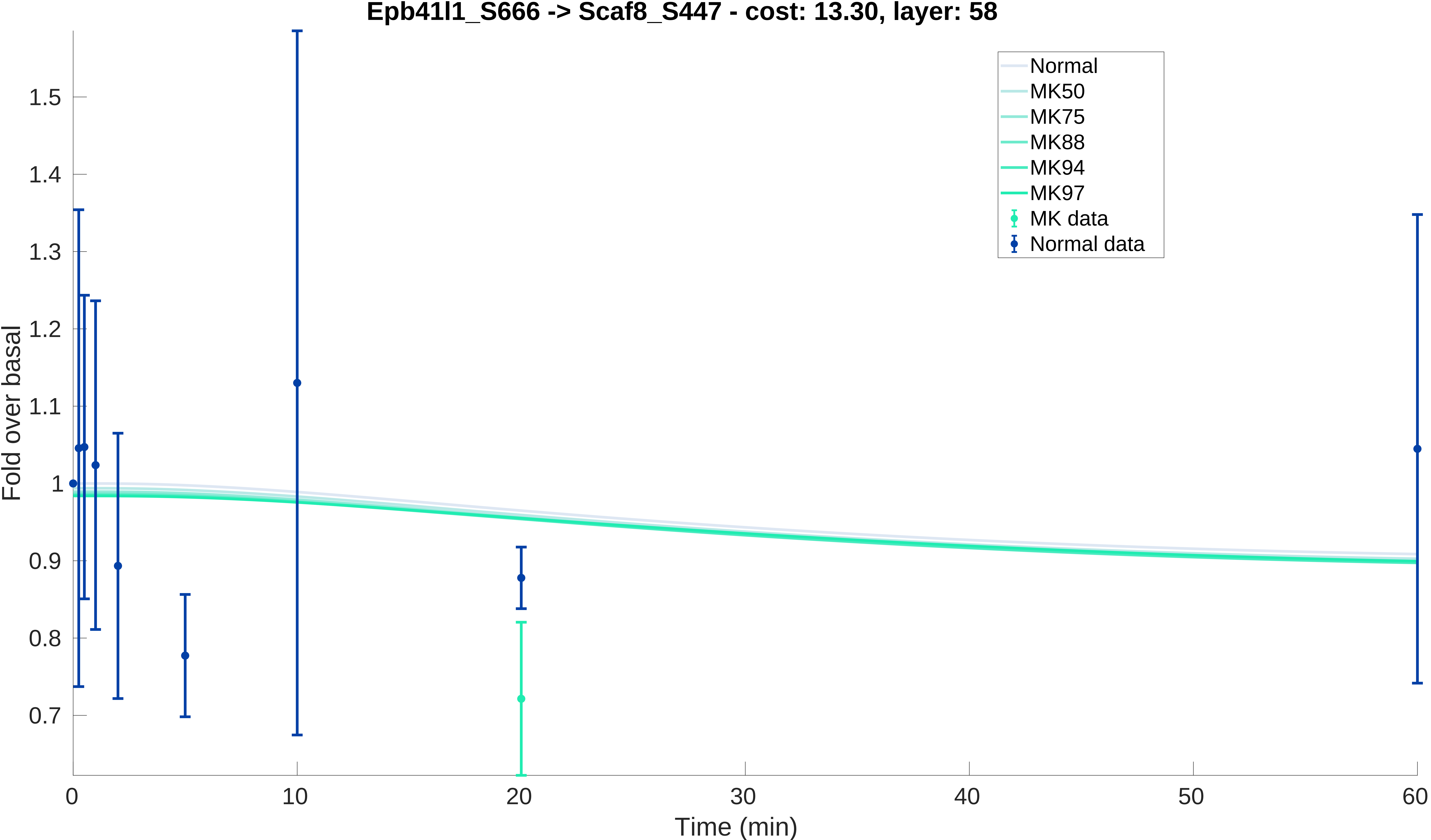

Epb41l1_S666 -> Scaf8_S447 - cost: 13.30, layer: 58
Fold over basal
Time (min)
1.5
1.4
1.3
1.2
1.1
1
0.9
0.8
0.7
0
10
20
30
40
50
60
Normal
MK50
MK75
MK88
MK94
MK97
MK data
Normal data

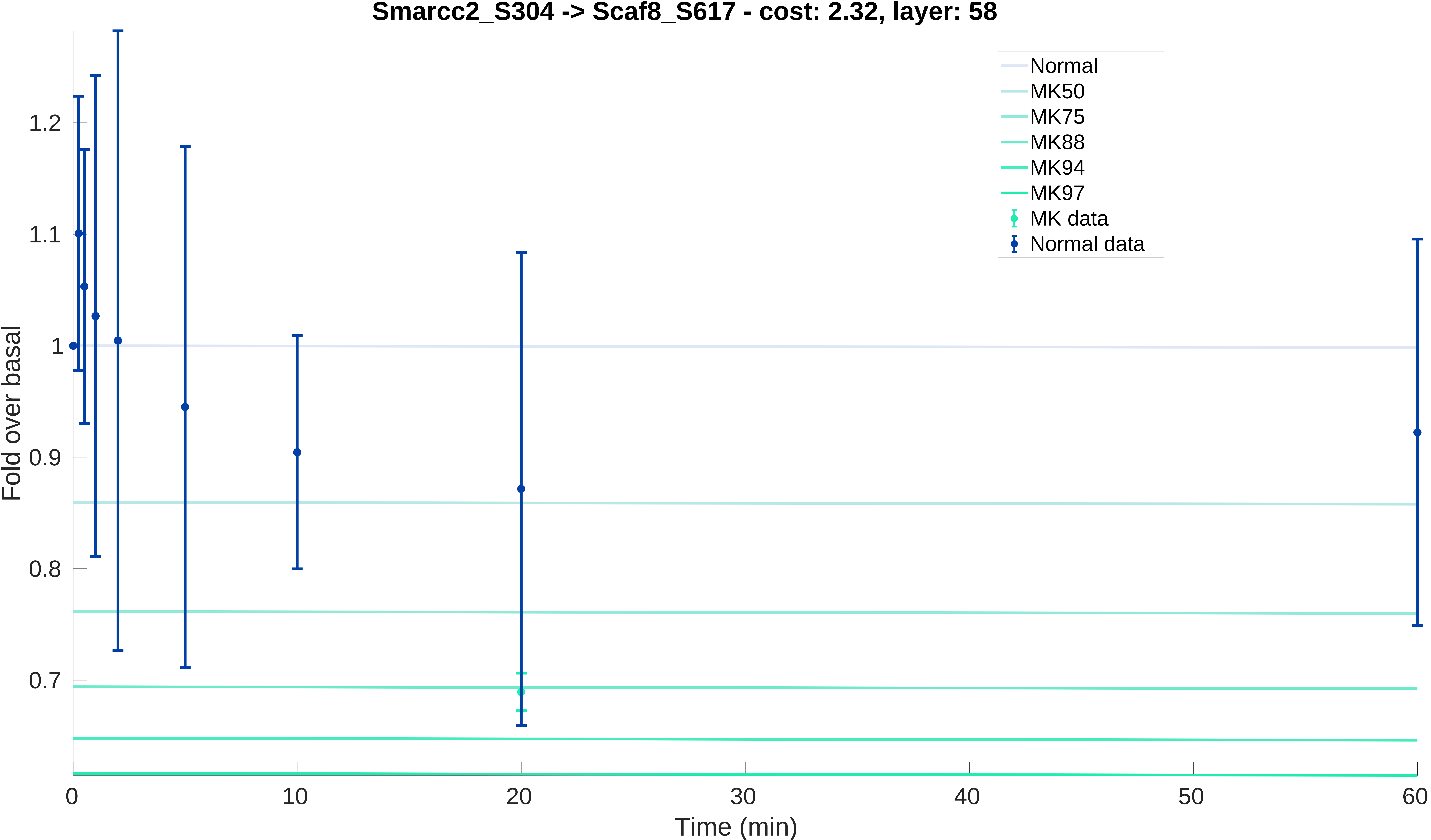

Smarcc2_S304 -> Scaf8_S617 - cost: 2.32, layer: 58
Fold over basal
Time (min)
0.7
0.8
0.9
1
1.1
1.2
0
10
20
30
40
50
60
Normal
MK50
MK75
MK88
MK94
MK97
MK data
Normal data

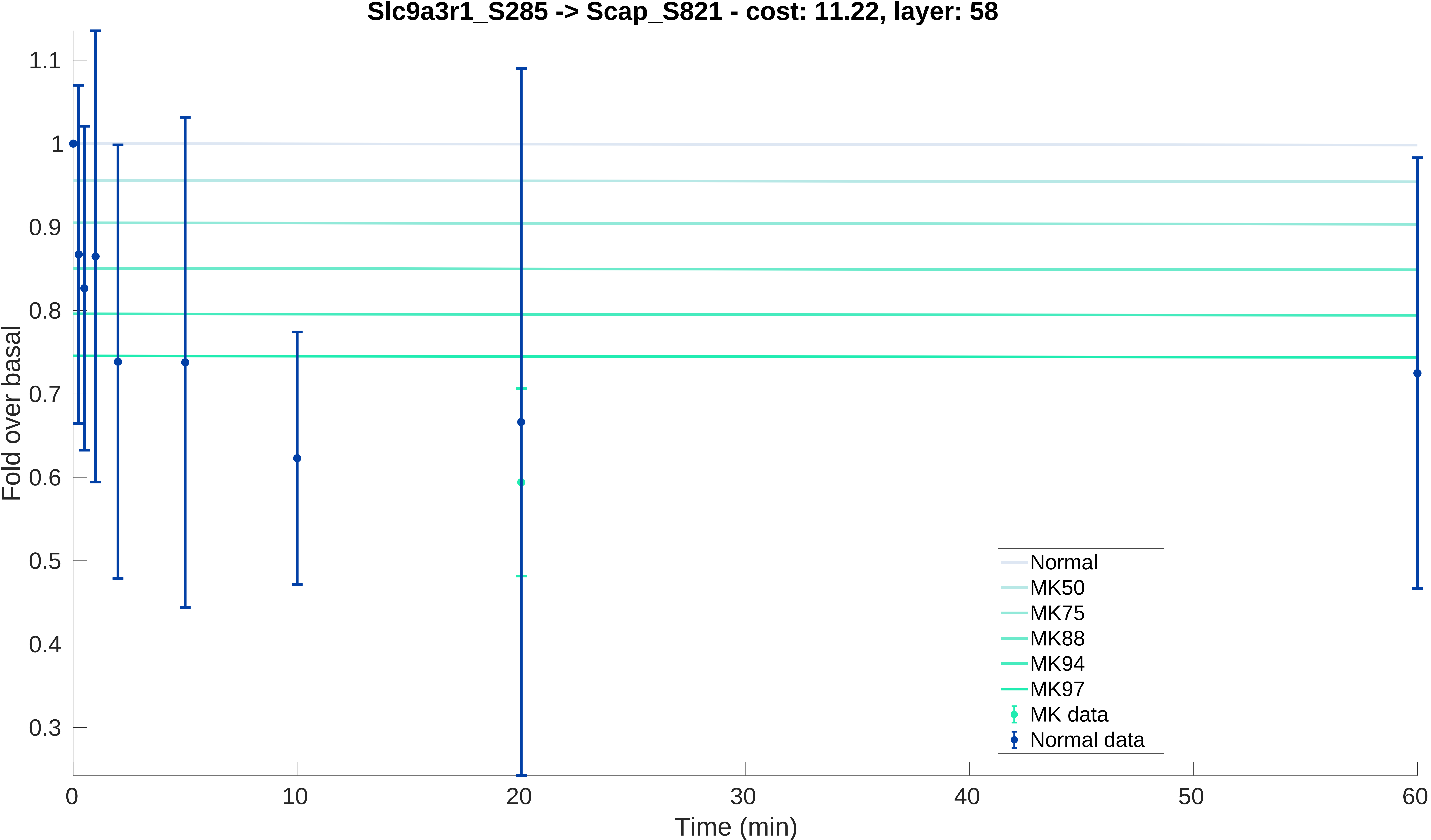

Slc9a3r1_S285 -> Scap_S821 - cost: 11.22, layer: 58
Fold over basal
Time (min)
0.3
0.4
0.5
0.6
0.7
0.8
0.9
1
1.1
0
10
20
30
40
50
60
Normal
MK50
MK75
MK88
MK94
MK97
MK data
Normal data

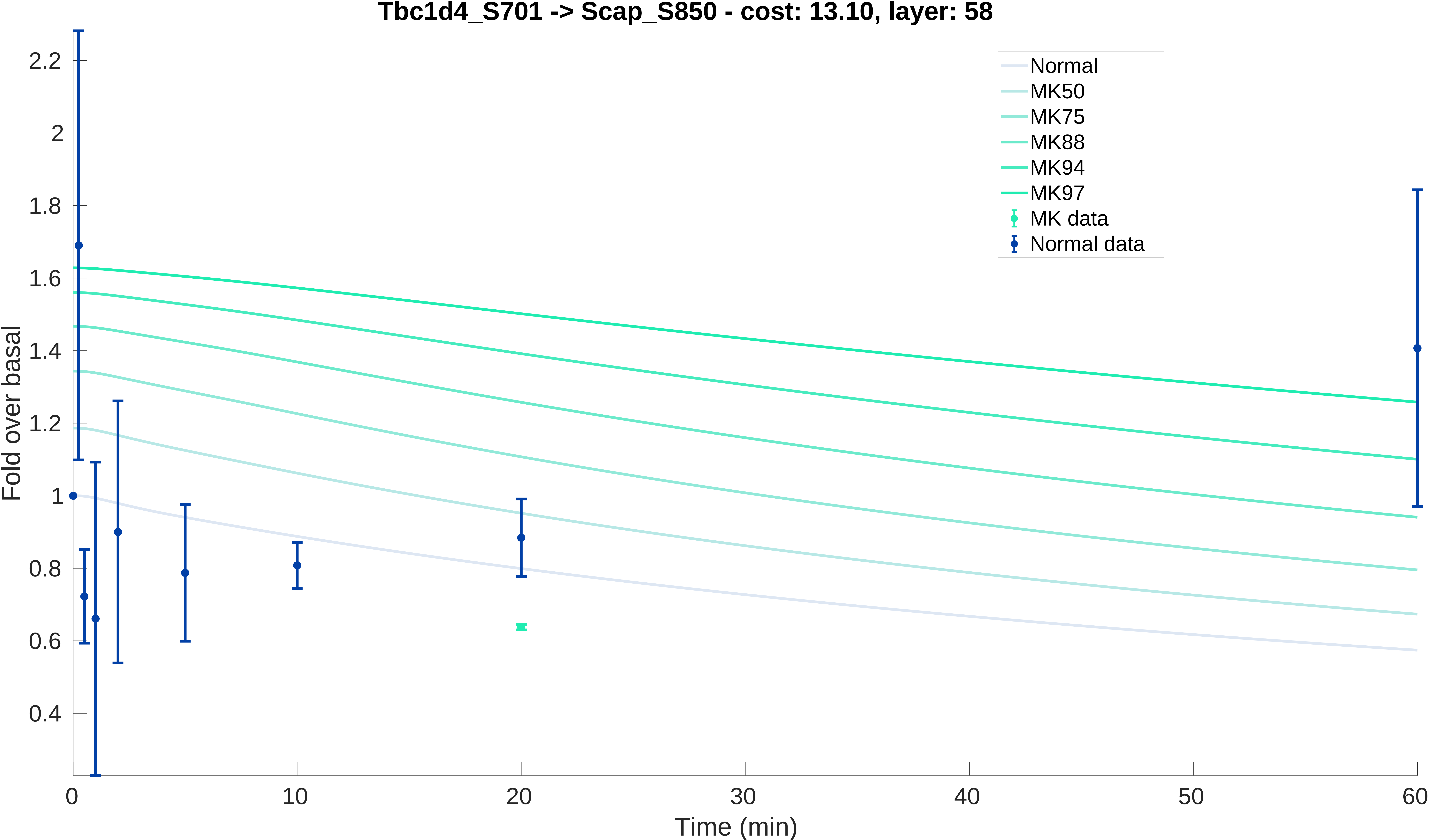

Tbc1d4_S701 -> Scap_S850 - cost: 13.10, layer: 58
Normal
MK50
MK75
MK88
MK94
MK97
MK data
Normal data
Fold over basal
Time (min)
0.4
0.6
0.8
1
1.2
1.4
1.6
1.8
2
2.2
0
10
20
30
40
50
60

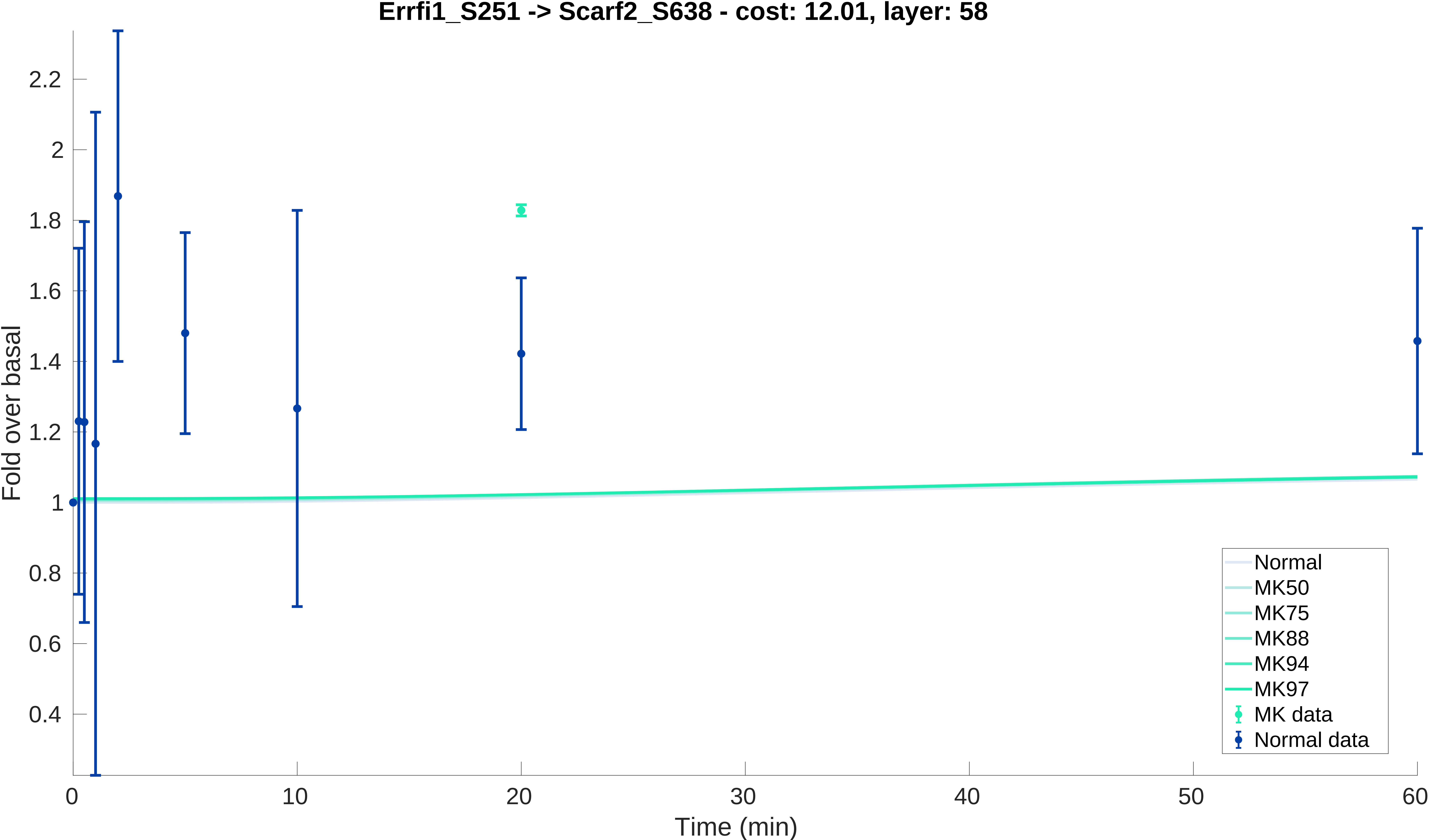

Errfi1_S251 -> Scarf2_S638 - cost: 12.01, layer: 58
Fold over basal
2.2
2
1.8
1.6
1.4
1.2
1
0.8
0.6
0.4
0
10
20
30
40
50
60
Time (min)
Normal
MK50
MK75
MK88
MK94
MK97
MK data
Normal data

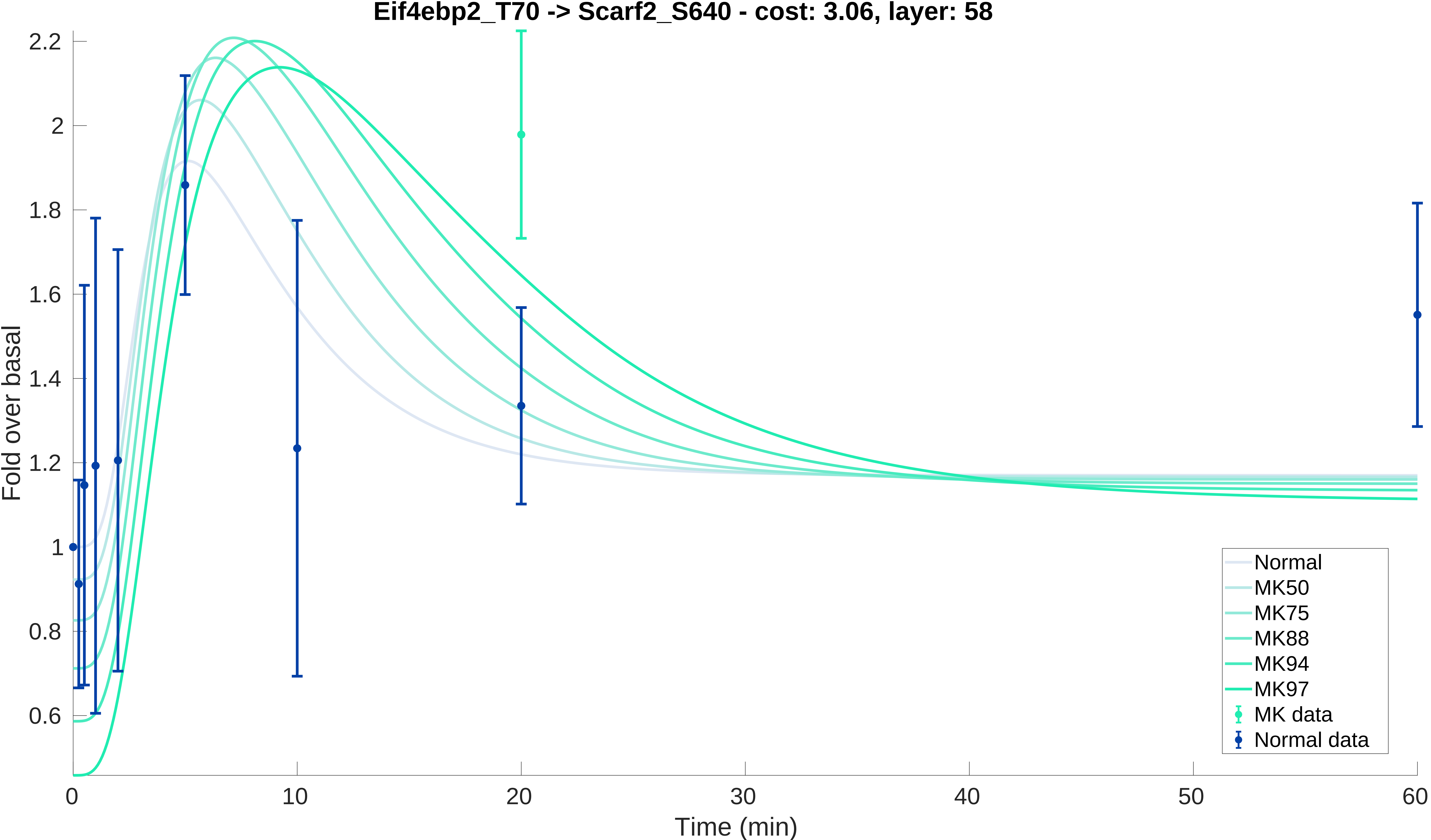
Eif4ebp2_T70 -> Scarf2_S640 - cost: 3.06, layer: 58
Fold over basal
Time (min)
0.6
0.8
1
1.2
1.4
1.6
1.8
2
2.2
0
10
20
30
40
50
60
Normal
MK50
MK75
MK88
MK94
MK97
MK data
Normal data

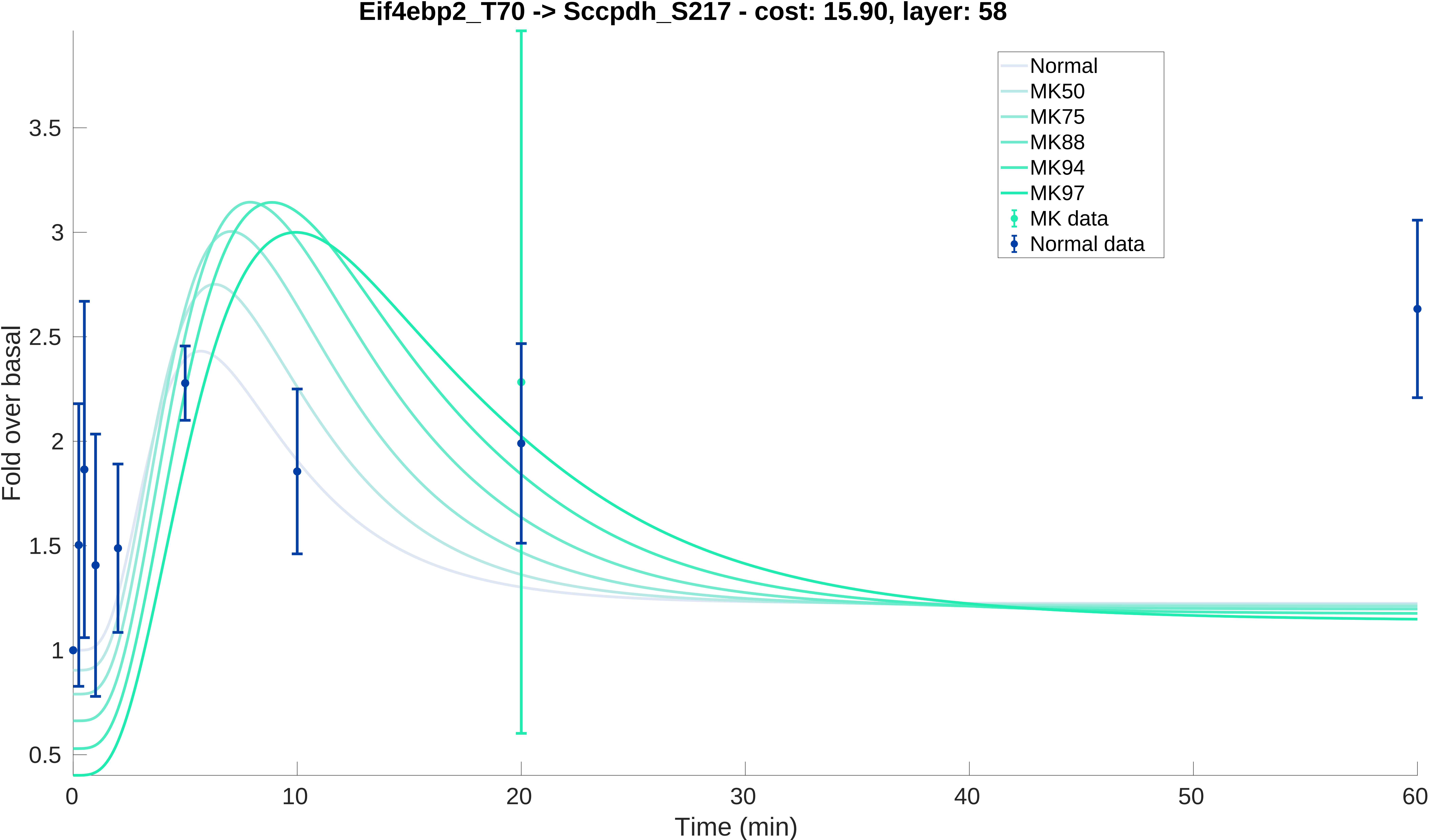

Eif4ebp2_T70 -> Sccpdh_S217 - cost: 15.90, layer: 58
Normal
MK50
MK75
MK88
MK94
MK97
MK data
Normal data
Fold over basal
3.5
3
2.5
2
1.5
1
0.5
0
10
20
30
40
50
60
Time (min)

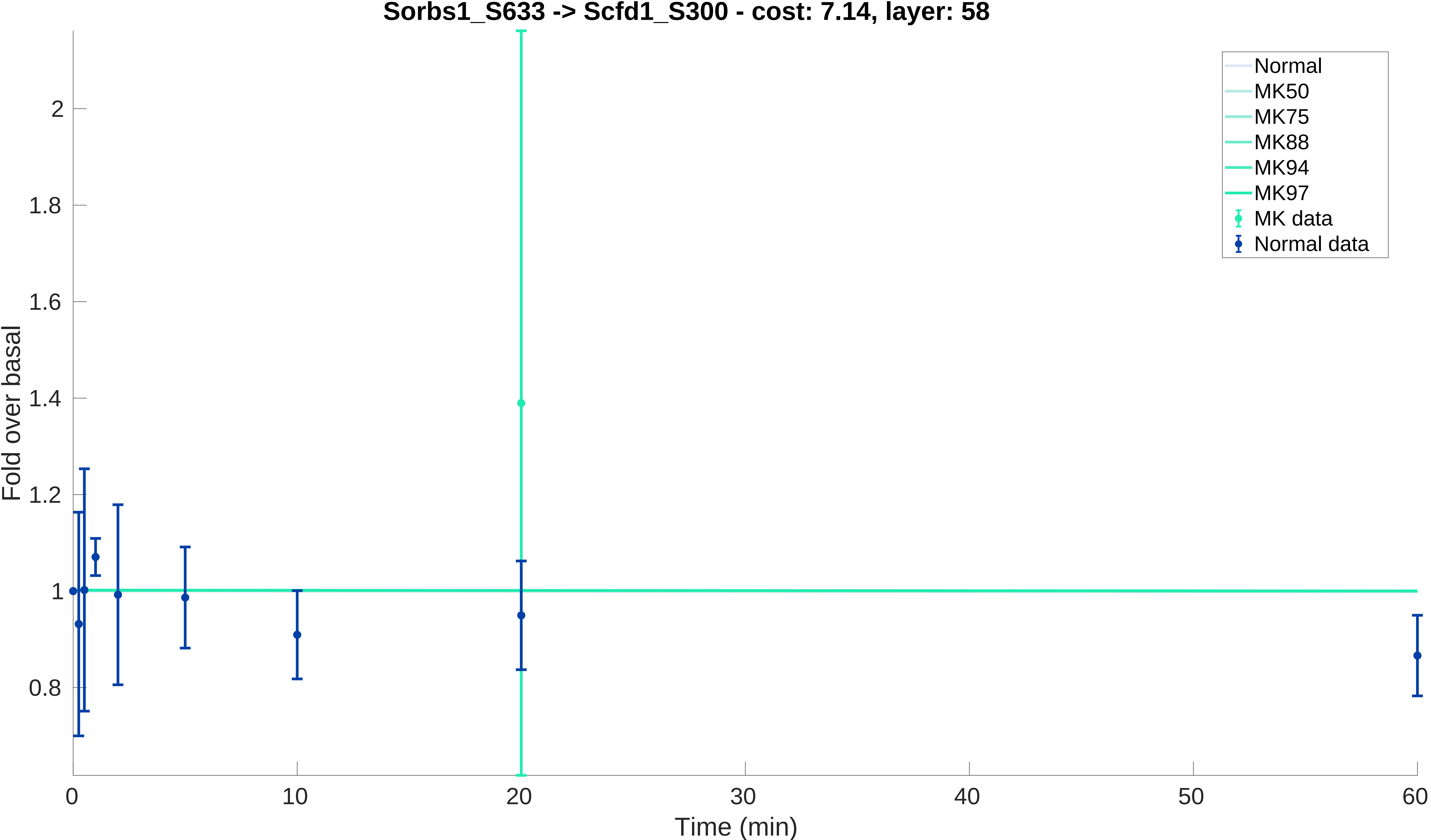
Sorbs1_S633 -> Scfd1_S300 - cost: 7.14, layer: 58
Normal
MK50
MK75
MK88
MK94
MK97
MK data
Normal data
Fold over basal
2
1.8
1.6
1.4
1.2
1
0.8
0
10
20
30
40
50
60
Time (min)

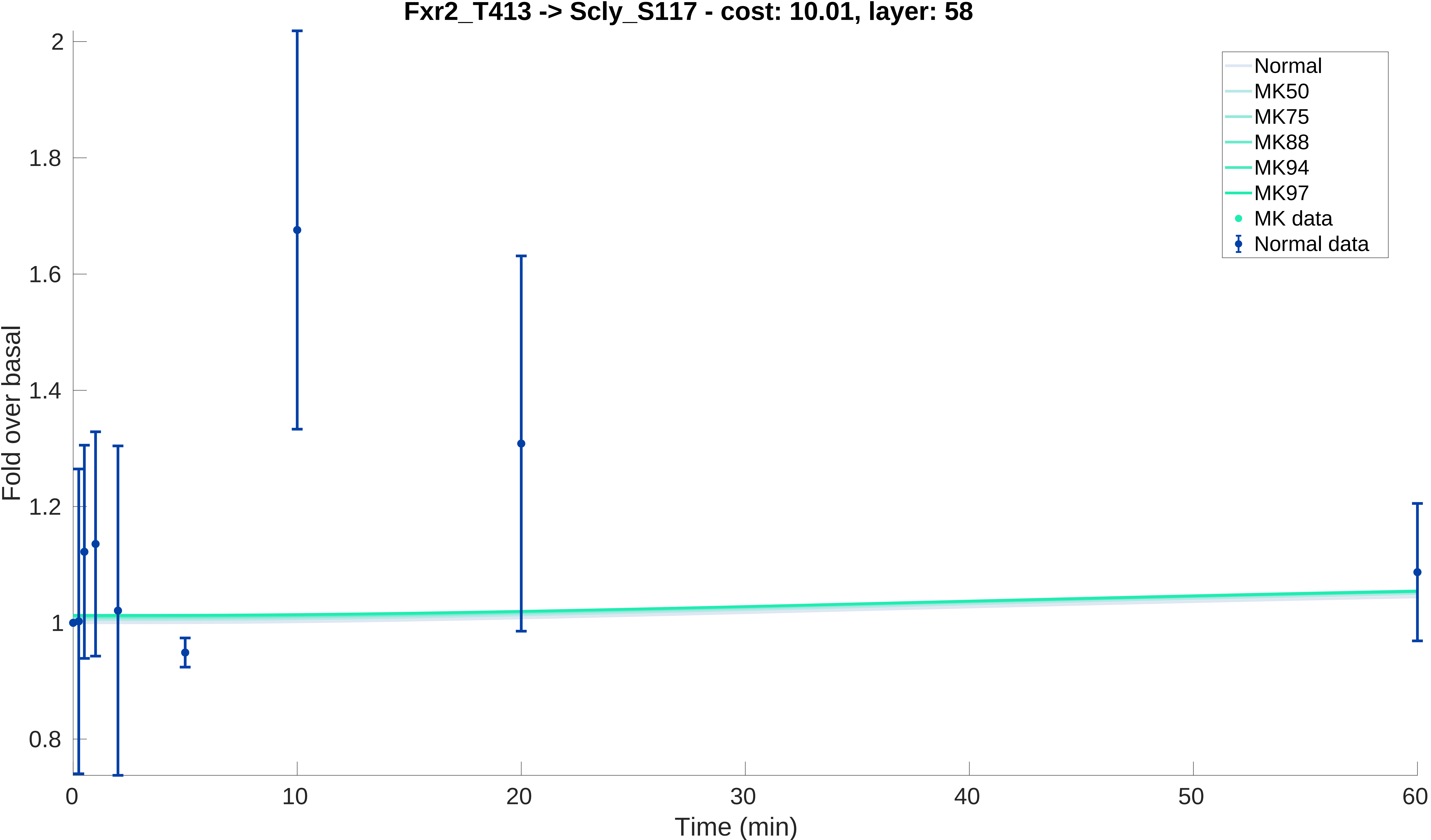
Fxr2_T413 -> Scly_S117 - cost: 10.01, layer: 58
Normal
MK50
MK75
MK88
MK94
MK97
MK data
Normal data
Fold over basal
2
1.8
1.6
1.4
1.2
1
0.8
0
10
20
30
40
50
60
Time (min)

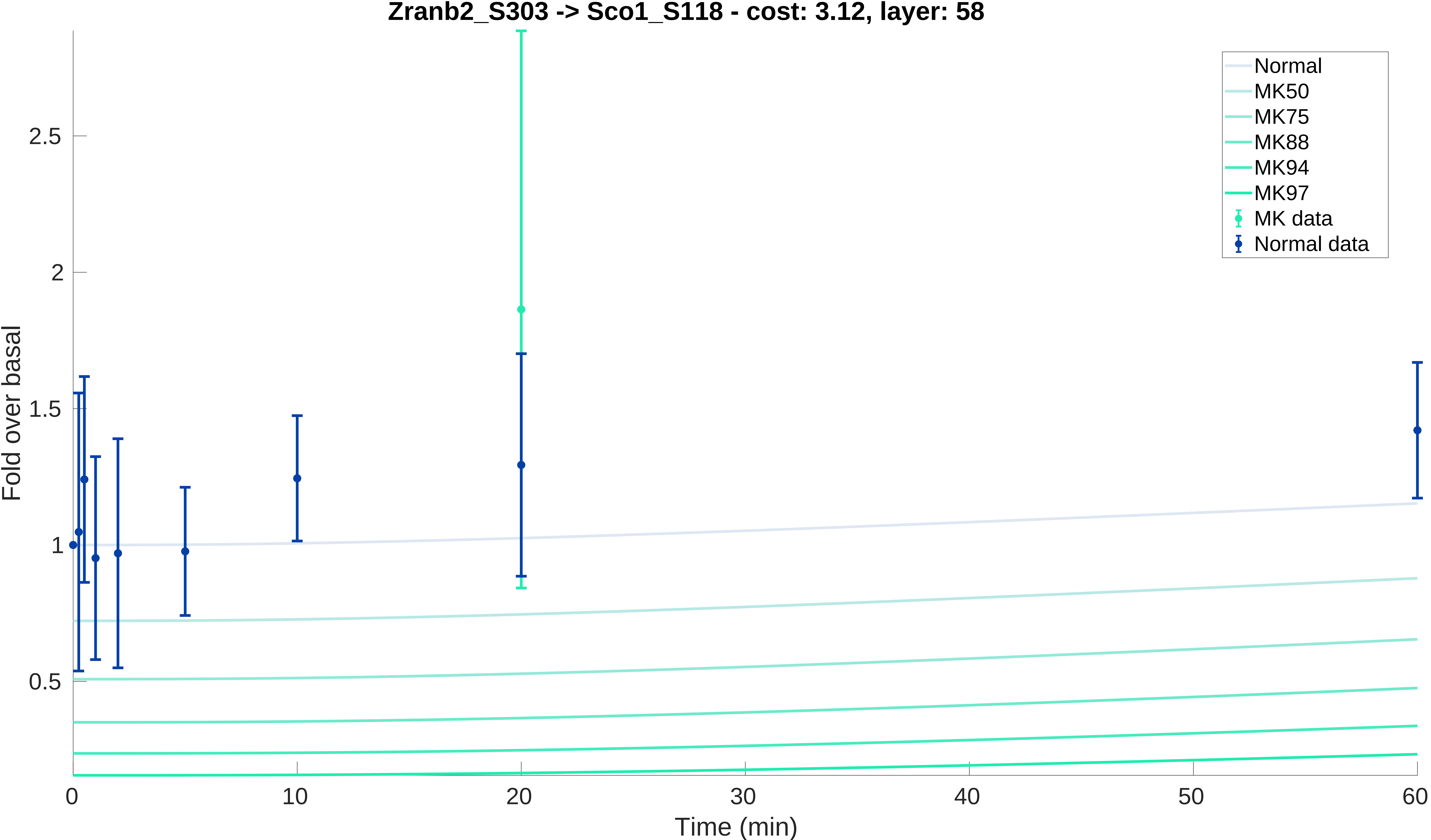

Zranb2_S303 -> Sco1_S118 - cost: 3.12, layer: 58
Fold over basal
Time (min)
0
10
20
30
40
50
60
0.5
1
1.5
2
2.5
Normal
MK50
MK75
MK88
MK94
MK97
MK data
Normal data

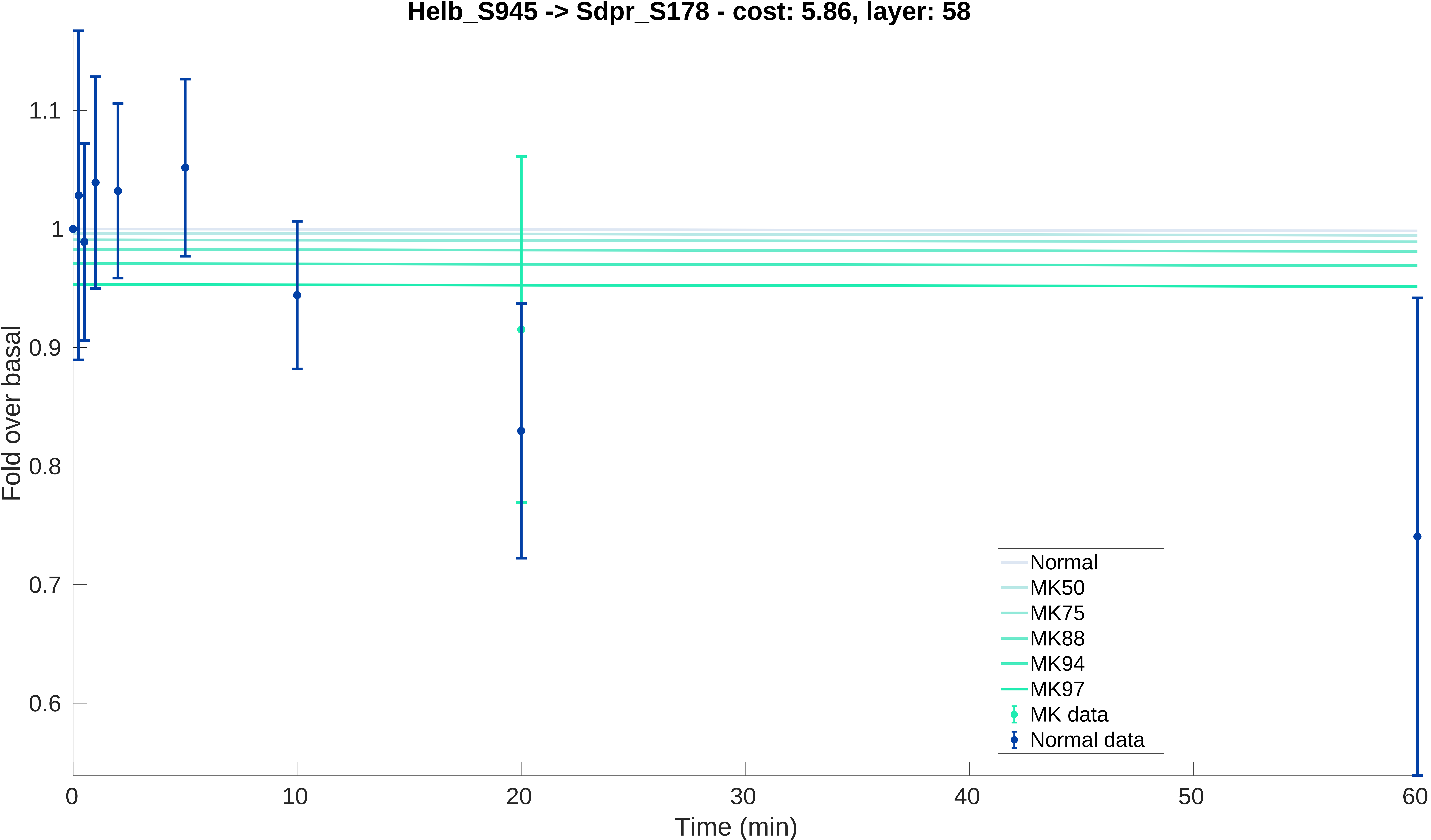

Helb_S945 -> Sdpr_S178 - cost: 5.86, layer: 58
Fold over basal
1.1
1
0.9
0.8
0.7
0.6
0
10
20
30
40
50
60
Time (min)
Normal
MK50
MK75
MK88
MK94
MK97
MK data
Normal data

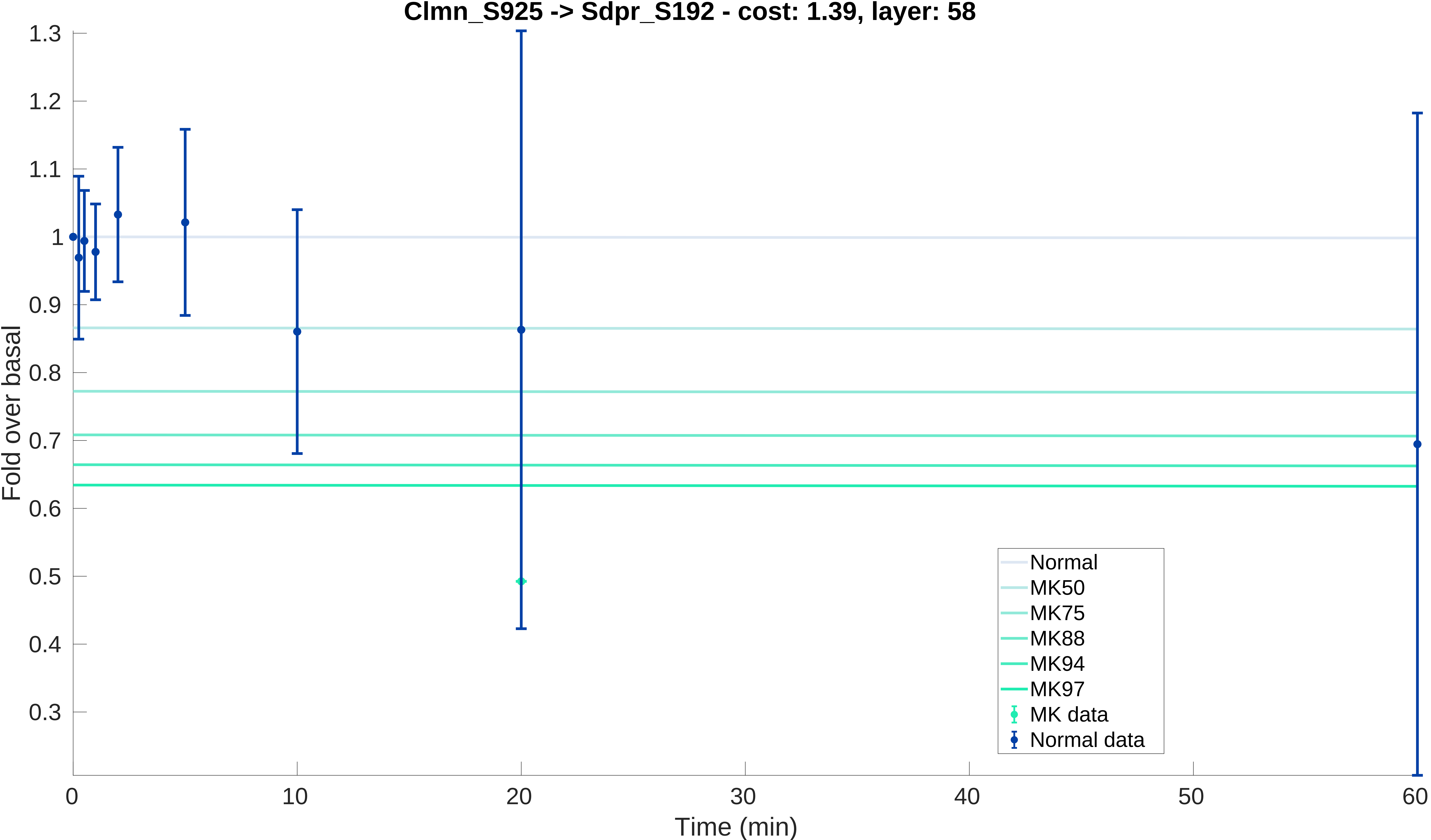

Clmn_S925 -> Sdpr_S192 - cost: 1.39, layer: 58
Fold over basal
Time (min)
1.3
1.2
1.1
1
0.9
0.8
0.7
0.6
0.5
0.4
0.3
0
10
20
30
40
50
60
Normal
MK50
MK75
MK88
MK94
MK97
MK data
Normal data

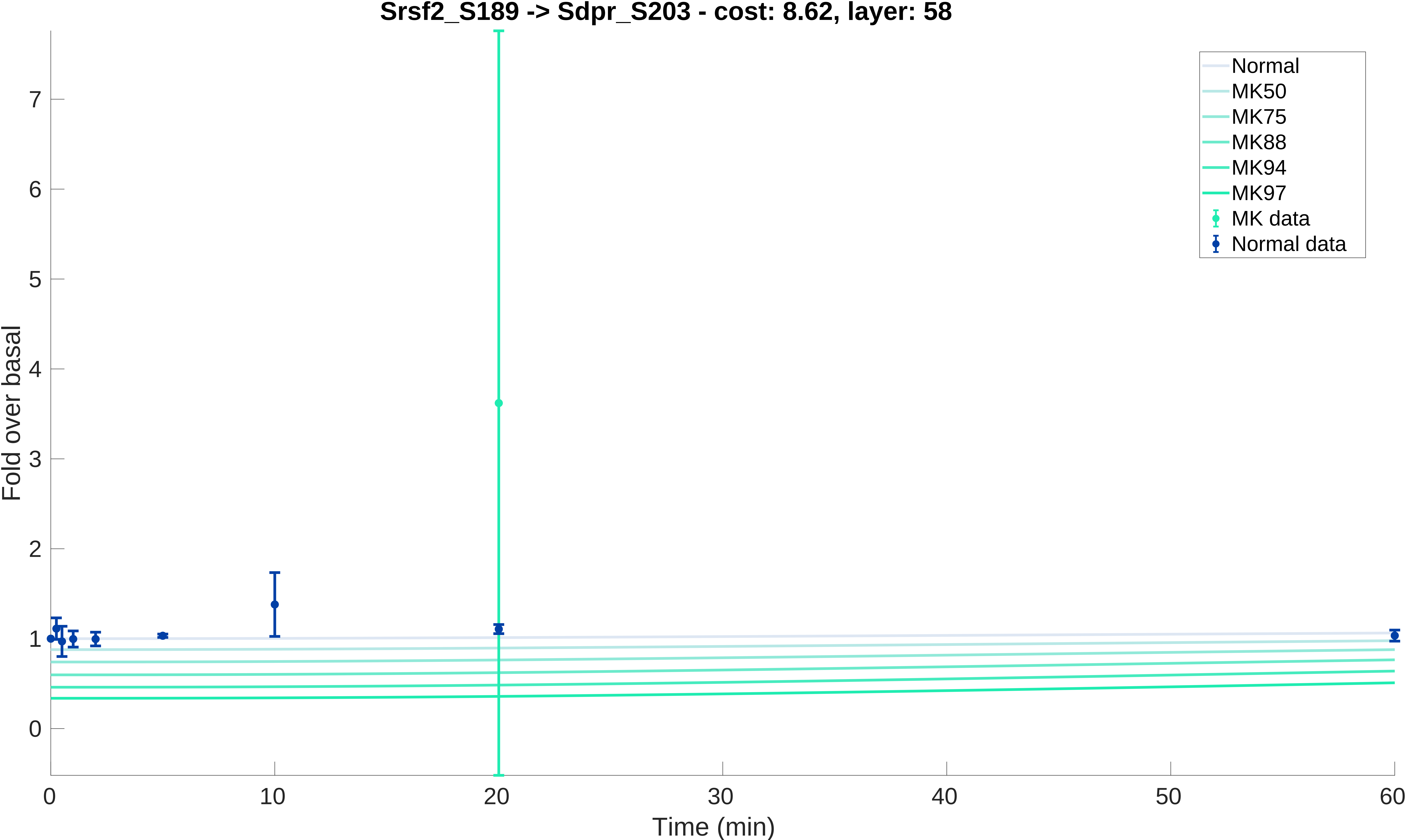

Srsf2_S189 -> Sdpr_S203 - cost: 8.62, layer: 58
Fold over basal
Time (min)
0
1
2
3
4
5
6
7
0
10
20
30
40
50
60
Normal
MK50
MK75
MK88
MK94
MK97
MK data
Normal data

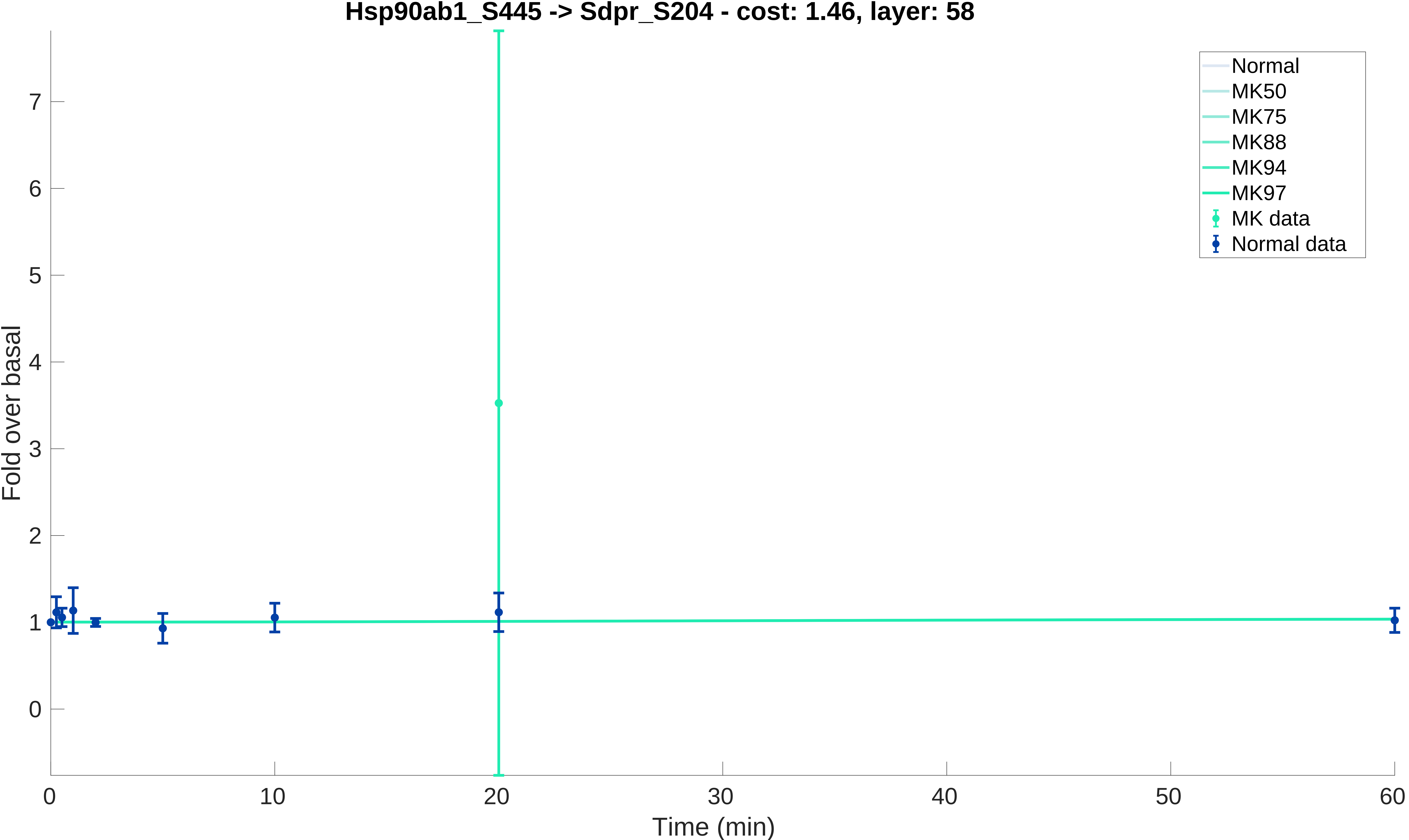

Hsp90ab1_S445 -> Sdpr_S204 - cost: 1.46, layer: 58
Fold over basal
Time (min)
0
1
2
3
4
5
6
7
0
10
20
30
40
50
60
Normal
MK50
MK75
MK88
MK94
MK97
MK data
Normal data

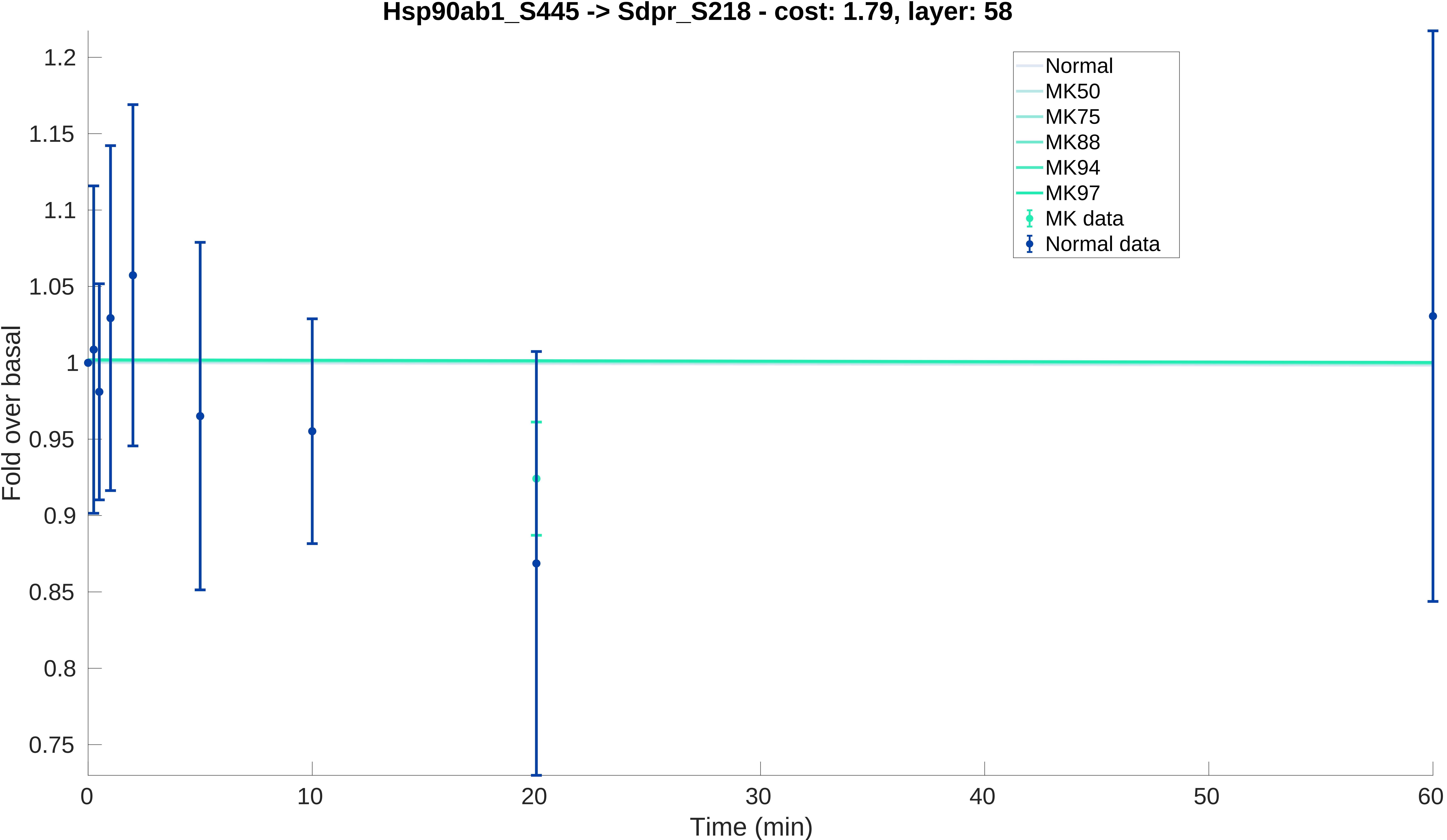

Hsp90ab1_S445 -> Sdpr_S218 - cost: 1.79, layer: 58
Normal
MK50
MK75
MK88
MK94
MK97
MK data
Normal data
Fold over basal
1.2
1.15
1.1
1.05
1
0.95
0.9
0.85
0.8
0.75
0
10
20
30
40
50
60
Time (min)

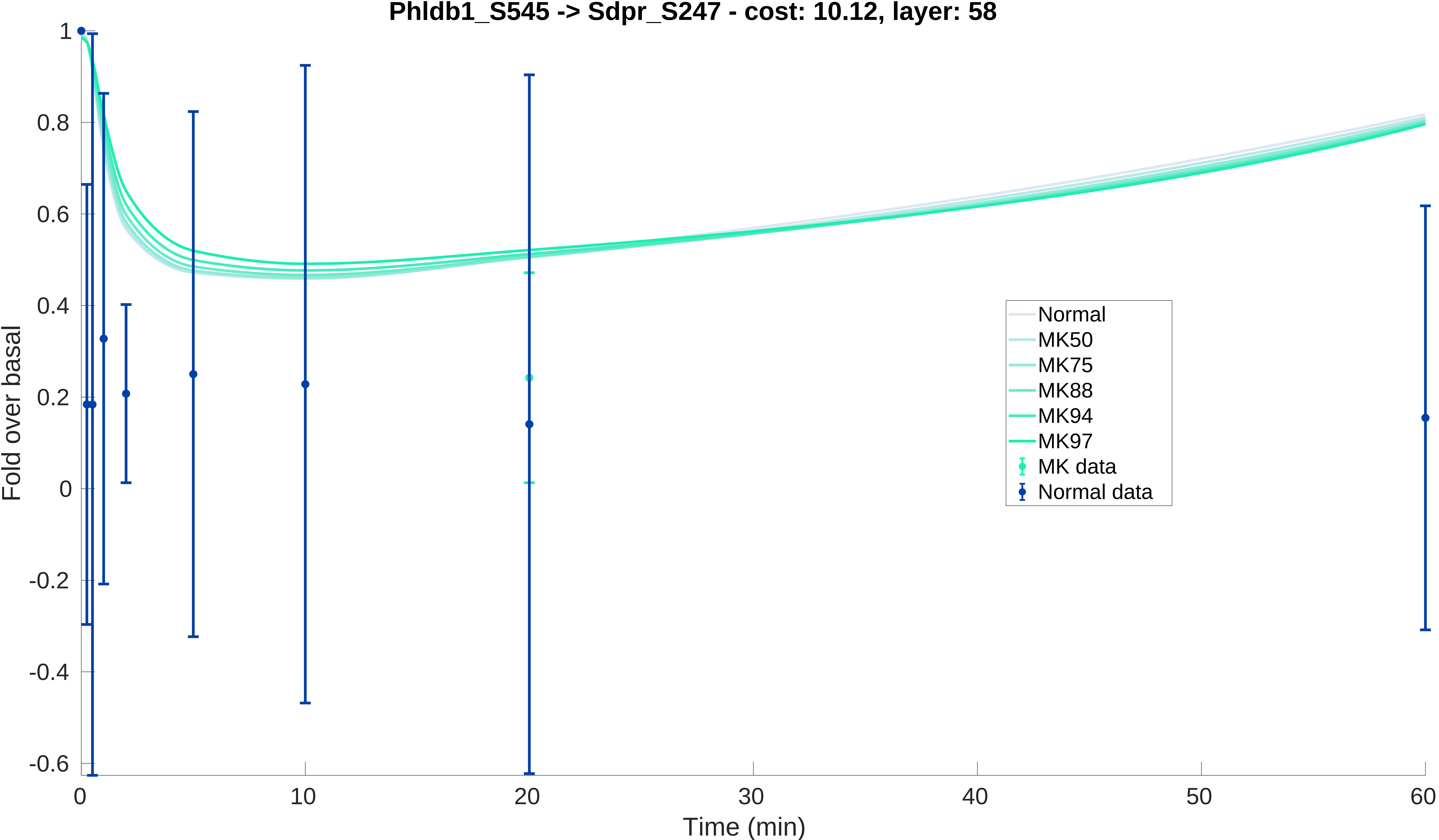

Phldb1_S545 -> Sdpr_S247 - cost: 10.12, layer: 58
Fold over basal
Time (min)
1
0.8
0.6
0.4
0.2
0
-0.2
-0.4
-0.6
0
10
20
30
40
50
60
Normal
MK50
MK75
MK88
MK94
MK97
MK data
Normal data

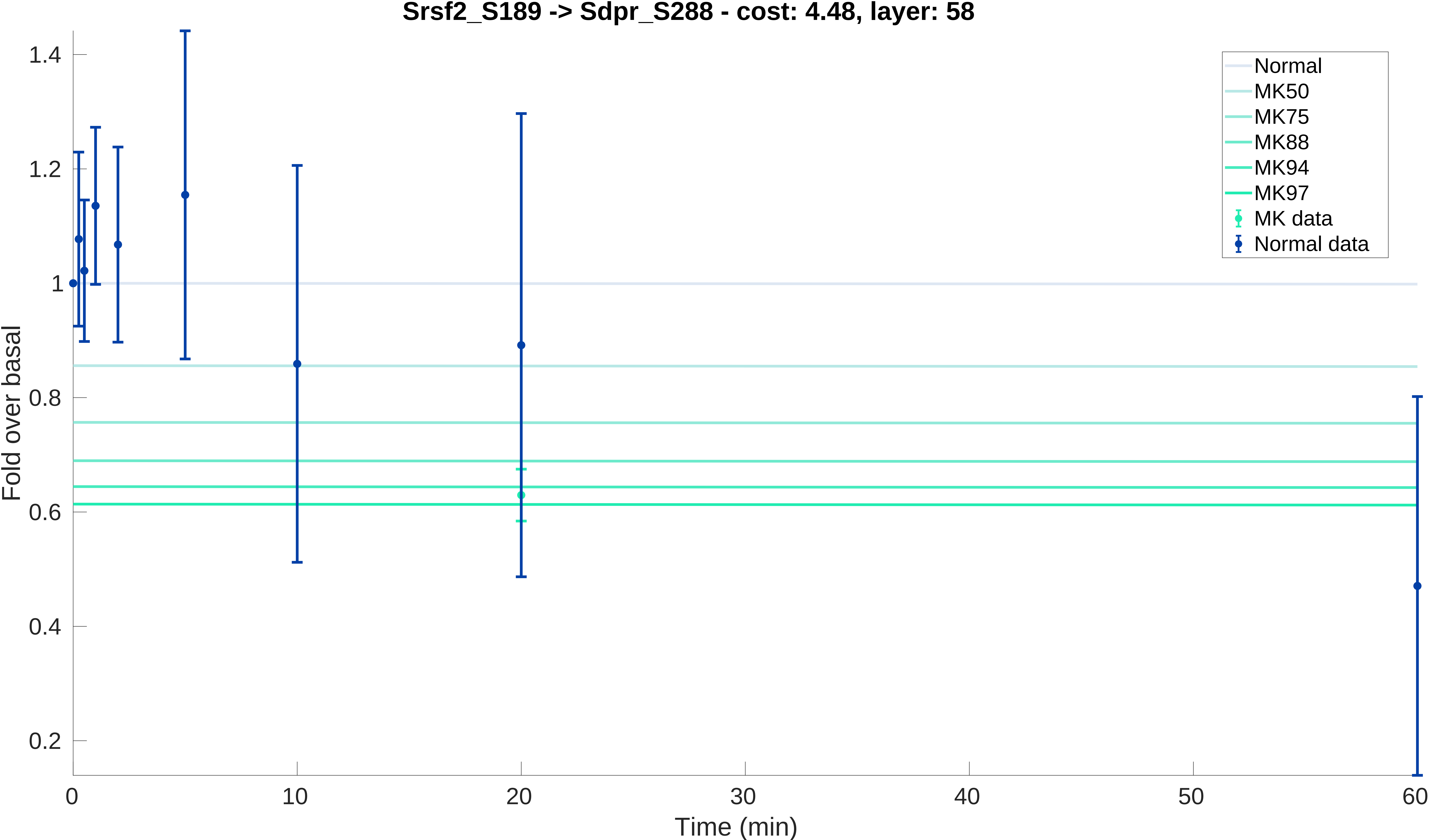
Srsf2_S189 -> Sdpr_S288 - cost: 4.48, layer: 58
Fold over basal
Time (min)
1.4
1.2
1
0.8
0.6
0.4
0.2
0
10
20
30
40
50
60
Normal
MK50
MK75
MK88
MK94
MK97
MK data
Normal data

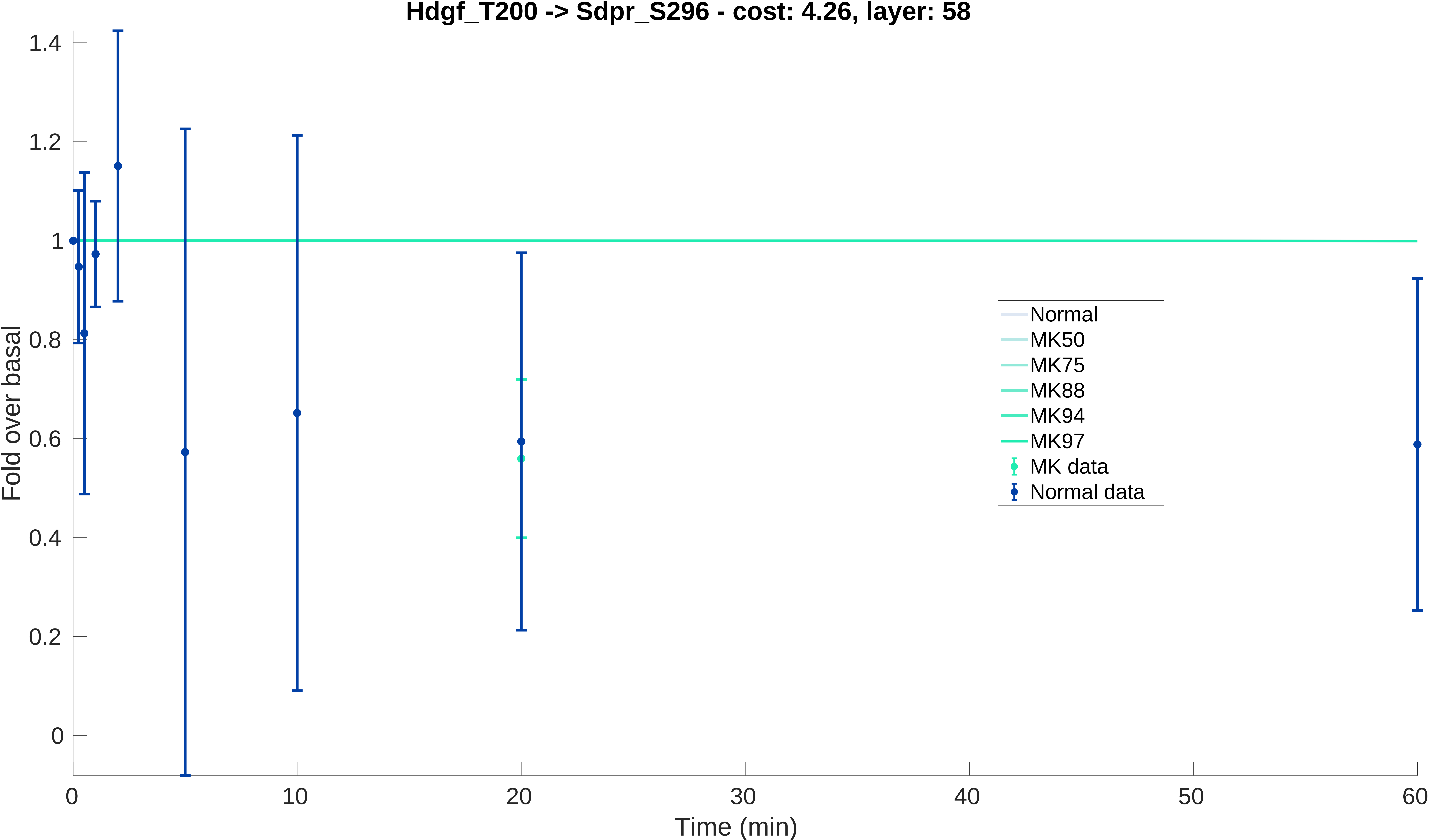

Hdgf_T200 -> Sdpr_S296 - cost: 4.26, layer: 58
1.4
1.2
1
0.8
0.6
0.4
0.2
0
Fold over basal
0
10
20
30
40
50
60
Time (min)
Normal
MK50
MK75
MK88
MK94
MK97
MK data
Normal data

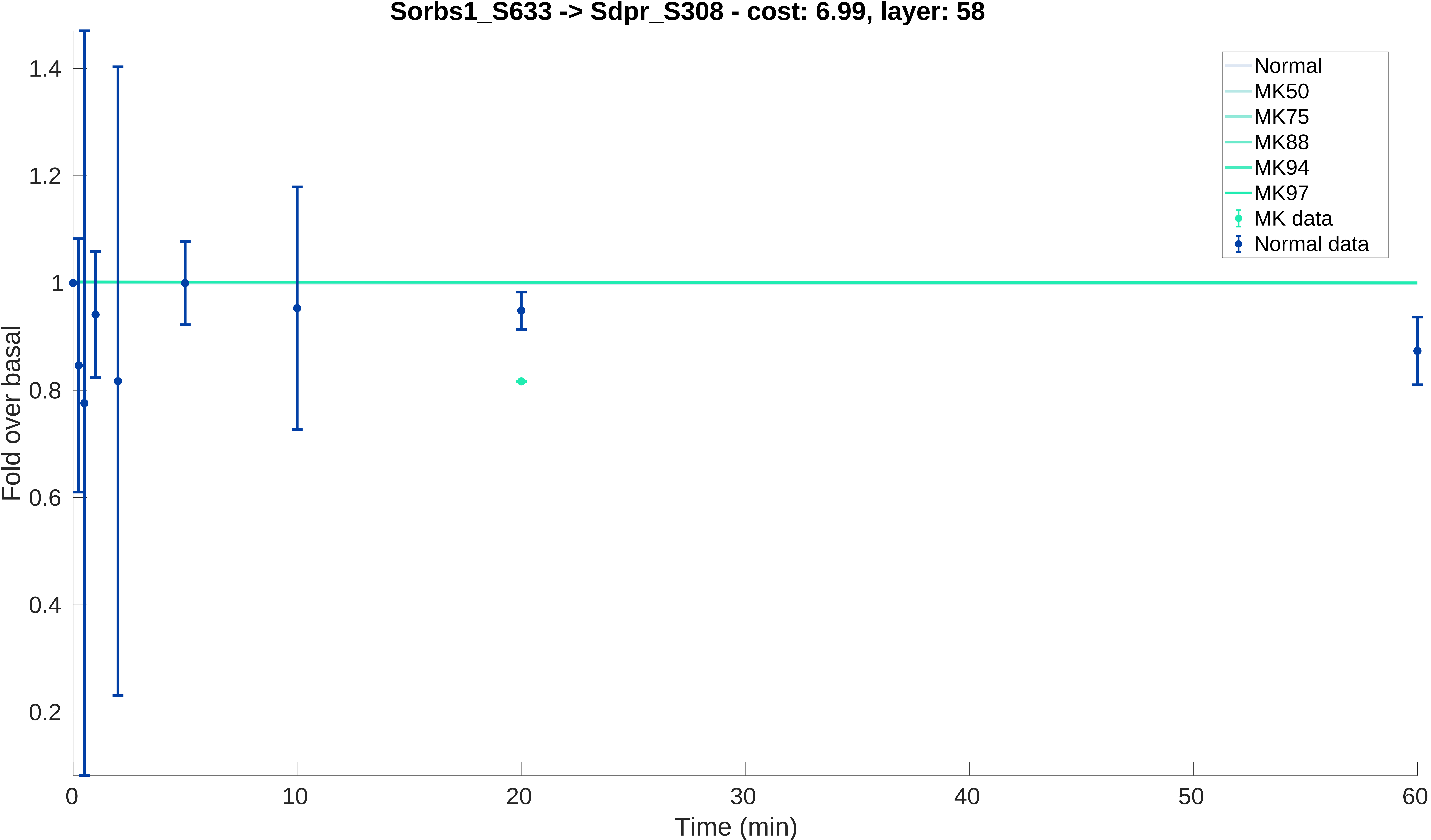

Sorbs1_S633 -> Sdpr_S308 - cost: 6.99, layer: 58
Fold over basal
Time (min)
0
10
20
30
40
50
60
0.2
0.4
0.6
0.8
1
1.2
1.4
Normal
MK50
MK75
MK88
MK94
MK97
MK data
Normal data

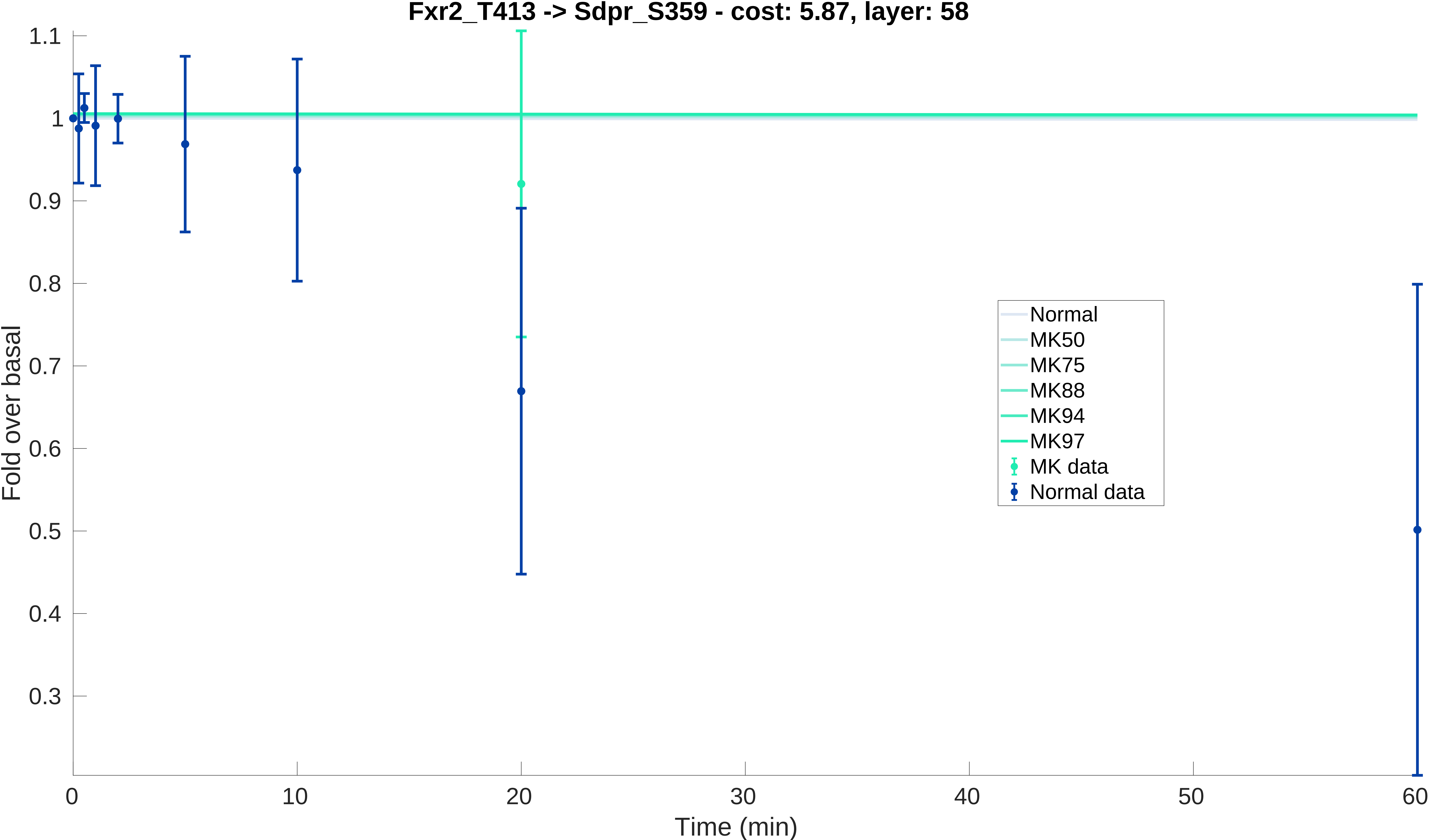

Fxr2_T413 -> Sdpr_S359 - cost: 5.87, layer: 58
Fold over basal
Time (min)
1.1
1
0.9
0.8
0.7
0.6
0.5
0.4
0.3
0
10
20
30
40
50
60
Normal
MK50
MK75
MK88
MK94
MK97
MK data
Normal data

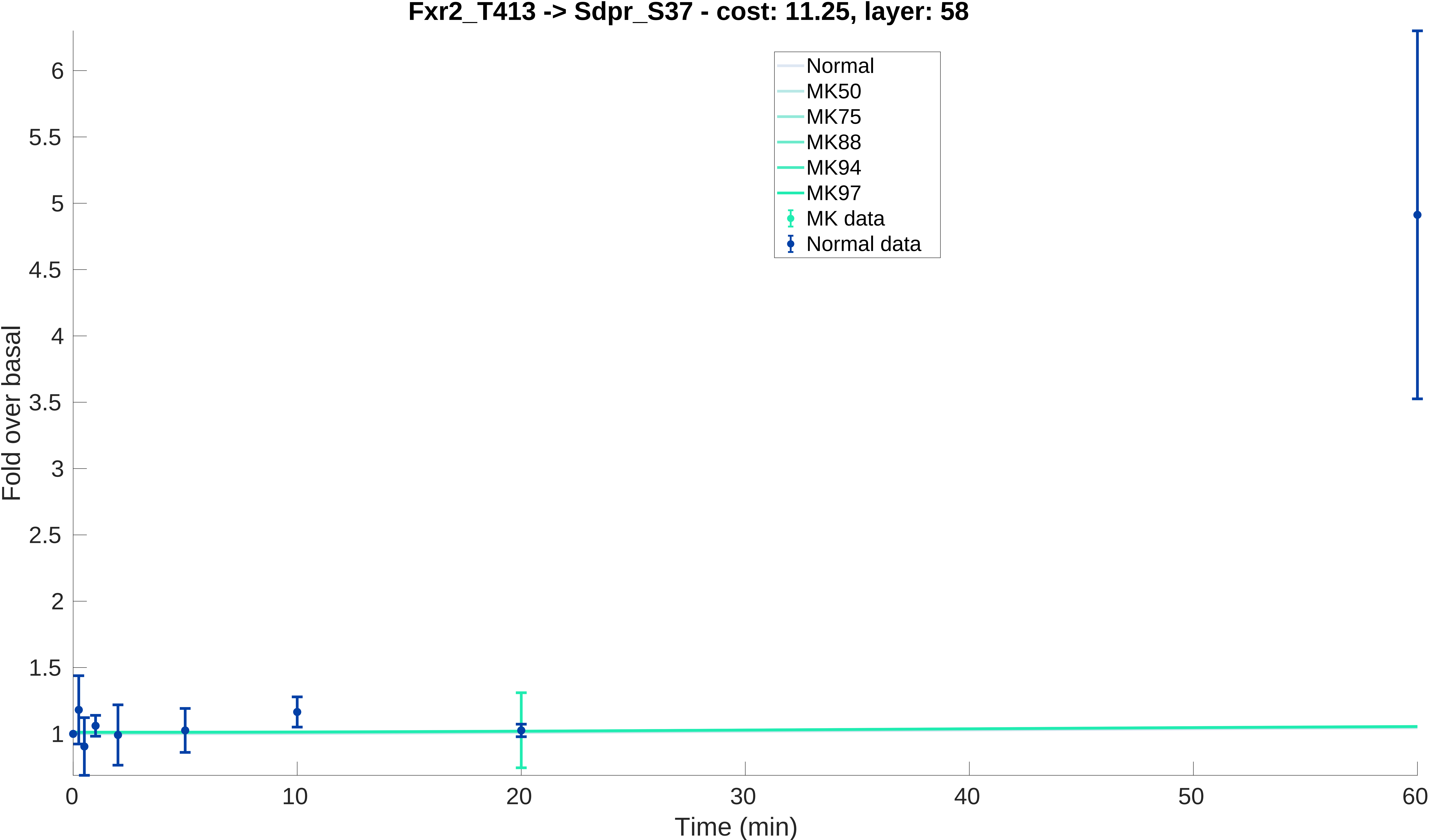
Fxr2_T413 -> Sdpr_S37 - cost: 11.25, layer: 58
Normal
MK50
MK75
MK88
MK94
MK97
MK data
Normal data
Fold over basal
Time (min)
0
10
20
30
40
50
60
1
1.5
2
2.5
3
3.5
4
4.5
5
5.5
6

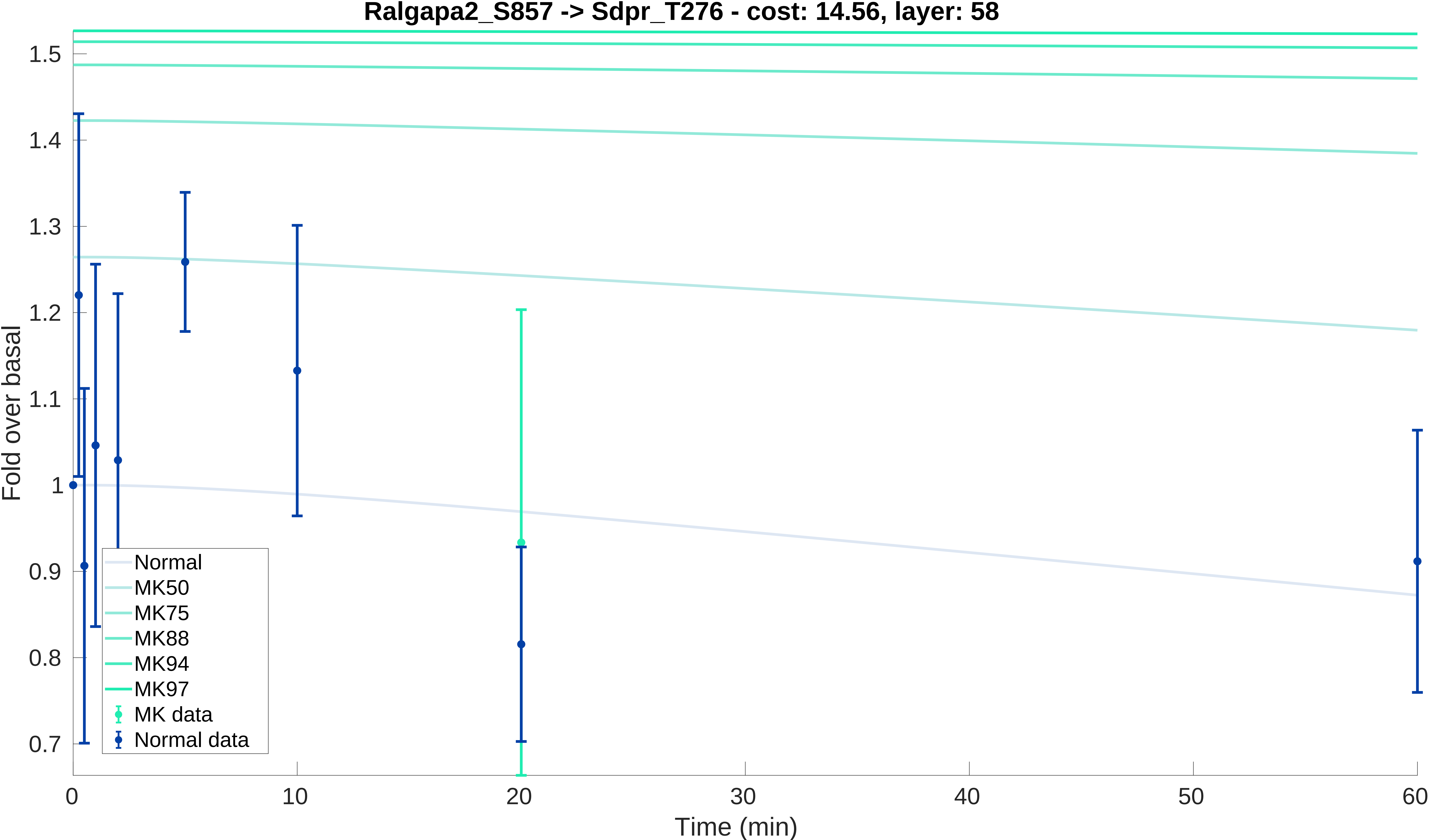

Ralgapa2_S857 -> Sdpr_T276 - cost: 14.56, layer: 58
Fold over basal
Time (min)
0
10
20
30
40
50
60
0.7
0.8
0.9
1
1.1
1.2
1.3
1.4
1.5
Normal
MK50
MK75
MK88
MK94
MK97
MK data
Normal data

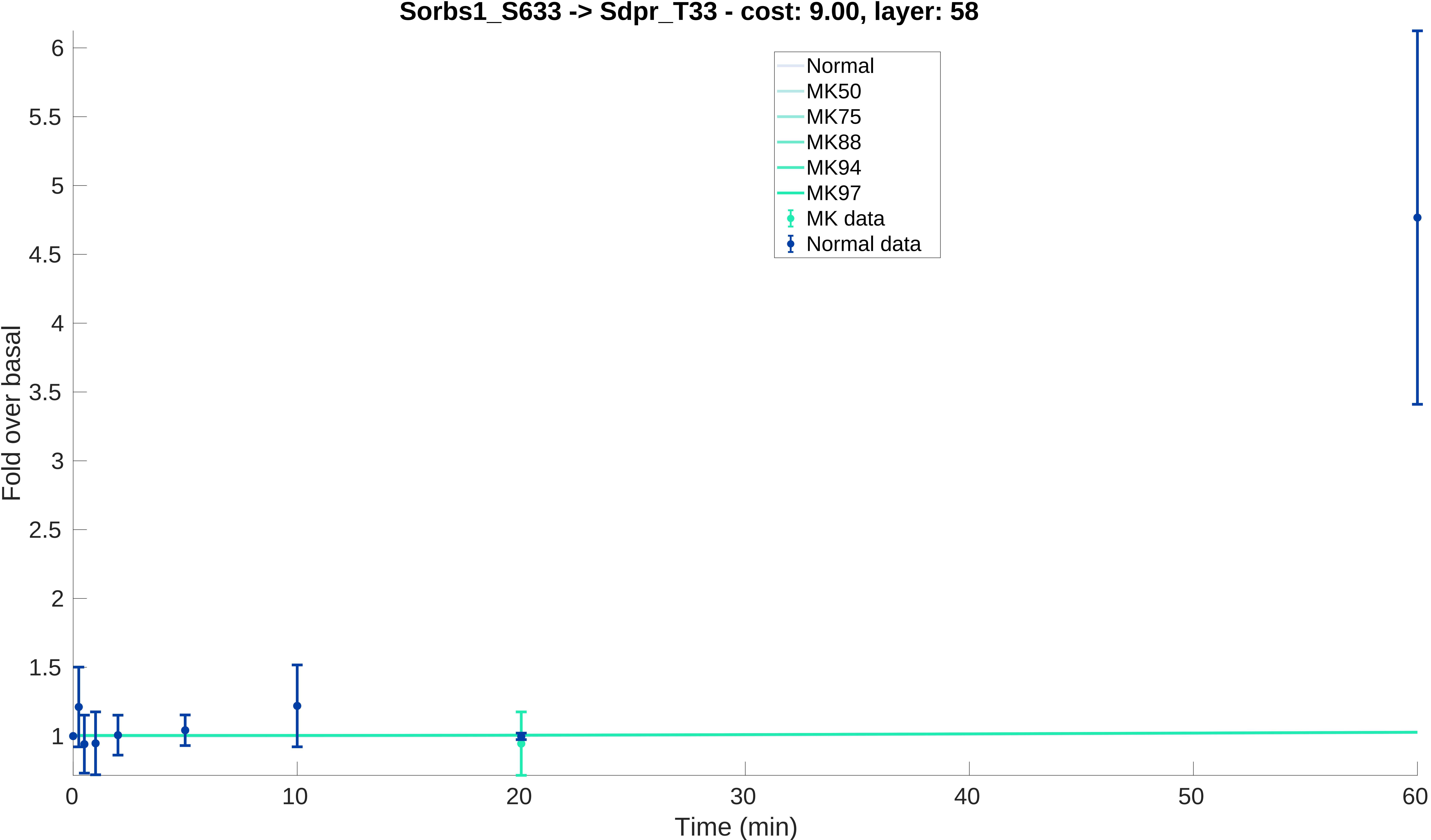
Sorbs1_S633 -> Sdpr_T33 - cost: 9.00, layer: 58
Fold over basal
Time (min)
0
10
20
30
40
50
60
1
1.5
2
2.5
3
3.5
4
4.5
5
5.5
6
Normal
MK50
MK75
MK88
MK94
MK97
MK data
Normal data

#### Pla2g4a_S434 -> Sec14l1_S578 - cost: 1.63, layer: 58

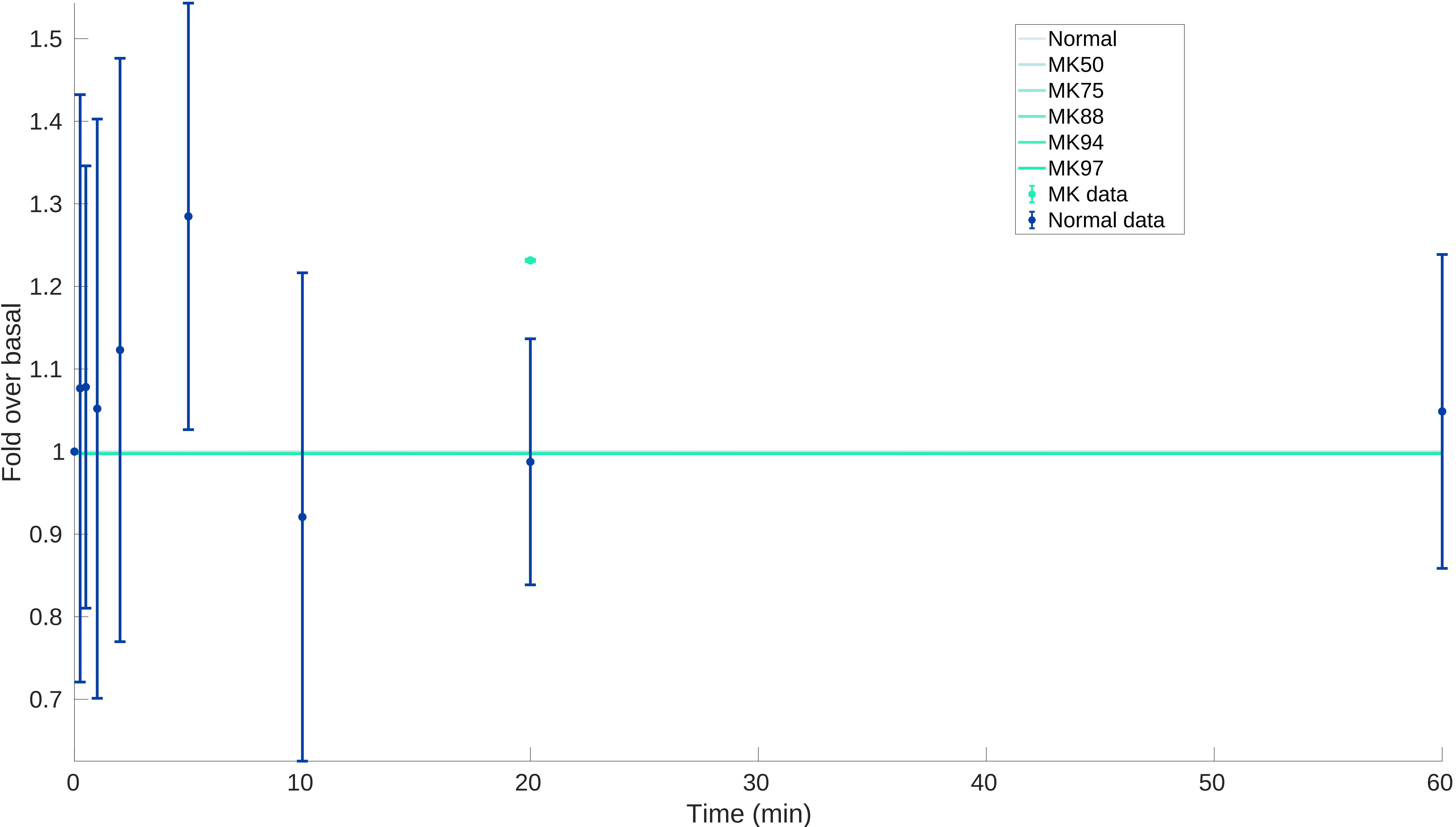

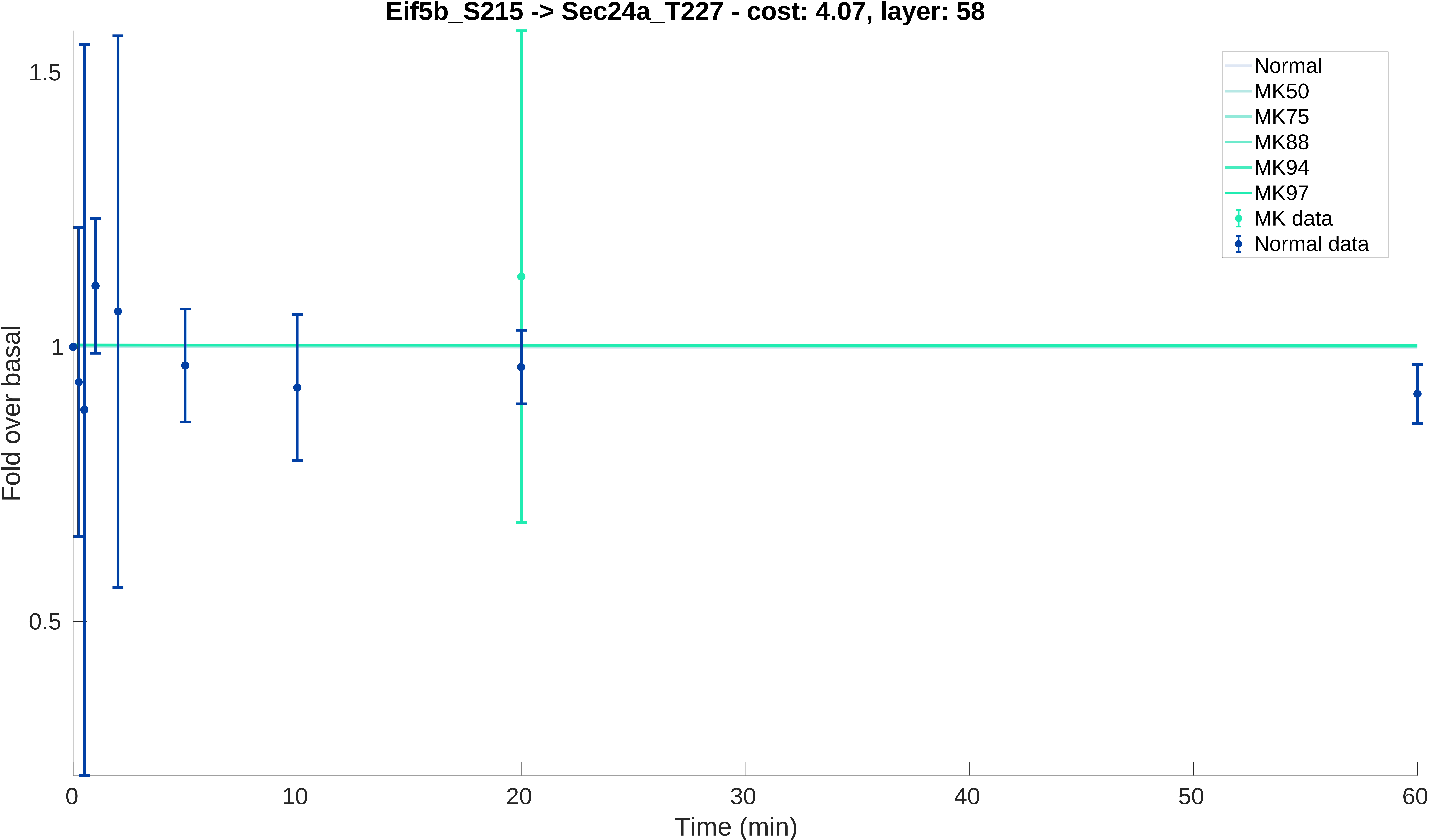

Eif5b_S215 -> Sec24a_T227 - cost: 4.07, layer: 58
Normal
MK50
MK75
MK88
MK94
MK97
MK data
Normal data
Fold over basal
1.5
1
0.5
0
10
20
30
40
50
60
Time (min)

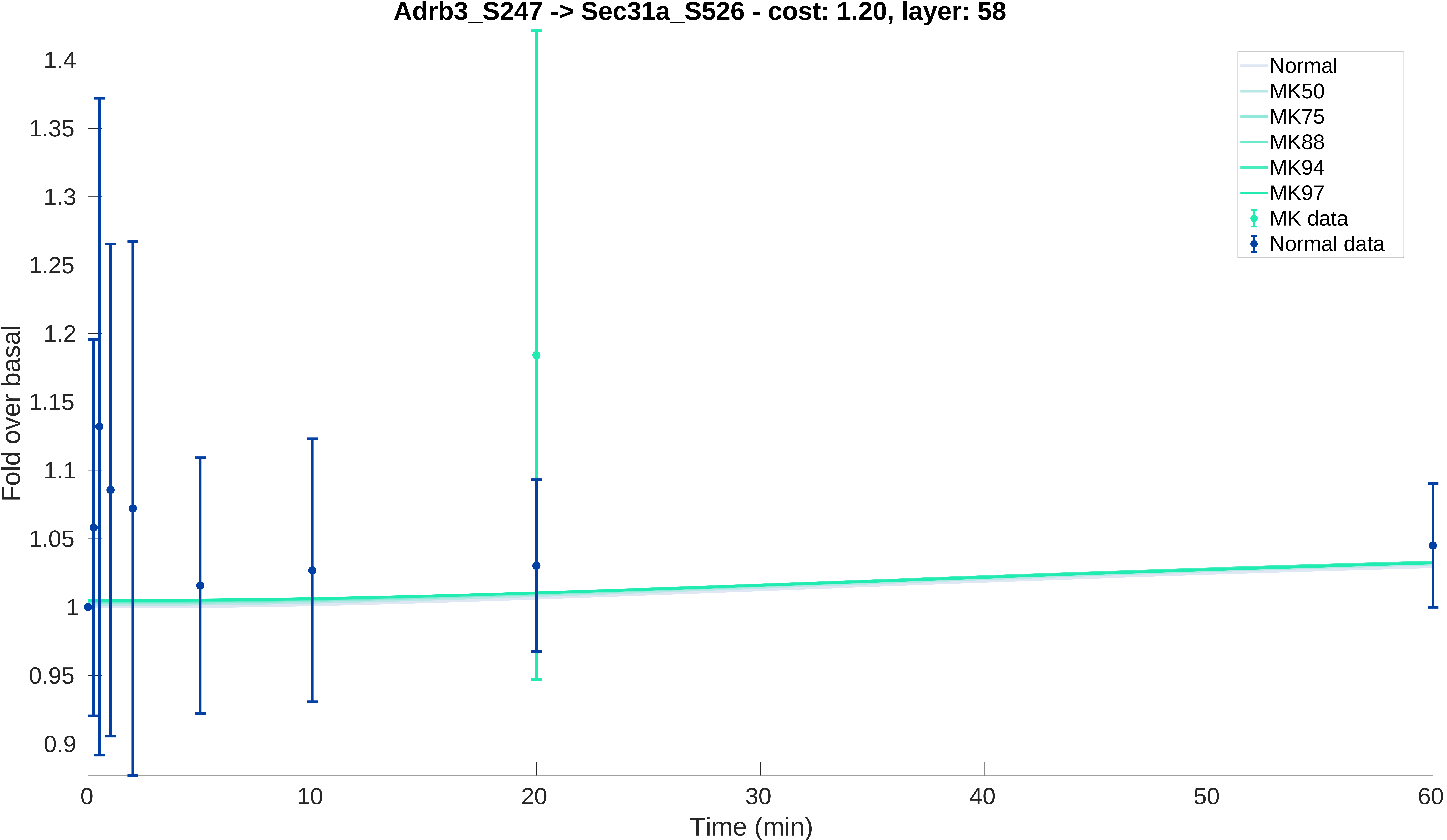

Adrb3_S247 -> Sec31a_S526 - cost: 1.20, layer: 58
Fold over basal
1.4
1.35
1.3
1.25
1.2
1.15
1.1
1.05
1
0.95
0.9
0
10
20
30
40
50
60
Time (min)
Normal
MK50
MK75
MK88
MK94
MK97
MK data
Normal data

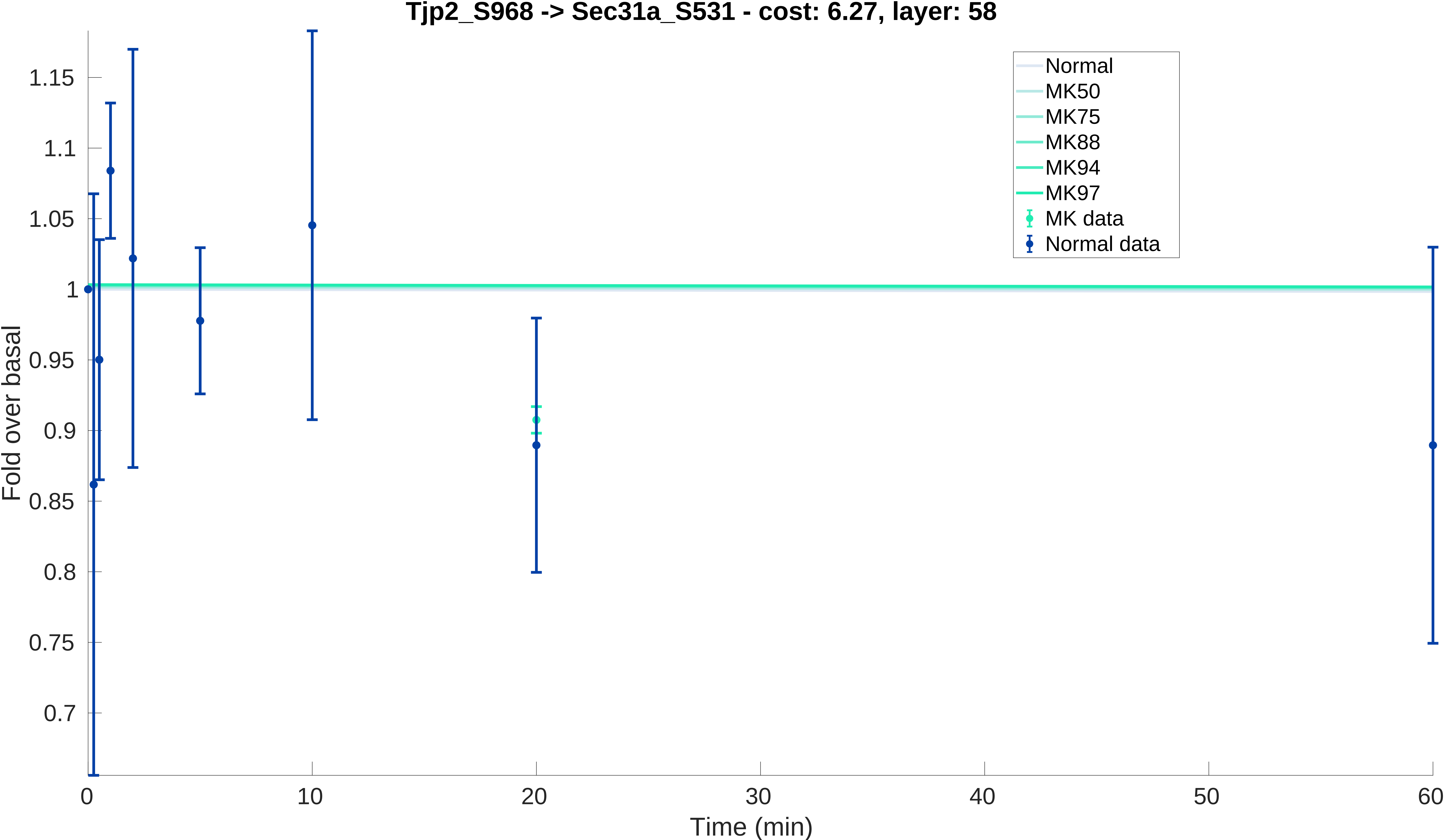

Tjp2_S968 -> Sec31a_S531 - cost: 6.27, layer: 58
Normal
MK50
MK75
MK88
MK94
MK97
MK data
Normal data
Fold over basal
1.15
1.1
1.05
1
0.95
0.9
0.85
0.8
0.75
0.7
0
10
20
30
40
50
60
Time (min)

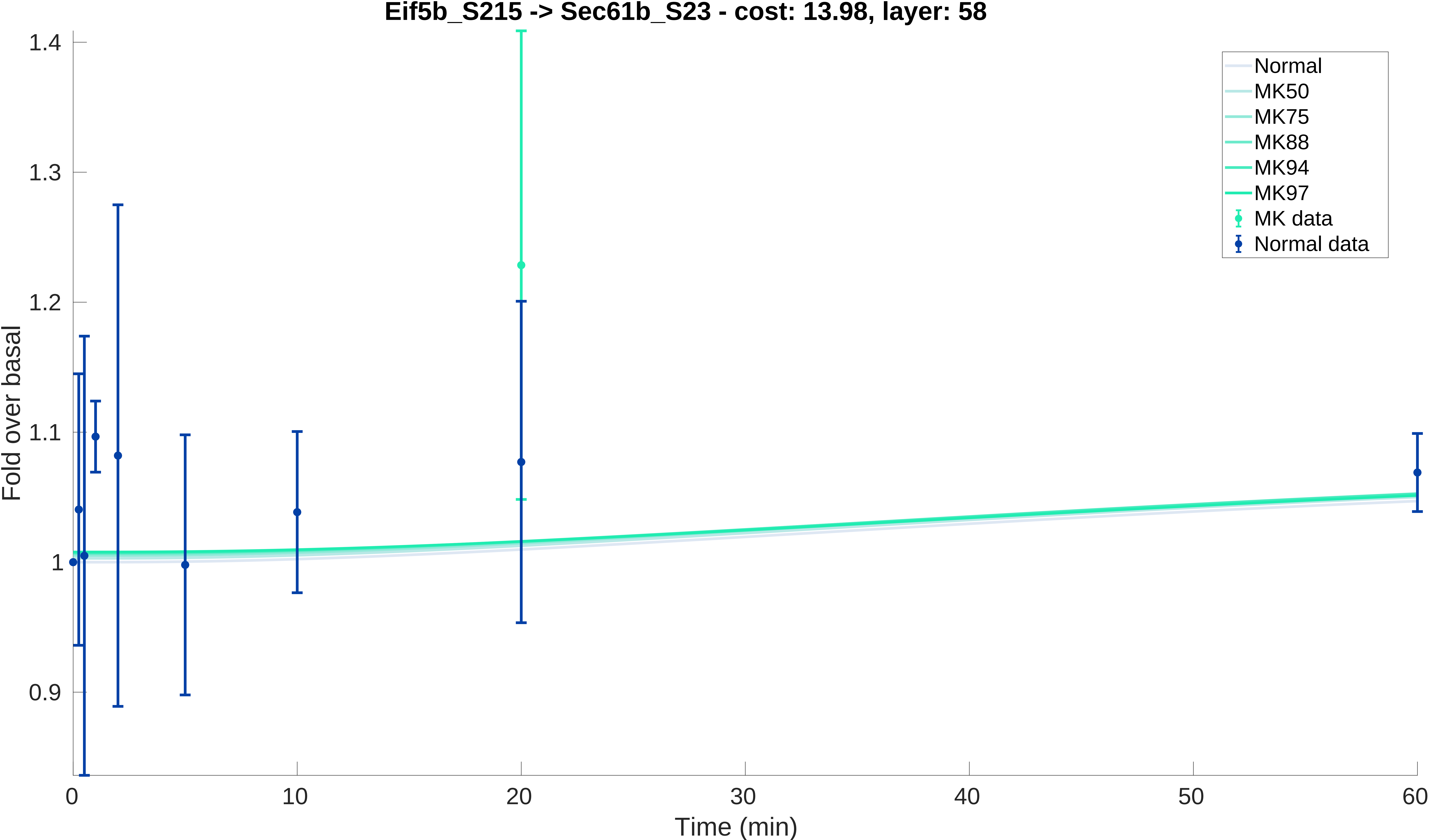

Eif5b_S215 -> Sec61b_S23 - cost: 13.98, layer: 58
Fold over basal
1.4
1.3
1.2
1.1
1
0.9
0
10
20
30
40
50
60
Time (min)
Normal
MK50
MK75
MK88
MK94
MK97
MK data
Normal data

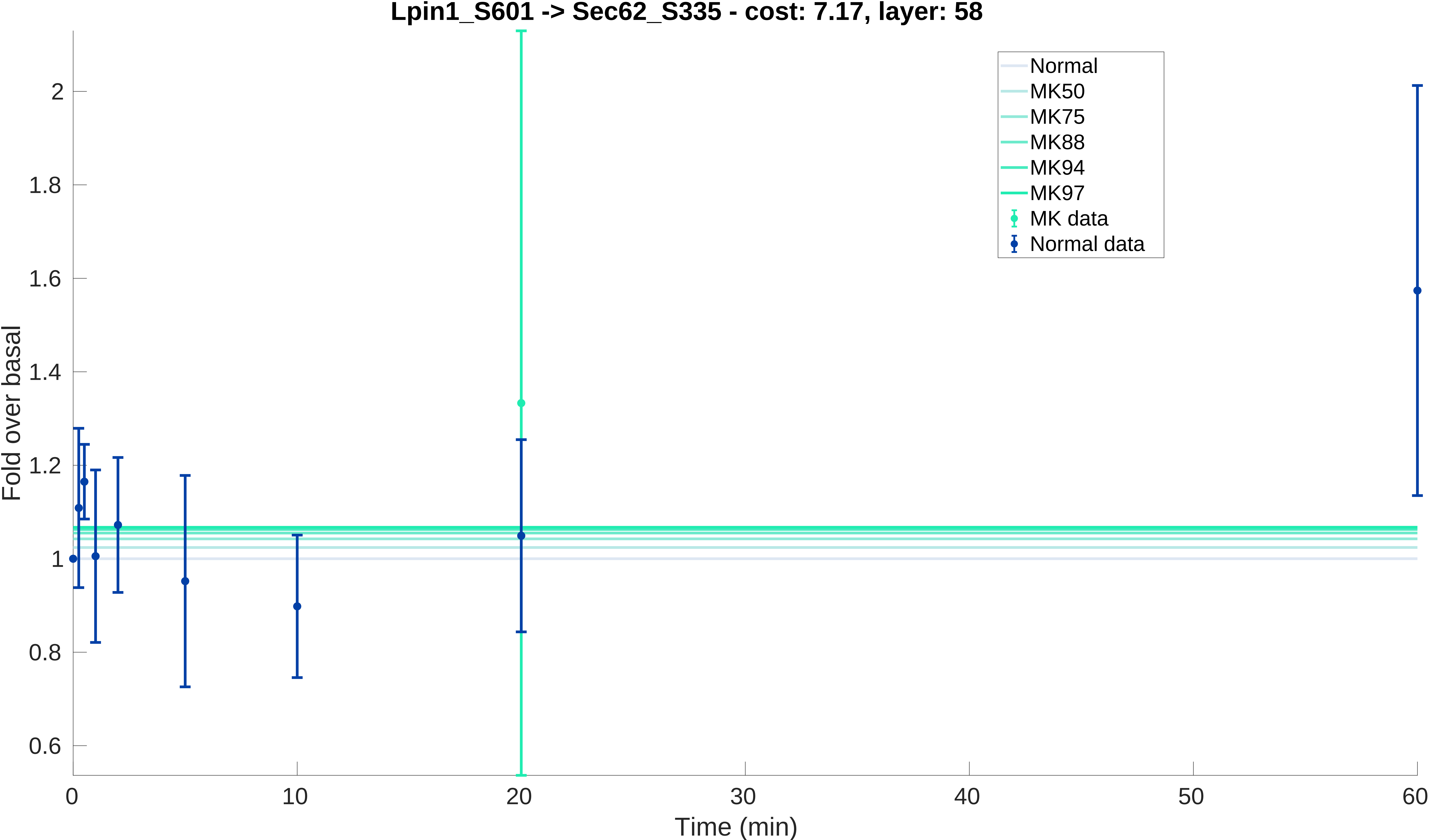

Lpin1_S601 -> Sec62_S335 - cost: 7.17, layer: 58
Normal
MK50
MK75
MK88
MK94
MK97
MK data
Normal data
Fold over basal
0.6
0.8
1
1.2
1.4
1.6
1.8
2
0
10
20
30
40
50
60
Time (min)

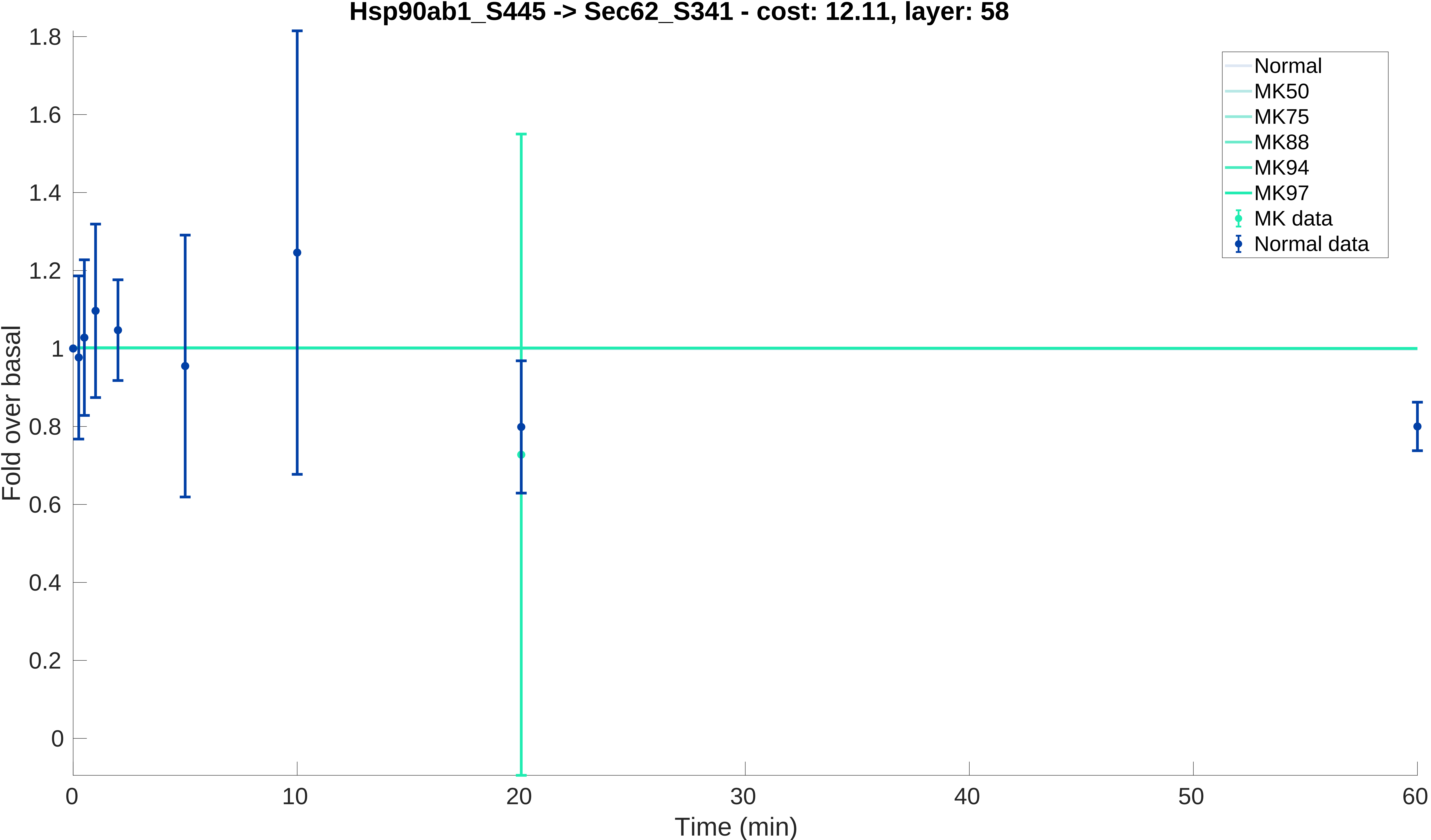

Hsp90ab1_S445 -> Sec62_S341 - cost: 12.11, layer: 58
Fold over basal
Time (min)
0
0.2
0.4
0.6
0.8
1
1.2
1.4
1.6
1.8
0
10
20
30
40
50
60
Normal
MK50
MK75
MK88
MK94
MK97
MK data
Normal data

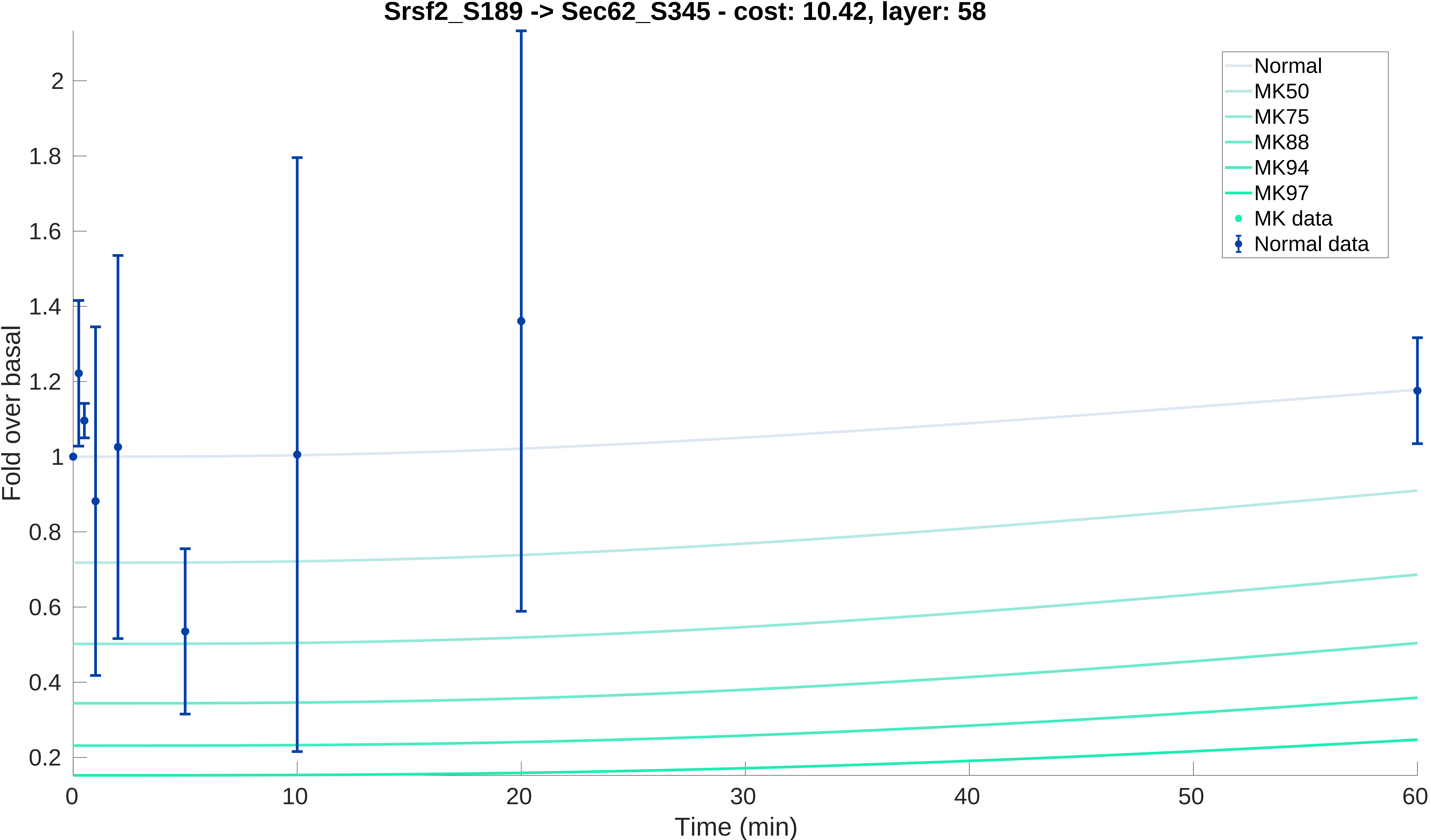

Srsf2_S189 -> Sec62_S345 - cost: 10.42, layer: 58
Fold over basal
Time (min)
0
10
20
30
40
50
60
0.2
0.4
0.6
0.8
1
1.2
1.4
1.6
1.8
2
Normal
MK50
MK75
MK88
MK94
MK97
MK data
Normal data

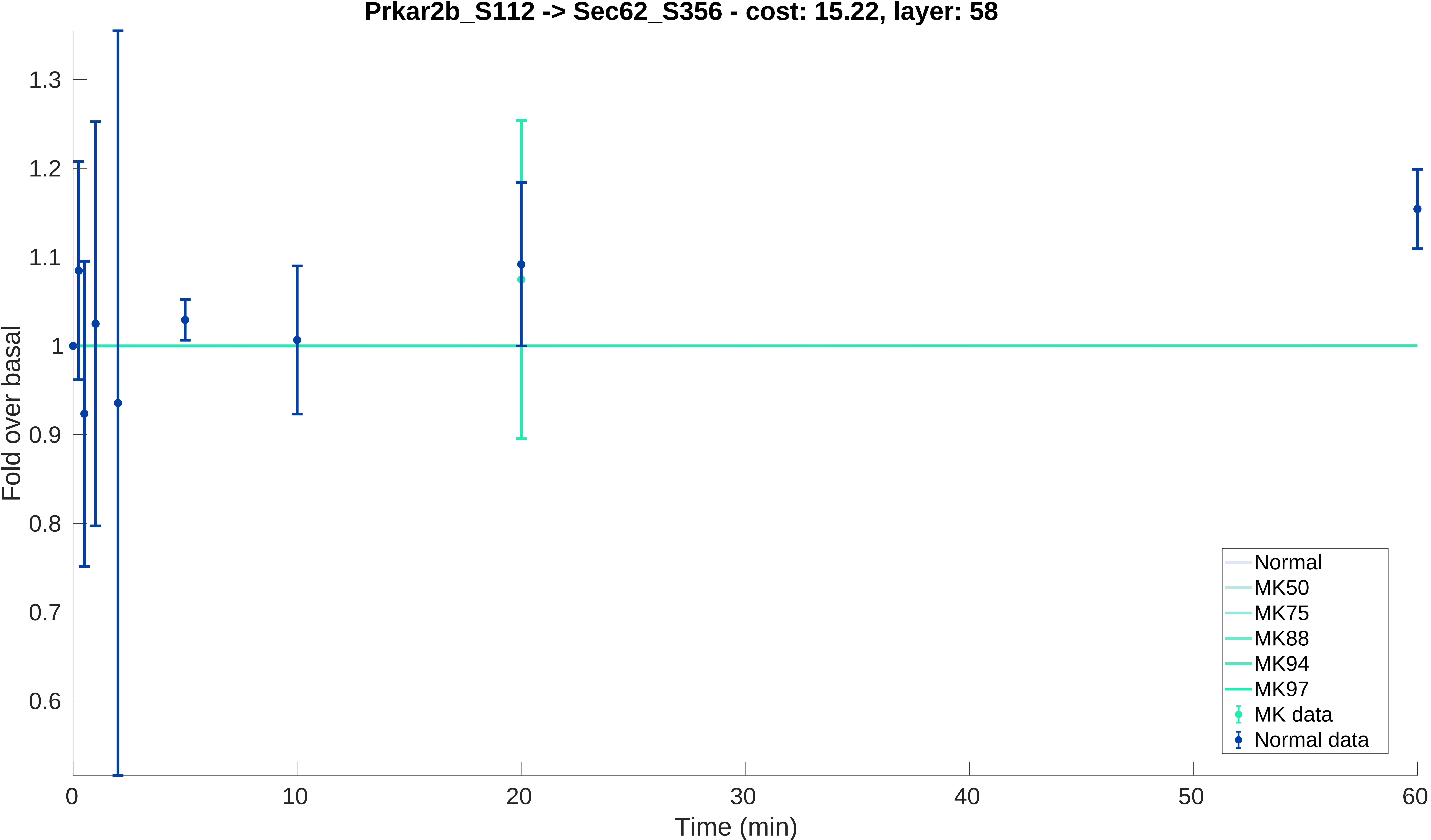
Prkar2b_S112 -> Sec62_S356 - cost: 15.22, layer: 58
Fold over basal
Time (min)
1.3
1.2
1.1
1
0.9
0.8
0.7
0.6
0
10
20
30
40
50
60
Normal
MK50
MK75
MK88
MK94
MK97
MK data
Normal data

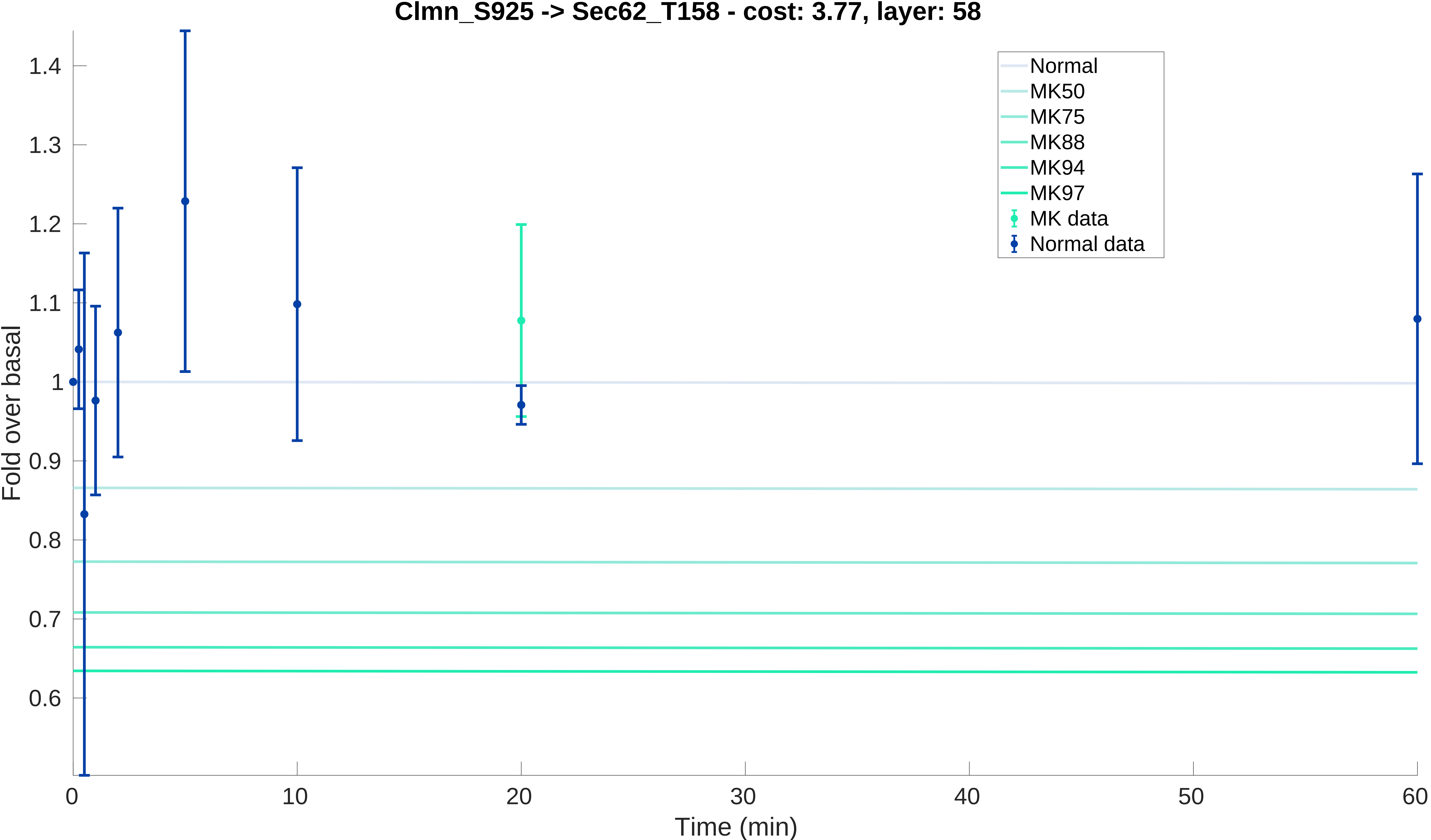

Clmn_S925 -> Sec62_T158 - cost: 3.77, layer: 58
Fold over basal
Time (min)
1.4
1.3
1.2
1.1
1
0.9
0.8
0.7
0.6
0
10
20
30
40
50
60
Normal
MK50
MK75
MK88
MK94
MK97
MK data
Normal data

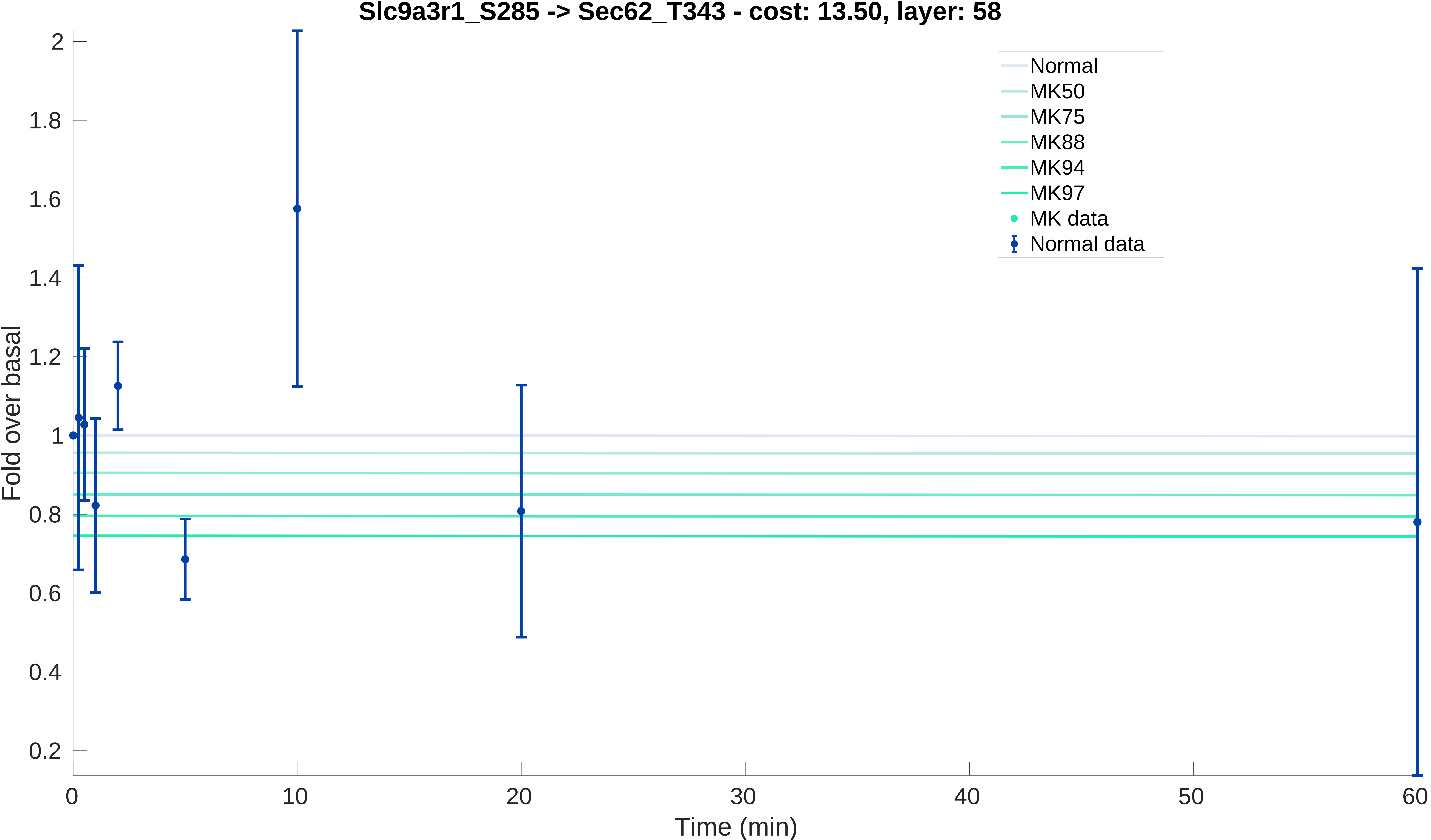

Slc9a3r1_S285 -> Sec62_T343 - cost: 13.50, layer: 58
Fold over basal
Time (min)
2
1.8
1.6
1.4
1.2
1
0.8
0.6
0.4
0.2
0
10
20
30
40
50
60
Normal
MK50
MK75
MK88
MK94
MK97
MK data
Normal data

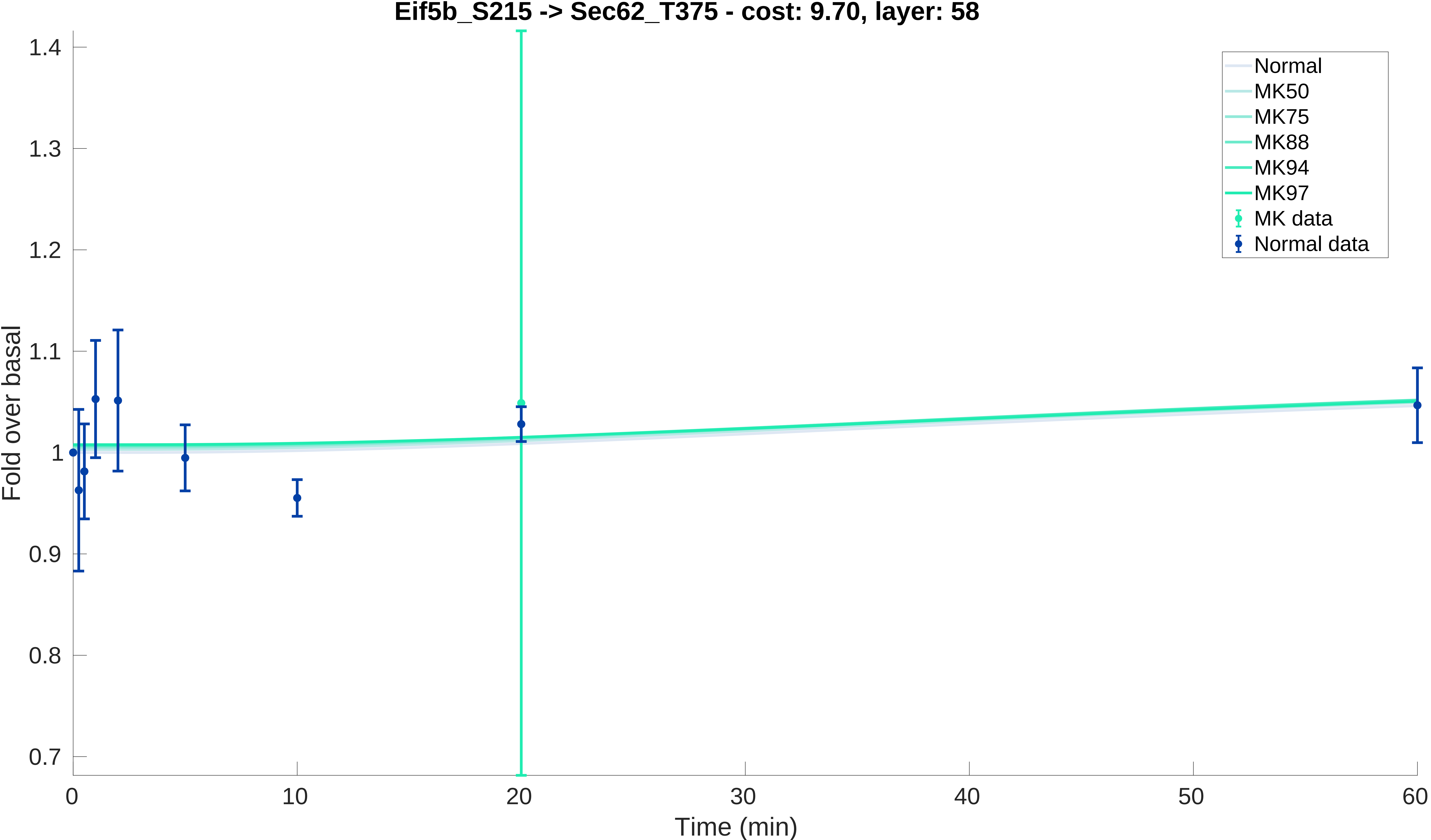

Eif5b_S215 -> Sec62_T375 - cost: 9.70, layer: 58
Fold over basal
1.4
1.3
1.2
1.1
1
0.9
0.8
0.7
0
10
20
30
40
50
60
Time (min)
Normal
MK50
MK75
MK88
MK94
MK97
MK data
Normal data

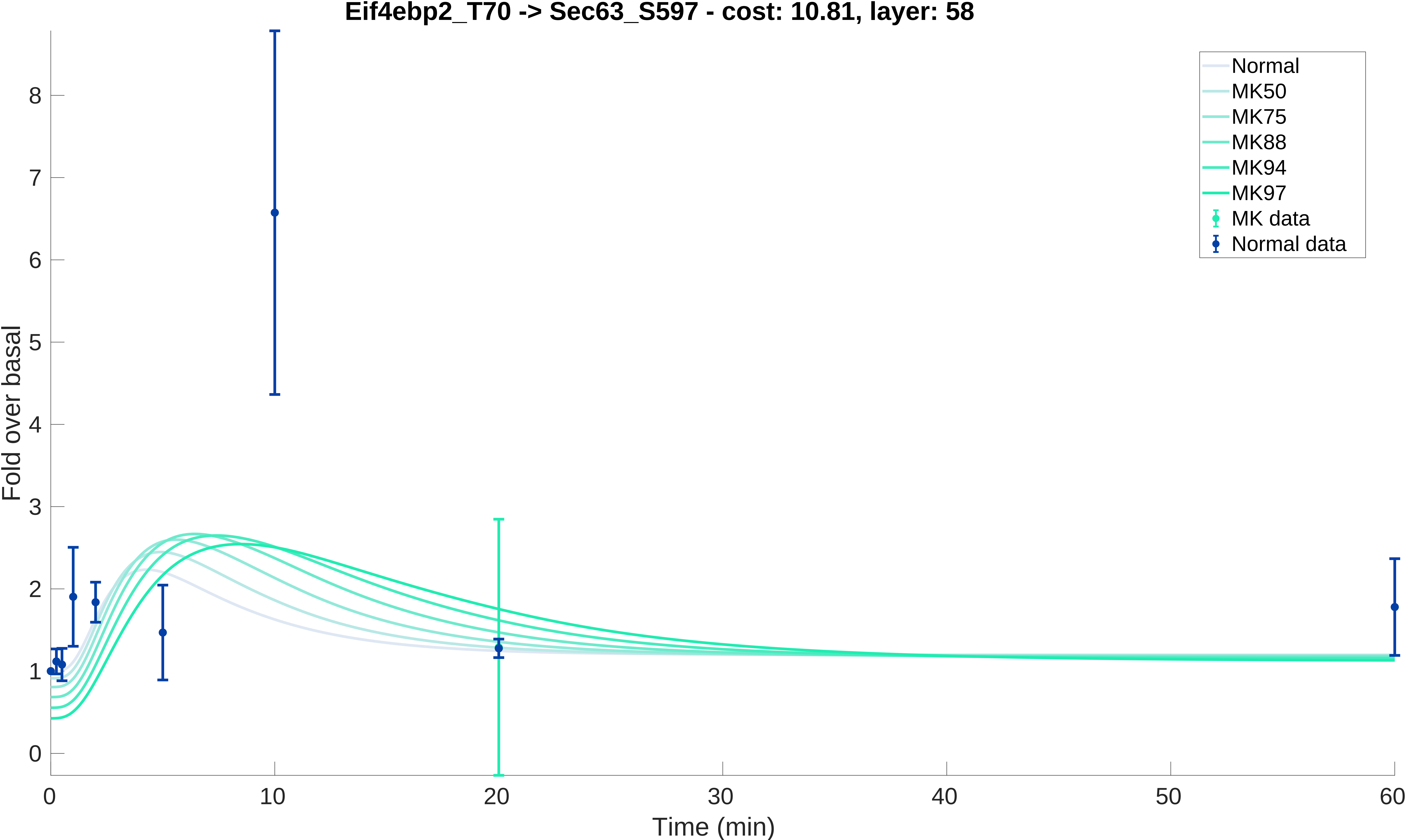
Eif4ebp2_T70 -> Sec63_S597 - cost: 10.81, layer: 58
Fold over basal
Time (min)
0
1
2
3
4
5
6
7
8
0
10
20
30
40
50
60
Normal
MK50
MK75
MK88
MK94
MK97
MK data
Normal data

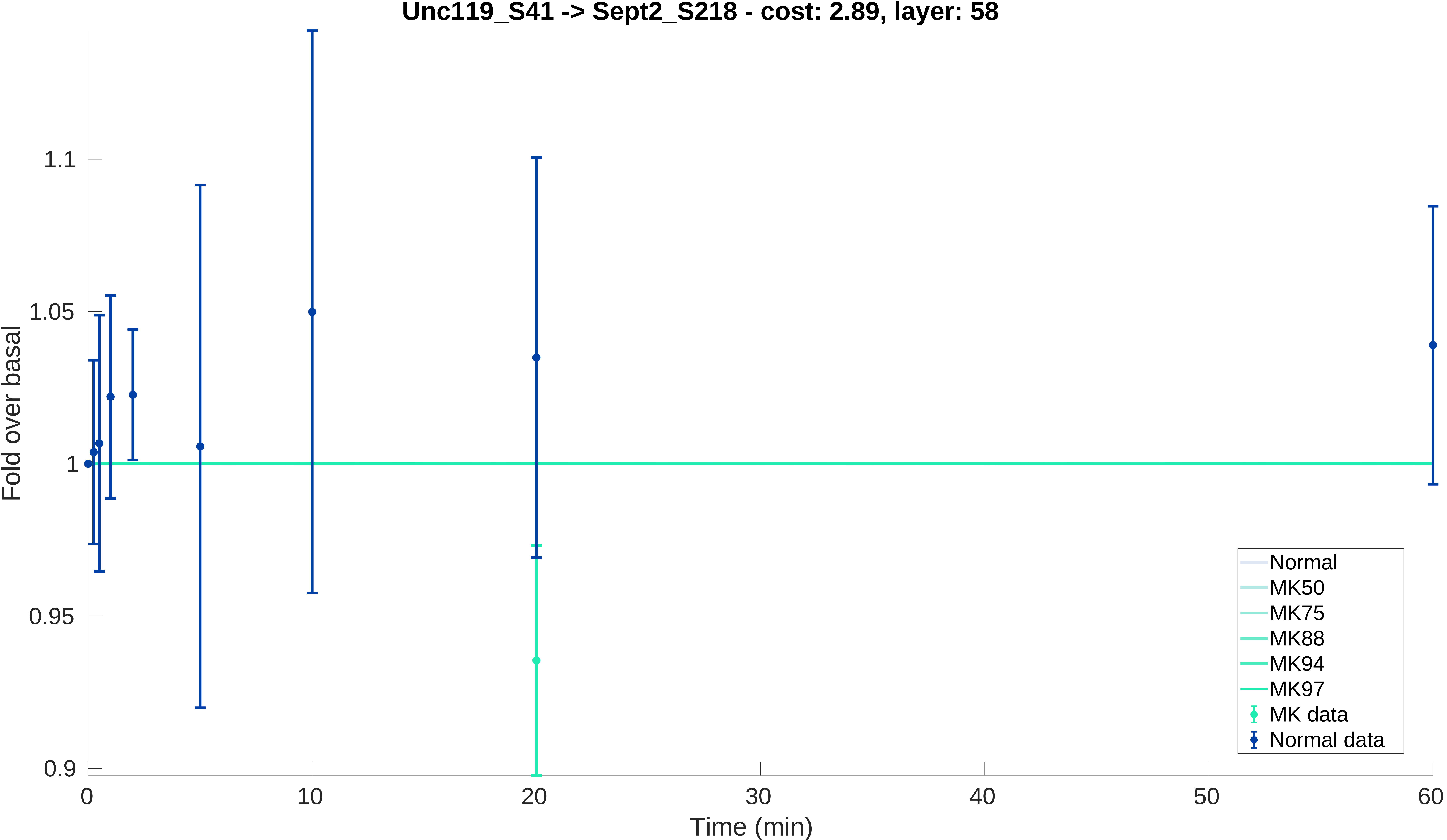
Unc119_S41 -> Sept2_S218 - cost: 2.89, layer: 58
Fold over basal
1.1
1.05
1
0.95
0.9
0
10
20
30
40
50
60
Time (min)
Normal
MK50
MK75
MK88
MK94
MK97
MK data
Normal data

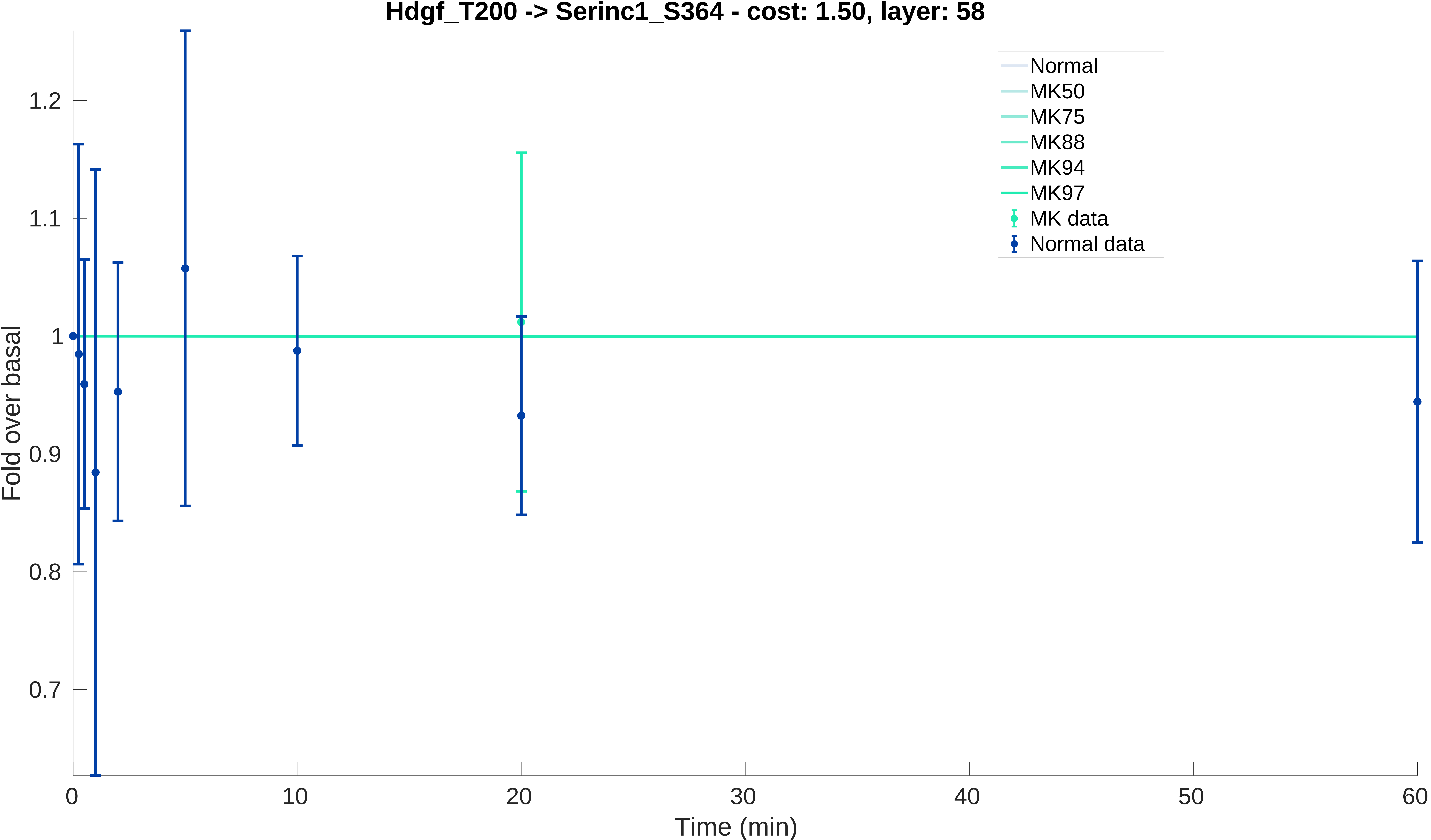

Hdgf_T200 -> Serinc1_S364 - cost: 1.50, layer: 58
Normal
MK50
MK75
MK88
MK94
MK97
MK data
Normal data
Fold over basal
1.2
1.1
1
0.9
0.8
0.7
0
10
20
30
40
50
60
Time (min)

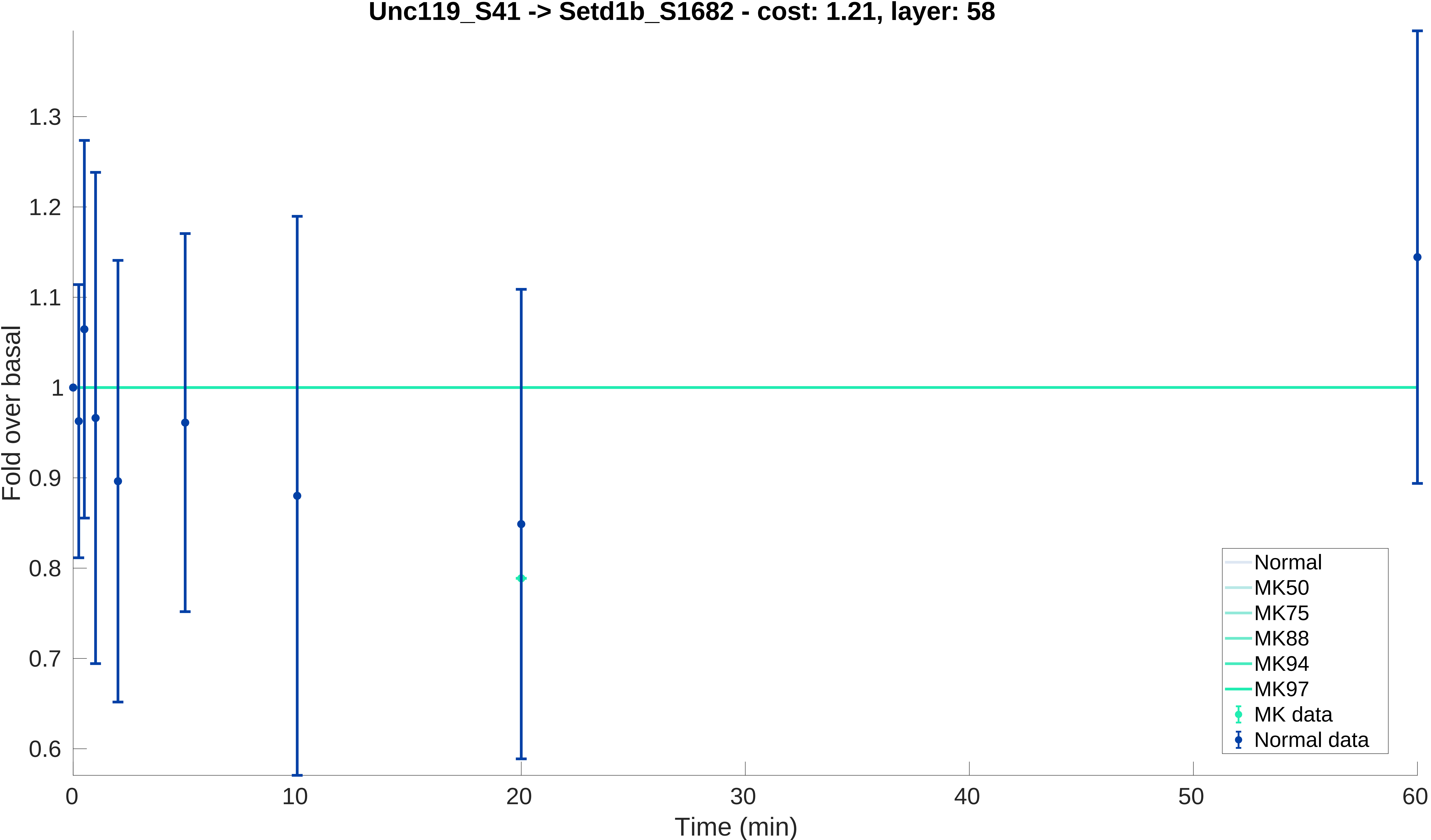

Unc119_S41 -> Setd1b_S1682 - cost: 1.21, layer: 58
Fold over basal
1.3
1.2
1.1
1
0.9
0.8
0.7
0.6
0
10
20
30
40
50
60
Time (min)
Normal
MK50
MK75
MK88
MK94
MK97
MK data
Normal data

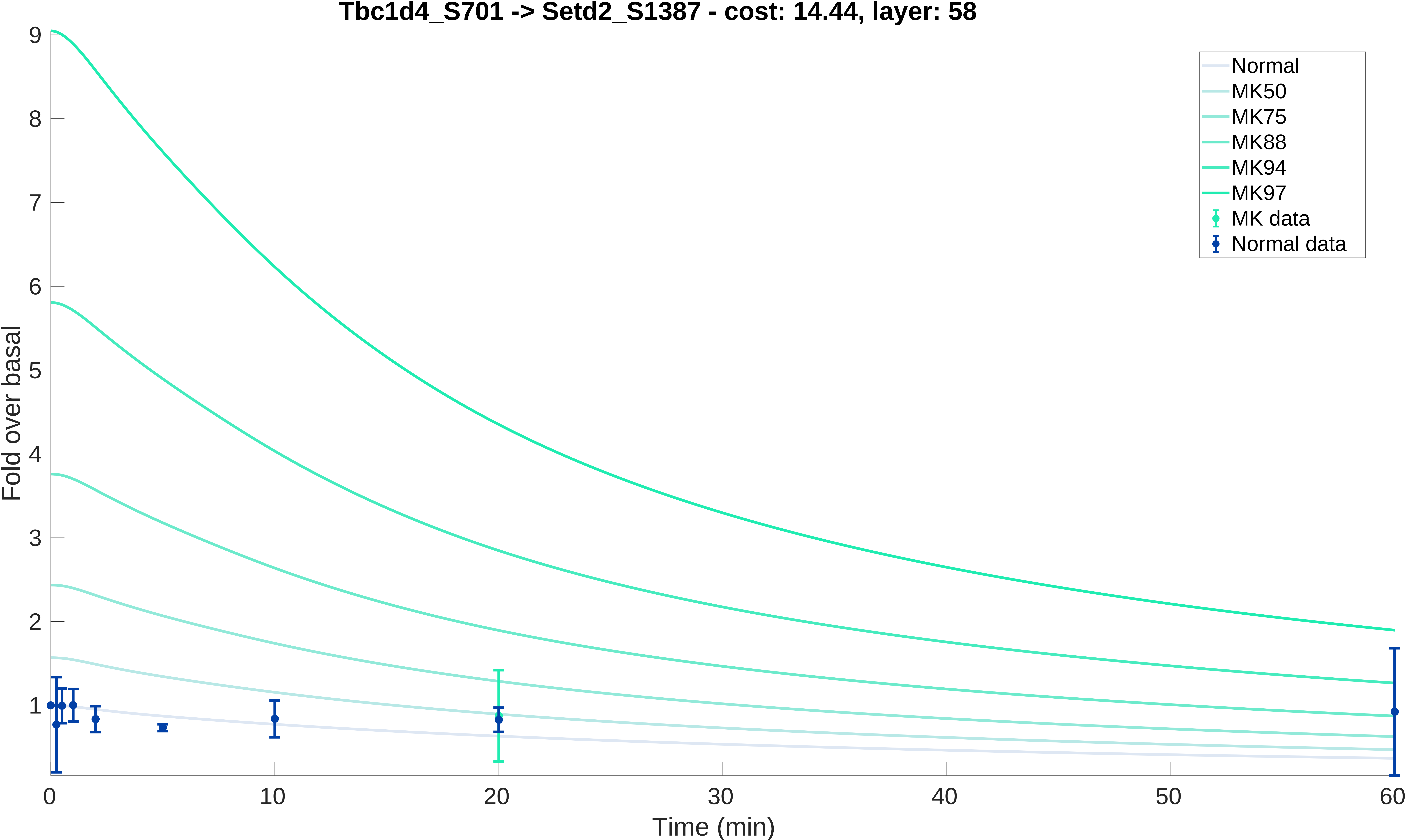
Tbc1d4_S701 -> Setd2_S1387 - cost: 14.44, layer: 58
Fold over basal
Time (min)
0
10
20
30
40
50
60
1
2
3
4
5
6
7
8
9
Normal
MK50
MK75
MK88
MK94
MK97
MK data
Normal data

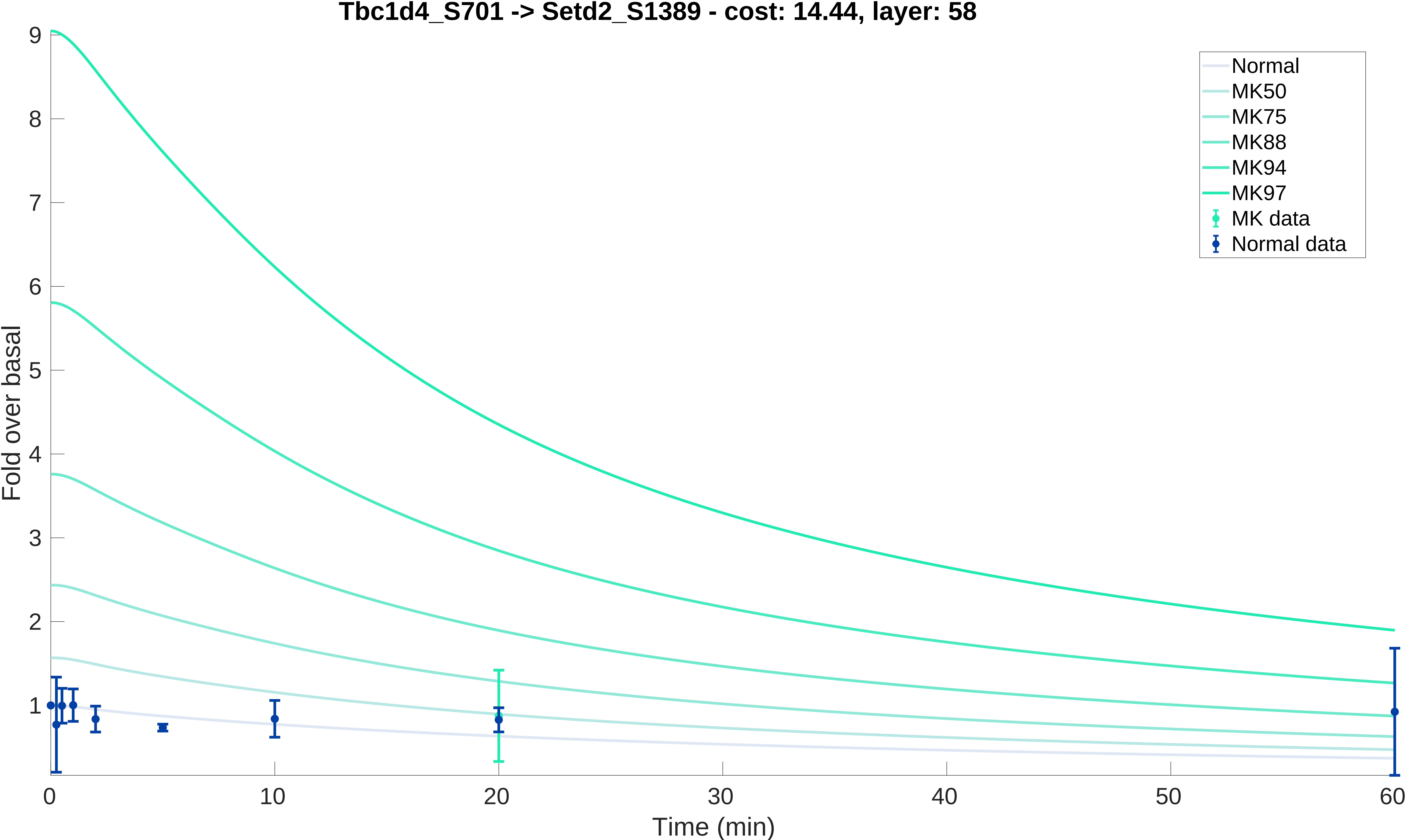

Tbc1d4_S701 -> Setd2_S1389 - cost: 14.44, layer: 58
Fold over basal
Time (min)
0
10
20
30
40
50
60
1
2
3
4
5
6
7
8
9
Normal
MK50
MK75
MK88
MK94
MK97
MK data
Normal data

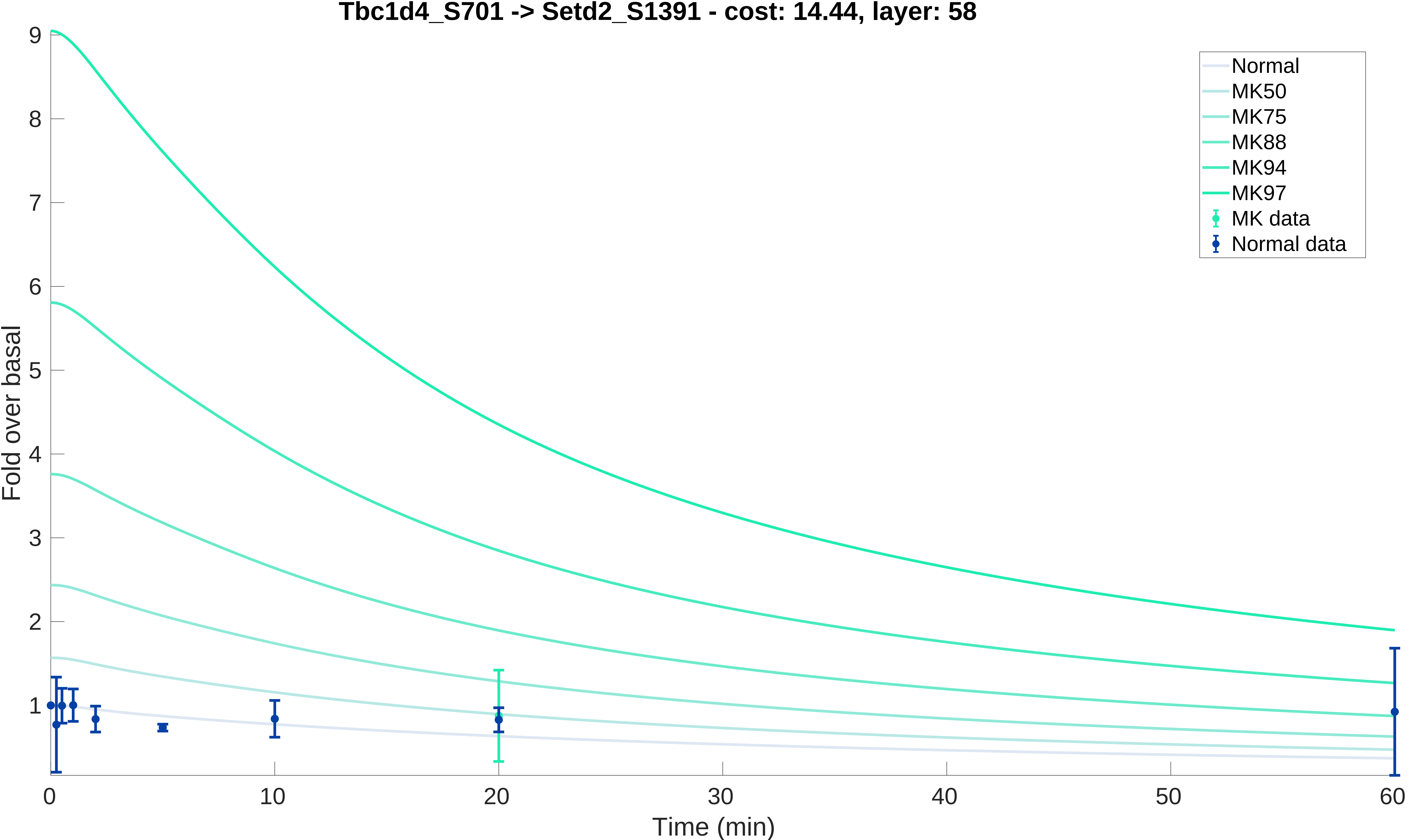

Tbc1d4_S701 -> Setd2_S1391 - cost: 14.44, layer: 58
Fold over basal
Time (min)
0
10
20
30
40
50
60
1
2
3
4
5
6
7
8
9
Normal
MK50
MK75
MK88
MK94
MK97
MK data
Normal data

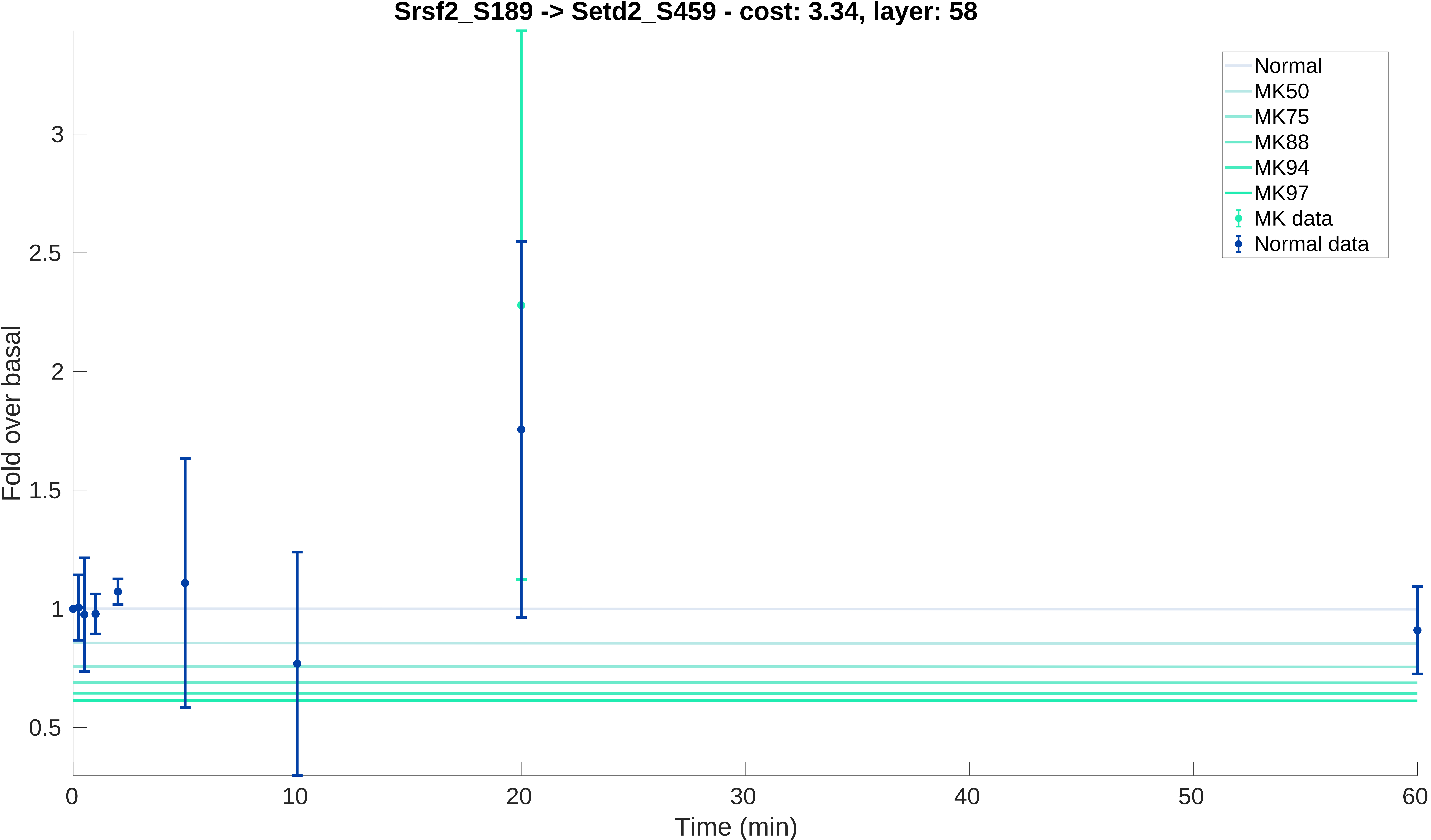

Srsf2_S189 -> Setd2_S459 - cost: 3.34, layer: 58
Fold over basal
Time (min)
0
10
20
30
40
50
60
0.5
1
1.5
2
2.5
3
Normal
MK50
MK75
MK88
MK94
MK97
MK data
Normal data

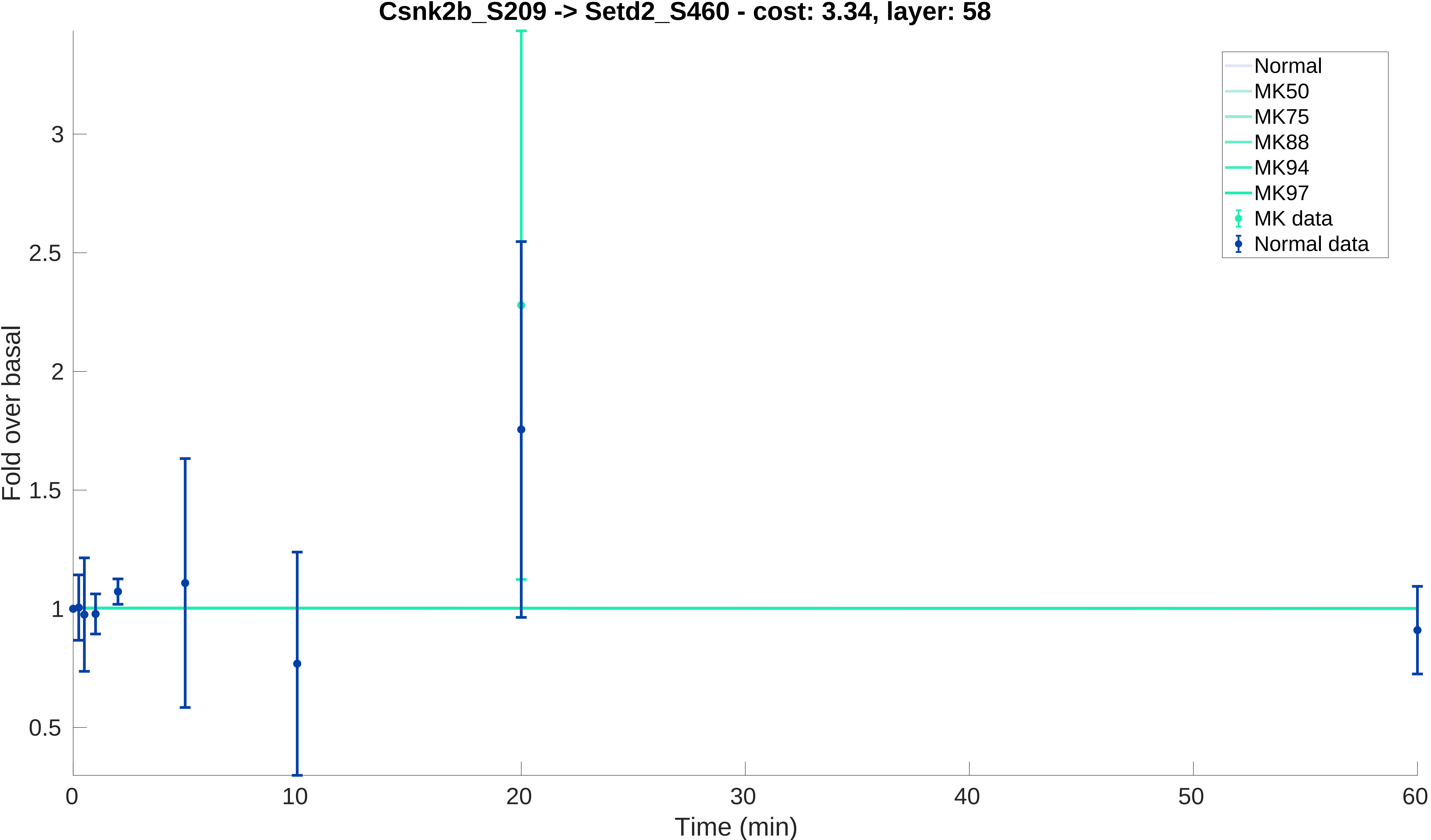

Csnk2b_S209 -> Setd2_S460 - cost: 3.34, layer: 58
Fold over basal
Time (min)
0.5
1
1.5
2
2.5
3
0
10
20
30
40
50
60
Normal
MK50
MK75
MK88
MK94
MK97
MK data
Normal data

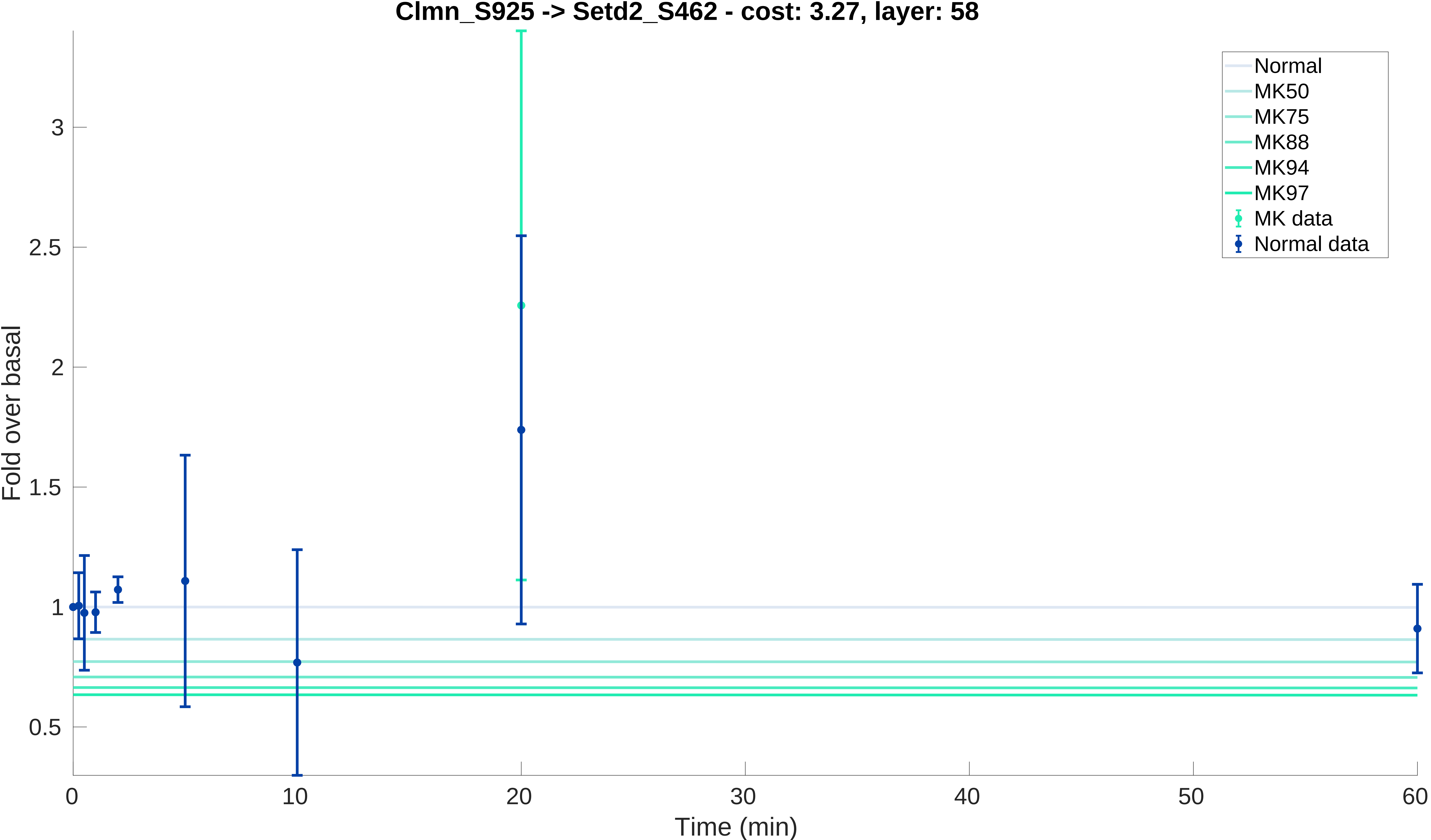

Clmn_S925 -> Setd2_S462 - cost: 3.27, layer: 58
Fold over basal
0.5
1
1.5
2
2.5
3
0
10
20
30
40
50
60
Time (min)
Normal
MK50
MK75
MK88
MK94
MK97
MK data
Normal data

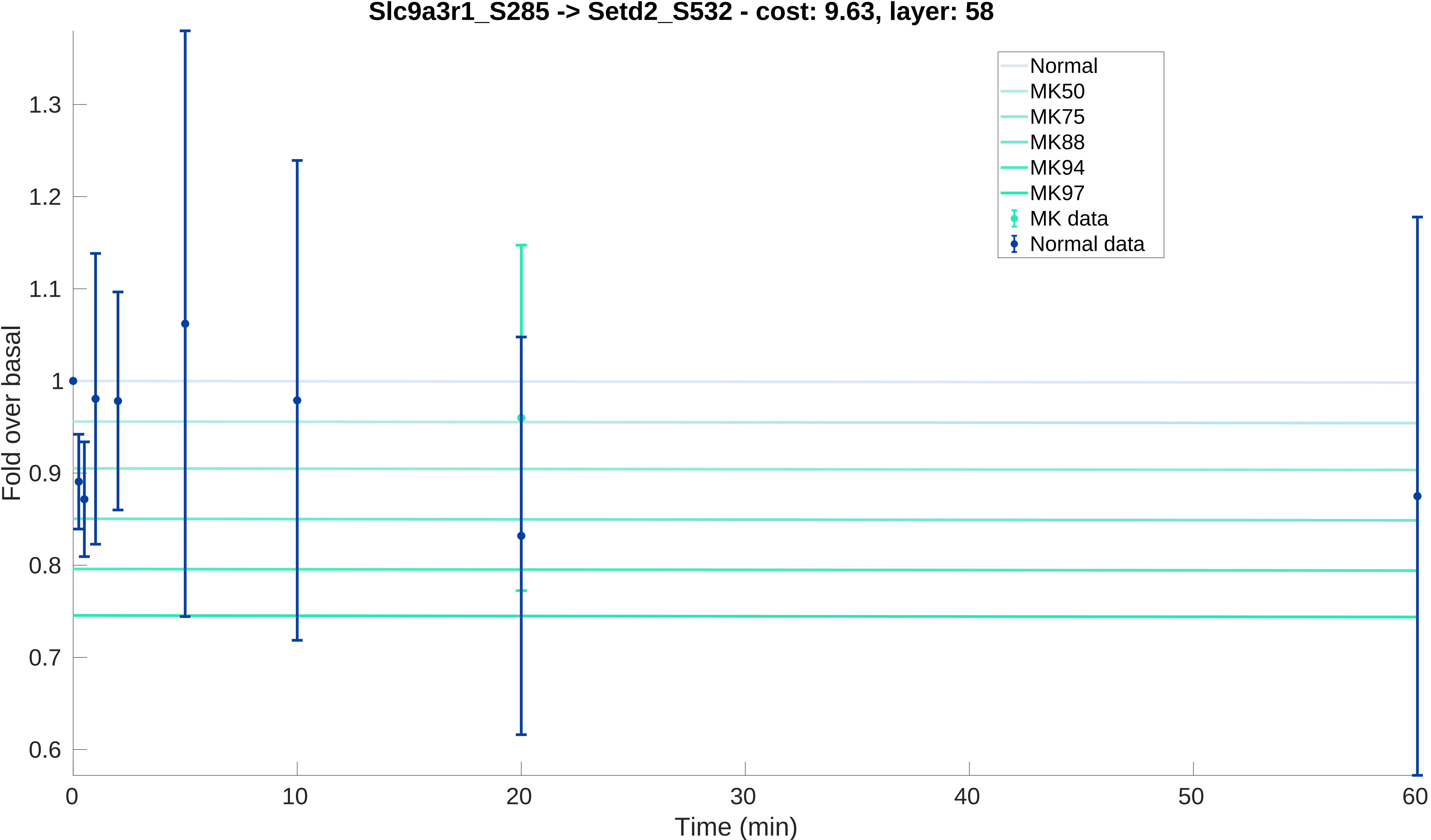

Slc9a3r1_S285 -> Setd2_S532 - cost: 9.63, layer: 58
Normal
MK50
MK75
MK88
MK94
MK97
MK data
Normal data
Fold over basal
Time (min)
0.6
0.7
0.8
0.9
1
1.1
1.2
1.3
0
10
20
30
40
50
60

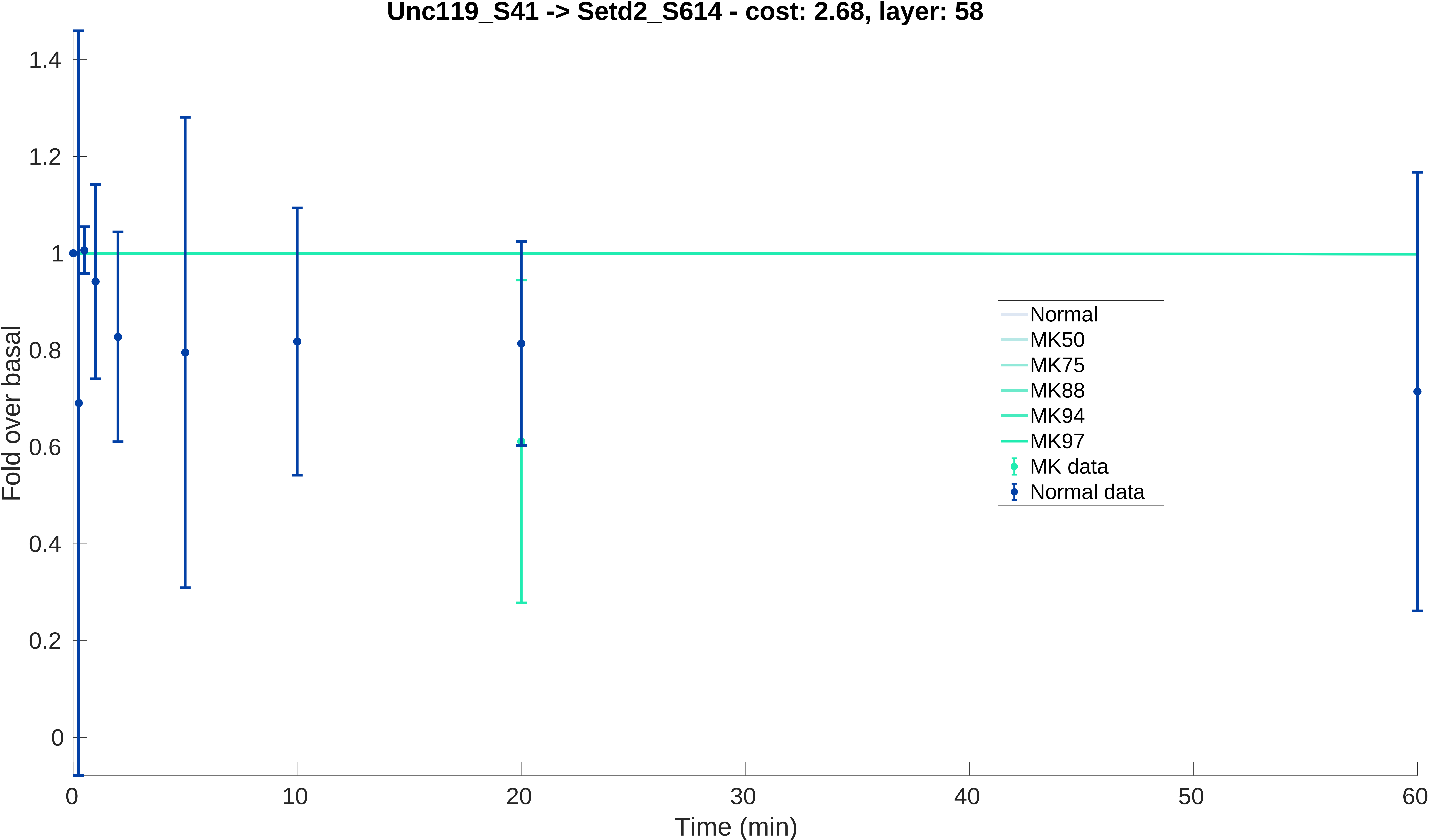
Unc119_S41 -> Setd2_S614 - cost: 2.68, layer: 58
Fold over basal
1.4
1.2
1
0.8
0.6
0.4
0.2
0
0
10
20
30
40
50
60
Time (min)
Normal
MK50
MK75
MK88
MK94
MK97
MK data
Normal data

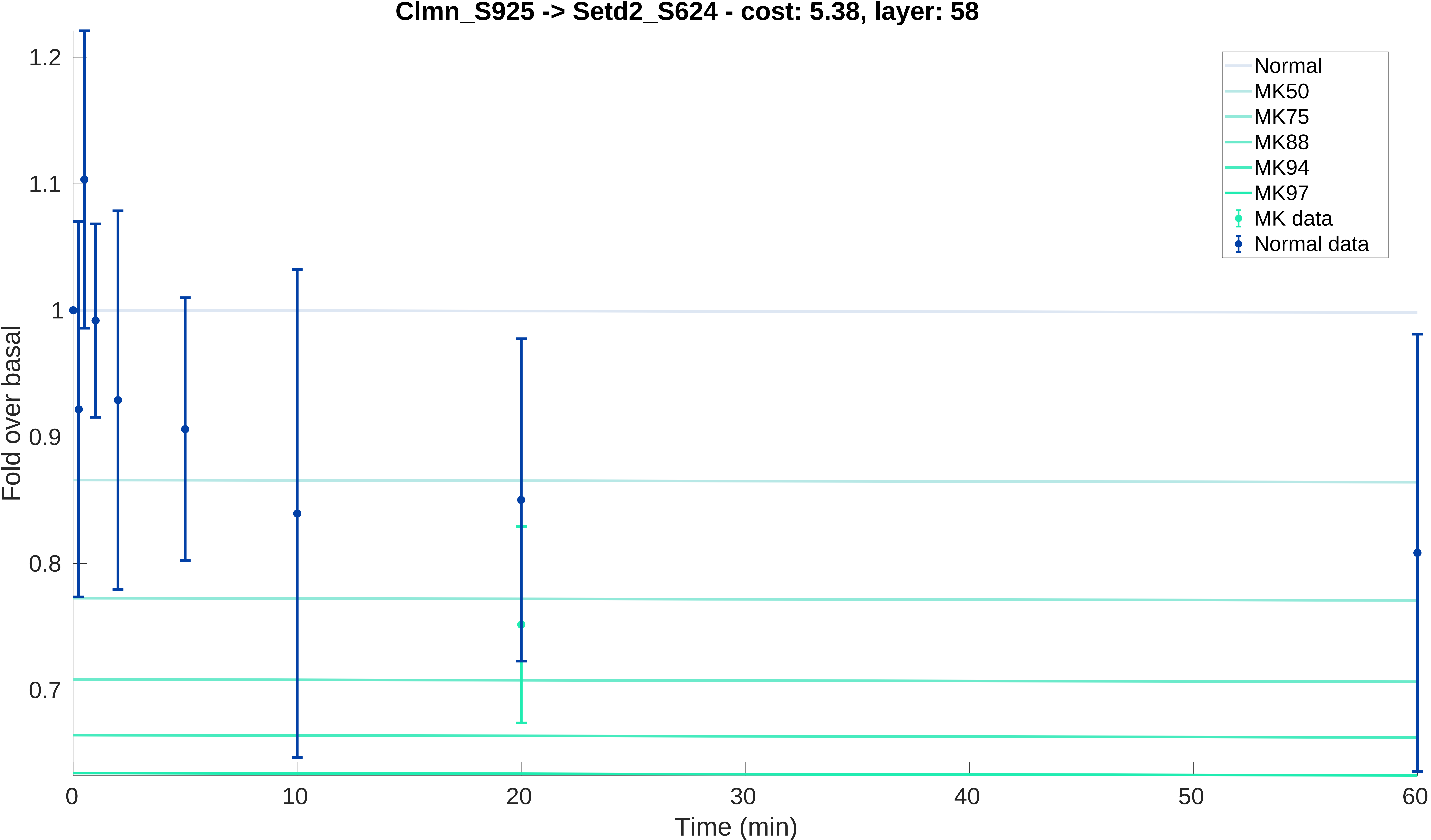

Clmn_S925 -> Setd2_S624 - cost: 5.38, layer: 58
Fold over basal
Time (min)
1.2
1.1
1
0.9
0.8
0.7
0
10
20
30
40
50
60
Normal
MK50
MK75
MK88
MK94
MK97
MK data
Normal data

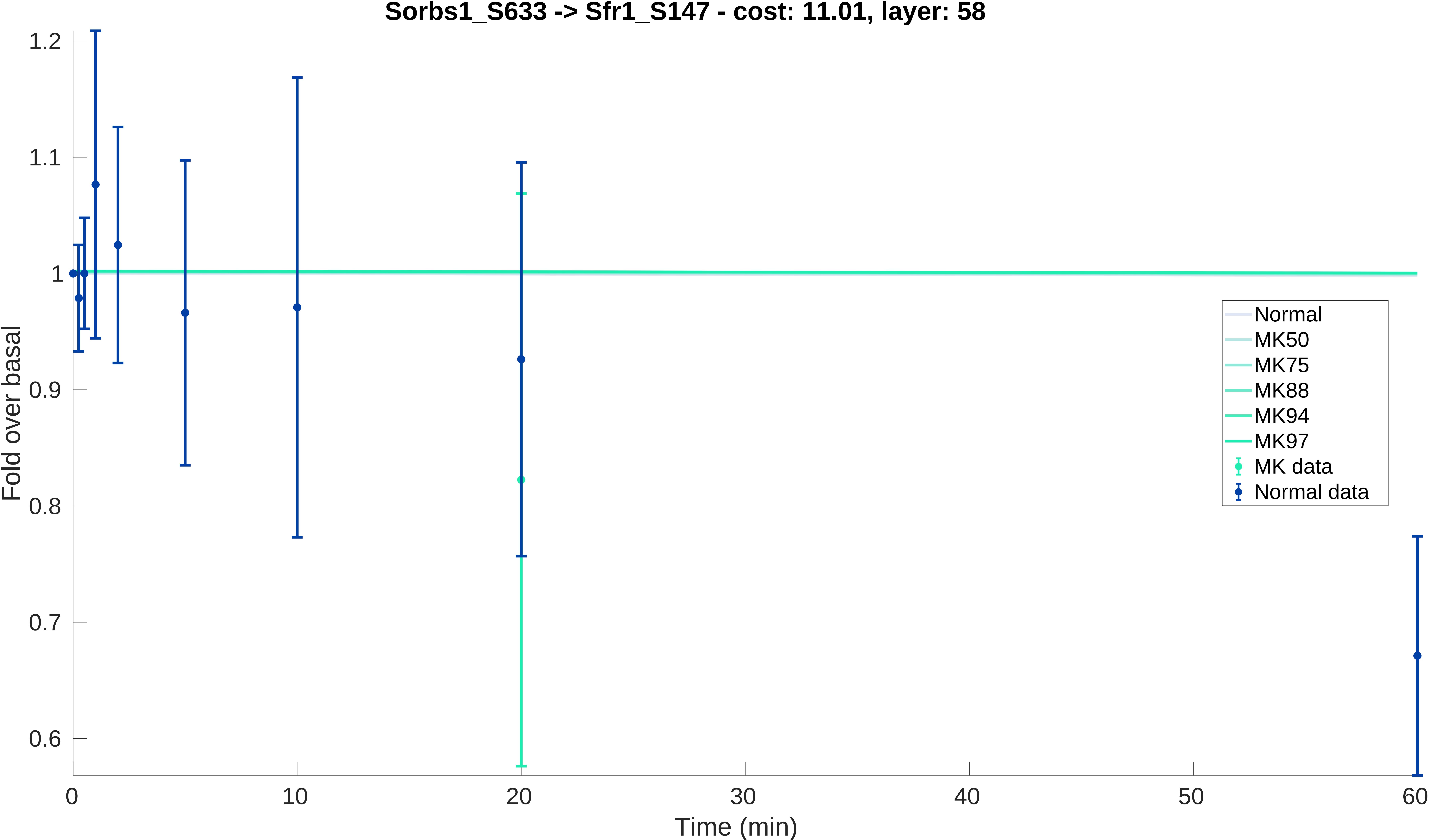

Sorbs1_S633 -> Sfr1_S147 - cost: 11.01, layer: 58
Fold over basal
Time (min)
1.2
1.1
1
0.9
0.8
0.7
0.6
0
10
20
30
40
50
60
Normal
MK50
MK75
MK88
MK94
MK97
MK data
Normal data

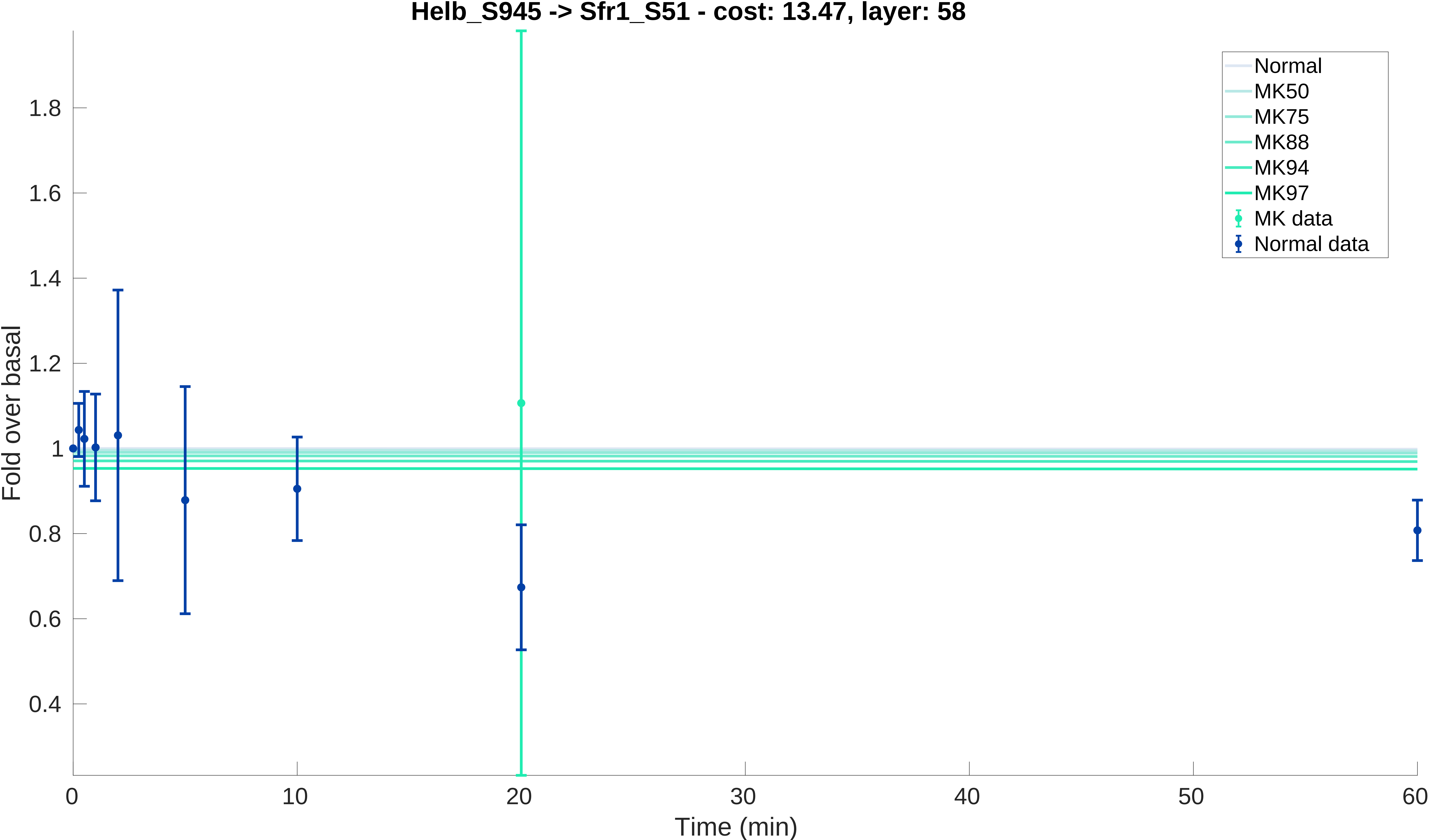
Helb_S945 -> Sfr1_S51 - cost: 13.47, layer: 58
Normal
MK50
MK75
MK88
MK94
MK97
MK data
Normal data
Fold over basal
1.8
1.6
1.4
1.2
1
0.8
0.6
0.4
0
10
20
30
40
50
60
Time (min)

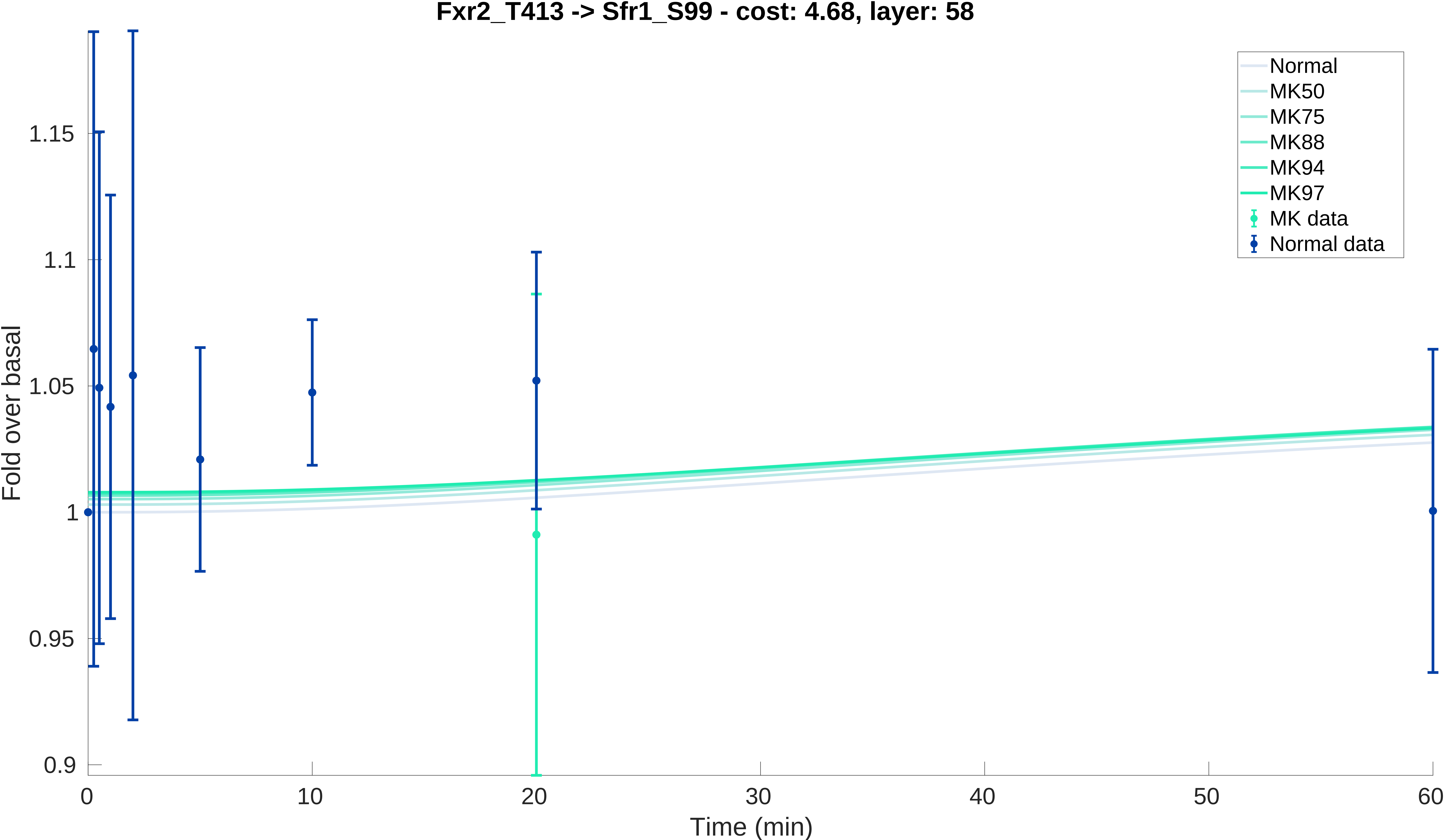

Fxr2_T413 -> Sfr1_S99 - cost: 4.68, layer: 58
Fold over basal
Time (min)
1.15
1.1
1.05
1
0.95
0.9
0
10
20
30
40
50
60
Normal
MK50
MK75
MK88
MK94
MK97
MK data
Normal data

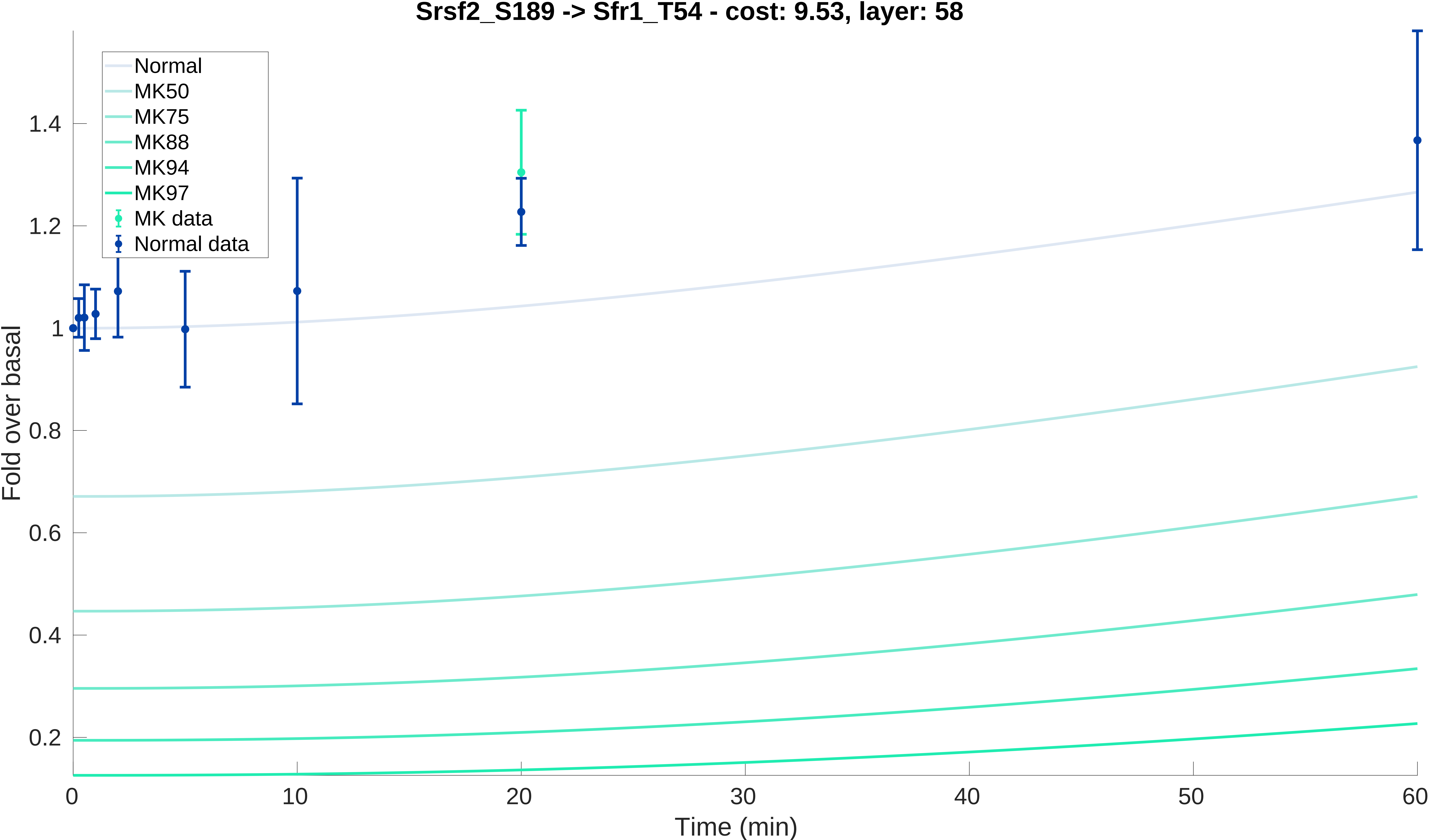

Srsf2_S189 -> Sfr1_T54 - cost: 9.53, layer: 58
Normal
MK50
MK75
MK88
MK94
MK97
MK data
Normal data
Fold over basal
Time (min)
1.4
1.2
1
0.8
0.6
0.4
0.2
0
10
20
30
40
50
60

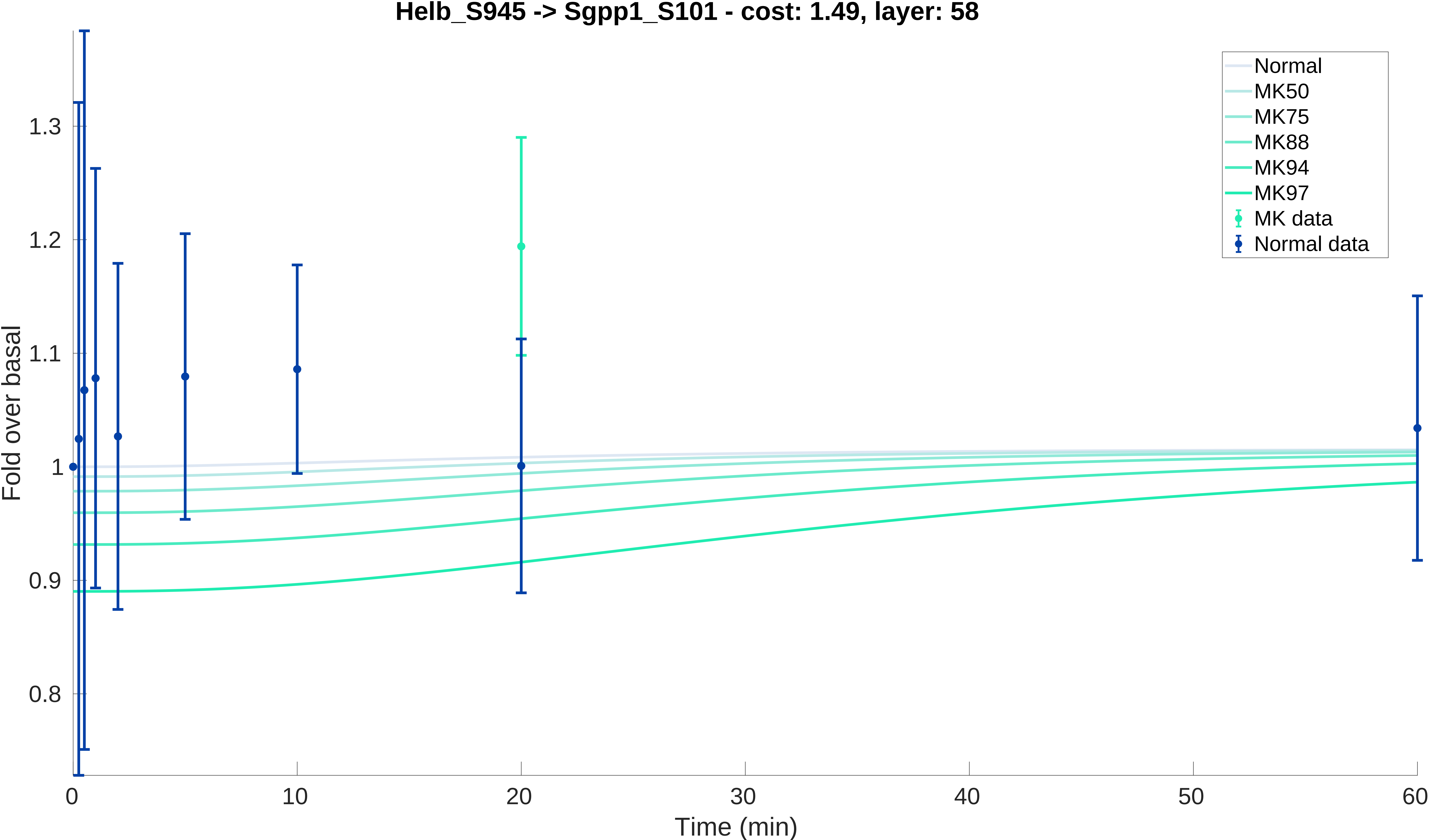
Helb_S945 -> Sgpp1_S101 - cost: 1.49, layer: 58
Fold over basal
1.3
1.2
1.1
1
0.9
0.8
0
10
20
30
40
50
60
Time (min)
Normal
MK50
MK75
MK88
MK94
MK97
MK data
Normal data

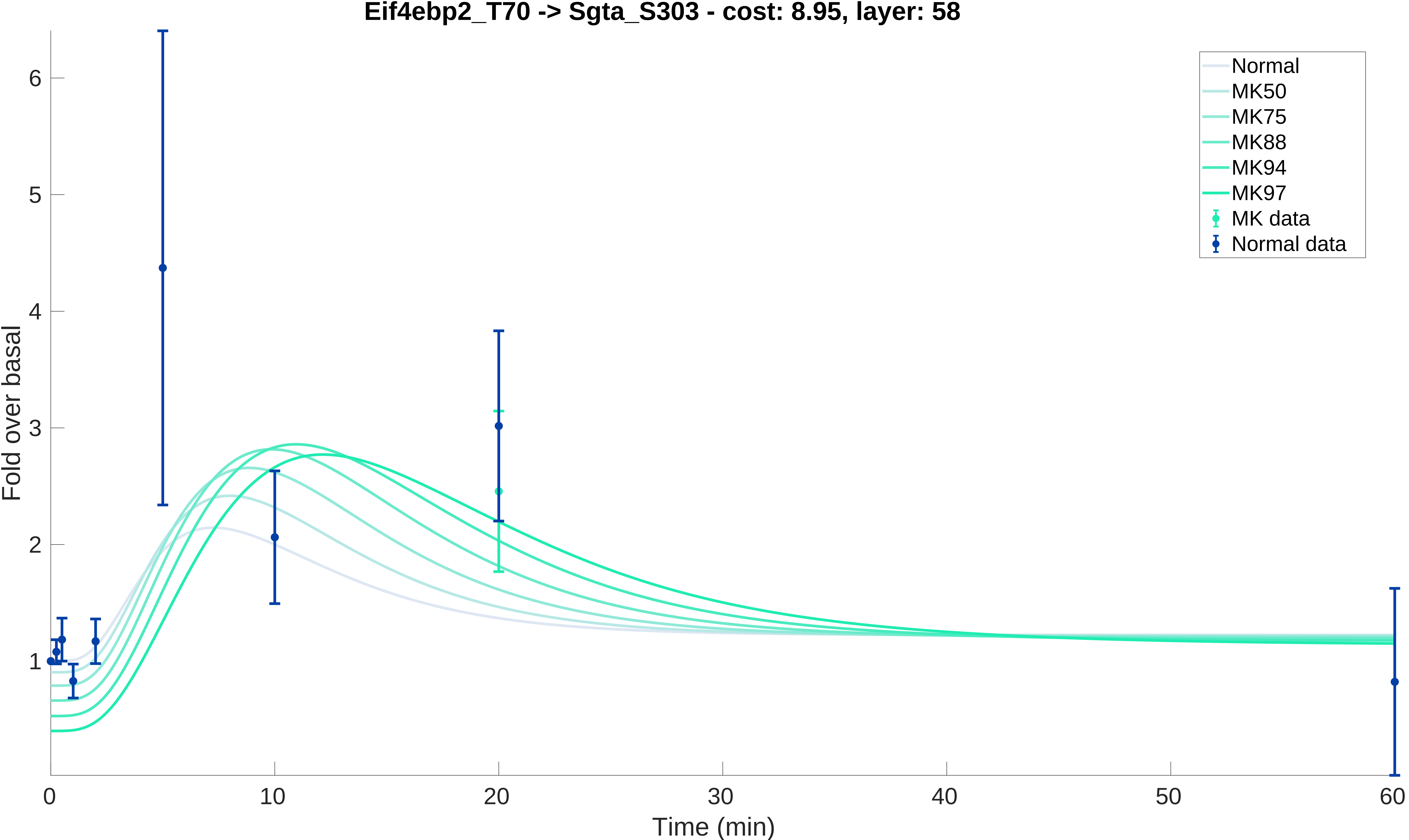
Eif4ebp2_T70 -> Sgta_S303 - cost: 8.95, layer: 58
Fold over basal
Time (min)
0
10
20
30
40
50
60
1
2
3
4
5
6
Normal
MK50
MK75
MK88
MK94
MK97
MK data
Normal data

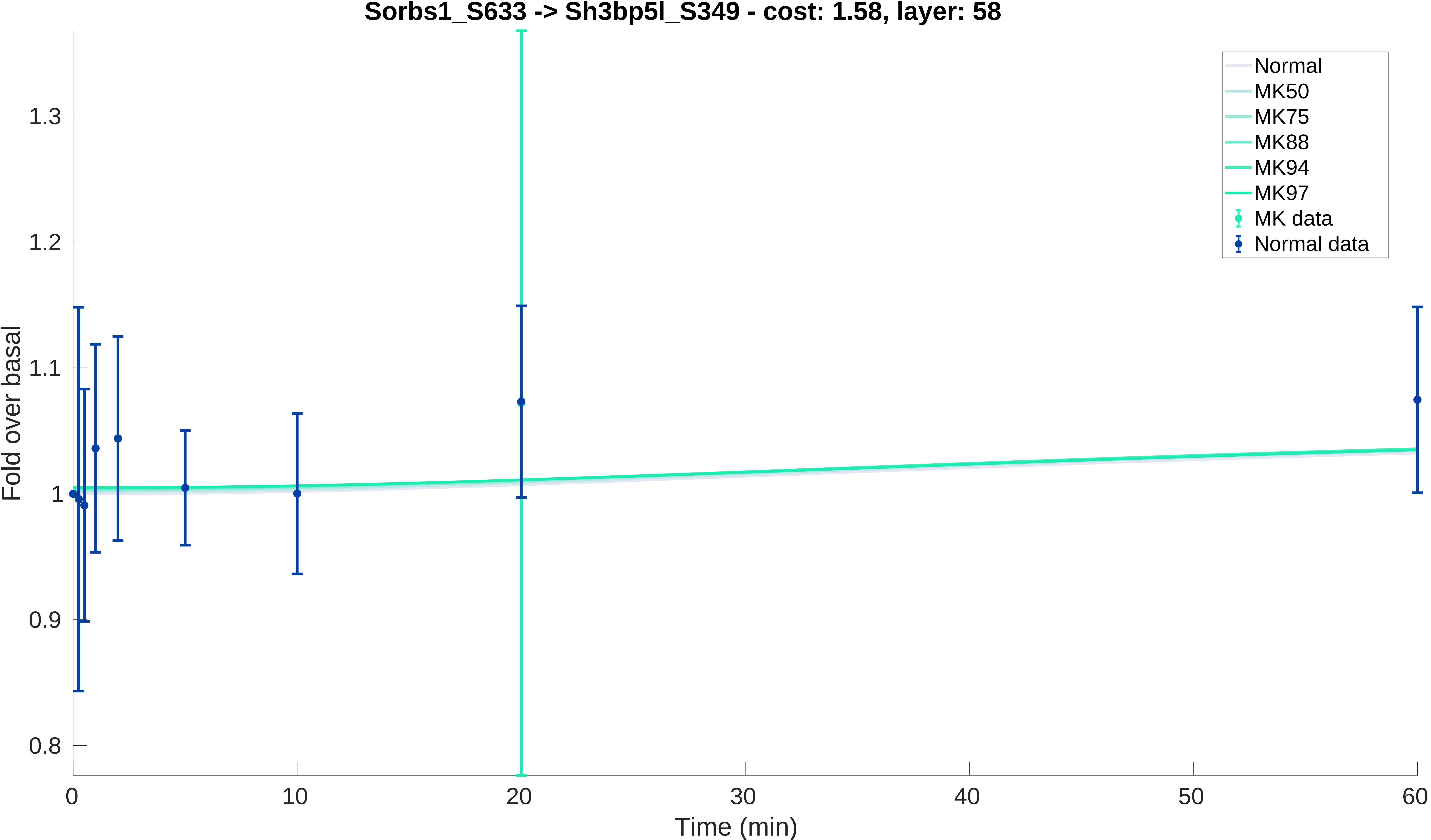

Sorbs1_S633 -> Sh3bp5l_S349 - cost: 1.58, layer: 58
Normal
MK50
MK75
MK88
MK94
MK97
MK data
Normal data
Fold over basal
1.3
1.2
1.1
1
0.9
0.8
0
10
20
30
40
50
60
Time (min)

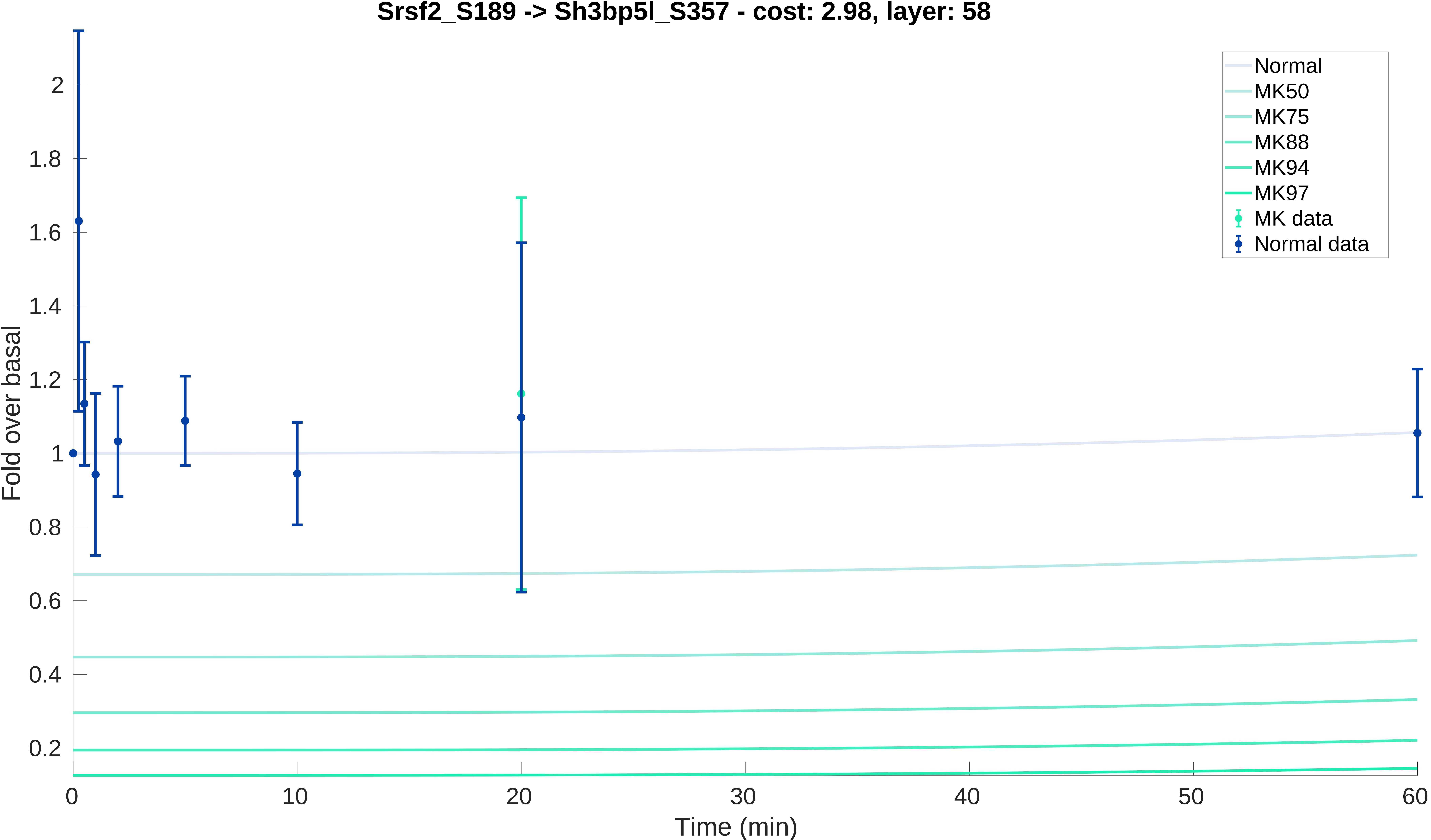

Srsf2_S189 -> Sh3bp5l_S357 - cost: 2.98, layer: 58
Fold over basal
Time (min)
0
10
20
30
40
50
60
0.2
0.4
0.6
0.8
1
1.2
1.4
1.6
1.8
2
Normal
MK50
MK75
MK88
MK94
MK97
MK data
Normal data

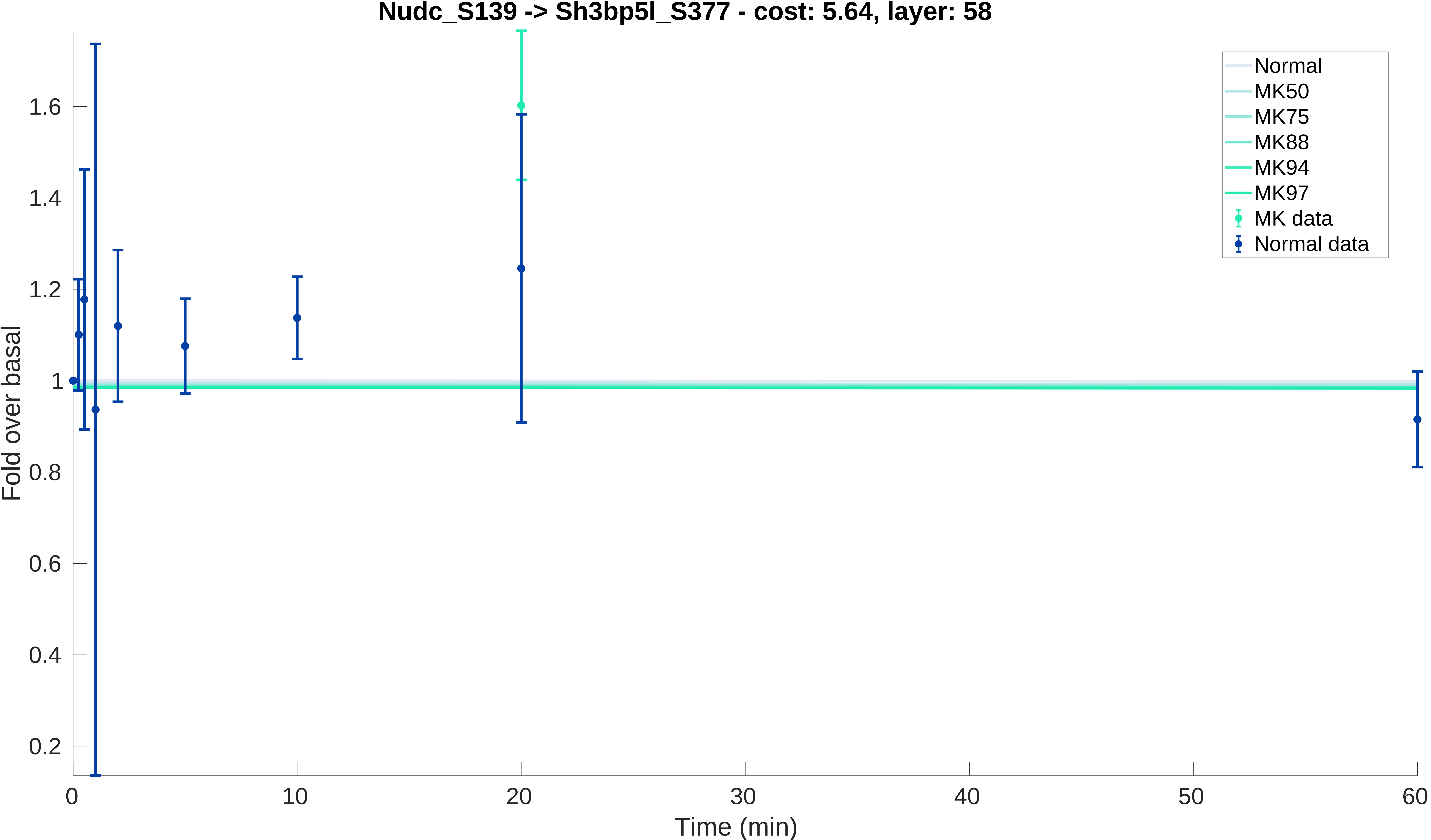

Nudc_S139 -> Sh3bp5l_S377 - cost: 5.64, layer: 58
Fold over basal
Time (min)
0
10
20
30
40
50
60
0.2
0.4
0.6
0.8
1
1.2
1.4
1.6
Normal
MK50
MK75
MK88
MK94
MK97
MK data
Normal data

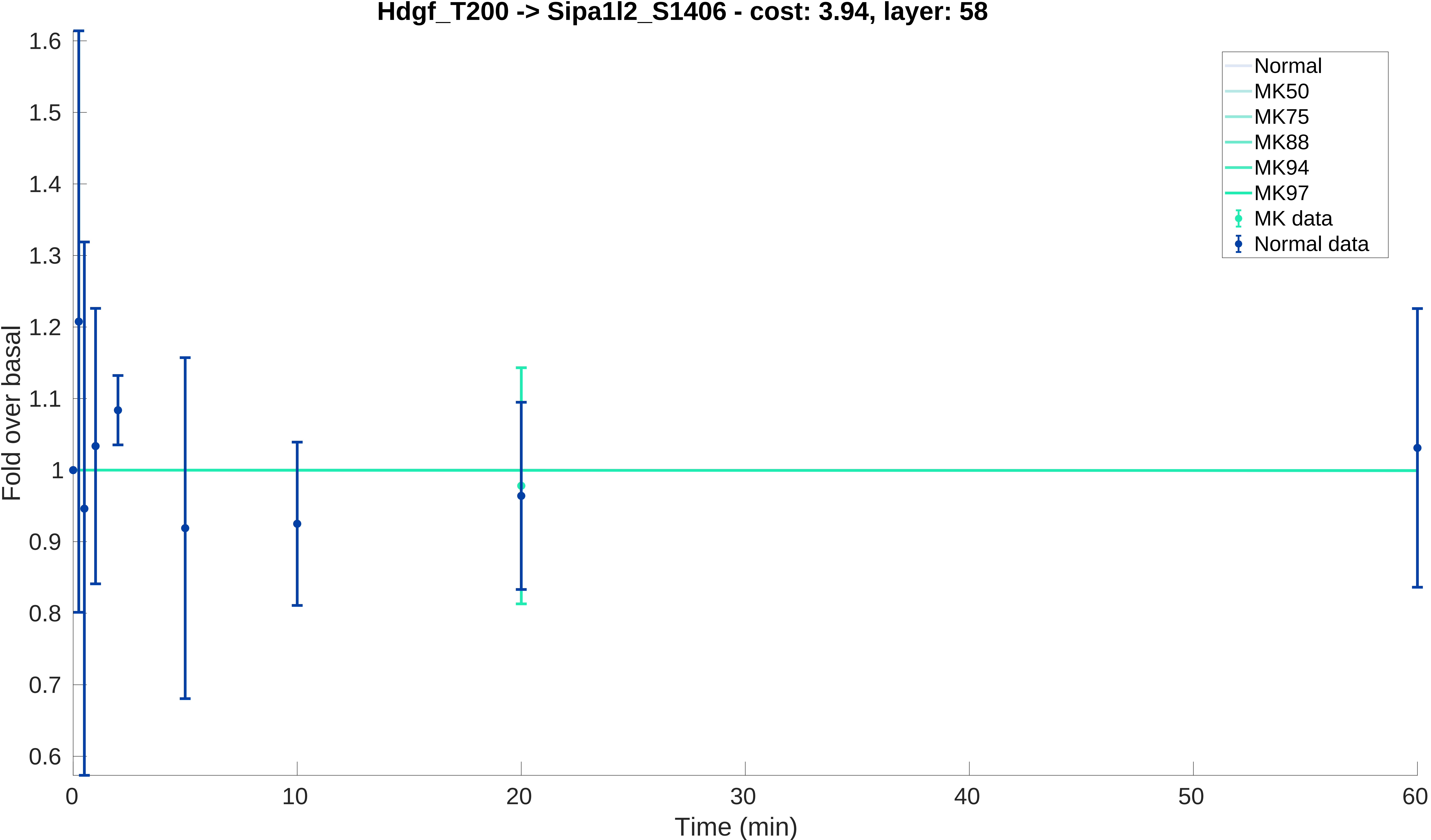

Hdgf_T200 -> Sipa1l2_S1406 - cost: 3.94, layer: 58
Fold over basal
Time (min)
Normal
MK50
MK75
MK88
MK94
MK97
MK data
Normal data
1.6
1.5
1.4
1.3
1.2
1.1
1
0.9
0.8
0.7
0.6
0
10
20
30
40
50
60

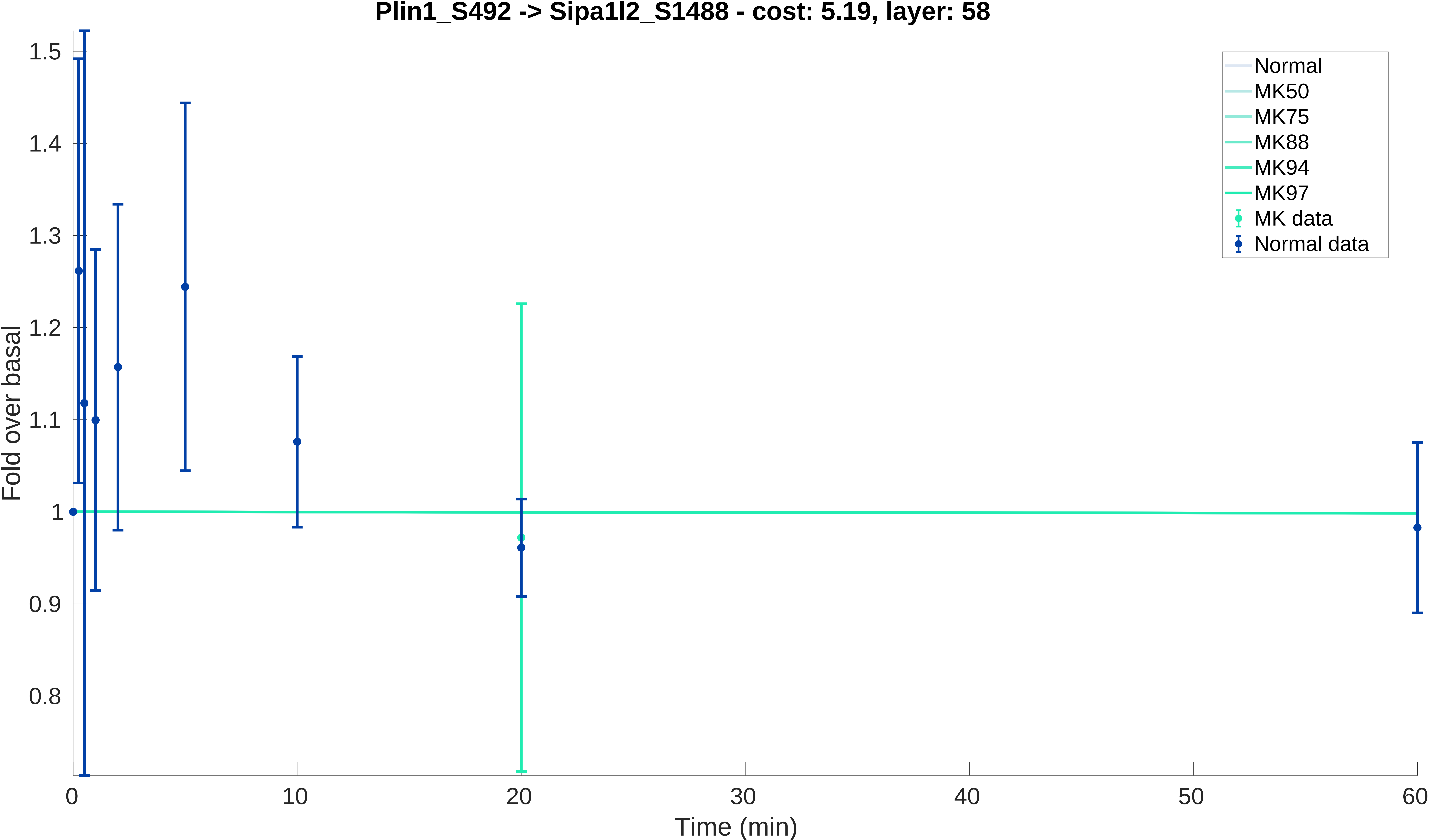
Plin1_S492 -> Sipa1l2_S1488 - cost: 5.19, layer: 58
Fold over basal
Time (min)
0
10
20
30
40
50
60
0.8
0.9
1
1.1
1.2
1.3
1.4
1.5
Normal
MK50
MK75
MK88
MK94
MK97
MK data
Normal data

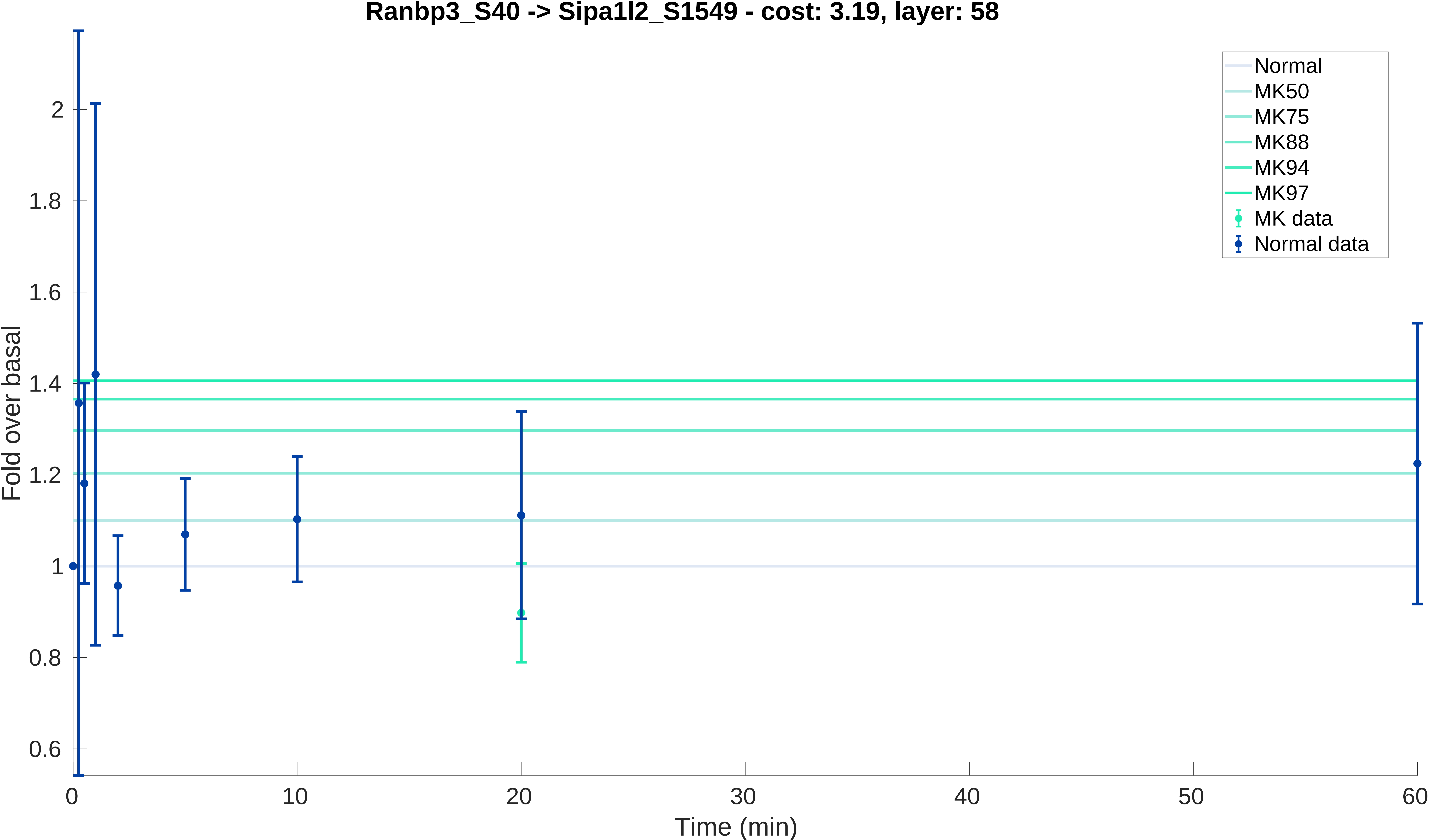
Ranbp3_S40 -> Sipa1l2_S1549 - cost: 3.19, layer: 58
Normal
MK50
MK75
MK88
MK94
MK97
MK data
Normal data
Fold over basal
2
1.8
1.6
1.4
1.2
1
0.8
0.6
0
10
20
30
40
50
60
Time (min)

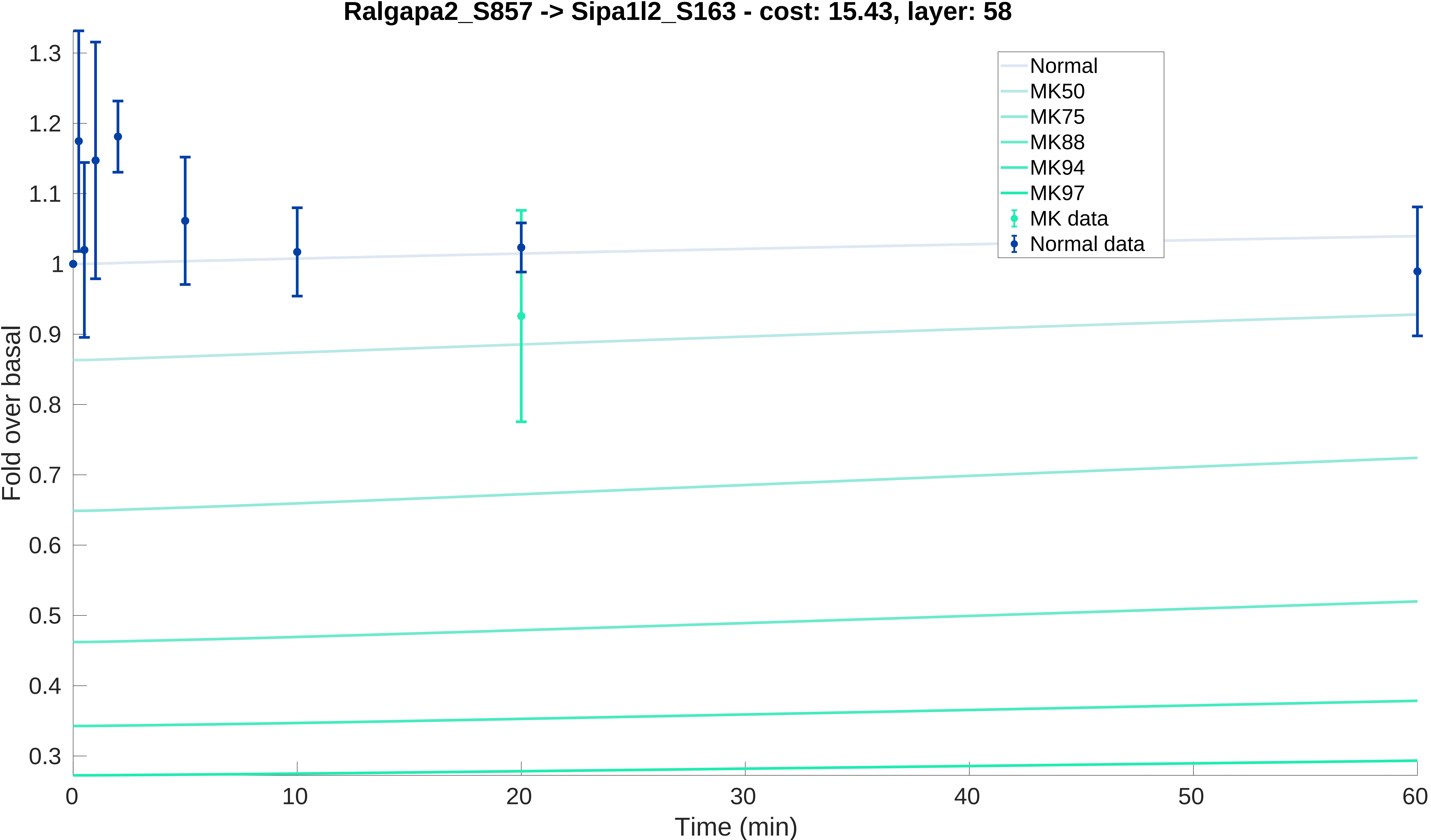

Ralgapa2_S857 -> Sipa1l2_S163 - cost: 15.43, layer: 58
Fold over basal
Time (min)
1.3
1.2
1.1
1
0.9
0.8
0.7
0.6
0.5
0.4
0.3
0
10
20
30
40
50
60
Normal
MK50
MK75
MK88
MK94
MK97
MK data
Normal data

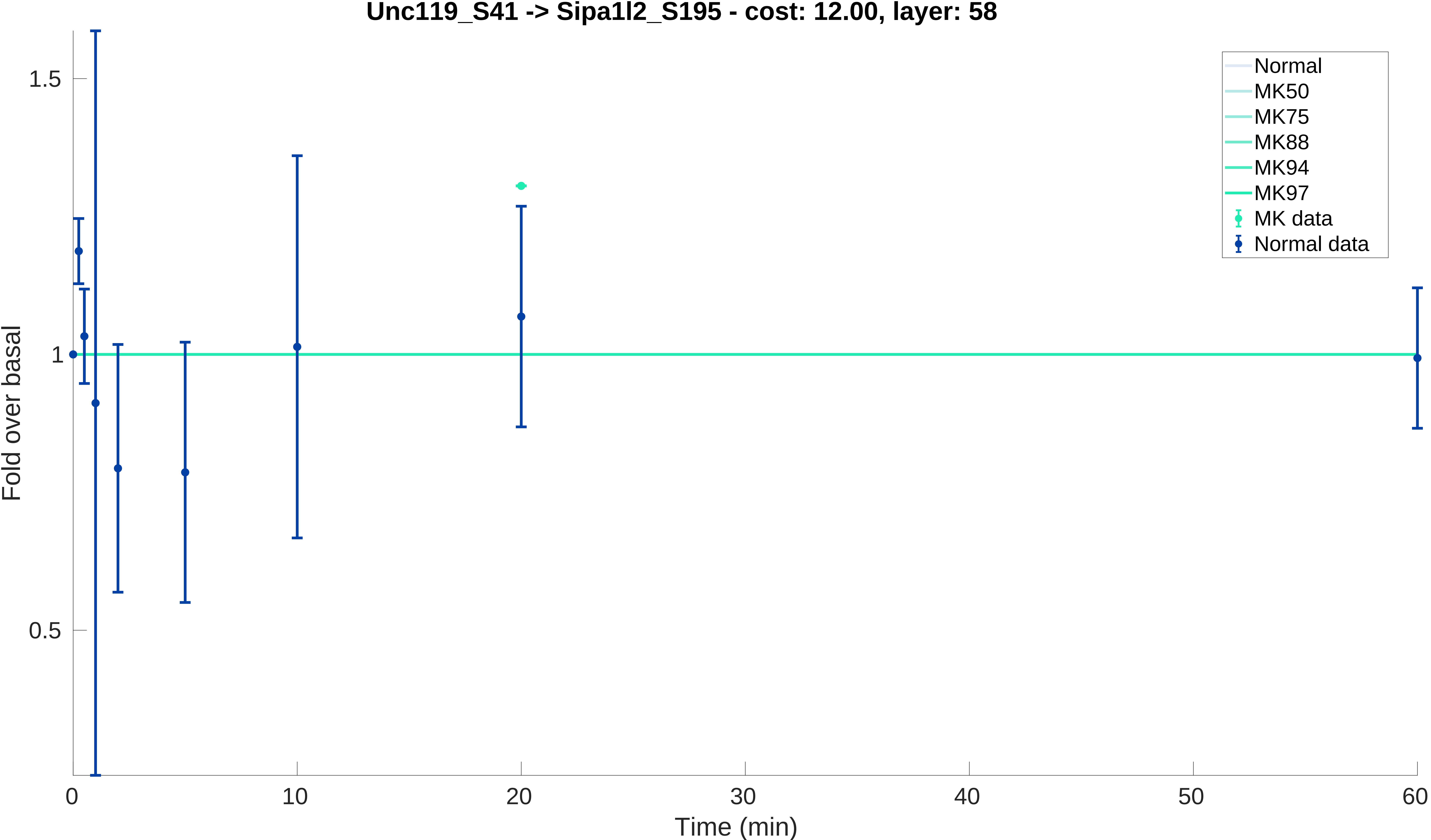

Unc119_S41 -> Sipa1l2_S195 - cost: 12.00, layer: 58
Fold over basal
Time (min)
1.5
1
0.5
0
10
20
30
40
50
60
Normal
MK50
MK75
MK88
MK94
MK97
MK data
Normal data

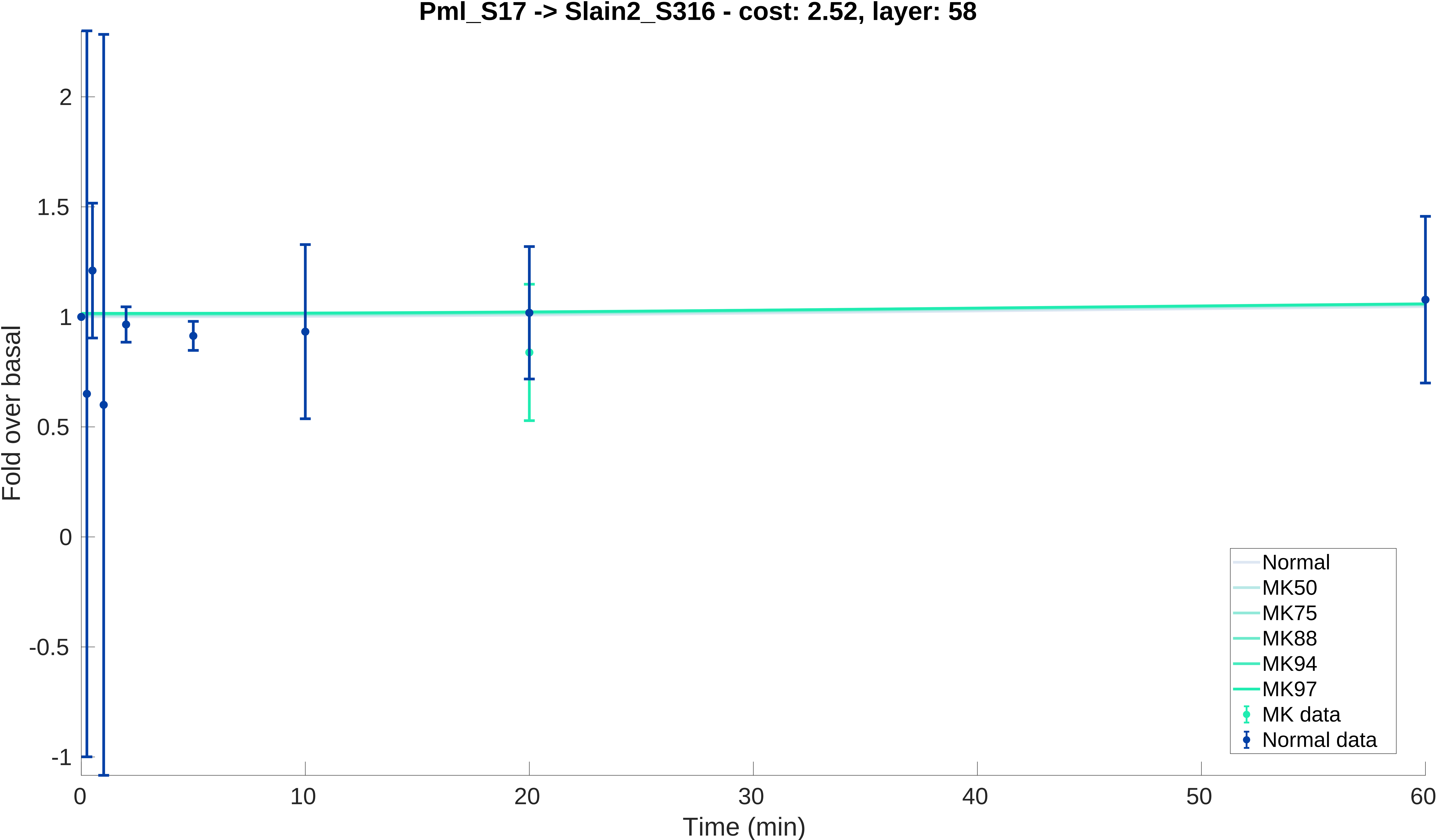

Pml_S17 -> Slain2_S316 - cost: 2.52, layer: 58
Fold over basal
Time (min)
2
1.5
1
0.5
0
-0.5
-1
0
10
20
30
40
50
60
Normal
MK50
MK75
MK88
MK94
MK97
MK data
Normal data

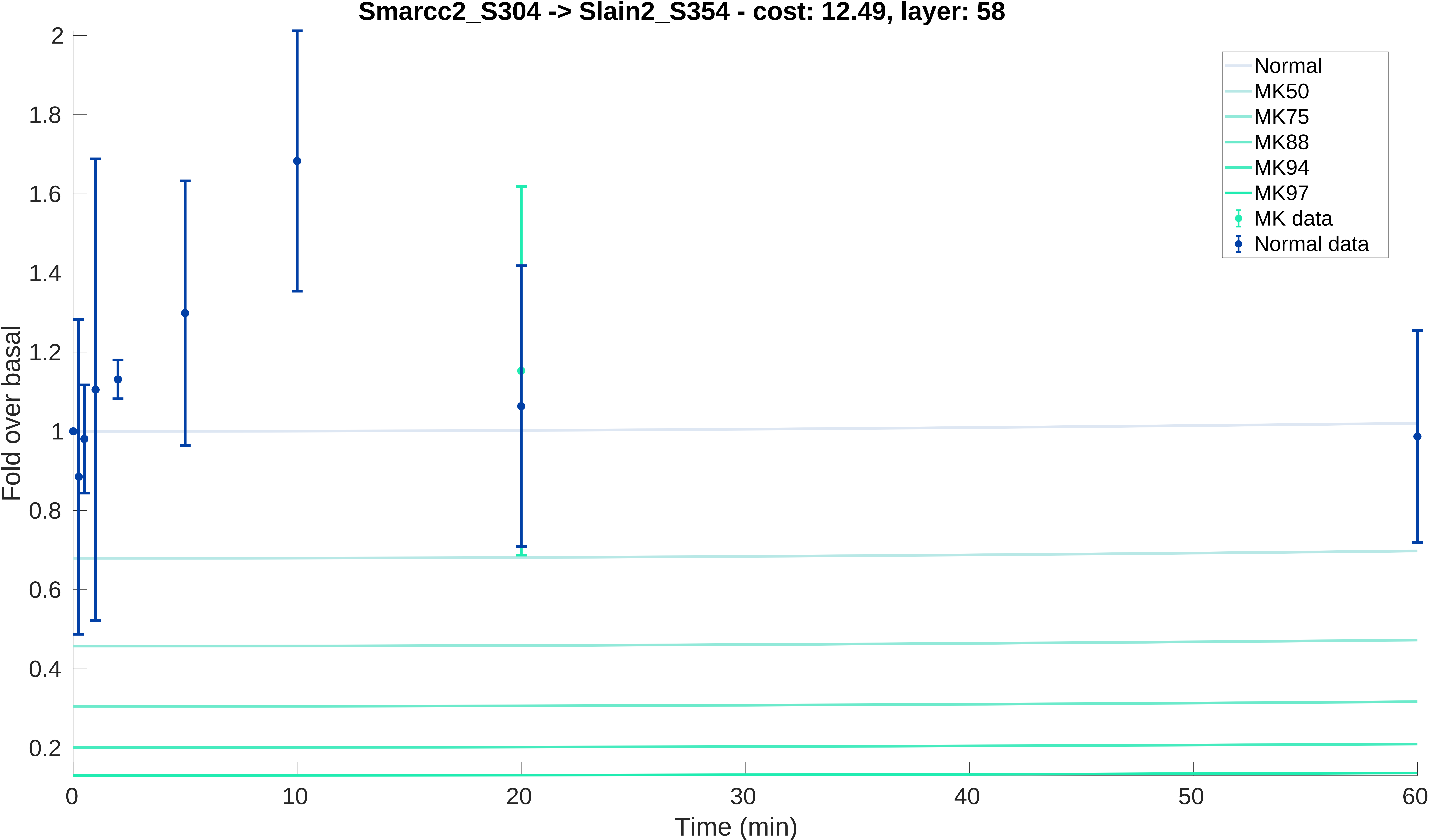

Smarcc2_S304 -> Slain2_S354 - cost: 12.49, layer: 58
Fold over basal
Time (min)
0
10
20
30
40
50
60
0.2
0.4
0.6
0.8
1
1.2
1.4
1.6
1.8
2
Normal
MK50
MK75
MK88
MK94
MK97
MK data
Normal data

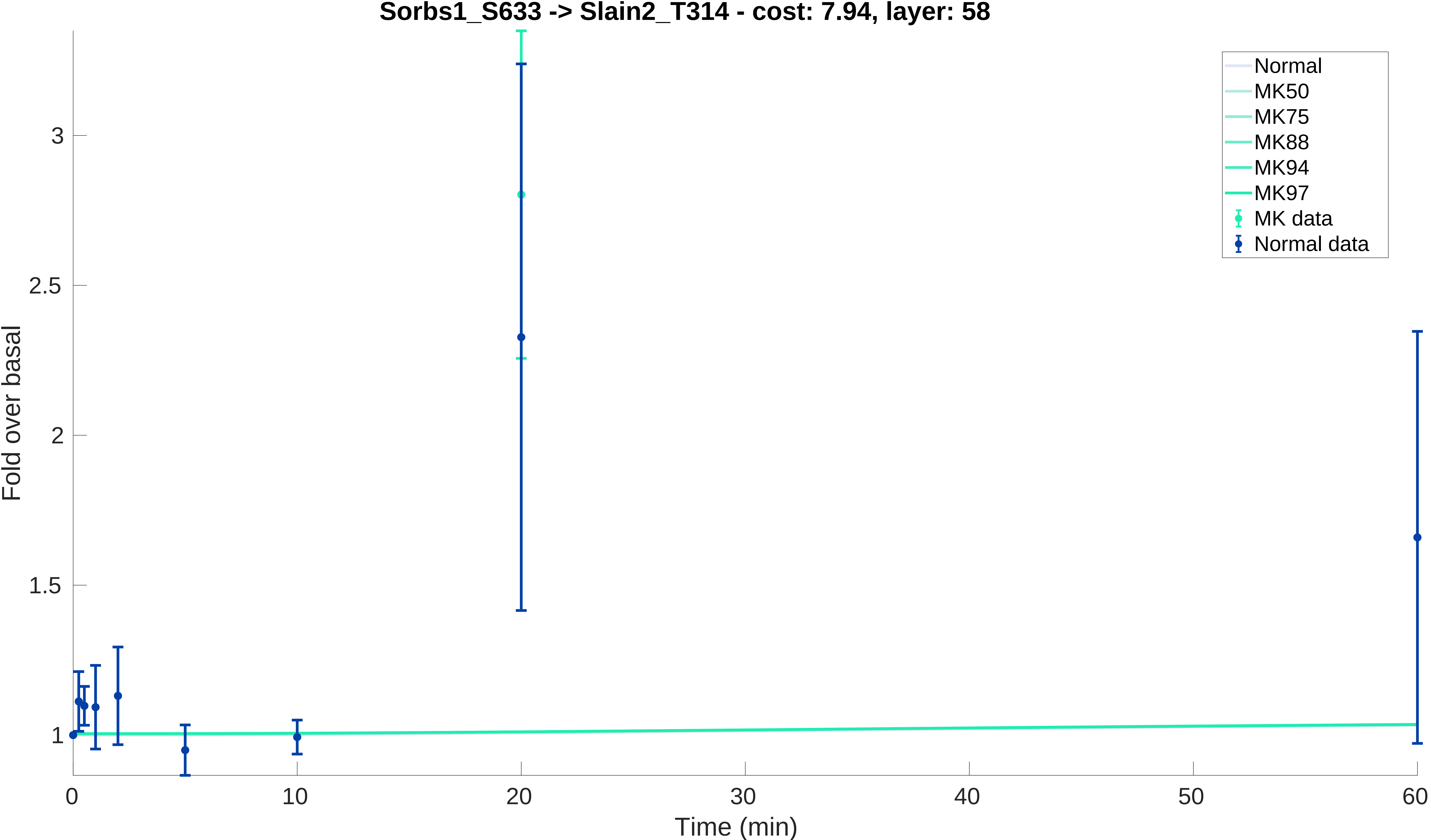

Sorbs1_S633 -> Slain2_T314 - cost: 7.94, layer: 58
Fold over basal
Time (min)
Normal
MK50
MK75
MK88
MK94
MK97
MK data
Normal data
1
1.5
2
2.5
3
0
10
20
30
40
50
60

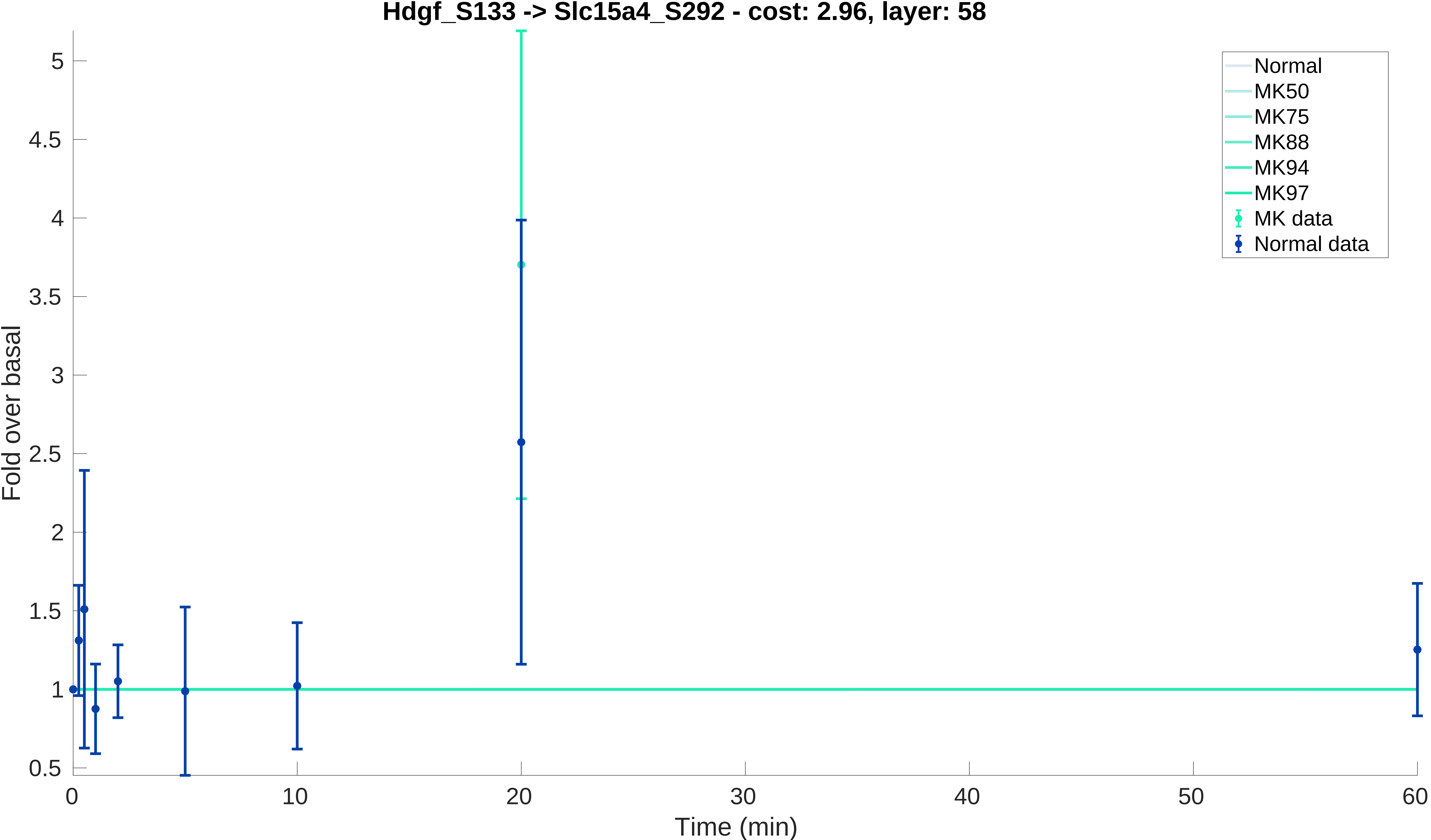
Hdgf_S133 -> Slc15a4_S292 - cost: 2.96, layer: 58
Fold over basal
Time (min)
0
10
20
30
40
50
60
0.5
1
1.5
2
2.5
3
3.5
4
4.5
5
Normal
MK50
MK75
MK88
MK94
MK97
MK data
Normal data

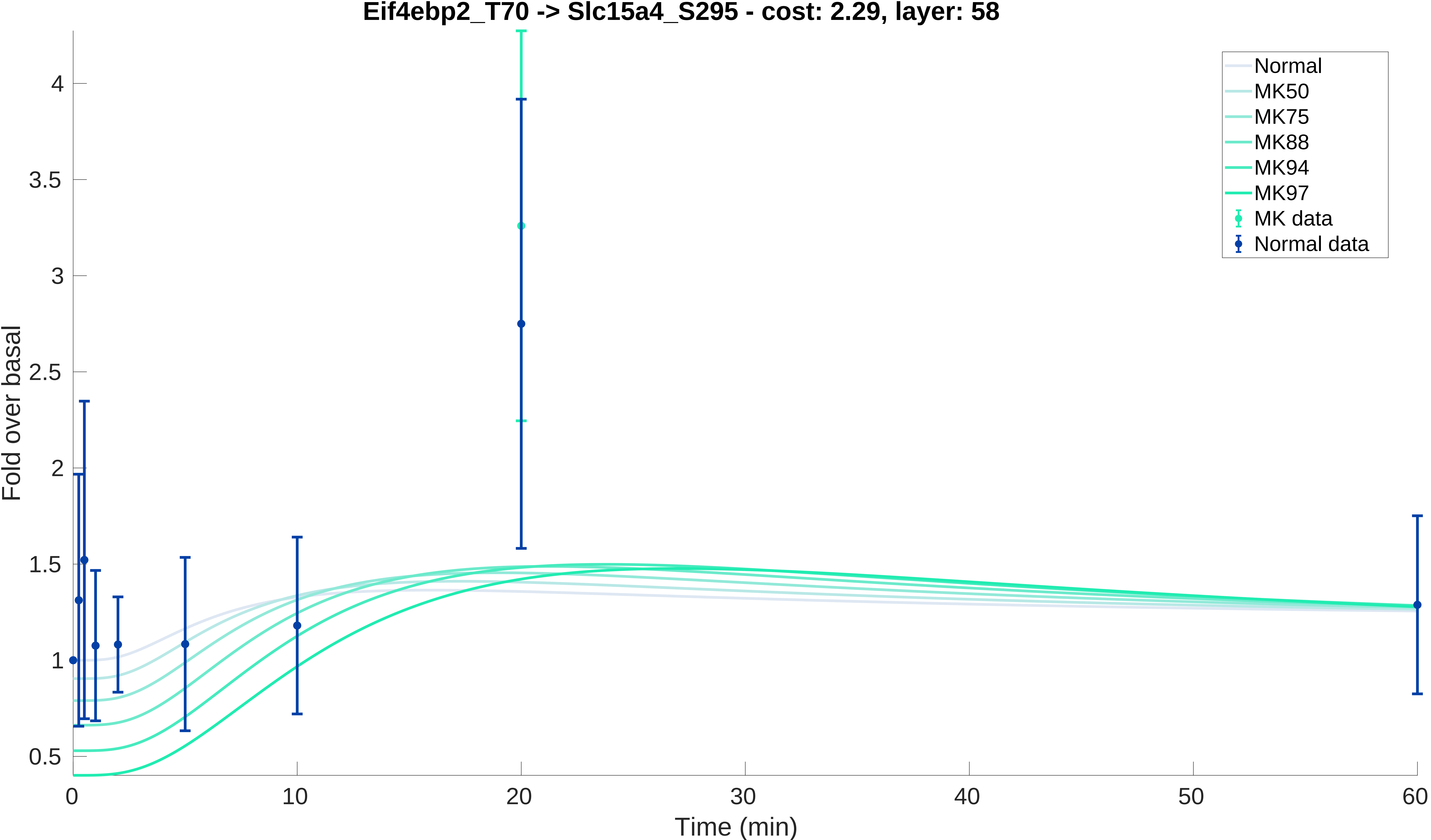

Eif4ebp2_T70 -> Slc15a4_S295 - cost: 2.29, layer: 58
Fold over basal
Time (min)
0.5
1
1.5
2
2.5
3
3.5
4
0
10
20
30
40
50
60
Normal
MK50
MK75
MK88
MK94
MK97
MK data
Normal data

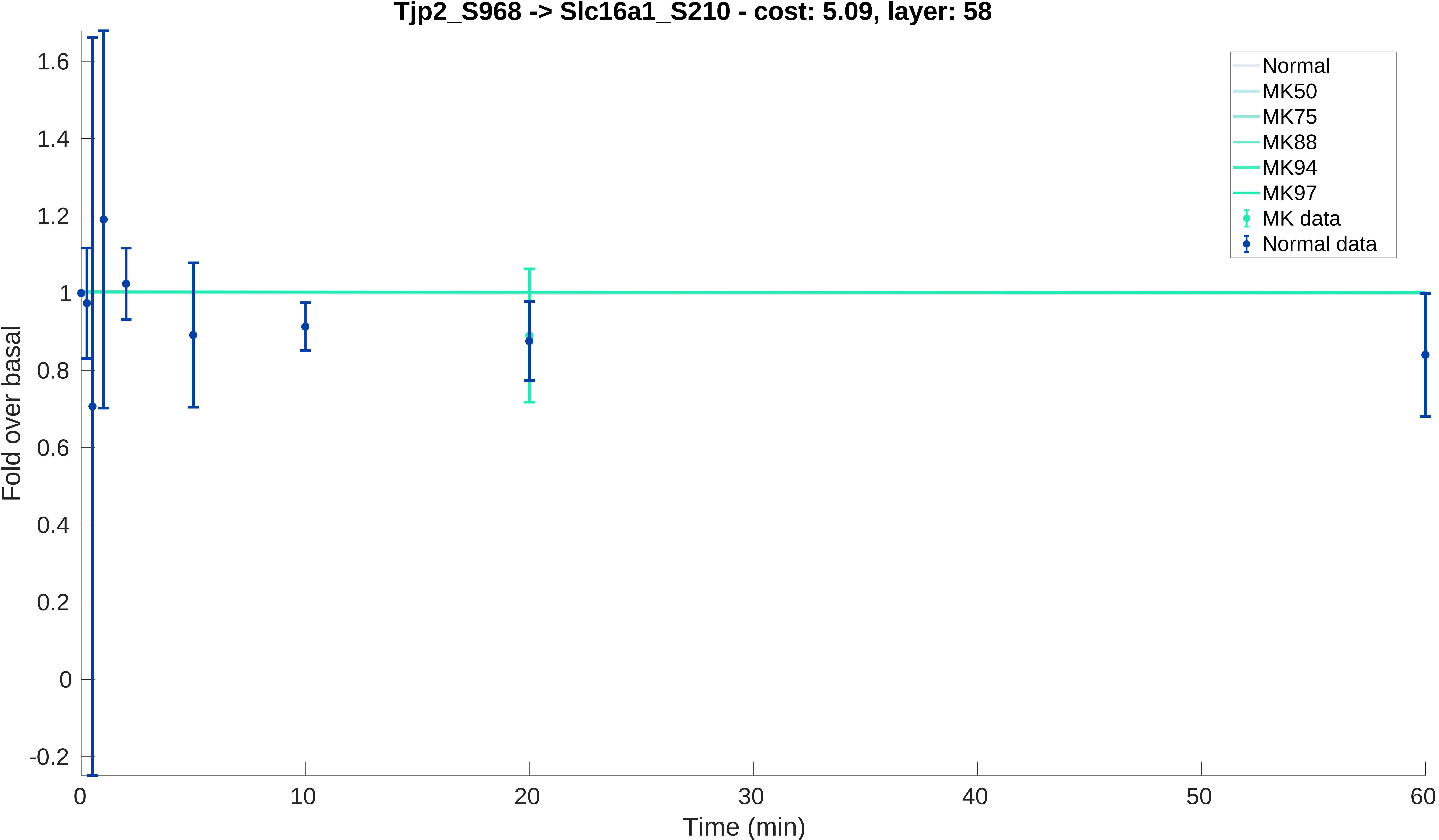

Tjp2_S968 -> Slc16a1_S210 - cost: 5.09, layer: 58
Fold over basal
Time (min)
1.6
1.4
1.2
1
0.8
0.6
0.4
0.2
0
-0.2
0
10
20
30
40
50
60
Normal
MK50
MK75
MK88
MK94
MK97
MK data
Normal data

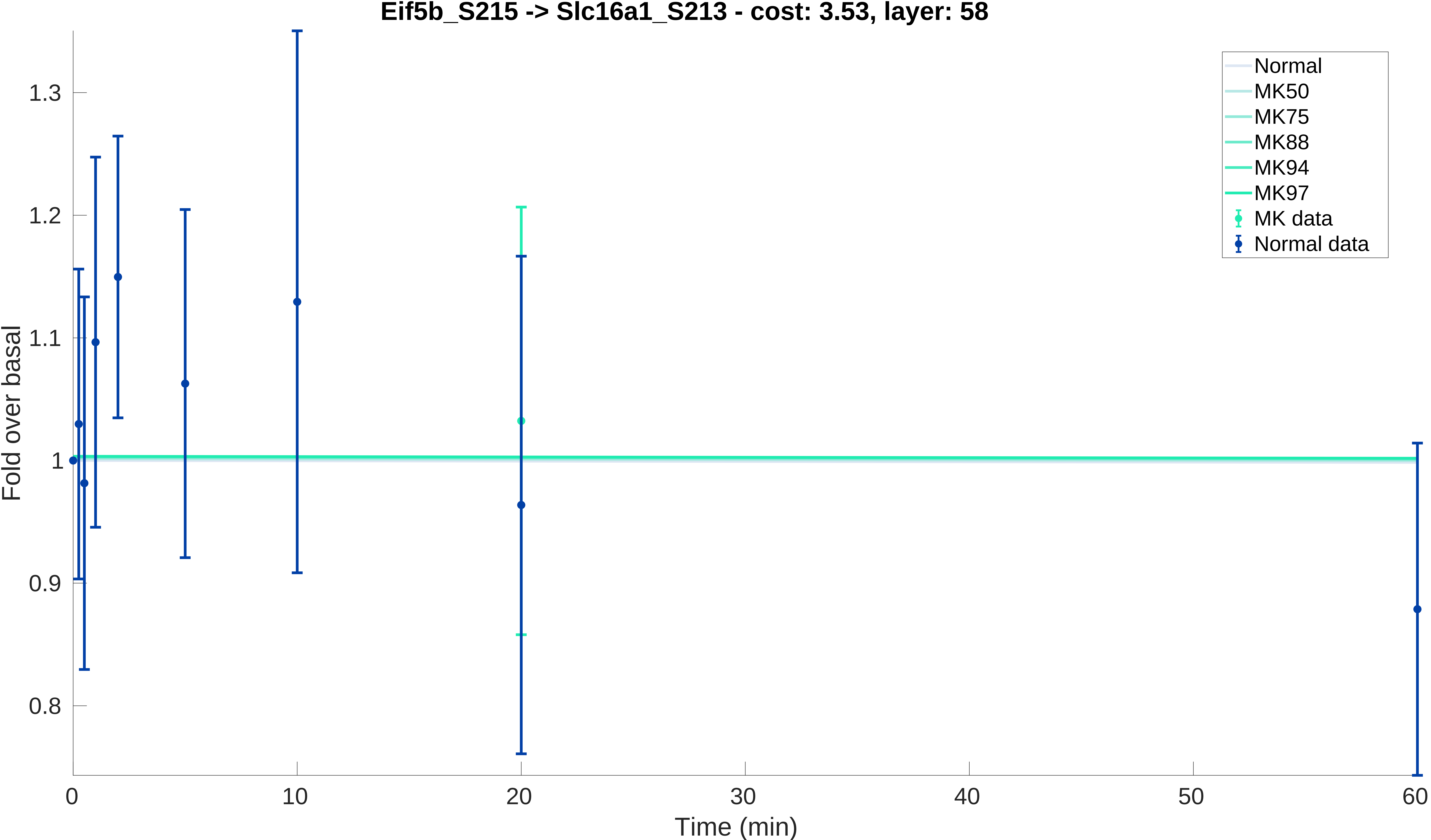

Eif5b_S215 -> Slc16a1_S213 - cost: 3.53, layer: 58
Fold over basal
Time (min)
1.3
1.2
1.1
1
0.9
0.8
0
10
20
30
40
50
60
Normal
MK50
MK75
MK88
MK94
MK97
MK data
Normal data

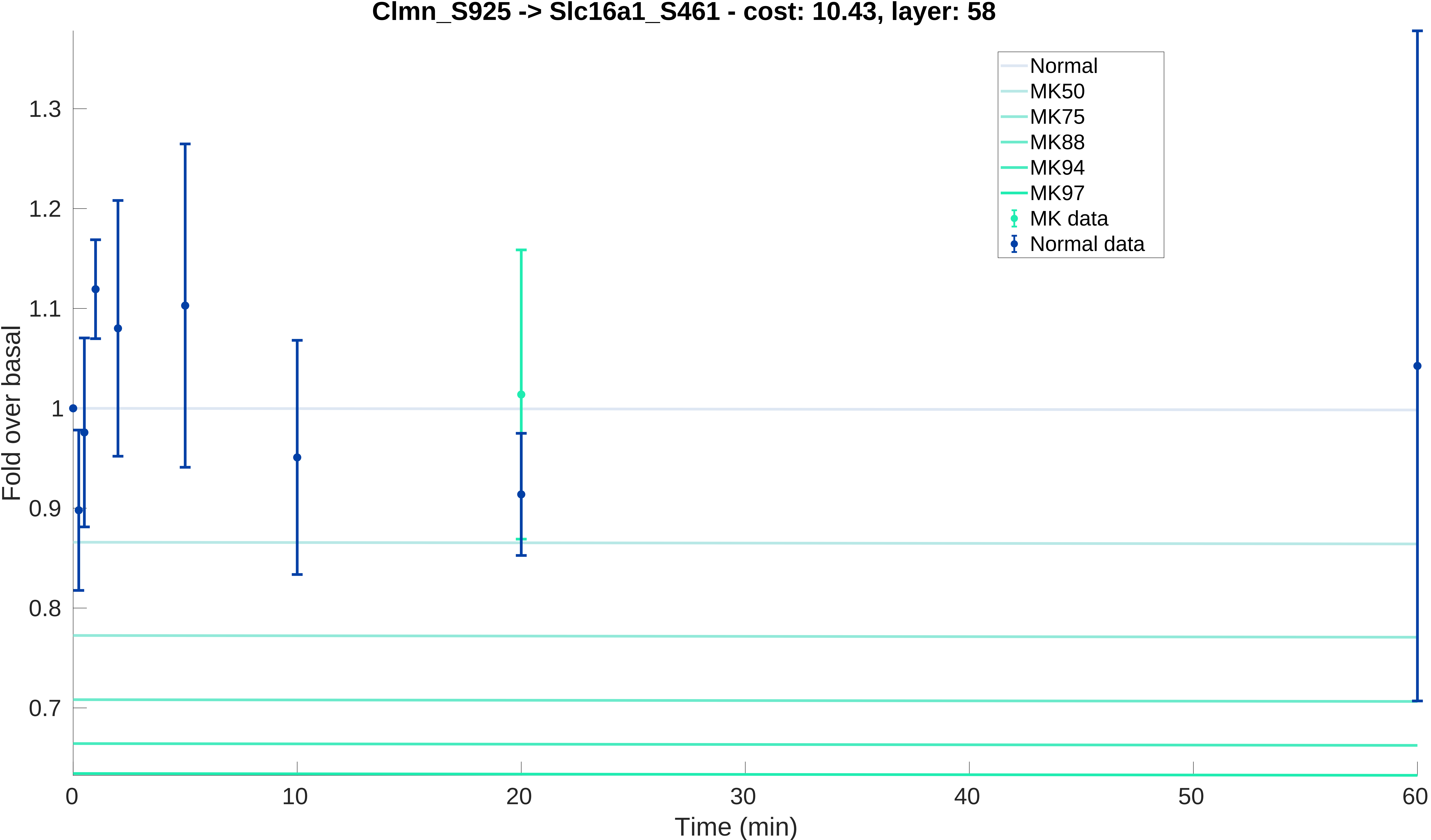

Clmn_S925 -> Slc16a1_S461 - cost: 10.43, layer: 58
Fold over basal
Time (min)
0
10
20
30
40
50
60
0.7
0.8
0.9
1
1.1
1.2
1.3
Normal
MK50
MK75
MK88
MK94
MK97
MK data
Normal data

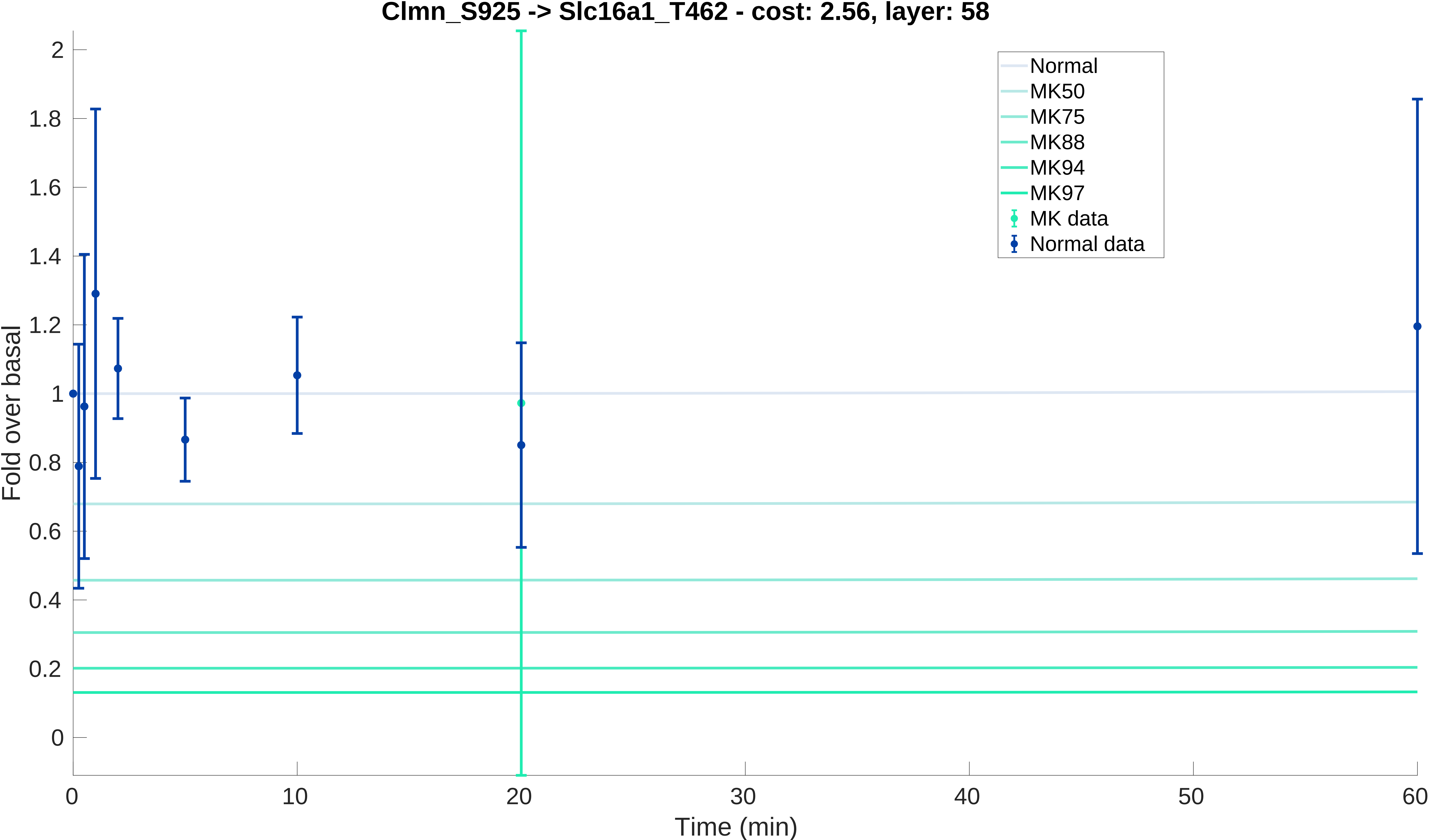

Clmn_S925 -> Slc16a1_T462 - cost: 2.56, layer: 58
Normal
MK50
MK75
MK88
MK94
MK97
MK data
Normal data
Fold over basal
Time (min)
0
0.2
0.4
0.6
0.8
1
1.2
1.4
1.6
1.8
2
0
10
20
30
40
50
60

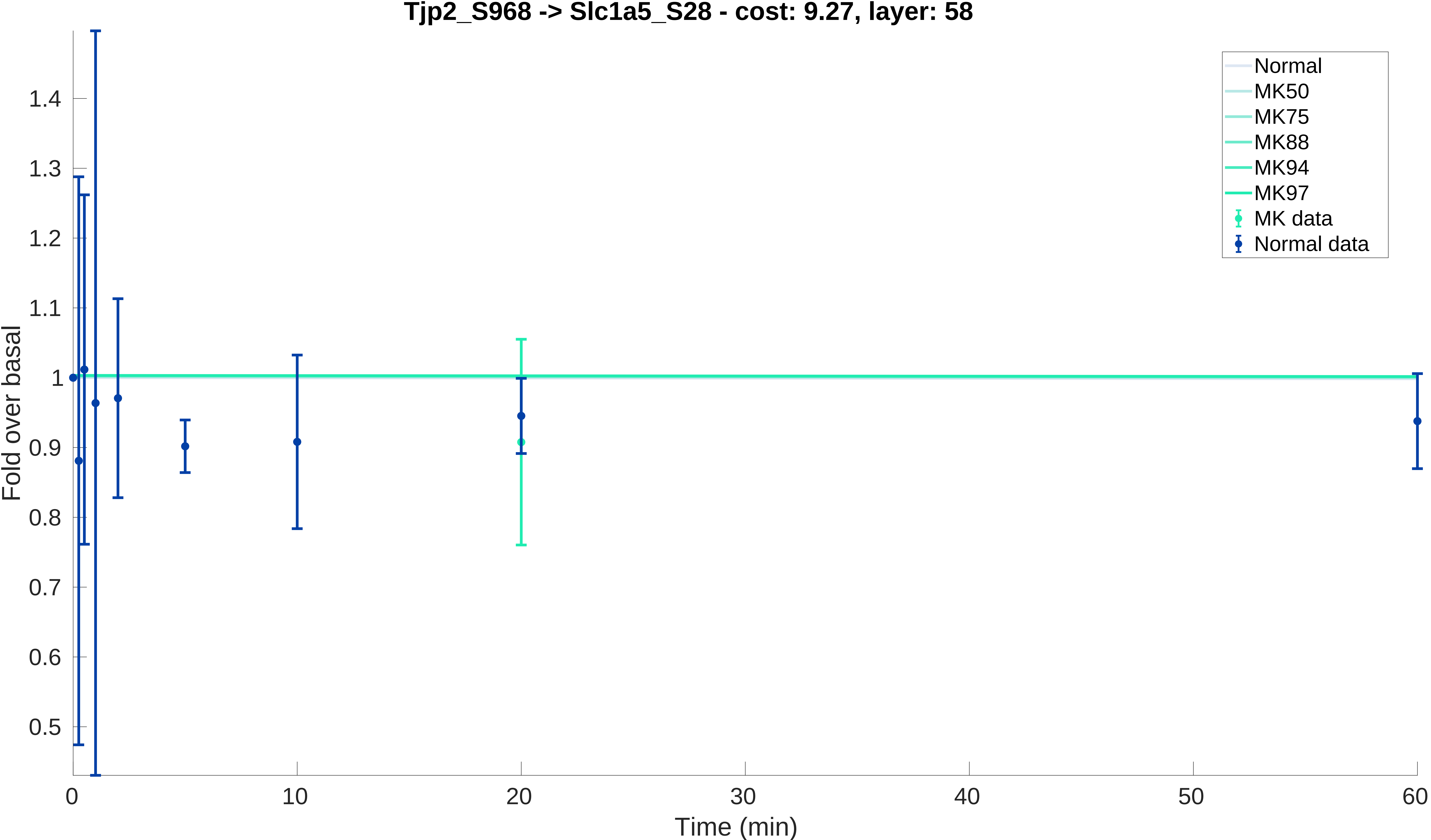

Tjp2_S968 -> Slc1a5_S28 - cost: 9.27, layer: 58
Normal
MK50
MK75
MK88
MK94
MK97
MK data
Normal data
Fold over basal
1.4
1.3
1.2
1.1
1
0.9
0.8
0.7
0.6
0.5
0
10
20
30
40
50
60
Time (min)

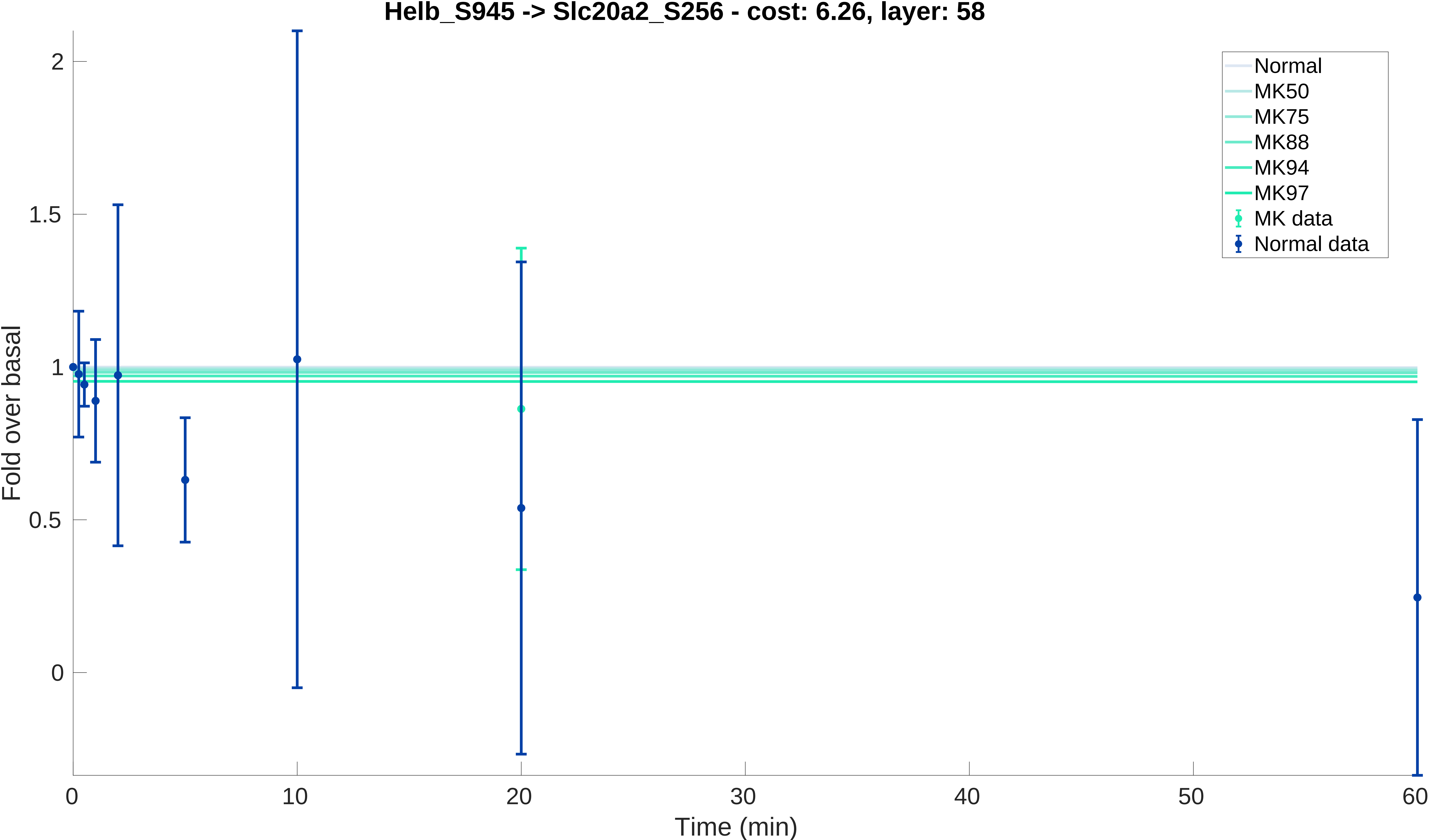
Helb_S945 -> Slc20a2_S256 - cost: 6.26, layer: 58
Fold over basal
Time (min)
0
10
20
30
40
50
60
0
0.5
1
1.5
2
Normal
MK50
MK75
MK88
MK94
MK97
MK data
Normal data

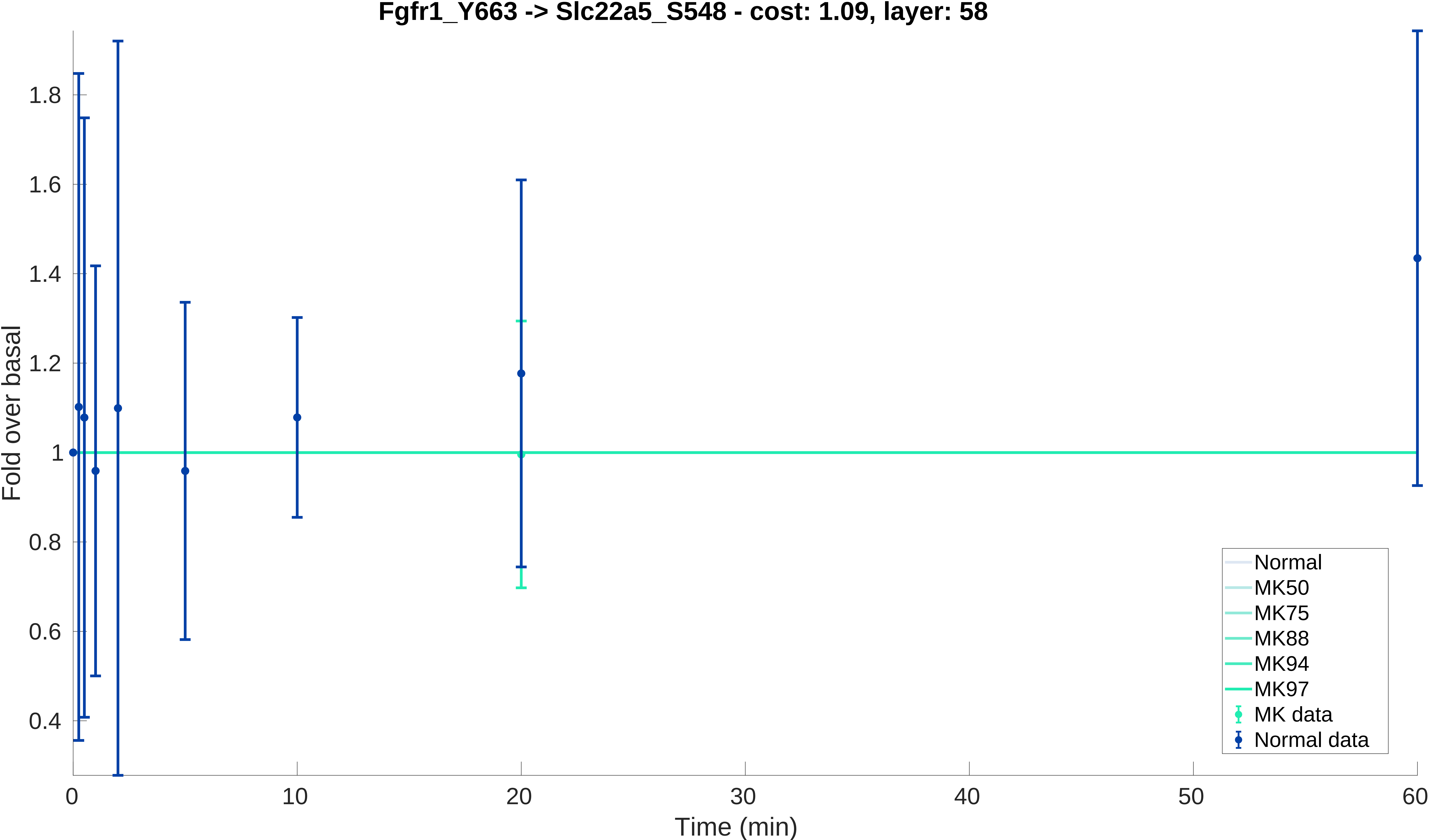
Fgfr1_Y663 -> Slc22a5_S548 - cost: 1.09, layer: 58
Fold over basal
1.8
1.6
1.4
1.2
1
0.8
0.6
0.4
0
10
20
30
40
50
60
Time (min)
Normal
MK50
MK75
MK88
MK94
MK97
MK data
Normal data

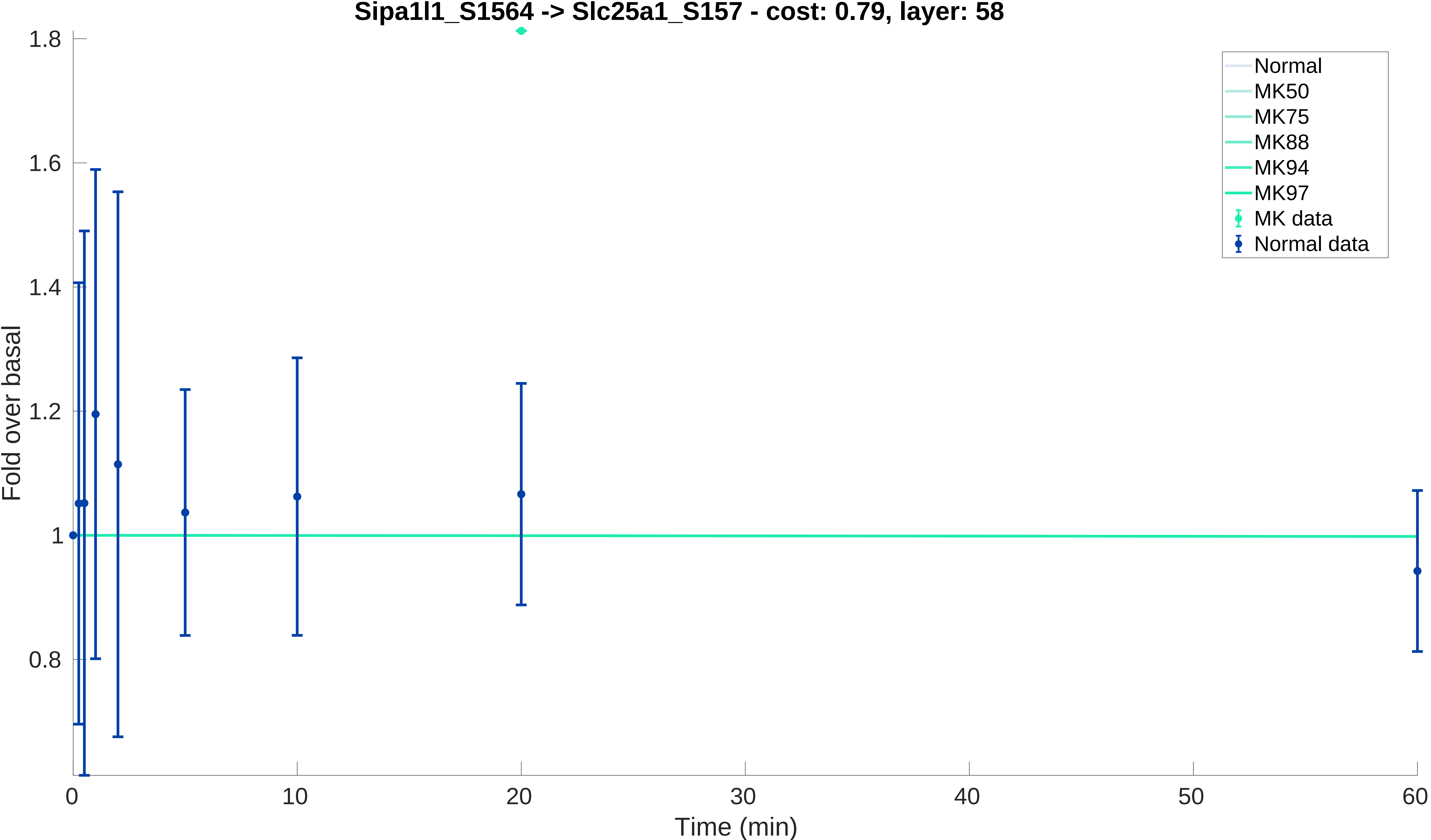

Sipa1l1_S1564 -> Slc25a1_S157 - cost: 0.79, layer: 58
Fold over basal
Time (min)
1.8
1.6
1.4
1.2
1
0.8
0
10
20
30
40
50
60
Normal
MK50
MK75
MK88
MK94
MK97
MK data
Normal data

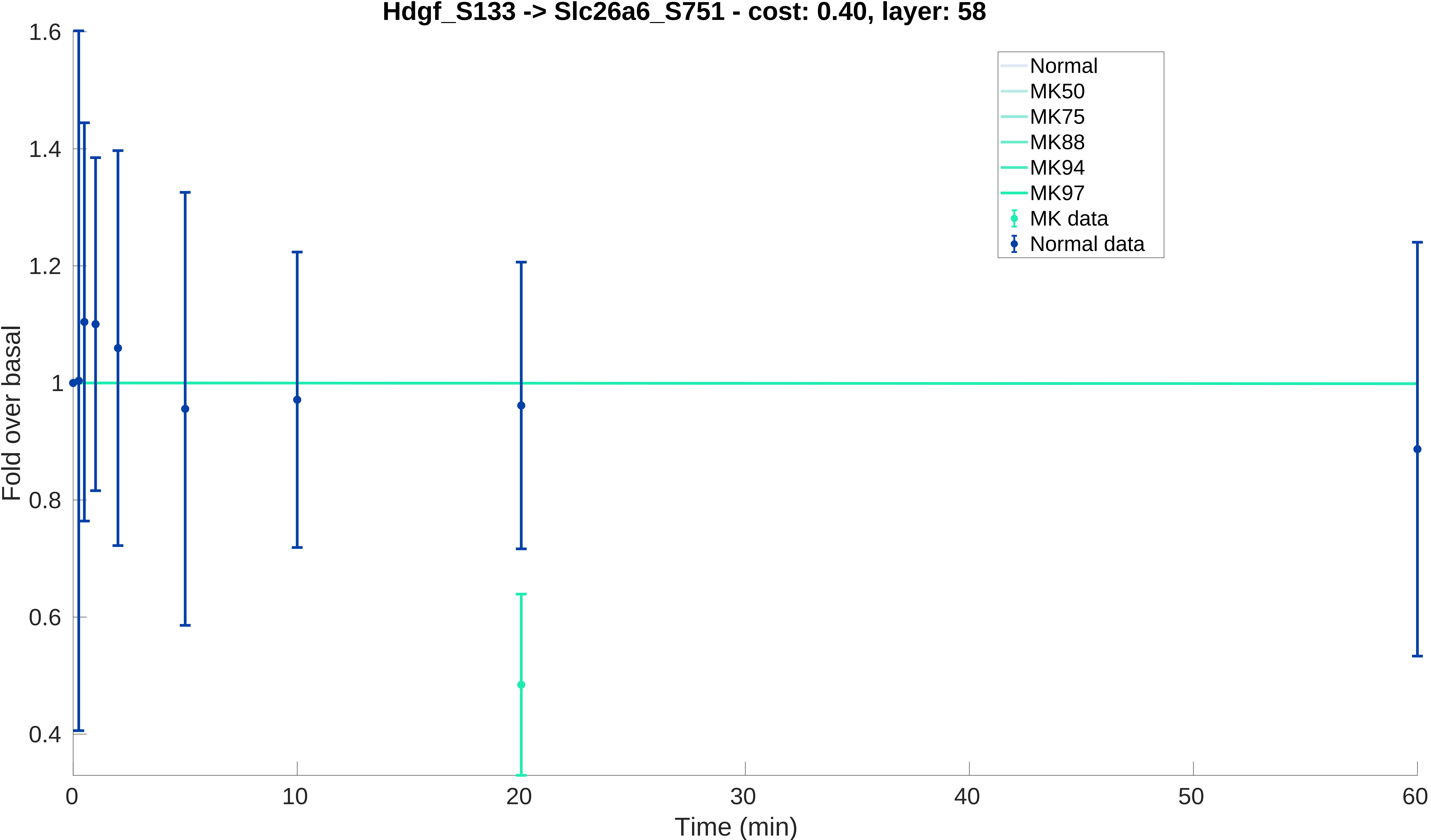

Hdgf_S133 -> Slc26a6_S751 - cost: 0.40, layer: 58
Fold over basal
Time (min)
1.6
1.4
1.2
1
0.8
0.6
0.4
0
10
20
30
40
50
60
Normal
MK50
MK75
MK88
MK94
MK97
MK data
Normal data

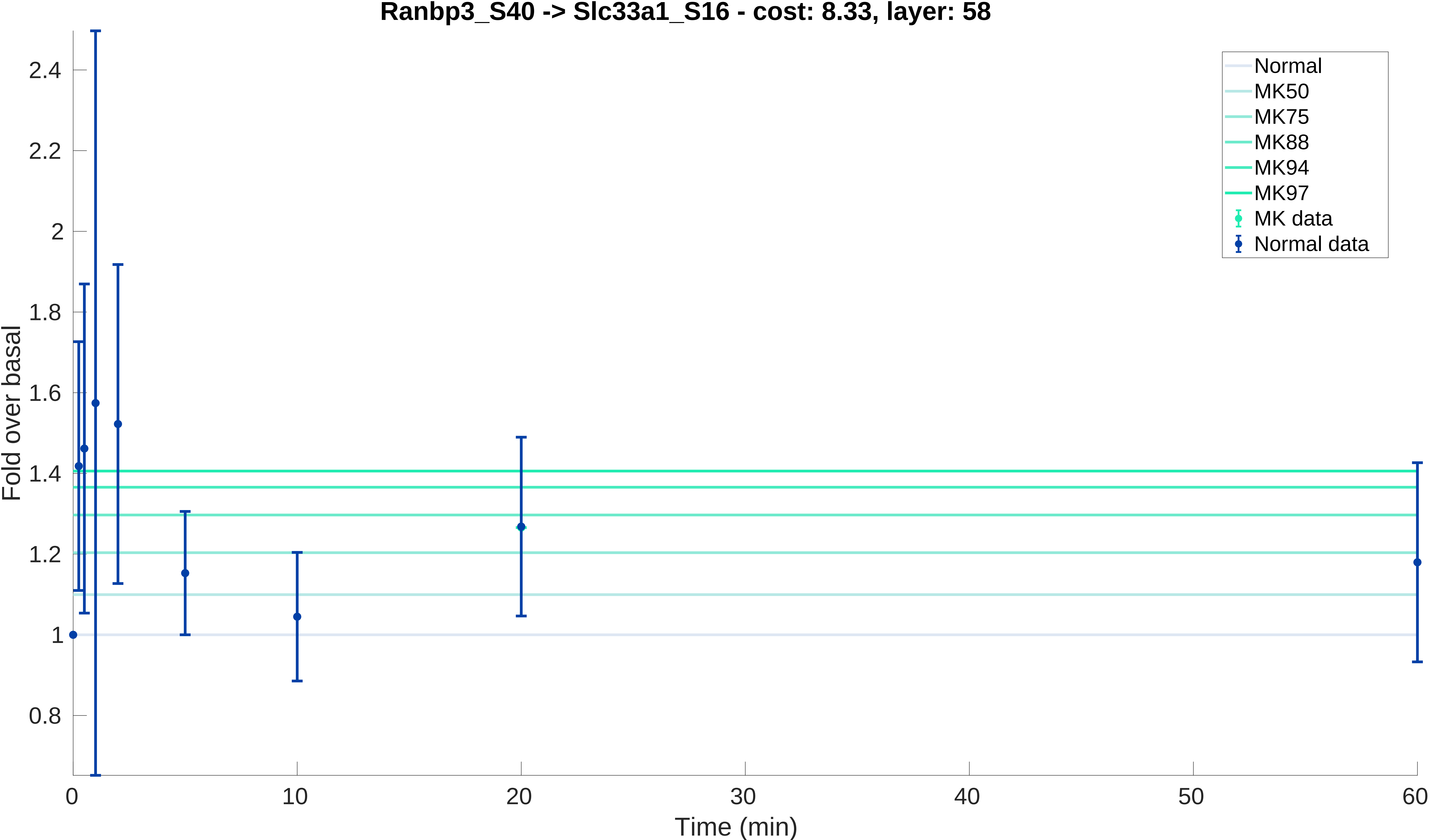
Ranbp3_S40 -> Slc33a1_S16 - cost: 8.33, layer: 58
Fold over basal
Time (min)
0.8
1
1.2
1.4
1.6
1.8
2
2.2
2.4
0
10
20
30
40
50
60
Normal
MK50
MK75
MK88
MK94
MK97
MK data
Normal data

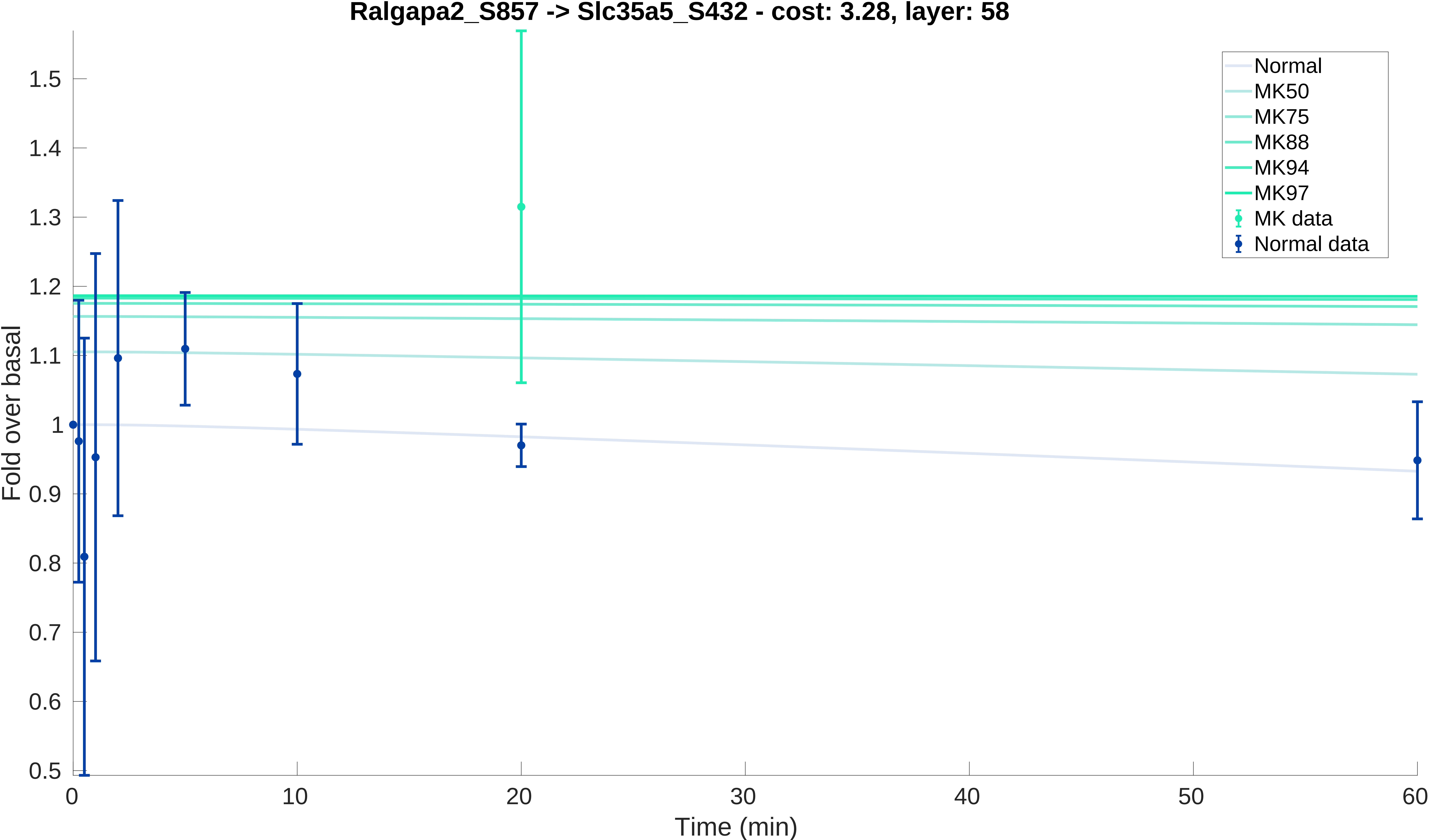

Ralgapa2_S857 -> Slc35a5_S432 - cost: 3.28, layer: 58
Fold over basal
Time (min)
1.5
1.4
1.3
1.2
1.1
1
0.9
0.8
0.7
0.6
0.5
0
10
20
30
40
50
60
Normal
MK50
MK75
MK88
MK94
MK97
MK data
Normal data

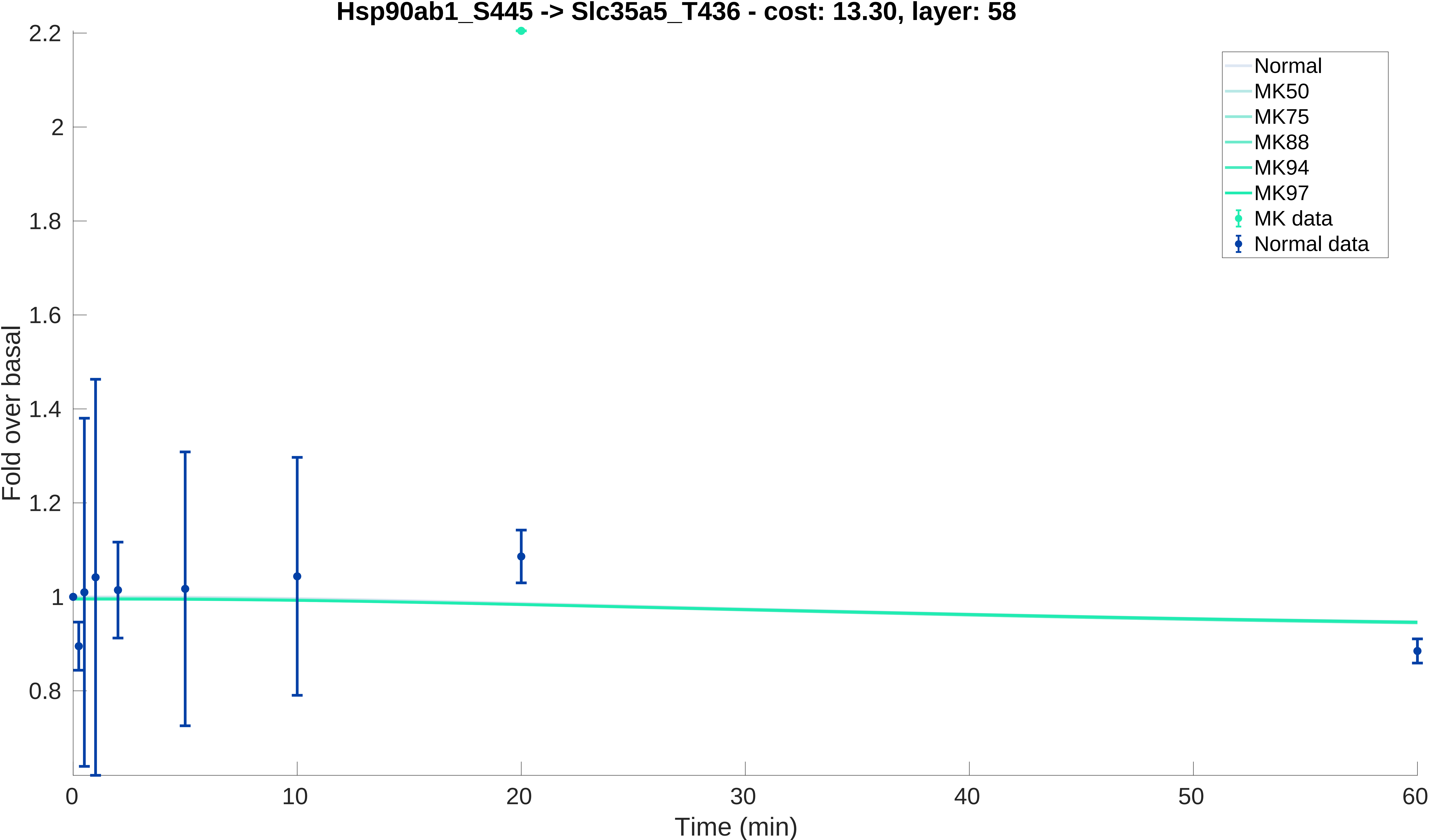

Hsp90ab1_S445 -> Slc35a5_T436 - cost: 13.30, layer: 58
Fold over basal
Time (min)
0.8
1
1.2
1.4
1.6
1.8
2
2.2
0
10
20
30
40
50
60
Normal
MK50
MK75
MK88
MK94
MK97
MK data
Normal data

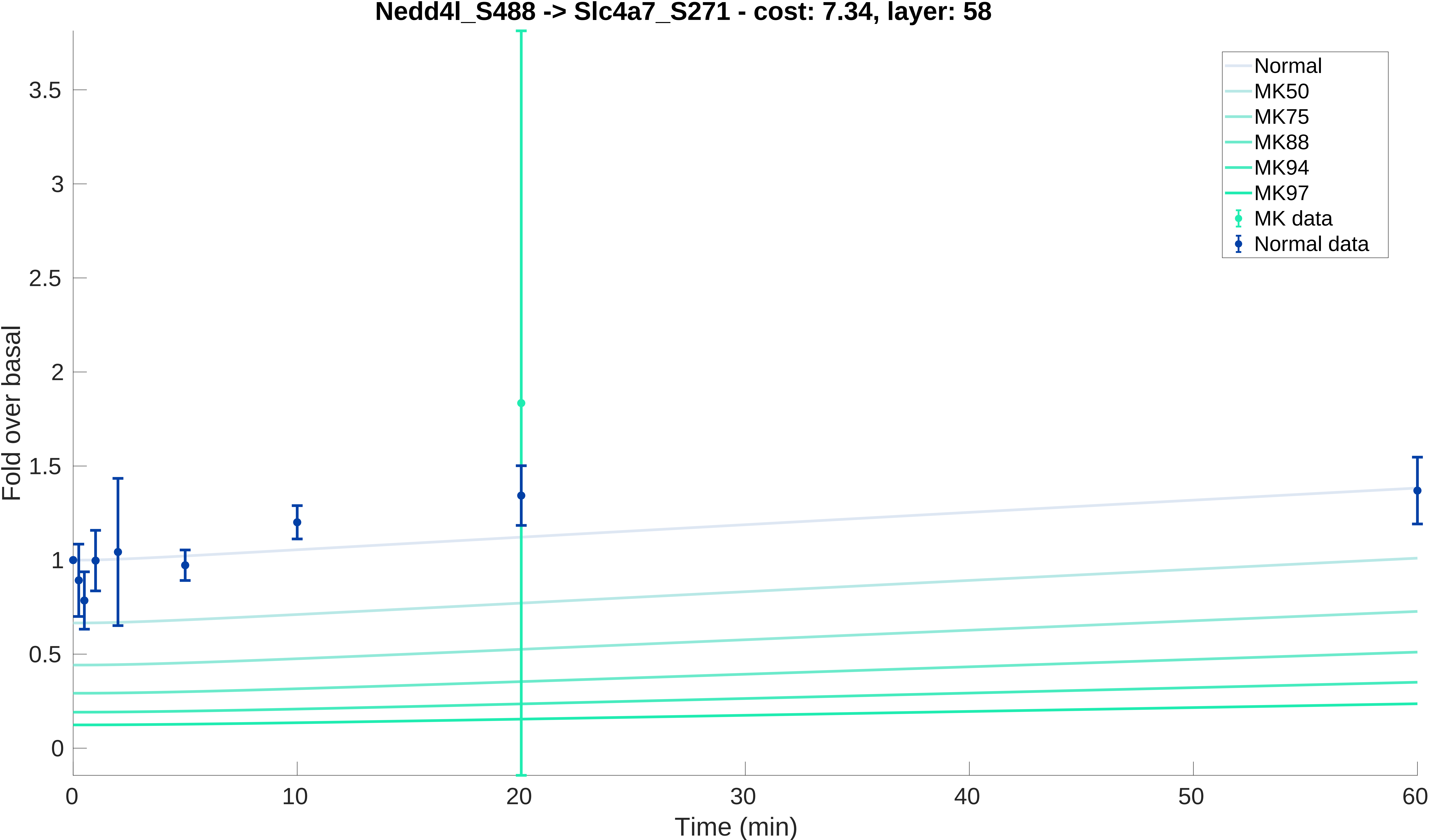

Nedd4l_S488 -> Slc4a7_S271 - cost: 7.34, layer: 58
Fold over basal
3.5
3
2.5
2
1.5
1
0.5
0
0
10
20
30
40
50
60
Time (min)
Normal
MK50
MK75
MK88
MK94
MK97
MK data
Normal data

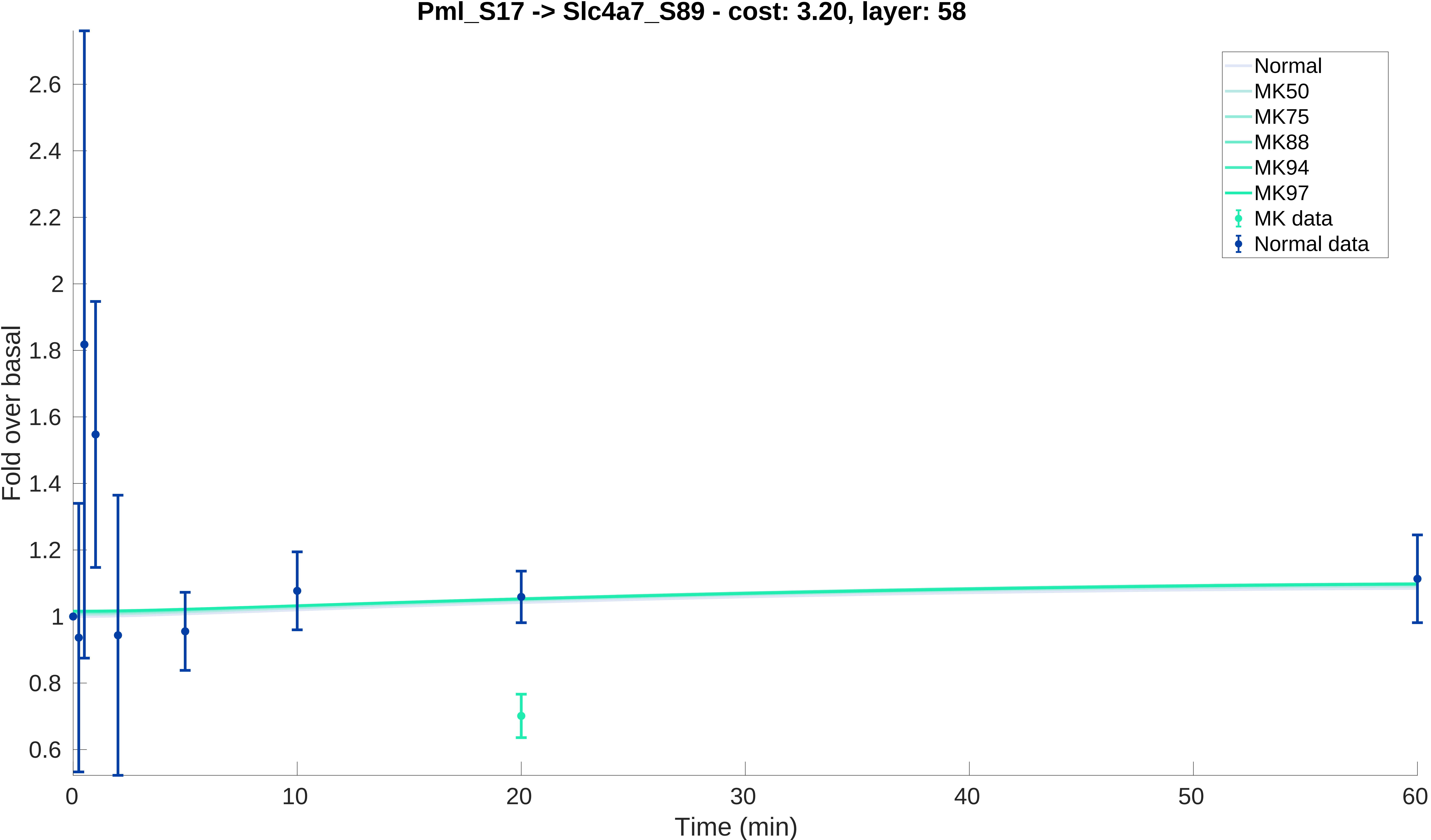

Pml_S17 -> Slc4a7_S89 - cost: 3.20, layer: 58
Fold over basal
Time (min)
2.6
2.4
2.2
2
1.8
1.6
1.4
1.2
1
0.8
0.6
0
10
20
30
40
50
60
Normal
MK50
MK75
MK88
MK94
MK97
MK data
Normal data

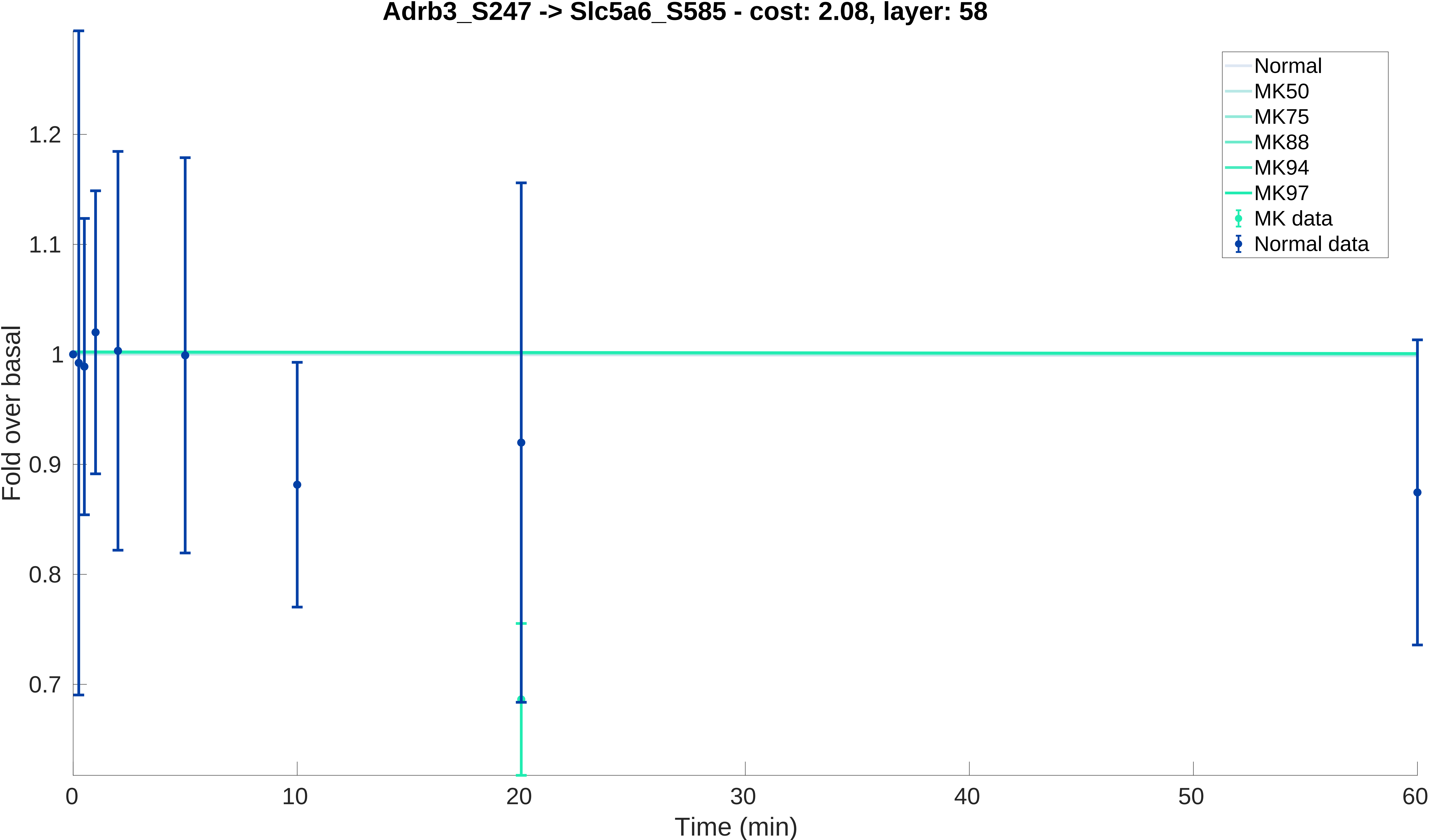

Adrb3_S247 -> Slc5a6_S585 - cost: 2.08, layer: 58
Normal
MK50
MK75
MK88
MK94
MK97
MK data
Normal data
Fold over basal
1.2
1.1
1
0.9
0.8
0.7
0
10
20
30
40
50
60
Time (min)

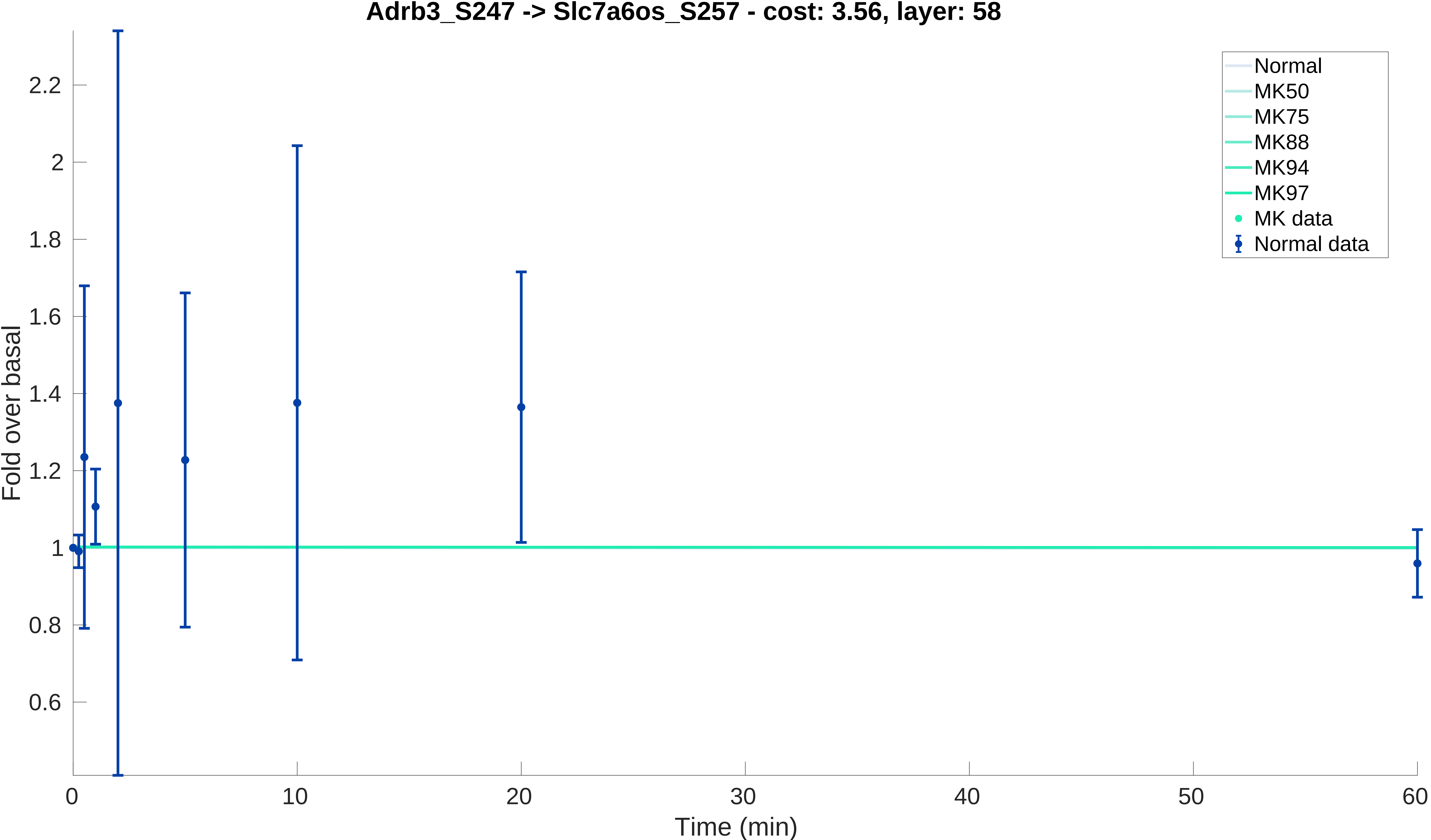
Adrb3_S247 -> Slc7a6os_S257 - cost: 3.56, layer: 58
Fold over basal
Time (min)
0.6
0.8
1
1.2
1.4
1.6
1.8
2
2.2
0
10
20
30
40
50
60
Normal
MK50
MK75
MK88
MK94
MK97
MK data
Normal data

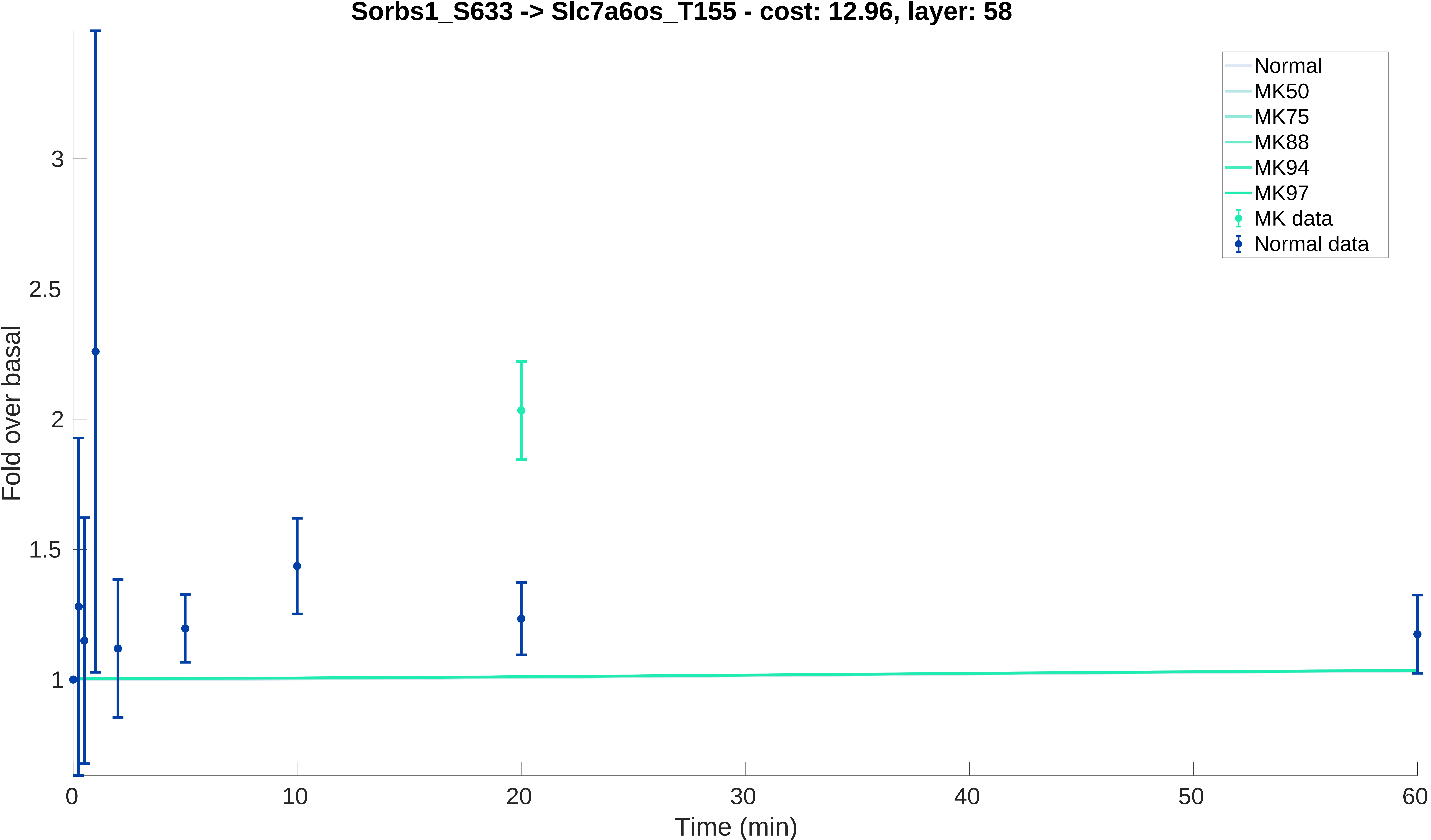

Sorbs1_S633 -> Slc7a6os_T155 - cost: 12.96, layer: 58
Fold over basal
Time (min)
0
10
20
30
40
50
60
1
1.5
2
2.5
3
Normal
MK50
MK75
MK88
MK94
MK97
MK data
Normal data

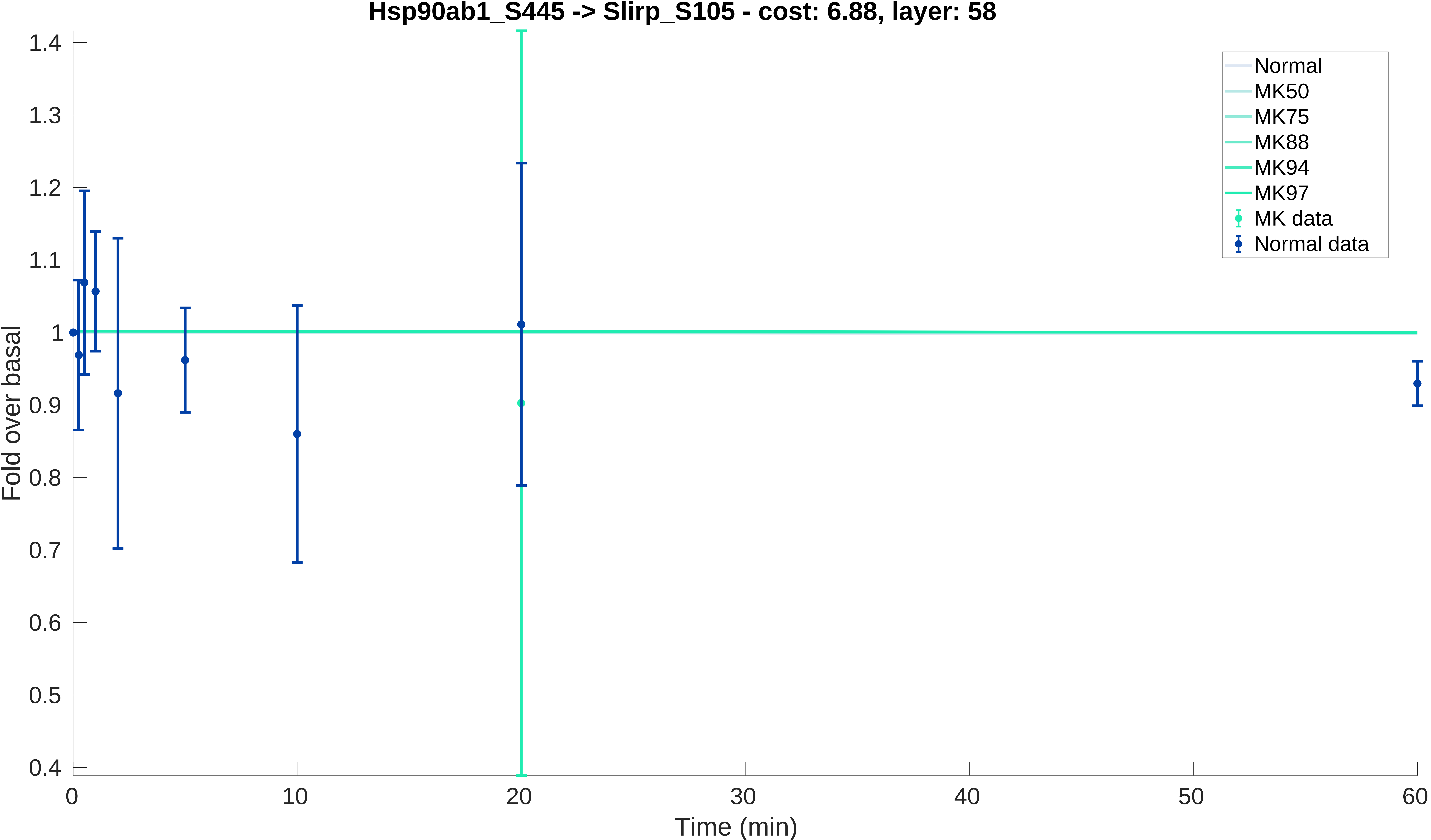

Hsp90ab1_S445 -> Slirp_S105 - cost: 6.88, layer: 58
Fold over basal
Time (min)
0.4
0.5
0.6
0.7
0.8
0.9
1
1.1
1.2
1.3
1.4
0
10
20
30
40
50
60
Normal
MK50
MK75
MK88
MK94
MK97
MK data
Normal data

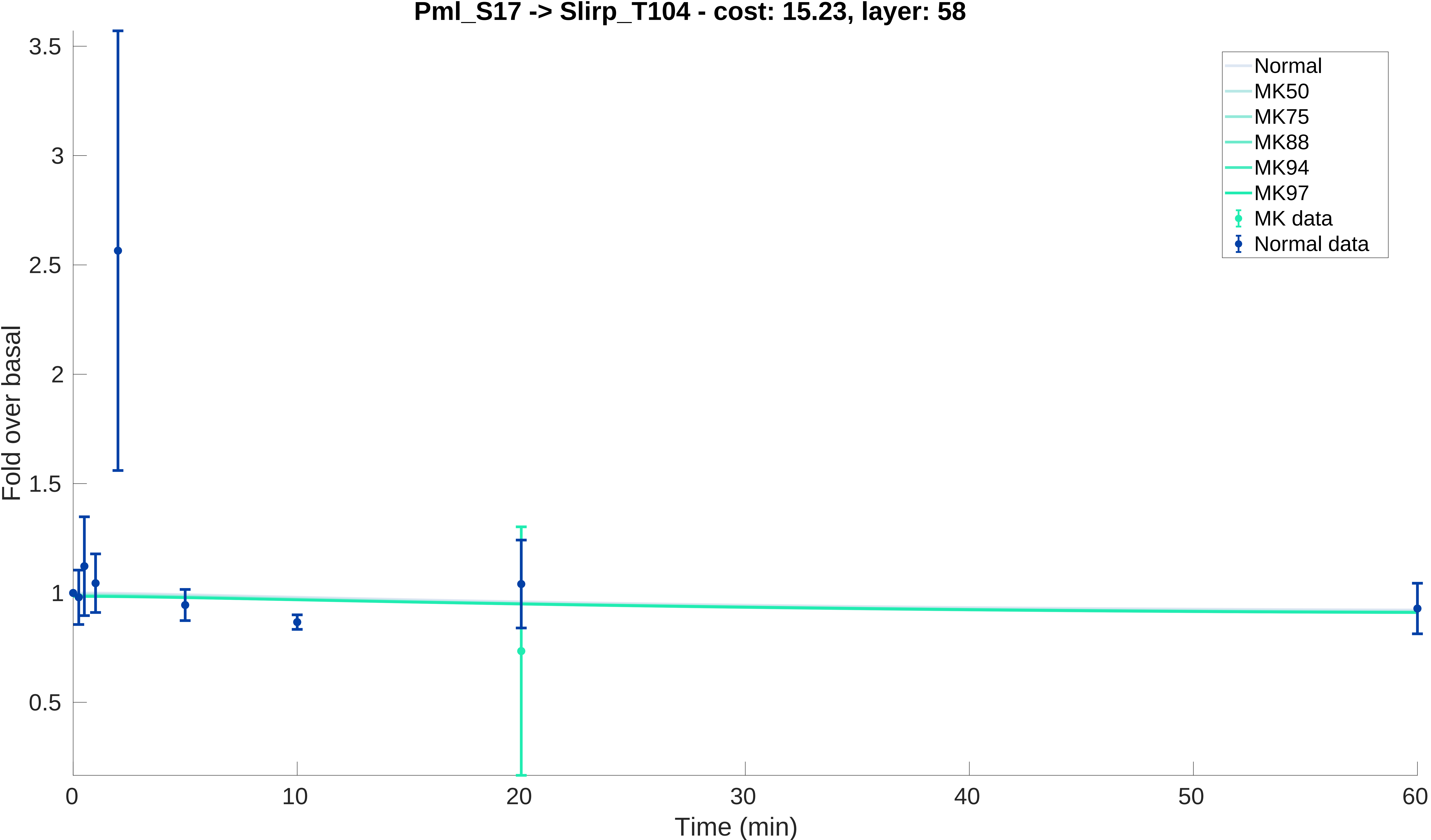

Pml_S17 -> Slirp_T104 - cost: 15.23, layer: 58
Fold over basal
Time (min)
0
10
20
30
40
50
60
0.5
1
1.5
2
2.5
3
3.5
Normal
MK50
MK75
MK88
MK94
MK97
MK data
Normal data

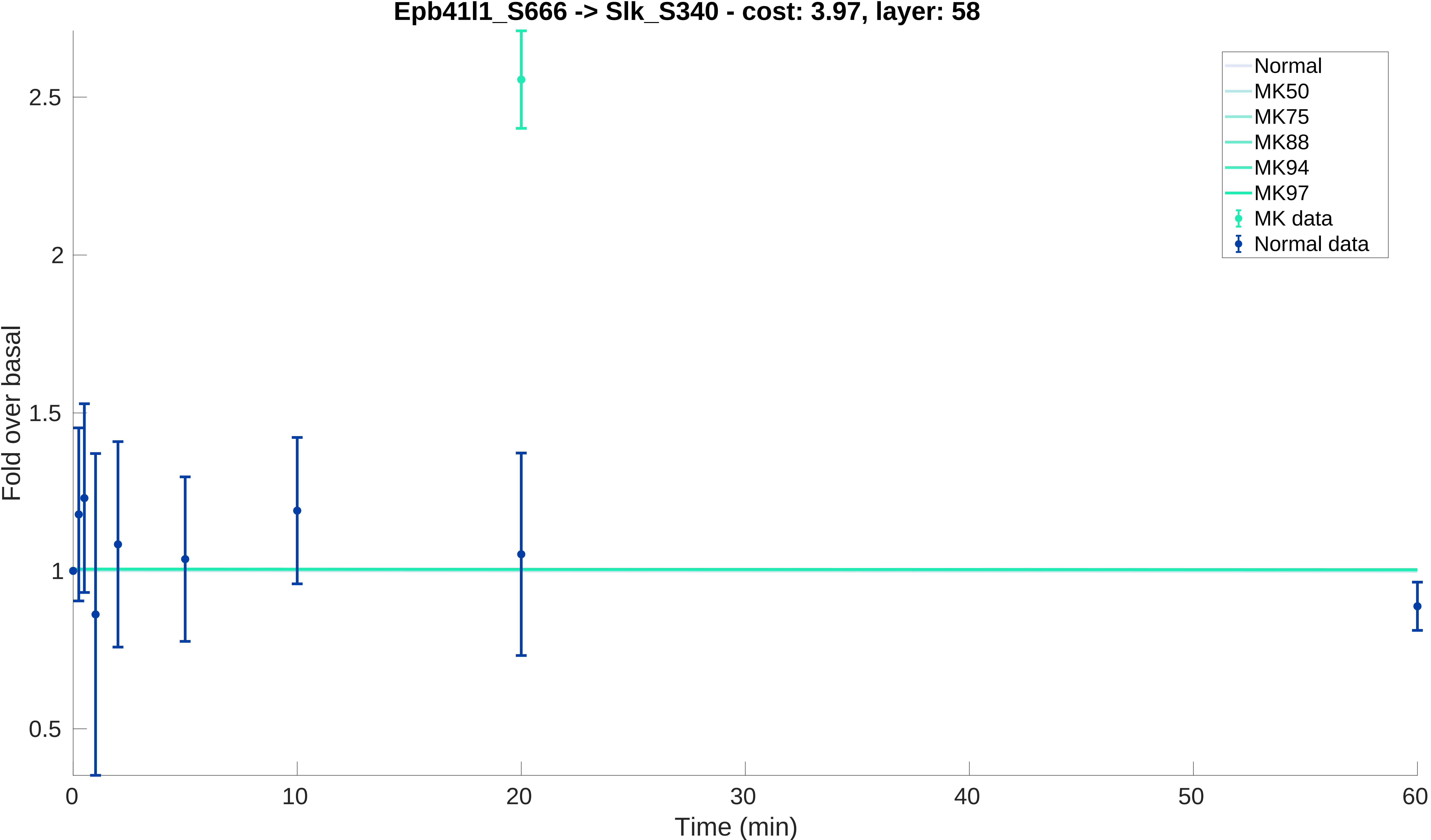

Epb41l1_S666 -> Slk_S340 - cost: 3.97, layer: 58
Normal
MK50
MK75
MK88
MK94
MK97
MK data
Normal data
Fold over basal
0.5
1
1.5
2
2.5
0
10
20
30
40
50
60
Time (min)

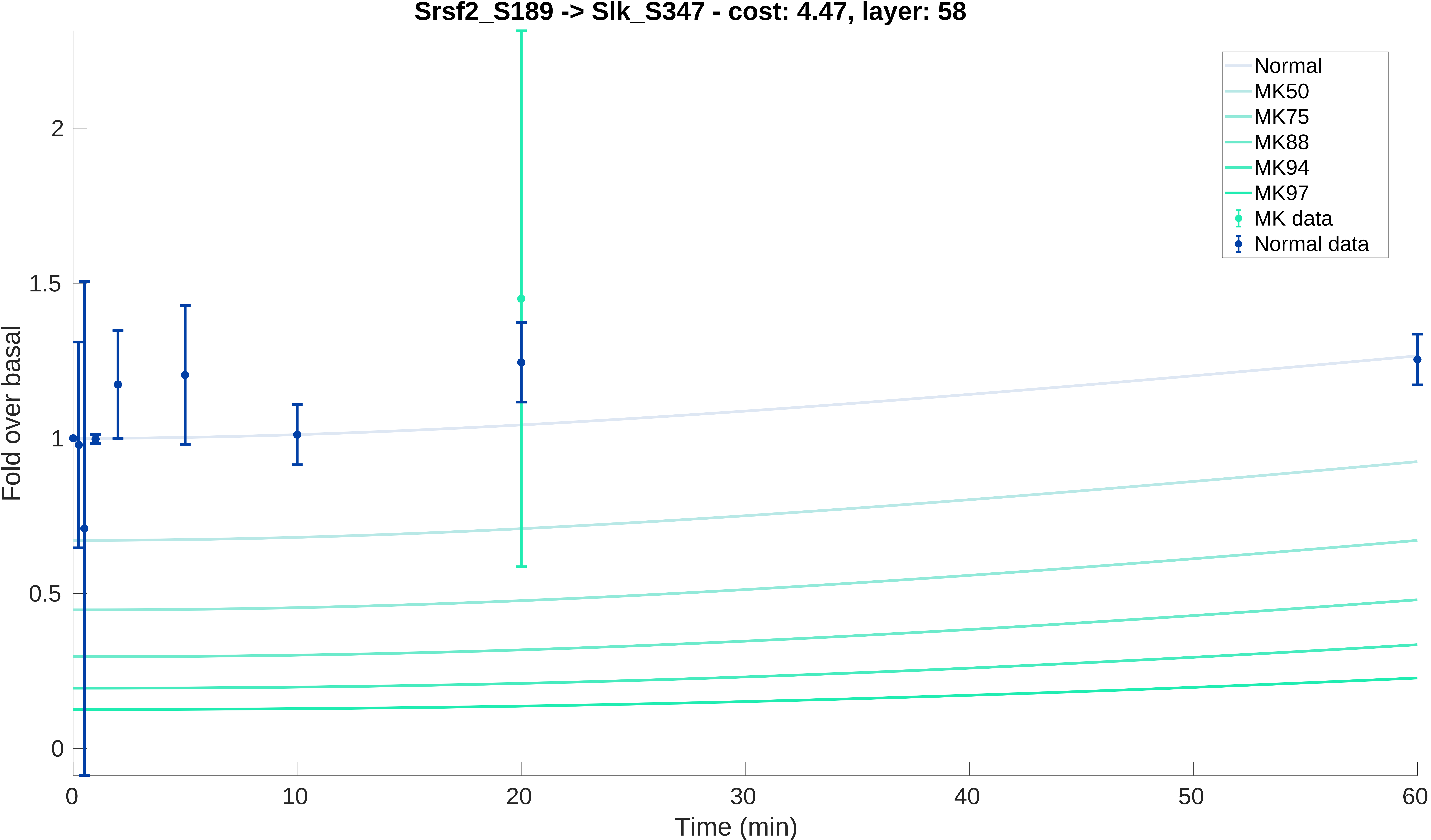
Srsf2_S189 -> Slk_S347 - cost: 4.47, layer: 58
Normal
MK50
MK75
MK88
MK94
MK97
MK data
Normal data
Fold over basal
0
0.5
1
1.5
2
0
10
20
30
40
50
60
Time (min)

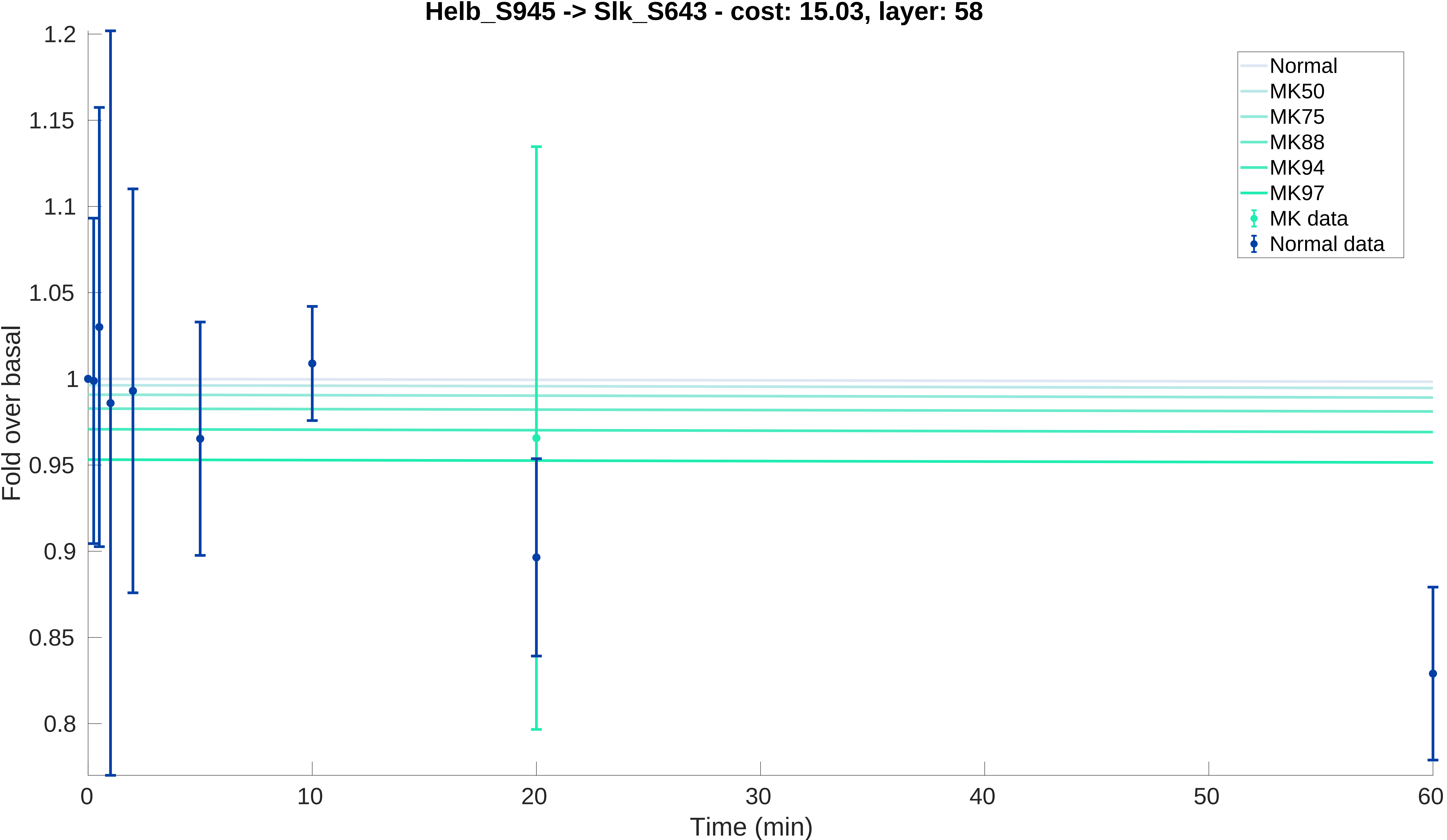

Helb_S945 -> Slk_S643 - cost: 15.03, layer: 58
Fold over basal
1.2
1.15
1.1
1.05
1
0.95
0.9
0.85
0.8
0
10
20
30
40
50
60
Time (min)
Normal
MK50
MK75
MK88
MK94
MK97
MK data
Normal data

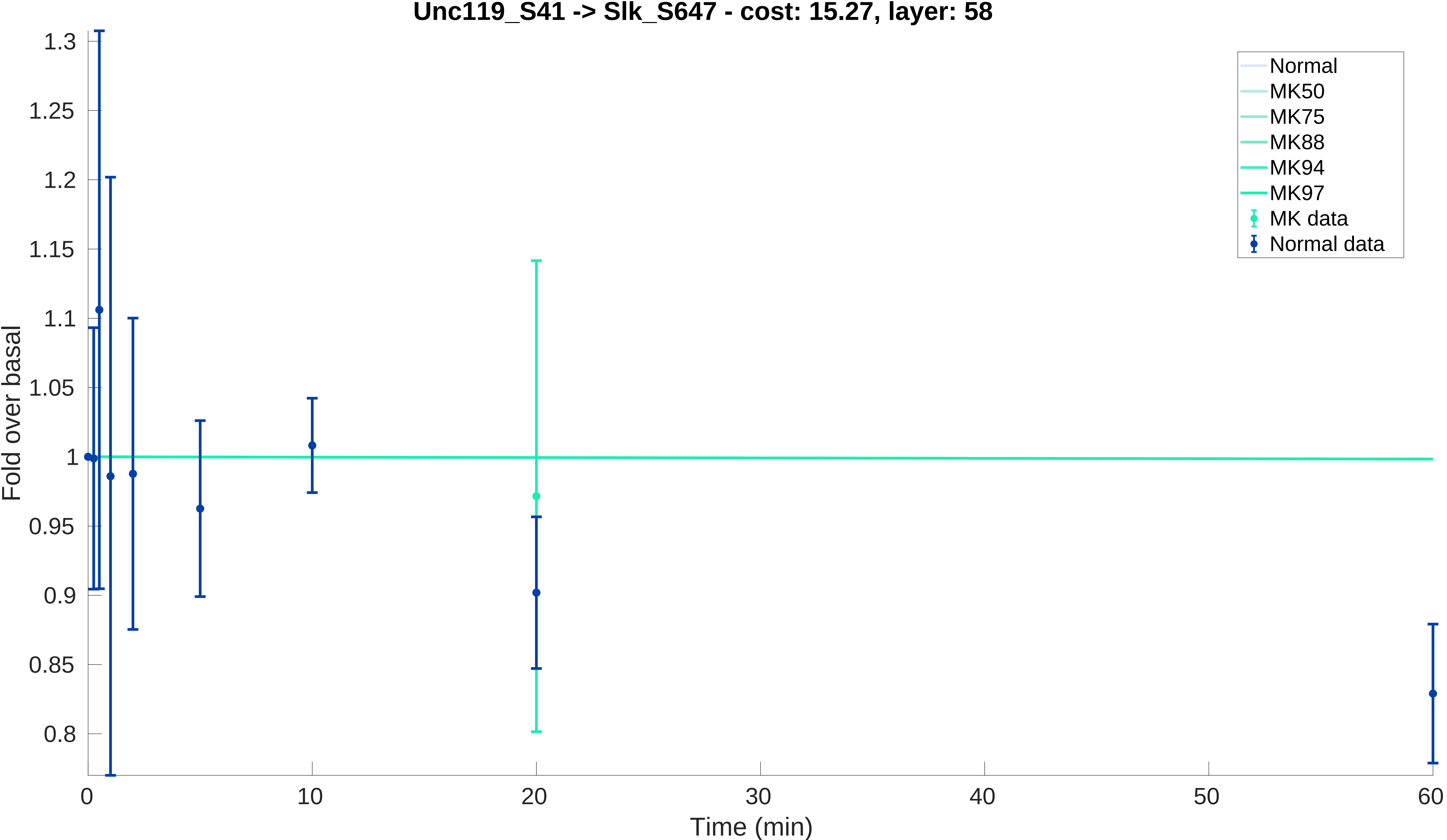

Unc119_S41 -> Slk_S647 - cost: 15.27, layer: 58
Fold over basal
1.3
1.25
1.2
1.15
1.1
1.05
1
0.95
0.9
0.85
0.8
0
10
20
30
40
50
60
Time (min)
Normal
MK50
MK75
MK88
MK94
MK97
MK data
Normal data

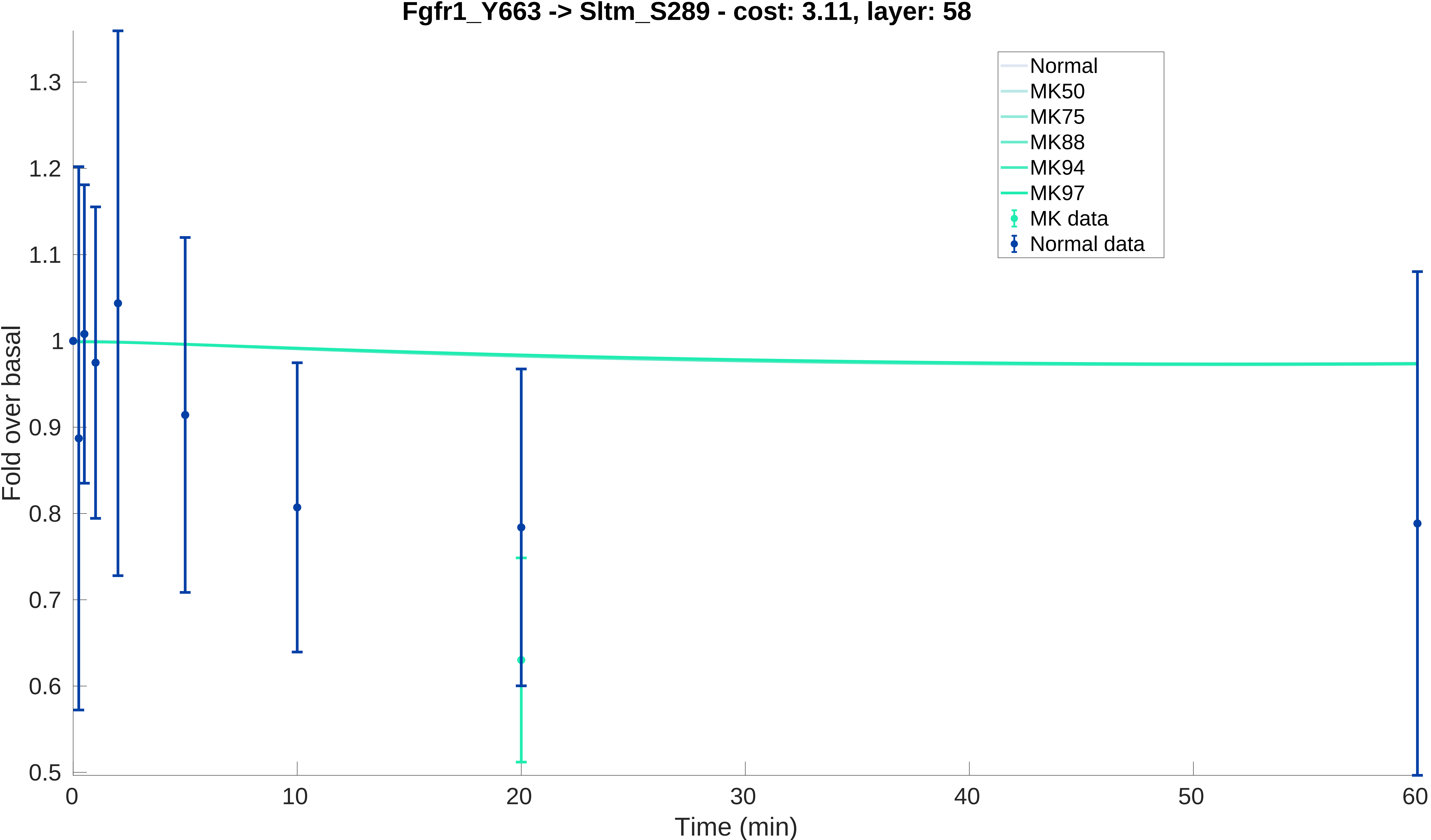

Fgfr1_Y663 -> Sltm_S289 - cost: 3.11, layer: 58
Normal
MK50
MK75
MK88
MK94
MK97
MK data
Normal data
Fold over basal
1.3
1.2
1.1
1
0.9
0.8
0.7
0.6
0.5
0
10
20
30
40
50
60
Time (min)

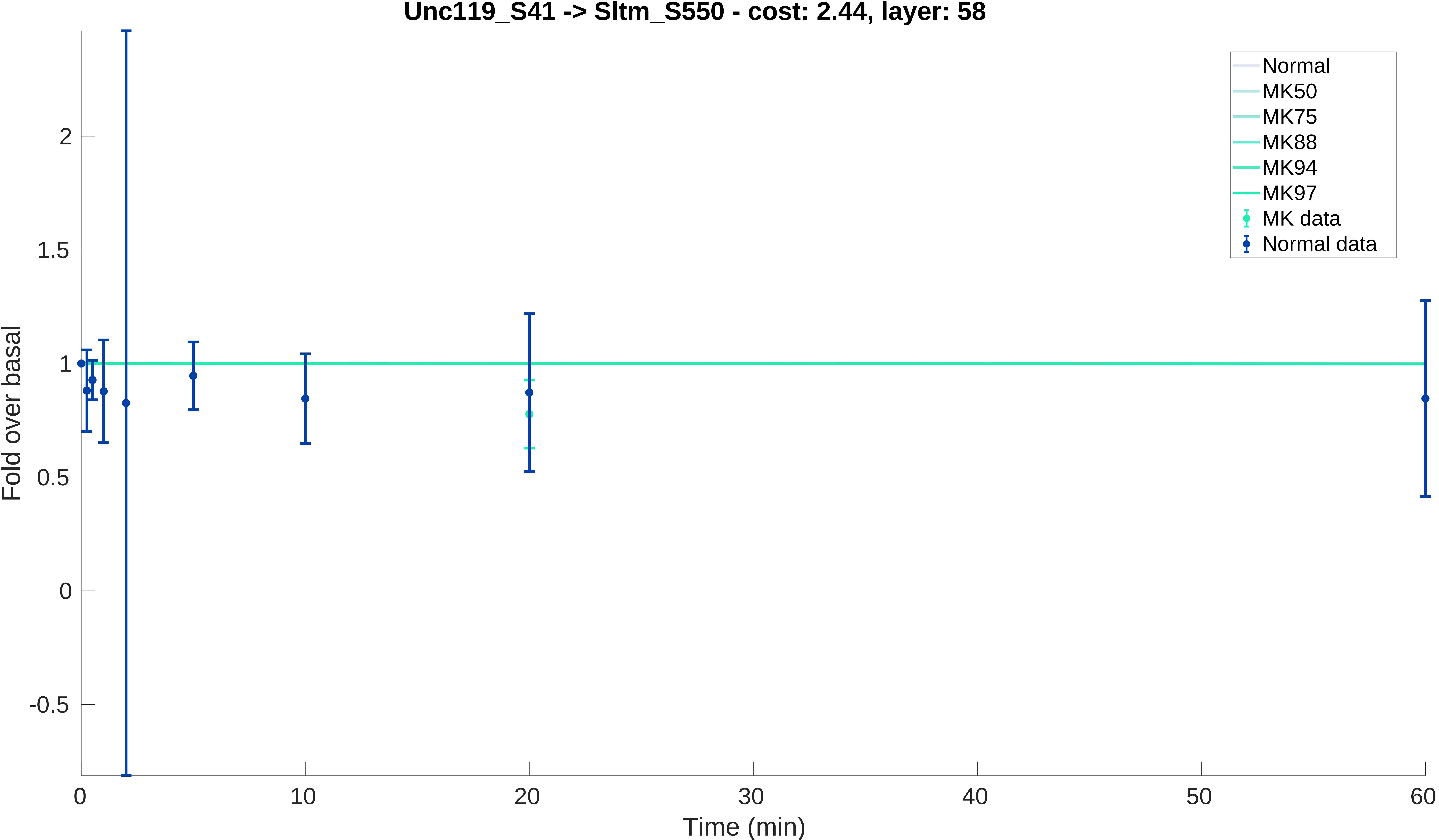

Unc119_S41 -> Sltm_S550 - cost: 2.44, layer: 58
Normal
MK50
MK75
MK88
MK94
MK97
MK data
Normal data
Fold over basal
2
1.5
1
0.5
0
-0.5
0
10
20
30
40
50
60
Time (min)

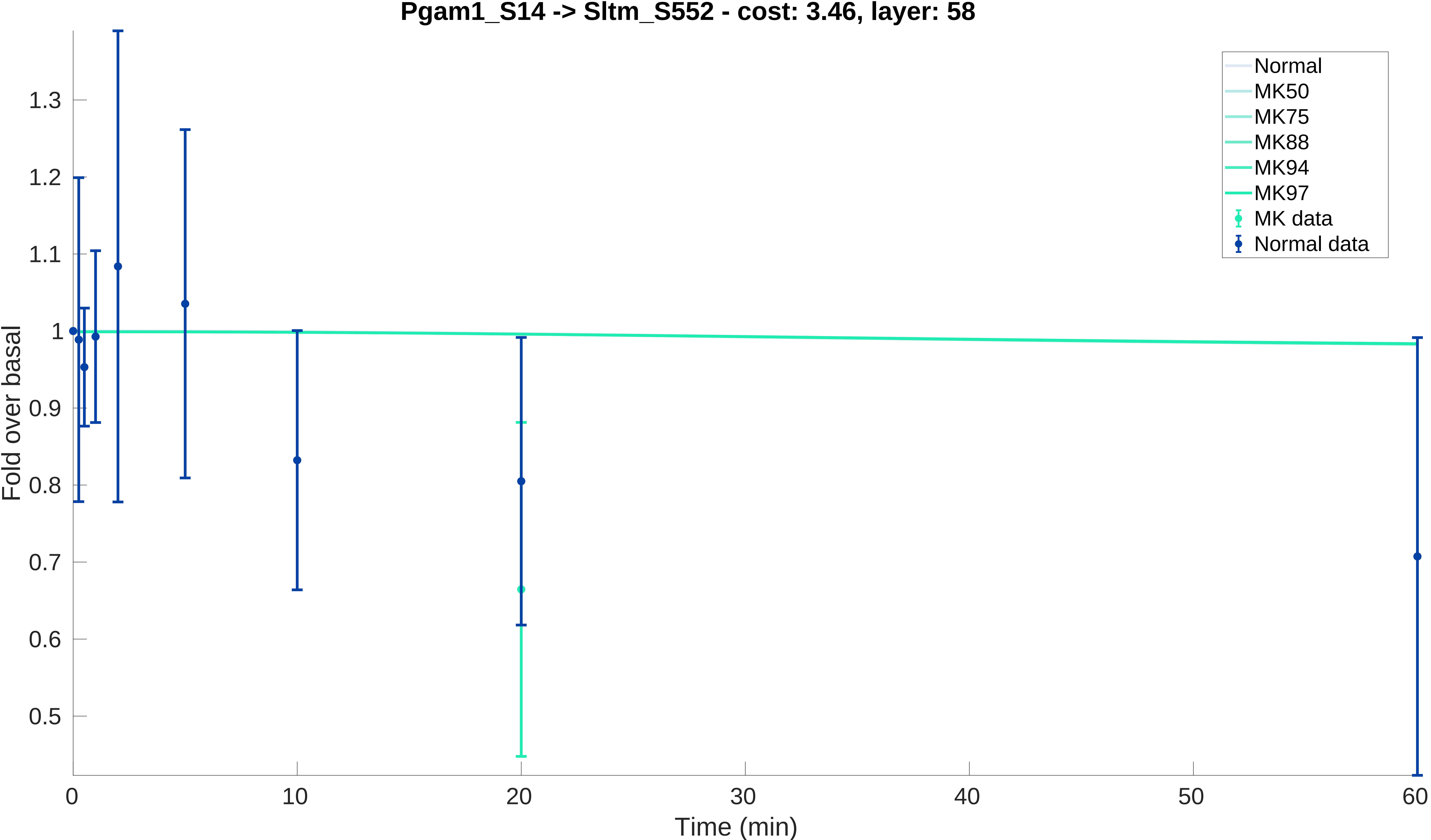

Pgam1_S14 -> Sltm_S552 - cost: 3.46, layer: 58
Normal
MK50
MK75
MK88
MK94
MK97
MK data
Normal data
Fold over basal
Time (min)
1.3
1.2
1.1
1
0.9
0.8
0.7
0.6
0.5
0
10
20
30
40
50
60

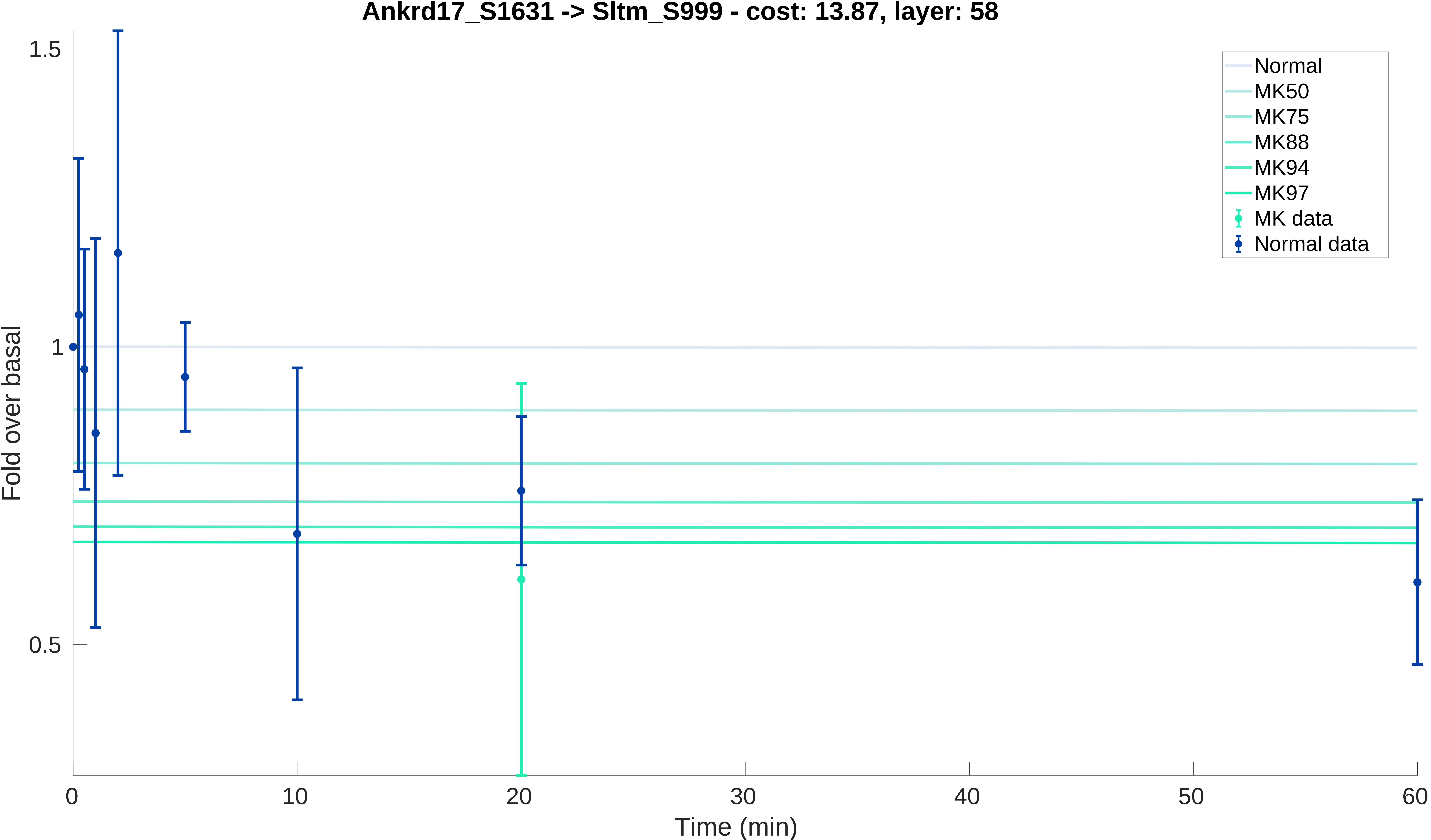

Ankrd17_S1631 -> Sltm_S999 - cost: 13.87, layer: 58
Fold over basal
1.5
1
0.5
Time (min)
0
10
20
30
40
50
60
Normal
MK50
MK75
MK88
MK94
MK97
MK data
Normal data

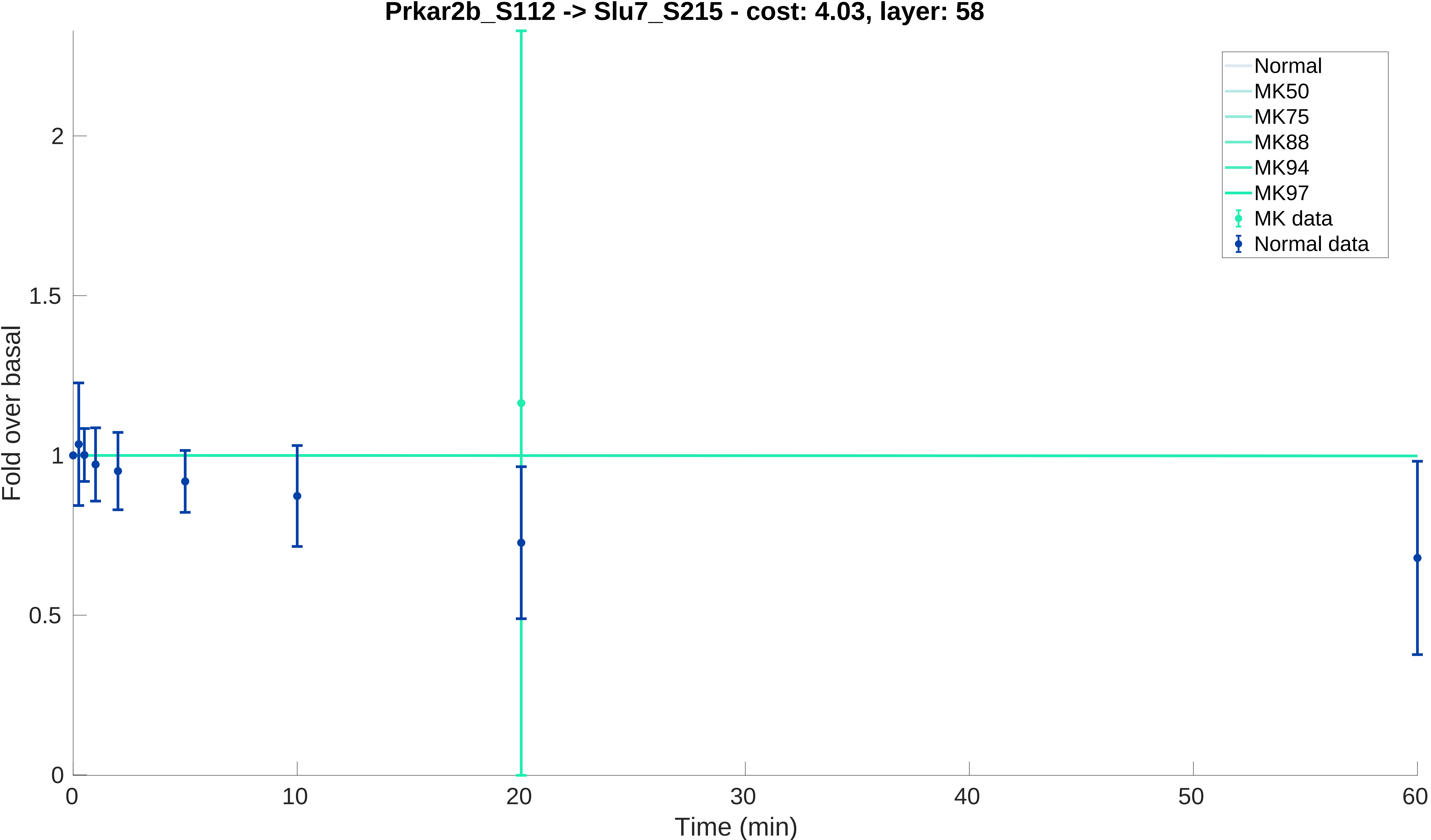

Prrkar2b_S112 -> Slu7_S215 - cost: 4.03, layer: 58
Fold over basal
Time (min)
0
0.5
1
1.5
2
0
10
20
30
40
50
60
Normal
MK50
MK75
MK88
MK94
MK97
MK data
Normal data

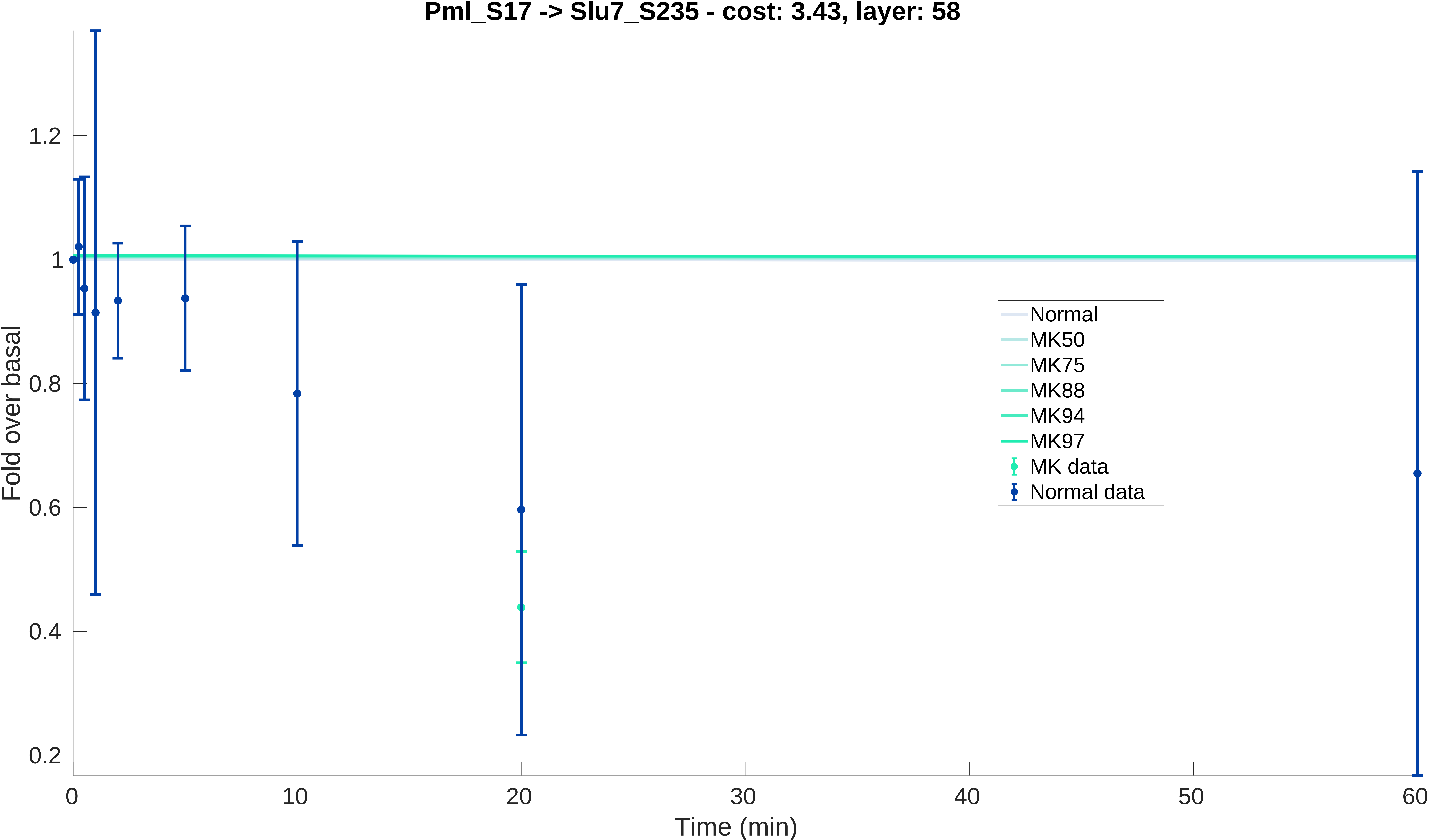

Pml_S17 -> Slu7_S235 - cost: 3.43, layer: 58
Fold over basal
Time (min)
0.2
0.4
0.6
0.8
1
1.2
0
10
20
30
40
50
60
Normal
MK50
MK75
MK88
MK94
MK97
MK data
Normal data

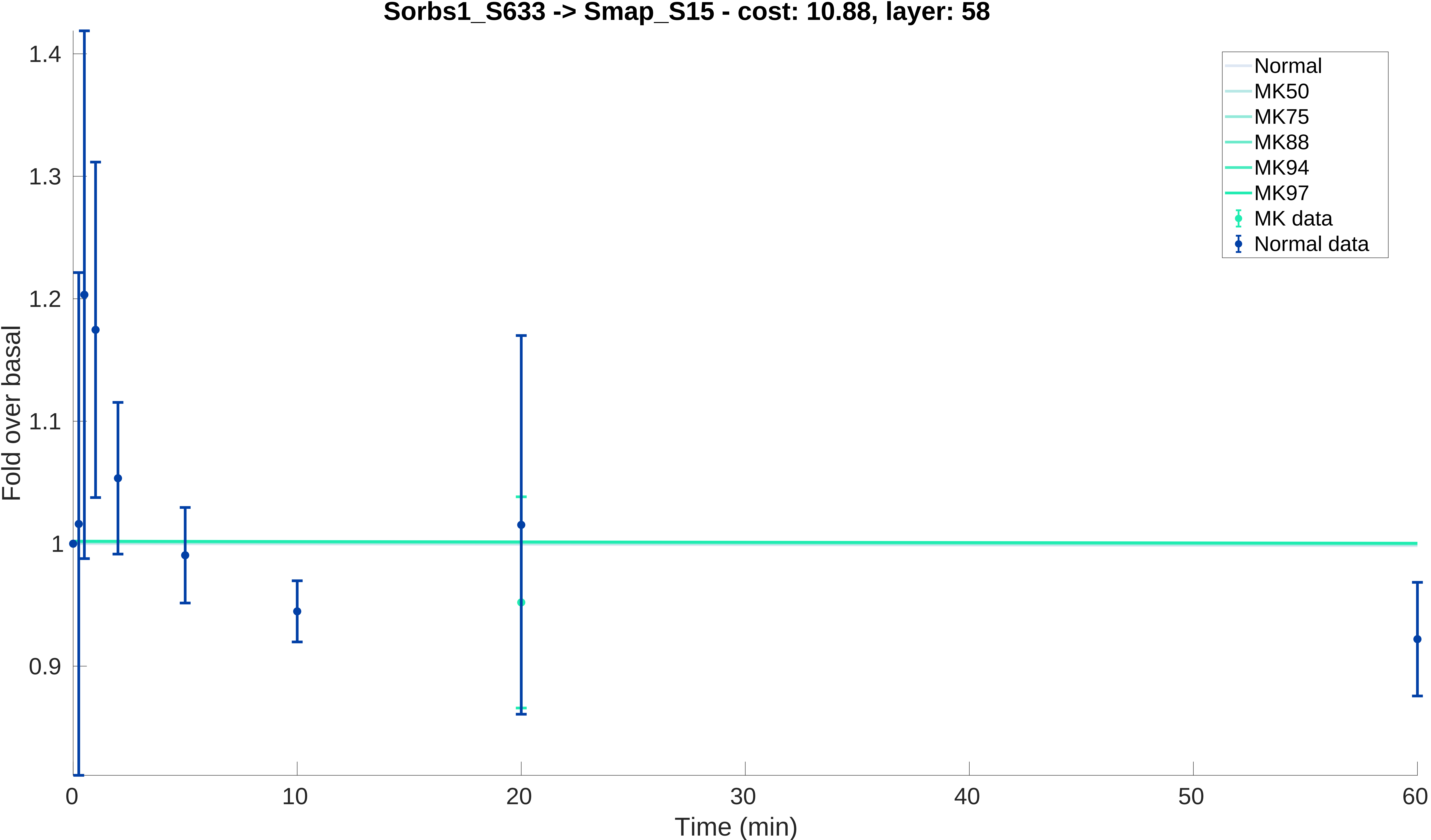

Sorbs1_S633 -> Smap_S15 - cost: 10.88, layer: 58
Fold over basal
1.4
1.3
1.2
1.1
1
0.9
0
10
20
30
40
50
60
Time (min)
Normal
MK50
MK75
MK88
MK94
MK97
MK data
Normal data

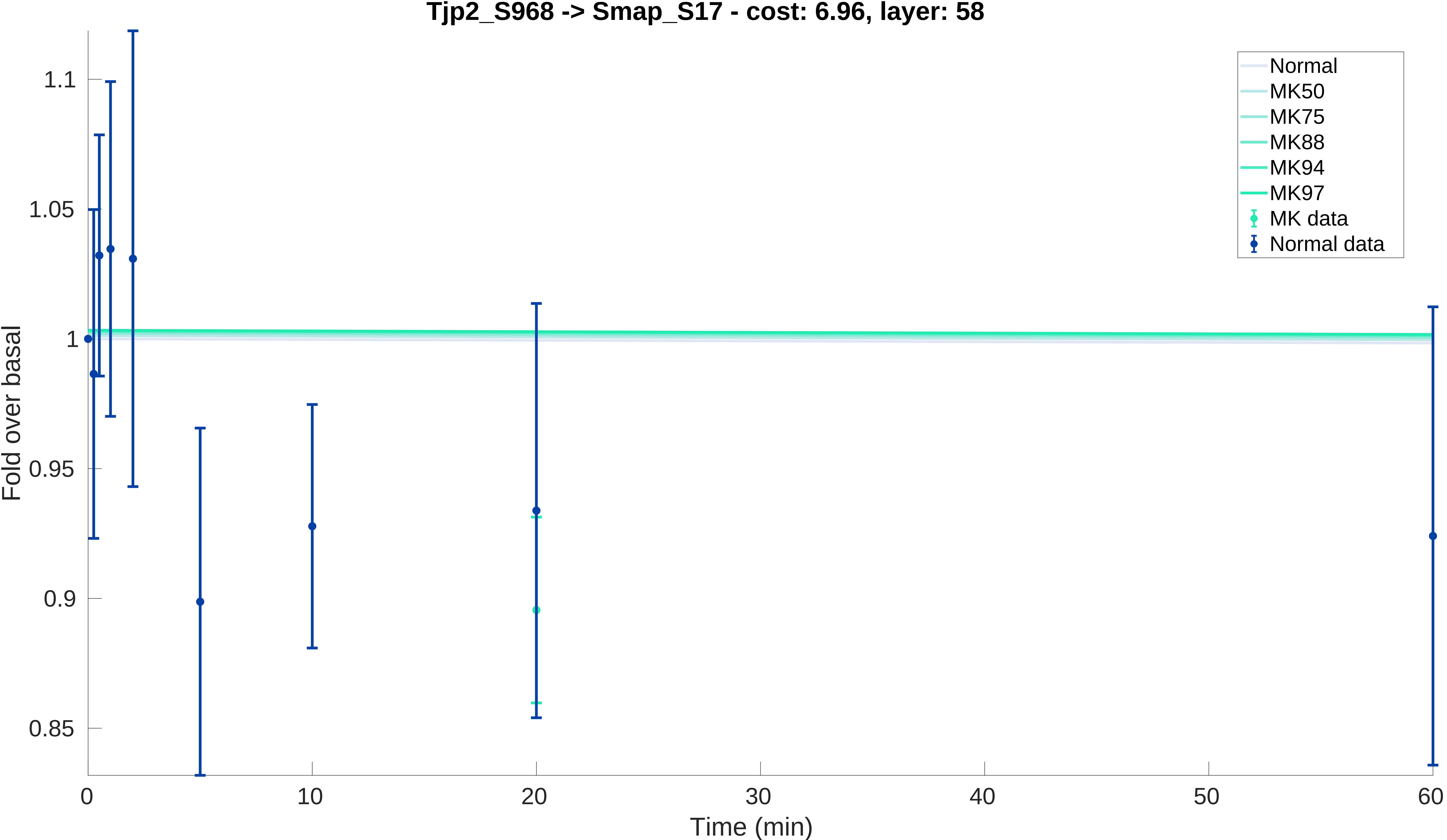
Tjp2_S968 -> Smap_S17 - cost: 6.96, layer: 58
Fold over basal
1.1
1.05
1
0.95
0.9
0.85
0
10
20
30
40
50
60
Time (min)
Normal
MK50
MK75
MK88
MK94
MK97
MK data
Normal data

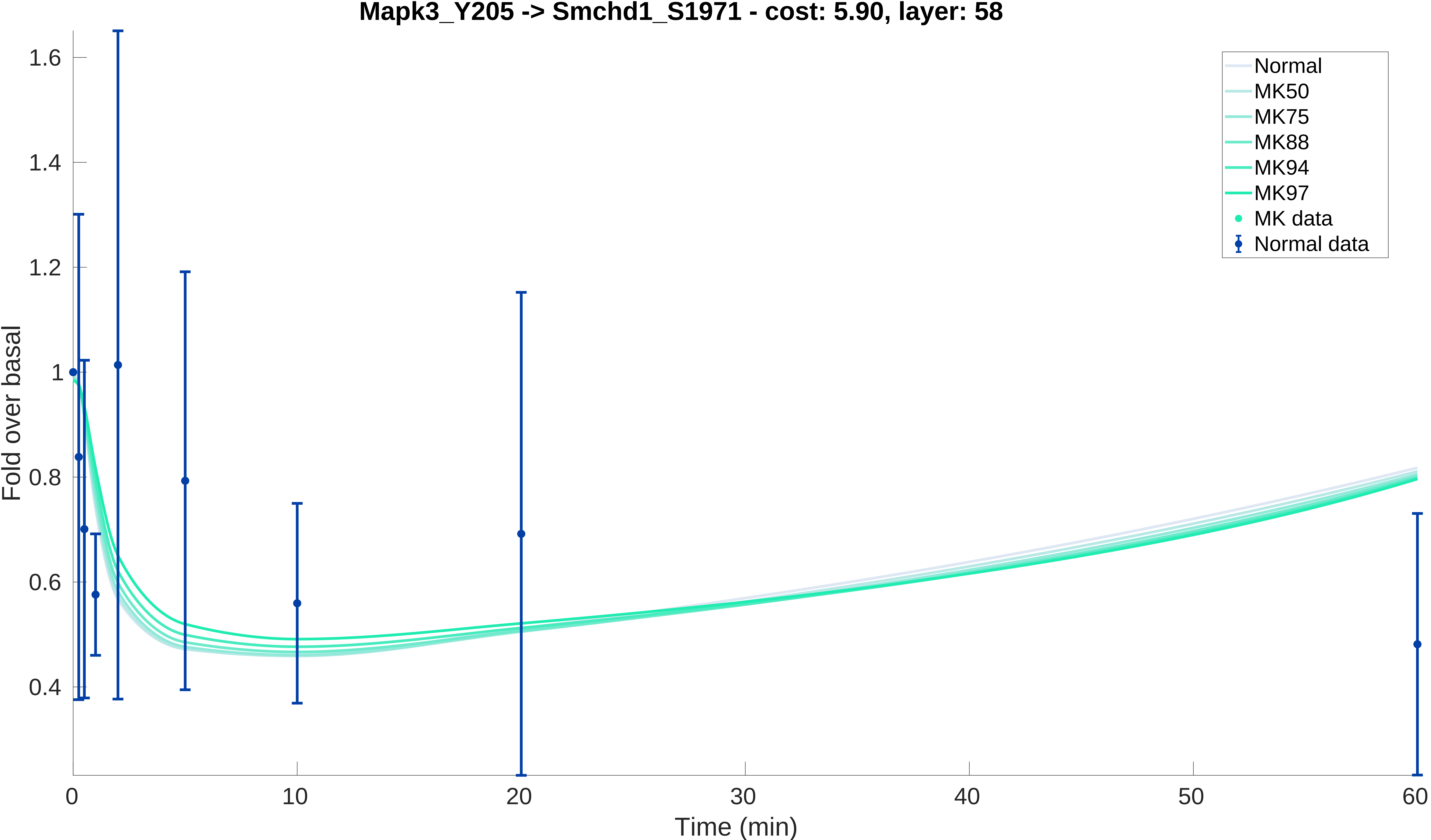

Mapk3_Y205 -> Smchd1_S1971 - cost: 5.90, layer: 58
Fold over basal
Time (min)
1.6
1.4
1.2
1
0.8
0.6
0.4
0
10
20
30
40
50
60
Normal
MK50
MK75
MK88
MK94
MK97
MK data
Normal data

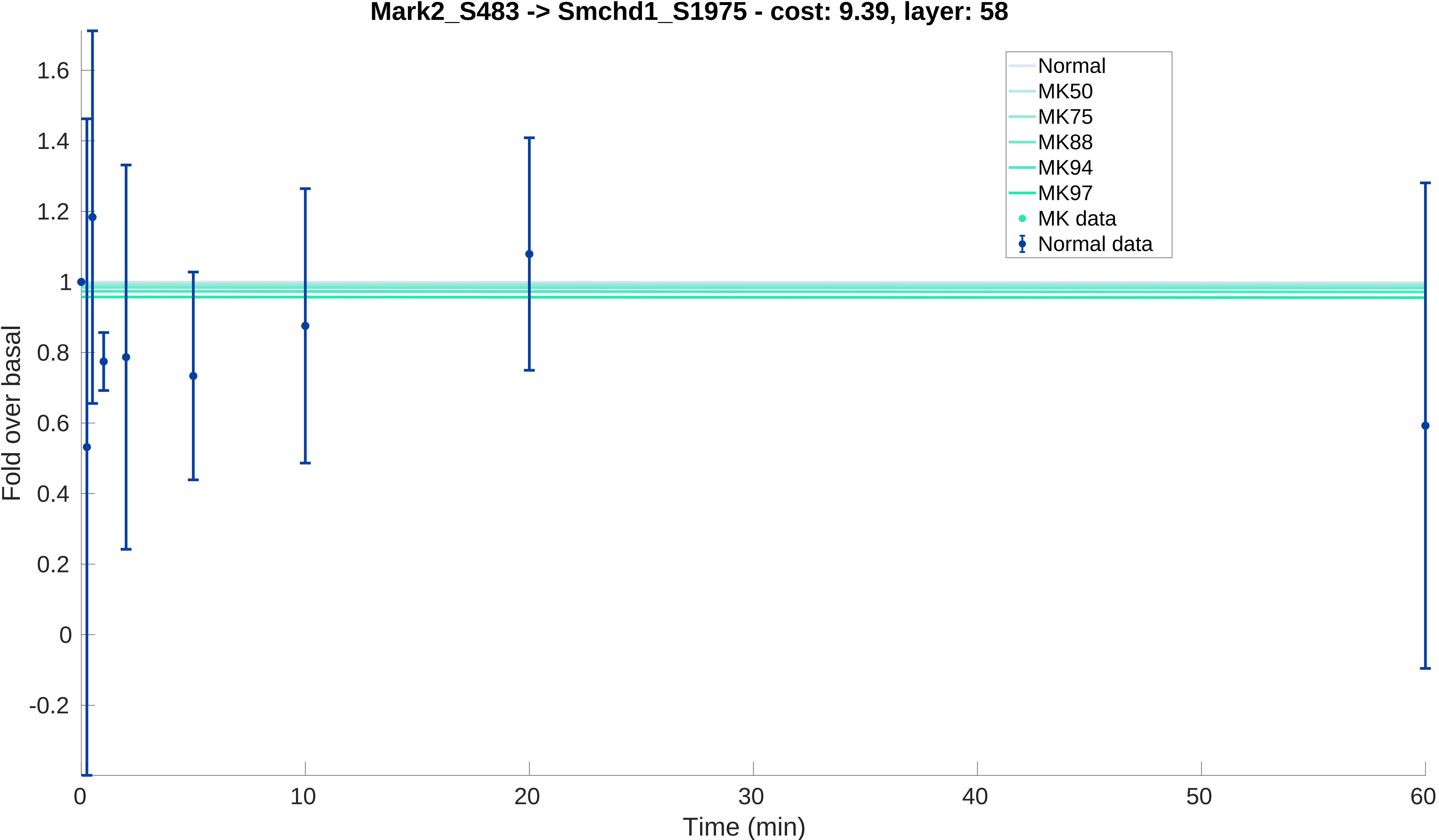

Mark2_S483 -> Smchd1_S1975 - cost: 9.39, layer: 58
Normal
MK50
MK75
MK88
MK94
MK97
MK data
Normal data
Fold over basal
1.6
1.4
1.2
1
0.8
0.6
0.4
0.2
0
-0.2
0
10
20
30
40
50
60
Time (min)

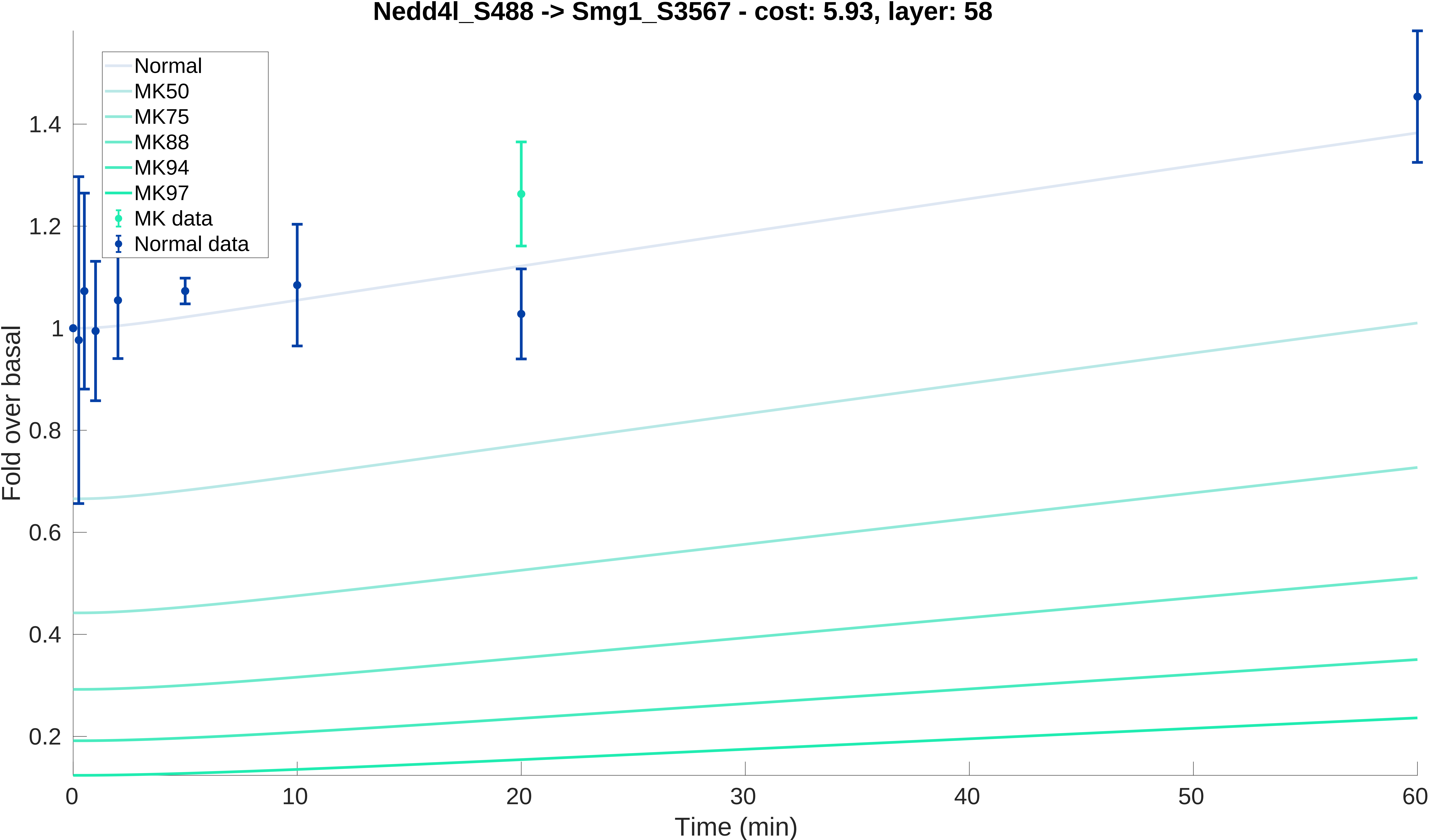

Nedd4l_S488 -> Smg1_S3567 - cost: 5.93, layer: 58
Normal
MK50
MK75
MK88
MK94
MK97
MK data
Normal data
Fold over basal
0.2
0.4
0.6
0.8
1
1.2
1.4
0
10
20
30
40
50
60
Time (min)

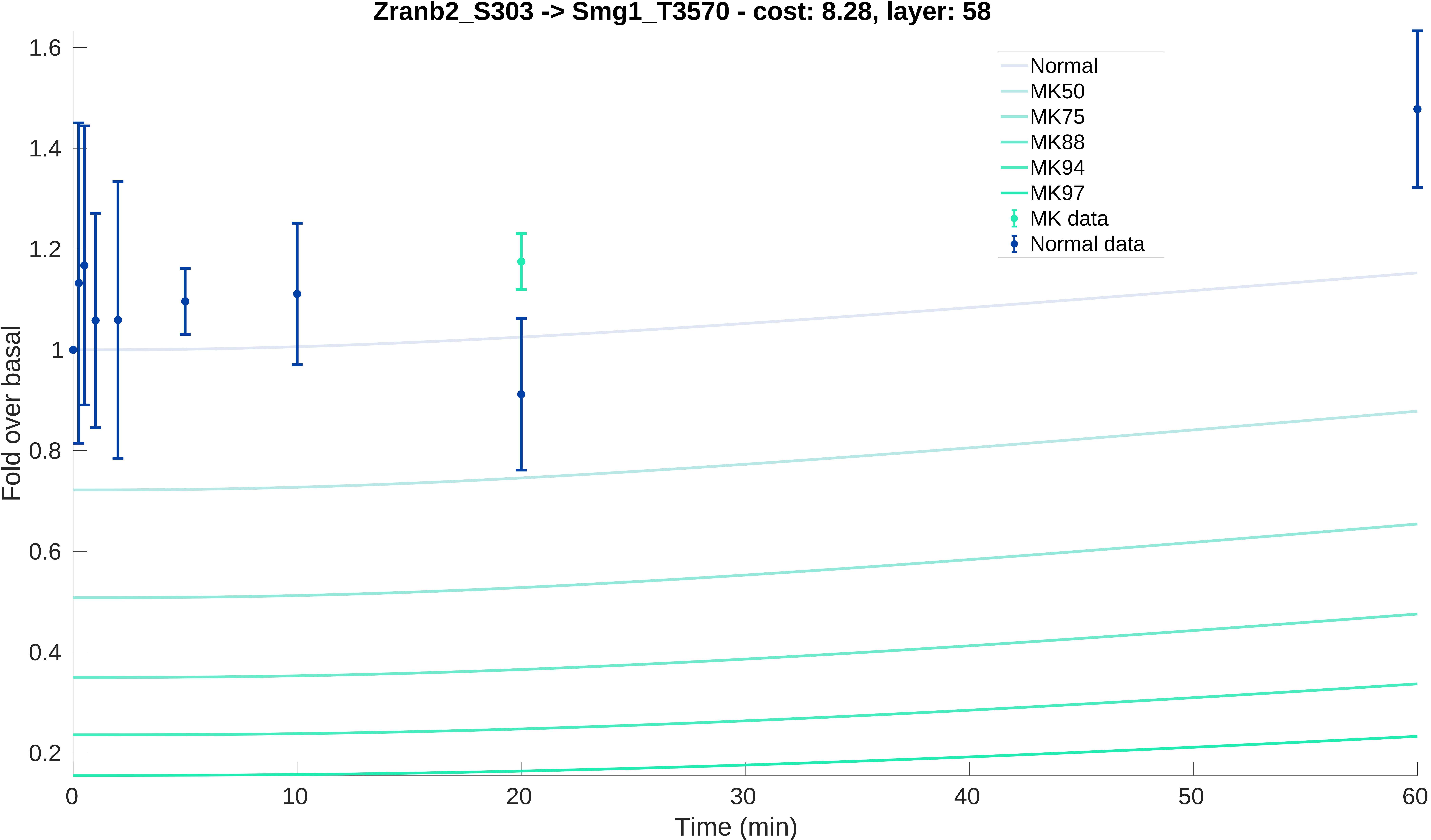
Zranb2_S303 -> Smg1_T3570 - cost: 8.28, layer: 58
Fold over basal
Time (min)
0
10
20
30
40
50
60
0.2
0.4
0.6
0.8
1
1.2
1.4
1.6
Normal
MK50
MK75
MK88
MK94
MK97
MK data
Normal data

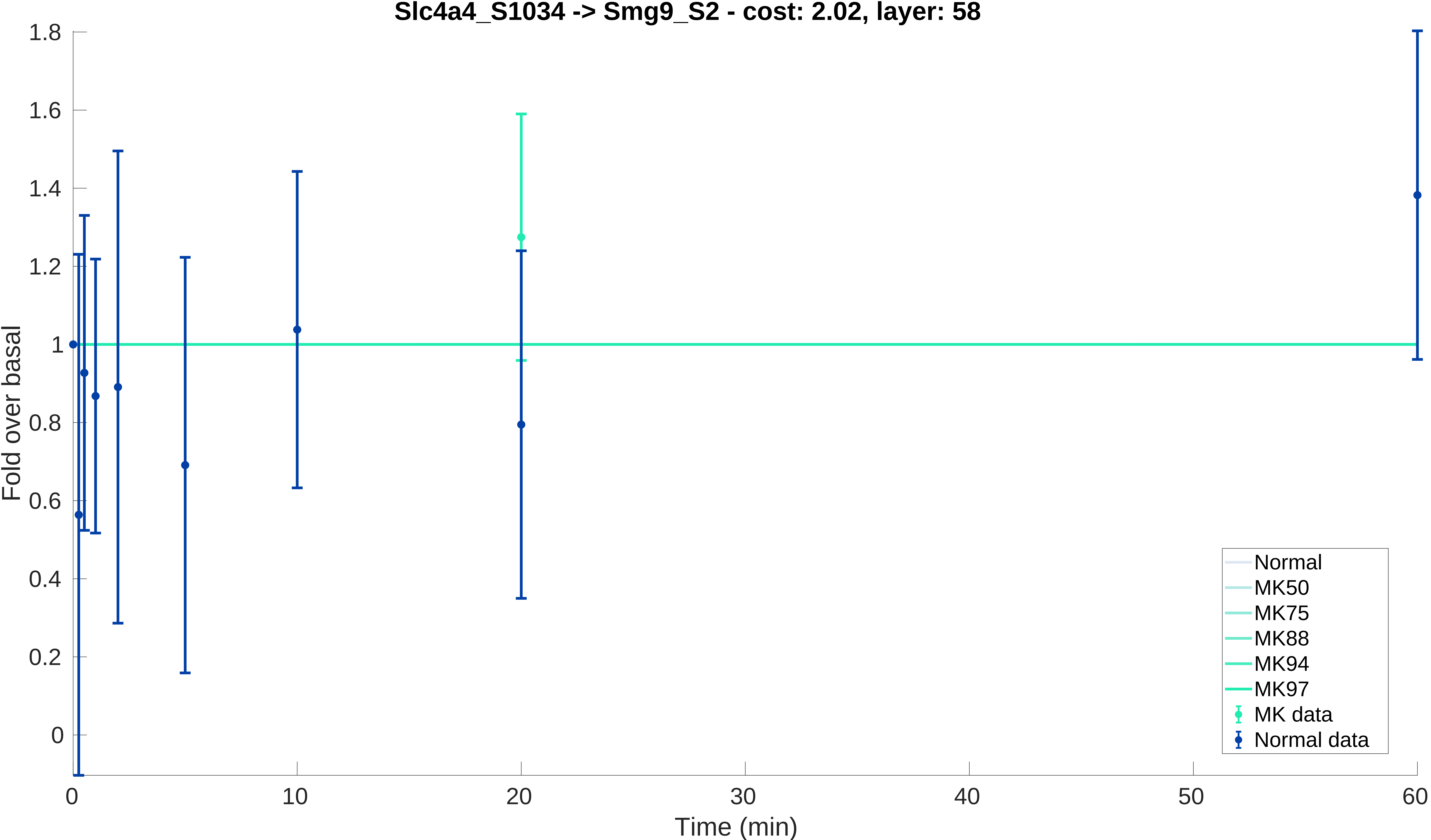
Slc4a4_S1034 -> Smg9_S2 - cost: 2.02, layer: 58
Fold over basal
Time (min)
1.8
1.6
1.4
1.2
1
0.8
0.6
0.4
0.2
0
10
20
30
40
50
60
Normal
MK50
MK75
MK88
MK94
MK97
MK data
Normal data

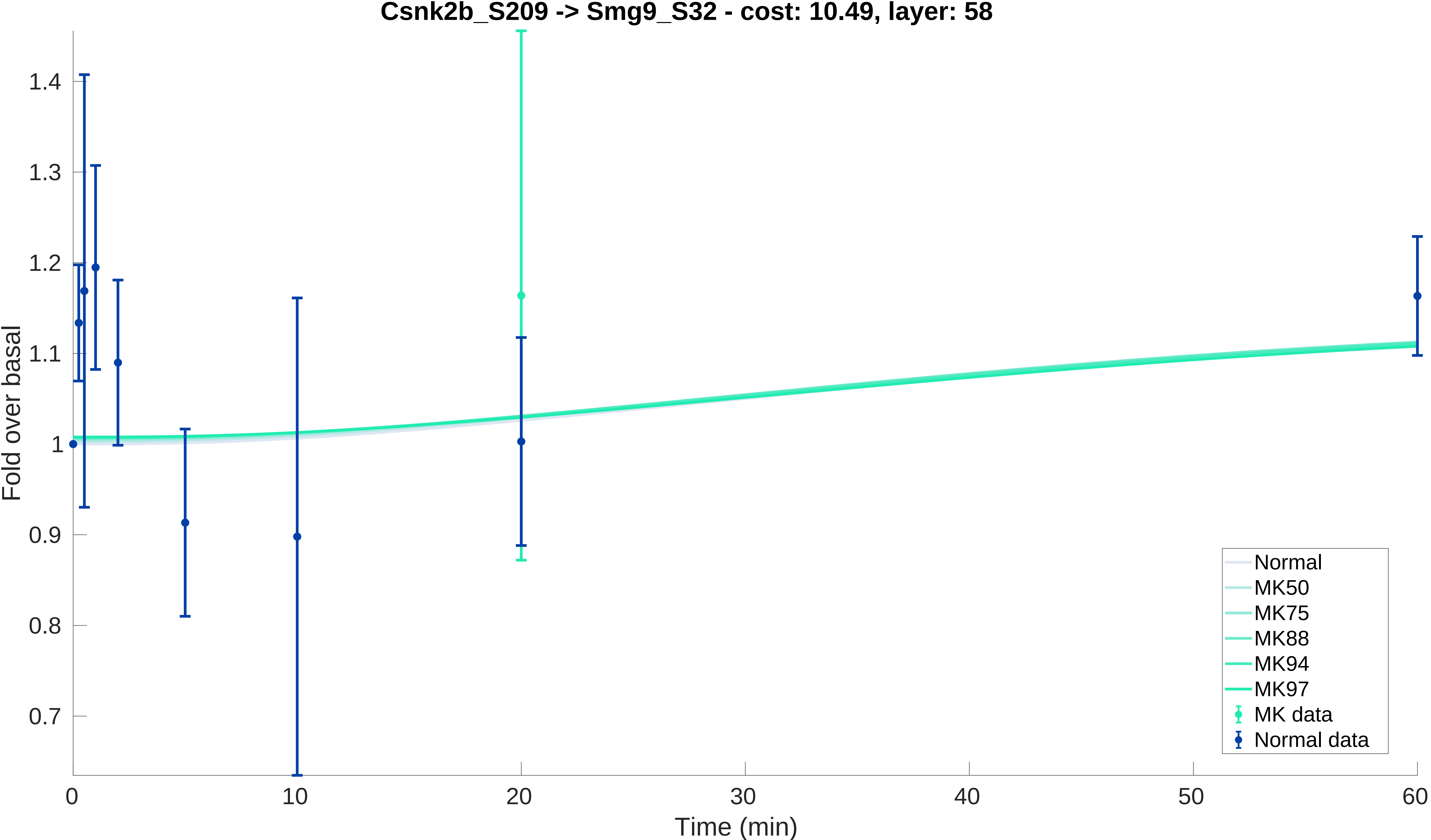

Csnk2b_S209 -> Smg9_S32 - cost: 10.49, layer: 58
Fold over basal
1.4
1.3
1.2
1.1
1
0.9
0.8
0.7
0
10
20
30
40
50
60
Time (min)
Normal
MK50
MK75
MK88
MK94
MK97
MK data
Normal data

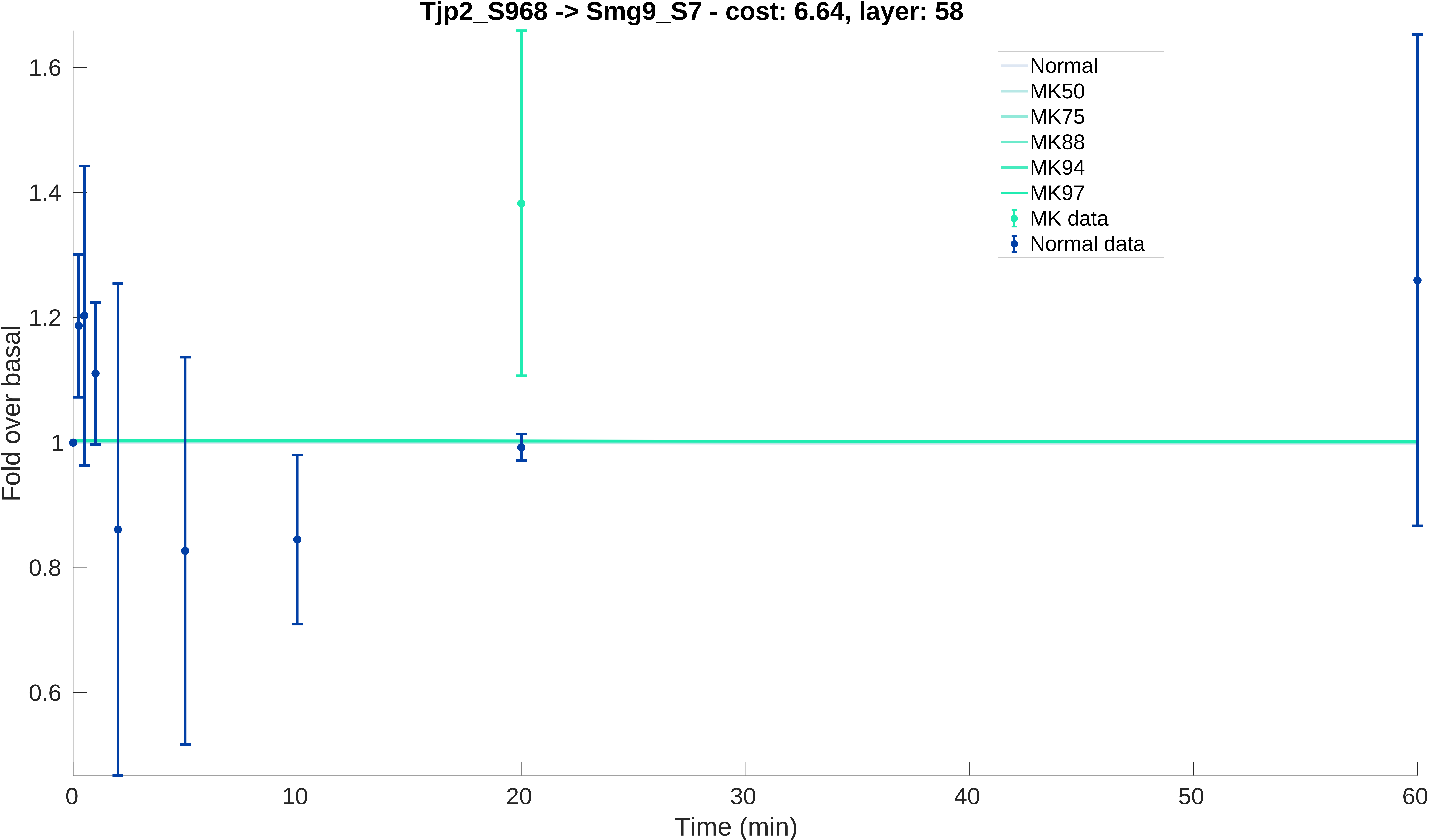

Tjp2_S968 -> Smg9_S7 - cost: 6.64, layer: 58
Normal
MK50
MK75
MK88
MK94
MK97
MK data
Normal data
Fold over basal
1.6
1.4
1.2
1
0.8
0.6
0
10
20
30
40
50
60
Time (min)

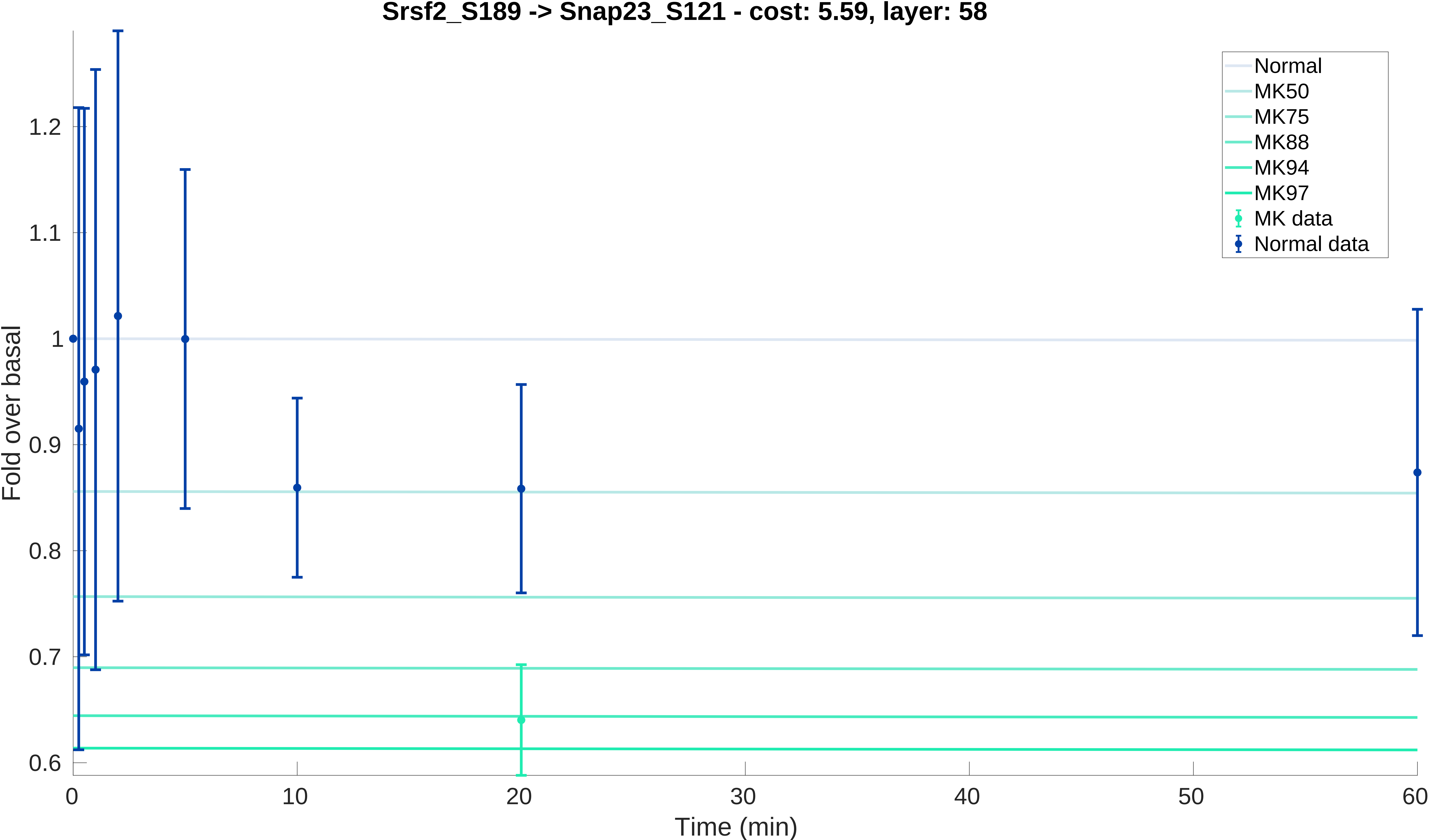

Srsf2_S189 -> Snap23_S121 - cost: 5.59, layer: 58
Fold over basal
Time (min)
0.6
0.7
0.8
0.9
1
1.1
1.2
0
10
20
30
40
50
60
Normal
MK50
MK75
MK88
MK94
MK97
MK data
Normal data

Sipa1l1_S1564 -> Snap23_S20 - cost: 12.31, layer: 58

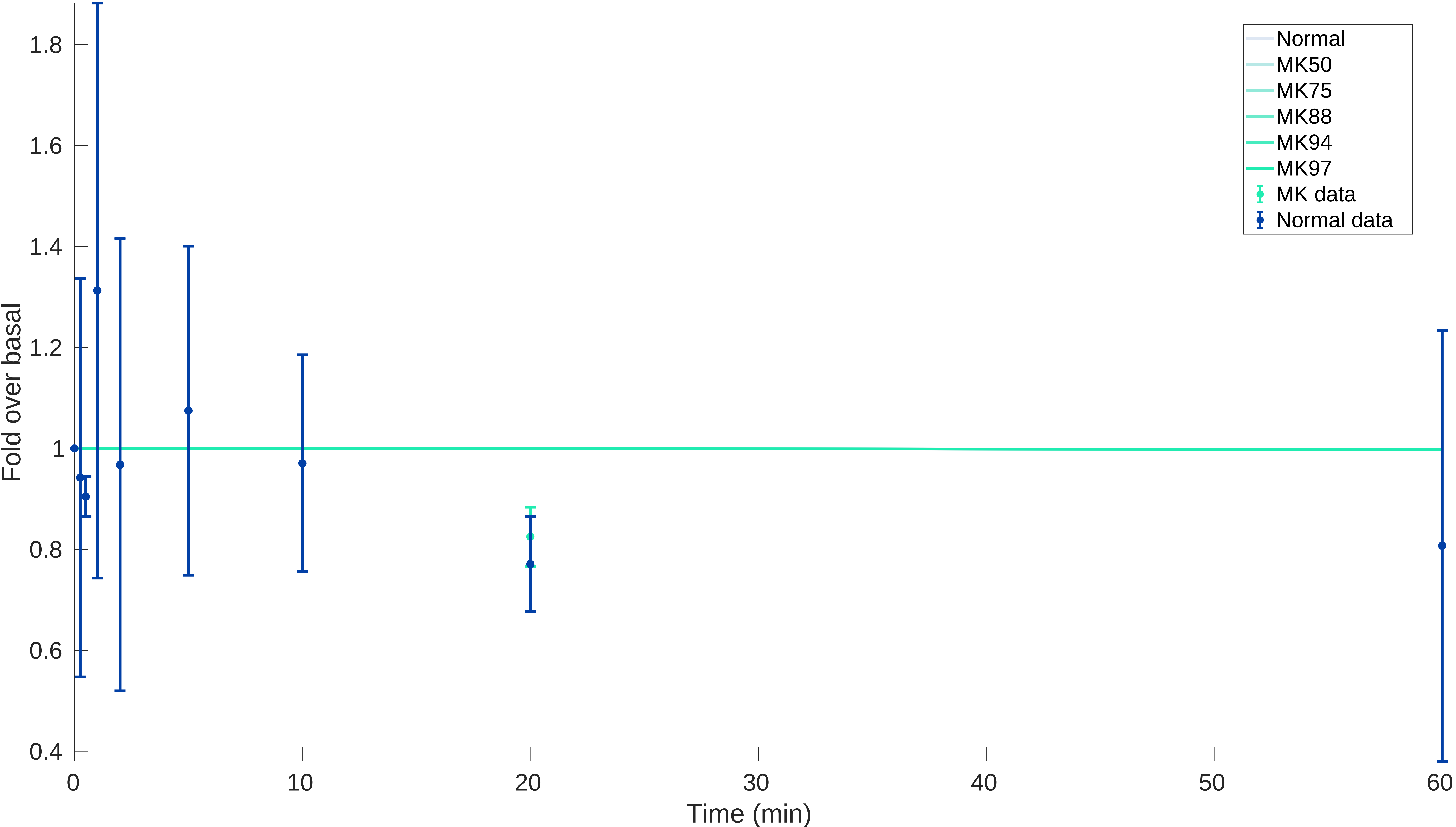

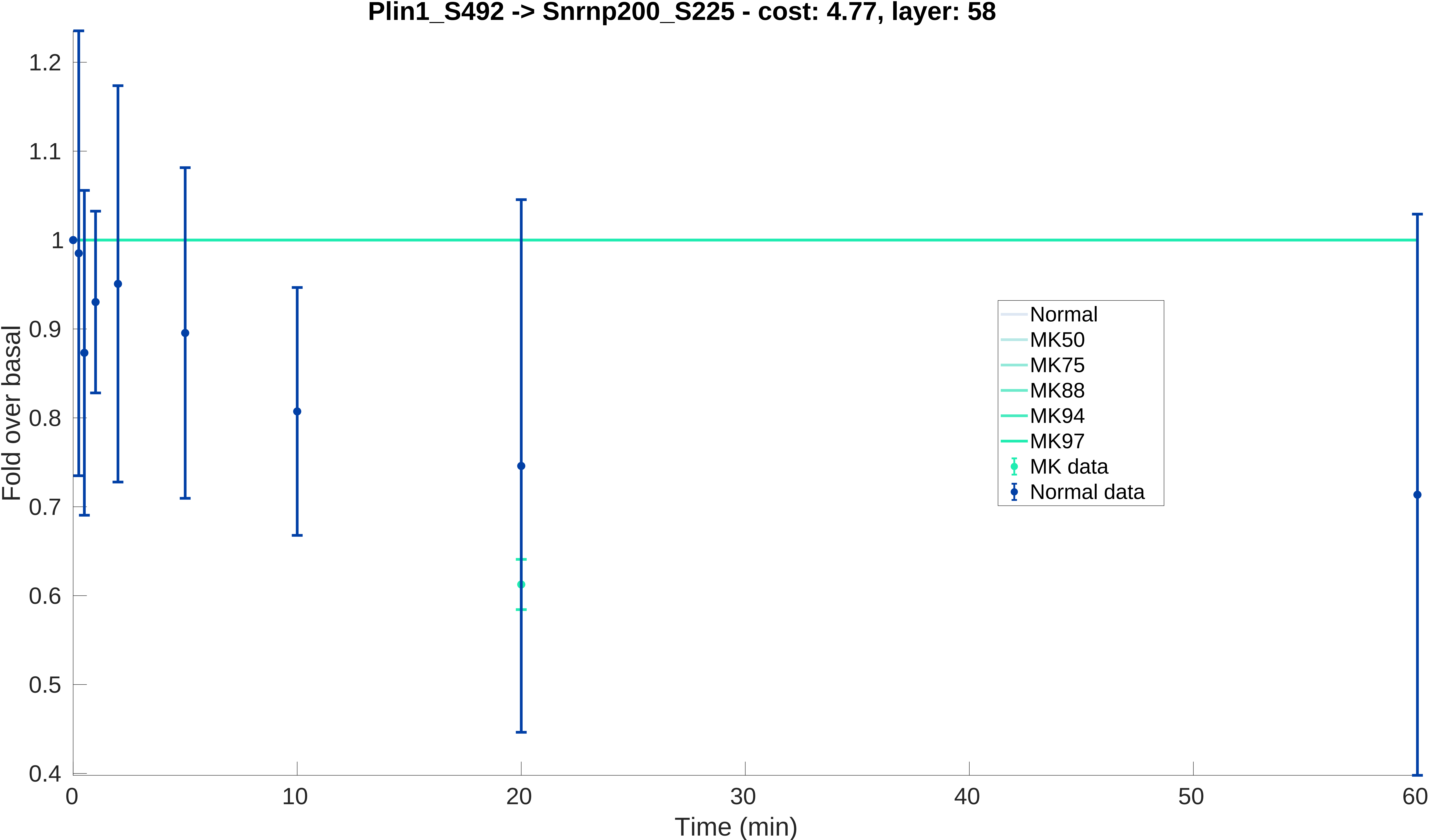

Plin1_S492 -> Snrnp200_S225 - cost: 4.77, layer: 58
Fold over basal
Time (min)
0.4
0.5
0.6
0.7
0.8
0.9
1
1.1
1.2
0
10
20
30
40
50
60
Normal
MK50
MK75
MK88
MK94
MK97
MK data
Normal data

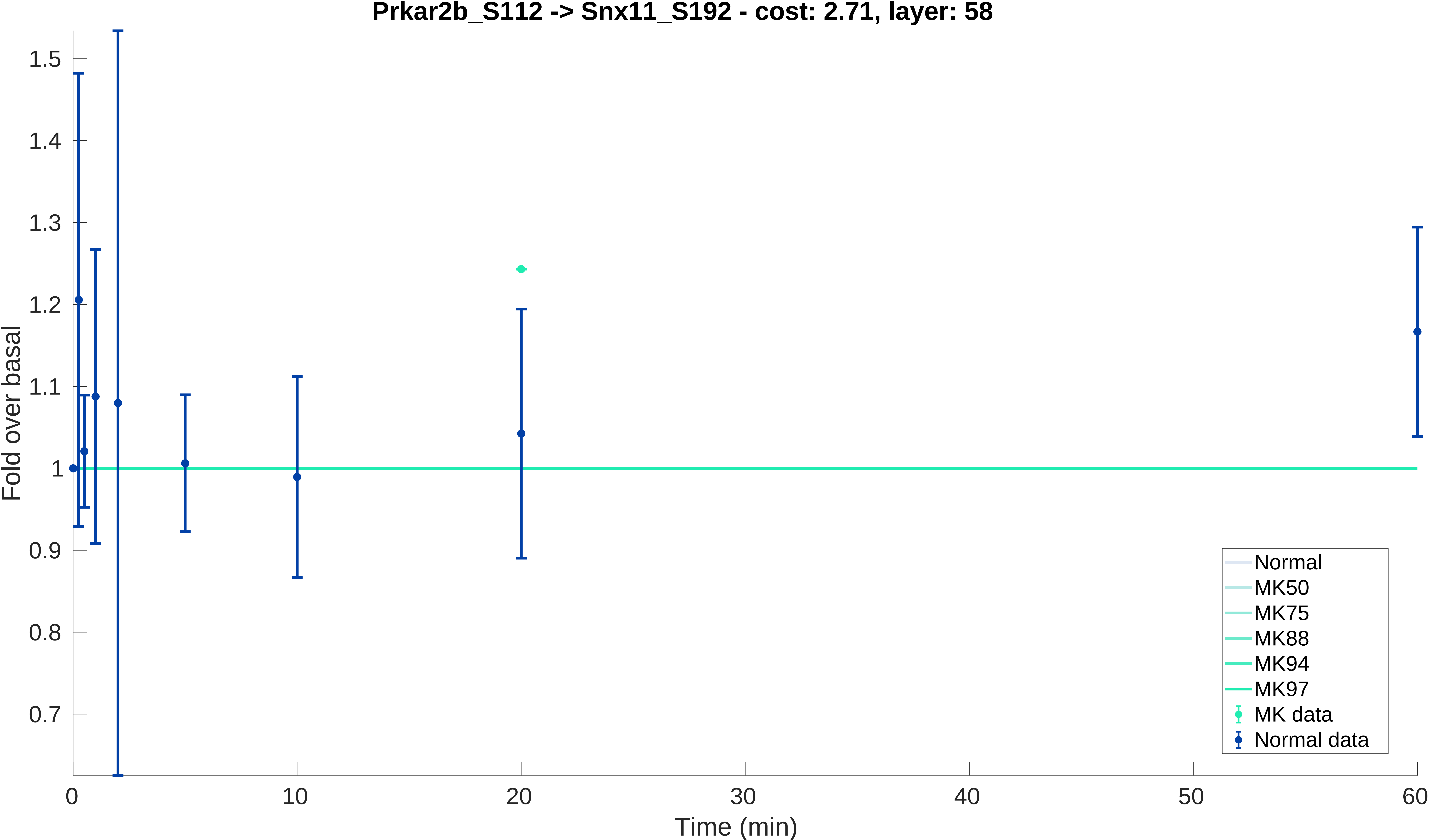

Prkar2b_S112 -> Snx11_S192 - cost: 2.71, layer: 58
Fold over basal
1.5
1.4
1.3
1.2
1.1
1
0.9
0.8
0.7
0
10
20
30
40
50
60
Time (min)
Normal
MK50
MK75
MK88
MK94
MK97
MK data
Normal data

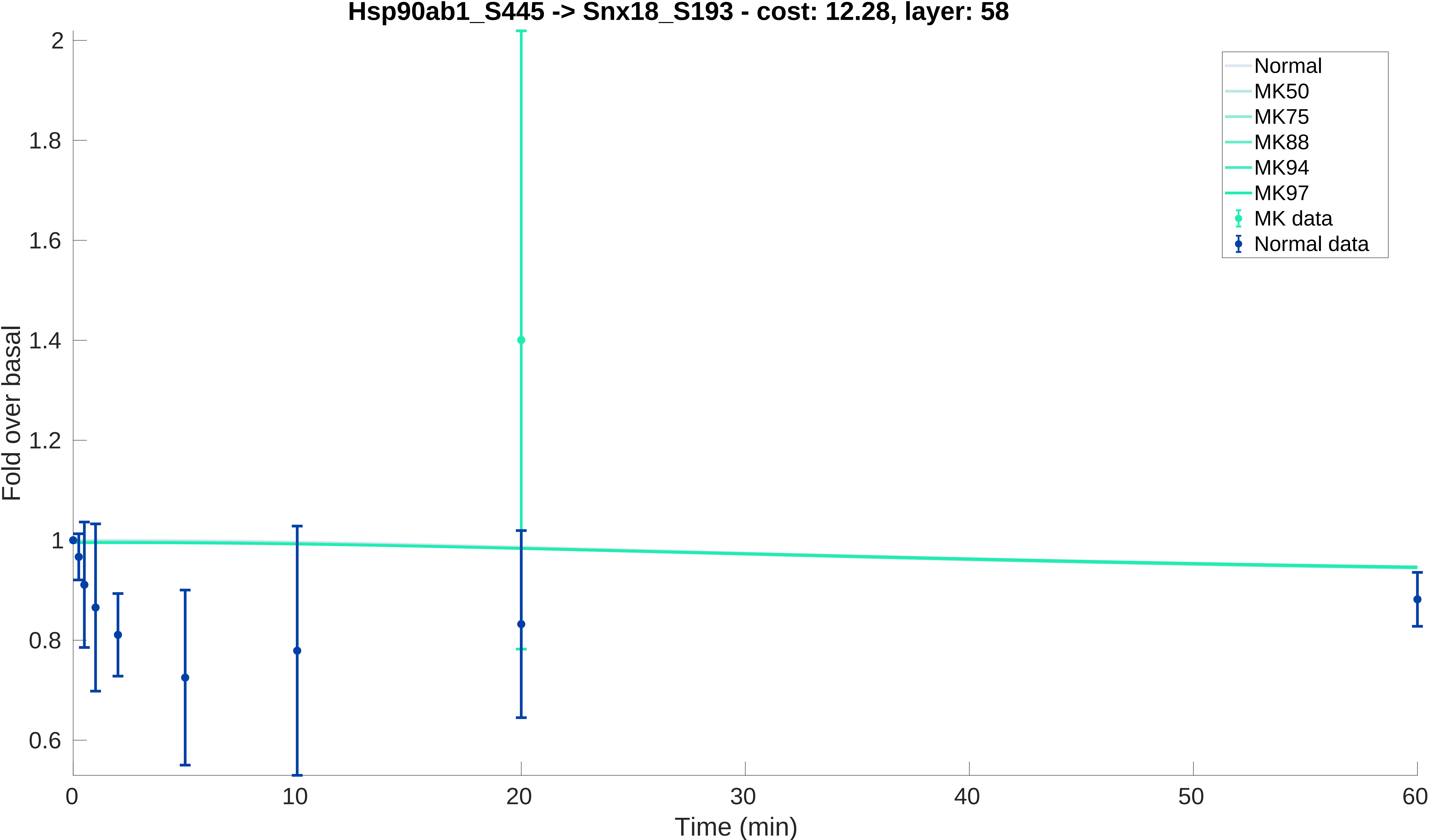

Hsp90ab1_S445 -> Snx18_S193 - cost: 12.28, layer: 58
Fold over basal
Time (min)
Normal
MK50
MK75
MK88
MK94
MK97
MK data
Normal data
0.6
0.8
1
1.2
1.4
1.6
1.8
2
0
10
20
30
40
50
60

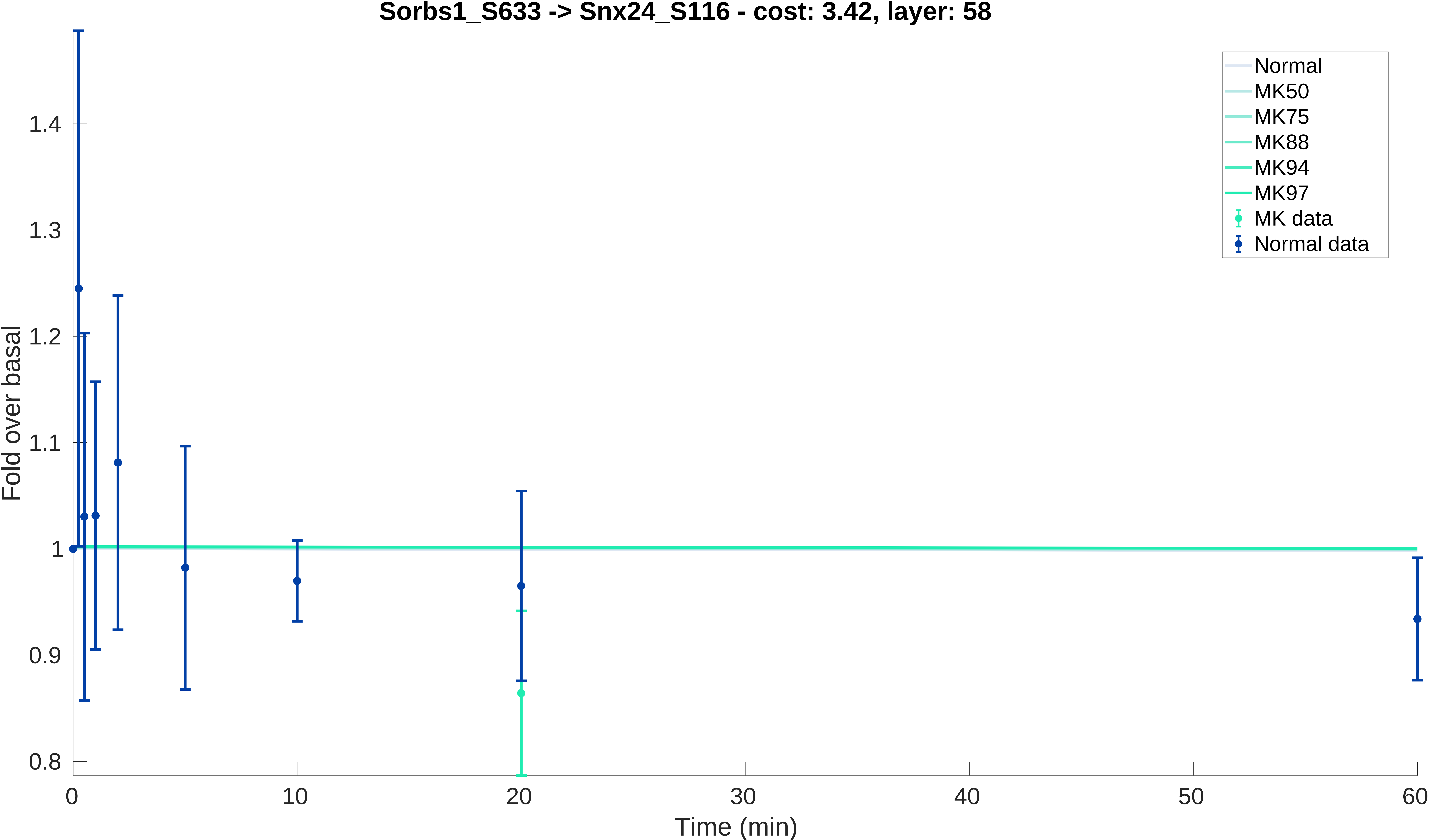
Sorbs1_S633 -> Snx24_S116 - cost: 3.42, layer: 58
Normal
MK50
MK75
MK88
MK94
MK97
MK data
Normal data
Fold over basal
1.4
1.3
1.2
1.1
1
0.9
0.8
0
10
20
30
40
50
60
Time (min)

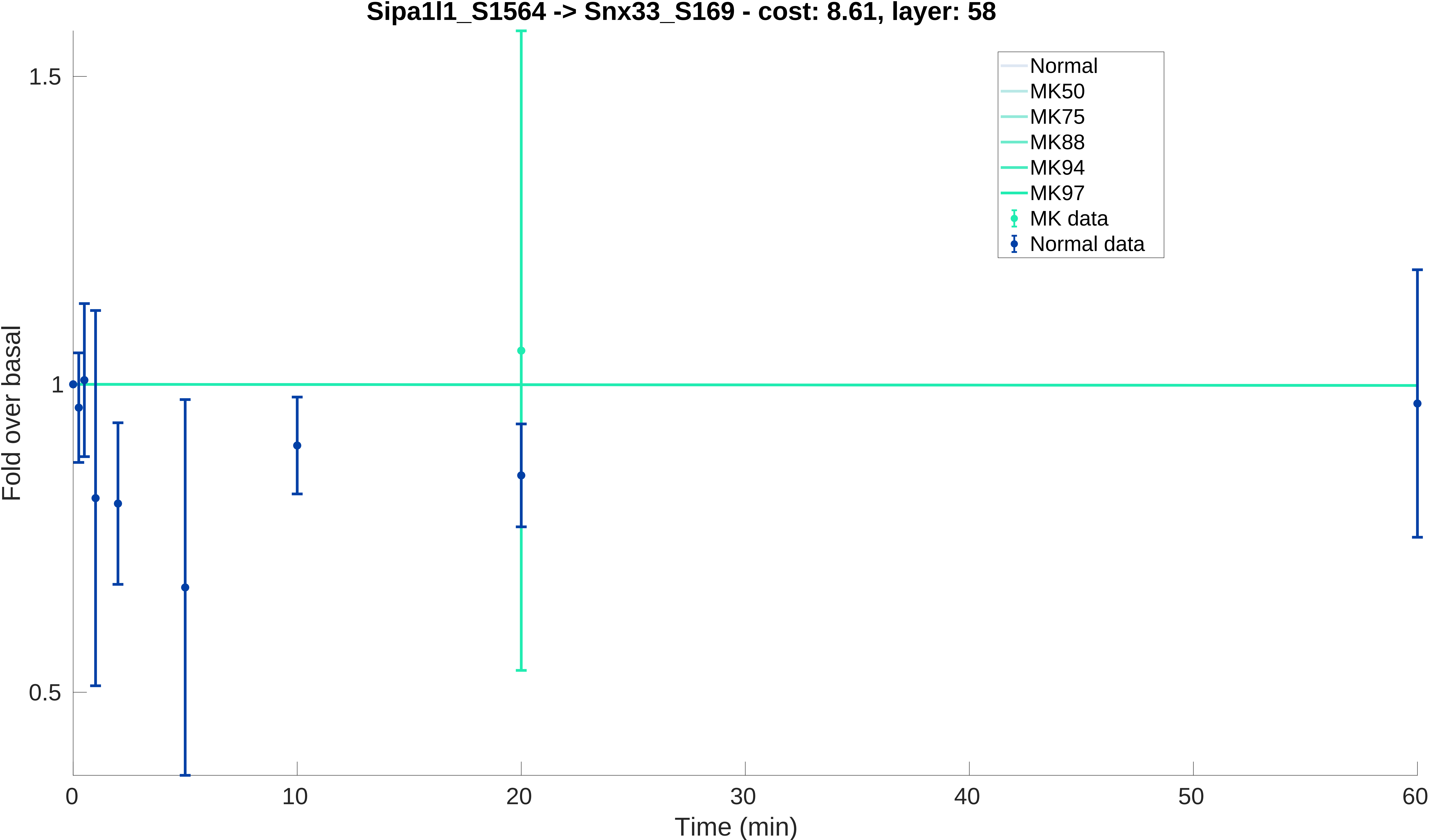

Sipa1l1_S1564 -> Snx33_S169 - cost: 8.61, layer: 58
Normal
MK50
MK75
MK88
MK94
MK97
MK data
Normal data
1.5
1
0.5
Fold over basal
0
10
20
30
40
50
60
Time (min)

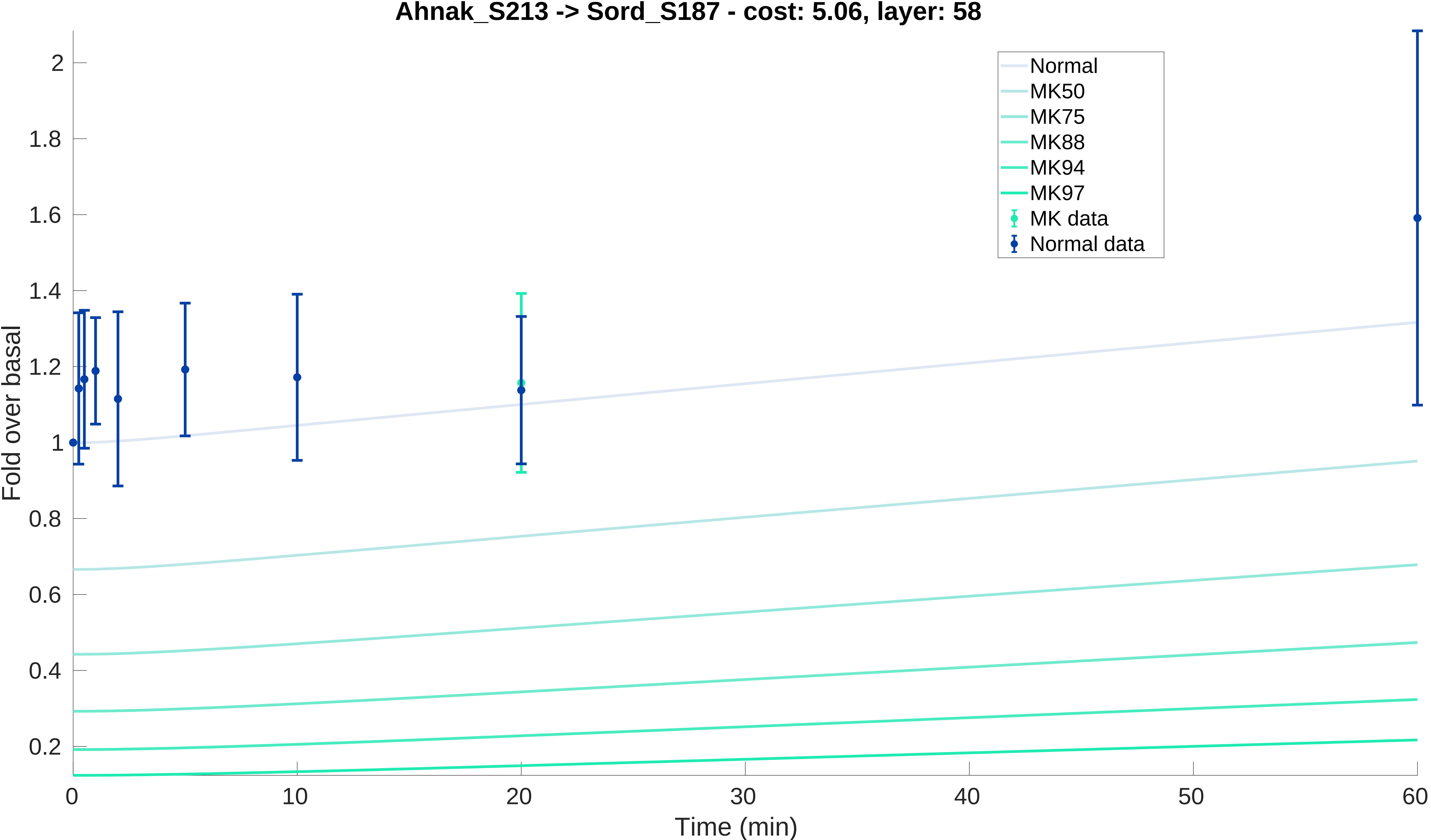
Ahnak_S213 -> Sord_S187 - cost: 5.06, layer: 58
Fold over basal
Time (min)
0
10
20
30
40
50
60
0.2
0.4
0.6
0.8
1
1.2
1.4
1.6
1.8
2
Normal
MK50
MK75
MK88
MK94
MK97
MK data
Normal data

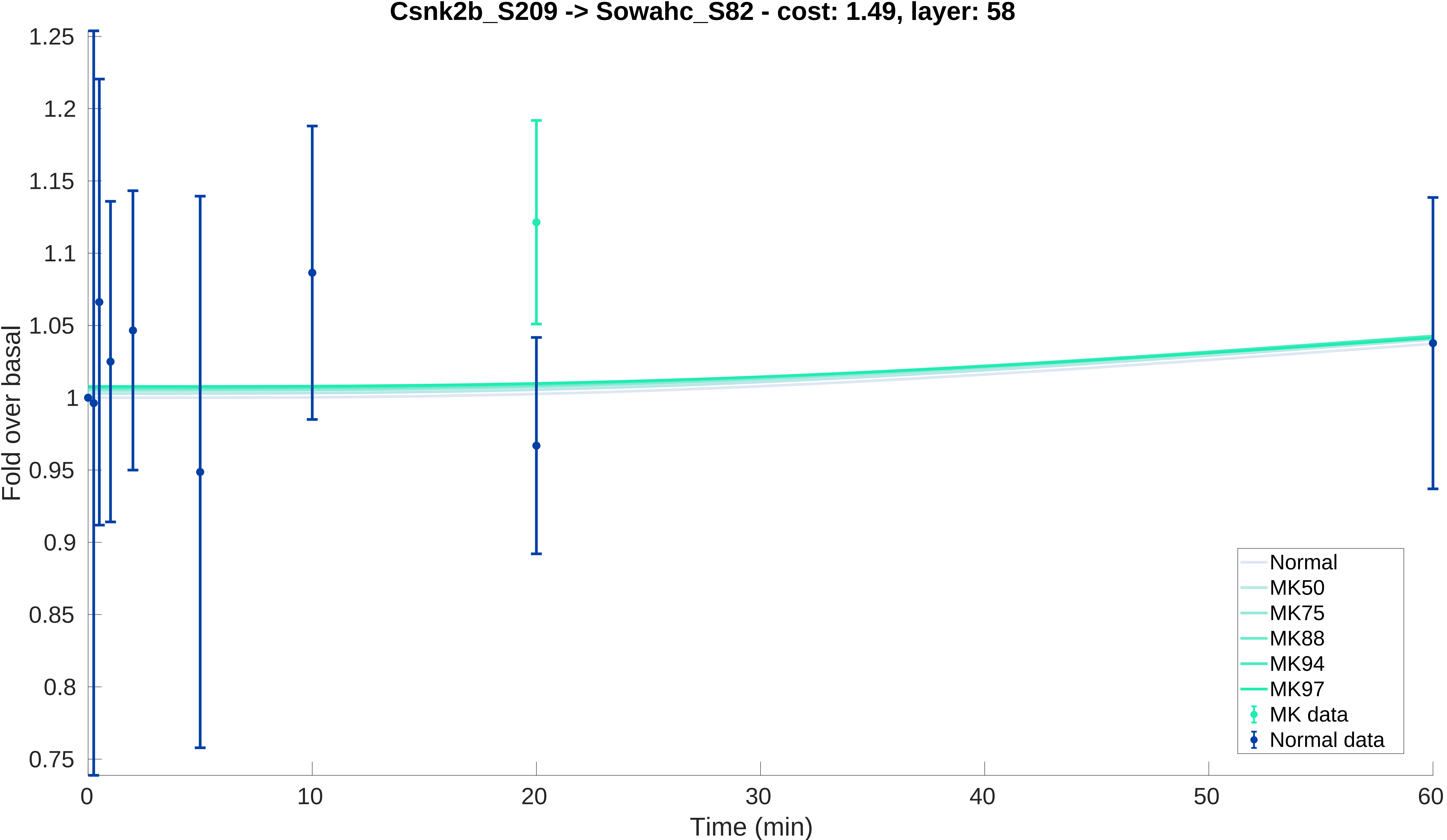

Csnk2b_S209 -> Sowahc_S82 - cost: 1.49, layer: 58
Fold over basal
Time (min)
1.25
1.2
1.15
1.1
1.05
1
0.95
0.9
0.85
0.8
0.75
0
10
20
30
40
50
60
Normal
MK50
MK75
MK88
MK94
MK97
MK data
Normal data

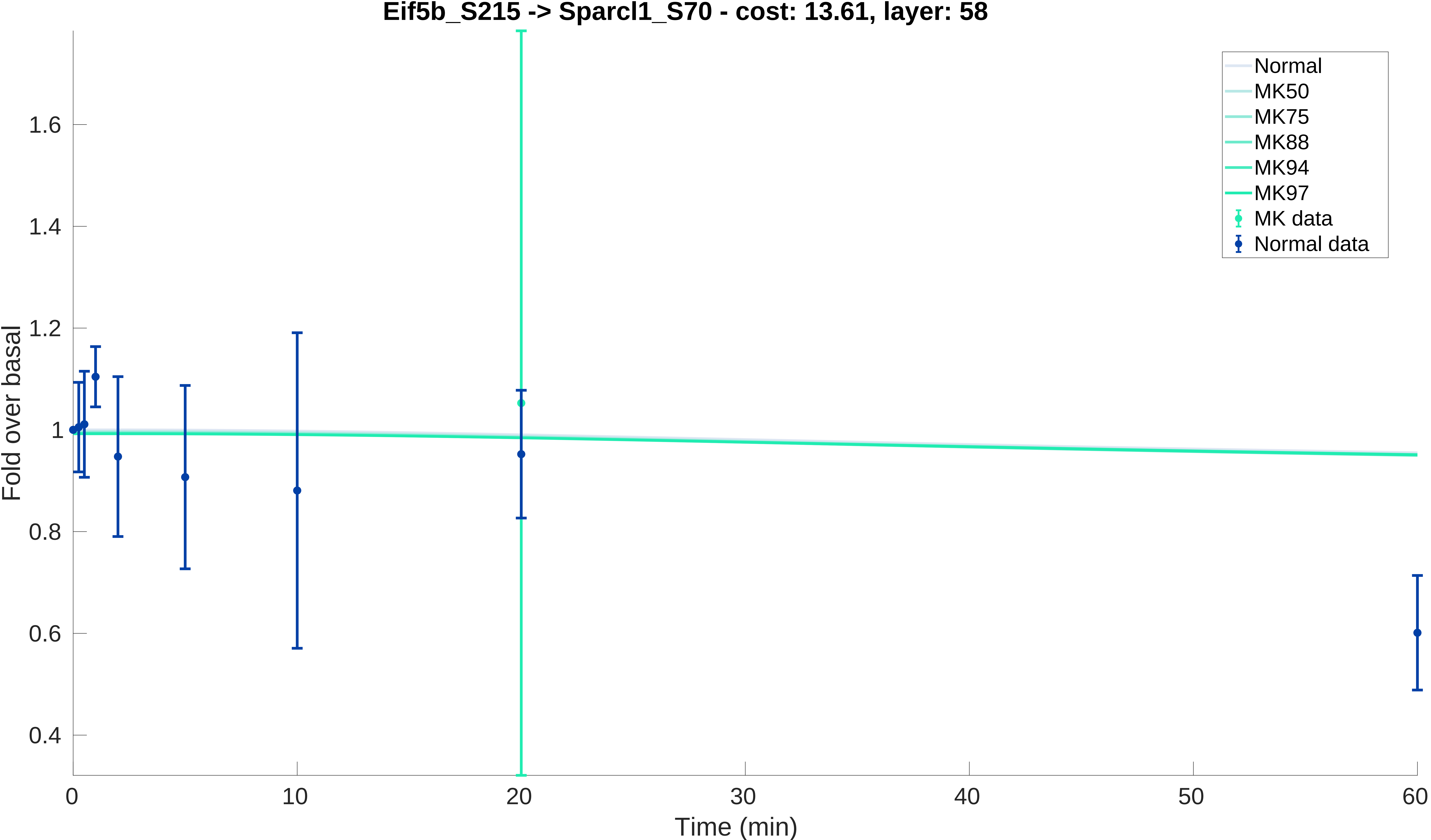

Eif5b_S215 -> Sparcl1_S70 - cost: 13.61, layer: 58
Normal
MK50
MK75
MK88
MK94
MK97
MK data
Normal data
Fold over basal
Time (min)
1.6
1.4
1.2
1
0.8
0.6
0.4
0
10
20
30
40
50
60

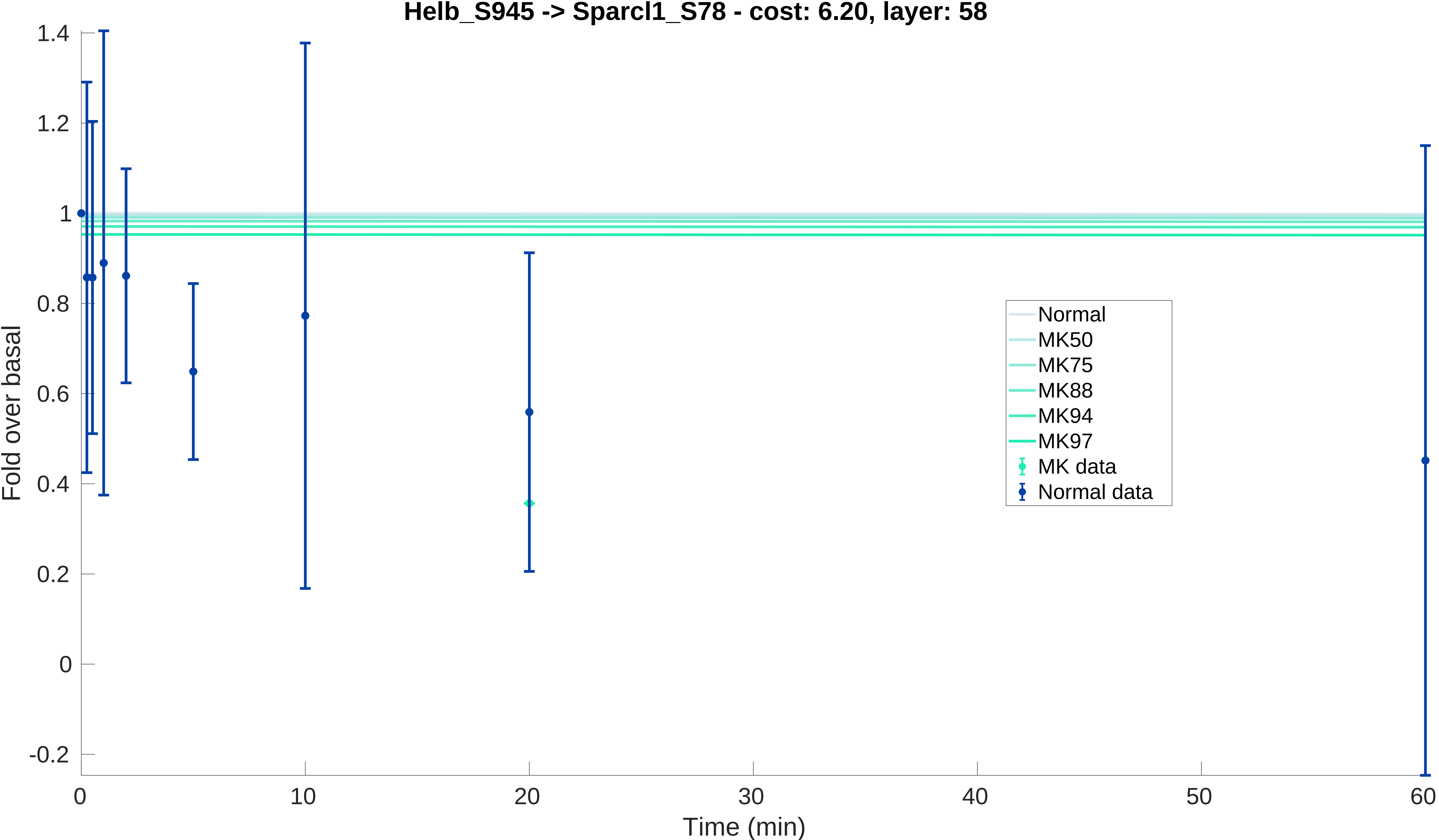

Helb_S945 -> Sparcl1_S78 - cost: 6.20, layer: 58
Fold over basal
Time (min)
1.4
1.2
1
0.8
0.6
0.4
0.2
0
-0.2
0
10
20
30
40
50
60
Normal
MK50
MK75
MK88
MK94
MK97
MK data
Normal data

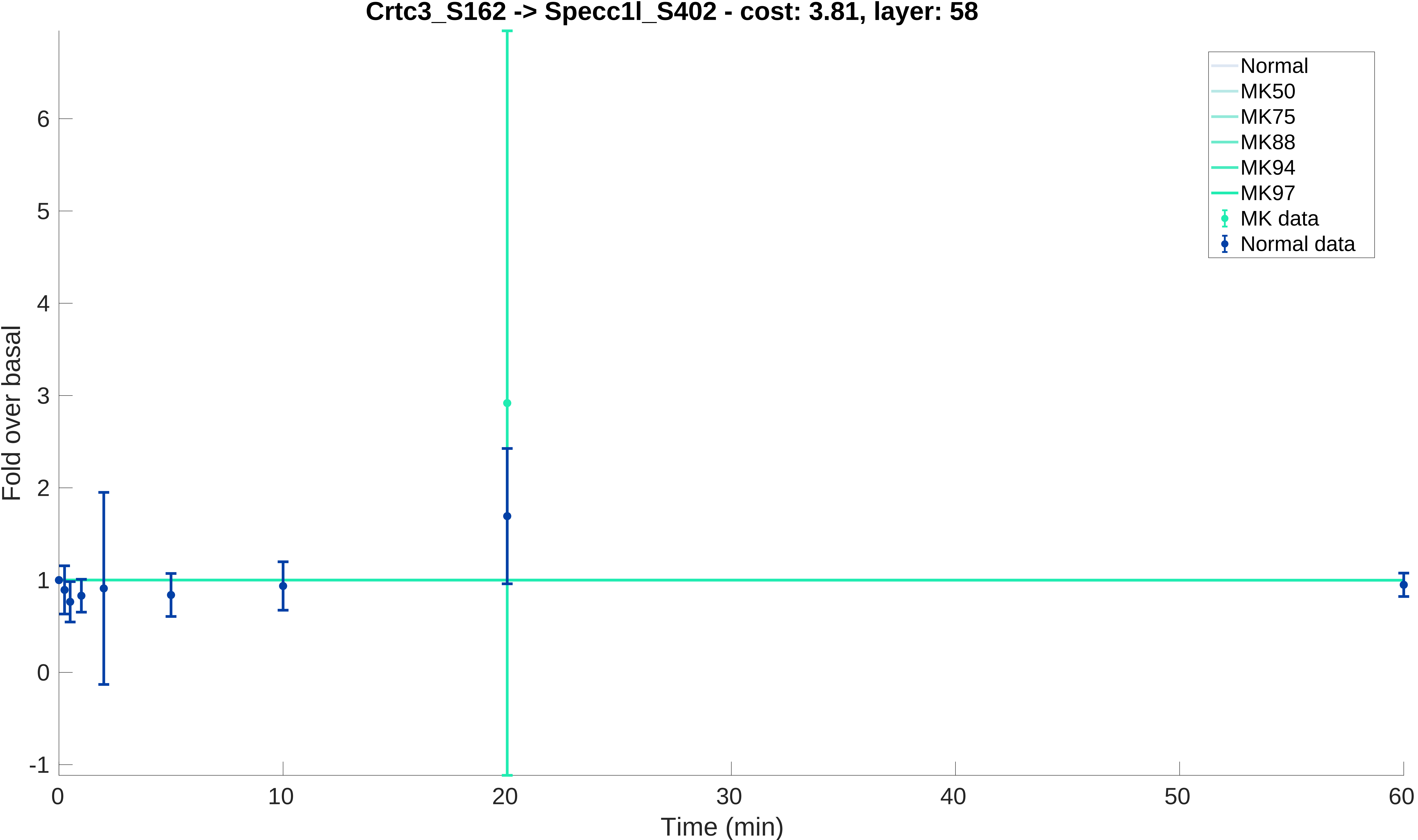

Crtc3_S162 -> Specc1l_S402 - cost: 3.81, layer: 58
Fold over basal
Time (min)
6
5
4
3
2
1
0
-1
0
10
20
30
40
50
60
Normal
MK50
MK75
MK88
MK94
MK97
MK data
Normal data

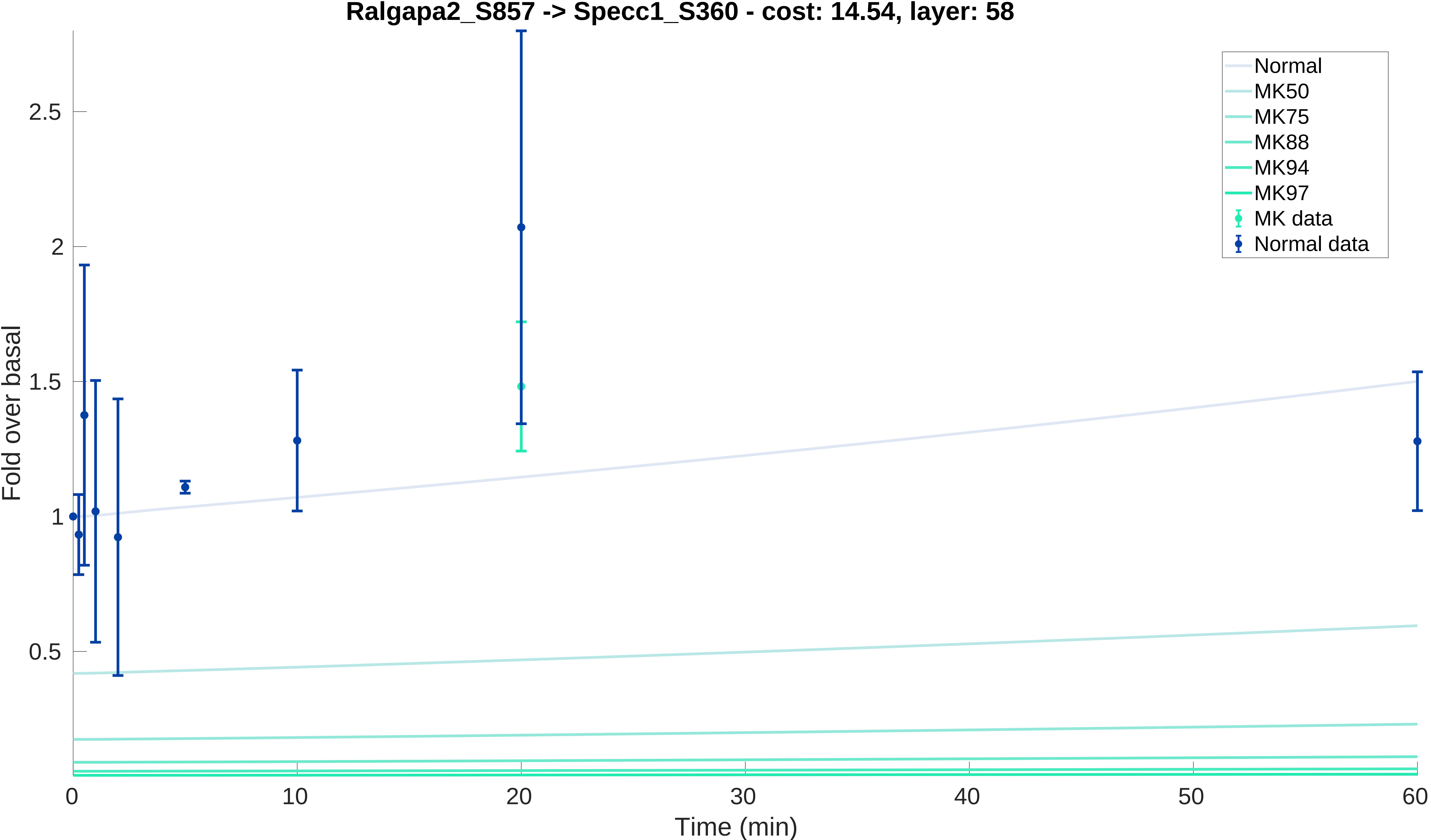

Ralgapa2_S857 -> Specc1_S360 - cost: 14.54, layer: 58
Fold over basal
0.5
1
1.5
2
2.5
0
10
20
30
40
50
60
Time (min)
Normal
MK50
MK75
MK88
MK94
MK97
MK data
Normal data

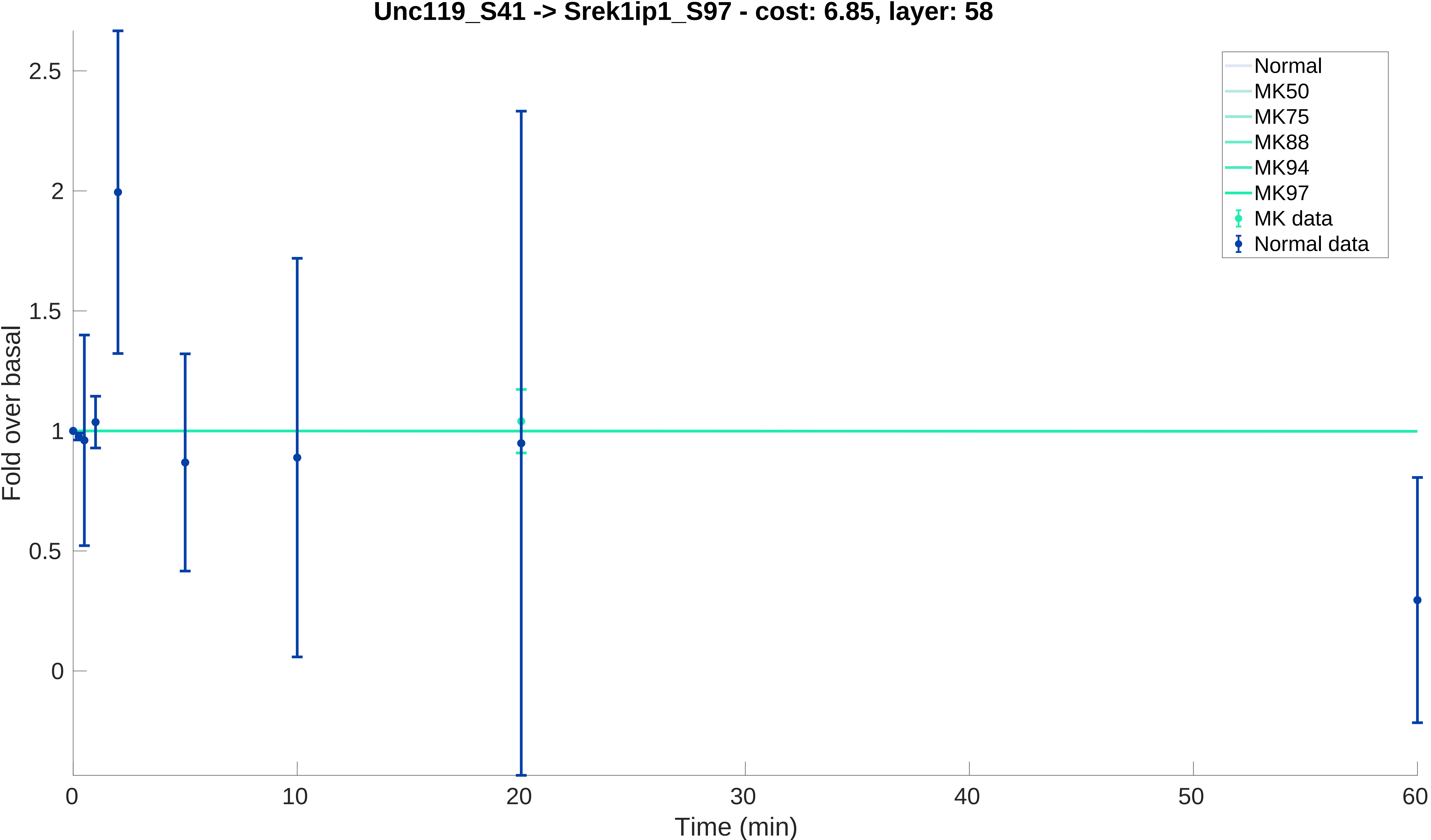

Unc119_S41 -> Srek1ip1_S97 - cost: 6.85, layer: 58
Fold over basal
Time (min)
2.5
2
1.5
1
0.5
0
0
10
20
30
40
50
60
Normal
MK50
MK75
MK88
MK94
MK97
MK data
Normal data

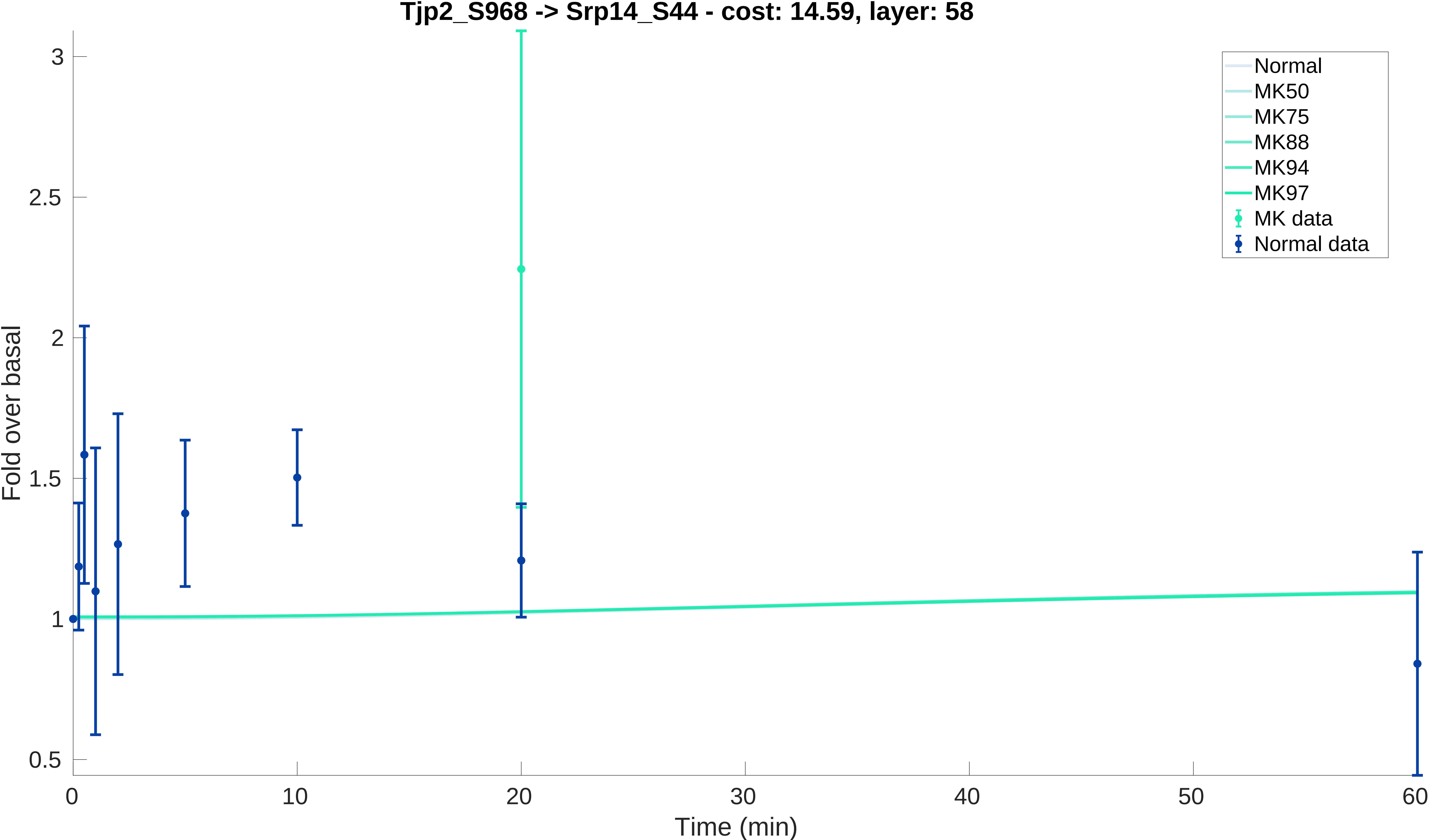

Tjp2_S968 -> Srp14_S44 - cost: 14.59, layer: 58
Fold over basal
Time (min)
0
10
20
30
40
50
60
0.5
1
1.5
2
2.5
3
Normal
MK50
MK75
MK88
MK94
MK97
MK data
Normal data

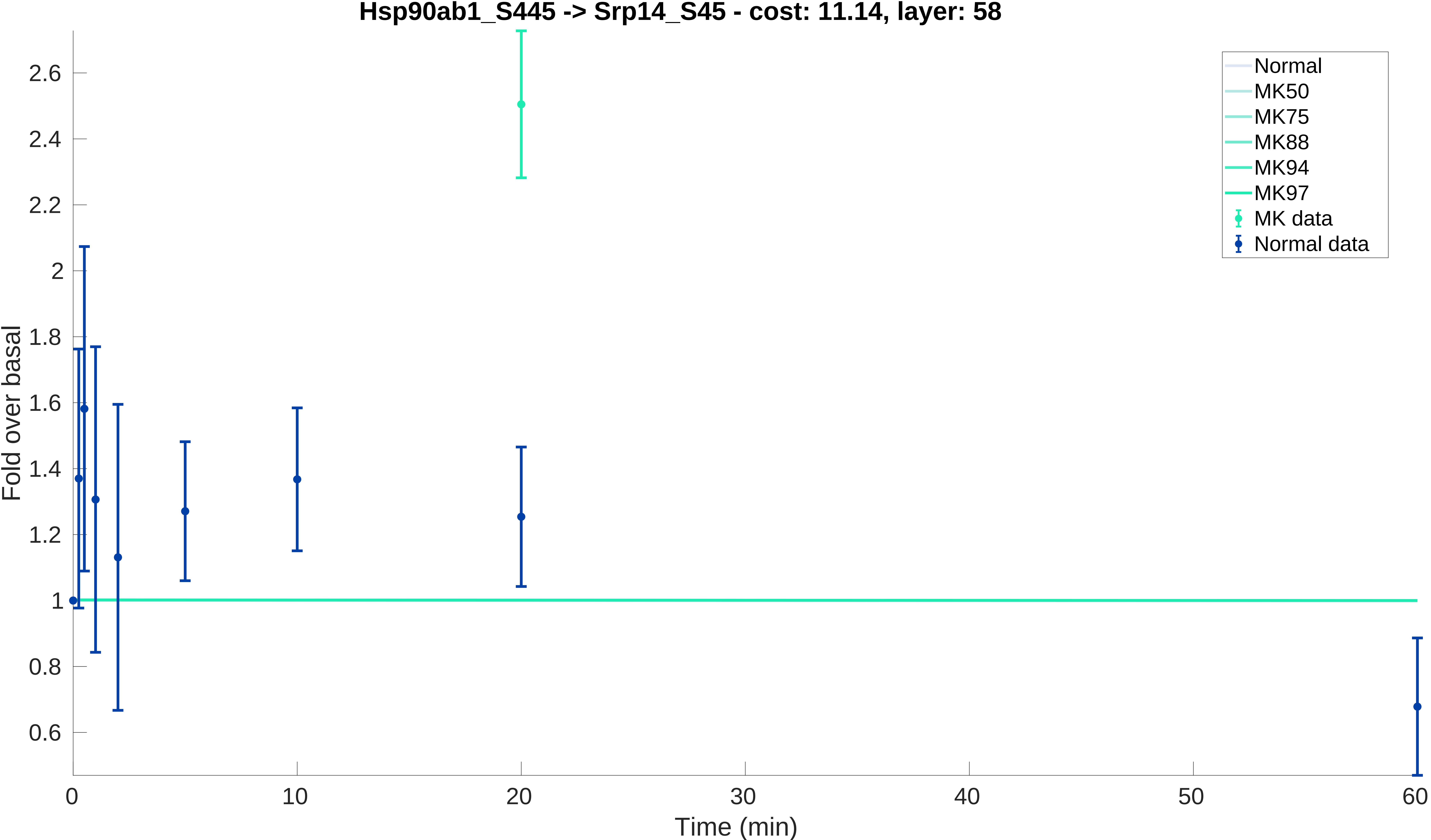
Hsp90ab1_S445 -> Srp14_S45 - cost: 11.14, layer: 58
Fold over basal
Time (min)
0.6
0.8
1
1.2
1.4
1.6
1.8
2
2.2
2.4
2.6
0
10
20
30
40
50
60
Normal
MK50
MK75
MK88
MK94
MK97
MK data
Normal data

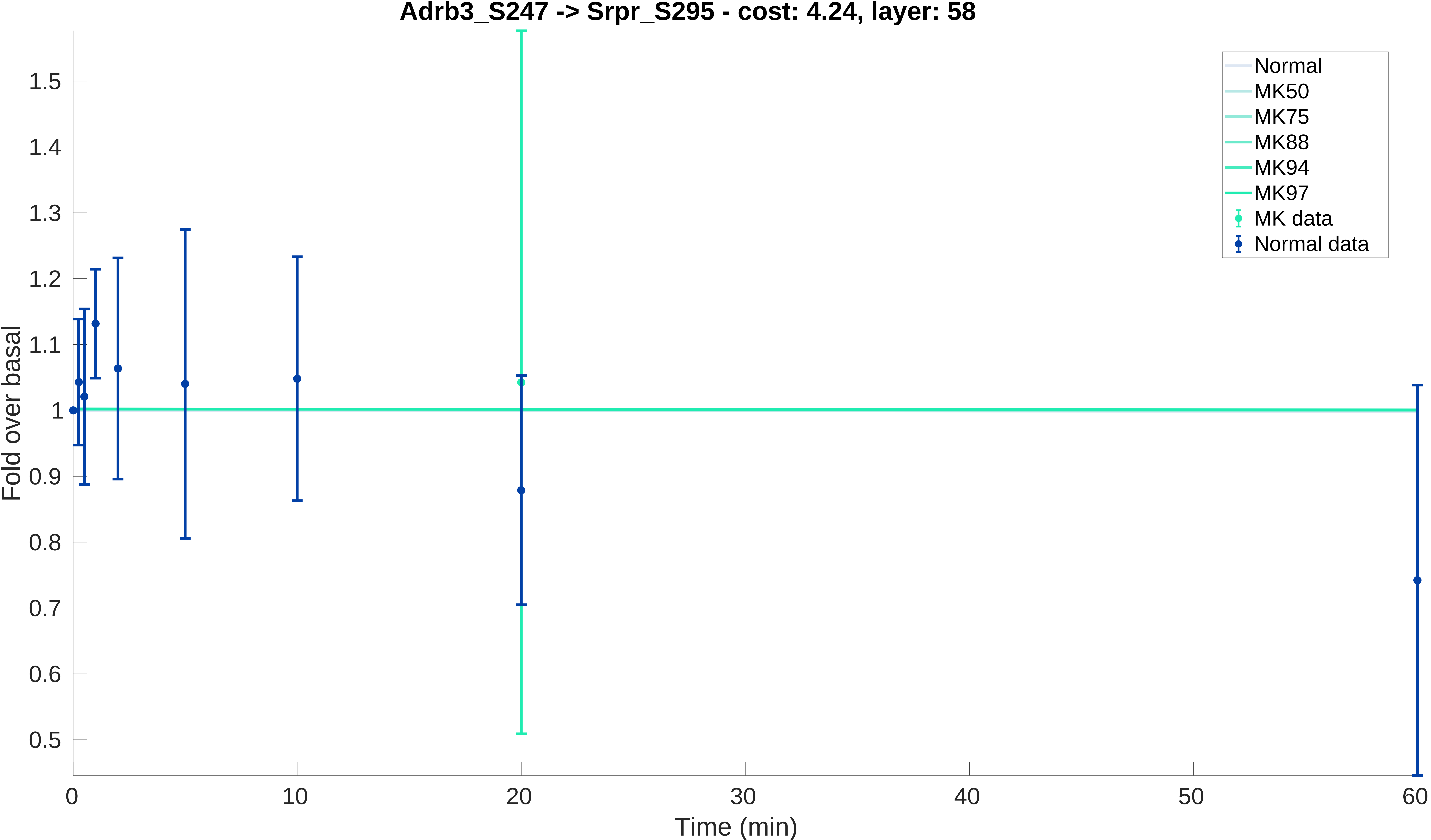

Adrb3_S247 -> Srpr_S295 - cost: 4.24, layer: 58
Normal
MK50
MK75
MK88
MK94
MK97
MK data
Normal data
Fold over basal
1.5
1.4
1.3
1.2
1.1
1
0.9
0.8
0.7
0.6
0.5
0
10
20
30
40
50
60
Time (min)

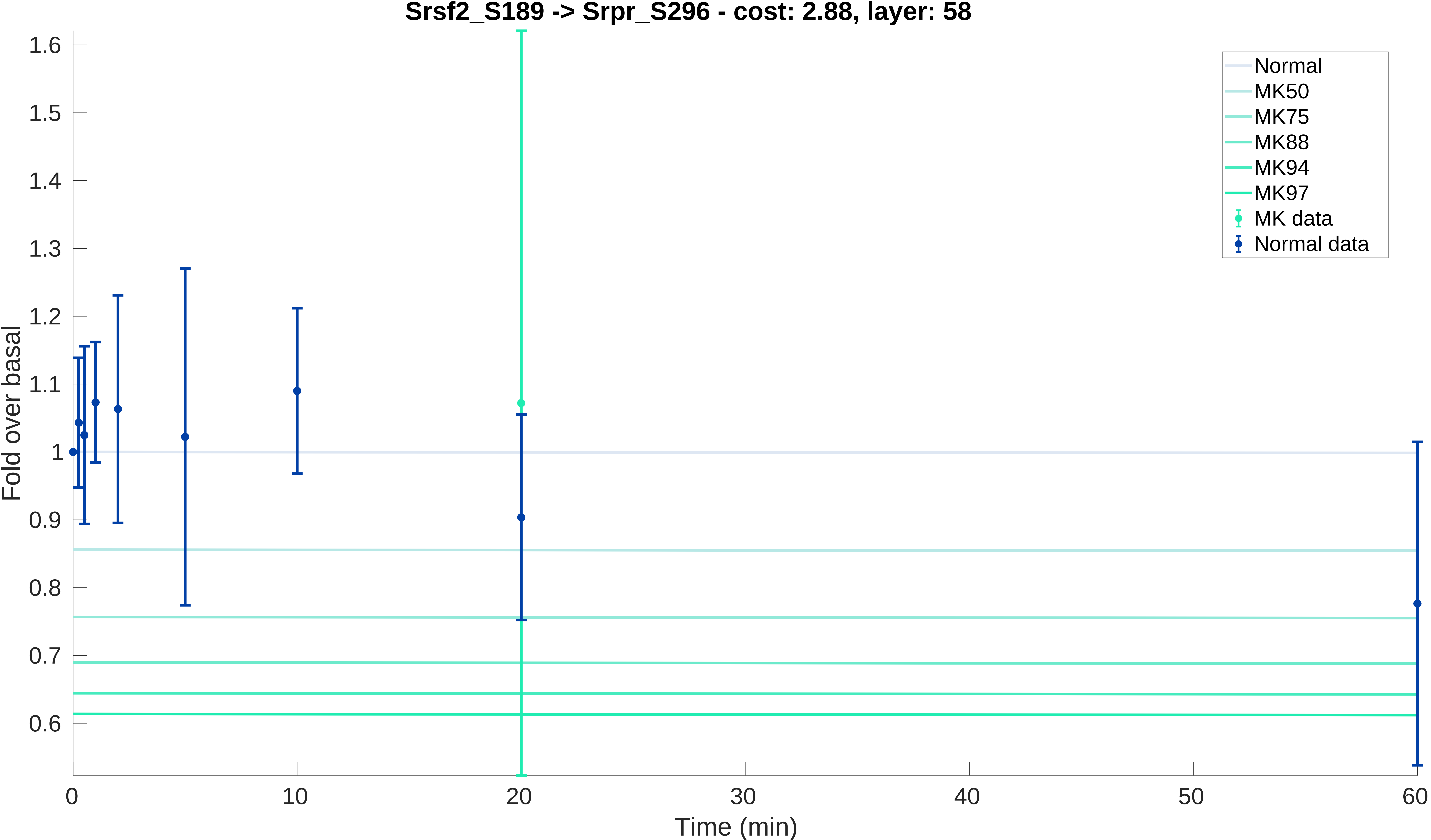

Srsf2_S189 -> Srpr_S296 - cost: 2.88, layer: 58
Fold over basal
Time (min)
1.6
1.5
1.4
1.3
1.2
1.1
1
0.9
0.8
0.7
0.6
0
10
20
30
40
50
60
Normal
MK50
MK75
MK88
MK94
MK97
MK data
Normal data

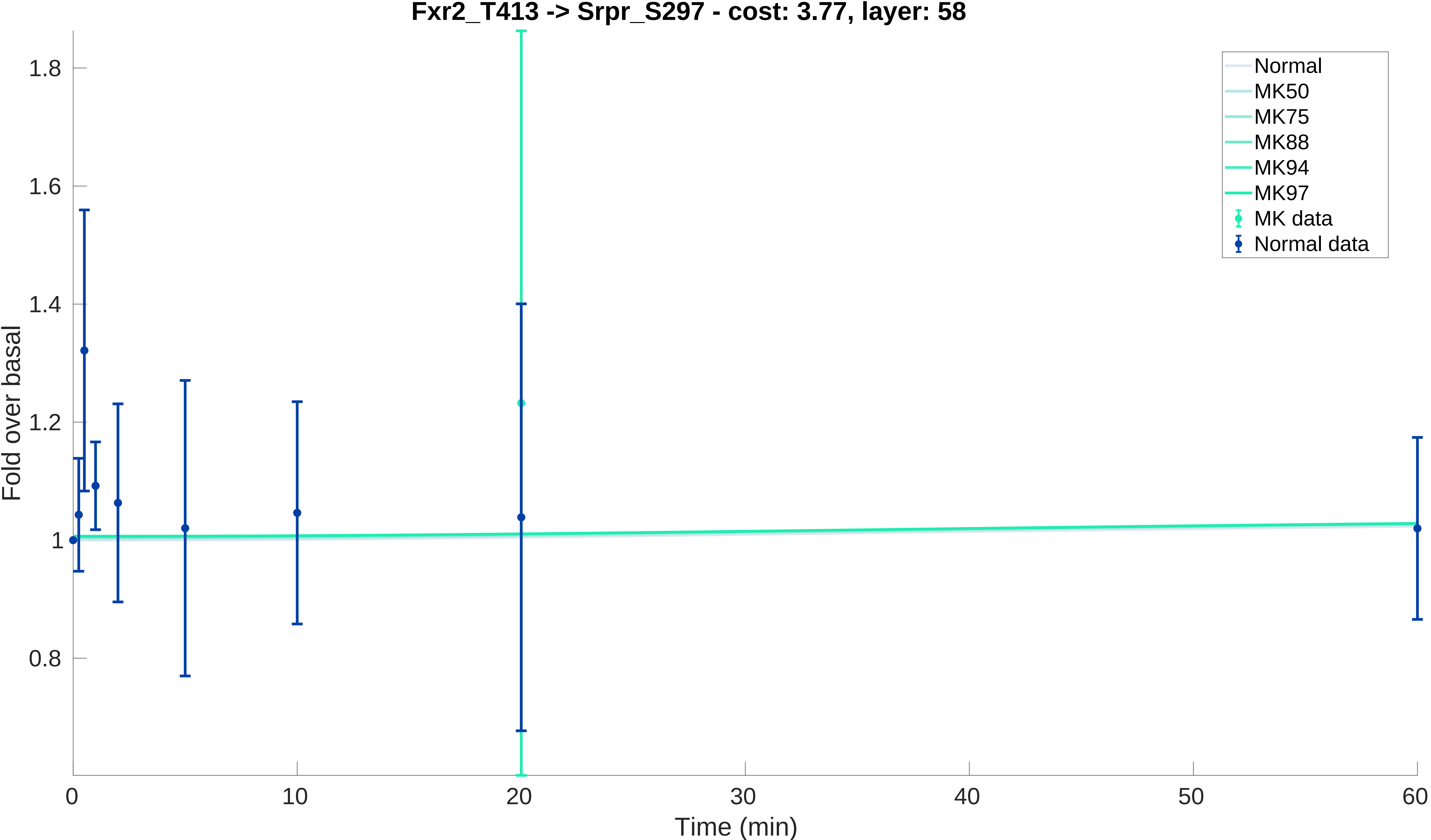
Fxr2_T413 -> Srpr_S297 - cost: 3.77, layer: 58
Fold over basal
Time (min)
1.8
1.6
1.4
1.2
1
0.8
0
10
20
30
40
50
60
Normal
MK50
MK75
MK88
MK94
MK97
MK data
Normal data

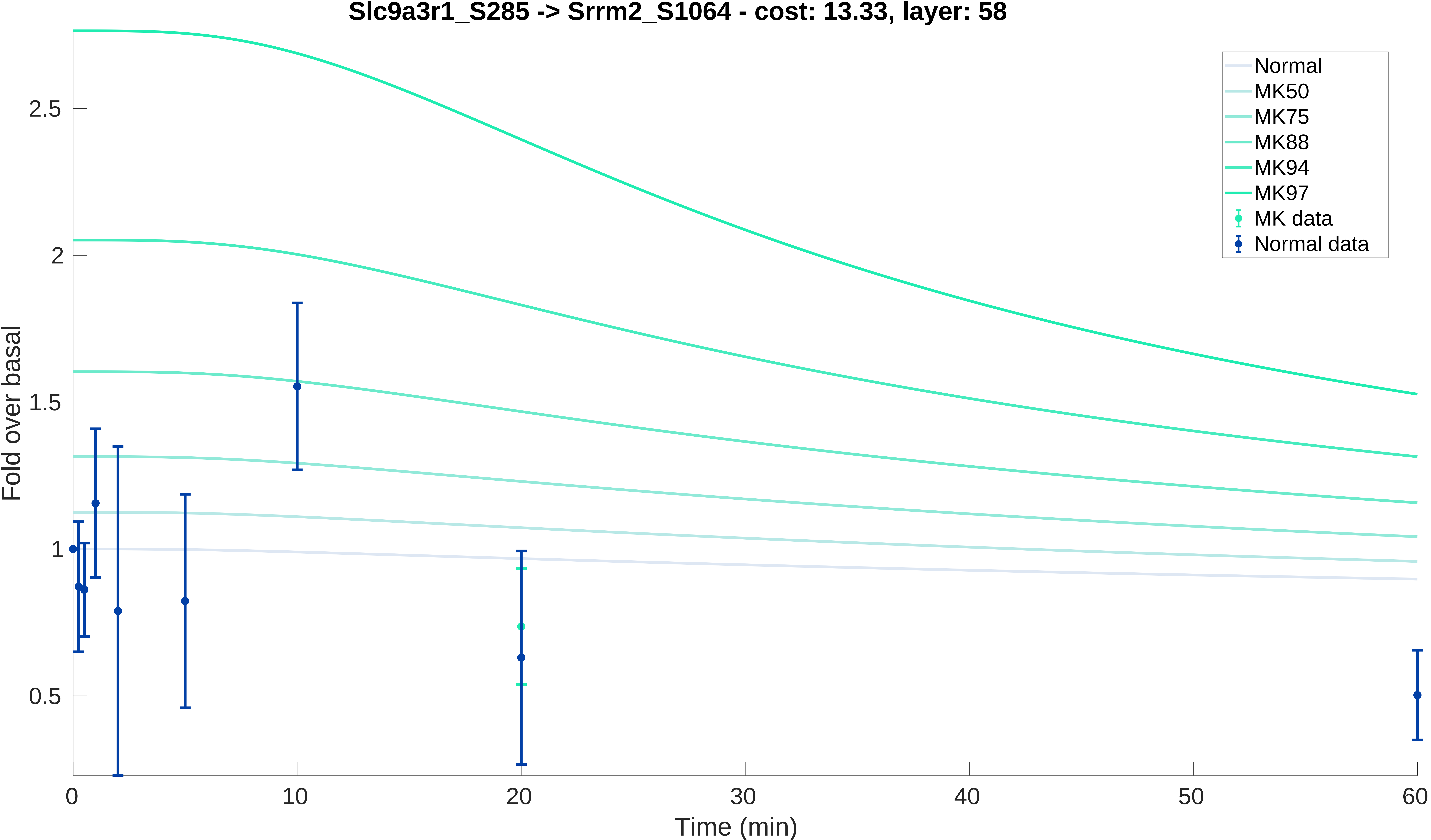
Slc9a3r1_S285 -> Srrm2_S1064 - cost: 13.33, layer: 58
Normal
MK50
MK75
MK88
MK94
MK97
MK data
Normal data
Fold over basal
Time (min)
0.5
1
1.5
2
2.5
0
10
20
30
40
50
60

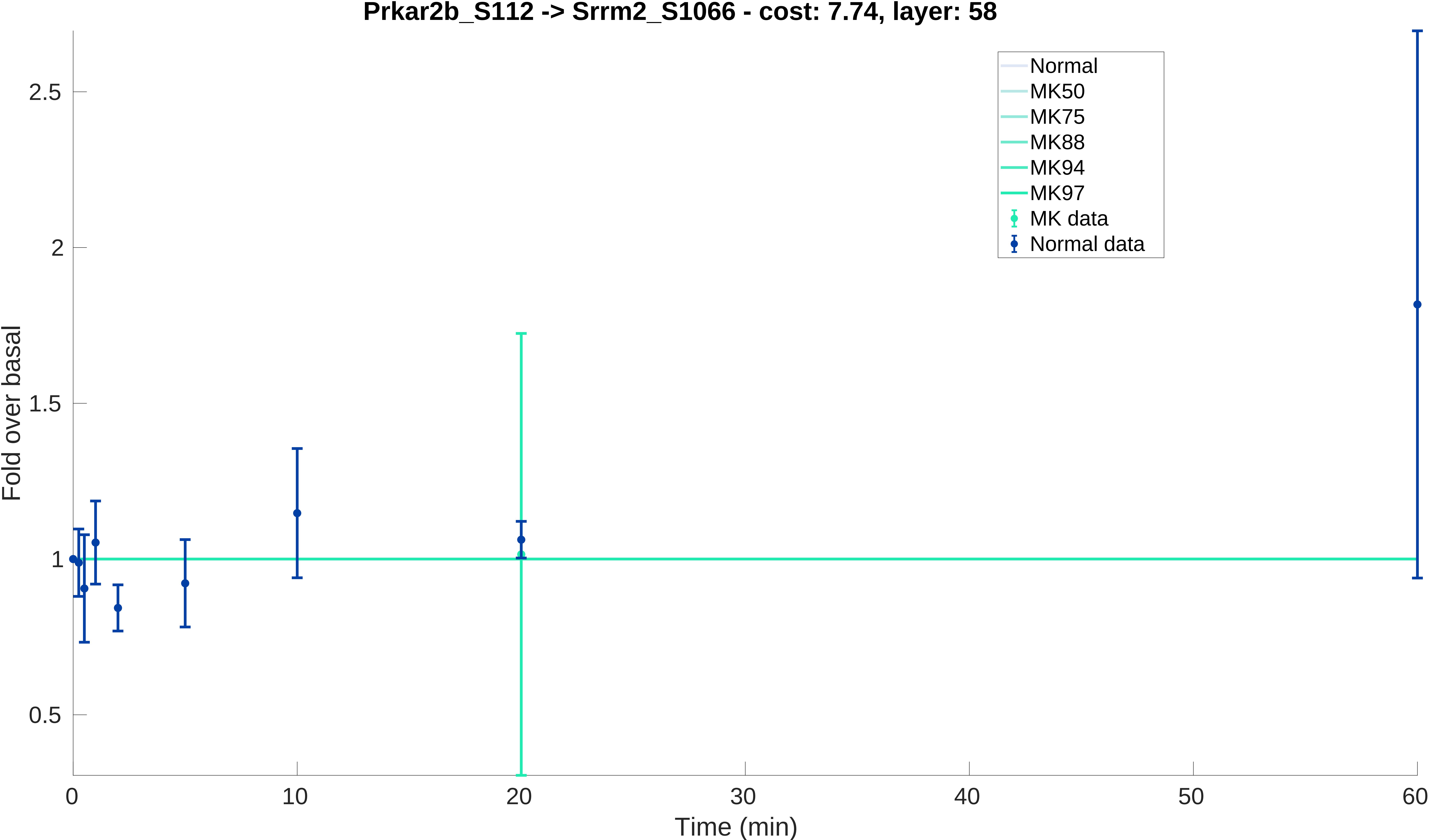
Prkar2b_S112 -> Srrm2_S1066 - cost: 7.74, layer: 58
Fold over basal
Time (min)
0.5
1
1.5
2
2.5
0
10
20
30
40
50
60
Normal
MK50
MK75
MK88
MK94
MK97
MK data
Normal data

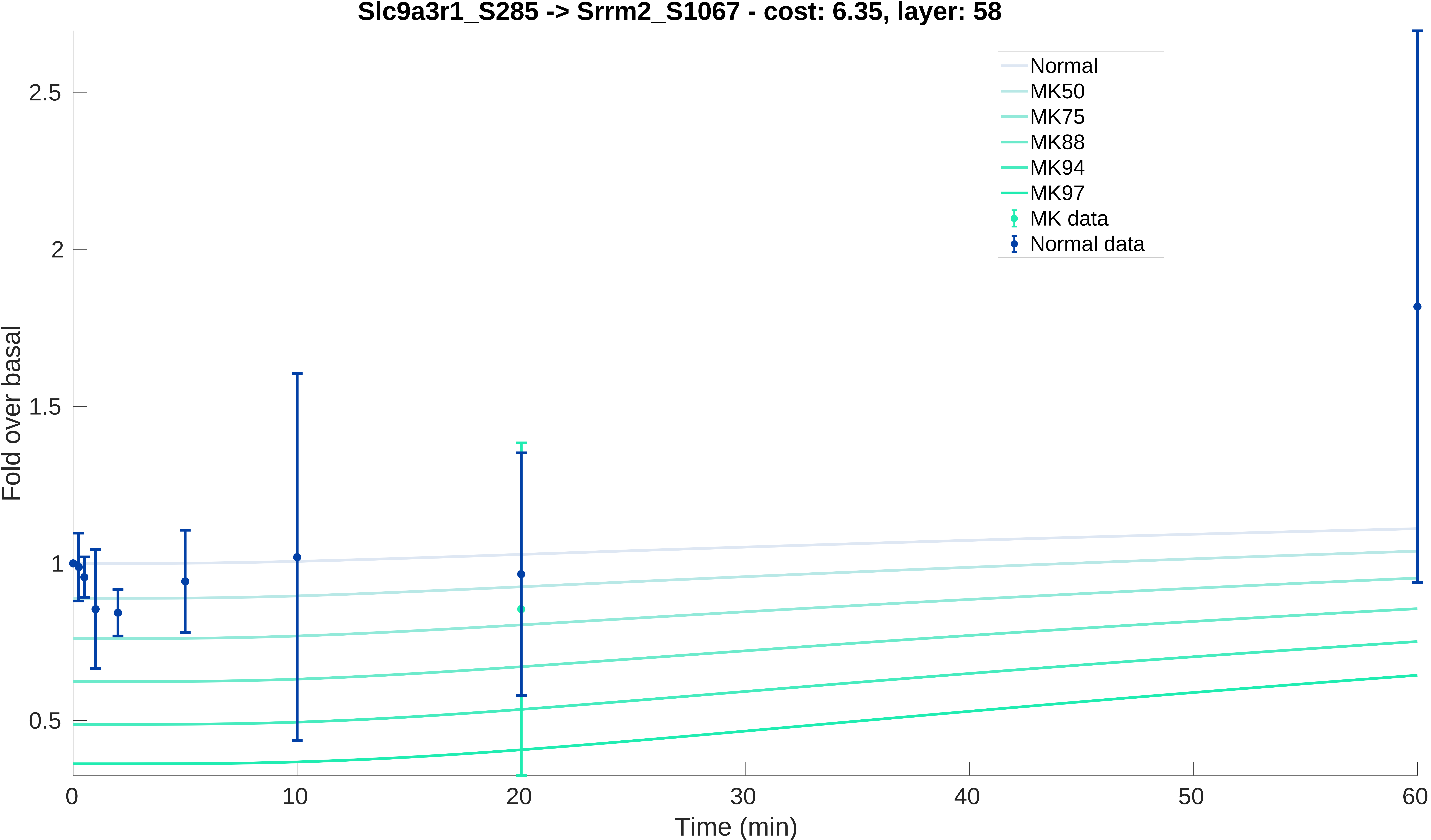
Slc9a3r1_S285 -> Srrm2_S1067 - cost: 6.35, layer: 58
Fold over basal
2.5
2
1.5
1
0.5
0
10
20
30
40
50
60
Time (min)
Normal
MK50
MK75
MK88
MK94
MK97
MK data
Normal data

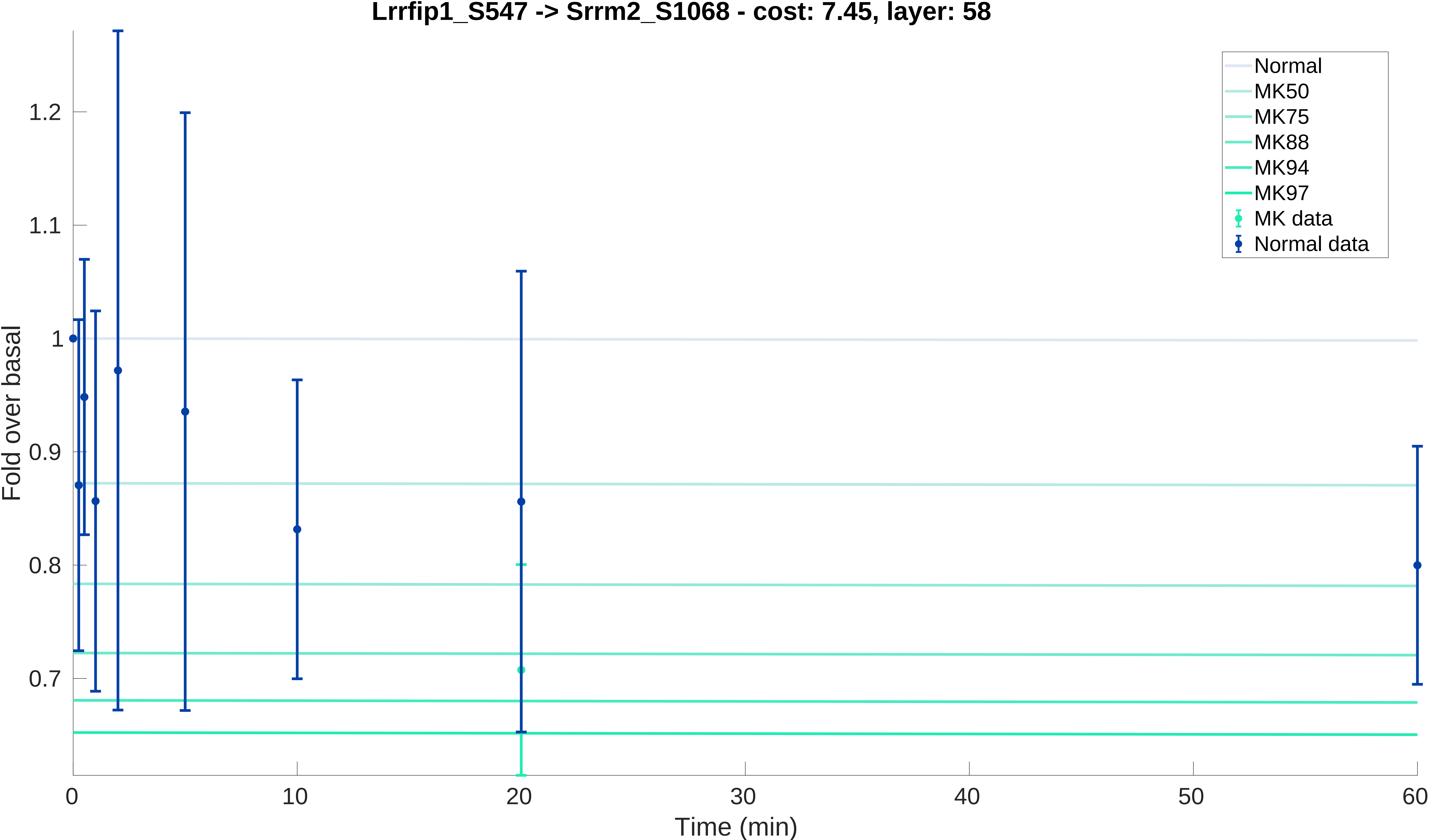

Lrrfip1_S547 -> Srrm2_S1068 - cost: 7.45, layer: 58
Fold over basal
Time (min)
0
10
20
30
40
50
60
0.7
0.8
0.9
1
1.1
1.2
Normal
MK50
MK75
MK88
MK94
MK97
MK data
Normal data

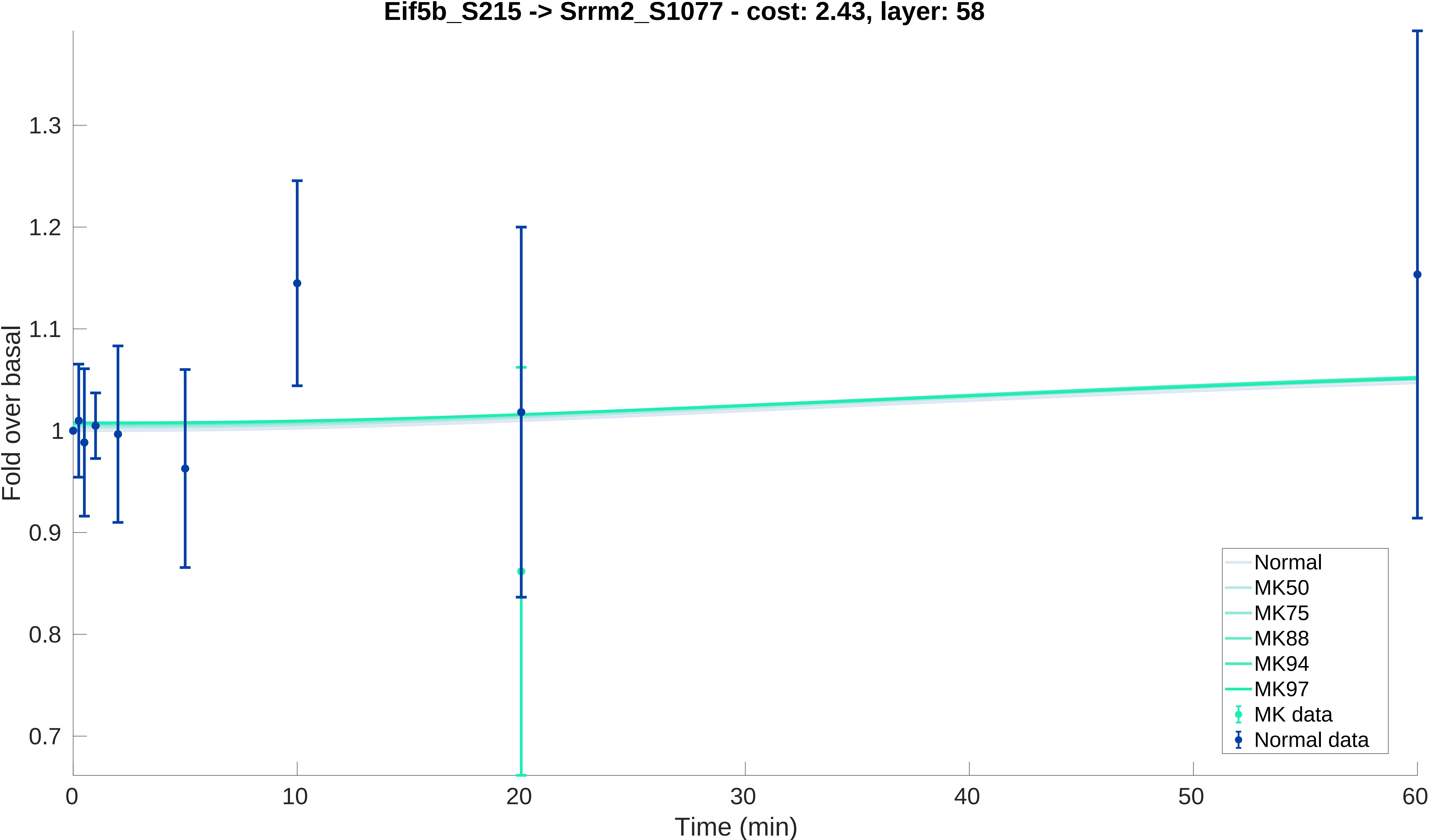

Eif5b_S215 -> Srrm2_S1077 - cost: 2.43, layer: 58
Fold over basal
Time (min)
0.7
0.8
0.9
1
1.1
1.2
1.3
0
10
20
30
40
50
60
Normal
MK50
MK75
MK88
MK94
MK97
MK data
Normal data

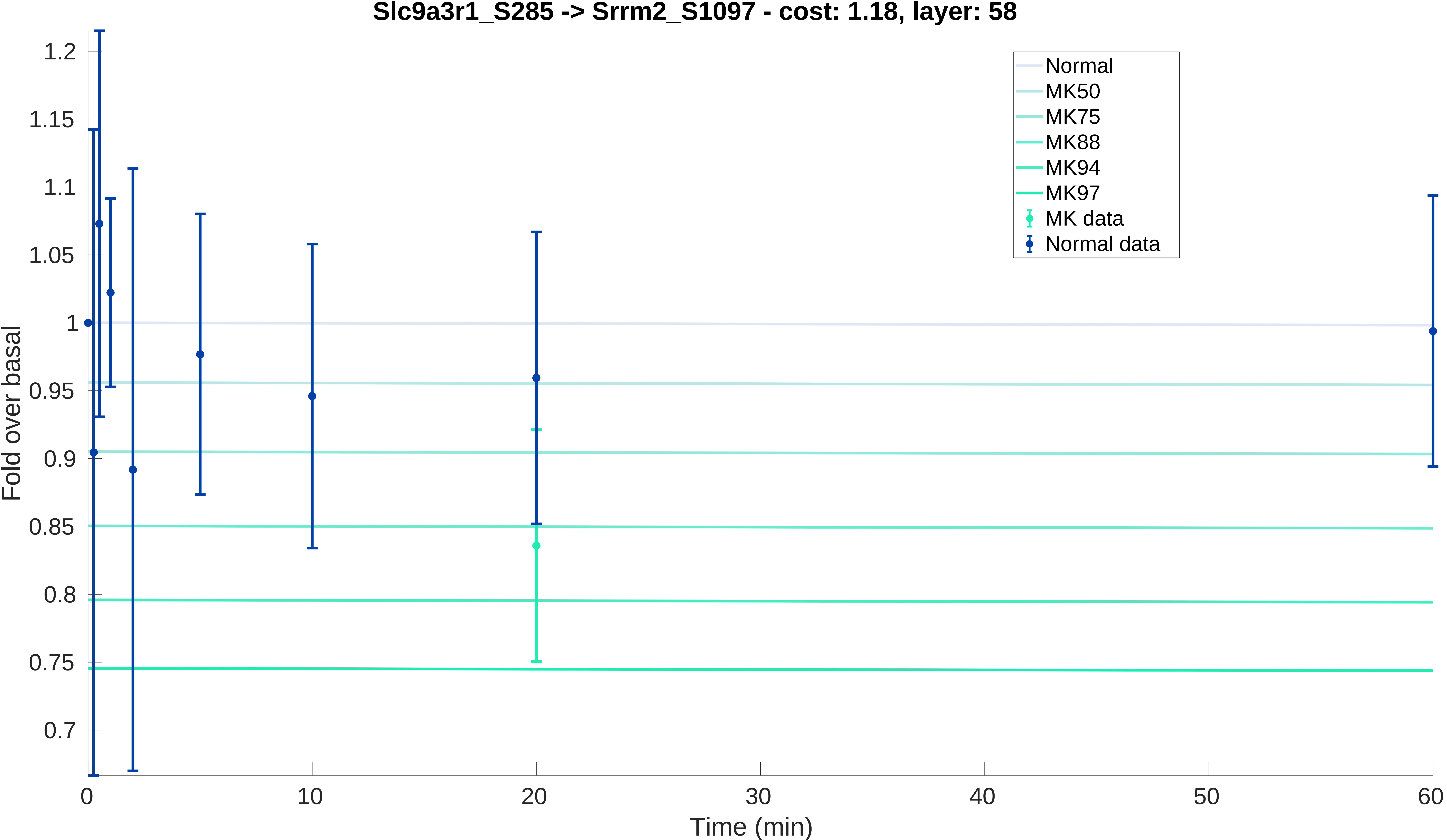

Slc9a3r1_S285 -> Srrm2_S1097 - cost: 1.18, layer: 58
Fold over basal
1.2
1.15
1.1
1.05
1
0.95
0.9
0.85
0.8
0.75
0.7
0
10
20
30
40
50
60
Time (min)
Normal
MK50
MK75
MK88
MK94
MK97
MK data
Normal data

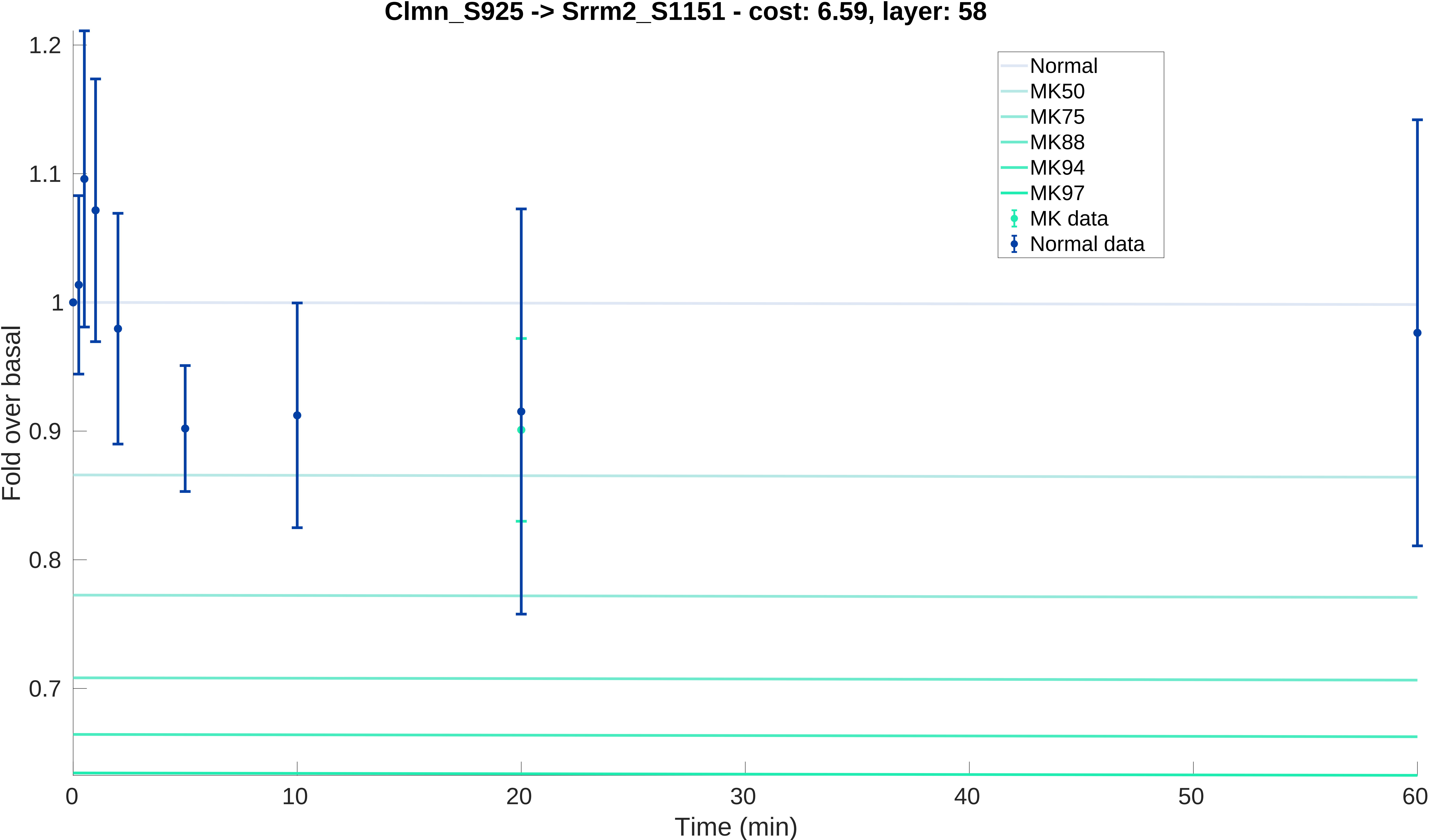

Clmn_S925 -> Srrm2_S1151 - cost: 6.59, layer: 58
Fold over basal
1.2
1.1
1
0.9
0.8
0.7
0
10
20
30
40
50
60
Time (min)
Normal
MK50
MK75
MK88
MK94
MK97
MK data
Normal data

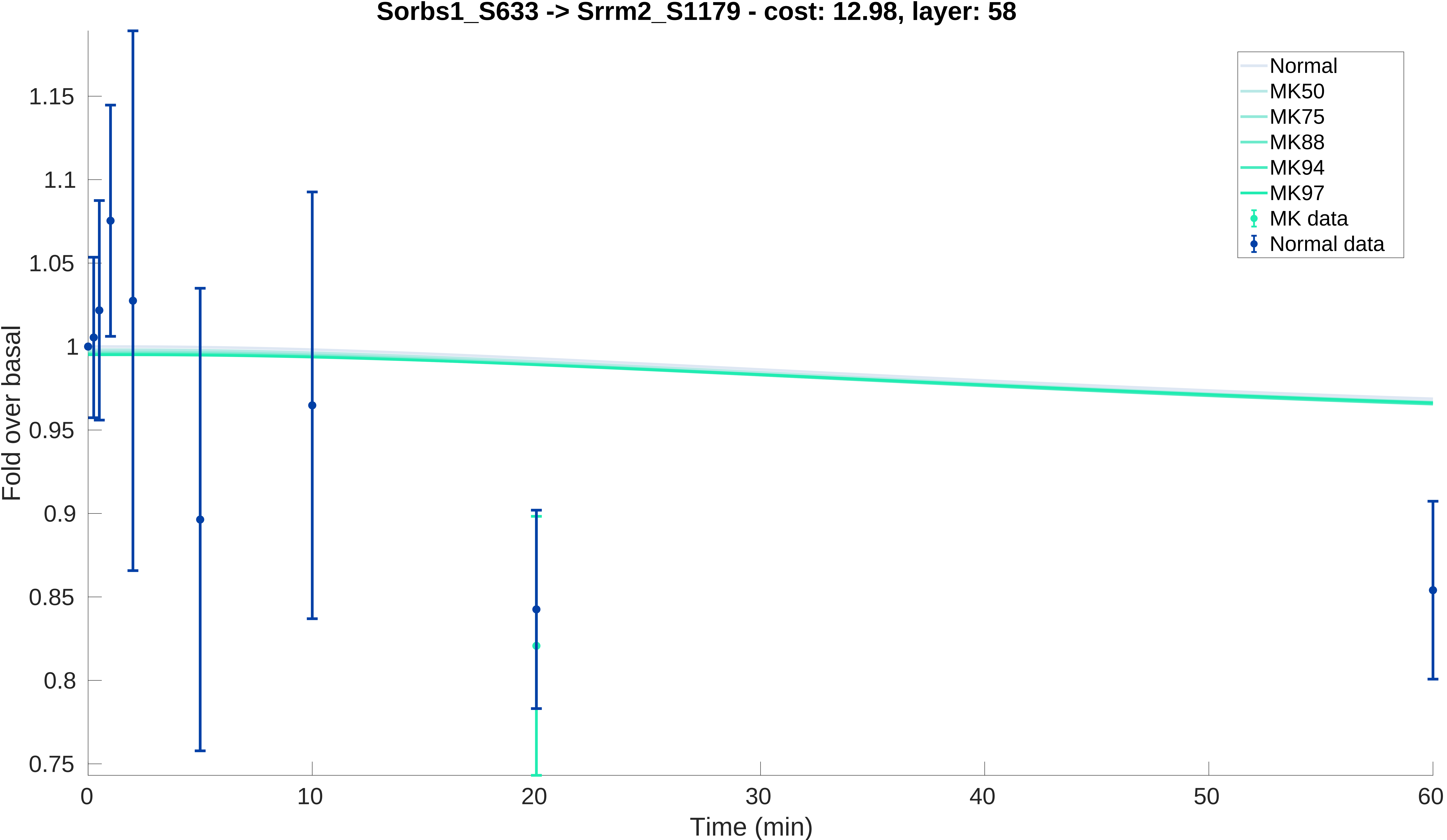

Sorbs1_S633 -> Srrm2_S1179 - cost: 12.98, layer: 58
Normal
MK50
MK75
MK88
MK94
MK97
MK data
Normal data
Fold over basal
1.15
1.1
1.05
1
0.95
0.9
0.85
0.8
0.75
0
10
20
30
40
50
60
Time (min)

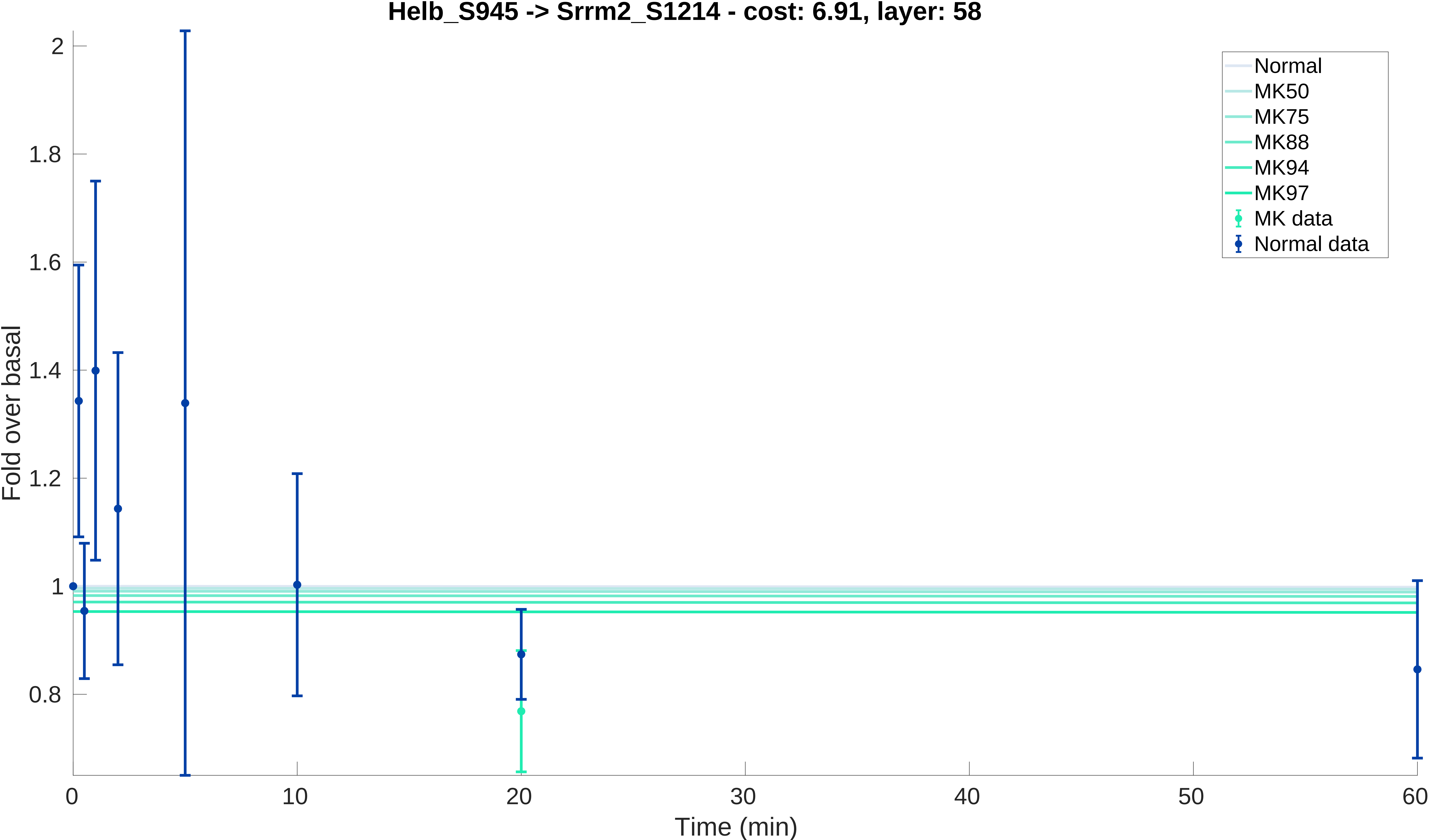

Helb_S945 -> Srrm2_S1214 - cost: 6.91, layer: 58
Fold over basal
Time (min)
0
10
20
30
40
50
60
0.8
1
1.2
1.4
1.6
1.8
2
Normal
MK50
MK75
MK88
MK94
MK97
MK data
Normal data

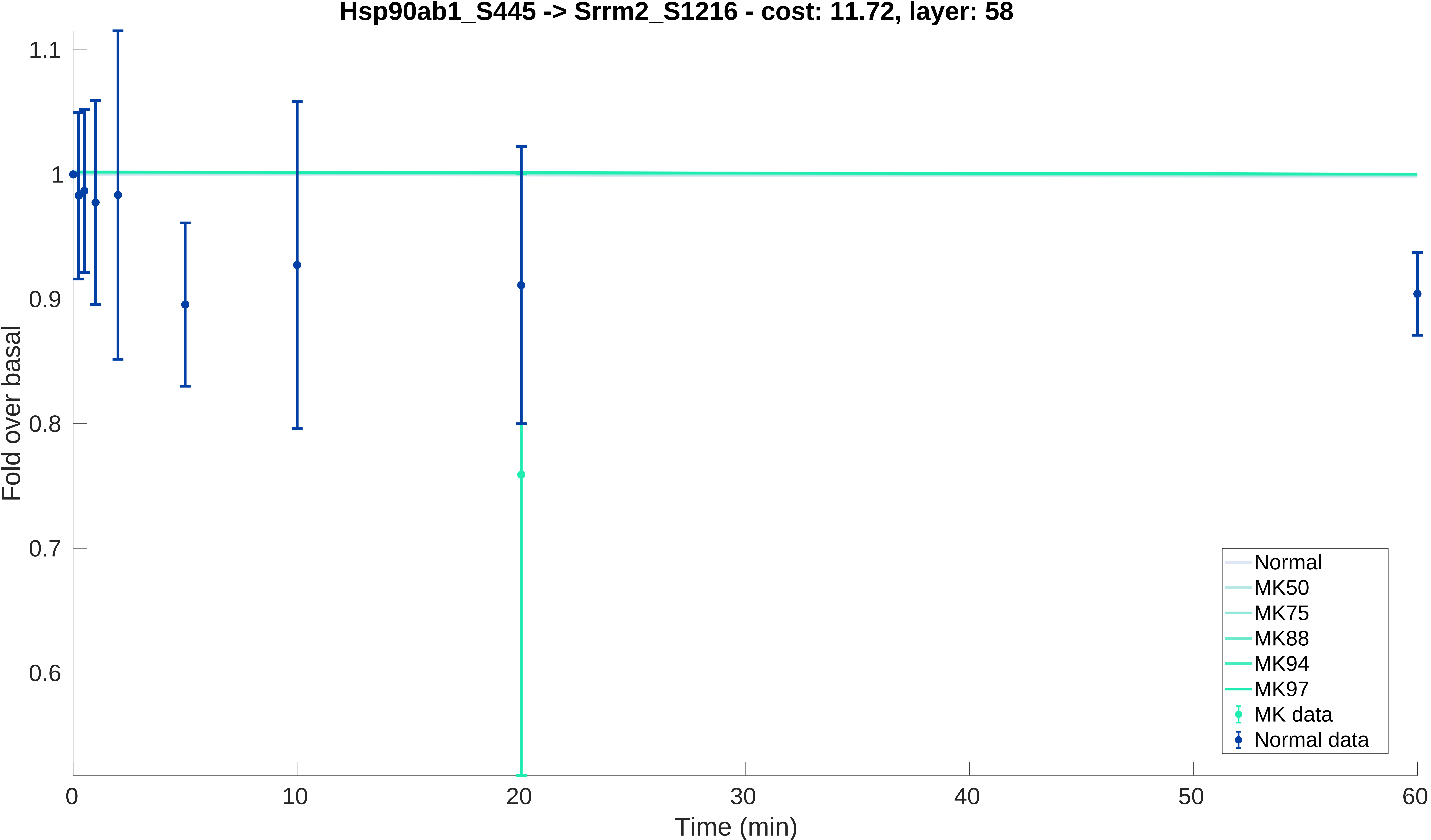
Hsp90ab1_S445 -> Srrm2_S1216 - cost: 11.72, layer: 58
Fold over basal
Time (min)
1.1
1
0.9
0.8
0.7
0.6
0
10
20
30
40
50
60
Normal
MK50
MK75
MK88
MK94
MK97
MK data
Normal data

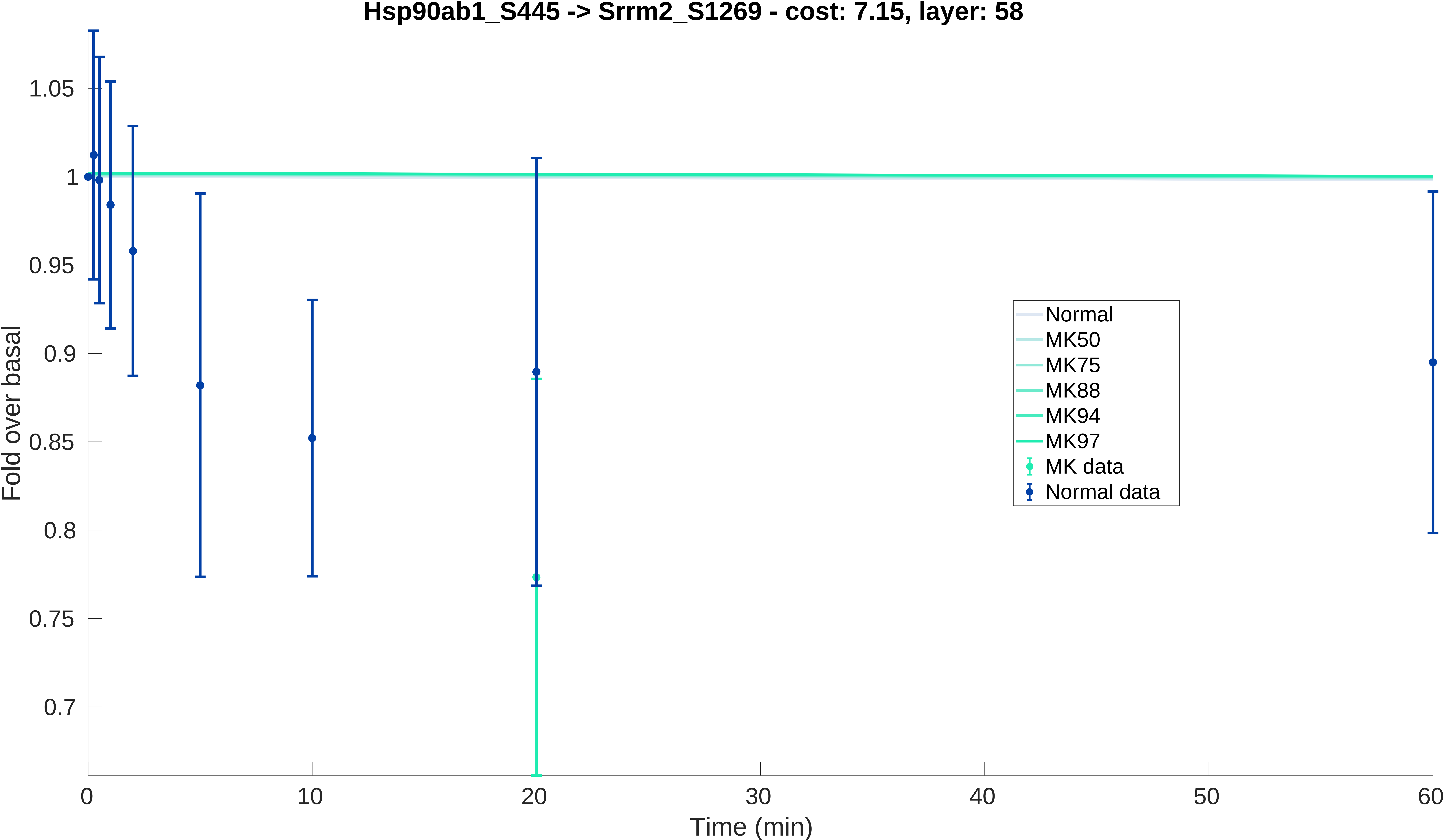

Hsp90ab1_S445 -> Srrm2_S1269 - cost: 7.15, layer: 58
Fold over basal
1.05
1
0.95
0.9
0.85
0.8
0.75
0.7
0
10
20
30
40
50
60
Time (min)
Normal
MK50
MK75
MK88
MK94
MK97
MK data
Normal data

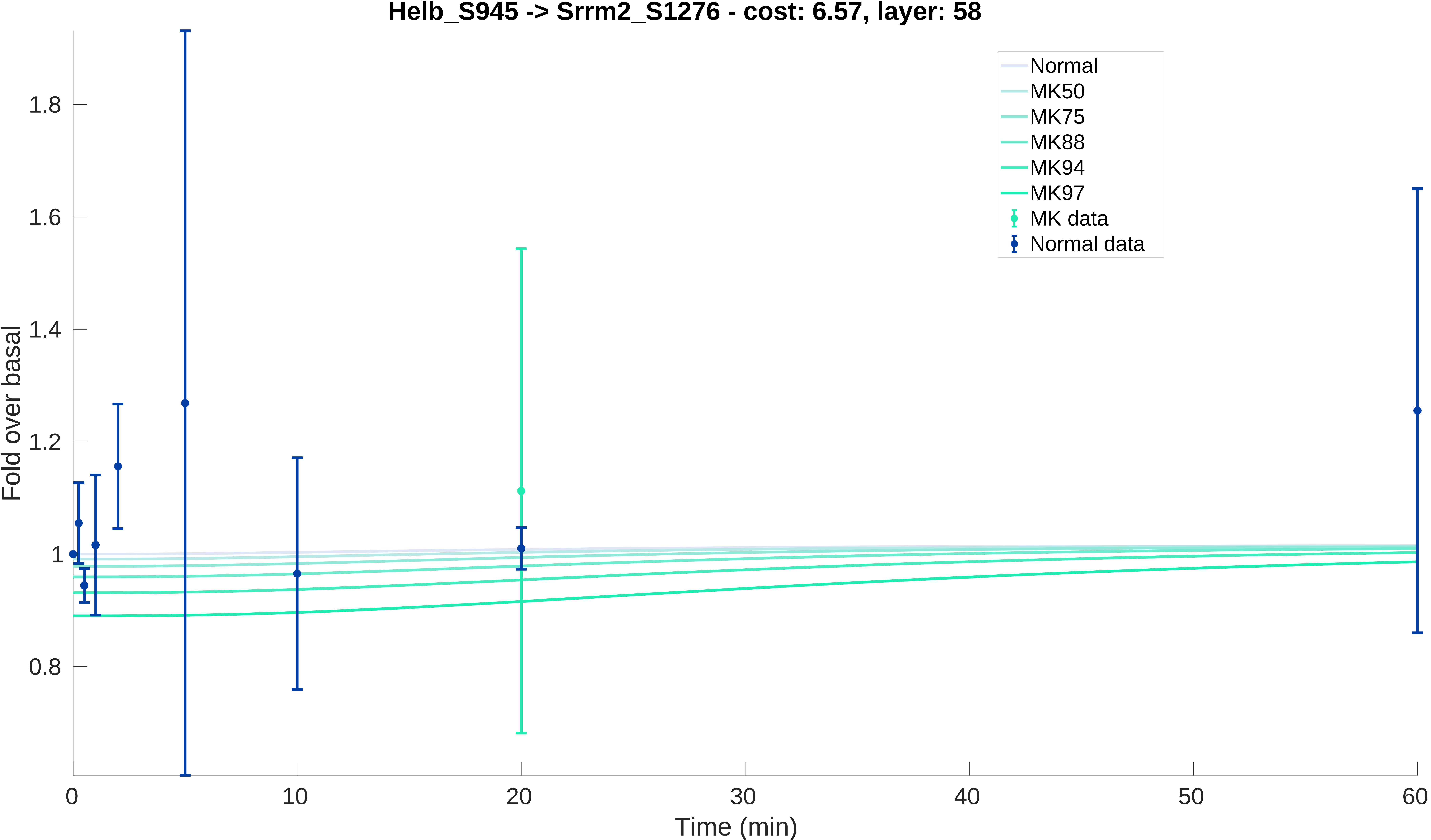

Helb_S945 -> Srrm2_S1276 - cost: 6.57, layer: 58
Normal
MK50
MK75
MK88
MK94
MK97
MK data
Normal data
Fold over basal
1.8
1.6
1.4
1.2
1
0.8
0
10
20
30
40
50
60
Time (min)

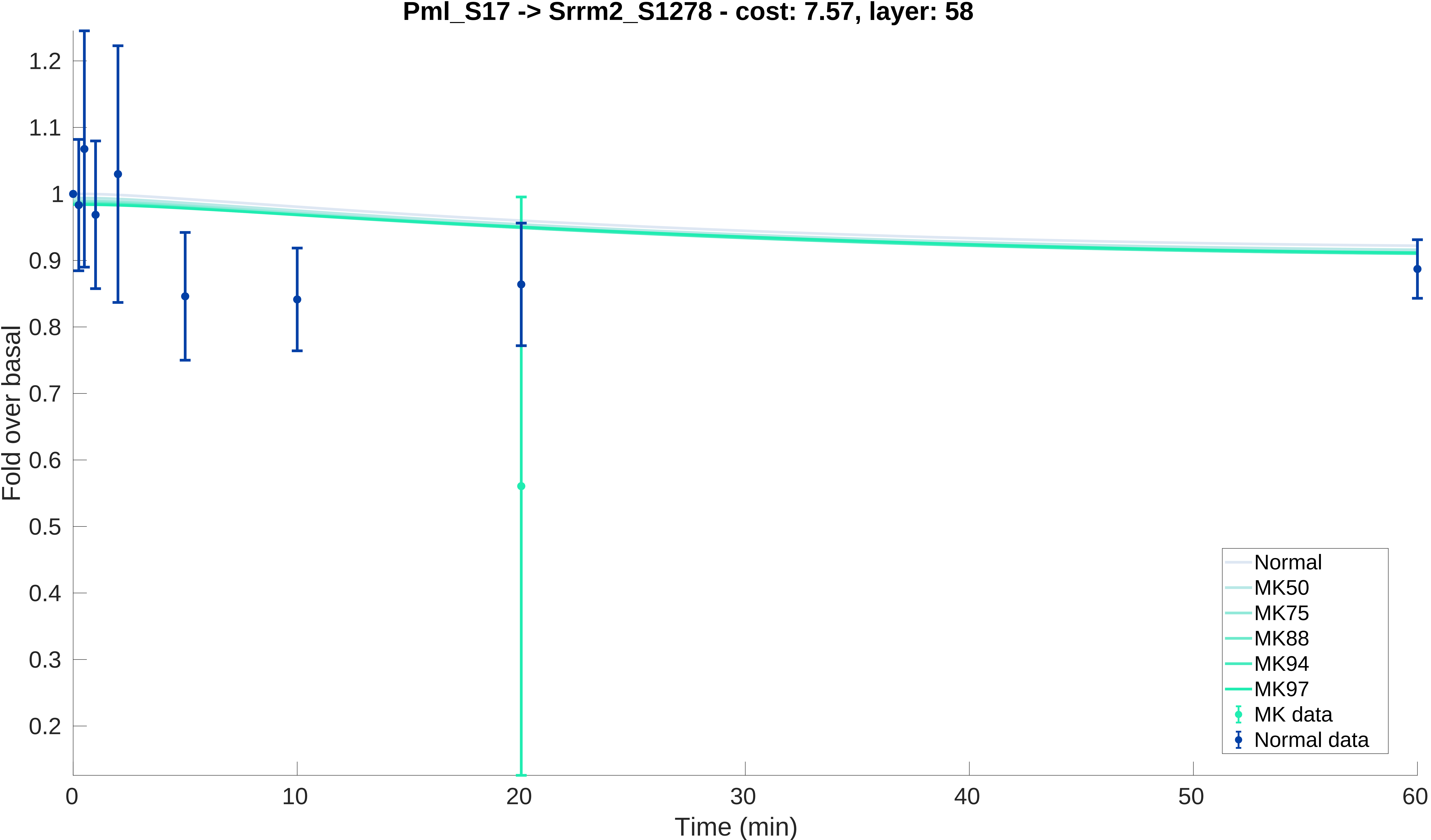

Pml_S17 -> Srrm2_S1278 - cost: 7.57, layer: 58
Fold over basal
Time (min)
0.2
0.3
0.4
0.5
0.6
0.7
0.8
0.9
1
1.1
1.2
0
10
20
30
40
50
60
Normal
MK50
MK75
MK88
MK94
MK97
MK data
Normal data

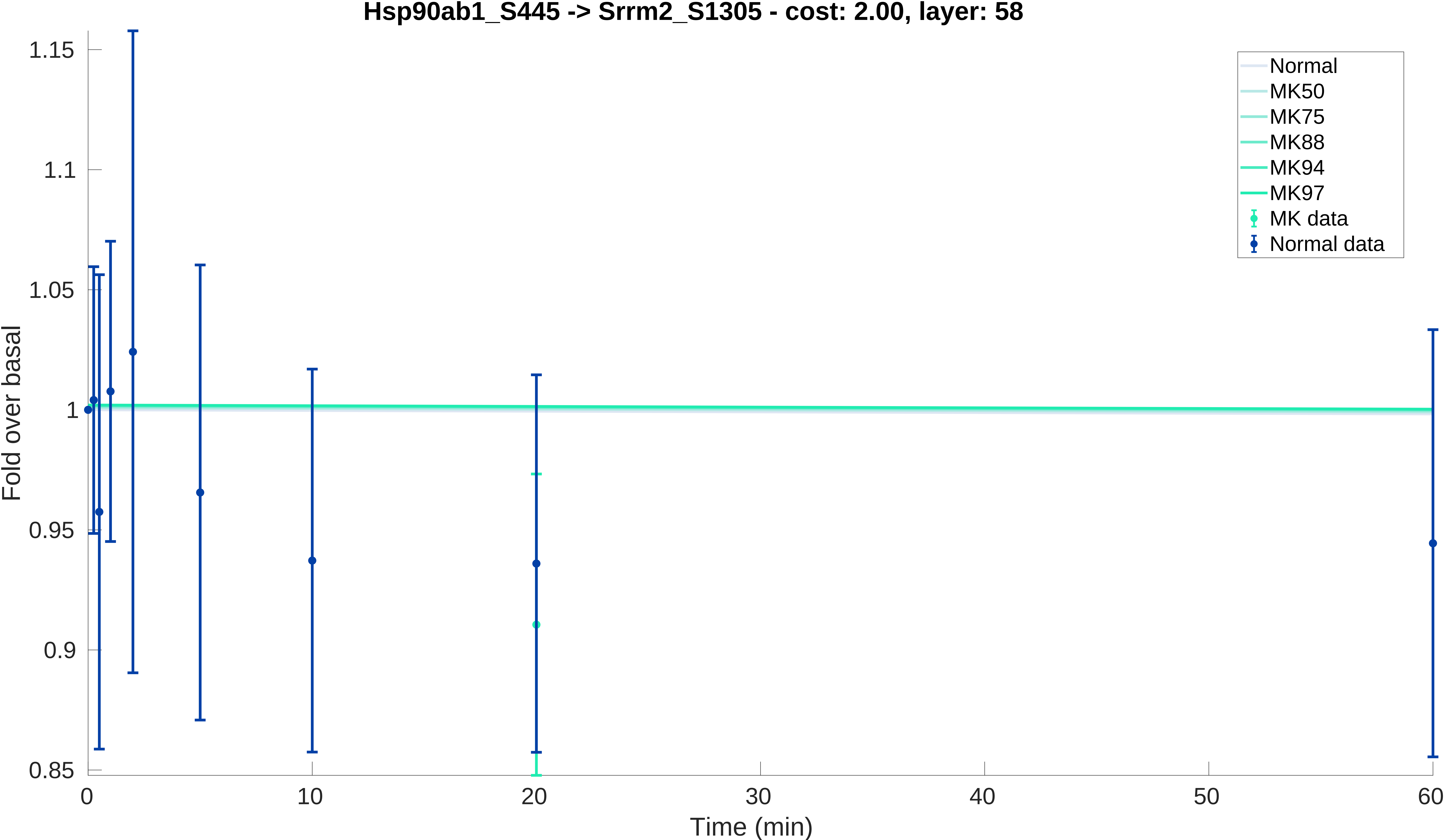

Hsp90ab1_S445 -> Srrm2_S1305 - cost: 2.00, layer: 58
Fold over basal
Time (min)
1.15
1.1
1.05
1
0.95
0.9
0.85
0
10
20
30
40
50
60
Normal
MK50
MK75
MK88
MK94
MK97
MK data
Normal data

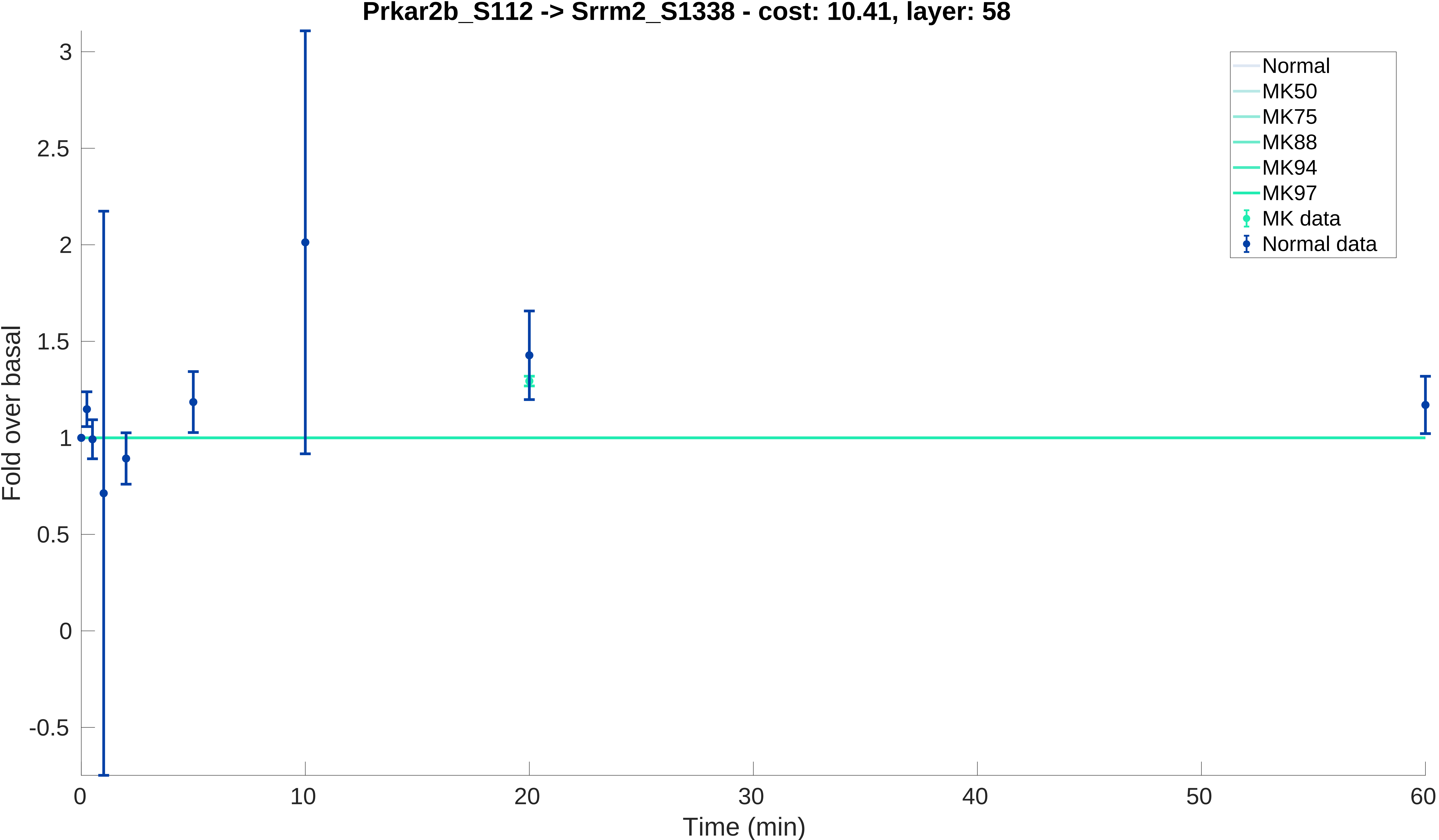

Prkar2b_S112 -> Srrm2_S1338 - cost: 10.41, layer: 58
Fold over basal
Time (min)
3
2.5
2
1.5
1
0.5
0
-0.5
0
10
20
30
40
50
60
Normal
MK50
MK75
MK88
MK94
MK97
MK data
Normal data

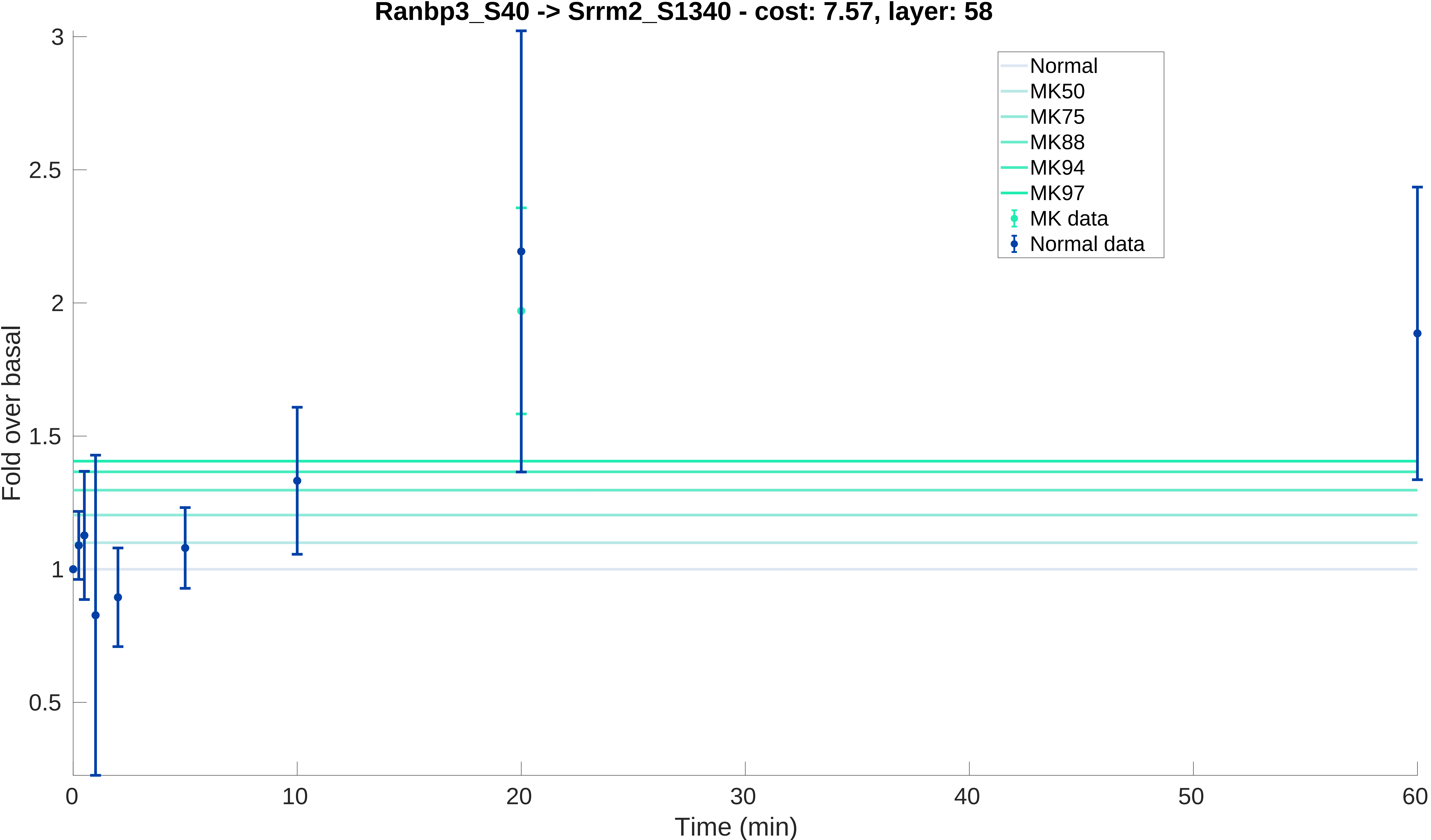

Ranbp3_S40 -> Srrm2_S1340 - cost: 7.57, layer: 58
Fold over basal
Time (min)
0
10
20
30
40
50
60
0.5
1
1.5
2
2.5
3
Normal
MK50
MK75
MK88
MK94
MK97
MK data
Normal data

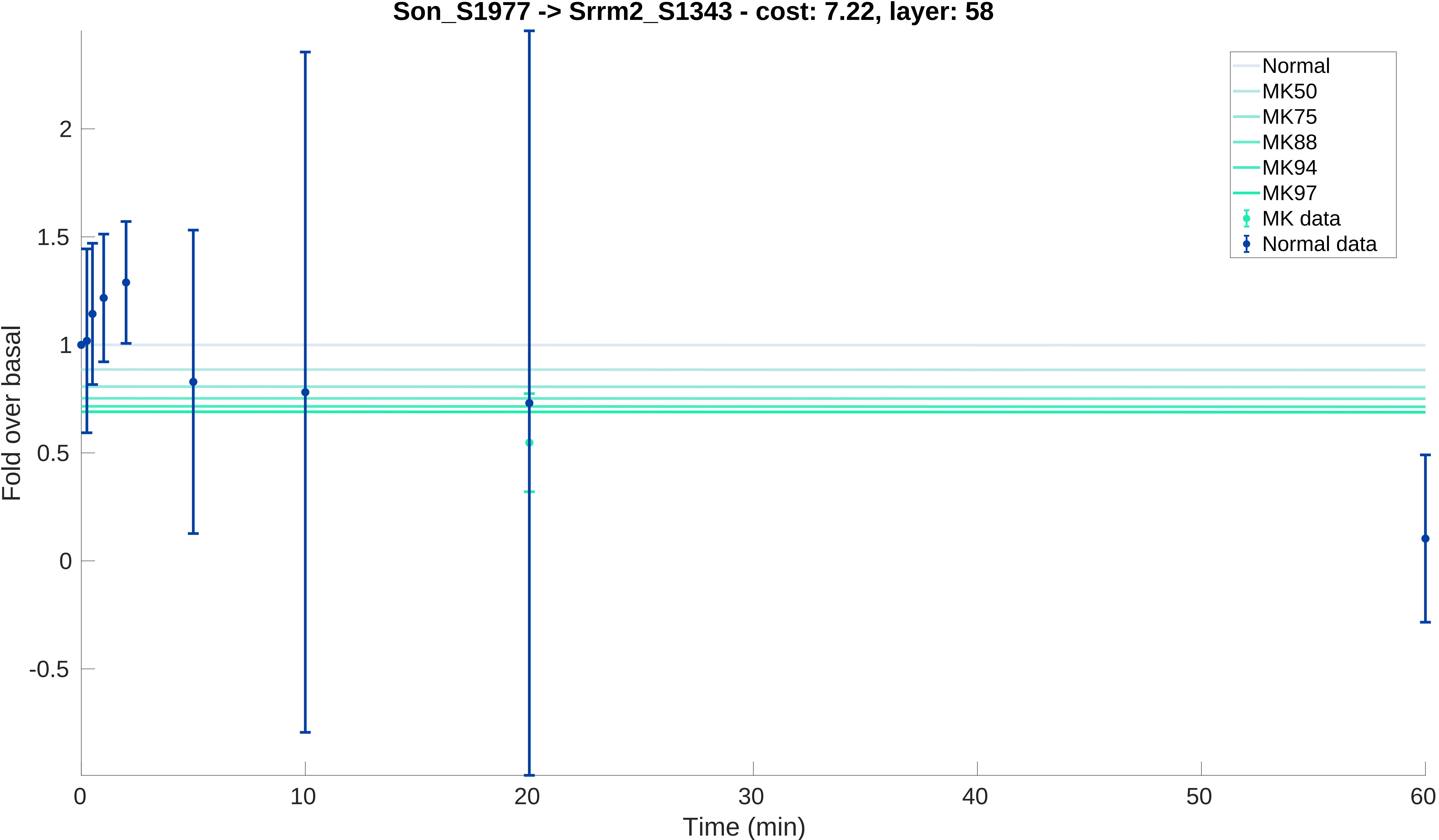

Son_S1977 -> Srrm2_S1343 - cost: 7.22, layer: 58
Normal
MK50
MK75
MK88
MK94
MK97
MK data
Normal data
Fold over basal
2
1.5
1
0.5
0
-0.5
0
10
20
30
40
50
60
Time (min)

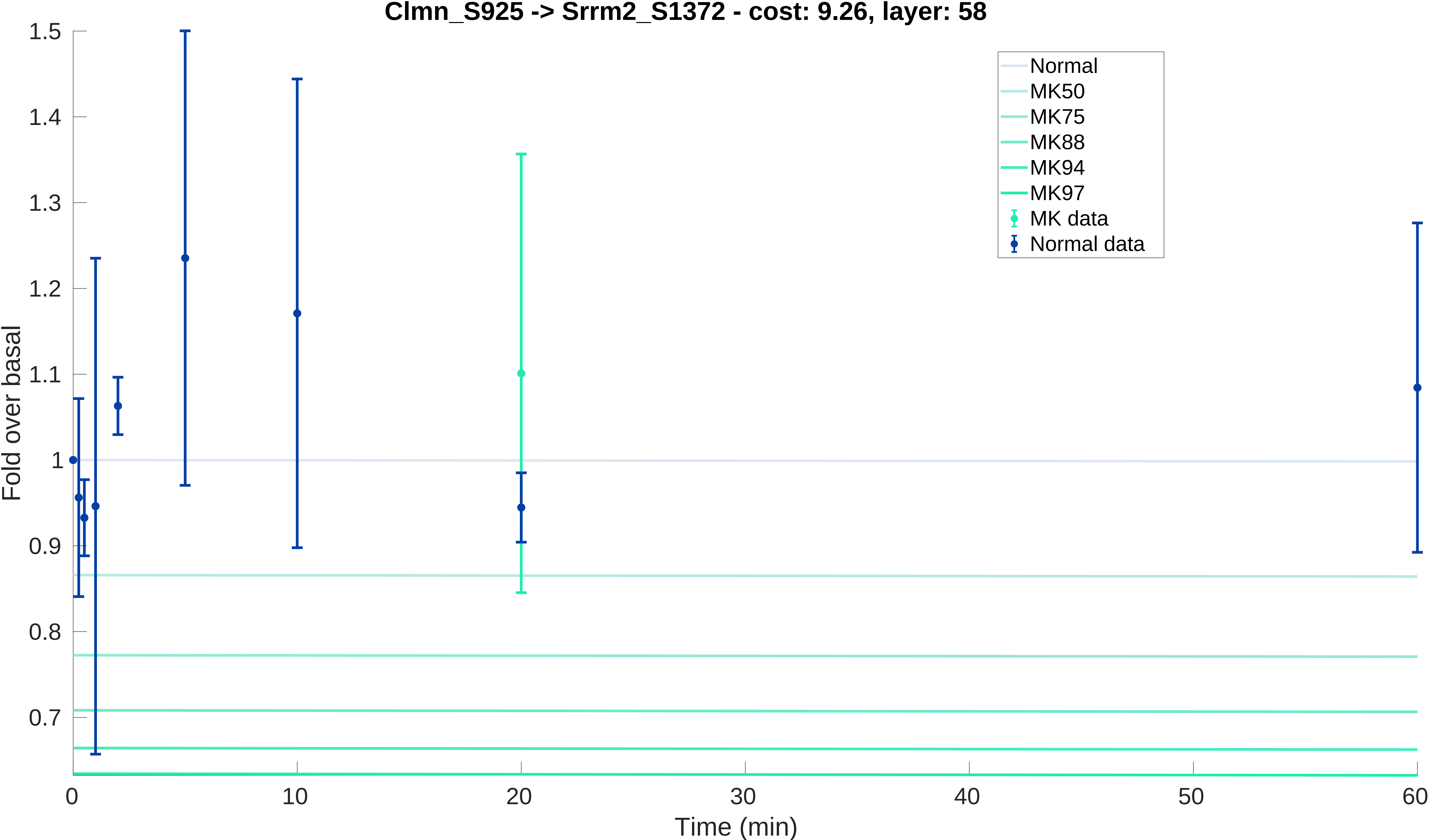

Clmn_S925 -> Srrm2_S1372 - cost: 9.26, layer: 58
Fold over basal
Time (min)
1.5
1.4
1.3
1.2
1.1
1
0.9
0.8
0.7
0
10
20
30
40
50
60
Normal
MK50
MK75
MK88
MK94
MK97
MK data
Normal data

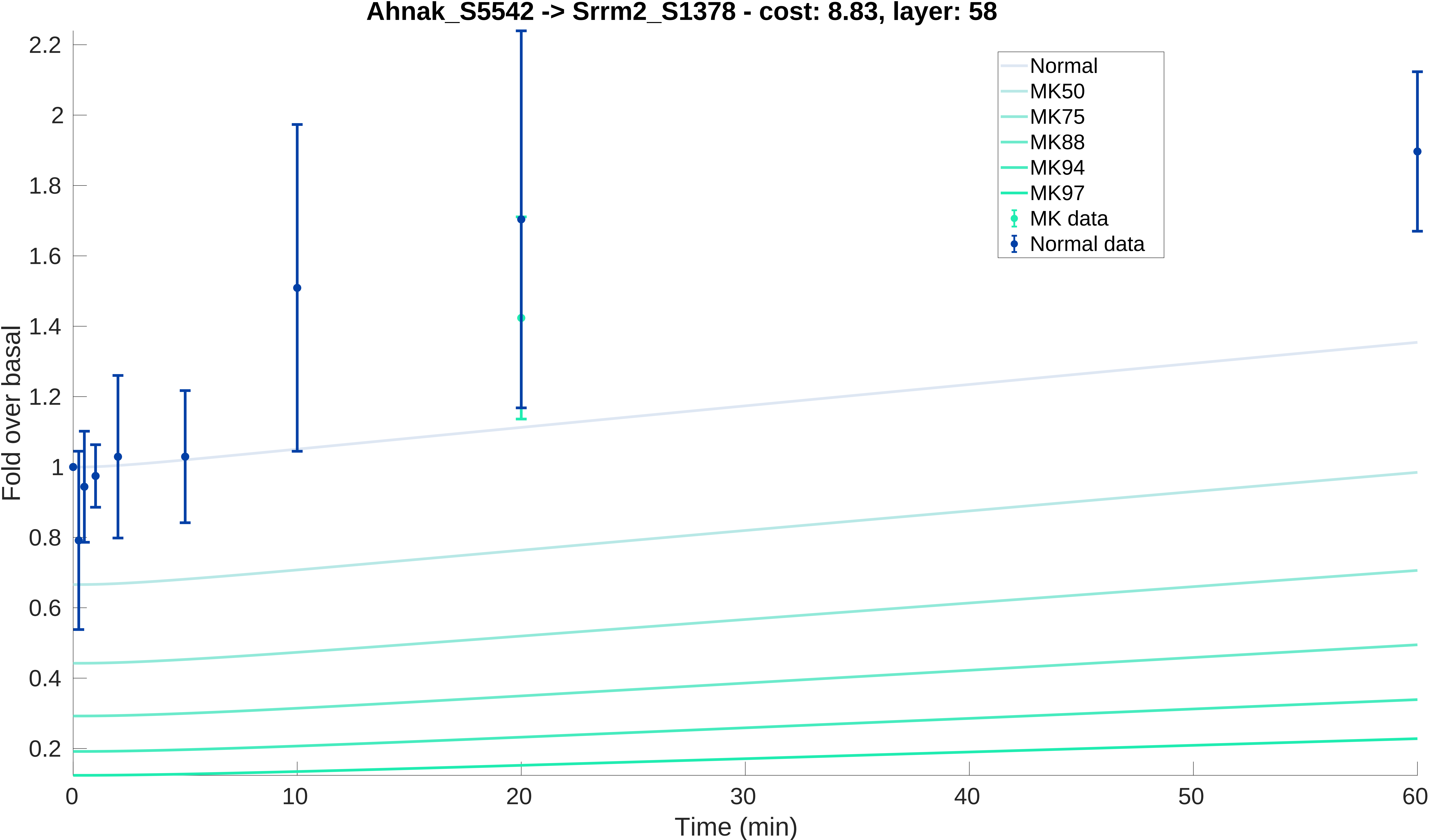

Ahnak_S5542 -> Srrm2_S1378 - cost: 8.83, layer: 58
Fold over basal
Time (min)
0
10
20
30
40
50
60
0.2
0.4
0.6
0.8
1
1.2
1.4
1.6
1.8
2
2.2
Normal
MK50
MK75
MK88
MK94
MK97
MK data
Normal data

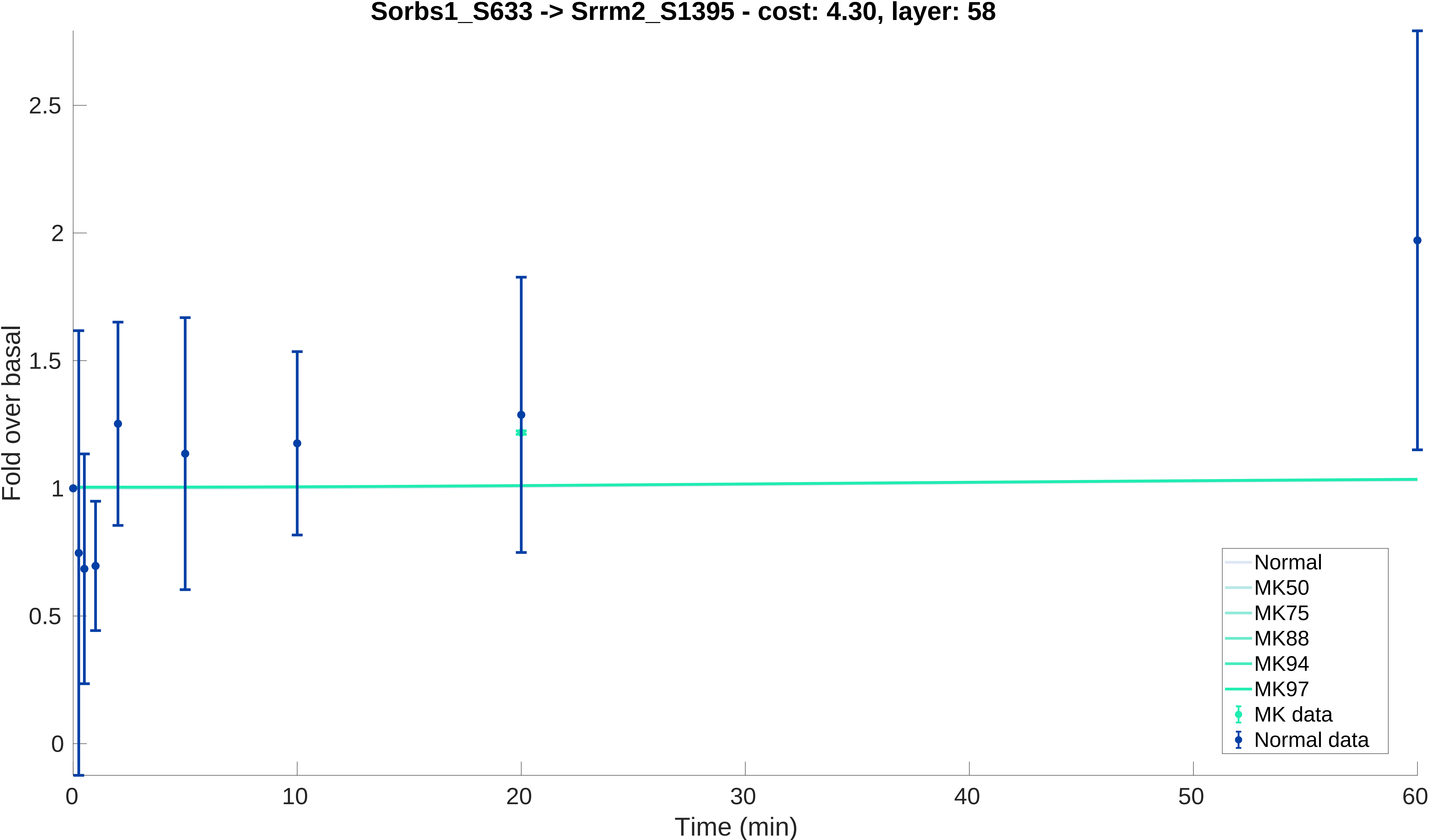

Sorbs1_S633 -> Srrm2_S1395 - cost: 4.30, layer: 58
Fold over basal
Time (min)
0
0.5
1
1.5
2
2.5
0
10
20
30
40
50
60
Normal
MK50
MK75
MK88
MK94
MK97
MK data
Normal data

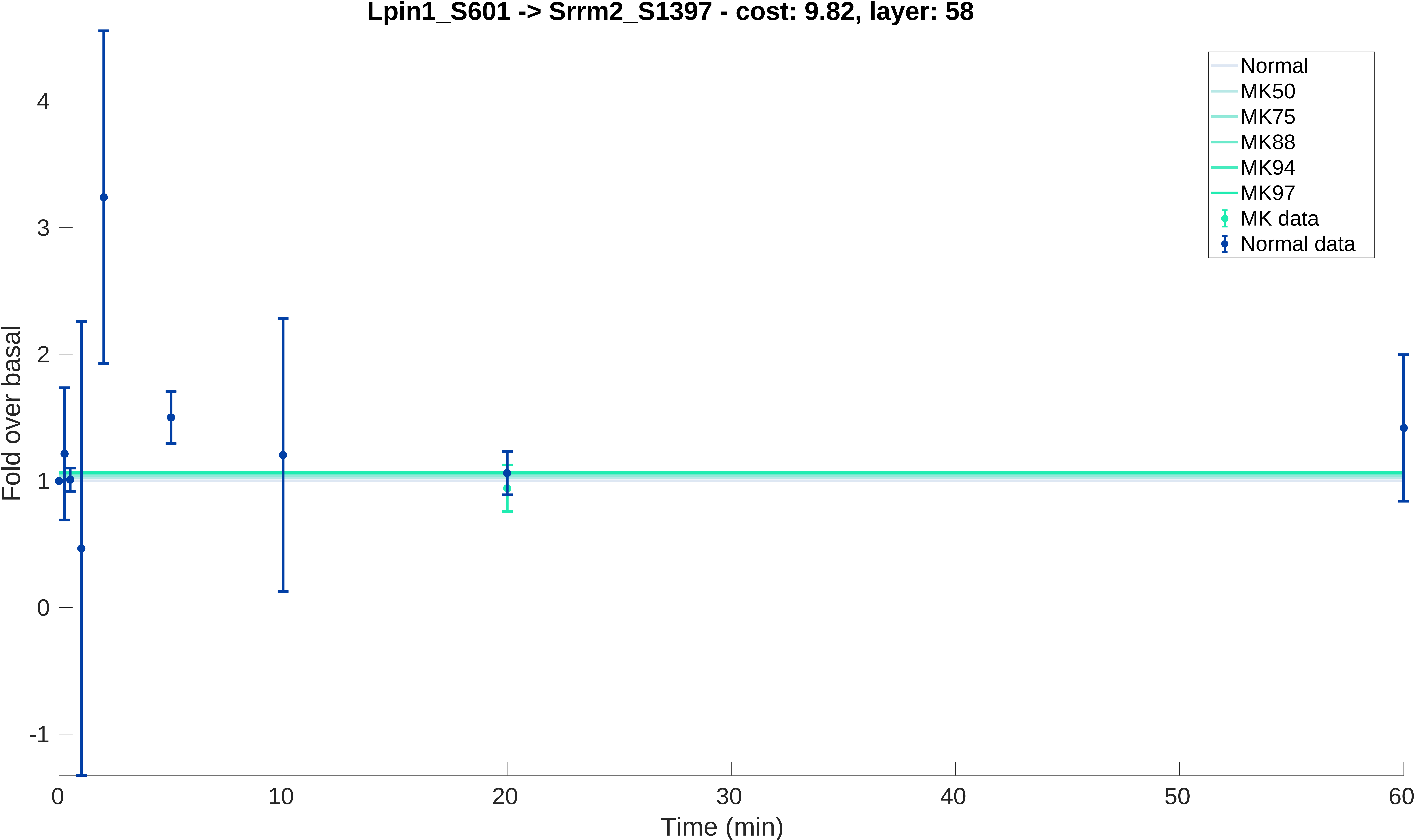

Lpin1_S601 -> Srrm2_S1397 - cost: 9.82, layer: 58
Normal
MK50
MK75
MK88
MK94
MK97
MK data
Normal data
Fold over basal
Time (min)
4
3
2
1
0
-1
0
10
20
30
40
50
60

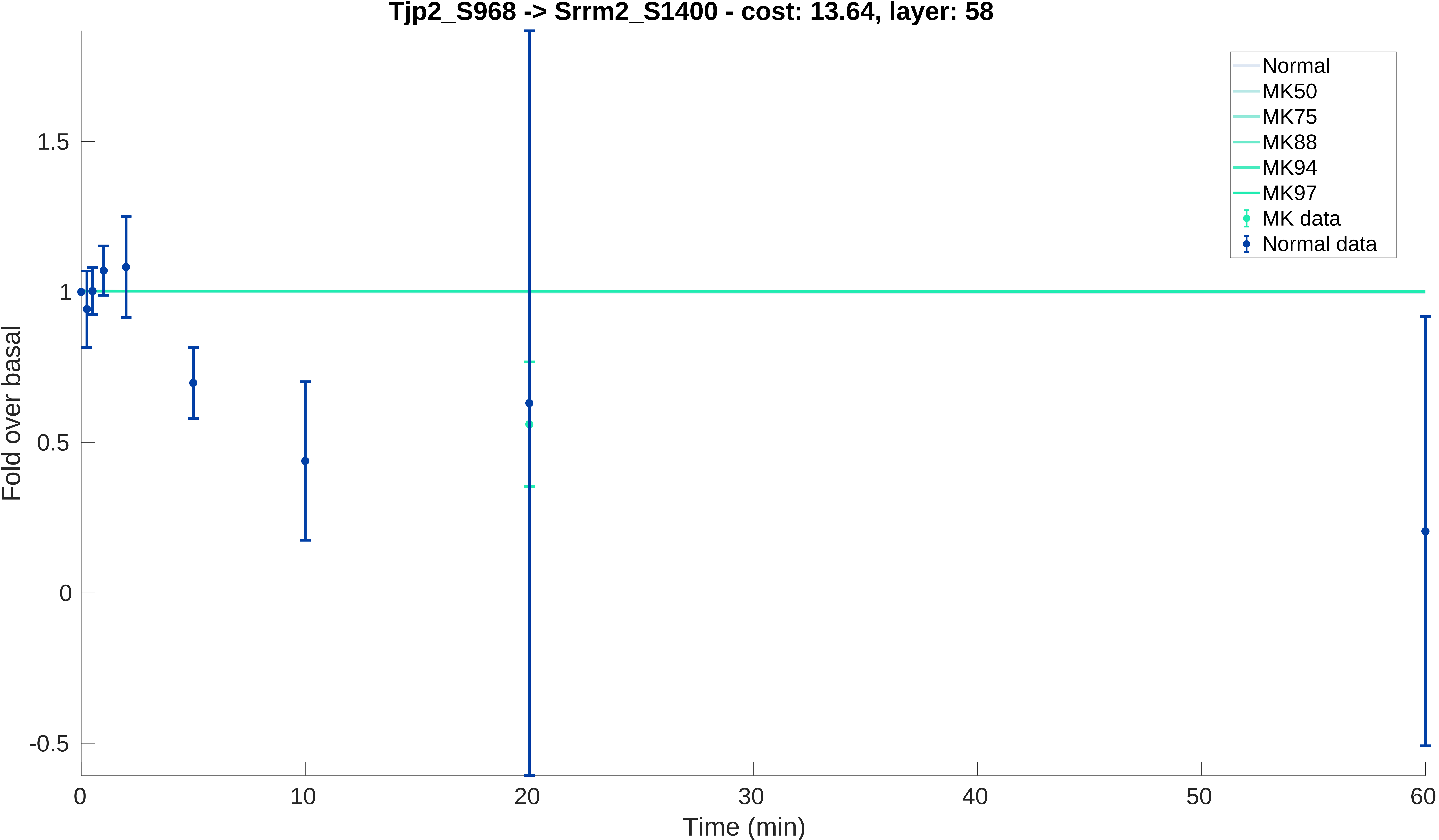

Tjp2_S968 -> Srrm2_S1400 - cost: 13.64, layer: 58
Fold over basal
Time (min)
1.5
1
0.5
0
-0.5
0
10
20
30
40
50
60
Normal
MK50
MK75
MK88
MK94
MK97
MK data
Normal data

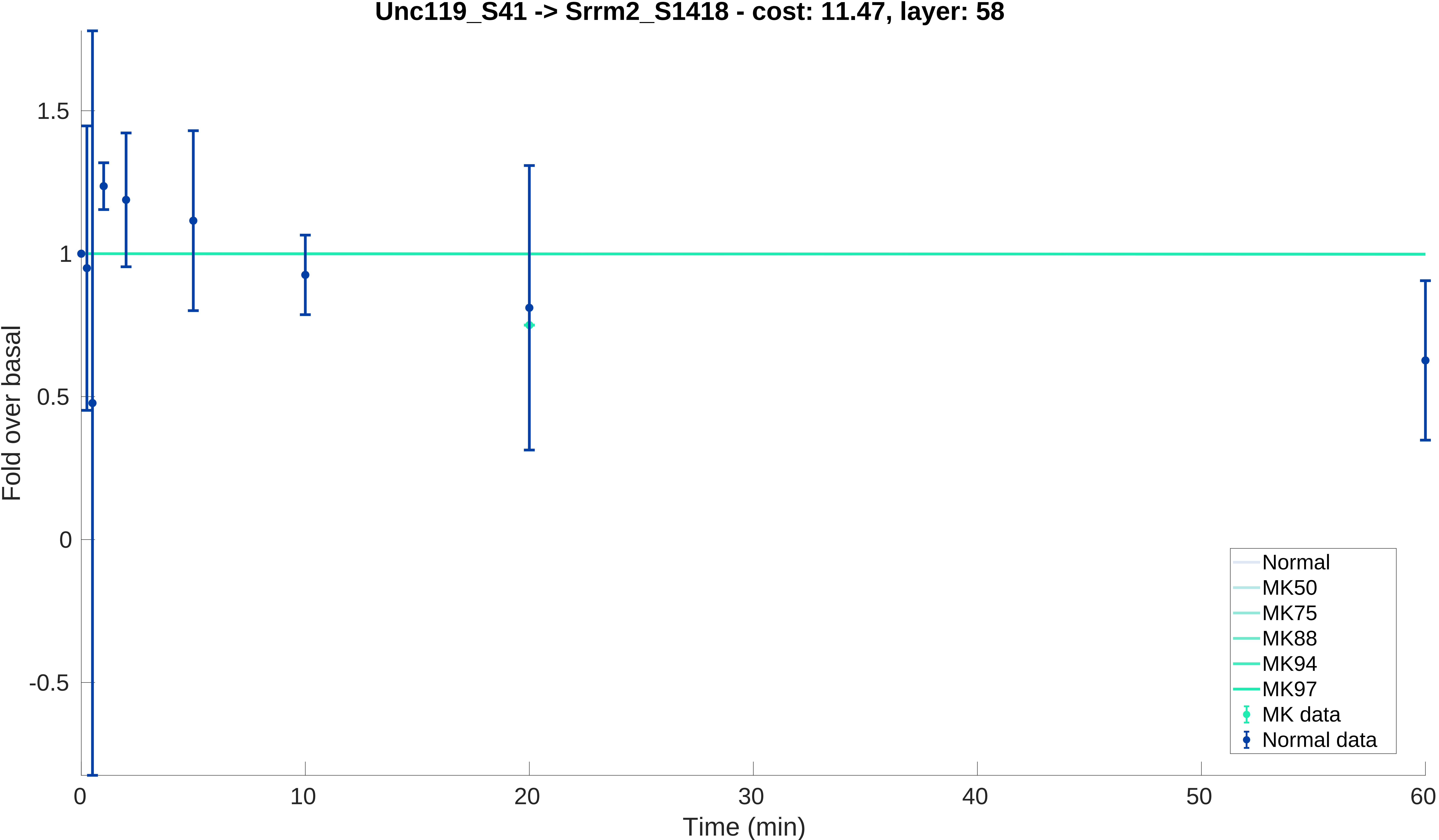

Unc119_S41 -> Srrm2_S1418 - cost: 11.47, layer: 58
Fold over basal
1.5
1
0.5
0
-0.5
0
10
20
30
40
50
60
Time (min)
Normal
MK50
MK75
MK88
MK94
MK97
MK data
Normal data

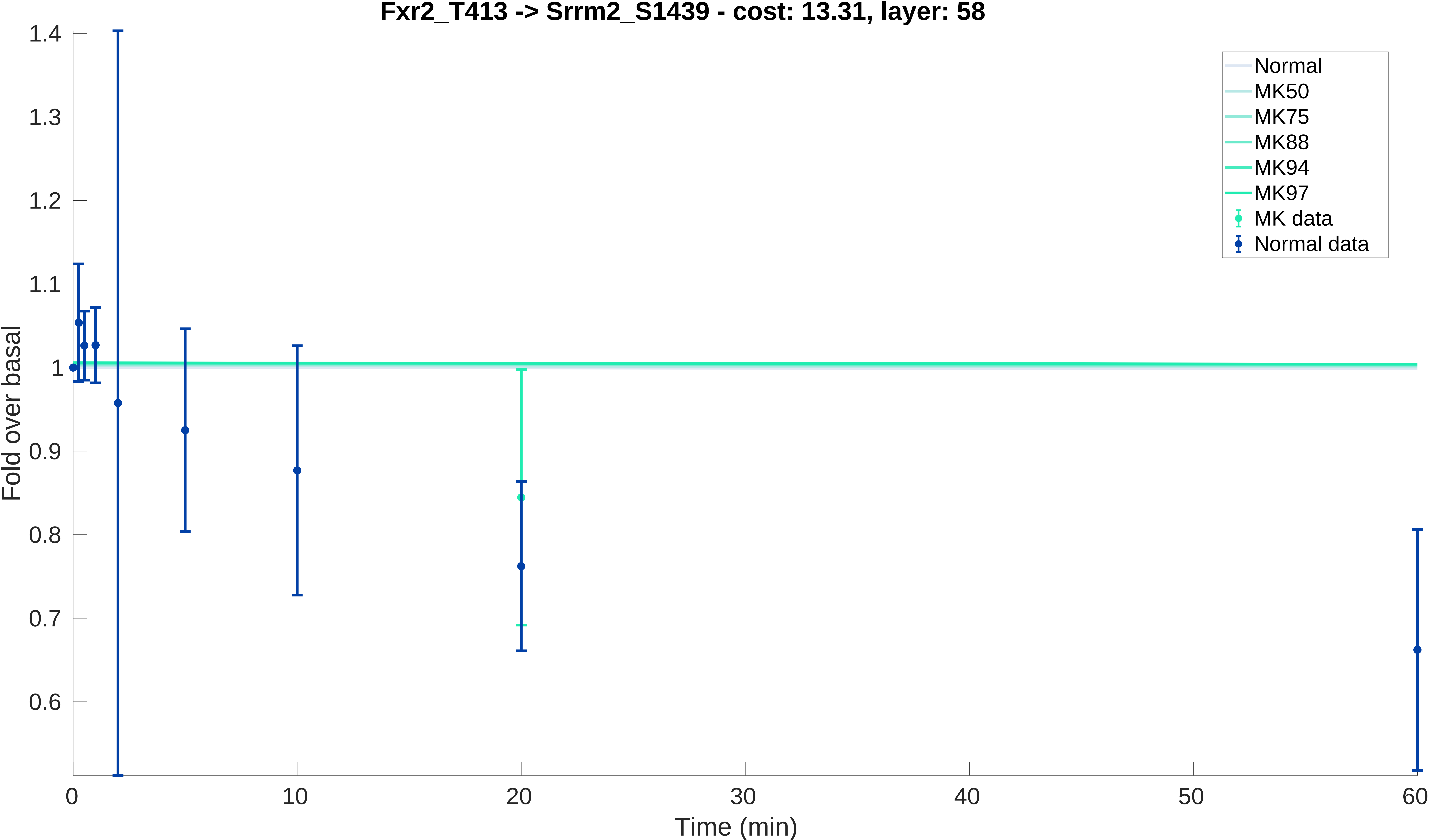

Fxr2_T413 -> Srrm2_S1439 - cost: 13.31, layer: 58
Fold over basal
Time (min)
1.4
1.3
1.2
1.1
1
0.9
0.8
0.7
0.6
0
10
20
30
40
50
60
Normal
MK50
MK75
MK88
MK94
MK97
MK data
Normal data

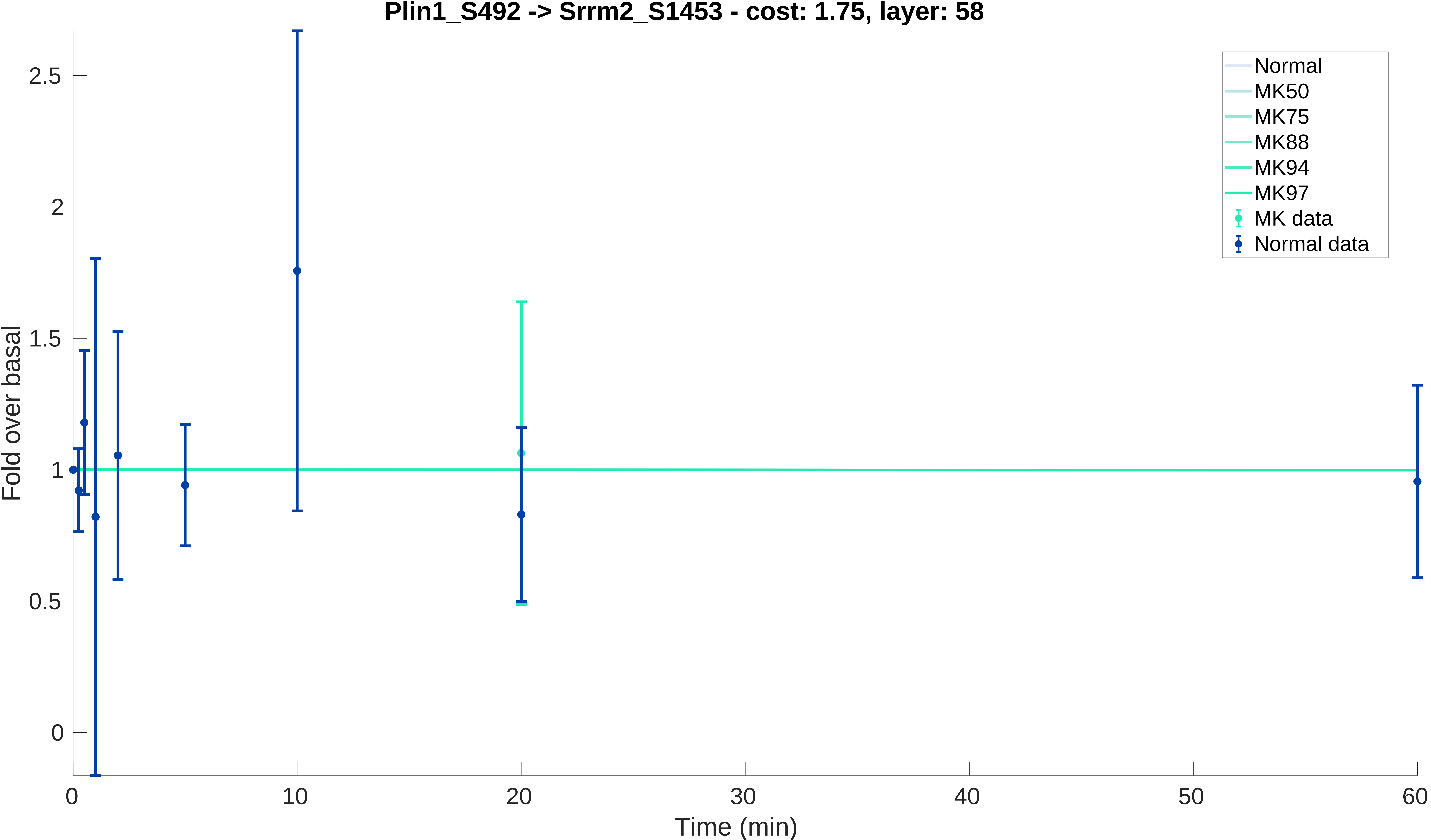

Plin1_S492 -> Srrm2_S1453 - cost: 1.75, layer: 58
Fold over basal
Time (min)
0
0.5
1
1.5
2
2.5
0
10
20
30
40
50
60
Normal
MK50
MK75
MK88
MK94
MK97
MK data
Normal data

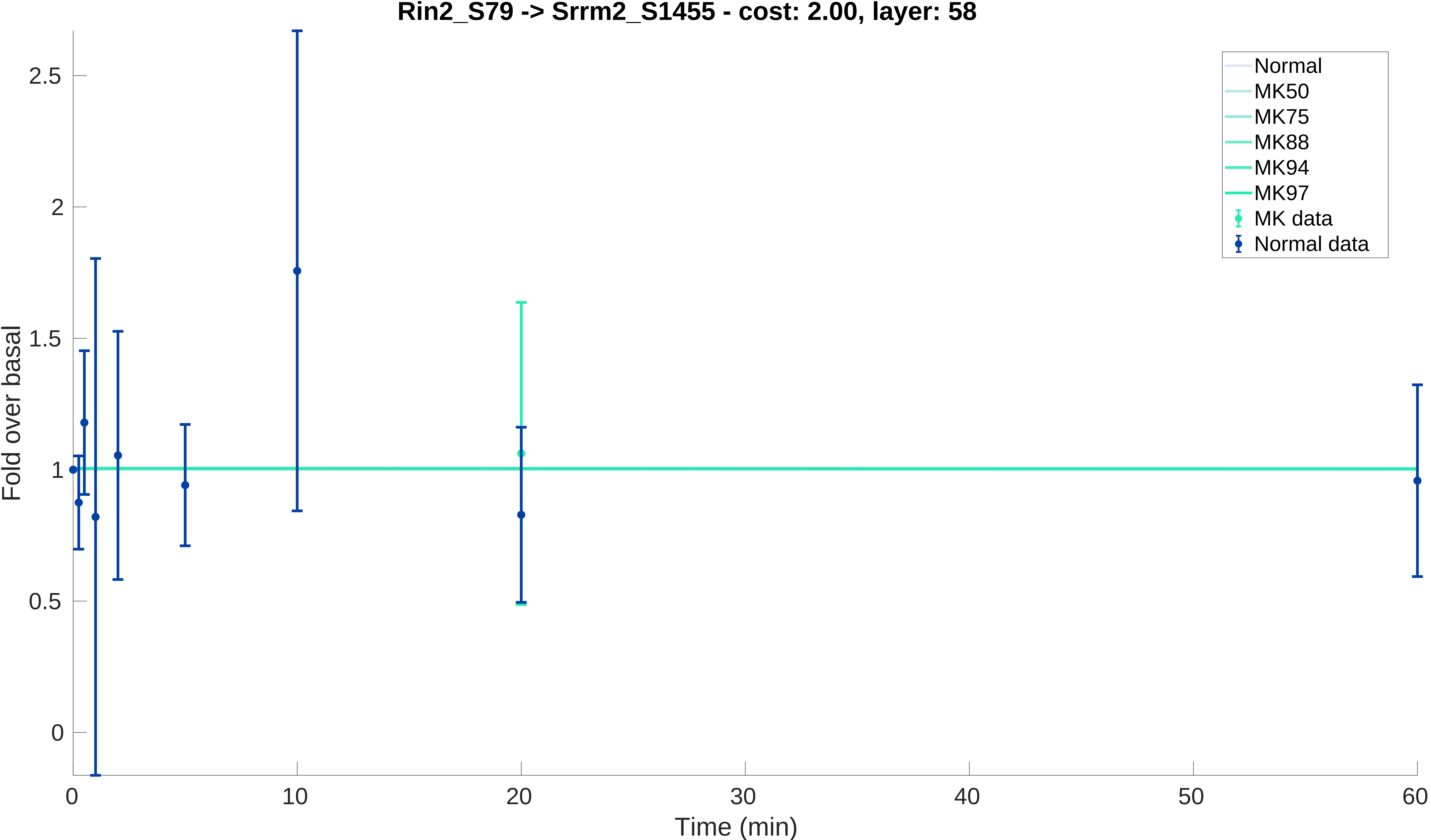
Rin2_S79 -> Srrm2_S1455 - cost: 2.00, layer: 58
Fold over basal
Time (min)
0
0.5
1
1.5
2
2.5
0
10
20
30
40
50
60
Normal
MK50
MK75
MK88
MK94
MK97
MK data
Normal data

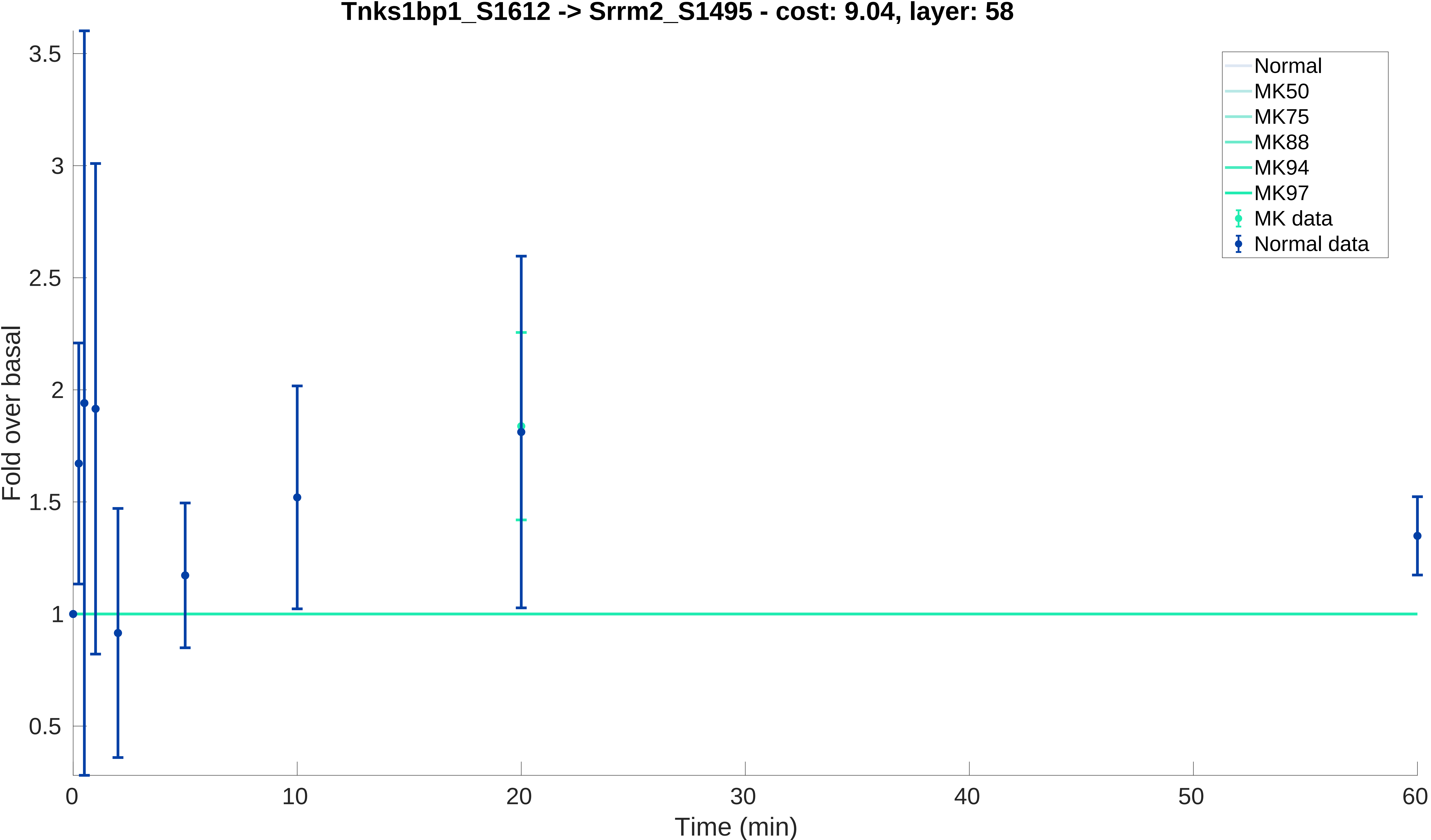

Tnks1bp1_S1612 -> Srrm2_S1495 - cost: 9.04, layer: 58
Fold over basal
Time (min)
0.5
1
1.5
2
2.5
3
3.5
0
10
20
30
40
50
60
Normal
MK50
MK75
MK88
MK94
MK97
MK data
Normal data

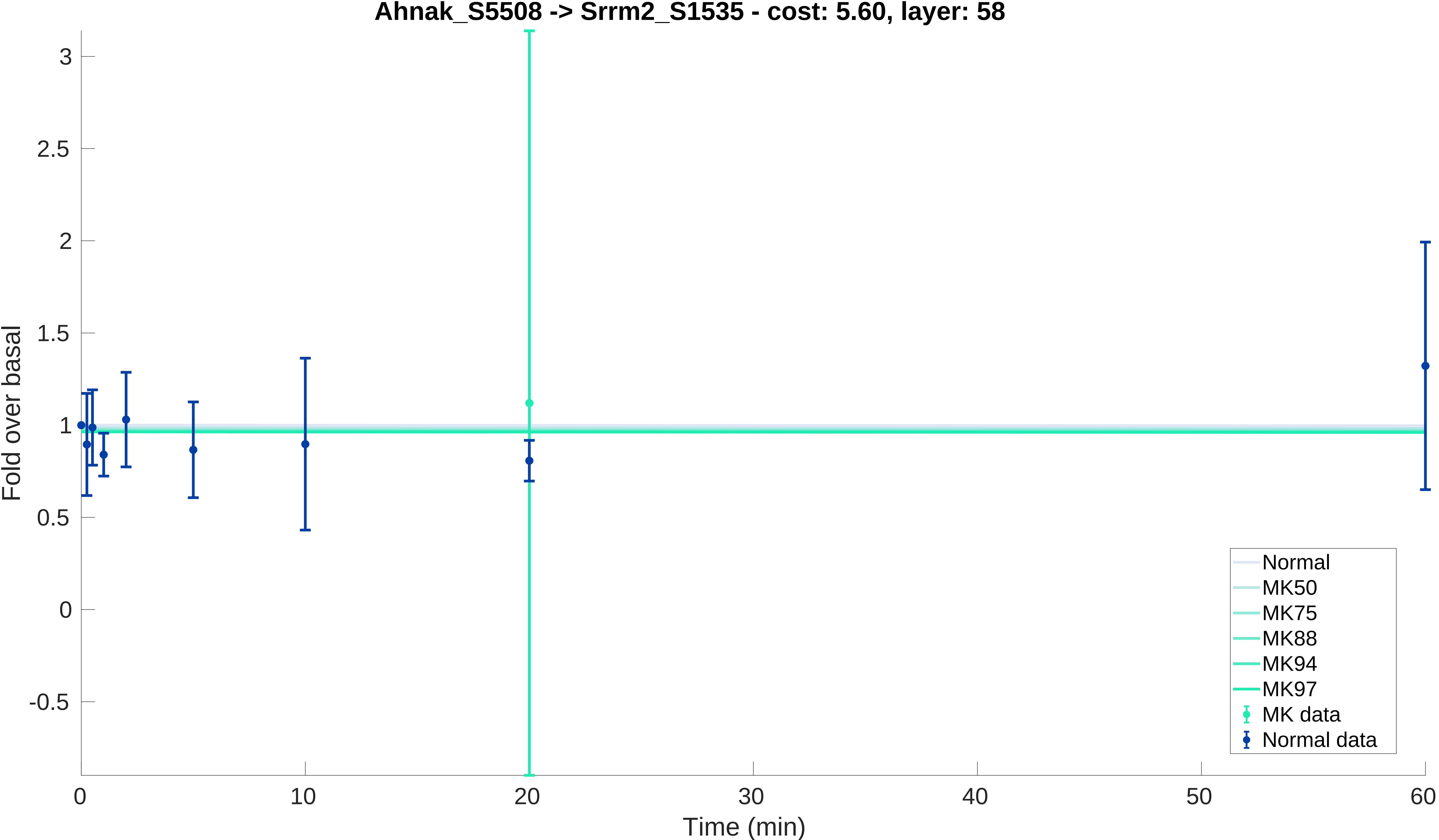

Ahnak_S5508 -> Srrm2_S1535 - cost: 5.60, layer: 58
Fold over basal
Time (min)
3
2.5
2
1.5
1
0.5
0
-0.5
0
10
20
30
40
50
60
Normal
MK50
MK75
MK88
MK94
MK97
MK data
Normal data

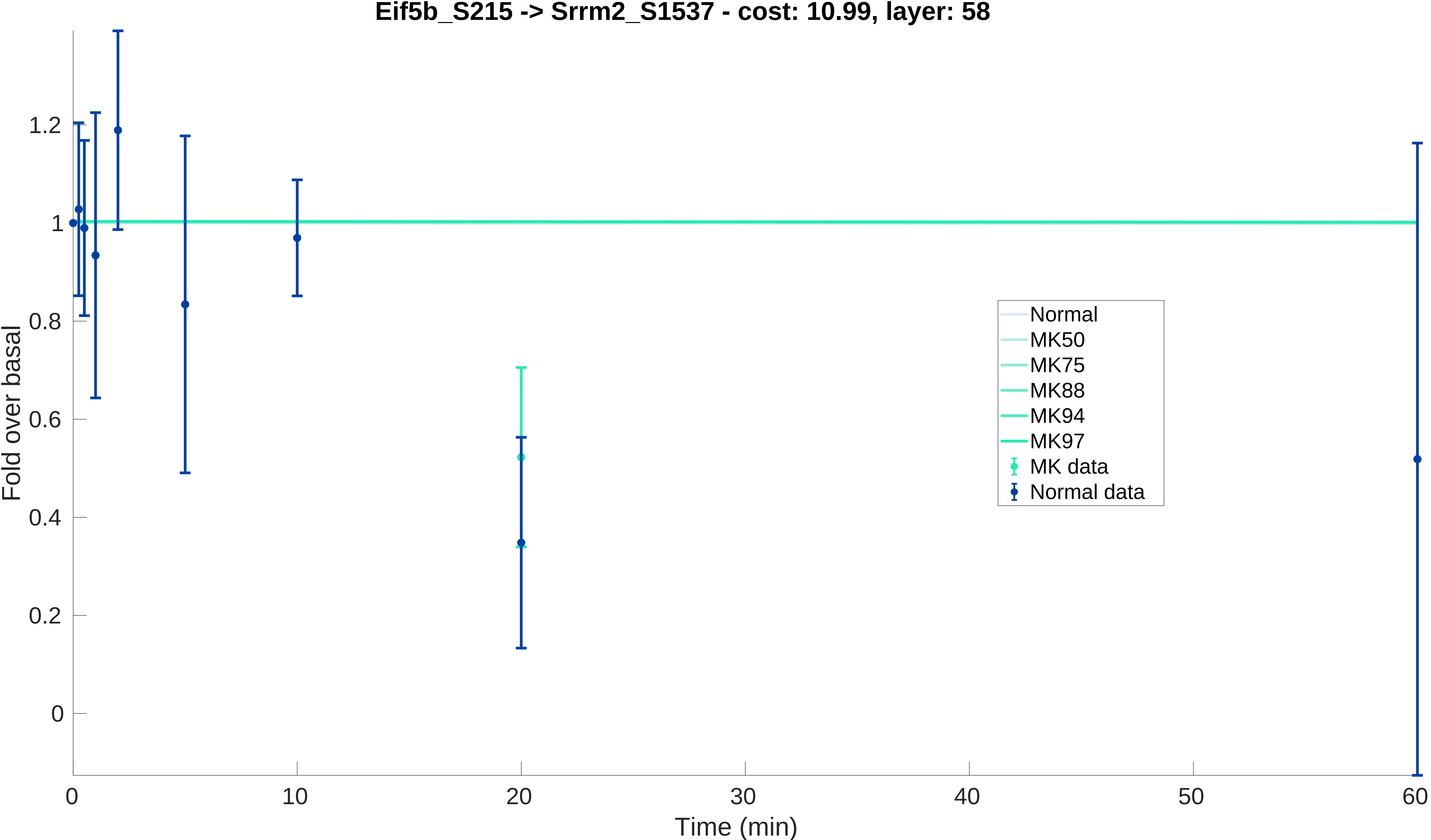

Eif5b_S215 -> Srrm2_S1537 - cost: 10.99, layer: 58
Fold over basal
Time (min)
1.2
1
0.8
0.6
0.4
0.2
0
0
10
20
30
40
50
60
Normal
MK50
MK75
MK88
MK94
MK97
MK data
Normal data

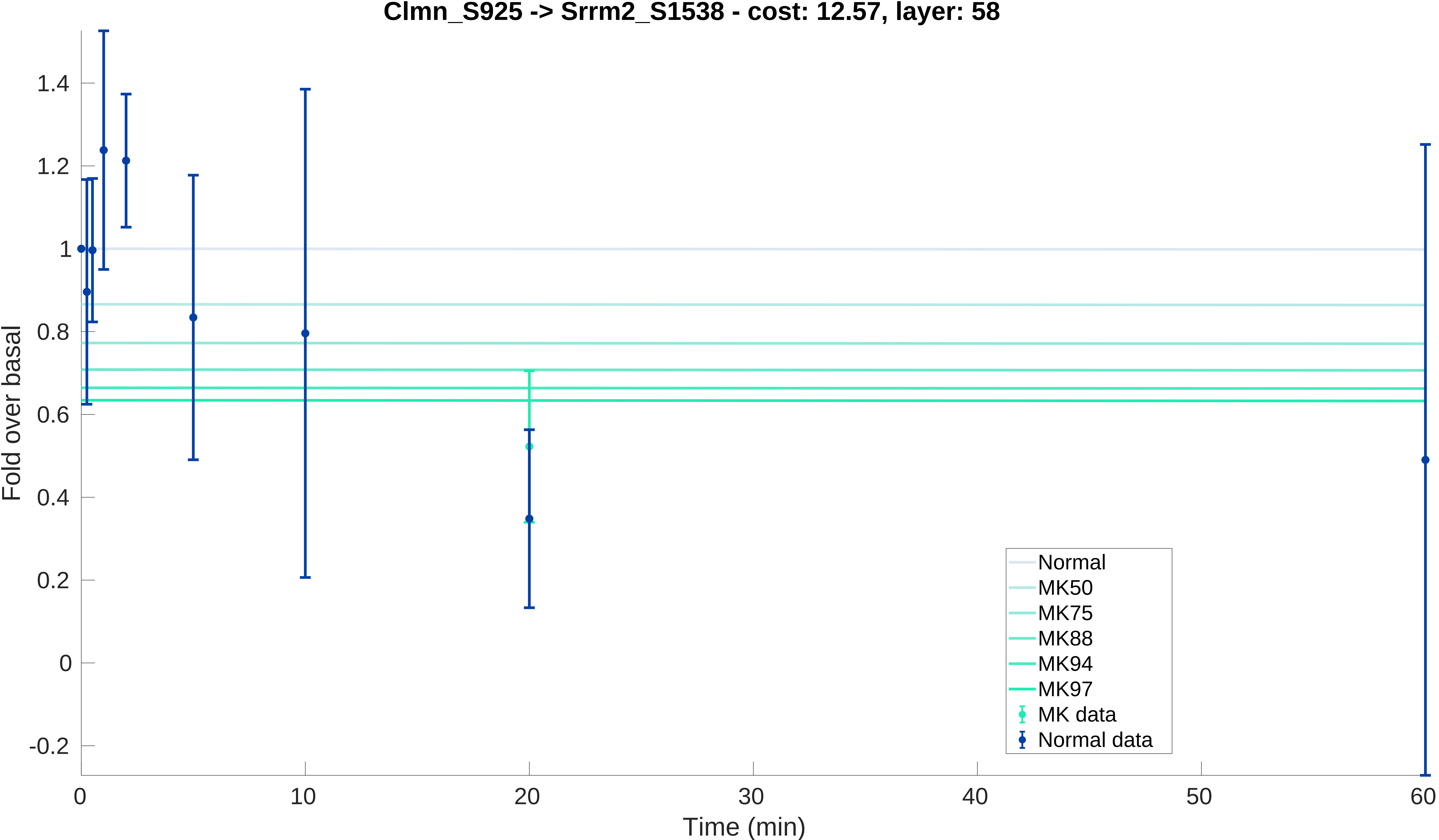

Clmn_S925 -> Srrm2_S1538 - cost: 12.57, layer: 58
Fold over basal
1.4
1.2
1
0.8
0.6
0.4
0.2
0
-0.2
0
10
20
30
40
50
60
Time (min)
Normal
MK50
MK75
MK88
MK94
MK97
MK data
Normal data

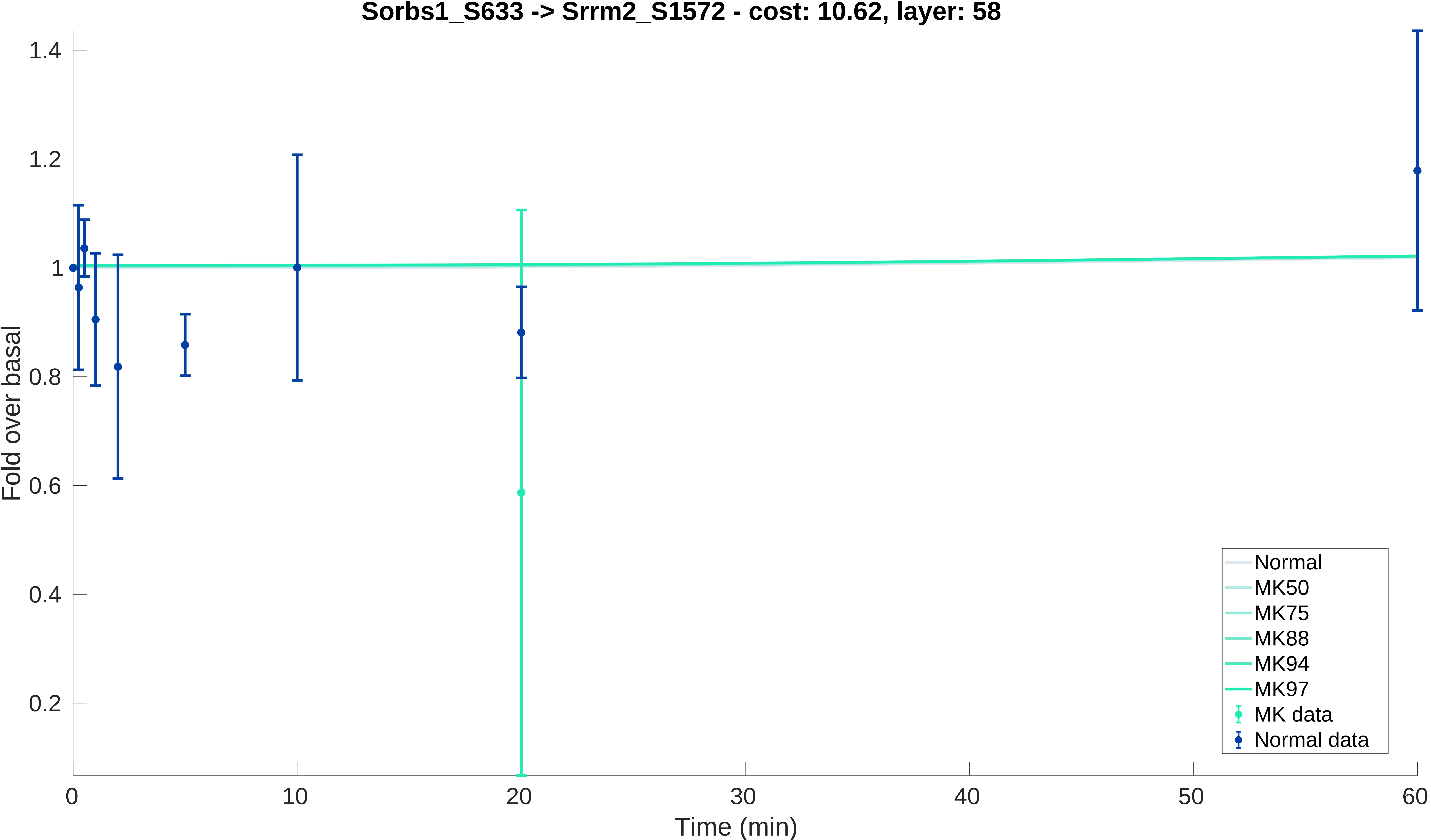

Sorbs1_S633 -> Srrm2_S1572 - cost: 10.62, layer: 58
Fold over basal
1.4
1.2
1
0.8
0.6
0.4
0.2
0
10
20
30
40
50
60
Time (min)
Normal
MK50
MK75
MK88
MK94
MK97
MK data
Normal data

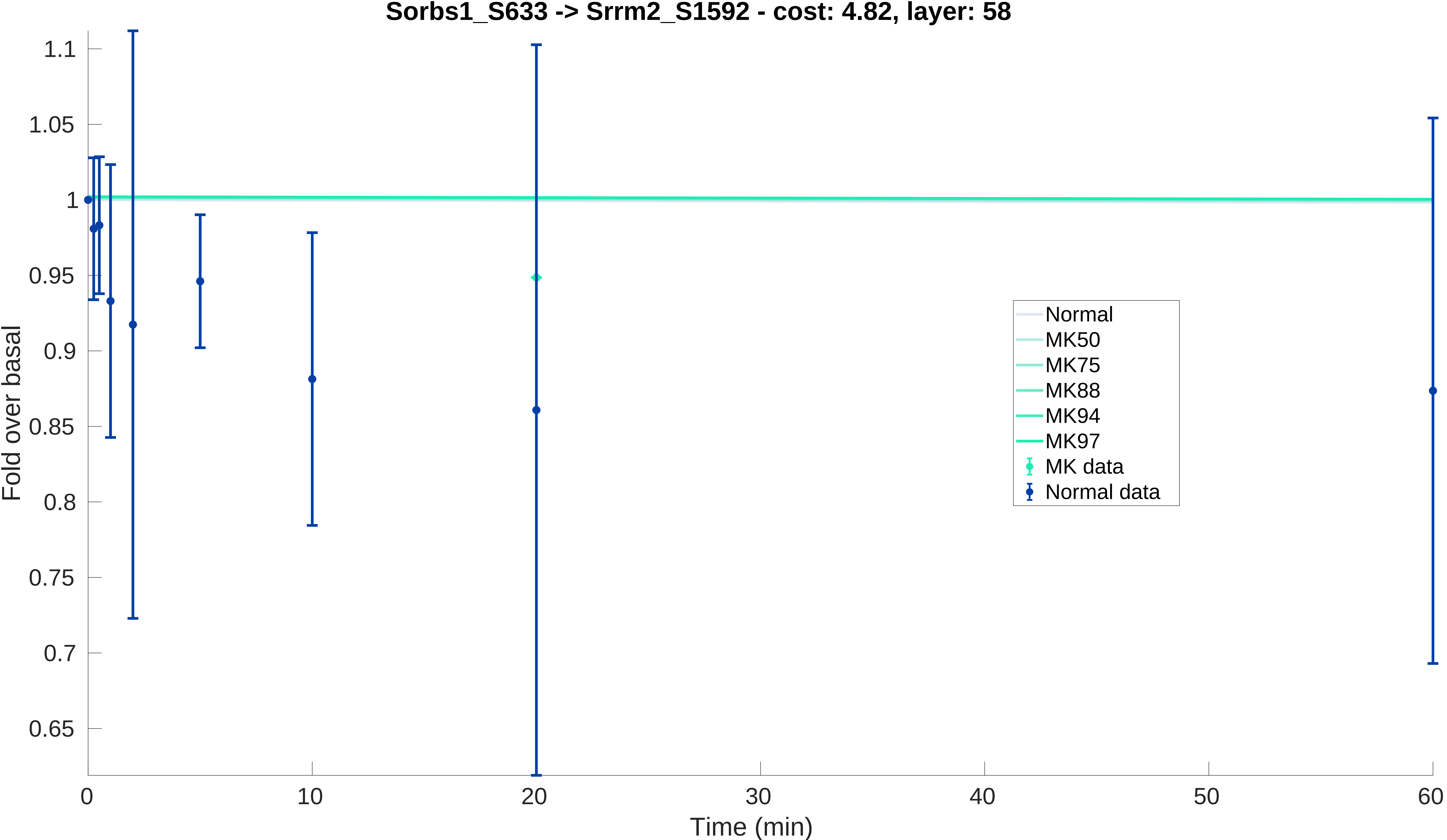

Sorbs1_S633 -> Srrm2_S1592 - cost: 4.82, layer: 58
Fold over basal
Time (min)
1.1
1.05
1
0.95
0.9
0.85
0.8
0.75
0.7
0.65
0
10
20
30
40
50
60
Normal
MK50
MK75
MK88
MK94
MK97
MK data
Normal data

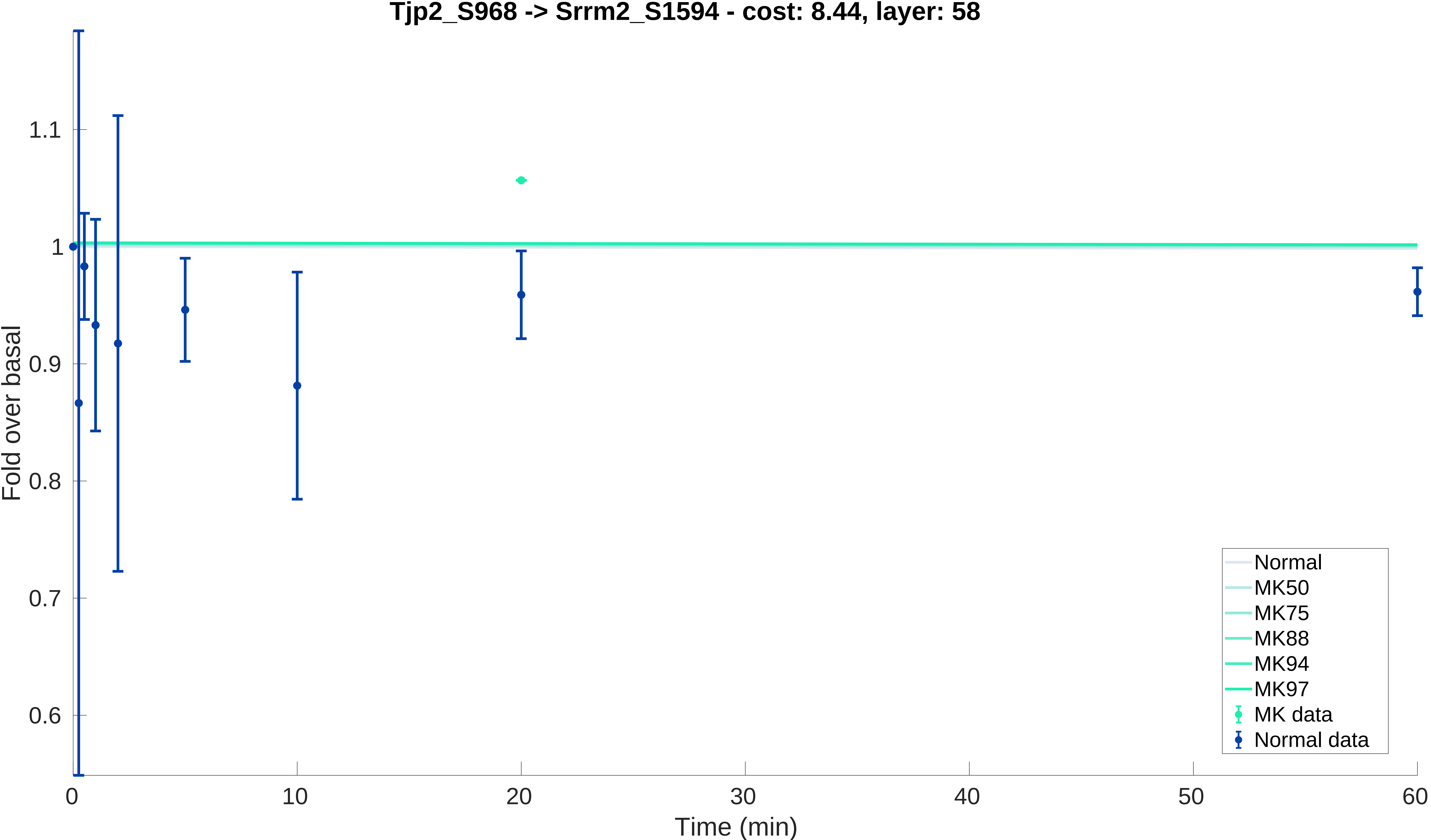
Tjp2_S968 -> Srrm2_S1594 - cost: 8.44, layer: 58
Fold over basal
1.1
1
0.9
0.8
0.7
0.6
0
10
20
30
40
50
60
Time (min)
Normal
MK50
MK75
MK88
MK94
MK97
MK data
Normal data

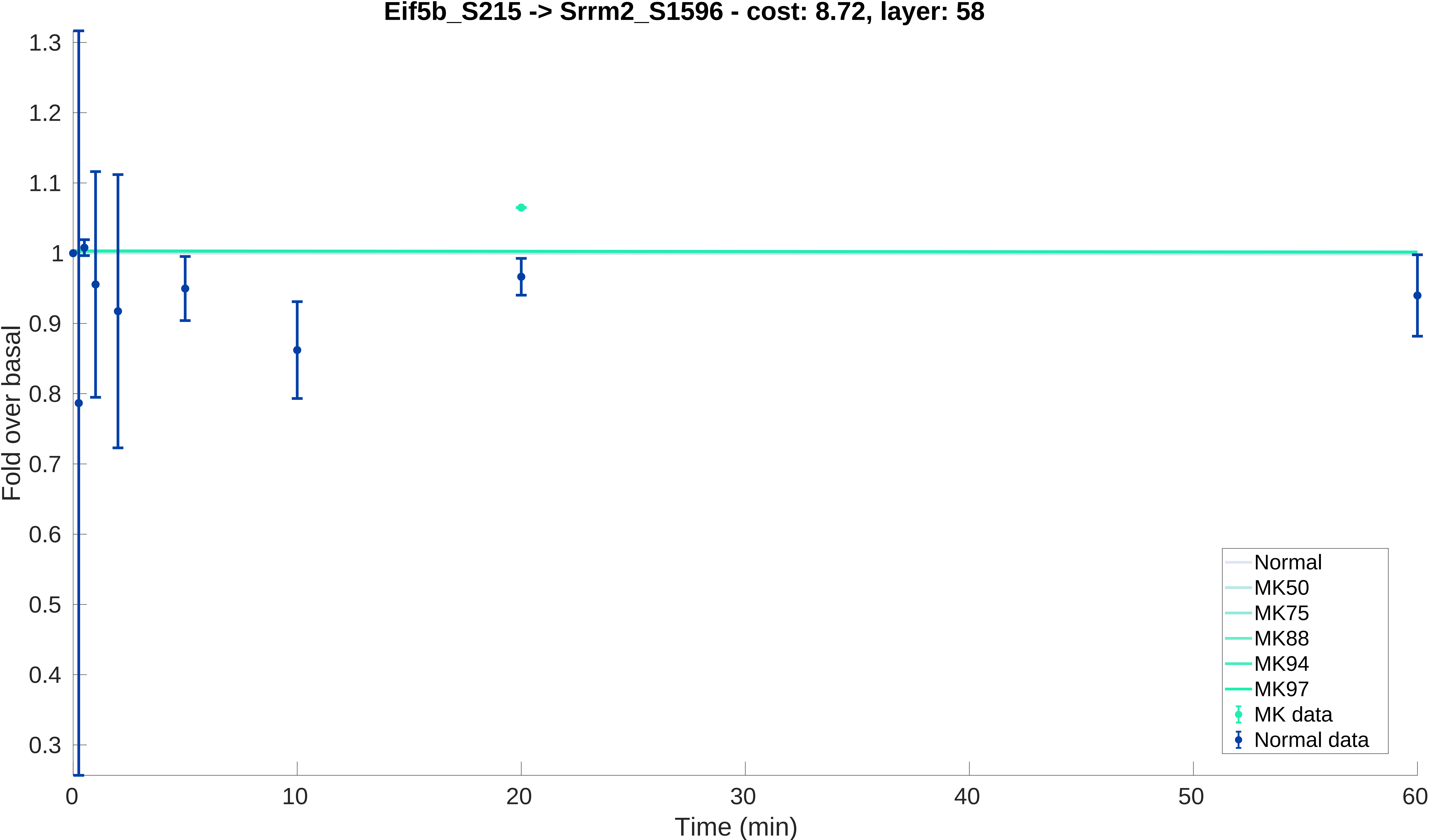

Eif5b_S215 -> Srrm2_S1596 - cost: 8.72, layer: 58
Fold over basal
Time (min)
1.3
1.2
1.1
1
0.9
0.8
0.7
0.6
0.5
0.4
0.3
0
10
20
30
40
50
60
Normal
MK50
MK75
MK88
MK94
MK97
MK data
Normal data

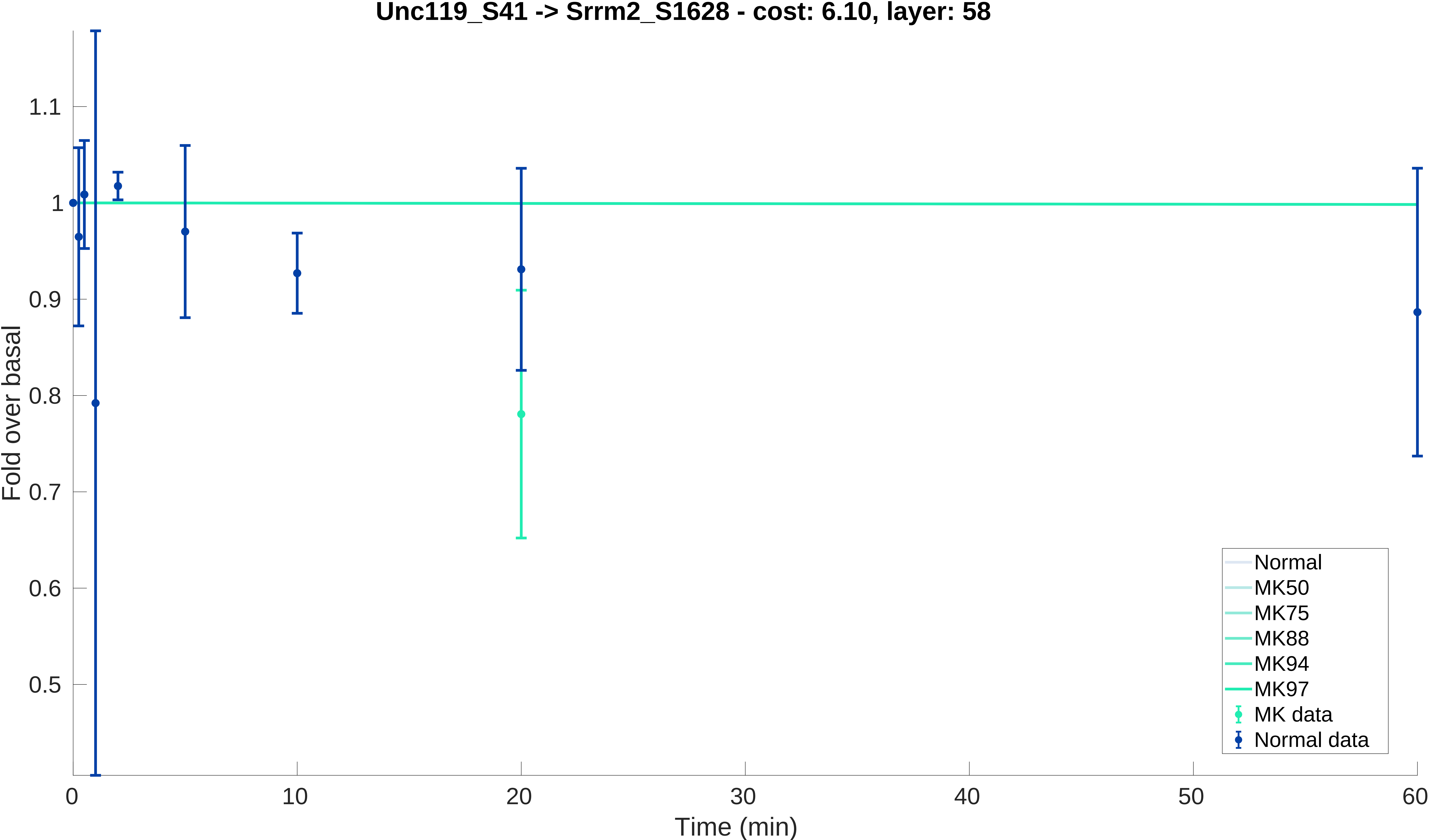
Unc119_S41 -> Srrm2_S1628 - cost: 6.10, layer: 58
Fold over basal
Time (min)
1.1
1
0.9
0.8
0.7
0.6
0.5
0
10
20
30
40
50
60
Normal
MK50
MK75
MK88
MK94
MK97
MK data
Normal data

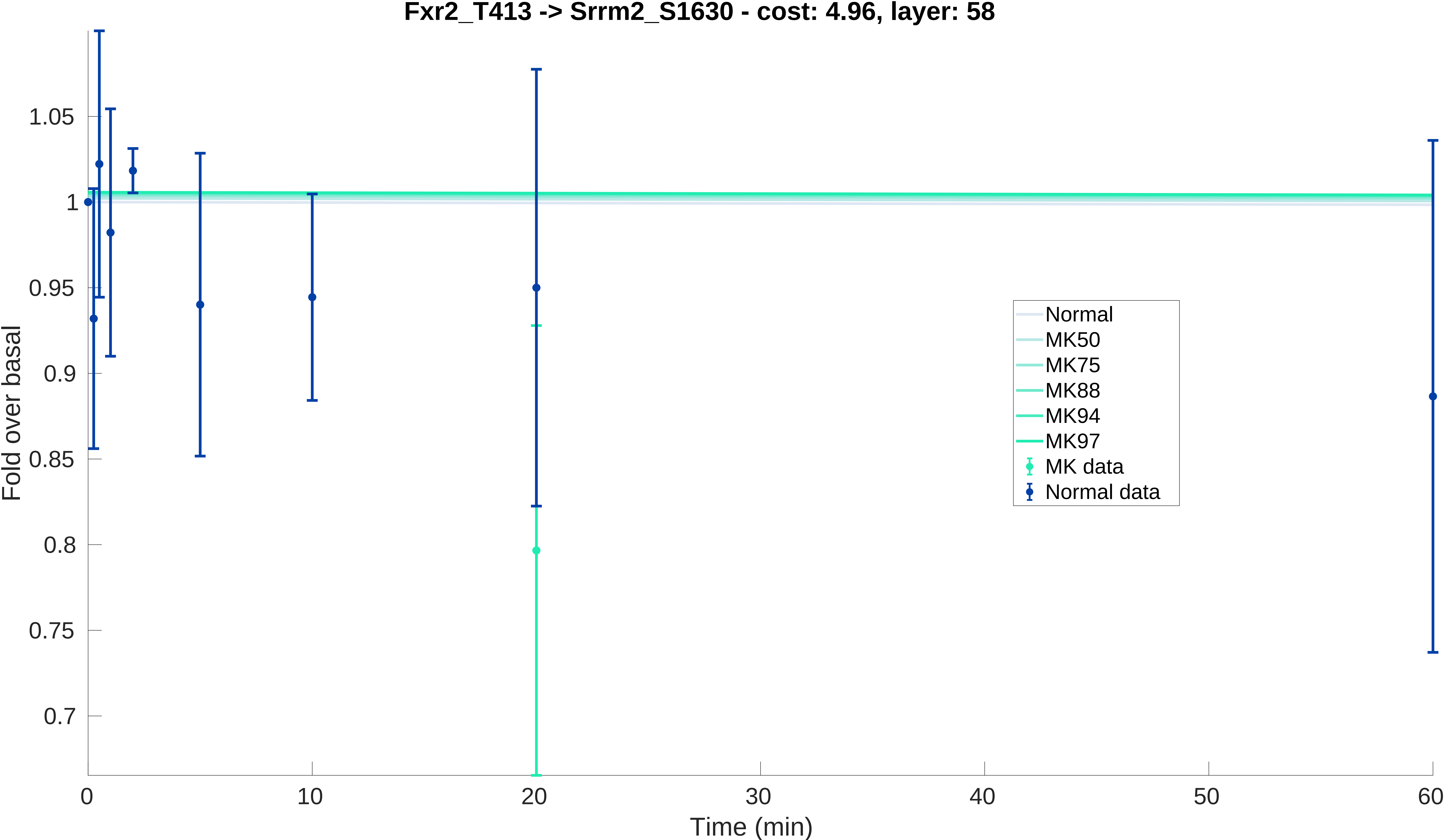

Fxr2_T413 -> Srrm2_S1630 - cost: 4.96, layer: 58
Fold over basal
1.05
1
0.95
0.9
0.85
0.8
0.75
0.7
0
10
20
30
40
50
60
Time (min)
Normal
MK50
MK75
MK88
MK94
MK97
MK data
Normal data

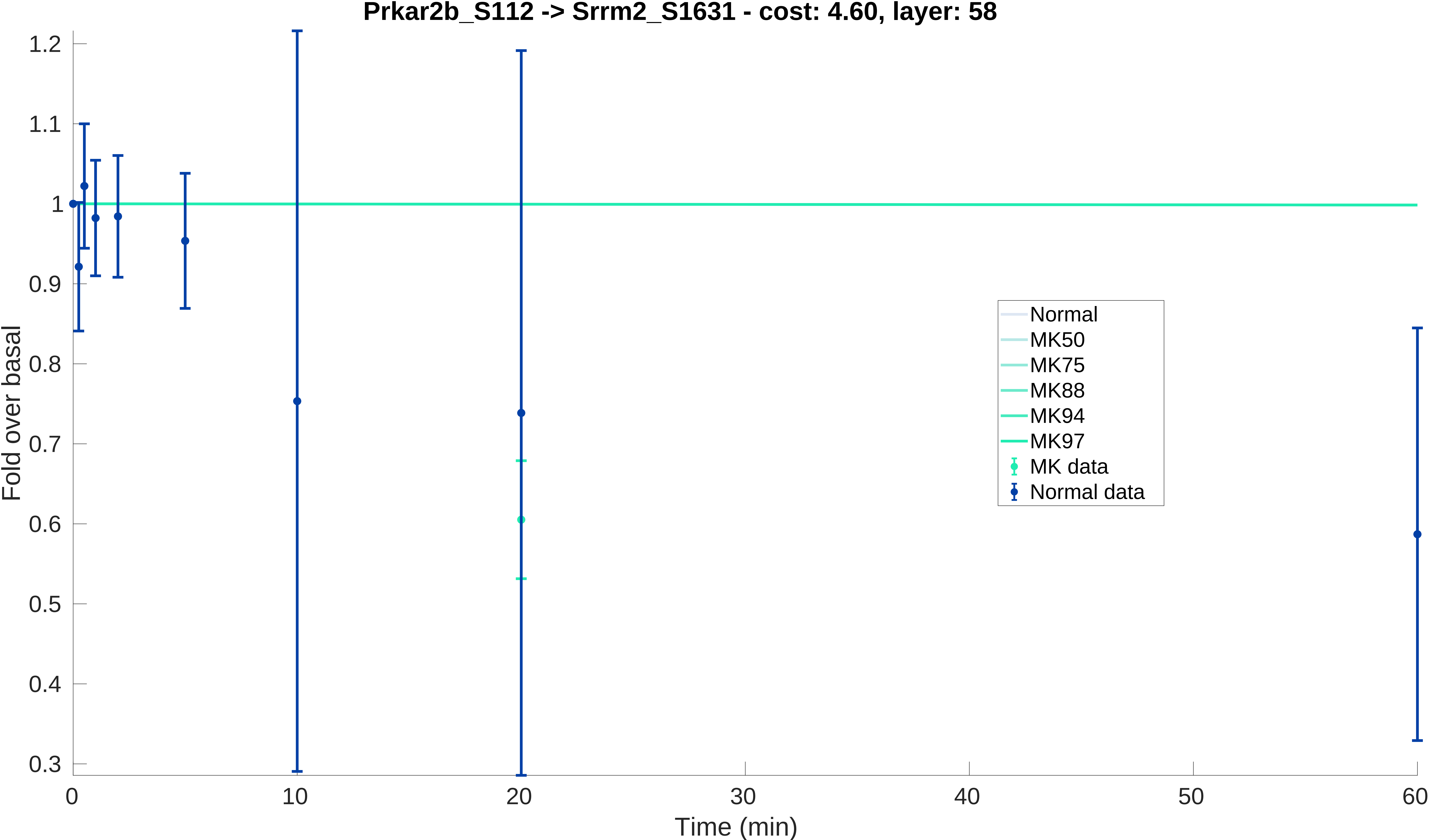

Prkar2b_S112 -> Srrm2_S1631 - cost: 4.60, layer: 58
Fold over basal
Time (min)
1.2
1.1
1
0.9
0.8
0.7
0.6
0.5
0.4
0.3
0
10
20
30
40
50
60
Normal
MK50
MK75
MK88
MK94
MK97
MK data
Normal data

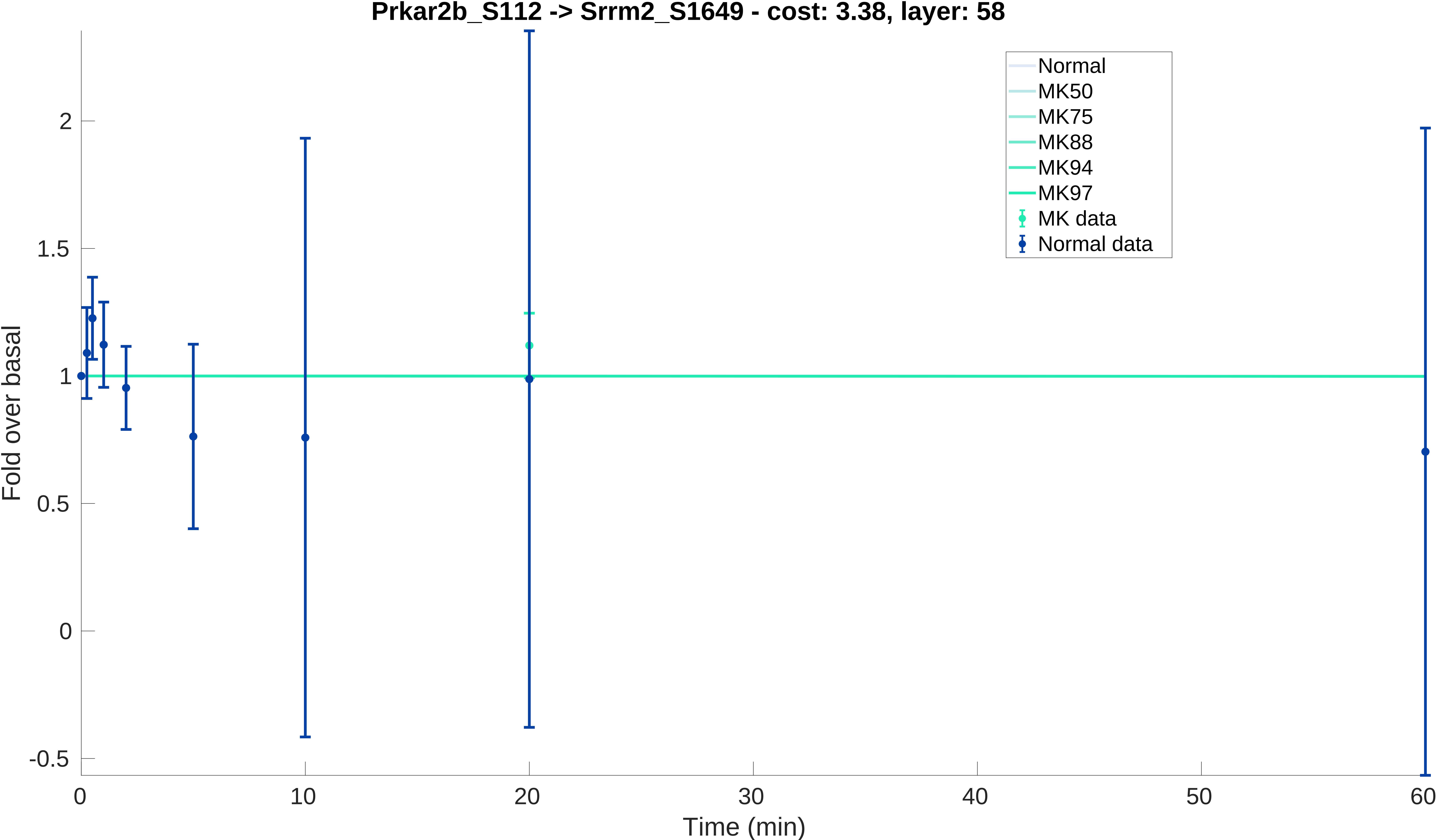

Prkar2b_S112 -> Srrm2_S1649 - cost: 3.38, layer: 58
Normal
MK50
MK75
MK88
MK94
MK97
MK data
Normal data
Fold over basal
Time (min)
2
1.5
1
0.5
0
-0.5
0
10
20
30
40
50
60

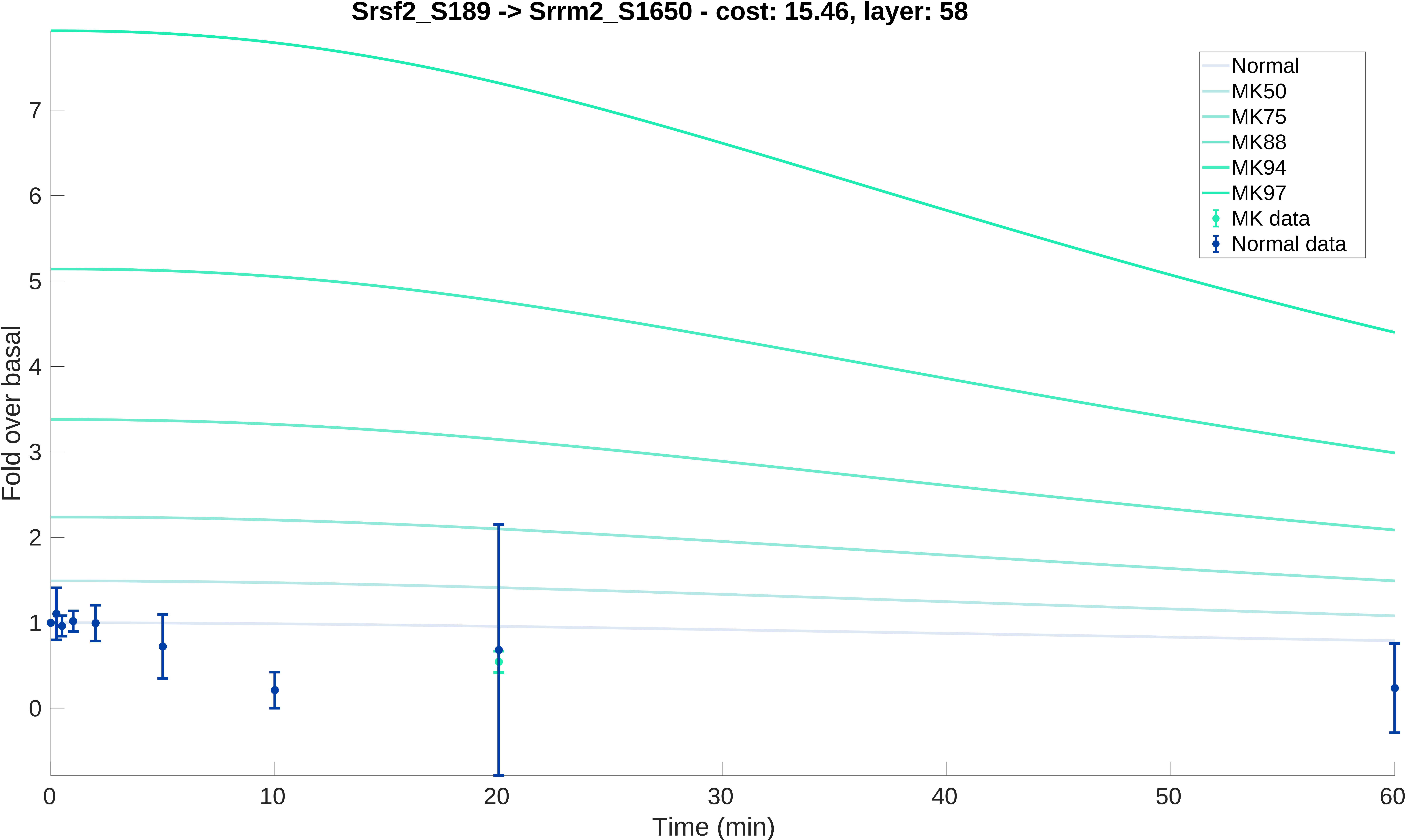
Srsf2_S189 -> Srrm2_S1650 - cost: 15.46, layer: 58
Fold over basal
Time (min)
Normal
MK50
MK75
MK88
MK94
MK97
MK data
Normal data
0
1
2
3
4
5
6
7
0
10
20
30
40
50
60

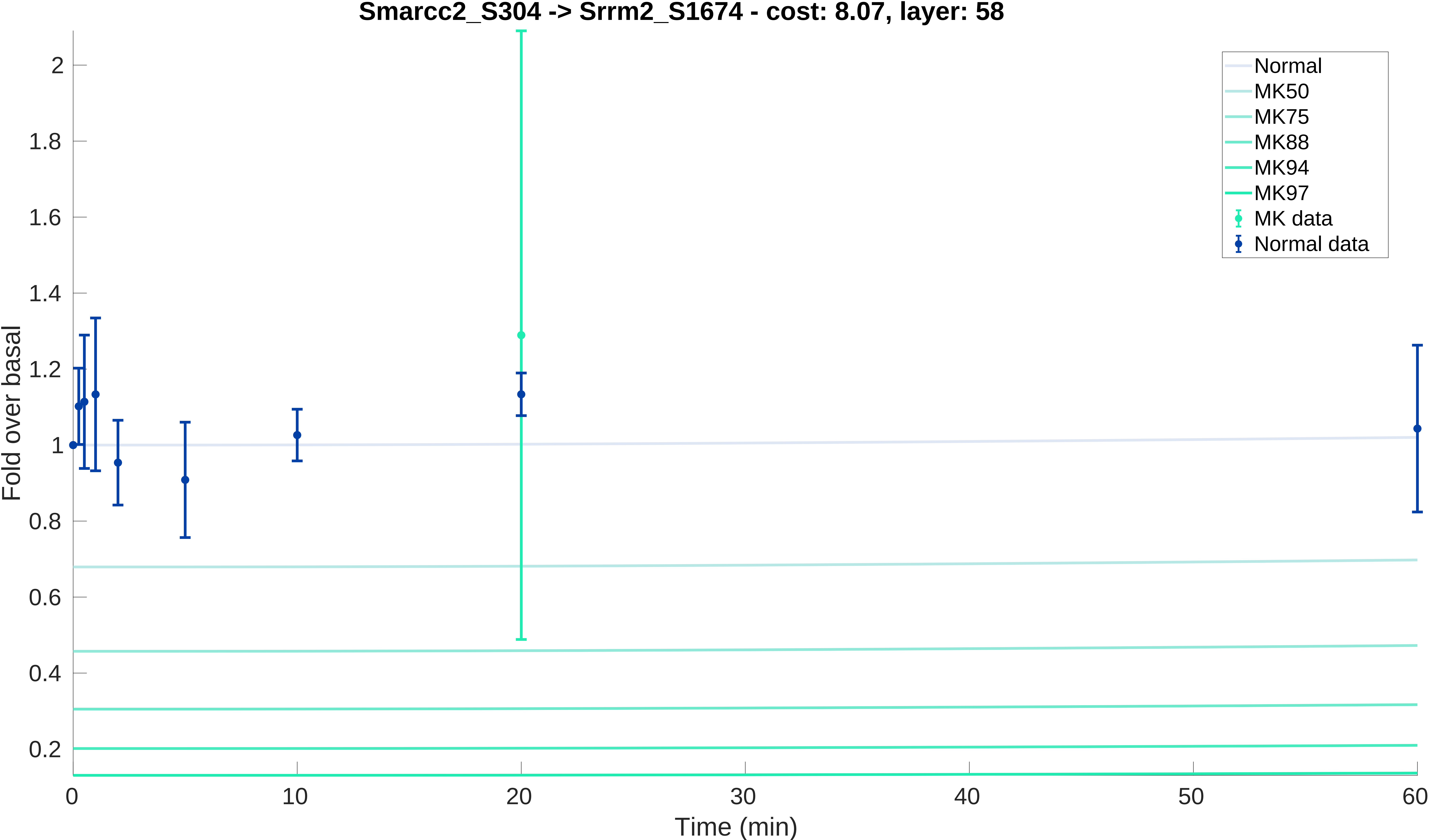

Smarcc2_S304 -> Srrm2_S1674 - cost: 8.07, layer: 58
Fold over basal
Time (min)
2
1.8
1.6
1.4
1.2
1
0.8
0.6
0.4
0.2
0
10
20
30
40
50
60
Normal
MK50
MK75
MK88
MK94
MK97
MK data
Normal data

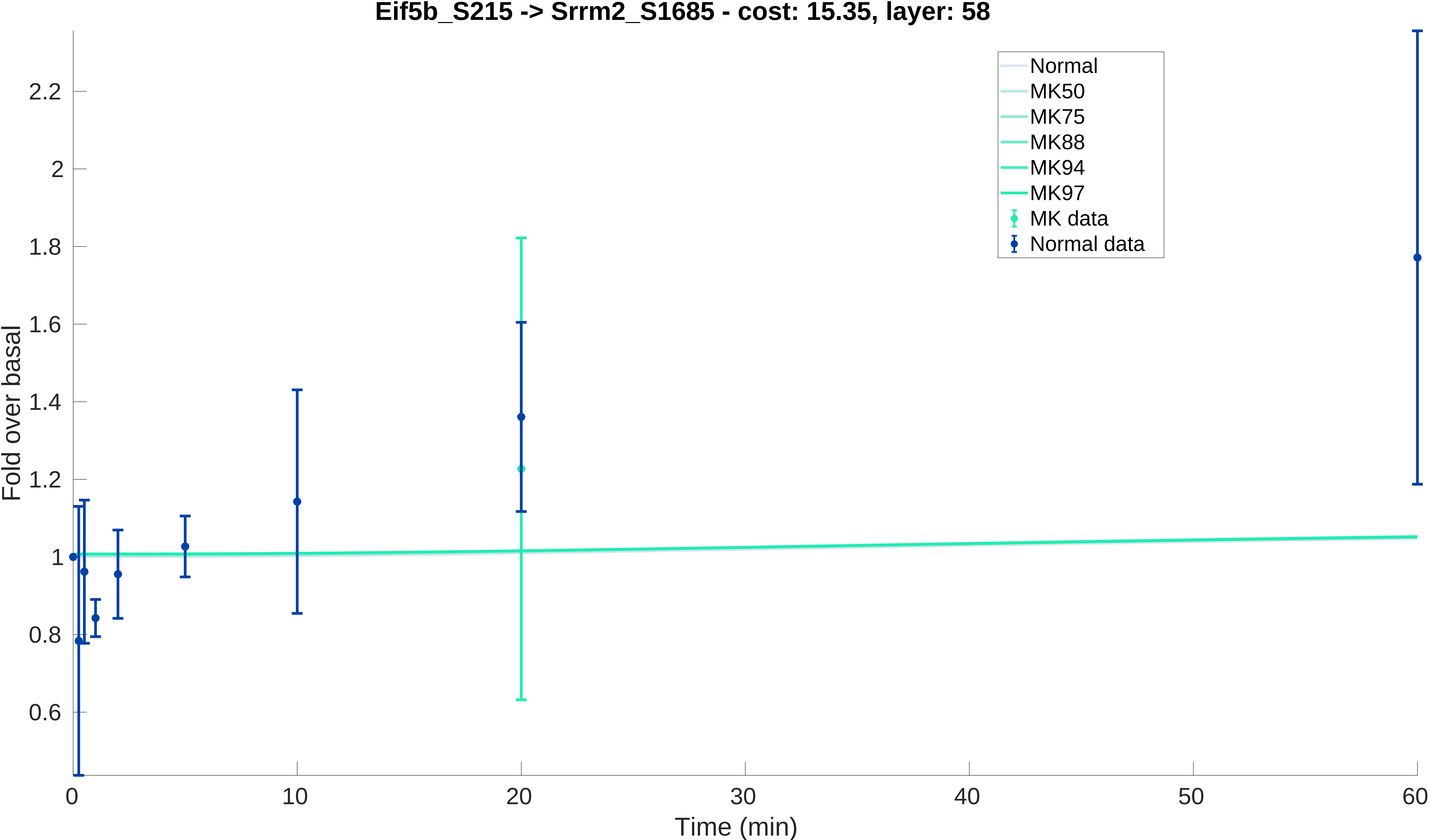

Eif5b_S215 -> Srrm2_S1685 - cost: 15.35, layer: 58
Fold over basal
Time (min)
Normal
MK50
MK75
MK88
MK94
MK97
MK data
Normal data
2.2
2
1.8
1.6
1.4
1.2
1
0.8
0.6
0
10
20
30
40
50
60

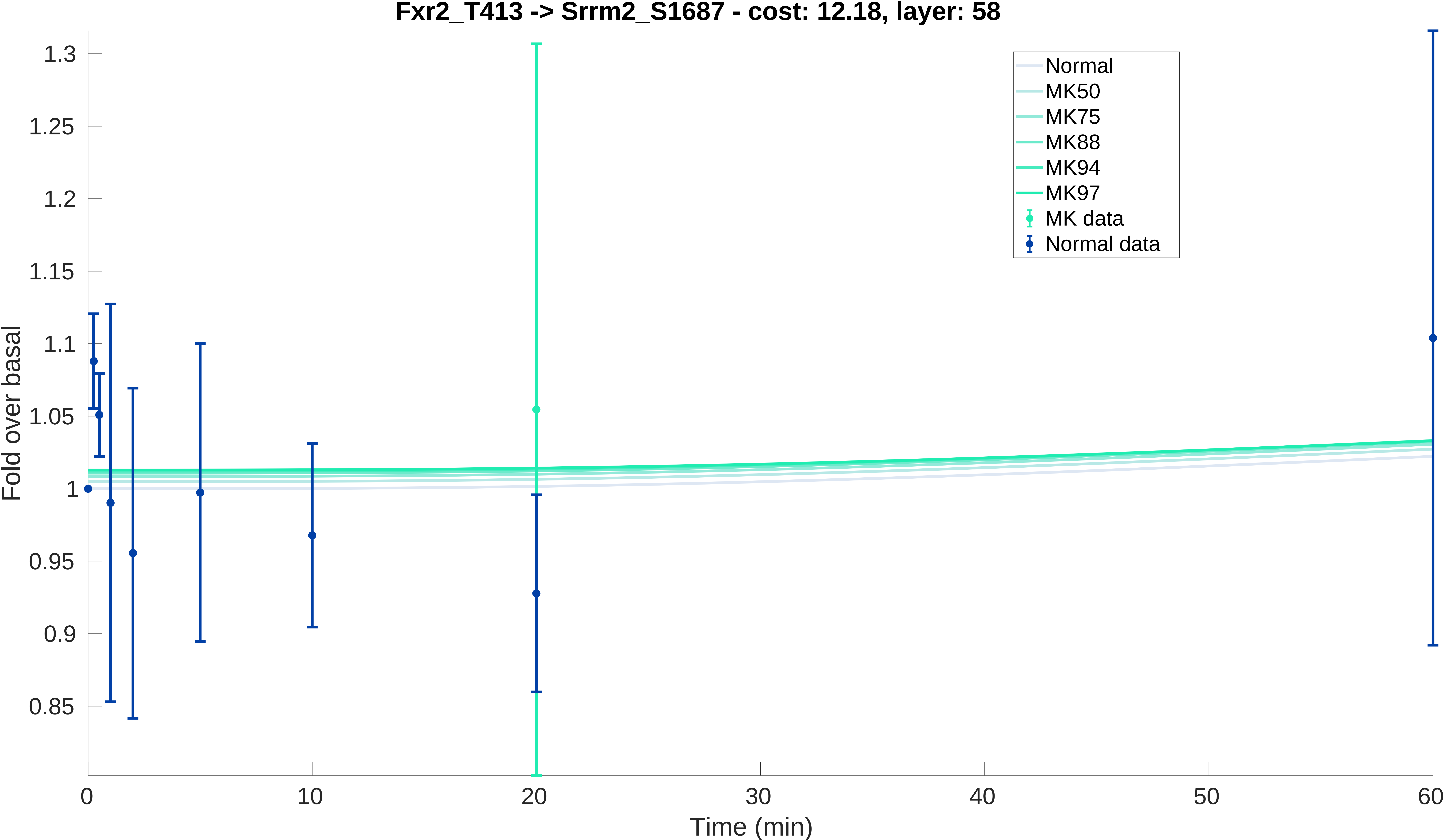

Fxr2_T413 -> Srrm2_S1687 - cost: 12.18, layer: 58
Normal
MK50
MK75
MK88
MK94
MK97
MK data
Normal data
Fold over basal
1.3
1.25
1.2
1.15
1.1
1.05
1
0.95
0.9
0.85
0
10
20
30
40
50
60
Time (min)

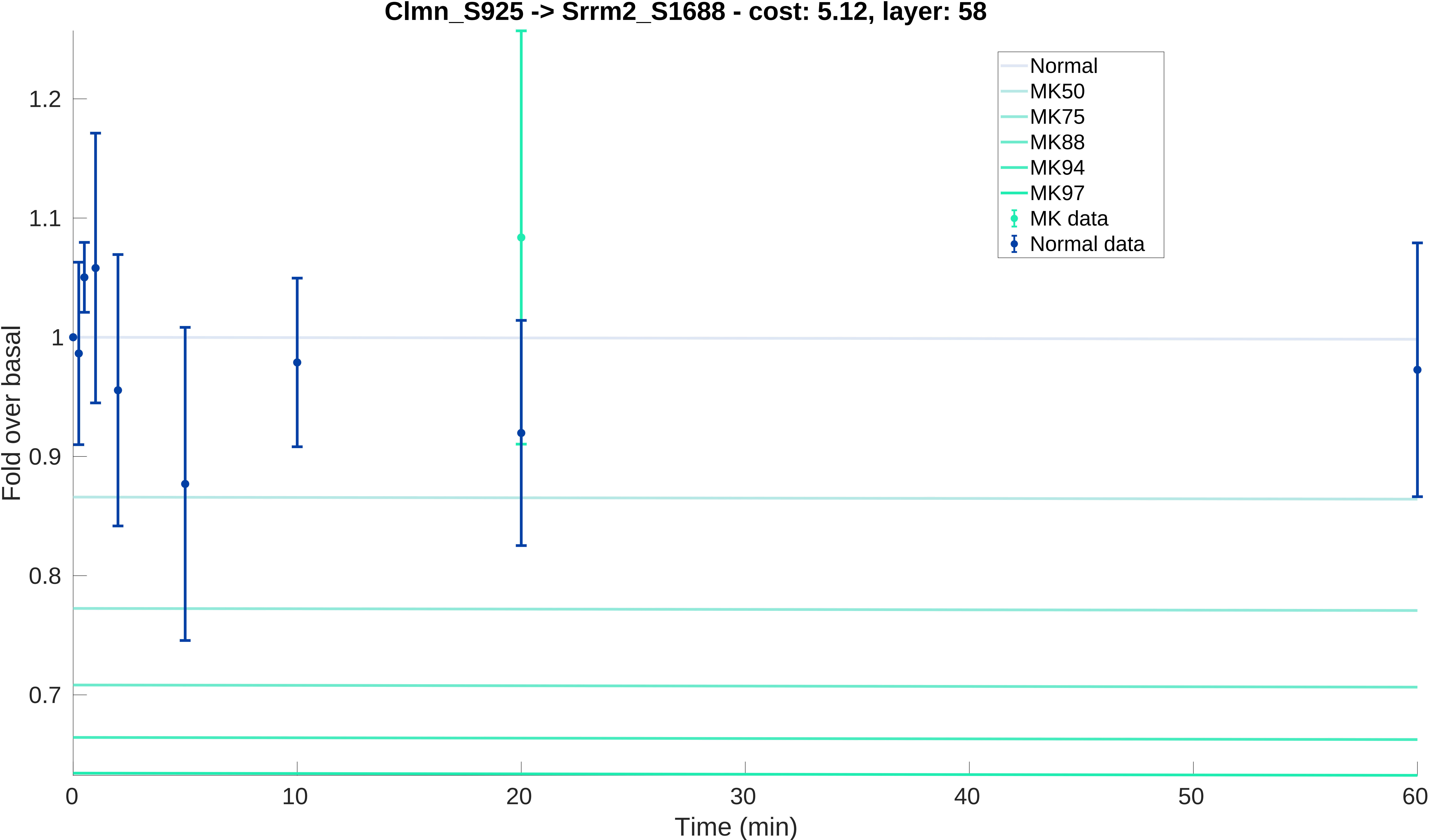
Clmn_S925 -> Srrm2_S1688 - cost: 5.12, layer: 58
Fold over basal
Time (min)
Normal
MK50
MK75
MK88
MK94
MK97
MK data
Normal data
1.2
1.1
1
0.9
0.8
0.7
0
10
20
30
40
50
60

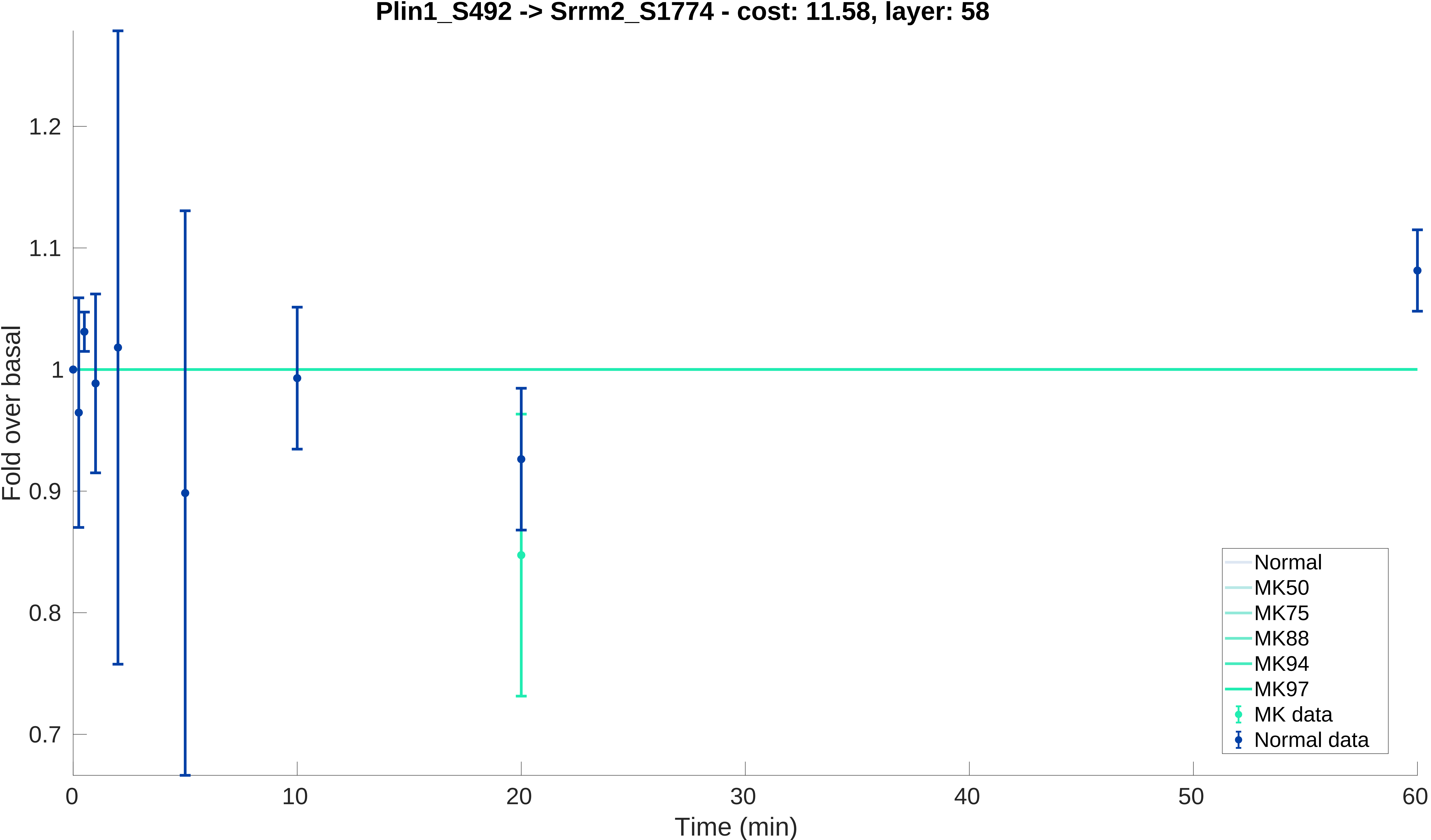

Plin1_S492 -> Srrm2_S1774 - cost: 11.58, layer: 58
Fold over basal
1.2
1.1
1
0.9
0.8
0.7
0
10
20
30
40
50
60
Time (min)
Normal
MK50
MK75
MK88
MK94
MK97
MK data
Normal data

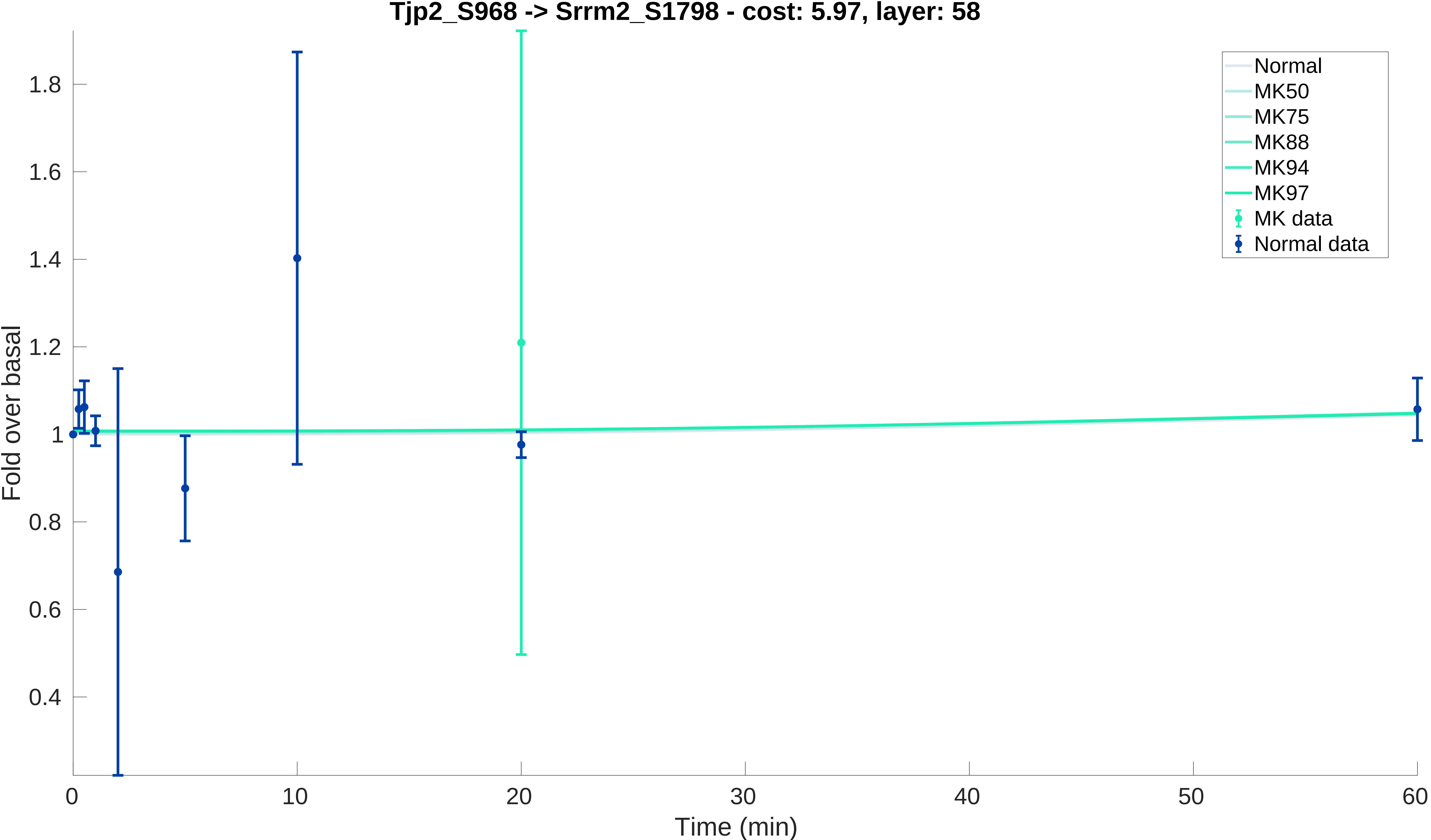
Tjp2_S968 -> Srrm2_S1798 - cost: 5.97, layer: 58
Fold over basal
Time (min)
Normal
MK50
MK75
MK88
MK94
MK97
MK data
Normal data
0.4
0.6
0.8
1
1.2
1.4
1.6
1.8
0
10
20
30
40
50
60

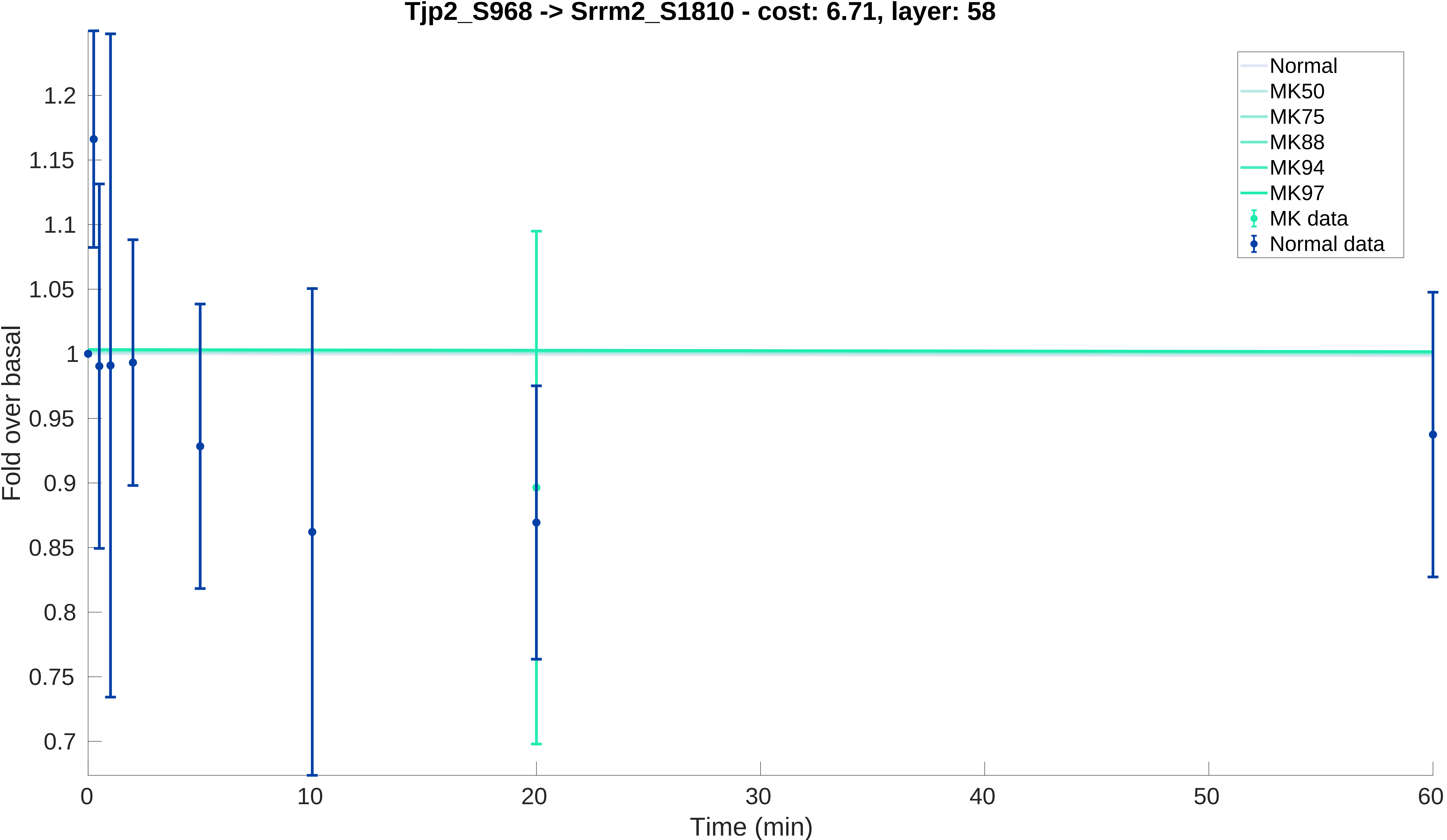

Tjp2_S968 -> Srrm2_S1810 - cost: 6.71, layer: 58
Normal
MK50
MK75
MK88
MK94
MK97
MK data
Normal data
Fold over basal
1.2
1.15
1.1
1.05
1
0.95
0.9
0.85
0.8
0.75
0.7
0
10
20
30
40
50
60
Time (min)

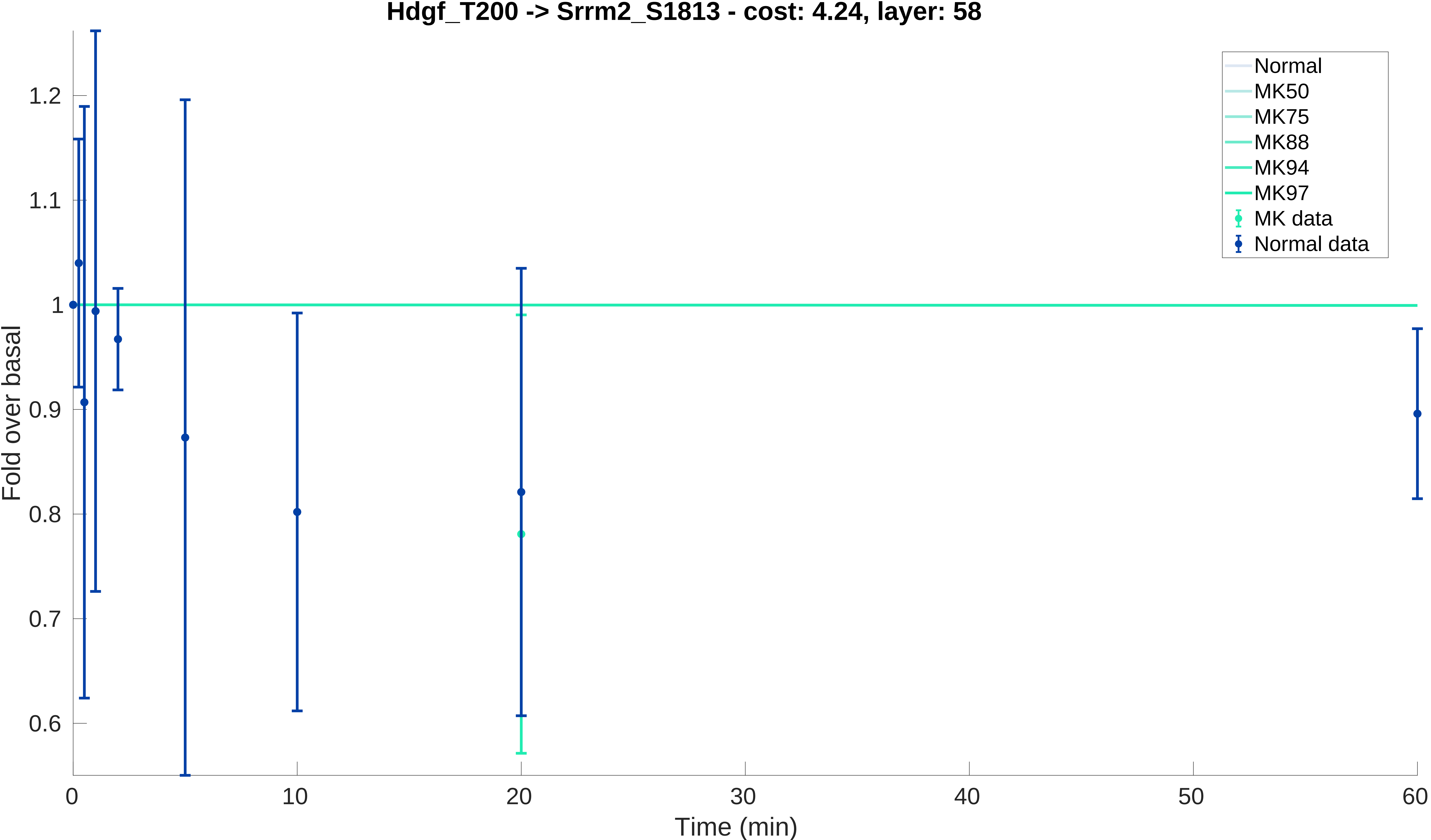

Hdgf_T200 -> Srrm2_S1813 - cost: 4.24, layer: 58
Normal
MK50
MK75
MK88
MK94
MK97
MK data
Normal data
Fold over basal
Time (min)
1.2
1.1
1
0.9
0.8
0.7
0.6
0
10
20
30
40
50
60

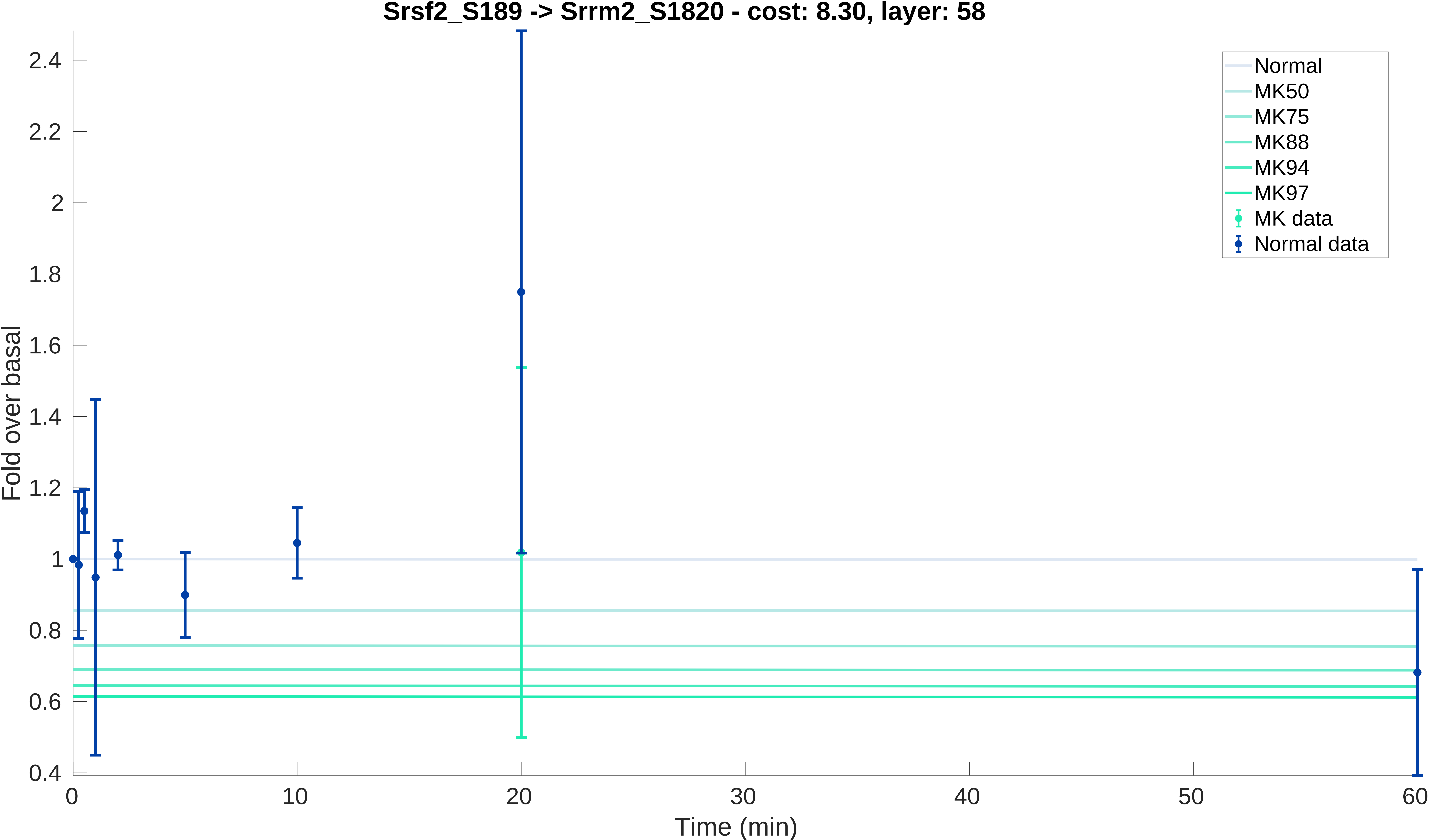
Srsf2_S189 -> Srrm2_S1820 - cost: 8.30, layer: 58
Fold over basal
Time (min)
2.4
2.2
2
1.8
1.6
1.4
1.2
1
0.8
0.6
0.4
0
10
20
30
40
50
60
Normal
MK50
MK75
MK88
MK94
MK97
MK data
Normal data

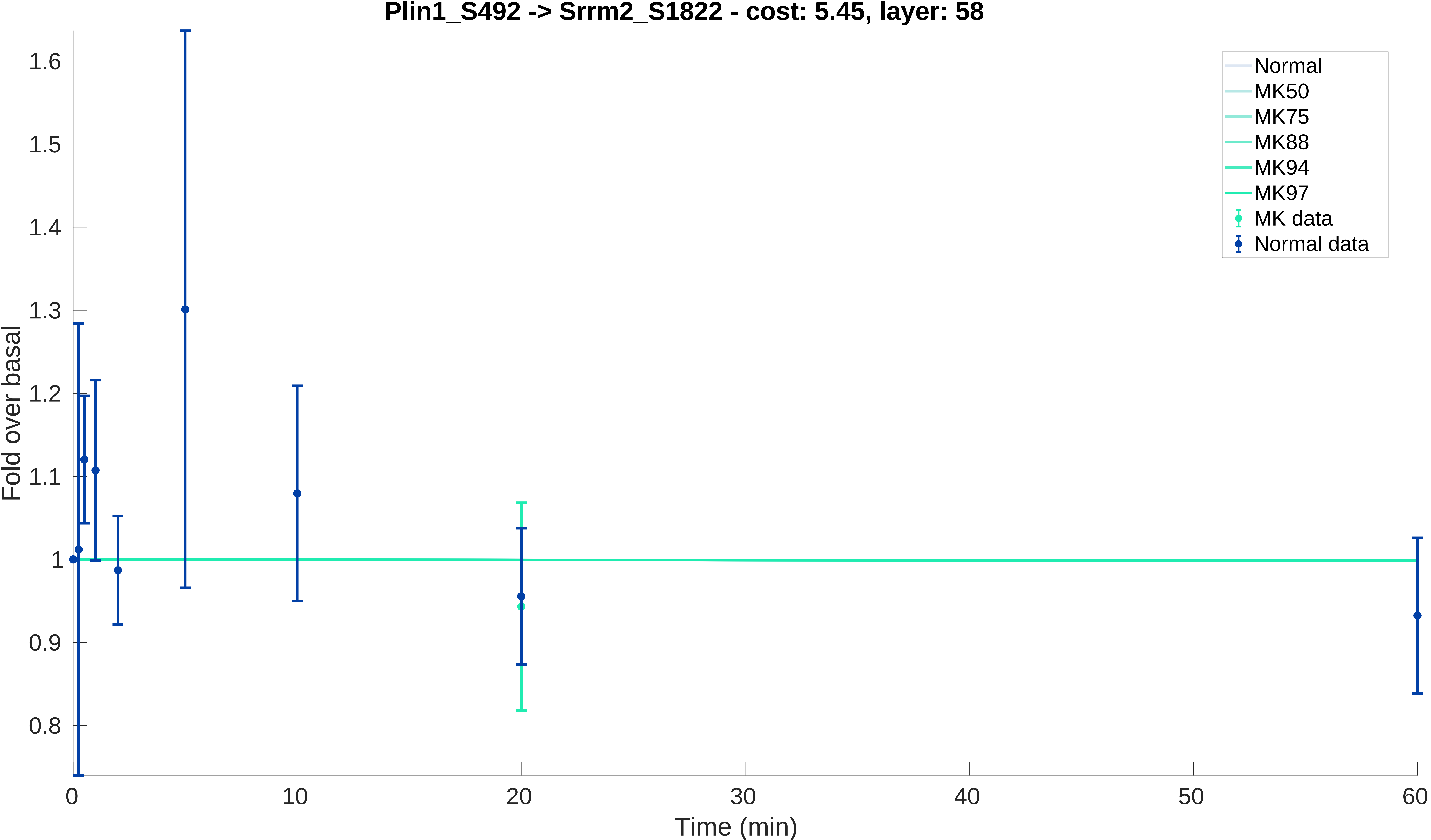

Plin1_S492 -> Srrm2_S1822 - cost: 5.45, layer: 58
Fold over basal
Time (min)
Normal
MK50
MK75
MK88
MK94
MK97
MK data
Normal data
1.6
1.5
1.4
1.3
1.2
1.1
1
0.9
0.8
0
10
20
30
40
50
60

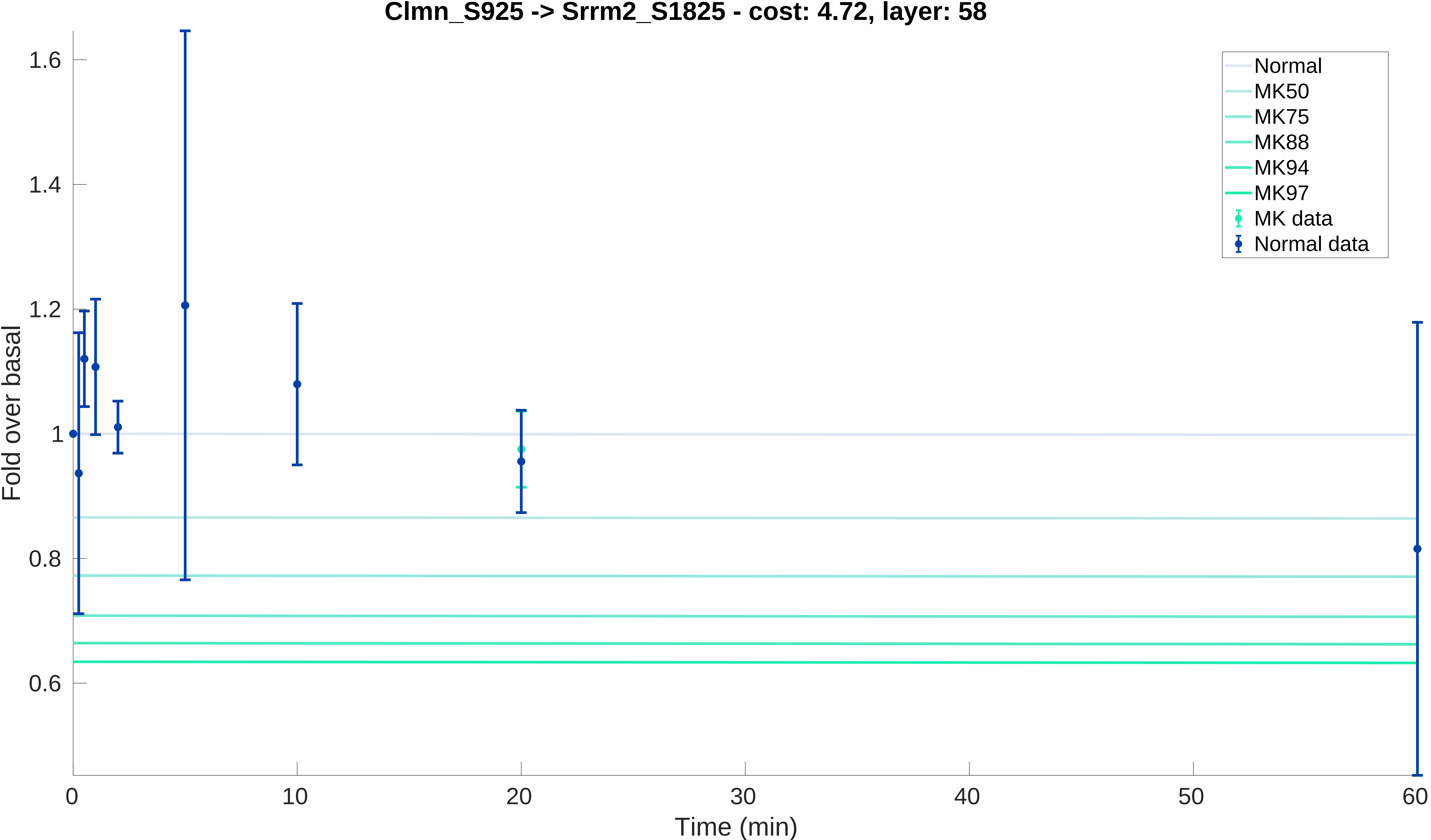
Clmn_S925 -> Srrm2_S1825 - cost: 4.72, layer: 58
Fold over basal
1.6
1.4
1.2
1
0.8
0.6
0
10
20
30
40
50
60
Time (min)
Normal
MK50
MK75
MK88
MK94
MK97
MK data
Normal data

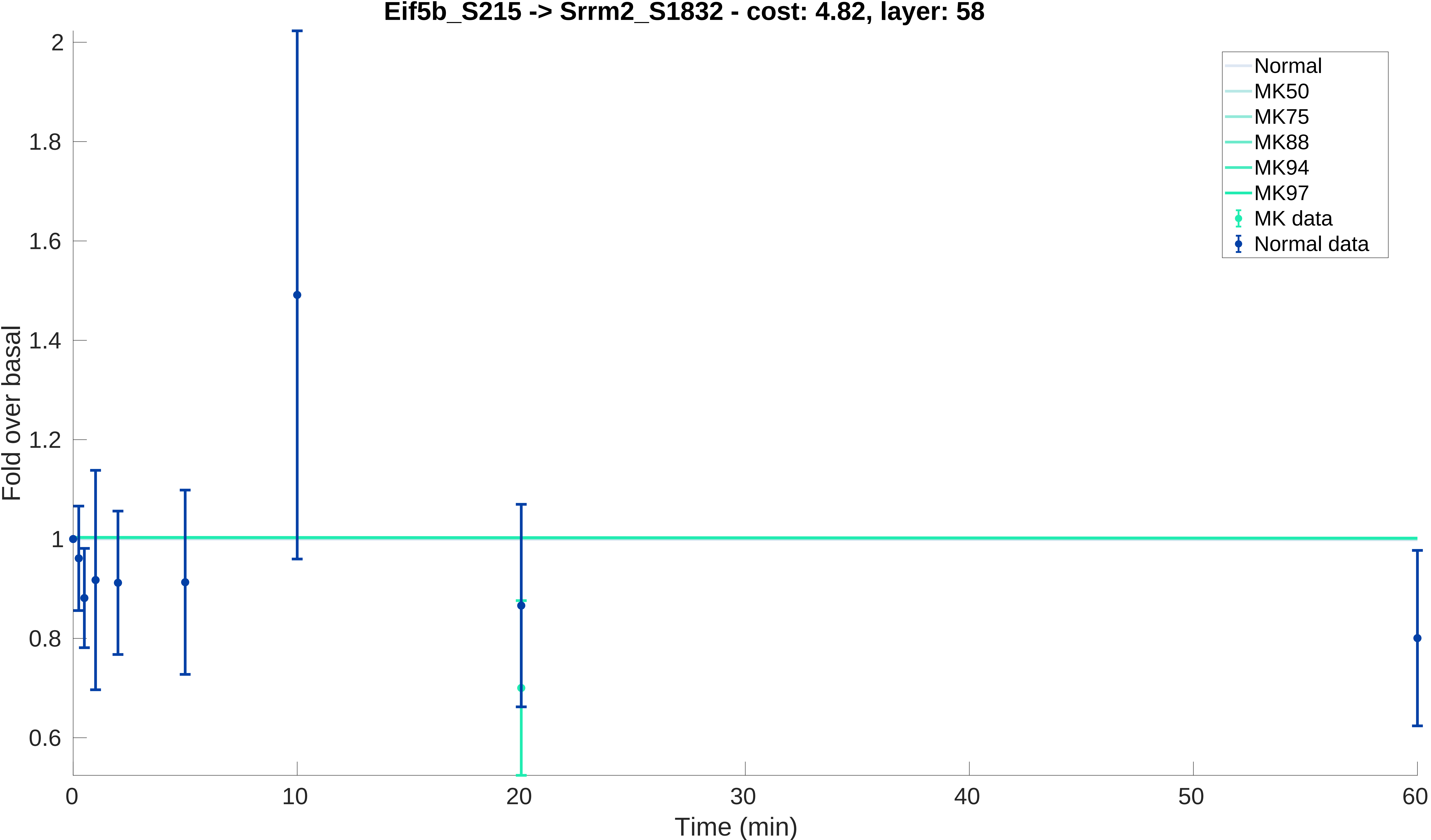
Eif5b_S215 -> Srrm2_S1832 - cost: 4.82, layer: 58
Fold over basal
Time (min)
2
1.8
1.6
1.4
1.2
1
0.8
0.6
0
10
20
30
40
50
60
Normal
MK50
MK75
MK88
MK94
MK97
MK data
Normal data

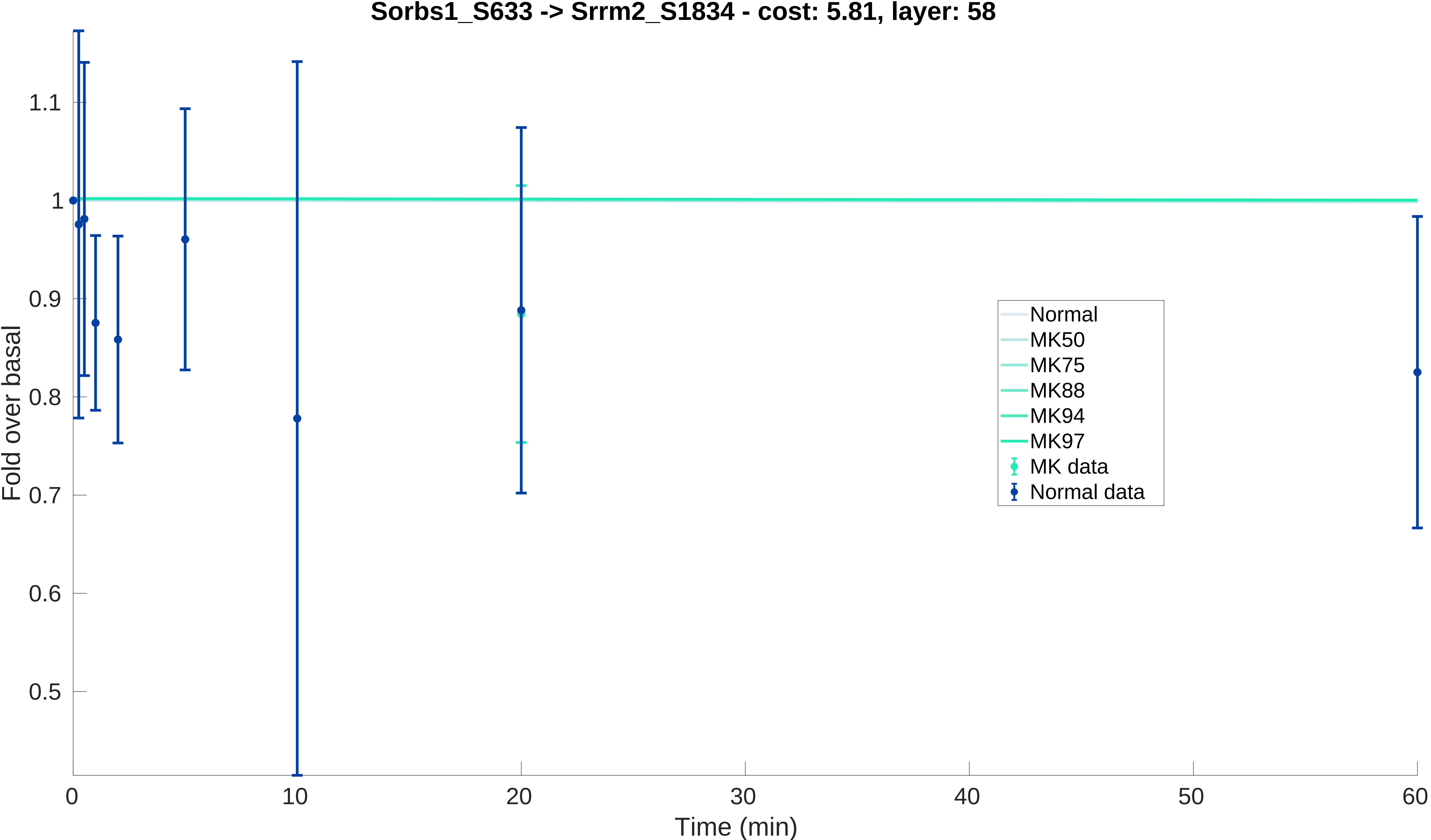

Sorbs1_S633 -> Srrm2_S1834 - cost: 5.81, layer: 58
Fold over basal
1.1
1
0.9
0.8
0.7
0.6
0.5
0
10
20
30
40
50
60
Time (min)
Normal
MK50
MK75
MK88
MK94
MK97
MK data
Normal data

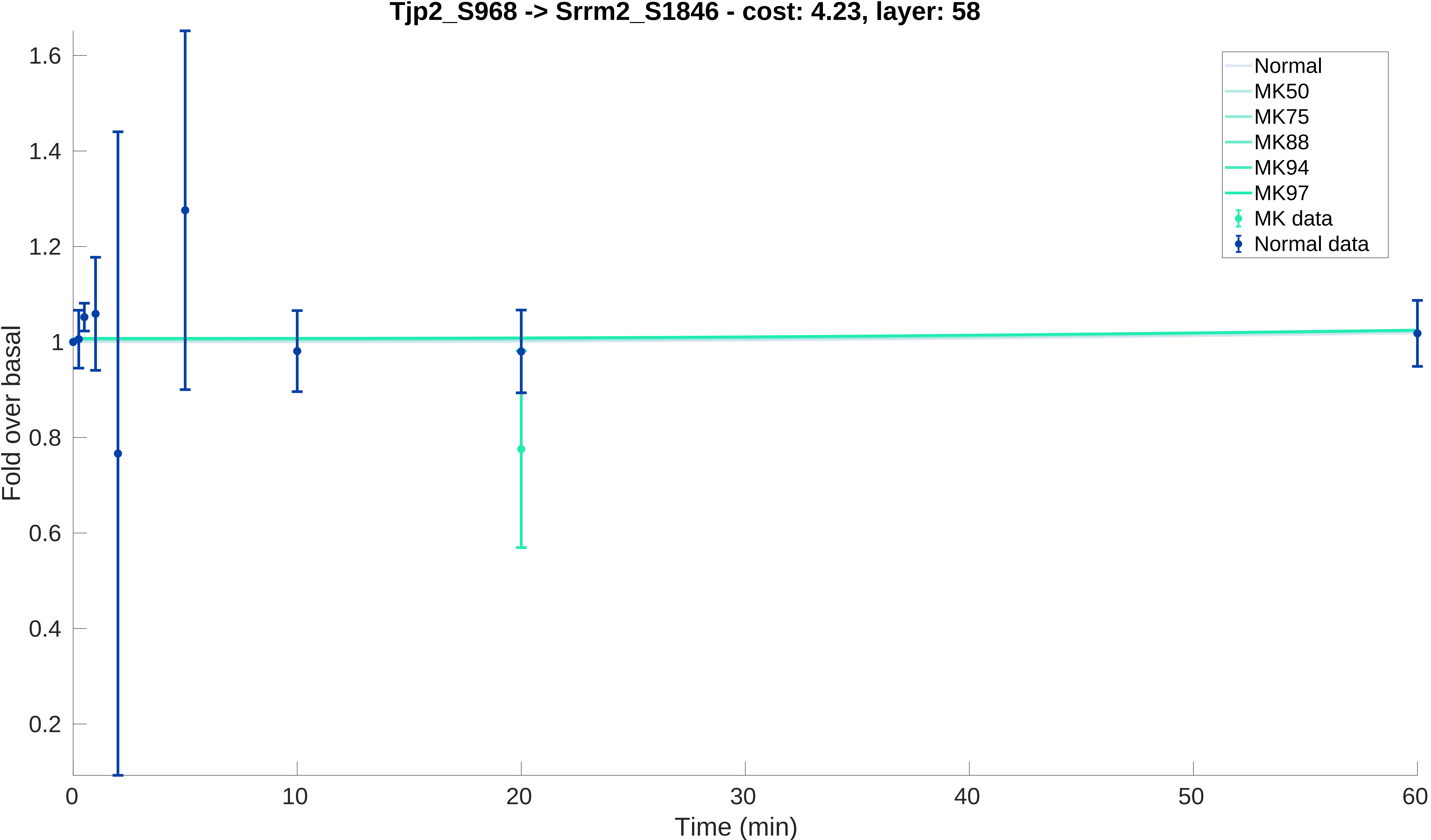

Tjp2_S968 -> Srrm2_S1846 - cost: 4.23, layer: 58
Fold over basal
Time (min)
1.6
1.4
1.2
1
0.8
0.6
0.4
0.2
0
10
20
30
40
50
60
Normal
MK50
MK75
MK88
MK94
MK97
MK data
Normal data

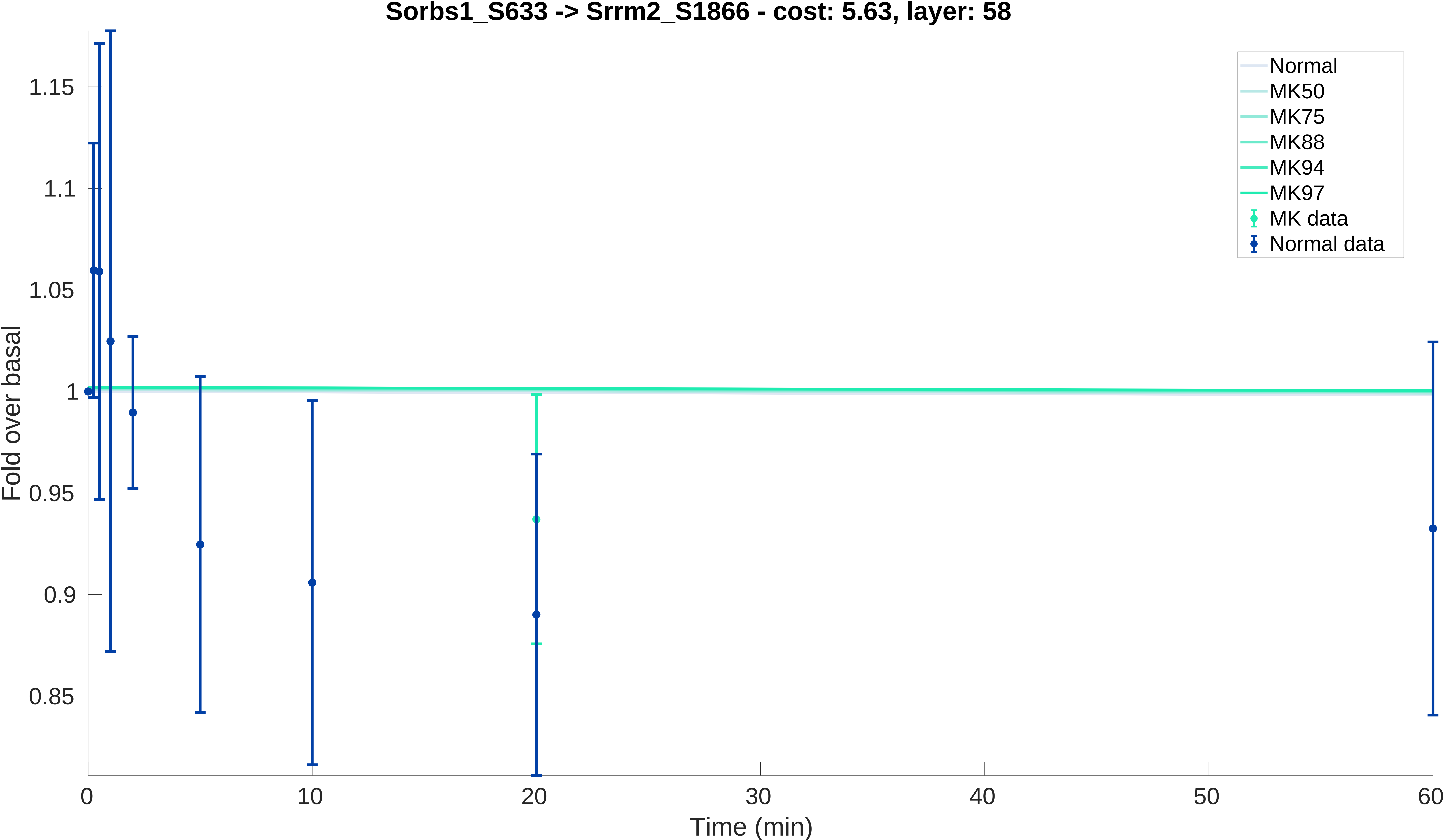
Sorbs1_S633 -> Srrm2_S1866 - cost: 5.63, layer: 58
Fold over basal
1.15
1.1
1.05
1
0.95
0.9
0.85
0
10
20
30
40
50
60
Time (min)
Normal
MK50
MK75
MK88
MK94
MK97
MK data
Normal data

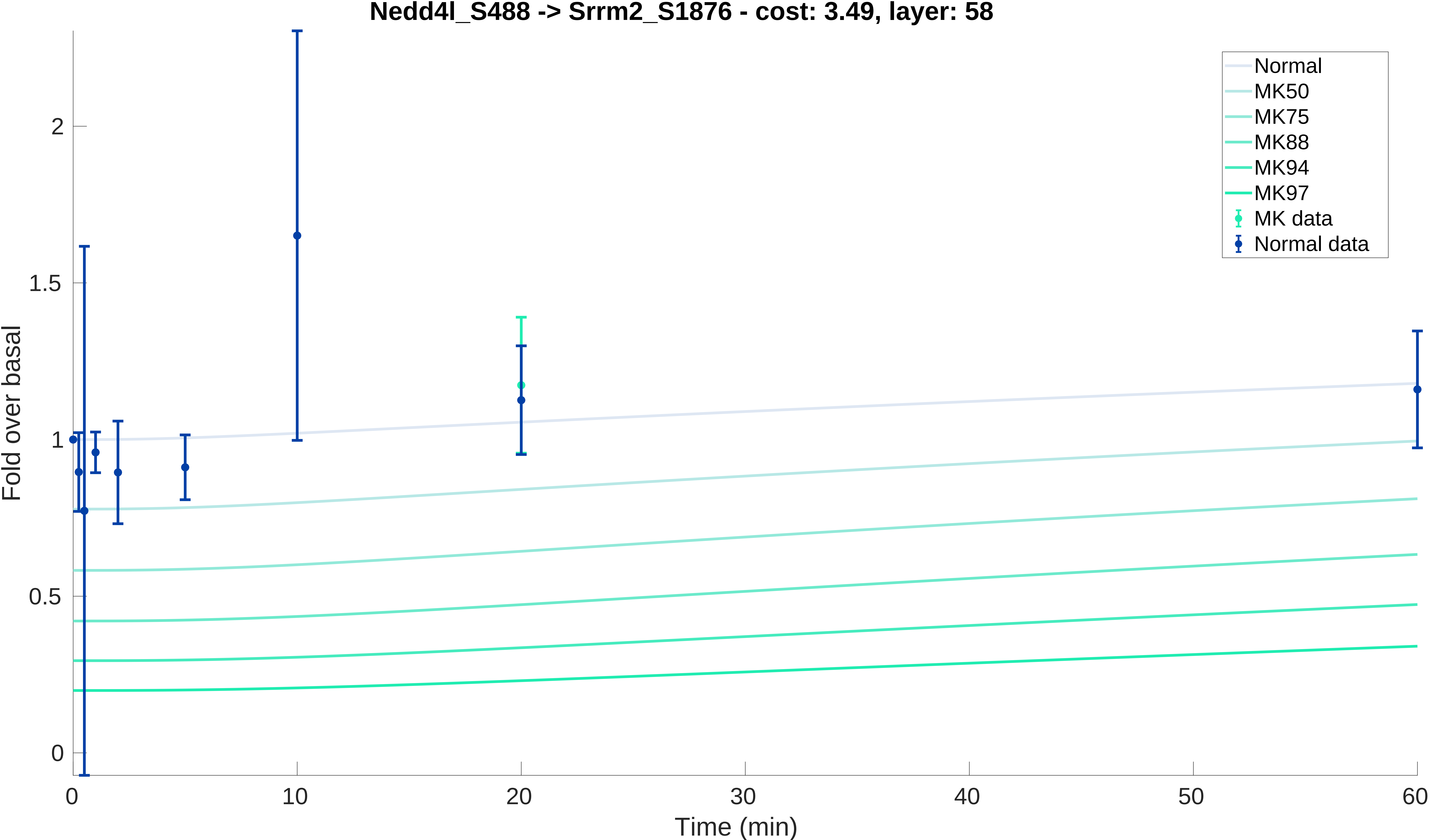

Nedd4l_S488 -> Srrm2_S1876 - cost: 3.49, layer: 58
Fold over basal
Time (min)
0
0.5
1
1.5
2
0
10
20
30
40
50
60
Normal
MK50
MK75
MK88
MK94
MK97
MK data
Normal data

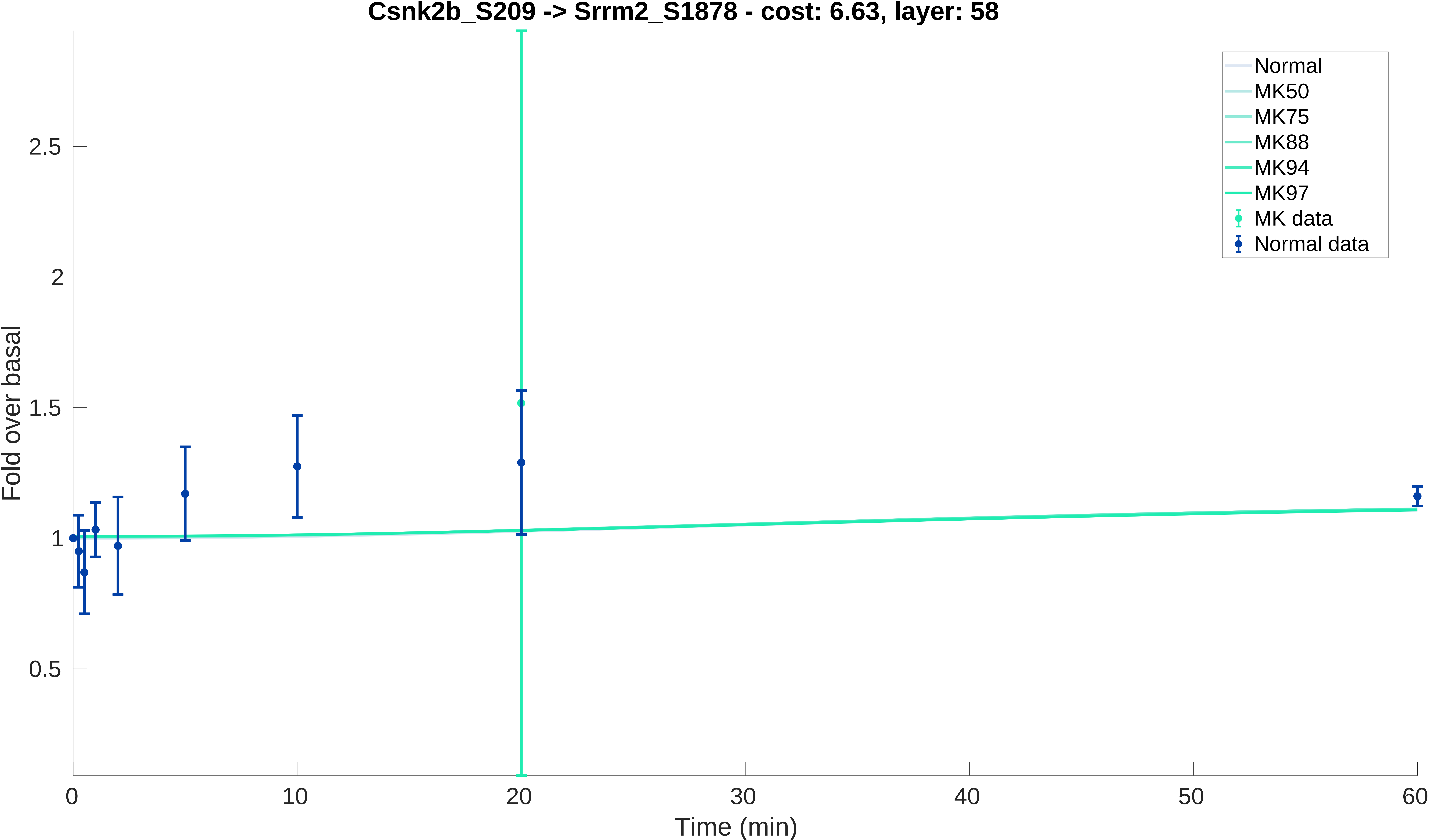
Csnk2b_S209 -> Srrm2_S1878 - cost: 6.63, layer: 58
Normal
MK50
MK75
MK88
MK94
MK97
MK data
Normal data
Fold over basal
0.5
1
1.5
2
2.5
0
10
20
30
40
50
60
Time (min)

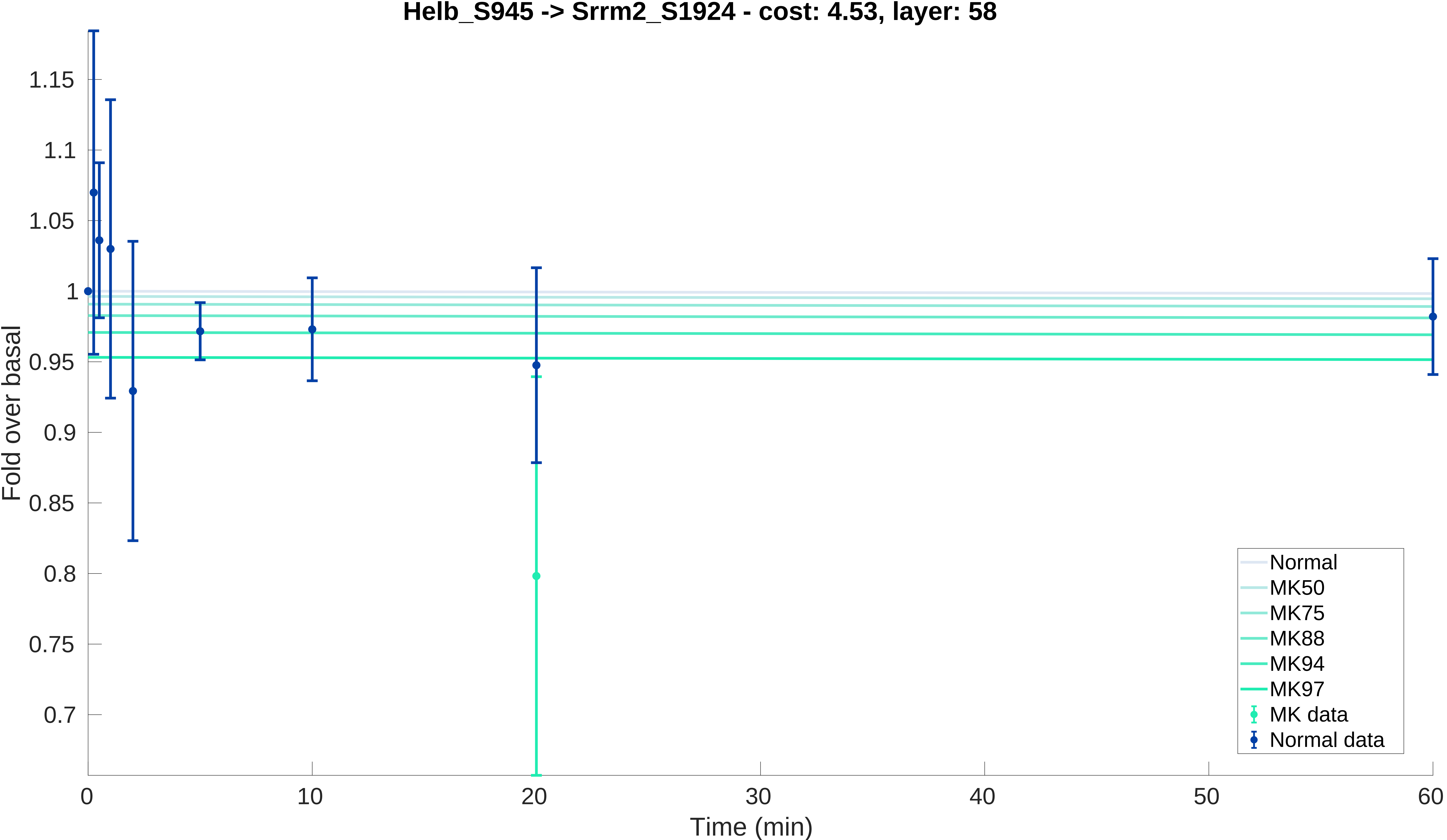

Helb_S945 -> Srrm2_S1924 - cost: 4.53, layer: 58
Fold over basal
1.15
1.1
1.05
1
0.95
0.9
0.85
0.8
0.75
0.7
0
10
20
30
40
50
60
Time (min)
Normal
MK50
MK75
MK88
MK94
MK97
MK data
Normal data

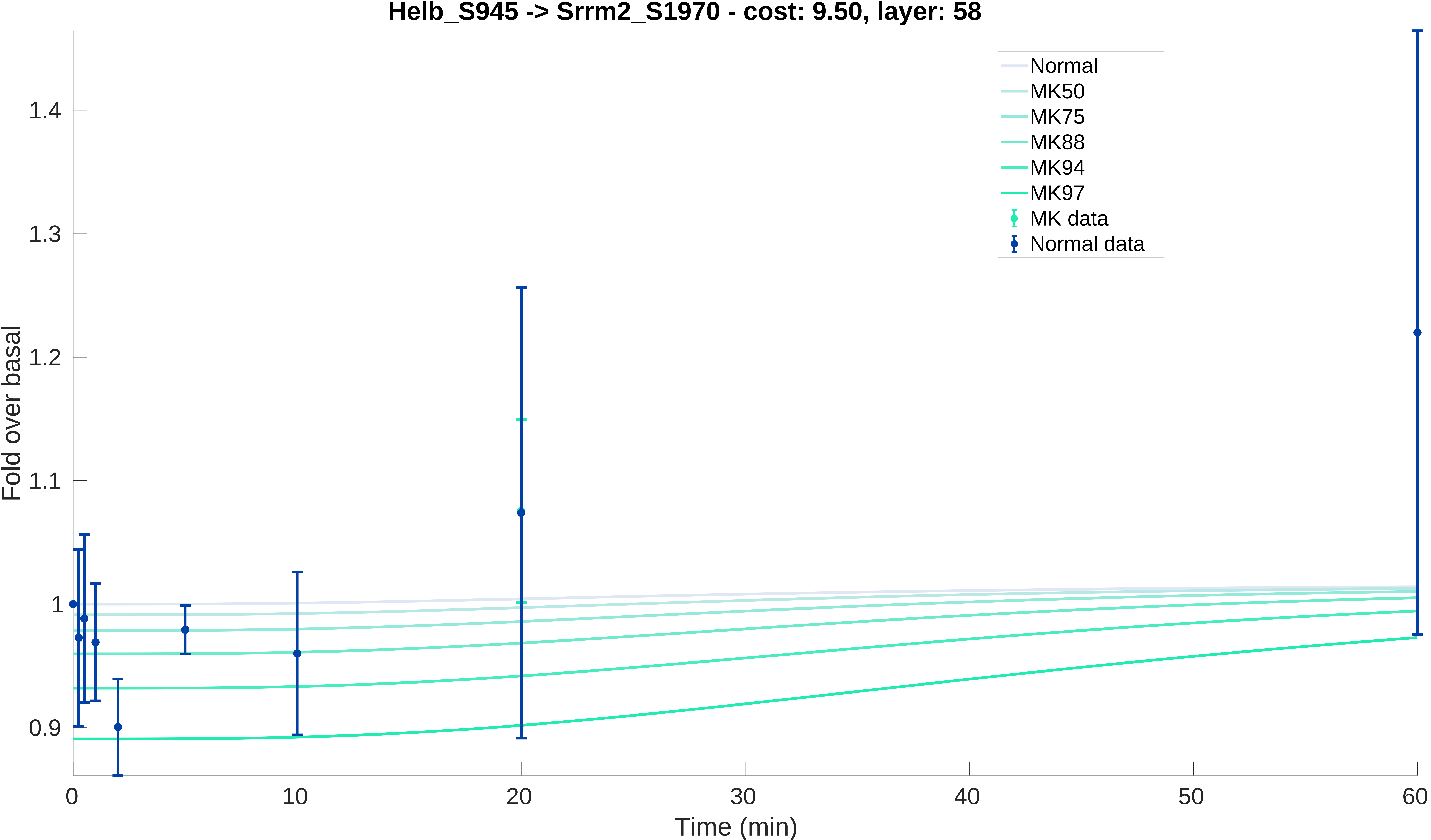
Helb_S945 -> Srrm2_S1970 - cost: 9.50, layer: 58
Normal
MK50
MK75
MK88
MK94
MK97
MK data
Normal data
Fold over basal
1.4
1.3
1.2
1.1
1
0.9
0
10
20
30
40
50
60
Time (min)

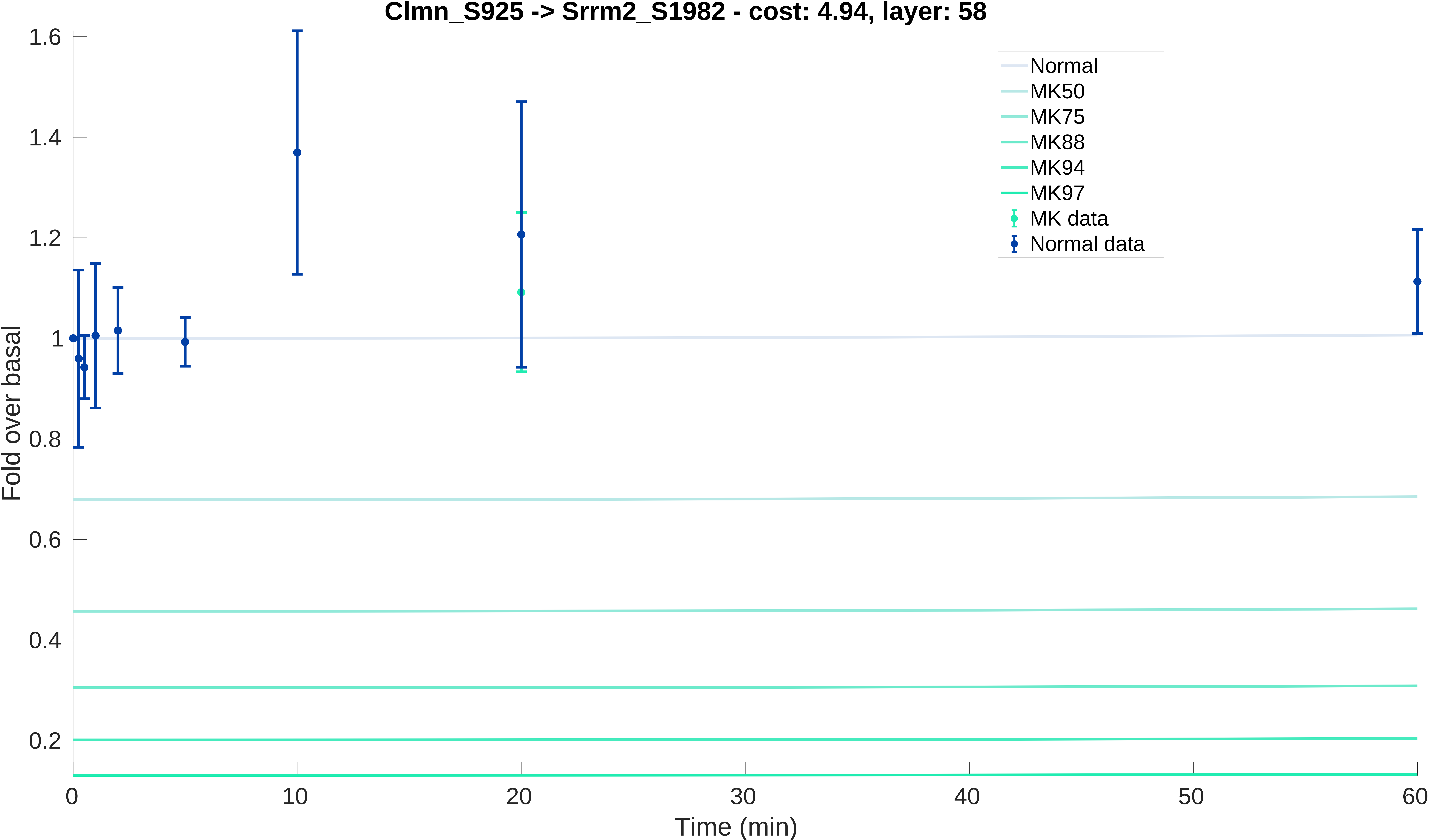

Clmn_S925 -> Srrm2_S1982 - cost: 4.94, layer: 58
Normal
MK50
MK75
MK88
MK94
MK97
MK data
Normal data
Fold over basal
Time (min)
0.2
0.4
0.6
0.8
1
1.2
1.4
1.6
0
10
20
30
40
50
60

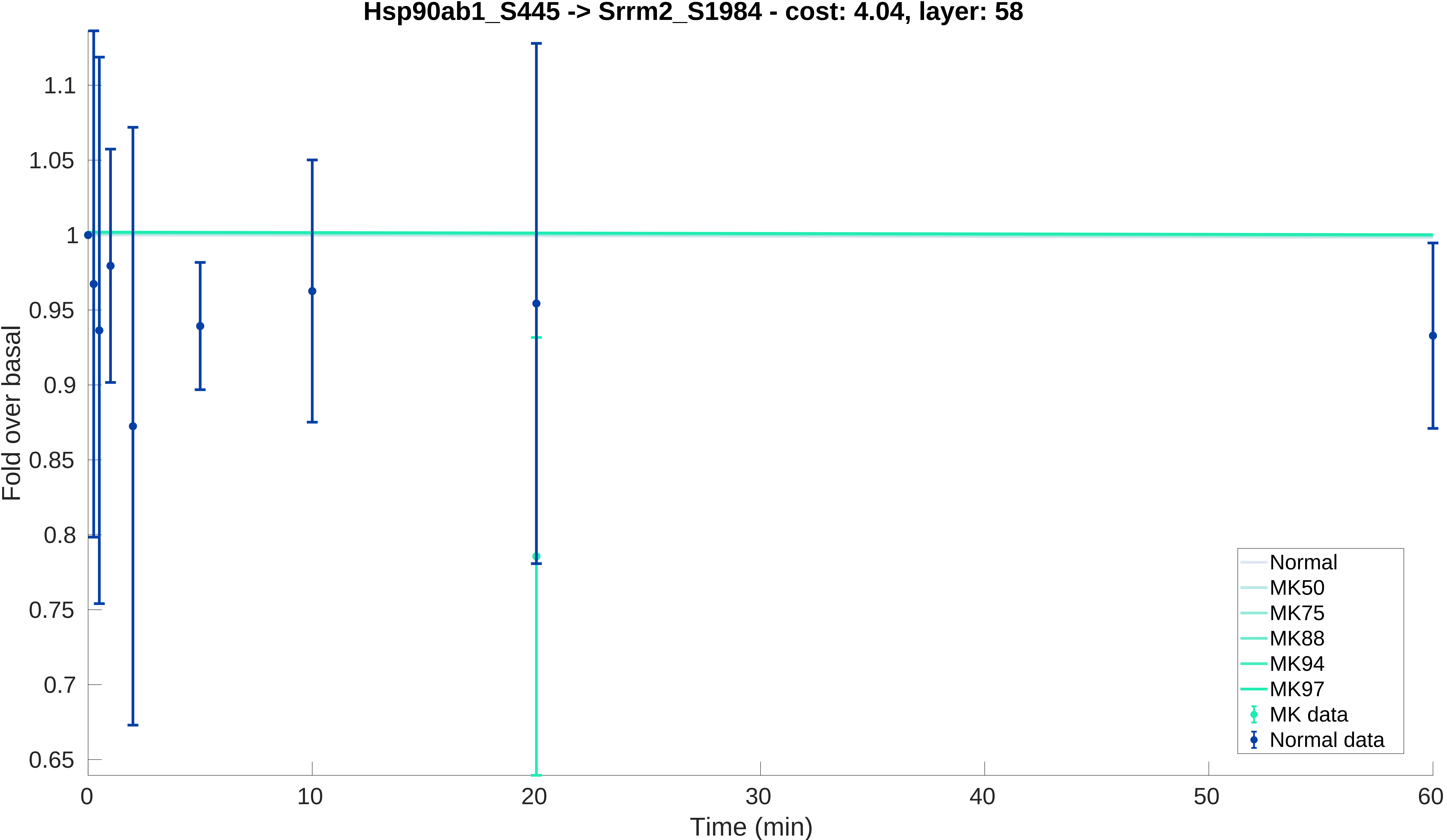

Hsp90ab1_S445 -> Srrm2_S1984 - cost: 4.04, layer: 58
Fold over basal
1.1
1.05
1
0.95
0.9
0.85
0.8
0.75
0.7
0.65
0
10
20
30
40
50
60
Time (min)
Normal
MK50
MK75
MK88
MK94
MK97
MK data
Normal data

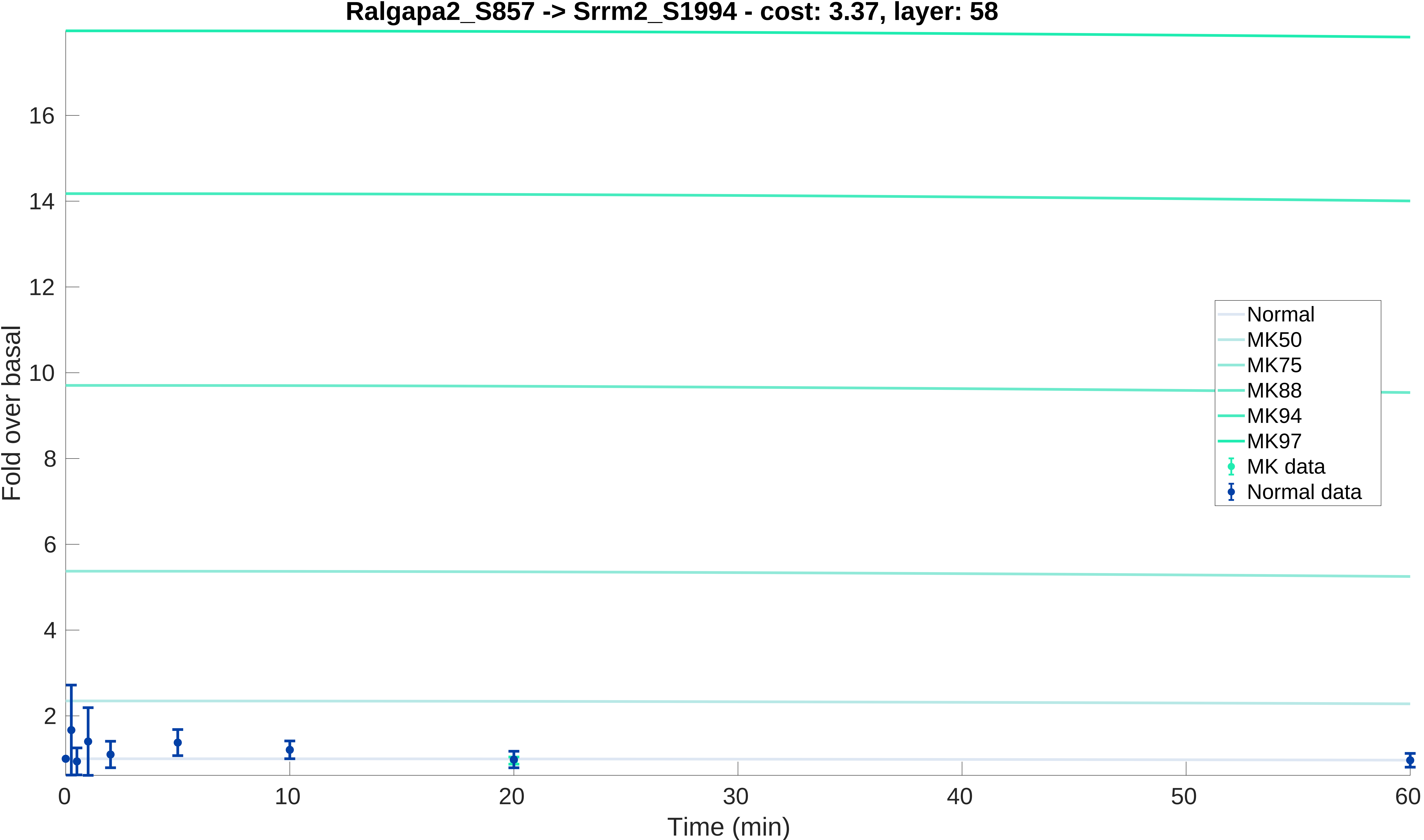

Ralgapa2_S857 -> Srrm2_S1994 - cost: 3.37, layer: 58
Fold over basal
Time (min)
0
10
20
30
40
50
60
2
4
6
8
10
12
14
16
Normal
MK50
MK75
MK88
MK94
MK97
MK data
Normal data

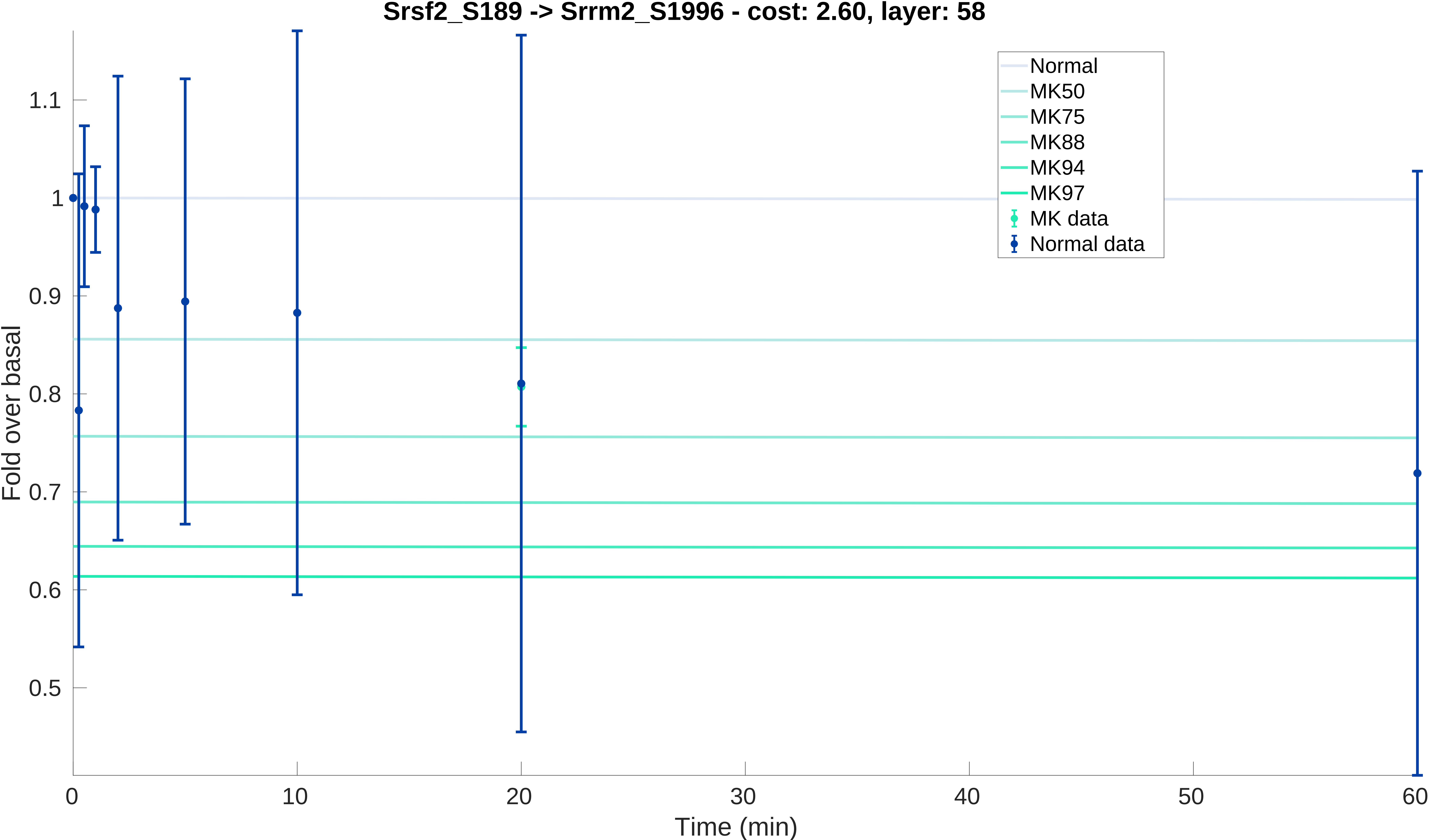

Srsf2_S189 -> Srrm2_S1996 - cost: 2.60, layer: 58
Fold over basal
Time (min)
Normal
MK50
MK75
MK88
MK94
MK97
MK data
Normal data
1.1
1
0.9
0.8
0.7
0.6
0.5
0
10
20
30
40
50
60

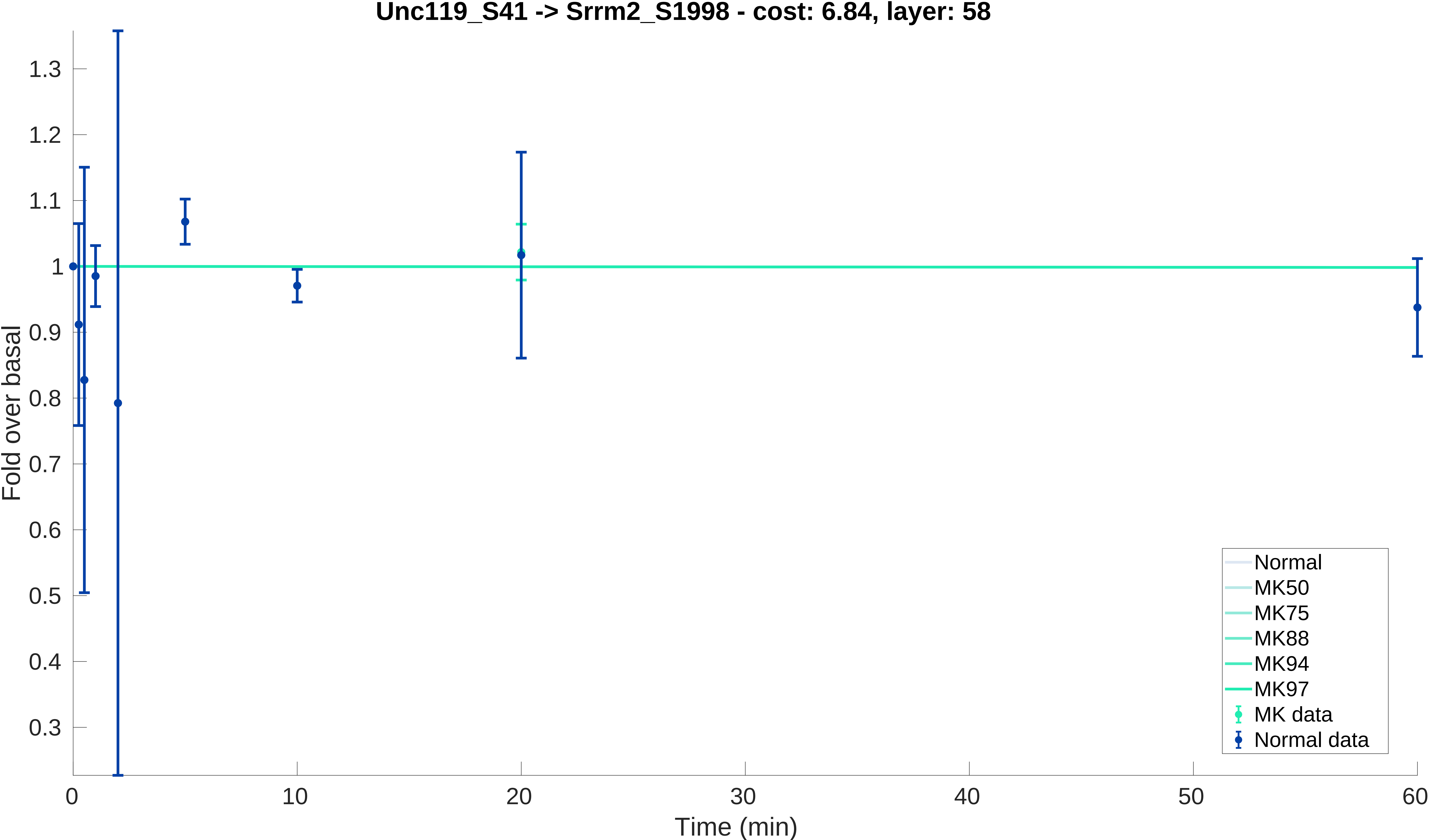

Unc119_S41 -> Srrm2_S1998 - cost: 6.84, layer: 58
Fold over basal
Time (min)
0
10
20
30
40
50
60
0.3
0.4
0.5
0.6
0.7
0.8
0.9
1
1.1
1.2
1.3
Normal
MK50
MK75
MK88
MK94
MK97
MK data
Normal data

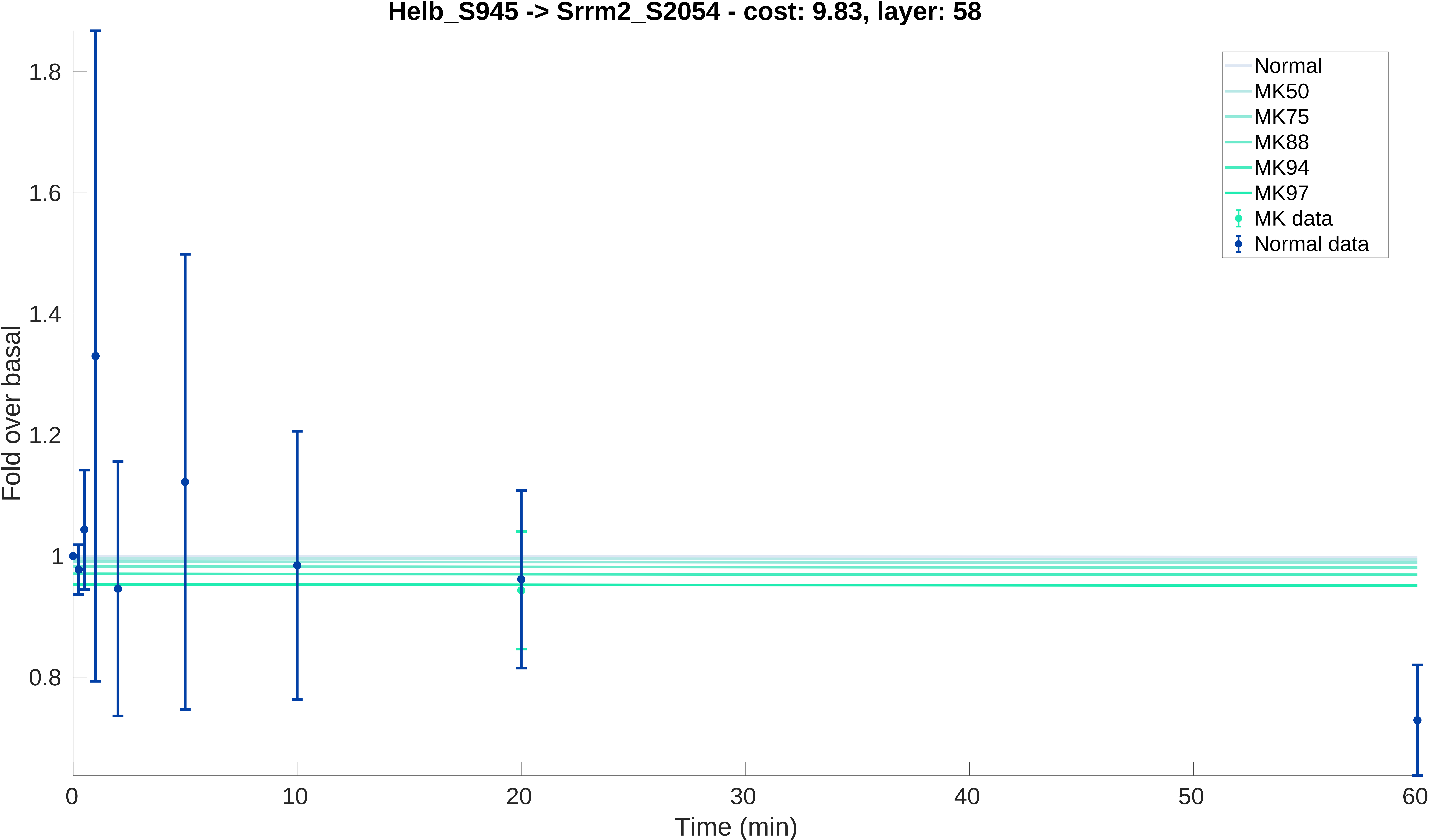

Helb_S945 -> Srrm2_S2054 - cost: 9.83, layer: 58
Fold over basal
Time (min)
1.8
1.6
1.4
1.2
1
0.8
0
10
20
30
40
50
60
Normal
MK50
MK75
MK88
MK94
MK97
MK data
Normal data

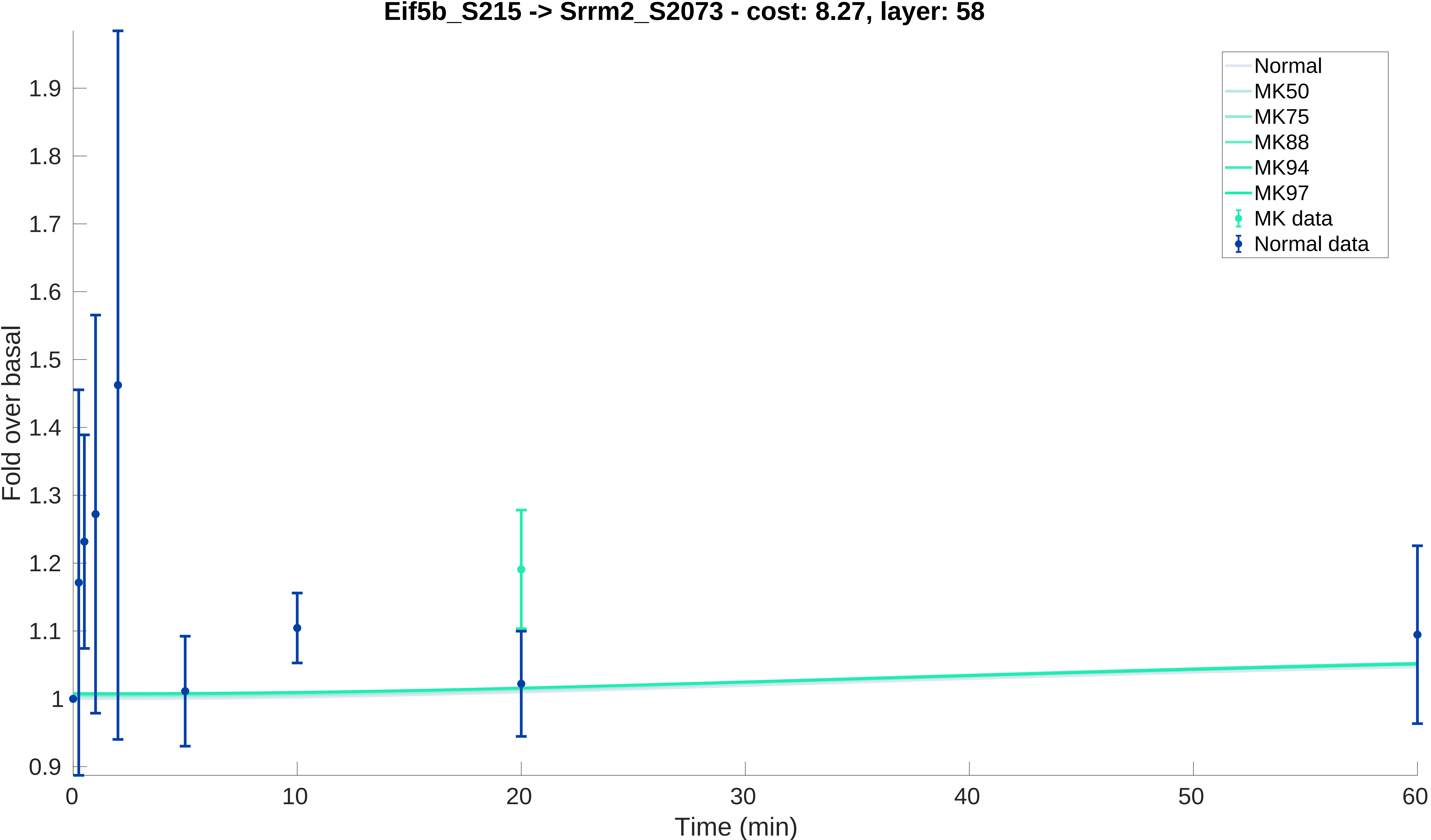

Eif5b_S215 -> Srrm2_S2073 - cost: 8.27, layer: 58
Fold over basal
1.9
1.8
1.7
1.6
1.5
1.4
1.3
1.2
1.1
1
0.9
0
10
20
30
40
50
60
Time (min)
Normal
MK50
MK75
MK88
MK94
MK97
MK data
Normal data

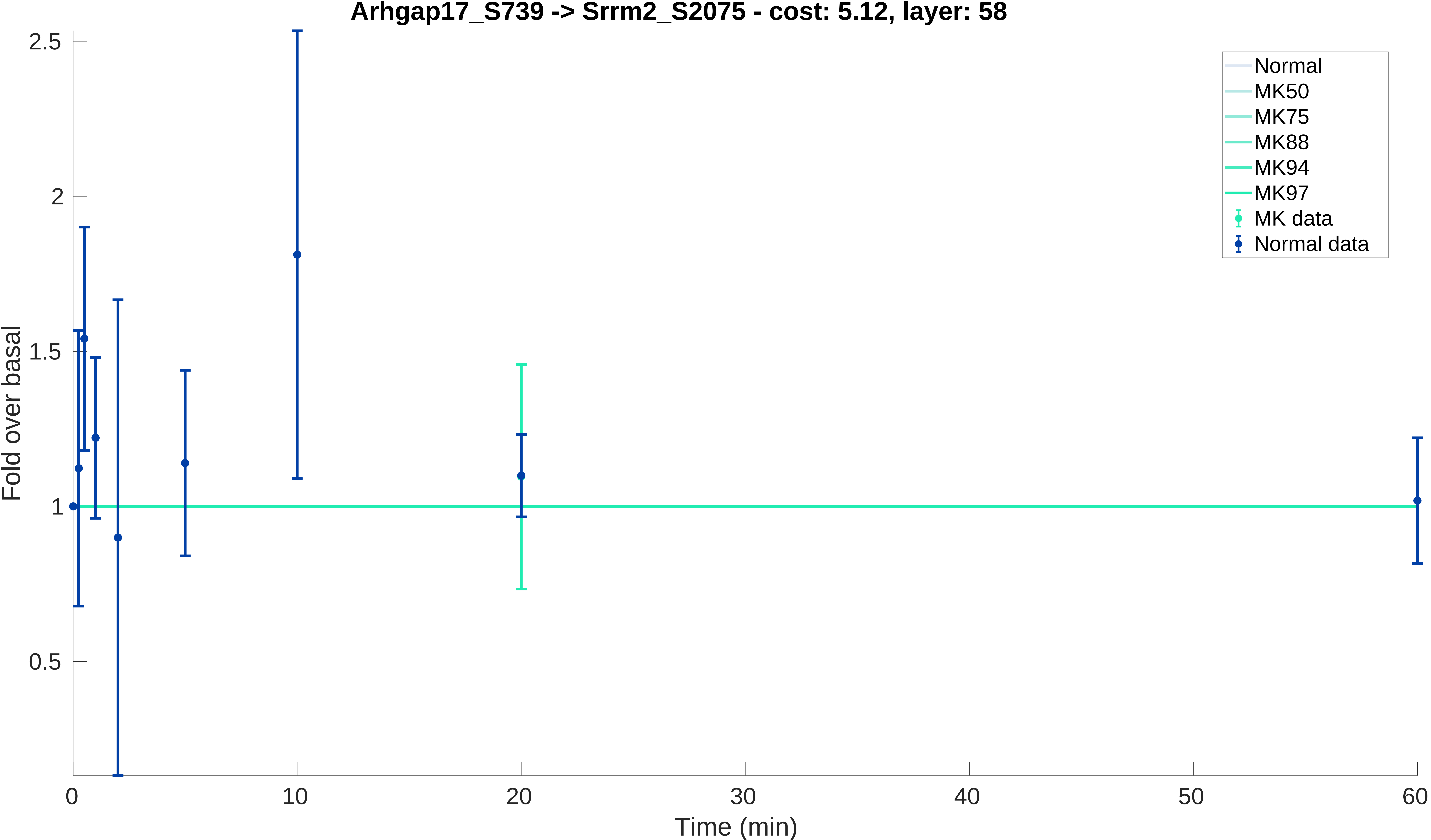

Arhgap17_S739 -> Srrm2_S2075 - cost: 5.12, layer: 58
Fold over basal
Time (min)
0
10
20
30
40
50
60
0.5
1
1.5
2
2.5
Normal
MK50
MK75
MK88
MK94
MK97
MK data
Normal data

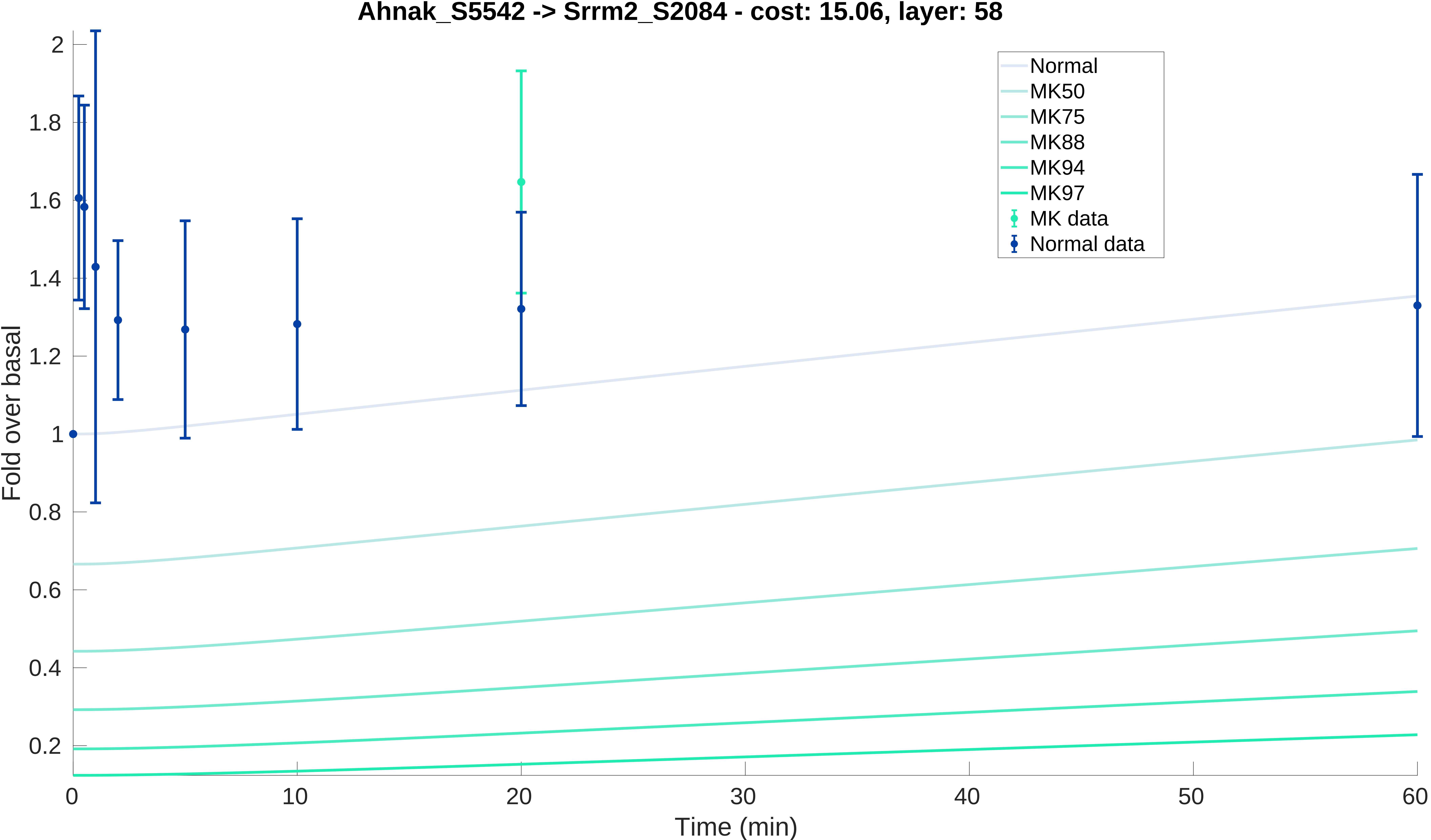

Ahnak_S5542 -> Srrm2_S2084 - cost: 15.06, layer: 58
Fold over basal
Time (min)
Normal
MK50
MK75
MK88
MK94
MK97
MK data
Normal data
2
1.8
1.6
1.4
1.2
1
0.8
0.6
0.4
0.2
0
10
20
30
40
50
60

#### Helb_S945 -> Srrm2_S2351 - cost: 10.34, layer: 58

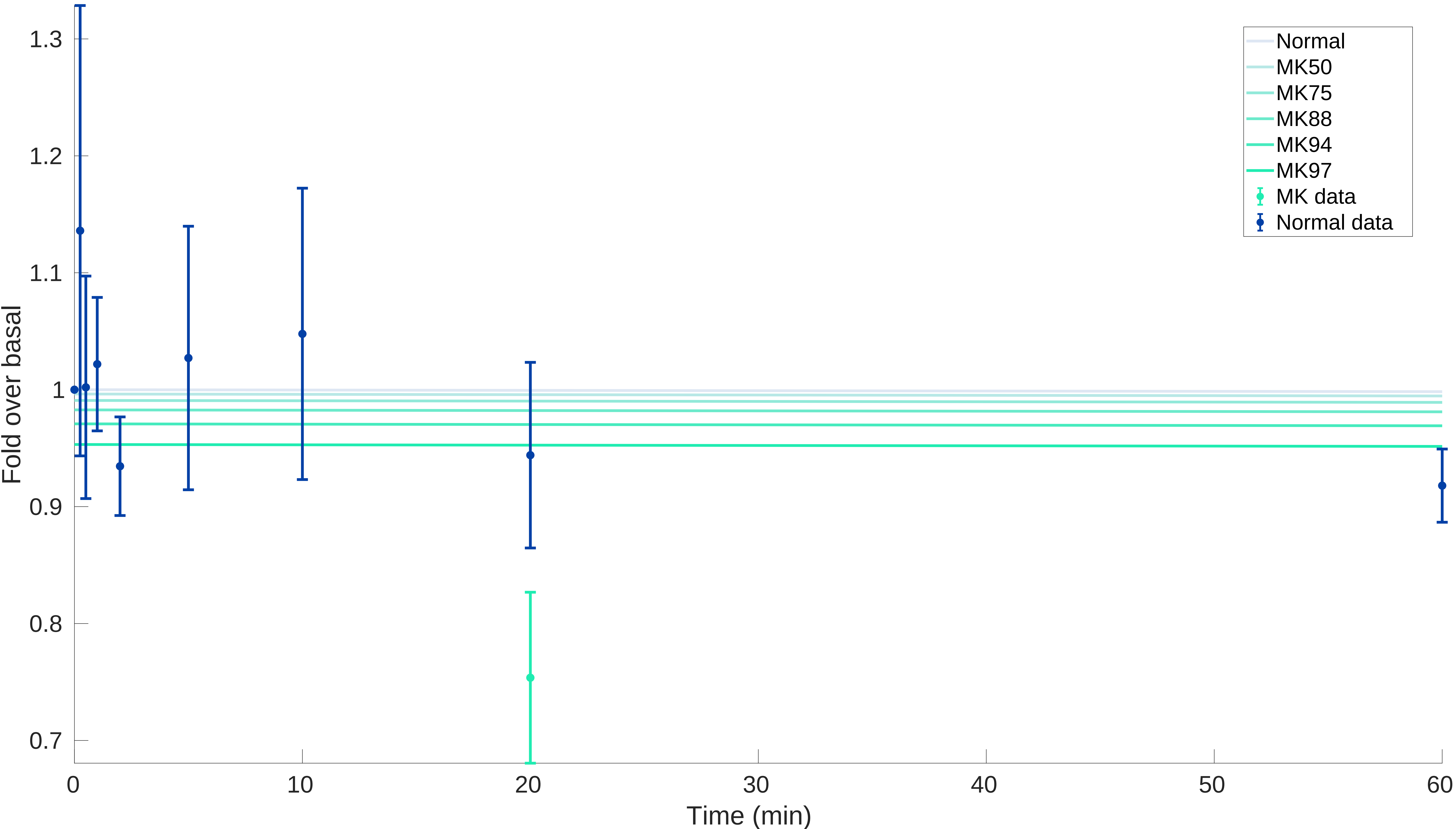

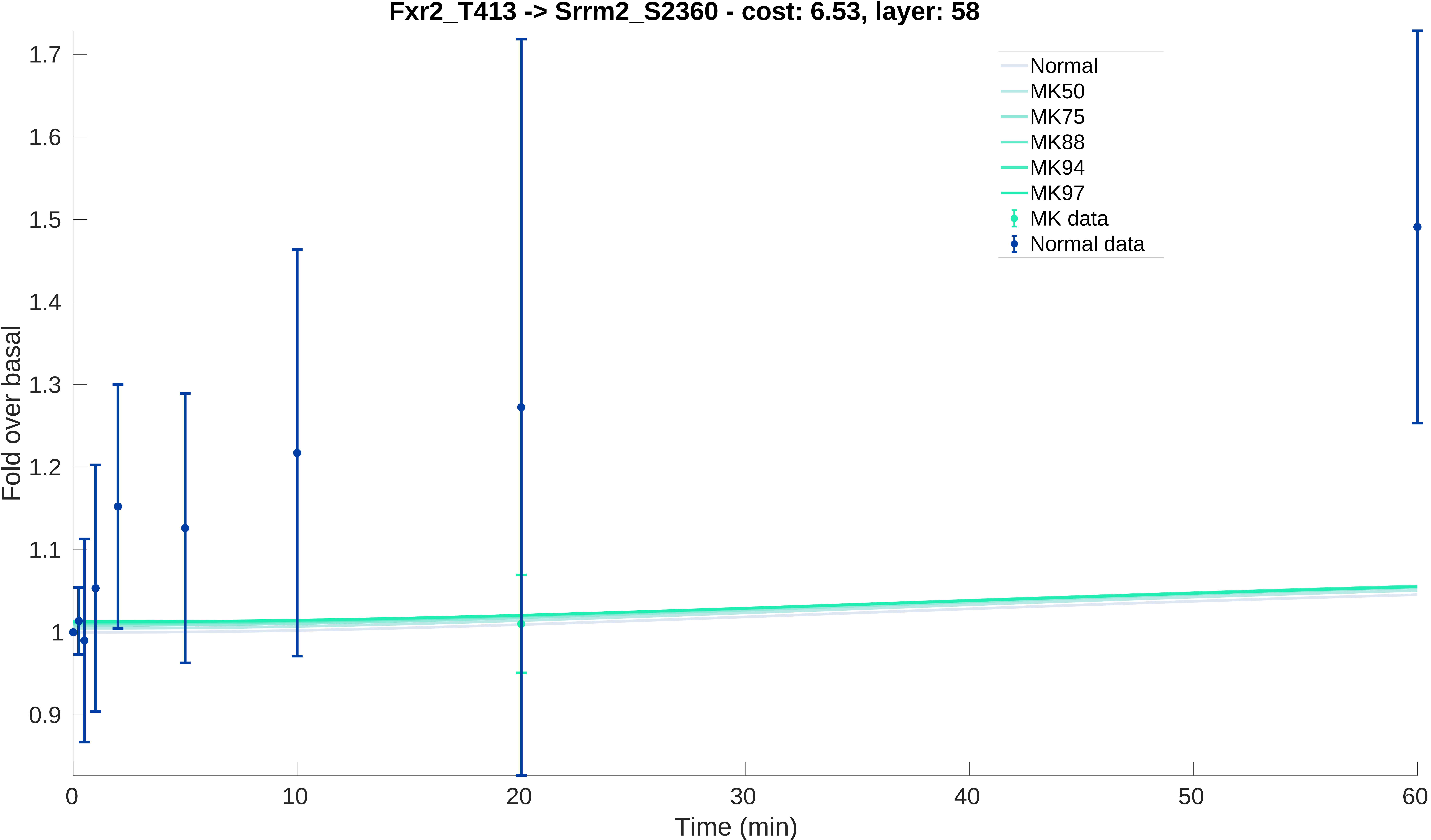

Fxr2_T413 -> Srrm2_S2360 - cost: 6.53, layer: 58
Fold over basal
Time (min)
0.9
1
1.1
1.2
1.3
1.4
1.5
1.6
1.7
0
10
20
30
40
50
60
Normal
MK50
MK75
MK88
MK94
MK97
MK data
Normal data

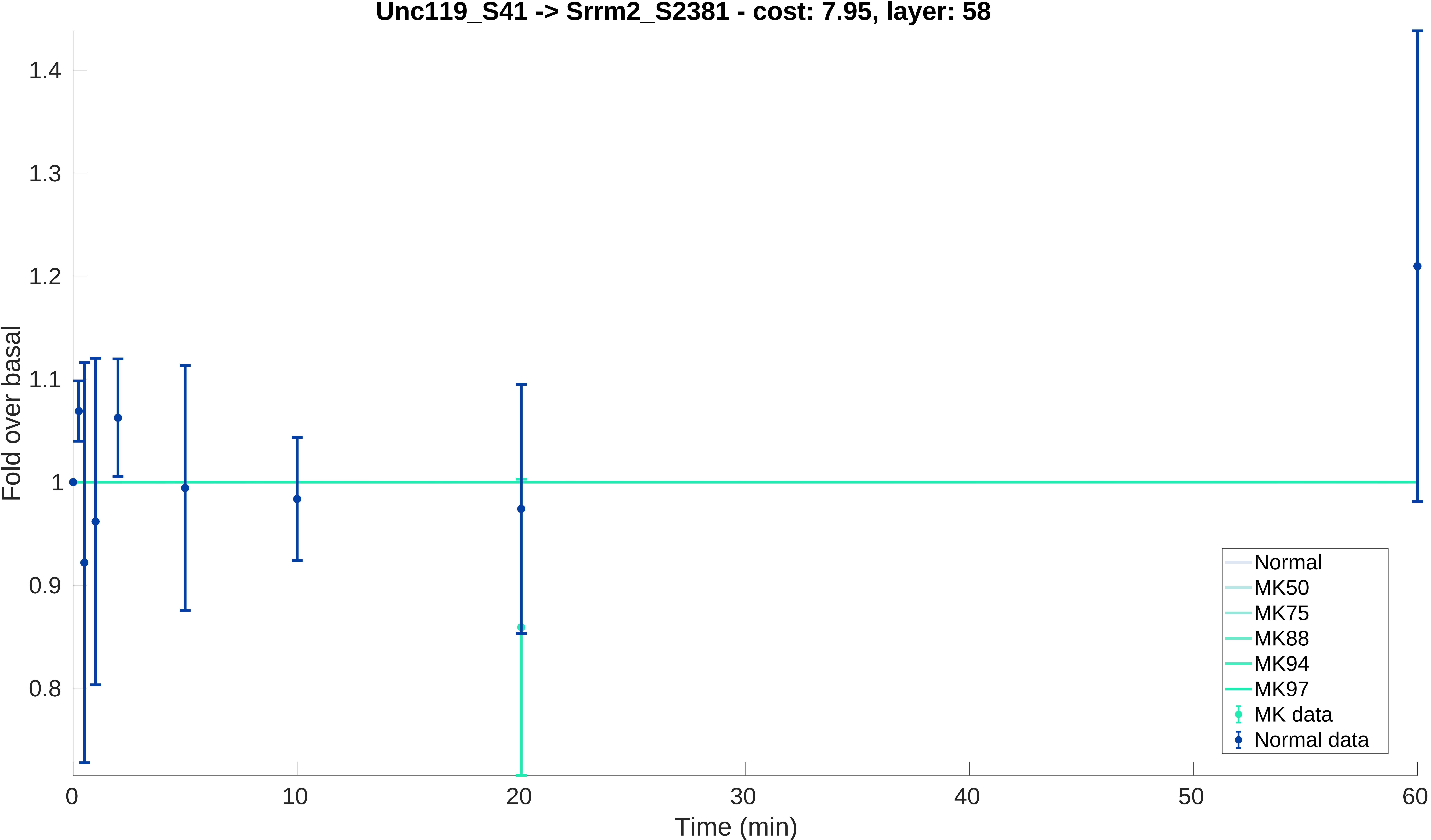

Unc119_S41 -> Srrm2_S2381 - cost: 7.95, layer: 58
Fold over basal
Time (min)
0.8
0.9
1
1.1
1.2
1.3
1.4
0
10
20
30
40
50
60
Normal
MK50
MK75
MK88
MK94
MK97
MK data
Normal data

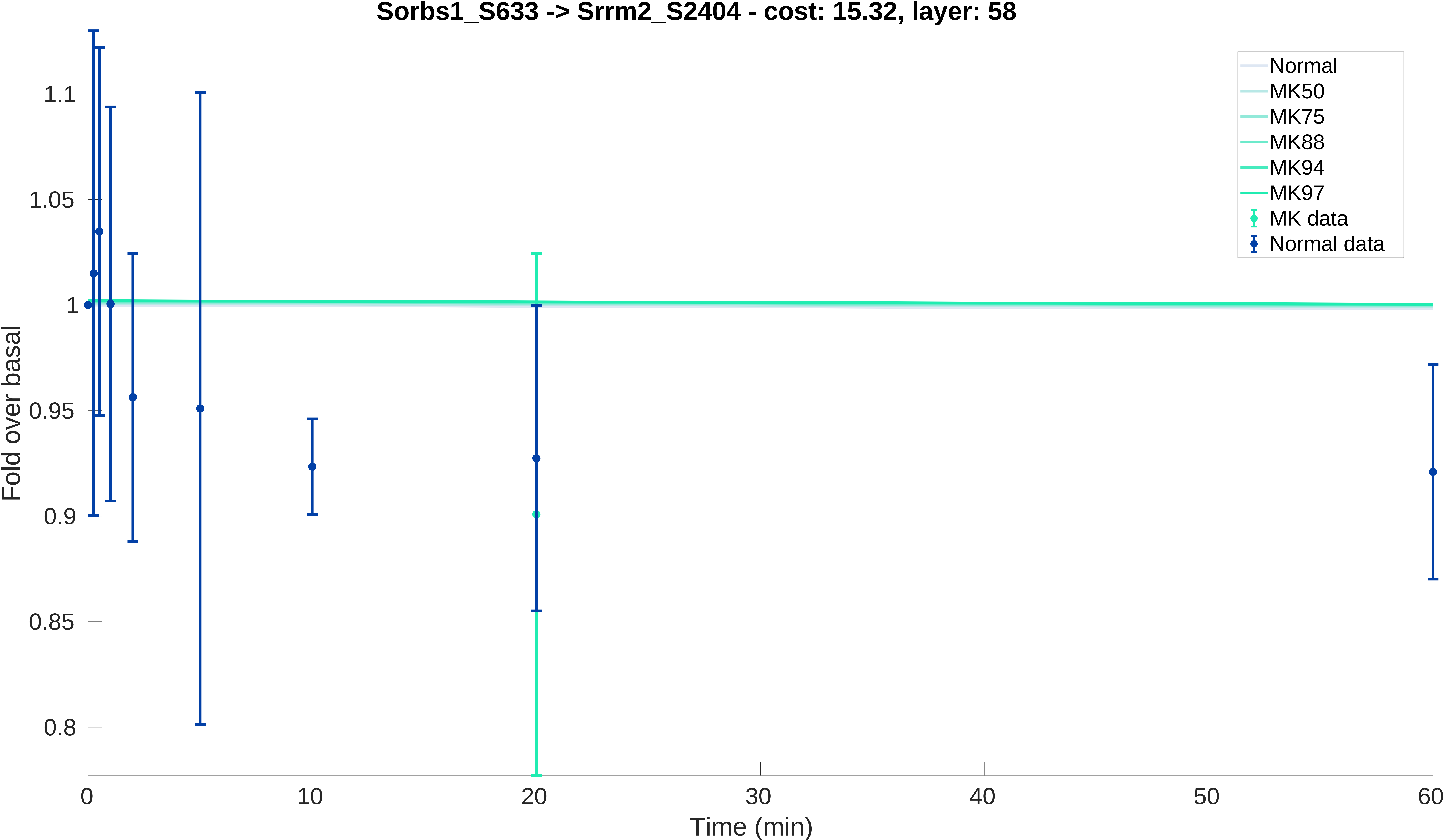
Sorbs1_S633 -> Srrm2_S2404 - cost: 15.32, layer: 58
Fold over basal
Time (min)
0.8
0.85
0.9
0.95
1
1.05
1.1
0
10
20
30
40
50
60
Normal
MK50
MK75
MK88
MK94
MK97
MK data
Normal data

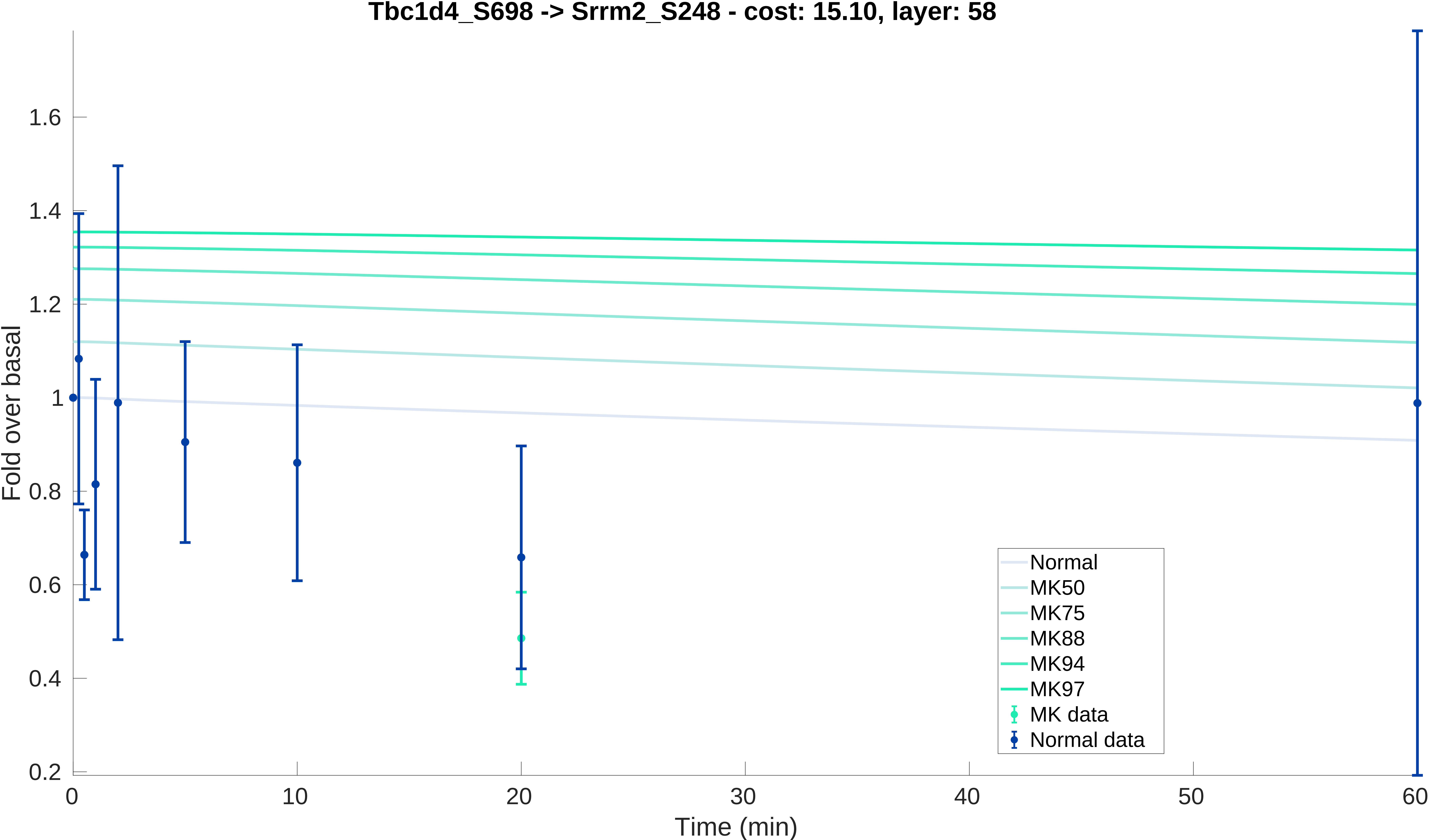

Tbc1d4_S698 -> Srrm2_S248 - cost: 15.10, layer: 58
Fold over basal
Time (min)
0
10
20
30
40
50
60
0.2
0.4
0.6
0.8
1
1.2
1.4
1.6
Normal
MK50
MK75
MK88
MK94
MK97
MK data
Normal data

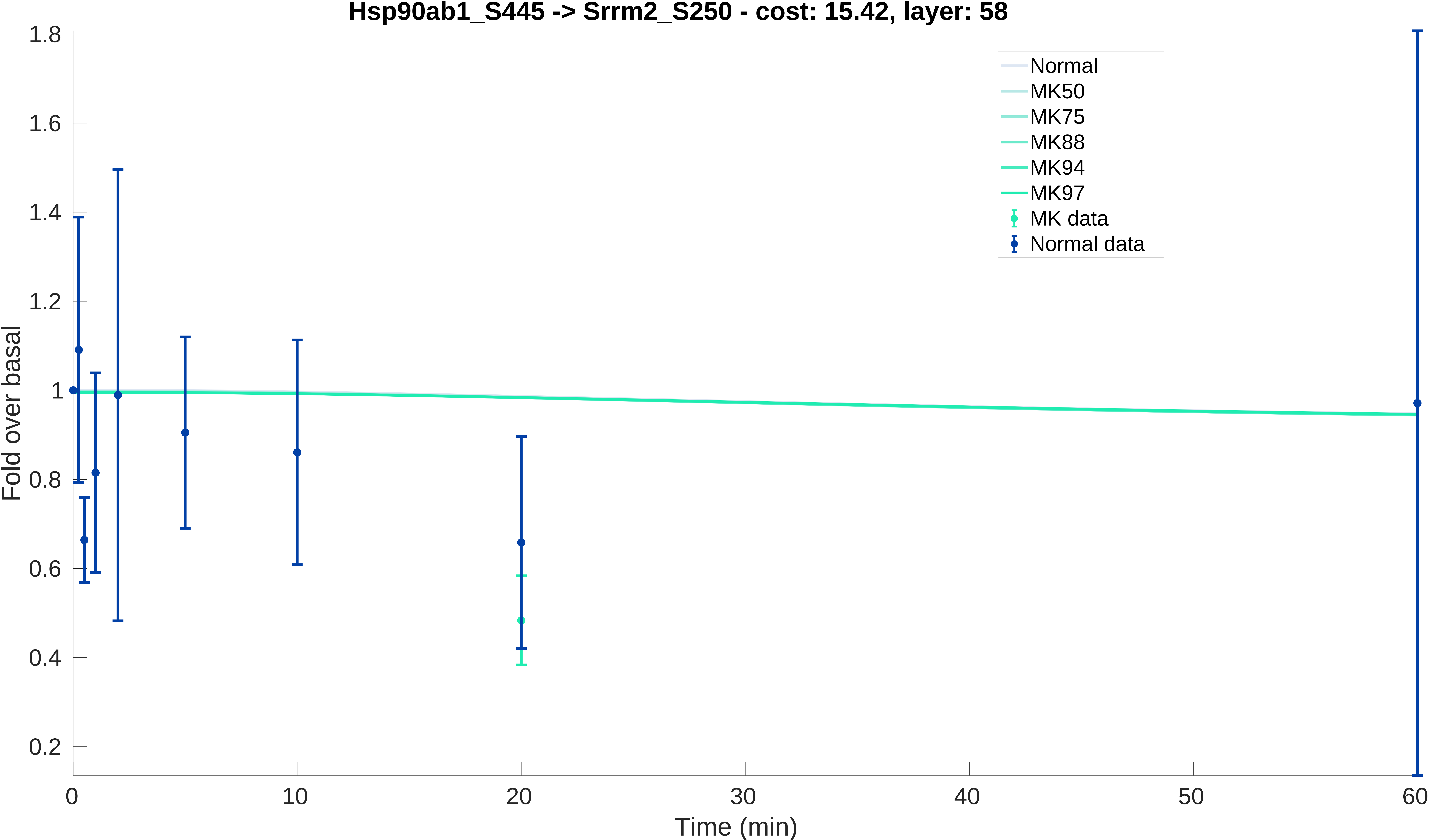

Hsp90ab1_S445 -> Srrm2_S250 - cost: 15.42, layer: 58
Fold over basal
Time (min)
1.8
1.6
1.4
1.2
1
0.8
0.6
0.4
0.2
0
10
20
30
40
50
60
Normal
MK50
MK75
MK88
MK94
MK97
MK data
Normal data

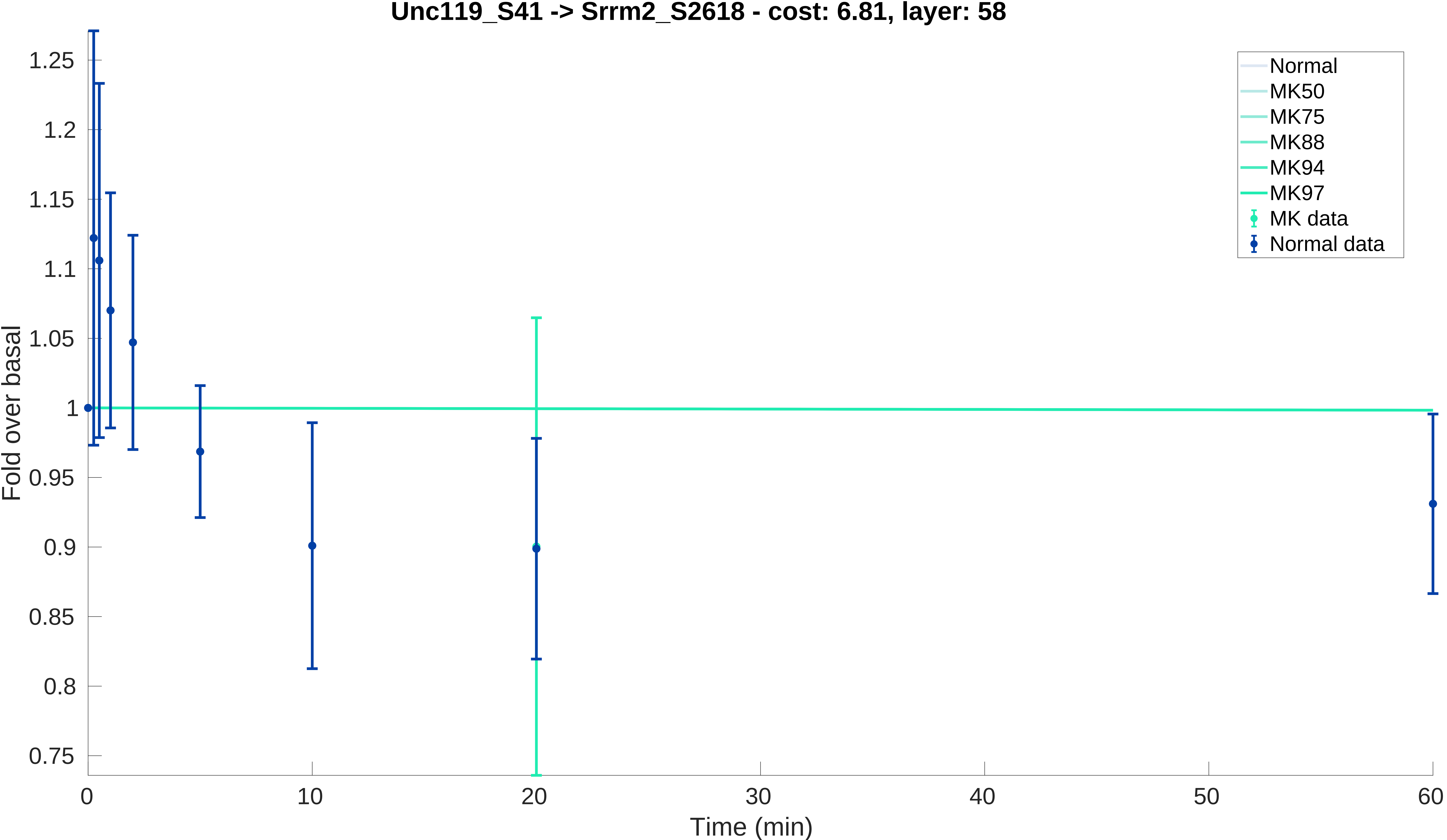

Unc119_S41 -> Srrm2_S2618 - cost: 6.81, layer: 58
Fold over basal
1.25
1.2
1.15
1.1
1.05
1
0.95
0.9
0.85
0.8
0.75
0
10
20
30
40
50
60
Time (min)
Normal
MK50
MK75
MK88
MK94
MK97
MK data
Normal data

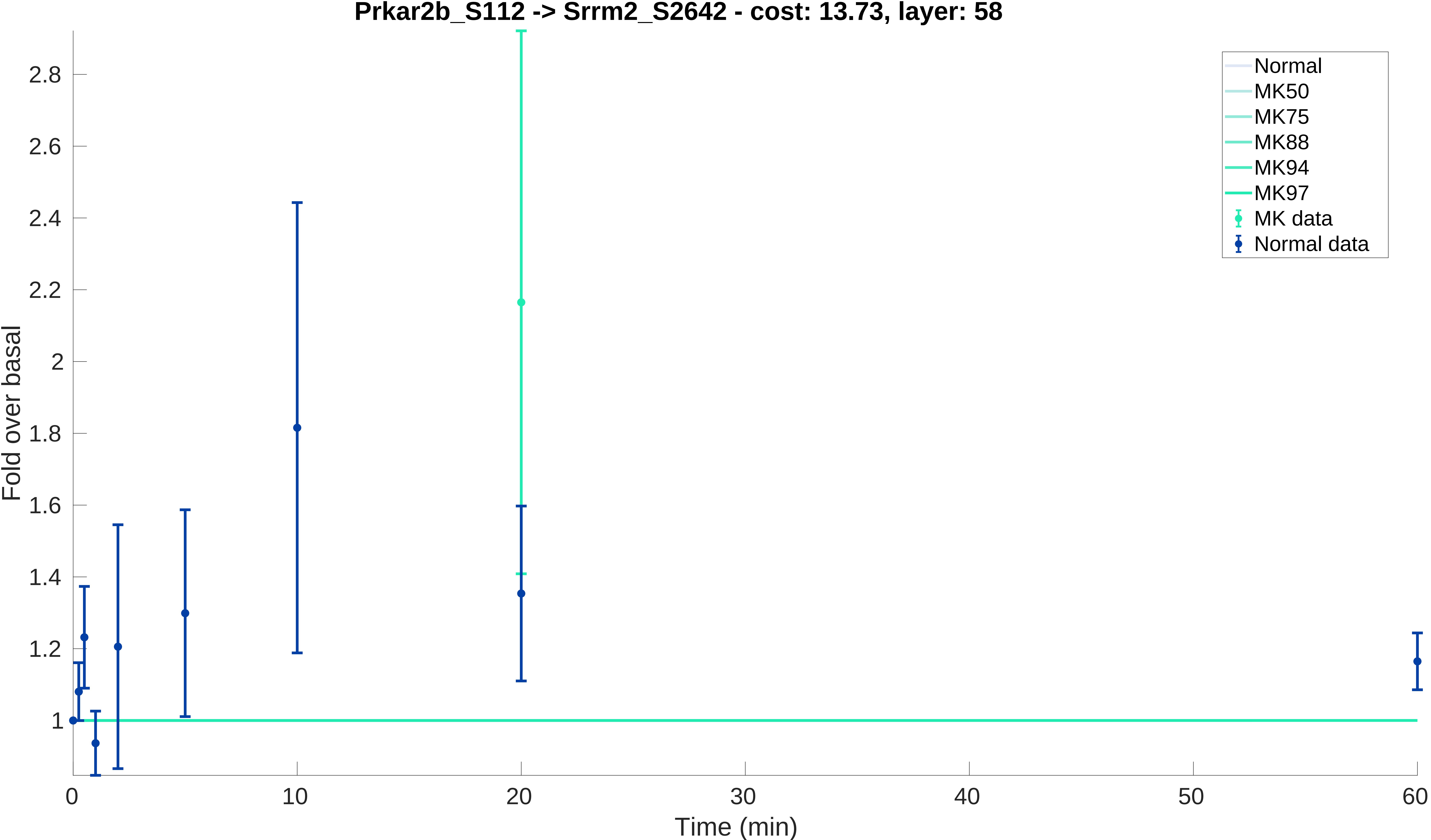

Prkar2b_S112 -> Srrm2_S2642 - cost: 13.73, layer: 58
Fold over basal
Time (min)
Normal
MK50
MK75
MK88
MK94
MK97
MK data
Normal data
2.8
2.6
2.4
2.2
2
1.8
1.6
1.4
1.2
1
0
10
20
30
40
50
60

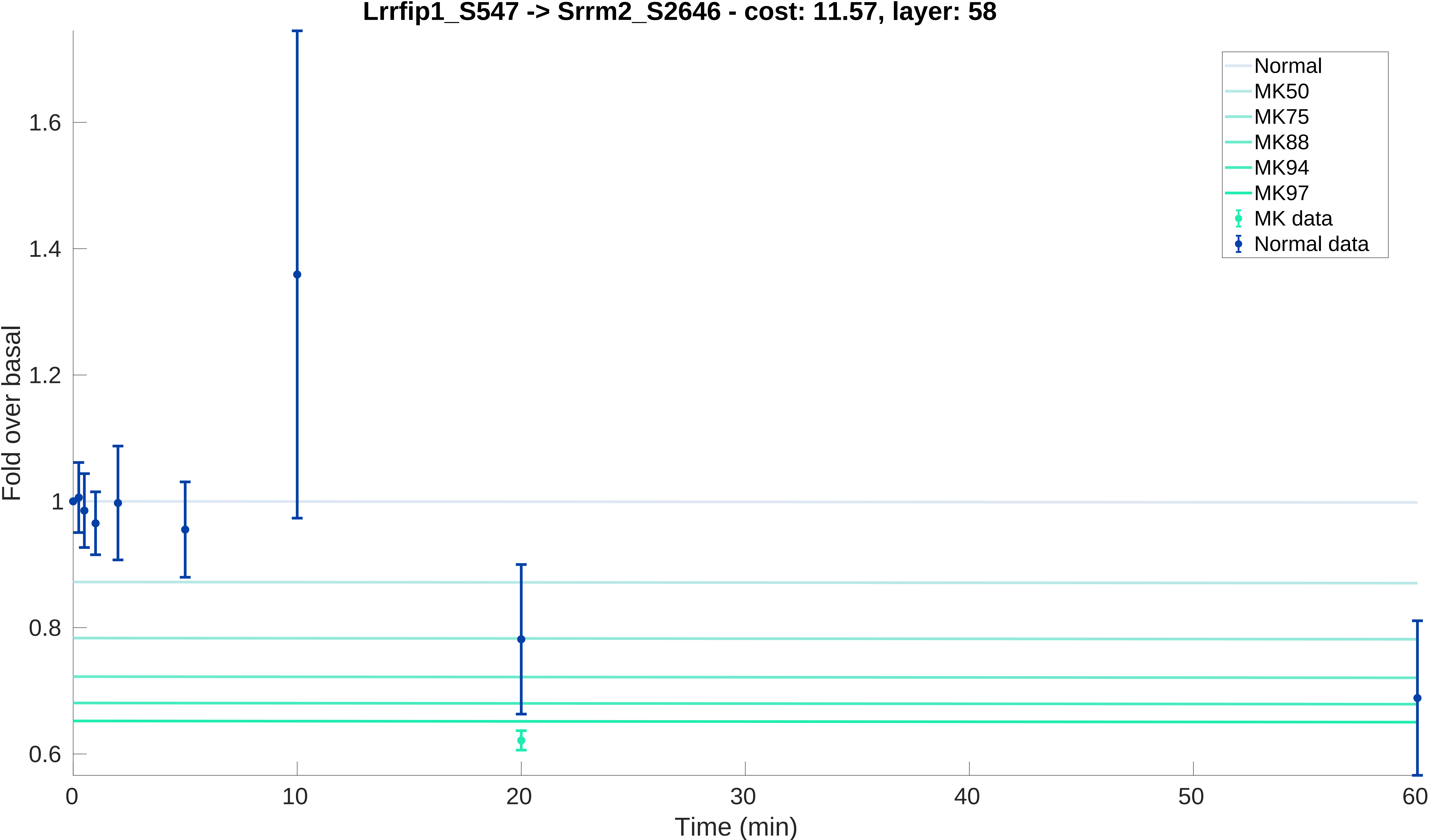

Lrrfip1_S547 -> Srrm2_S2646 - cost: 11.57, layer: 58
Fold over basal
Time (min)
0.6
0.8
1
1.2
1.4
1.6
0
10
20
30
40
50
60
Normal
MK50
MK75
MK88
MK94
MK97
MK data
Normal data

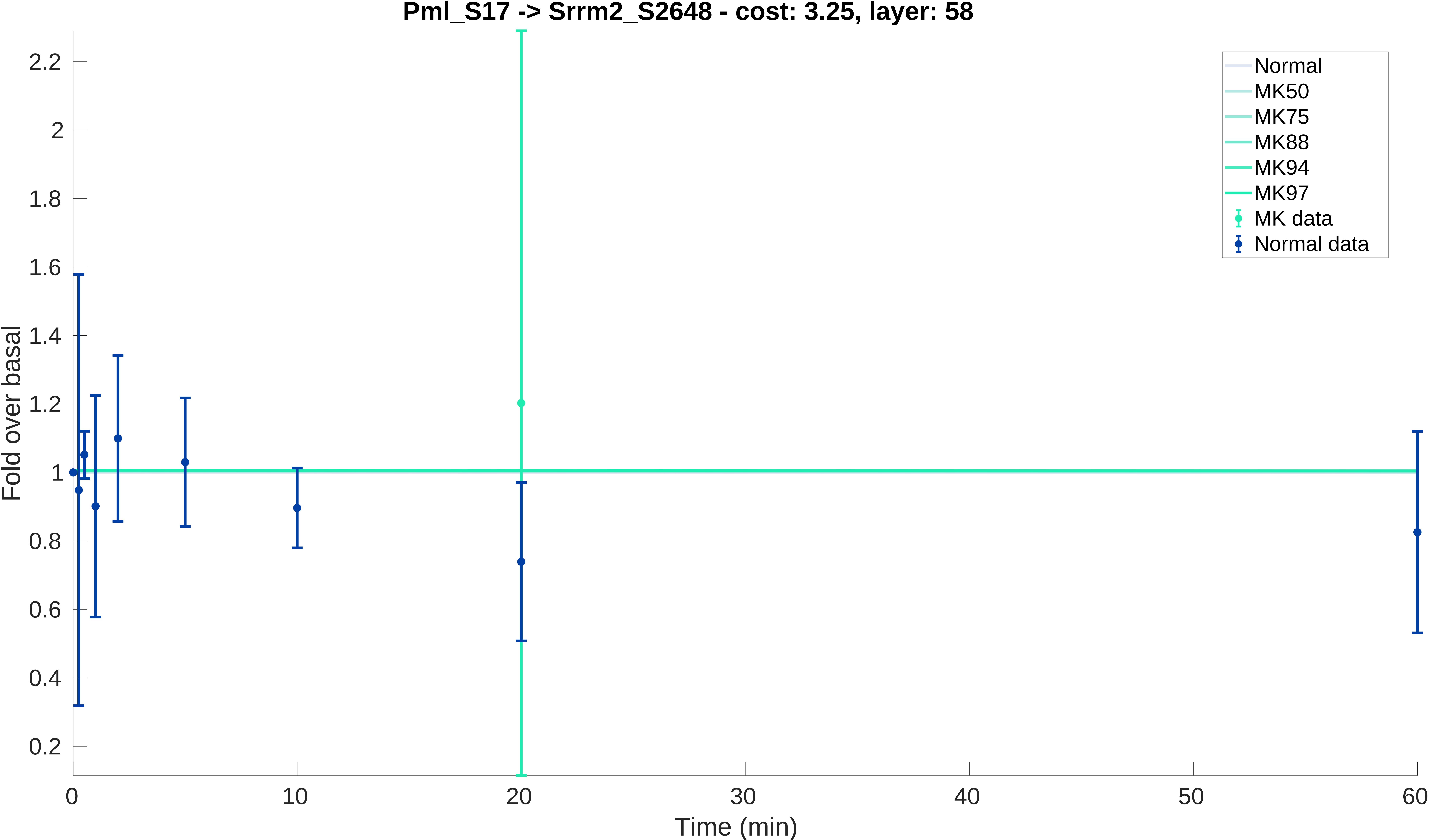
Pml_S17 -> Srrm2_S2648 - cost: 3.25, layer: 58
Fold over basal
Time (min)
0.2
0.4
0.6
0.8
1
1.2
1.4
1.6
1.8
2
2.2
0
10
20
30
40
50
60
Normal
MK50
MK75
MK88
MK94
MK97
MK data
Normal data

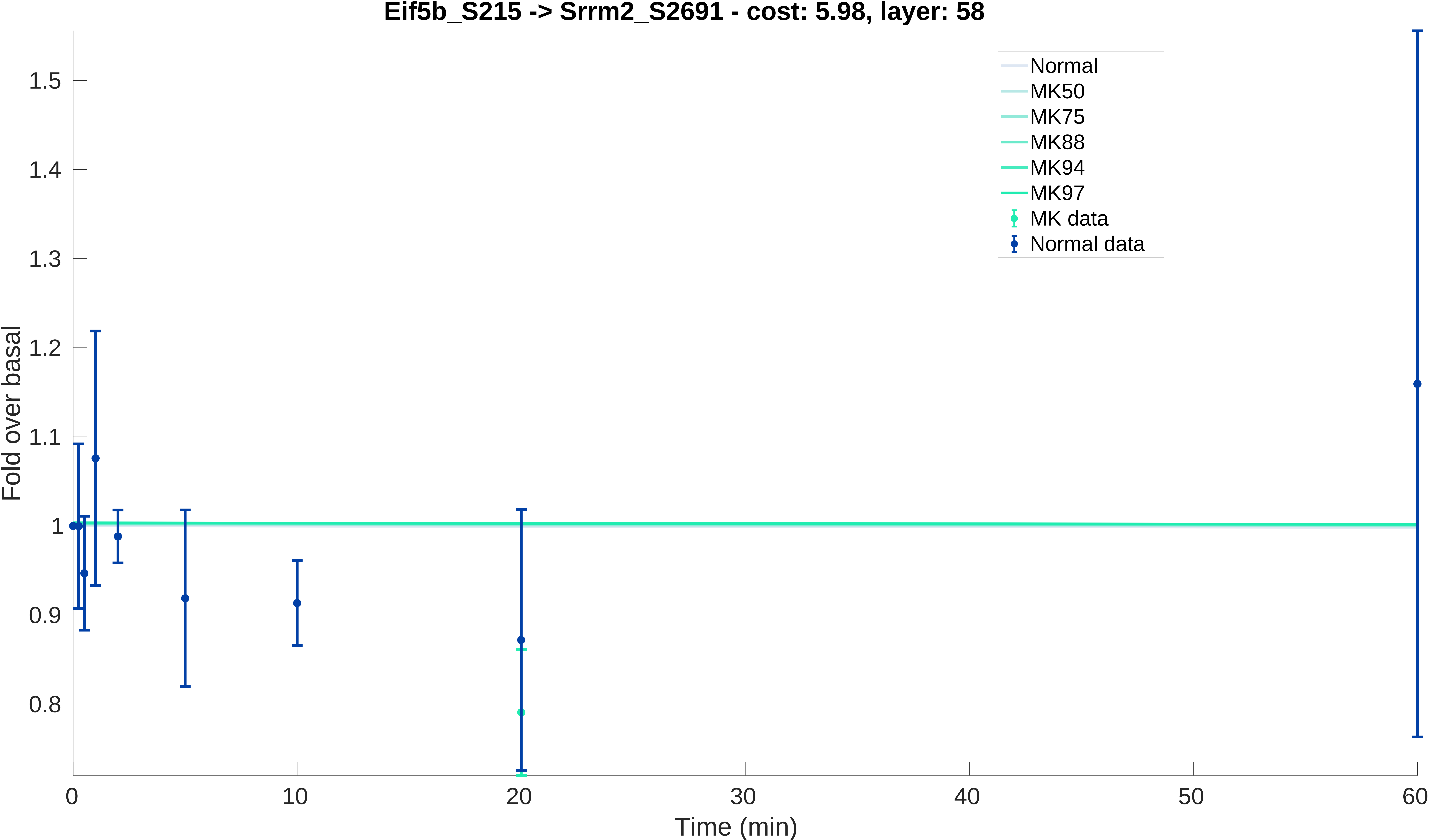
Eif5b_S215 -> Srrm2_S2691 - cost: 5.98, layer: 58
Fold over basal
1.5
1.4
1.3
1.2
1.1
1
0.9
0.8
0
10
20
30
40
50
60
Time (min)
Normal
MK50
MK75
MK88
MK94
MK97
MK data
Normal data

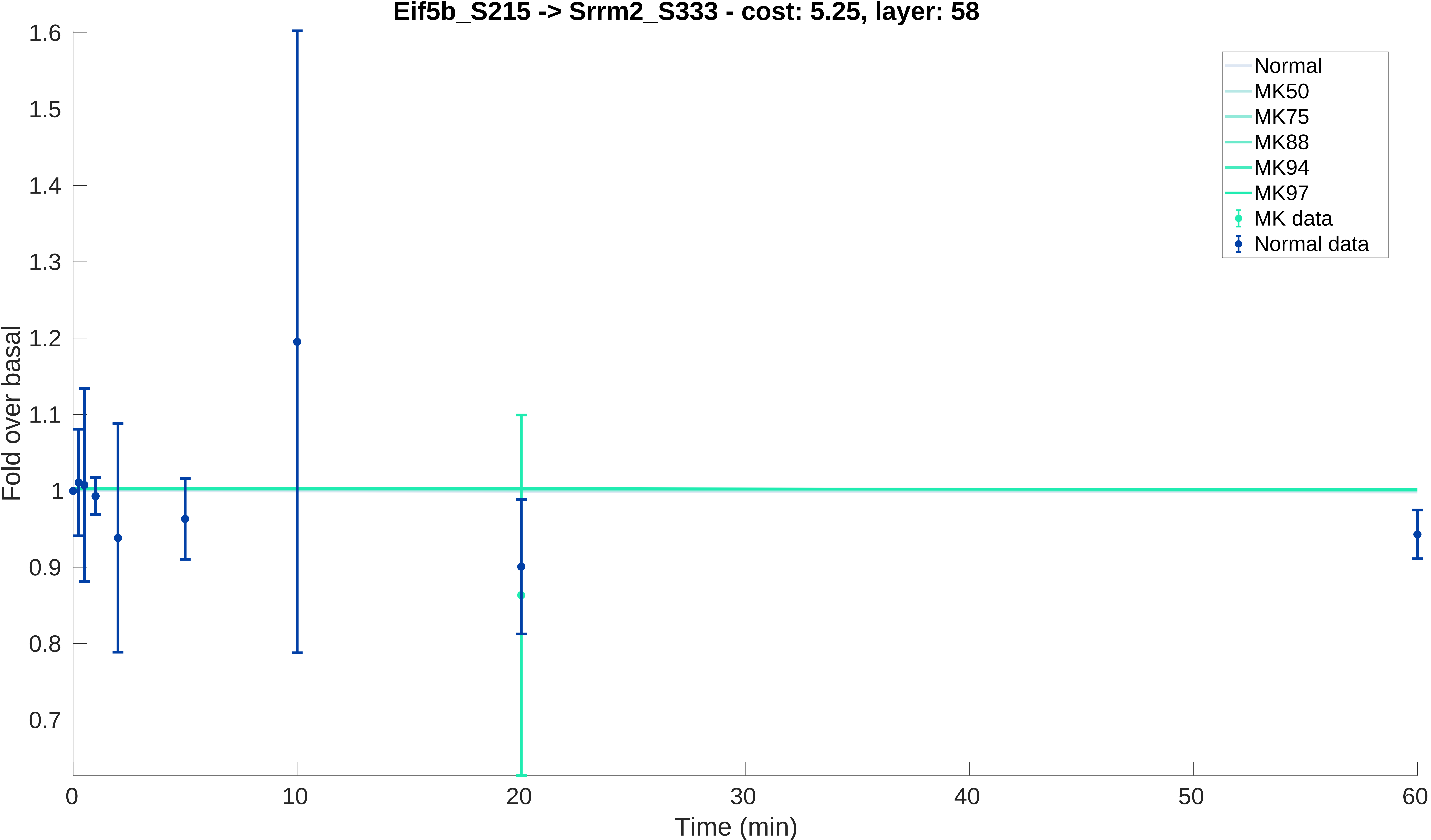
Eif5b_S215 -> Srrm2_S333 - cost: 5.25, layer: 58
Fold over basal
Time (min)
1.6
1.5
1.4
1.3
1.2
1.1
1
0.9
0.8
0.7
0
10
20
30
40
50
60
Normal
MK50
MK75
MK88
MK94
MK97
MK data
Normal data

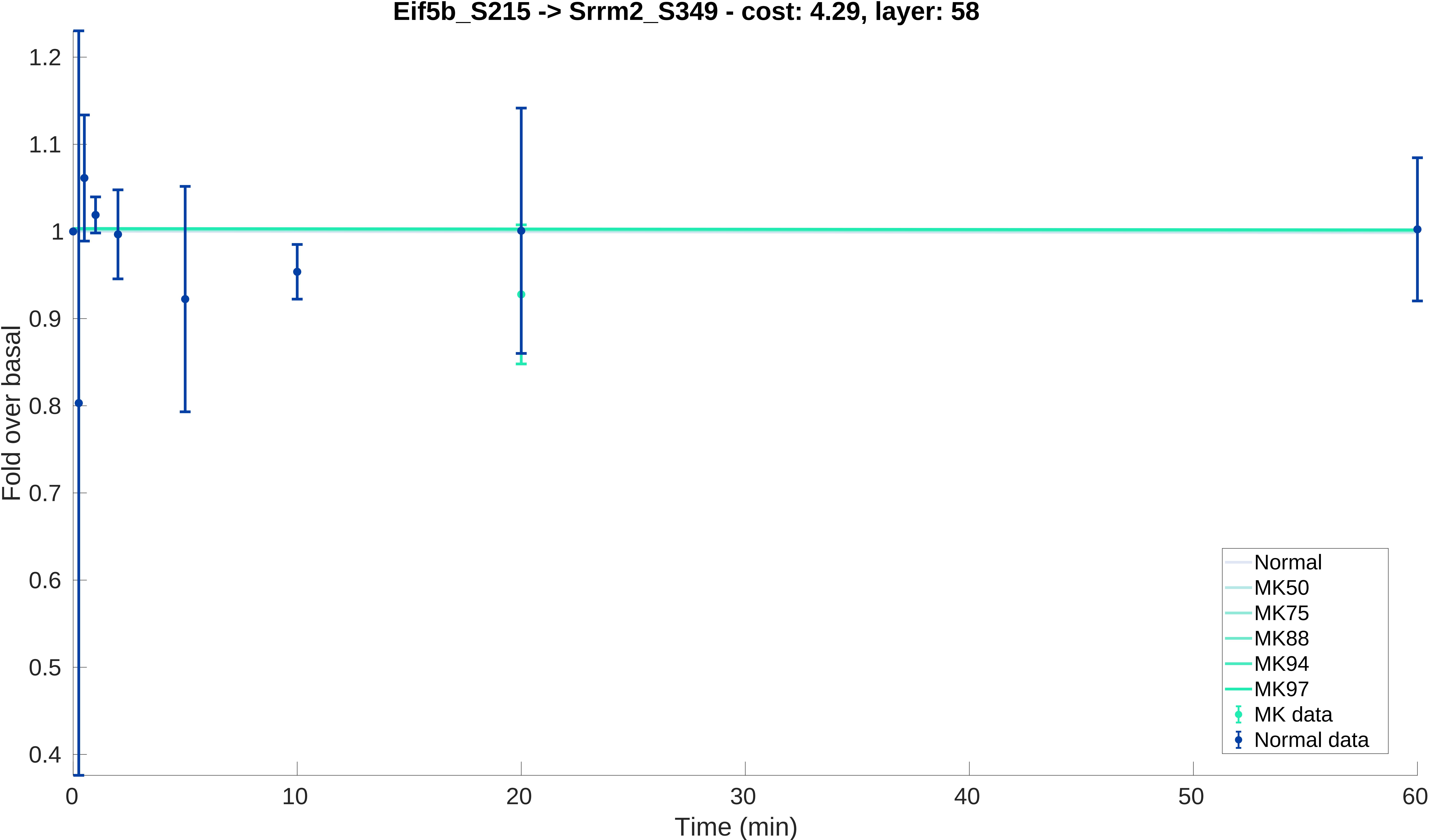

Eif5b_S215 -> Srrm2_S349 - cost: 4.29, layer: 58
Fold over basal
Time (min)
1.2
1.1
1
0.9
0.8
0.7
0.6
0.5
0.4
0
10
20
30
40
50
60
Normal
MK50
MK75
MK88
MK94
MK97
MK data
Normal data

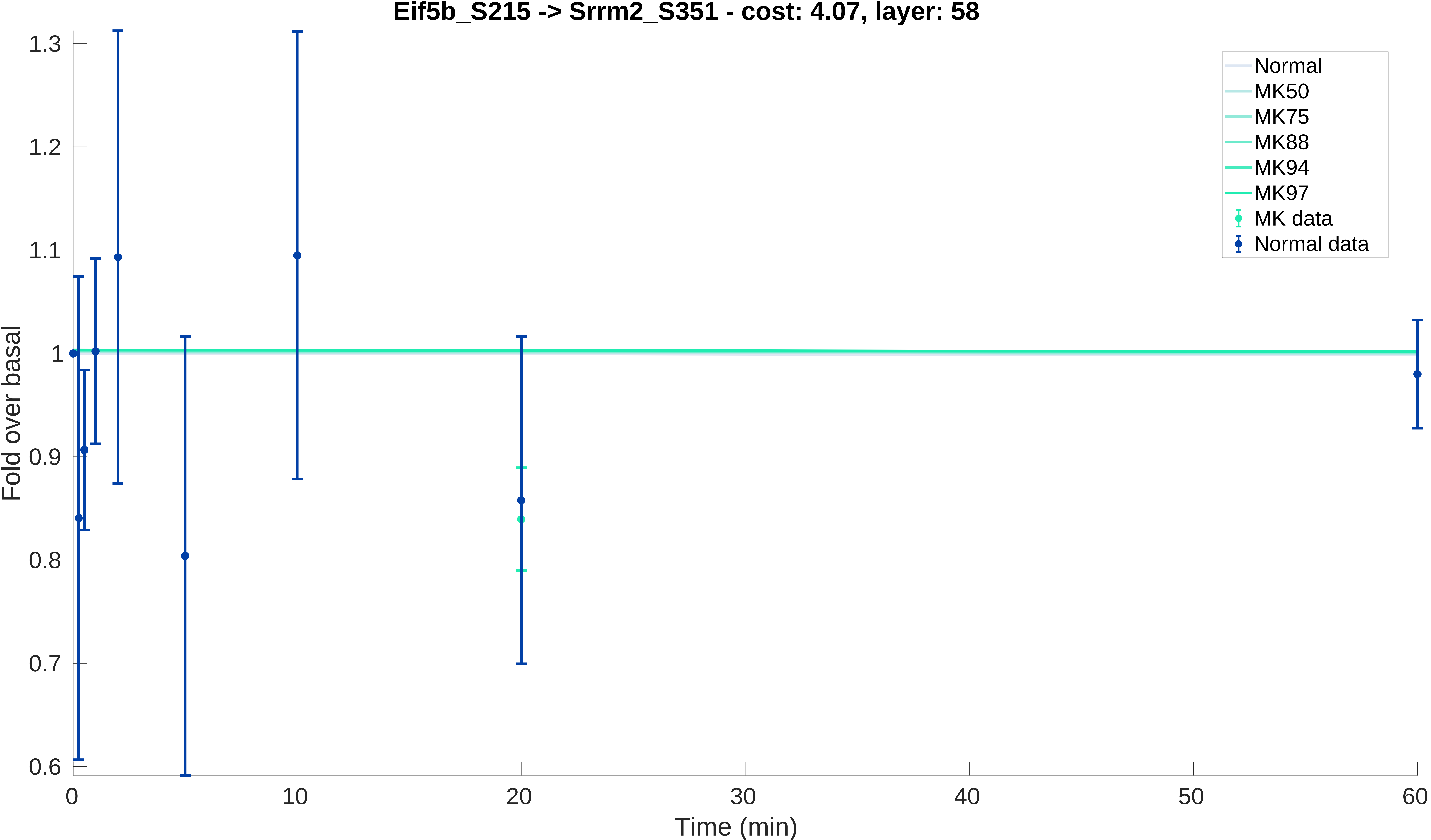
Eif5b_S215 -> Srrm2_S351 - cost: 4.07, layer: 58
Fold over basal
Time (min)
Normal
MK50
MK75
MK88
MK94
MK97
MK data
Normal data
1.3
1.2
1.1
1
0.9
0.8
0.7
0.6
0
10
20
30
40
50
60

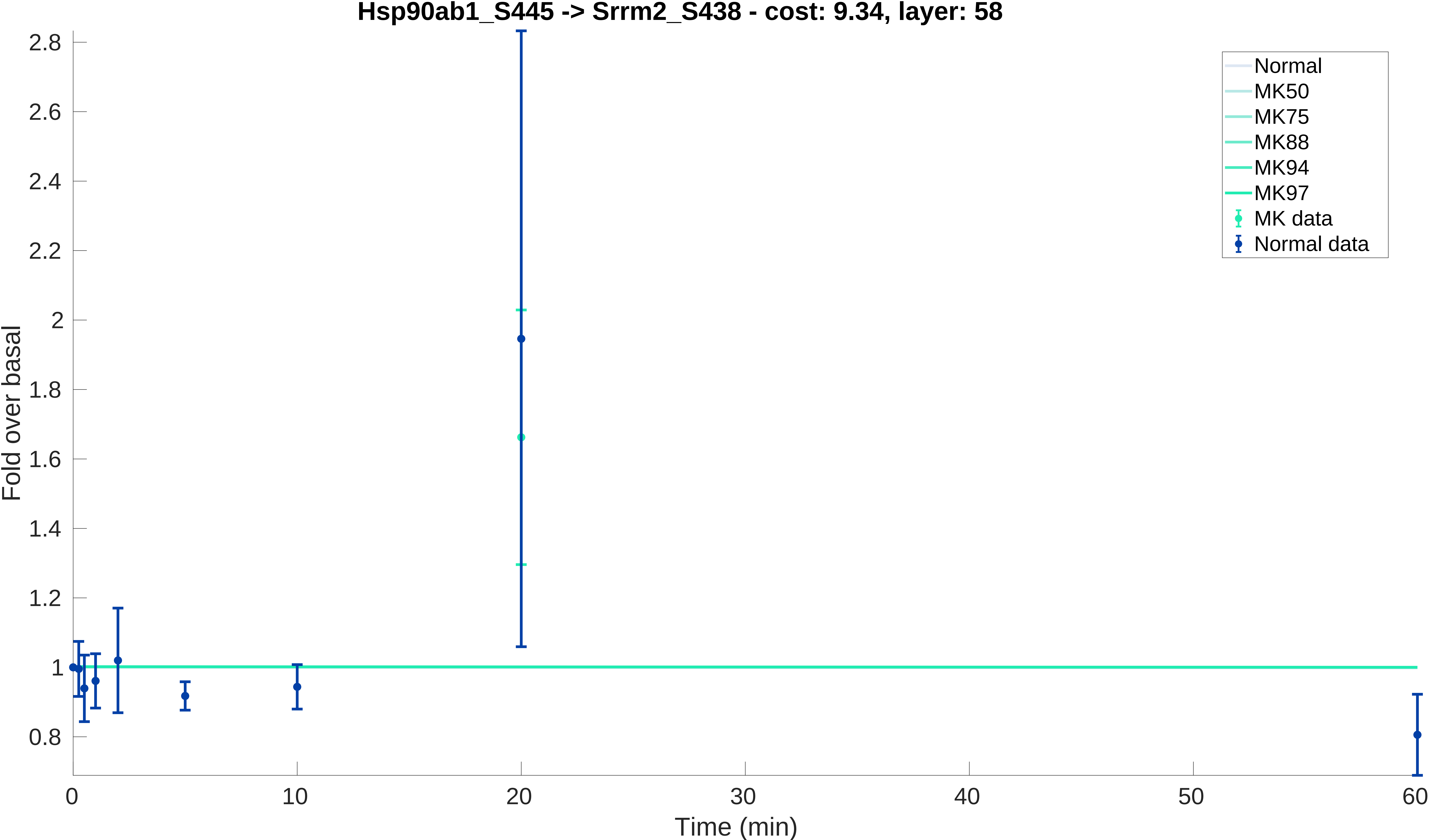

Hsp90ab1_S445 -> Srrm2_S438 - cost: 9.34, layer: 58
Fold over basal
Time (min)
0.8
1
1.2
1.4
1.6
1.8
2
2.2
2.4
2.6
2.8
0
10
20
30
40
50
60
Normal
MK50
MK75
MK88
MK94
MK97
MK data
Normal data

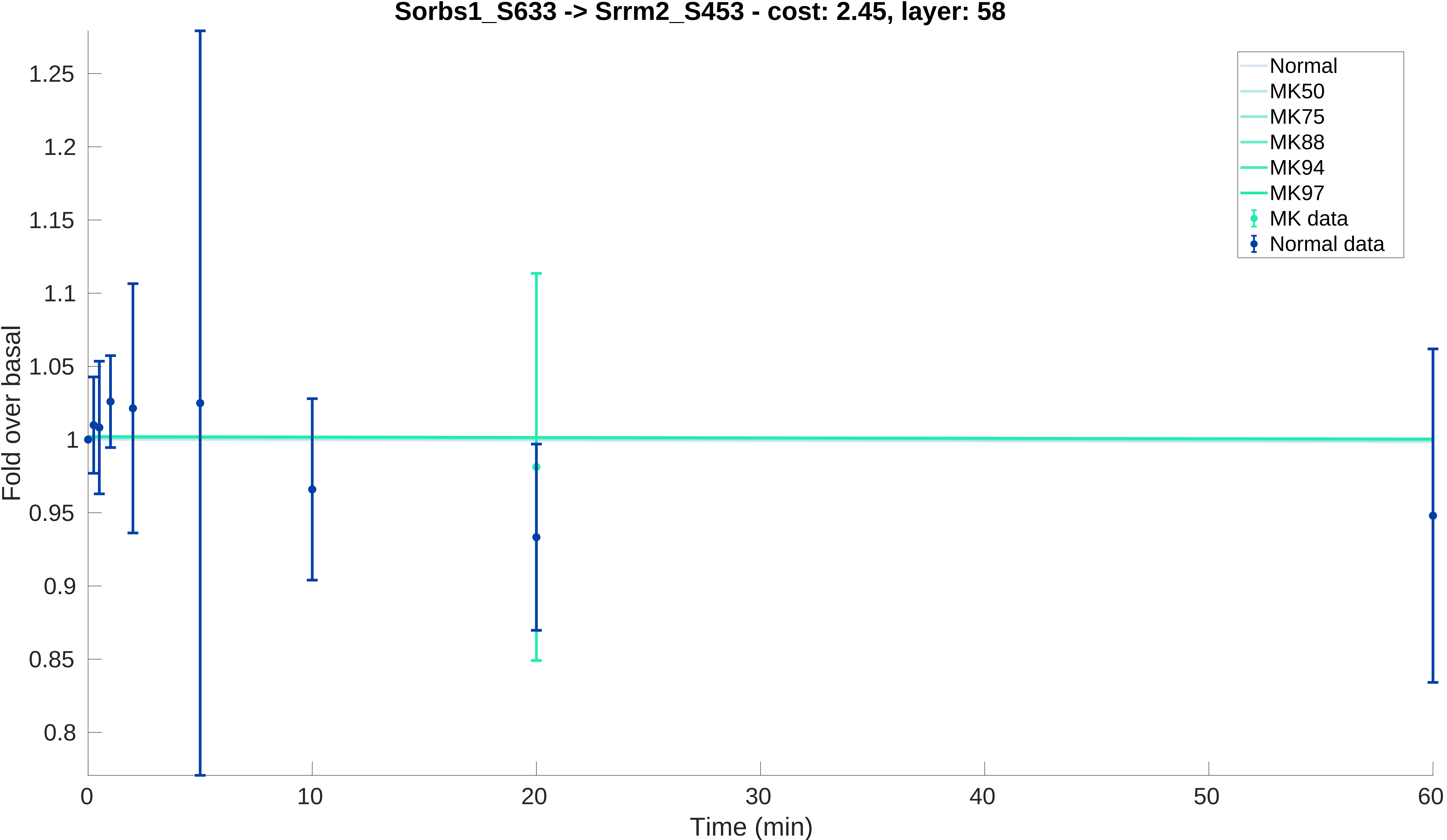

Sorbs1_S633 -> Srrm2_S453 - cost: 2.45, layer: 58
Normal
MK50
MK75
MK88
MK94
MK97
MK data
Normal data
Fold over basal
1.25
1.2
1.15
1.1
1.05
1
0.95
0.9
0.85
0.8
0
10
20
30
40
50
60
Time (min)

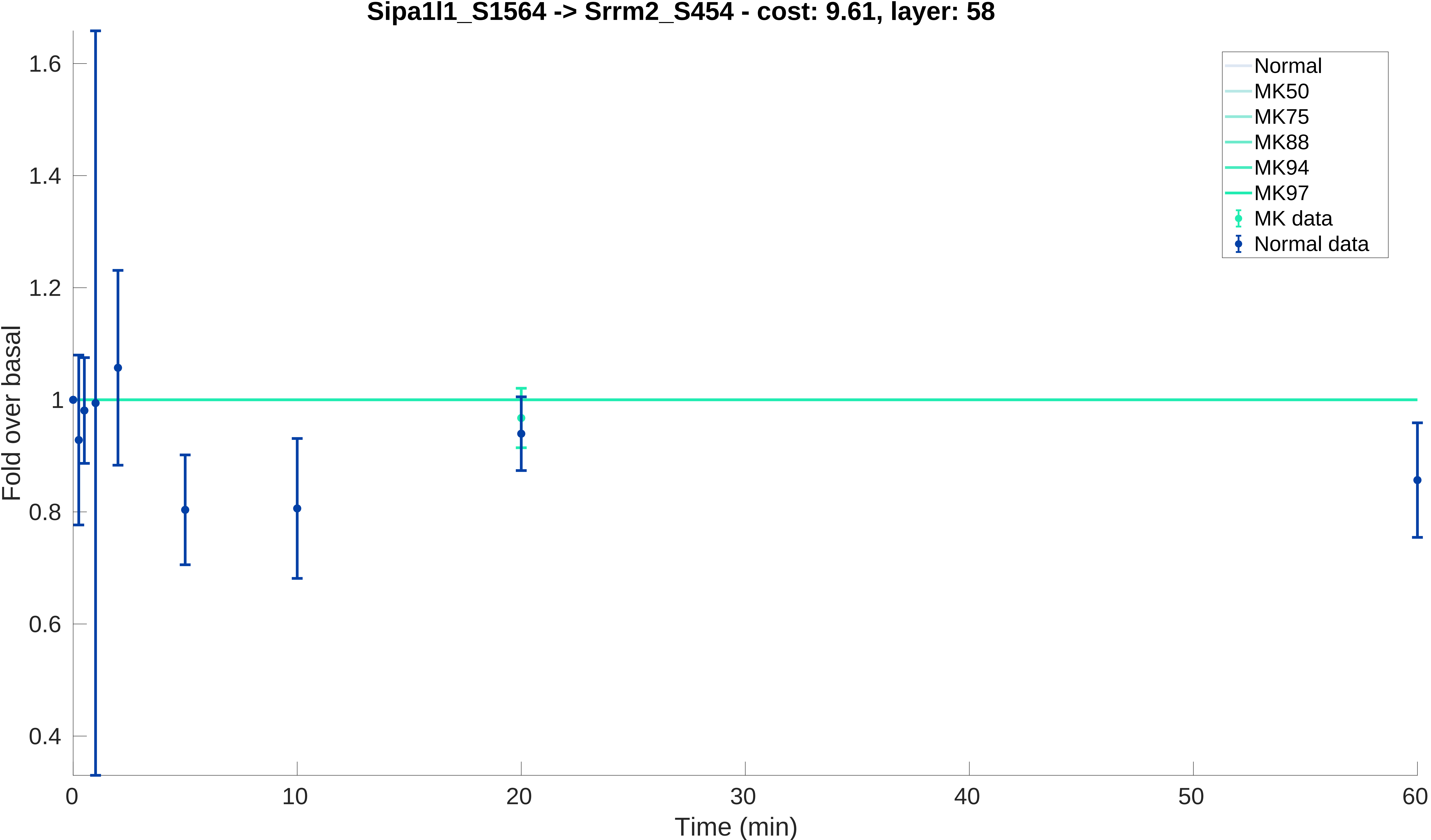
Sipa1l1_S1564 -> Srrm2_S454 - cost: 9.61, layer: 58
Fold over basal
Time (min)
1.6
1.4
1.2
1
0.8
0.6
0.4
0
10
20
30
40
50
60
Normal
MK50
MK75
MK88
MK94
MK97
MK data
Normal data

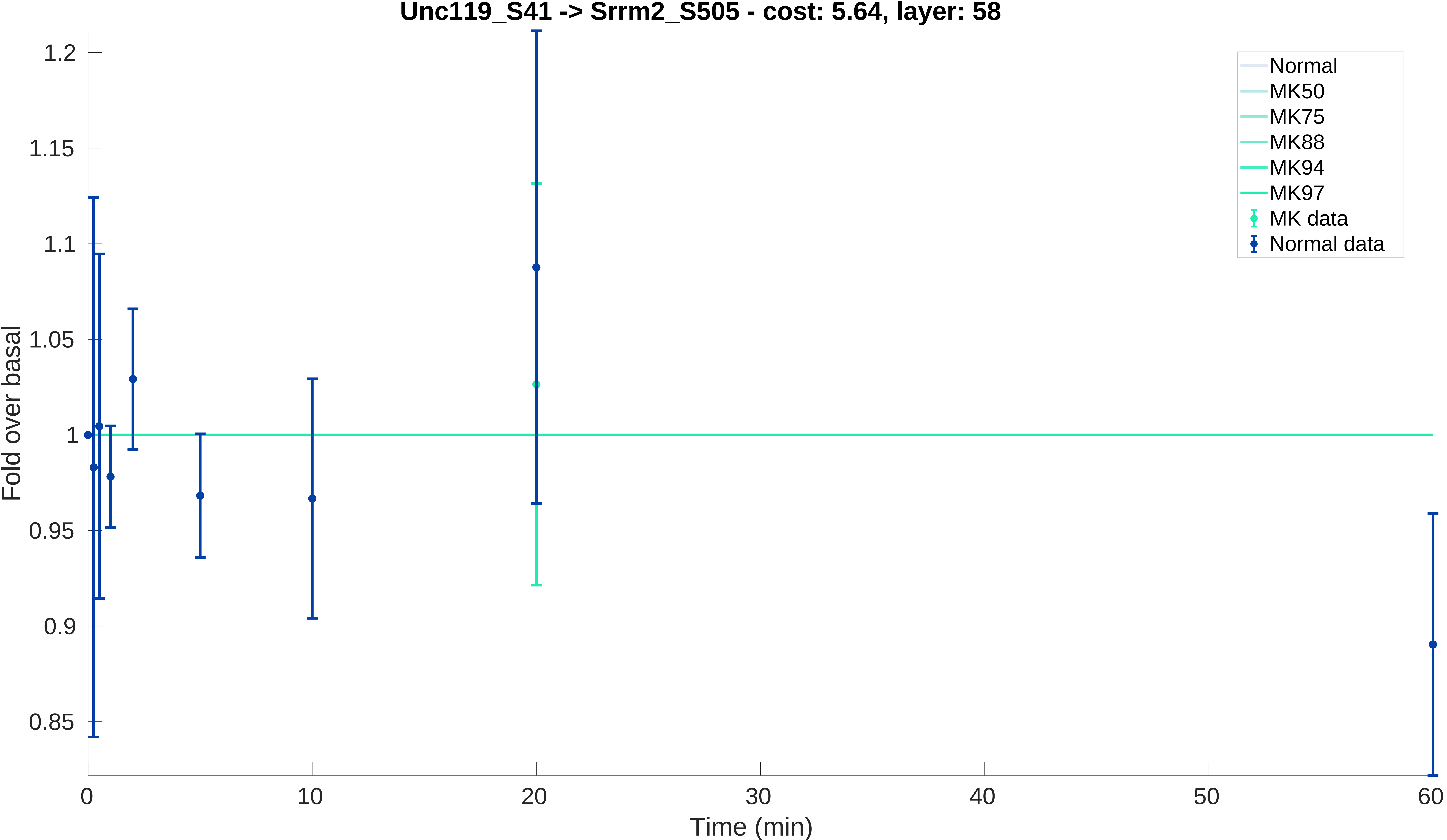

Unc119_S41 -> Srrm2_S505 - cost: 5.64, layer: 58
Fold over basal
Time (min)
1.2
1.15
1.1
1.05
1
0.95
0.9
0.85
0
10
20
30
40
50
60
Normal
MK50
MK75
MK88
MK94
MK97
MK data
Normal data

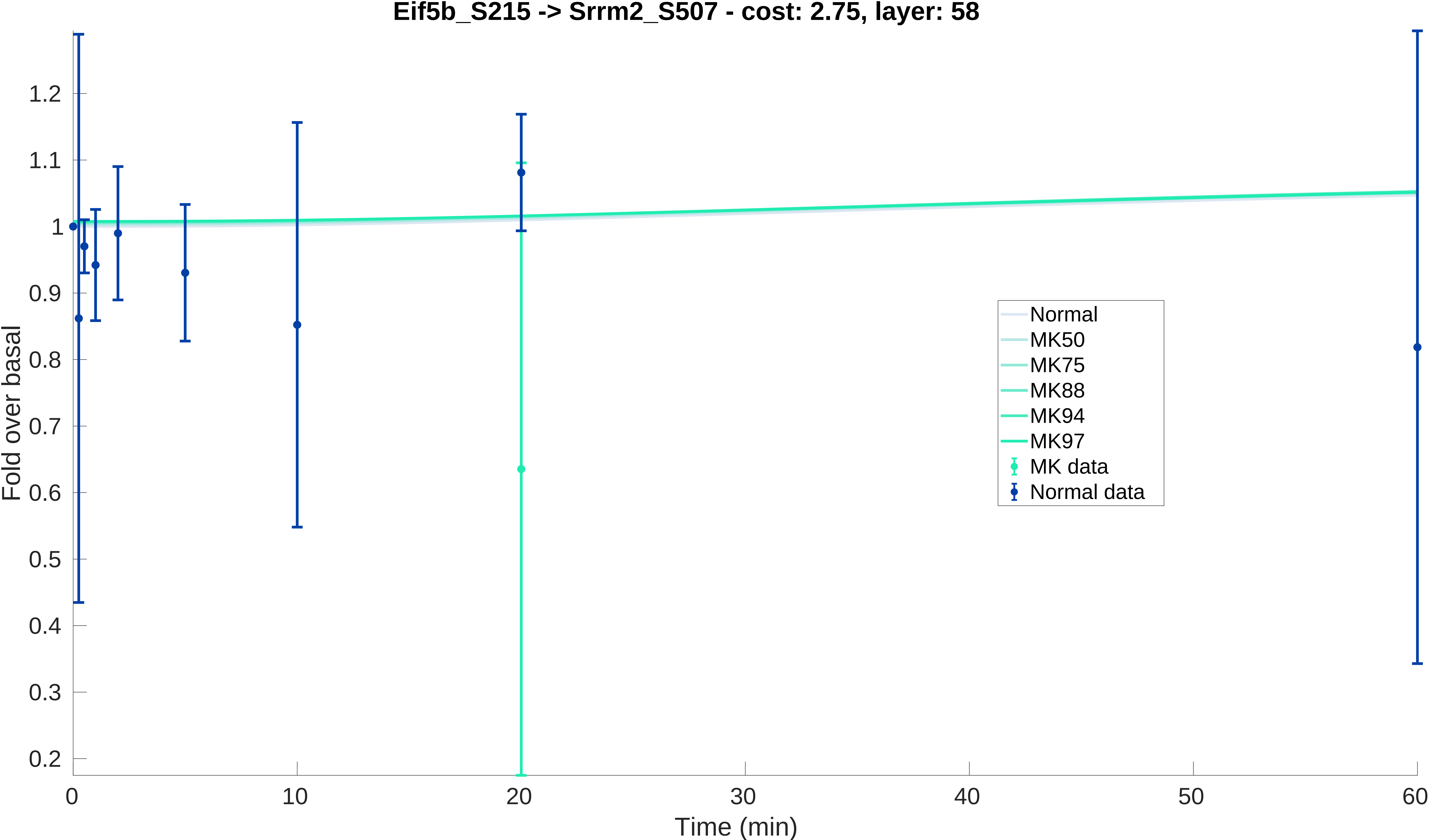

Eif5b_S215 -> Srrm2_S507 - cost: 2.75, layer: 58
Fold over basal
Time (min)
Normal
MK50
MK75
MK88
MK94
MK97
MK data
Normal data
0
10
20
30
40
50
60
0.2
0.3
0.4
0.5
0.6
0.7
0.8
0.9
1
1.1
1.2

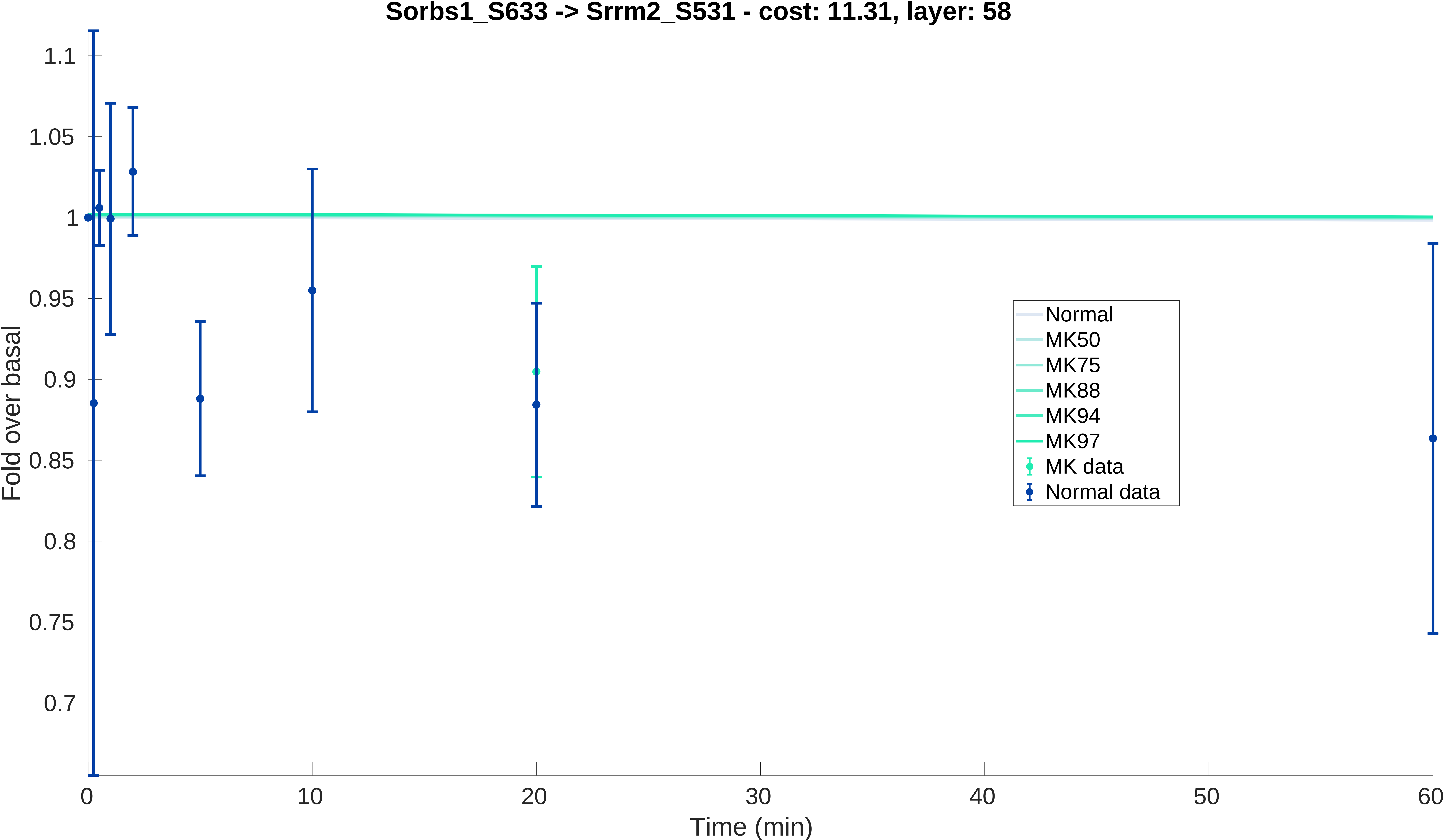

Sorbs1_S633 -> Srrm2_S531 - cost: 11.31, layer: 58
Fold over basal
Time (min)
1.1
1.05
1
0.95
0.9
0.85
0.8
0.75
0.7
0
10
20
30
40
50
60
Normal
MK50
MK75
MK88
MK94
MK97
MK data
Normal data

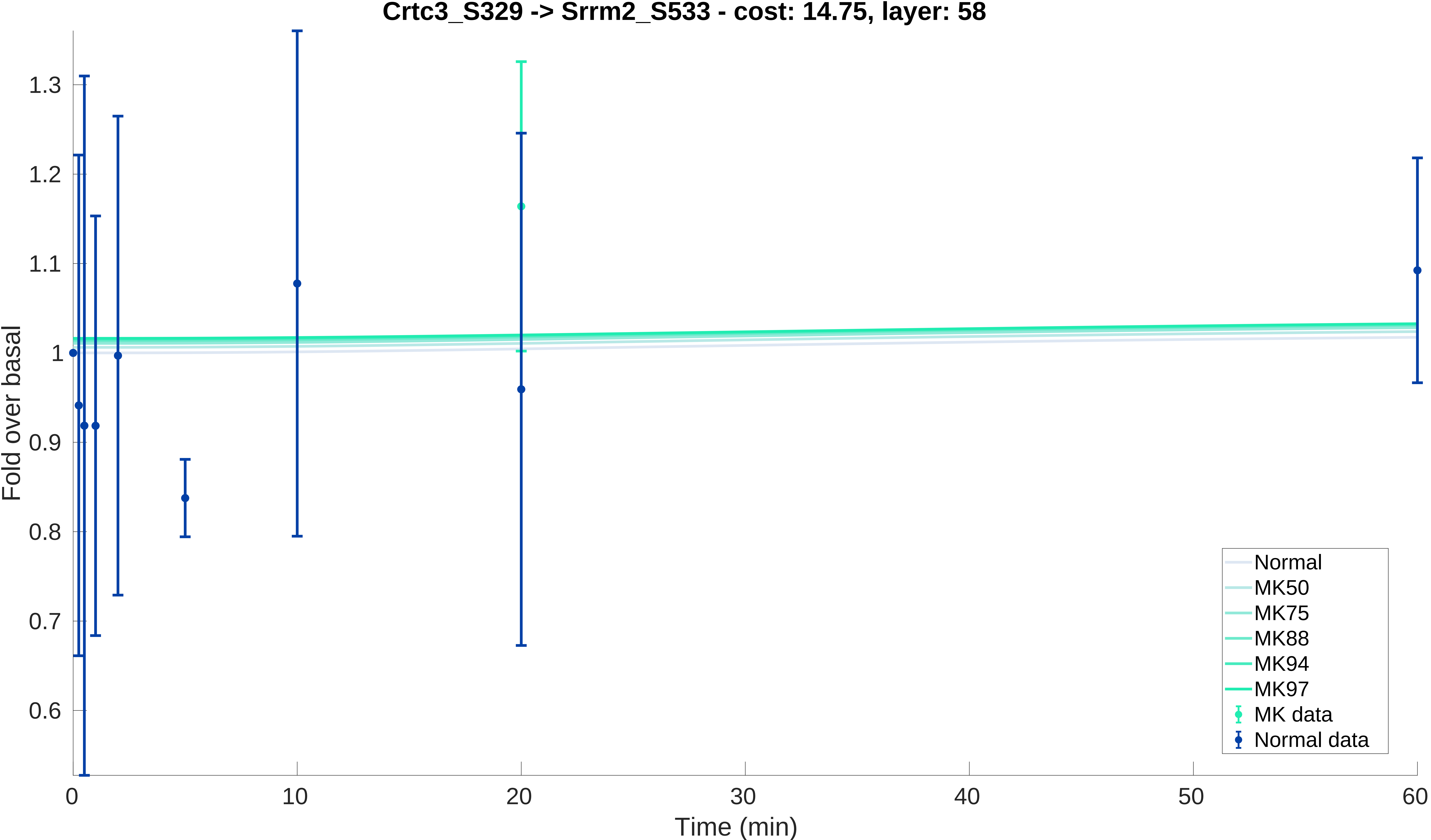

Crtc3_S329 -> Srrm2_S533 - cost: 14.75, layer: 58
Fold over basal
1.3
1.2
1.1
1
0.9
0.8
0.7
0.6
0
10
20
30
40
50
60
Time (min)
Normal
MK50
MK75
MK88
MK94
MK97
MK data
Normal data

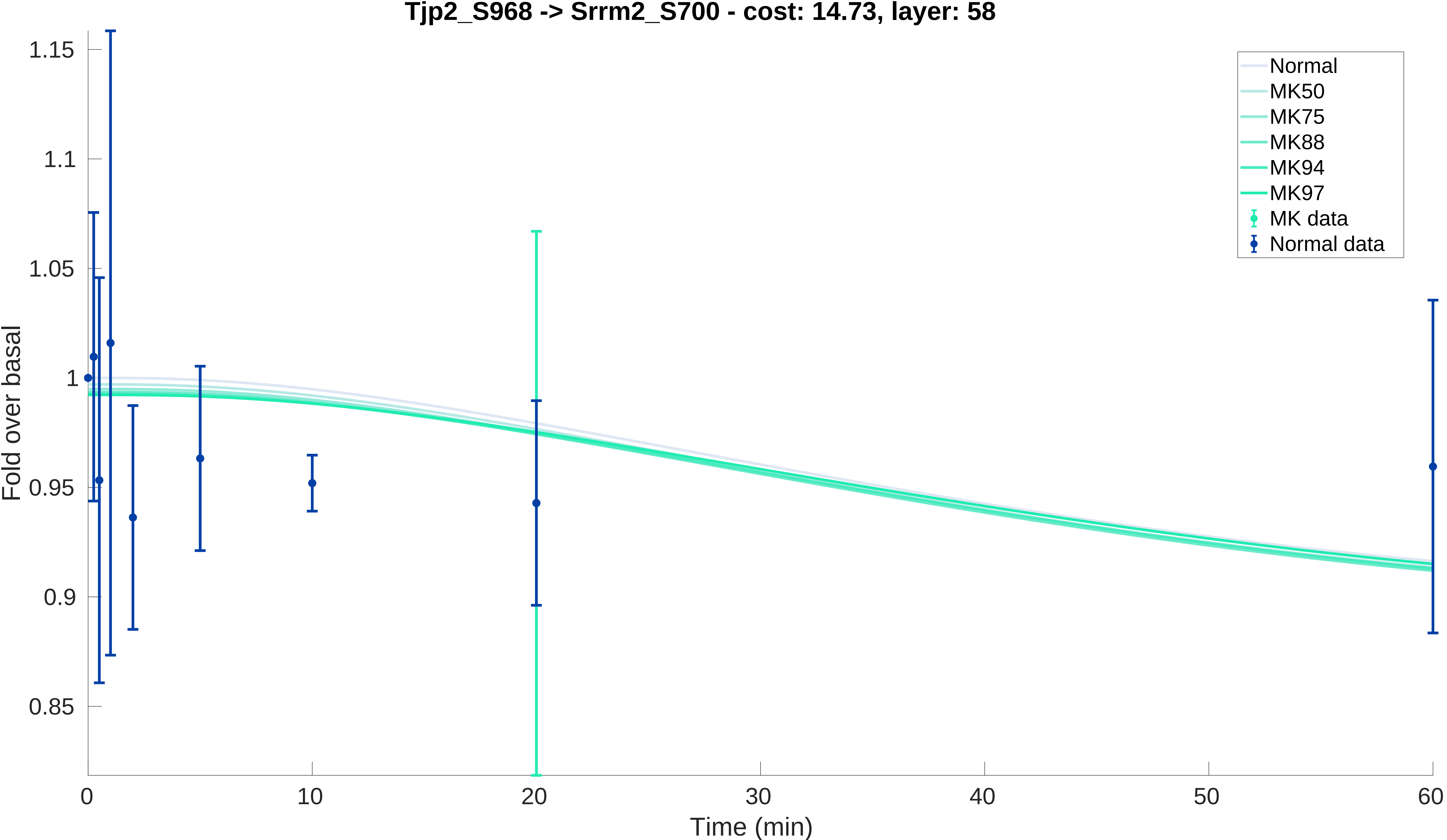

Tjp2_S968 -> Srrm2_S700 - cost: 14.73, layer: 58
Fold over basal
1.15
1.1
1.05
1
0.95
0.9
0.85
0
10
20
30
40
50
60
Time (min)
Normal
MK50
MK75
MK88
MK94
MK97
MK data
Normal data

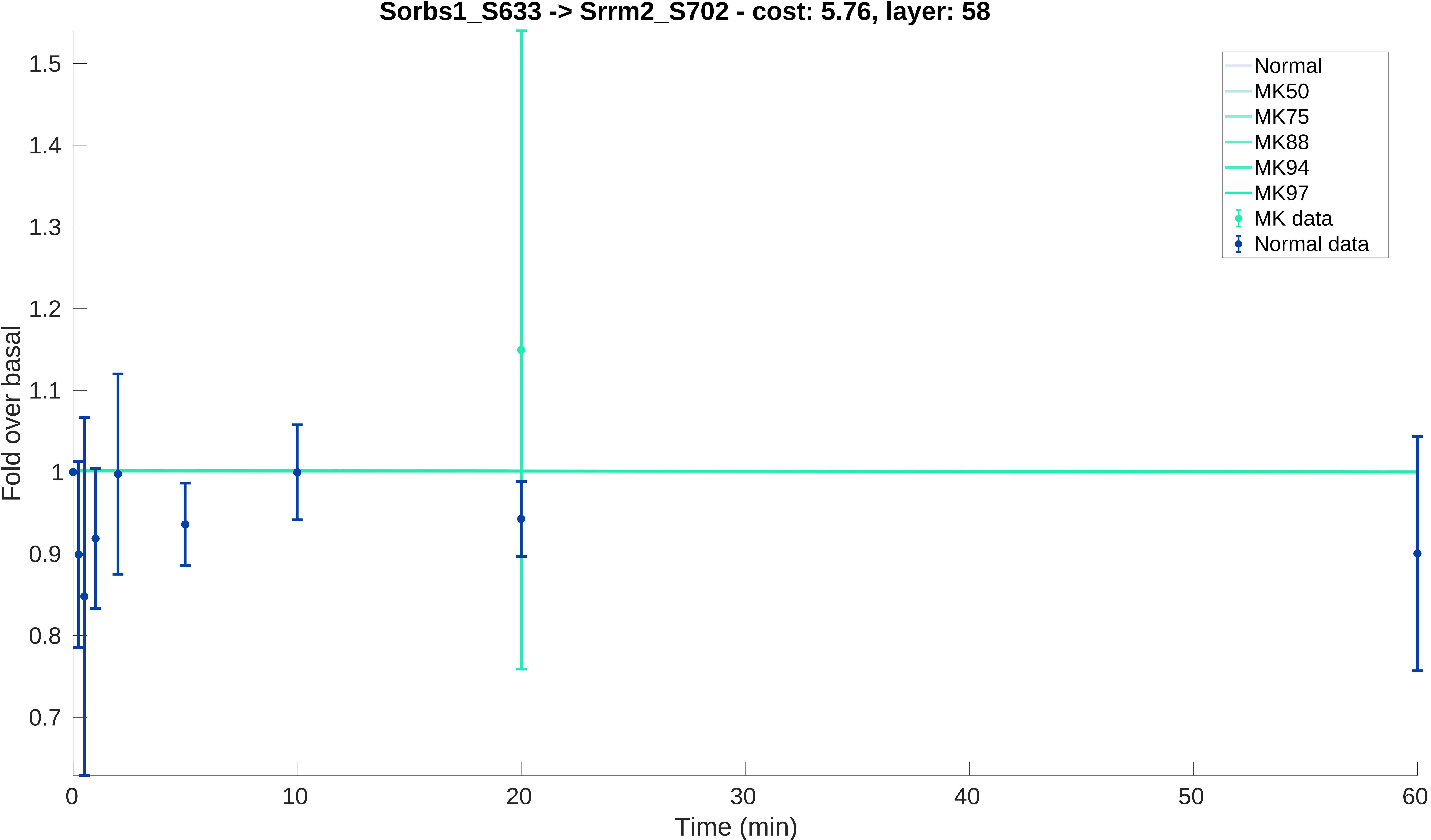

Sorbs1_S633 -> Srrm2_S702 - cost: 5.76, layer: 58
Fold over basal
Time (min)
1.5
1.4
1.3
1.2
1.1
1
0.9
0.8
0.7
0
10
20
30
40
50
60
Normal
MK50
MK75
MK88
MK94
MK97
MK data
Normal data

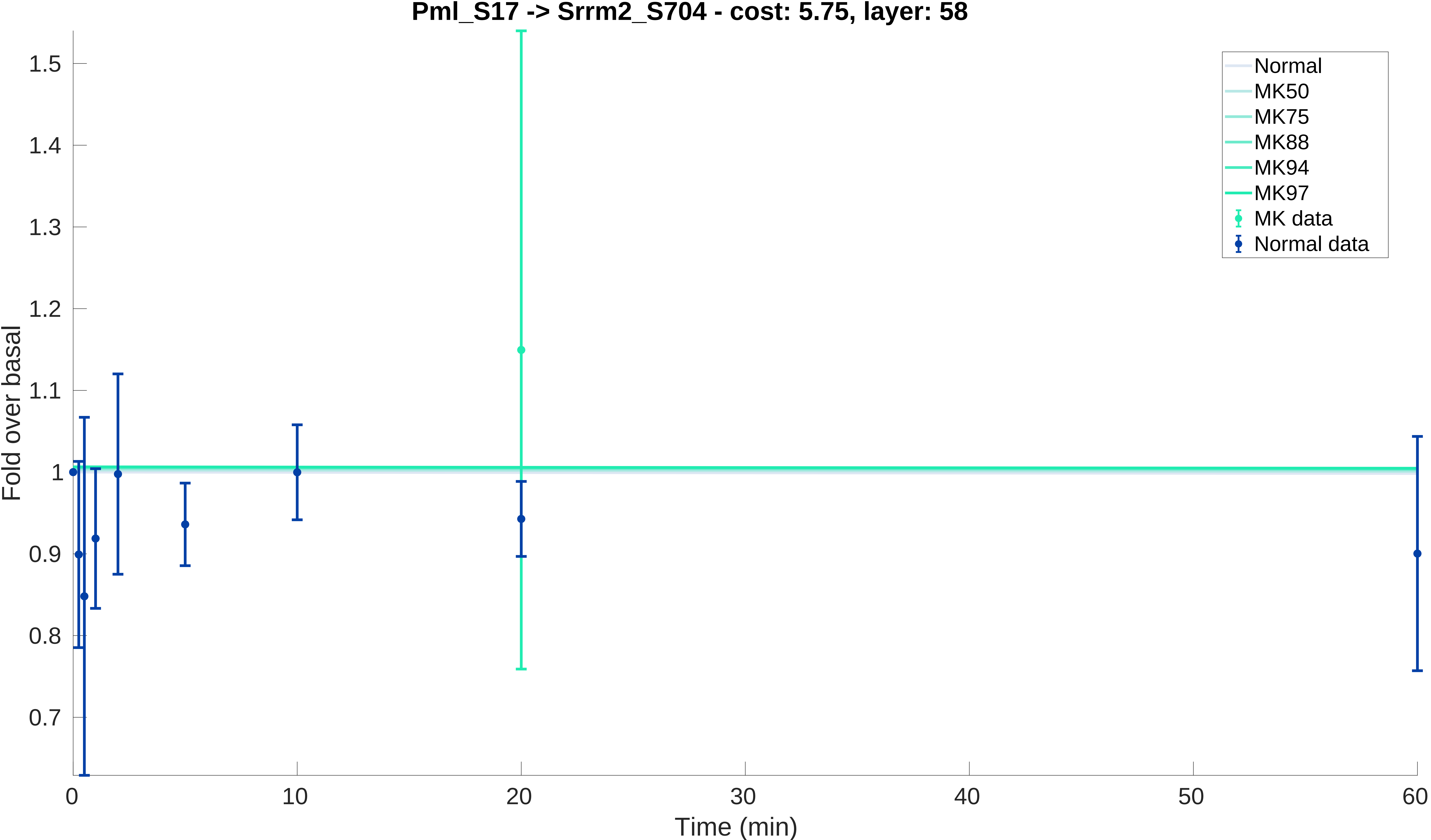

Pml_S17 -> Srrm2_S704 - cost: 5.75, layer: 58
Fold over basal
Time (min)
0
10
20
30
40
50
60
0.7
0.8
0.9
1
1.1
1.2
1.3
1.4
1.5
Normal
MK50
MK75
MK88
MK94
MK97
MK data
Normal data

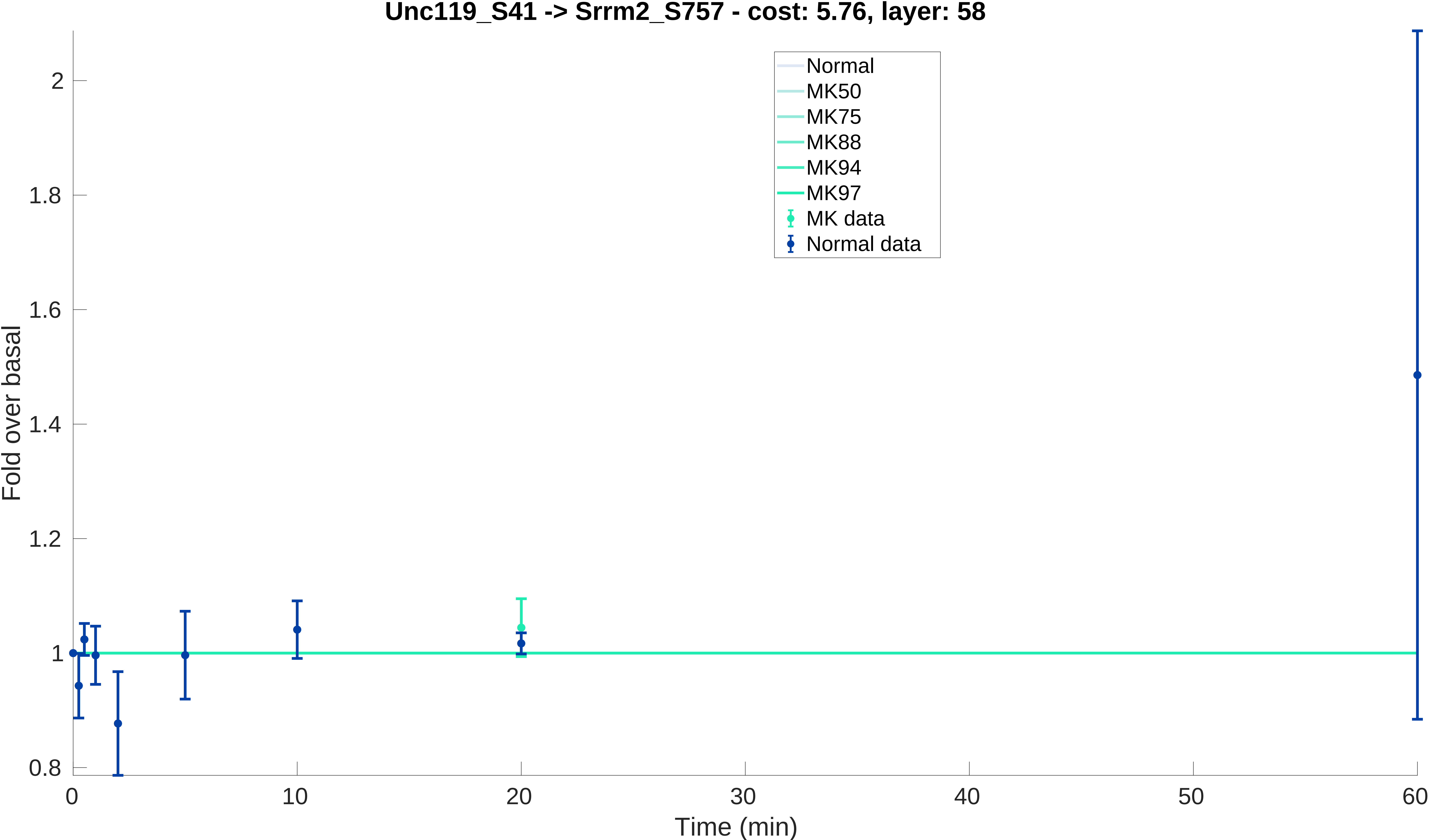

Unc119_S41 -> Srrm2_S757 - cost: 5.76, layer: 58
Normal
MK50
MK75
MK88
MK94
MK97
MK data
Normal data
Fold over basal
Time (min)
0.8
1
1.2
1.4
1.6
1.8
2
0
10
20
30
40
50
60

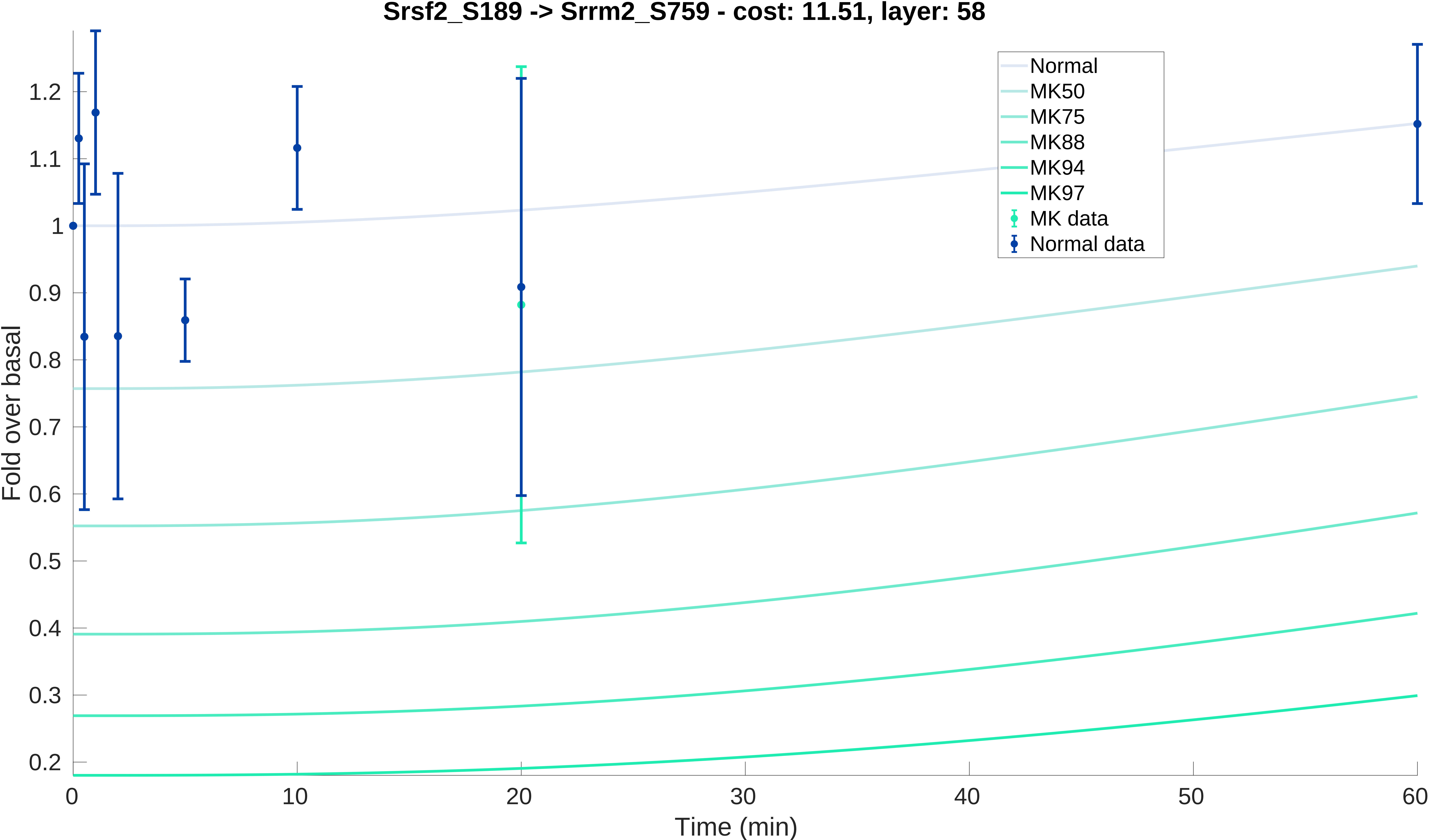

Srsf2_S189 -> Srrm2_S759 - cost: 11.51, layer: 58
Normal
MK50
MK75
MK88
MK94
MK97
MK data
Normal data
Fold over basal
1.2
1.1
1
0.9
0.8
0.7
0.6
0.5
0.4
0.3
0.2
0
10
20
30
40
50
60
Time (min)

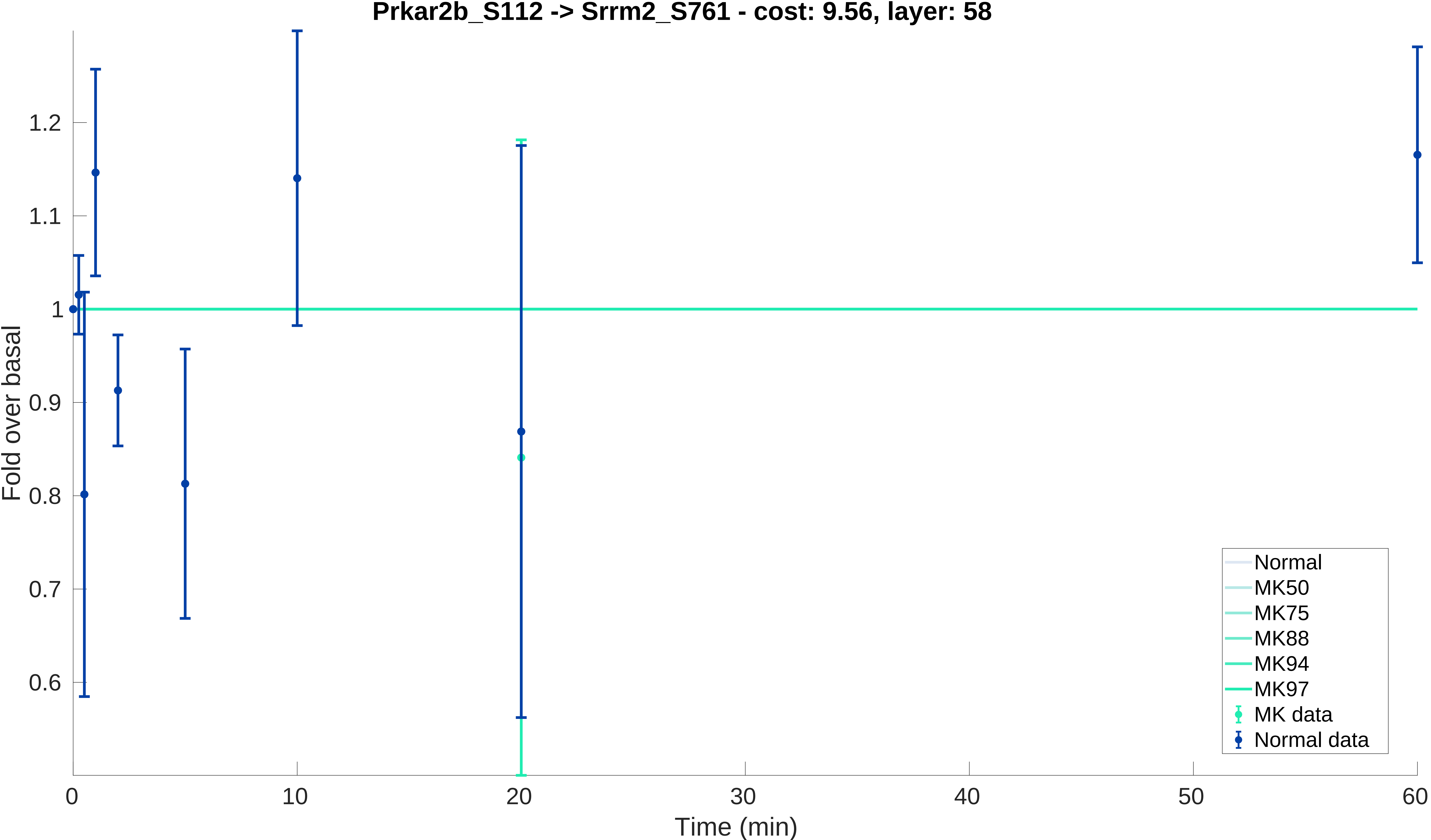

Prkar2b_S112 -> Srrm2_S761 - cost: 9.56, layer: 58
Fold over basal
Time (min)
1.2
1.1
1
0.9
0.8
0.7
0.6
0
10
20
30
40
50
60
Normal
MK50
MK75
MK88
MK94
MK97
MK data
Normal data

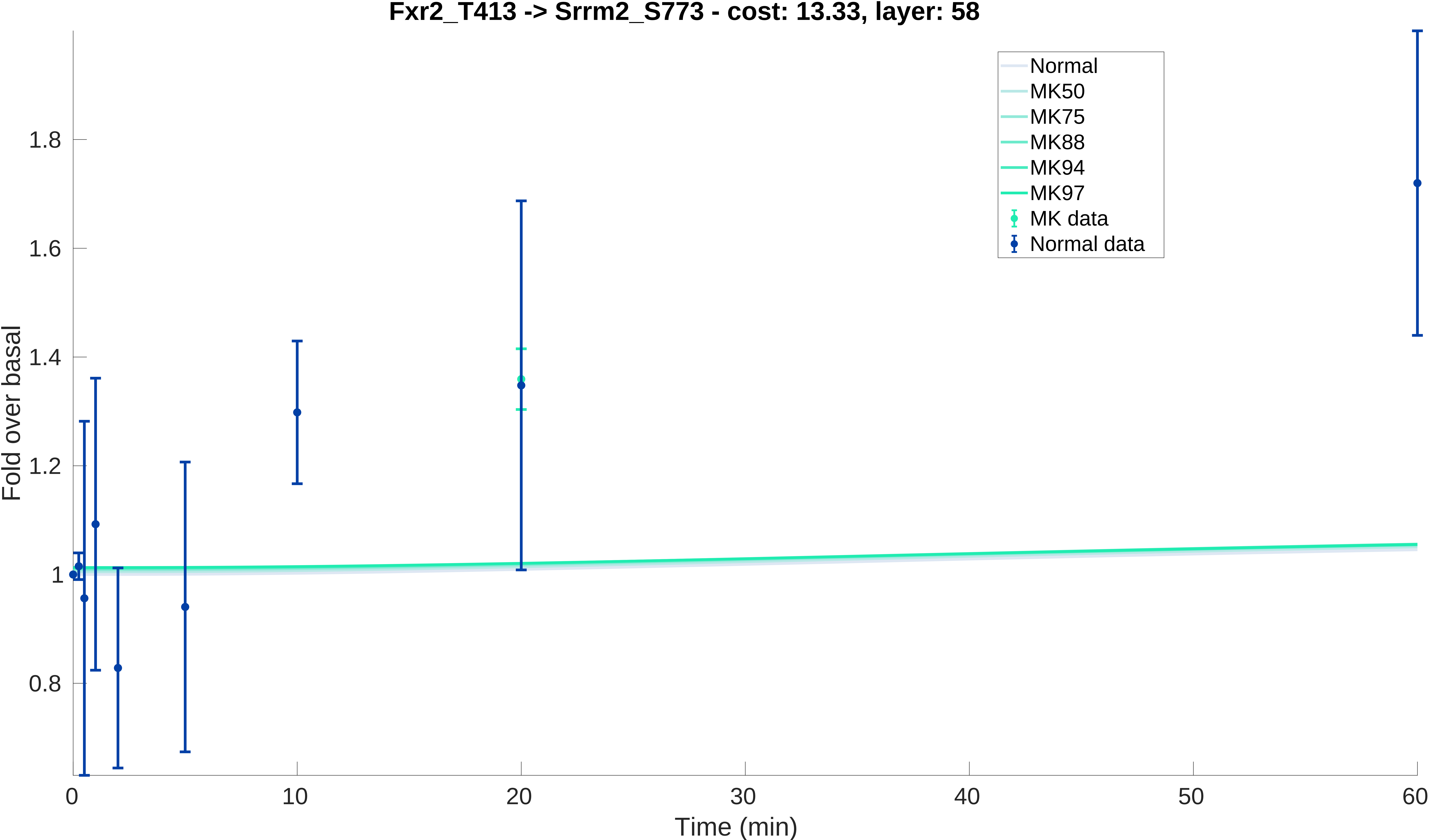

Fxr2_T413 -> Srrm2_S773 - cost: 13.33, layer: 58
Normal
MK50
MK75
MK88
MK94
MK97
MK data
Normal data
Fold over basal
1.8
1.6
1.4
1.2
1
0.8
0
10
20
30
40
50
60
Time (min)

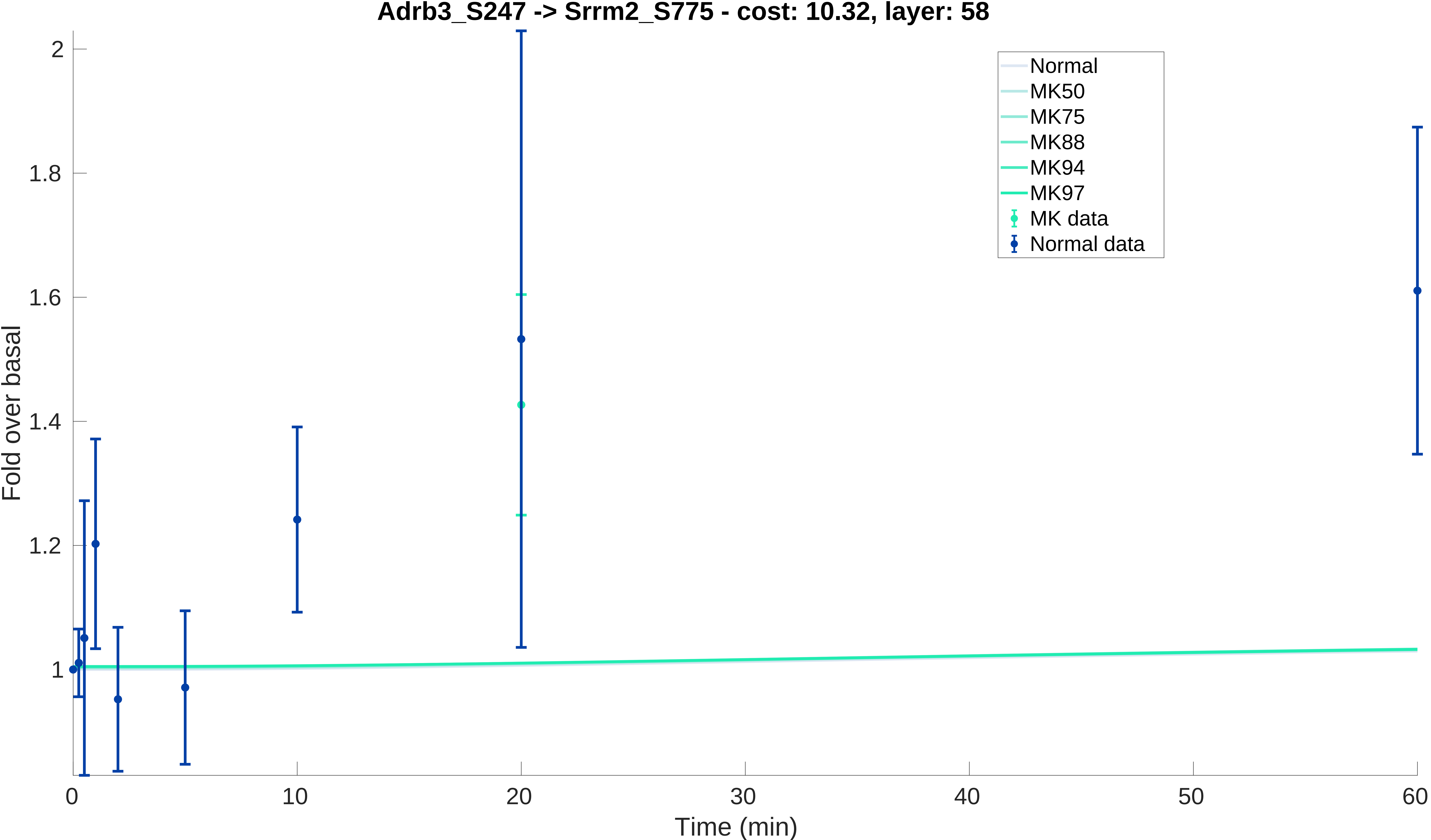
Adrb3_S247 -> Srrm2_S775 - cost: 10.32, layer: 58
Fold over basal
Time (min)
2
1.8
1.6
1.4
1.2
1
0
10
20
30
40
50
60
Normal
MK50
MK75
MK88
MK94
MK97
MK data
Normal data

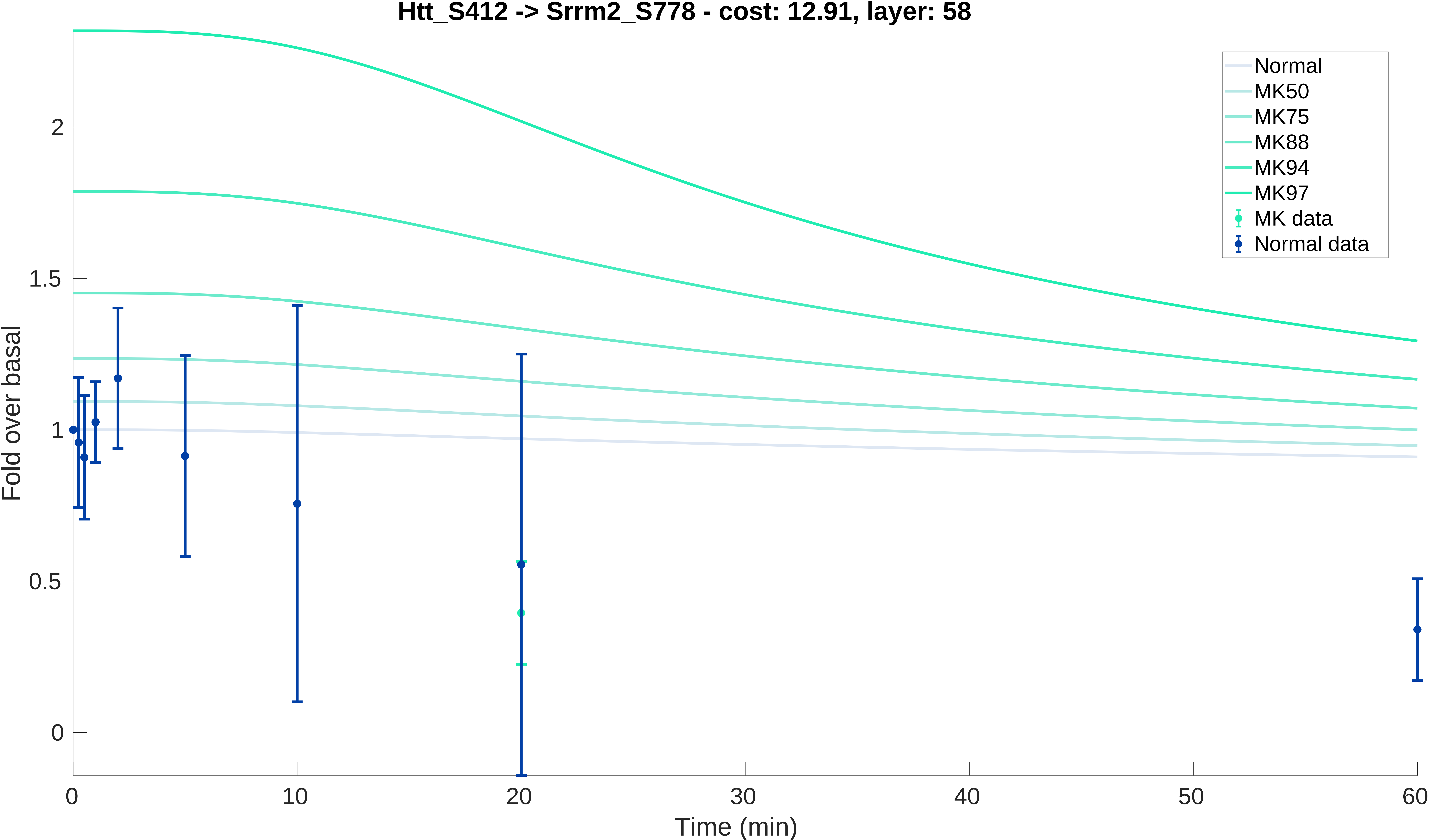
Htt_S412 -> Srrm2_S778 - cost: 12.91, layer: 58
Normal
MK50
MK75
MK88
MK94
MK97
MK data
Normal data
Fold over basal
Time (min)
0
0.5
1
1.5
2
0
10
20
30
40
50
60

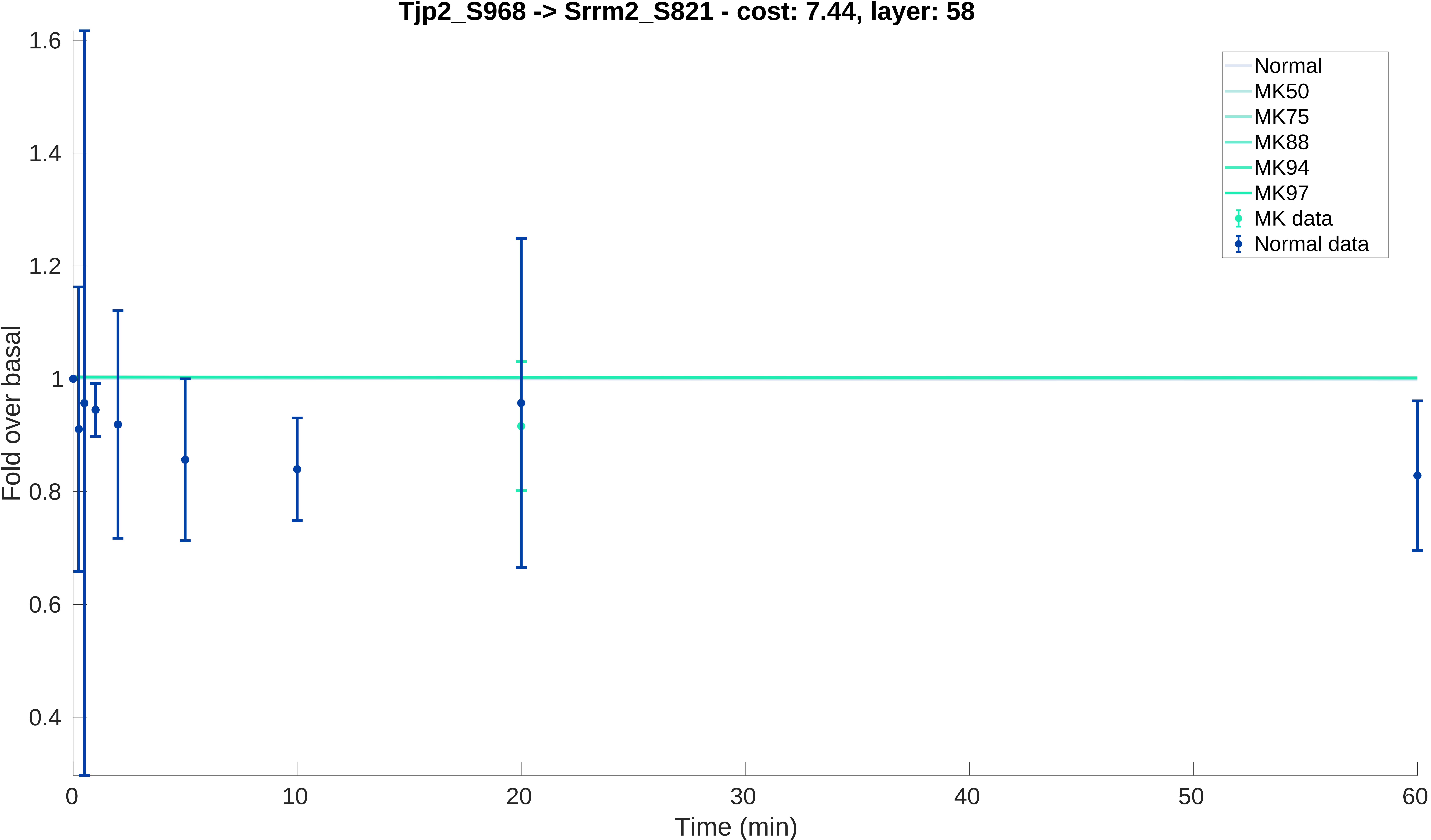

Tjp2_S968 -> Srrm2_S821 - cost: 7.44, layer: 58
Fold over basal
Time (min)
0.4
0.6
0.8
1
1.2
1.4
1.6
0
10
20
30
40
50
60
Normal
MK50
MK75
MK88
MK94
MK97
MK data
Normal data

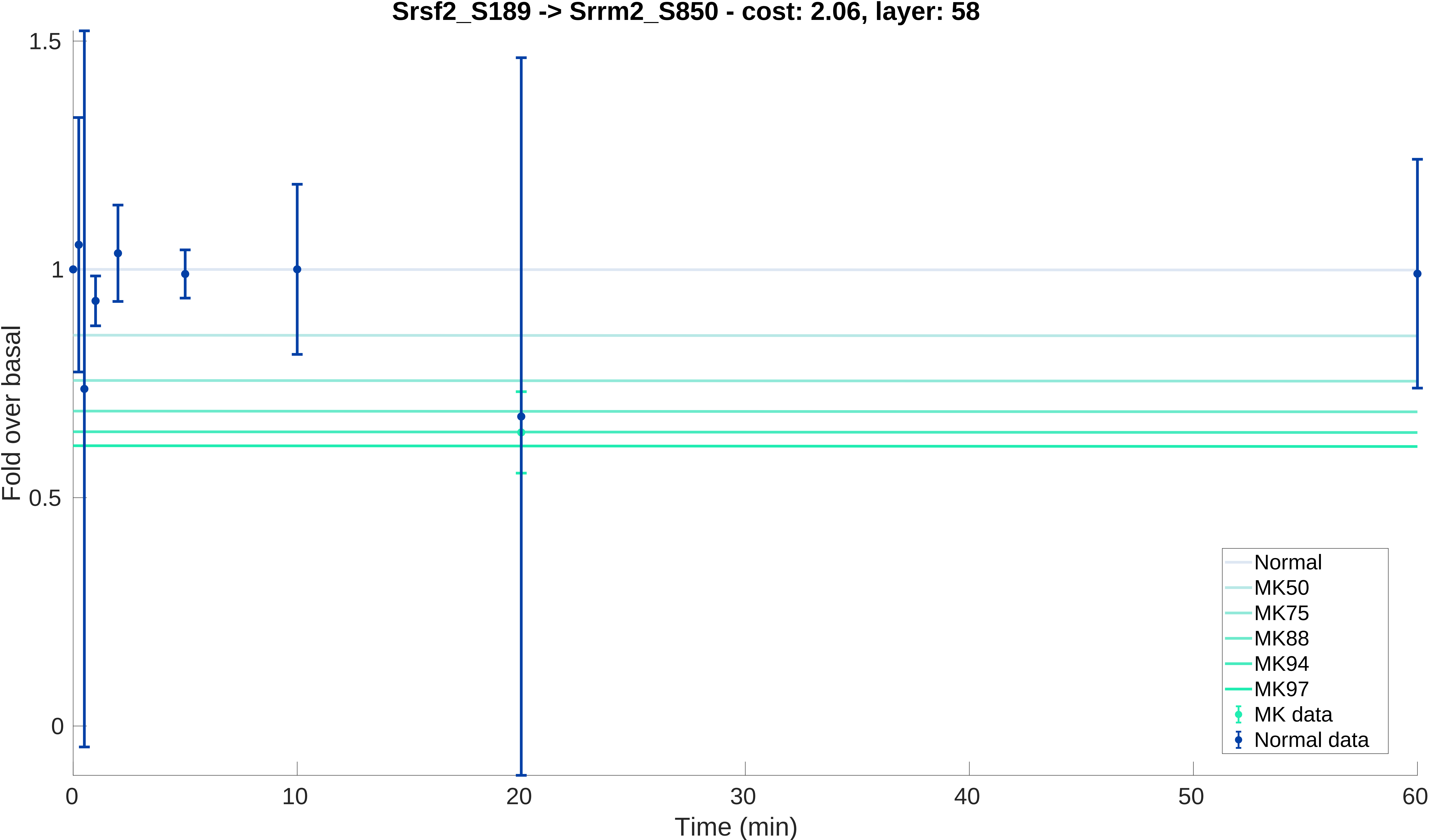

Srsf2_S189 -> Srrm2_S850 - cost: 2.06, layer: 58
Fold over basal
Time (min)
0
0.5
1
1.5
0
10
20
30
40
50
60
Normal
MK50
MK75
MK88
MK94
MK97
MK data
Normal data

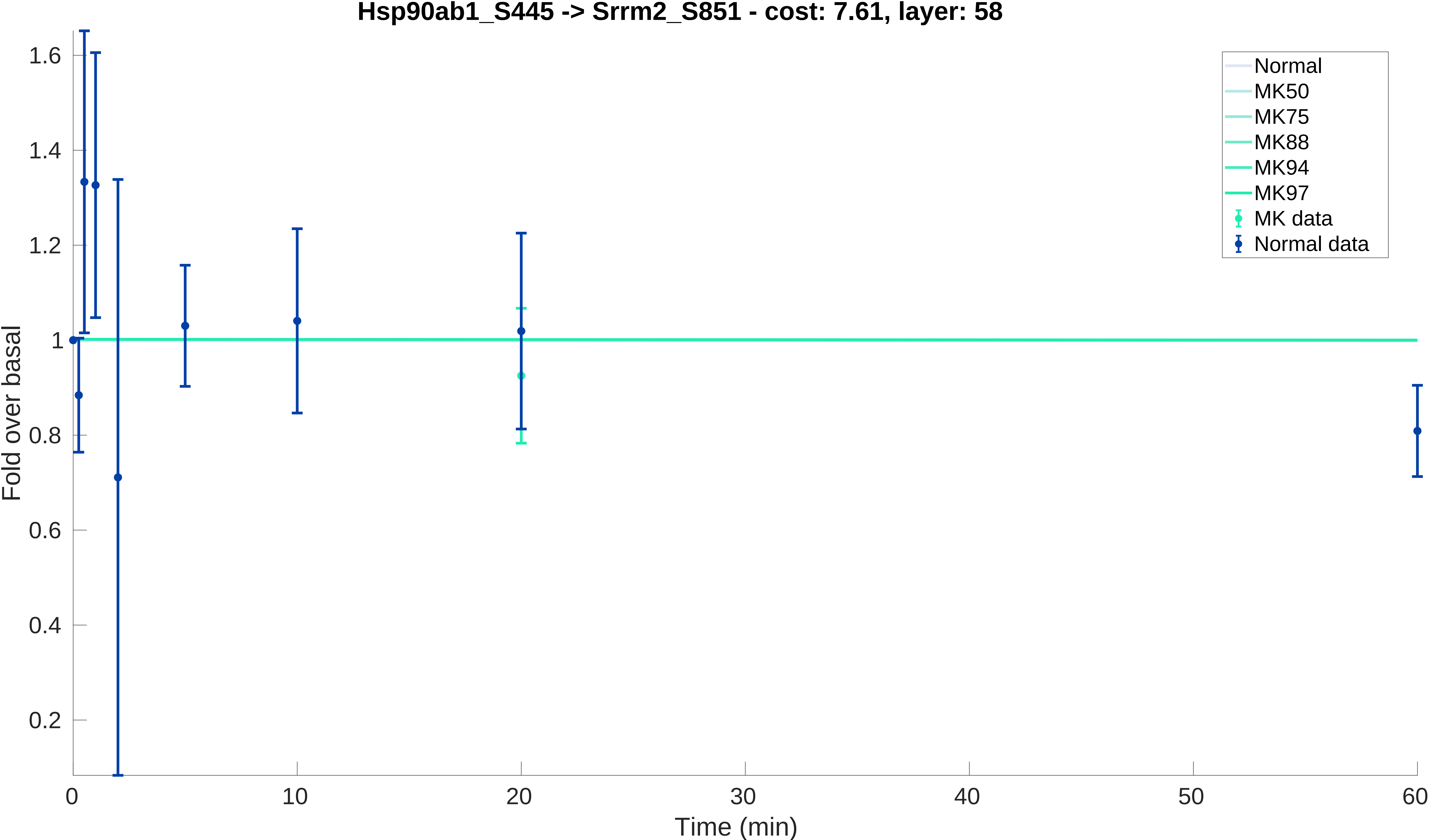

Hsp90ab1_S445 -> Srrm2_S851 - cost: 7.61, layer: 58
Fold over basal
Time (min)
0
10
20
30
40
50
60
0.2
0.4
0.6
0.8
1
1.2
1.4
1.6
Normal
MK50
MK75
MK88
MK94
MK97
MK data
Normal data

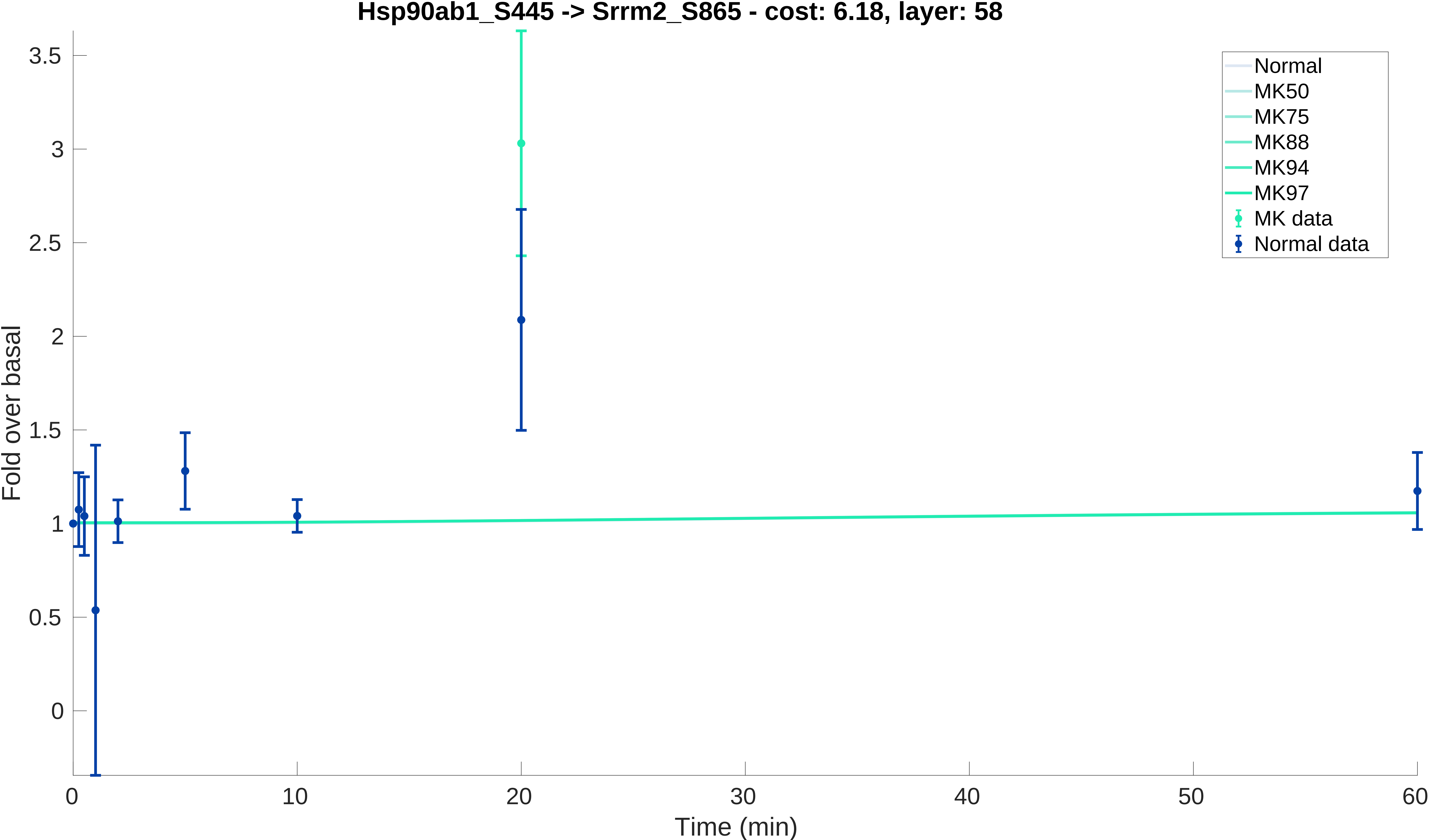

Hsp90ab1_S445 -> Srrm2_S865 - cost: 6.18, layer: 58
Fold over basal
Time (min)
3.5
3
2.5
2
1.5
1
0.5
0
0
10
20
30
40
50
60
Normal
MK50
MK75
MK88
MK94
MK97
MK data
Normal data

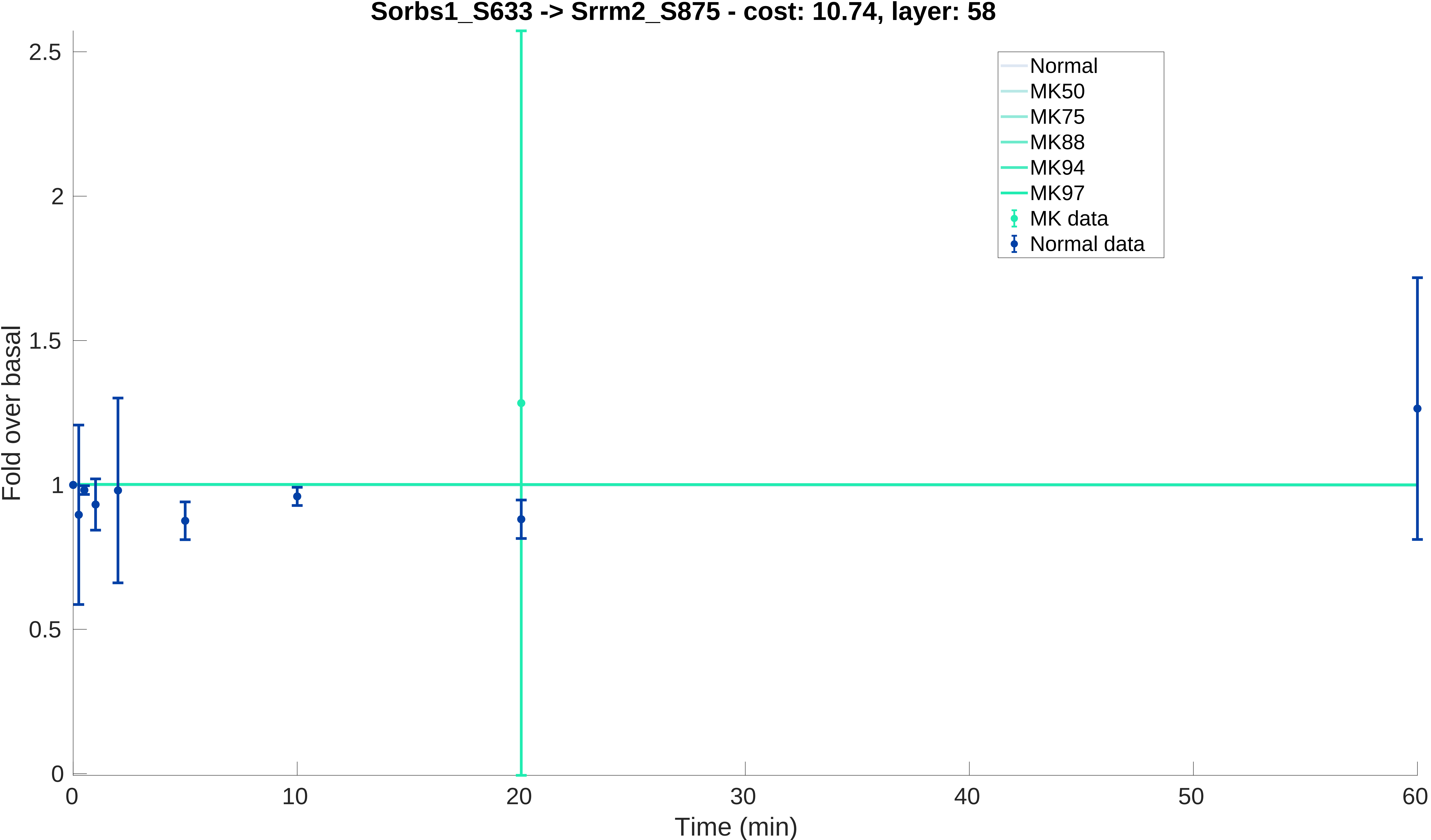

Sorbs1_S633 -> Srrm2_S875 - cost: 10.74, layer: 58
Fold over basal
Time (min)
0
0.5
1
1.5
2
2.5
0
10
20
30
40
50
60
Normal
MK50
MK75
MK88
MK94
MK97
MK data
Normal data

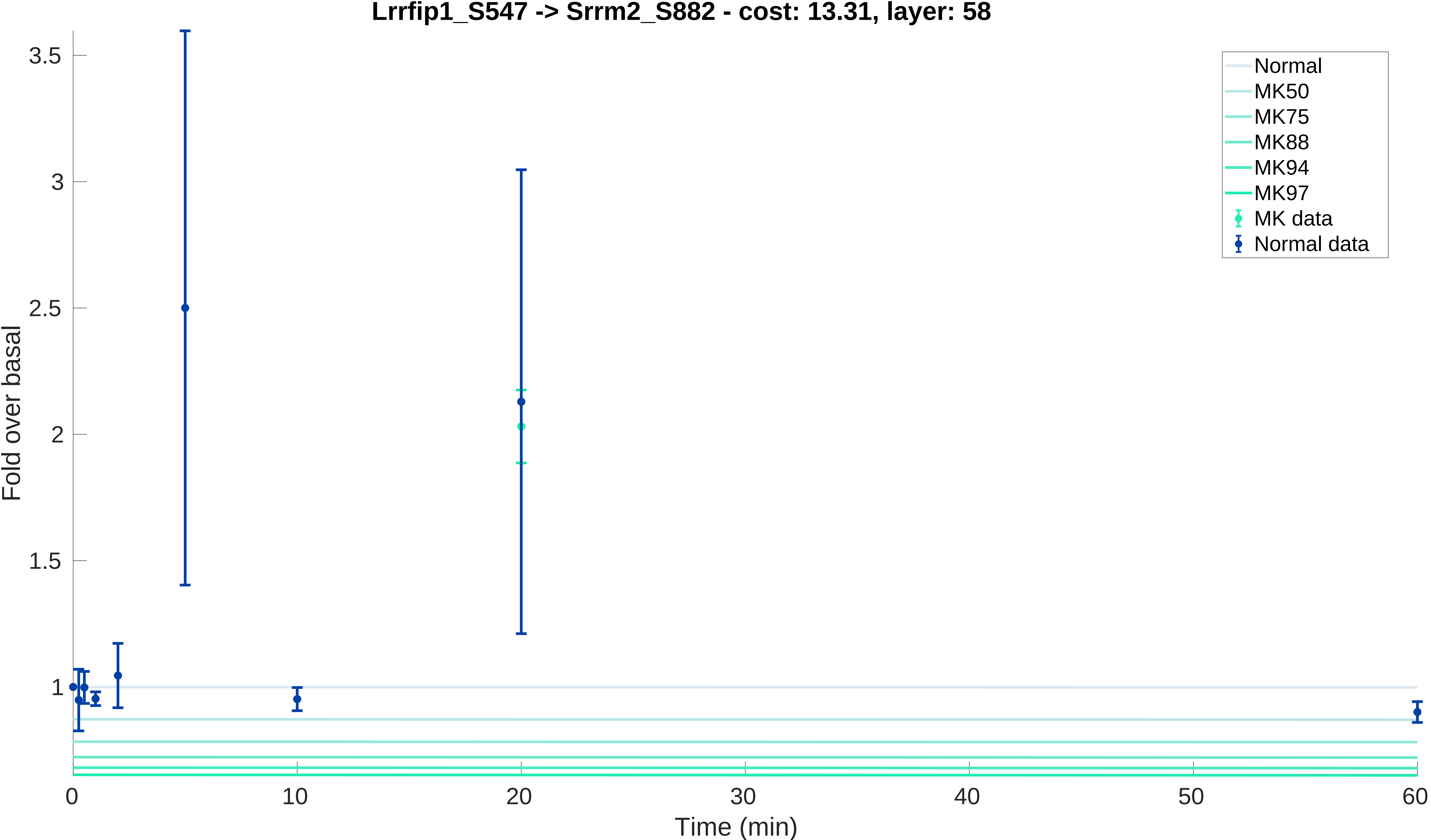

Lrrfip1_S547 -> Srrm2_S882 - cost: 13.31, layer: 58
Fold over basal
3.5
3
2.5
2
1.5
1
0
10
20
30
40
50
60
Time (min)
Normal
MK50
MK75
MK88
MK94
MK97
MK data
Normal data

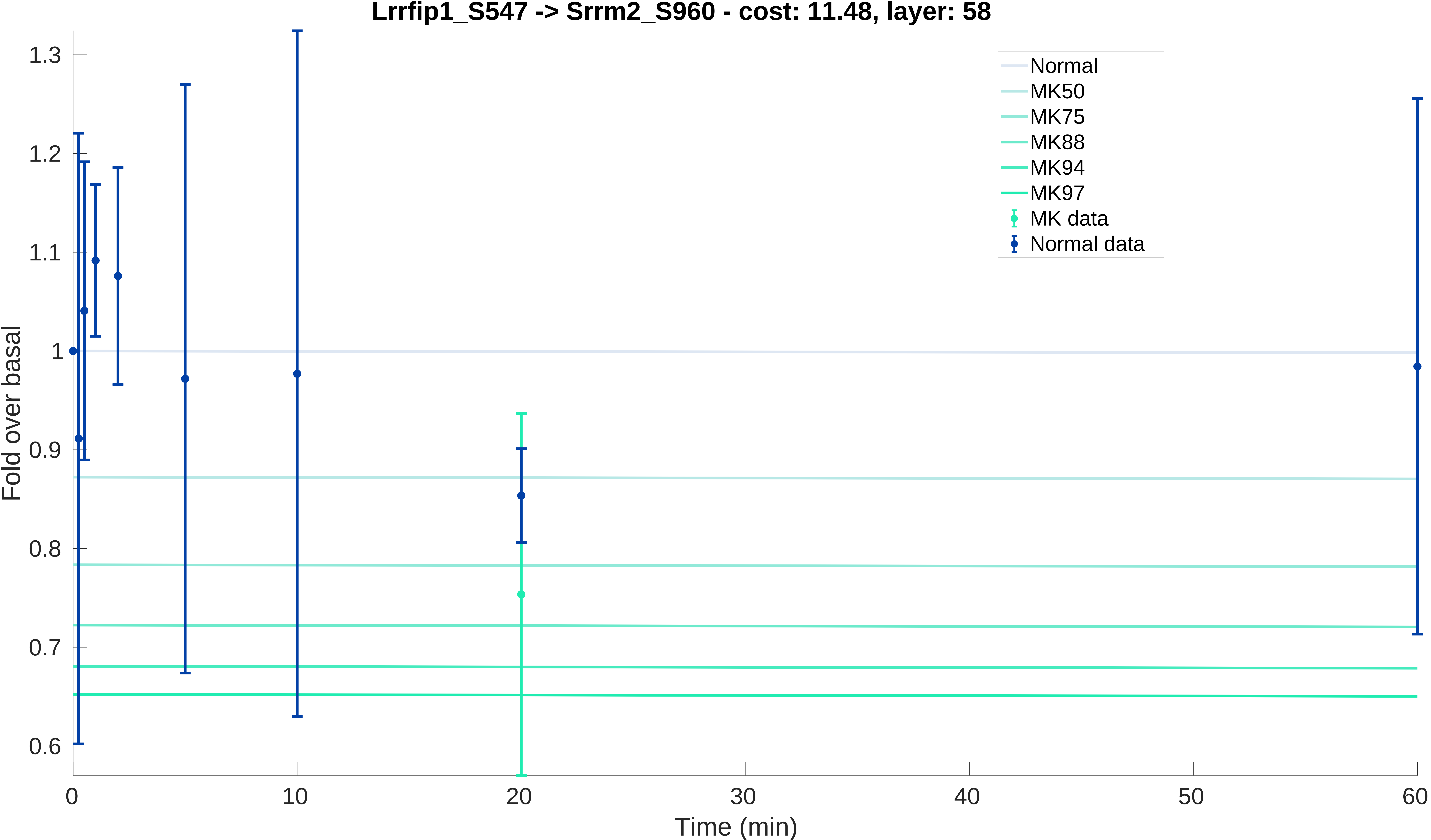

Lrrfip1_S547 -> Srrm2_S960 - cost: 11.48, layer: 58
Fold over basal
Time (min)
Normal
MK50
MK75
MK88
MK94
MK97
MK data
Normal data
1.3
1.2
1.1
1
0.9
0.8
0.7
0.6
0
10
20
30
40
50
60

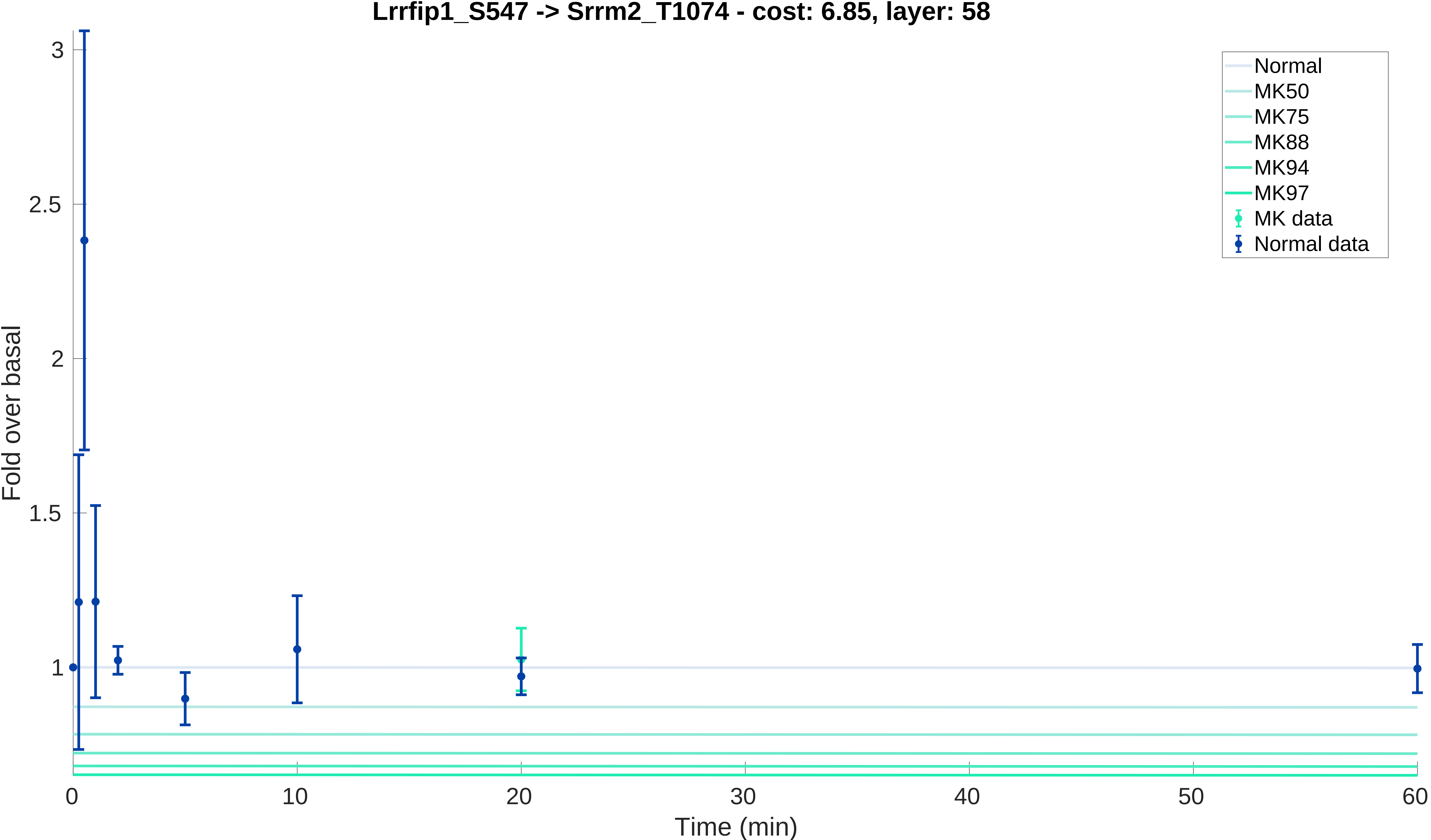

Lrrfip1_S547 -> Srrm2_T1074 - cost: 6.85, layer: 58
Fold over basal
Time (min)
3
2.5
2
1.5
1
0
10
20
30
40
50
60
Normal
MK50
MK75
MK88
MK94
MK97
MK data
Normal data

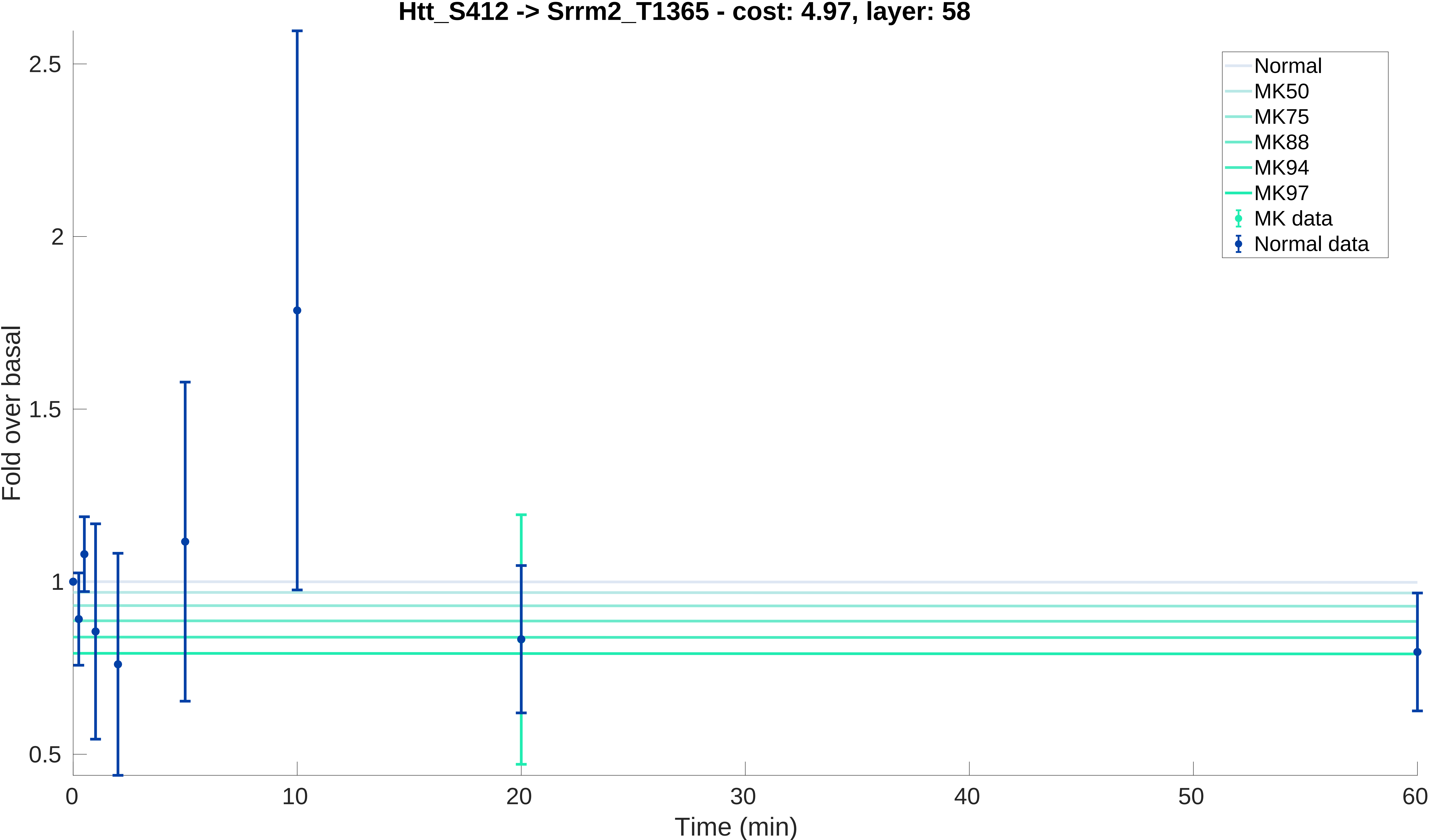
Htt_S412 -> Srrm2_T1365 - cost: 4.97, layer: 58
Fold over basal
Time (min)
0.5
1
1.5
2
2.5
0
10
20
30
40
50
60
Normal
MK50
MK75
MK88
MK94
MK97
MK data
Normal data

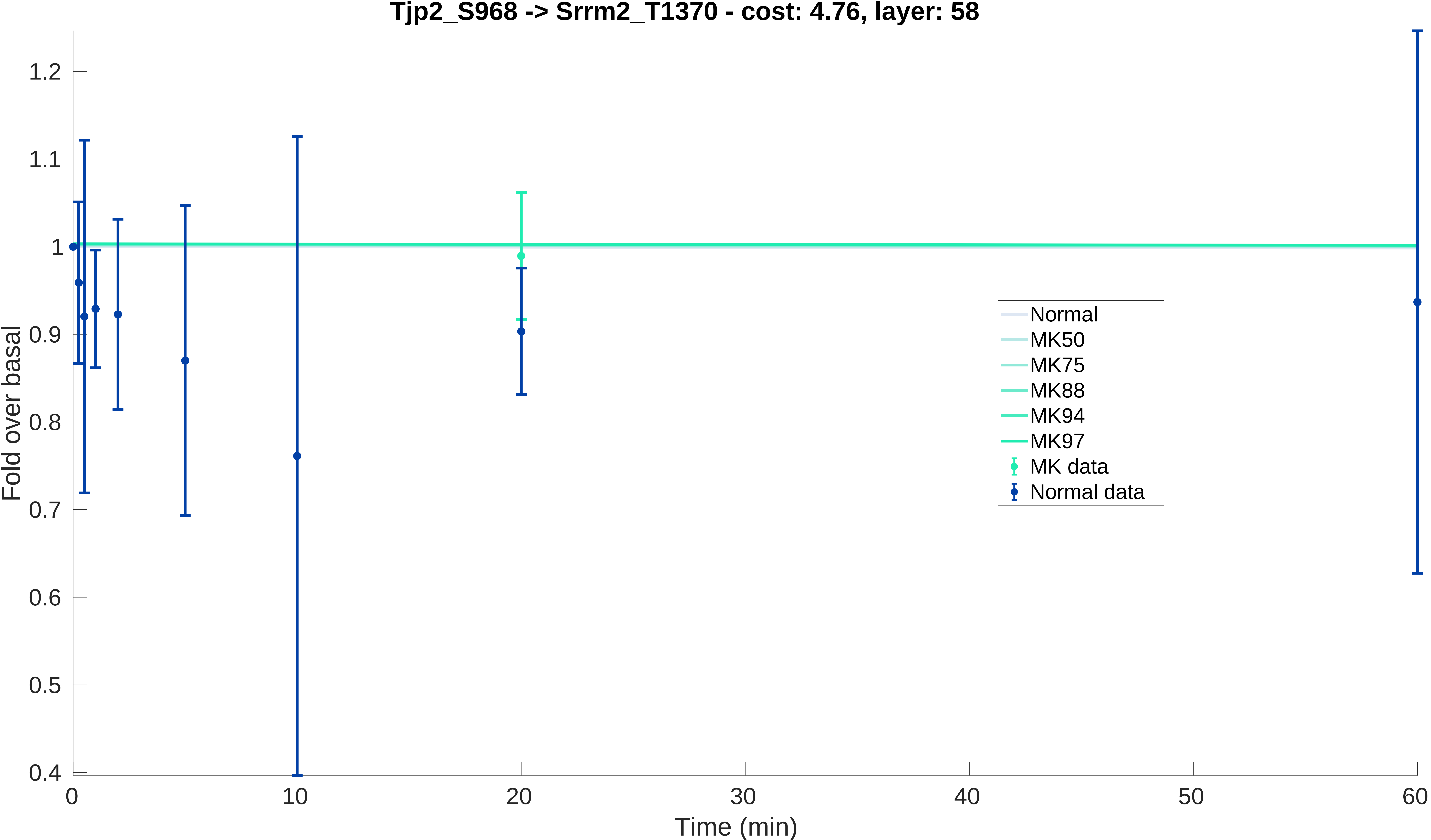

Tjp2_S968 -> Srrm2_T1370 - cost: 4.76, layer: 58
Fold over basal
Time (min)
1.2
1.1
1
0.9
0.8
0.7
0.6
0.5
0.4
0
10
20
30
40
50
60
Normal
MK50
MK75
MK88
MK94
MK97
MK data
Normal data

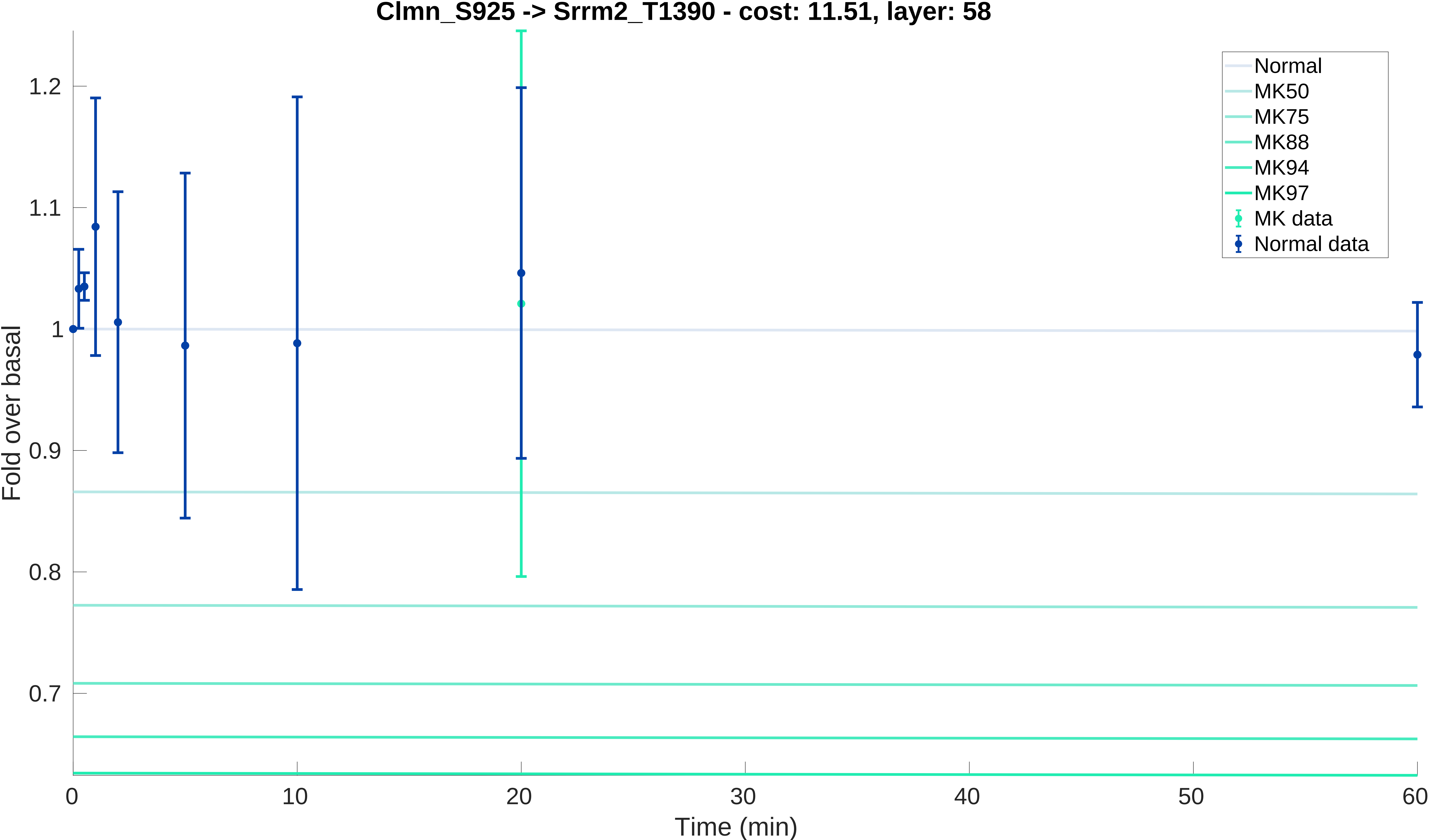

Clmn_S925 -> Srrm2_T1390 - cost: 11.51, layer: 58
Fold over basal
Time (min)
1.2
1.1
1
0.9
0.8
0.7
0
10
20
30
40
50
60
Normal
MK50
MK75
MK88
MK94
MK97
MK data
Normal data

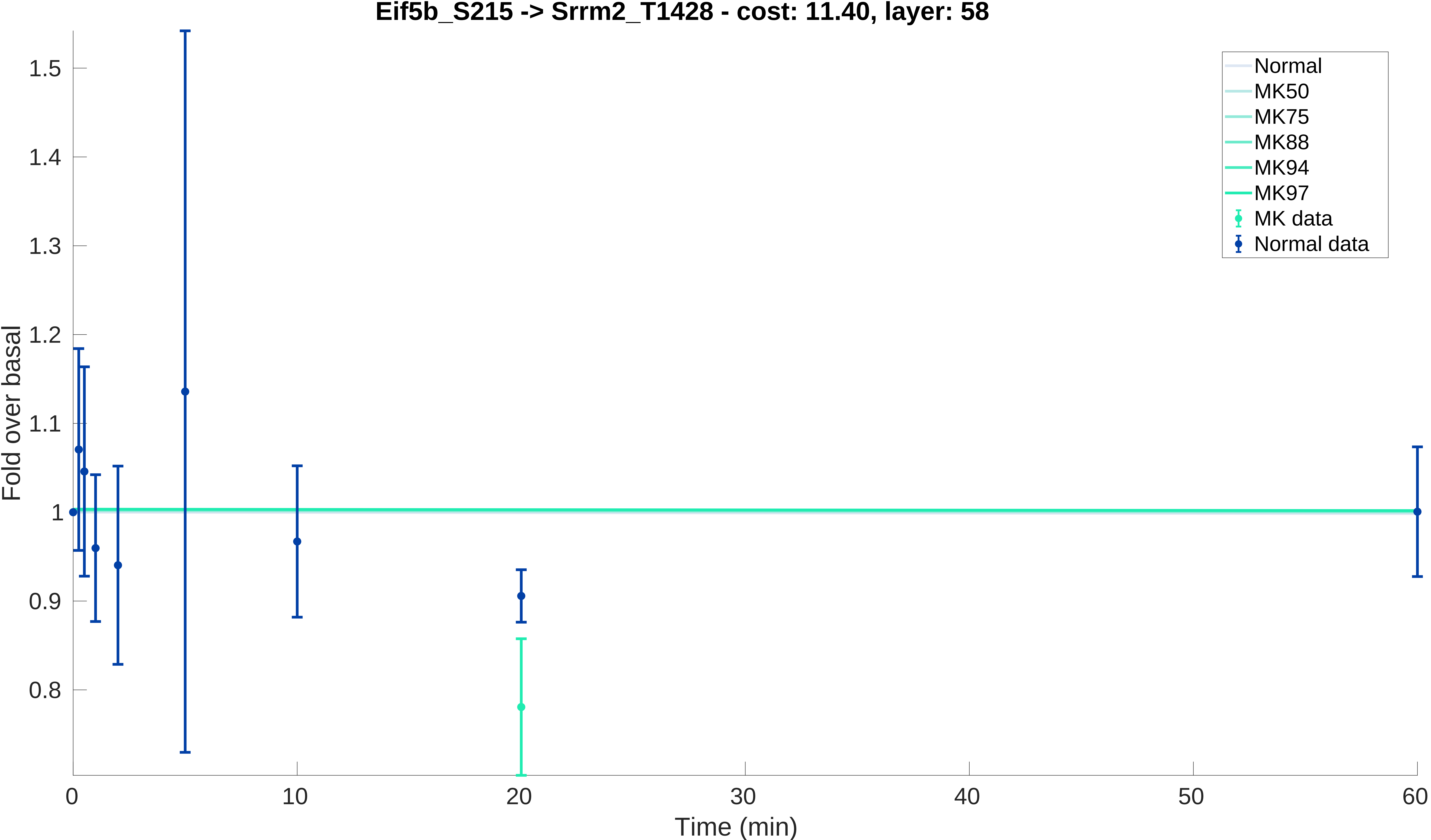

Eif5b_S215 -> Srrm2_T1428 - cost: 11.40, layer: 58
Fold over basal
Time (min)
1.5
1.4
1.3
1.2
1.1
1
0.9
0.8
0
10
20
30
40
50
60
Normal
MK50
MK75
MK88
MK94
MK97
MK data
Normal data

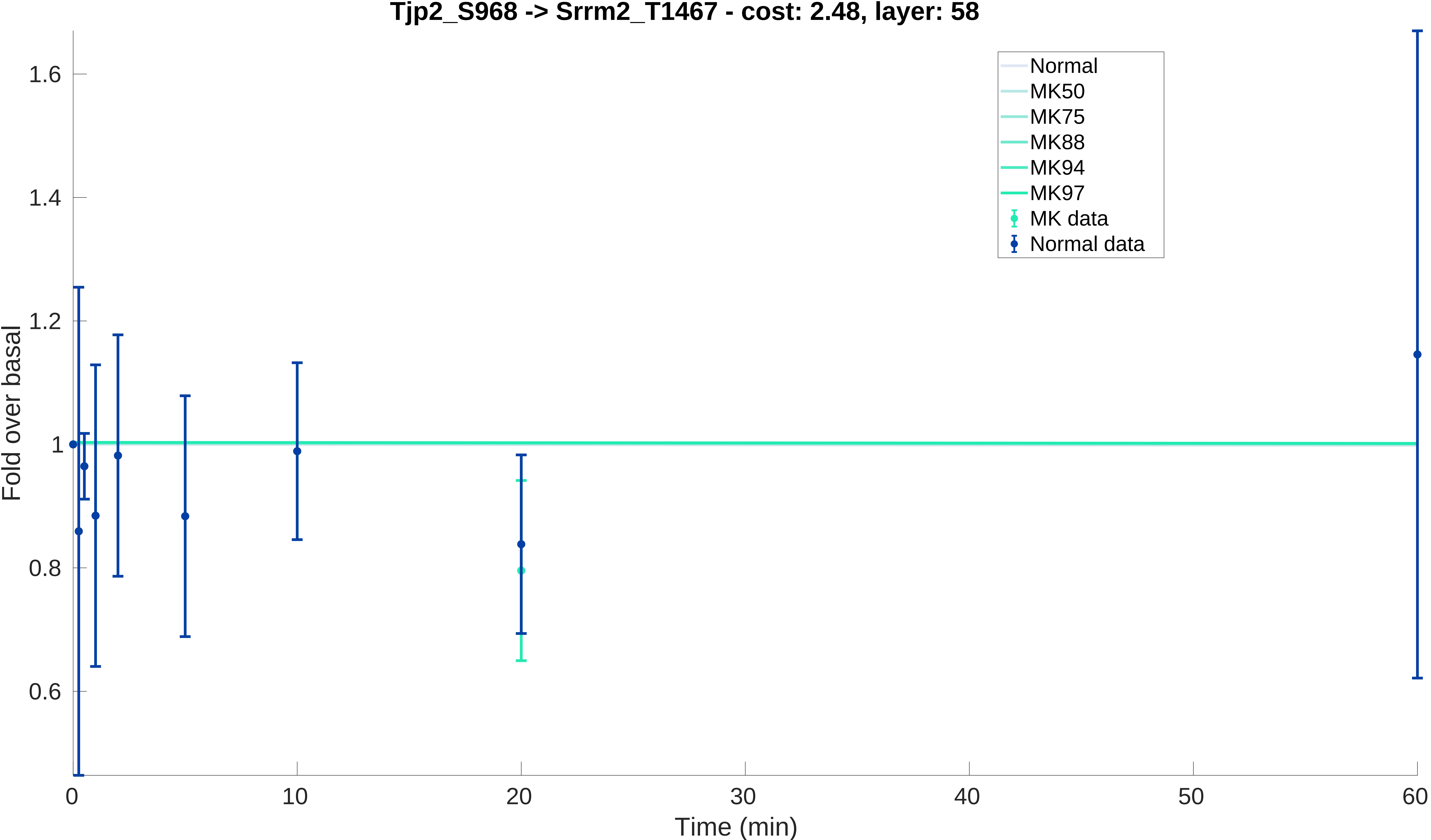

Tjp2_S968 -> Srrm2_T1467 - cost: 2.48, layer: 58
Fold over basal
Time (min)
0
10
20
30
40
50
60
0.6
0.8
1
1.2
1.4
1.6
Normal
MK50
MK75
MK88
MK94
MK97
MK data
Normal data

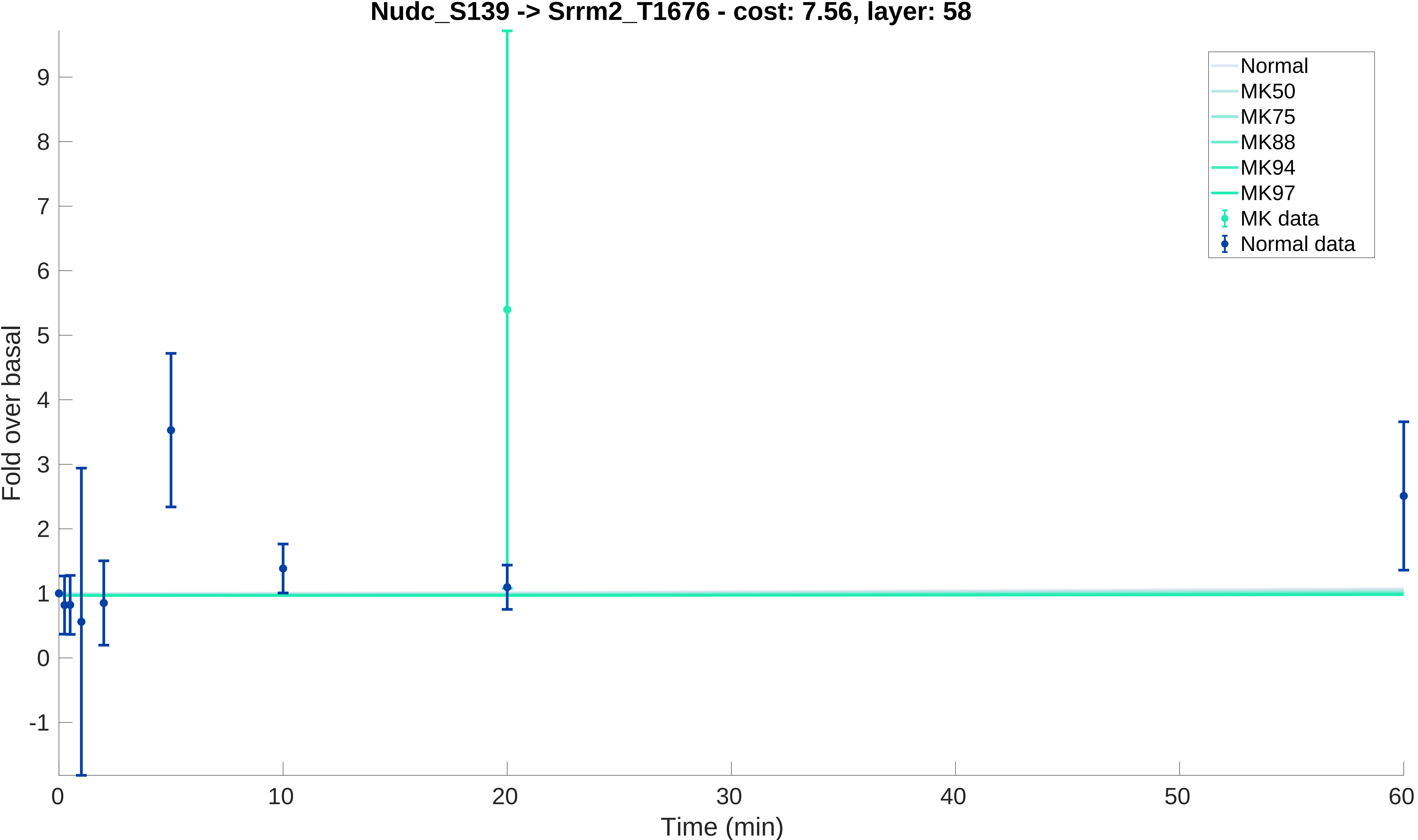

Nudc_S139 -> Srrm2_T1676 - cost: 7.56, layer: 58
Fold over basal
Time (min)
9
8
7
6
5
4
3
2
1
0
-1
0
10
20
30
40
50
60
Normal
MK50
MK75
MK88
MK94
MK97
MK data
Normal data

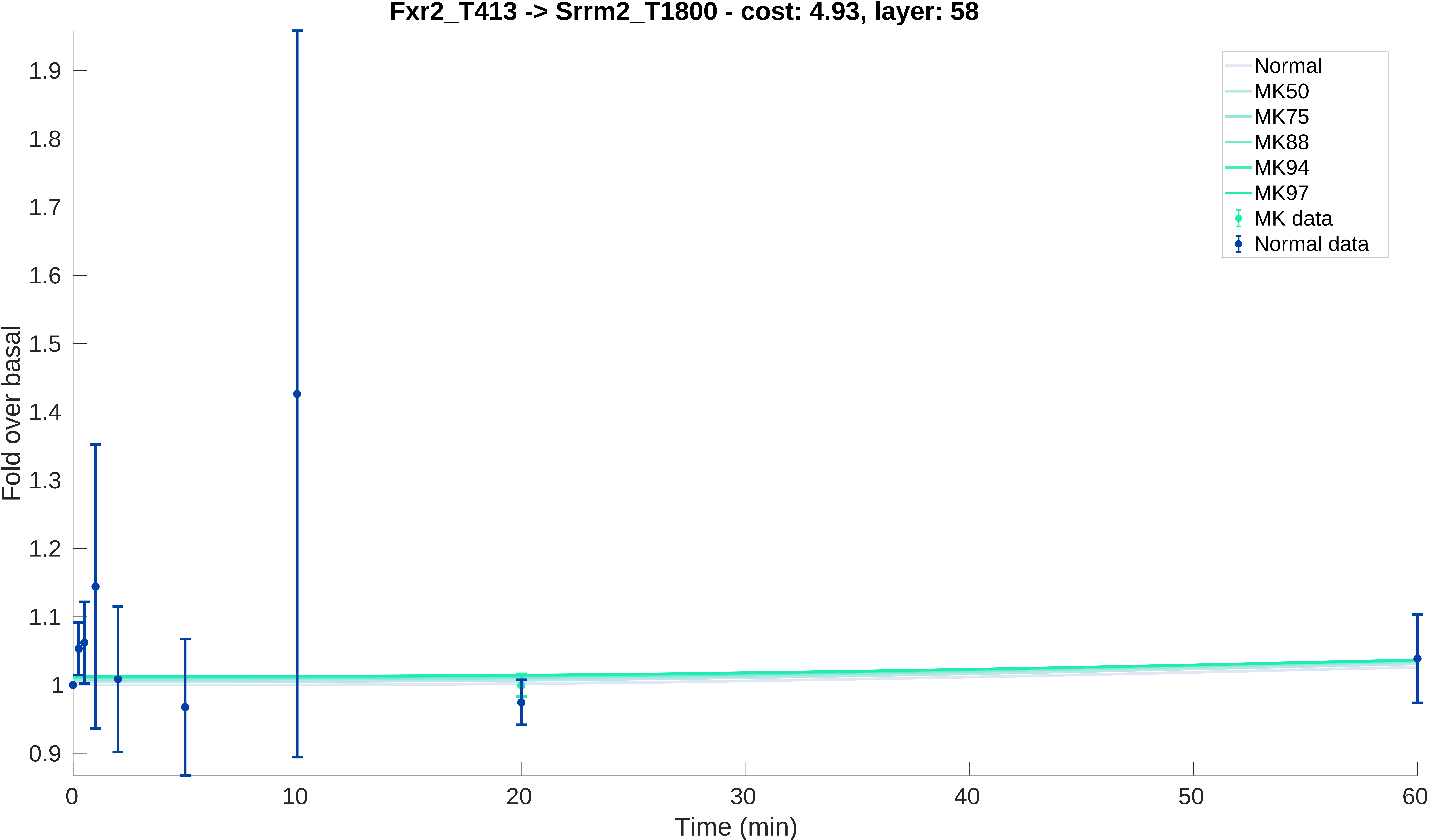

Fxr2_T413 -> Srrm2_T1800 - cost: 4.93, layer: 58
Fold over basal
Time (min)
Normal
MK50
MK75
MK88
MK94
MK97
MK data
Normal data
0.9
1
1.1
1.2
1.3
1.4
1.5
1.6
1.7
1.8
1.9
0
10
20
30
40
50
60

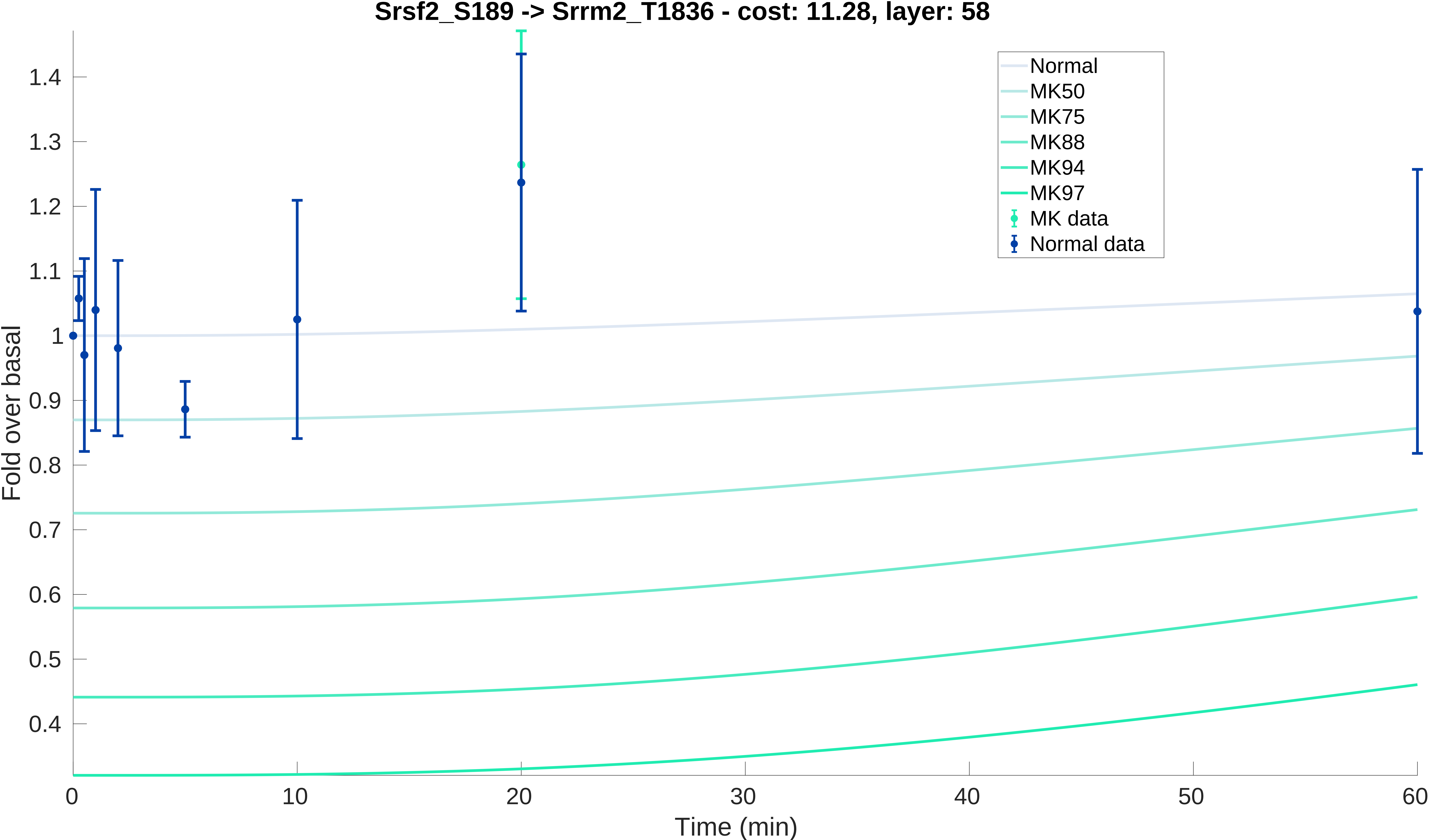

Srsf2_S189 -> Srrm2_T1836 - cost: 11.28, layer: 58
Fold over basal
Time (min)
1.4
1.3
1.2
1.1
1
0.9
0.8
0.7
0.6
0.5
0.4
0
10
20
30
40
50
60
Normal
MK50
MK75
MK88
MK94
MK97
MK data
Normal data

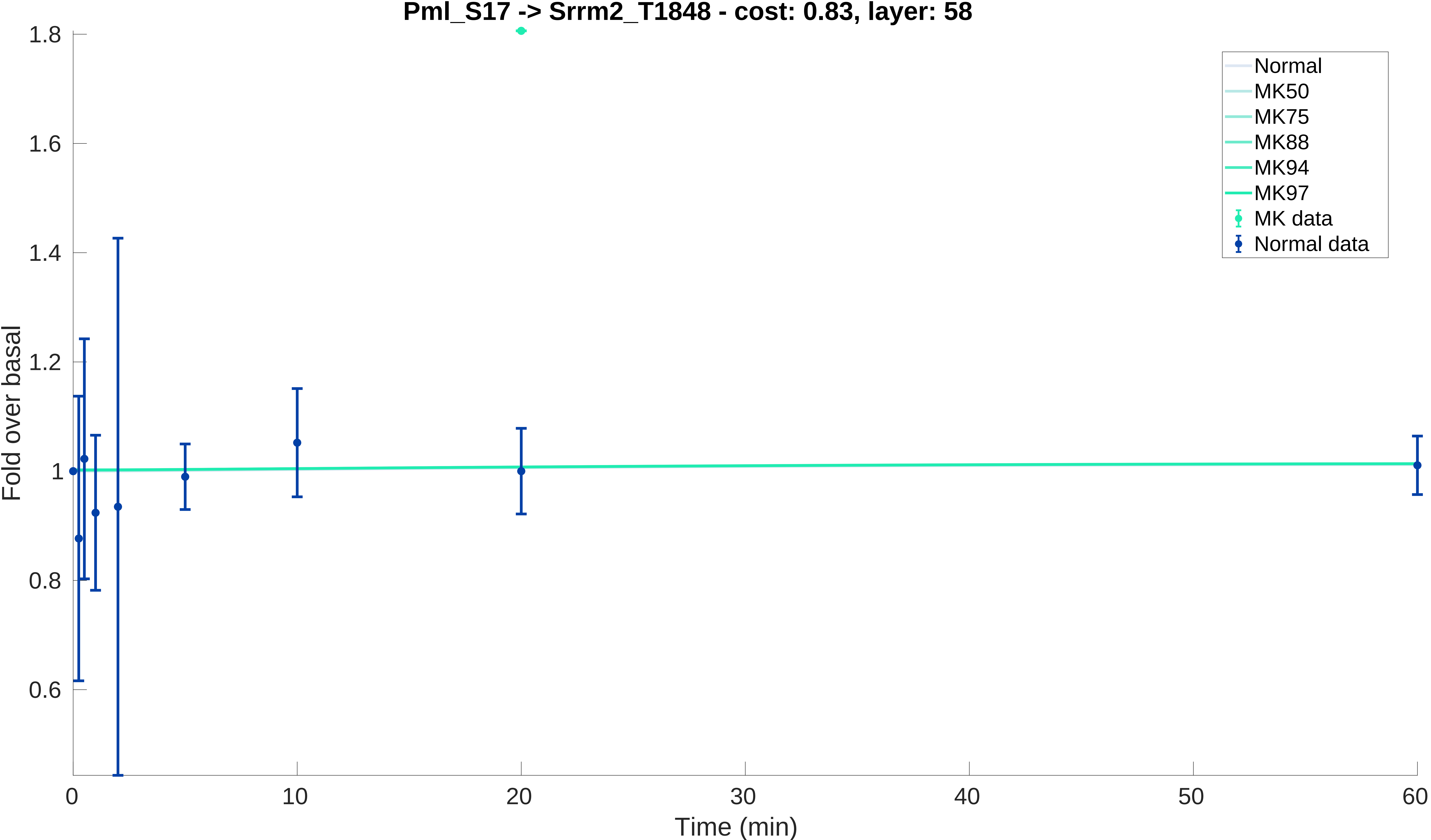

Pml_S17 -> Srrm2_T1848 - cost: 0.83, layer: 58
Fold over basal
Time (min)
1.8
1.6
1.4
1.2
1
0.8
0.6
0
10
20
30
40
50
60
Normal
MK50
MK75
MK88
MK94
MK97
MK data
Normal data

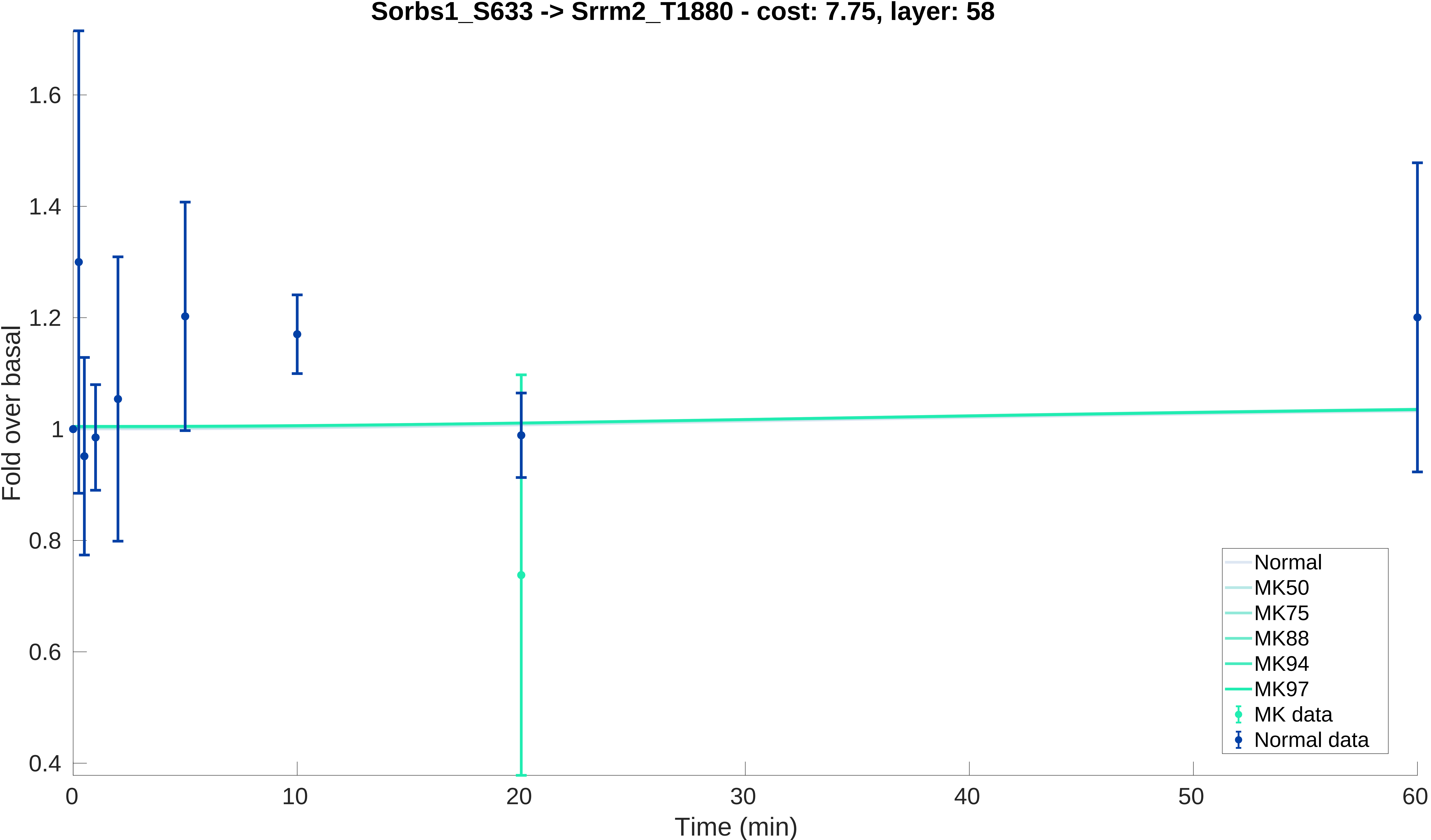

Sorbs1_S633 -> Srrm2_T1880 - cost: 7.75, layer: 58
Fold over basal
Time (min)
0.4
0.6
0.8
1
1.2
1.4
1.6
0
10
20
30
40
50
60
Normal
MK50
MK75
MK88
MK94
MK97
MK data
Normal data

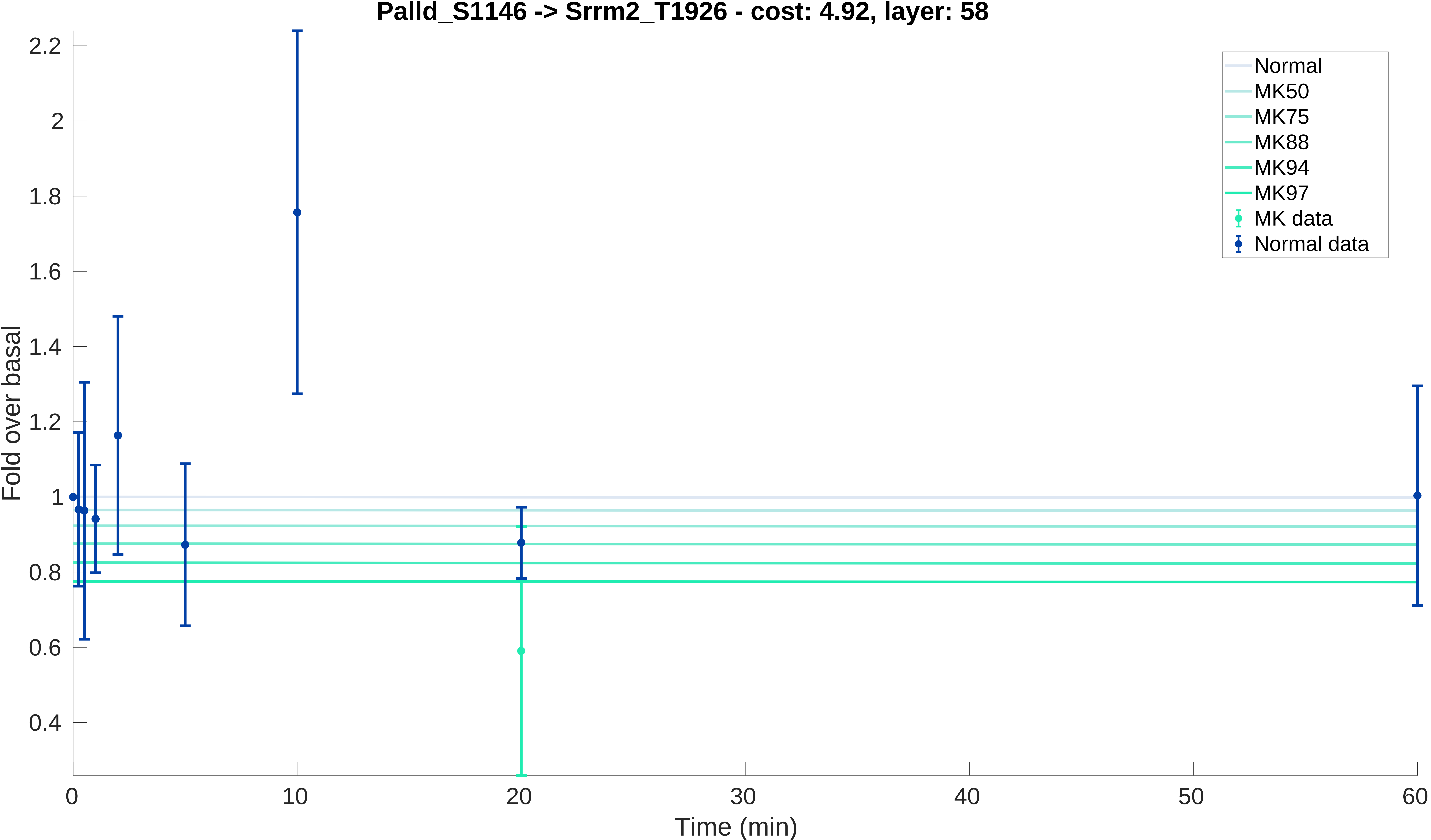
Palld_S1146 -> Srrm2_T1926 - cost: 4.92, layer: 58
Fold over basal
Time (min)
2.2
2
1.8
1.6
1.4
1.2
1
0.8
0.6
0.4
0
10
20
30
40
50
60
Normal
MK50
MK75
MK88
MK94
MK97
MK data
Normal data

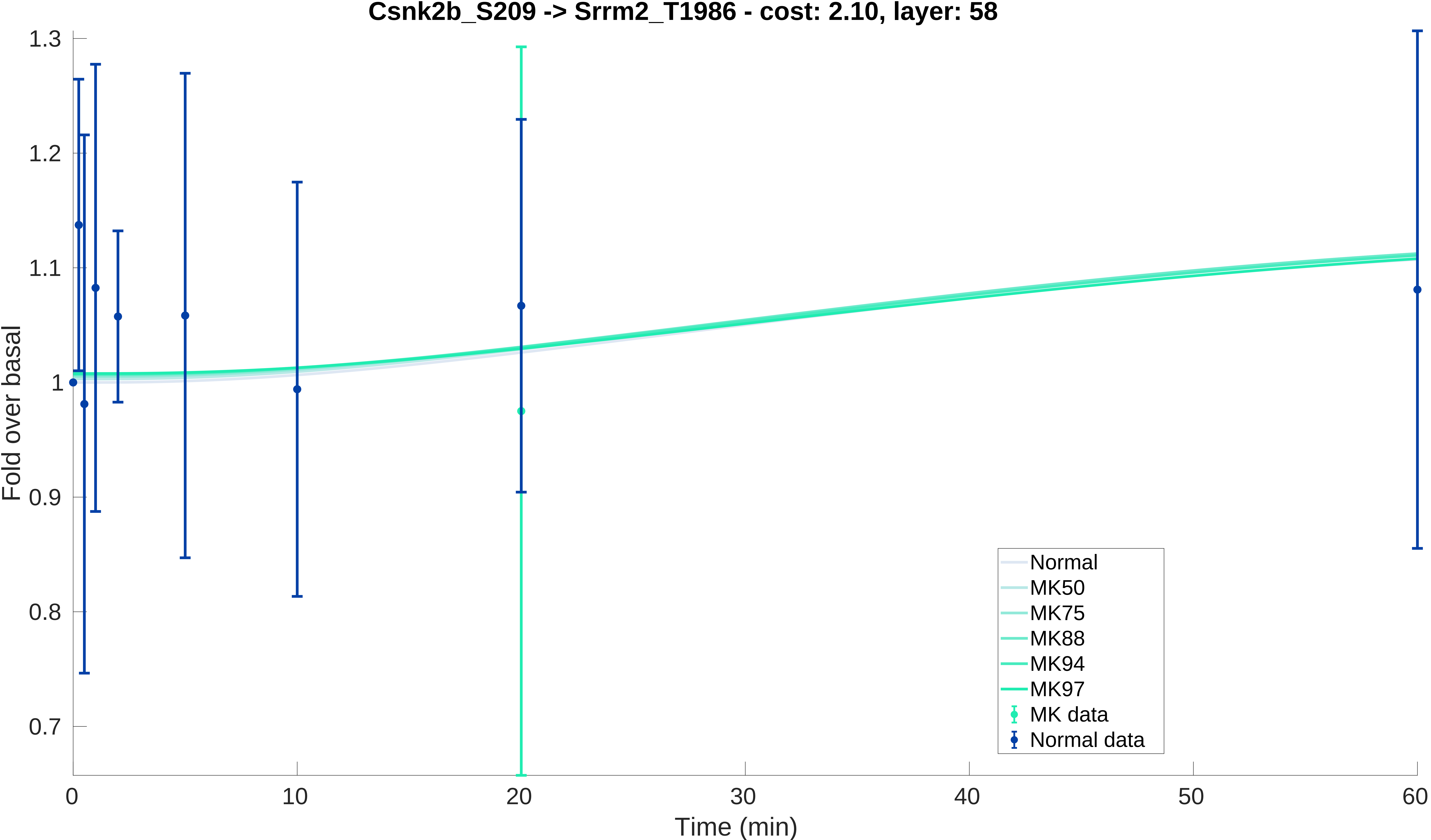
Csnk2b_S209 -> Srrm2_T1986 - cost: 2.10, layer: 58
Fold over basal
1.3
1.2
1.1
1
0.9
0.8
0.7
0
10
20
30
40
50
60
Time (min)
Normal
MK50
MK75
MK88
MK94
MK97
MK data
Normal data

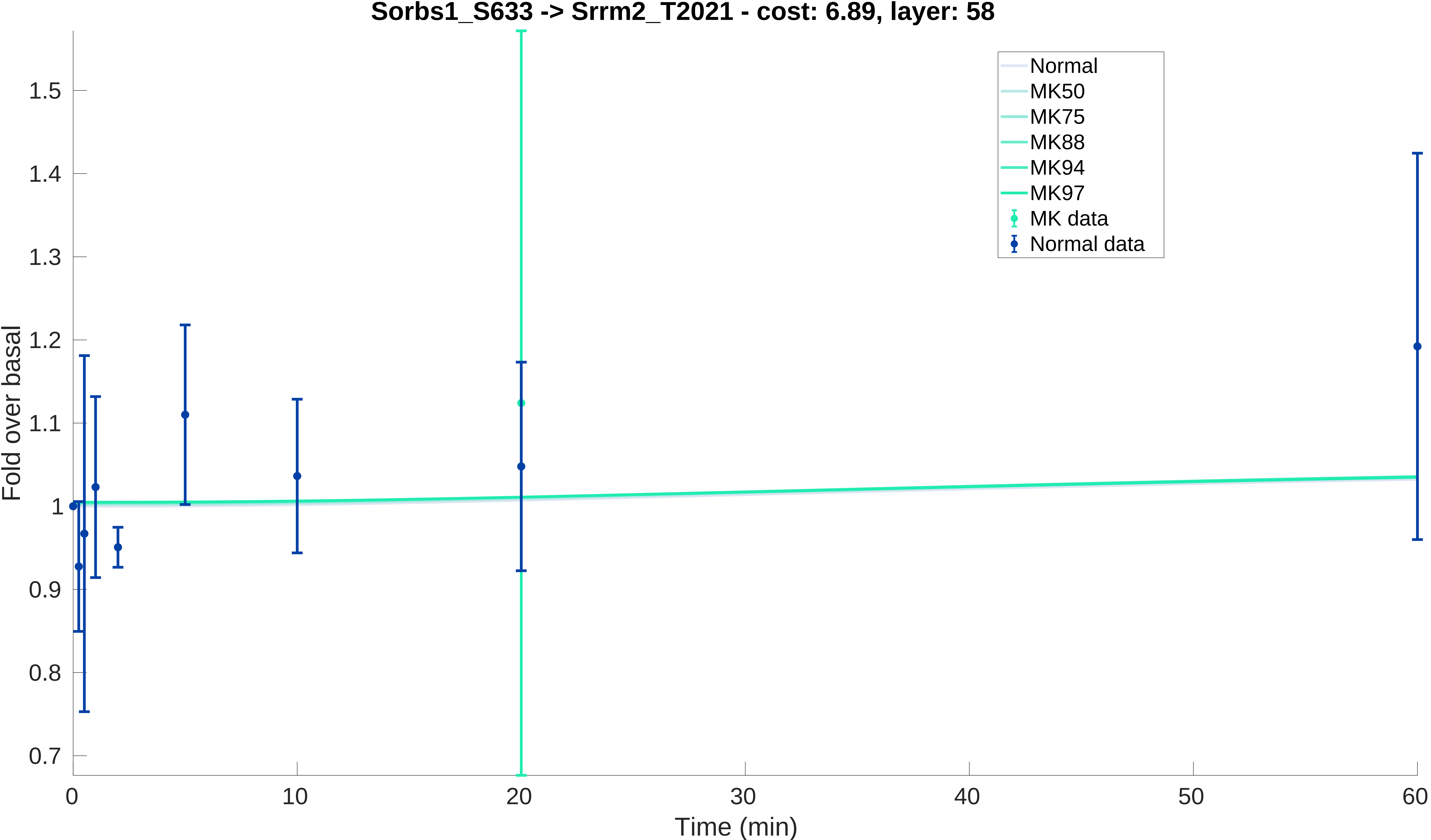

Sorbs1_S633 -> Srrm2_T2021 - cost: 6.89, layer: 58
Normal
MK50
MK75
MK88
MK94
MK97
MK data
Normal data
Fold over basal
1.5
1.4
1.3
1.2
1.1
1
0.9
0.8
0.7
0
10
20
30
40
50
60
Time (min)

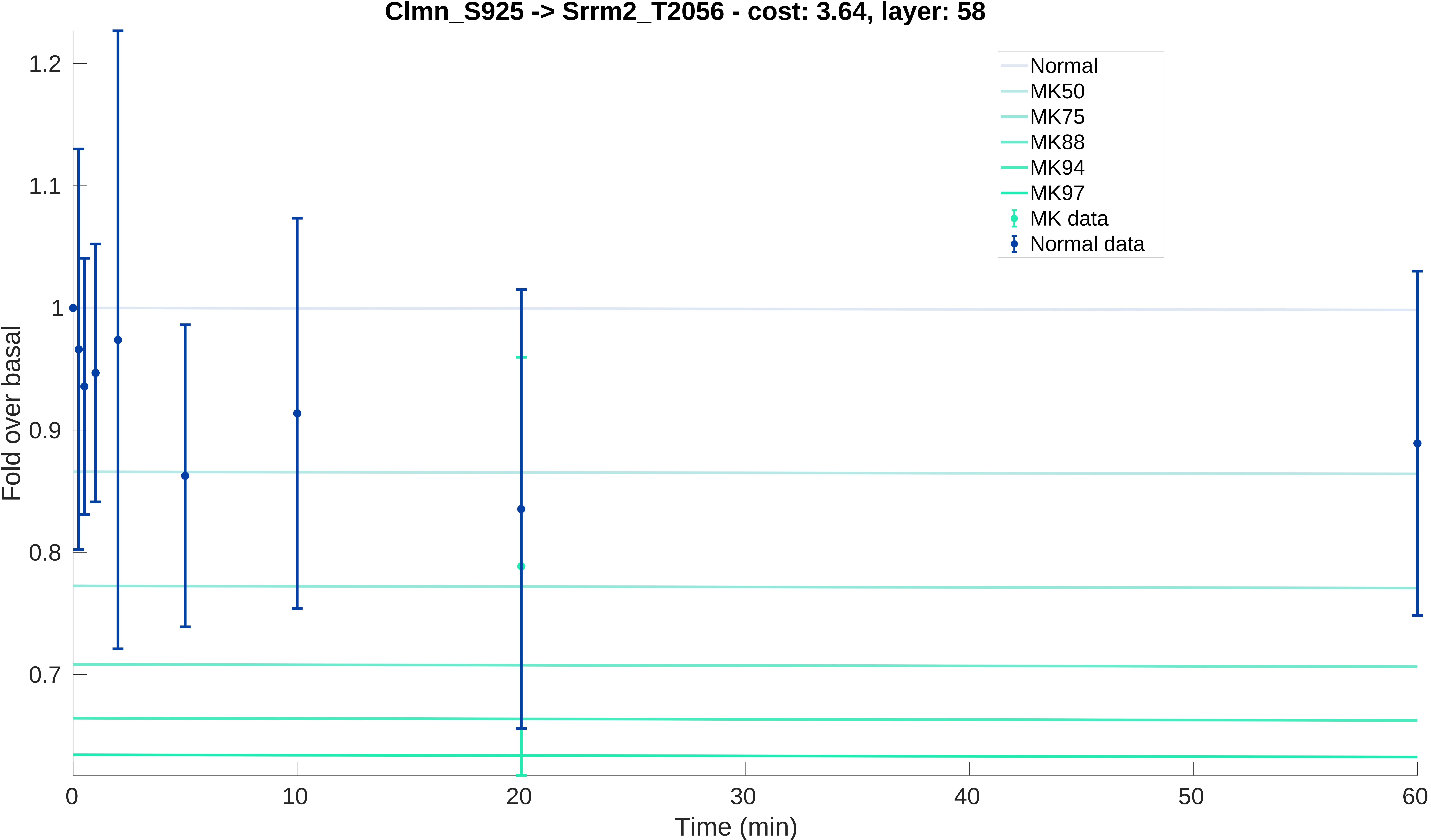

Clmn_S925 -> Srrm2_T2056 - cost: 3.64, layer: 58
Fold over basal
Time (min)
1.2
1.1
1
0.9
0.8
0.7
0
10
20
30
40
50
60
Normal
MK50
MK75
MK88
MK94
MK97
MK data
Normal data

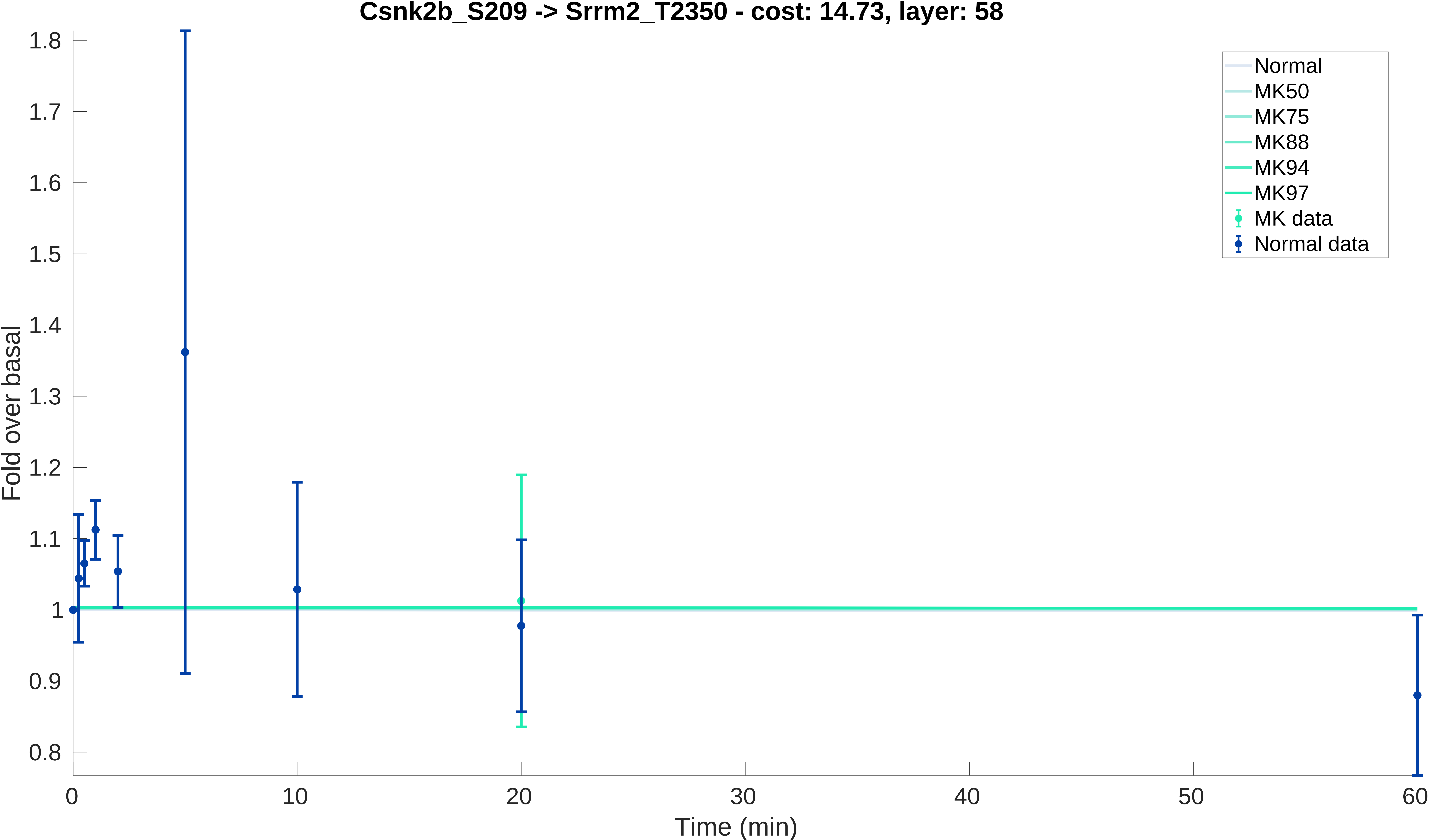

Csnk2b_S209 -> Srrm2_T2350 - cost: 14.73, layer: 58
Fold over basal
Time (min)
Normal
MK50
MK75
MK88
MK94
MK97
MK data
Normal data
0.8
0.9
1
1.1
1.2
1.3
1.4
1.5
1.6
1.7
1.8
0
10
20
30
40
50
60

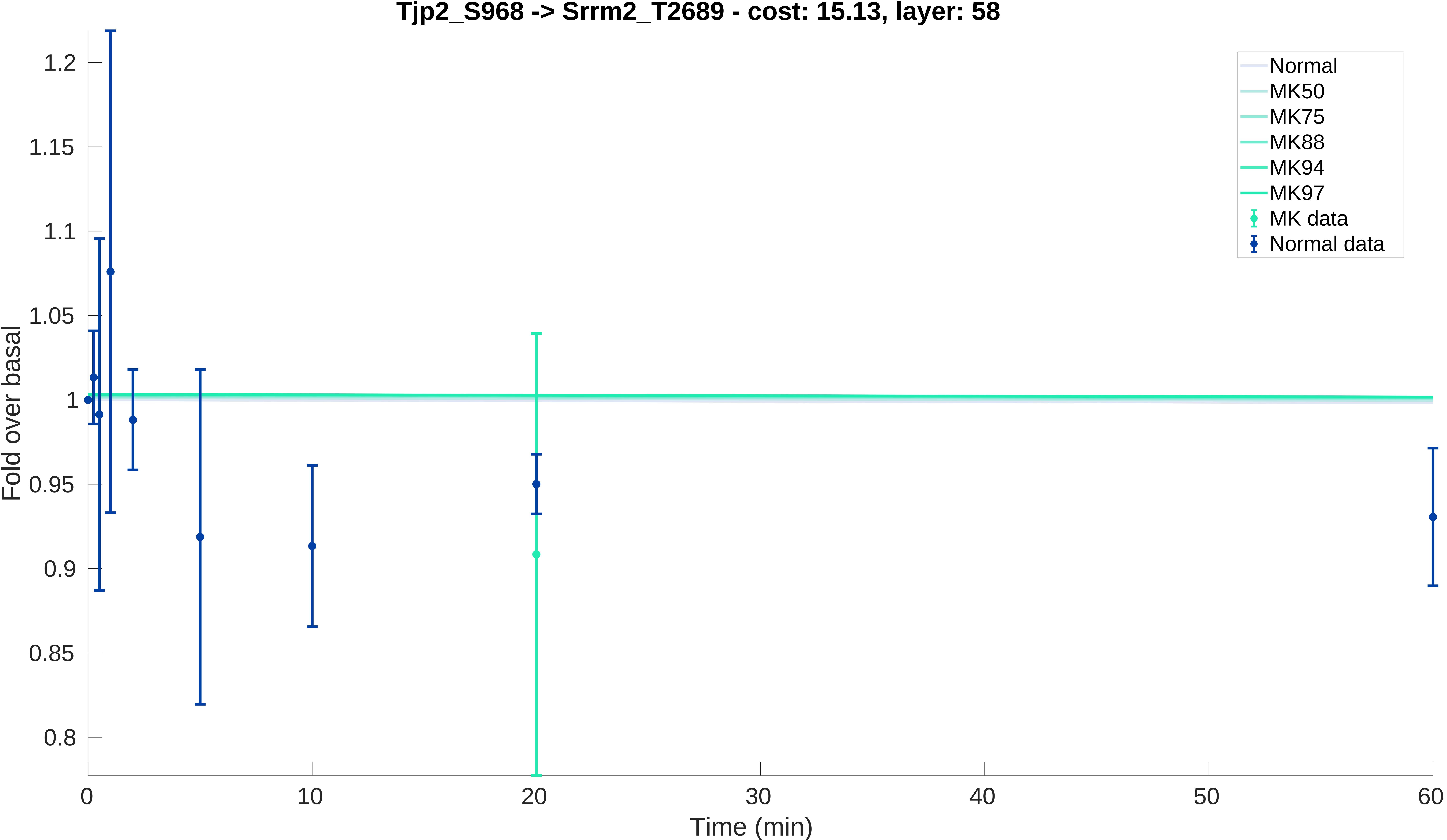

Tjp2_S968 -> Srrm2_T2689 - cost: 15.13, layer: 58
Fold over basal
Time (min)
1.2
1.15
1.1
1.05
1
0.95
0.9
0.85
0.8
0
10
20
30
40
50
60
Normal
MK50
MK75
MK88
MK94
MK97
MK data
Normal data

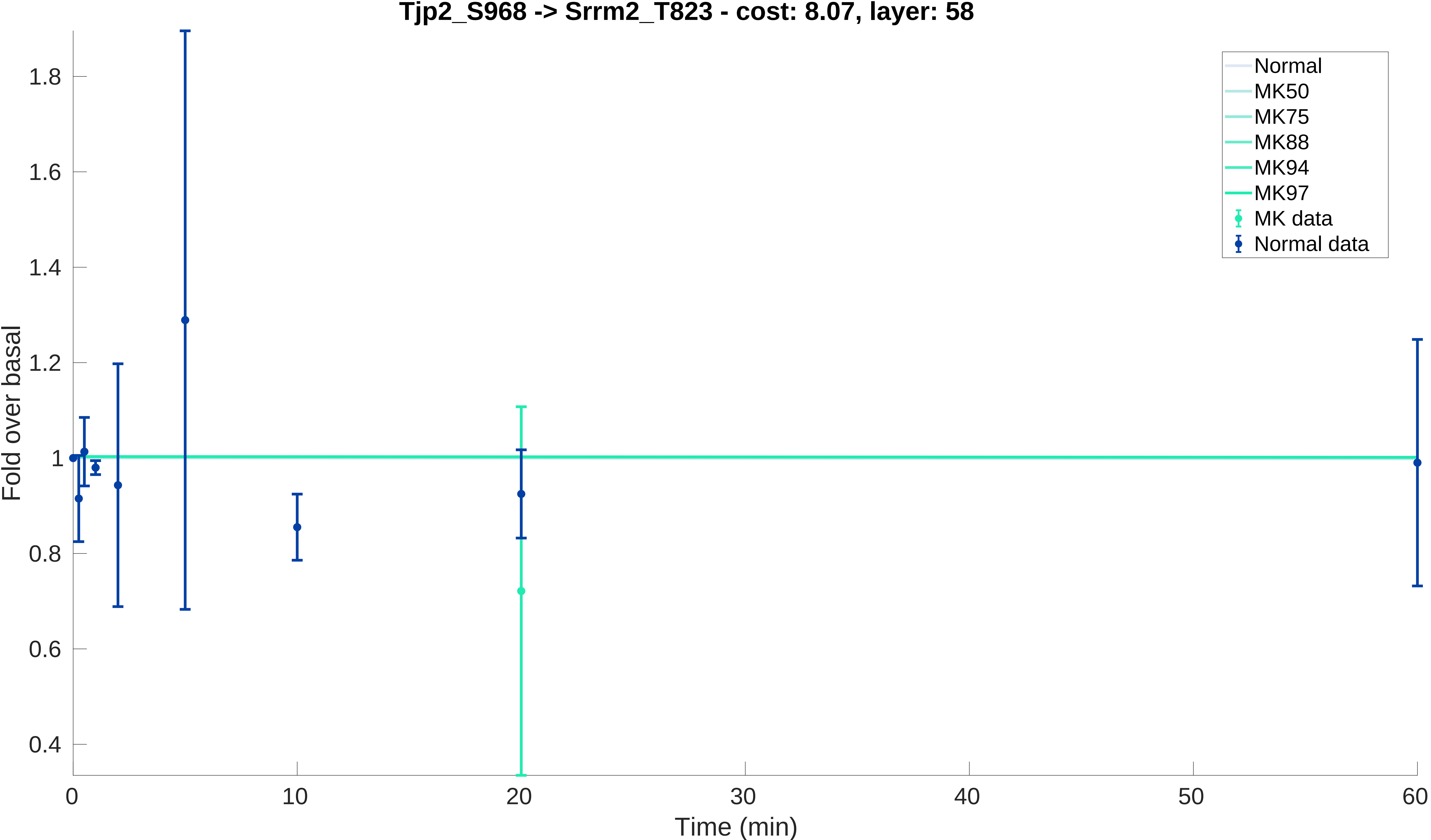
Tjp2_S968 -> Srrm2_T823 - cost: 8.07, layer: 58
Fold over basal
Time (min)
0.4
0.6
0.8
1
1.2
1.4
1.6
1.8
0
10
20
30
40
50
60
Normal
MK50
MK75
MK88
MK94
MK97
MK data
Normal data

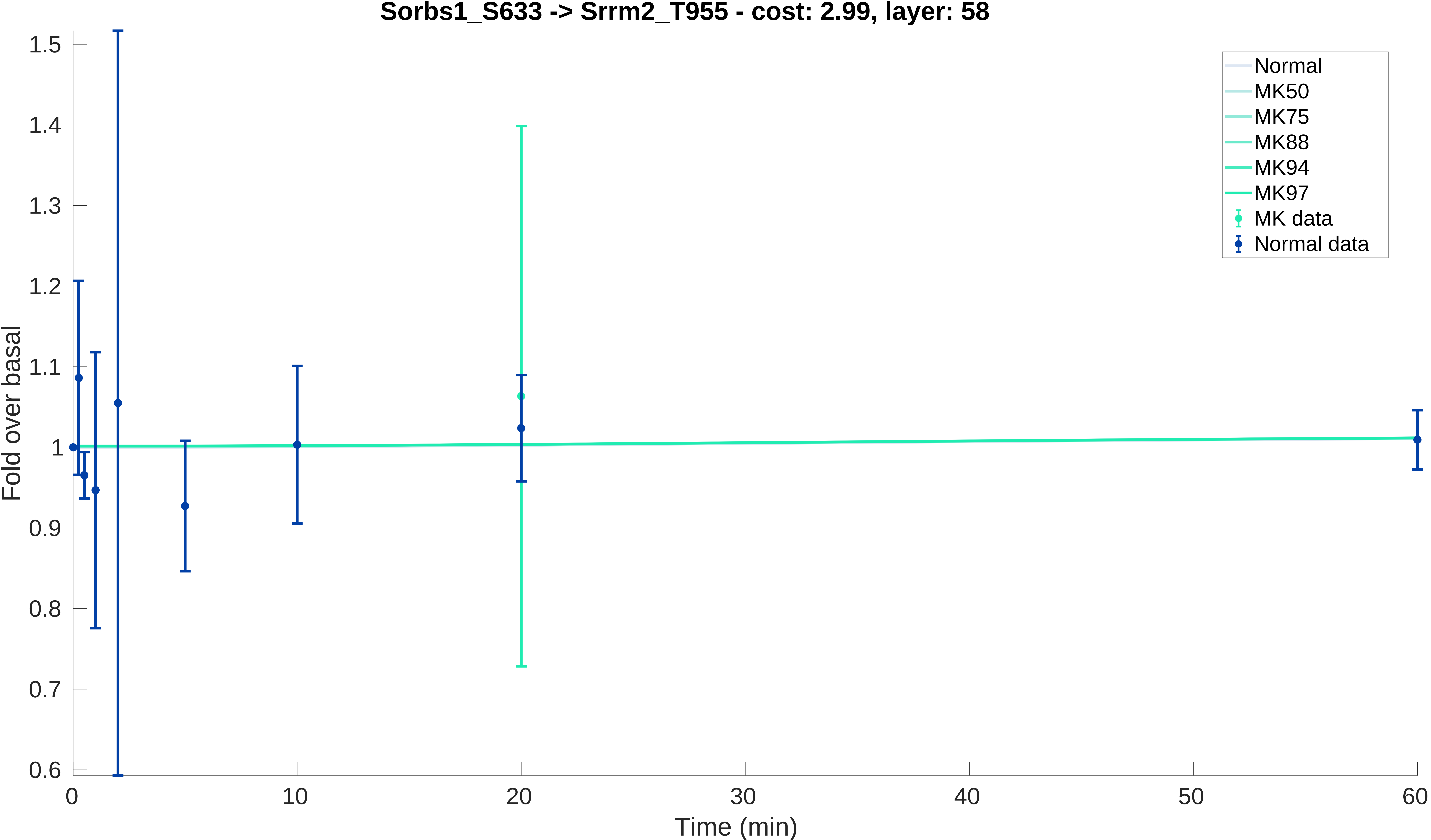

Sorbs1_S633 -> Srrm2_T955 - cost: 2.99, layer: 58
Fold over basal
Time (min)
0.6
0.7
0.8
0.9
1
1.1
1.2
1.3
1.4
1.5
0
10
20
30
40
50
60
Normal
MK50
MK75
MK88
MK94
MK97
MK data
Normal data

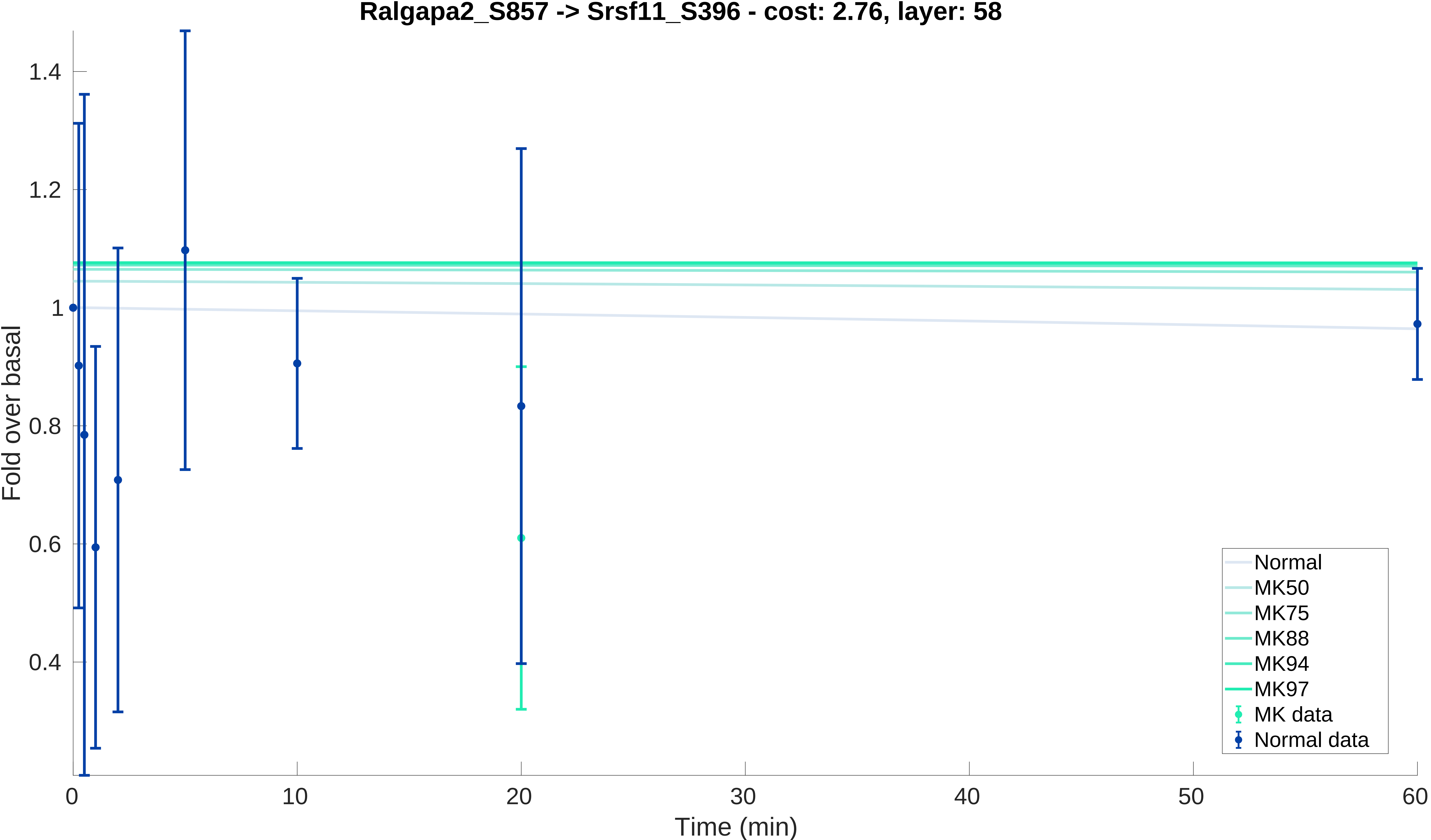
Ralgapa2_S857 -> Srsf11_S396 - cost: 2.76, layer: 58
Fold over basal
1.4
1.2
1
0.8
0.6
0.4
0
10
20
30
40
50
60
Time (min)
Normal
MK50
MK75
MK88
MK94
MK97
MK data
Normal data

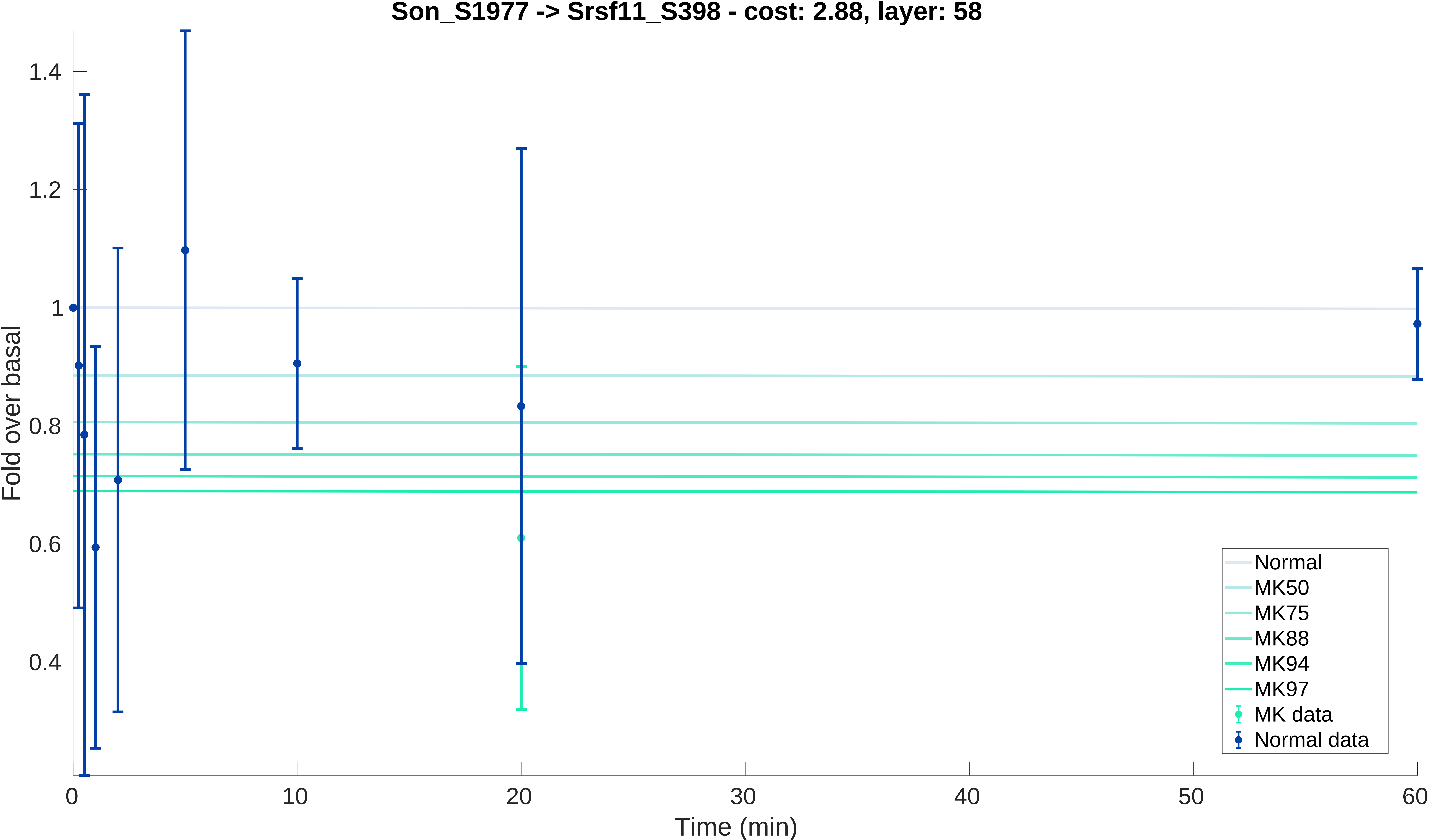

Son_S1977 -> Srsf11_S398 - cost: 2.88, layer: 58
Fold over basal
1.4
1.2
1
0.8
0.6
0.4
0
10
20
30
40
50
60
Time (min)
Normal
MK50
MK75
MK88
MK94
MK97
MK data
Normal data

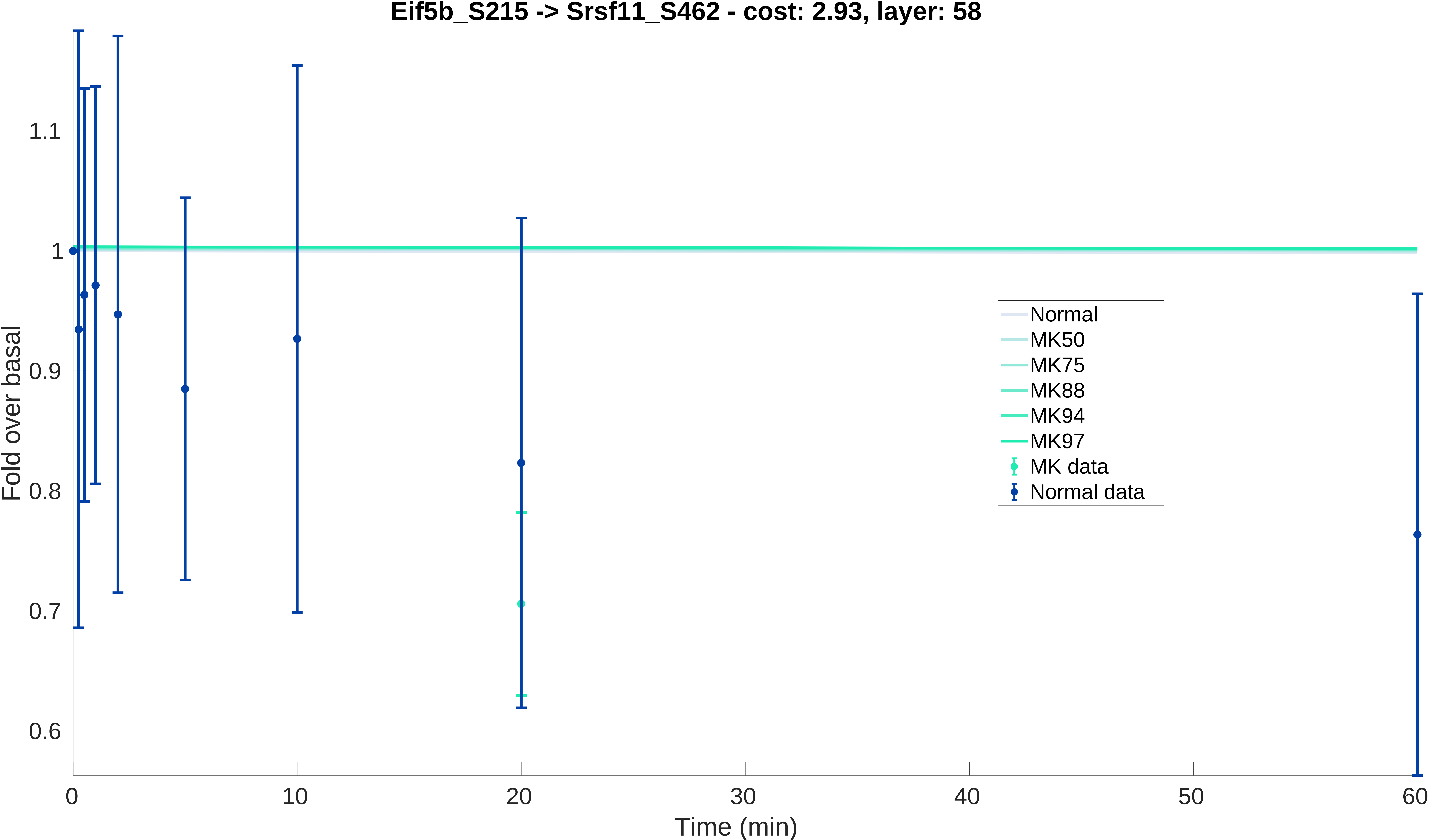

Eif5b_S215 -> Srsf11_S462 - cost: 2.93, layer: 58
Fold over basal
Time (min)
0.6
0.7
0.8
0.9
1
1.1
0
10
20
30
40
50
60
Normal
MK50
MK75
MK88
MK94
MK97
MK data
Normal data

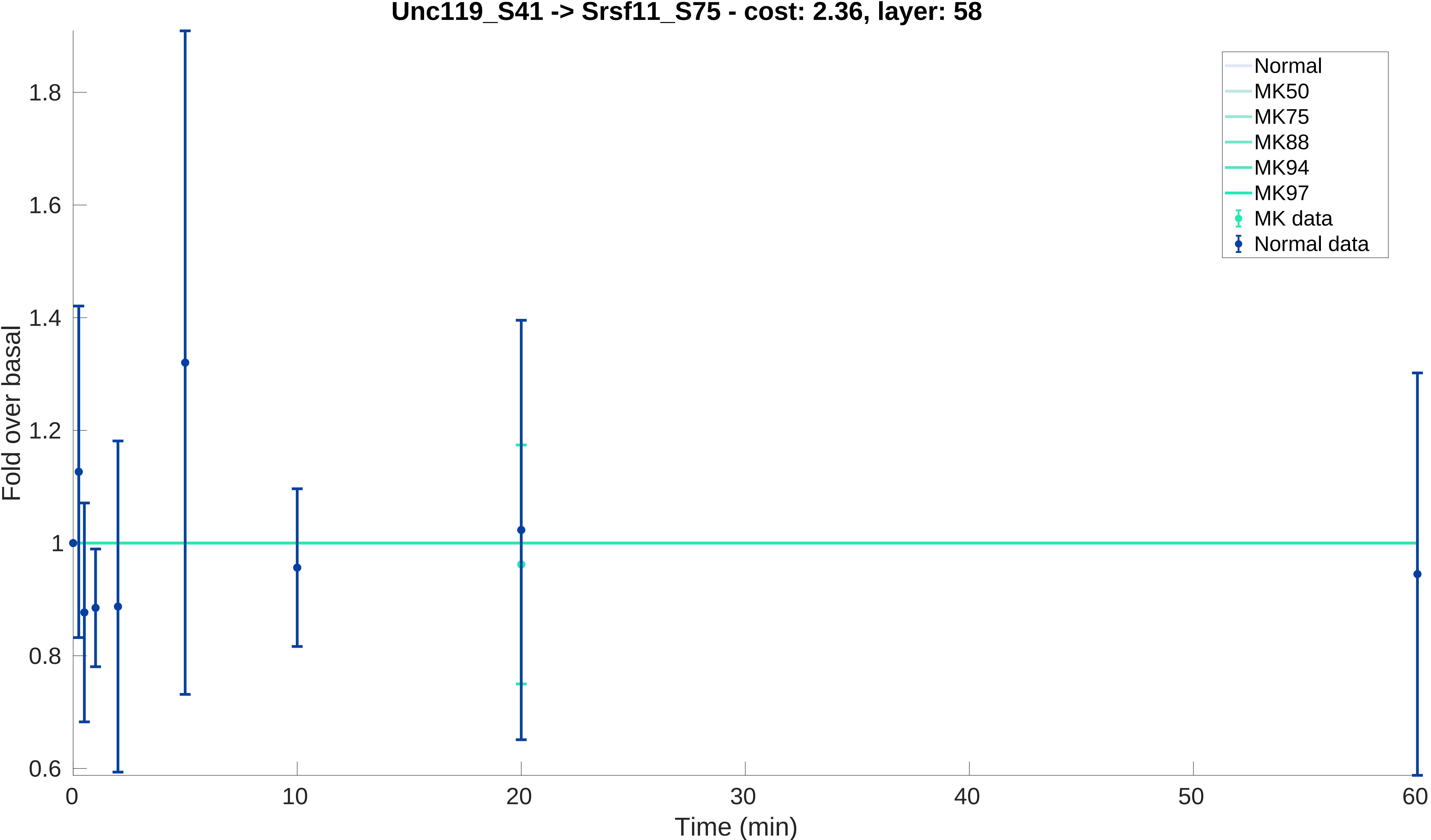

Unc119_S41 -> Srsf11_S75 - cost: 2.36, layer: 58
Fold over basal
Time (min)
0.6
0.8
1
1.2
1.4
1.6
1.8
0
10
20
30
40
50
60
Normal
MK50
MK75
MK88
MK94
MK97
MK data
Normal data

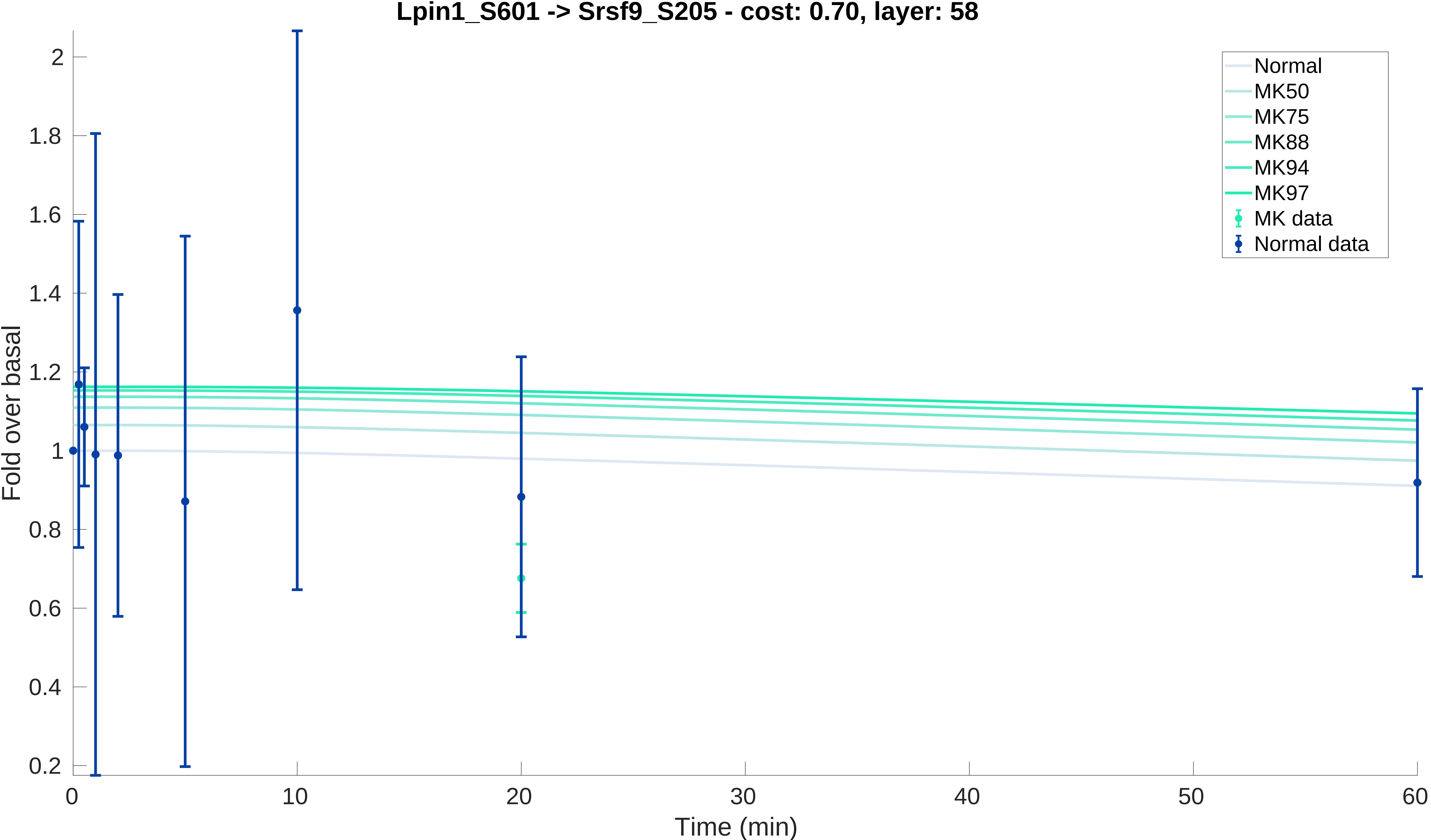

Lpin1_S601 -> Srsf9_S205 - cost: 0.70, layer: 58
Normal
MK50
MK75
MK88
MK94
MK97
MK data
Normal data
Fold over basal
Time (min)
0.2
0.4
0.6
0.8
1
1.2
1.4
1.6
1.8
2
0
10
20
30
40
50
60

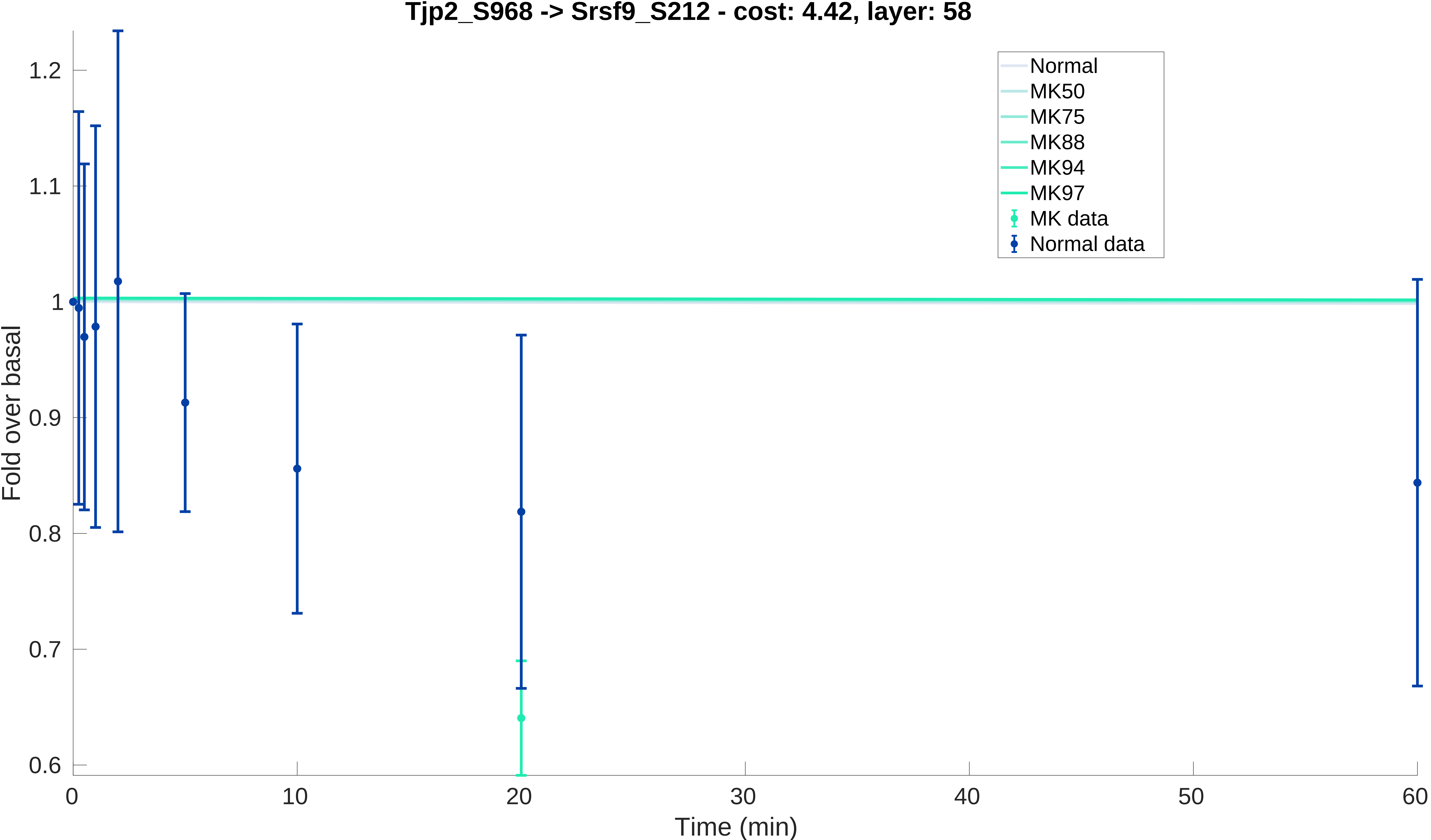

Tjp2_S968 -> Srsf9_S212 - cost: 4.42, layer: 58
Normal
MK50
MK75
MK88
MK94
MK97
MK data
Normal data
Fold over basal
Time (min)
1.2
1.1
1
0.9
0.8
0.7
0.6
0
10
20
30
40
50
60

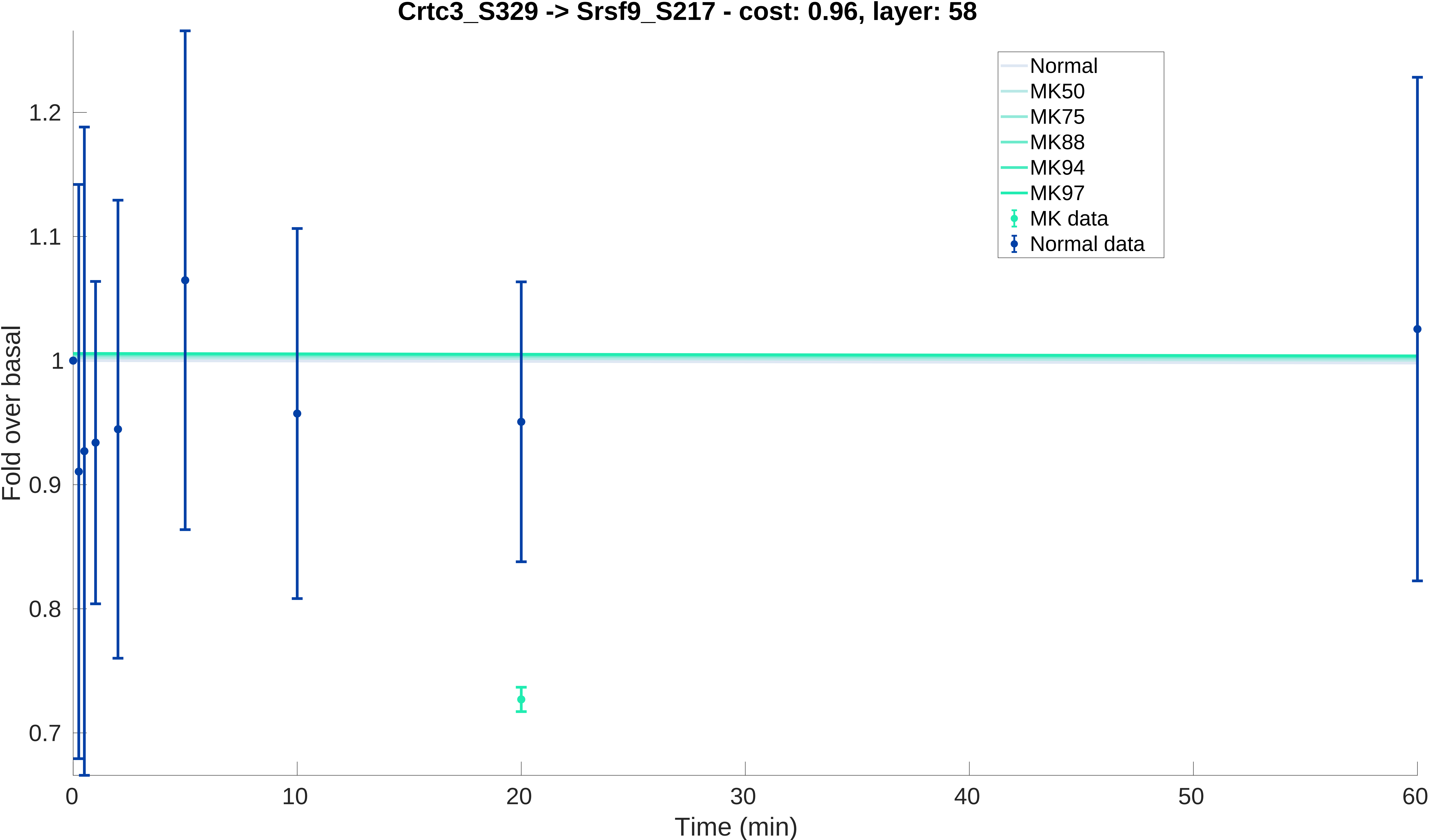

Crtc3_S329 -> Srsf9_S217 - cost: 0.96, layer: 58
Fold over basal
Time (min)
0.7
0.8
0.9
1
1.1
1.2
0
10
20
30
40
50
60
Normal
MK50
MK75
MK88
MK94
MK97
MK data
Normal data

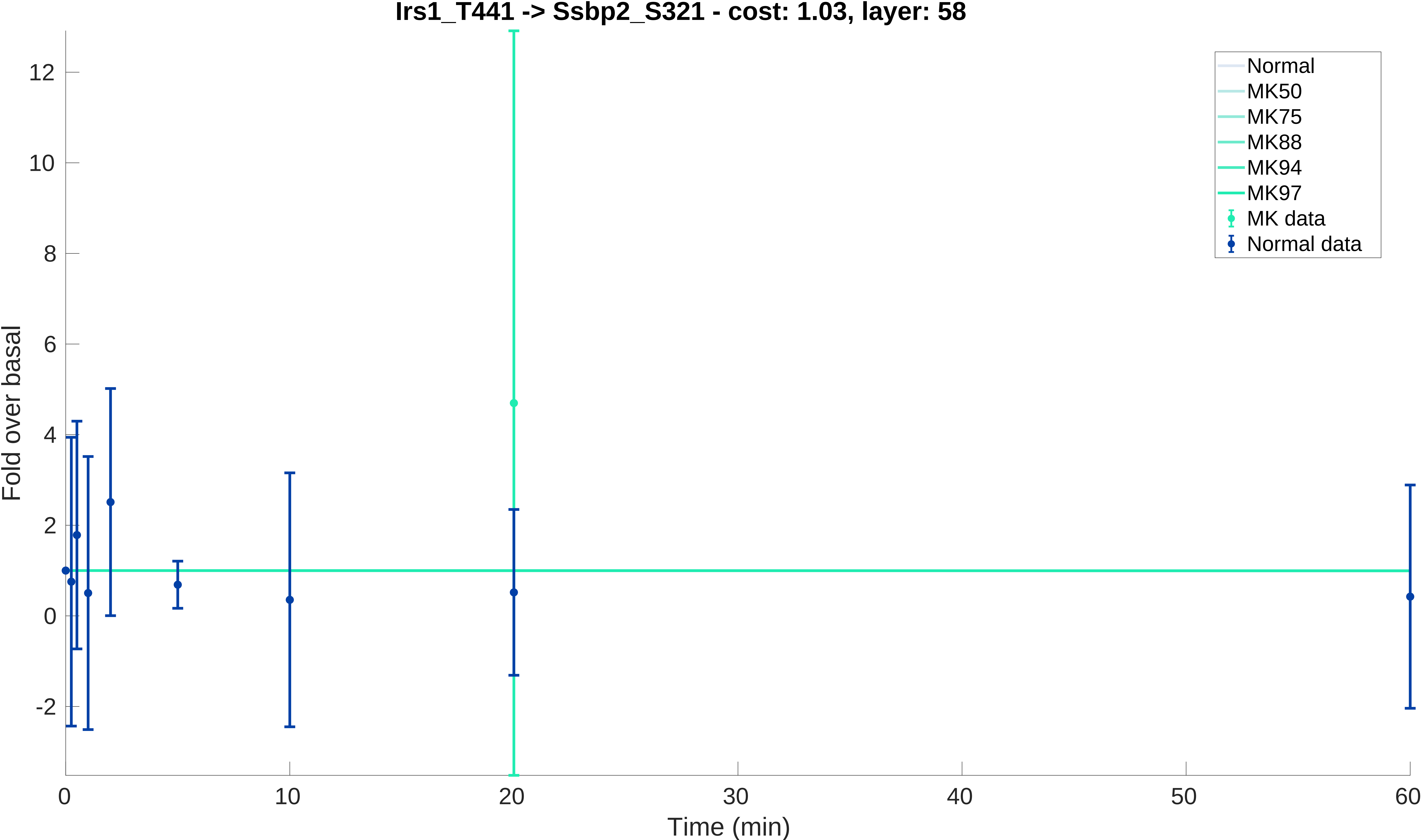
Irs1_T441 -> Ssbp2_S321 - cost: 1.03, layer: 58
Fold over basal
Time (min)
Normal
MK50
MK75
MK88
MK94
MK97
MK data
Normal data
12
10
8
6
4
2
0
-2
0
10
20
30
40
50
60

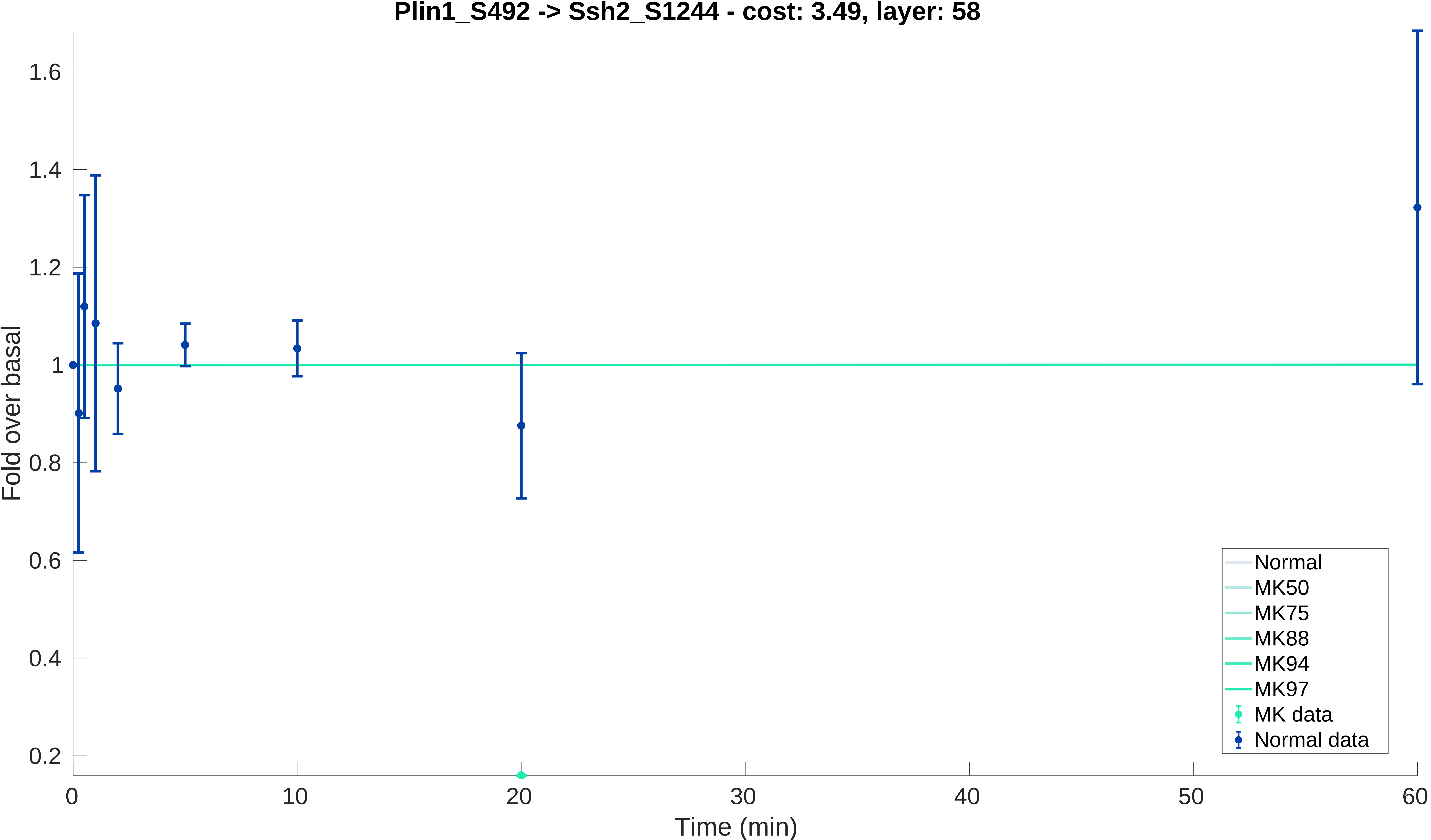

Plin1_S492 -> Ssh2_S1244 - cost: 3.49, layer: 58
Fold over basal
1.6
1.4
1.2
1
0.8
0.6
0.4
0.2
0
10
20
30
40
50
60
Time (min)
Normal
MK50
MK75
MK88
MK94
MK97
MK data
Normal data

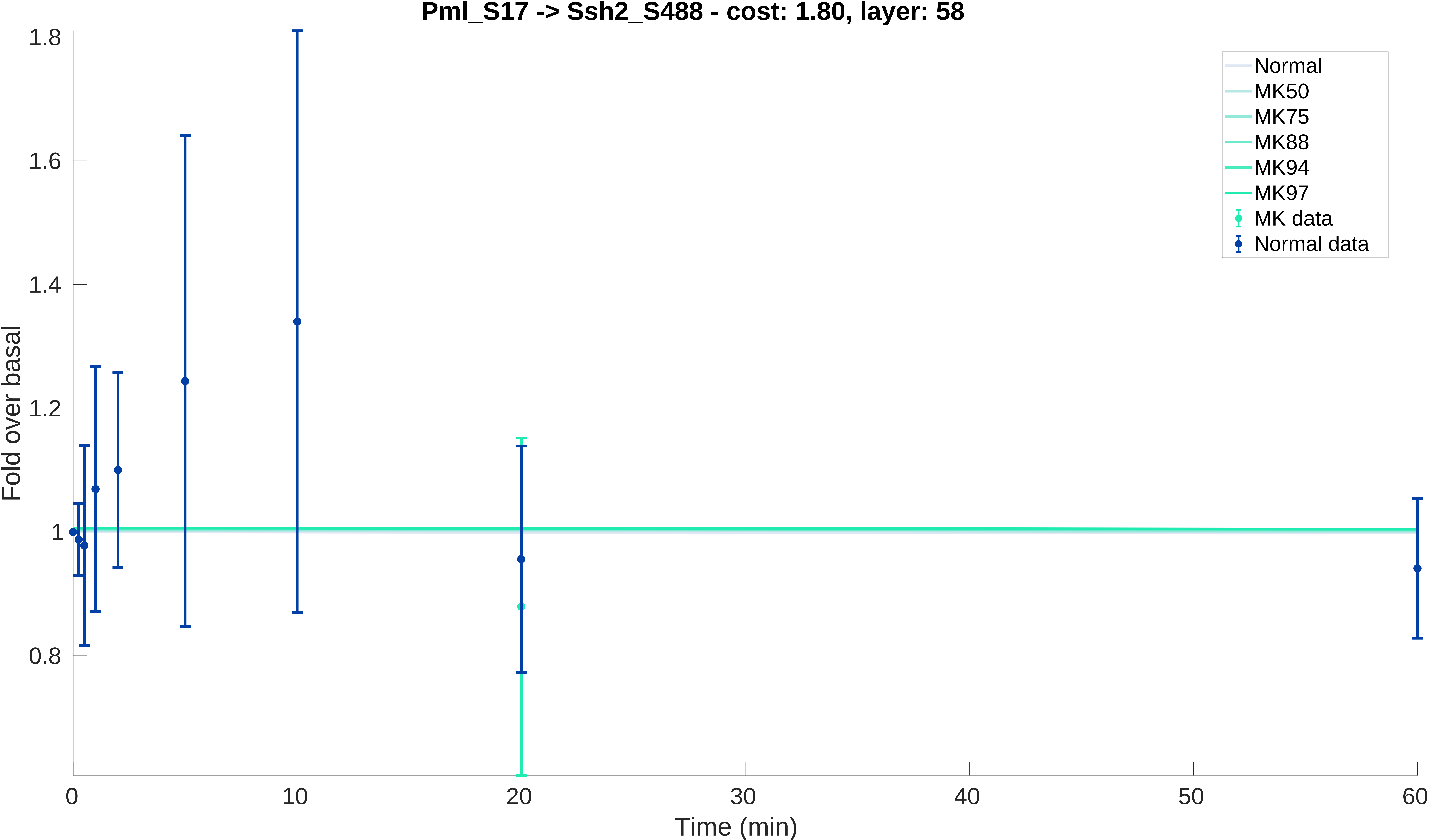

Pml_S17 -> Ssh2_S488 - cost: 1.80, layer: 58
Fold over basal
Time (min)
0
10
20
30
40
50
60
0.8
1
1.2
1.4
1.6
1.8
Normal
MK50
MK75
MK88
MK94
MK97
MK data
Normal data

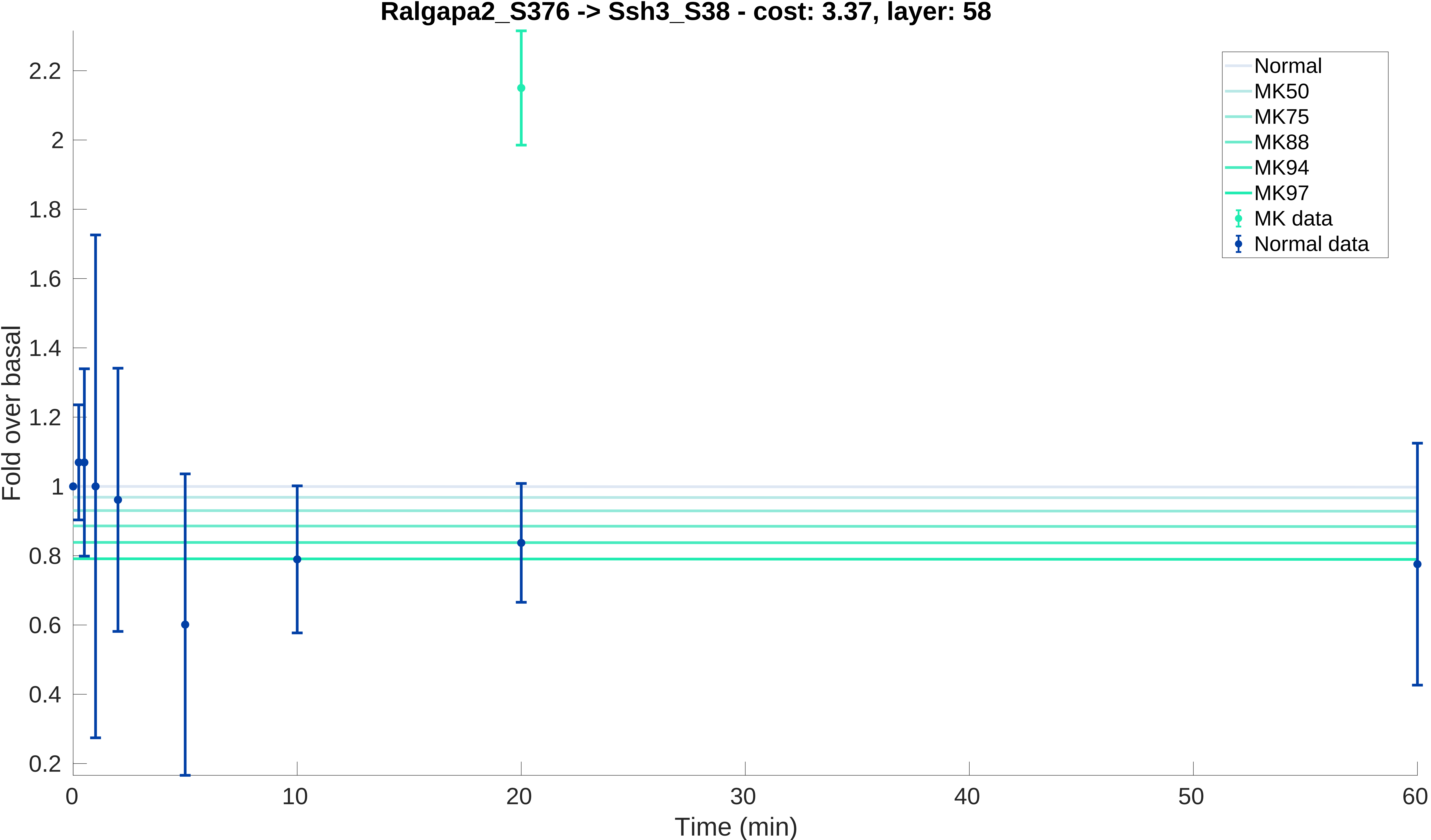
Ralgapa2_S376 -> Ssh3_S38 - cost: 3.37, layer: 58
Fold over basal
Time (min)
0.2
0.4
0.6
0.8
1
1.2
1.4
1.6
1.8
2
2.2
0
10
20
30
40
50
60
Normal
MK50
MK75
MK88
MK94
MK97
MK data
Normal data

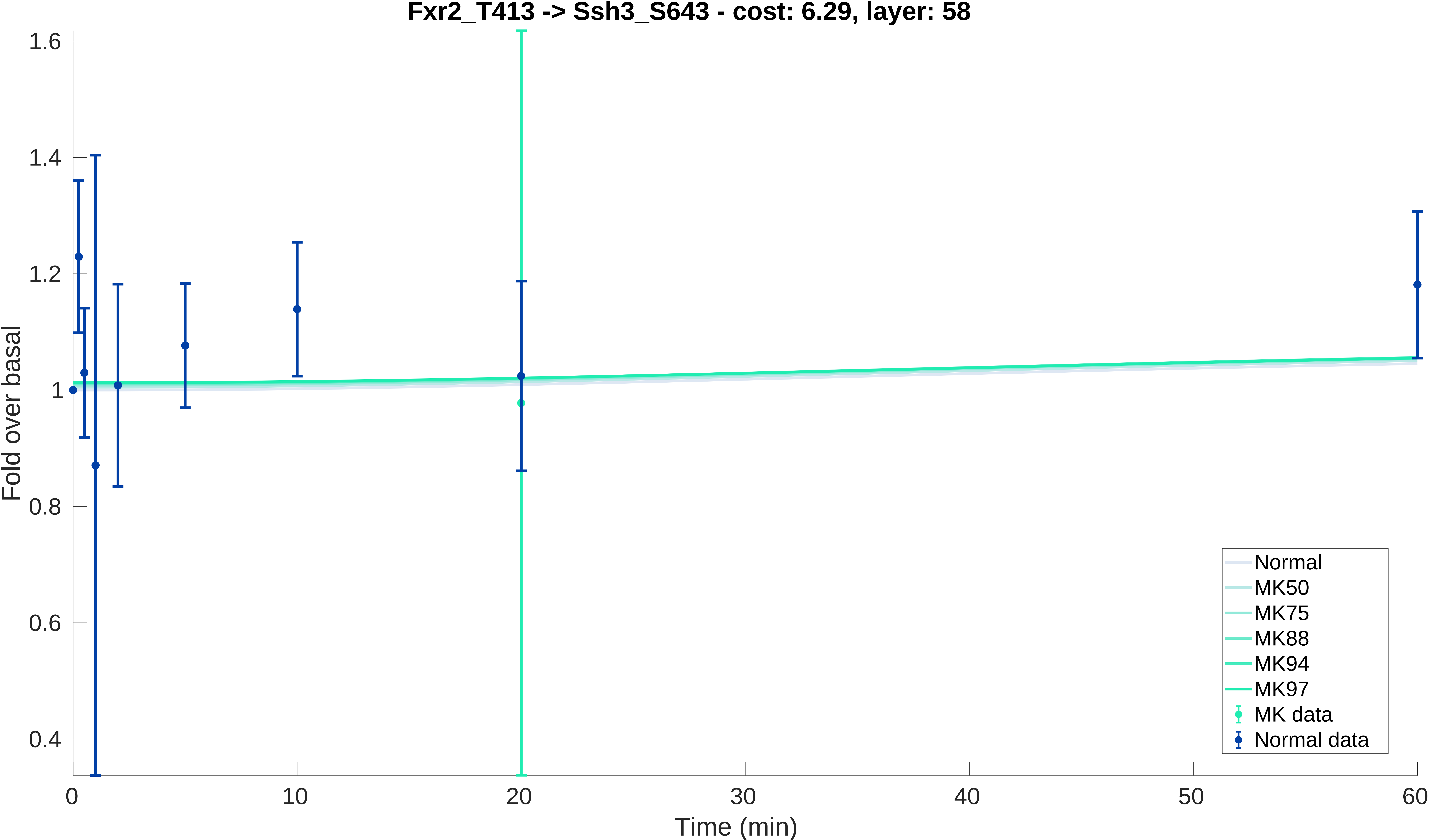

Fxr2_T413 -> Ssh3_S643 - cost: 6.29, layer: 58
Fold over basal
Time (min)
0.4
0.6
0.8
1
1.2
1.4
1.6
0
10
20
30
40
50
60
Normal
MK50
MK75
MK88
MK94
MK97
MK data
Normal data

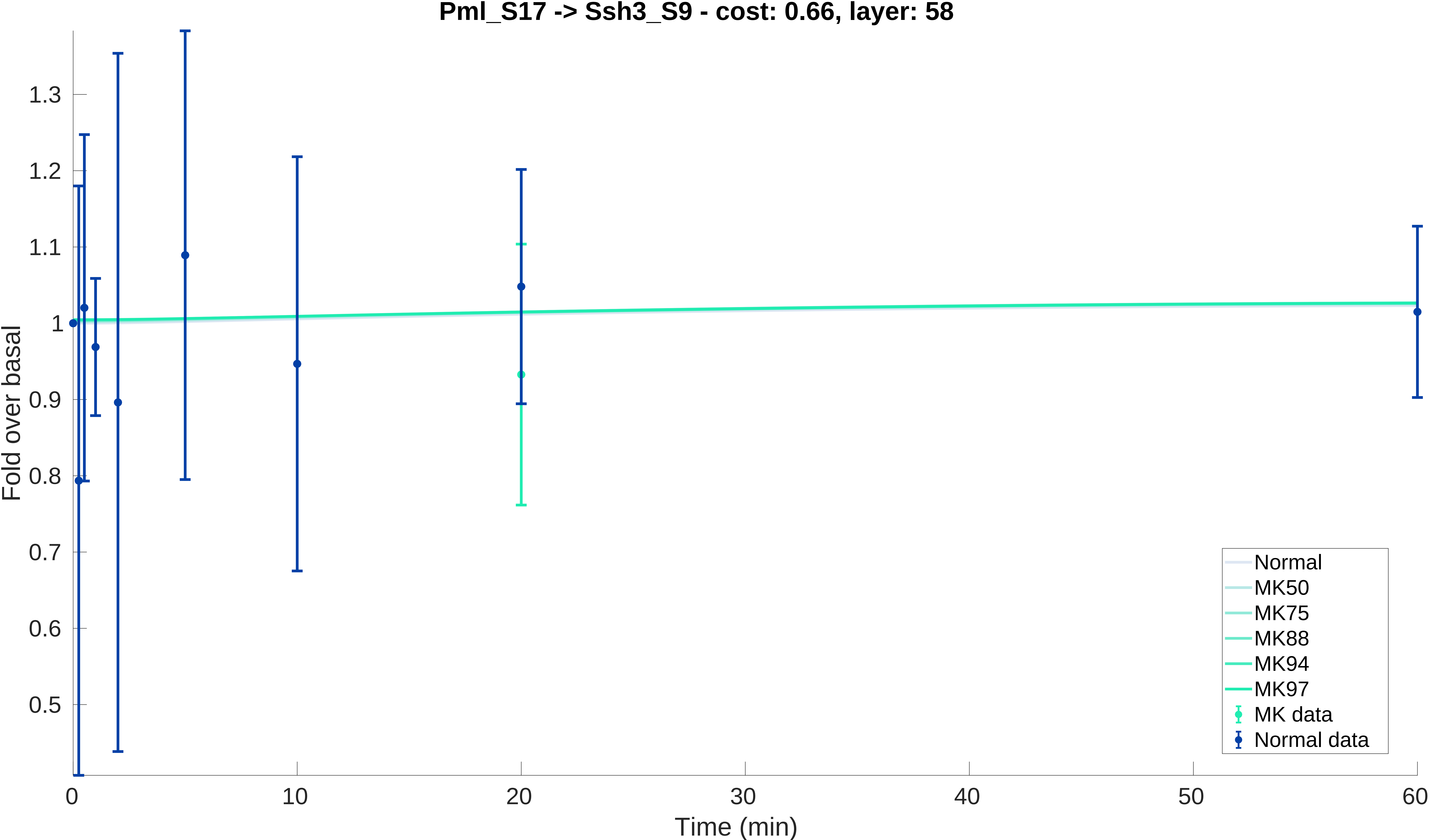

Pml_S17 -> Ssh3_S9 - cost: 0.66, layer: 58
Fold over basal
1.3
1.2
1.1
1
0.9
0.8
0.7
0.6
0.5
0
10
20
30
40
50
60
Time (min)
Normal
MK50
MK75
MK88
MK94
MK97
MK data
Normal data

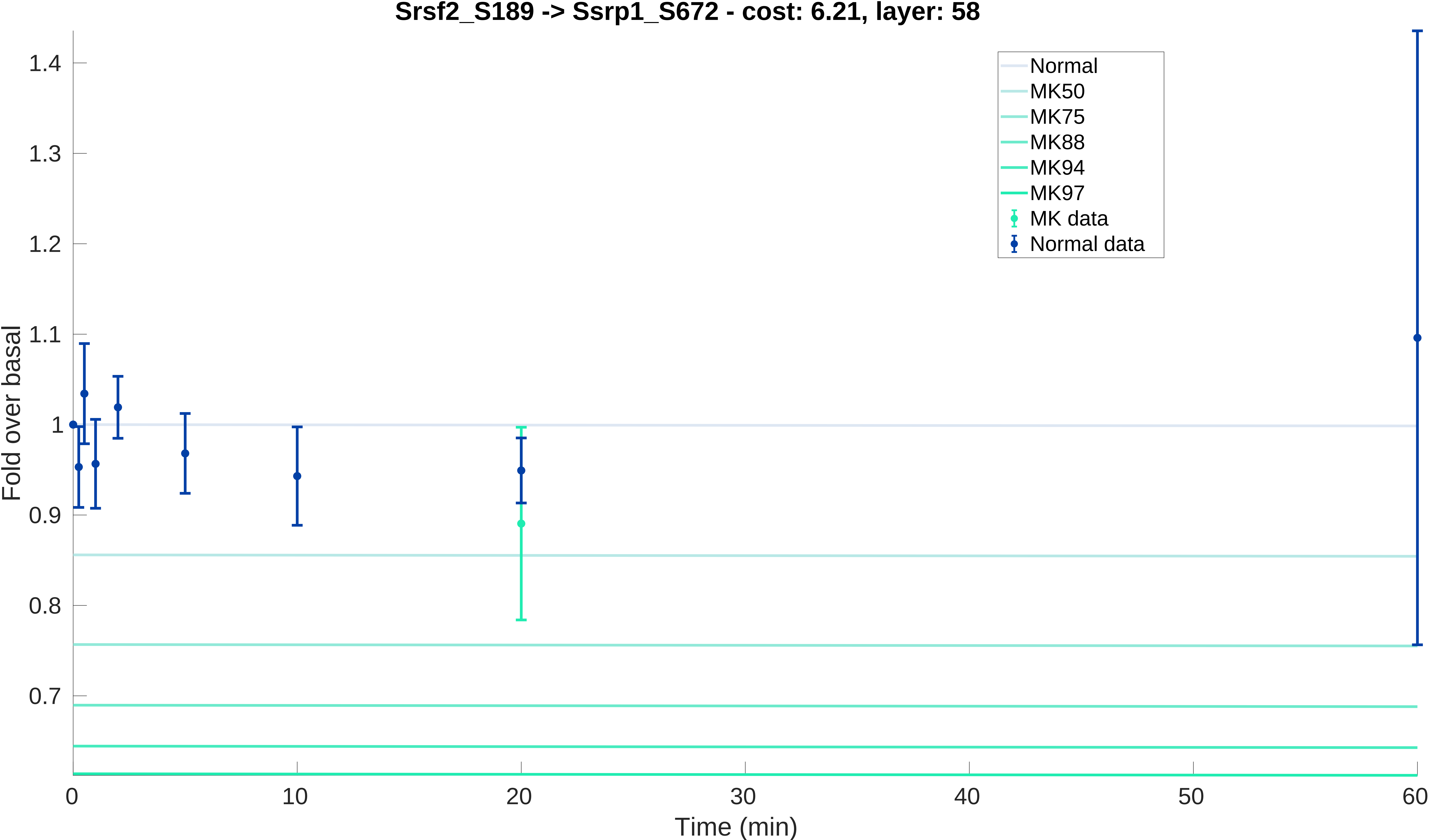

Srsf2_S189 -> Ssrp1_S672 - cost: 6.21, layer: 58
Fold over basal
1.4
1.3
1.2
1.1
1
0.9
0.8
0.7
0
10
20
30
40
50
60
Time (min)
Normal
MK50
MK75
MK88
MK94
MK97
MK data
Normal data

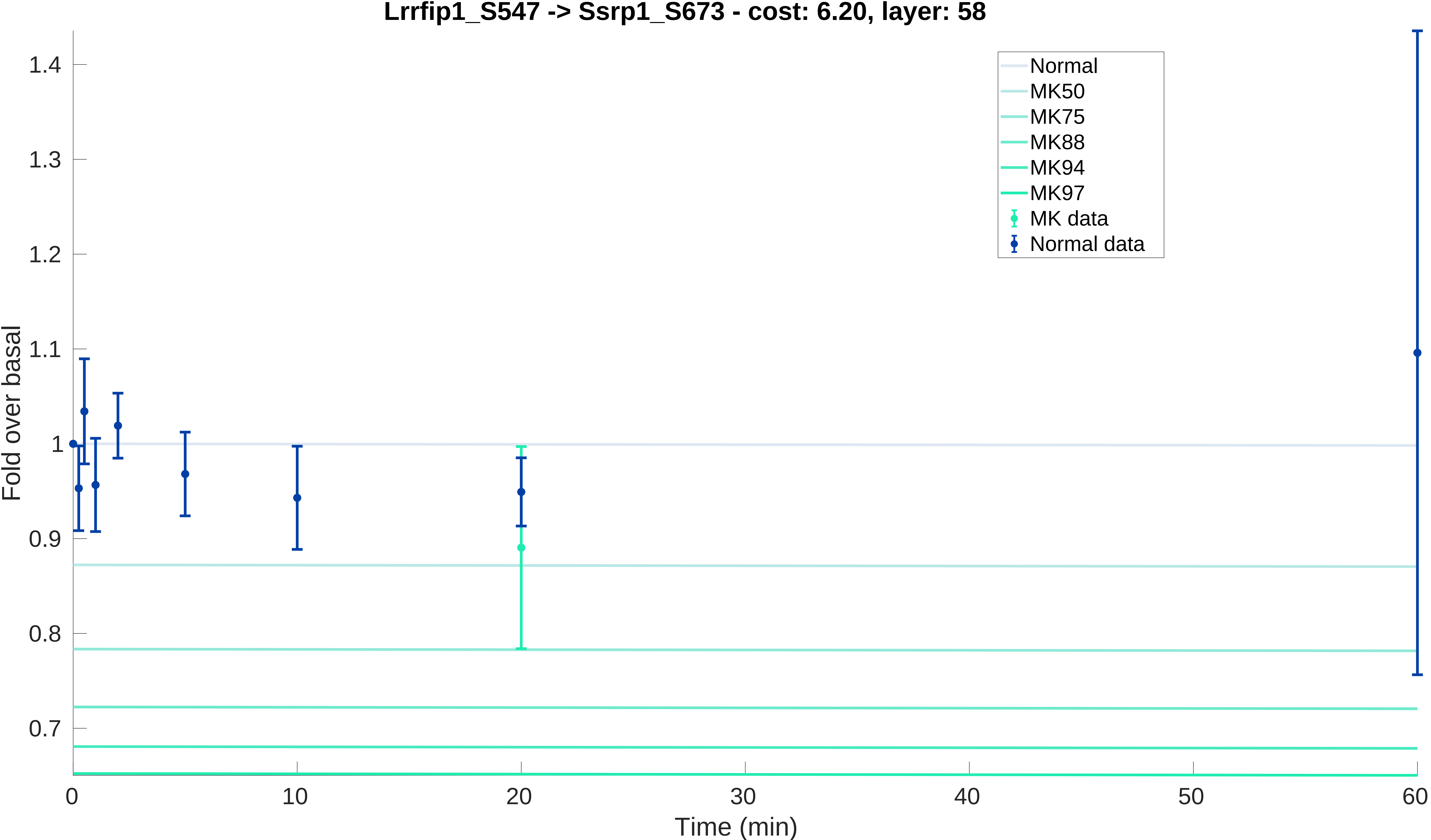

Lrrfip1_S547 -> Ssrp1_S673 - cost: 6.20, layer: 58
Fold over basal
1.4
1.3
1.2
1.1
1
0.9
0.8
0.7
0
10
20
30
40
50
60
Time (min)
Normal
MK50
MK75
MK88
MK94
MK97
MK data
Normal data

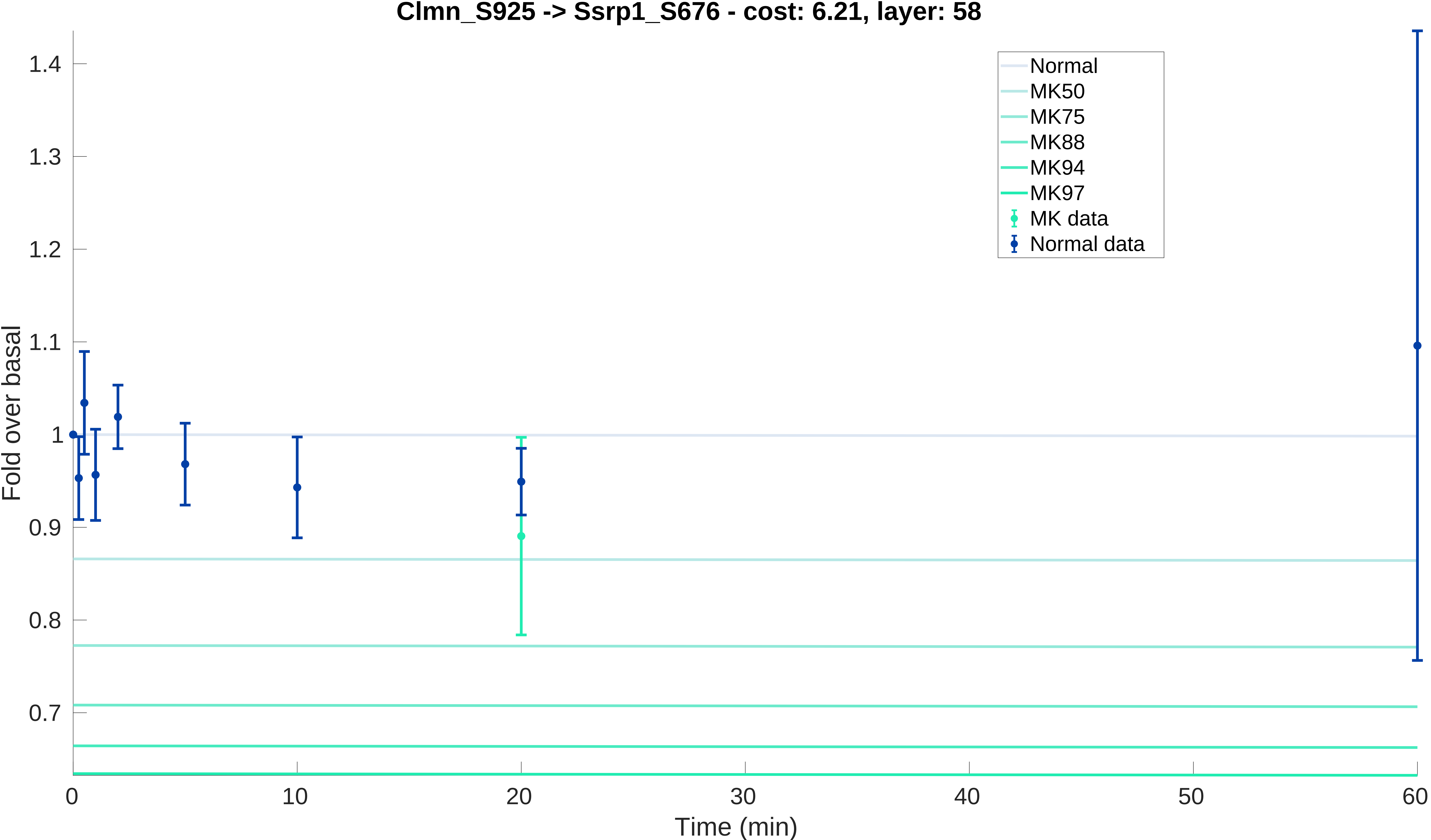

Clmn_S925 -> Ssrp1_S676 - cost: 6.21, layer: 58
Normal
MK50
MK75
MK88
MK94
MK97
MK data
Normal data
Fold over basal
1.4
1.3
1.2
1.1
1
0.9
0.8
0.7
0
10
20
30
40
50
60
Time (min)

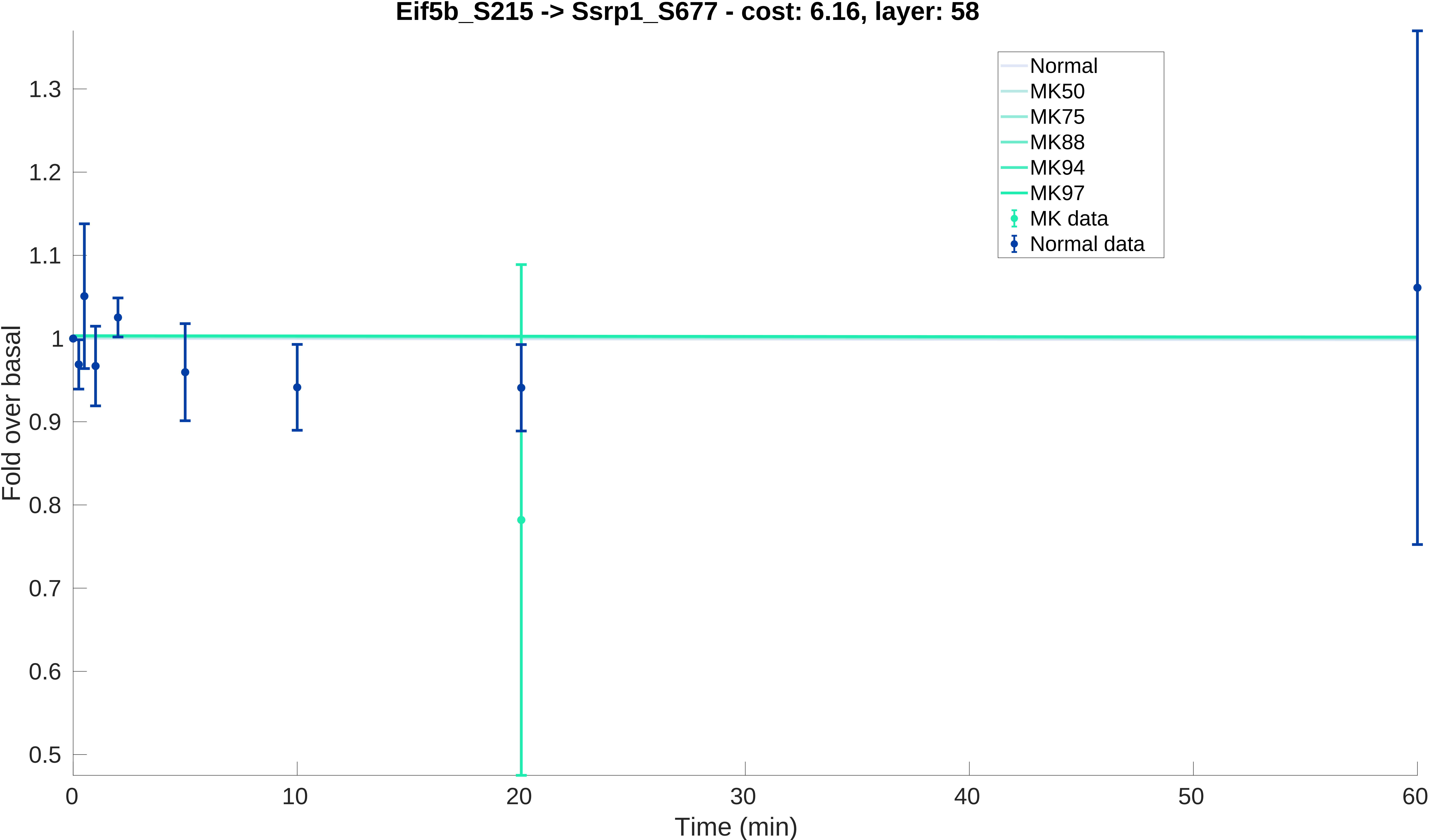

Eif5b_S215 -> Ssrp1_S677 - cost: 6.16, layer: 58
Normal
MK50
MK75
MK88
MK94
MK97
MK data
Normal data
Fold over basal
1.3
1.2
1.1
1
0.9
0.8
0.7
0.6
0.5
0
10
20
30
40
50
60
Time (min)

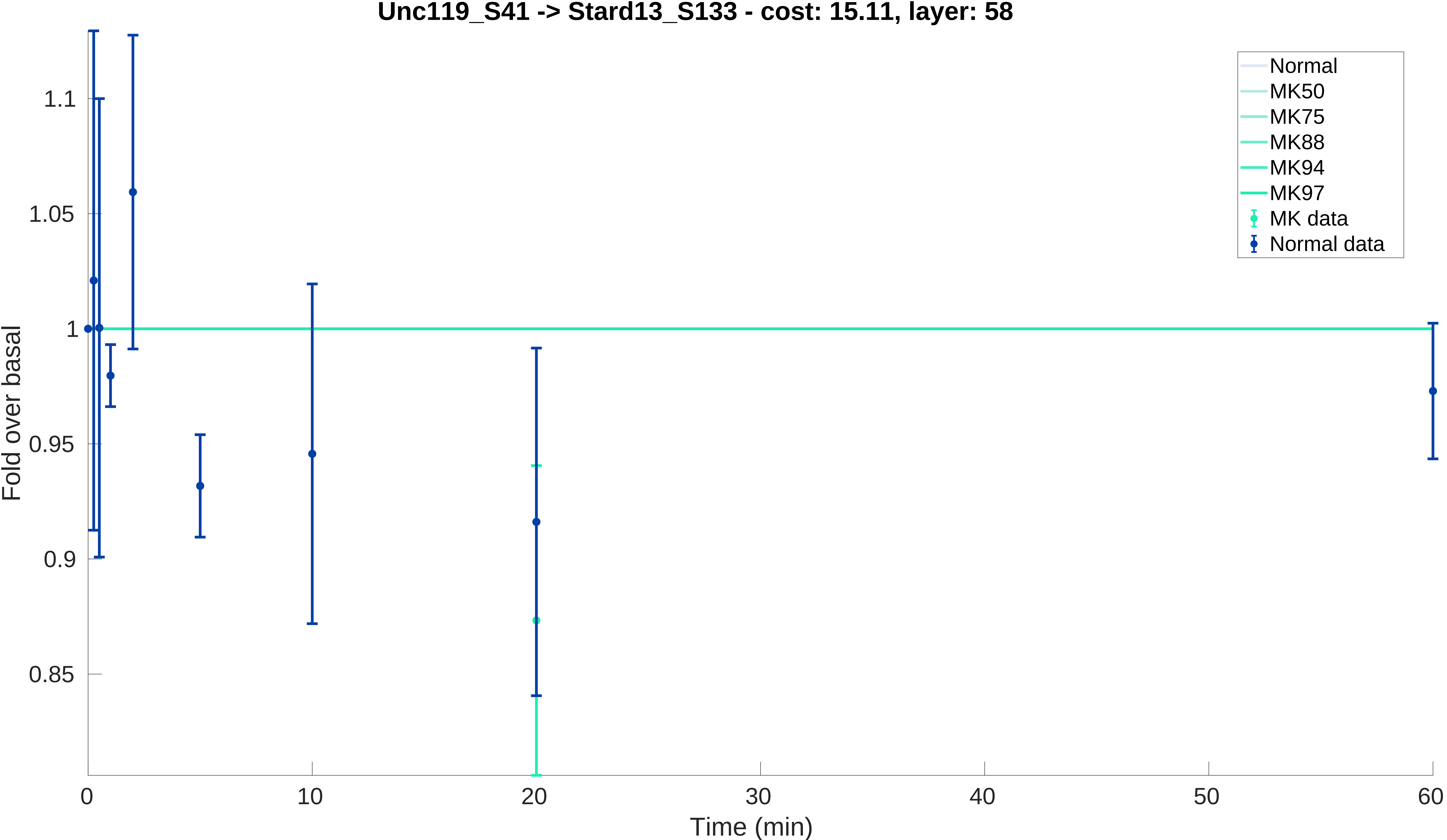

Unc119_S41 -> Stard13_S133 - cost: 15.11, layer: 58
Normal
MK50
MK75
MK88
MK94
MK97
MK data
Normal data
Fold over basal
1.1
1.05
1
0.95
0.9
0.85
0
10
20
30
40
50
60
Time (min)

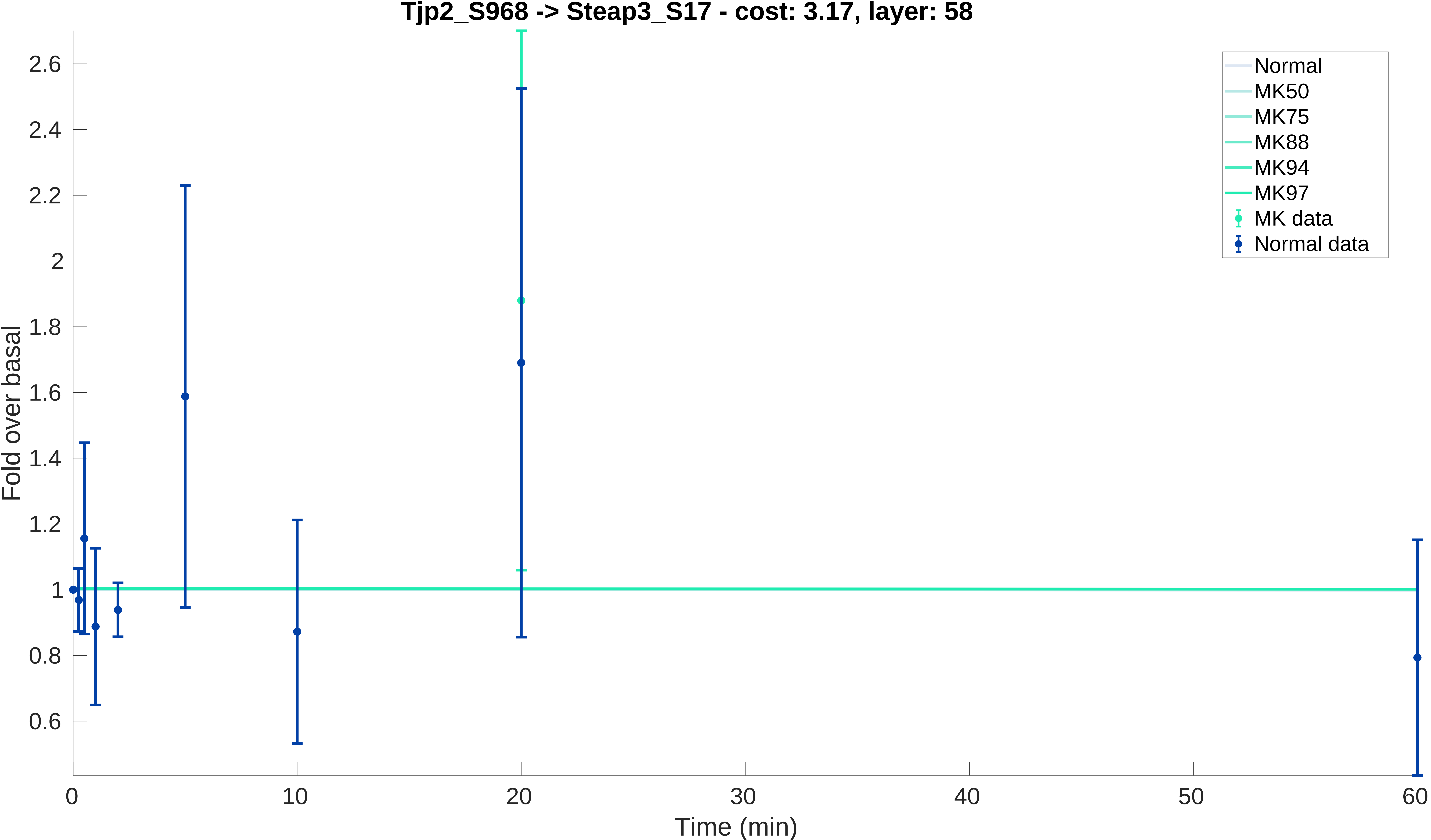

Tjp2_S968 -> Steap3_S17 - cost: 3.17, layer: 58
Fold over basal
Time (min)
0.6
0.8
1
1.2
1.4
1.6
1.8
2
2.2
2.4
2.6
0
10
20
30
40
50
60
Normal
MK50
MK75
MK88
MK94
MK97
MK data
Normal data

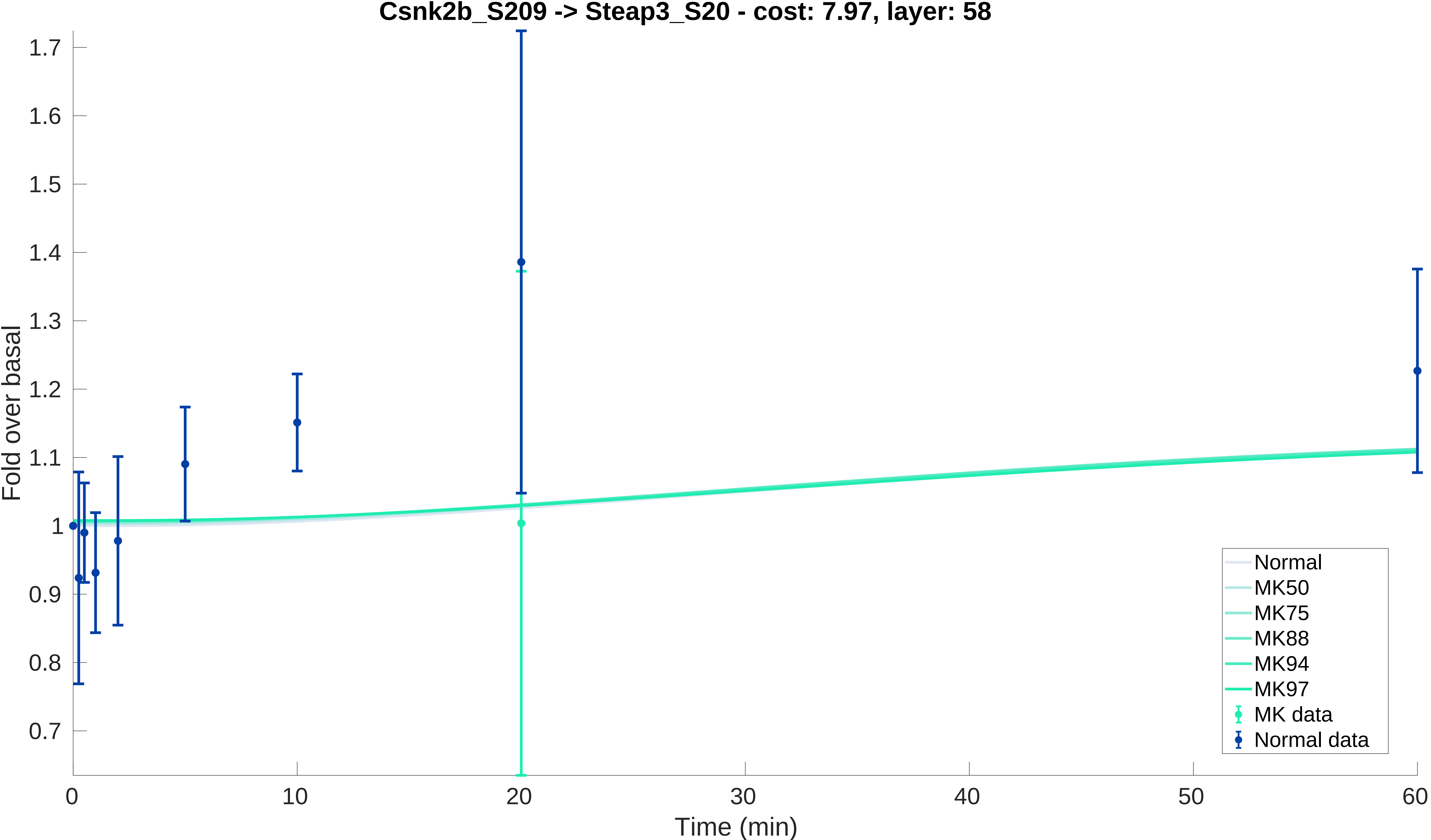

Csnk2b_S209 -> Steap3_S20 - cost: 7.97, layer: 58
Fold over basal
Time (min)
1.7
1.6
1.5
1.4
1.3
1.2
1.1
1
0.9
0.8
0.7
0
10
20
30
40
50
60
Normal
MK50
MK75
MK88
MK94
MK97
MK data
Normal data

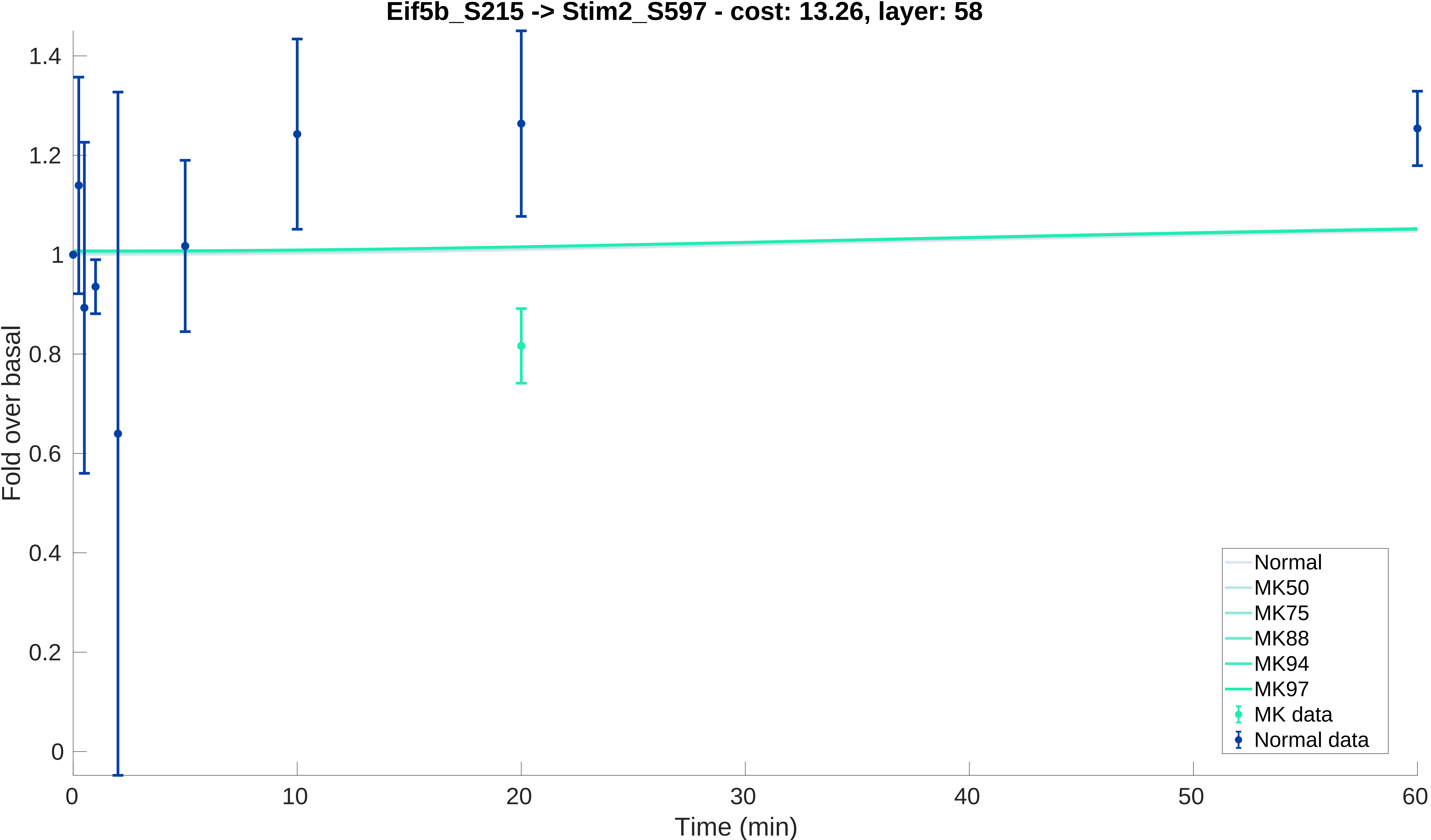
Eif5b_S215 -> Stim2_S597 - cost: 13.26, layer: 58
Fold over basal
Time (min)
0
0.2
0.4
0.6
0.8
1
1.2
1.4
0
10
20
30
40
50
60
Normal
MK50
MK75
MK88
MK94
MK97
MK data
Normal data

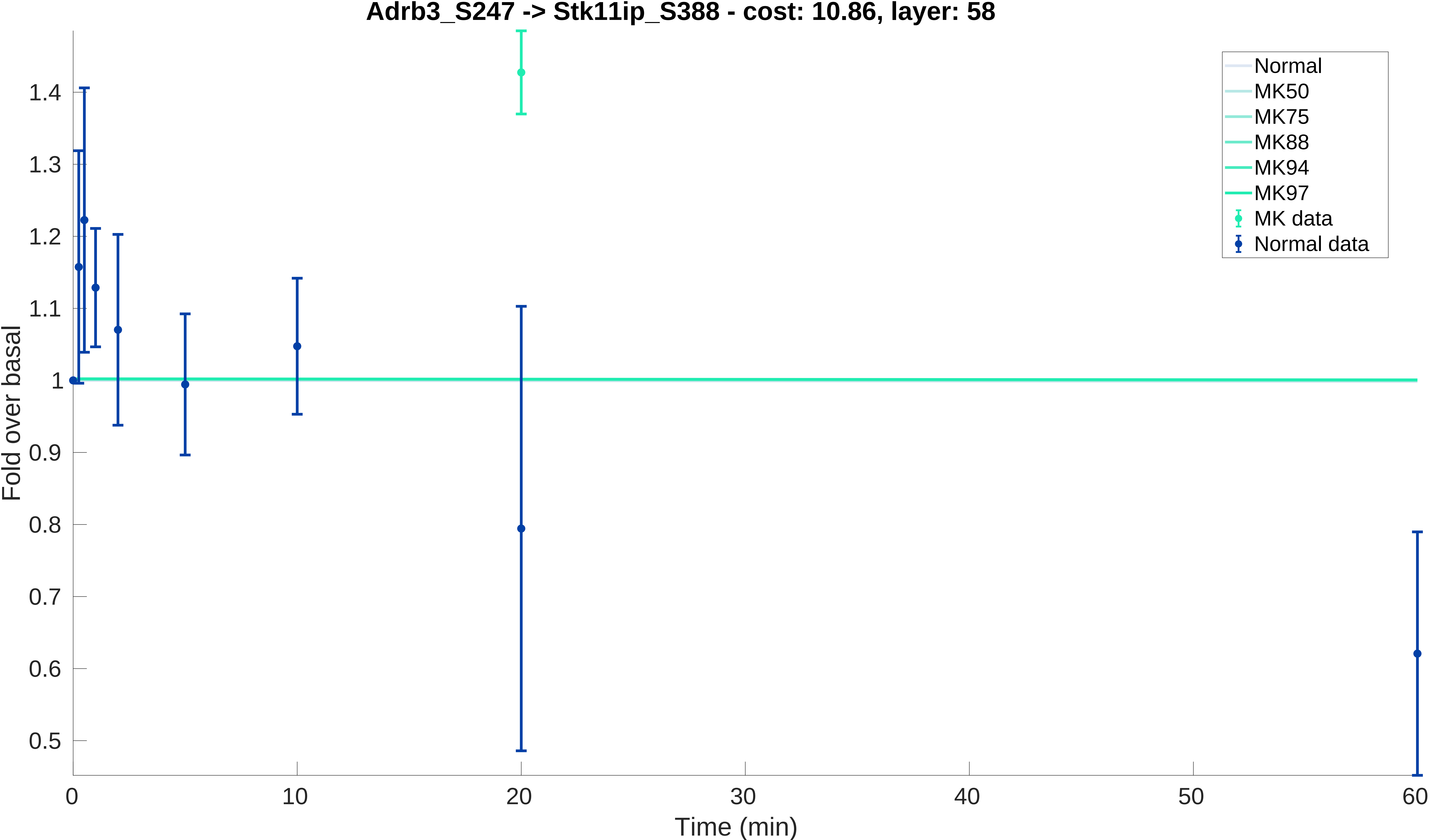

Adrb3_S247 -> Stk11ip_S388 - cost: 10.86, layer: 58
Normal
MK50
MK75
MK88
MK94
MK97
MK data
Normal data
Fold over basal
Time (min)
1.4
1.3
1.2
1.1
1
0.9
0.8
0.7
0.6
0.5
0
10
20
30
40
50
60

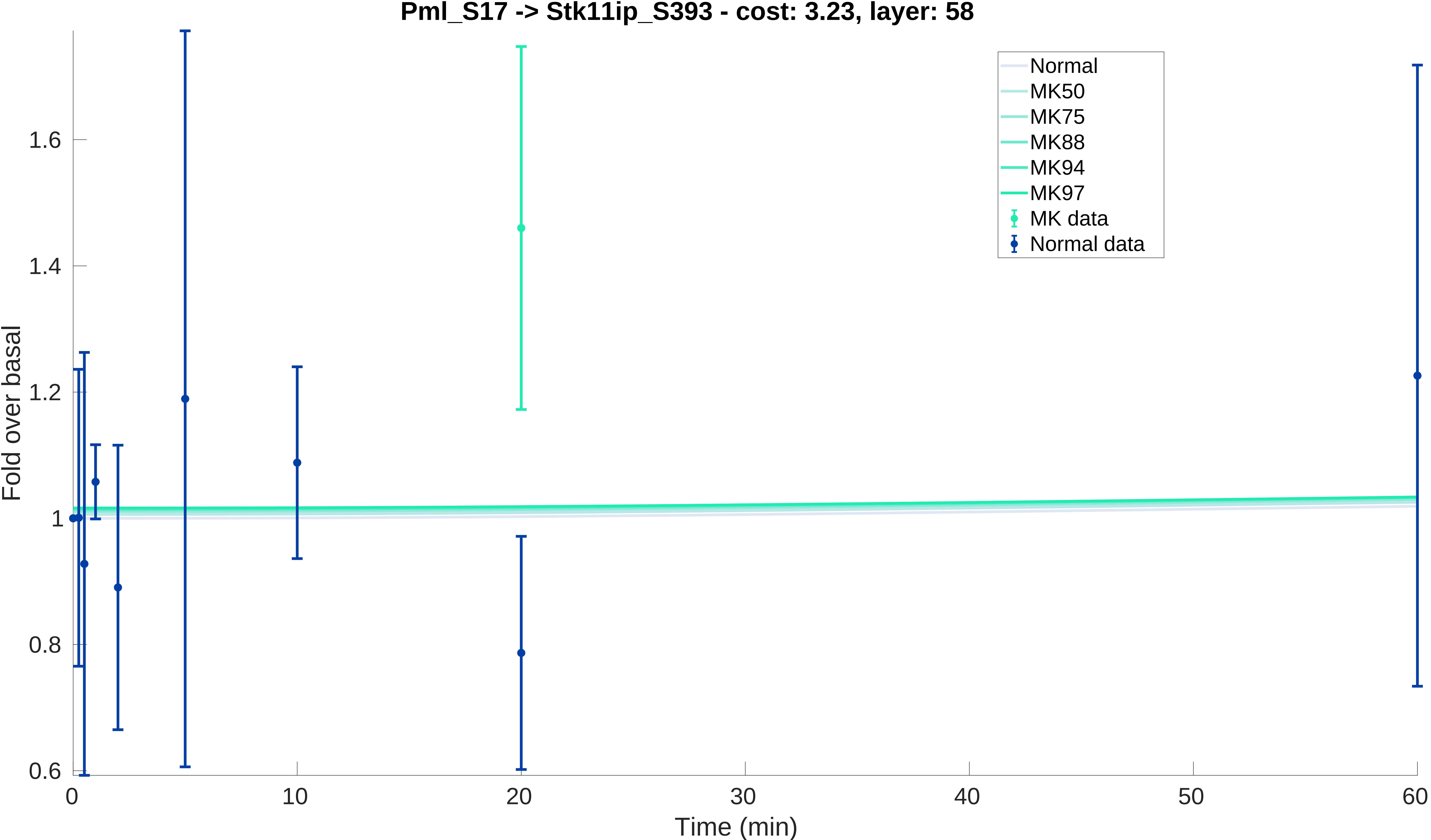

Pml_S17 -> Stk11ip_S393 - cost: 3.23, layer: 58
Normal
MK50
MK75
MK88
MK94
MK97
MK data
Normal data
Fold over basal
1.6
1.4
1.2
1
0.8
0.6
0
10
20
30
40
50
60
Time (min)

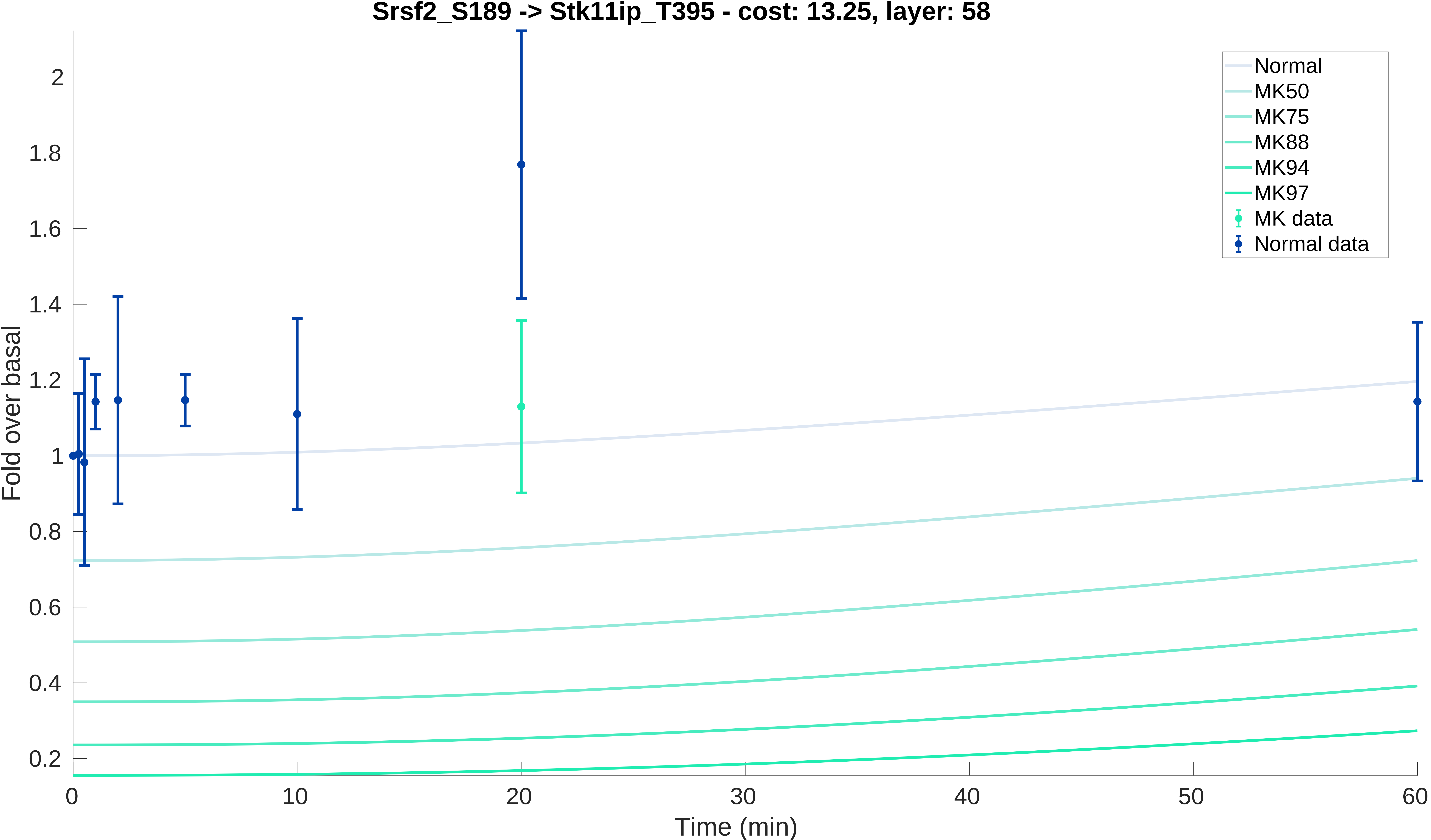

Srsf2_S189 -> Stk11ip_T395 - cost: 13.25, layer: 58
Fold over basal
Time (min)
Normal
MK50
MK75
MK88
MK94
MK97
MK data
Normal data
0.2
0.4
0.6
0.8
1
1.2
1.4
1.6
1.8
2
0
10
20
30
40
50
60

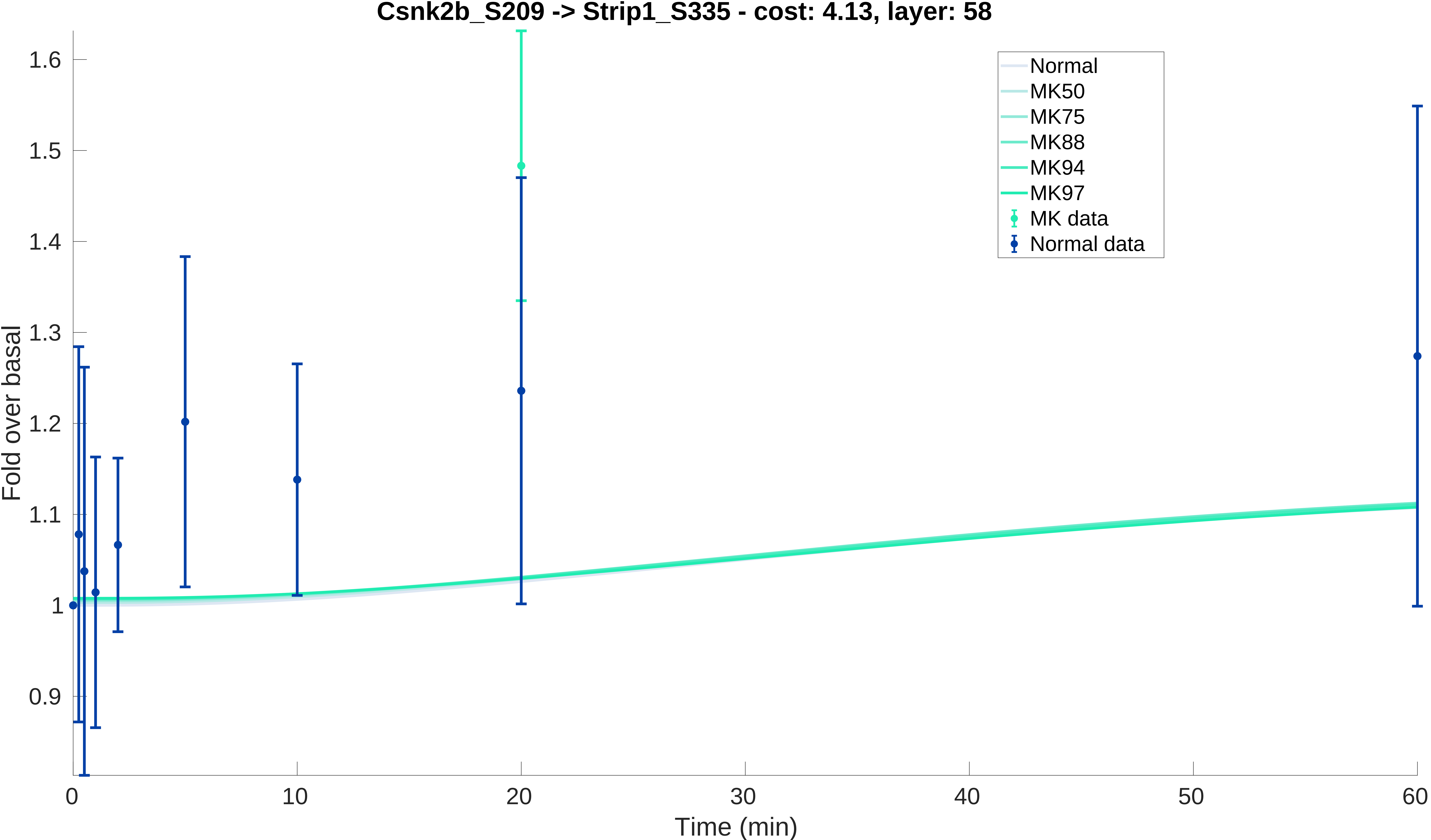

Csnk2b_S209 -> Strip1_S335 - cost: 4.13, layer: 58
Normal
MK50
MK75
MK88
MK94
MK97
MK data
Normal data
Fold over basal
Time (min)
1.6
1.5
1.4
1.3
1.2
1.1
1
0.9
0
10
20
30
40
50
60

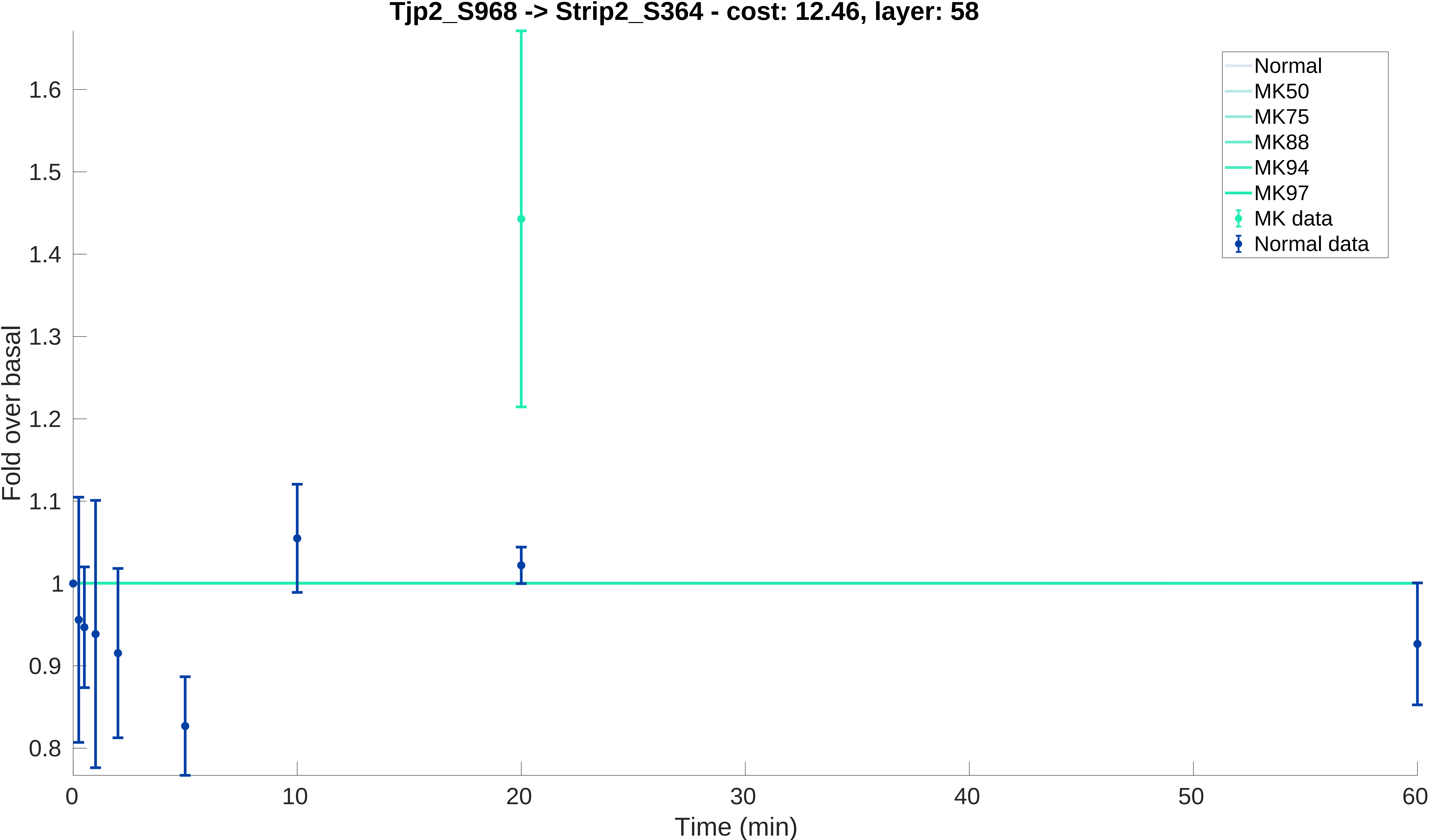

Tjp2_S968 -> Strip2_S364 - cost: 12.46, layer: 58
Fold over basal
Time (min)
0
10
20
30
40
50
60
0.8
0.9
1
1.1
1.2
1.3
1.4
1.5
1.6
Normal
MK50
MK75
MK88
MK94
MK97
MK data
Normal data

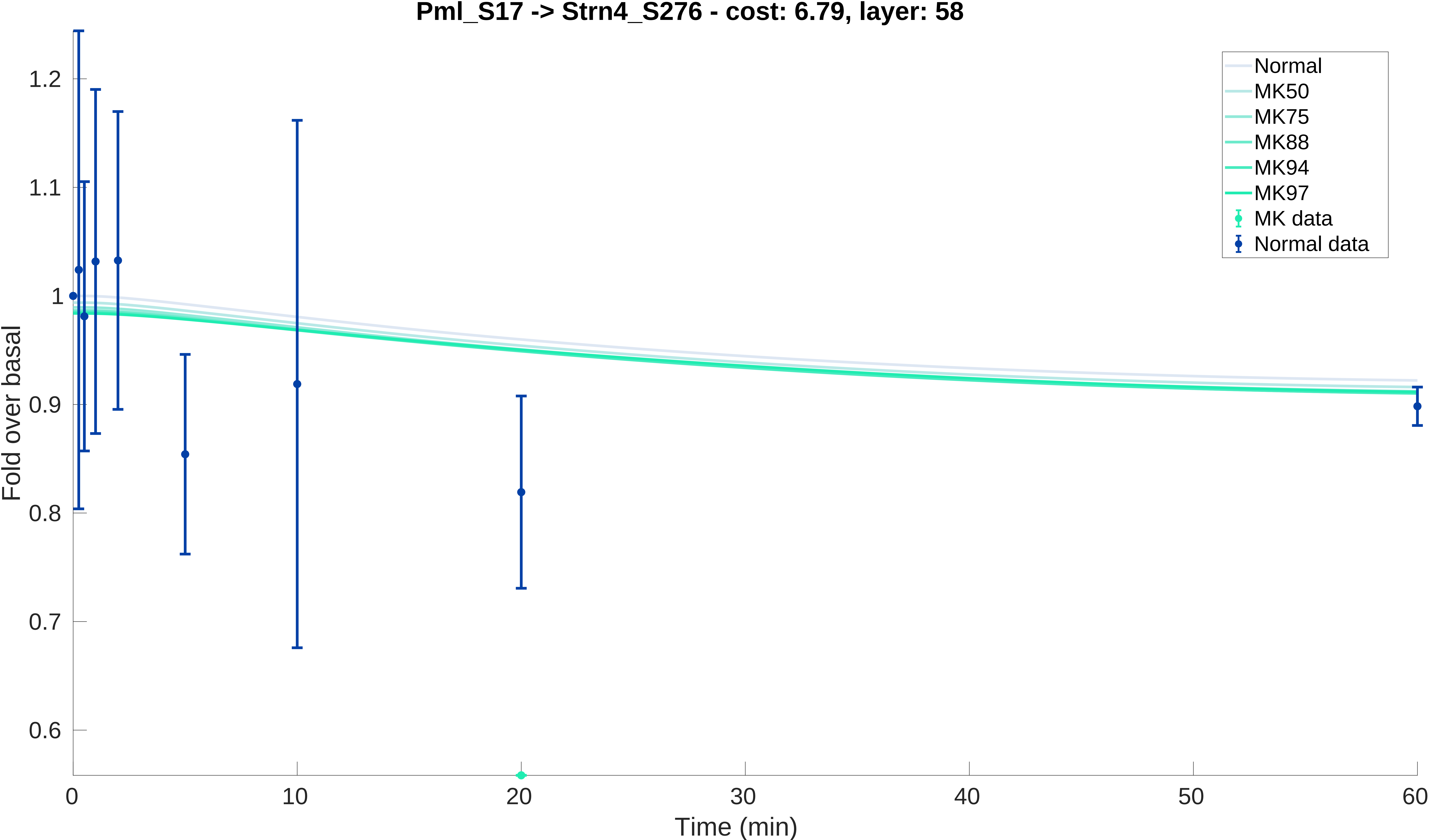

Pml_S17 -> Strn4_S276 - cost: 6.79, layer: 58
Fold over basal
Time (min)
1.2
1.1
1
0.9
0.8
0.7
0.6
0
10
20
30
40
50
60
Normal
MK50
MK75
MK88
MK94
MK97
MK data
Normal data

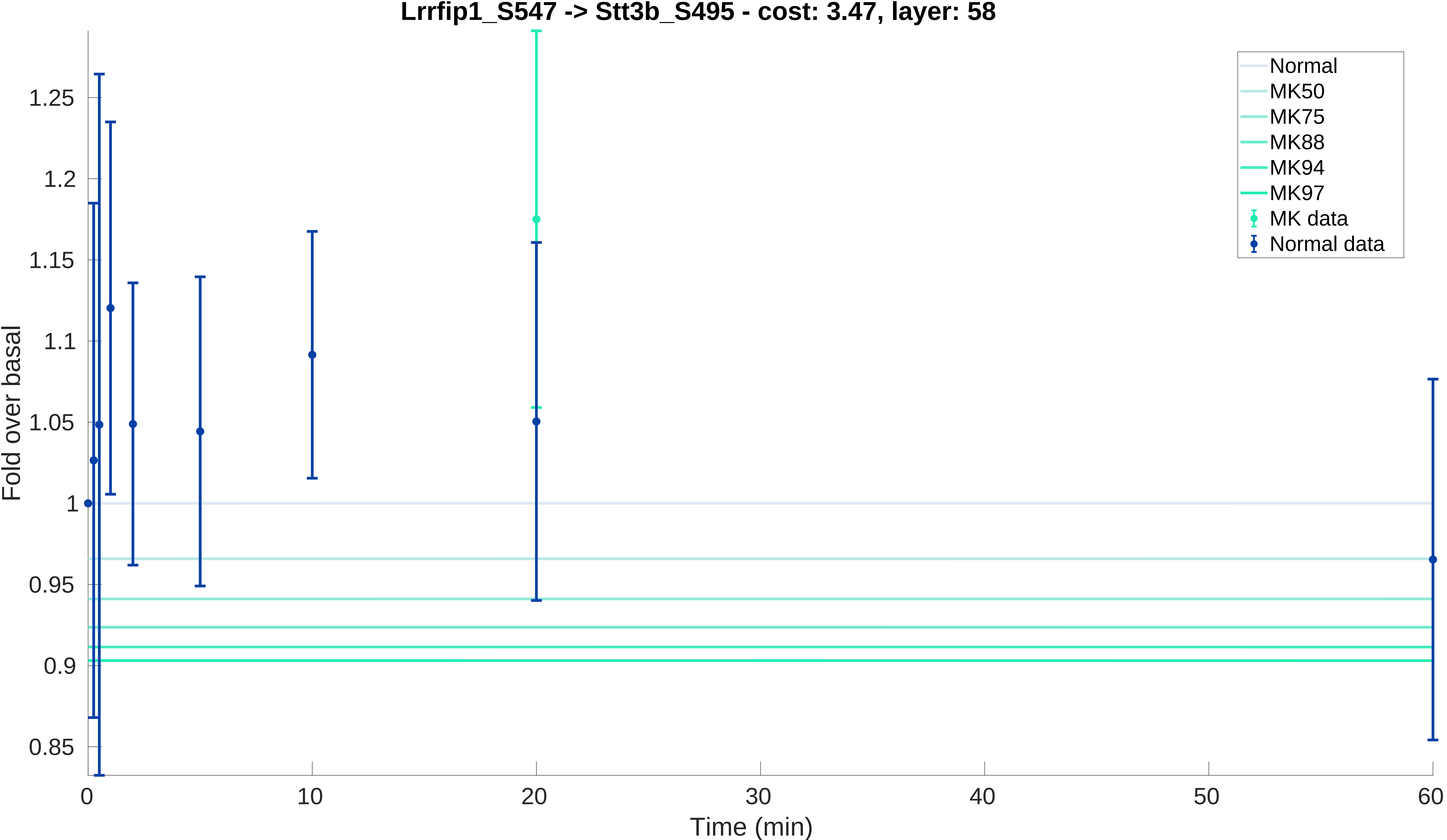

Lrrfip1_S547 -> Stt3b_S495 - cost: 3.47, layer: 58
Fold over basal
1.25
1.2
1.15
1.1
1.05
1
0.95
0.9
0.85
0
10
20
30
40
50
60
Time (min)
Normal
MK50
MK75
MK88
MK94
MK97
MK data
Normal data

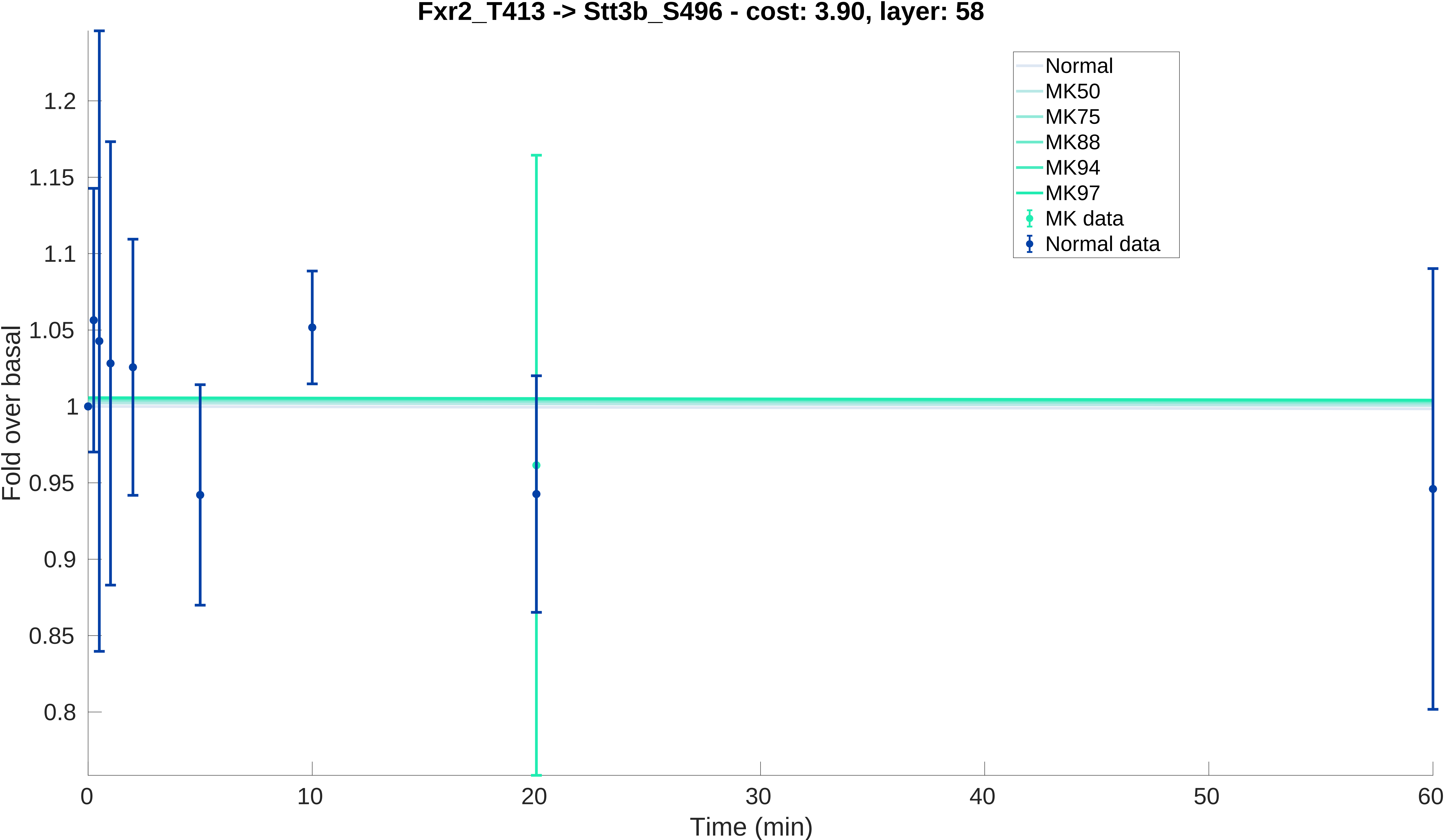

Fxr2_T413 -> Stt3b_S496 - cost: 3.90, layer: 58
Normal
MK50
MK75
MK88
MK94
MK97
MK data
Normal data
Fold over basal
1.2
1.15
1.1
1.05
1
0.95
0.9
0.85
0.8
0
10
20
30
40
50
60
Time (min)

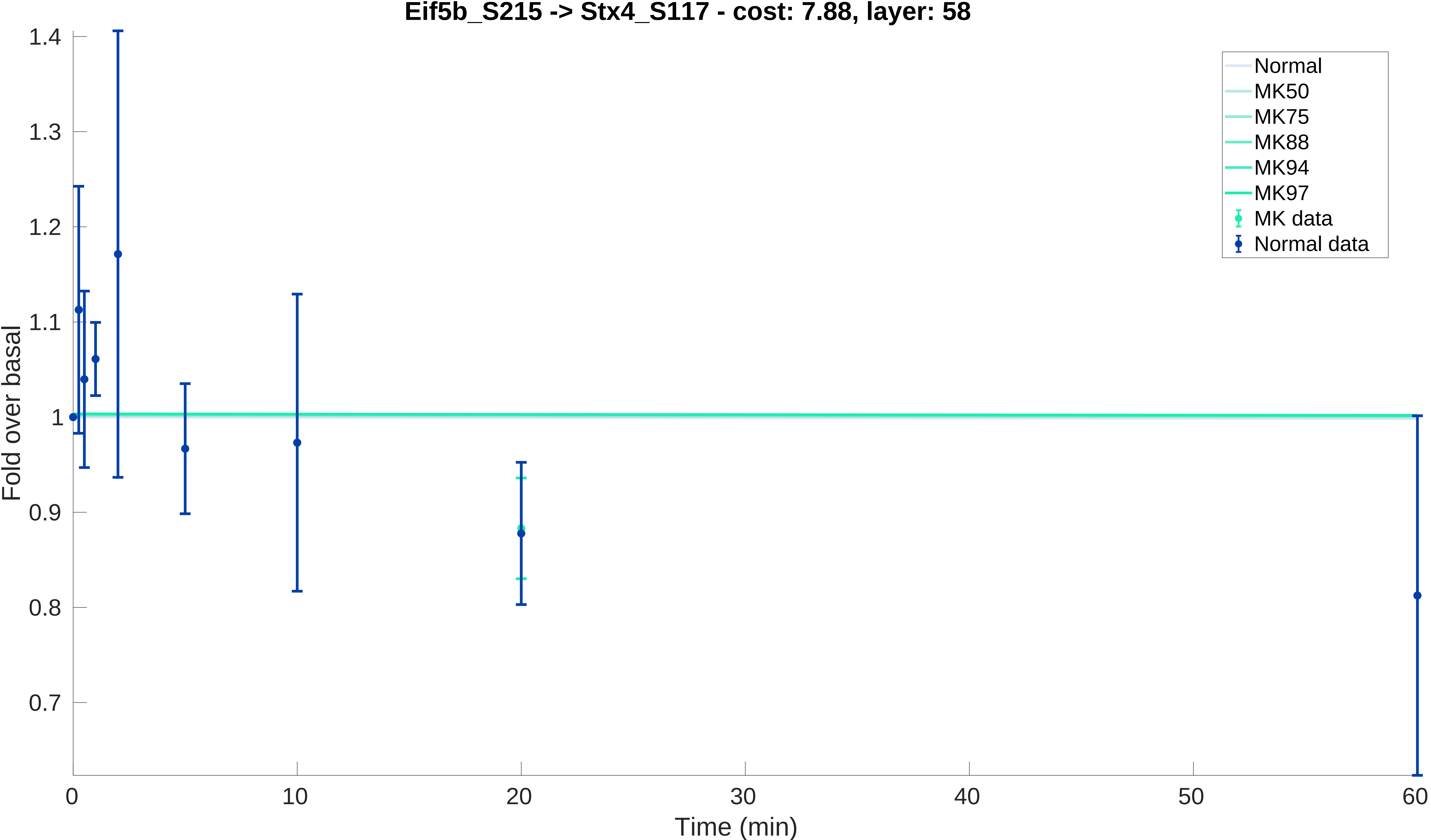
Eif5b_S215 -> Stx4_S117 - cost: 7.88, layer: 58
Fold over basal
Time (min)
1.4
1.3
1.2
1.1
1
0.9
0.8
0.7
0
10
20
30
40
50
60
Normal
MK50
MK75
MK88
MK94
MK97
MK data
Normal data

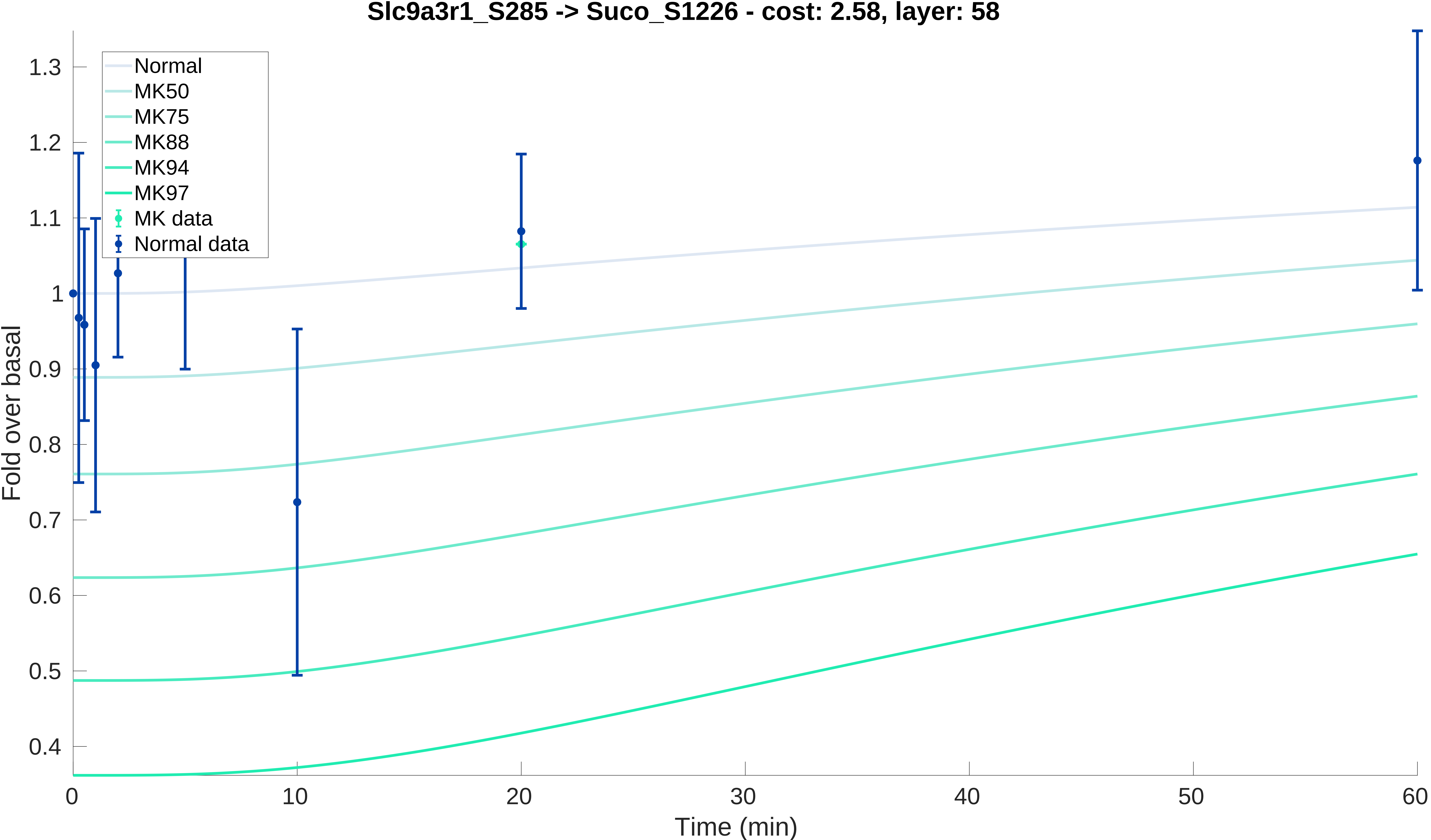

Slc9a3r1_S285 -> Suco_S1226 - cost: 2.58, layer: 58
Normal
MK50
MK75
MK88
MK94
MK97
MK data
Normal data
Fold over basal
Time (min)
1.3
1.2
1.1
1
0.9
0.8
0.7
0.6
0.5
0.4
0
10
20
30
40
50
60

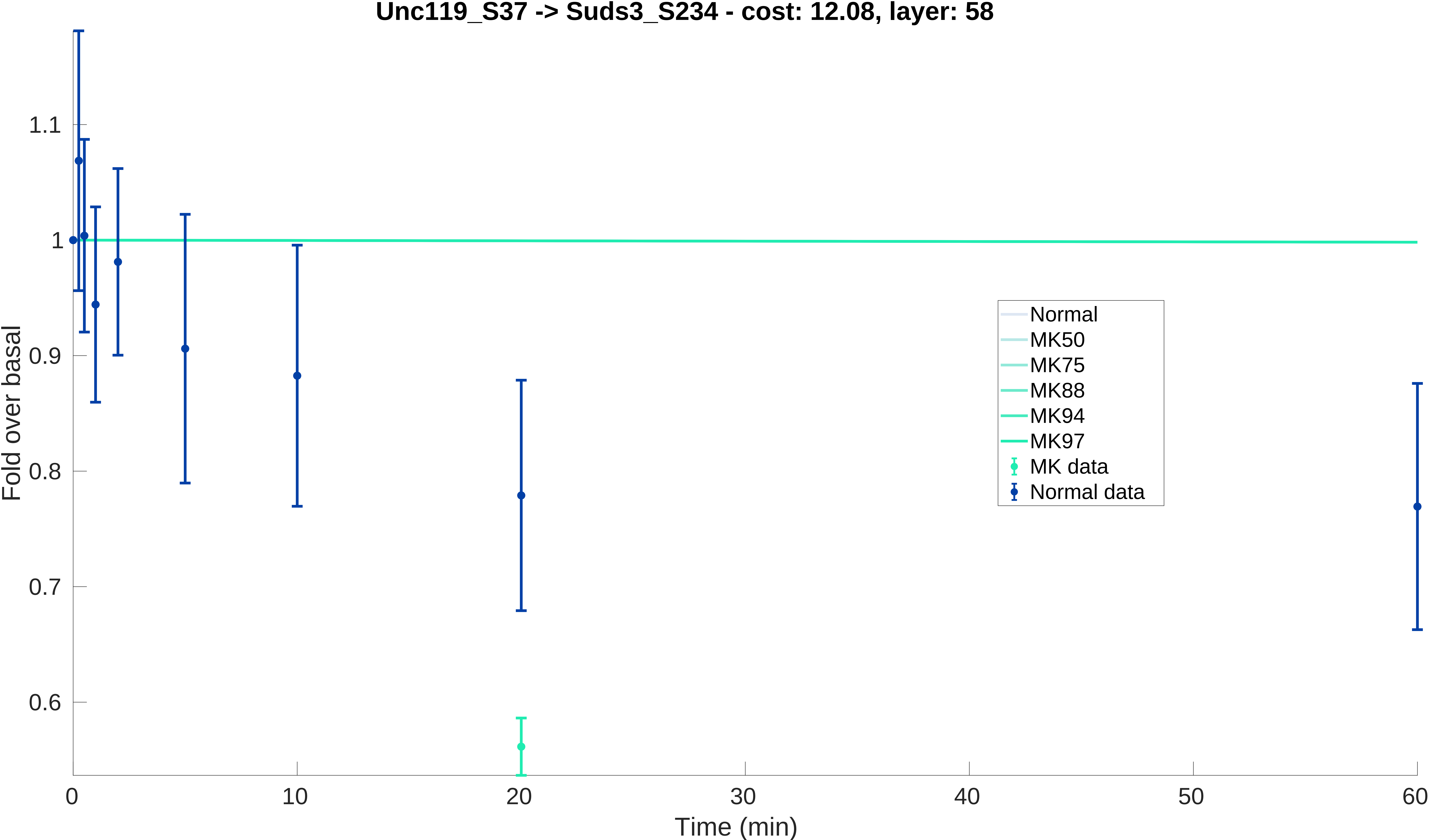

Unc119_S37 -> Suds3_S234 - cost: 12.08, layer: 58
Fold over basal
1.1
1
0.9
0.8
0.7
0.6
0
10
20
30
40
50
60
Time (min)
Normal
MK50
MK75
MK88
MK94
MK97
MK data
Normal data

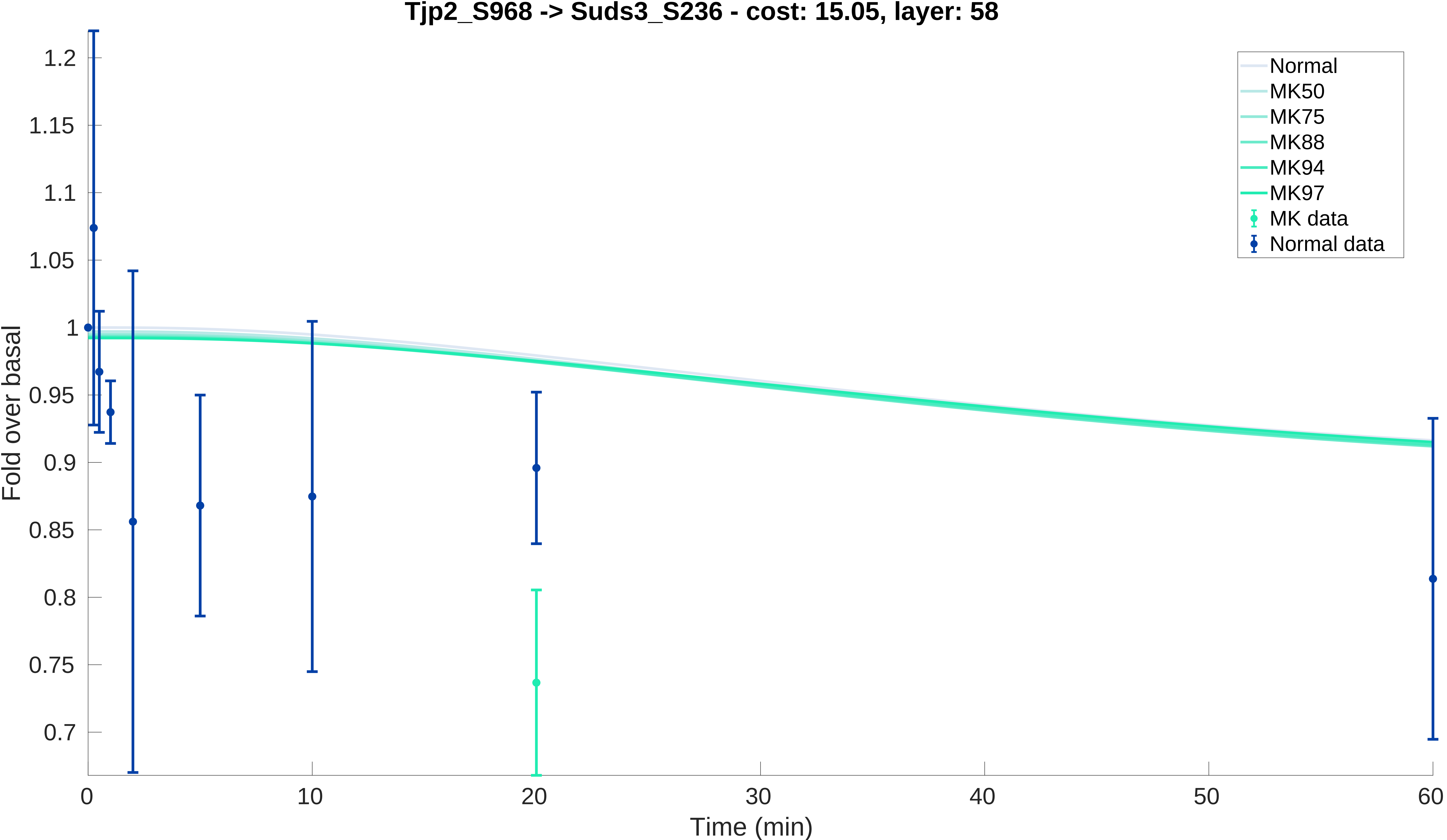

Tjp2_S968 -> Suds3_S236 - cost: 15.05, layer: 58
Fold over basal
Time (min)
1.2
1.15
1.1
1.05
1
0.95
0.9
0.85
0.8
0.75
0.7
0
10
20
30
40
50
60
Normal
MK50
MK75
MK88
MK94
MK97
MK data
Normal data

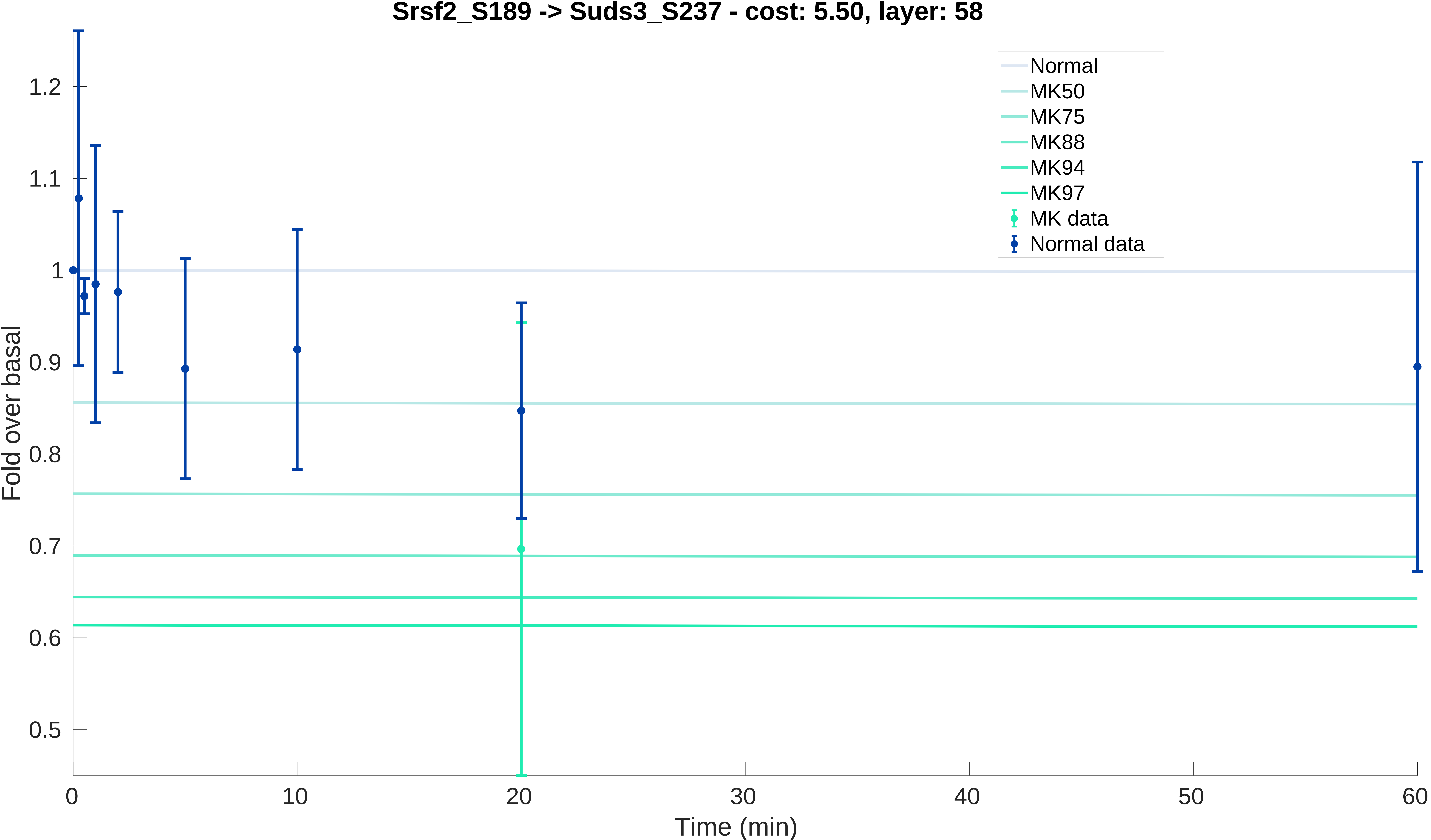
Srsf2_S189 -> Suds3_S237 - cost: 5.50, layer: 58
Normal
MK50
MK75
MK88
MK94
MK97
MK data
Normal data
Fold over basal
1.2
1.1
1
0.9
0.8
0.7
0.6
0.5
0
10
20
30
40
50
60
Time (min)

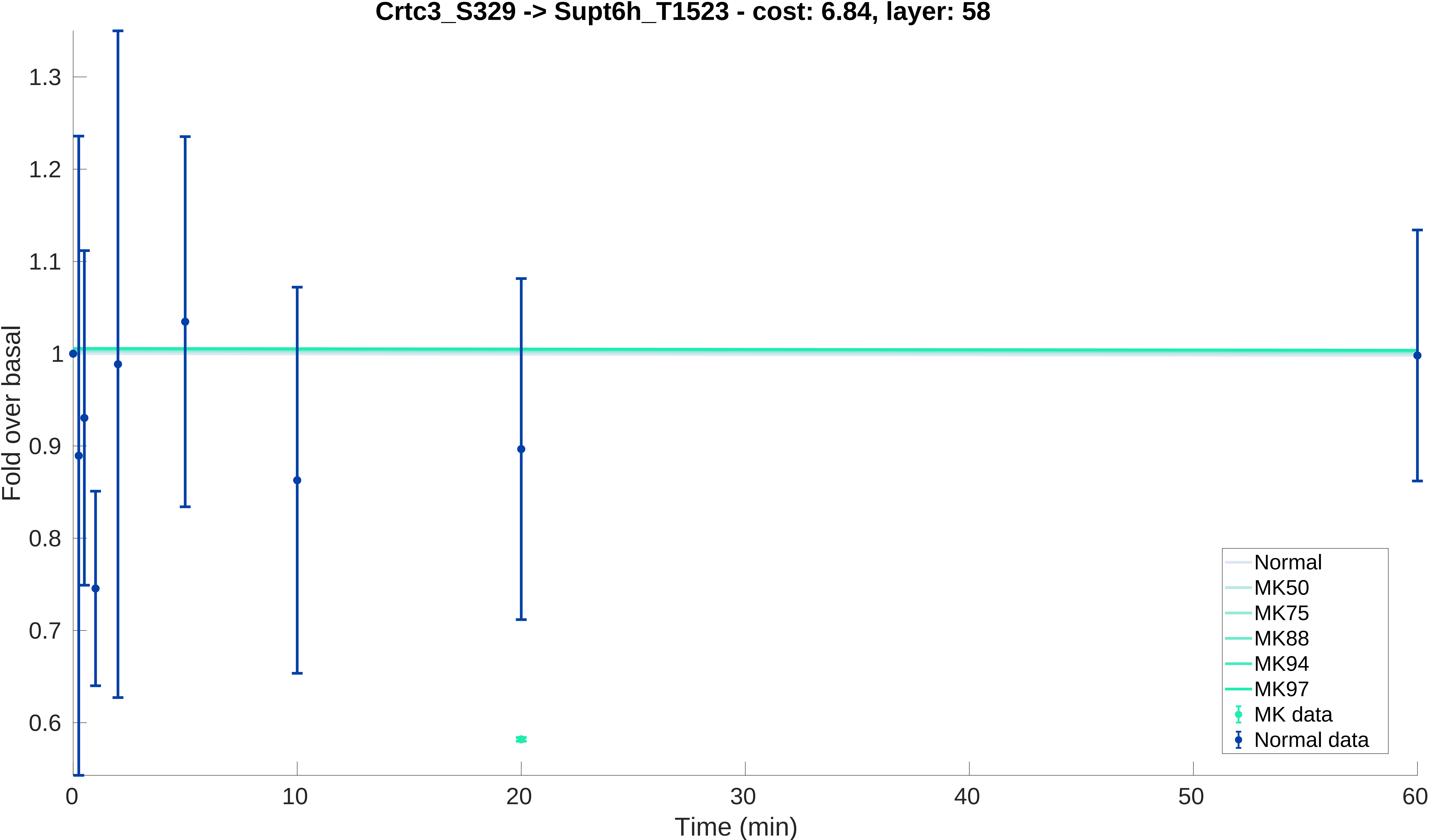

Crtc3_S329 -> Supt6h_T1523 - cost: 6.84, layer: 58
Fold over basal
Time (min)
1.3
1.2
1.1
1
0.9
0.8
0.7
0.6
0
10
20
30
40
50
60
Normal
MK50
MK75
MK88
MK94
MK97
MK data
Normal data

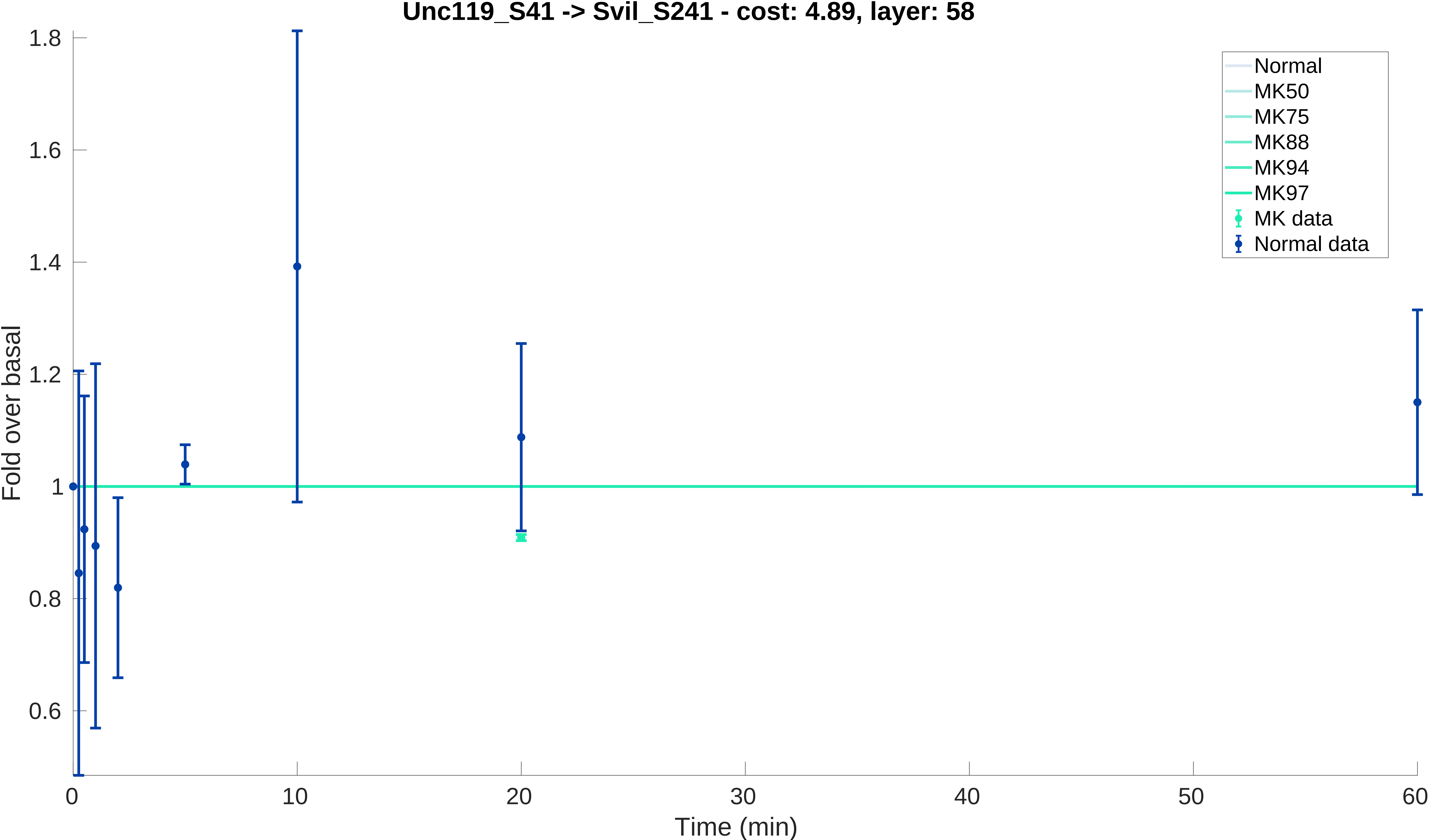

Unc119_S41 -> Svil_S241 - cost: 4.89, layer: 58
Fold over basal
Time (min)
1.8
1.6
1.4
1.2
1
0.8
0.6
0
10
20
30
40
50
60
Normal
MK50
MK75
MK88
MK94
MK97
MK data
Normal data

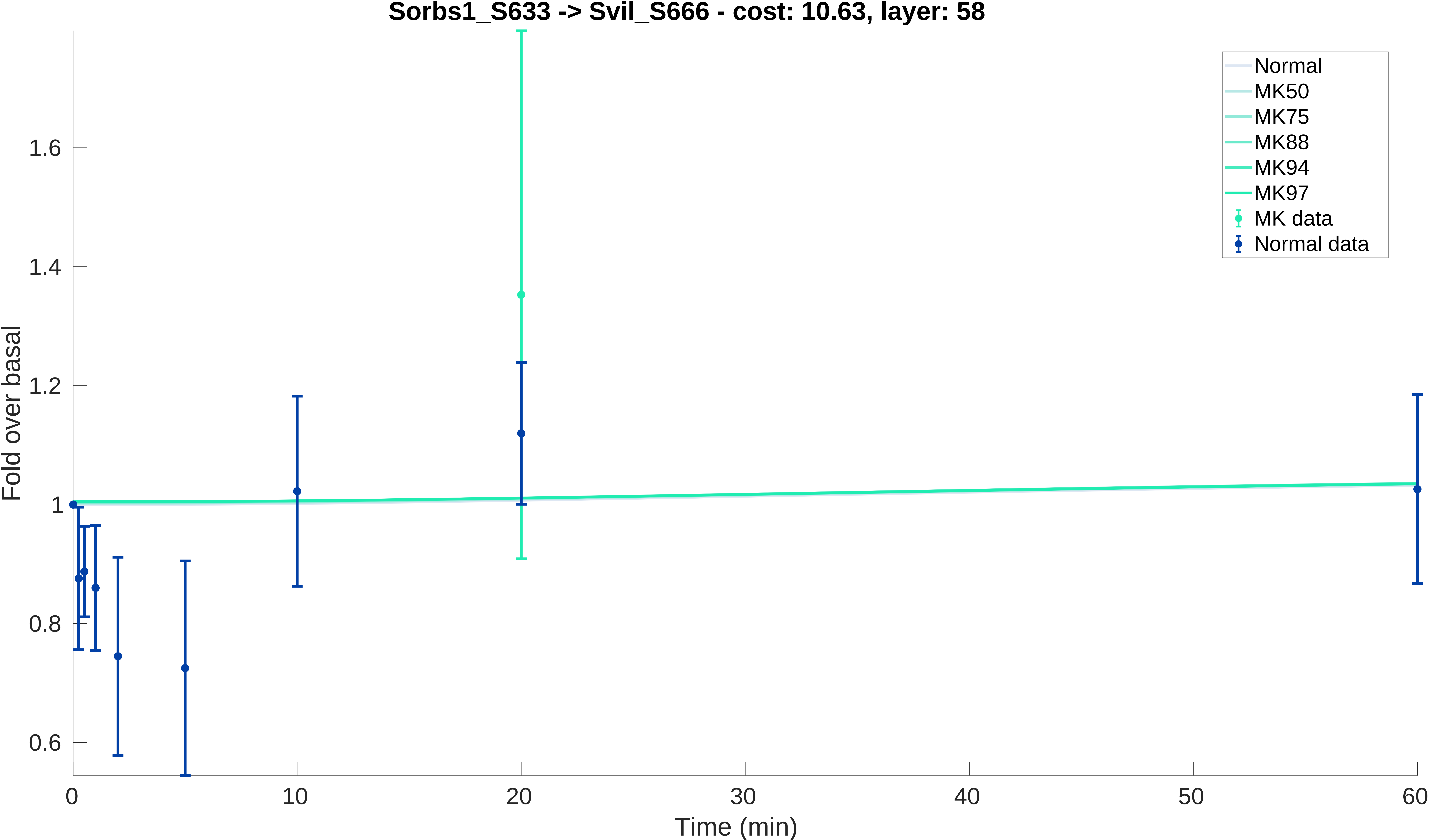
Sorbs1_S633 -> Svil_S666 - cost: 10.63, layer: 58
Normal
MK50
MK75
MK88
MK94
MK97
MK data
Normal data
Fold over basal
Time (min)
0.6
0.8
1
1.2
1.4
1.6
0
10
20
30
40
50
60

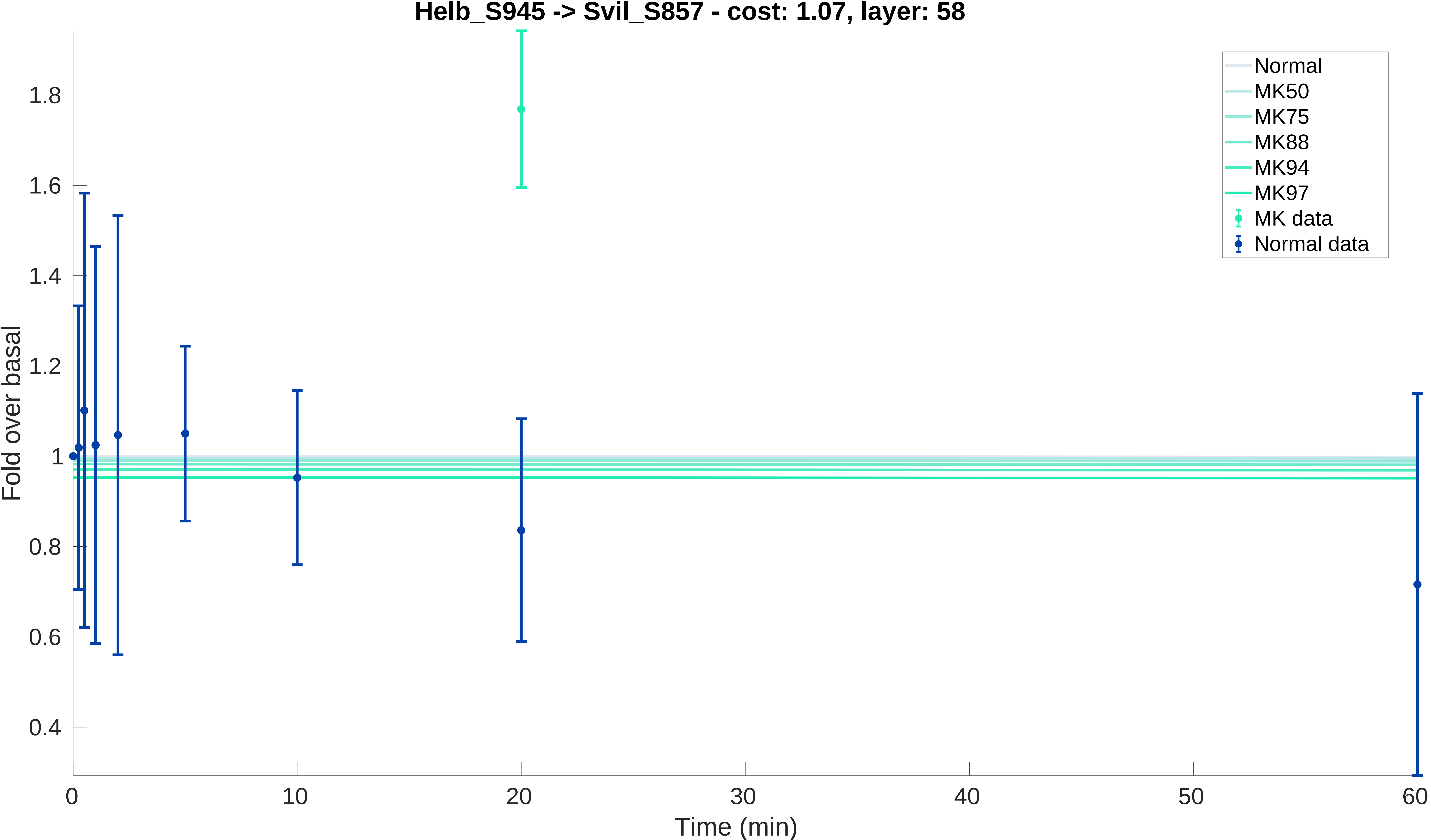

Helb_S945 -> Svil_S857 - cost: 1.07, layer: 58
Fold over basal
Time (min)
0
10
20
30
40
50
60
0.4
0.6
0.8
1
1.2
1.4
1.6
1.8
Normal
MK50
MK75
MK88
MK94
MK97
MK data
Normal data

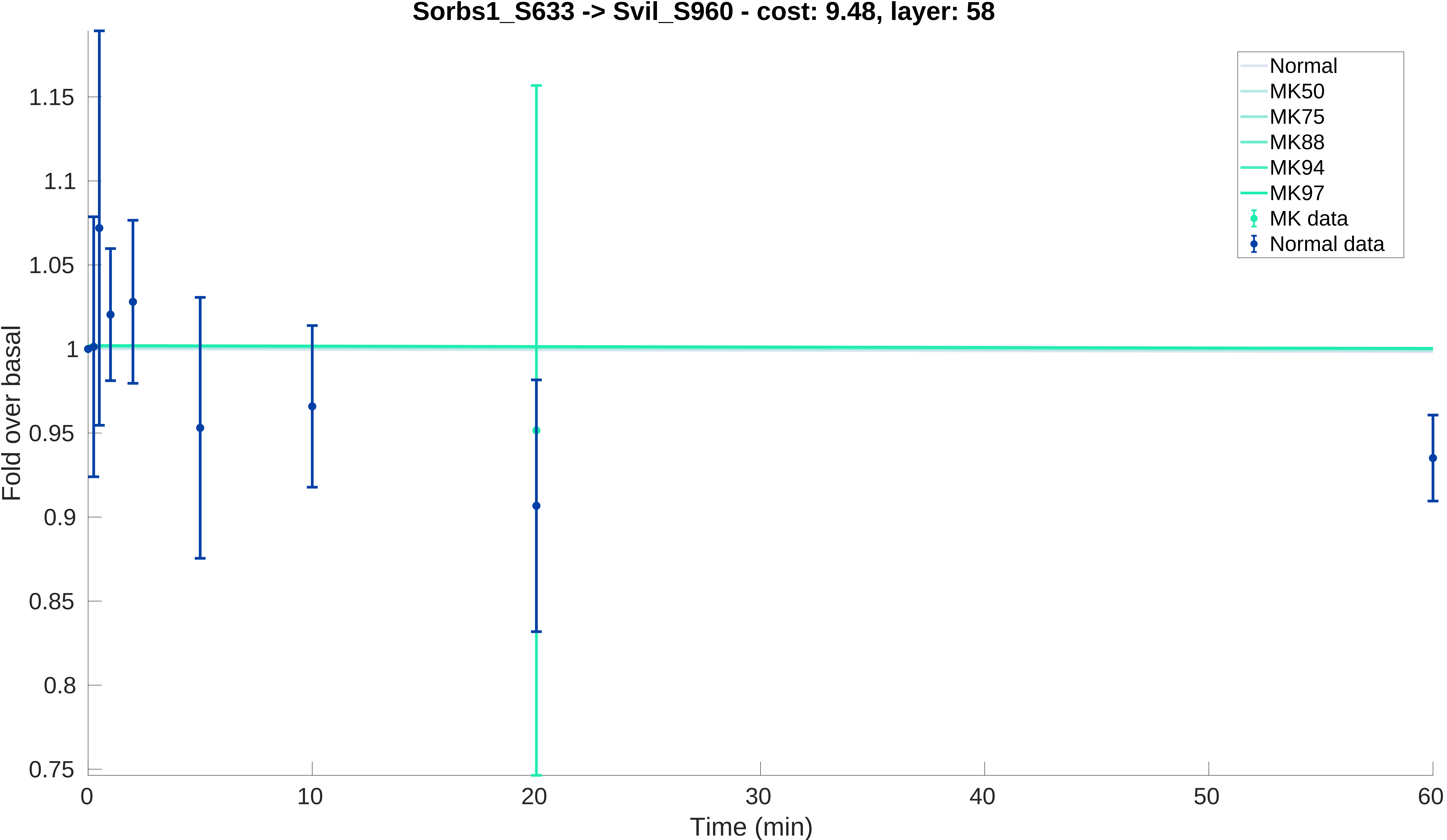

Sorbs1_S633 -> Svil_S960 - cost: 9.48, layer: 58
Normal
MK50
MK75
MK88
MK94
MK97
MK data
Normal data
Fold over basal
1.15
1.1
1.05
1
0.95
0.9
0.85
0.8
0.75
0
10
20
30
40
50
60
Time (min)

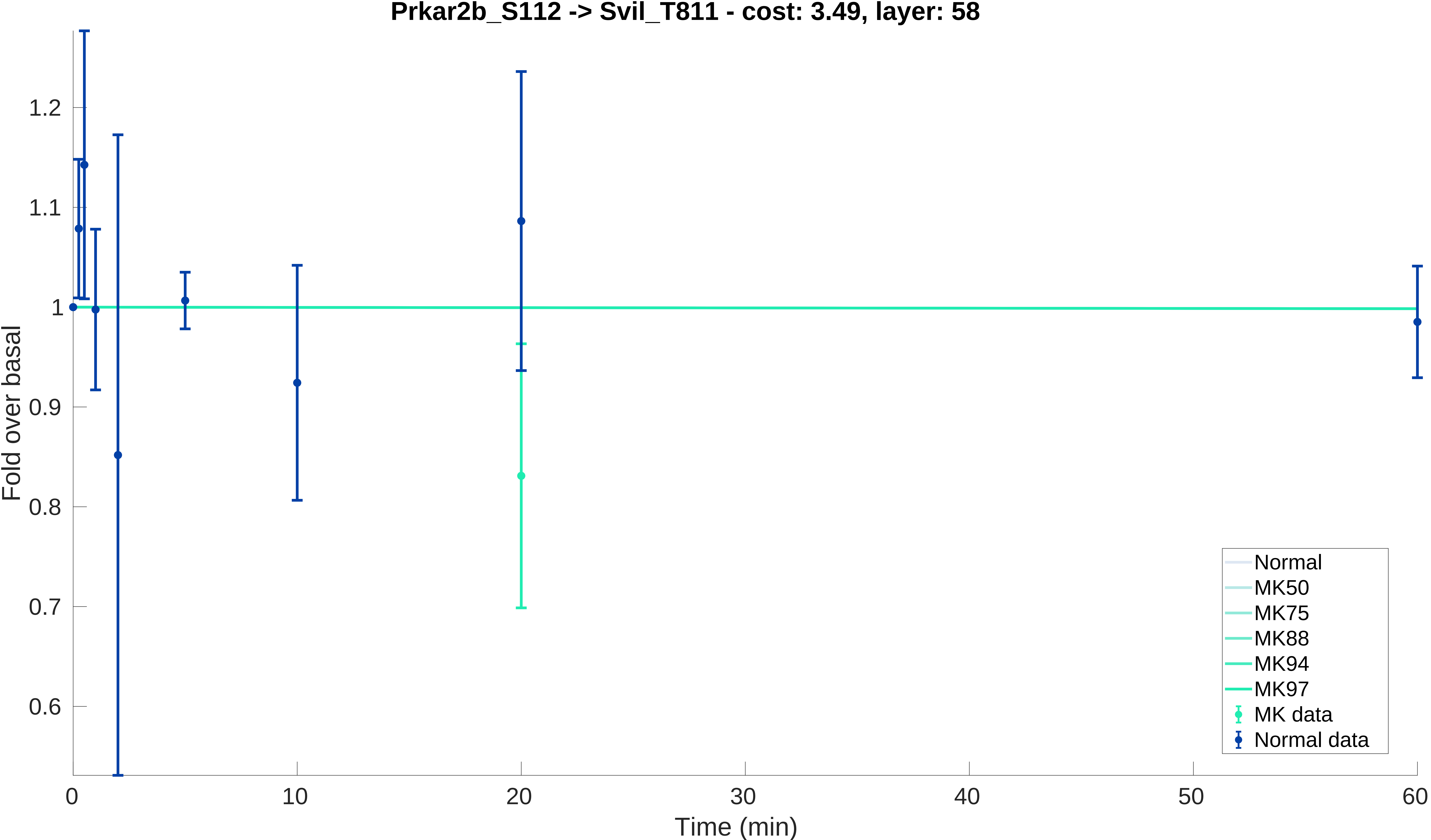

Prkar2b_S112 -> Svil_T811 - cost: 3.49, layer: 58
Fold over basal
1.2
1.1
1
0.9
0.8
0.7
0.6
0
10
20
30
40
50
60
Time (min)
Normal
MK50
MK75
MK88
MK94
MK97
MK data
Normal data

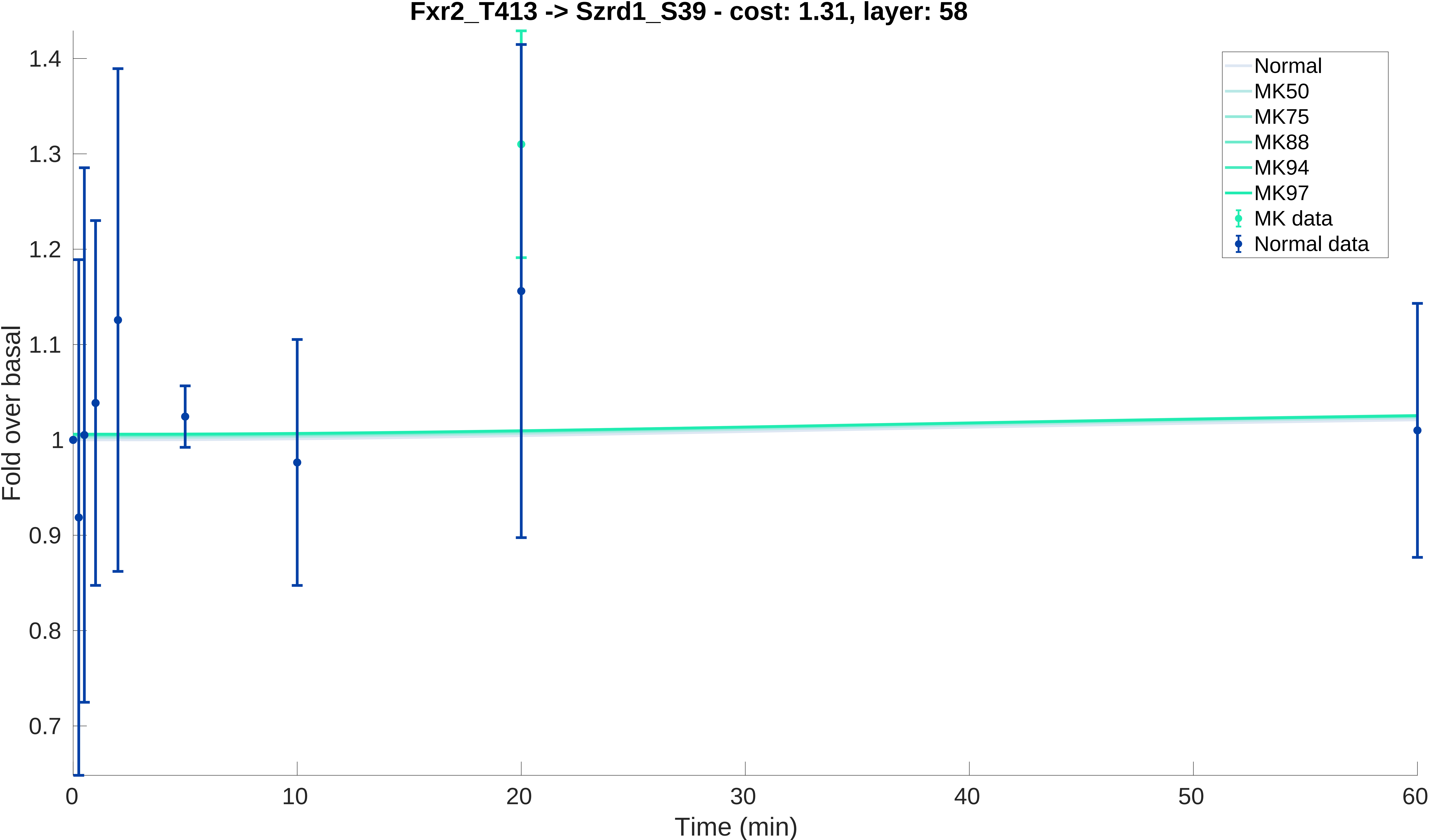

Fxr2_T413 -> Szrd1_S39 - cost: 1.31, layer: 58
Fold over basal
Time (min)
0.7
0.8
0.9
1
1.1
1.2
1.3
1.4
0
10
20
30
40
50
60
Normal
MK50
MK75
MK88
MK94
MK97
MK data
Normal data

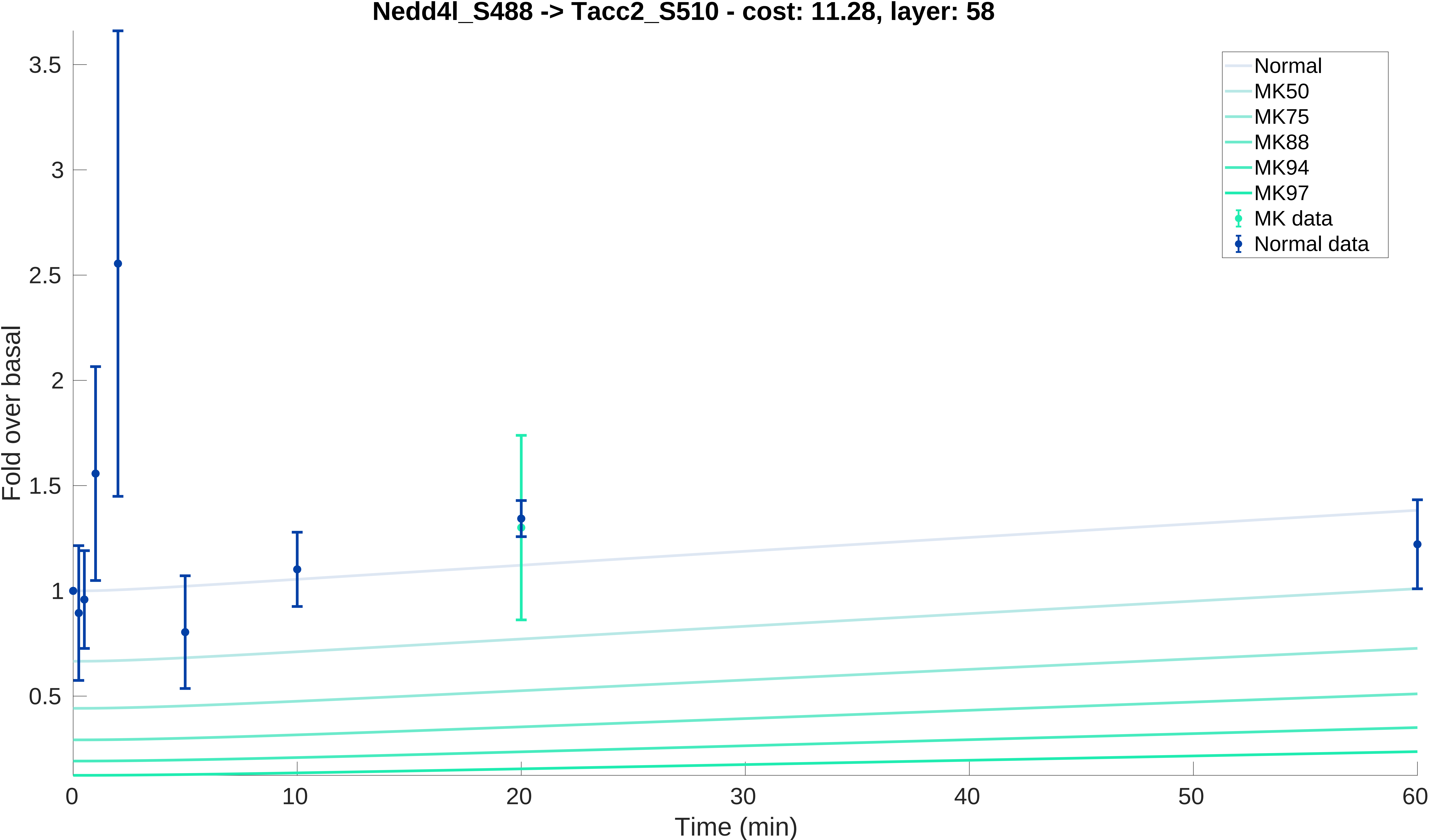
Nedd4l_S488 -> Tacc2_S510 - cost: 11.28, layer: 58
Fold over basal
Time (min)
0.5
1
1.5
2
2.5
3
3.5
0
10
20
30
40
50
60
Normal
MK50
MK75
MK88
MK94
MK97
MK data
Normal data

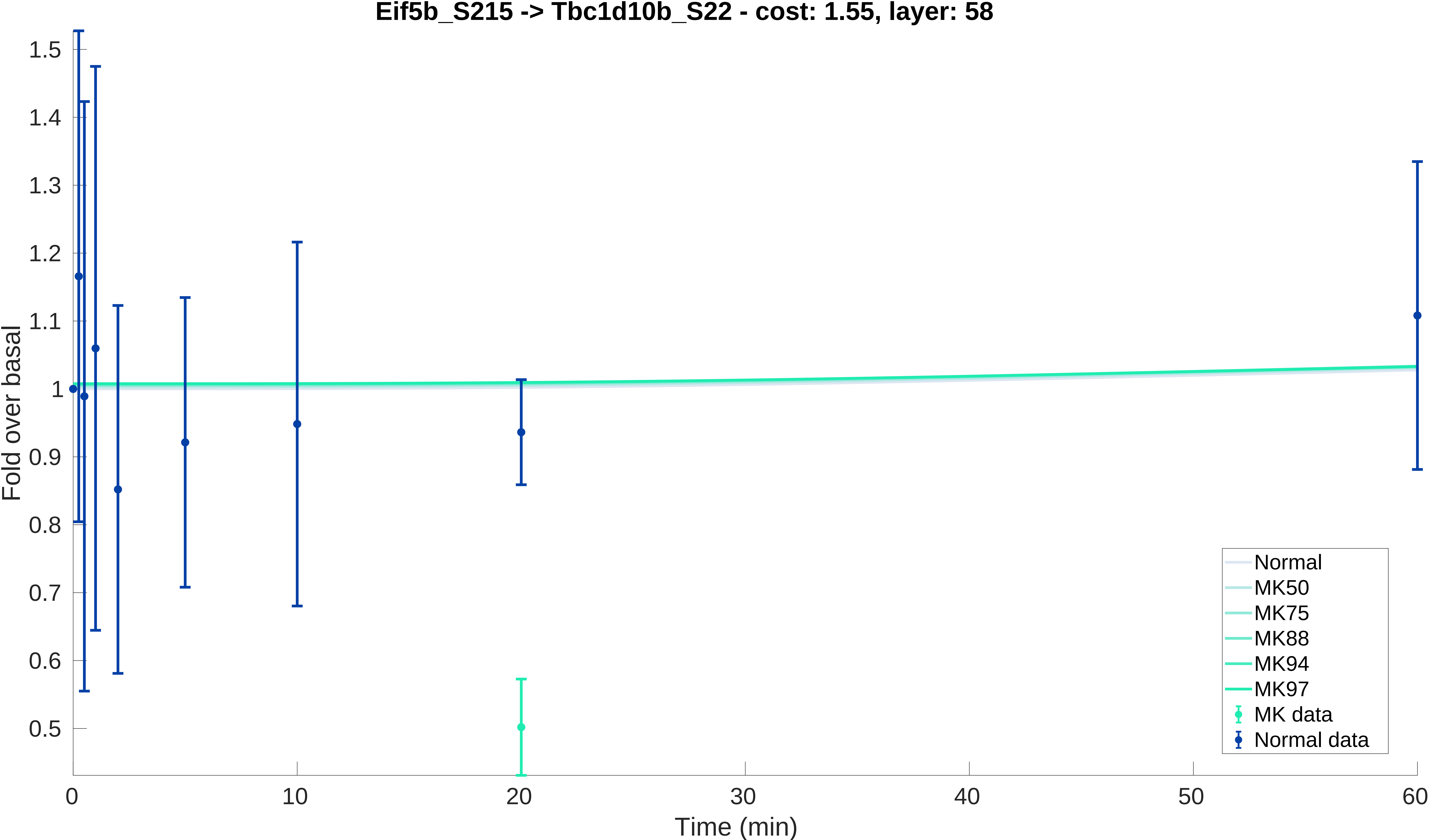
Eif5b_S215 -> Tbc1d10b_S22 - cost: 1.55, layer: 58
Fold over basal
Time (min)
1.5
1.4
1.3
1.2
1.1
1
0.9
0.8
0.7
0.6
0.5
0
10
20
30
40
50
60
Normal
MK50
MK75
MK88
MK94
MK97
MK data
Normal data

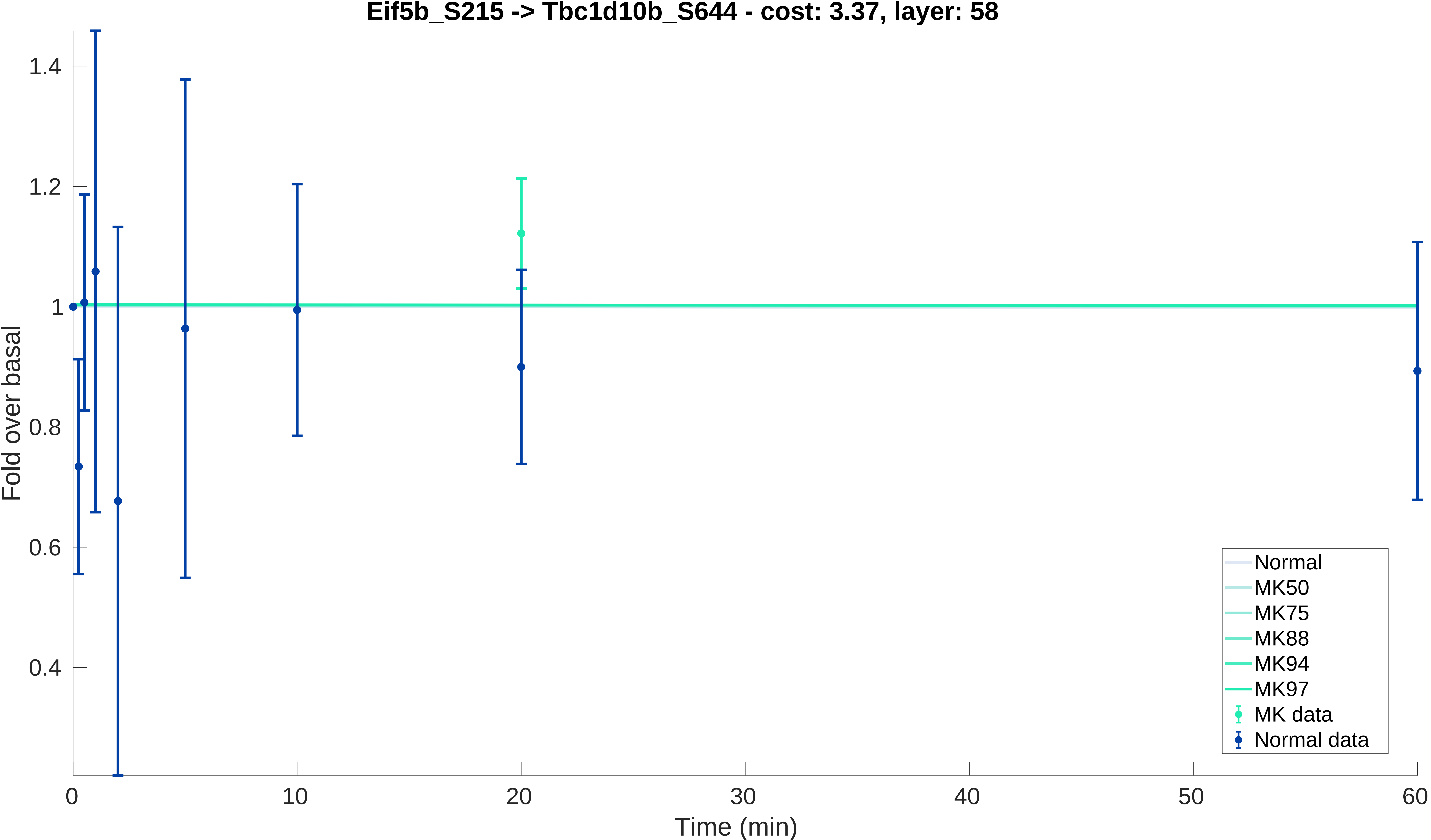

Eif5b_S215 -> Tbc1d10b_S644 - cost: 3.37, layer: 58
Fold over basal
1.4
1.2
1
0.8
0.6
0.4
0
10
20
30
40
50
60
Time (min)
Normal
MK50
MK75
MK88
MK94
MK97
MK data
Normal data

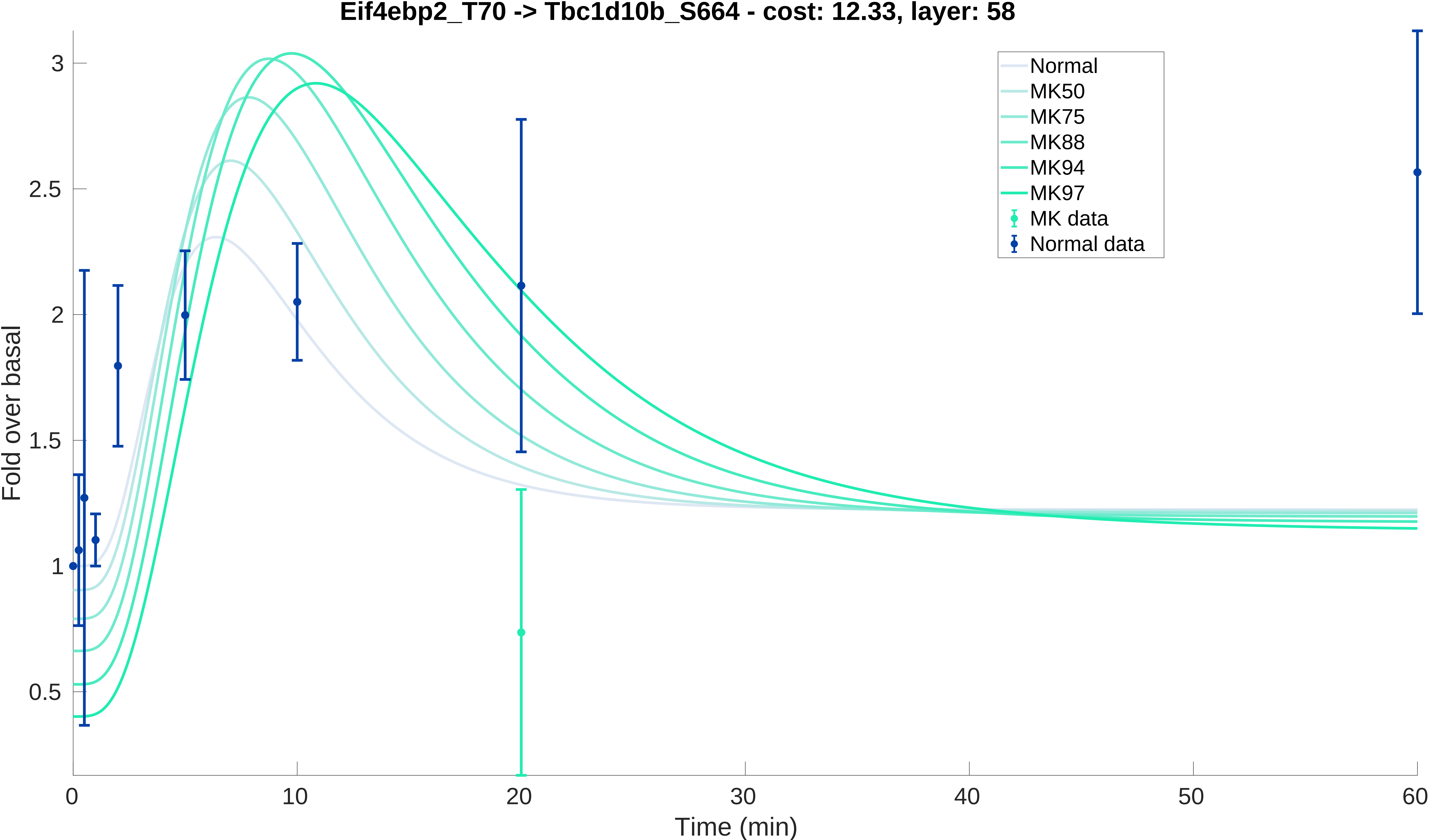

Eif4ebp2_T70 -> Tbc1d10b_S664 - cost: 12.33, layer: 58
Fold over basal
Time (min)
0
10
20
30
40
50
60
0.5
1
1.5
2
2.5
3
Normal
MK50
MK75
MK88
MK94
MK97
MK data
Normal data

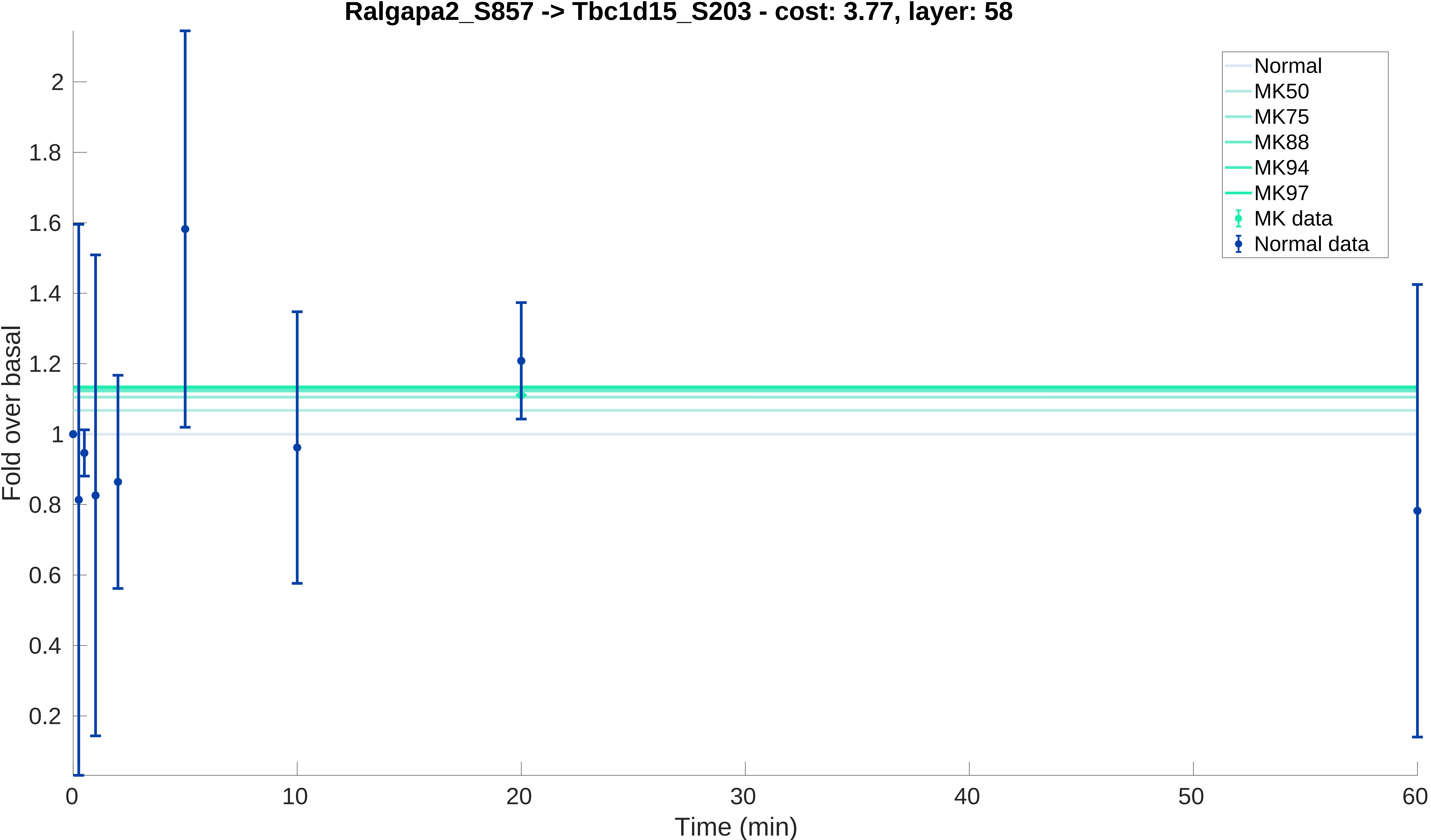

Ralgapa2_S857 -> Tbc1d15_S203 - cost: 3.77, layer: 58
Fold over basal
Time (min)
0.2
0.4
0.6
0.8
1
1.2
1.4
1.6
1.8
2
0
10
20
30
40
50
60
Normal
MK50
MK75
MK88
MK94
MK97
MK data
Normal data

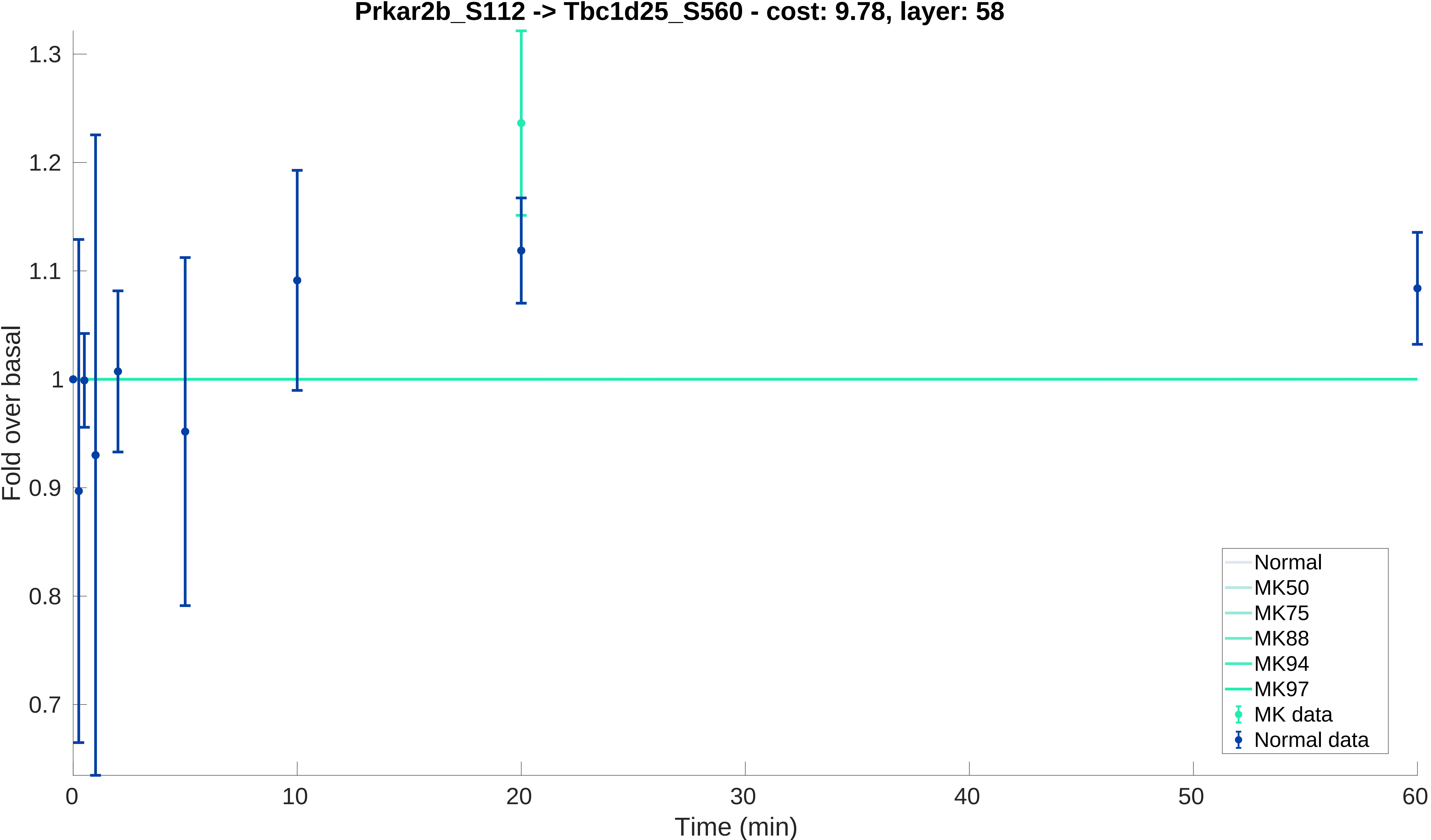

Prkar2b_S112 -> Tbc1d25_S560 - cost: 9.78, layer: 58
Fold over basal
1.3
1.2
1.1
1
0.9
0.8
0.7
0
10
20
30
40
50
60
Time (min)
Normal
MK50
MK75
MK88
MK94
MK97
MK data
Normal data

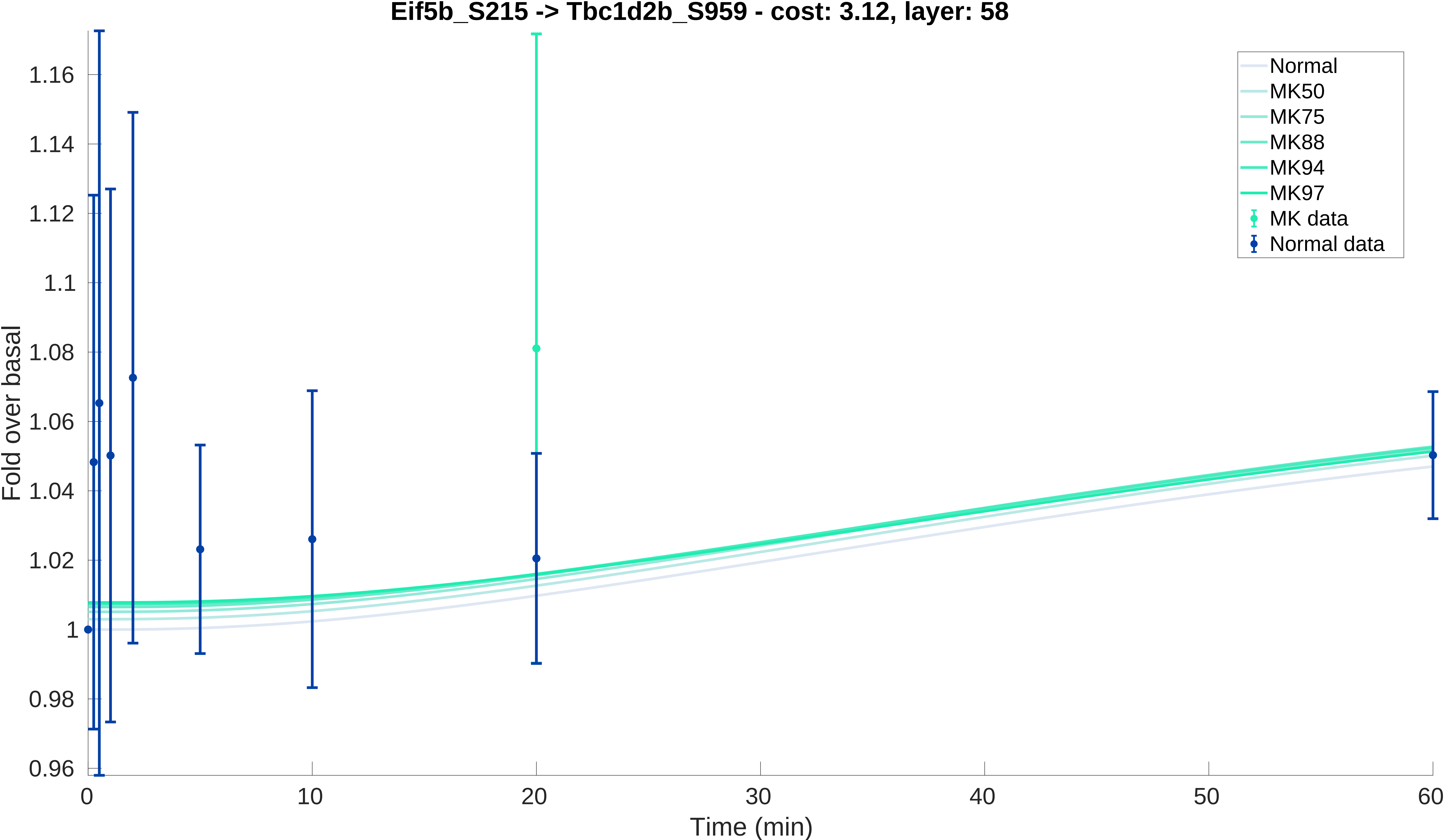

Eif5b_S215 -> Tbc1d2b_S959 - cost: 3.12, layer: 58
Fold over basal
1.16
1.14
1.12
1.1
1.08
1.06
1.04
1.02
1
0.98
0.96
Time (min)
0
10
20
30
40
50
60
Normal
MK50
MK75
MK88
MK94
MK97
MK data
Normal data

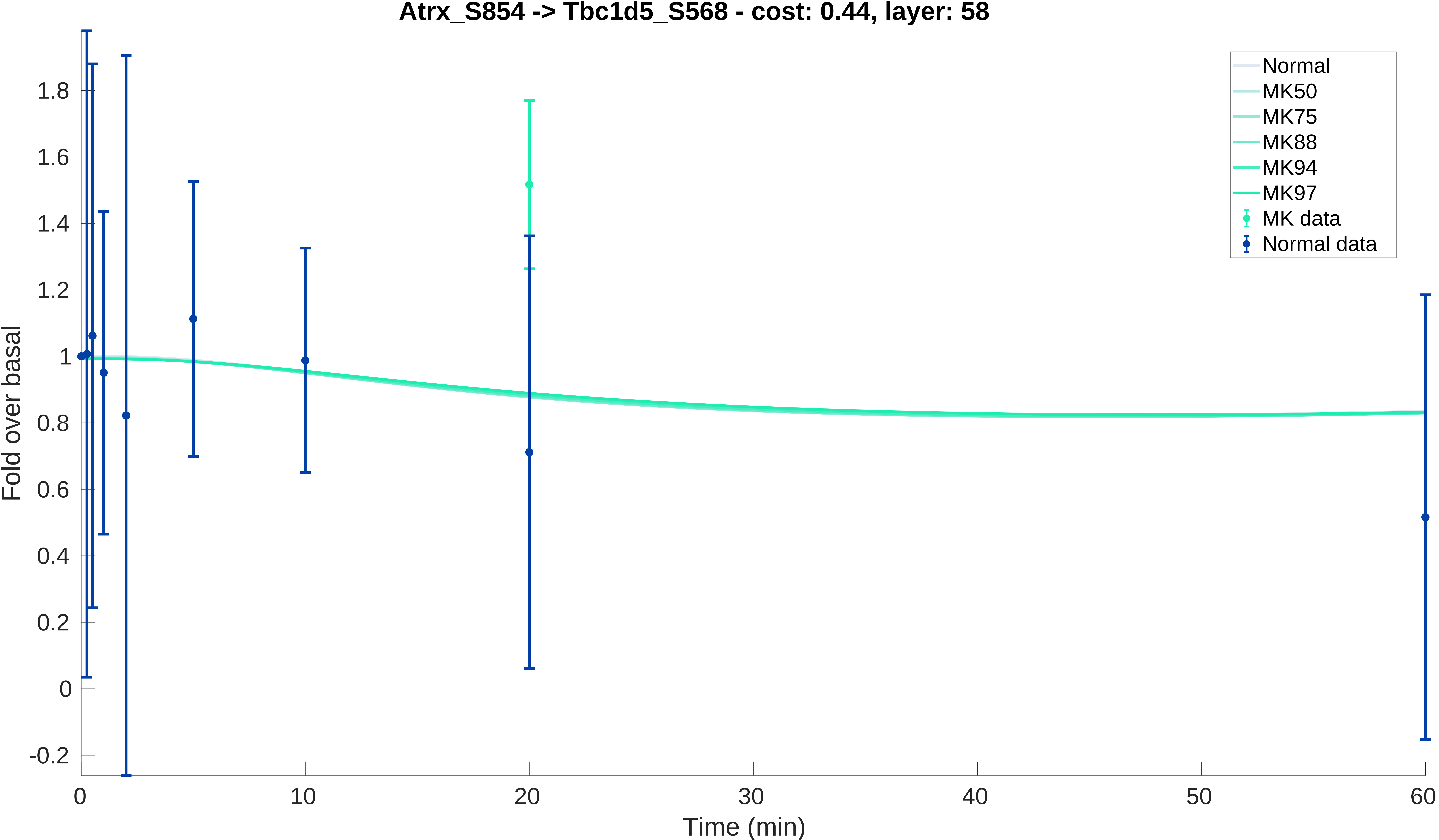

Atrx_S854 -> Tbc1d5_S568 - cost: 0.44, layer: 58
Normal
MK50
MK75
MK88
MK94
MK97
MK data
Normal data
Fold over basal
1.8
1.6
1.4
1.2
1
0.8
0.6
0.4
0.2
0
-0.2
0
10
20
30
40
50
60
Time (min)

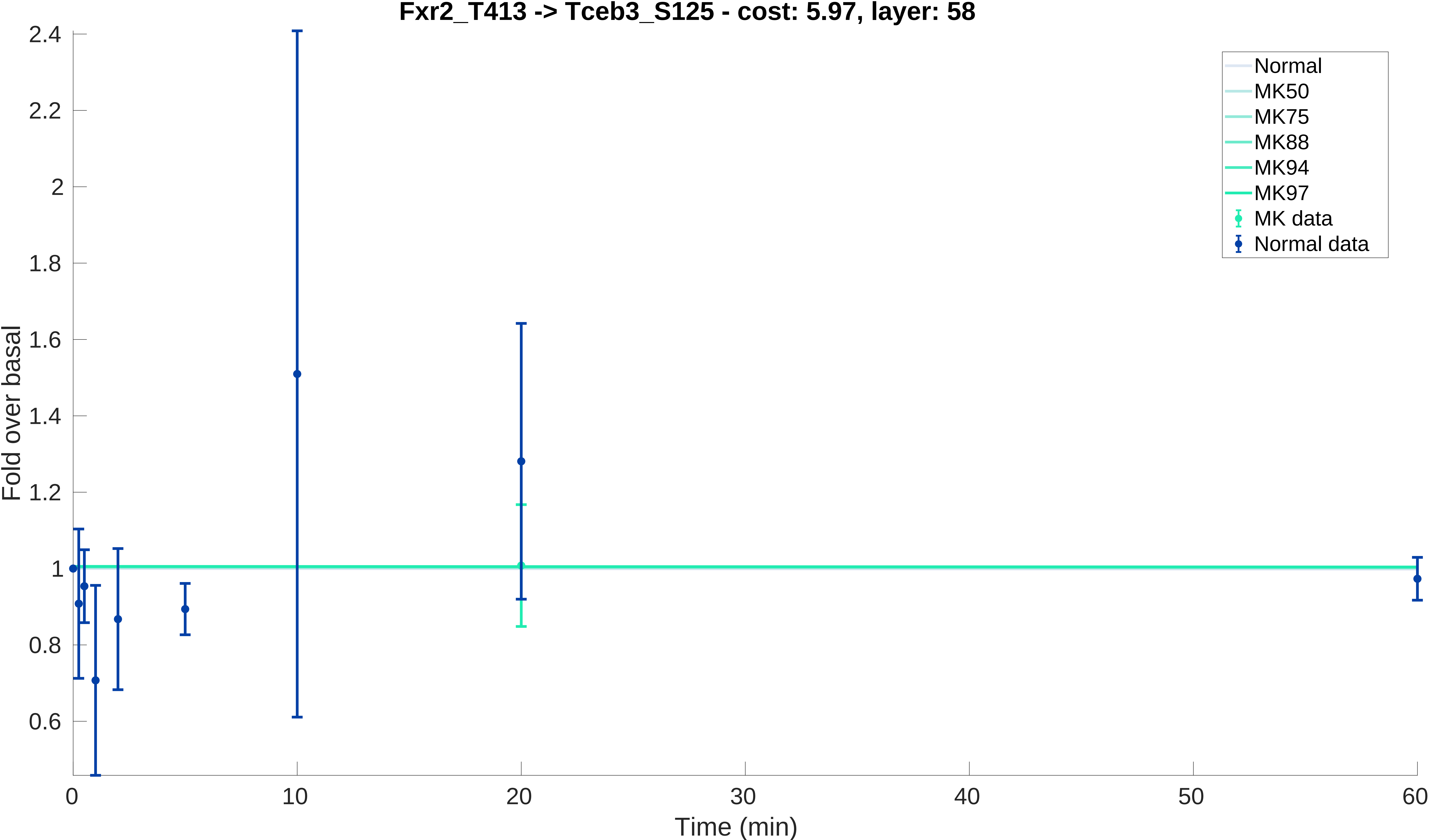

Fxr2_T413 -> Tceb3_S125 - cost: 5.97, layer: 58
Fold over basal
Time (min)
0.6
0.8
1
1.2
1.4
1.6
1.8
2
2.2
2.4
0
10
20
30
40
50
60
Normal
MK50
MK75
MK88
MK94
MK97
MK data
Normal data

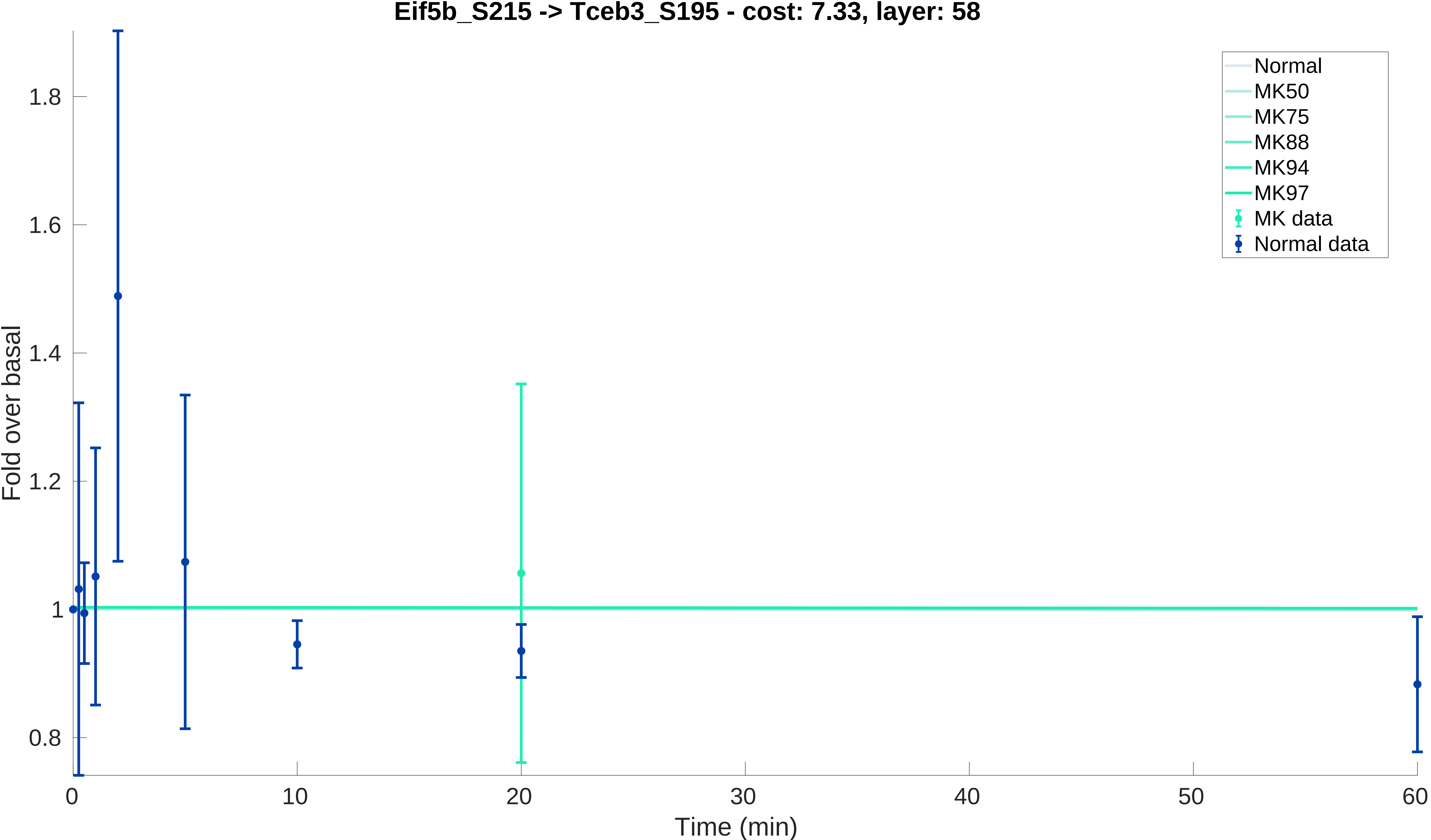

Eif5b_S215 -> Tceb3_S195 - cost: 7.33, layer: 58
Normal
MK50
MK75
MK88
MK94
MK97
MK data
Normal data
Fold over basal
1.8
1.6
1.4
1.2
1
0.8
0
10
20
30
40
50
60
Time (min)

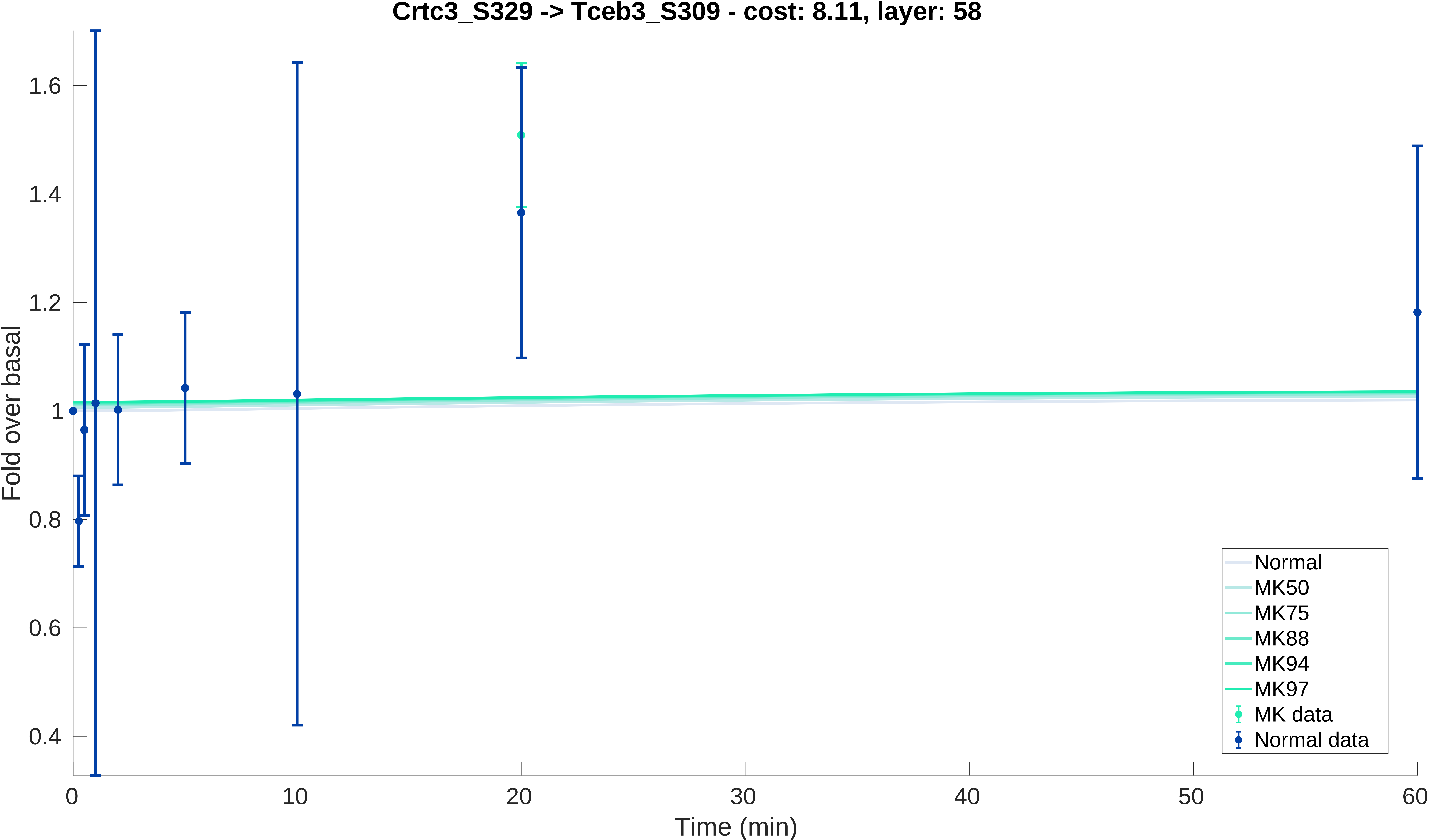

Crtc3_S329 -> Tceb3_S309 - cost: 8.11, layer: 58
Fold over basal
Time (min)
0.4
0.6
0.8
1
1.2
1.4
1.6
0
10
20
30
40
50
60
Normal
MK50
MK75
MK88
MK94
MK97
MK data
Normal data

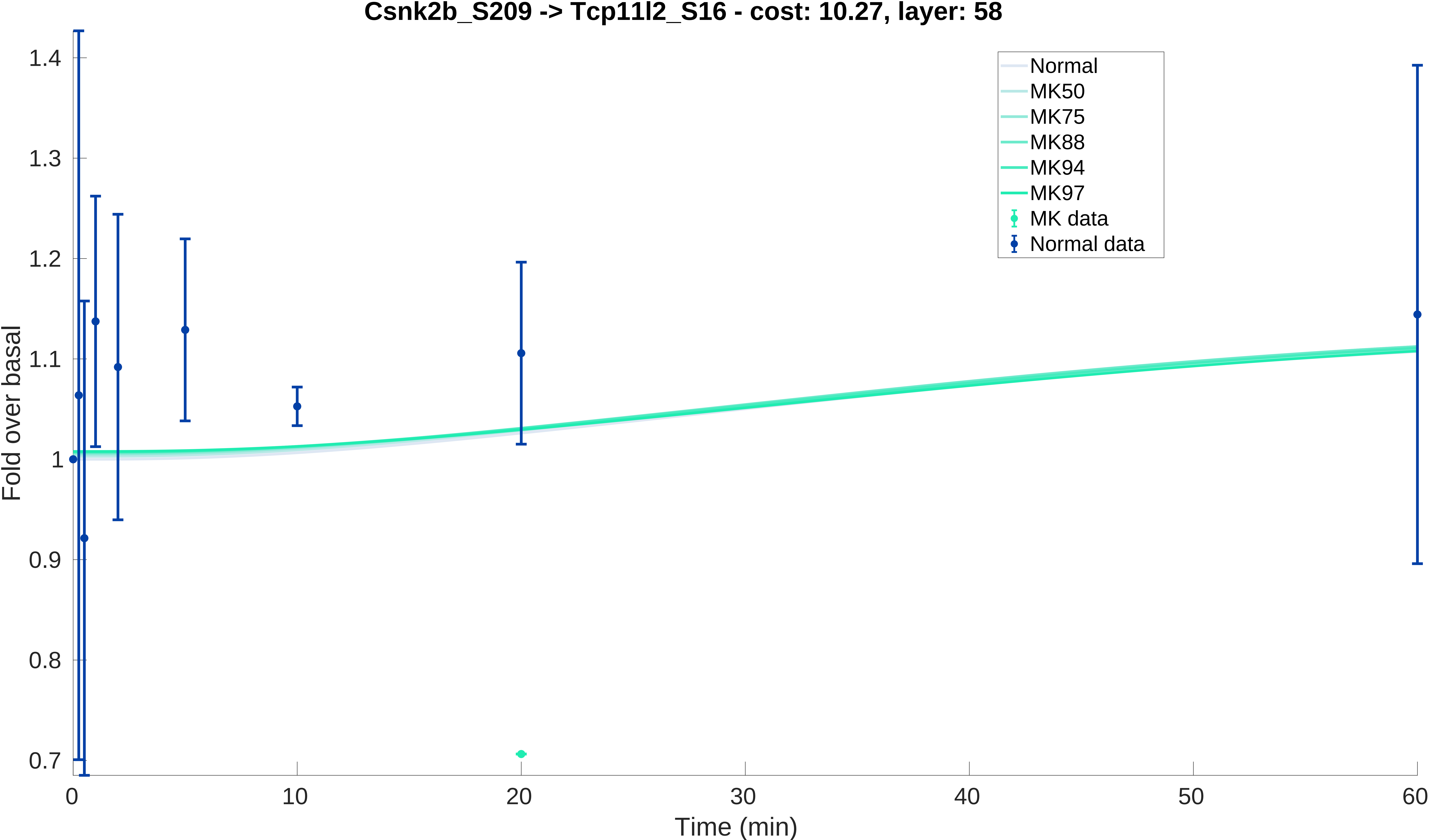

Csnk2b_S209 -> Tcp11l2_S16 - cost: 10.27, layer: 58
Normal
MK50
MK75
MK88
MK94
MK97
MK data
Normal data
Fold over basal
Time (min)
1.4
1.3
1.2
1.1
1
0.9
0.8
0.7
0
10
20
30
40
50
60

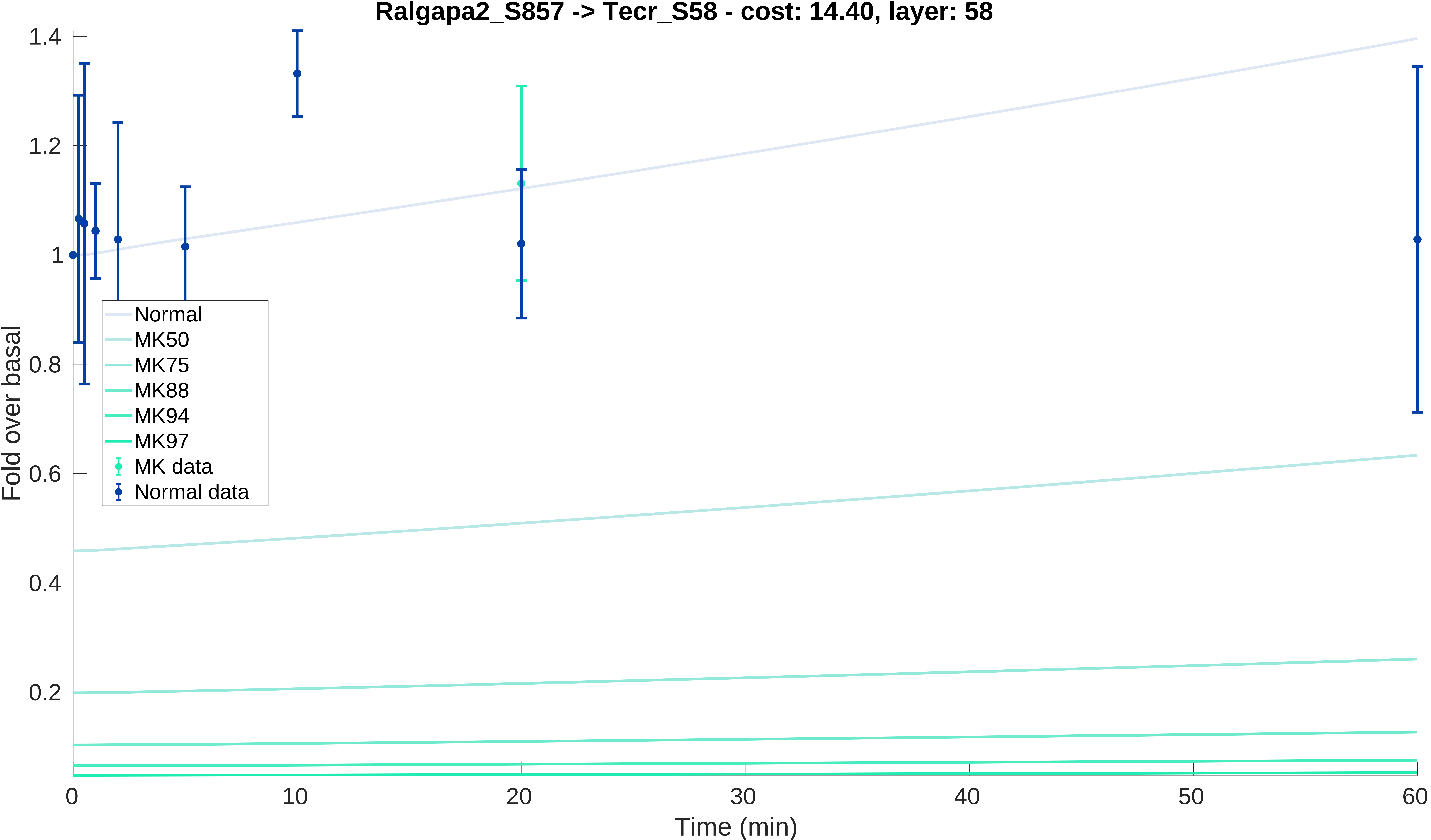

Ralgapa2_S857 -> Tecr_S58 - cost: 14.40, layer: 58
Fold over basal
Time (min)
1.4
1.2
1
0.8
0.6
0.4
0.2
0
10
20
30
40
50
60
Normal
MK50
MK75
MK88
MK94
MK97
MK data
Normal data

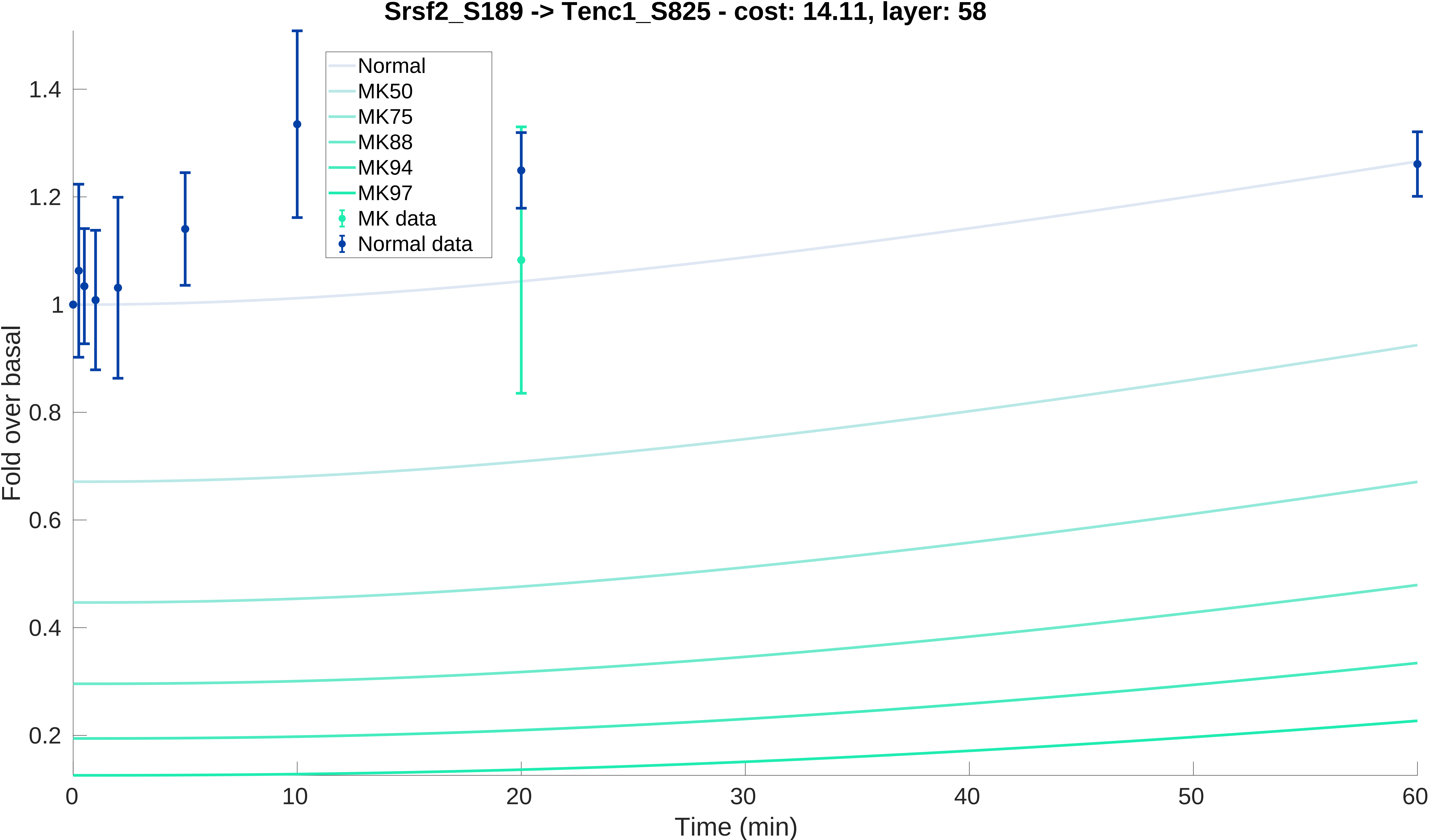

Srsf2_S189 -> Tenc1_S825 - cost: 14.11, layer: 58
Normal
MK50
MK75
MK88
MK94
MK97
MK data
Normal data
Fold over basal
Time (min)
1.4
1.2
1
0.8
0.6
0.4
0.2
0
10
20
30
40
50
60

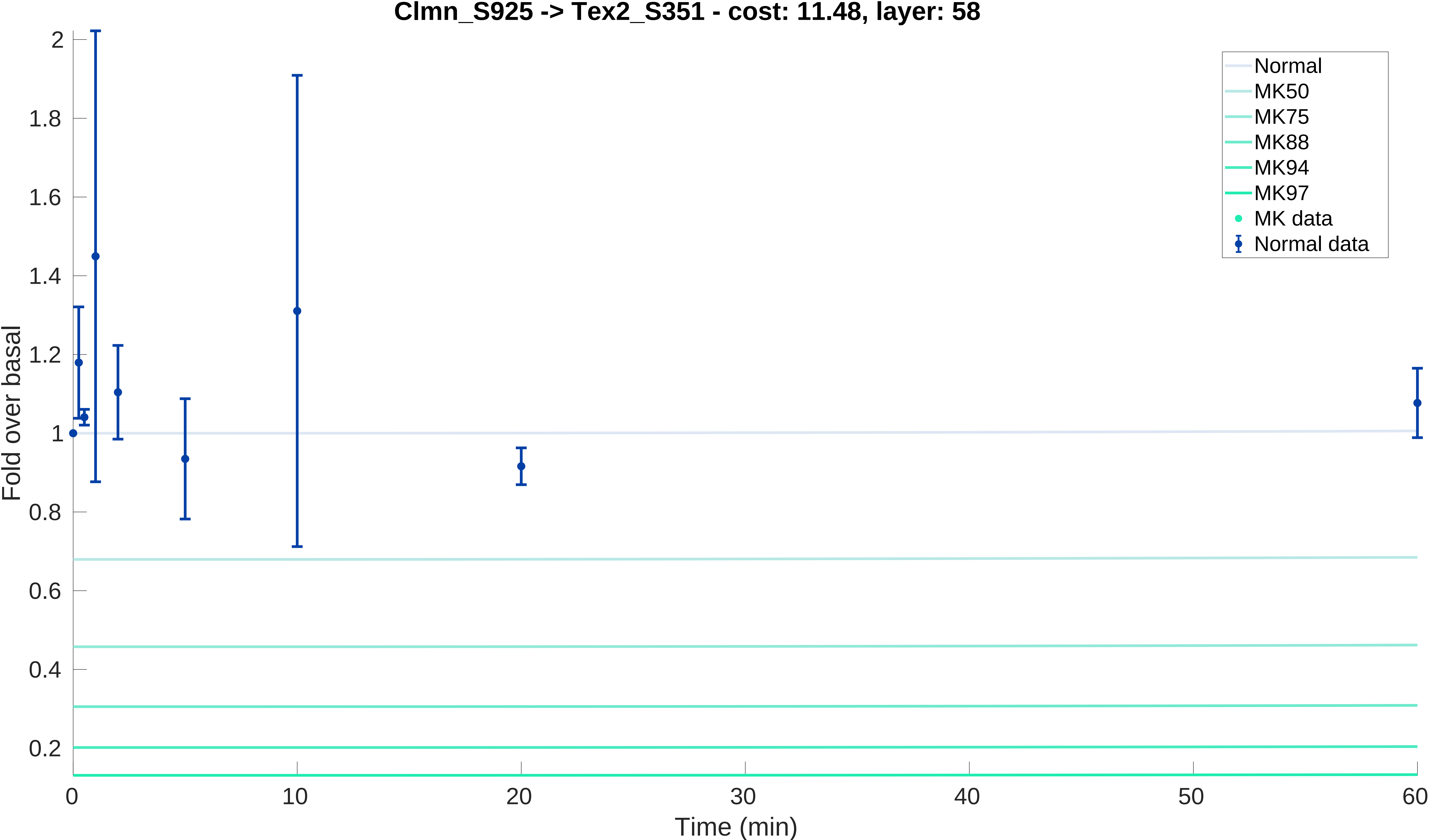

Clmn_S925 -> Tex2_S351 - cost: 11.48, layer: 58
Fold over basal
Time (min)
0
10
20
30
40
50
60
0.2
0.4
0.6
0.8
1
1.2
1.4
1.6
1.8
2
Normal
MK50
MK75
MK88
MK94
MK97
MK data
Normal data

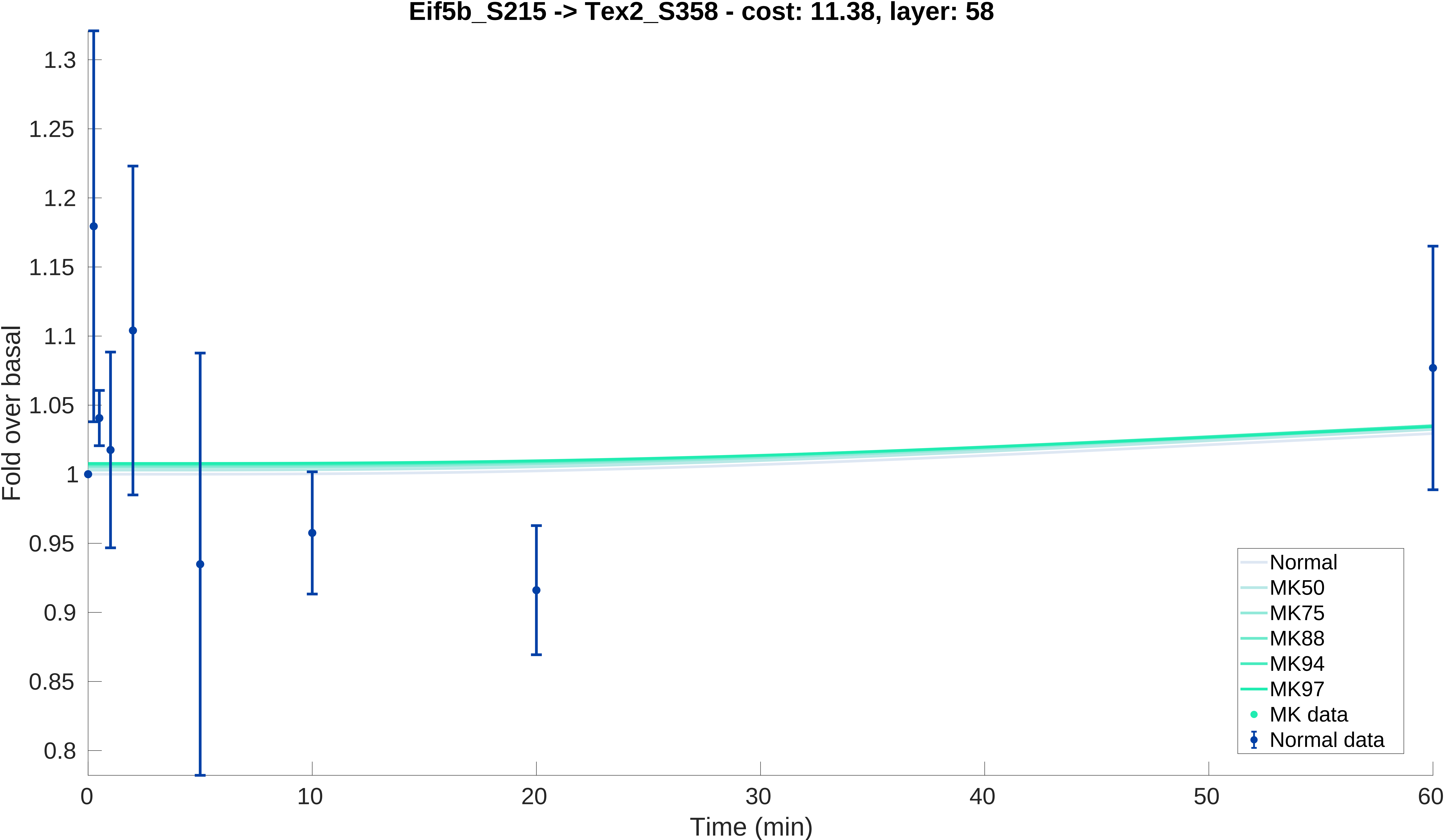

Eif5b_S215 -> Tex2_S358 - cost: 11.38, layer: 58
Fold over basal
1.3
1.25
1.2
1.15
1.1
1.05
1
0.95
0.9
0.85
0.8
0
10
20
30
40
50
60
Time (min)
Normal
MK50
MK75
MK88
MK94
MK97
MK data
Normal data

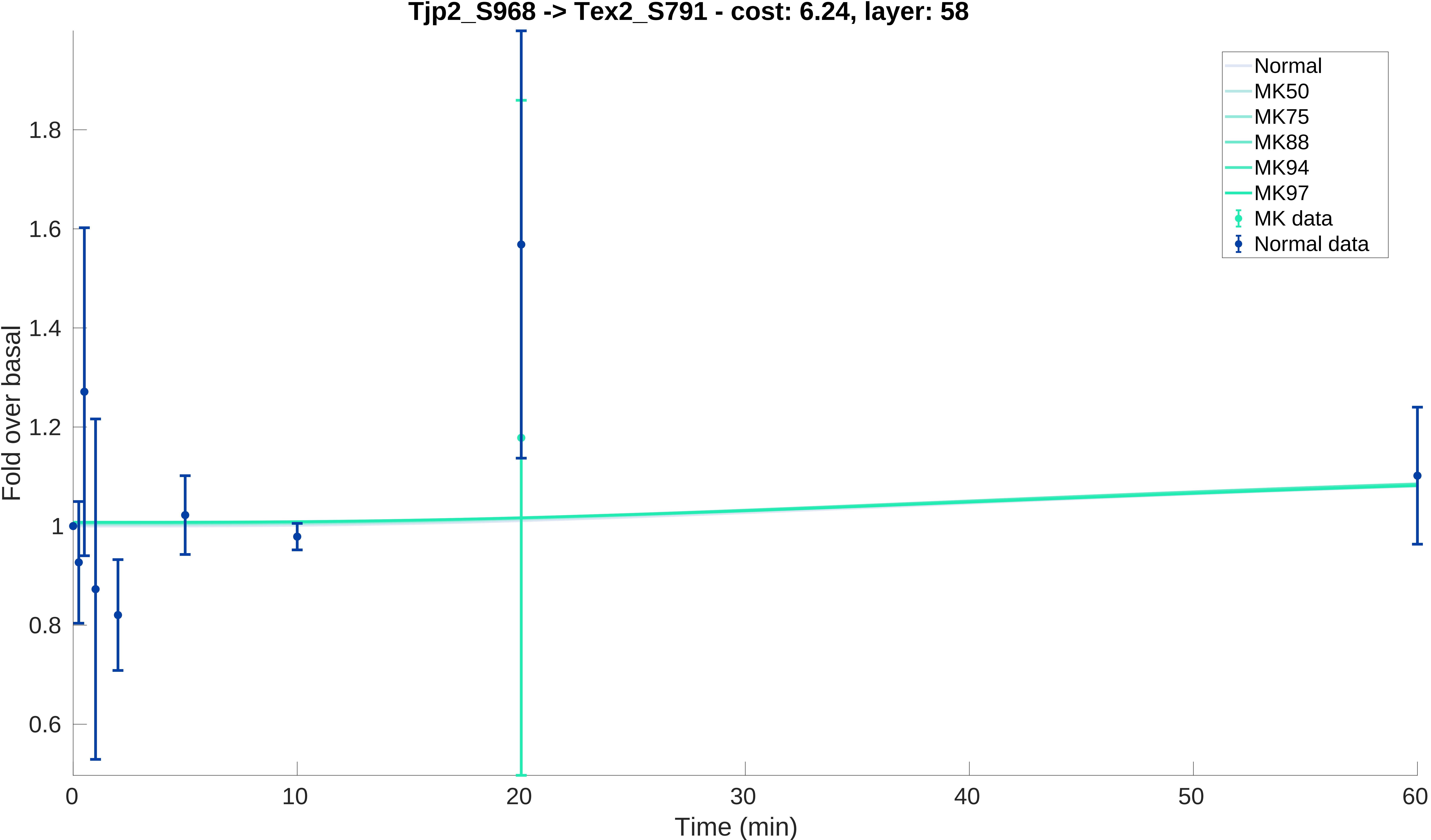

Tjp2_S968 -> Tex2_S791 - cost: 6.24, layer: 58
Fold over basal
Time (min)
0.6
0.8
1
1.2
1.4
1.6
1.8
0
10
20
30
40
50
60
Normal
MK50
MK75
MK88
MK94
MK97
MK data
Normal data

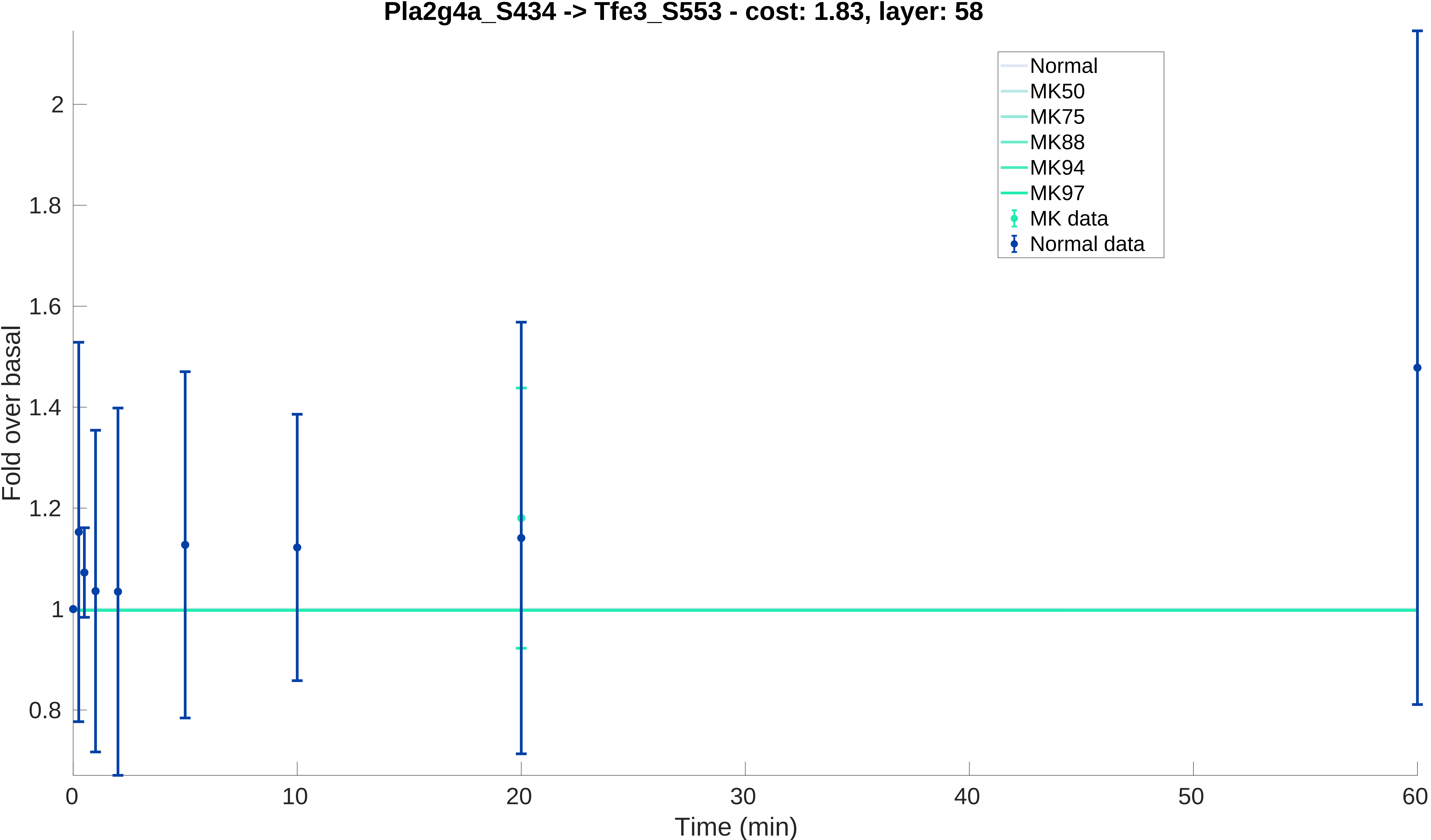

Pla2g4a_S434 -> Tfe3_S553 - cost: 1.83, layer: 58
Normal
MK50
MK75
MK88
MK94
MK97
MK data
Normal data
Fold over basal
2
1.8
1.6
1.4
1.2
1
0.8
0
10
20
30
40
50
60
Time (min)

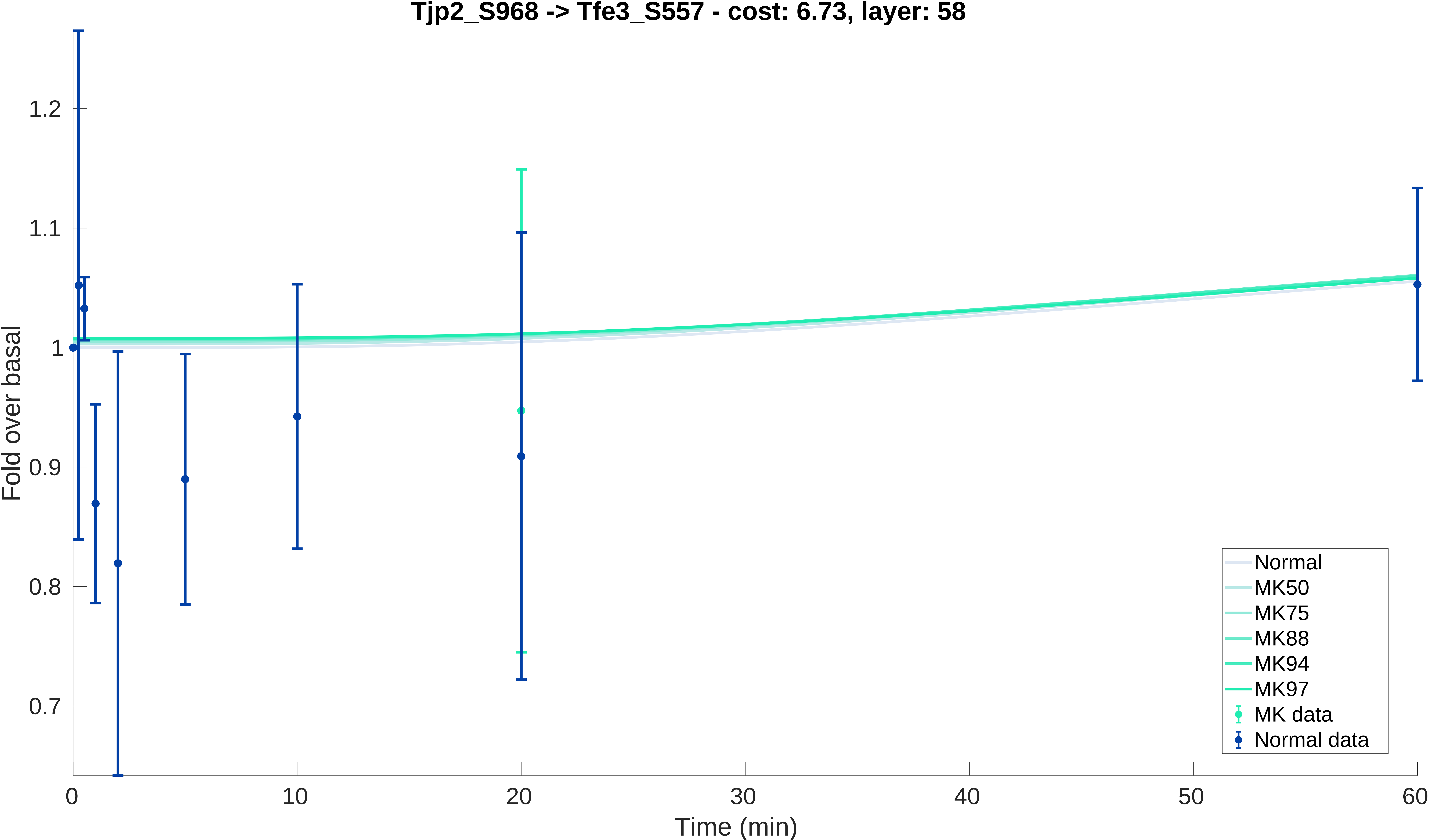

Tjp2_S968 -> Tfe3_S557 - cost: 6.73, layer: 58
Fold over basal
Time (min)
1.2
1.1
1
0.9
0.8
0.7
0
10
20
30
40
50
60
Normal
MK50
MK75
MK88
MK94
MK97
MK data
Normal data

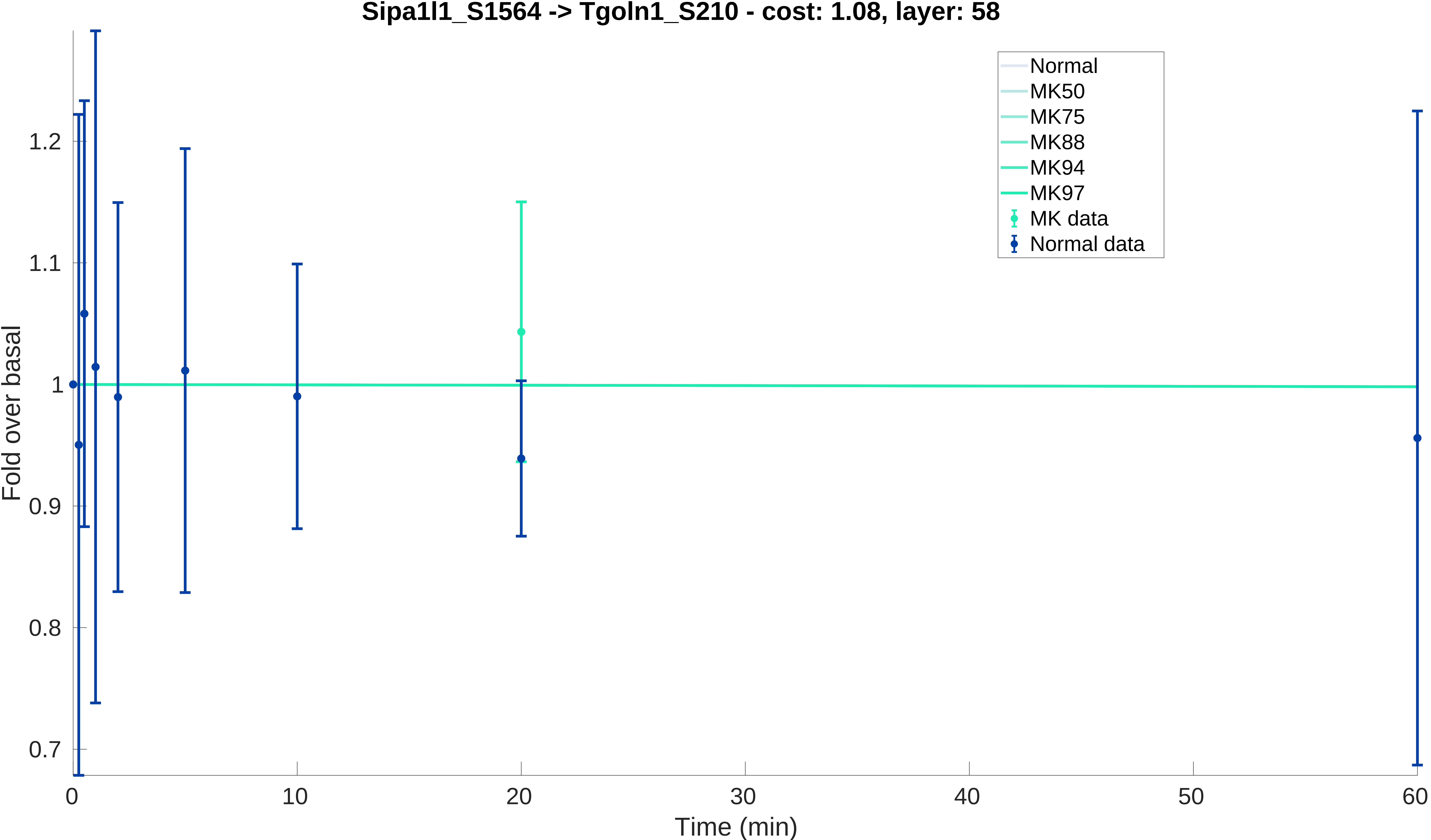

Sipa1l1_S1564 -> Tgoln1_S210 - cost: 1.08, layer: 58
Normal
MK50
MK75
MK88
MK94
MK97
MK data
Normal data
Fold over basal
1.2
1.1
1
0.9
0.8
0.7
0
10
20
30
40
50
60
Time (min)

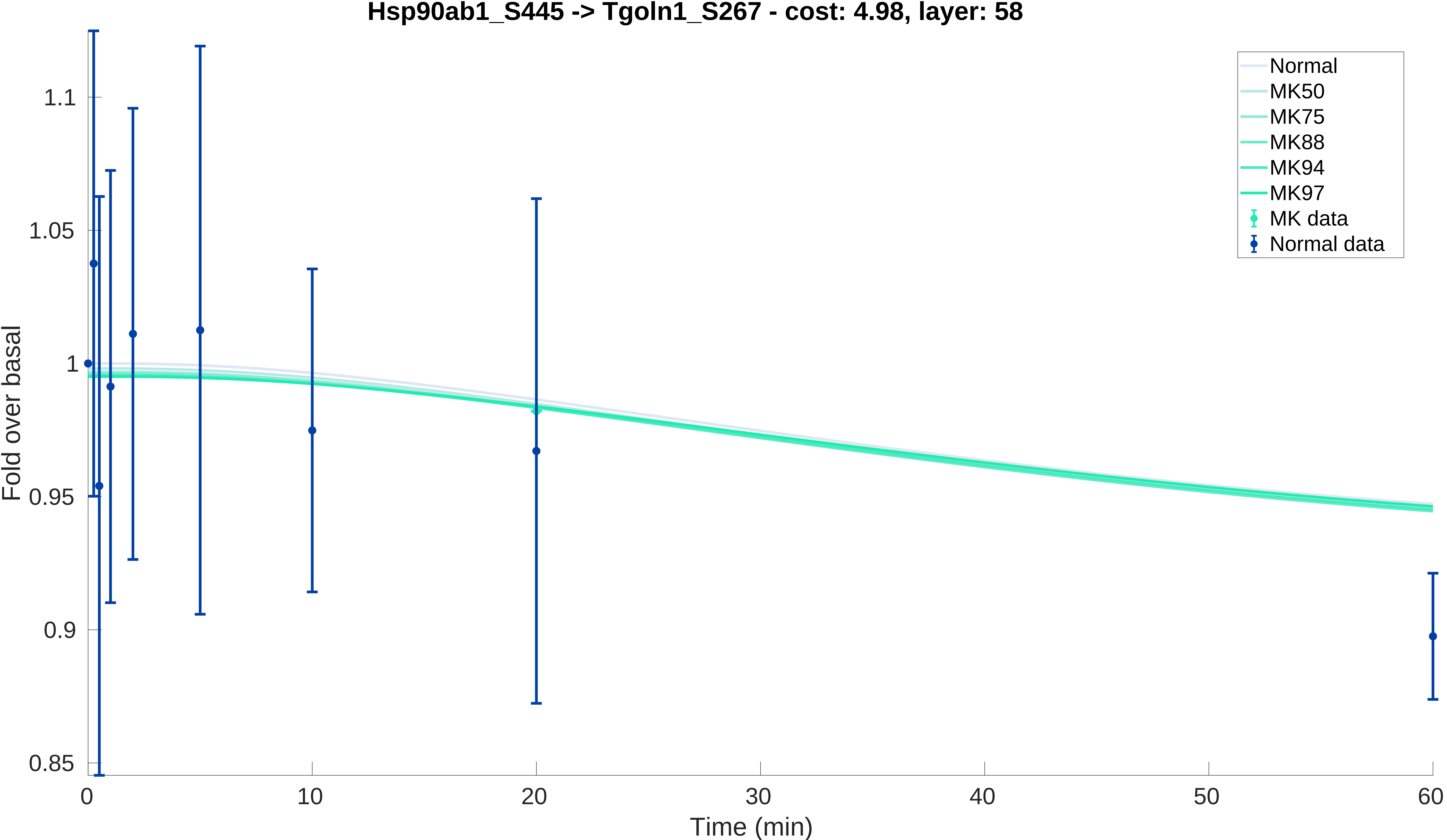

Hsp90ab1_S445 -> Tgoln1_S267 - cost: 4.98, layer: 58
Fold over basal
Time (min)
1.1
1.05
1
0.95
0.9
0.85
0
10
20
30
40
50
60
Normal
MK50
MK75
MK88
MK94
MK97
MK data
Normal data

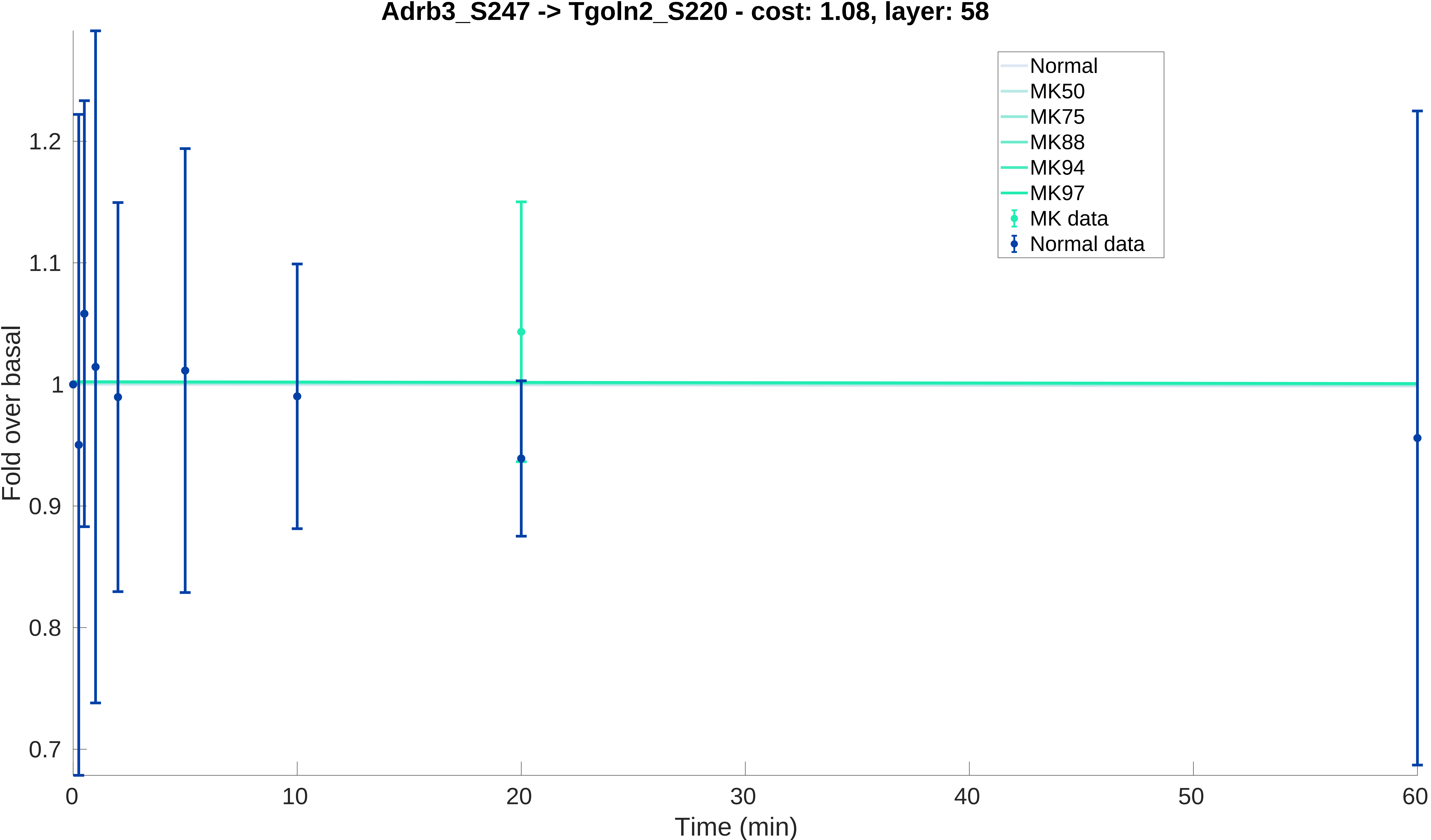

Adrb3_S247 -> Tgoln2_S220 - cost: 1.08, layer: 58
Normal
MK50
MK75
MK88
MK94
MK97
MK data
Normal data
Fold over basal
1.2
1.1
1
0.9
0.8
0.7
0
10
20
30
40
50
60
Time (min)

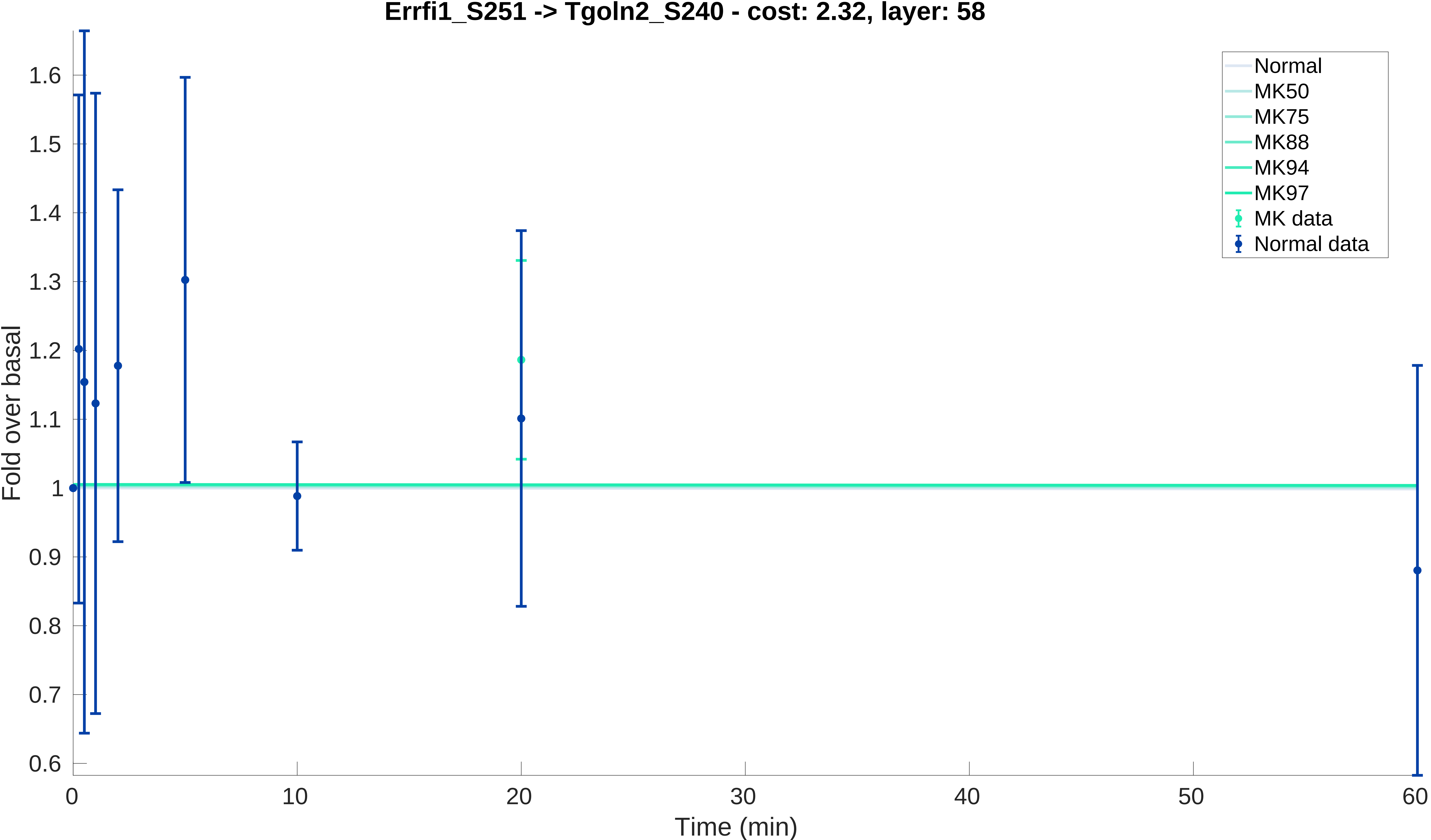

Errfi1_S251 -> Tgoln2_S240 - cost: 2.32, layer: 58
Fold over basal
Time (min)
1.6
1.5
1.4
1.3
1.2
1.1
1
0.9
0.8
0.7
0.6
0
10
20
30
40
50
60
Normal
MK50
MK75
MK88
MK94
MK97
MK data
Normal data

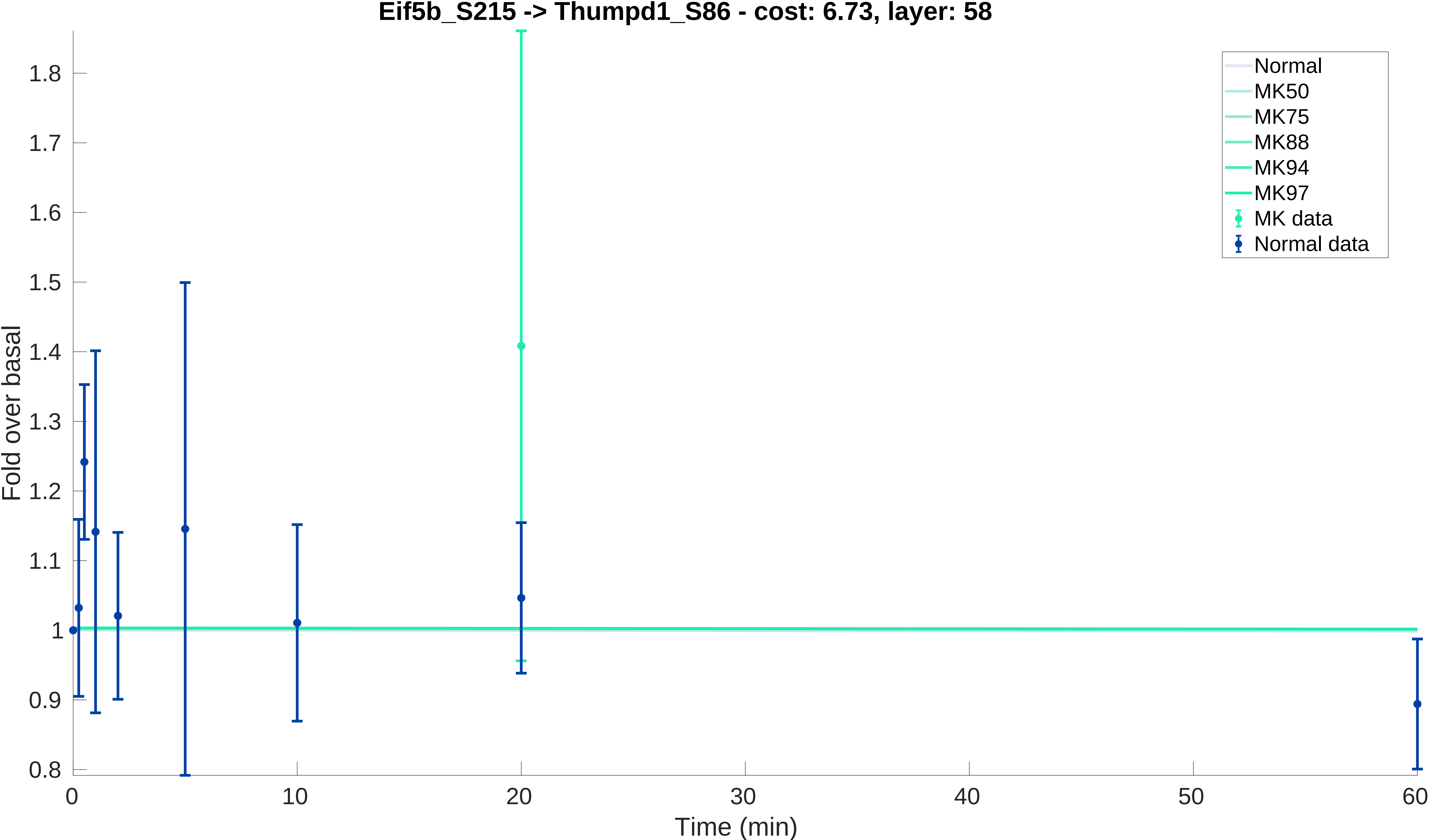

Eif5b_S215 -> Thumpd1_S86 - cost: 6.73, layer: 58
Fold over basal
Time (min)
0.8
0.9
1
1.1
1.2
1.3
1.4
1.5
1.6
1.7
1.8
0
10
20
30
40
50
60
Normal
MK50
MK75
MK88
MK94
MK97
MK data
Normal data

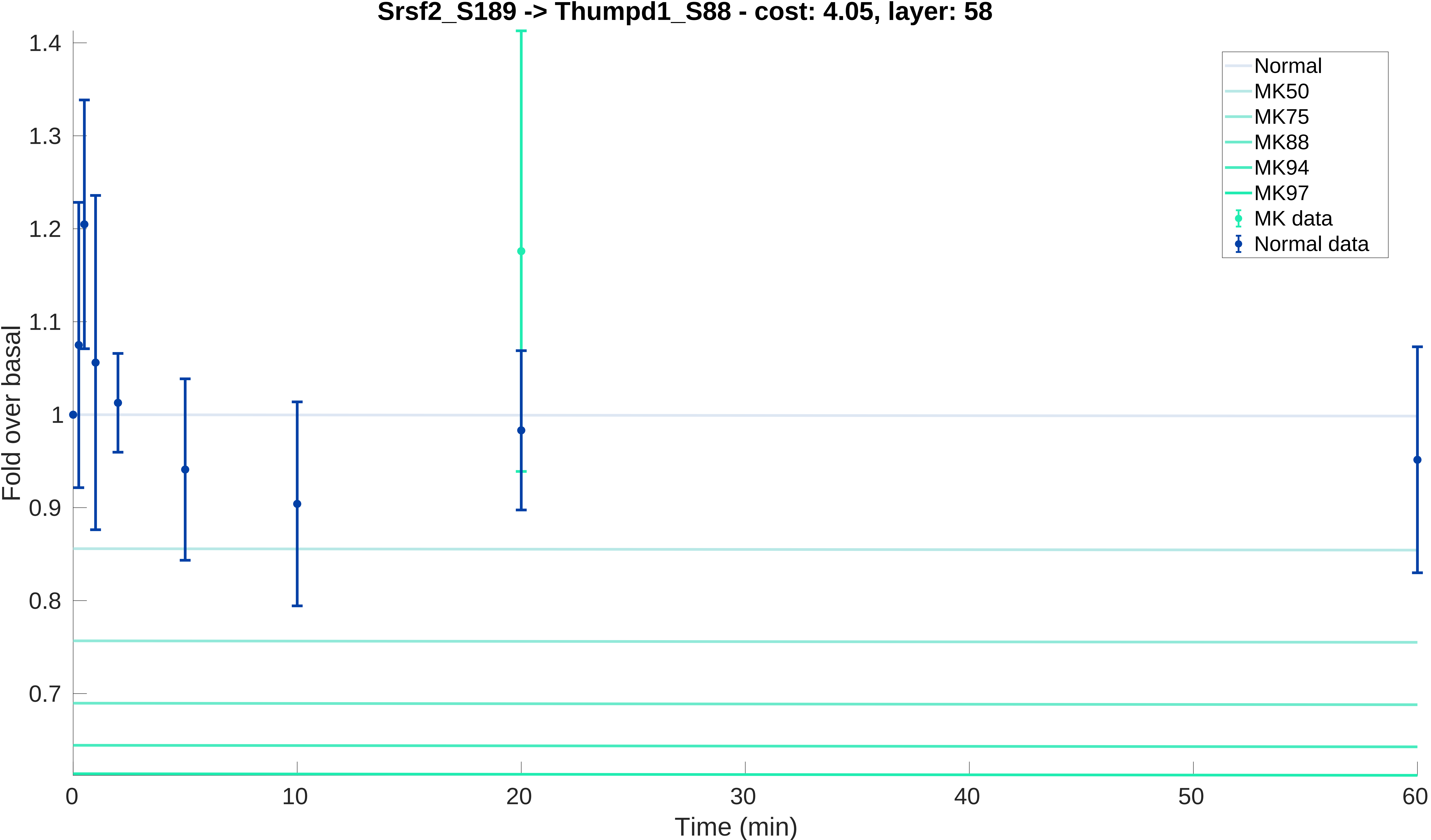
Srsf2_S189 -> Thumpd1_S88 - cost: 4.05, layer: 58
Fold over basal
Time (min)
1.4
1.3
1.2
1.1
1
0.9
0.8
0.7
0
10
20
30
40
50
60
Normal
MK50
MK75
MK88
MK94
MK97
MK data
Normal data

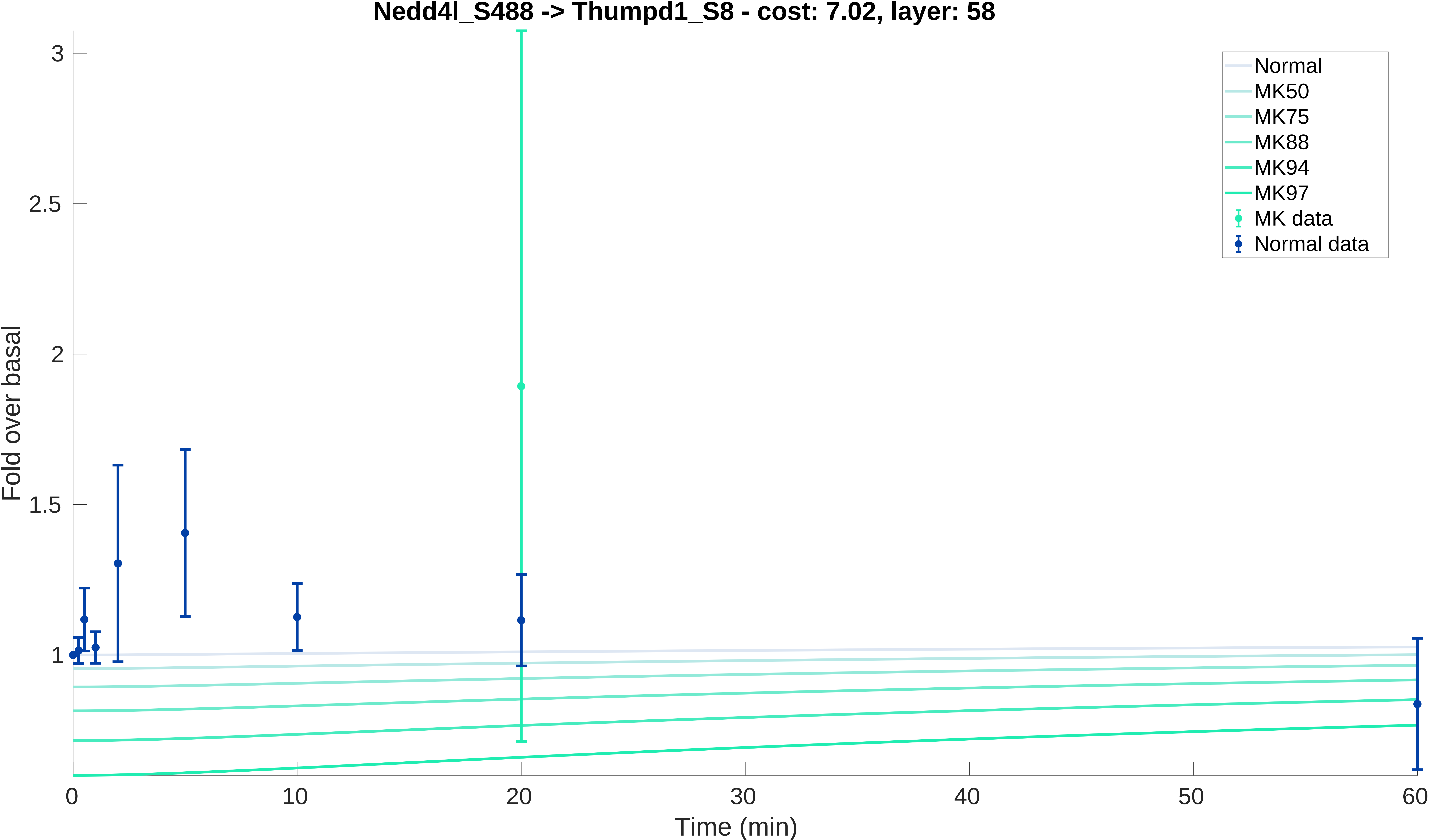

Nedd4l_S488 -> Thumpd1_S8 - cost: 7.02, layer: 58
Fold over basal
Time (min)
3
2.5
2
1.5
1
0
10
20
30
40
50
60
Normal
MK50
MK75
MK88
MK94
MK97
MK data
Normal data

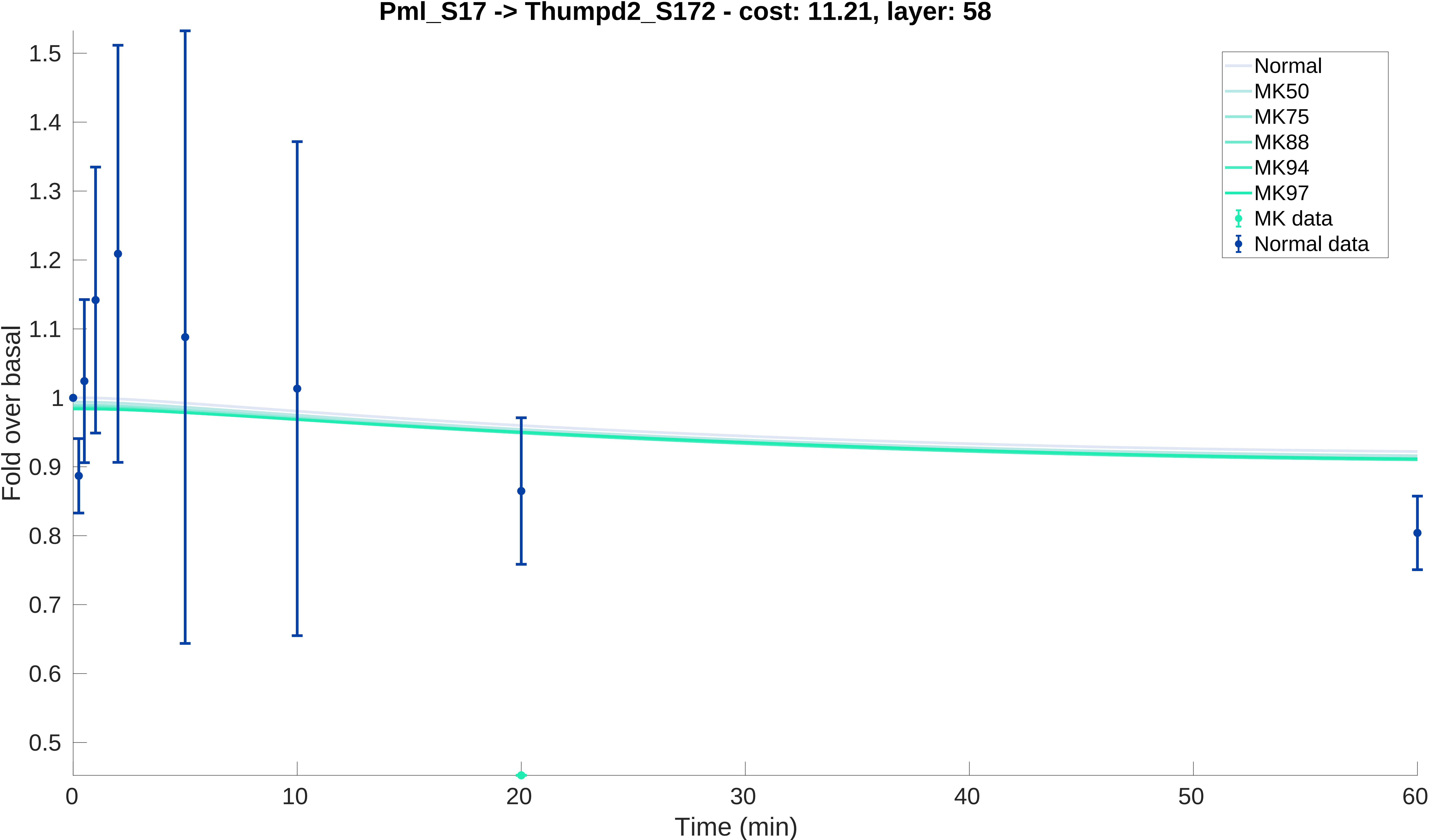

Pml_S17 -> Thumpd2_S172 - cost: 11.21, layer: 58
Fold over basal
Time (min)
1.5
1.4
1.3
1.2
1.1
1
0.9
0.8
0.7
0.6
0.5
0
10
20
30
40
50
60
Normal
MK50
MK75
MK88
MK94
MK97
MK data
Normal data

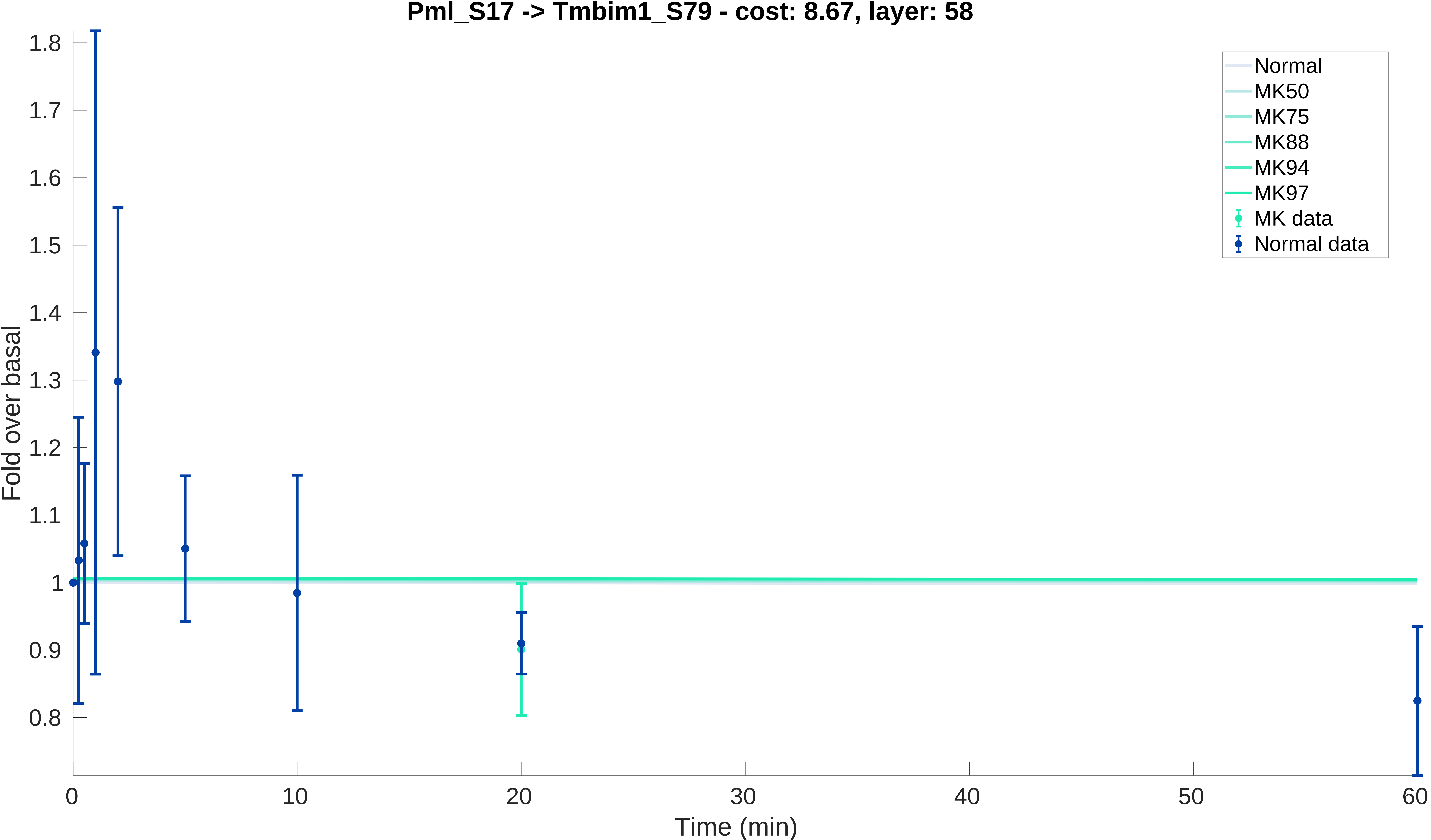

Pml_S17 -> Tmbim1_S79 - cost: 8.67, layer: 58
Fold over basal
Time (min)
1.8
1.7
1.6
1.5
1.4
1.3
1.2
1.1
1
0.9
0.8
0
10
20
30
40
50
60
Normal
MK50
MK75
MK88
MK94
MK97
MK data
Normal data

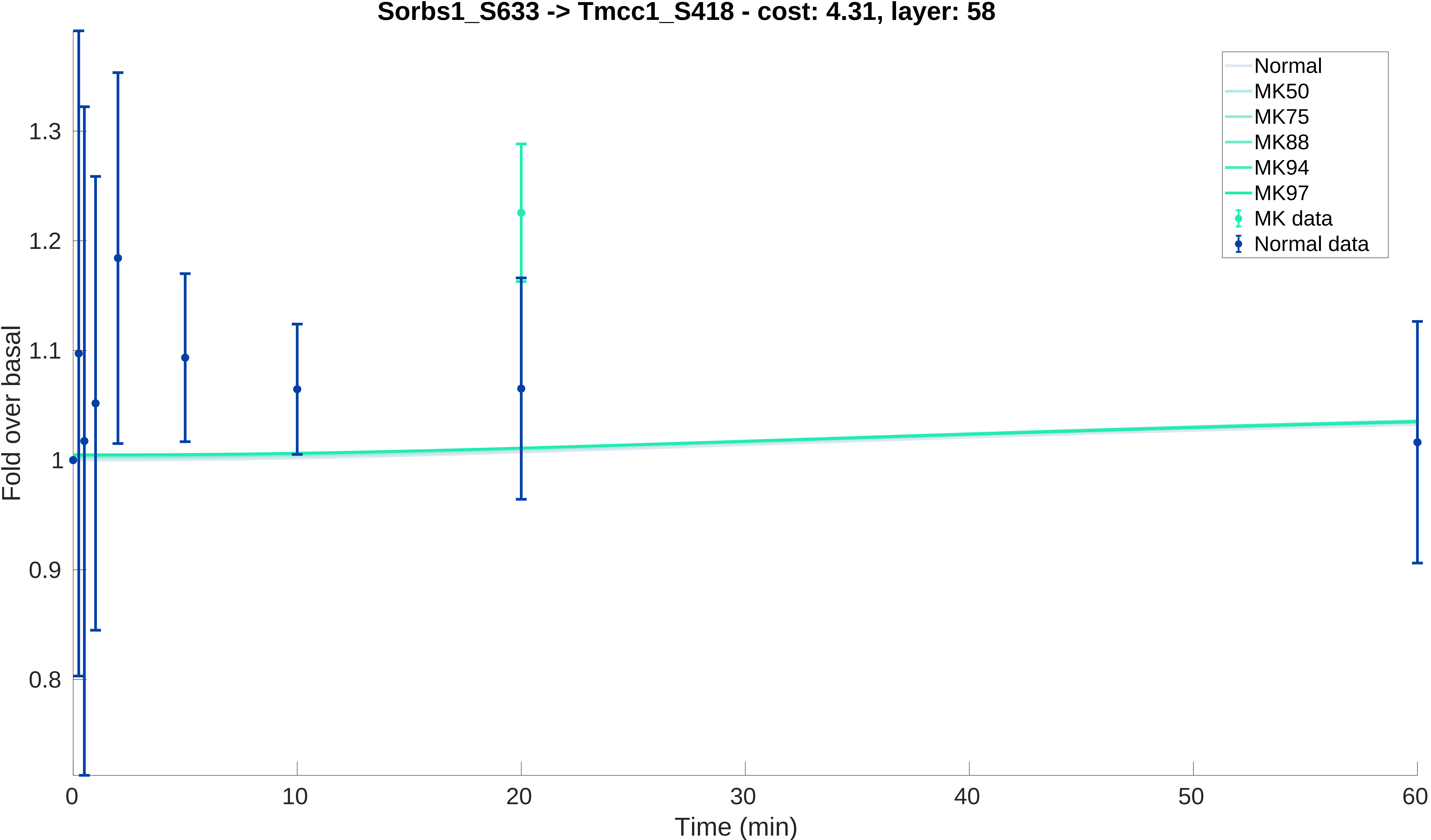

Sorbs1_S633 -> Tmcc1_S418 - cost: 4.31, layer: 58
Fold over basal
Time (min)
1.3
1.2
1.1
1
0.9
0.8
0
10
20
30
40
50
60
Normal
MK50
MK75
MK88
MK94
MK97
MK data
Normal data

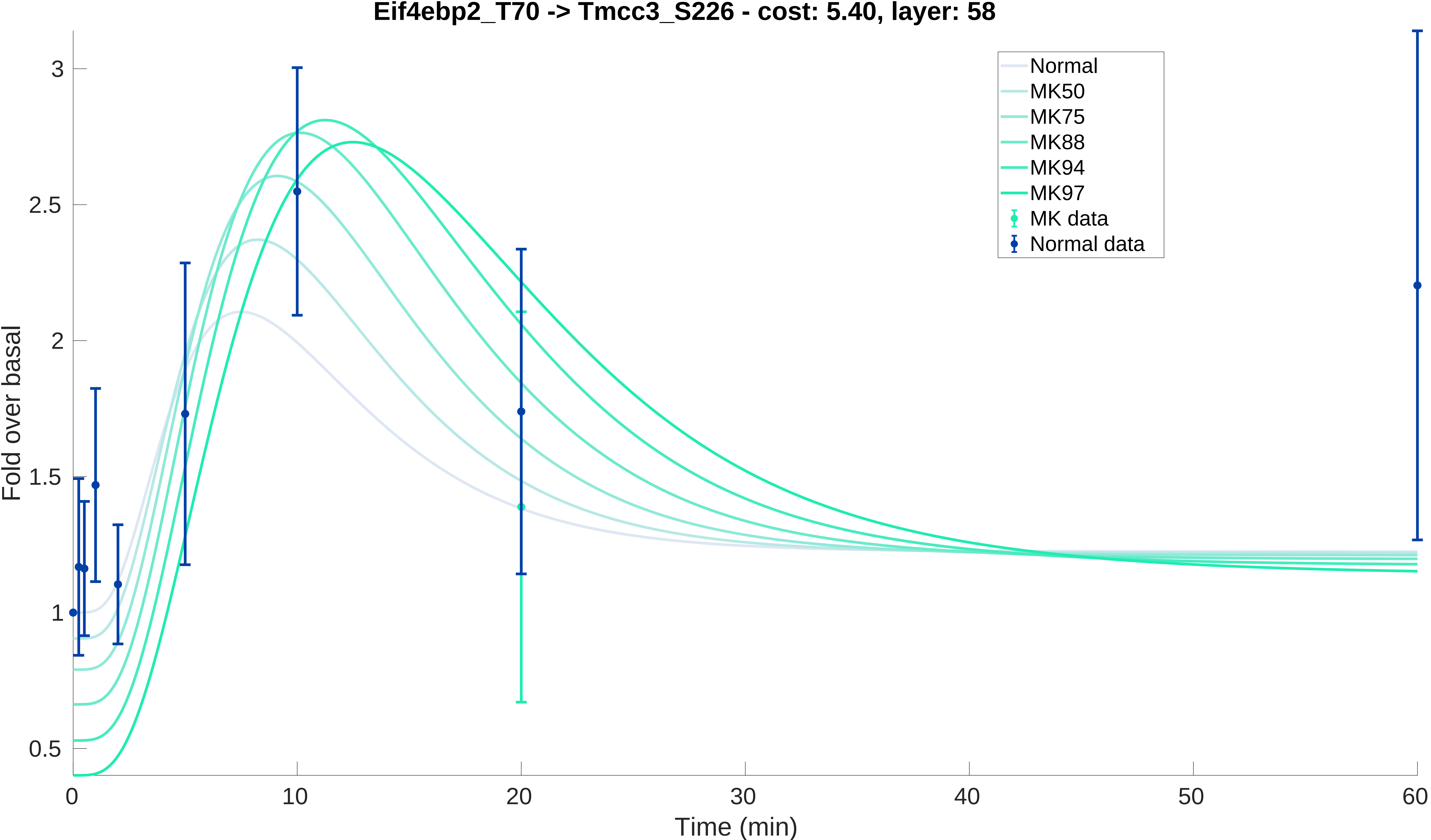

Eif4ebp2_T70 -> Tmcc3_S226 - cost: 5.40, layer: 58
Fold over basal
Time (min)
0.5
1
1.5
2
2.5
3
0
10
20
30
40
50
60
Normal
MK50
MK75
MK88
MK94
MK97
MK data
Normal data

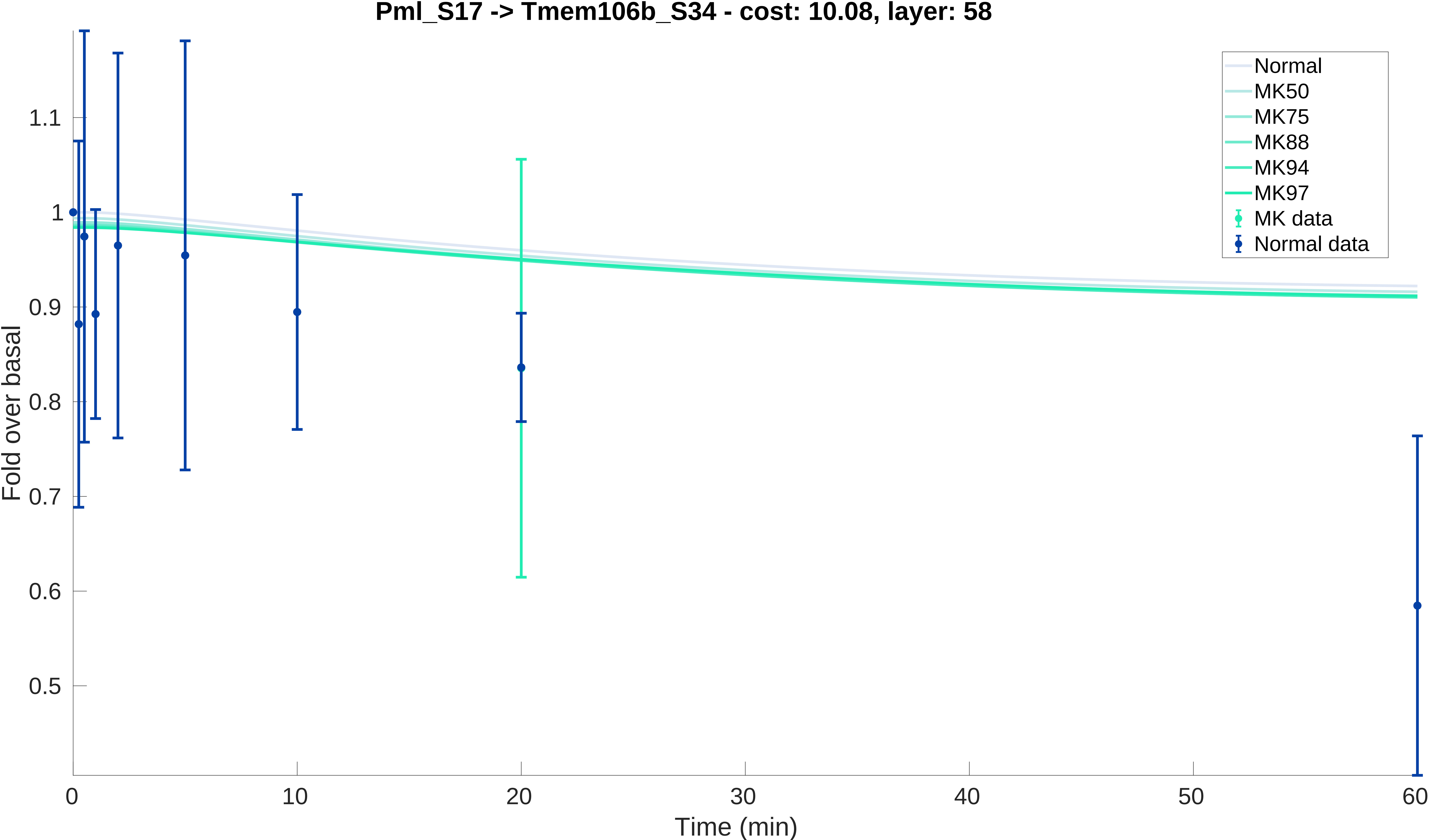
Pml_S17 -> Tmem106b_S34 - cost: 10.08, layer: 58
Fold over basal
Time (min)
0.5
0.6
0.7
0.8
0.9
1
1.1
0
10
20
30
40
50
60
Normal
MK50
MK75
MK88
MK94
MK97
MK data
Normal data

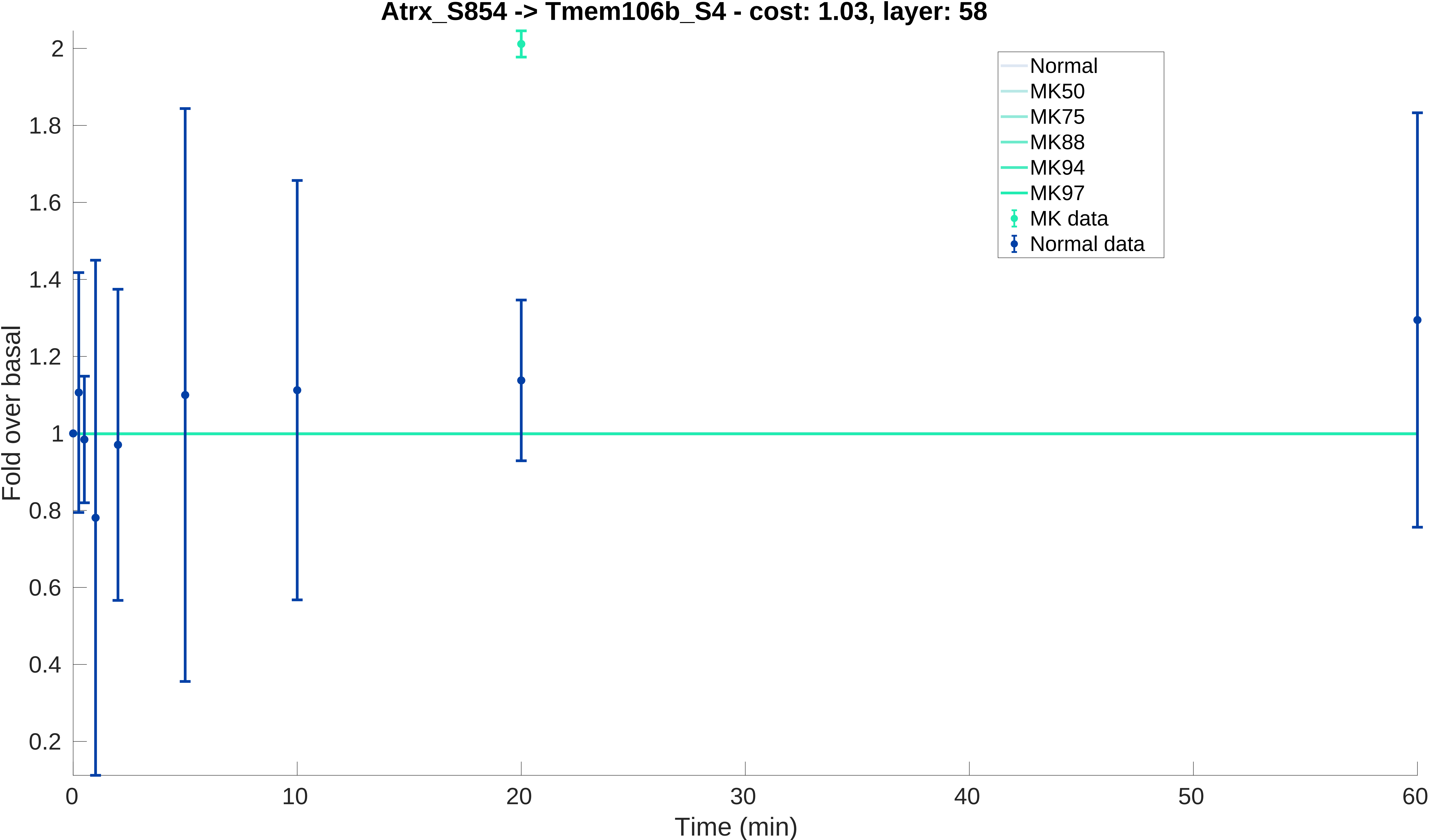

Atrx_S854 -> Tmem106b_S4 - cost: 1.03, layer: 58
Normal
MK50
MK75
MK88
MK94
MK97
MK data
Normal data
Fold over basal
Time (min)
0
10
20
30
40
50
60
0.2
0.4
0.6
0.8
1
1.2
1.4
1.6
1.8
2

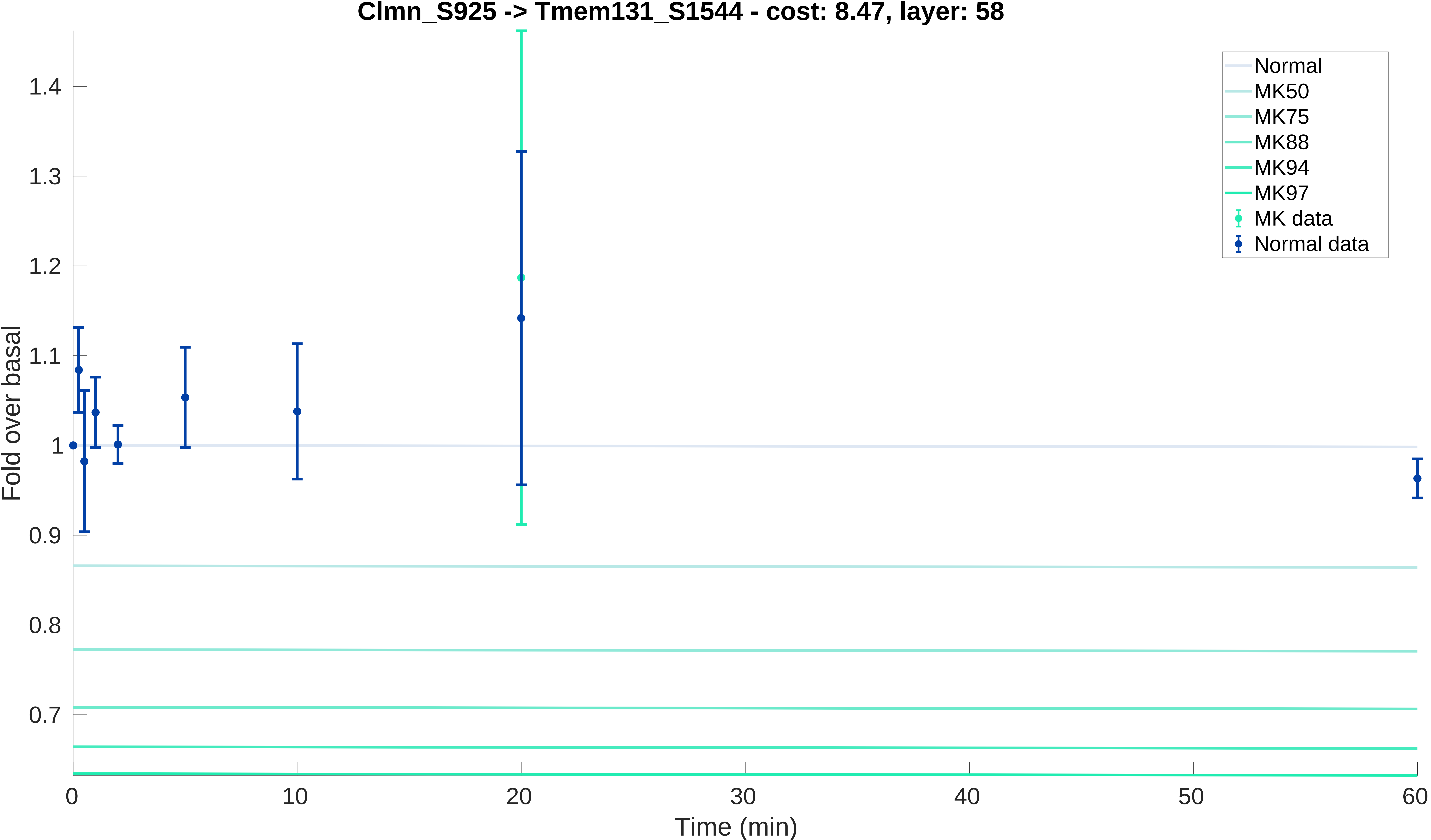
Clmn_S925 -> Tmem131_S1544 - cost: 8.47, layer: 58
Fold over basal
Time (min)
1.4
1.3
1.2
1.1
1
0.9
0.8
0.7
0
10
20
30
40
50
60
Normal
MK50
MK75
MK88
MK94
MK97
MK data
Normal data

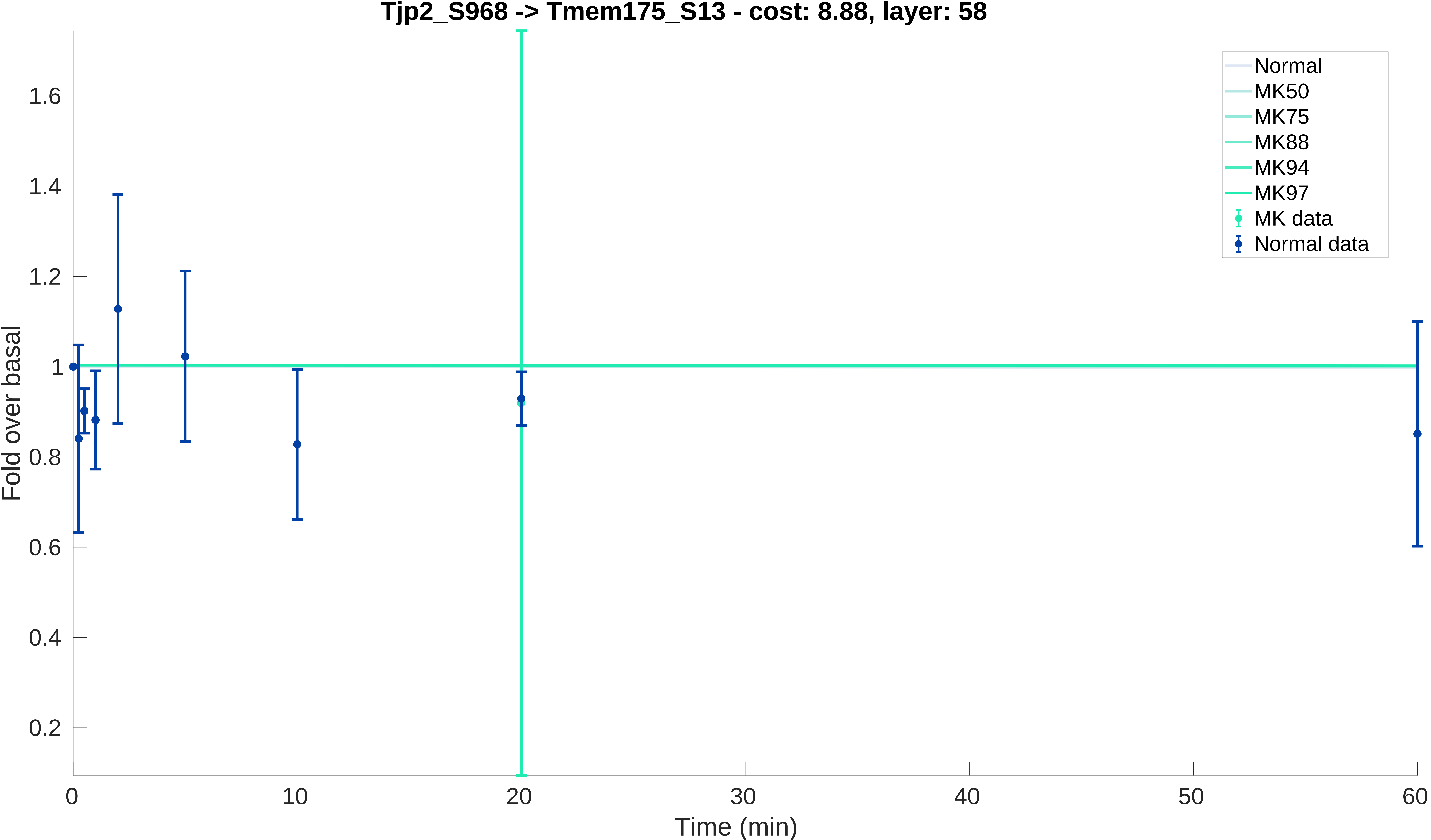
Tjp2_S968 -> Tmem175_S13 - cost: 8.88, layer: 58
Fold over basal
Time (min)
0.2
0.4
0.6
0.8
1
1.2
1.4
1.6
0
10
20
30
40
50
60
Normal
MK50
MK75
MK88
MK94
MK97
MK data
Normal data

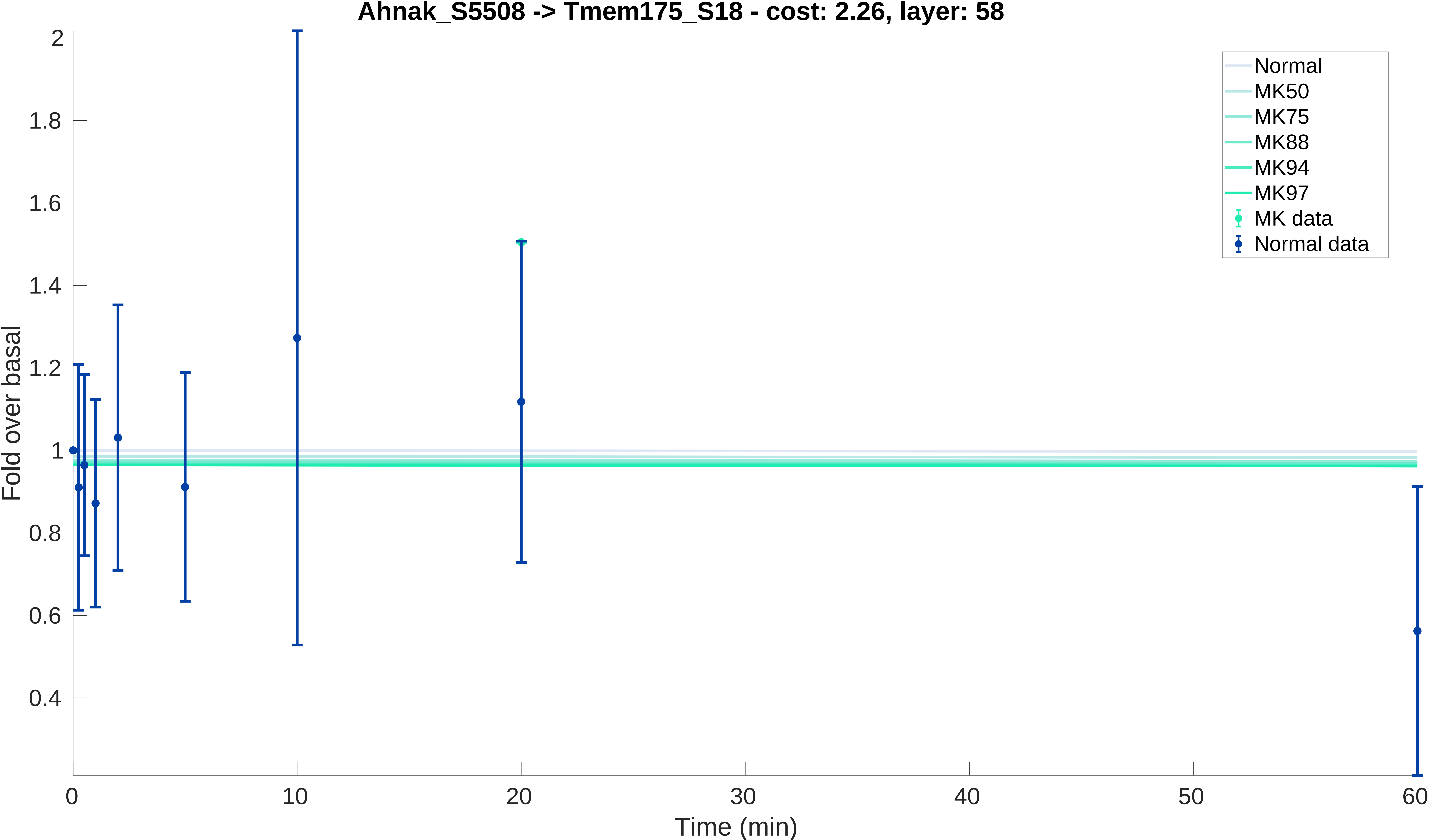

Ahnak_S5508 -> Tmem175_S18 - cost: 2.26, layer: 58
Fold over basal
Time (min)
0.4
0.6
0.8
1
1.2
1.4
1.6
1.8
2
0
10
20
30
40
50
60
Normal
MK50
MK75
MK88
MK94
MK97
MK data
Normal data

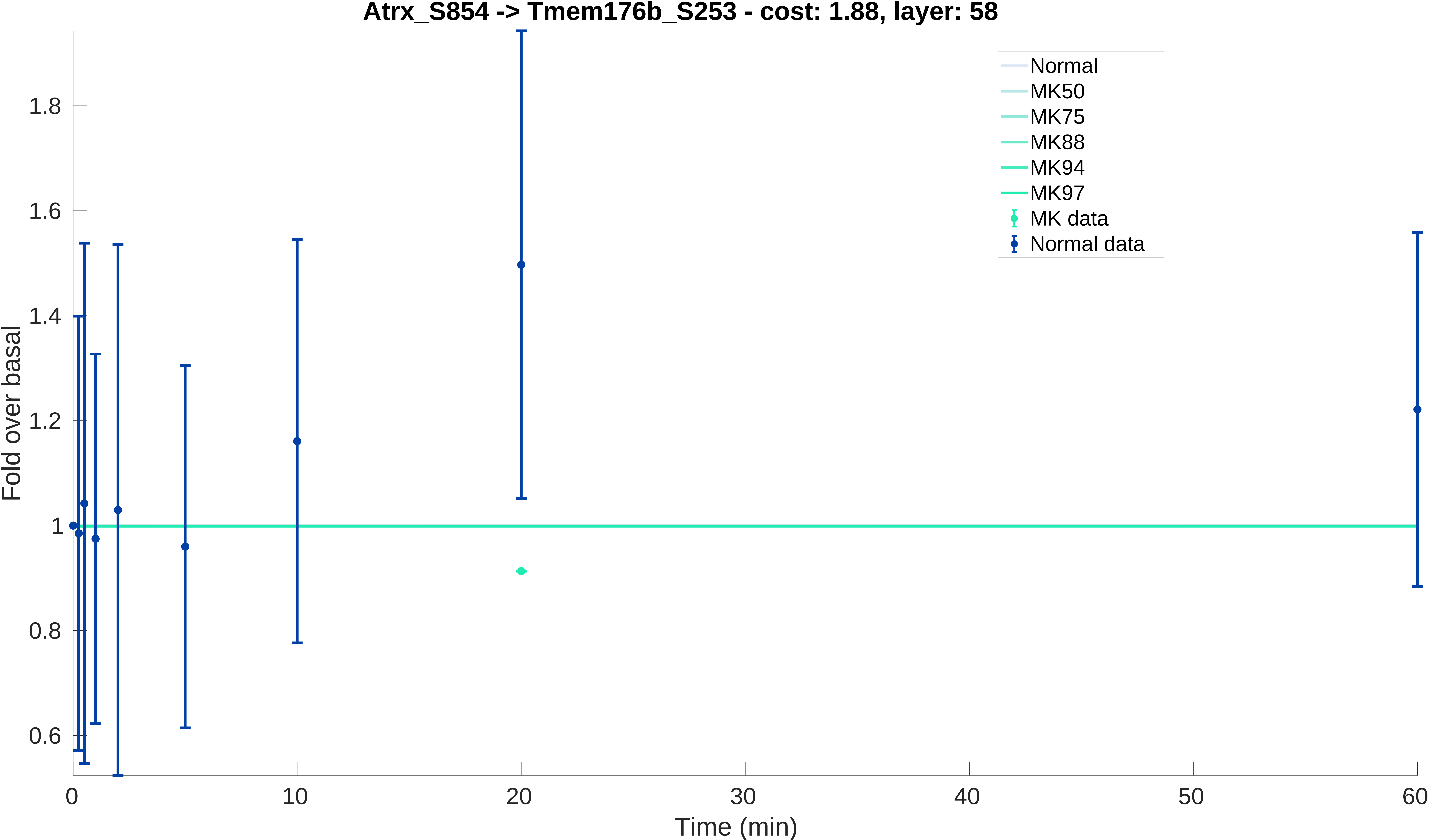

Atrx_S854 -> Tmem176b_S253 - cost: 1.88, layer: 58
Normal
MK50
MK75
MK88
MK94
MK97
MK data
Normal data
Fold over basal
1.8
1.6
1.4
1.2
1
0.8
0.6
0
10
20
30
40
50
60
Time (min)

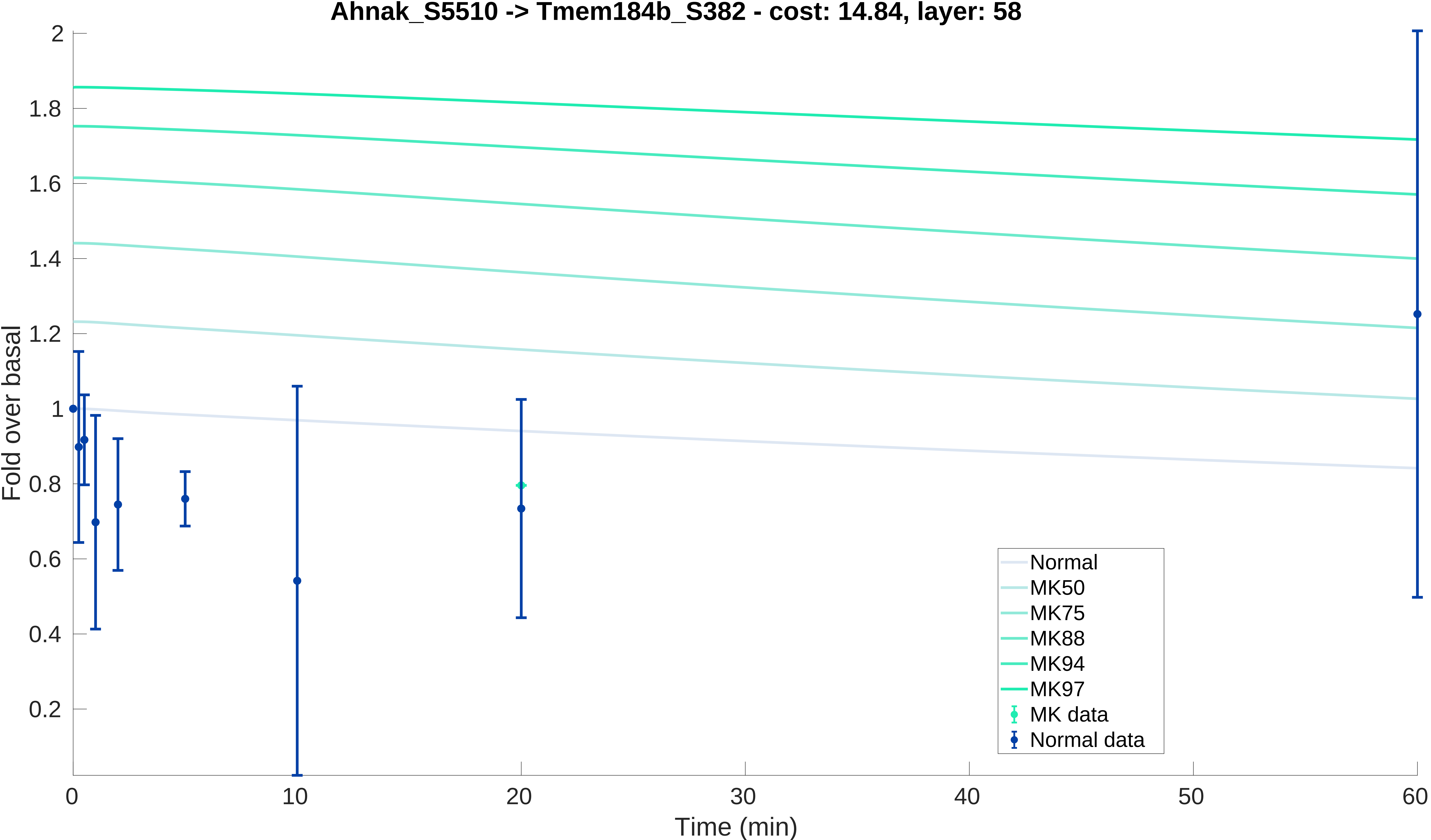

Ahnak_S5510 -> Tmem184b_S382 - cost: 14.84, layer: 58
Fold over basal
Time (min)
0
10
20
30
40
50
60
0.2
0.4
0.6
0.8
1
1.2
1.4
1.6
1.8
2
Normal
MK50
MK75
MK88
MK94
MK97
MK data
Normal data

#### Ccdc88a_S237 -> Tmem184b_S384 - cost: 5.48, layer: 58

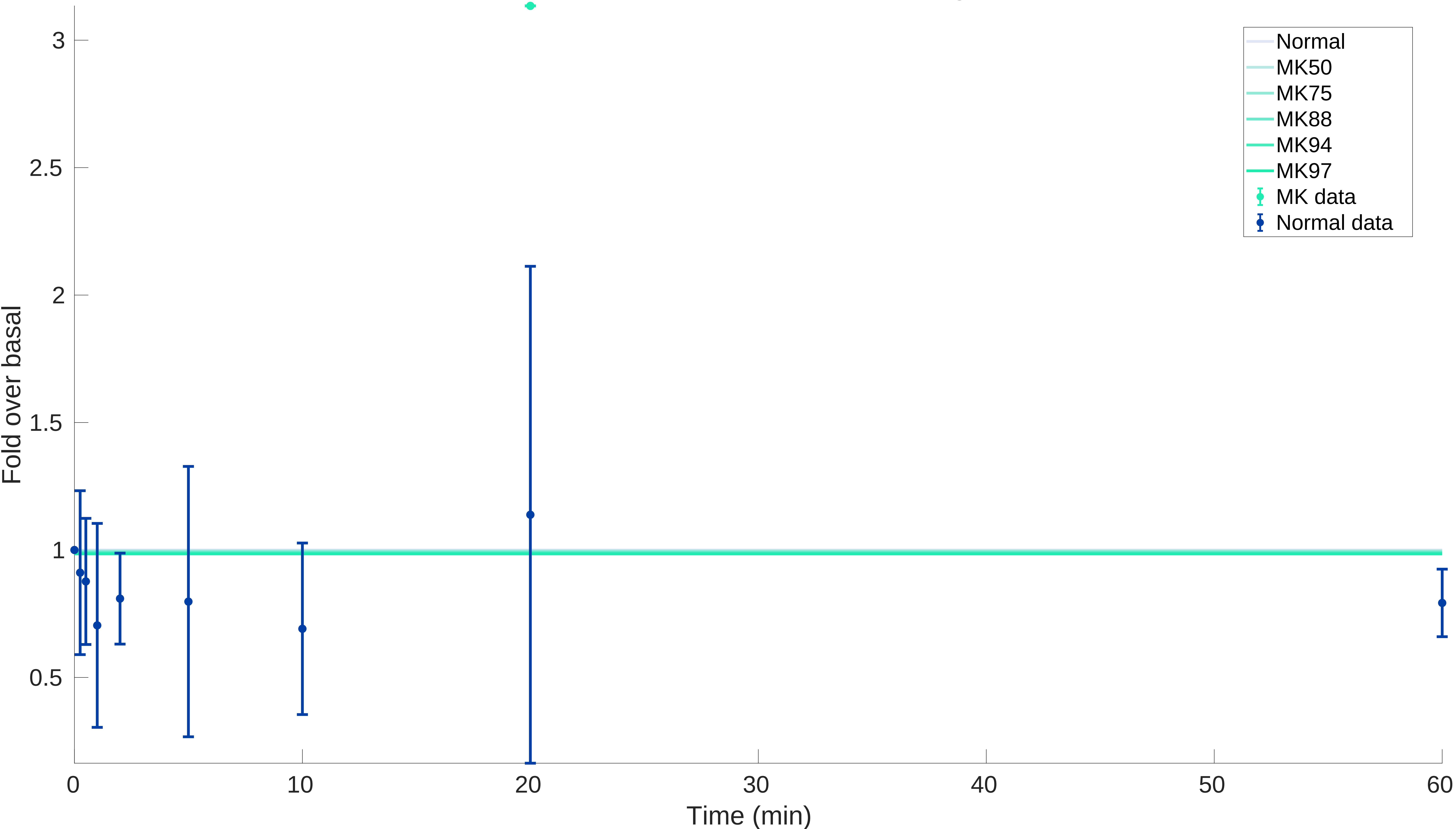

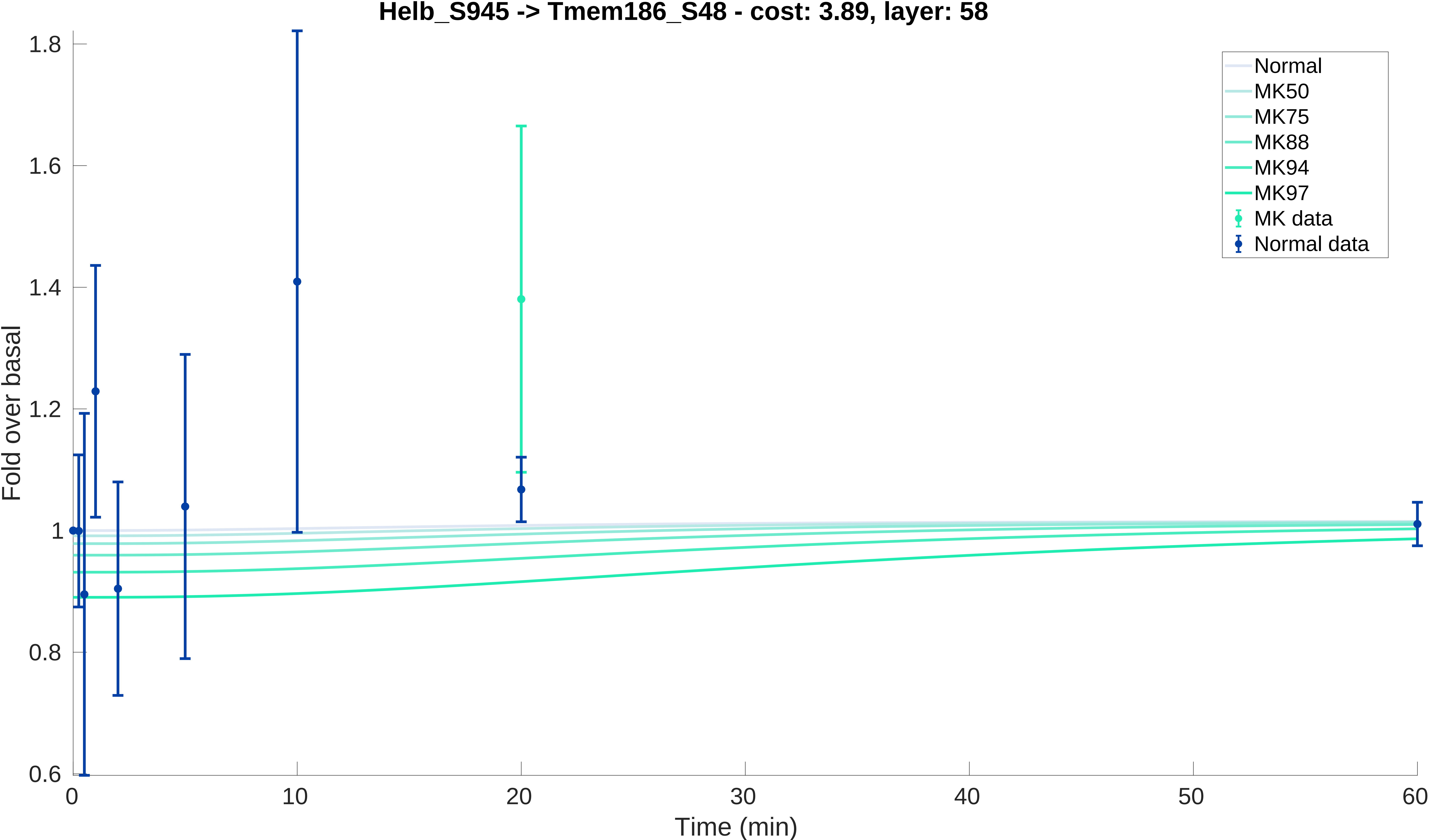

Helb_S945 -> Tmem186_S48 - cost: 3.89, layer: 58
Fold over basal
Time (min)
1.8
1.6
1.4
1.2
1
0.8
0.6
0
10
20
30
40
50
60
Normal
MK50
MK75
MK88
MK94
MK97
MK data
Normal data

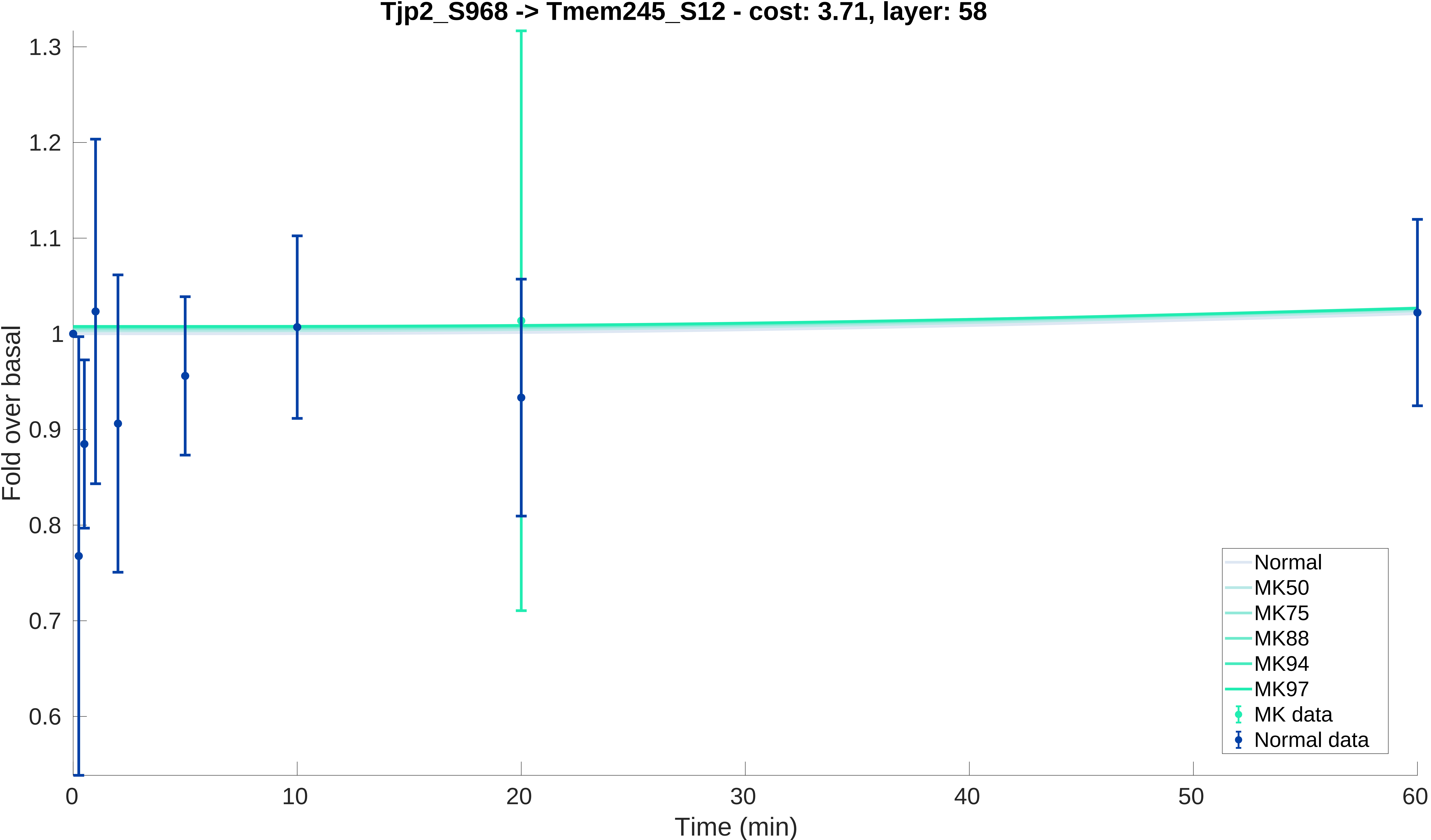

Tjp2_S968 -> Tmem245_S12 - cost: 3.71, layer: 58
Fold over basal
Time (min)
1.3
1.2
1.1
1
0.9
0.8
0.7
0.6
0
10
20
30
40
50
60
Normal
MK50
MK75
MK88
MK94
MK97
MK data
Normal data

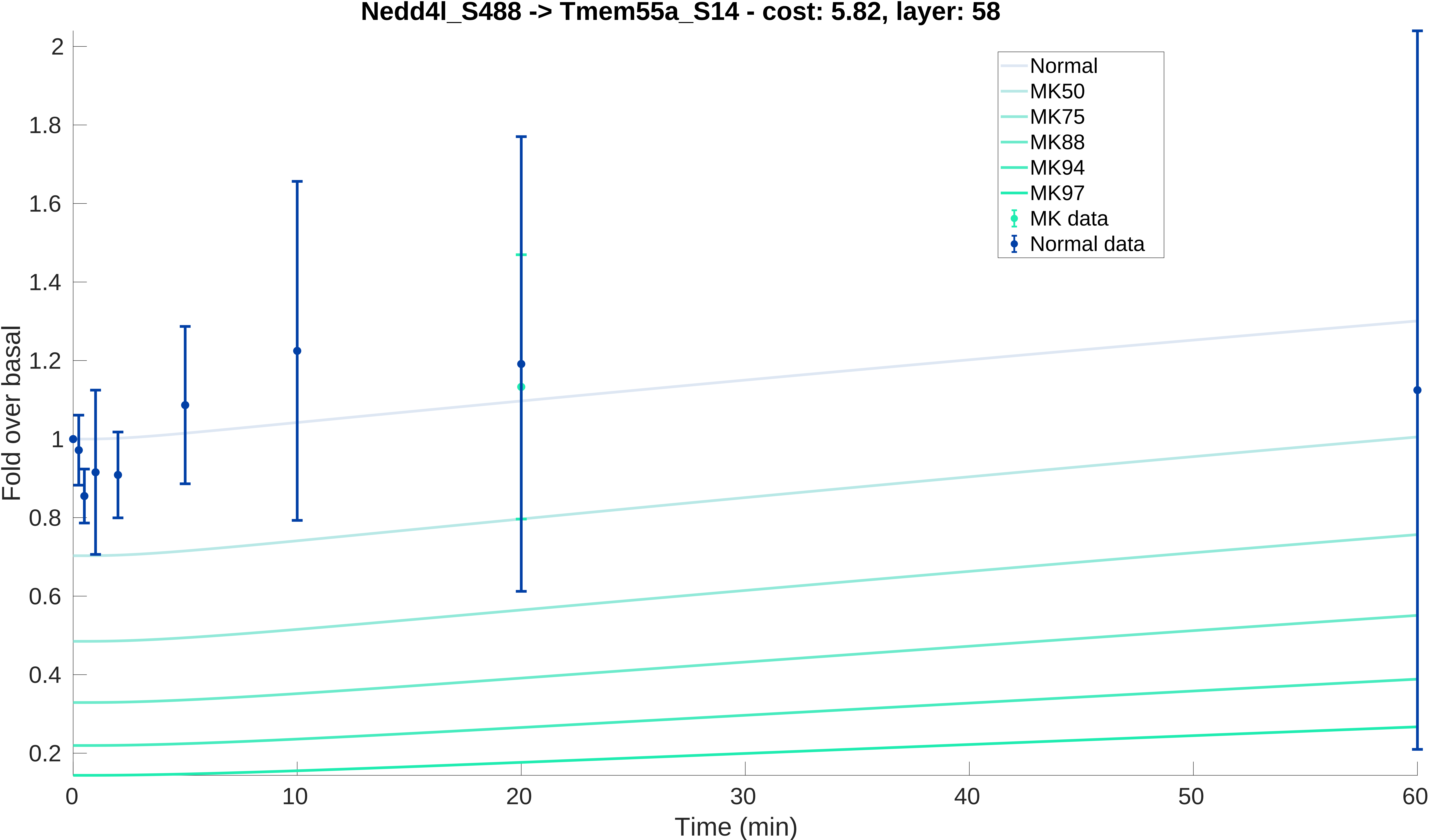
Nedd4l_S488 -> Tmem55a_S14 - cost: 5.82, layer: 58
Fold over basal
Time (min)
0
10
20
30
40
50
60
0.2
0.4
0.6
0.8
1
1.2
1.4
1.6
1.8
2
Normal
MK50
MK75
MK88
MK94
MK97
MK data
Normal data

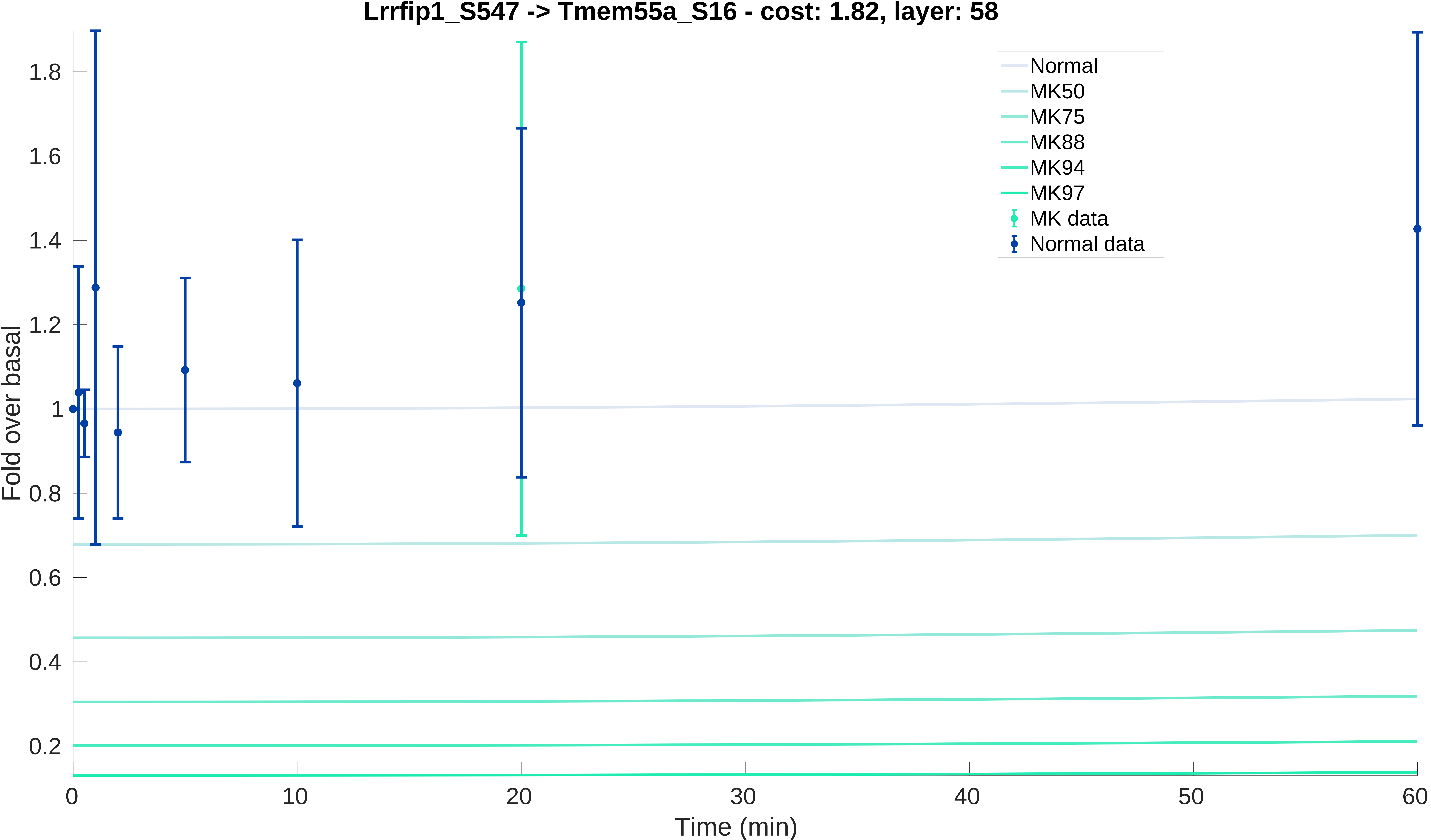

Lrrfip1_S547 -> Tmem55a_S16 - cost: 1.82, layer: 58
Fold over basal
Time (min)
0
10
20
30
40
50
60
0.2
0.4
0.6
0.8
1
1.2
1.4
1.6
1.8
Normal
MK50
MK75
MK88
MK94
MK97
MK data
Normal data

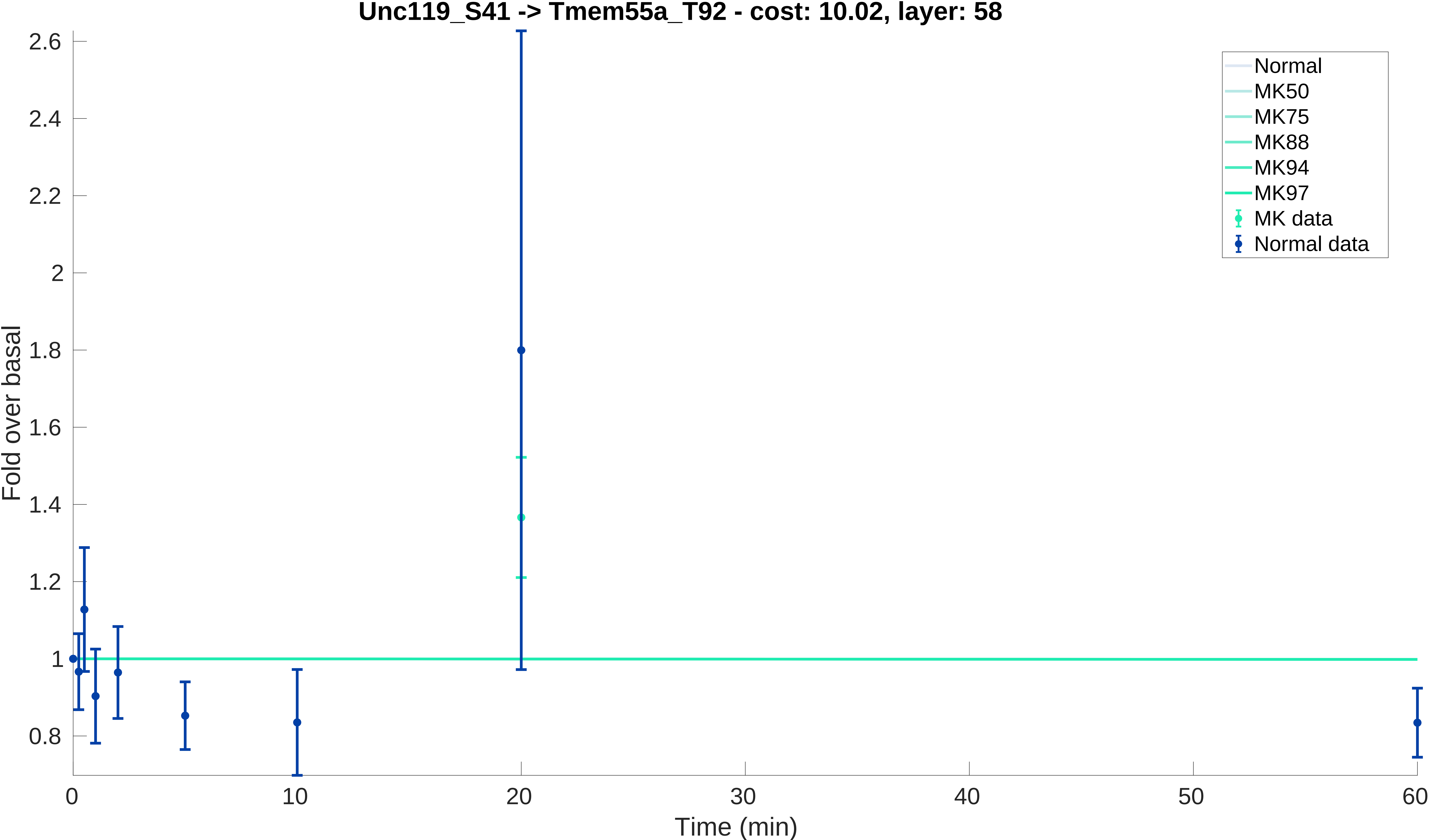
Unc119_S41 -> Tmem55a_T92 - cost: 10.02, layer: 58
Fold over basal
Time (min)
Normal
MK50
MK75
MK88
MK94
MK97
MK data
Normal data
0.8
1
1.2
1.4
1.6
1.8
2
2.2
2.4
2.6
0
10
20
30
40
50
60

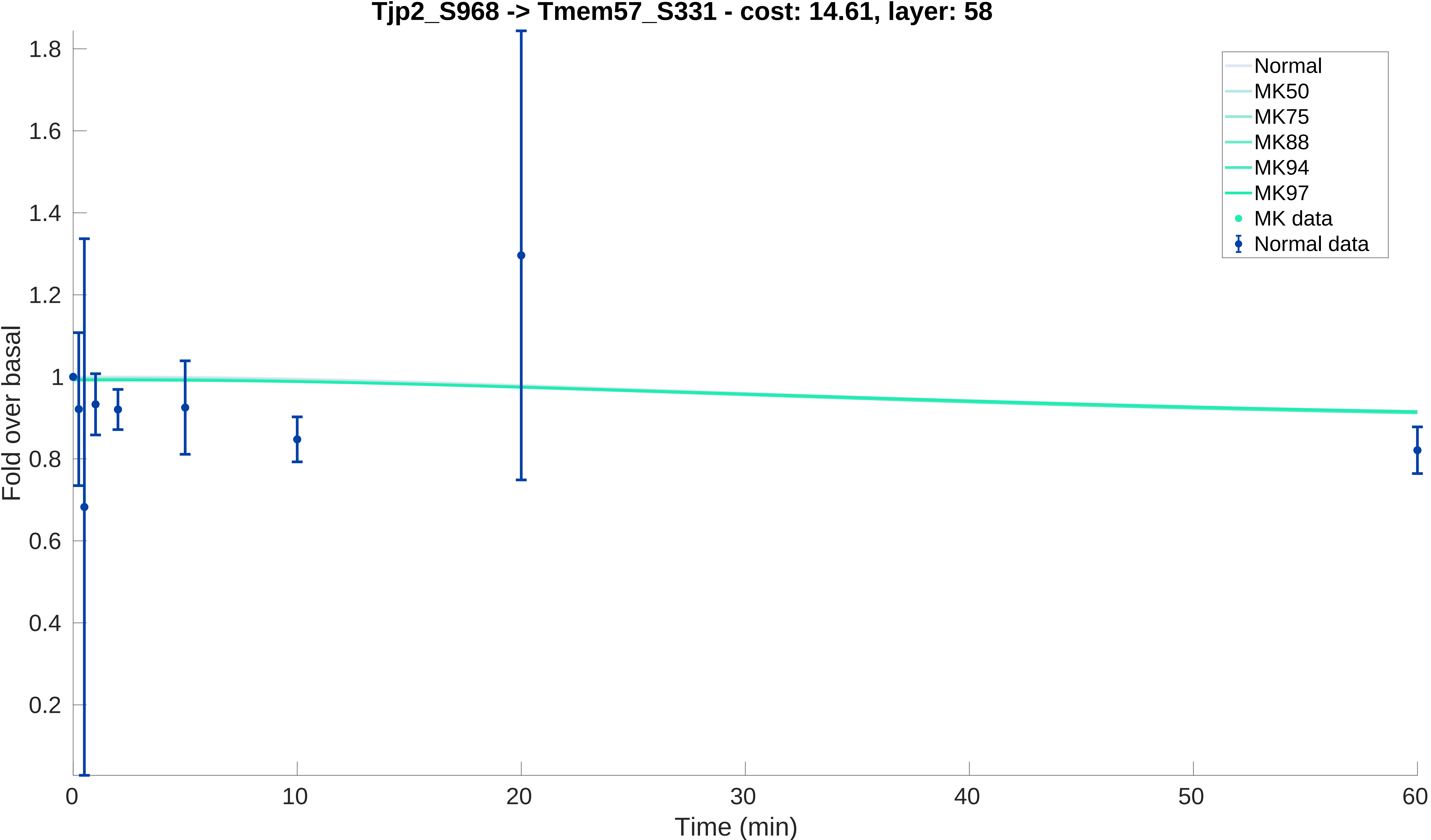

Tjp2_S968 -> Tmem57_S331 - cost: 14.61, layer: 58
Fold over basal
Time (min)
0
10
20
30
40
50
60
0.2
0.4
0.6
0.8
1
1.2
1.4
1.6
1.8
Normal
MK50
MK75
MK88
MK94
MK97
MK data
Normal data

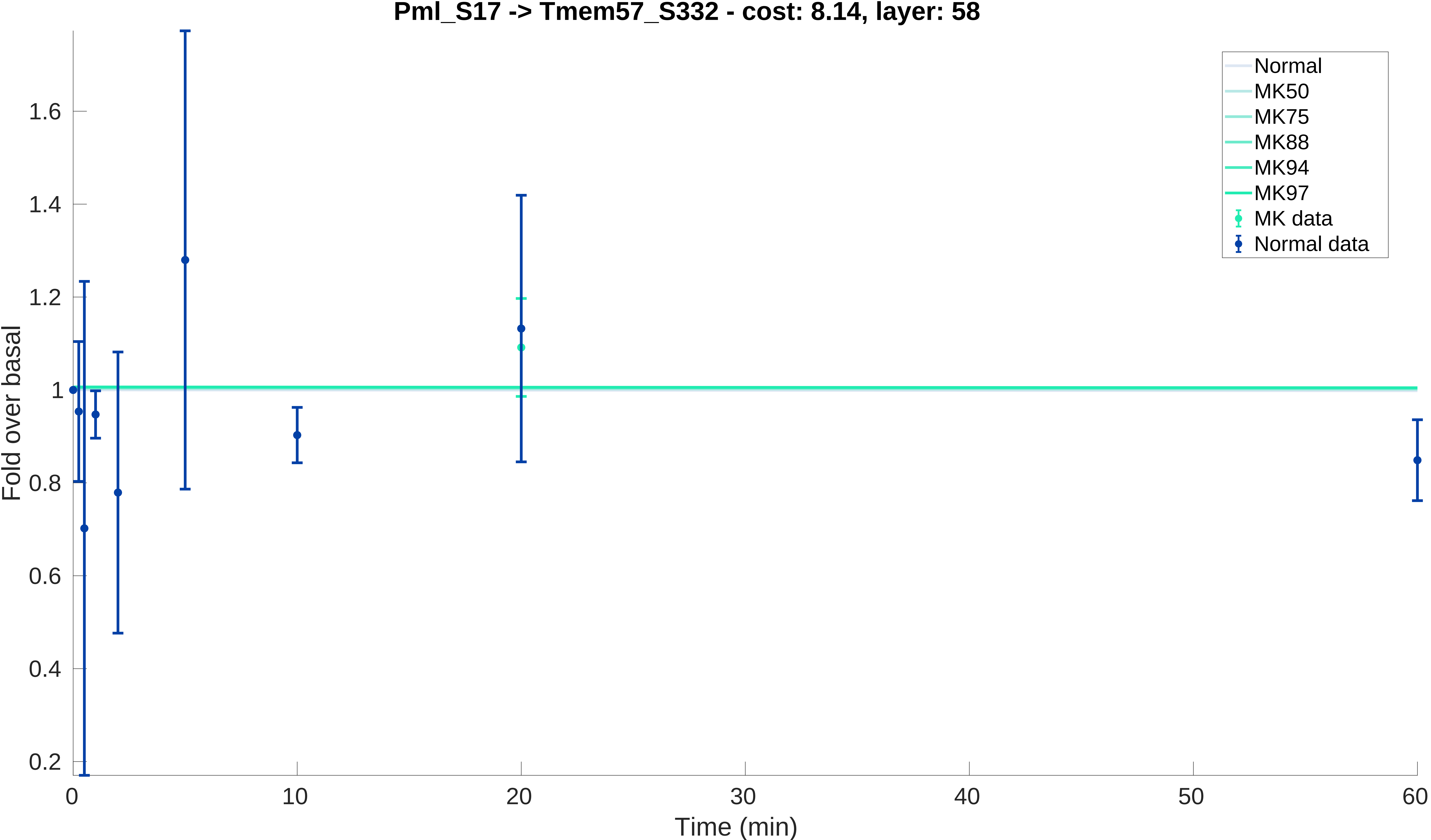

Pml_S17 -> Tmem57_S332 - cost: 8.14, layer: 58
Normal
MK50
MK75
MK88
MK94
MK97
MK data
Normal data
Fold over basal
Time (min)
0.2
0.4
0.6
0.8
1
1.2
1.4
1.6
0
10
20
30
40
50
60

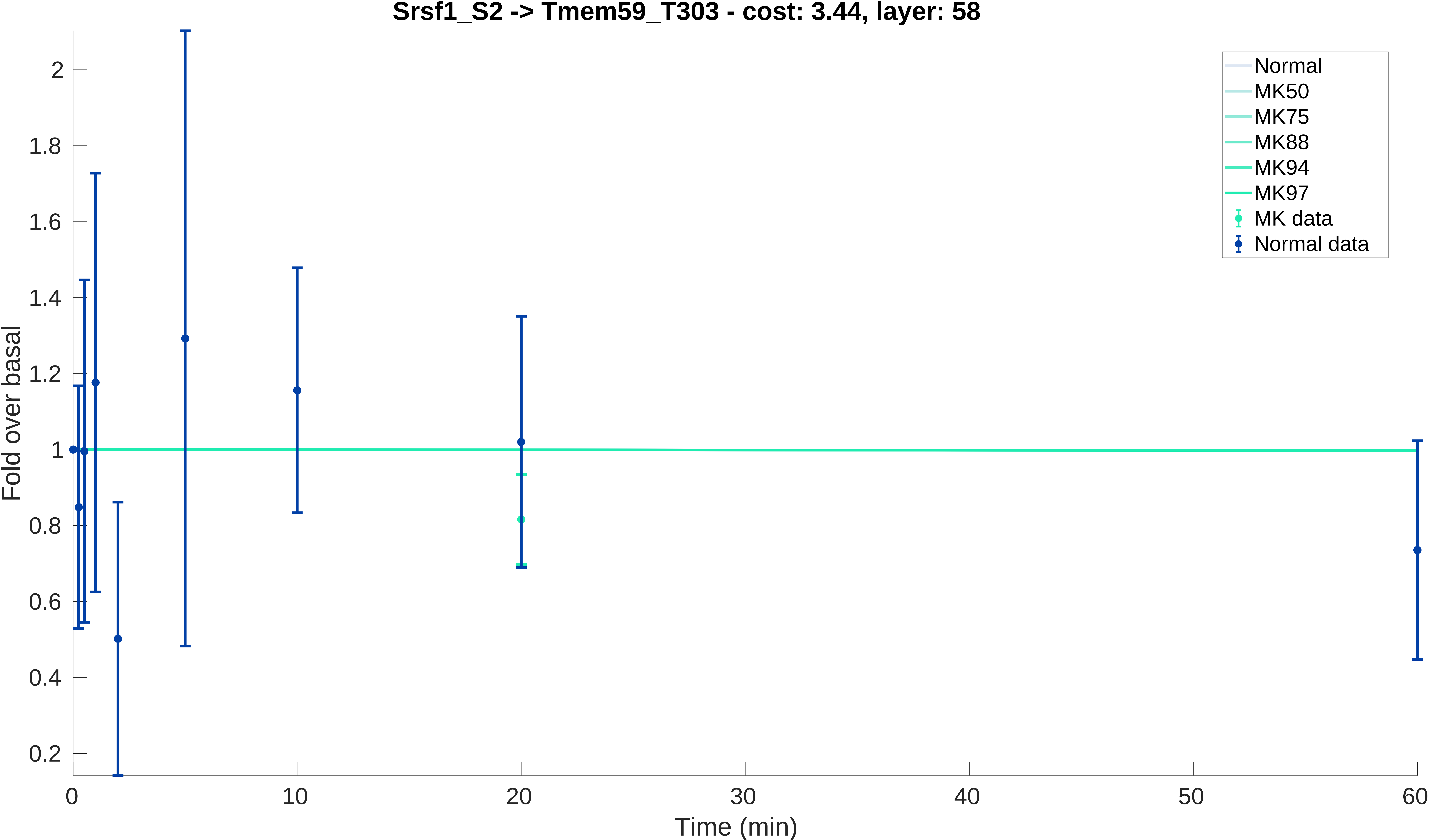

Srsf1_S2 -> Tmem59_T303 - cost: 3.44, layer: 58
Normal
MK50
MK75
MK88
MK94
MK97
MK data
Normal data
Fold over basal
Time (min)
0.2
0.4
0.6
0.8
1
1.2
1.4
1.6
1.8
2
0
10
20
30
40
50
60

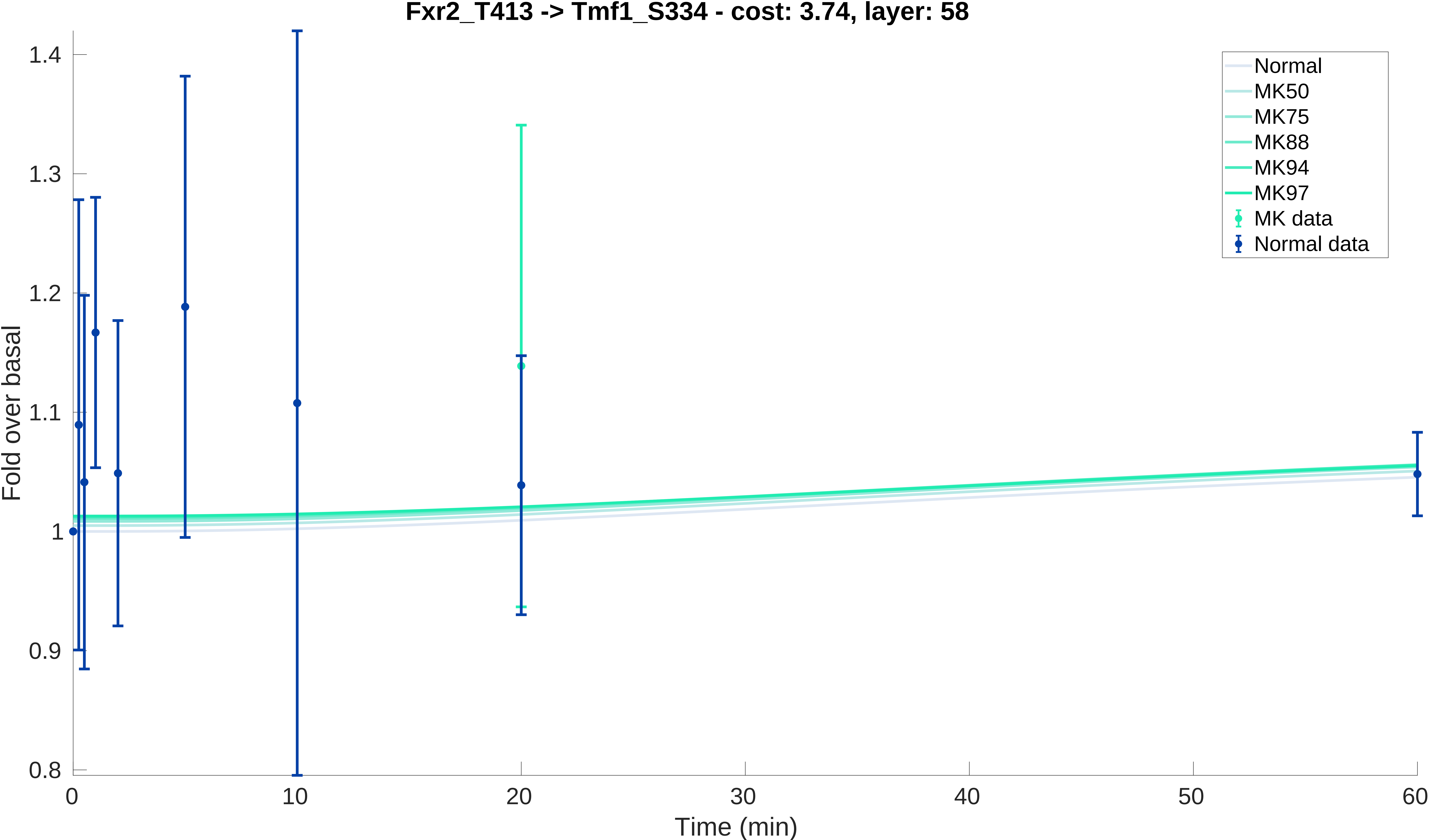

Fxr2_T413 -> Tmf1_S334 - cost: 3.74, layer: 58
Fold over basal
Time (min)
0.8
0.9
1
1.1
1.2
1.3
1.4
0
10
20
30
40
50
60
Normal
MK50
MK75
MK88
MK94
MK97
MK data
Normal data

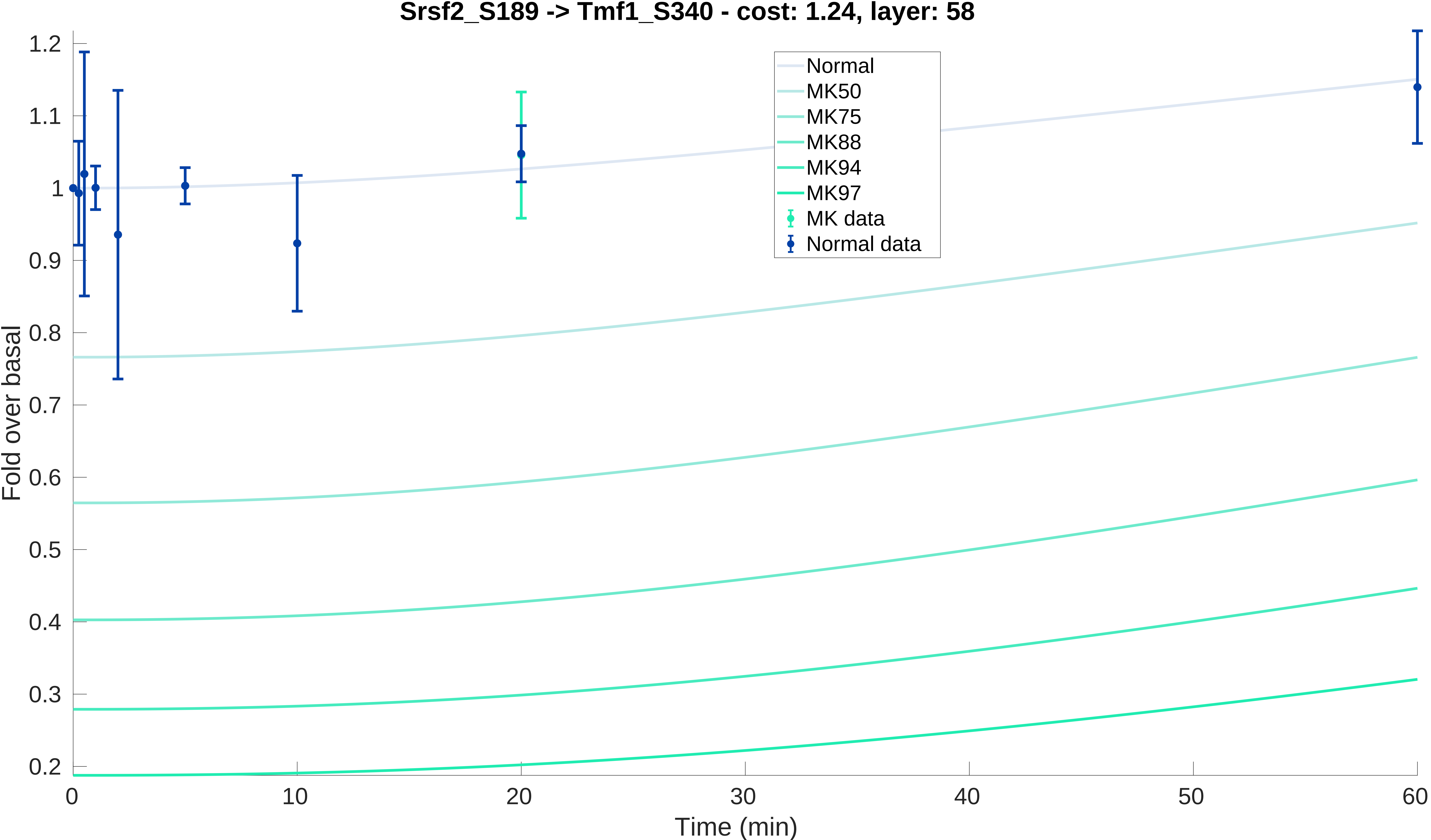
Srsf2_S189 -> Tmf1_S340 - cost: 1.24, layer: 58
Fold over basal
1.2
1.1
1
0.9
0.8
0.7
0.6
0.5
0.4
0.3
0.2
0
10
20
30
40
50
60
Time (min)
Normal
MK50
MK75
MK88
MK94
MK97
MK data
Normal data

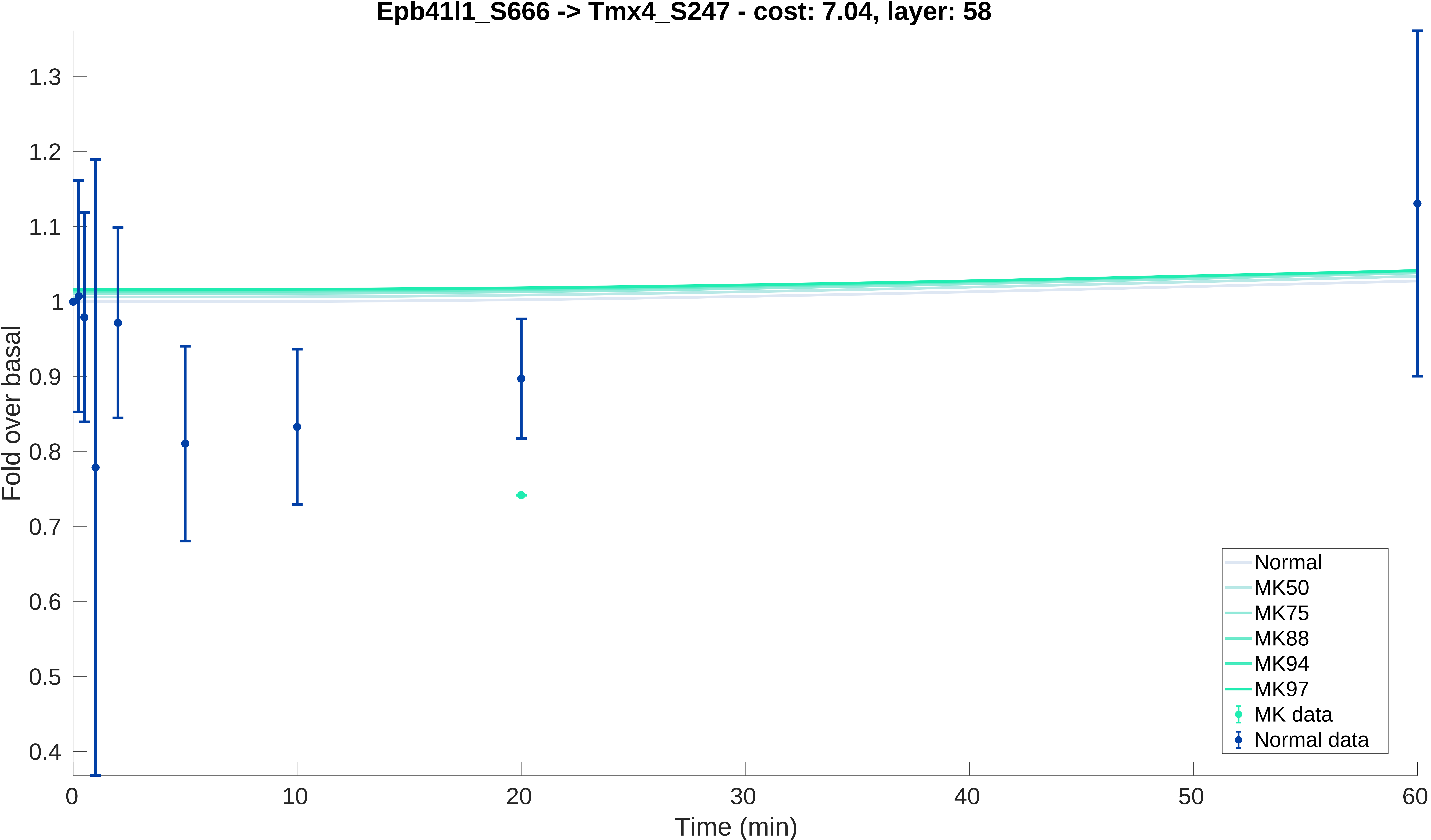

Epb41l1_S666 -> Tmx4_S247 - cost: 7.04, layer: 58
Fold over basal
Time (min)
0
10
20
30
40
50
60
0.4
0.5
0.6
0.7
0.8
0.9
1
1.1
1.2
1.3
Normal
MK50
MK75
MK88
MK94
MK97
MK data
Normal data

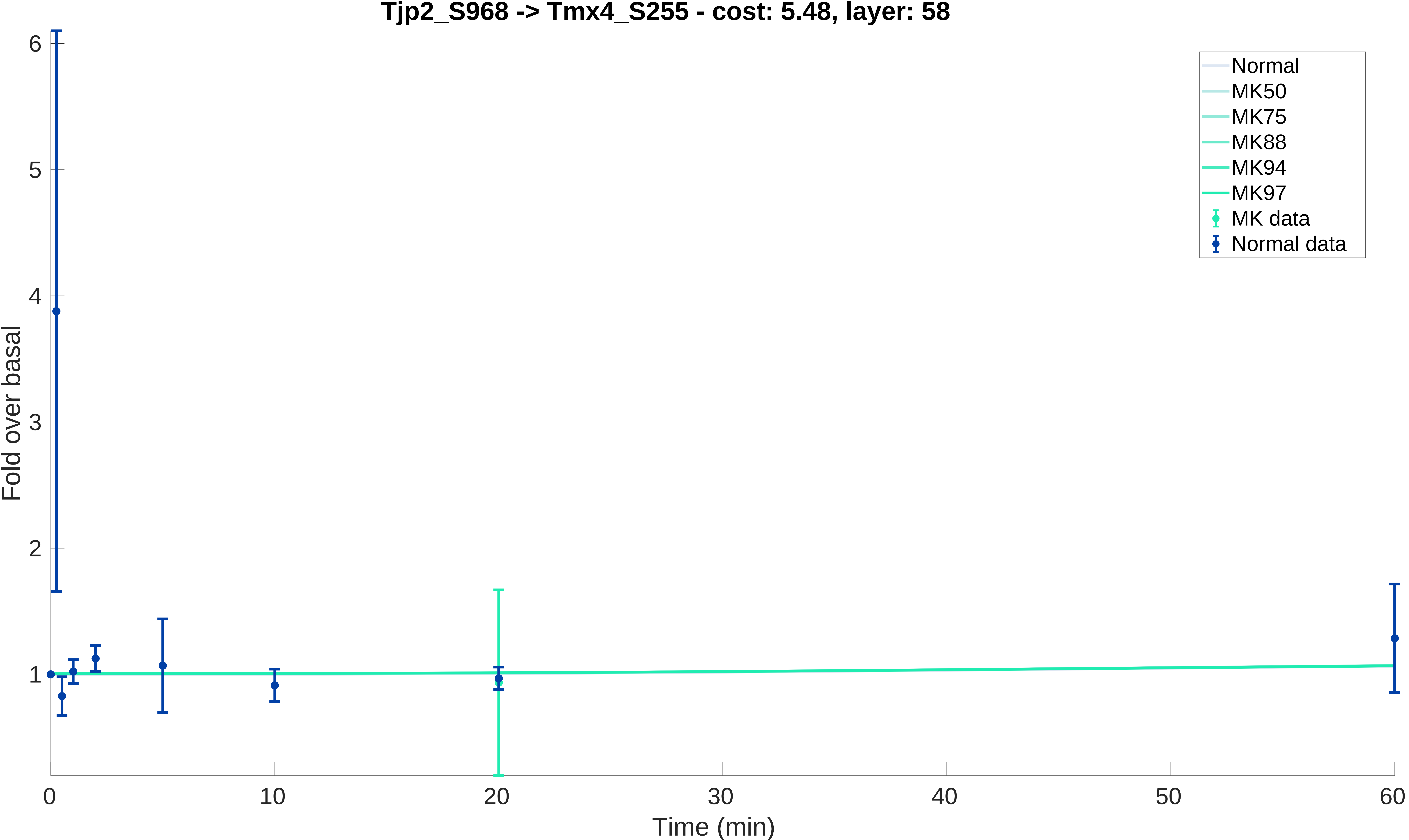

Tjp2_S968 -> Tmx4_S255 - cost: 5.48, layer: 58
Fold over basal
Time (min)
0
10
20
30
40
50
60
1
2
3
4
5
6
Normal
MK50
MK75
MK88
MK94
MK97
MK data
Normal data

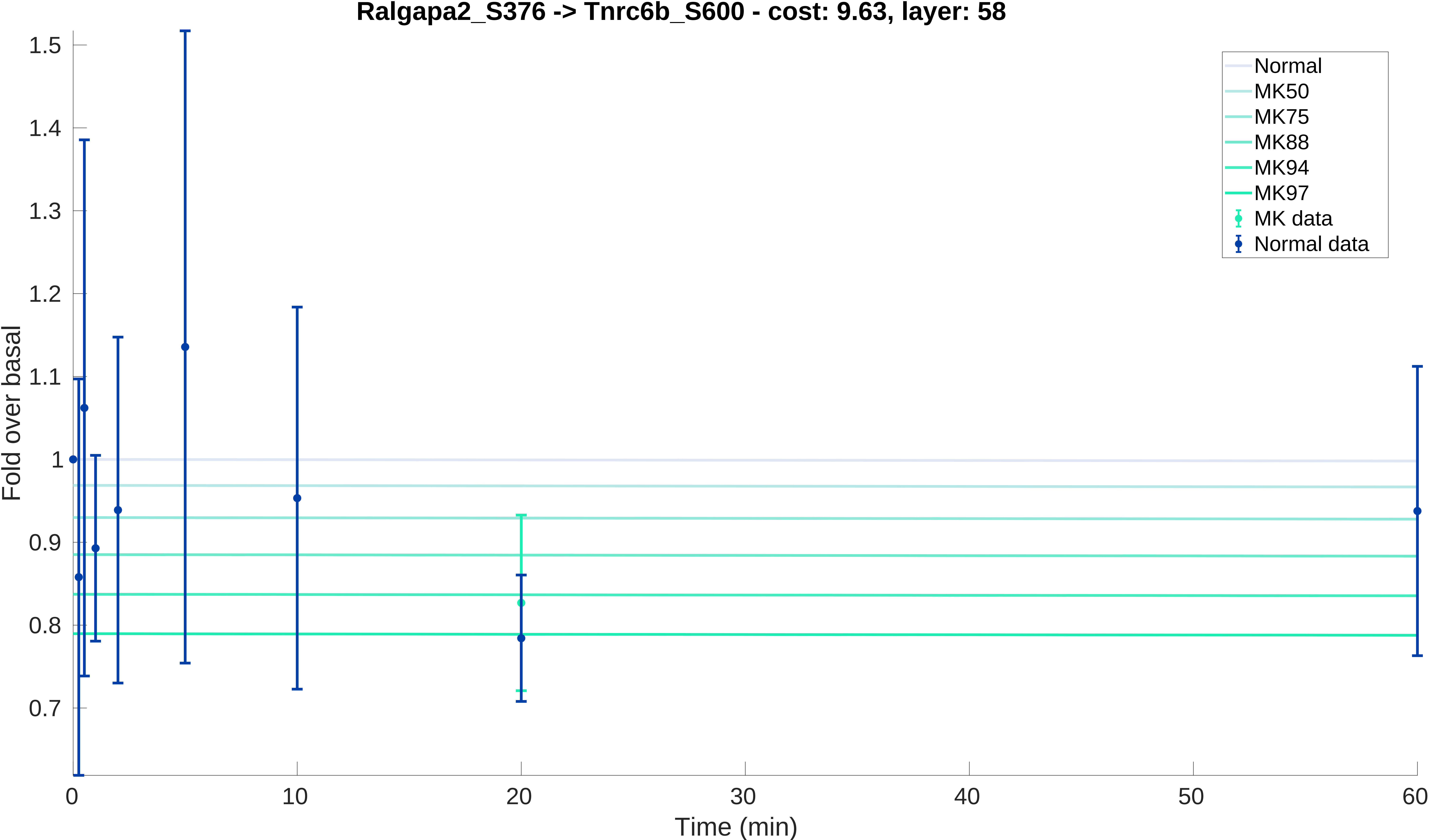

Ralgapa2_S376 -> Tnrc6b_S600 - cost: 9.63, layer: 58
Fold over basal
Time (min)
1.5
1.4
1.3
1.2
1.1
1
0.9
0.8
0.7
0
10
20
30
40
50
60
Normal
MK50
MK75
MK88
MK94
MK97
MK data
Normal data

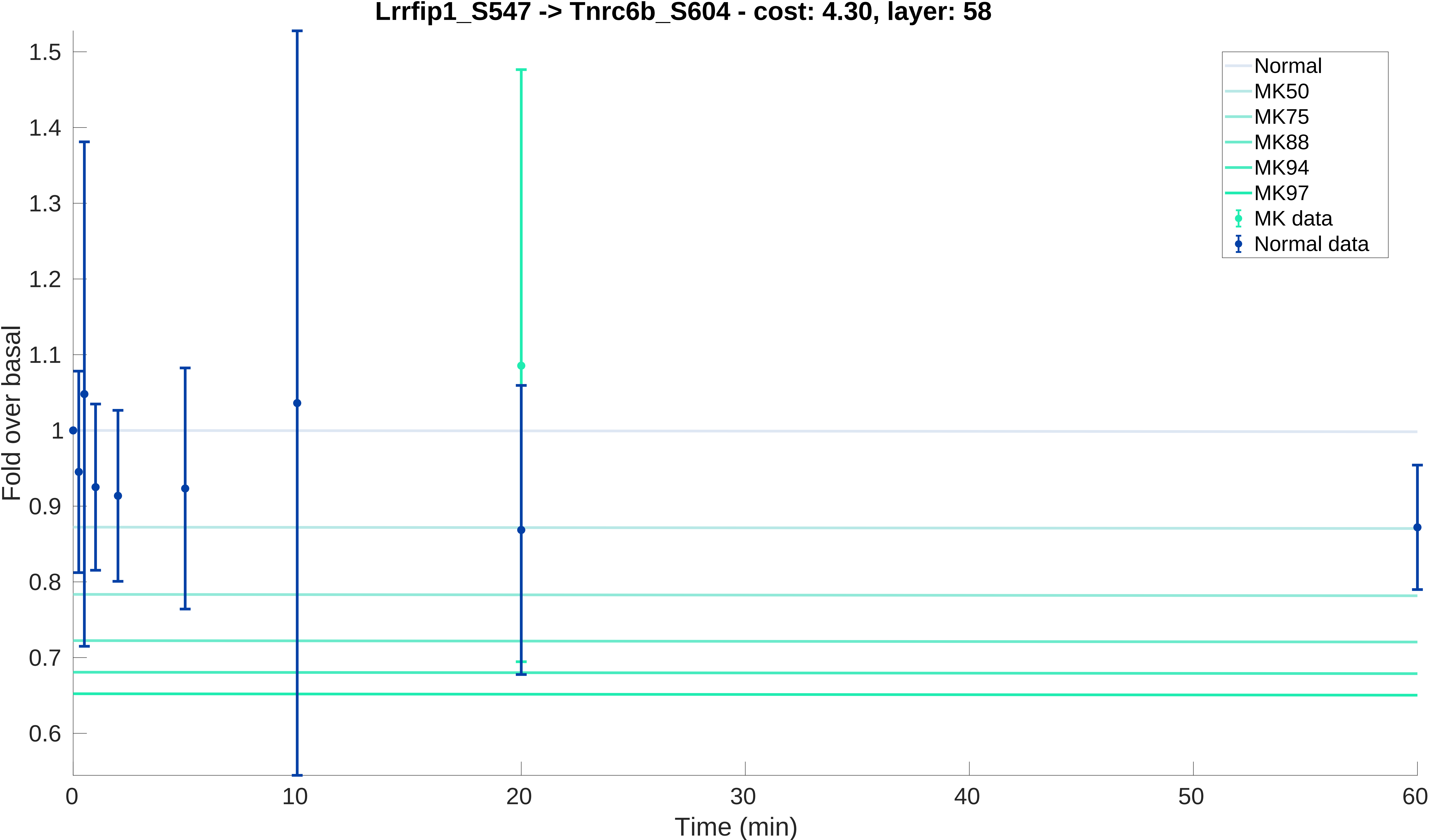

Lrrfip1_S547 -> Tnrc6b_S604 - cost: 4.30, layer: 58
Fold over basal
Time (min)
0.6
0.7
0.8
0.9
1
1.1
1.2
1.3
1.4
1.5
0
10
20
30
40
50
60
Normal
MK50
MK75
MK88
MK94
MK97
MK data
Normal data

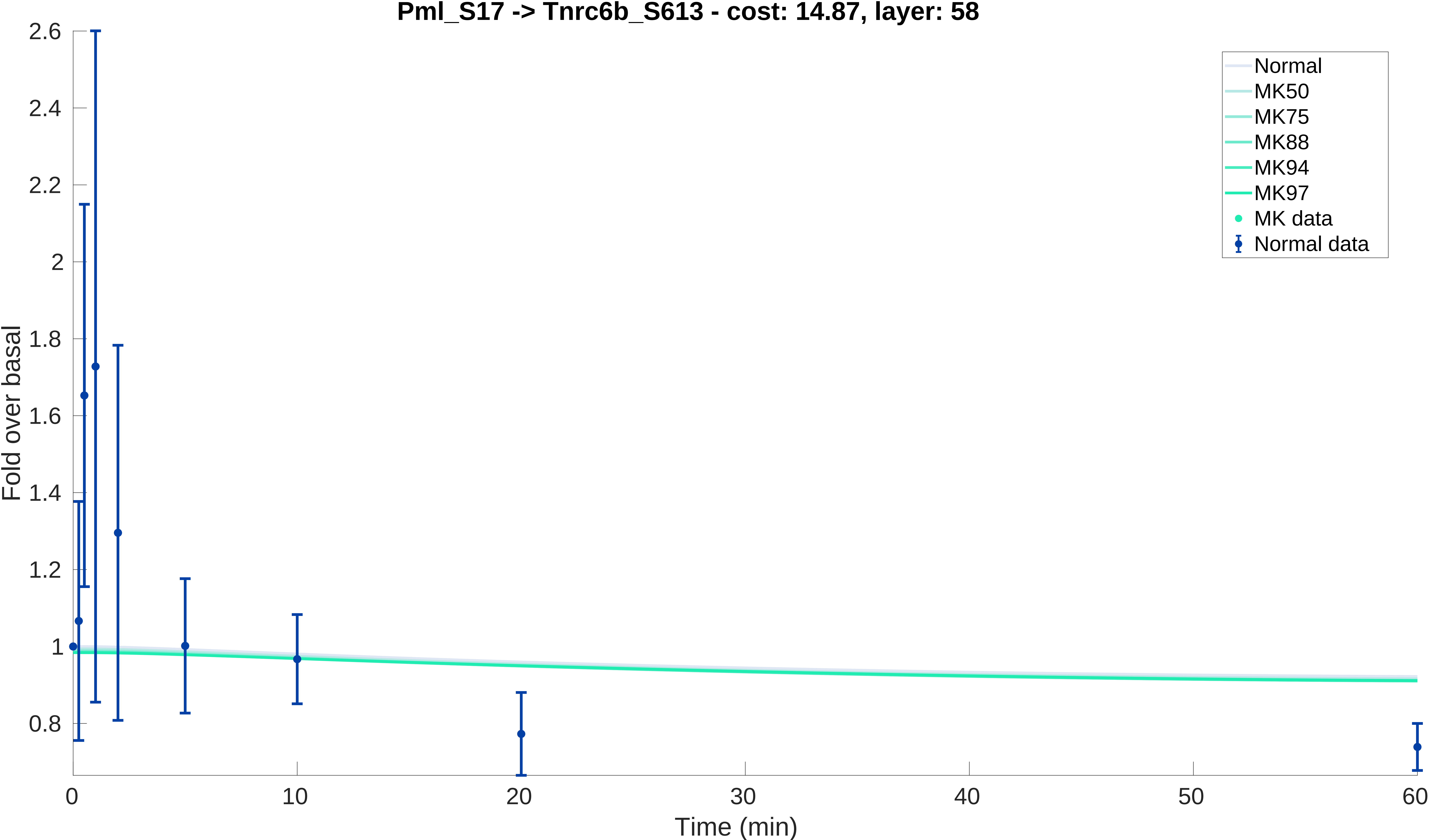

Pml_S17 -> Tnrc6b_S613 - cost: 14.87, layer: 58
Fold over basal
Time (min)
2.6
2.4
2.2
2
1.8
1.6
1.4
1.2
1
0.8
0
10
20
30
40
50
60
Normal
MK50
MK75
MK88
MK94
MK97
MK data
Normal data

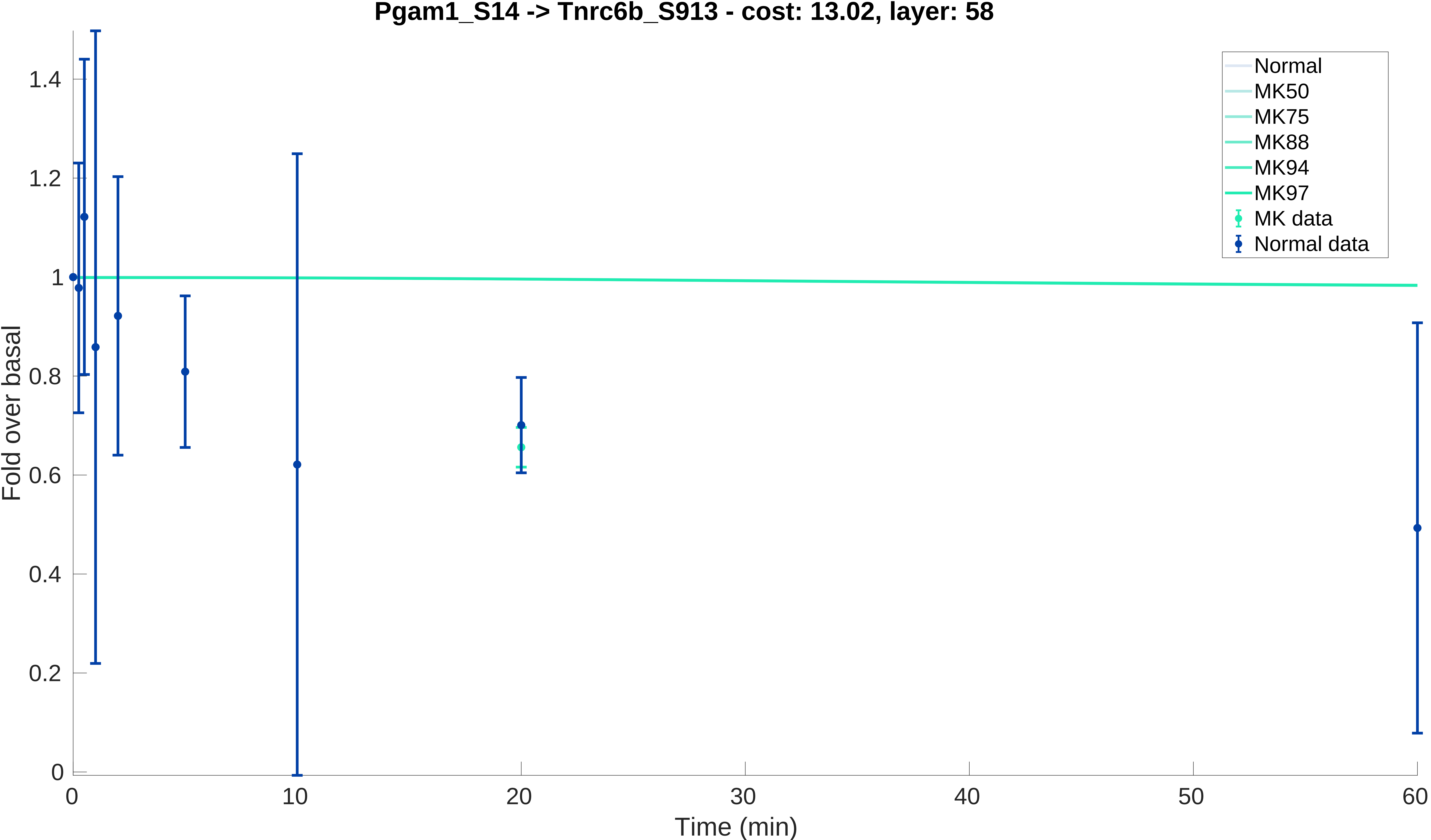

Pgam1_S14 -> Tnrc6b_S913 - cost: 13.02, layer: 58
Fold over basal
Time (min)
0
0.2
0.4
0.6
0.8
1
1.2
1.4
0
10
20
30
40
50
60
Normal
MK50
MK75
MK88
MK94
MK97
MK data
Normal data

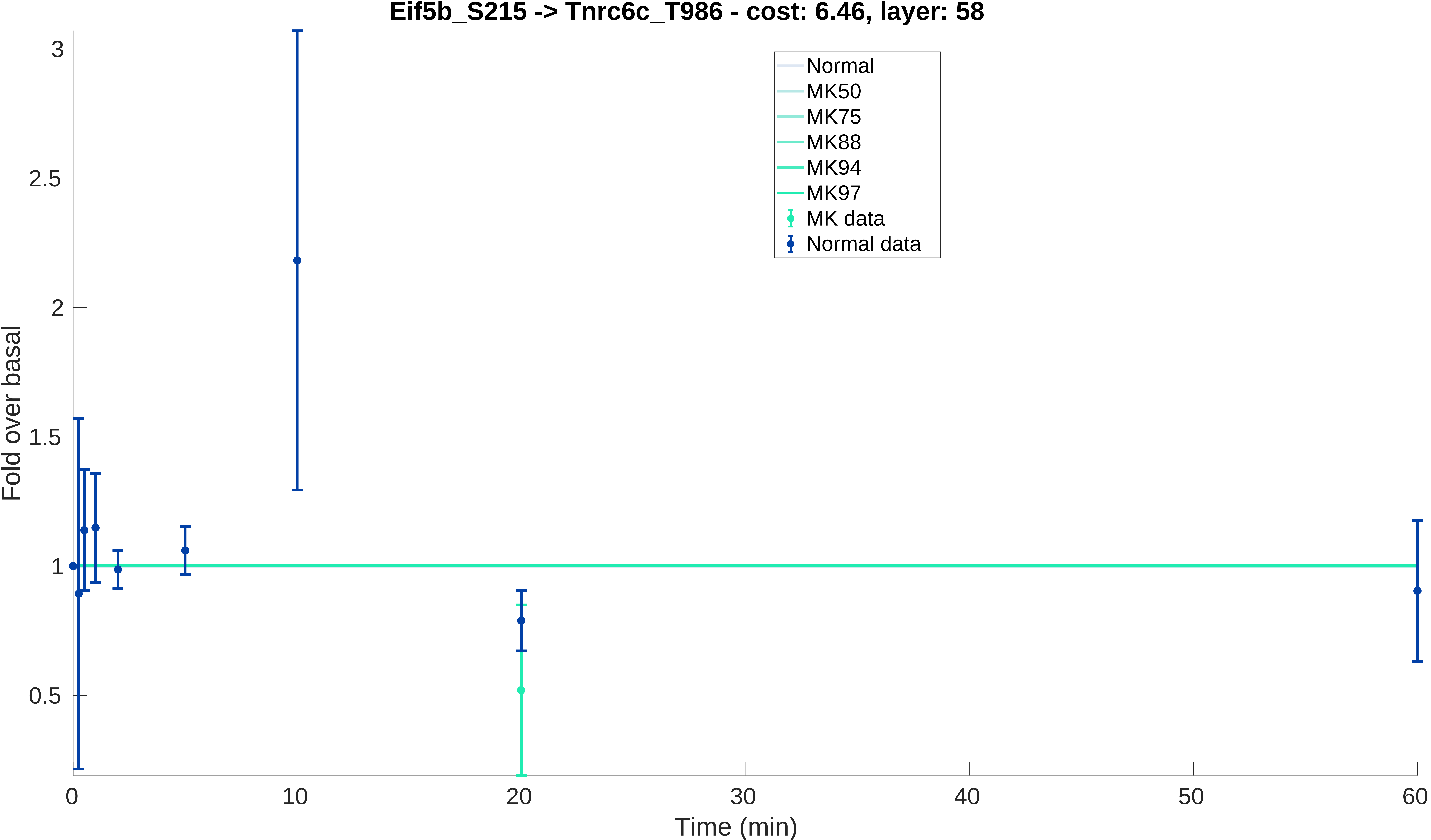

Eif5b_S215 -> Tnrc6c_T986 - cost: 6.46, layer: 58
Normal
MK50
MK75
MK88
MK94
MK97
MK data
Normal data
Fold over basal
0.5
1
1.5
2
2.5
3
0
10
20
30
40
50
60
Time (min)

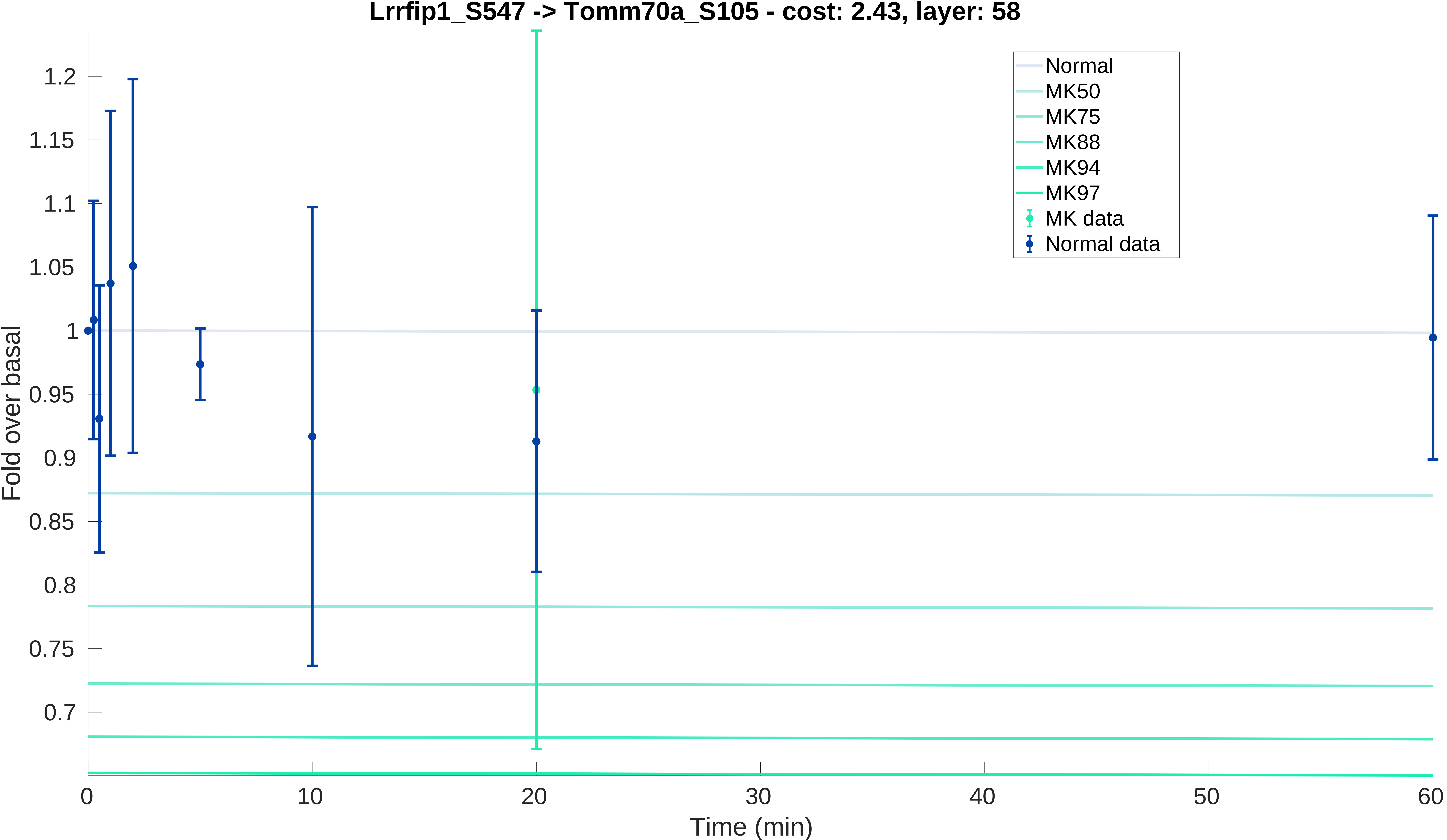

Lrrfip1_S547 -> Tomm70a_S105 - cost: 2.43, layer: 58
Fold over basal
Time (min)
1.2
1.15
1.1
1.05
1
0.95
0.9
0.85
0.8
0.75
0.7
0
10
20
30
40
50
60
Normal
MK50
MK75
MK88
MK94
MK97
MK data
Normal data

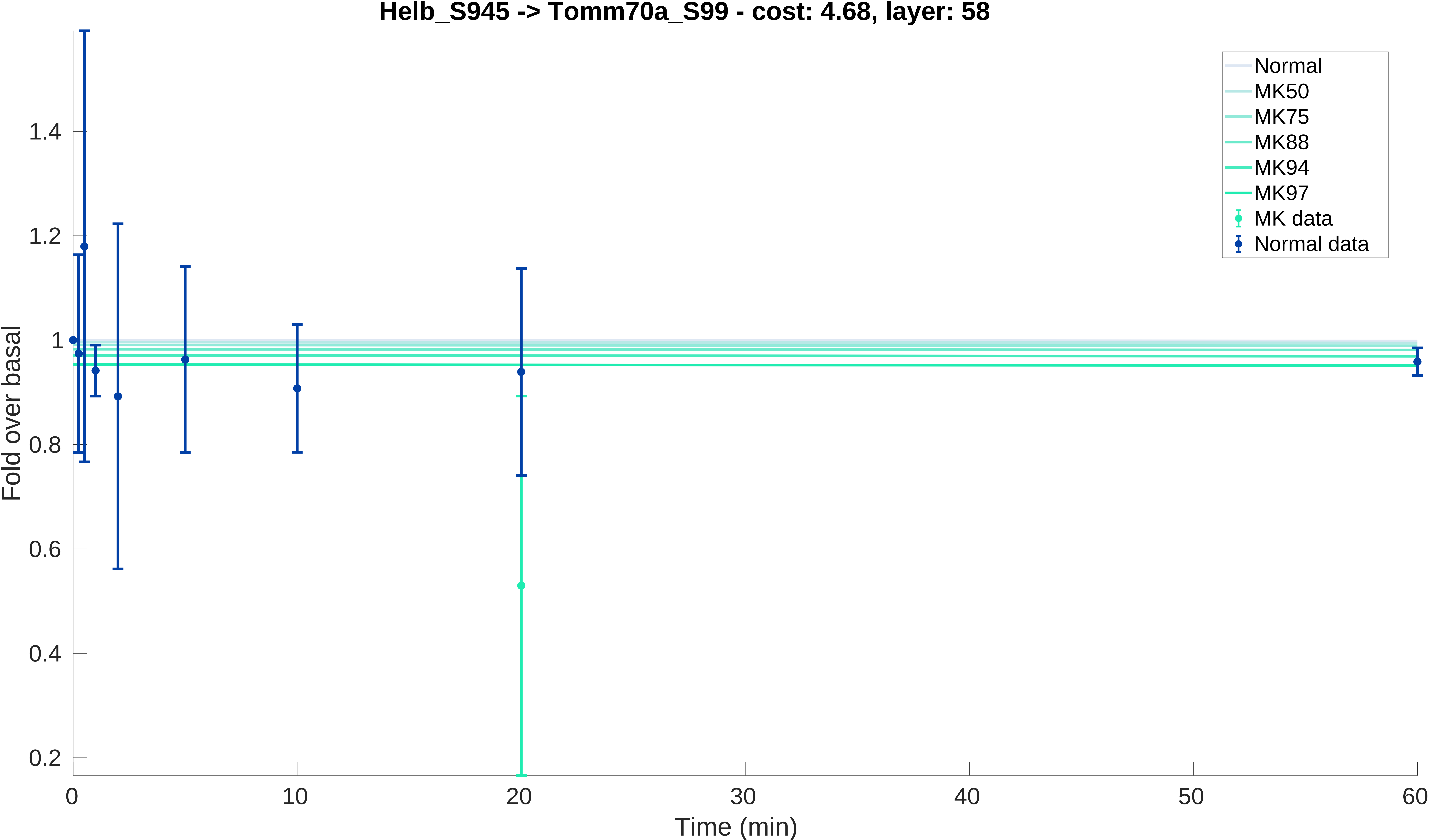

Helb_S945 -> Tomm70a_S99 - cost: 4.68, layer: 58
Normal
MK50
MK75
MK88
MK94
MK97
MK data
Normal data
Fold over basal
1.4
1.2
1
0.8
0.6
0.4
0.2
0
10
20
30
40
50
60
Time (min)

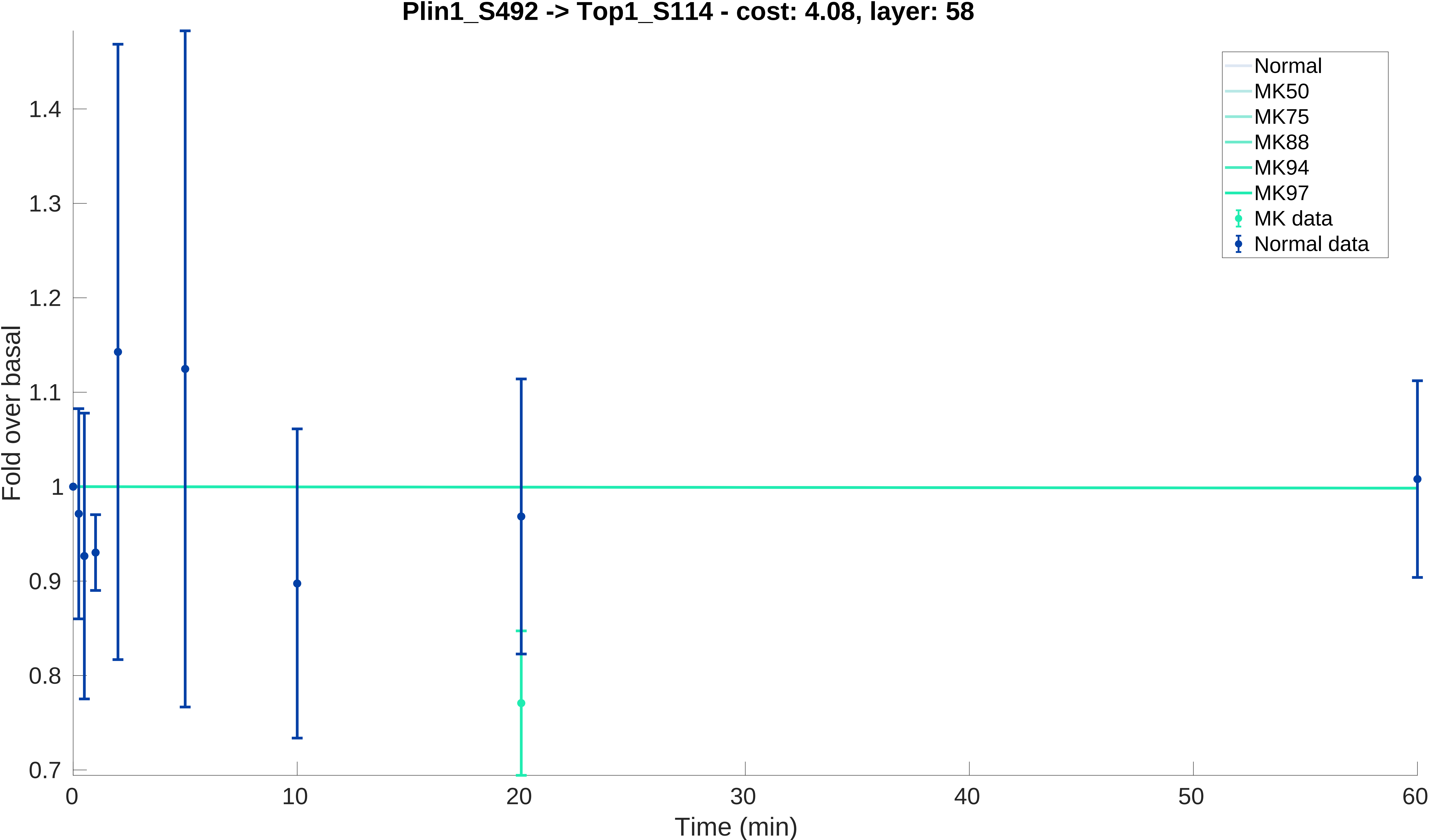
Plin1_S492 -> Top1_S114 - cost: 4.08, layer: 58
Normal
MK50
MK75
MK88
MK94
MK97
MK data
Normal data
Fold over basal
Time (min)
0.7
0.8
0.9
1
1.1
1.2
1.3
1.4
0
10
20
30
40
50
60

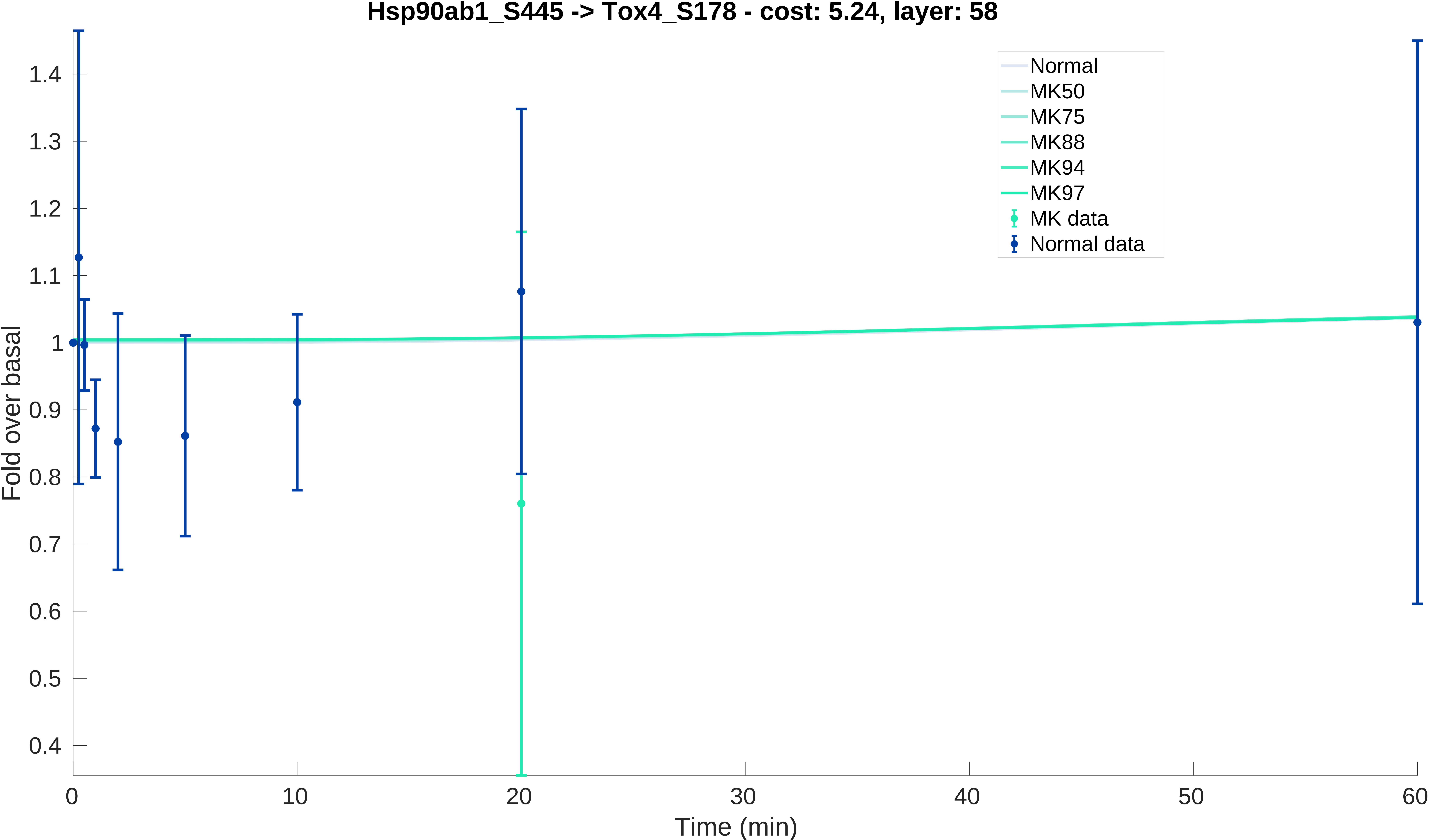

Hsp90ab1_S445 -> Tox4_S178 - cost: 5.24, layer: 58
Normal
MK50
MK75
MK88
MK94
MK97
MK data
Normal data
Fold over basal
1.4
1.3
1.2
1.1
1
0.9
0.8
0.7
0.6
0.5
0.4
0
10
20
30
40
50
60
Time (min)

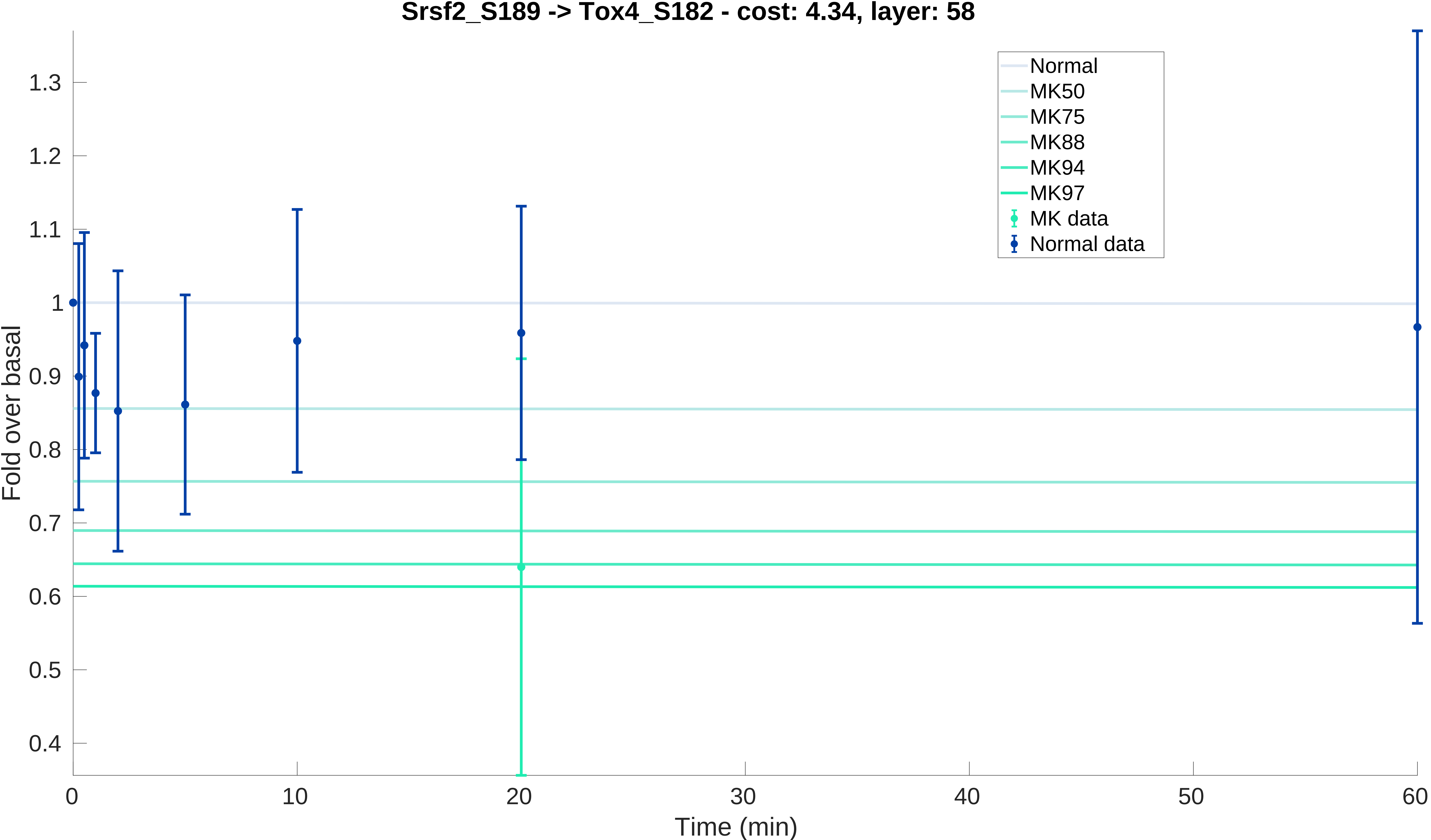

Srsf2_S189 -> Tox4_S182 - cost: 4.34, layer: 58
Normal
MK50
MK75
MK88
MK94
MK97
MK data
Normal data
Fold over basal
Time (min)
1.3
1.2
1.1
1
0.9
0.8
0.7
0.6
0.5
0.4
0
10
20
30
40
50
60

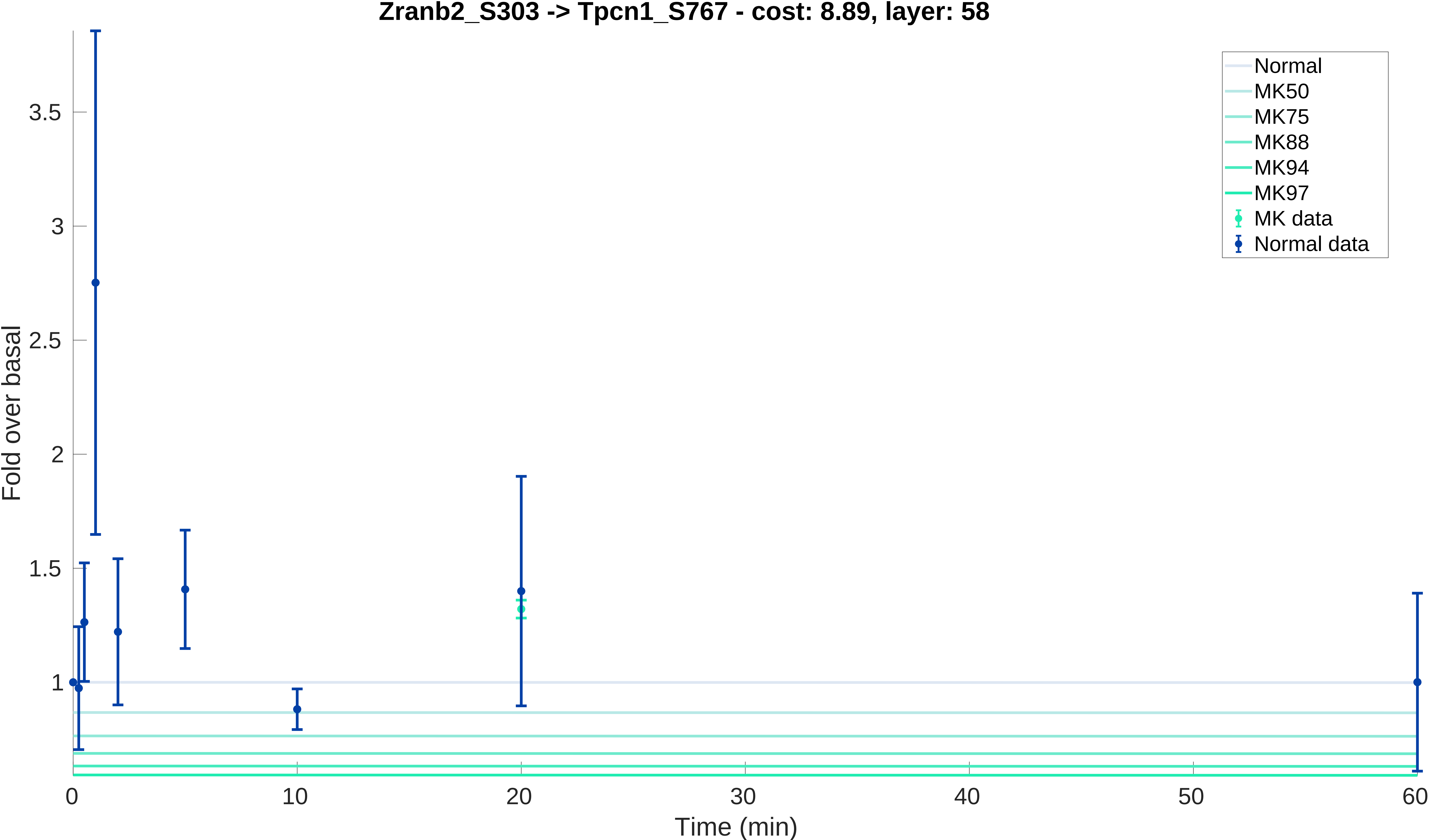

Zranb2_S303 -> Tpcn1_S767 - cost: 8.89, layer: 58
Normal
MK50
MK75
MK88
MK94
MK97
MK data
Normal data
Fold over basal
3.5
3
2.5
2
1.5
1
0
10
20
30
40
50
60
Time (min)

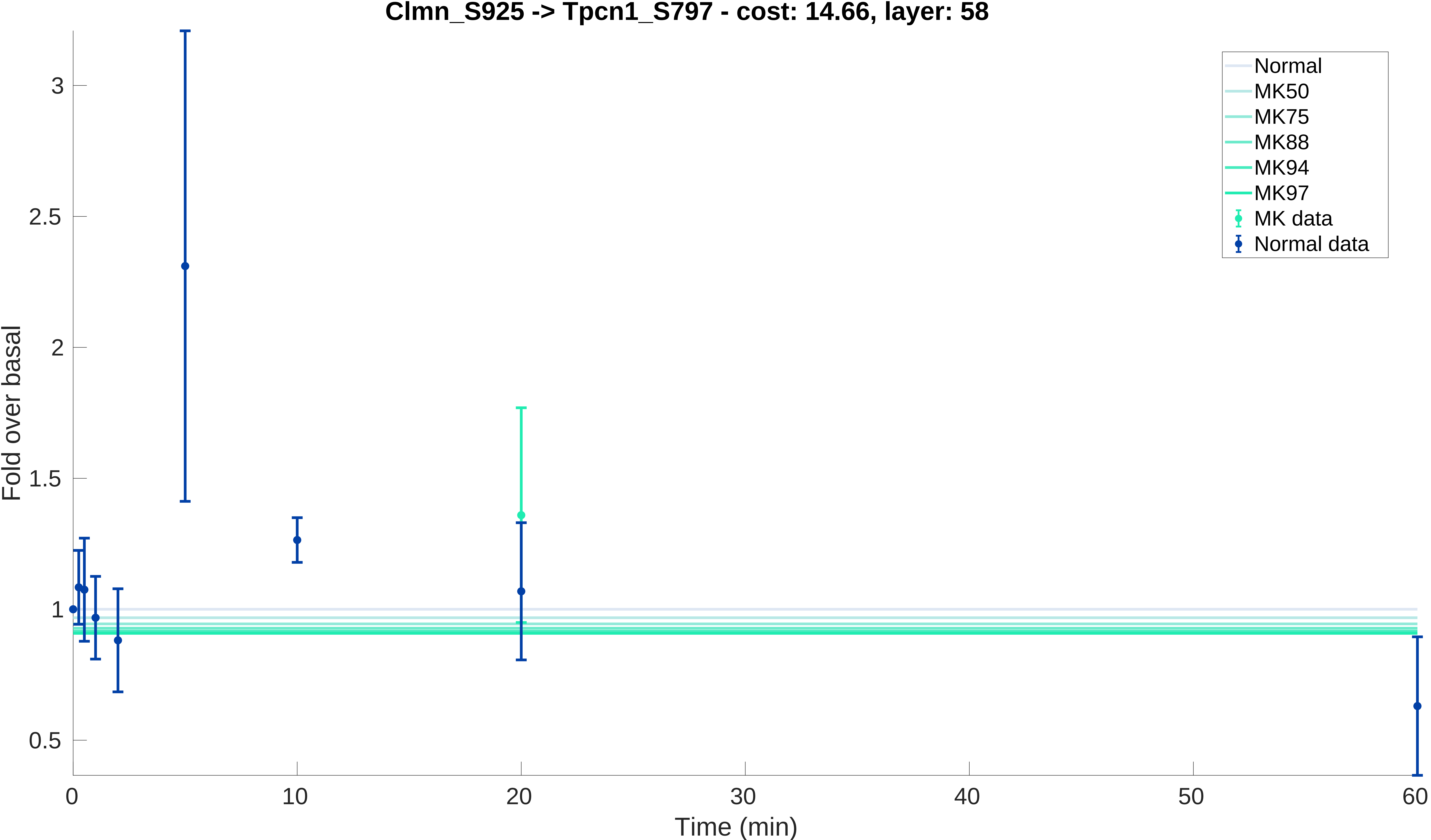

Clmn_S925 -> Tpcn1_S797 - cost: 14.66, layer: 58
Fold over basal
3
2.5
2
1.5
1
0.5
0
10
20
30
40
50
60
Time (min)
Normal
MK50
MK75
MK88
MK94
MK97
MK data
Normal data

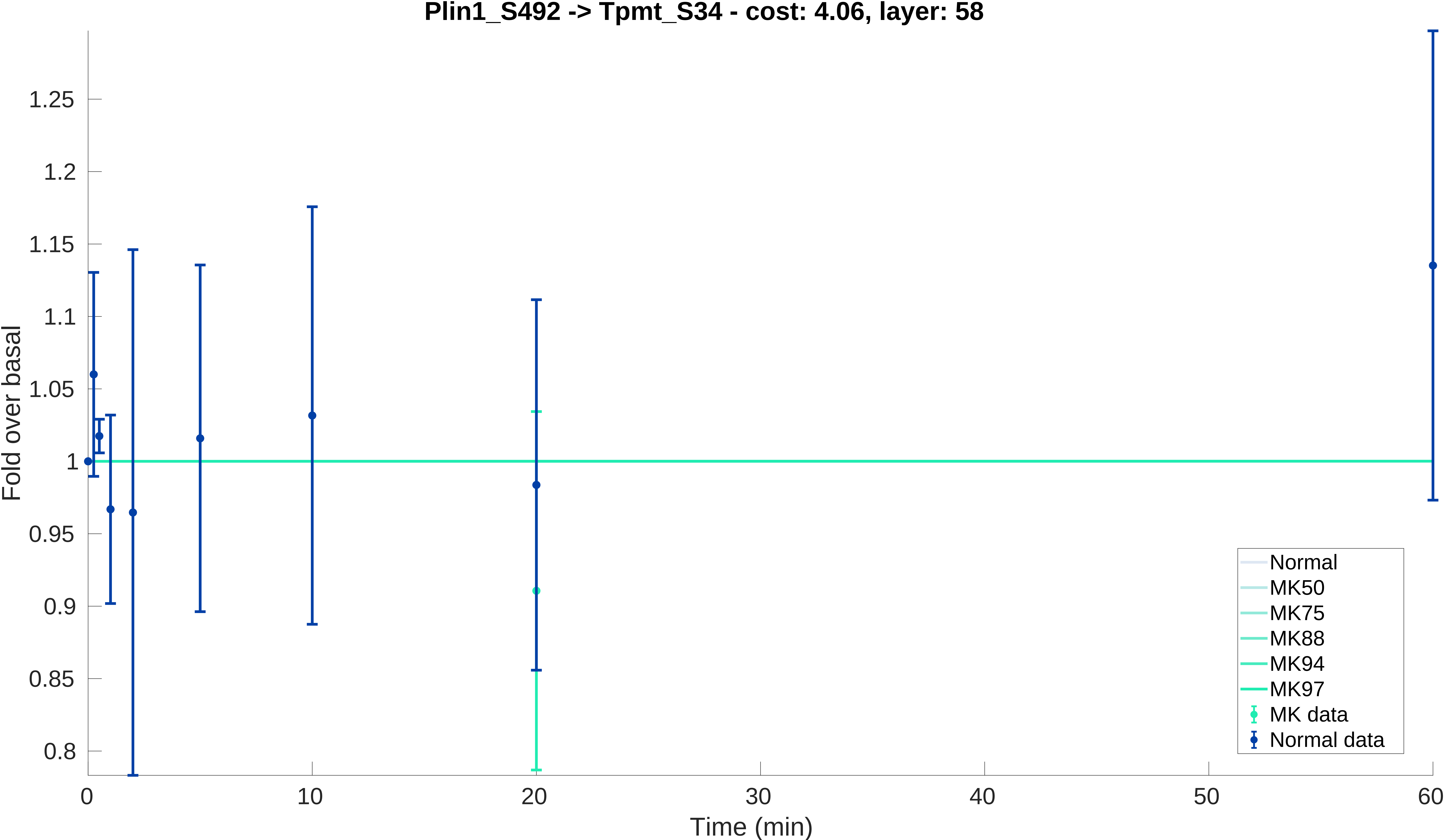
Plin1_S492 -> Tpmt_S34 - cost: 4.06, layer: 58
Fold over basal
1.25
1.2
1.15
1.1
1.05
1
0.95
0.9
0.85
0.8
0
10
20
30
40
50
60
Time (min)
Normal
MK50
MK75
MK88
MK94
MK97
MK data
Normal data

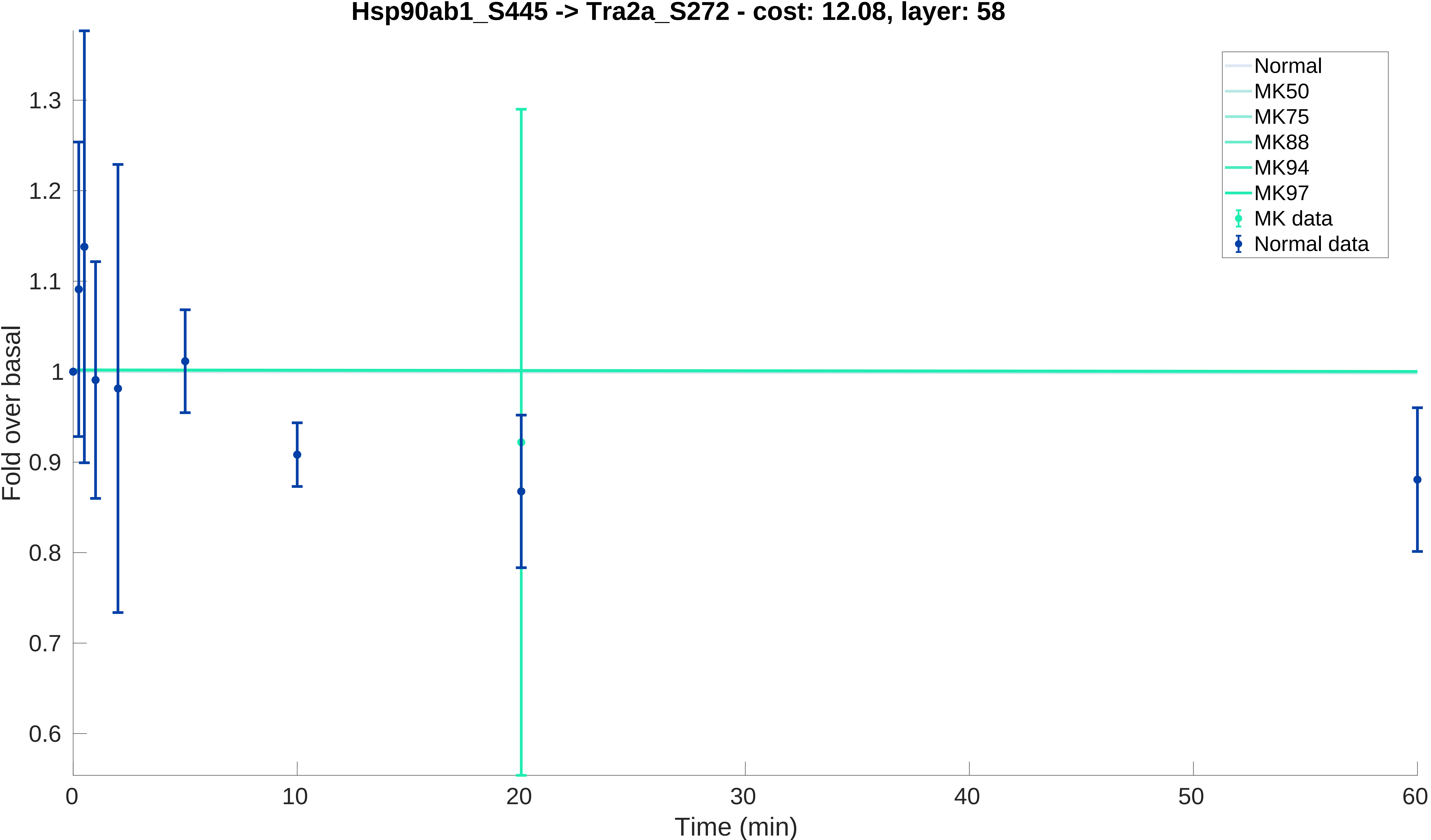
Hsp90ab1_S445 -> Tra2a_S272 - cost: 12.08, layer: 58
Normal
MK50
MK75
MK88
MK94
MK97
MK data
Normal data
Fold over basal
1.3
1.2
1.1
1
0.9
0.8
0.7
0.6
0
10
20
30
40
50
60
Time (min)

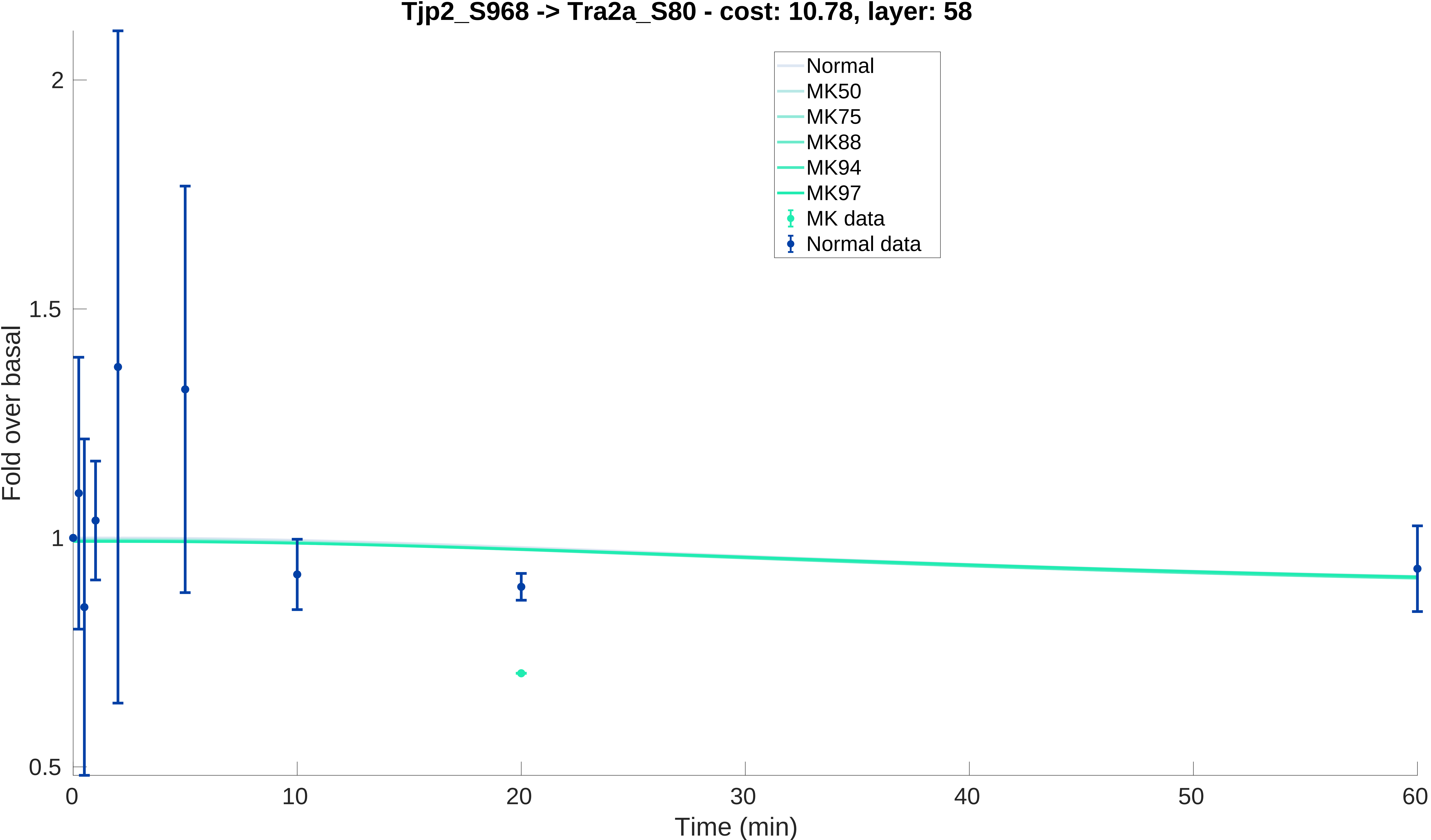

Tjp2_S968 -> Tra2a_S80 - cost: 10.78, layer: 58
Normal
MK50
MK75
MK88
MK94
MK97
MK data
Normal data
Fold over basal
Time (min)
0.5
1
1.5
2
0
10
20
30
40
50
60

#### Sorbs1_S633 -> Tra2a_S82 - cost: 7.25, layer: 58

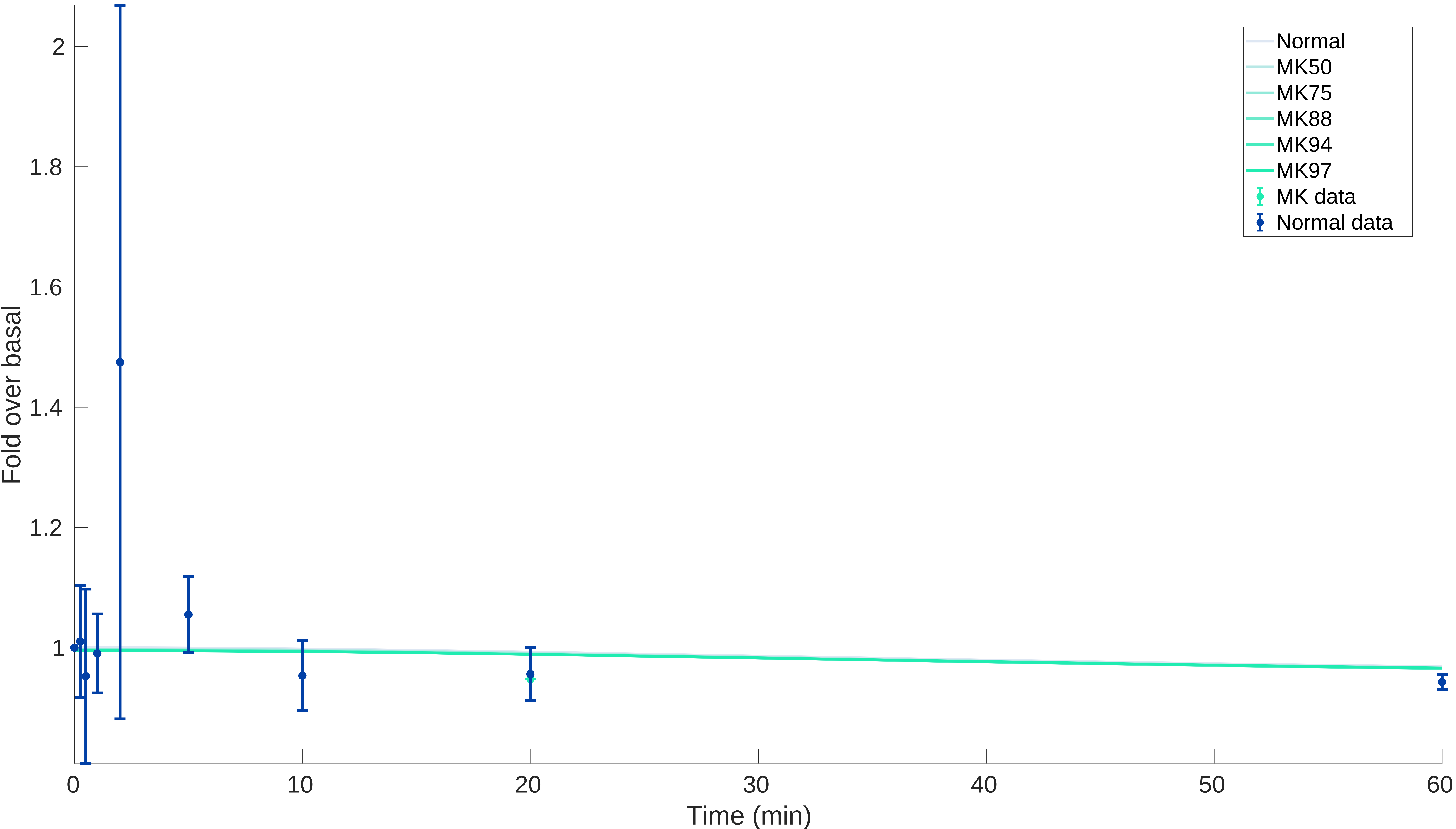

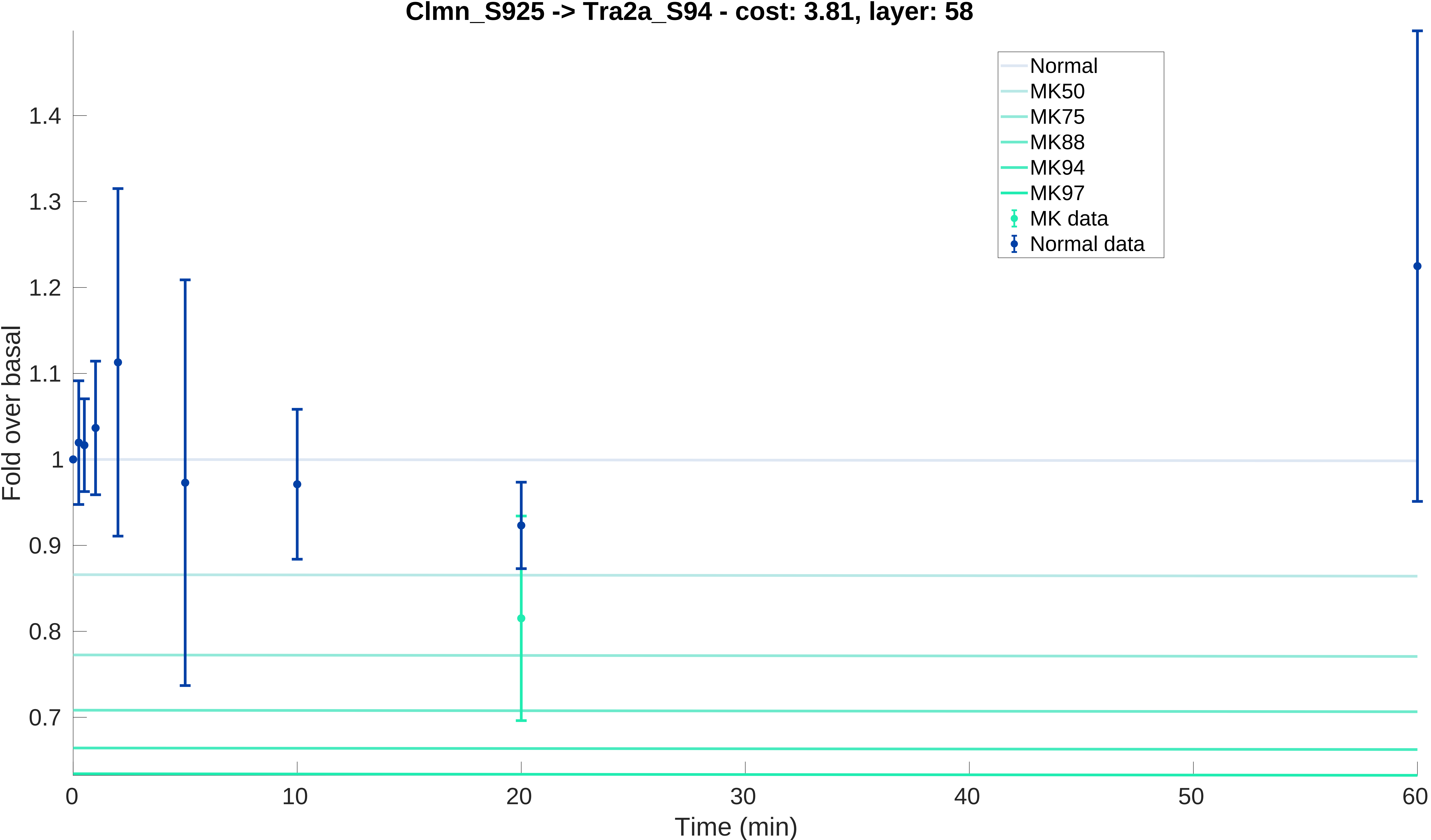

Clmn_S925 -> Tra2a_S94 - cost: 3.81, layer: 58
Normal
MK50
MK75
MK88
MK94
MK97
MK data
Normal data
1.4
1.3
1.2
1.1
1
0.9
0.8
0.7
Fold over basal
0
10
20
30
40
50
60
Time (min)

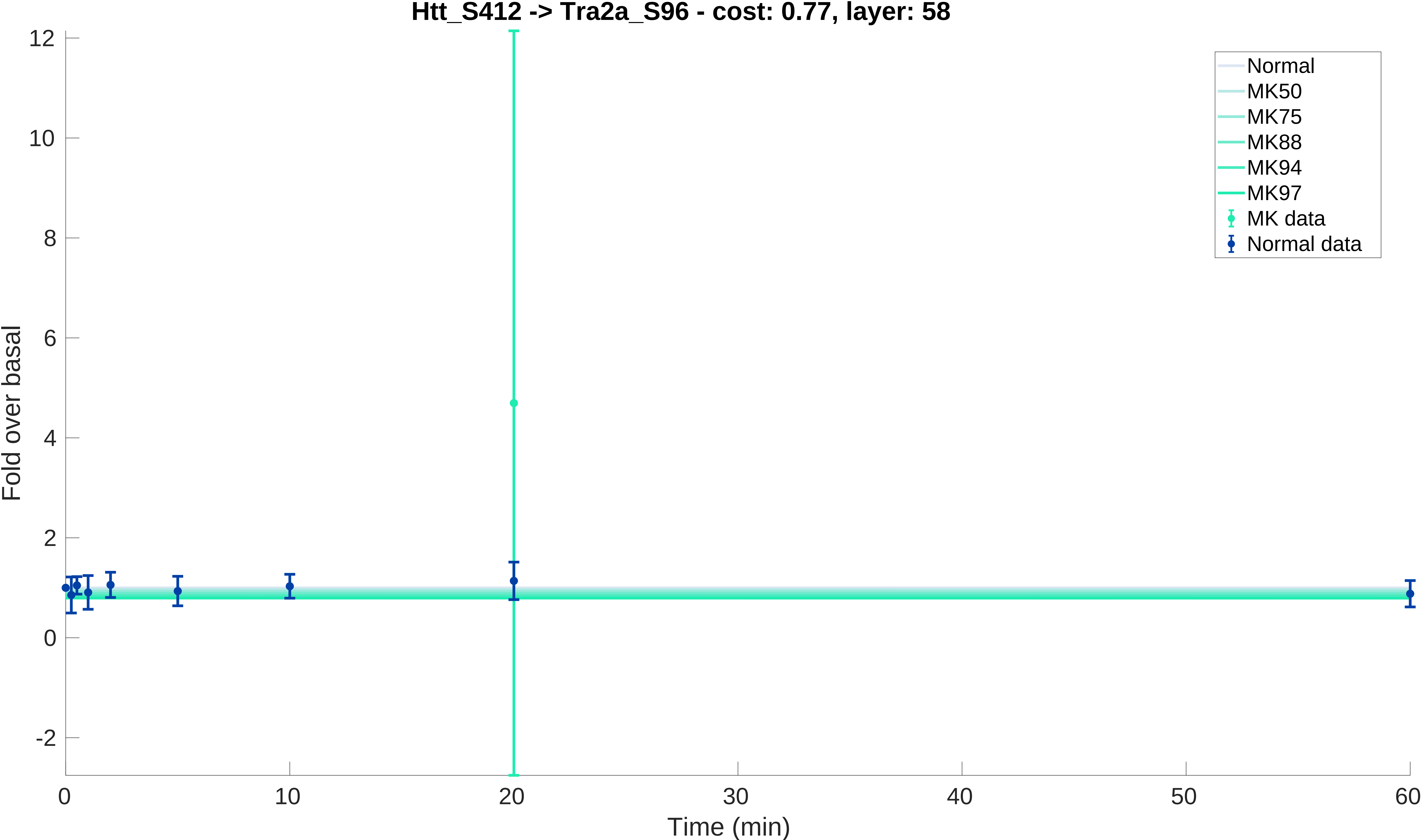

Htt_S412 -> Tra2a_S96 - cost: 0.77, layer: 58
Fold over basal
Time (min)
12
10
8
6
4
2
0
-2
0
10
20
30
40
50
60
Normal
MK50
MK75
MK88
MK94
MK97
MK data
Normal data

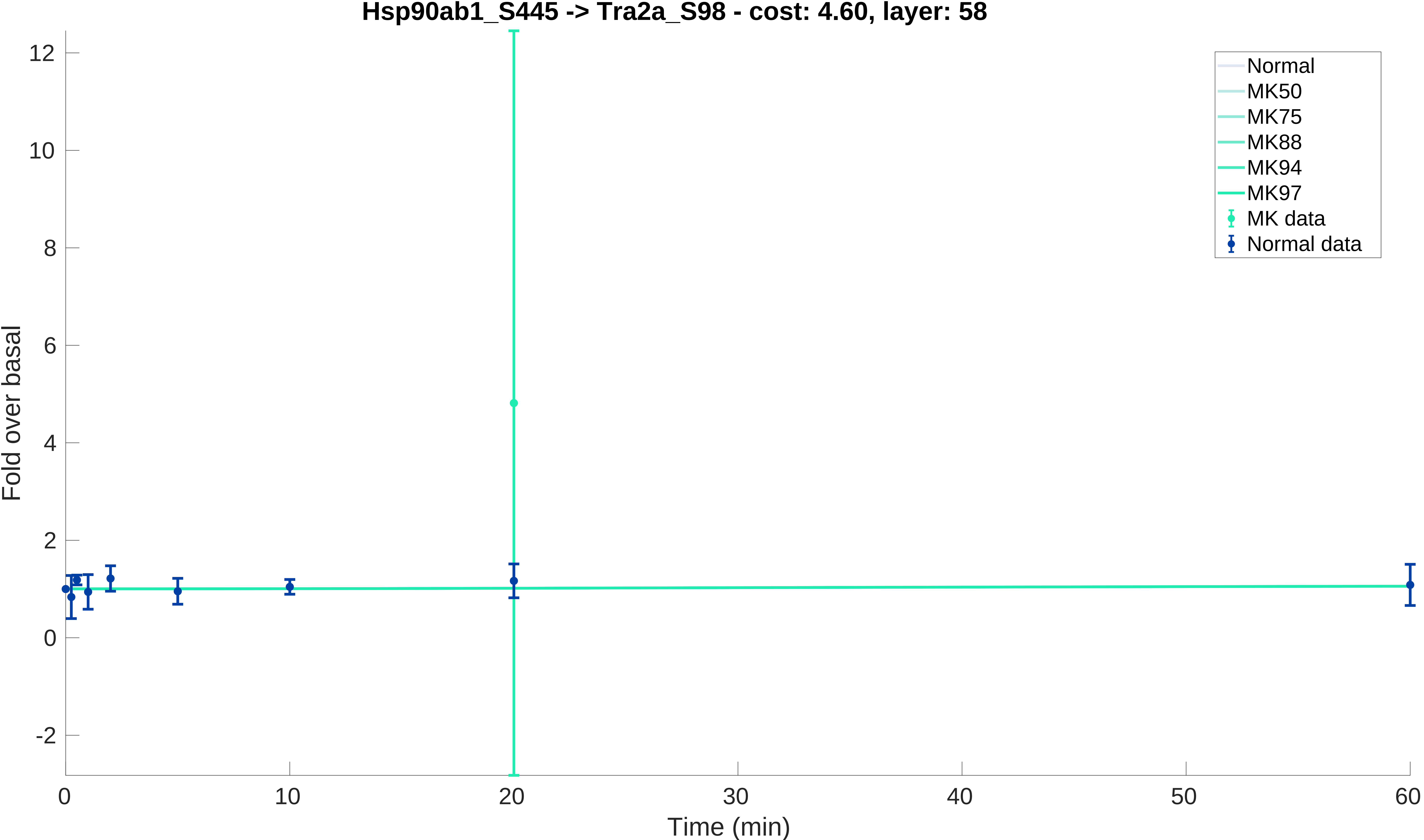

Hsp90ab1_S445 -> Tra2a_S98 - cost: 4.60, layer: 58
Fold over basal
Time (min)
12
10
8
6
4
2
0
-2
0
10
20
30
40
50
60
Normal
MK50
MK75
MK88
MK94
MK97
MK data
Normal data

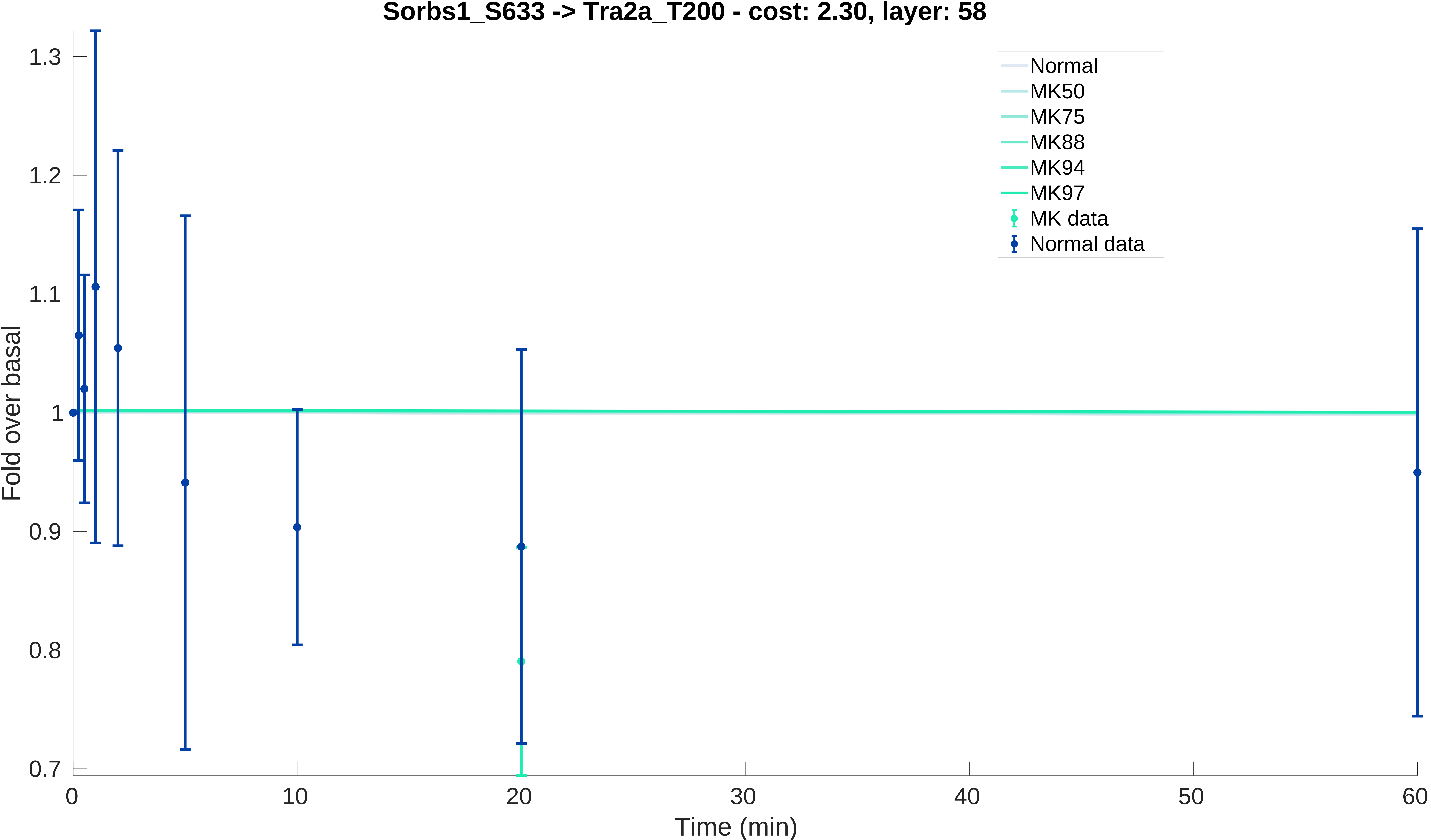

Sorbs1_S633 -> Tra2a_T200 - cost: 2.30, layer: 58
Fold over basal
Time (min)
0
10
20
30
40
50
60
0.7
0.8
0.9
1
1.1
1.2
1.3
Normal
MK50
MK75
MK88
MK94
MK97
MK data
Normal data

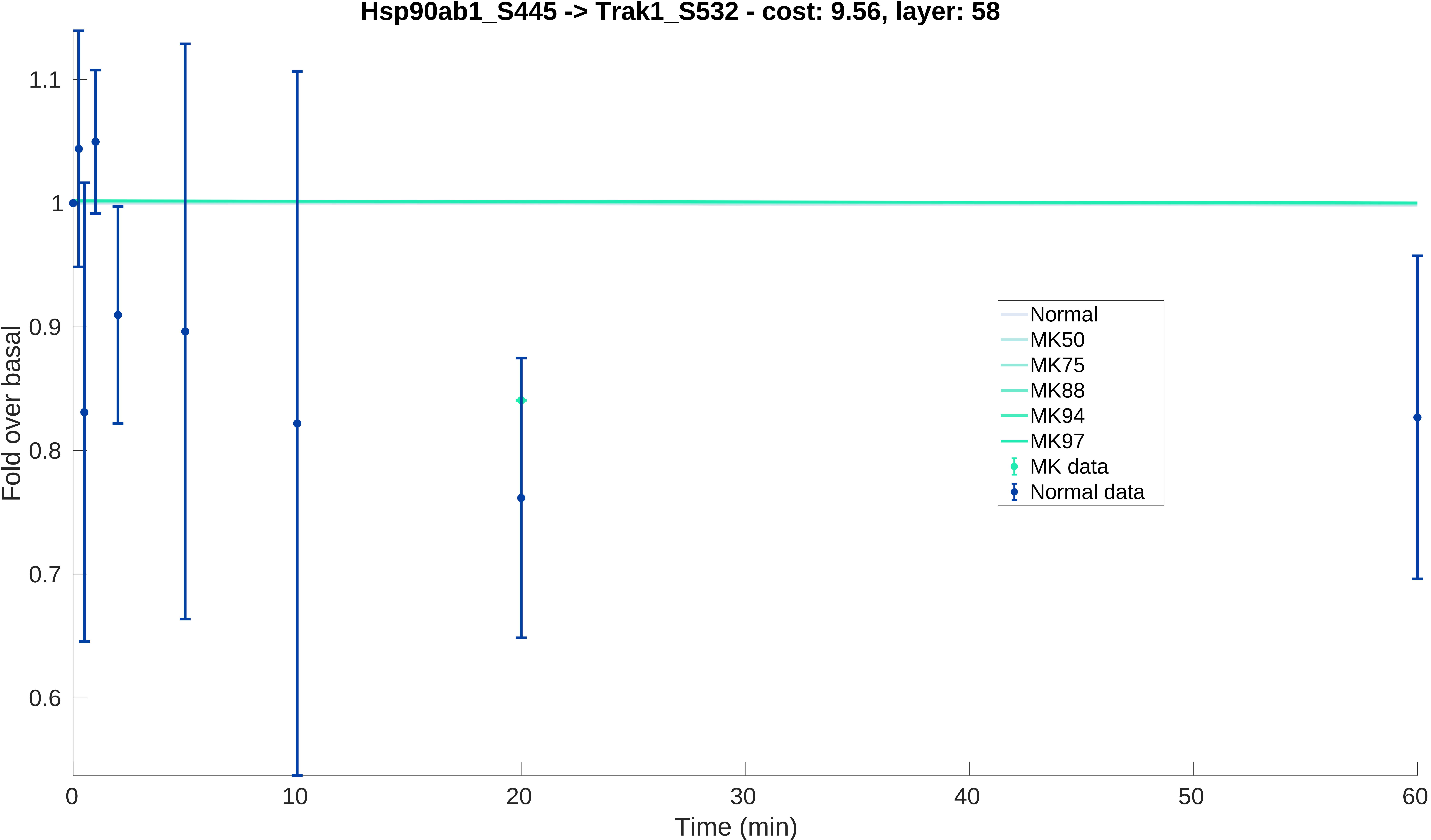

Hsp90ab1_S445 -> Trak1_S532 - cost: 9.56, layer: 58
Fold over basal
1.1
1
0.9
0.8
0.7
0.6
0
10
20
30
40
50
60
Time (min)
Normal
MK50
MK75
MK88
MK94
MK97
MK data
Normal data

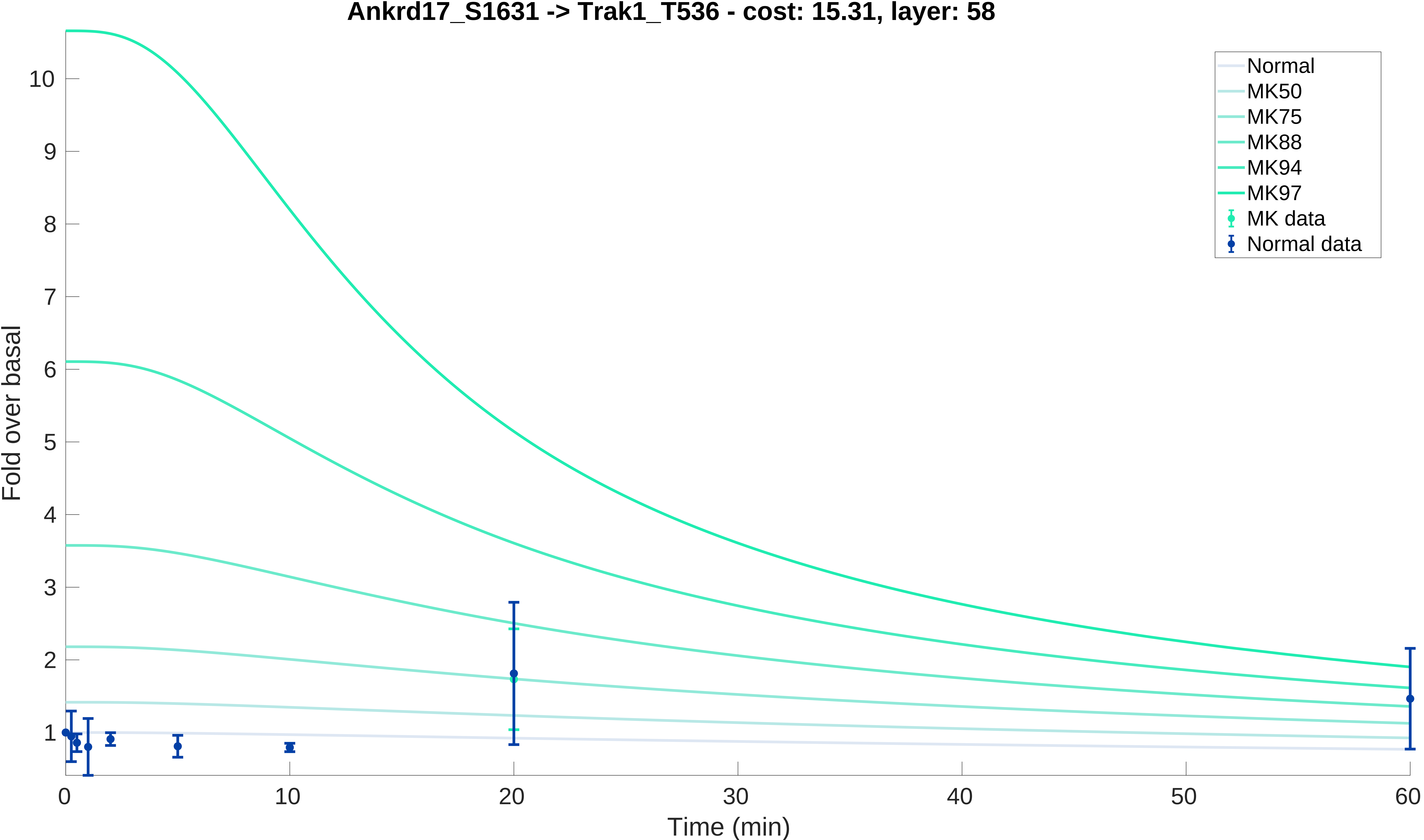
Ankrd17_S1631 -> Trak1_T536 - cost: 15.31, layer: 58
Fold over basal
Time (min)
Normal
MK50
MK75
MK88
MK94
MK97
MK data
Normal data
1
2
3
4
5
6
7
8
9
10
0
10
20
30
40
50
60

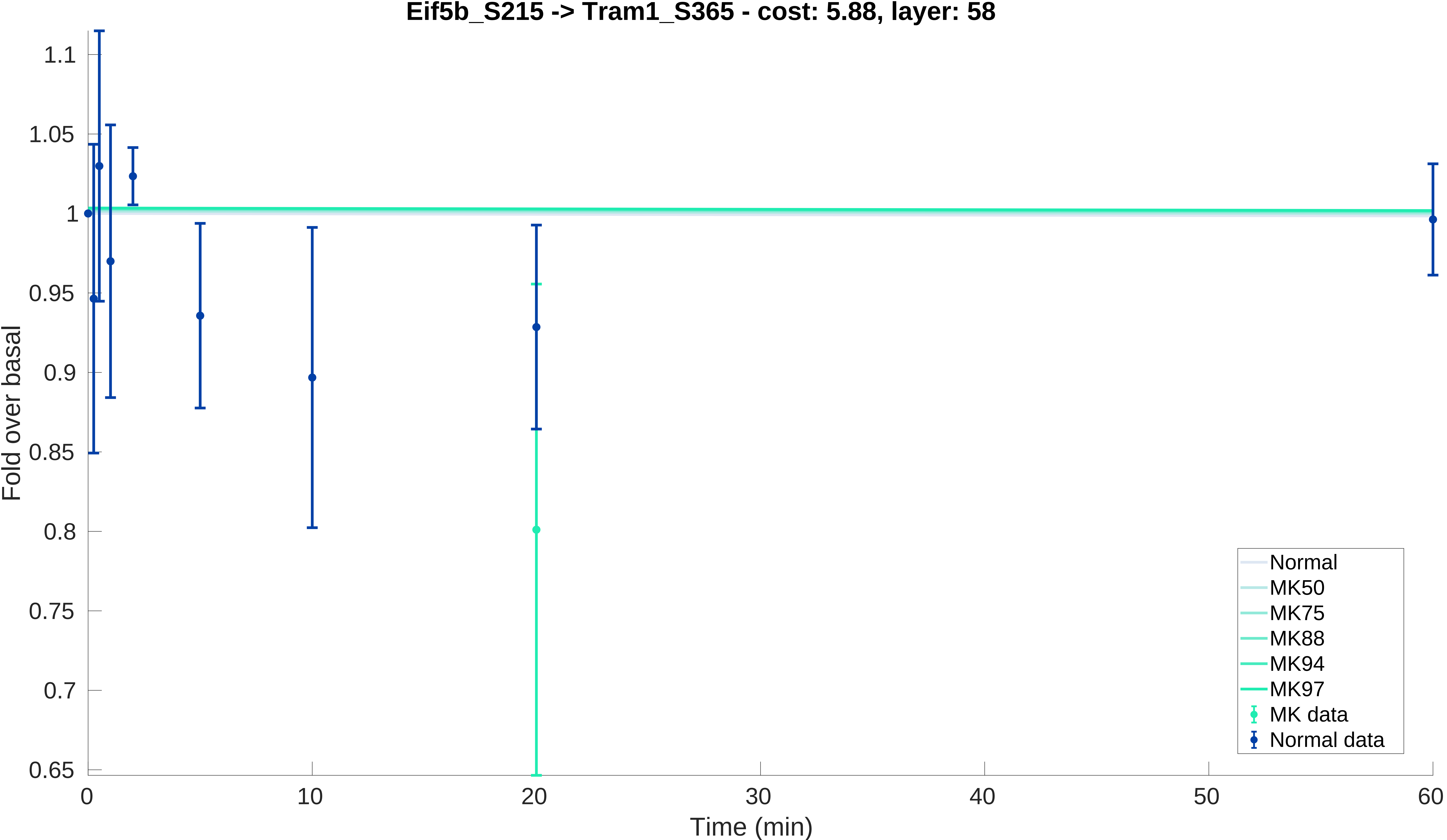

Eif5b_S215 -> Tram1_S365 - cost: 5.88, layer: 58
Fold over basal
1.1
1.05
1
0.95
0.9
0.85
0.8
0.75
0.7
0.65
0
10
20
30
40
50
60
Time (min)
Normal
MK50
MK75
MK88
MK94
MK97
MK data
Normal data

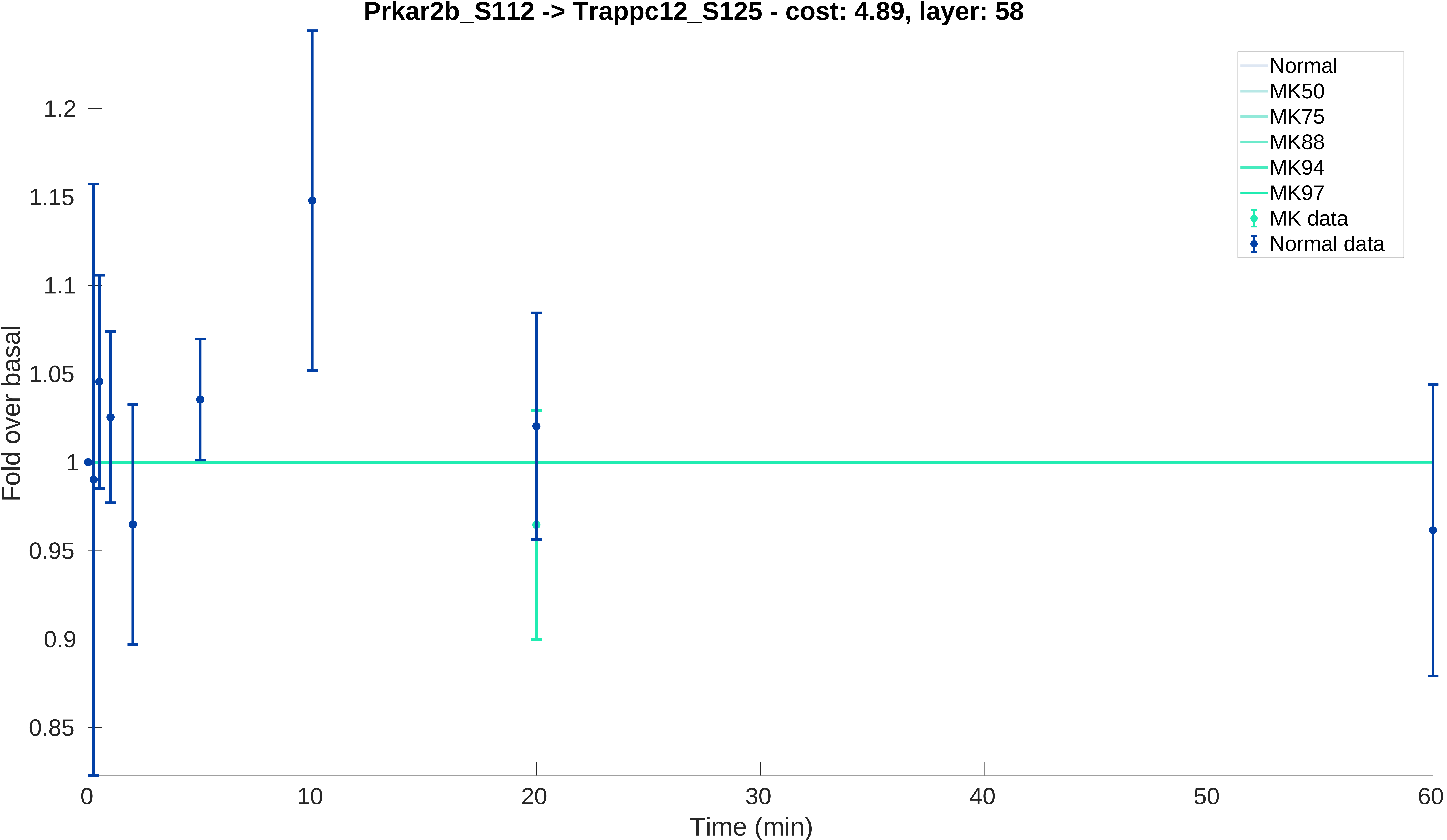

Prkar2b_S112 -> Trappc12_S125 - cost: 4.89, layer: 58
Normal
MK50
MK75
MK88
MK94
MK97
MK data
Normal data
Fold over basal
Time (min)
1.2
1.15
1.1
1.05
1
0.95
0.9
0.85
0
10
20
30
40
50
60

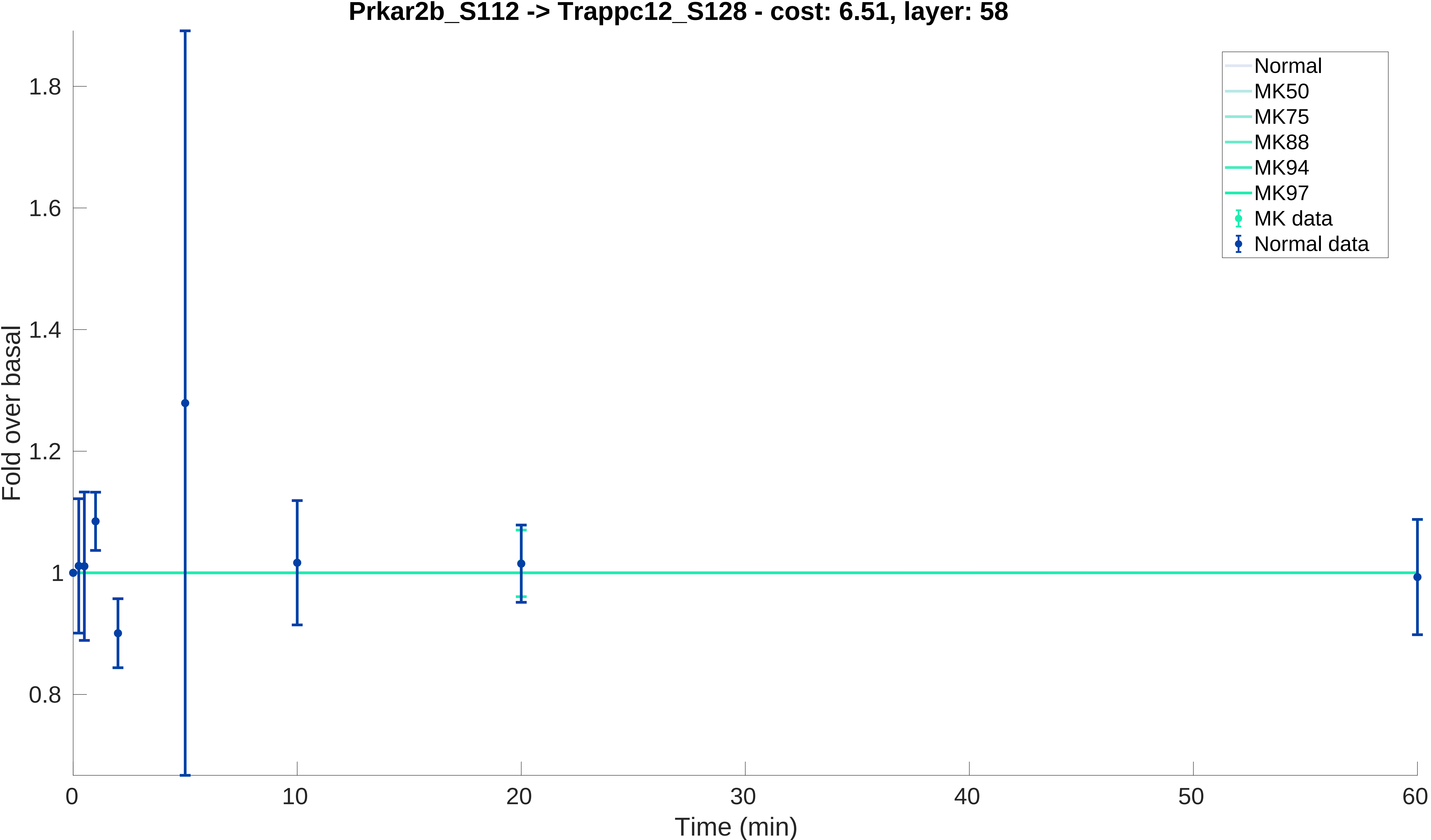

Prkar2b_S112 -> Trappc12_S128 - cost: 6.51, layer: 58
Fold over basal
1.8
1.6
1.4
1.2
1
0.8
0
10
20
30
40
50
60
Time (min)
Normal
MK50
MK75
MK88
MK94
MK97
MK data
Normal data

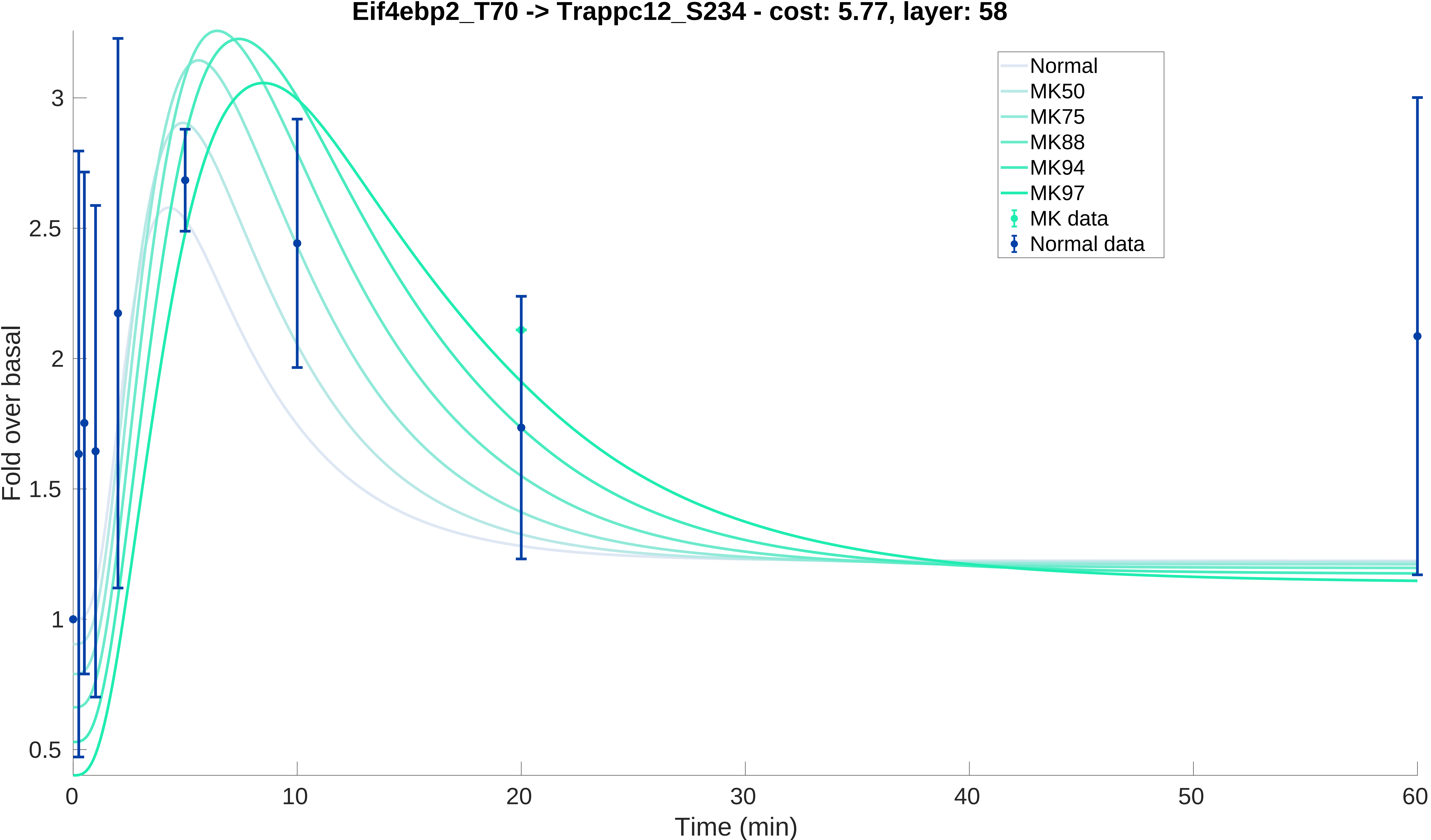

Eif4ebp2_T70 -> Trappc12_S234 - cost: 5.77, layer: 58
Fold over basal
Time (min)
0.5
1
1.5
2
2.5
3
0
10
20
30
40
50
60
Normal
MK50
MK75
MK88
MK94
MK97
MK data
Normal data

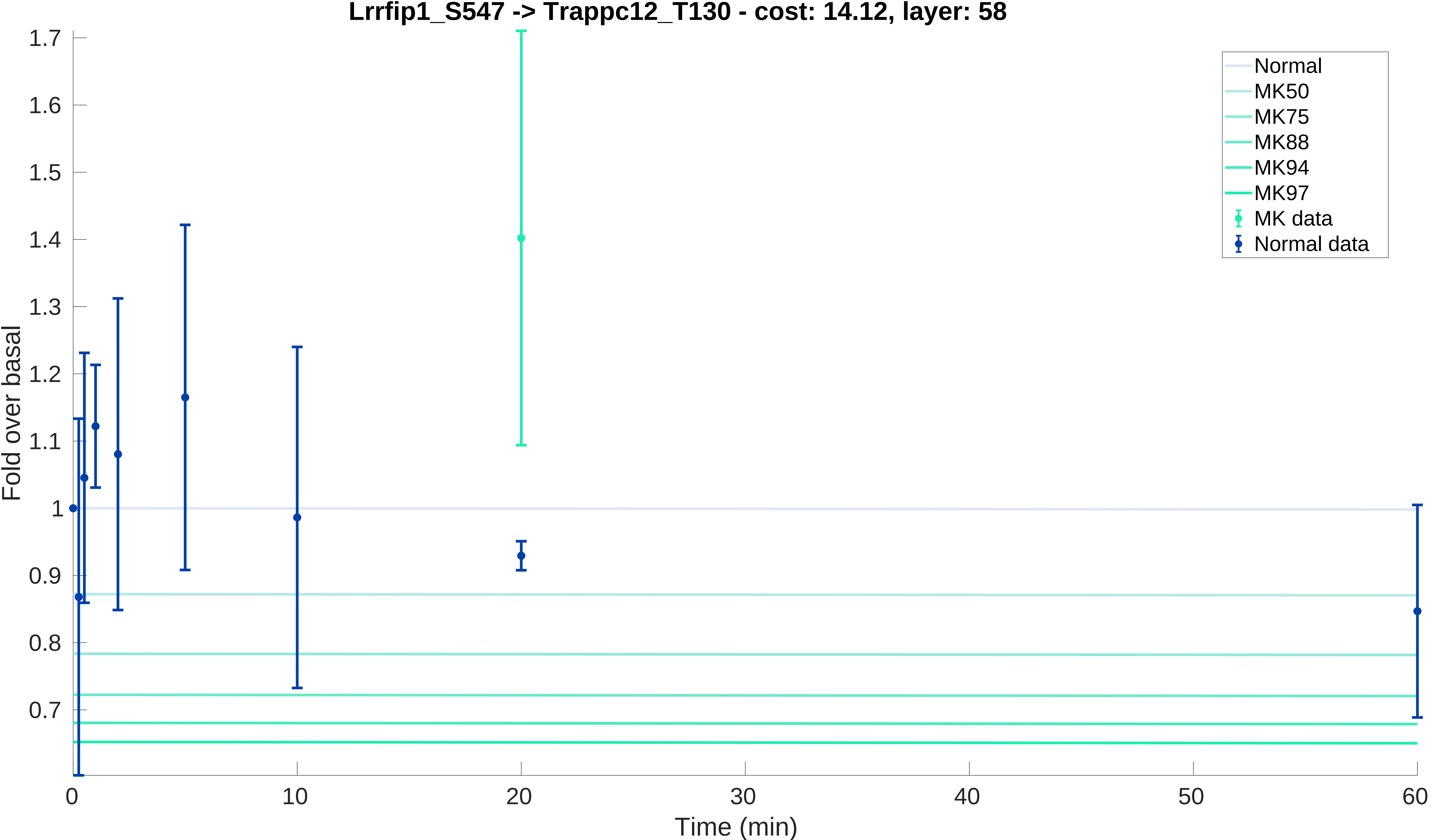

Lrrfip1_S547 -> Trappc12_T130 - cost: 14.12, layer: 58
Fold over basal
Time (min)
1.7
1.6
1.5
1.4
1.3
1.2
1.1
1
0.9
0.8
0.7
0
10
20
30
40
50
60
Normal
MK50
MK75
MK88
MK94
MK97
MK data
Normal data

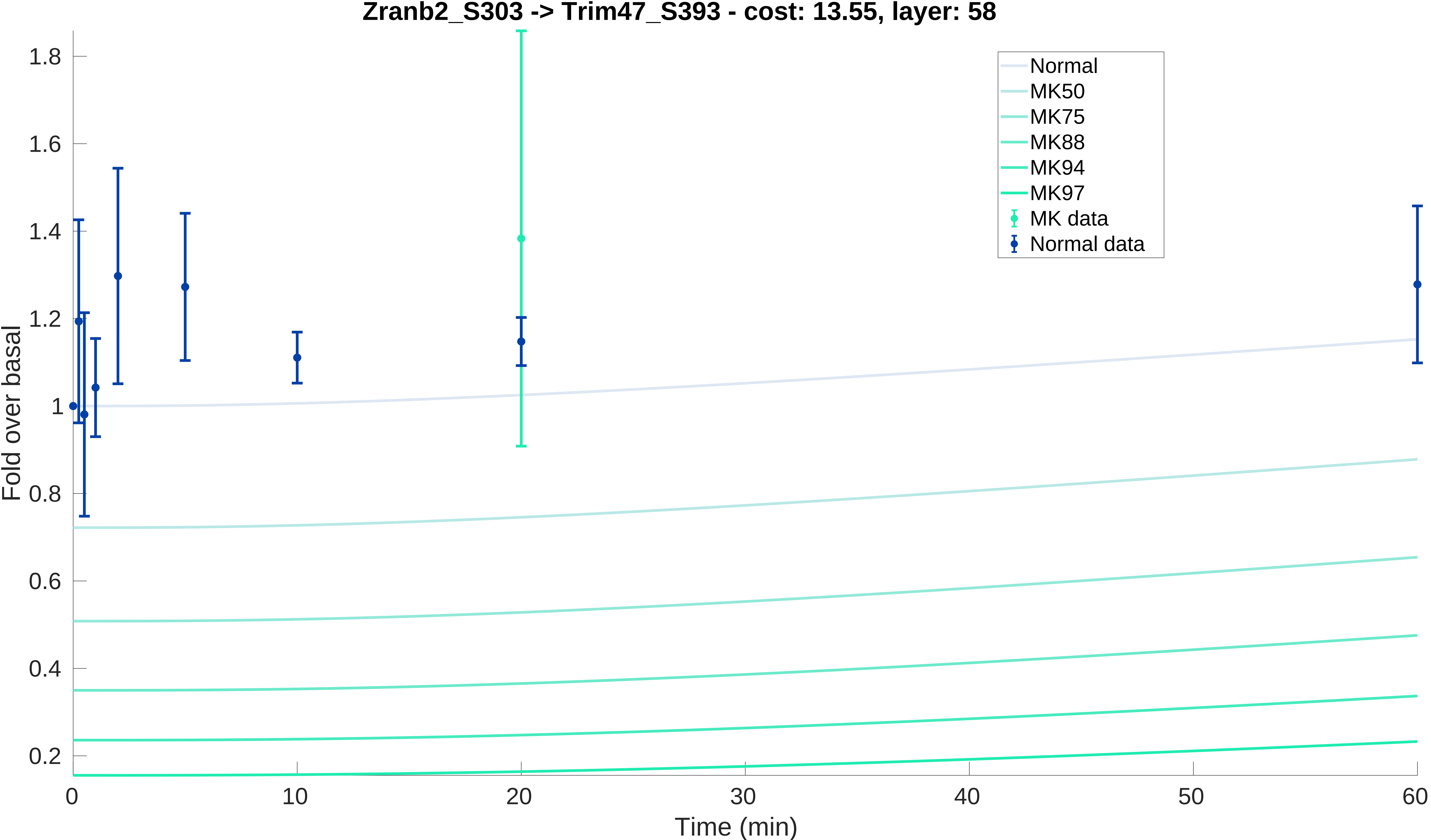

Zranb2_S303 -> Trim47_S393 - cost: 13.55, layer: 58
Fold over basal
Time (min)
0
10
20
30
40
50
60
0.2
0.4
0.6
0.8
1
1.2
1.4
1.6
1.8
Normal
MK50
MK75
MK88
MK94
MK97
MK data
Normal data

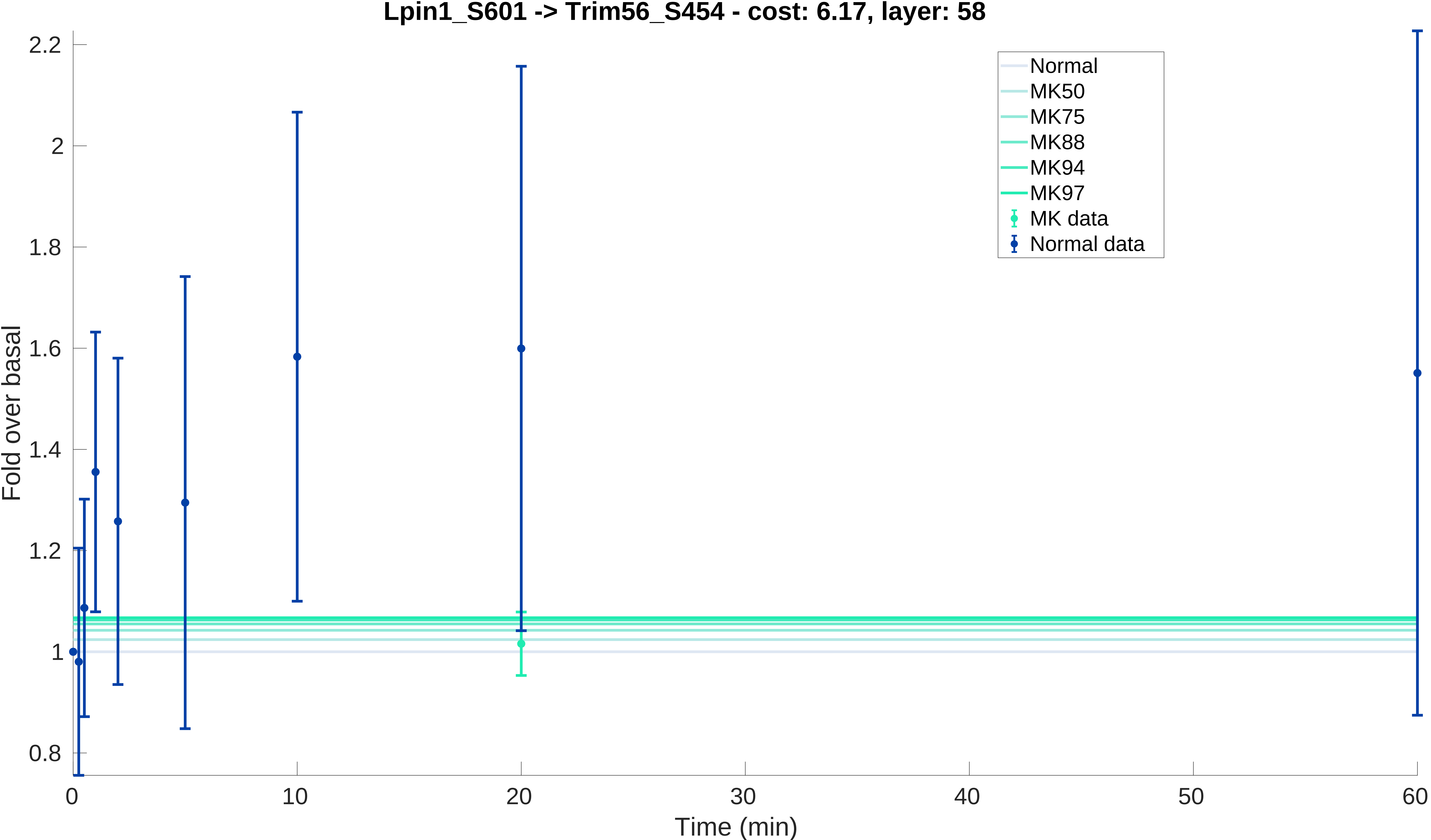

Lpin1_S601 -> Trim56_S454 - cost: 6.17, layer: 58
Normal
MK50
MK75
MK88
MK94
MK97
MK data
Normal data
Fold over basal
Time (min)
2.2
2
1.8
1.6
1.4
1.2
1
0.8
0
10
20
30
40
50
60

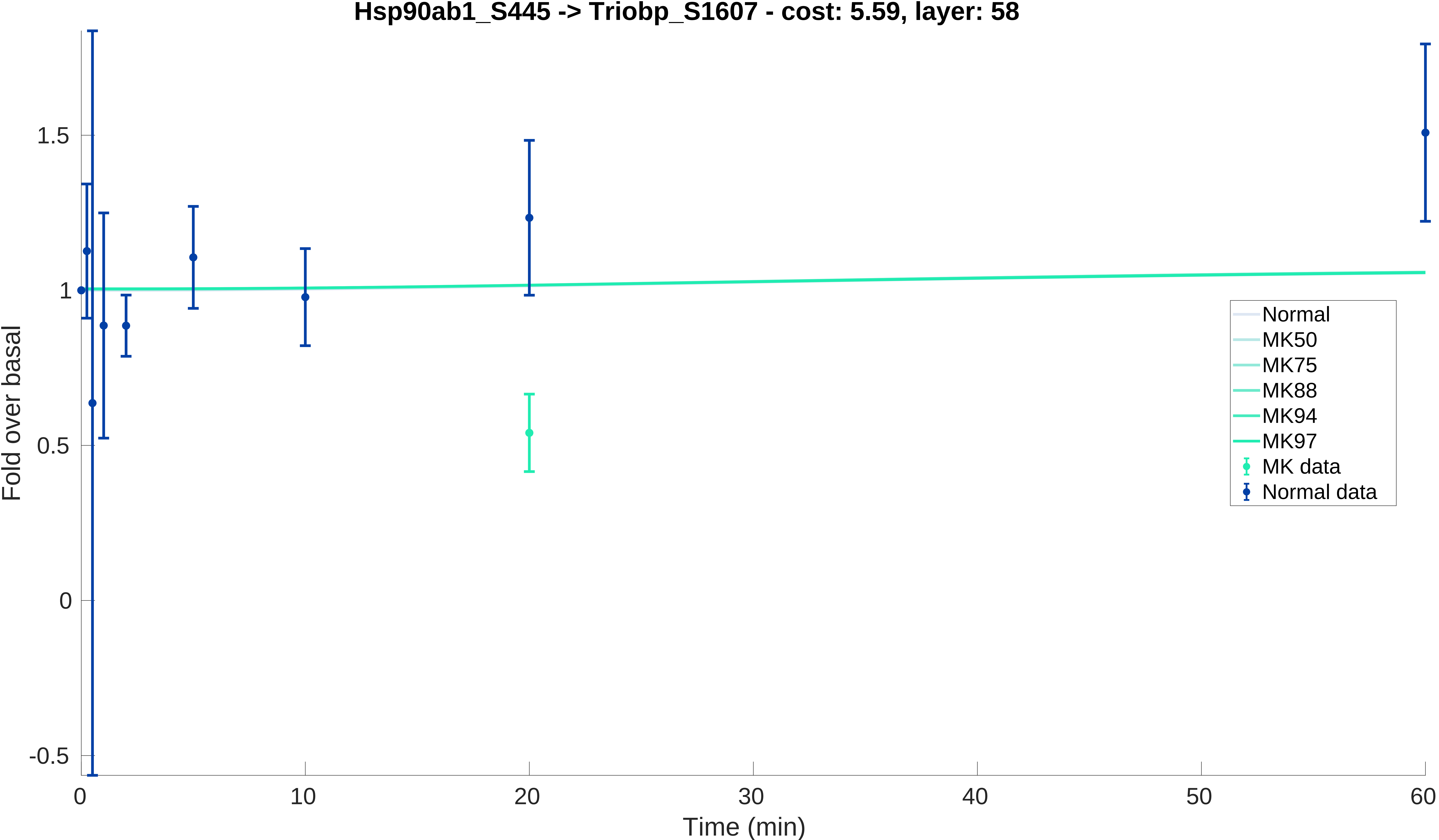

Hsp90ab1_S445 -> Triobp_S1607 - cost: 5.59, layer: 58
Fold over basal
Time (min)
1.5
1
0.5
0
-0.5
0
10
20
30
40
50
60
Normal
MK50
MK75
MK88
MK94
MK97
MK data
Normal data

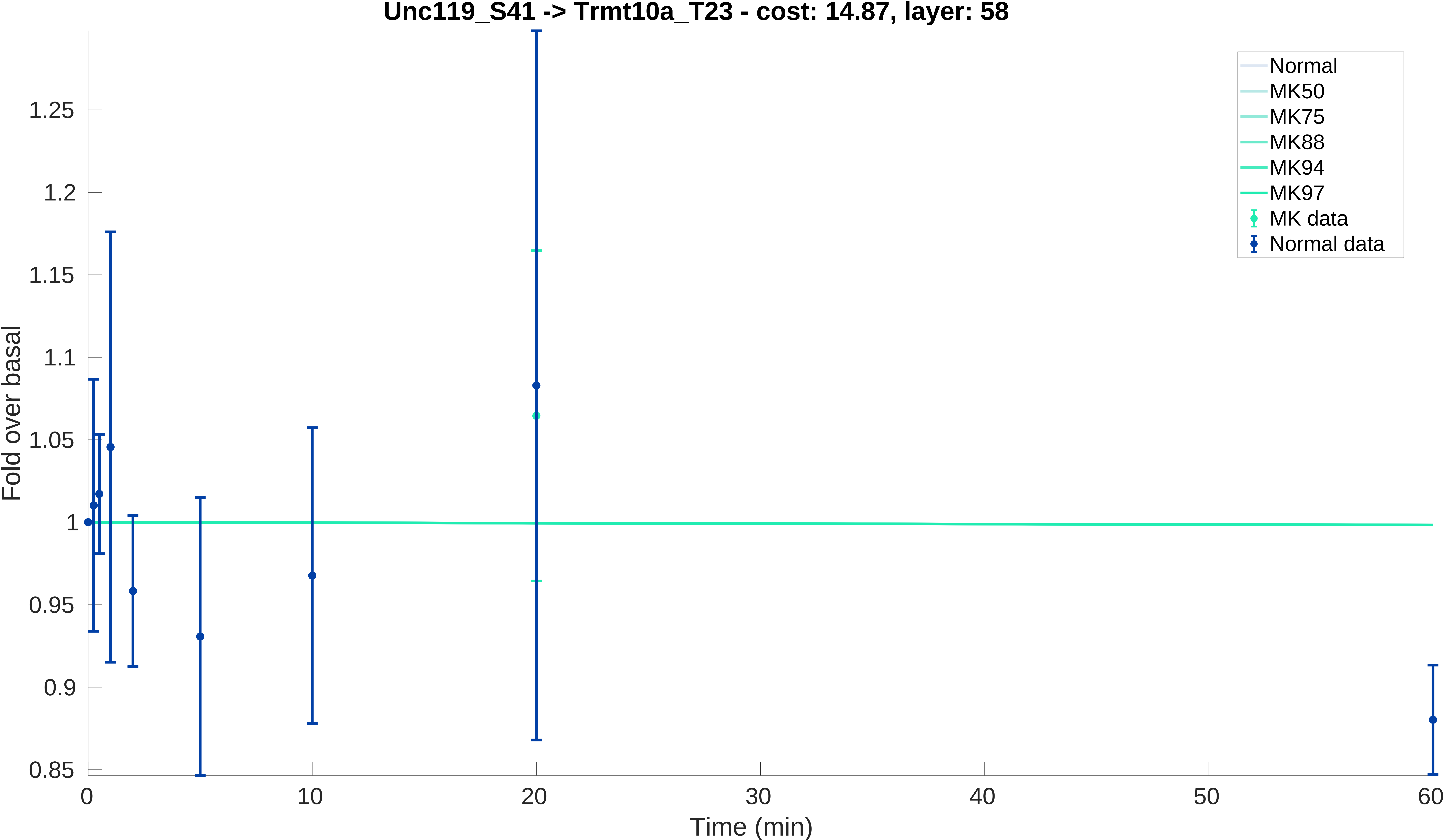

Unc119_S41 -> Trmt10a_T23 - cost: 14.87, layer: 58
Fold over basal
Time (min)
1.25
1.2
1.15
1.1
1.05
1
0.95
0.9
0.85
0
10
20
30
40
50
60
Normal
MK50
MK75
MK88
MK94
MK97
MK data
Normal data

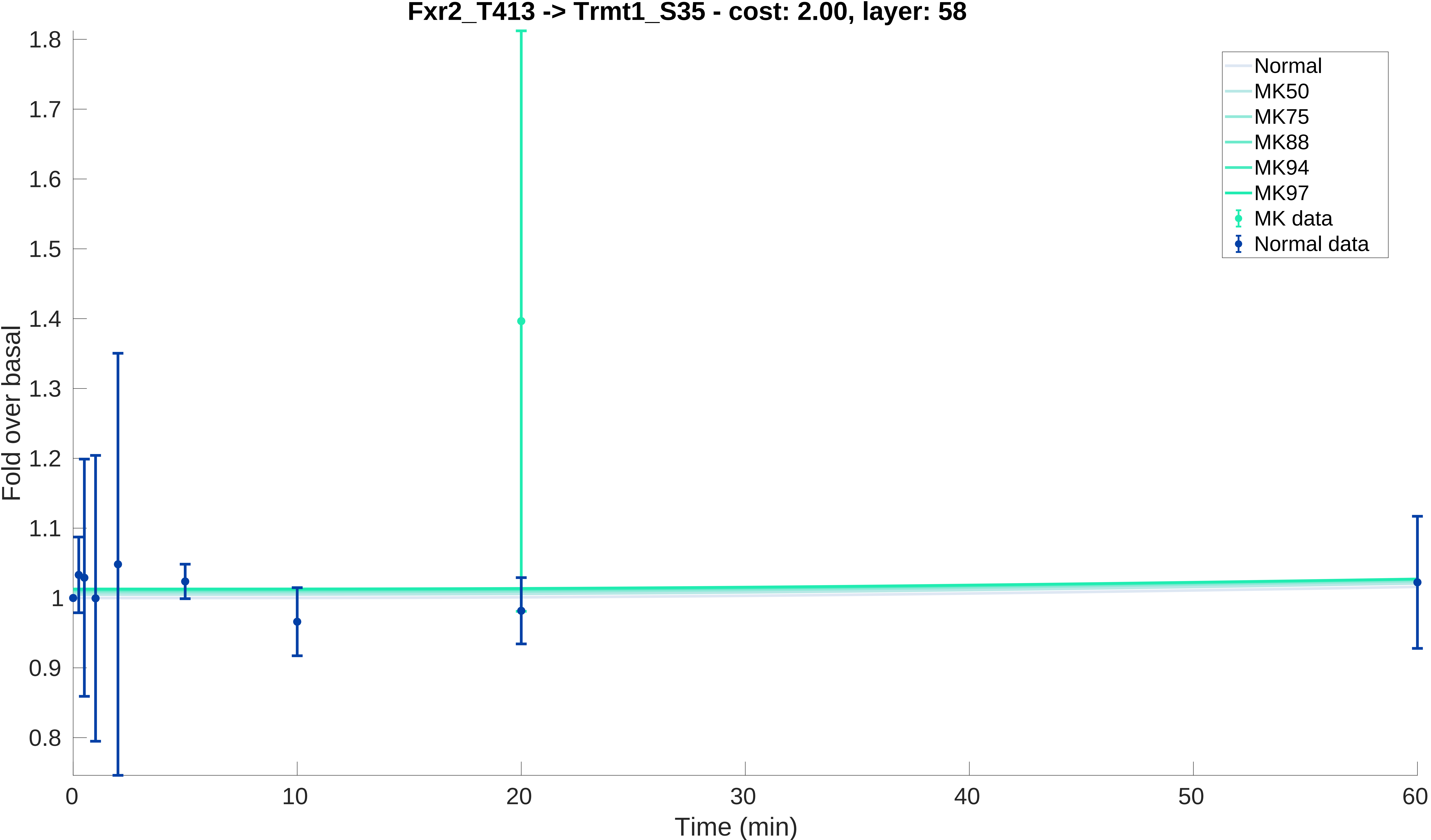

Fxr2_T413 -> Trmt1_S35 - cost: 2.00, layer: 58
Fold over basal
Time (min)
0.8
0.9
1
1.1
1.2
1.3
1.4
1.5
1.6
1.7
1.8
0
10
20
30
40
50
60
Normal
MK50
MK75
MK88
MK94
MK97
MK data
Normal data

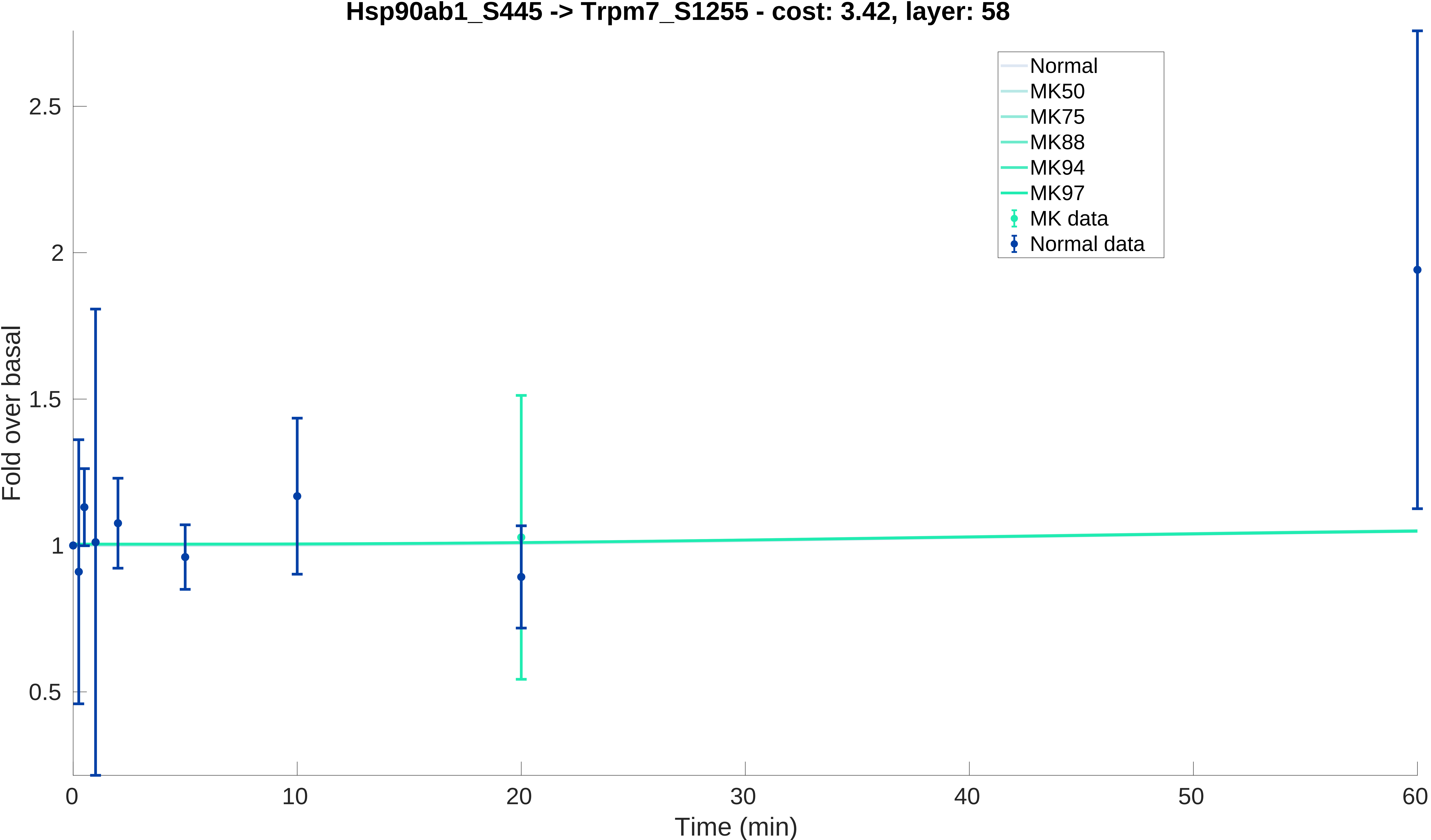

Hsp90ab1_S445 -> Trpm7_S1255 - cost: 3.42, layer: 58
Fold over basal
Time (min)
2.5
2
1.5
1
0.5
0
10
20
30
40
50
60
Normal
MK50
MK75
MK88
MK94
MK97
MK data
Normal data

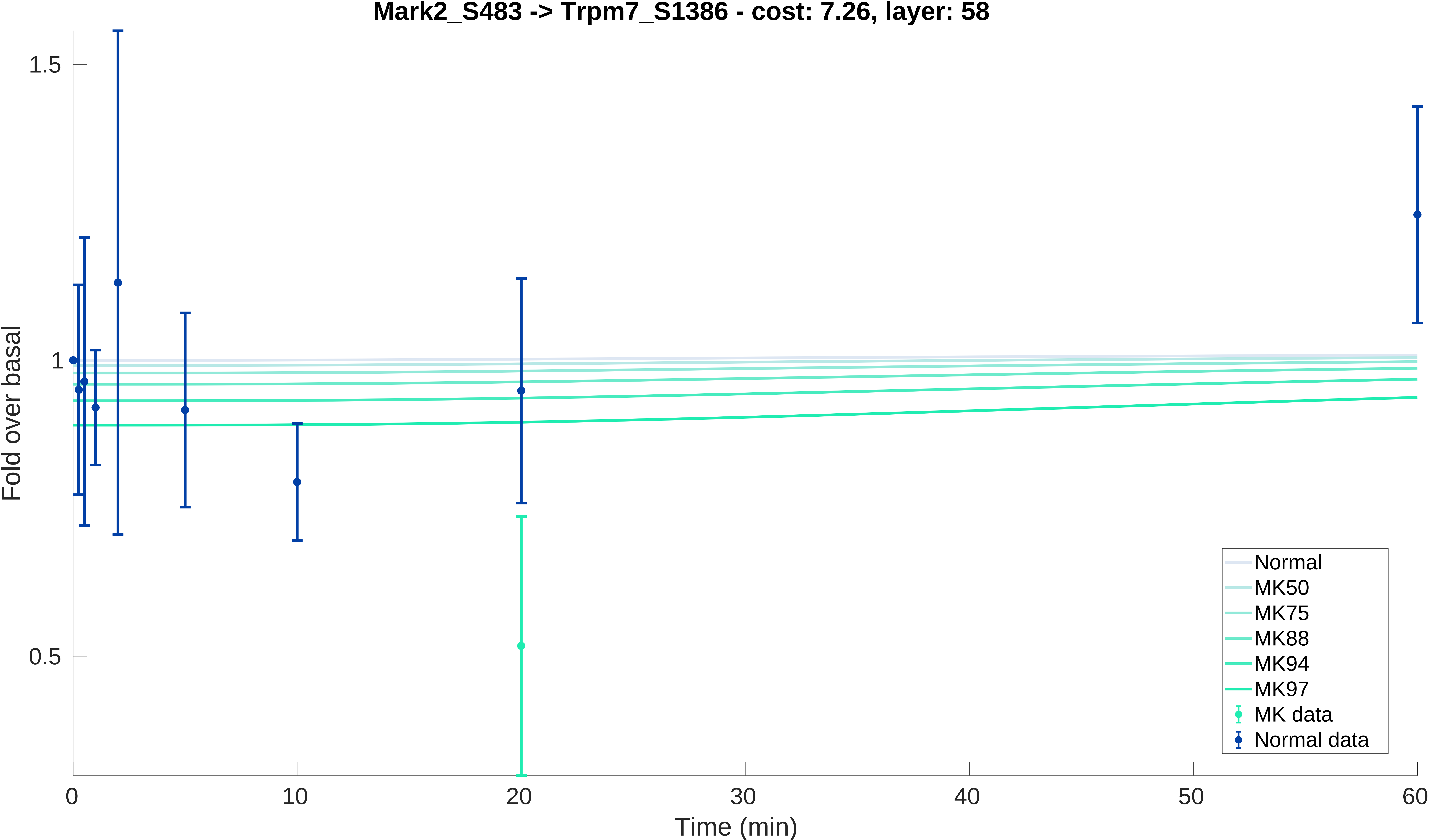

Mark2_S483 -> Trpm7_S1386 - cost: 7.26, layer: 58
1.5
1
0.5
Fold over basal
0
10
20
30
40
50
60
Time (min)
Normal
MK50
MK75
MK88
MK94
MK97
MK data
Normal data

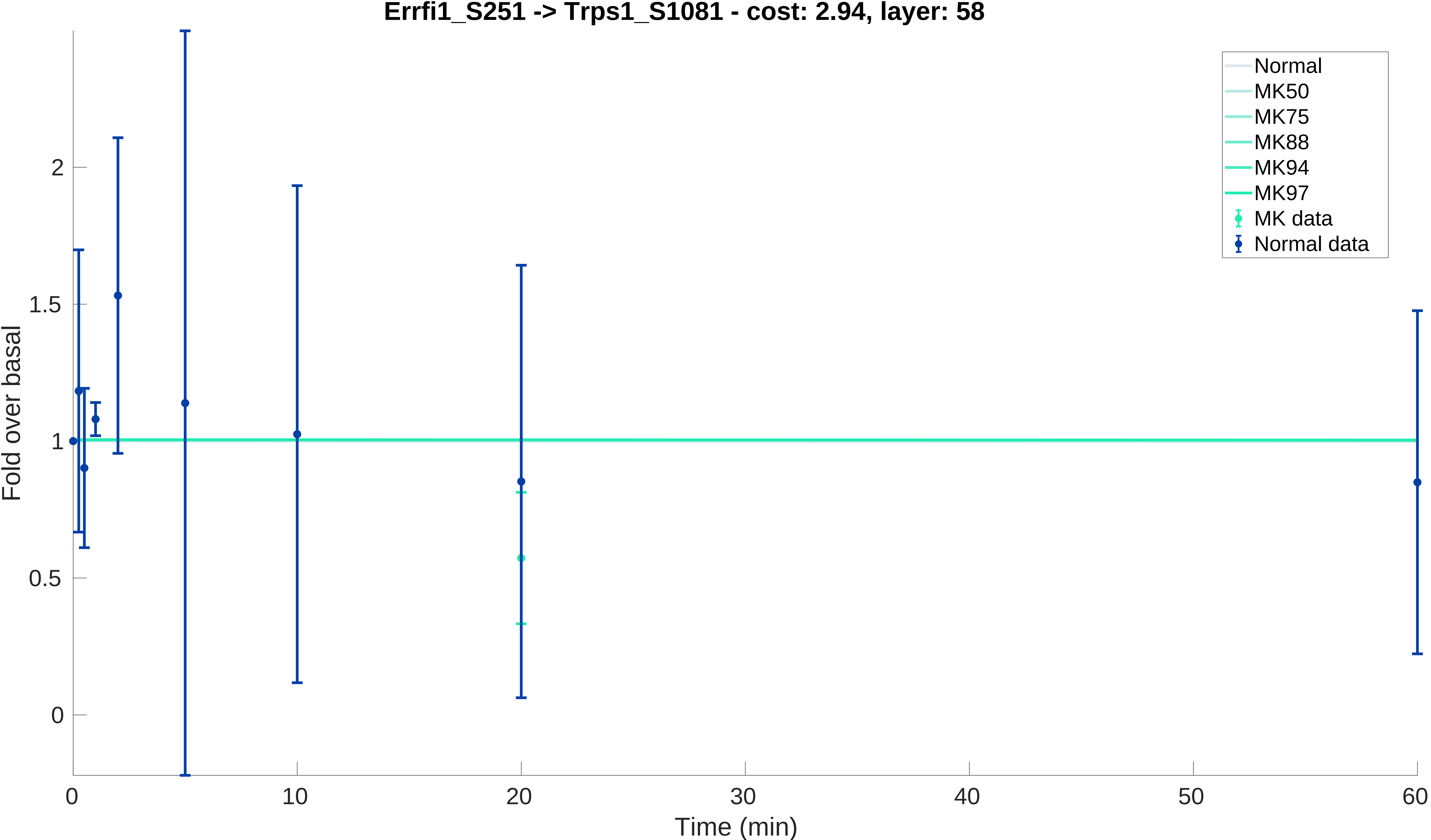

Errfi1_S251 -> Trps1_S1081 - cost: 2.94, layer: 58
Normal
MK50
MK75
MK88
MK94
MK97
MK data
Normal data
Fold over basal
0
0.5
1
1.5
2
0
10
20
30
40
50
60
Time (min)

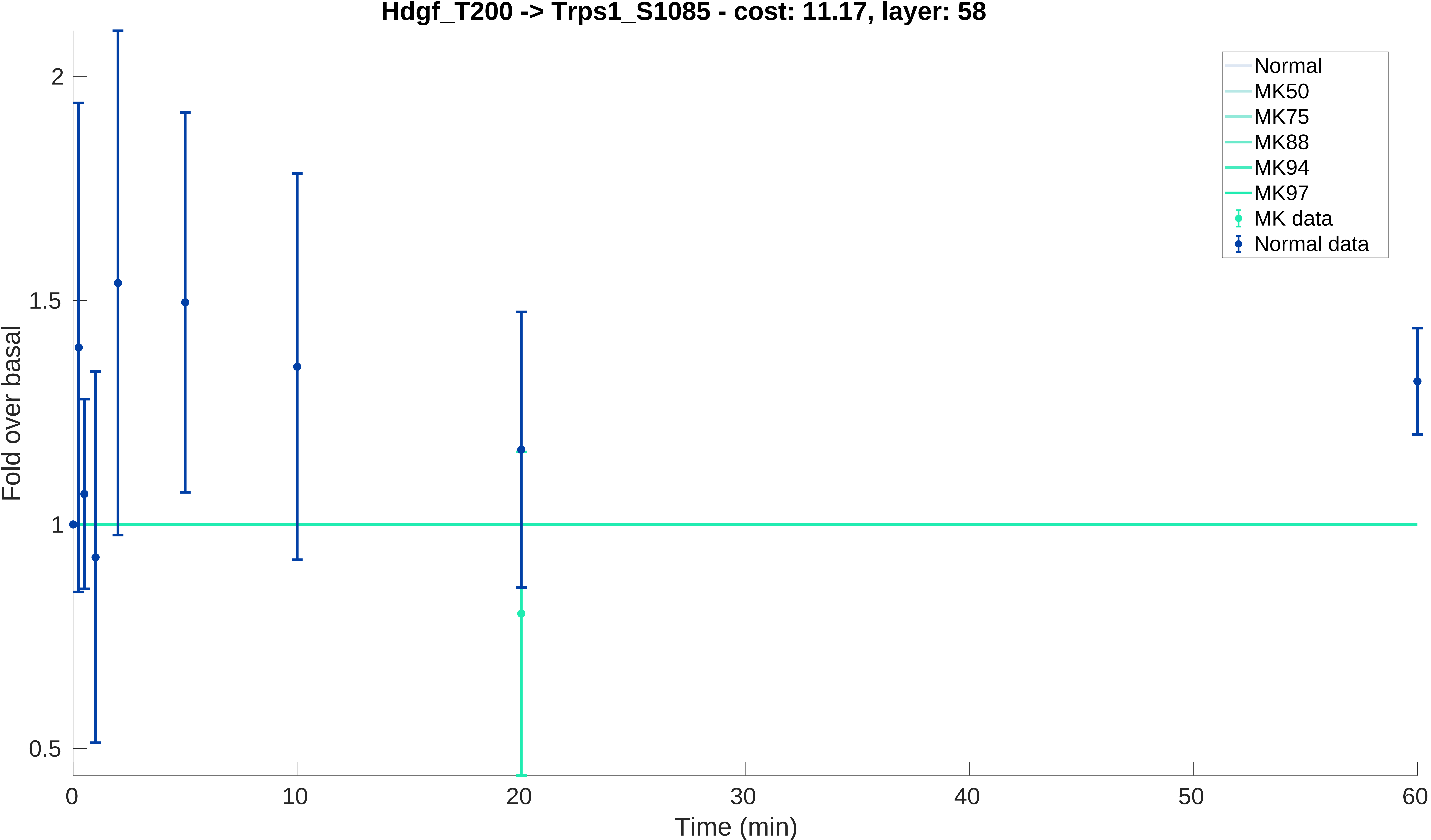

Hdgf_T200 -> Trps1_S1085 - cost: 11.17, layer: 58
Fold over basal
Time (min)
0.5
1
1.5
2
0
10
20
30
40
50
60
Normal
MK50
MK75
MK88
MK94
MK97
MK data
Normal data

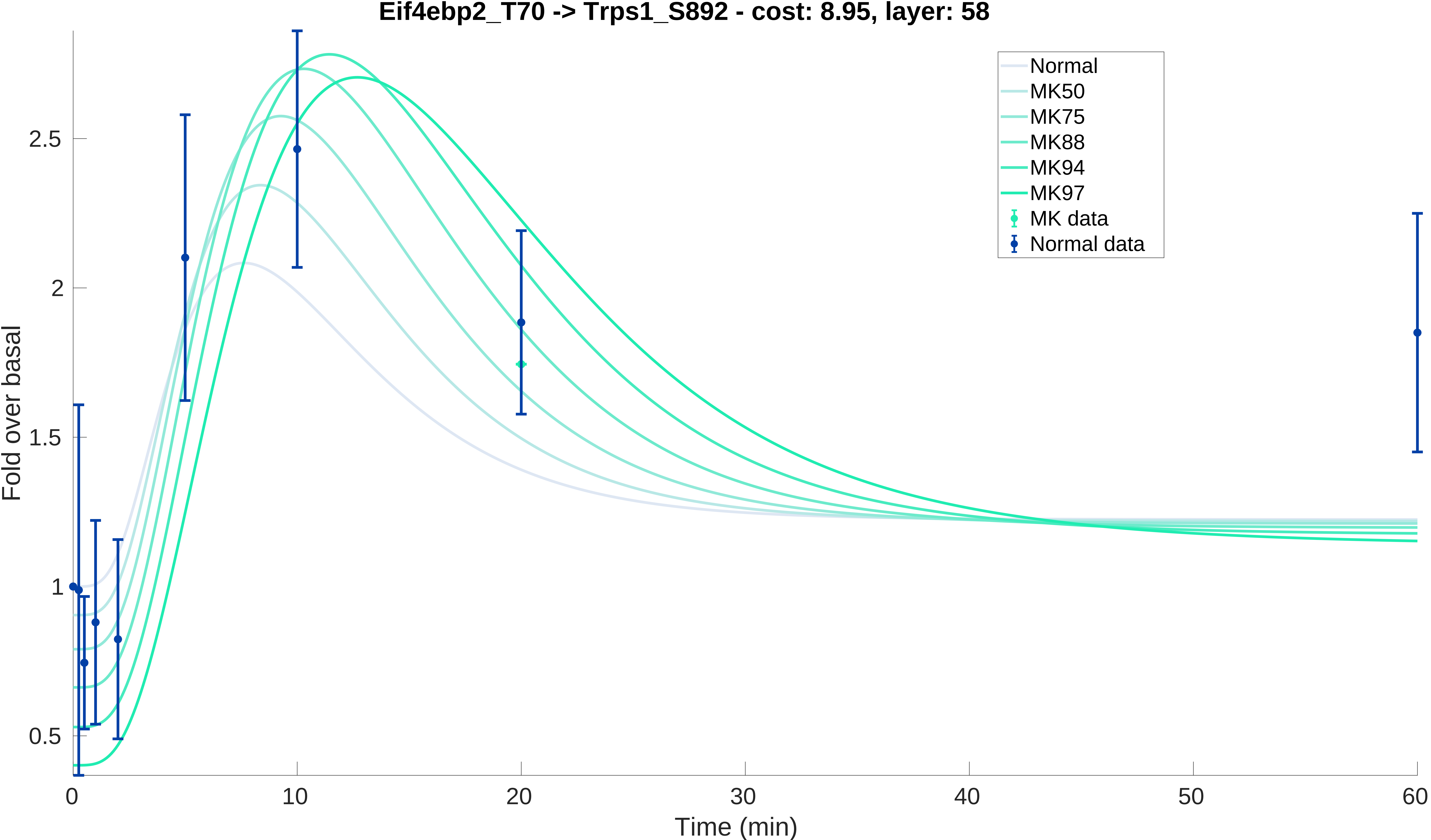

Eif4ebp2_T70 -> Trps1_S892 - cost: 8.95, layer: 58
Fold over basal
Time (min)
0.5
1
1.5
2
2.5
0
10
20
30
40
50
60
Normal
MK50
MK75
MK88
MK94
MK97
MK data
Normal data

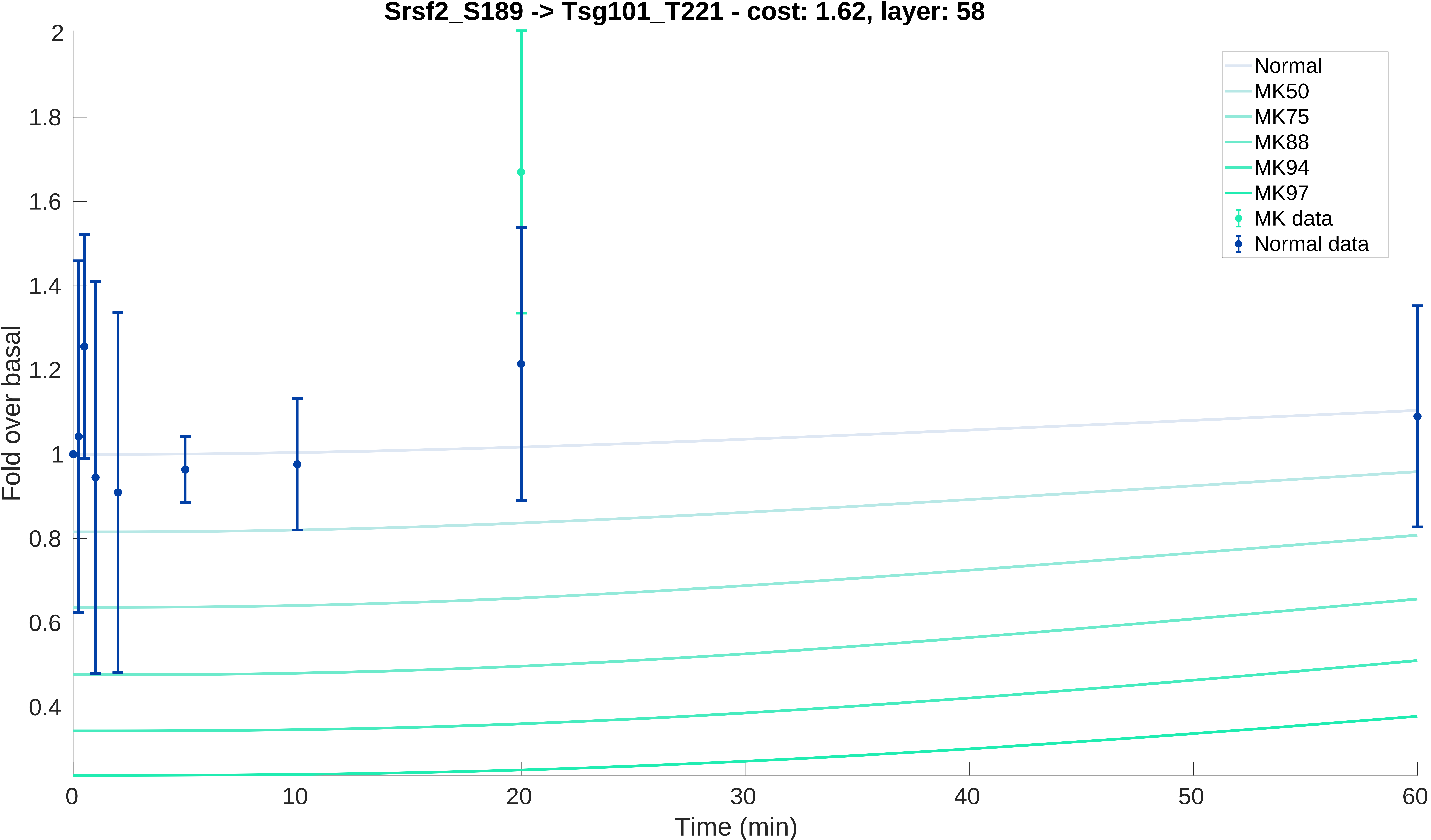
Srsf2_S189 -> Tsg101_T221 - cost: 1.62, layer: 58
Fold over basal
Time (min)
0
10
20
30
40
50
60
0.4
0.6
0.8
1
1.2
1.4
1.6
1.8
2
Normal
MK50
MK75
MK88
MK94
MK97
MK data
Normal data

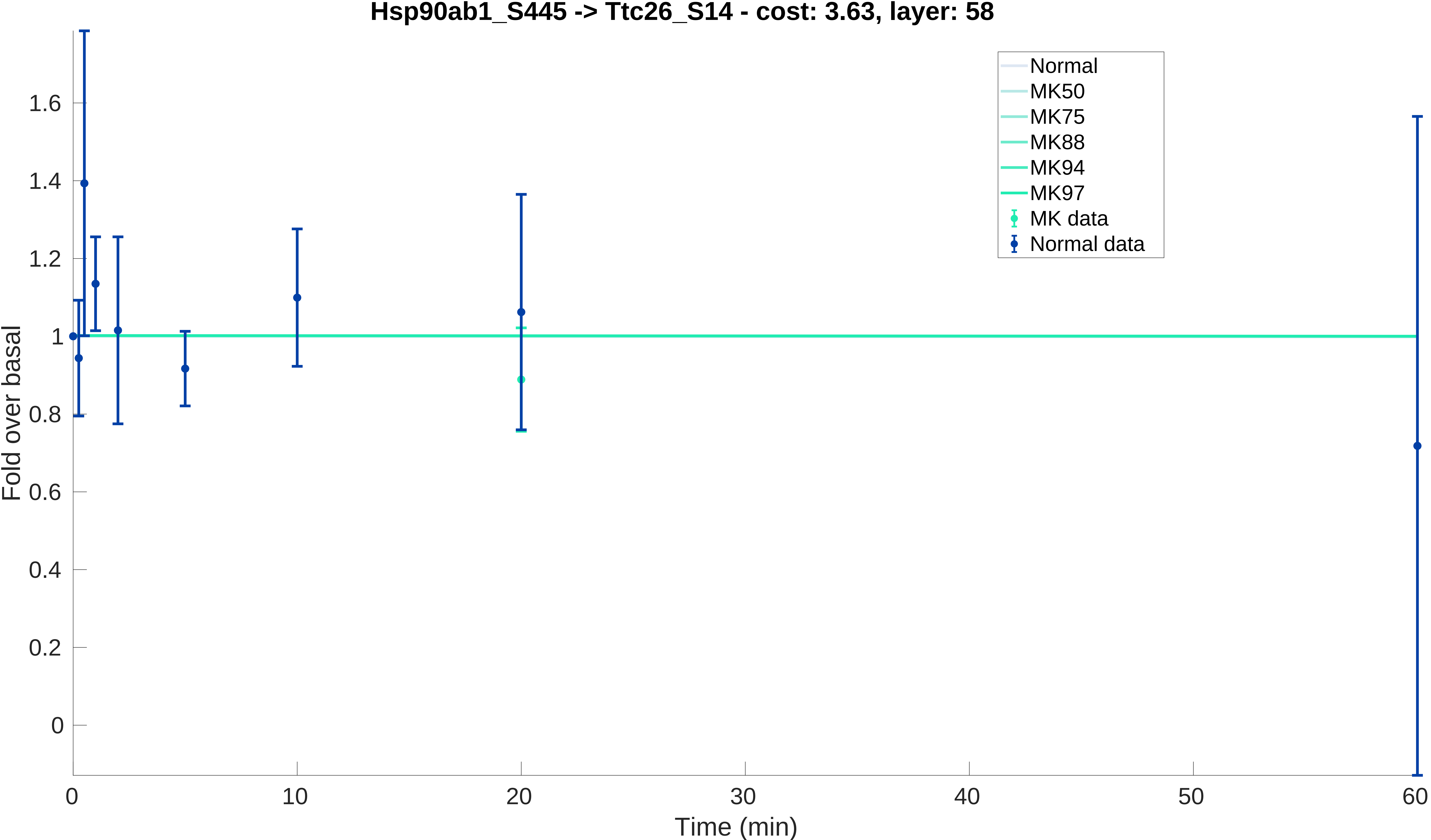
Hsp90ab1_S445 -> Ttc26_S14 - cost: 3.63, layer: 58
Fold over basal
Time (min)
0
0.2
0.4
0.6
0.8
1
1.2
1.4
1.6
0
10
20
30
40
50
60
Normal
MK50
MK75
MK88
MK94
MK97
MK data
Normal data

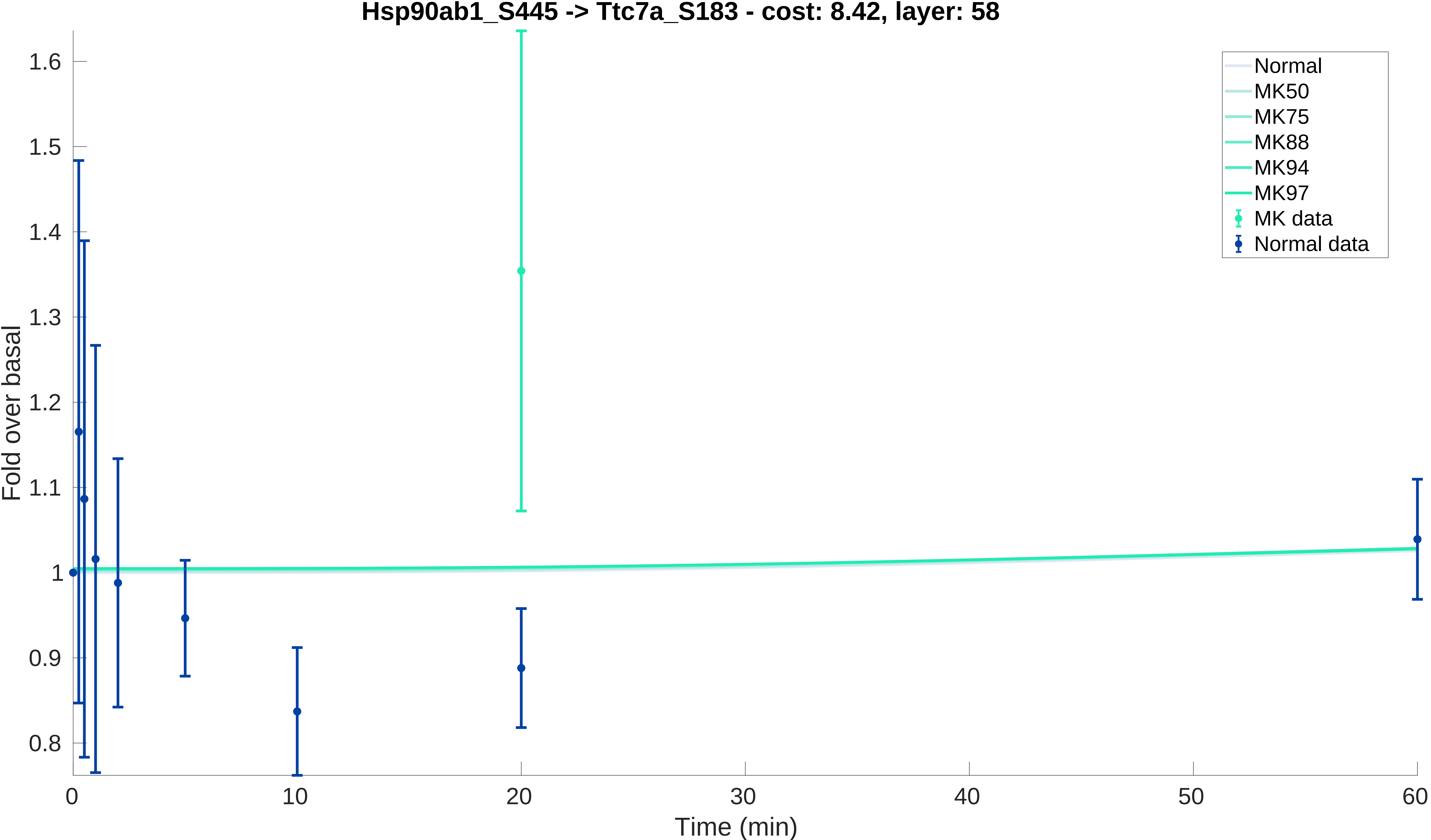

Hsp90ab1_S445 -> Ttc7a_S183 - cost: 8.42, layer: 58
Fold over basal
Time (min)
0.8
0.9
1
1.1
1.2
1.3
1.4
1.5
1.6
0
10
20
30
40
50
60
Normal
MK50
MK75
MK88
MK94
MK97
MK data
Normal data

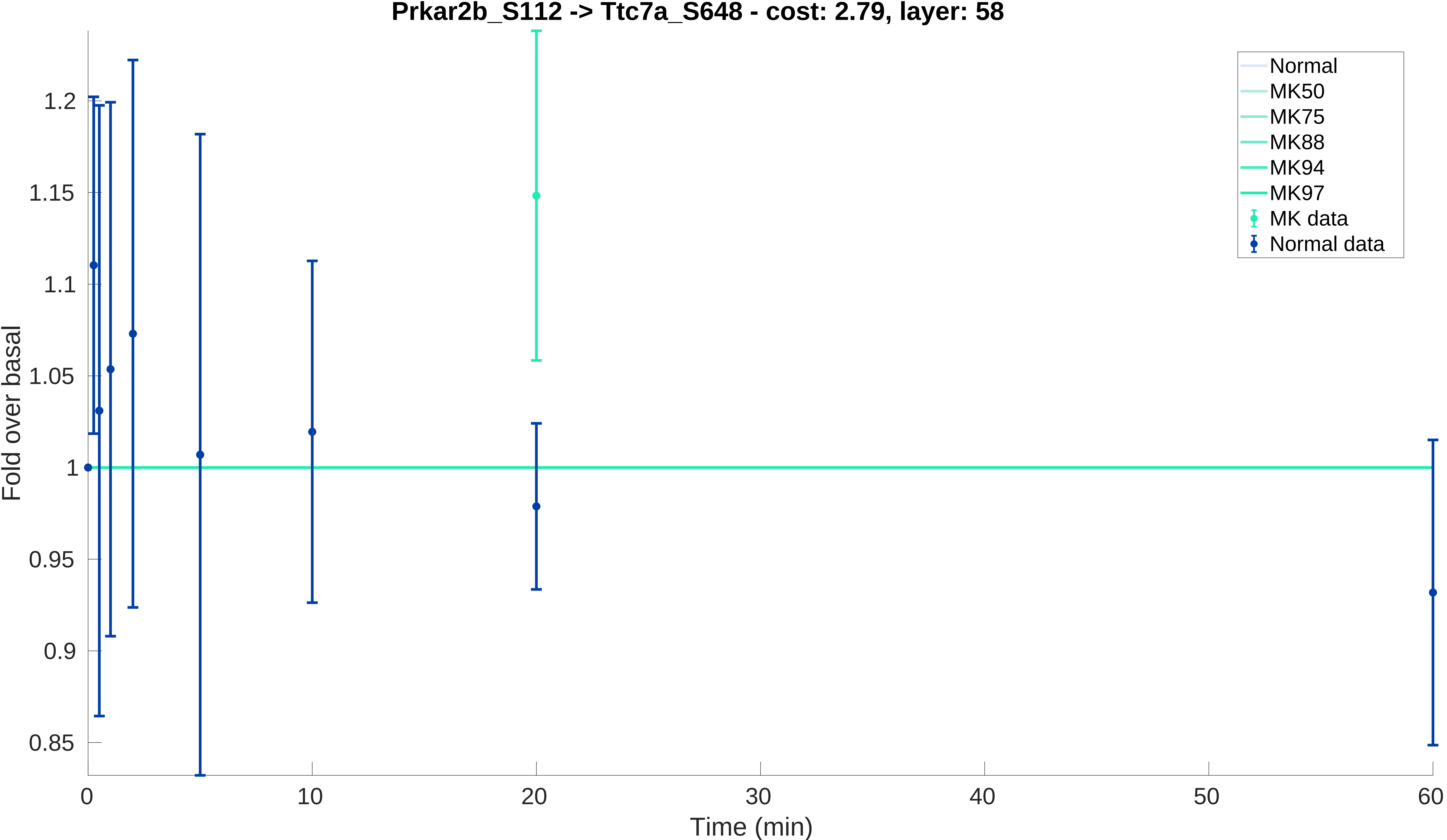

Prkar2b_S112 -> Ttc7a_S648 - cost: 2.79, layer: 58
Fold over basal
1.2
1.15
1.1
1.05
1
0.95
0.9
0.85
0
10
20
30
40
50
60
Time (min)
Normal
MK50
MK75
MK88
MK94
MK97
MK data
Normal data

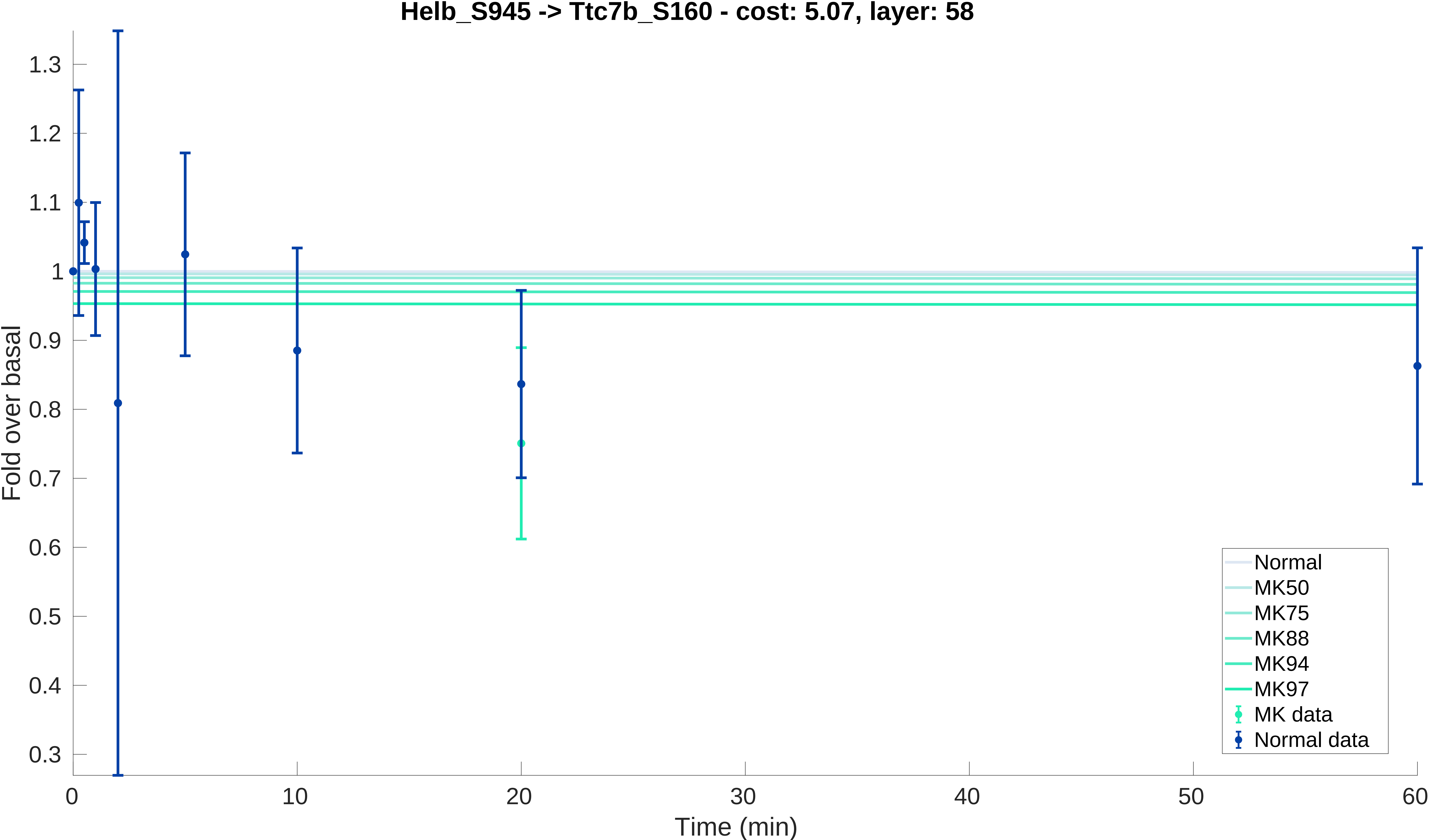

Helb_S945 -> Ttc7b_S160 - cost: 5.07, layer: 58
Fold over basal
1.3
1.2
1.1
1
0.9
0.8
0.7
0.6
0.5
0.4
0.3
0
10
20
30
40
50
60
Time (min)
Normal
MK50
MK75
MK88
MK94
MK97
MK data
Normal data

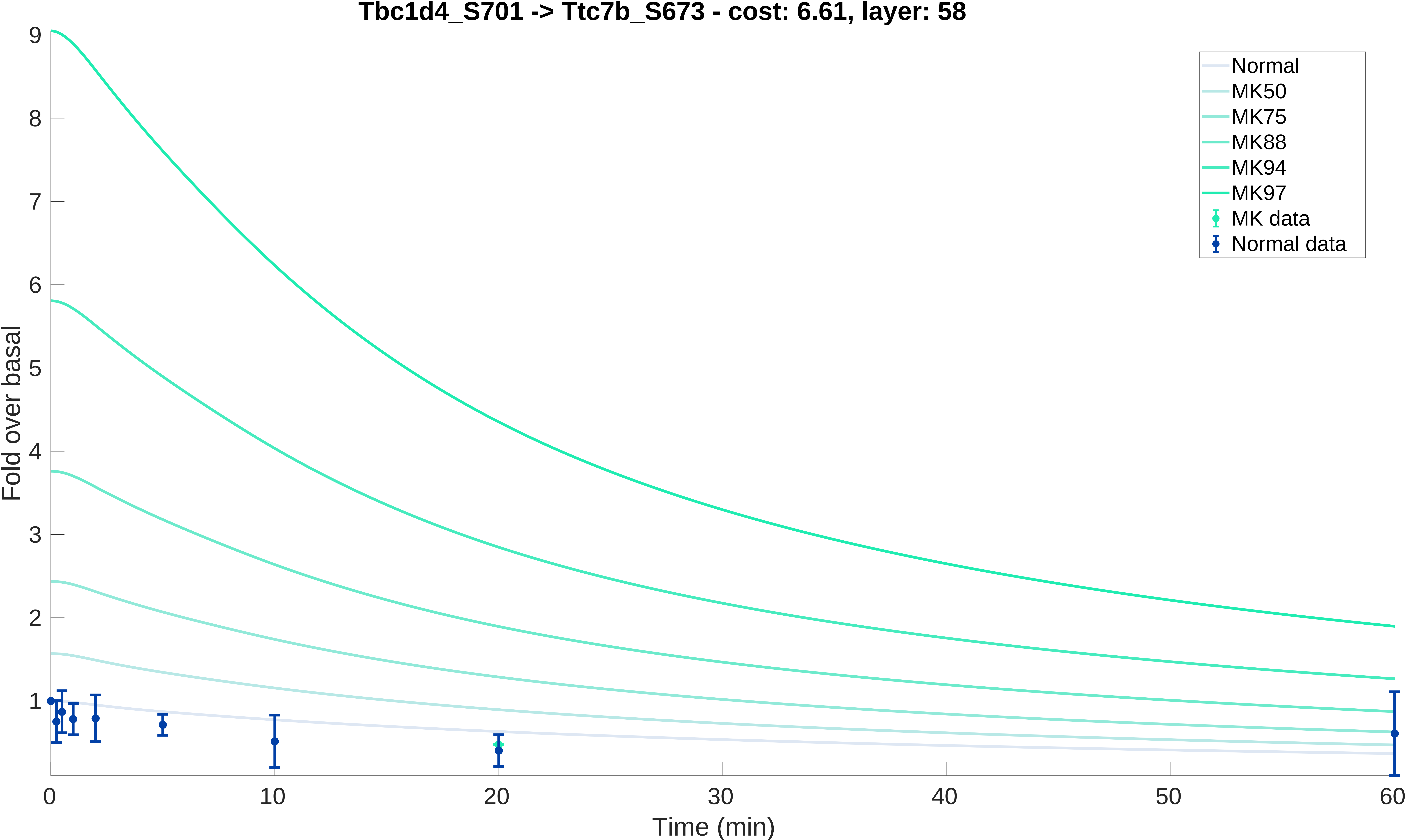

Tbc1d4_S701 -> Ttc7b_S673 - cost: 6.61, layer: 58
Fold over basal
Time (min)
Normal
MK50
MK75
MK88
MK94
MK97
MK data
Normal data
0
10
20
30
40
50
60
1
2
3
4
5
6
7
8
9

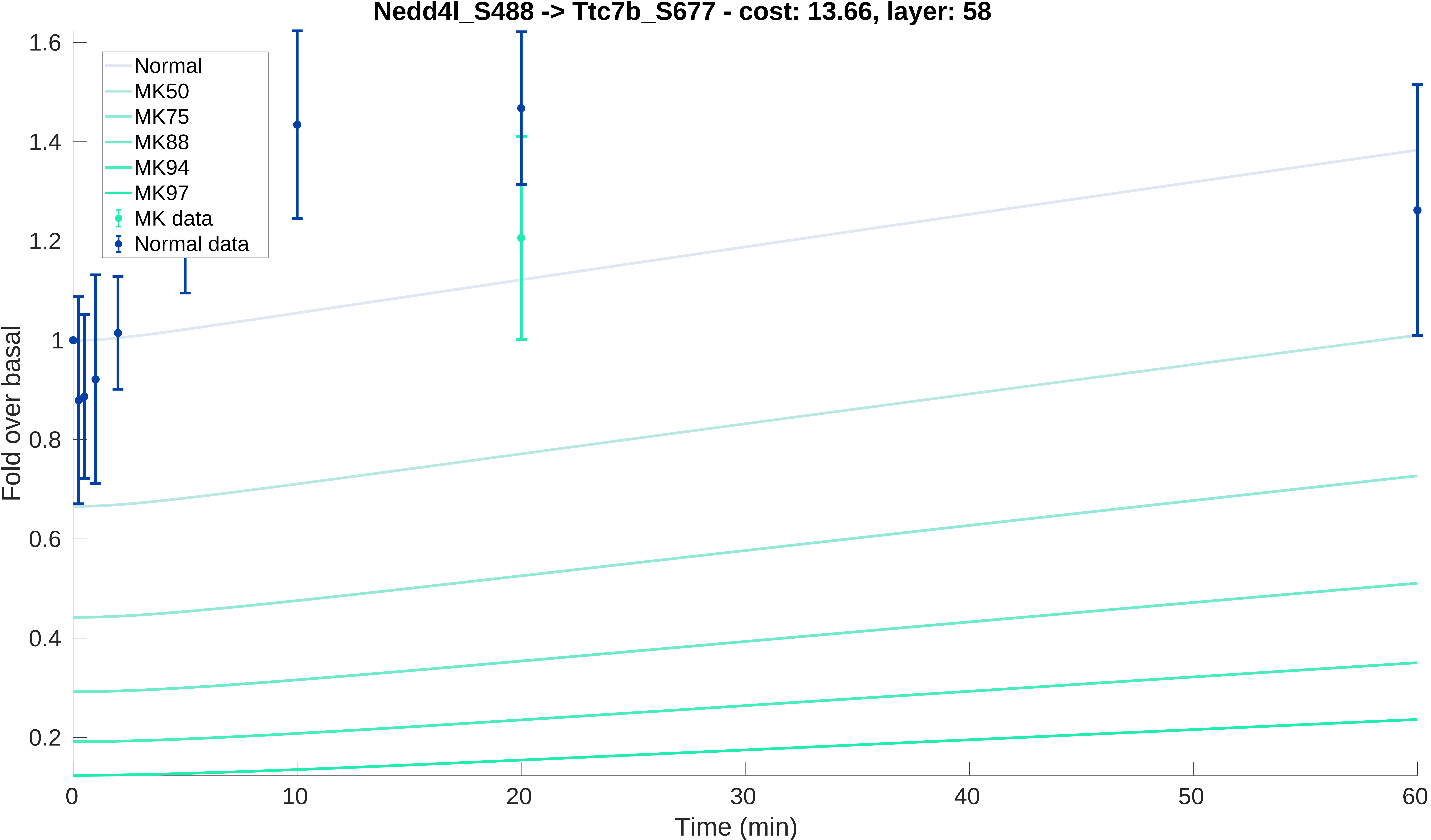

Nedd4l_S488 -> Ttc7b_S677 - cost: 13.66, layer: 58
Normal
MK50
MK75
MK88
MK94
MK97
MK data
Normal data
Fold over basal
Time (min)
1.6
1.4
1.2
1
0.8
0.6
0.4
0.2
0
10
20
30
40
50
60

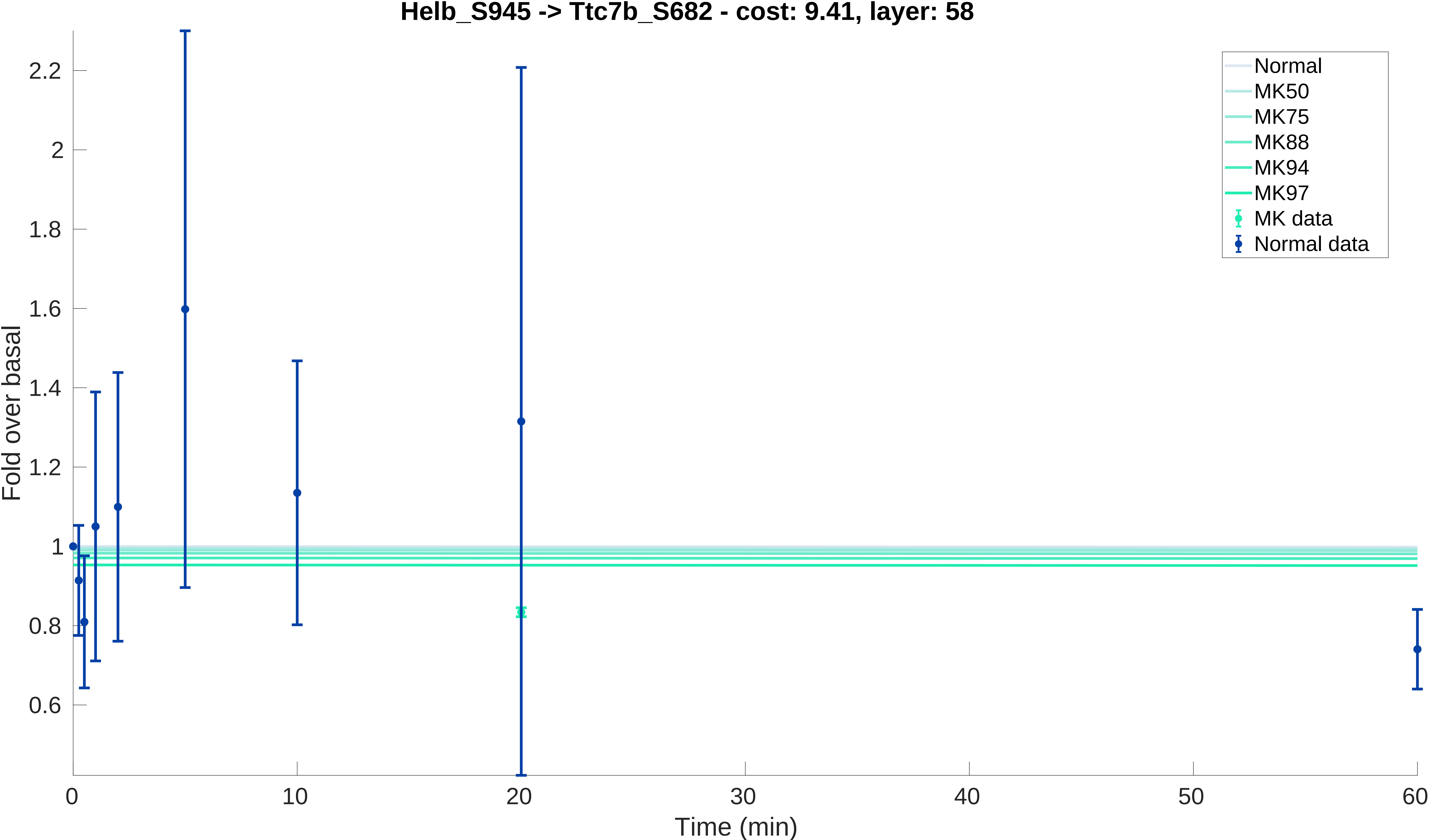

Helb_S945 -> Ttc7b_S682 - cost: 9.41, layer: 58
Normal
MK50
MK75
MK88
MK94
MK97
MK data
Normal data
Fold over basal
Time (min)
0.6
0.8
1
1.2
1.4
1.6
1.8
2
2.2
0
10
20
30
40
50
60

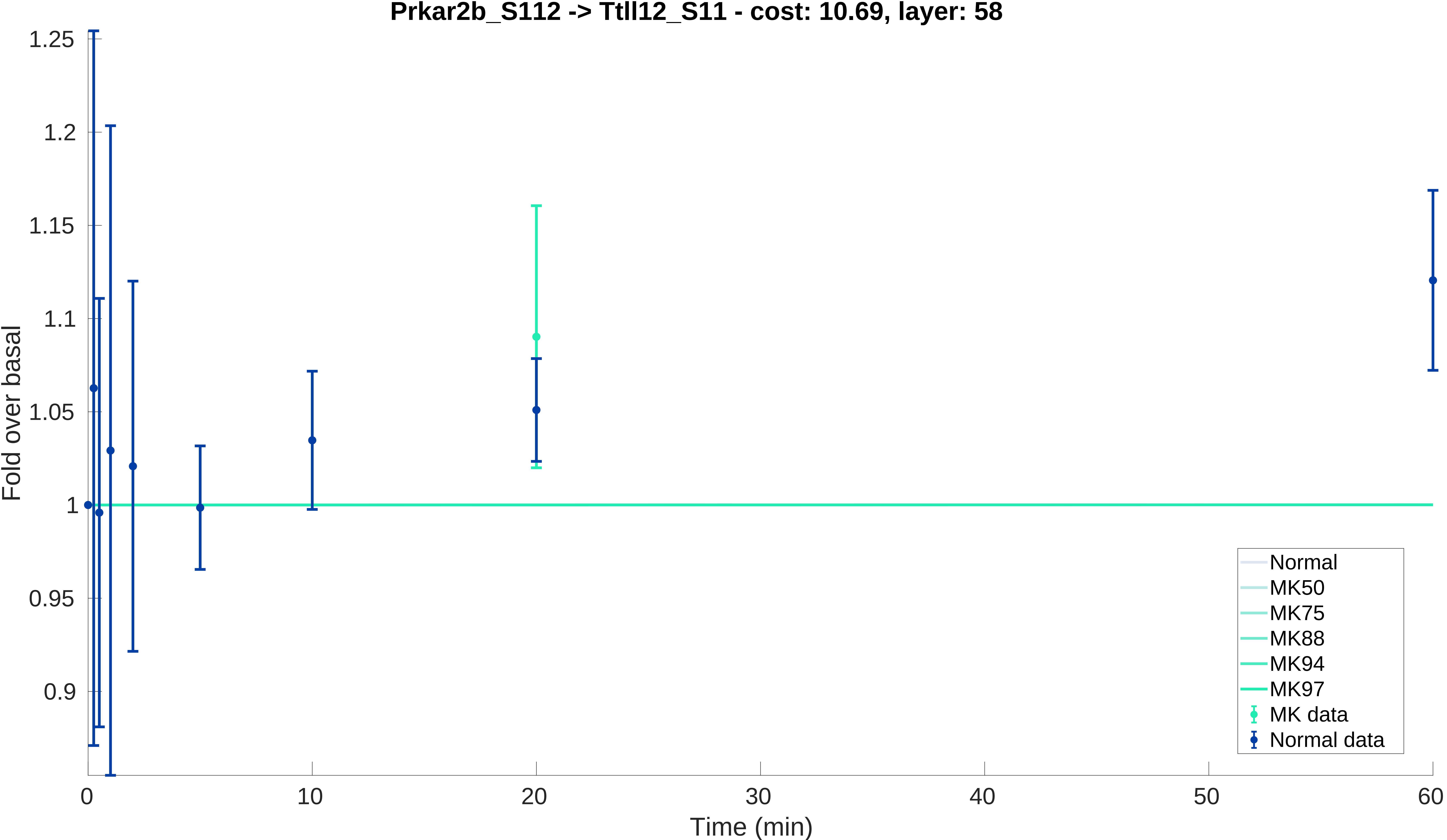

Prkar2b_S112 -> Ttll12_S11 - cost: 10.69, layer: 58
Fold over basal
1.25
1.2
1.15
1.1
1.05
1
0.95
0.9
0
10
20
30
40
50
60
Time (min)
Normal
MK50
MK75
MK88
MK94
MK97
MK data
Normal data

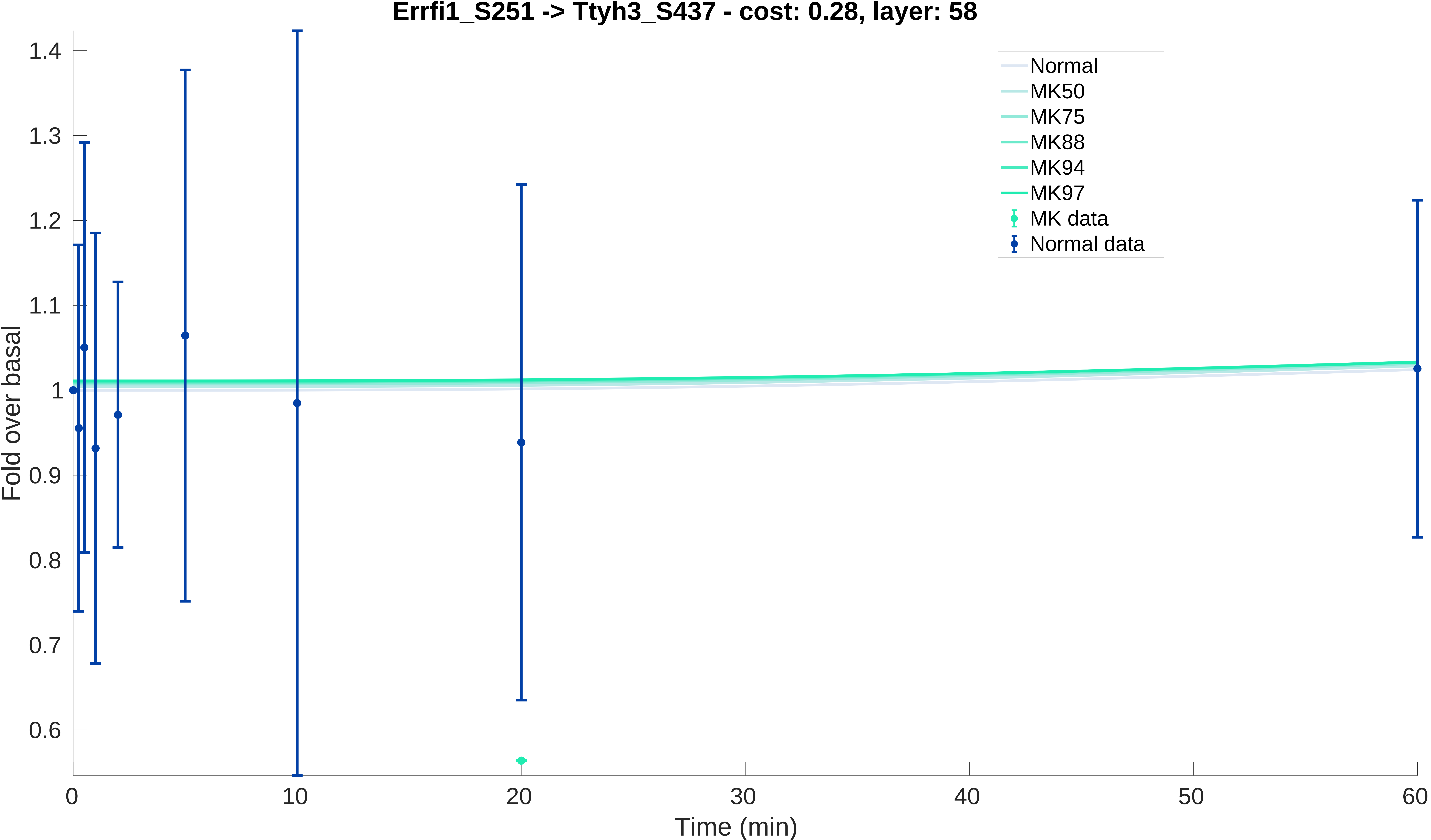

Errfi1_S251 -> Ttyh3_S437 - cost: 0.28, layer: 58
Fold over basal
1.4
1.3
1.2
1.1
1
0.9
0.8
0.7
0.6
Time (min)
0
10
20
30
40
50
60
Normal
MK50
MK75
MK88
MK94
MK97
MK data
Normal data

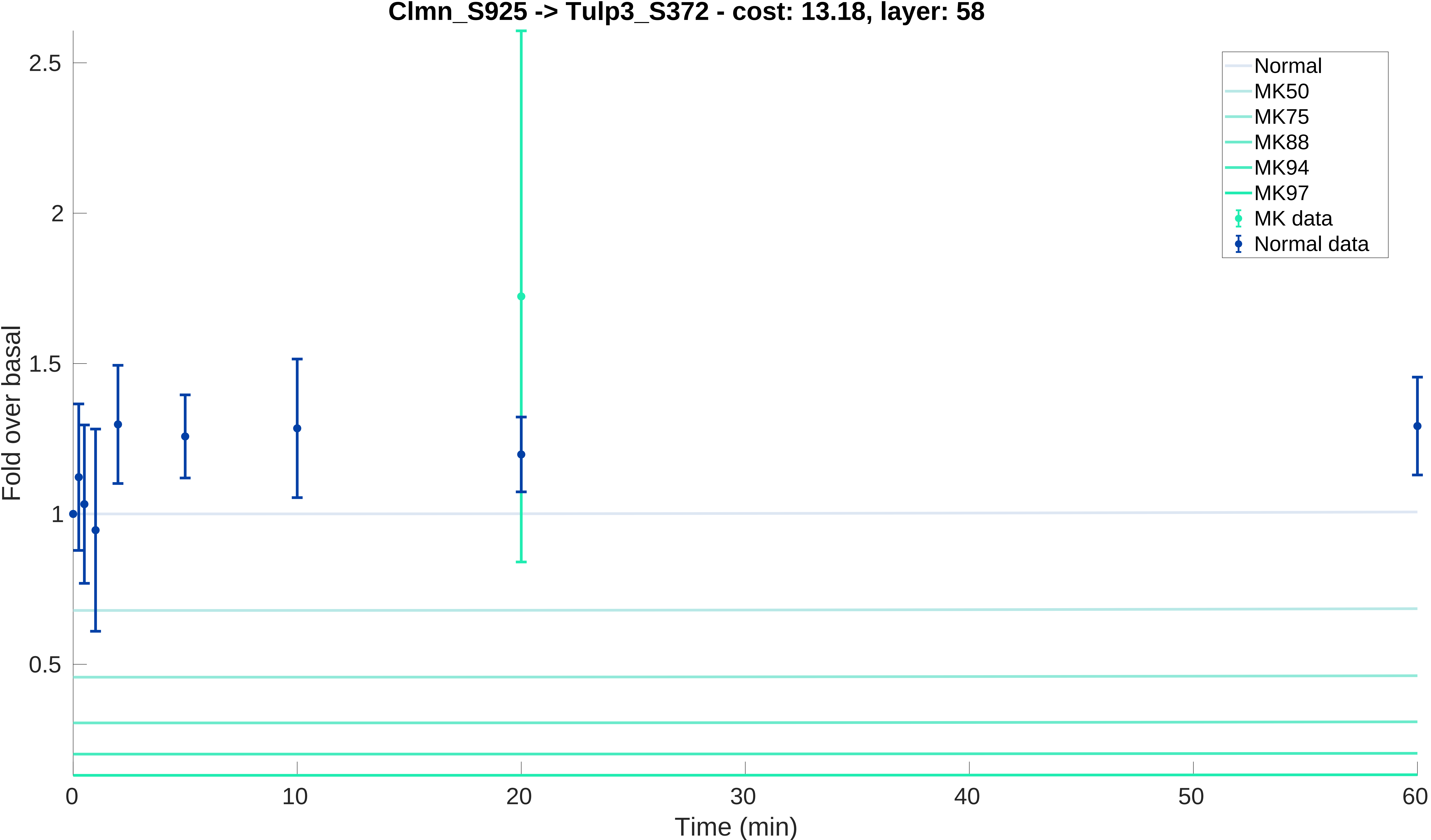
Clmn_S925 -> Tulp3_S372 - cost: 13.18, layer: 58
Fold over basal
2.5
2
1.5
1
0.5
0
10
20
30
40
50
60
Time (min)
Normal
MK50
MK75
MK88
MK94
MK97
MK data
Normal data

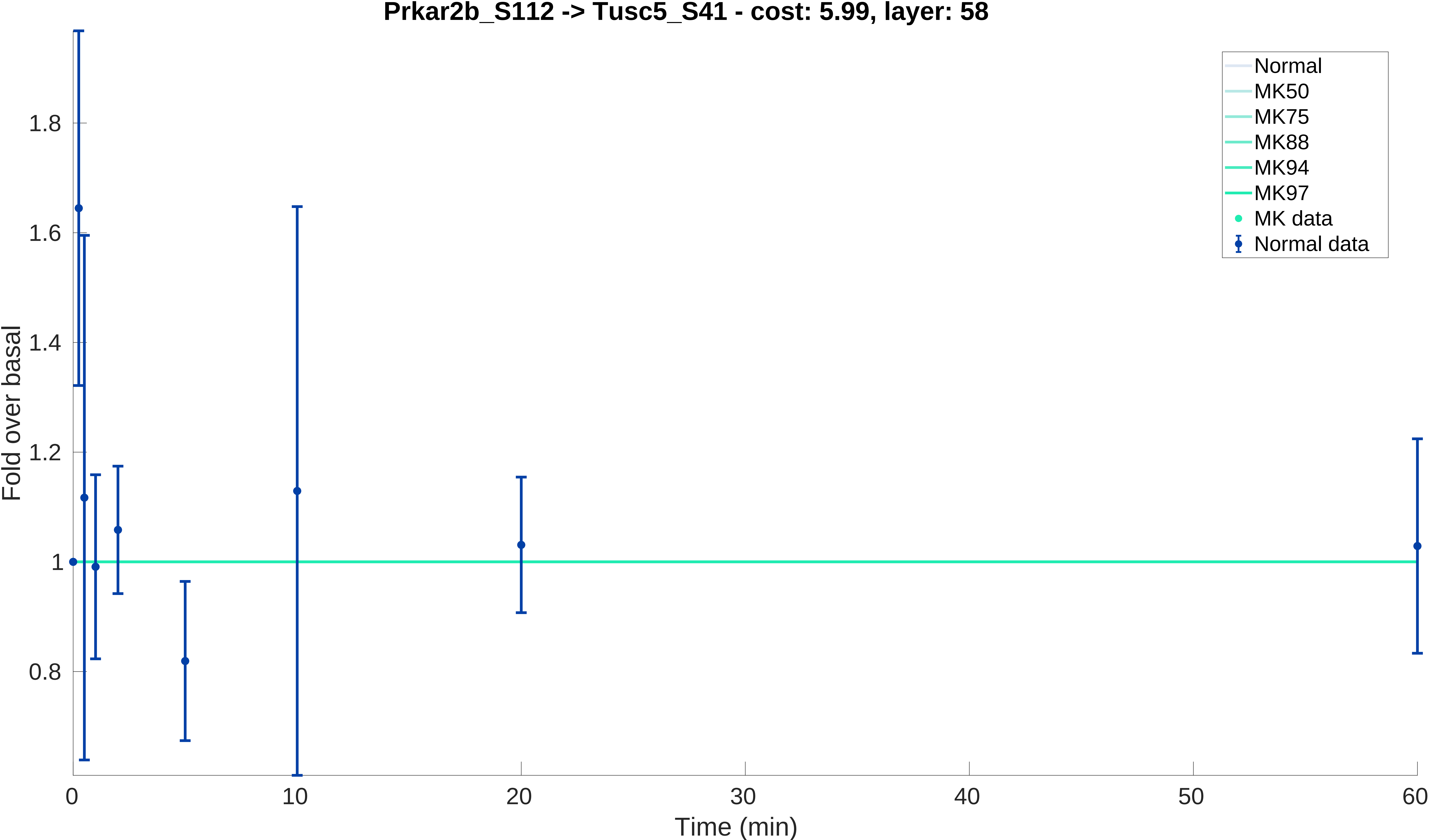

Prkar2b_S112 -> Tusc5_S41 - cost: 5.99, layer: 58
Normal
MK50
MK75
MK88
MK94
MK97
MK data
Normal data
Fold over basal
1.8
1.6
1.4
1.2
1
0.8
0
10
20
30
40
50
60
Time (min)

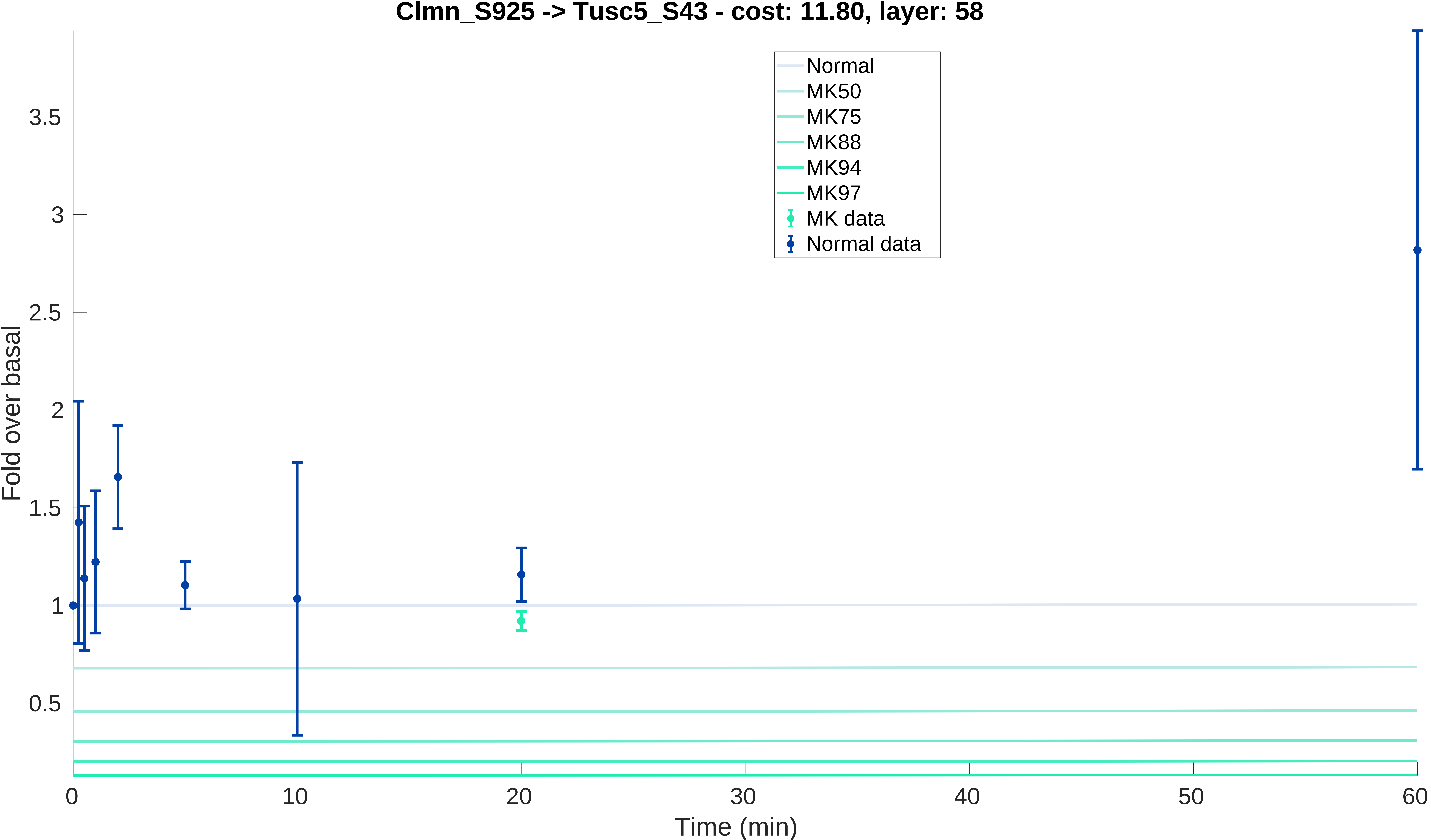

Clmn_S925 -> Tusc5_S43 - cost: 11.80, layer: 58
Normal
MK50
MK75
MK88
MK94
MK97
MK data
Normal data
Fold over basal
3.5
3
2.5
2
1.5
1
0.5
0
10
20
30
40
50
60
Time (min)

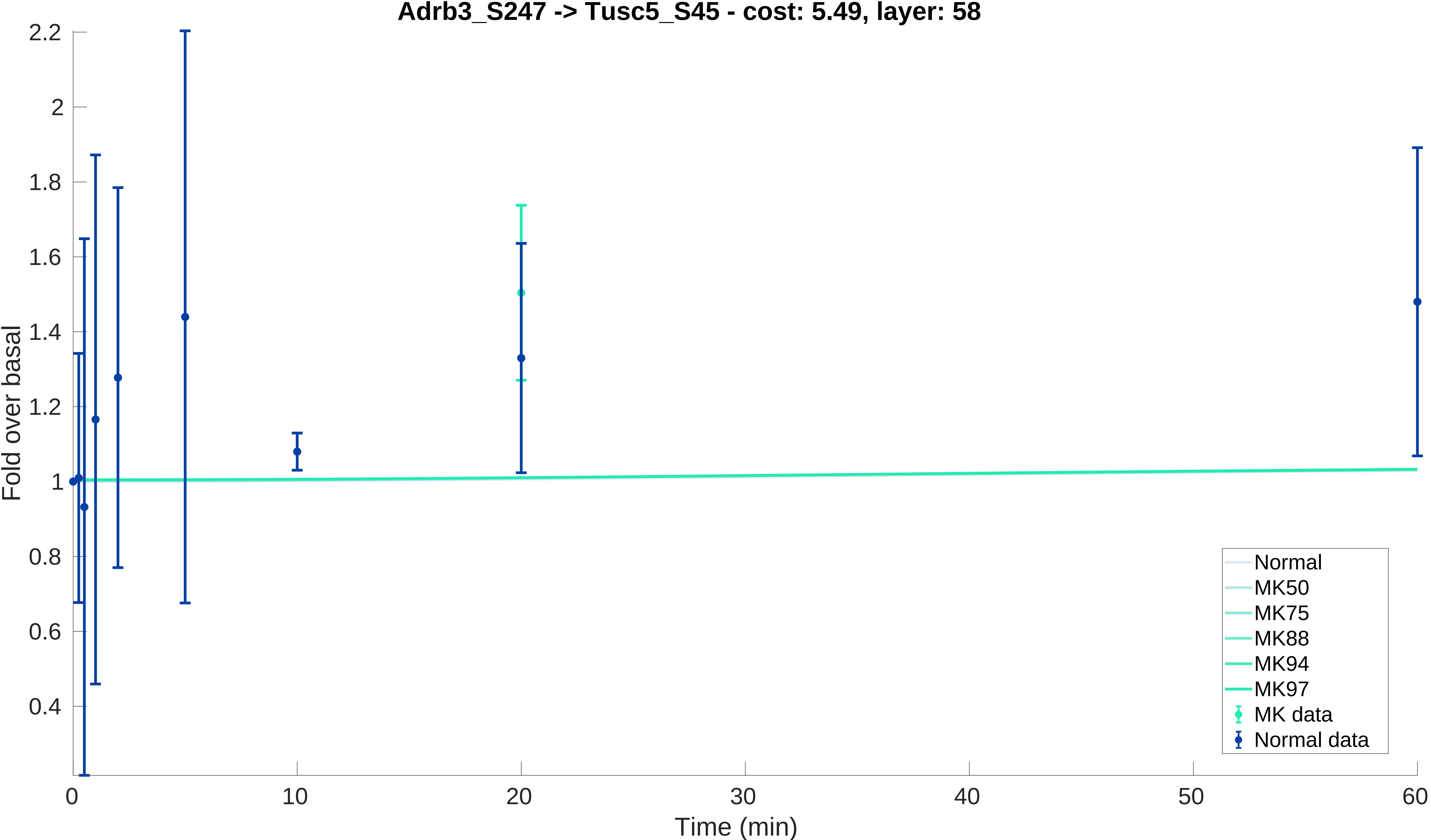

Adrb3_S247 -> Tusc5_S45 - cost: 5.49, layer: 58
Fold over basal
Time (min)
2.2
2
1.8
1.6
1.4
1.2
1
0.8
0.6
0.4
0
10
20
30
40
50
60
Normal
MK50
MK75
MK88
MK94
MK97
MK data
Normal data

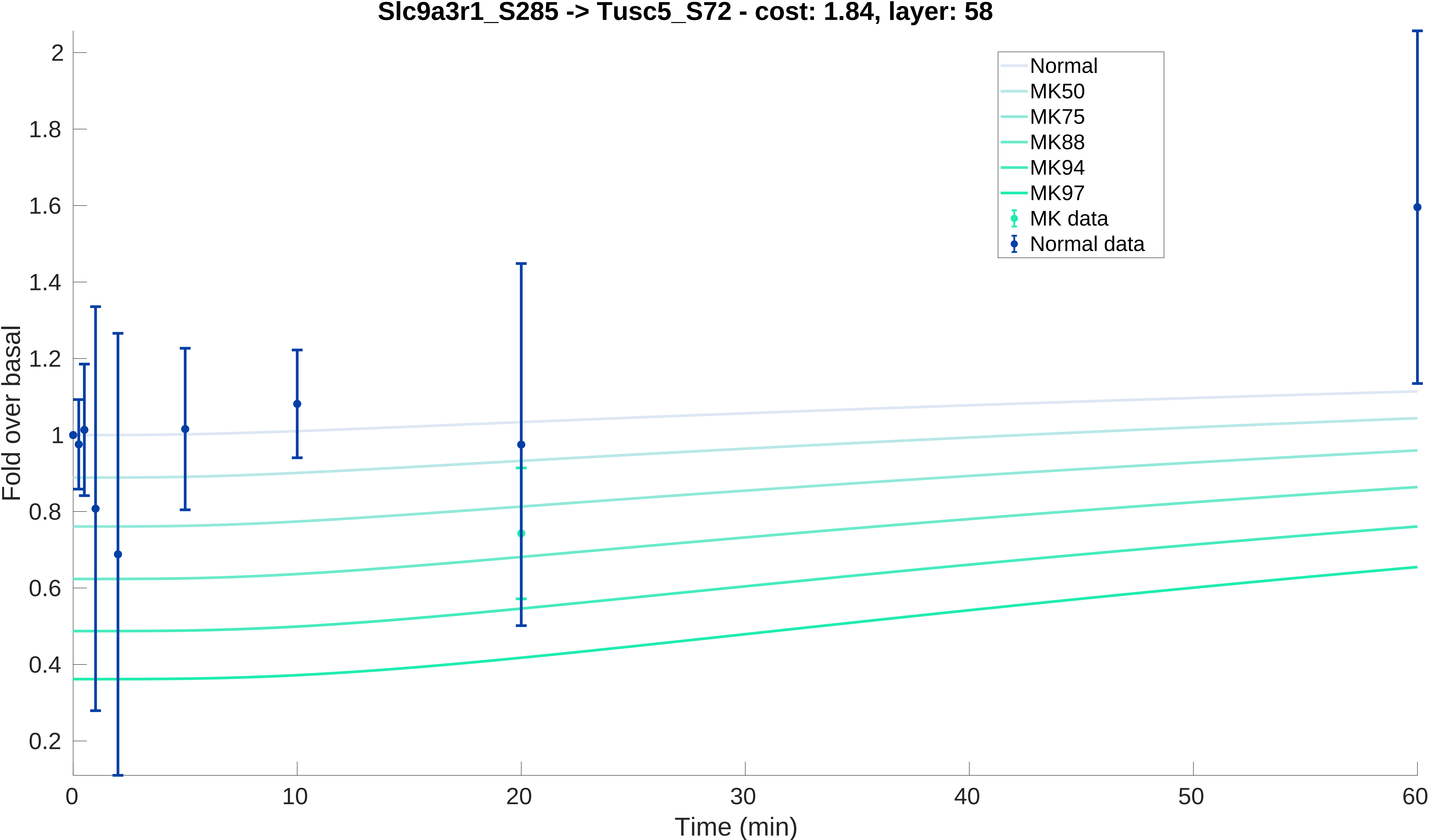

Slc9a3r1_S285 -> Tusc5_S72 - cost: 1.84, layer: 58
Fold over basal
Time (min)
0
10
20
30
40
50
60
0.2
0.4
0.6
0.8
1
1.2
1.4
1.6
1.8
2
Normal
MK50
MK75
MK88
MK94
MK97
MK data
Normal data

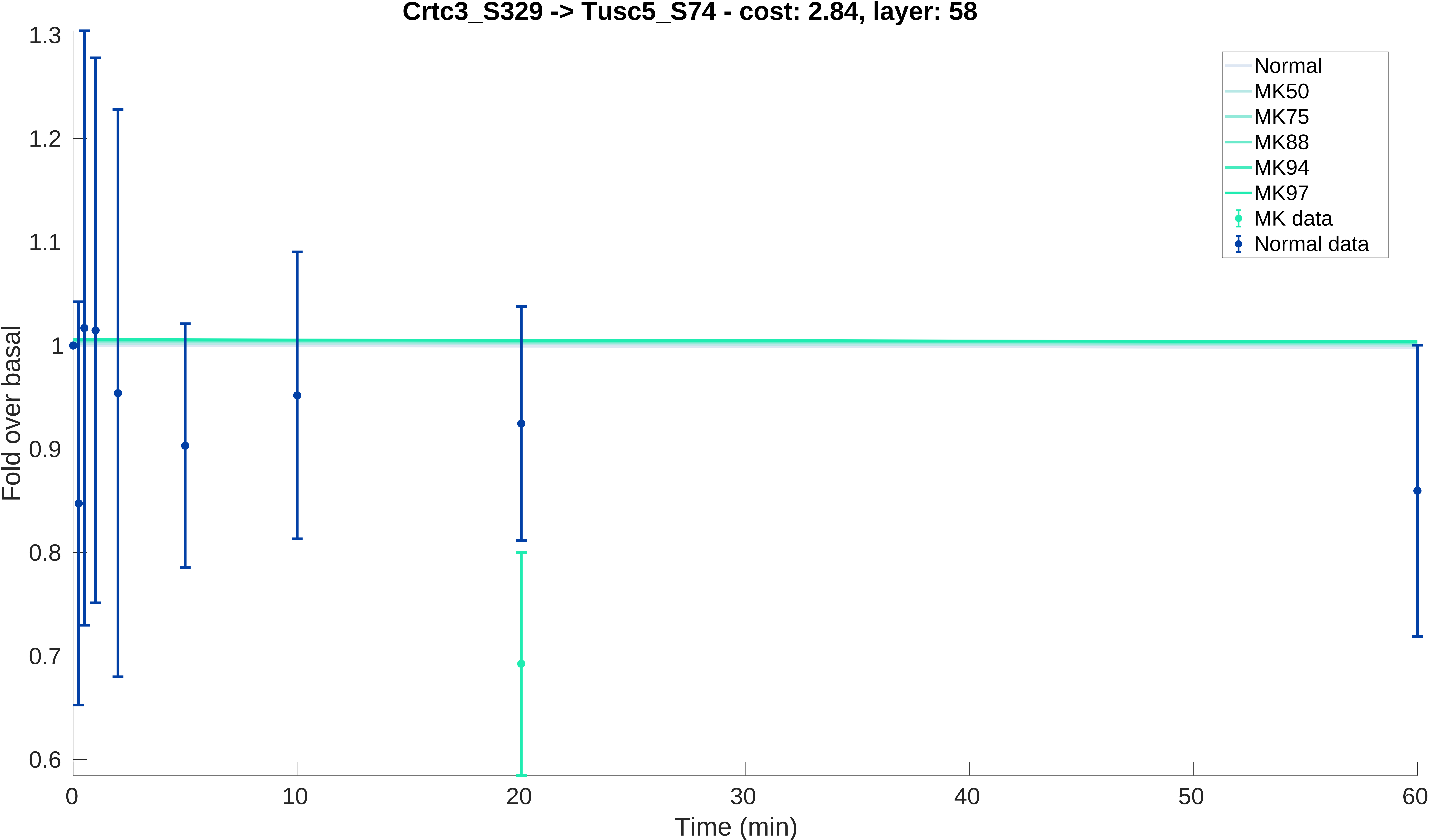

Crtc3_S329 -> Tusc5_S74 - cost: 2.84, layer: 58
Normal
MK50
MK75
MK88
MK94
MK97
MK data
Normal data
Fold over basal
Time (min)
1.3
1.2
1.1
1
0.9
0.8
0.7
0.6
0
10
20
30
40
50
60

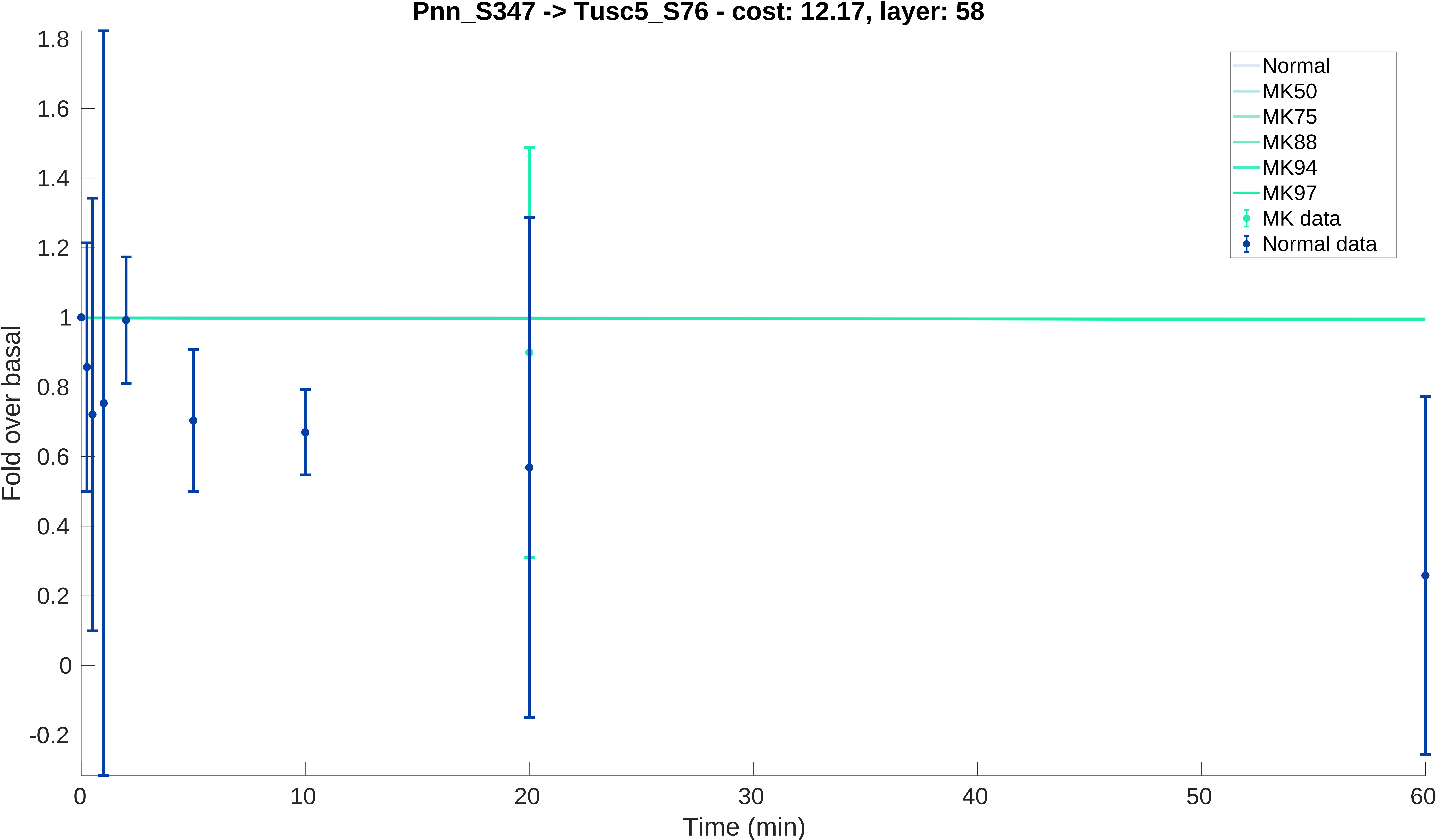

Pnn_S347 -> Tusc5_S76 - cost: 12.17, layer: 58
Fold over basal
Time (min)
1.8
1.6
1.4
1.2
1
0.8
0.6
0.4
0.2
0
-0.2
0
10
20
30
40
50
60
Normal
MK50
MK75
MK88
MK94
MK97
MK data
Normal data

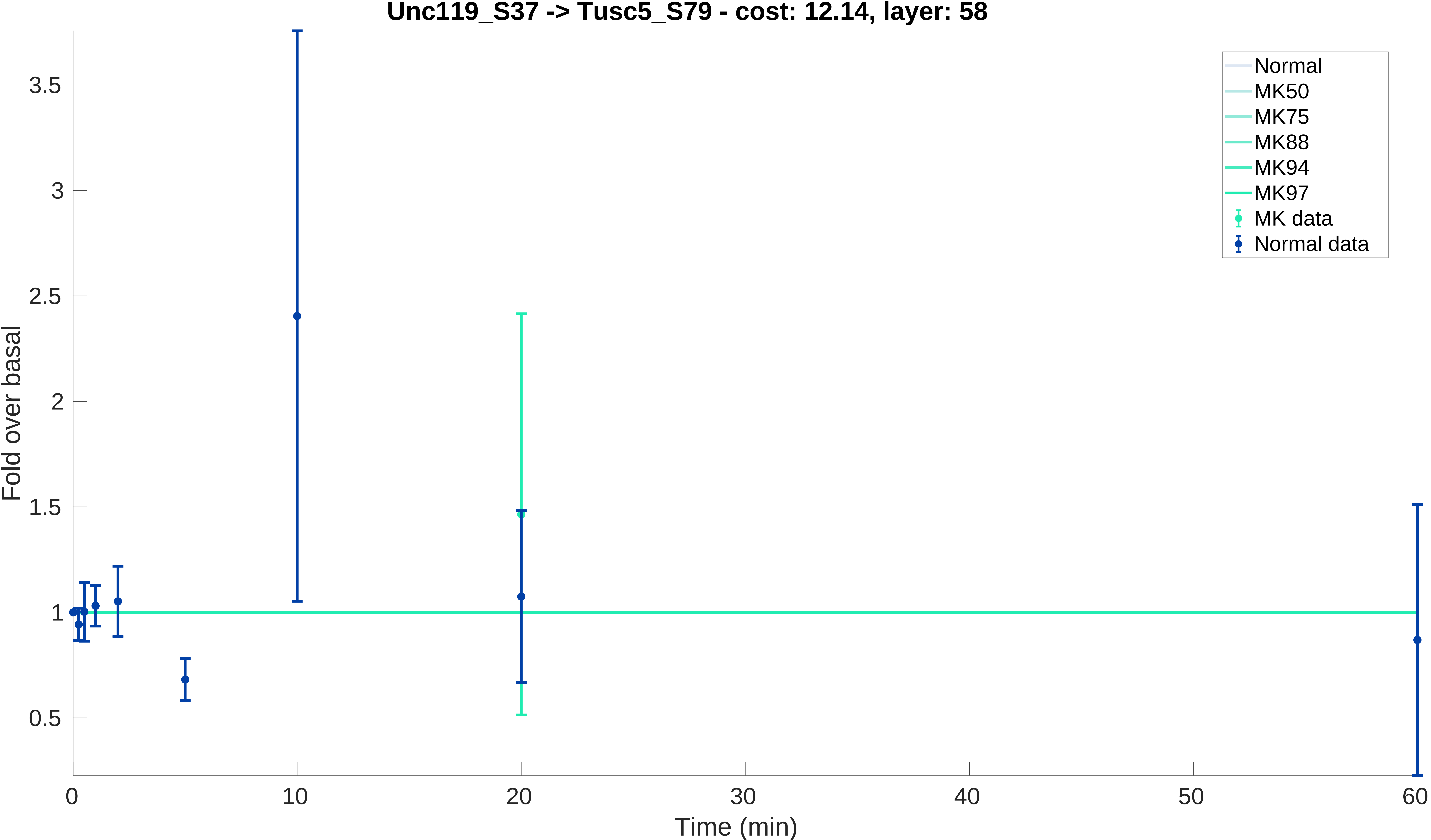

Unc119_S37 -> Tusc5_S79 - cost: 12.14, layer: 58
Normal
MK50
MK75
MK88
MK94
MK97
MK data
Normal data
Fold over basal
3.5
3
2.5
2
1.5
1
0.5
0
10
20
30
40
50
60
Time (min)

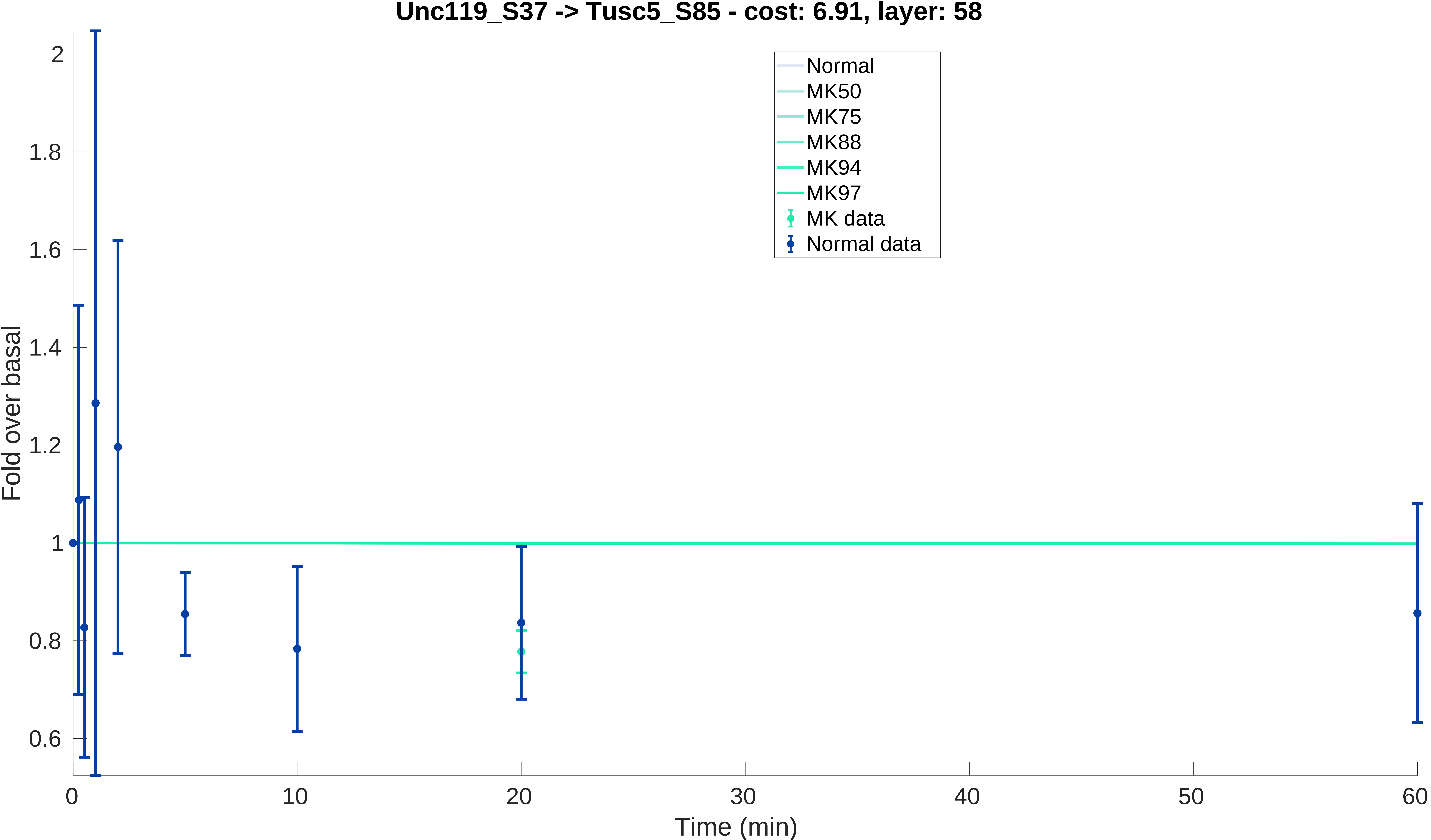

Unc119_S37 -> Tusc5_S85 - cost: 6.91, layer: 58
Normal
MK50
MK75
MK88
MK94
MK97
MK data
Normal data
Fold over basal
2
1.8
1.6
1.4
1.2
1
0.8
0.6
0
10
20
30
40
50
60
Time (min)

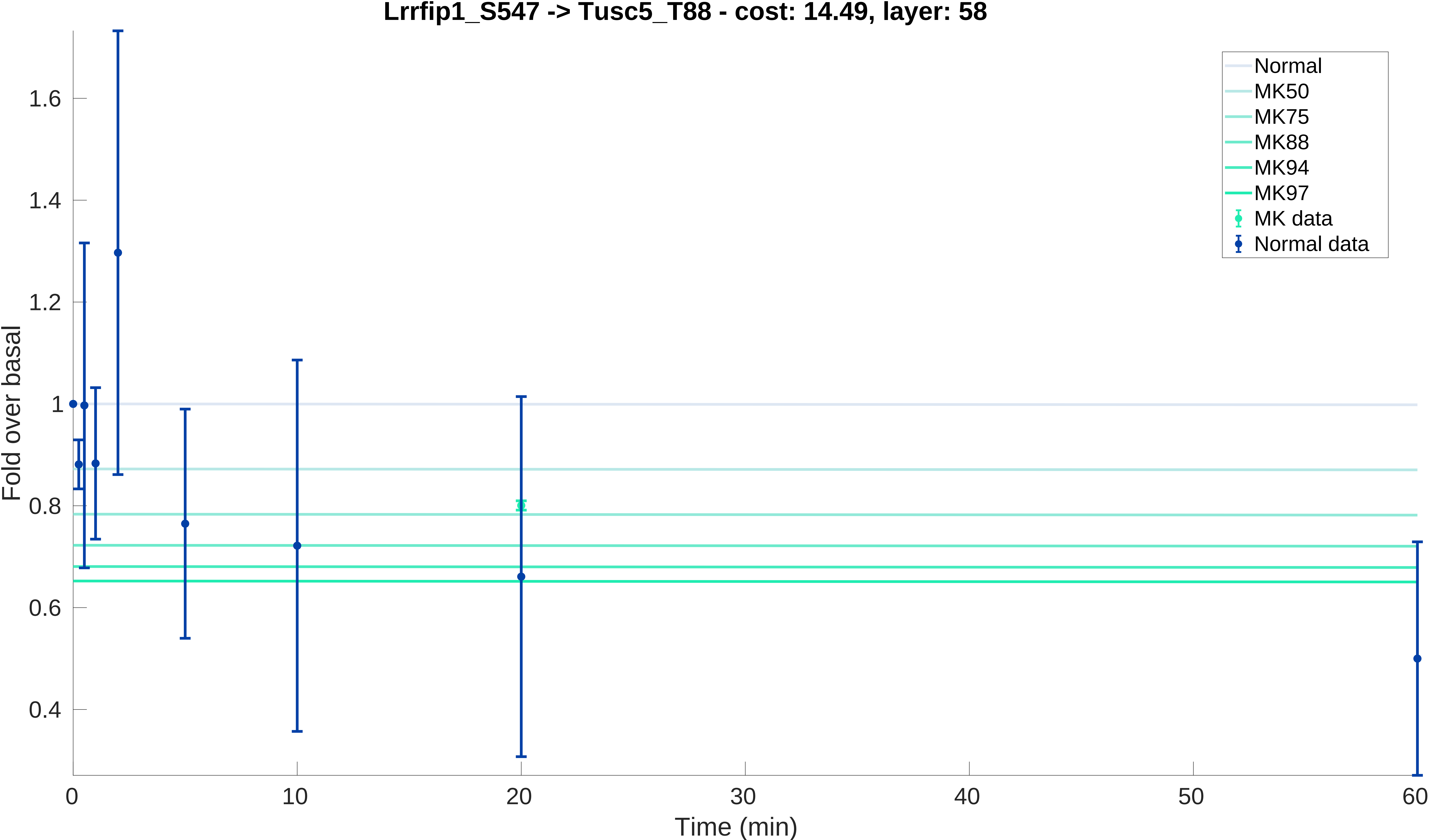

Lrrfip1_S547 -> Tusc5_T88 - cost: 14.49, layer: 58
Fold over basal
1.6
1.4
1.2
1
0.8
0.6
0.4
0
10
20
30
40
50
60
Time (min)
Normal
MK50
MK75
MK88
MK94
MK97
MK data
Normal data

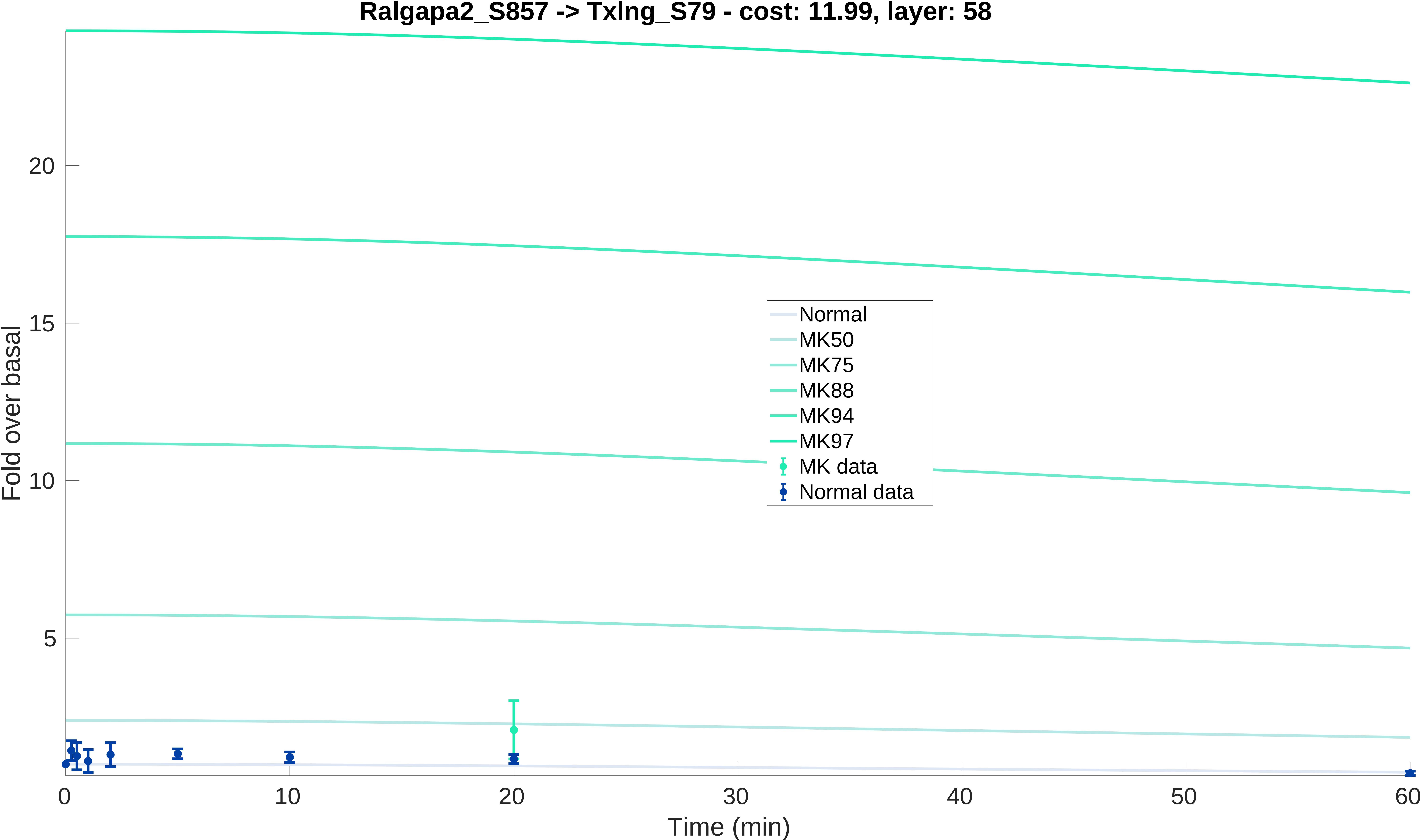
Ralgapa2_S857 -> Txlng_S79 - cost: 11.99, layer: 58
Fold over basal
20
15
10
5
0
10
20
30
40
50
60
Time (min)
Normal
MK50
MK75
MK88
MK94
MK97
MK data
Normal data

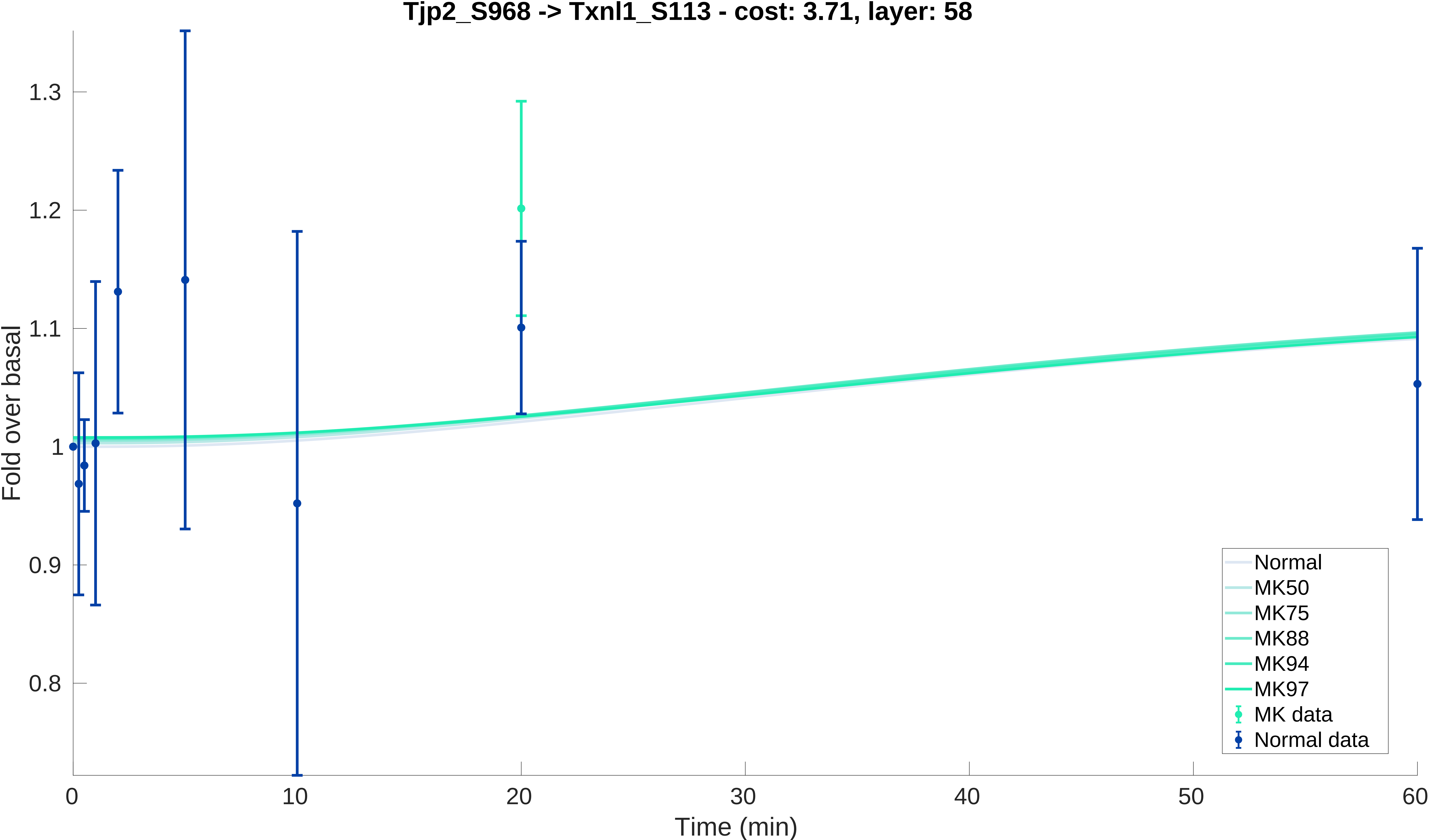

Tjp2_S968 -> Txnl1_S113 - cost: 3.71, layer: 58
Fold over basal
1.3
1.2
1.1
1
0.9
0.8
0
10
20
30
40
50
60
Time (min)
Normal
MK50
MK75
MK88
MK94
MK97
MK data
Normal data

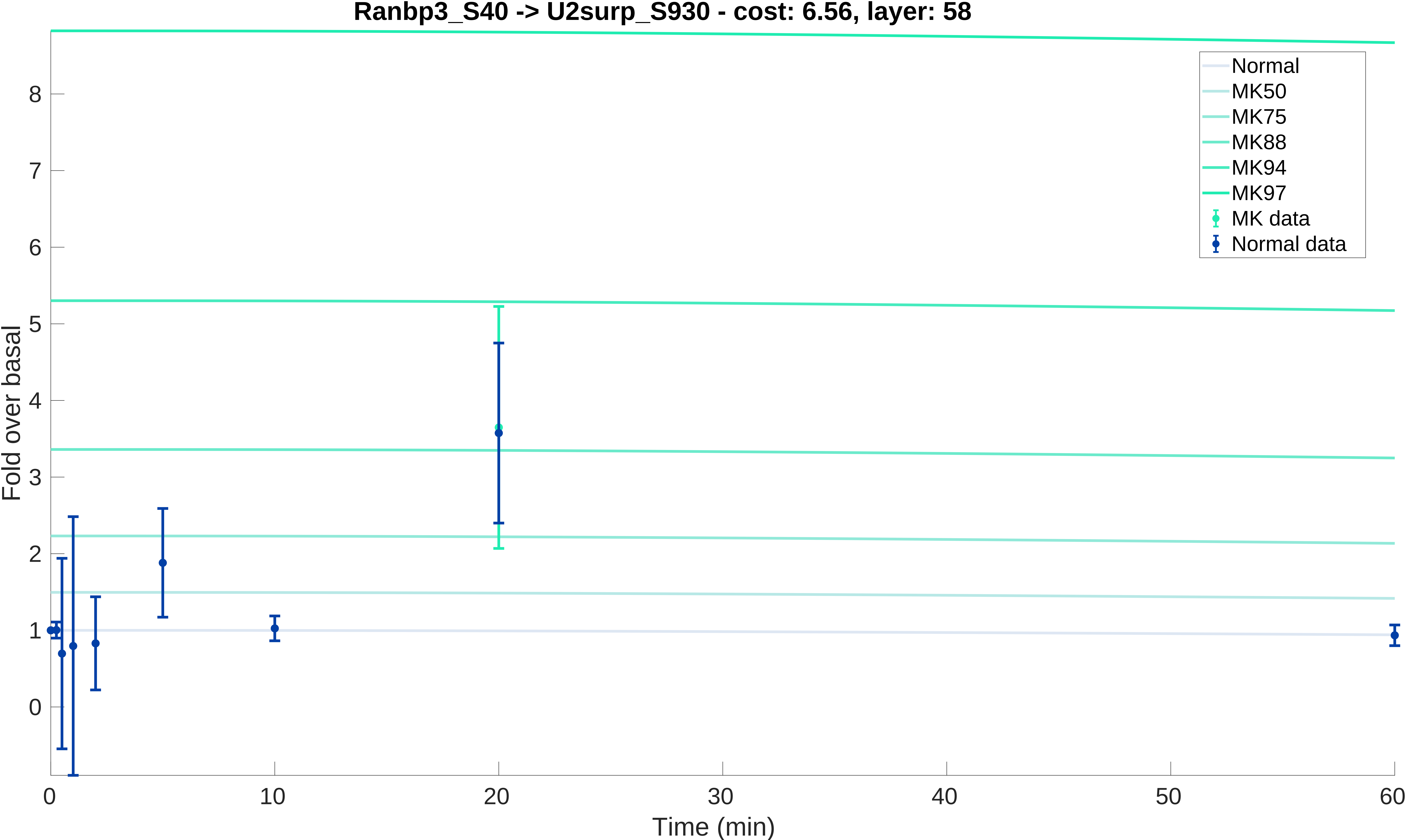

Ranbp3_S40 -> U2surp_S930 - cost: 6.56, layer: 58
Normal
MK50
MK75
MK88
MK94
MK97
MK data
Normal data
Fold over basal
Time (min)
0
10
20
30
40
50
60
8
7
6
5
4
3
2
1
0

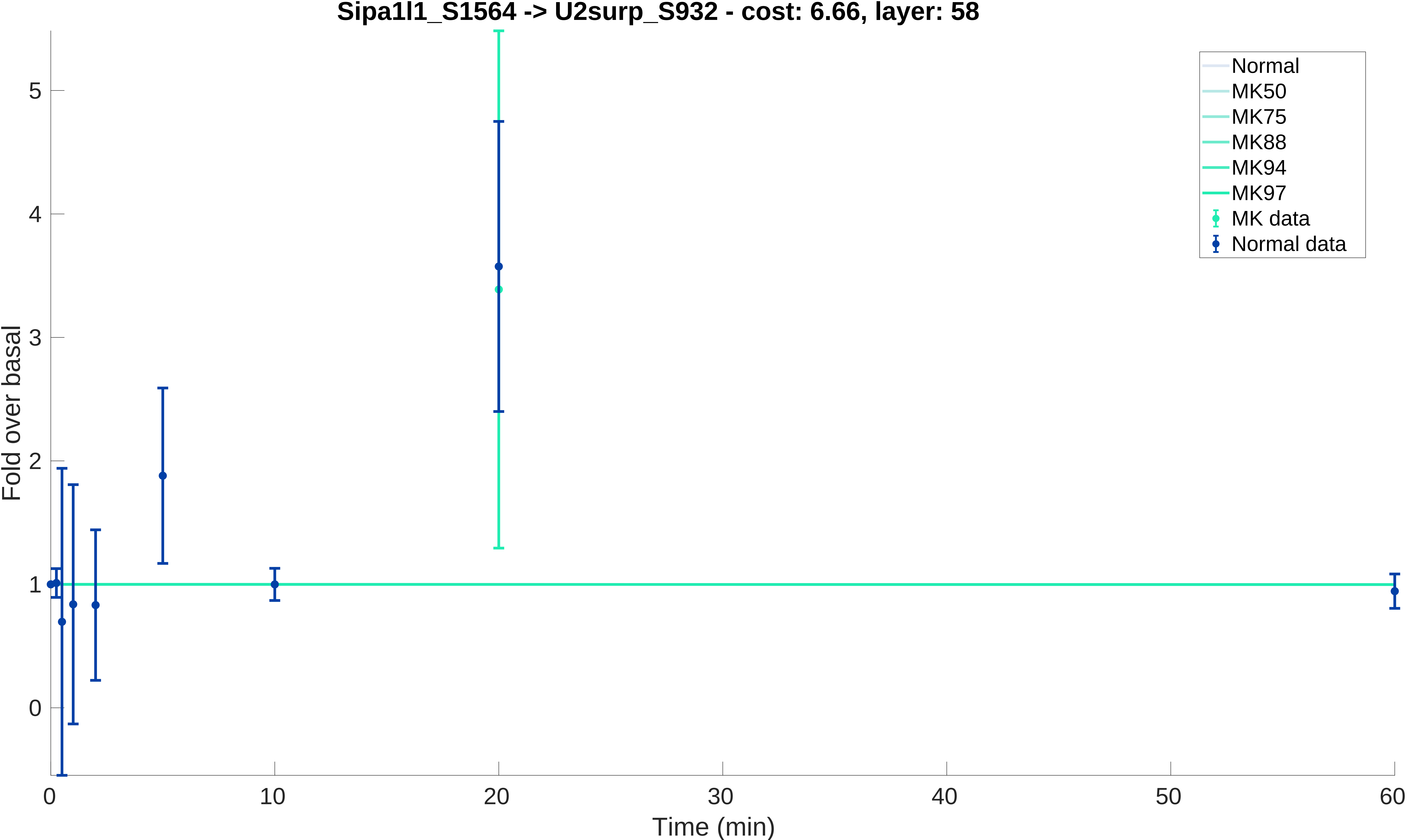

Sipa1l1_S1564 -> U2surp_S932 - cost: 6.66, layer: 58
Fold over basal
Time (min)
0
1
2
3
4
5
0
10
20
30
40
50
60
Normal
MK50
MK75
MK88
MK94
MK97
MK data
Normal data

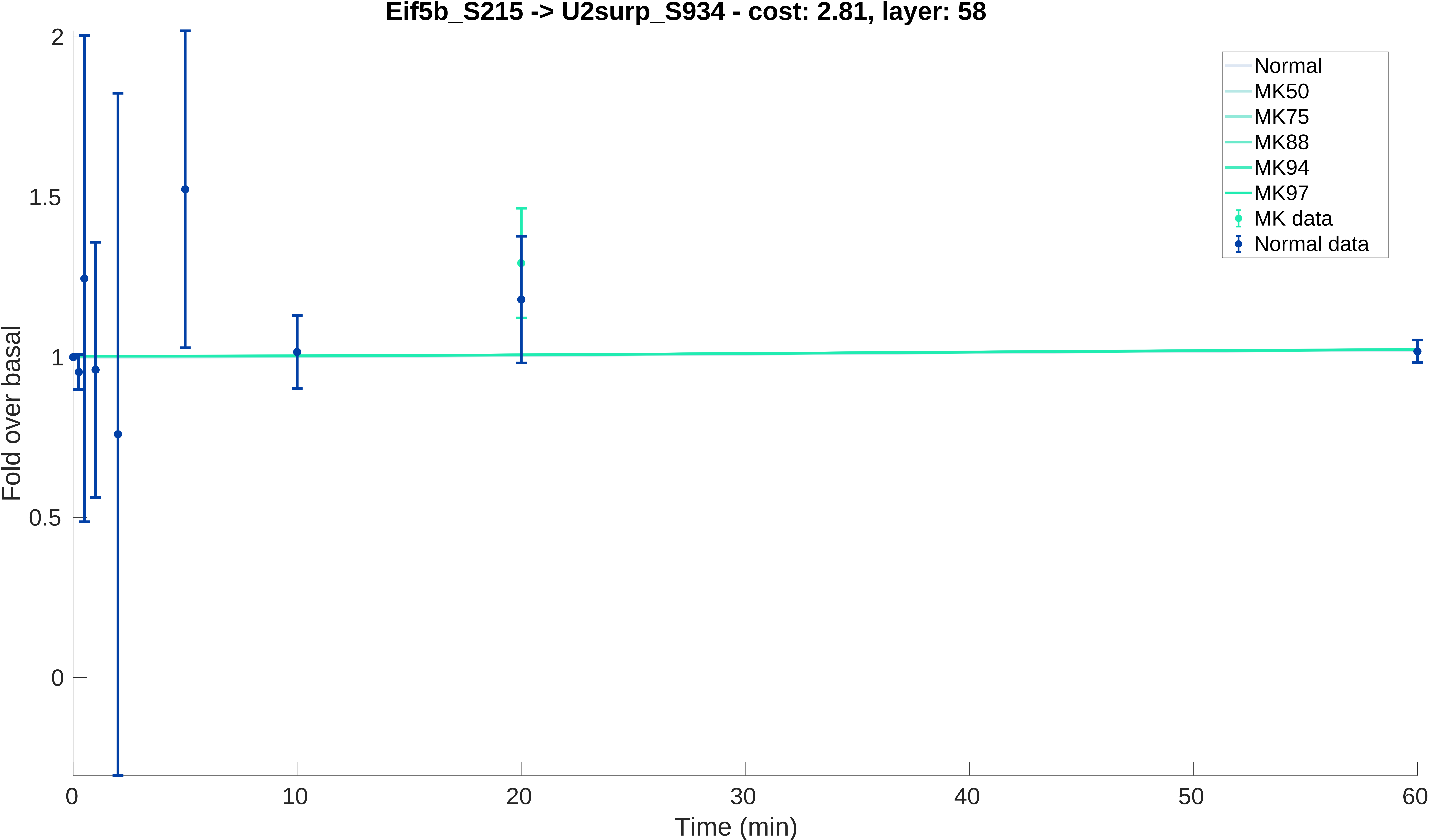
Eif5b_S215 -> U2surp_S934 - cost: 2.81, layer: 58
Fold over basal
Time (min)
0
10
20
30
40
50
60
2
1.5
1
0.5
0
Normal
MK50
MK75
MK88
MK94
MK97
MK data
Normal data

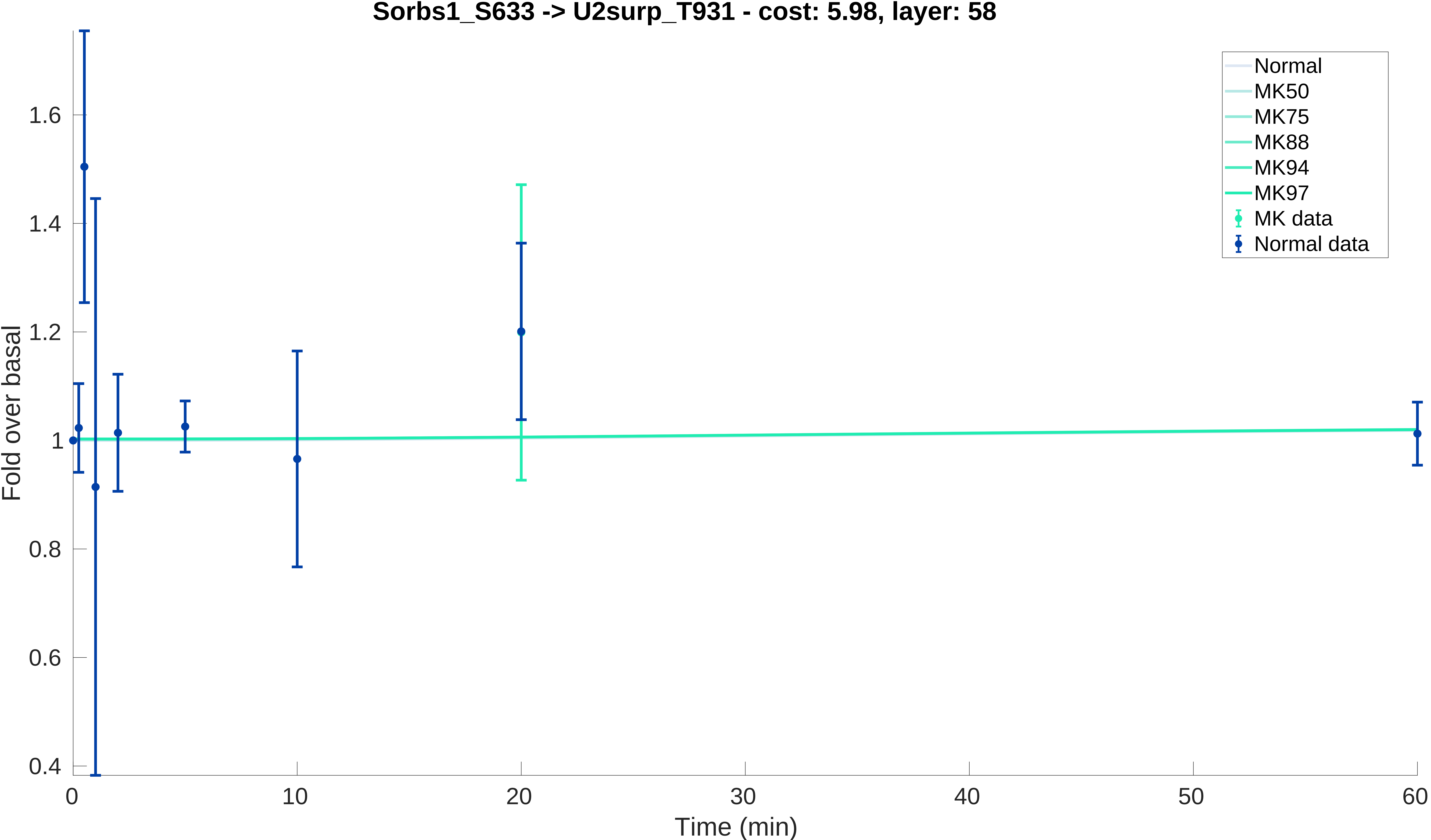

Sorbs1_S633 -> U2surp_T931 - cost: 5.98, layer: 58
Normal
MK50
MK75
MK88
MK94
MK97
MK data
Normal data
Fold over basal
1.6
1.4
1.2
1
0.8
0.6
0.4
0
10
20
30
40
50
60
Time (min)

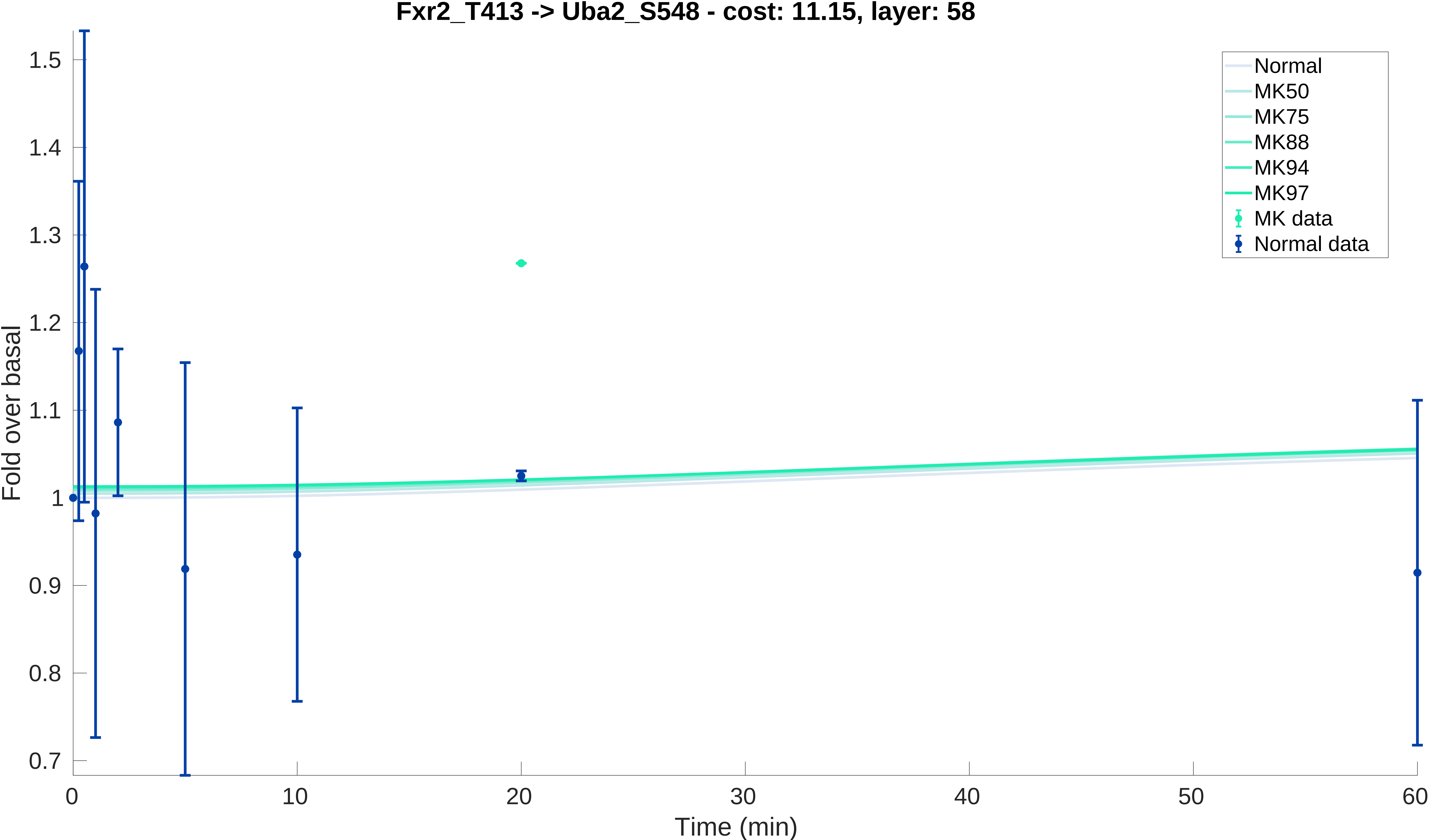

Fxr2_T413 -> Uba2_S548 - cost: 11.15, layer: 58
Fold over basal
Time (min)
0
10
20
30
40
50
60
0.7
0.8
0.9
1
1.1
1.2
1.3
1.4
1.5
Normal
MK50
MK75
MK88
MK94
MK97
MK data
Normal data

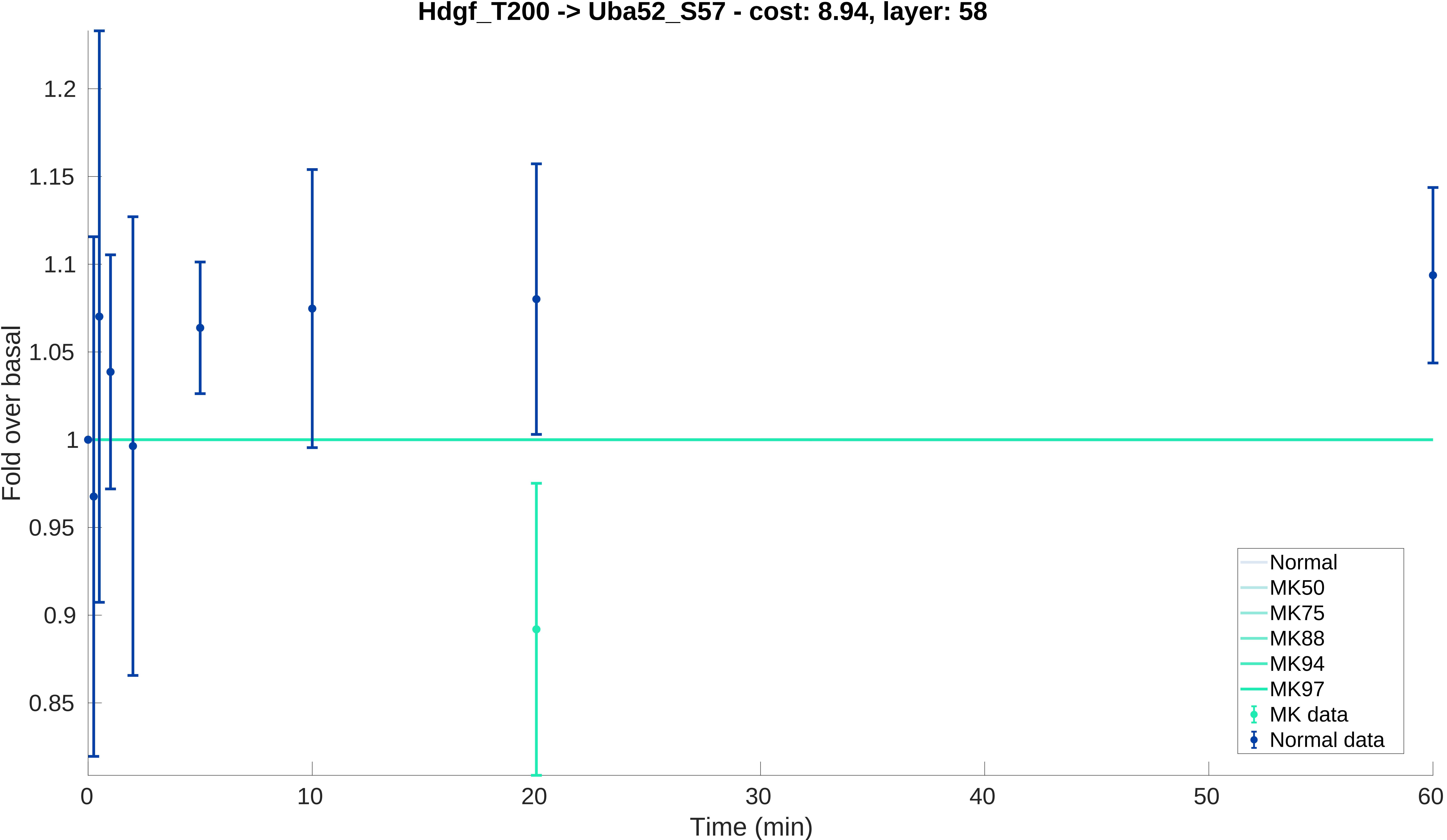

Hdgf_T200 -> Uba52_S57 - cost: 8.94, layer: 58
Fold over basal
1.2
1.15
1.1
1.05
1
0.95
0.9
0.85
0
10
20
30
40
50
60
Time (min)
Normal
MK50
MK75
MK88
MK94
MK97
MK data
Normal data

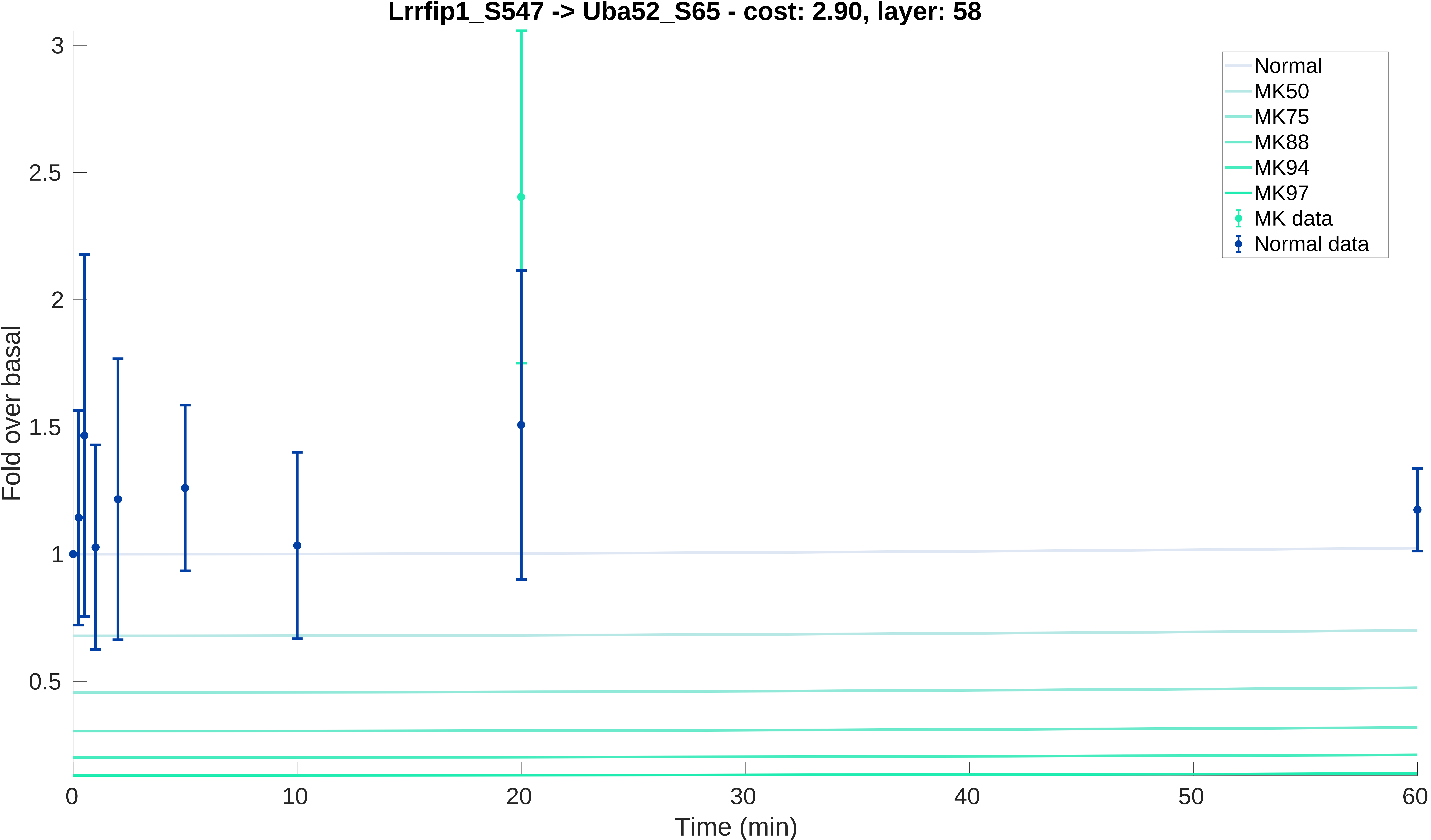

Lrrfip1_S547 -> Uba52_S65 - cost: 2.90, layer: 58
Fold over basal
Time (min)
0.5
1
1.5
2
2.5
3
0
10
20
30
40
50
60
Normal
MK50
MK75
MK88
MK94
MK97
MK data
Normal data

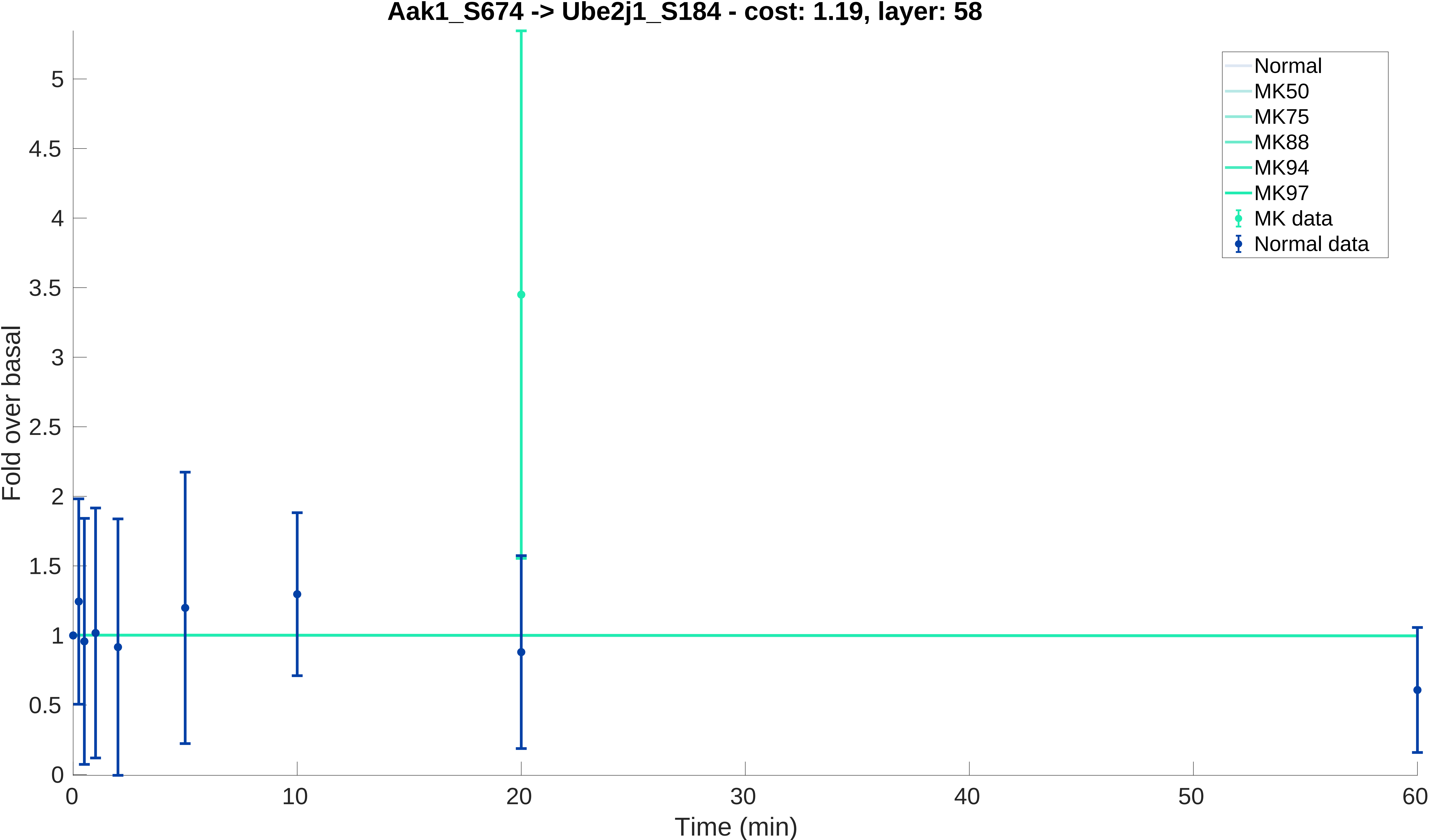

Aak1_S674 -> Ube2j1_S184 - cost: 1.19, layer: 58
Normal
MK50
MK75
MK88
MK94
MK97
MK data
Normal data
Fold over basal
Time (min)
0
0.5
1
1.5
2
2.5
3
3.5
4
4.5
5
0
10
20
30
40
50
60

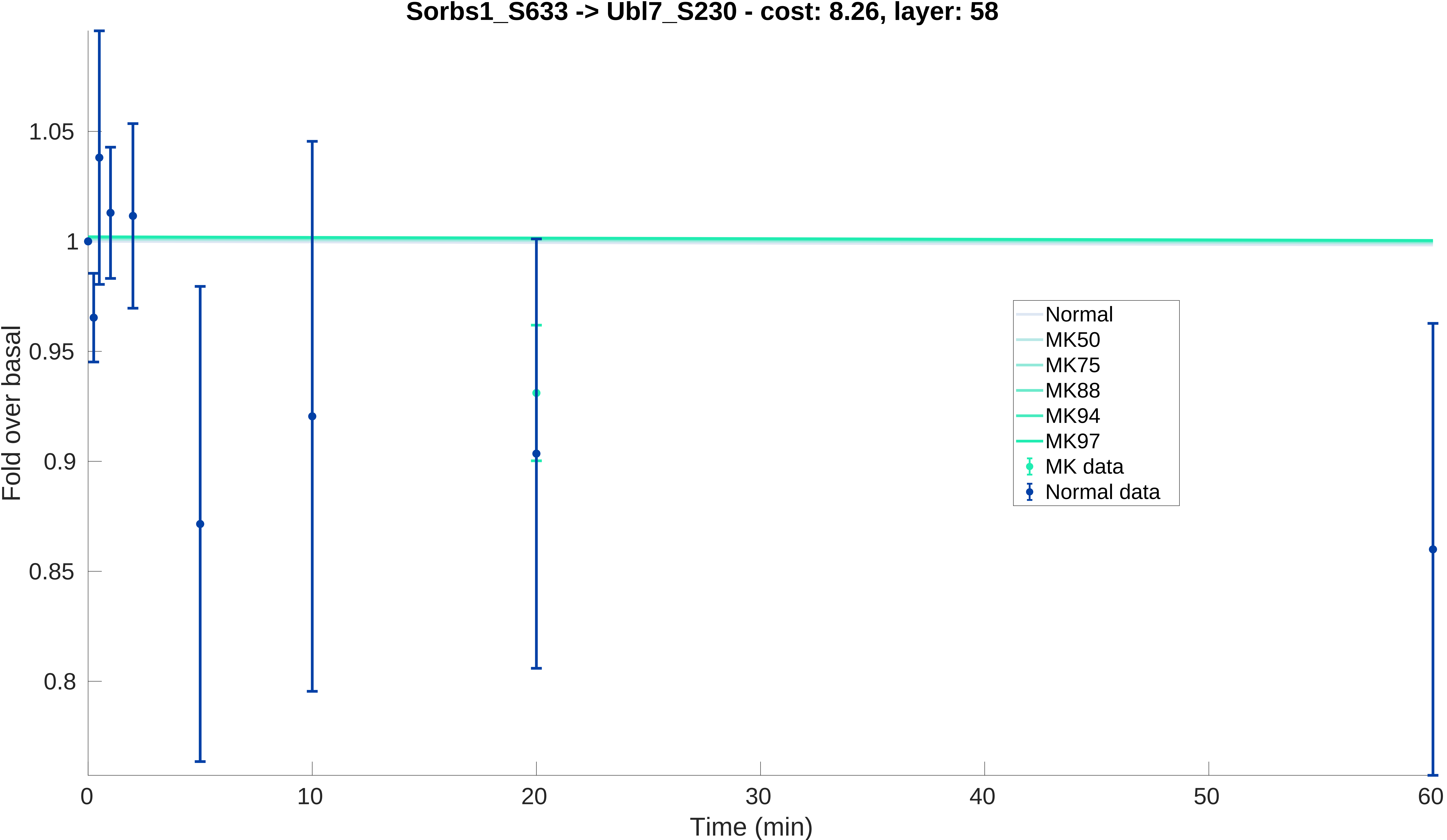

Sorbs1_S633 -> Ubl7_S230 - cost: 8.26, layer: 58
Fold over basal
1.05
1
0.95
0.9
0.85
0.8
0
10
20
30
40
50
60
Time (min)
Normal
MK50
MK75
MK88
MK94
MK97
MK data
Normal data

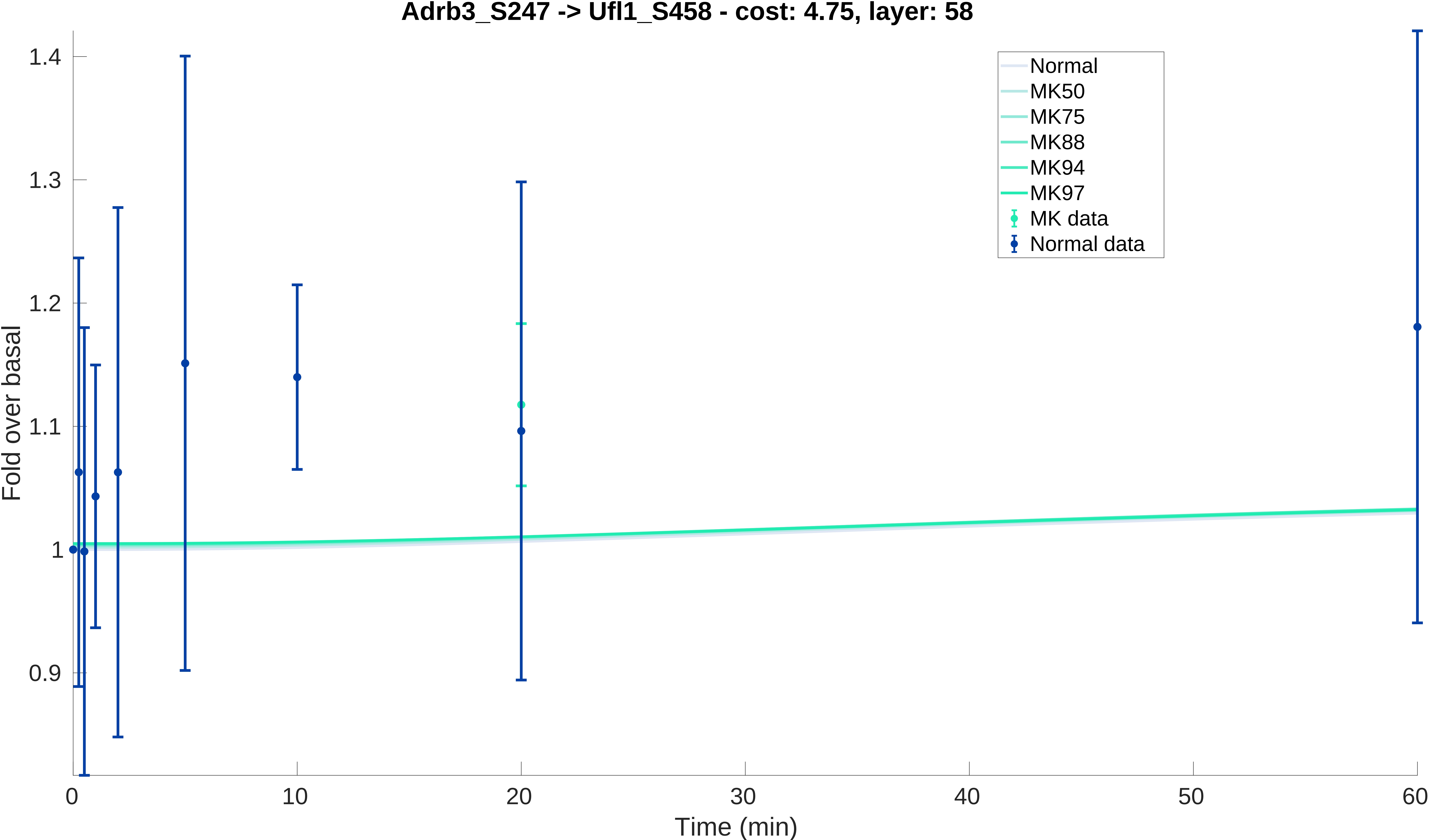

Adrb3_S247 -> Ufl1_S458 - cost: 4.75, layer: 58
Fold over basal
Time (min)
1.4
1.3
1.2
1.1
1
0.9
0
10
20
30
40
50
60
Normal
MK50
MK75
MK88
MK94
MK97
MK data
Normal data

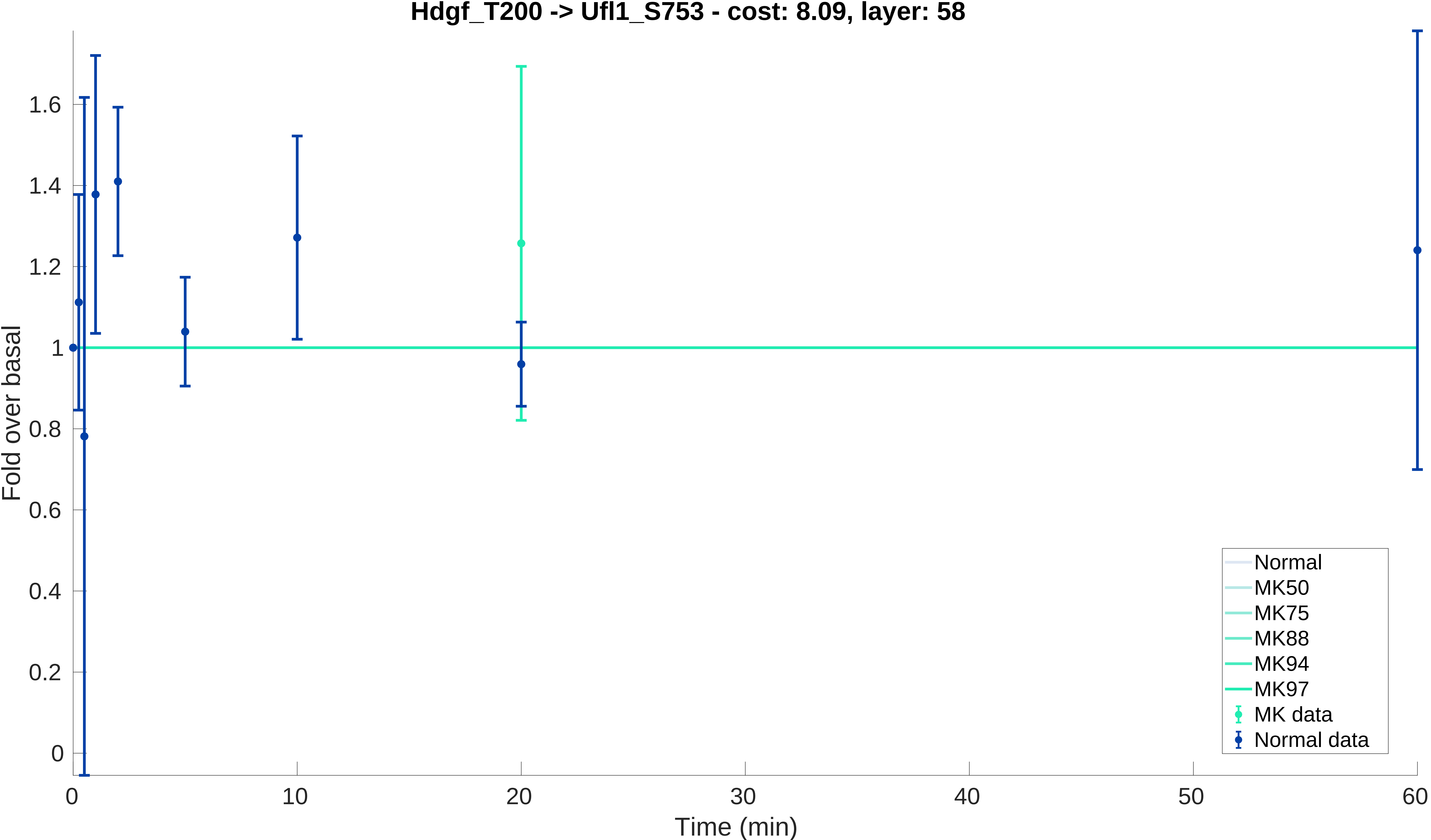

Hdgf_T200 -> Ufl1_S753 - cost: 8.09, layer: 58
Fold over basal
Time (min)
0
0.2
0.4
0.6
0.8
1
1.2
1.4
1.6
0
10
20
30
40
50
60
Normal
MK50
MK75
MK88
MK94
MK97
MK data
Normal data

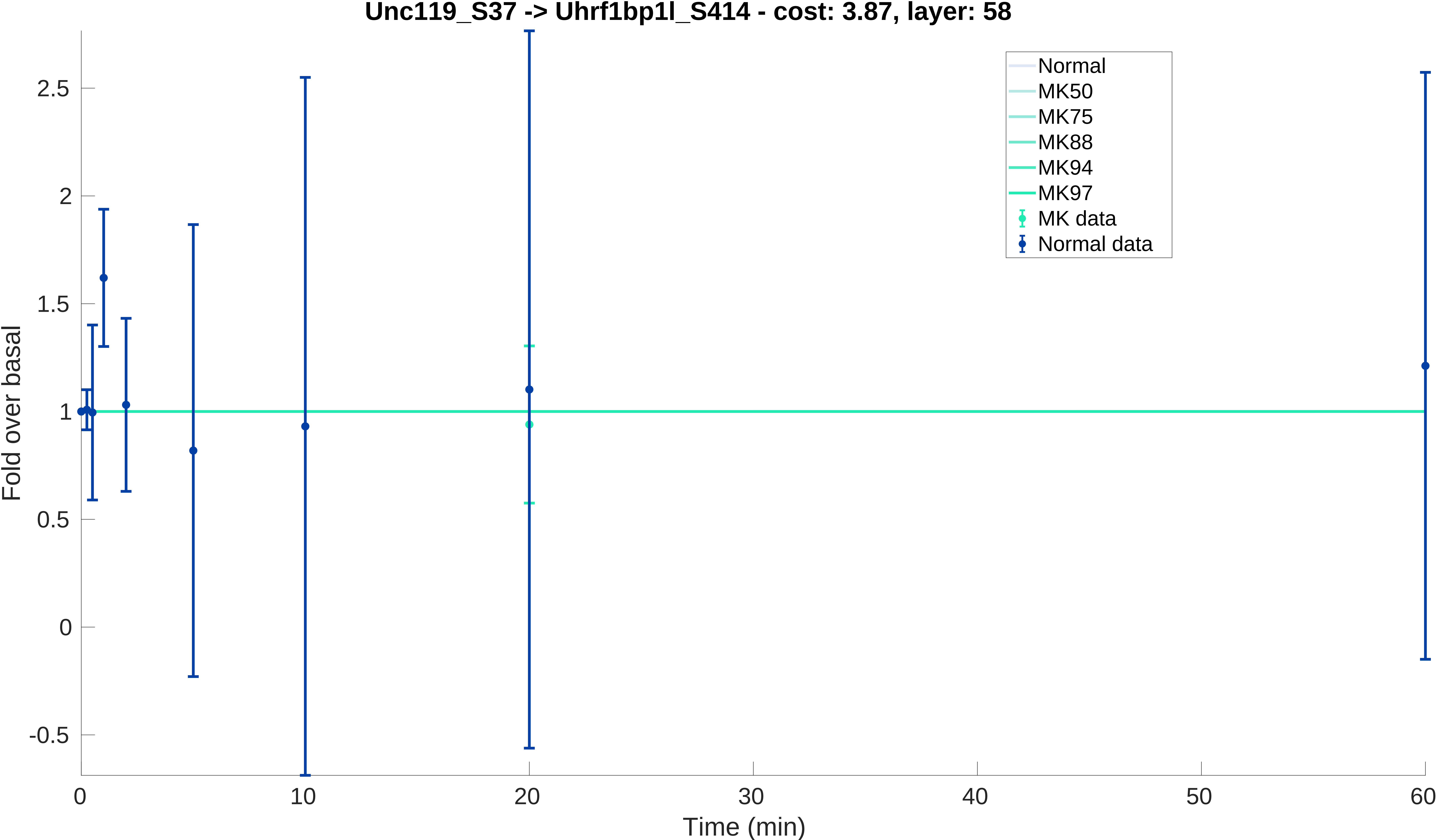

Unc119_S37 -> Uhrf1bp1l_S414 - cost: 3.87, layer: 58
Fold over basal
Time (min)
Normal
MK50
MK75
MK88
MK94
MK97
MK data
Normal data

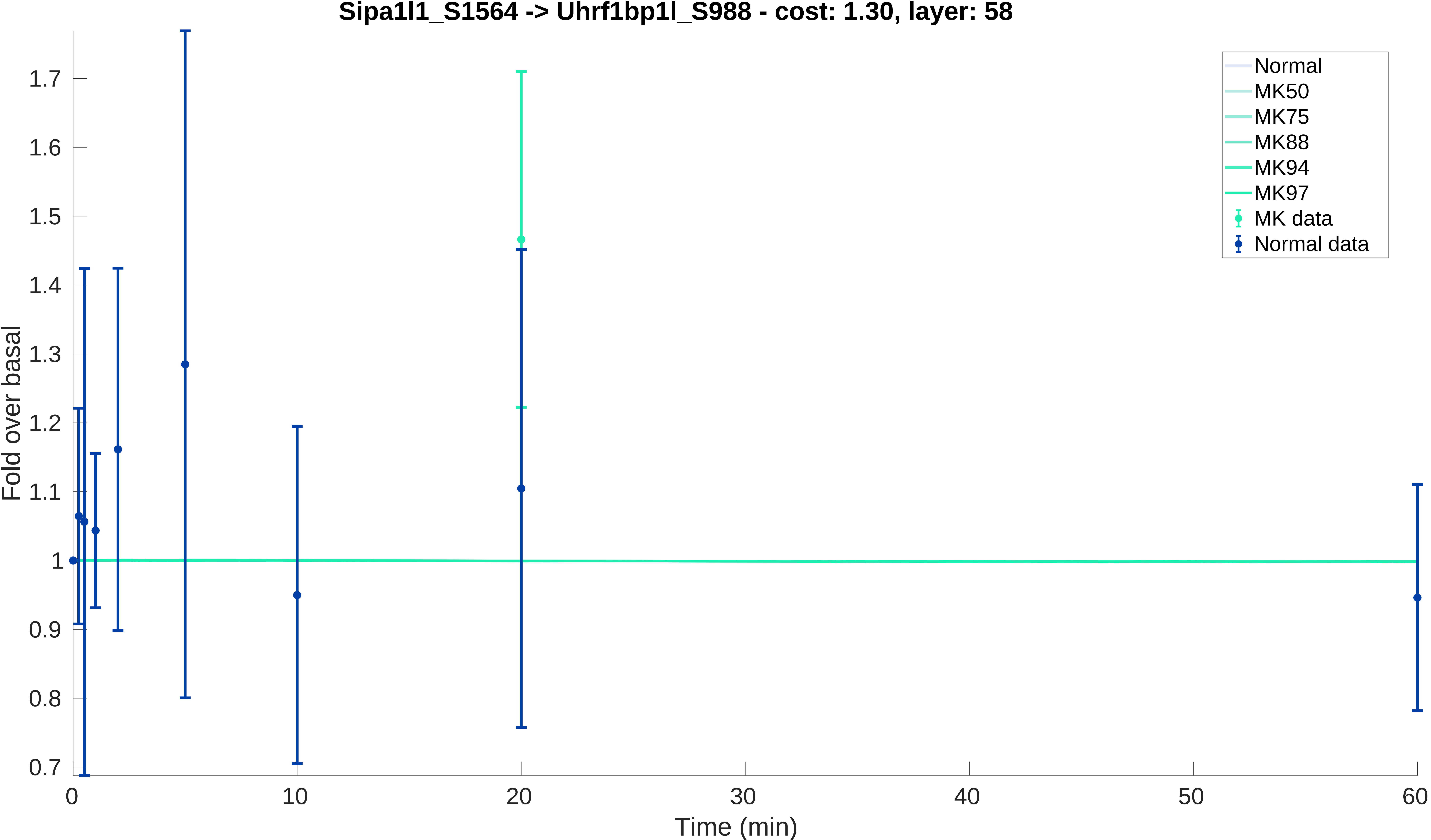

Sipa1l1_S1564 -> Uhrf1bp1l_S988 - cost: 1.30, layer: 58
Normal
MK50
MK75
MK88
MK94
MK97
MK data
Normal data
Fold over basal
Time (min)
1.7
1.6
1.5
1.4
1.3
1.2
1.1
1
0.9
0.8
0.7
0
10
20
30
40
50
60

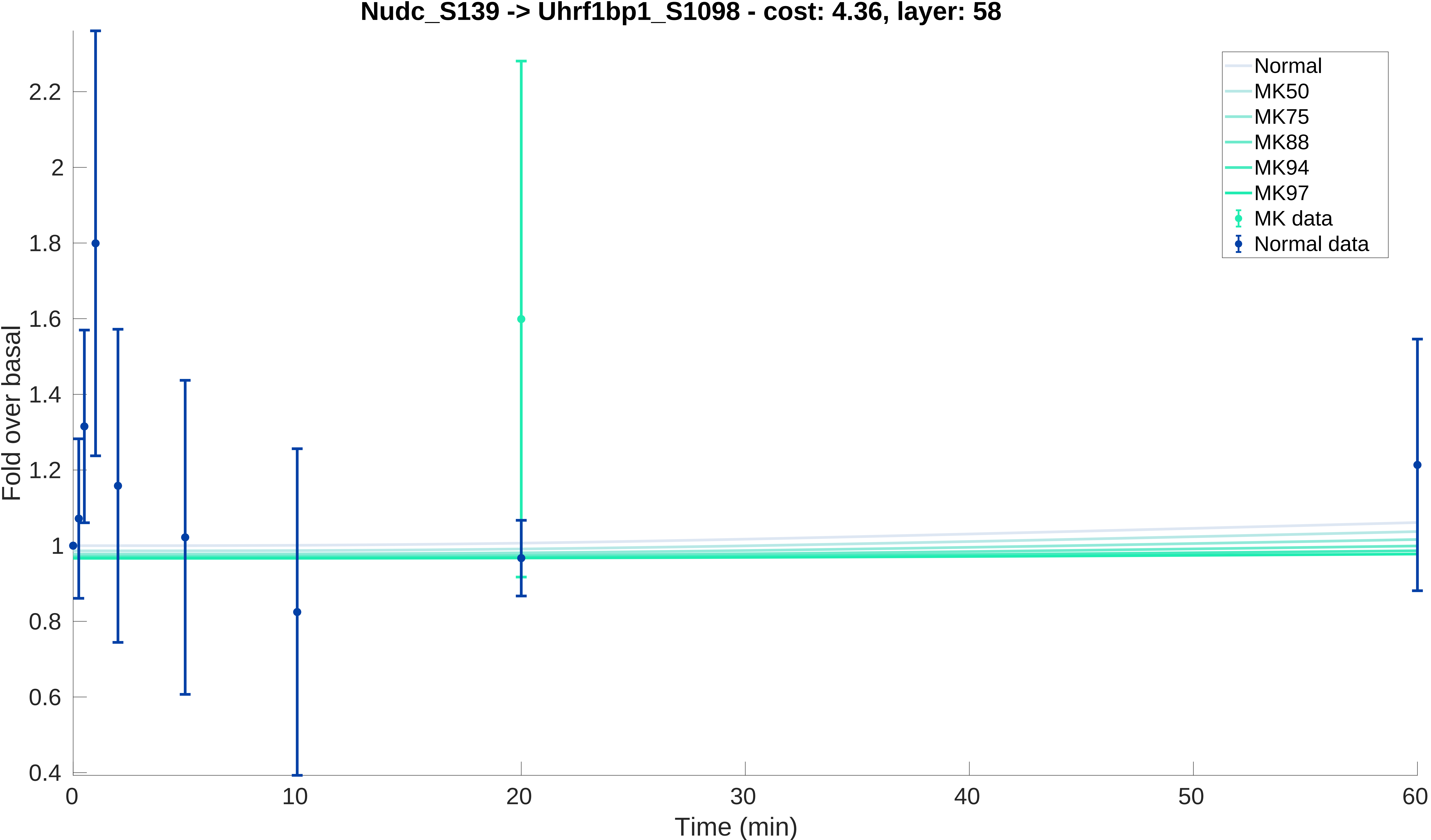

Nudc_S139 -> Uhrf1bp1_S1098 - cost: 4.36, layer: 58
Normal
MK50
MK75
MK88
MK94
MK97
MK data
Normal data
Fold over basal
Time (min)
0.4
0.6
0.8
1
1.2
1.4
1.6
1.8
2
2.2
0
10
20
30
40
50
60

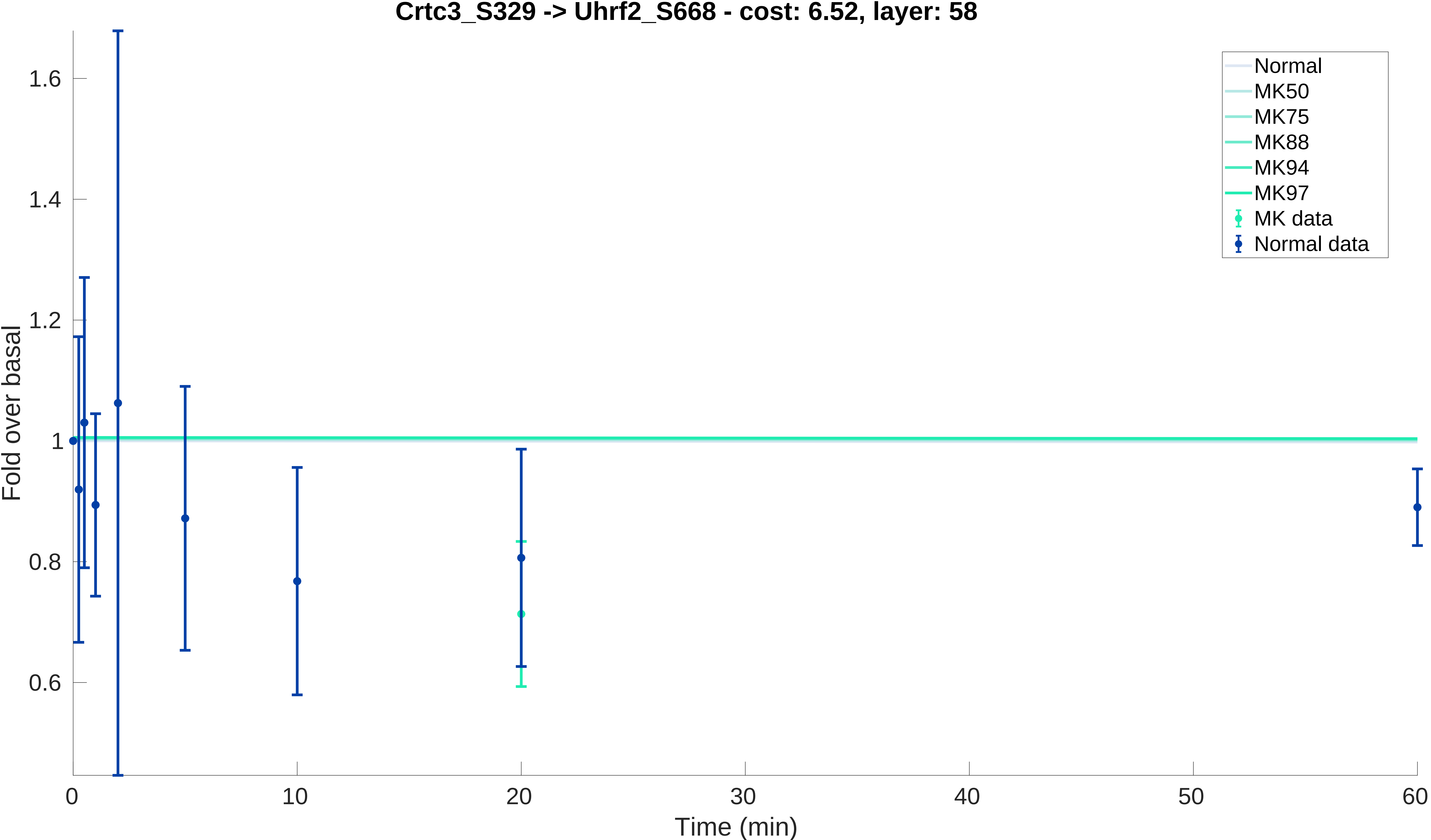
Crtc3_S329 -> Uhrf2_S668 - cost: 6.52, layer: 58
Normal
MK50
MK75
MK88
MK94
MK97
MK data
Normal data
Fold over basal
1.6
1.4
1.2
1
0.8
0.6
0
10
20
30
40
50
60
Time (min)

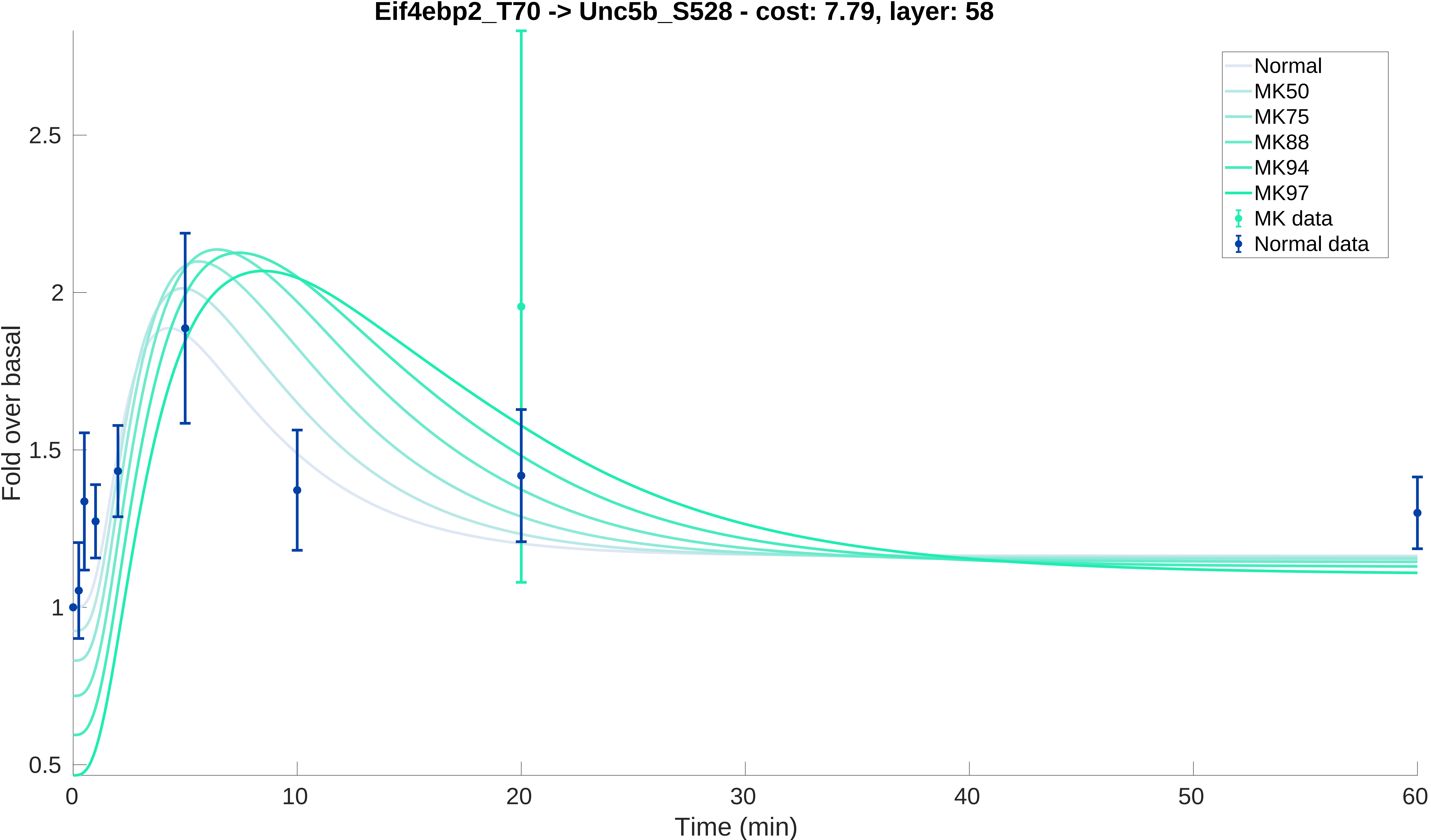

Eif4ebp2_T70 -> Unc5b_S528 - cost: 7.79, layer: 58
Fold over basal
Time (min)
0.5
1
1.5
2
2.5
0
10
20
30
40
50
60
Normal
MK50
MK75
MK88
MK94
MK97
MK data
Normal data

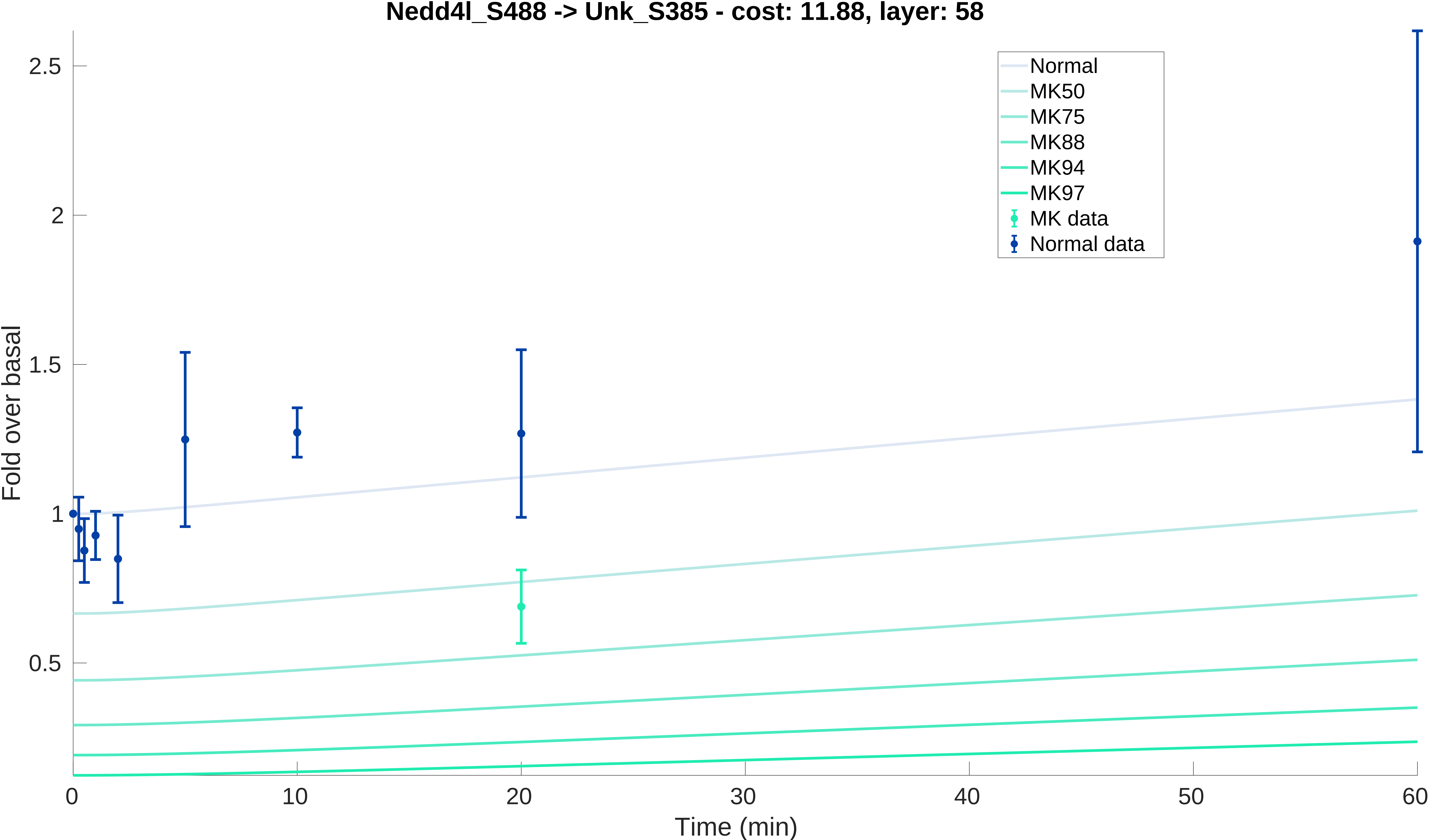

Nedd4l_S488 -> Unk_S385 - cost: 11.88, layer: 58
Normal
MK50
MK75
MK88
MK94
MK97
MK data
Normal data
Fold over basal
2.5
2
1.5
1
0.5
0
10
20
30
40
50
60
Time (min)

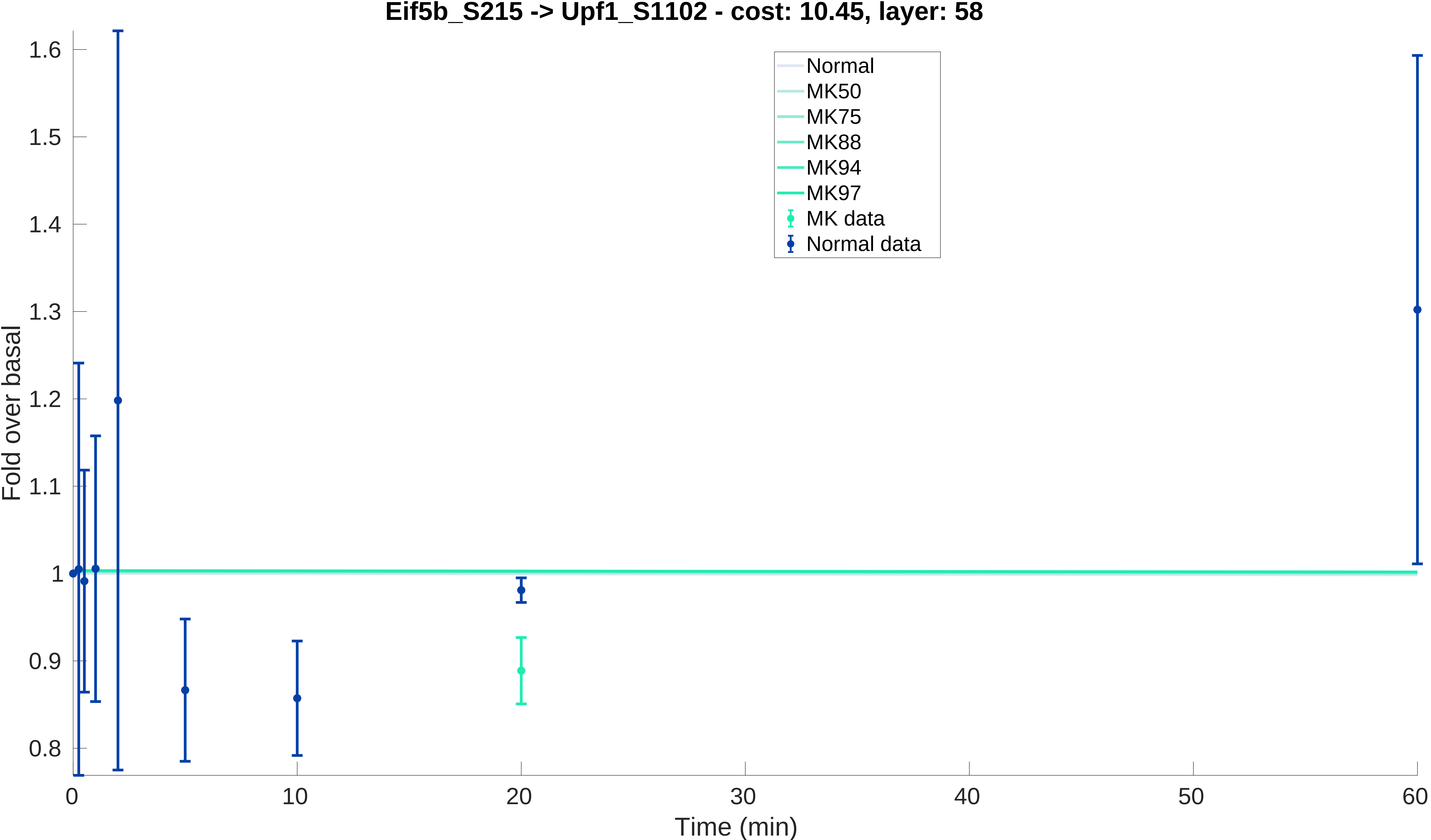

Eif5b_S215 -> Upf1_S1102 - cost: 10.45, layer: 58
Normal
MK50
MK75
MK88
MK94
MK97
MK data
Normal data
Fold over basal
1.6
1.5
1.4
1.3
1.2
1.1
1
0.9
0.8
0
10
20
30
40
50
60
Time (min)

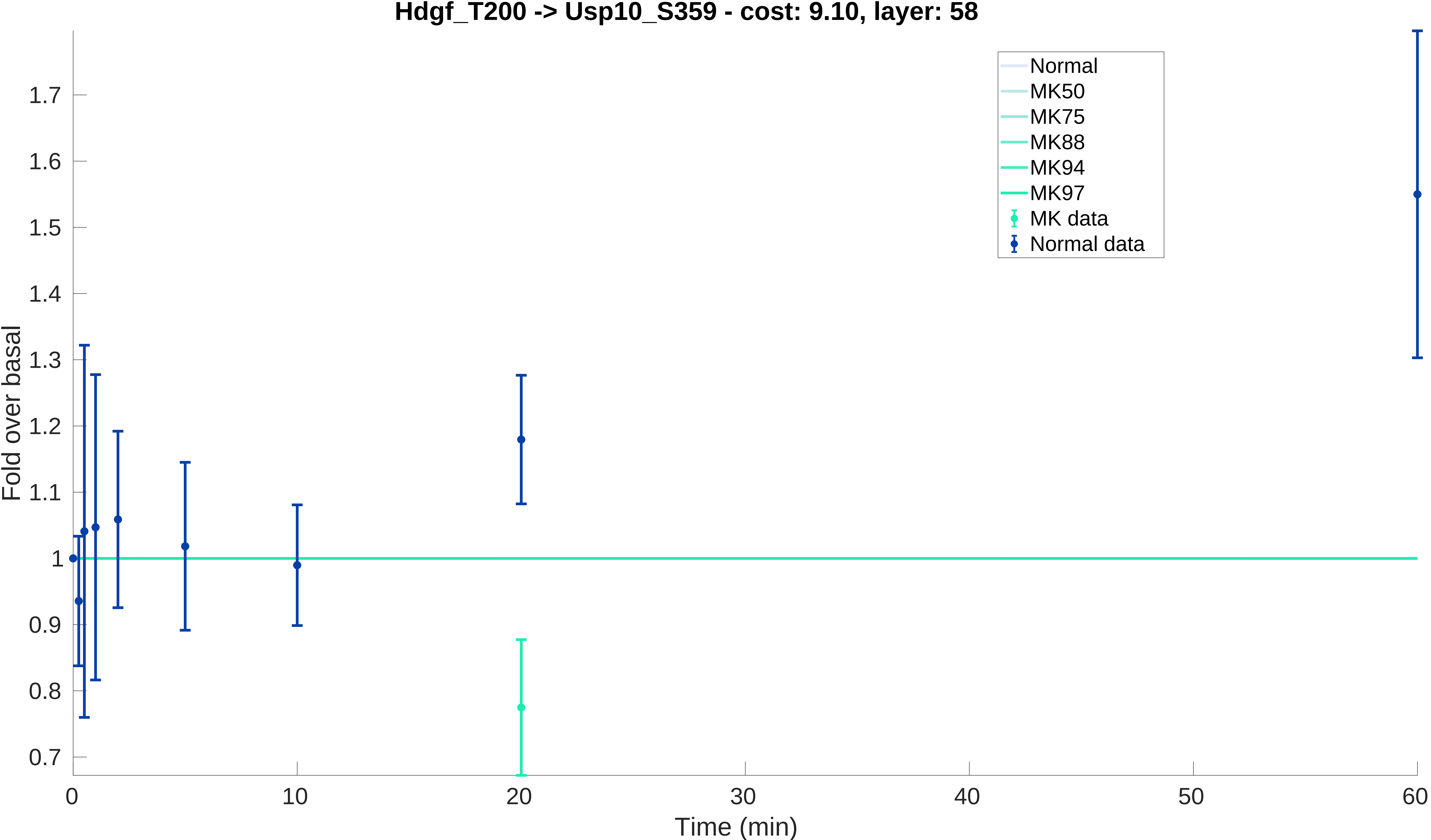

Hdgf_T200 -> Usp10_S359 - cost: 9.10, layer: 58
Normal
MK50
MK75
MK88
MK94
MK97
MK data
Normal data
Fold over basal
Time (min)
0.7
0.8
0.9
1
1.1
1.2
1.3
1.4
1.5
1.6
1.7
0
10
20
30
40
50
60

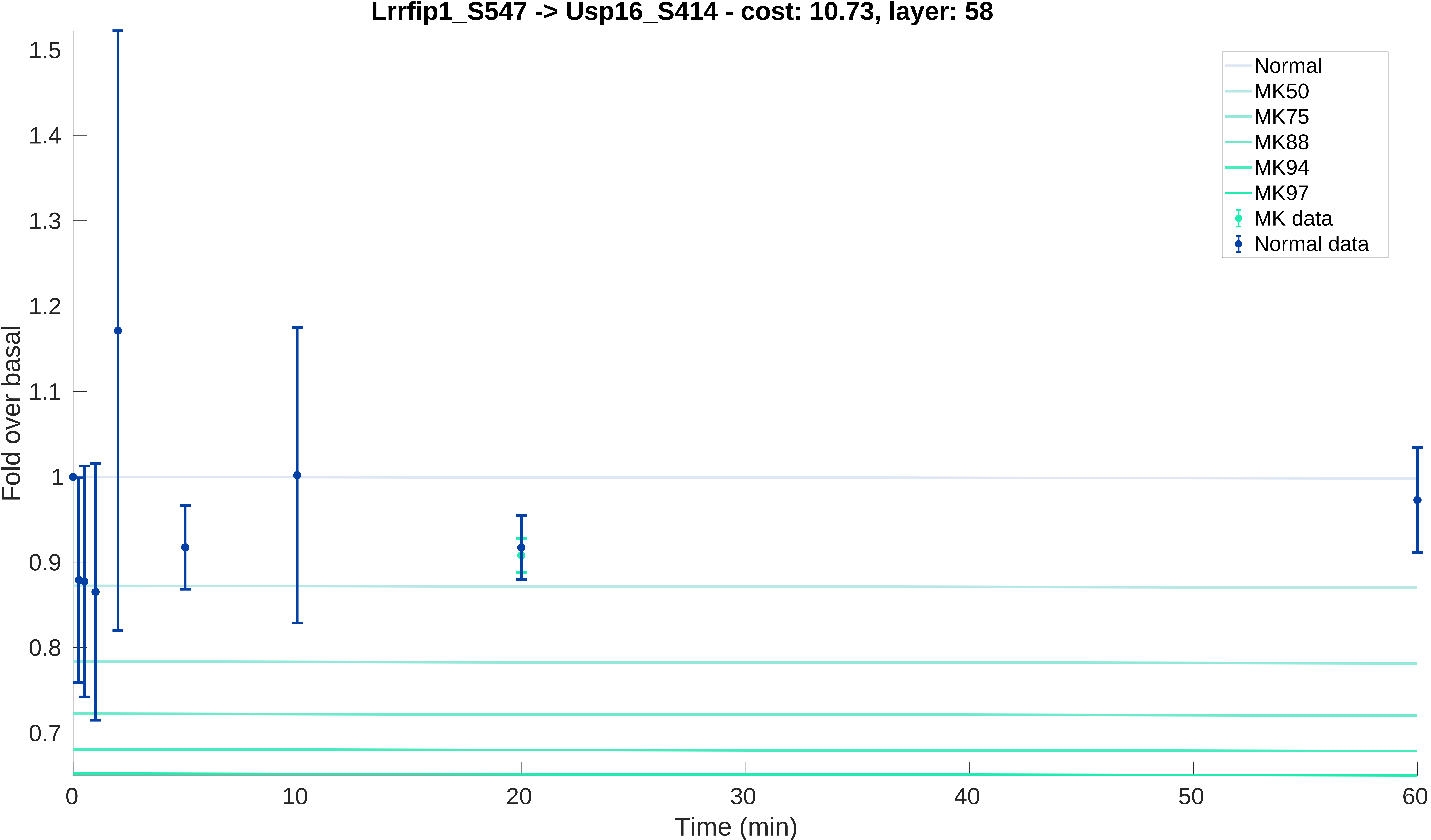

Lrrfip1_S547 -> Usp16_S414 - cost: 10.73, layer: 58
Fold over basal
Time (min)
1.5
1.4
1.3
1.2
1.1
1
0.9
0.8
0.7
0
10
20
30
40
50
60
Normal
MK50
MK75
MK88
MK94
MK97
MK data
Normal data

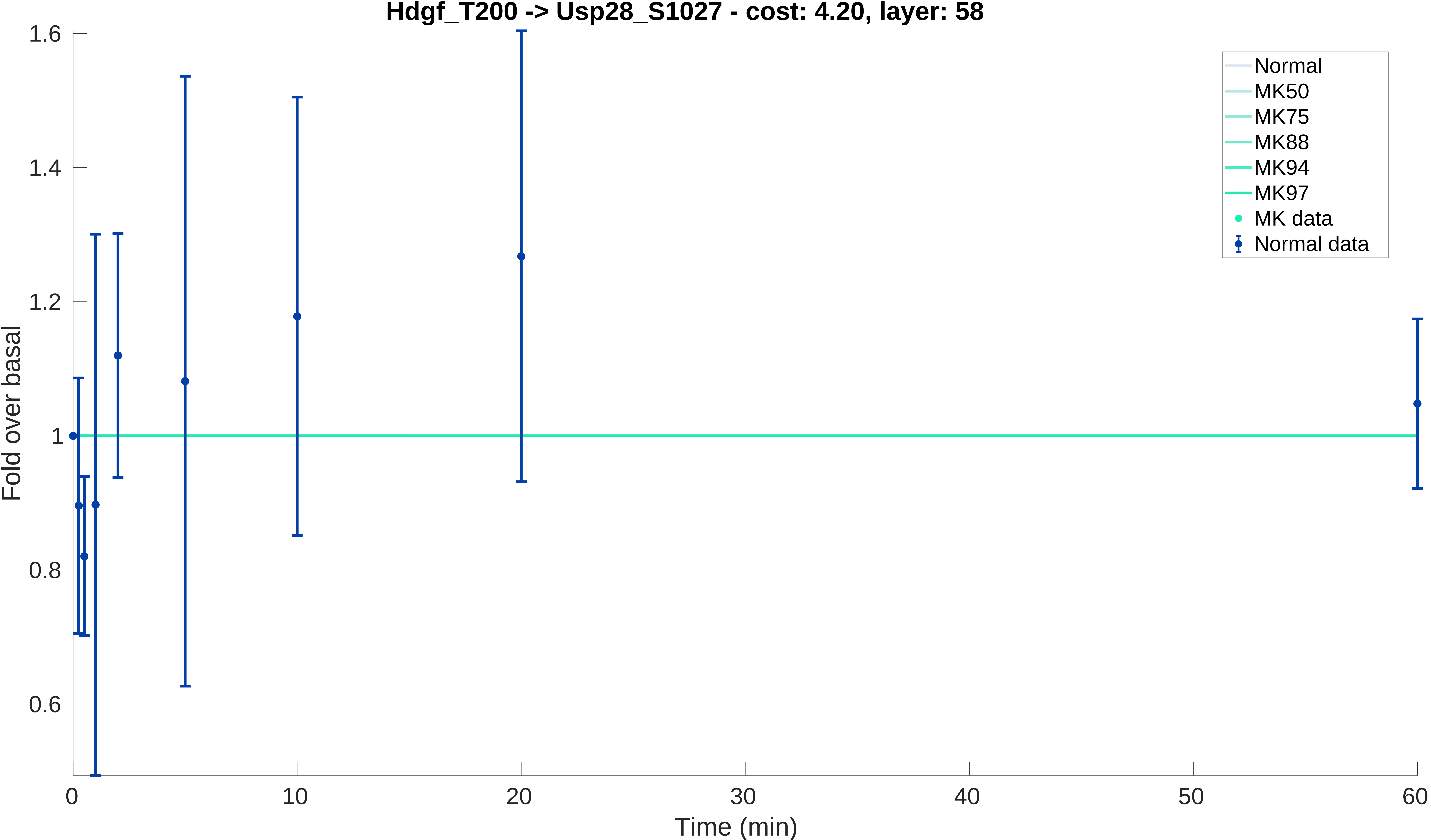

Hdgf_T200 -> Usp28_S1027 - cost: 4.20, layer: 58
Fold over basal
Time (min)
1.6
1.4
1.2
1
0.8
0.6
0
10
20
30
40
50
60
Normal
MK50
MK75
MK88
MK94
MK97
MK data
Normal data

#### Sorbs1_S633 -> Usp39_S46 - cost: 13.74, layer: 58

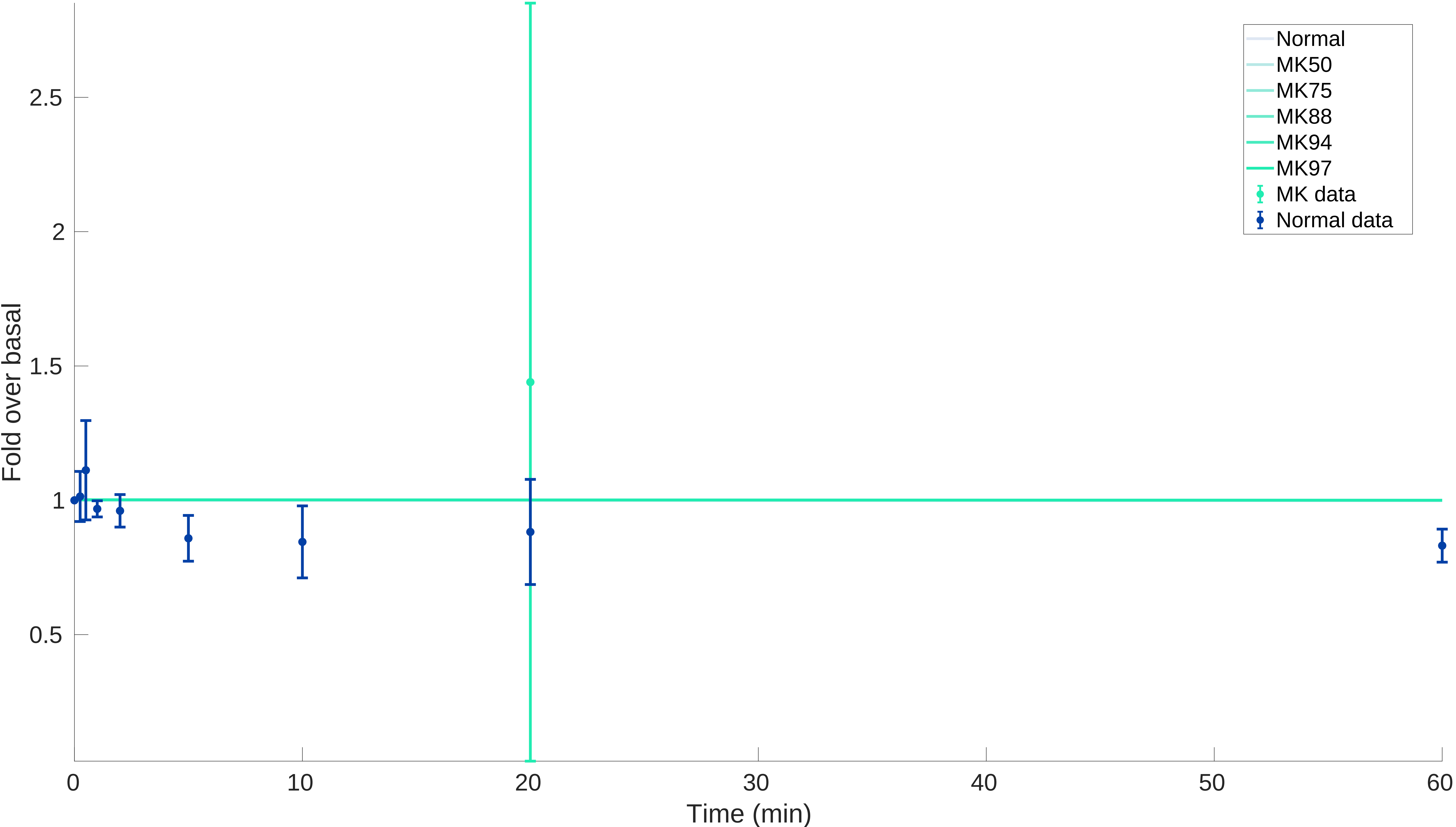

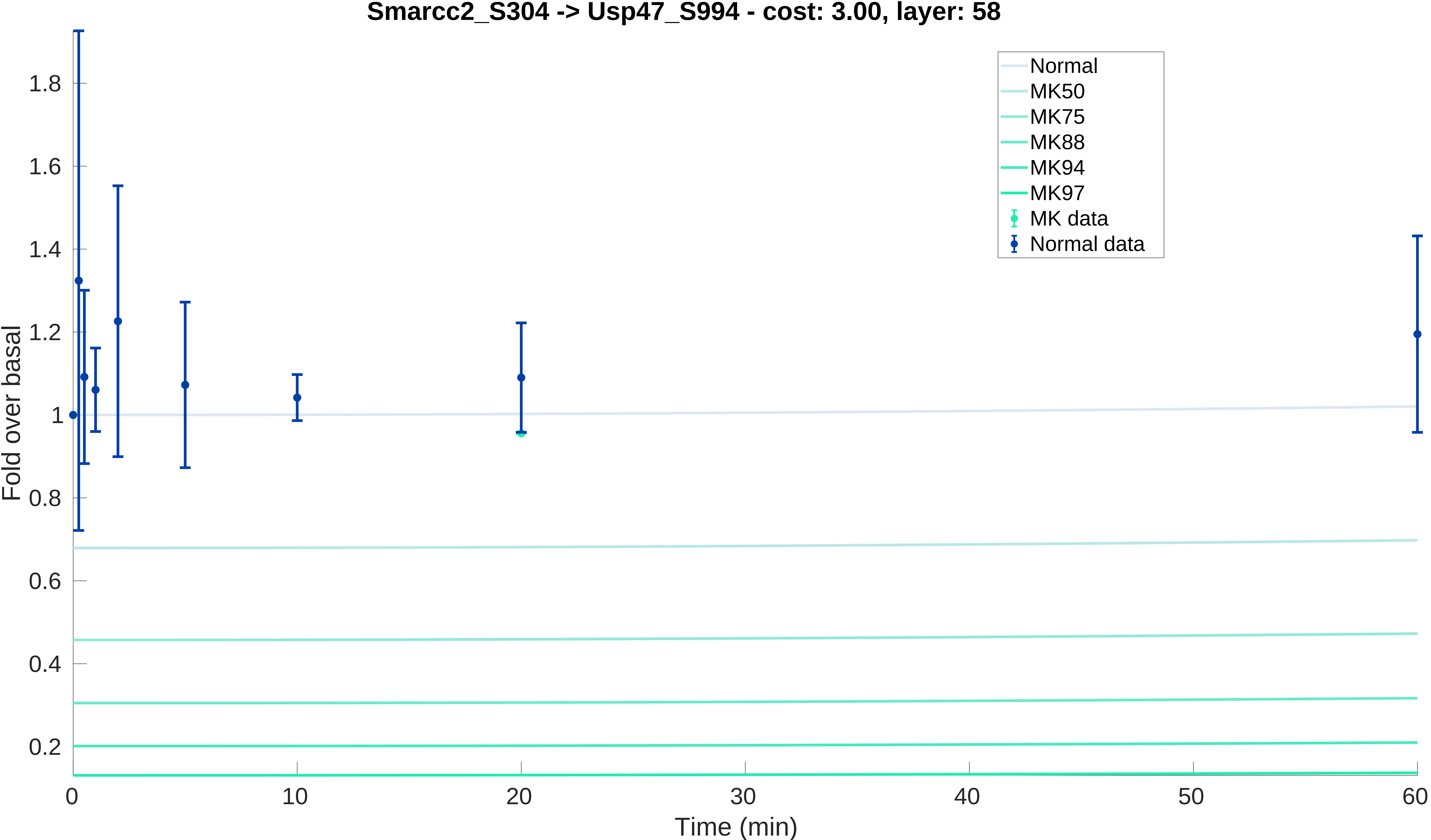

Smarcc2_S304 -> Usp47_S994 - cost: 3.00, layer: 58
Normal
MK50
MK75
MK88
MK94
MK97
MK data
Normal data
Fold over basal
1.8
1.6
1.4
1.2
1
0.8
0.6
0.4
0.2
0
10
20
30
40
50
60
Time (min)

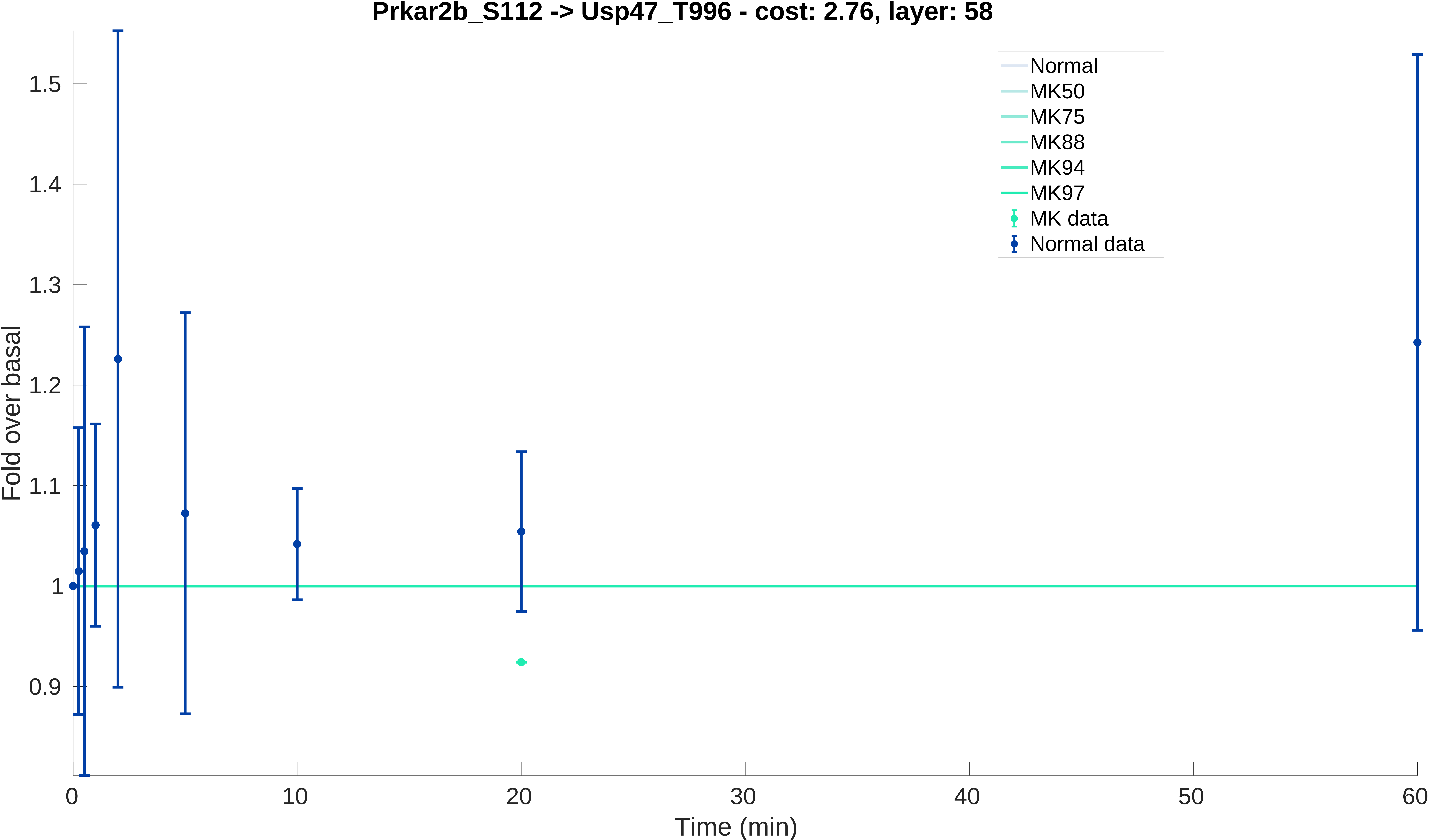

Prkar2b_S112 -> Usp47_T996 - cost: 2.76, layer: 58
Fold over basal
Time (min)
0.9
1
1.1
1.2
1.3
1.4
1.5
0
10
20
30
40
50
60
Normal
MK50
MK75
MK88
MK94
MK97
MK data
Normal data

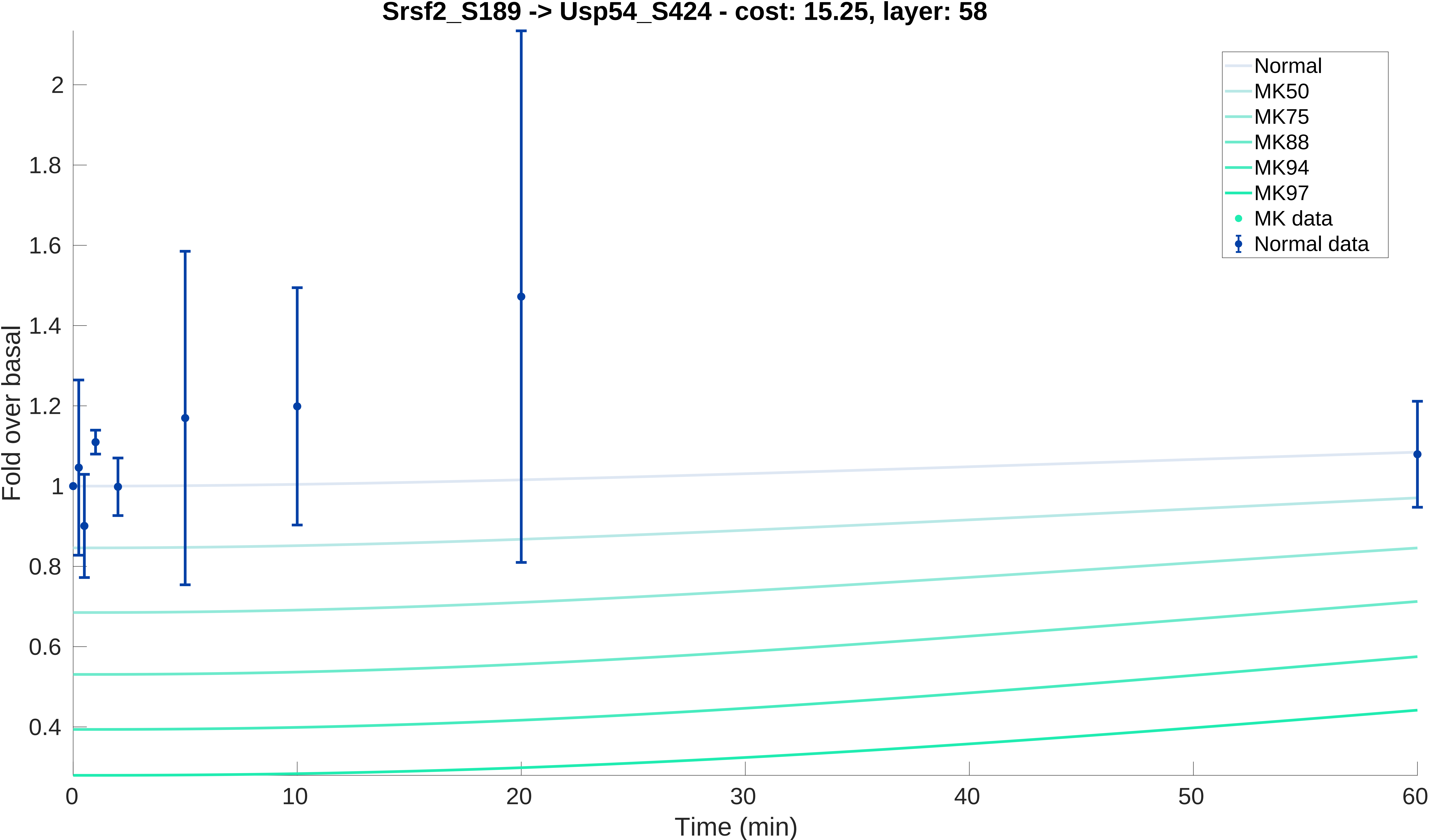
Srsf2_S189 -> Usp54_S424 - cost: 15.25, layer: 58
Fold over basal
Time (min)
0.4
0.6
0.8
1
1.2
1.4
1.6
1.8
2
0
10
20
30
40
50
60
Normal
MK50
MK75
MK88
MK94
MK97
MK data
Normal data

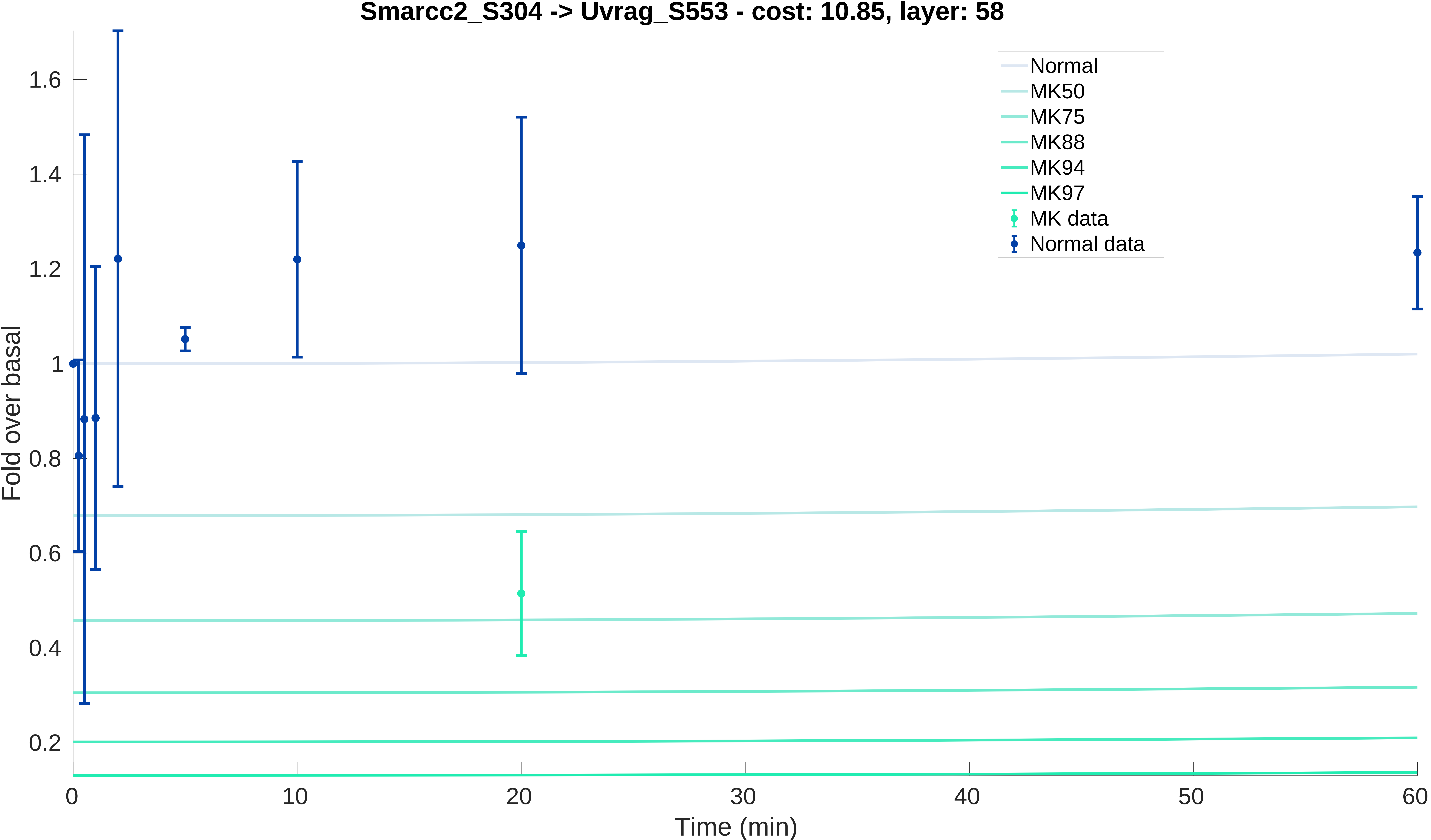

Smarcc2_S304 -> Uvrag_S553 - cost: 10.85, layer: 58
Fold over basal
1.6
1.4
1.2
1
0.8
0.6
0.4
0.2
0
10
20
30
40
50
60
Time (min)
Normal
MK50
MK75
MK88
MK94
MK97
MK data
Normal data

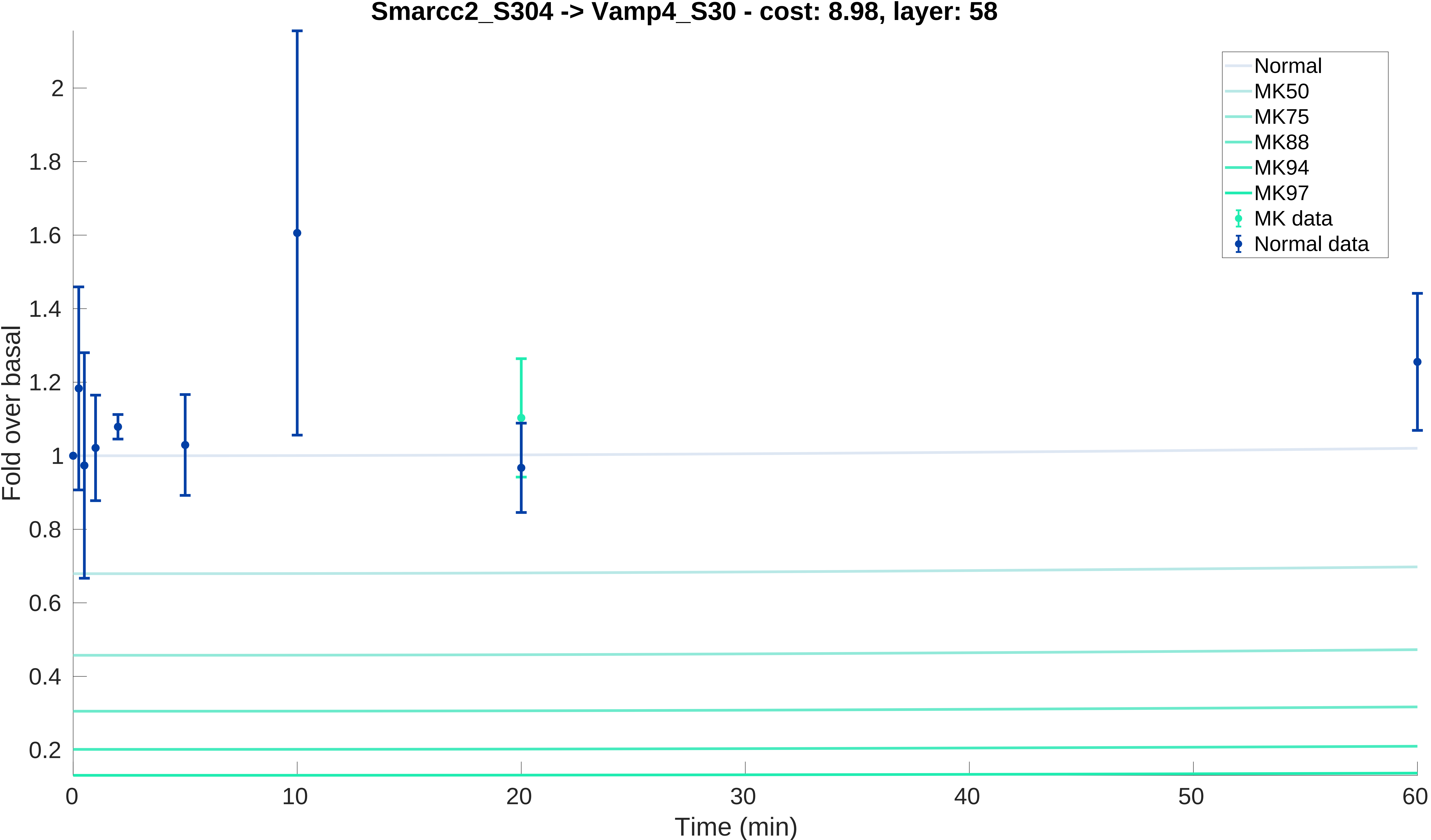
Smarcc2_S304 -> Vamp4_S30 - cost: 8.98, layer: 58
Fold over basal
Time (min)
0.2
0.4
0.6
0.8
1
1.2
1.4
1.6
1.8
2
0
10
20
30
40
50
60
Normal
MK50
MK75
MK88
MK94
MK97
MK data
Normal data

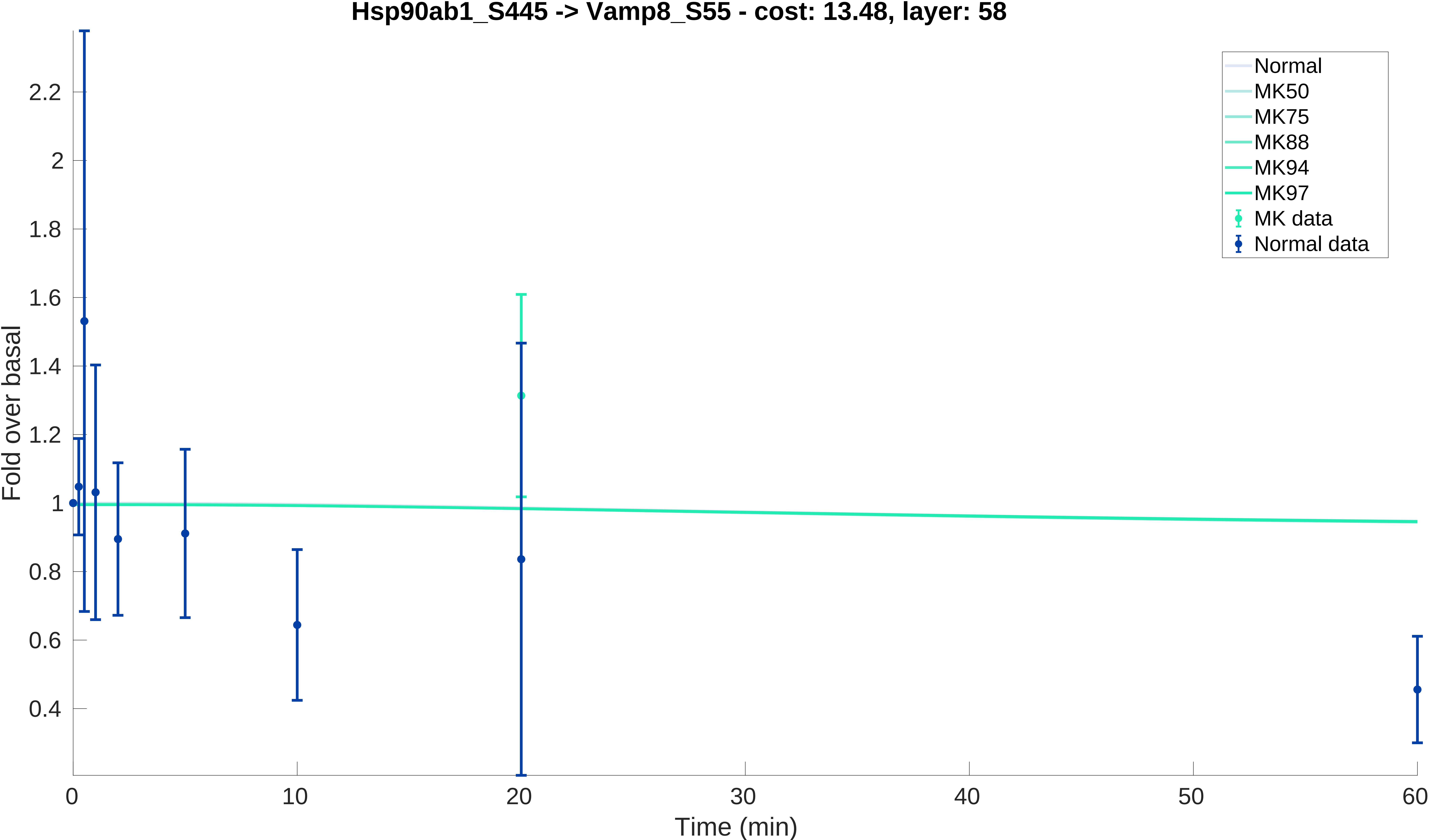

Hsp90ab1_S445 -> Vamp8_S55 - cost: 13.48, layer: 58
Fold over basal
Time (min)
0.4
0.6
0.8
1
1.2
1.4
1.6
1.8
2
2.2
0
10
20
30
40
50
60
Normal
MK50
MK75
MK88
MK94
MK97
MK data
Normal data

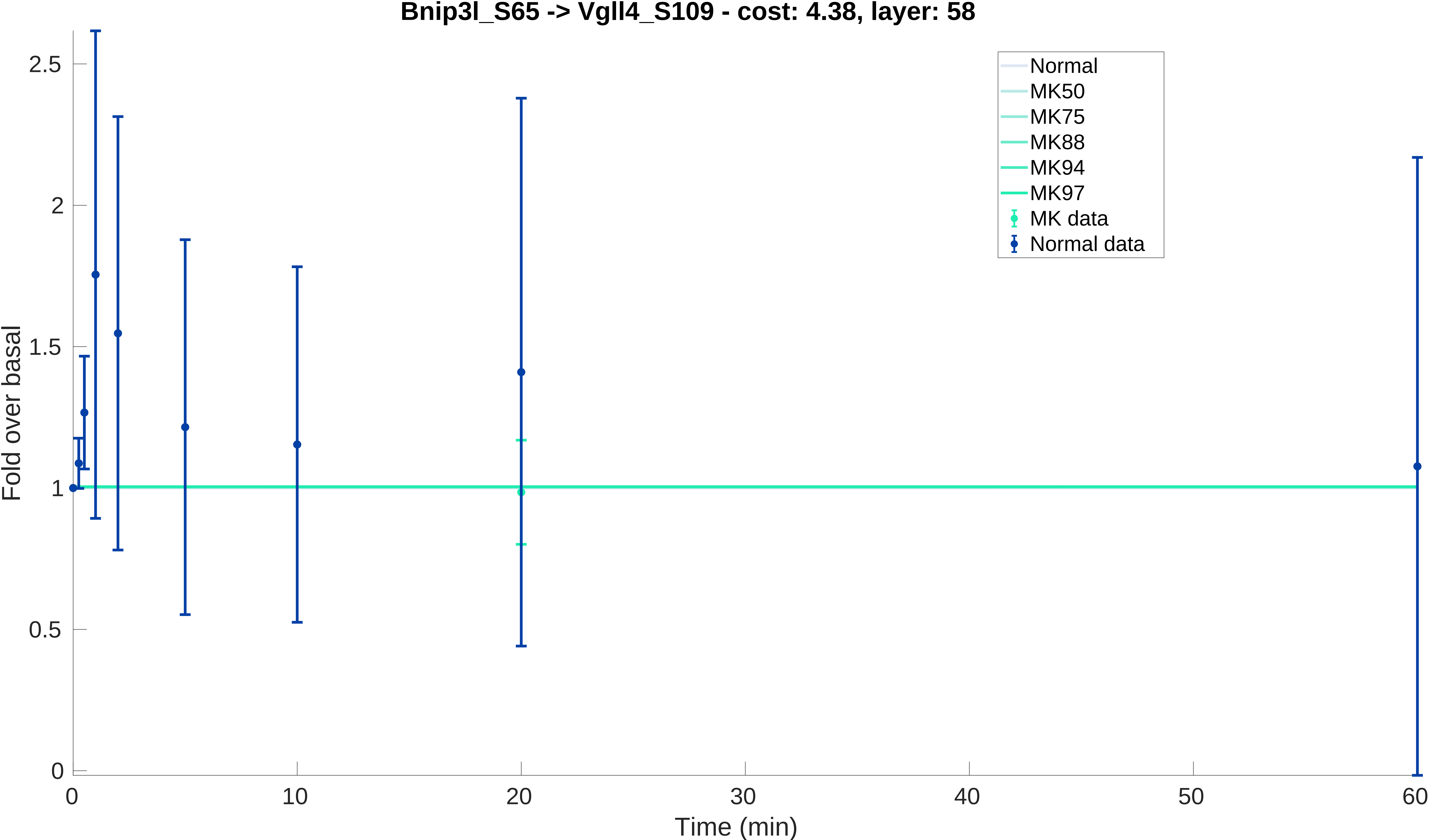

Bnip3l_S65 -> Vgll4_S109 - cost: 4.38, layer: 58
Fold over basal
Time (min)
0
0.5
1
1.5
2
2.5
0
10
20
30
40
50
60
Normal
MK50
MK75
MK88
MK94
MK97
MK data
Normal data

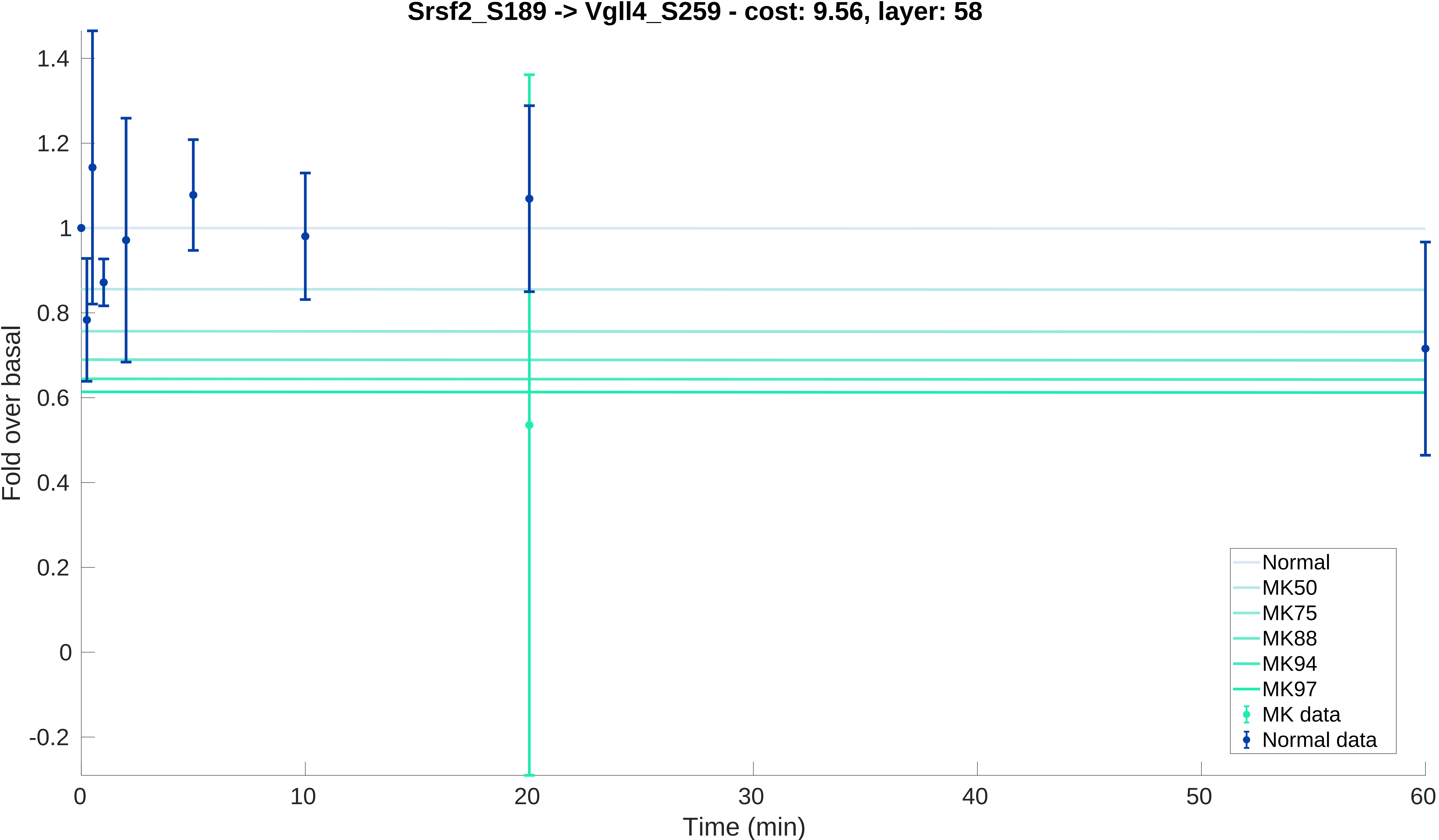
Srsf2_S189 -> Vgll4_S259 - cost: 9.56, layer: 58
Fold over basal
Time (min)
1.4
1.2
1
0.8
0.6
0.4
0.2
0
-0.2
0
10
20
30
40
50
60
Normal
MK50
MK75
MK88
MK94
MK97
MK data
Normal data

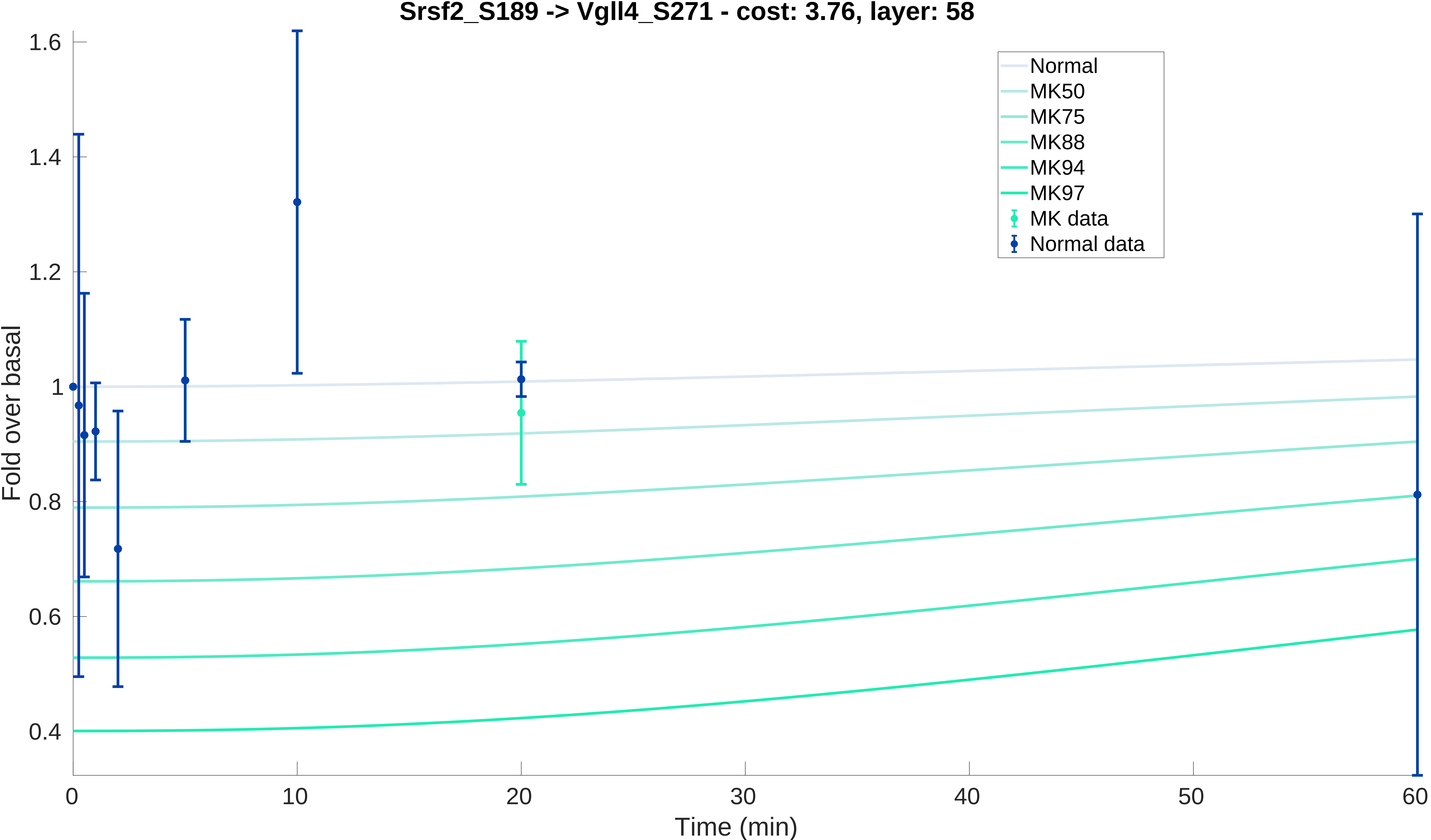

Srsf2_S189 -> Vgll4_S271 - cost: 3.76, layer: 58
Fold over basal
Time (min)
1.6
1.4
1.2
1
0.8
0.6
0.4
0
10
20
30
40
50
60
Normal
MK50
MK75
MK88
MK94
MK97
MK data
Normal data

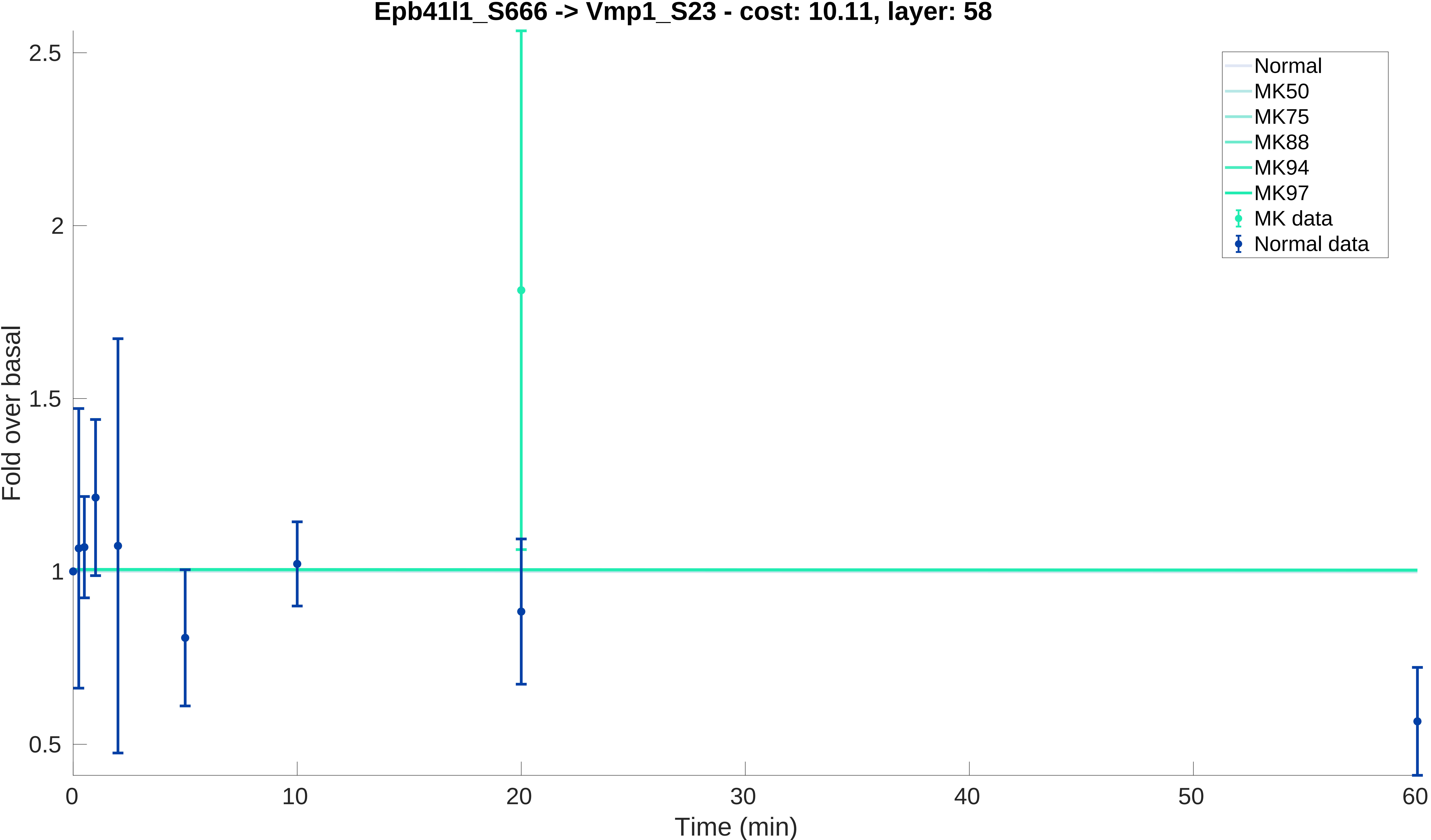

Epb41l1_S666 -> Vmp1_S23 - cost: 10.11, layer: 58
Fold over basal
Time (min)
2.5
2
1.5
1
0.5
0
10
20
30
40
50
60
Normal
MK50
MK75
MK88
MK94
MK97
MK data
Normal data

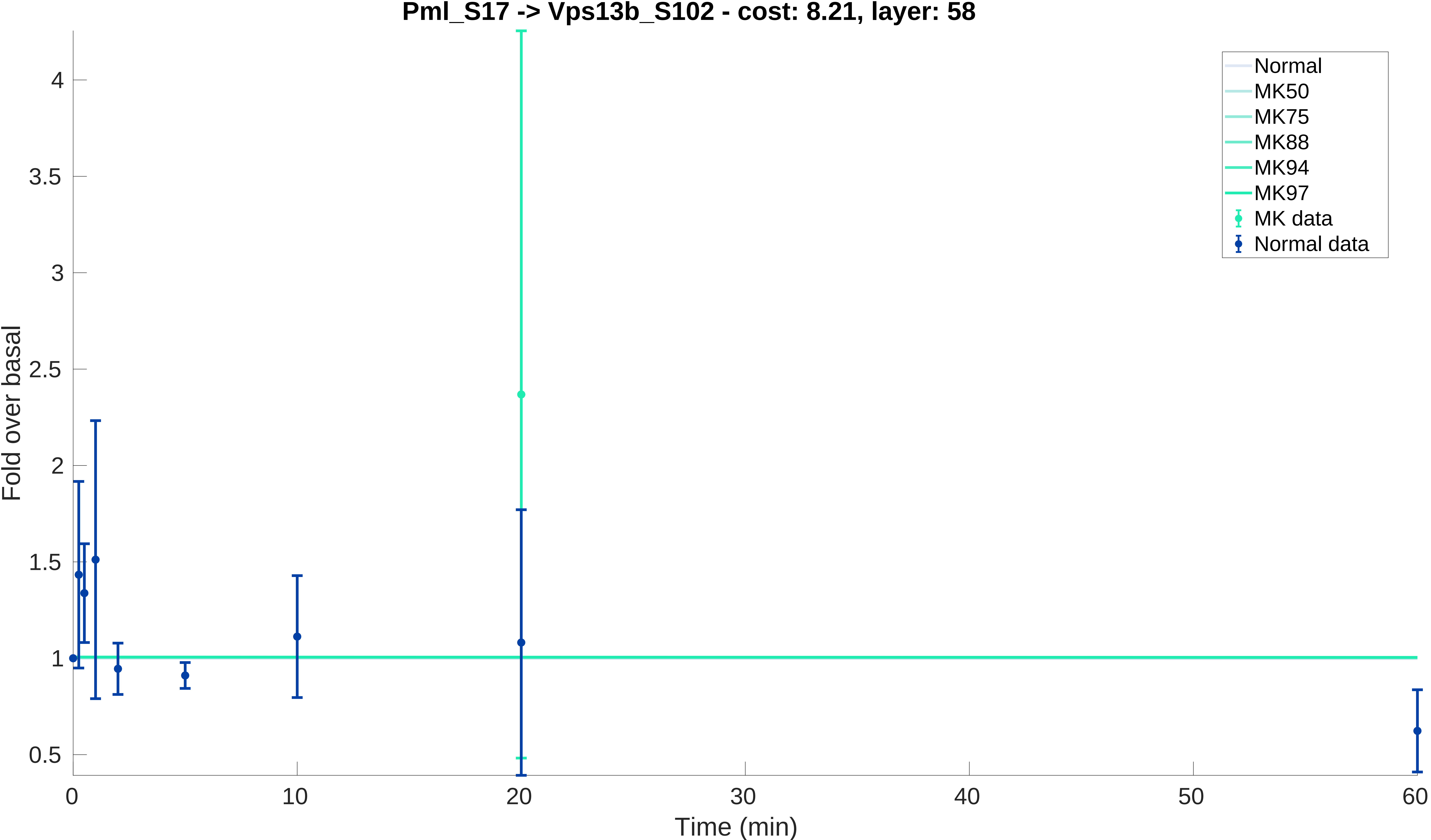

Pml_S17 -> Vps13b_S102 - cost: 8.21, layer: 58
Fold over basal
Time (min)
0
10
20
30
40
50
60
0.5
1
1.5
2
2.5
3
3.5
4
Normal
MK50
MK75
MK88
MK94
MK97
MK data
Normal data

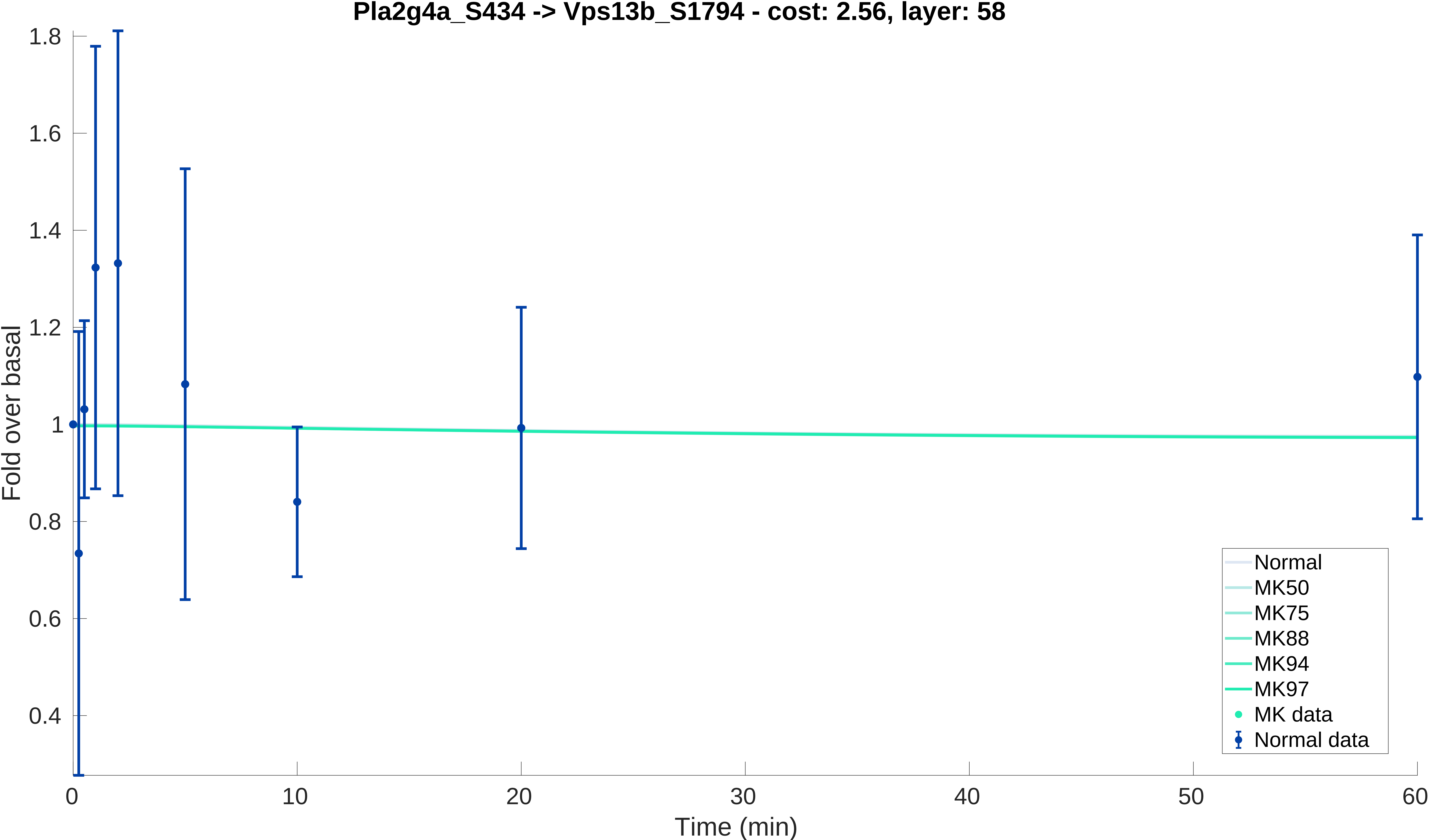

Pla2g4a_S434 -> Vps13b_S1794 - cost: 2.56, layer: 58
Fold over basal
Time (min)
1.8
1.6
1.4
1.2
1
0.8
0.6
0.4
0
10
20
30
40
50
60
Normal
MK50
MK75
MK88
MK94
MK97
MK data
Normal data

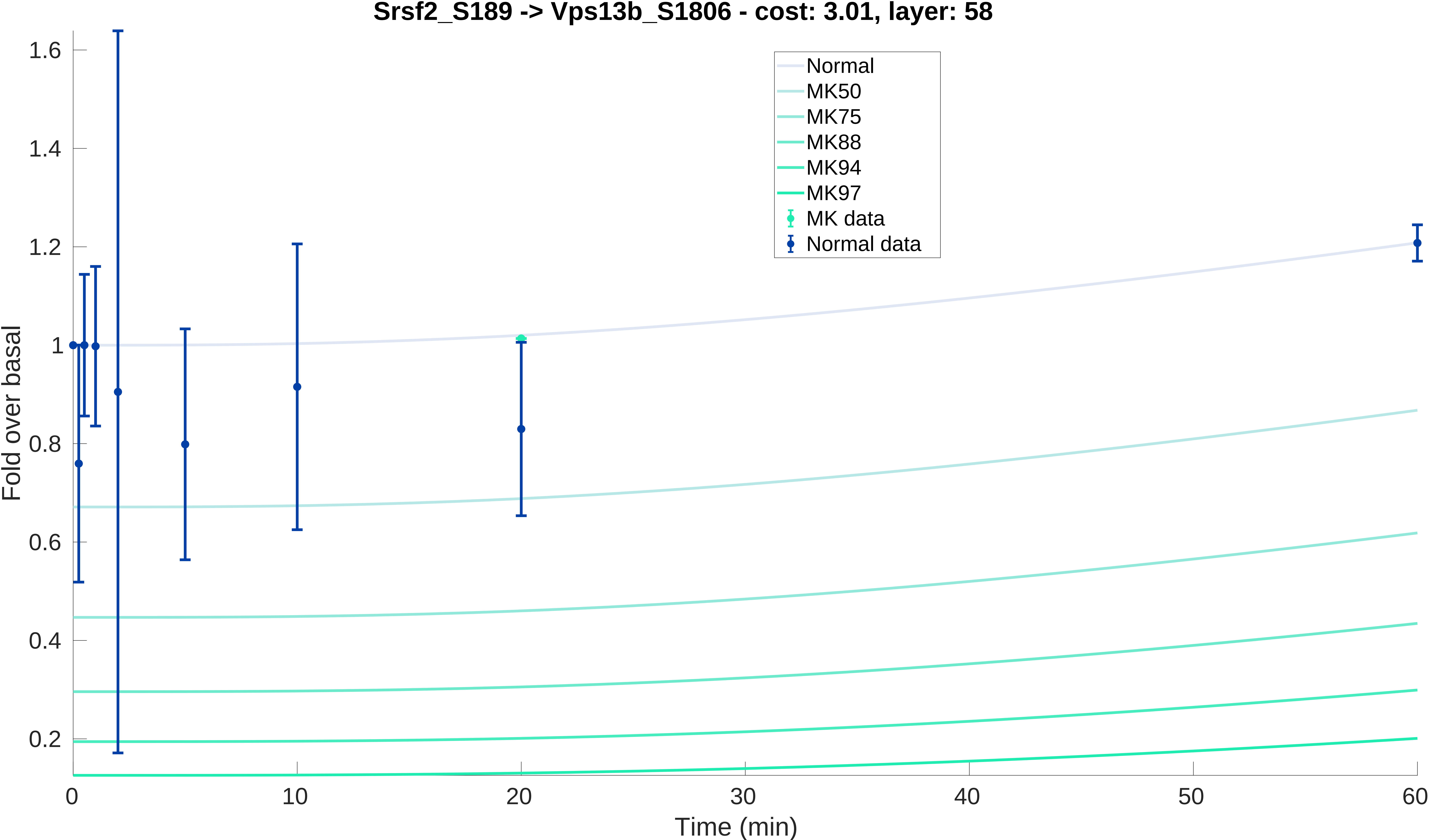

Srsf2_S189 -> Vps13b_S1806 - cost: 3.01, layer: 58
Normal
MK50
MK75
MK88
MK94
MK97
MK data
Normal data
Fold over basal
Time (min)
1.6
1.4
1.2
1
0.8
0.6
0.4
0.2
0
10
20
30
40
50
60

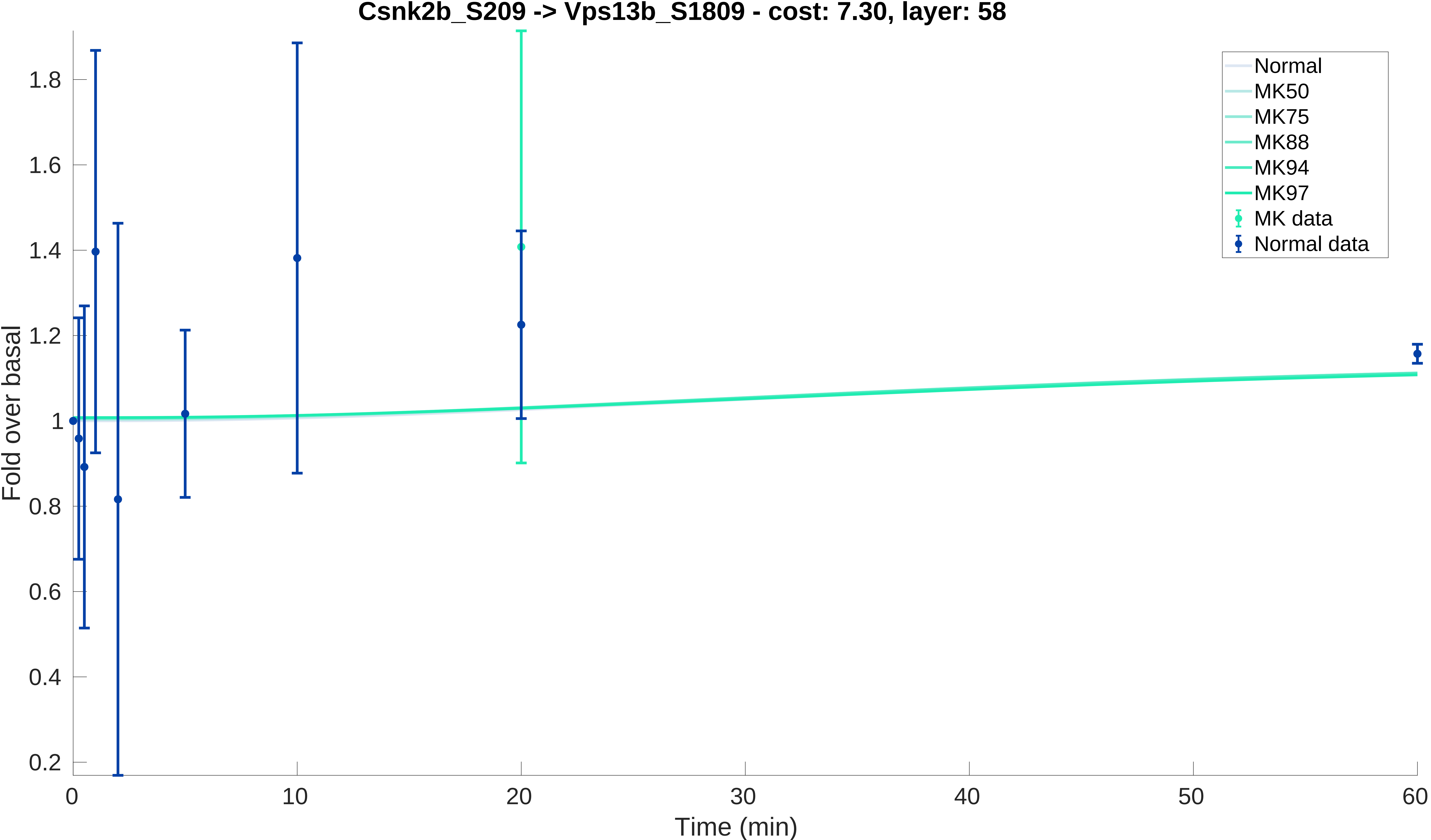

Csnk2b_S209 -> Vps13b_S1809 - cost: 7.30, layer: 58
Fold over basal
Time (min)
0.2
0.4
0.6
0.8
1
1.2
1.4
1.6
1.8
0
10
20
30
40
50
60
Normal
MK50
MK75
MK88
MK94
MK97
MK data
Normal data

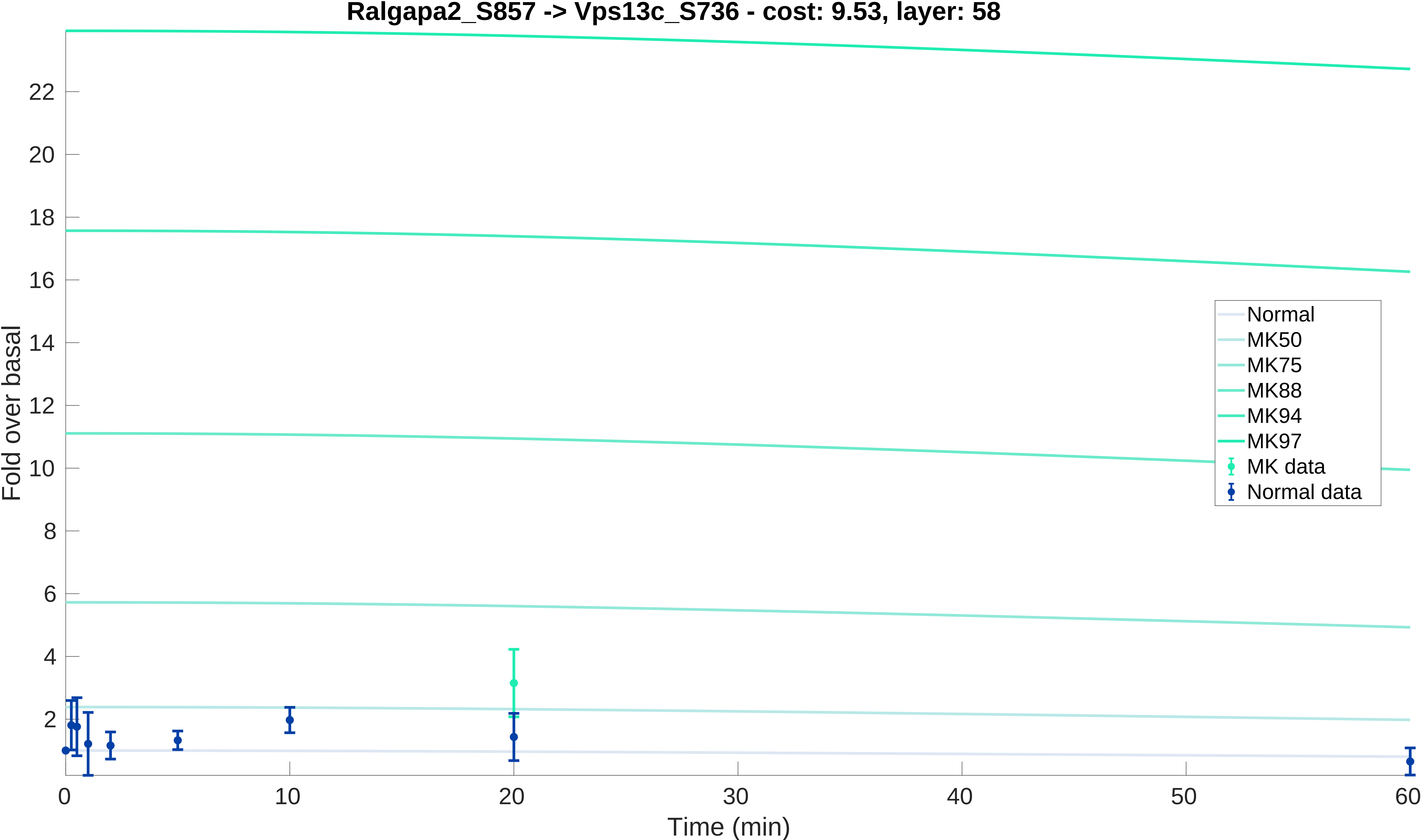

Ralgapa2_S857 -> Vps13c_S736 - cost: 9.53, layer: 58
Fold over basal
Time (min)
22
20
18
16
14
12
10
8
6
4
2
0
10
20
30
40
50
60
Normal
MK50
MK75
MK88
MK94
MK97
MK data
Normal data

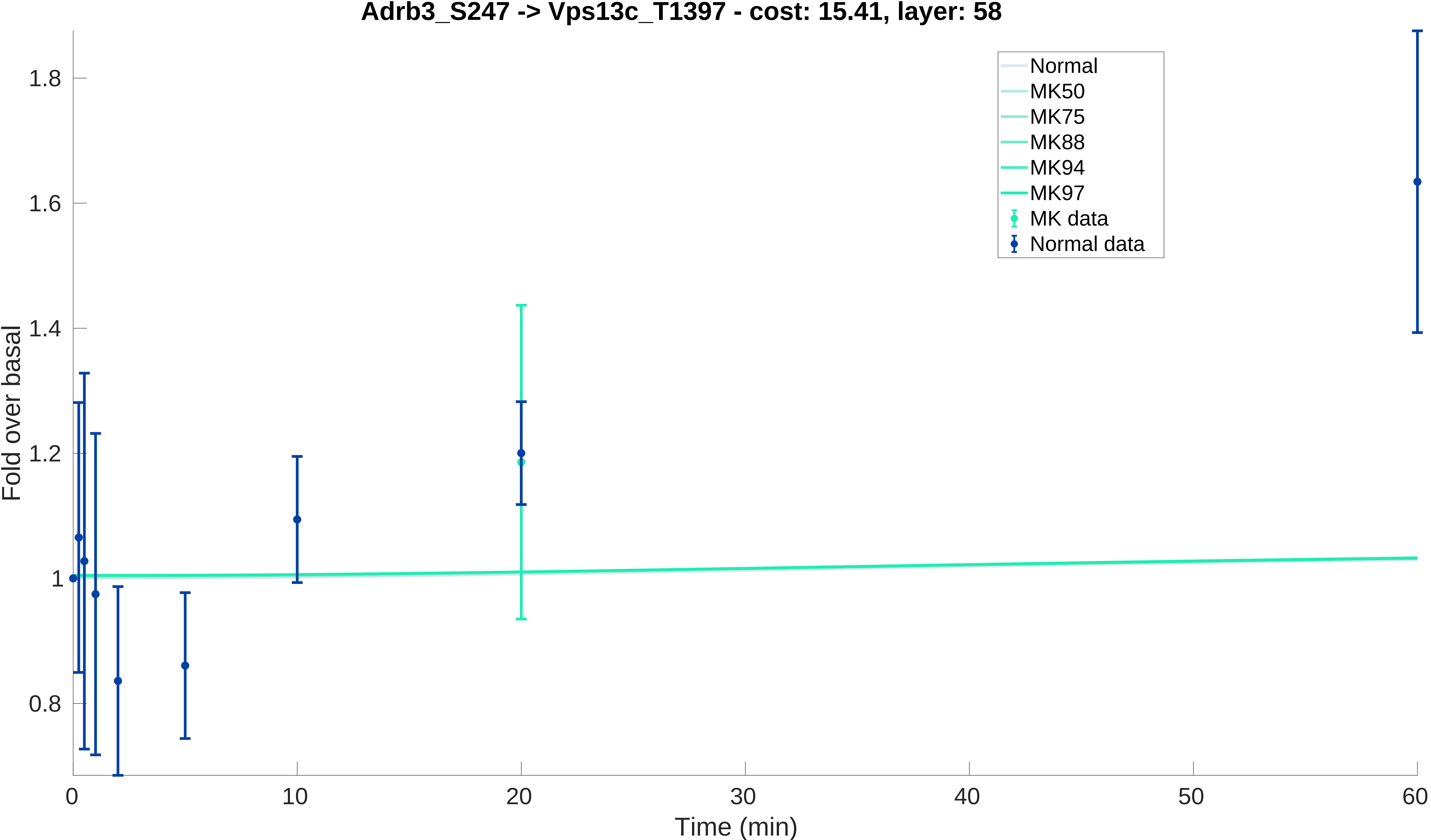
Adrb3_S247 -> Vps13c_T1397 - cost: 15.41, layer: 58
Fold over basal
Time (min)
1.8
1.6
1.4
1.2
1
0.8
0
10
20
30
40
50
60
Normal
MK50
MK75
MK88
MK94
MK97
MK data
Normal data

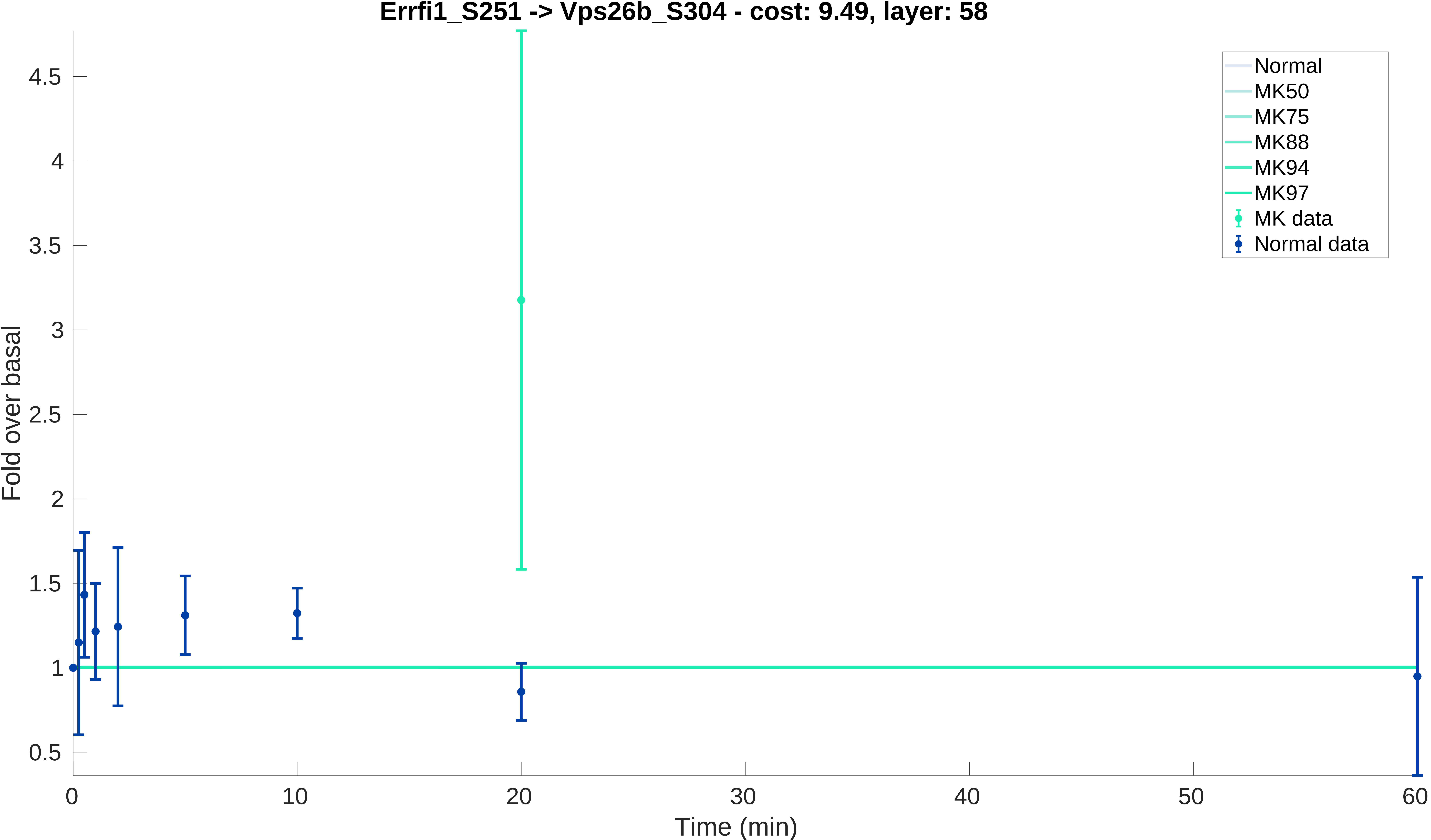

Errfi1_S251 -> Vps26b_S304 - cost: 9.49, layer: 58
Fold over basal
Time (min)
4.5
4
3.5
3
2.5
2
1.5
1
0.5
0
10
20
30
40
50
60
Normal
MK50
MK75
MK88
MK94
MK97
MK data
Normal data

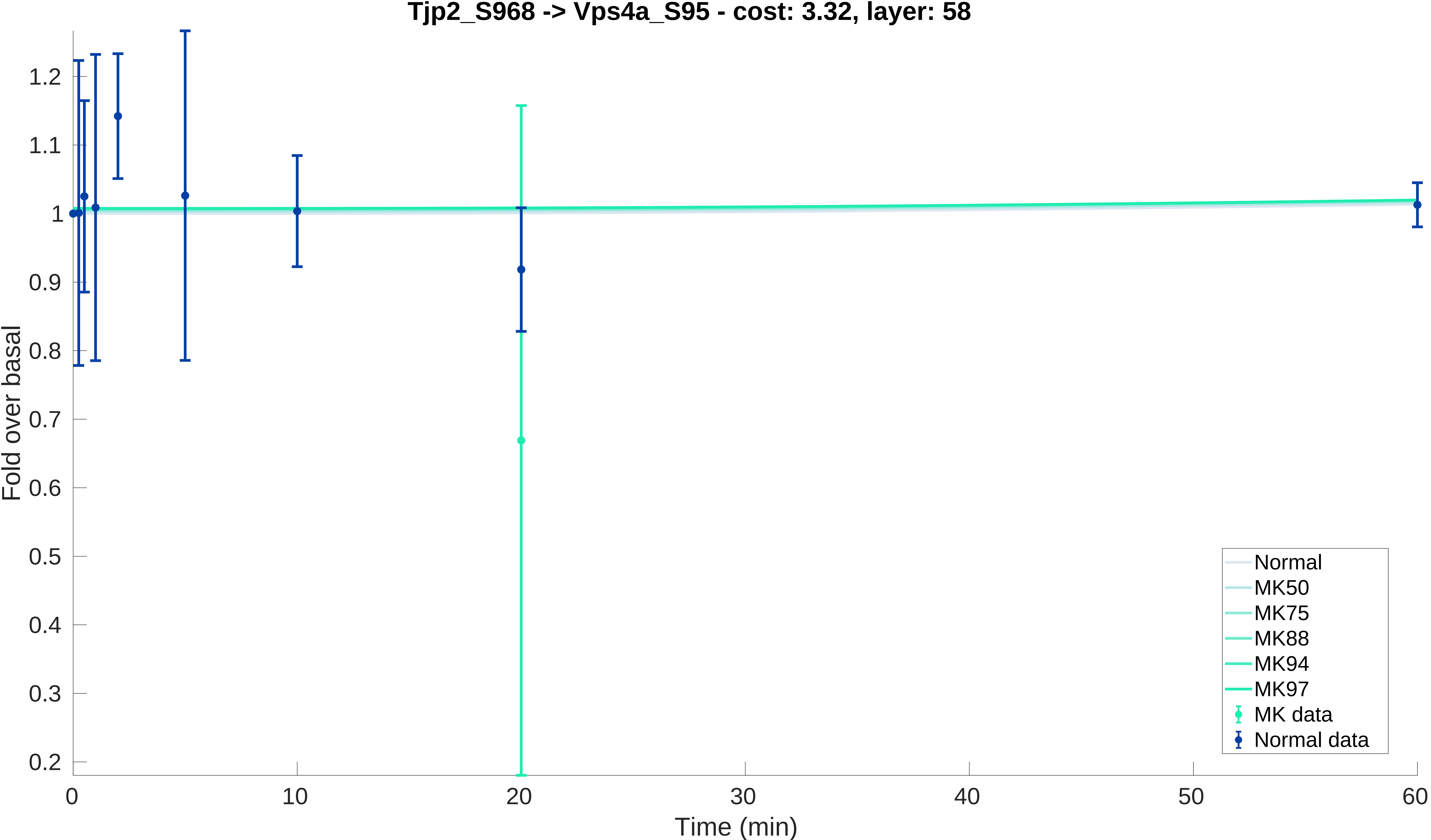

Tjp2_S968 -> Vps4a_S95 - cost: 3.32, layer: 58
Fold over basal
Time (min)
0
10
20
30
40
50
60
0.2
0.3
0.4
0.5
0.6
0.7
0.8
0.9
1
1.1
1.2
Normal
MK50
MK75
MK88
MK94
MK97
MK data
Normal data

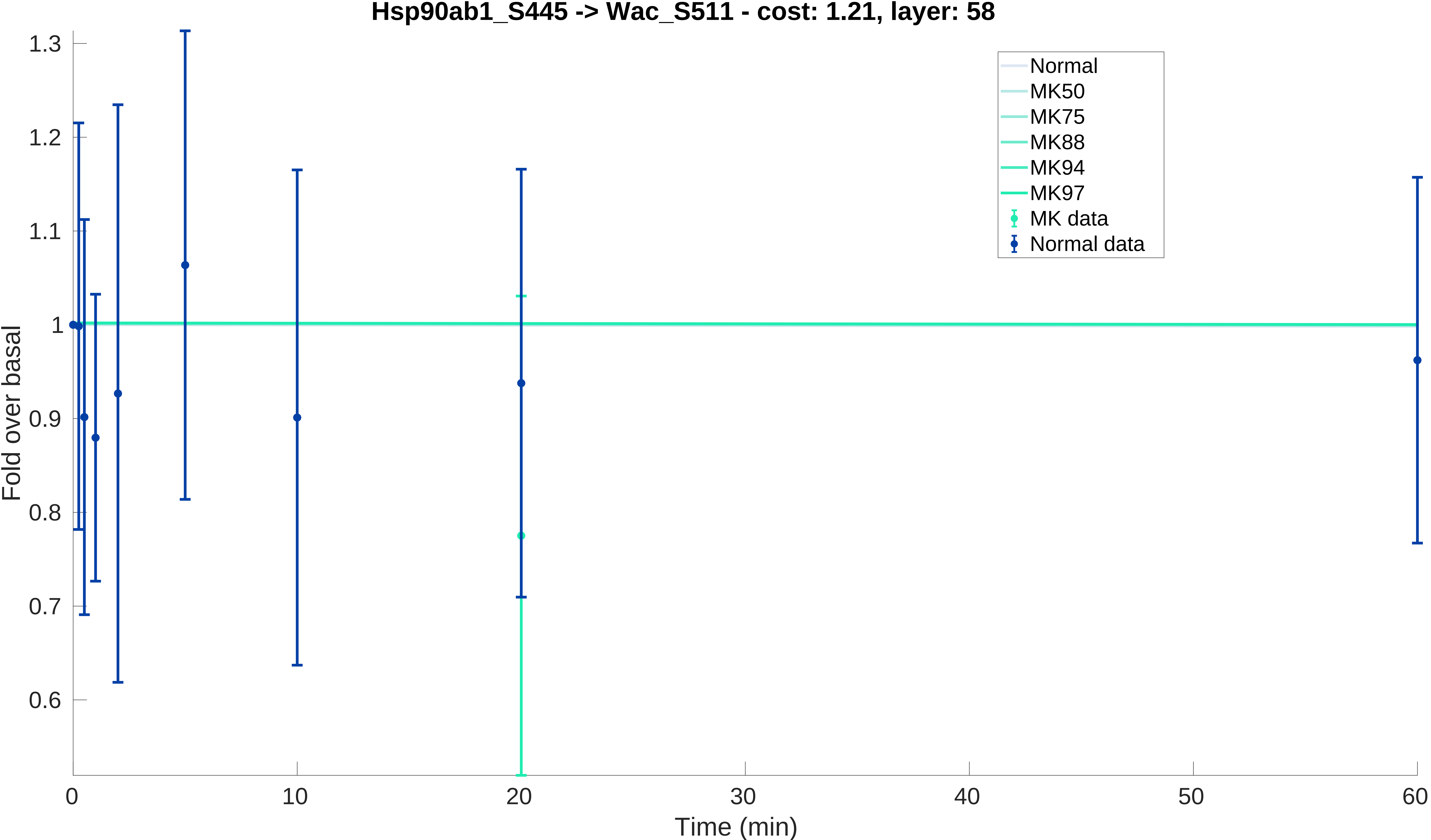

Hsp90ab1_S445 -> Wac_S511 - cost: 1.21, layer: 58
Fold over basal
1.3
1.2
1.1
1
0.9
0.8
0.7
0.6
0
10
20
30
40
50
60
Time (min)
Normal
MK50
MK75
MK88
MK94
MK97
MK data
Normal data

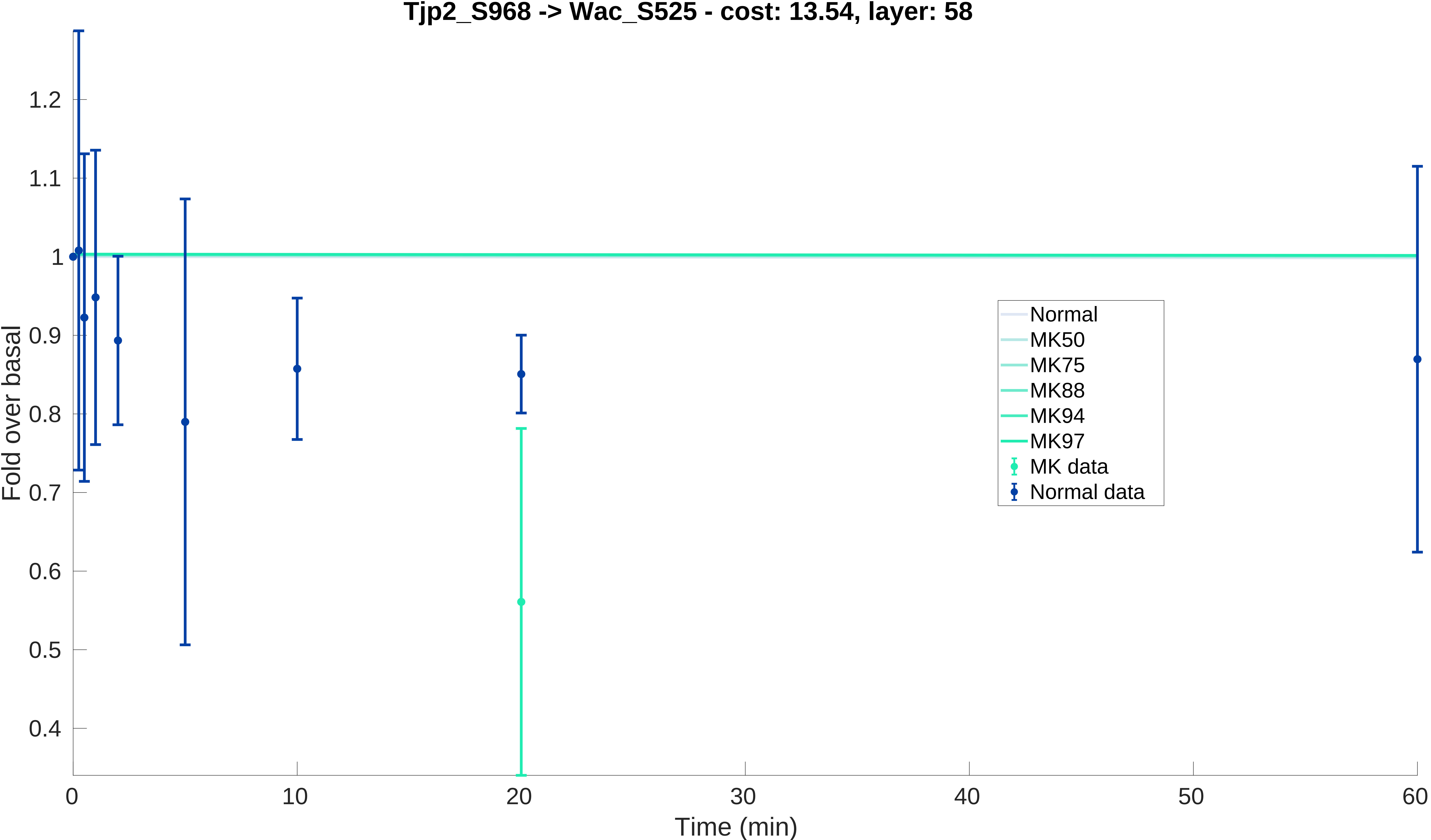

Tjp2_S968 -> Wac_S525 - cost: 13.54, layer: 58
Fold over basal
1.2
1.1
1
0.9
0.8
0.7
0.6
0.5
0.4
0
10
20
30
40
50
60
Time (min)
Normal
MK50
MK75
MK88
MK94
MK97
MK data
Normal data

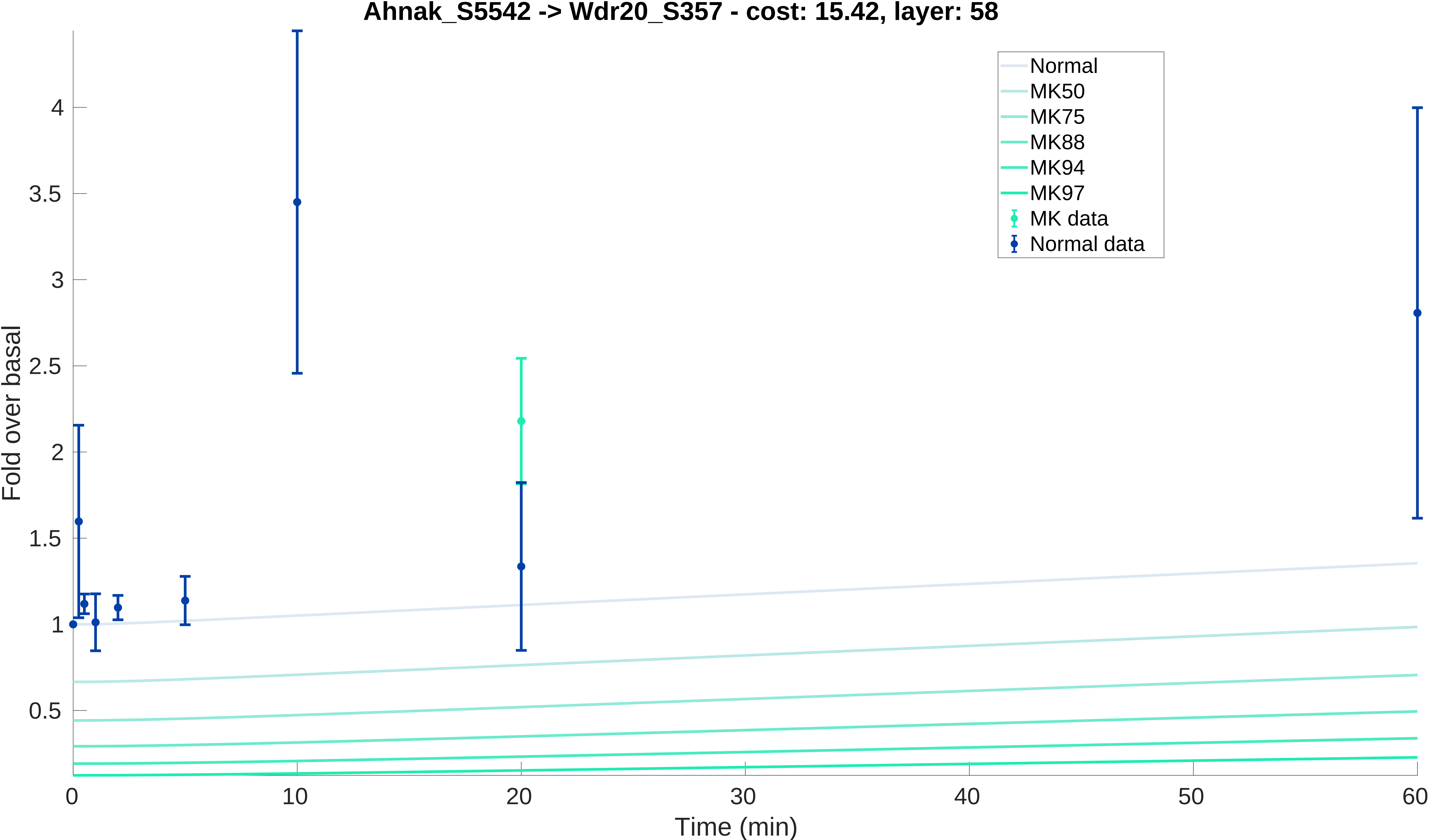

Ahnak_S5542 -> Wdr20_S357 - cost: 15.42, layer: 58
Normal
MK50
MK75
MK88
MK94
MK97
MK data
Normal data
Fold over basal
Time (min)
0
10
20
30
40
50
60
0.5
1
1.5
2
2.5
3
3.5
4

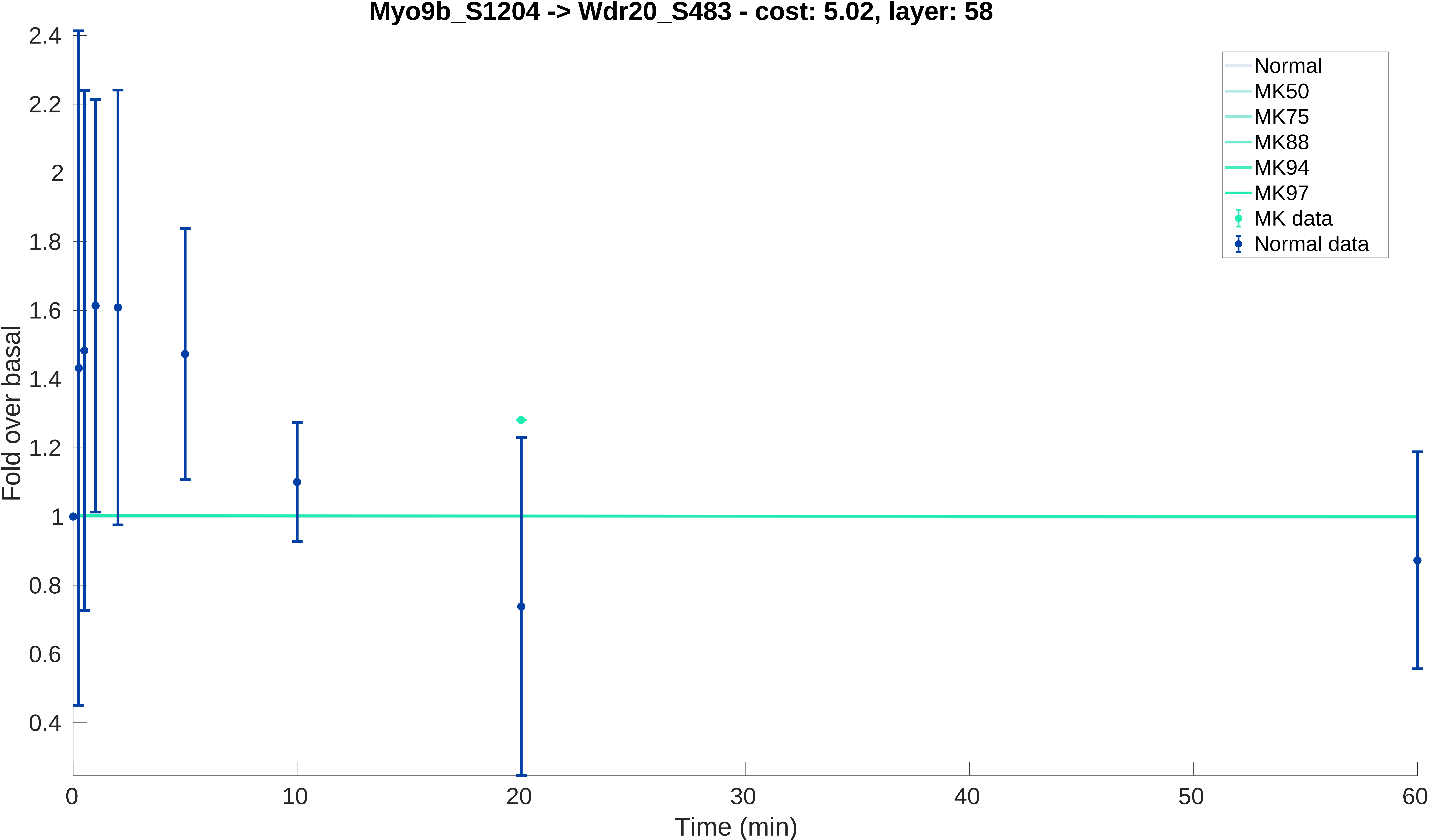

Myo9b_S1204 -> Wdr20_S483 - cost: 5.02, layer: 58
Fold over basal
2.4
2.2
2
1.8
1.6
1.4
1.2
1
0.8
0.6
0.4
0
10
20
30
40
50
60
Time (min)
Normal
MK50
MK75
MK88
MK94
MK97
MK data
Normal data

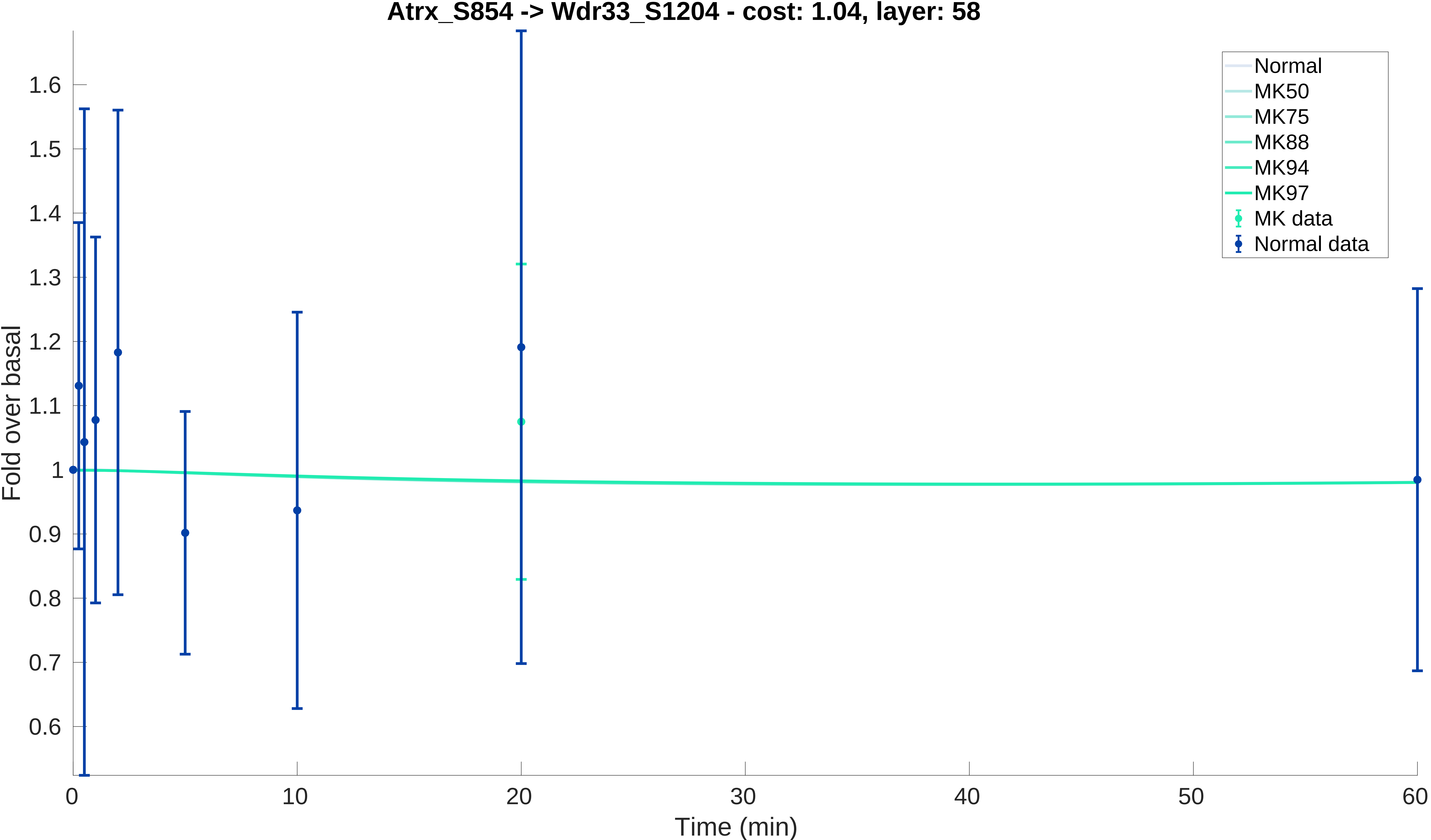
Atrx_S854 -> Wdr33_S1204 - cost: 1.04, layer: 58
Fold over basal
Time (min)
Normal
MK50
MK75
MK88
MK94
MK97
MK data
Normal data
0
10
20
30
40
50
60
0.6
0.7
0.8
0.9
1
1.1
1.2
1.3
1.4
1.5
1.6

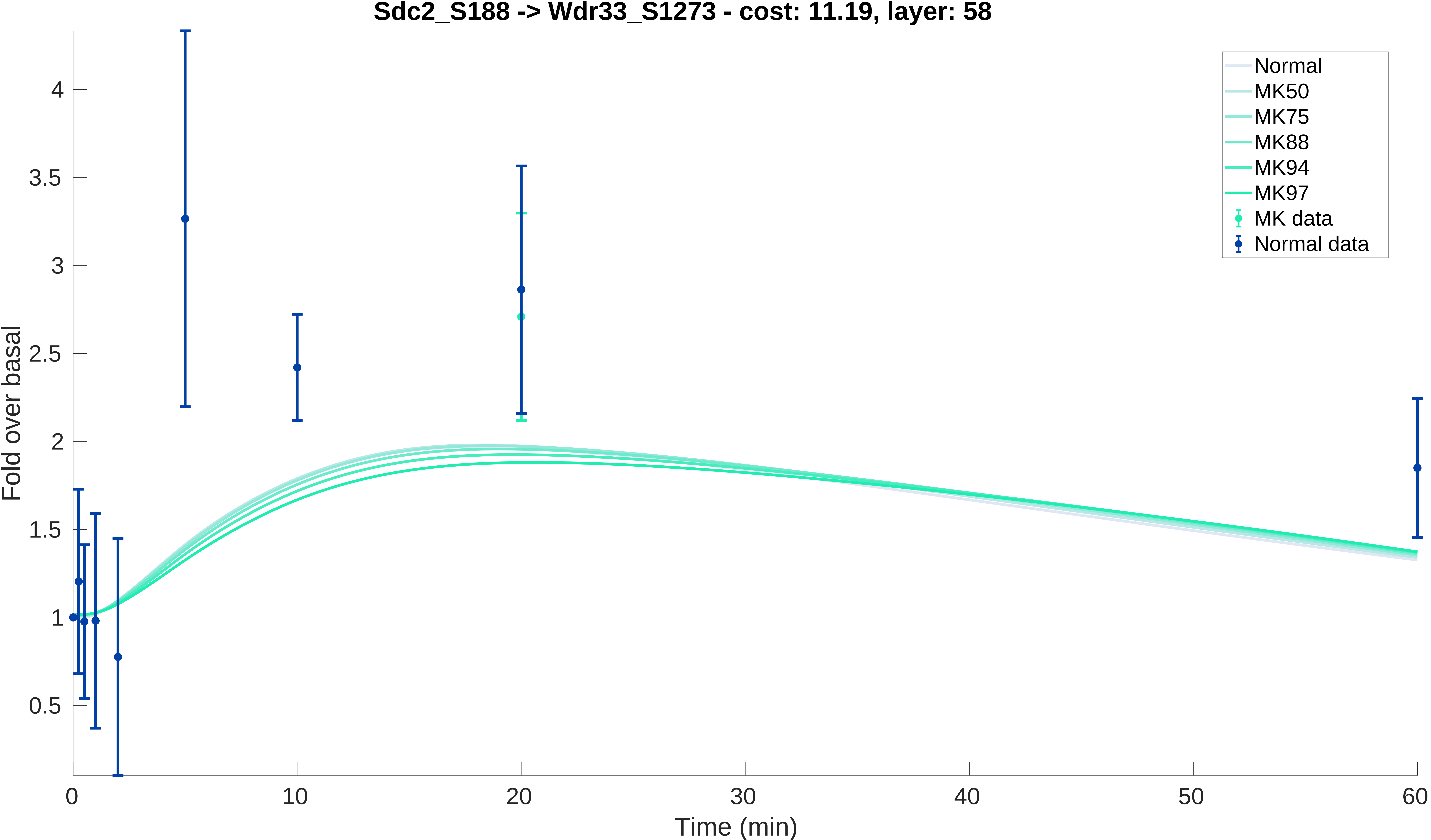

Sdc2_S188 -> Wdr33_S1273 - cost: 11.19, layer: 58
Fold over basal
Time (min)
0
10
20
30
40
50
60
0.5
1
1.5
2
2.5
3
3.5
4
Normal
MK50
MK75
MK88
MK94
MK97
MK data
Normal data

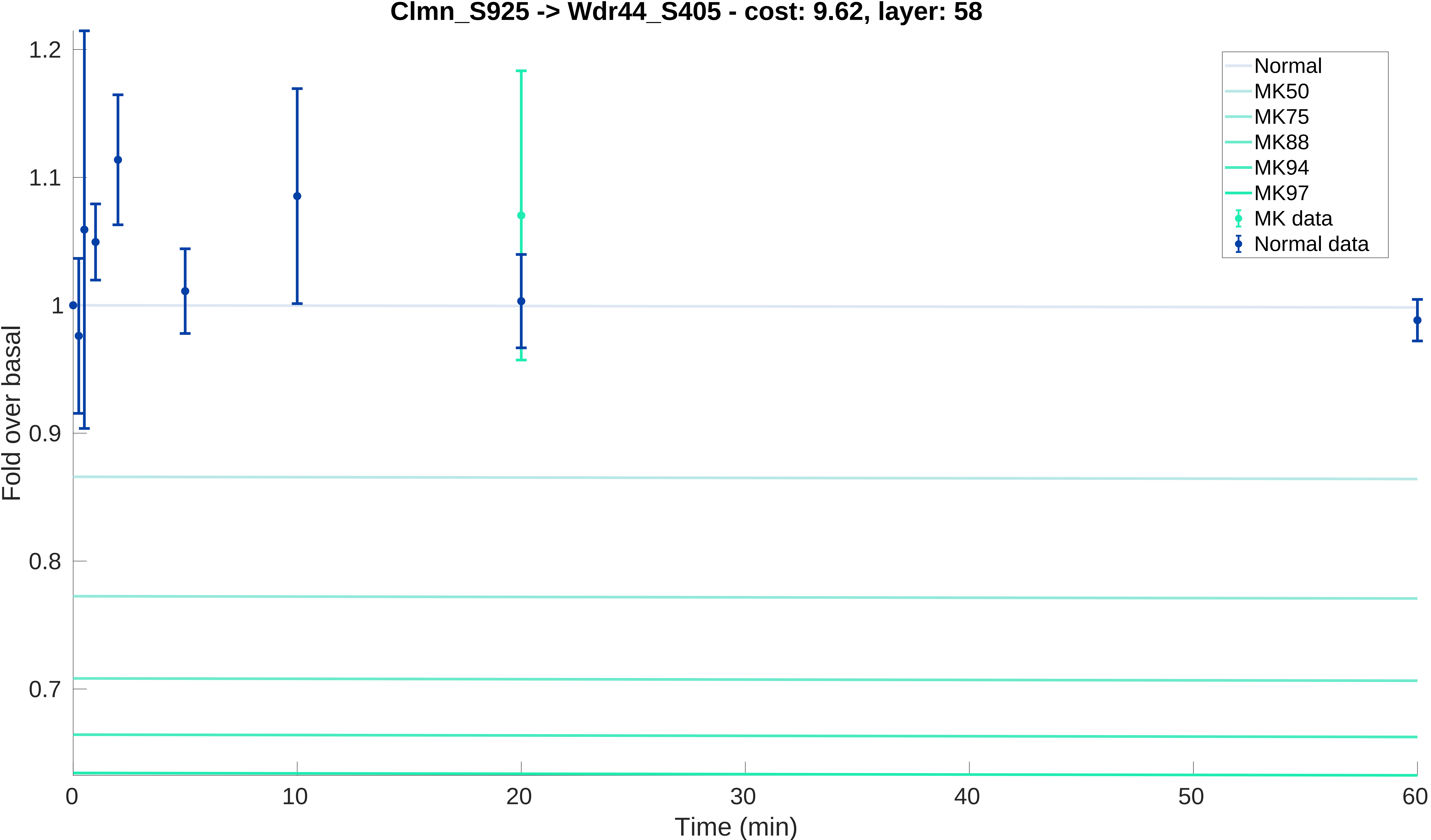

Clmn_S925 -> Wdr44_S405 - cost: 9.62, layer: 58
Fold over basal
Time (min)
1.2
1.1
1
0.9
0.8
0.7
0
10
20
30
40
50
60
Normal
MK50
MK75
MK88
MK94
MK97
MK data
Normal data

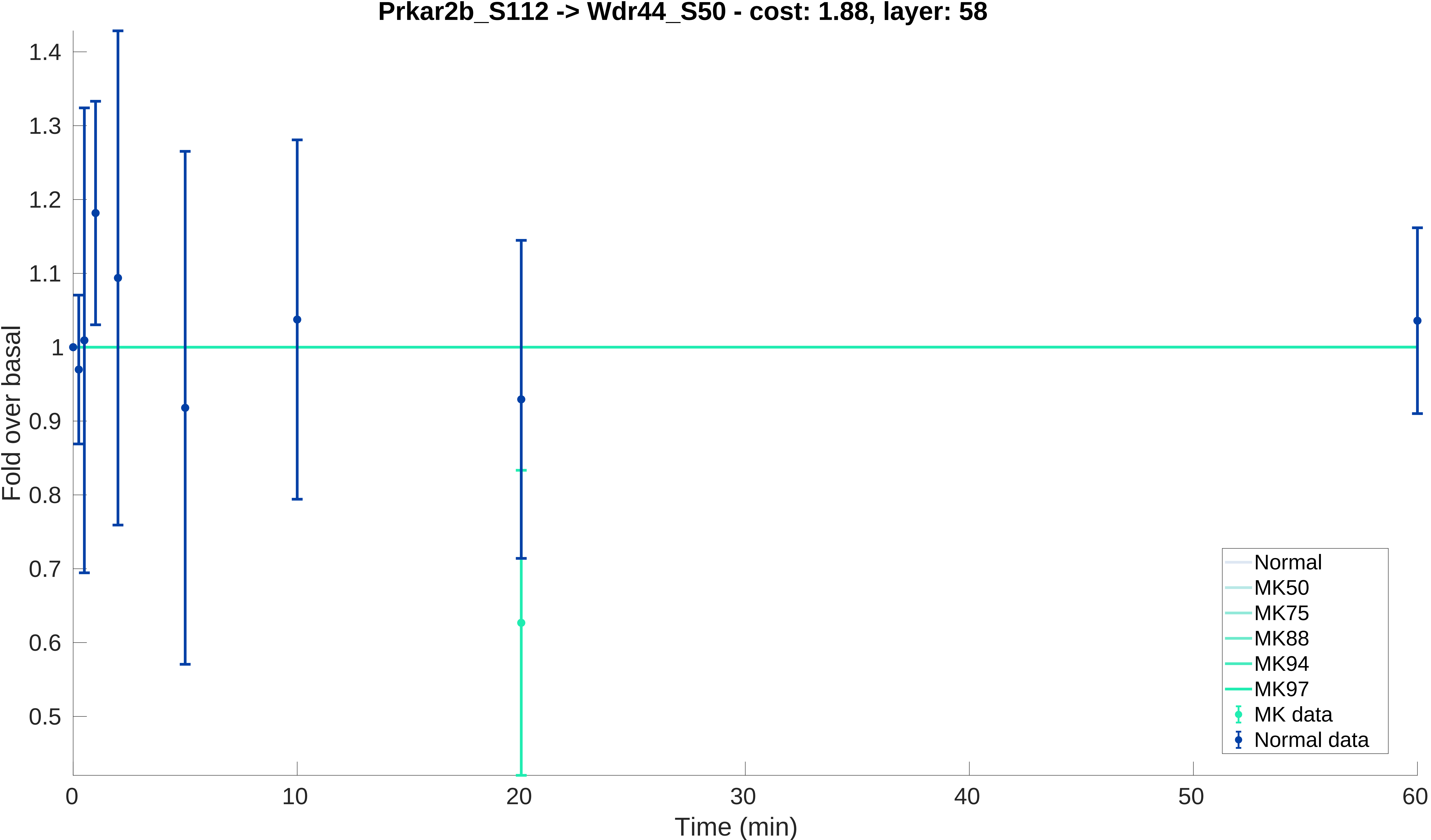

Prkar2b_S112 -> Wdr44_S50 - cost: 1.88, layer: 58
Fold over basal
Time (min)
1.4
1.3
1.2
1.1
1
0.9
0.8
0.7
0.6
0.5
0
10
20
30
40
50
60
Normal
MK50
MK75
MK88
MK94
MK97
MK data
Normal data

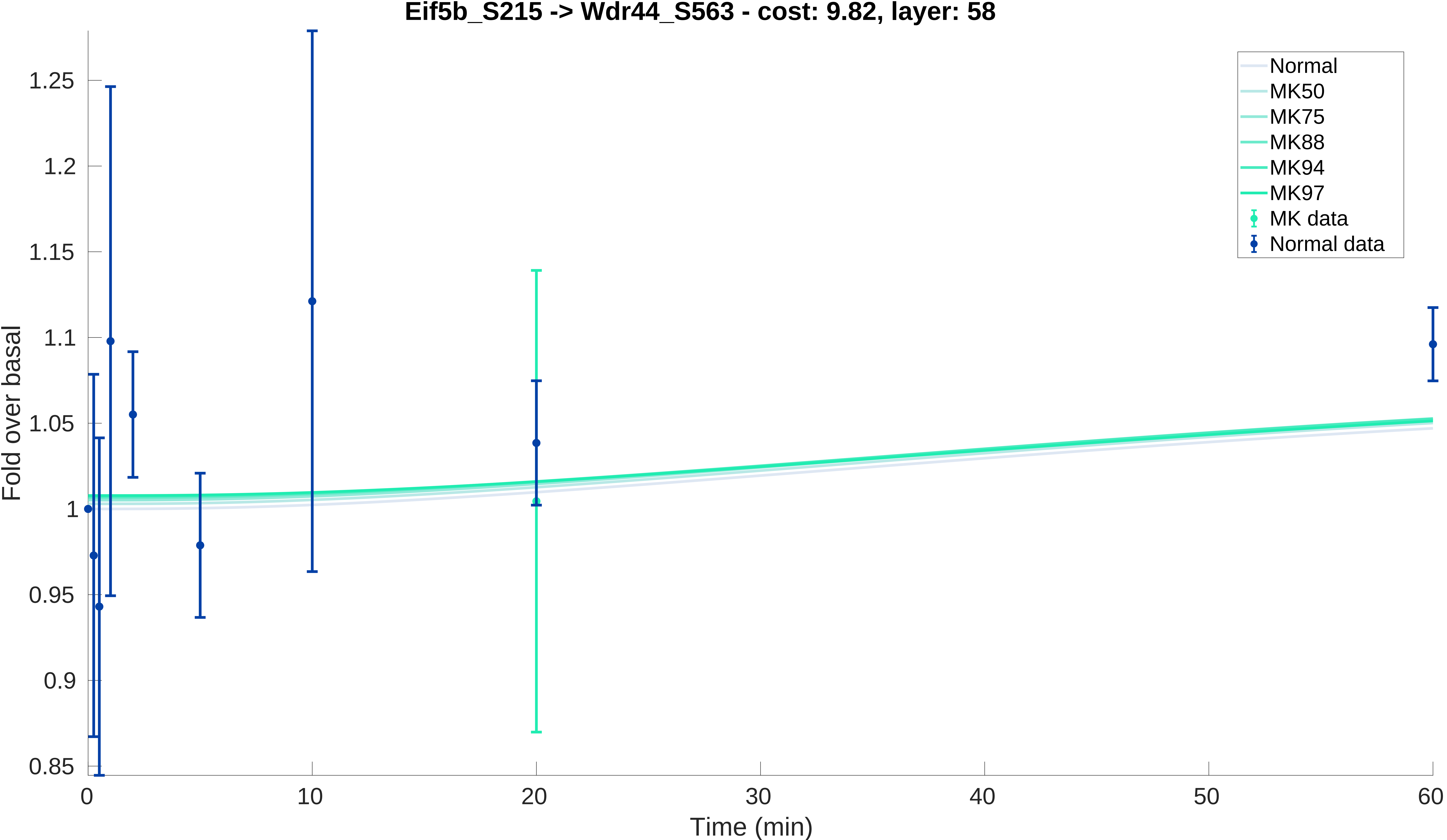

Eif5b_S215 -> Wdr44_S563 - cost: 9.82, layer: 58
Normal
MK50
MK75
MK88
MK94
MK97
MK data
Normal data
Fold over basal
1.25
1.2
1.15
1.1
1.05
1
0.95
0.9
0.85
0
10
20
30
40
50
60
Time (min)

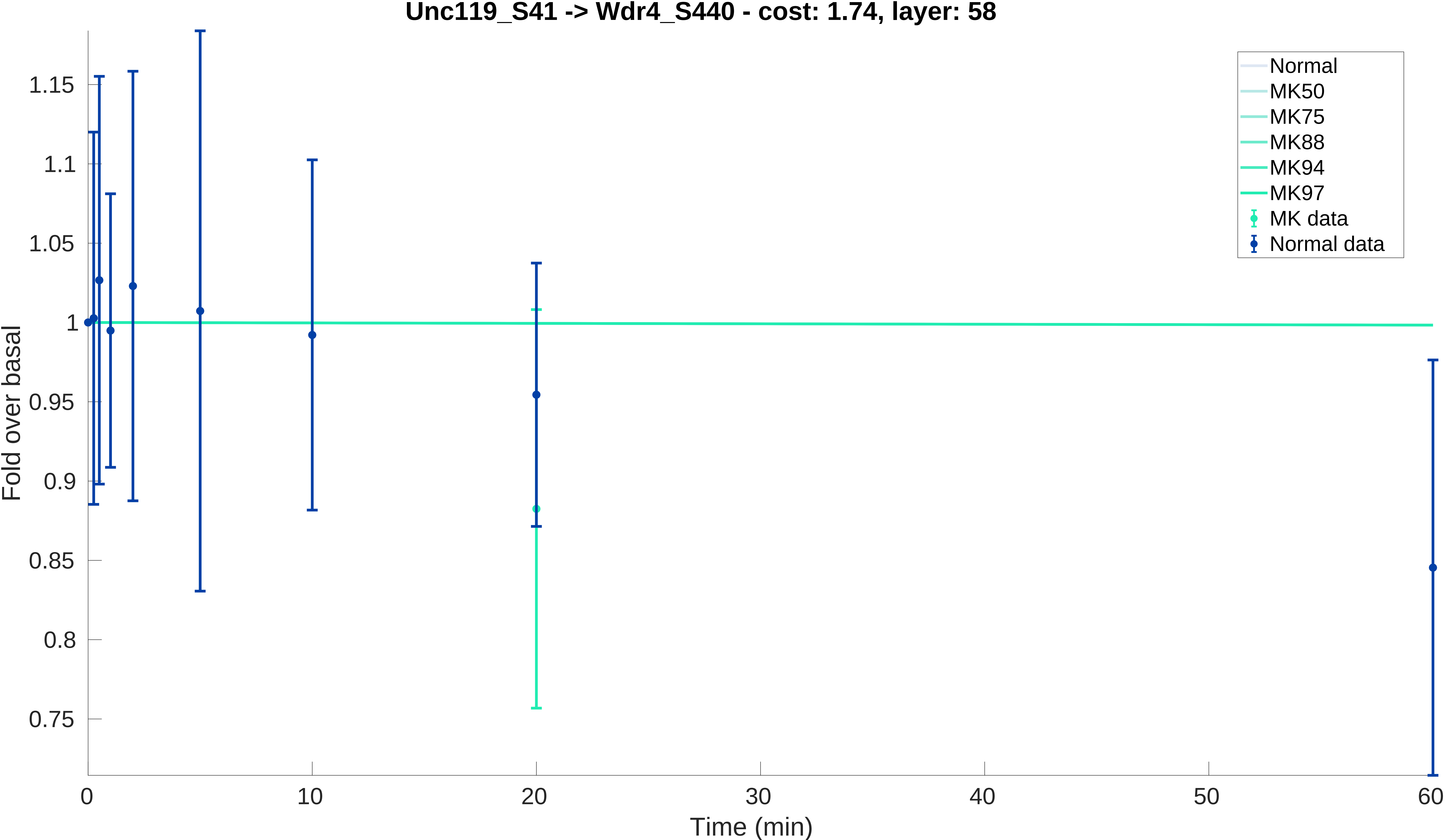

Unc119_S41 -> Wdr4_S440 - cost: 1.74, layer: 58
Normal
MK50
MK75
MK88
MK94
MK97
MK data
Normal data
Fold over basal
Time (min)
1.15
1.1
1.05
1
0.95
0.9
0.85
0.8
0.75
0
10
20
30
40
50
60

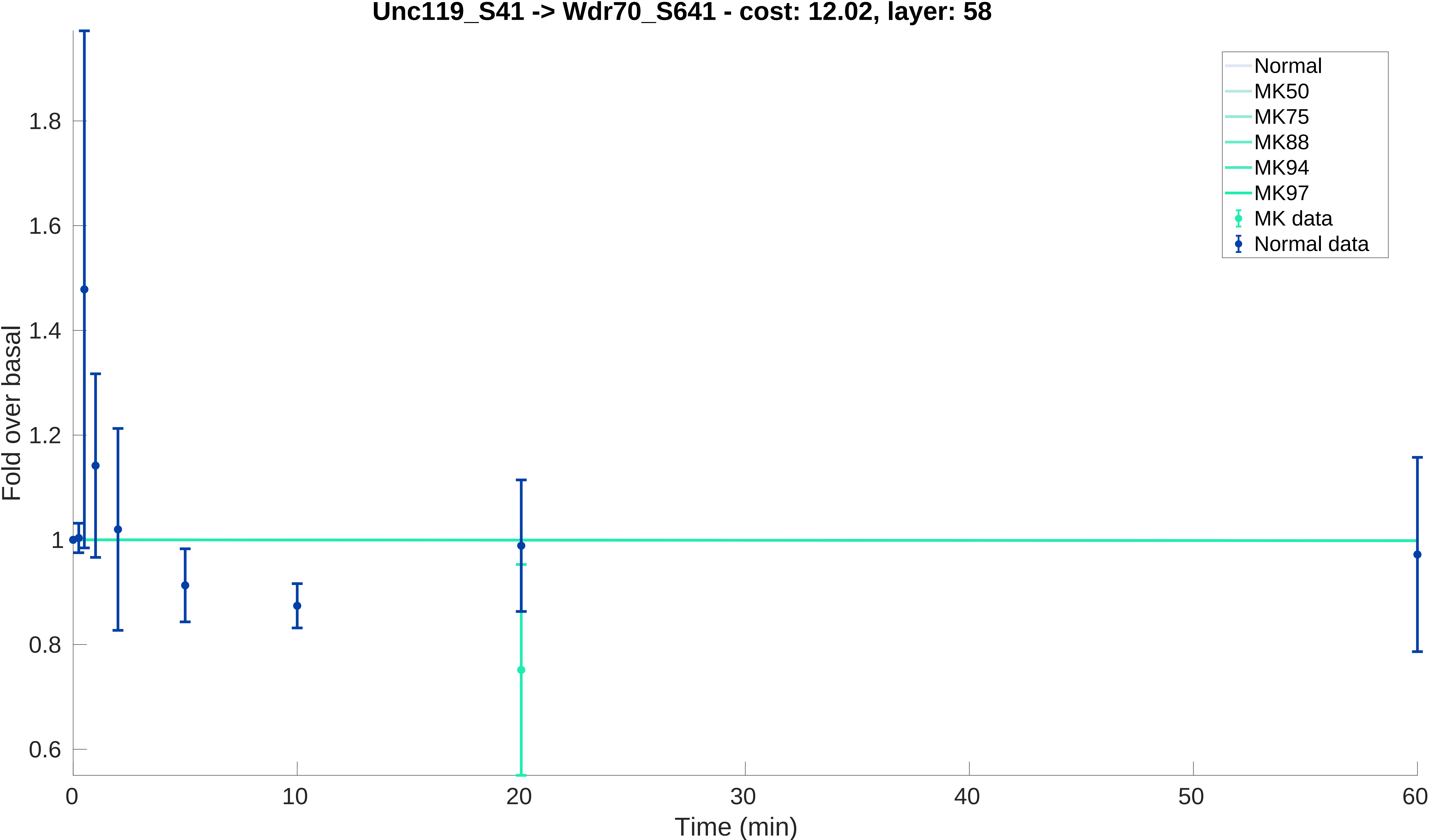

Unc119_S41 -> Wdr70_S641 - cost: 12.02, layer: 58
Normal
MK50
MK75
MK88
MK94
MK97
MK data
Normal data
Fold over basal
1.8
1.6
1.4
1.2
1
0.8
0.6
0
10
20
30
40
50
60
Time (min)

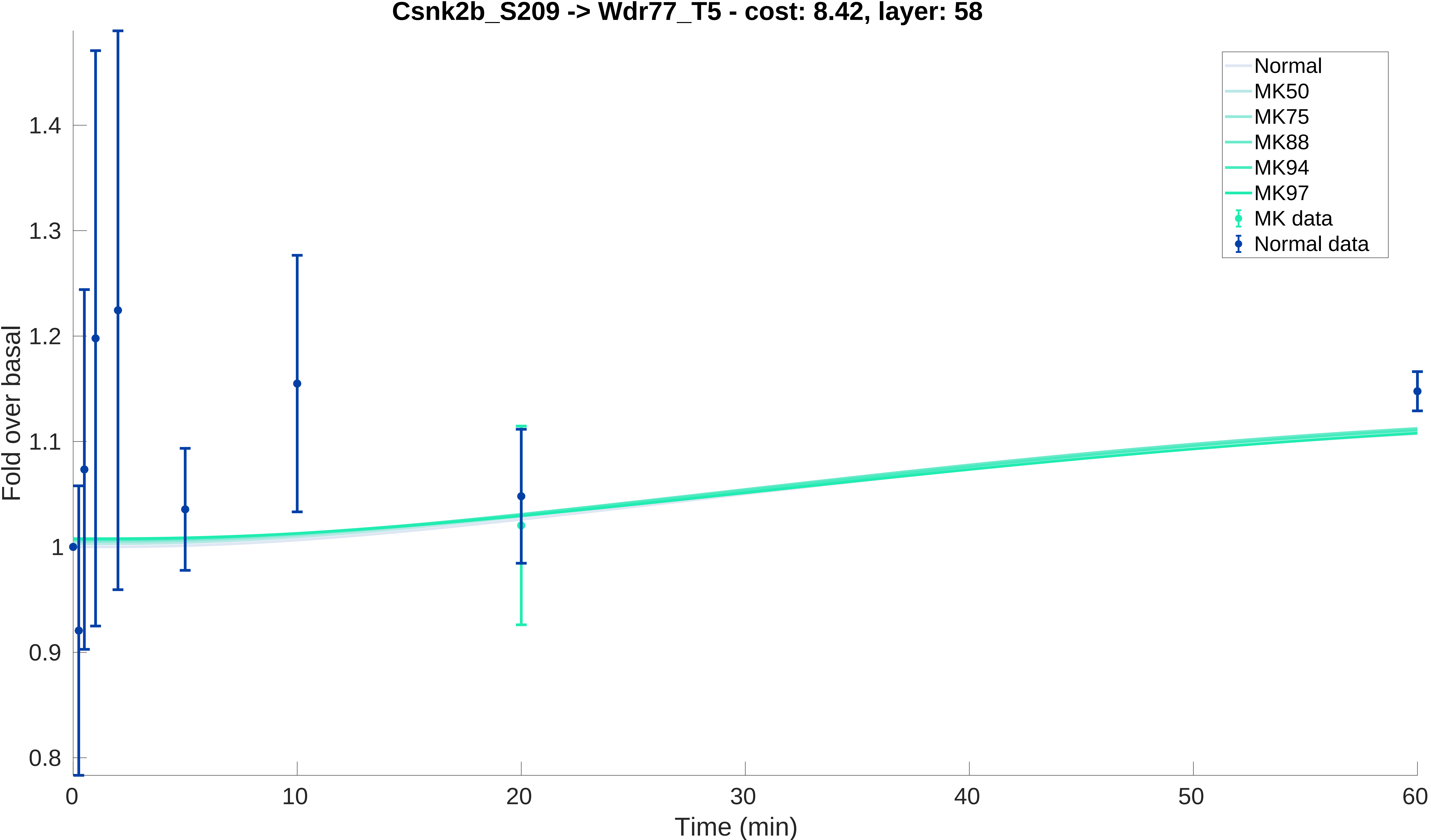

Csnk2b_S209 -> Wdr77_T5 - cost: 8.42, layer: 58
Fold over basal
Time (min)
1.4
1.3
1.2
1.1
1
0.9
0.8
0
10
20
30
40
50
60
Normal
MK50
MK75
MK88
MK94
MK97
MK data
Normal data

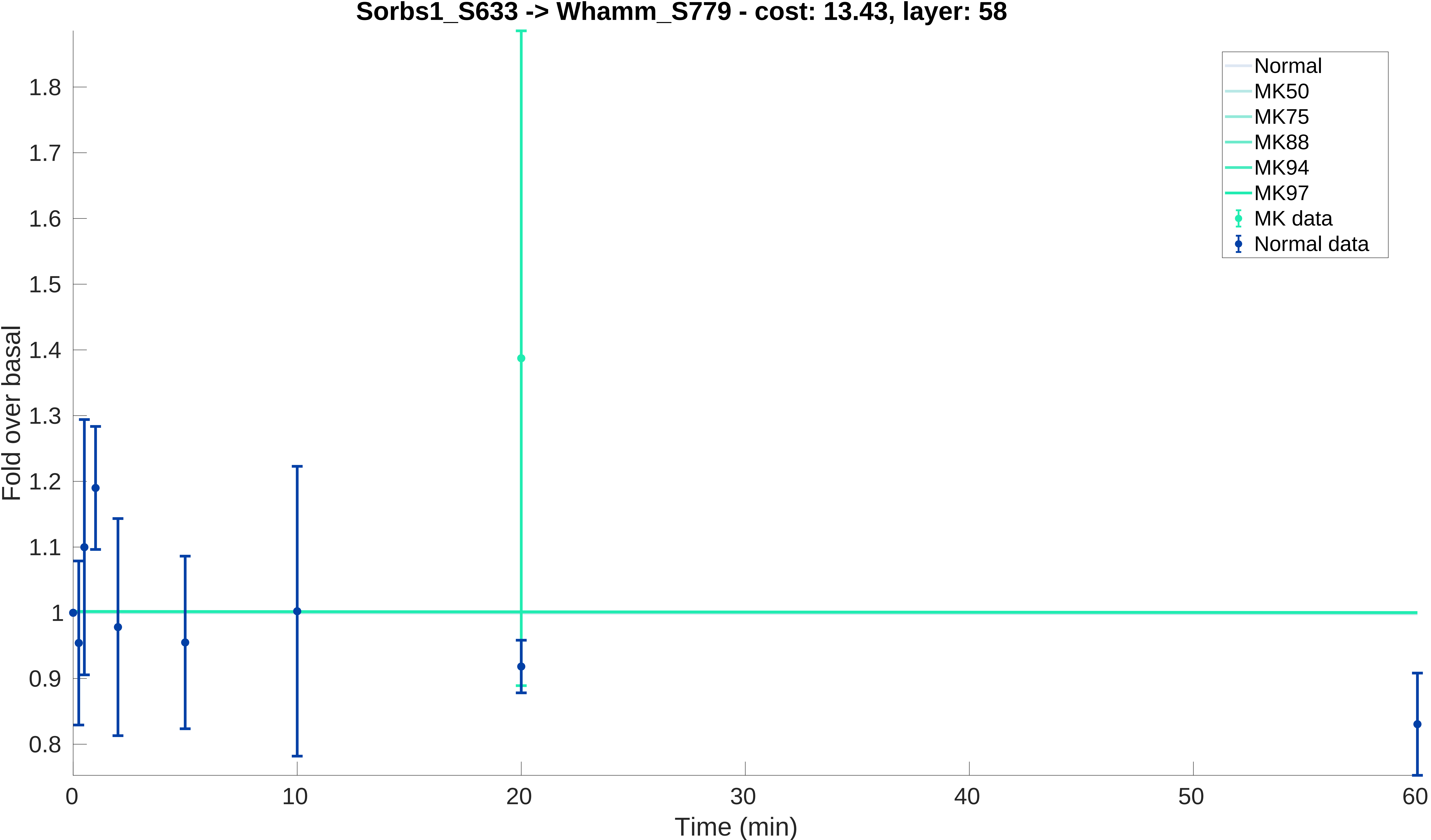
Sorbs1_S633 -> Whamm_S779 - cost: 13.43, layer: 58
Fold over basal
Time (min)
0
10
20
30
40
50
60
0.8
0.9
1
1.1
1.2
1.3
1.4
1.5
1.6
1.7
1.8
Normal
MK50
MK75
MK88
MK94
MK97
MK data
Normal data

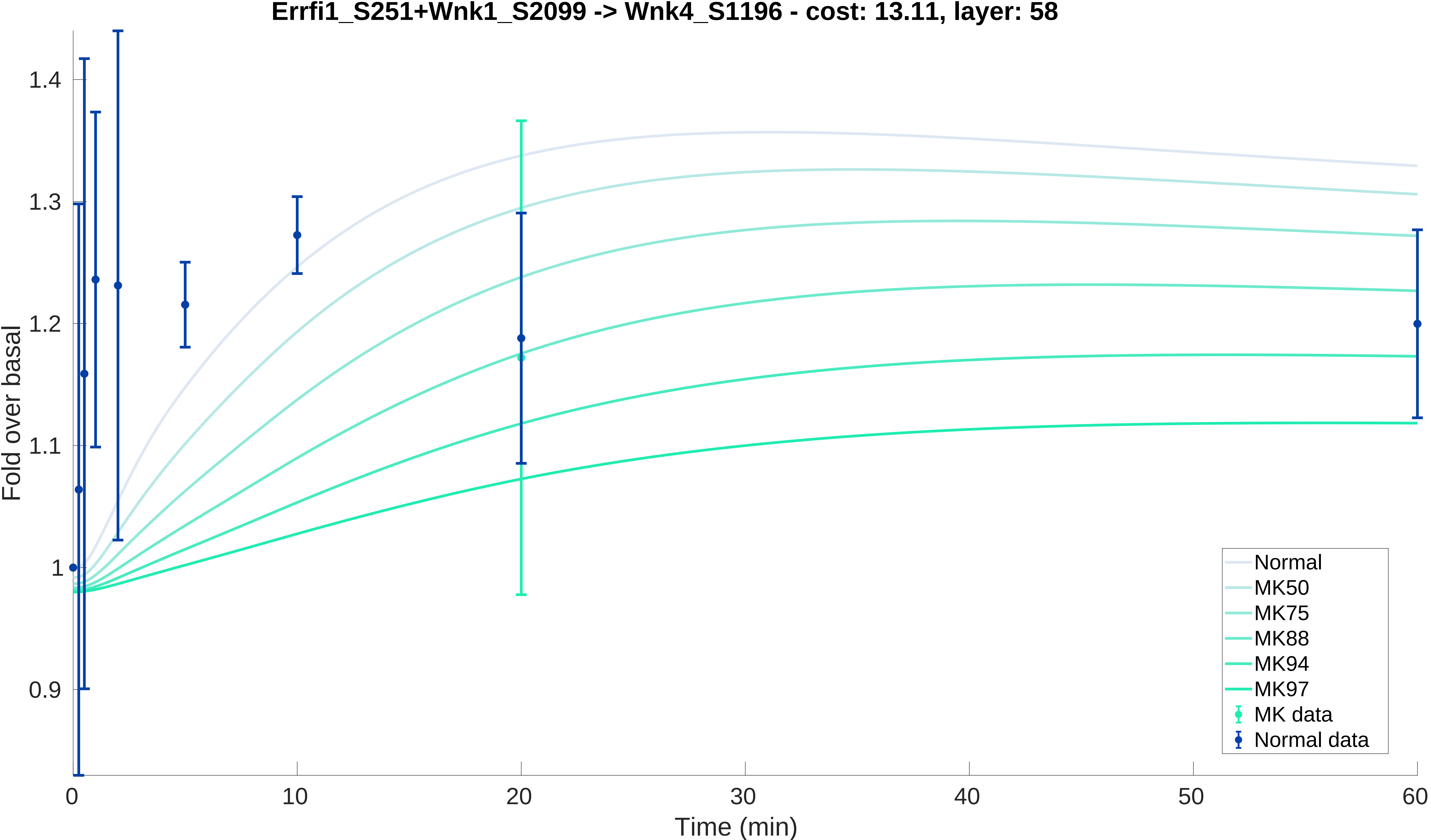

Errfi1_S251+Wnk1_S2099 -> Wnk4_S1196 - cost: 13.11, layer: 58
Fold over basal
1.4
1.3
1.2
1.1
1
0.9
0
10
20
30
40
50
60
Time (min)
Normal
MK50
MK75
MK88
MK94
MK97
MK data
Normal data

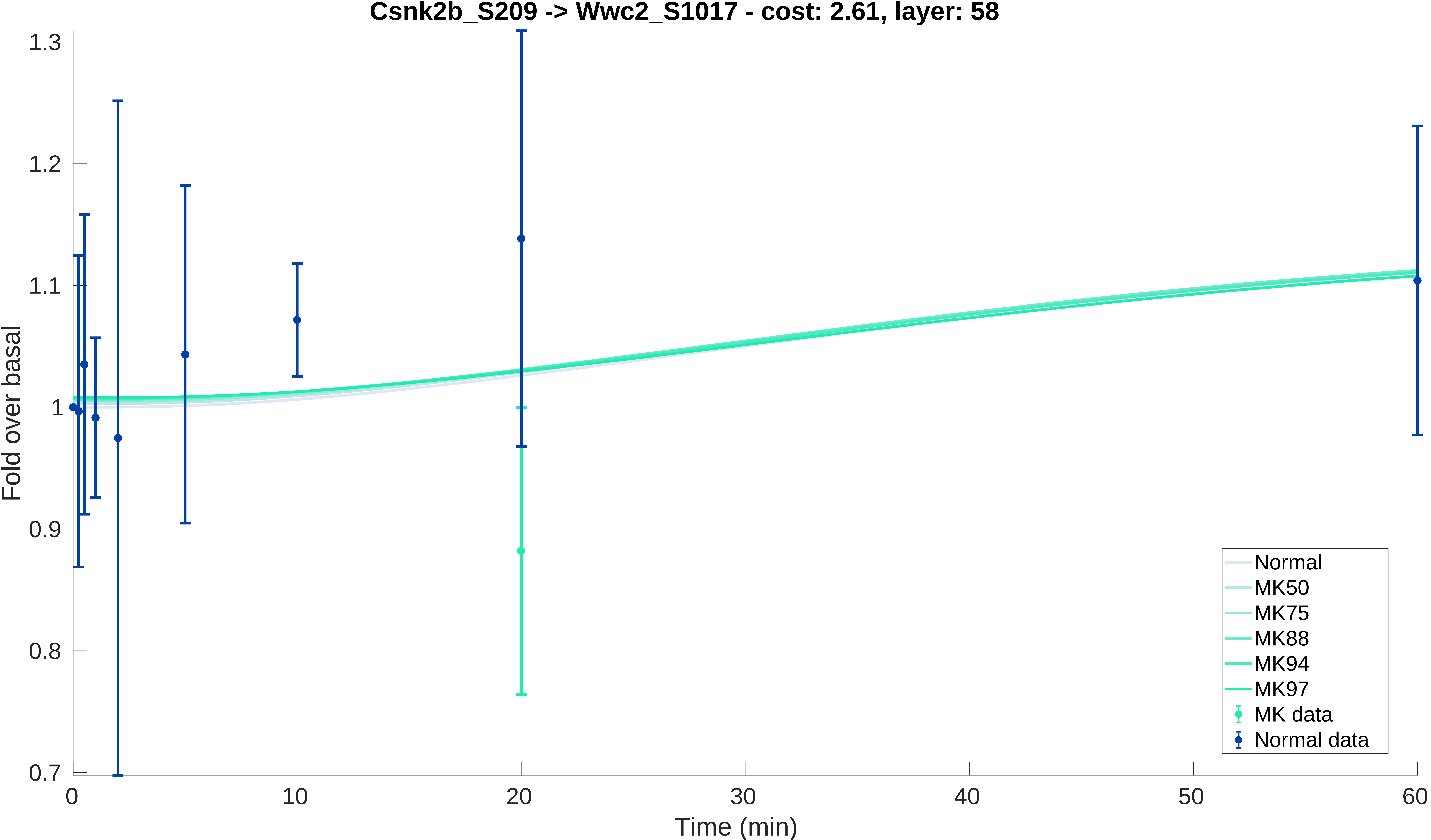

Csnk2b_S209 -> Wwc2_S1017 - cost: 2.61, layer: 58
Fold over basal
Time (min)
1.3
1.2
1.1
1
0.9
0.8
0.7
0
10
20
30
40
50
60
Normal
MK50
MK75
MK88
MK94
MK97
MK data
Normal data

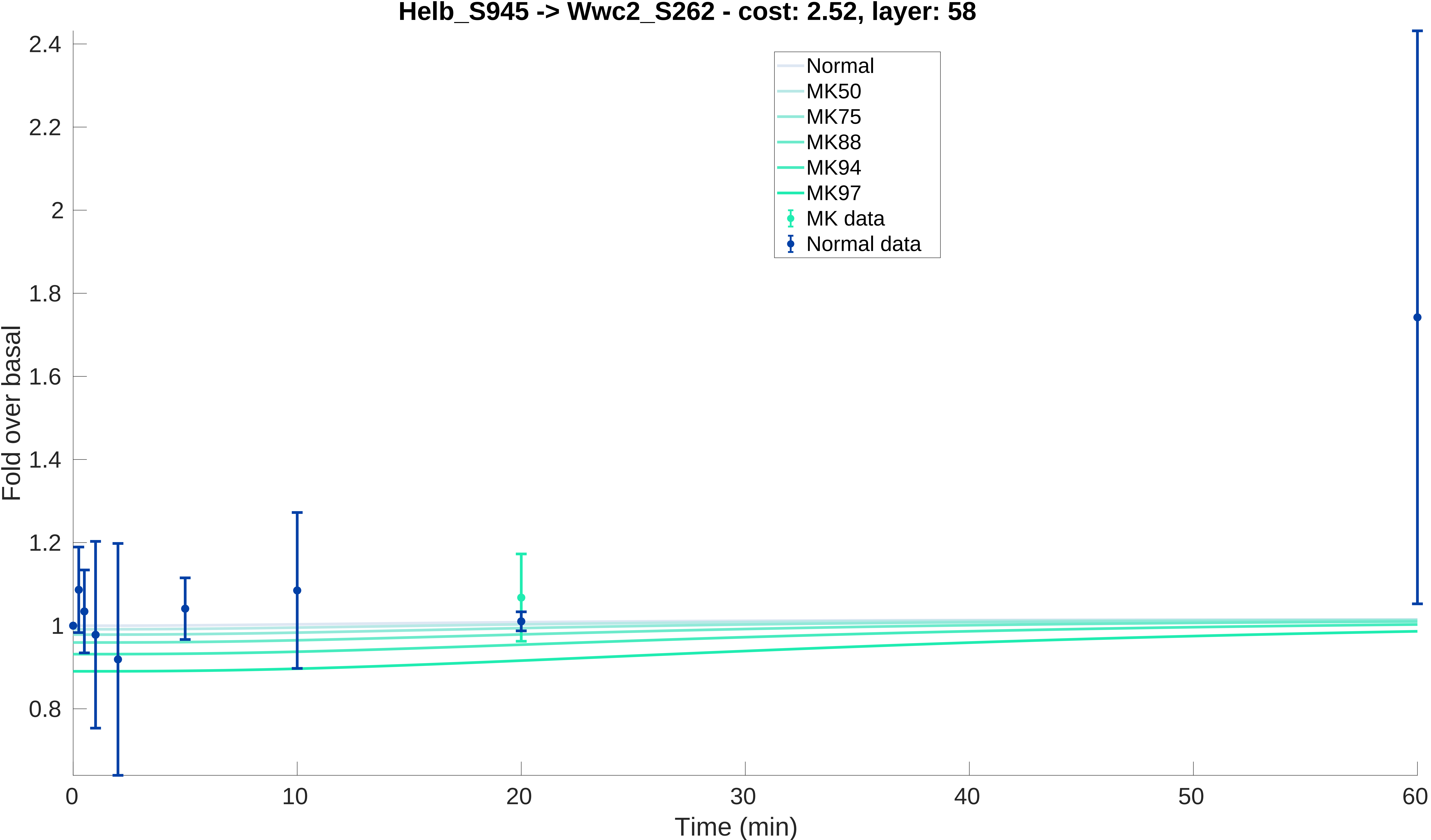

Helb_S945 -> Wwc2_S262 - cost: 2.52, layer: 58
Normal
MK50
MK75
MK88
MK94
MK97
MK data
Normal data
Fold over basal
2.4
2.2
2
1.8
1.6
1.4
1.2
1
0.8
0
10
20
30
40
50
60
Time (min)

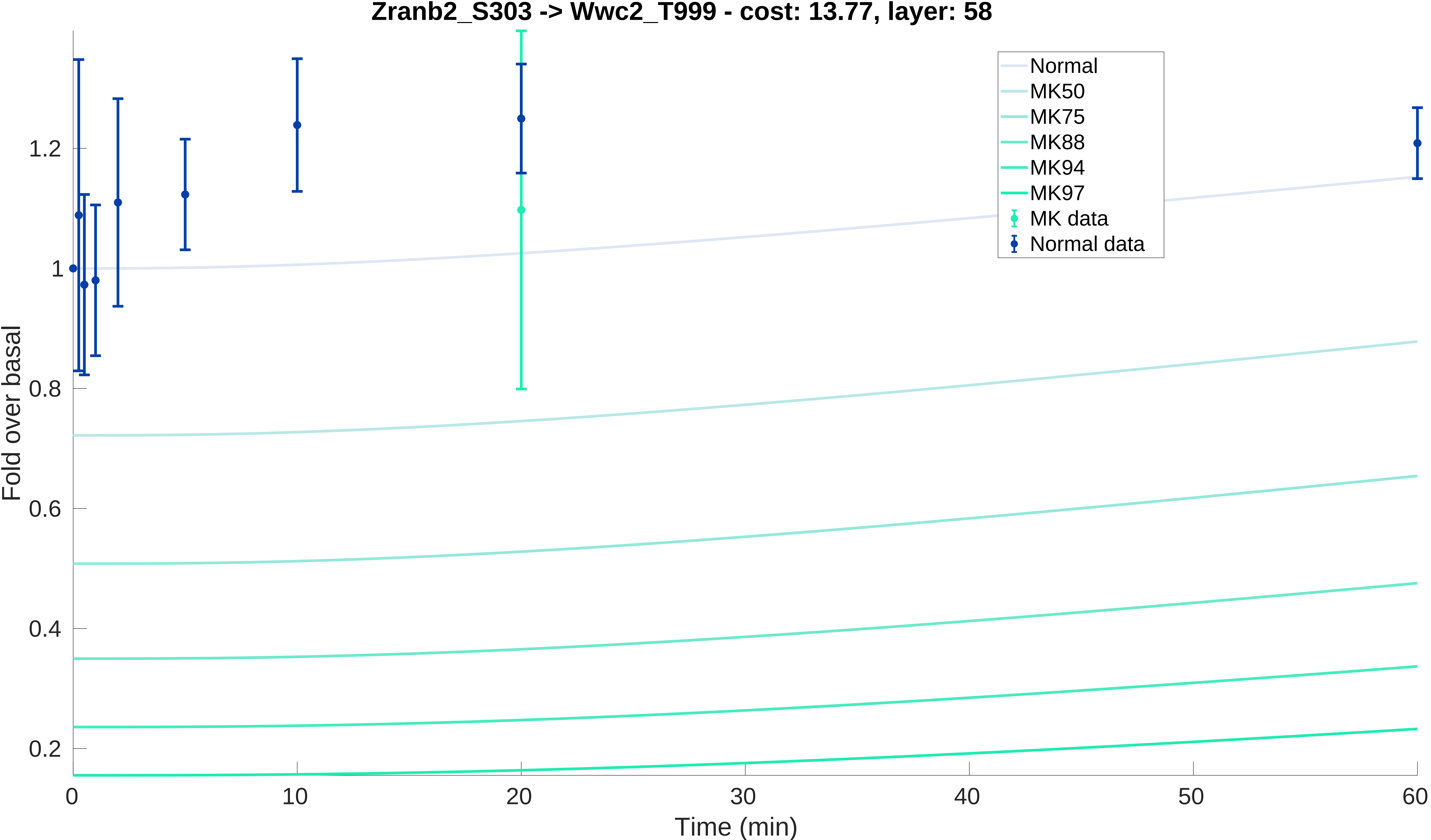

Zranb2_S303 -> Wwc2_T999 - cost: 13.77, layer: 58
Normal
MK50
MK75
MK88
MK94
MK97
MK data
Normal data
Fold over basal
1.2
1
0.8
0.6
0.4
0.2
0
10
20
30
40
50
60
Time (min)

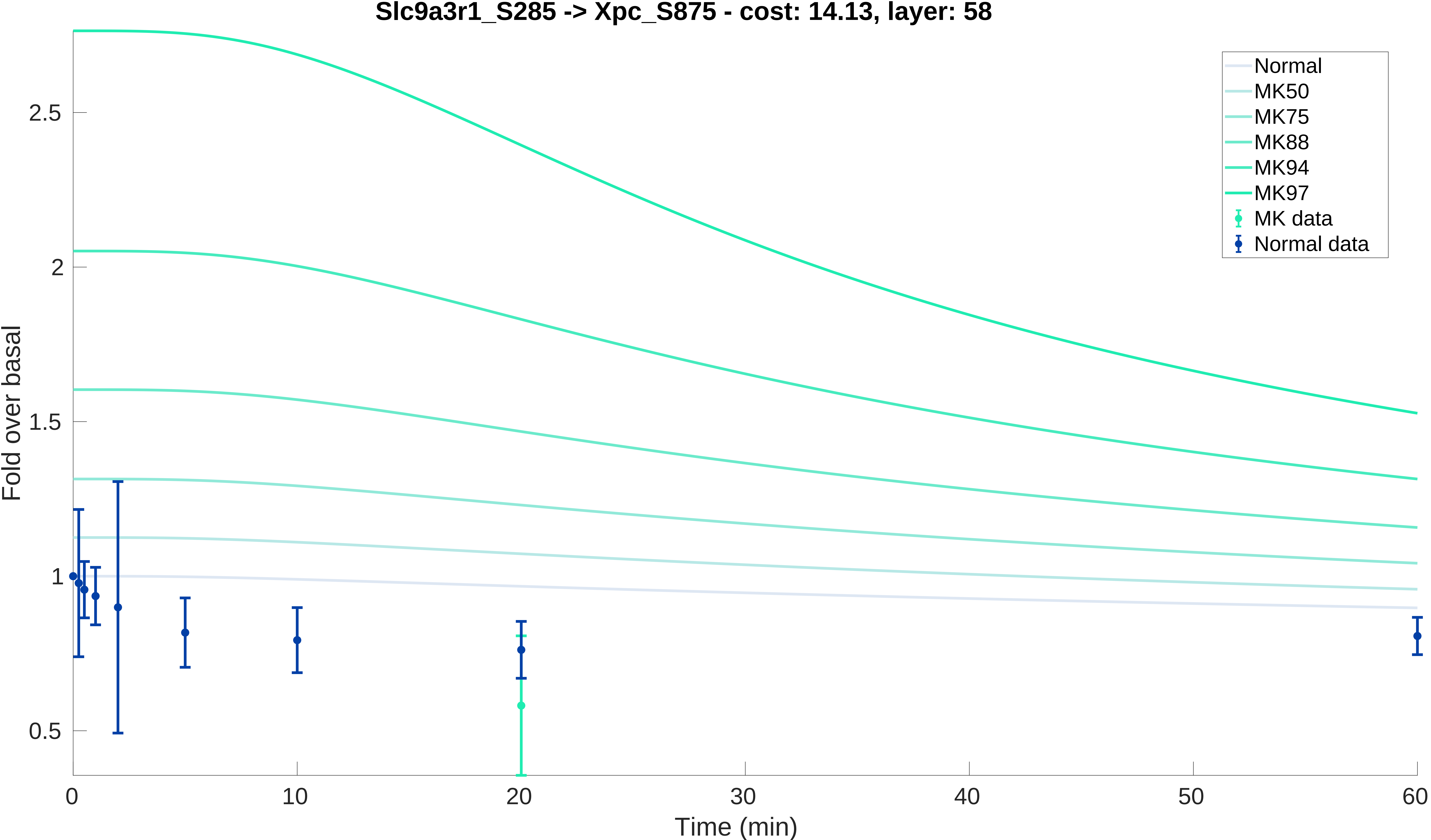
Slc9a3r1_S285 -> Xpc_S875 - cost: 14.13, layer: 58
Normal
MK50
MK75
MK88
MK94
MK97
MK data
Normal data
Fold over basal
Time (min)
0.5
1
1.5
2
2.5
0
10
20
30
40
50
60

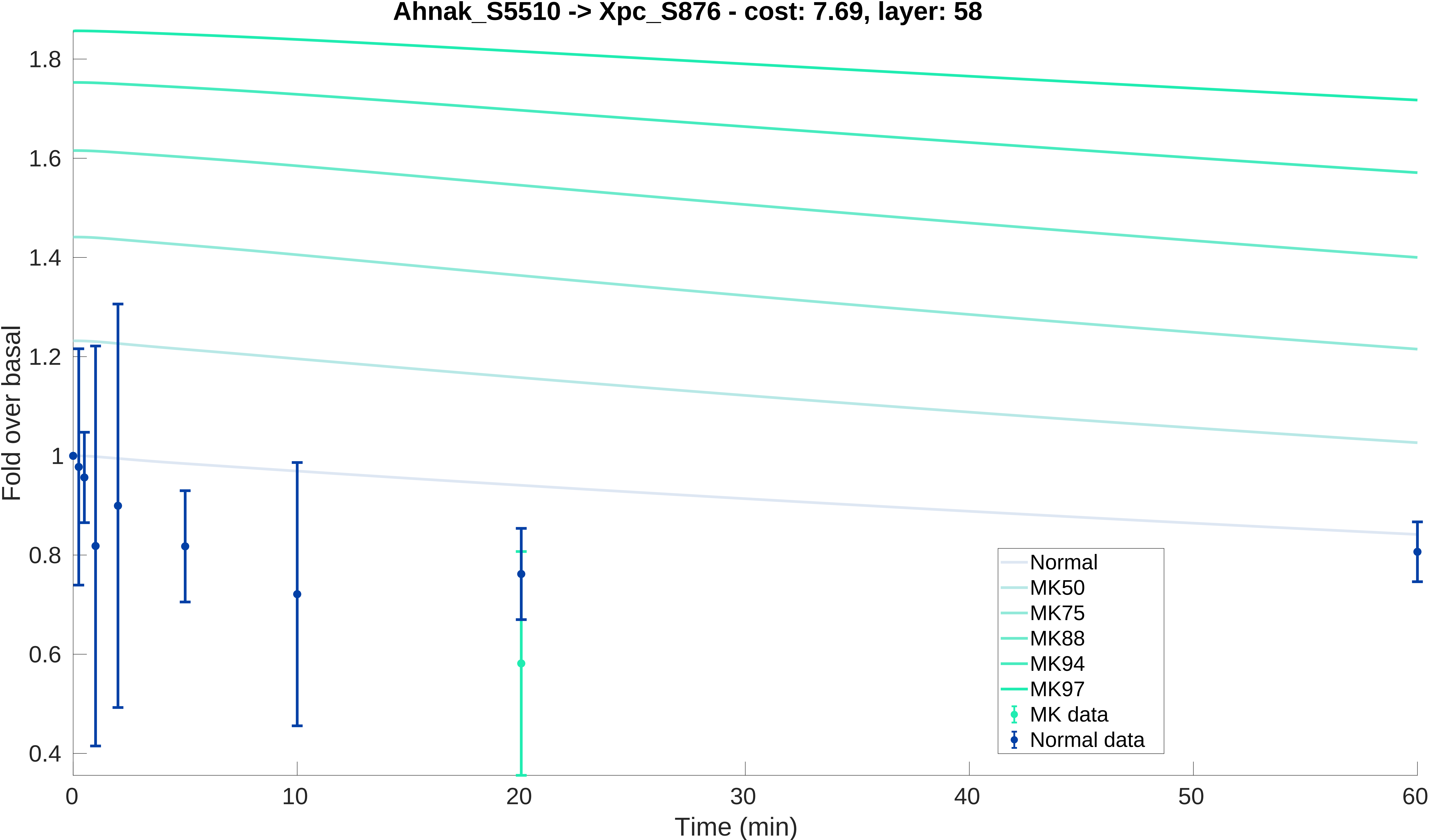

Ahnak_S5510 -> Xpc_S876 - cost: 7.69, layer: 58
Fold over basal
Time (min)
1.8
1.6
1.4
1.2
1
0.8
0.6
0.4
0
10
20
30
40
50
60
Normal
MK50
MK75
MK88
MK94
MK97
MK data
Normal data

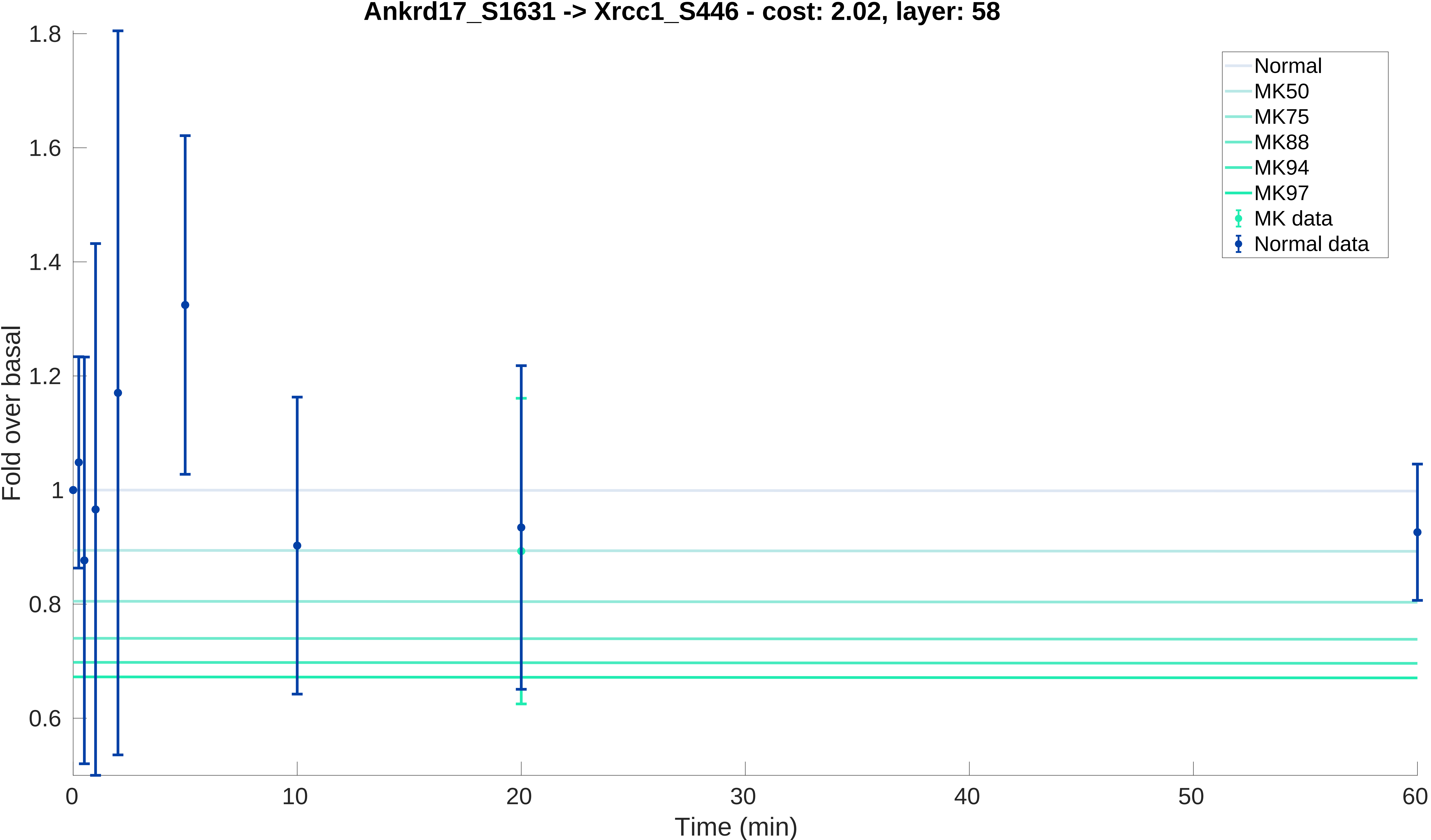

Ankrd17_S1631 -> Xrcc1_S446 - cost: 2.02, layer: 58
Fold over basal
Time (min)
Normal
MK50
MK75
MK88
MK94
MK97
MK data
Normal data
1.8
1.6
1.4
1.2
1
0.8
0.6
0
10
20
30
40
50
60

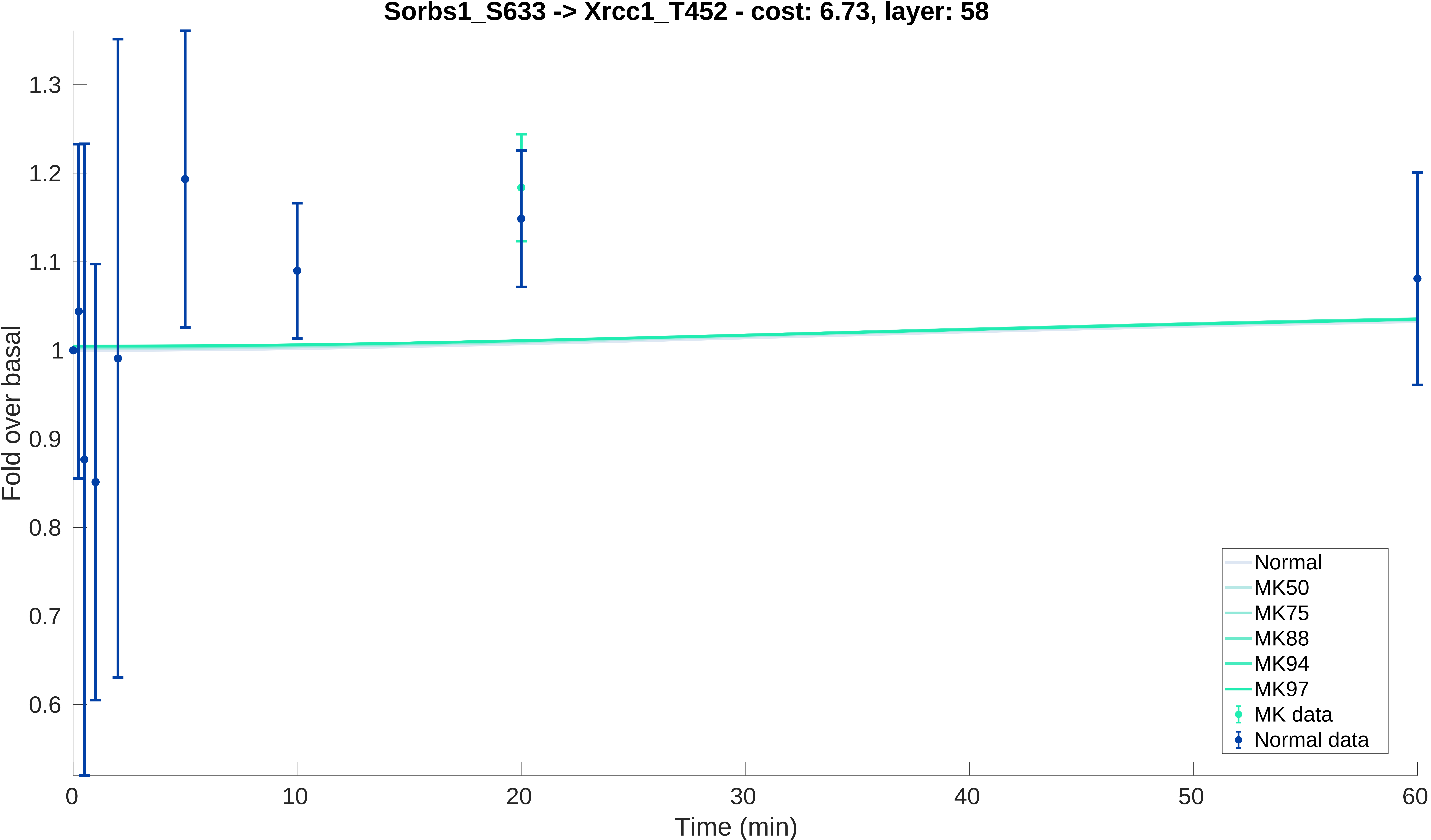

Sorbs1_S633 -> Xrcc1_T452 - cost: 6.73, layer: 58
Fold over basal
Time (min)
1.3
1.2
1.1
1
0.9
0.8
0.7
0.6
0
10
20
30
40
50
60
Normal
MK50
MK75
MK88
MK94
MK97
MK data
Normal data

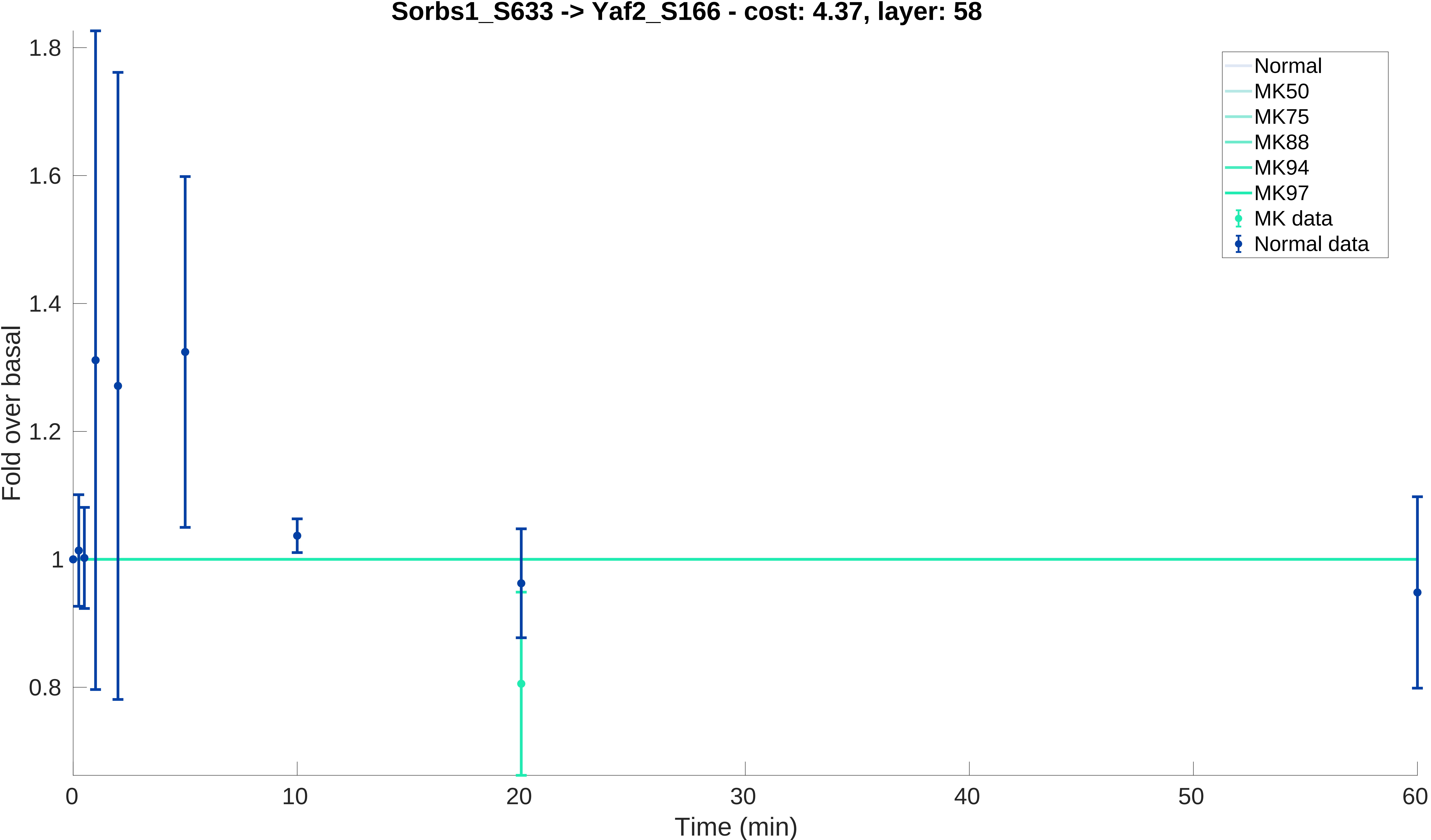

Sorbs1_S633 -> Yaf2_S166 - cost: 4.37, layer: 58
Fold over basal
Time (min)
1.8
1.6
1.4
1.2
1
0.8
0
10
20
30
40
50
60
Normal
MK50
MK75
MK88
MK94
MK97
MK data
Normal data

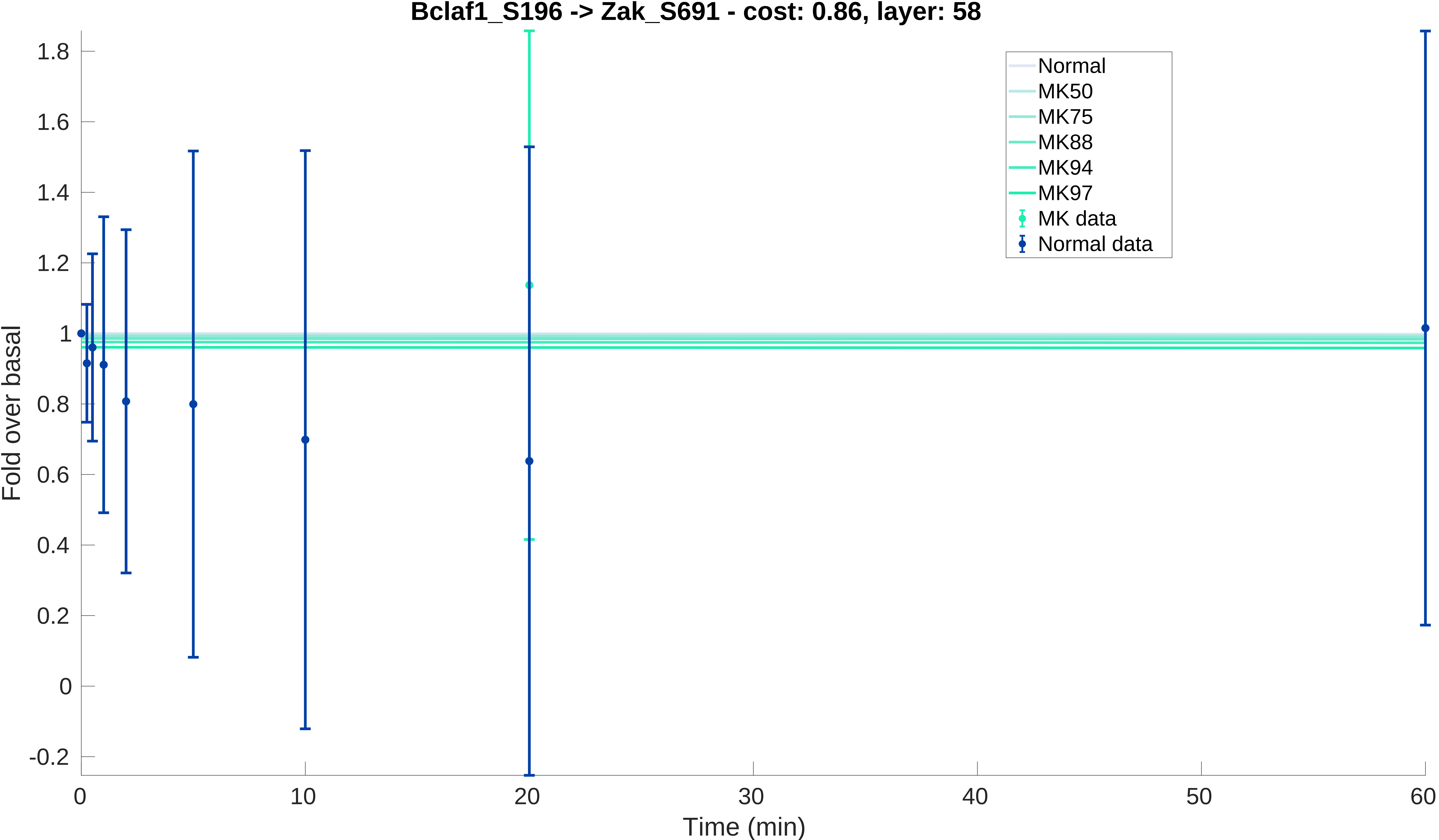

Bclaf1_S196 -> Zak_S691 - cost: 0.86, layer: 58
Normal
MK50
MK75
MK88
MK94
MK97
MK data
Normal data
Fold over basal
Time (min)
1.8
1.6
1.4
1.2
1
0.8
0.6
0.4
0.2
0
-0.2
0
10
20
30
40
50
60

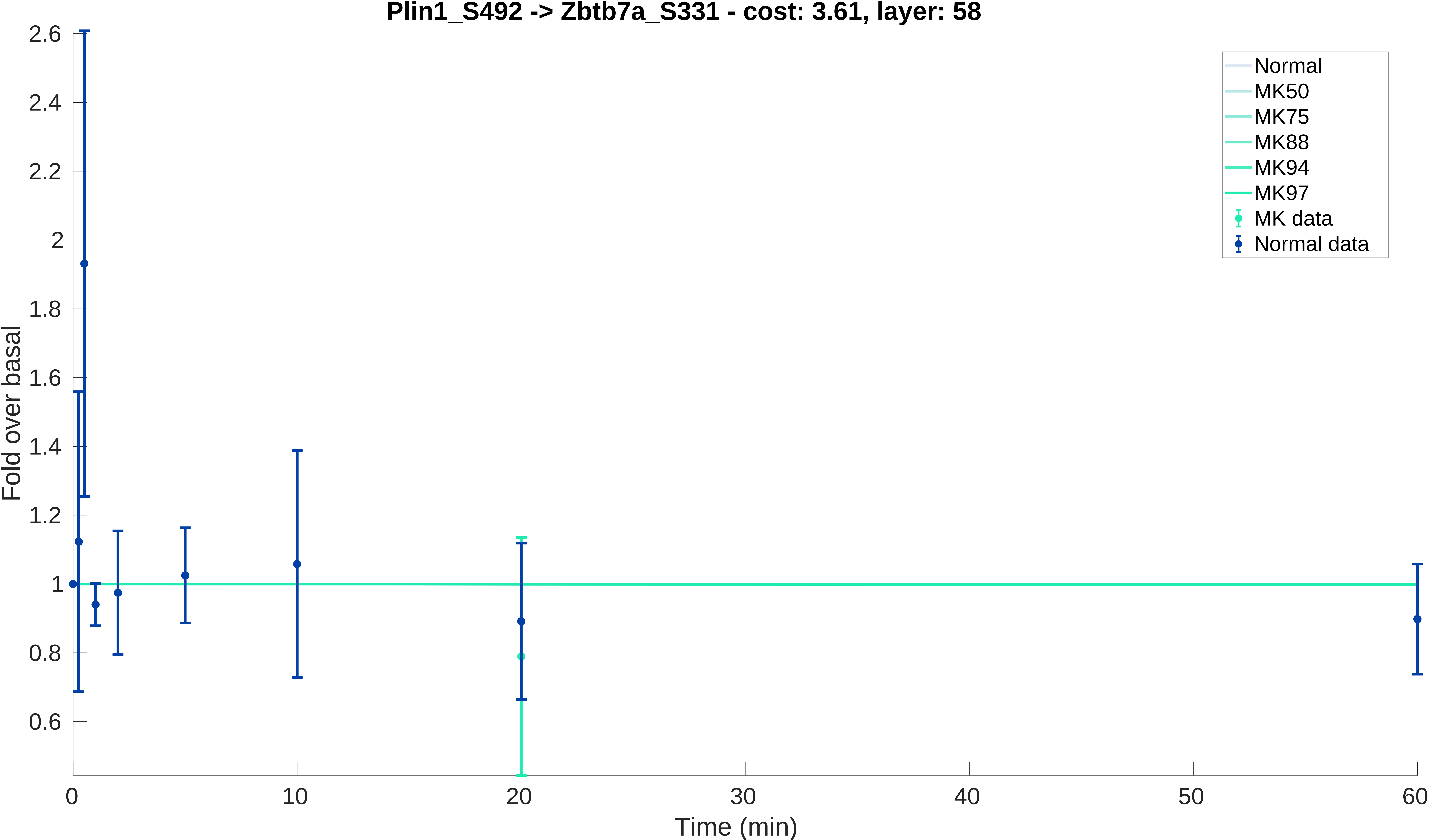

Plin1_S492 -> Zbtb7a_S331 - cost: 3.61, layer: 58
Fold over basal
Time (min)
2.6
2.4
2.2
2
1.8
1.6
1.4
1.2
1
0.8
0.6
0
10
20
30
40
50
60
Normal
MK50
MK75
MK88
MK94
MK97
MK data
Normal data

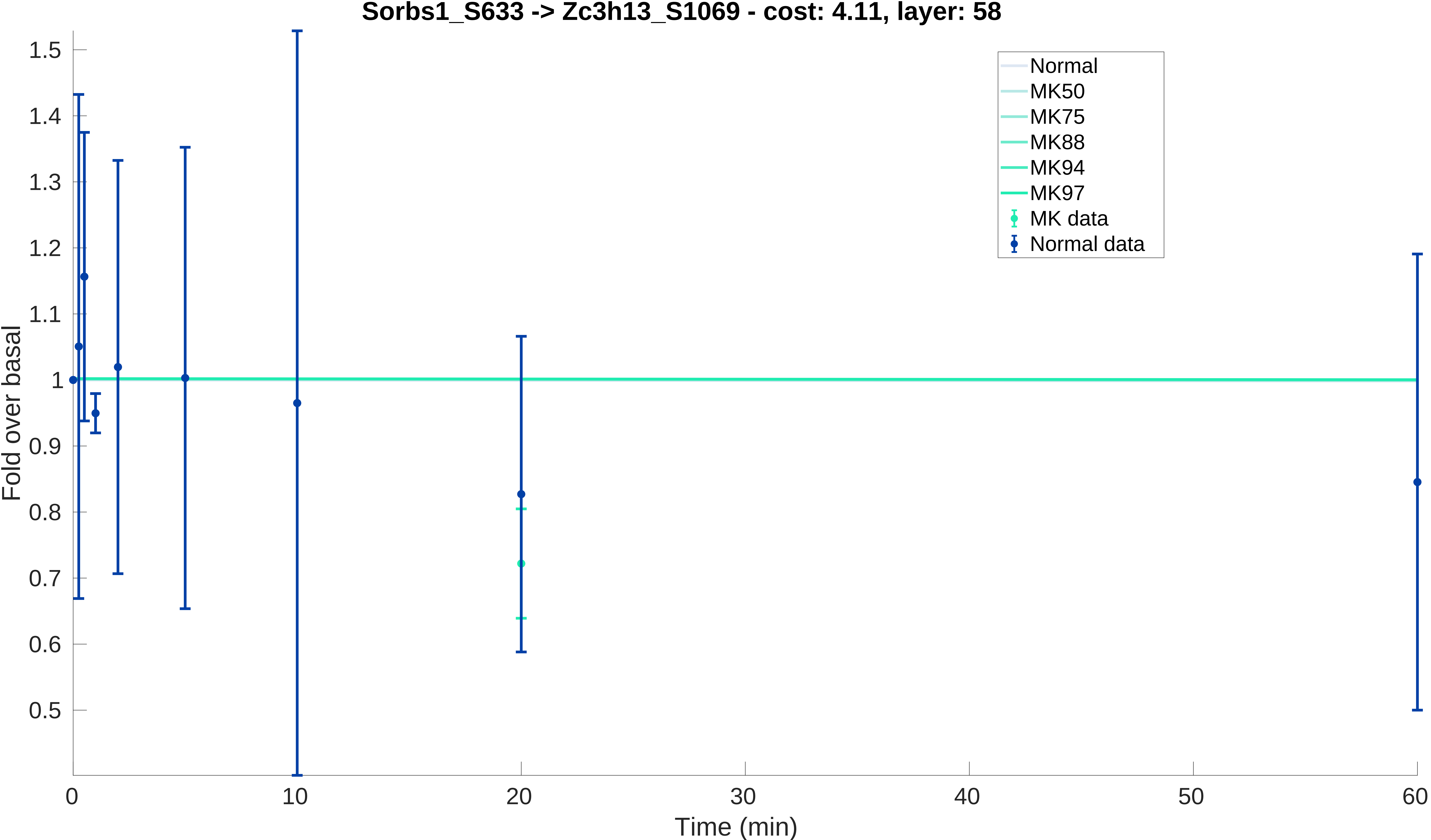

Sorbs1_S633 -> Zc3h13_S1069 - cost: 4.11, layer: 58
Normal
MK50
MK75
MK88
MK94
MK97
MK data
Normal data
Fold over basal
1.5
1.4
1.3
1.2
1.1
1
0.9
0.8
0.7
0.6
0.5
0
10
20
30
40
50
60
Time (min)

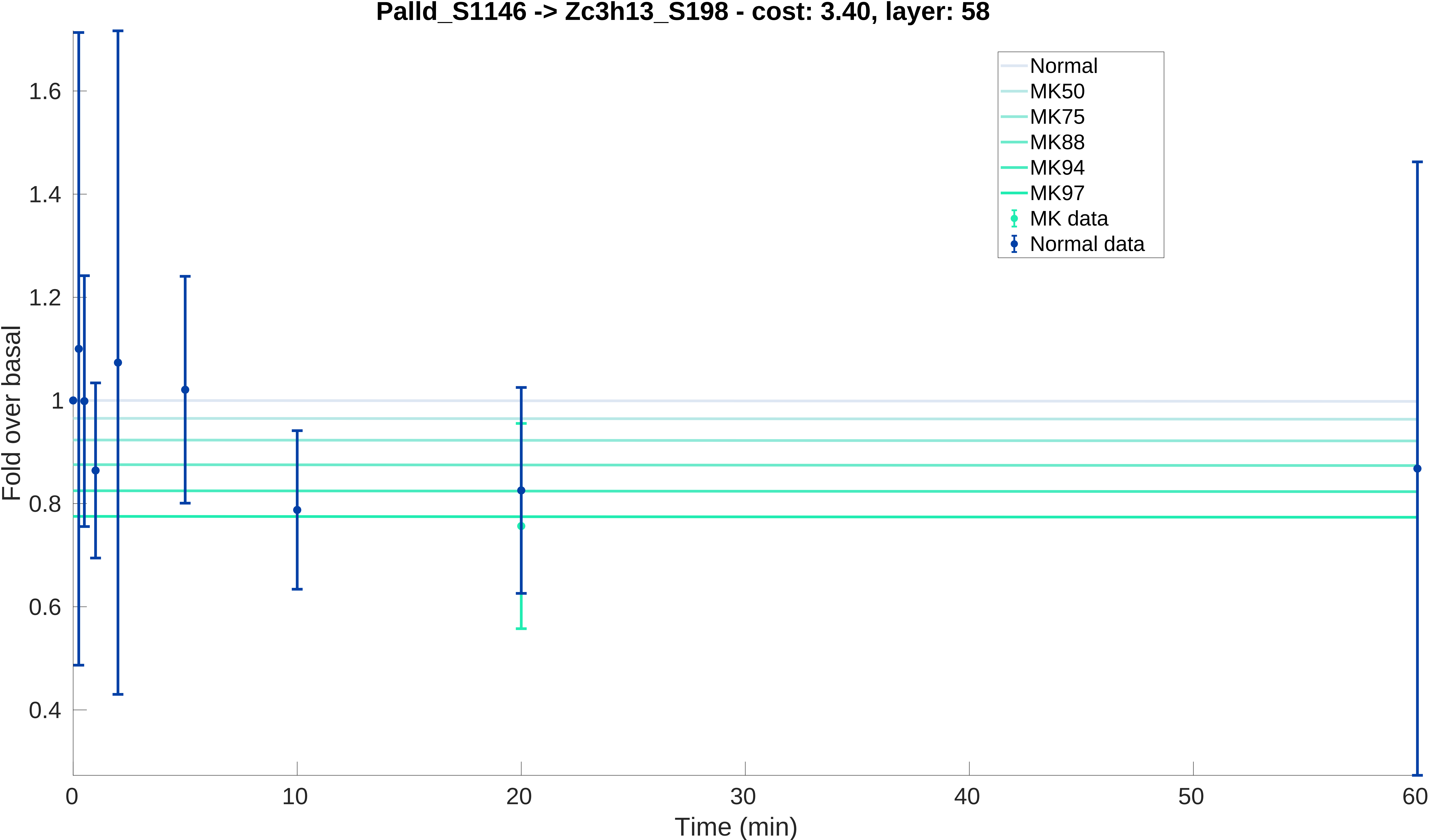
Palld_S1146 -> Zc3h13_S198 - cost: 3.40, layer: 58
Normal
MK50
MK75
MK88
MK94
MK97
MK data
Normal data
Fold over basal
Time (min)
0.4
0.6
0.8
1
1.2
1.4
1.6
0
10
20
30
40
50
60

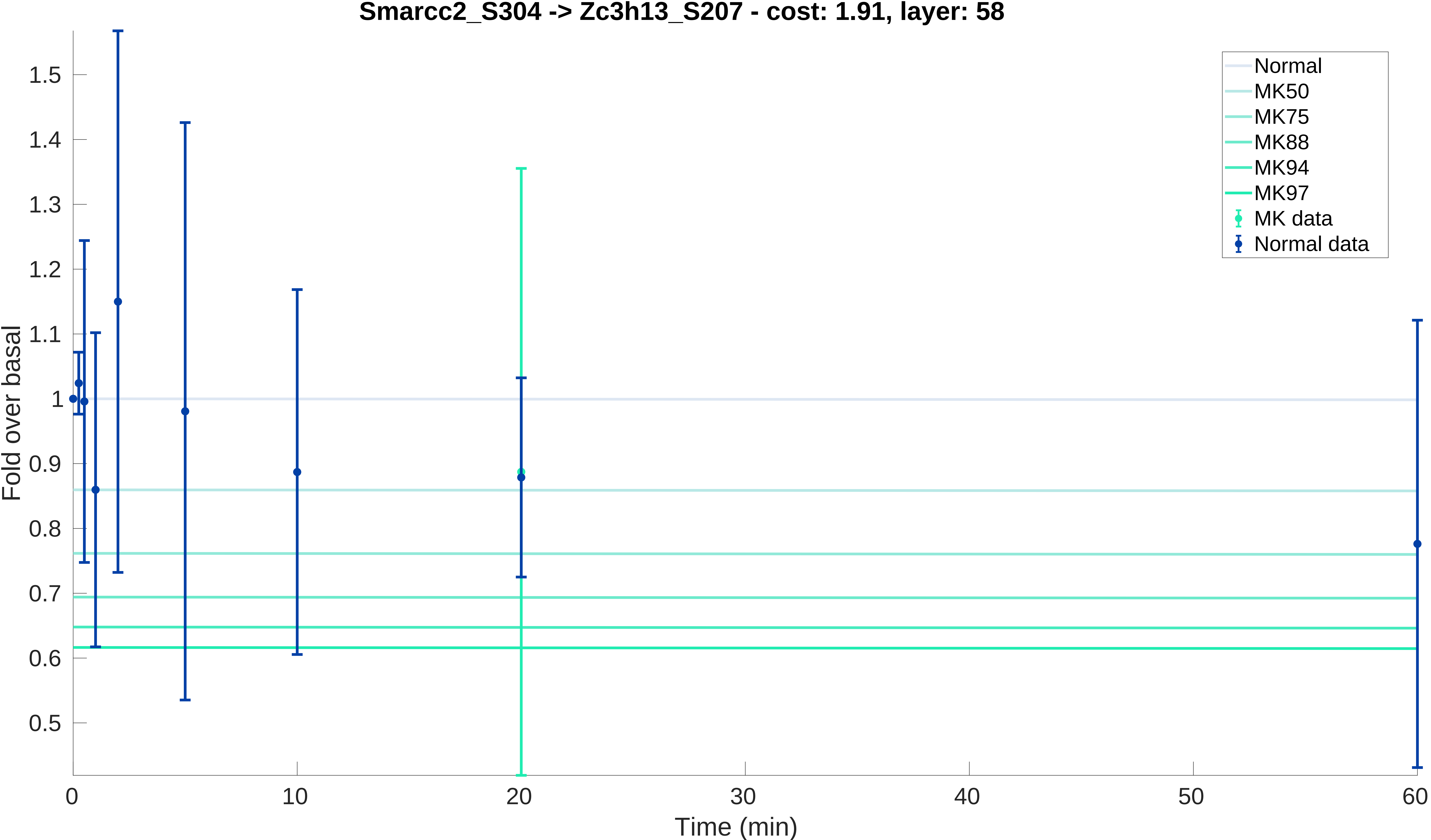

Smarcc2_S304 -> Zc3h13_S207 - cost: 1.91, layer: 58
Fold over basal
1.5
1.4
1.3
1.2
1.1
1
0.9
0.8
0.7
0.6
0.5
0
10
20
30
40
50
60
Time (min)
Normal
MK50
MK75
MK88
MK94
MK97
MK data
Normal data

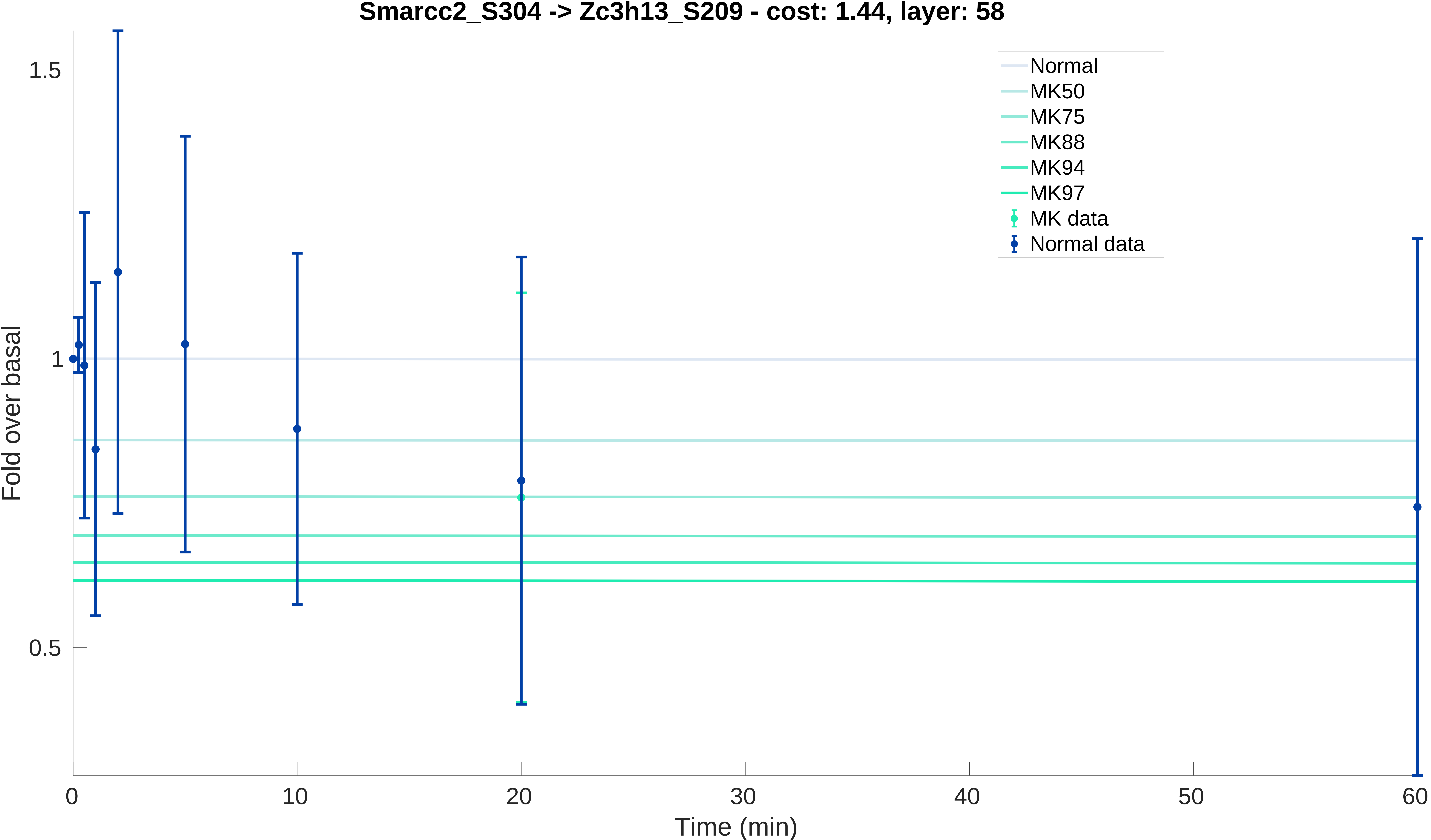

Smarcc2_S304 -> Zc3h13_S209 - cost: 1.44, layer: 58
Normal
MK50
MK75
MK88
MK94
MK97
MK data
Normal data
1.5
1
0.5
Fold over basal
0
10
20
30
40
50
60
Time (min)

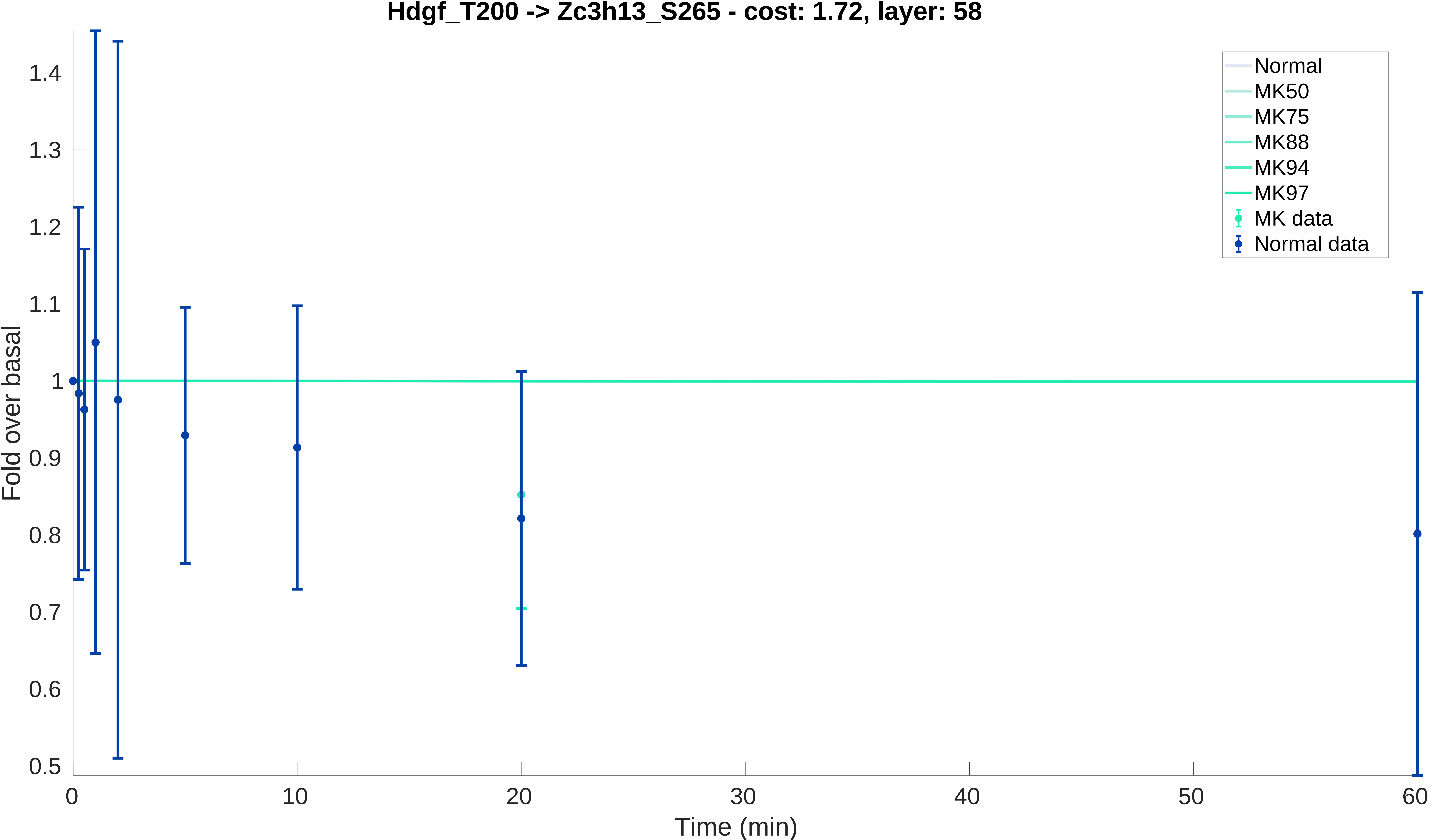

Hdgf_T200 -> Zc3h13_S265 - cost: 1.72, layer: 58
Fold over basal
Time (min)
0
10
20
30
40
50
60
0.5
0.6
0.7
0.8
0.9
1
1.1
1.2
1.3
1.4
Normal
MK50
MK75
MK88
MK94
MK97
MK data
Normal data

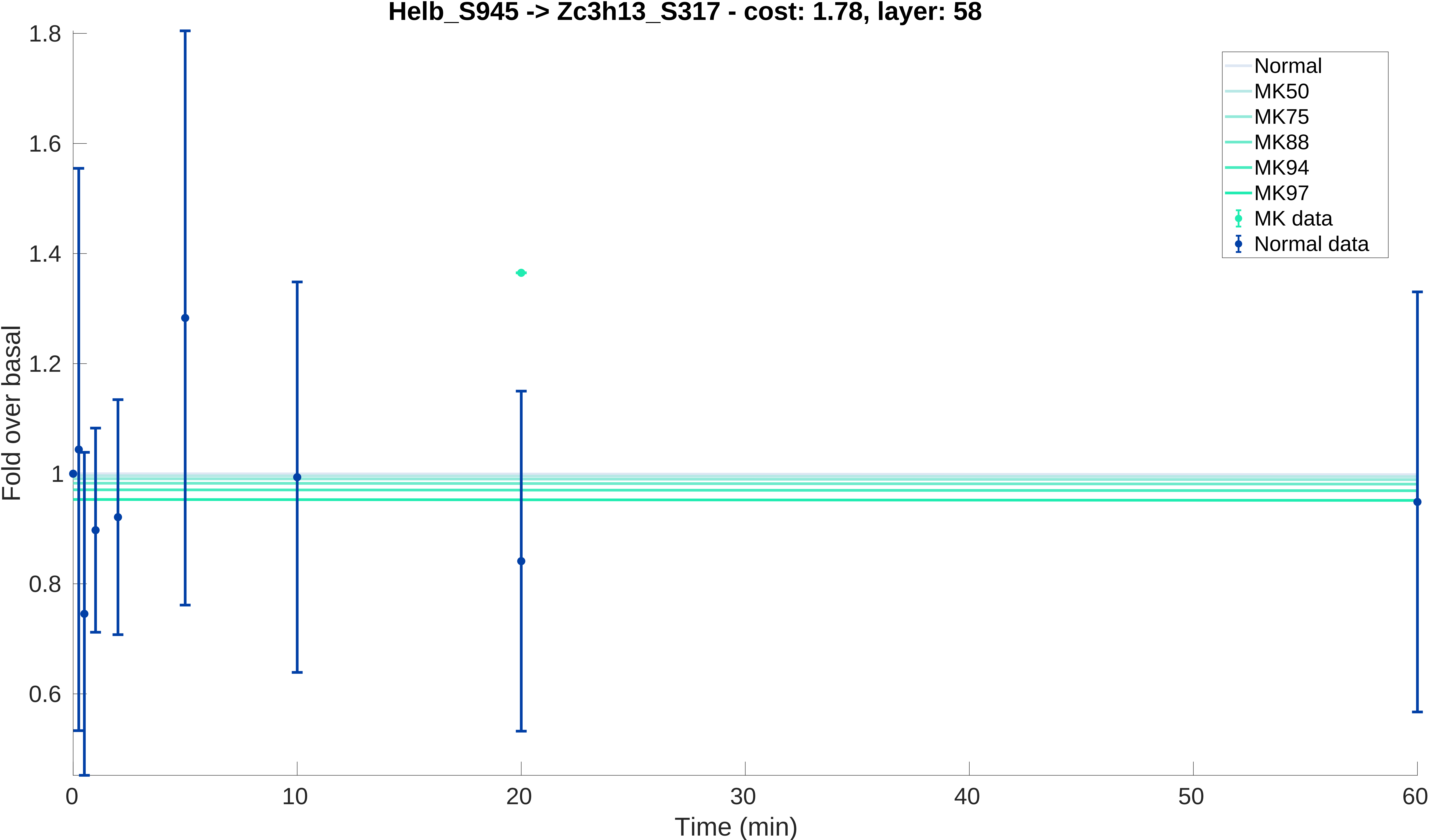

Helb_S945 -> Zc3h13_S317 - cost: 1.78, layer: 58
Fold over basal
Time (min)
1.8
1.6
1.4
1.2
1
0.8
0.6
0
10
20
30
40
50
60
Normal
MK50
MK75
MK88
MK94
MK97
MK data
Normal data

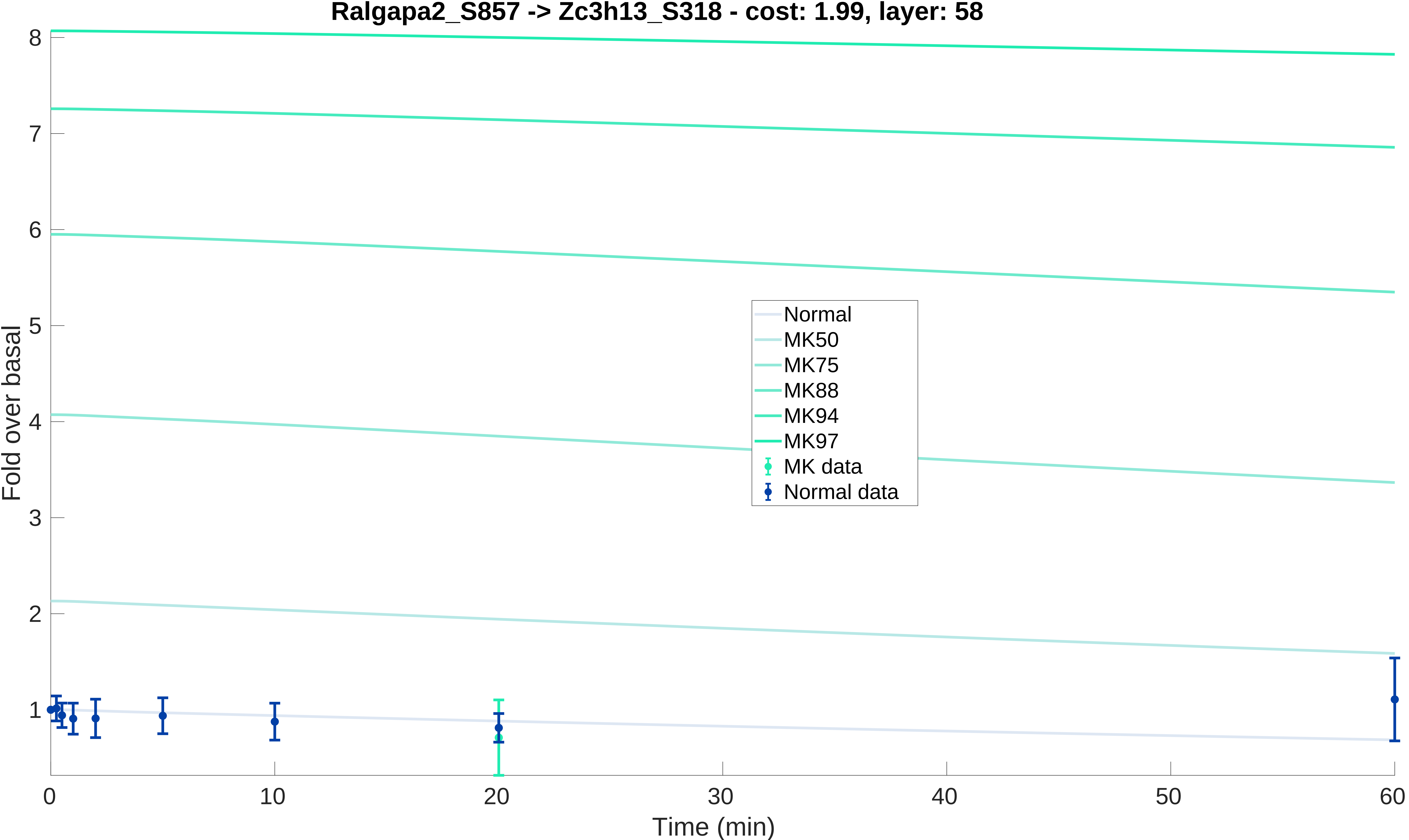

Ralgapa2_S857 -> Zc3h13_S318 - cost: 1.99, layer: 58
Fold over basal
Time (min)
0
10
20
30
40
50
60
1
2
3
4
5
6
7
8
Normal
MK50
MK75
MK88
MK94
MK97
MK data
Normal data

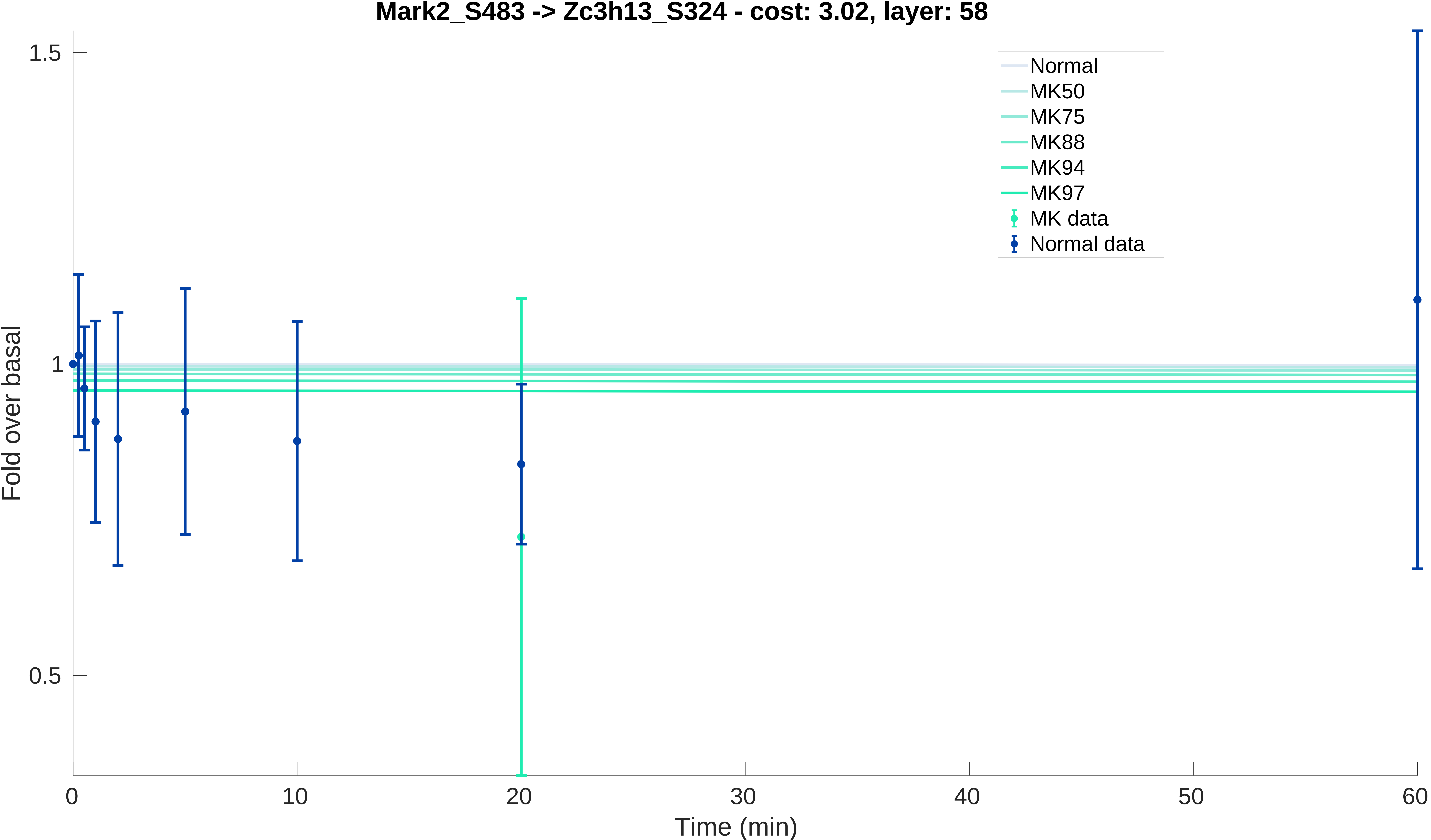

Mark2_S483 -> Zc3h13_S324 - cost: 3.02, layer: 58
Normal
MK50
MK75
MK88
MK94
MK97
MK data
Normal data
Fold over basal
1.5
1
0.5
0
10
20
30
40
50
60
Time (min)

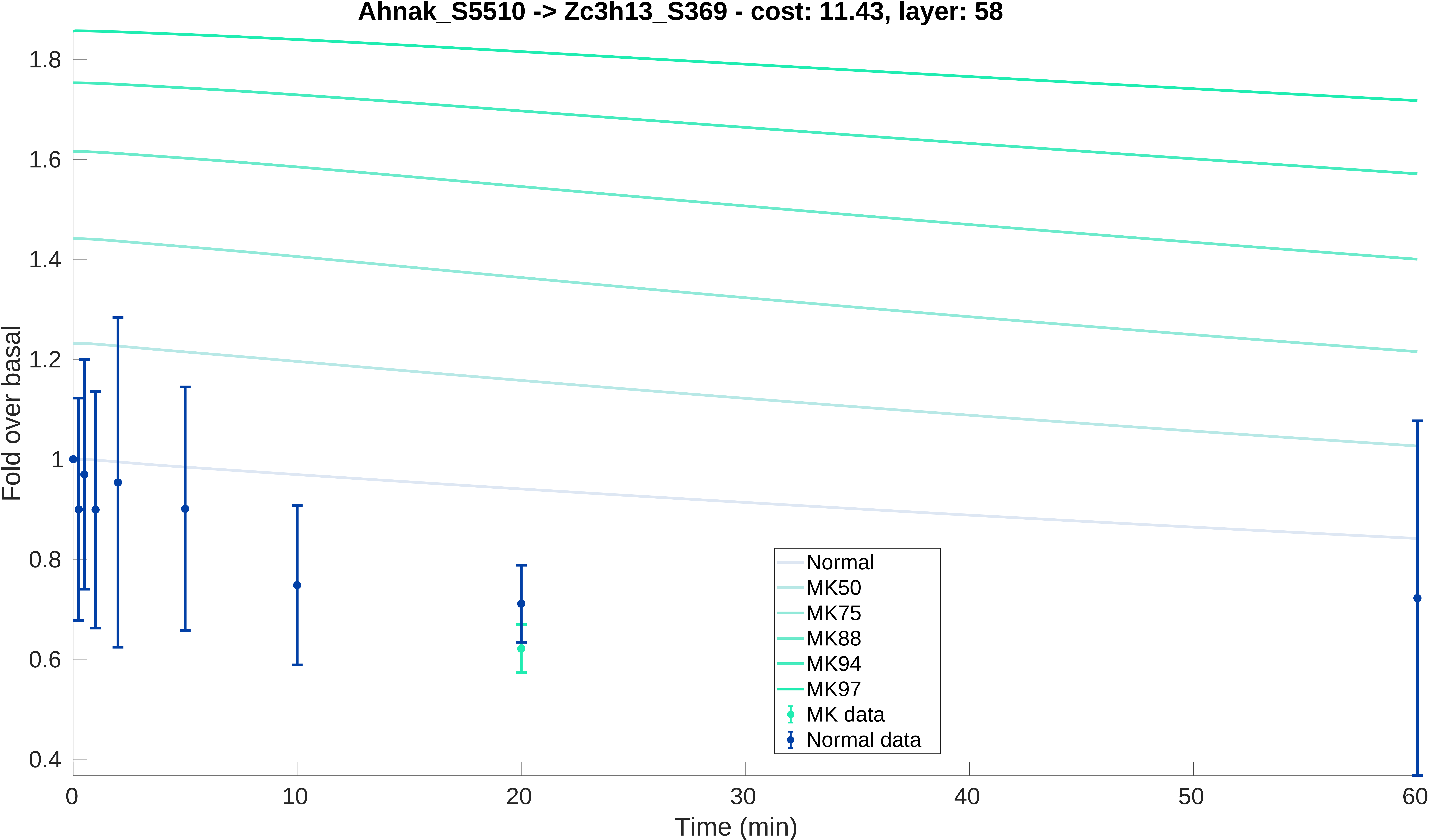
Ahnak_S5510 -> Zc3h13_S369 - cost: 11.43, layer: 58
Fold over basal
Time (min)
0.4
0.6
0.8
1
1.2
1.4
1.6
1.8
0
10
20
30
40
50
60
Normal
MK50
MK75
MK88
MK94
MK97
MK data
Normal data

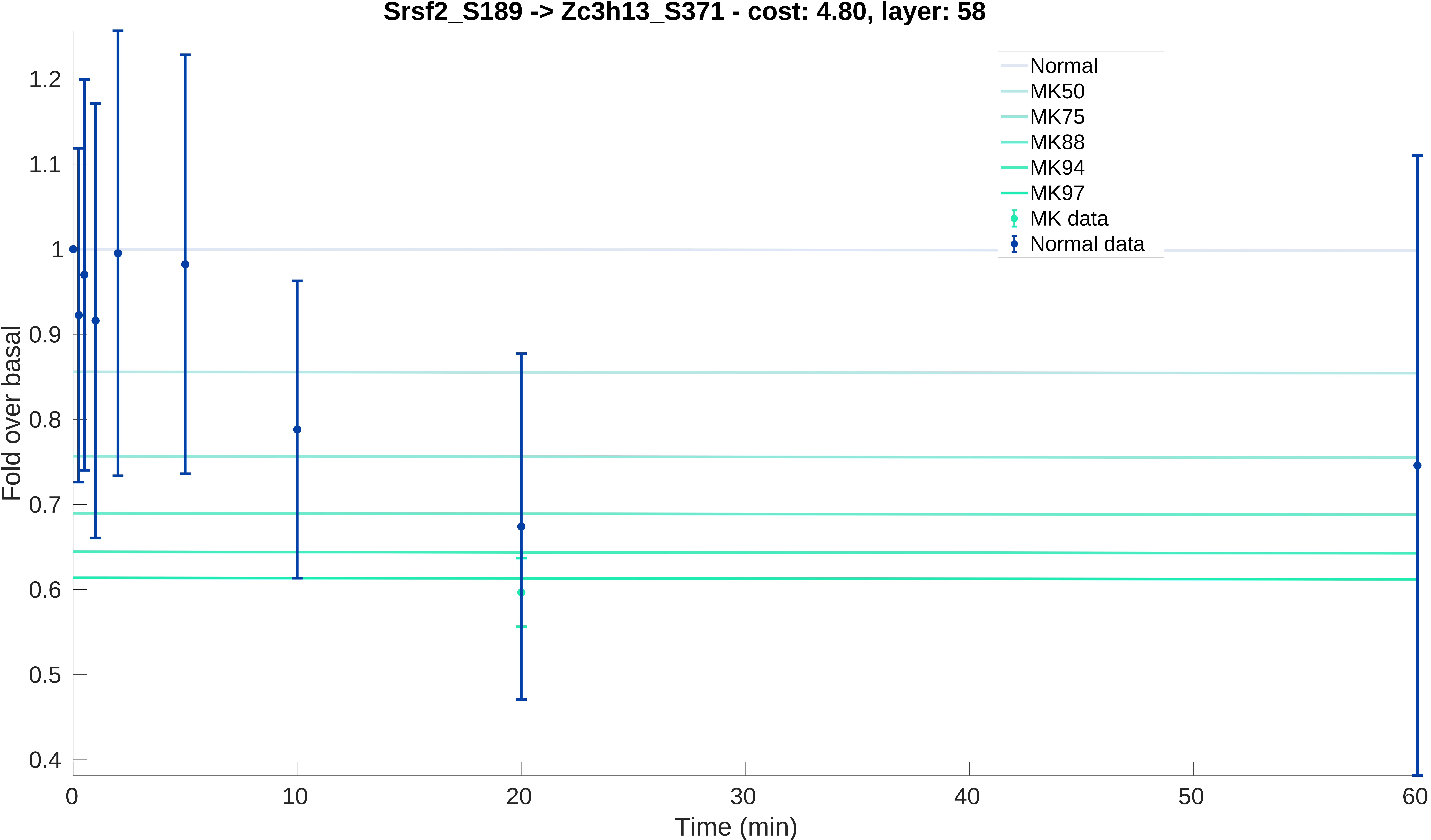

Srsf2_S189 -> Zc3h13_S371 - cost: 4.80, layer: 58
Fold over basal
Time (min)
1.2
1.1
1
0.9
0.8
0.7
0.6
0.5
0.4
0
10
20
30
40
50
60
Normal
MK50
MK75
MK88
MK94
MK97
MK data
Normal data

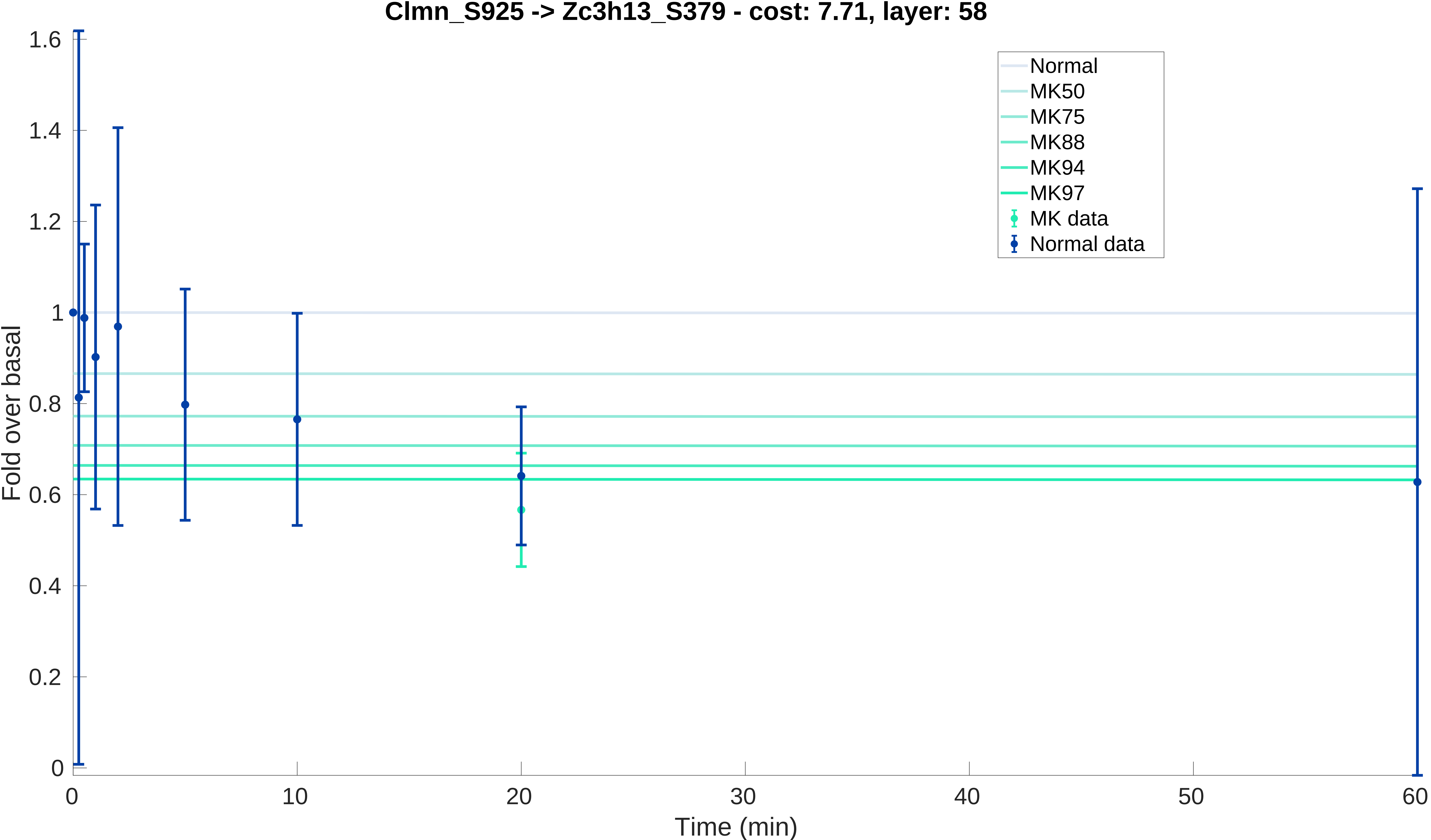
Clmn_S925 -> Zc3h13_S379 - cost: 7.71, layer: 58
Normal
MK50
MK75
MK88
MK94
MK97
MK data
Normal data
Fold over basal
0
0.2
0.4
0.6
0.8
1
1.2
1.4
1.6
Time (min)
0
10
20
30
40
50
60

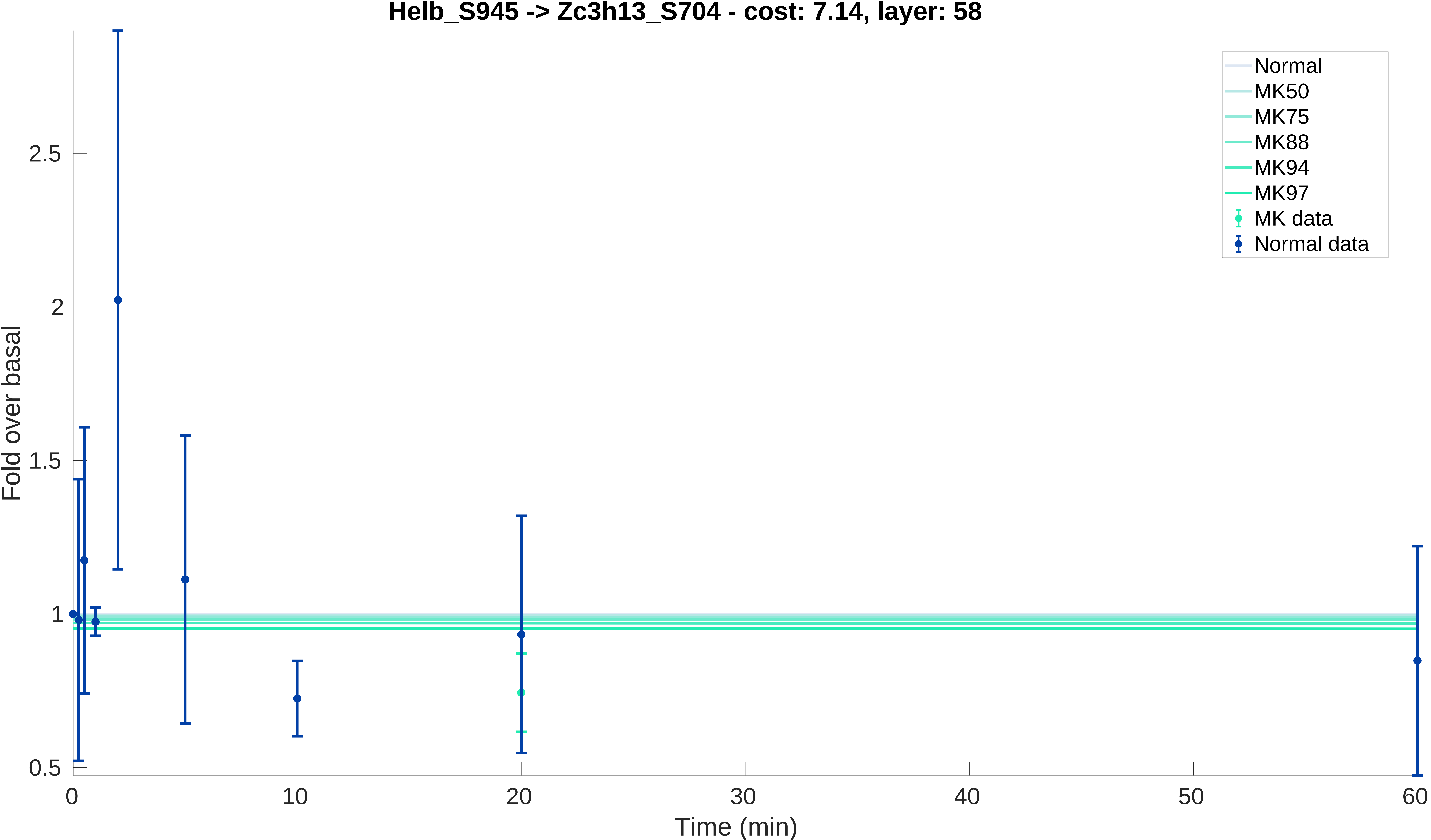

Helb_S945 -> Zc3h13_S704 - cost: 7.14, layer: 58
Fold over basal
Time (min)
Normal
MK50
MK75
MK88
MK94
MK97
MK data
Normal data
0.5
1
1.5
2
2.5
0
10
20
30
40
50
60

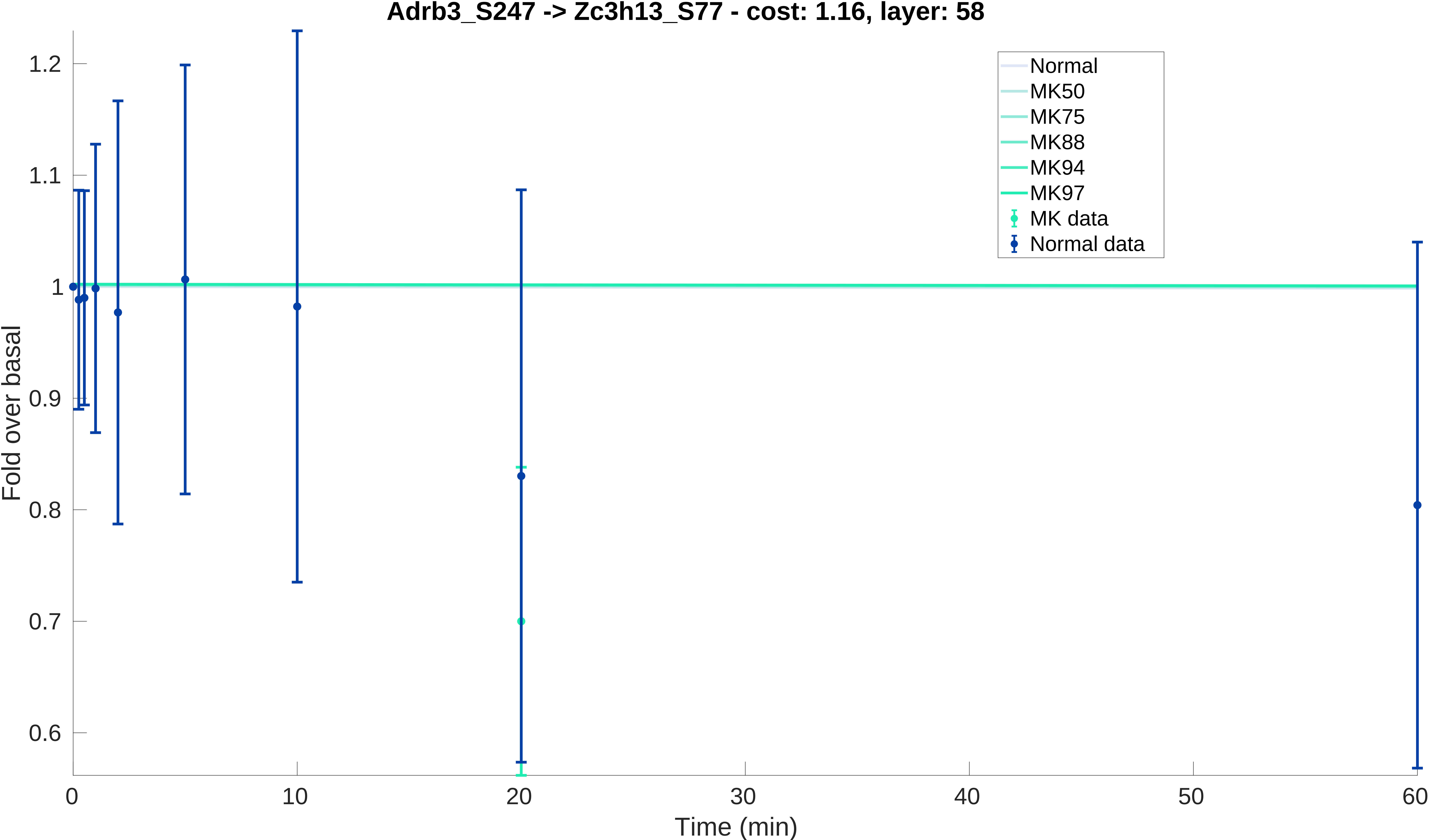

Adrb3_S247 -> Zc3h13_S77 - cost: 1.16, layer: 58
Fold over basal
Time (min)
Normal
MK50
MK75
MK88
MK94
MK97
MK data
Normal data
1.2
1.1
1
0.9
0.8
0.7
0.6
0
10
20
30
40
50
60

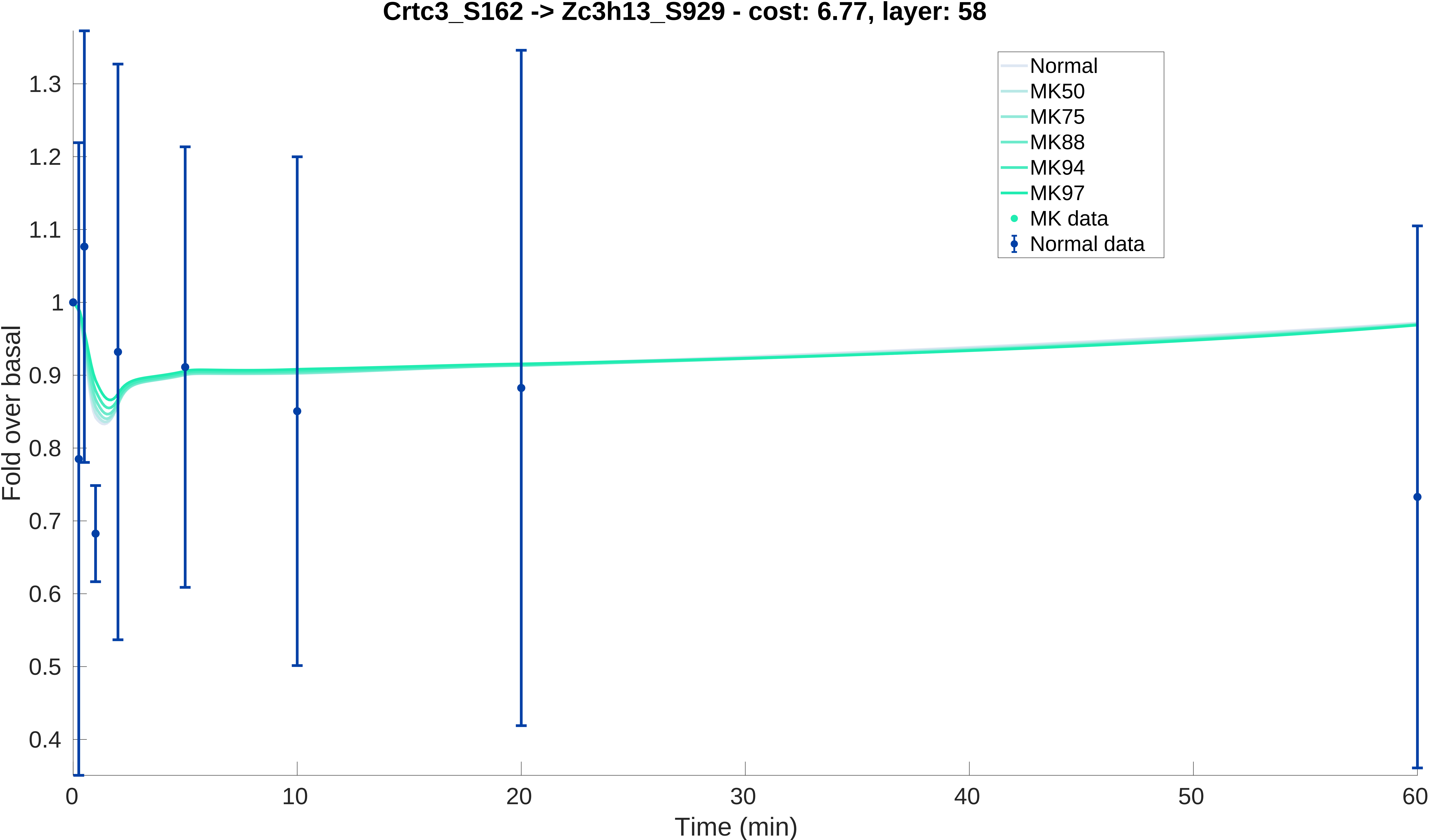

Crtc3_S162 -> Zc3h13_S929 - cost: 6.77, layer: 58
Fold over basal
Time (min)
Normal
MK50
MK75
MK88
MK94
MK97
MK data
Normal data
0.4
0.5
0.6
0.7
0.8
0.9
1
1.1
1.2
1.3
0
10
20
30
40
50
60

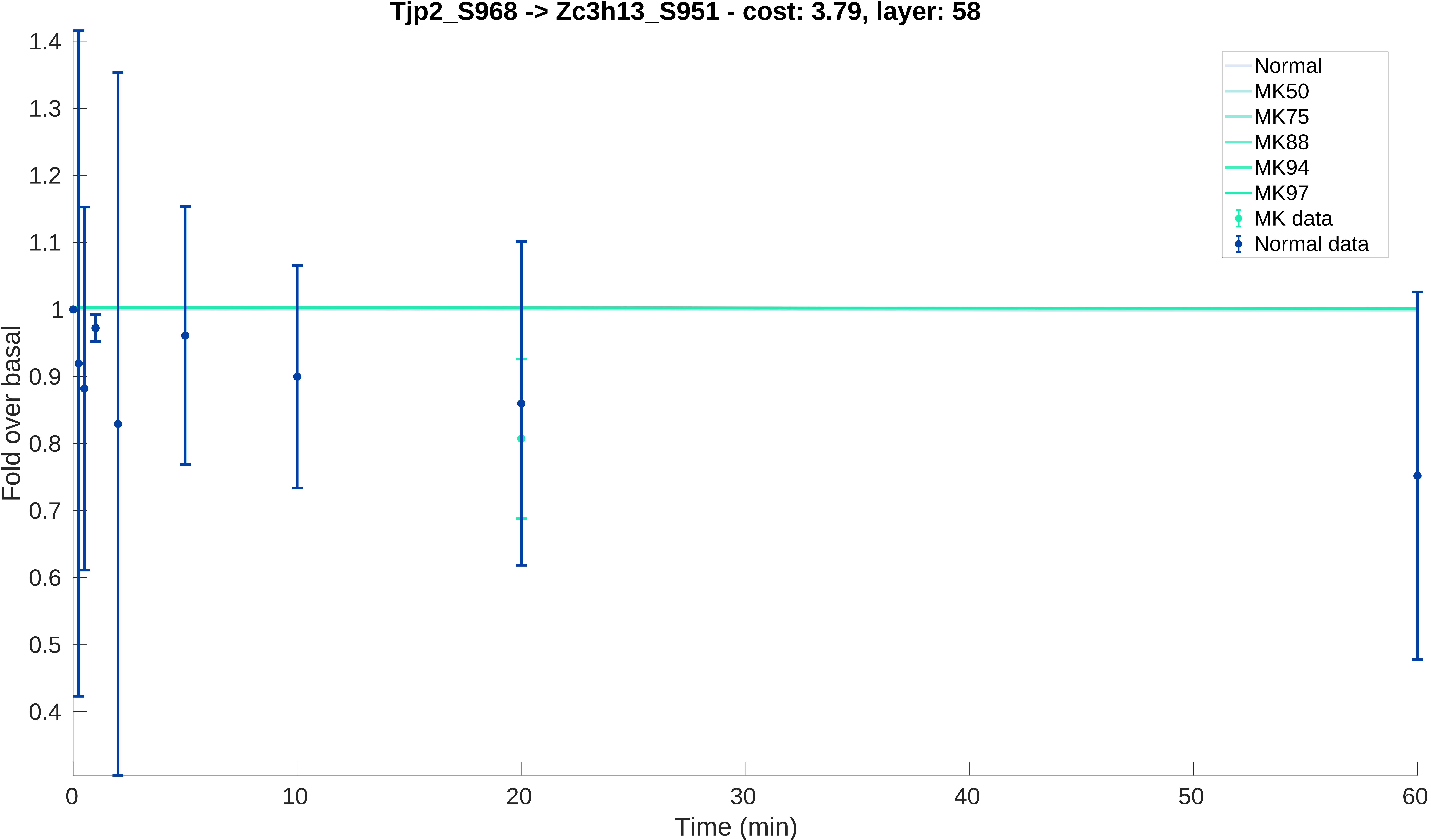

Tjp2_S968 -> Zc3h13_S951 - cost: 3.79, layer: 58
Fold over basal
Time (min)
1.4
1.3
1.2
1.1
1
0.9
0.8
0.7
0.6
0.5
0.4
0
10
20
30
40
50
60
Normal
MK50
MK75
MK88
MK94
MK97
MK data
Normal data

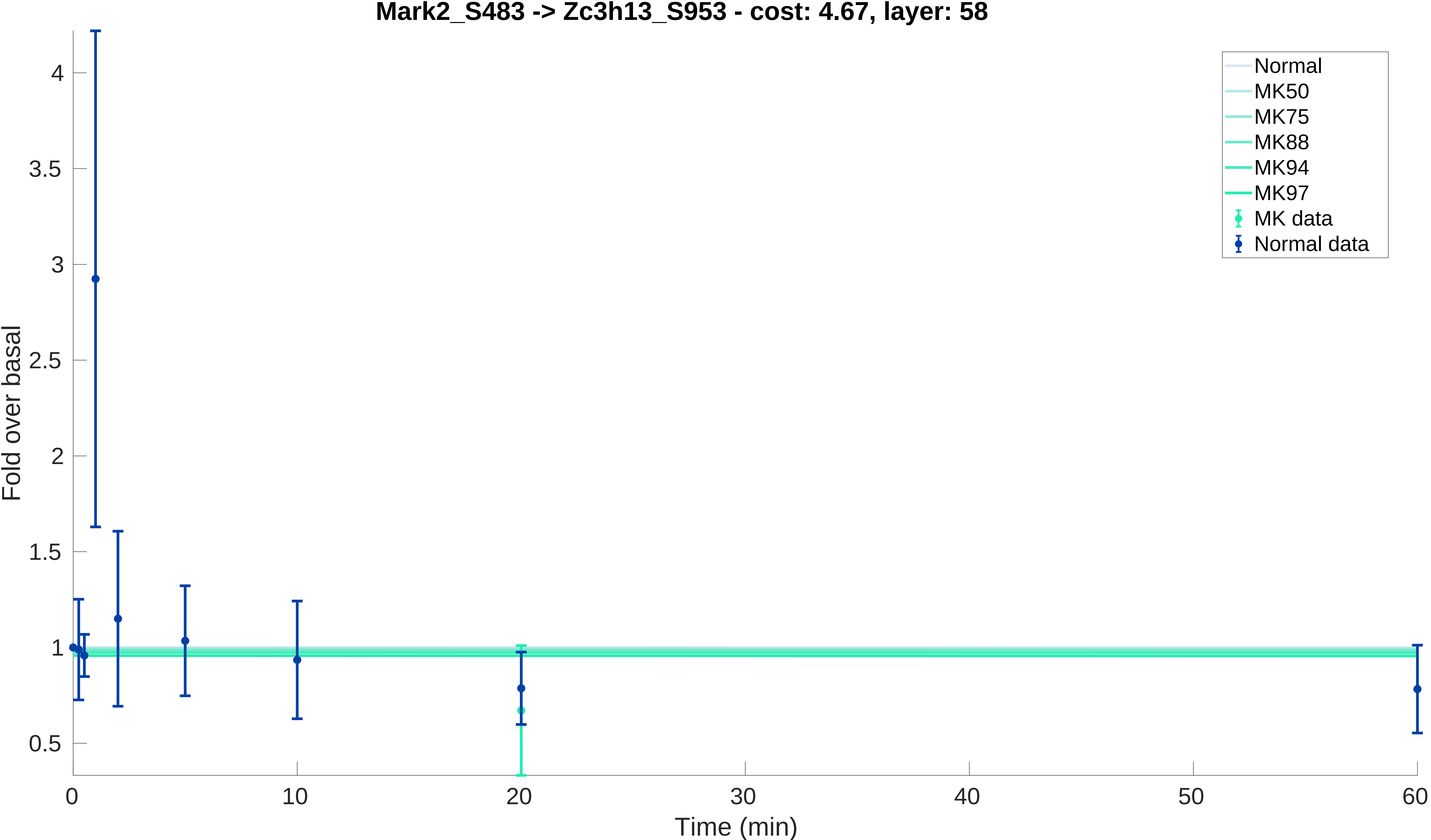

Mark2_S483 -> Zc3h13_S953 - cost: 4.67, layer: 58
Fold over basal
Time (min)
0.5
1
1.5
2
2.5
3
3.5
4
0
10
20
30
40
50
60
Normal
MK50
MK75
MK88
MK94
MK97
MK data
Normal data

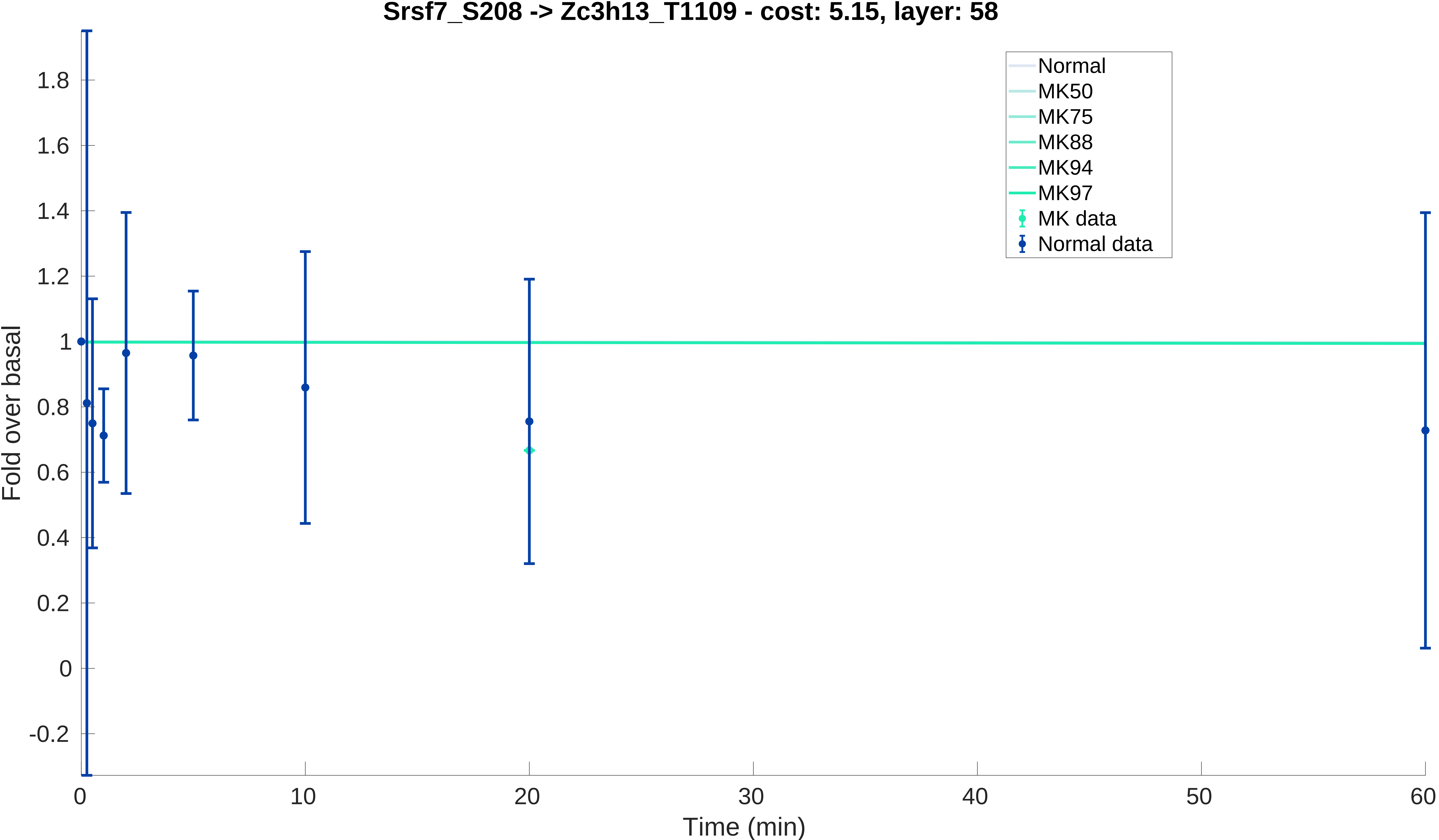

Srsf7_S208 -> Zc3h13_T1109 - cost: 5.15, layer: 58
Normal
MK50
MK75
MK88
MK94
MK97
MK data
Normal data
Fold over basal
Time (min)
1.8
1.6
1.4
1.2
1
0.8
0.6
0.4
0.2
0
-0.2
0
10
20
30
40
50
60

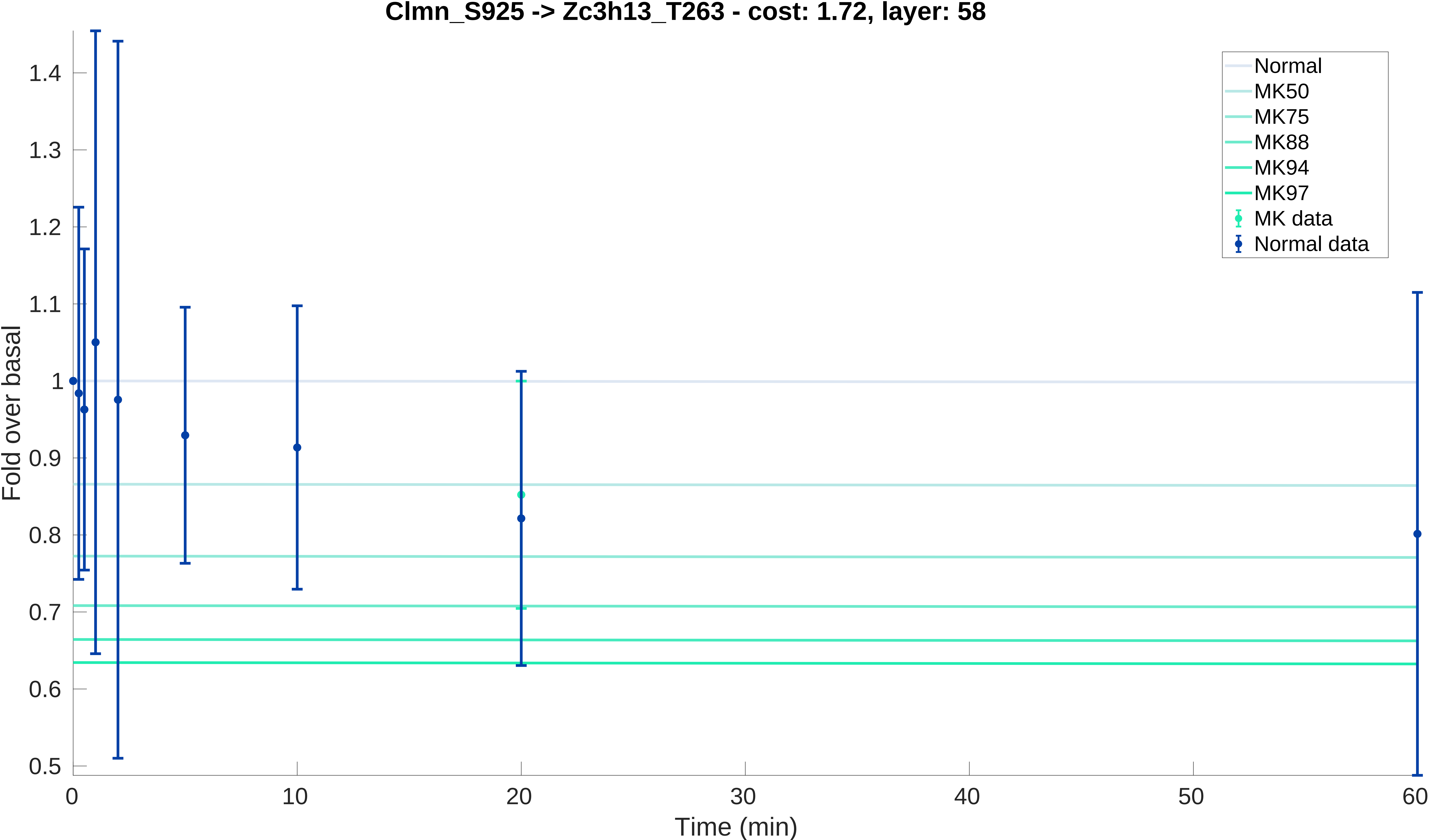

Clmn_S925 -> Zc3h13_T263 - cost: 1.72, layer: 58
Normal
MK50
MK75
MK88
MK94
MK97
MK data
Normal data
Fold over basal
1.4
1.3
1.2
1.1
1
0.9
0.8
0.7
0.6
0.5
0
10
20
30
40
50
60
Time (min)

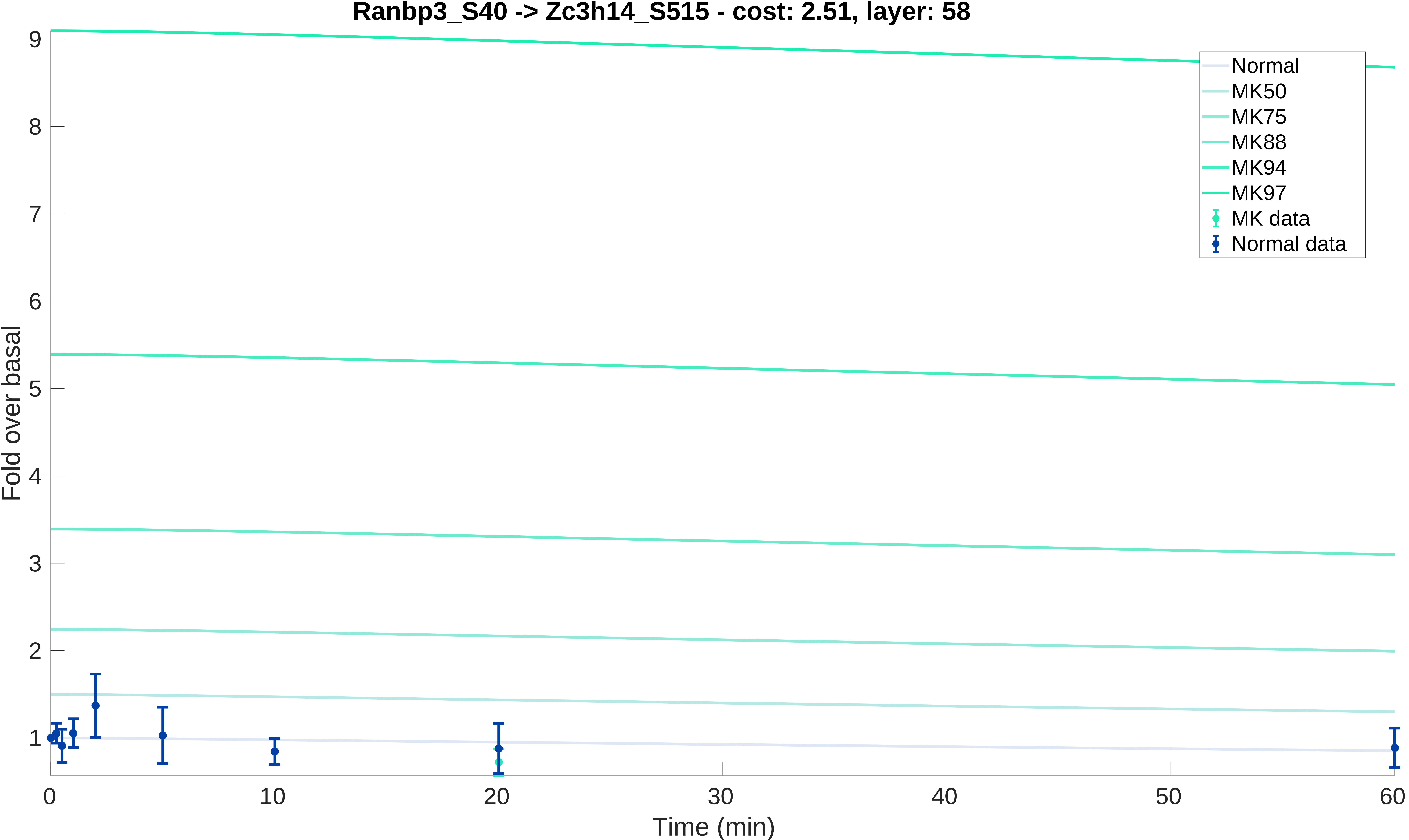

Ranbp3_S40 -> Zc3h14_S515 - cost: 2.51, layer: 58
Fold over basal
Time (min)
0
10
20
30
40
50
60
1
2
3
4
5
6
7
8
9
Normal
MK50
MK75
MK88
MK94
MK97
MK data
Normal data

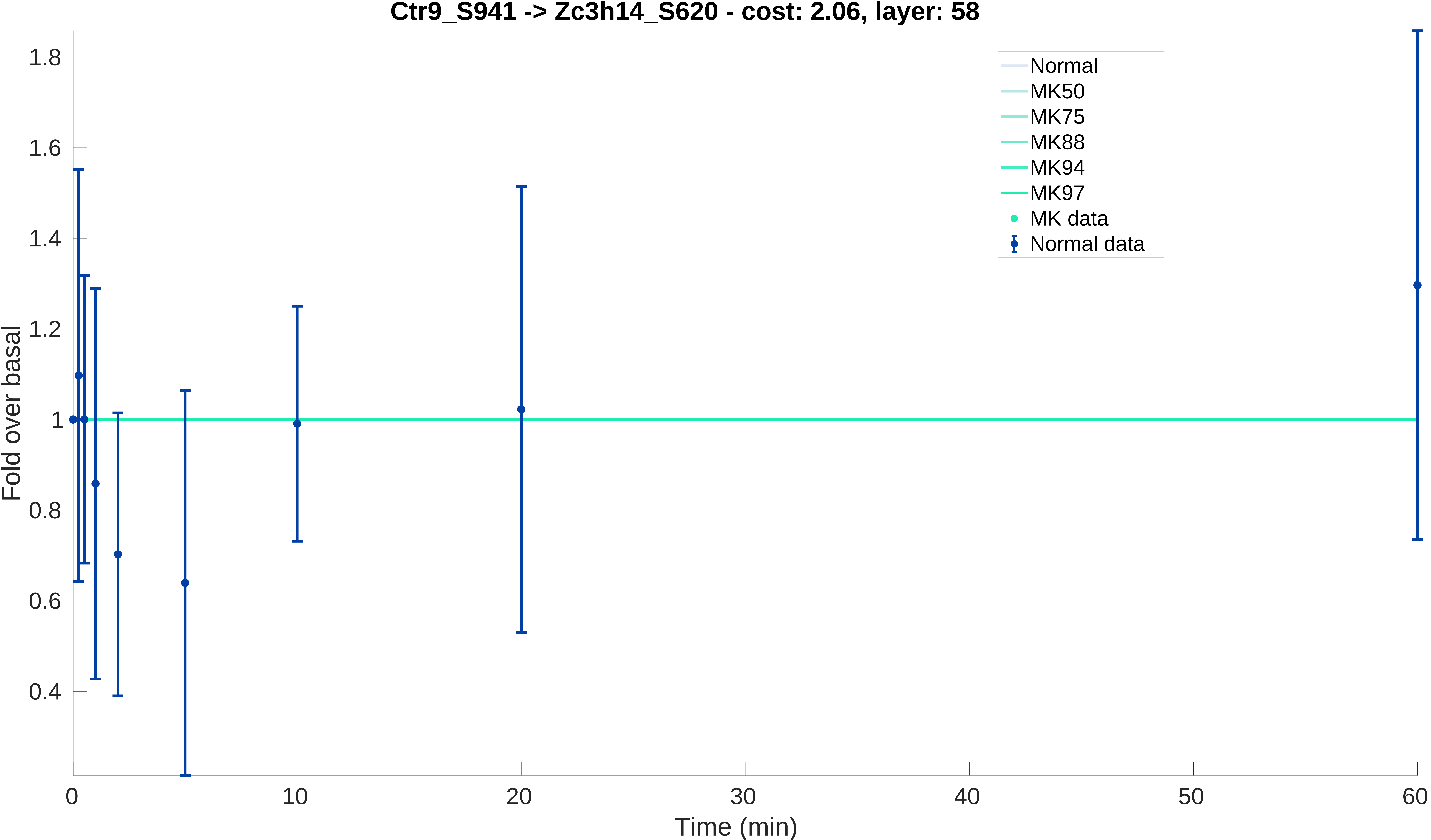
Ctr9_S941 -> Zc3h14_S620 - cost: 2.06, layer: 58
Fold over basal
Time (min)
0
10
20
30
40
50
60
0.4
0.6
0.8
1
1.2
1.4
1.6
1.8
Normal
MK50
MK75
MK88
MK94
MK97
MK data
Normal data

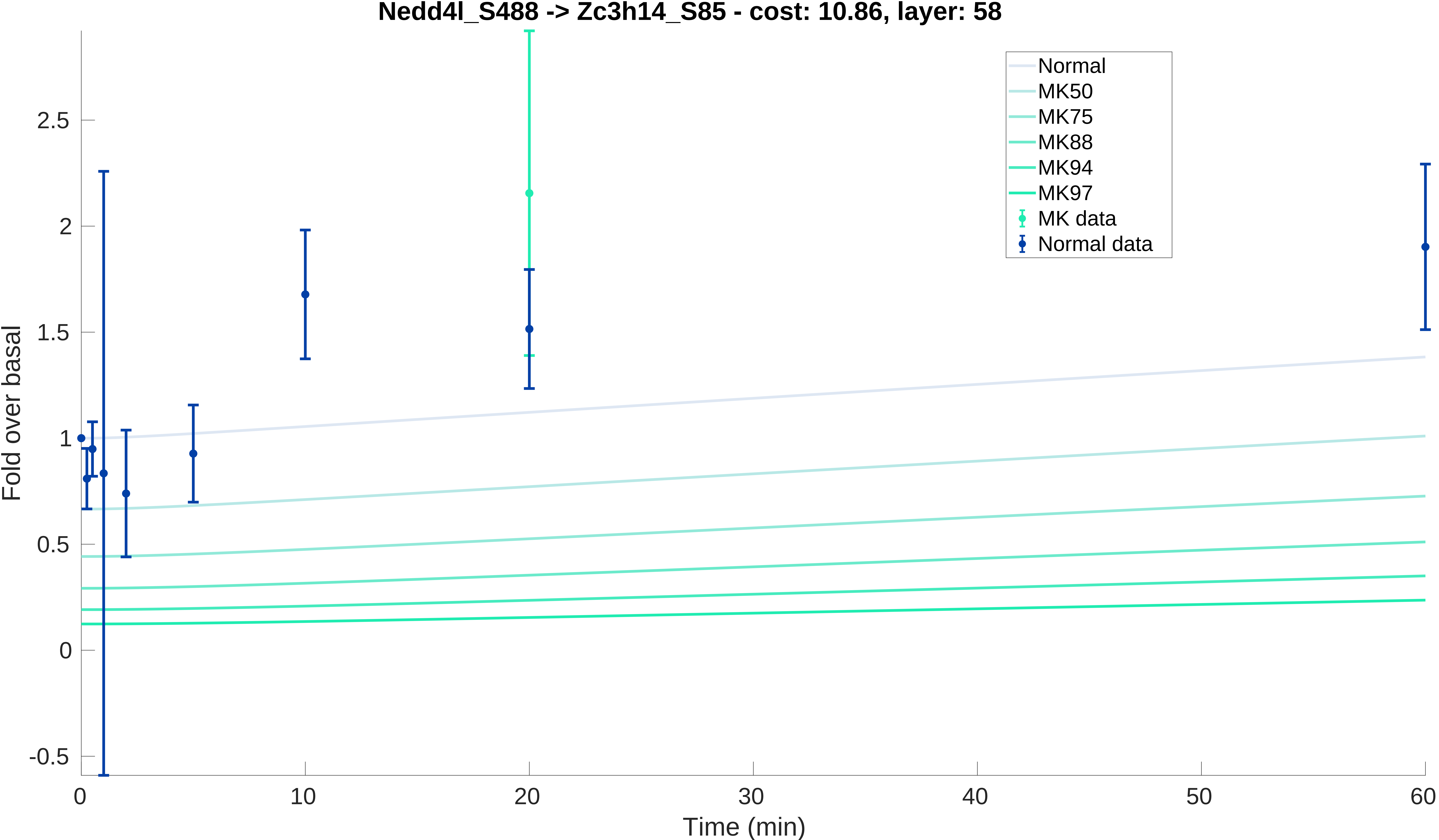

Nedd4l_S488 -> Zc3h14_S85 - cost: 10.86, layer: 58
Fold over basal
Time (min)
Normal
MK50
MK75
MK88
MK94
MK97
MK data
Normal data
2.5
2
1.5
1
0.5
0
-0.5
0
10
20
30
40
50
60

Eif5b_S215 -> Zc3h18_S169 - cost: 11.23, layer: 58

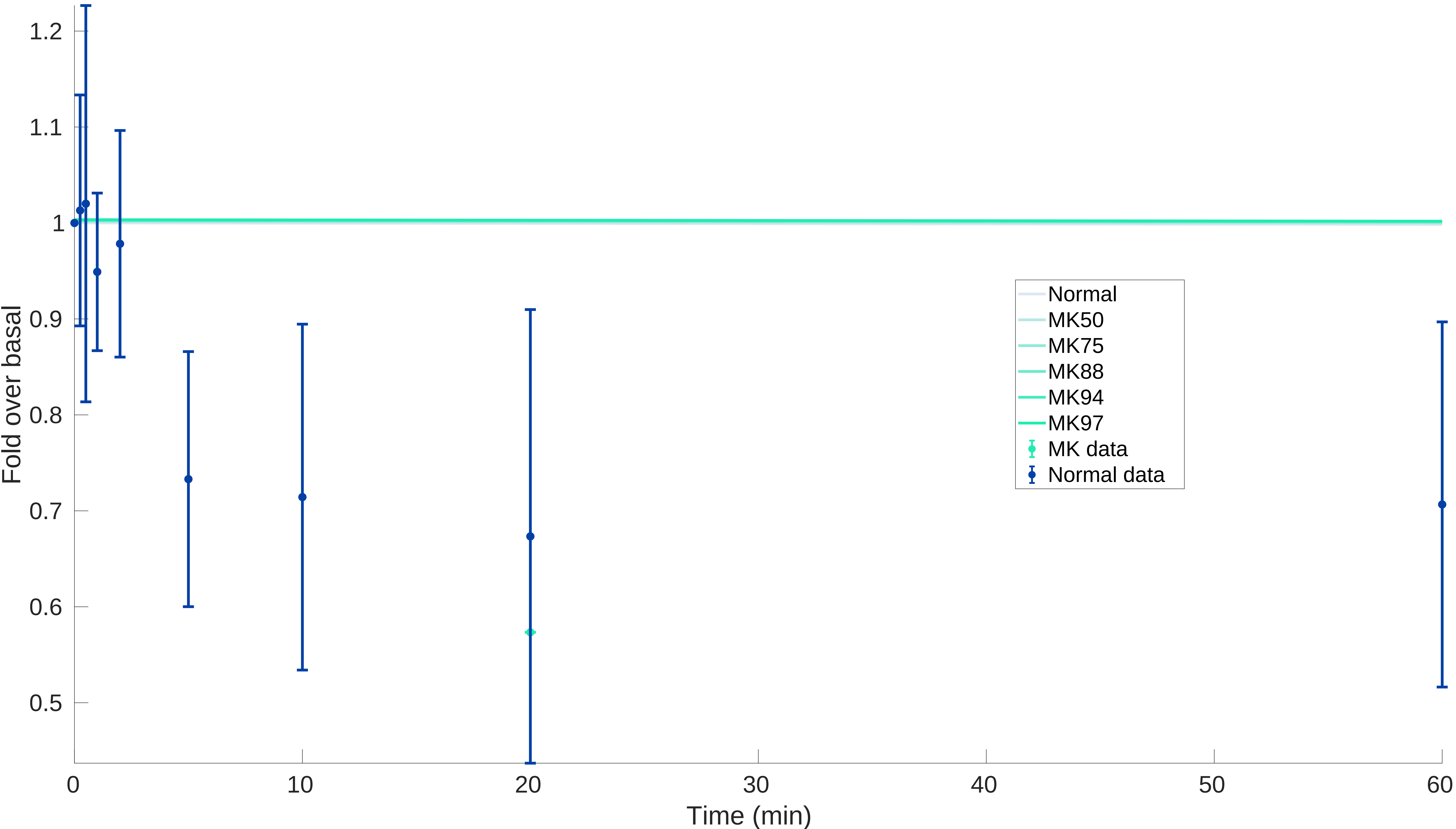

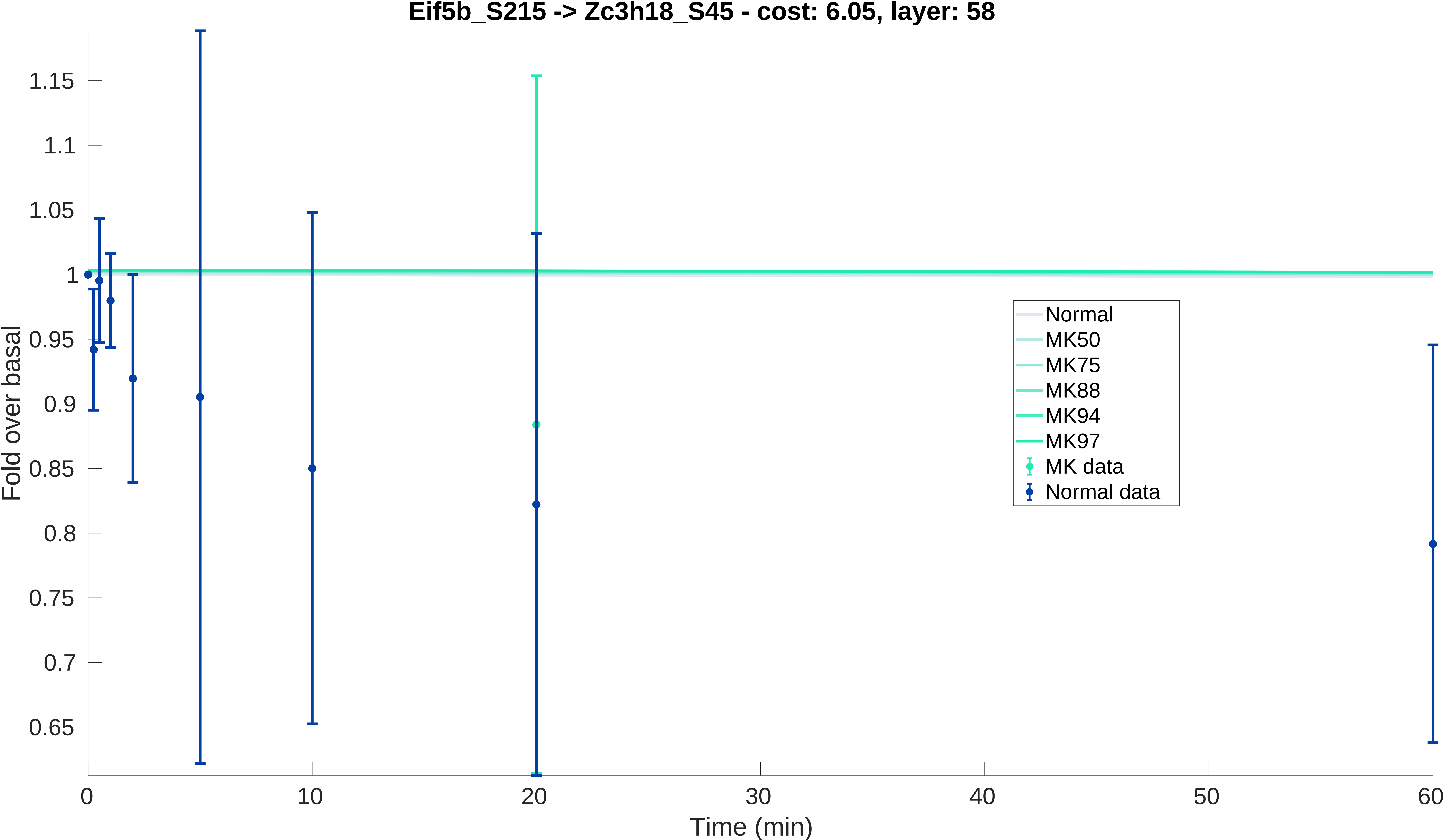

Eif5b_S215 -> Zc3h18_S45 - cost: 6.05, layer: 58
Fold over basal
1.15
1.1
1.05
1
0.95
0.9
0.85
0.8
0.75
0.7
0.65
0
10
20
30
40
50
60
Time (min)
Normal
MK50
MK75
MK88
MK94
MK97
MK data
Normal data

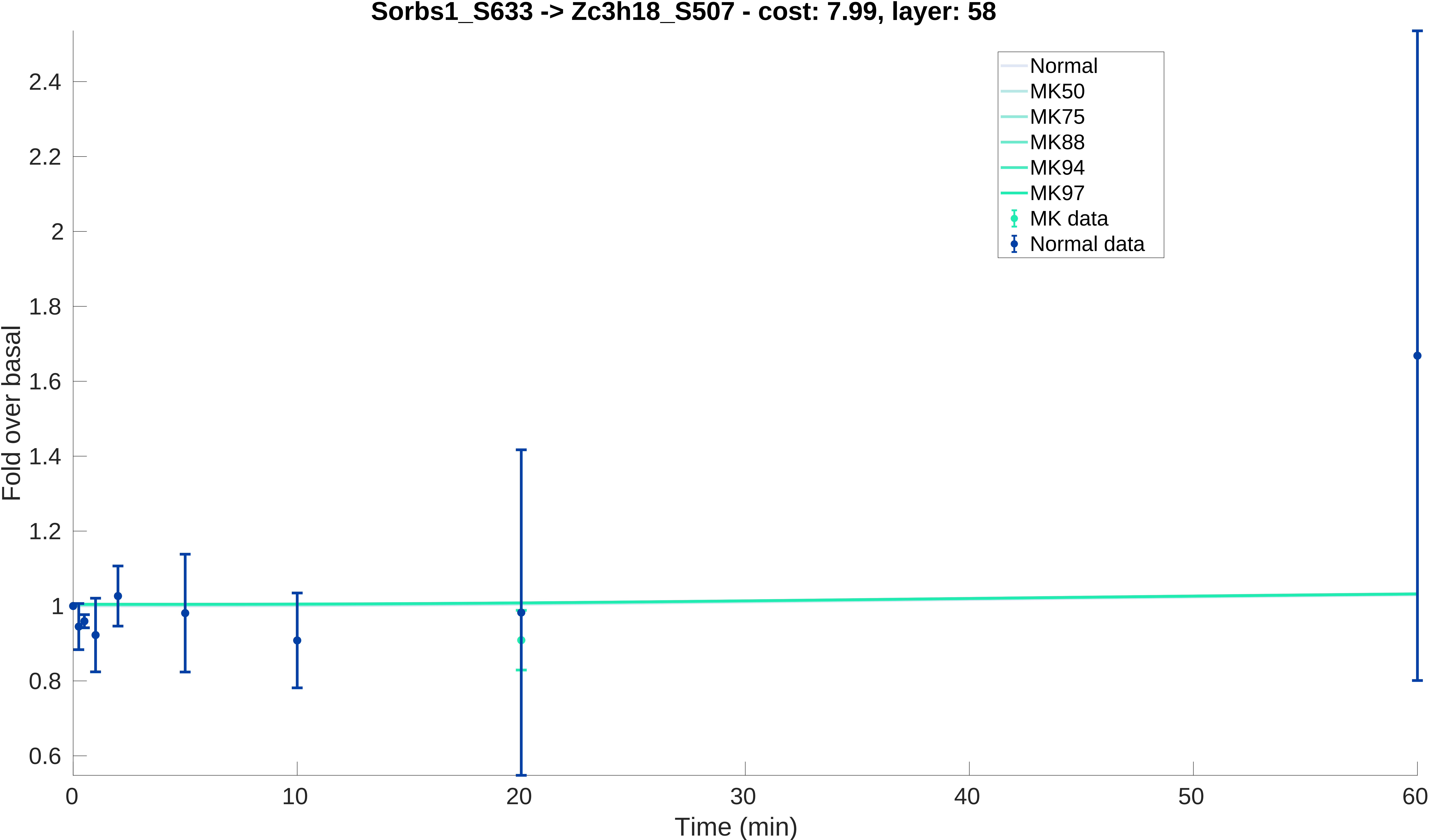
Sorbs1_S633 -> Zc3h18_S507 - cost: 7.99, layer: 58
Fold over basal
2.4
2.2
2
1.8
1.6
1.4
1.2
1
0.8
0.6
0
10
20
30
40
50
60
Time (min)
Normal
MK50
MK75
MK88
MK94
MK97
MK data
Normal data

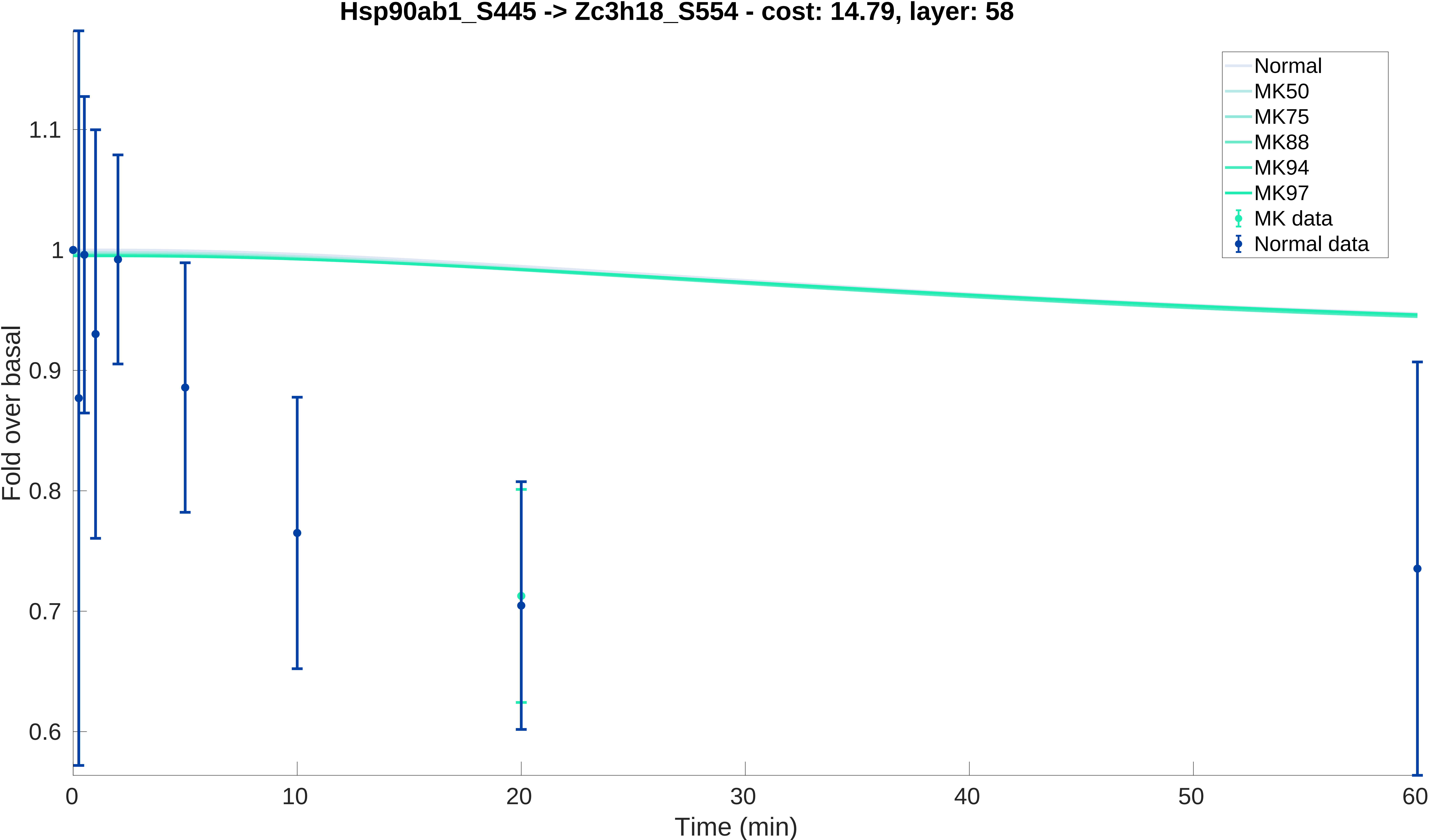

Hsp90ab1_S445 -> Zc3h18_S554 - cost: 14.79, layer: 58
Fold over basal
Time (min)
0
10
20
30
40
50
60
0.6
0.7
0.8
0.9
1
1.1
Normal
MK50
MK75
MK88
MK94
MK97
MK data
Normal data

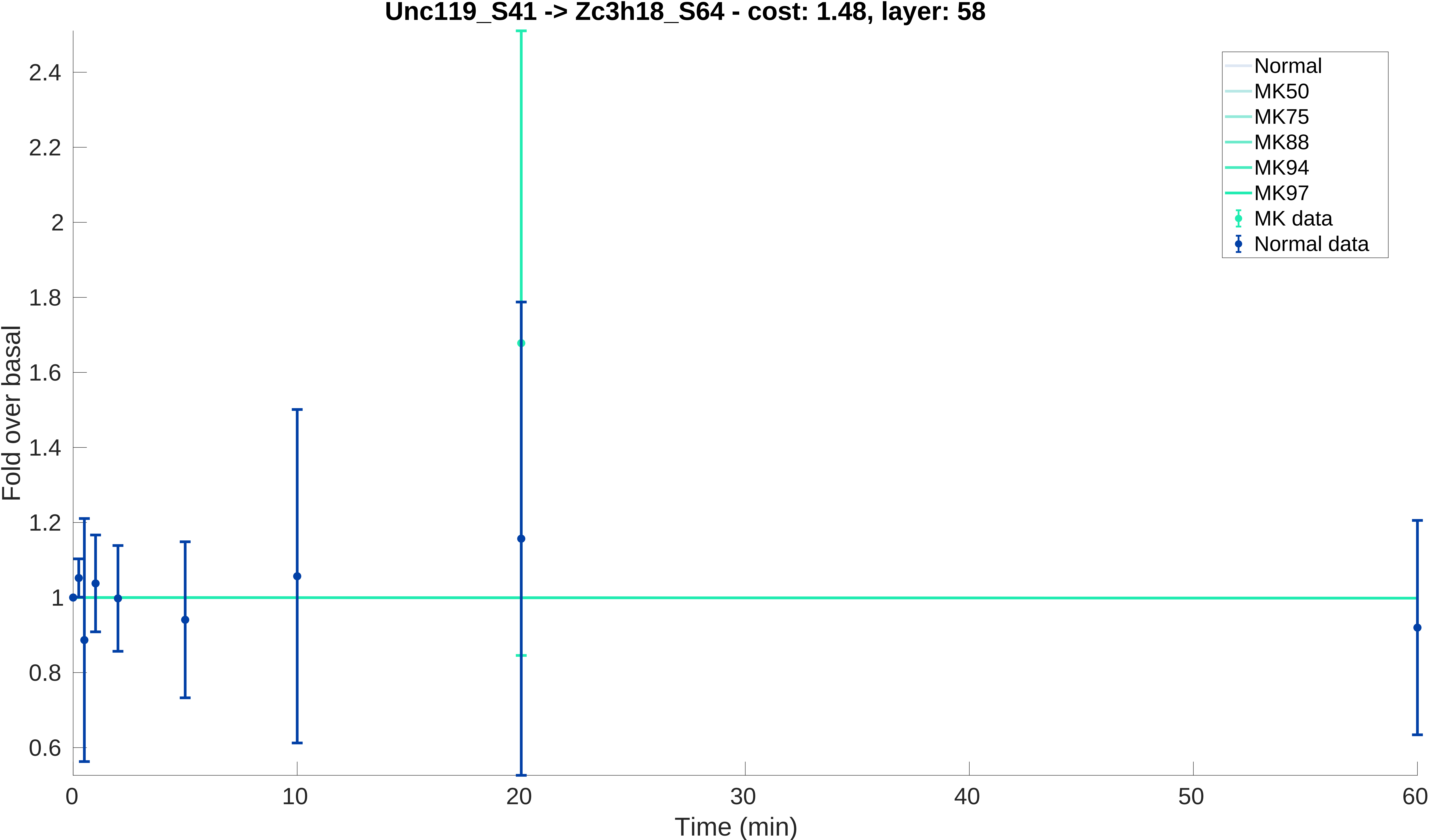
Unc119_S41 -> Zc3h18_S64 - cost: 1.48, layer: 58
Fold over basal
Time (min)
0.6
0.8
1
1.2
1.4
1.6
1.8
2
2.2
2.4
0
10
20
30
40
50
60
Normal
MK50
MK75
MK88
MK94
MK97
MK data
Normal data

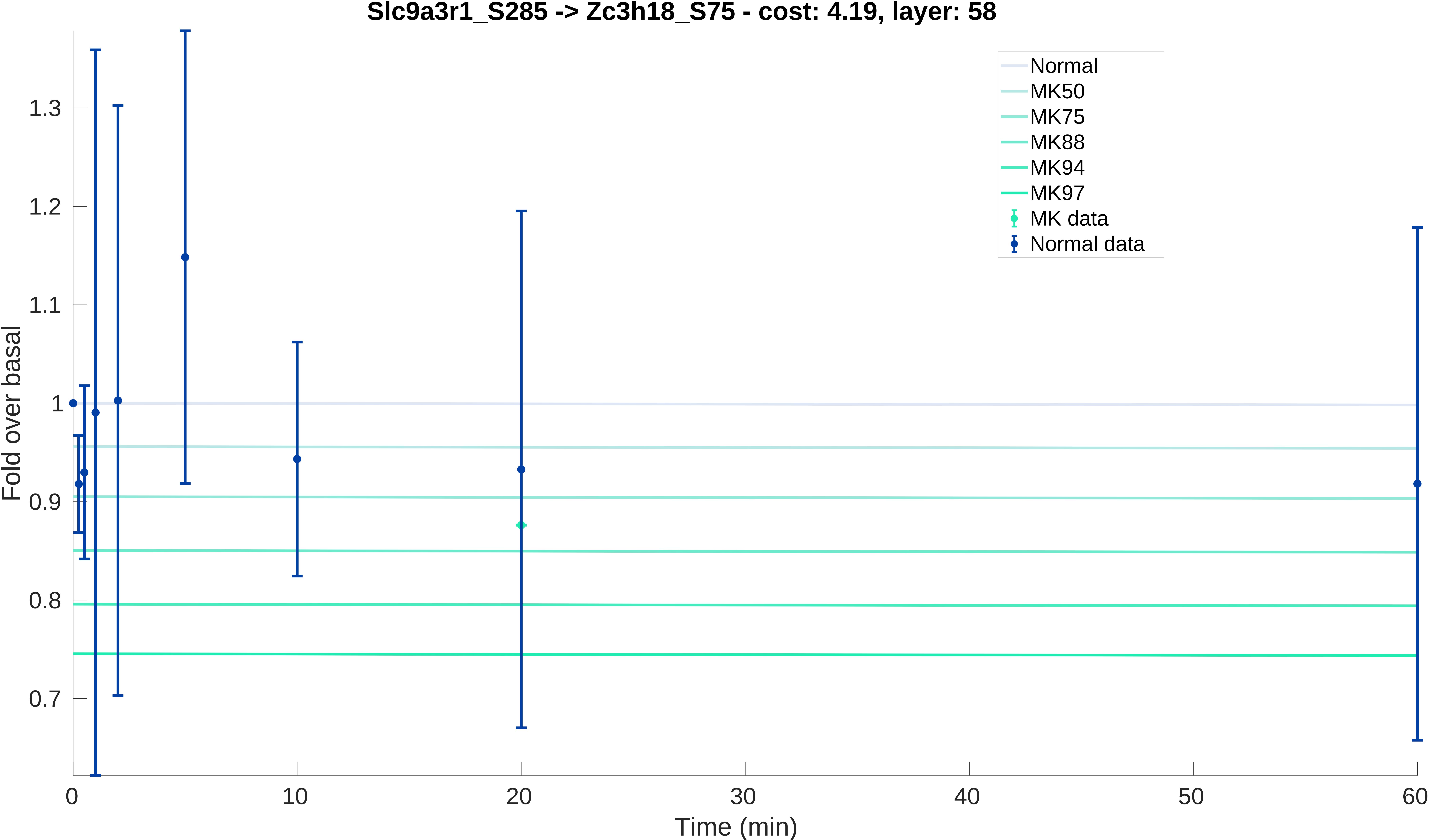

Slc9a3r1_S285 -> Zc3h18_S75 - cost: 4.19, layer: 58
Normal
MK50
MK75
MK88
MK94
MK97
MK data
Normal data
Fold over basal
1.3
1.2
1.1
1
0.9
0.8
0.7
0
10
20
30
40
50
60
Time (min)

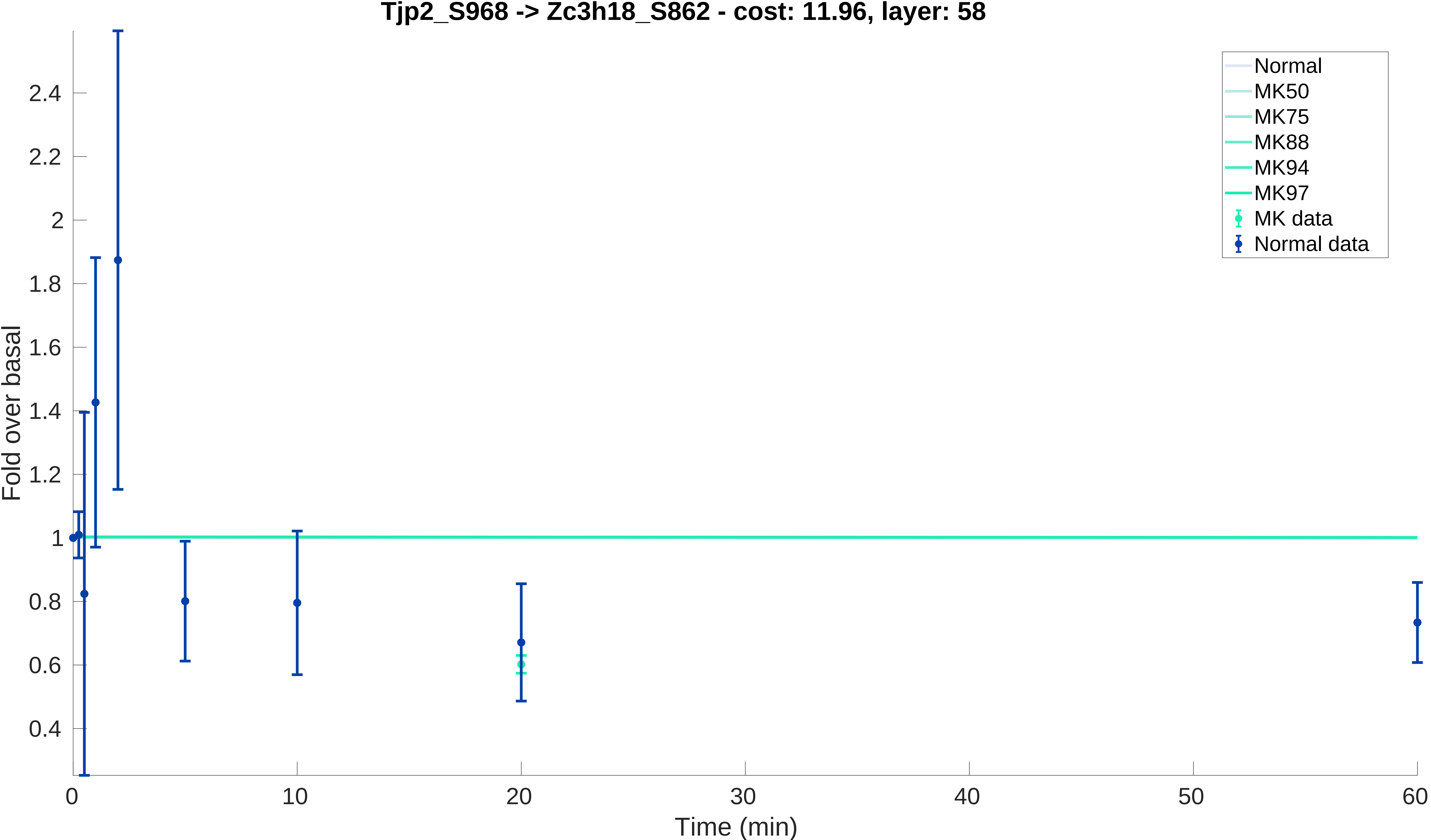
Tjp2_S968 -> Zc3h18_S862 - cost: 11.96, layer: 58
Fold over basal
Time (min)
0
10
20
30
40
50
60
0.4
0.6
0.8
1
1.2
1.4
1.6
1.8
2
2.2
2.4
Normal
MK50
MK75
MK88
MK94
MK97
MK data
Normal data

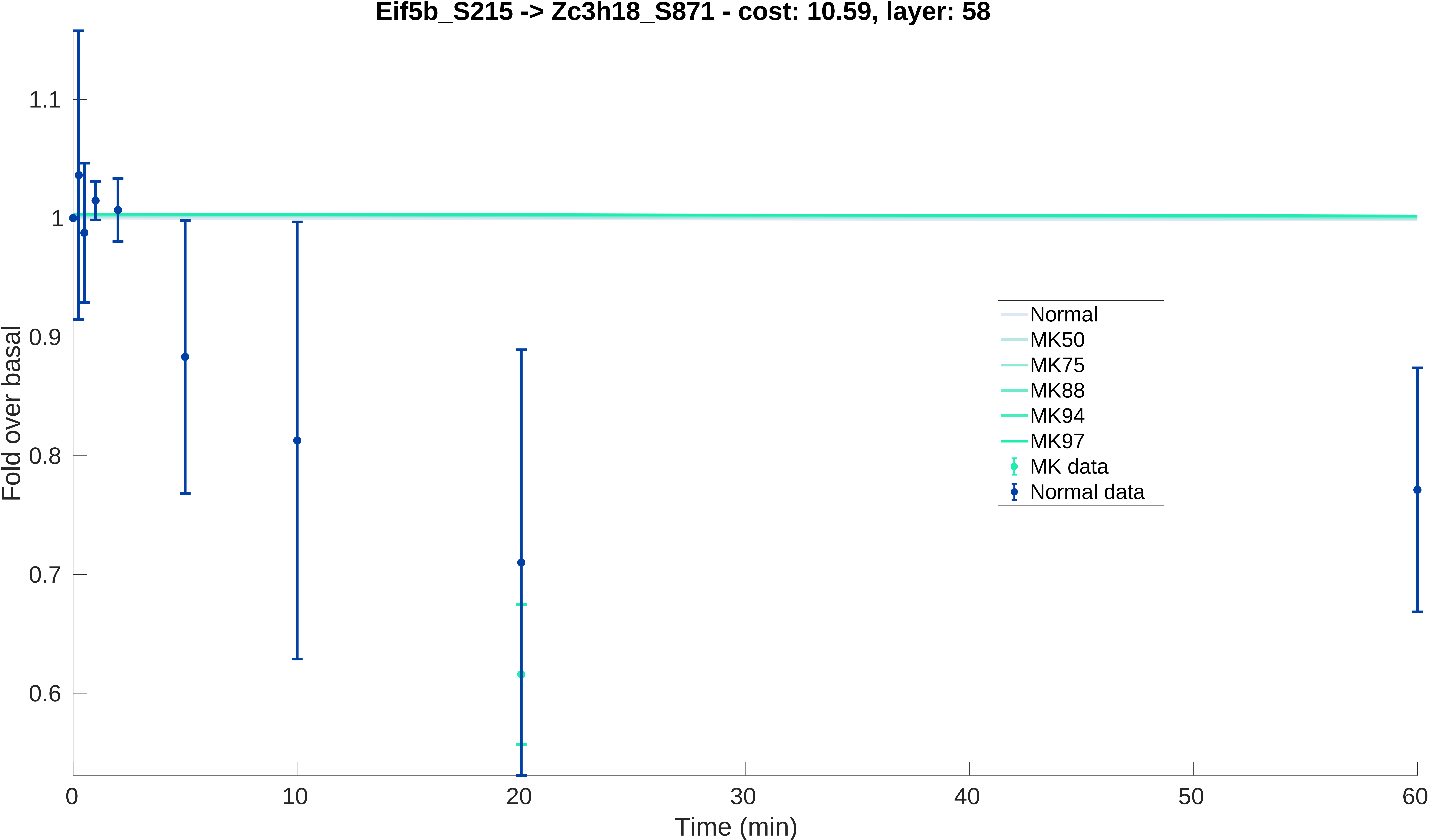

Eif5b_S215 -> Zc3h18_S871 - cost: 10.59, layer: 58
Fold over basal
1.1
1
0.9
0.8
0.7
0.6
0
10
20
30
40
50
60
Time (min)
Normal
MK50
MK75
MK88
MK94
MK97
MK data
Normal data

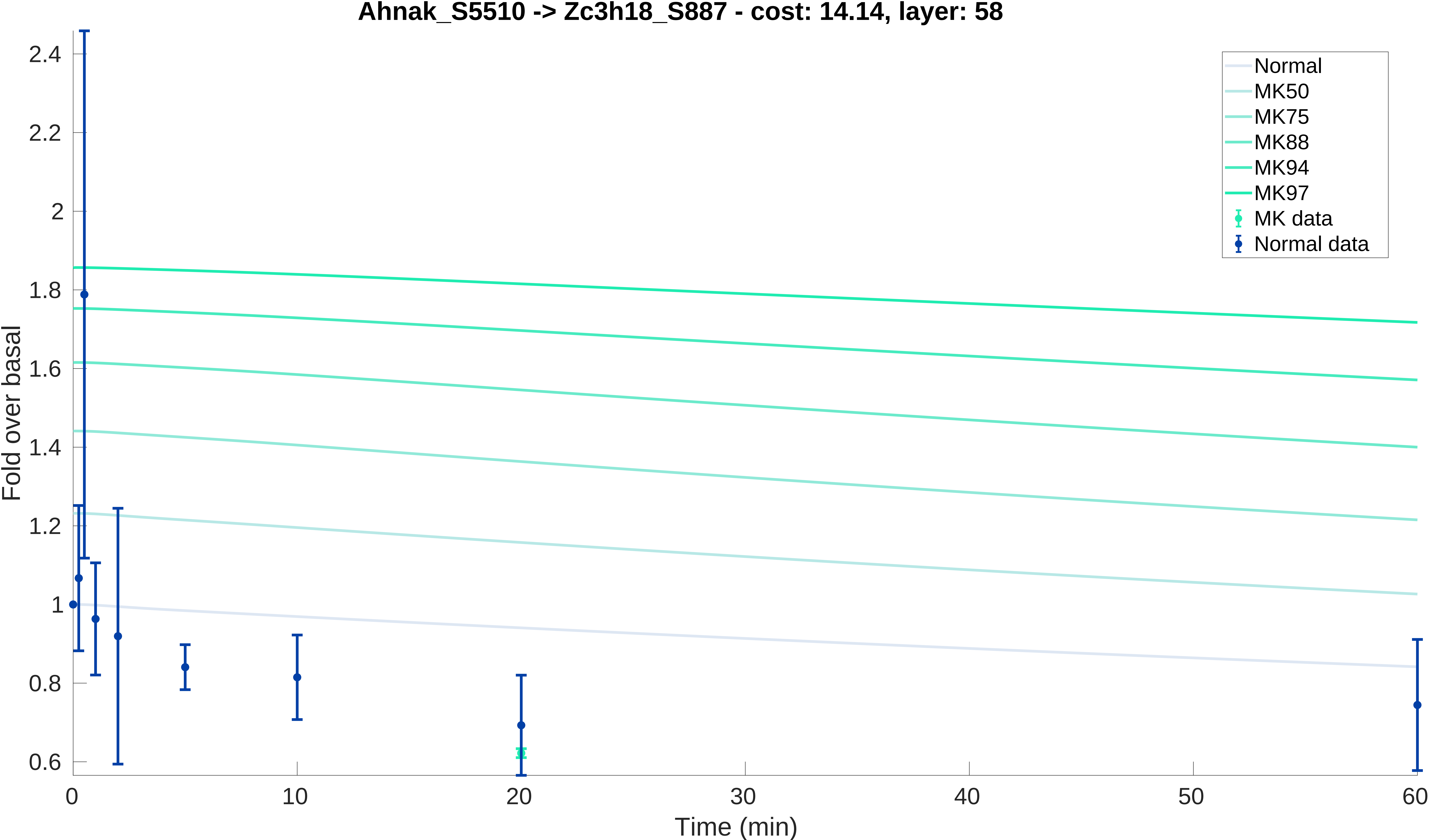

Ahnak_S5510 -> Zc3h18_S887 - cost: 14.14, layer: 58
Fold over basal
Time (min)
0.6
0.8
1
1.2
1.4
1.6
1.8
2
2.2
2.4
0
10
20
30
40
50
60
Normal
MK50
MK75
MK88
MK94
MK97
MK data
Normal data

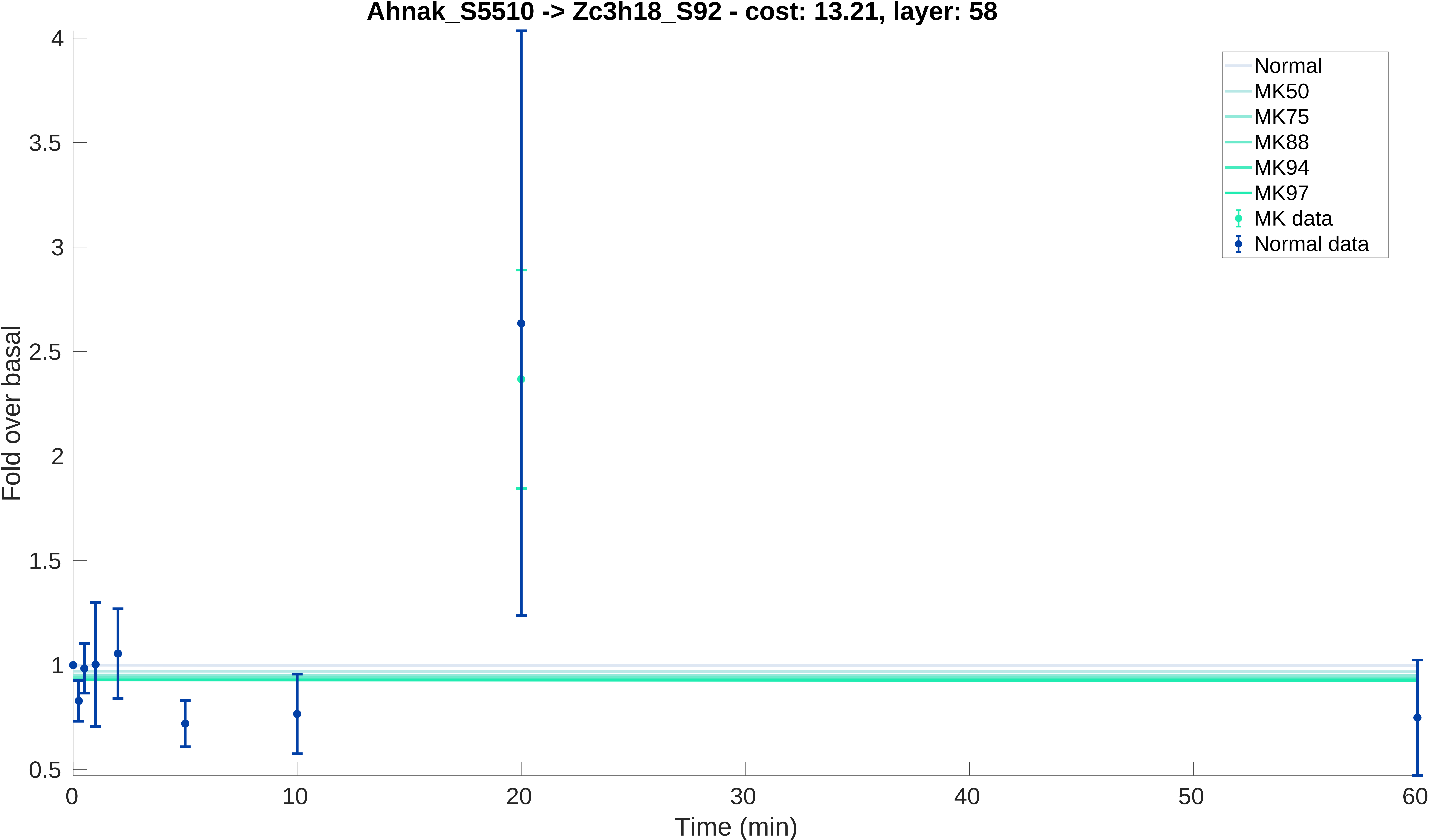

Ahnak_S5510 -> Zc3h18_S92 - cost: 13.21, layer: 58
Fold over basal
Time (min)
0.5
1
1.5
2
2.5
3
3.5
4
0
10
20
30
40
50
60
Normal
MK50
MK75
MK88
MK94
MK97
MK data
Normal data

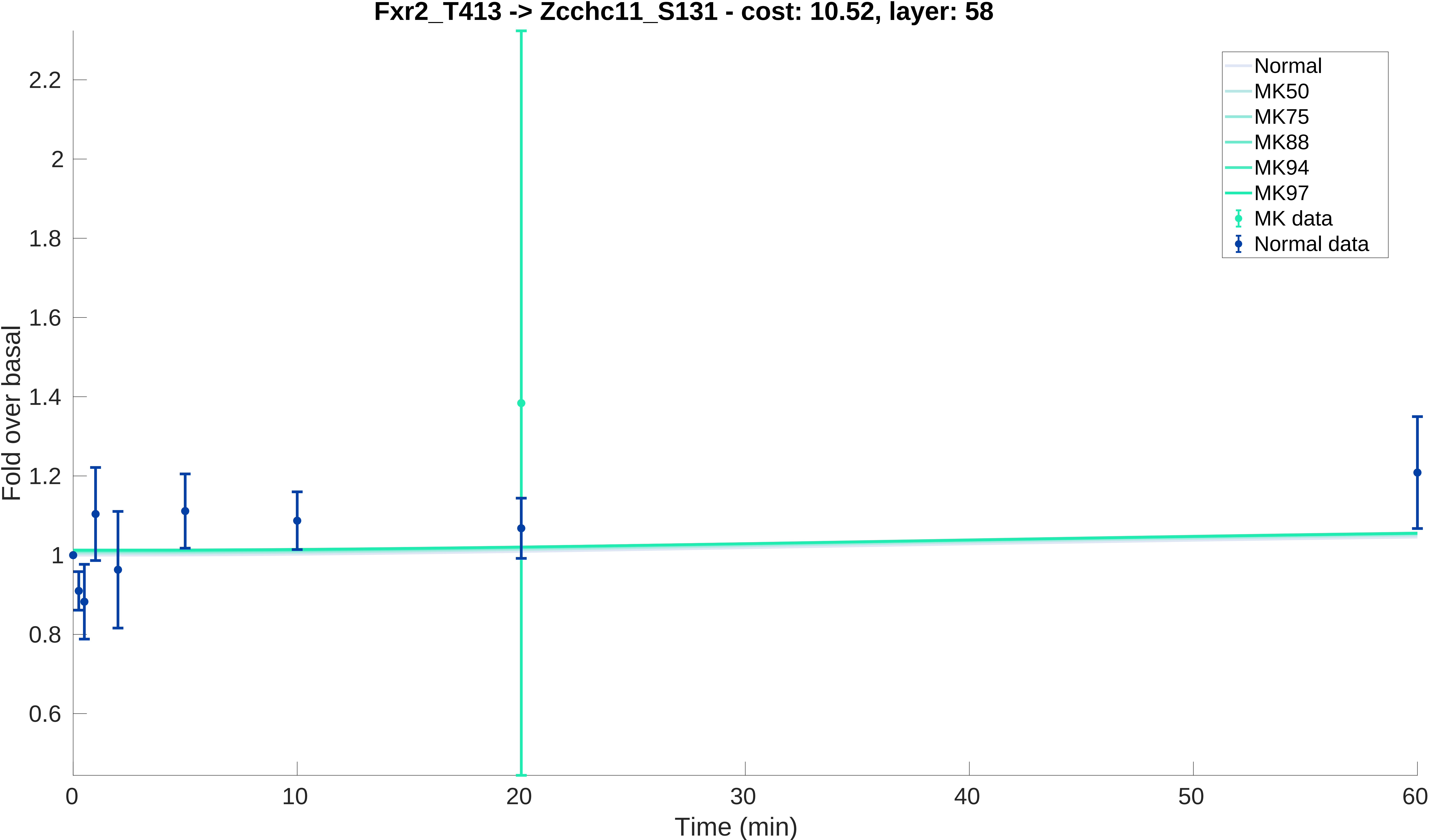

Fxr2_T413 -> Zcchc11_S131 - cost: 10.52, layer: 58
Fold over basal
Time (min)
0.6
0.8
1
1.2
1.4
1.6
1.8
2
2.2
0
10
20
30
40
50
60
Normal
MK50
MK75
MK88
MK94
MK97
MK data
Normal data

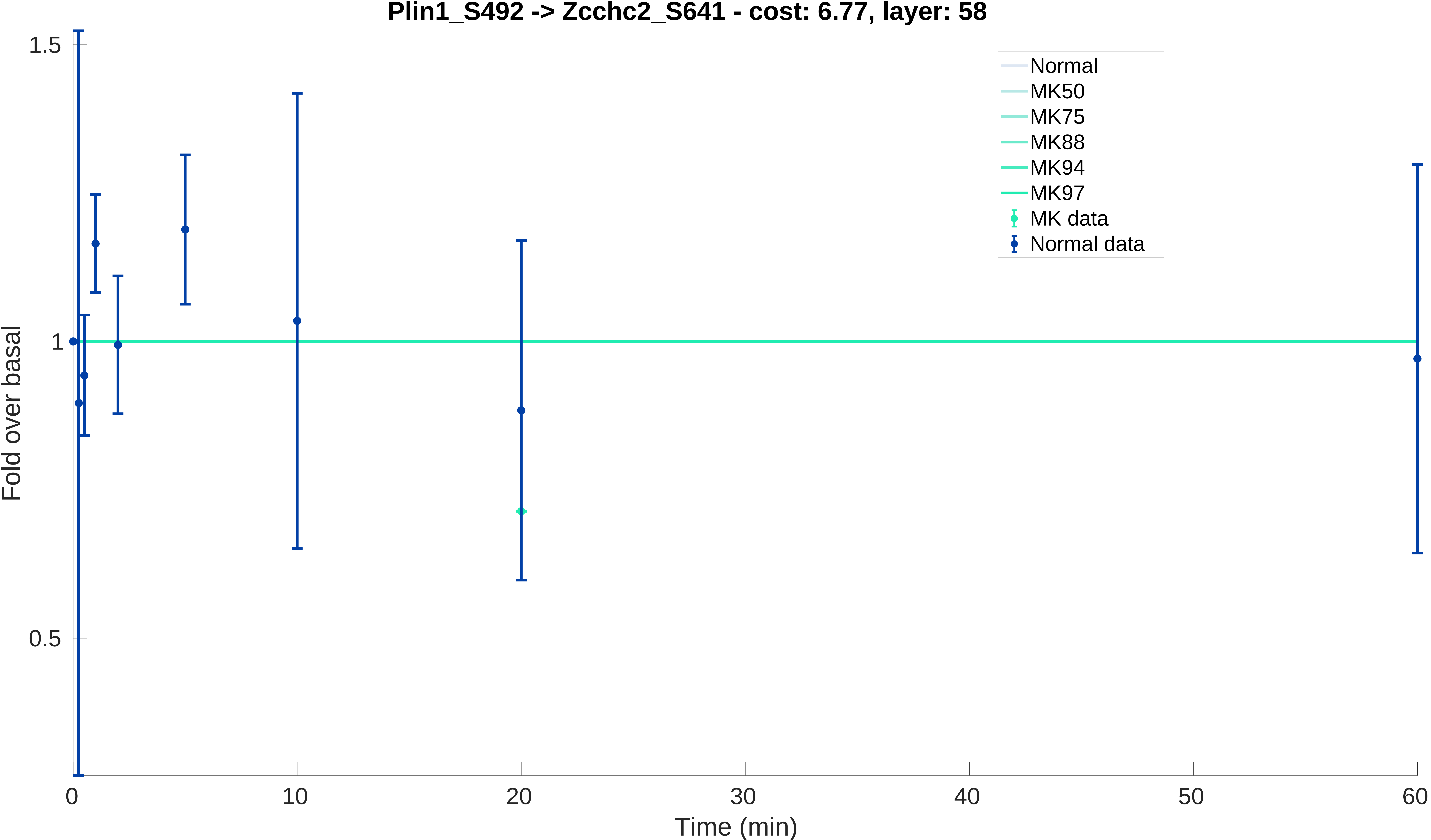

Plin1_S492 -> Zcchc2_S641 - cost: 6.77, layer: 58
Fold over basal
1.5
1
0.5
Time (min)
0
10
20
30
40
50
60
Normal
MK50
MK75
MK88
MK94
MK97
MK data
Normal data

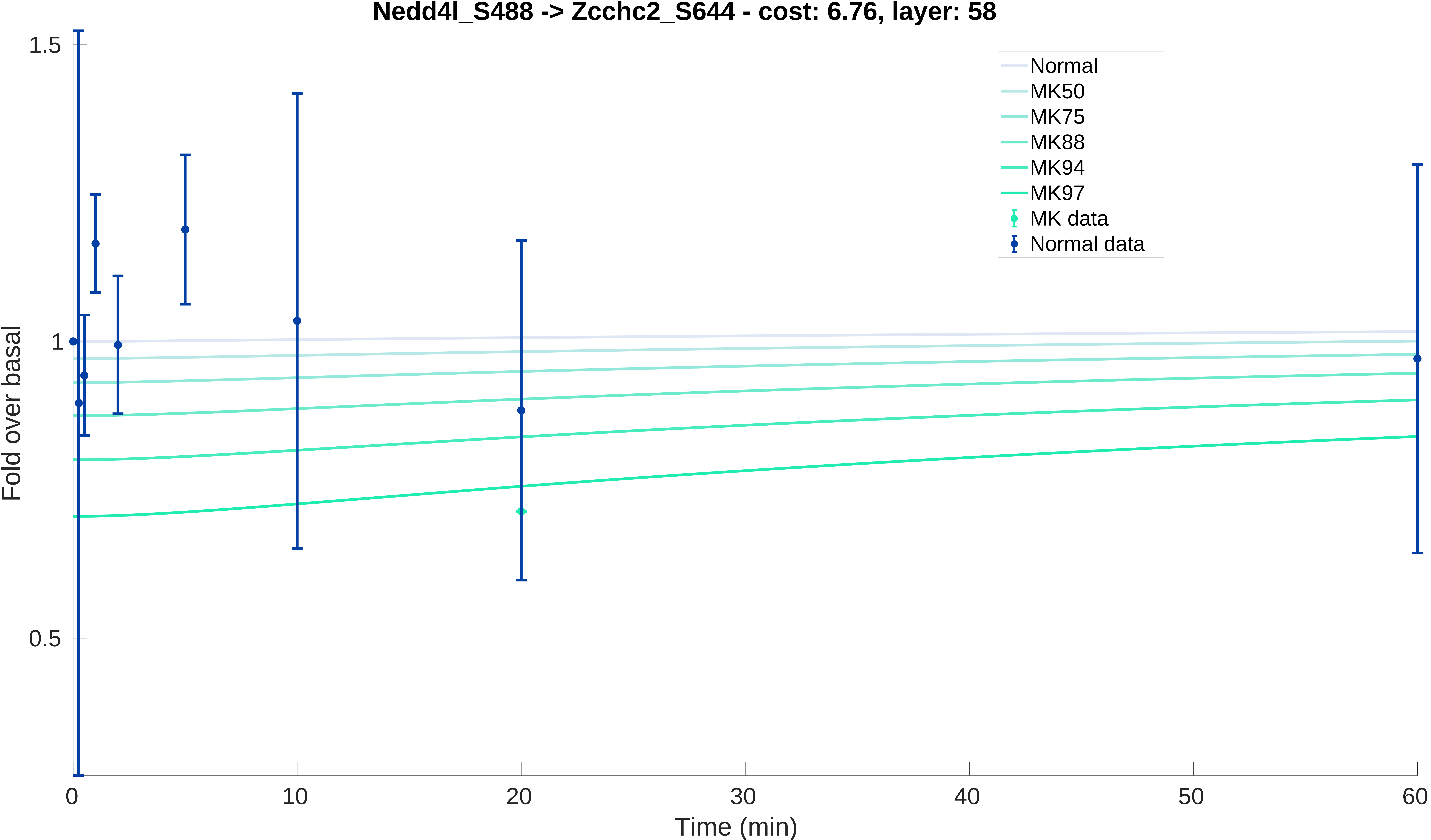

Nedd4l_S488 -> Zcchc2_S644 - cost: 6.76, layer: 58
Fold over basal
1.5
1
0.5
0
10
20
30
40
50
60
Time (min)
Normal
MK50
MK75
MK88
MK94
MK97
MK data
Normal data

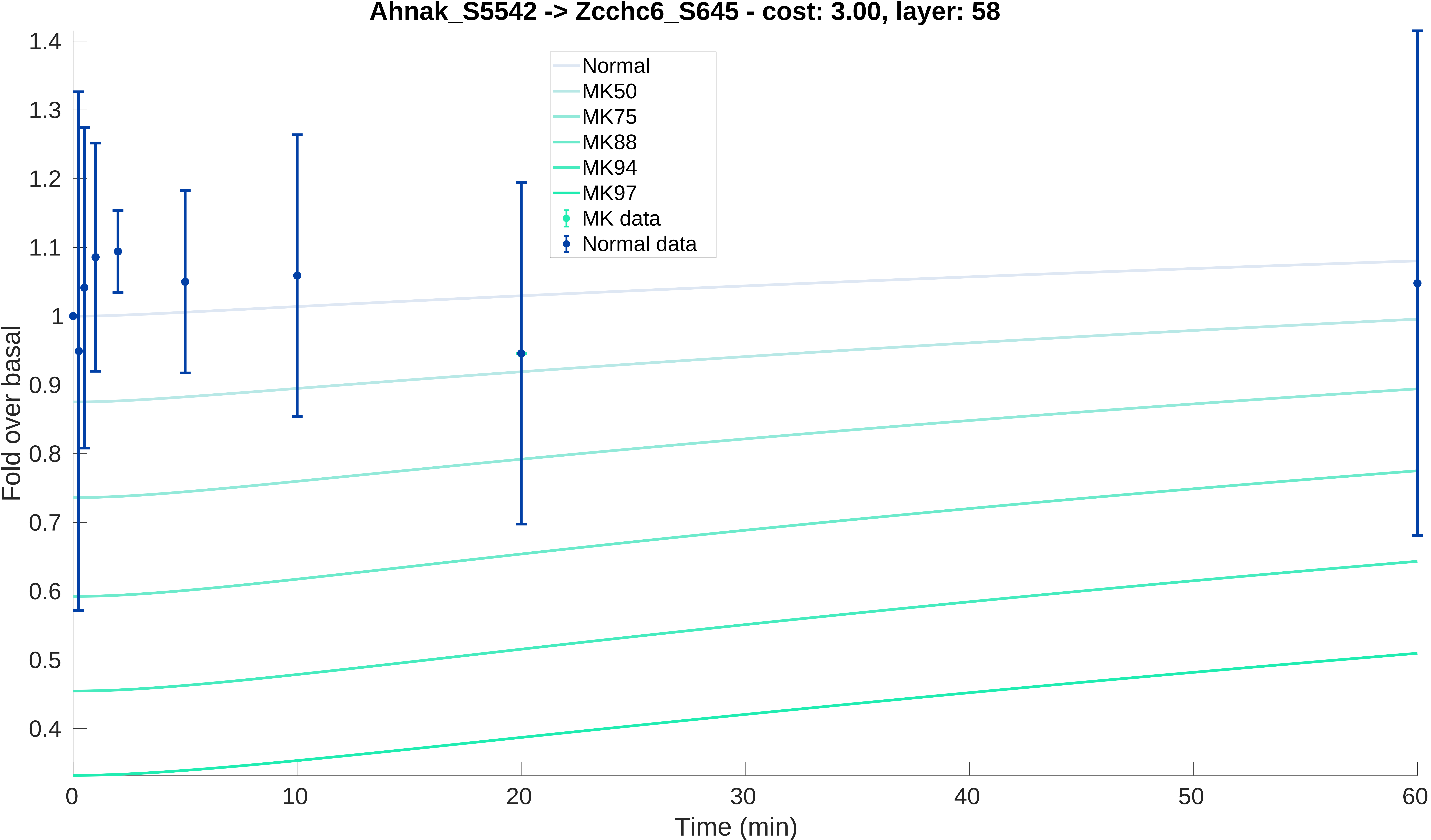

Ahnak_S5542 -> Zcchc6_S645 - cost: 3.00, layer: 58
Fold over basal
Time (min)
Normal
MK50
MK75
MK88
MK94
MK97
MK data
Normal data
1.4
1.3
1.2
1.1
1
0.9
0.8
0.7
0.6
0.5
0.4
0
10
20
30
40
50
60

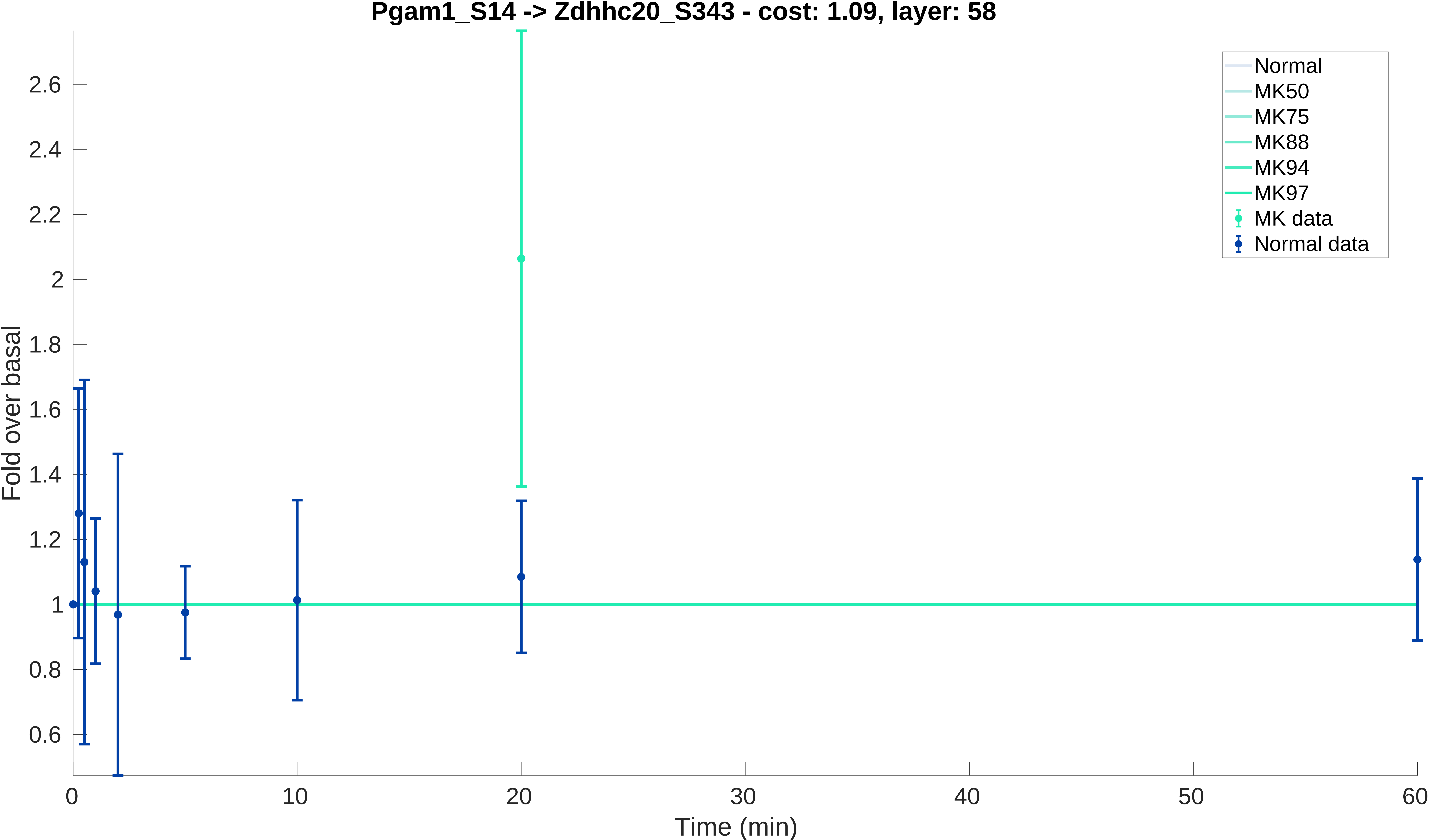

Pgam1_S14 -> Zdhhc20_S343 - cost: 1.09, layer: 58
Normal
MK50
MK75
MK88
MK94
MK97
MK data
Normal data
Fold over basal
Time (min)
0.6
0.8
1
1.2
1.4
1.6
1.8
2
2.2
2.4
2.6
0
10
20
30
40
50
60

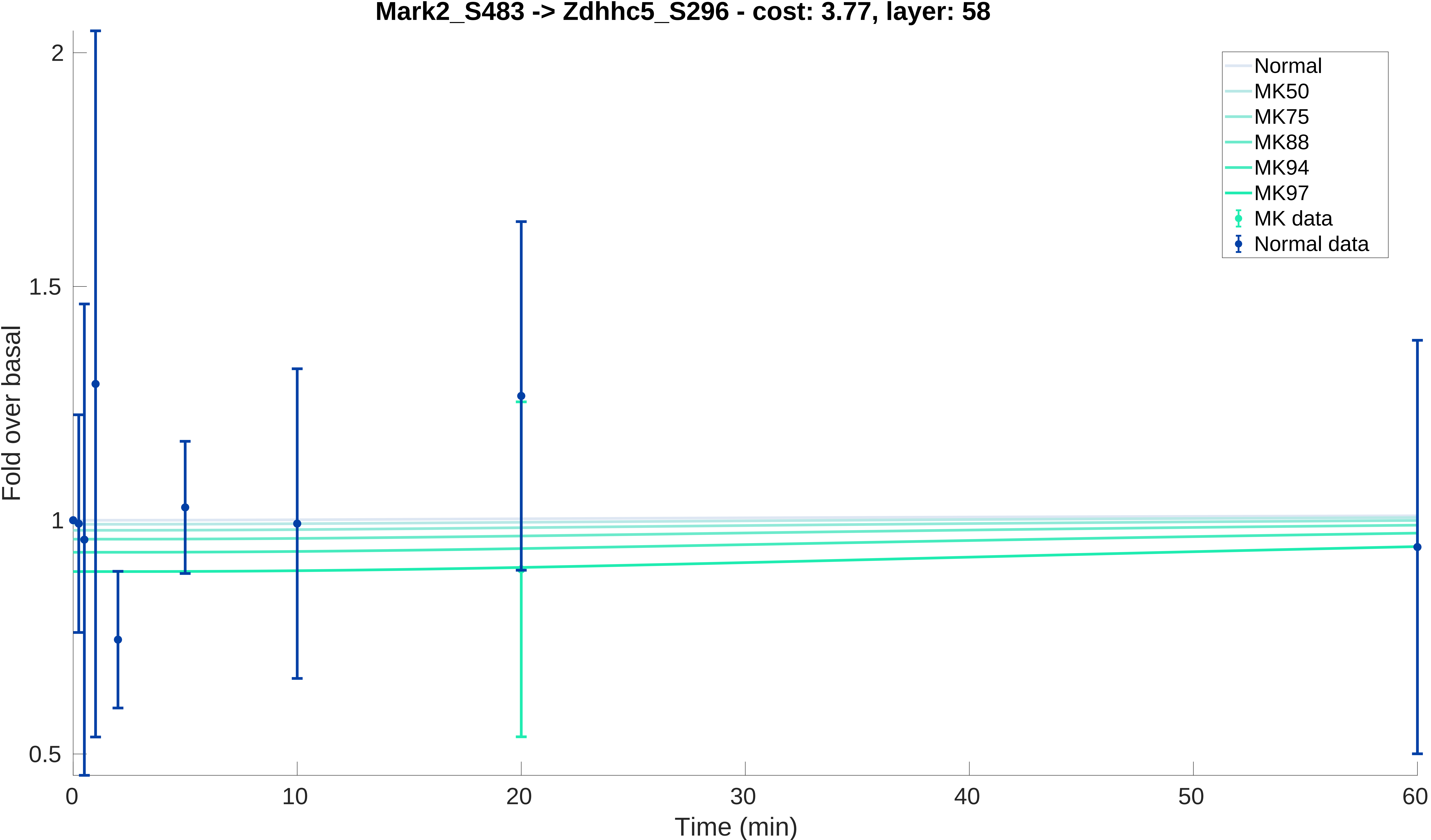

Mark2_S483 -> Zdhhc5_S296 - cost: 3.77, layer: 58
Fold over basal
Time (min)
0.5
1
1.5
2
0
10
20
30
40
50
60
Normal
MK50
MK75
MK88
MK94
MK97
MK data
Normal data

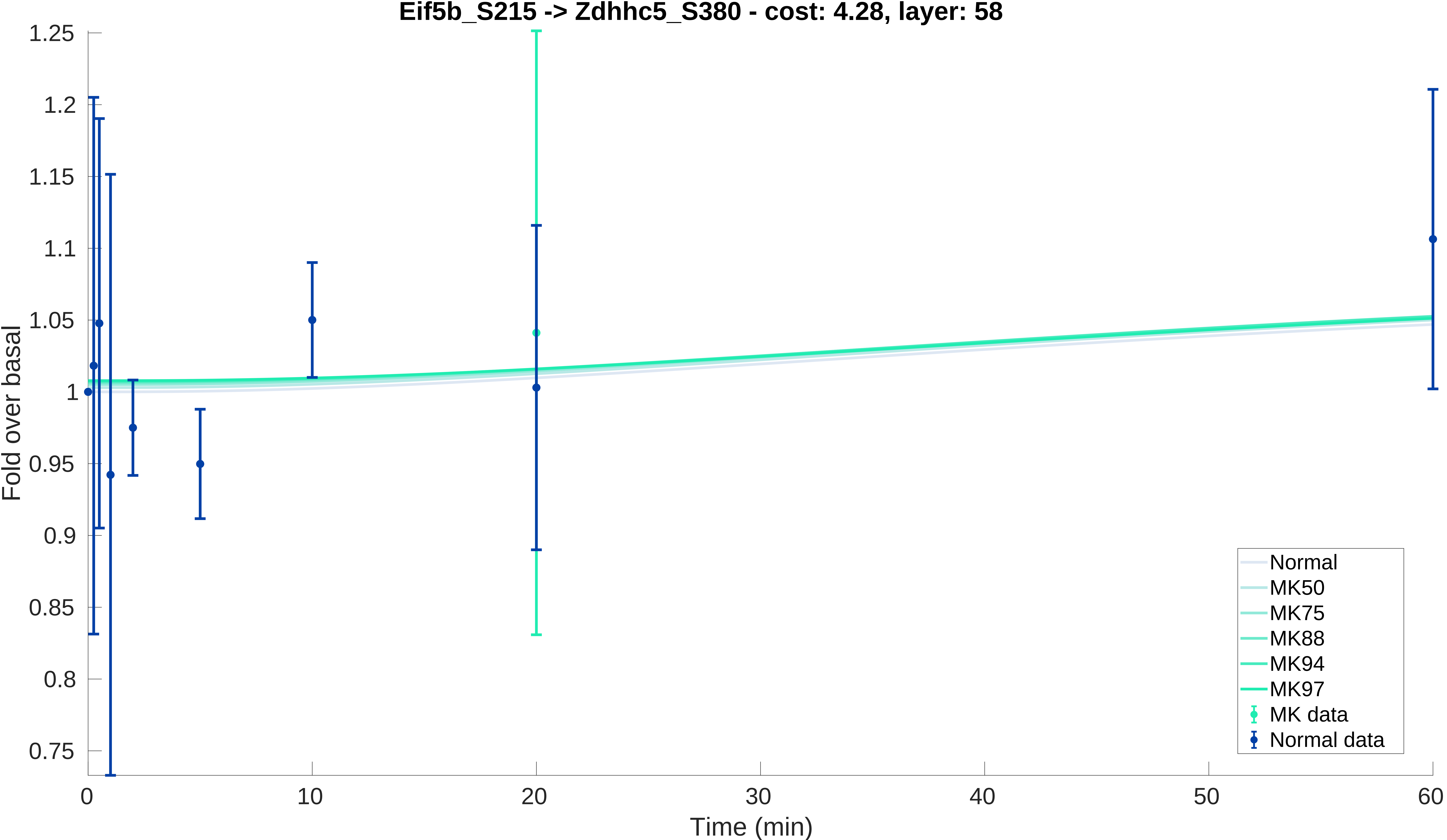

Eif5b_S215 -> Zdhhc5_S380 - cost: 4.28, layer: 58
Fold over basal
Time (min)
1.25
1.2
1.15
1.1
1.05
1
0.95
0.9
0.85
0.8
0.75
0
10
20
30
40
50
60
Normal
MK50
MK75
MK88
MK94
MK97
MK data
Normal data

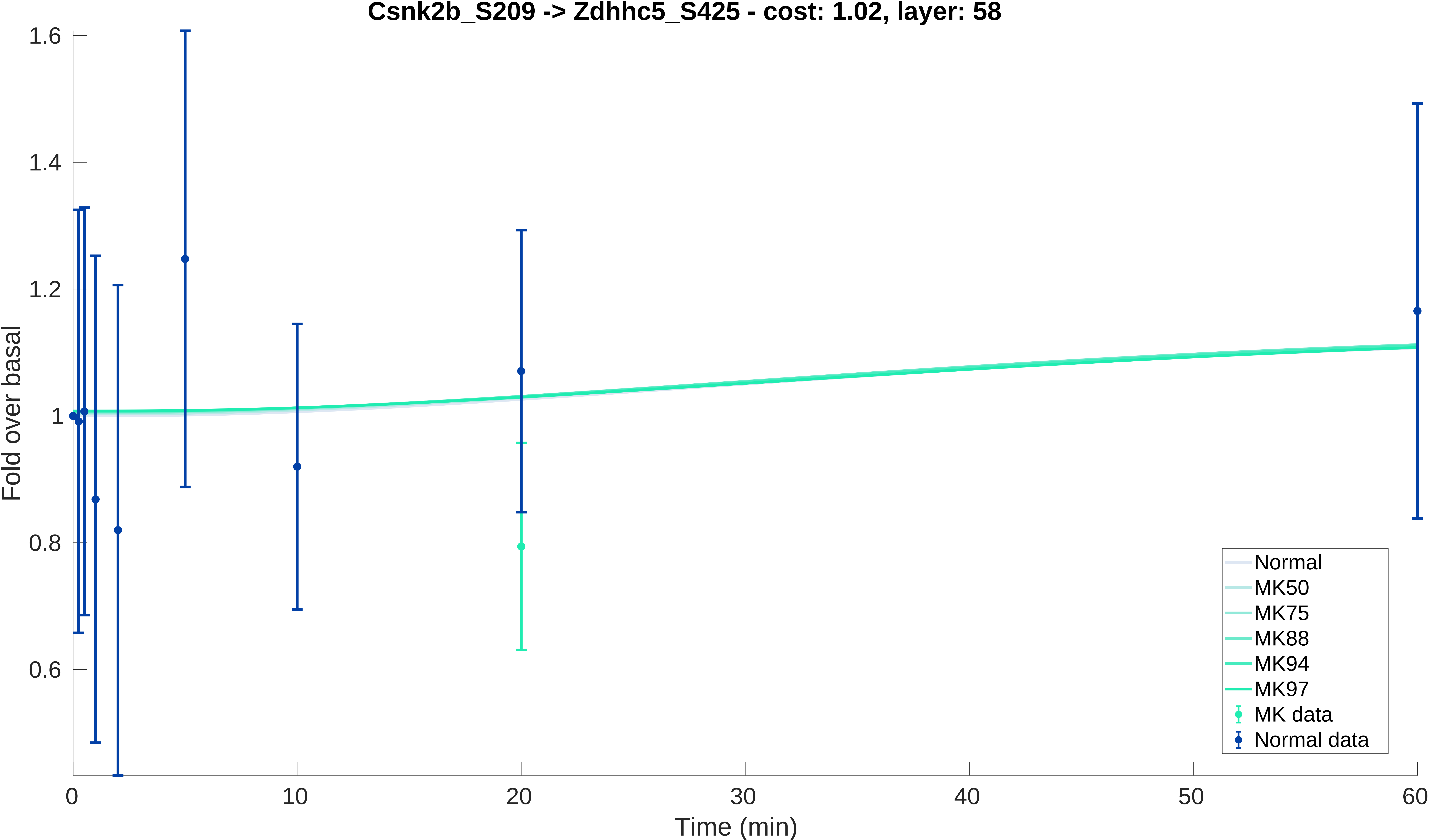

Csnk2b_S209 -> Zdhhc5_S425 - cost: 1.02, layer: 58
Fold over basal
Time (min)
1.6
1.4
1.2
1
0.8
0.6
0
10
20
30
40
50
60
Normal
MK50
MK75
MK88
MK94
MK97
MK data
Normal data

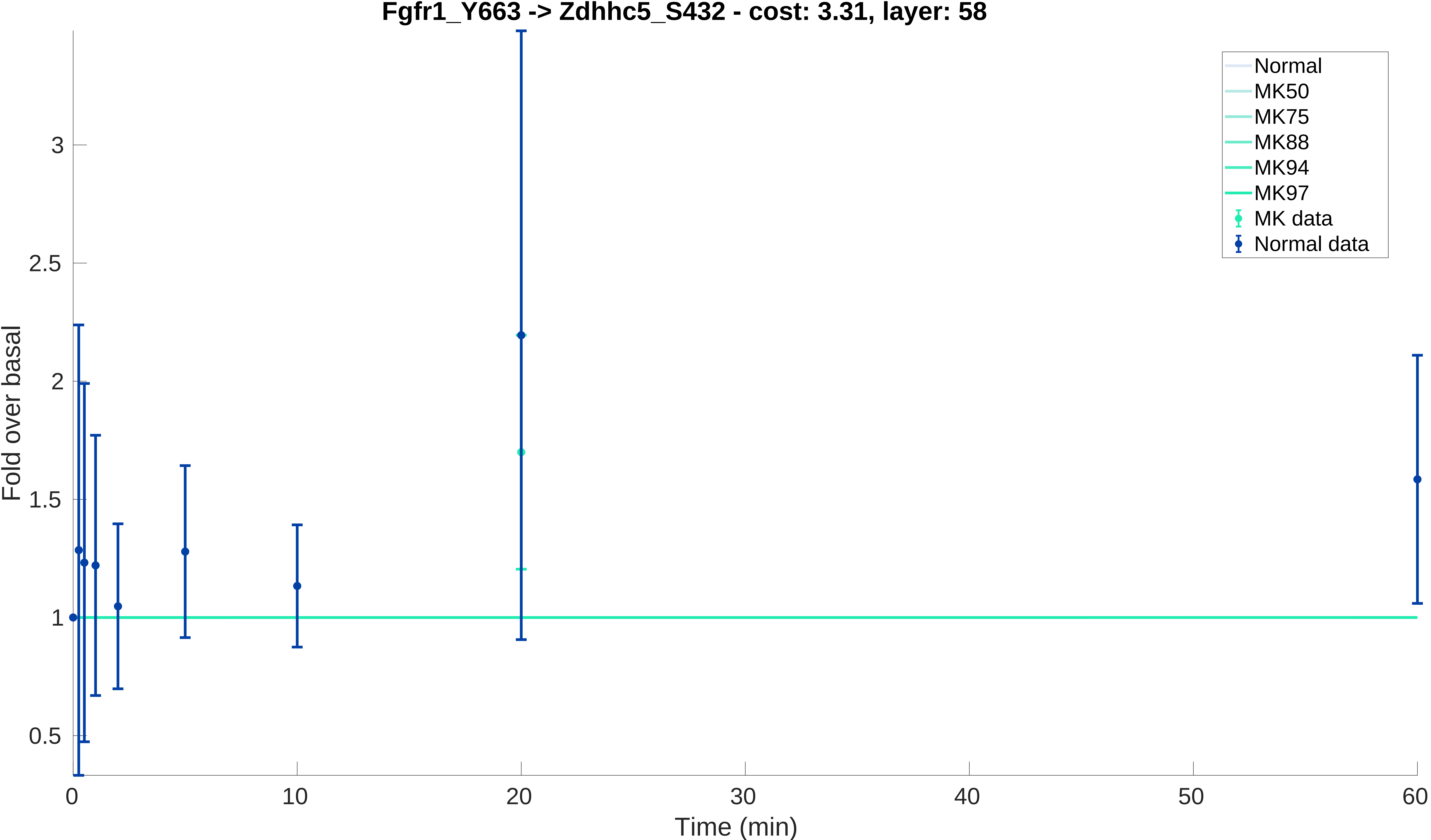

Fgfr1_Y663 -> Zdhhc5_S432 - cost: 3.31, layer: 58
Normal
MK50
MK75
MK88
MK94
MK97
MK data
Normal data
Fold over basal
3
2.5
2
1.5
1
0.5
0
10
20
30
40
50
60
Time (min)

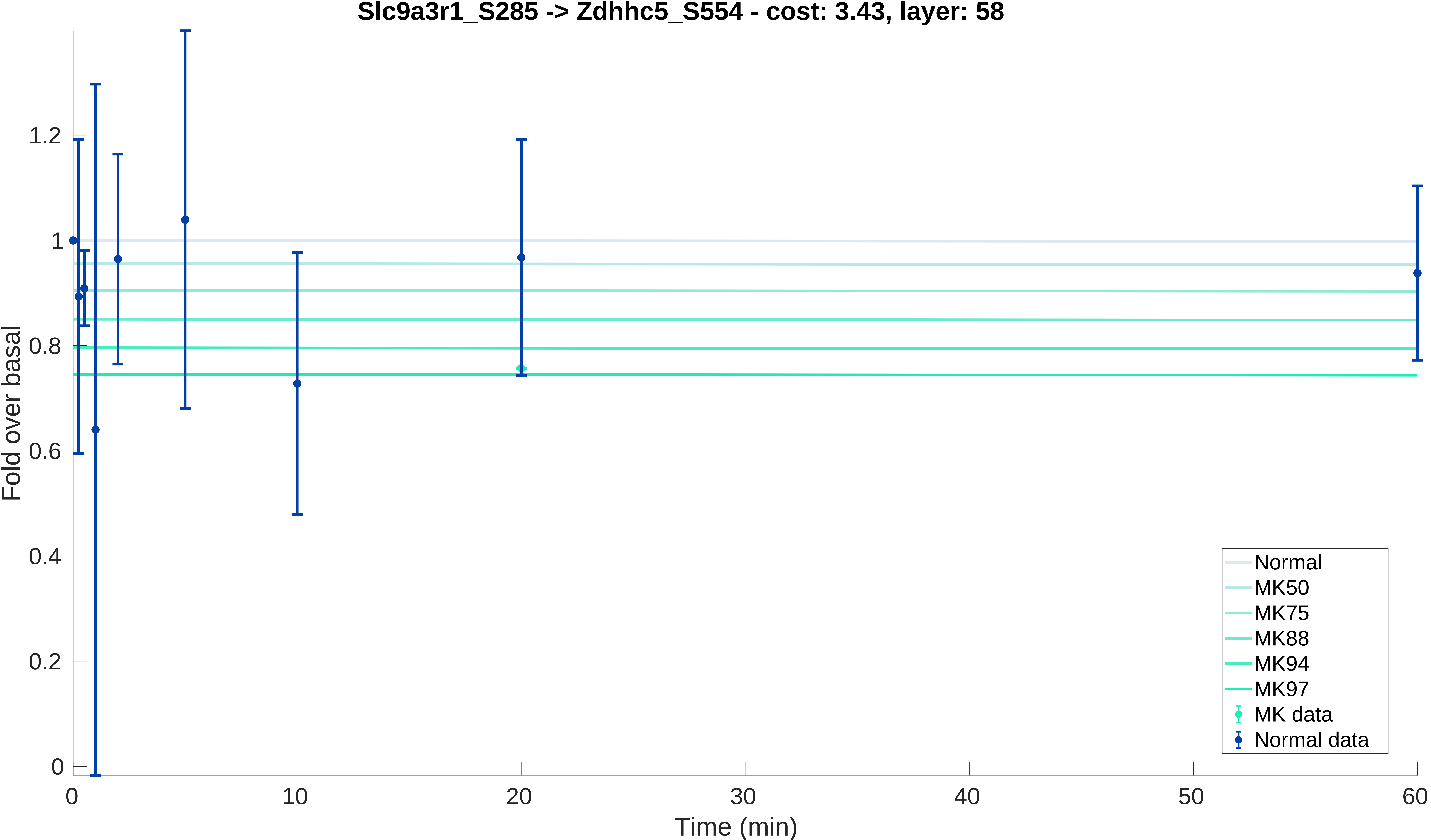

Slc9a3r1_S285 -> Zdhhc5_S554 - cost: 3.43, layer: 58
Fold over basal
Time (min)
0
0.2
0.4
0.6
0.8
1
1.2
0
10
20
30
40
50
60
Normal
MK50
MK75
MK88
MK94
MK97
MK data
Normal data

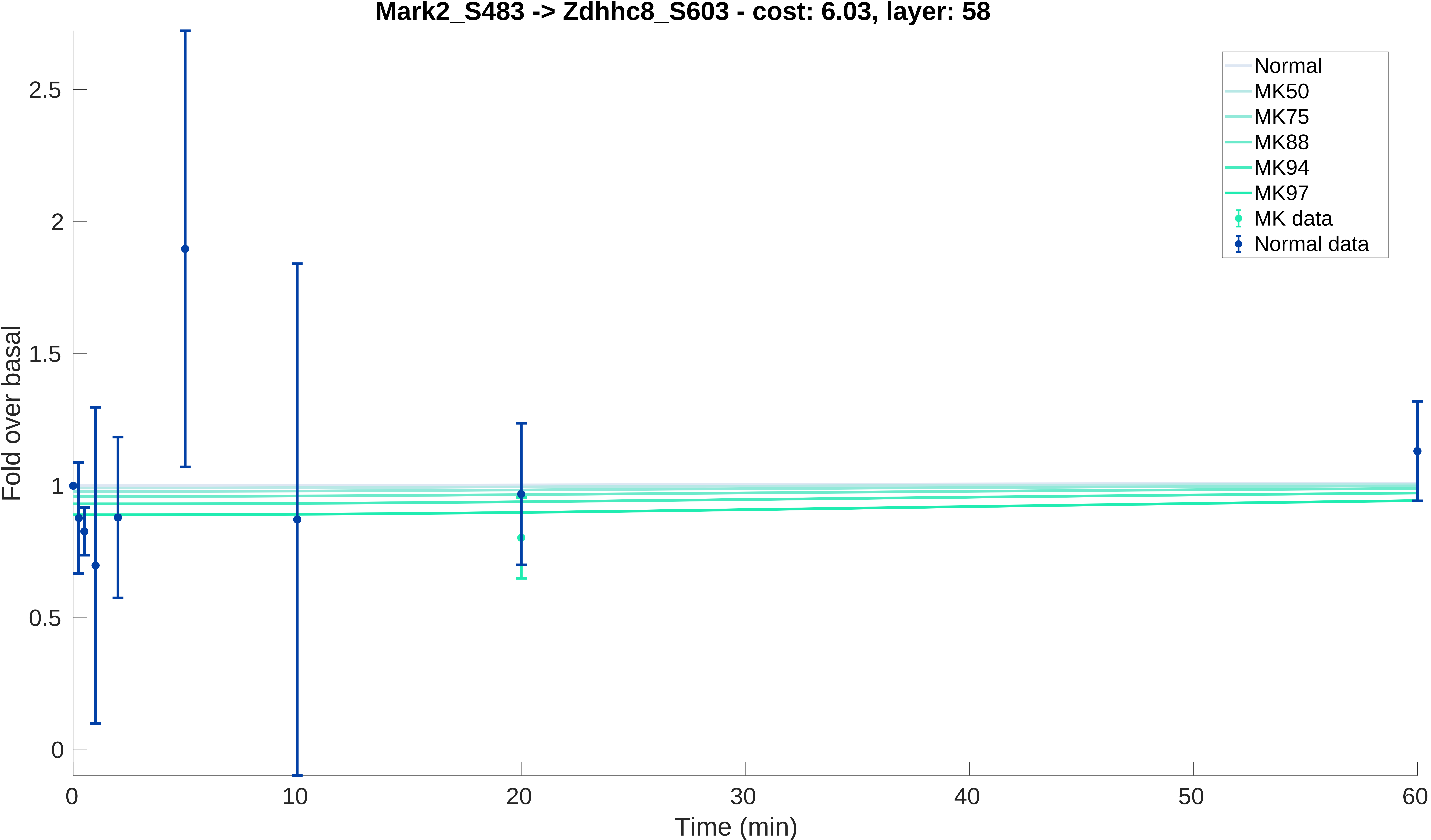

Mark2_S483 -> Zdhhc8_S603 - cost: 6.03, layer: 58
Normal
MK50
MK75
MK88
MK94
MK97
MK data
Normal data
Fold over basal
Time (min)
0
0.5
1
1.5
2
2.5
0
10
20
30
40
50
60

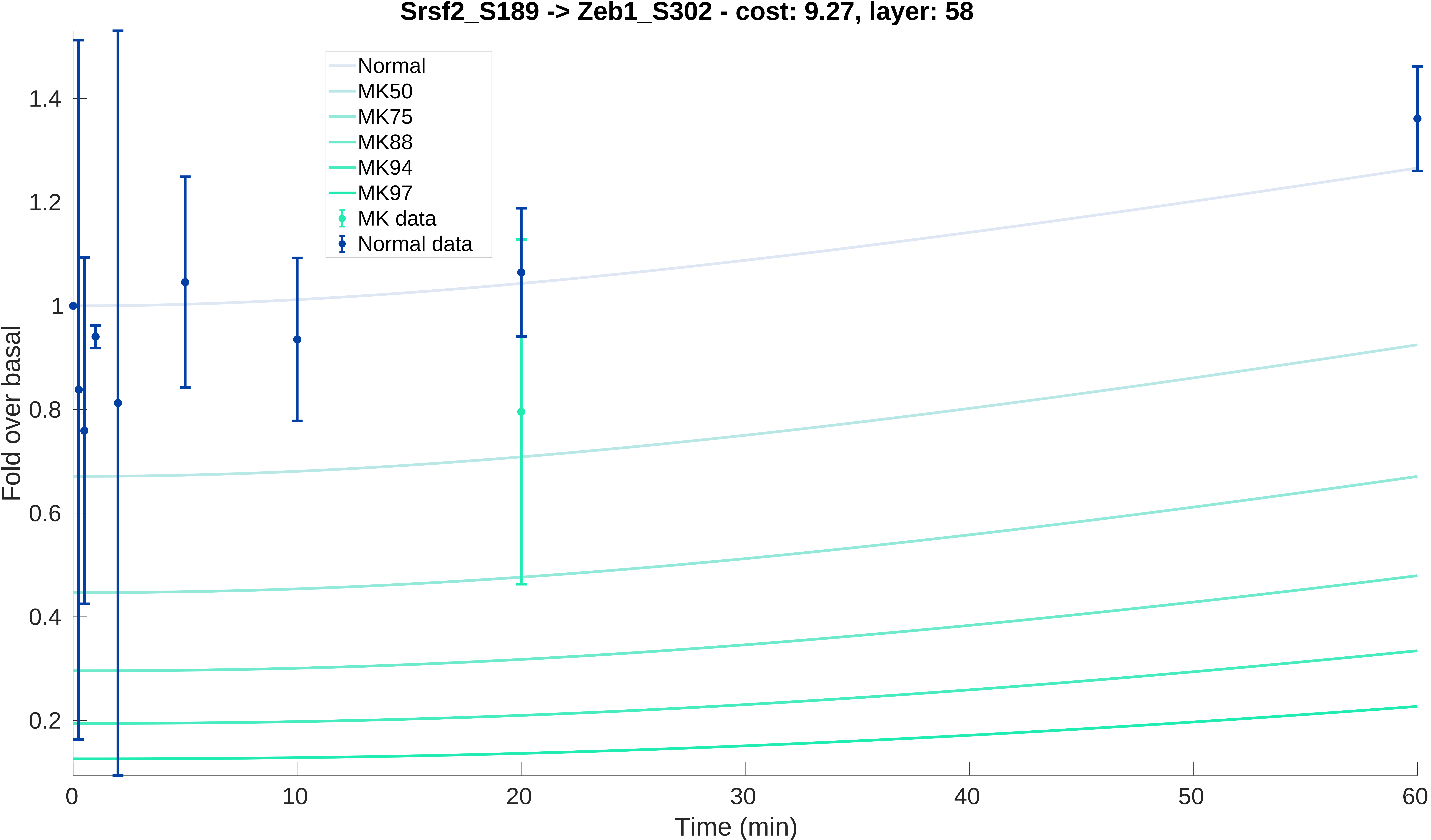

Srsf2_S189 -> Zeb1_S302 - cost: 9.27, layer: 58
Normal
MK50
MK75
MK88
MK94
MK97
MK data
Normal data
Fold over basal
1.4
1.2
1
0.8
0.6
0.4
0.2
0
10
20
30
40
50
60
Time (min)

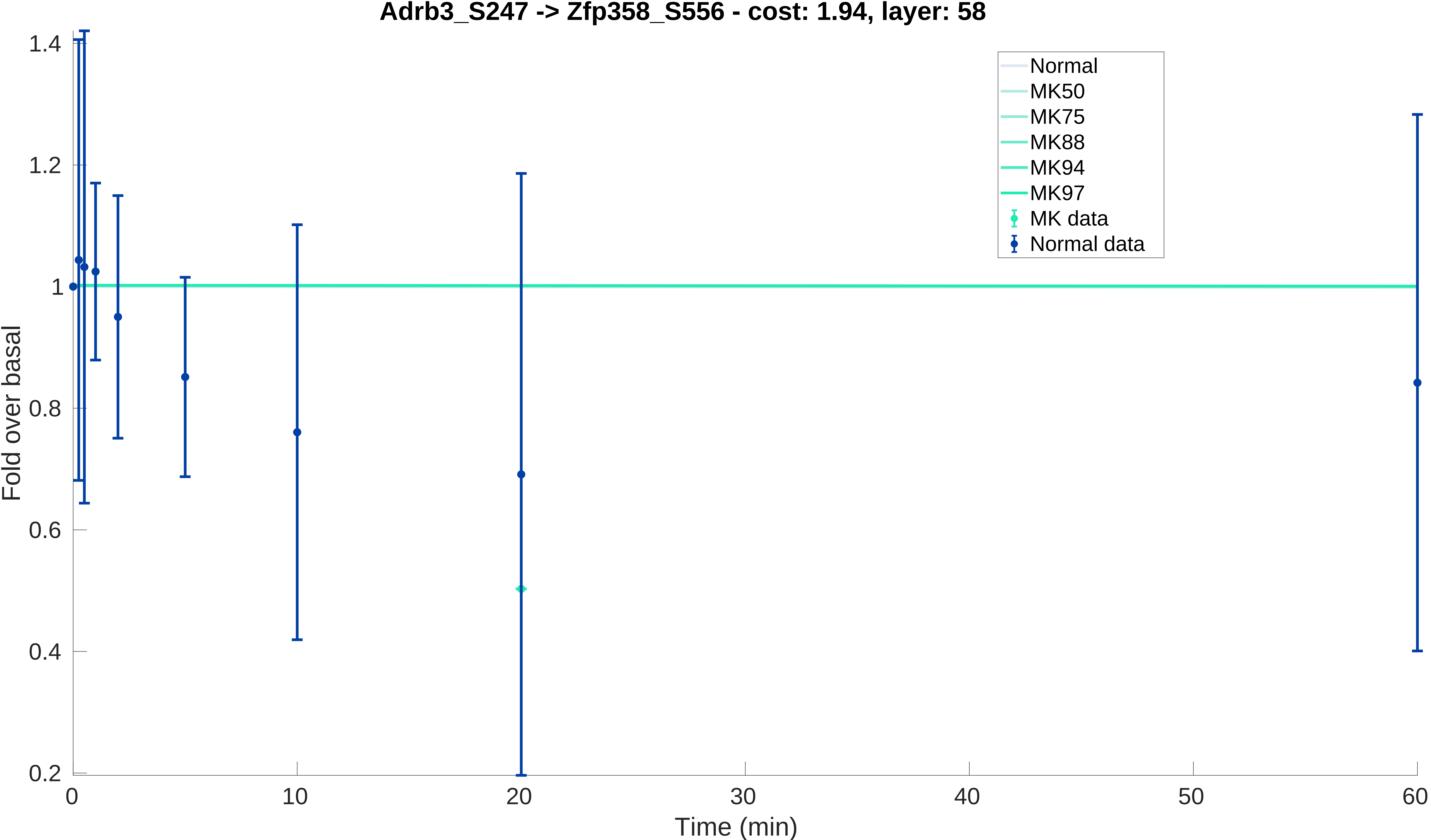

Adrb3_S247 -> Zfp358_S556 - cost: 1.94, layer: 58
Normal
MK50
MK75
MK88
MK94
MK97
MK data
Normal data
Fold over basal
Time (min)
0.2
0.4
0.6
0.8
1
1.2
1.4
0
10
20
30
40
50
60

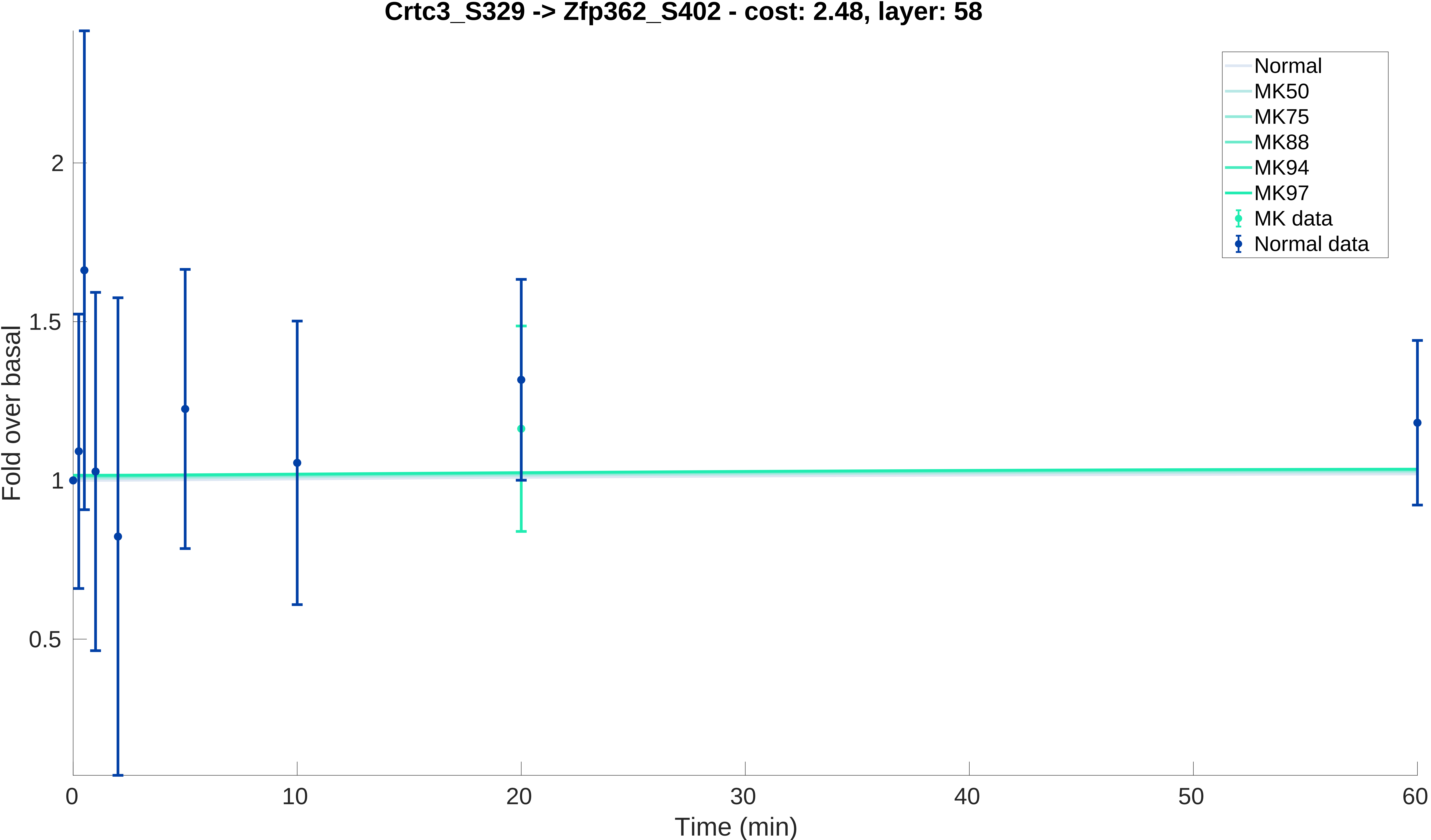

Crtc3_S329 -> Zfp362_S402 - cost: 2.48, layer: 58
Normal
MK50
MK75
MK88
MK94
MK97
MK data
Normal data
Fold over basal
2
1.5
1
0.5
0
10
20
30
40
50
60
Time (min)

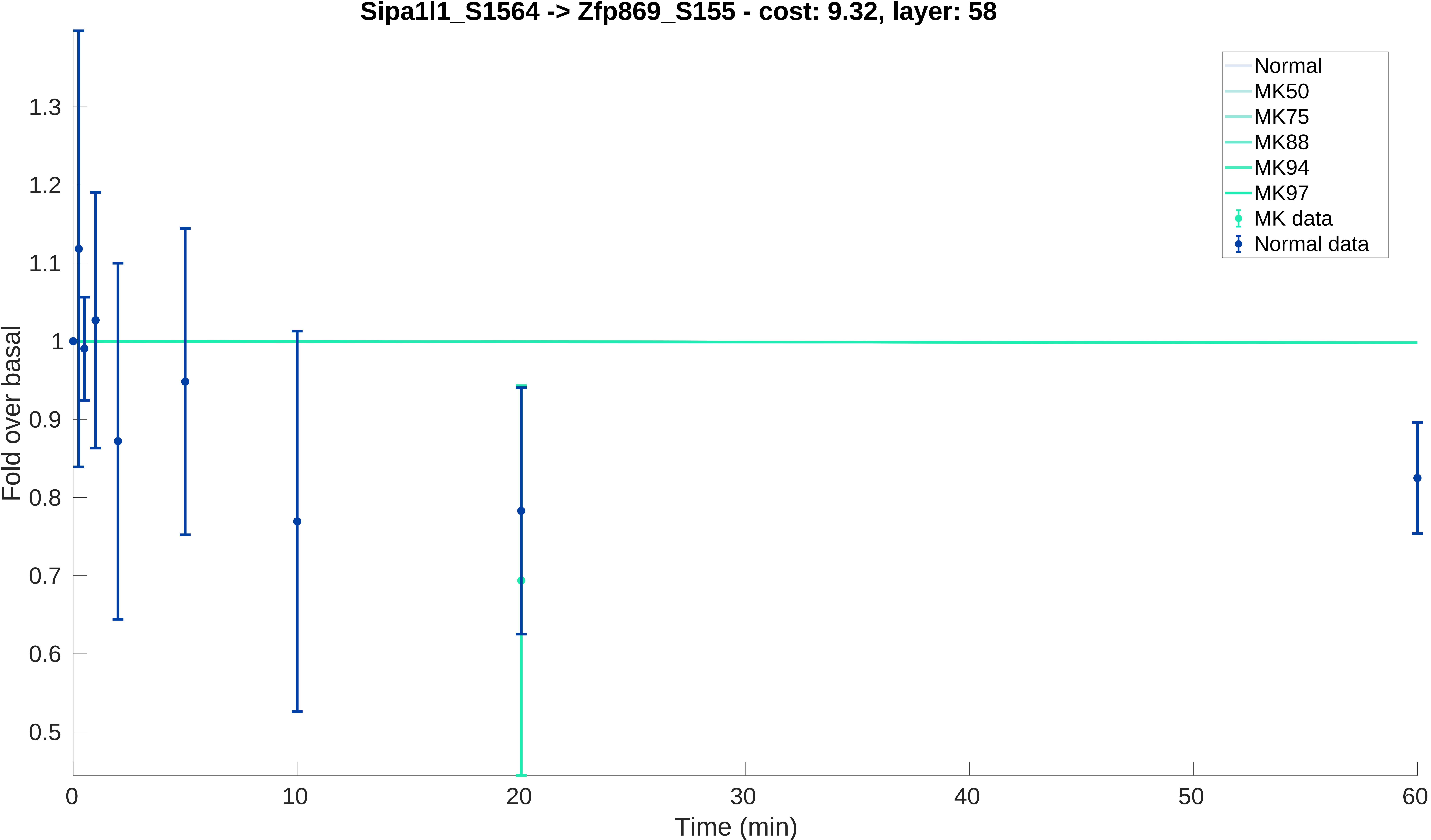

Sipa1l1_S1564 -> Zfp869_S155 - cost: 9.32, layer: 58
Fold over basal
Time (min)
1.3
1.2
1.1
1
0.9
0.8
0.7
0.6
0.5
0
10
20
30
40
50
60
Normal
MK50
MK75
MK88
MK94
MK97
MK data
Normal data

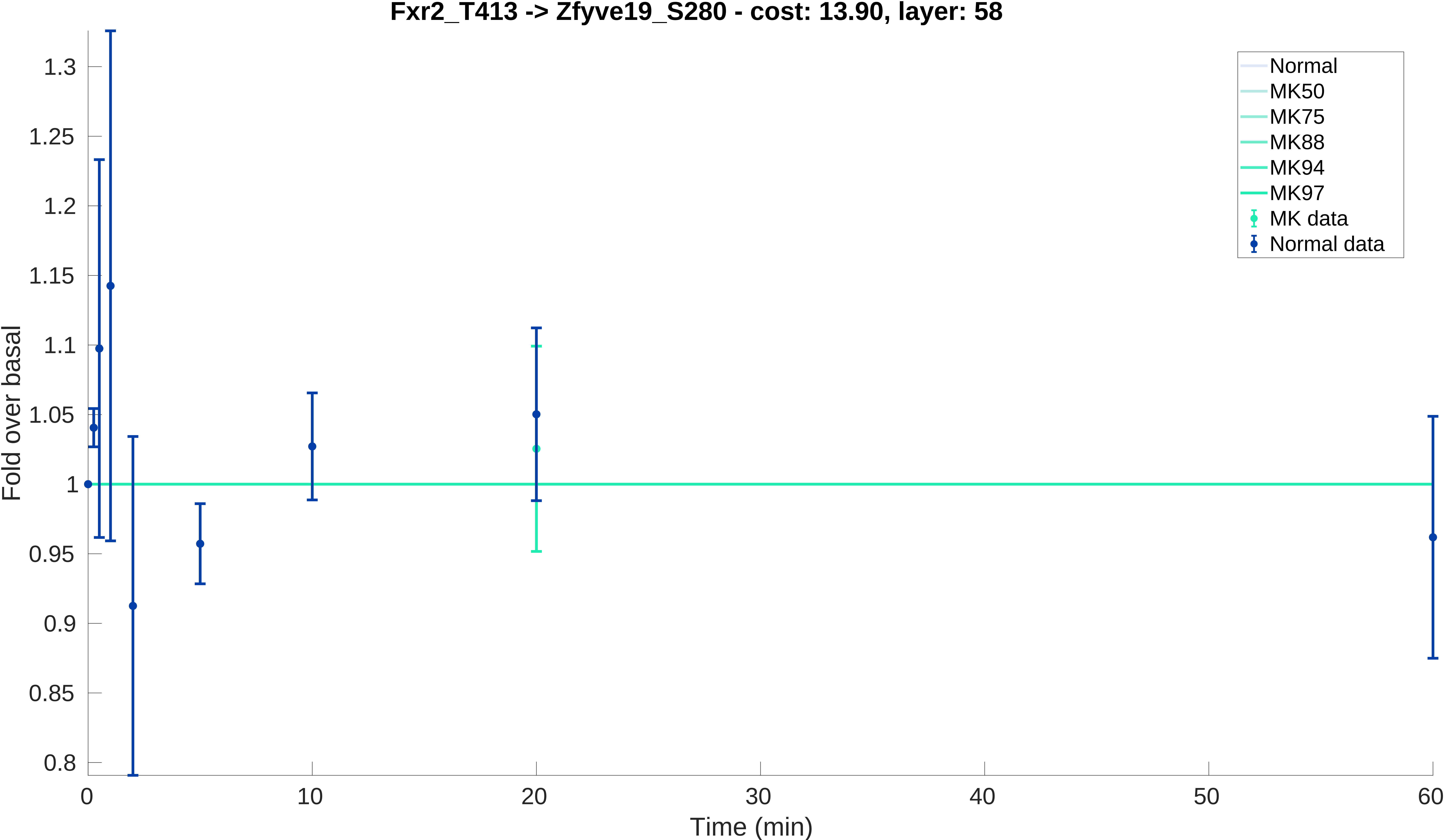
Fxr2_T413 -> Zfyve19_S280 - cost: 13.90, layer: 58
Fold over basal
Time (min)
1.3
1.25
1.2
1.15
1.1
1.05
1
0.95
0.9
0.85
0.8
0
10
20
30
40
50
60
Normal
MK50
MK75
MK88
MK94
MK97
MK data
Normal data

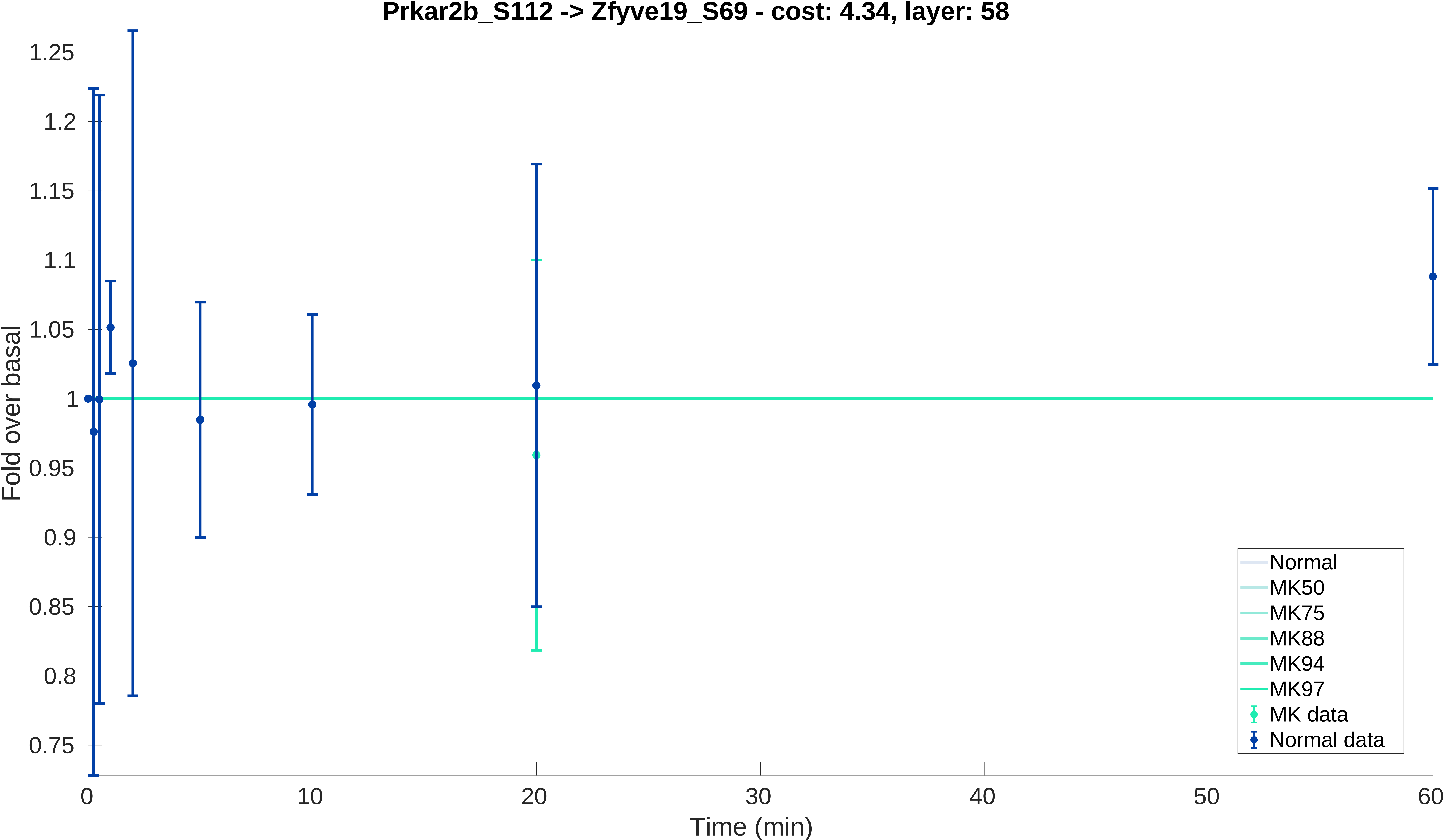

Prkar2b_S112 -> Zfyve19_S69 - cost: 4.34, layer: 58
Fold over basal
Time (min)
1.25
1.2
1.15
1.1
1.05
1
0.95
0.9
0.85
0.8
0.75
0
10
20
30
40
50
60
Normal
MK50
MK75
MK88
MK94
MK97
MK data
Normal data

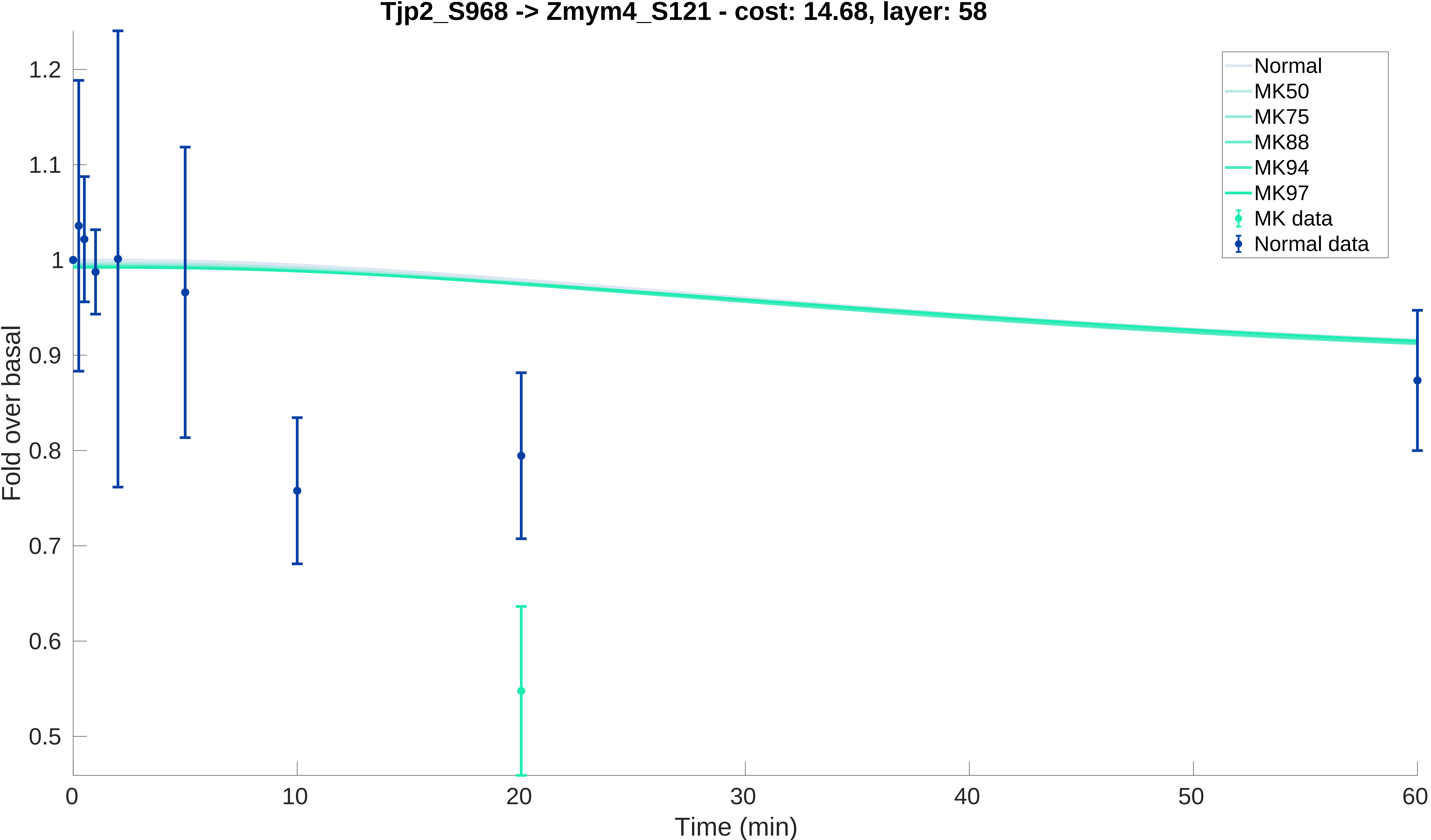

Tjp2_S968 -> Zmym4_S121 - cost: 14.68, layer: 58
Normal
MK50
MK75
MK88
MK94
MK97
MK data
Normal data
Fold over basal
Time (min)
1.2
1.1
1
0.9
0.8
0.7
0.6
0.5
0
10
20
30
40
50
60

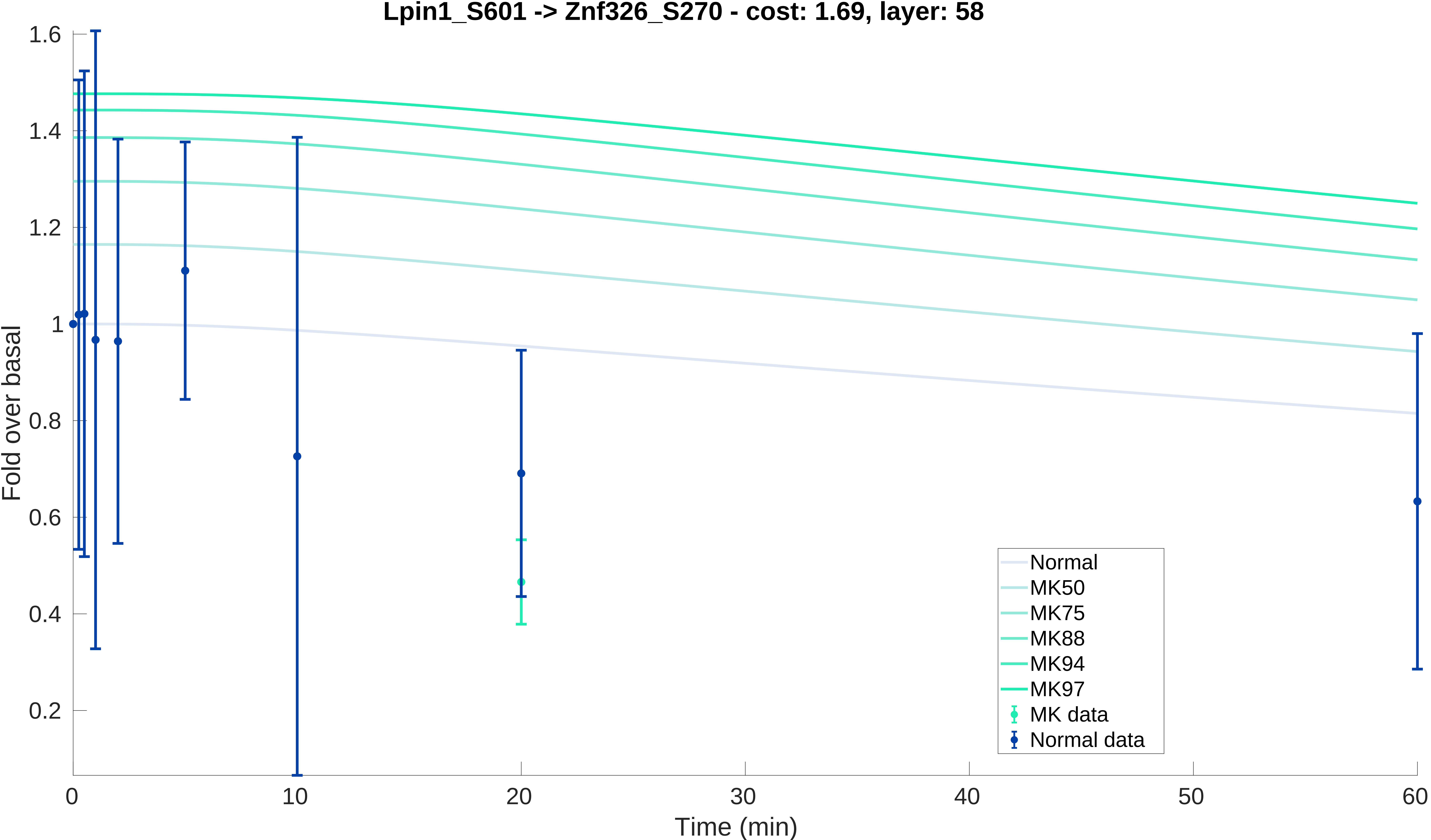

Lpin1_S601 -> Znf326_S270 - cost: 1.69, layer: 58
1.6
1.4
1.2
1
0.8
0.6
0.4
0.2
Fold over basal
0
10
20
30
40
50
60
Time (min)
Normal
MK50
MK75
MK88
MK94
MK97
MK data
Normal data

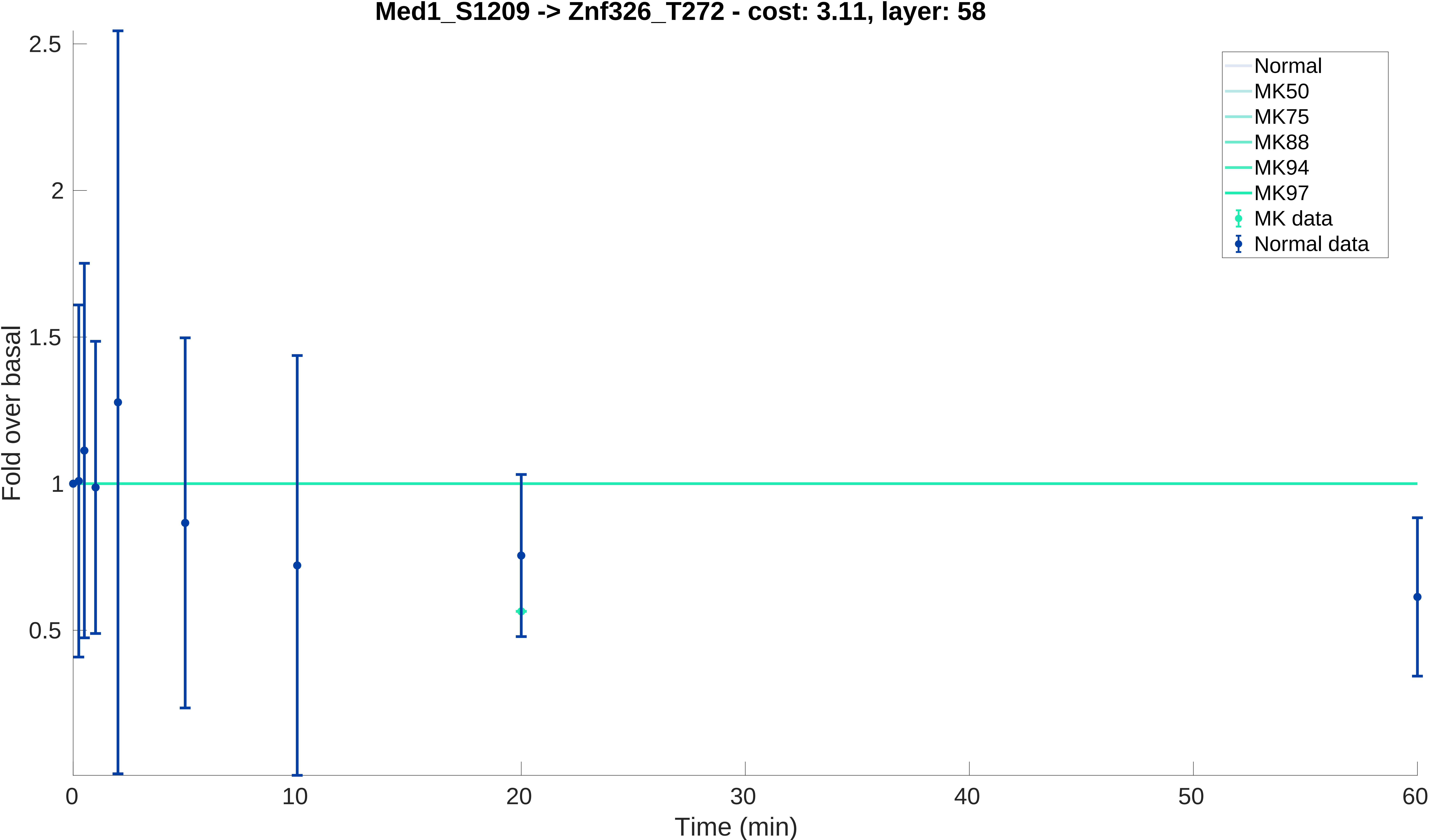
Med1_S1209 -> Znf326_T272 - cost: 3.11, layer: 58
Fold over basal
Time (min)
0
10
20
30
40
50
60
0.5
1
1.5
2
2.5
Normal
MK50
MK75
MK88
MK94
MK97
MK data
Normal data

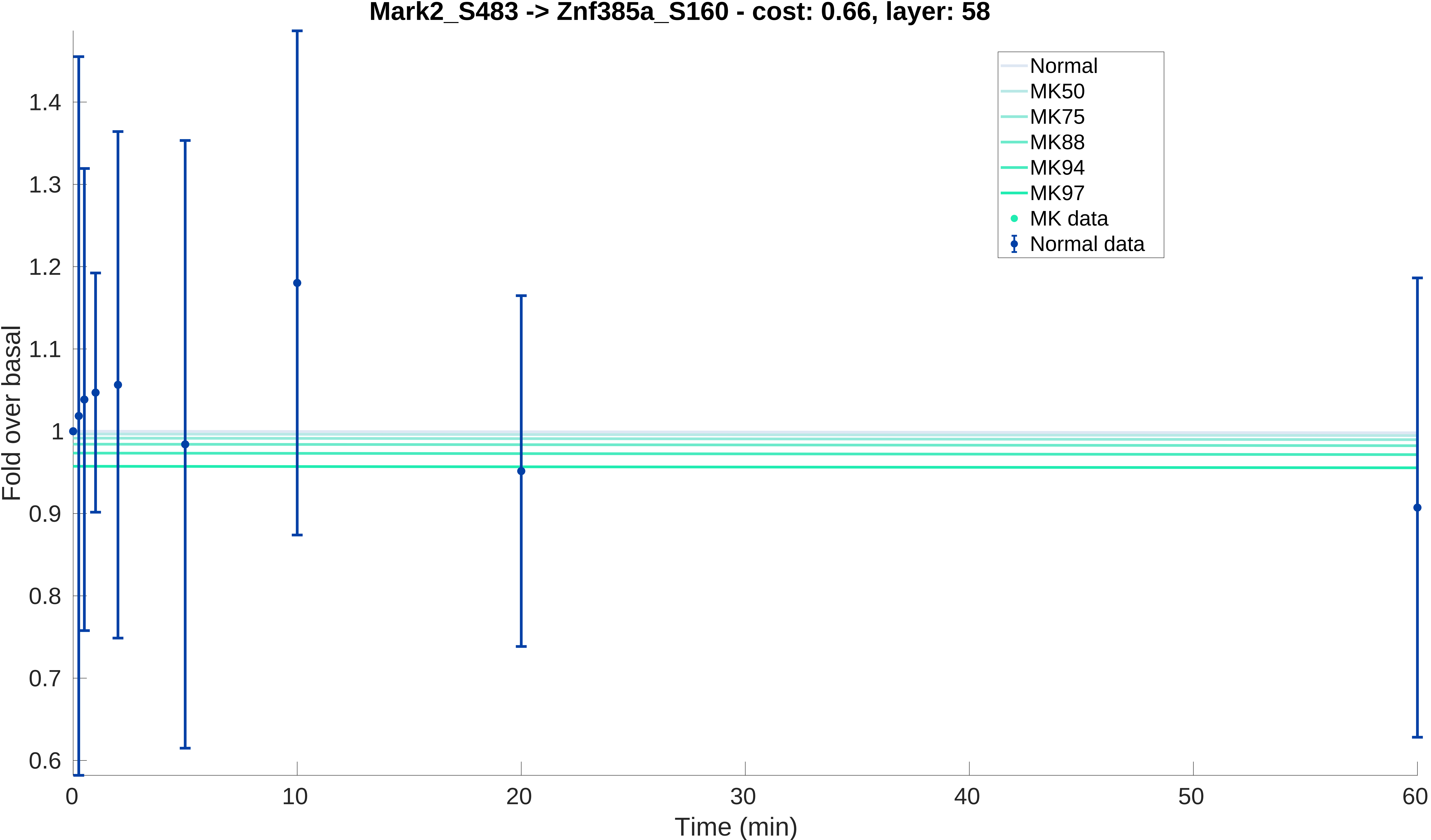
Mark2_S483 -> Znf385a_S160 - cost: 0.66, layer: 58
Fold over basal
Time (min)
0.6
0.7
0.8
0.9
1
1.1
1.2
1.3
1.4
0
10
20
30
40
50
60
Normal
MK50
MK75
MK88
MK94
MK97
MK data
Normal data

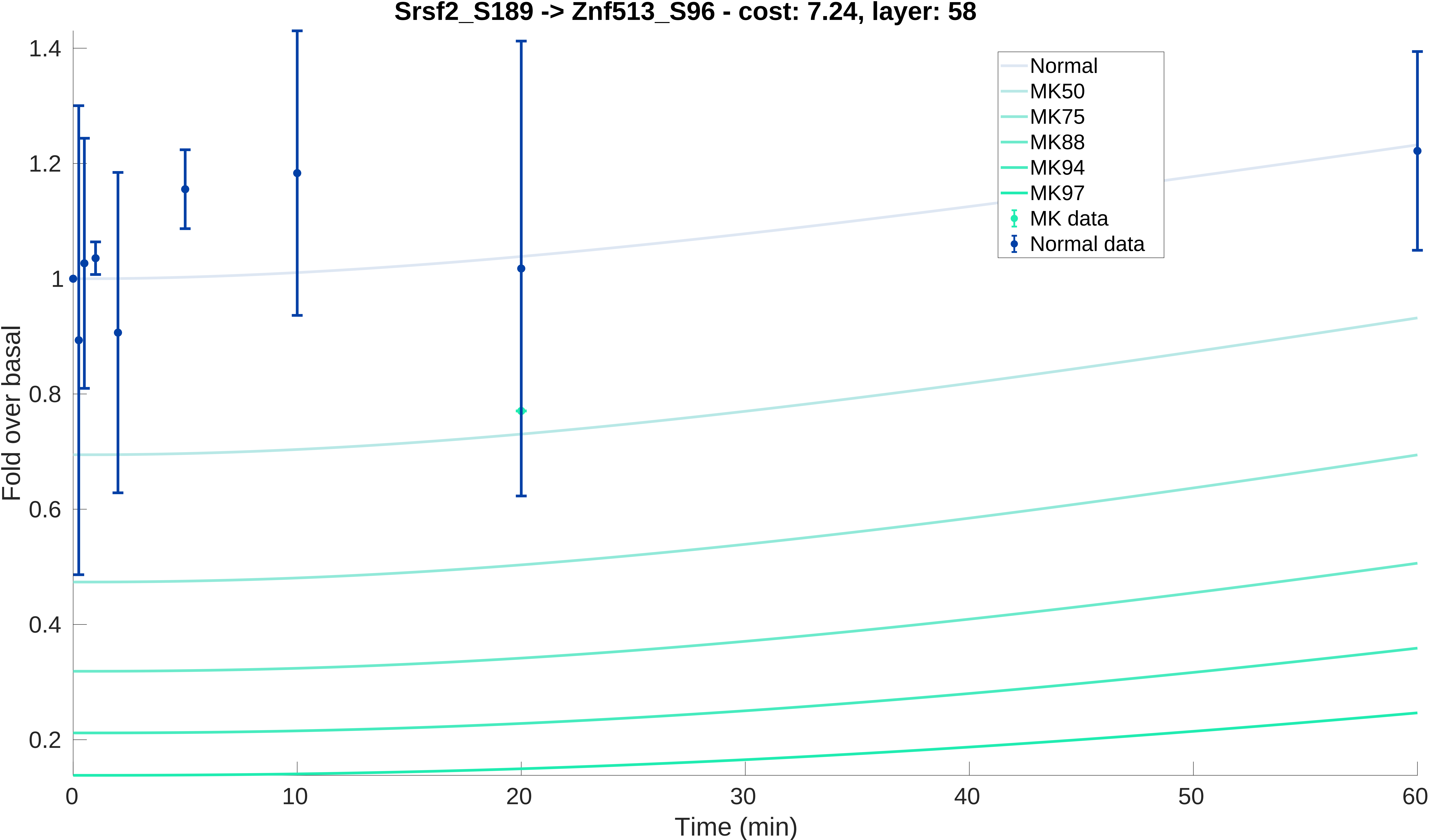

Srsf2_S189 -> Znf513_S96 - cost: 7.24, layer: 58
Fold over basal
Time (min)
0
10
20
30
40
50
60
0.2
0.4
0.6
0.8
1
1.2
1.4
Normal
MK50
MK75
MK88
MK94
MK97
MK data
Normal data

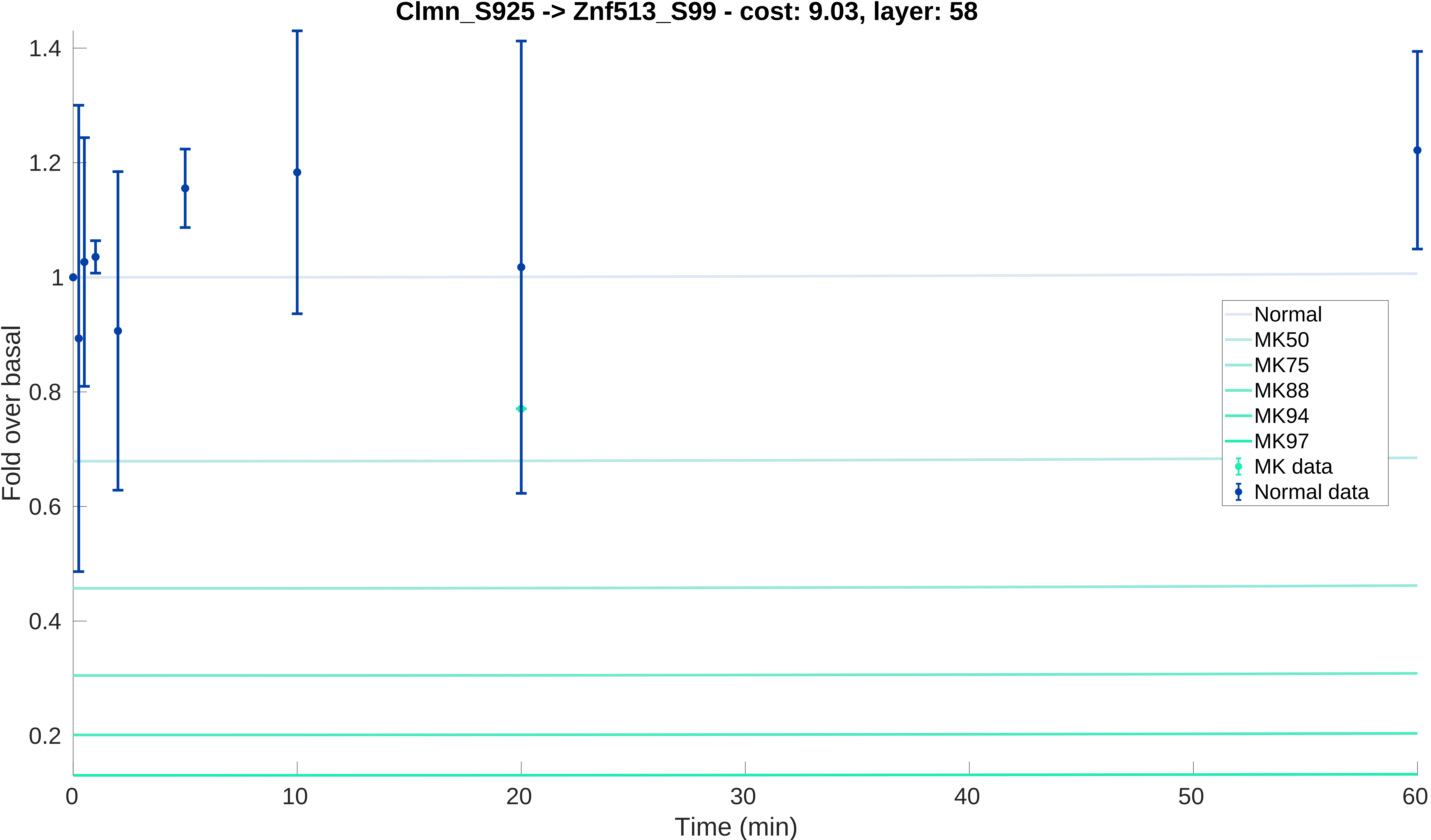

Clmn_S925 -> Znf513_S99 - cost: 9.03, layer: 58
Fold over basal
1.4
1.2
1
0.8
0.6
0.4
0.2
0
10
20
30
40
50
60
Time (min)
Normal
MK50
MK75
MK88
MK94
MK97
MK data
Normal data

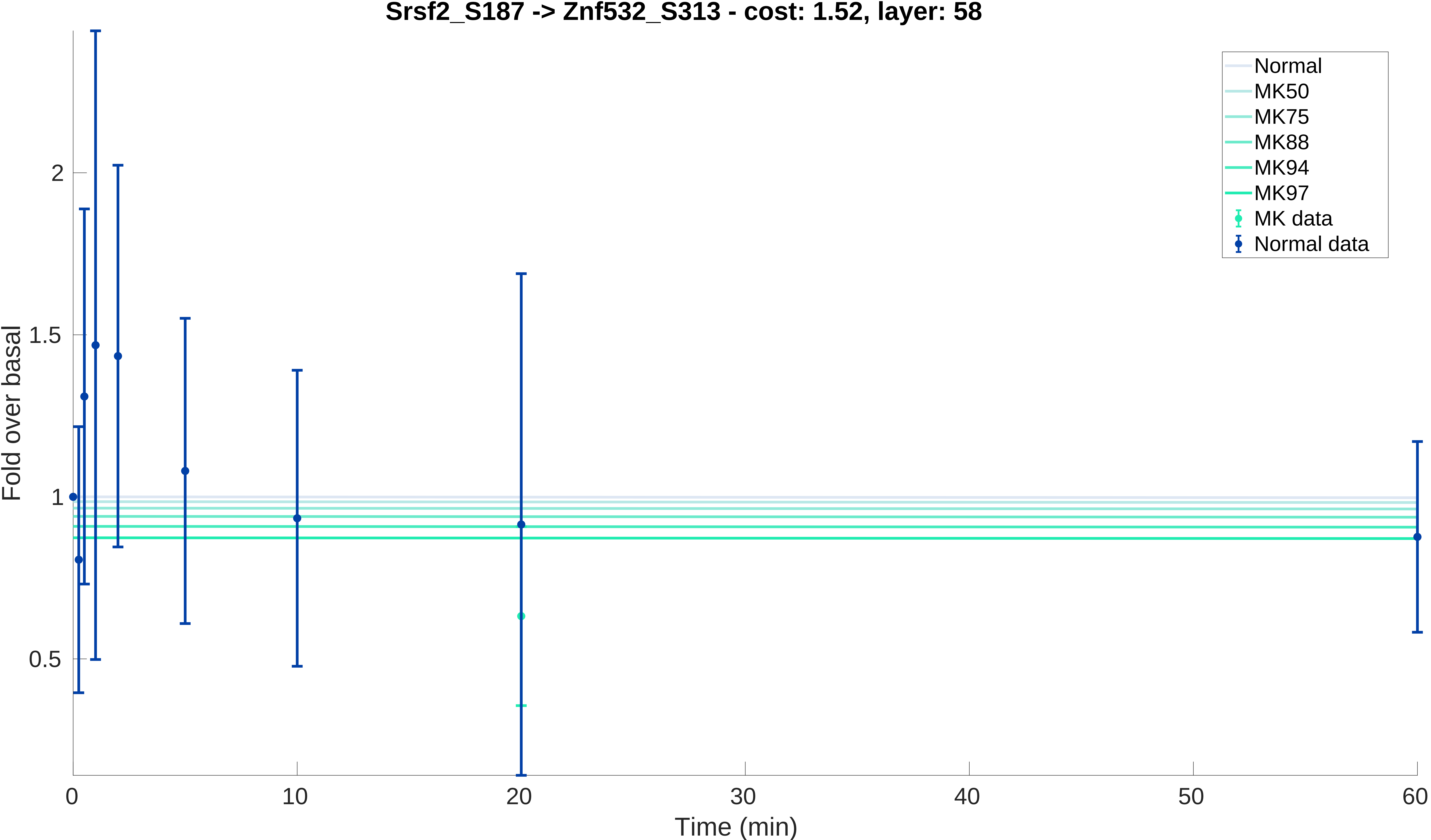

Srsf2_S187 -> Znf532_S313 - cost: 1.52, layer: 58
Normal
MK50
MK75
MK88
MK94
MK97
MK data
Normal data
Fold over basal
0.5
1
1.5
2
0
10
20
30
40
50
60
Time (min)

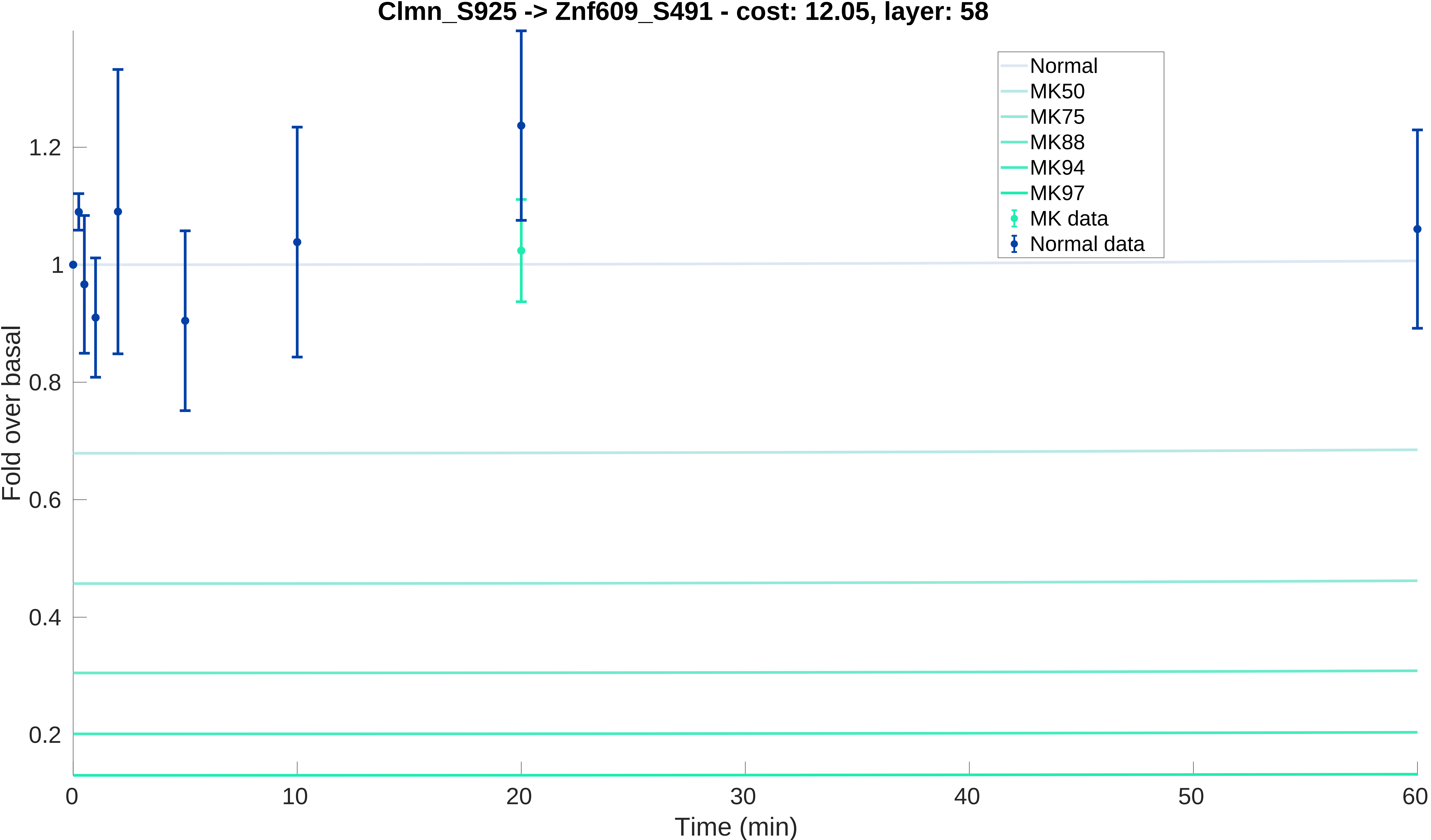

Clmn_S925 -> Znf609_S491 - cost: 12.05, layer: 58
Fold over basal
Time (min)
0
10
20
30
40
50
60
0.2
0.4
0.6
0.8
1
1.2
Normal
MK50
MK75
MK88
MK94
MK97
MK data
Normal data

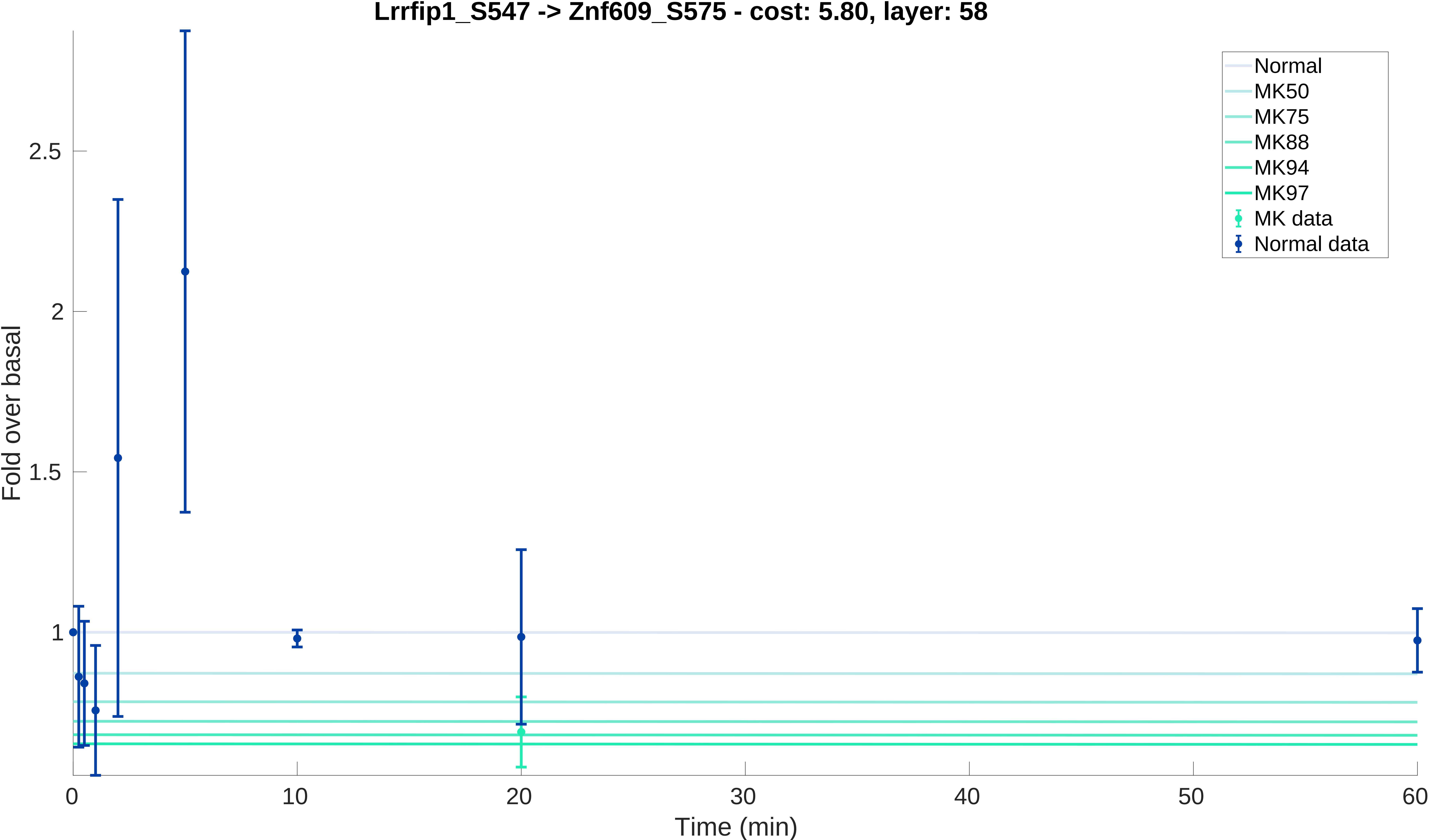
Lrrfip1_S547 -> Znf609_S575 - cost: 5.80, layer: 58
Fold over basal
Time (min)
2.5
2
1.5
1
0
10
20
30
40
50
60
Normal
MK50
MK75
MK88
MK94
MK97
MK data
Normal data

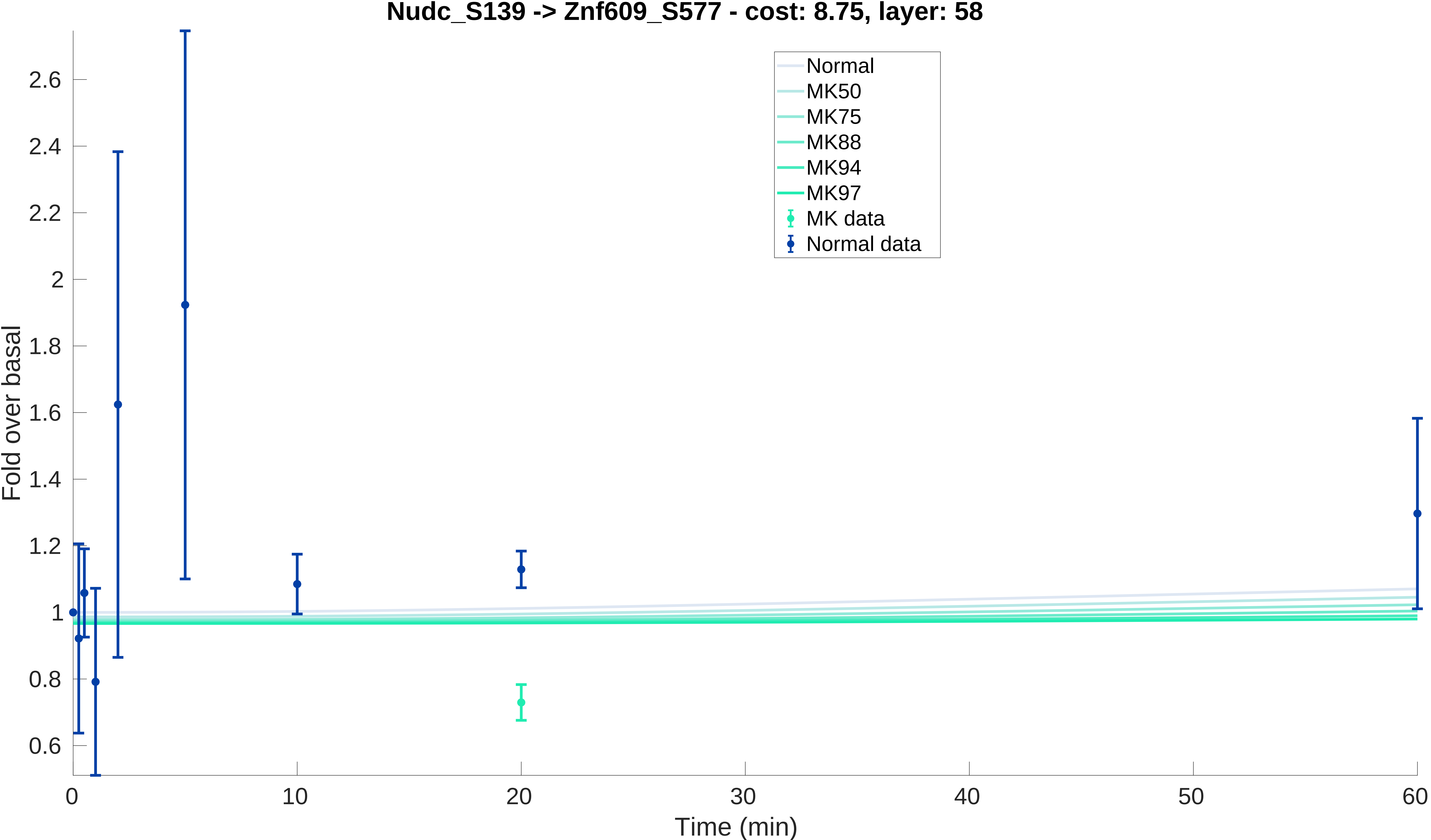
Nudc_S139 -> Znf609_S577 - cost: 8.75, layer: 58
Normal
MK50
MK75
MK88
MK94
MK97
MK data
Normal data
Fold over basal
Time (min)
0.6
0.8
1
1.2
1.4
1.6
1.8
2
2.2
2.4
2.6
0
10
20
30
40
50
60

Hsp90ab1_S445 -> Znf638_S1569 - cost: 3.36, layer: 58

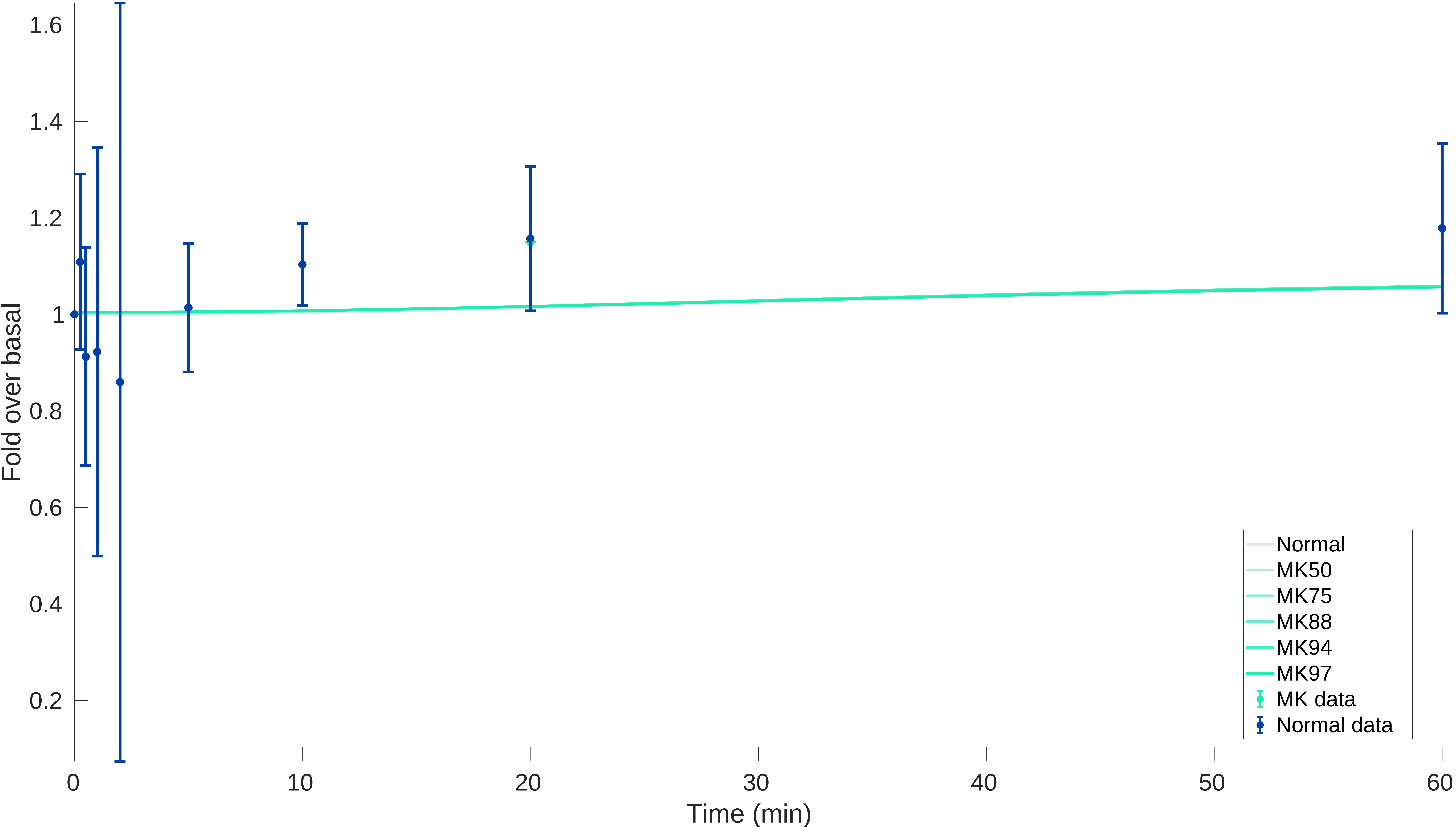

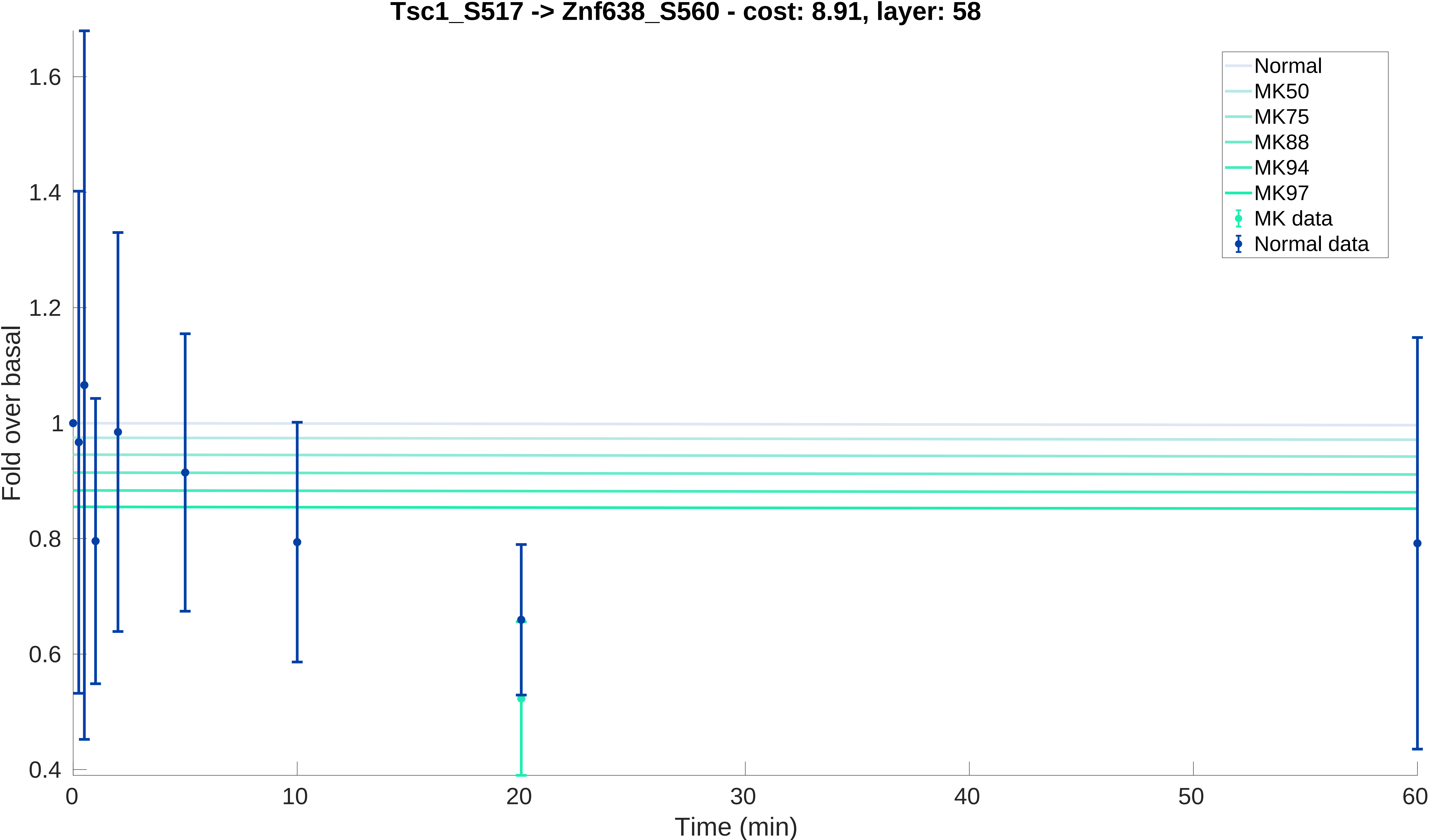

Tsc1_S517 -> Znf638_S560 - cost: 8.91, layer: 58
Fold over basal
Time (min)
1.6
1.4
1.2
1
0.8
0.6
0.4
0
10
20
30
40
50
60
Normal
MK50
MK75
MK88
MK94
MK97
MK data
Normal data

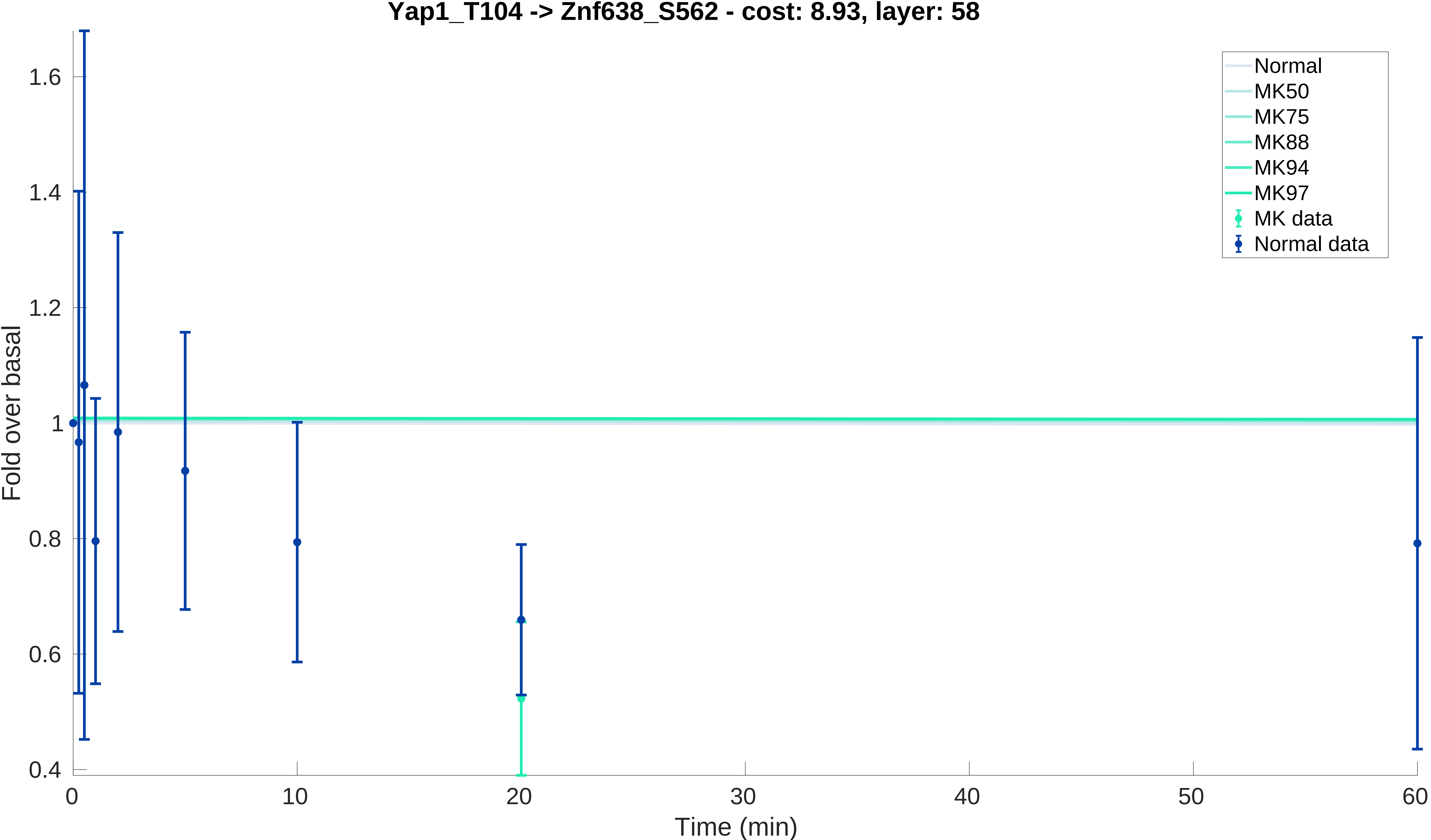

Yap1_T104 -> Znf638_S562 - cost: 8.93, layer: 58
Normal
MK50
MK75
MK88
MK94
MK97
MK data
Normal data
Fold over basal
1.6
1.4
1.2
1
0.8
0.6
0.4
0
10
20
30
40
50
60
Time (min)

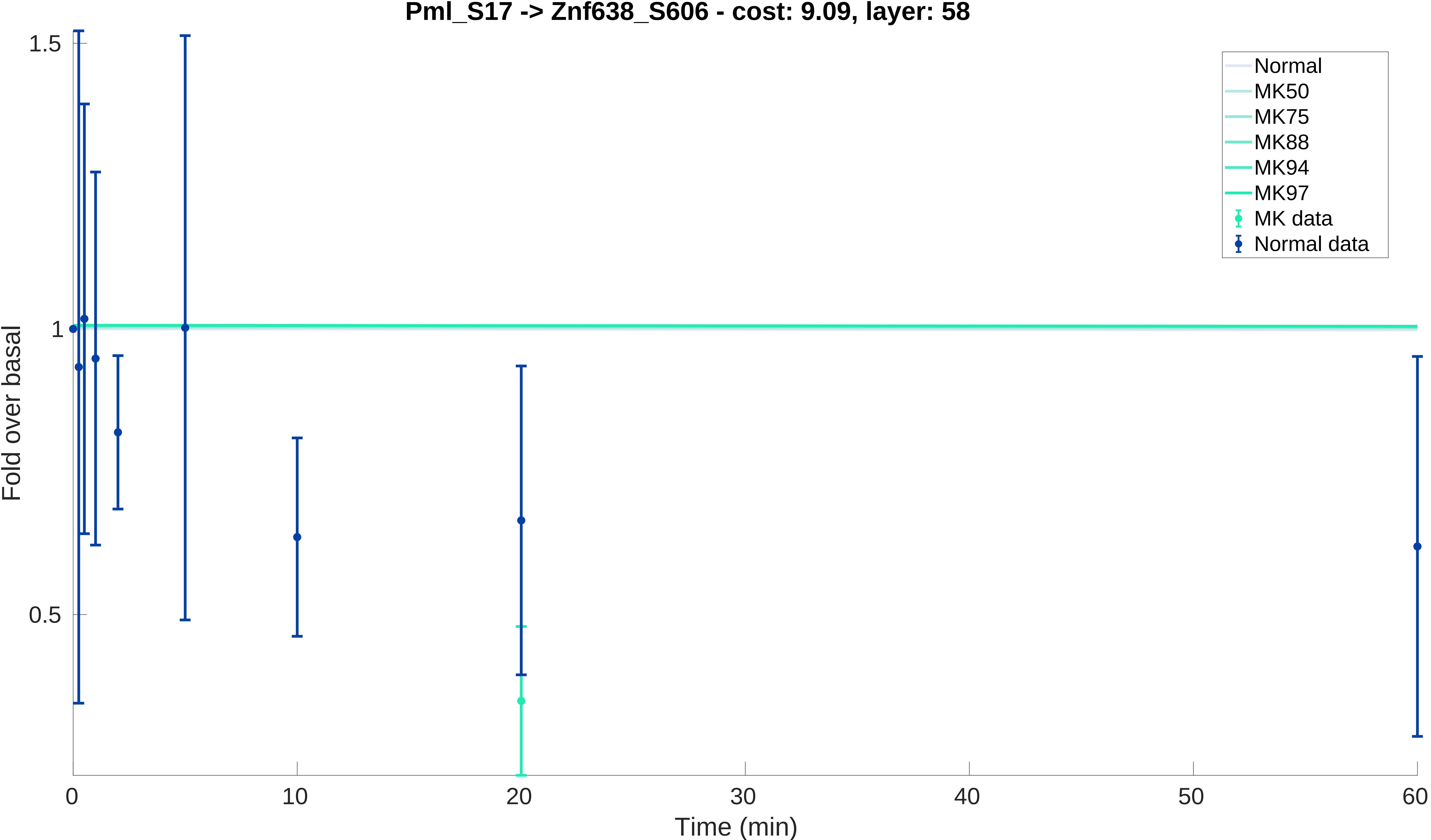

Pml_S17 -> Znf638_S606 - cost: 9.09, layer: 58
Fold over basal
1.5
1
0.5
0
10
20
30
40
50
60
Time (min)
Normal
MK50
MK75
MK88
MK94
MK97
MK data
Normal data

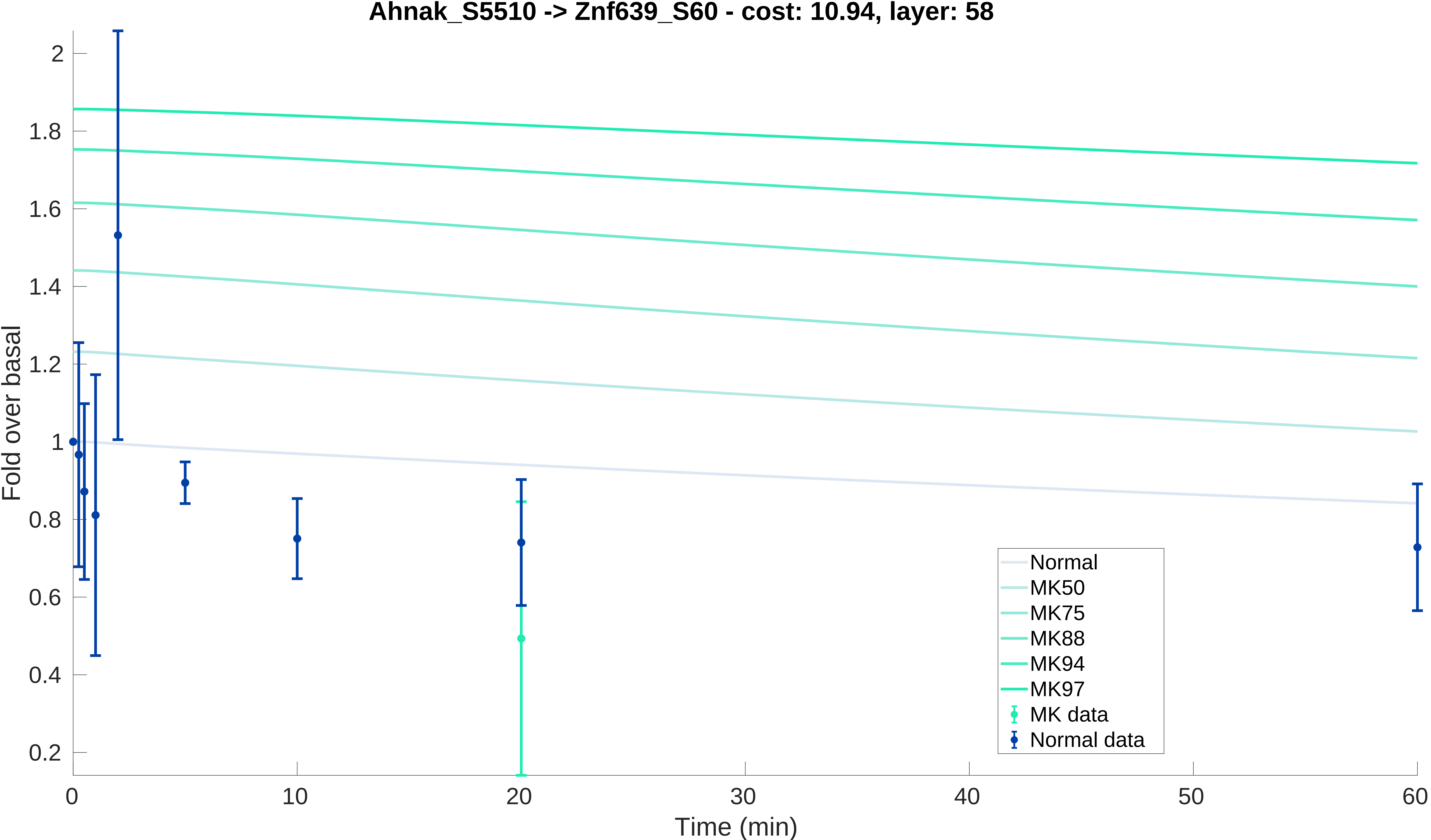

Ahnak_S5510 -> Znf639_S60 - cost: 10.94, layer: 58
Fold over basal
Time (min)
0
10
20
30
40
50
60
0.2
0.4
0.6
0.8
1
1.2
1.4
1.6
1.8
2
Normal
MK50
MK75
MK88
MK94
MK97
MK data
Normal data

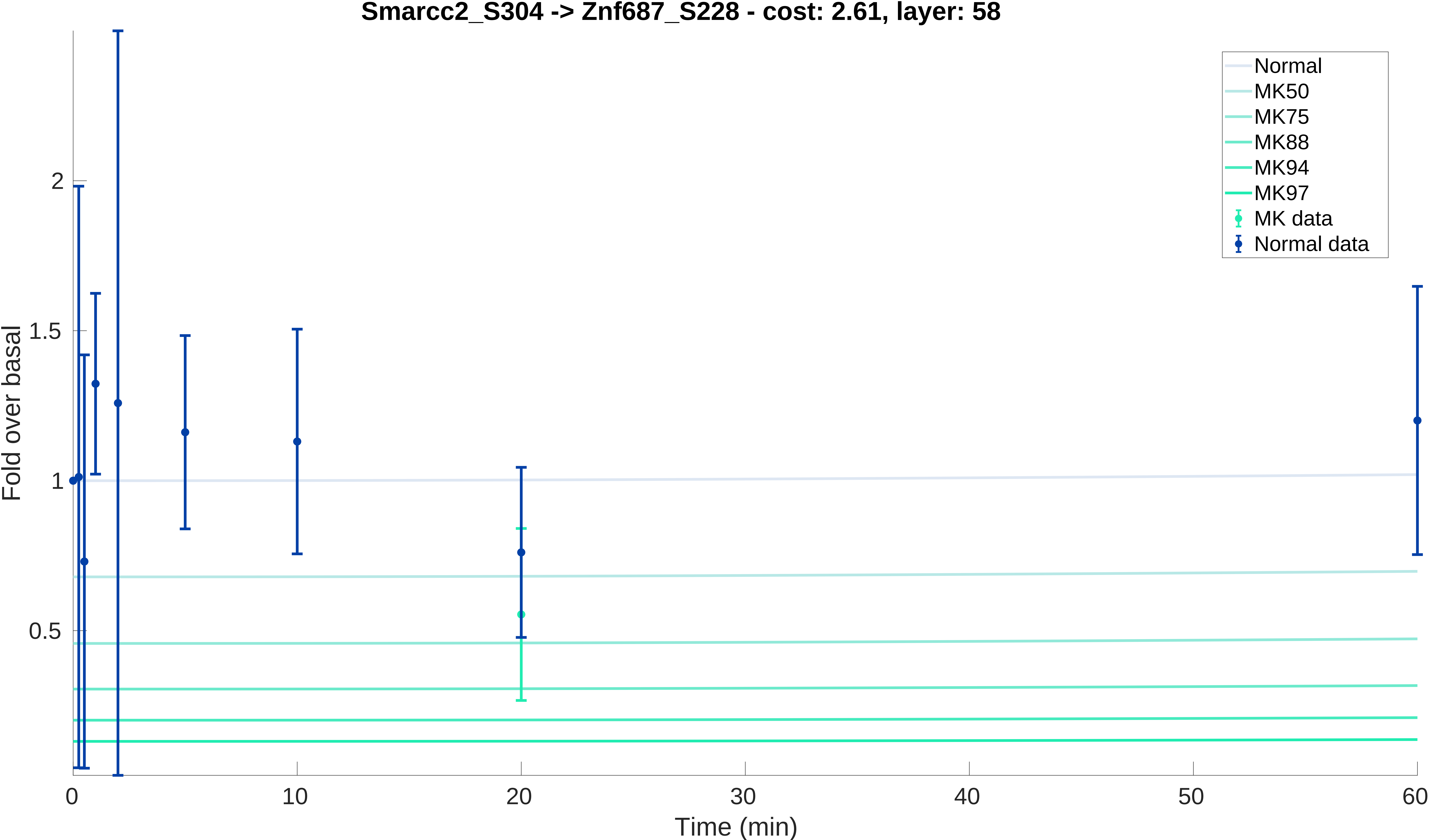

Smarcc2_S304 -> Znf687_S228 - cost: 2.61, layer: 58
Normal
MK50
MK75
MK88
MK94
MK97
MK data
Normal data
Fold over basal
2
1.5
1
0.5
0
10
20
30
40
50
60
Time (min)

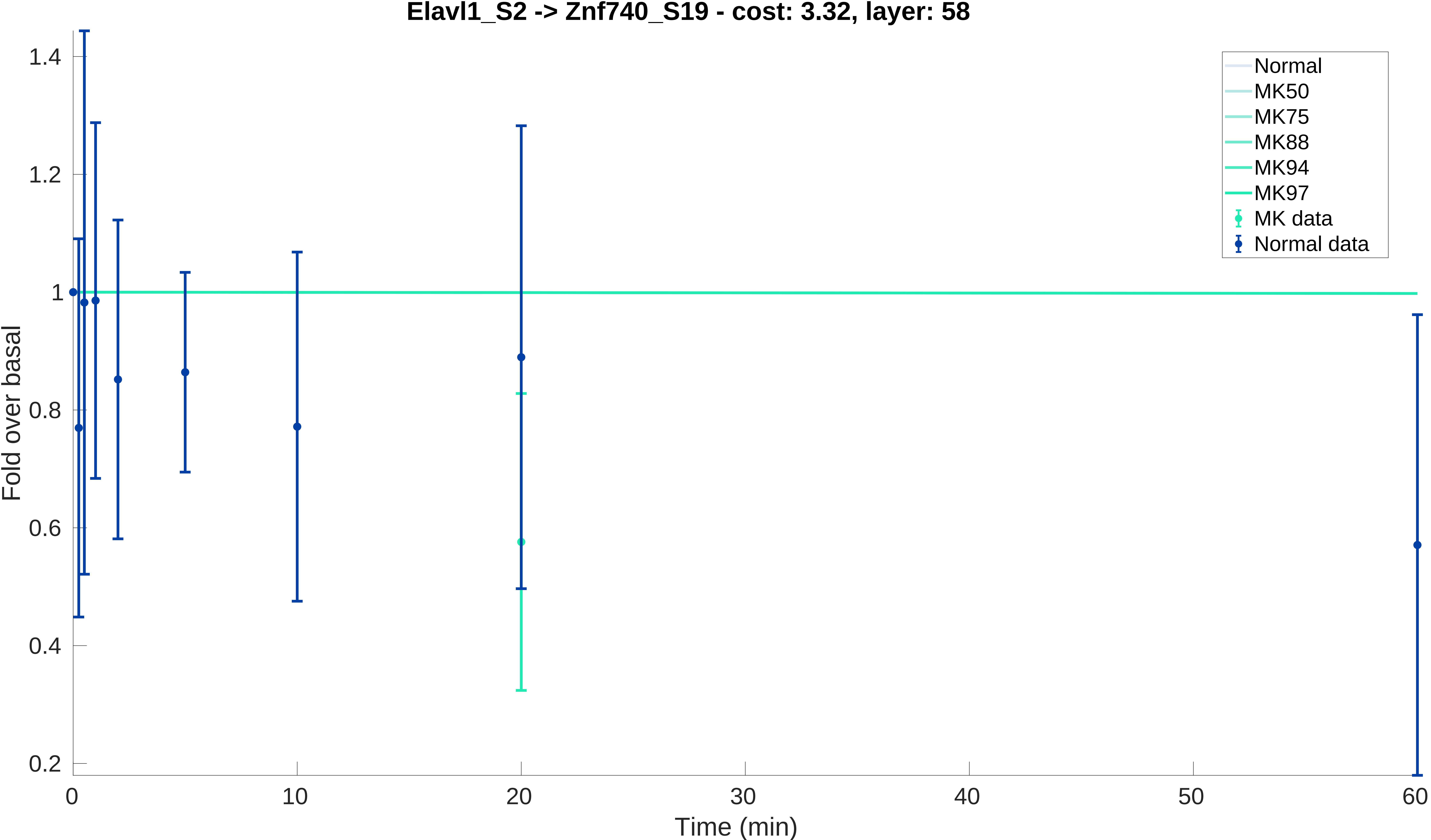

Elavl1_S2 -> Znf740_S19 - cost: 3.32, layer: 58
Normal
MK50
MK75
MK88
MK94
MK97
MK data
Normal data
Fold over basal
Time (min)
1.4
1.2
1
0.8
0.6
0.4
0.2
0
10
20
30
40
50
60

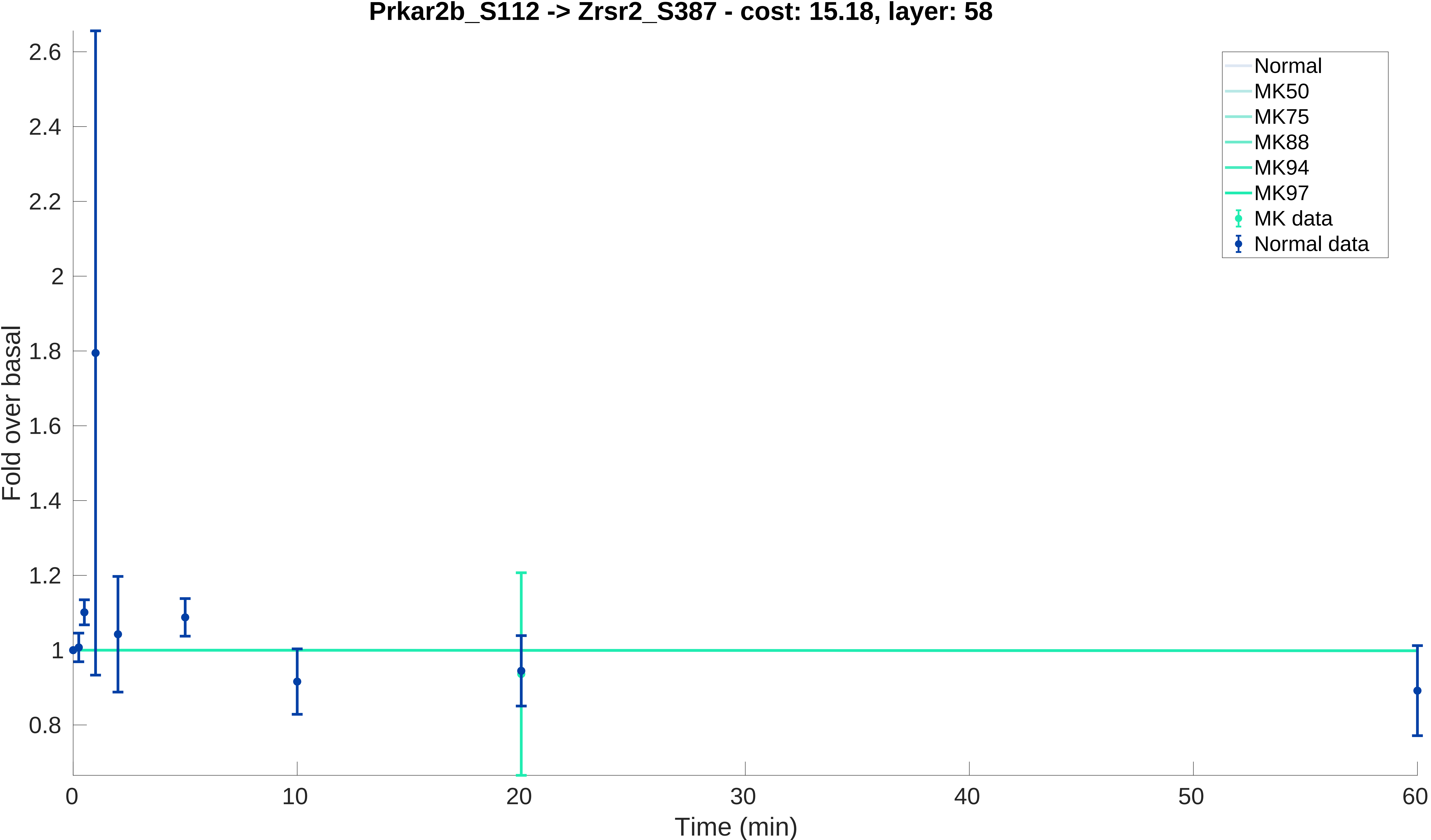

Prkar2b_S112 -> Zrsr2_S387 - cost: 15.18, layer: 58
Fold over basal
Time (min)
2.6
2.4
2.2
2
1.8
1.6
1.4
1.2
1
0.8
0
10
20
30
40
50
60
Normal
MK50
MK75
MK88
MK94
MK97
MK data
Normal data

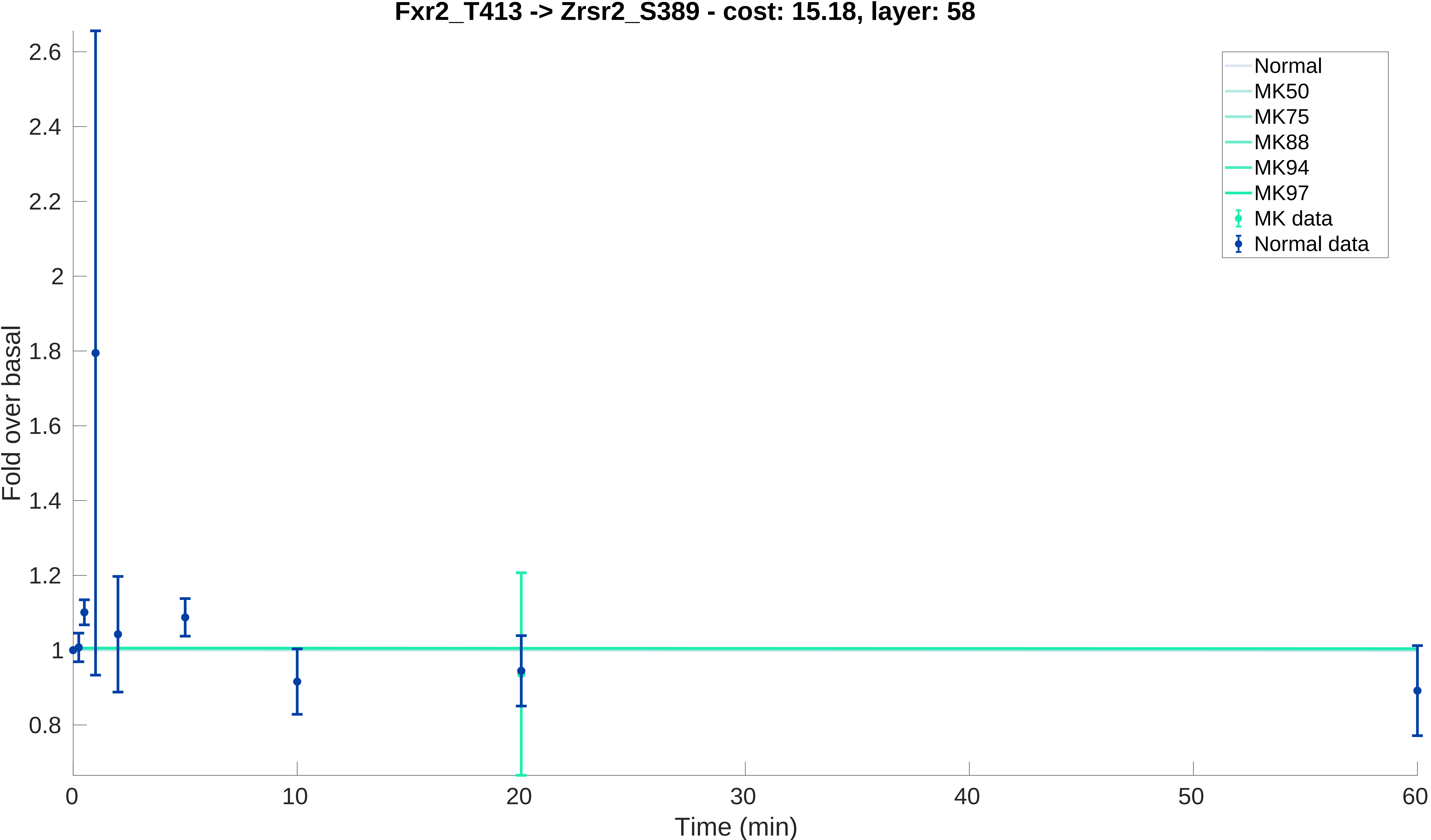

Fxr2_T413 -> Zrsr2_S389 - cost: 15.18, layer: 58
Fold over basal
Time (min)
Normal
MK50
MK75
MK88
MK94
MK97
MK data
Normal data
2.6
2.4
2.2
2
1.8
1.6
1.4
1.2
1
0.8
0
10
20
30
40
50
60

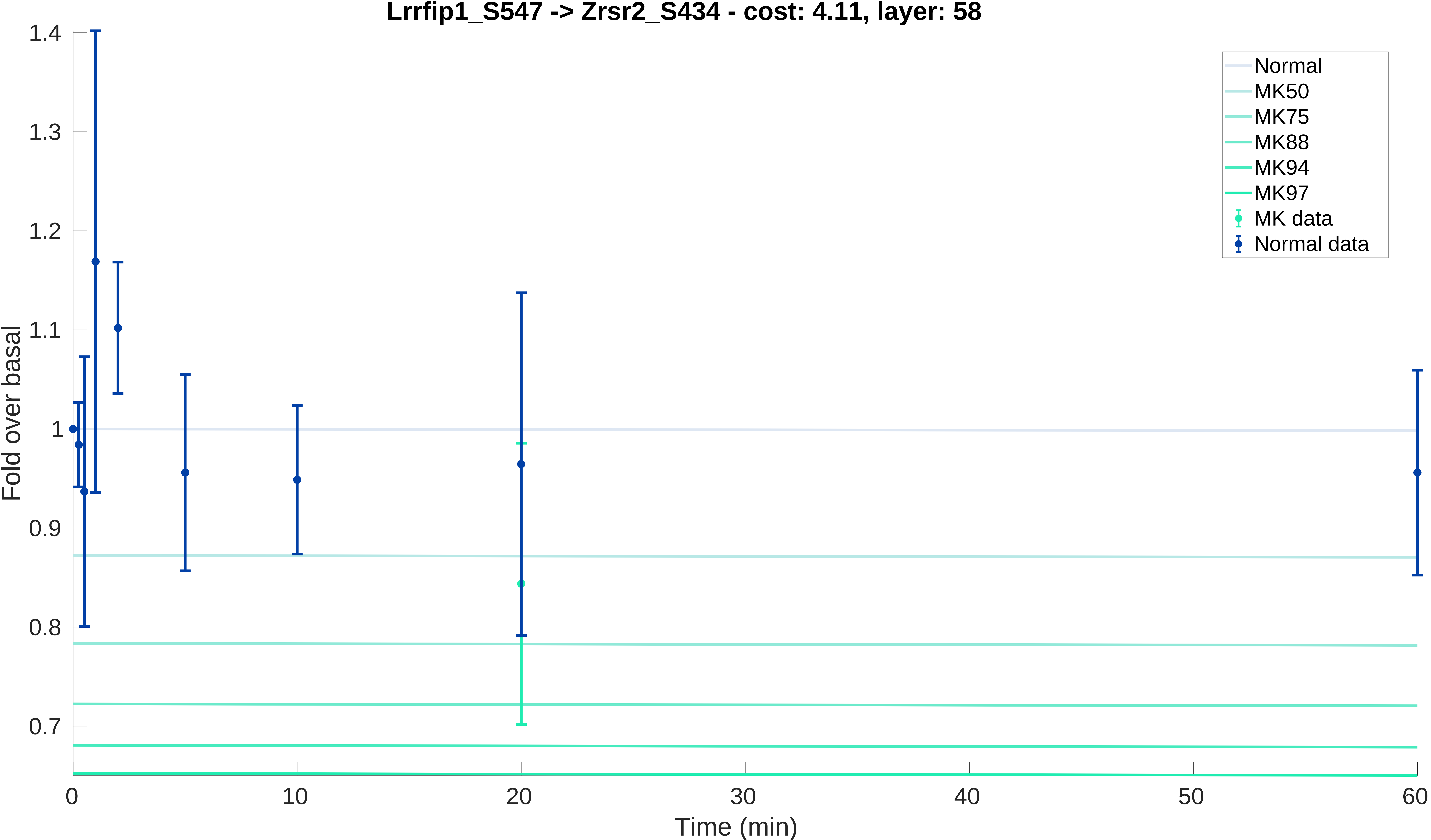

Lrrfip1_S547 -> Zrsr2_S434 - cost: 4.11, layer: 58
Fold over basal
1.4
1.3
1.2
1.1
1
0.9
0.8
0.7
0
10
20
30
40
50
60
Time (min)
Normal
MK50
MK75
MK88
MK94
MK97
MK data
Normal data

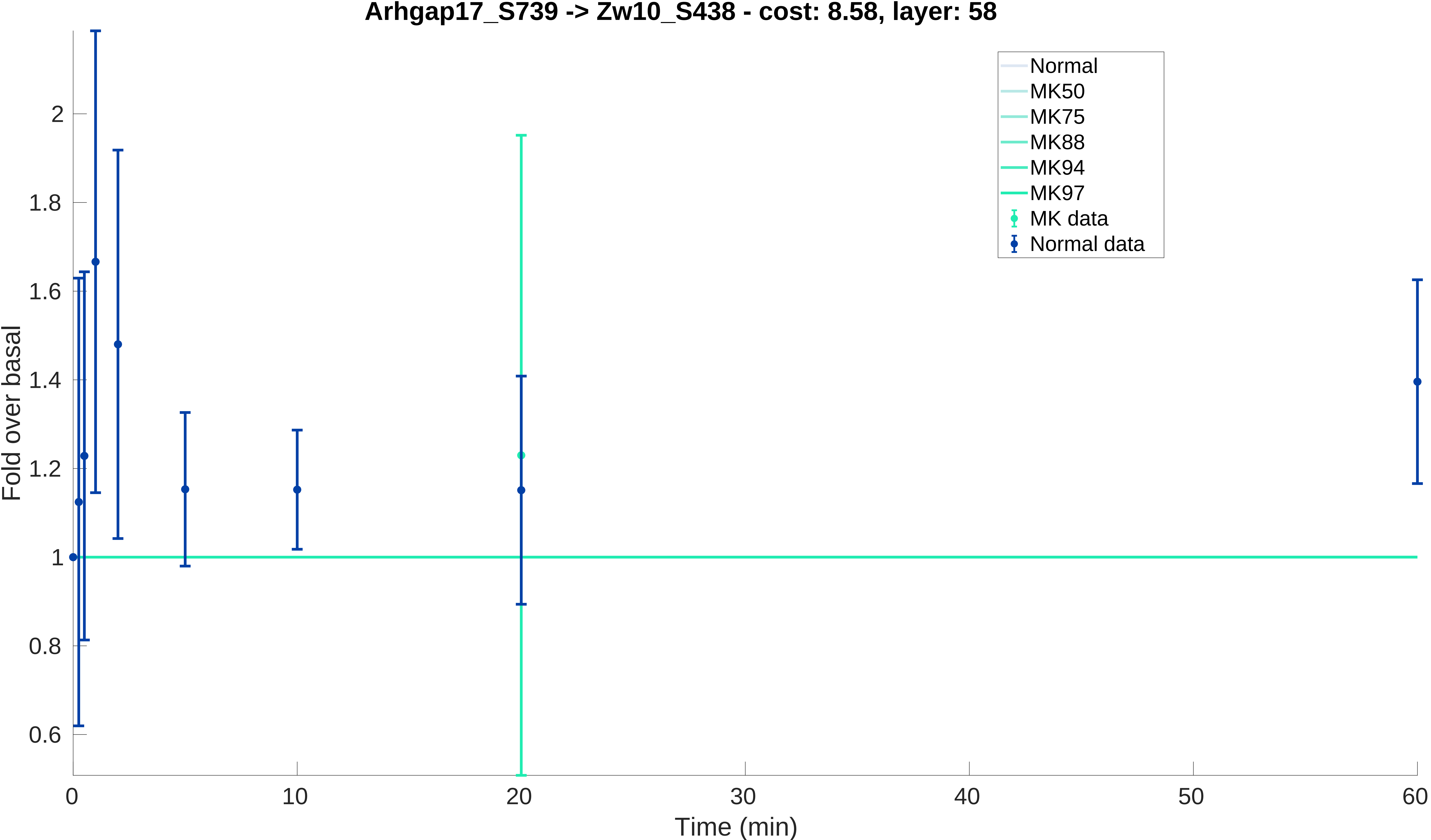

Arhgap17_S739 -> Zw10_S438 - cost: 8.58, layer: 58
Normal
MK50
MK75
MK88
MK94
MK97
MK data
Normal data
Fold over basal
0.6
0.8
1
1.2
1.4
1.6
1.8
2
0
10
20
30
40
50
60
Time (min)

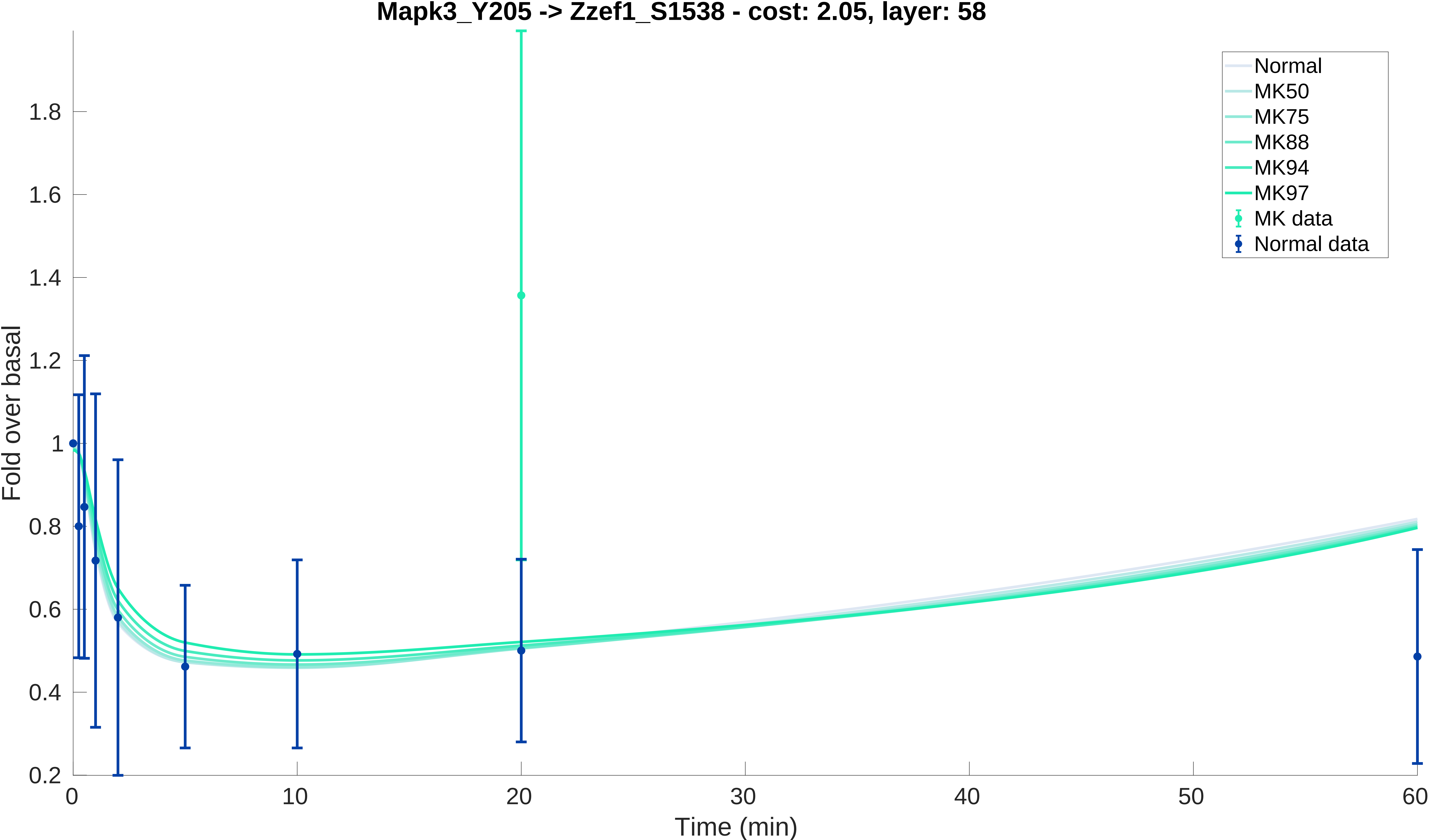

Mapk3_Y205 -> Zzef1_S1538 - cost: 2.05, layer: 58
Fold over basal
Time (min)
0
10
20
30
40
50
60
0.2
0.4
0.6
0.8
1
1.2
1.4
1.6
1.8
Normal
MK50
MK75
MK88
MK94
MK97
MK data
Normal data

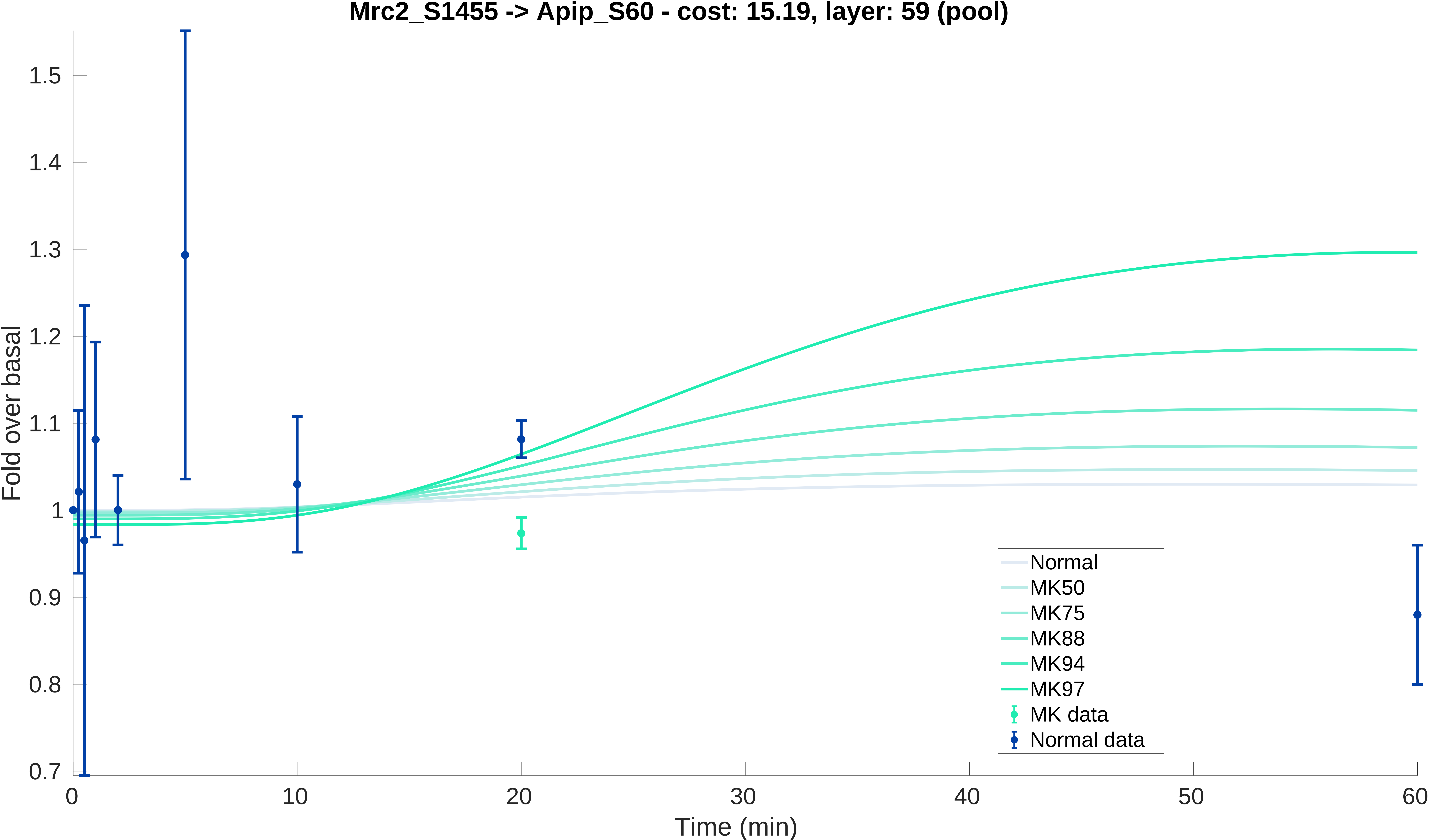

Mrc2_S1455 -> Apip_S60 - cost: 15.19, layer: 59 (pool)
Fold over basal
1.5
1.4
1.3
1.2
1.1
1
0.9
0.8
0.7
0
10
20
30
40
50
60
Time (min)
Normal
MK50
MK75
MK88
MK94
MK97
MK data
Normal data

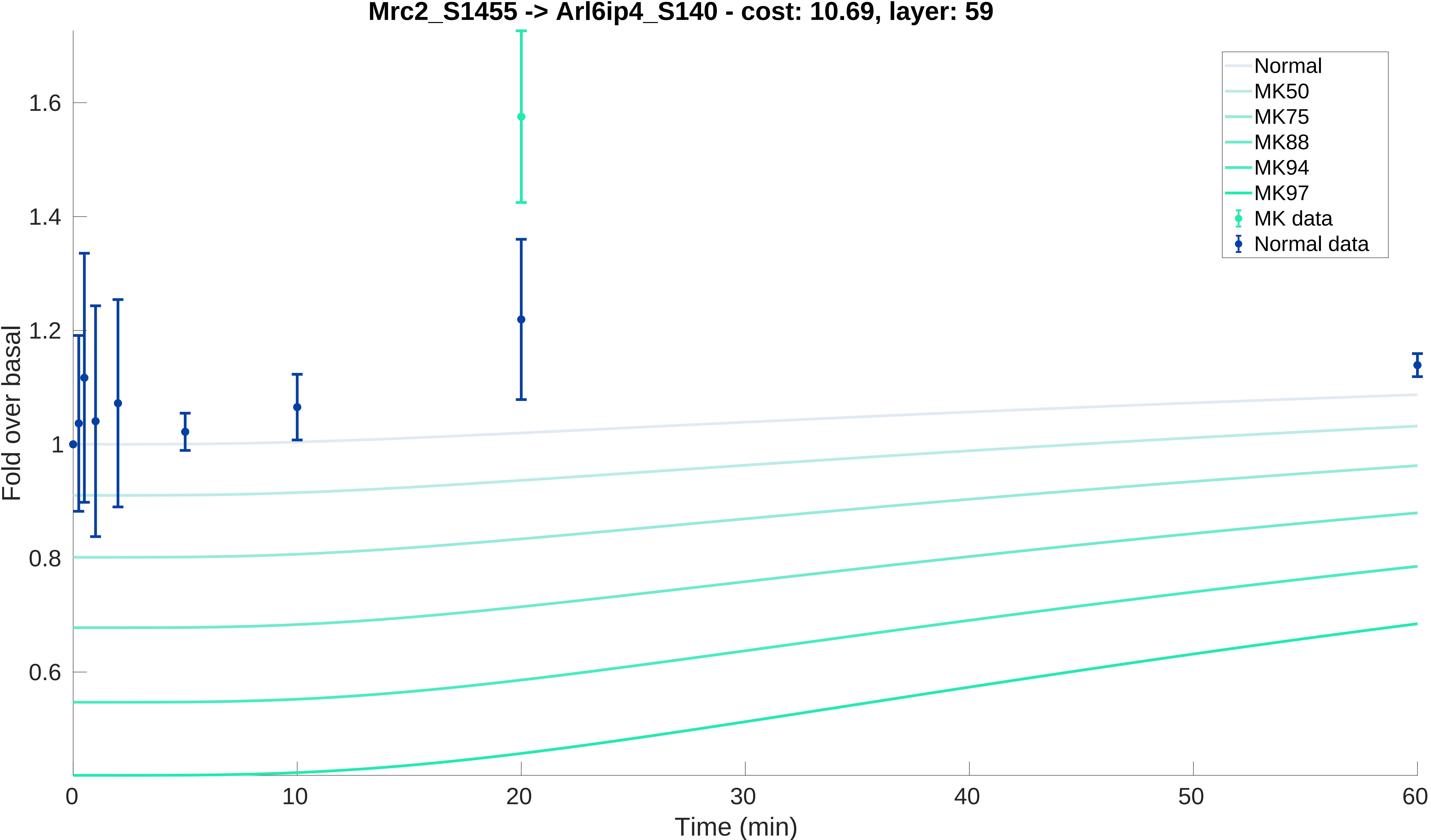

Mrc2_S1455 -> Arl6ip4_S140 - cost: 10.69, layer: 59
Fold over basal
1.6
1.4
1.2
1
0.8
0.6
0
10
20
30
40
50
60
Time (min)
Normal
MK50
MK75
MK88
MK94
MK97
MK data
Normal data

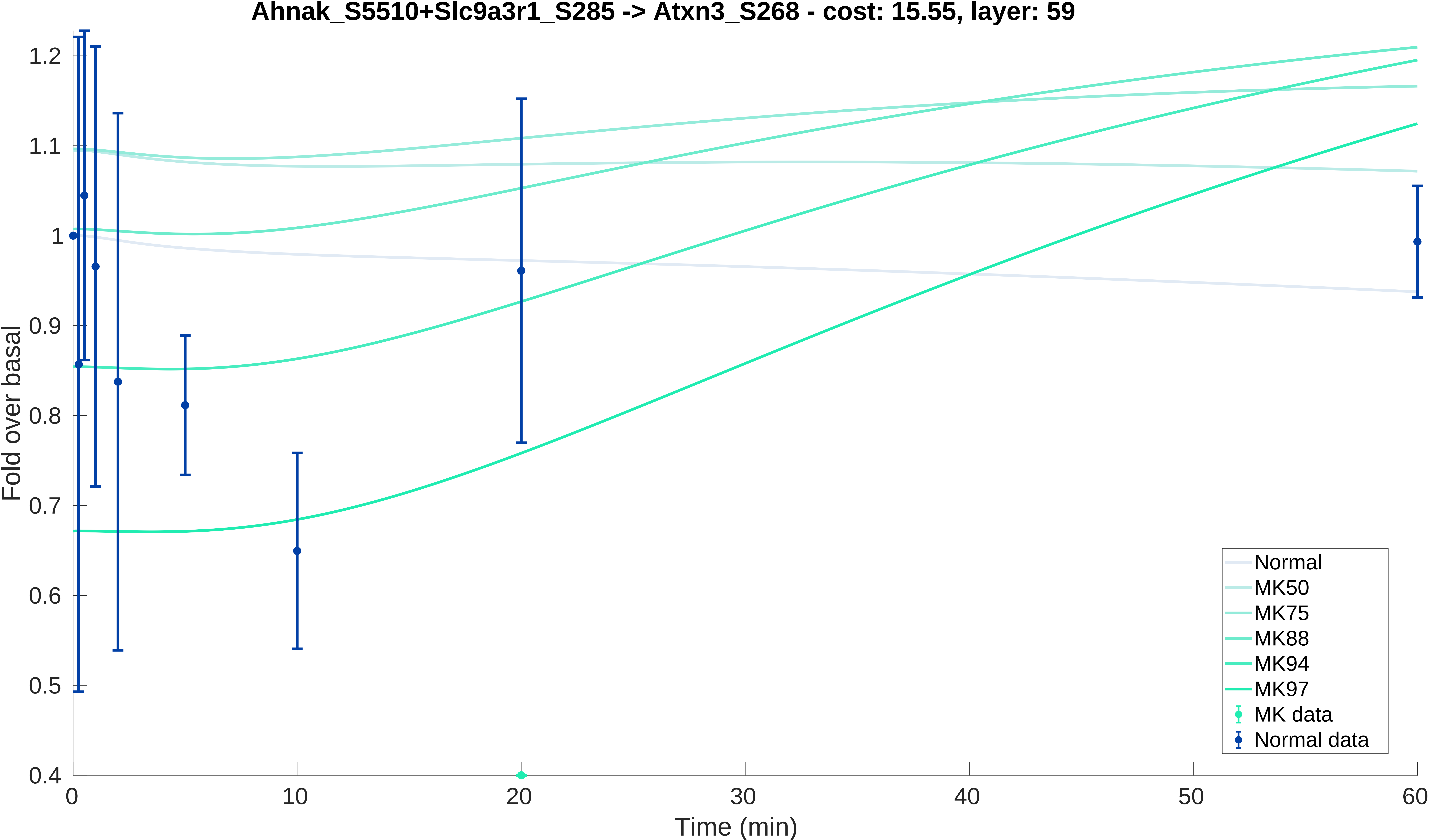

Ahnak_S5510+Slc9a3r1_S285 -> Atxn3_S268 - cost: 15.55, layer: 59
Fold over basal
Time (min)
0.4
0.5
0.6
0.7
0.8
0.9
1
1.1
1.2
0
10
20
30
40
50
60
Normal
MK50
MK75
MK88
MK94
MK97
MK data
Normal data

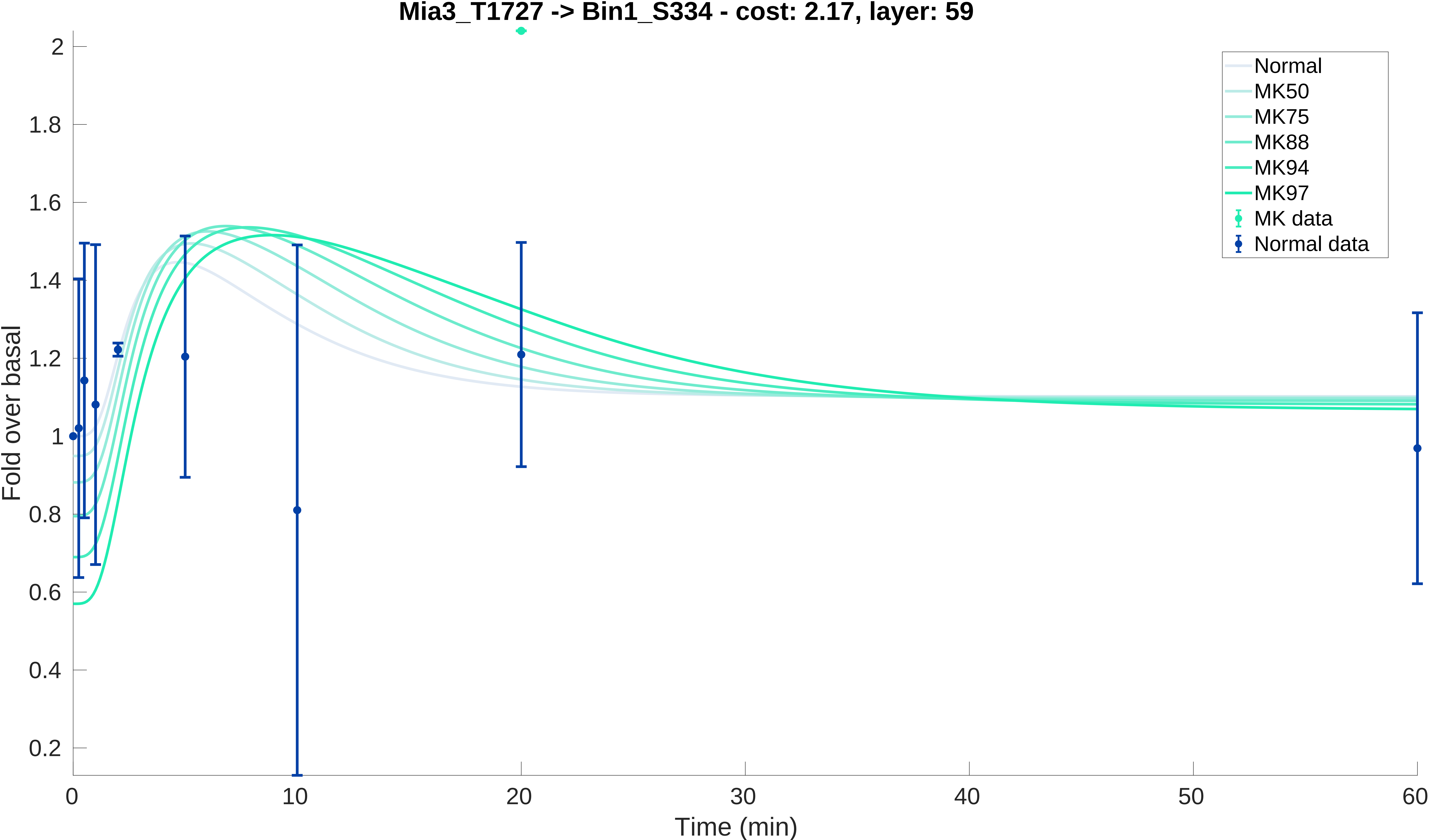

Mia3_T1727 -> Bin1_S334 - cost: 2.17, layer: 59
Fold over basal
Time (min)
0
10
20
30
40
50
60
0.2
0.4
0.6
0.8
1
1.2
1.4
1.6
1.8
2
Normal
MK50
MK75
MK88
MK94
MK97
MK data
Normal data

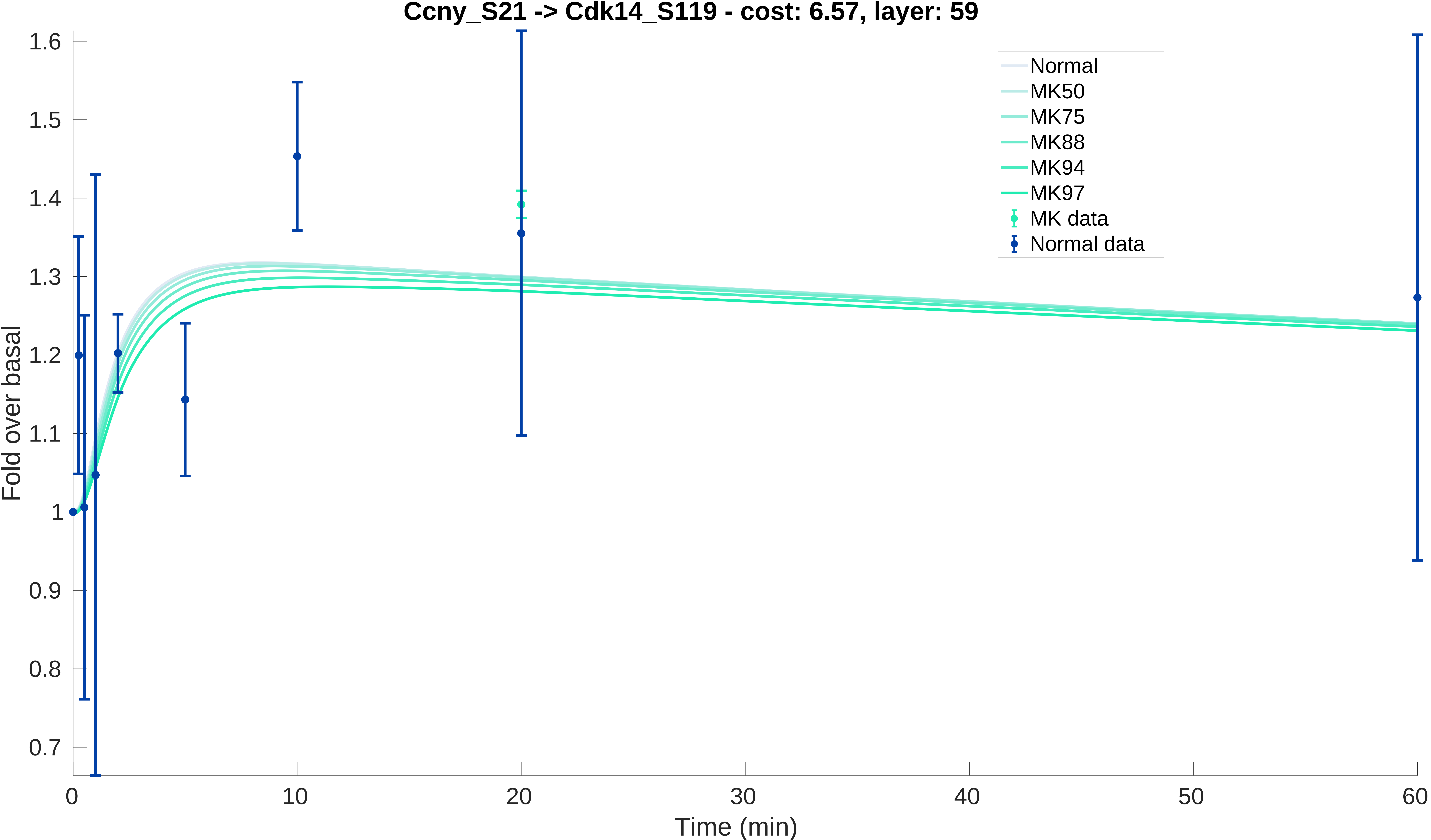

Ccny_S21 -> Cdk14_S119 - cost: 6.57, layer: 59
Fold over basal
Time (min)
1.6
1.5
1.4
1.3
1.2
1.1
1
0.9
0.8
0.7
0
10
20
30
40
50
60
Normal
MK50
MK75
MK88
MK94
MK97
MK data
Normal data

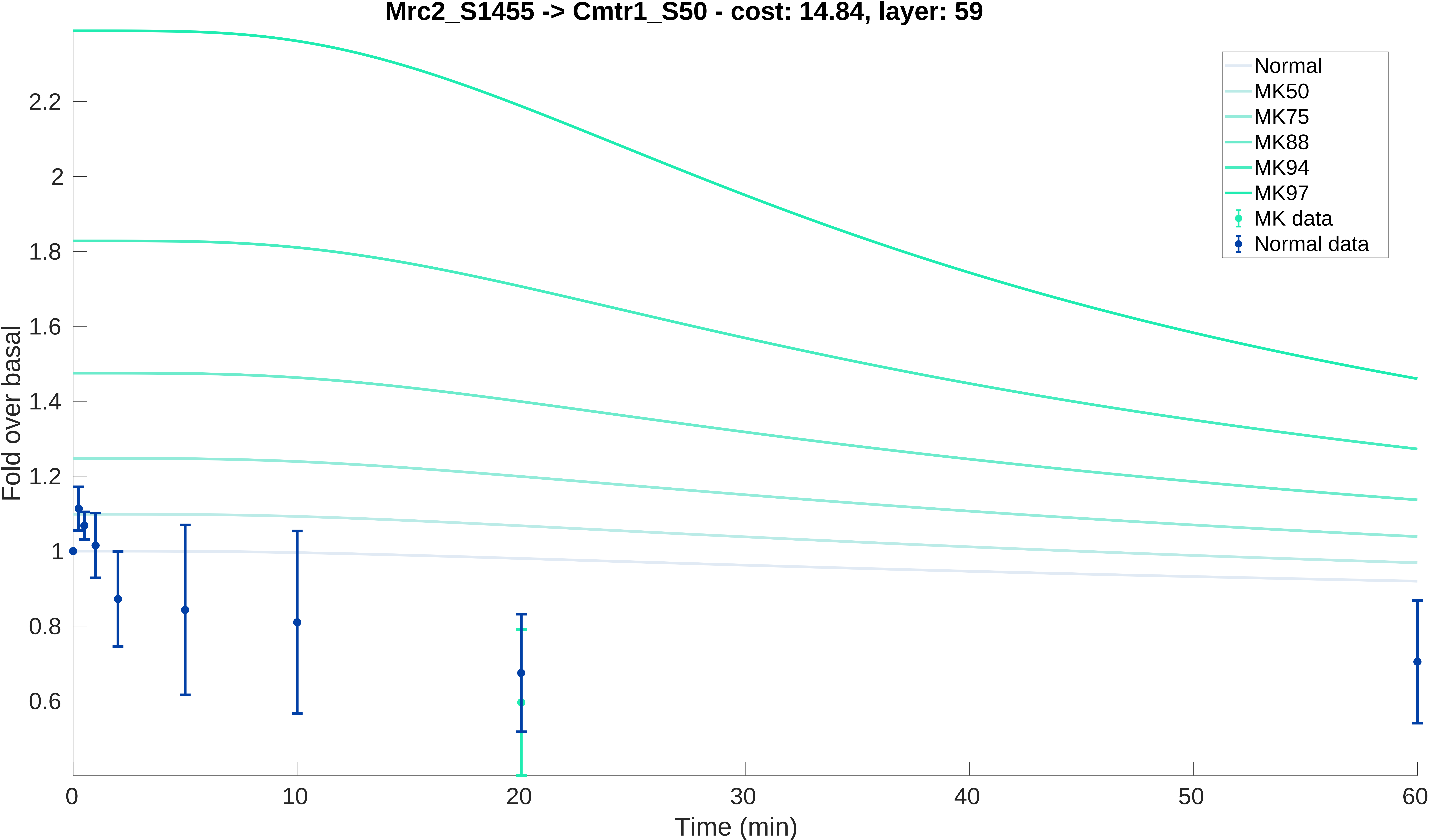

Mrc2_S1455 -> Cmtr1_S50 - cost: 14.84, layer: 59
Fold over basal
Time (min)
2.2
2
1.8
1.6
1.4
1.2
1
0.8
0.6
0
10
20
30
40
50
60
Normal
MK50
MK75
MK88
MK94
MK97
MK data
Normal data

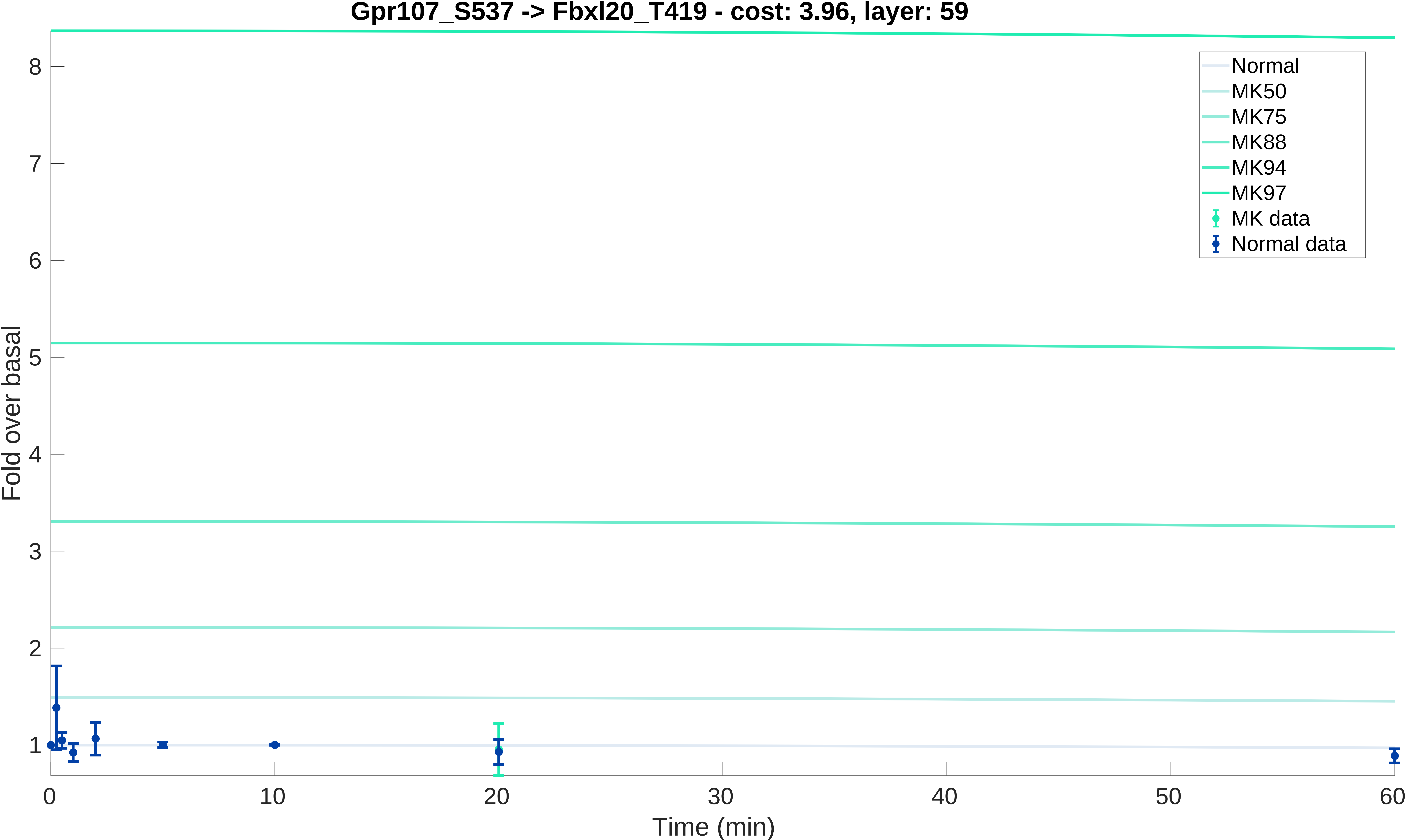
Gpr107_S537 -> Fbxl20_T419 - cost: 3.96, layer: 59
Fold over basal
Time (min)
Normal
MK50
MK75
MK88
MK94
MK97
MK data
Normal data
0
10
20
30
40
50
60
1
2
3
4
5
6
7
8

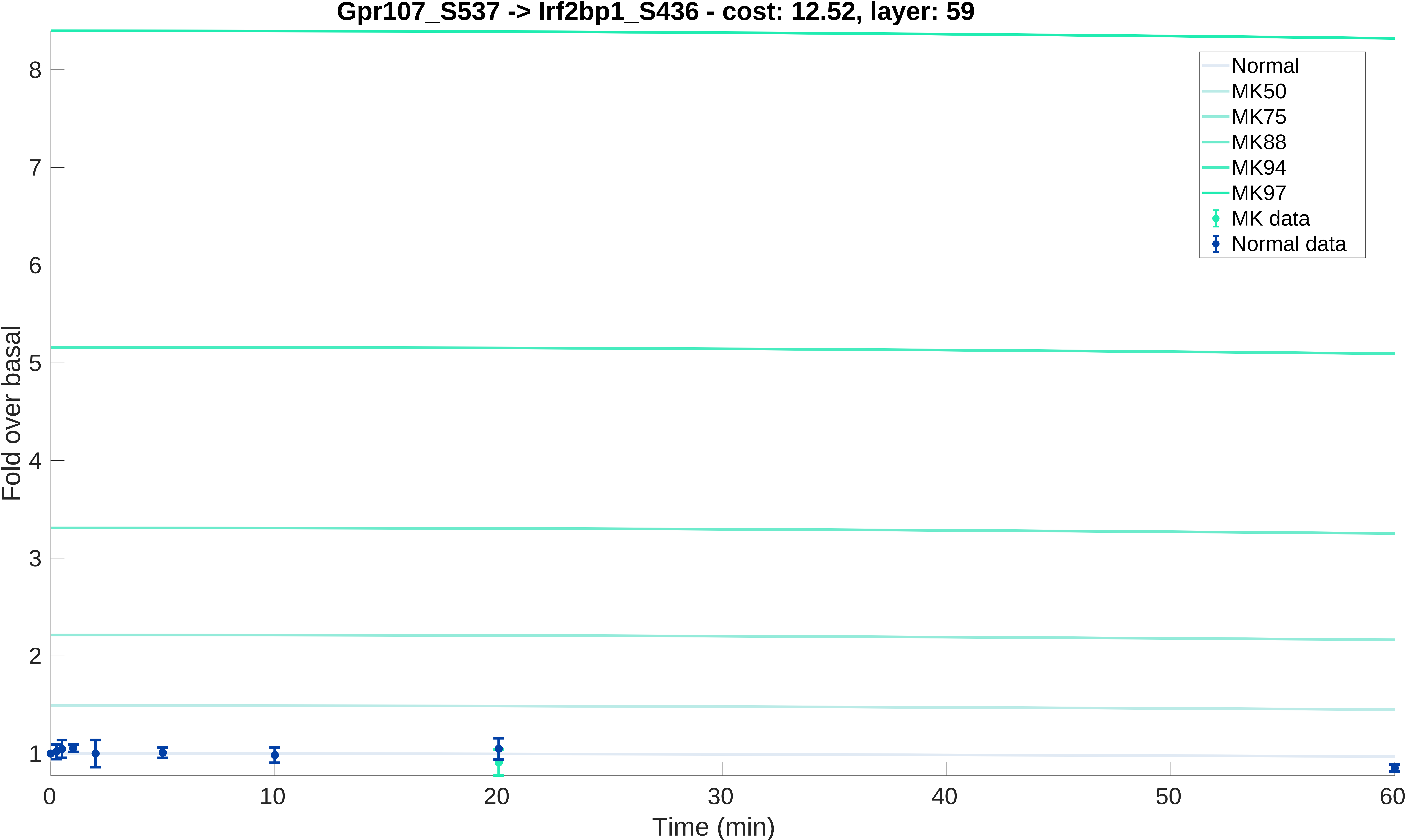

Gpr107_S537 -> Irf2bp1_S436 - cost: 12.52, layer: 59
Fold over basal
Time (min)
Normal
MK50
MK75
MK88
MK94
MK97
MK data
Normal data
1
2
3
4
5
6
7
8
0
10
20
30
40
50
60

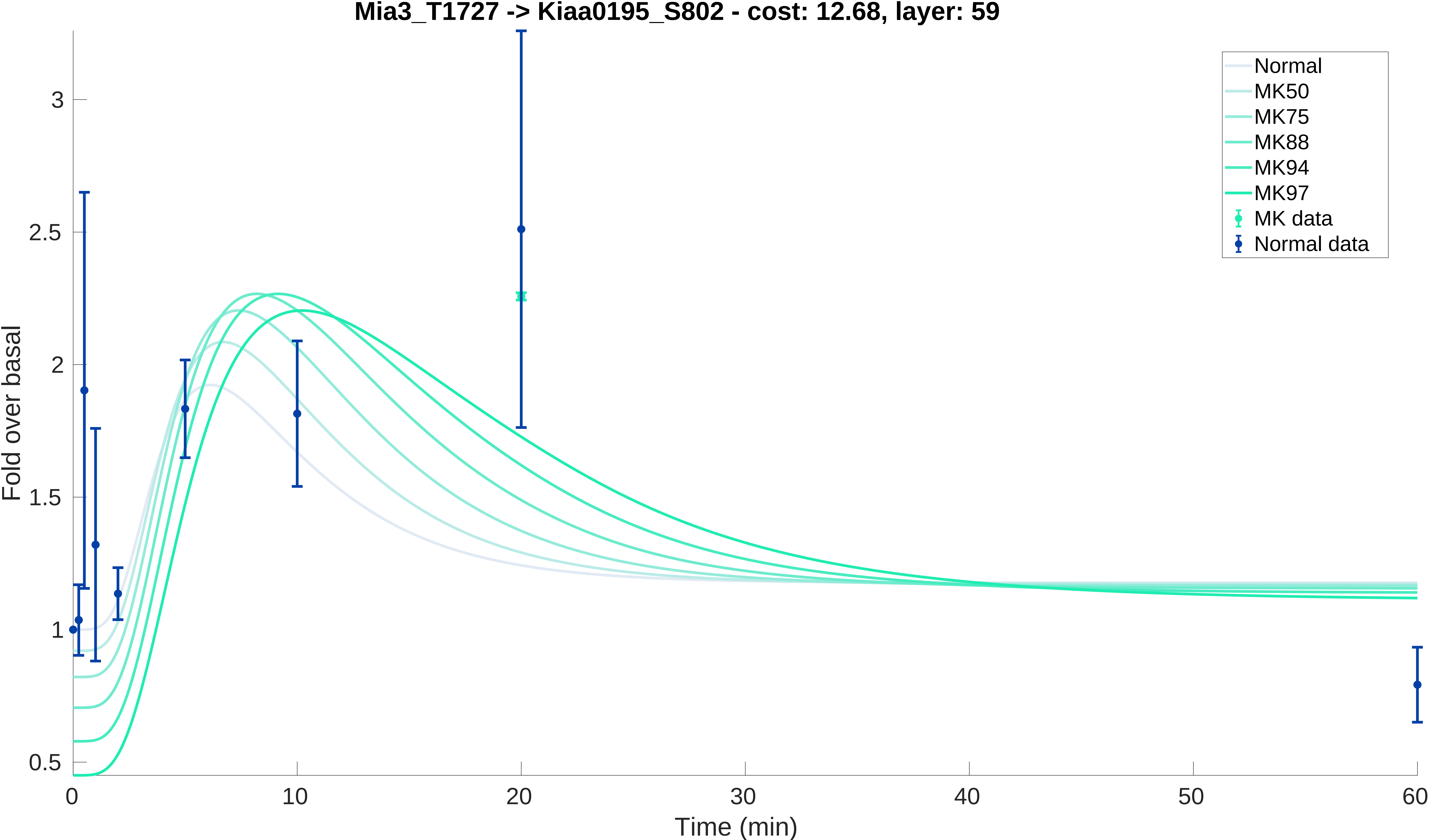

Mia3_T1727 -> Kiaa0195_S802 - cost: 12.68, layer: 59
Normal
MK50
MK75
MK88
MK94
MK97
MK data
Normal data
Fold over basal
0.5
1
1.5
2
2.5
3
0
10
20
30
40
50
60
Time (min)

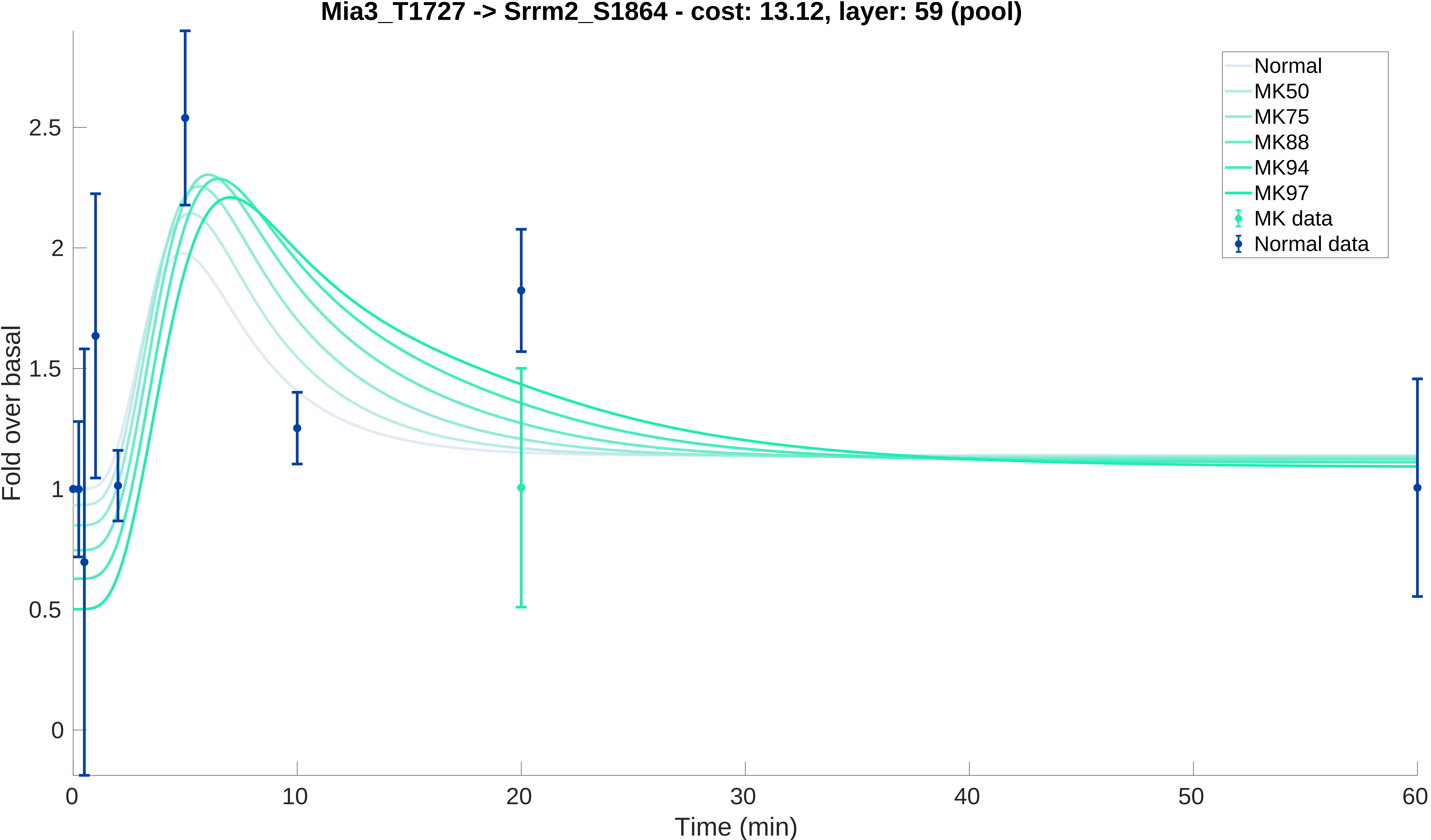
Mia3_T1727 -> Srrm2_S1864 - cost: 13.12, layer: 59 (pool)
Normal
MK50
MK75
MK88
MK94
MK97
MK data
Normal data
Fold over basal
Time (min)
0
0.5
1
1.5
2
2.5
0
10
20
30
40
50
60

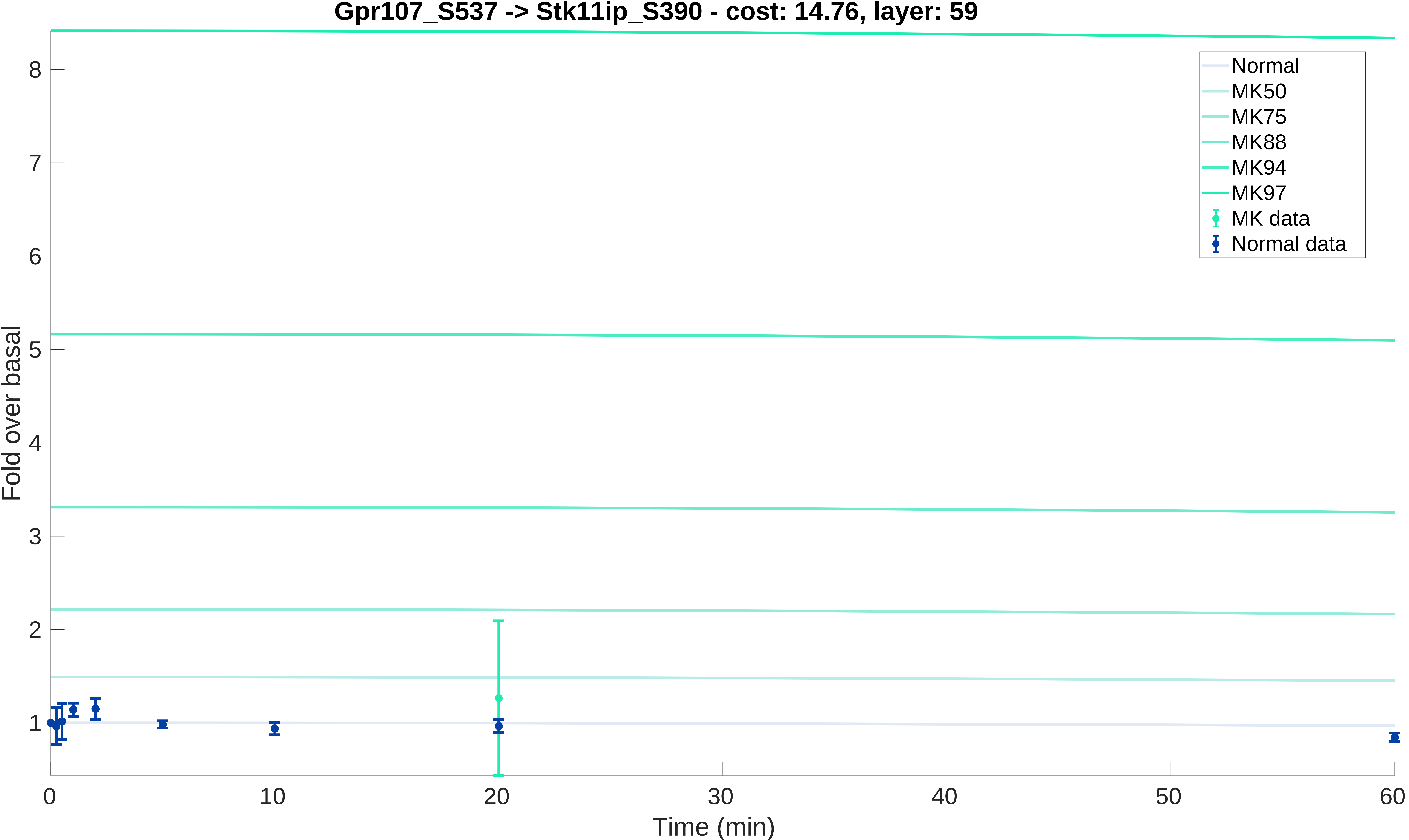
Gpr107_S537 -> Stk11ip_S390 - cost: 14.76, layer: 59
Fold over basal
Time (min)
0
10
20
30
40
50
60
1
2
3
4
5
6
7
8
Normal
MK50
MK75
MK88
MK94
MK97
MK data
Normal data

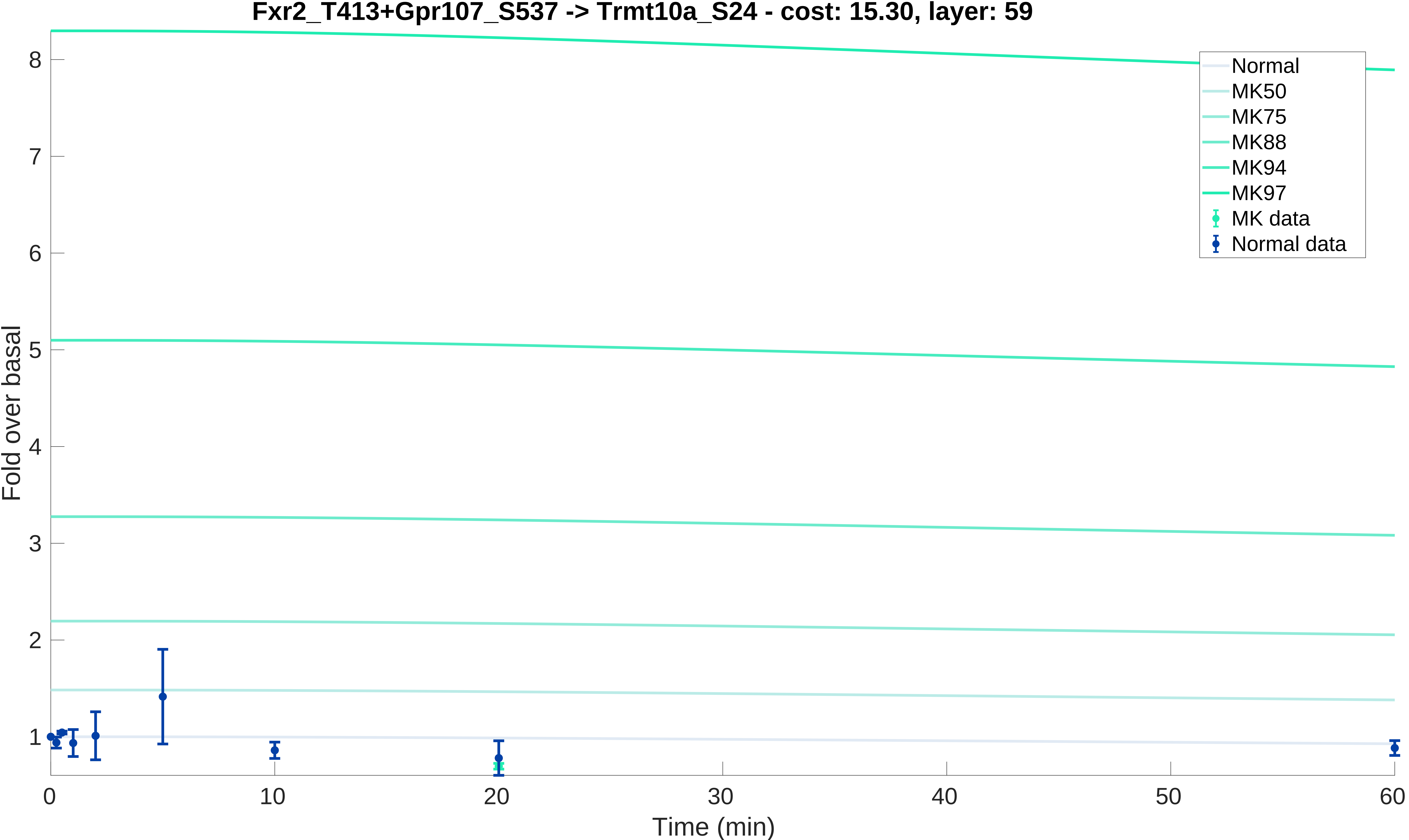

Fxr2_T413+Gpr107_S537 -> Trmt10a_S24 - cost: 15.30, layer: 59
Fold over basal
Time (min)
0
10
20
30
40
50
60
1
2
3
4
5
6
7
8
Normal
MK50
MK75
MK88
MK94
MK97
MK data
Normal data

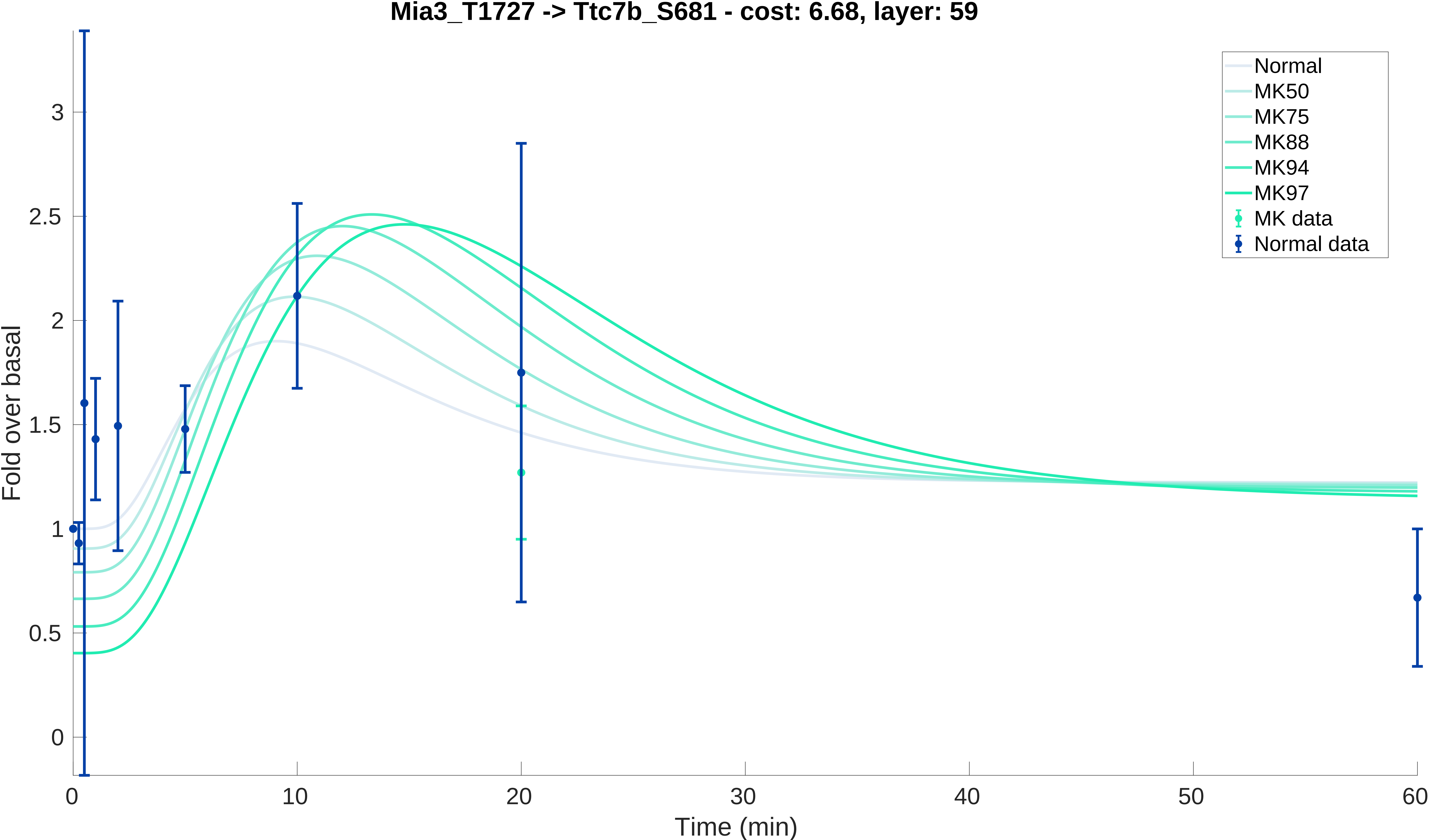

Mia3_T1727 -> Ttc7b_S681 - cost: 6.68, layer: 59
Normal
MK50
MK75
MK88
MK94
MK97
MK data
Normal data
Fold over basal
Time (min)
0
0.5
1
1.5
2
2.5
3
0
10
20
30
40
50
60

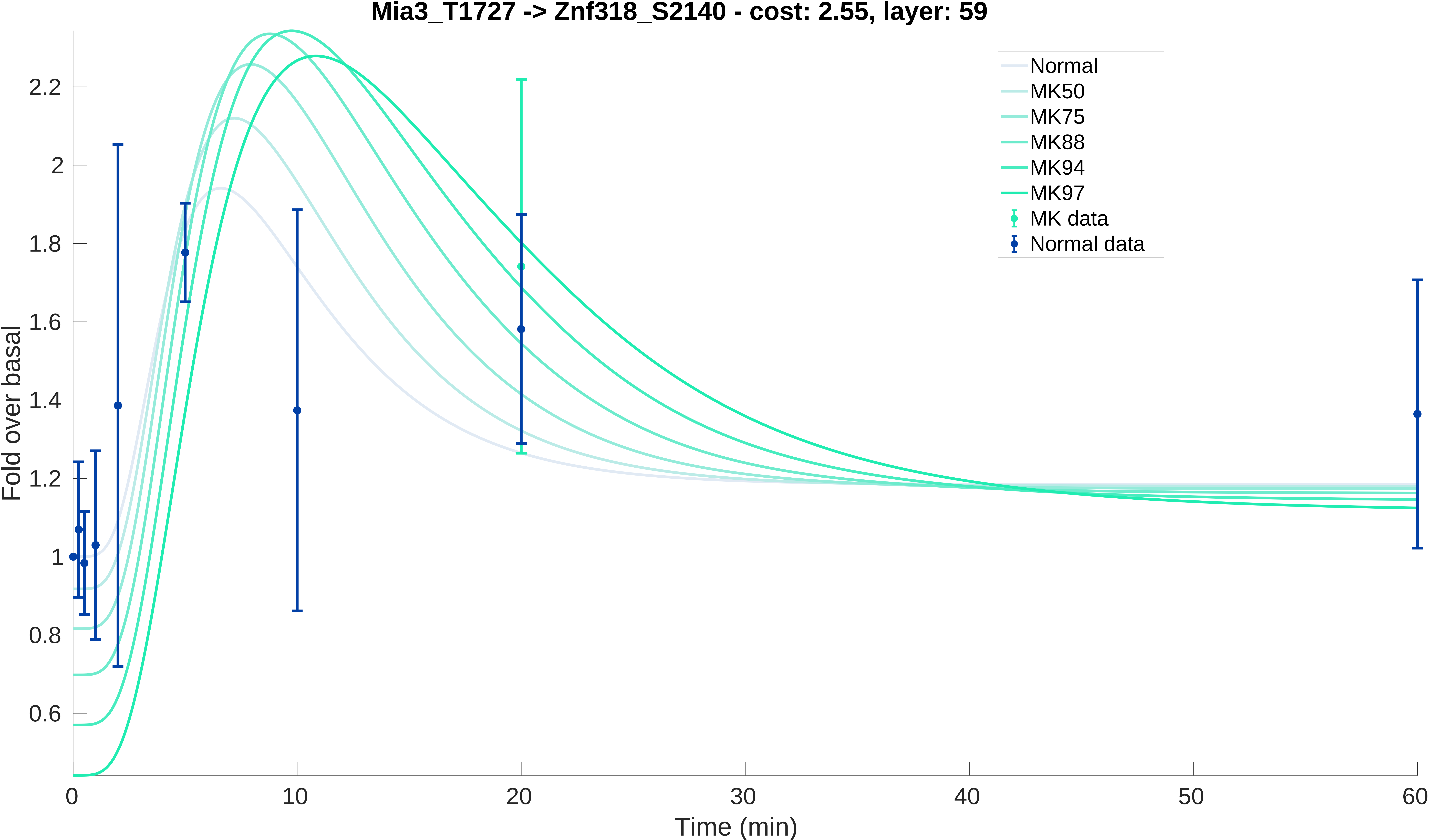
Mia3_T1727 -> Znf318_S2140 - cost: 2.55, layer: 59
Fold over basal
Time (min)
0.6
0.8
1
1.2
1.4
1.6
1.8
2
2.2
0
10
20
30
40
50
60
Normal
MK50
MK75
MK88
MK94
MK97
MK data
Normal data

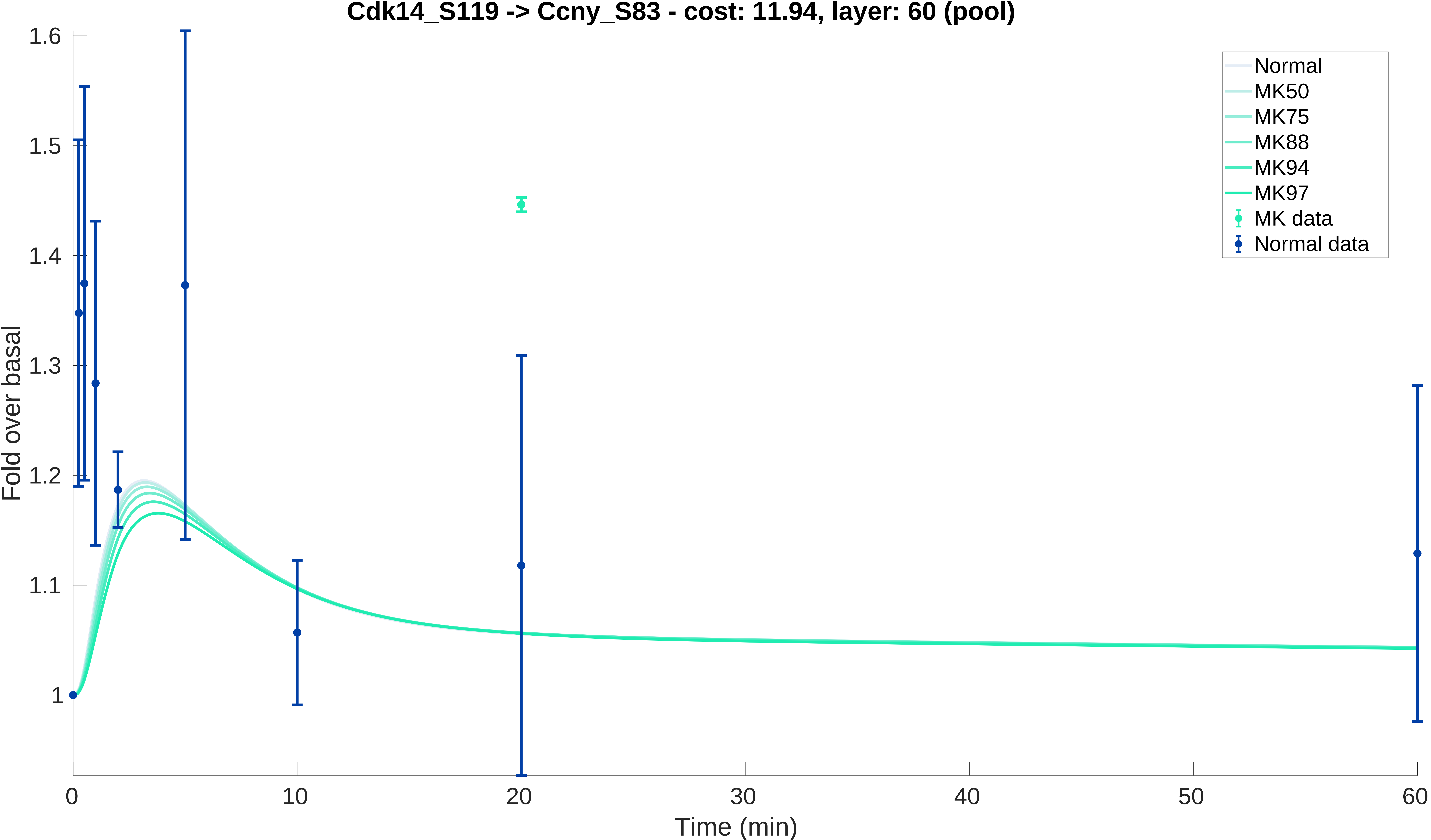

Cdk14_S119 -> Ccny_S83 - cost: 11.94, layer: 60 (pool)
Fold over basal
Time (min)
1.6
1.5
1.4
1.3
1.2
1.1
1
0
10
20
30
40
50
60
Normal
MK50
MK75
MK88
MK94
MK97
MK data
Normal data

### 3 Insulin stimulation, normal and with the PKB inhibitor

The following pages give all simulations of insulin stimulation normally and with the PKB inhibitor MK2206.

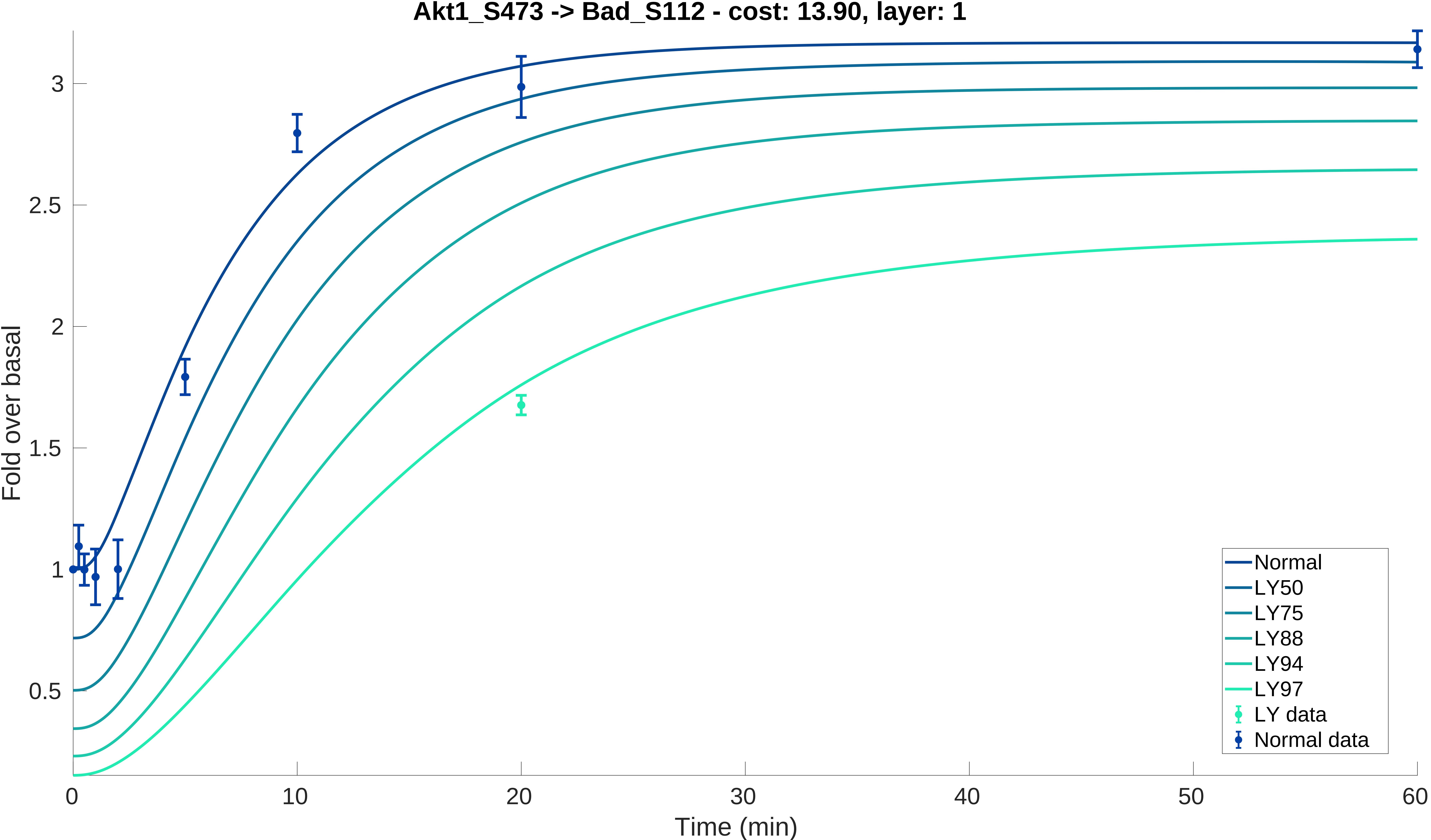

Akt1_S473 -> Bad_S112 - cost: 13.90, layer: 1
Fold over basal
Time (min)
0
10
20
30
40
50
60
0.5
1
1.5
2
2.5
3
Normal
LY50
LY75
LY88
LY94
LY97
LY data
Normal data

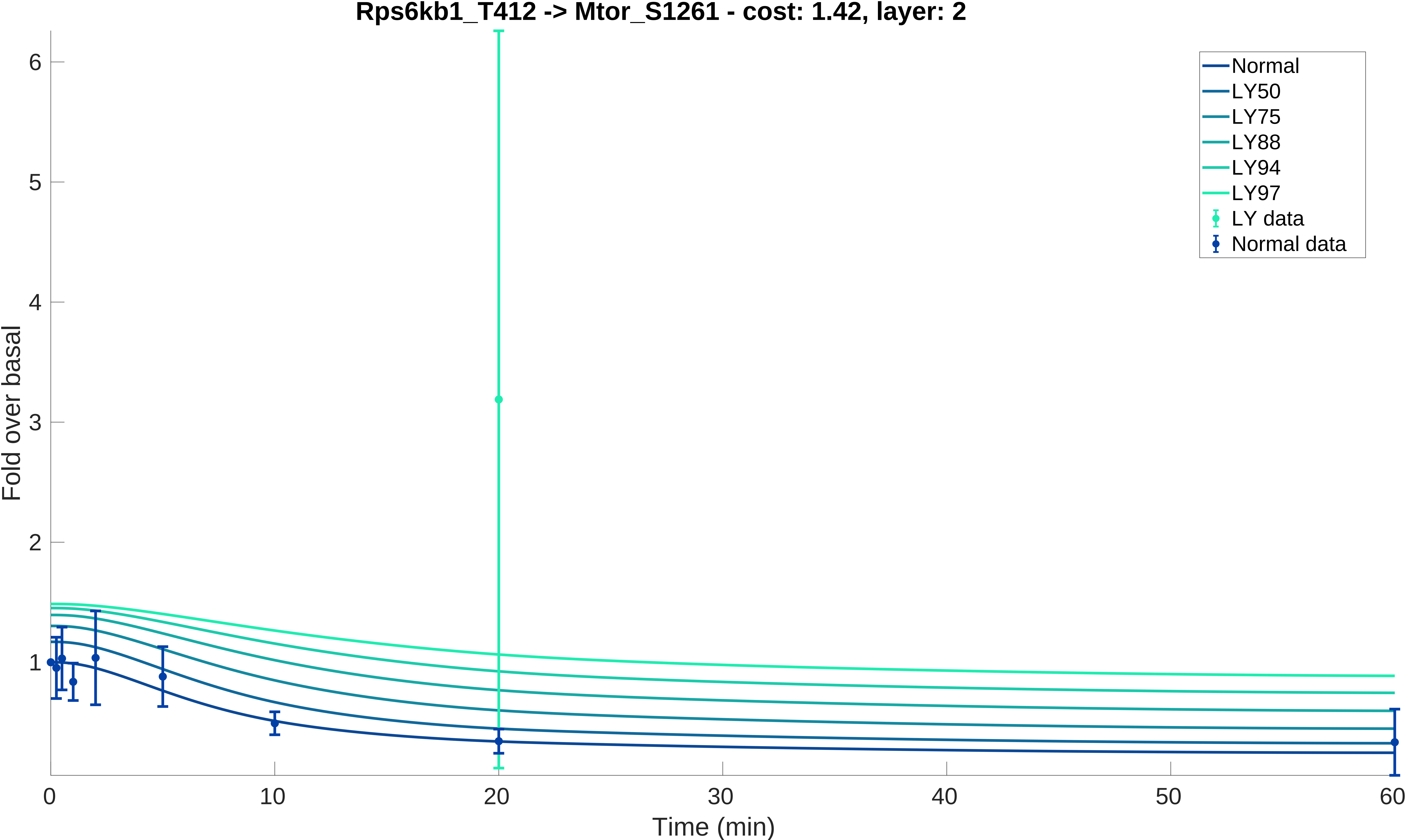

Rps6kb1_T412 -> Mtor_S1261 - cost: 1.42, layer: 2
Fold over basal
Time (min)
0
10
20
30
40
50
60
1
2
3
4
5
6
Normal
LY50
LY75
LY88
LY94
LY97
LY data
Normal data

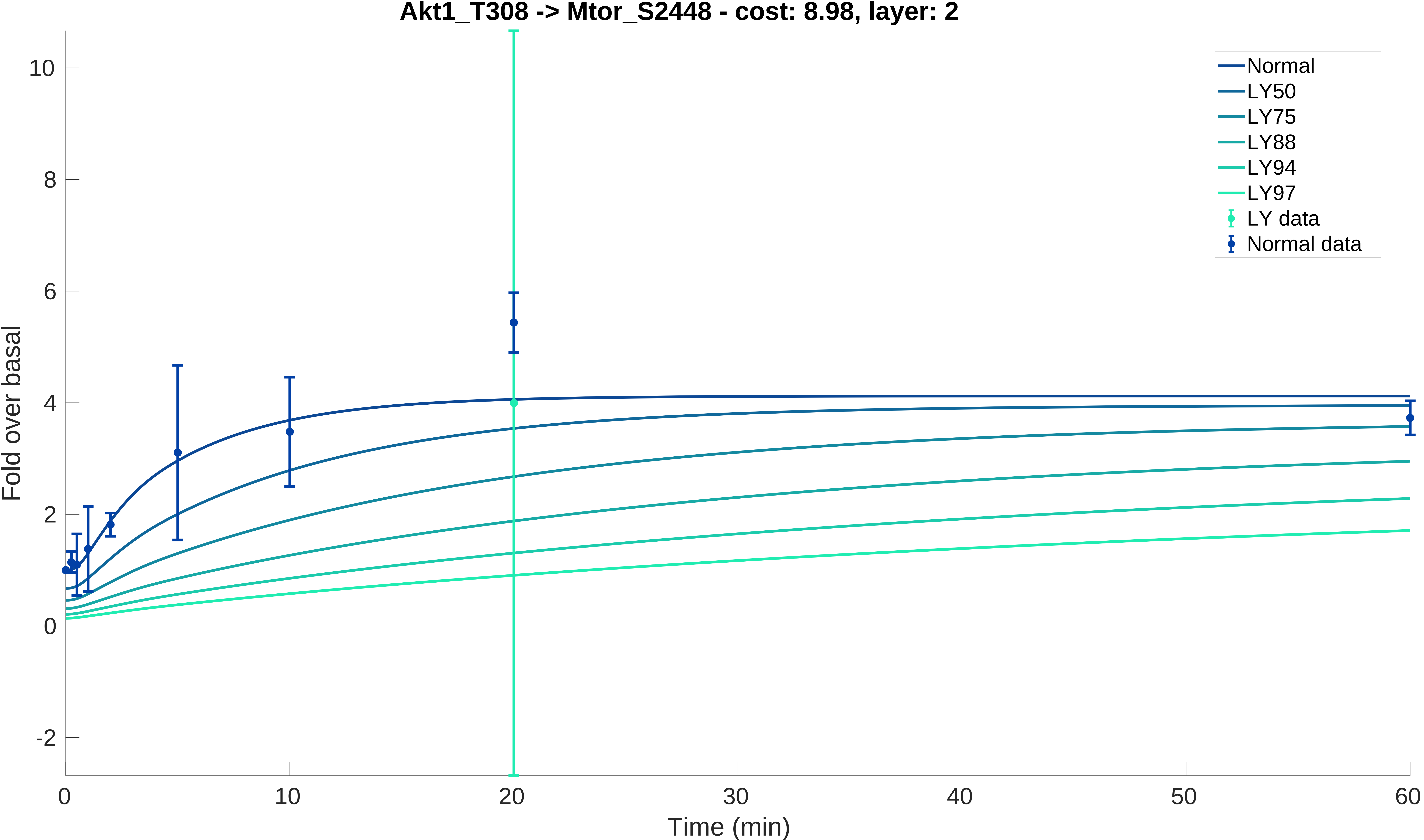

Akt1_T308 -> Mtor_S2448 - cost: 8.98, layer: 2
Fold over basal
Time (min)
-2
0
2
4
6
8
10
0
10
20
30
40
50
60
Normal
LY50
LY75
LY88
LY94
LY97
LY data
Normal data

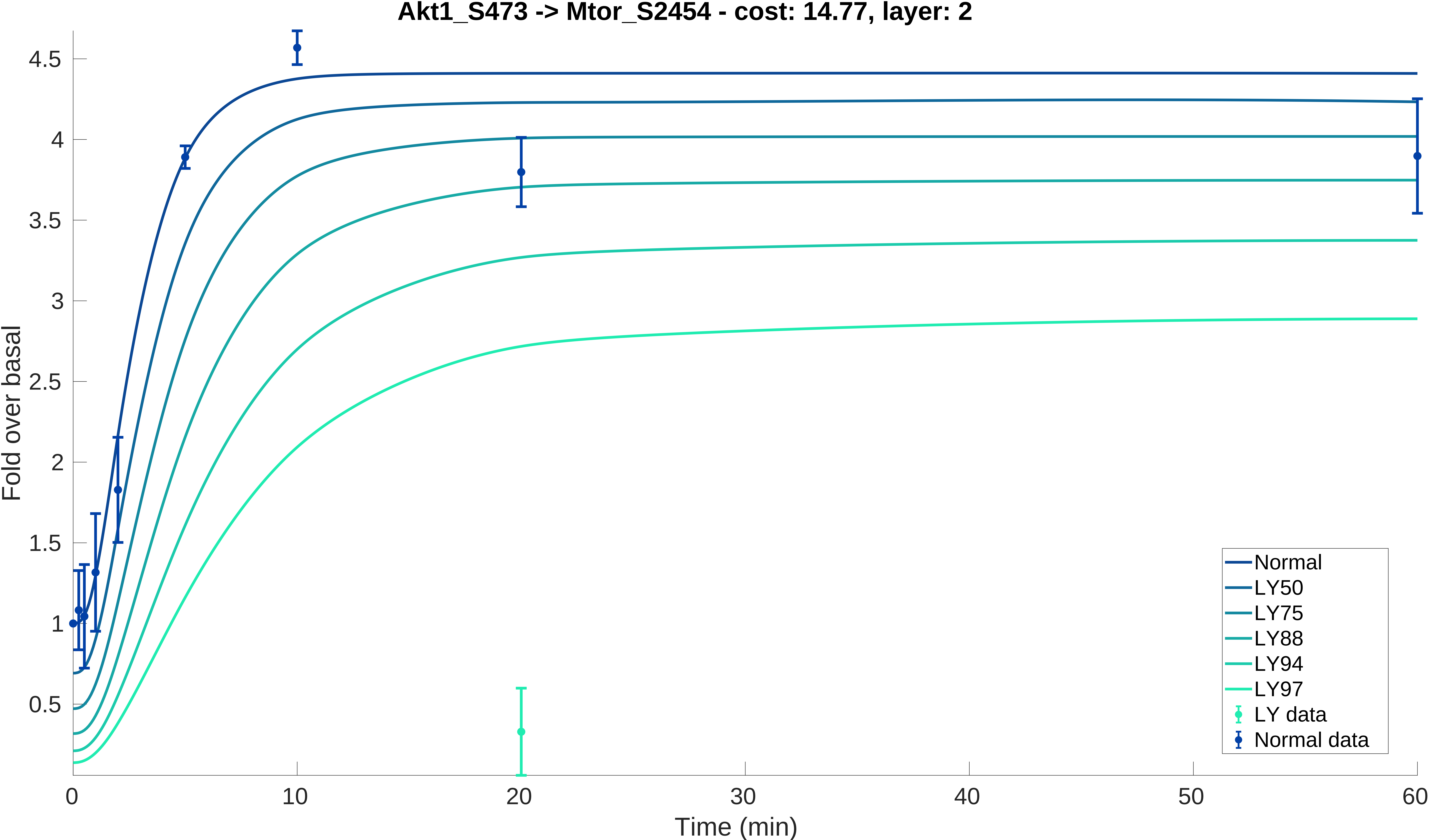

Akt1_S473 -> Mtor_S2454 - cost: 14.77, layer: 2
Fold over basal
0.5
1
1.5
2
2.5
3
3.5
4
4.5
0
10
20
30
40
50
60
Time (min)
Normal
LY50
LY75
LY88
LY94
LY97
LY data
Normal data

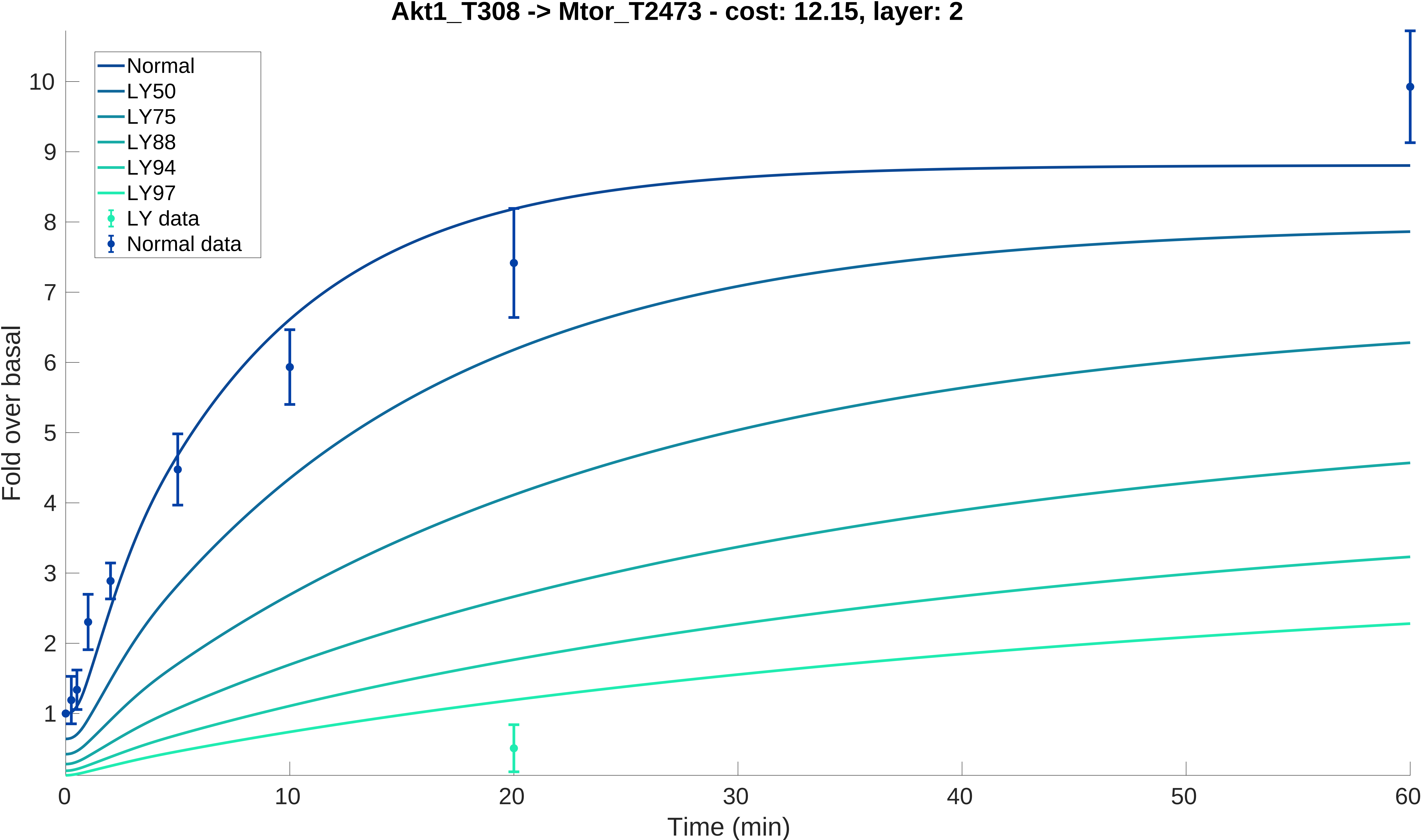

Akt1_T308 -> Mtor_T2473 - cost: 12.15, layer: 2
Normal
LY50
LY75
LY88
LY94
LY97
LY data
Normal data
Fold over basal
Time (min)
1
2
3
4
5
6
7
8
9
10
0
10
20
30
40
50
60

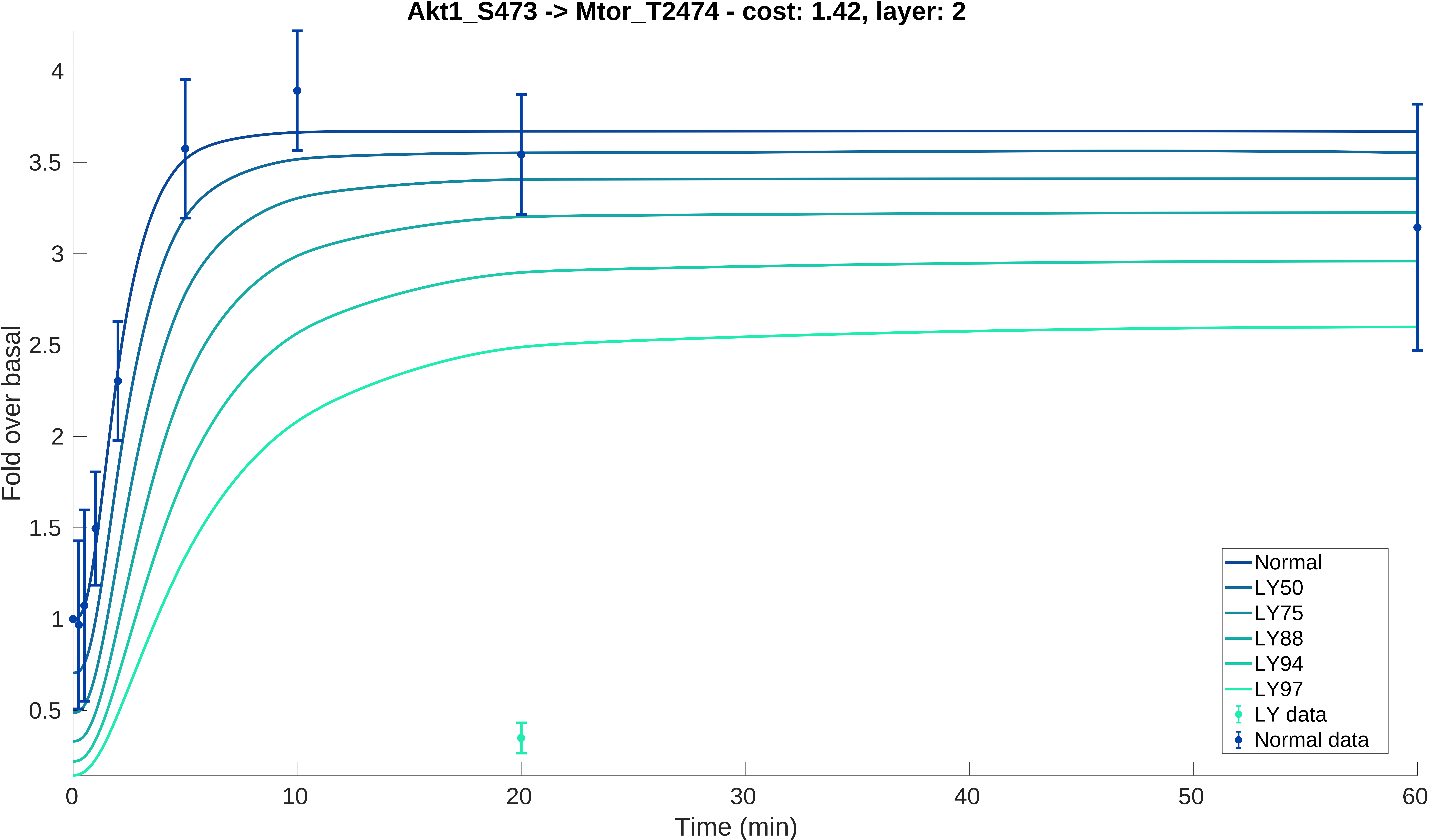

Akt1_S473 -> Mtor_T2474 - cost: 1.42, layer: 2
Fold over basal
Time (min)
0.5
1
1.5
2
2.5
3
3.5
4
0
10
20
30
40
50
60
Normal
LY50
LY75
LY88
LY94
LY97
LY data
Normal data

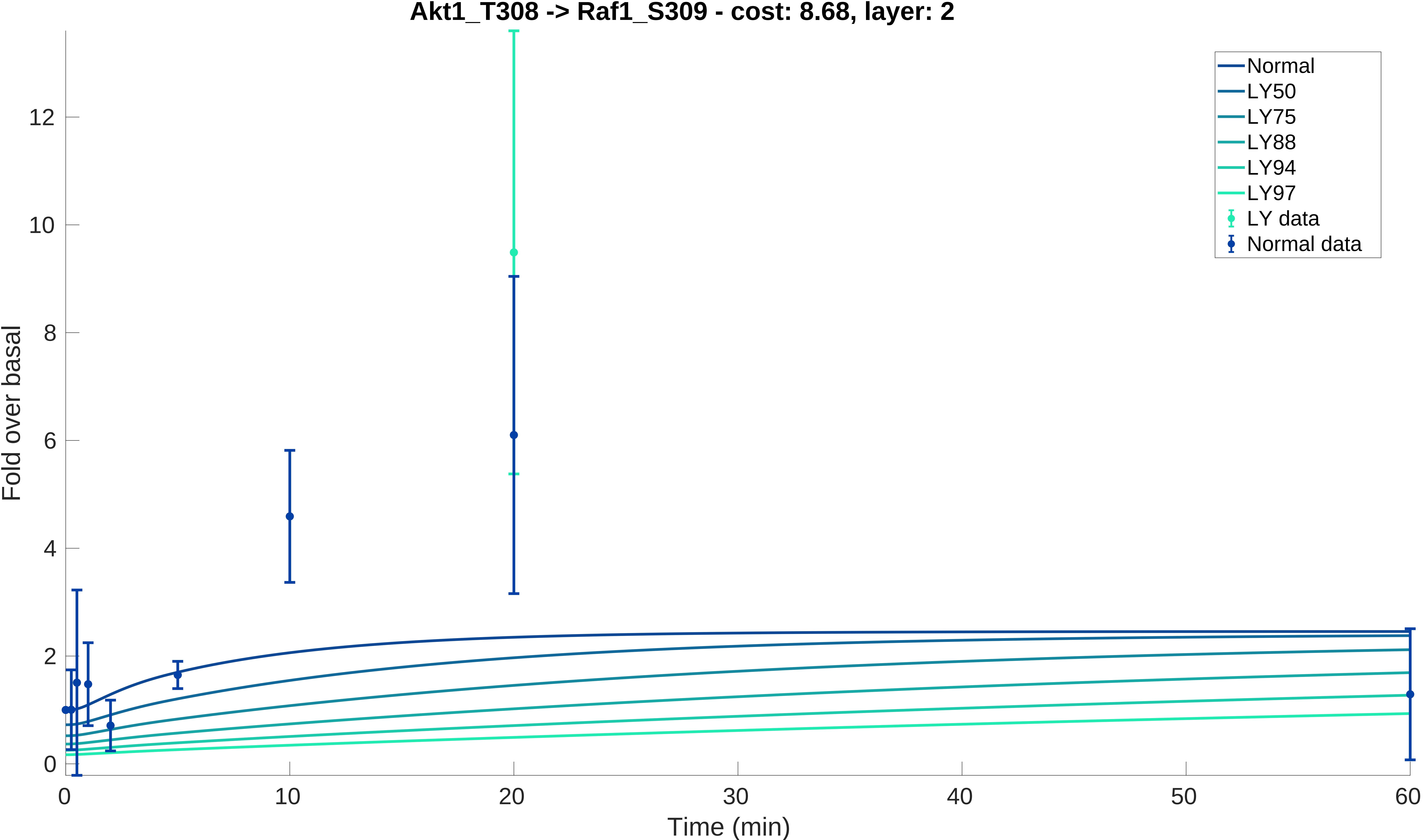

Akt1_T308 -> Raf1_S309 - cost: 8.68, layer: 2
Normal
LY50
LY75
LY88
LY94
LY97
LY data
Normal data
Fold over basal
Time (min)
0
2
4
6
8
10
12
0
10
20
30
40
50
60

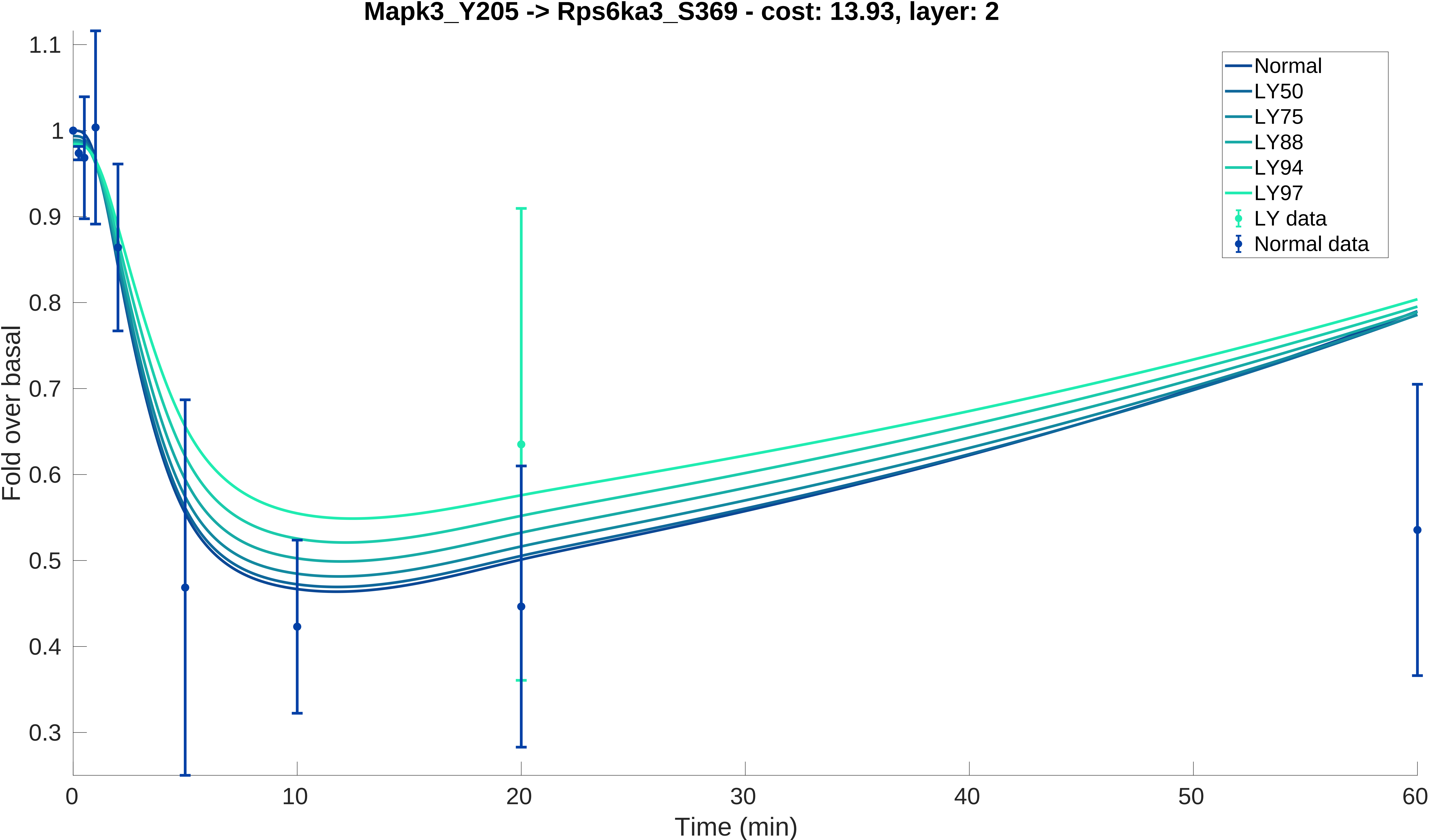

Mapk3_Y205 -> Rps6ka3_S369 - cost: 13.93, layer: 2
Fold over basal
Time (min)
1.1
1
0.9
0.8
0.7
0.6
0.5
0.4
0.3
0
10
20
30
40
50
60
Normal
LY50
LY75
LY88
LY94
LY97
LY data
Normal data

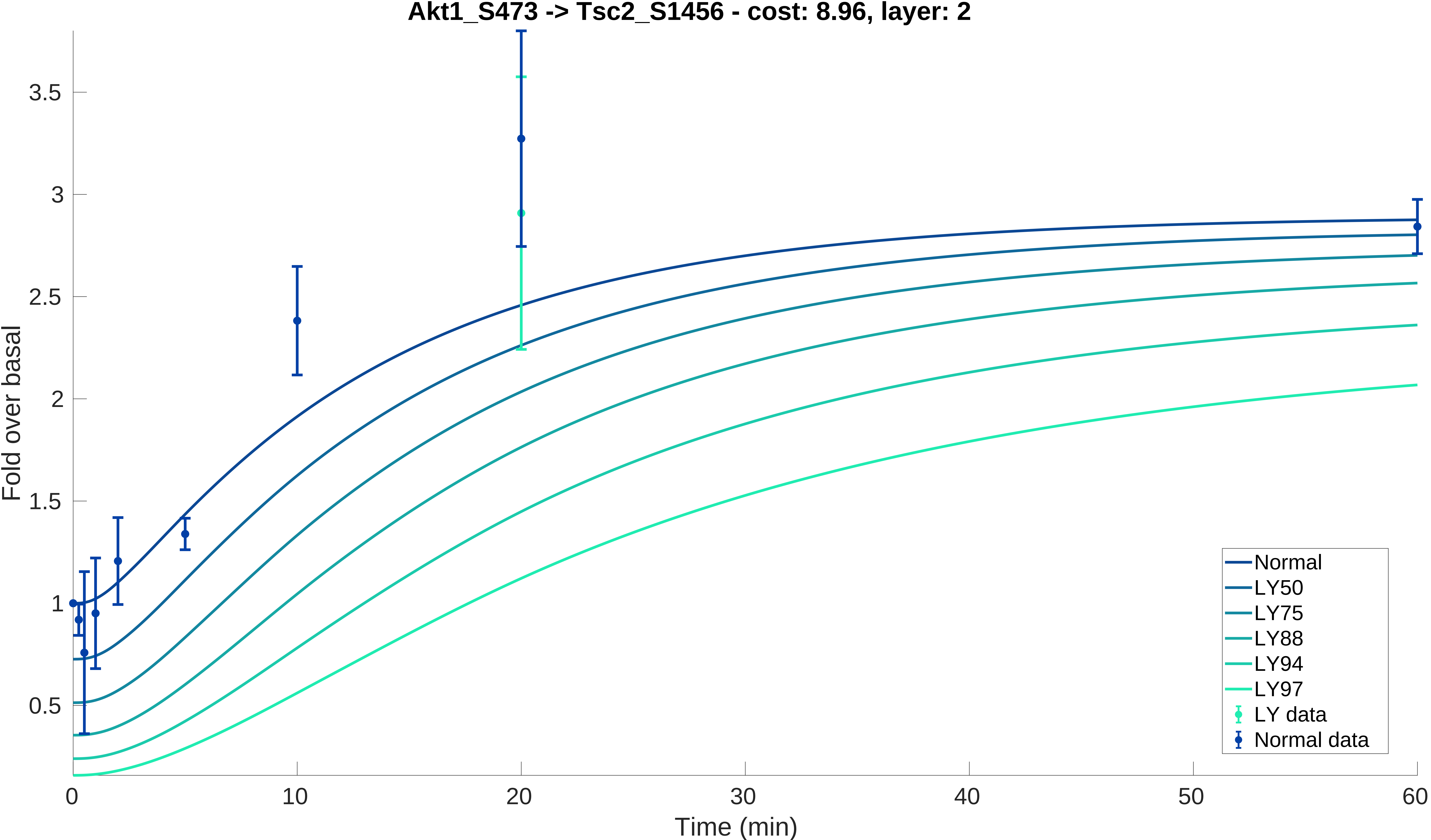

Akt1_S473 -> Tsc2_S1456 - cost: 8.96, layer: 2
Fold over basal
Time (min)
0.5
1
1.5
2
2.5
3
3.5
0
10
20
30
40
50
60
Normal
LY50
LY75
LY88
LY94
LY97
LY data
Normal data

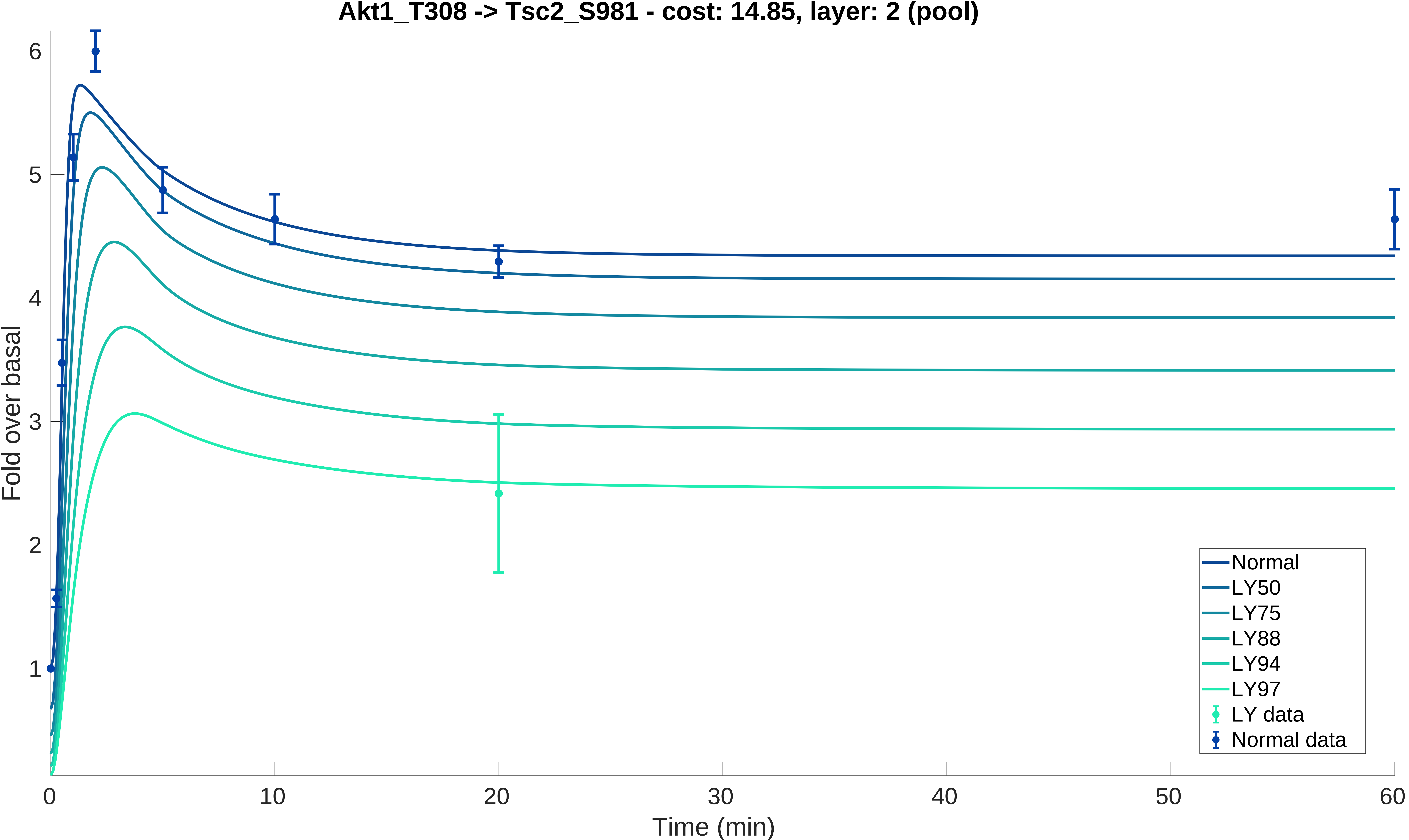

Akt1_T308 -> Tsc2_S981 - cost: 14.85, layer: 2 (pool)
Fold over basal
Time (min)
0
10
20
30
40
50
60
1
2
3
4
5
6
Normal
LY50
LY75
LY88
LY94
LY97
LY data
Normal data

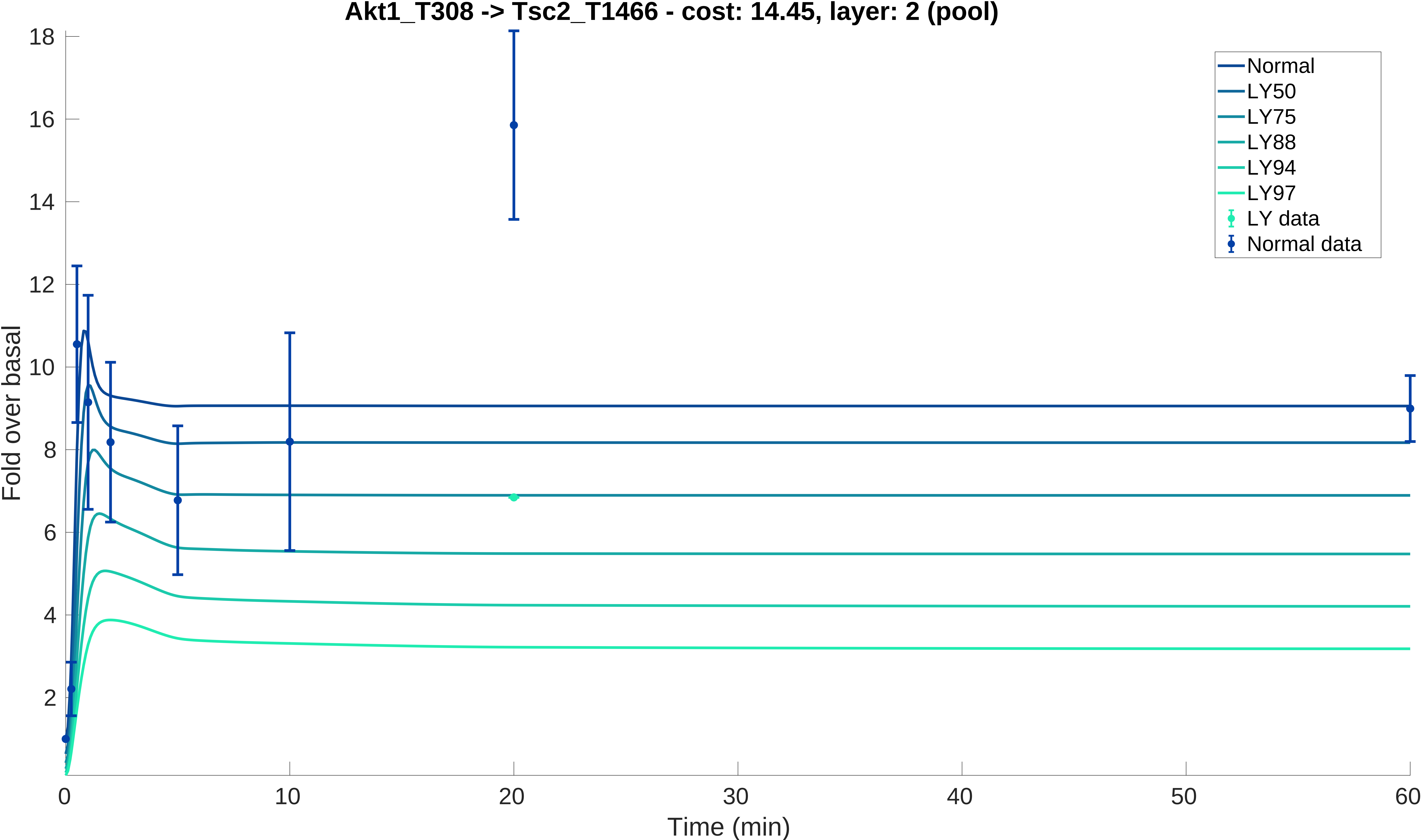
Akt1_T308 -> Tsc2_T1466 - cost: 14.45, layer: 2 (pool)
Fold over basal
Time (min)
0
10
20
30
40
50
60
2
4
6
8
10
12
14
16
18
Normal
LY50
LY75
LY88
LY94
LY97
LY data
Normal data

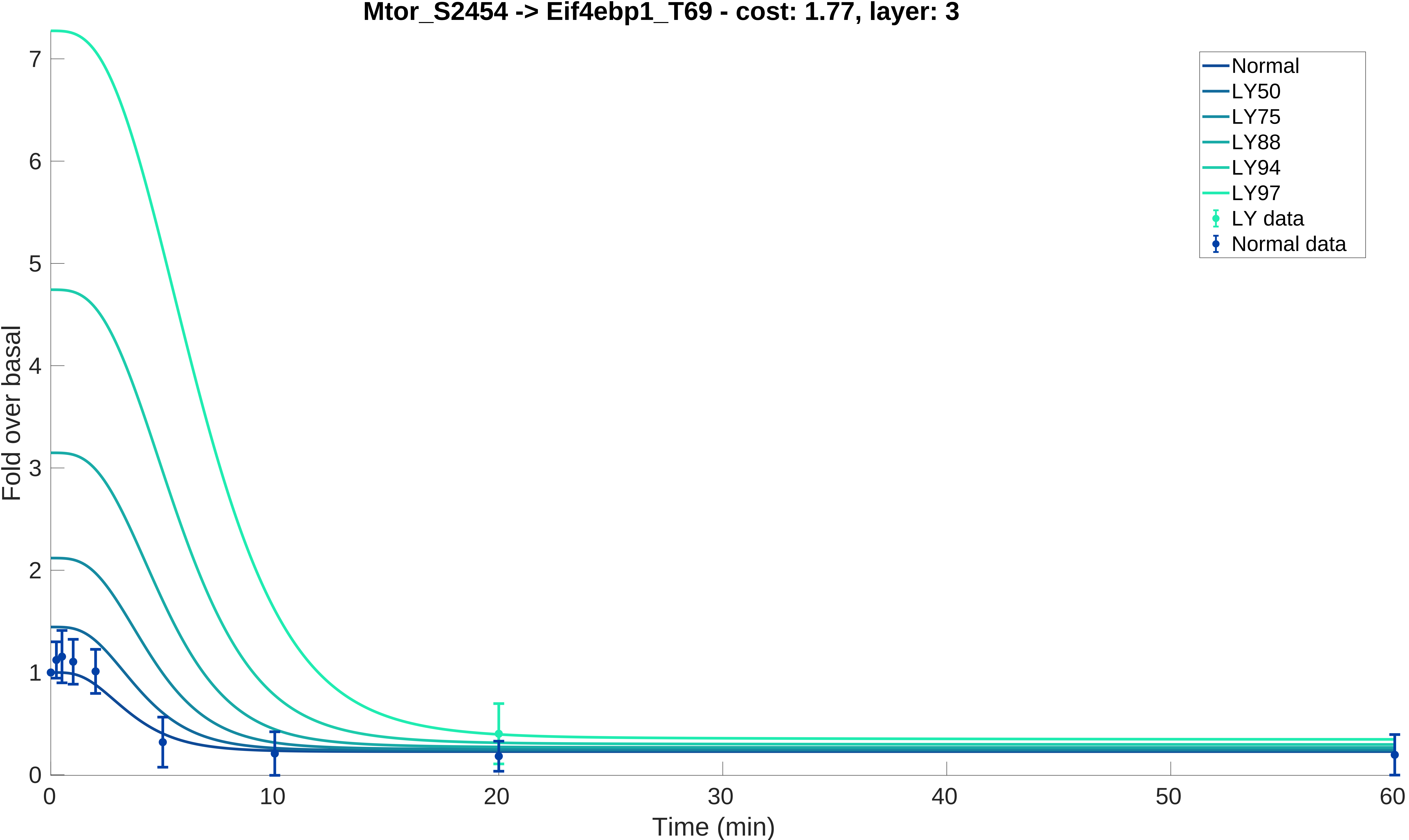
Mtor_S2454 -> Eif4ebp1_T69 - cost: 1.77, layer: 3
Fold over basal
Time (min)
0
1
2
3
4
5
6
7
0
10
20
30
40
50
60
Normal
LY50
LY75
LY88
LY94
LY97
LY data
Normal data

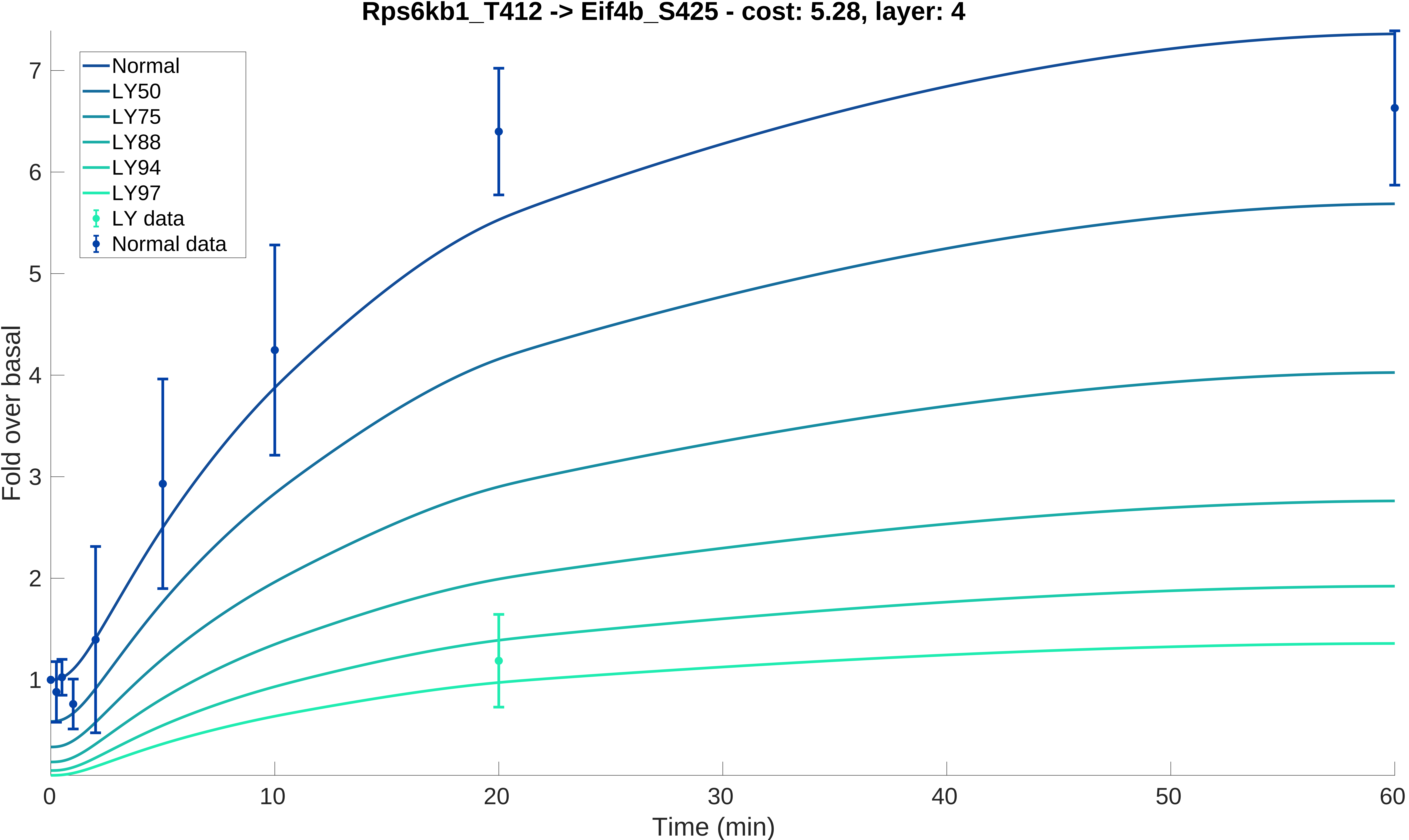

Rps6kb1_T412 -> Eif4b_S425 - cost: 5.28, layer: 4
Normal
LY50
LY75
LY88
LY94
LY97
LY data
Normal data
Fold over basal
1
2
3
4
5
6
7
0
10
20
30
40
50
60
Time (min)

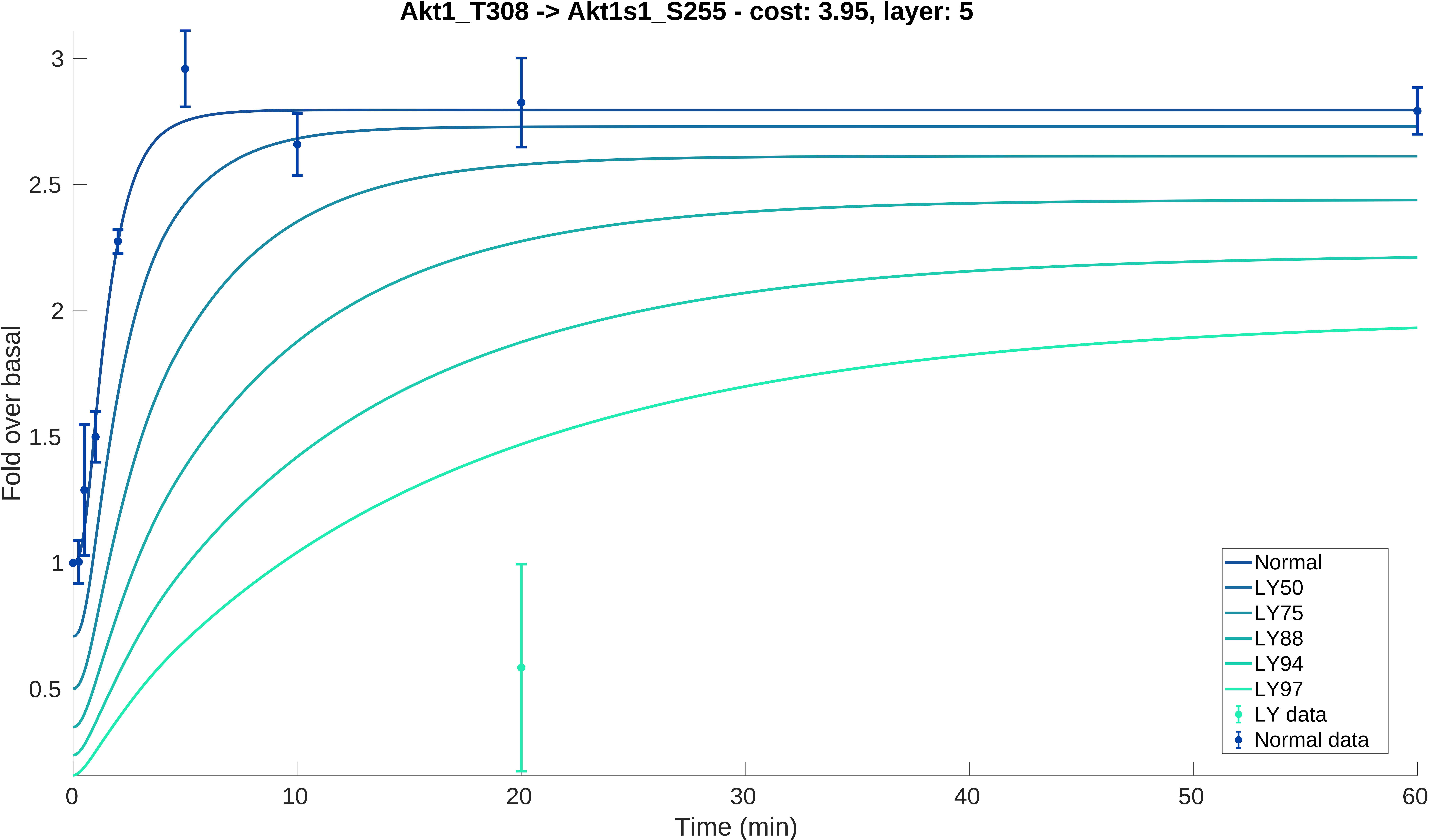
Akt1_T308 -> Akt1s1_S255 - cost: 3.95, layer: 5
Fold over basal
Time (min)
0.5
1
1.5
2
2.5
3
0
10
20
30
40
50
60
Normal
LY50
LY75
LY88
LY94
LY97
LY data
Normal data

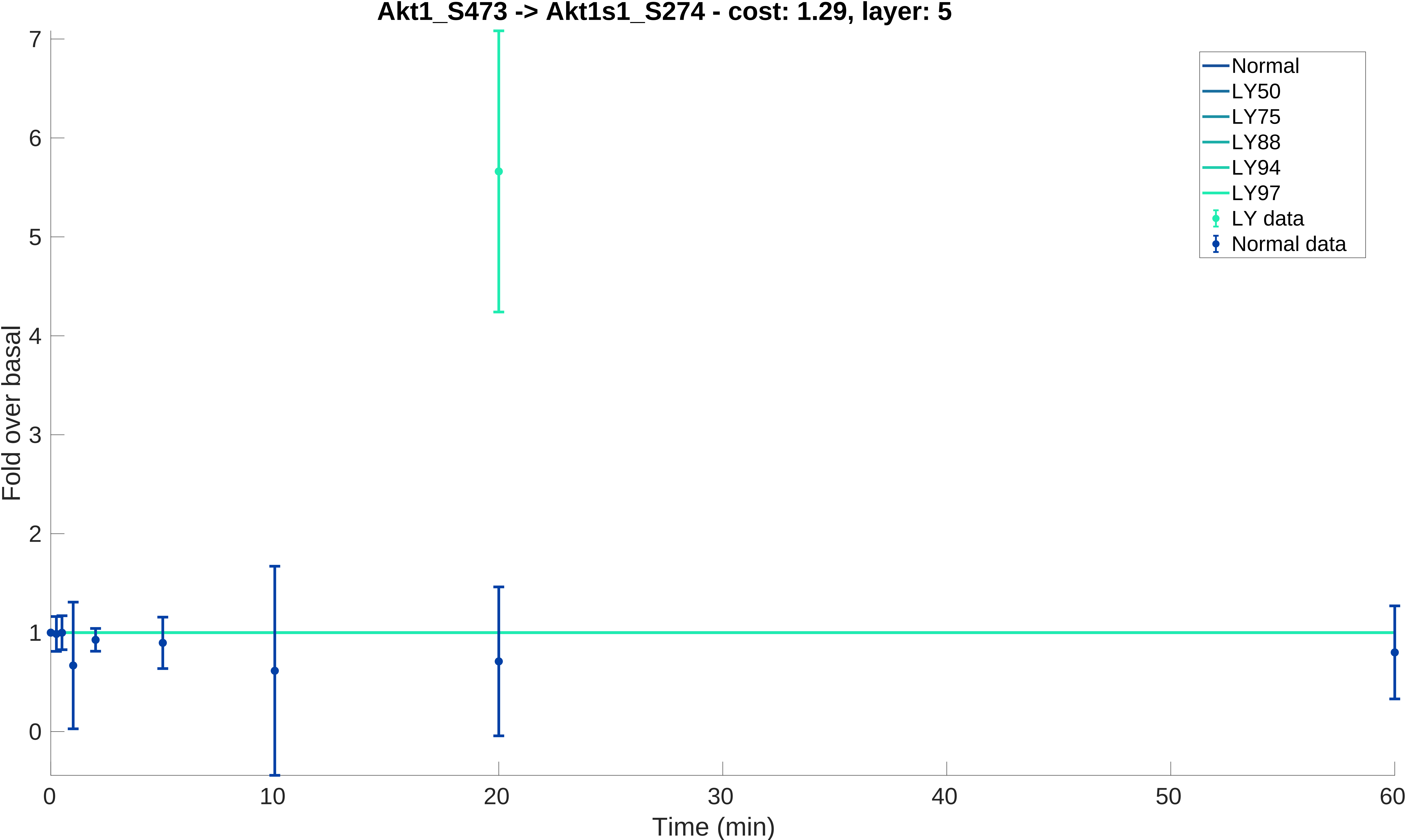

Akt1_S473 -> Akt1s1_S274 - cost: 1.29, layer: 5
Fold over basal
Time (min)
0
1
2
3
4
5
6
7
0
10
20
30
40
50
60
Normal
LY50
LY75
LY88
LY94
LY97
LY data
Normal data

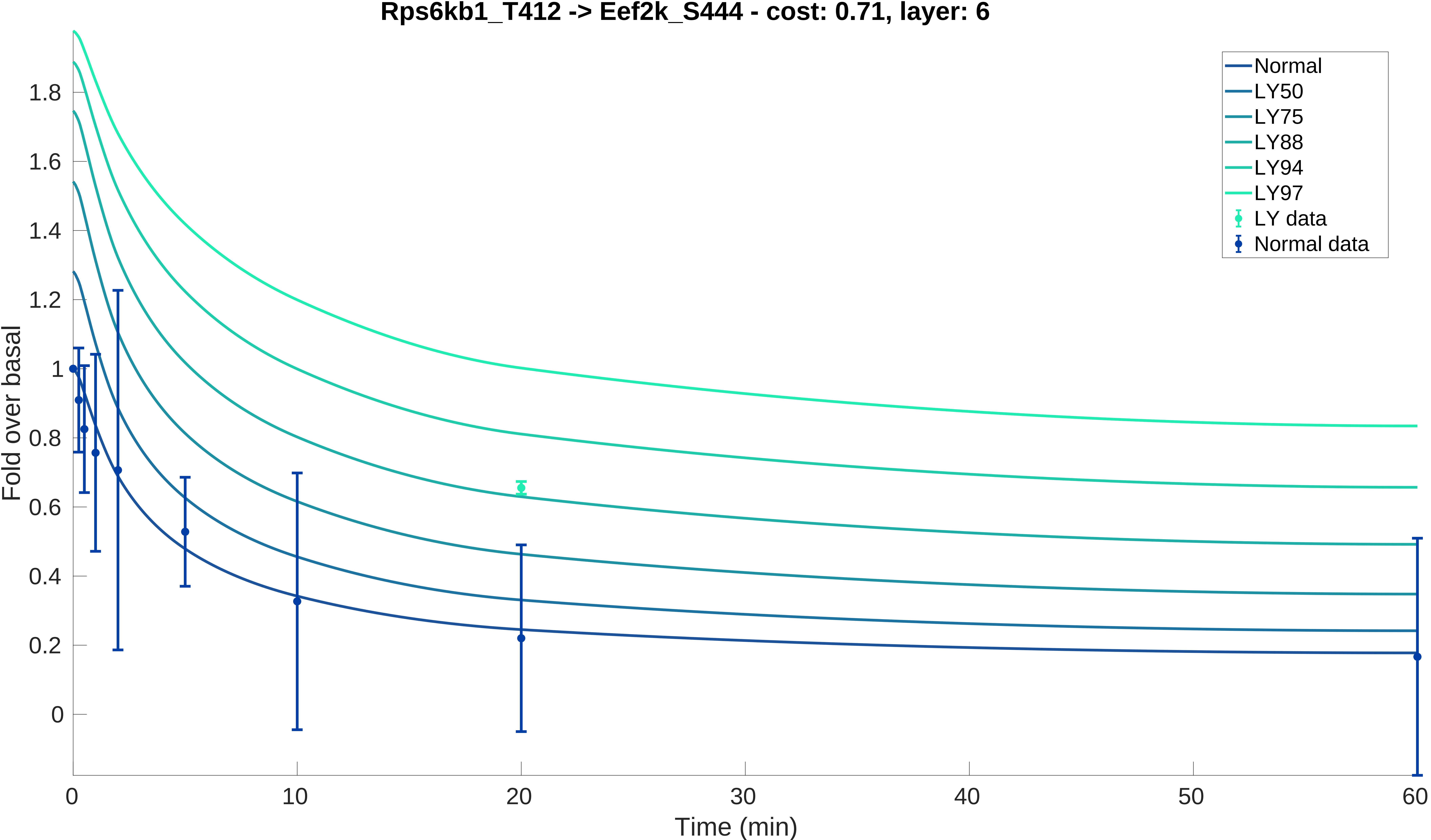
Rps6kb1_T412 -> Eef2k_S444 - cost: 0.71, layer: 6
Fold over basal
Time (min)
1.8
1.6
1.4
1.2
1
0.8
0.6
0.4
0.2
0
0
10
20
30
40
50
60
Normal
LY50
LY75
LY88
LY94
LY97
LY data
Normal data

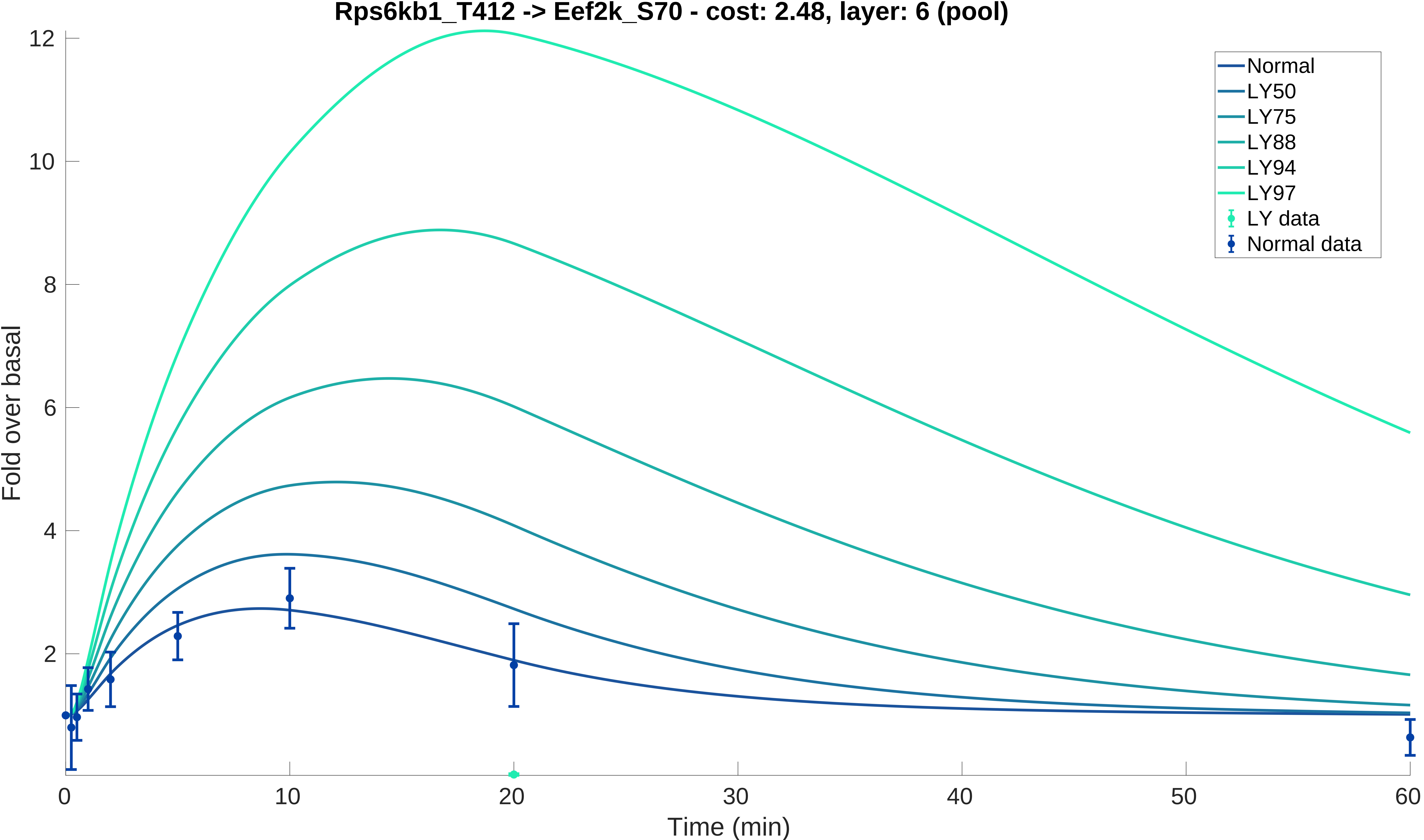
Rps6kb1_T412 -> Eef2k_S70 - cost: 2.48, layer: 6 (pool)
Fold over basal
Time (min)
0
10
20
30
40
50
60
2
4
6
8
10
12
Normal
LY50
LY75
LY88
LY94
LY97
LY data
Normal data

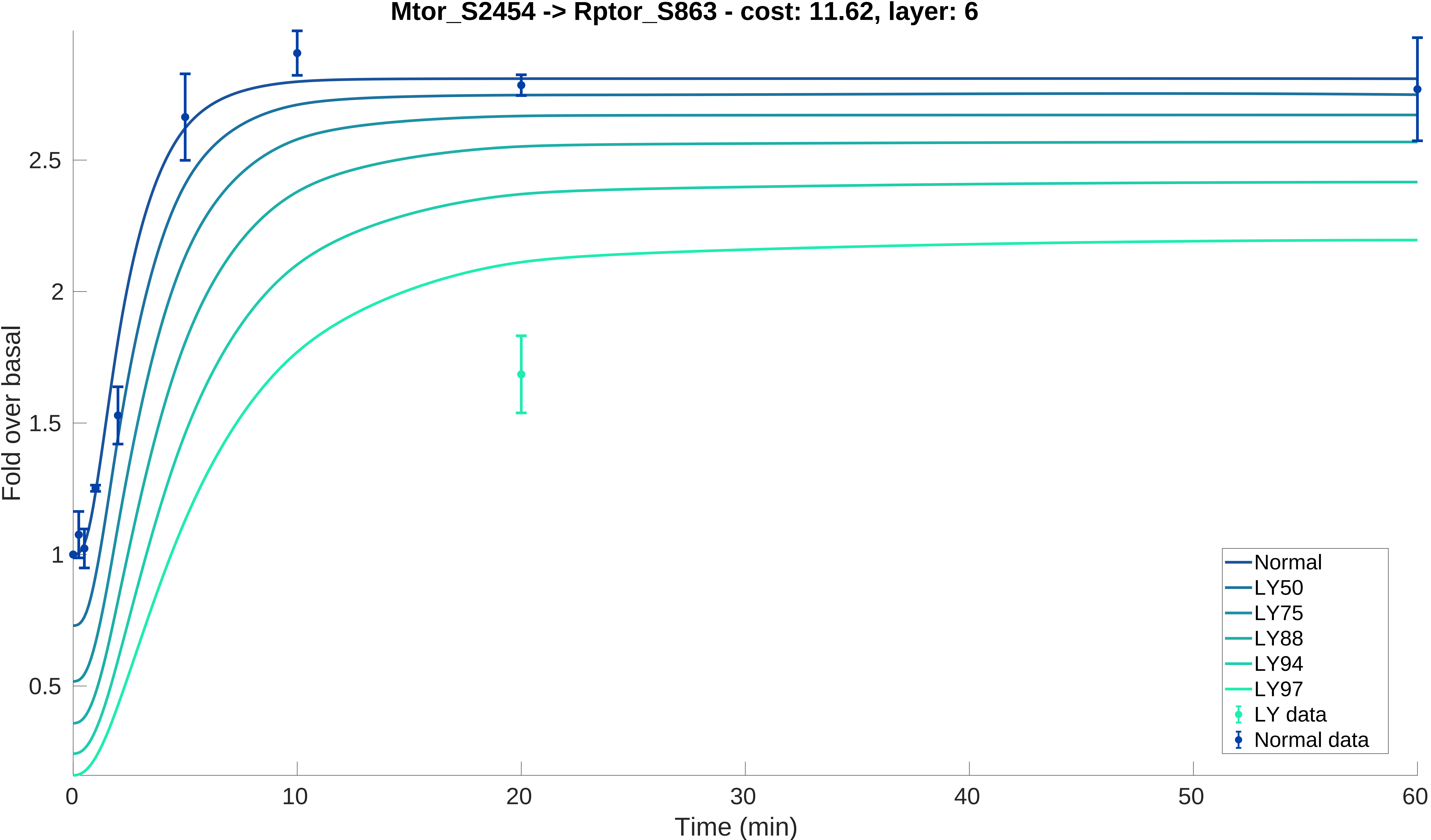

Mtor_S2454 -> Rptor_S863 - cost: 11.62, layer: 6
Fold over basal
Time (min)
2.5
2
1.5
1
0.5
0
10
20
30
40
50
60
Normal
LY50
LY75
LY88
LY94
LY97
LY data
Normal data

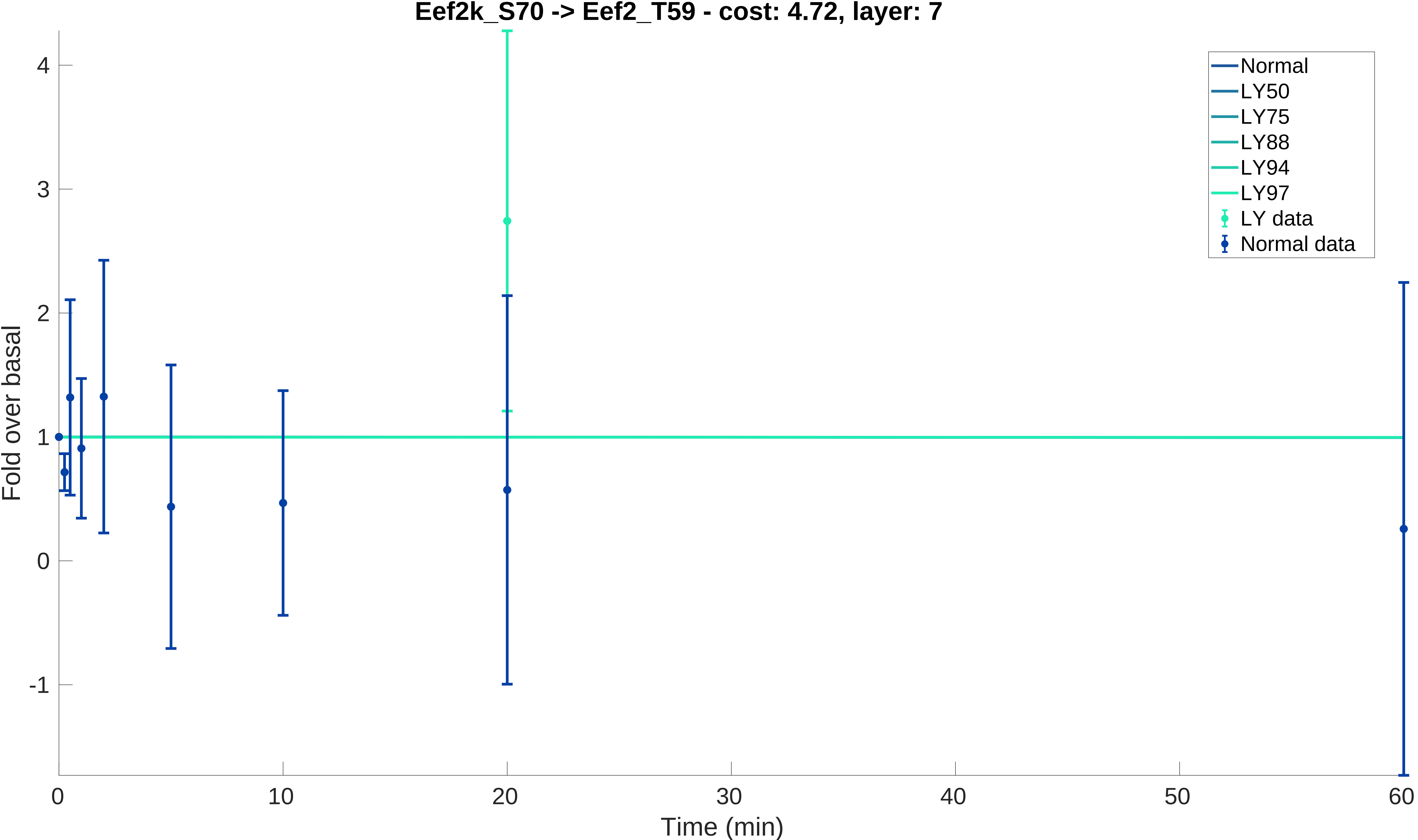

Eef2k_S70 -> Eef2_T59 - cost: 4.72, layer: 7
Fold over basal
Time (min)
4
3
2
1
0
-1
0
10
20
30
40
50
60
Normal
LY50
LY75
LY88
LY94
LY97
LY data
Normal data

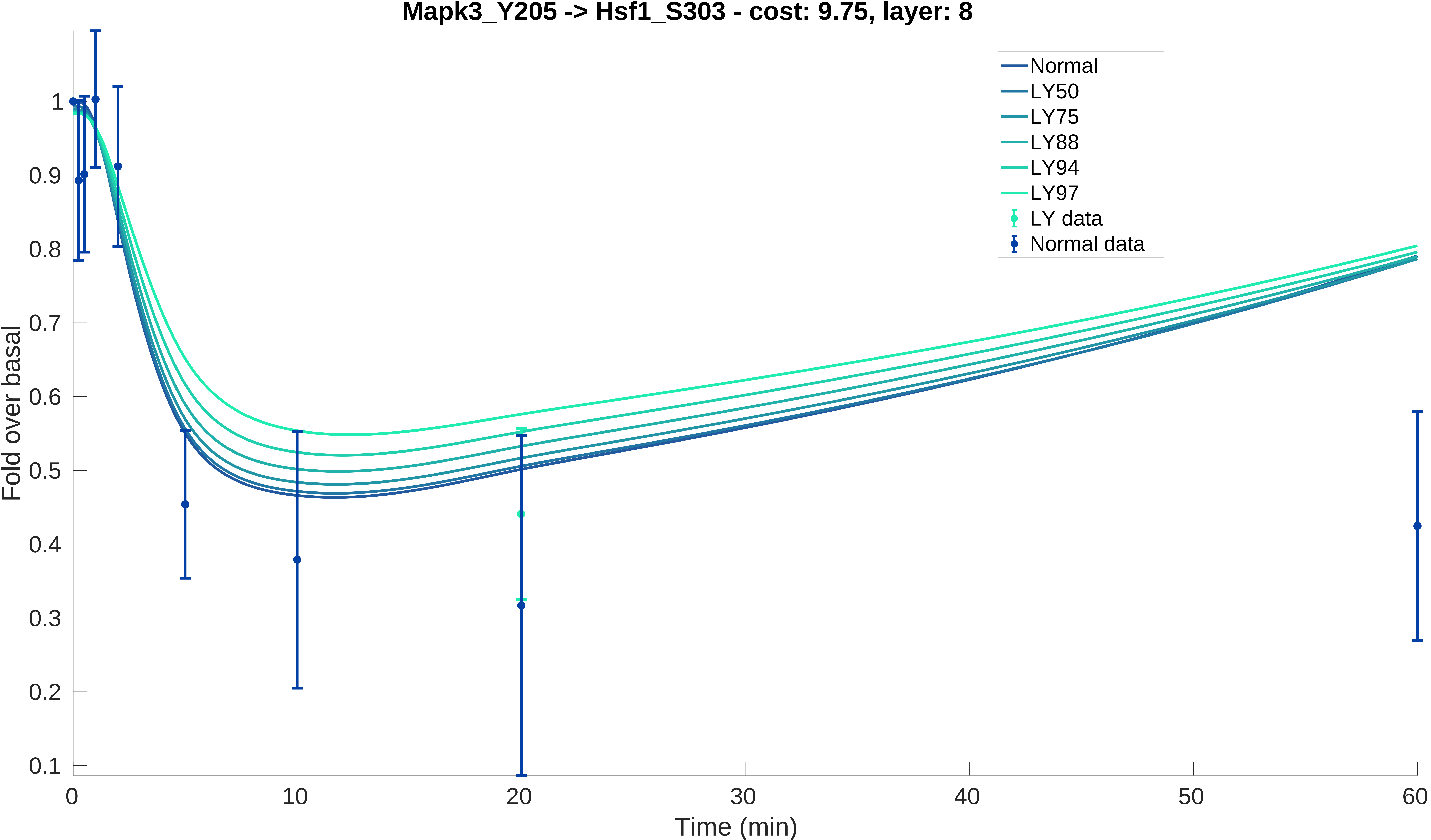
Mapk3_Y205 -> Hsf1_S303 - cost: 9.75, layer: 8
Normal
LY50
LY75
LY88
LY94
LY97
LY data
Normal data
Fold over basal
Time (min)
0.1
0.2
0.3
0.4
0.5
0.6
0.7
0.8
0.9
1
0
10
20
30
40
50
60

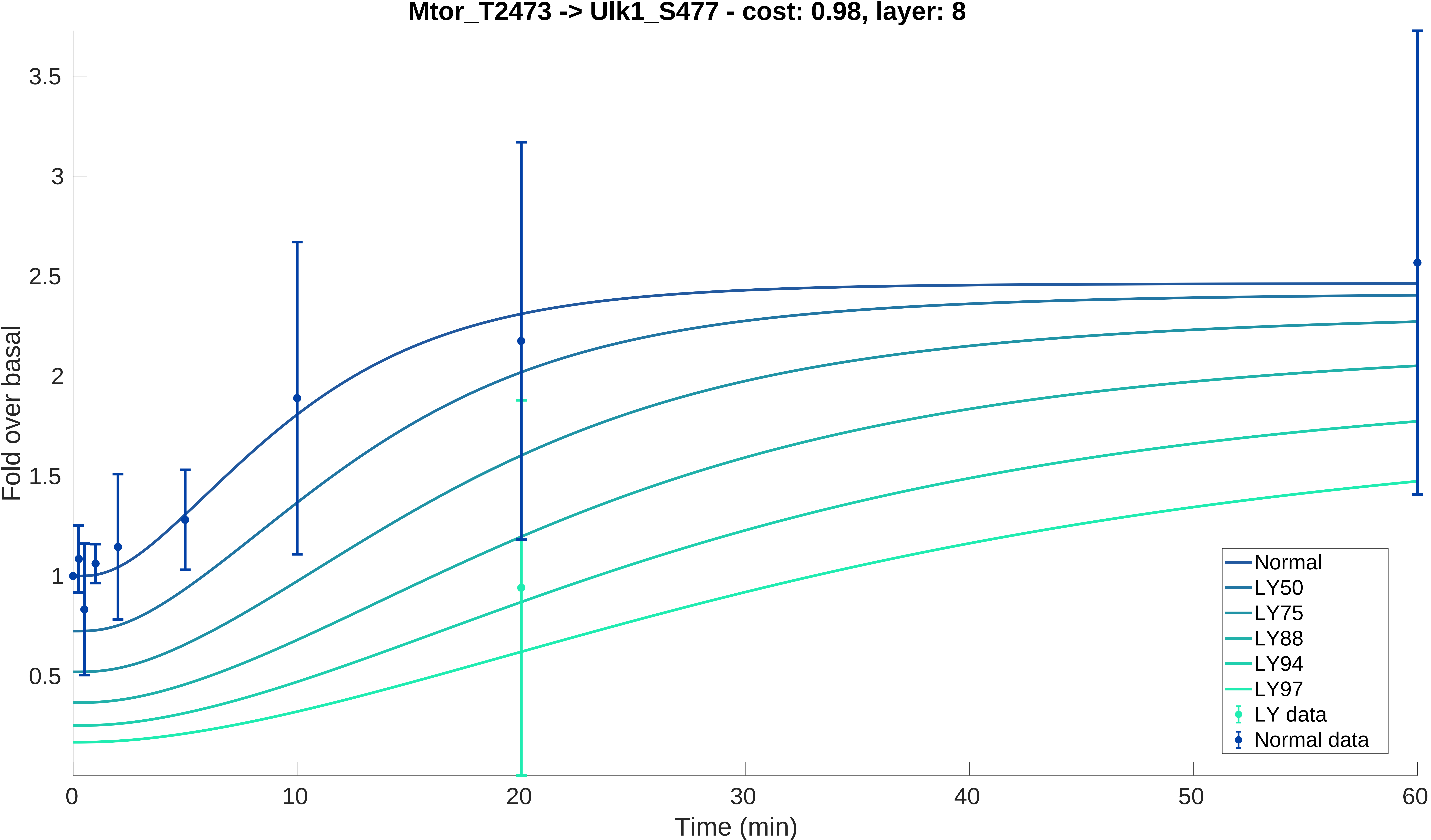

Mtor_T2473 -> Ulk1_S477 - cost: 0.98, layer: 8
Fold over basal
3.5
3
2.5
2
1.5
1
0.5
0
10
20
30
40
50
60
Time (min)
Normal
LY50
LY75
LY88
LY94
LY97
LY data
Normal data

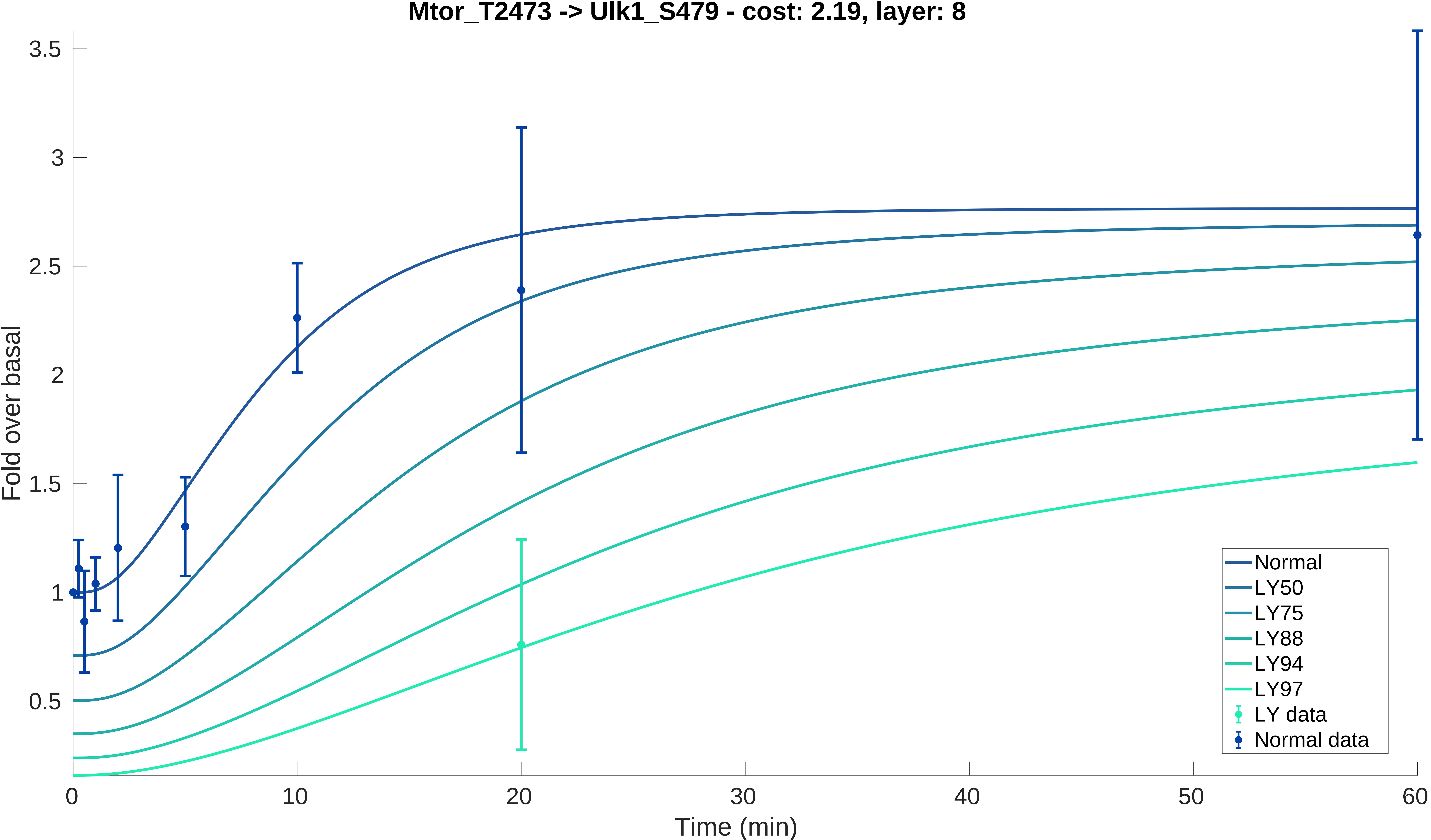

Mtor_T2473 -> Ulk1_S479 - cost: 2.19, layer: 8
Fold over basal
Time (min)
0.5
1
1.5
2
2.5
3
3.5
0
10
20
30
40
50
60
Normal
LY50
LY75
LY88
LY94
LY97
LY data
Normal data

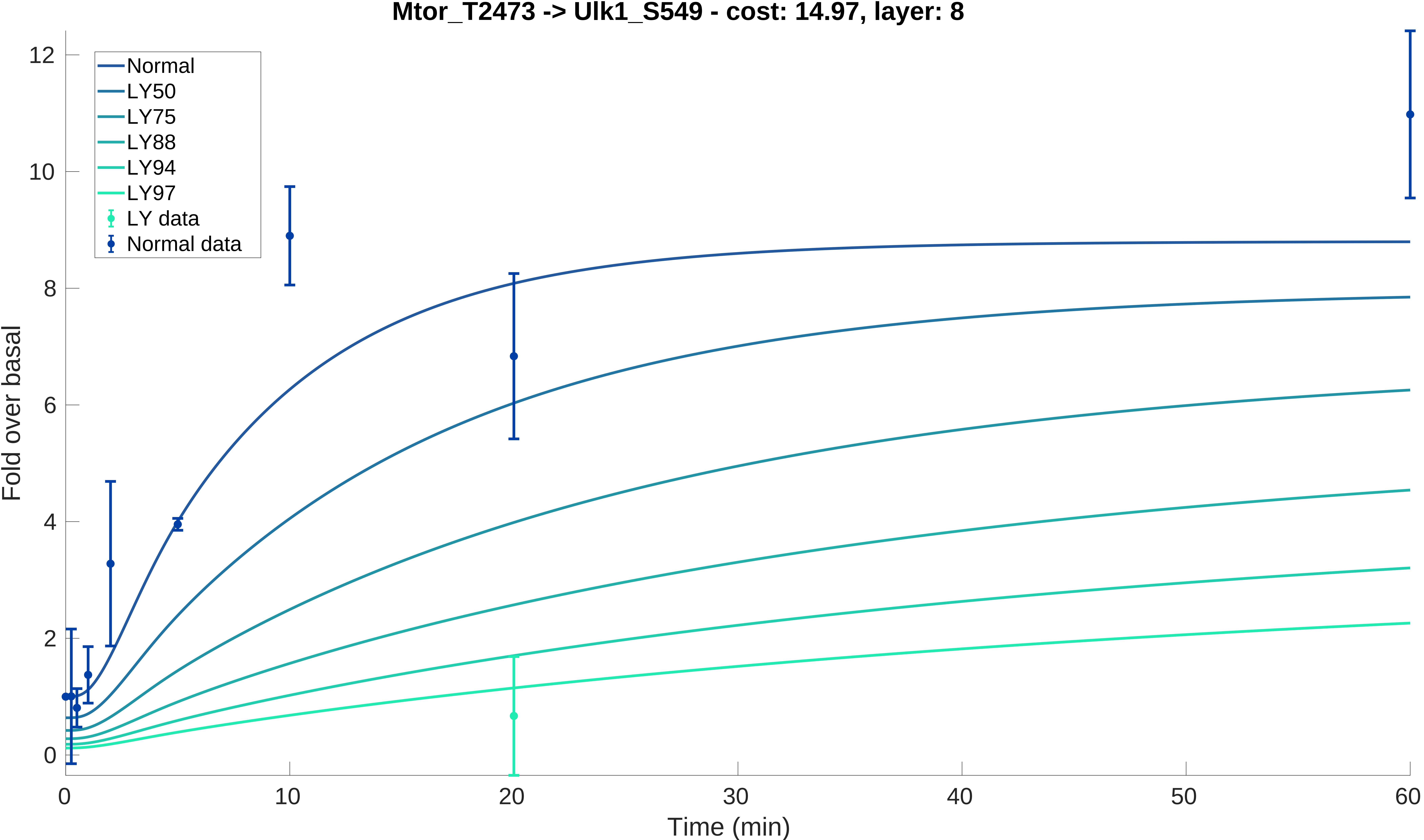

Mtor_T2473 -> Ulk1_S549 - cost: 14.97, layer: 8
Normal
LY50
LY75
LY88
LY94
LY97
LY data
Normal data
Fold over basal
Time (min)
0
2
4
6
8
10
12
0
10
20
30
40
50
60

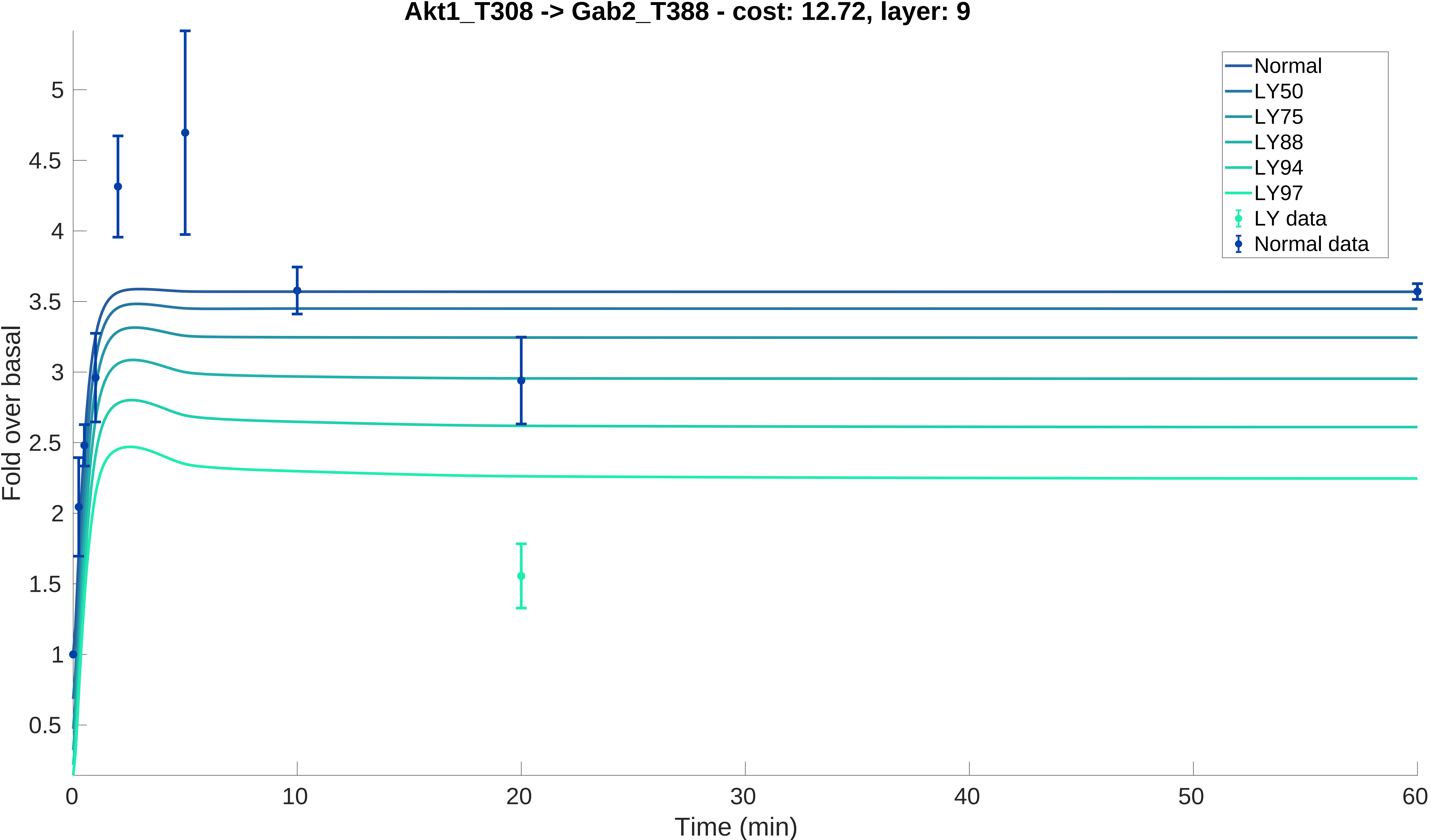

Akt1_T308 -> Gab2_T388 - cost: 12.72, layer: 9
Normal
LY50
LY75
LY88
LY94
LY97
LY data
Normal data
Fold over basal
5
4.5
4
3.5
3
2.5
2
1.5
1
0.5
0
10
20
30
40
50
60
Time (min)

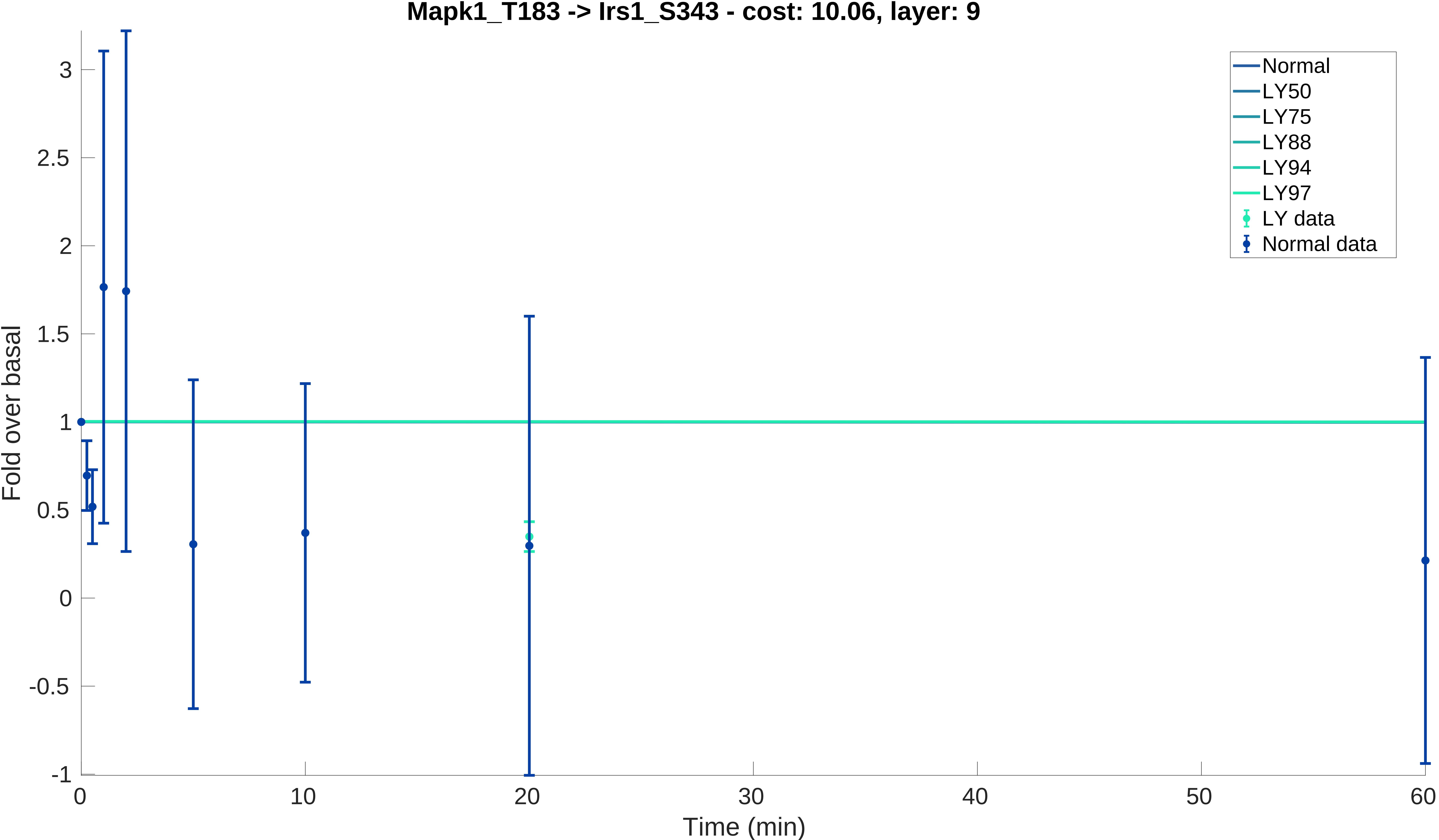

Mapk1_T183 -> Irs1_S343 - cost: 10.06, layer: 9
Normal
LY50
LY75
LY88
LY94
LY97
LY data
Normal data
Fold over basal
Time (min)
3
2.5
2
1.5
1
0.5
0
-0.5
-1
0
10
20
30
40
50
60

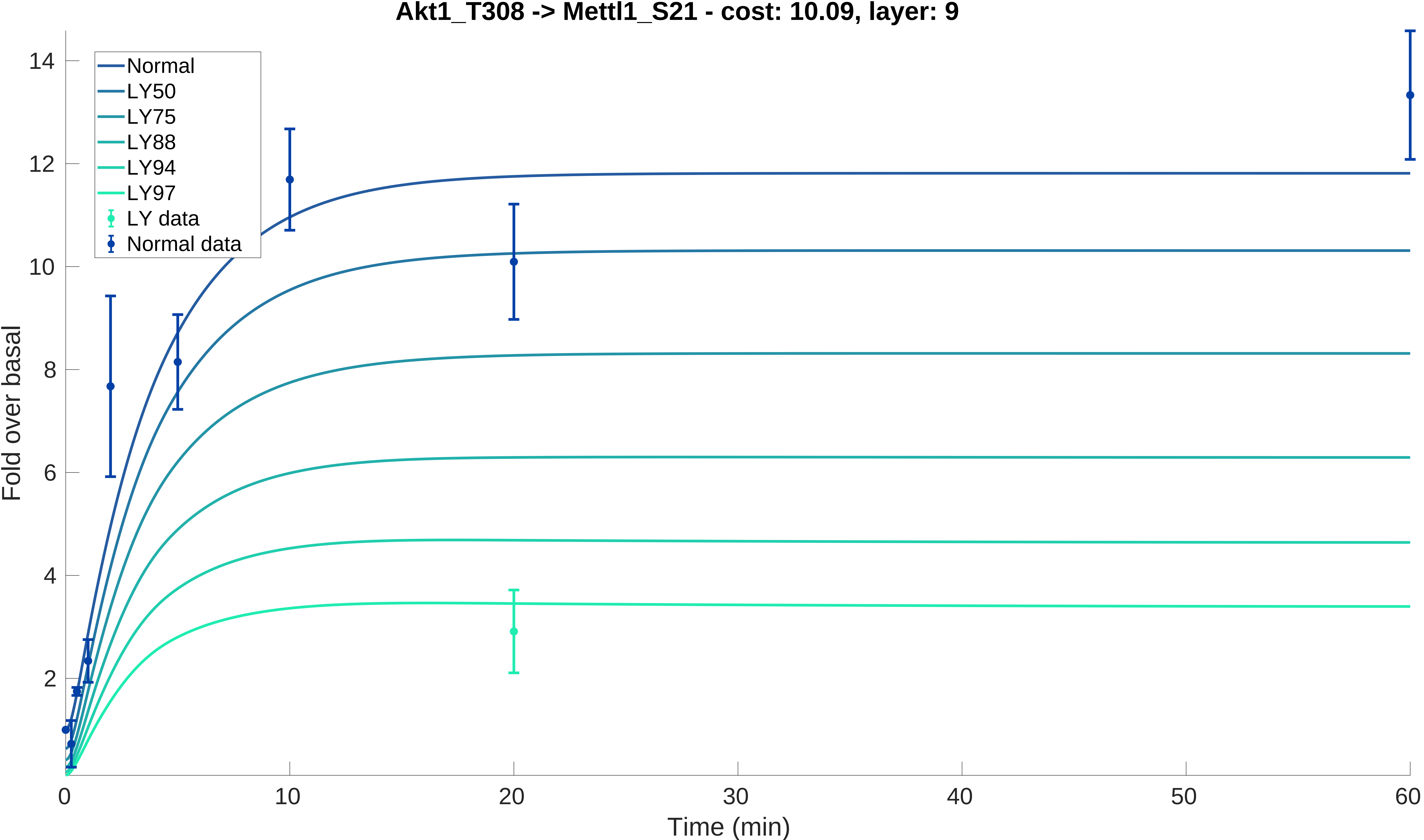

Akt1_T308 -> Mettl1_S21 - cost: 10.09, layer: 9
Normal
LY50
LY75
LY88
LY94
LY97
LY data
Normal data
Fold over basal
Time (min)
0
10
20
30
40
50
60
2
4
6
8
10
12
14

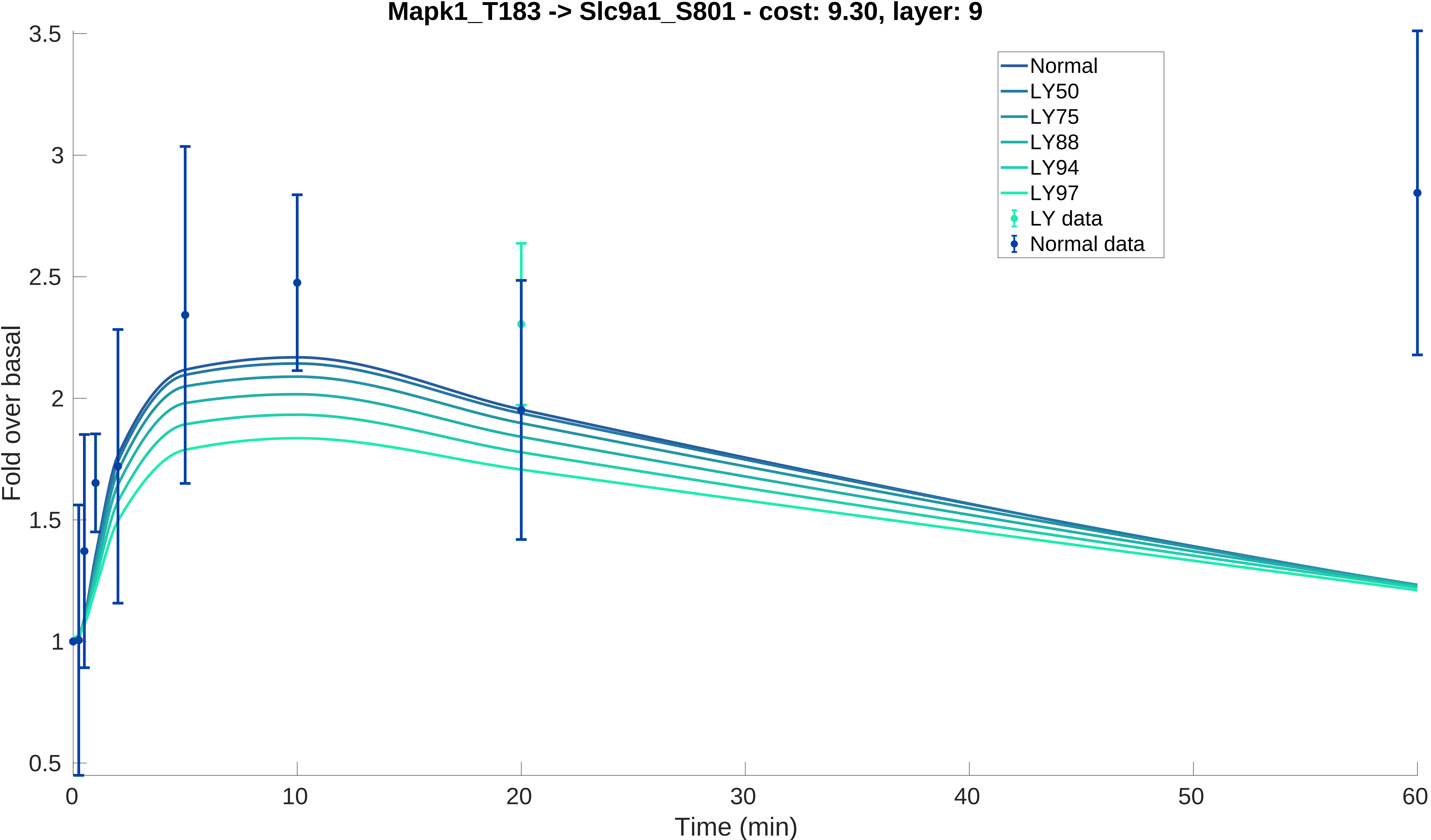

Mapk1_T183 -> Slc9a1_S801 - cost: 9.30, layer: 9
Fold over basal
Time (min)
3.5
3
2.5
2
1.5
1
0.5
0
10
20
30
40
50
60
Normal
LY50
LY75
LY88
LY94
LY97
LY data
Normal data

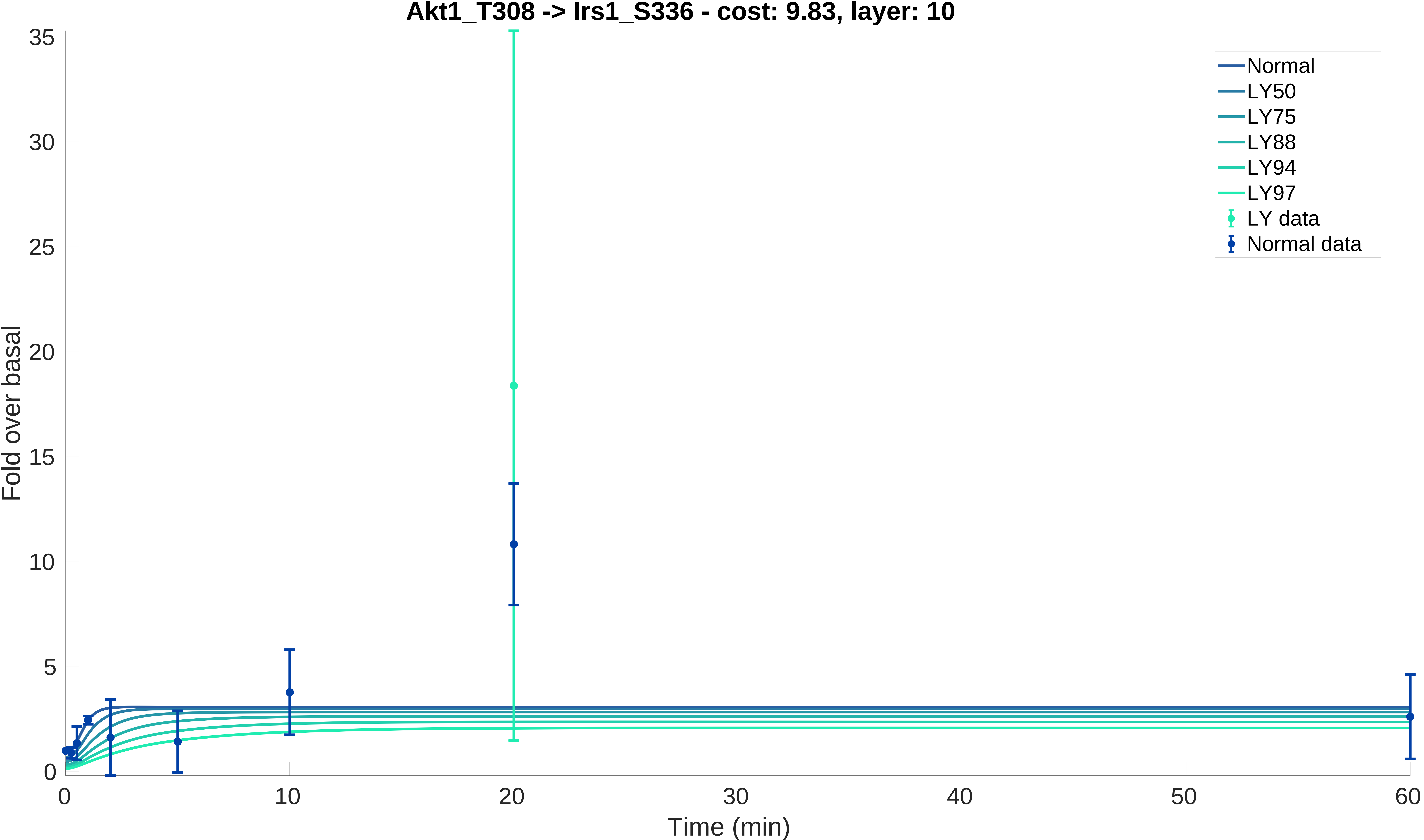

Akt1_T308 -> Irs1_S336 - cost: 9.83, layer: 10
Fold over basal
Time (min)
0
5
10
15
20
25
30
35
0
10
20
30
40
50
60
Normal
LY50
LY75
LY88
LY94
LY97
LY data
Normal data

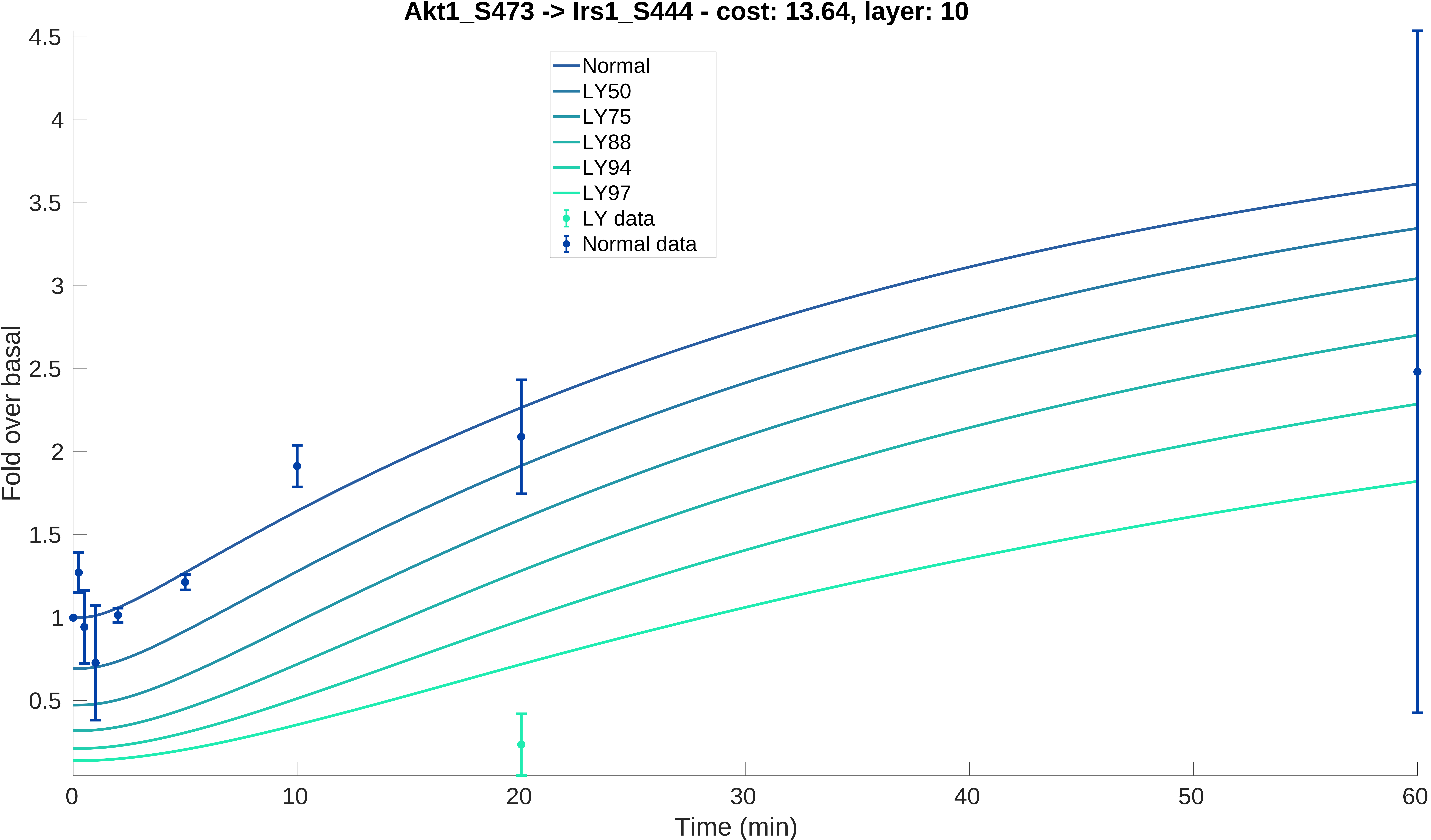
Akt1_S473 -> Irs1_S444 - cost: 13.64, layer: 10
Normal
LY50
LY75
LY88
LY94
LY97
LY data
Normal data
Fold over basal
Time (min)
4.5
4
3.5
3
2.5
2
1.5
1
0.5
0
10
20
30
40
50
60

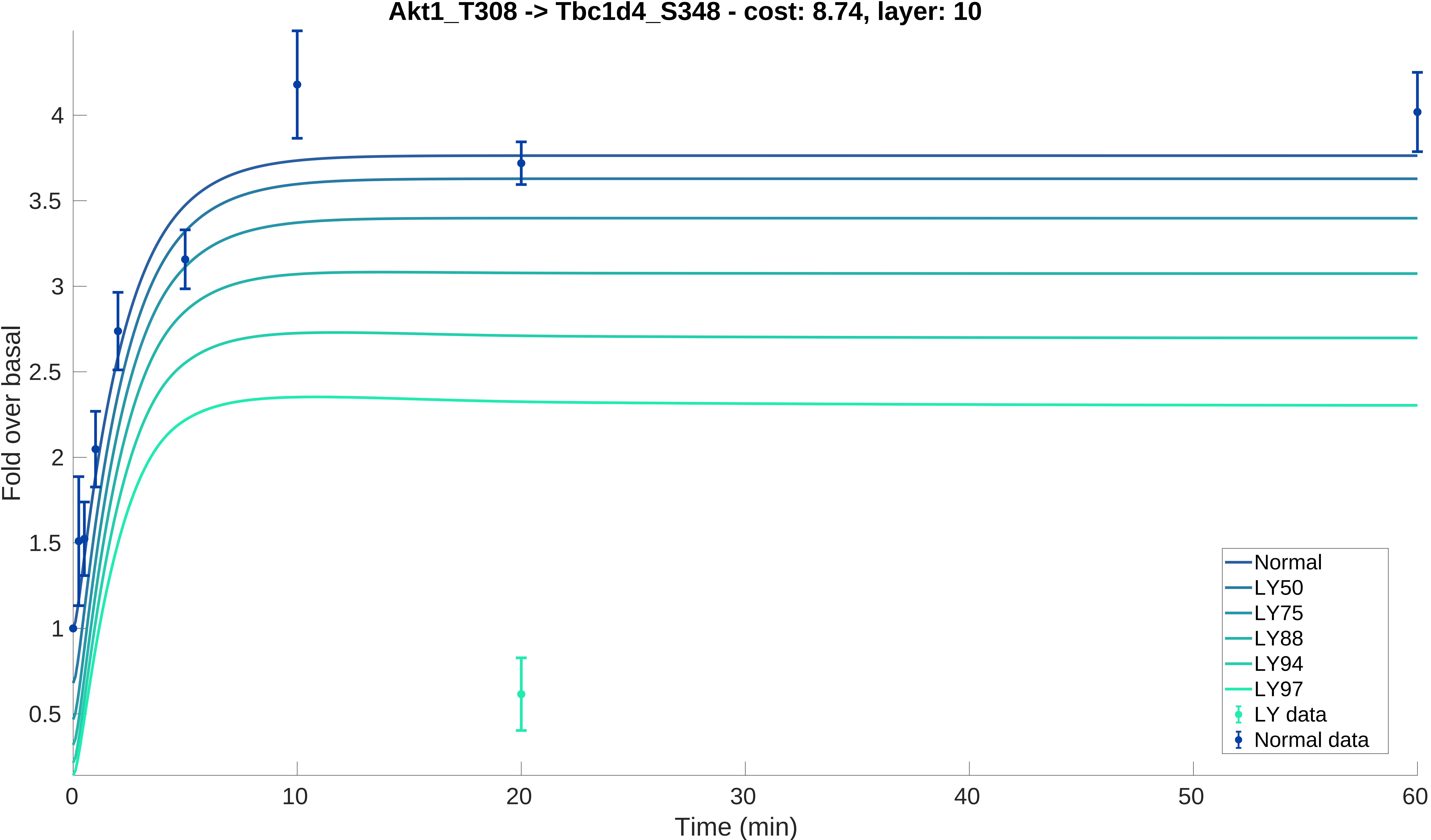

Akt1_T308 -> Tbc1d4_S348 - cost: 8.74, layer: 10
Fold over basal
Time (min)
0.5
1
1.5
2
2.5
3
3.5
4
0
10
20
30
40
50
60
Normal
LY50
LY75
LY88
LY94
LY97
LY data
Normal data

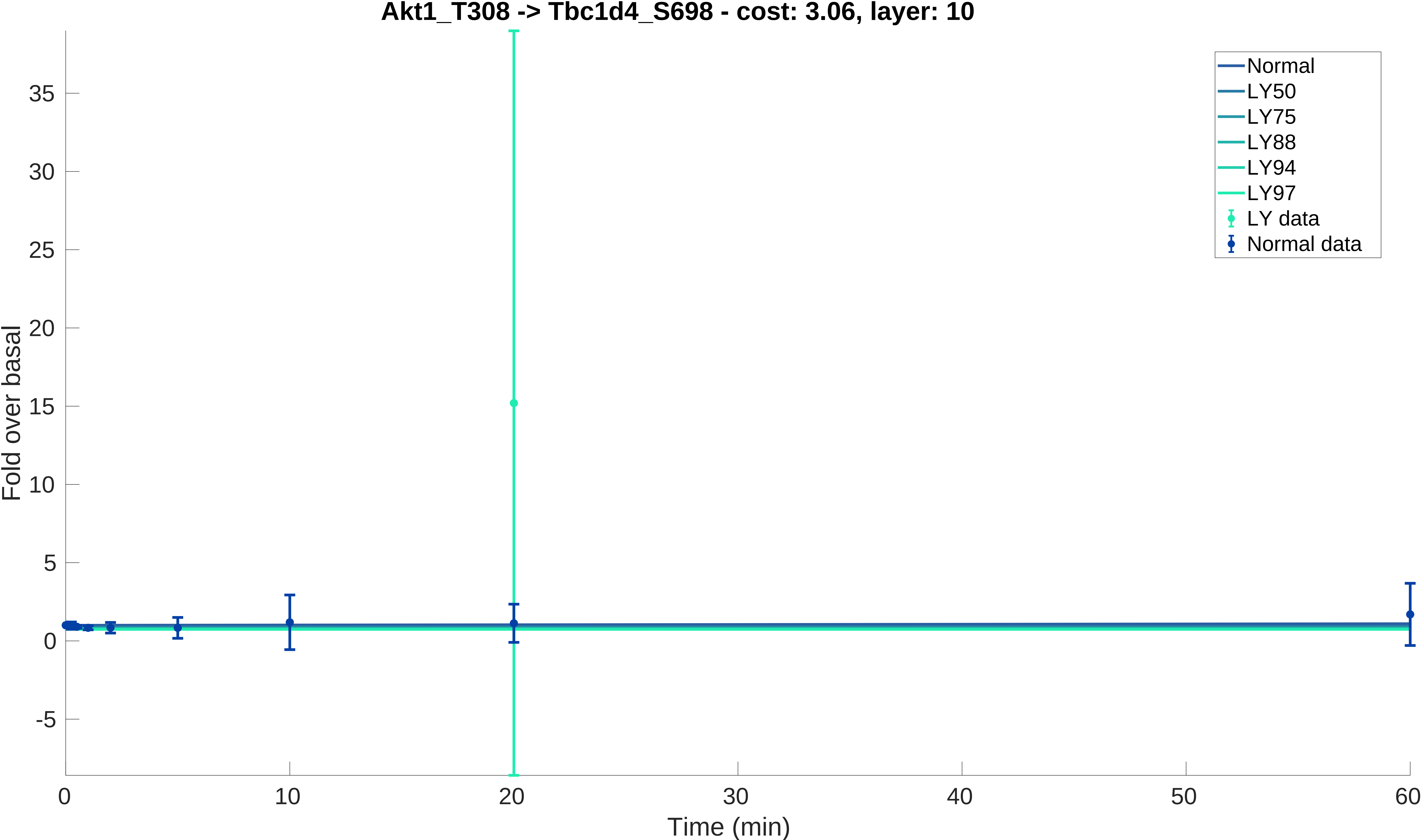
Akt1_T308 -> Tbc1d4_S698 - cost: 3.06, layer: 10
Fold over basal
Time (min)
35
30
25
20
15
10
5
0
-5
0
10
20
30
40
50
60
Normal
LY50
LY75
LY88
LY94
LY97
LY data
Normal data

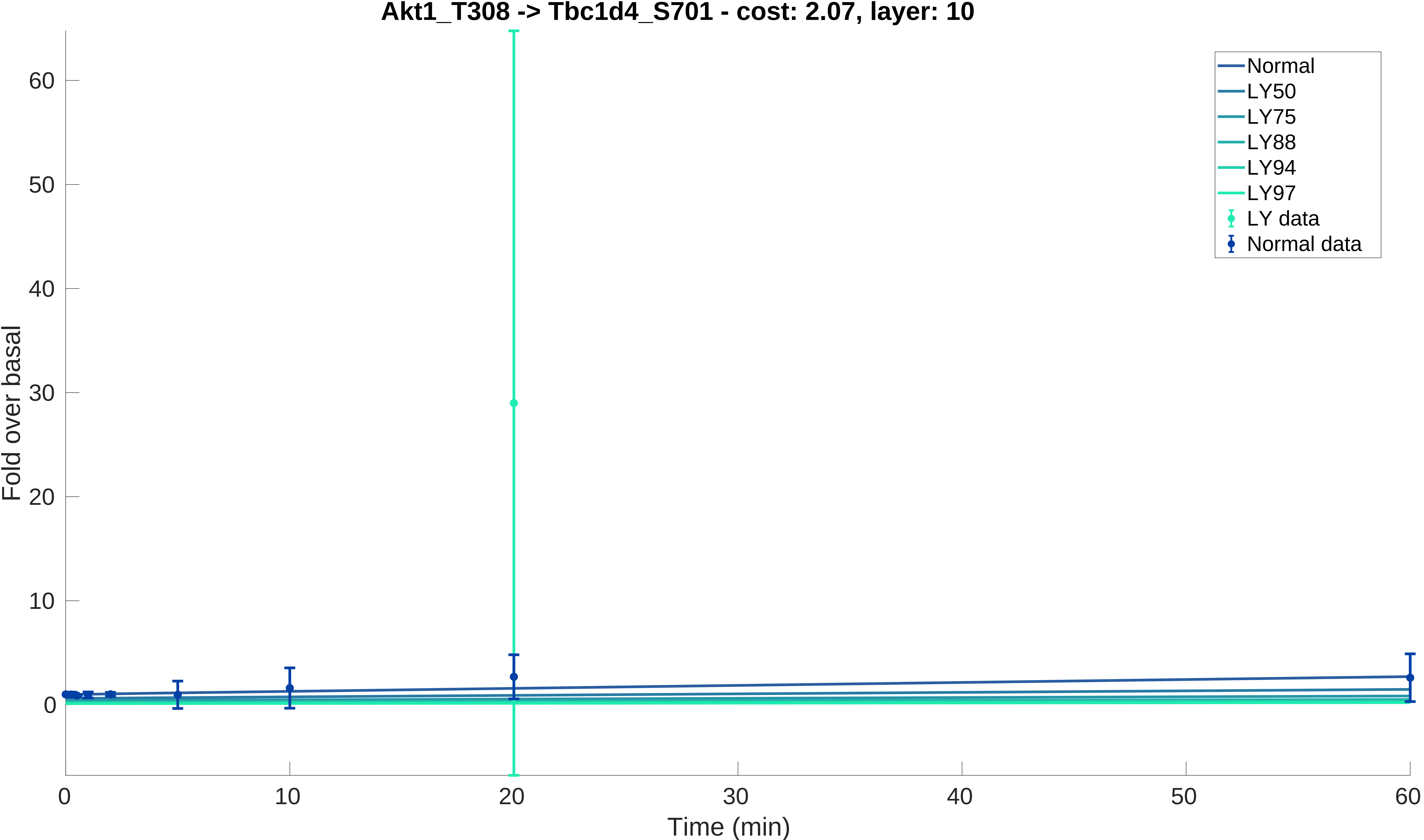

Akt1_T308 -> Tbc1d4_S701 - cost: 2.07, layer: 10
Fold over basal
60
50
40
30
20
10
0
0
10
20
30
40
50
60
Time (min)
Normal
LY50
LY75
LY88
LY94
LY97
LY data
Normal data

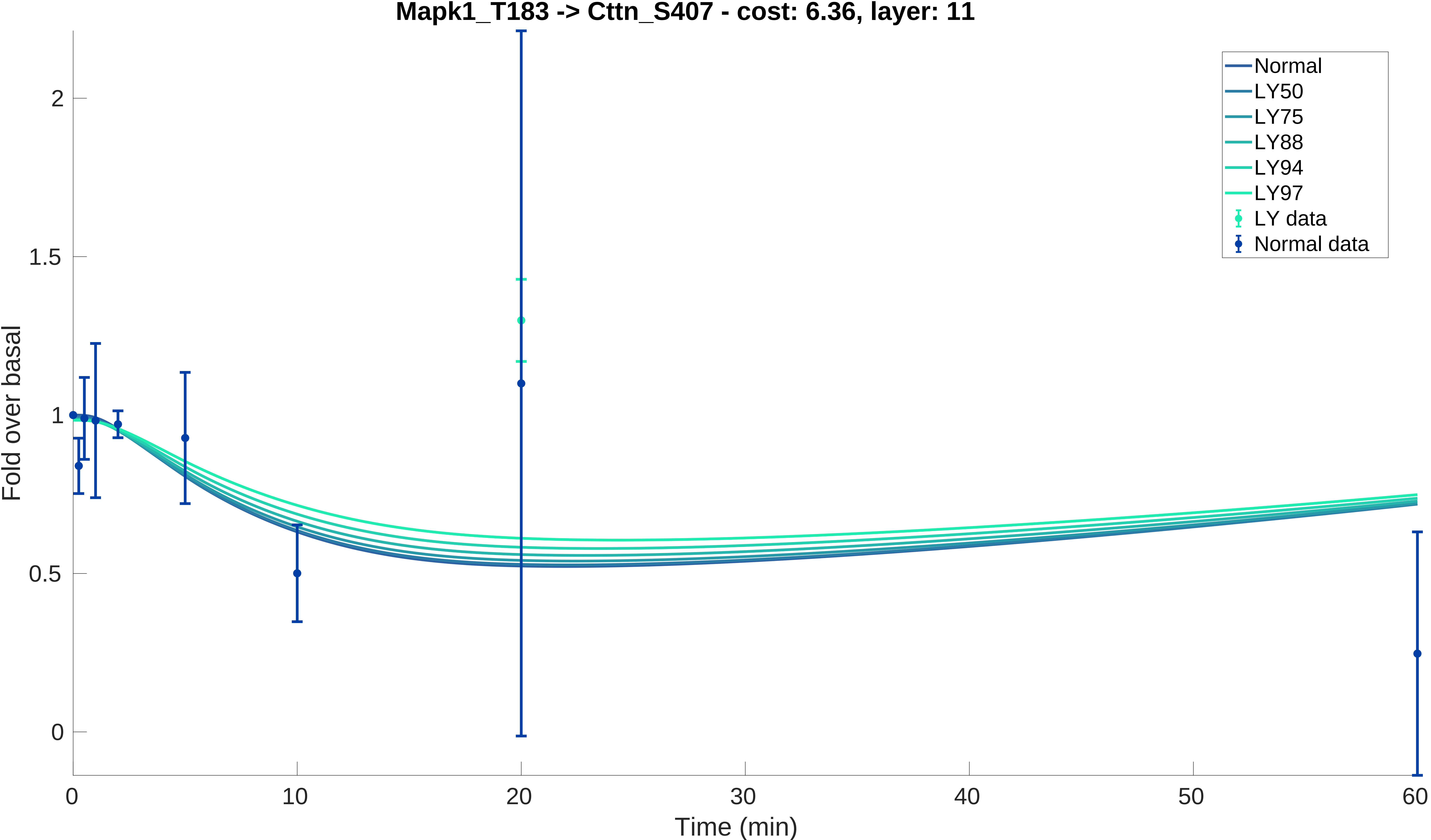
Mapk1_T183 -> Cttn_S407 - cost: 6.36, layer: 11
Normal
LY50
LY75
LY88
LY94
LY97
LY data
Normal data
Fold over basal
Time (min)
0
0.5
1
1.5
2
0
10
20
30
40
50
60

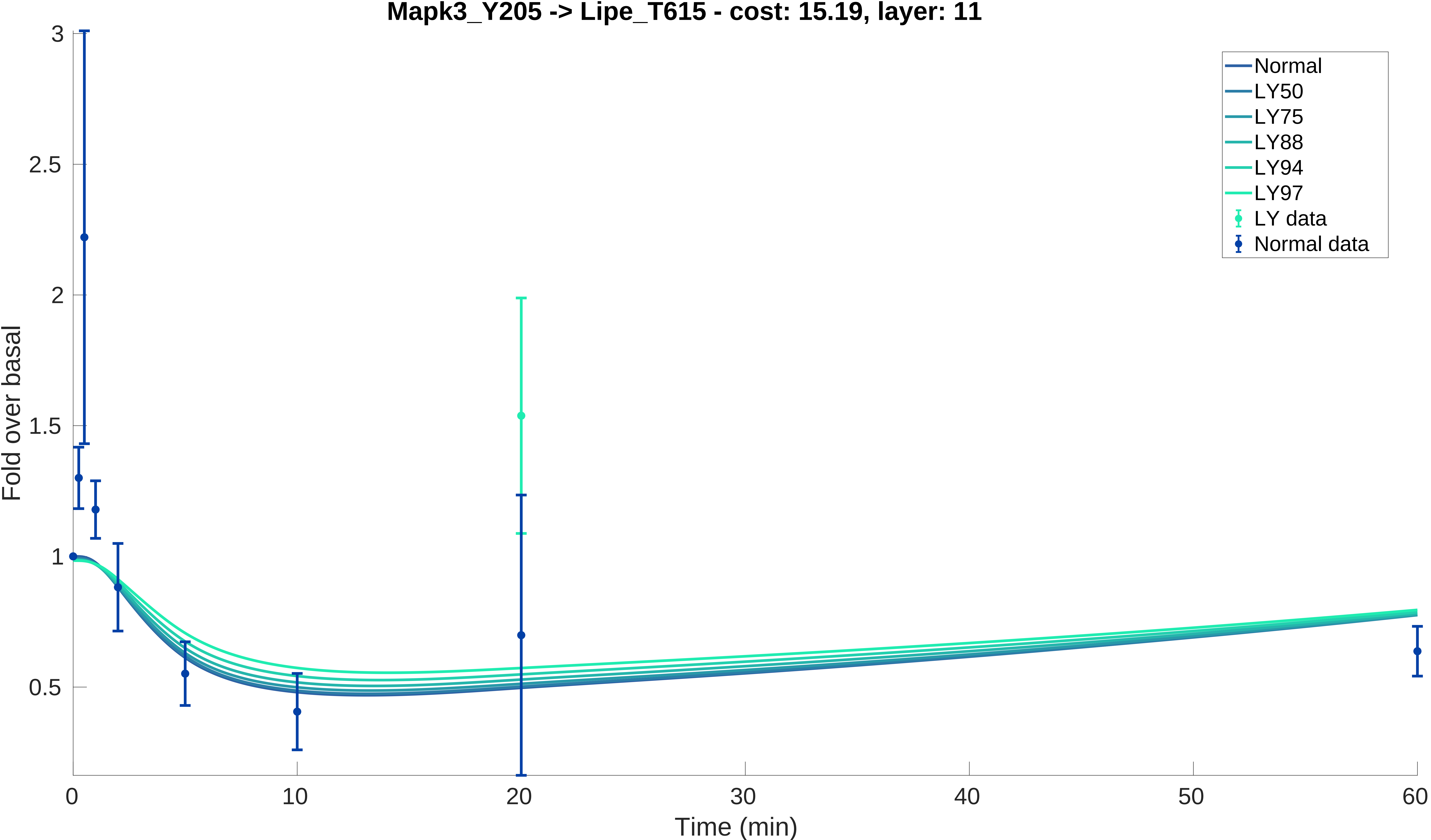

Mapk3_Y205 -> Lipe_T615 - cost: 15.19, layer: 11
Fold over basal
Time (min)
3
2.5
2
1.5
1
0.5
0
10
20
30
40
50
60
Normal
LY50
LY75
LY88
LY94
LY97
LY data
Normal data

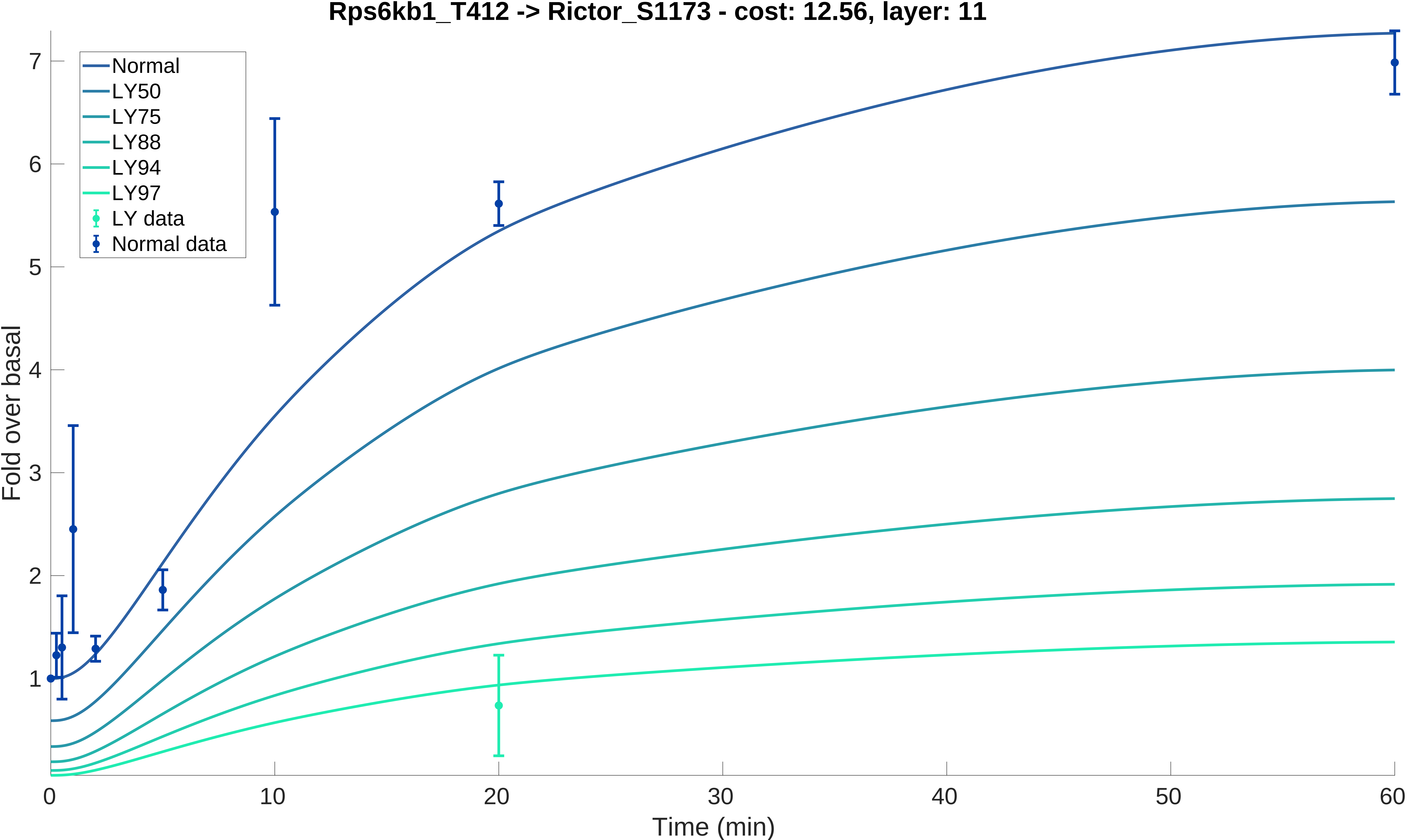

Rps6kb1_T412 -> Rictor_S1173 - cost: 12.56, layer: 11
Normal
LY50
LY75
LY88
LY94
LY97
LY data
Normal data
Fold over basal
Time (min)
0
10
20
30
40
50
60
1
2
3
4
5
6
7

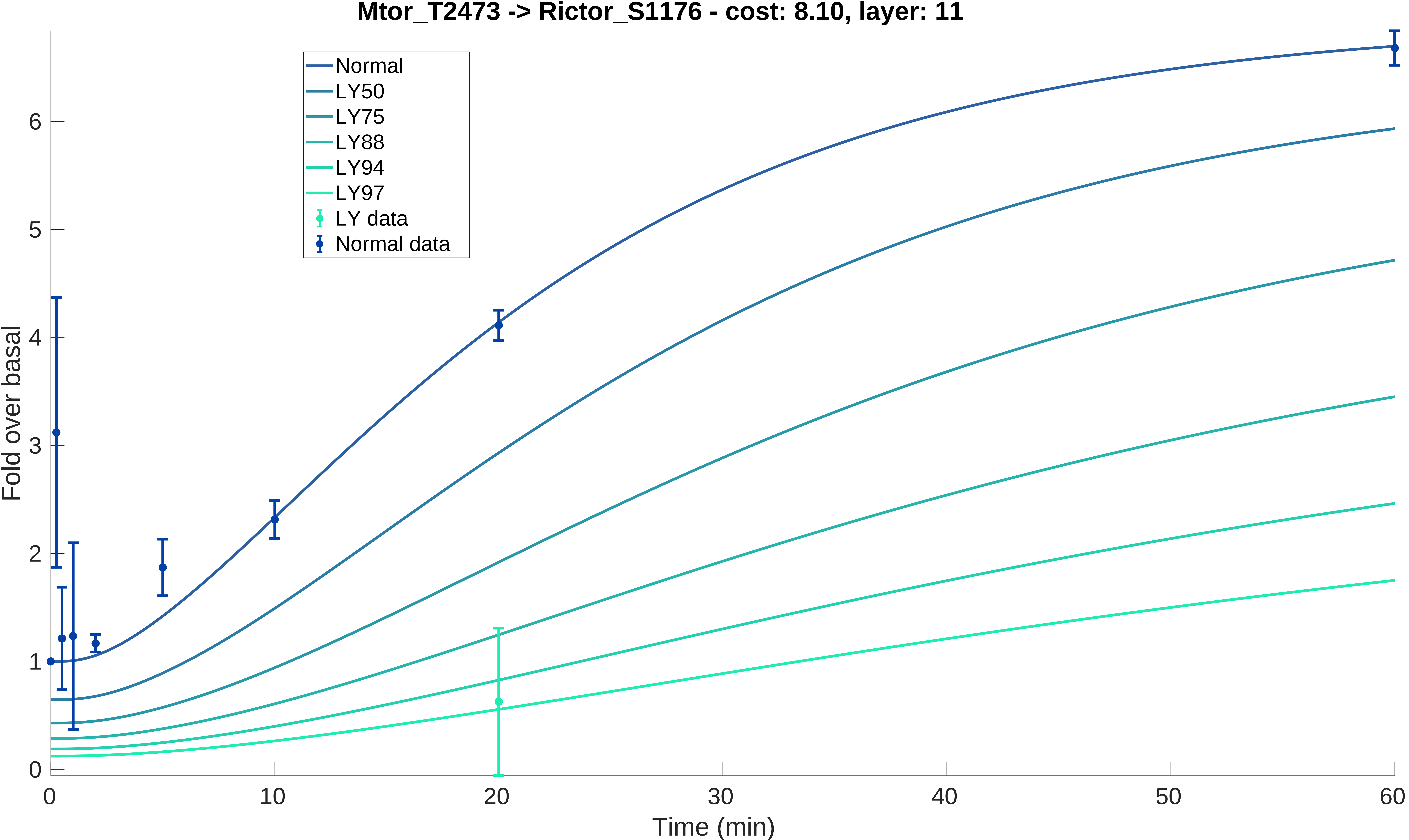

Mtor_T2473 -> Rictor_S1176 - cost: 8.10, layer: 11
Normal
LY50
LY75
LY88
LY94
LY97
LY data
Normal data
Fold over basal
0
1
2
3
4
5
6
0
10
20
30
40
50
60
Time (min)

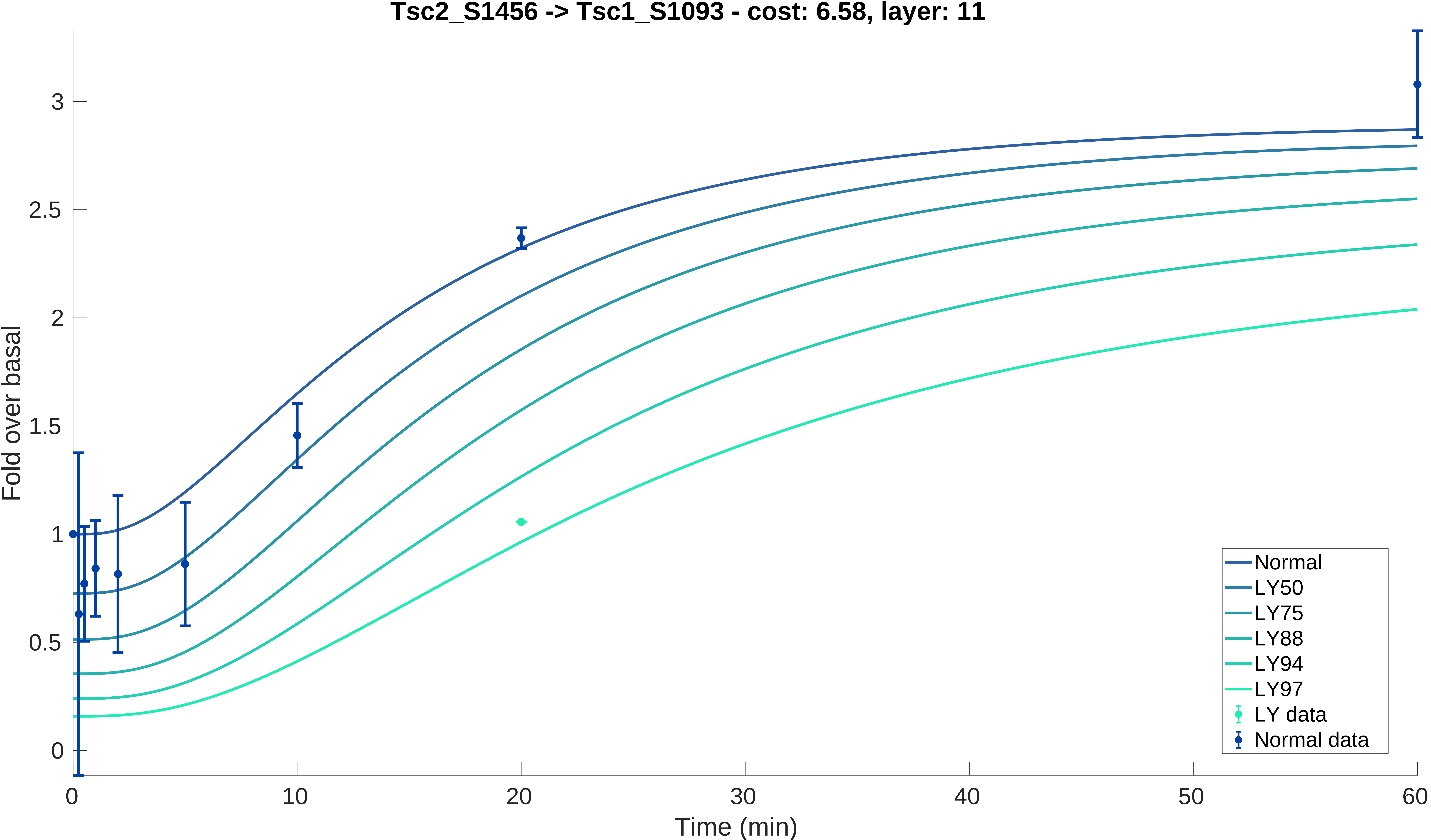

Tsc2_S1456 -> Tsc1_S1093 - cost: 6.58, layer: 11
Fold over basal
Time (min)
0
0.5
1
1.5
2
2.5
3
0
10
20
30
40
50
60
Normal
LY50
LY75
LY88
LY94
LY97
LY data
Normal data

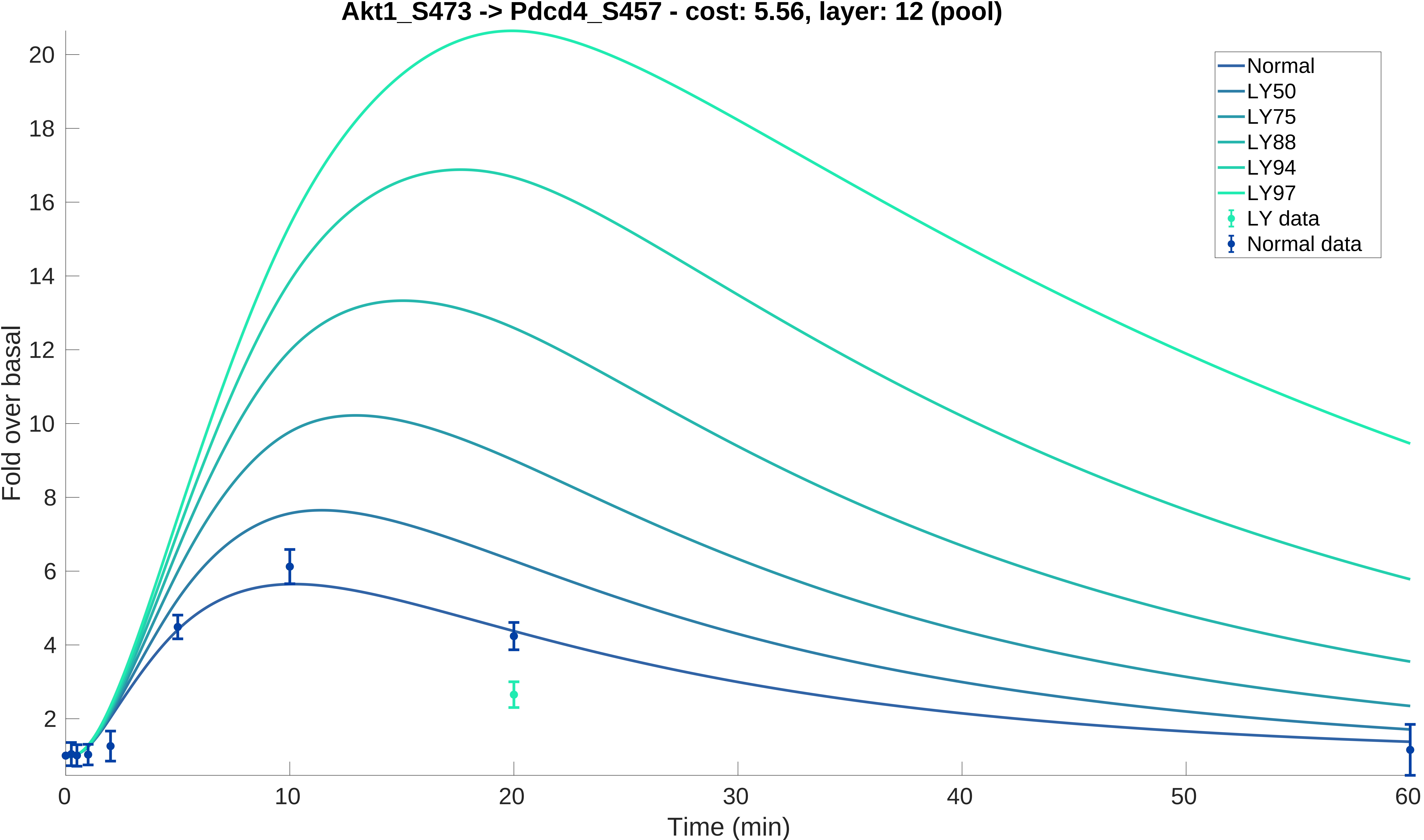

Akt1_S473 -> Pdcd4_S457 - cost: 5.56, layer: 12 (pool)
Fold over basal
Time (min)
0
10
20
30
40
50
60
2
4
6
8
10
12
14
16
18
20
Normal
LY50
LY75
LY88
LY94
LY97
LY data
Normal data

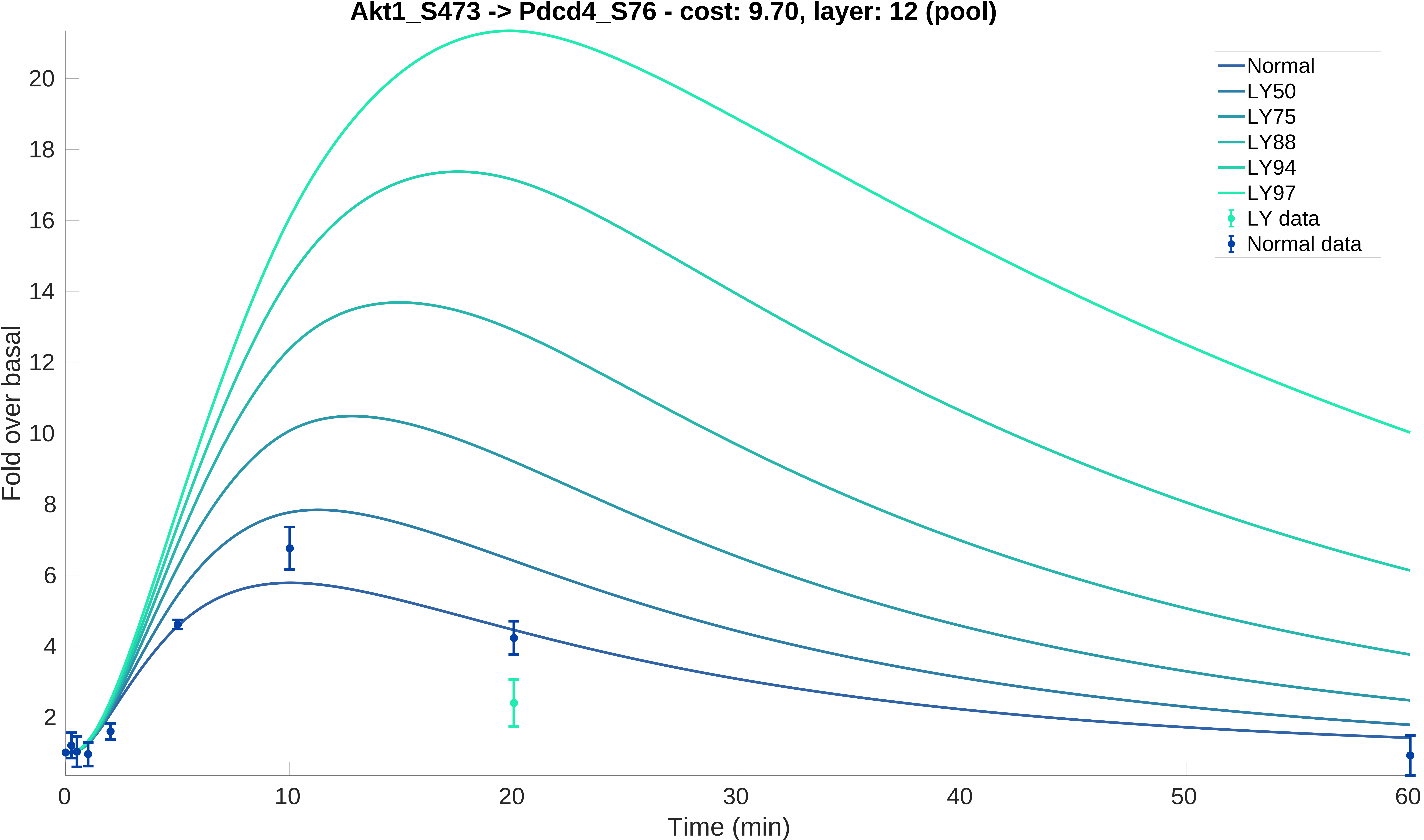
Akt1_S473 -> Pdcd4_S76 - cost: 9.70, layer: 12 (pool)
Normal
LY50
LY75
LY88
LY94
LY97
LY data
Normal data
Fold over basal
Time (min)
0
10
20
30
40
50
60
2
4
6
8
10
12
14
16
18
20

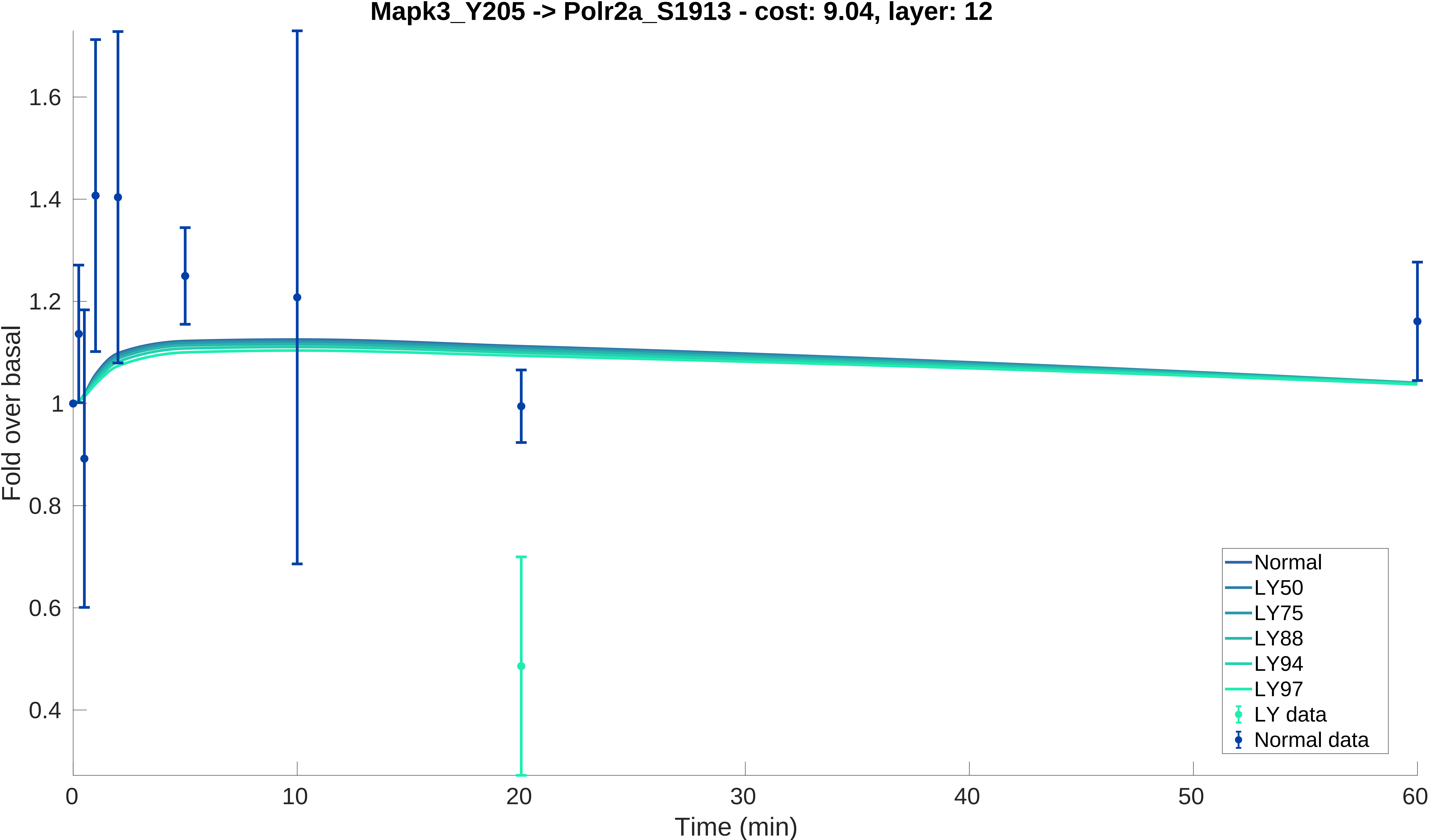

Mapk3_Y205 -> Polr2a_S1913 - cost: 9.04, layer: 12
Fold over basal
1.6
1.4
1.2
1
0.8
0.6
0.4
0
10
20
30
40
50
60
Time (min)
Normal
LY50
LY75
LY88
LY94
LY97
LY data
Normal data

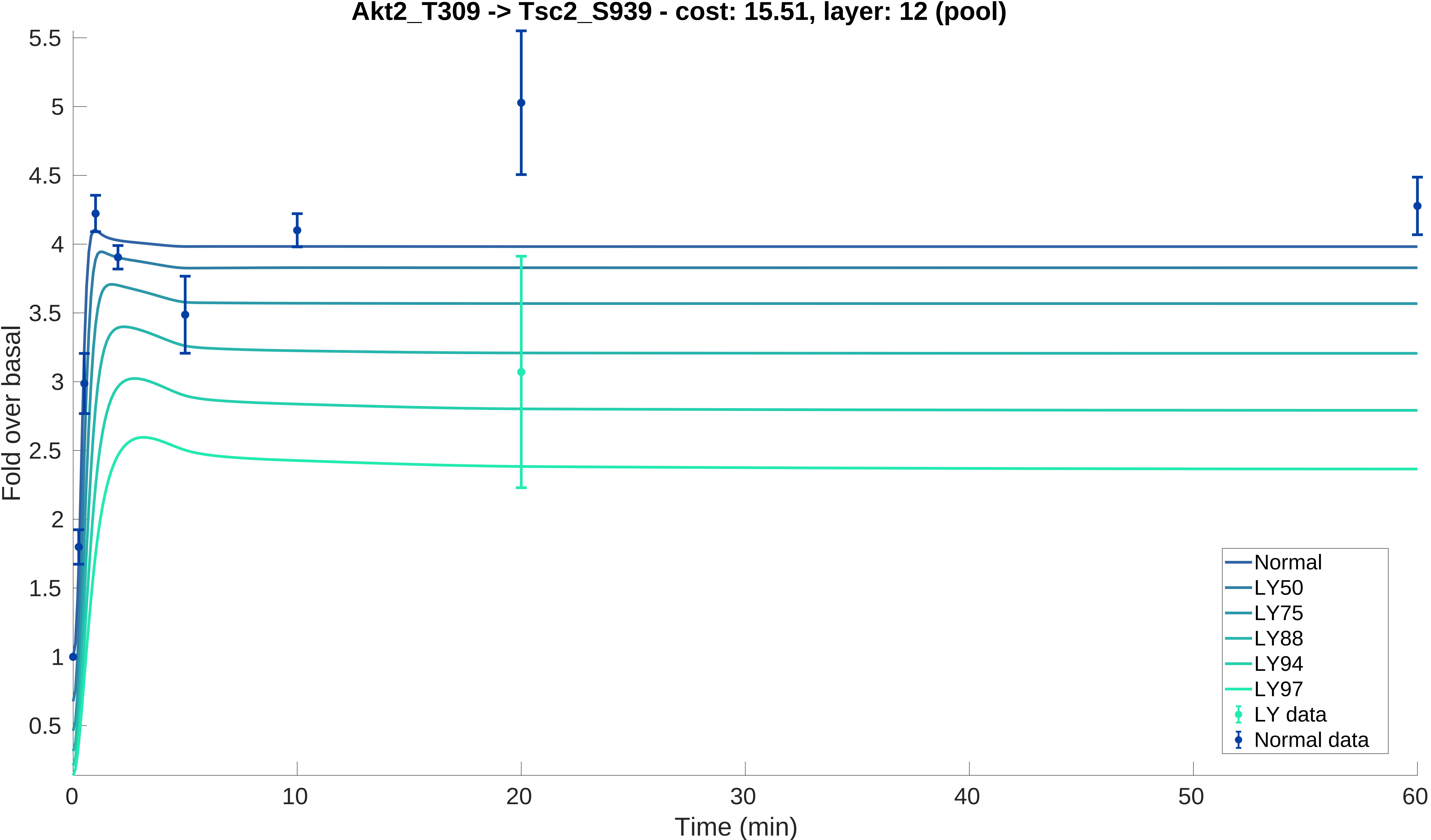
Akt2_T309 -> Tsc2_S939 - cost: 15.51, layer: 12 (pool)
Fold over basal
Time (min)
0
10
20
30
40
50
60
0.5
1
1.5
2
2.5
3
3.5
4
4.5
5
5.5
Normal
LY50
LY75
LY88
LY94
LY97
LY data
Normal data

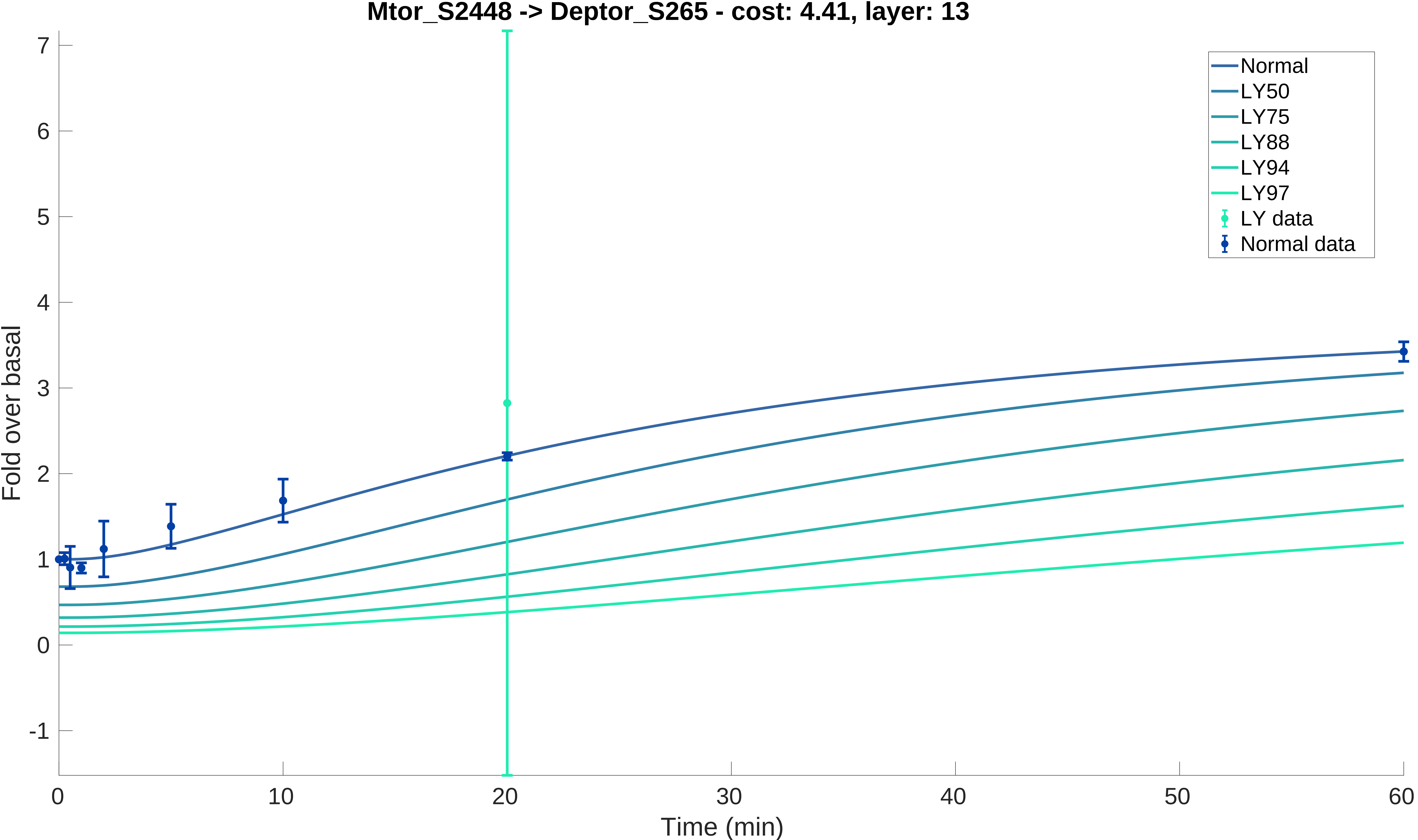
Mtor_S2448 -> Deptor_S265 - cost: 4.41, layer: 13
Normal
LY50
LY75
LY88
LY94
LY97
LY data
Normal data
Fold over basal
Time (min)
7
6
5
4
3
2
1
0
-1
0
10
20
30
40
50
60

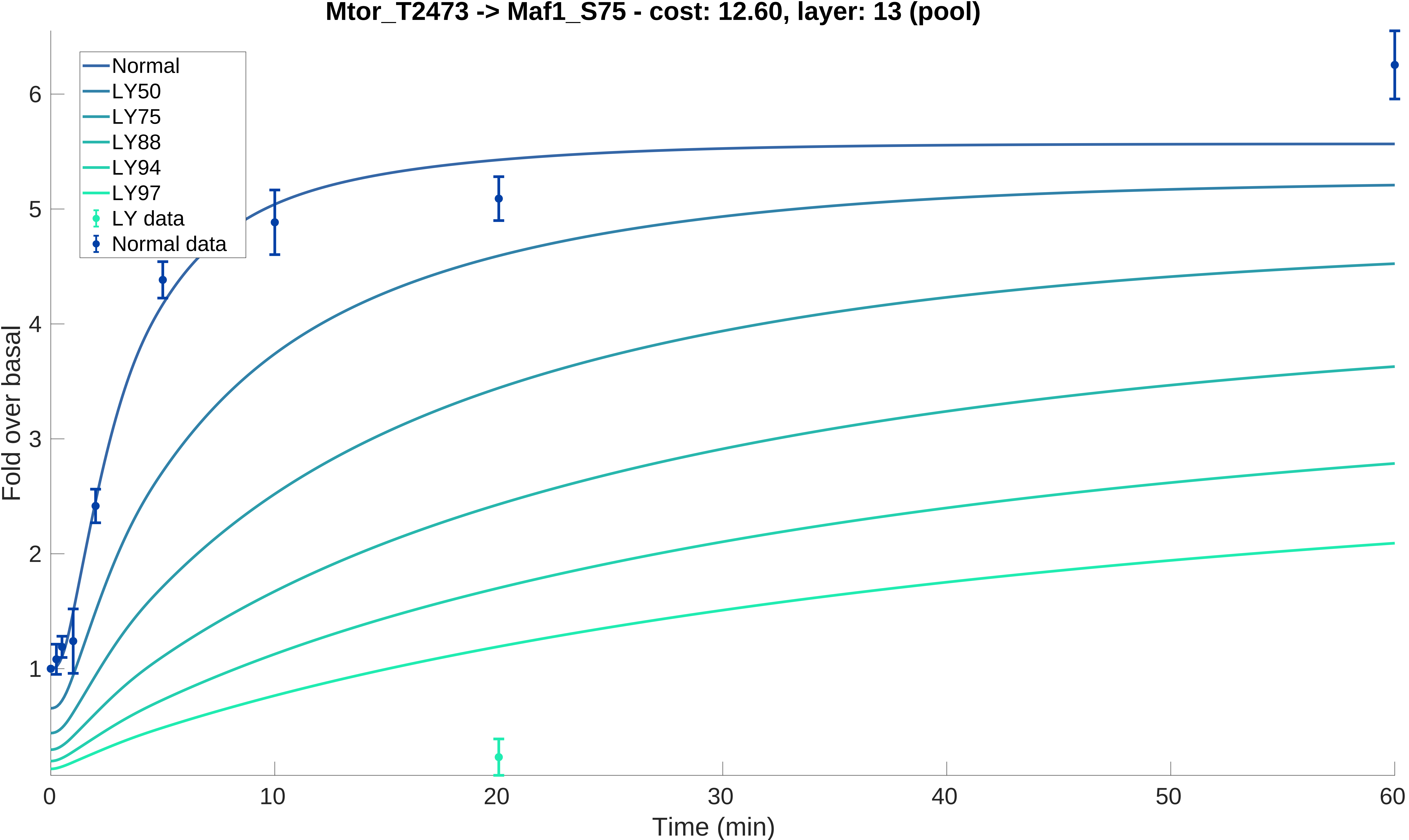

Mtor_T2473 -> Maf1_S75 - cost: 12.60, layer: 13 (pool)
Normal
LY50
LY75
LY88
LY94
LY97
LY data
Normal data
Fold over basal
Time (min)
0
10
20
30
40
50
60
1
2
3
4
5
6

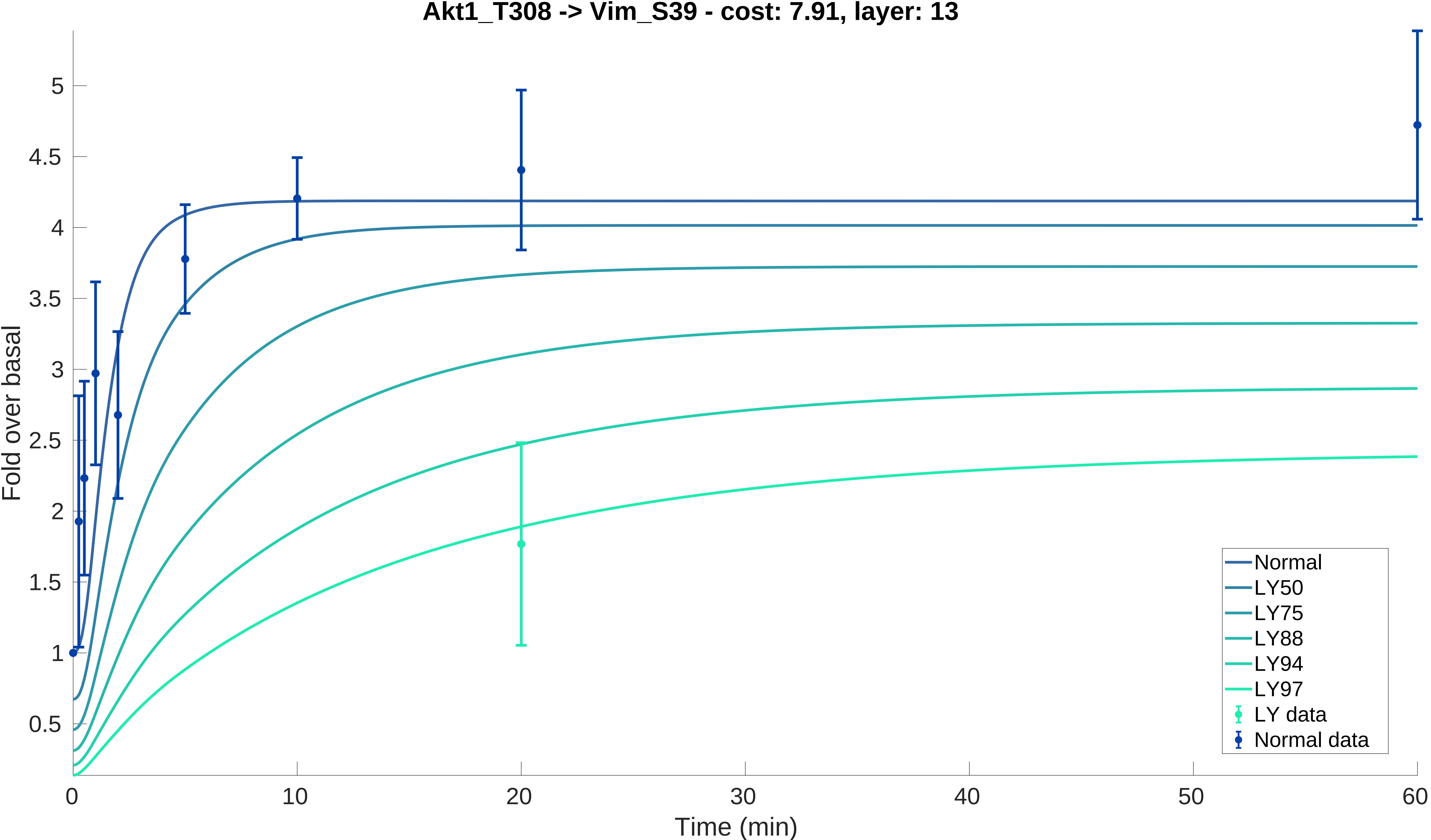

Akt1_T308 -> Vim_S39 - cost: 7.91, layer: 13
Fold over basal
Time (min)
0
10
20
30
40
50
60
0.5
1
1.5
2
2.5
3
3.5
4
4.5
5
Normal
LY50
LY75
LY88
LY94
LY97
LY data
Normal data

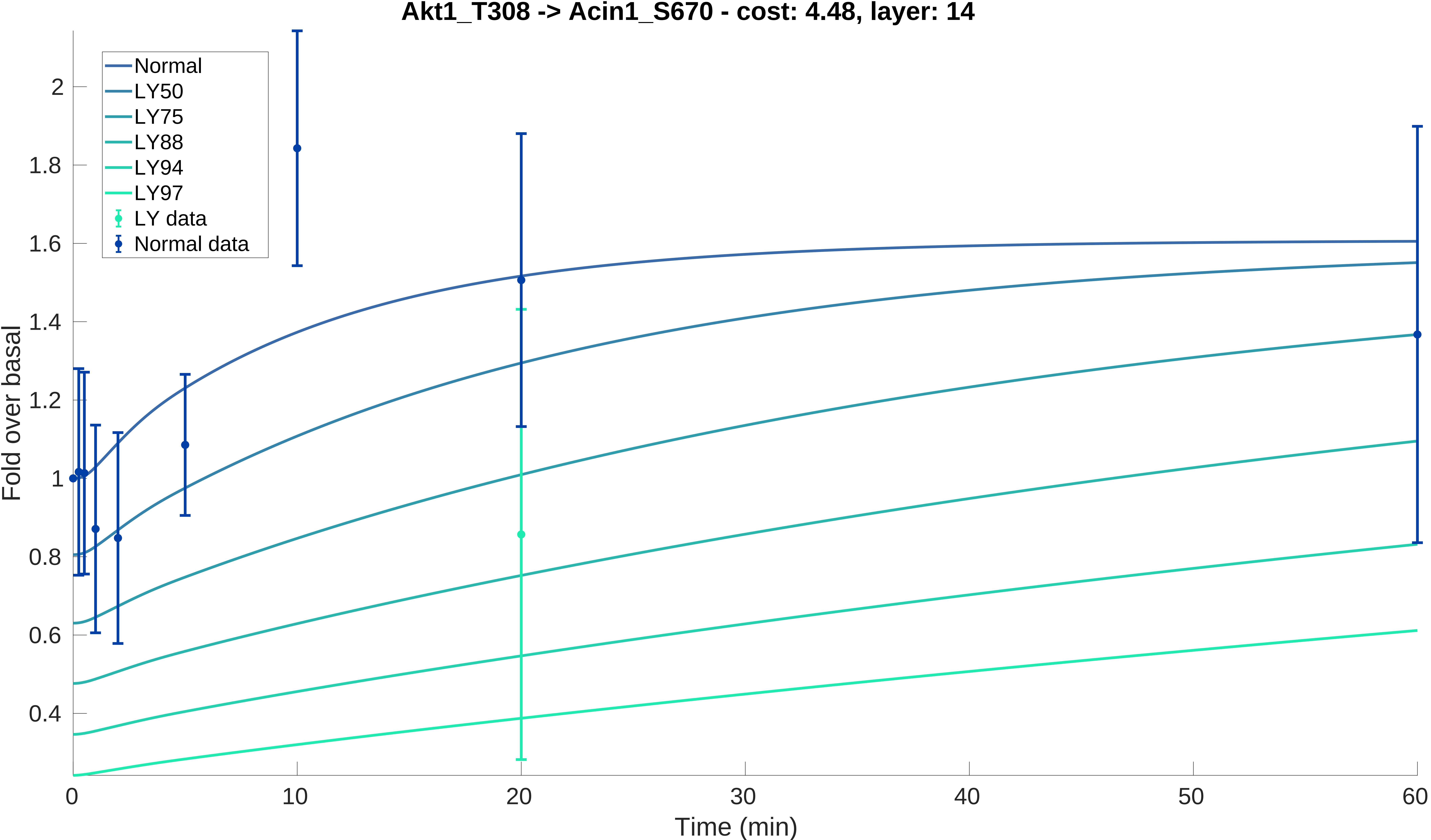

Akt1_T308 -> Acin1_S670 - cost: 4.48, layer: 14
Normal
LY50
LY75
LY88
LY94
LY97
LY data
Normal data
Fold over basal
0.4
0.6
0.8
1
1.2
1.4
1.6
1.8
2
0
10
20
30
40
50
60
Time (min)

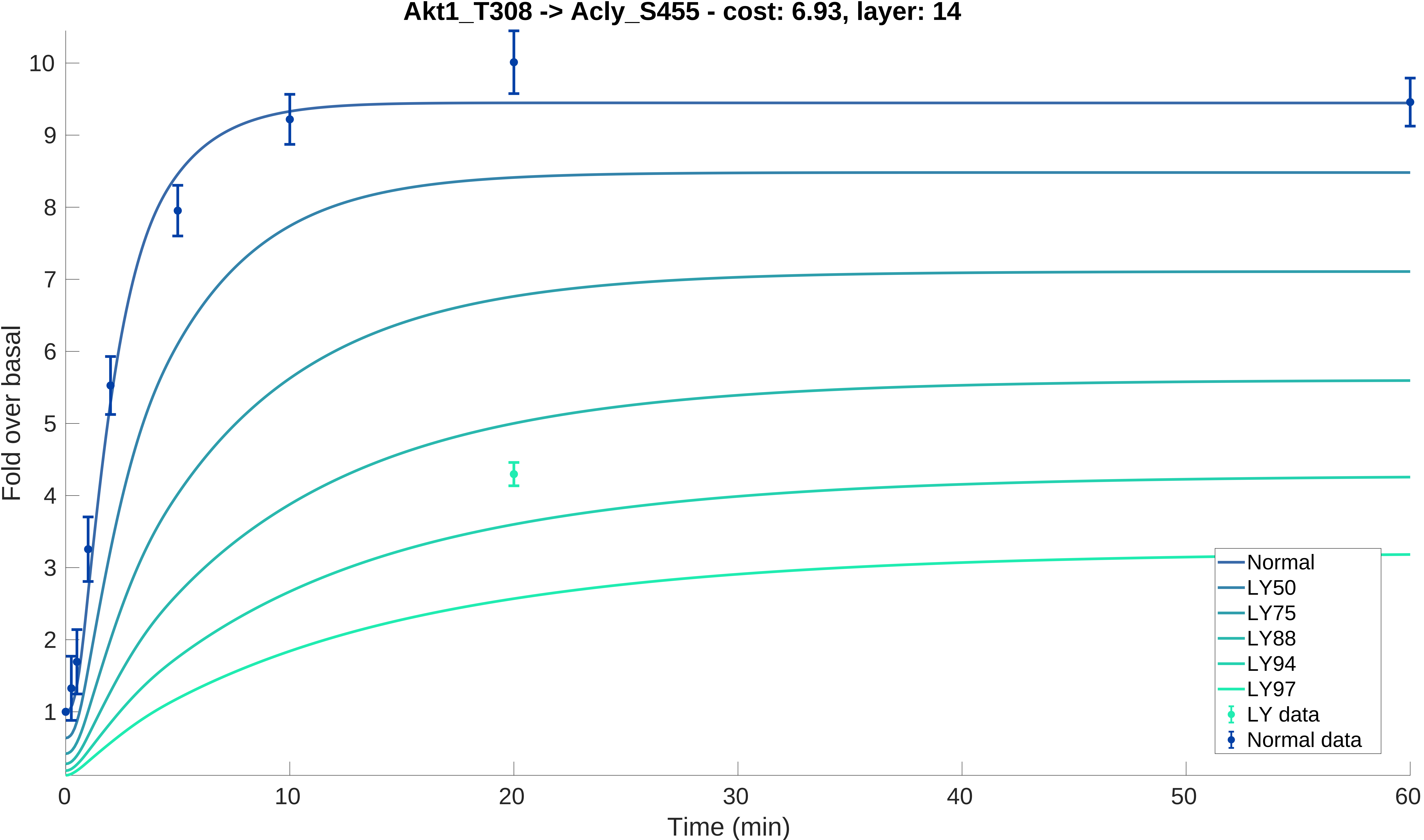

Akt1_T308 -> Acly_S455 - cost: 6.93, layer: 14
Fold over basal
Time (min)
10
9
8
7
6
5
4
3
2
1
0
10
20
30
40
50
60
Normal
LY50
LY75
LY88
LY94
LY97
LY data
Normal data

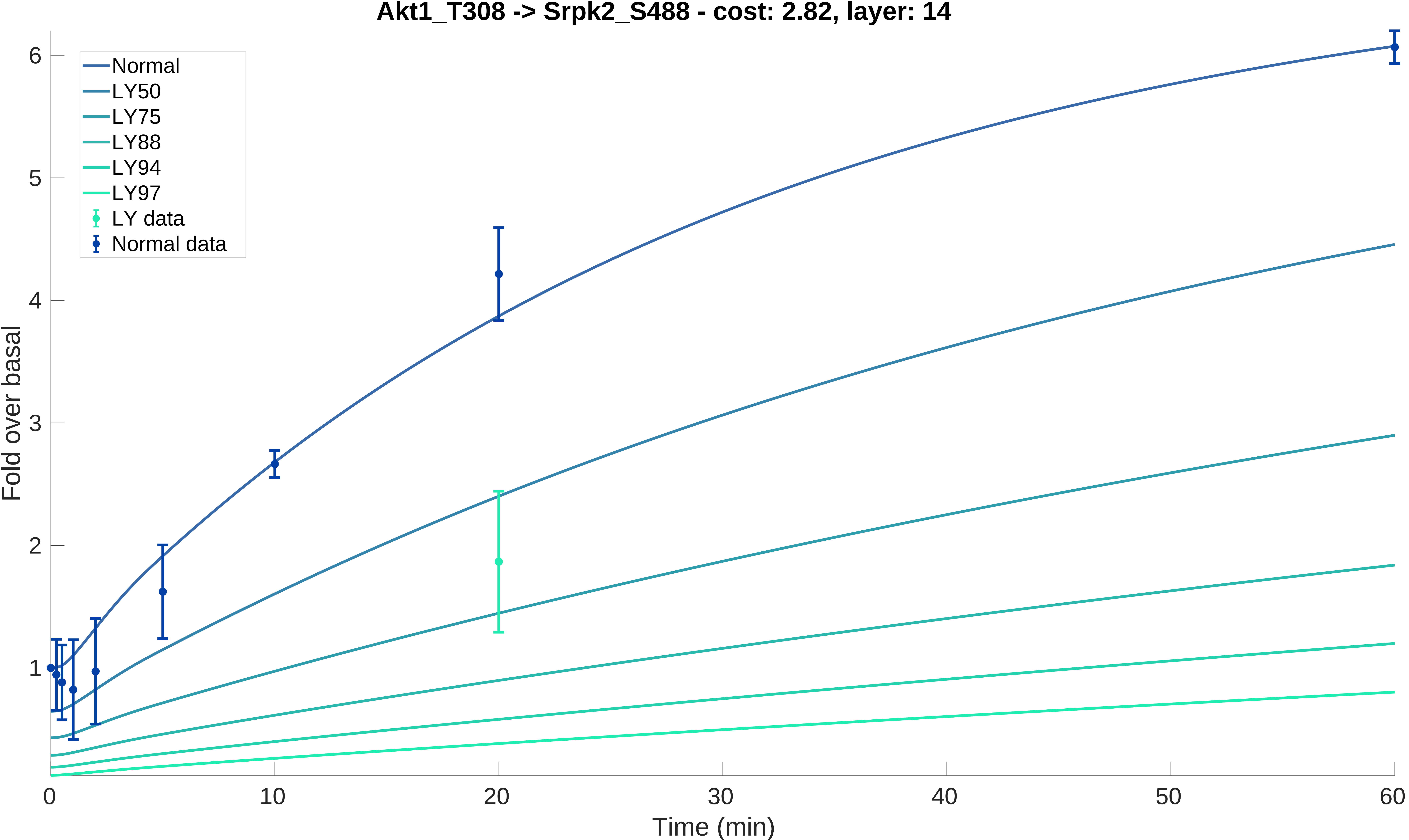
Akt1_T308 -> Srpk2_S488 - cost: 2.82, layer: 14
Normal
LY50
LY75
LY88
LY94
LY97
LY data
Normal data
Fold over basal
Time (min)
0
10
20
30
40
50
60
1
2
3
4
5
6

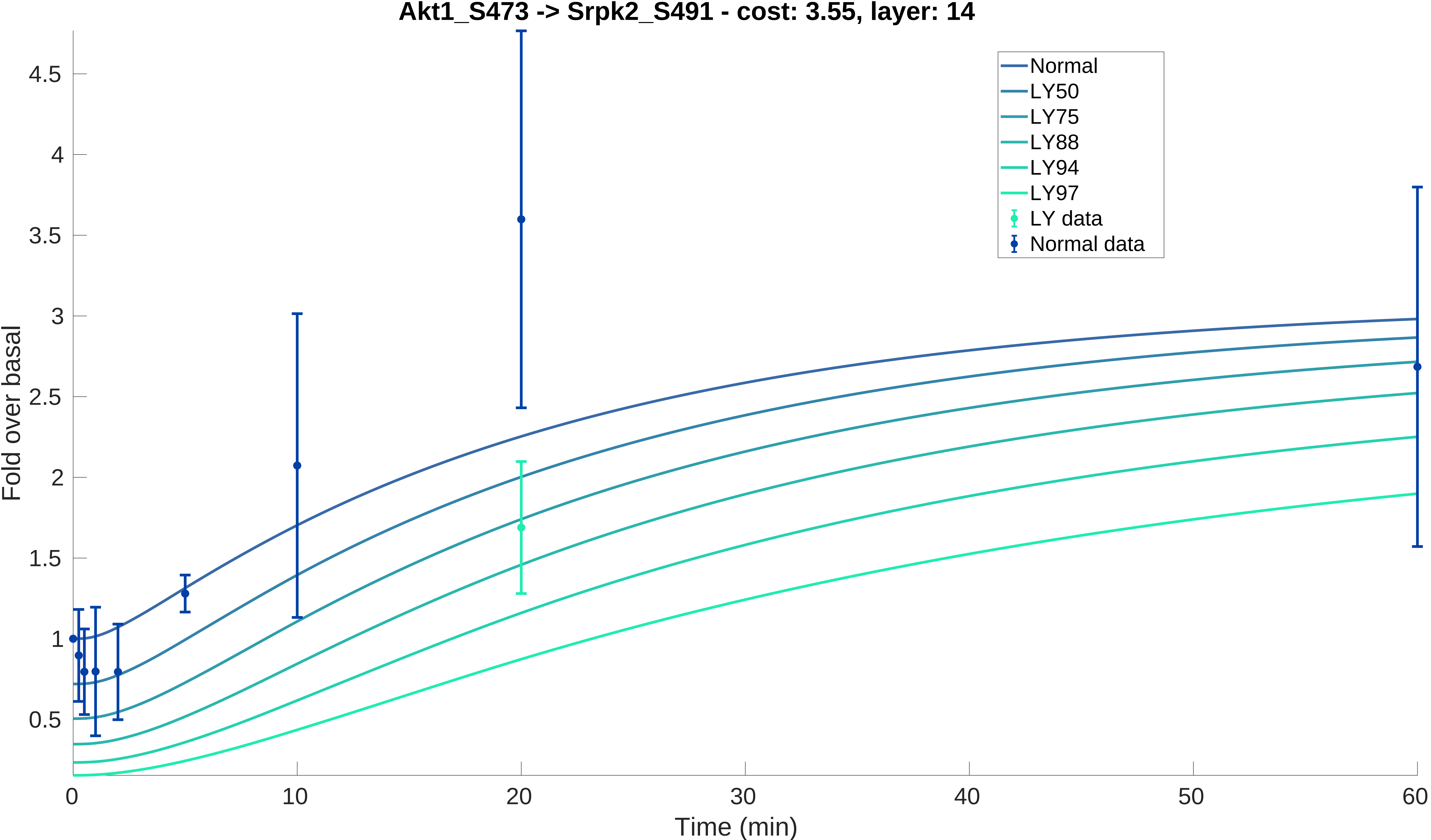
Akt1_S473 -> Srpk2_S491 - cost: 3.55, layer: 14
Normal
LY50
LY75
LY88
LY94
LY97
LY data
Normal data
Fold over basal
Time (min)
0.5
1
1.5
2
2.5
3
3.5
4
4.5
0
10
20
30
40
50
60

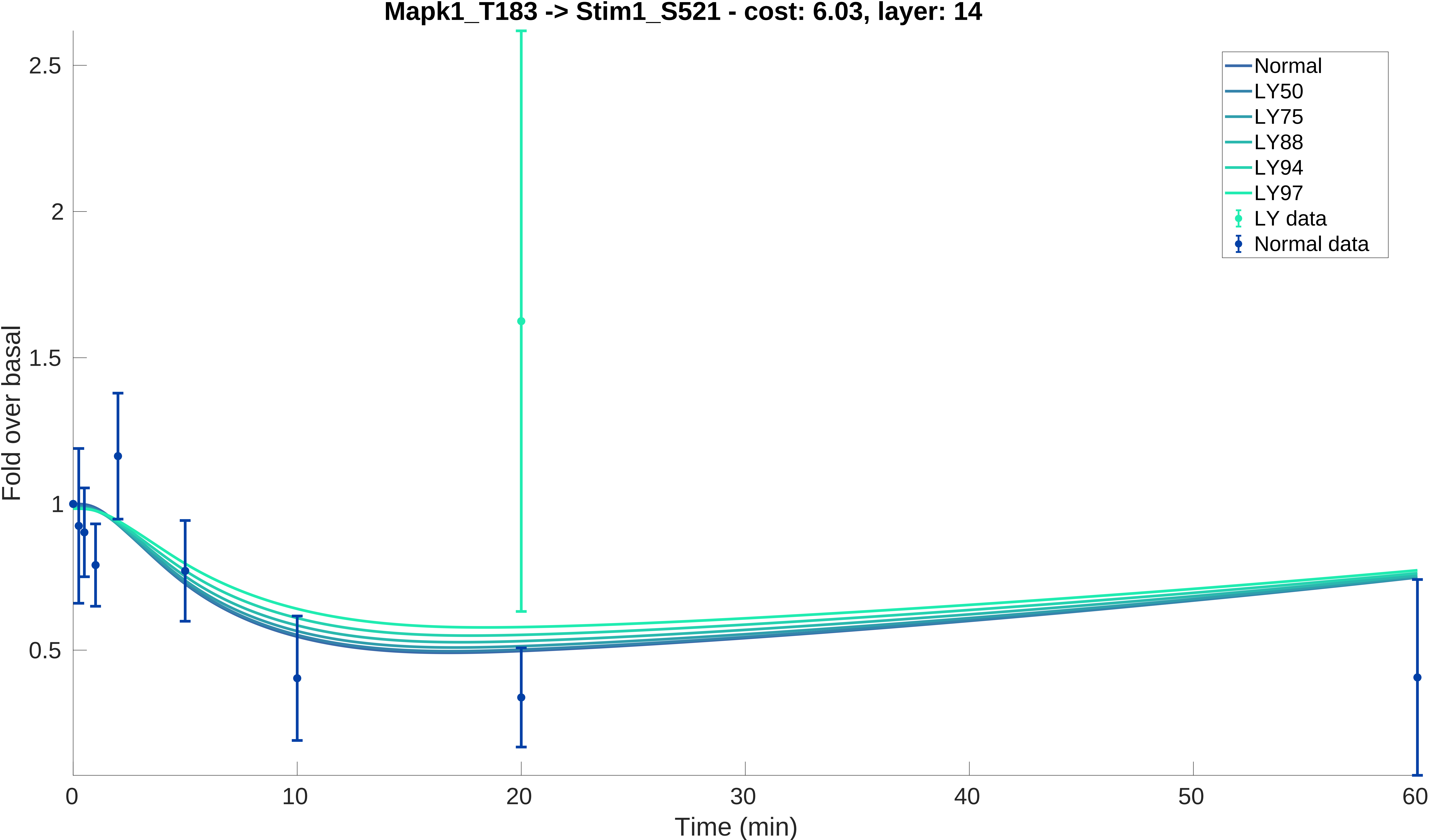

Mapk1_T183 -> Stim1_S521 - cost: 6.03, layer: 14
Normal
LY50
LY75
LY88
LY94
LY97
LY data
Normal data
Fold over basal
Time (min)
0.5
1
1.5
2
2.5
0
10
20
30
40
50
60

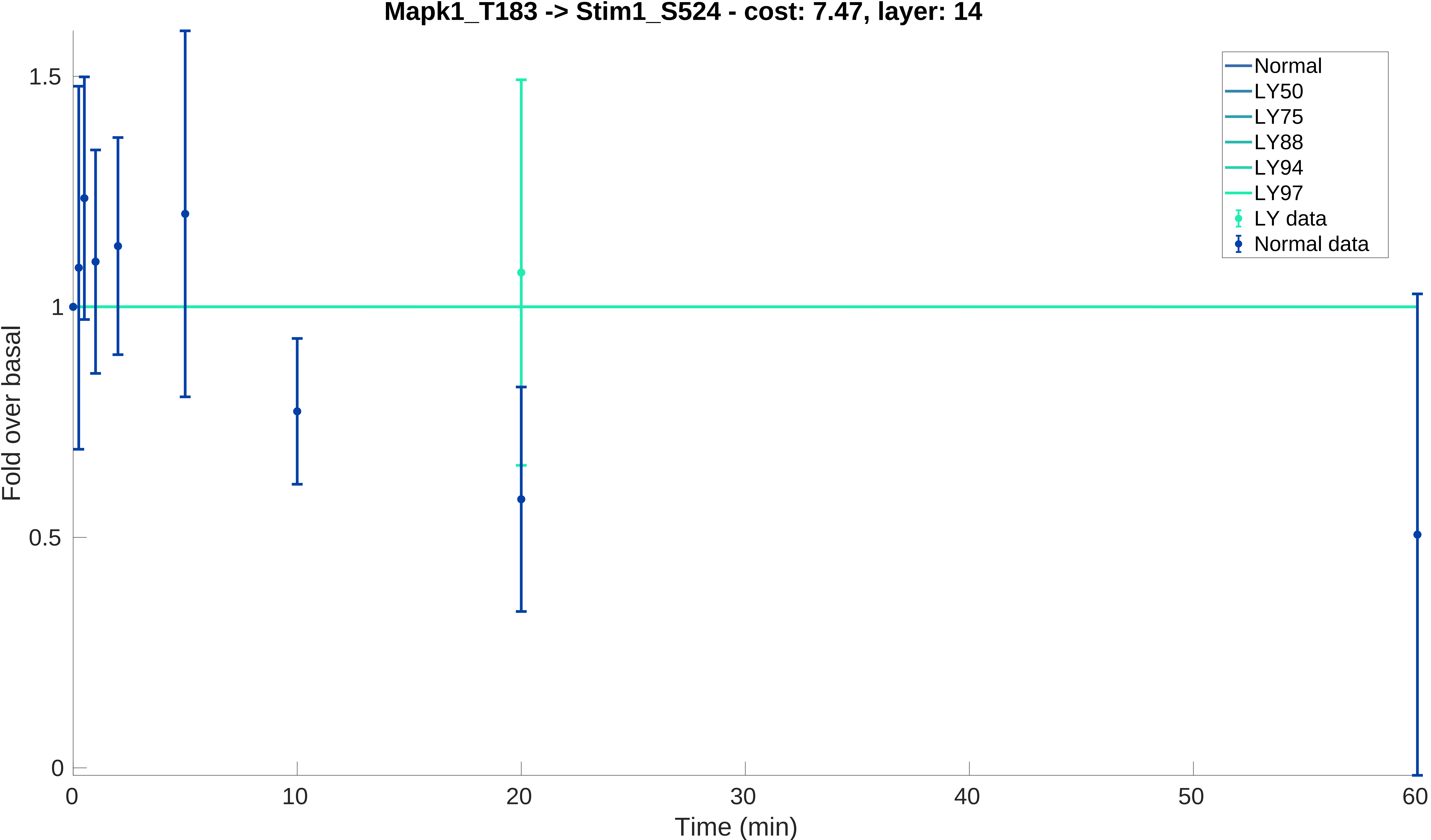
Mapk1_T183 -> Stim1_S524 - cost: 7.47, layer: 14
Fold over basal
Time (min)
0
0.5
1
1.5
0
10
20
30
40
50
60
Normal
LY50
LY75
LY88
LY94
LY97
LY data
Normal data

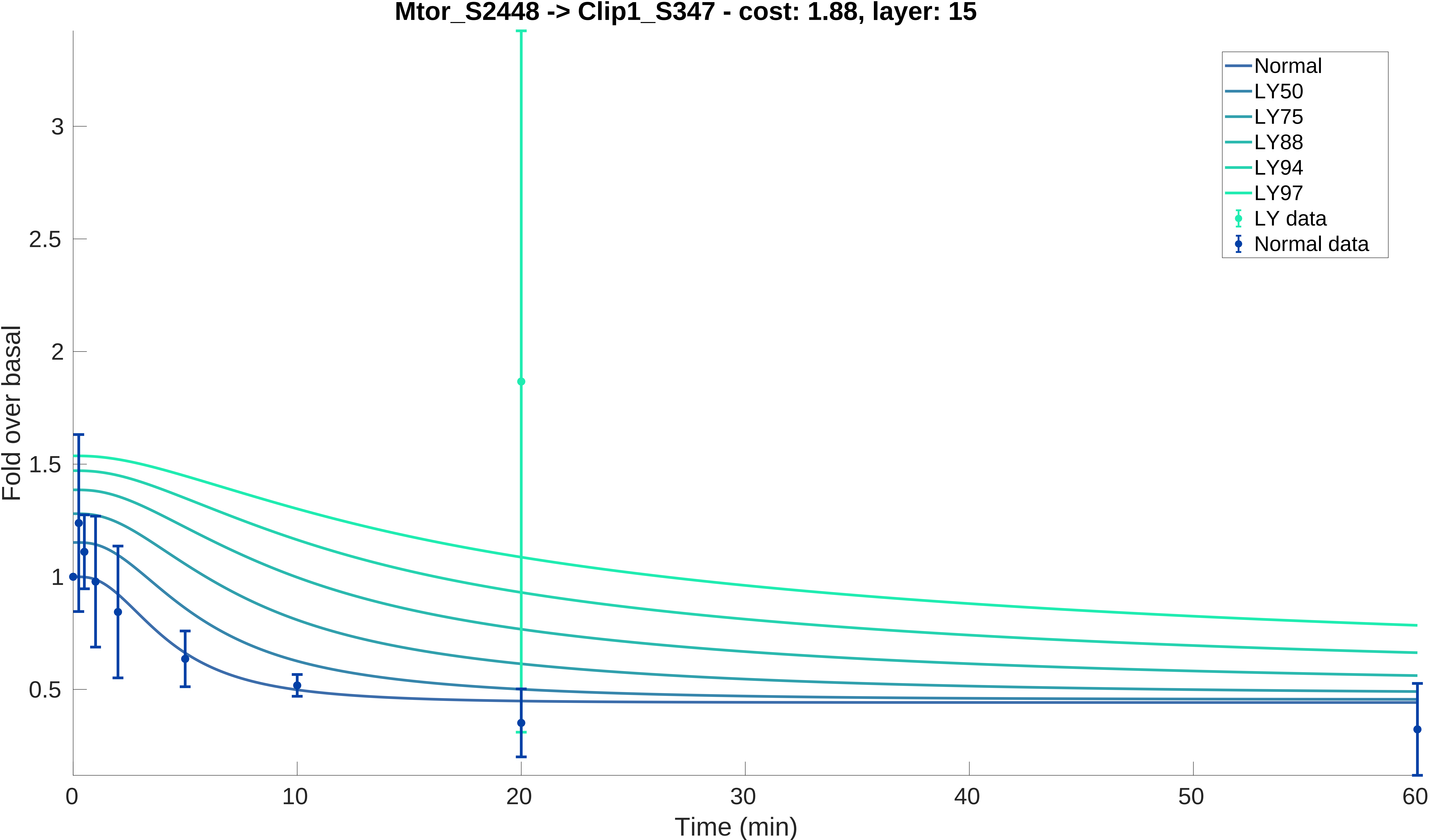
Mtor_S2448 -> Clip1_S347 - cost: 1.88, layer: 15
Normal
LY50
LY75
LY88
LY94
LY97
LY data
Normal data
Fold over basal
3
2.5
2
1.5
1
0.5
0
10
20
30
40
50
60
Time (min)

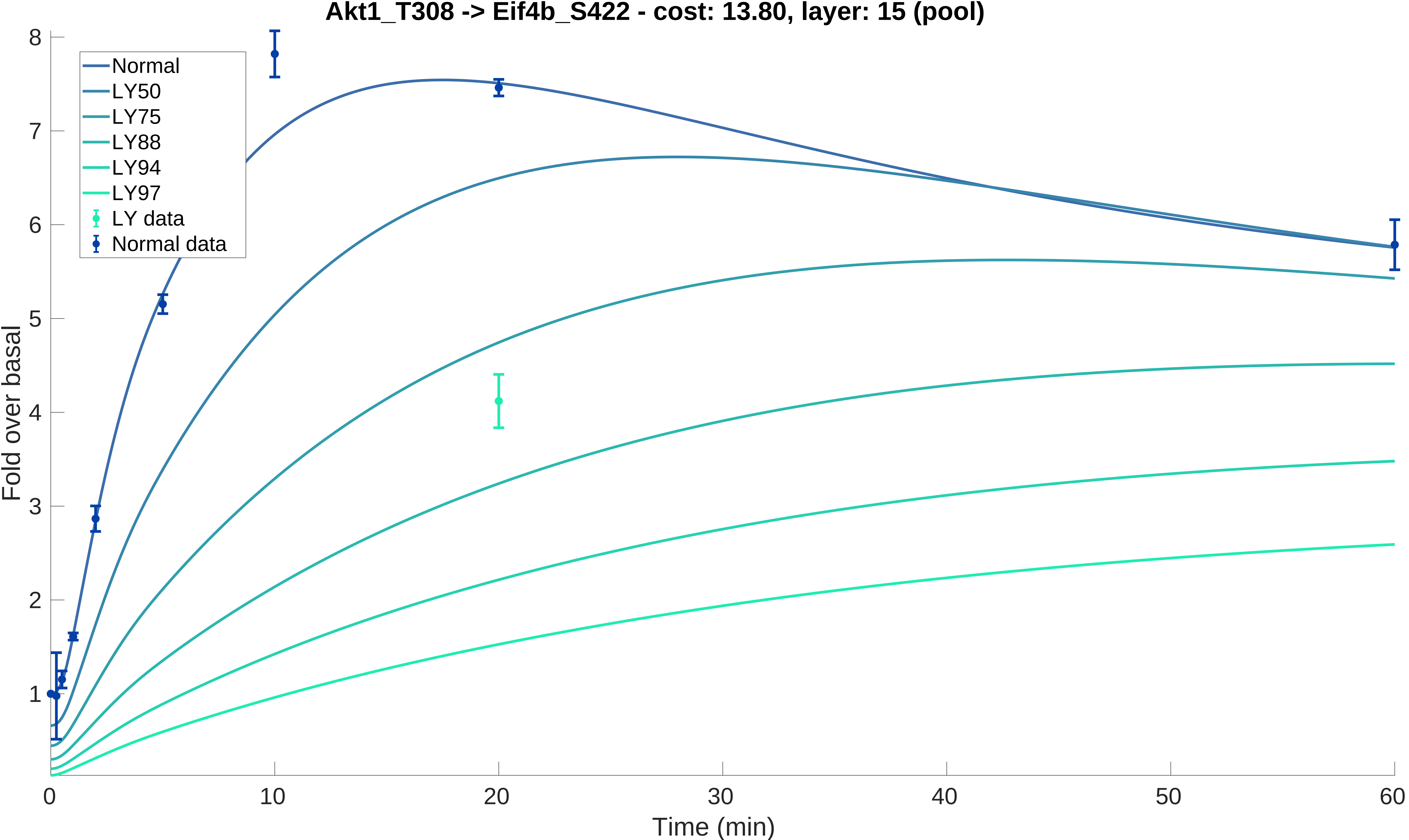

Akt1_T308 -> Eif4b_S422 - cost: 13.80, layer: 15 (pool)
Normal
LY50
LY75
LY88
LY94
LY97
LY data
Normal data
Fold over basal
Time (min)
8
7
6
5
4
3
2
1
0
10
20
30
40
50
60

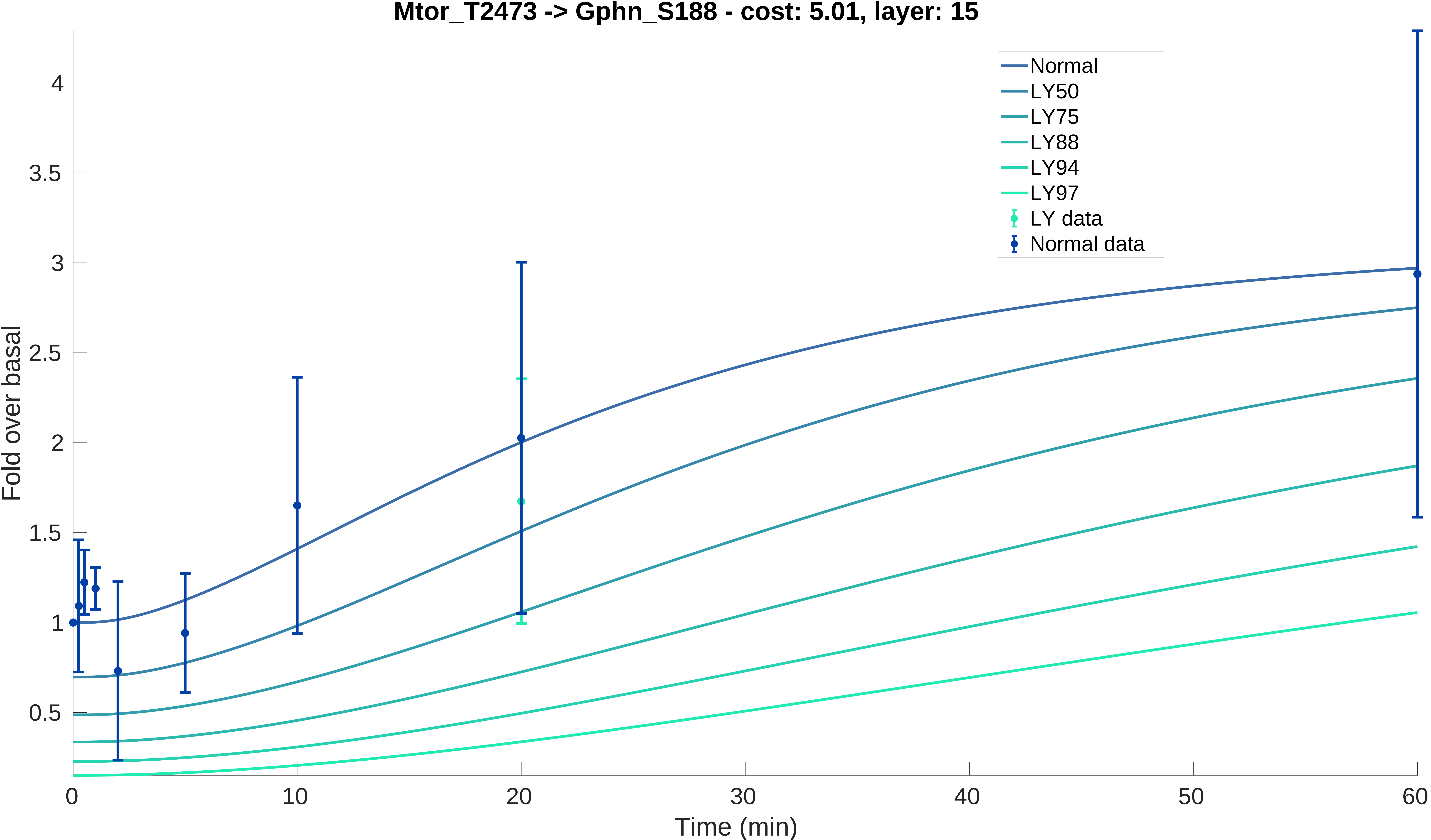

Mtor_T2473 -> Gphn_S188 - cost: 5.01, layer: 15
Fold over basal
Time (min)
0
10
20
30
40
50
60
0.5
1
1.5
2
2.5
3
3.5
4
Normal
LY50
LY75
LY88
LY94
LY97
LY data
Normal data

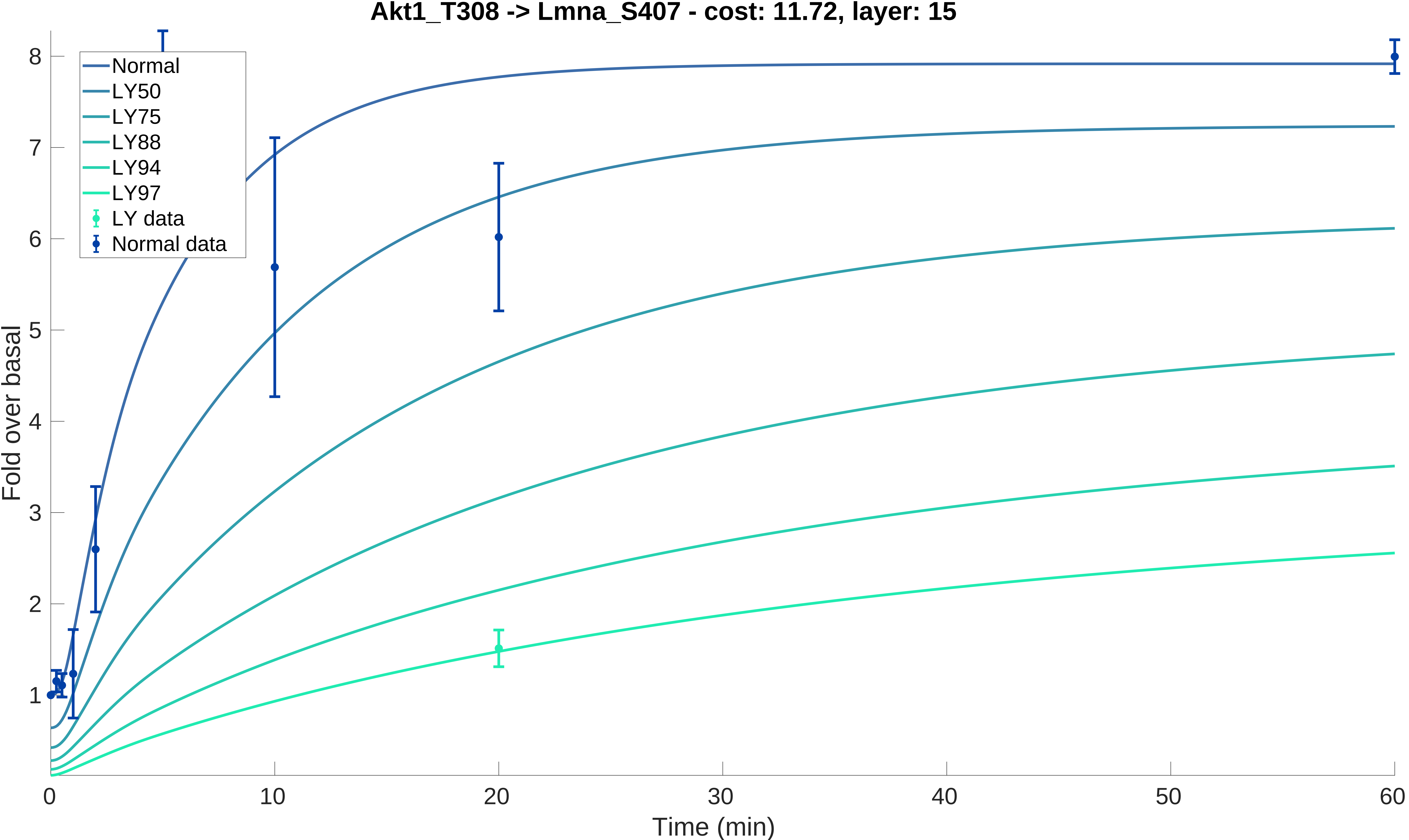

Akt1_T308 -> Lmna_S407 - cost: 11.72, layer: 15
Normal
LY50
LY75
LY88
LY94
LY97
LY data
Normal data
Fold over basal
Time (min)
0
10
20
30
40
50
60
1
2
3
4
5
6
7
8

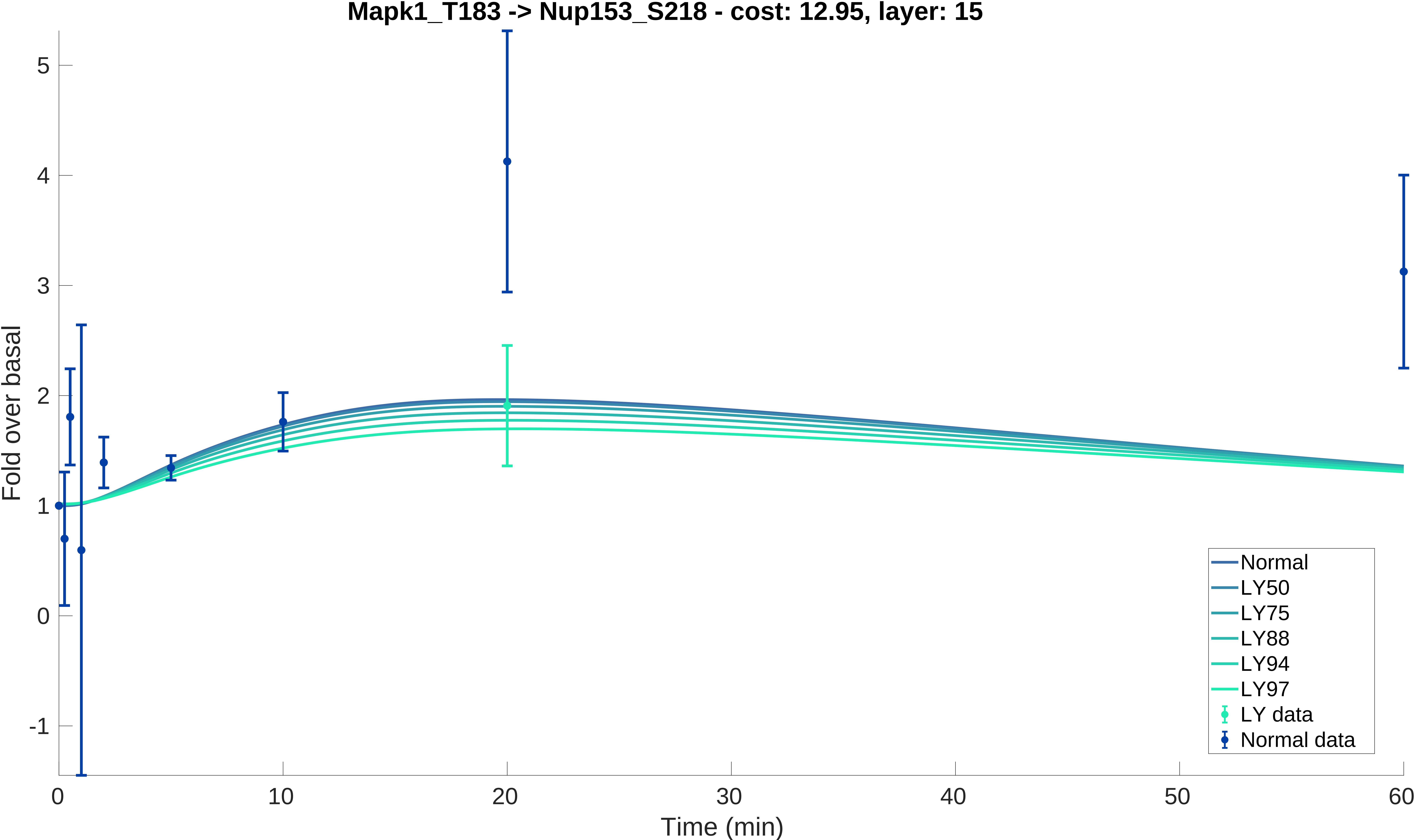

Mapk1_T183 -> Nup153_S218 - cost: 12.95, layer: 15
Fold over basal
Time (min)
5
4
3
2
1
0
-1
0
10
20
30
40
50
60
Normal
LY50
LY75
LY88
LY94
LY97
LY data
Normal data

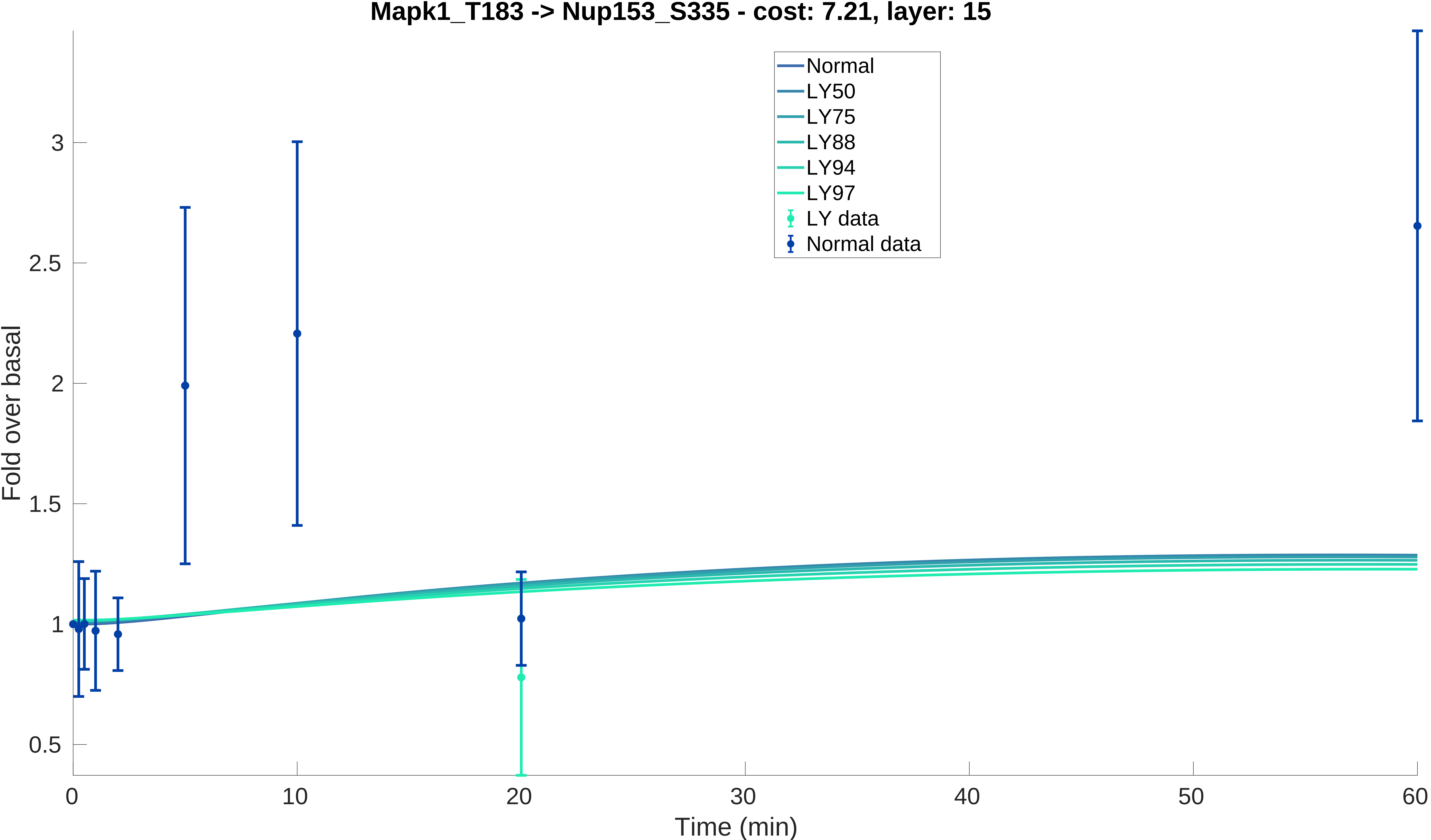

Mapk1_T183 -> Nup153_S335 - cost: 7.21, layer: 15
Normal
LY50
LY75
LY88
LY94
LY97
LY data
Normal data
Fold over basal
0.5
1
1.5
2
2.5
3
0
10
20
30
40
50
60
Time (min)

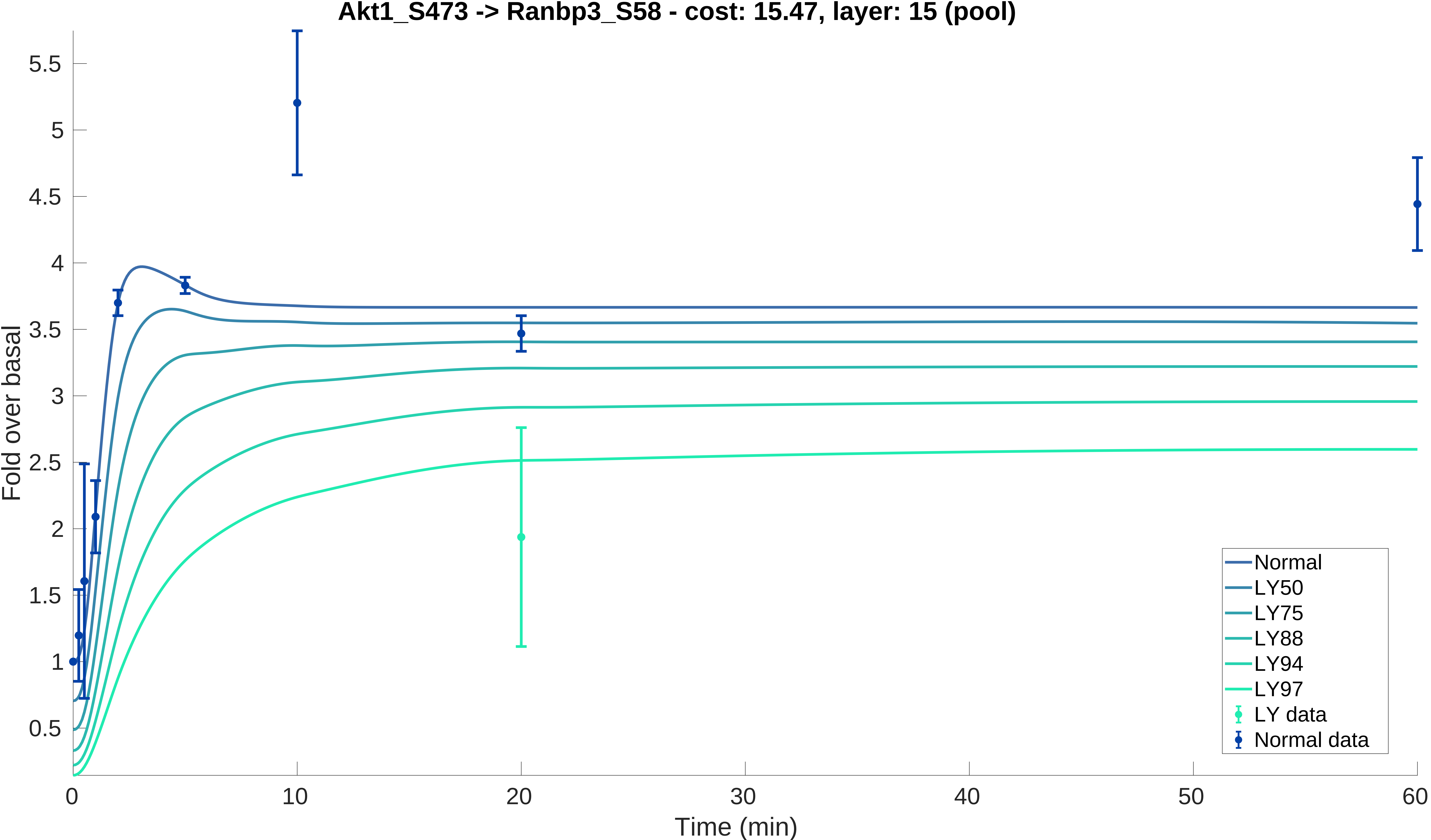

Akt1_S473 -> Ranbp3_S58 - cost: 15.47, layer: 15 (pool)
Fold over basal
5.5
5
4.5
4
3.5
3
2.5
2
1.5
1
0.5
0
10
20
30
40
50
60
Time (min)
Normal
LY50
LY75
LY88
LY94
LY97
LY data
Normal data

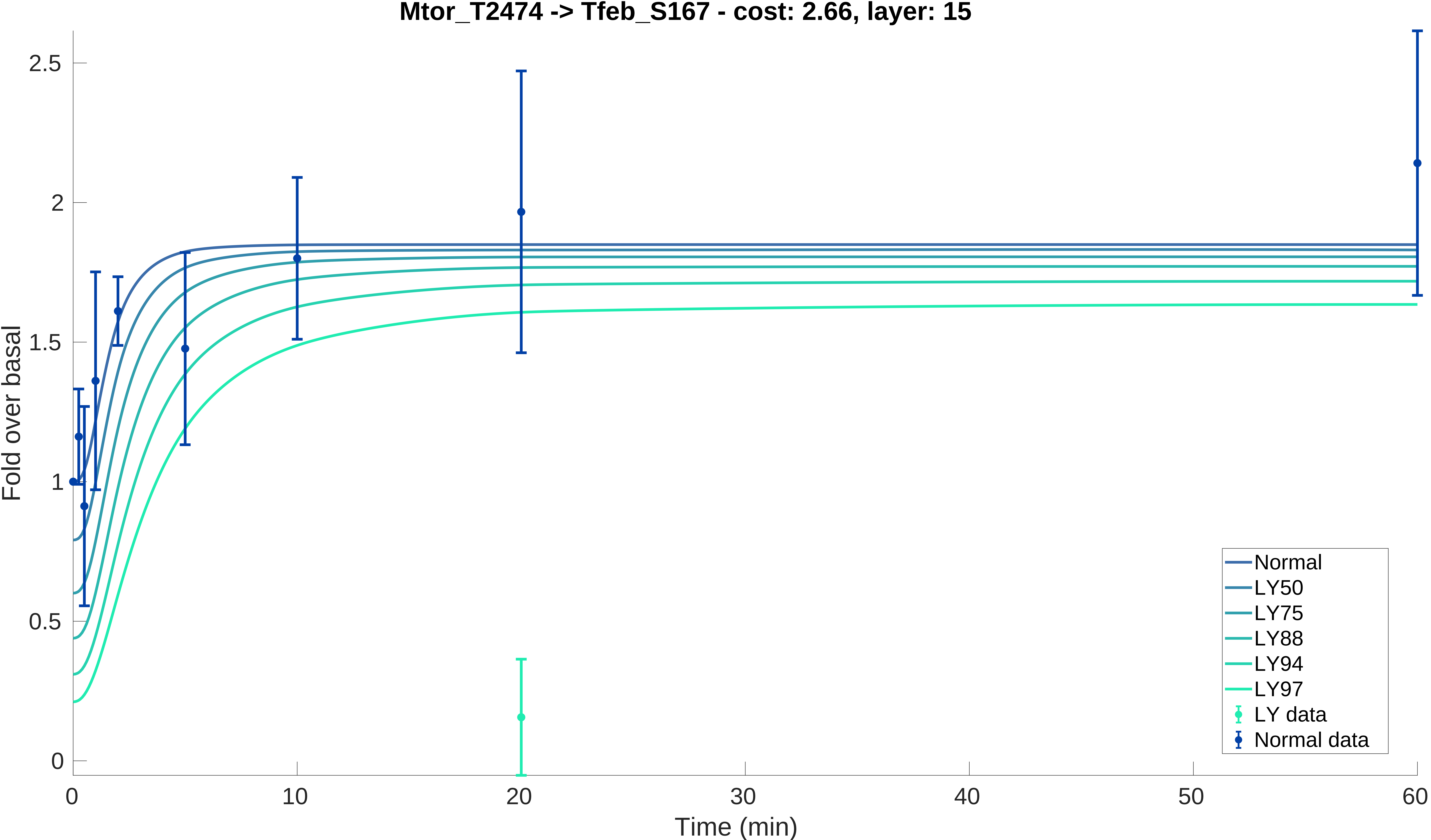

Mtor_T2474 -> Tfeb_S167 - cost: 2.66, layer: 15
Fold over basal
Time (min)
0
0.5
1
1.5
2
2.5
0
10
20
30
40
50
60
Normal
LY50
LY75
LY88
LY94
LY97
LY data
Normal data

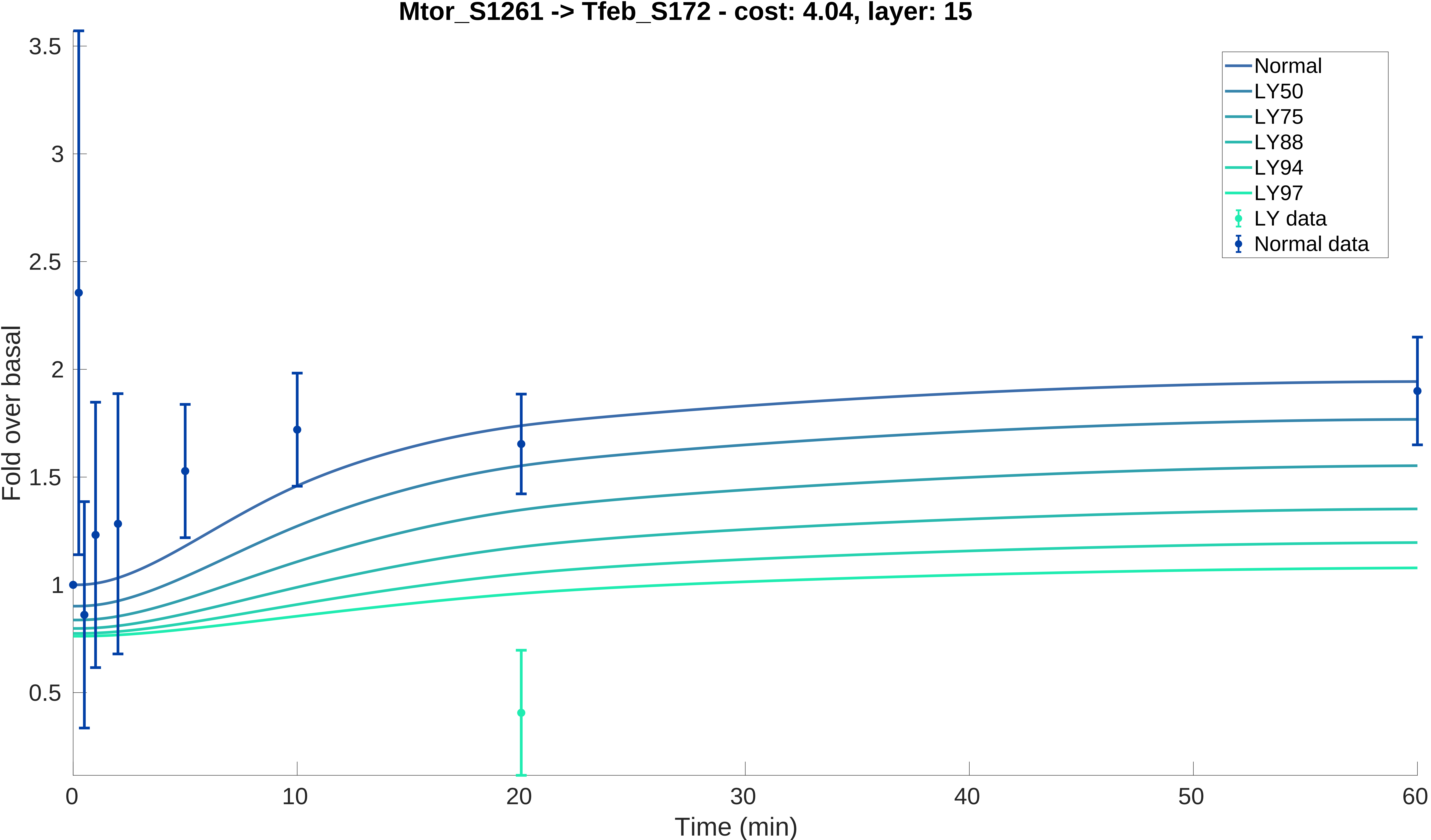

Mtor_S1261 -> Tfeb_S172 - cost: 4.04, layer: 15
Fold over basal
Time (min)
0.5
1
1.5
2
2.5
3
3.5
0
10
20
30
40
50
60
Normal
LY50
LY75
LY88
LY94
LY97
LY data
Normal data

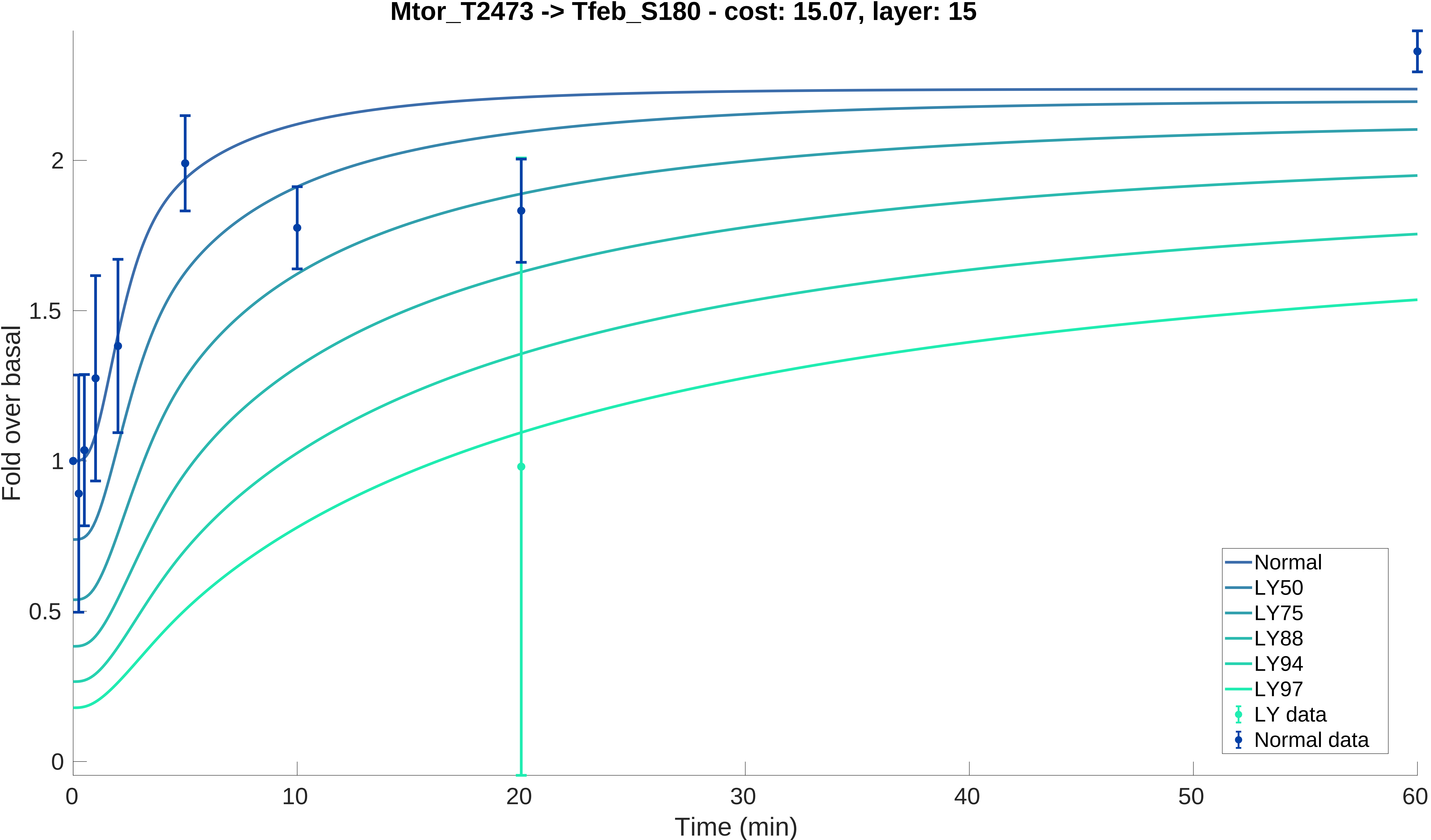

Mtor_T2473 -> Tfeb_S180 - cost: 15.07, layer: 15
Fold over basal
Time (min)
0
0.5
1
1.5
2
0
10
20
30
40
50
60
Normal
LY50
LY75
LY88
LY94
LY97
LY data
Normal data

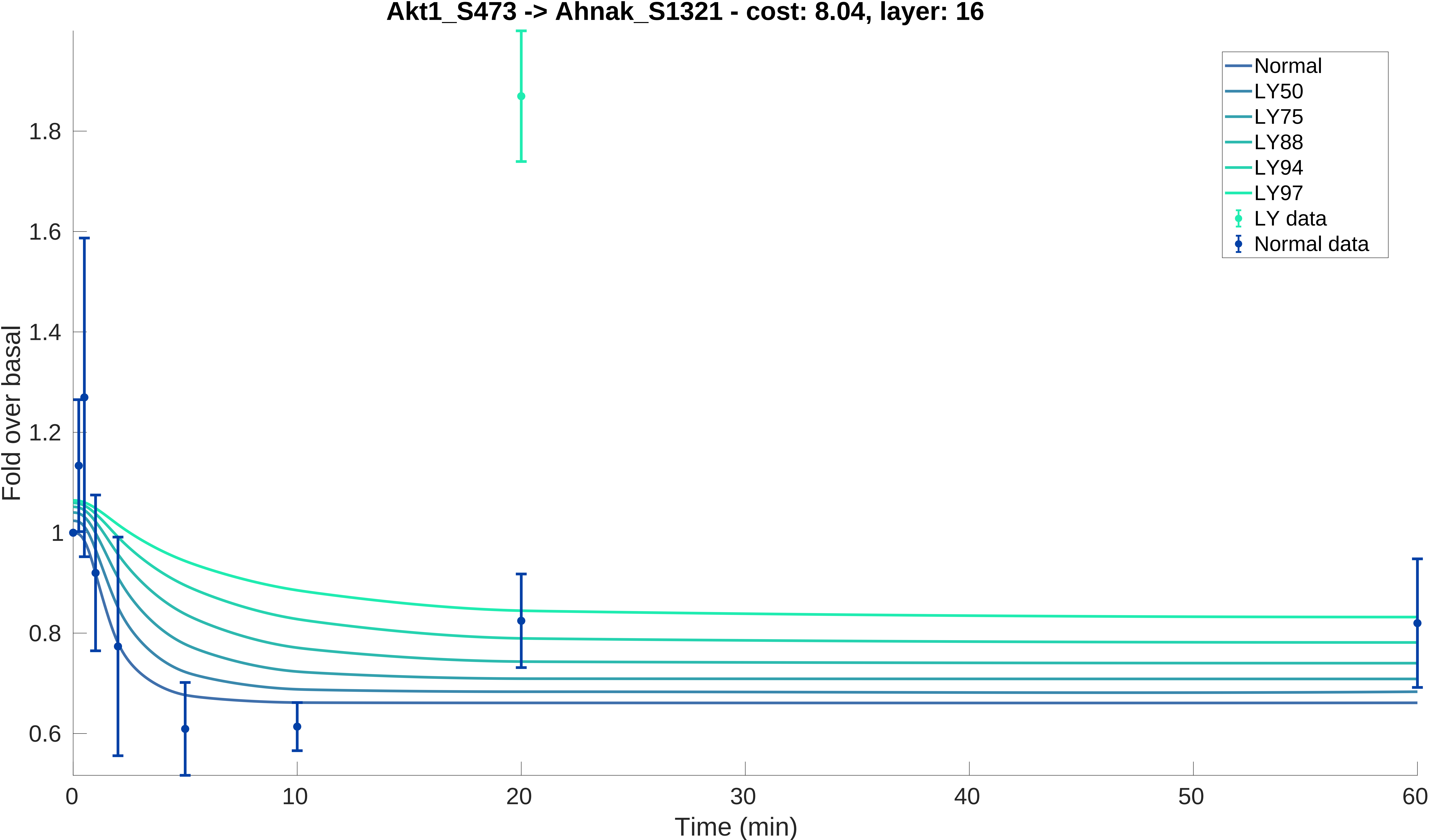

Akt1_S473 -> Ahnak_S1321 - cost: 8.04, layer: 16
Fold over basal
Time (min)
1.8
1.6
1.4
1.2
1
0.8
0.6
0
10
20
30
40
50
60
Normal
LY50
LY75
LY88
LY94
LY97
LY data
Normal data

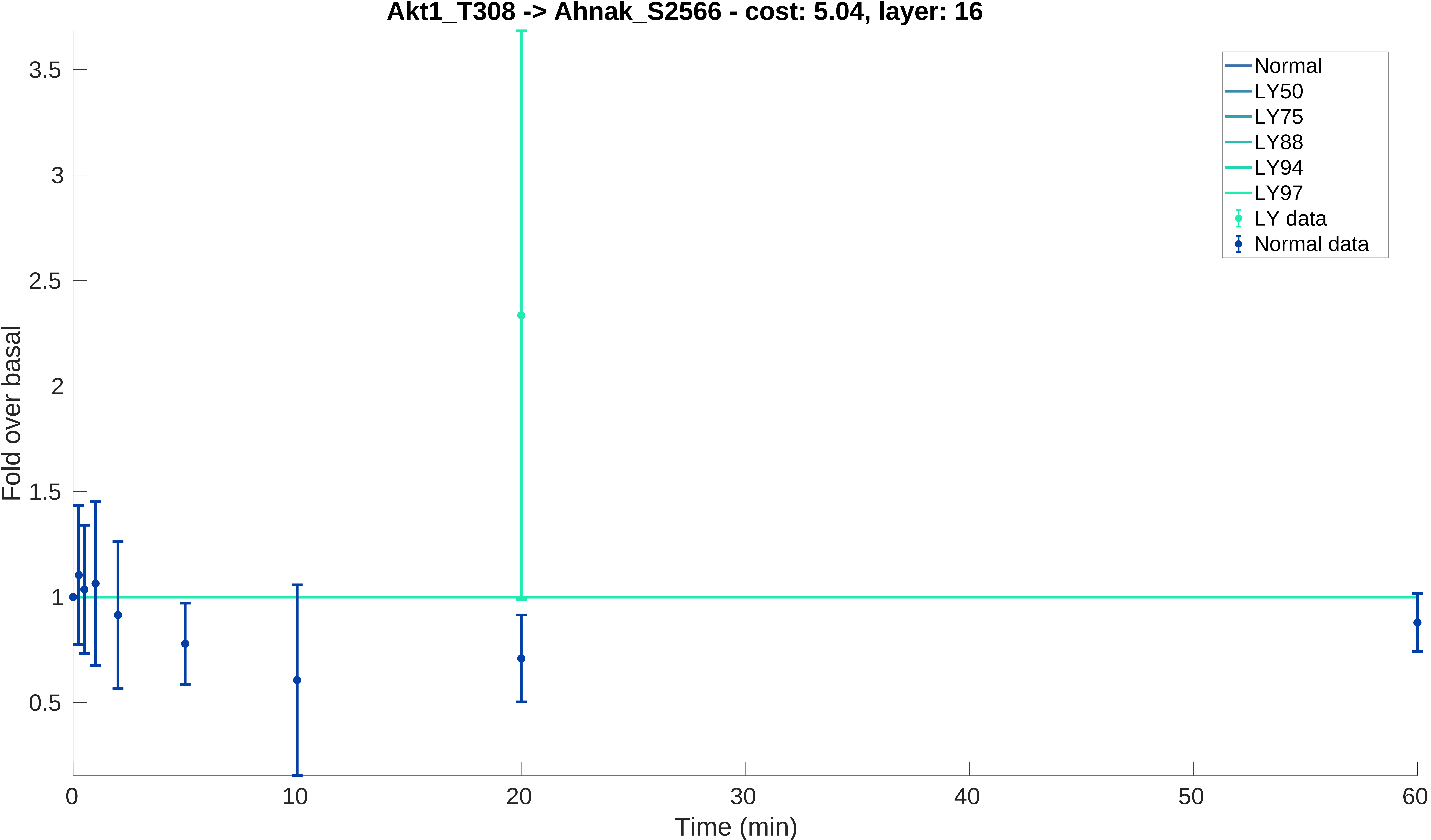
Akt1_T308 -> Ahnak_S2566 - cost: 5.04, layer: 16
Fold over basal
Time (min)
0.5
1
1.5
2
2.5
3
3.5
0
10
20
30
40
50
60
Normal
LY50
LY75
LY88
LY94
LY97
LY data
Normal data

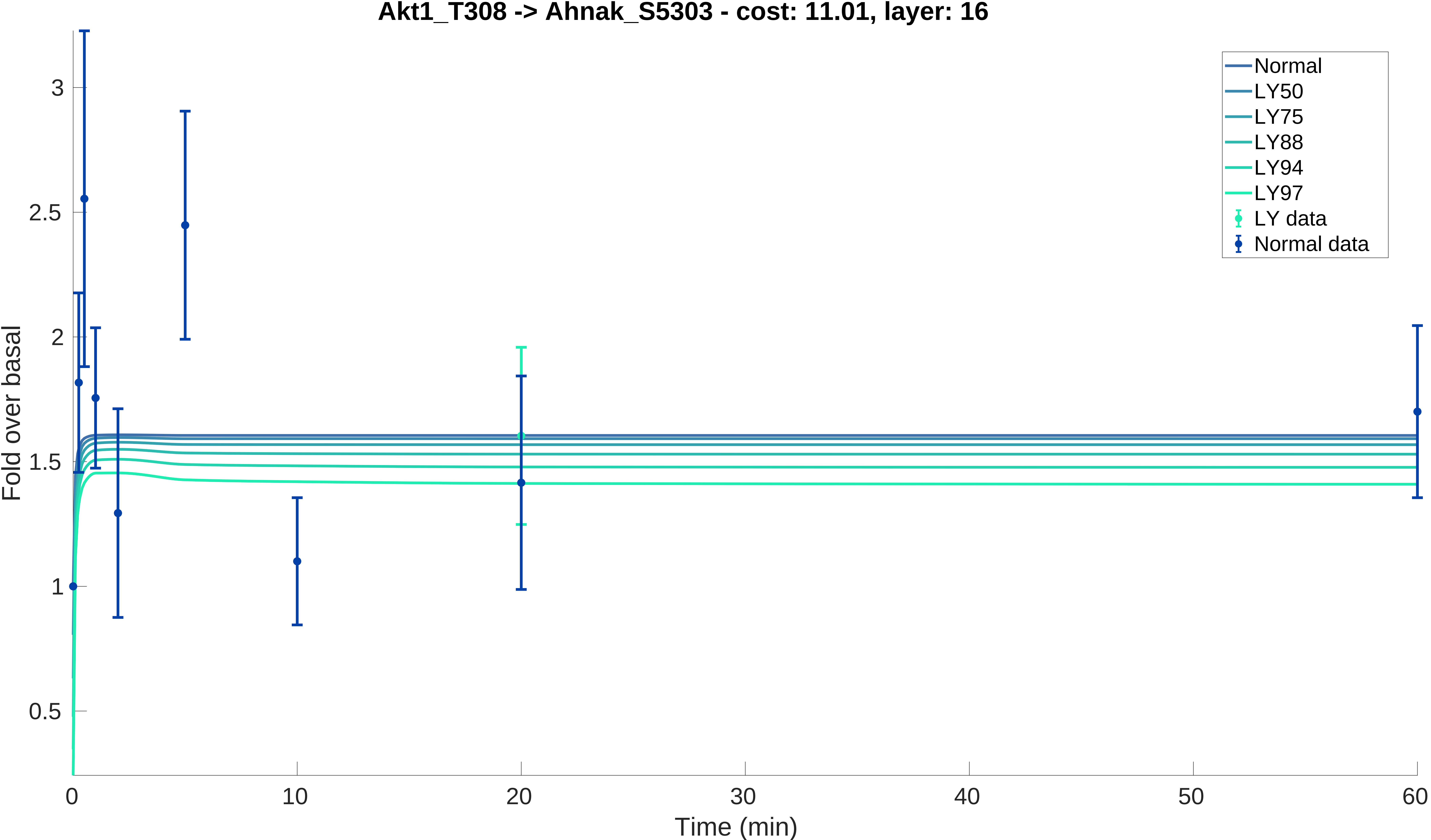

Akt1_T308 -> Ahnak_S5303 - cost: 11.01, layer: 16
Normal
LY50
LY75
LY88
LY94
LY97
LY data
Normal data
Fold over basal
3
2.5
2
1.5
1
0.5
0
10
20
30
40
50
60
Time (min)

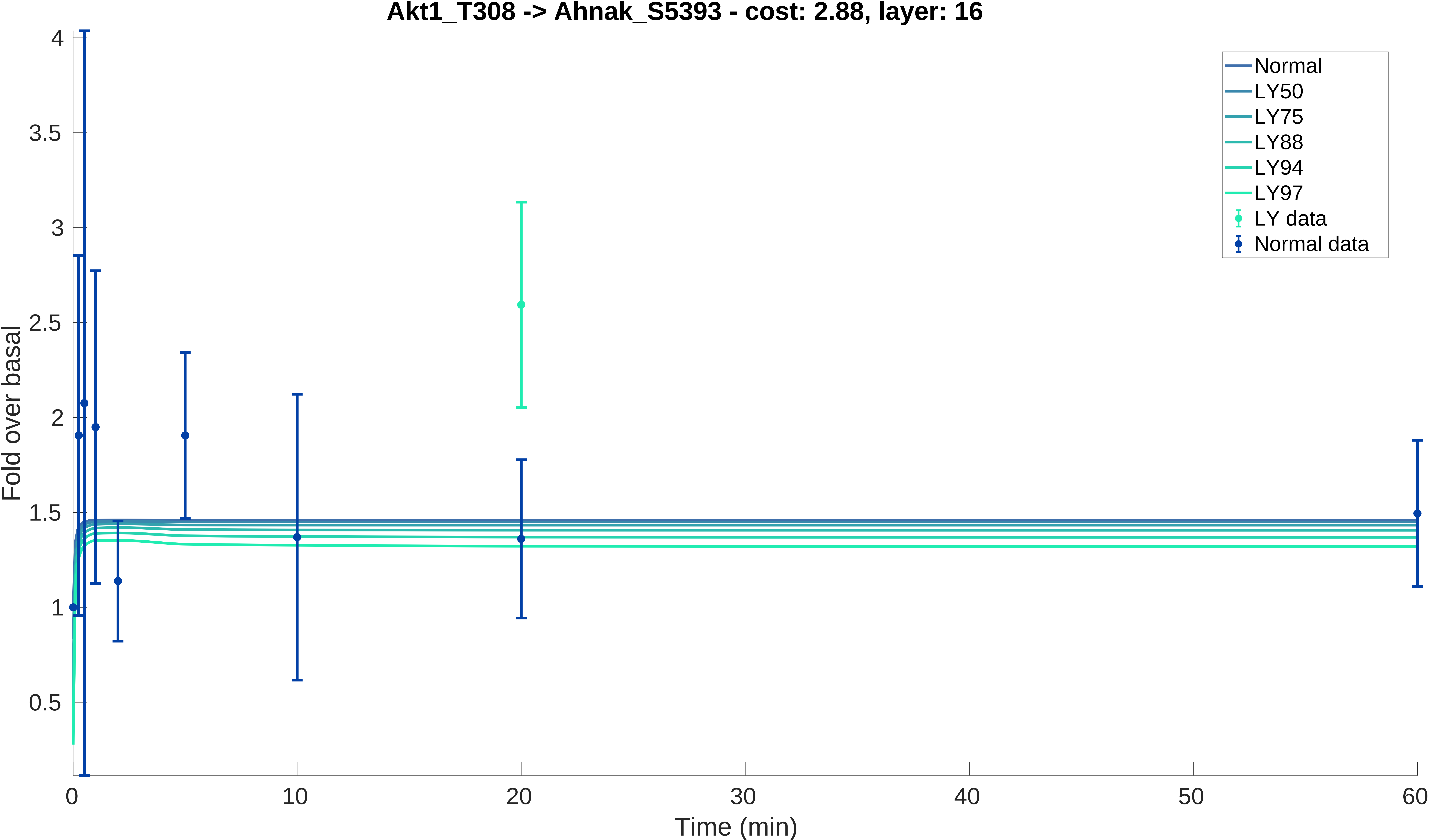

Akt1_T308 -> Ahnak_S5393 - cost: 2.88, layer: 16
Fold over basal
Time (min)
Normal
LY50
LY75
LY88
LY94
LY97
LY data
Normal data
0
10
20
30
40
50
60
0.5
1
1.5
2
2.5
3
3.5
4

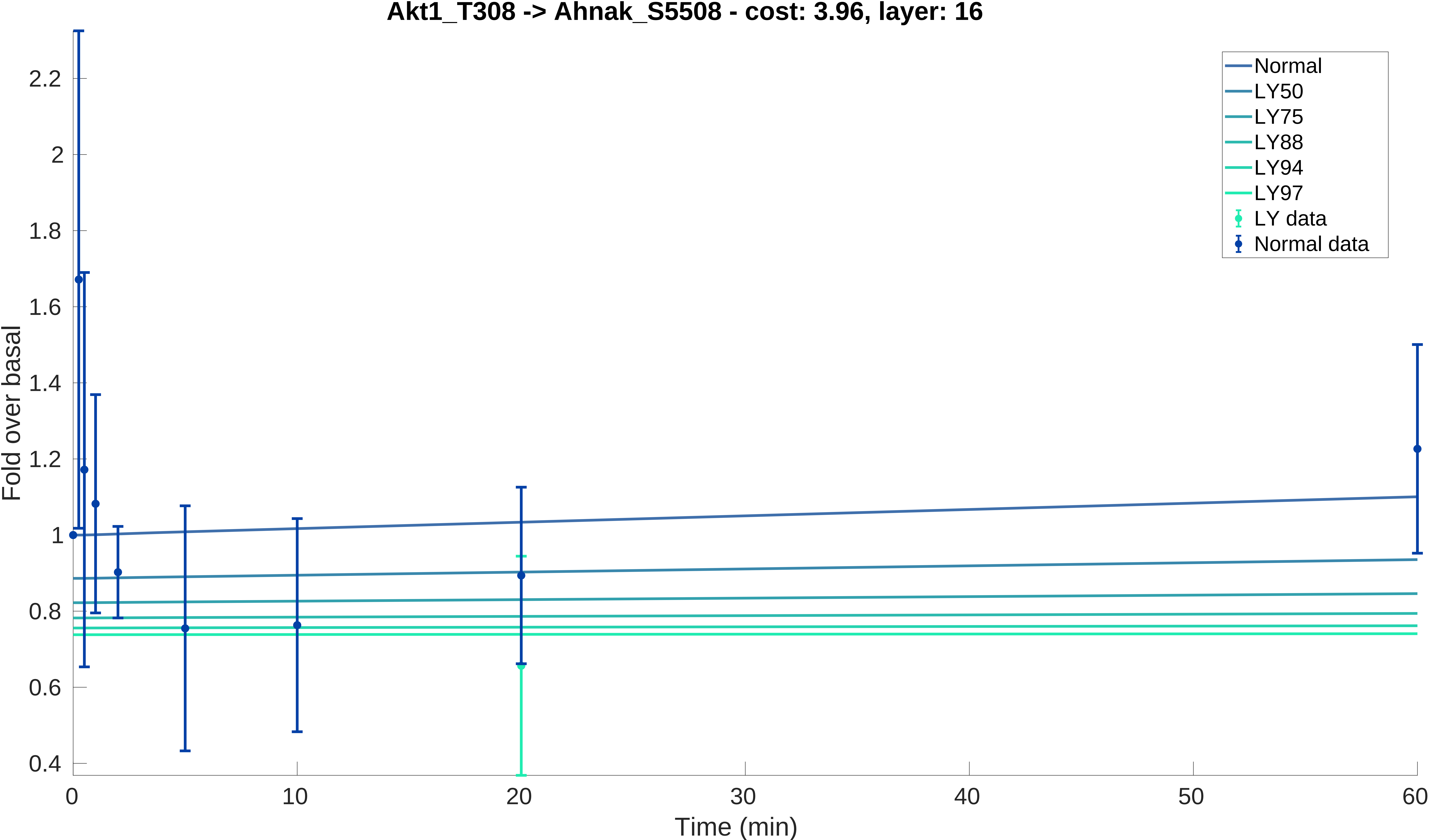

Akt1_T308 -> Ahnak_S5508 - cost: 3.96, layer: 16
Fold over basal
Time (min)
Normal
LY50
LY75
LY88
LY94
LY97
LY data
Normal data
0.4
0.6
0.8
1
1.2
1.4
1.6
1.8
2
2.2
0
10
20
30
40
50
60

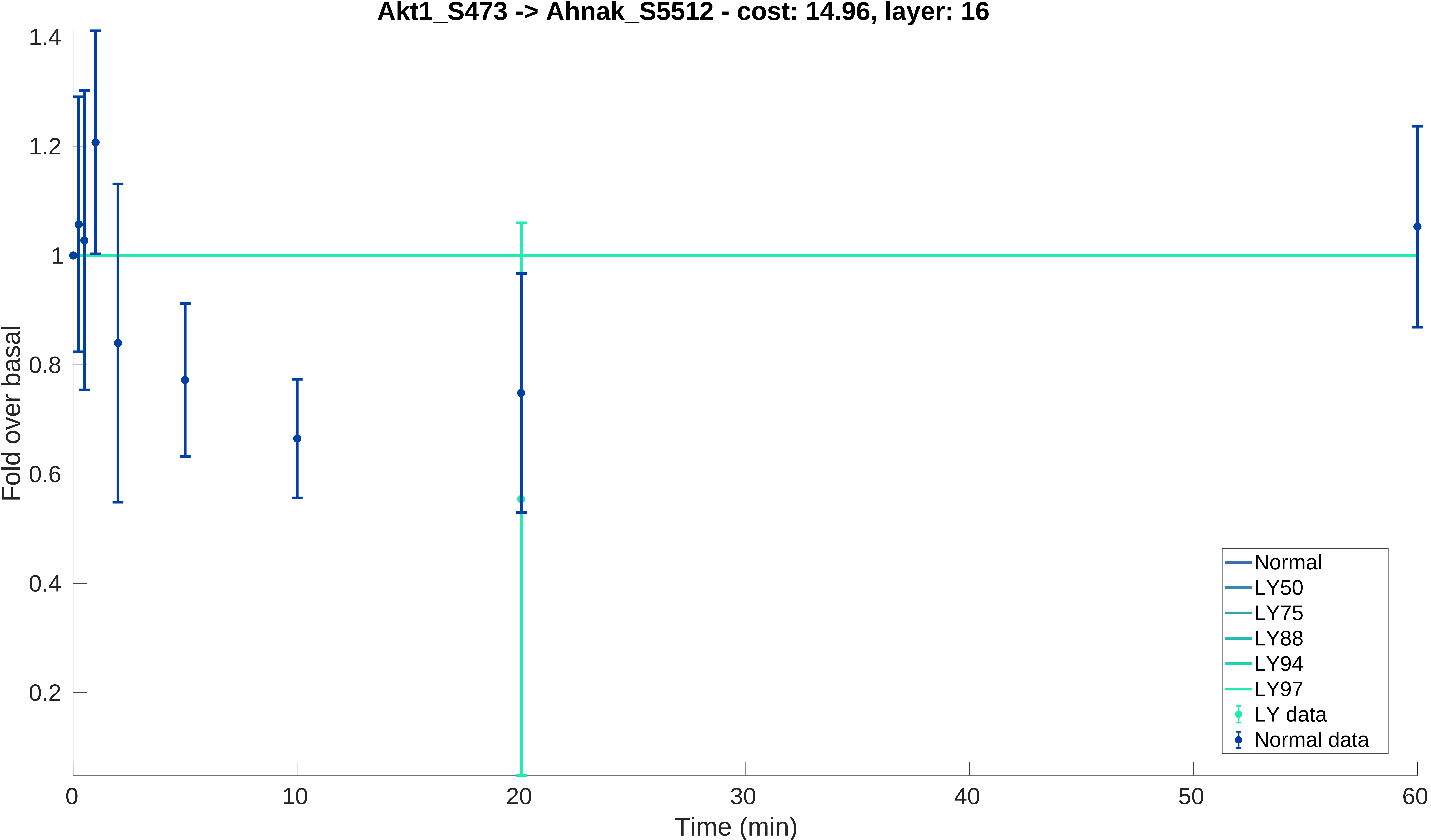

Akt1_S473 -> Ahnak_S5512 - cost: 14.96, layer: 16
Fold over basal
Time (min)
0.2
0.4
0.6
0.8
1
1.2
1.4
0
10
20
30
40
50
60
Normal
LY50
LY75
LY88
LY94
LY97
LY data
Normal data

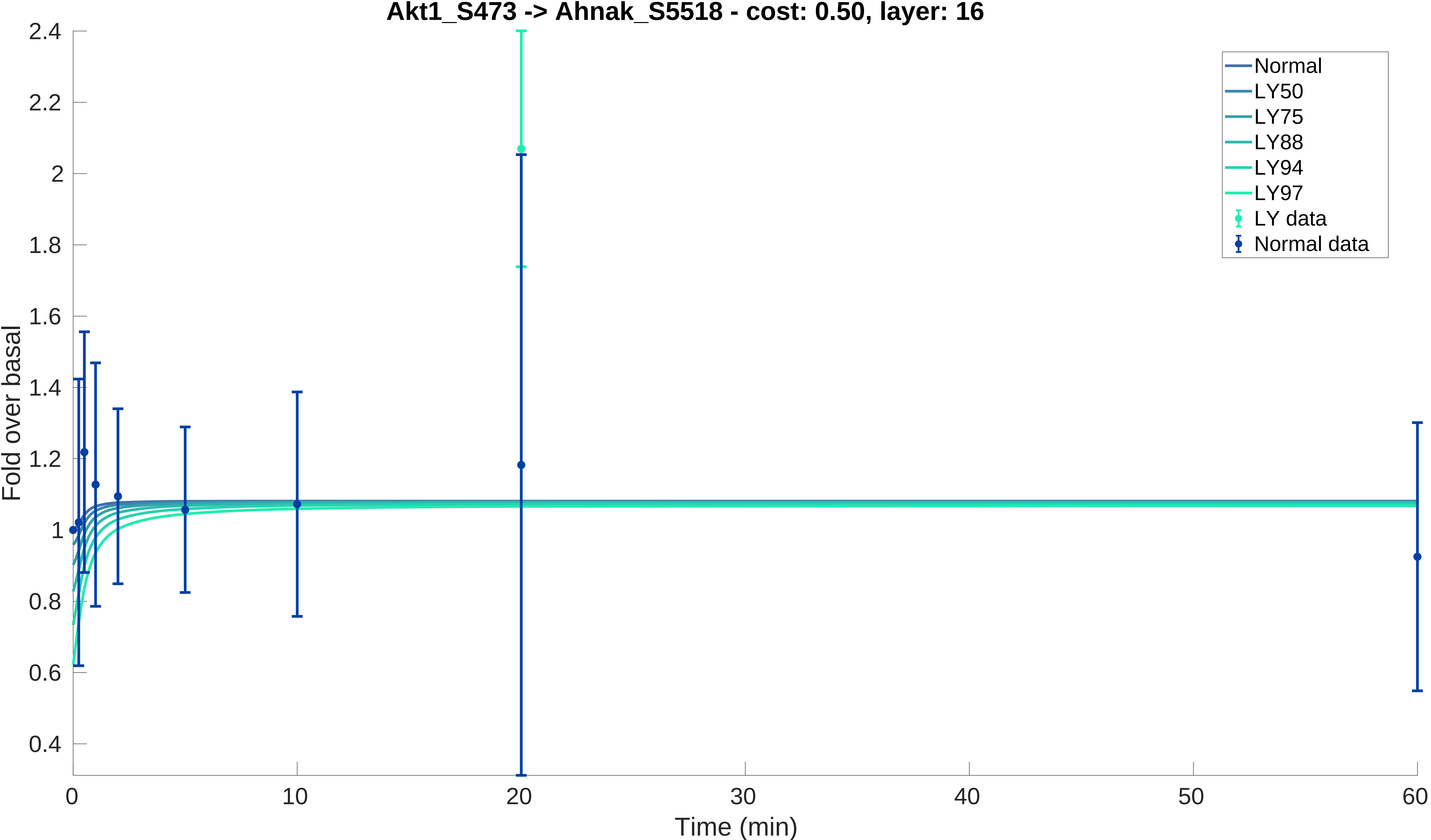

Akt1_S473 -> Ahnak_S5518 - cost: 0.50, layer: 16
Fold over basal
Time (min)
2.4
2.2
2
1.8
1.6
1.4
1.2
1
0.8
0.6
0.4
0
10
20
30
40
50
60
Normal
LY50
LY75
LY88
LY94
LY97
LY data
Normal data

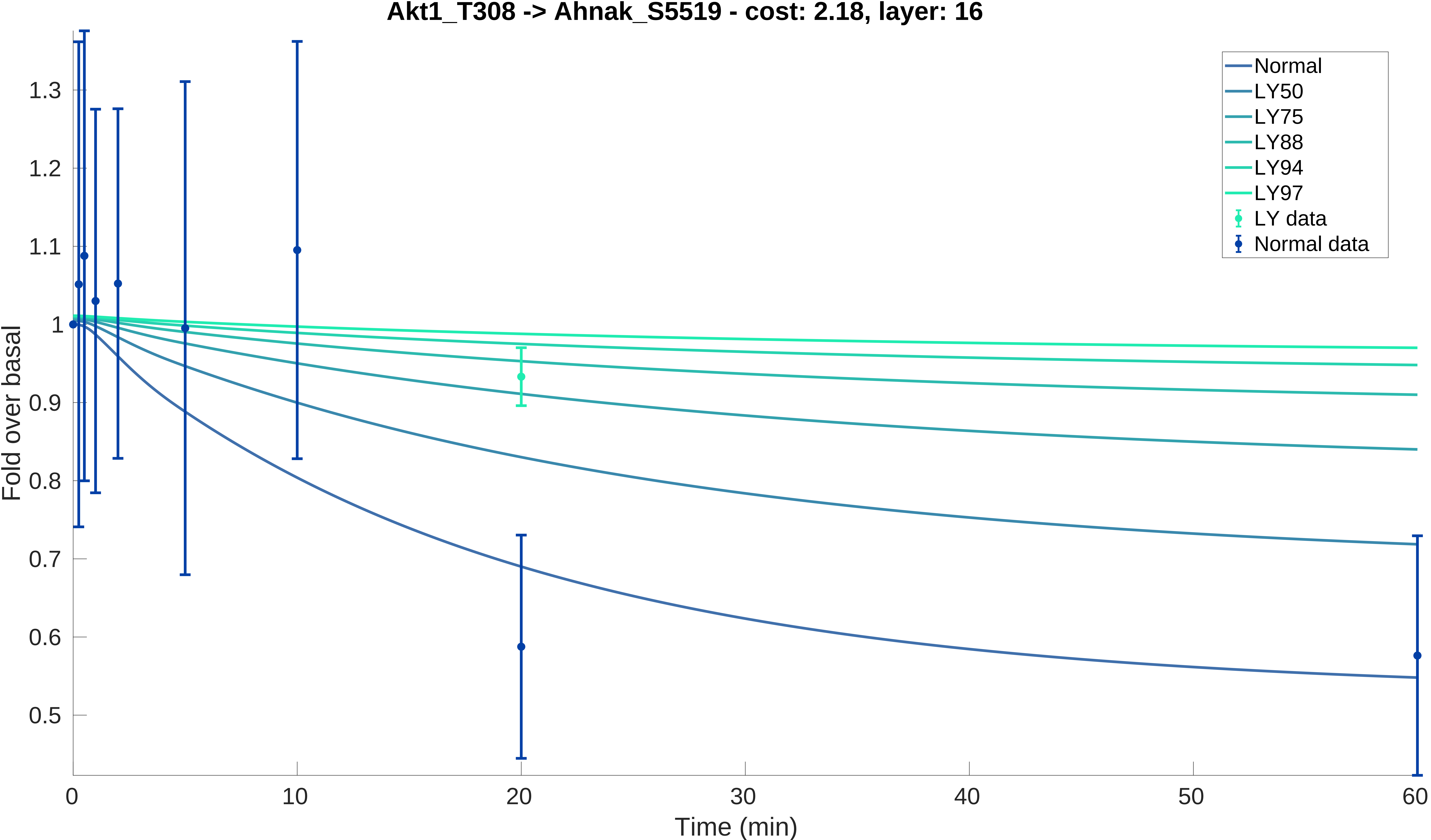
Akt1_T308 -> Ahnak_S5519 - cost: 2.18, layer: 16
Normal
LY50
LY75
LY88
LY94
LY97
LY data
Normal data
Fold over basal
1.3
1.2
1.1
1
0.9
0.8
0.7
0.6
0.5
0
10
20
30
40
50
60
Time (min)

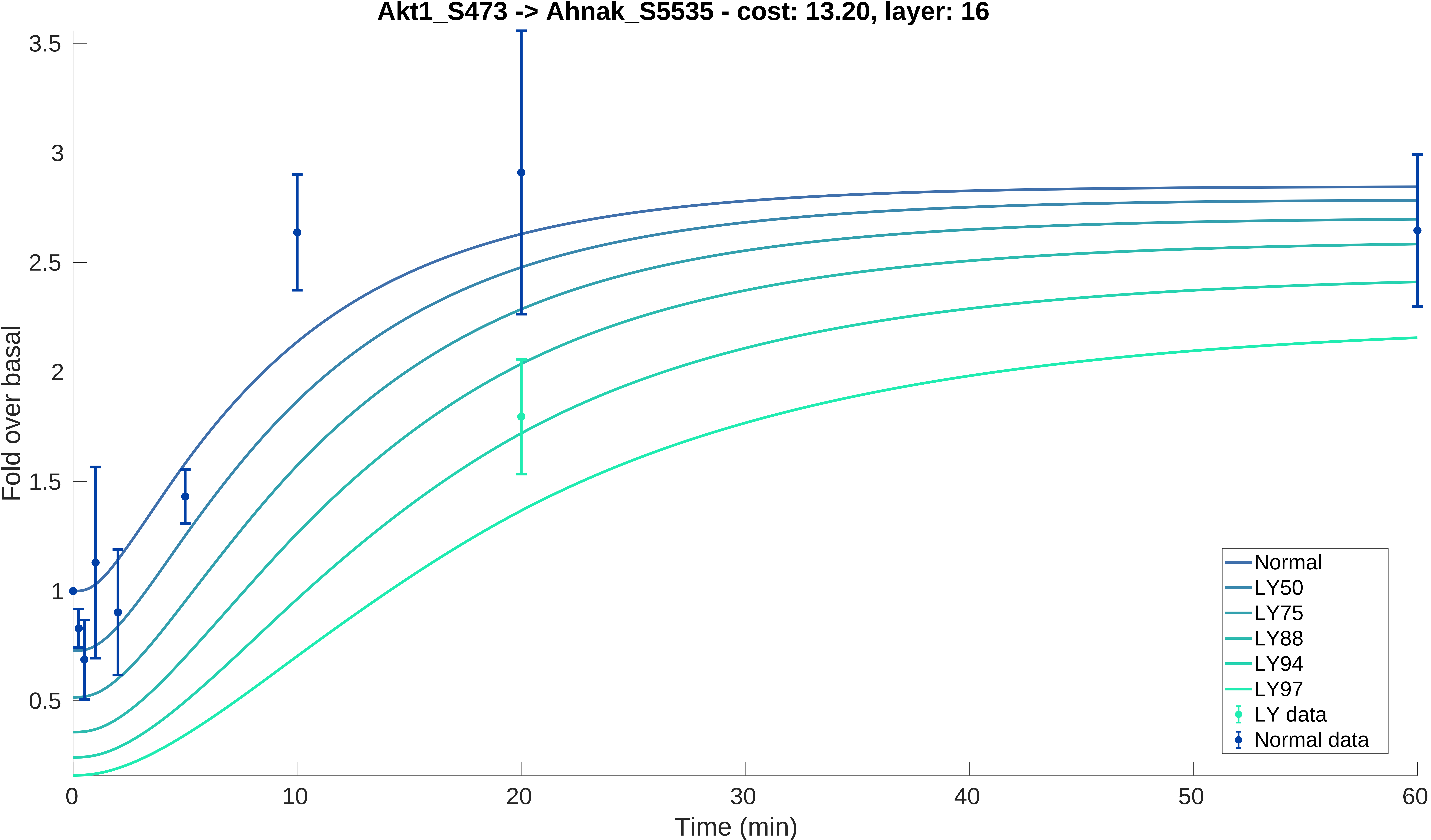

Akt1_S473 -> Ahnak_S5535 - cost: 13.20, layer: 16
Fold over basal
Time (min)
0.5
1
1.5
2
2.5
3
3.5
0
10
20
30
40
50
60
Normal
LY50
LY75
LY88
LY94
LY97
LY data
Normal data

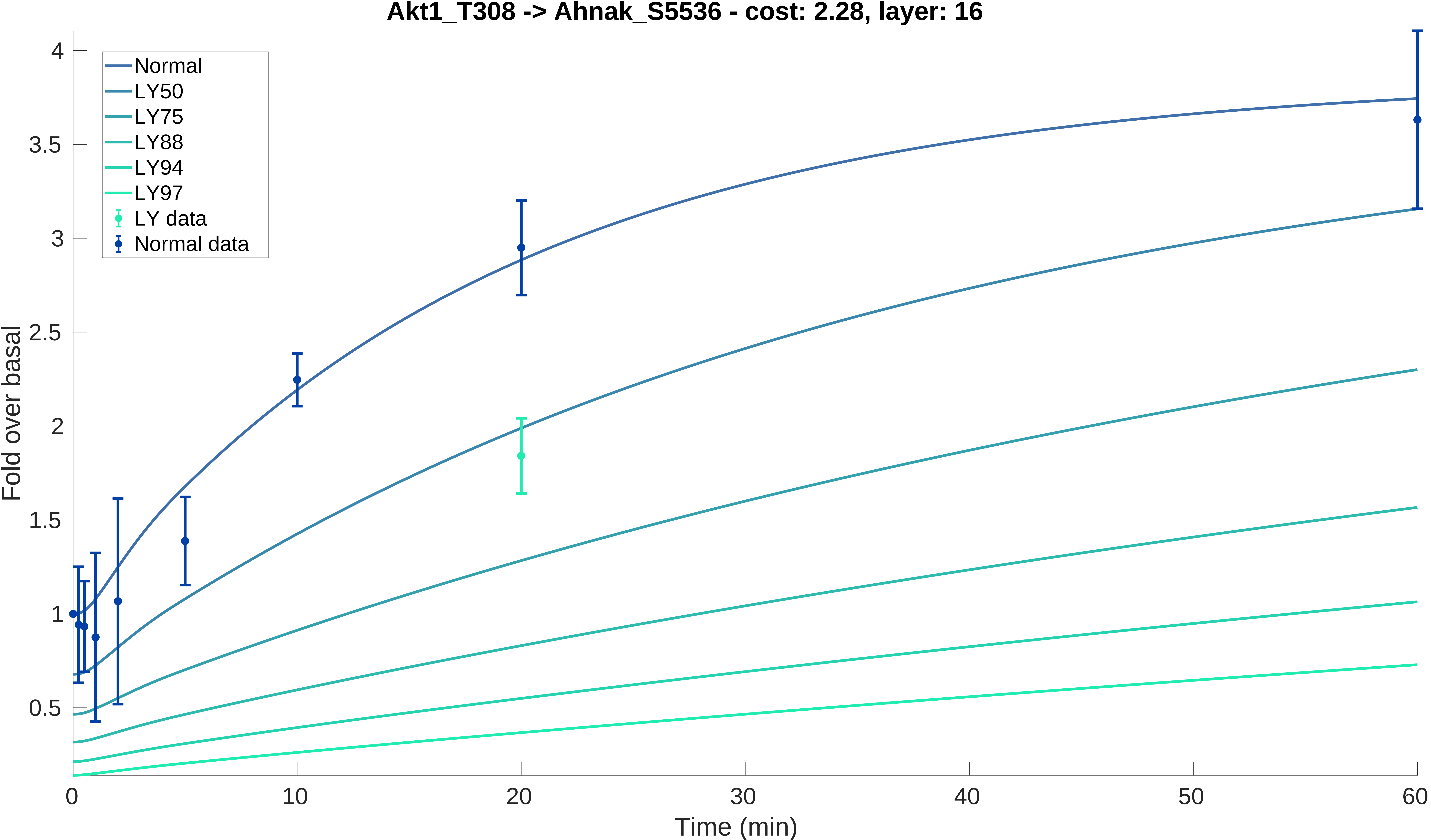

Akt1_T308 -> Ahnak_S5536 - cost: 2.28, layer: 16
Normal
LY50
LY75
LY88
LY94
LY97
LY data
Normal data
Fold over basal
Time (min)
0
10
20
30
40
50
60
0.5
1
1.5
2
2.5
3
3.5
4

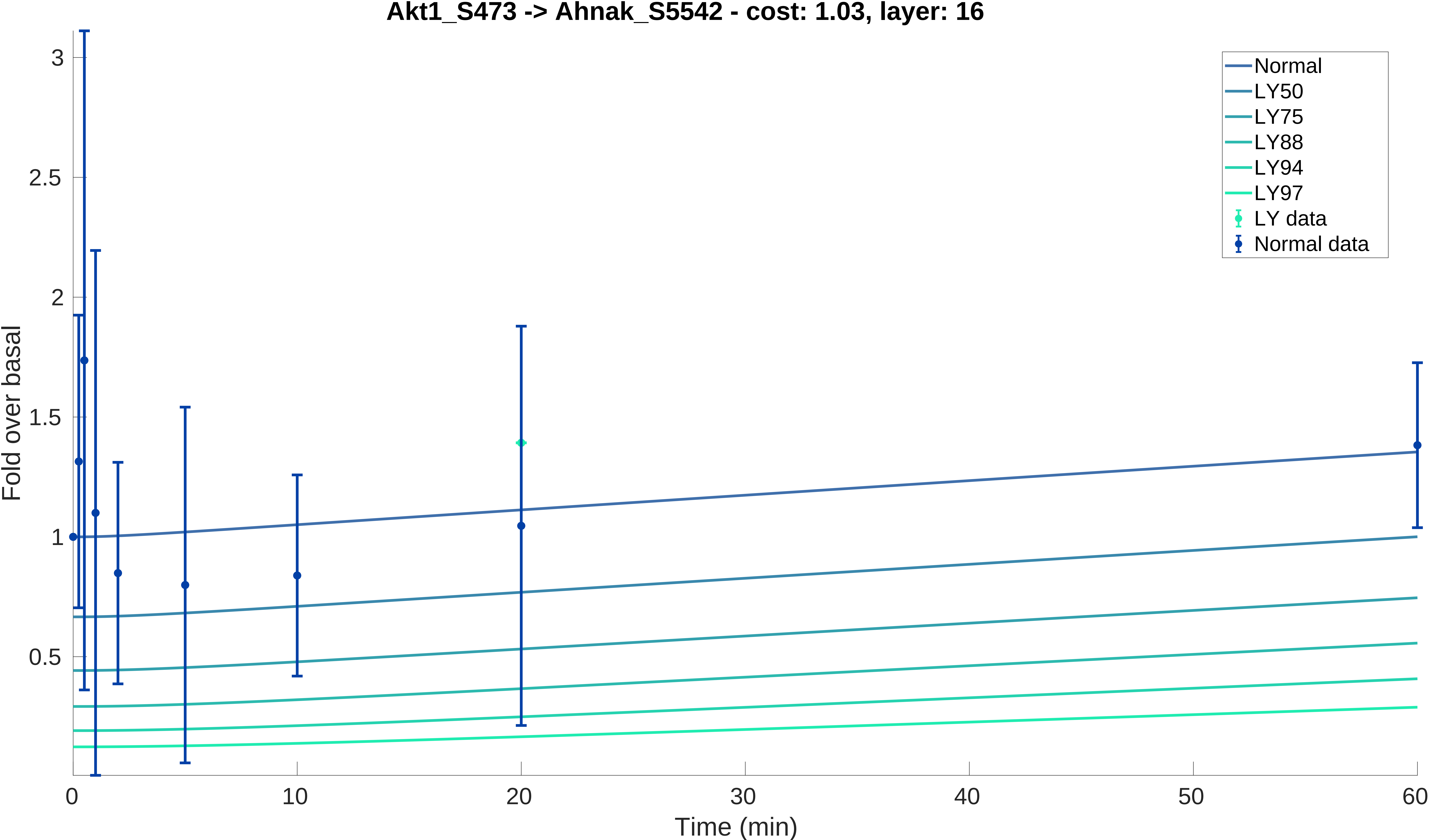

Akt1_S473 -> Ahnak_S5542 - cost: 1.03, layer: 16
Fold over basal
Time (min)
Normal
LY50
LY75
LY88
LY94
LY97
LY data
Normal data
0
0.5
1
1.5
2
2.5
3
10
20
30
40
50
60

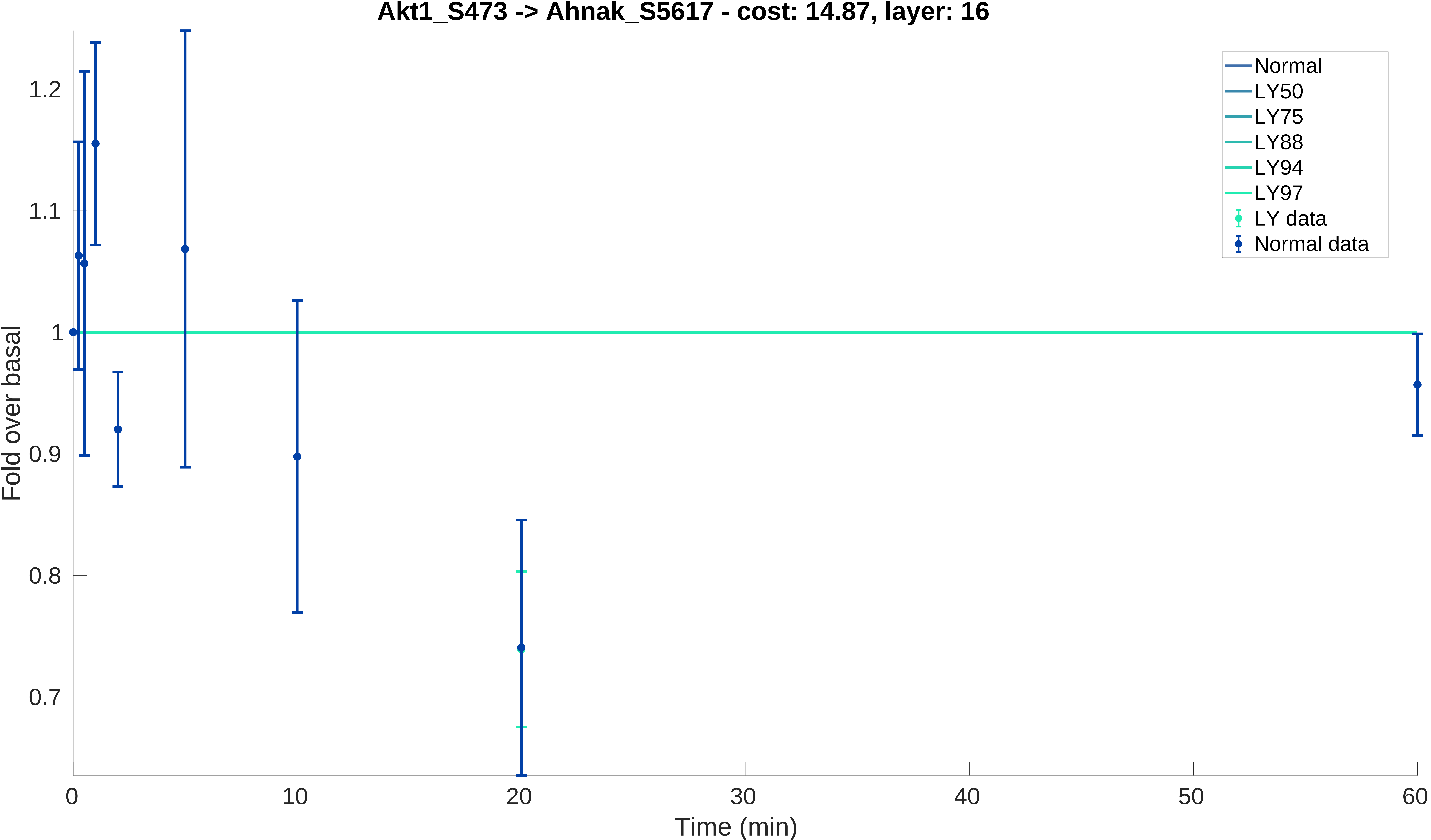
Akt1_S473 -> Ahnak_S5617 - cost: 14.87, layer: 16
Fold over basal
Time (min)
0.7
0.8
0.9
1
1.1
1.2
0
10
20
30
40
50
60
Normal
LY50
LY75
LY88
LY94
LY97
LY data
Normal data

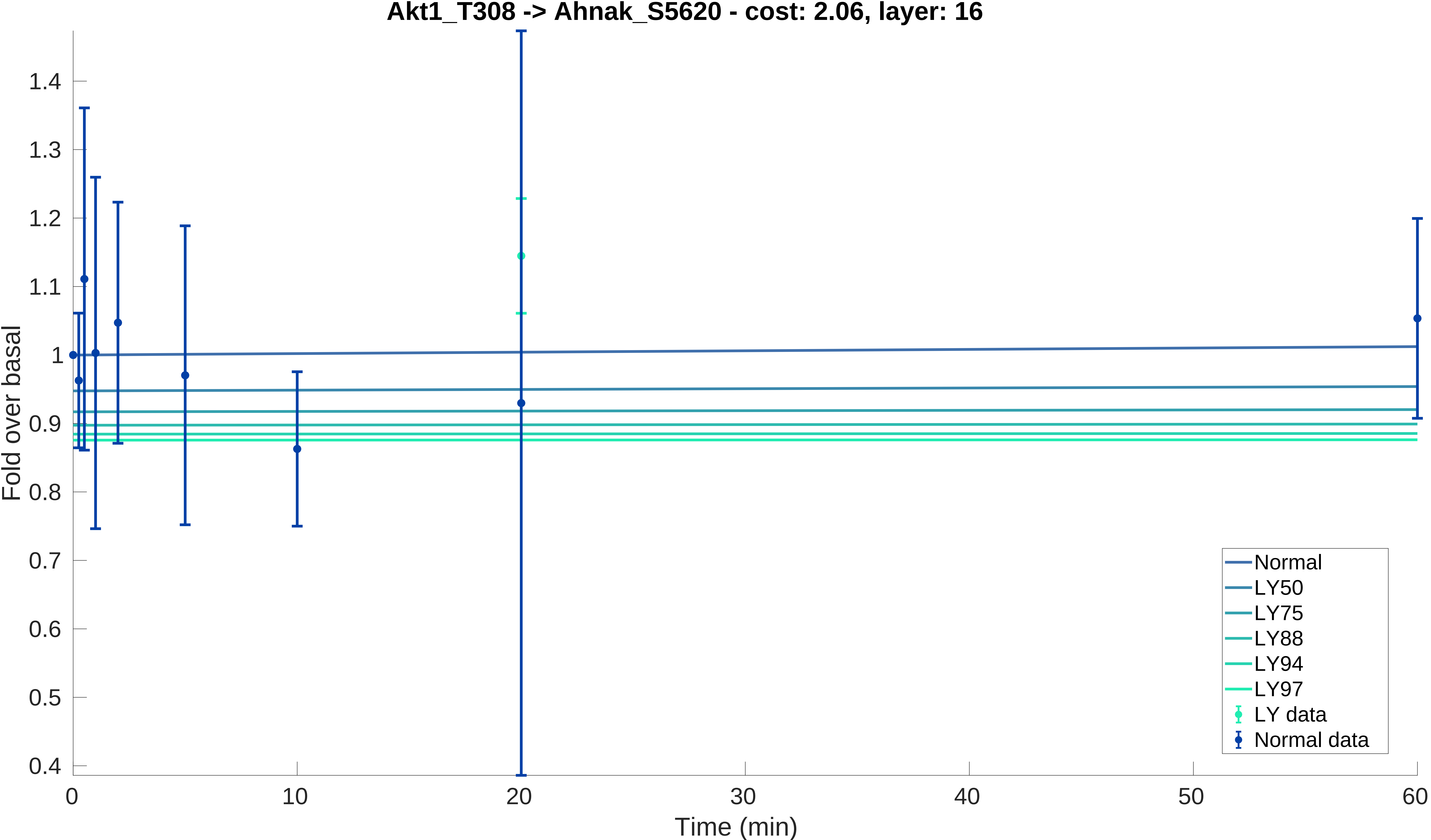

Akt1_T308 -> Ahnak_S5620 - cost: 2.06, layer: 16
Fold over basal
1.4
1.3
1.2
1.1
1
0.9
0.8
0.7
0.6
0.5
0.4
0
10
20
30
40
50
60
Time (min)
Normal
LY50
LY75
LY88
LY94
LY97
LY data
Normal data

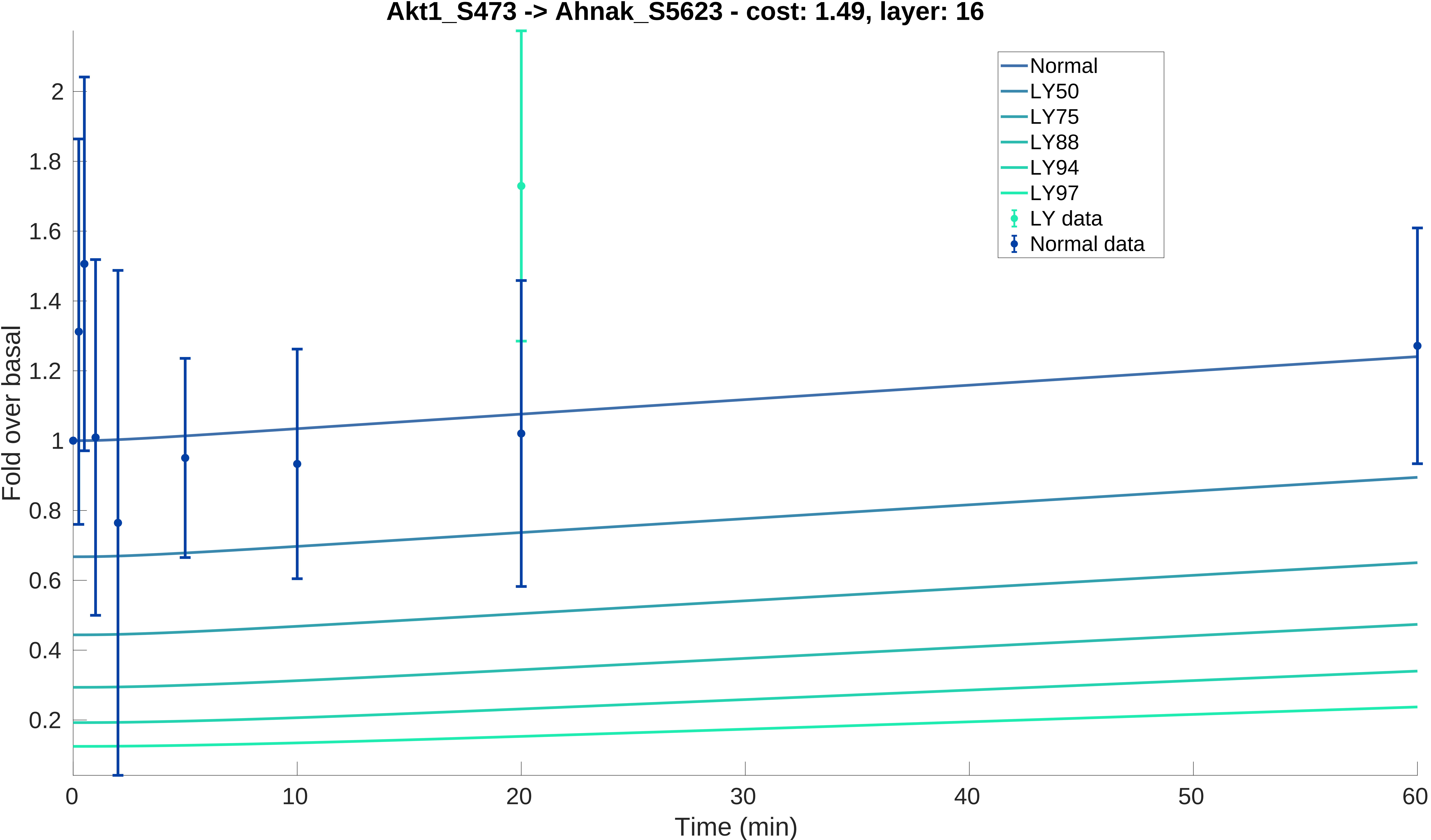

Akt1_S473 -> Ahnak_S5623 - cost: 1.49, layer: 16
Normal
LY50
LY75
LY88
LY94
LY97
LY data
Normal data
Fold over basal
Time (min)
0
10
20
30
40
50
60
0.2
0.4
0.6
0.8
1
1.2
1.4
1.6
1.8
2

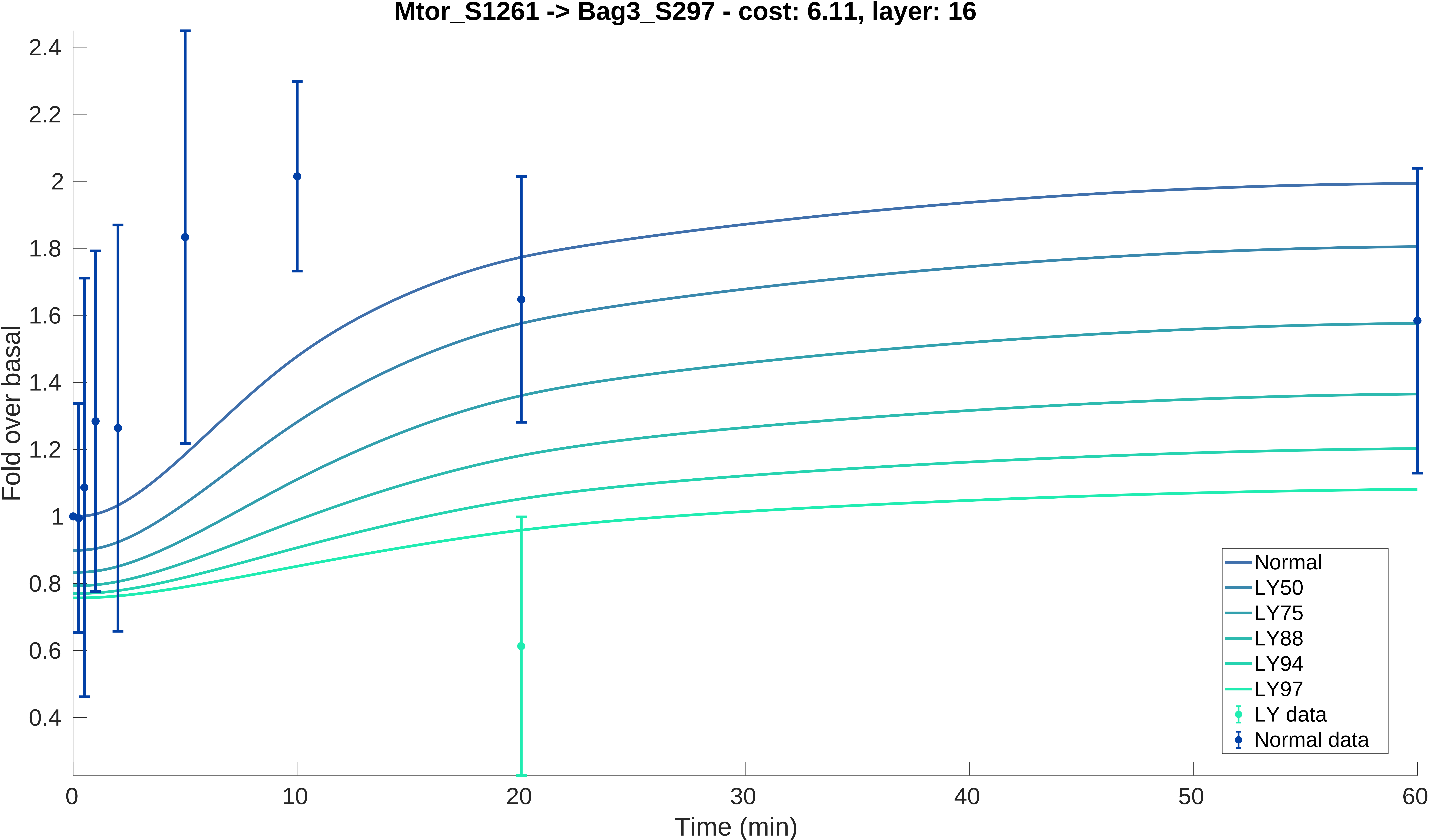

Mtor_S1261 -> Bag3_S297 - cost: 6.11, layer: 16
Fold over basal
2.4
2.2
2
1.8
1.6
1.4
1.2
1
0.8
0.6
0.4
0
10
20
30
40
50
60
Time (min)
Normal
LY50
LY75
LY88
LY94
LY97
LY data
Normal data

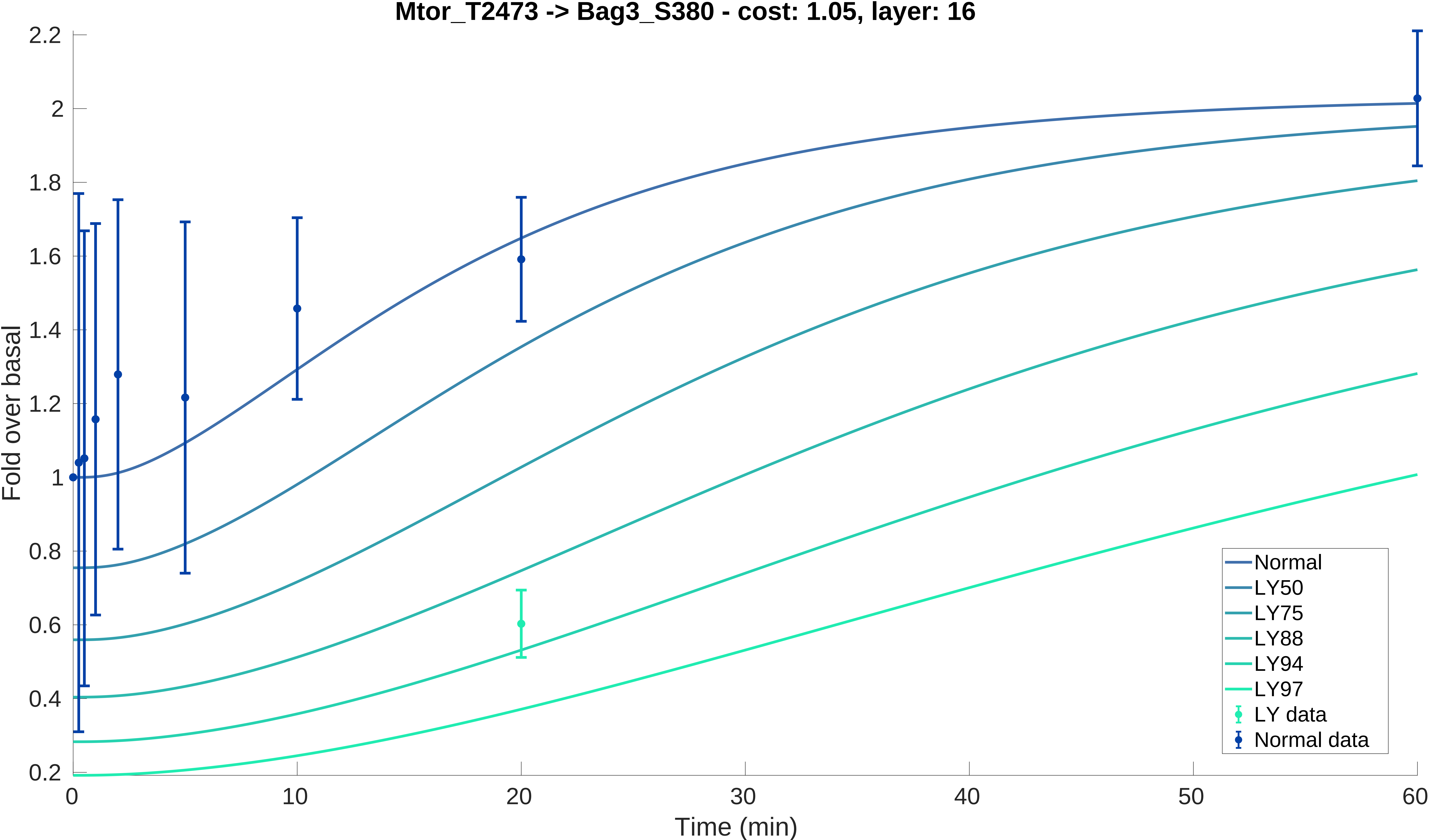

Mtor_T2473 -> Bag3_S380 - cost: 1.05, layer: 16
Fold over basal
Time (min)
0.2
0.4
0.6
0.8
1
1.2
1.4
1.6
1.8
2
2.2
0
10
20
30
40
50
60
Normal
LY50
LY75
LY88
LY94
LY97
LY data
Normal data

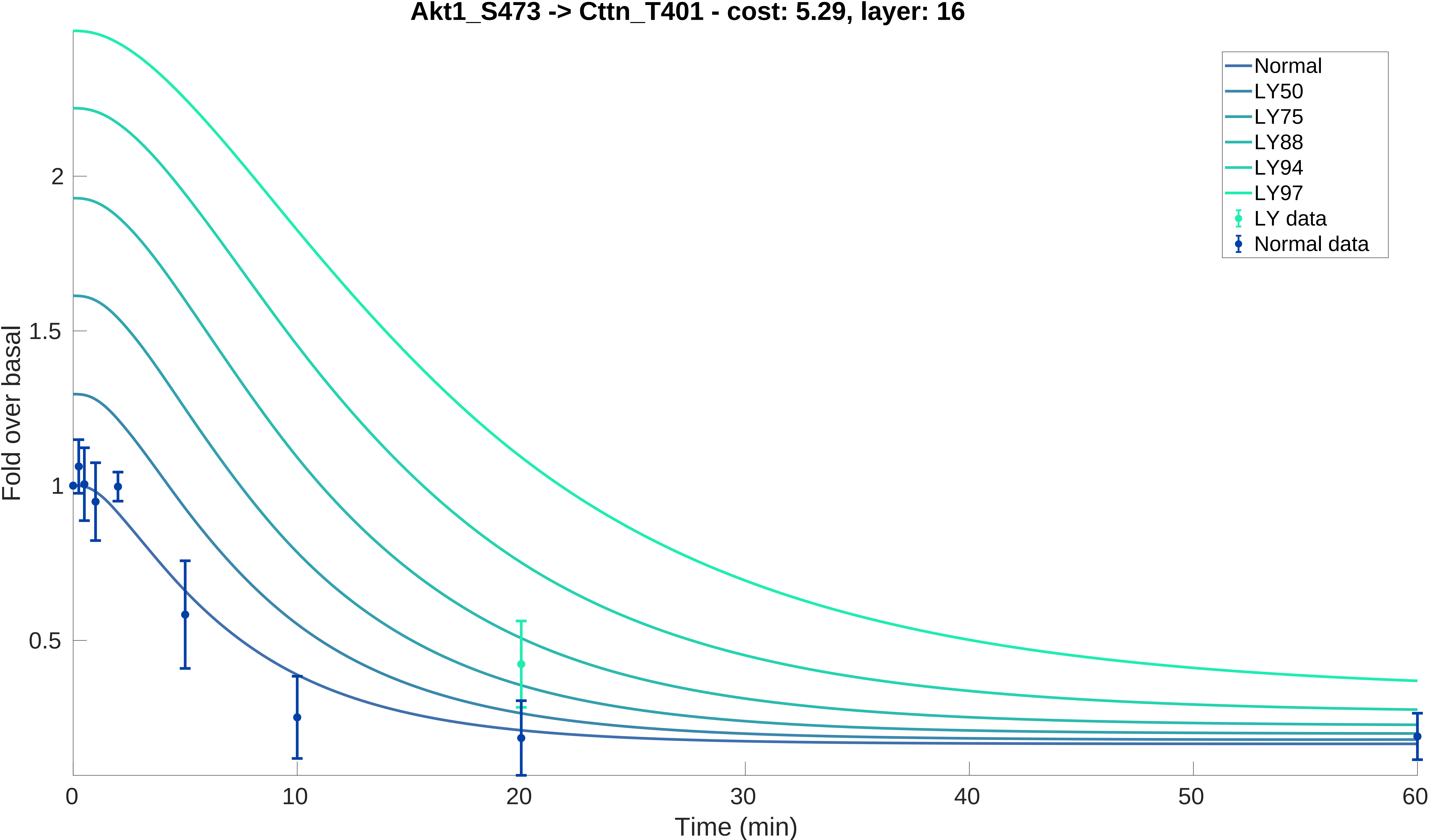

Akt1_S473 -> Cttn_T401 - cost: 5.29, layer: 16
Normal
LY50
LY75
LY88
LY94
LY97
LY data
Normal data
Fold over basal
2
1.5
1
0.5
0
10
20
30
40
50
60
Time (min)

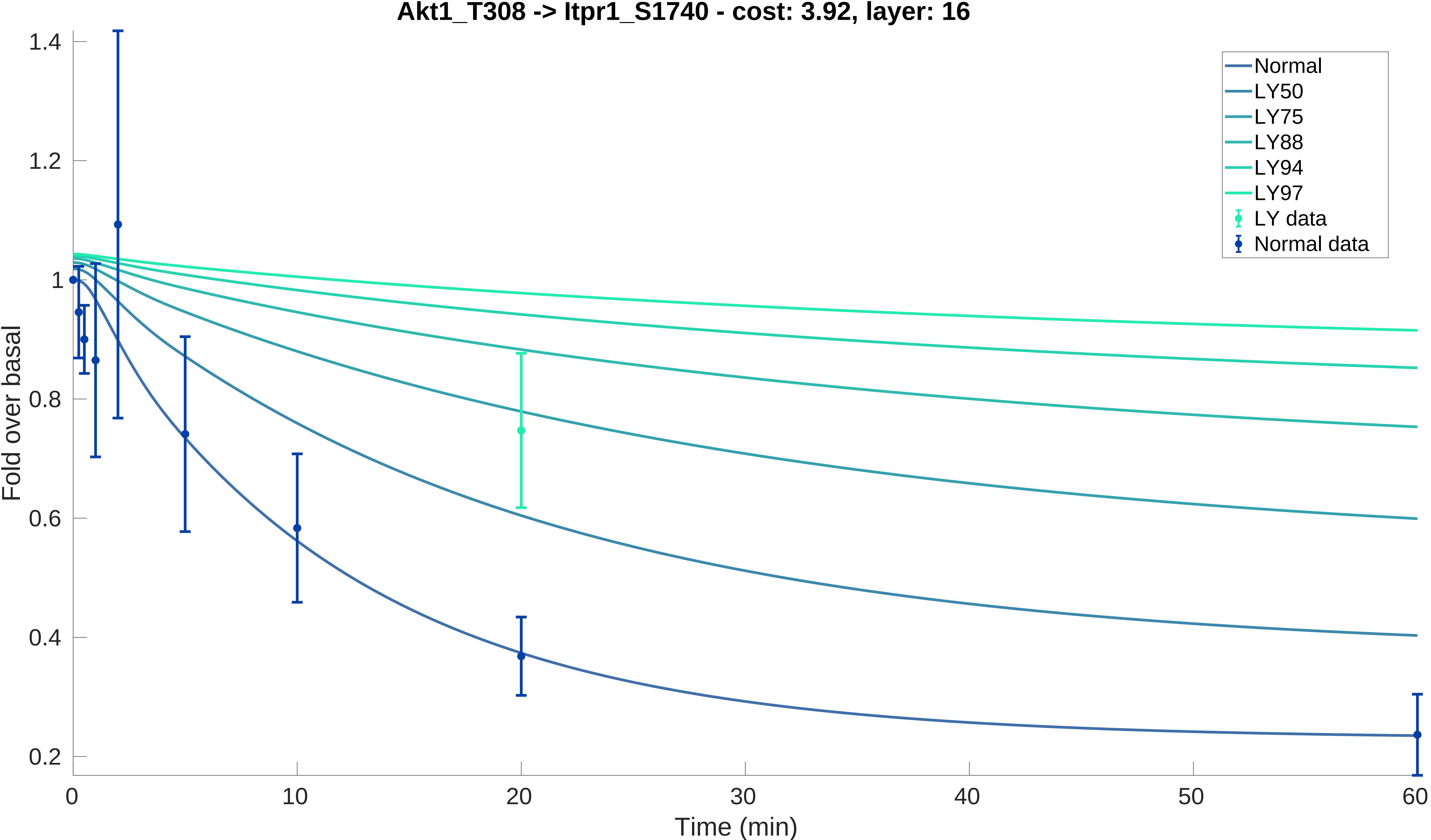

Akt1_T308 -> Itpr1_S1740 - cost: 3.92, layer: 16
Fold over basal
Time (min)
0.2
0.4
0.6
0.8
1
1.2
1.4
0
10
20
30
40
50
60
Normal
LY50
LY75
LY88
LY94
LY97
LY data
Normal data

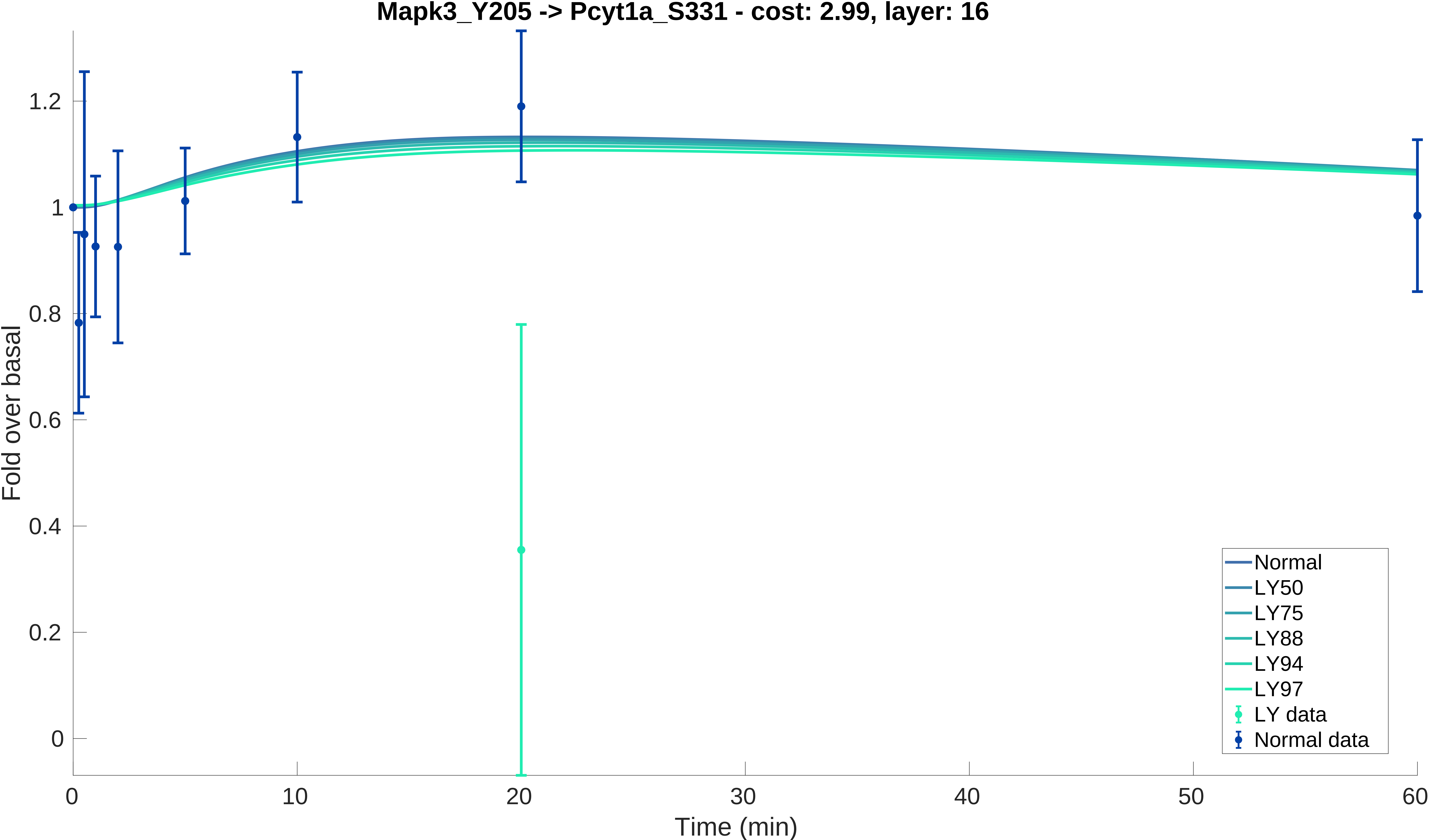

Mapk3_Y205 -> Pcyt1a_S331 - cost: 2.99, layer: 16
Fold over basal
Time (min)
0
0.2
0.4
0.6
0.8
1
1.2
0
10
20
30
40
50
60
Normal
LY50
LY75
LY88
LY94
LY97
LY data
Normal data

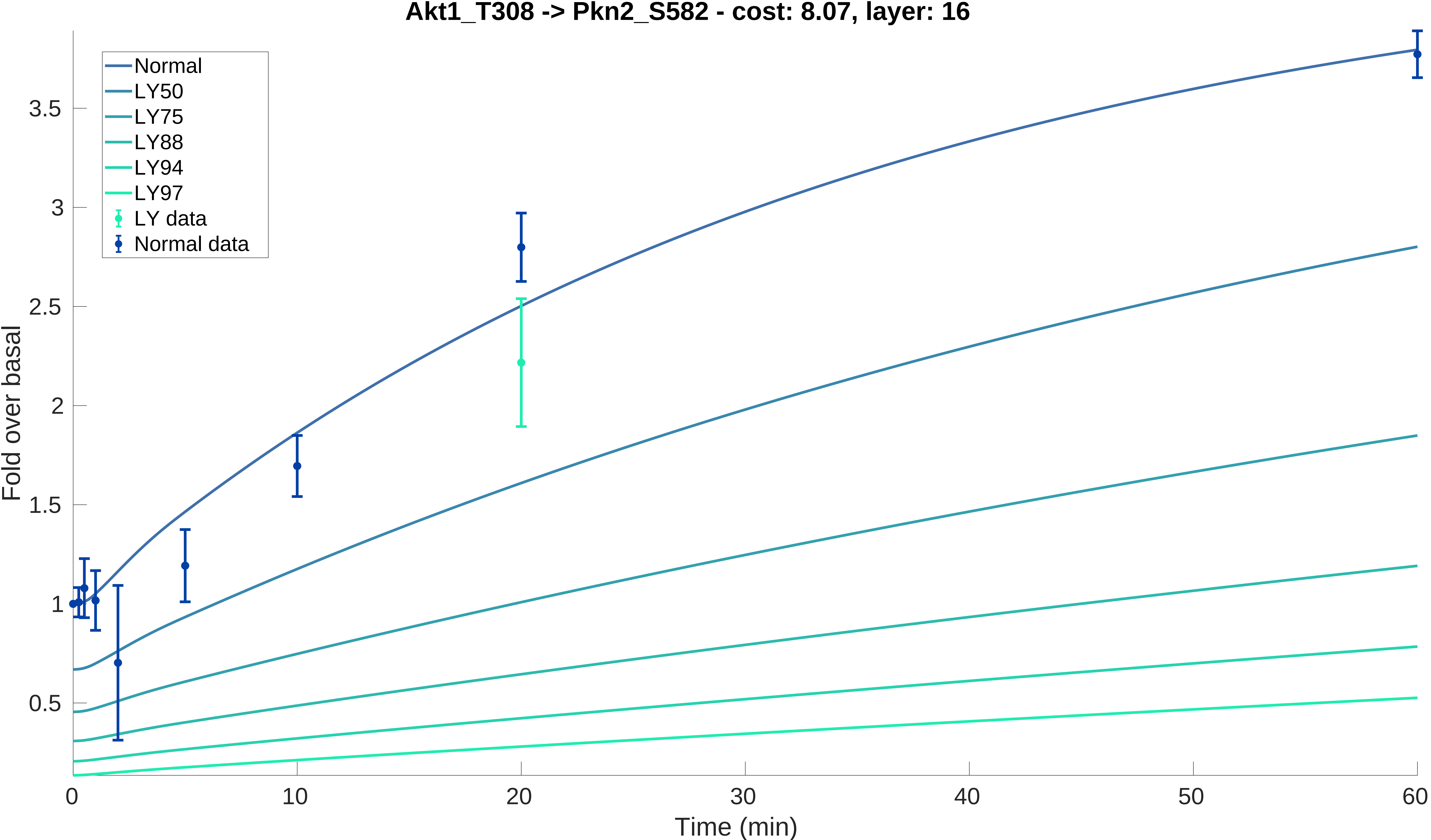

Akt1_T308 -> Pkn2_S582 - cost: 8.07, layer: 16
Normal
LY50
LY75
LY88
LY94
LY97
LY data
Normal data
Fold over basal
3.5
3
2.5
2
1.5
1
0.5
0
10
20
30
40
50
60
Time (min)

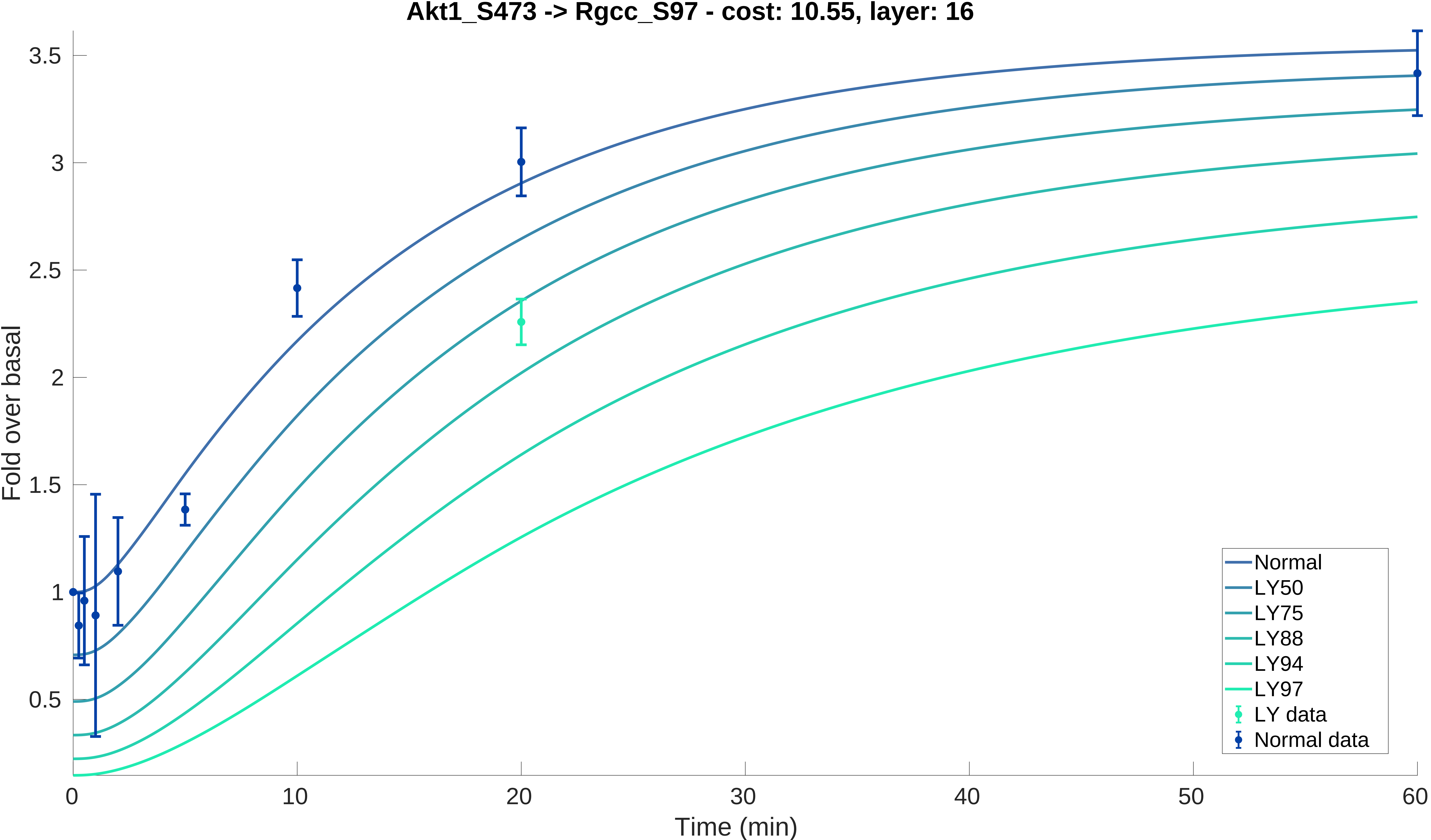

Akt1_S473 -> Rgcc_S97 - cost: 10.55, layer: 16
Fold over basal
0.5
1
1.5
2
2.5
3
3.5
0
10
20
30
40
50
60
Time (min)
Normal
LY50
LY75
LY88
LY94
LY97
LY data
Normal data

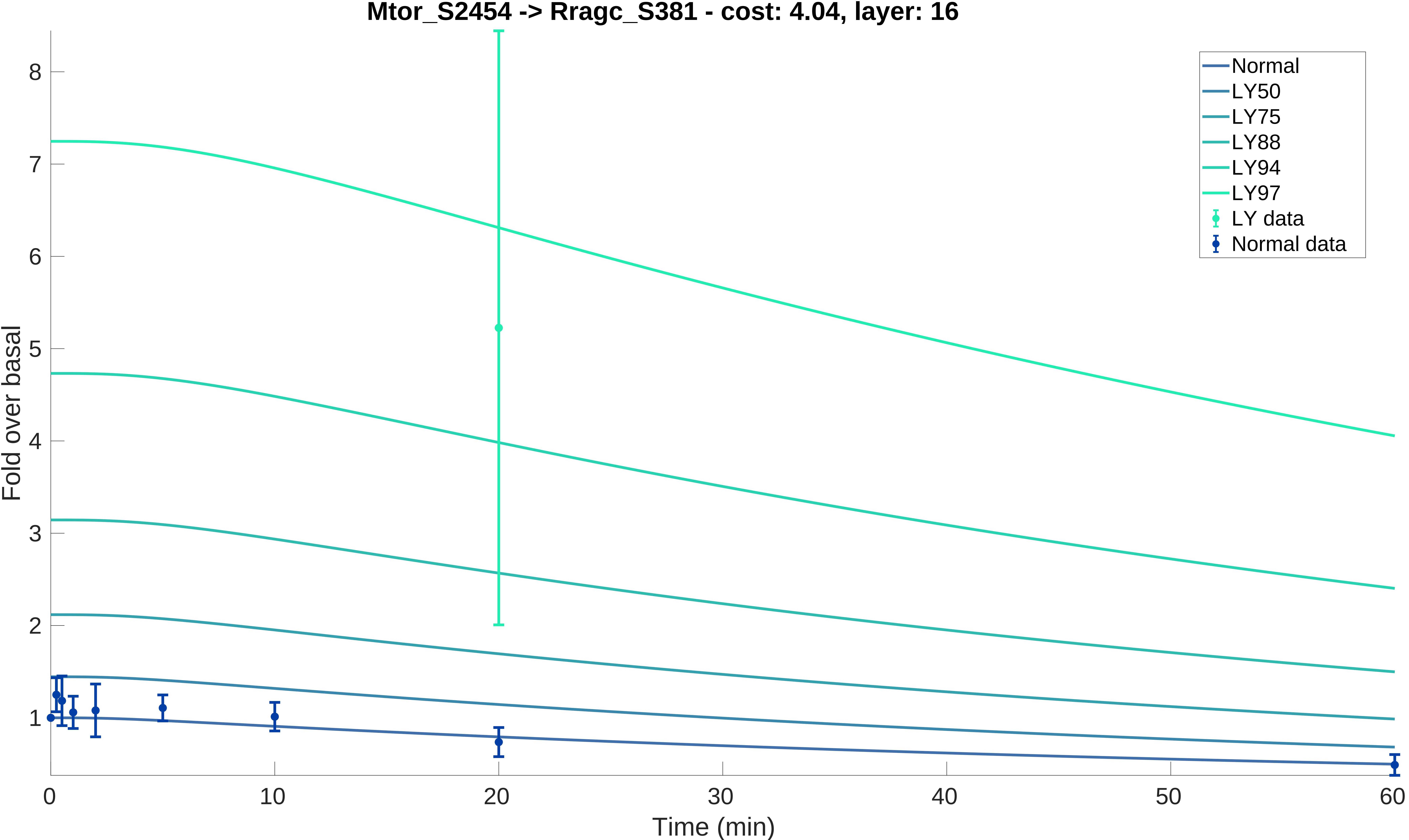
Mtor_S2454 -> Rragc_S381 - cost: 4.04, layer: 16
Fold over basal
Time (min)
0
10
20
30
40
50
60
1
2
3
4
5
6
7
8
Normal
LY50
LY75
LY88
LY94
LY97
LY data
Normal data

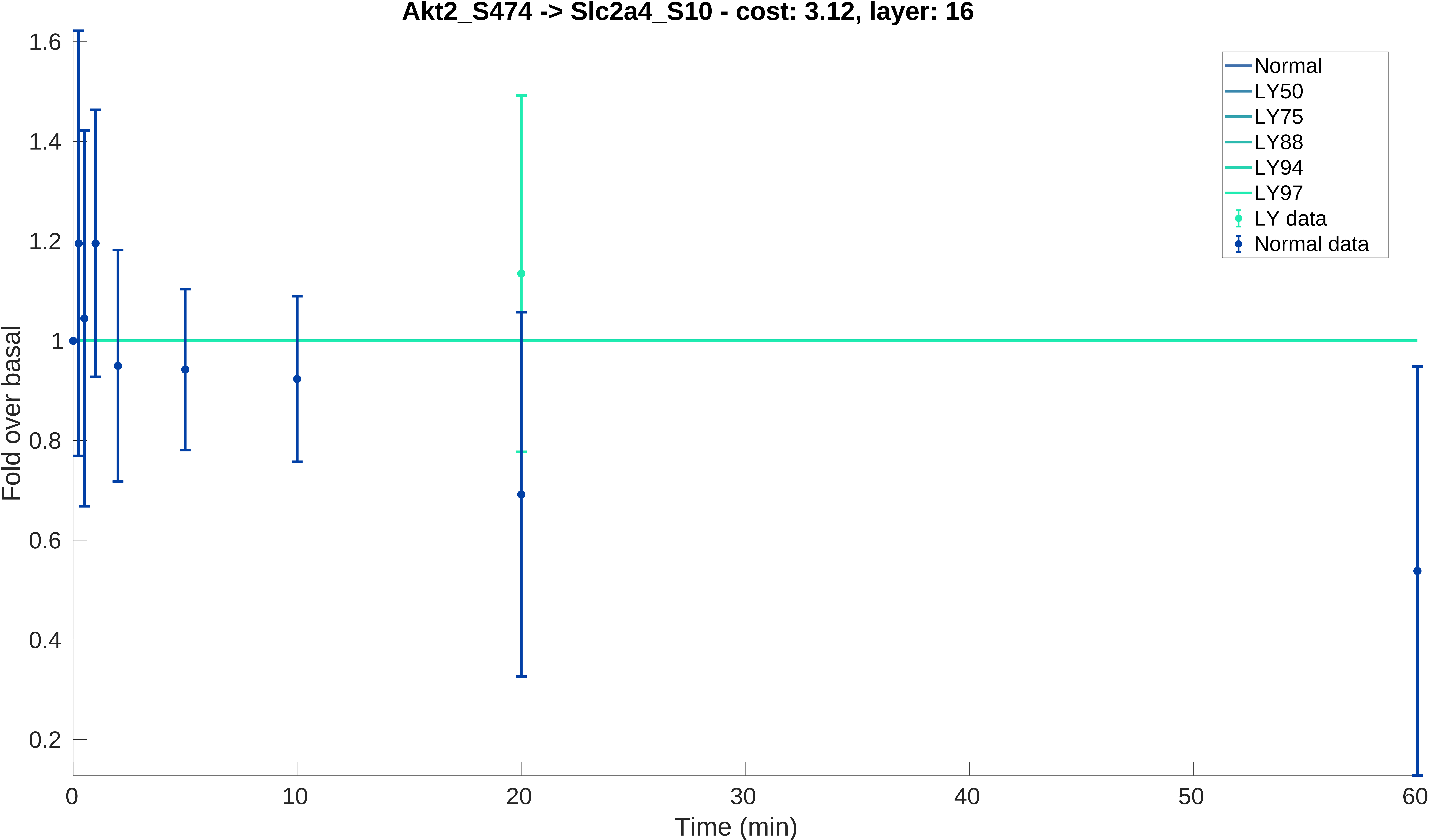

Akt2_S474 -> Slc2a4_S10 - cost: 3.12, layer: 16
Fold over basal
1.6
1.4
1.2
1
0.8
0.6
0.4
0.2
0
10
20
30
40
50
60
Time (min)
Normal
LY50
LY75
LY88
LY94
LY97
LY data
Normal data

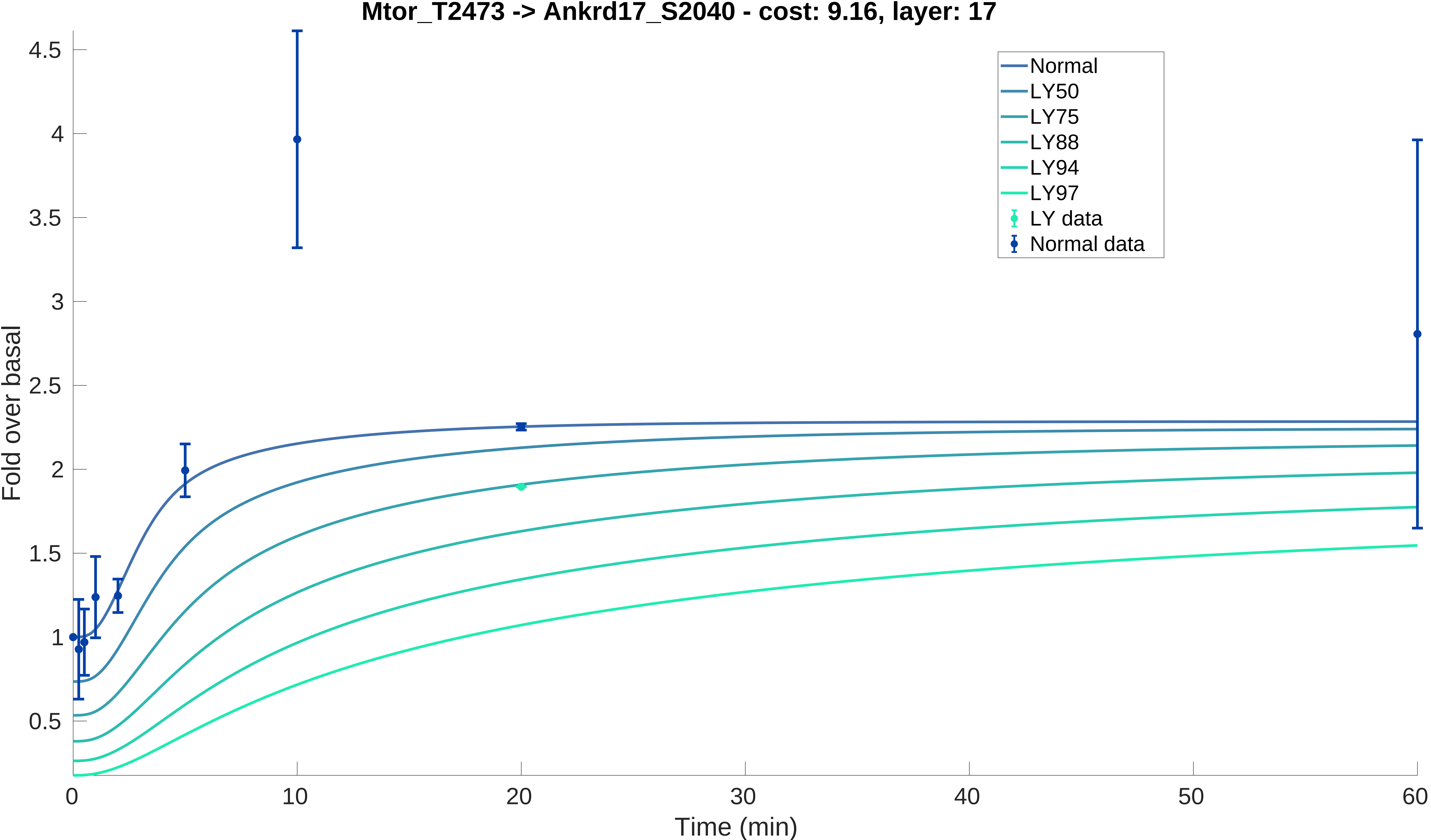

Mtor_T2473 -> Ankrd17_S2040 - cost: 9.16, layer: 17
Fold over basal
Time (min)
0
10
20
30
40
50
60
0.5
1
1.5
2
2.5
3
3.5
4
4.5
Normal
LY50
LY75
LY88
LY94
LY97
LY data
Normal data

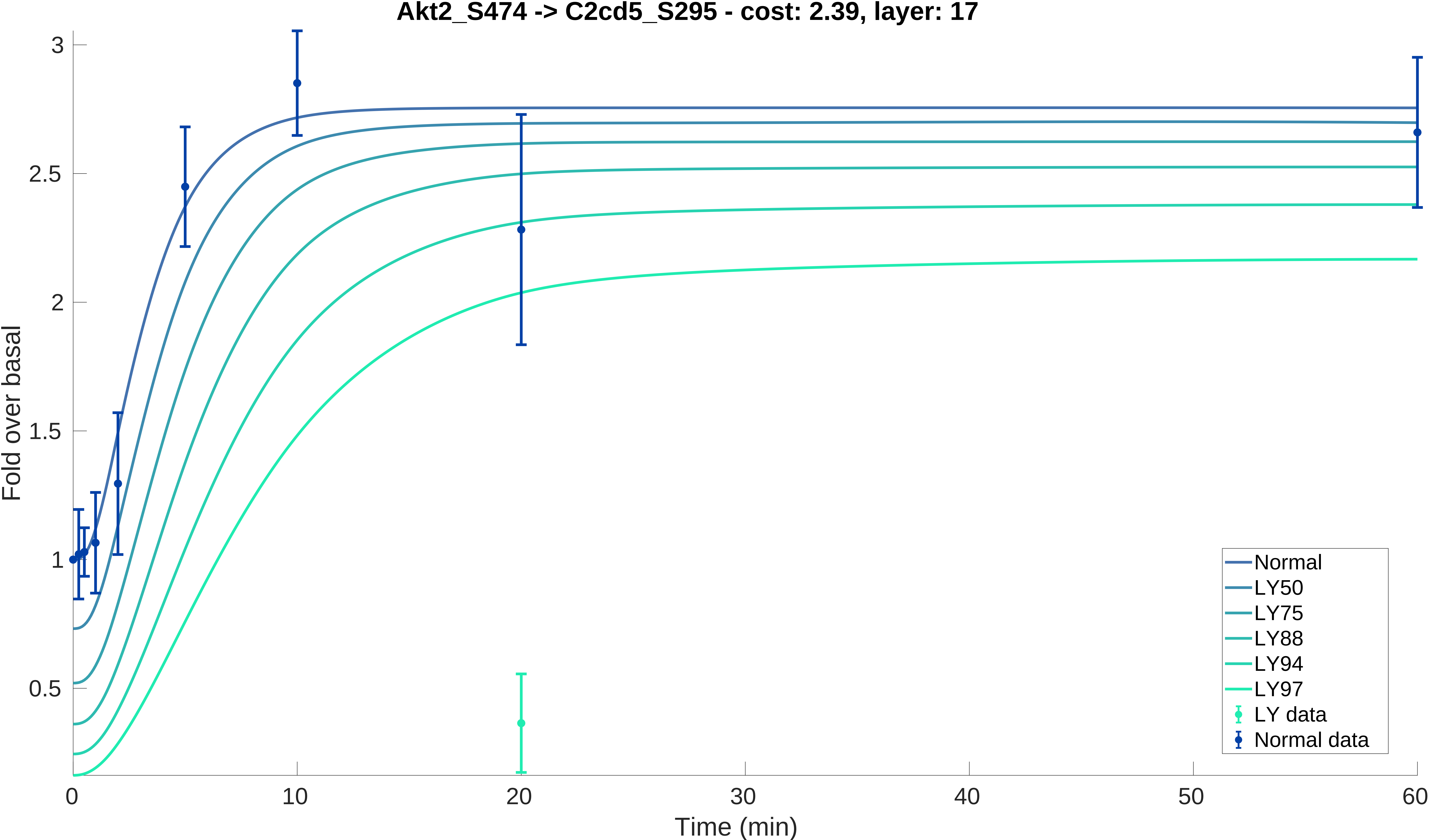
Akt2_S474 -> C2cd5_S295 - cost: 2.39, layer: 17
Fold over basal
Time (min)
0
10
20
30
40
50
60
0.5
1
1.5
2
2.5
3
Normal
LY50
LY75
LY88
LY94
LY97
LY data
Normal data

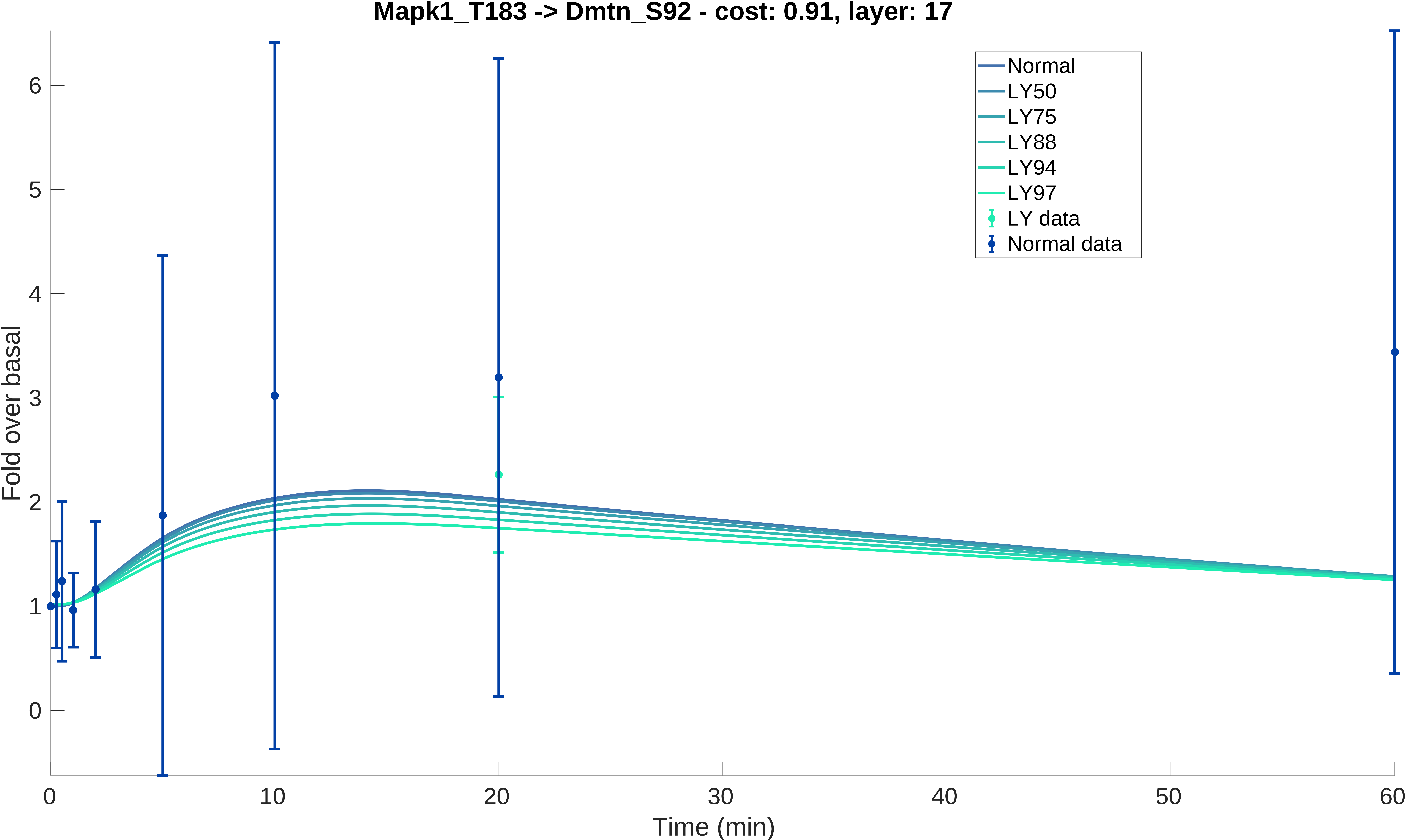
Mapk1_T183 -> Dmtn_S92 - cost: 0.91, layer: 17
Fold over basal
Time (min)
Normal
LY50
LY75
LY88
LY94
LY97
LY data
Normal data
0
1
2
3
4
5
6
0
10
20
30
40
50
60

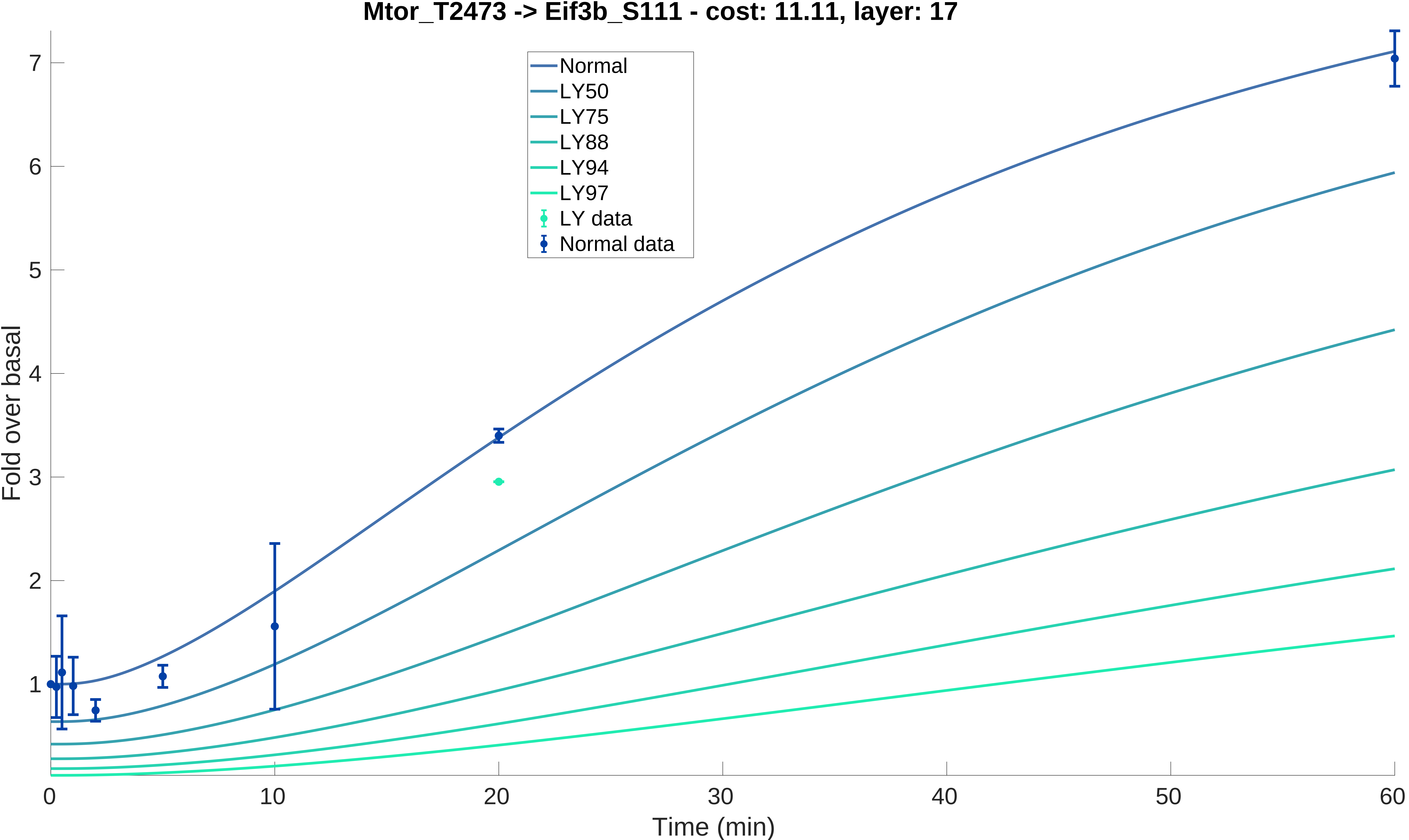

Mtor_T2473 -> Eif3b_S111 - cost: 11.11, layer: 17
Normal
LY50
LY75
LY88
LY94
LY97
LY data
Normal data
Fold over basal
Time (min)
0
10
20
30
40
50
60
1
2
3
4
5
6
7

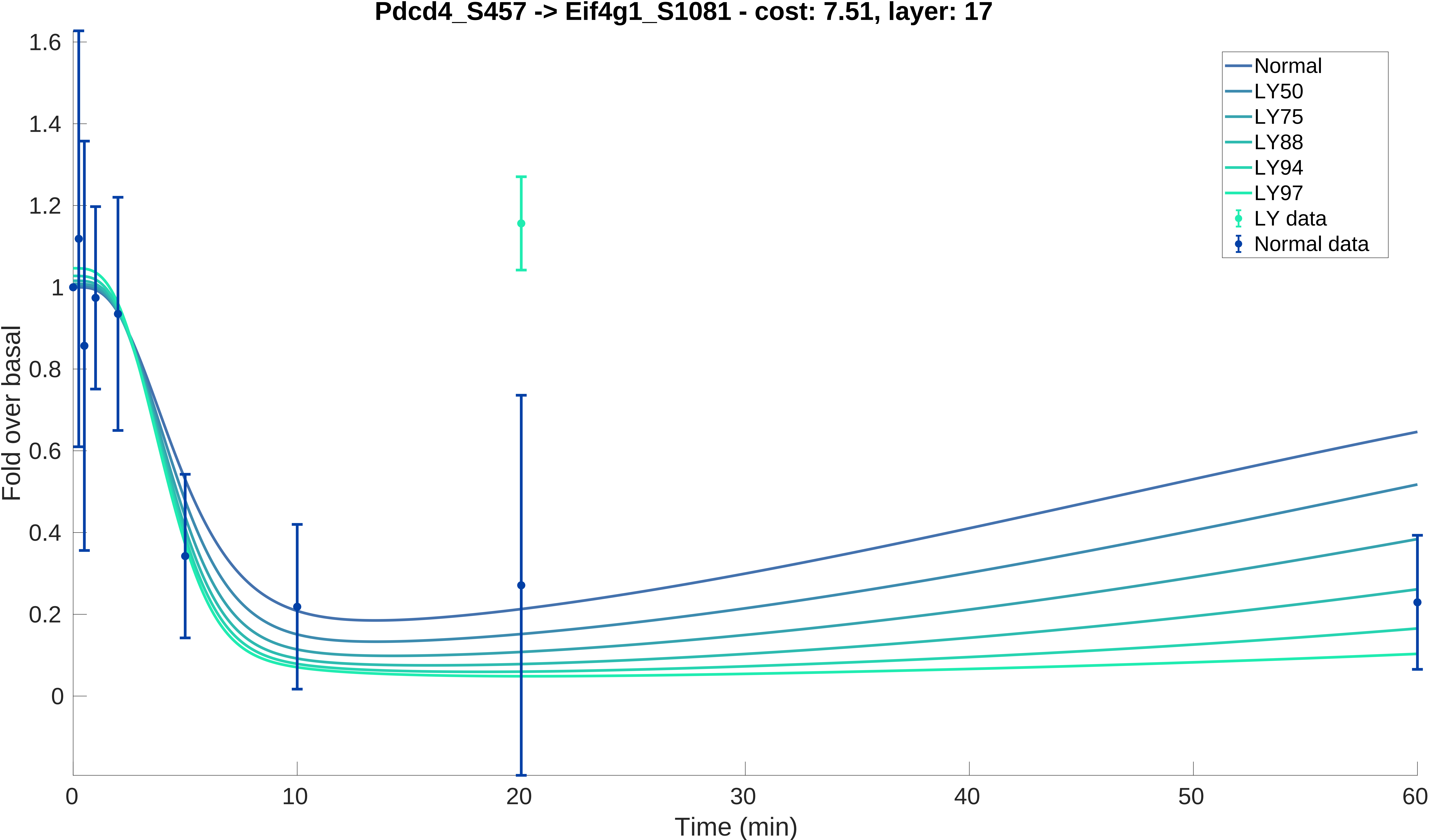

Pdcd4_S457 -> Eif4g1_S1081 - cost: 7.51, layer: 17
Fold over basal
Time (min)
1.6
1.4
1.2
1
0.8
0.6
0.4
0.2
0
0
10
20
30
40
50
60
Normal
LY50
LY75
LY88
LY94
LY97
LY data
Normal data

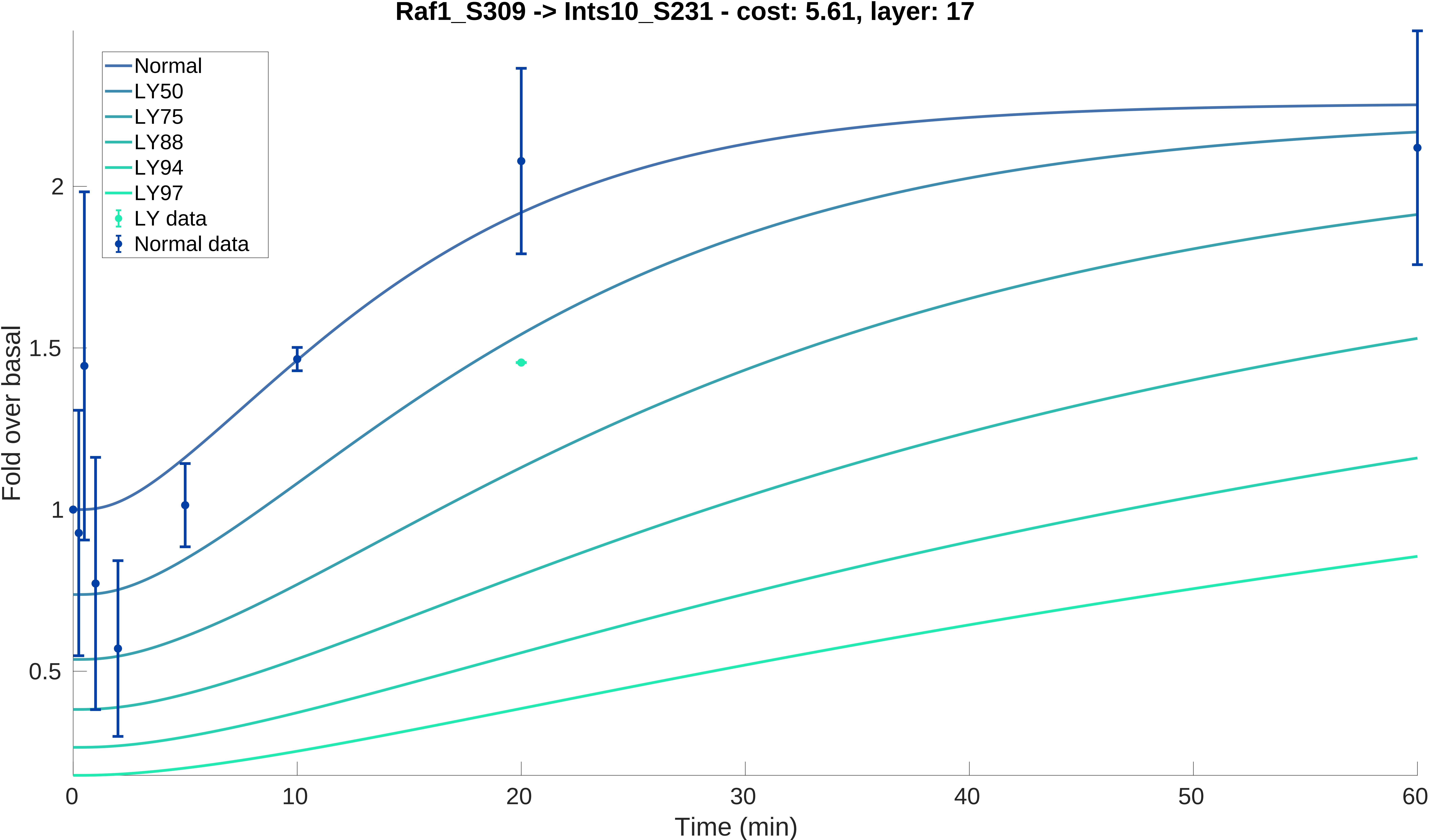

Raf1_S309 -> Ints10_S231 - cost: 5.61, layer: 17
Normal
LY50
LY75
LY88
LY94
LY97
LY data
Normal data
Fold over basal
0.5
1
1.5
2
0
10
20
30
40
50
60
Time (min)

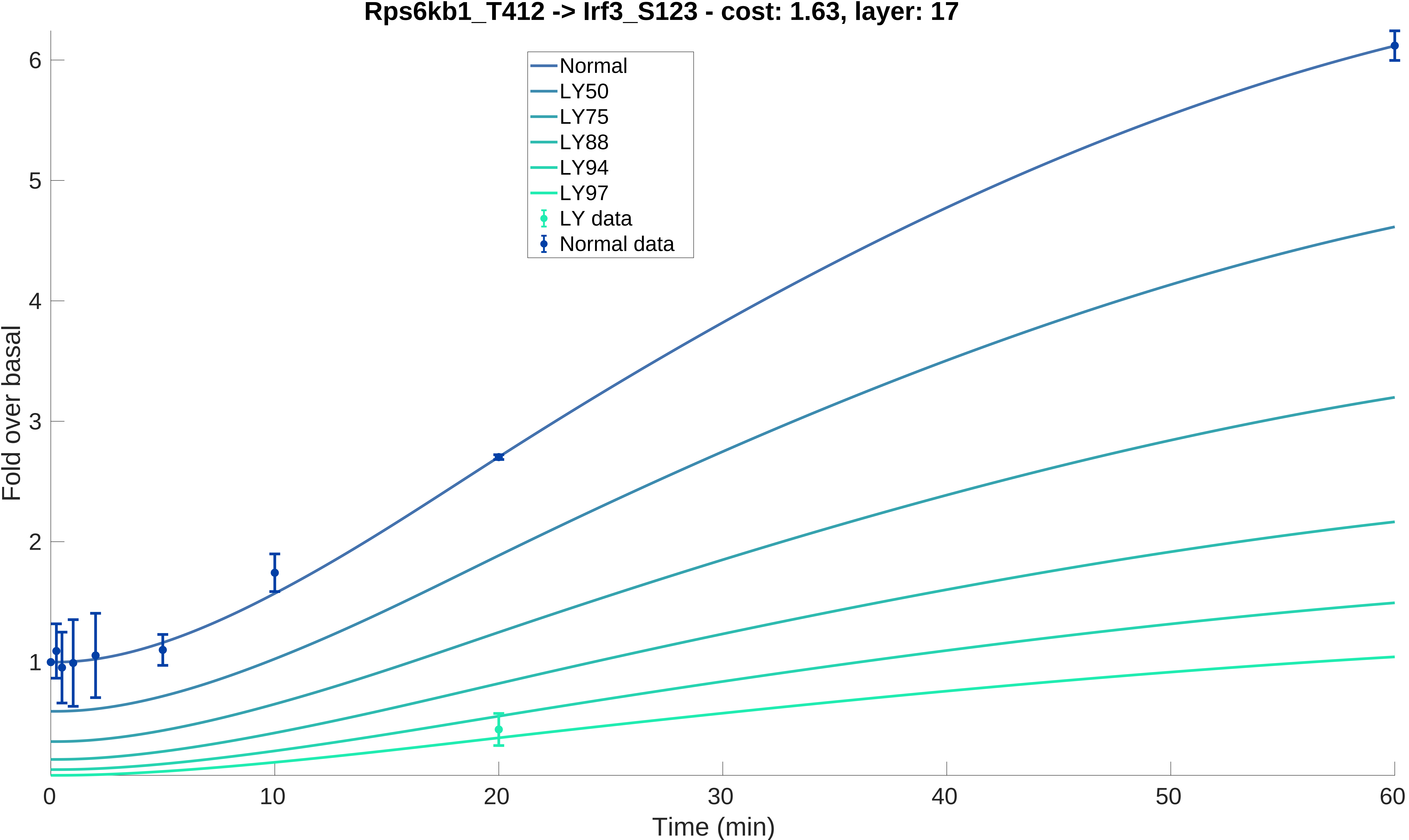
Rps6kb1_T412 -> Irf3_S123 - cost: 1.63, layer: 17
Normal
LY50
LY75
LY88
LY94
LY97
LY data
Normal data
Fold over basal
Time (min)
1
2
3
4
5
6
0
10
20
30
40
50
60

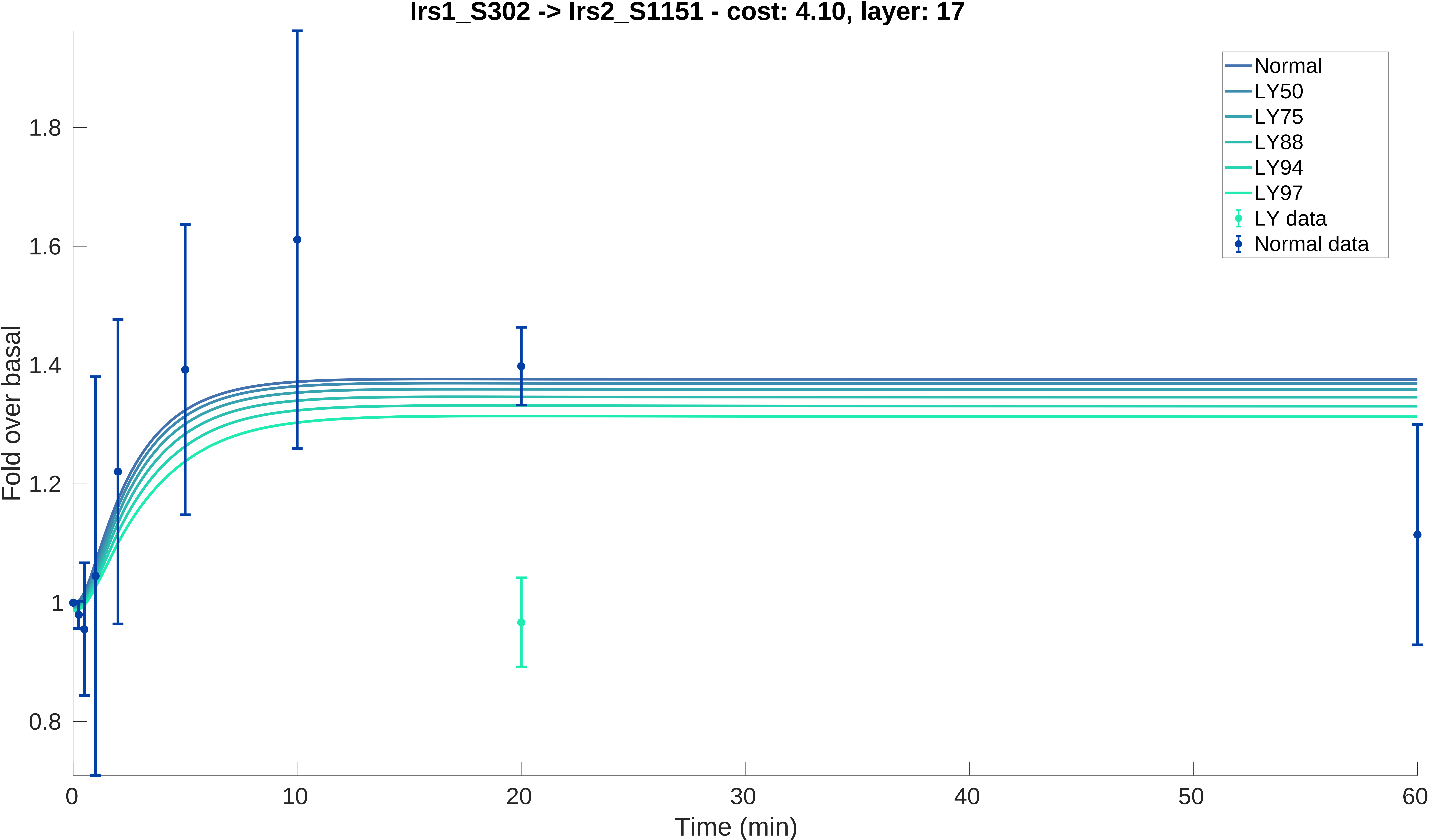

Irs1_S302 -> Irs2_S1151 - cost: 4.10, layer: 17
Normal
LY50
LY75
LY88
LY94
LY97
LY data
Normal data
Fold over basal
Time (min)
0.8
1
1.2
1.4
1.6
1.8
0
10
20
30
40
50
60

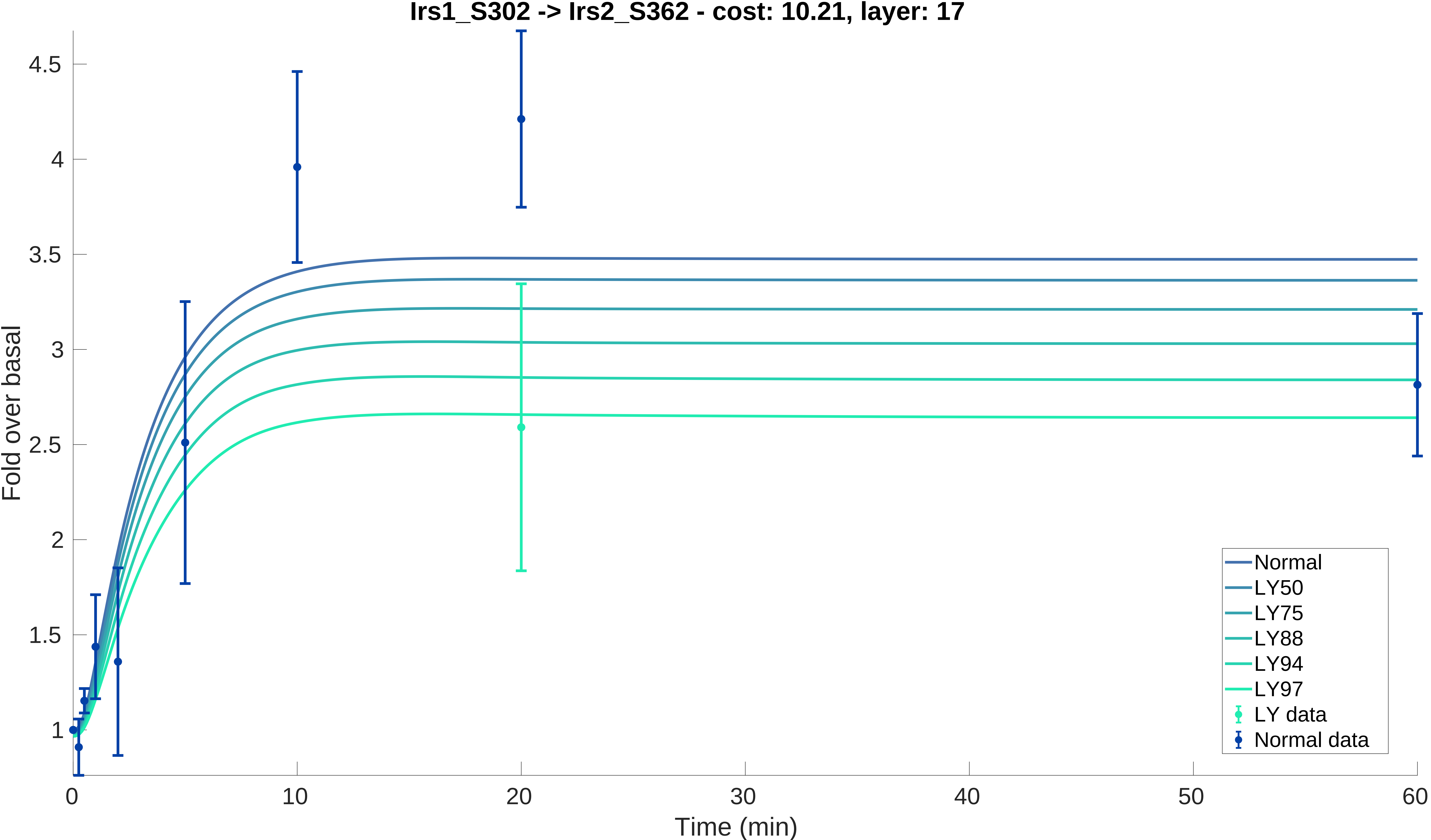

Irs1_S302 -> Irs2_S362 - cost: 10.21, layer: 17
Fold over basal
Time (min)
4.5
4
3.5
3
2.5
2
1.5
1
0
10
20
30
40
50
60
Normal
LY50
LY75
LY88
LY94
LY97
LY data
Normal data

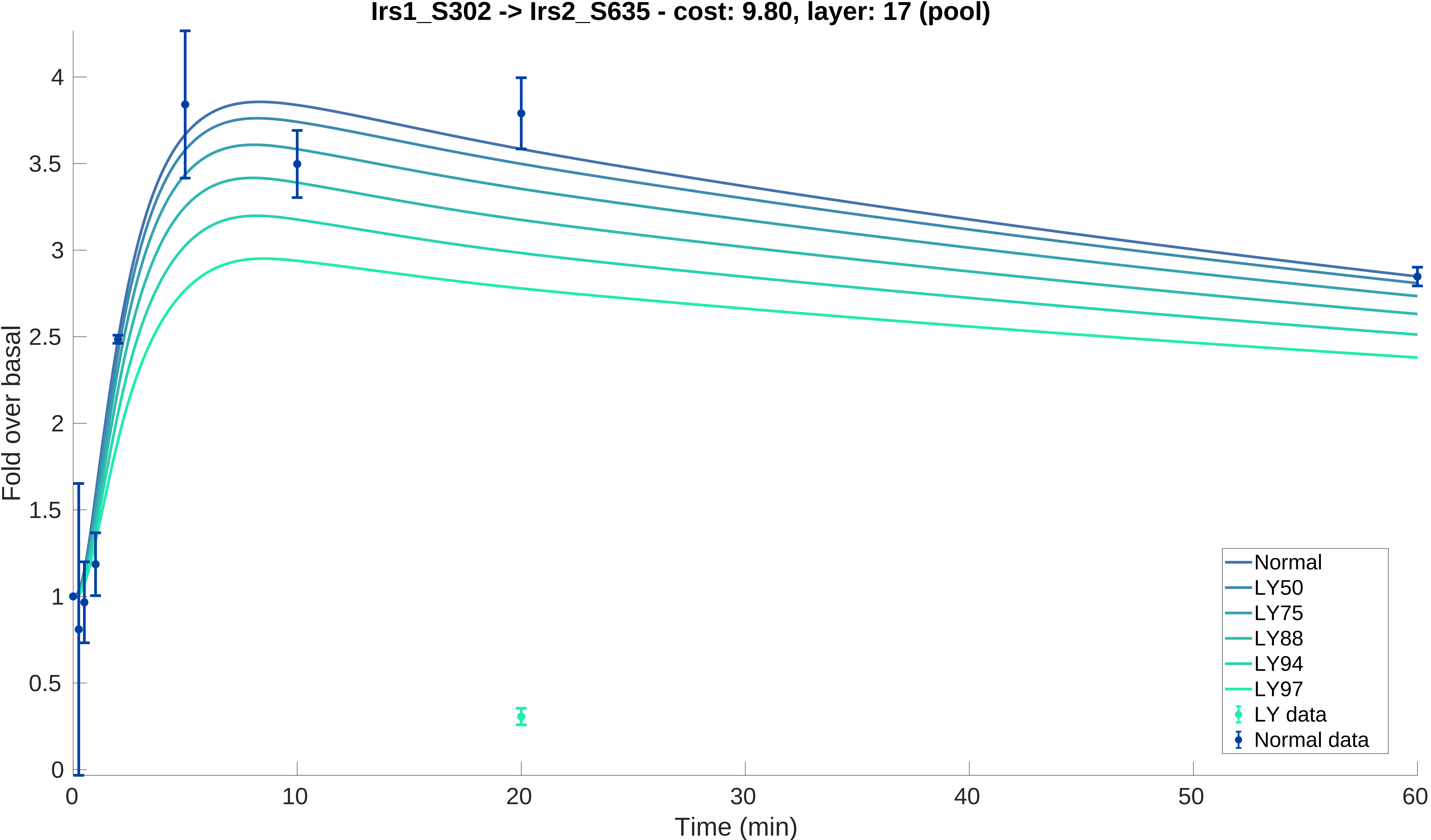

Irs1_S302 -> Irs2_S635 - cost: 9.80, layer: 17 (pool)
Fold over basal
Time (min)
0
0.5
1
1.5
2
2.5
3
3.5
4
0
10
20
30
40
50
60
Normal
LY50
LY75
LY88
LY94
LY97
LY data
Normal data

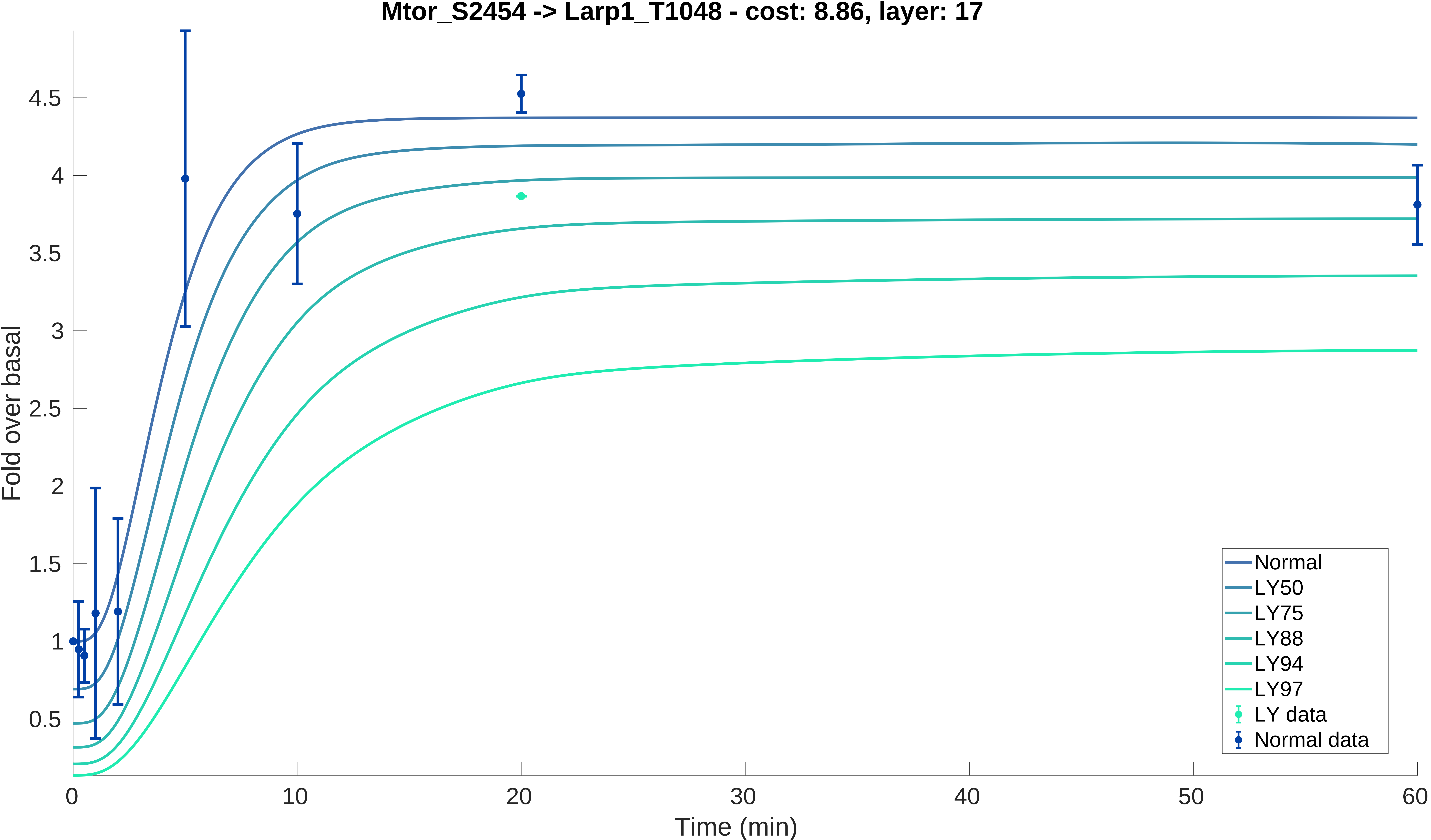

Mtor_S2454 -> Larp1_T1048 - cost: 8.86, layer: 17
Fold over basal
4.5
4
3.5
3
2.5
2
1.5
1
0.5
0
10
20
30
40
50
60
Time (min)
Normal
LY50
LY75
LY88
LY94
LY97
LY data
Normal data

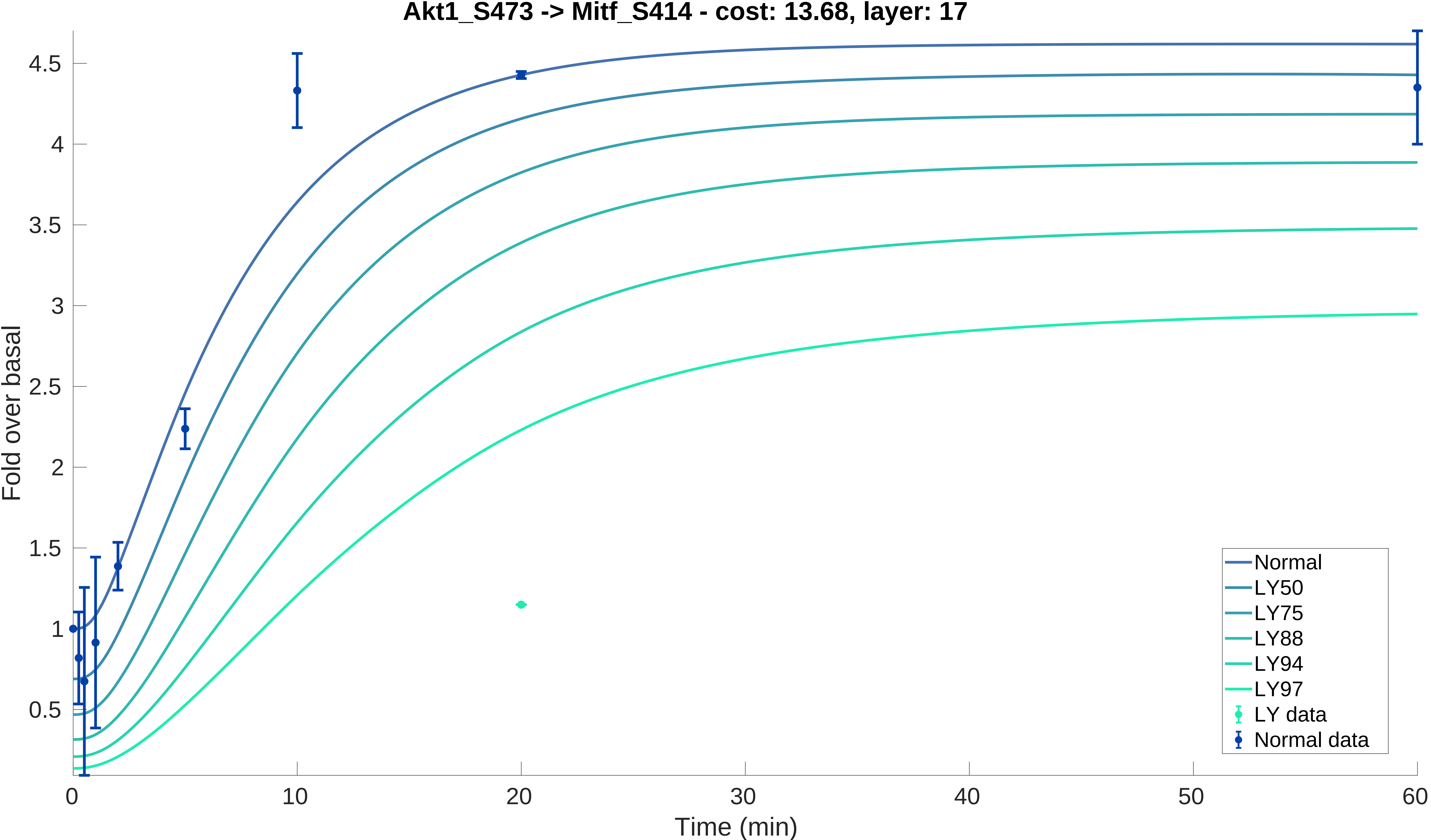

Akt1_S473 -> Mitf_S414 - cost: 13.68, layer: 17
Fold over basal
Time (min)
0.5
1
1.5
2
2.5
3
3.5
4
4.5
0
10
20
30
40
50
60
Normal
LY50
LY75
LY88
LY94
LY97
LY data
Normal data

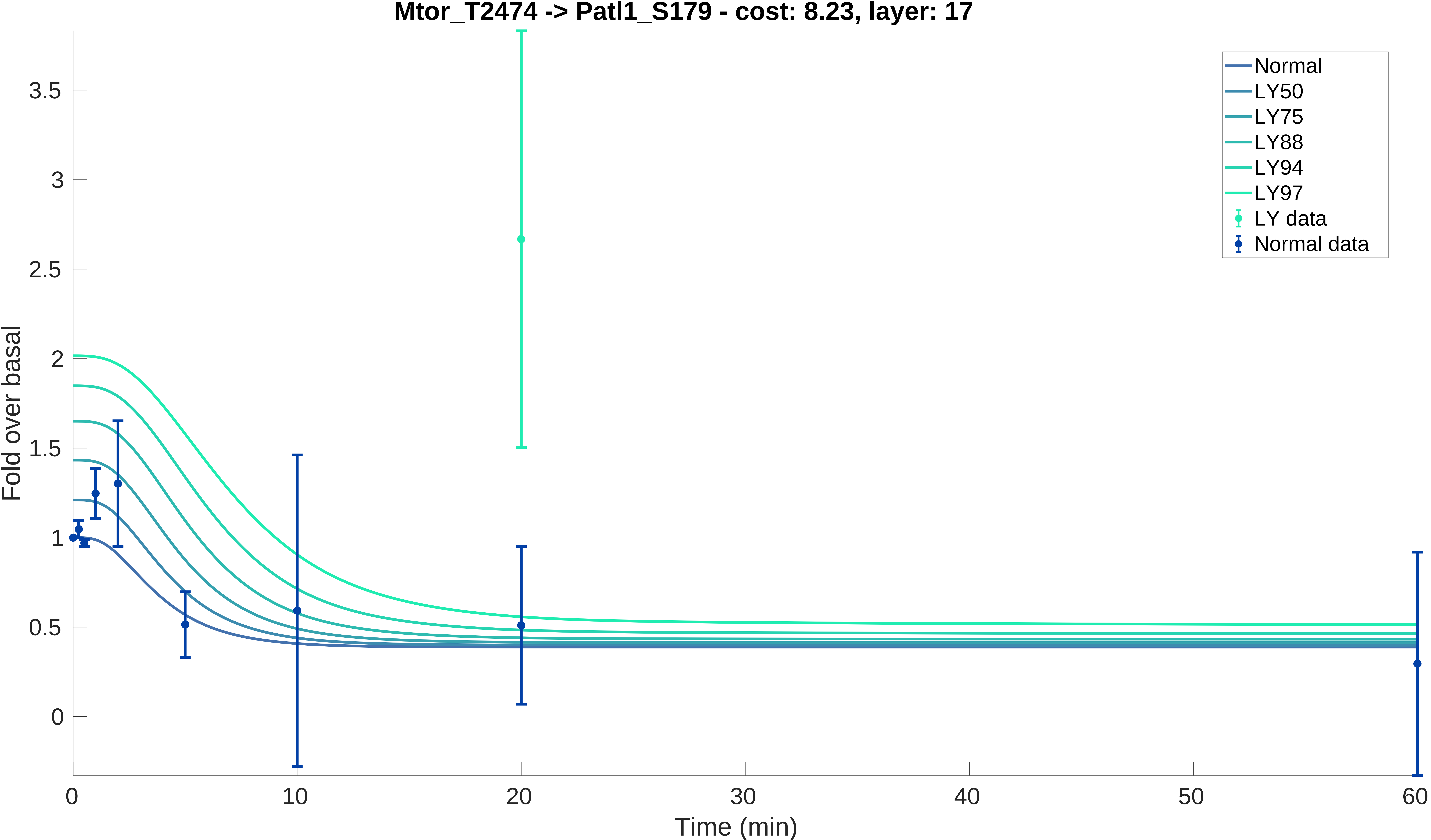

Mtor_T2474 -> Patl1_S179 - cost: 8.23, layer: 17
Fold over basal
Time (min)
Normal
LY50
LY75
LY88
LY94
LY97
LY data
Normal data
3.5
3
2.5
2
1.5
1
0.5
0
0
10
20
30
40
50
60

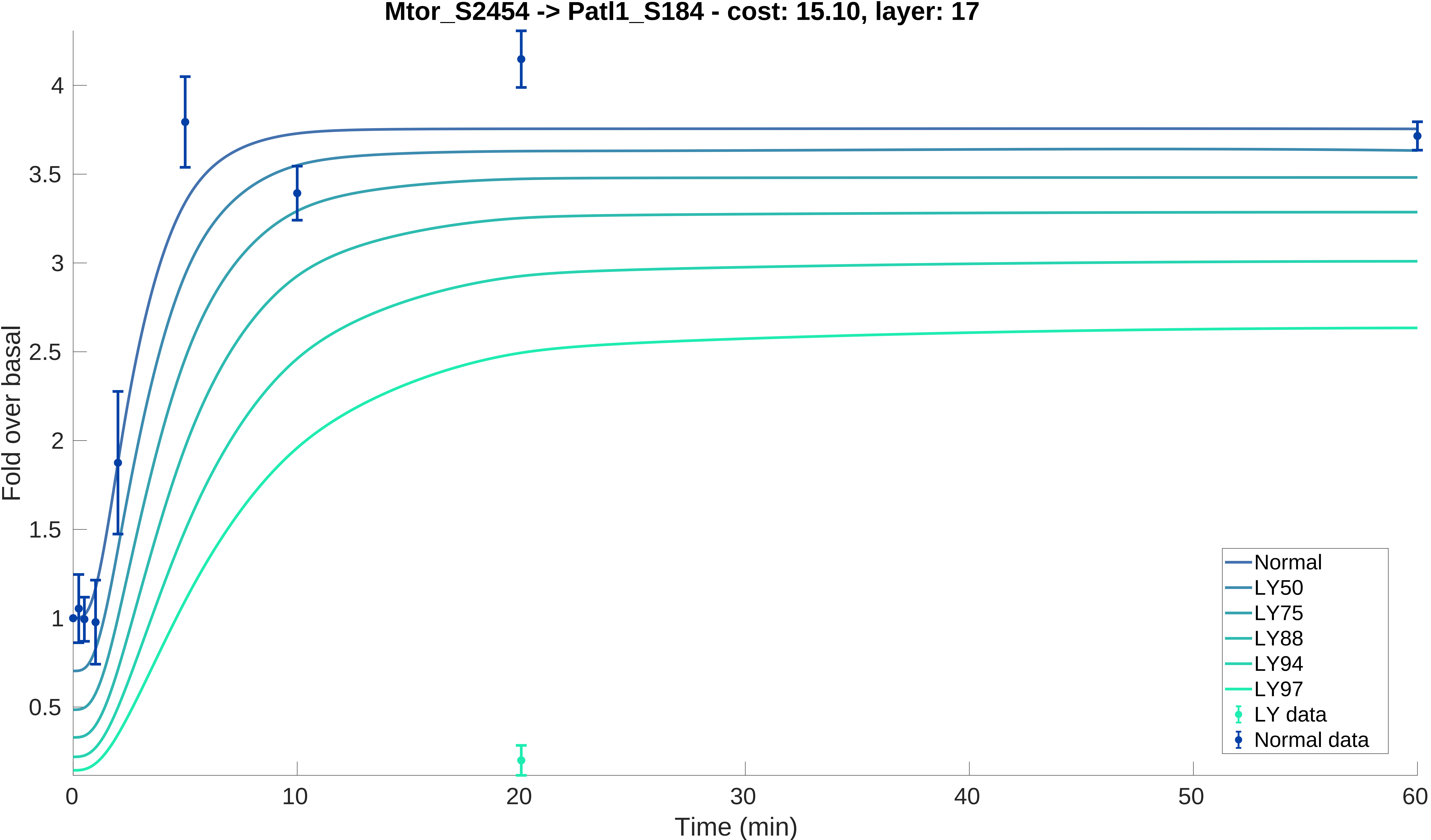

Mtor_S2454 -> Patl1_S184 - cost: 15.10, layer: 17
Fold over basal
Time (min)
0.5
1
1.5
2
2.5
3
3.5
4
0
10
20
30
40
50
60
Normal
LY50
LY75
LY88
LY94
LY97
LY data
Normal data

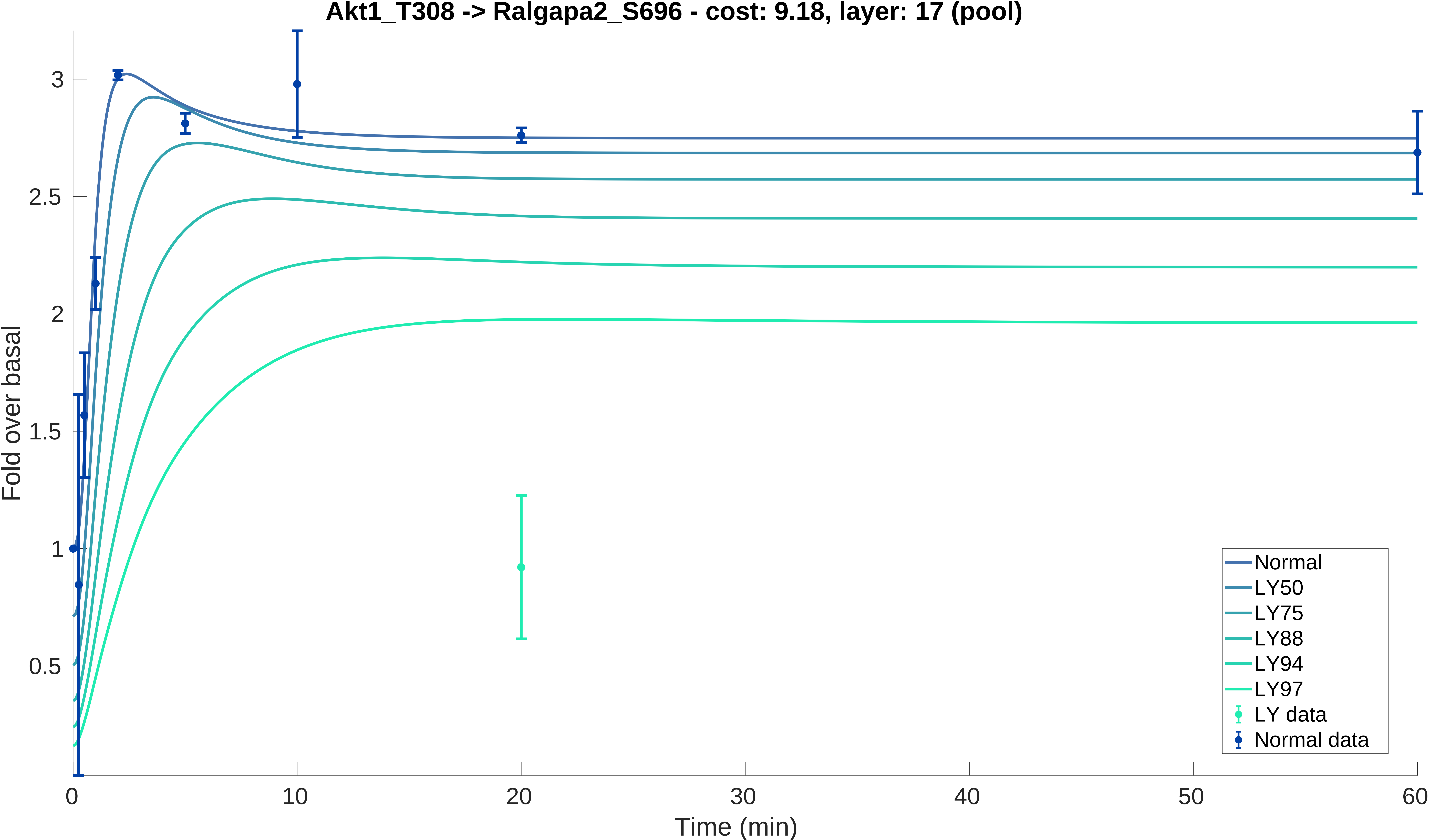
Akt1_T308 -> Ralgapa2_S696 - cost: 9.18, layer: 17 (pool)
Fold over basal
Time (min)
0
10
20
30
40
50
60
0.5
1
1.5
2
2.5
3
Normal
LY50
LY75
LY88
LY94
LY97
LY data
Normal data

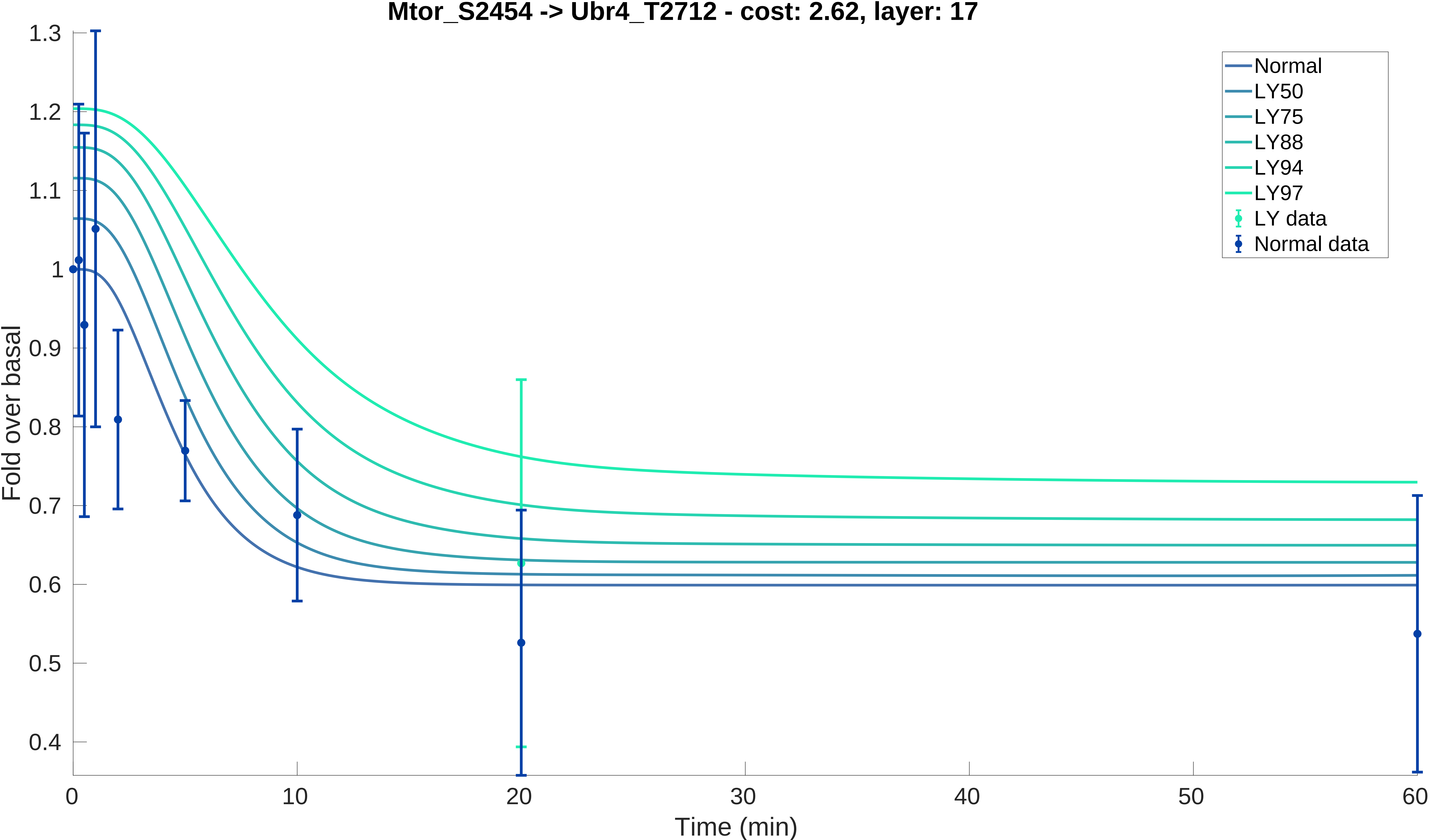

Mtor_S2454 -> Ubr4_T2712 - cost: 2.62, layer: 17
Fold over basal
Time (min)
1.3
1.2
1.1
1
0.9
0.8
0.7
0.6
0.5
0.4
0
10
20
30
40
50
60
Normal
LY50
LY75
LY88
LY94
LY97
LY data
Normal data

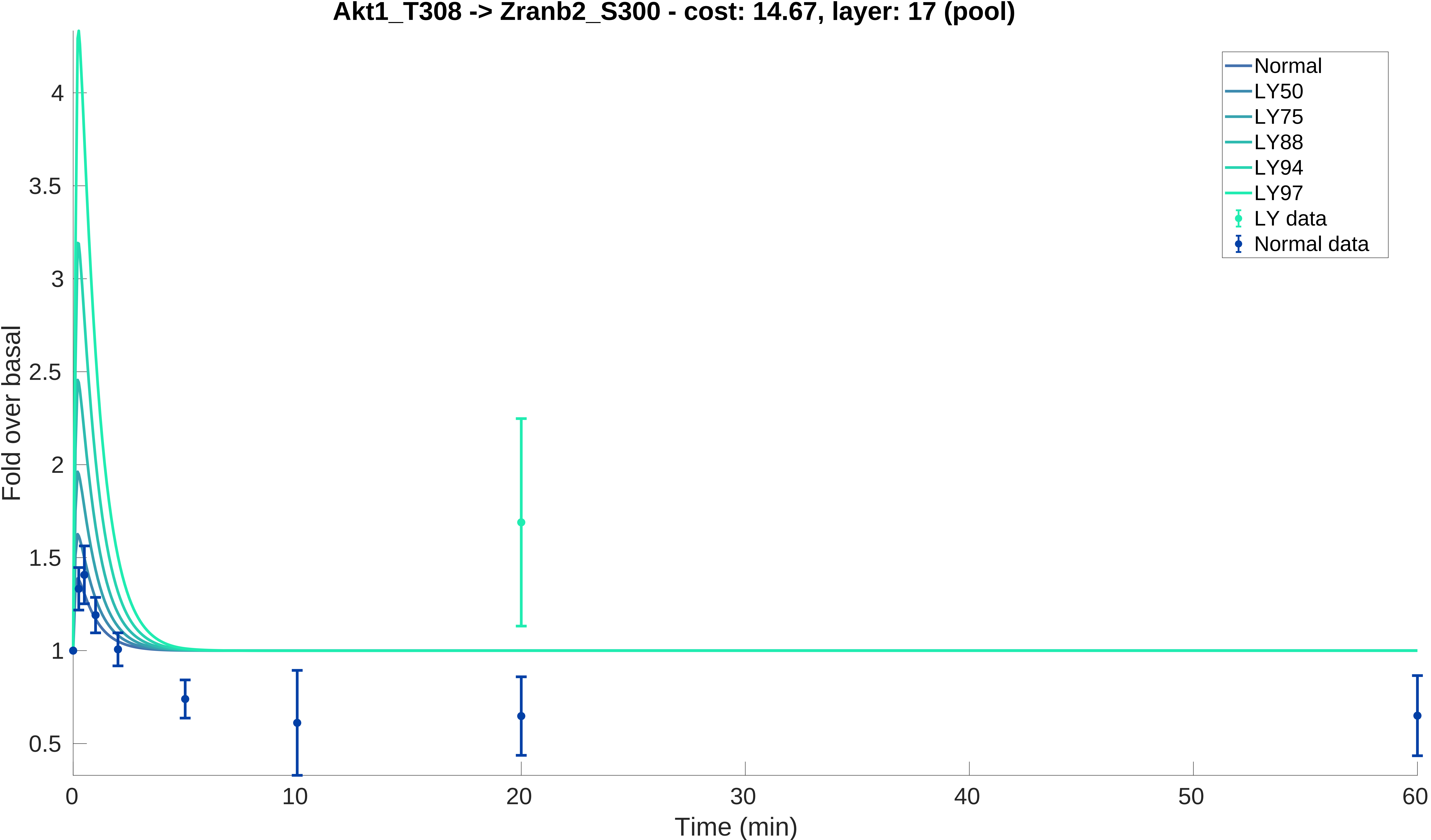

Akt1_T308 -> Zranb2_S300 - cost: 14.67, layer: 17 (pool)
Fold over basal
Time (min)
0.5
1
1.5
2
2.5
3
3.5
4
0
10
20
30
40
50
60
Normal
LY50
LY75
LY88
LY94
LY97
LY data
Normal data

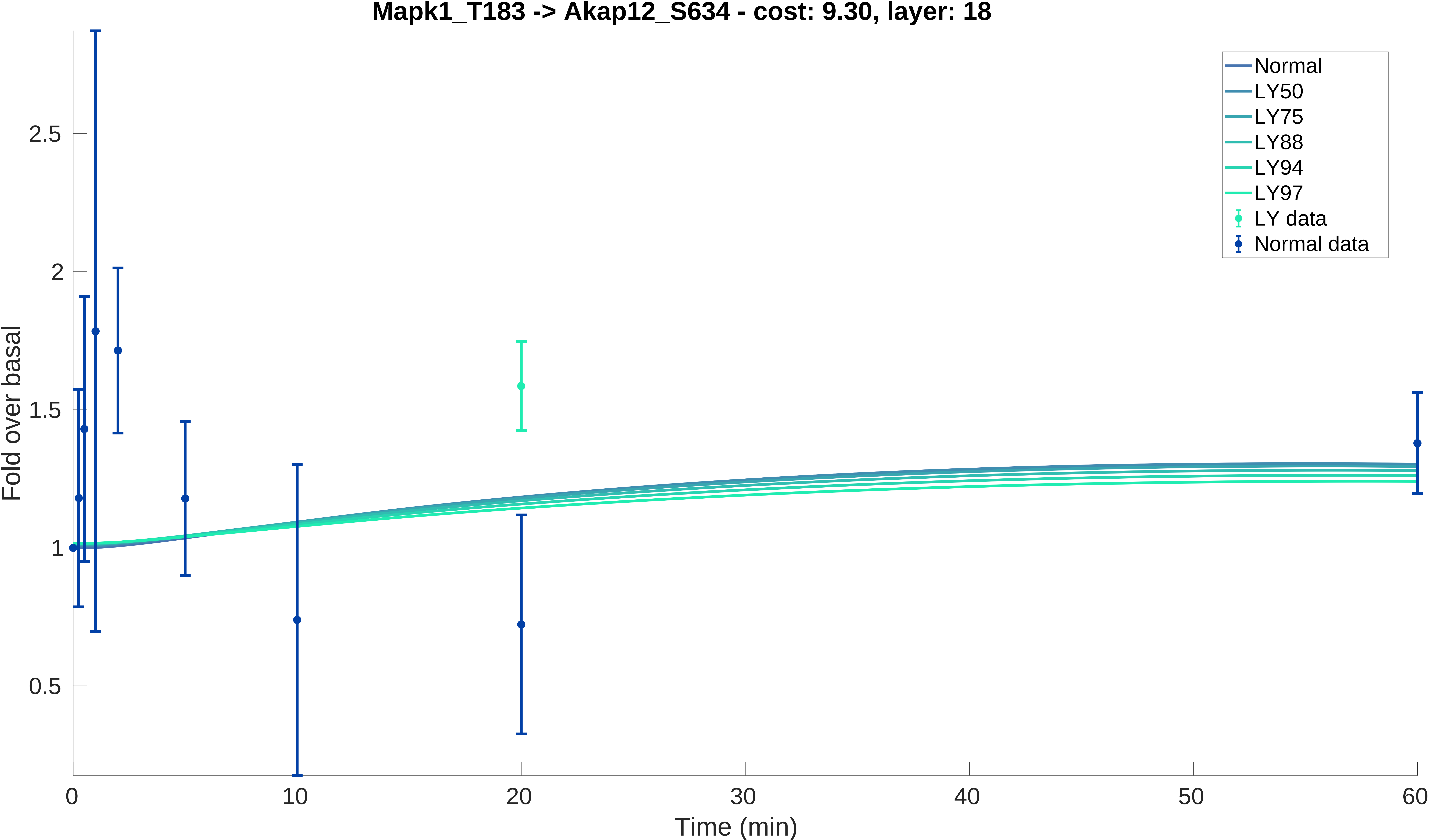

Mapk1_T183 -> Akap12_S634 - cost: 9.30, layer: 18
Normal
LY50
LY75
LY88
LY94
LY97
LY data
Normal data
Fold over basal
2.5
2
1.5
1
0.5
0
10
20
30
40
50
60
Time (min)

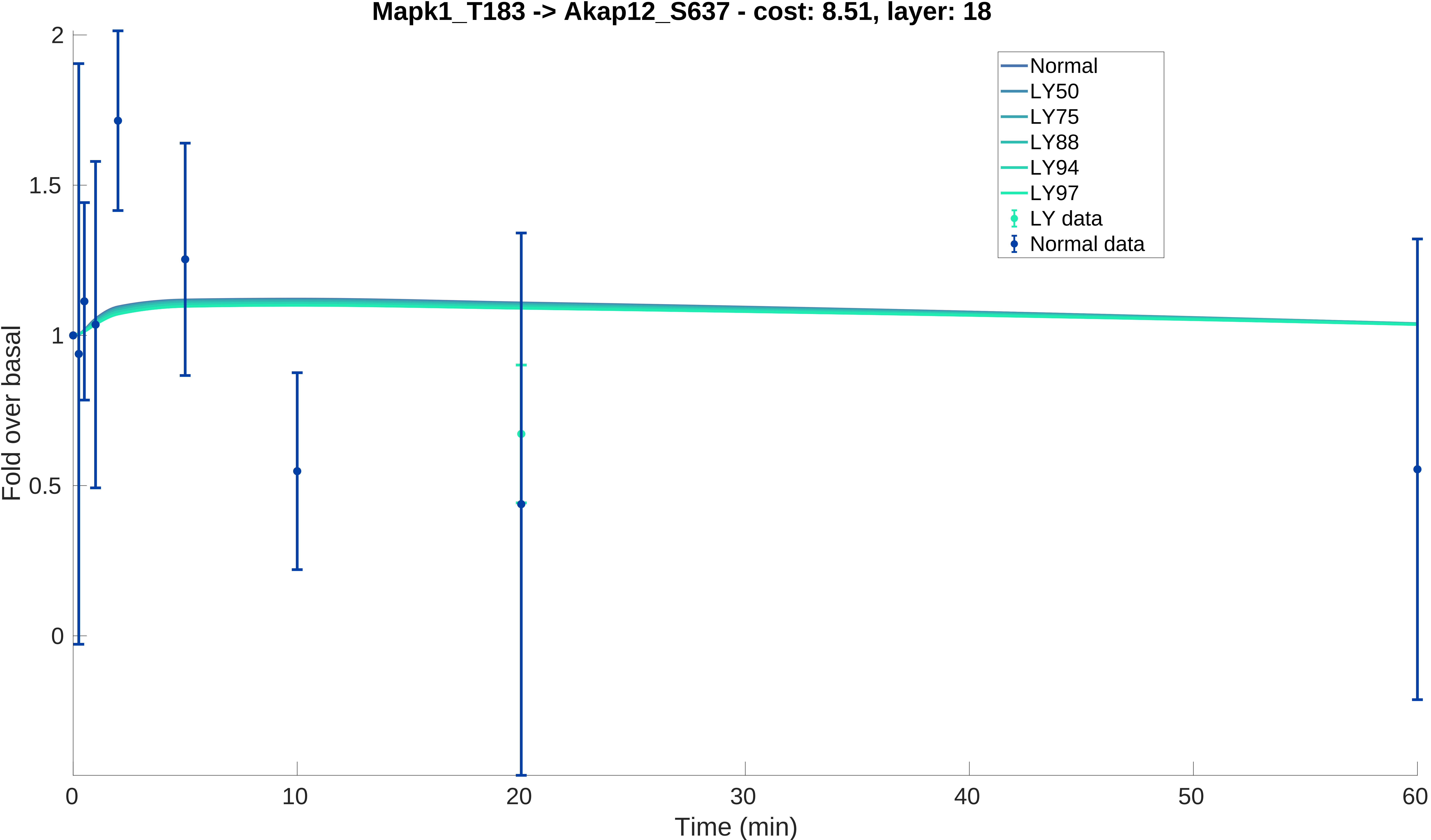

Mapk1_T183 -> Akap12_S637 - cost: 8.51, layer: 18
Normal
LY50
LY75
LY88
LY94
LY97
LY data
Normal data
Fold over basal
Time (min)
0
0.5
1
1.5
2
0
10
20
30
40
50
60

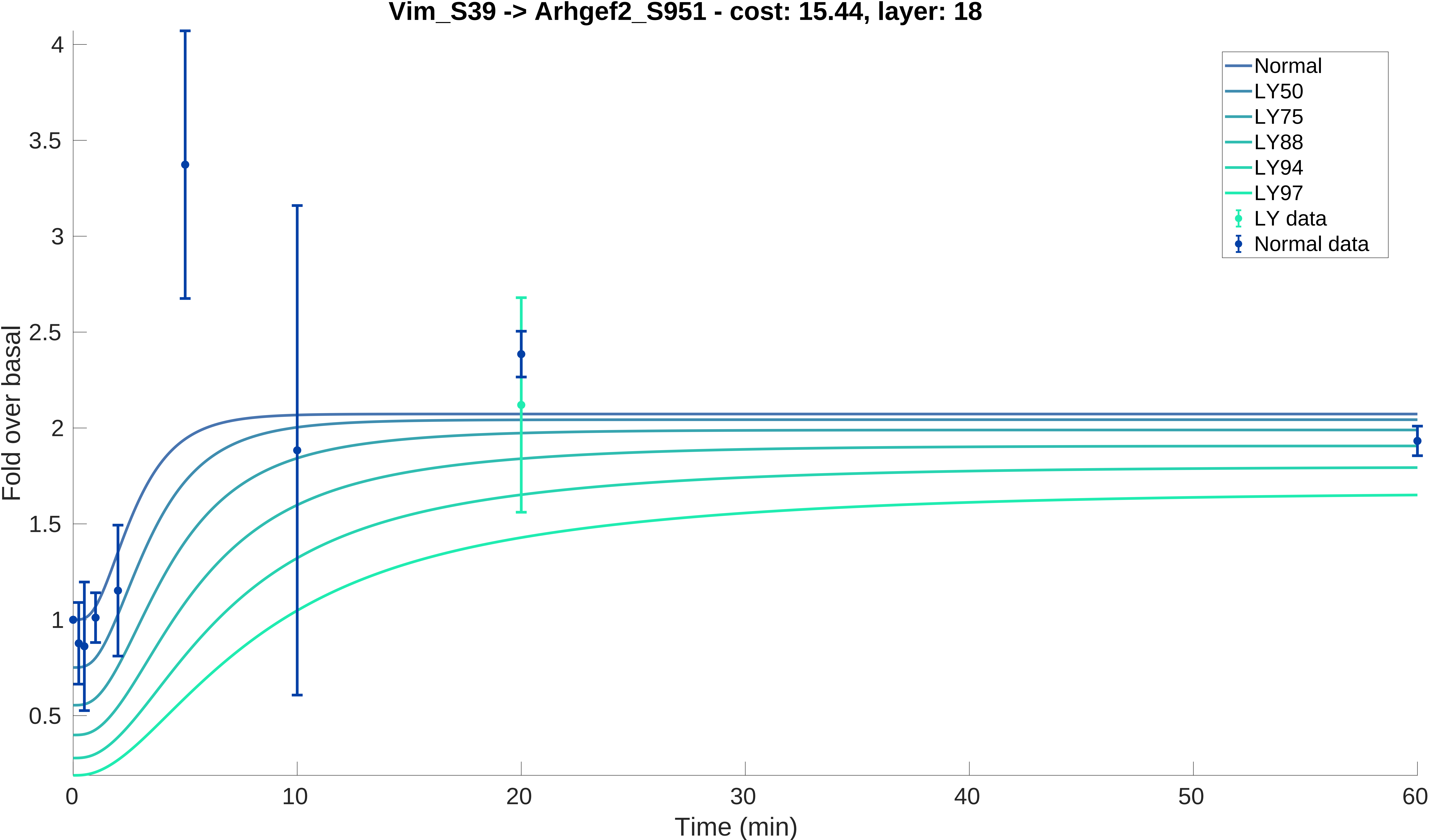

Vim_S39 -> Arhgef2_S951 - cost: 15.44, layer: 18
Normal
LY50
LY75
LY88
LY94
LY97
LY data
Normal data
Fold over basal
Time (min)
0.5
1
1.5
2
2.5
3
3.5
4
0
10
20
30
40
50
60

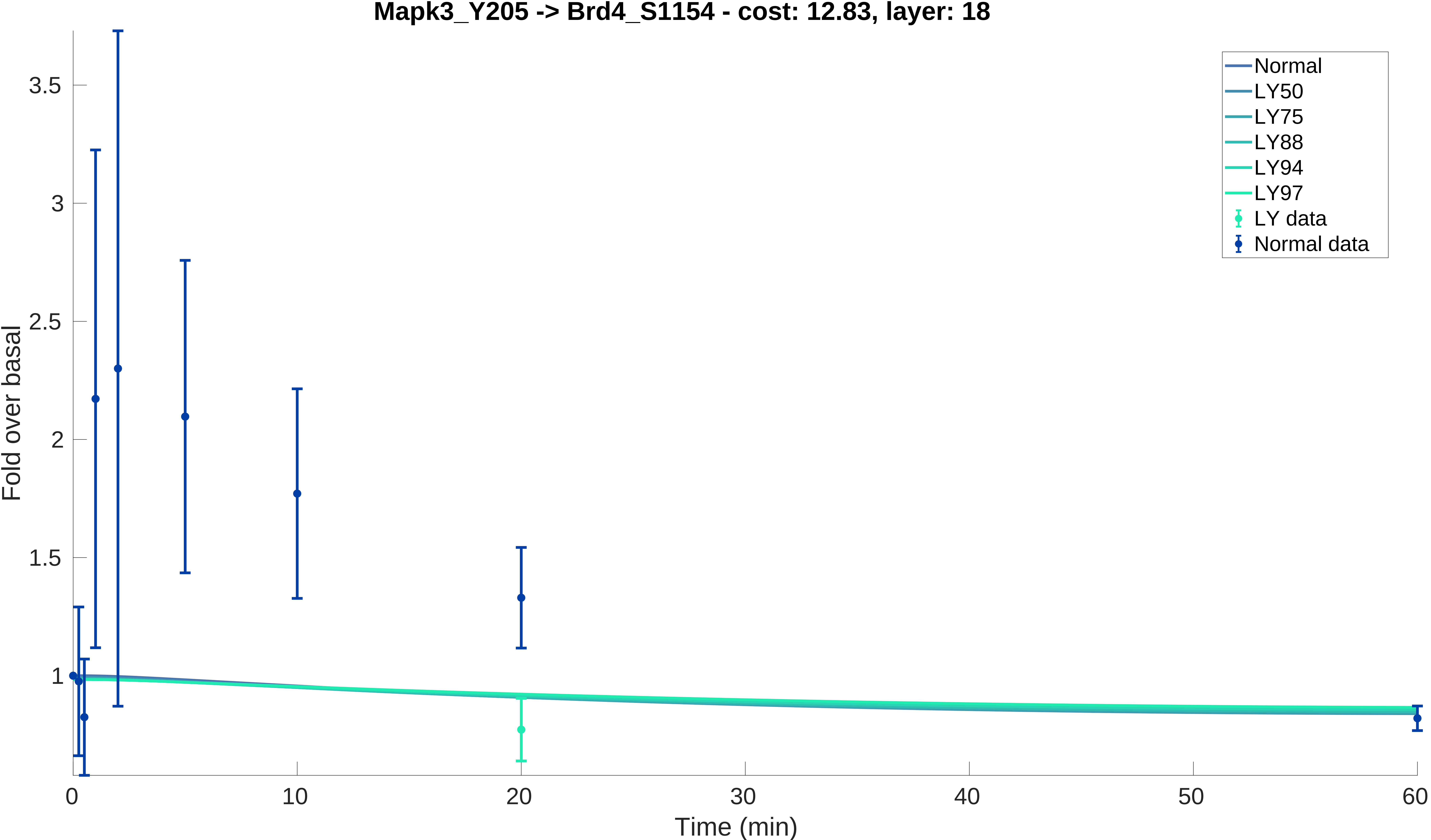

Mapk3_Y205 -> Brd4_S1154 - cost: 12.83, layer: 18
Fold over basal
Time (min)
3.5
3
2.5
2
1.5
1
0
10
20
30
40
50
60
Normal
LY50
LY75
LY88
LY94
LY97
LY data
Normal data

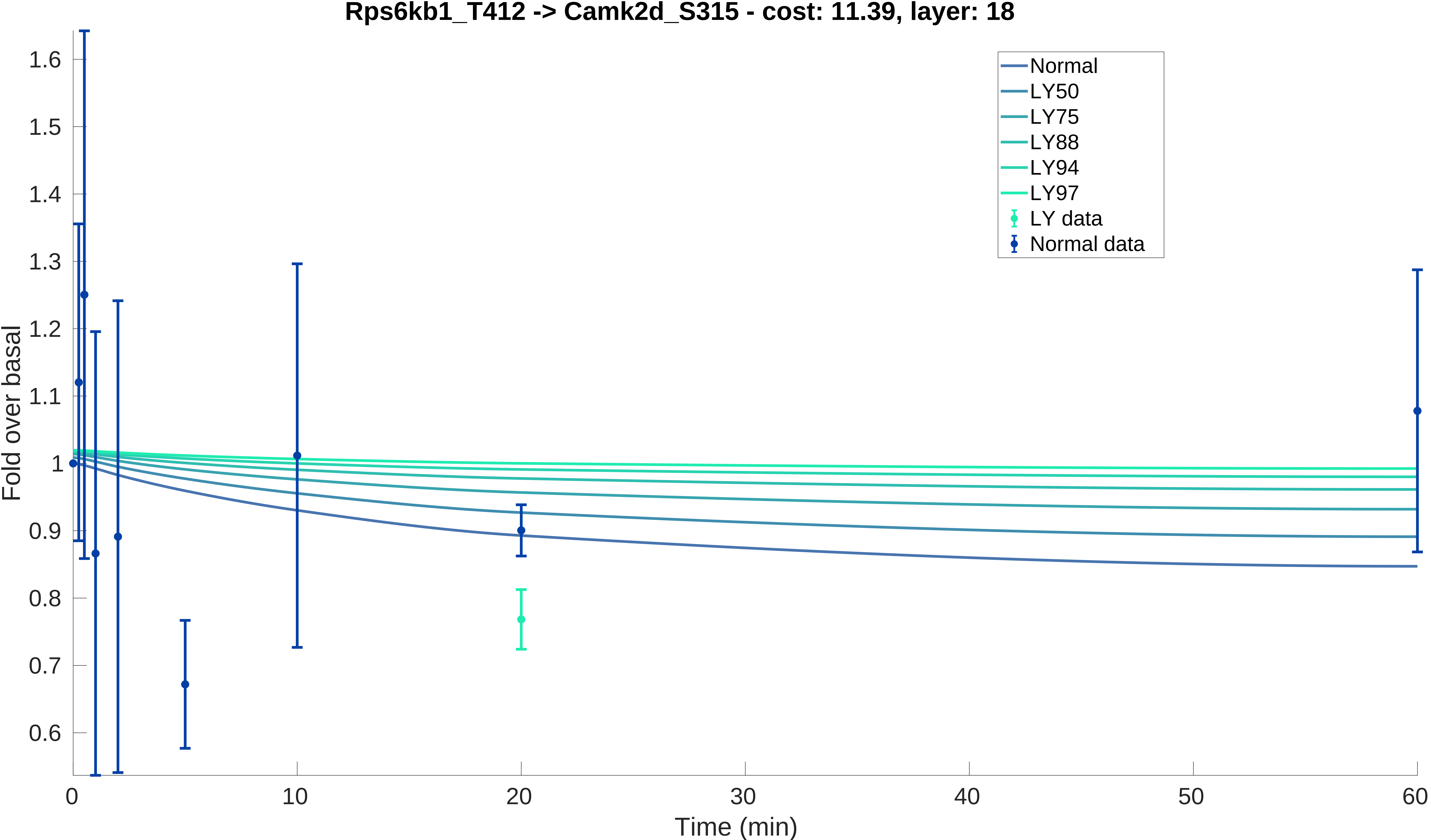
Rps6kb1_T412 -> Camk2d_S315 - cost: 11.39, layer: 18
Fold over basal
Time (min)
0
10
20
30
40
50
60
0.6
0.7
0.8
0.9
1
1.1
1.2
1.3
1.4
1.5
1.6
Normal
LY50
LY75
LY88
LY94
LY97
LY data
Normal data

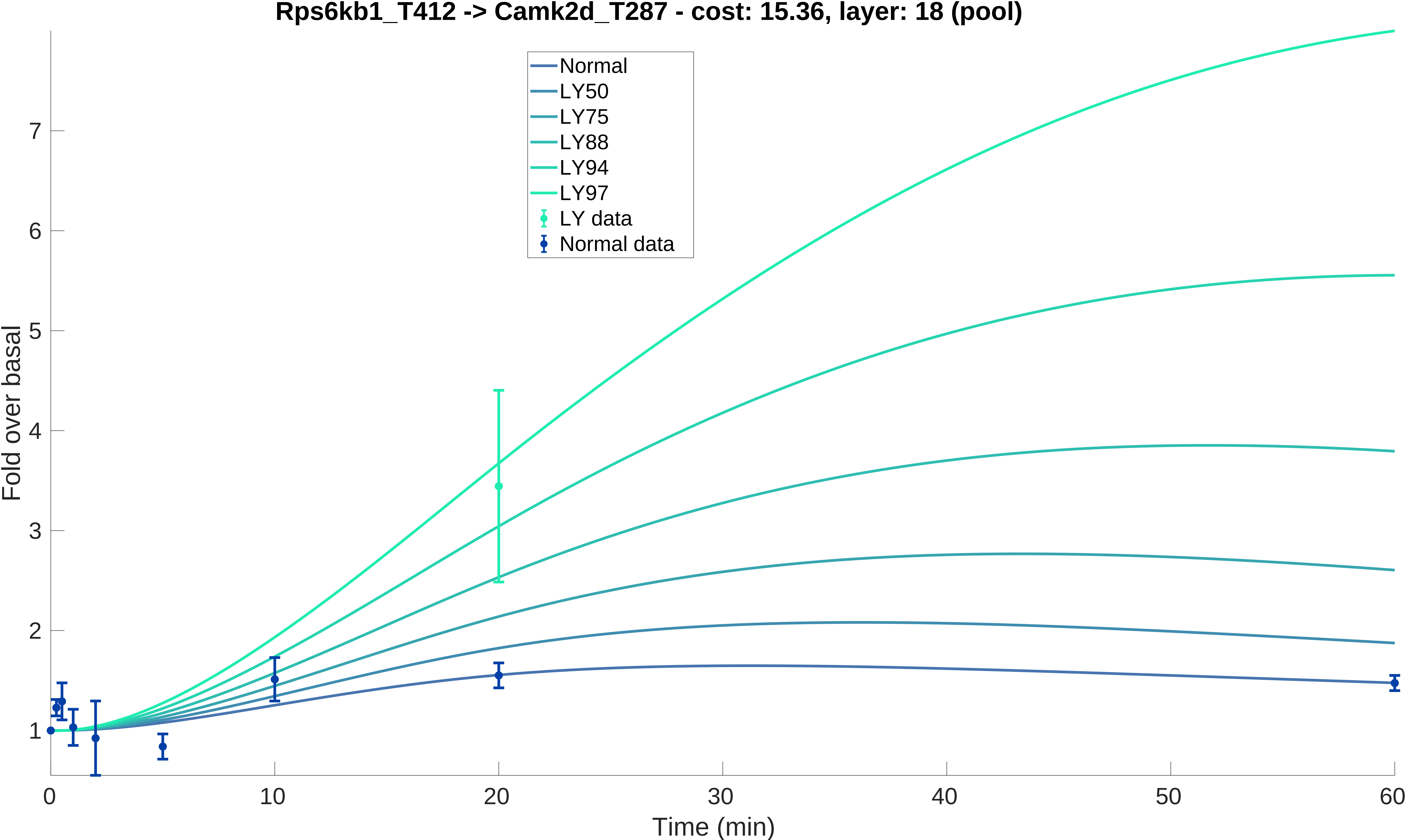

Rps6kb1_T412 -> Camk2d_T287 - cost: 15.36, layer: 18 (pool)
Normal
LY50
LY75
LY88
LY94
LY97
LY data
Normal data
Fold over basal
Time (min)
1
2
3
4
5
6
7
0
10
20
30
40
50
60

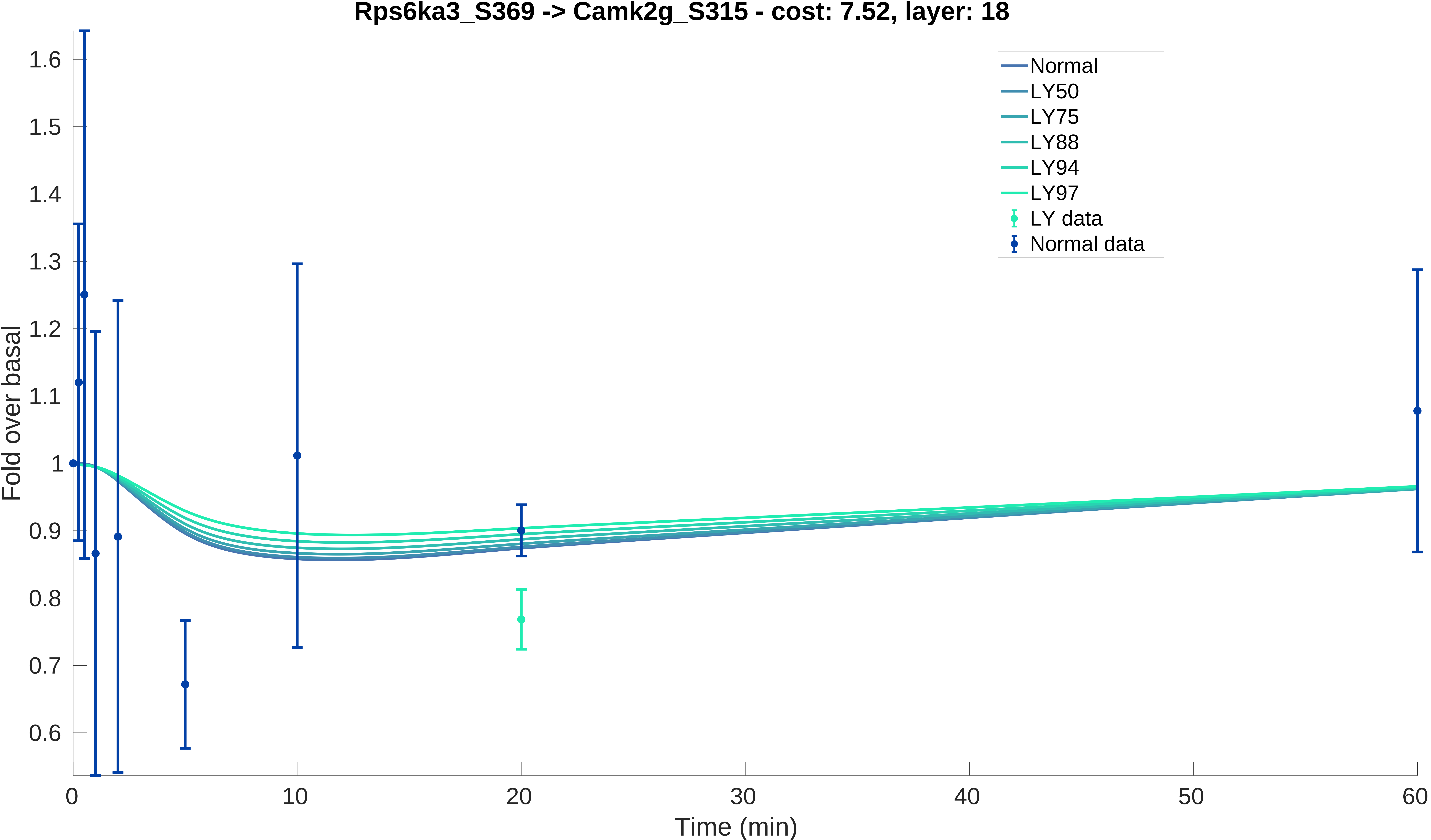
Rps6ka3_S369 -> Camk2g_S315 - cost: 7.52, layer: 18
Fold over basal
Time (min)
Normal
LY50
LY75
LY88
LY94
LY97
LY data
Normal data
1.6
1.5
1.4
1.3
1.2
1.1
1
0.9
0.8
0.7
0.6
0
10
20
30
40
50
60

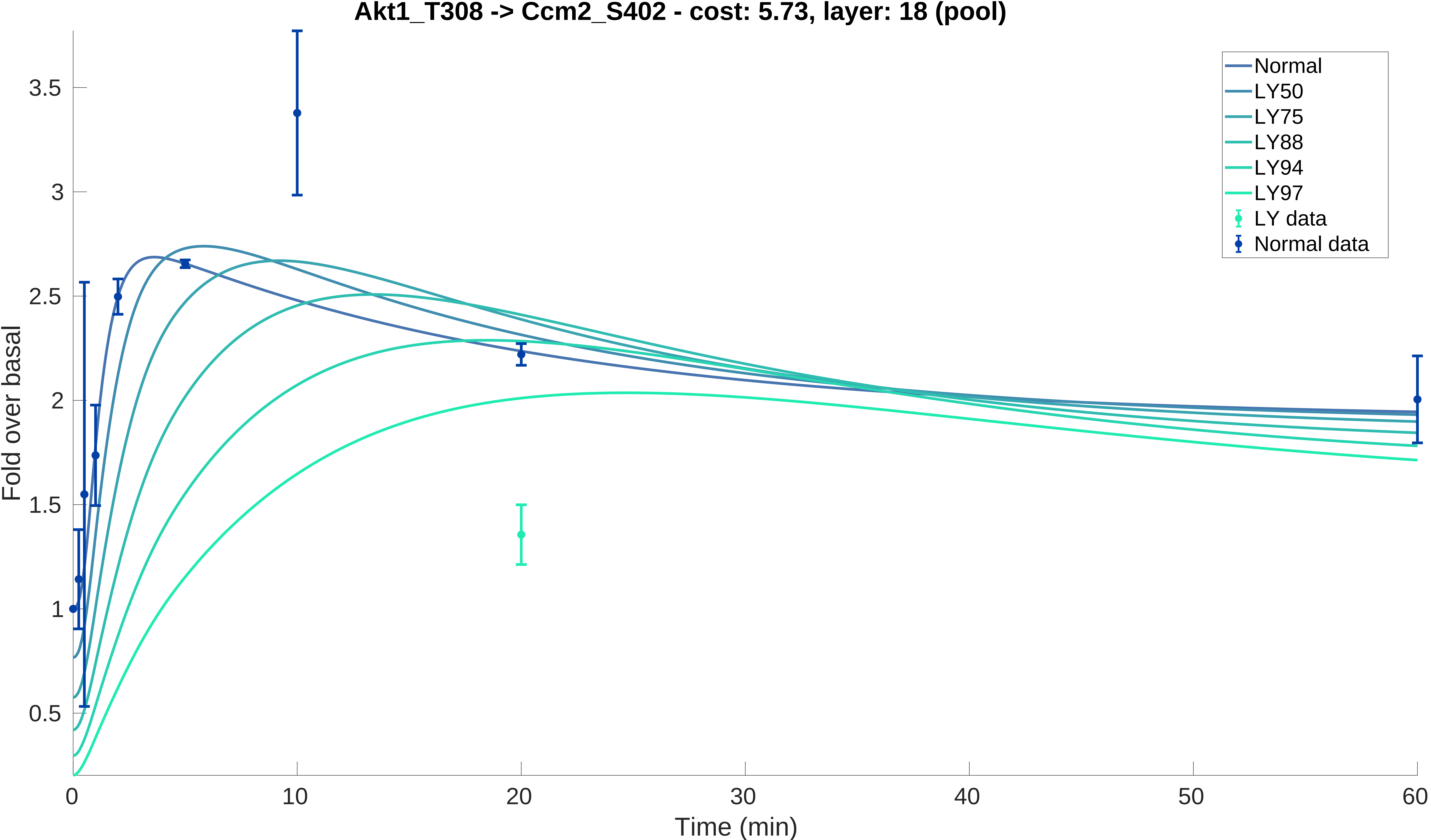

Akt1_T308 -> Ccm2_S402 - cost: 5.73, layer: 18 (pool)
Fold over basal
Time (min)
0.5
1
1.5
2
2.5
3
3.5
0
10
20
30
40
50
60
Normal
LY50
LY75
LY88
LY94
LY97
LY data
Normal data

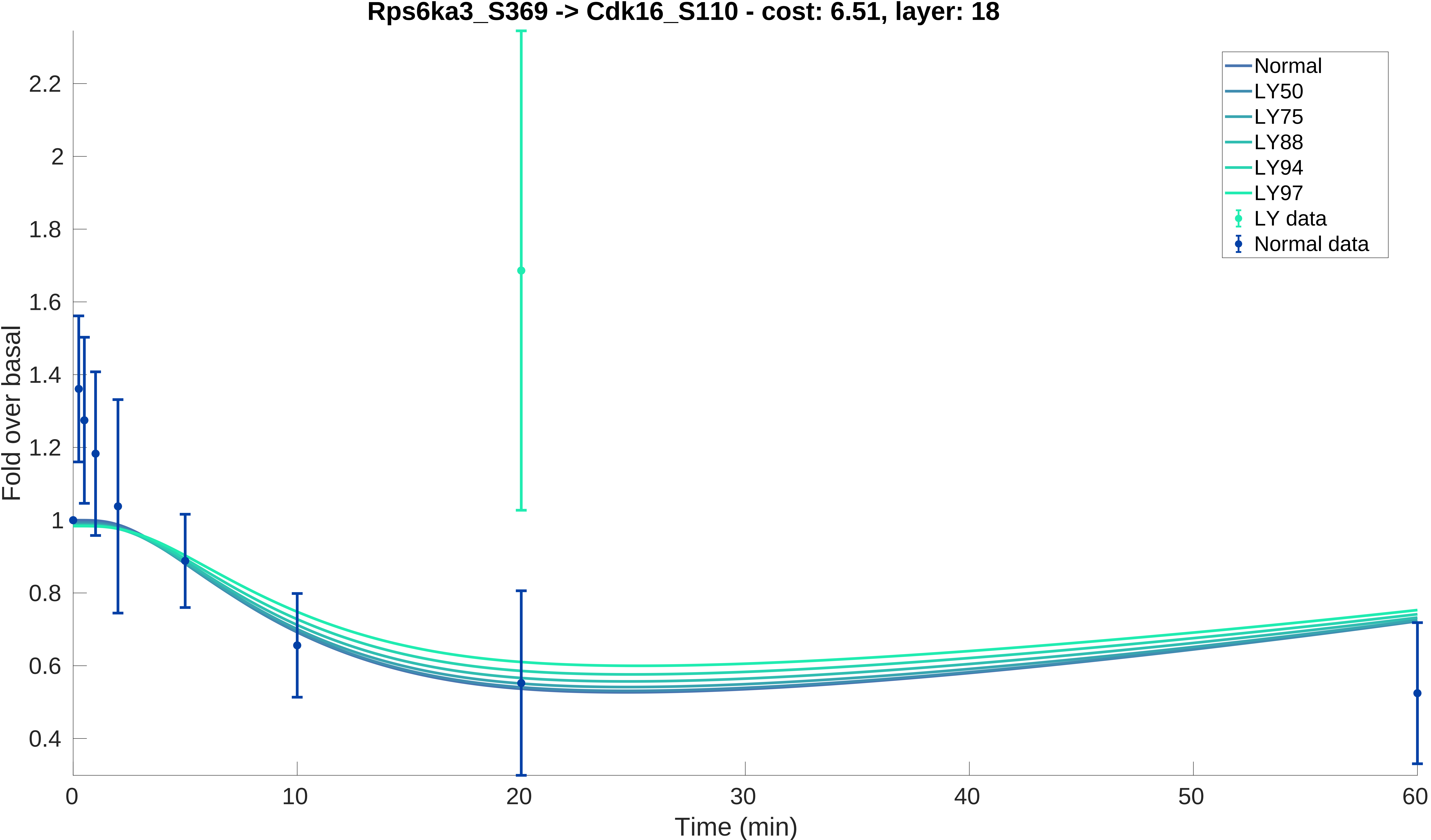

Rps6ka3_S369 -> Cdk16_S110 - cost: 6.51, layer: 18
Fold over basal
Time (min)
0
10
20
30
40
50
60
0.4
0.6
0.8
1
1.2
1.4
1.6
1.8
2
2.2
Normal
LY50
LY75
LY88
LY94
LY97
LY data
Normal data

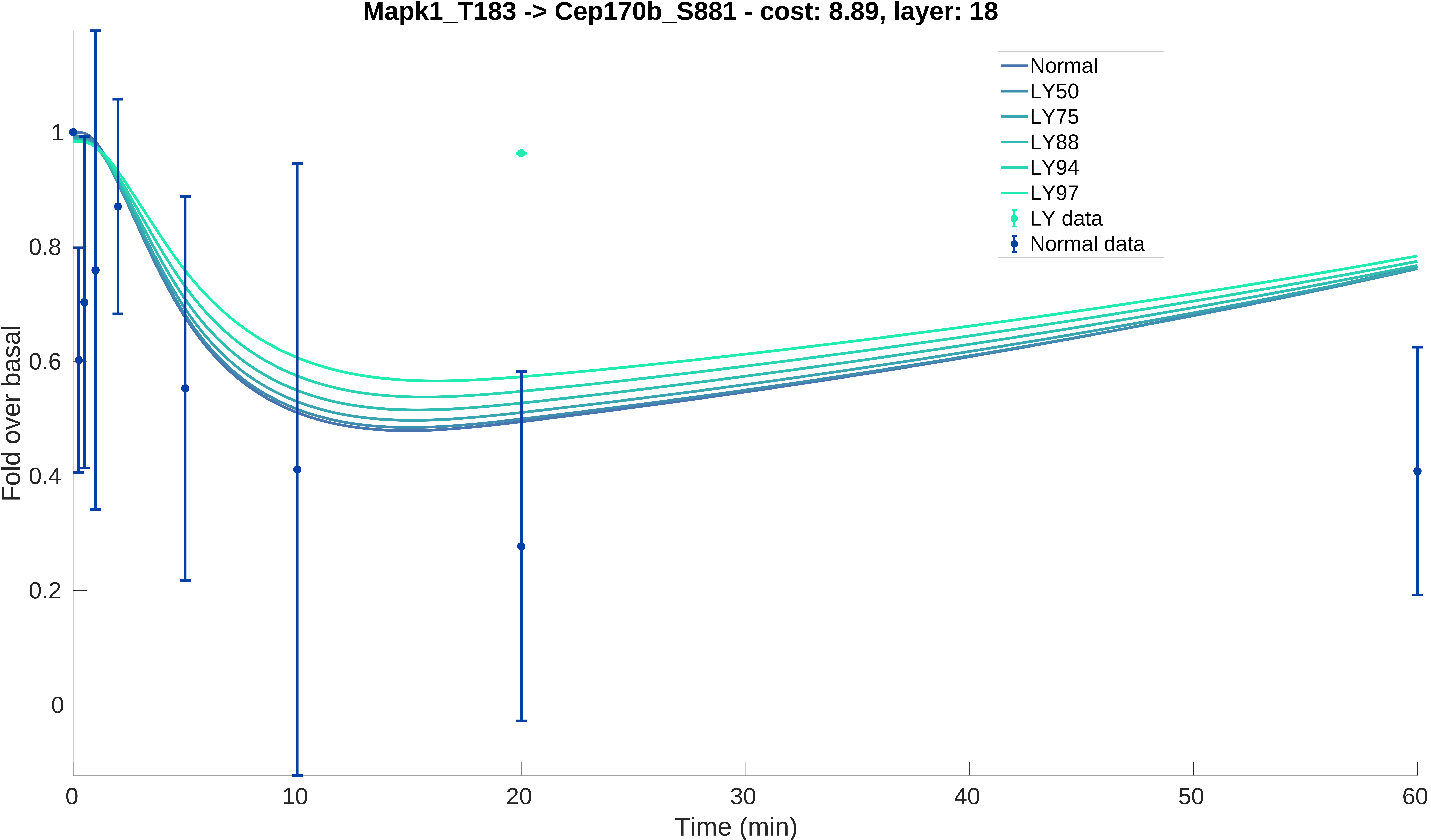

Mapk1_T183 -> Cep170b_S881 - cost: 8.89, layer: 18
Normal
LY50
LY75
LY88
LY94
LY97
LY data
Normal data
Fold over basal
0
0.2
0.4
0.6
0.8
1
0
10
20
30
40
50
60
Time (min)

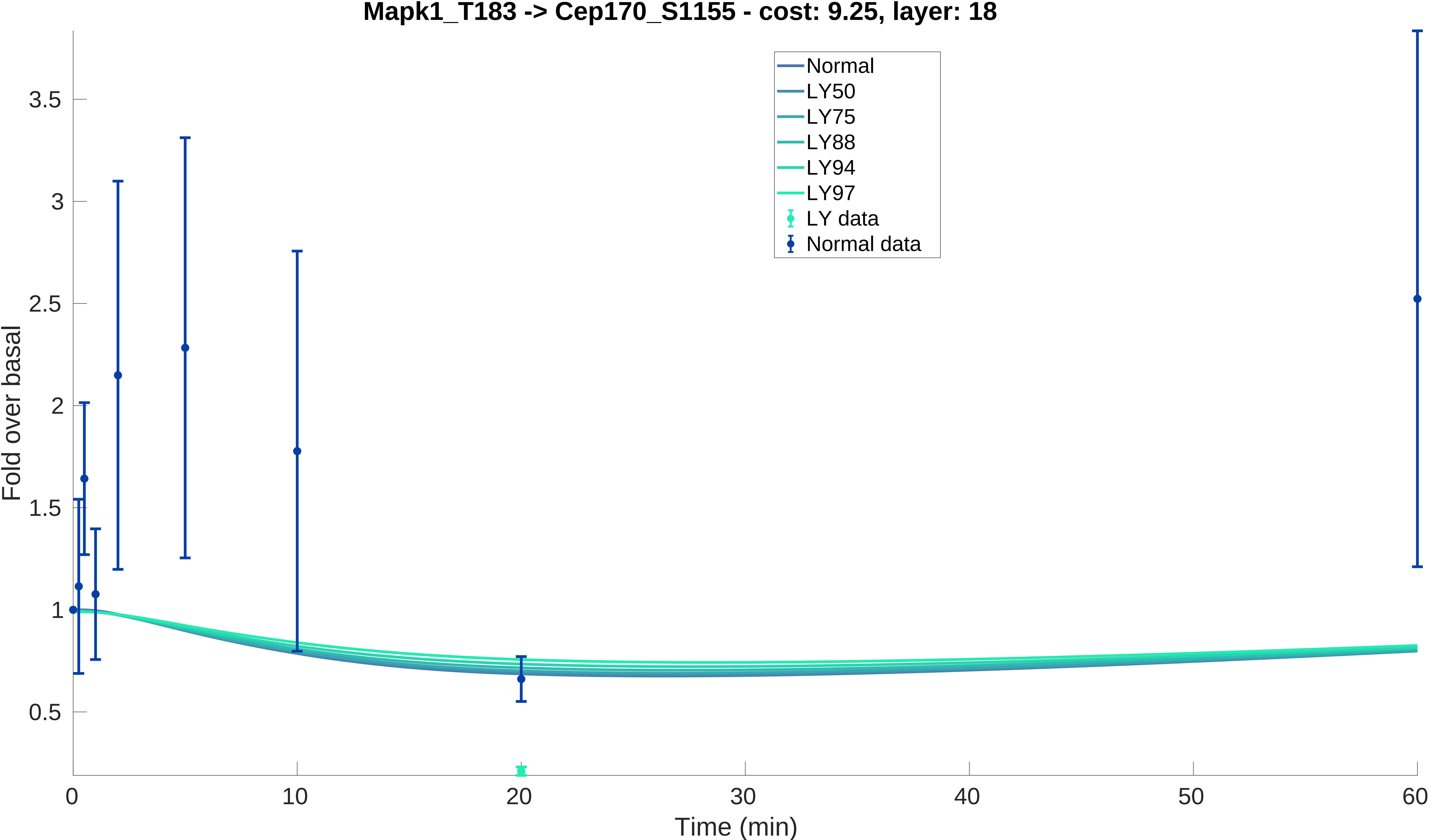
Mapk1_T183 -> Cep170_S1155 - cost: 9.25, layer: 18
Normal
LY50
LY75
LY88
LY94
LY97
LY data
Normal data
Fold over basal
3.5
3
2.5
2
1.5
1
0.5
0
10
20
30
40
50
60
Time (min)

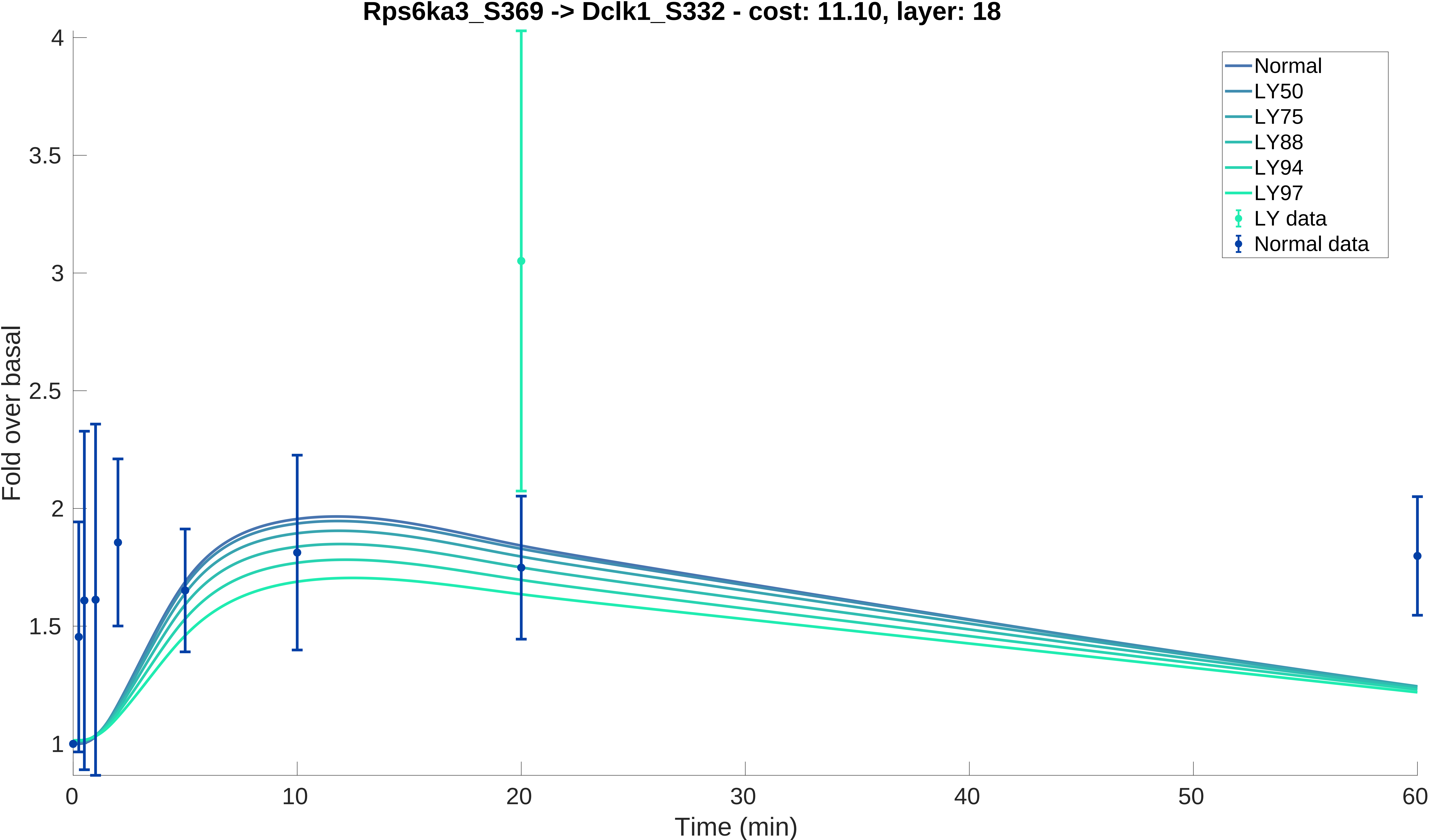

Rps6ka3_S369 -> Dclk1_S332 - cost: 11.10, layer: 18
Fold over basal
Time (min)
Normal
LY50
LY75
LY88
LY94
LY97
LY data
Normal data
1
1.5
2
2.5
3
3.5
4
0
10
20
30
40
50
60

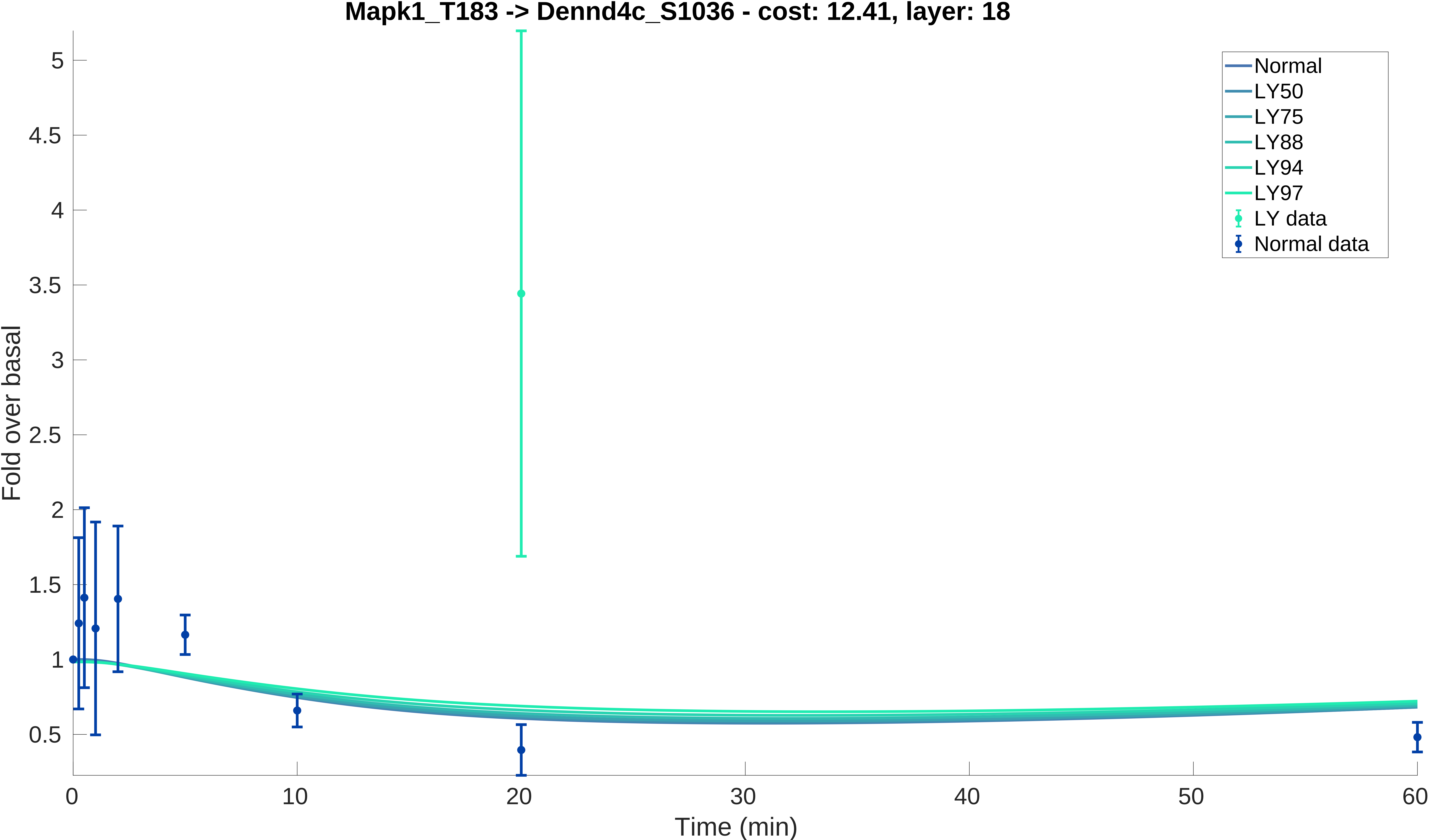

Mapk1_T183 -> Dennd4c_S1036 - cost: 12.41, layer: 18
Fold over basal
Time (min)
0.5
1
1.5
2
2.5
3
3.5
4
4.5
5
0
10
20
30
40
50
60
Normal
LY50
LY75
LY88
LY94
LY97
LY data
Normal data

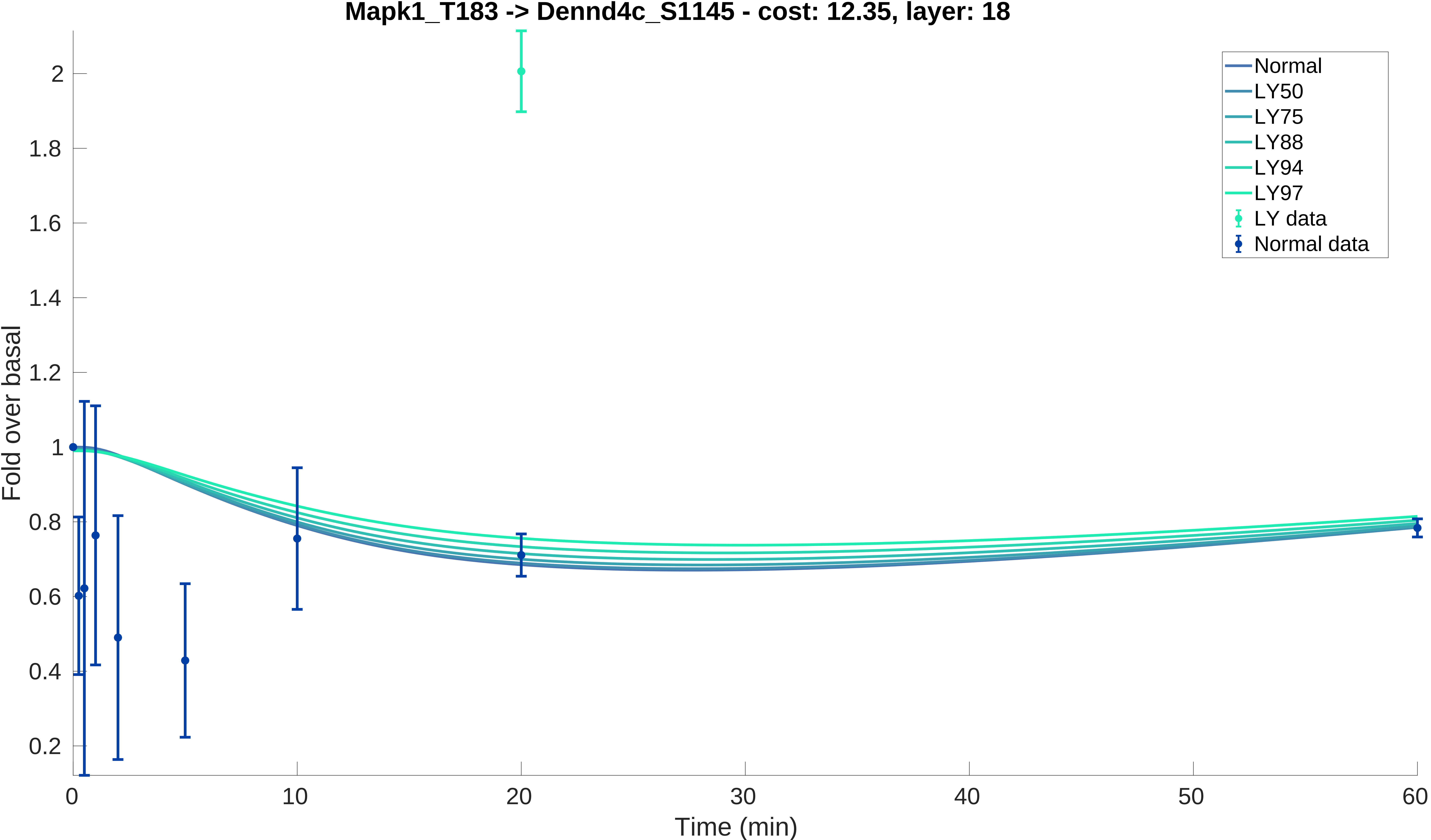

Mapk1_T183 -> Dennd4c_S1145 - cost: 12.35, layer: 18
Normal
LY50
LY75
LY88
LY94
LY97
LY data
Normal data
Fold over basal
Time (min)
0
10
20
30
40
50
60
0.2
0.4
0.6
0.8
1
1.2
1.4
1.6
1.8
2

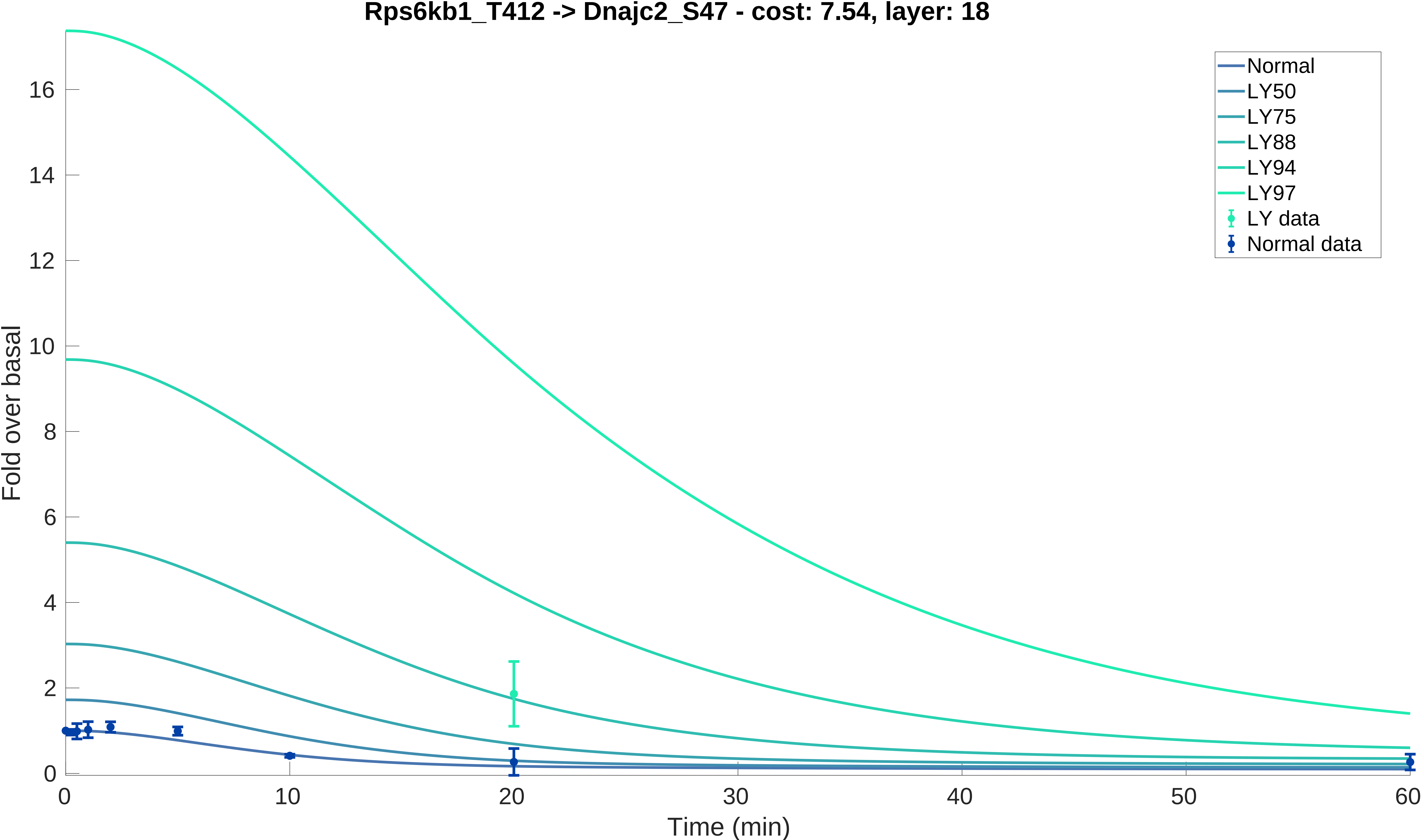

Rps6kb1_T412 -> Dnajc2_S47 - cost: 7.54, layer: 18
Normal
LY50
LY75
LY88
LY94
LY97
LY data
Normal data
Fold over basal
0
2
4
6
8
10
12
14
16
0
10
20
30
40
50
60
Time (min)

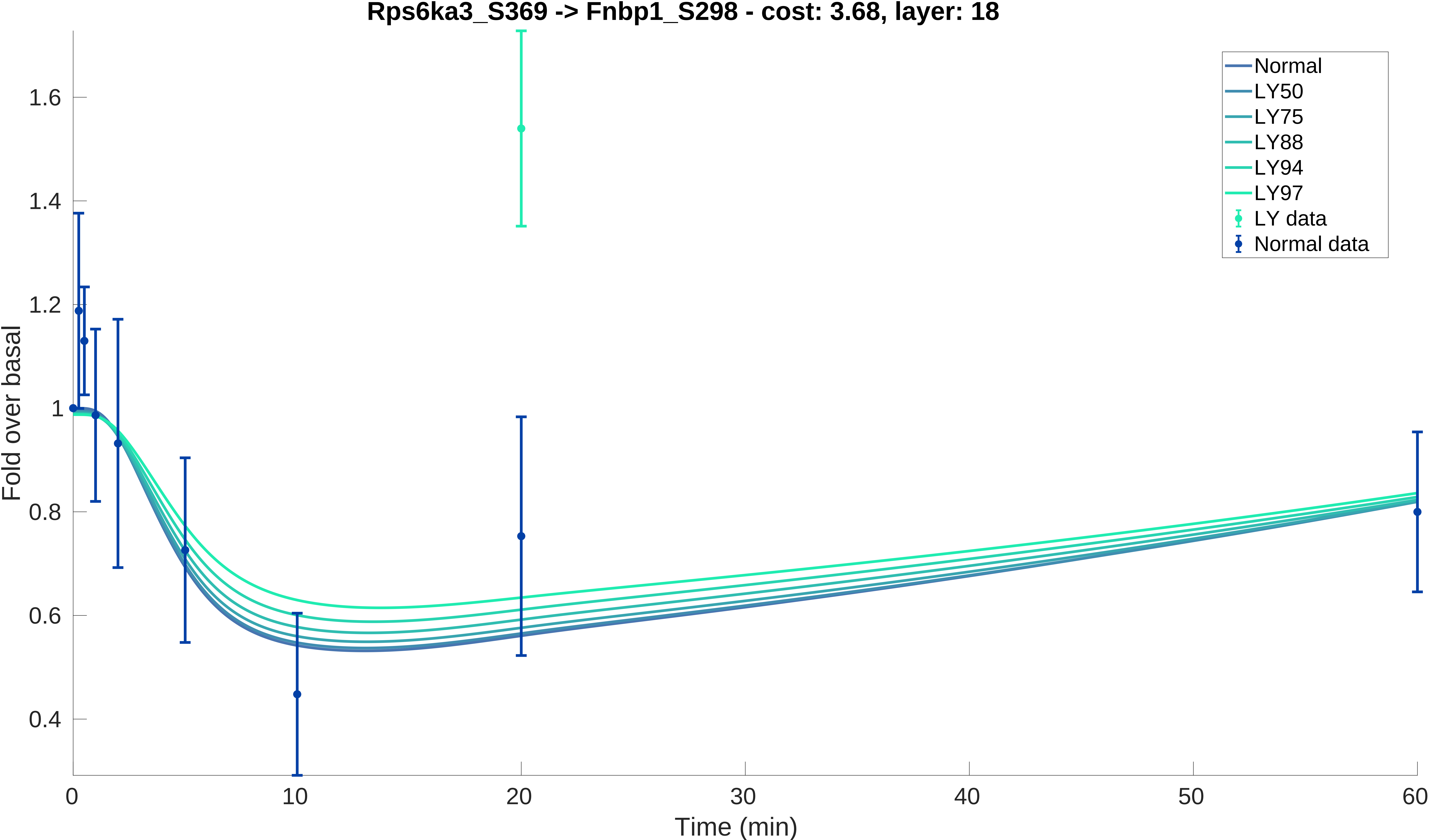
Rps6ka3_S369 -> Fnbp1_S298 - cost: 3.68, layer: 18
Fold over basal
1.6
1.4
1.2
1
0.8
0.6
0.4
0
10
20
30
40
50
60
Time (min)
Normal
LY50
LY75
LY88
LY94
LY97
LY data
Normal data

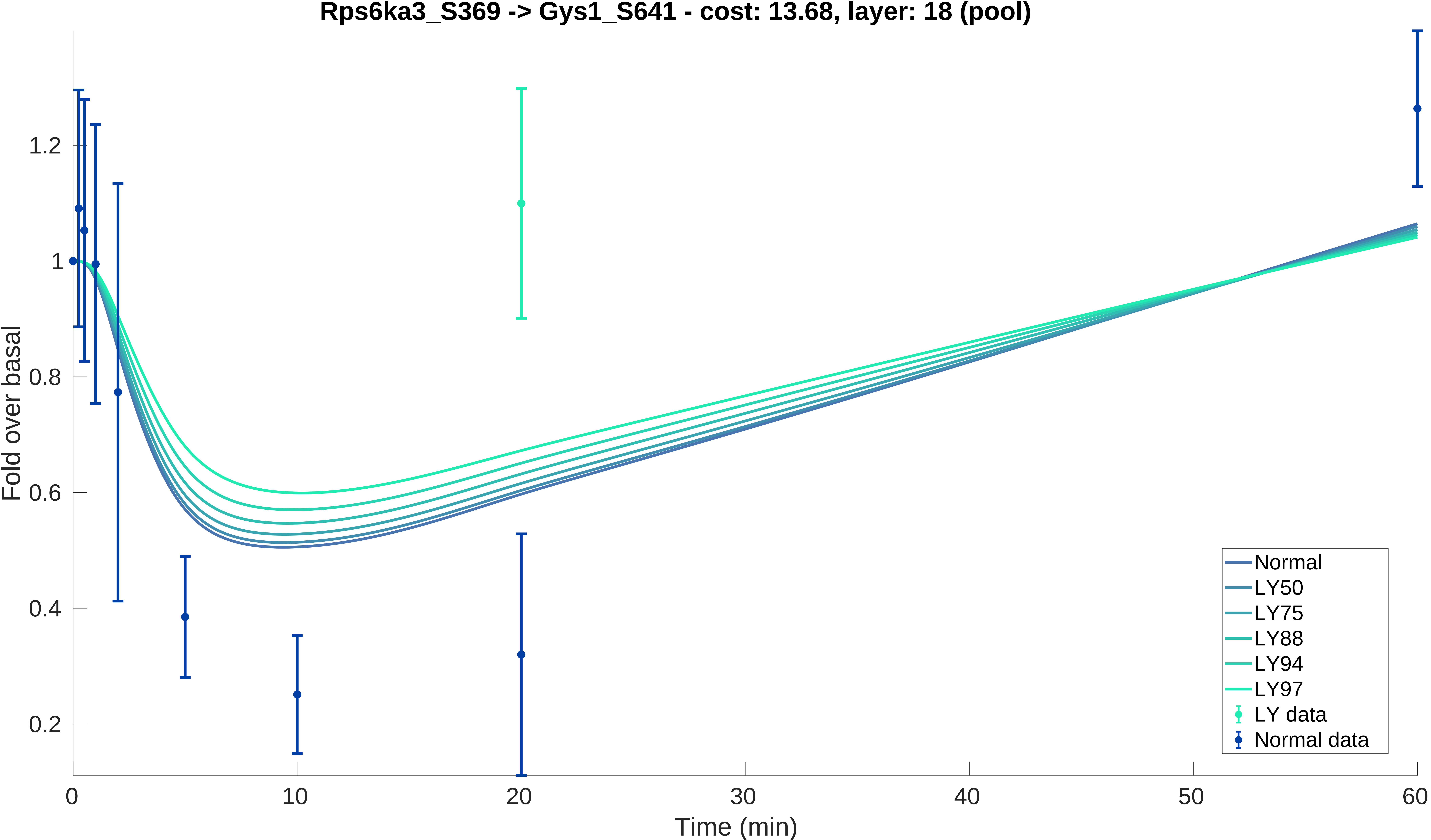
Rps6ka3_S369 -> Gys1_S641 - cost: 13.68, layer: 18 (pool)
Fold over basal
1.2
1
0.8
0.6
0.4
0.2
0
10
20
30
40
50
60
Time (min)
Normal
LY50
LY75
LY88
LY94
LY97
LY data
Normal data

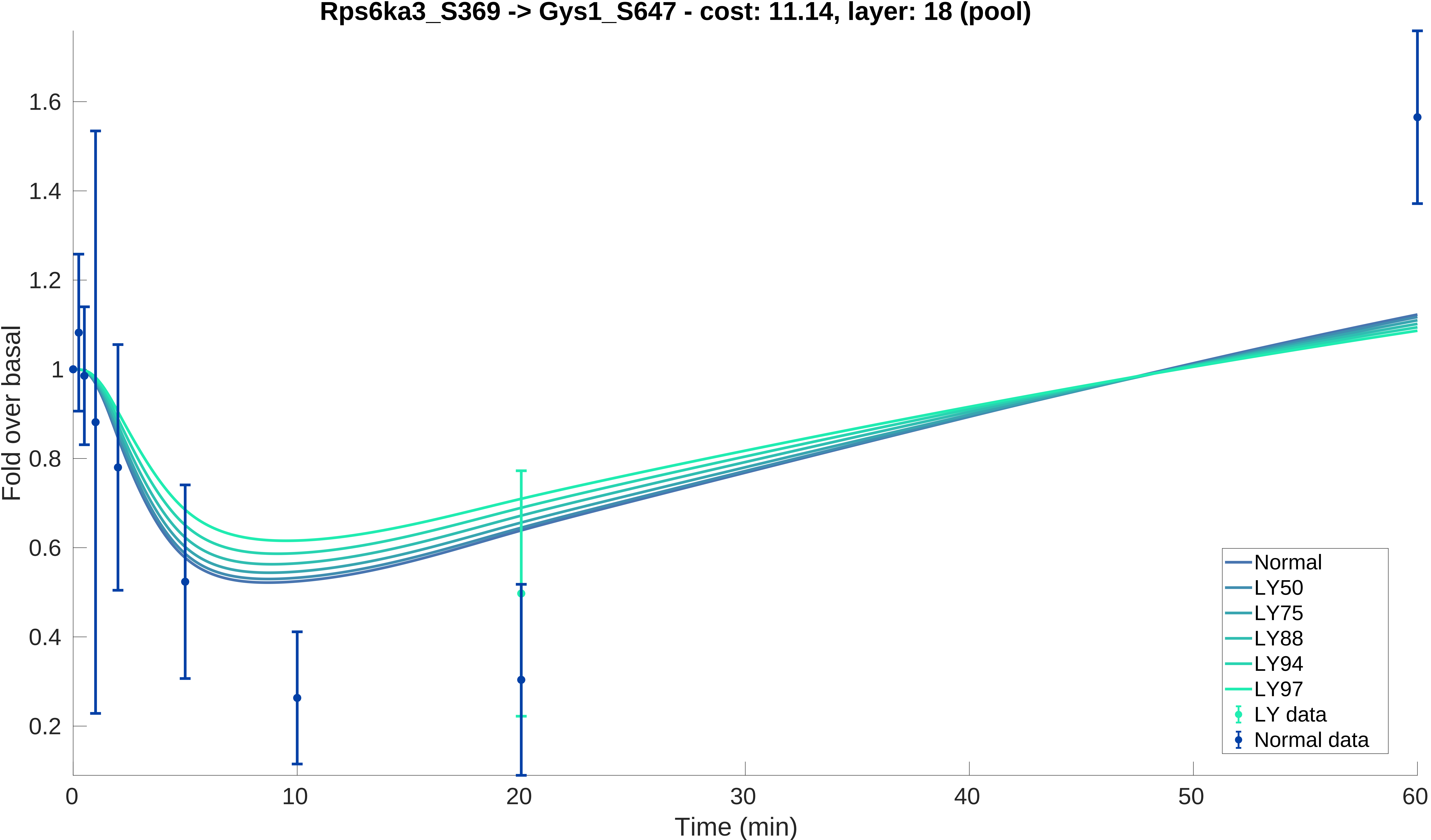
Rps6ka3_S369 -> Gys1_S647 - cost: 11.14, layer: 18 (pool)
Fold over basal
1.6
1.4
1.2
1
0.8
0.6
0.4
0.2
0
10
20
30
40
50
60
Time (min)
Normal
LY50
LY75
LY88
LY94
LY97
LY data
Normal data

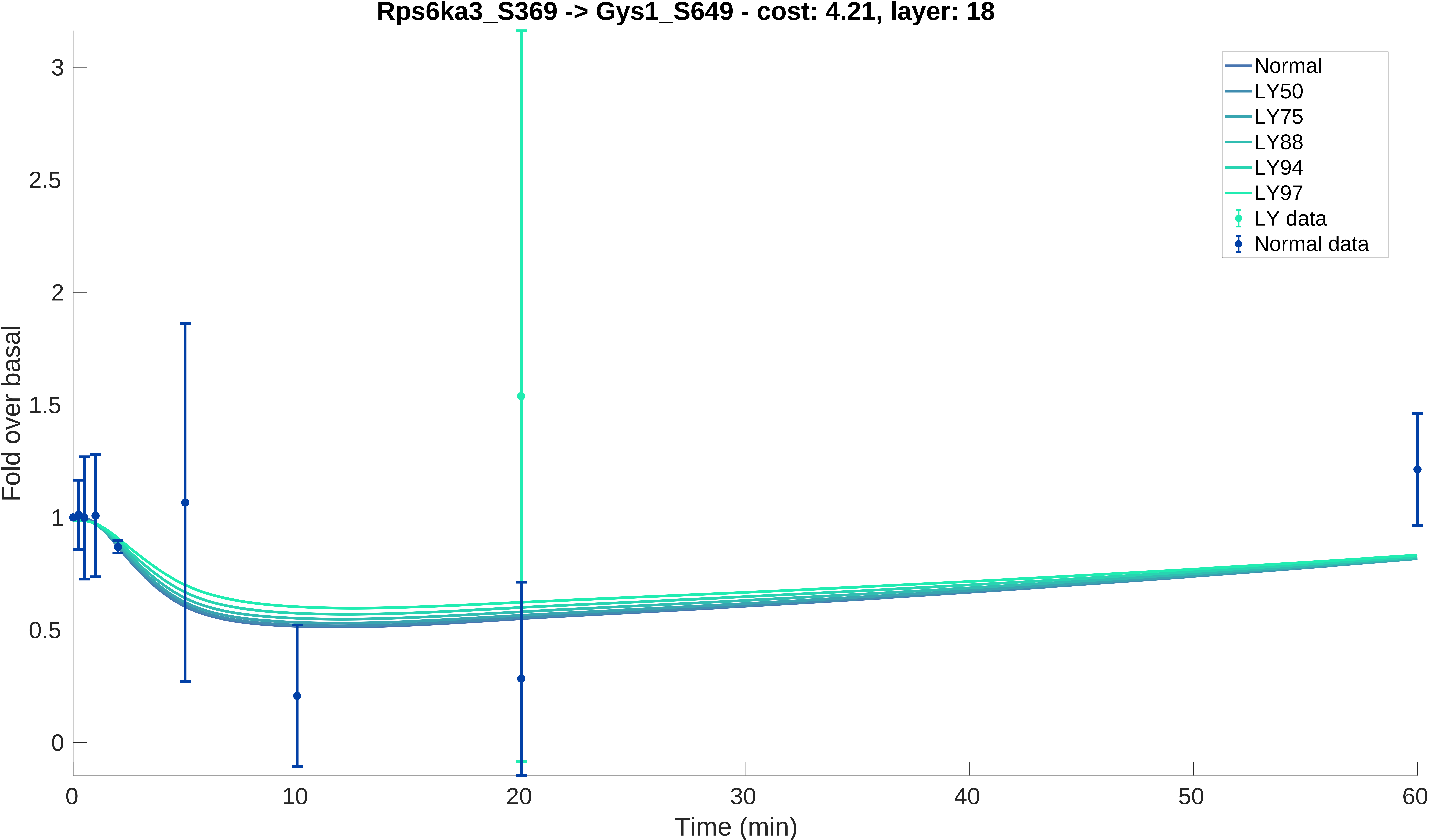

Rps6ka3_S369 -> Gys1_S649 - cost: 4.21, layer: 18
Fold over basal
Time (min)
0
0.5
1
1.5
2
2.5
3
0
10
20
30
40
50
60
Normal
LY50
LY75
LY88
LY94
LY97
LY data
Normal data

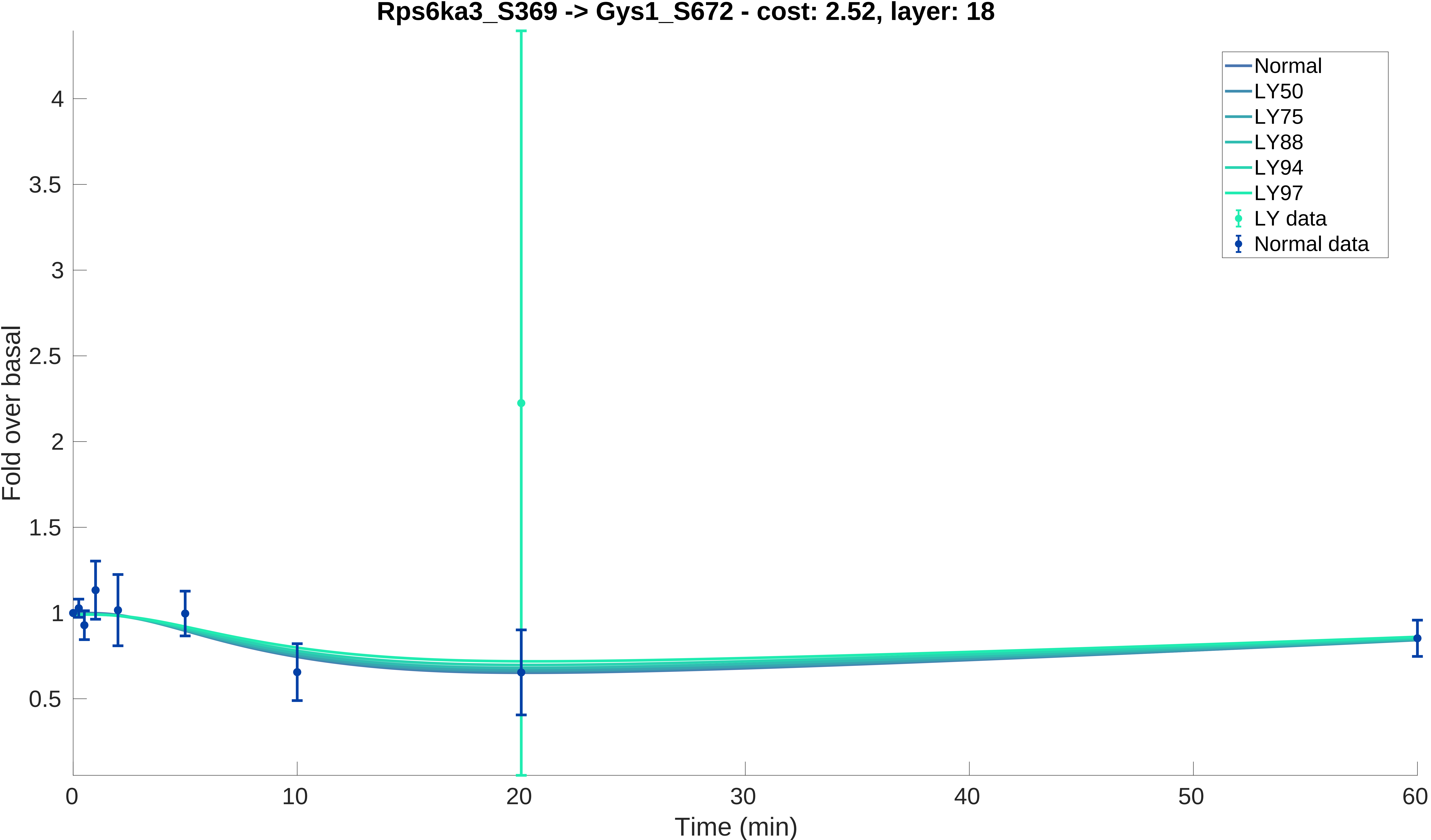
Rps6ka3_S369 -> Gys1_S672 - cost: 2.52, layer: 18
Fold over basal
4
3.5
3
2.5
2
1.5
1
0.5
0
10
20
30
40
50
60
Time (min)
Normal
LY50
LY75
LY88
LY94
LY97
LY data
Normal data

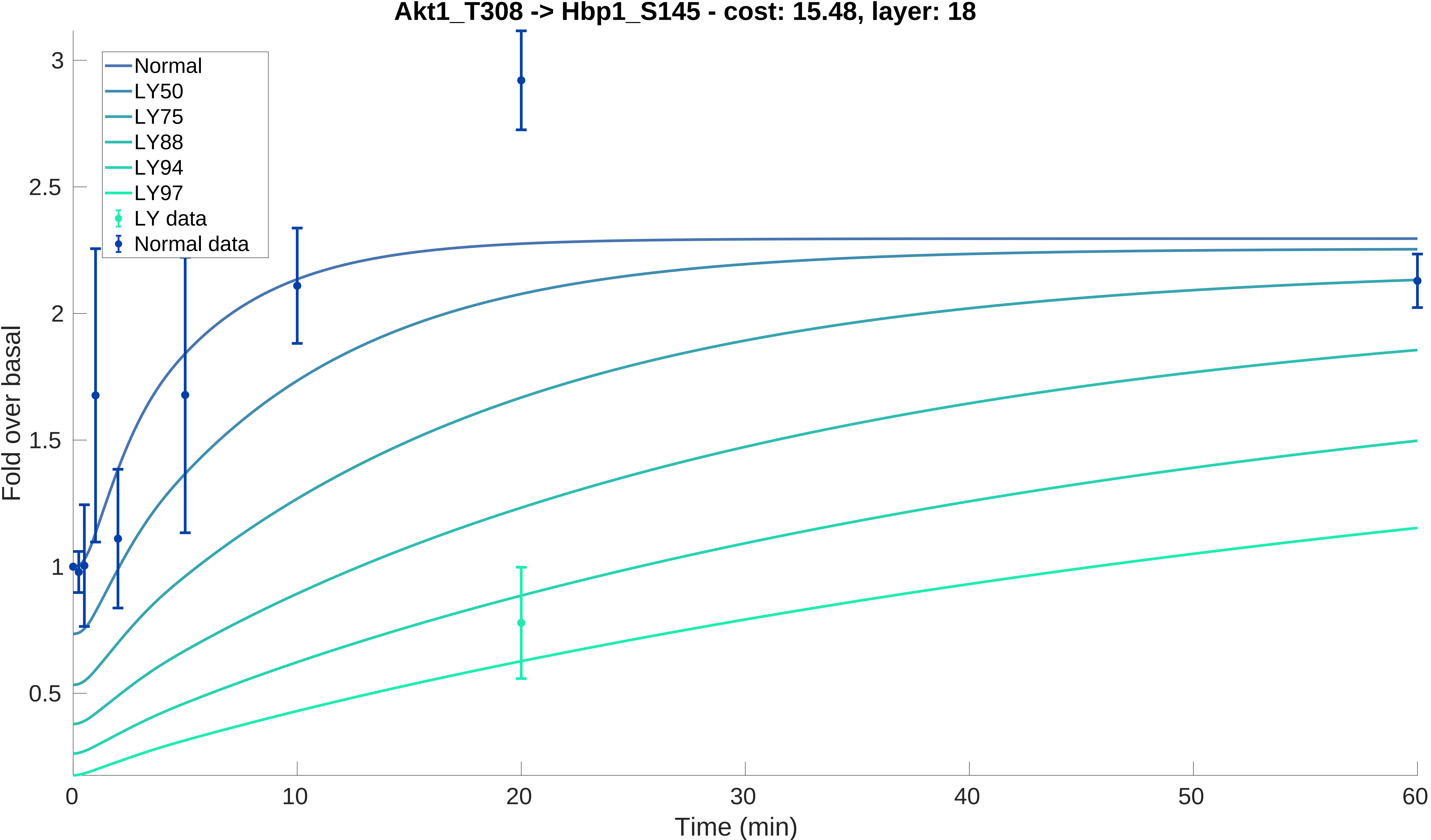

Akt1_T308 -> Hbp1_S145 - cost: 15.48, layer: 18
Normal
LY50
LY75
LY88
LY94
LY97
LY data
Normal data
Fold over basal
Time (min)
0.5
1
1.5
2
2.5
3
0
10
20
30
40
50
60

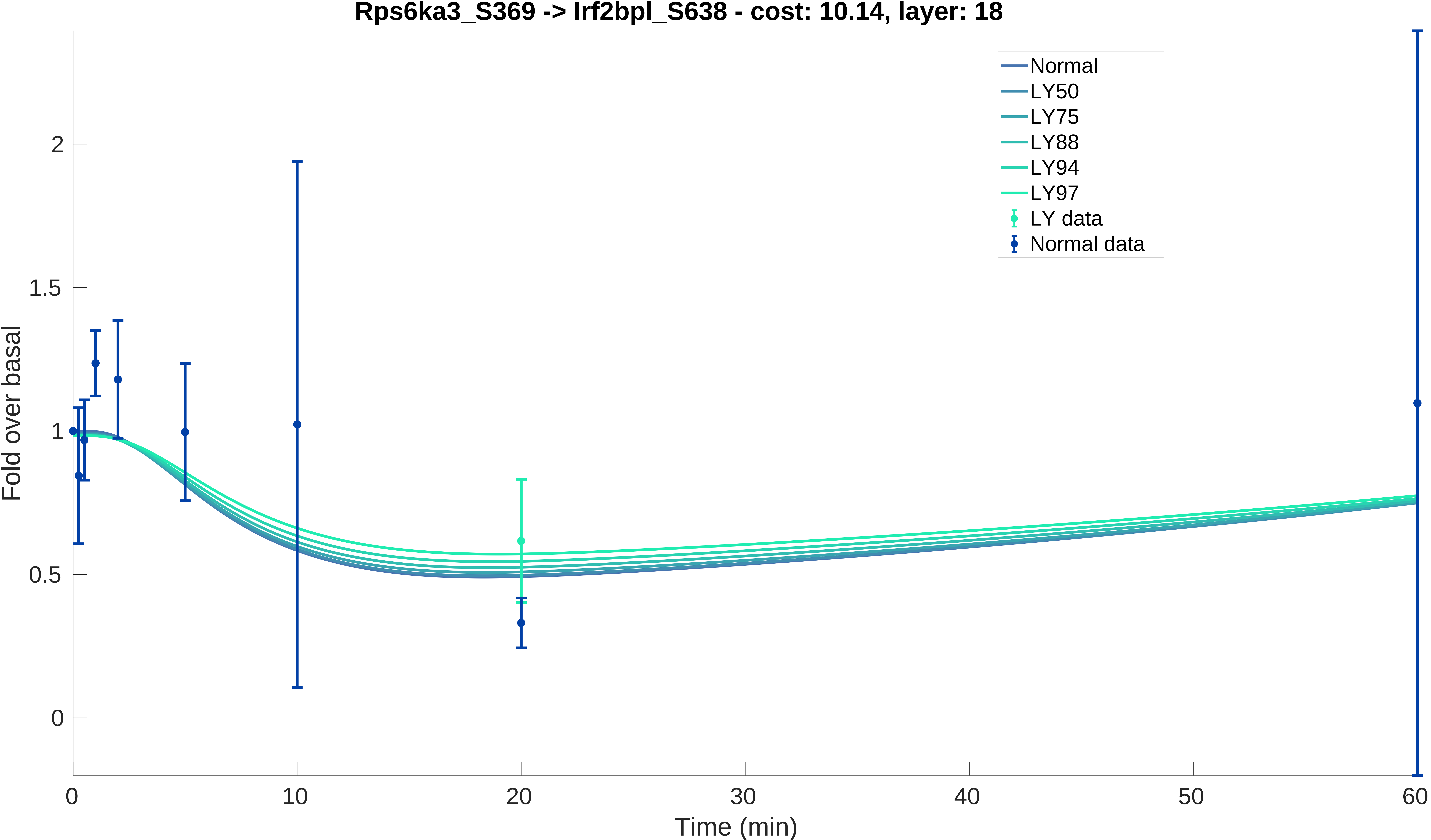

Rps6ka3_S369 -> Irf2bpl_S638 - cost: 10.14, layer: 18
Normal
LY50
LY75
LY88
LY94
LY97
LY data
Normal data
Fold over basal
Time (min)
0
0.5
1
1.5
2
0
10
20
30
40
50
60

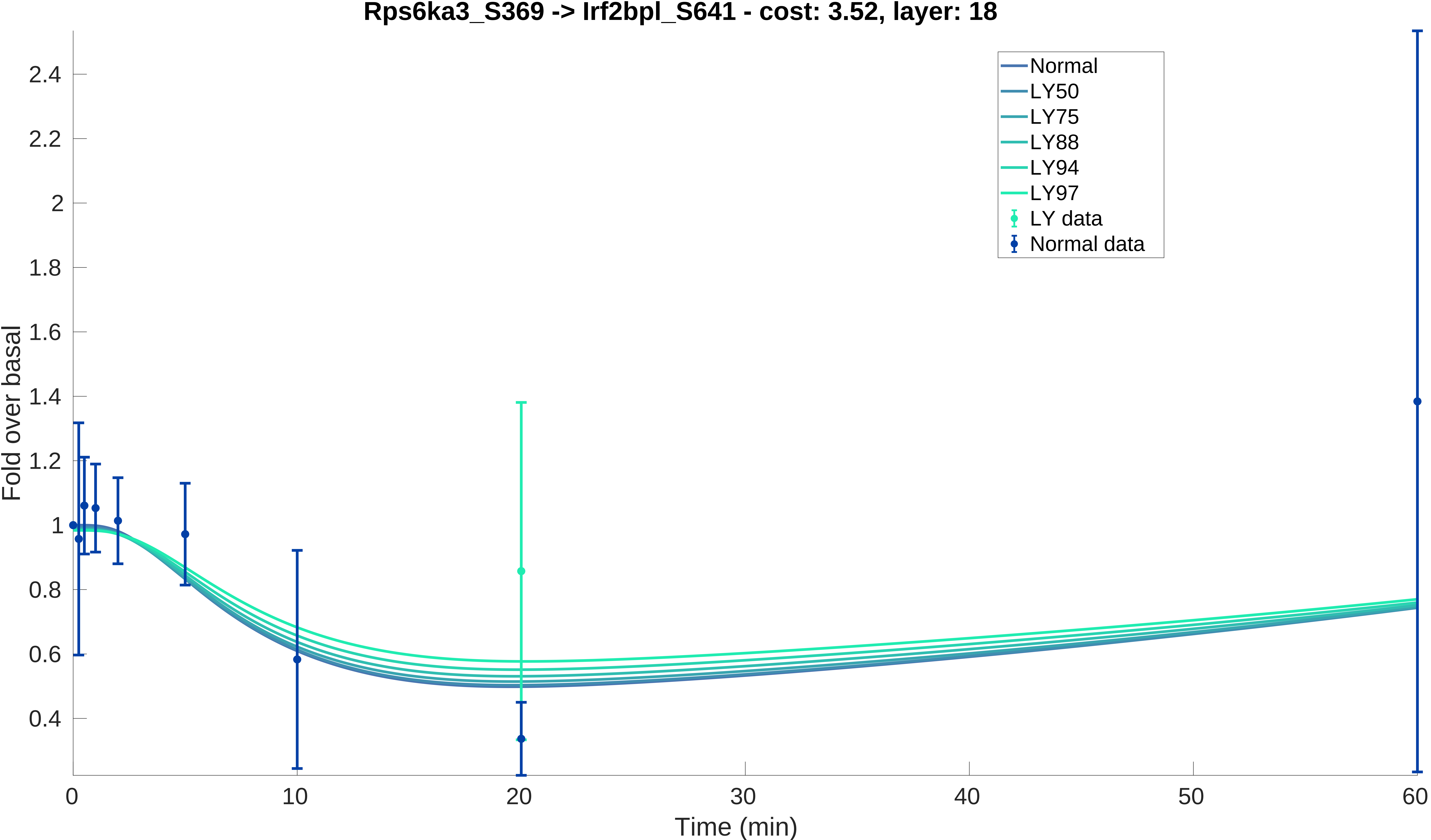

Rps6ka3_S369 -> Irf2bpl_S641 - cost: 3.52, layer: 18
Normal
LY50
LY75
LY88
LY94
LY97
LY data
Normal data
Fold over basal
Time (min)
2.4
2.2
2
1.8
1.6
1.4
1.2
1
0.8
0.6
0.4
0
10
20
30
40
50
60

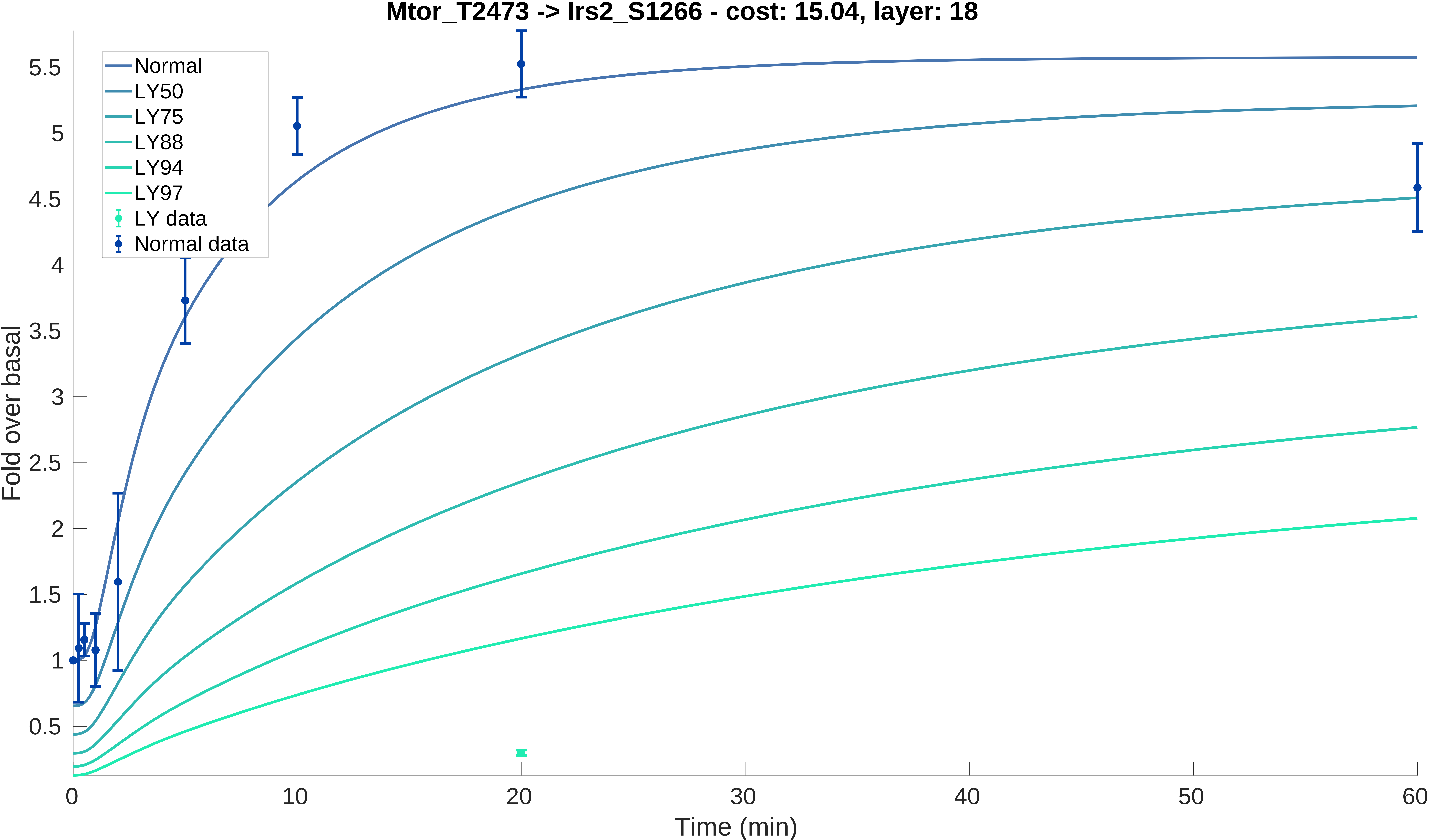

Mtor_T2473 -> Irs2_S1266 - cost: 15.04, layer: 18
Normal
LY50
LY75
LY88
LY94
LY97
LY data
Normal data
Fold over basal
Time (min)
0
10
20
30
40
50
60
0.5
1
1.5
2
2.5
3
3.5
4
4.5
5
5.5

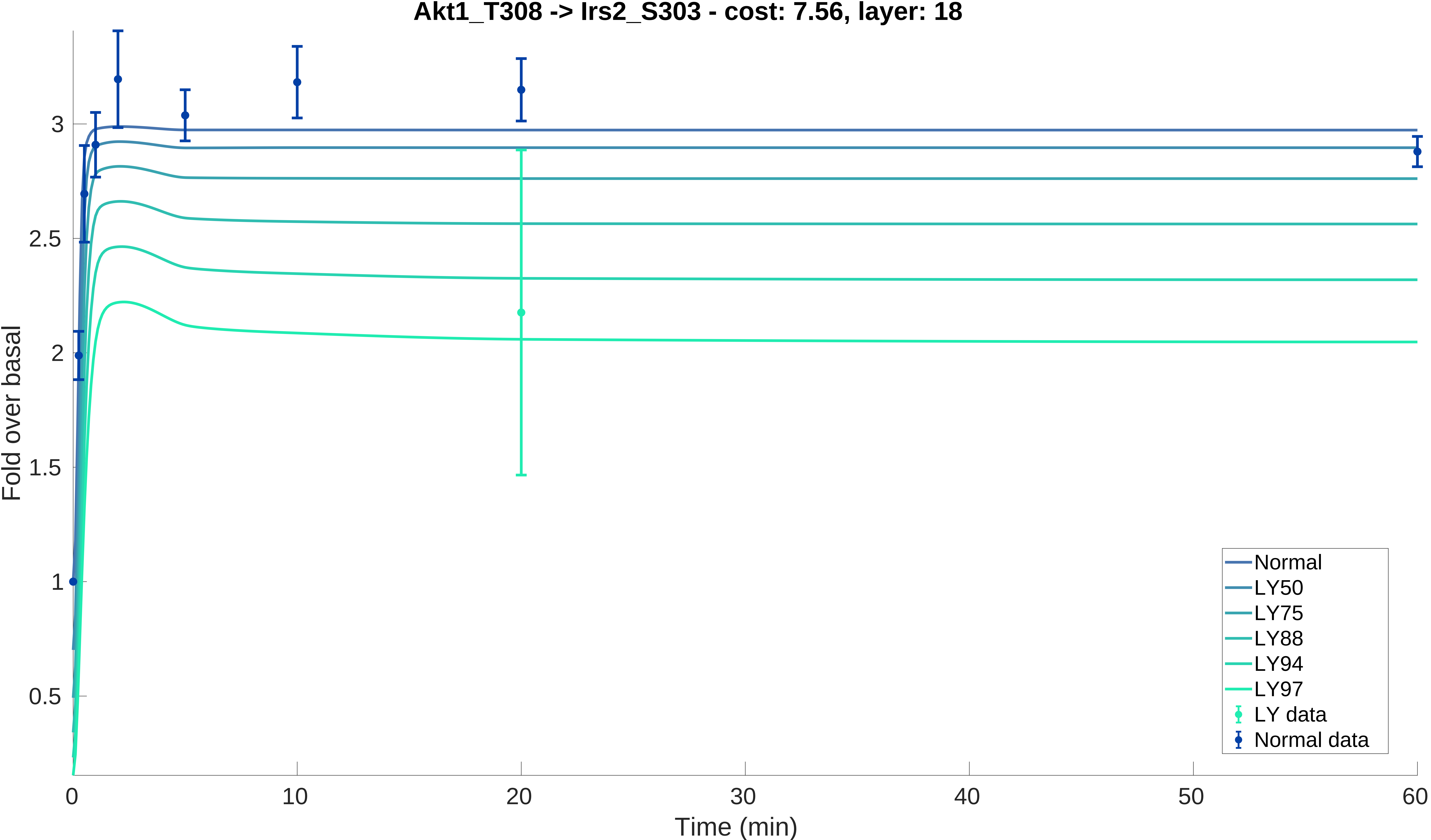

Akt1_T308 -> Irs2_S303 - cost: 7.56, layer: 18
Fold over basal
3
2.5
2
1.5
1
0.5
0
10
20
30
40
50
60
Time (min)
Normal
LY50
LY75
LY88
LY94
LY97
LY data
Normal data

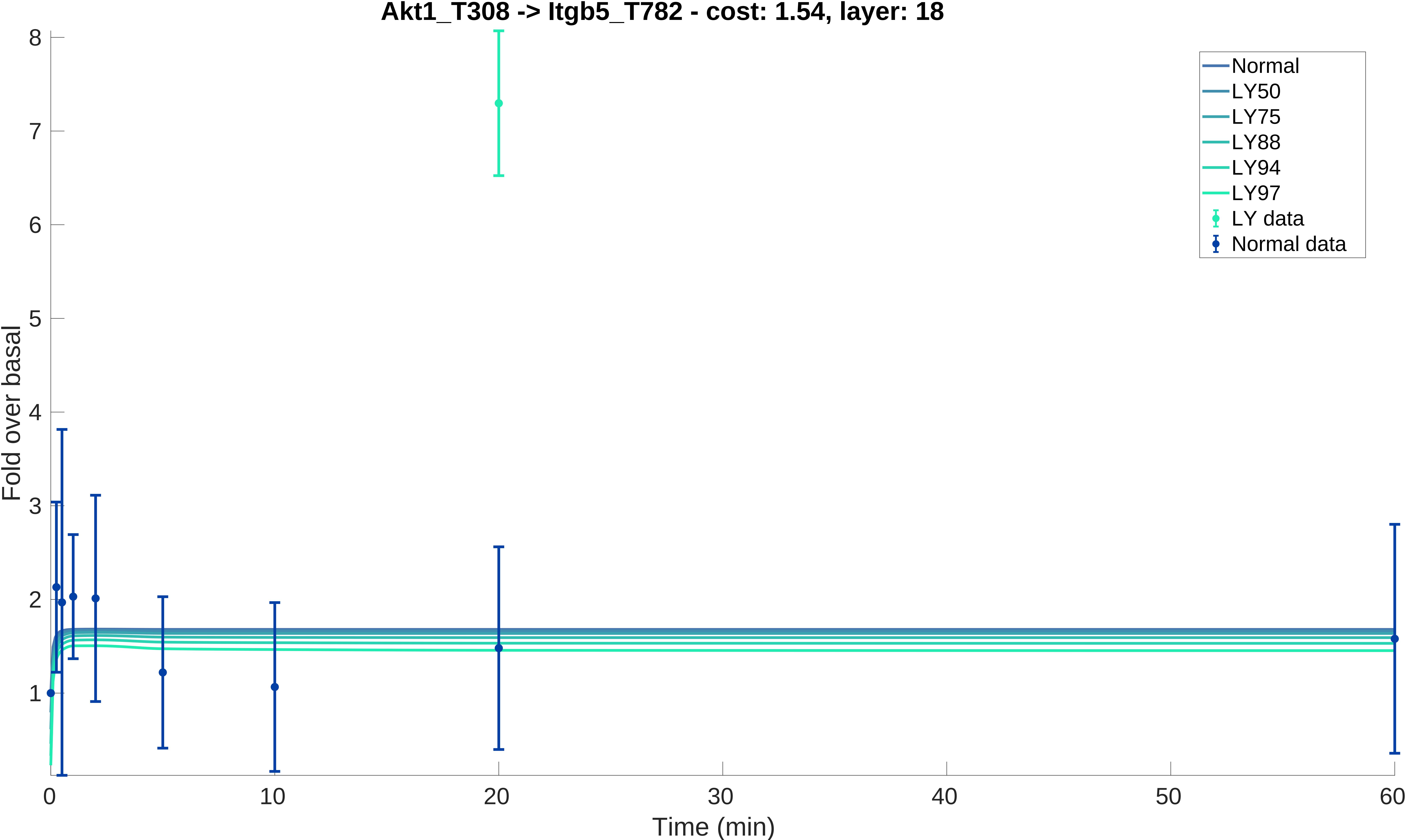

Akt1_T308 -> Itgb5_T782 - cost: 1.54, layer: 18
Fold over basal
Time (min)
0
10
20
30
40
50
60
1
2
3
4
5
6
7
8
Normal
LY50
LY75
LY88
LY94
LY97
LY data
Normal data

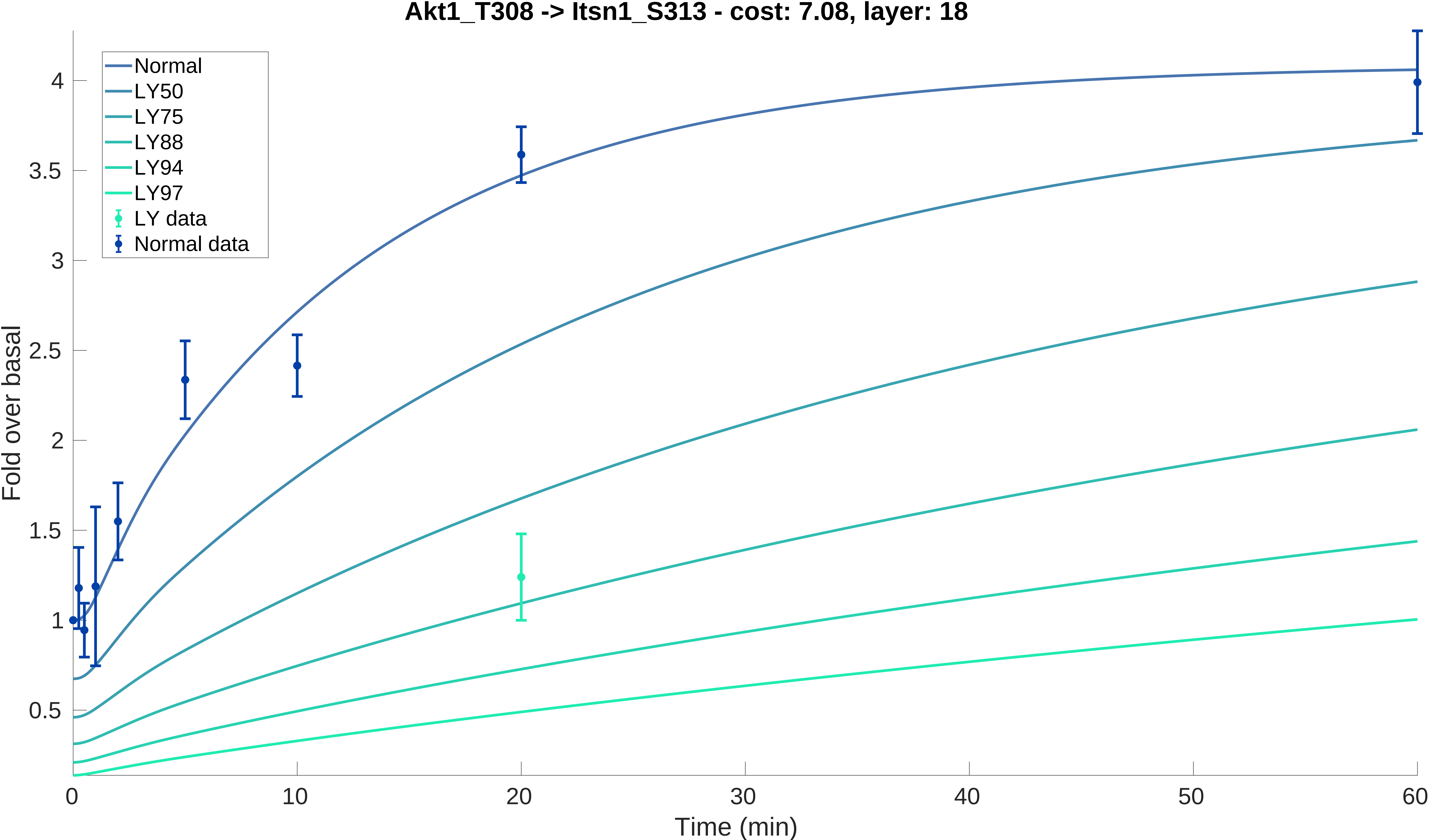

Akt1_T308 -> Itsn1_S313 - cost: 7.08, layer: 18
Normal
LY50
LY75
LY88
LY94
LY97
LY data
Normal data
Fold over basal
Time (min)
0
10
20
30
40
50
60
0.5
1
1.5
2
2.5
3
3.5
4

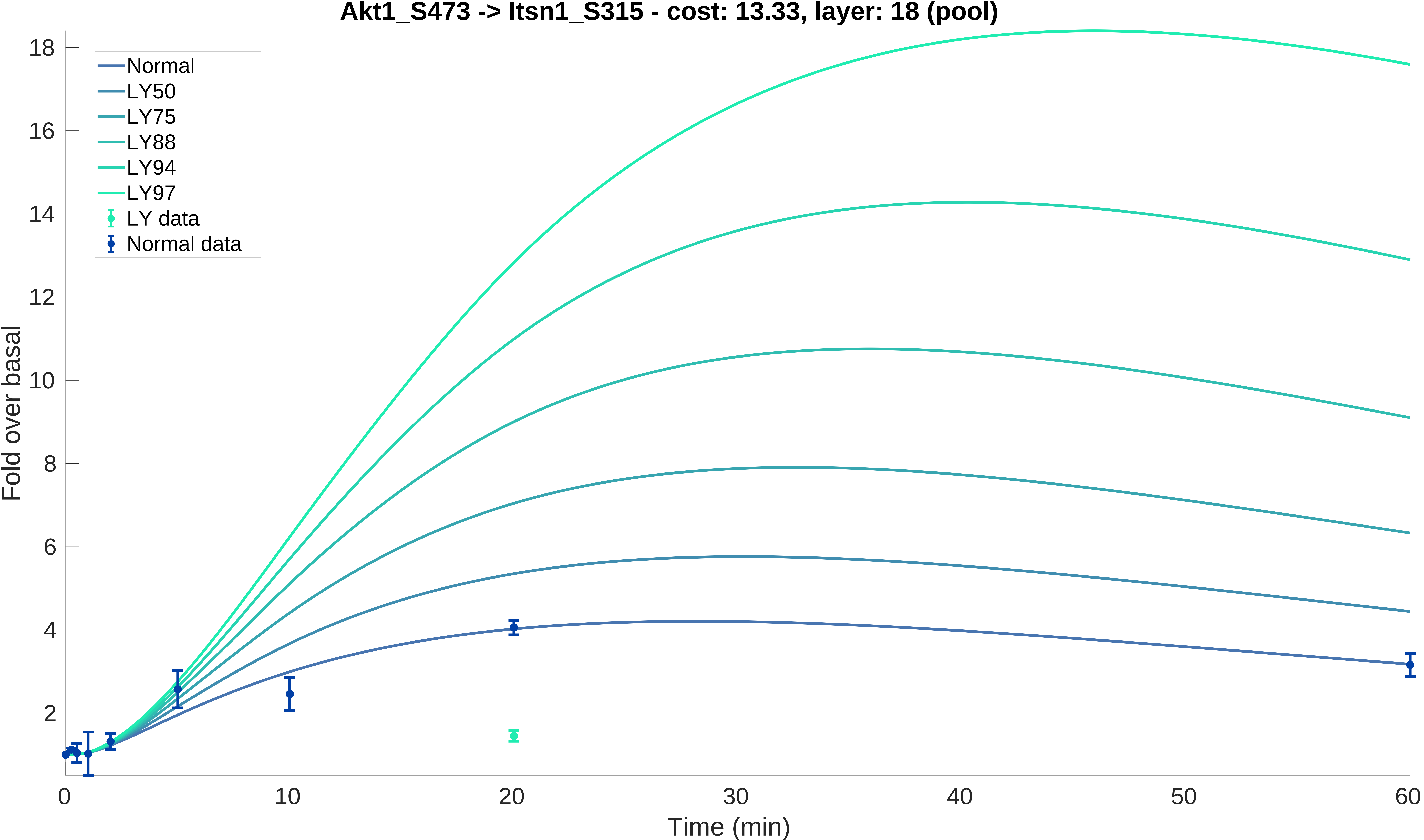
Akt1_S473 -> Itsn1_S315 - cost: 13.33, layer: 18 (pool)
Normal
LY50
LY75
LY88
LY94
LY97
LY data
Normal data
Fold over basal
Time (min)
0
10
20
30
40
50
60
2
4
6
8
10
12
14
16
18

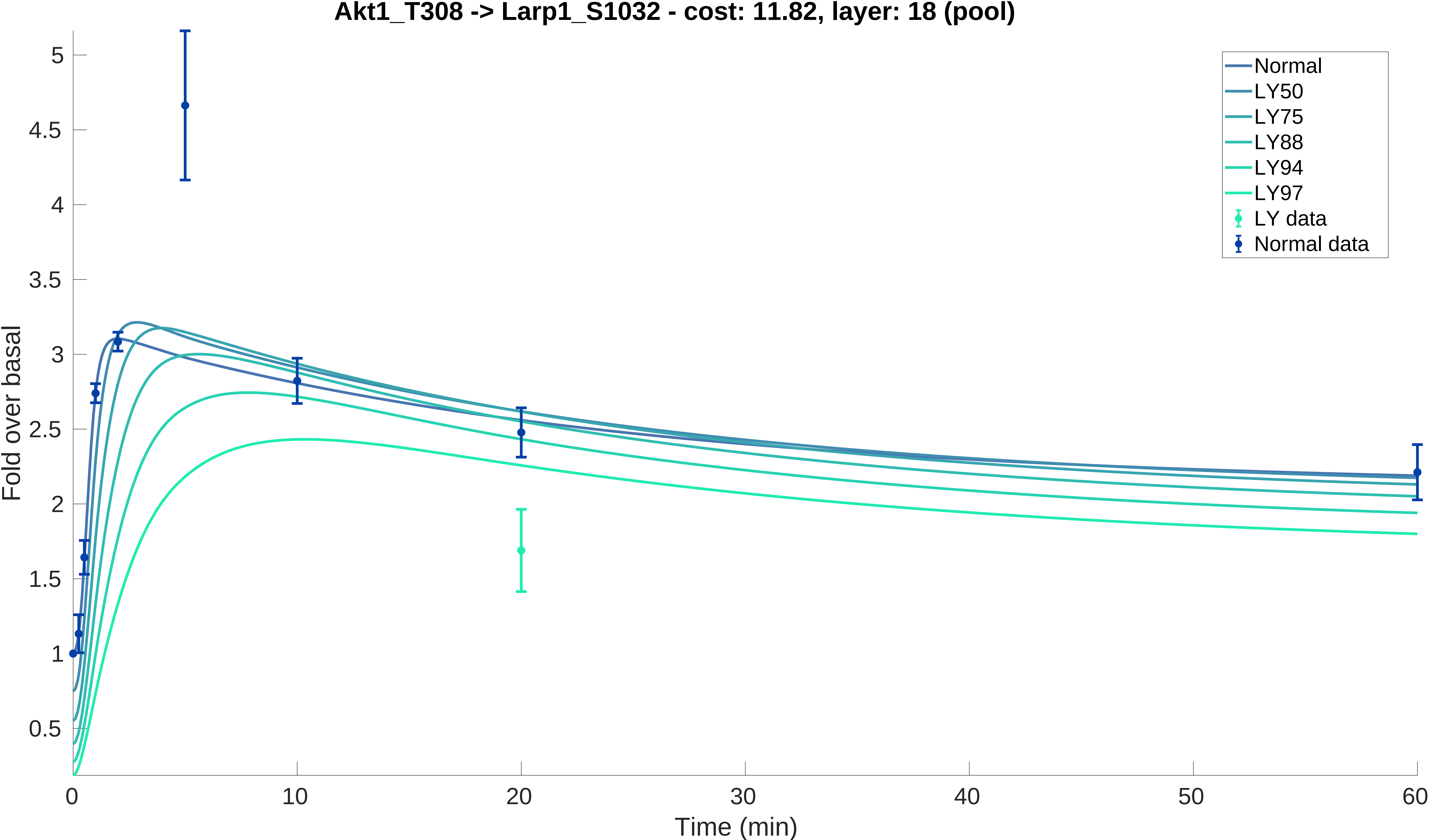

Akt1_T308 -> Larp1_S1032 - cost: 11.82, layer: 18 (pool)
Fold over basal
Time (min)
0.5
1
1.5
2
2.5
3
3.5
4
4.5
5
0
10
20
30
40
50
60
Normal
LY50
LY75
LY88
LY94
LY97
LY data
Normal data

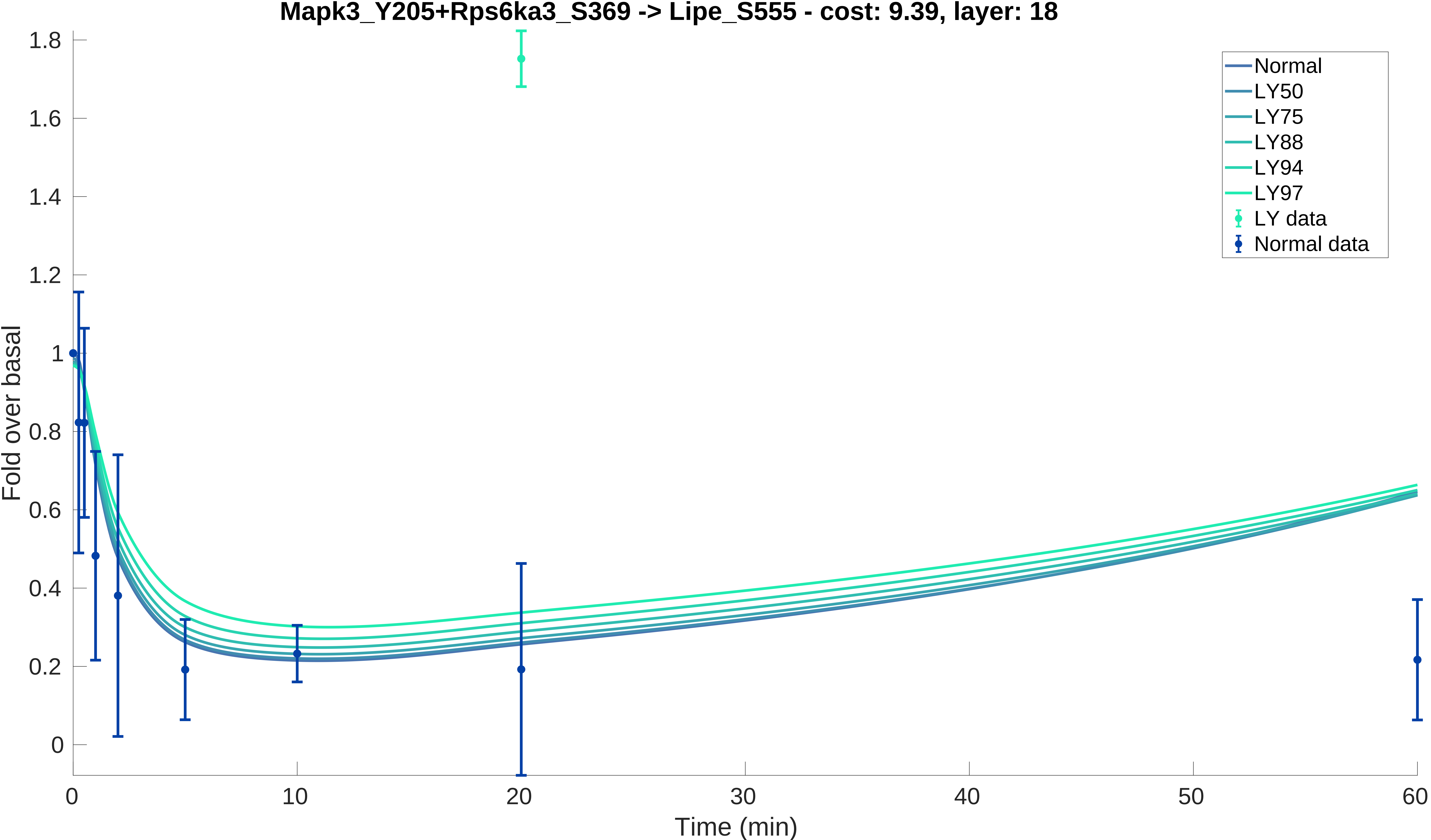
Mapk3_Y205+Rps6ka3_S369 -> Lipe_S555 - cost: 9.39, layer: 18
Fold over basal
Time (min)
0
0.2
0.4
0.6
0.8
1
1.2
1.4
1.6
1.8
0
10
20
30
40
50
60
Normal
LY50
LY75
LY88
LY94
LY97
LY data
Normal data

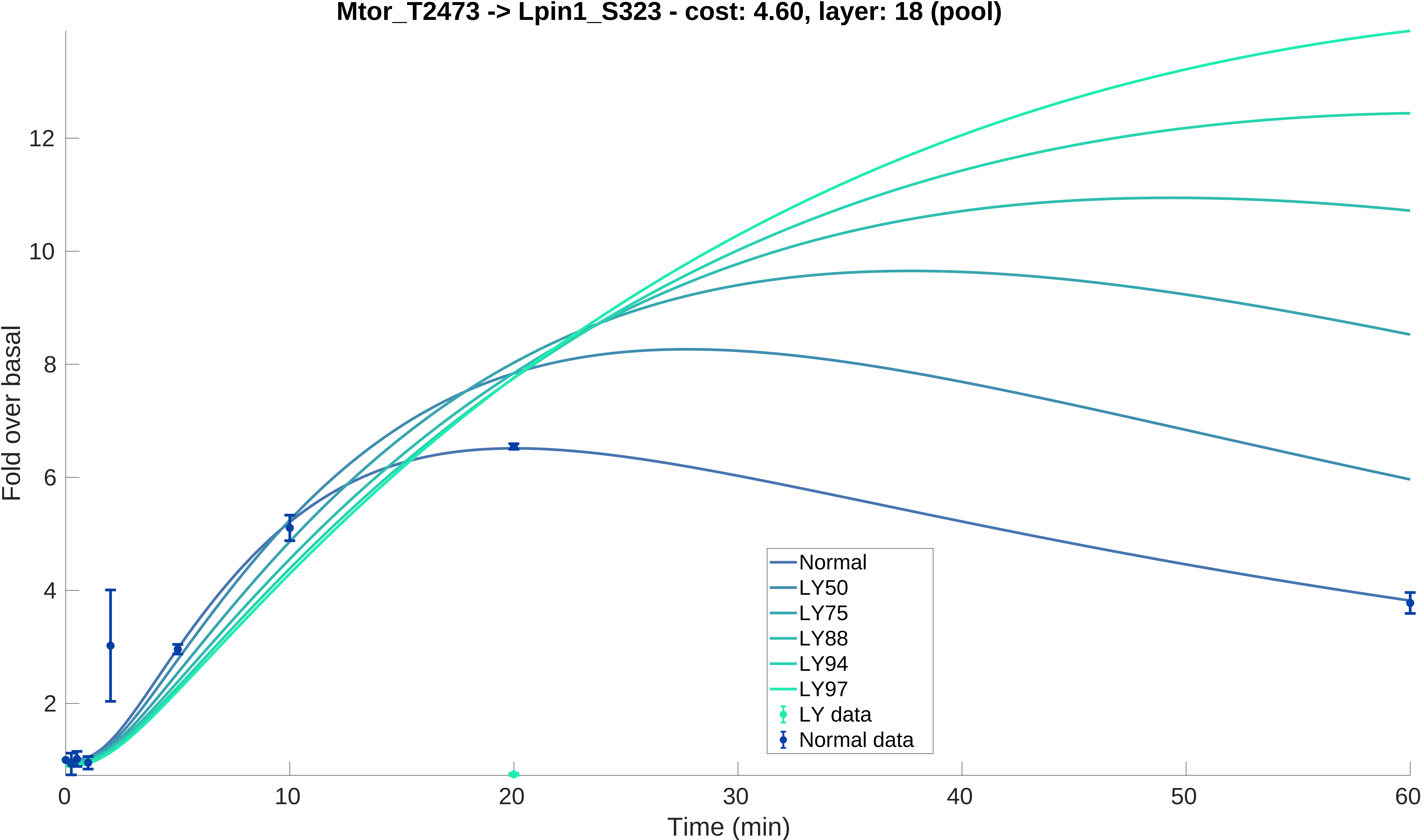

Mtor_T2473 -> Lpin1_S323 - cost: 4.60, layer: 18 (pool)
Fold over basal
Time (min)
0
10
20
30
40
50
60
2
4
6
8
10
12
Normal
LY50
LY75
LY88
LY94
LY97
LY data
Normal data

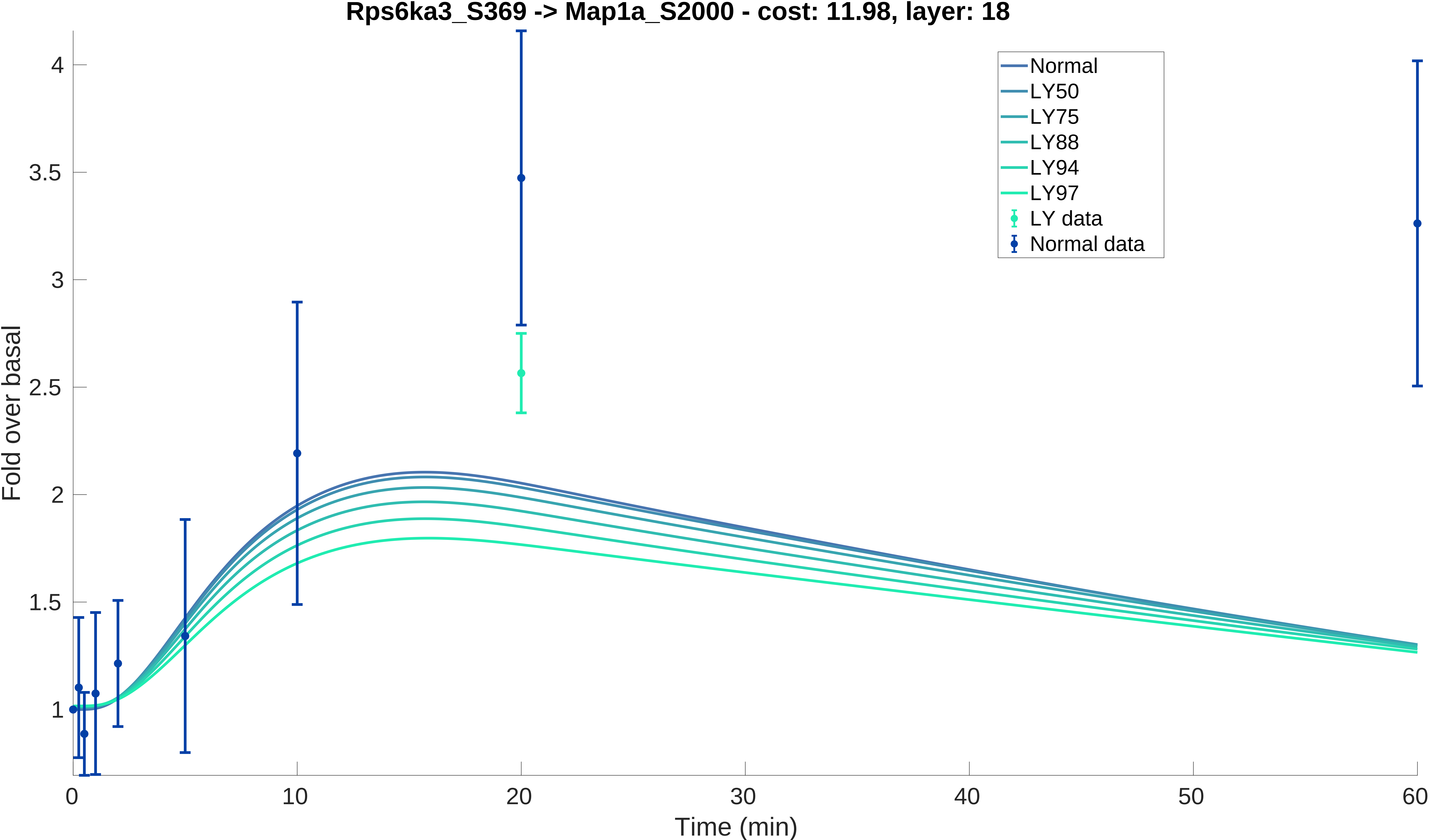
Rps6ka3_S369 -> Map1a_S2000 - cost: 11.98, layer: 18
Fold over basal
Time (min)
Normal
LY50
LY75
LY88
LY94
LY97
LY data
Normal data
4
3.5
3
2.5
2
1.5
1
0
10
20
30
40
50
60

#### Akt1_T308 -> Map2_S1787 - cost: 0.55, layer: 18

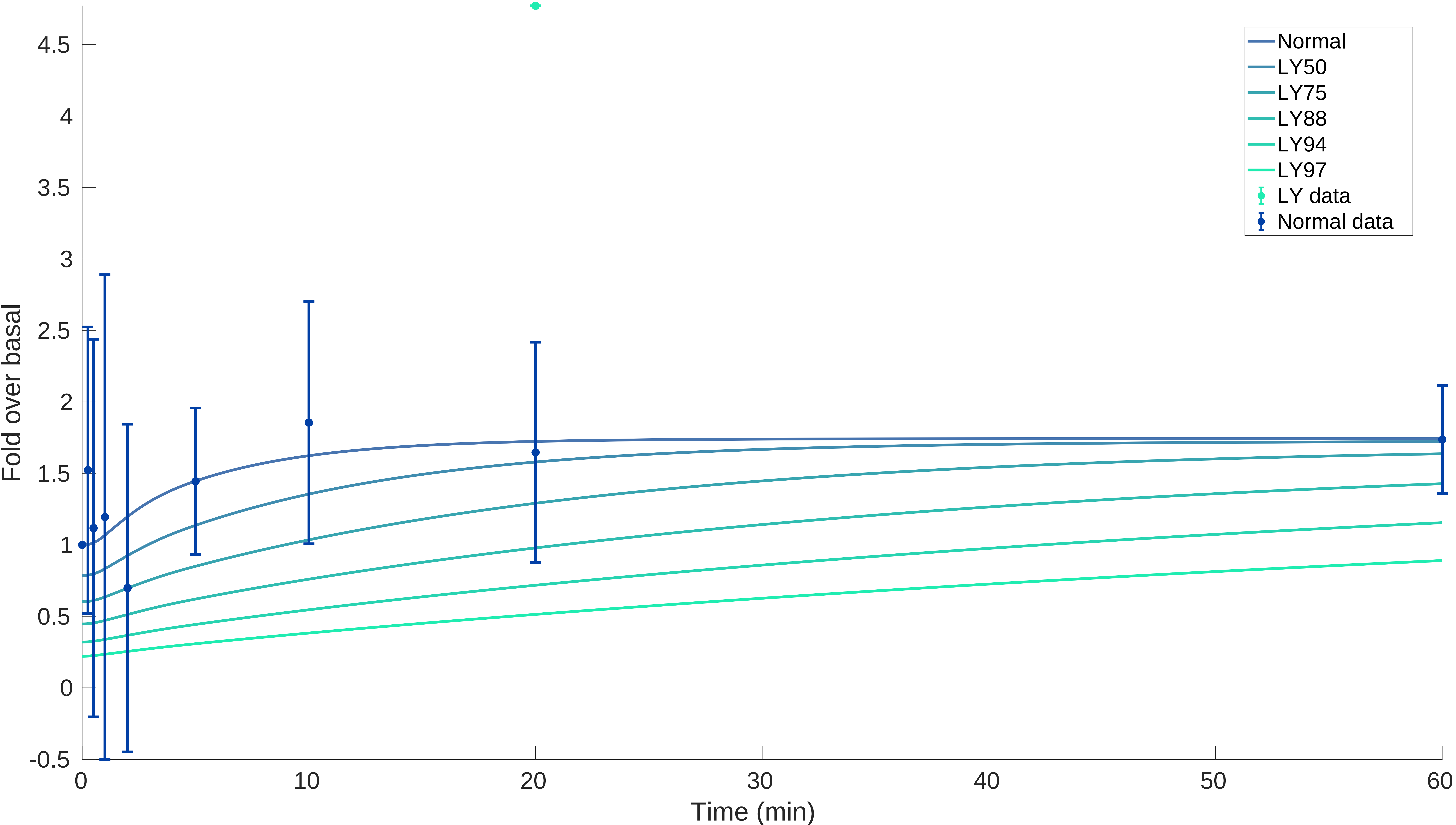

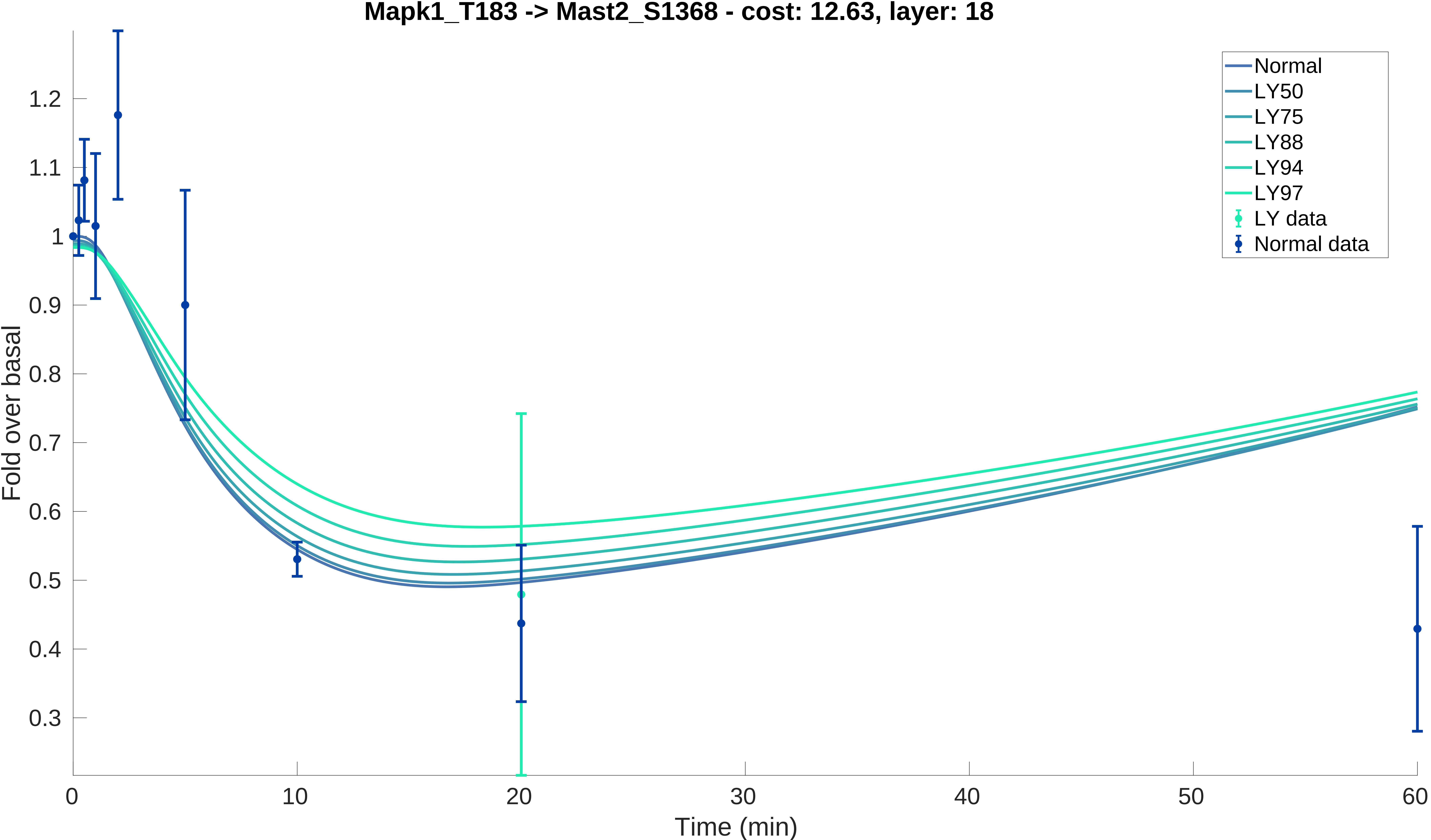

Mapk1_T183 -> Mast2_S1368 - cost: 12.63, layer: 18
Normal
LY50
LY75
LY88
LY94
LY97
LY data
Normal data
Fold over basal
Time (min)
1.2
1.1
1
0.9
0.8
0.7
0.6
0.5
0.4
0.3
0
10
20
30
40
50
60

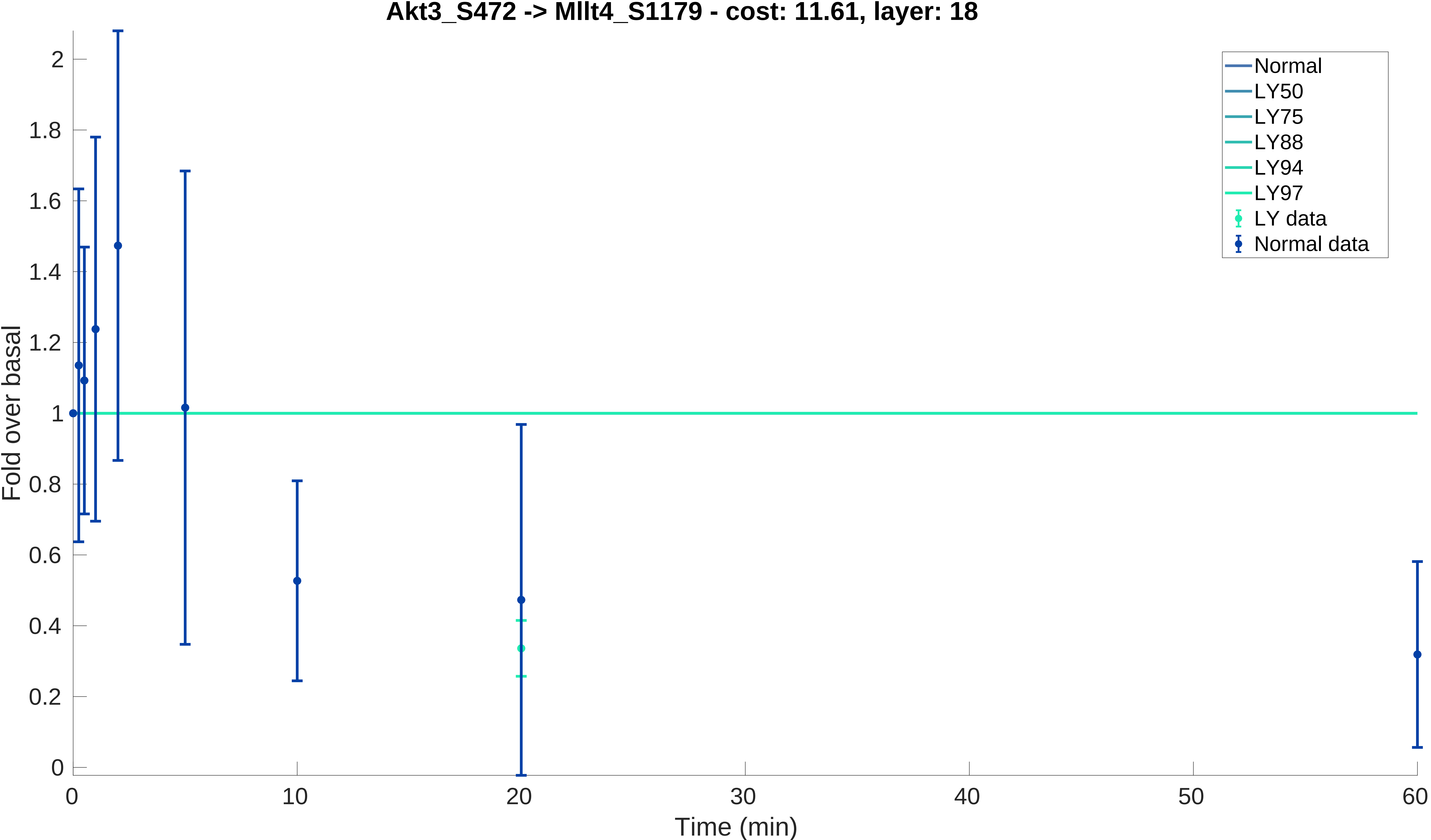

Akt3_S472 -> Mllt4_S1179 - cost: 11.61, layer: 18
Fold over basal
Time (min)
0
0.2
0.4
0.6
0.8
1
1.2
1.4
1.6
1.8
2
0
10
20
30
40
50
60
Normal
LY50
LY75
LY88
LY94
LY97
LY data
Normal data

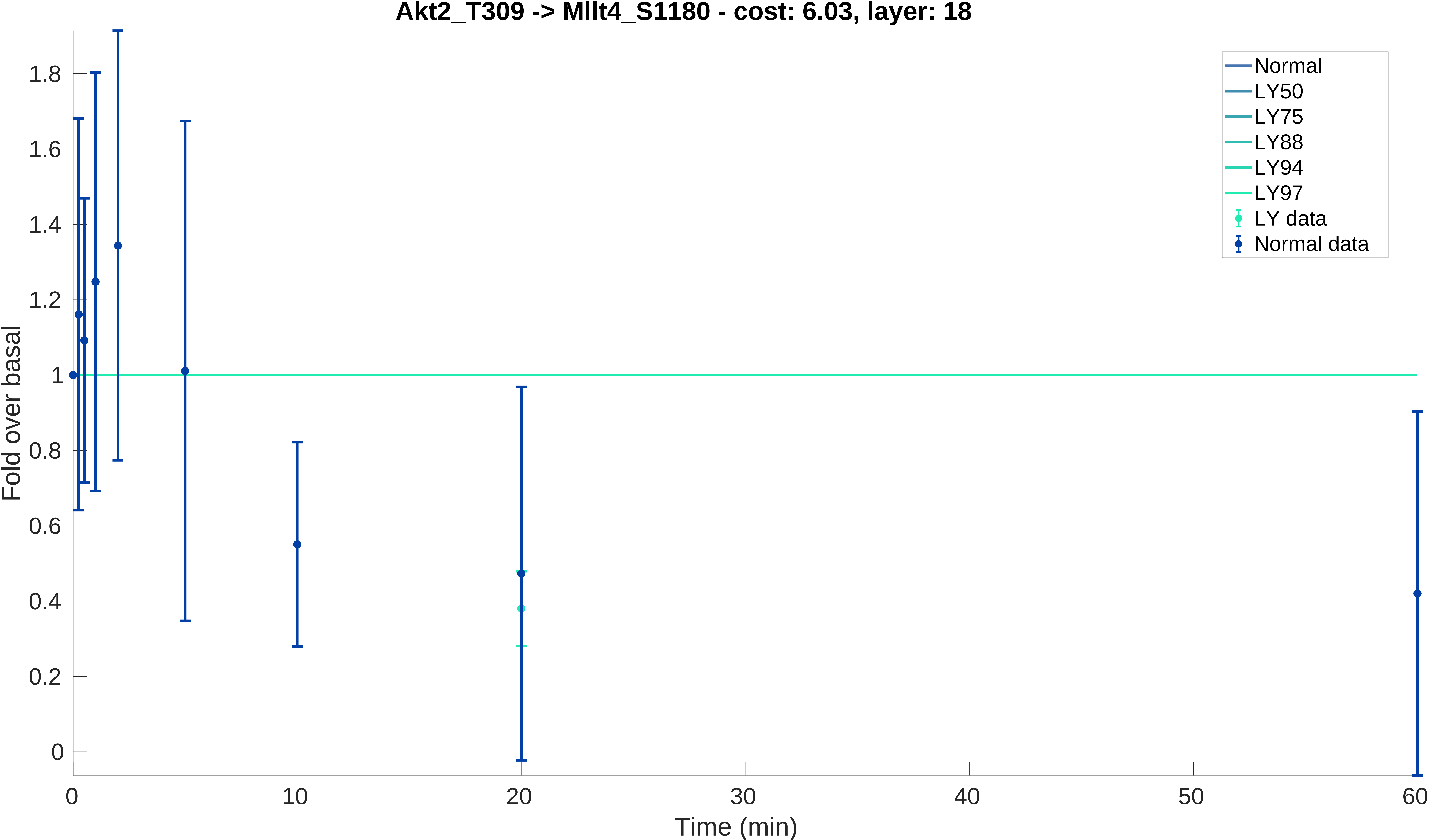

Akt2_T309 -> Mllt4_S1180 - cost: 6.03, layer: 18
Normal
LY50
LY75
LY88
LY94
LY97
LY data
Normal data
Fold over basal
Time (min)
0
0.2
0.4
0.6
0.8
1
1.2
1.4
1.6
1.8
0
10
20
30
40
50
60

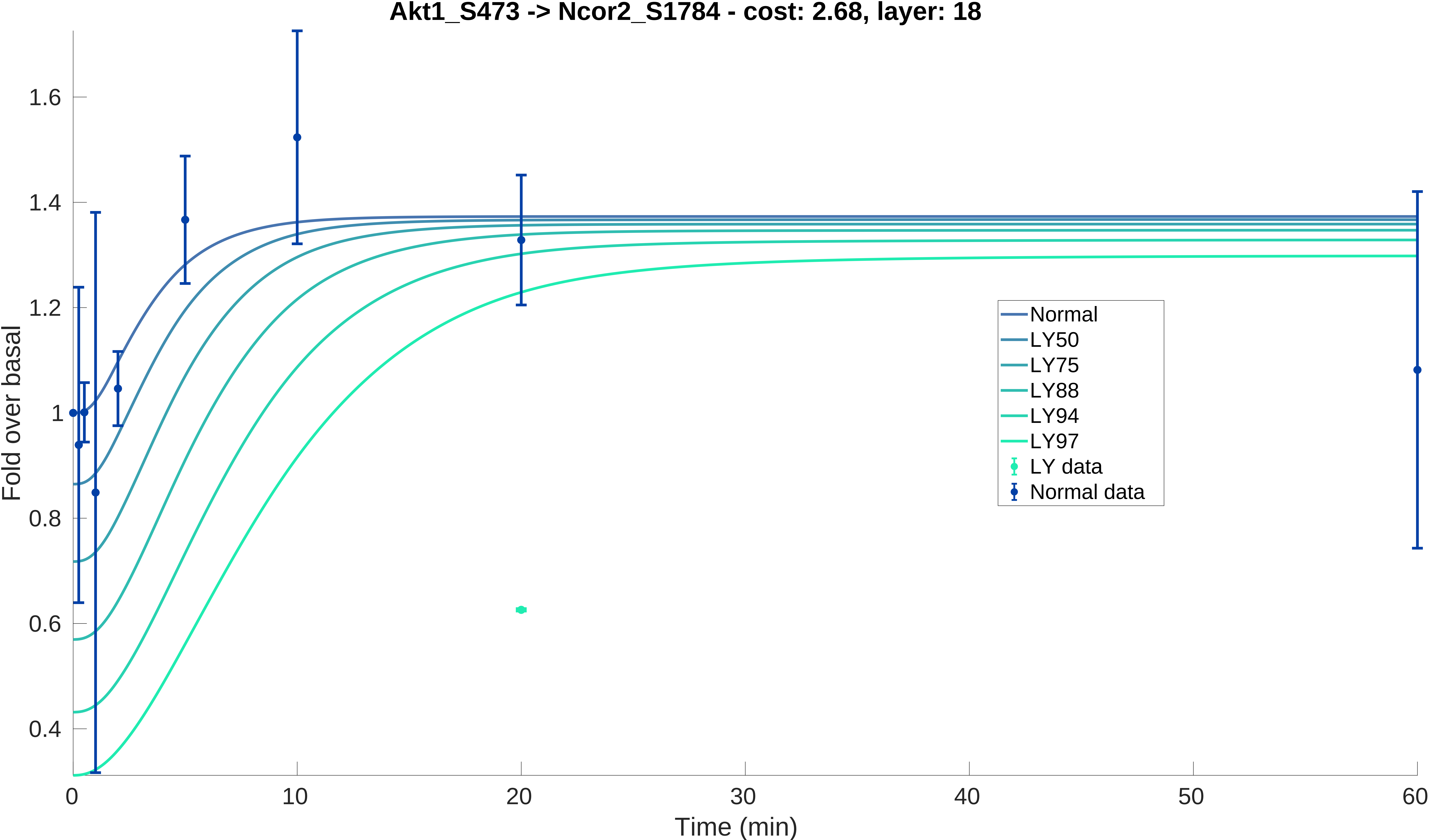

Akt1_S473 -> Ncor2_S1784 - cost: 2.68, layer: 18
Fold over basal
Time (min)
0.4
0.6
0.8
1
1.2
1.4
1.6
0
10
20
30
40
50
60
Normal
LY50
LY75
LY88
LY94
LY97
LY data
Normal data

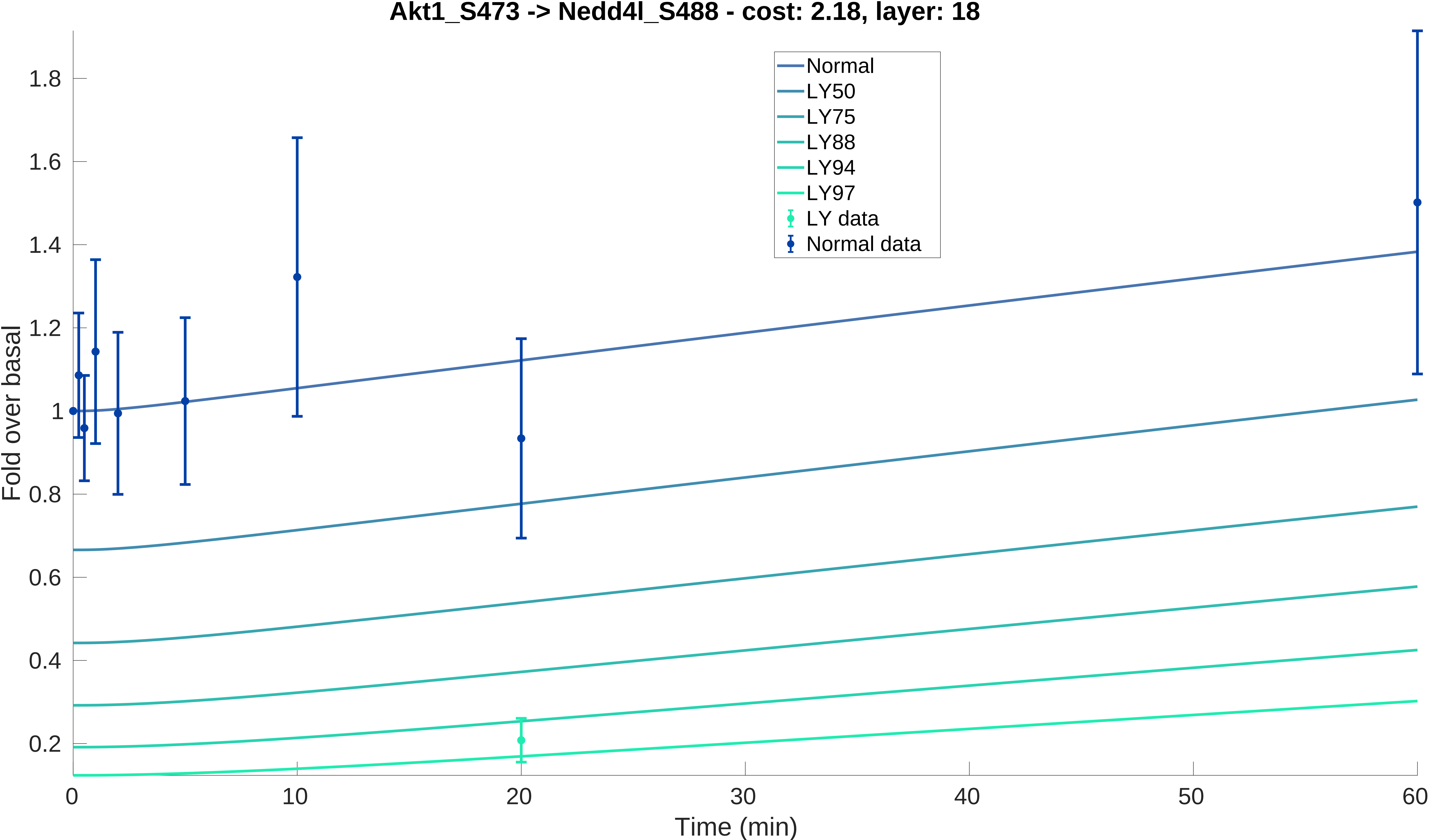

Akt1_S473 -> Nedd4l_S488 - cost: 2.18, layer: 18
Fold over basal
Time (min)
1.8
1.6
1.4
1.2
1
0.8
0.6
0.4
0.2
0
10
20
30
40
50
60
Normal
LY50
LY75
LY88
LY94
LY97
LY data
Normal data

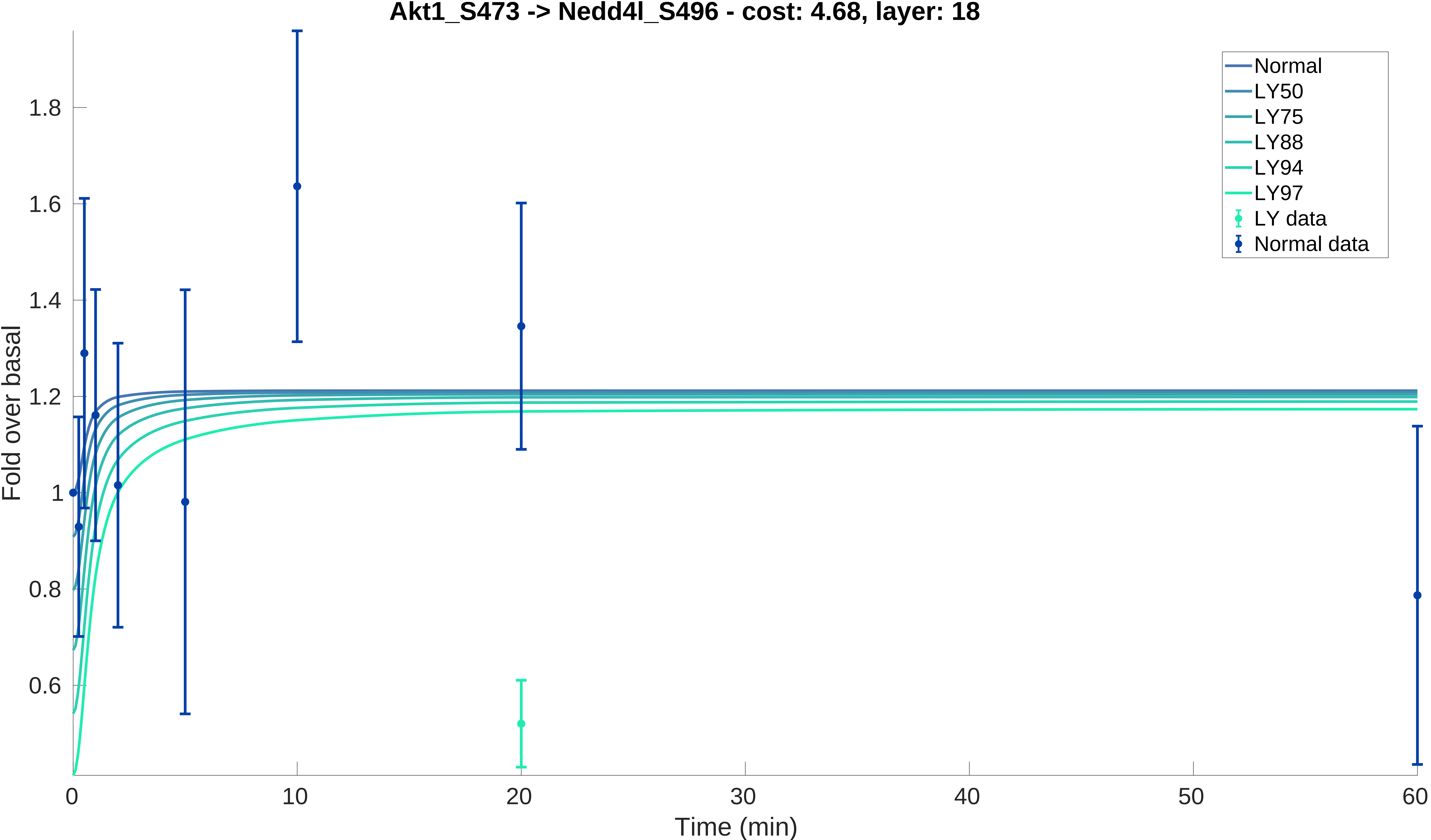
Akt1_S473 -> Nedd4l_S496 - cost: 4.68, layer: 18
Fold over basal
1.8
1.6
1.4
1.2
1
0.8
0.6
0
10
20
30
40
50
60
Time (min)
Normal
LY50
LY75
LY88
LY94
LY97
LY data
Normal data

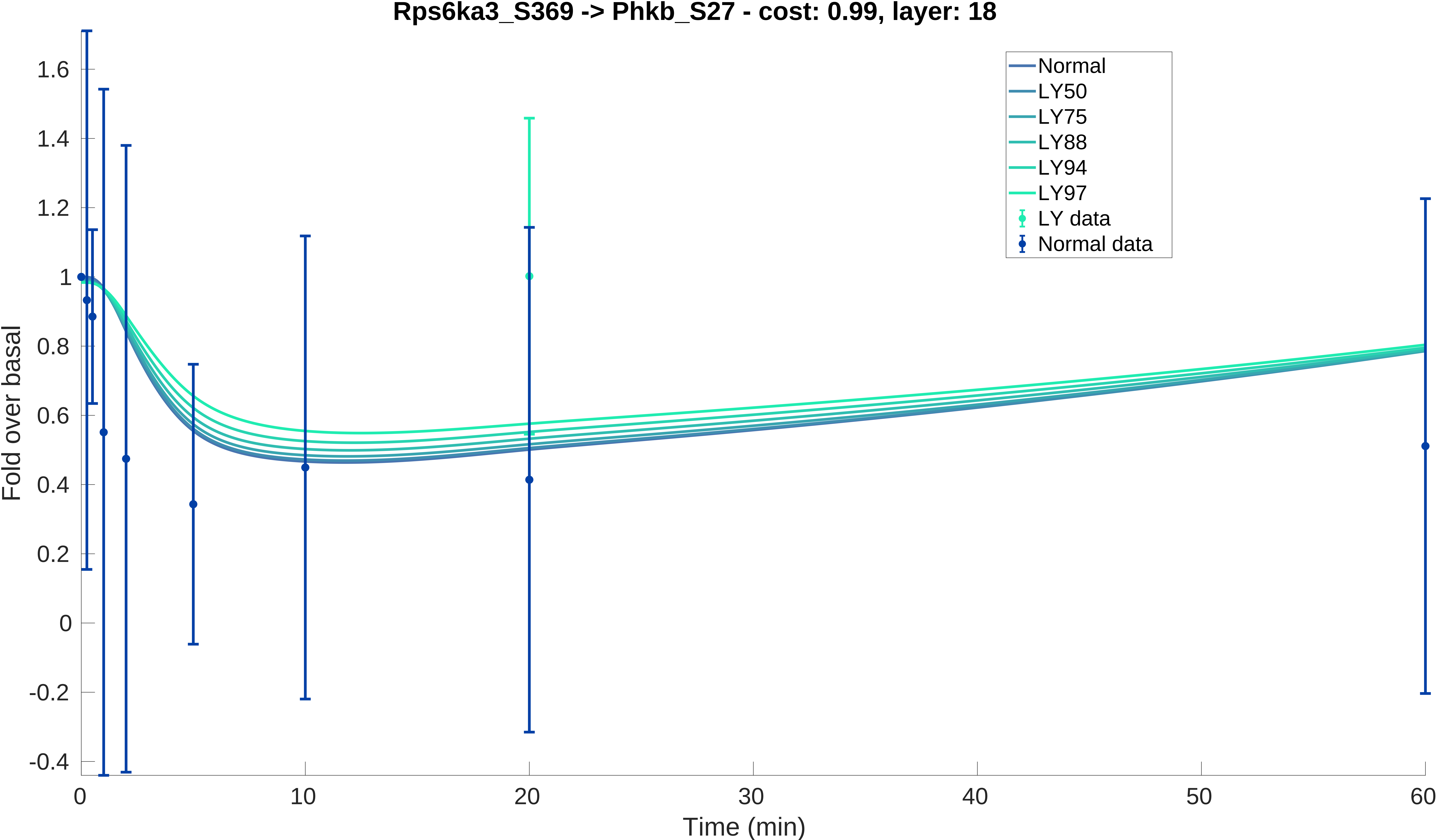

Rps6ka3_S369 -> Phkb_S27 - cost: 0.99, layer: 18
Normal
LY50
LY75
LY88
LY94
LY97
LY data
Normal data
Fold over basal
Time (min)
1.6
1.4
1.2
1
0.8
0.6
0.4
0.2
0
-0.2
-0.4
0
10
20
30
40
50
60

#### Rps6ka3_S369 -> Plec_T3889 - cost: 9.99, layer: 18

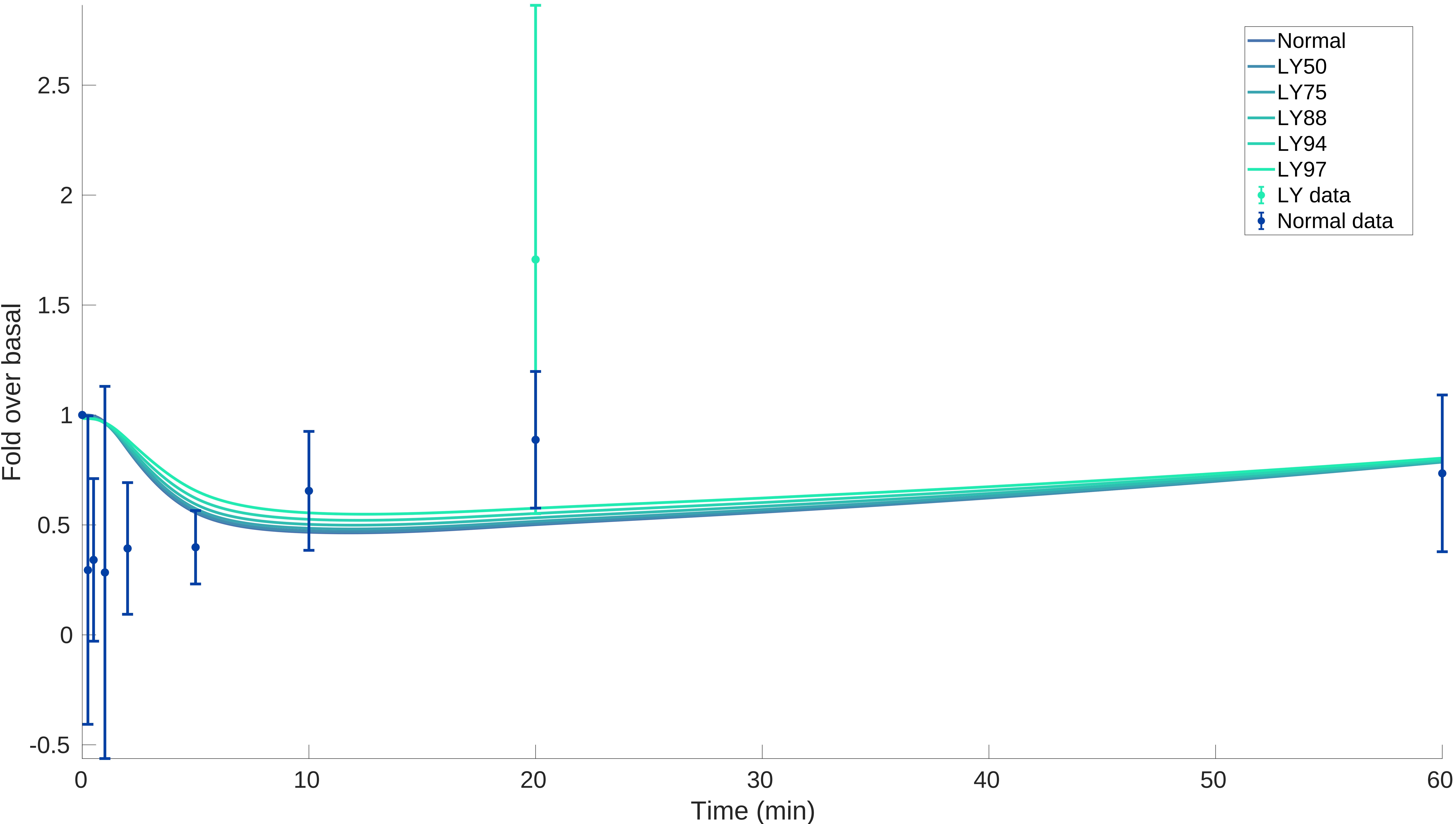

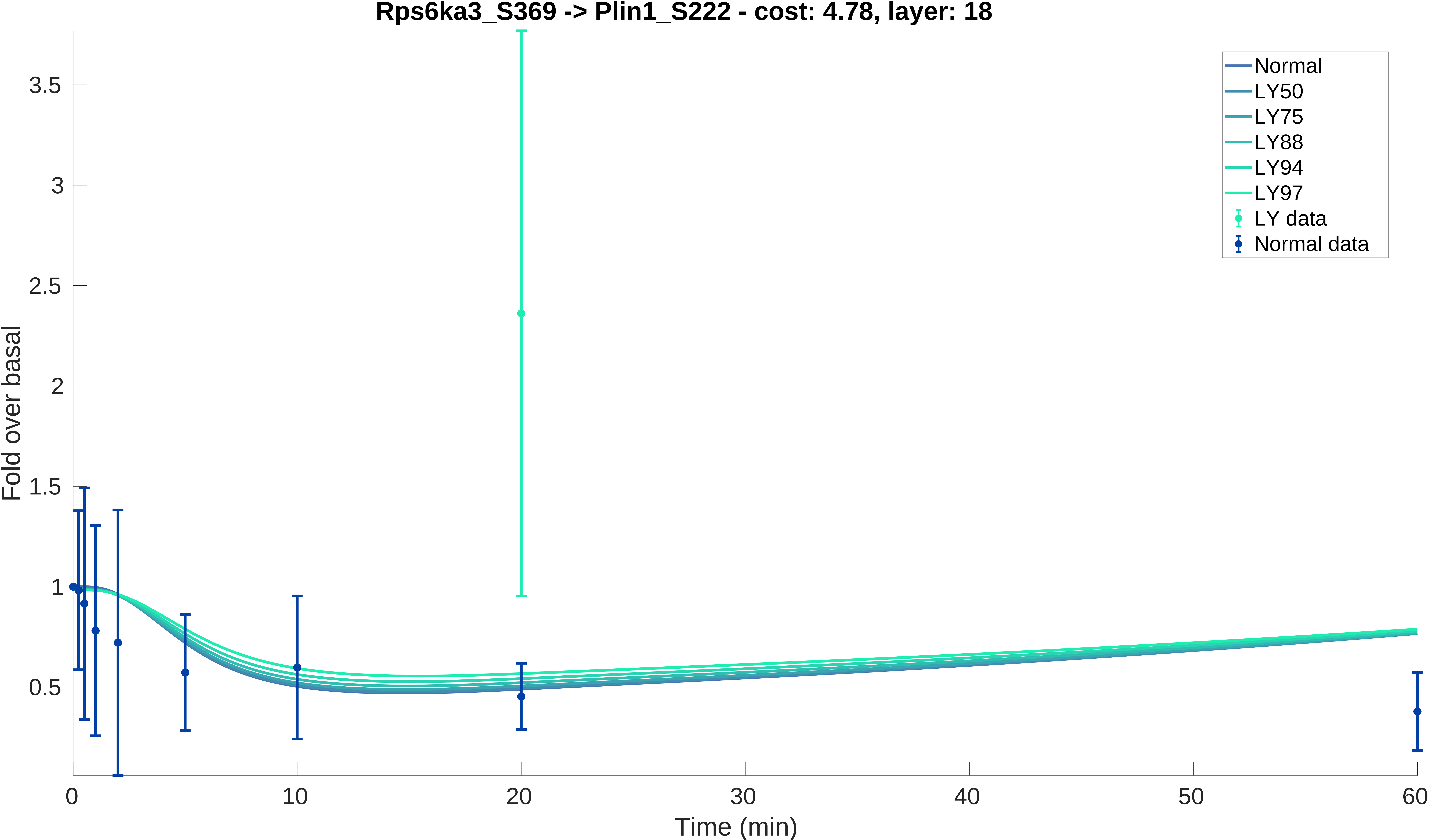
Rps6ka3_S369 -> Plin1_S222 - cost: 4.78, layer: 18
Fold over basal
Time (min)
3.5
3
2.5
2
1.5
1
0.5
0
10
20
30
40
50
60
Normal
LY50
LY75
LY88
LY94
LY97
LY data
Normal data

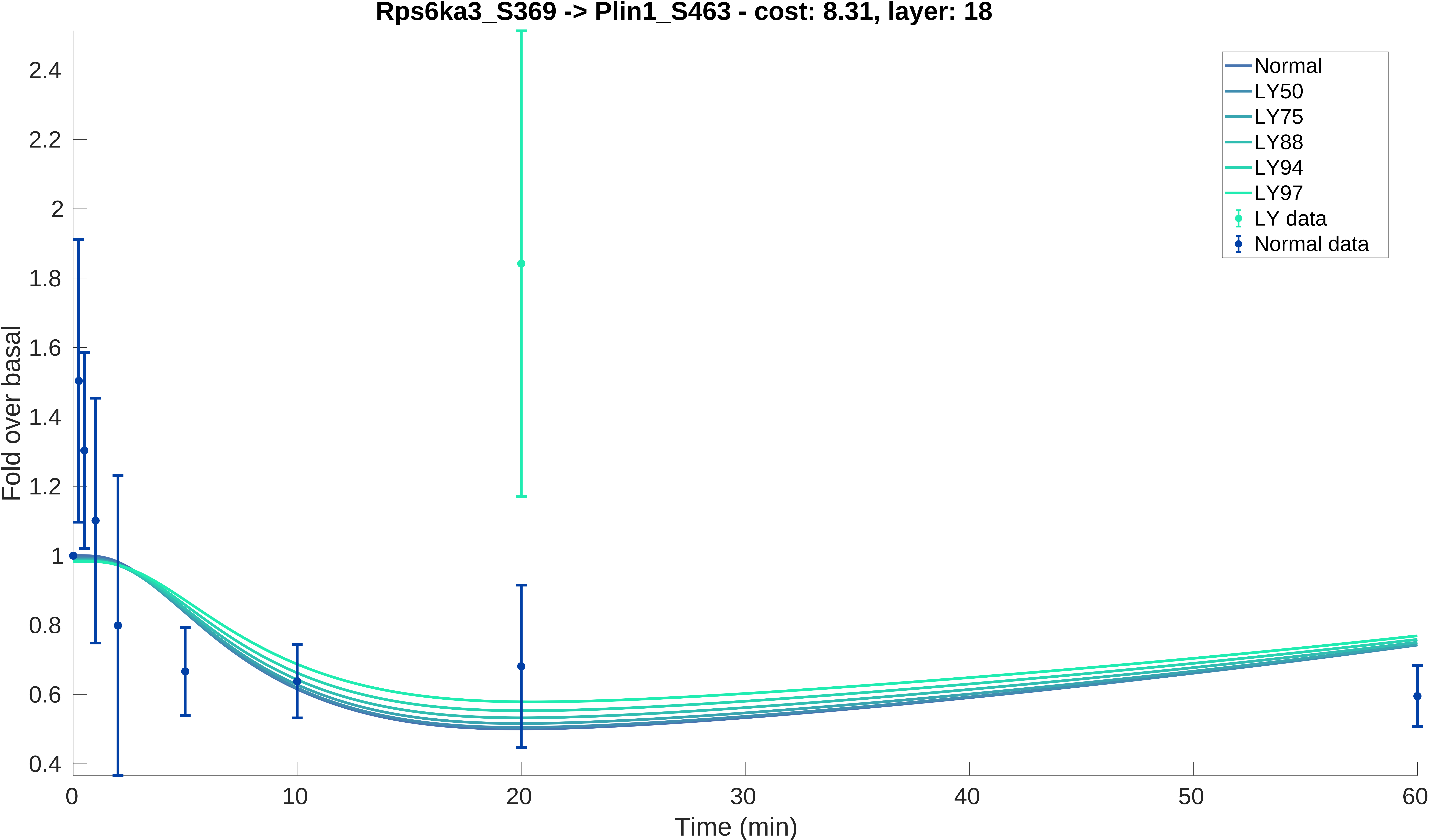

Rps6ka3_S369 -> Plin1_S463 - cost: 8.31, layer: 18
Fold over basal
Time (min)
Normal
LY50
LY75
LY88
LY94
LY97
LY data
Normal data
2.4
2.2
2
1.8
1.6
1.4
1.2
1
0.8
0.6
0.4
0
10
20
30
40
50
60

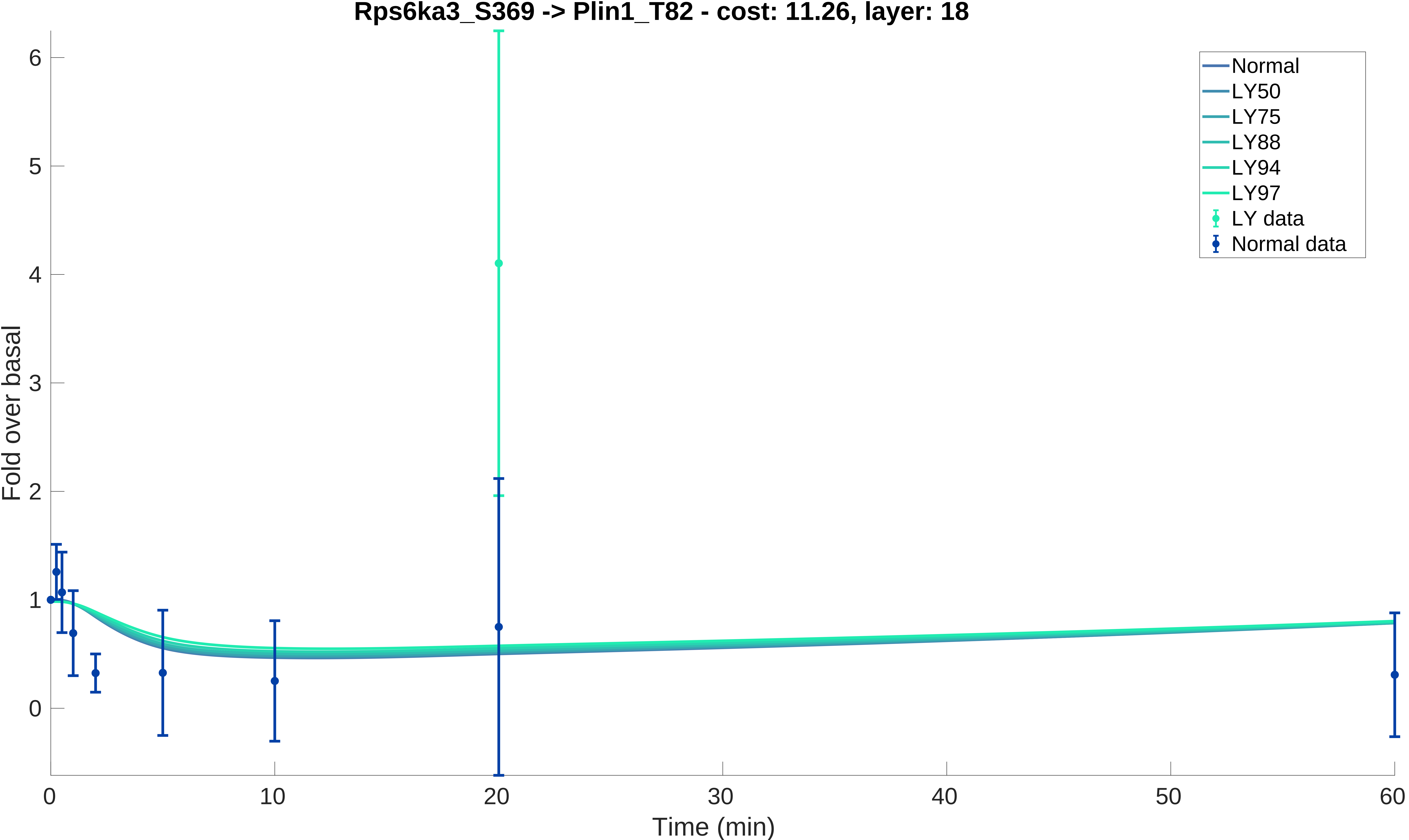
Rps6ka3_S369 -> Plin1_T82 - cost: 11.26, layer: 18
Fold over basal
Time (min)
0
1
2
3
4
5
6
0
10
20
30
40
50
60
Normal
LY50
LY75
LY88
LY94
LY97
LY data
Normal data

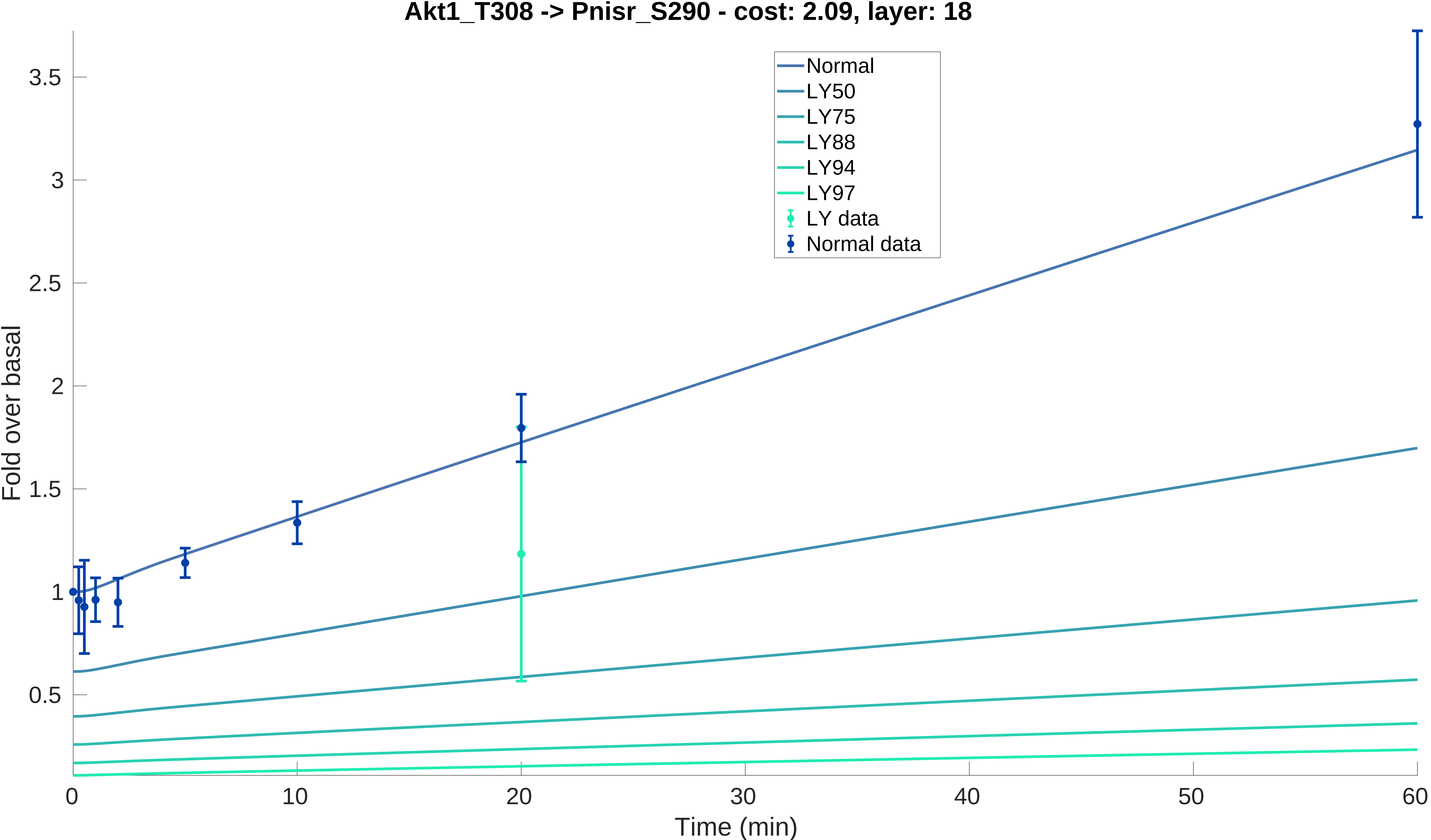

Akt1_T308 -> Pnisr_S290 - cost: 2.09, layer: 18
Normal
LY50
LY75
LY88
LY94
LY97
LY data
Normal data
Fold over basal
3.5
3
2.5
2
1.5
1
0.5
0
10
20
30
40
50
60
Time (min)

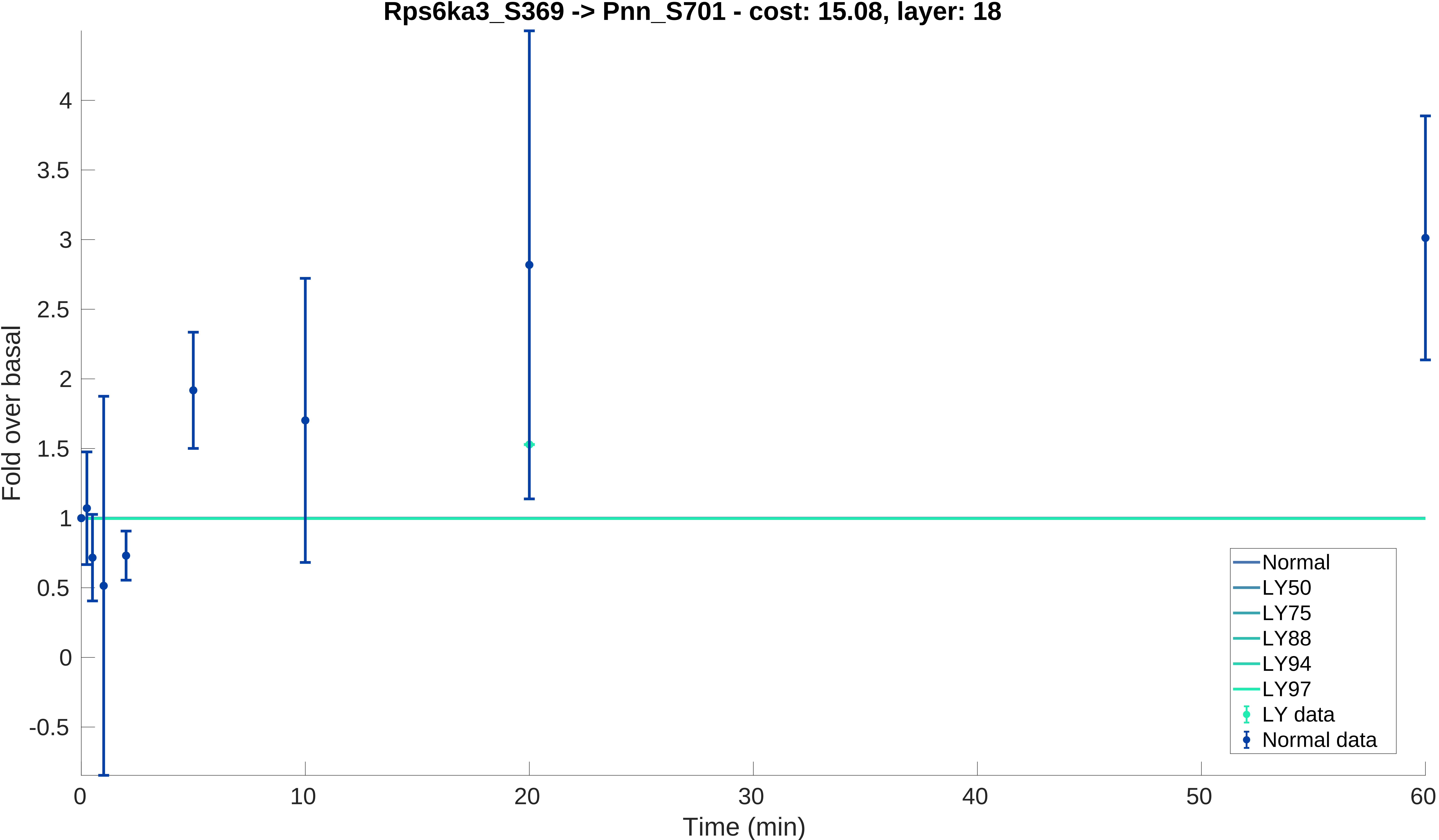
Rps6ka3_S369 -> Pnn_S701 - cost: 15.08, layer: 18
Fold over basal
4
3.5
3
2.5
2
1.5
1
0.5
0
-0.5
0
10
20
30
40
50
60
Time (min)
Normal
LY50
LY75
LY88
LY94
LY97
LY data
Normal data

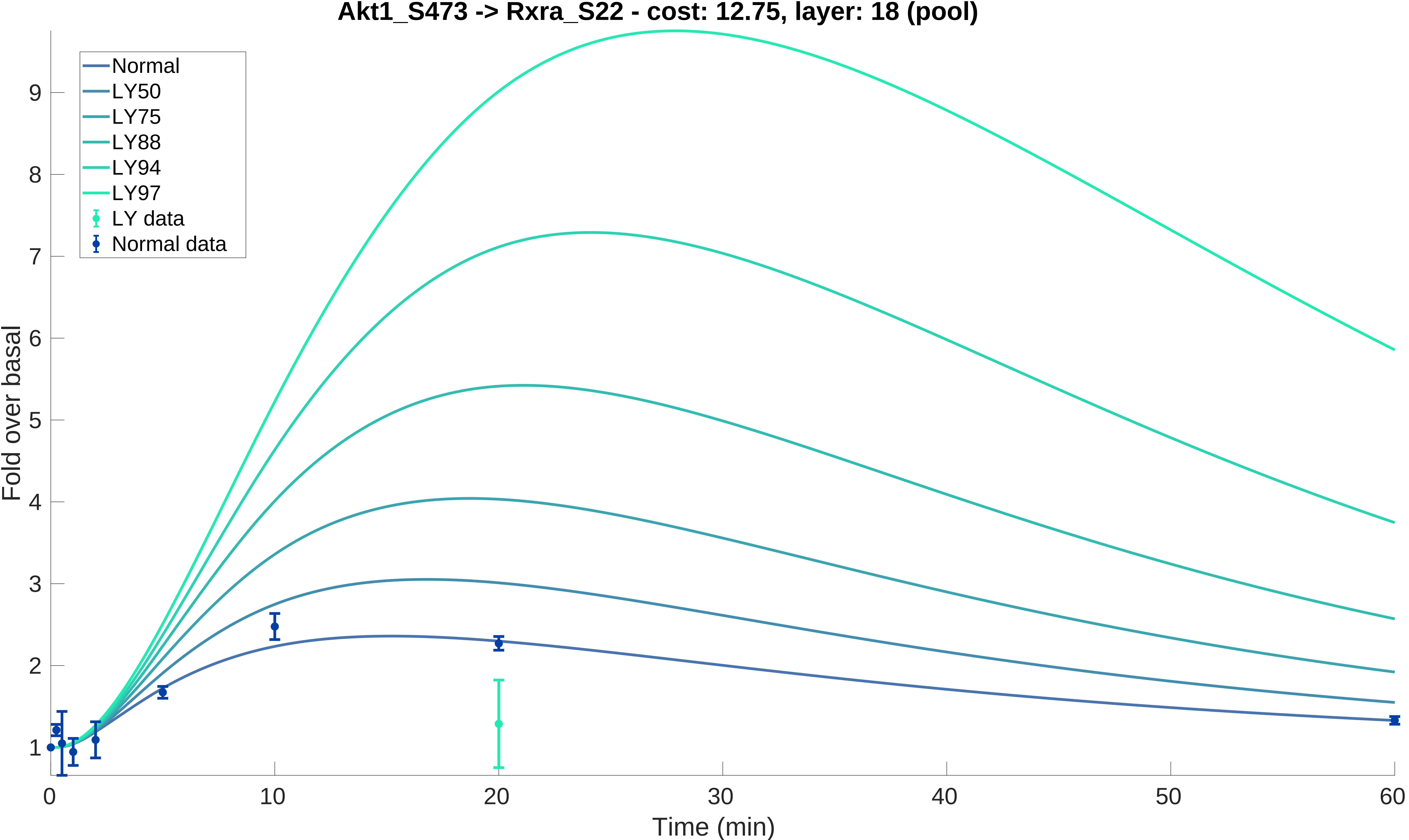
Akt1_S473 -> Rxra_S22 - cost: 12.75, layer: 18 (pool)
Normal
LY50
LY75
LY88
LY94
LY97
LY data
Normal data
Fold over basal
Time (min)
1
2
3
4
5
6
7
8
9
0
10
20
30
40
50
60

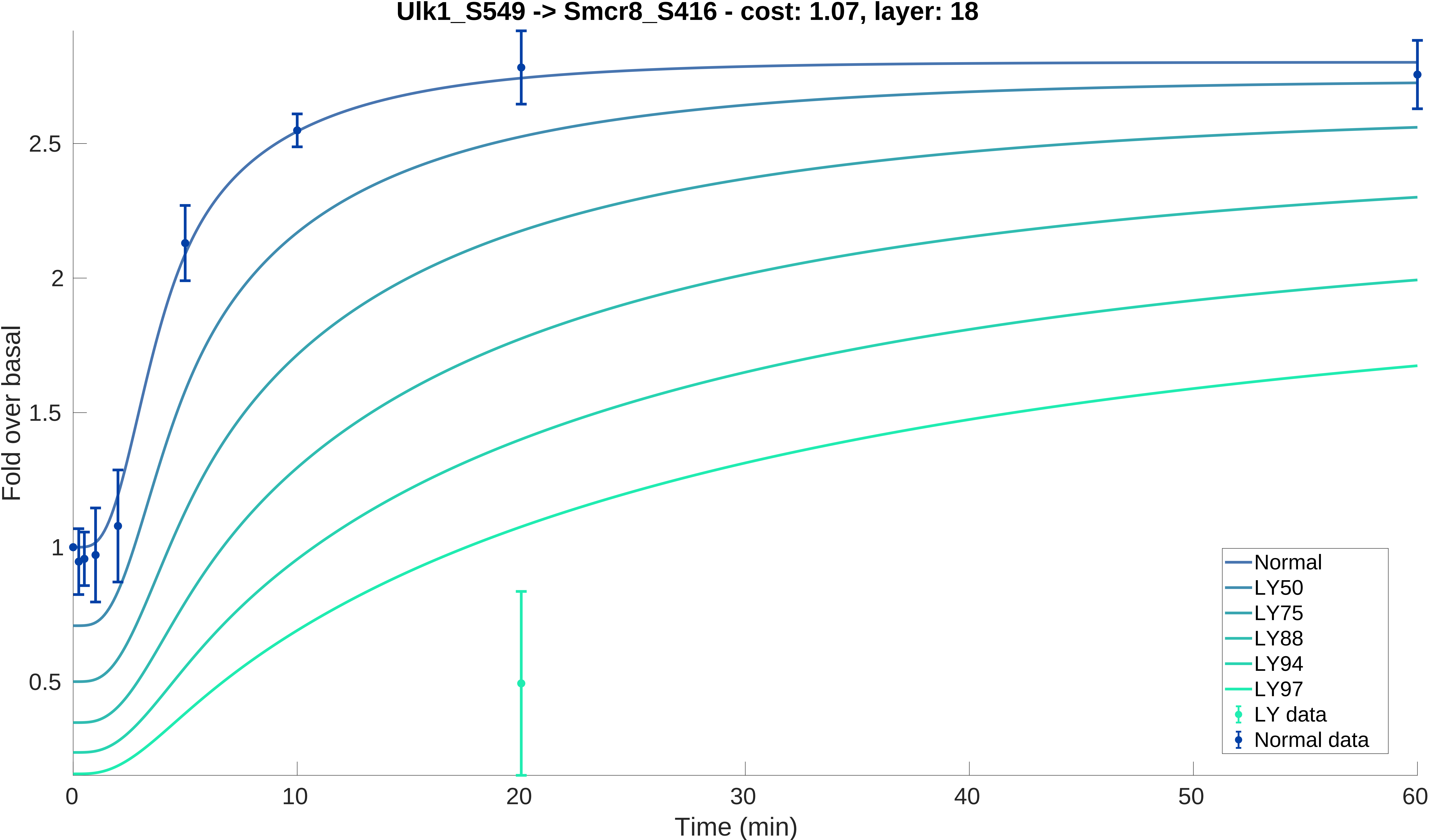

Ulk1_S549 -> Smcr8_S416 - cost: 1.07, layer: 18
Fold over basal
Time (min)
0.5
1
1.5
2
2.5
0
10
20
30
40
50
60
Normal
LY50
LY75
LY88
LY94
LY97
LY data
Normal data

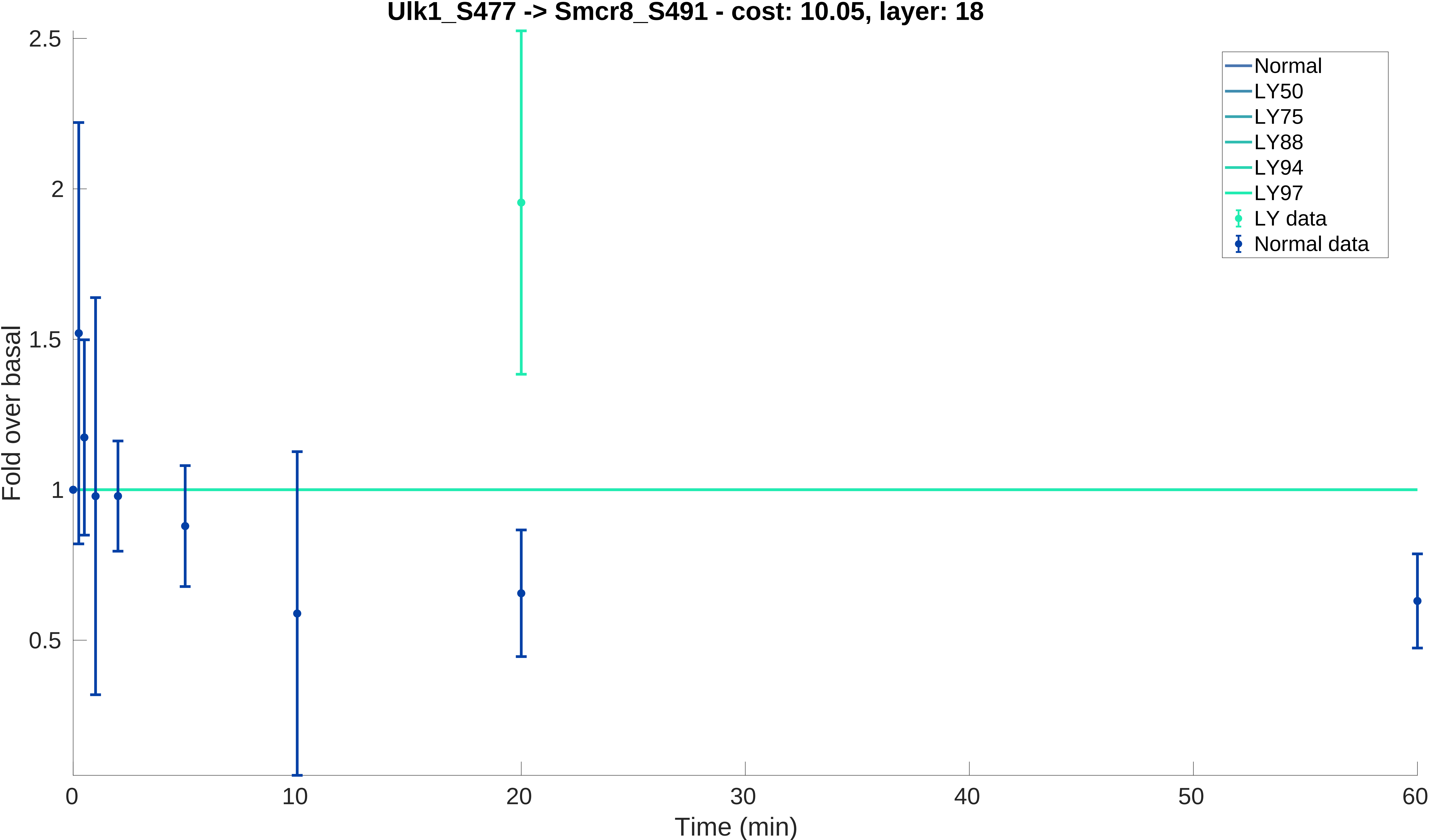

Ulk1_S477 -> Smcr8_S491 - cost: 10.05, layer: 18
Fold over basal
Time (min)
0.5
1
1.5
2
2.5
0
10
20
30
40
50
60
Normal
LY50
LY75
LY88
LY94
LY97
LY data
Normal data

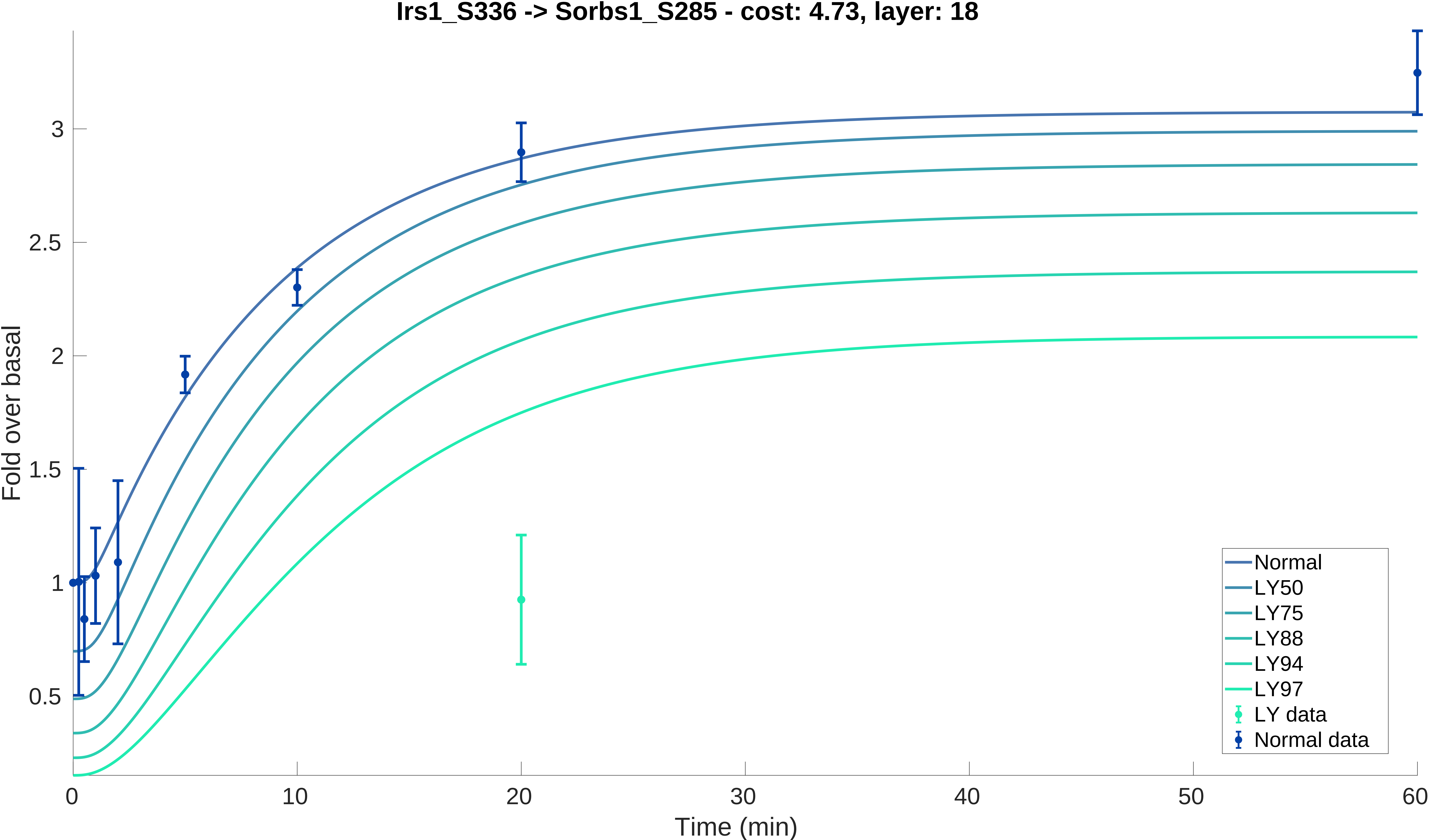

Irs1_S336 -> Sorbs1_S285 - cost: 4.73, layer: 18
Fold over basal
3
2.5
2
1.5
1
0.5
0
10
20
30
40
50
60
Time (min)
Normal
LY50
LY75
LY88
LY94
LY97
LY data
Normal data

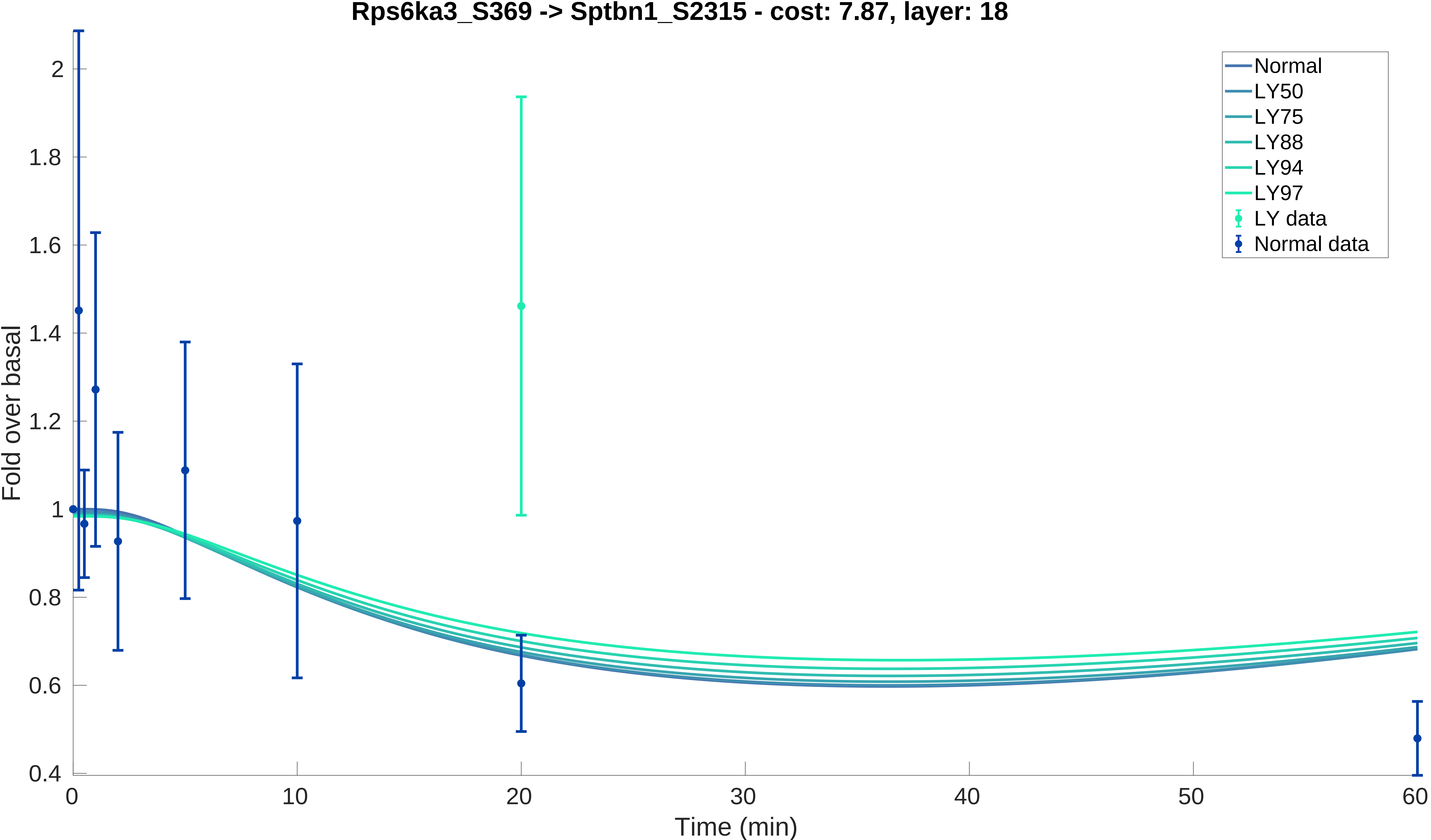

Rps6ka3_S369 -> Sptbn1_S2315 - cost: 7.87, layer: 18
Fold over basal
Time (min)
0
10
20
30
40
50
60
0.4
0.6
0.8
1
1.2
1.4
1.6
1.8
2
Normal
LY50
LY75
LY88
LY94
LY97
LY data
Normal data

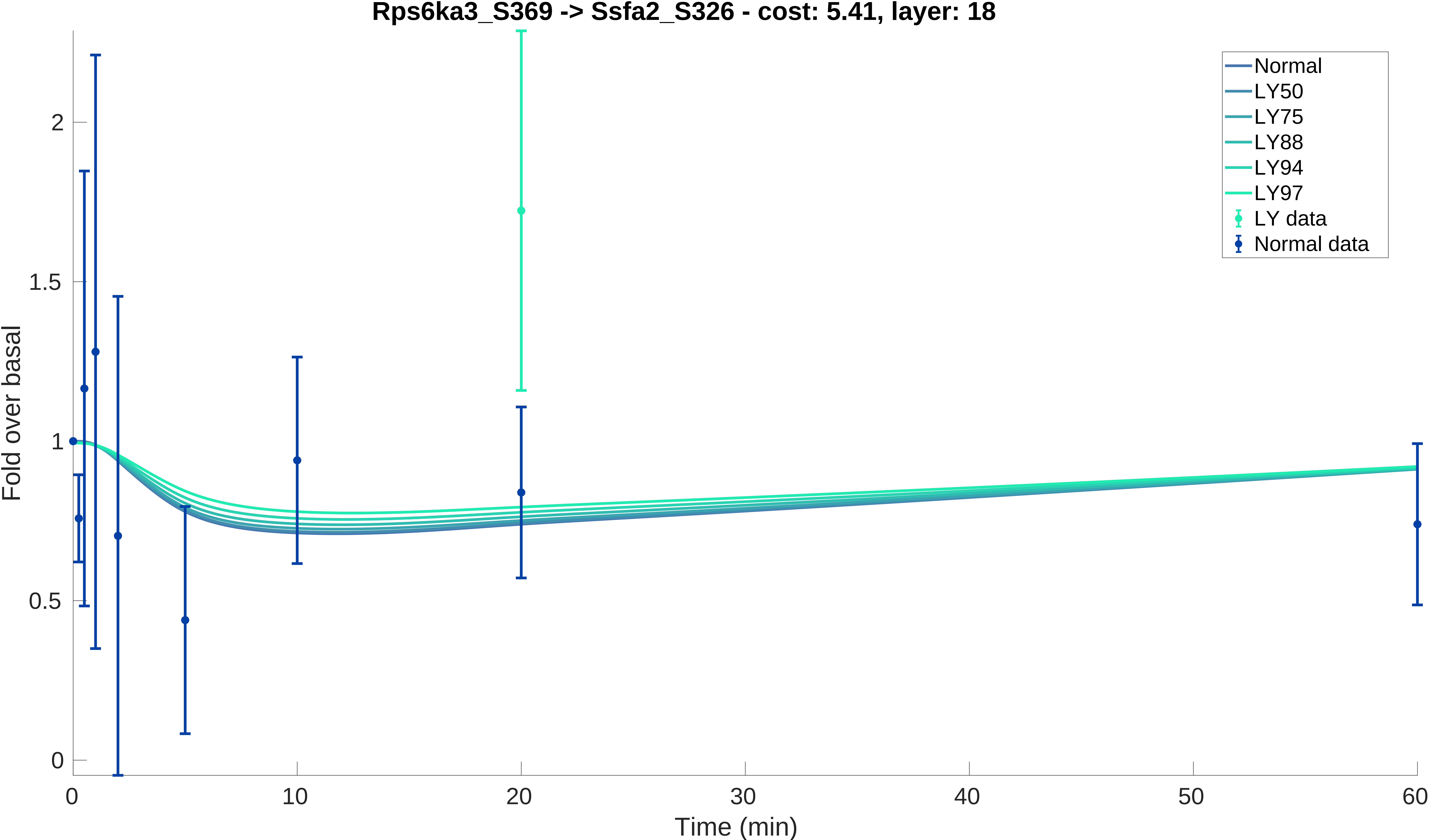

Rps6ka3_S369 -> Ssfa2_S326 - cost: 5.41, layer: 18
Normal
LY50
LY75
LY88
LY94
LY97
LY data
Normal data
Fold over basal
0
0.5
1
1.5
2
0
10
20
30
40
50
60
Time (min)

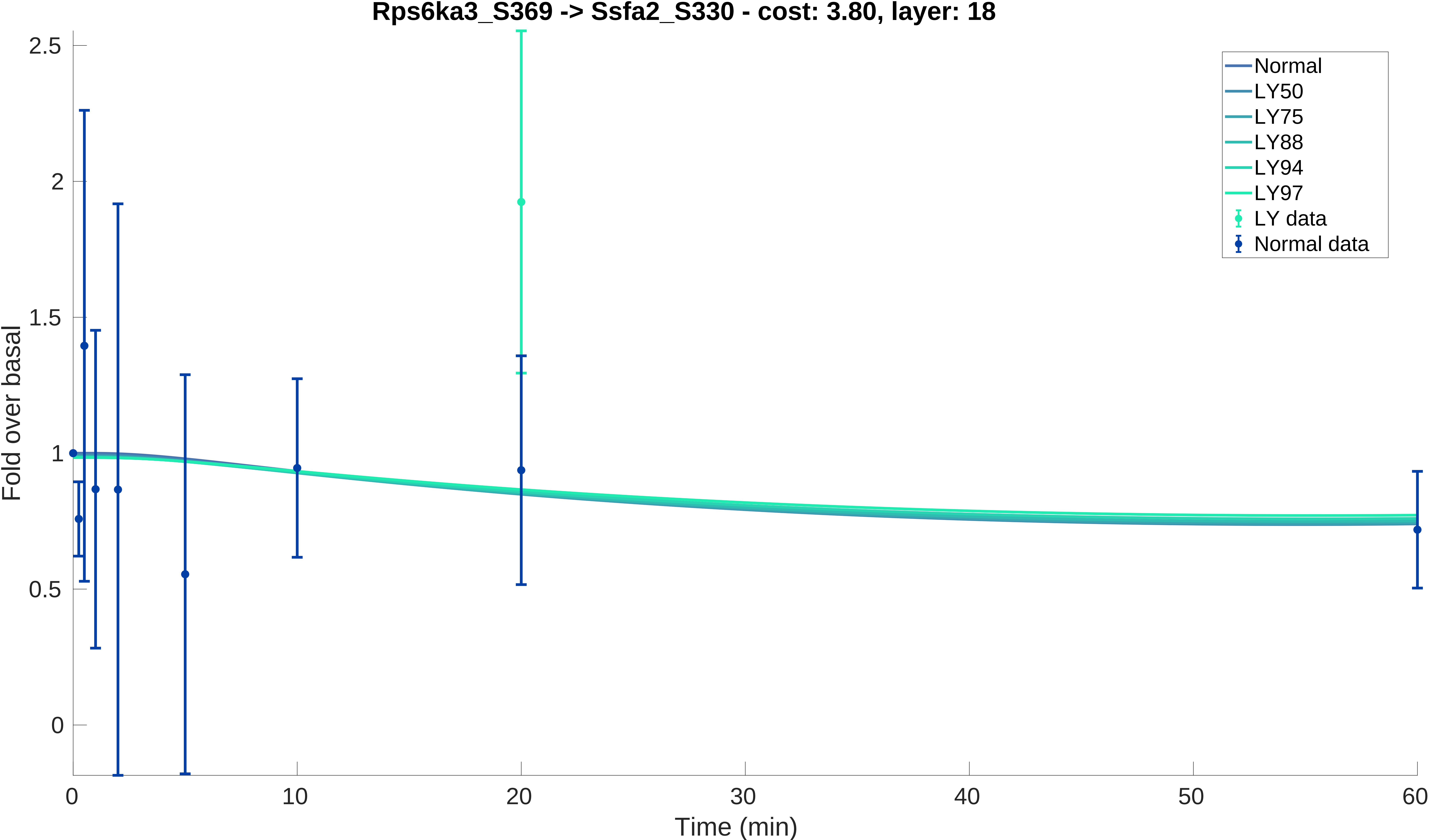

Rps6ka3_S369 -> Ssfa2_S330 - cost: 3.80, layer: 18
Fold over basal
Time (min)
0
0.5
1
1.5
2
2.5
0
10
20
30
40
50
60
Normal
LY50
LY75
LY88
LY94
LY97
LY data
Normal data

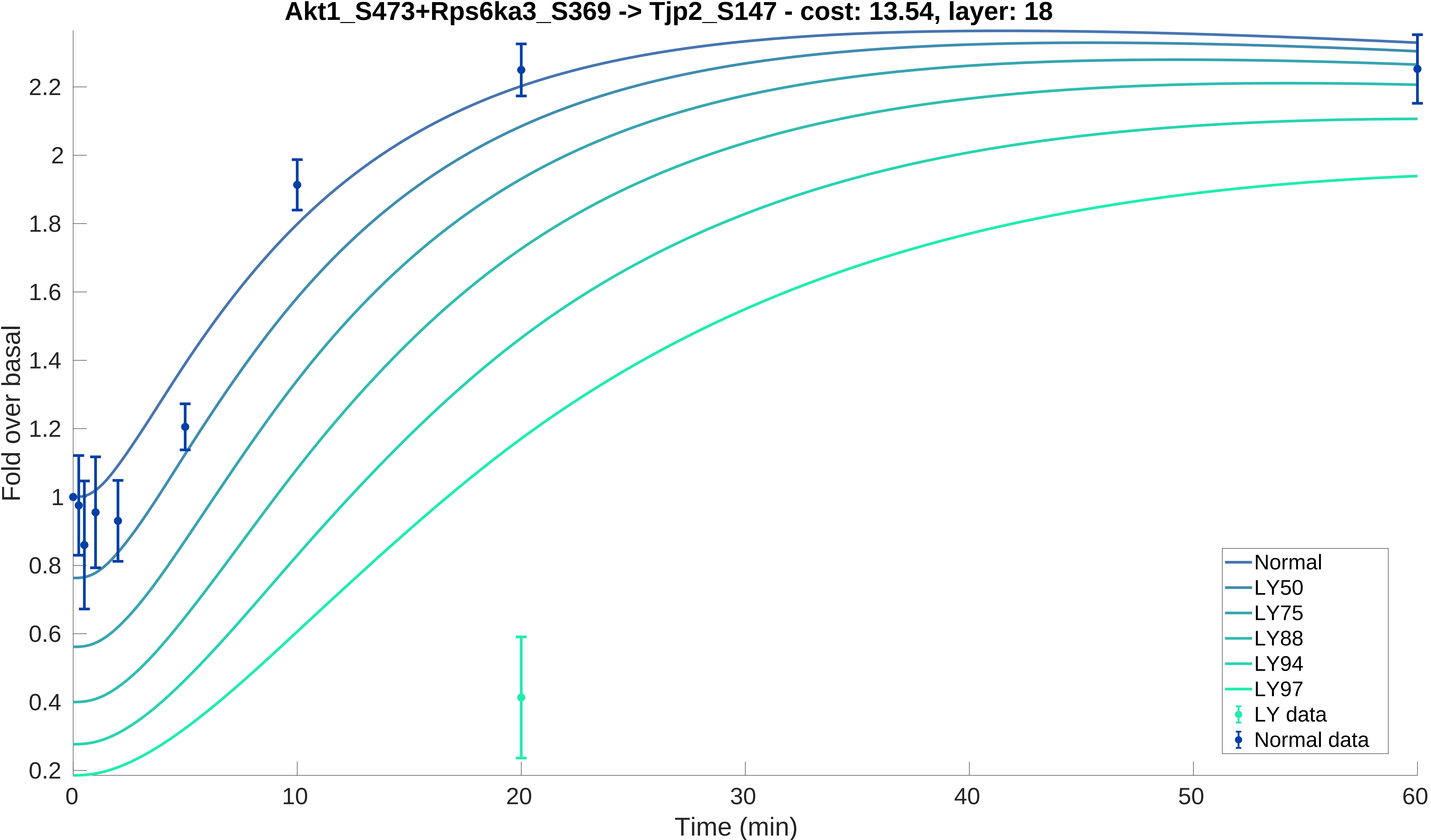

Akt1_S473+Rps6ka3_S369 -> Tjp2_S147 - cost: 13.54, layer: 18
Fold over basal
Time (min)
0.2
0.4
0.6
0.8
1
1.2
1.4
1.6
1.8
2
2.2
0
10
20
30
40
50
60
Normal
LY50
LY75
LY88
LY94
LY97
LY data
Normal data

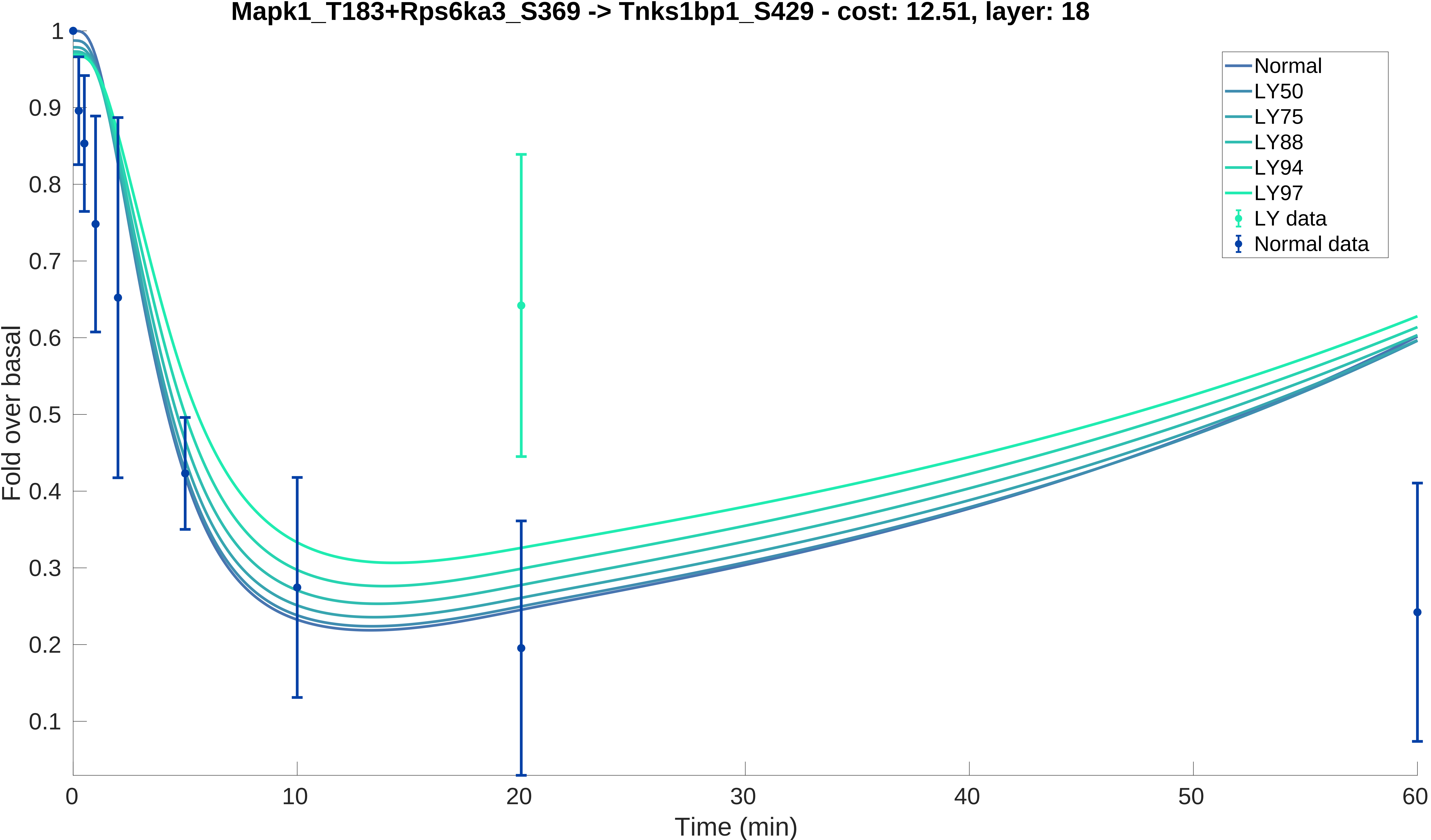
Mapk1_T183+Rps6ka3_S369 -> Tnks1bp1_S429 - cost: 12.51, layer: 18
Normal
LY50
LY75
LY88
LY94
LY97
LY data
Normal data
Fold over basal
Time (min)
1
0.9
0.8
0.7
0.6
0.5
0.4
0.3
0.2
0.1
0
10
20
30
40
50
60

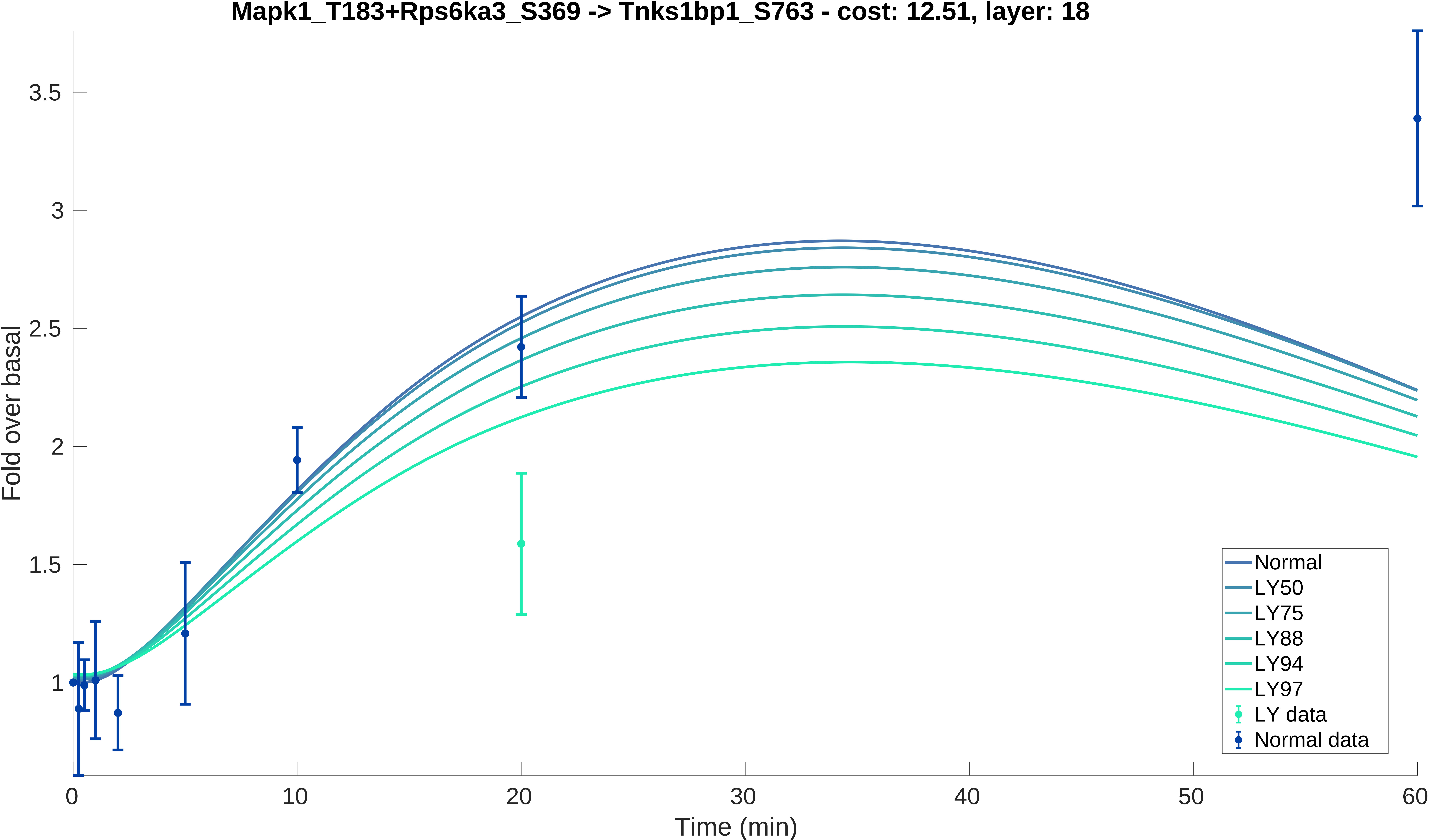

Mapk1_T183+Rps6ka3_S369 -> Tnks1bp1_S763 - cost: 12.51, layer: 18
Fold over basal
3.5
3
2.5
2
1.5
1
0
10
20
30
40
50
60
Time (min)
Normal
LY50
LY75
LY88
LY94
LY97
LY data
Normal data

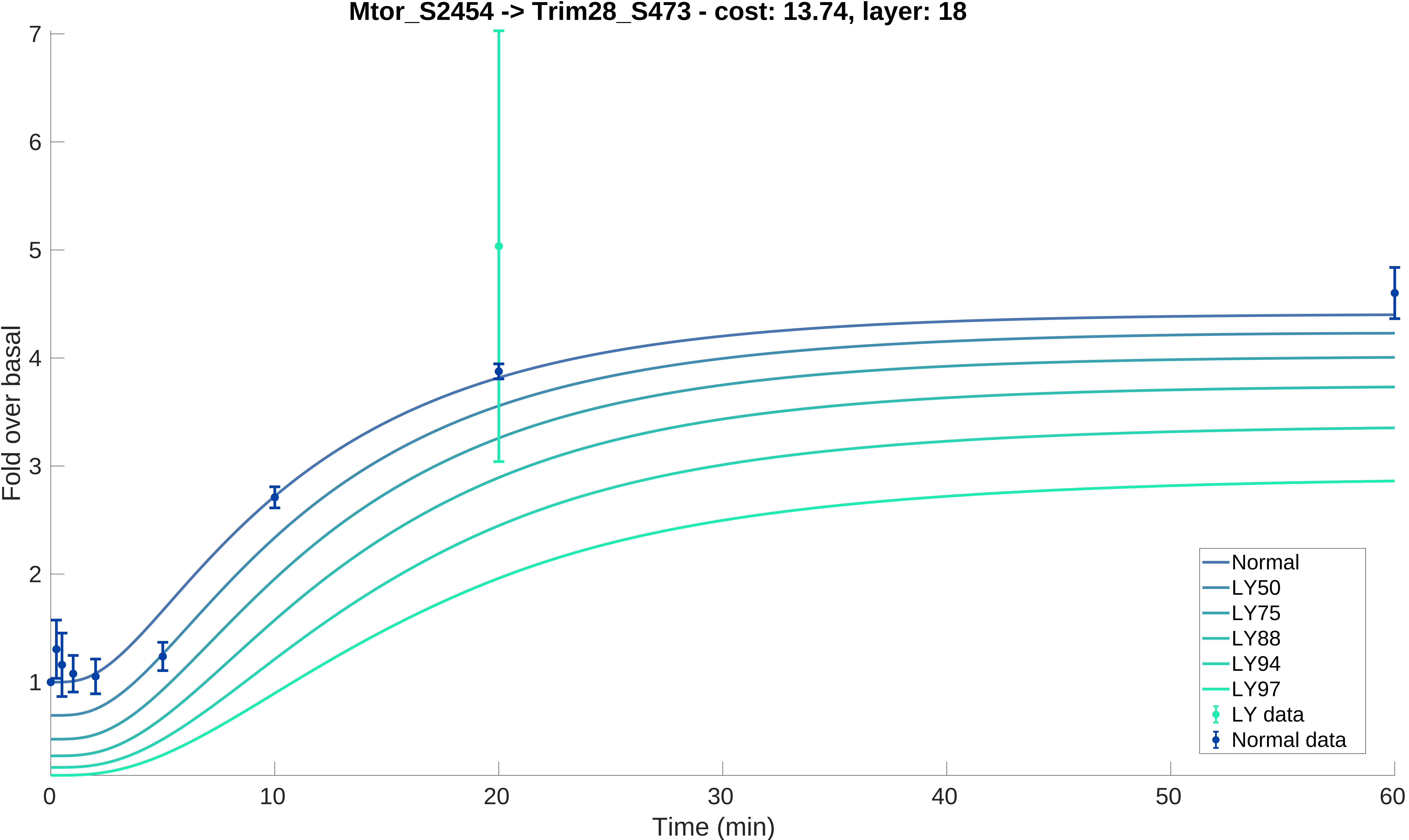

Mtor_S2454 -> Trim28_S473 - cost: 13.74, layer: 18
Fold over basal
Time (min)
0
10
20
30
40
50
60
1
2
3
4
5
6
7
Normal
LY50
LY75
LY88
LY94
LY97
LY data
Normal data

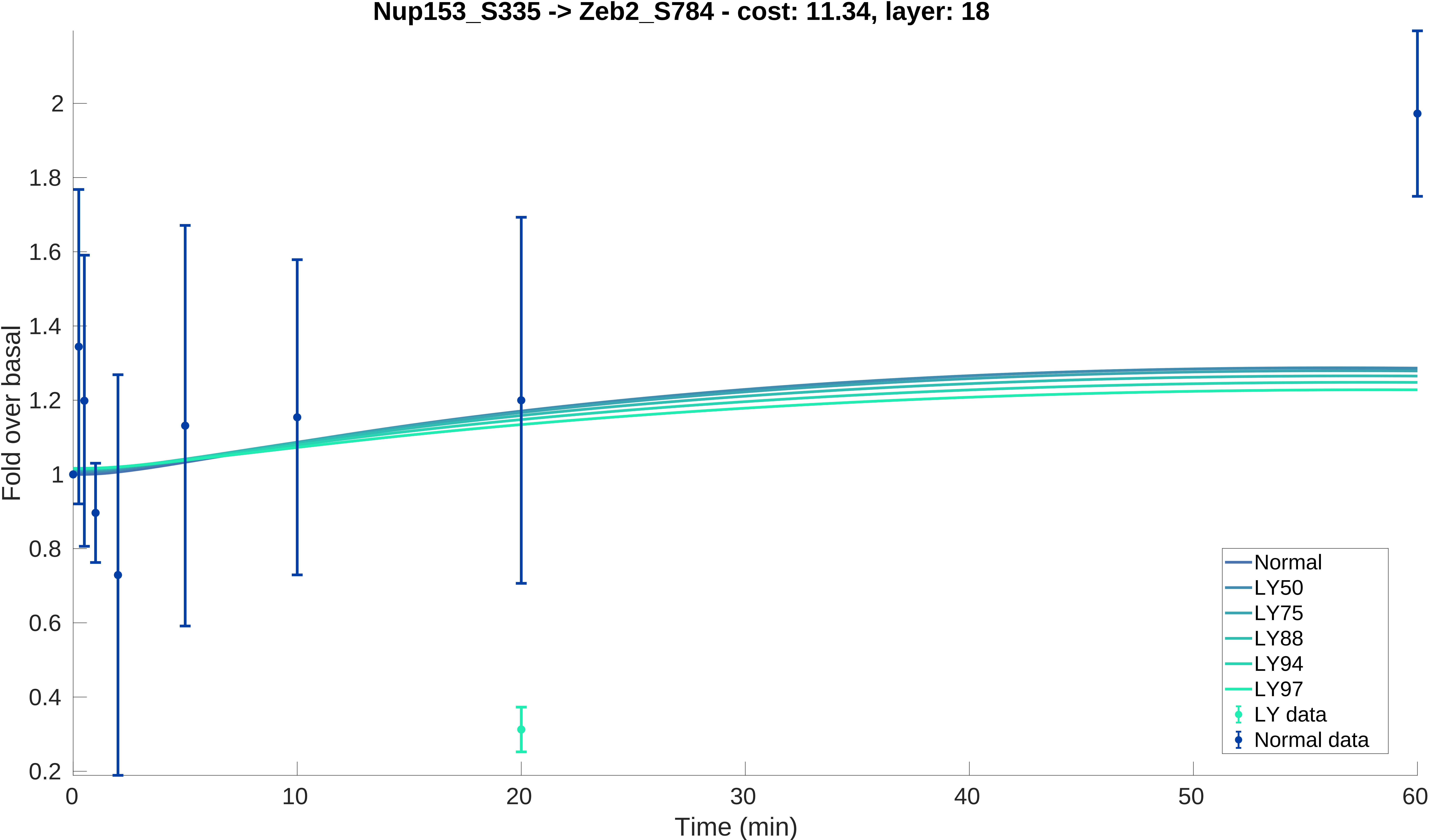
Nup153_S335 -> Zeb2_S784 - cost: 11.34, layer: 18
Fold over basal
Time (min)
0
10
20
30
40
50
60
0.2
0.4
0.6
0.8
1
1.2
1.4
1.6
1.8
2
Normal
LY50
LY75
LY88
LY94
LY97
LY data
Normal data

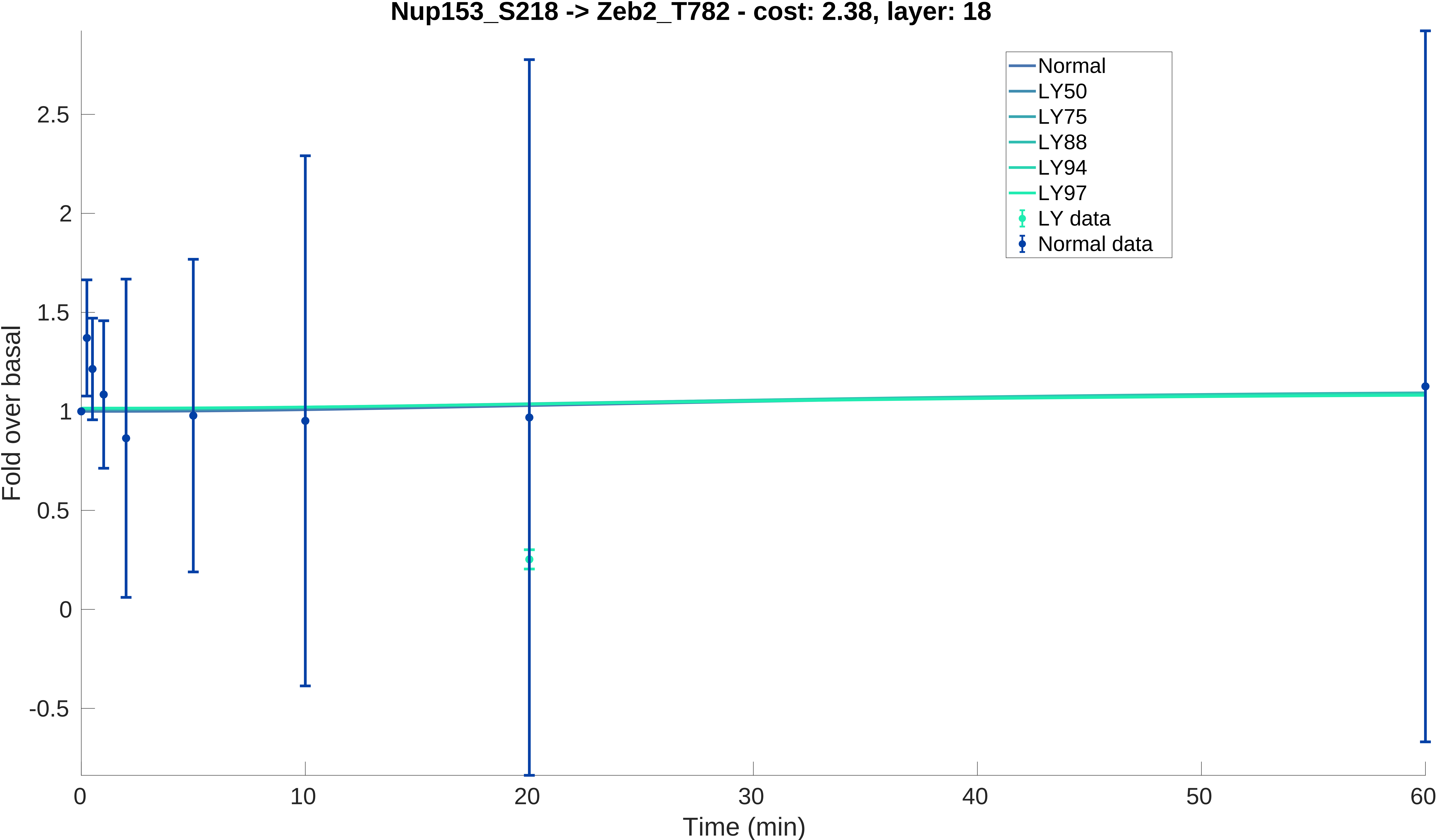
Nup153_S218 -> Zeb2_T782 - cost: 2.38, layer: 18
Normal
LY50
LY75
LY88
LY94
LY97
LY data
Normal data
Fold over basal
2.5
2
1.5
1
0.5
0
-0.5
0
10
20
30
40
50
60
Time (min)

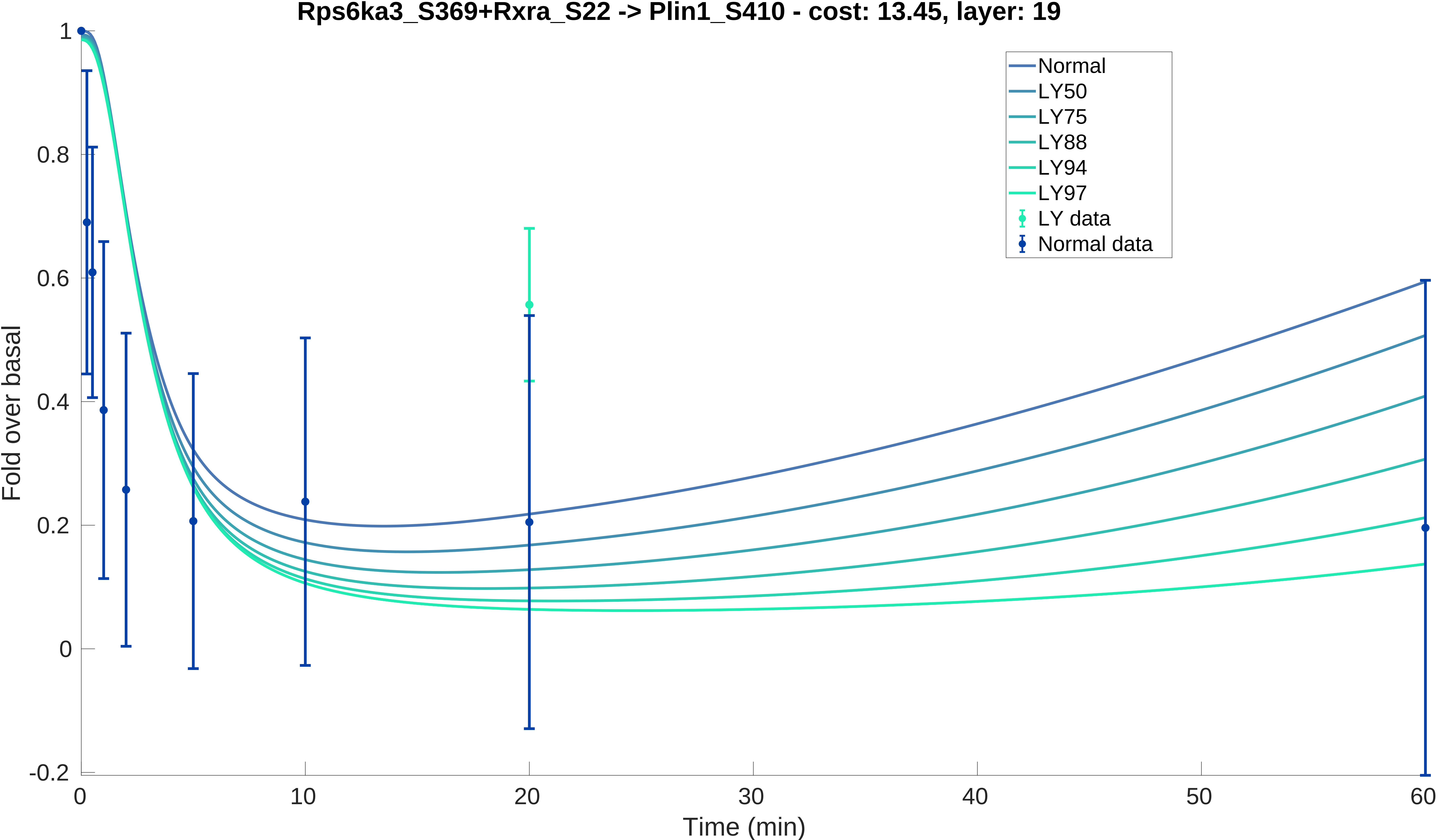

Rps6ka3_S369+Rxra_S22 -> Plin1_S410 - cost: 13.45, layer: 19
Fold over basal
Time (min)
1
0.8
0.6
0.4
0.2
0
-0.2
0
10
20
30
40
50
60
Normal
LY50
LY75
LY88
LY94
LY97
LY data
Normal data

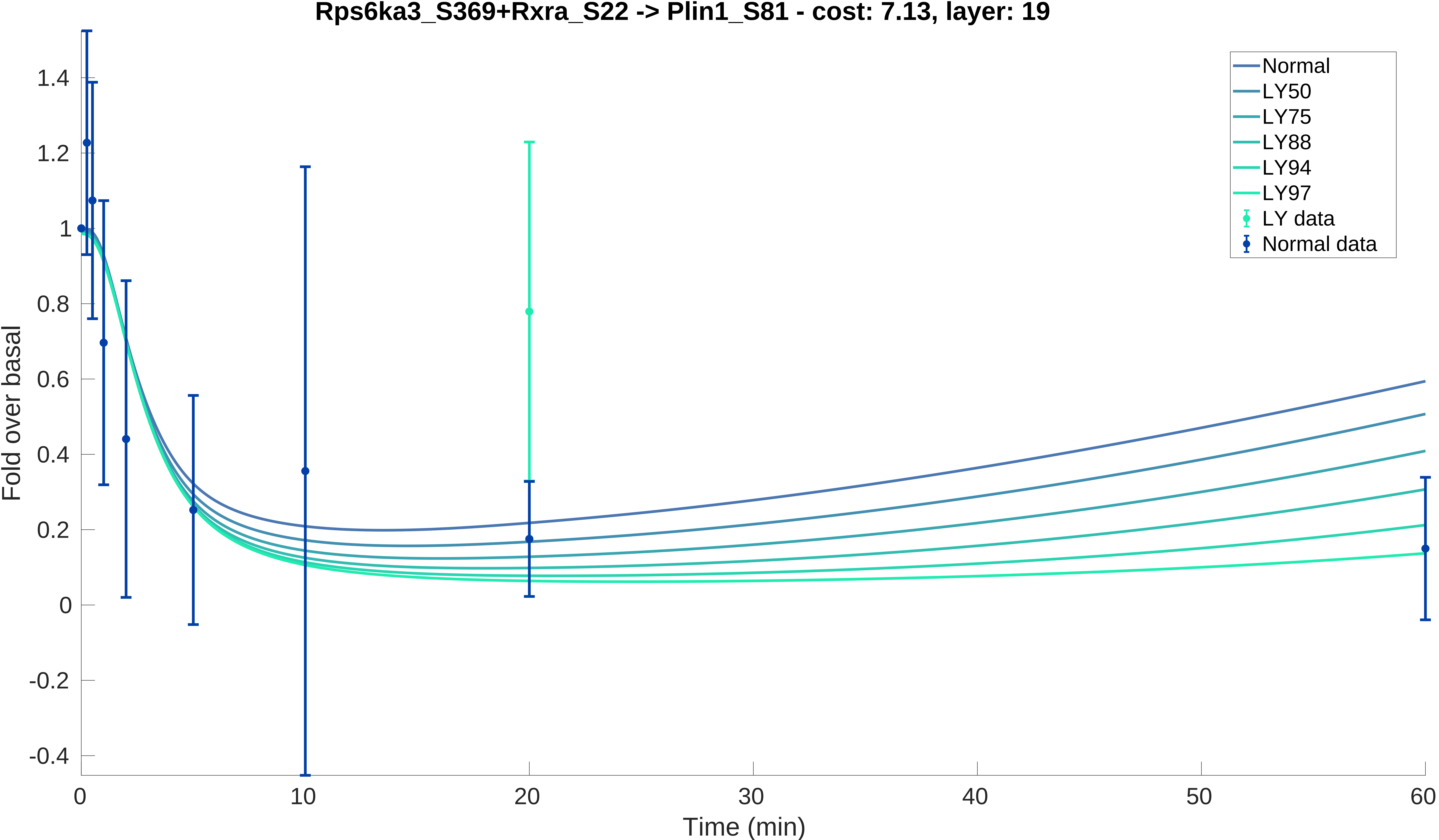

Rps6ka3_S369+Rxra_S22 -> Plin1_S81 - cost: 7.13, layer: 19
Fold over basal
Time (min)
Normal
LY50
LY75
LY88
LY94
LY97
LY data
Normal data
1.4
1.2
1
0.8
0.6
0.4
0.2
0
-0.2
-0.4
0
10
20
30
40
50
60

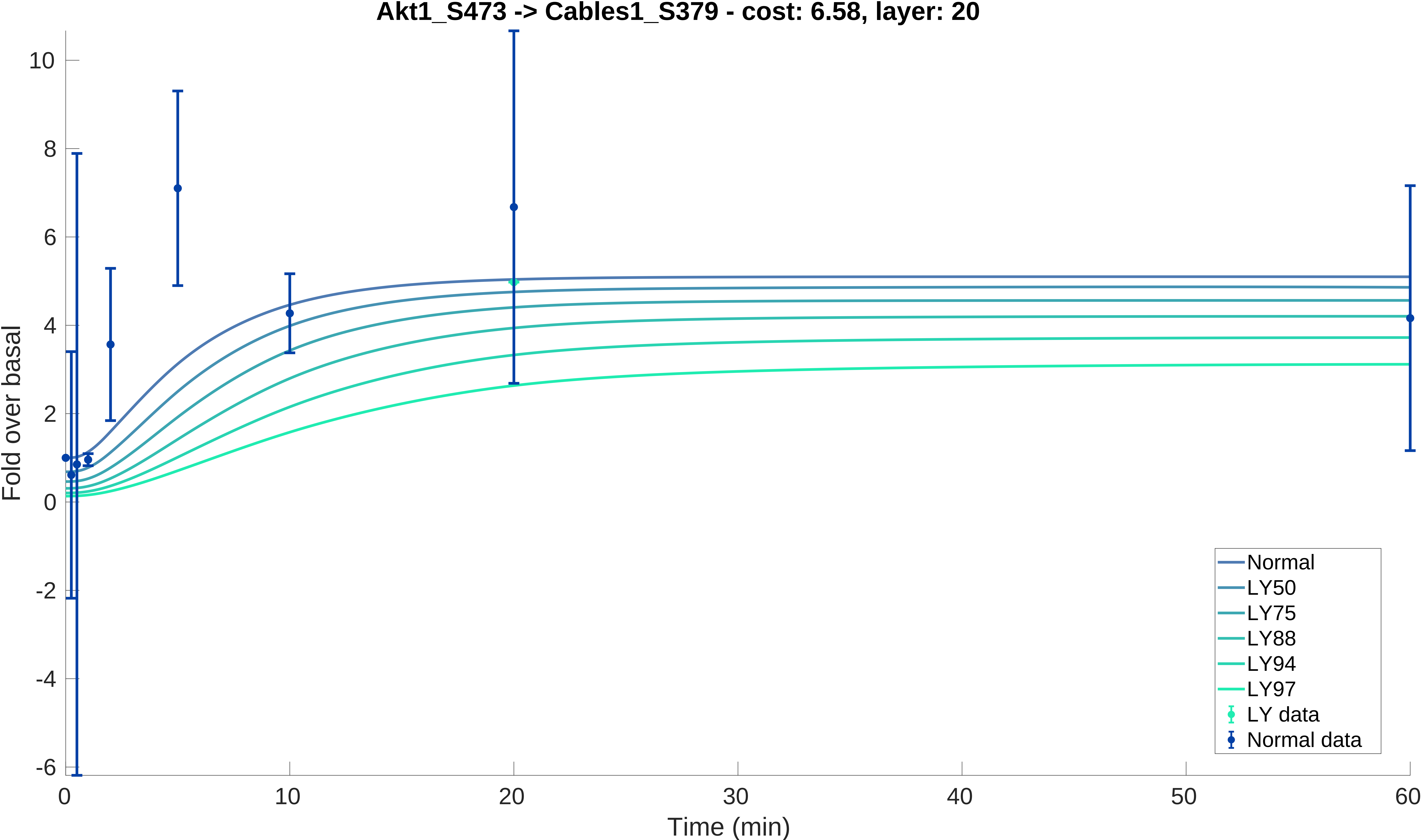

Akt1_S473 -> Cables1_S379 - cost: 6.58, layer: 20
Fold over basal
Time (min)
10
8
6
4
2
0
-2
-4
-6
0
10
20
30
40
50
60
Normal
LY50
LY75
LY88
LY94
LY97
LY data
Normal data

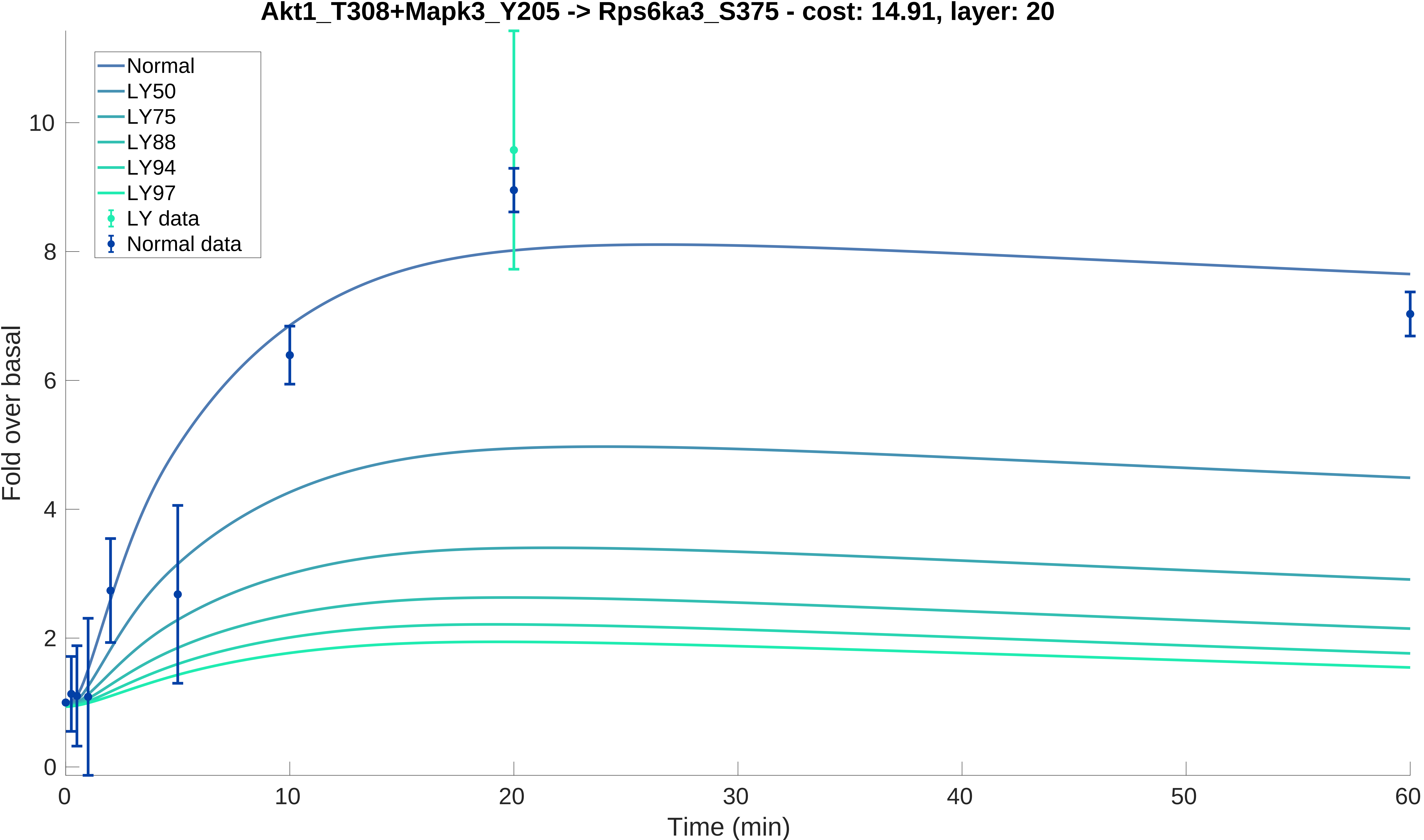

Akt1_T308+Mapk3_Y205 -> Rps6ka3_S375 - cost: 14.91, layer: 20
Normal
LY50
LY75
LY88
LY94
LY97
LY data
Normal data
Fold over basal
Time (min)
0
2
4
6
8
10
0
10
20
30
40
50
60

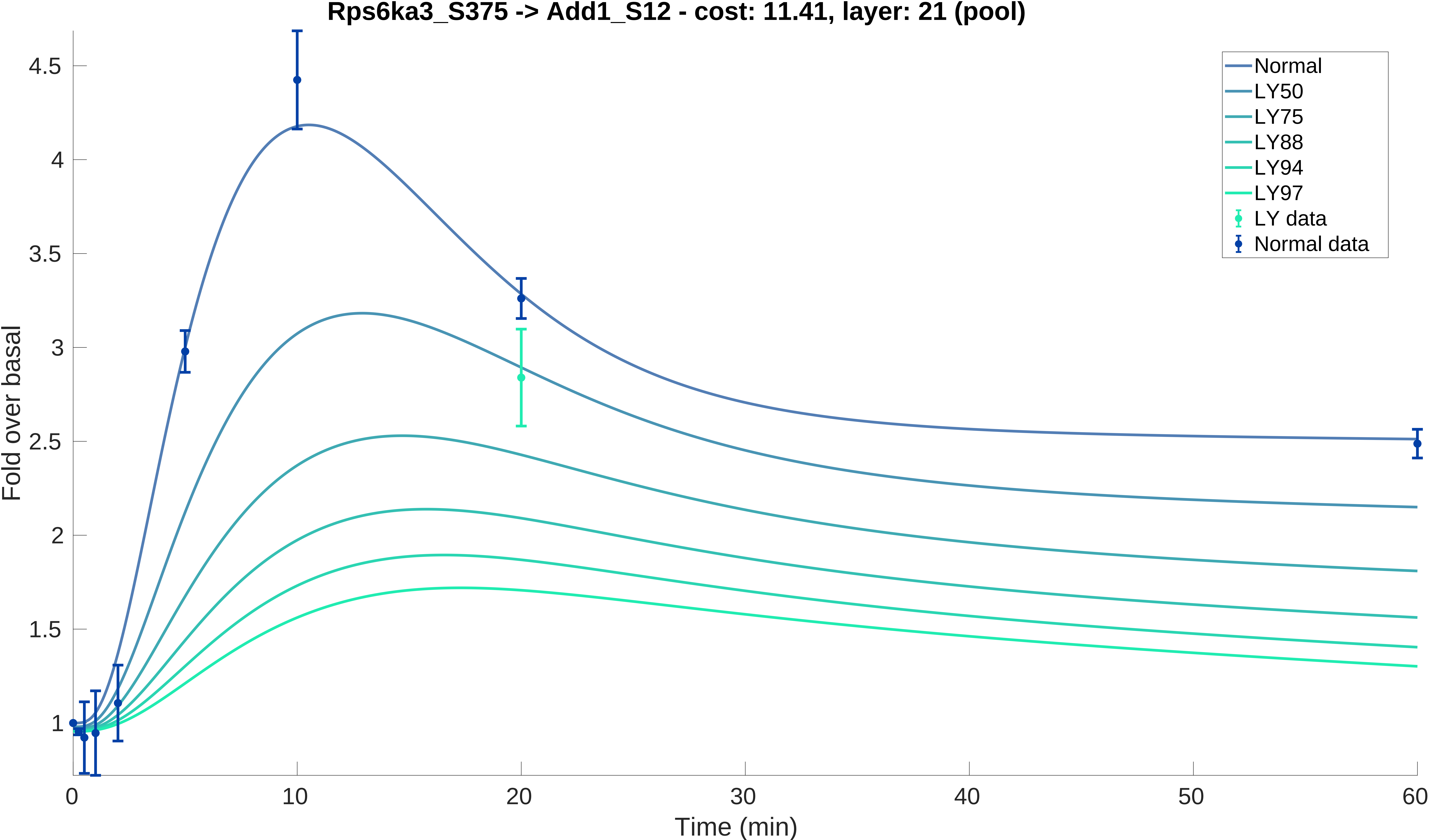

Rps6ka3_S375 -> Add1_S12 - cost: 11.41, layer: 21 (pool)
Fold over basal
Time (min)
4.5
4
3.5
3
2.5
2
1.5
1
0
10
20
30
40
50
60
Normal
LY50
LY75
LY88
LY94
LY97
LY data
Normal data

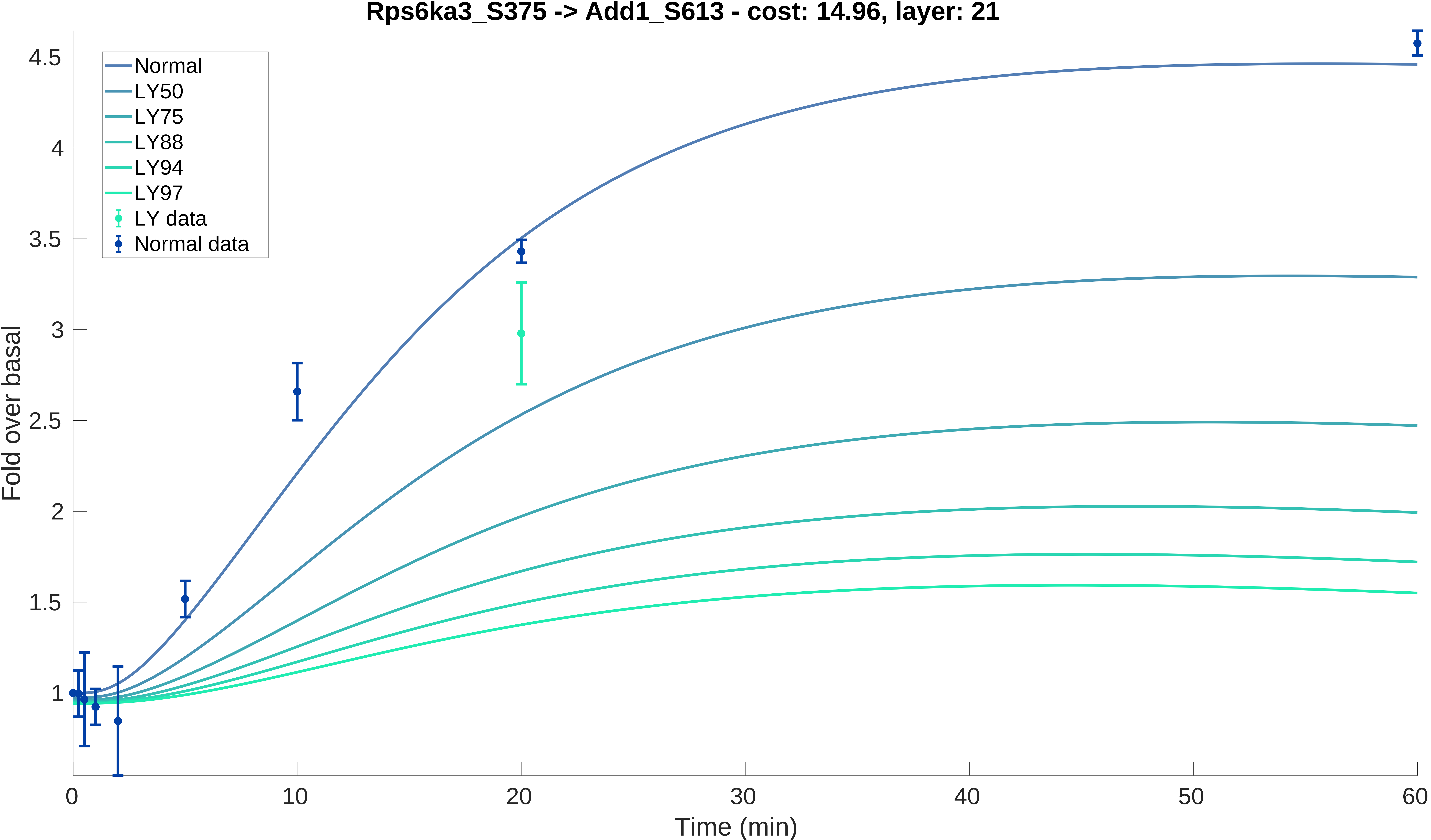
Rps6ka3_S375 -> Add1_S613 - cost: 14.96, layer: 21
Normal
LY50
LY75
LY88
LY94
LY97
LY data
Normal data
Fold over basal
Time (min)
4.5
4
3.5
3
2.5
2
1.5
1
0
10
20
30
40
50
60

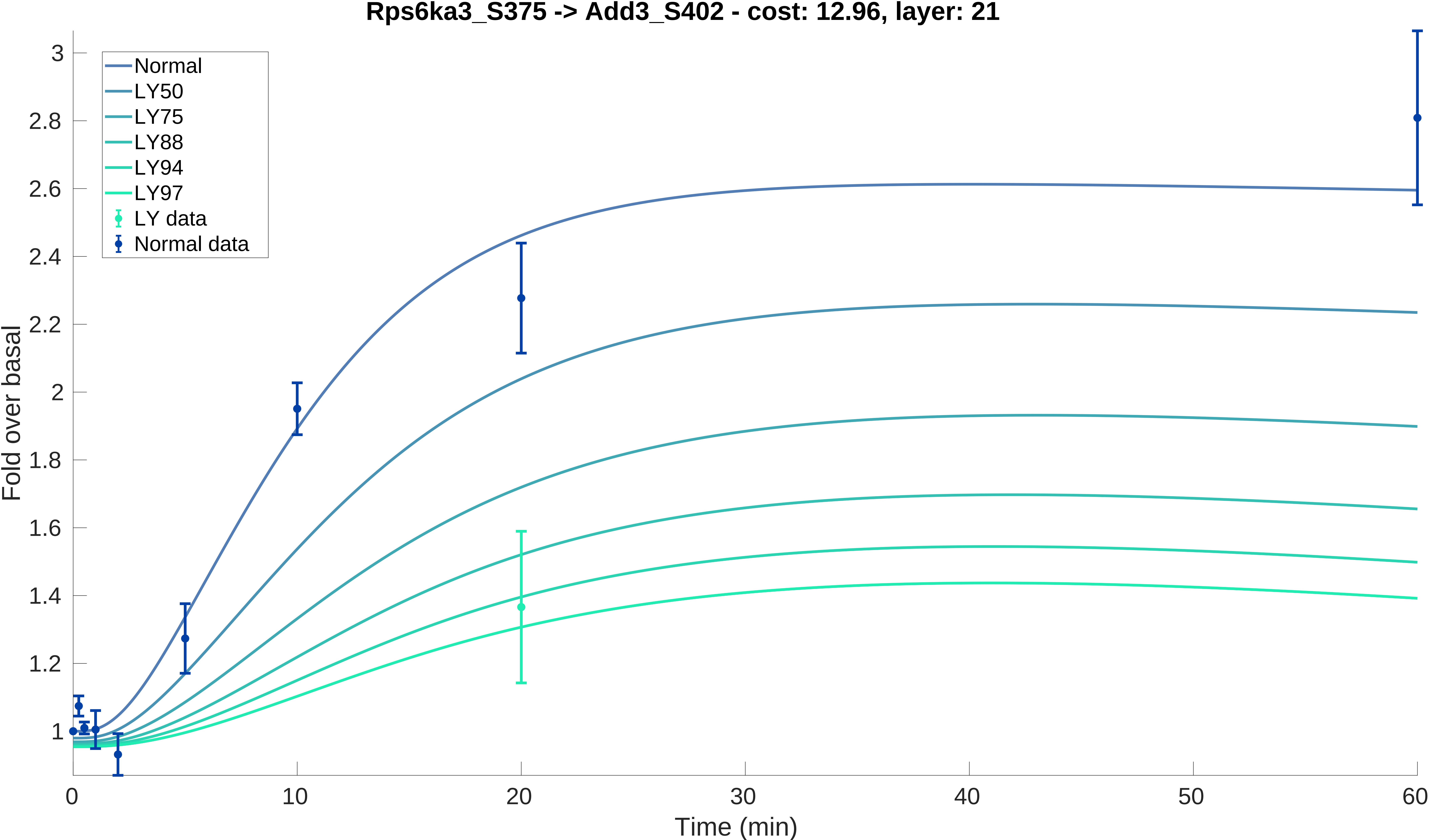

Rps6ka3_S375 -> Add3_S402 - cost: 12.96, layer: 21
Normal
LY50
LY75
LY88
LY94
LY97
LY data
Normal data
Fold over basal
3
2.8
2.6
2.4
2.2
2
1.8
1.6
1.4
1.2
1
0
10
20
30
40
50
60
Time (min)

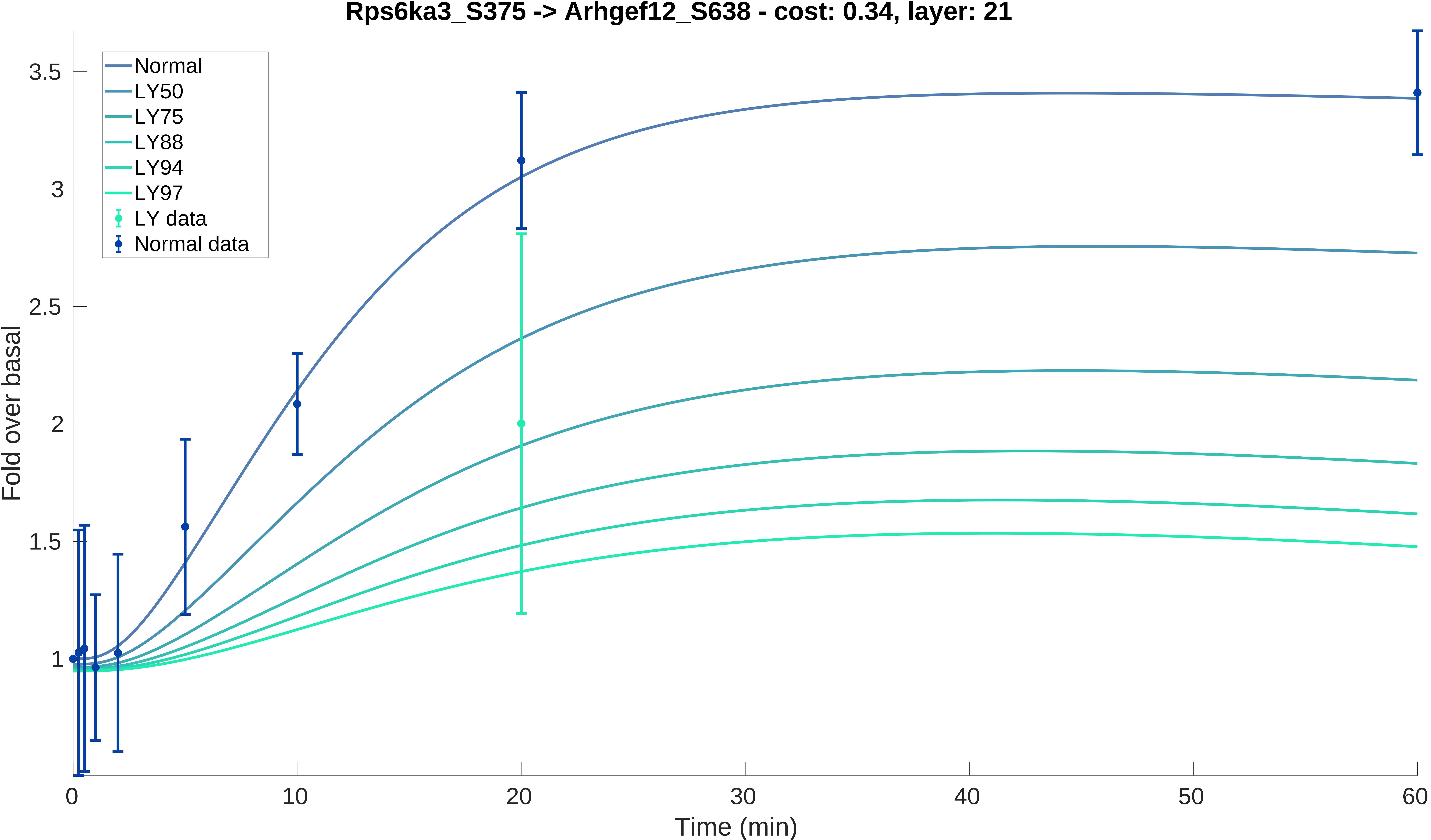

Rps6ka3_S375 -> Arhgef12_S638 - cost: 0.34, layer: 21
Normal
LY50
LY75
LY88
LY94
LY97
LY data
Normal data
Fold over basal
3.5
3
2.5
2
1.5
1
0
10
20
30
40
50
60
Time (min)

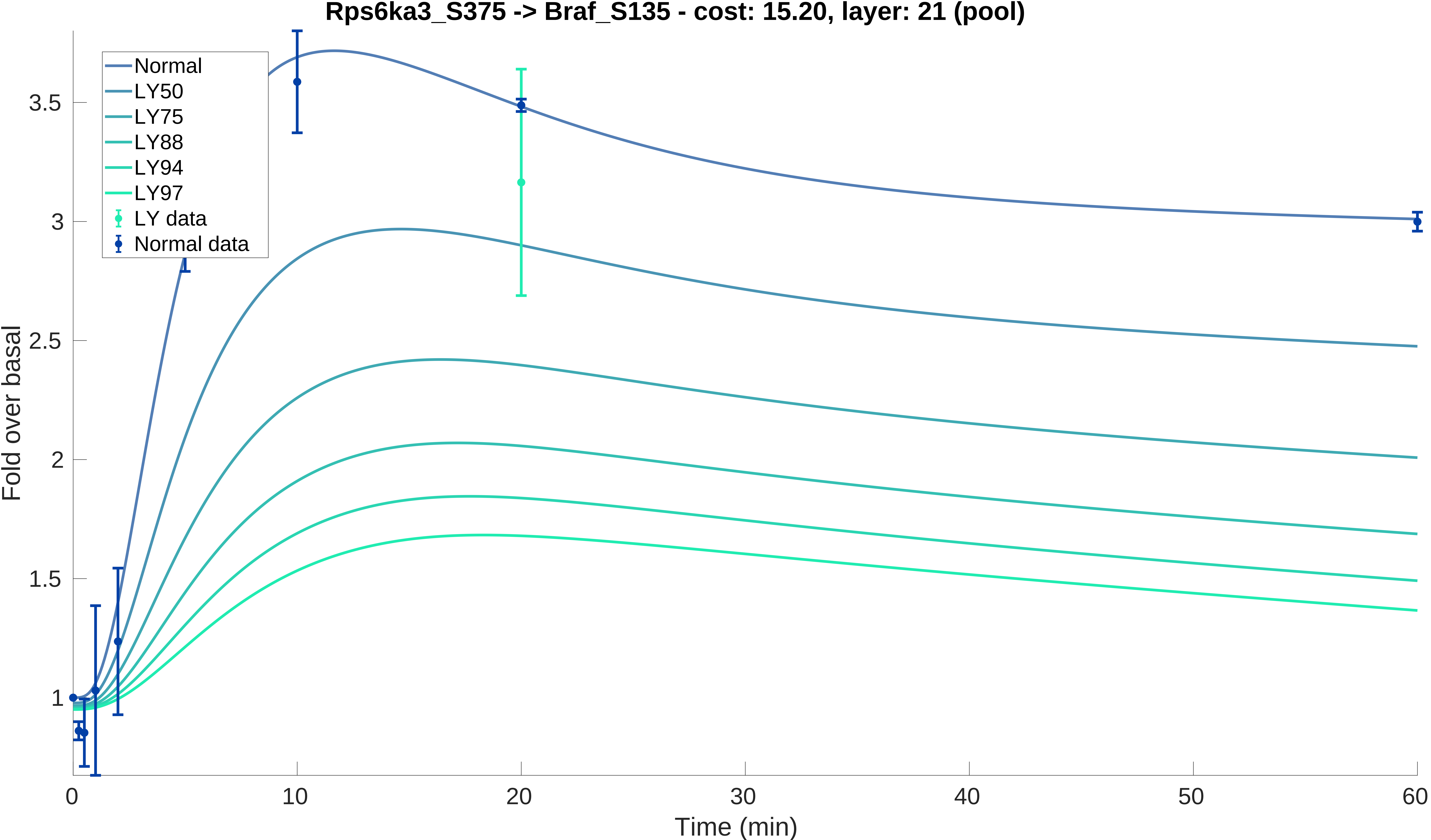

Rps6ka3_S375 -> Braf_S135 - cost: 15.20, layer: 21 (pool)
Normal
LY50
LY75
LY88
LY94
LY97
LY data
Normal data
Fold over basal
3.5
3
2.5
2
1.5
1
0
10
20
30
40
50
60
Time (min)

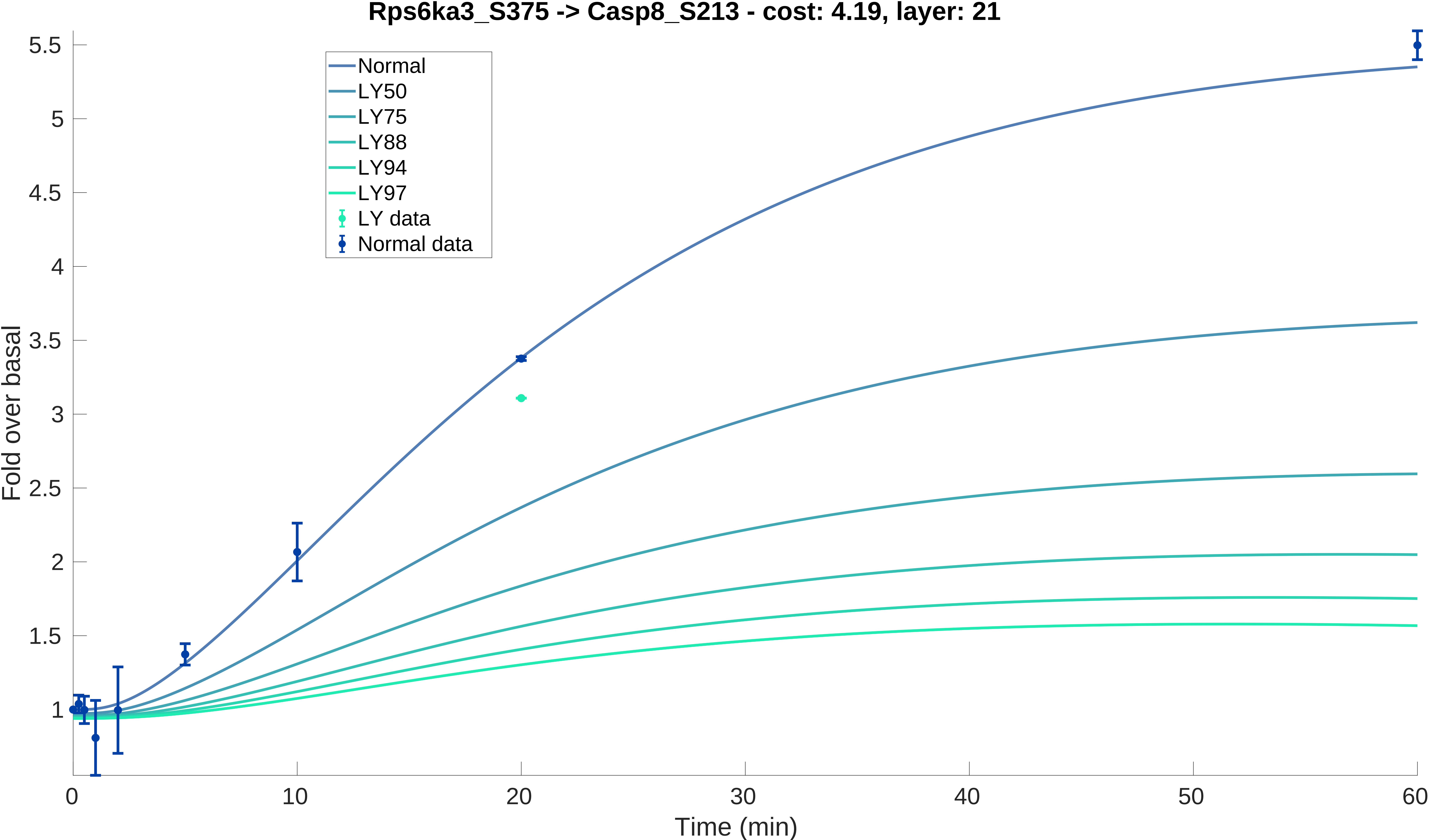

Rps6ka3_S375 -> Casp8_S213 - cost: 4.19, layer: 21
Normal
LY50
LY75
LY88
LY94
LY97
LY data
Normal data
Fold over basal
Time (min)
5.5
5
4.5
4
3.5
3
2.5
2
1.5
1
0
10
20
30
40
50
60

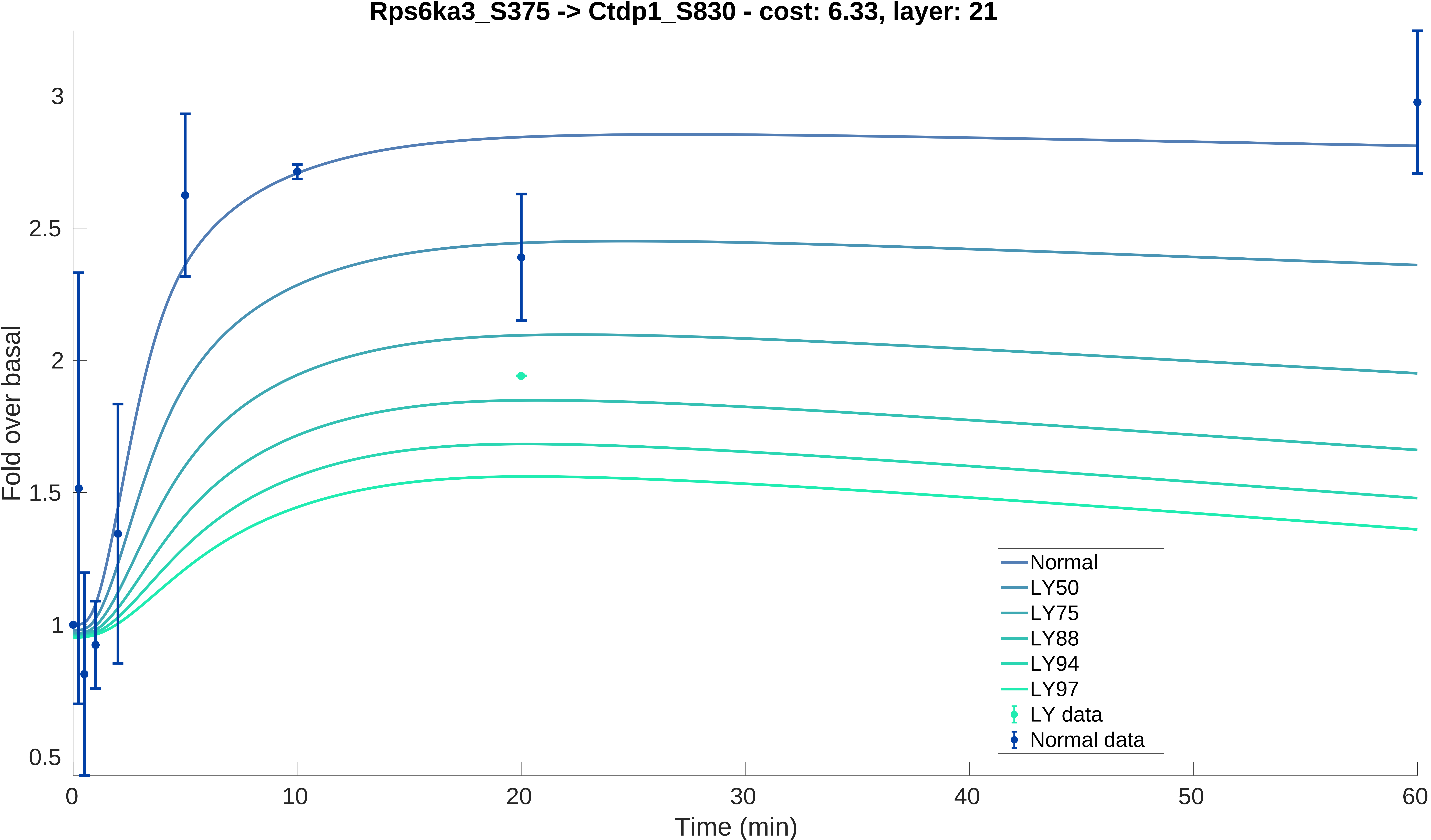

Rps6ka3_S375 -> Ctdp1_S830 - cost: 6.33, layer: 21
Fold over basal
Time (min)
0.5
1
1.5
2
2.5
3
0
10
20
30
40
50
60
Normal
LY50
LY75
LY88
LY94
LY97
LY data
Normal data

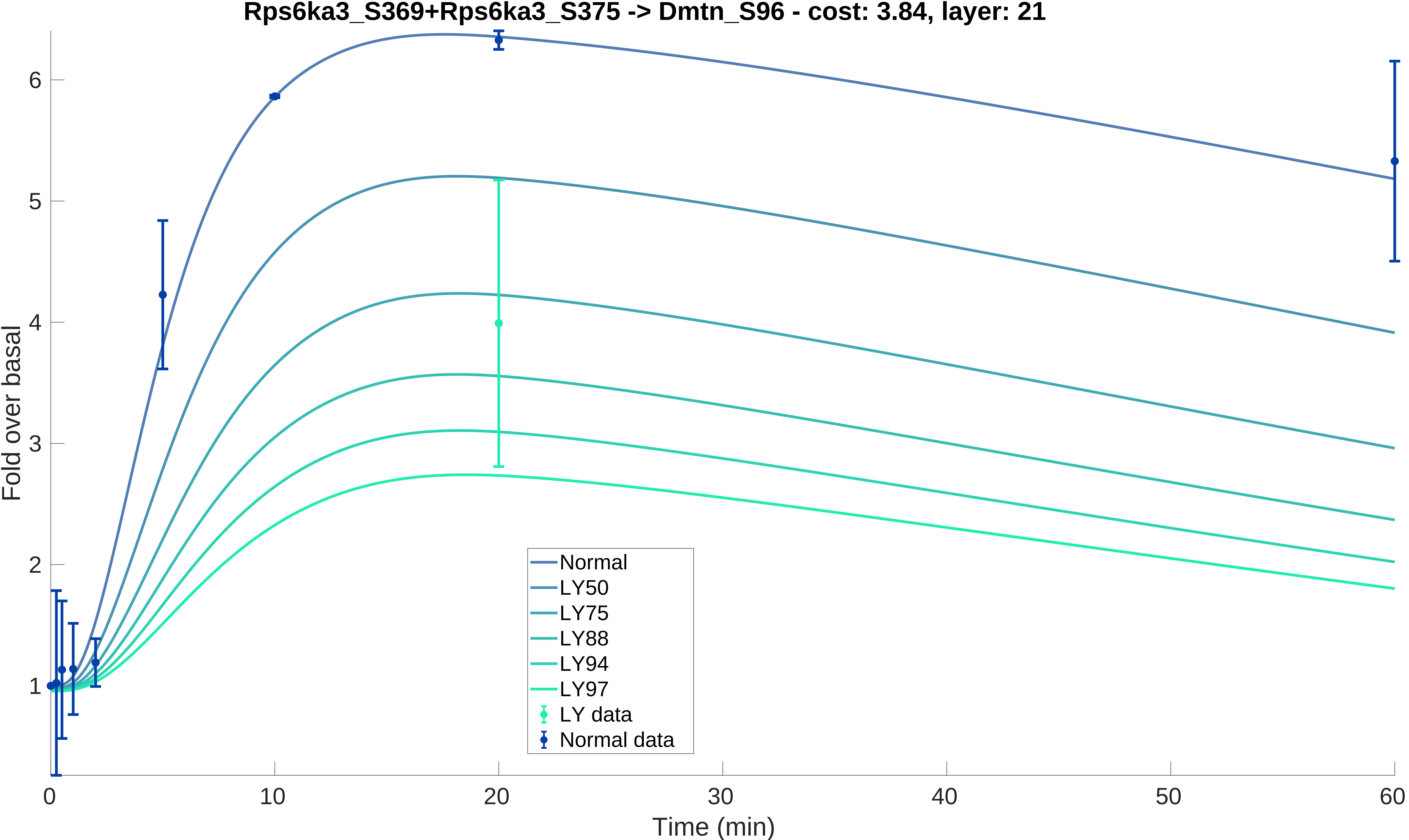

Rps6ka3_S369+Rps6ka3_S375 -> Dmtn_S96 - cost: 3.84, layer: 21
Fold over basal
Time (min)
0
10
20
30
40
50
60
1
2
3
4
5
6
Normal
LY50
LY75
LY88
LY94
LY97
LY data
Normal data

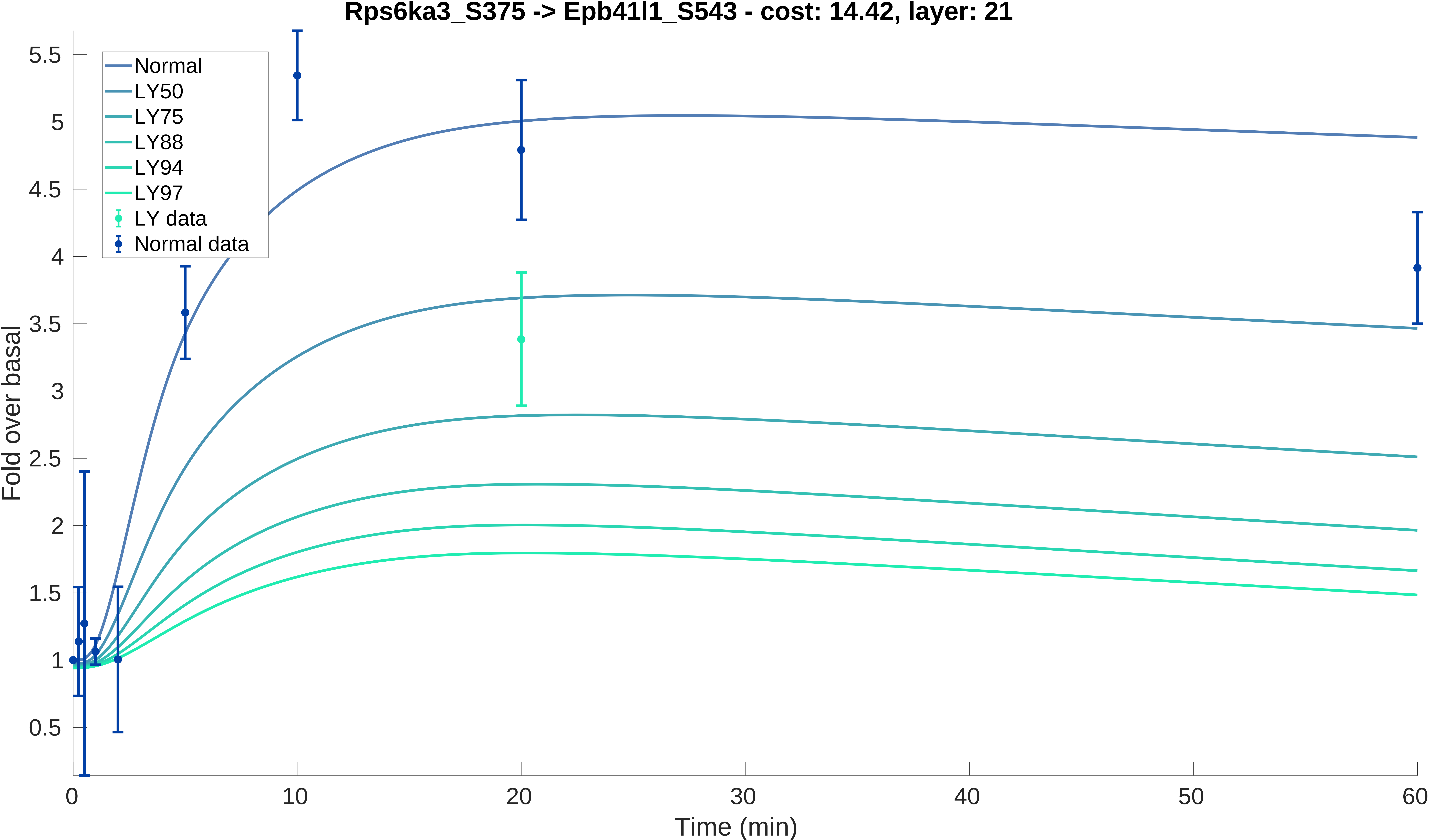

Rps6ka3_S375 -> Epb41l1_S543 - cost: 14.42, layer: 21
Normal
LY50
LY75
LY88
LY94
LY97
LY data
Normal data
Fold over basal
Time (min)
0.5
1
1.5
2
2.5
3
3.5
4
4.5
5
5.5
0
10
20
30
40
50
60

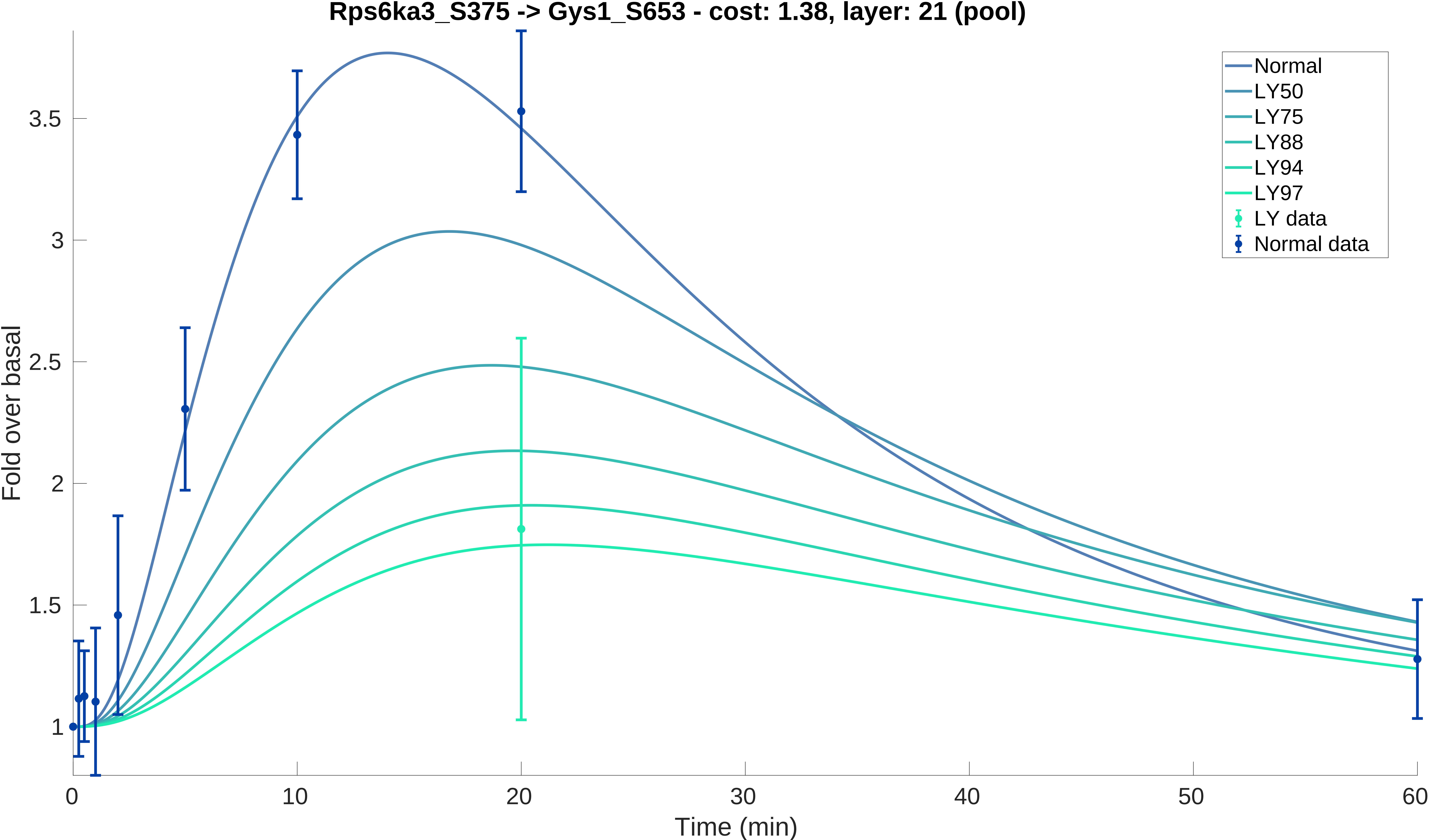

Rps6ka3_S375 -> Gys1_S653 - cost: 1.38, layer: 21 (pool)
Fold over basal
Time (min)
0
10
20
30
40
50
60
1
1.5
2
2.5
3
3.5
Normal
LY50
LY75
LY88
LY94
LY97
LY data
Normal data

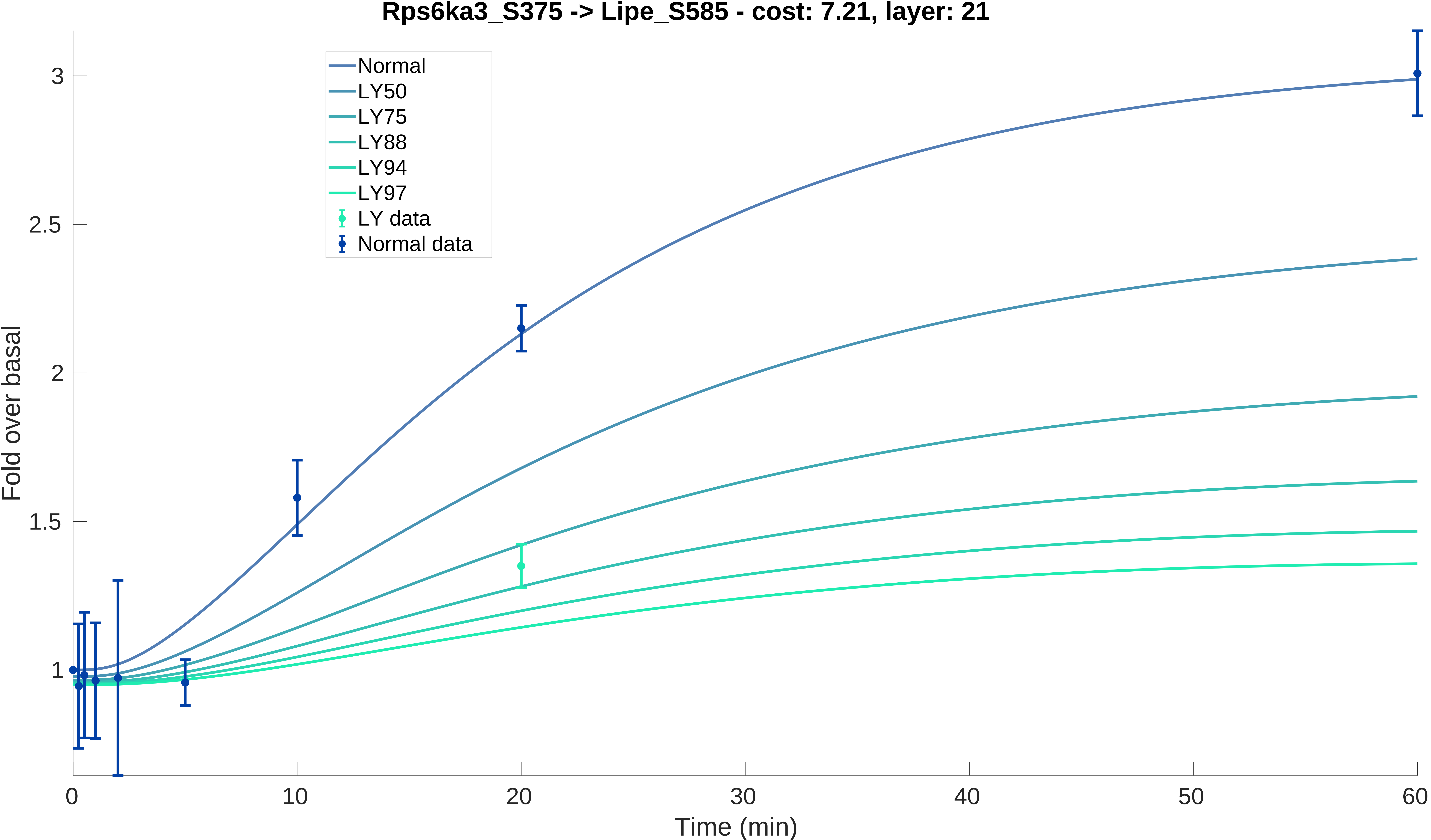
Rps6ka3_S375 -> Lipe_S585 - cost: 7.21, layer: 21
Normal
LY50
LY75
LY88
LY94
LY97
LY data
Normal data
Fold over basal
Time (min)
3
2.5
2
1.5
1
0
10
20
30
40
50
60

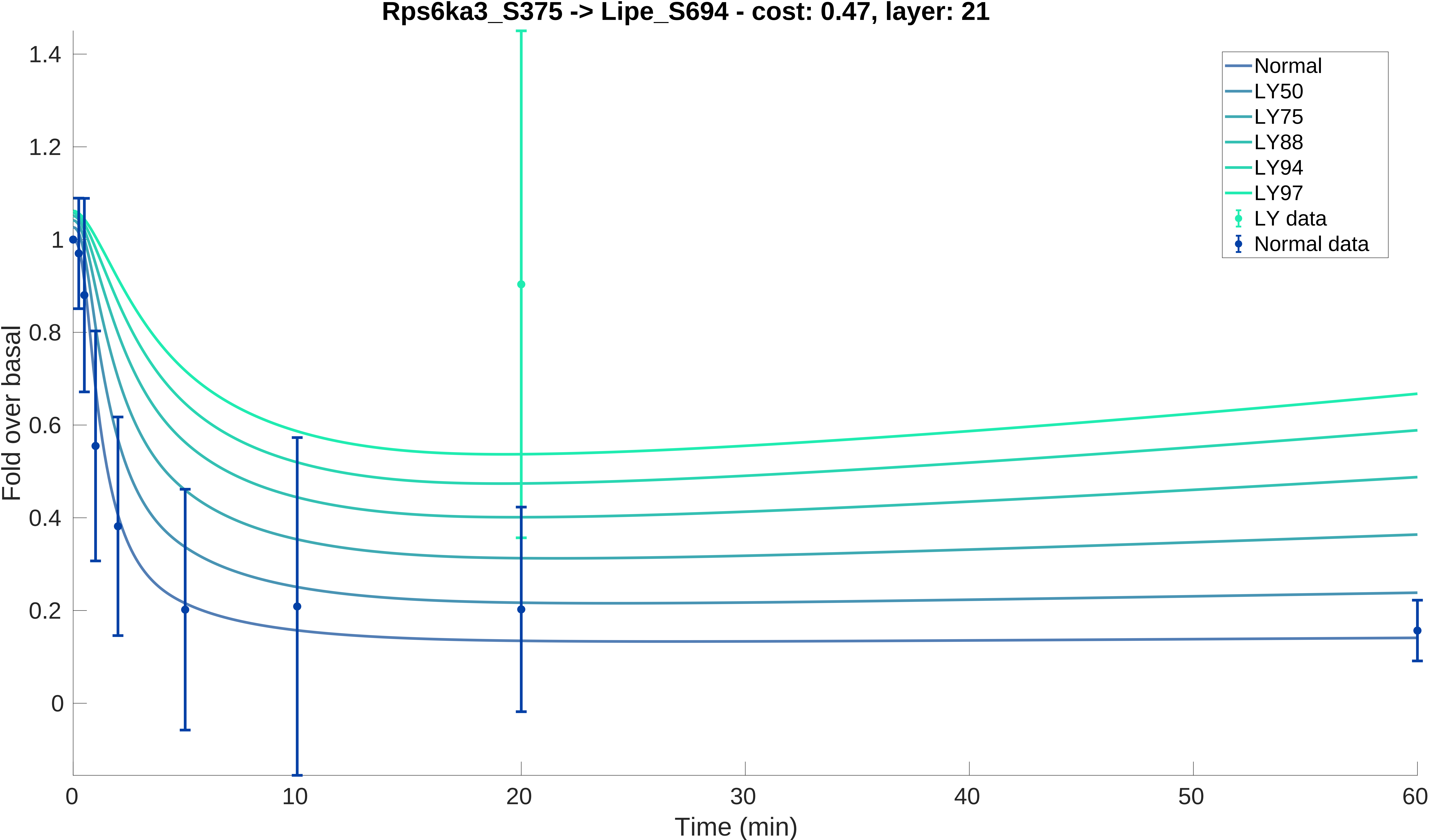

Rps6ka3_S375 -> Lipe_S694 - cost: 0.47, layer: 21
Normal
LY50
LY75
LY88
LY94
LY97
LY data
Normal data
Fold over basal
Time (min)
0
10
20
30
40
50
60
0
0.2
0.4
0.6
0.8
1
1.2
1.4

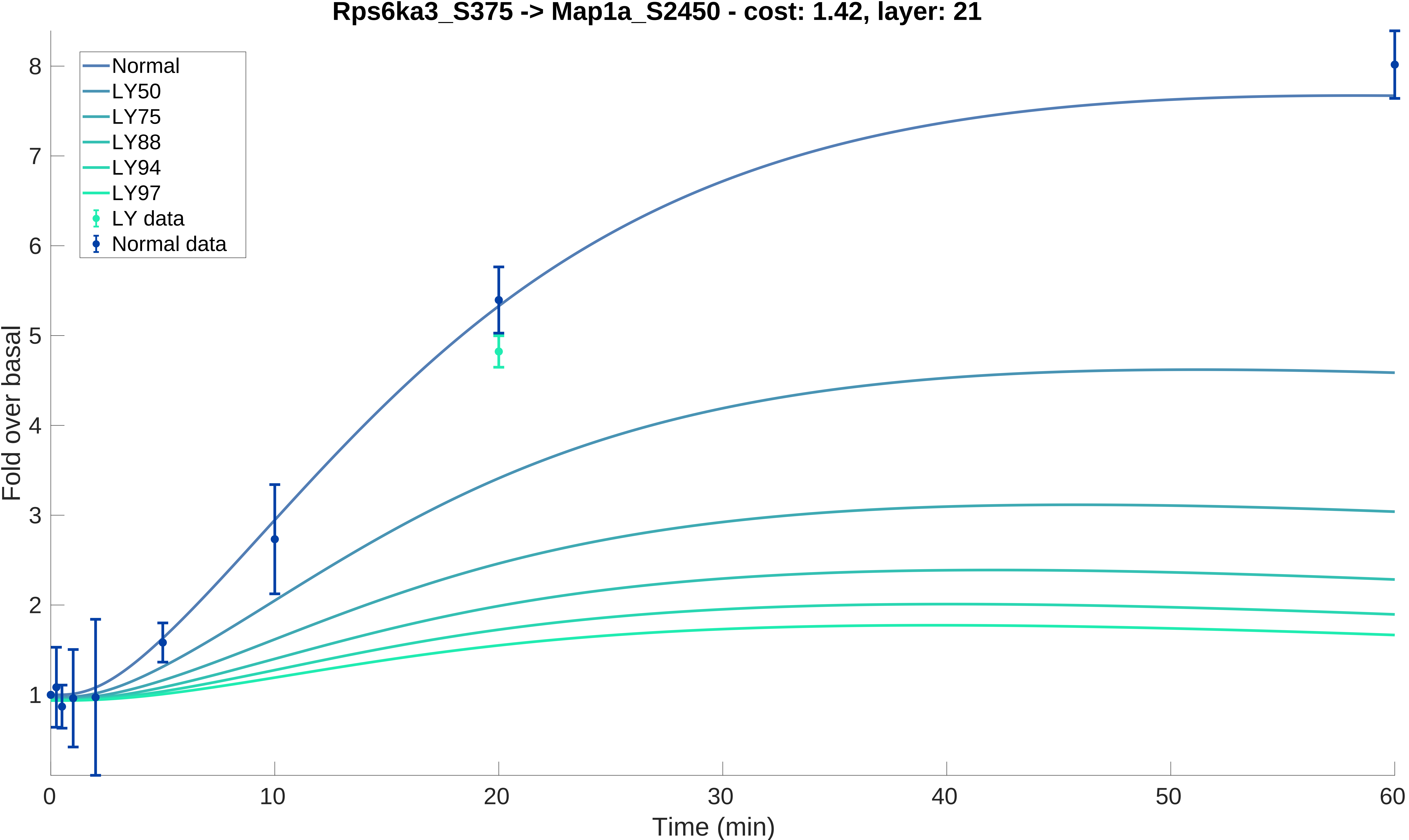

Rps6ka3_S375 -> Map1a_S2450 - cost: 1.42, layer: 21
Normal
LY50
LY75
LY88
LY94
LY97
LY data
Normal data
Fold over basal
1
2
3
4
5
6
7
8
0
10
20
30
40
50
60
Time (min)

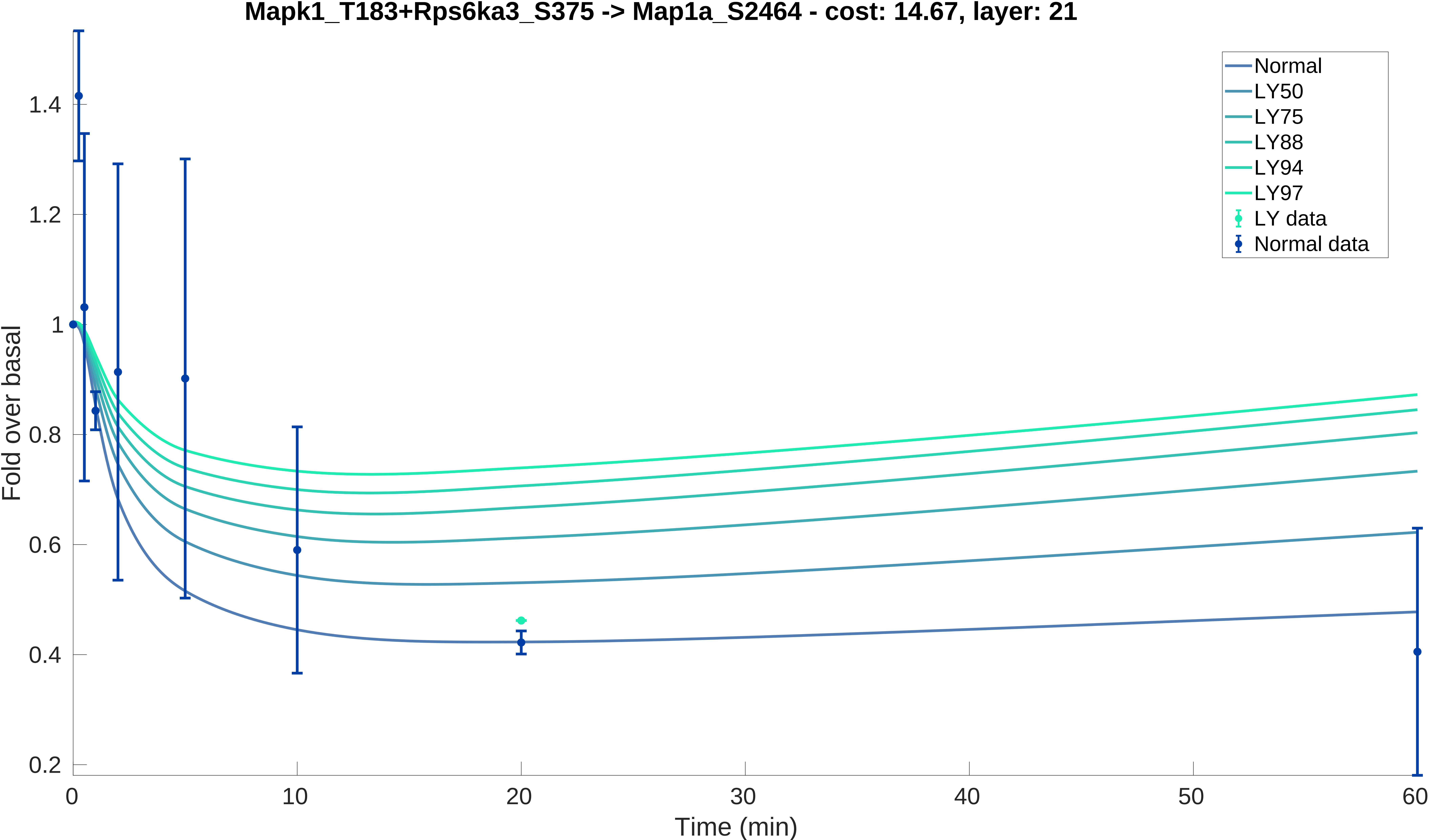

Mapk1_T183+Rps6ka3_S375 -> Map1a_S2464 - cost: 14.67, layer: 21
Fold over basal
Time (min)
0.2
0.4
0.6
0.8
1
1.2
1.4
0
10
20
30
40
50
60
Normal
LY50
LY75
LY88
LY94
LY97
LY data
Normal data

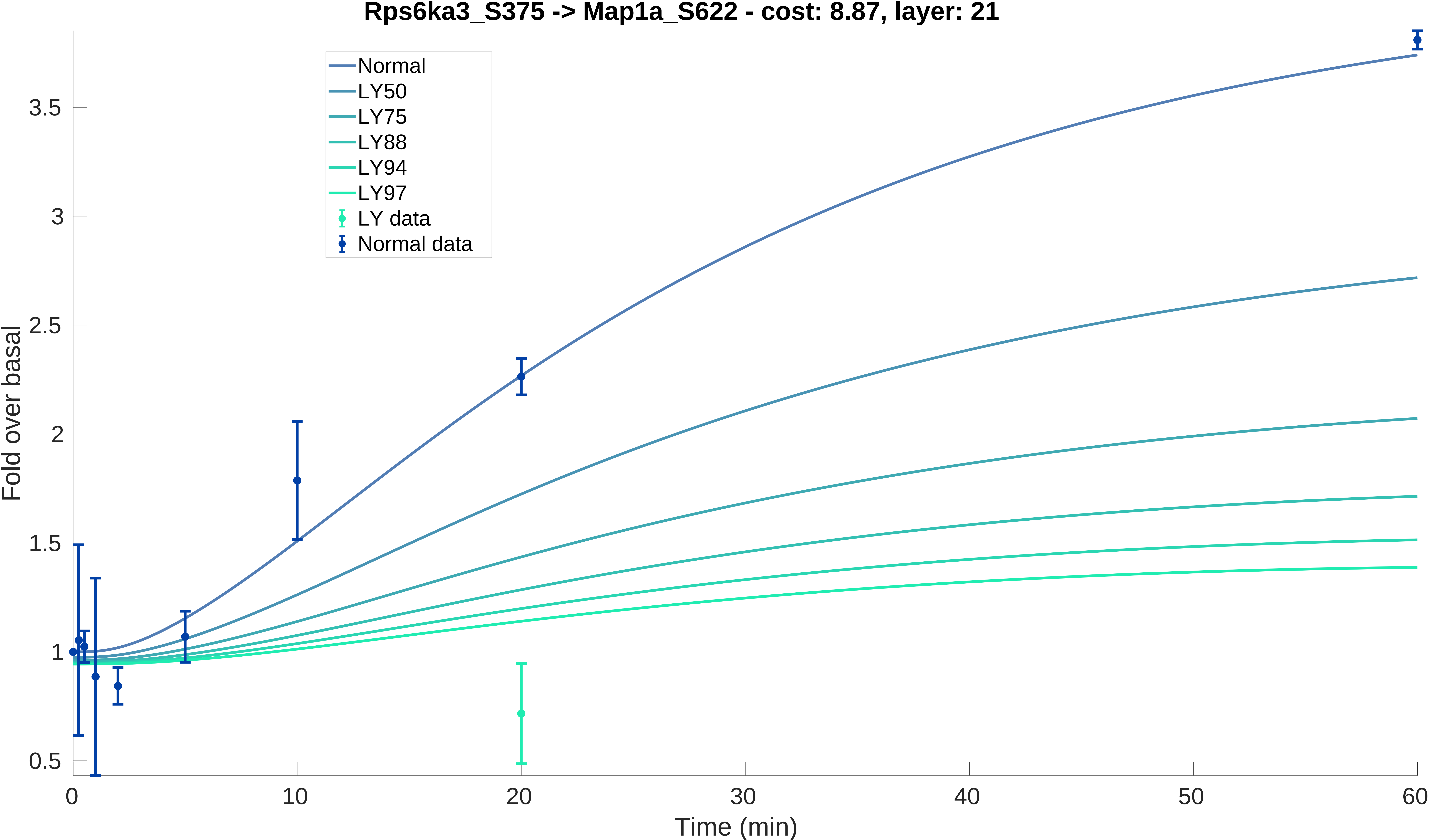
Rps6ka3_S375 -> Map1a_S622 - cost: 8.87, layer: 21
Normal
LY50
LY75
LY88
LY94
LY97
LY data
Normal data
Fold over basal
0.5
1
1.5
2
2.5
3
3.5
0
10
20
30
40
50
60
Time (min)

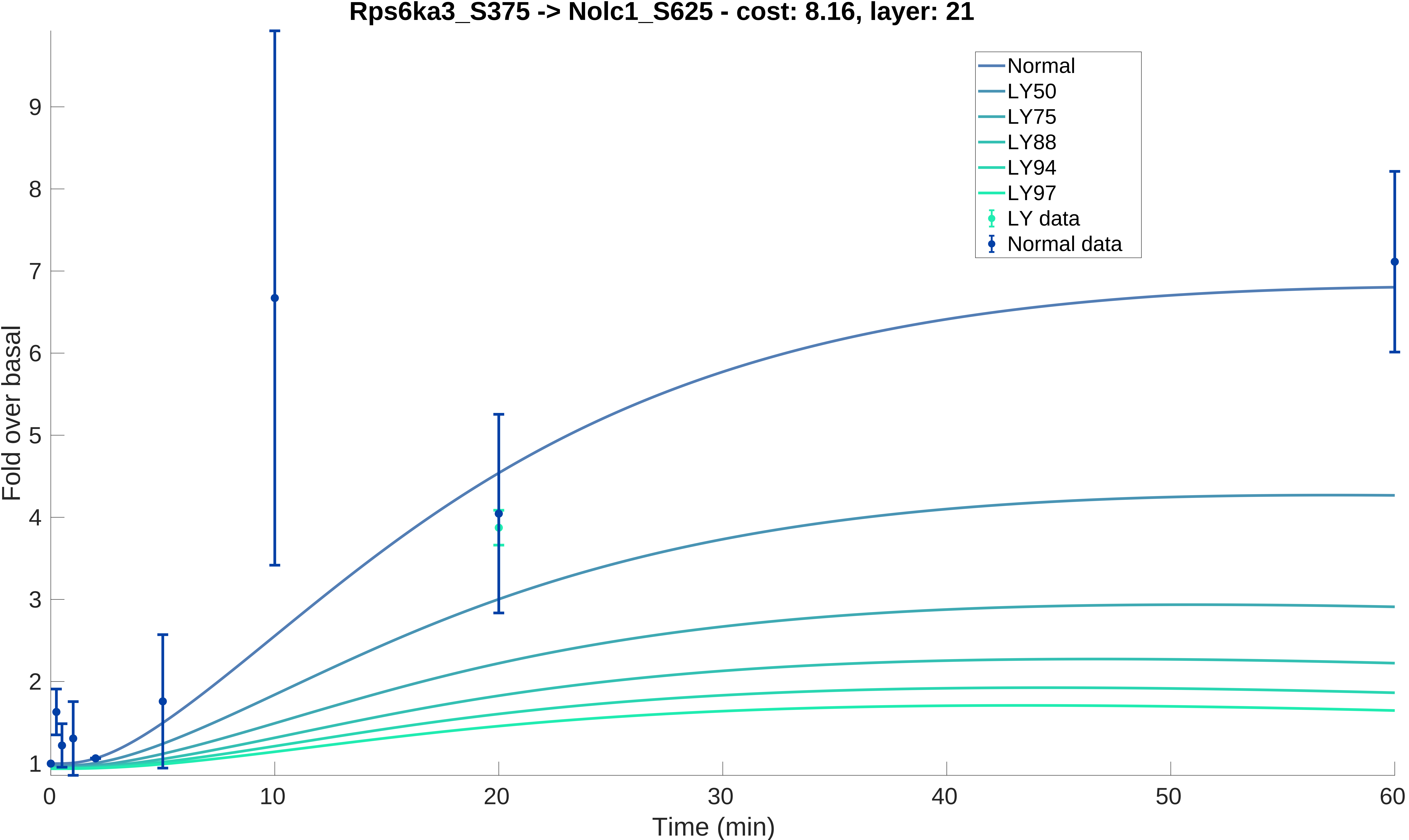

Rps6ka3_S375 -> Nolc1_S625 - cost: 8.16, layer: 21
Fold over basal
Time (min)
1
2
3
4
5
6
7
8
9
0
10
20
30
40
50
60
Normal
LY50
LY75
LY88
LY94
LY97
LY data
Normal data

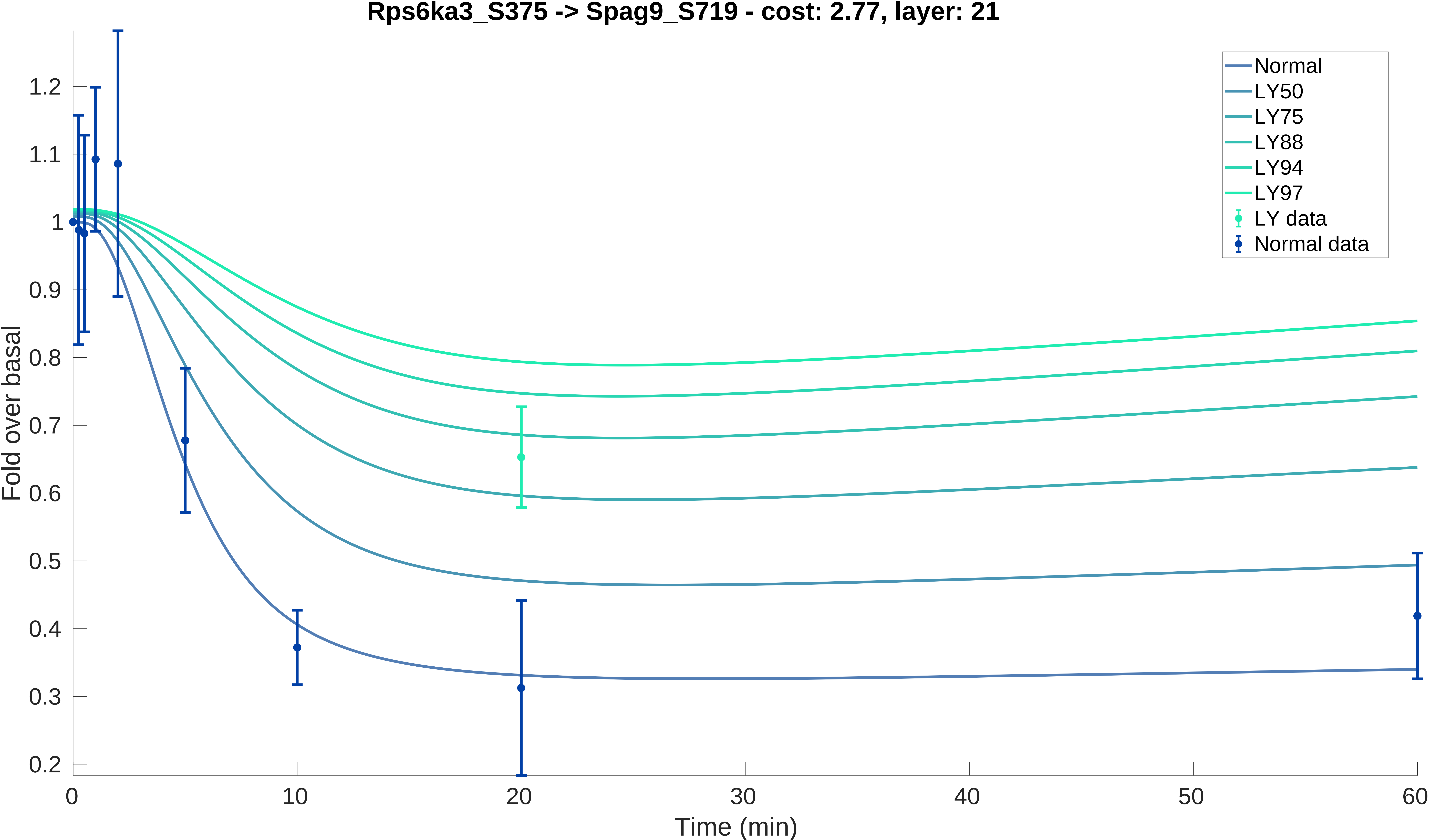
Rps6ka3_S375 -> Spag9_S719 - cost: 2.77, layer: 21
Fold over basal
Time (min)
0
10
20
30
40
50
60
0.2
0.3
0.4
0.5
0.6
0.7
0.8
0.9
1
1.1
1.2
Normal
LY50
LY75
LY88
LY94
LY97
LY data
Normal data

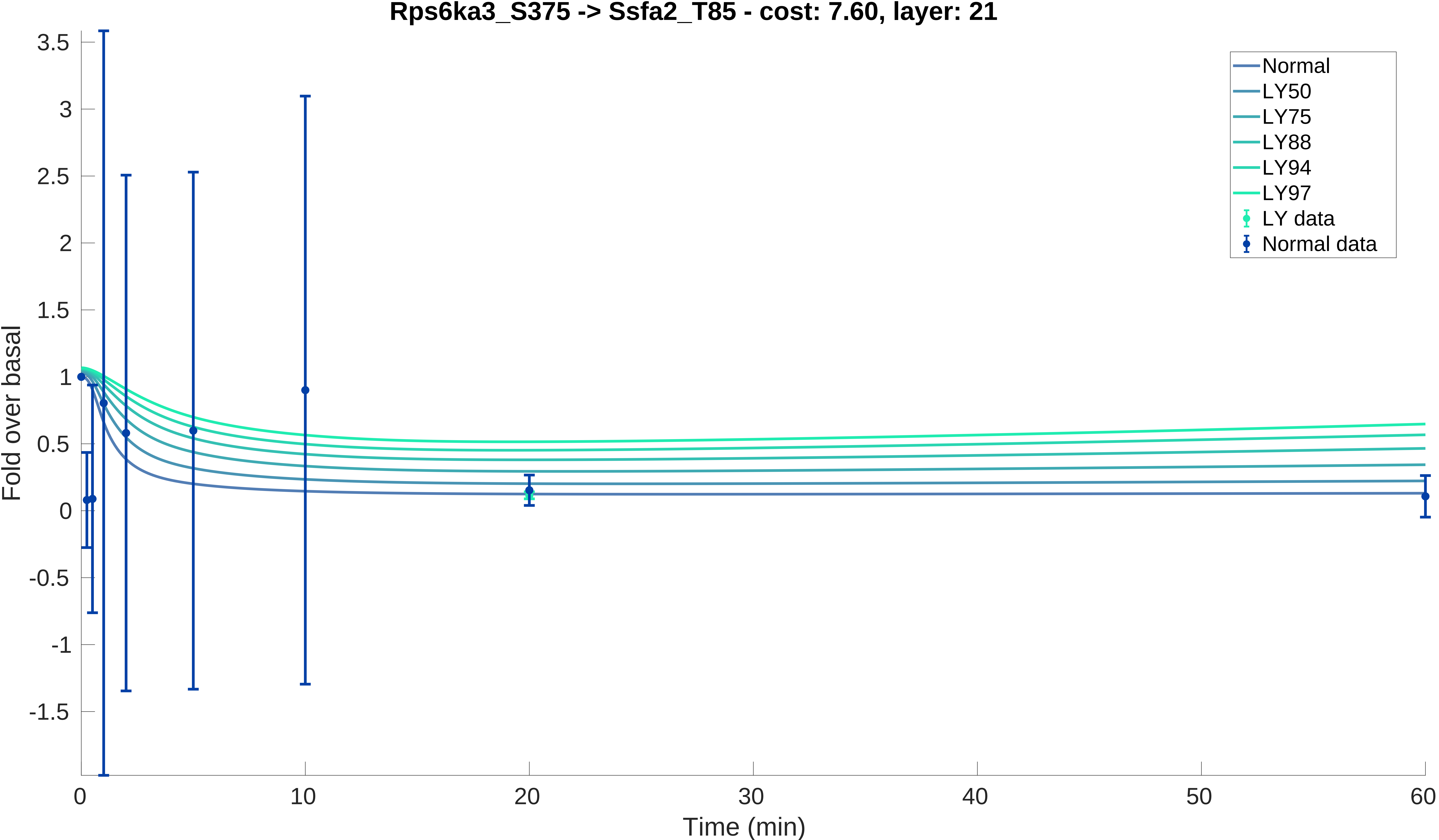

Rps6ka3_S375 -> Ssfa2_T85 - cost: 7.60, layer: 21
3.5
3
2.5
2
1.5
1
0.5
0
-0.5
-1
-1.5
Fold over basal
0
10
20
30
40
50
60
Time (min)
Normal
LY50
LY75
LY88
LY94
LY97
LY data
Normal data

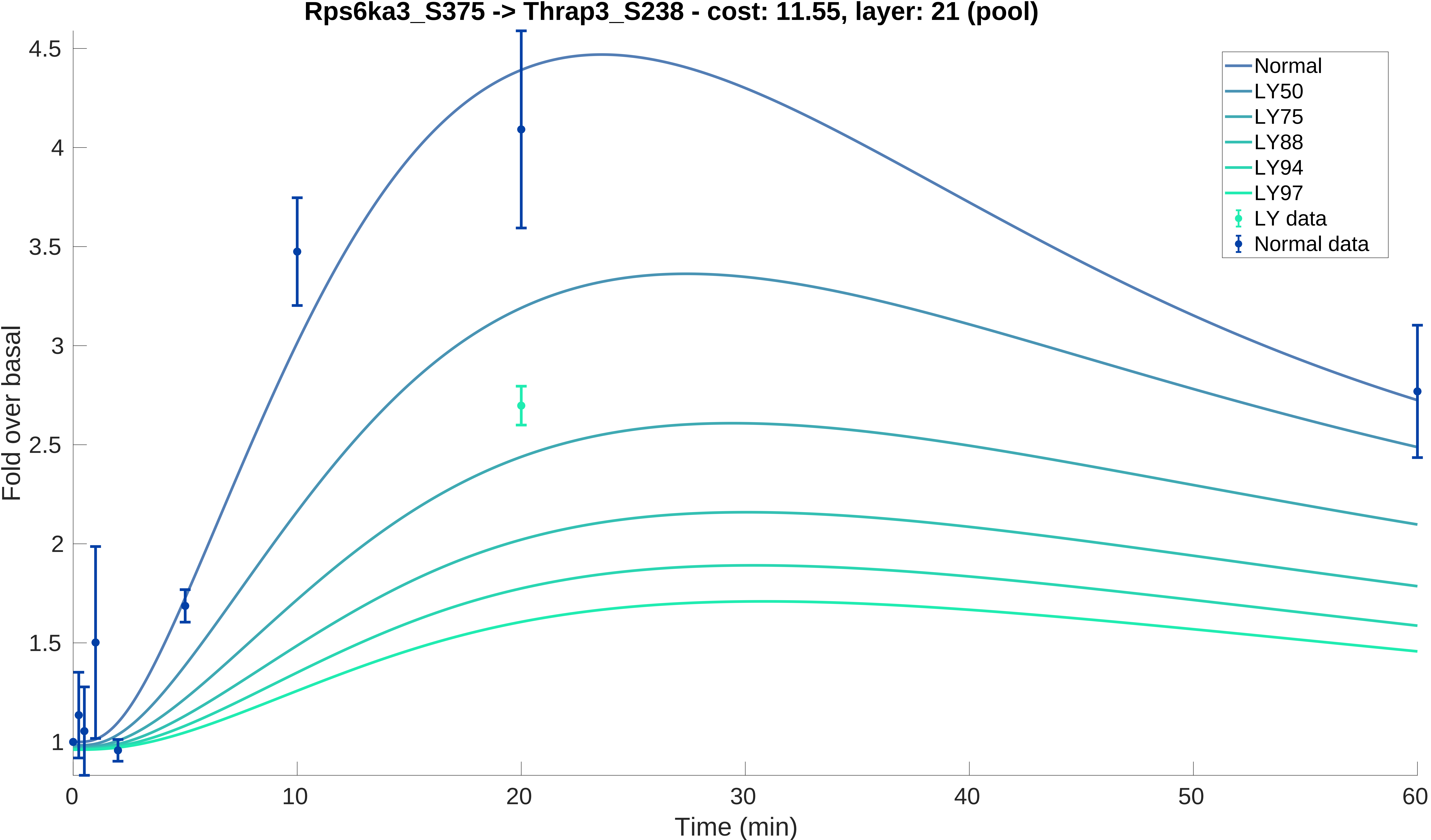

Rps6ka3_S375 -> Thrap3_S238 - cost: 11.55, layer: 21 (pool)
Fold over basal
Time (min)
Normal
LY50
LY75
LY88
LY94
LY97
LY data
Normal data
4.5
4
3.5
3
2.5
2
1.5
1
0
10
20
30
40
50
60

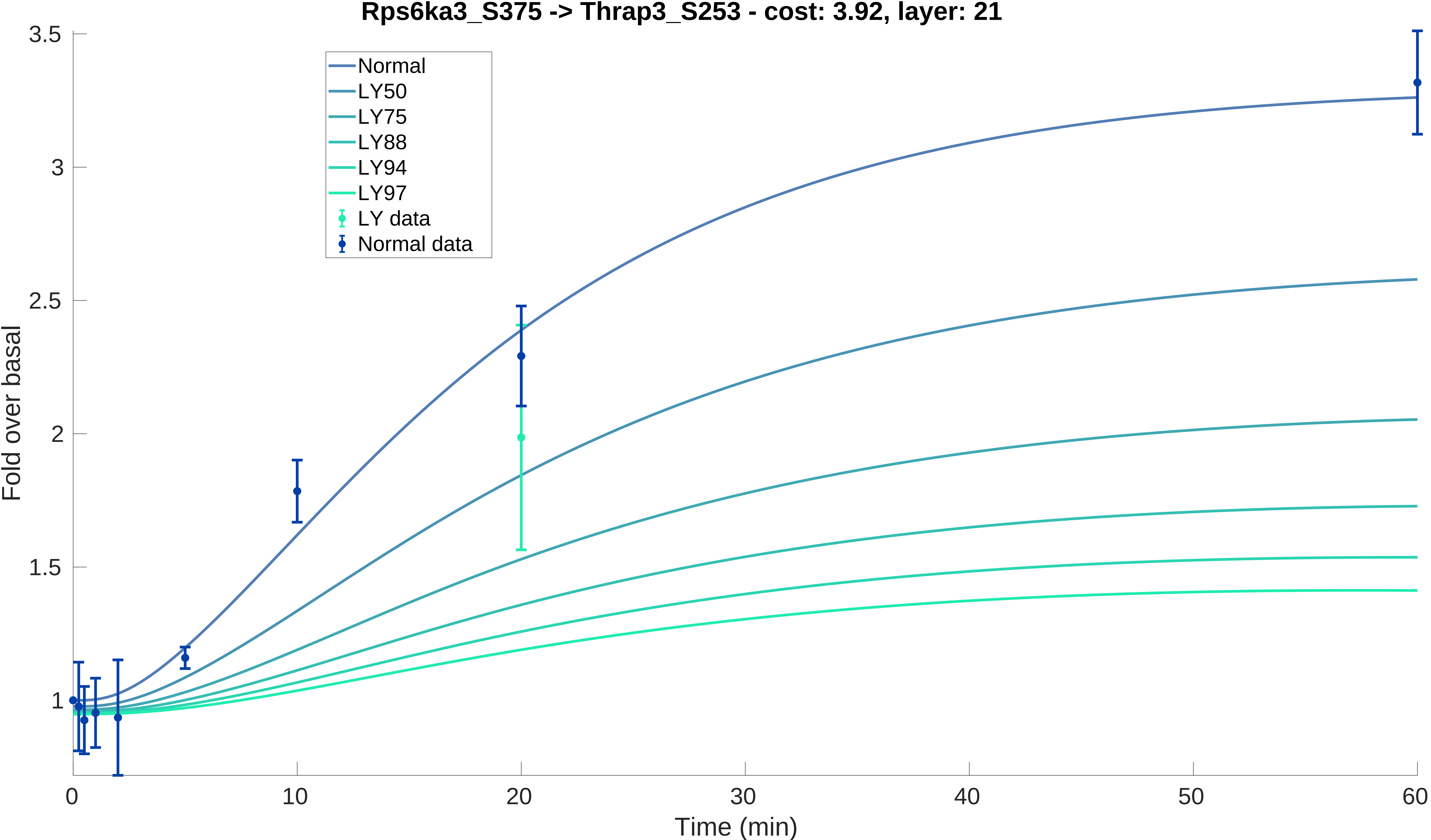

Rps6ka3_S375 -> Thrap3_S253 - cost: 3.92, layer: 21
Normal
LY50
LY75
LY88
LY94
LY97
LY data
Normal data
Fold over basal
Time (min)
3.5
3
2.5
2
1.5
1
0
10
20
30
40
50
60

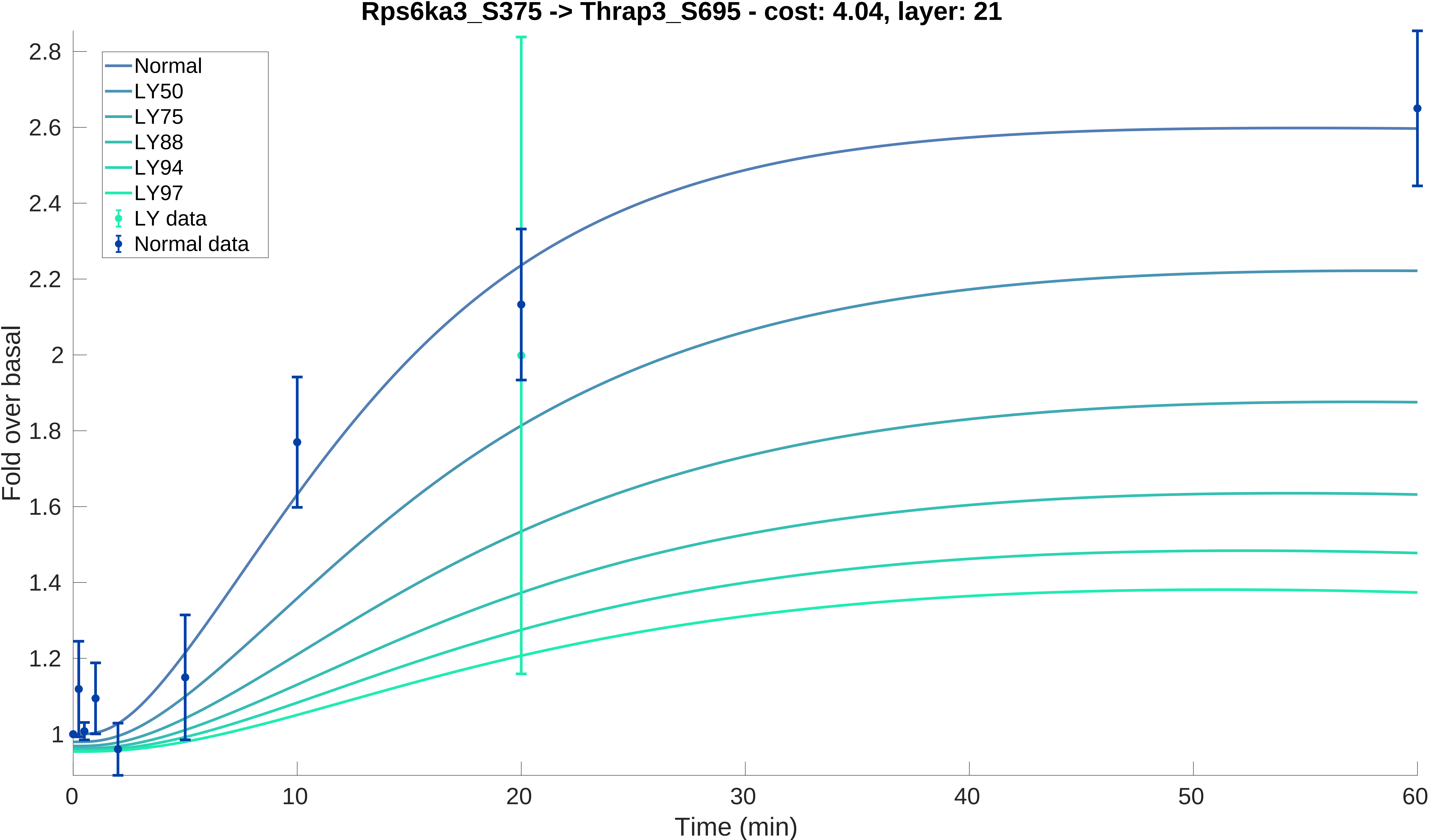

Rps6ka3_S375 -> Thrap3_S695 - cost: 4.04, layer: 21
Normal
LY50
LY75
LY88
LY94
LY97
LY data
Normal data
Fold over basal
Time (min)
2.8
2.6
2.4
2.2
2
1.8
1.6
1.4
1.2
1
0
10
20
30
40
50
60

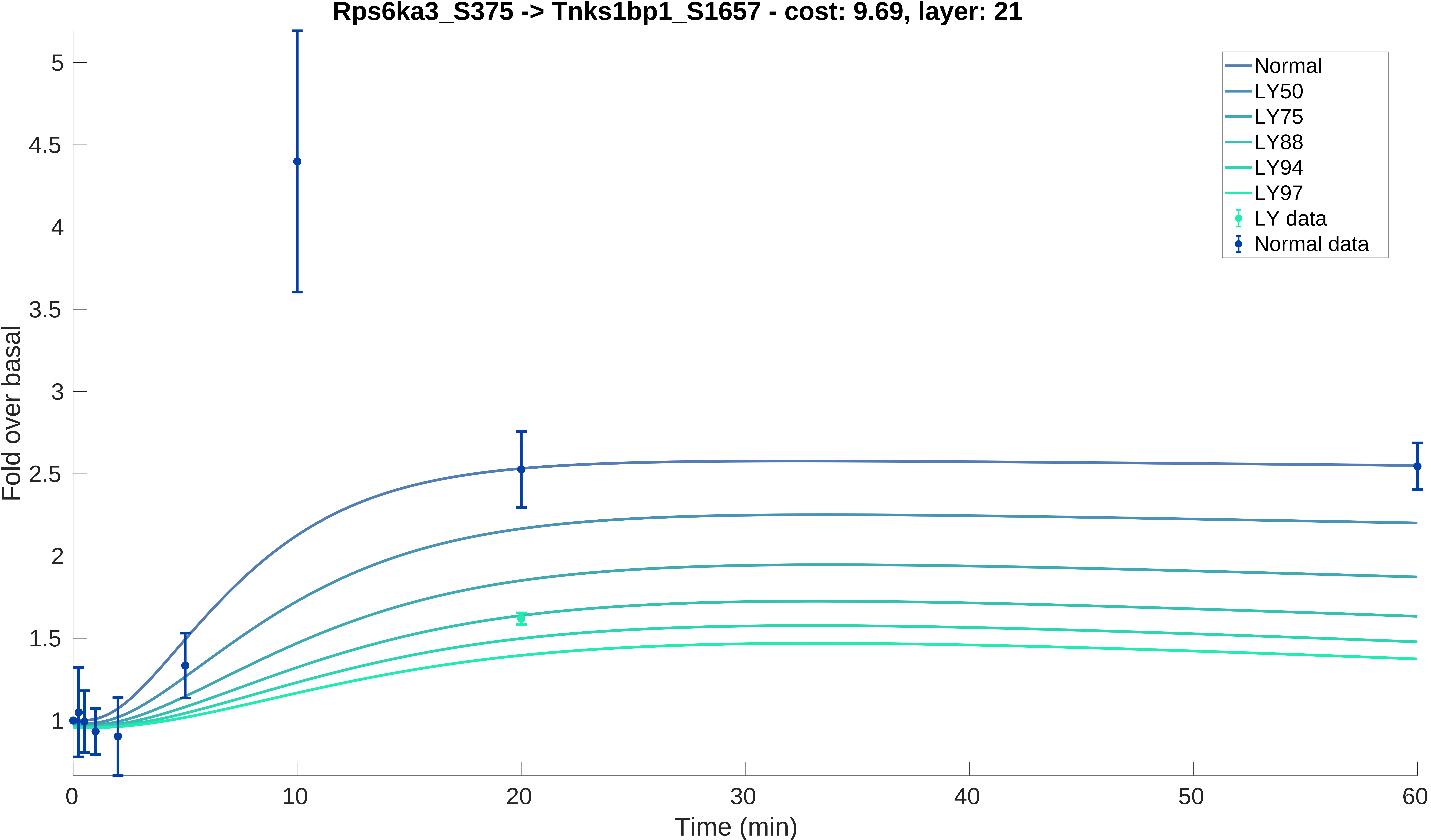
Rps6ka3_S375 -> Tnks1bp1_S1657 - cost: 9.69, layer: 21
Fold over basal
Time (min)
0
10
20
30
40
50
60
1
1.5
2
2.5
3
3.5
4
4.5
5
Normal
LY50
LY75
LY88
LY94
LY97
LY data
Normal data

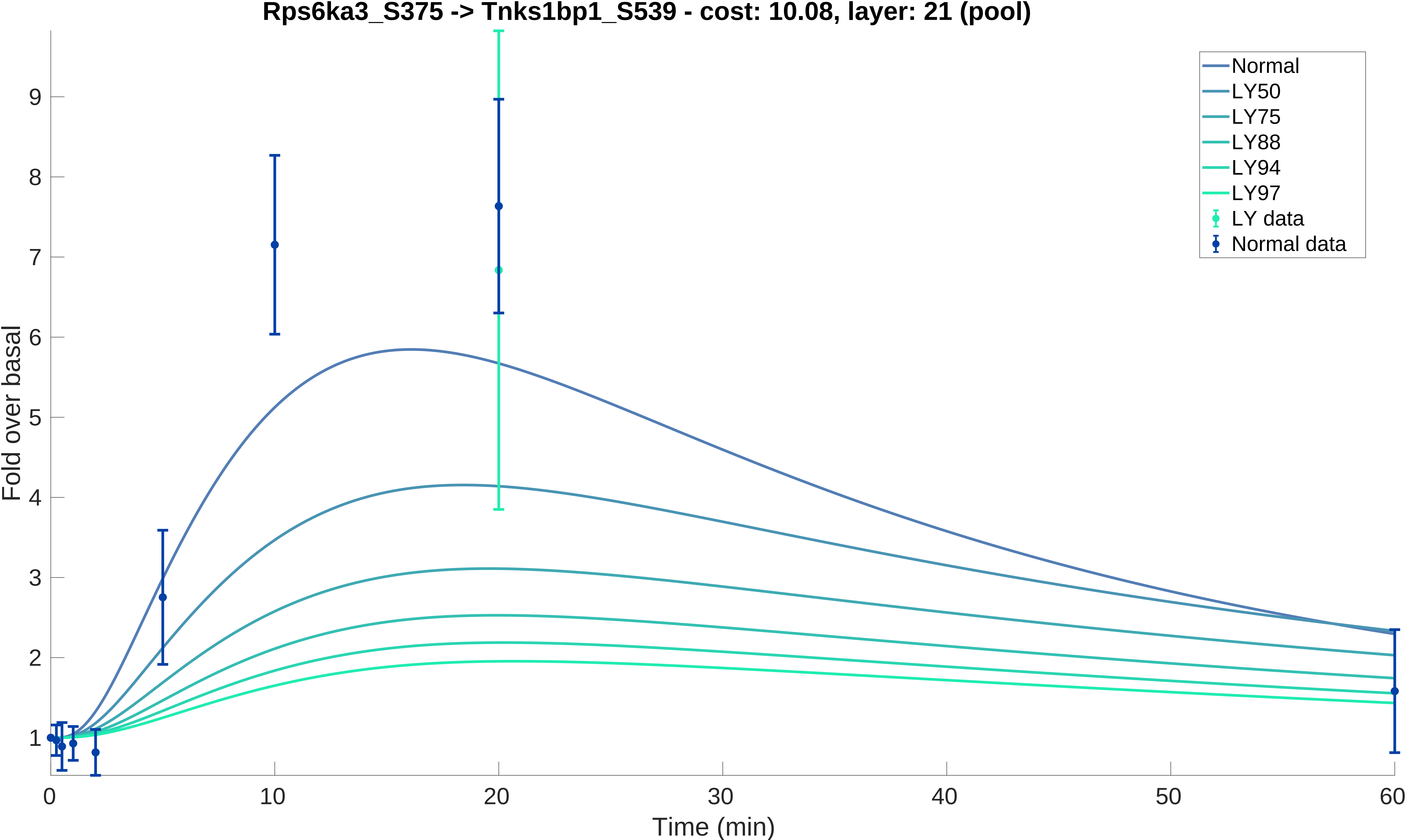

Rps6ka3_S375 -> Tnks1bp1_S539 - cost: 10.08, layer: 21 (pool)
Normal
LY50
LY75
LY88
LY94
LY97
LY data
Normal data
Fold over basal
Time (min)
0
10
20
30
40
50
60
1
2
3
4
5
6
7
8
9

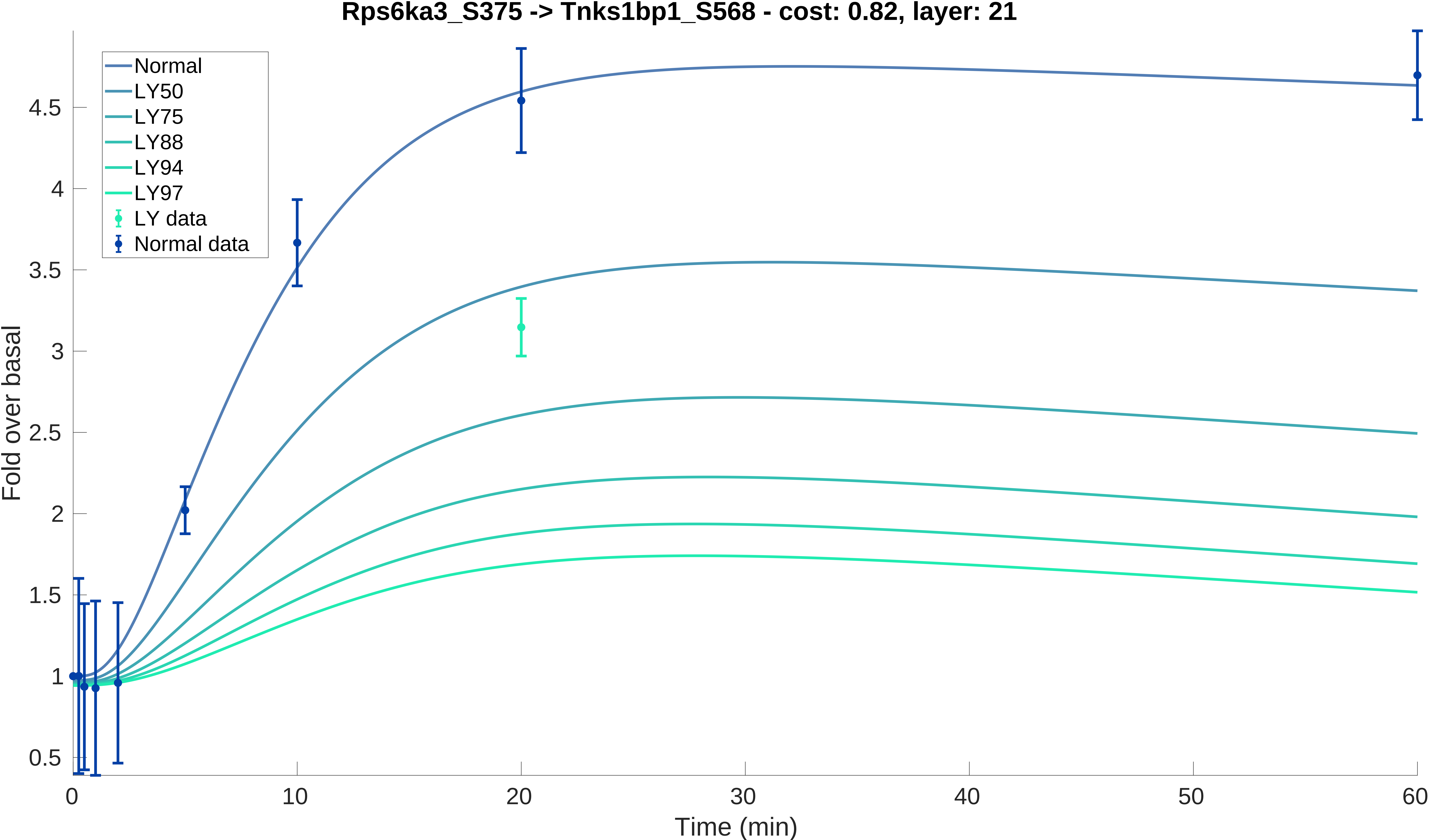

Rps6ka3_S375 -> Tnks1bp1_S568 - cost: 0.82, layer: 21
Normal
LY50
LY75
LY88
LY94
LY97
LY data
Normal data
Fold over basal
4.5
4
3.5
3
2.5
2
1.5
1
0.5
0
10
20
30
40
50
60
Time (min)

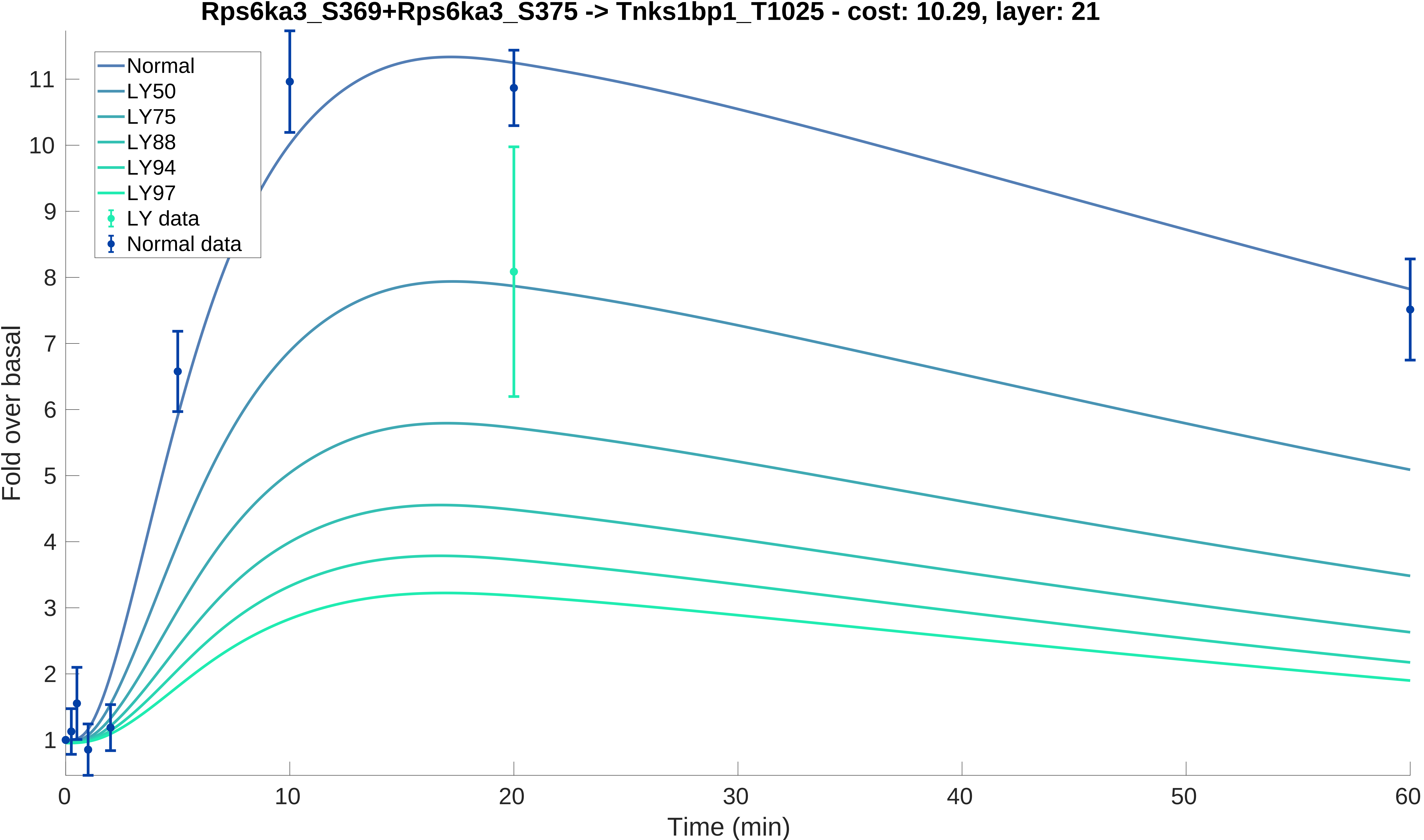
Rps6ka3_S369+Rps6ka3_S375 -> Tnks1bp1_T1025 - cost: 10.29, layer: 21
Normal
LY50
LY75
LY88
LY94
LY97
LY data
Normal data
Fold over basal
Time (min)
0
10
20
30
40
50
60
1
2
3
4
5
6
7
8
9
10
11

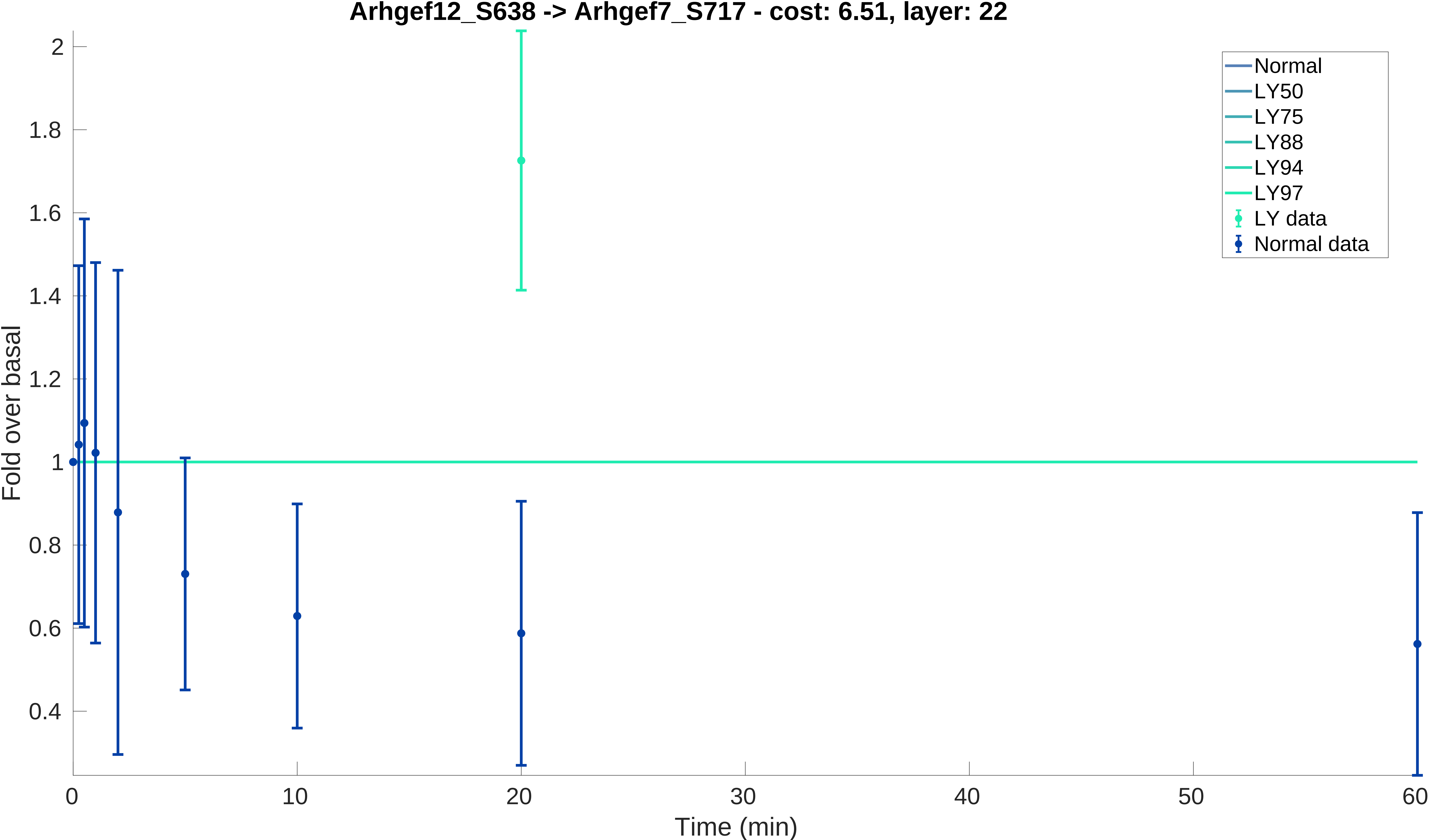

Arhgef12_S638 -> Arhgef7_S717 - cost: 6.51, layer: 22
Fold over basal
Time (min)
0
10
20
30
40
50
60
0.4
0.6
0.8
1
1.2
1.4
1.6
1.8
2
Normal
LY50
LY75
LY88
LY94
LY97
LY data
Normal data

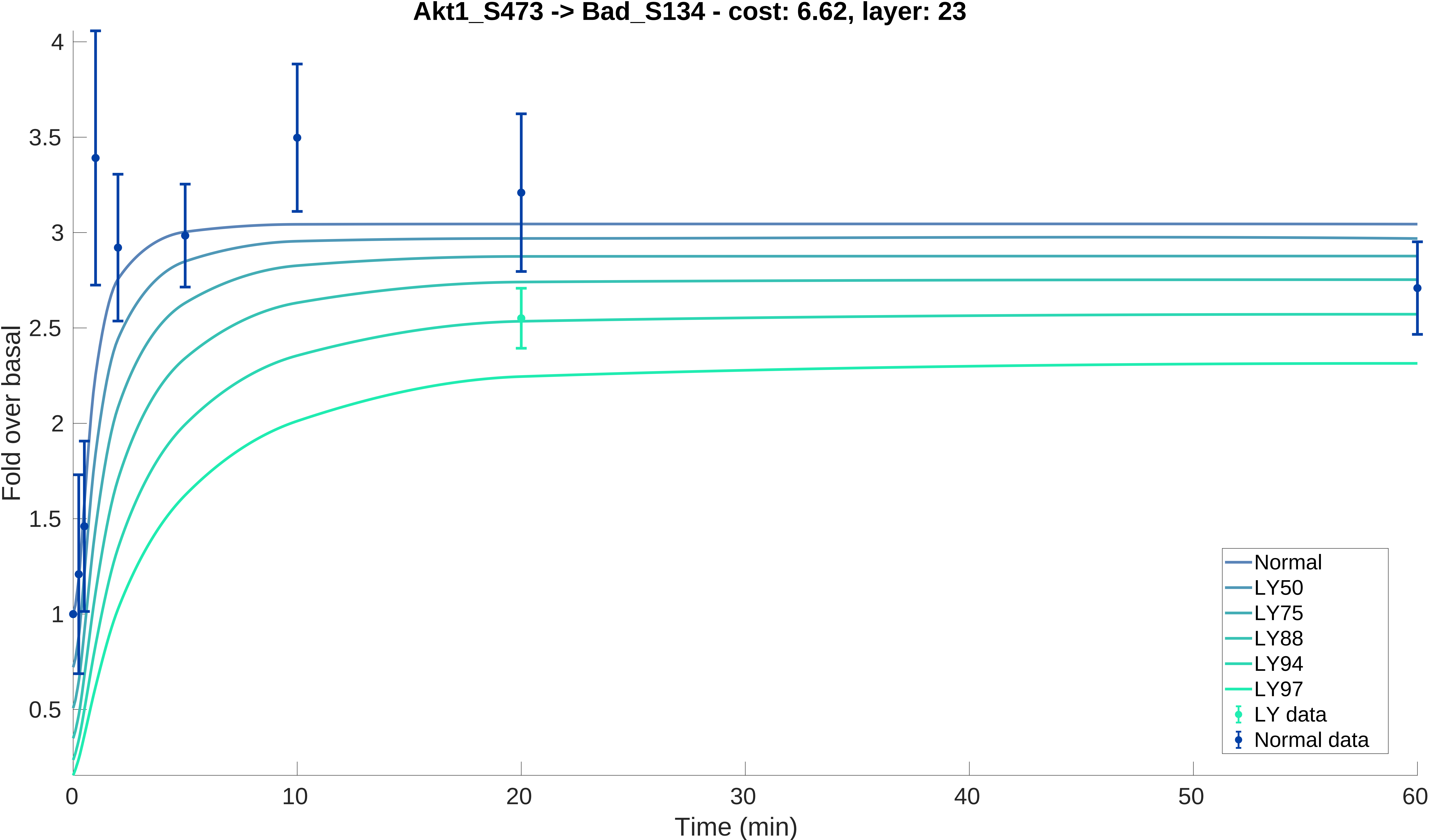
Akt1_S473 -> Bad_S134 - cost: 6.62, layer: 23
Fold over basal
Time (min)
0.5
1
1.5
2
2.5
3
3.5
4
0
10
20
30
40
50
60
Normal
LY50
LY75
LY88
LY94
LY97
LY data
Normal data

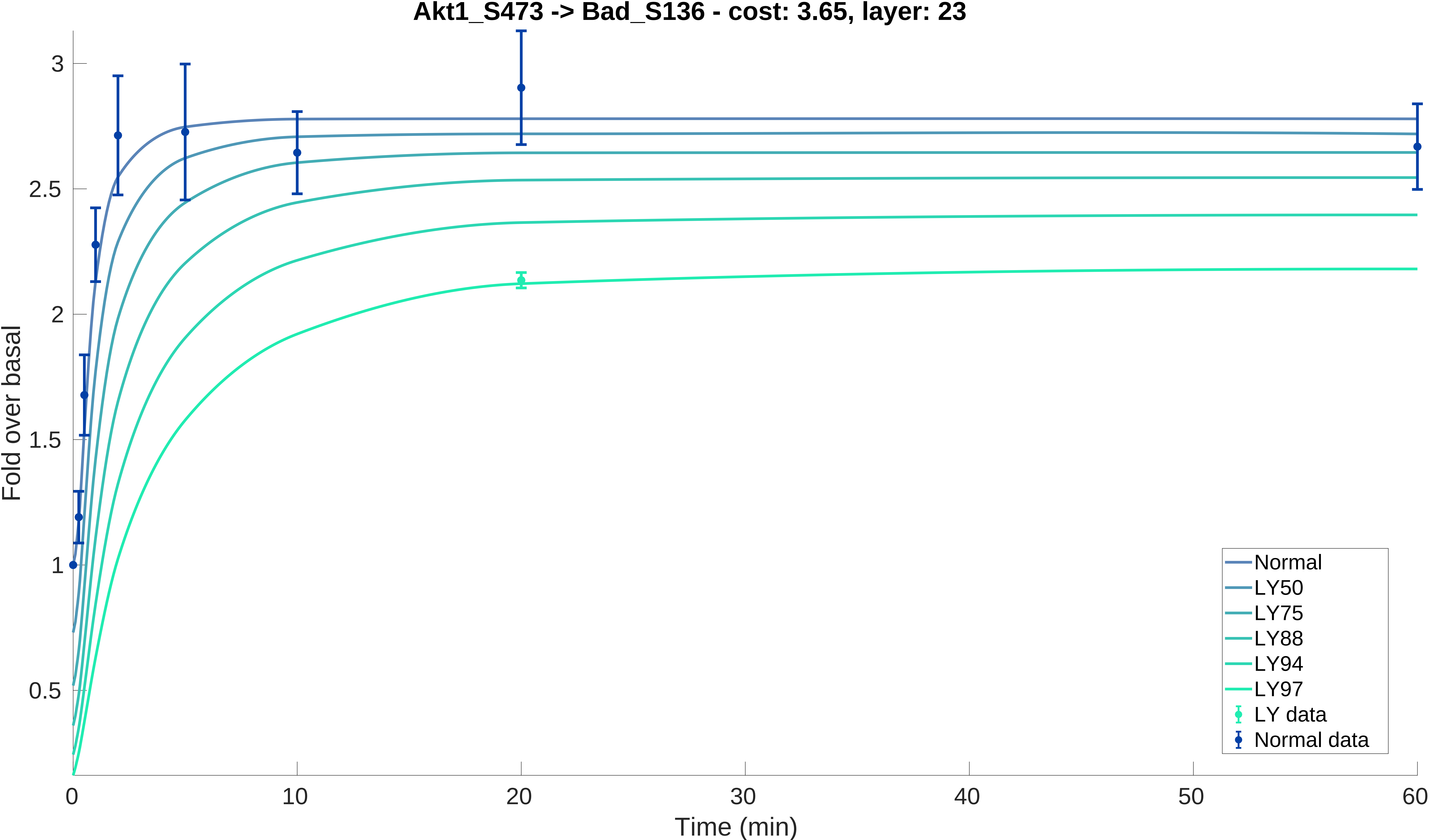
Akt1_S473 -> Bad_S136 - cost: 3.65, layer: 23
Fold over basal
Time (min)
0.5
1
1.5
2
2.5
3
0
10
20
30
40
50
60
Normal
LY50
LY75
LY88
LY94
LY97
LY data
Normal data

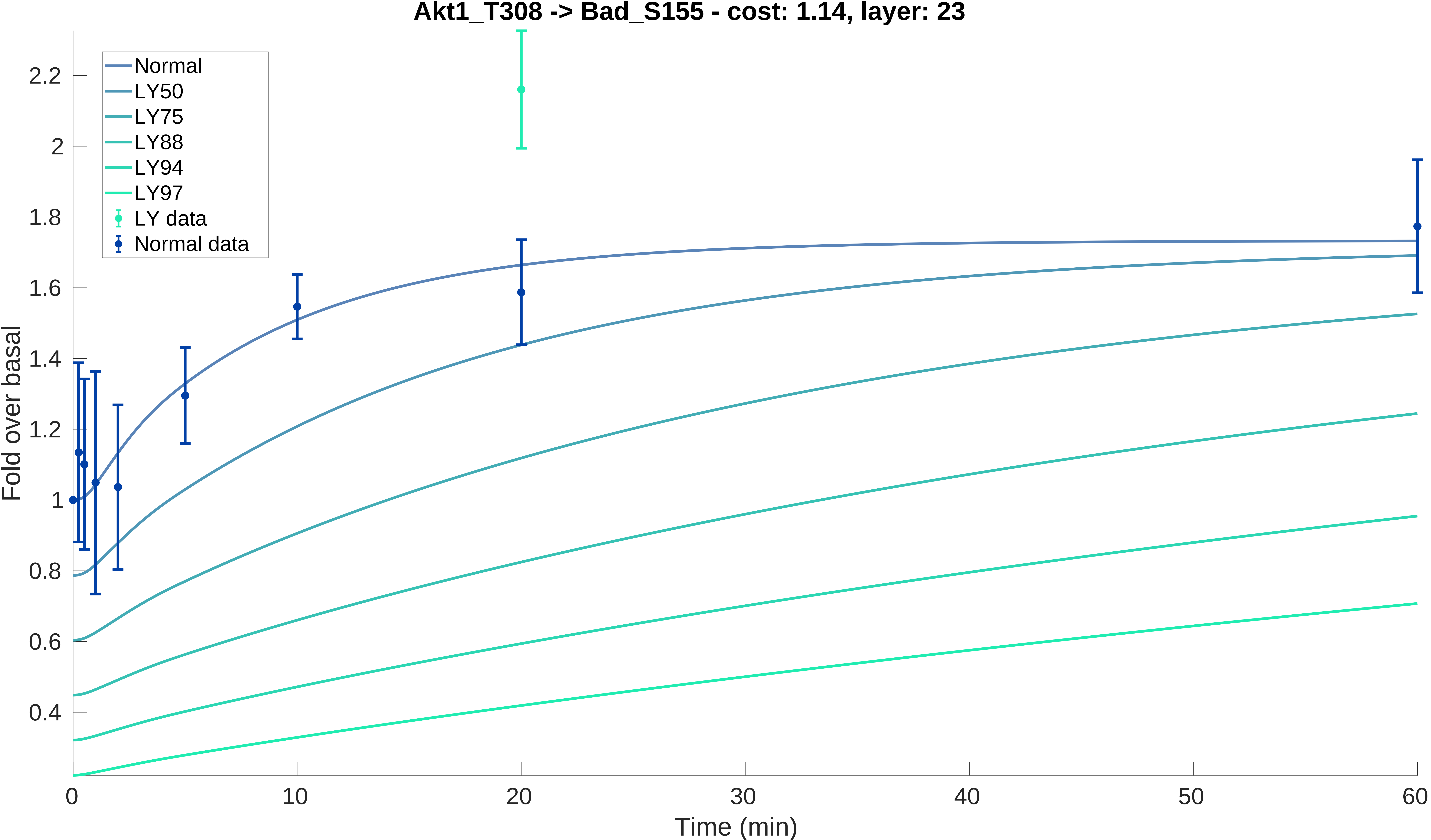

Akt1_T308 -> Bad_S155 - cost: 1.14, layer: 23
Normal
LY50
LY75
LY88
LY94
LY97
LY data
Normal data
Fold over basal
Time (min)
2.2
2
1.8
1.6
1.4
1.2
1
0.8
0.6
0.4
0
10
20
30
40
50
60

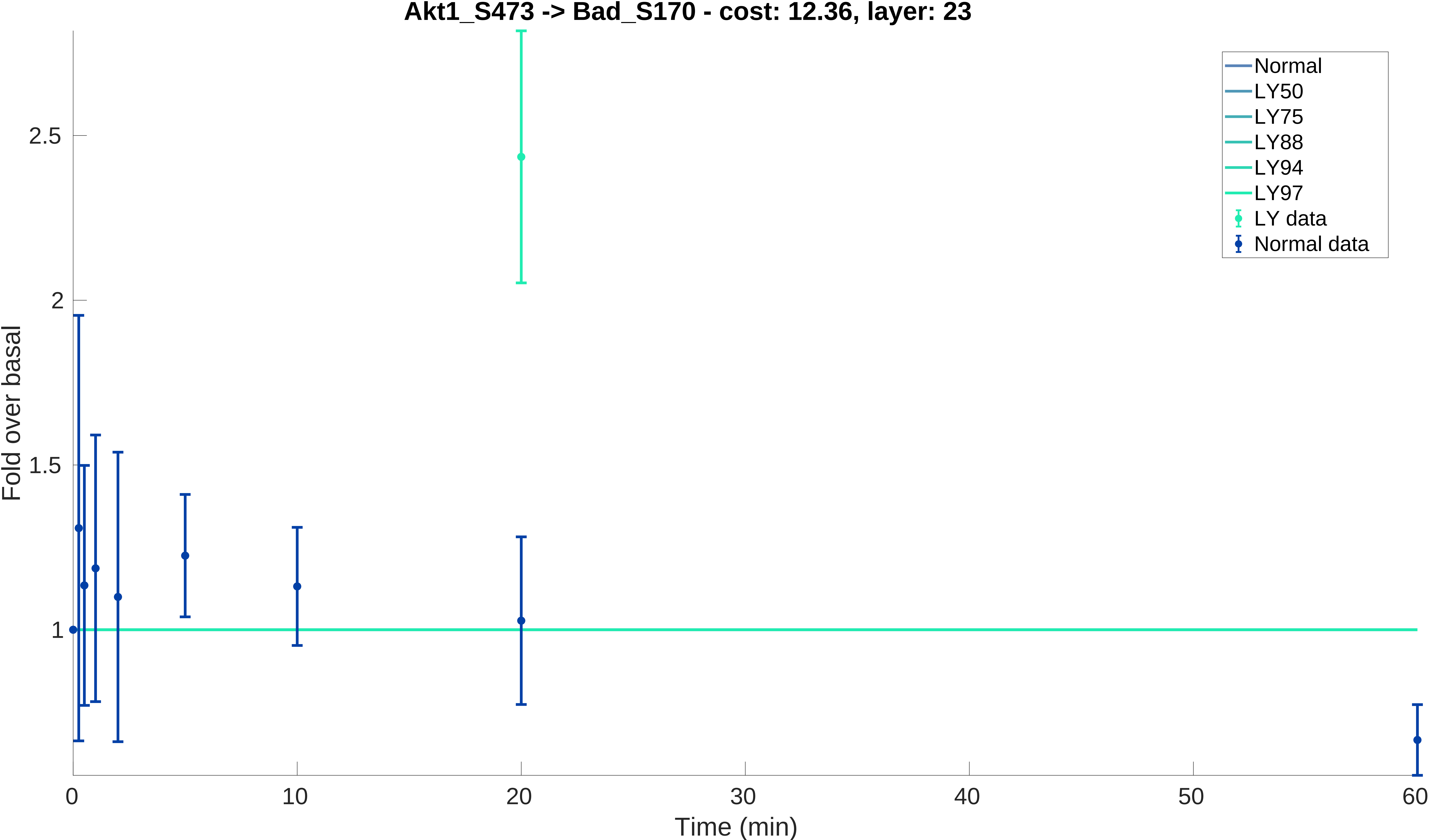

Akt1_S473 -> Bad_S170 - cost: 12.36, layer: 23
Normal
LY50
LY75
LY88
LY94
LY97
LY data
Normal data
Fold over basal
2.5
2
1.5
1
0
10
20
30
40
50
60
Time (min)

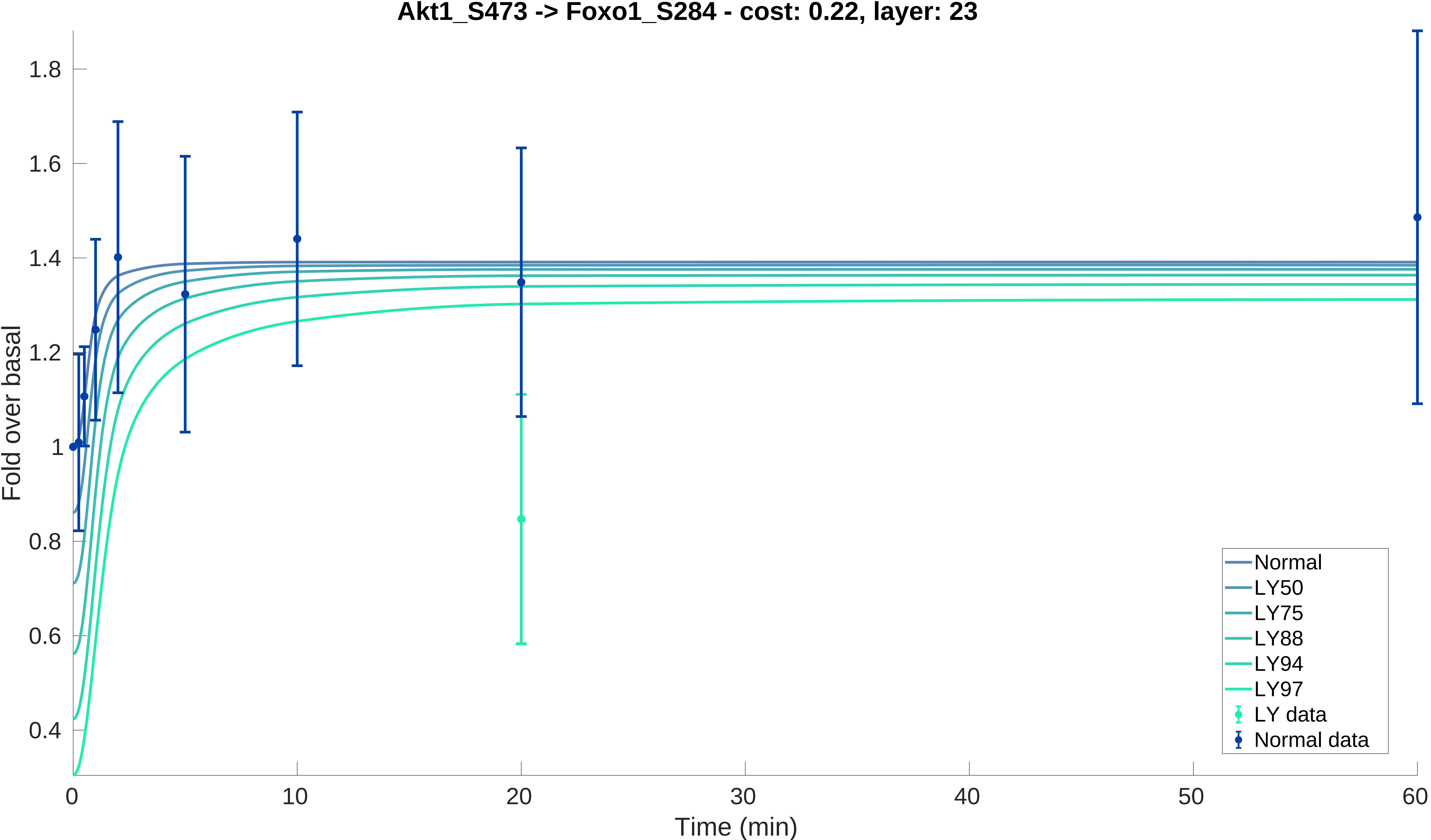

Akt1_S473 -> Foxo1_S284 - cost: 0.22, layer: 23
Fold over basal
0.4
0.6
0.8
1
1.2
1.4
1.6
1.8
0
10
20
30
40
50
60
Time (min)
Normal
LY50
LY75
LY88
LY94
LY97
LY data
Normal data

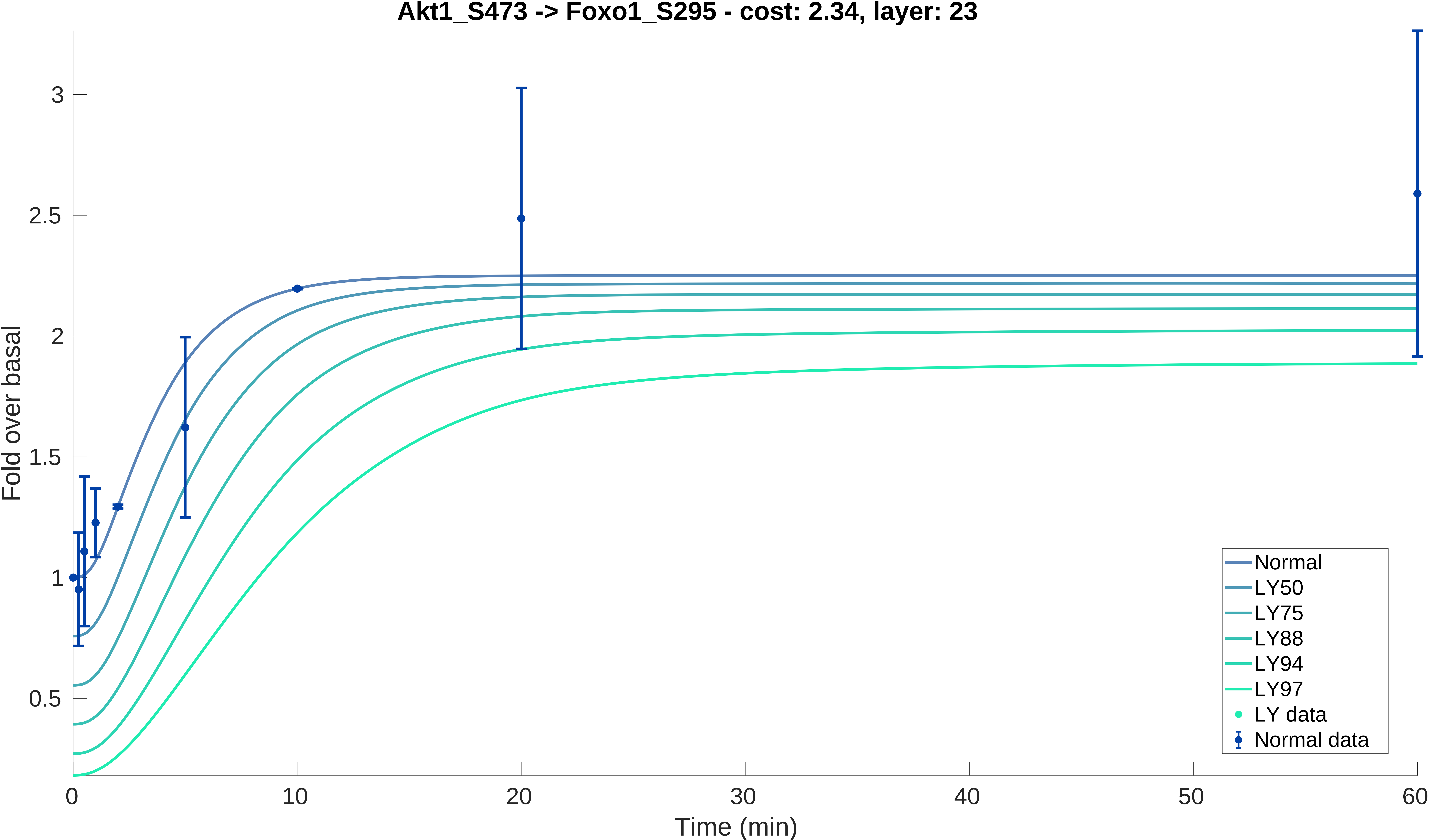

Akt1_S473 -> Foxo1_S295 - cost: 2.34, layer: 23
Fold over basal
Time (min)
0.5
1
1.5
2
2.5
3
0
10
20
30
40
50
60
Normal
LY50
LY75
LY88
LY94
LY97
LY data
Normal data

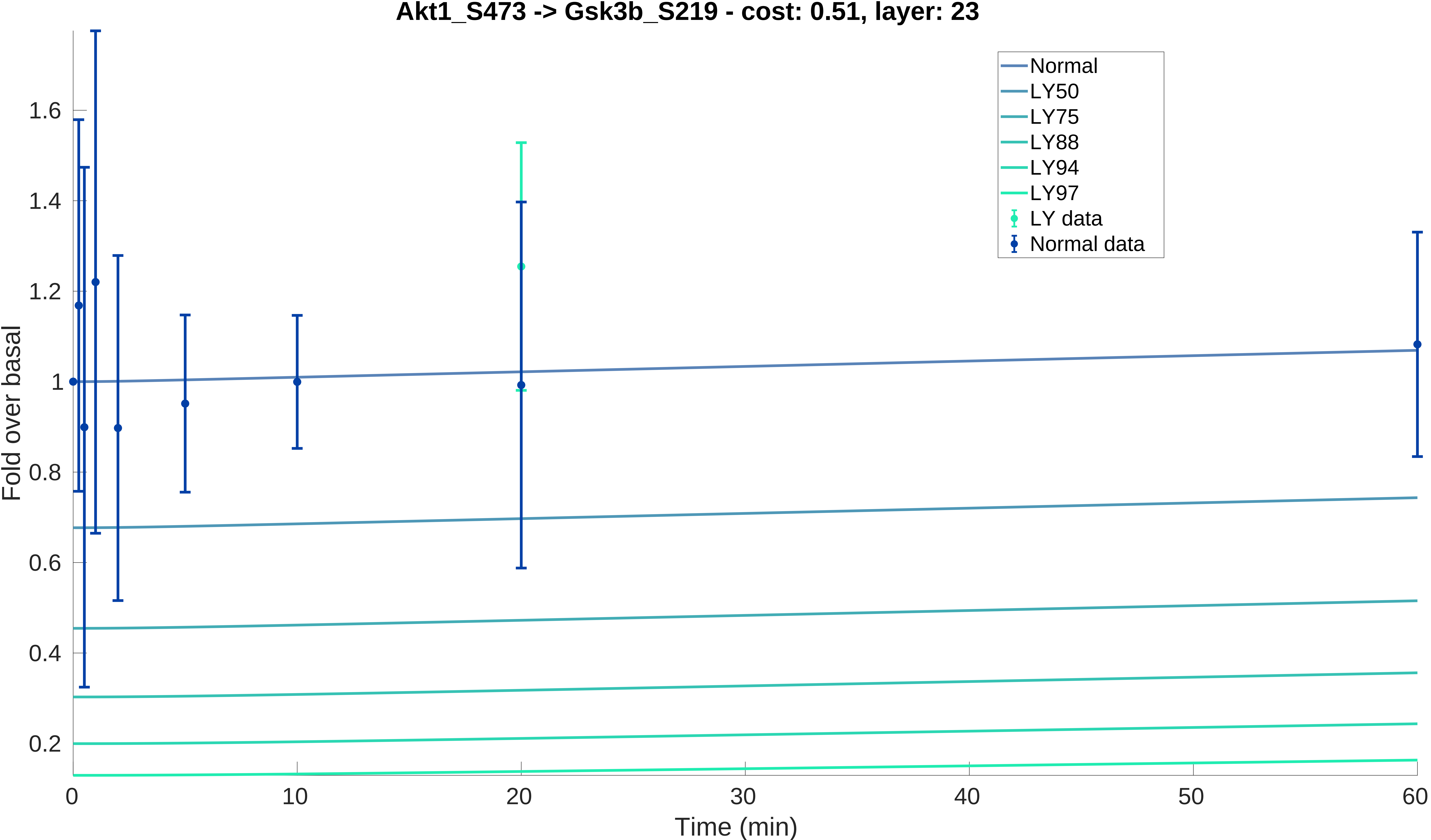

Akt1_S473 -> Gsk3b_S219 - cost: 0.51, layer: 23
Normal
LY50
LY75
LY88
LY94
LY97
LY data
Normal data
Fold over basal
Time (min)
0
10
20
30
40
50
60
0.2
0.4
0.6
0.8
1
1.2
1.4
1.6

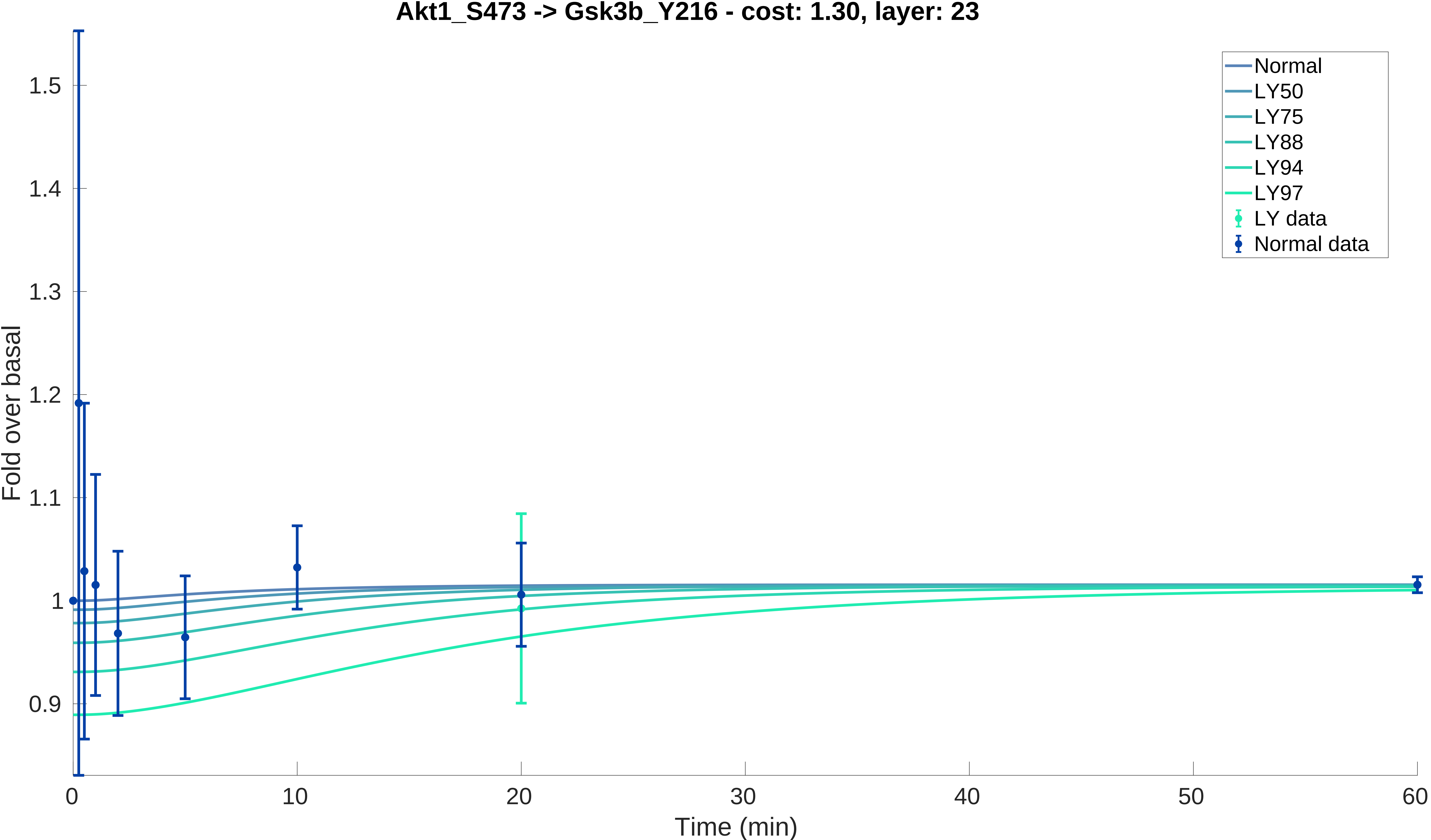

Akt1_S473 -> Gsk3b_Y216 - cost: 1.30, layer: 23
Fold over basal
Time (min)
0.9
1
1.1
1.2
1.3
1.4
1.5
0
10
20
30
40
50
60
Normal
LY50
LY75
LY88
LY94
LY97
LY data
Normal data

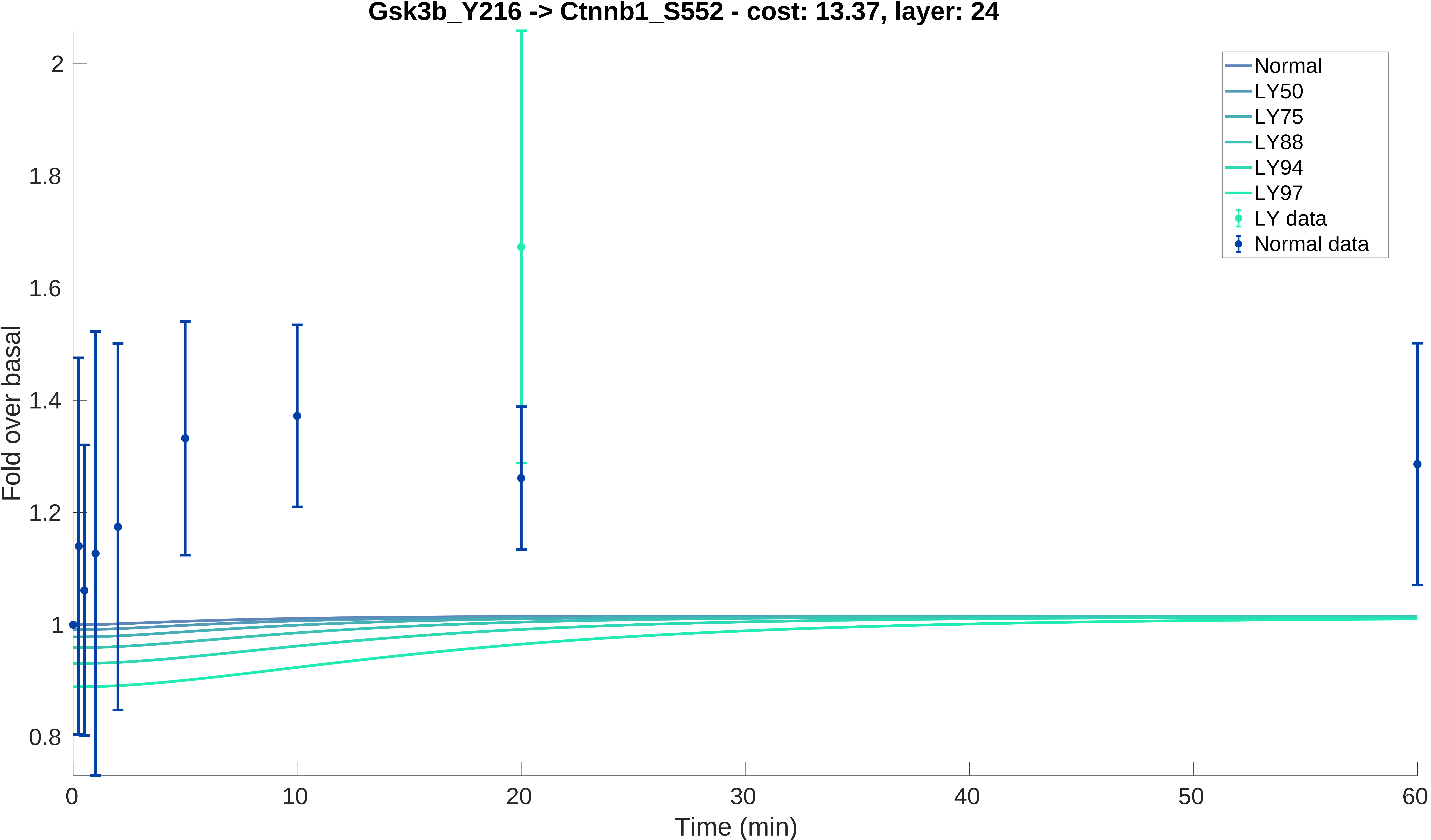
Gsk3b_Y216 -> Ctnnb1_S552 - cost: 13.37, layer: 24
Fold over basal
Time (min)
0.8
1
1.2
1.4
1.6
1.8
2
0
10
20
30
40
50
60
Normal
LY50
LY75
LY88
LY94
LY97
LY data
Normal data

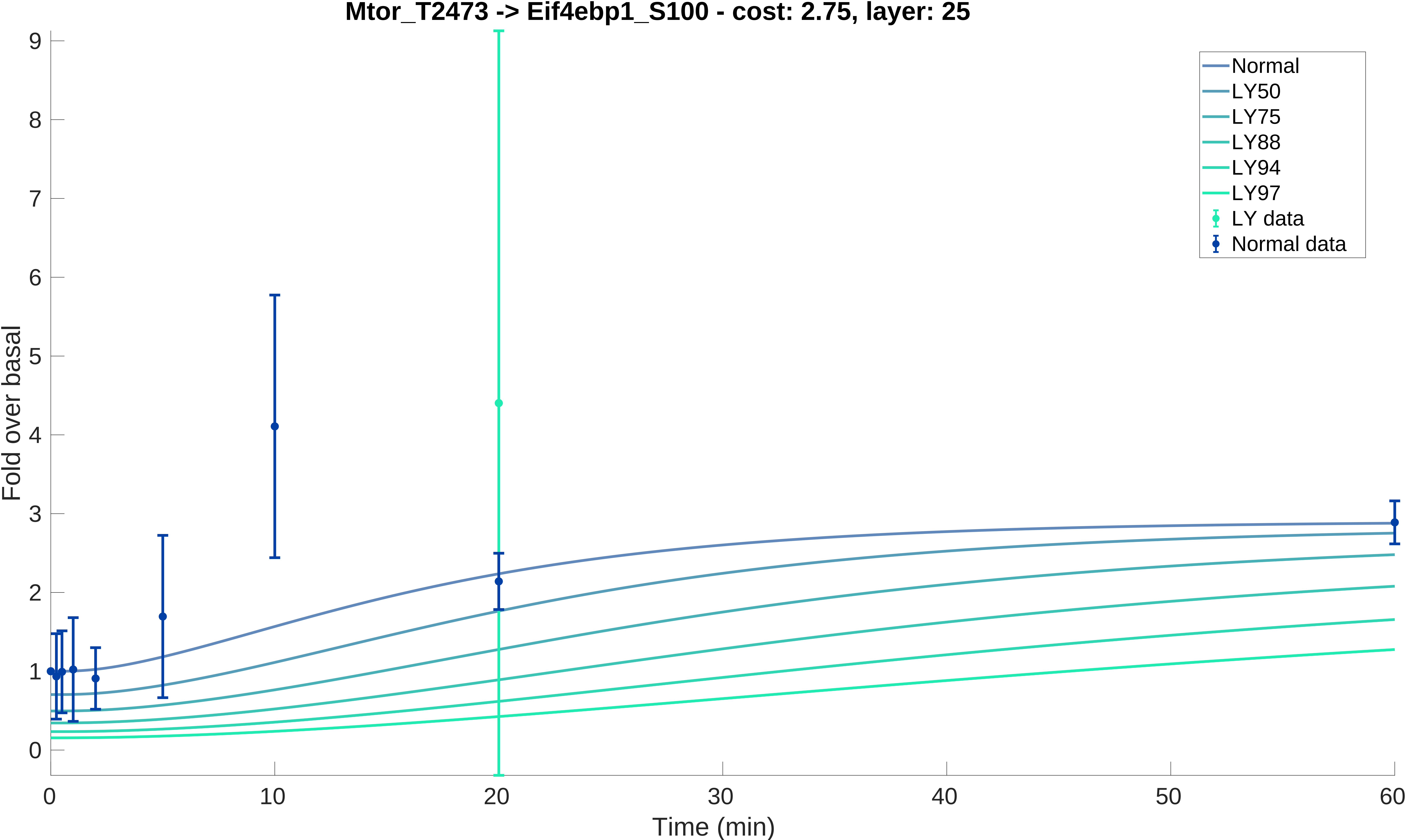

Mtor_T2473 -> Eif4ebp1_S100 - cost: 2.75, layer: 25
Fold over basal
Time (min)
0
1
2
3
4
5
6
7
8
9
0
10
20
30
40
50
60
Normal
LY50
LY75
LY88
LY94
LY97
LY data
Normal data

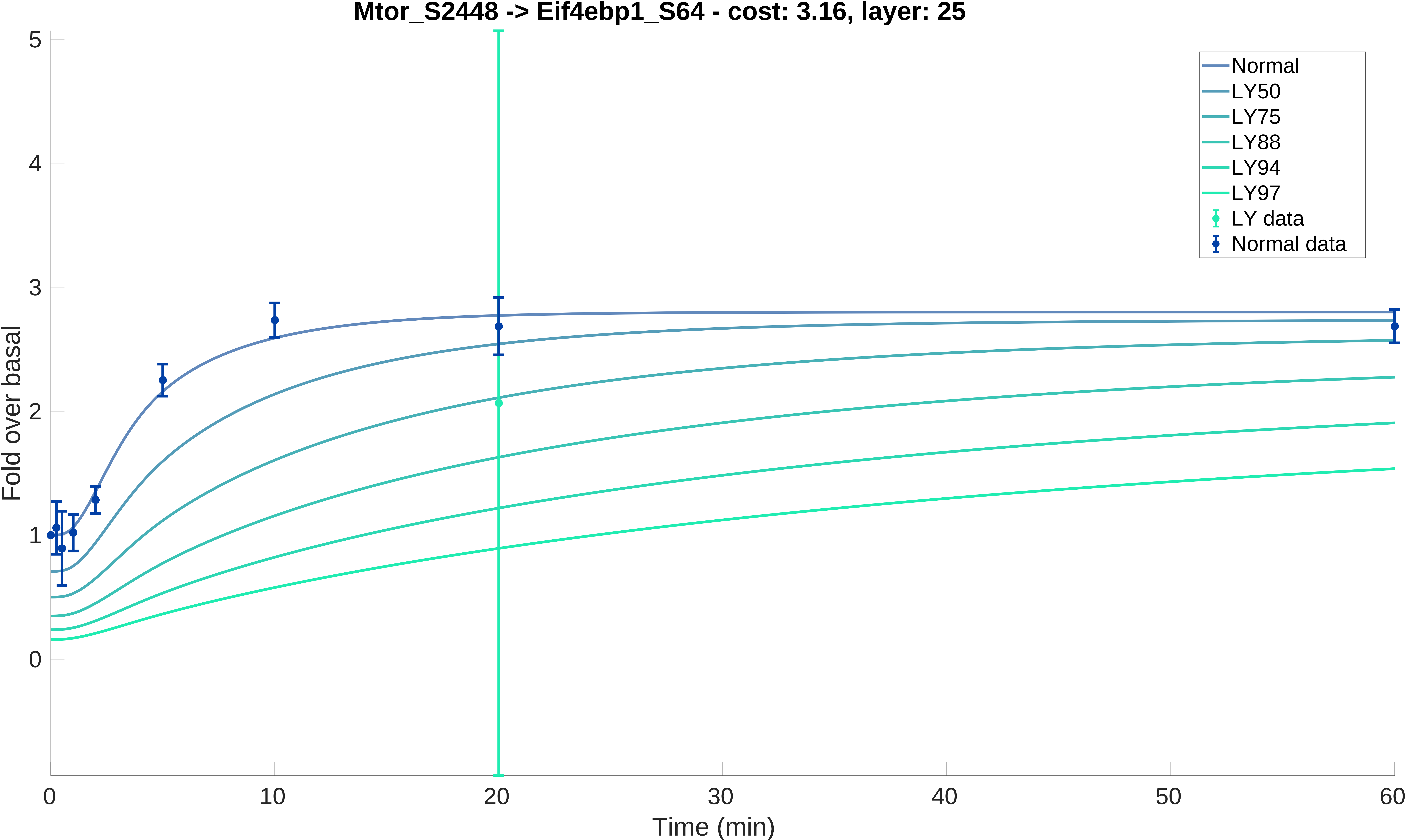

Mtor_S2448 -> Eif4ebp1_S64 - cost: 3.16, layer: 25
Fold over basal
Time (min)
0
10
20
30
40
50
60
5
4
3
2
1
0
Normal
LY50
LY75
LY88
LY94
LY97
LY data
Normal data

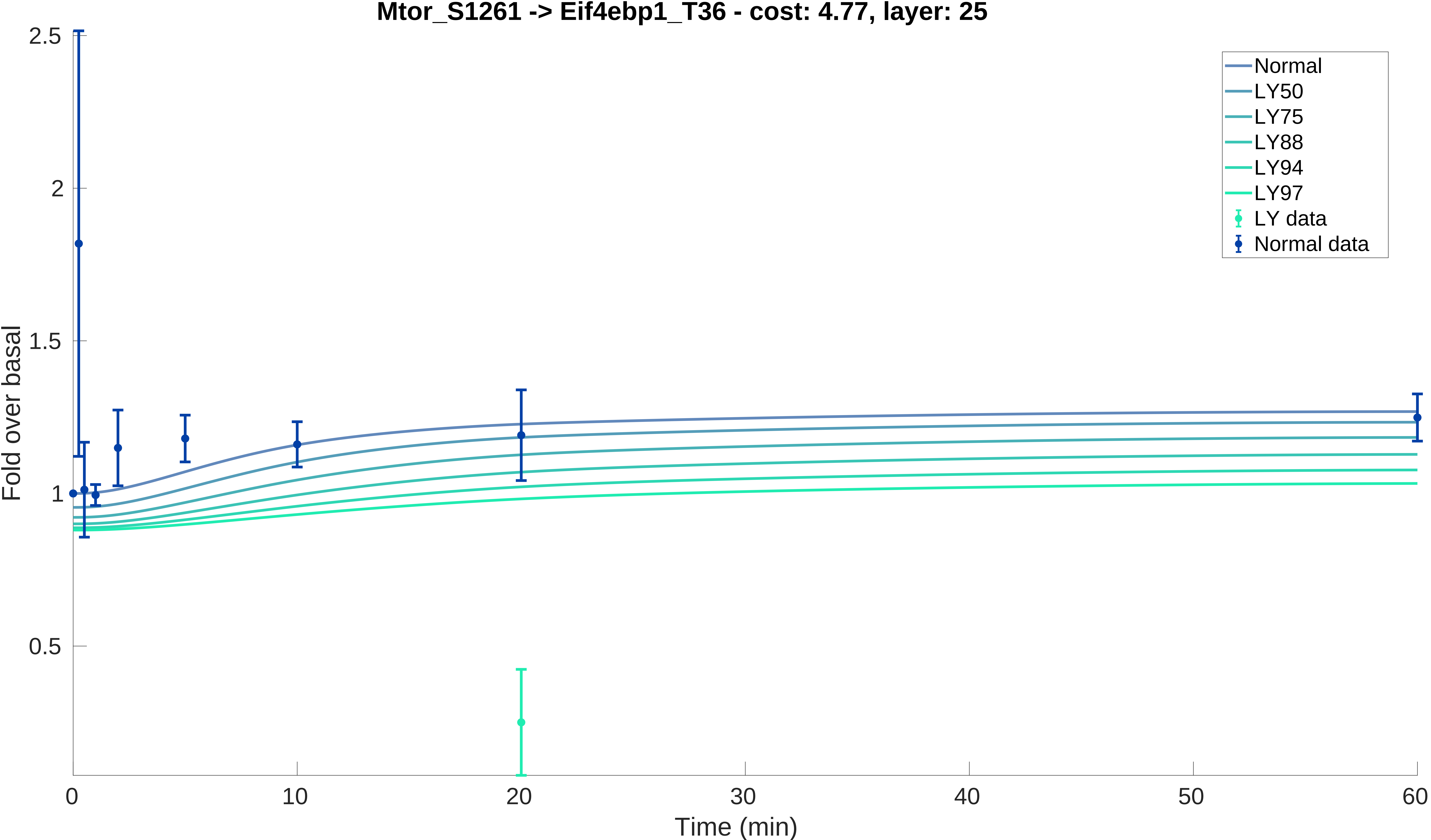
Mtor_S1261 -> Eif4ebp1_T36 - cost: 4.77, layer: 25
Fold over basal
2.5
2
1.5
1
0.5
0
10
20
30
40
50
60
Time (min)
Normal
LY50
LY75
LY88
LY94
LY97
LY data
Normal data

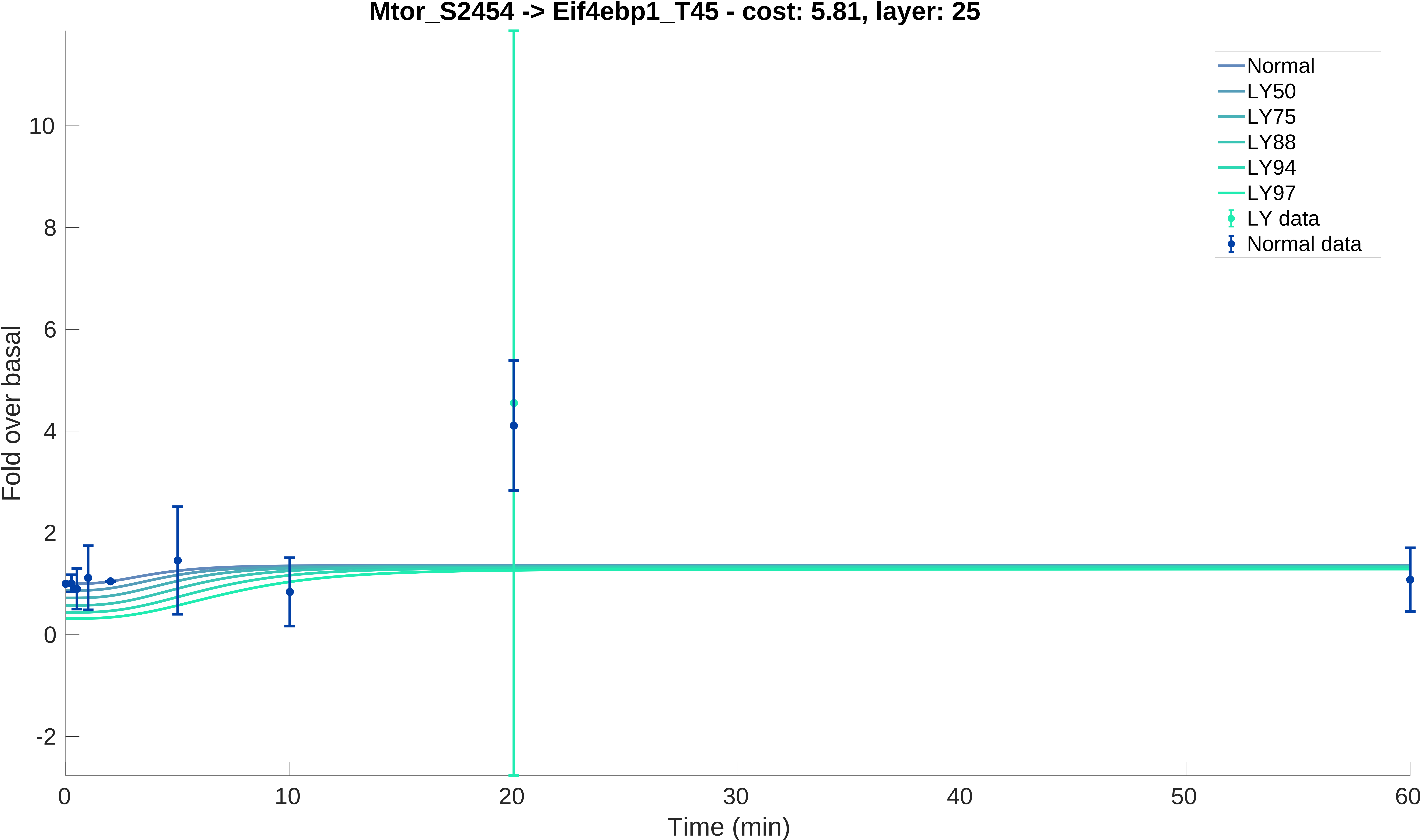
Mtor_S2454 -> Eif4ebp1_T45 - cost: 5.81, layer: 25
Normal
LY50
LY75
LY88
LY94
LY97
LY data
Normal data
Fold over basal
10
8
6
4
2
0
-2
0
10
20
30
40
50
60
Time (min)

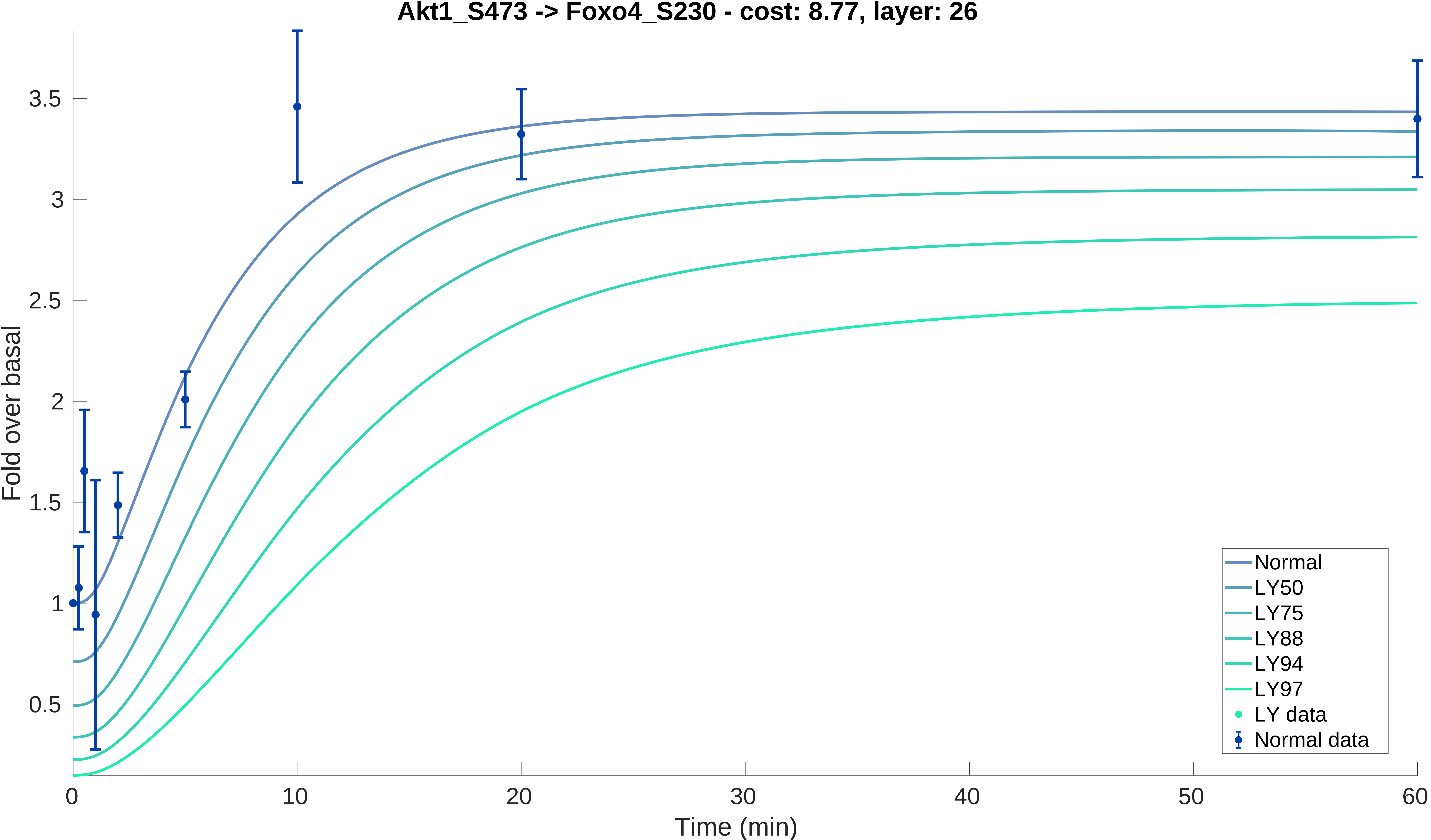

Akt1_S473 -> Foxo4_S230 - cost: 8.77, layer: 26
Fold over basal
Time (min)
0.5
1
1.5
2
2.5
3
3.5
0
10
20
30
40
50
60
Normal
LY50
LY75
LY88
LY94
LY97
LY data
Normal data

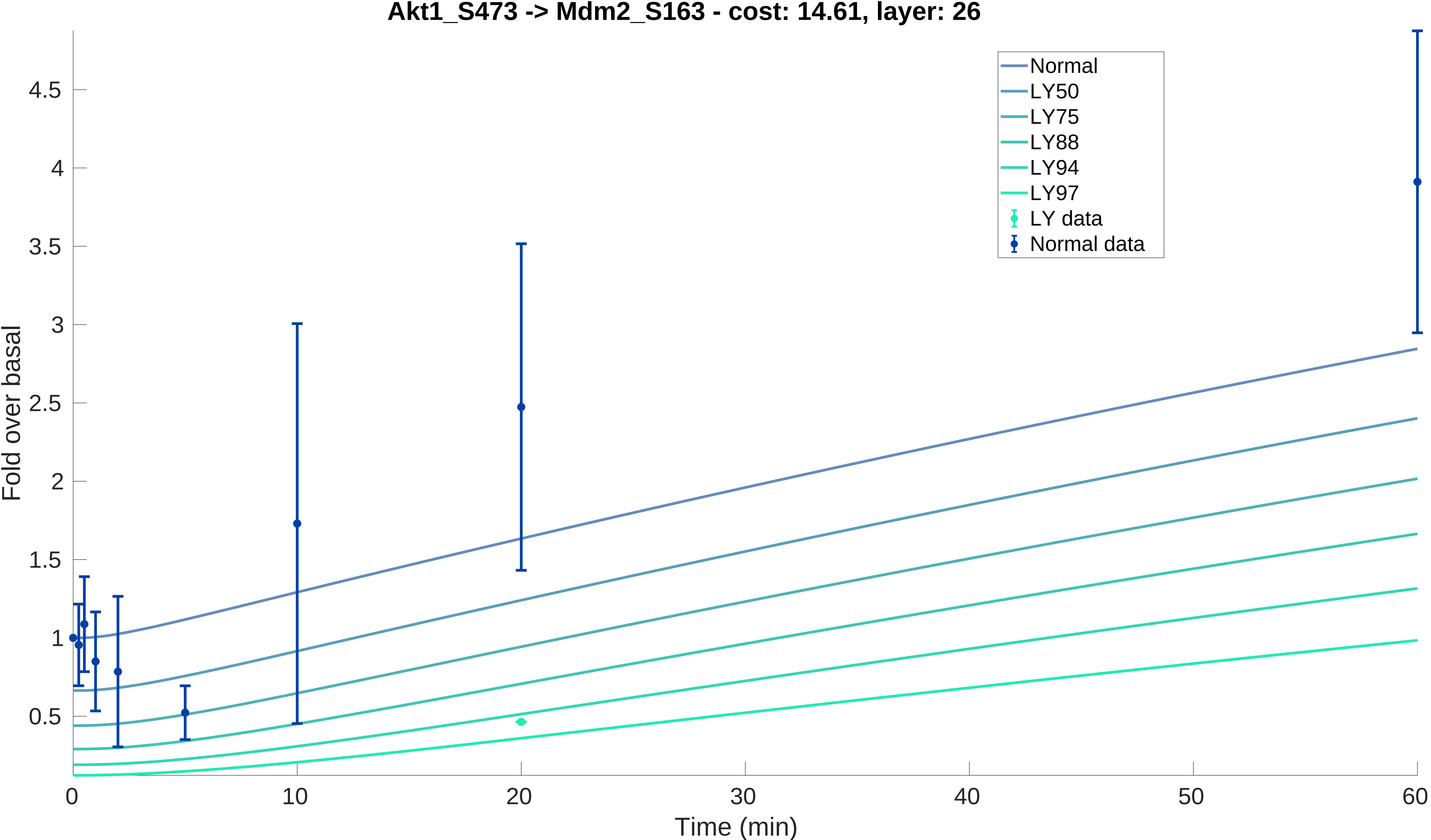

Akt1_S473 -> Mdm2_S163 - cost: 14.61, layer: 26
Fold over basal
Time (min)
0.5
1
1.5
2
2.5
3
3.5
4
4.5
0
10
20
30
40
50
60
Normal
LY50
LY75
LY88
LY94
LY97
LY data
Normal data

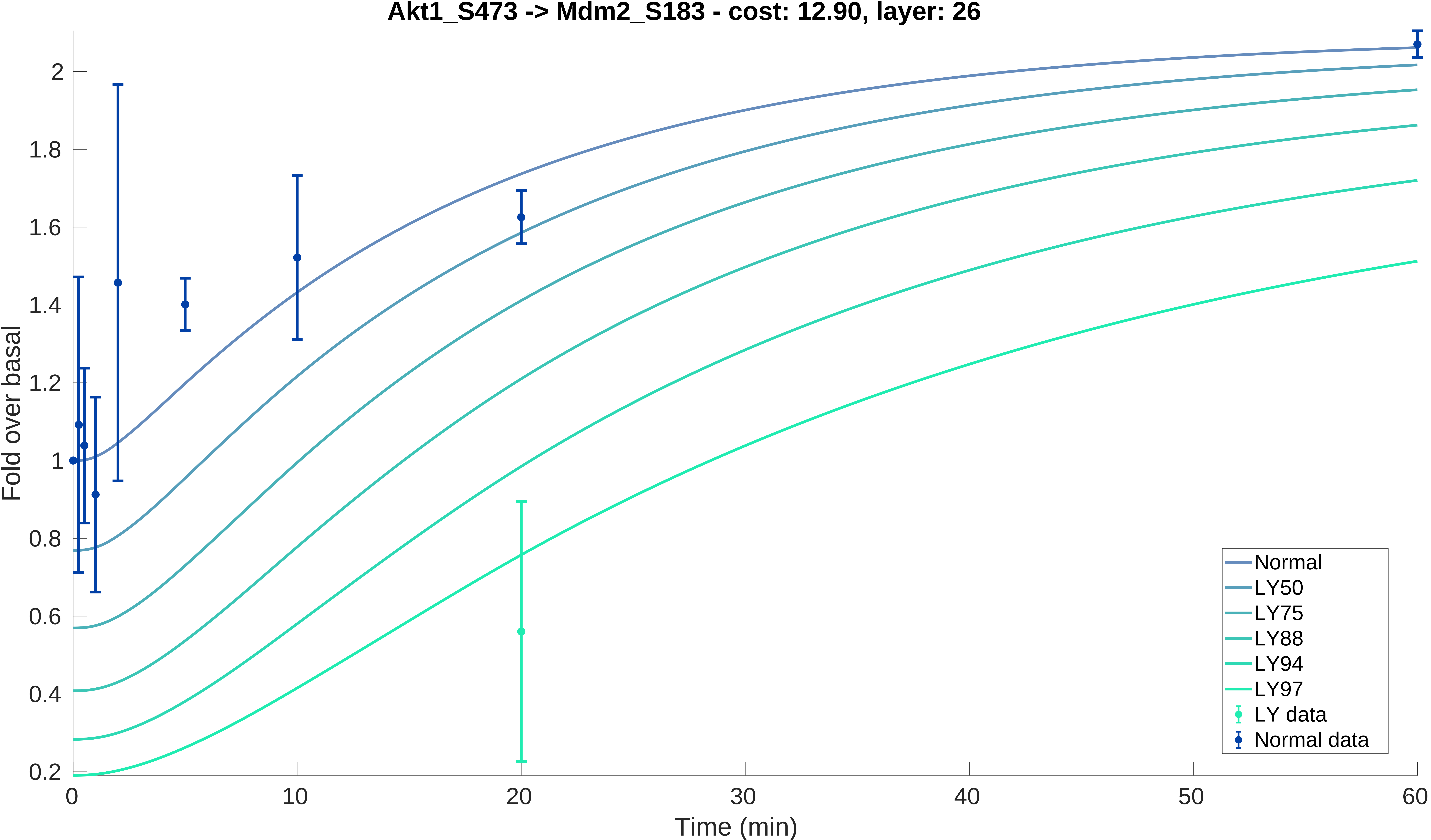

Akt1_S473 -> Mdm2_S183 - cost: 12.90, layer: 26
Fold over basal
Time (min)
0
10
20
30
40
50
60
0.2
0.4
0.6
0.8
1
1.2
1.4
1.6
1.8
2
Normal
LY50
LY75
LY88
LY94
LY97
LY data
Normal data

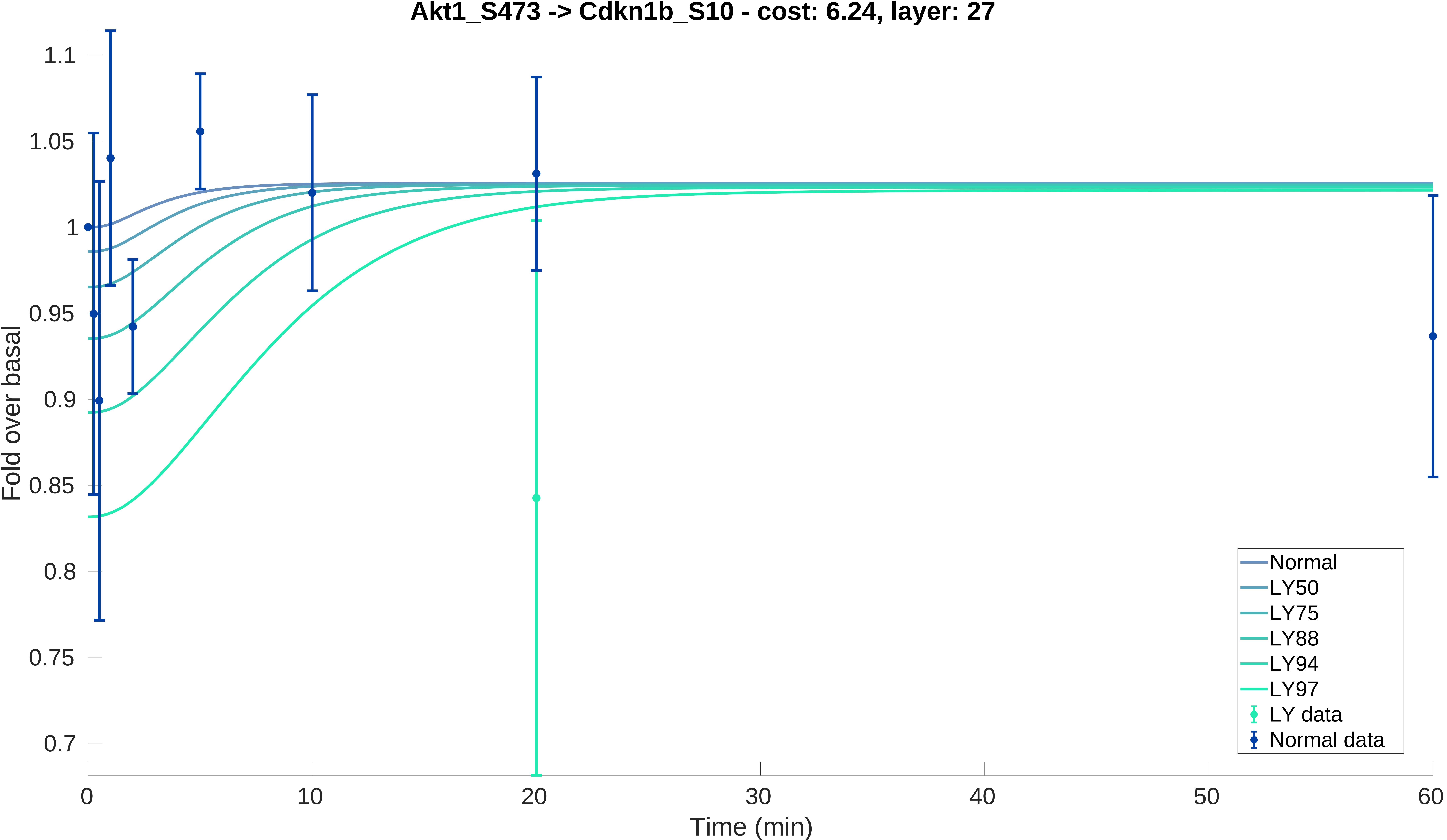

Akt1_S473 -> Cdkn1b_S10 - cost: 6.24, layer: 27
Fold over basal
Time (min)
1.1
1.05
1
0.95
0.9
0.85
0.8
0.75
0.7
0
10
20
30
40
50
60
Normal
LY50
LY75
LY88
LY94
LY97
LY data
Normal data

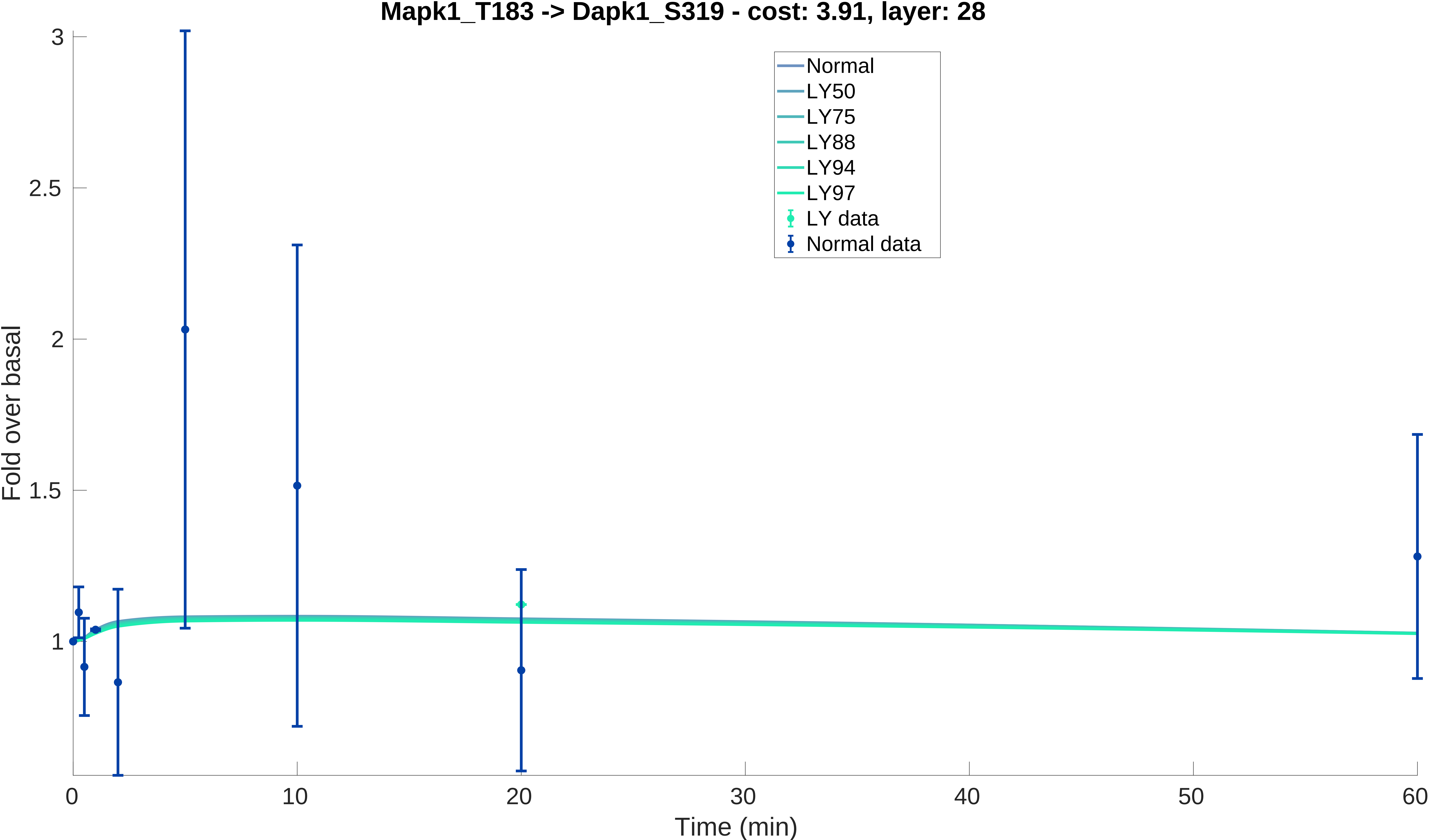

Mapk1_T183 -> Dapk1_S319 - cost: 3.91, layer: 28
Normal
LY50
LY75
LY88
LY94
LY97
LY data
Normal data
Fold over basal
Time (min)
0
10
20
30
40
50
60
1
1.5
2
2.5
3

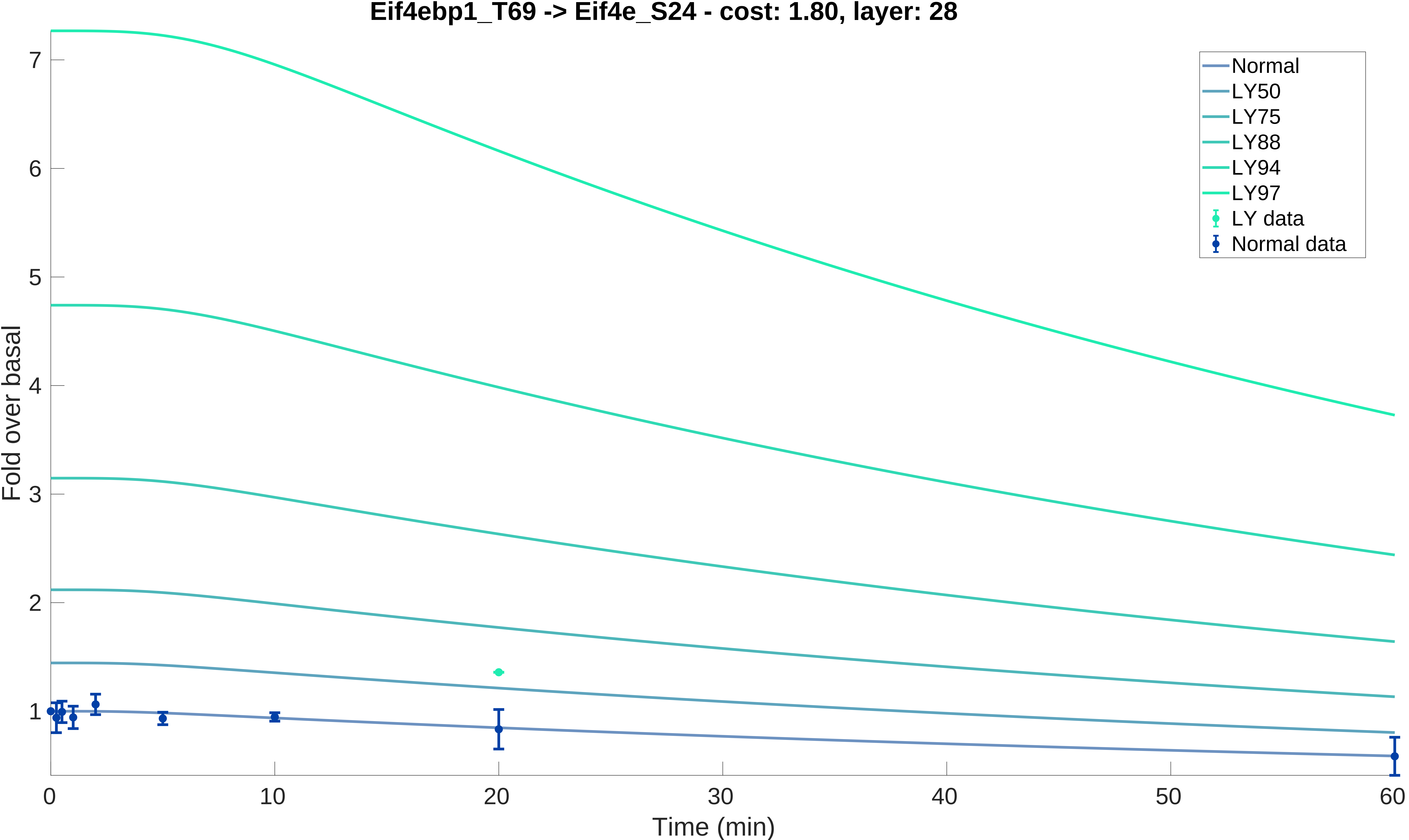
Eif4ebp1_T69 -> Eif4e_S24 - cost: 1.80, layer: 28
Fold over basal
Time (min)
Normal
LY50
LY75
LY88
LY94
LY97
LY data
Normal data
0
10
20
30
40
50
60
1
2
3
4
5
6
7

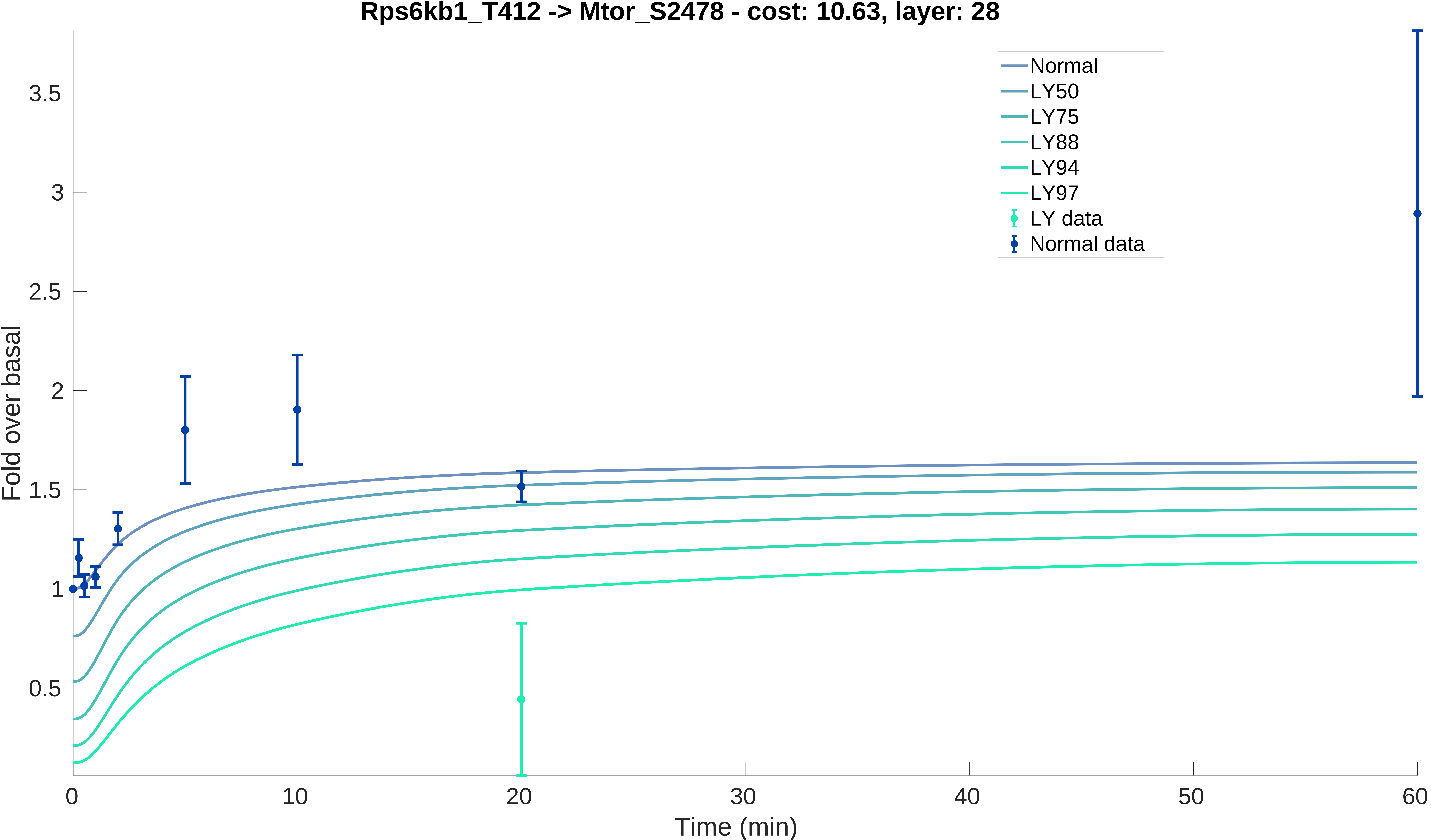
Rps6kb1_T412 -> Mtor_S2478 - cost: 10.63, layer: 28
Normal
LY50
LY75
LY88
LY94
LY97
LY data
Normal data
Fold over basal
0.5
1
1.5
2
2.5
3
3.5
0
10
20
30
40
50
60
Time (min)

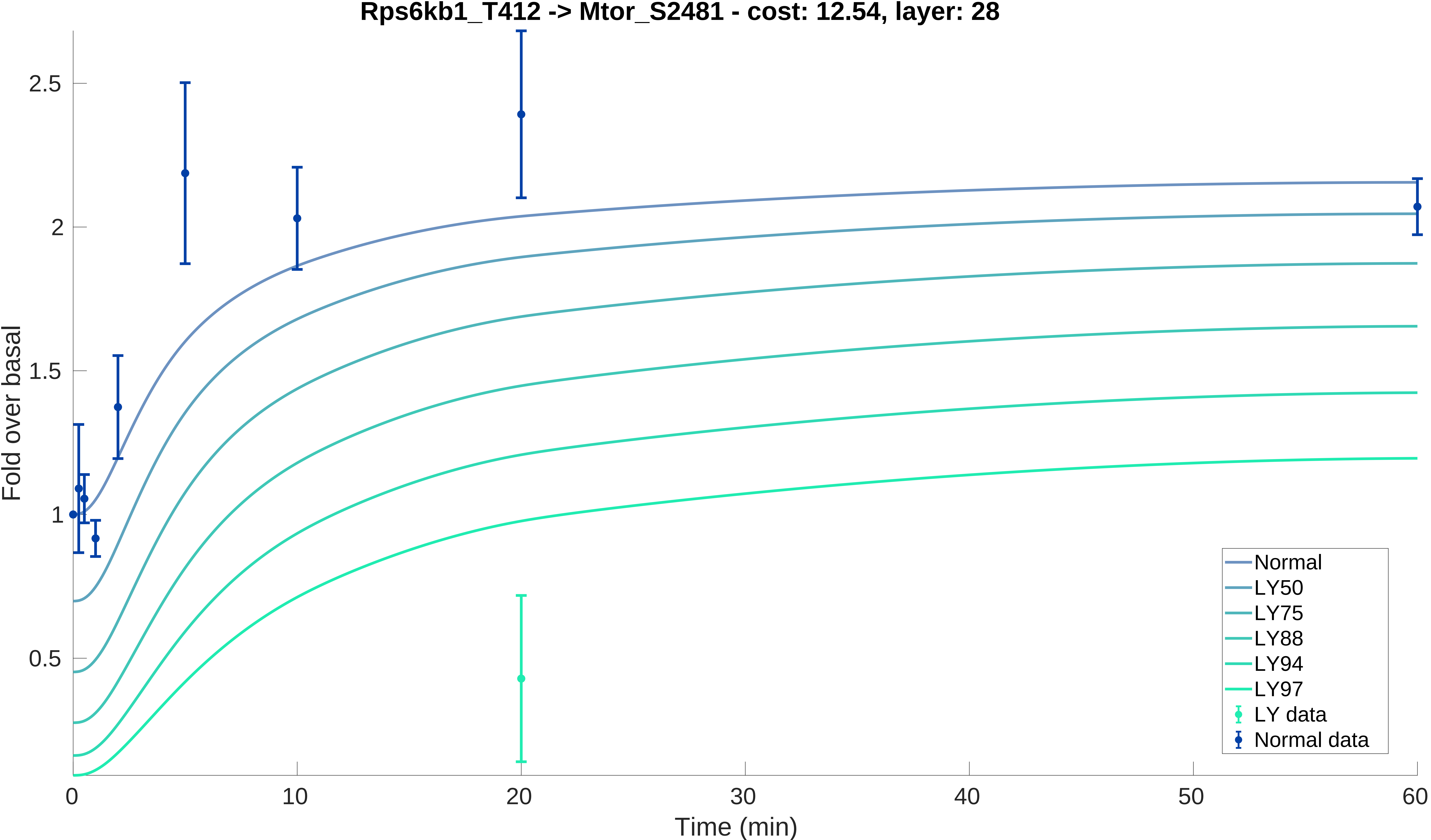

Rps6kb1_T412 -> Mtor_S2481 - cost: 12.54, layer: 28
Fold over basal
2.5
2
1.5
1
0.5
0
10
20
30
40
50
60
Time (min)
Normal
LY50
LY75
LY88
LY94
LY97
LY data
Normal data

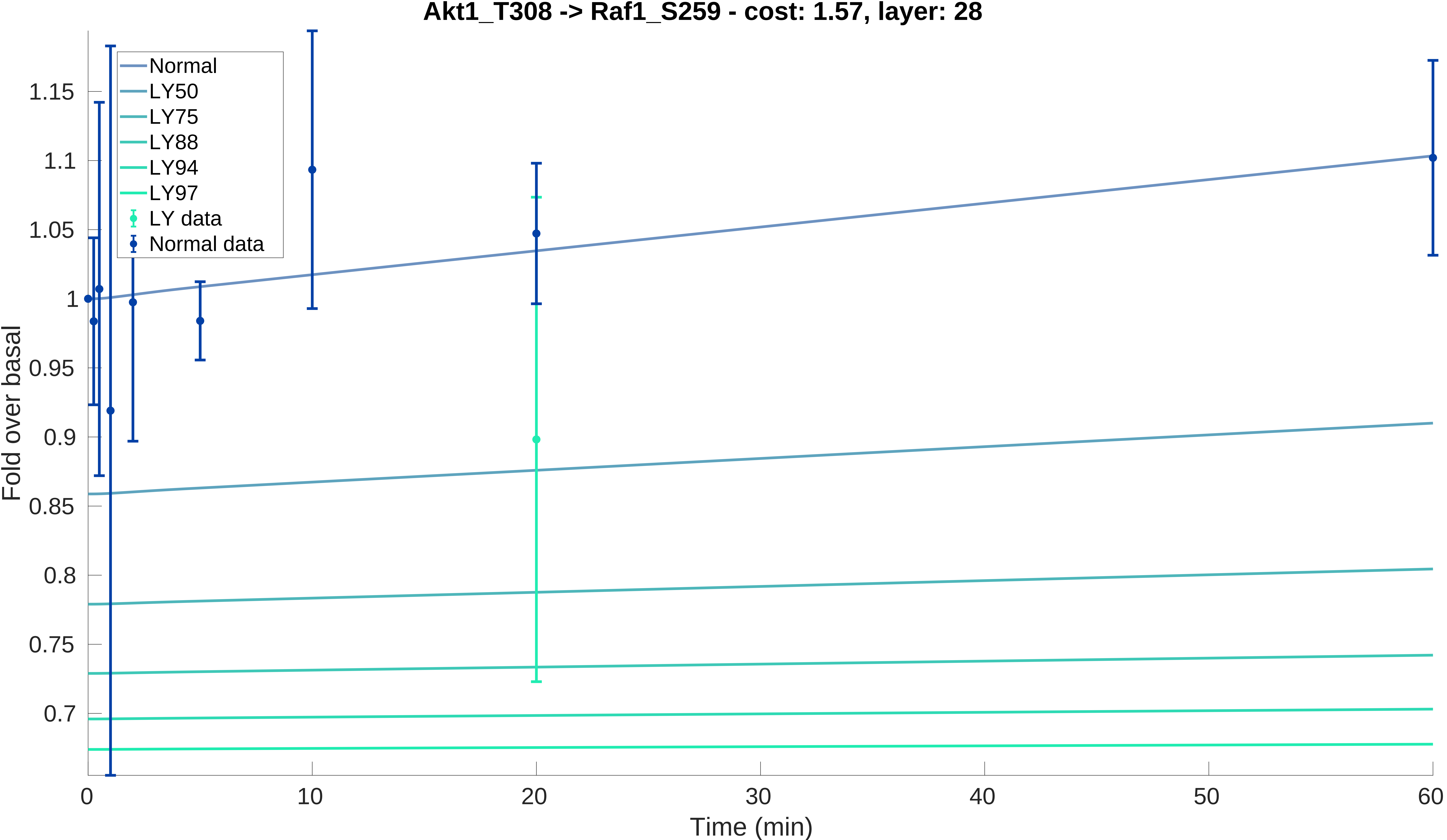

Akt1_T308 -> Raf1_S259 - cost: 1.57, layer: 28
Normal
LY50
LY75
LY88
LY94
LY97
LY data
Normal data
Fold over basal
1.15
1.1
1.05
1
0.95
0.9
0.85
0.8
0.75
0.7
0
10
20
30
40
50
60
Time (min)

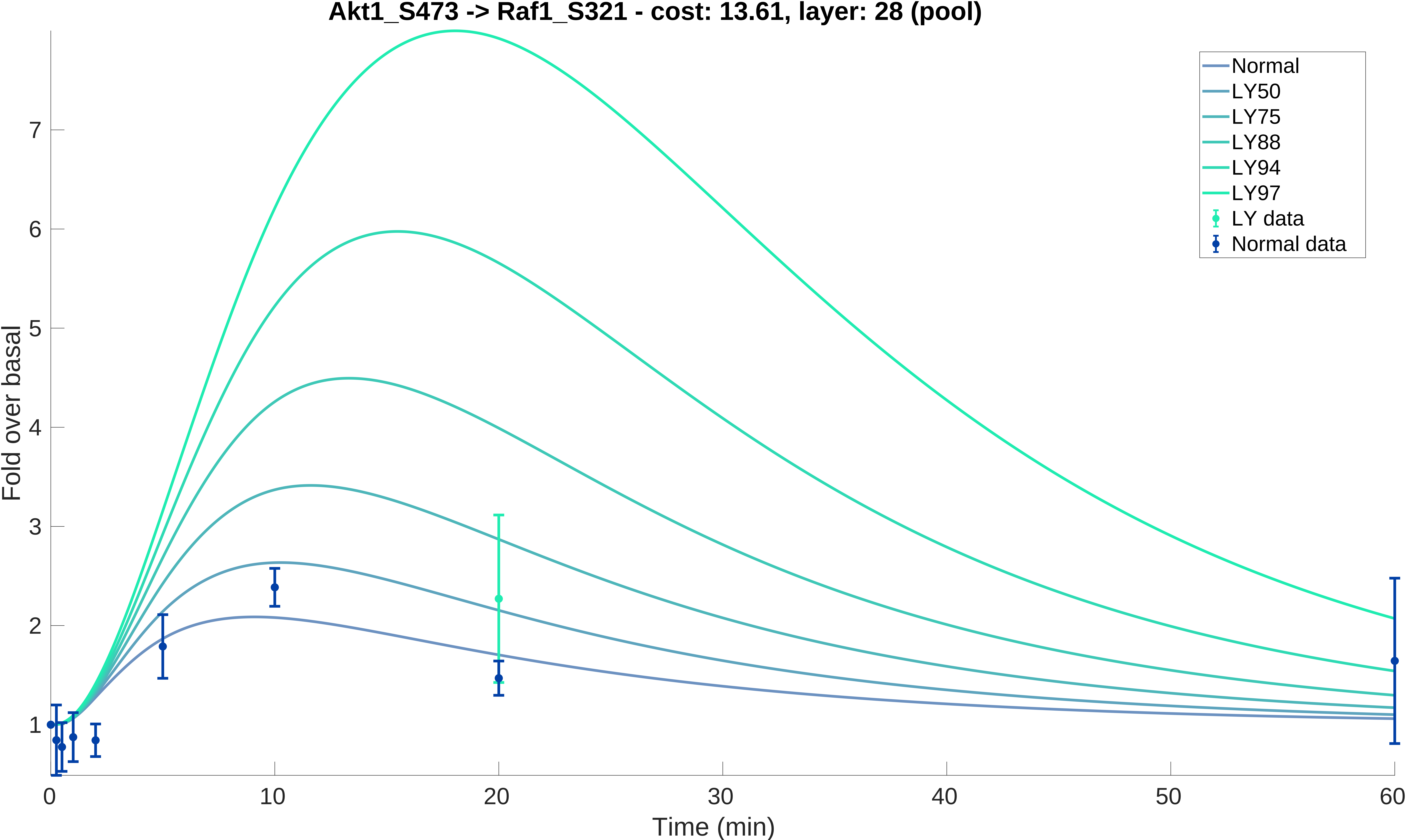
Akt1_S473 -> Raf1_S321 - cost: 13.61, layer: 28 (pool)
Fold over basal
Time (min)
0
10
20
30
40
50
60
1
2
3
4
5
6
7
Normal
LY50
LY75
LY88
LY94
LY97
LY data
Normal data

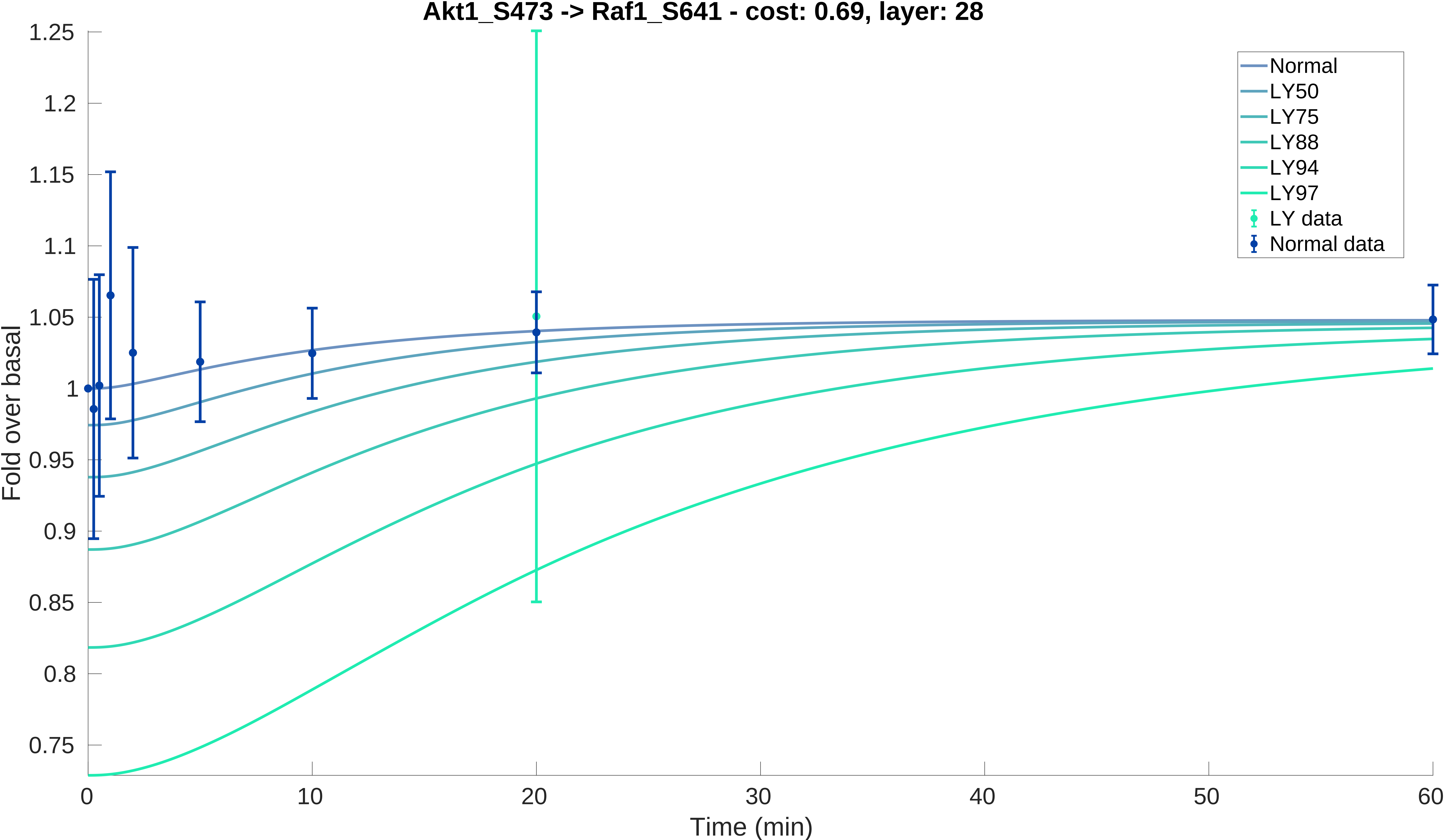
Akt1_S473 -> Raf1_S641 - cost: 0.69, layer: 28
1.25
1.2
1.15
1.1
1.05
1
0.95
0.9
0.85
0.8
0.75
Fold over basal
0
10
20
30
40
50
60
Time (min)
Normal
LY50
LY75
LY88
LY94
LY97
LY data
Normal data

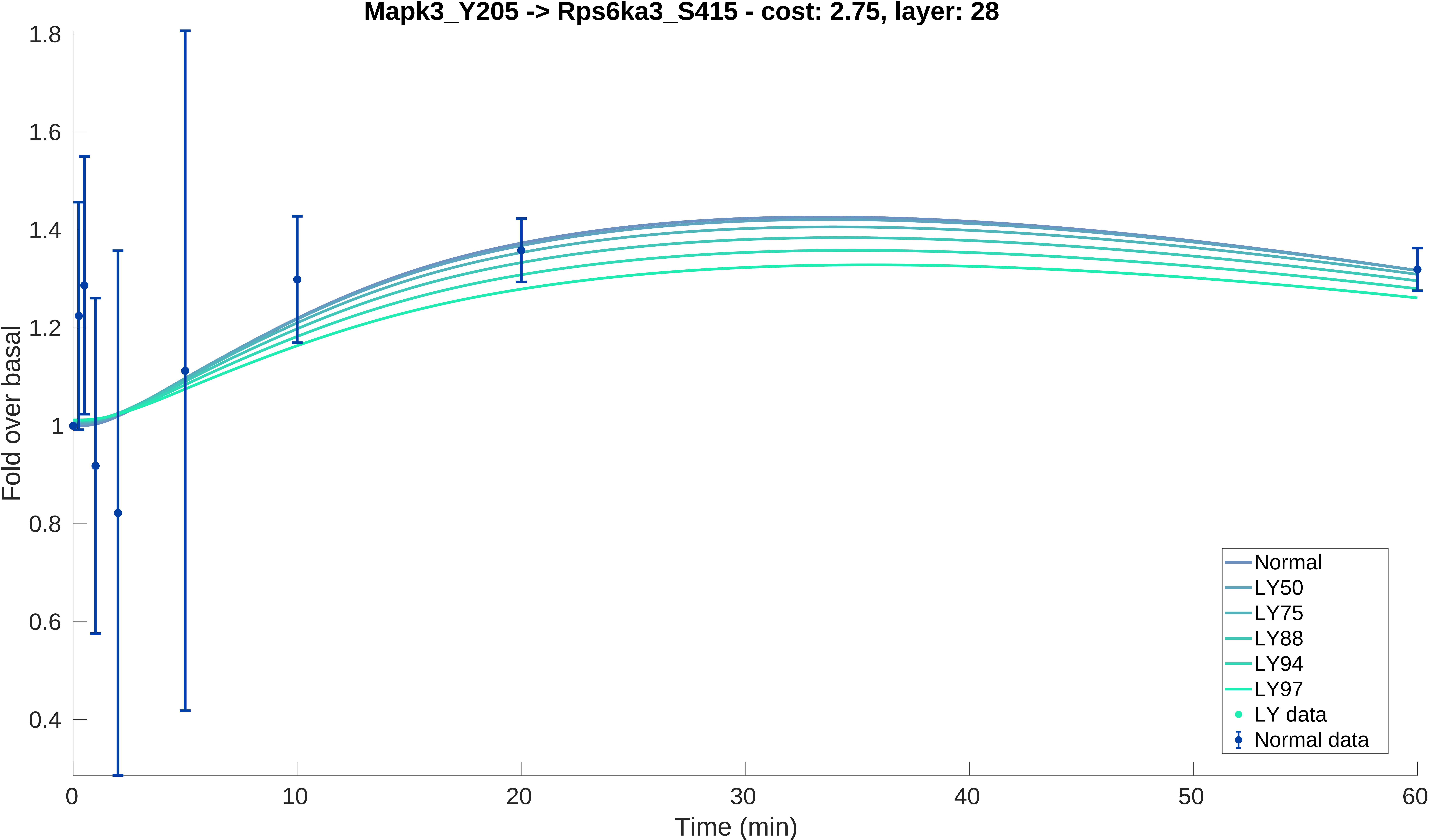
Mapk3_Y205 -> Rps6ka3_S415 - cost: 2.75, layer: 28
1.8
1.6
1.4
1.2
1
0.8
0.6
0.4
Fold over basal
0
10
20
30
40
50
60
Time (min)
Normal
LY50
LY75
LY88
LY94
LY97
LY data
Normal data

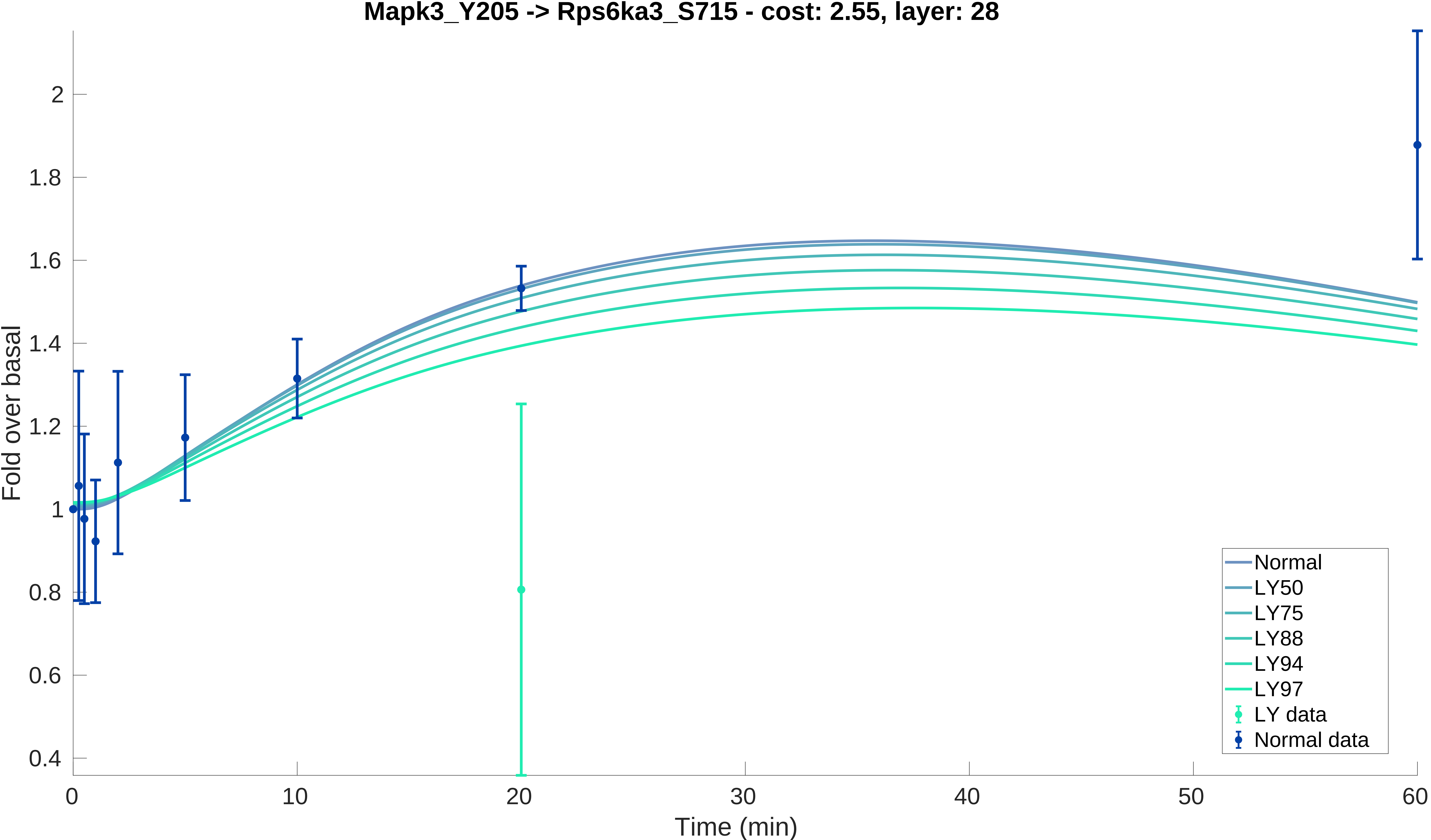

Mapk3_Y205 -> Rps6ka3_S715 - cost: 2.55, layer: 28
Fold over basal
2
1.8
1.6
1.4
1.2
1
0.8
0.6
0.4
0
10
20
30
40
50
60
Time (min)
Normal
LY50
LY75
LY88
LY94
LY97
LY data
Normal data

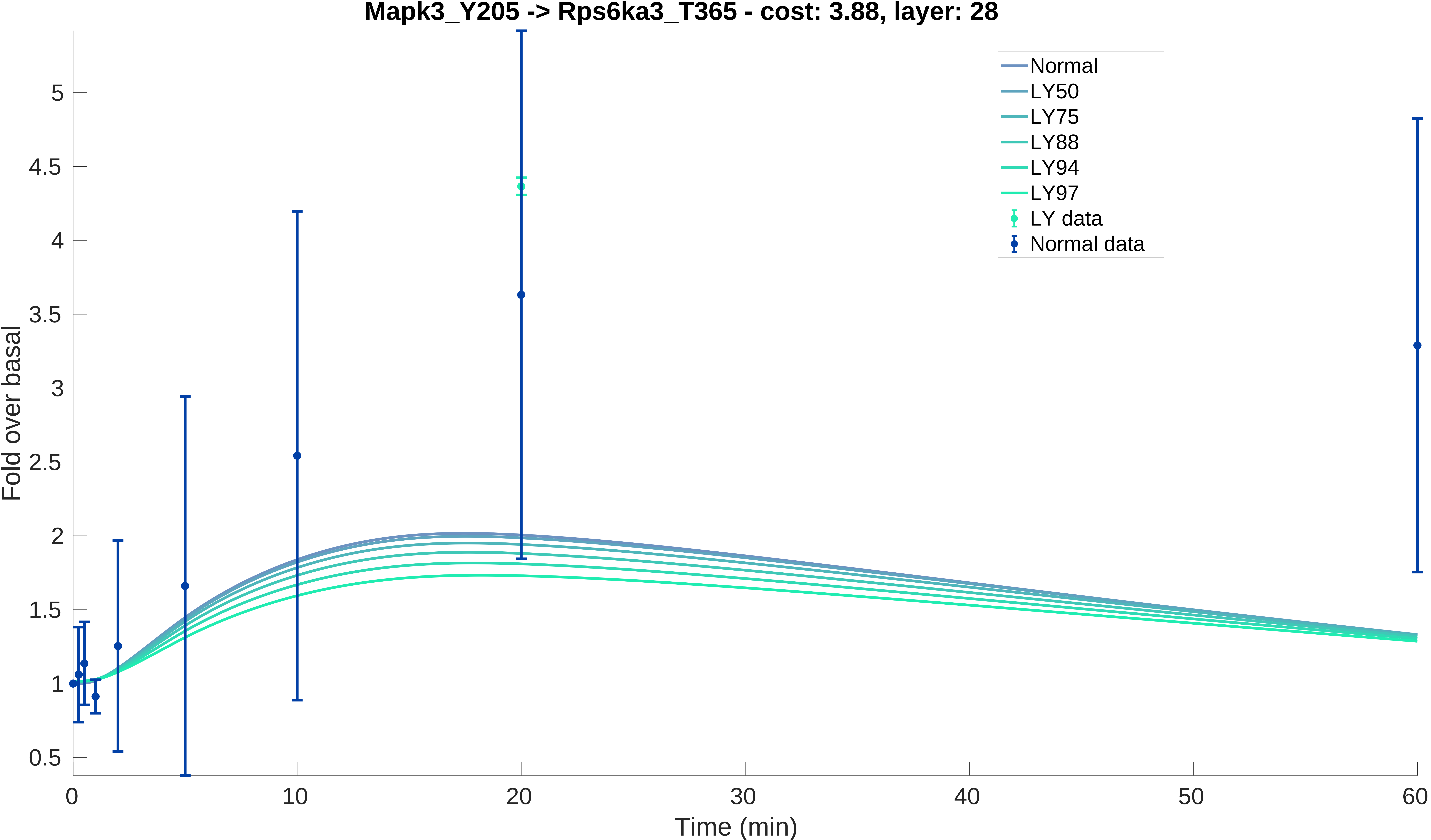

Mapk3_Y205 -> Rps6ka3_T365 - cost: 3.88, layer: 28
Normal
LY50
LY75
LY88
LY94
LY97
LY data
Normal data
Fold over basal
Time (min)
0.5
1
1.5
2
2.5
3
3.5
4
4.5
5
0
10
20
30
40
50
60

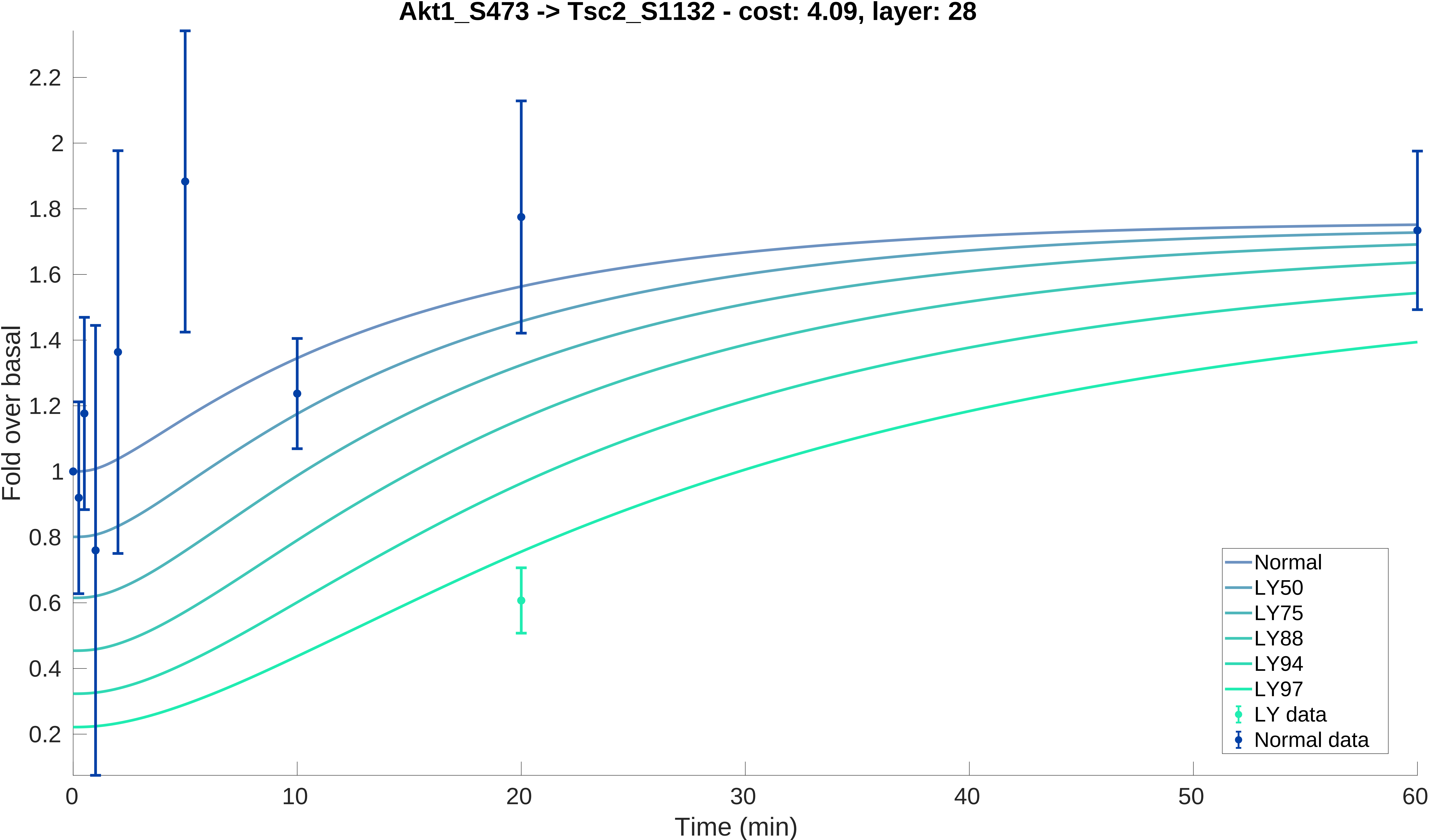

Akt1_S473 -> Tsc2_S1132 - cost: 4.09, layer: 28
Fold over basal
2.2
2
1.8
1.6
1.4
1.2
1
0.8
0.6
0.4
0.2
0
10
20
30
40
50
60
Time (min)
Normal
LY50
LY75
LY88
LY94
LY97
LY data
Normal data

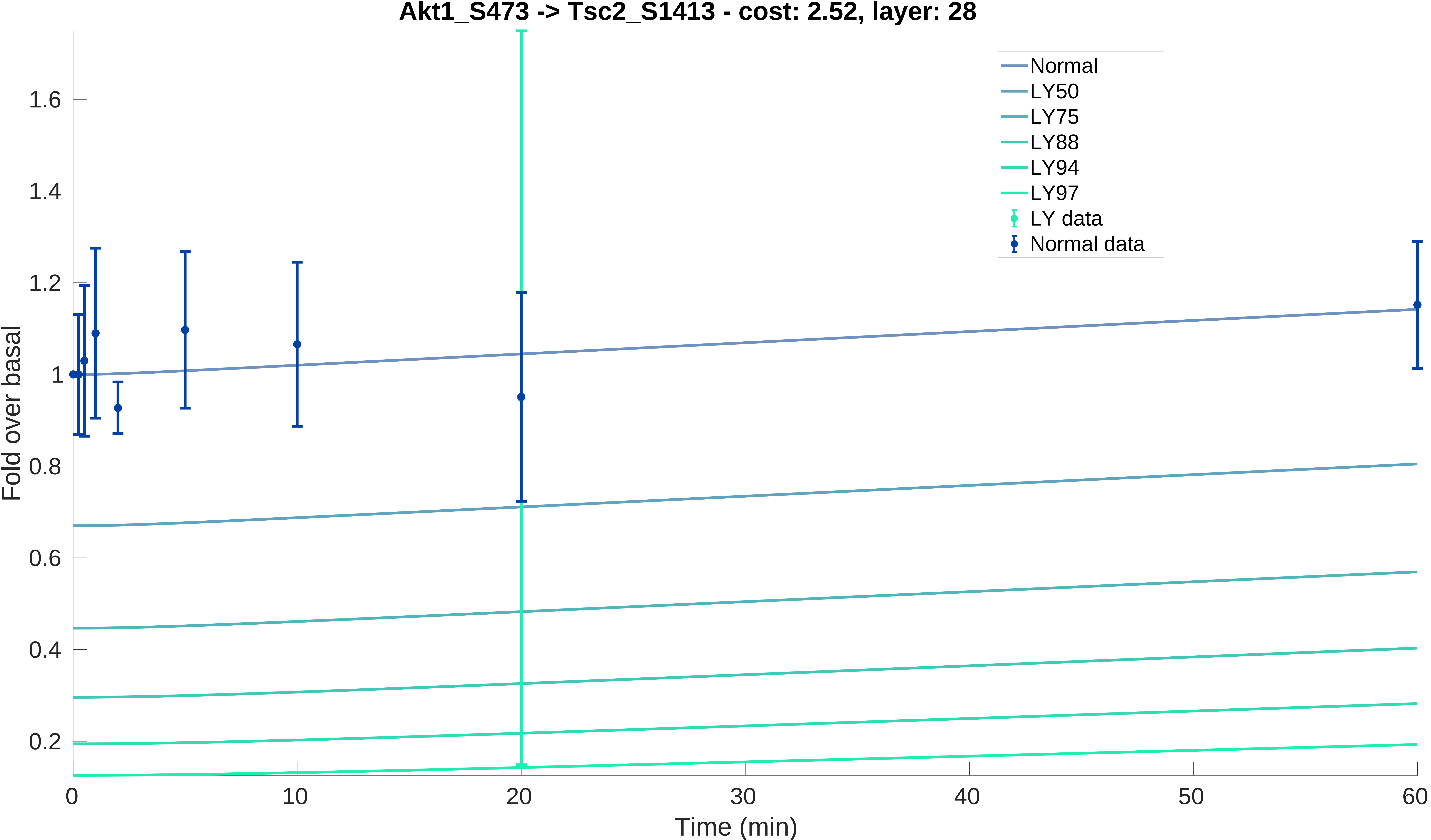

Akt1_S473 -> Tsc2_S1413 - cost: 2.52, layer: 28
Normal
LY50
LY75
LY88
LY94
LY97
LY data
Normal data
Fold over basal
Time (min)
1.6
1.4
1.2
1
0.8
0.6
0.4
0.2
0
10
20
30
40
50
60

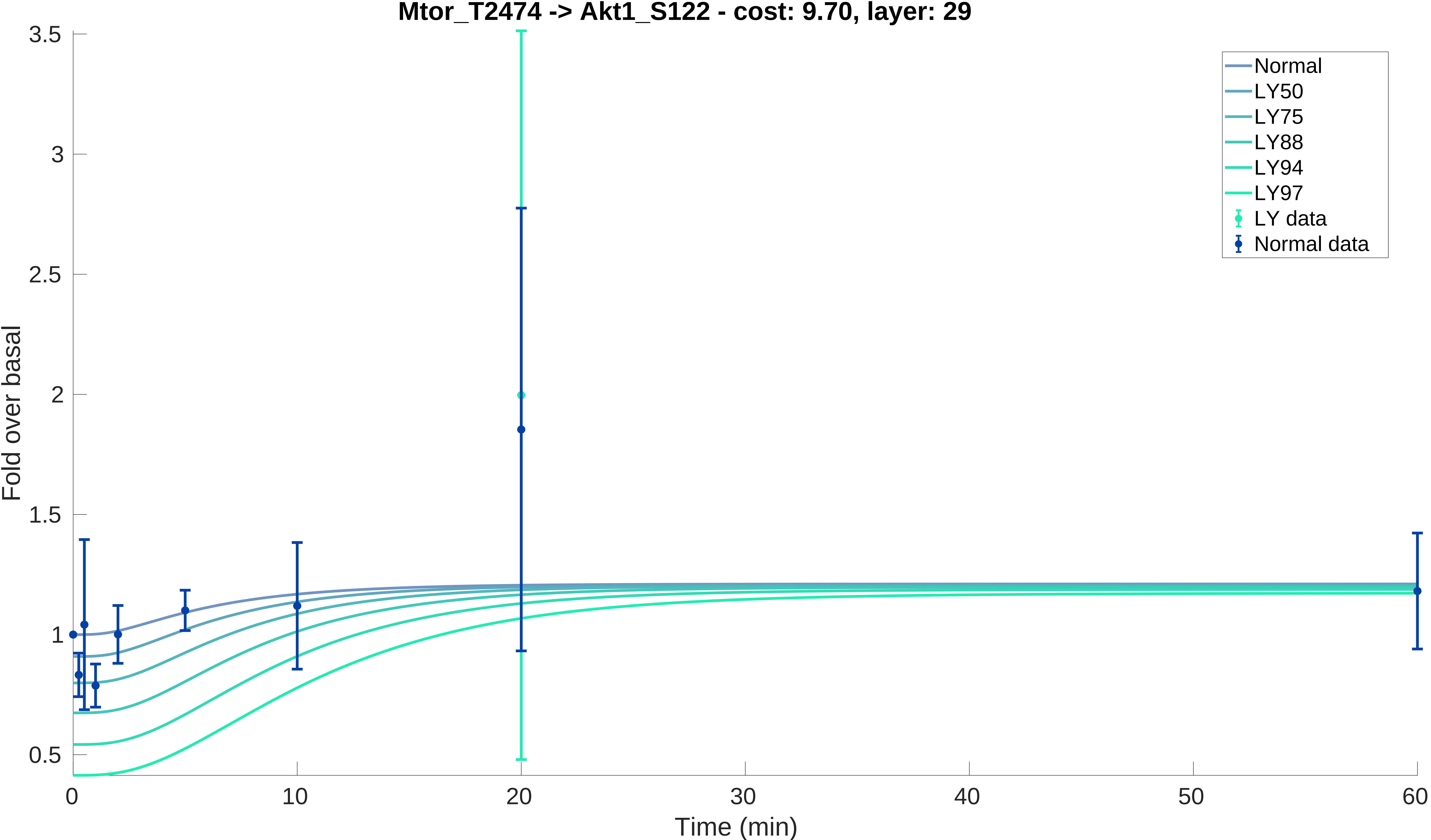

Mtor_T2474 -> Akt1_S122 - cost: 9.70, layer: 29
Fold over basal
Time (min)
Normal
LY50
LY75
LY88
LY94
LY97
LY data
Normal data
0.5
1
1.5
2
2.5
3
3.5
0
10
20
30
40
50
60

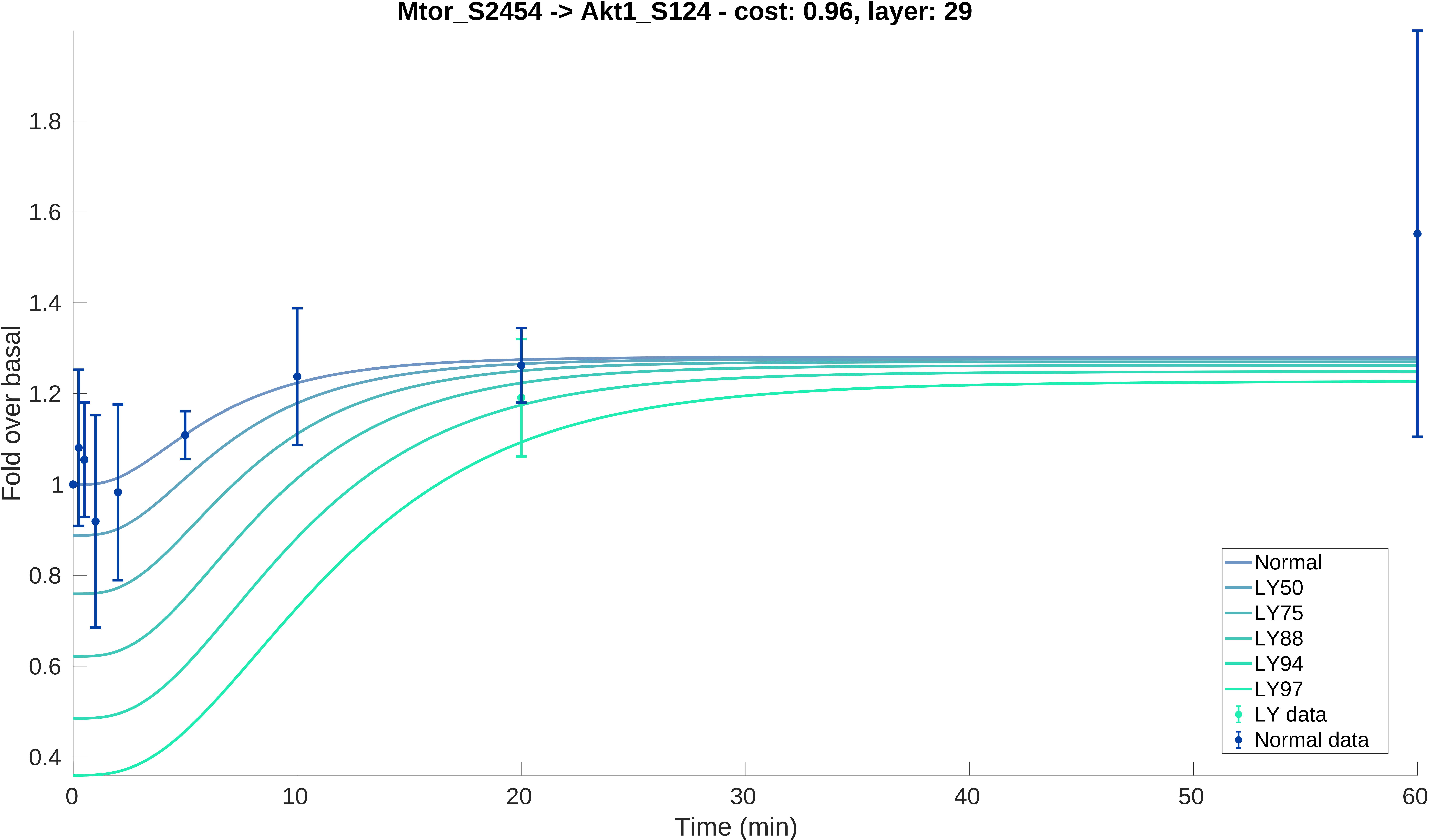

Mtor_S2454 -> Akt1_S124 - cost: 0.96, layer: 29
Fold over basal
1.8
1.6
1.4
1.2
1
0.8
0.6
0.4
0
10
20
30
40
50
60
Time (min)
Normal
LY50
LY75
LY88
LY94
LY97
LY data
Normal data

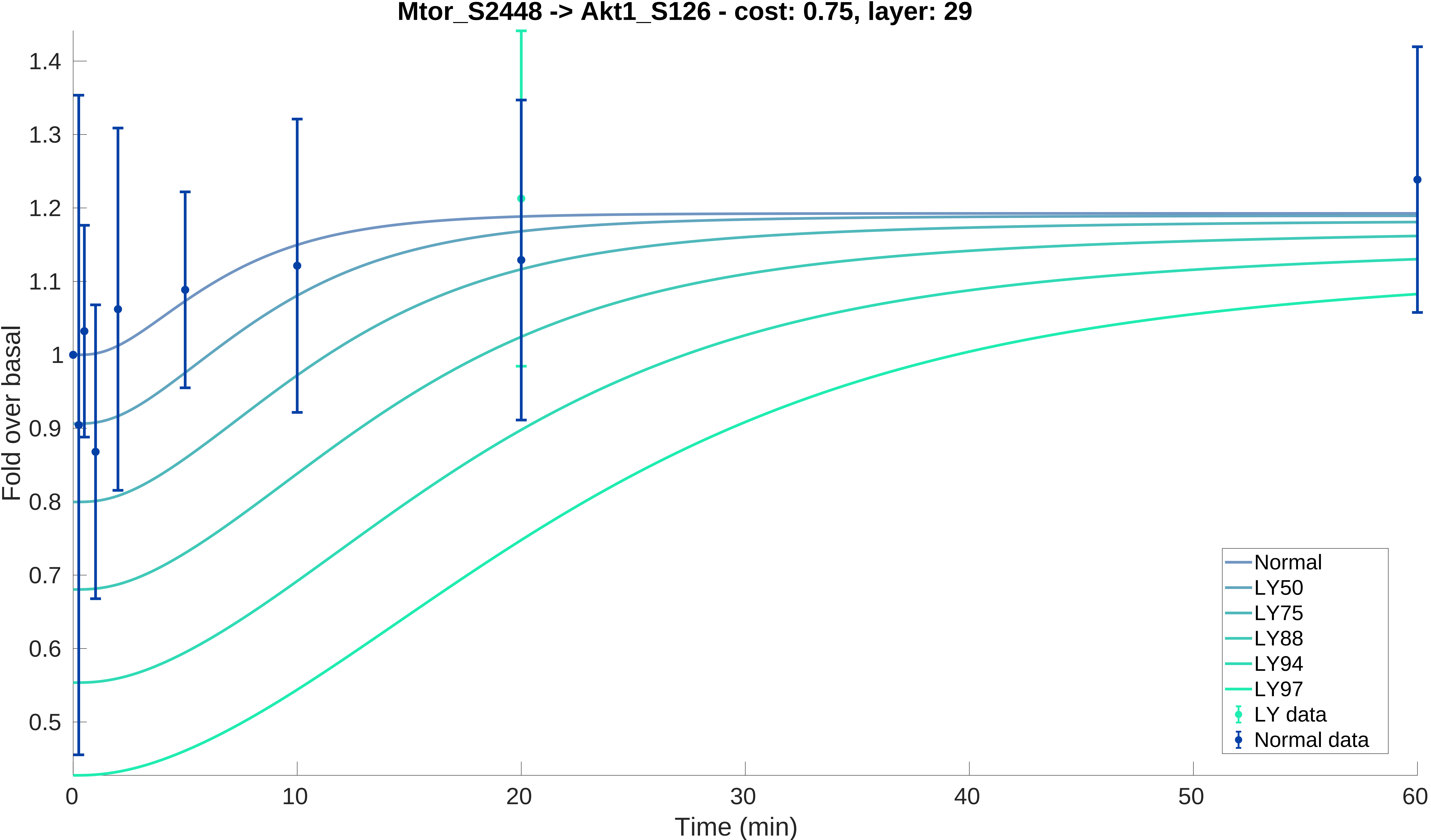

Mtor_S2448 -> Akt1_S126 - cost: 0.75, layer: 29
Fold over basal
Time (min)
1.4
1.3
1.2
1.1
1
0.9
0.8
0.7
0.6
0.5
0
10
20
30
40
50
60
Normal
LY50
LY75
LY88
LY94
LY97
LY data
Normal data

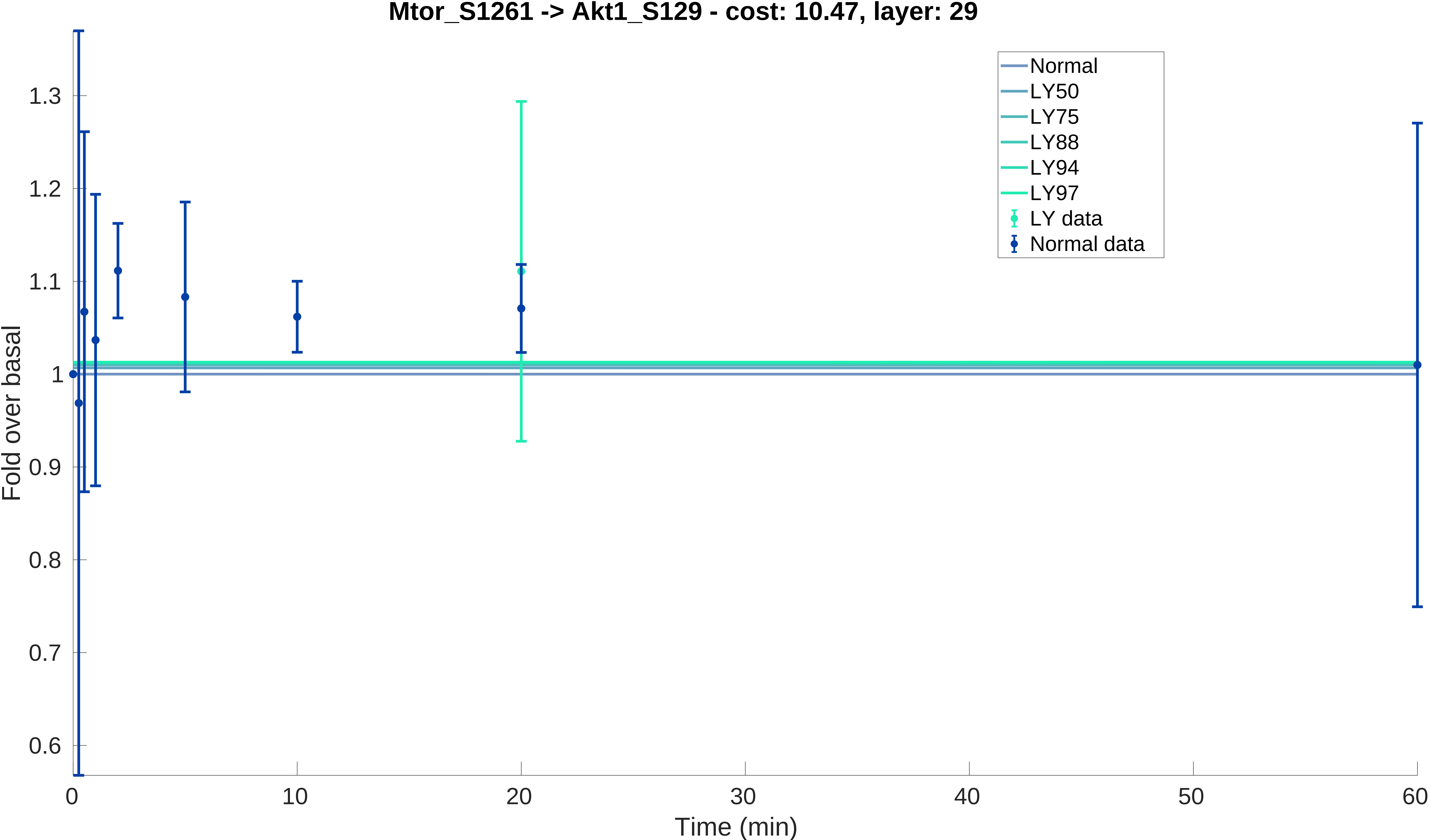

Mtor_S1261 -> Akt1_S129 - cost: 10.47, layer: 29
Normal
LY50
LY75
LY88
LY94
LY97
LY data
Normal data
Fold over basal
1.3
1.2
1.1
1
0.9
0.8
0.7
0.6
0
10
20
30
40
50
60
Time (min)

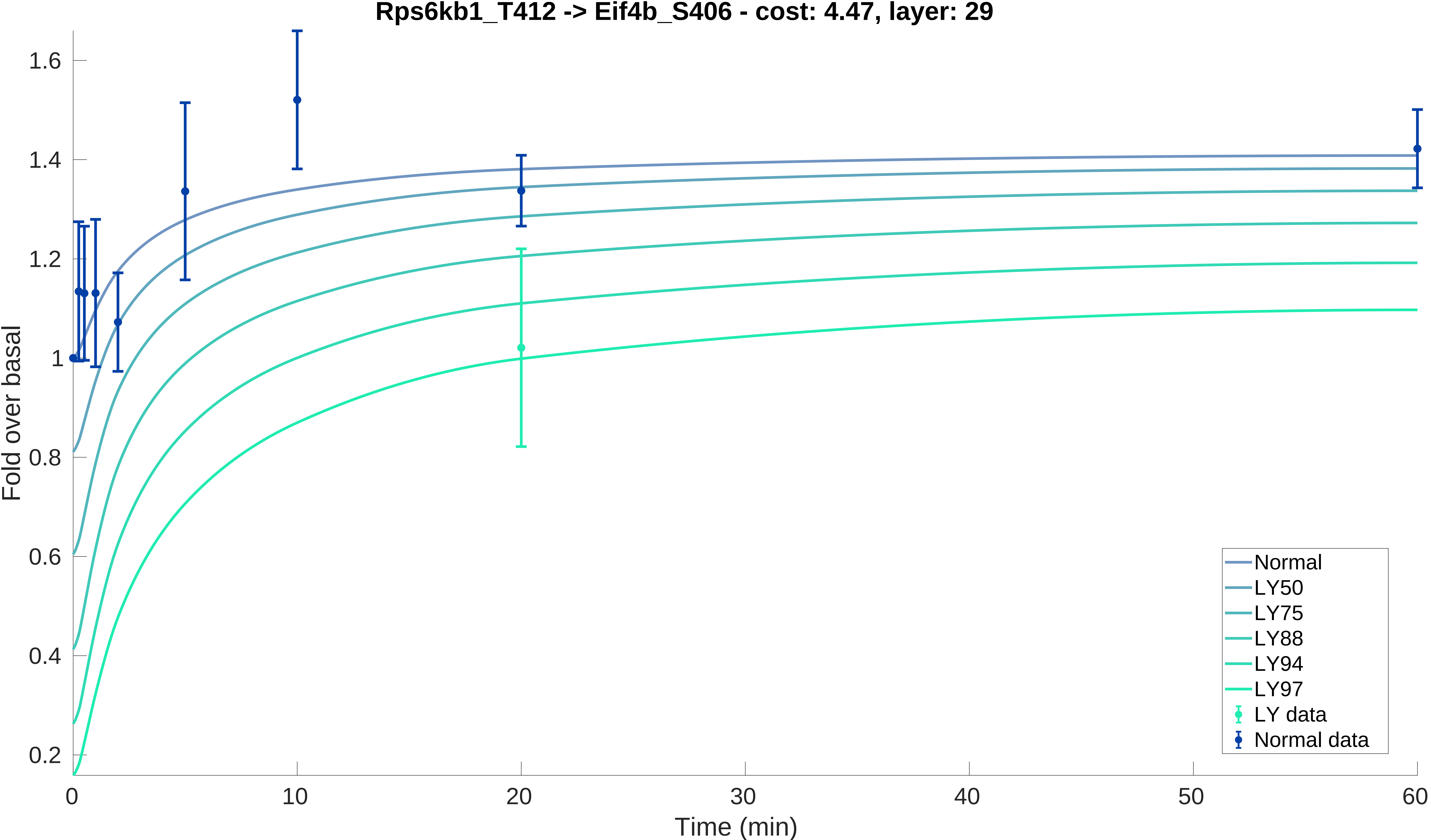
Rps6kb1_T412 -> Eif4b_S406 - cost: 4.47, layer: 29
Fold over basal
1.6
1.4
1.2
1
0.8
0.6
0.4
0.2
0
10
20
30
40
50
60
Time (min)
Normal
LY50
LY75
LY88
LY94
LY97
LY data
Normal data

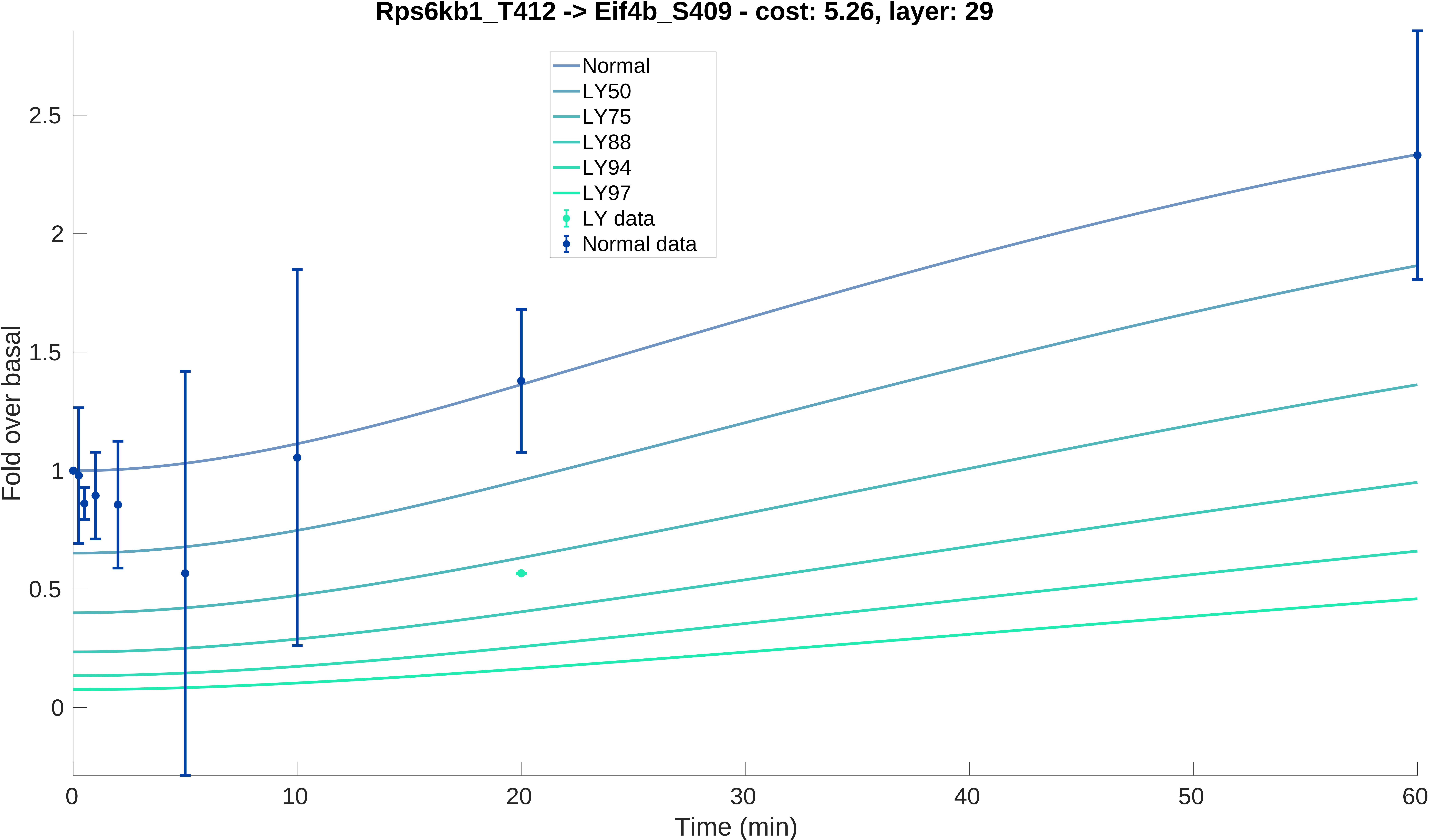
Rps6kb1_T412 -> Eif4b_S409 - cost: 5.26, layer: 29
Normal
LY50
LY75
LY88
LY94
LY97
LY data
Normal data
Fold over basal
Time (min)
2.5
2
1.5
1
0.5
0
0
10
20
30
40
50
60

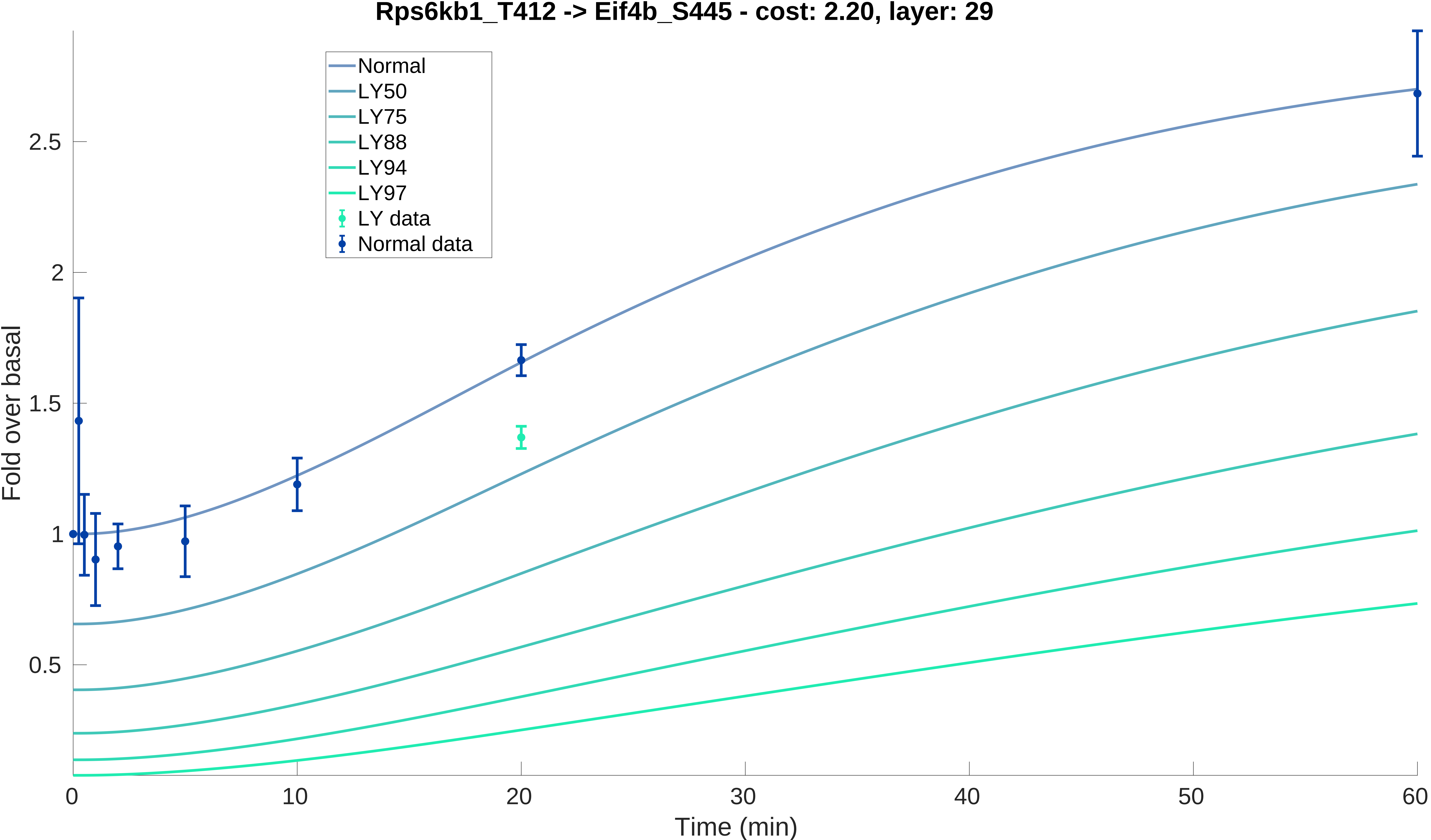
Rps6kb1_T412 -> Eif4b_S445 - cost: 2.20, layer: 29
Normal
LY50
LY75
LY88
LY94
LY97
LY data
Normal data
Fold over basal
2.5
2
1.5
1
0.5
0
10
20
30
40
50
60
Time (min)

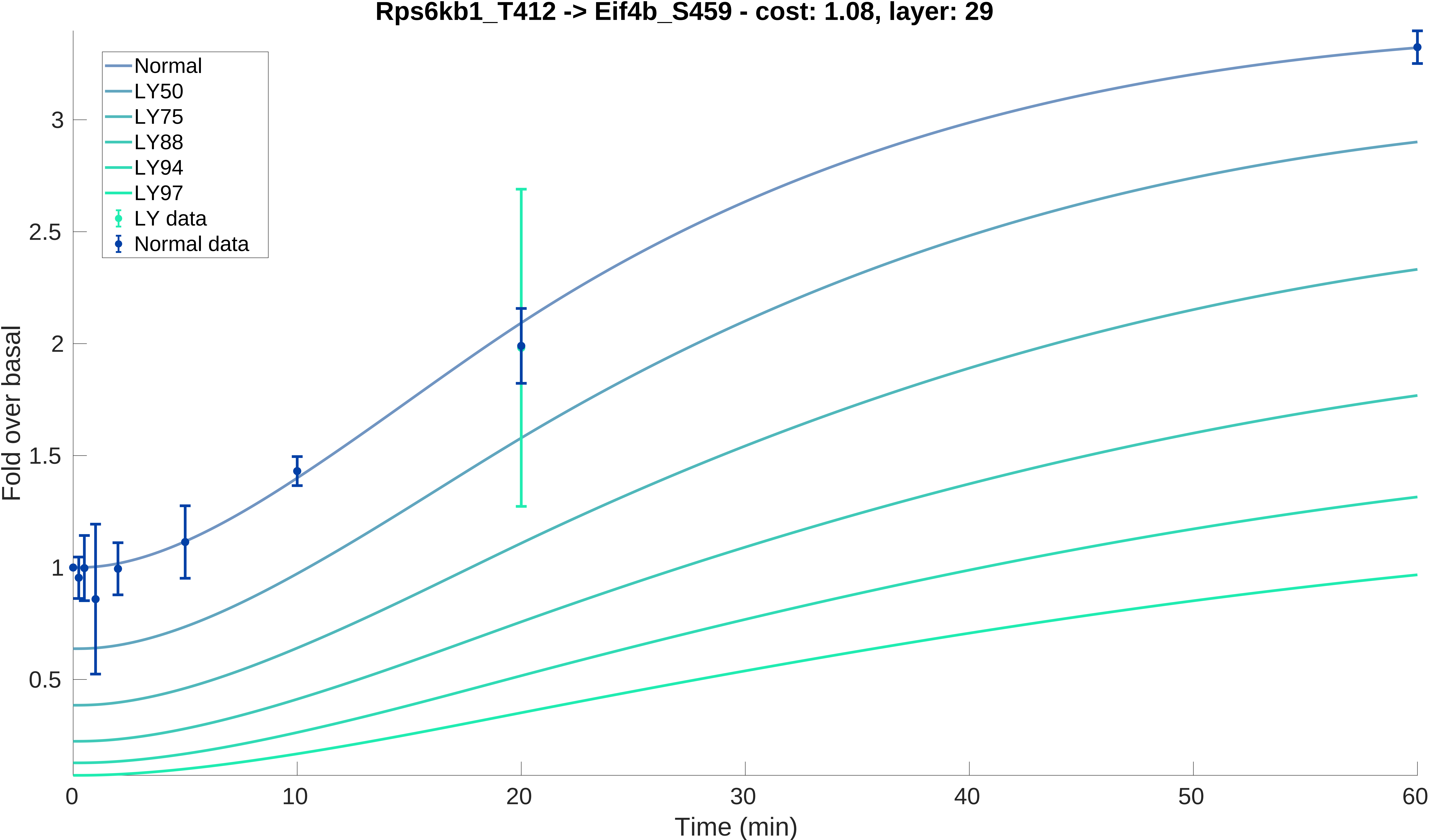

Rps6kb1_T412 -> Eif4b_S459 - cost: 1.08, layer: 29
Normal
LY50
LY75
LY88
LY94
LY97
LY data
Normal data
Fold over basal
Time (min)
0
10
20
30
40
50
60
0.5
1
1.5
2
2.5
3

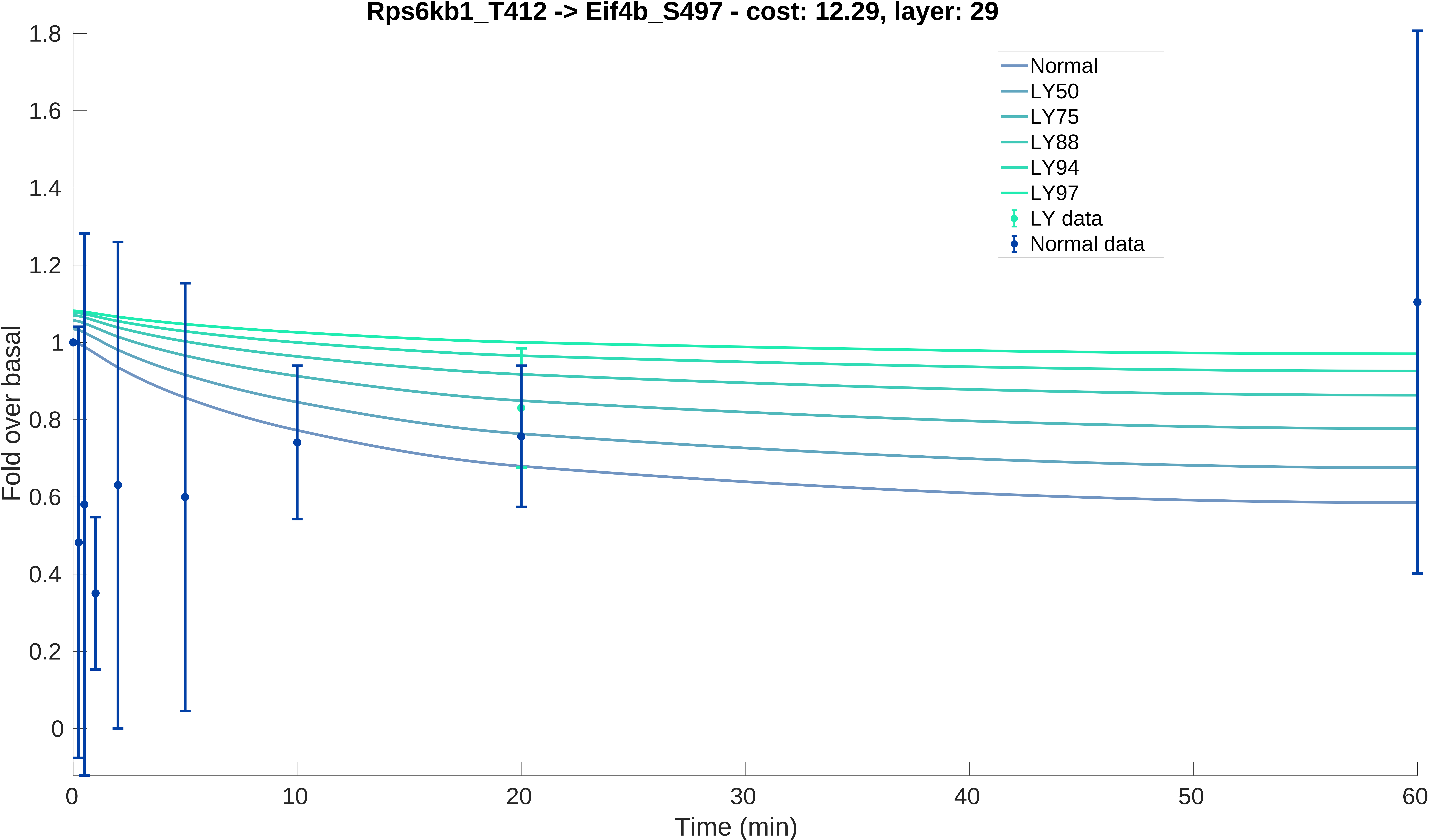

Rps6kb1_T412 -> Eif4b_S497 - cost: 12.29, layer: 29
Fold over basal
Time (min)
Normal
LY50
LY75
LY88
LY94
LY97
LY data
Normal data
1.8
1.6
1.4
1.2
1
0.8
0.6
0.4
0.2
0
0
10
20
30
40
50
60

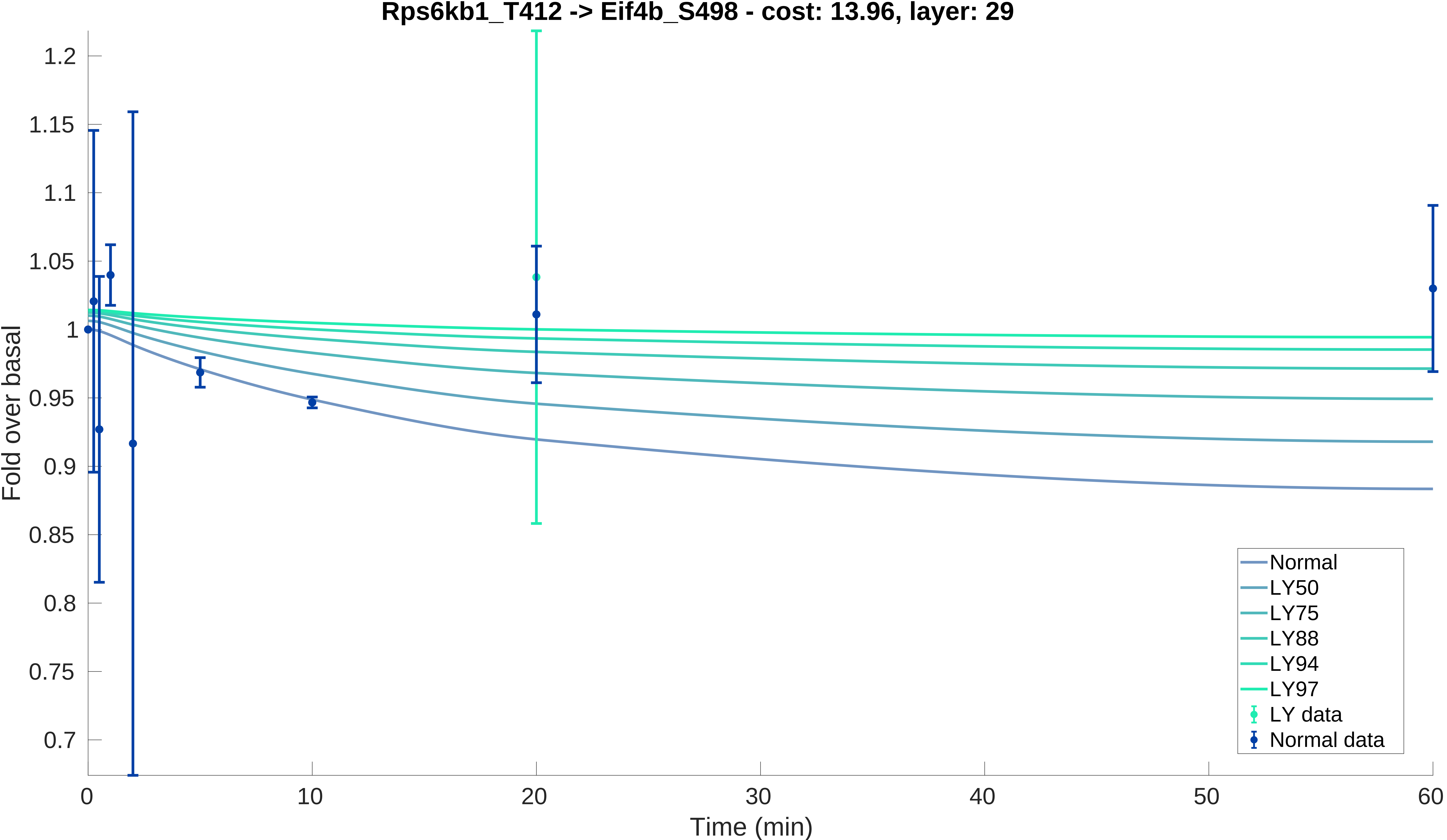

Rps6kb1_T412 -> Eif4b_S498 - cost: 13.96, layer: 29
Fold over basal
Time (min)
1.2
1.15
1.1
1.05
1
0.95
0.9
0.85
0.8
0.75
0.7
0
10
20
30
40
50
60
Normal
LY50
LY75
LY88
LY94
LY97
LY data
Normal data

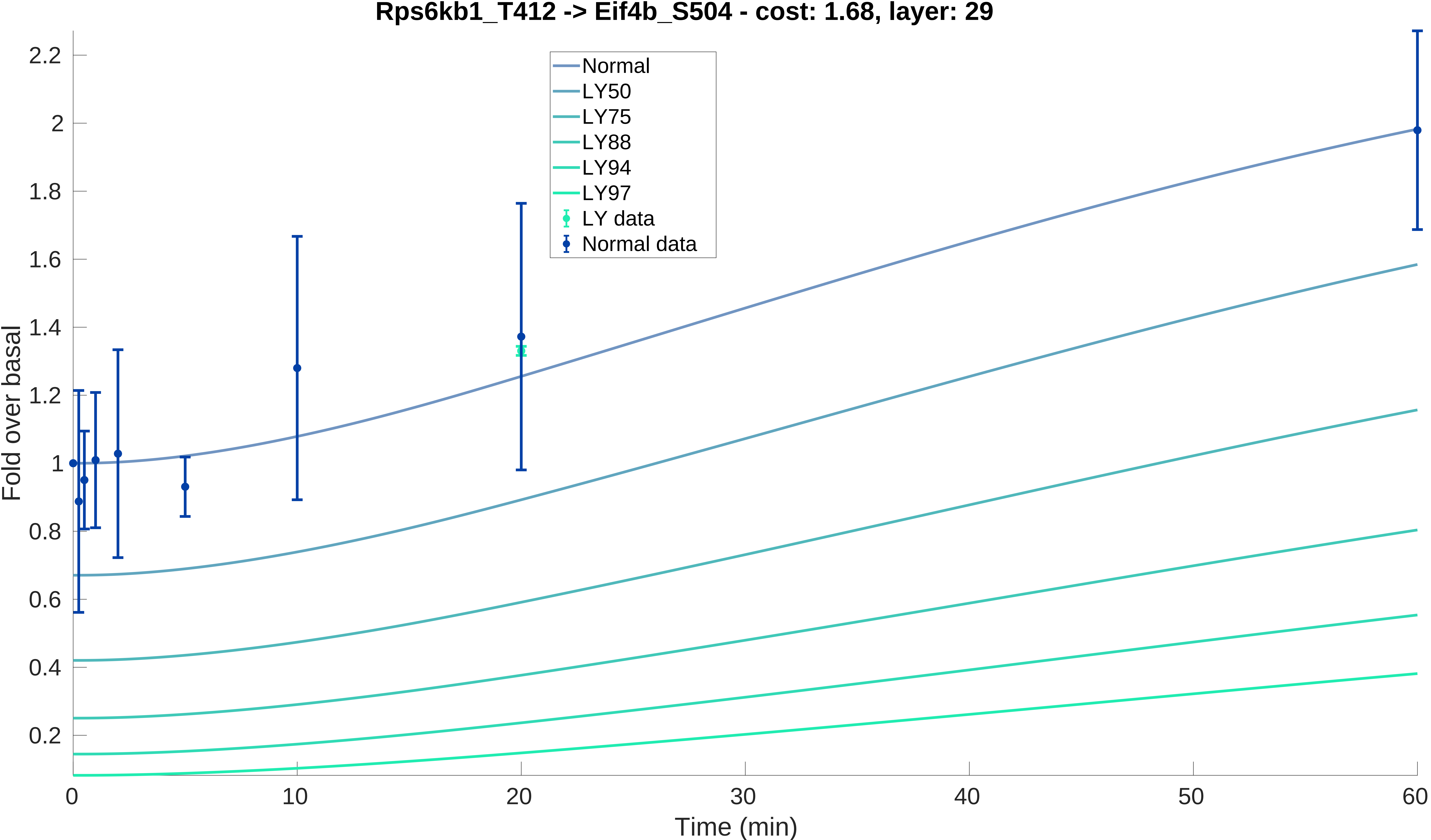
Rps6kb1_T412 -> Eif4b_S504 - cost: 1.68, layer: 29
Normal
LY50
LY75
LY88
LY94
LY97
LY data
Normal data
Fold over basal
Time (min)
0
10
20
30
40
50
60
0.2
0.4
0.6
0.8
1
1.2
1.4
1.6
1.8
2
2.2

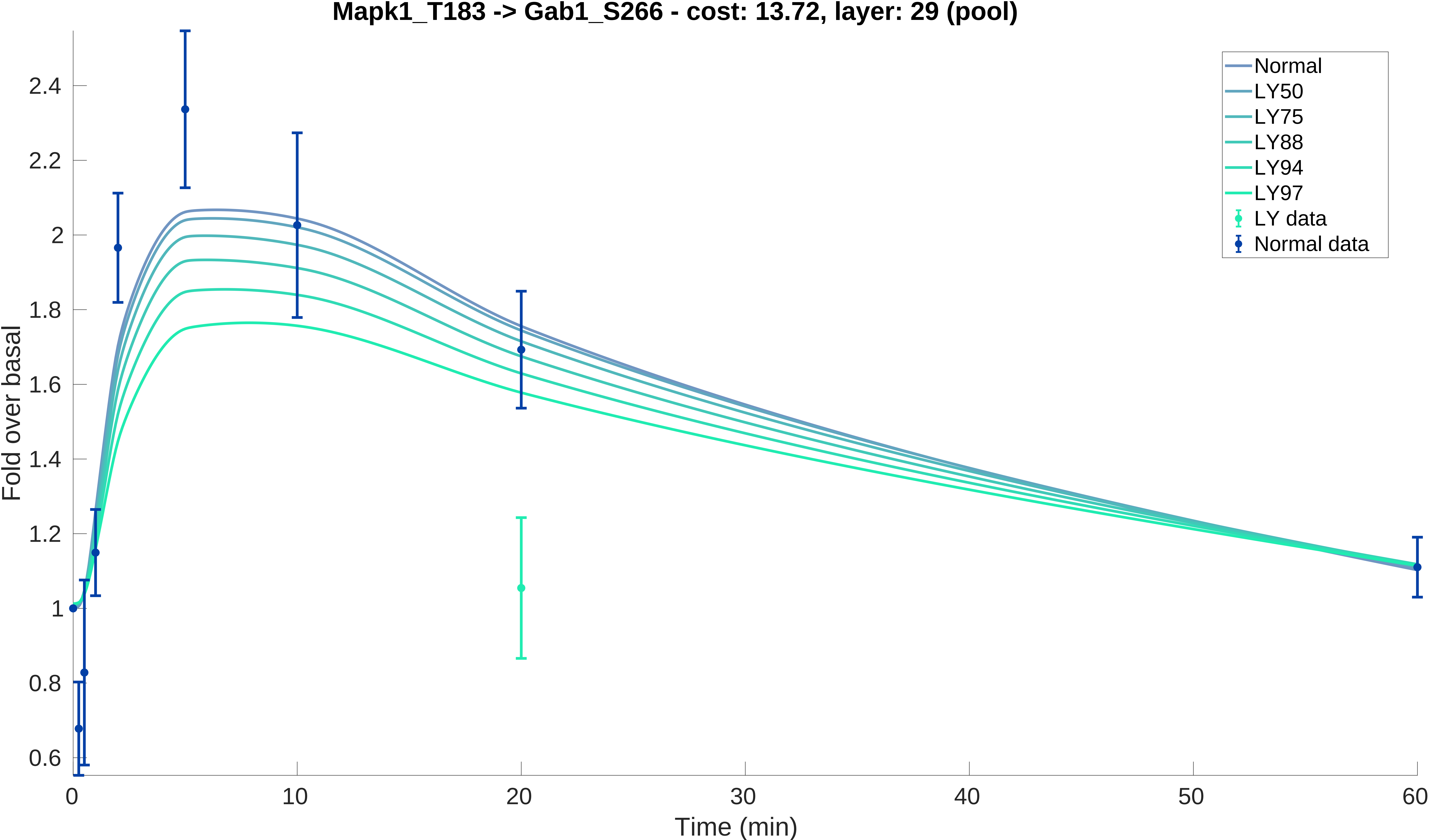

Mapk1_T183 -> Gab1_S266 - cost: 13.72, layer: 29 (pool)
Fold over basal
Time (min)
Normal
LY50
LY75
LY88
LY94
LY97
LY data
Normal data
0.6
0.8
1
1.2
1.4
1.6
1.8
2
2.2
2.4
0
10
20
30
40
50
60

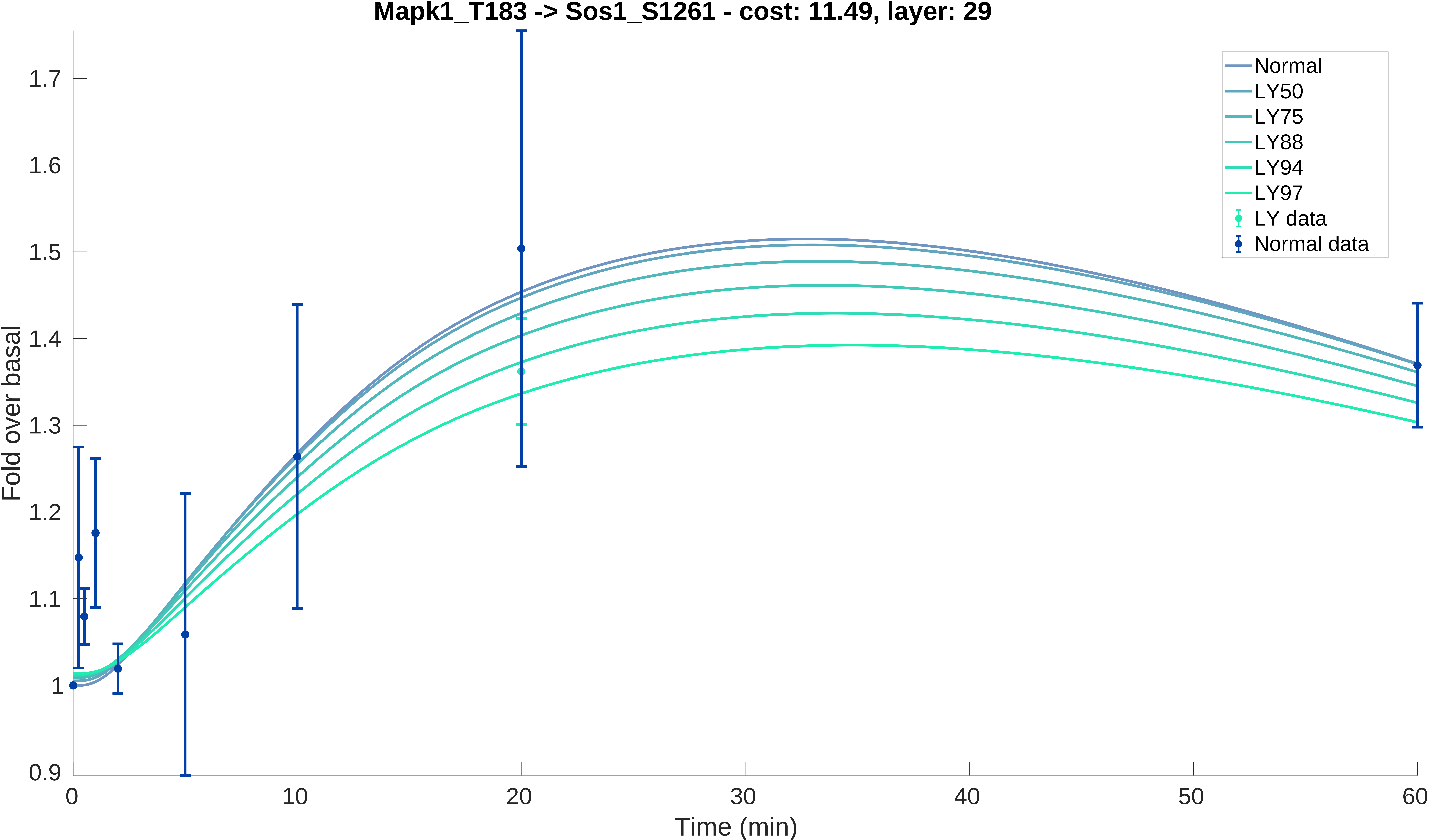

Mapk1_T183 -> Sos1_S1261 - cost: 11.49, layer: 29
Normal
LY50
LY75
LY88
LY94
LY97
LY data
Normal data
Fold over basal
1.7
1.6
1.5
1.4
1.3
1.2
1.1
1
0.9
0
10
20
30
40
50
60
Time (min)

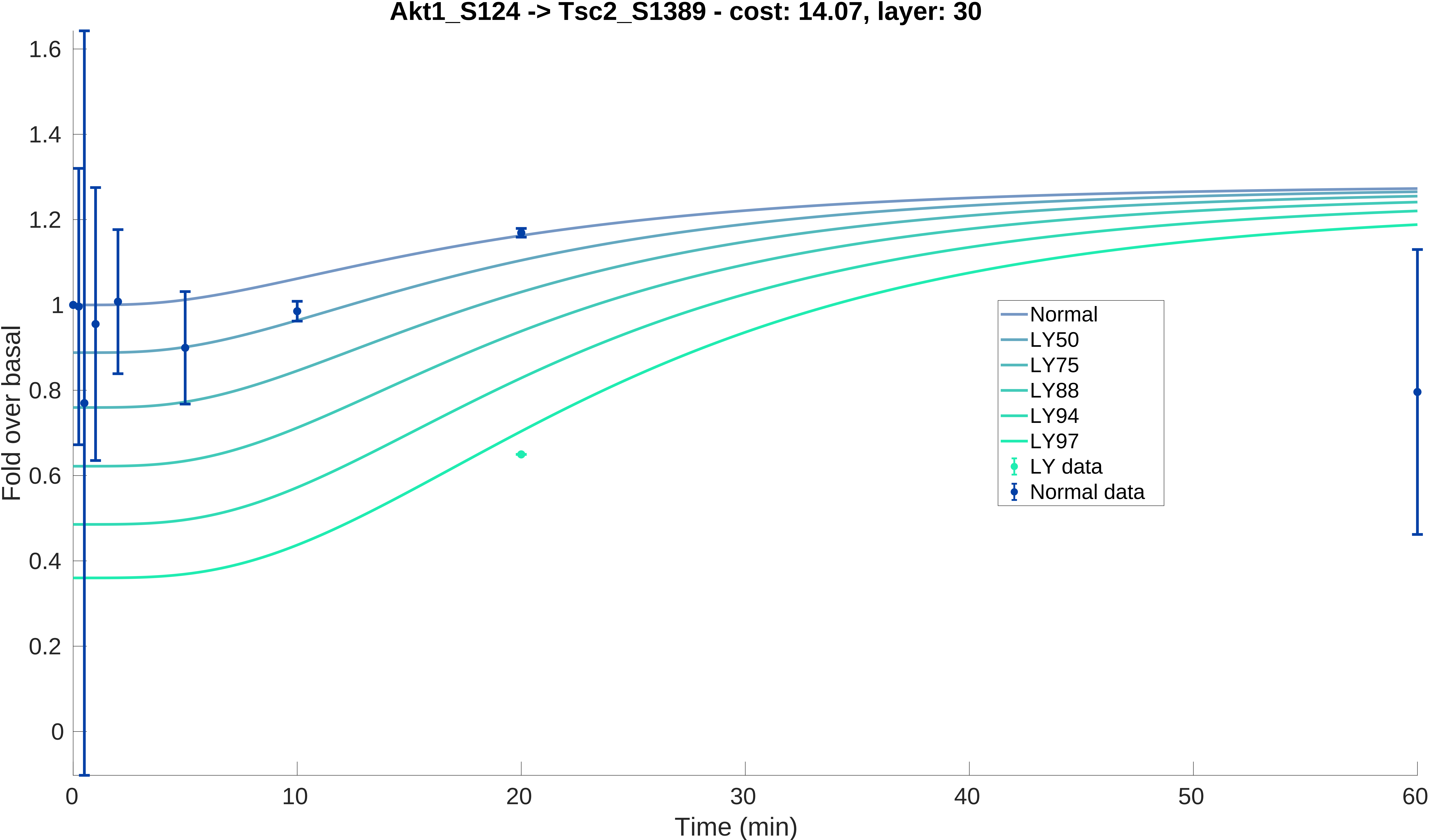

Akt1_S124 -> Tsc2_S1389 - cost: 14.07, layer: 30
Fold over basal
1.6
1.4
1.2
1
0.8
0.6
0.4
0.2
0
Time (min)
0
10
20
30
40
50
60
Normal
LY50
LY75
LY88
LY94
LY97
LY data
Normal data

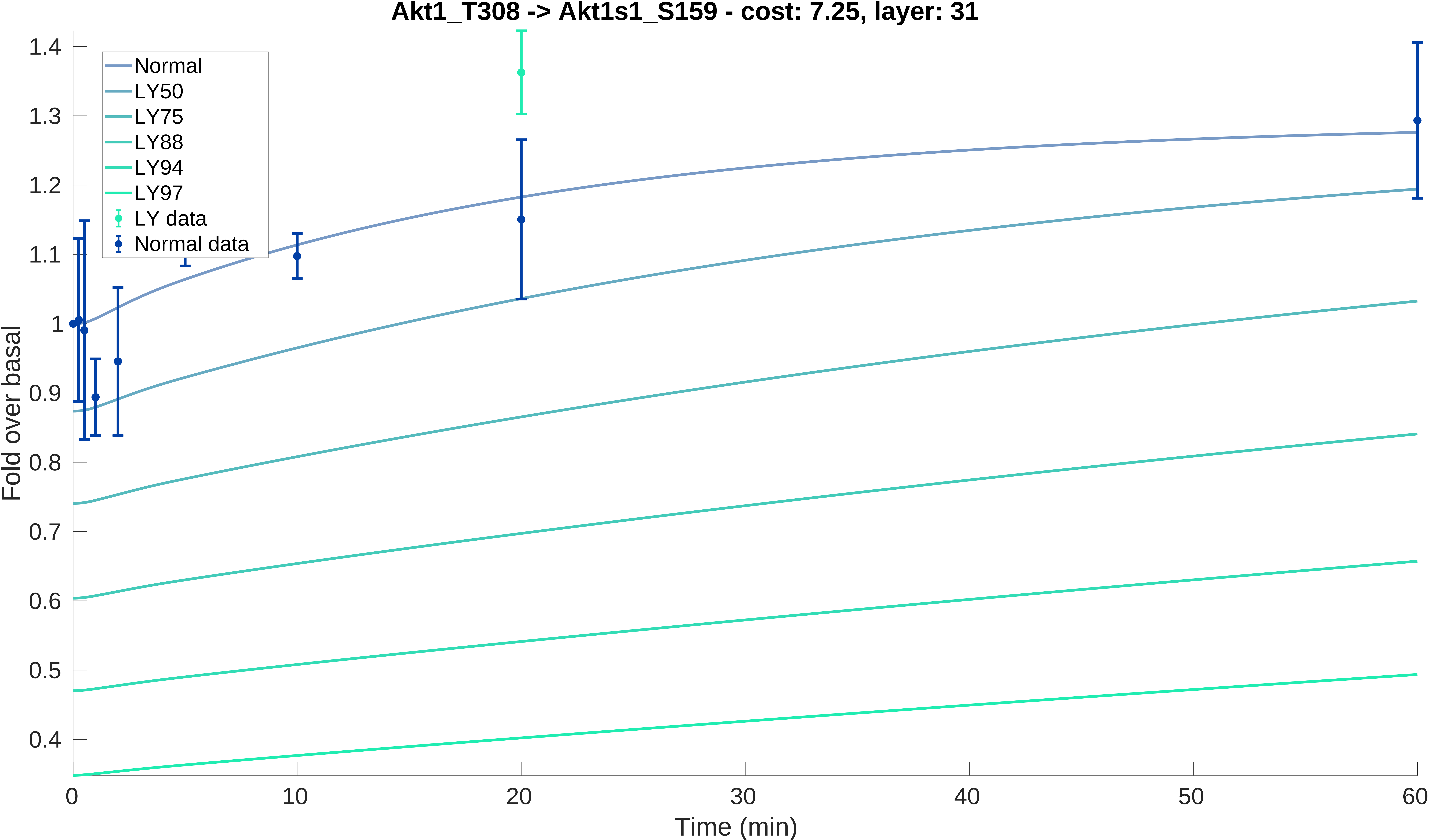

Akt1_T308 -> Akt1s1_S159 - cost: 7.25, layer: 31
Fold over basal
Time (min)
Normal
LY50
LY75
LY88
LY94
LY97
LY data
Normal data
1.4
1.3
1.2
1.1
1
0.9
0.8
0.7
0.6
0.5
0.4
0
10
20
30
40
50
60

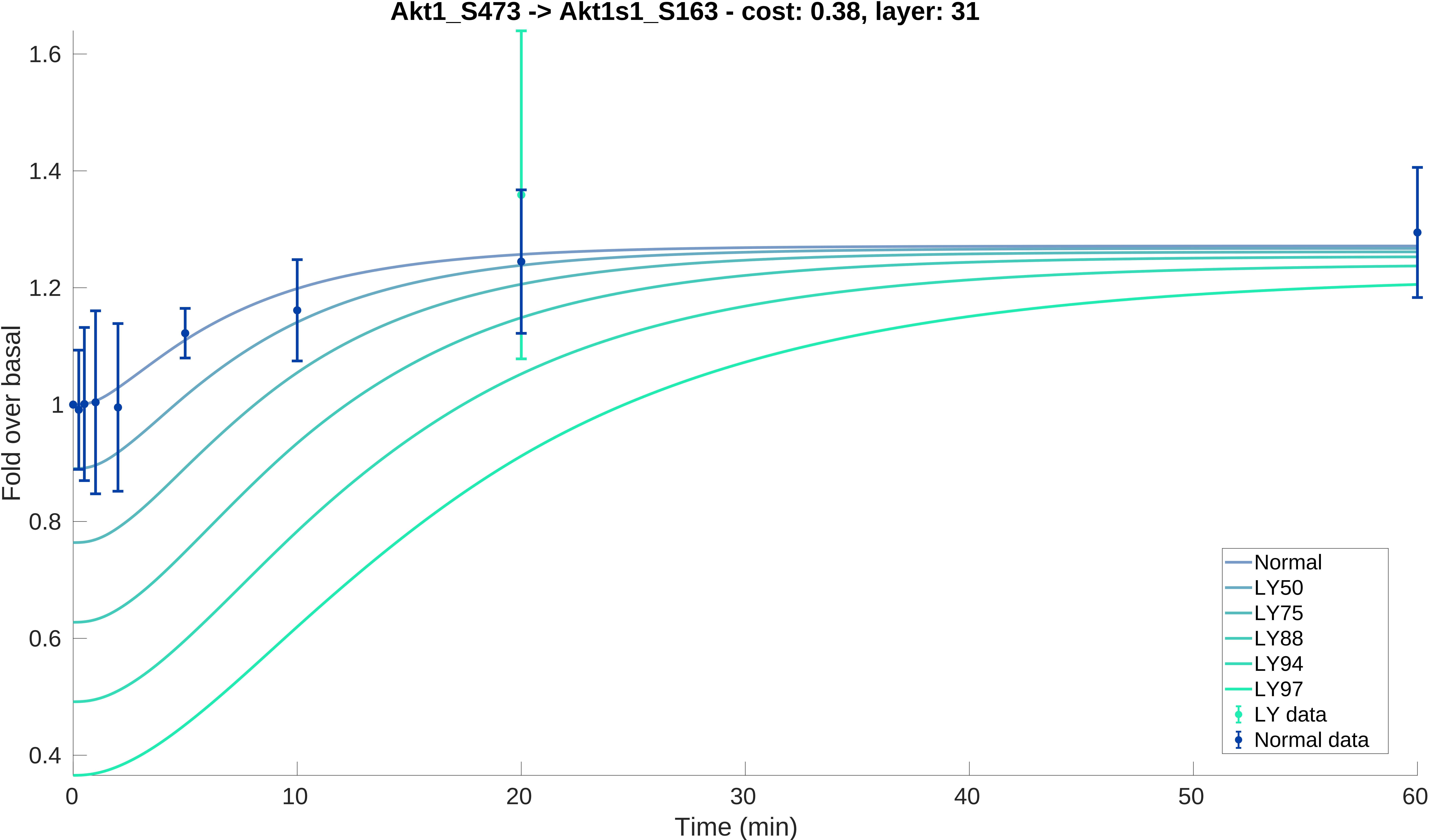
Akt1_S473 -> Akt1s1_S163 - cost: 0.38, layer: 31
Fold over basal
Time (min)
1.6
1.4
1.2
1
0.8
0.6
0.4
0
10
20
30
40
50
60
Normal
LY50
LY75
LY88
LY94
LY97
LY data
Normal data

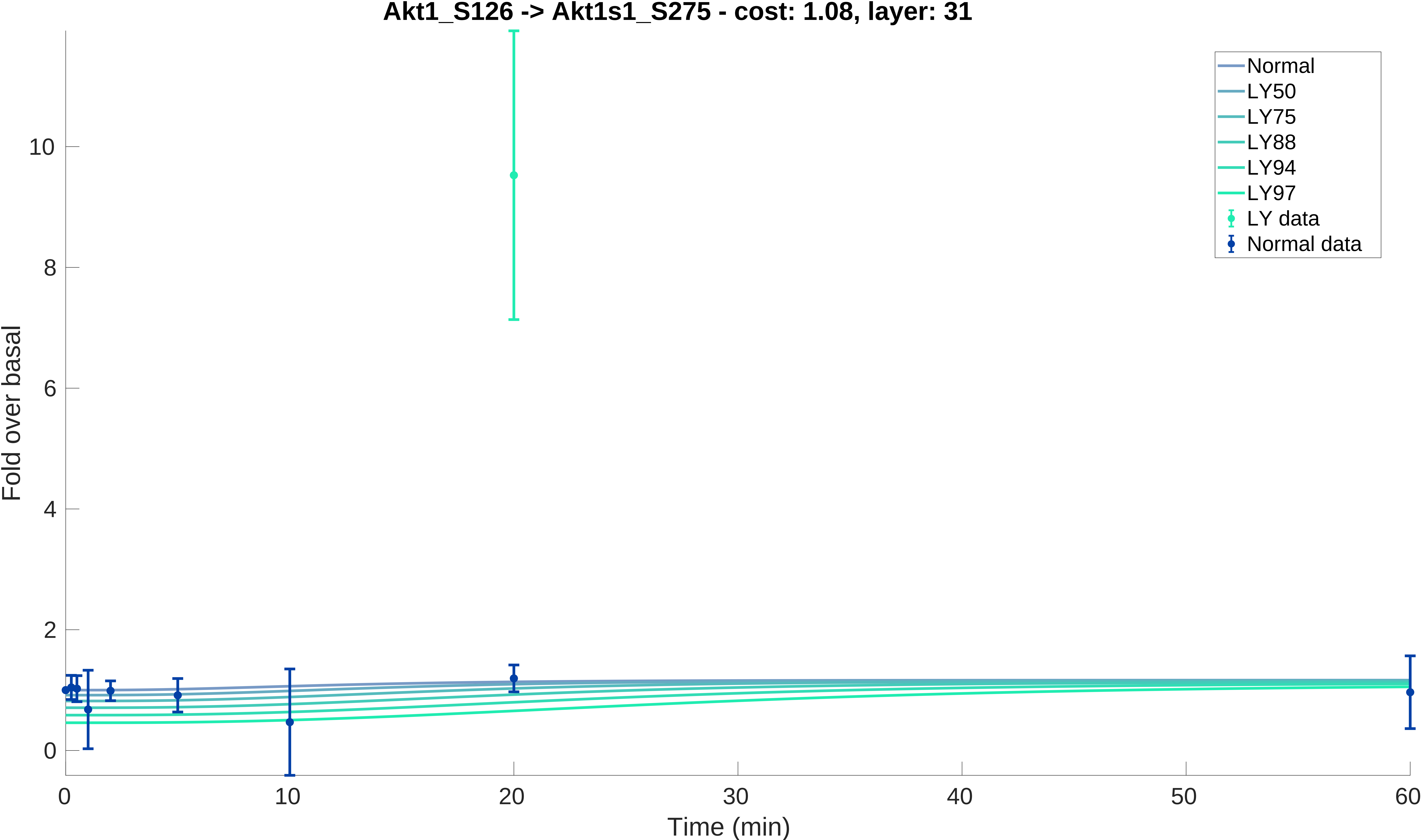

Akt1_S126 -> Akt1s1_S275 - cost: 1.08, layer: 31
Fold over basal
Time (min)
0
2
4
6
8
10
0
10
20
30
40
50
60
Normal
LY50
LY75
LY88
LY94
LY97
LY data
Normal data

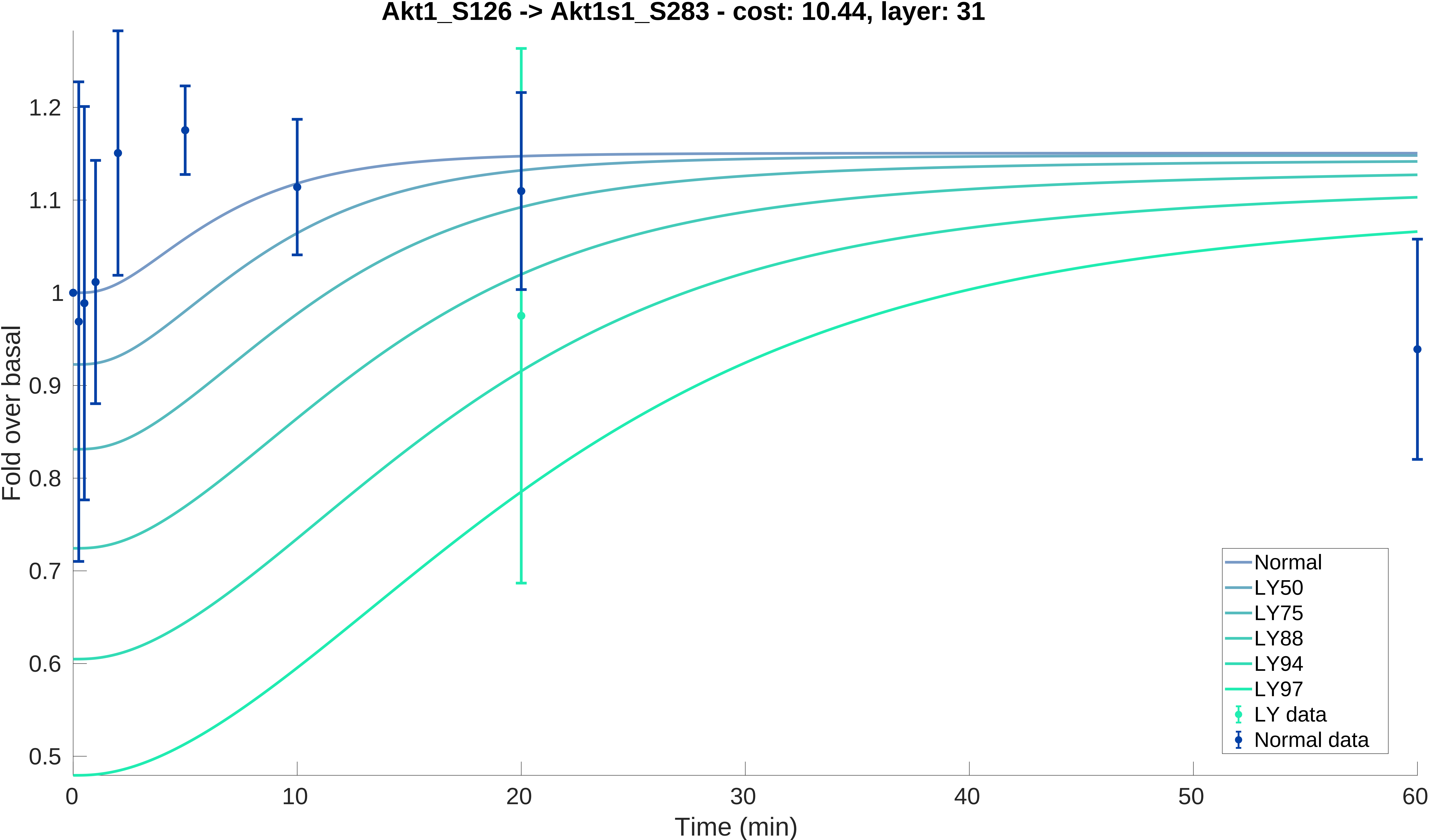
Akt1_S126 -> Akt1s1_S283 - cost: 10.44, layer: 31
Fold over basal
Time (min)
0.5
0.6
0.7
0.8
0.9
1
1.1
1.2
0
10
20
30
40
50
60
Normal
LY50
LY75
LY88
LY94
LY97
LY data
Normal data

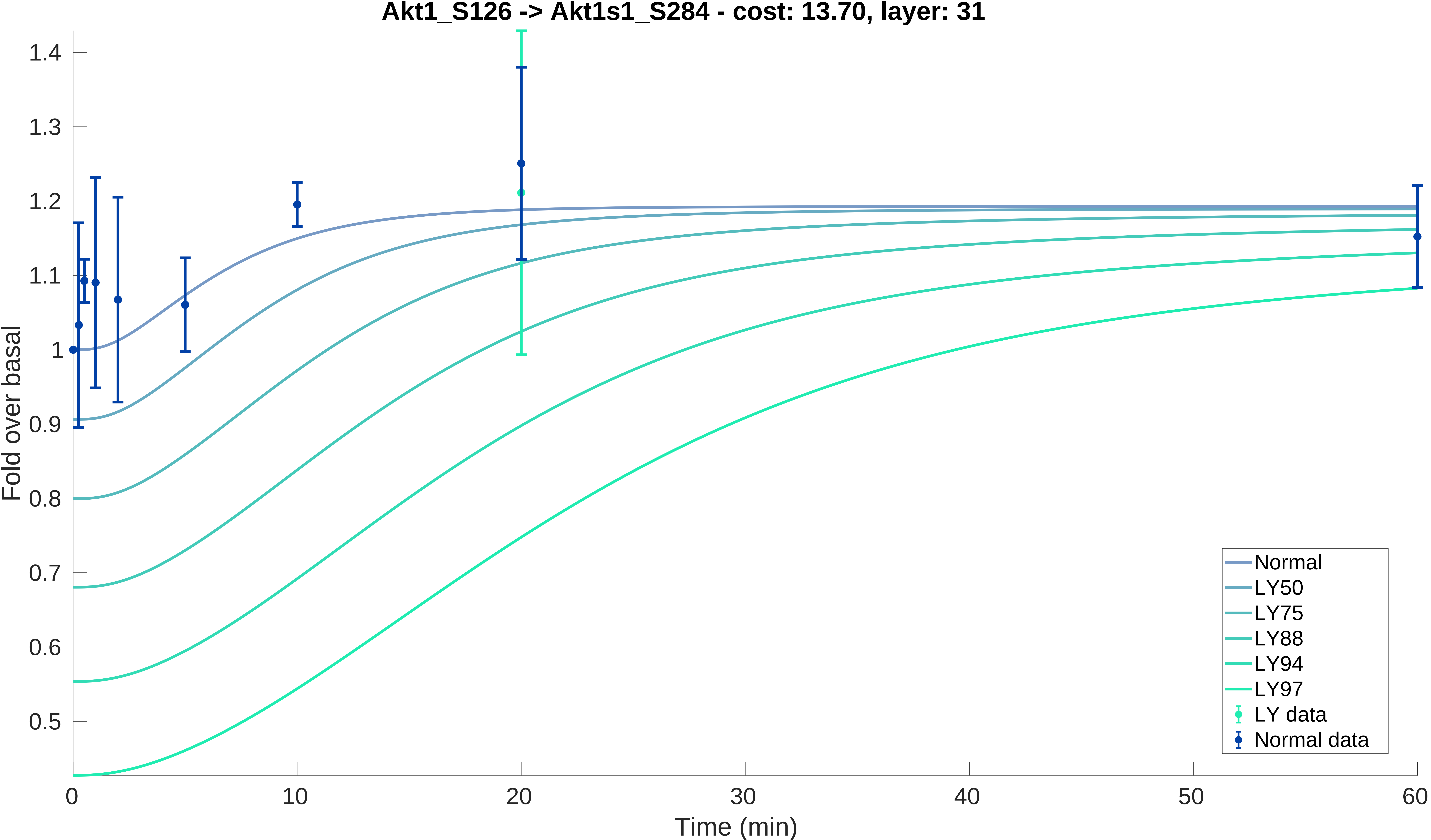
Akt1_S126 -> Akt1s1_S284 - cost: 13.70, layer: 31
Fold over basal
Time (min)
1.4
1.3
1.2
1.1
1
0.9
0.8
0.7
0.6
0.5
0
10
20
30
40
50
60
Normal
LY50
LY75
LY88
LY94
LY97
LY data
Normal data

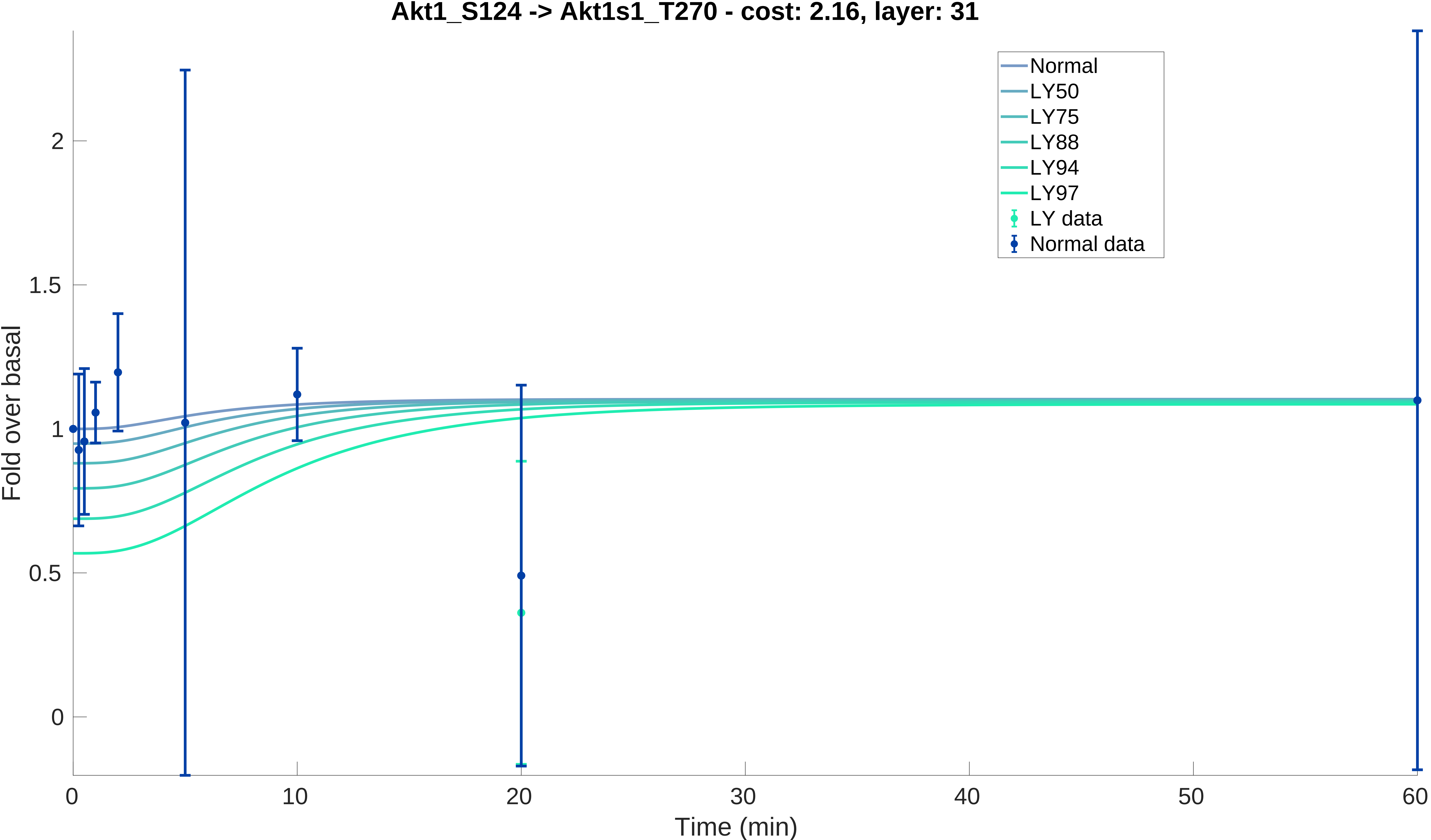

Akt1_S124 -> Akt1s1_T270 - cost: 2.16, layer: 31
Normal
LY50
LY75
LY88
LY94
LY97
LY data
Normal data
Fold over basal
Time (min)
0
0.5
1
1.5
2
0
10
20
30
40
50
60

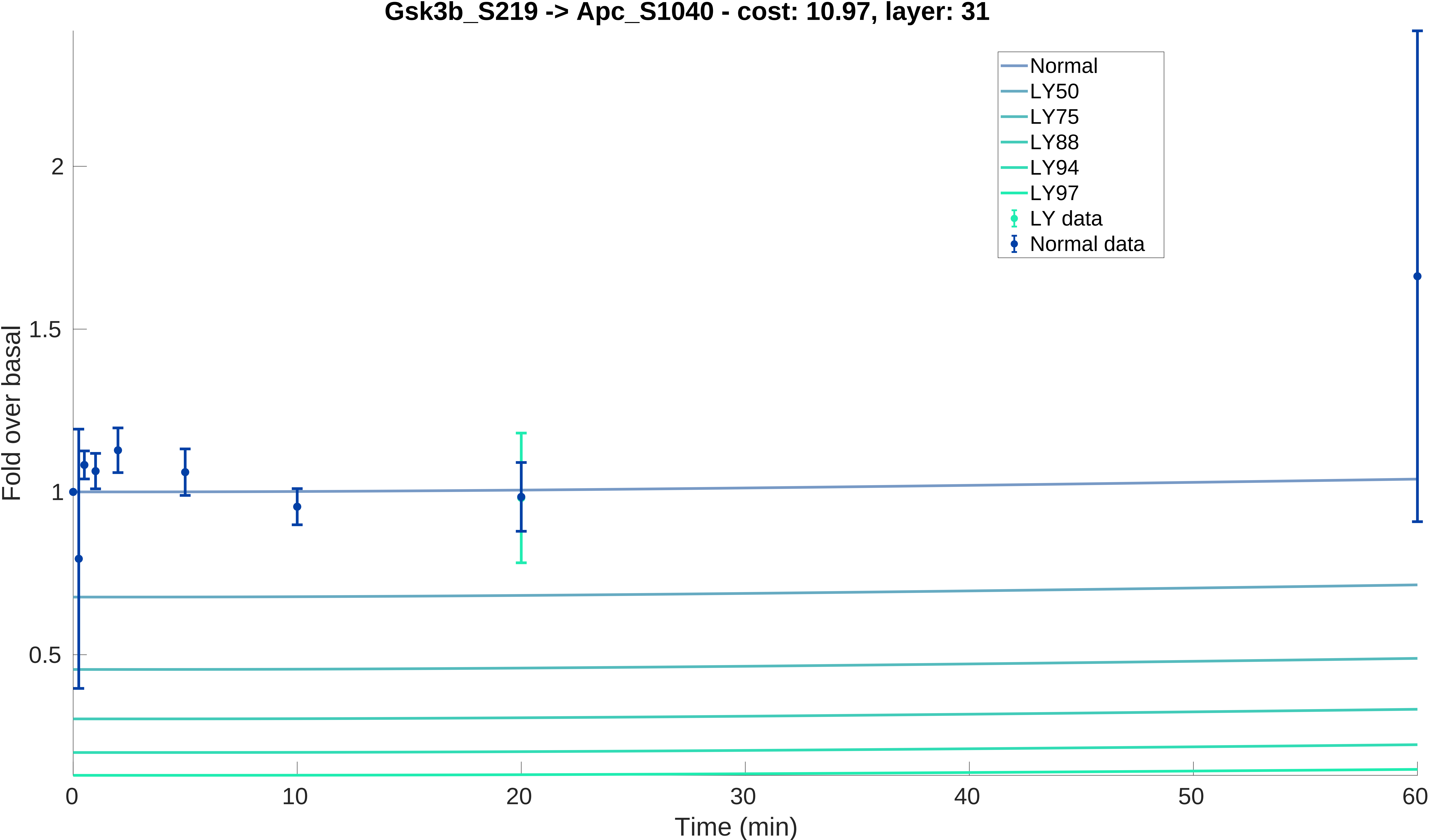

Gsk3b_S219 -> Apc_S1040 - cost: 10.97, layer: 31
Normal
LY50
LY75
LY88
LY94
LY97
LY data
Normal data
Fold over basal
2
1.5
1
0.5
0
10
20
30
40
50
60
Time (min)

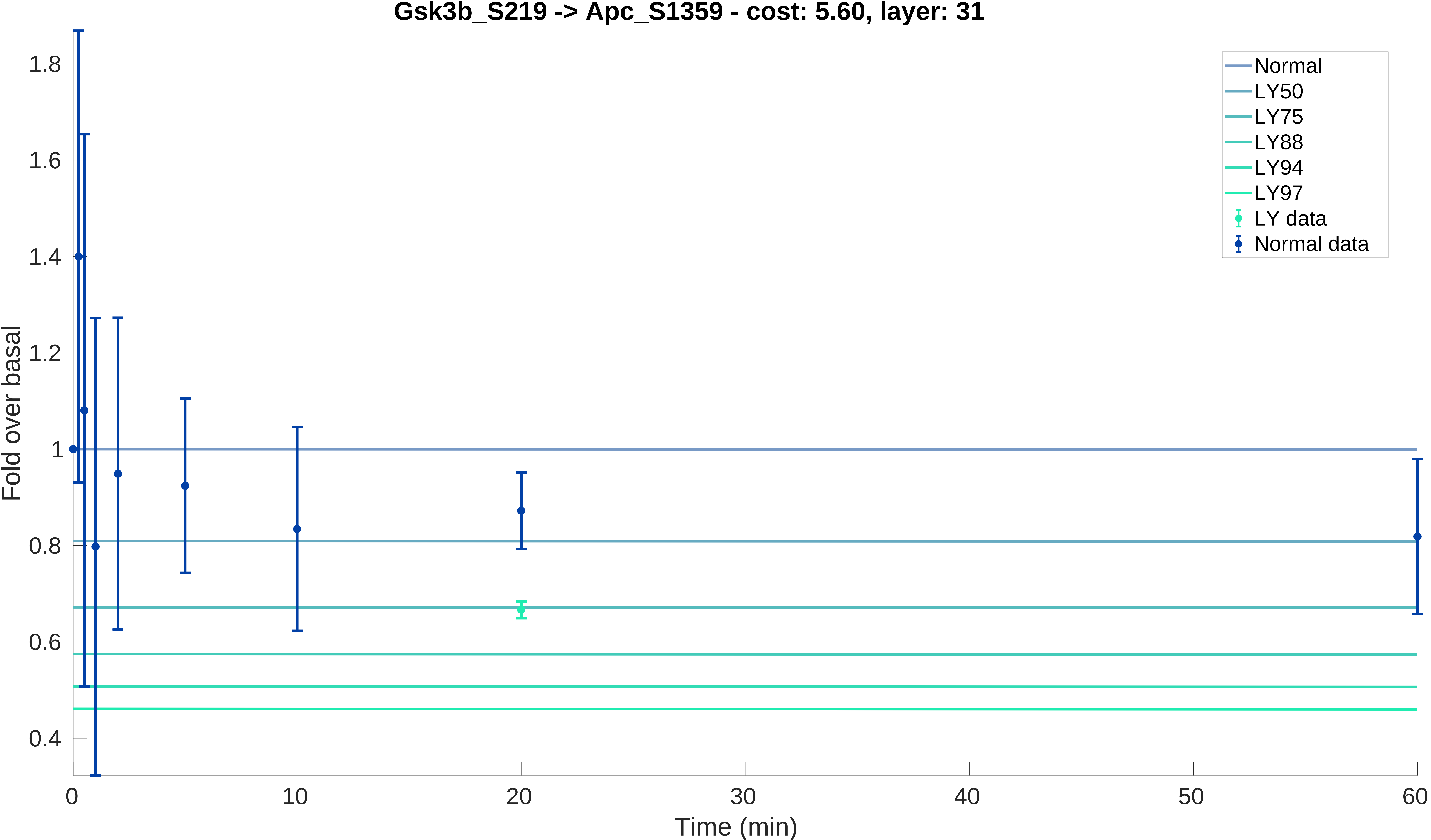

Gsk3b_S219 -> Apc_S1359 - cost: 5.60, layer: 31
Fold over basal
Time (min)
0.4
0.6
0.8
1
1.2
1.4
1.6
1.8
0
10
20
30
40
50
60
Normal
LY50
LY75
LY88
LY94
LY97
LY data
Normal data

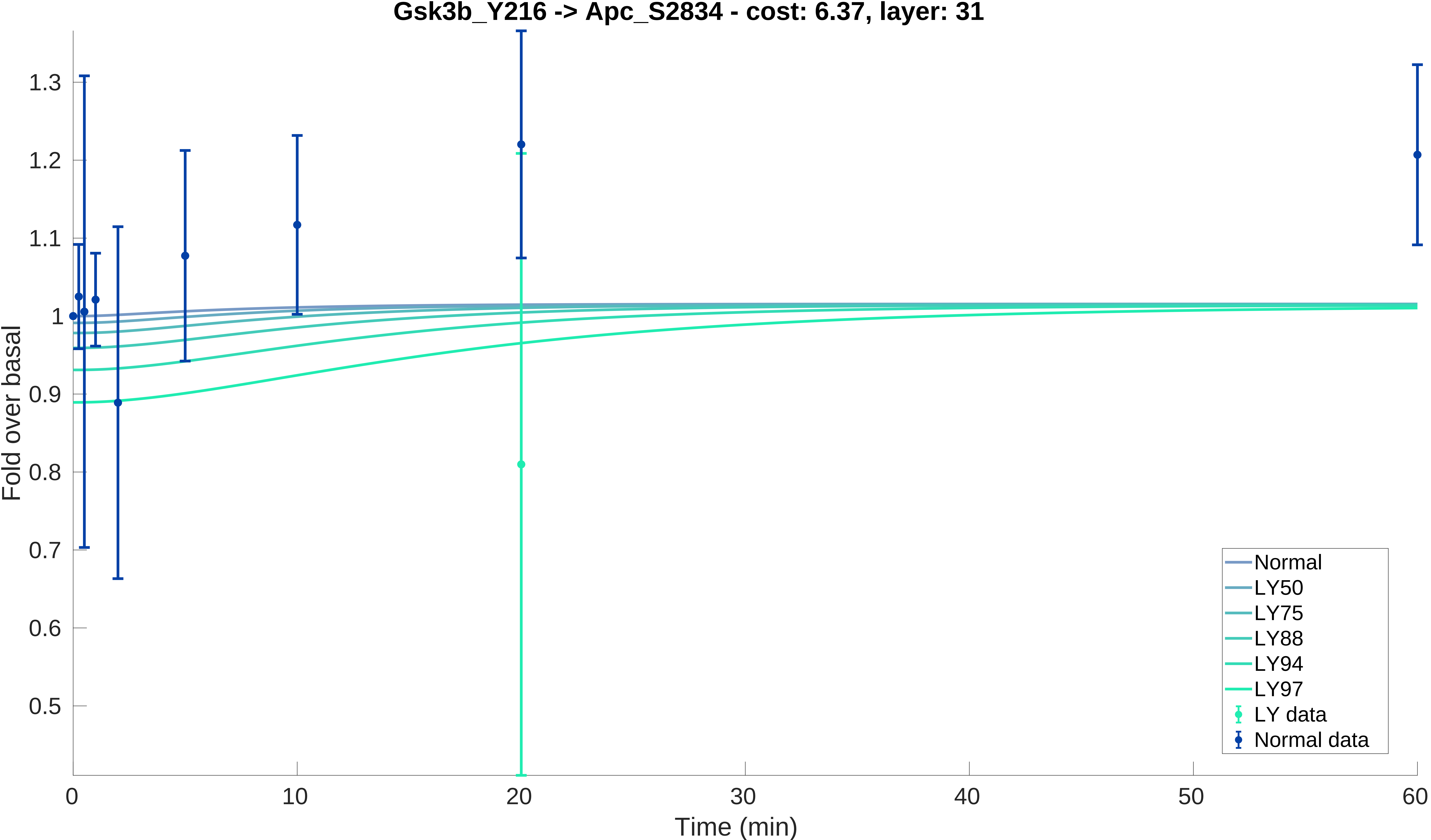

Gsk3b_Y216 -> Apc_S2834 - cost: 6.37, layer: 31
Fold over basal
Time (min)
1.3
1.2
1.1
1
0.9
0.8
0.7
0.6
0.5
0
10
20
30
40
50
60
Normal
LY50
LY75
LY88
LY94
LY97
LY data
Normal data

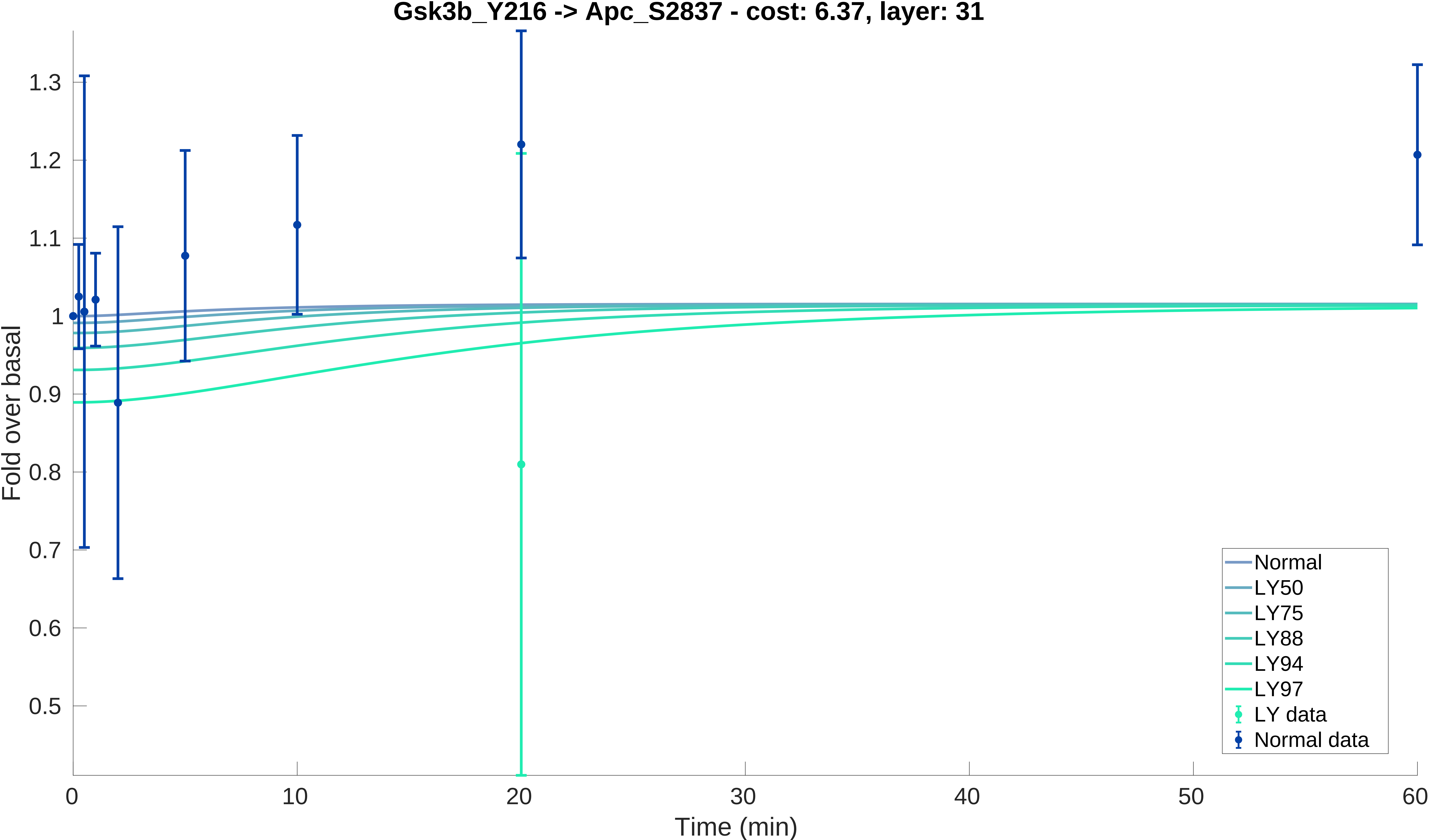
Gsk3b_Y216 -> Apc_S2837 - cost: 6.37, layer: 31
Fold over basal
1.3
1.2
1.1
1
0.9
0.8
0.7
0.6
0.5
0
10
20
30
40
50
60
Time (min)
Normal
LY50
LY75
LY88
LY94
LY97
LY data
Normal data

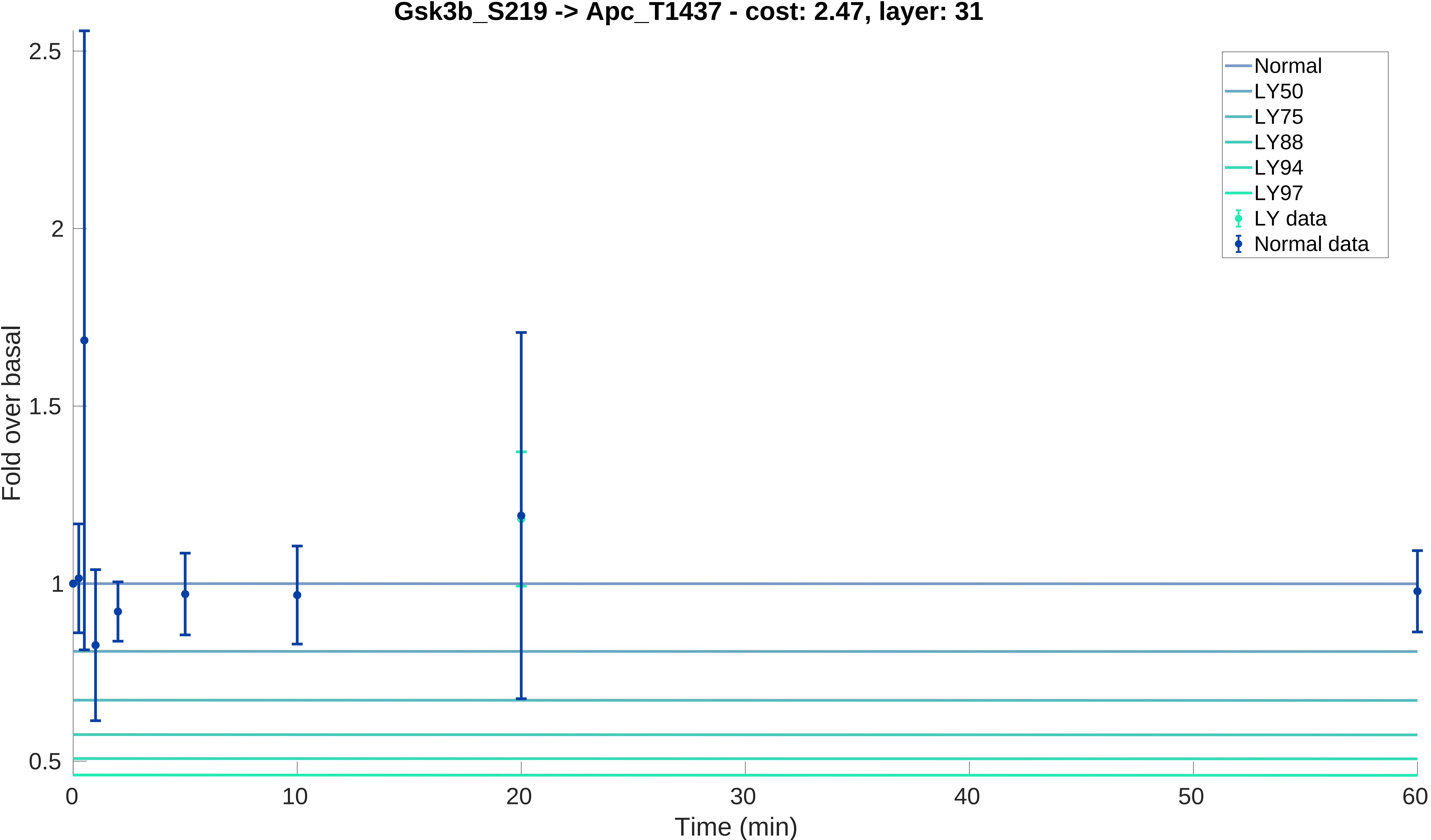

Gsk3b_S219 -> Apc_T1437 - cost: 2.47, layer: 31
Fold over basal
Time (min)
2.5
2
1.5
1
0.5
0
10
20
30
40
50
60
Normal
LY50
LY75
LY88
LY94
LY97
LY data
Normal data

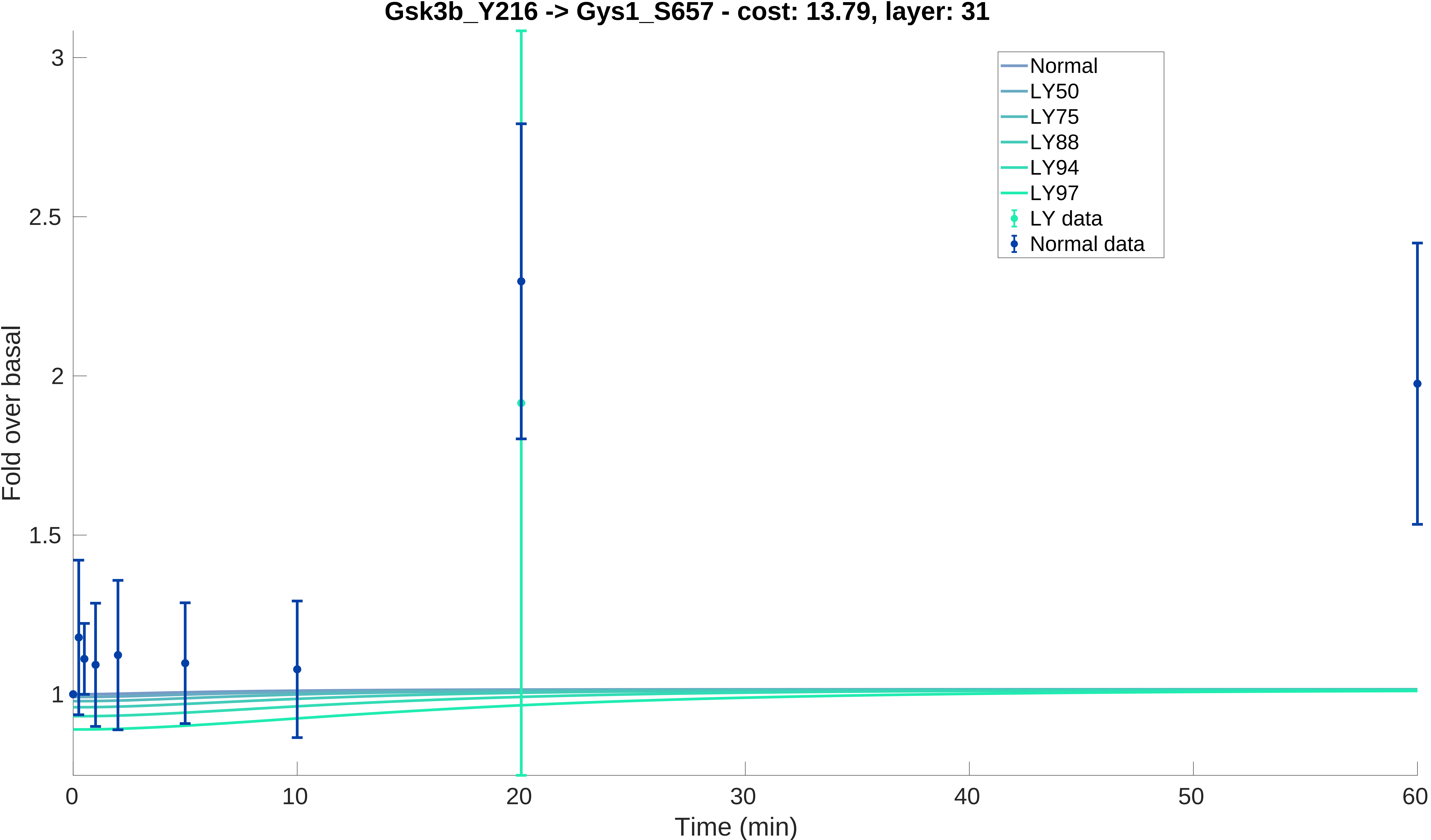

Gsk3b_Y216 -> Gys1_S657 - cost: 13.79, layer: 31
Fold over basal
Time (min)
3
2.5
2
1.5
1
0
10
20
30
40
50
60
Normal
LY50
LY75
LY88
LY94
LY97
LY data
Normal data

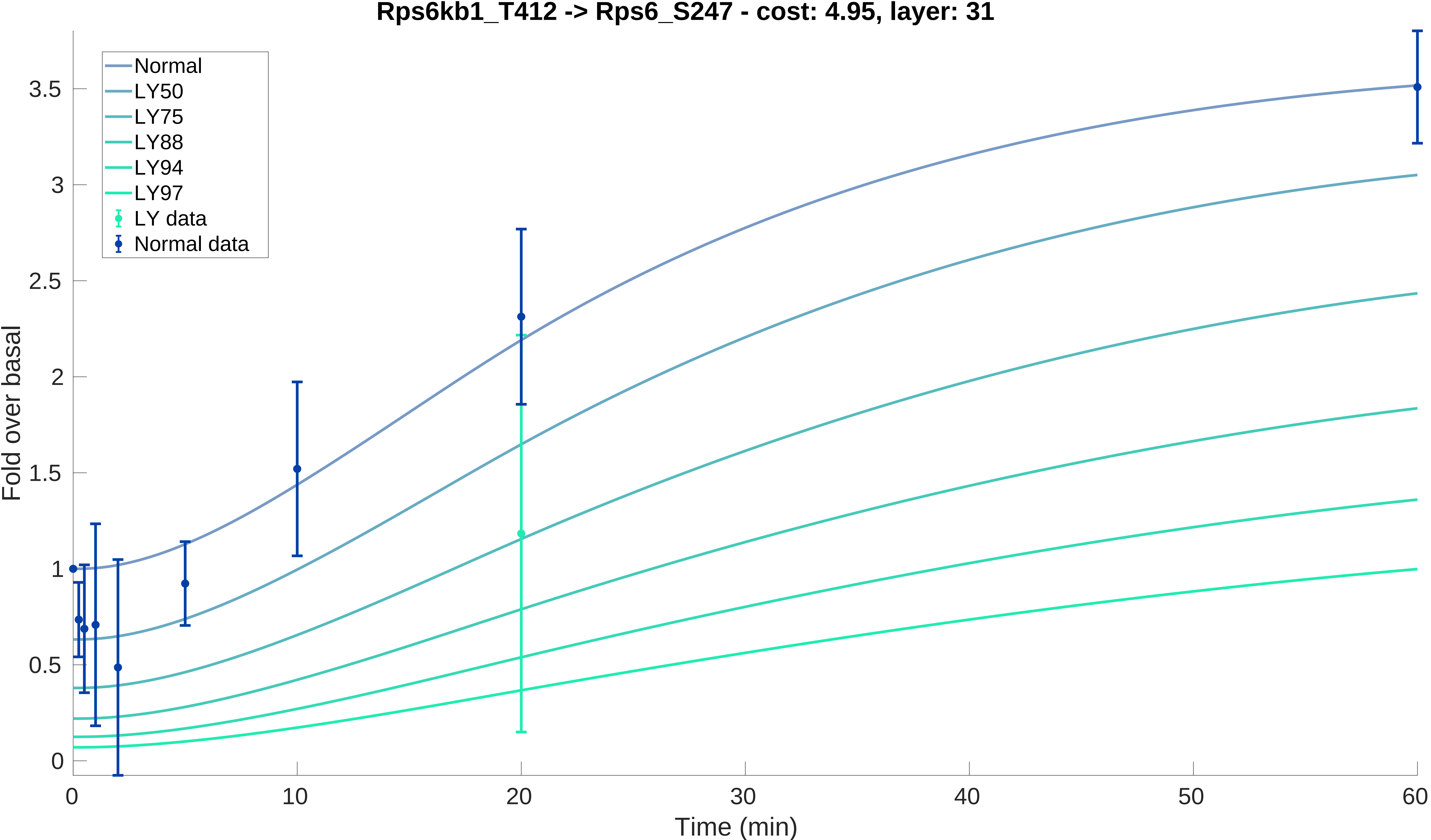

Rps6kb1_T412 -> Rps6_S247 - cost: 4.95, layer: 31
Normal
LY50
LY75
LY88
LY94
LY97
LY data
Normal data
Fold over basal
Time (min)
0
0.5
1
1.5
2
2.5
3
3.5
0
10
20
30
40
50
60

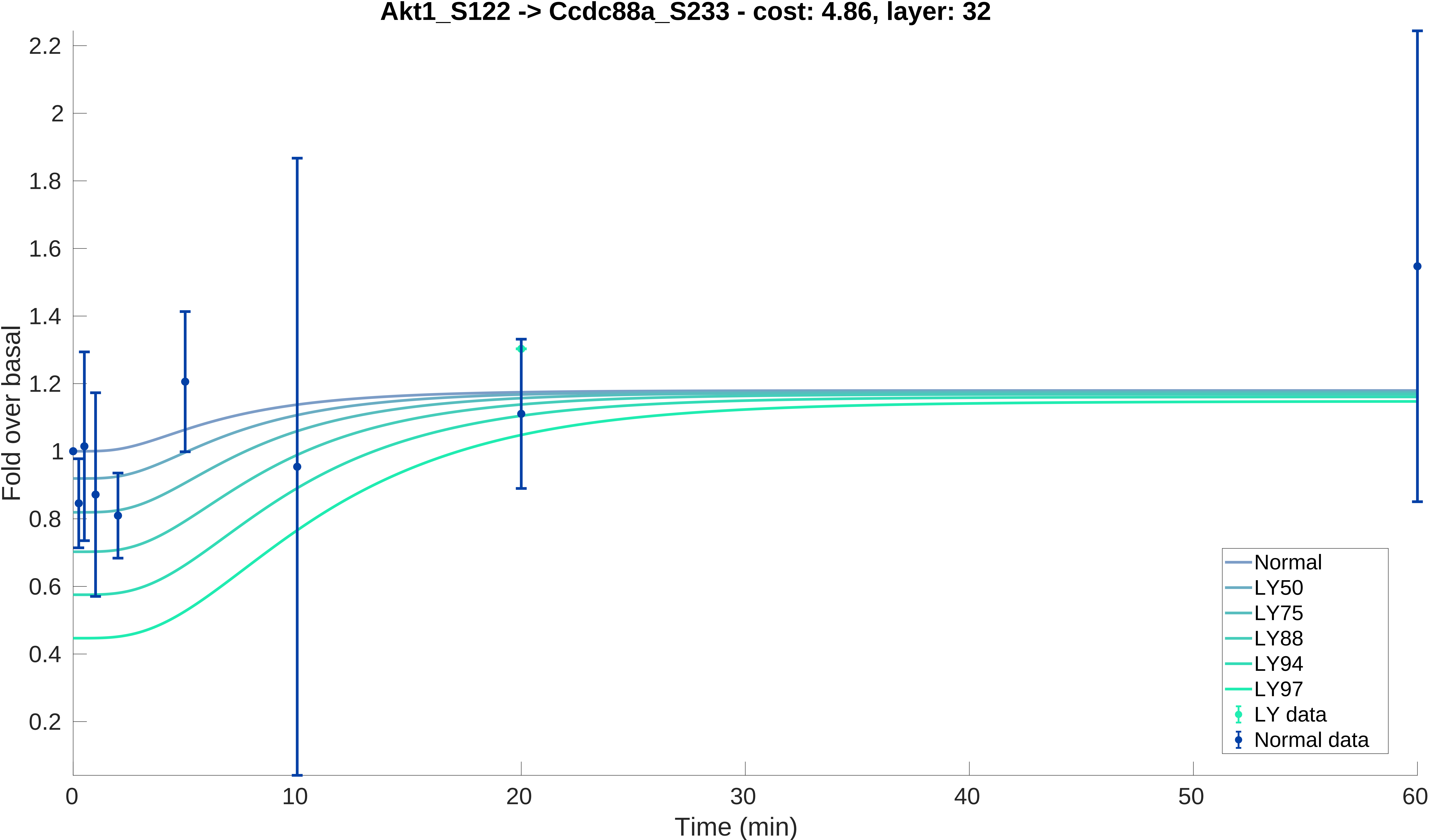

Akt1_S122 -> Ccdc88a_S233 - cost: 4.86, layer: 32
Fold over basal
Time (min)
0.2
0.4
0.6
0.8
1
1.2
1.4
1.6
1.8
2
2.2
0
10
20
30
40
50
60
Normal
LY50
LY75
LY88
LY94
LY97
LY data
Normal data

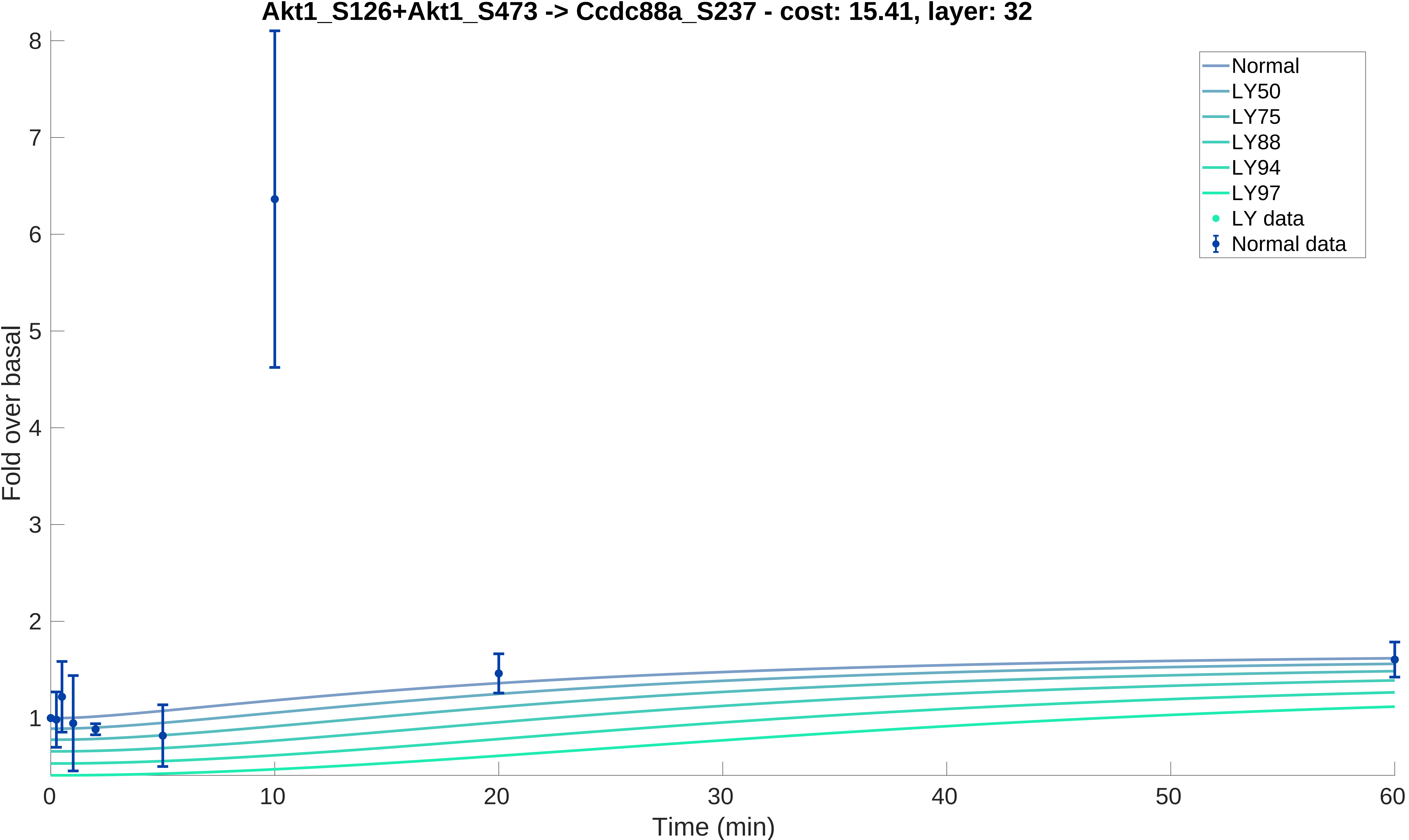

Akt1_S126+Akt1_S473 -> Ccdc88a_S237 - cost: 15.41, layer: 32
Fold over basal
1
2
3
4
5
6
7
8
0
10
20
30
40
50
60
Time (min)
Normal
LY50
LY75
LY88
LY94
LY97
LY data
Normal data

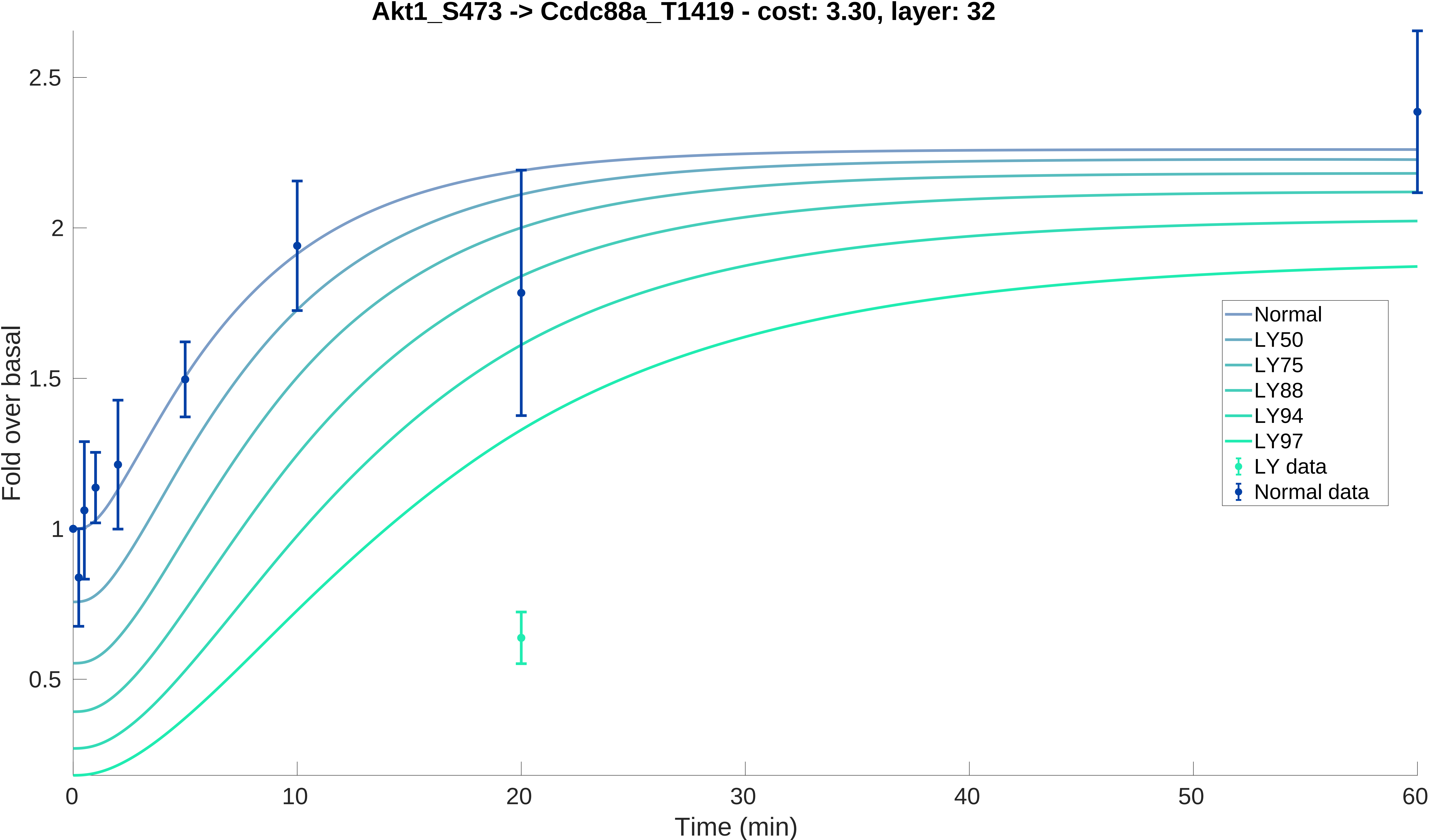

Akt1_S473 -> Ccdc88a_T1419 - cost: 3.30, layer: 32
Fold over basal
Time (min)
0.5
1
1.5
2
2.5
0
10
20
30
40
50
60
Normal
LY50
LY75
LY88
LY94
LY97
LY data
Normal data

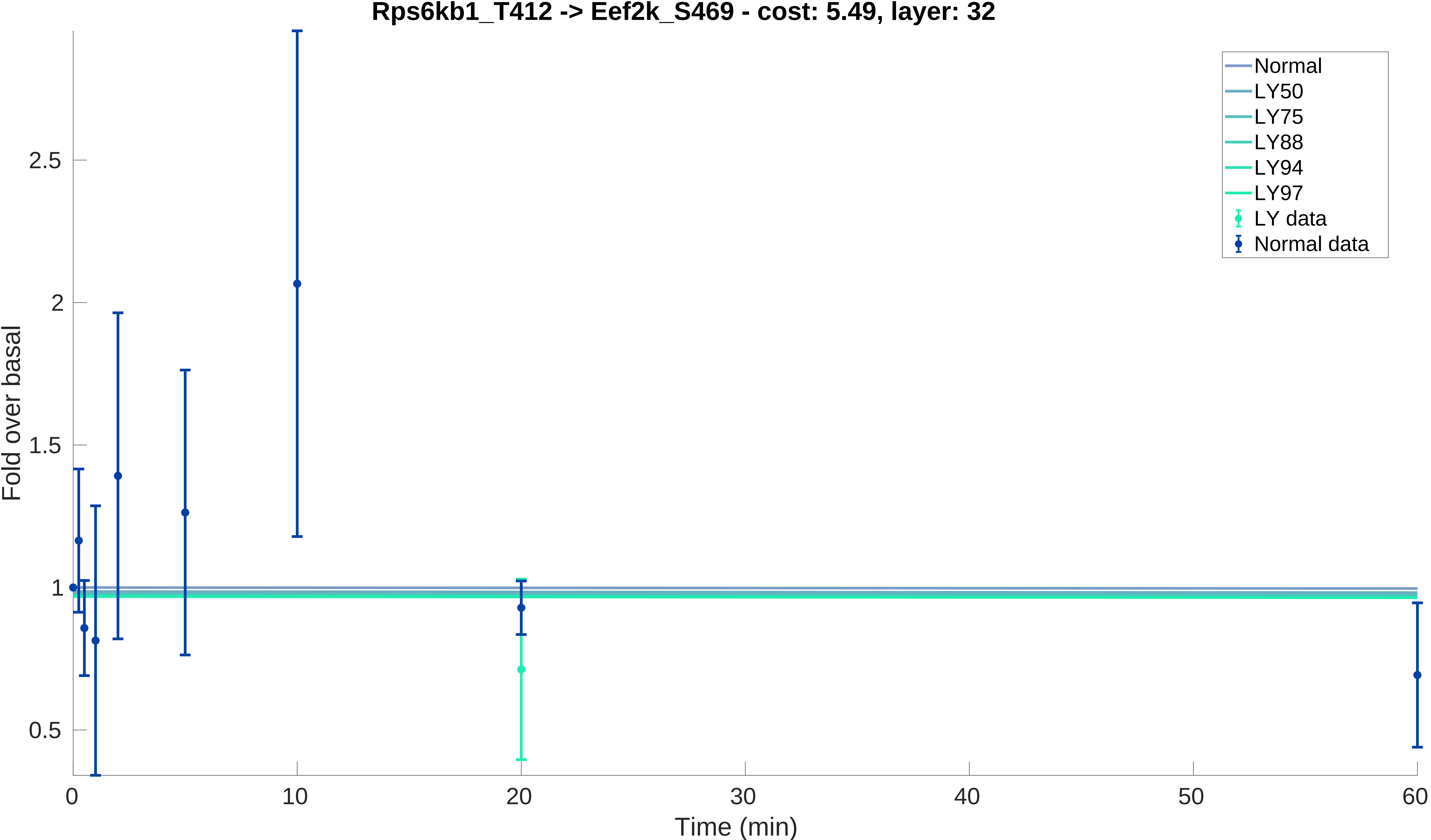

Rps6kb1_T412 -> Eef2k_S469 - cost: 5.49, layer: 32
Fold over basal
Time (min)
0.5
1
1.5
2
2.5
0
10
20
30
40
50
60
Normal
LY50
LY75
LY88
LY94
LY97
LY data
Normal data

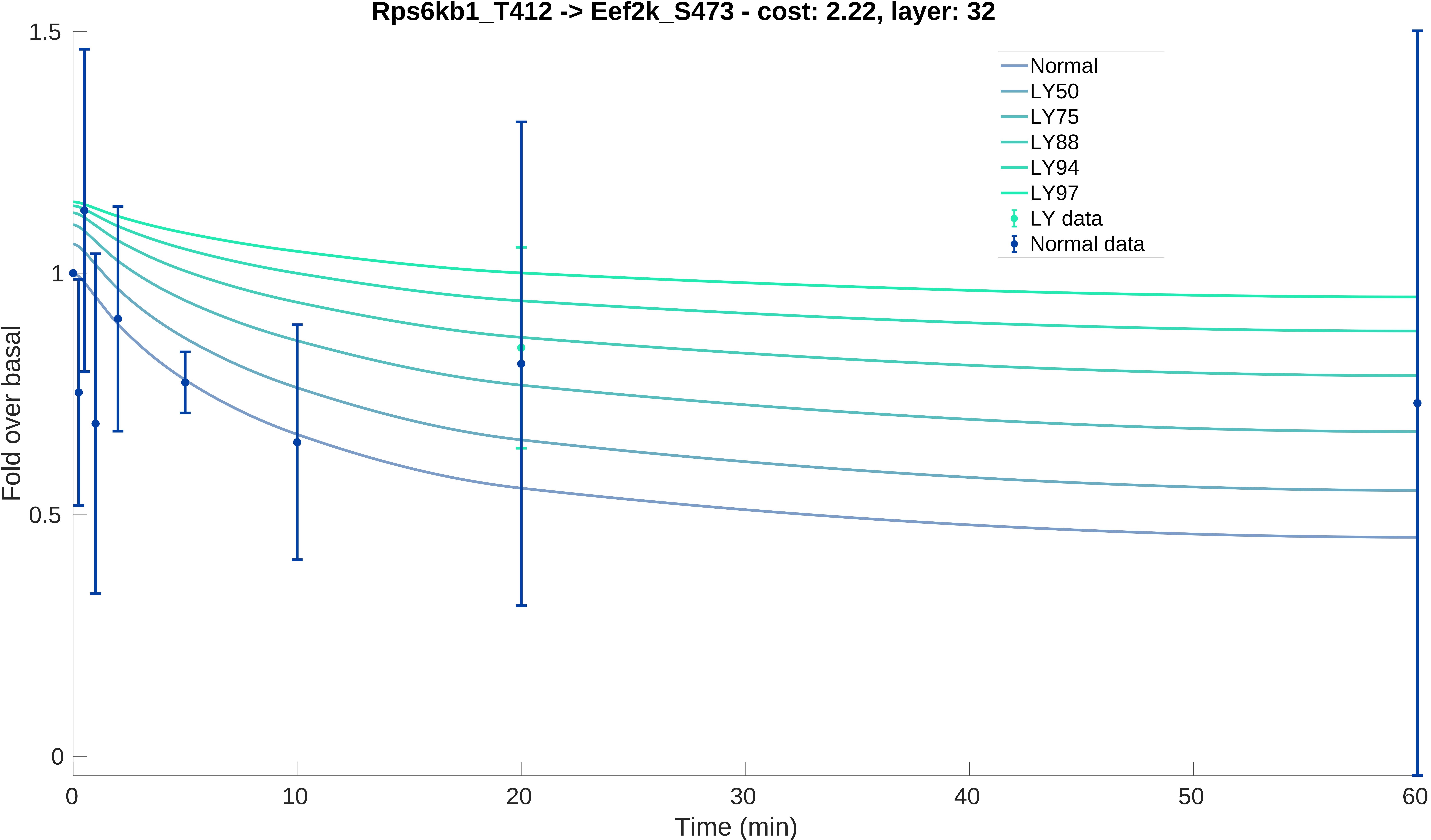
Rps6kb1_T412 -> Eef2k_S473 - cost: 2.22, layer: 32
1.5
1
0.5
0
Fold over basal
0
10
20
30
40
50
60
Time (min)
Normal
LY50
LY75
LY88
LY94
LY97
LY data
Normal data

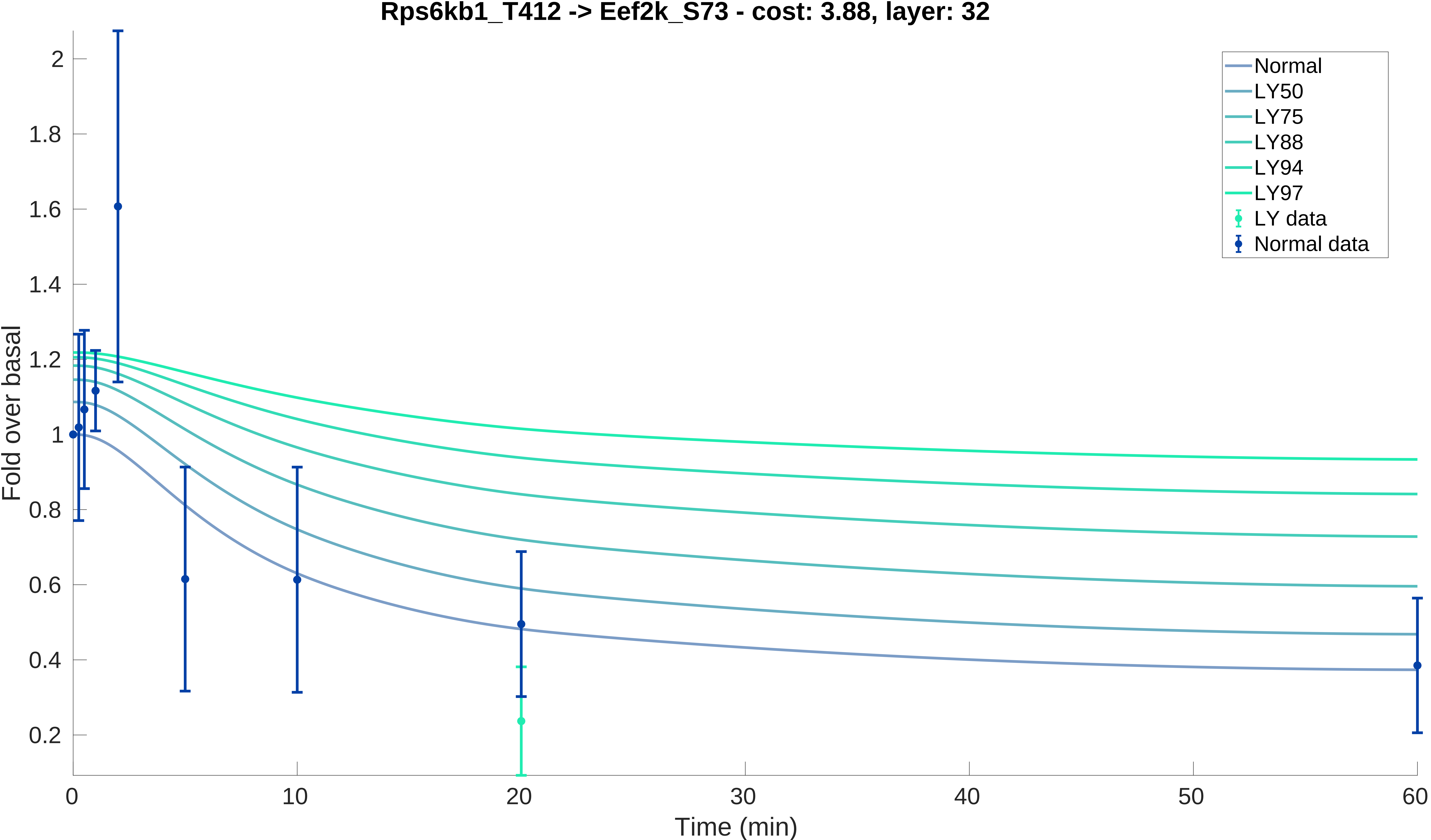

Rps6kb1_T412 -> Eef2k_S73 - cost: 3.88, layer: 32
Normal
LY50
LY75
LY88
LY94
LY97
LY data
Normal data
Fold over basal
Time (min)
0
10
20
30
40
50
60
0.2
0.4
0.6
0.8
1
1.2
1.4
1.6
1.8
2

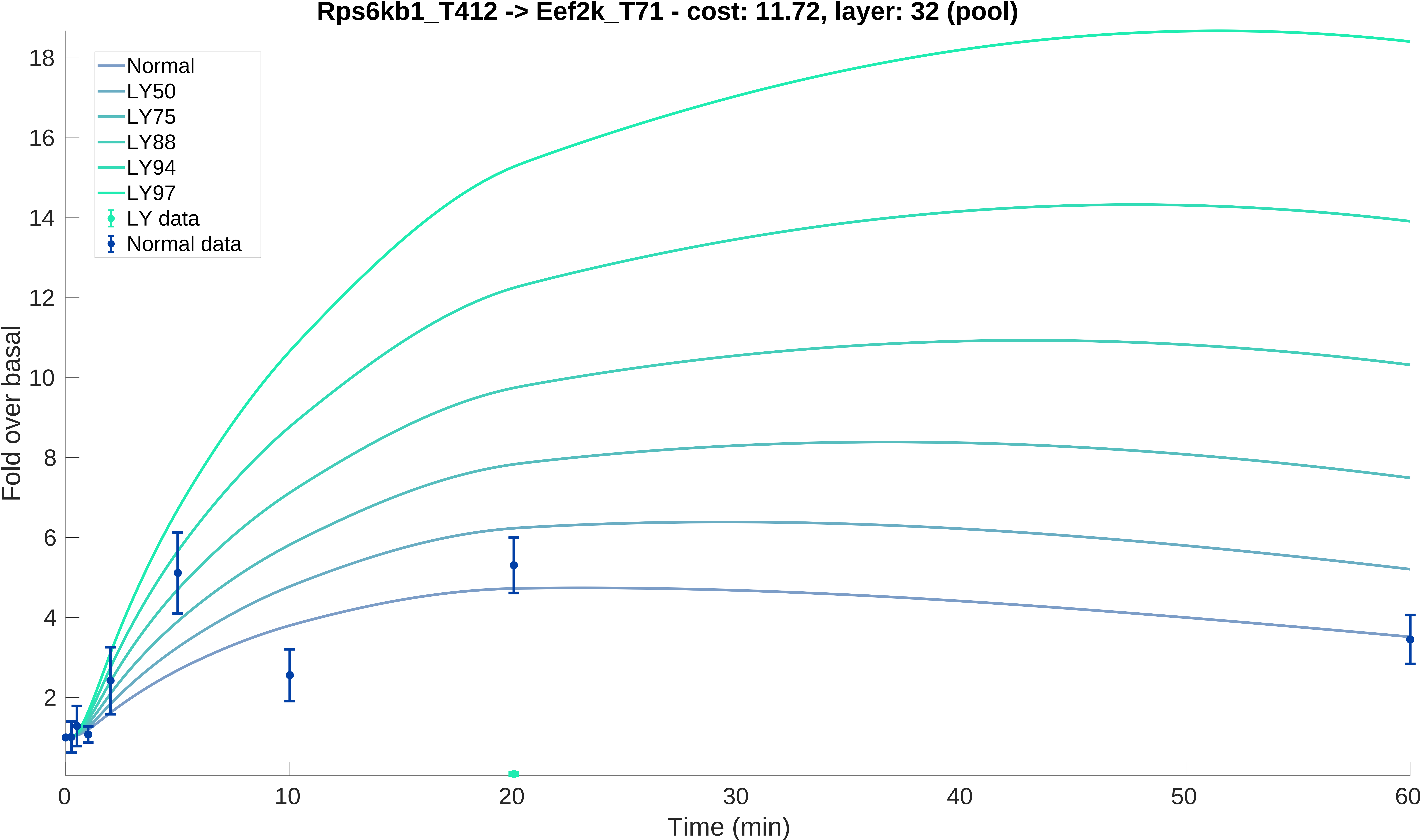

Rps6kb1_T412 -> Eef2k_T71 - cost: 11.72, layer: 32 (pool)
Normal
LY50
LY75
LY88
LY94
LY97
LY data
Normal data
Fold over basal
Time (min)
0
10
20
30
40
50
60
2
4
6
8
10
12
14
16
18

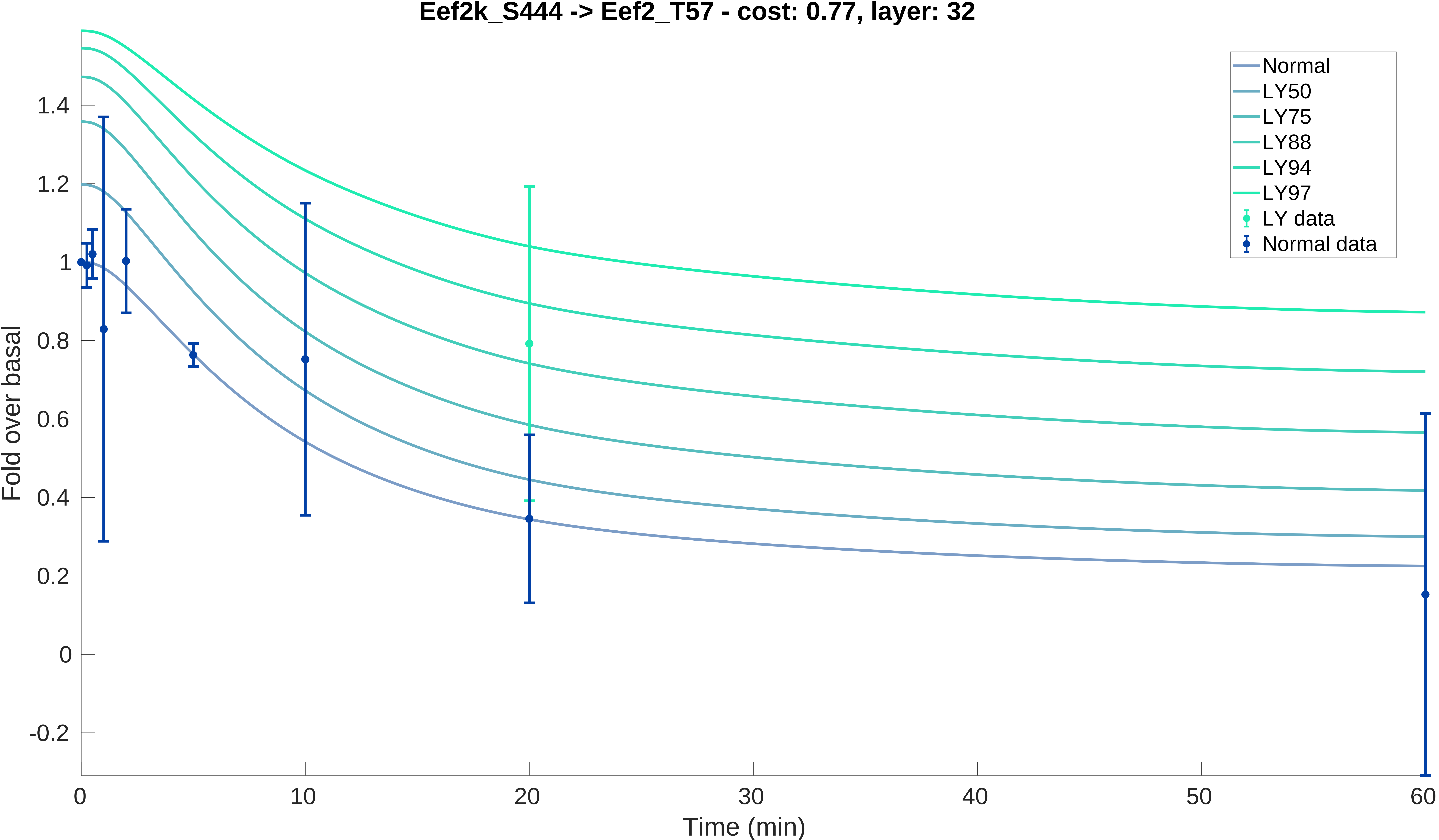

Eef2k_S444 -> Eef2_T57 - cost: 0.77, layer: 32
Normal
LY50
LY75
LY88
LY94
LY97
LY data
Normal data
Fold over basal
1.4
1.2
1
0.8
0.6
0.4
0.2
0
-0.2
0
10
20
30
40
50
60
Time (min)

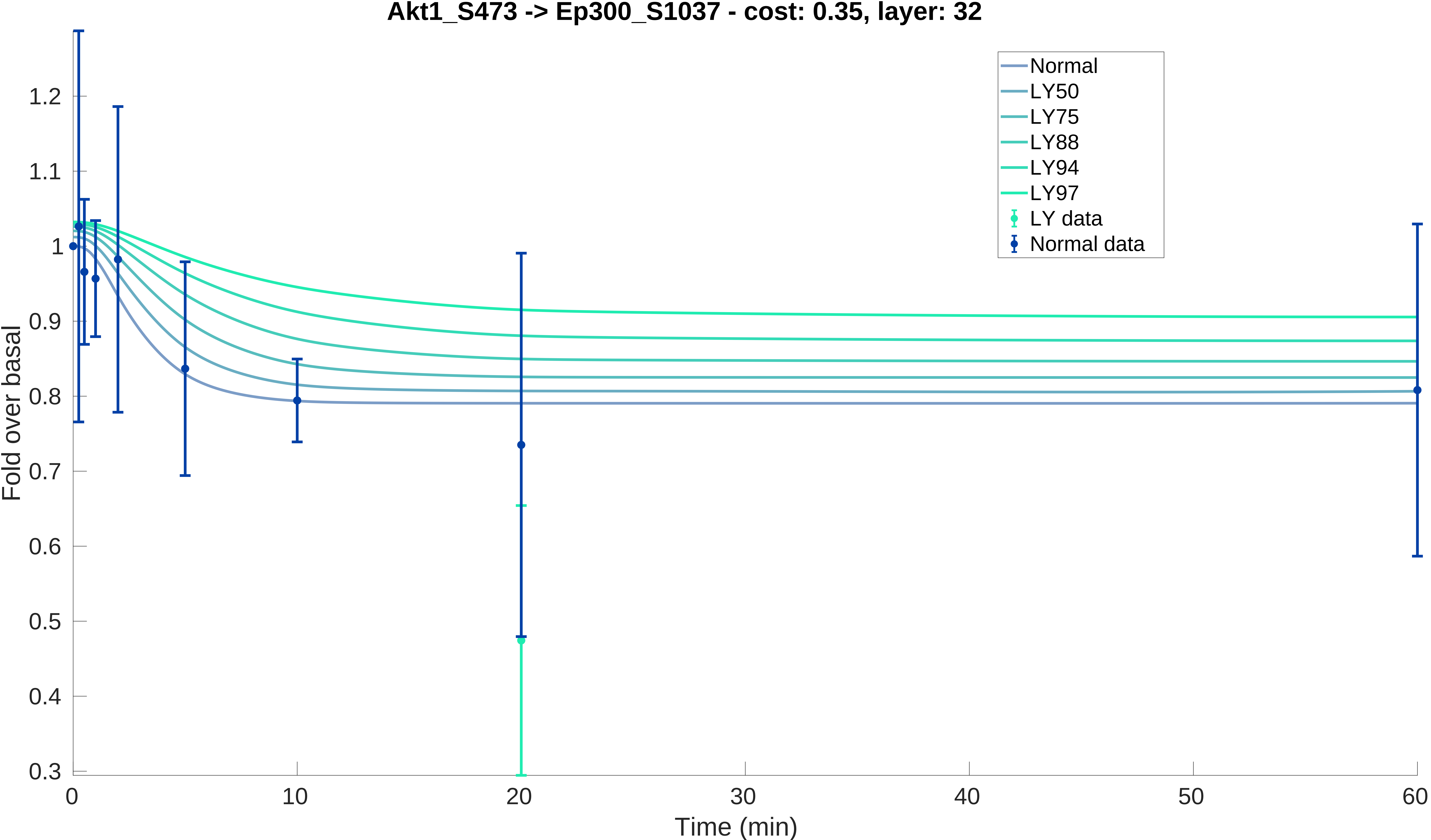
Akt1_S473 -> Ep300_S1037 - cost: 0.35, layer: 32
Normal
LY50
LY75
LY88
LY94
LY97
LY data
Normal data
Fold over basal
1.2
1.1
1
0.9
0.8
0.7
0.6
0.5
0.4
0.3
0
10
20
30
40
50
60
Time (min)

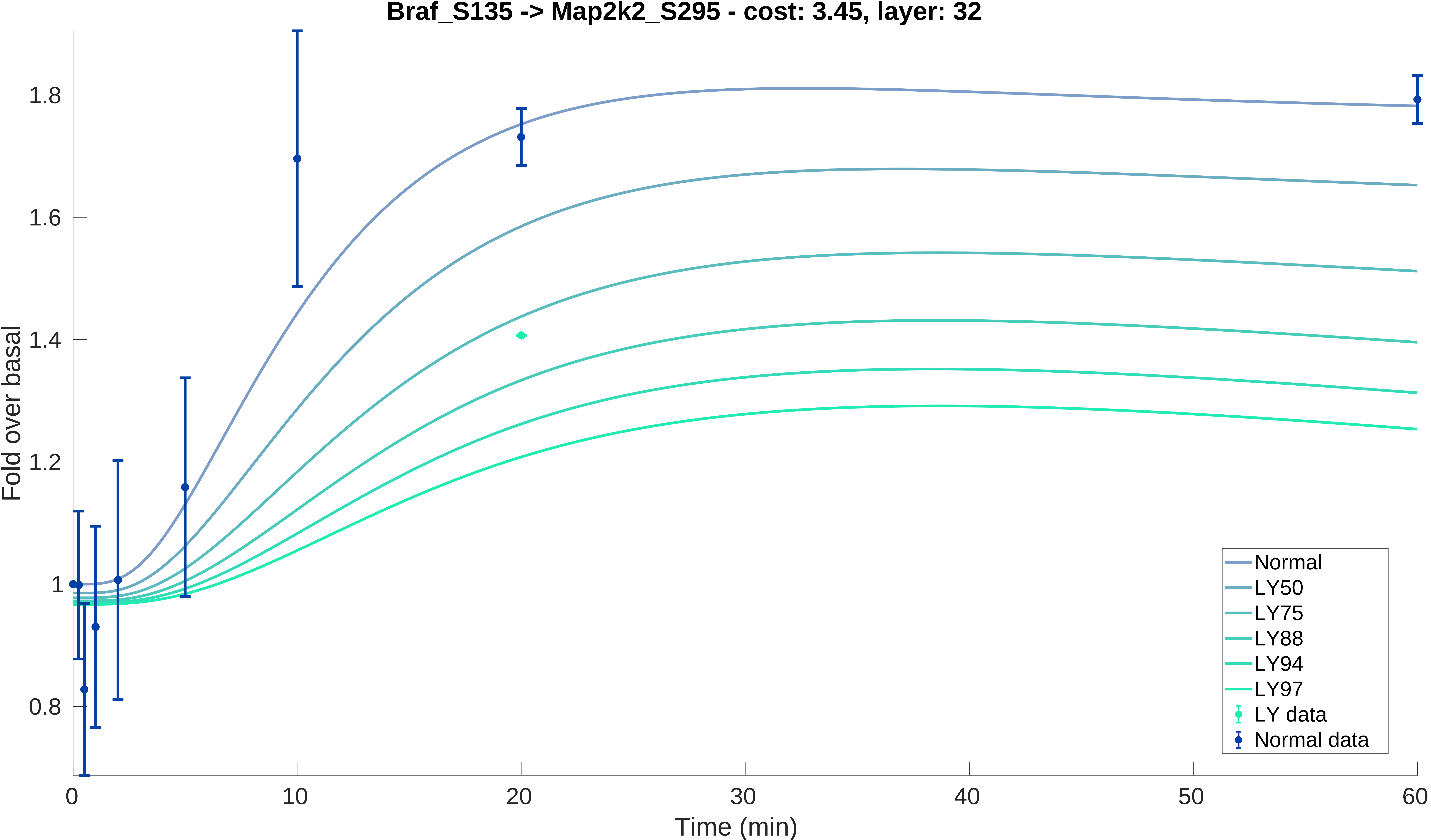

Braf_S135 -> Map2k2_S295 - cost: 3.45, layer: 32
Fold over basal
1.8
1.6
1.4
1.2
1
0.8
0
10
20
30
40
50
60
Time (min)
Normal
LY50
LY75
LY88
LY94
LY97
LY data
Normal data

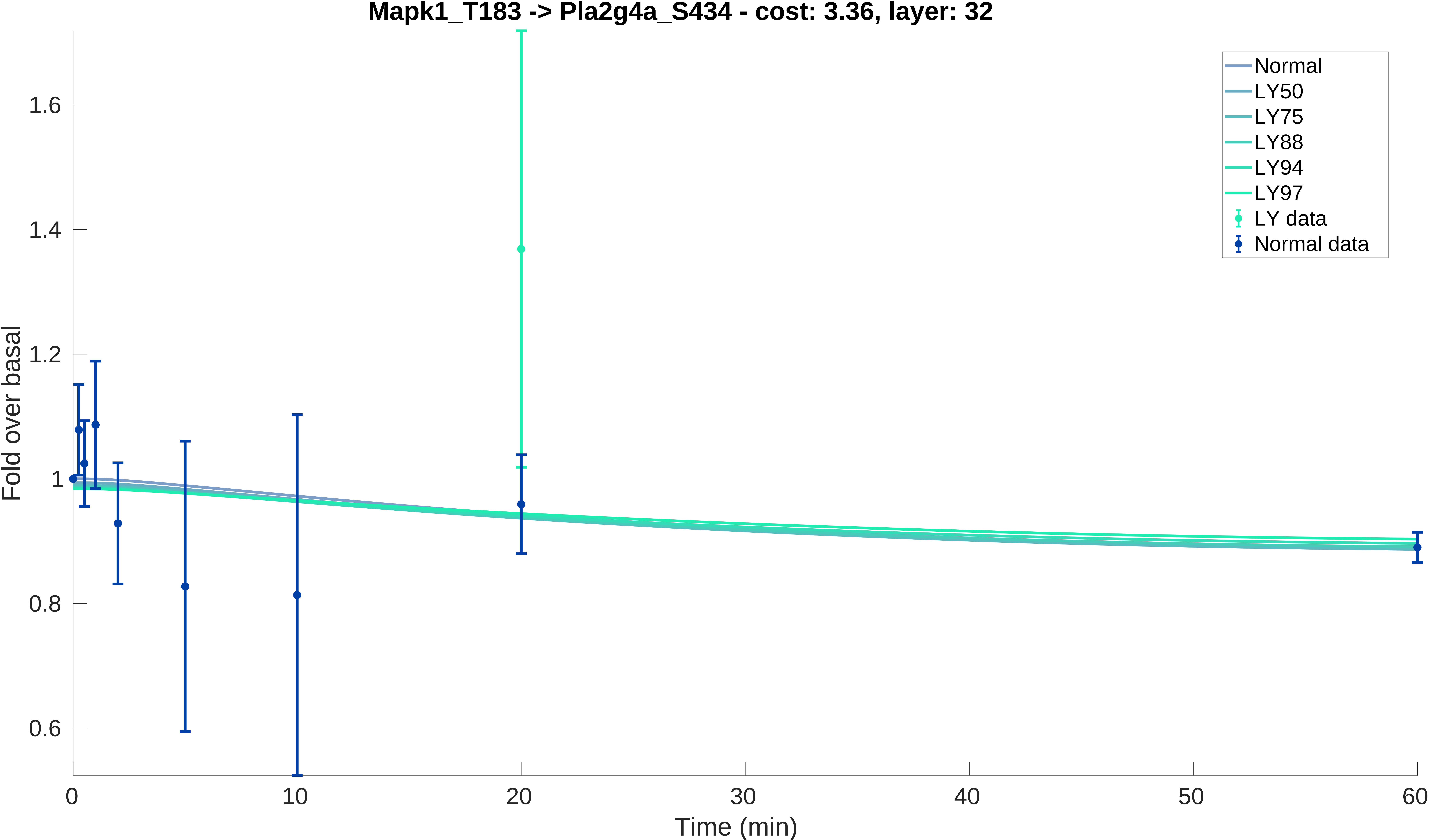

Mapk1_T183 -> Pla2g4a_S434 - cost: 3.36, layer: 32
Fold over basal
Time (min)
1.6
1.4
1.2
1
0.8
0.6
0
10
20
30
40
50
60
Normal
LY50
LY75
LY88
LY94
LY97
LY data
Normal data

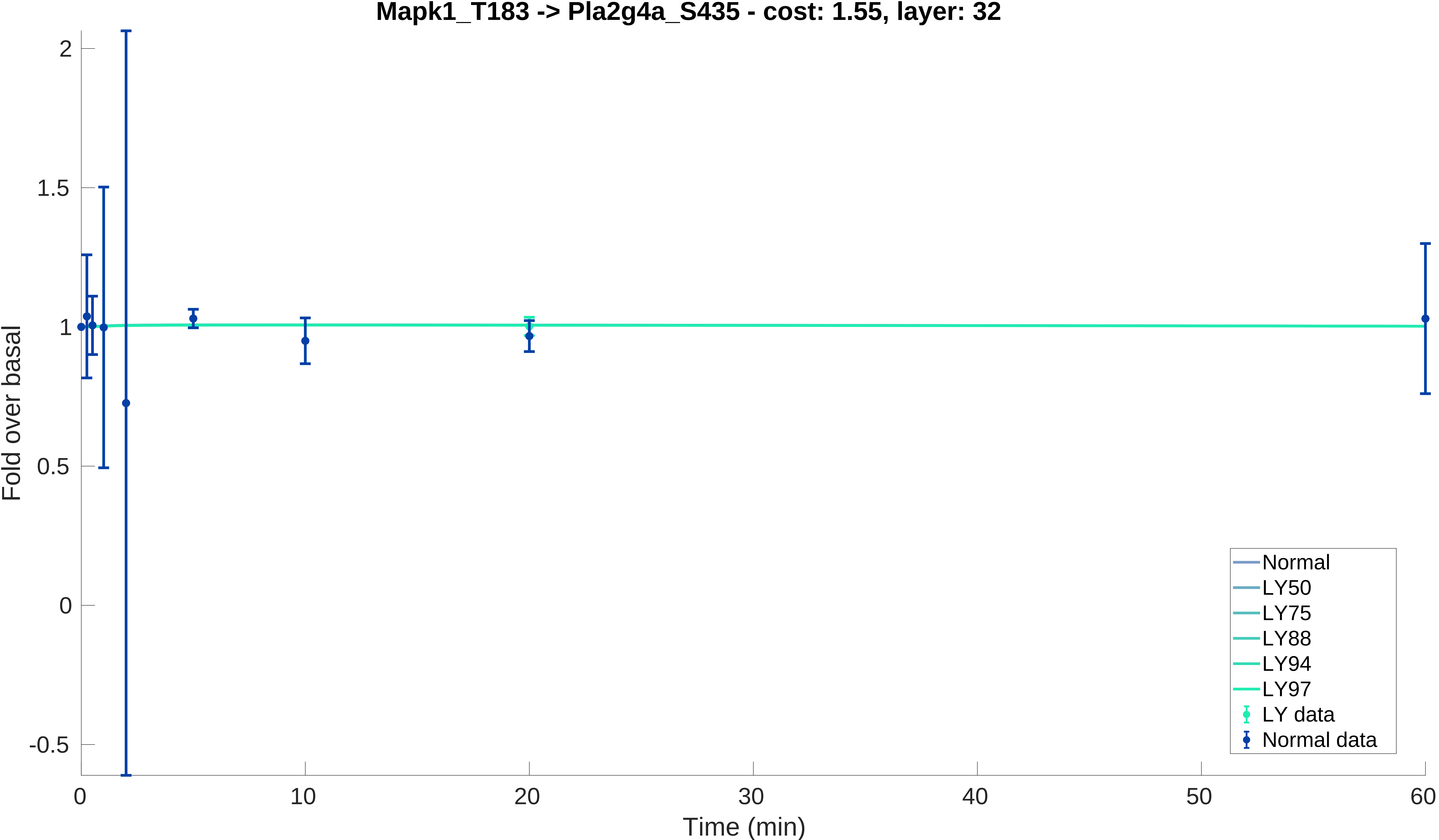

Mapk1_T183 -> Pla2g4a_S435 - cost: 1.55, layer: 32
Fold over basal
Time (min)
2
1.5
1
0.5
0
-0.5
0
10
20
30
40
50
60
Normal
LY50
LY75
LY88
LY94
LY97
LY data
Normal data

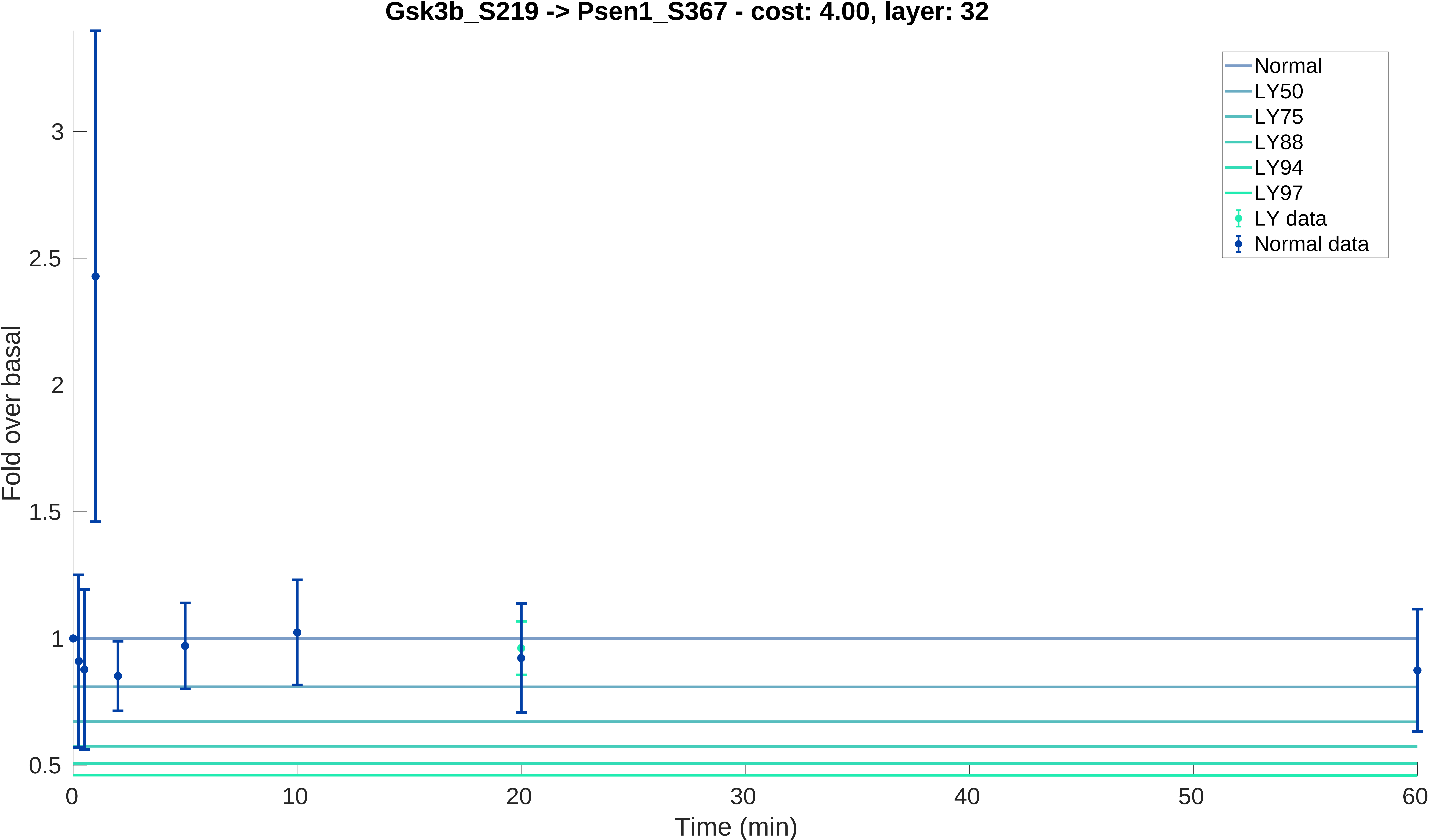

Gsk3b_S219 -> Psen1_S367 - cost: 4.00, layer: 32
Normal
LY50
LY75
LY88
LY94
LY97
LY data
Normal data
Fold over basal
3
2.5
2
1.5
1
0.5
0
10
20
30
40
50
60
Time (min)

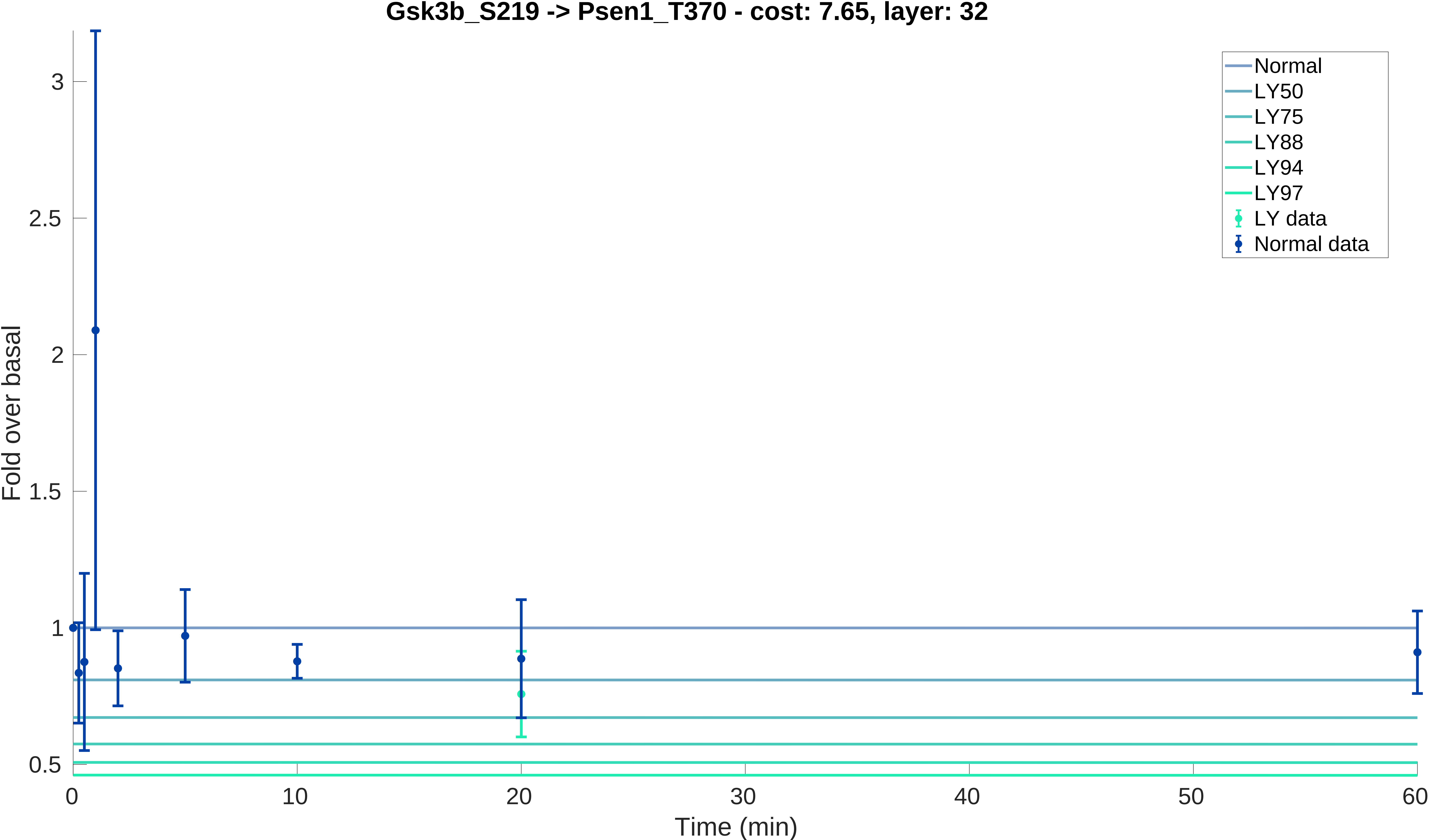

Gsk3b_S219 -> Psen1_T370 - cost: 7.65, layer: 32
Fold over basal
3
2.5
2
1.5
1
0.5
0
10
20
30
40
50
60
Time (min)
Normal
LY50
LY75
LY88
LY94
LY97
LY data
Normal data

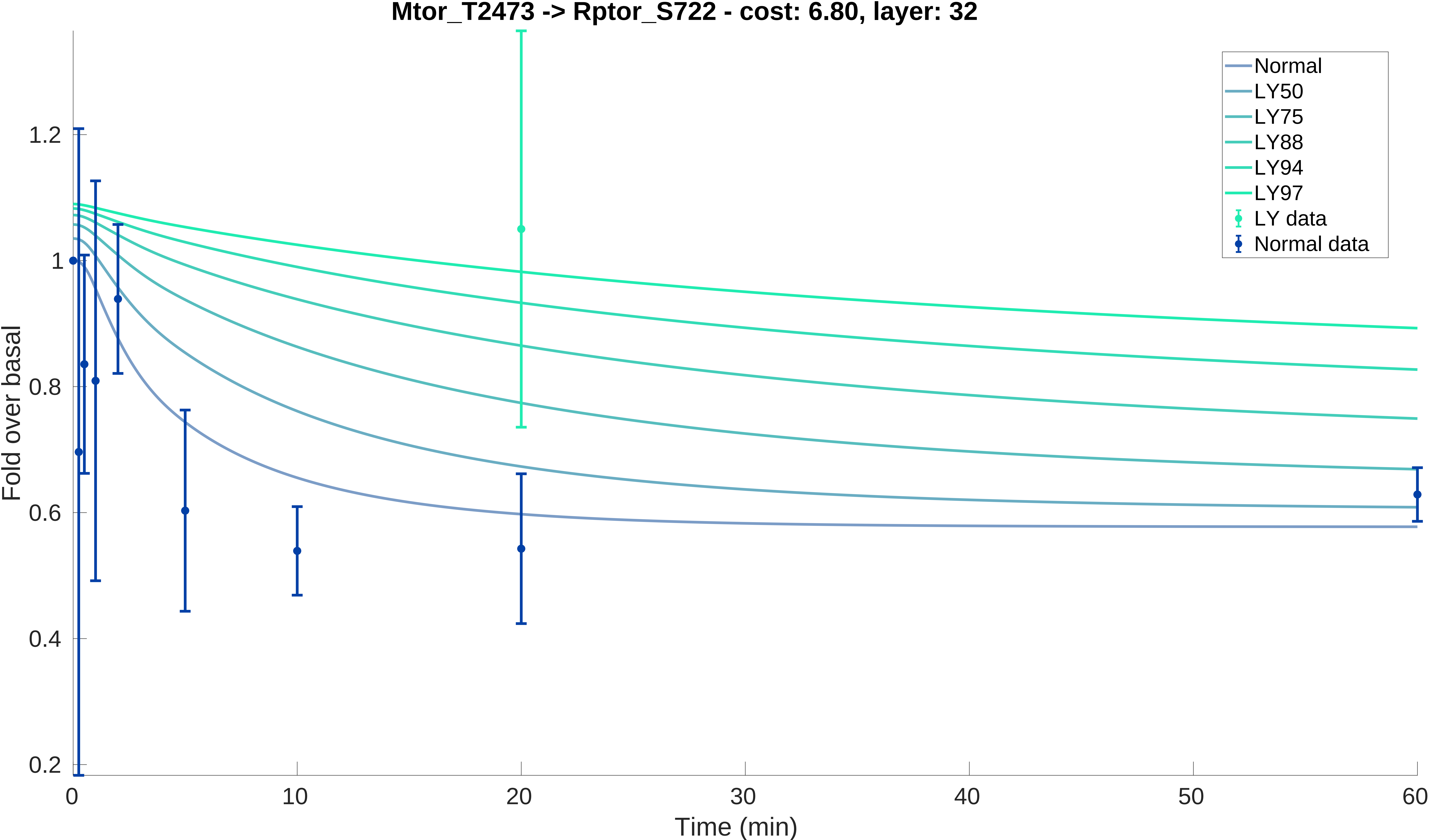

Mtor_T2473 -> Rptor_S722 - cost: 6.80, layer: 32
Fold over basal
Time (min)
0.2
0.4
0.6
0.8
1
1.2
0
10
20
30
40
50
60
Normal
LY50
LY75
LY88
LY94
LY97
LY data
Normal data

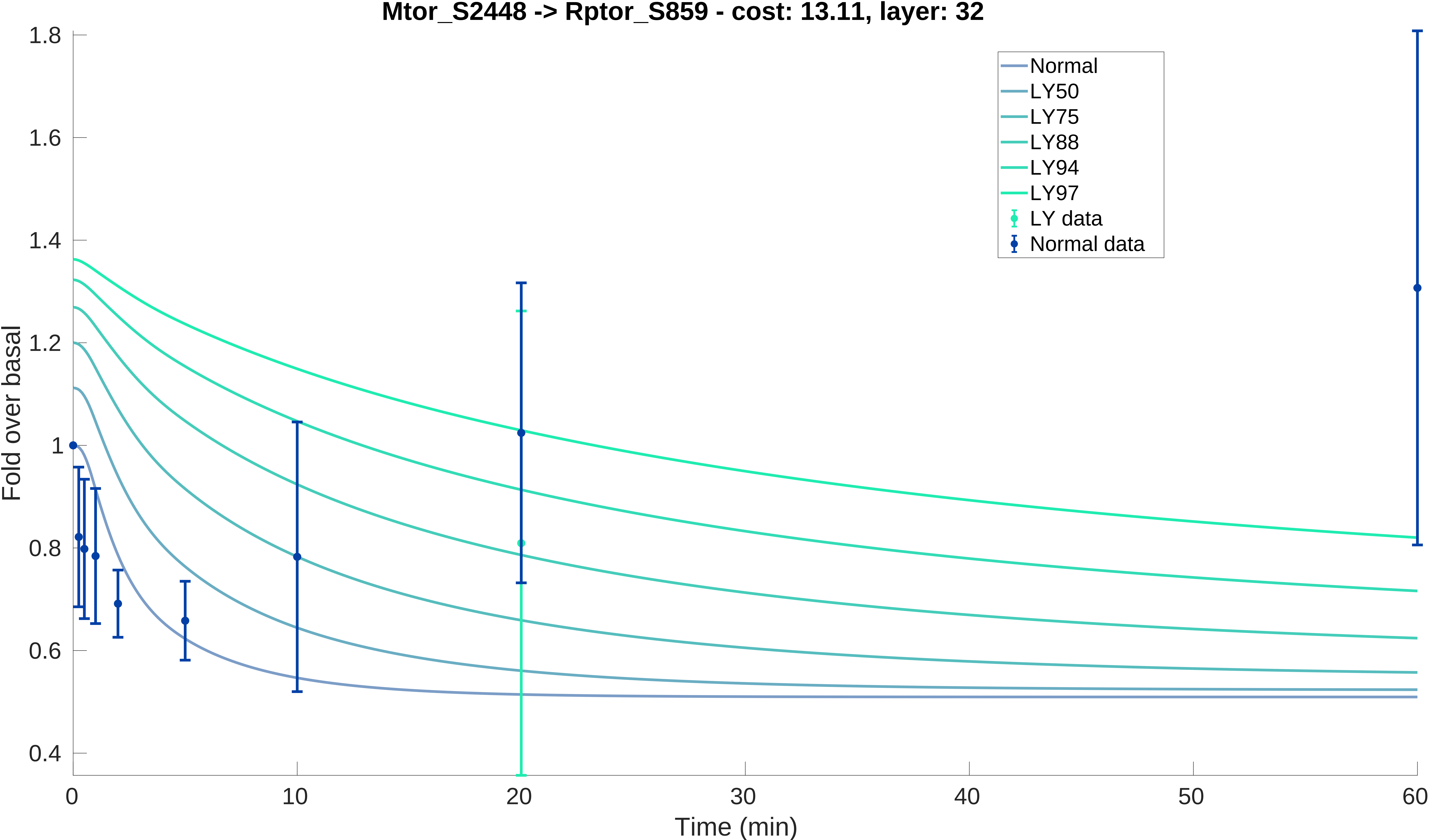
Mtor_S2448 -> Rptor_S859 - cost: 13.11, layer: 32
Fold over basal
Time (min)
1.8
1.6
1.4
1.2
1
0.8
0.6
0.4
0
10
20
30
40
50
60
Normal
LY50
LY75
LY88
LY94
LY97
LY data
Normal data

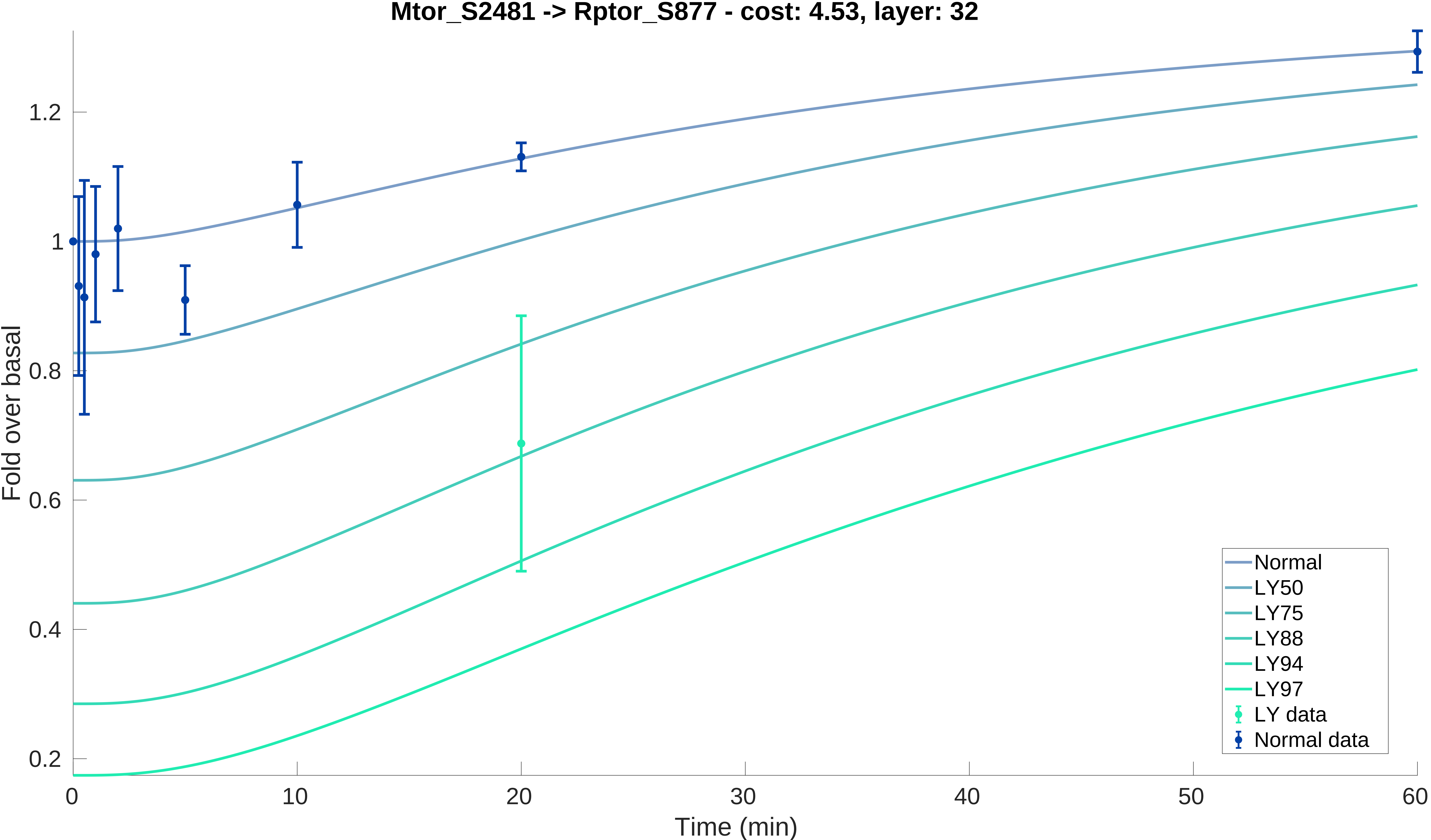

Mtor_S2481 -> Rptor_S877 - cost: 4.53, layer: 32
Fold over basal
Time (min)
0.2
0.4
0.6
0.8
1
1.2
0
10
20
30
40
50
60
Normal
LY50
LY75
LY88
LY94
LY97
LY data
Normal data

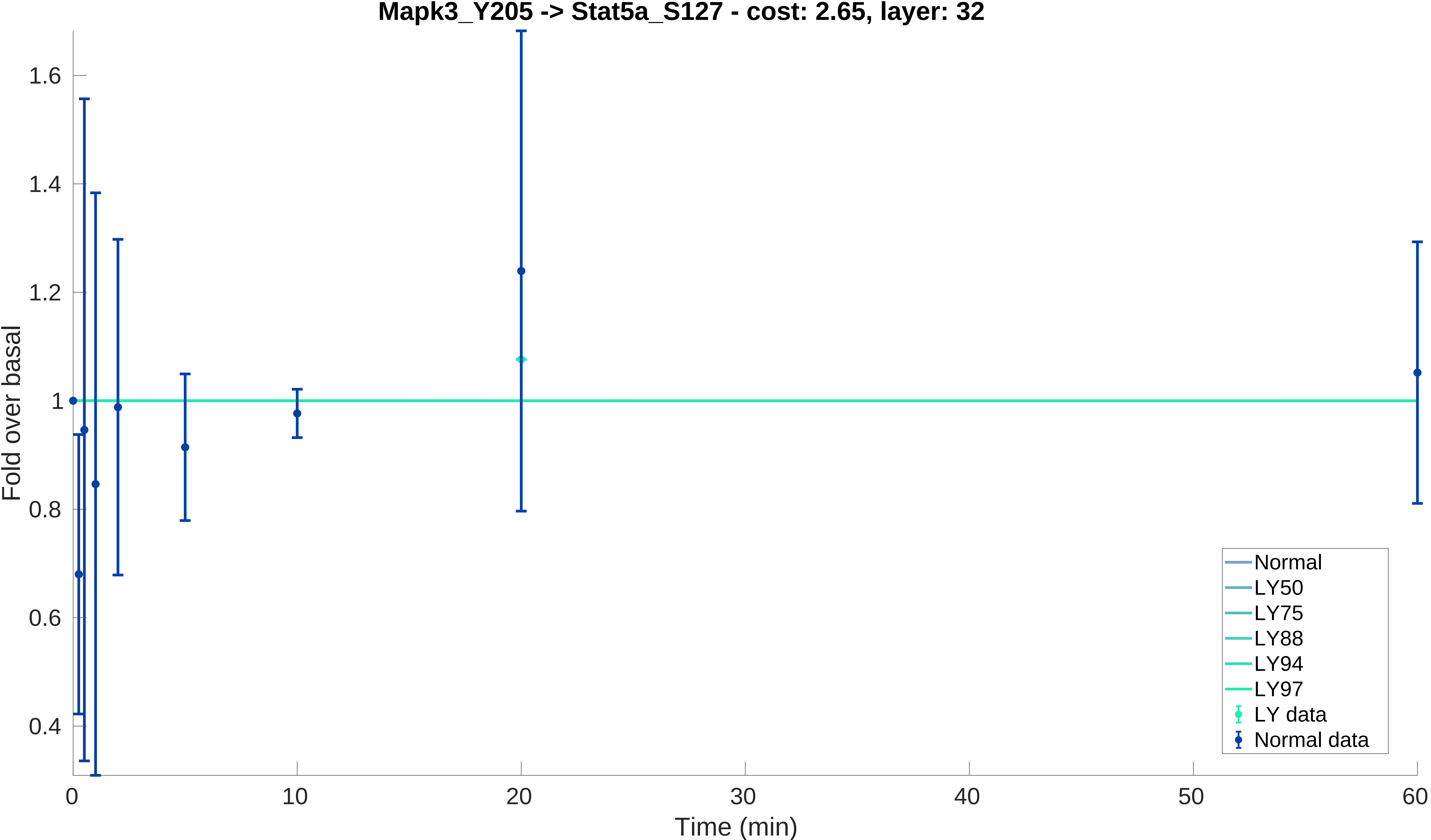

Mapk3_Y205 -> Stat5a_S127 - cost: 2.65, layer: 32
Fold over basal
1.6
1.4
1.2
1
0.8
0.6
0.4
0
10
20
30
40
50
60
Time (min)
Normal
LY50
LY75
LY88
LY94
LY97
LY data
Normal data

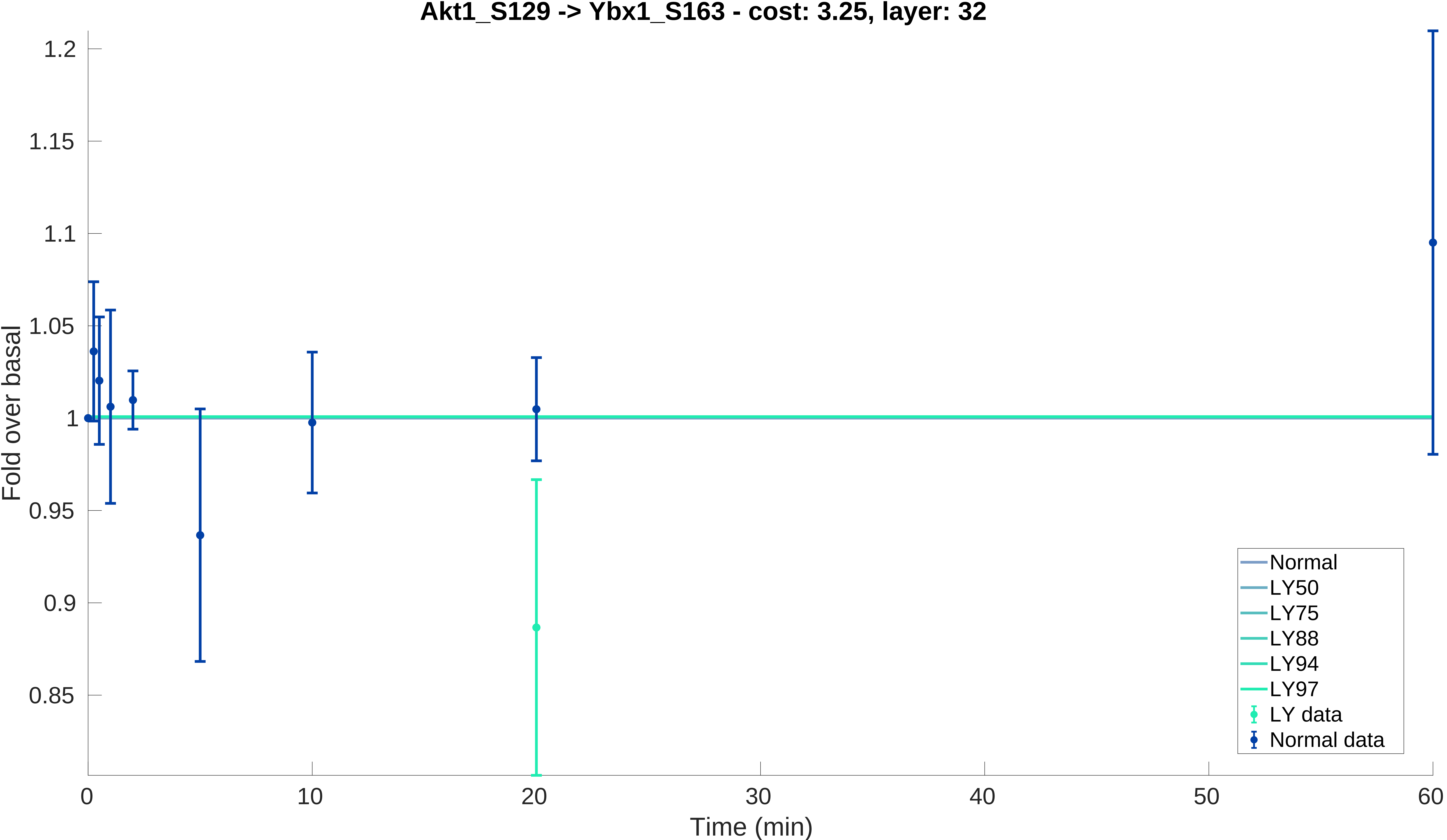

Akt1_S129 -> Ybx1_S163 - cost: 3.25, layer: 32
Fold over basal
1.2
1.15
1.1
1.05
1
0.95
0.9
0.85
0
10
20
30
40
50
60
Time (min)
Normal
LY50
LY75
LY88
LY94
LY97
LY data
Normal data

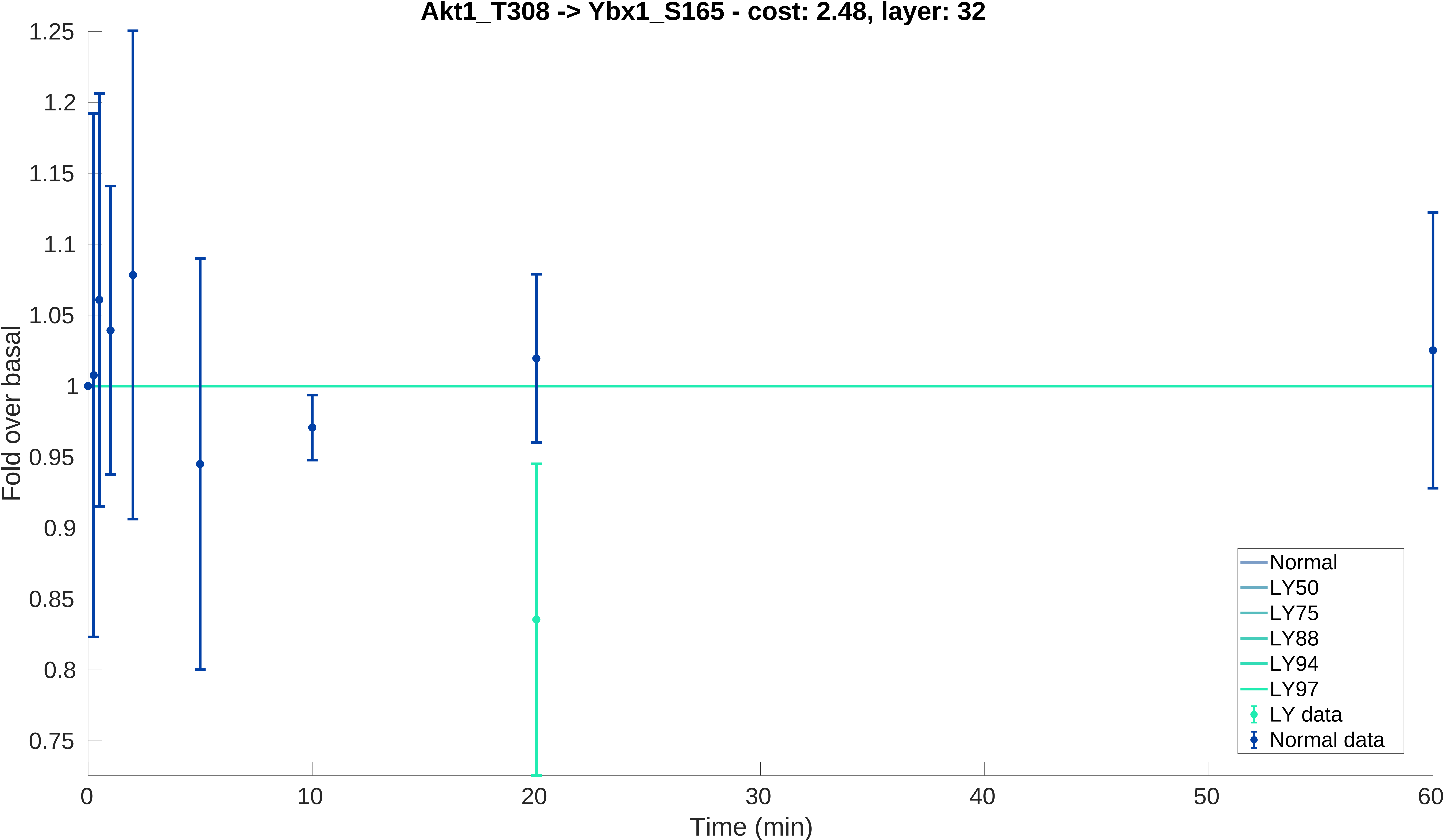

Akt1_T308 -> Ybx1_S165 - cost: 2.48, layer: 32
Fold over basal
Time (min)
1.25
1.2
1.15
1.1
1.05
1
0.95
0.9
0.85
0.8
0.75
0
10
20
30
40
50
60
Normal
LY50
LY75
LY88
LY94
LY97
LY data
Normal data

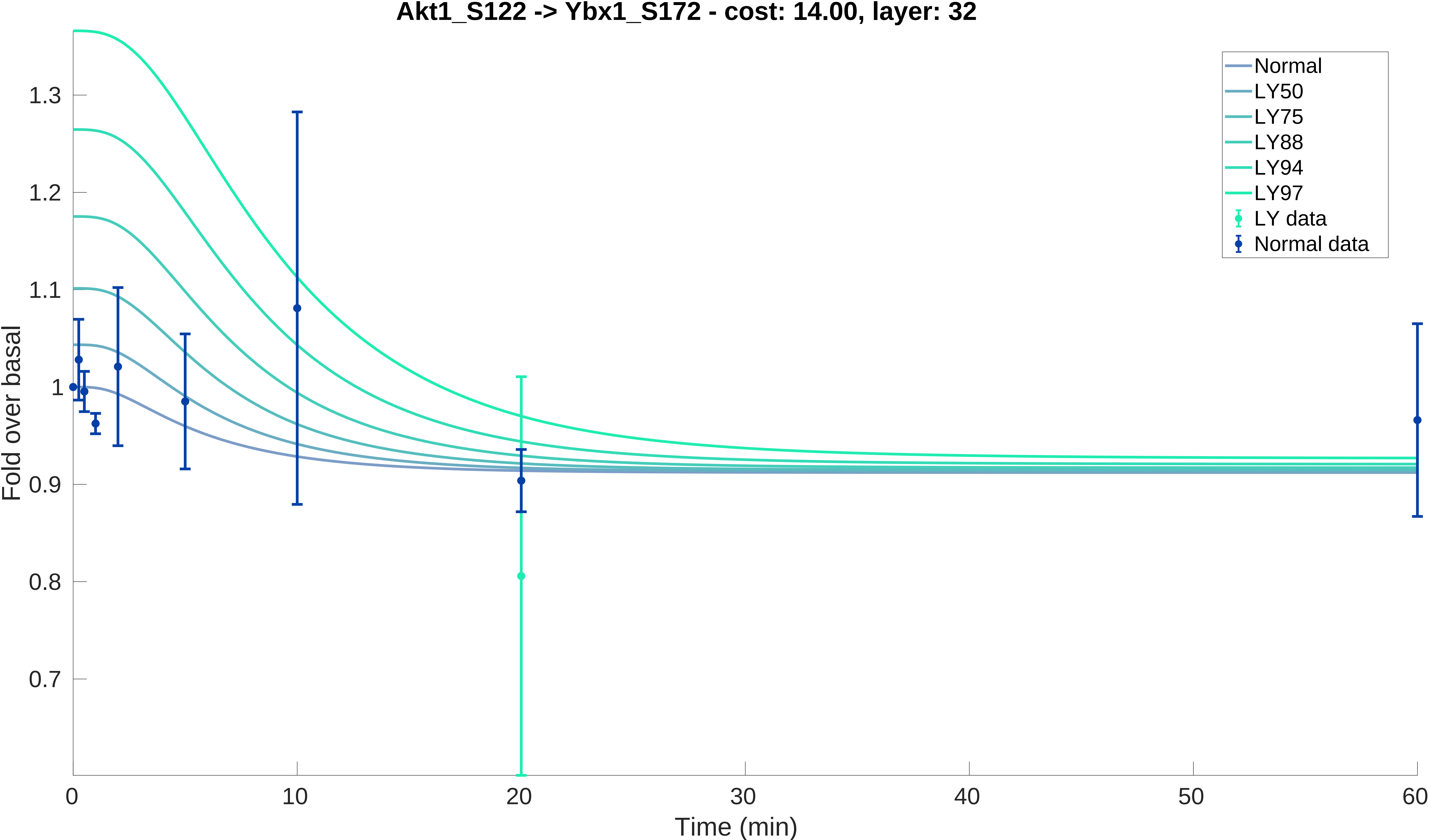

Akt1_S122 -> Ybx1_S172 - cost: 14.00, layer: 32
Normal
LY50
LY75
LY88
LY94
LY97
LY data
Normal data
Fold over basal
1.3
1.2
1.1
1
0.9
0.8
0.7
0
10
20
30
40
50
60
Time (min)

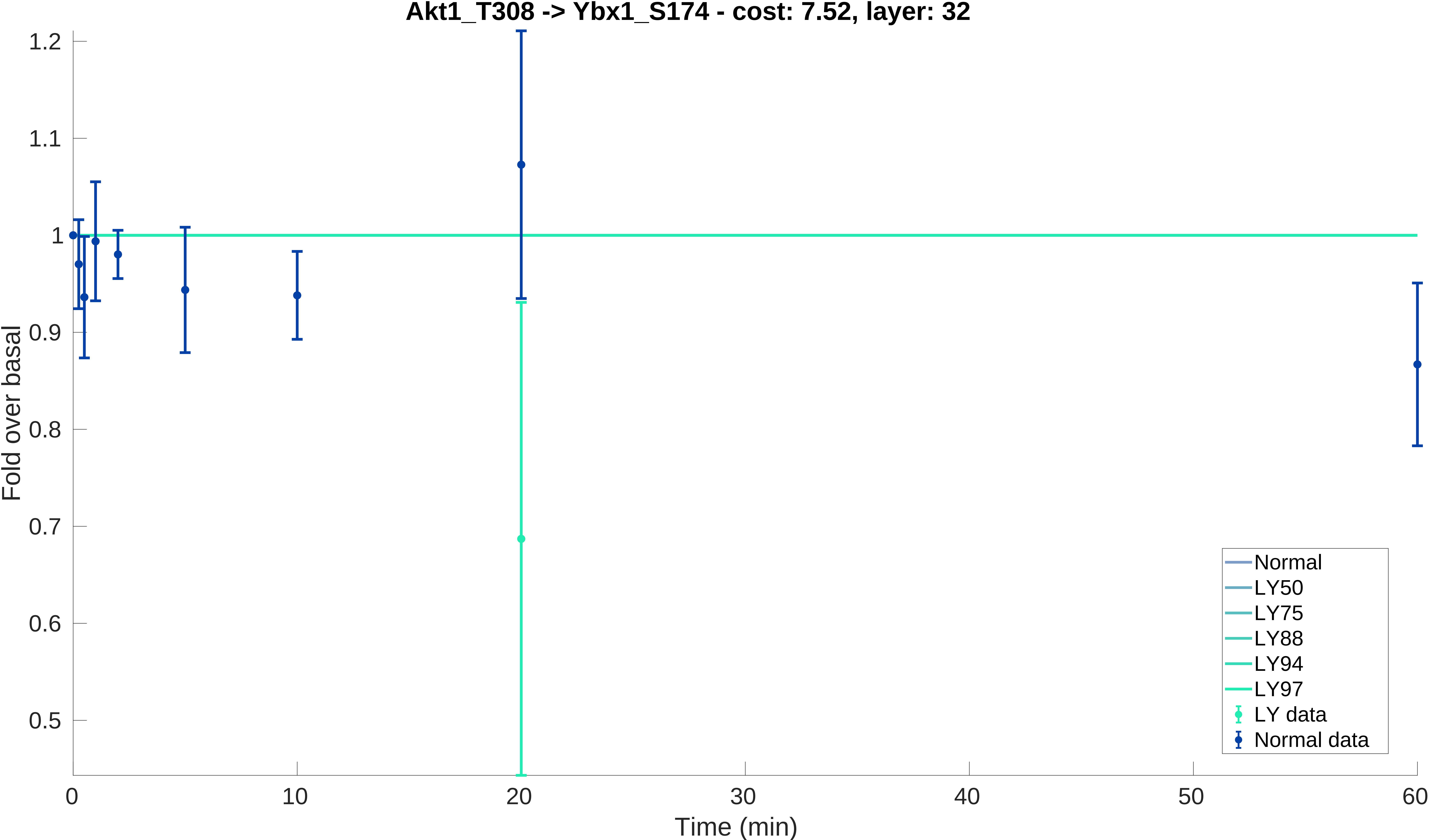
Akt1_T308 -> Ybx1_S174 - cost: 7.52, layer: 32
1.2
1.1
1
0.9
0.8
0.7
0.6
0.5
Fold over basal
0
10
20
30
40
50
60
Time (min)
Normal
LY50
LY75
LY88
LY94
LY97
LY data
Normal data

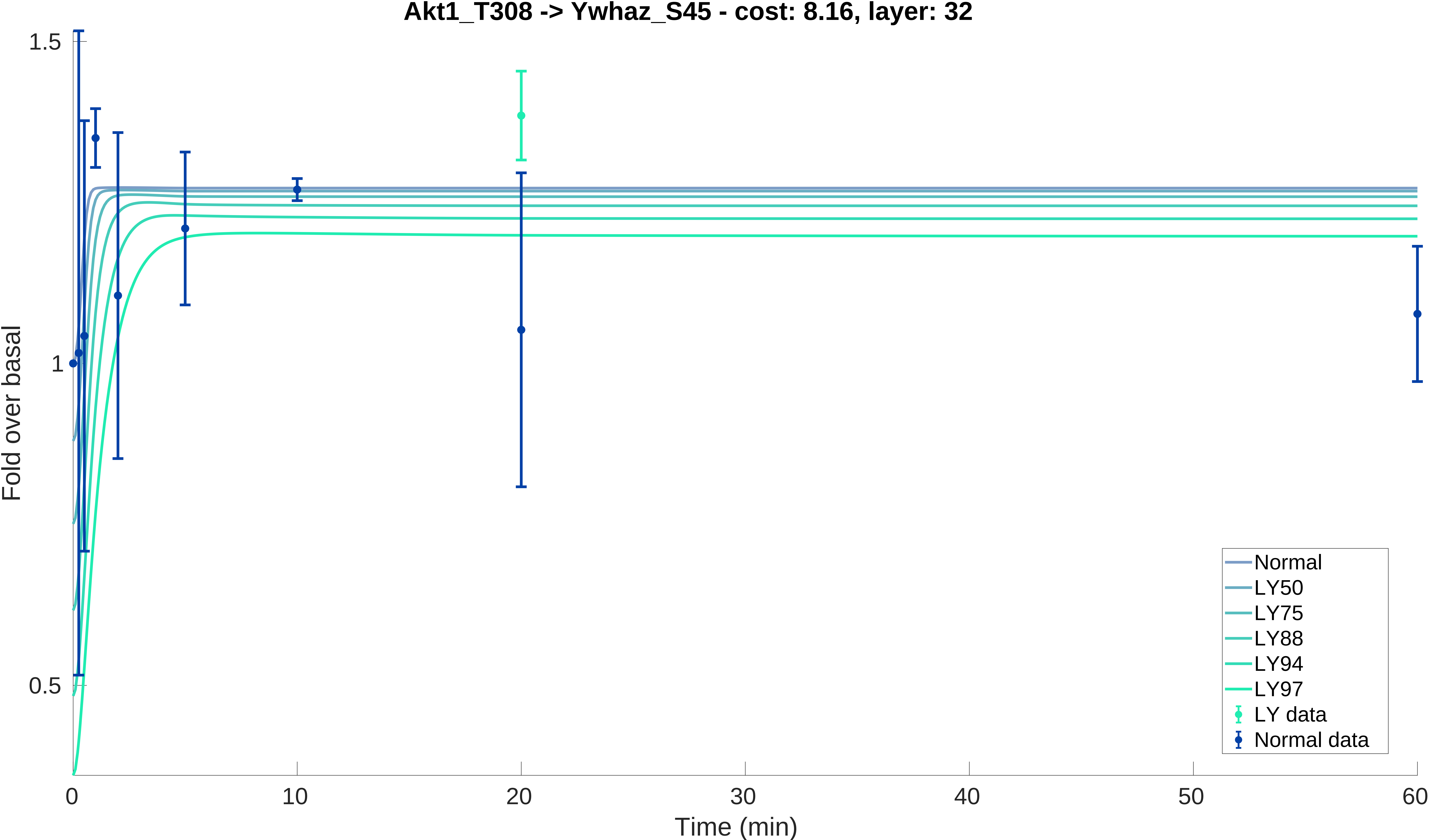

Akt1_T308 -> Ywhaz_S45 - cost: 8.16, layer: 32
Fold over basal
1.5
1
0.5
0
10
20
30
40
50
60
Time (min)
Normal
LY50
LY75
LY88
LY94
LY97
LY data
Normal data

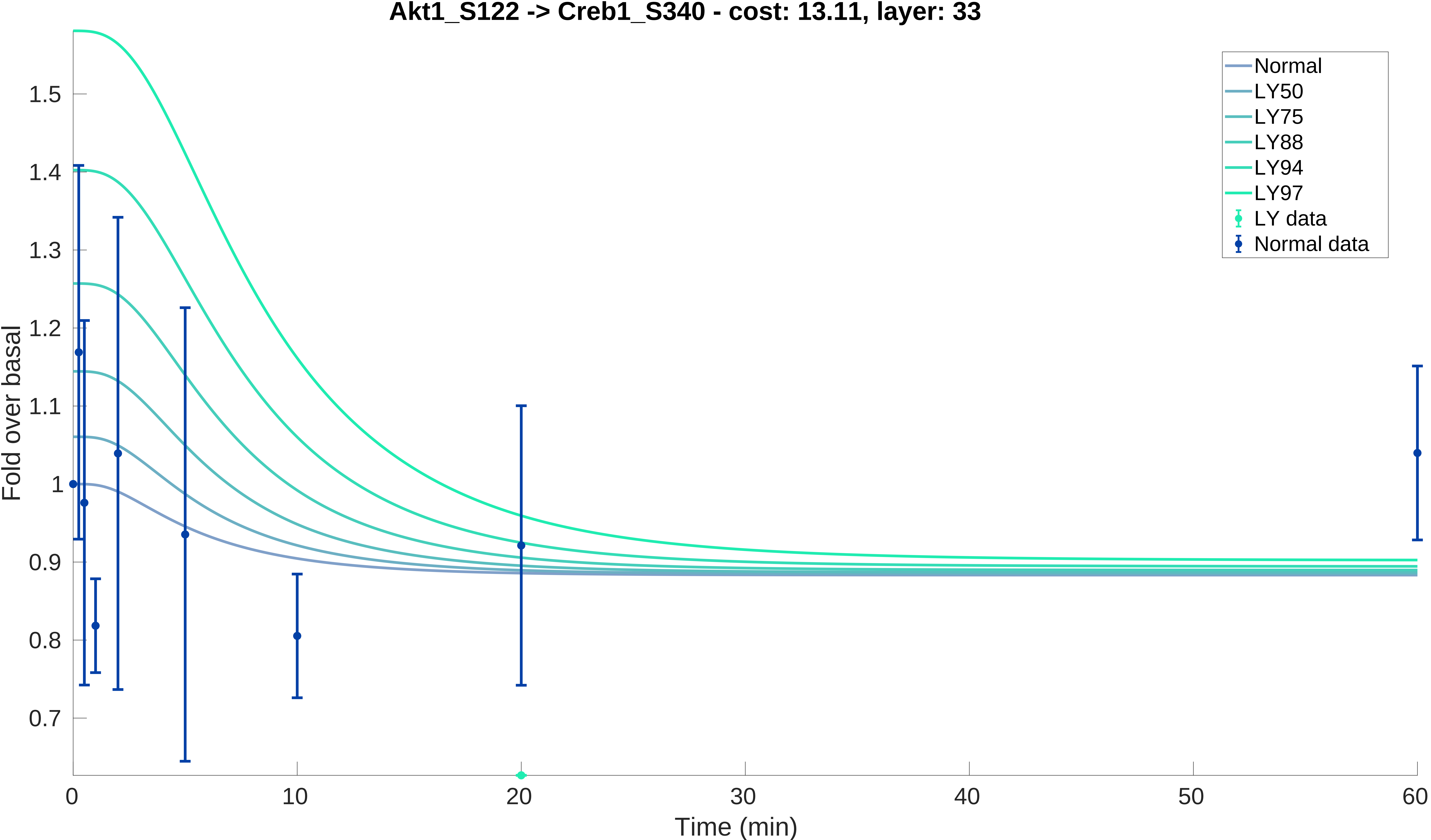

Akt1_S122 -> Creb1_S340 - cost: 13.11, layer: 33
Fold over basal
Time (min)
1.5
1.4
1.3
1.2
1.1
1
0.9
0.8
0.7
0
10
20
30
40
50
60
Normal
LY50
LY75
LY88
LY94
LY97
LY data
Normal data

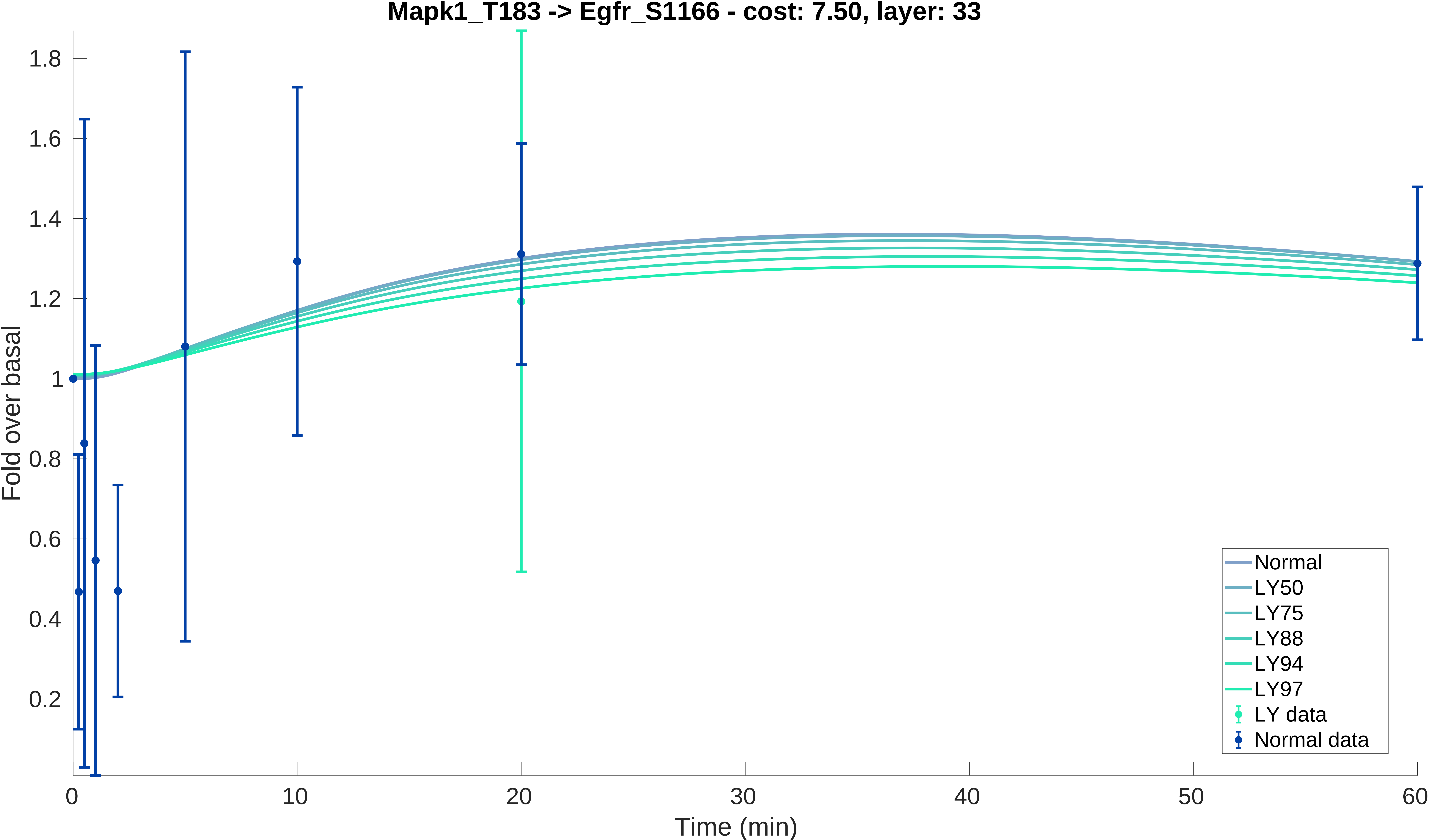
Mapk1_T183 -> Egfr_S1166 - cost: 7.50, layer: 33
Fold over basal
Time (min)
0
10
20
30
40
50
60
0.2
0.4
0.6
0.8
1
1.2
1.4
1.6
1.8
Normal
LY50
LY75
LY88
LY94
LY97
LY data
Normal data

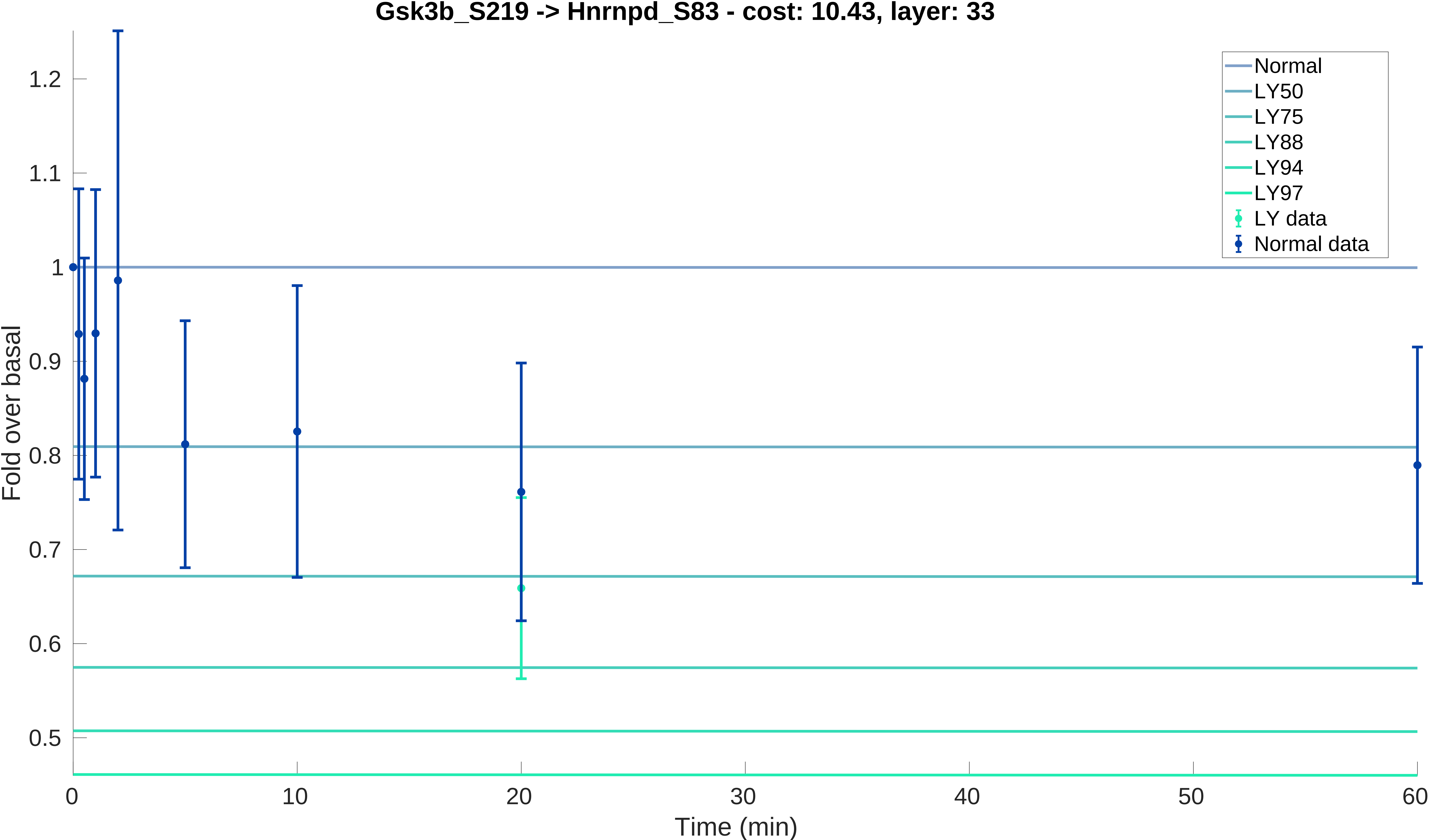

Gsk3b_S219 -> Hnrnpd_S83 - cost: 10.43, layer: 33
Normal
LY50
LY75
LY88
LY94
LY97
LY data
Normal data
Fold over basal
1.2
1.1
1
0.9
0.8
0.7
0.6
0.5
0
10
20
30
40
50
60
Time (min)

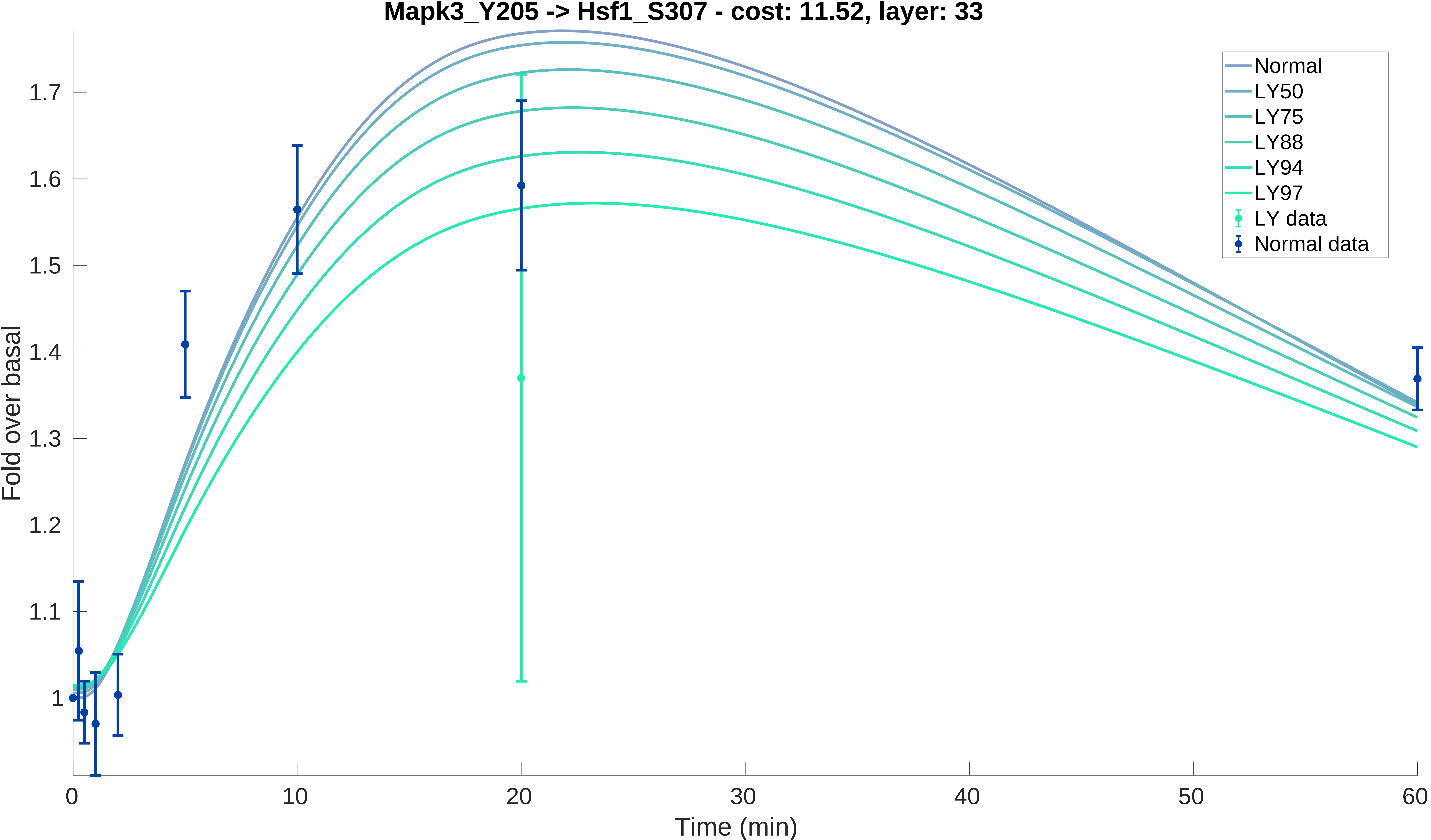

Mapk3_Y205 -> Hsf1_S307 - cost: 11.52, layer: 33
Fold over basal
Time (min)
1.7
1.6
1.5
1.4
1.3
1.2
1.1
1
0
10
20
30
40
50
60
Normal
LY50
LY75
LY88
LY94
LY97
LY data
Normal data

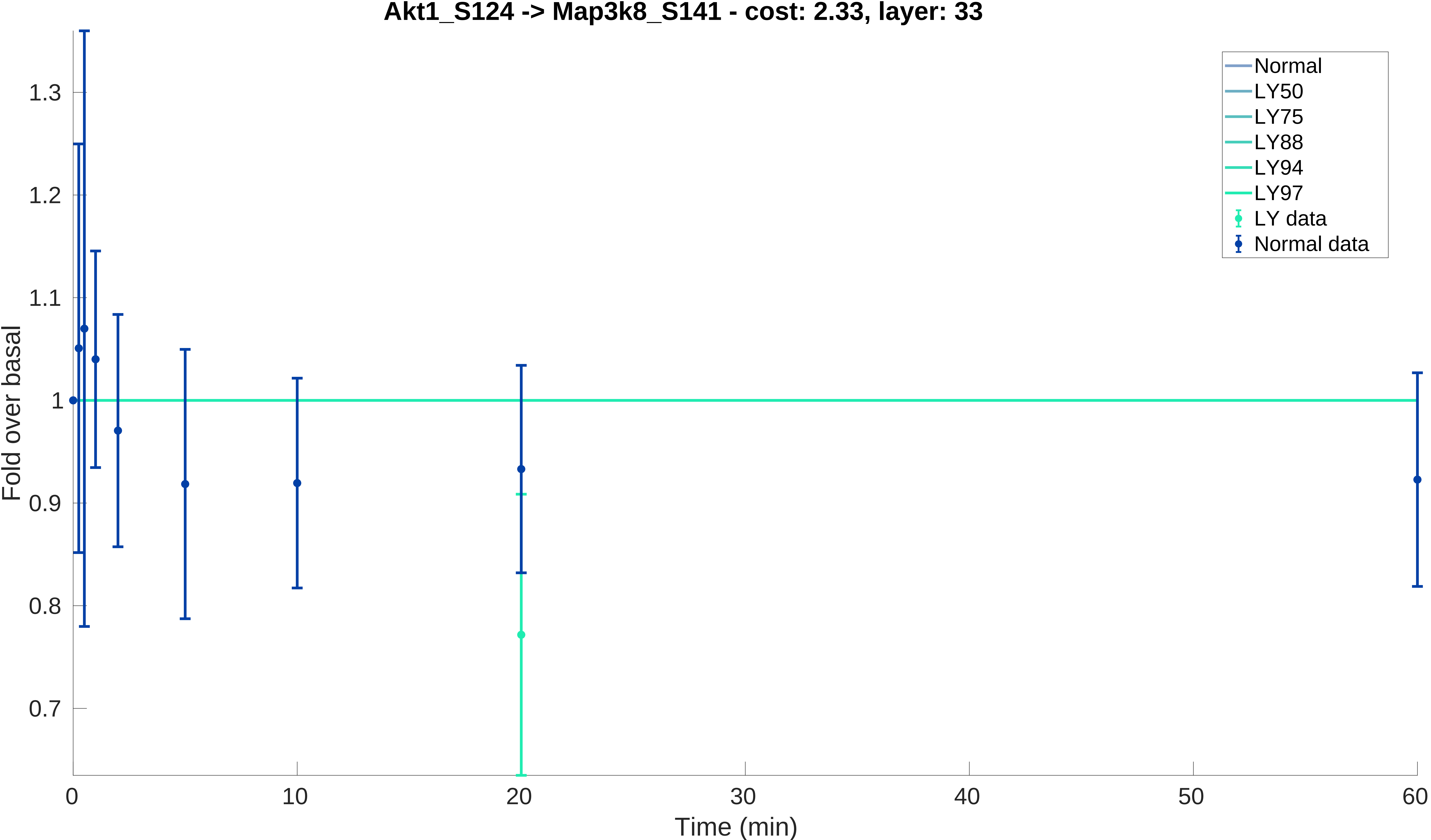

Akt1_S124 -> Map3k8_S141 - cost: 2.33, layer: 33
Fold over basal
Time (min)
Normal
LY50
LY75
LY88
LY94
LY97
LY data
Normal data
1.3
1.2
1.1
1
0.9
0.8
0.7
0
10
20
30
40
50
60

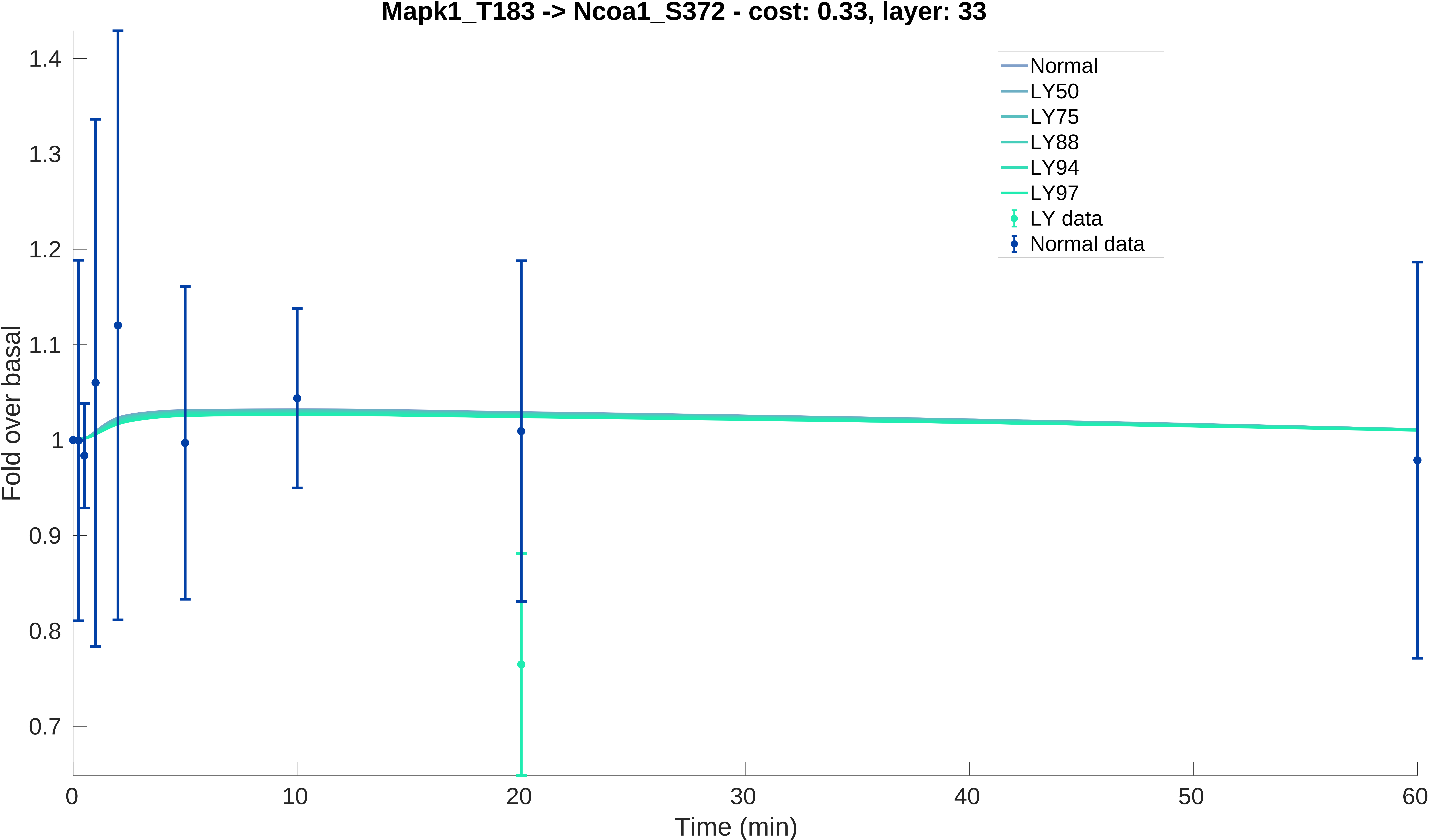

Mapk1_T183 -> Ncoa1_S372 - cost: 0.33, layer: 33
Normal
LY50
LY75
LY88
LY94
LY97
LY data
Normal data
Fold over basal
1.4
1.3
1.2
1.1
1
0.9
0.8
0.7
0
10
20
30
40
50
60
Time (min)

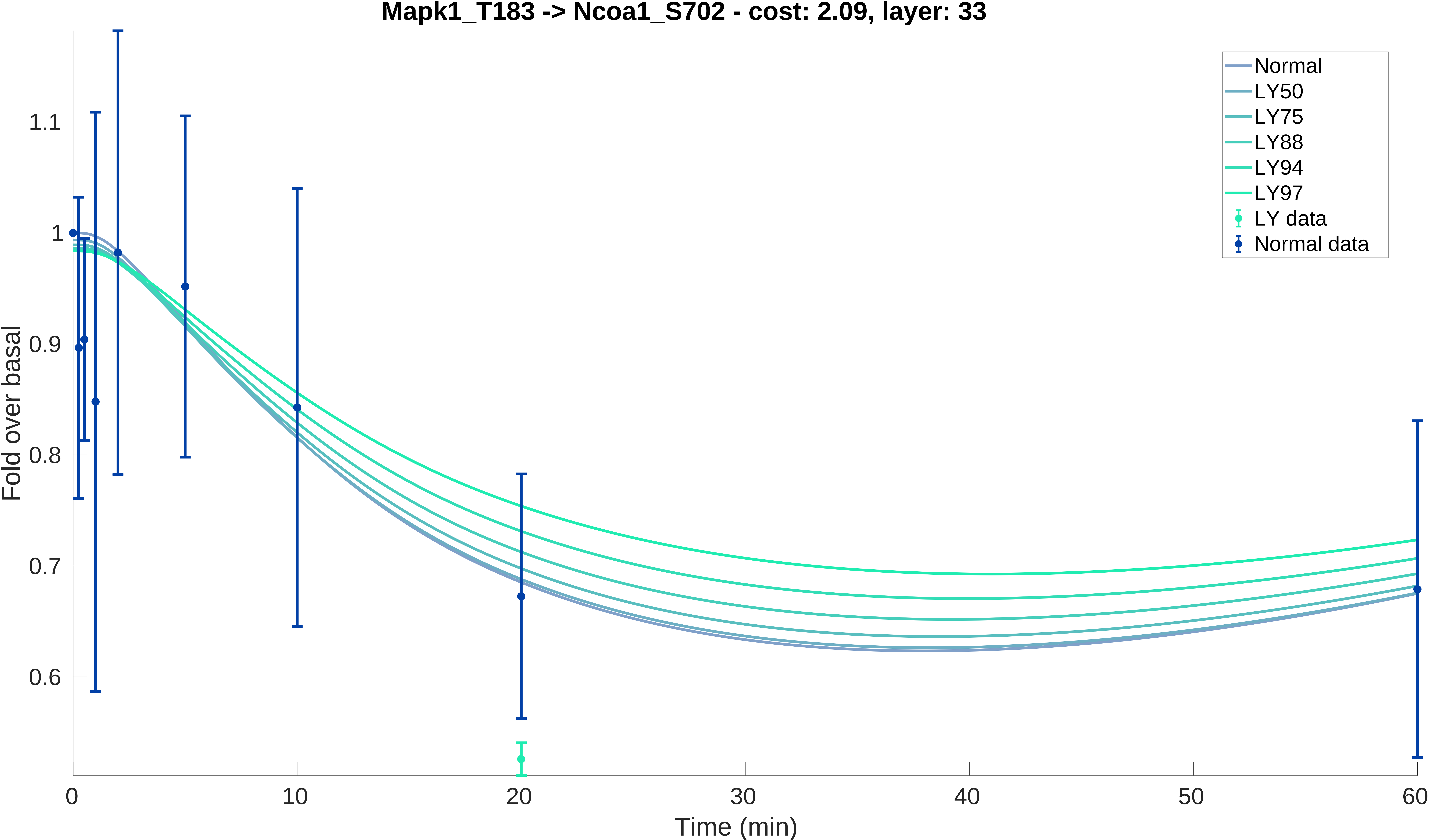

Mapk1_T183 -> Ncoa1_S702 - cost: 2.09, layer: 33
Normal
LY50
LY75
LY88
LY94
LY97
LY data
Normal data
Fold over basal
1.1
1
0.9
0.8
0.7
0.6
0
10
20
30
40
50
60
Time (min)

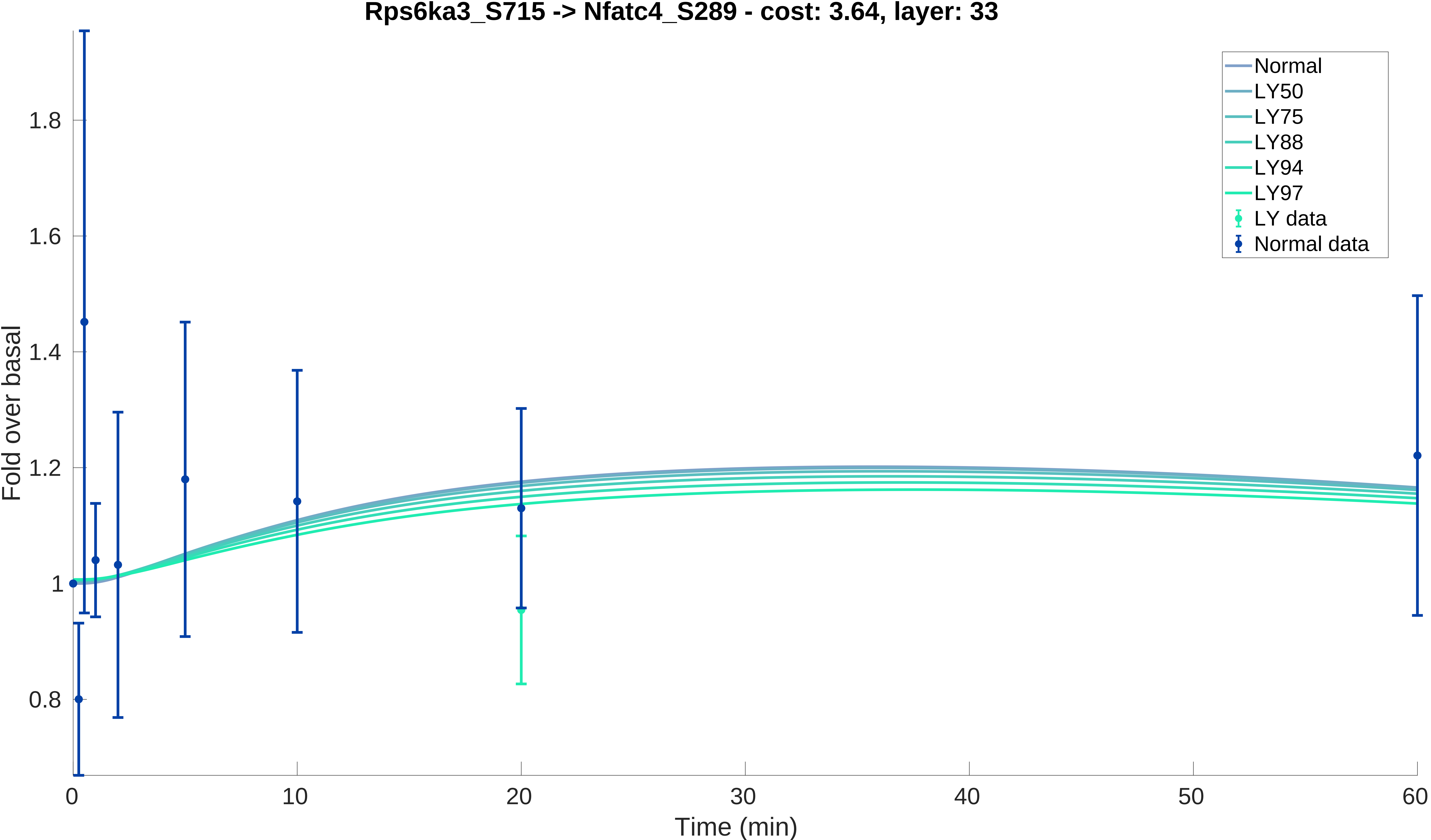

Rps6ka3_S715 -> Nfatc4_S289 - cost: 3.64, layer: 33
Normal
LY50
LY75
LY88
LY94
LY97
LY data
Normal data
Fold over basal
1.8
1.6
1.4
1.2
1
0.8
0
10
20
30
40
50
60
Time (min)

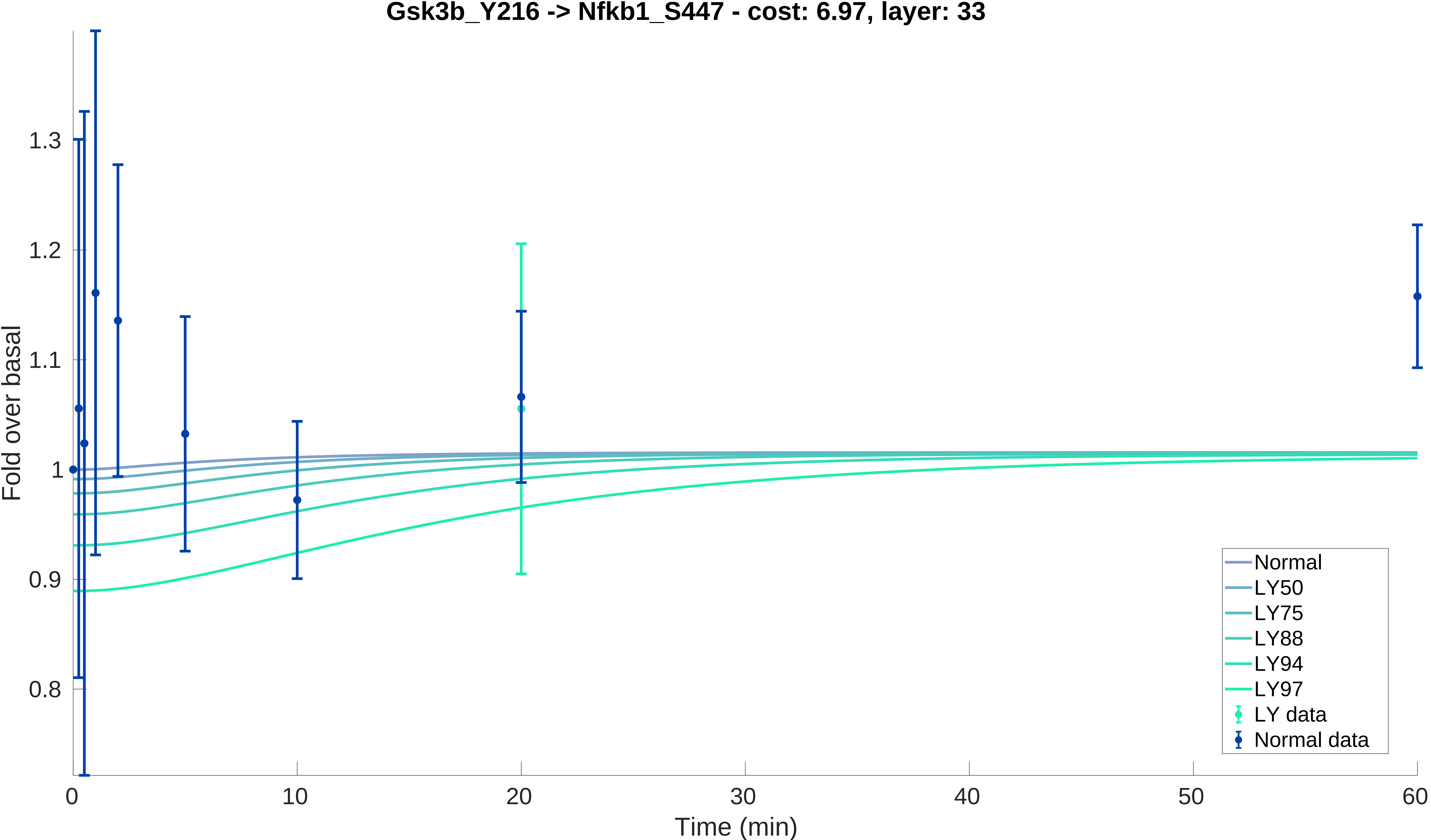

Gsk3b_Y216 -> Nfkb1_S447 - cost: 6.97, layer: 33
Fold over basal
1.3
1.2
1.1
1
0.9
0.8
0
10
20
30
40
50
60
Time (min)
Normal
LY50
LY75
LY88
LY94
LY97
LY data
Normal data

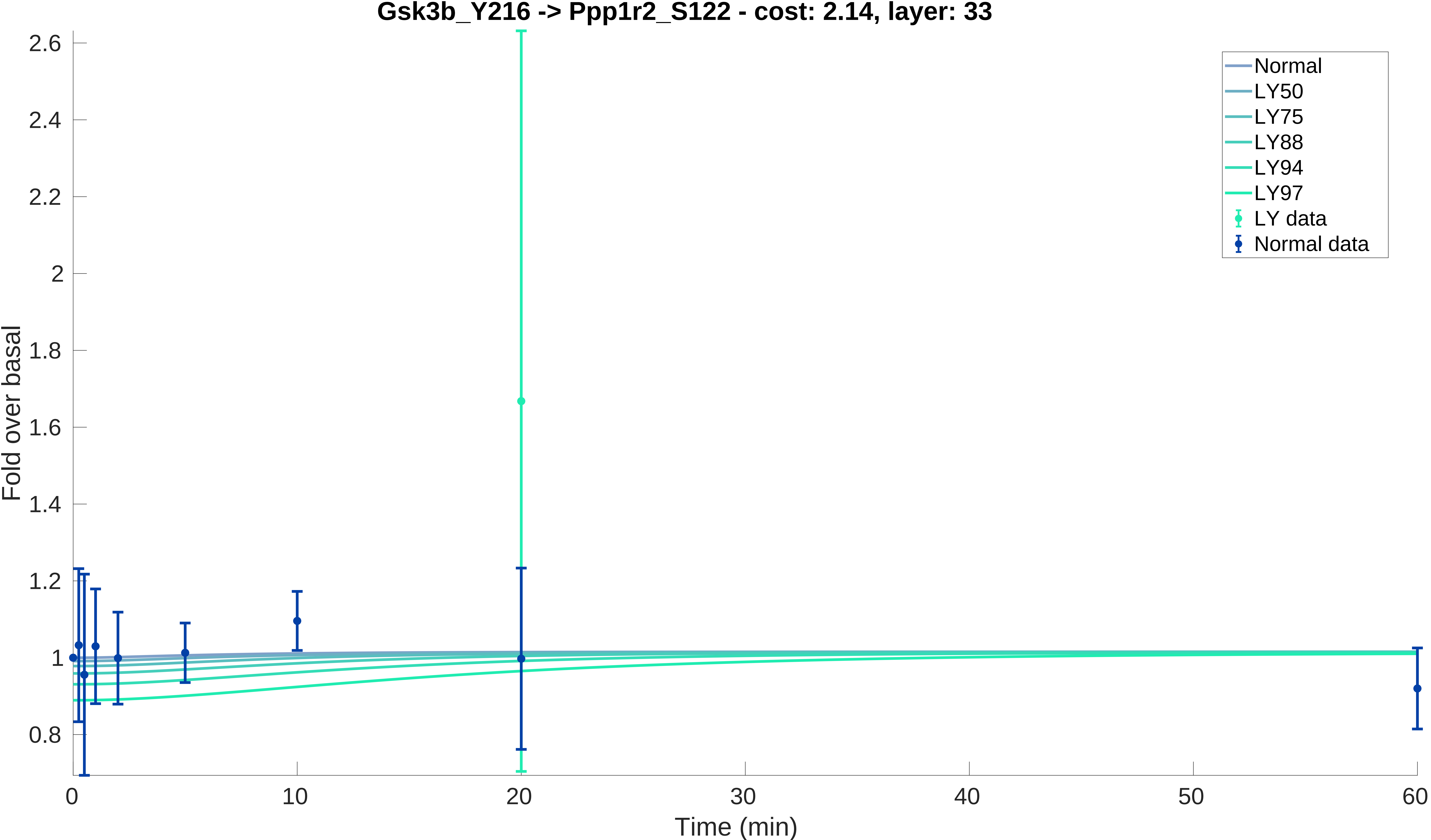

Gsk3b_Y216 -> Ppp1r2_S122 - cost: 2.14, layer: 33
Fold over basal
Time (min)
2.6
2.4
2.2
2
1.8
1.6
1.4
1.2
1
0.8
0
10
20
30
40
50
60
Normal
LY50
LY75
LY88
LY94
LY97
LY data
Normal data

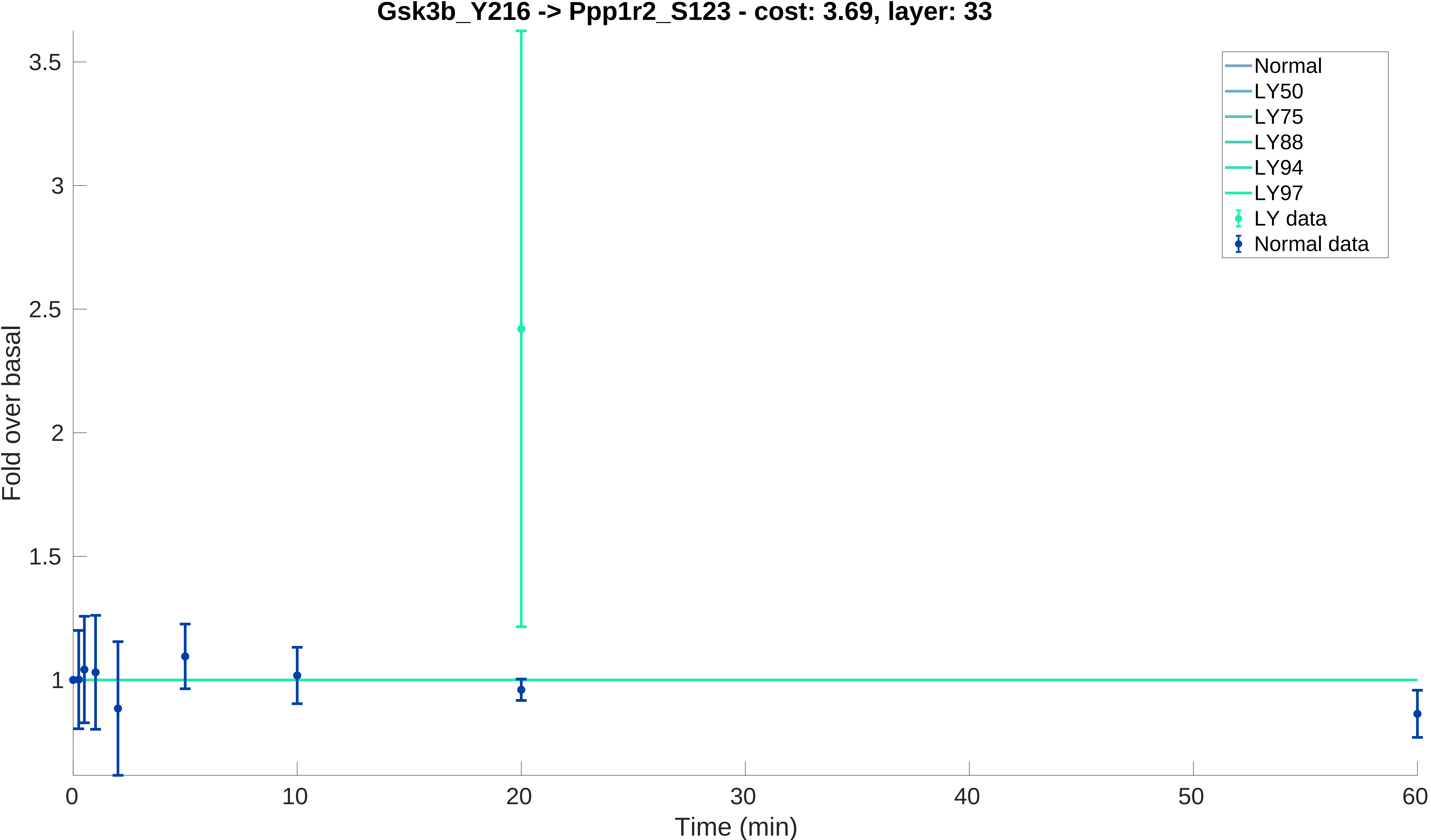

Gsk3b_Y216 -> Ppp1r2_S123 - cost: 3.69, layer: 33
Fold over basal
3.5
3
2.5
2
1.5
1
0
10
20
30
40
50
60
Time (min)
Normal
LY50
LY75
LY88
LY94
LY97
LY data
Normal data

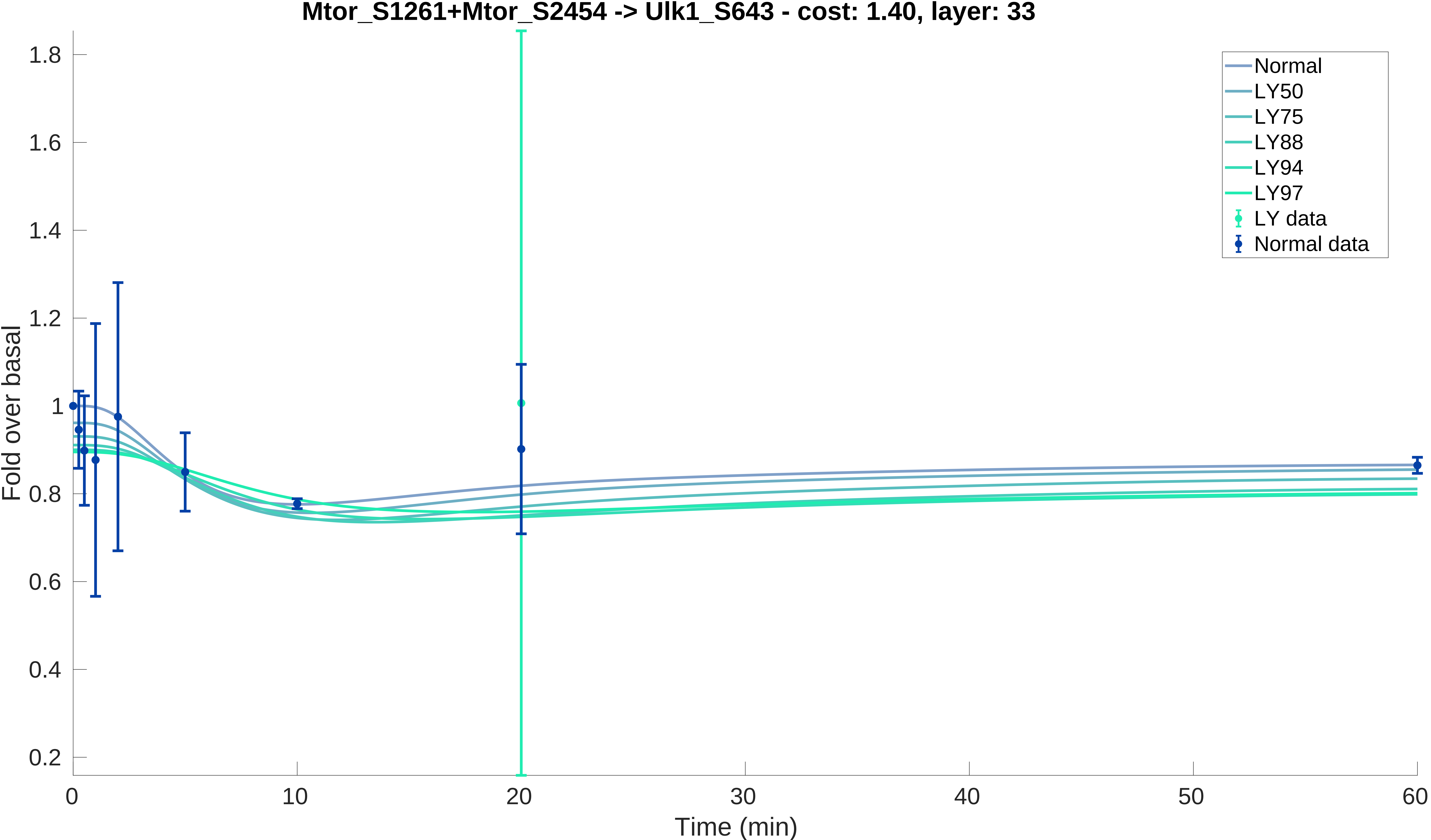

Mtor_S1261+Mtor_S2454 -> Ulk1_S643 - cost: 1.40, layer: 33
Fold over basal
Time (min)
0
10
20
30
40
50
60
0.2
0.4
0.6
0.8
1
1.2
1.4
1.6
1.8
Normal
LY50
LY75
LY88
LY94
LY97
LY data
Normal data

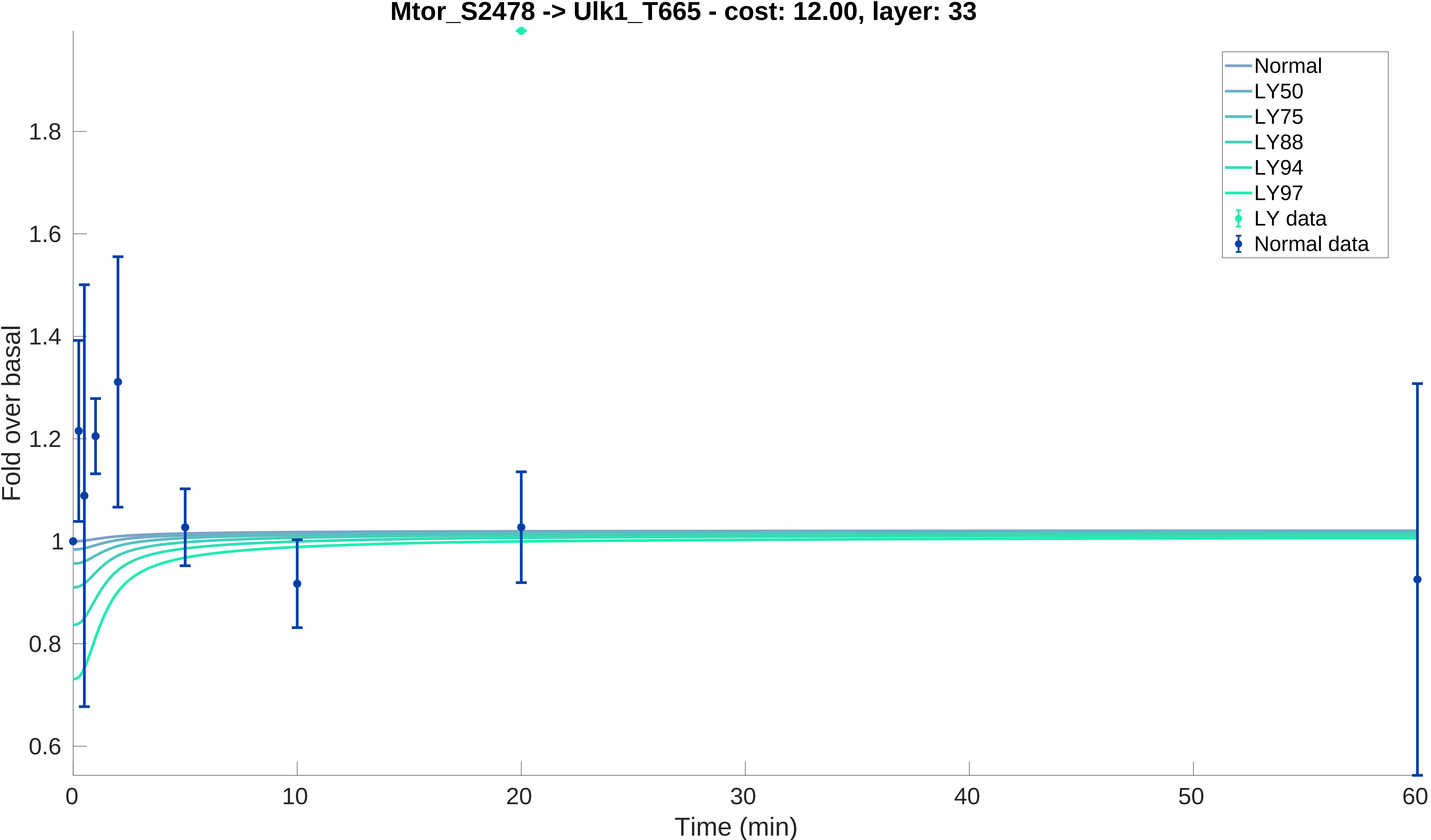
Mtor_S2478 -> Ulk1_T665 - cost: 12.00, layer: 33
Normal
LY50
LY75
LY88
LY94
LY97
LY data
Normal data
Fold over basal
1.8
1.6
1.4
1.2
1
0.8
0.6
0
10
20
30
40
50
60
Time (min)

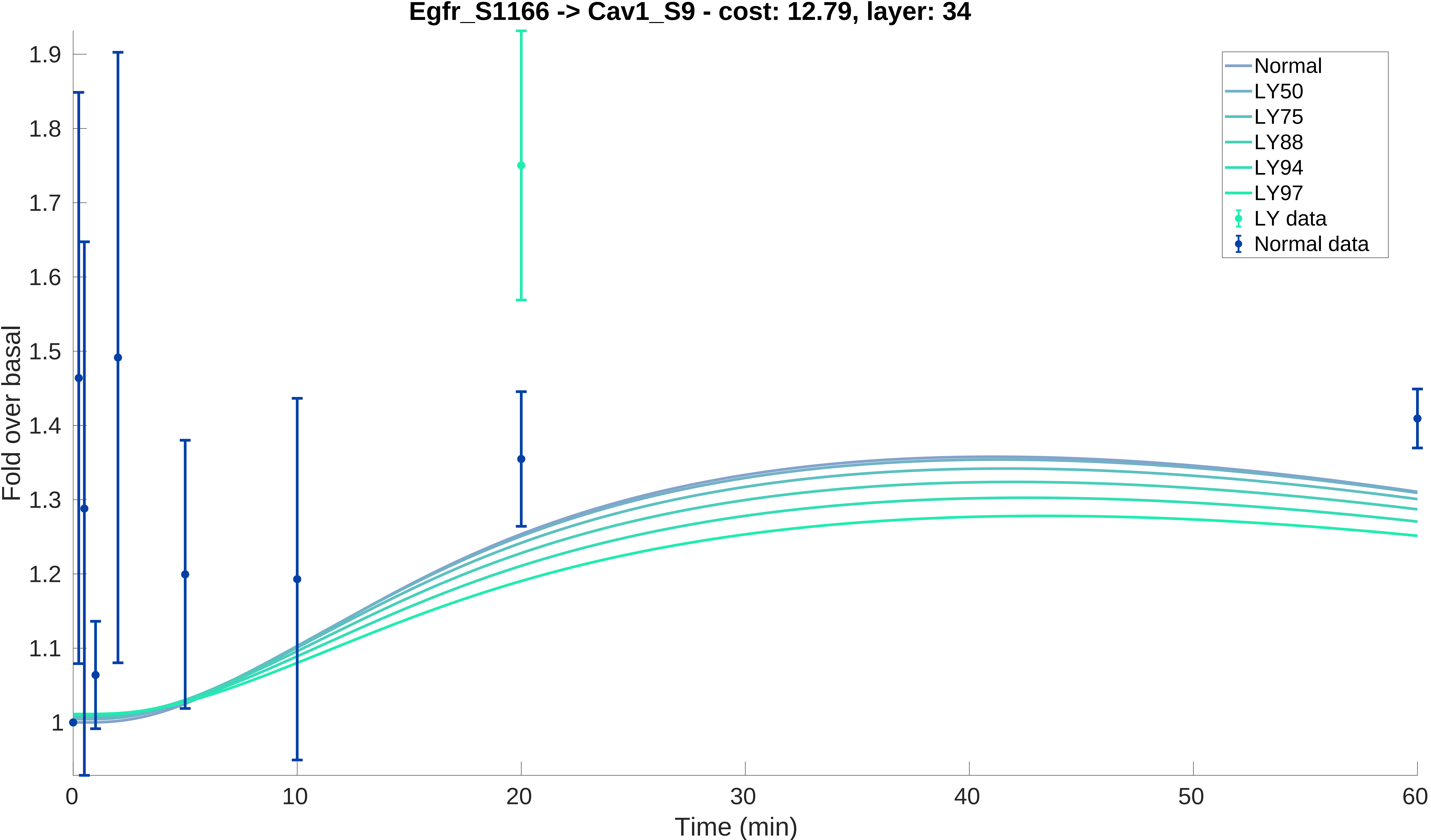
Egfr_S1166 -> Cav1_S9 - cost: 12.79, layer: 34
Fold over basal
Time (min)
1.9
1.8
1.7
1.6
1.5
1.4
1.3
1.2
1.1
1
0
10
20
30
40
50
60
Normal
LY50
LY75
LY88
LY94
LY97
LY data
Normal data

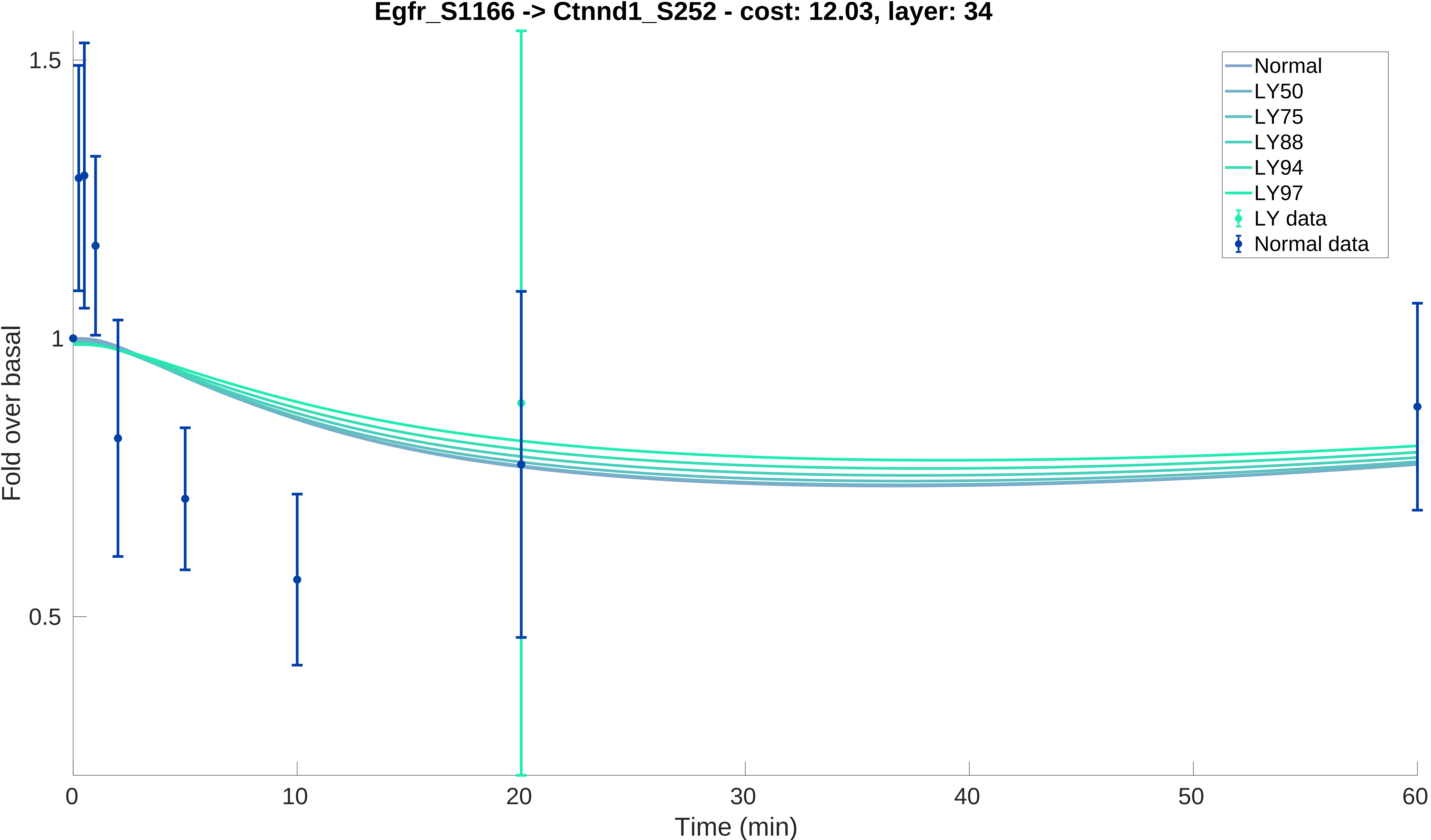

Egfr_S1166 -> Ctnnd1_S252 - cost: 12.03, layer: 34
Normal
LY50
LY75
LY88
LY94
LY97
LY data
Normal data
Fold over basal
Time (min)
1.5
1
0.5
0
10
20
30
40
50
60

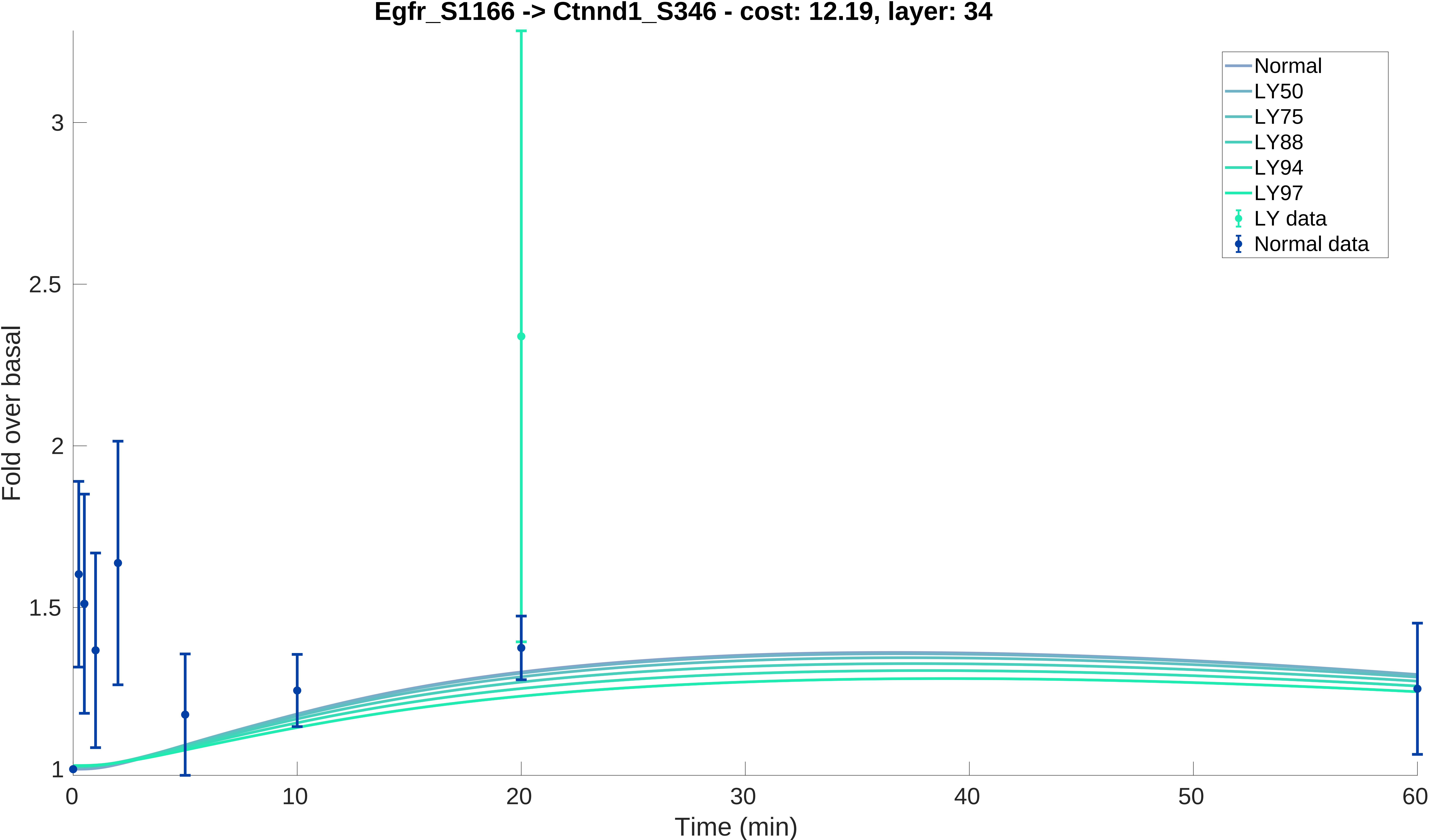

Egfr_S1166 -> Ctnnd1_S346 - cost: 12.19, layer: 34
Normal
LY50
LY75
LY88
LY94
LY97
LY data
Normal data
Fold over basal
3
2.5
2
1.5
1
0
10
20
30
40
50
60
Time (min)

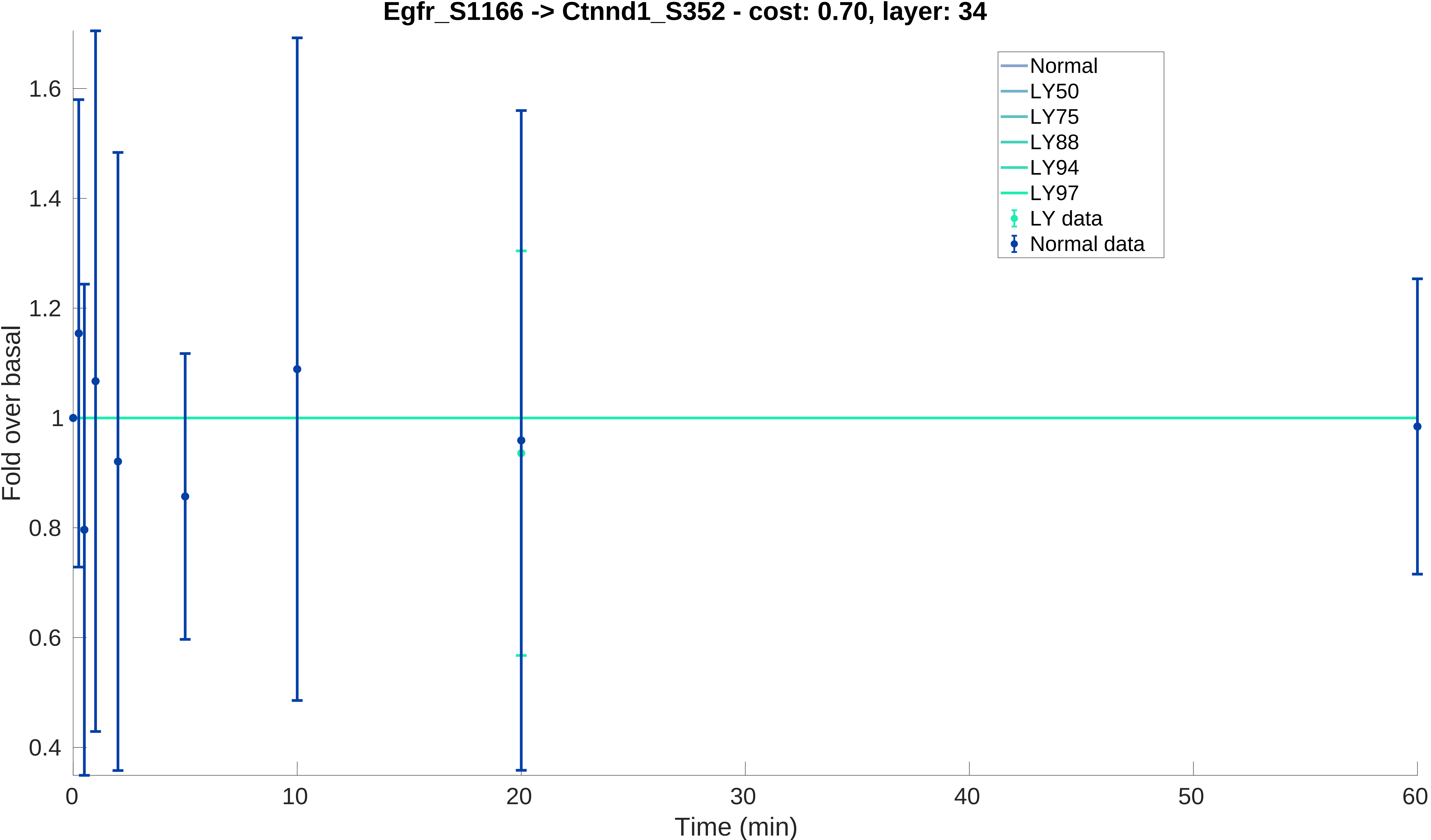
Egfr_S1166 -> Ctnnd1_S352 - cost: 0.70, layer: 34
Normal
LY50
LY75
LY88
LY94
LY97
LY data
Normal data
Fold over basal
Time (min)
0.4
0.6
0.8
1
1.2
1.4
1.6
0
10
20
30
40
50
60

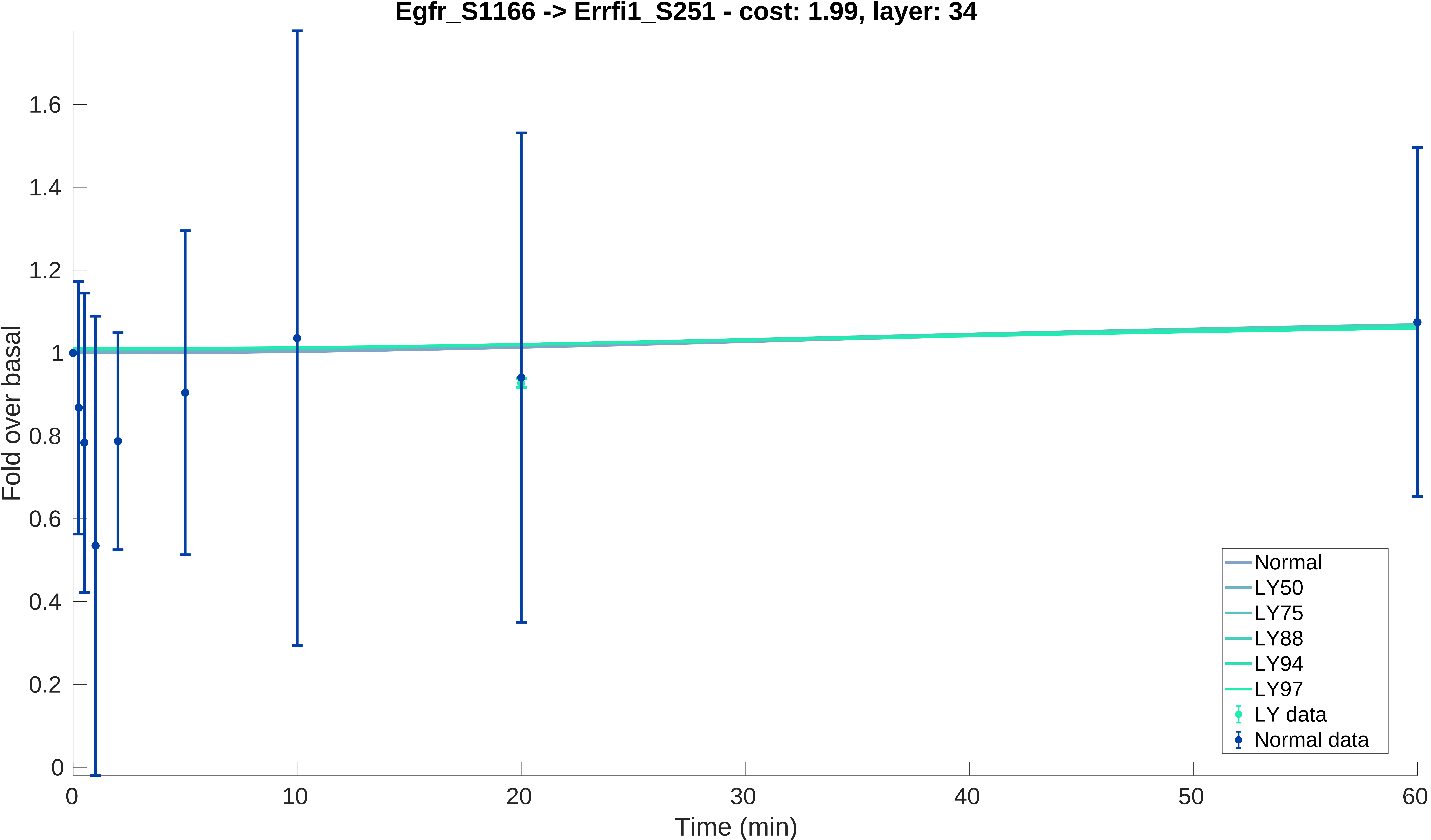
Egfr_S1166 -> Errfi1_S251 - cost: 1.99, layer: 34
Fold over basal
Time (min)
0
0.2
0.4
0.6
0.8
1
1.2
1.4
1.6
0
10
20
30
40
50
60
Normal
LY50
LY75
LY88
LY94
LY97
LY data
Normal data

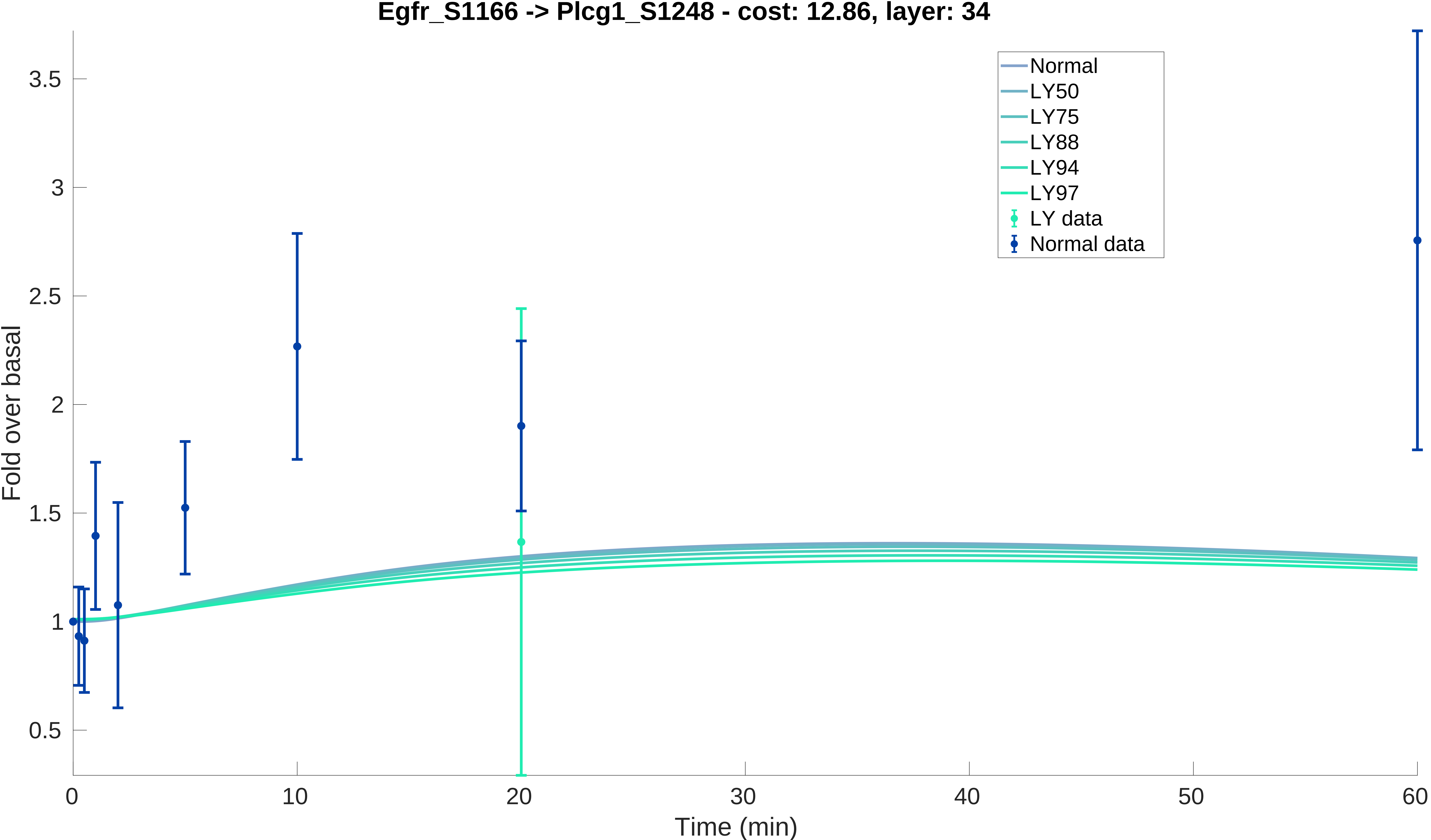
Egfr_S1166 -> Plcg1_S1248 - cost: 12.86, layer: 34
Fold over basal
3.5
3
2.5
2
1.5
1
0.5
0
10
20
30
40
50
60
Time (min)
Normal
LY50
LY75
LY88
LY94
LY97
LY data
Normal data

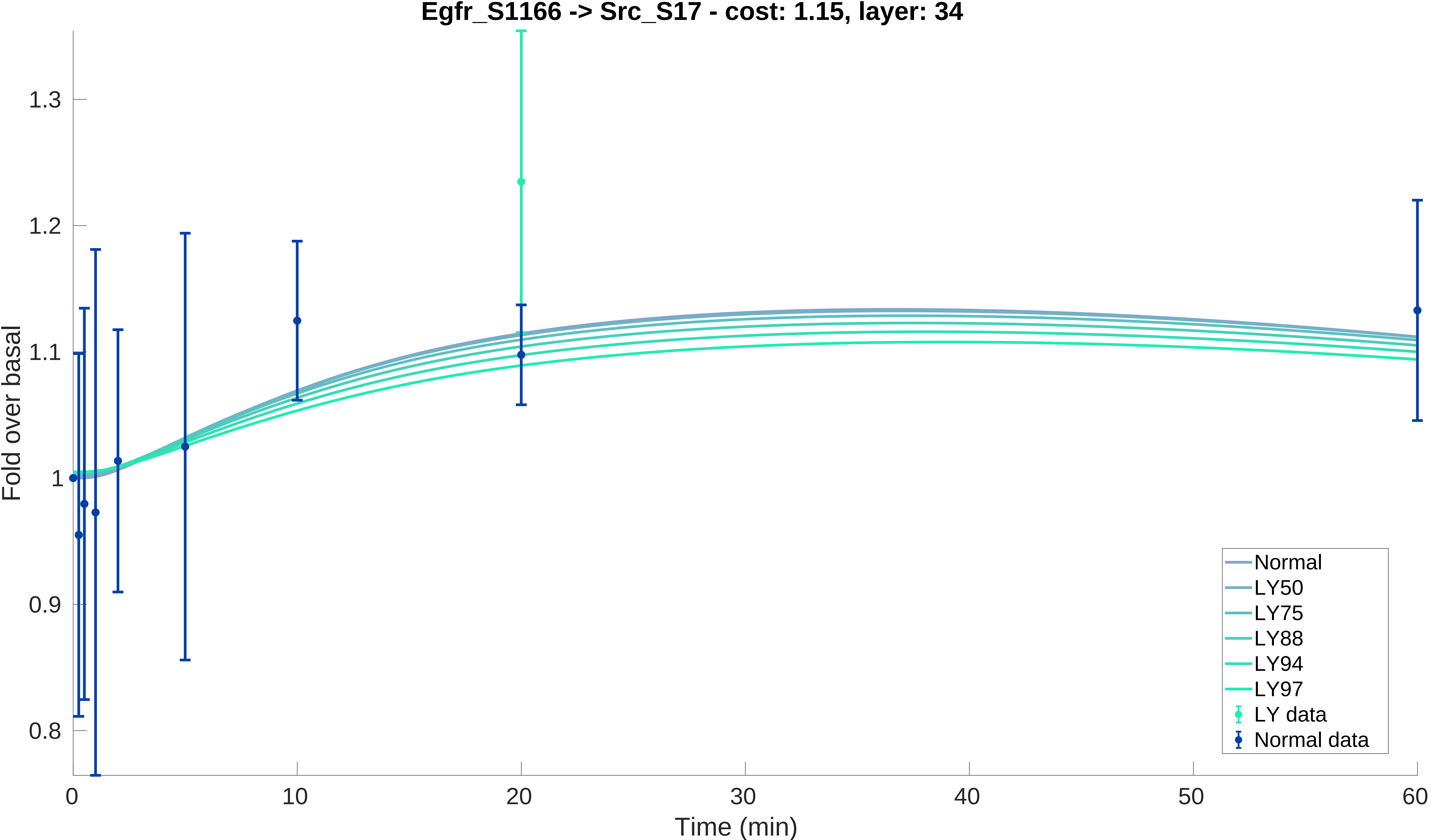

Egfr_S1166 -> Src_S17 - cost: 1.15, layer: 34
Fold over basal
1.3
1.2
1.1
1
0.9
0.8
0
10
20
30
40
50
60
Time (min)
Normal
LY50
LY75
LY88
LY94
LY97
LY data
Normal data

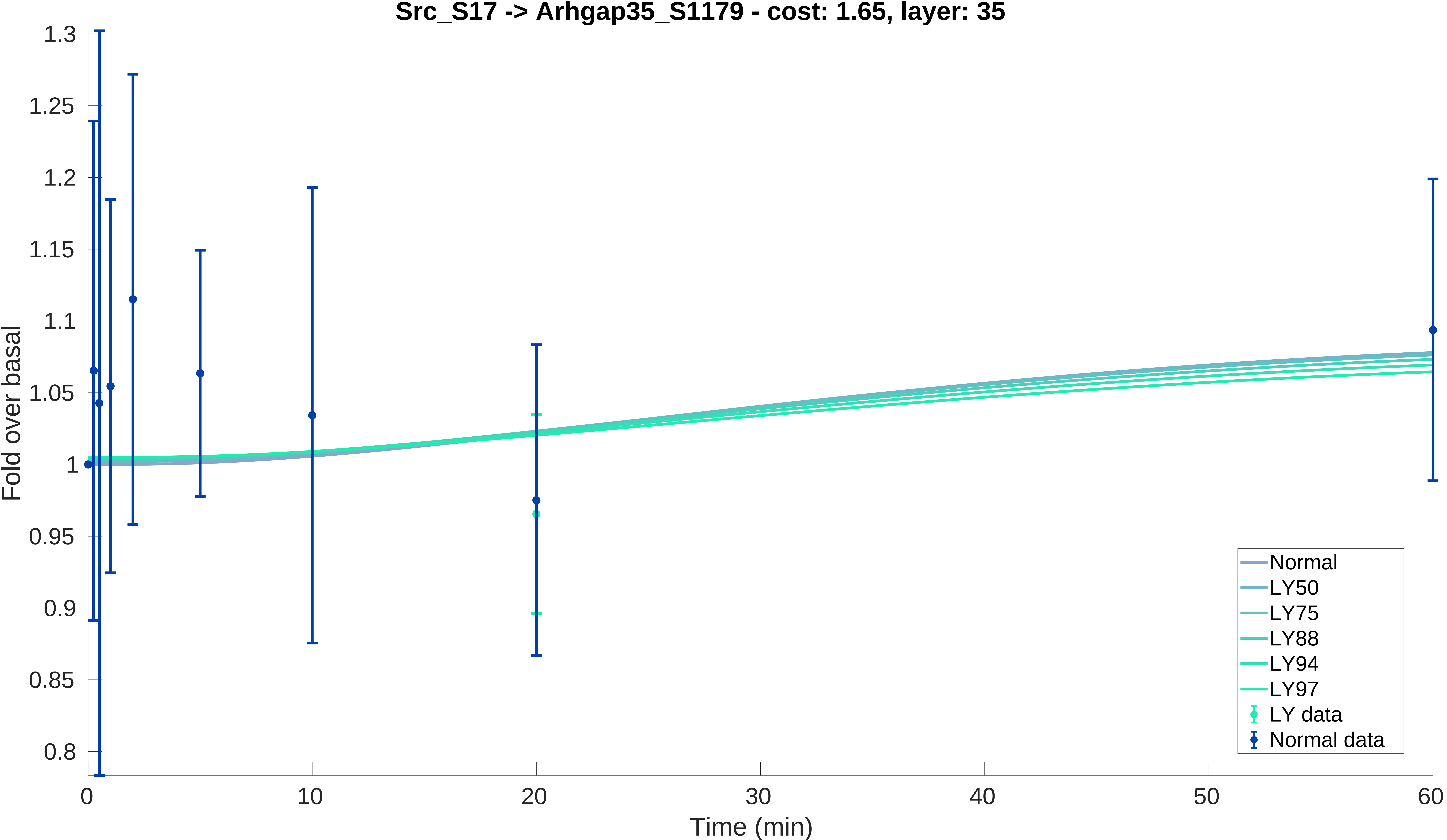

Src_S17 -> Arhgap35_S1179 - cost: 1.65, layer: 35
Fold over basal
1.3
1.25
1.2
1.15
1.1
1.05
1
0.95
0.9
0.85
0.8
0
10
20
30
40
50
60
Time (min)
Normal
LY50
LY75
LY88
LY94
LY97
LY data
Normal data

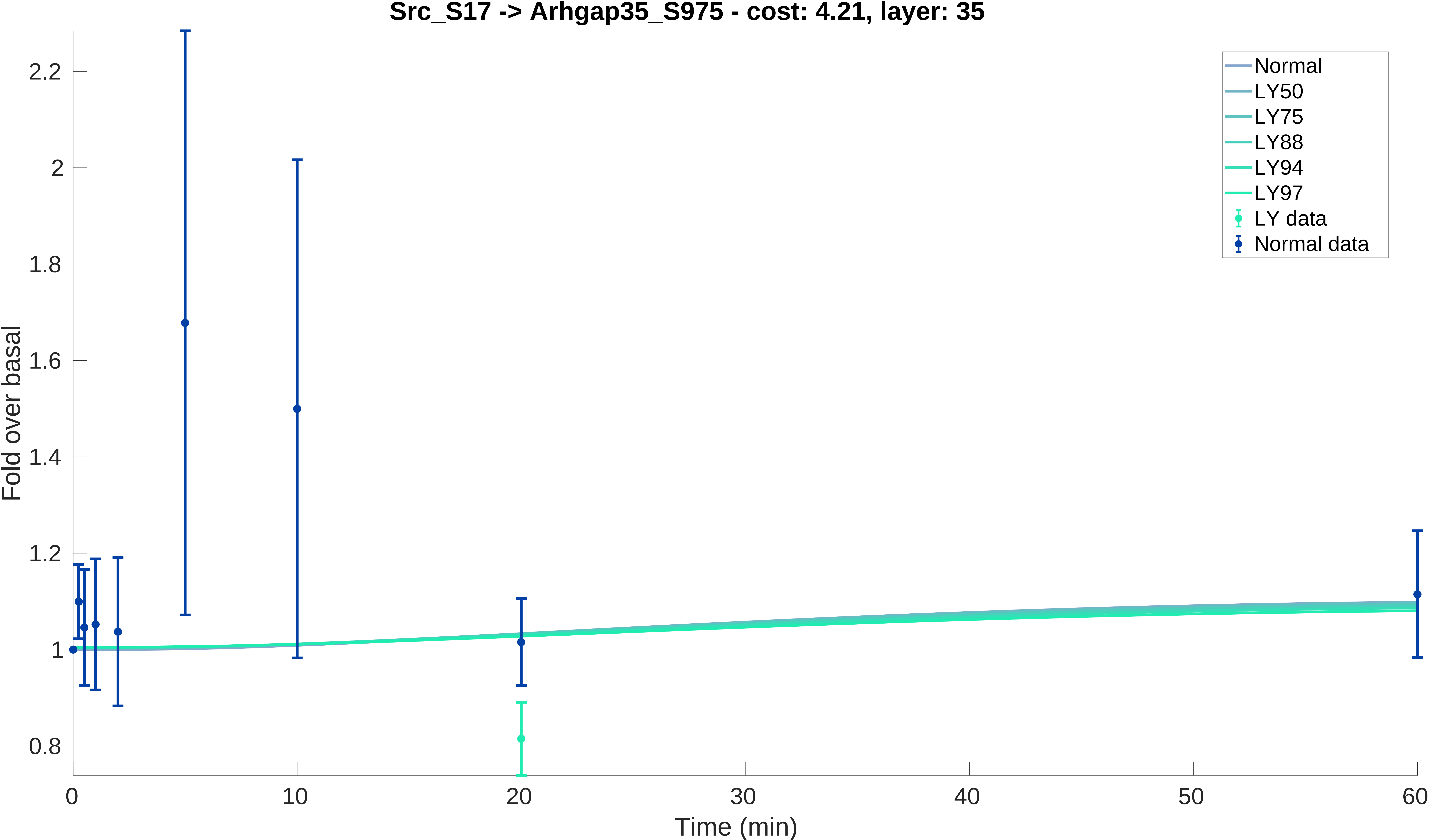
Src_S17 -> Arhgap35_S975 - cost: 4.21, layer: 35
Fold over basal
Time (min)
0.8
1
1.2
1.4
1.6
1.8
2
2.2
0
10
20
30
40
50
60
Normal
LY50
LY75
LY88
LY94
LY97
LY data
Normal data

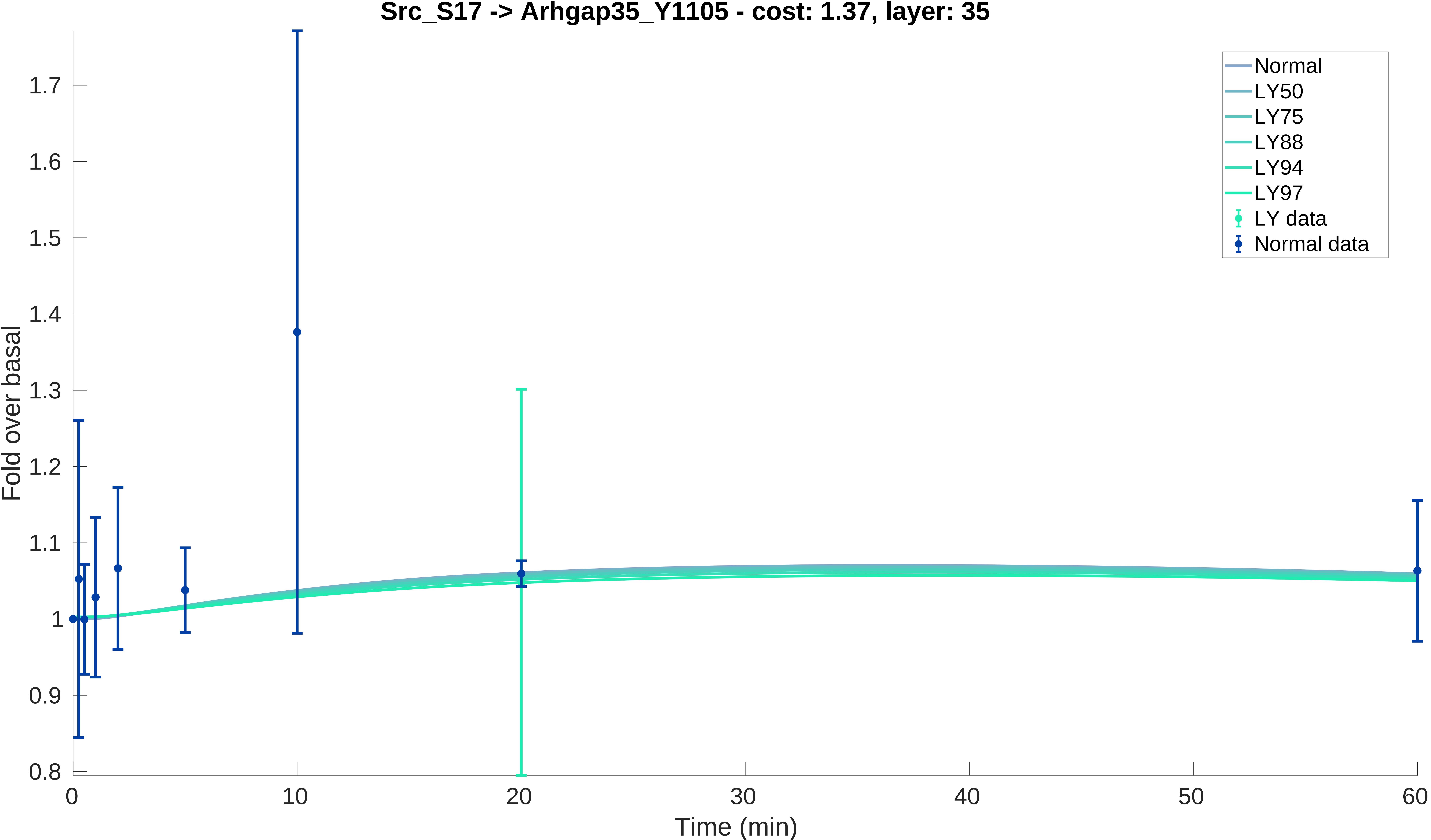

Src_S17 -> Arhgap35_Y1105 - cost: 1.37, layer: 35
Normal
LY50
LY75
LY88
LY94
LY97
LY data
Normal data
Fold over basal
Time (min)
0.8
0.9
1
1.1
1.2
1.3
1.4
1.5
1.6
1.7
0
10
20
30
40
50
60

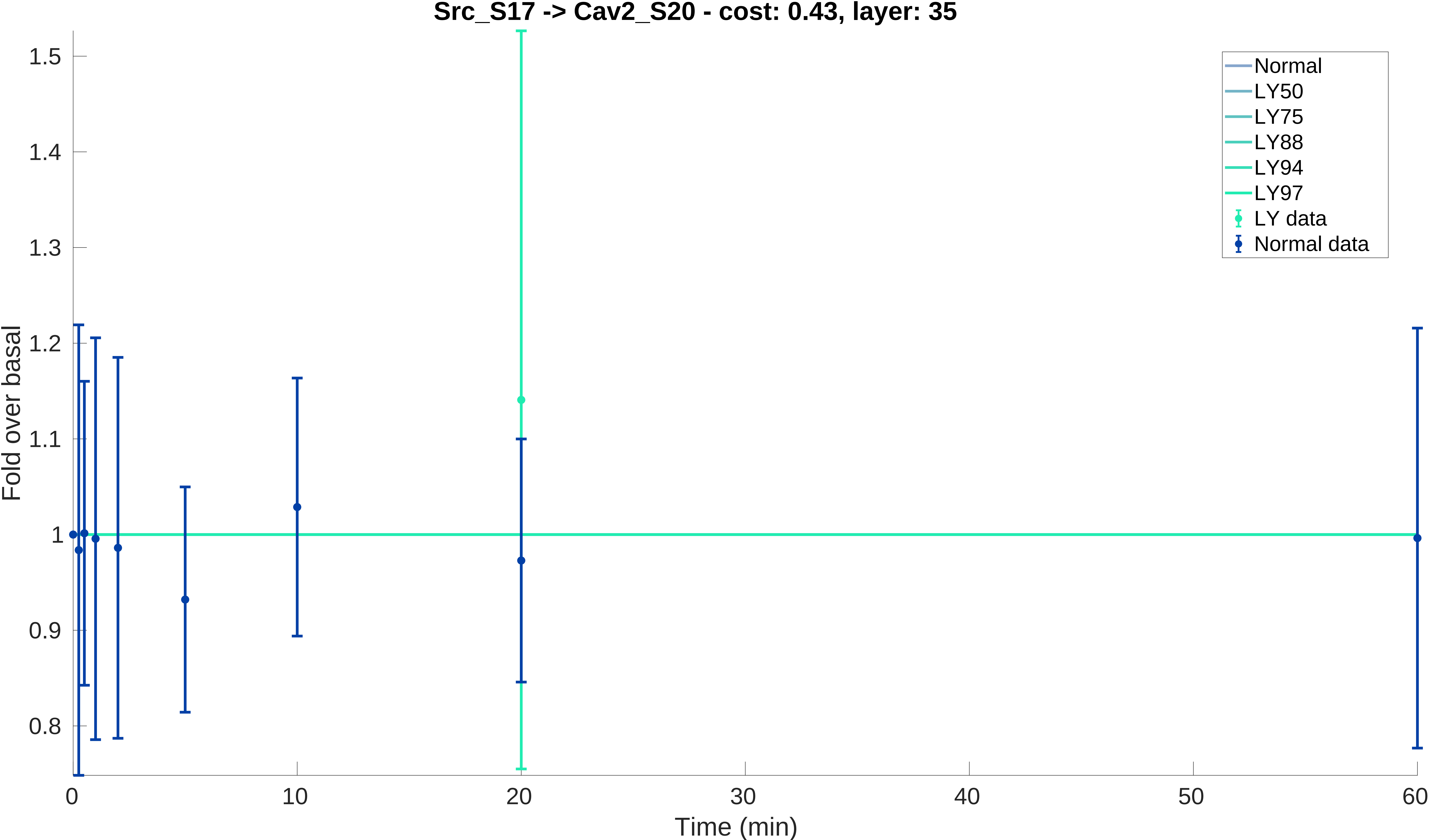

Src_S17 -> Cav2_S20 - cost: 0.43, layer: 35
Normal
LY50
LY75
LY88
LY94
LY97
LY data
Normal data
Fold over basal
1.5
1.4
1.3
1.2
1.1
1
0.9
0.8
0
10
20
30
40
50
60
Time (min)

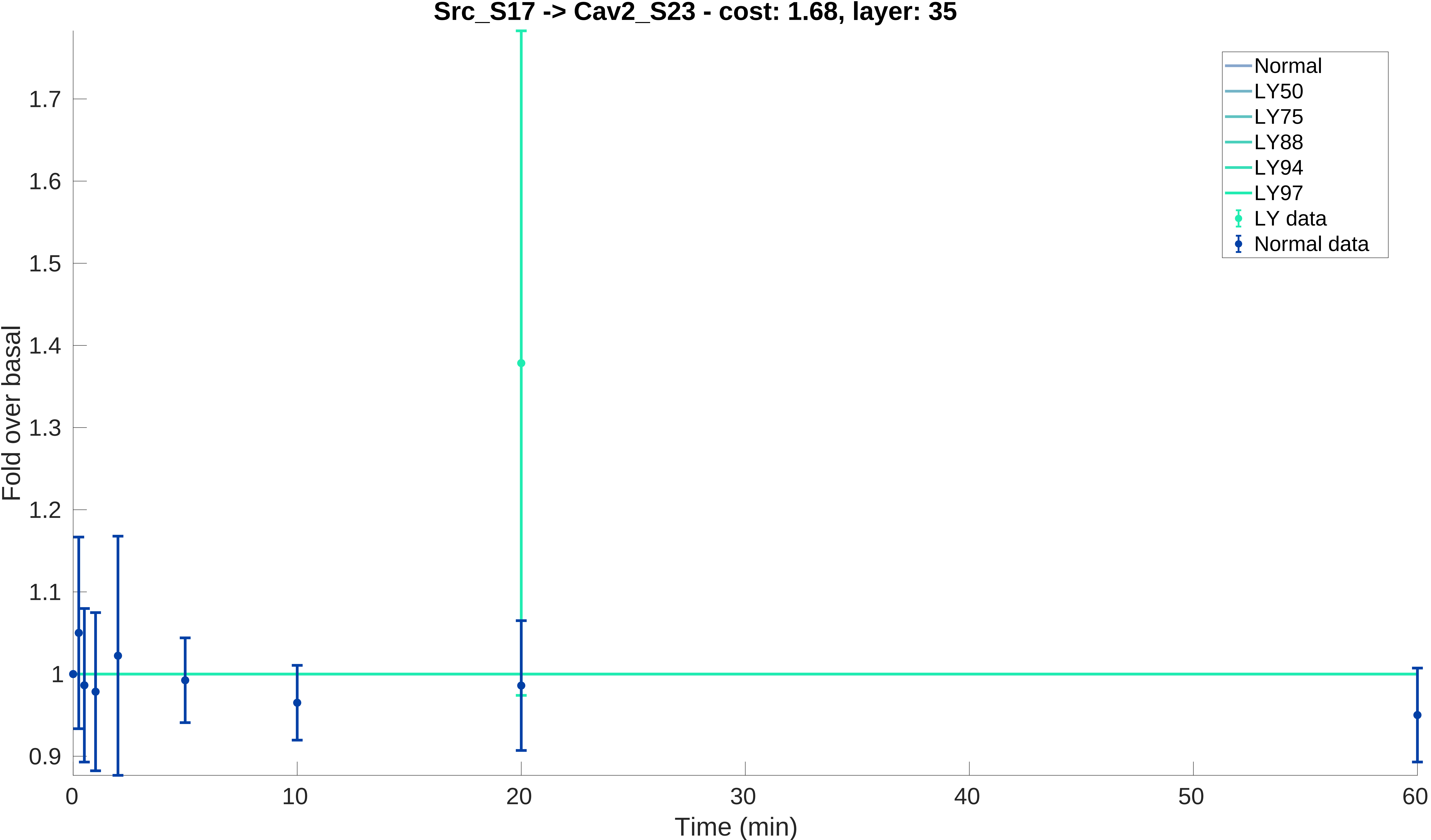

Src_S17 -> Cav2_S23 - cost: 1.68, layer: 35
Fold over basal
Time (min)
0.9
1
1.1
1.2
1.3
1.4
1.5
1.6
1.7
0
10
20
30
40
50
60
Normal
LY50
LY75
LY88
LY94
LY97
LY data
Normal data

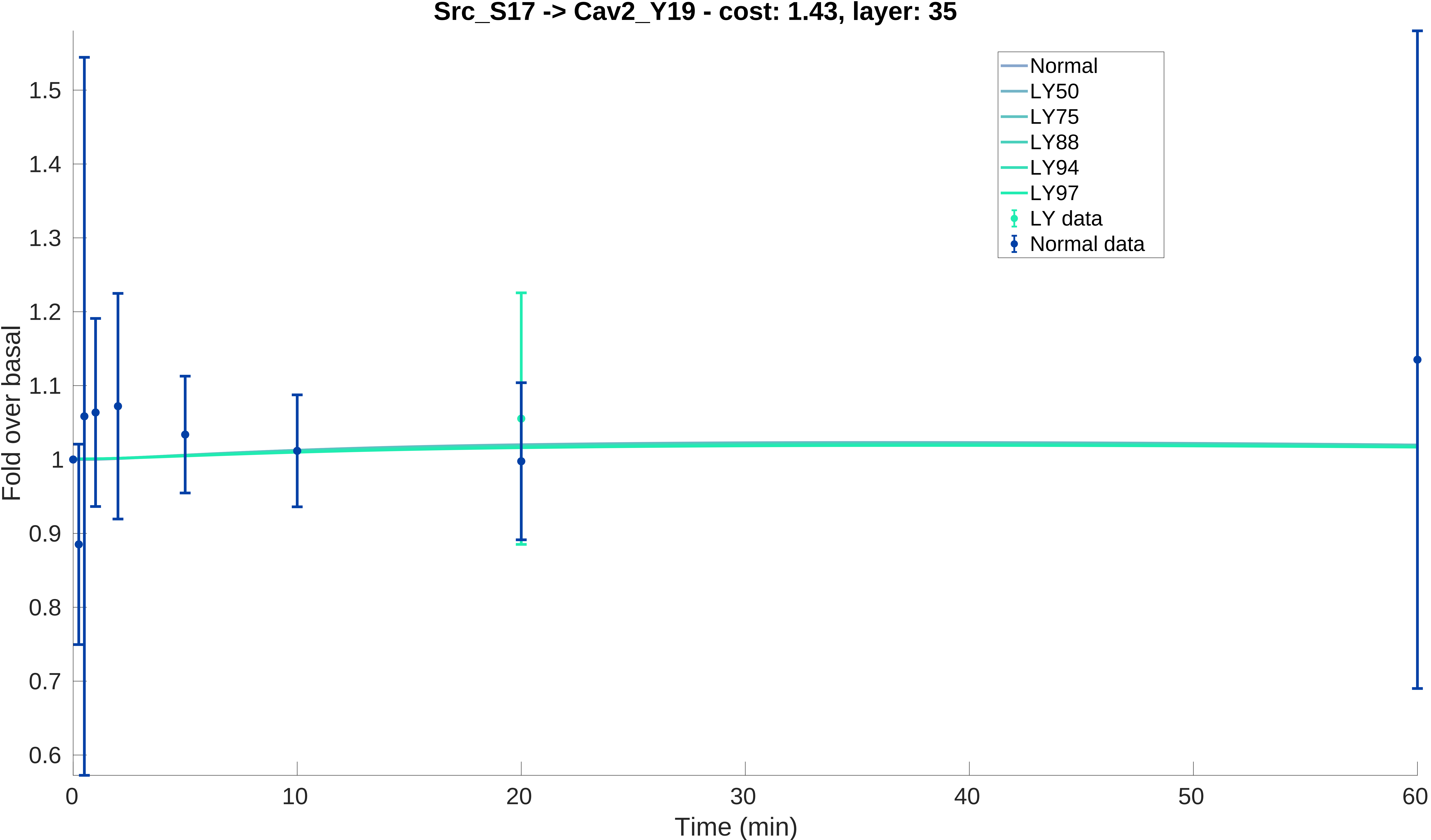

Src_S17 -> Cav2_Y19 - cost: 1.43, layer: 35
Fold over basal
Time (min)
Normal
LY50
LY75
LY88
LY94
LY97
LY data
Normal data
0.6
0.7
0.8
0.9
1
1.1
1.2
1.3
1.4
1.5
0
10
20
30
40
50
60

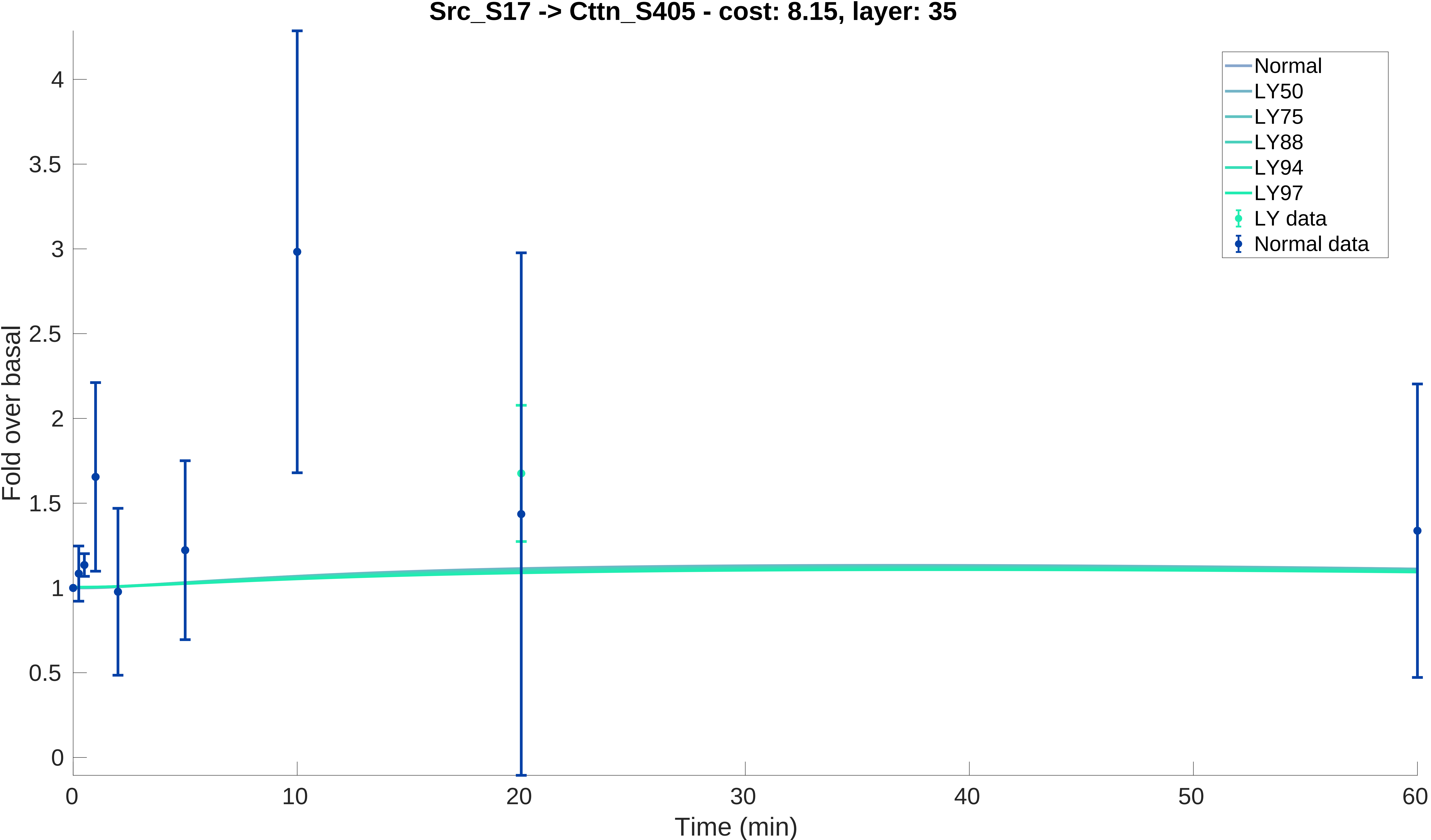

Src_S17 -> Cttn_S405 - cost: 8.15, layer: 35
Normal
LY50
LY75
LY88
LY94
LY97
LY data
Normal data
Fold over basal
4
3.5
3
2.5
2
1.5
1
0.5
0
0
10
20
30
40
50
60
Time (min)

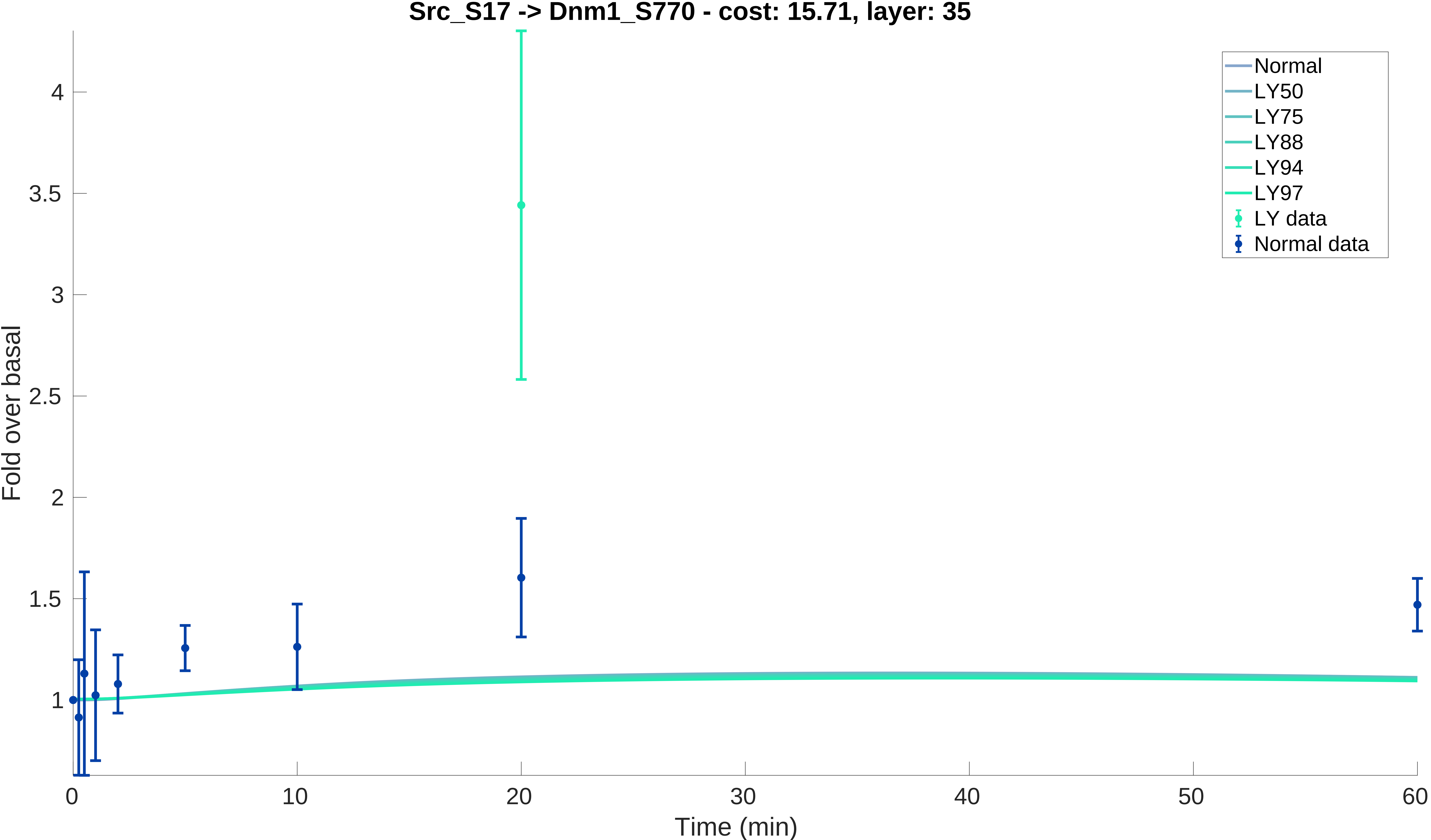

Src_S17 -> Dnm1_S770 - cost: 15.71, layer: 35
Fold over basal
Time (min)
0
10
20
30
40
50
60
1
1.5
2
2.5
3
3.5
4
Normal
LY50
LY75
LY88
LY94
LY97
LY data
Normal data

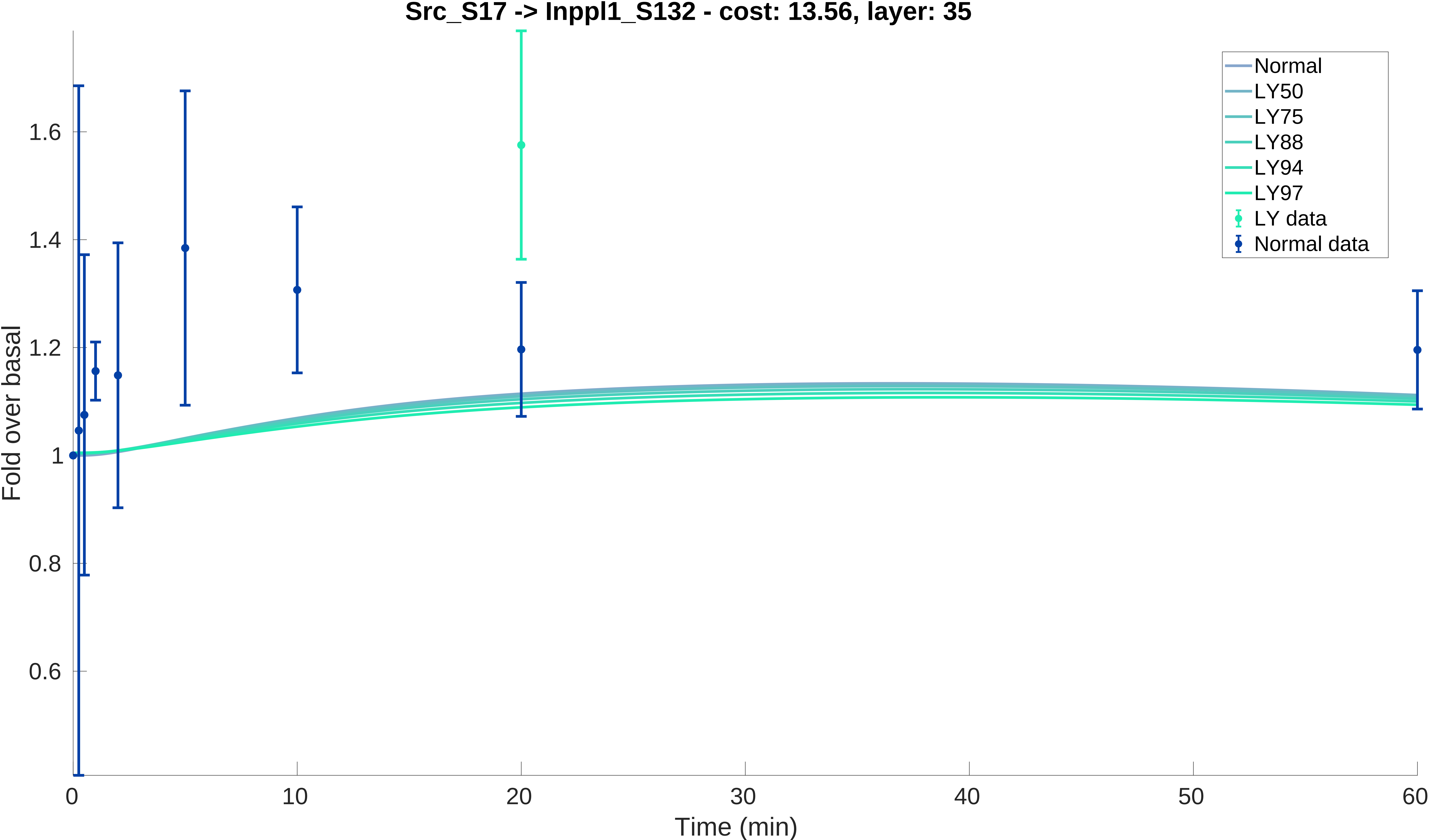

Src_S17 -> Inppl1_S132 - cost: 13.56, layer: 35
Normal
LY50
LY75
LY88
LY94
LY97
LY data
Normal data
Fold over basal
1.6
1.4
1.2
1
0.8
0.6
0
10
20
30
40
50
60
Time (min)

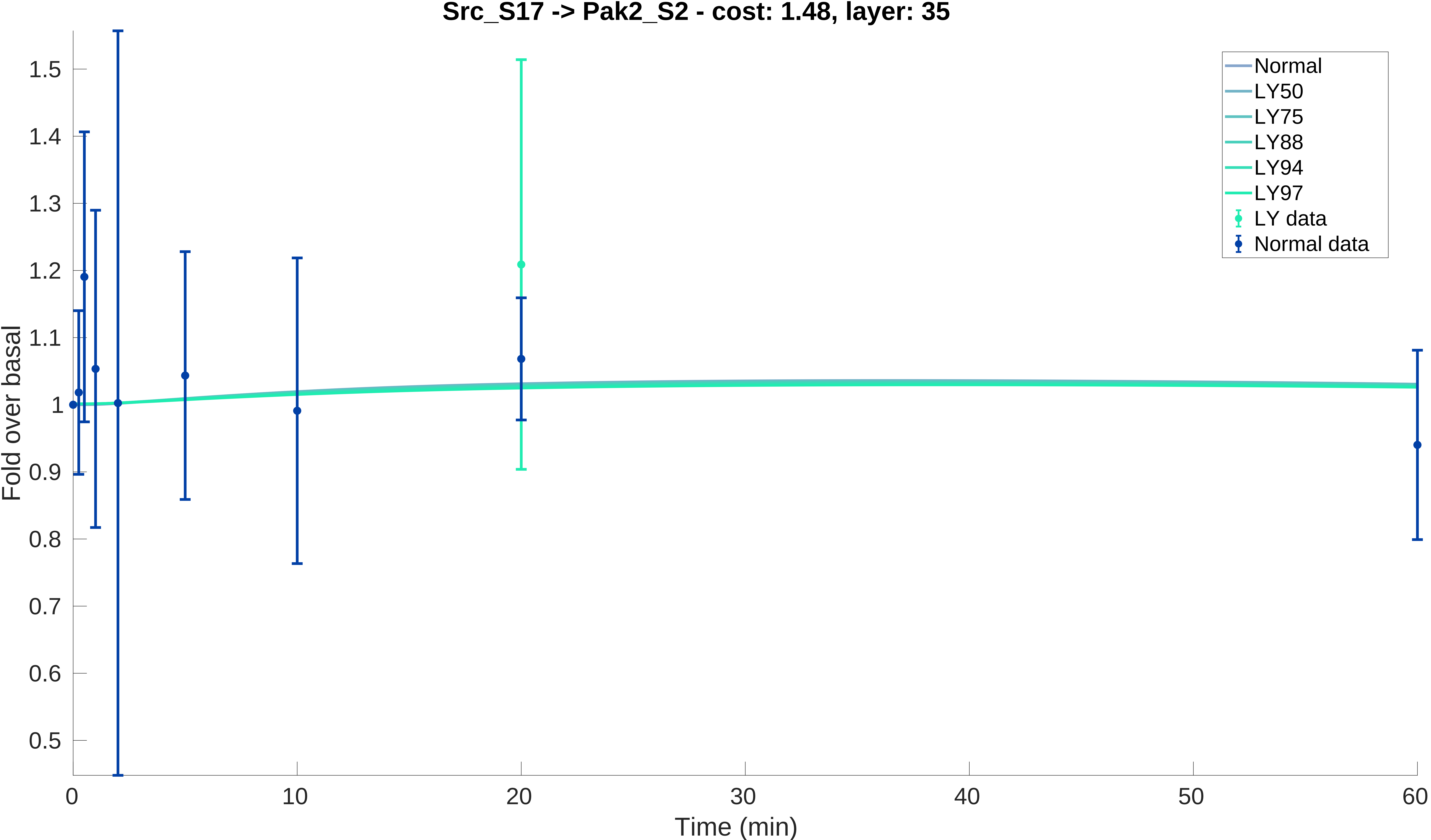

Src_S17 -> Pak2_S2 - cost: 1.48, layer: 35
Normal
LY50
LY75
LY88
LY94
LY97
LY data
Normal data
Fold over basal
1.5
1.4
1.3
1.2
1.1
1
0.9
0.8
0.7
0.6
0.5
0
10
20
30
40
50
60
Time (min)

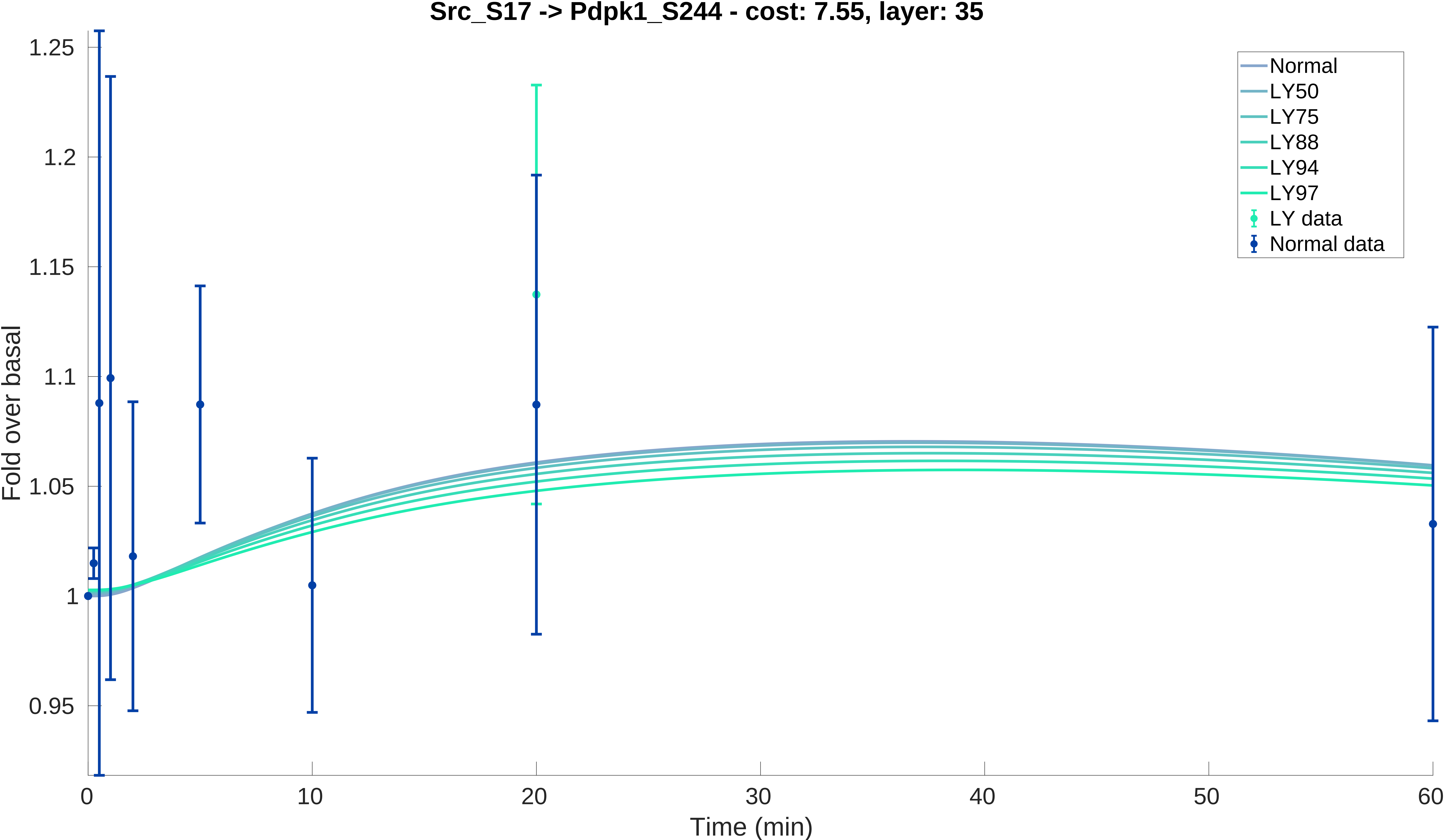

Src_S17 -> Pdpk1_S244 - cost: 7.55, layer: 35
Fold over basal
Time (min)
1.25
1.2
1.15
1.1
1.05
1
0.95
0
10
20
30
40
50
60
Normal
LY50
LY75
LY88
LY94
LY97
LY data
Normal data

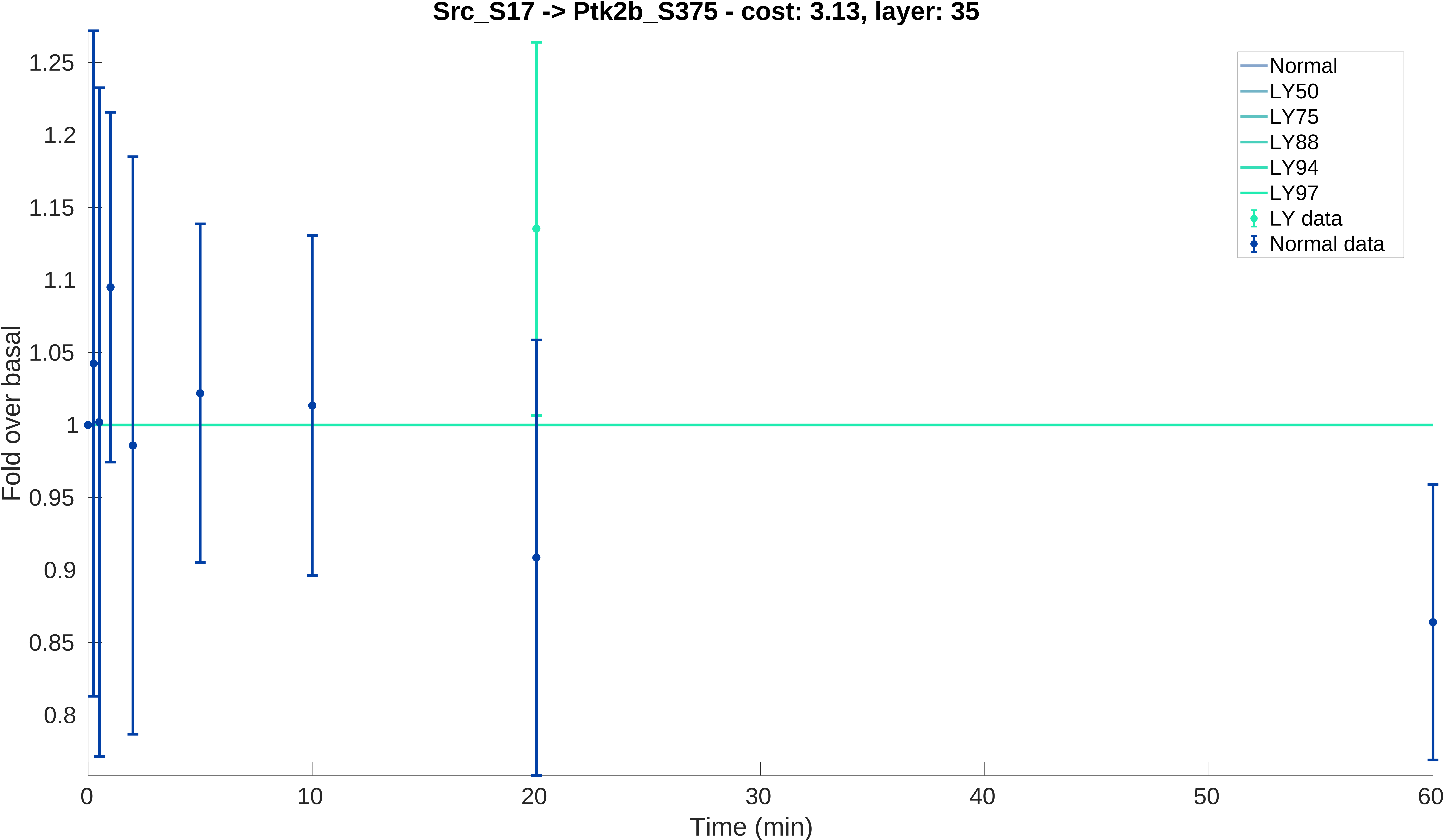

Src_S17 -> Ptk2b_S375 - cost: 3.13, layer: 35
Fold over basal
1.25
1.2
1.15
1.1
1.05
1
0.95
0.9
0.85
0.8
0
10
20
30
40
50
60
Time (min)
Normal
LY50
LY75
LY88
LY94
LY97
LY data
Normal data

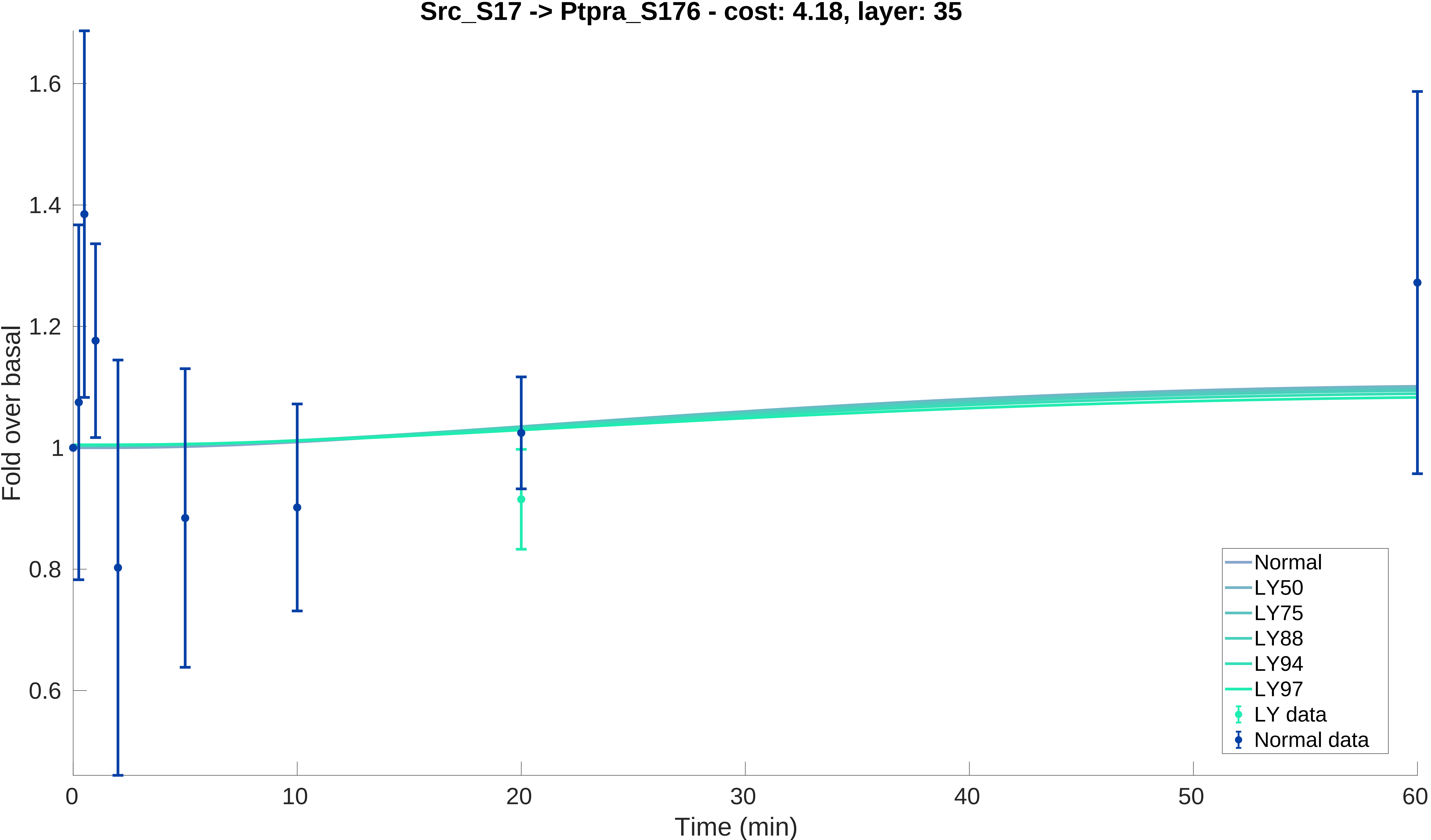

Src_S17 -> Ptpra_S176 - cost: 4.18, layer: 35
Fold over basal
1.6
1.4
1.2
1
0.8
0.6
0
10
20
30
40
50
60
Time (min)
Normal
LY50
LY75
LY88
LY94
LY97
LY data
Normal data

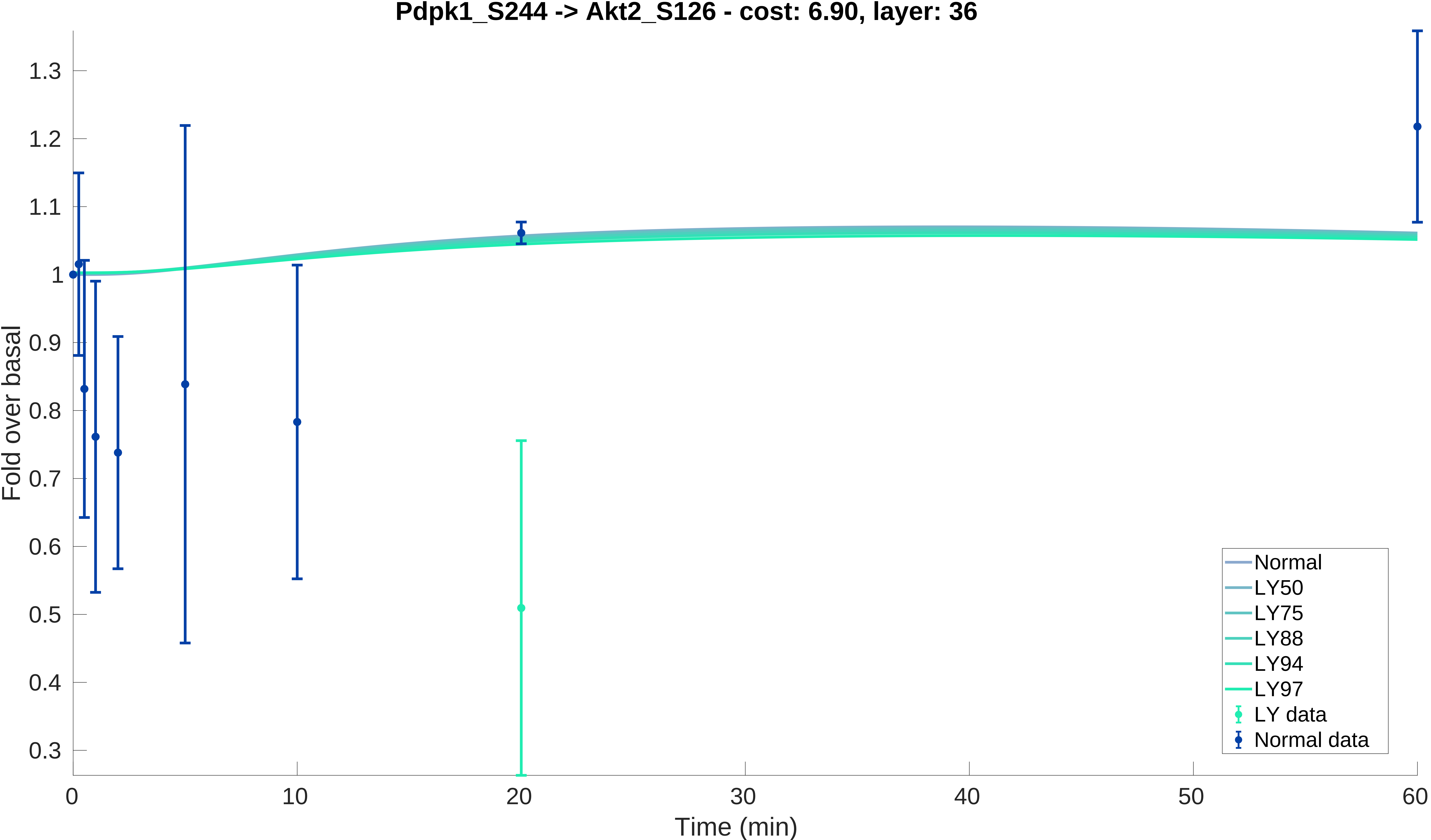

Pdpk1_S244 -> Akt2_S126 - cost: 6.90, layer: 36
Fold over basal
Time (min)
0
10
20
30
40
50
60
0.3
0.4
0.5
0.6
0.7
0.8
0.9
1
1.1
1.2
1.3
Normal
LY50
LY75
LY88
LY94
LY97
LY data
Normal data

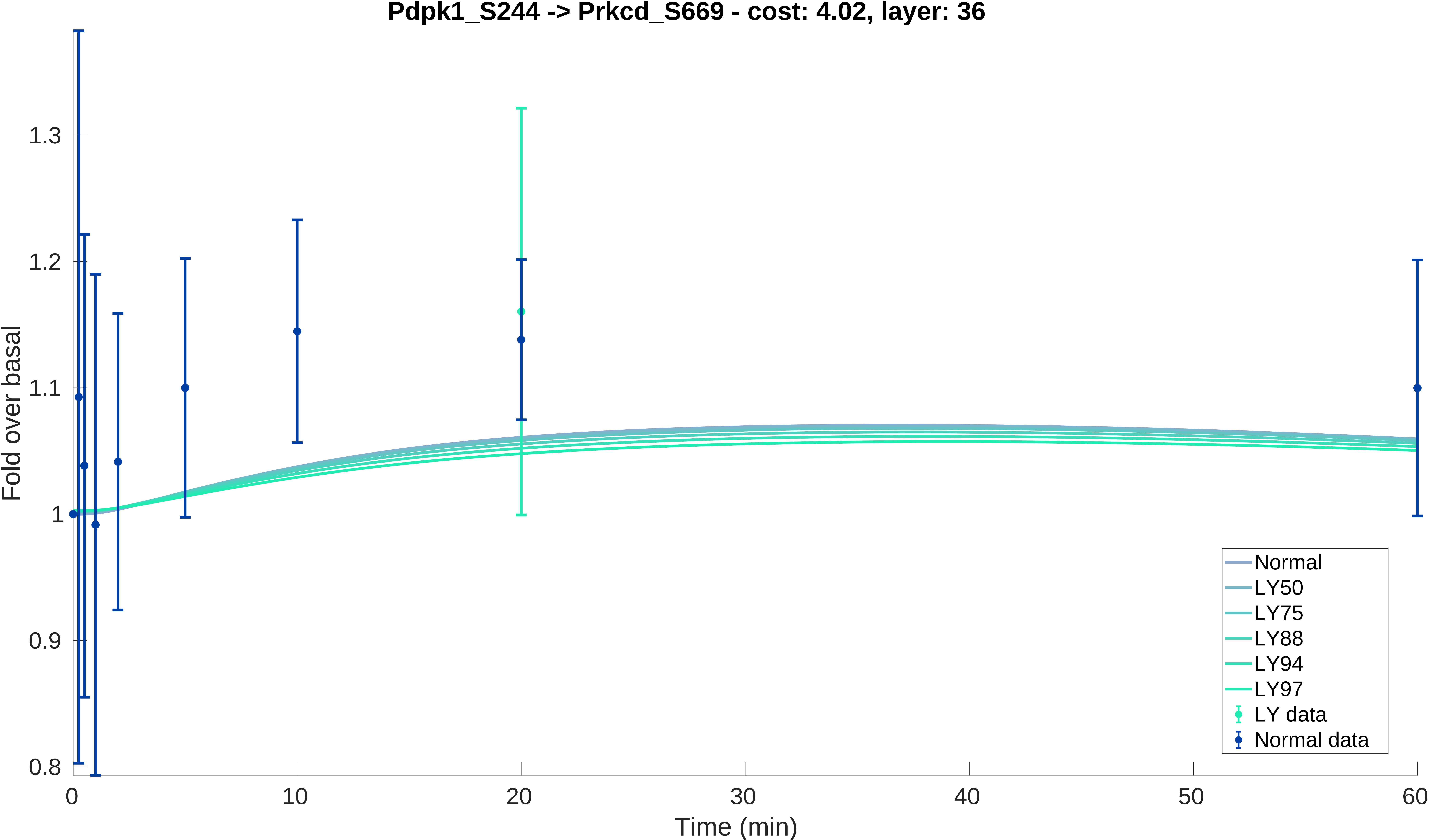

Pdpk1_S244 -> Prkcd_S669 - cost: 4.02, layer: 36
Fold over basal
1.3
1.2
1.1
1
0.9
0.8
0
10
20
30
40
50
60
Time (min)
Normal
LY50
LY75
LY88
LY94
LY97
LY data
Normal data

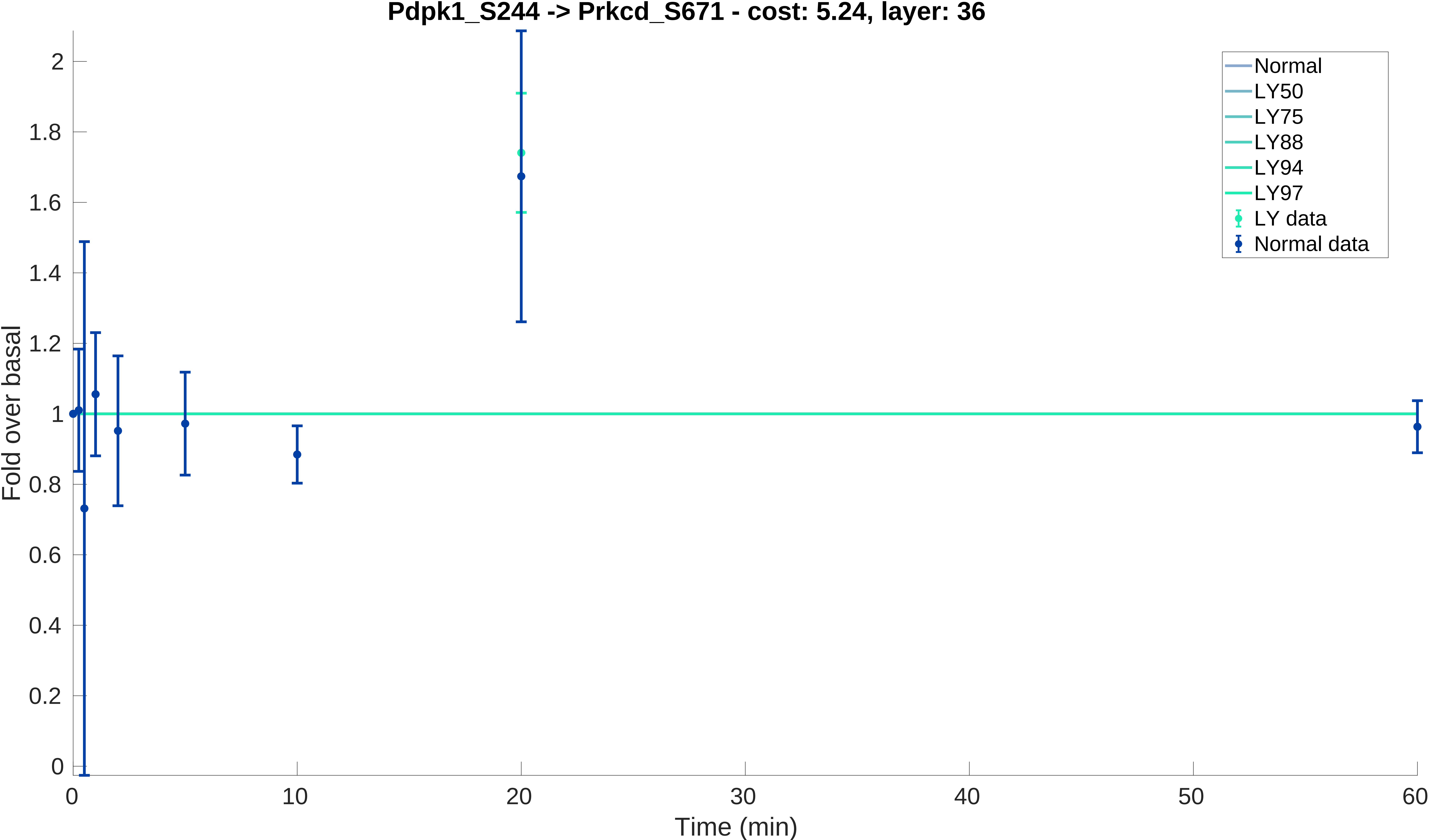

Pdpk1_S244 -> Prkcd_S671 - cost: 5.24, layer: 36
Fold over basal
Time (min)
0
0.2
0.4
0.6
0.8
1
1.2
1.4
1.6
1.8
2
0
10
20
30
40
50
60
Normal
LY50
LY75
LY88
LY94
LY97
LY data
Normal data

#### Pdpk1_S244 -> Prkcd_S688 - cost: 5.53, layer: 36

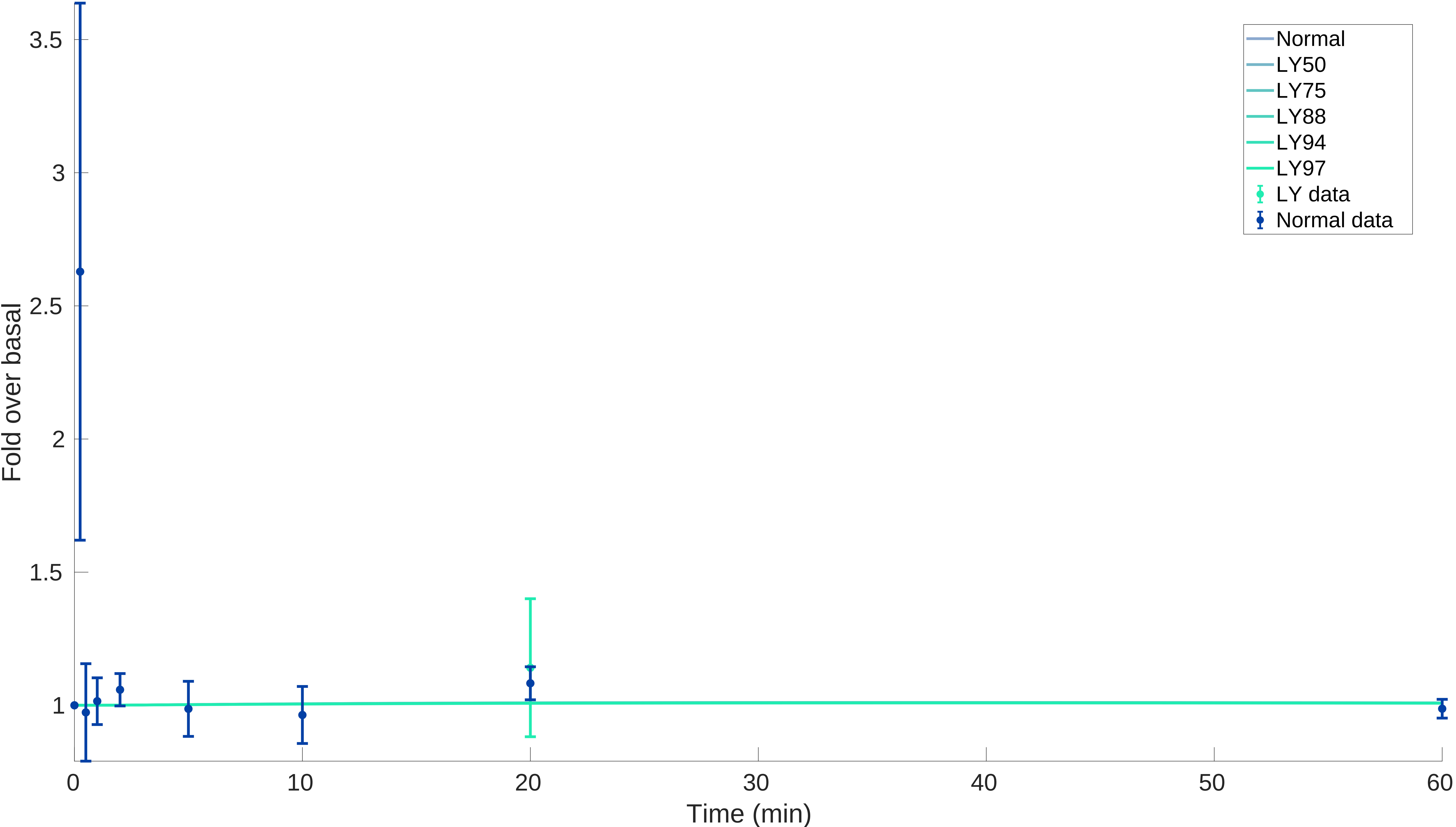

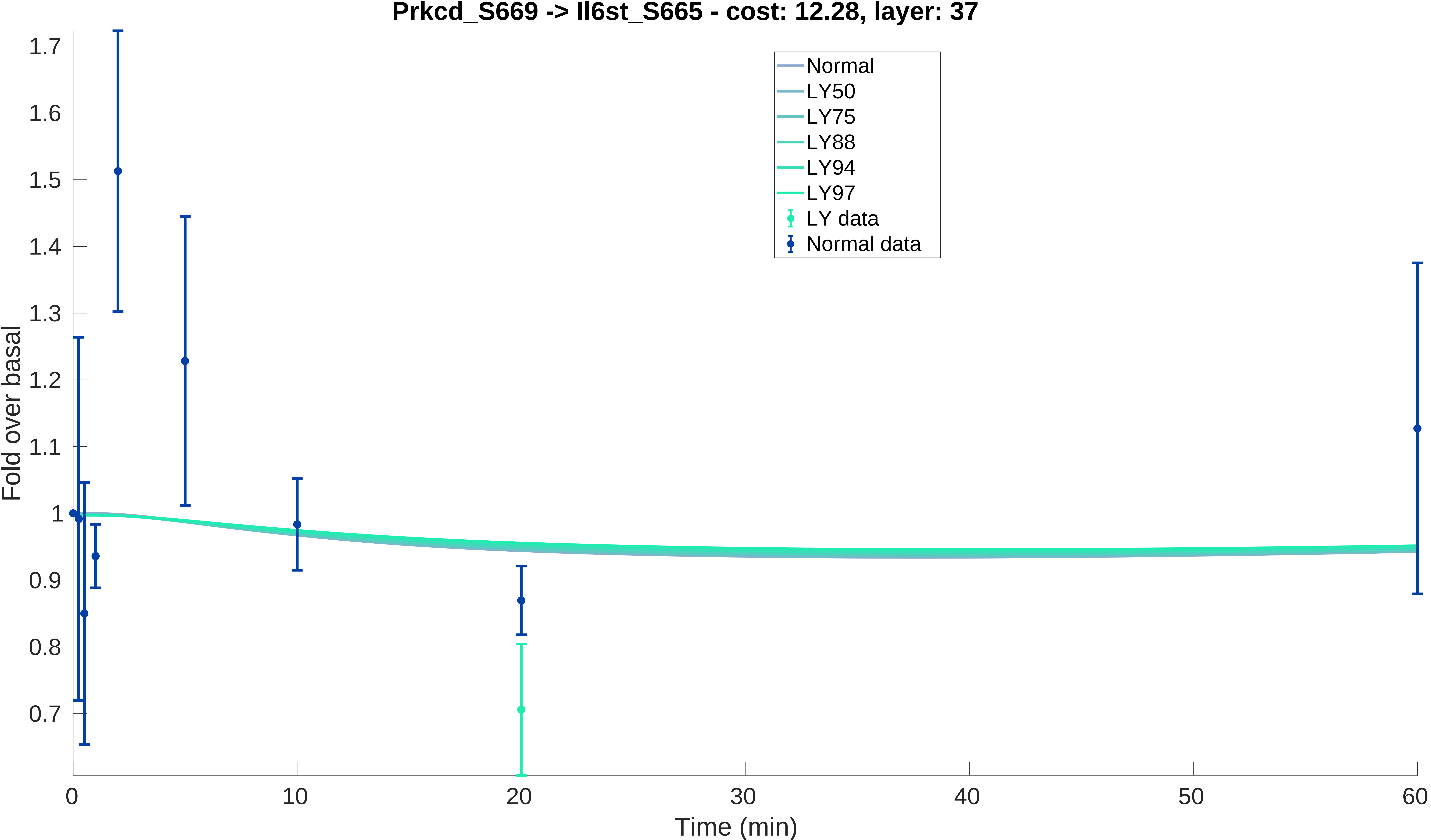
Prkcd_S669 -> Il6st_S665 - cost: 12.28, layer: 37
Fold over basal
Time (min)
Normal
LY50
LY75
LY88
LY94
LY97
LY data
Normal data
1.7
1.6
1.5
1.4
1.3
1.2
1.1
1
0.9
0.8
0.7
0
10
20
30
40
50
60

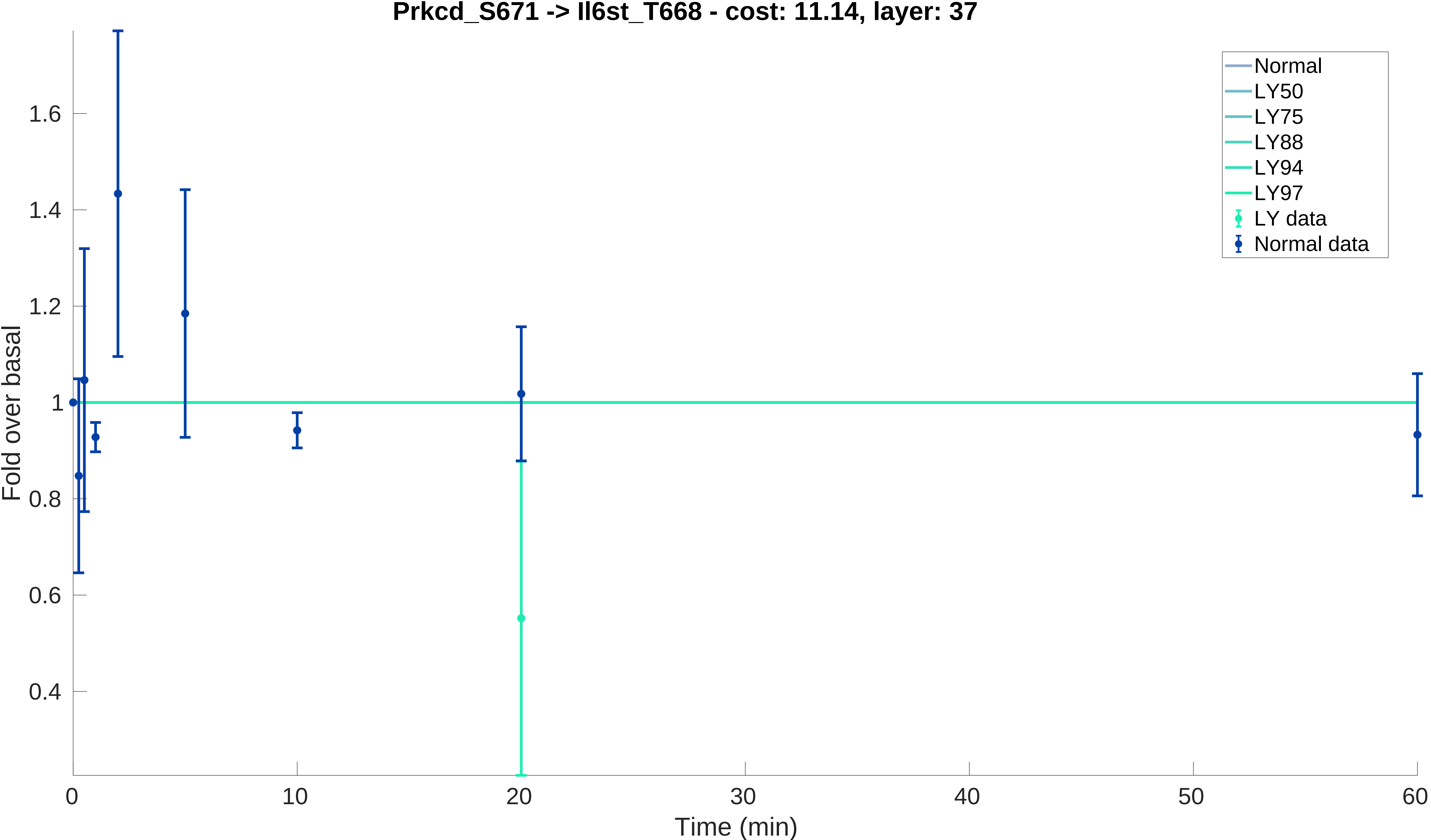

Prkcd_S671 -> Il6st_T668 - cost: 11.14, layer: 37
Fold over basal
Time (min)
0
10
20
30
40
50
60
0.4
0.6
0.8
1
1.2
1.4
1.6
Normal
LY50
LY75
LY88
LY94
LY97
LY data
Normal data

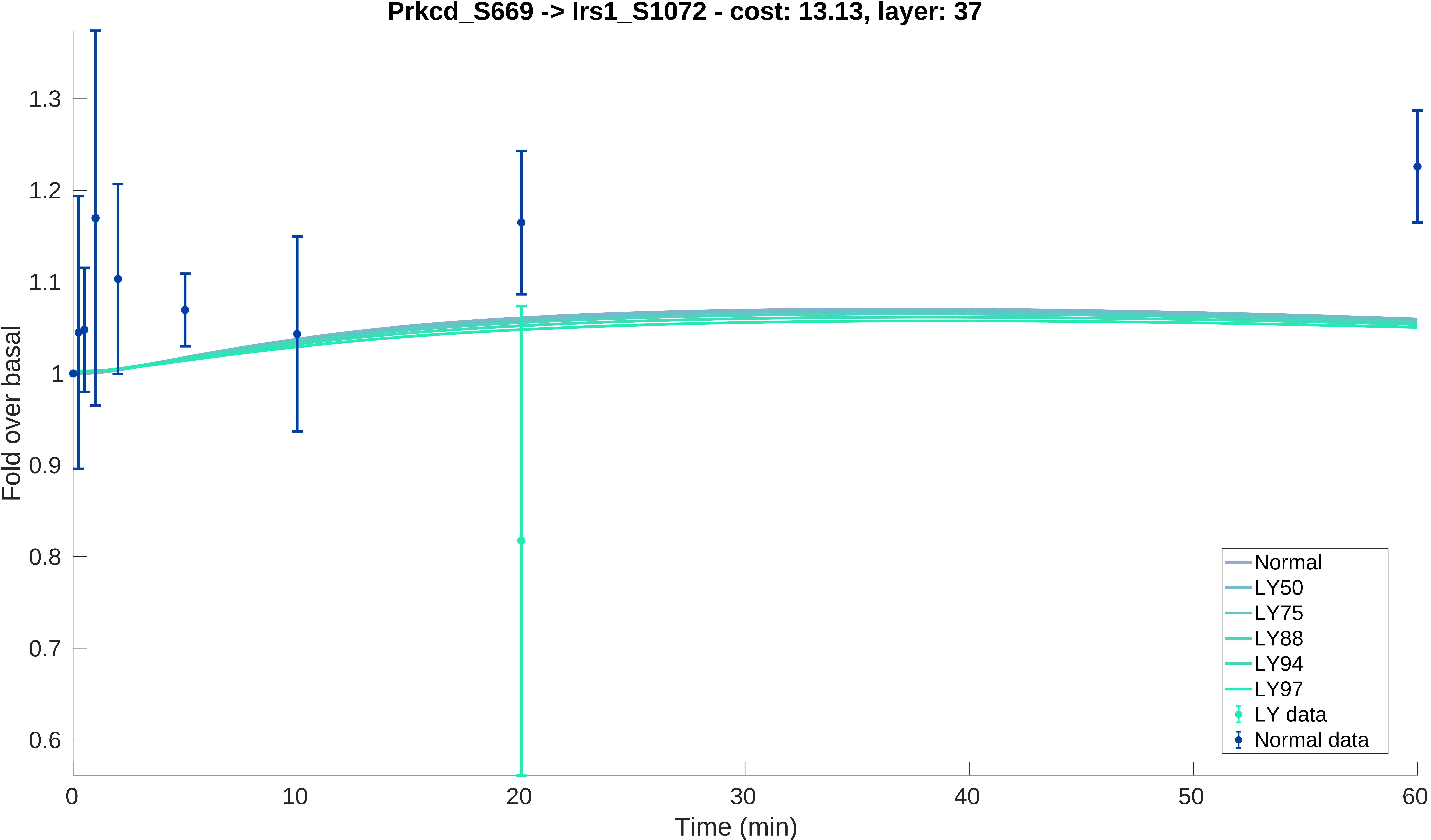

Prkcd_S669 -> Irs1_S1072 - cost: 13.13, layer: 37
Fold over basal
1.3
1.2
1.1
1
0.9
0.8
0.7
0.6
0
10
20
30
40
50
60
Time (min)
Normal
LY50
LY75
LY88
LY94
LY97
LY data
Normal data

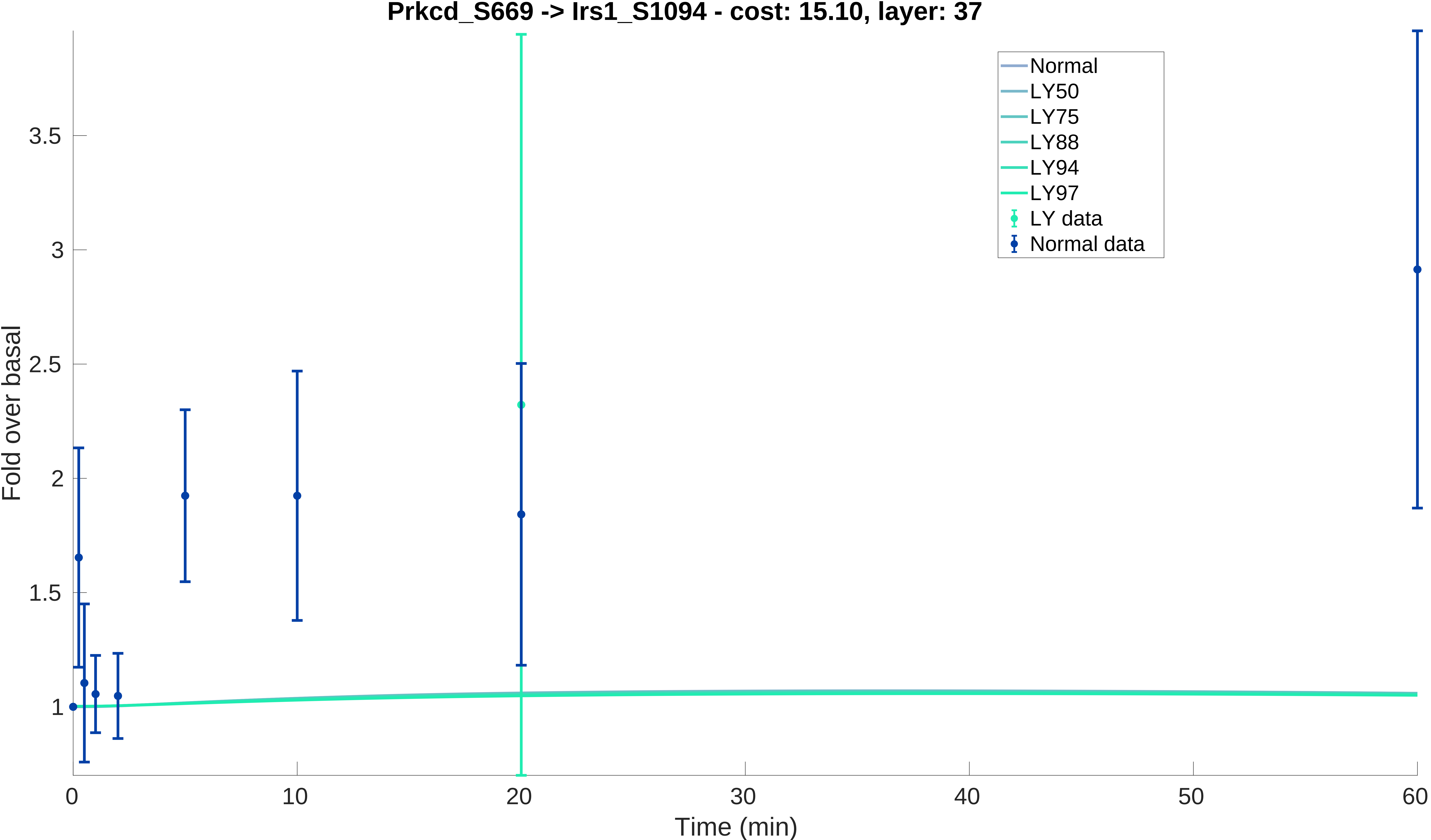

Prkcd_S669 -> Irs1_S1094 - cost: 15.10, layer: 37
Normal
LY50
LY75
LY88
LY94
LY97
LY data
Normal data
Fold over basal
Time (min)
3.5
3
2.5
2
1.5
1
0
10
20
30
40
50
60

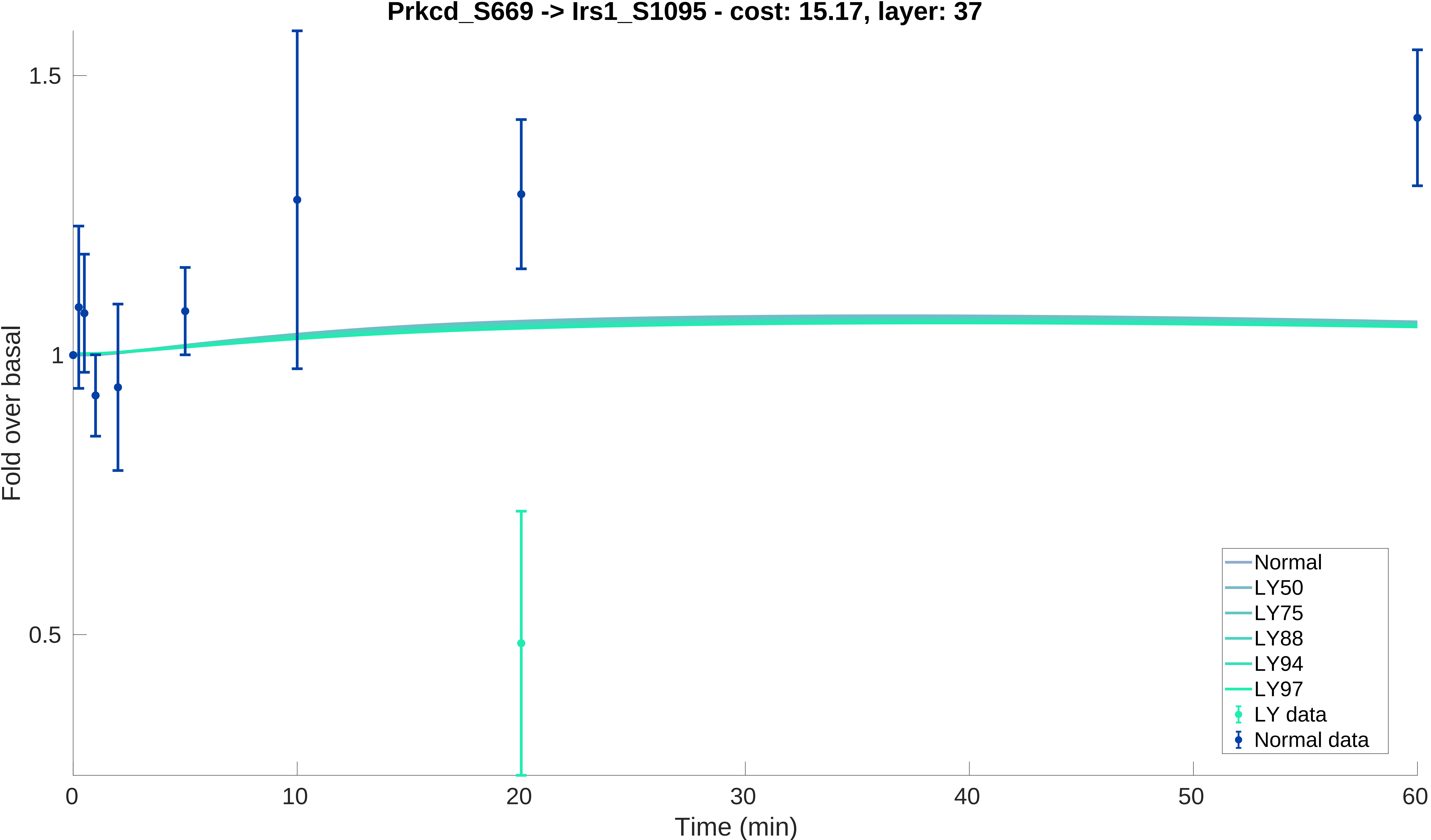

Prkcd_S669 -> Irs1_S1095 - cost: 15.17, layer: 37
Fold over basal
1.5
1
0.5
0
10
20
30
40
50
60
Time (min)
Normal
LY50
LY75
LY88
LY94
LY97
LY data
Normal data

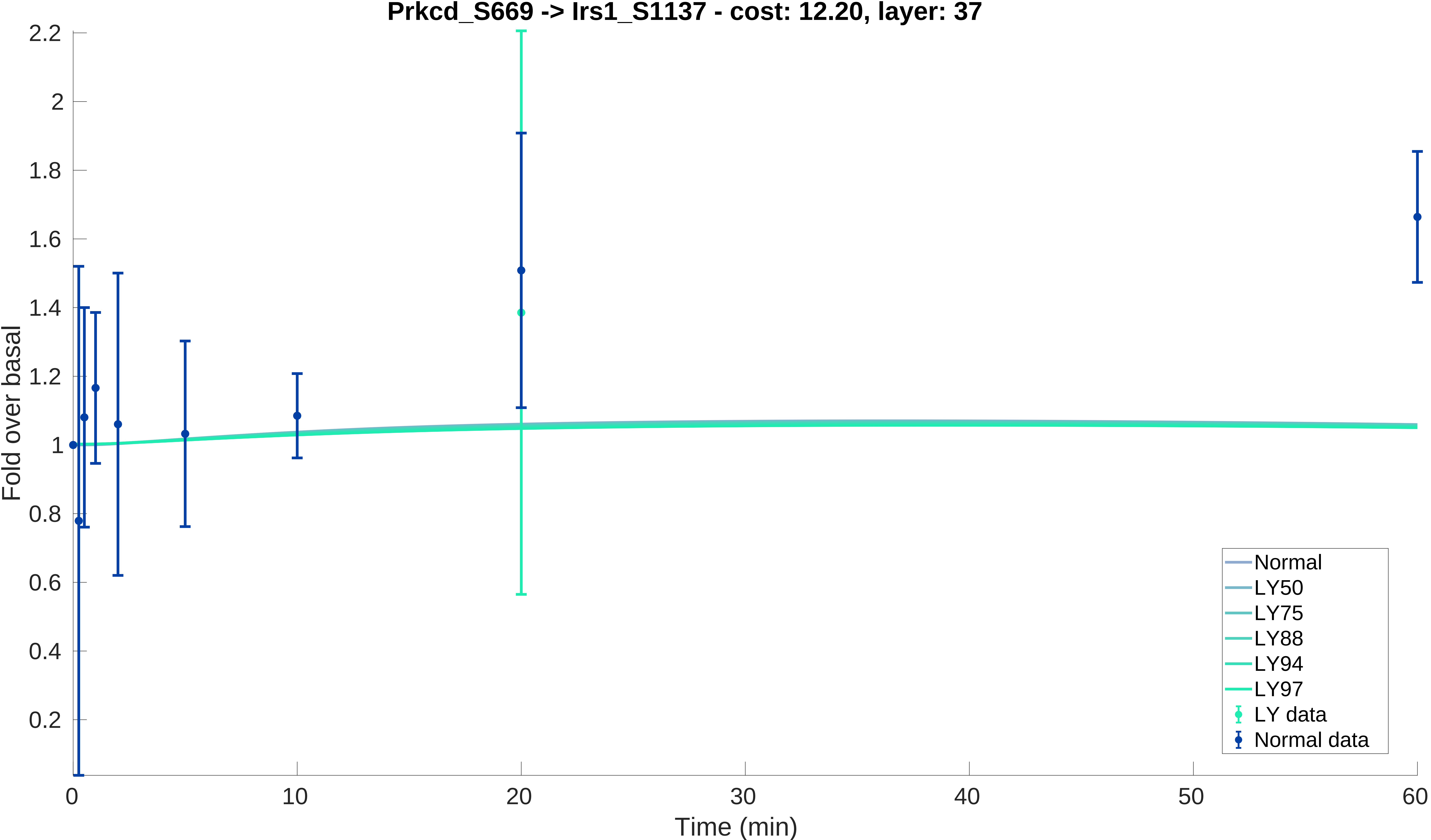

Prkcd_S669 -> Irs1_S1137 - cost: 12.20, layer: 37
Fold over basal
Time (min)
2.2
2
1.8
1.6
1.4
1.2
1
0.8
0.6
0.4
0.2
0
10
20
30
40
50
60
Normal
LY50
LY75
LY88
LY94
LY97
LY data
Normal data

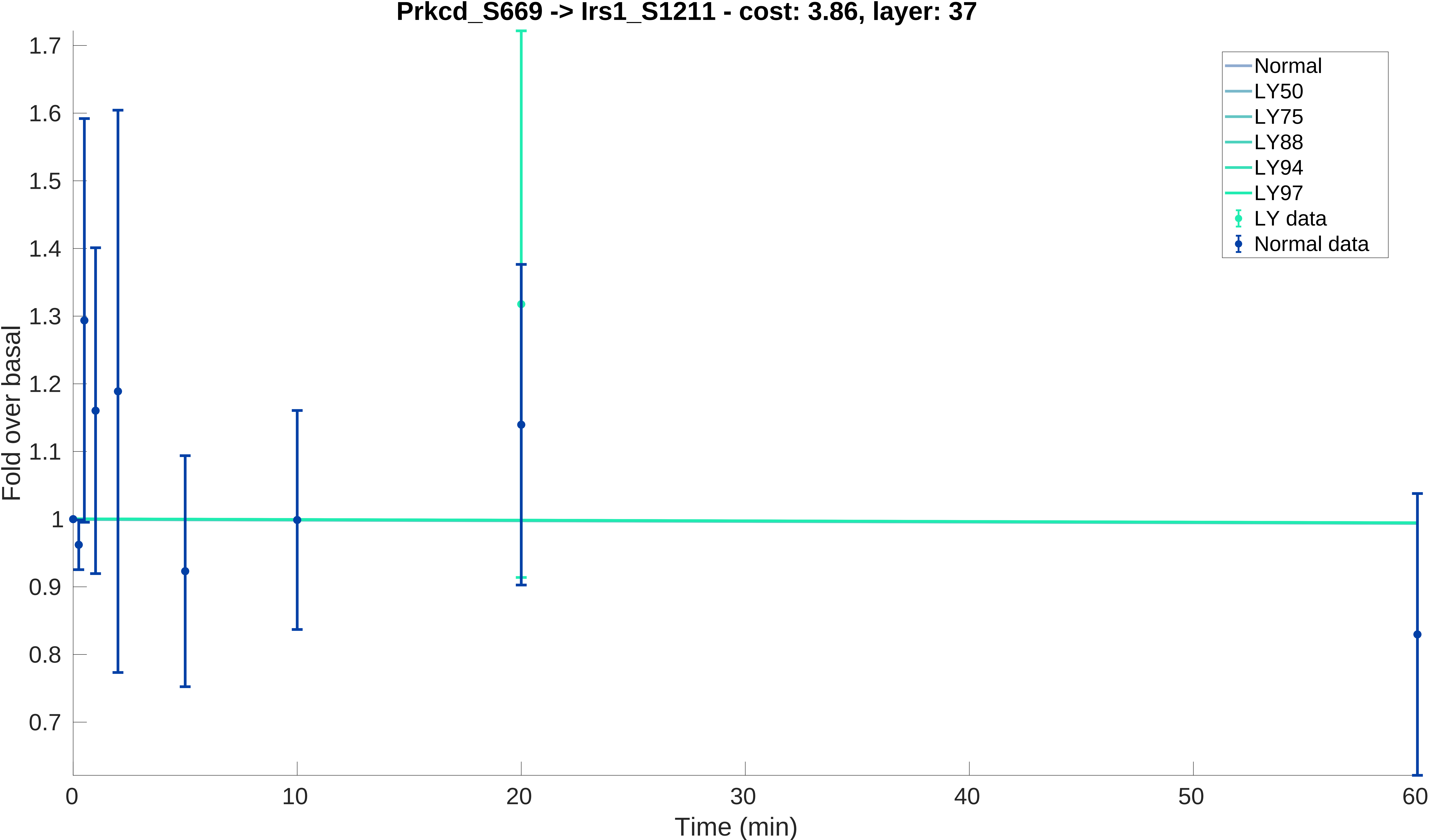

Prkcd_S669 -> Irs1_S1211 - cost: 3.86, layer: 37
Fold over basal
Time (min)
1.7
1.6
1.5
1.4
1.3
1.2
1.1
1
0.9
0.8
0.7
0
10
20
30
40
50
60
Normal
LY50
LY75
LY88
LY94
LY97
LY data
Normal data

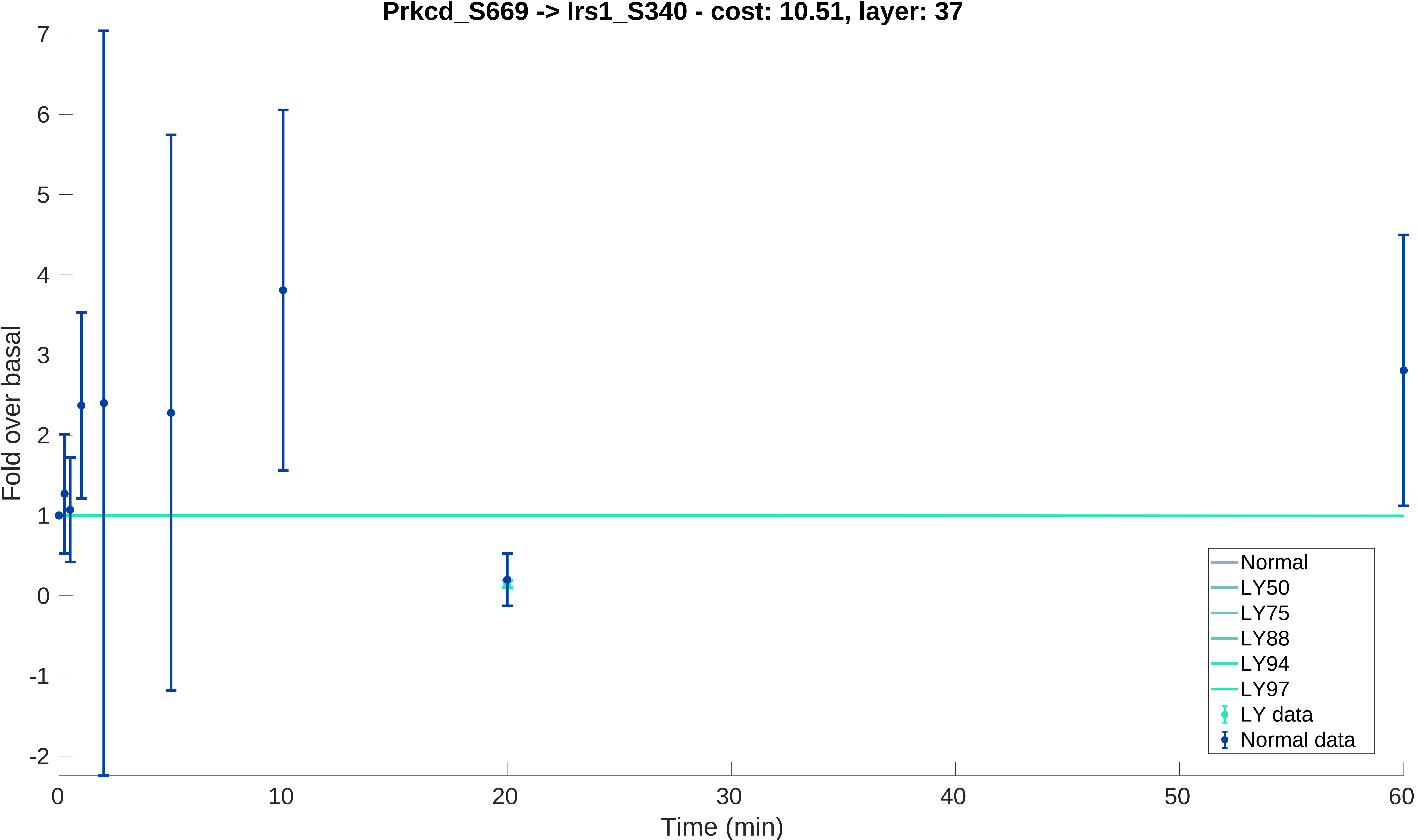

Prkcd_S669 -> Irs1_S340 - cost: 10.51, layer: 37
Fold over basal
7
6
5
4
3
2
1
0
-1
-2
0
10
20
30
40
50
60
Time (min)
Normal
LY50
LY75
LY88
LY94
LY97
LY data
Normal data

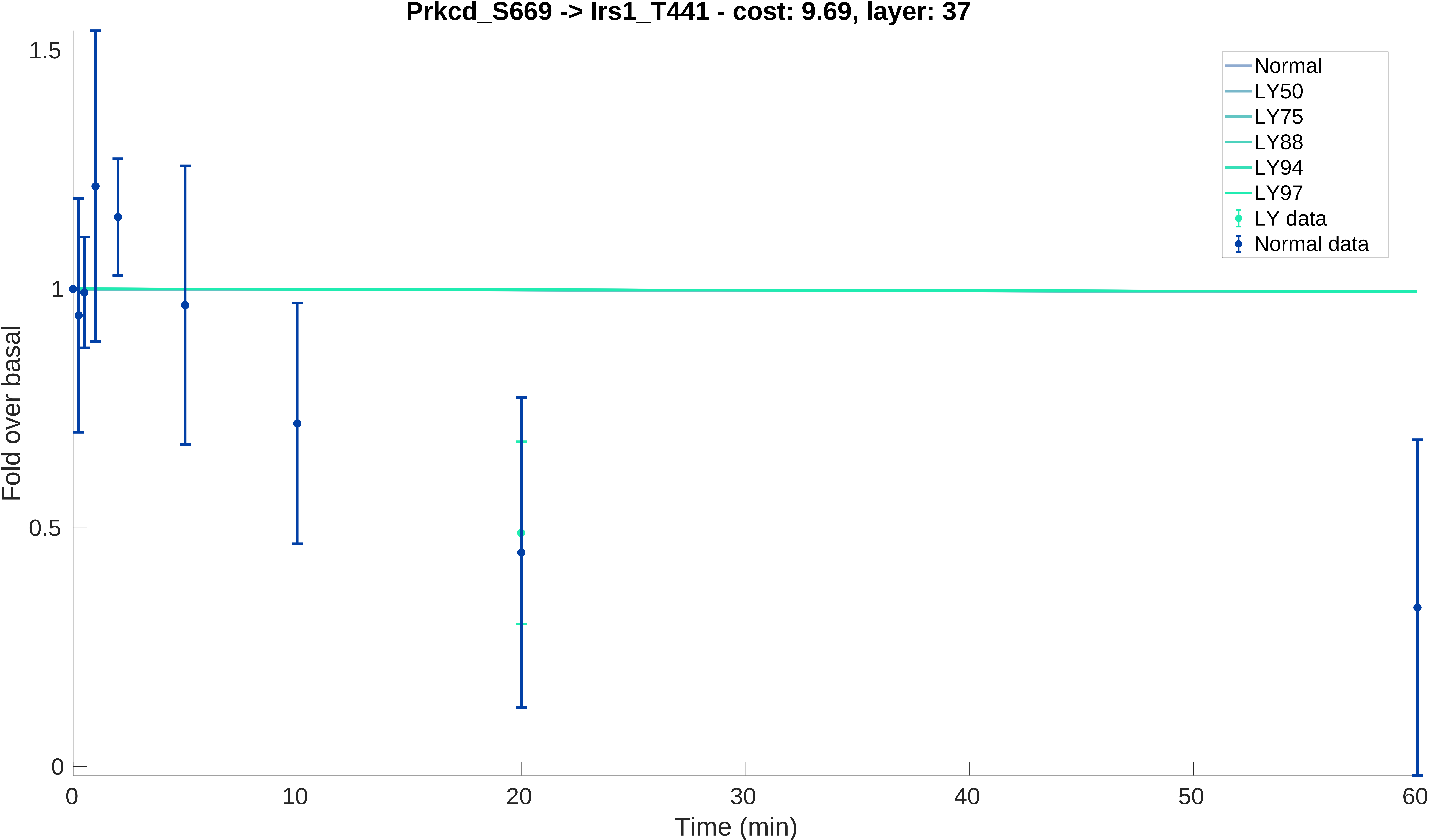

Prkcd_S669 -> Irs1_T441 - cost: 9.69, layer: 37
Fold over basal
Time (min)
0
0.5
1
1.5
0
10
20
30
40
50
60
Normal
LY50
LY75
LY88
LY94
LY97
LY data
Normal data

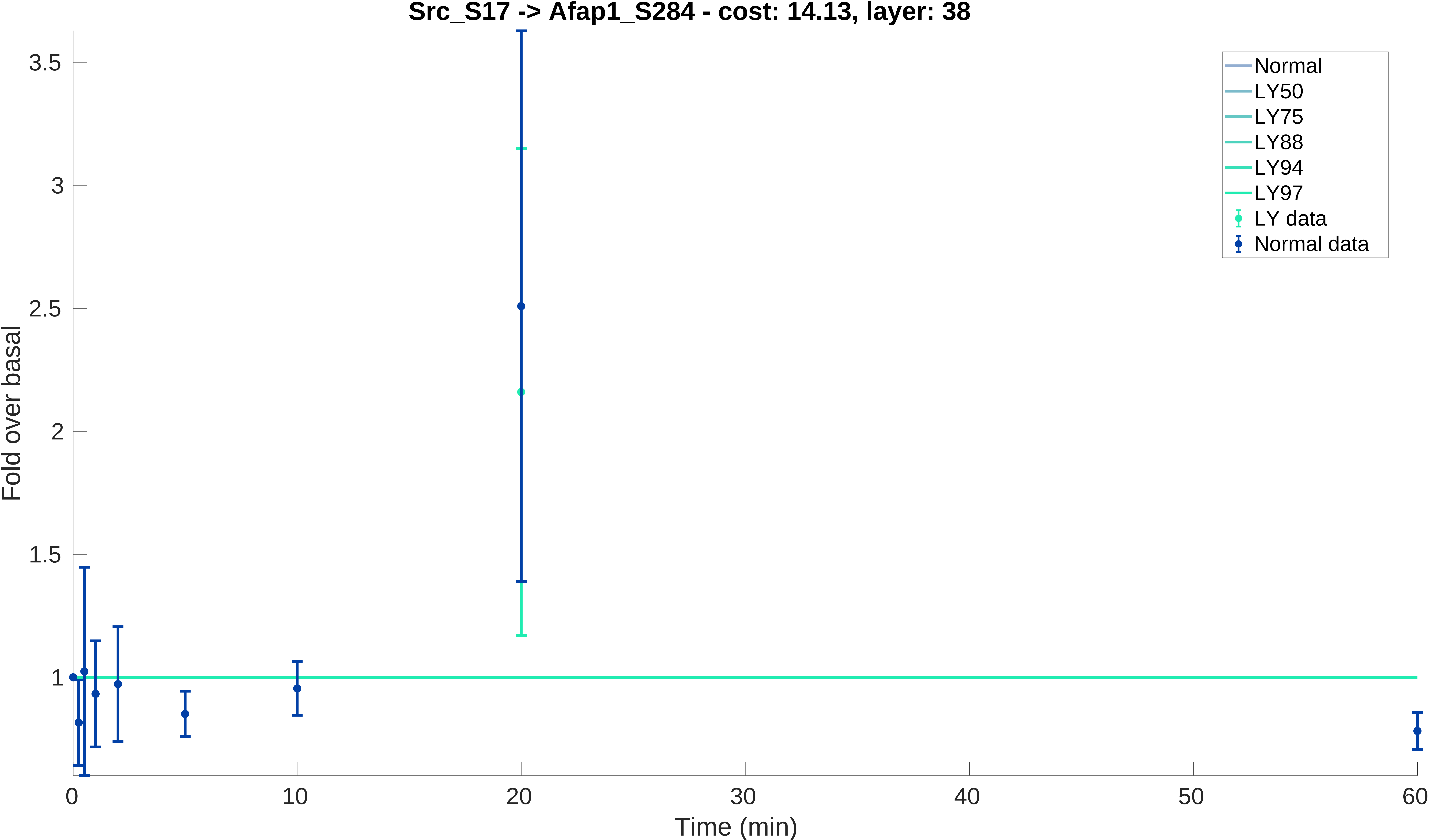

Src_S17 -> Afap1_S284 - cost: 14.13, layer: 38
Fold over basal
Time (min)
3.5
3
2.5
2
1.5
1
0
10
20
30
40
50
60
Normal
LY50
LY75
LY88
LY94
LY97
LY data
Normal data

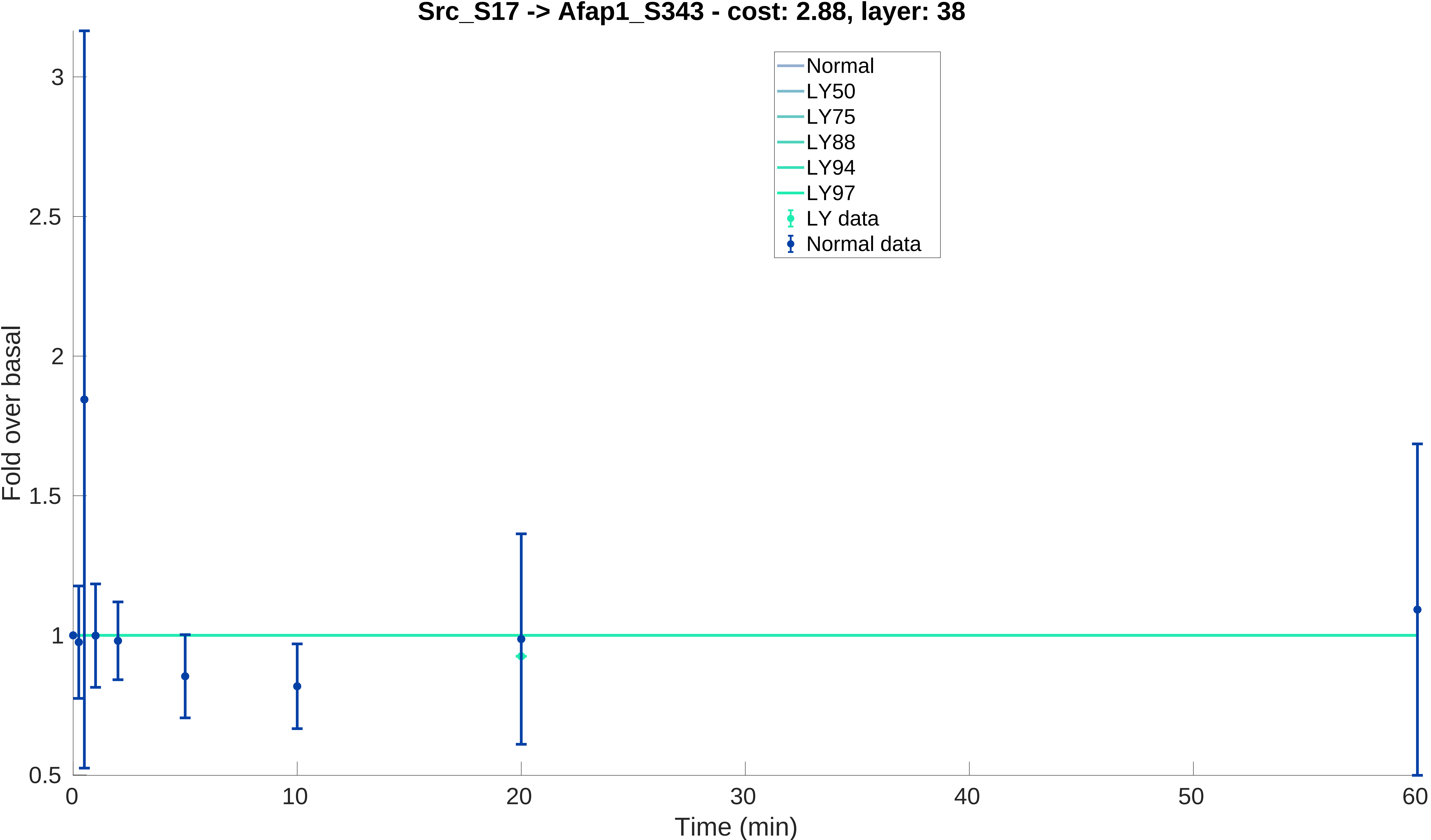

Src_S17 -> Afap1_S343 - cost: 2.88, layer: 38
Normal
LY50
LY75
LY88
LY94
LY97
LY data
Normal data
Fold over basal
3
2.5
2
1.5
1
0.5
0
10
20
30
40
50
60
Time (min)

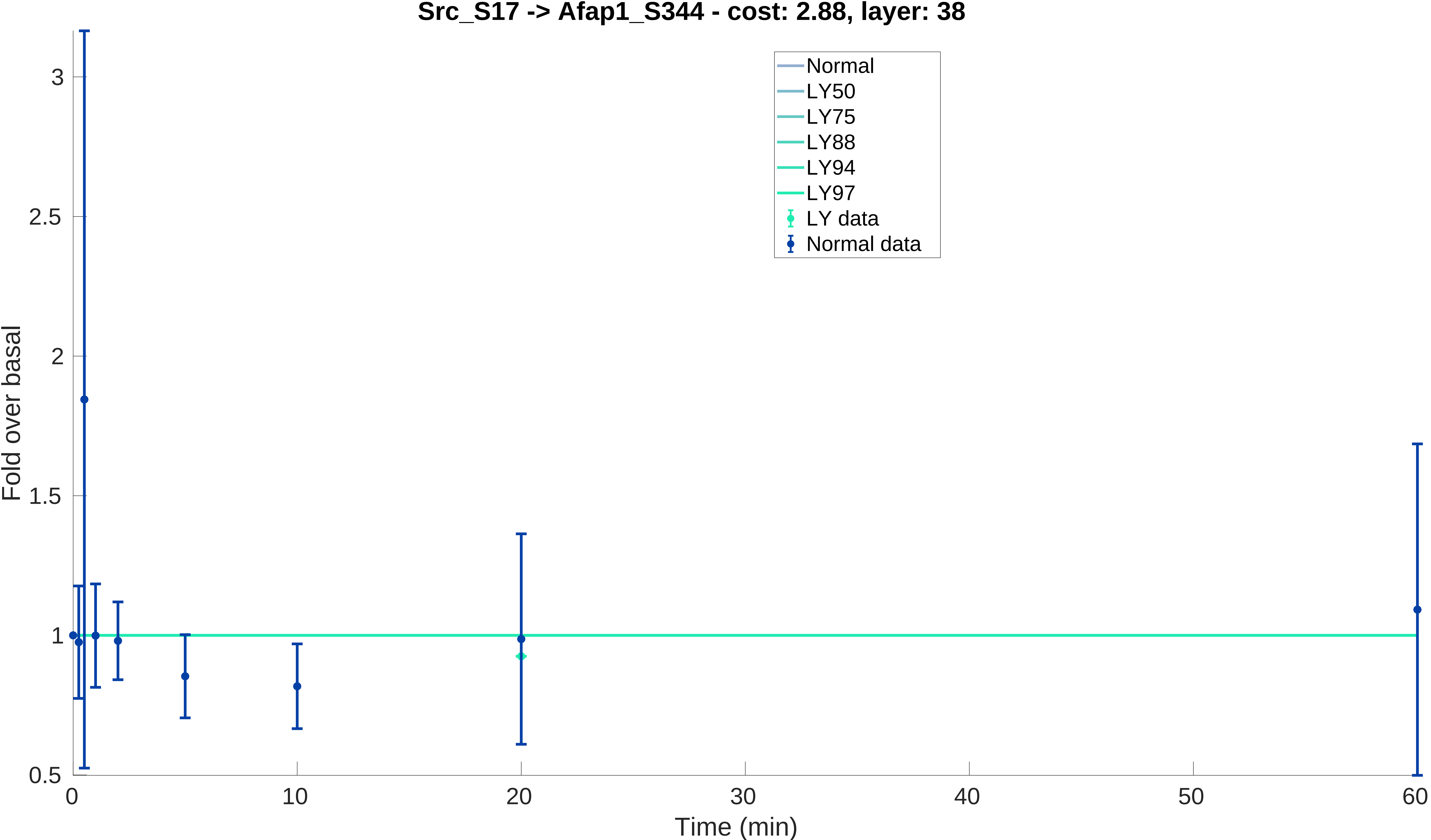

Src_S17 -> Afap1_S344 - cost: 2.88, layer: 38
Normal
LY50
LY75
LY88
LY94
LY97
LY data
Normal data
Fold over basal
Time (min)
0.5
1
1.5
2
2.5
3
0
10
20
30
40
50
60

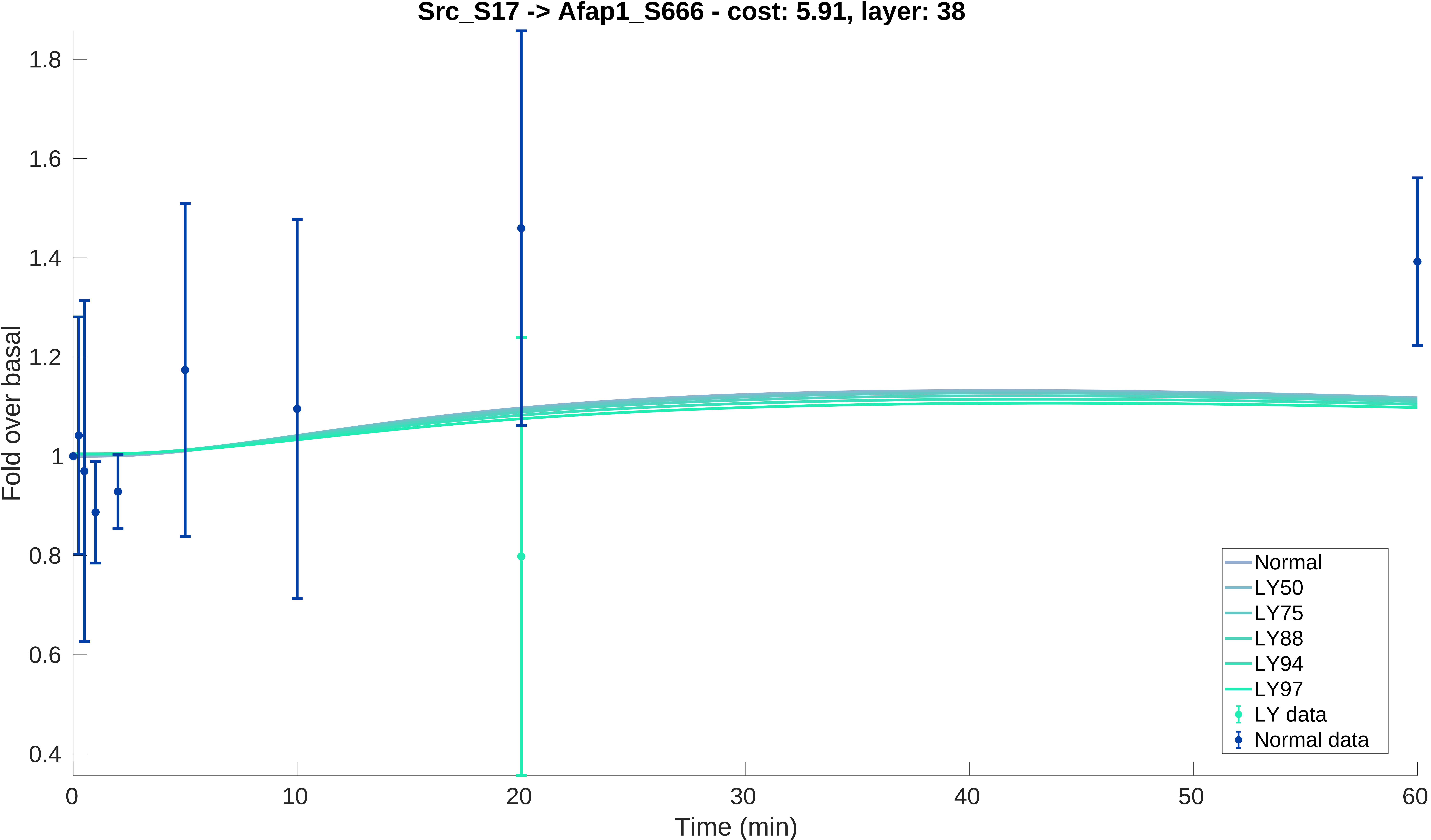

Src_S17 -> Afap1_S666 - cost: 5.91, layer: 38
Fold over basal
Time (min)
0.4
0.6
0.8
1
1.2
1.4
1.6
1.8
0
10
20
30
40
50
60
Normal
LY50
LY75
LY88
LY94
LY97
LY data
Normal data

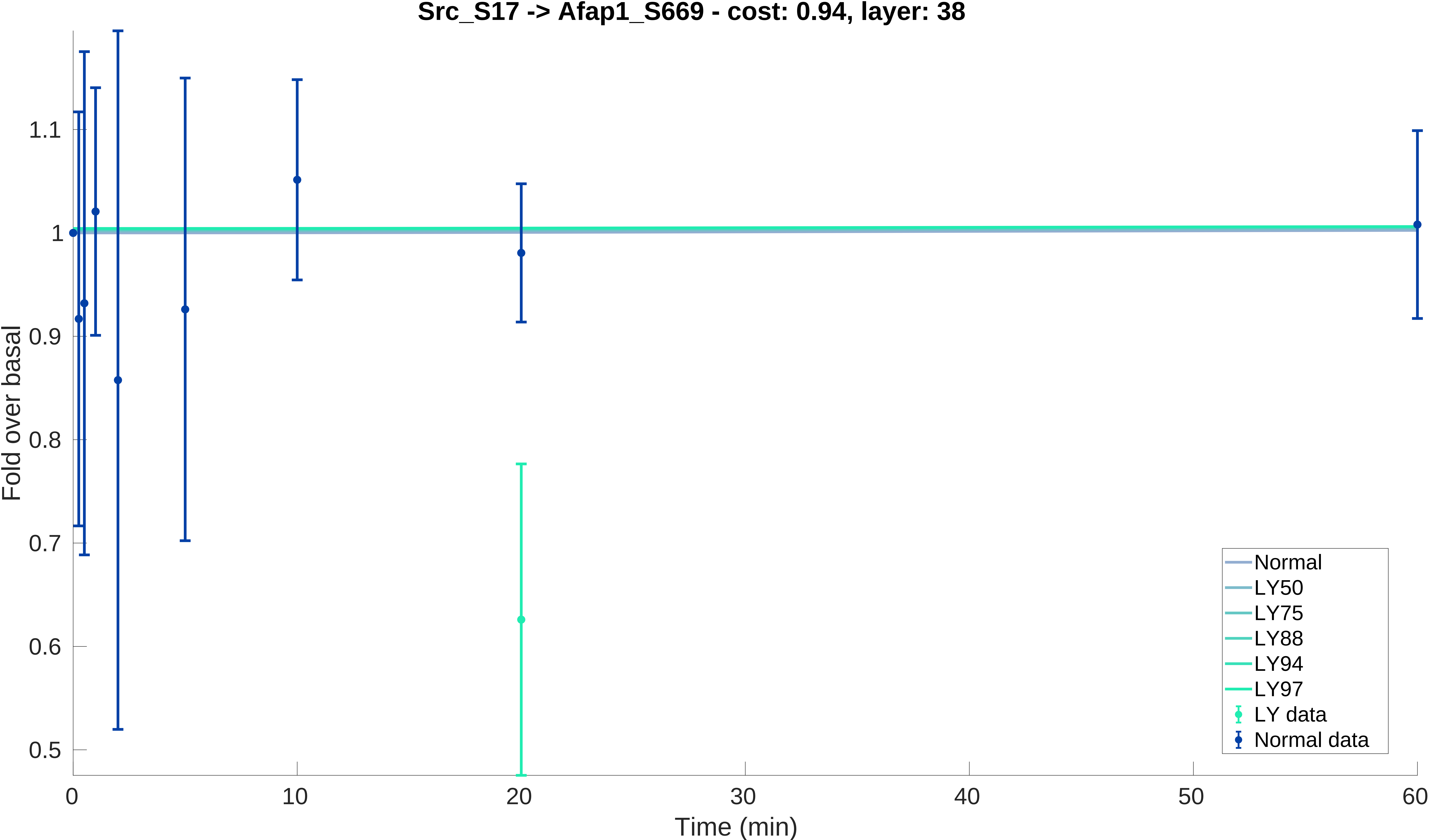

Src_S17 -> Afap1_S669 - cost: 0.94, layer: 38
Fold over basal
1.1
1
0.9
0.8
0.7
0.6
0.5
0
10
20
30
40
50
60
Time (min)
Normal
LY50
LY75
LY88
LY94
LY97
LY data
Normal data

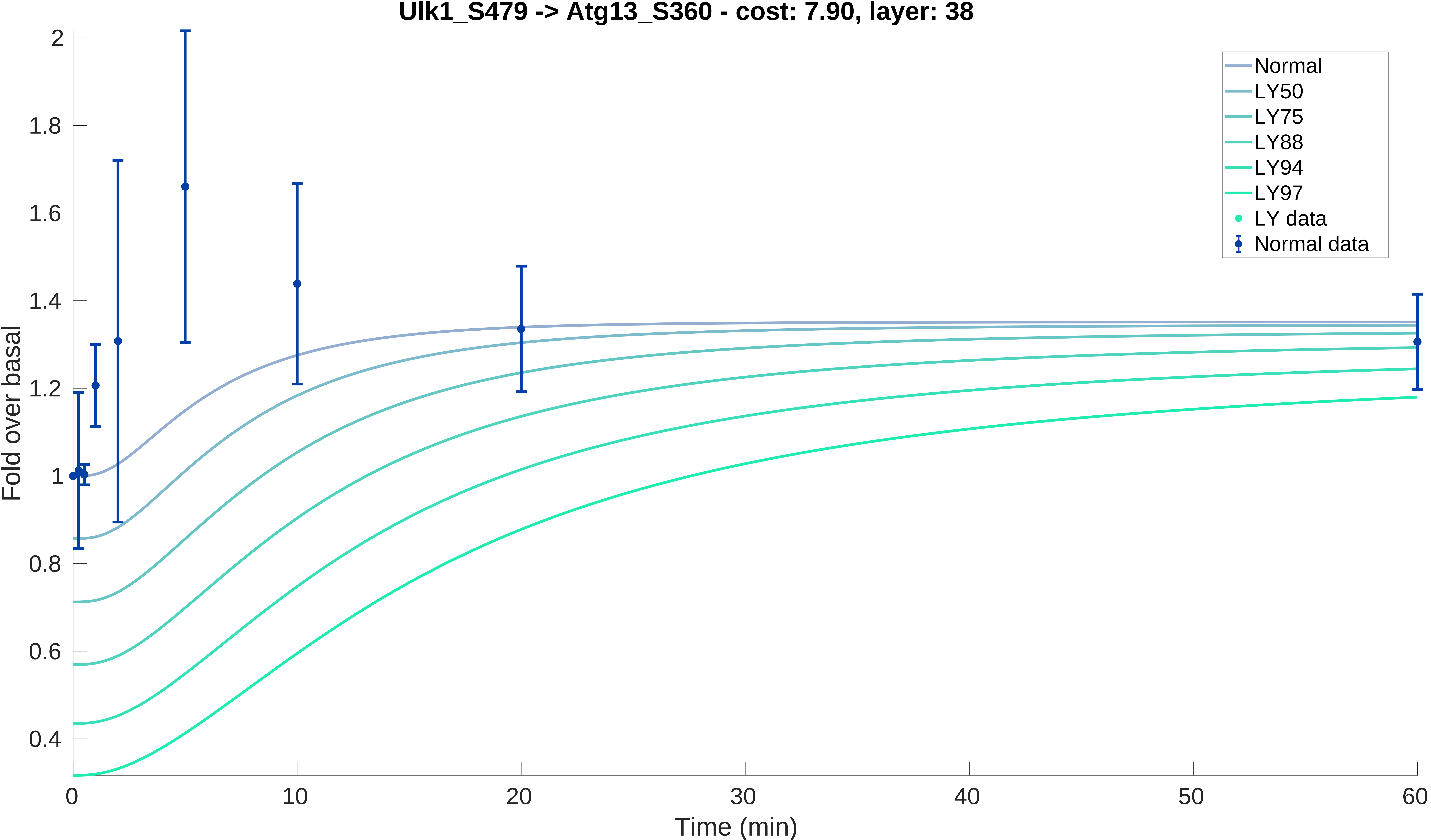
Ulk1_S479 -> Atg13_S360 - cost: 7.90, layer: 38
Fold over basal
Time (min)
0
10
20
30
40
50
60
0.4
0.6
0.8
1
1.2
1.4
1.6
1.8
2
Normal
LY50
LY75
LY88
LY94
LY97
LY data
Normal data

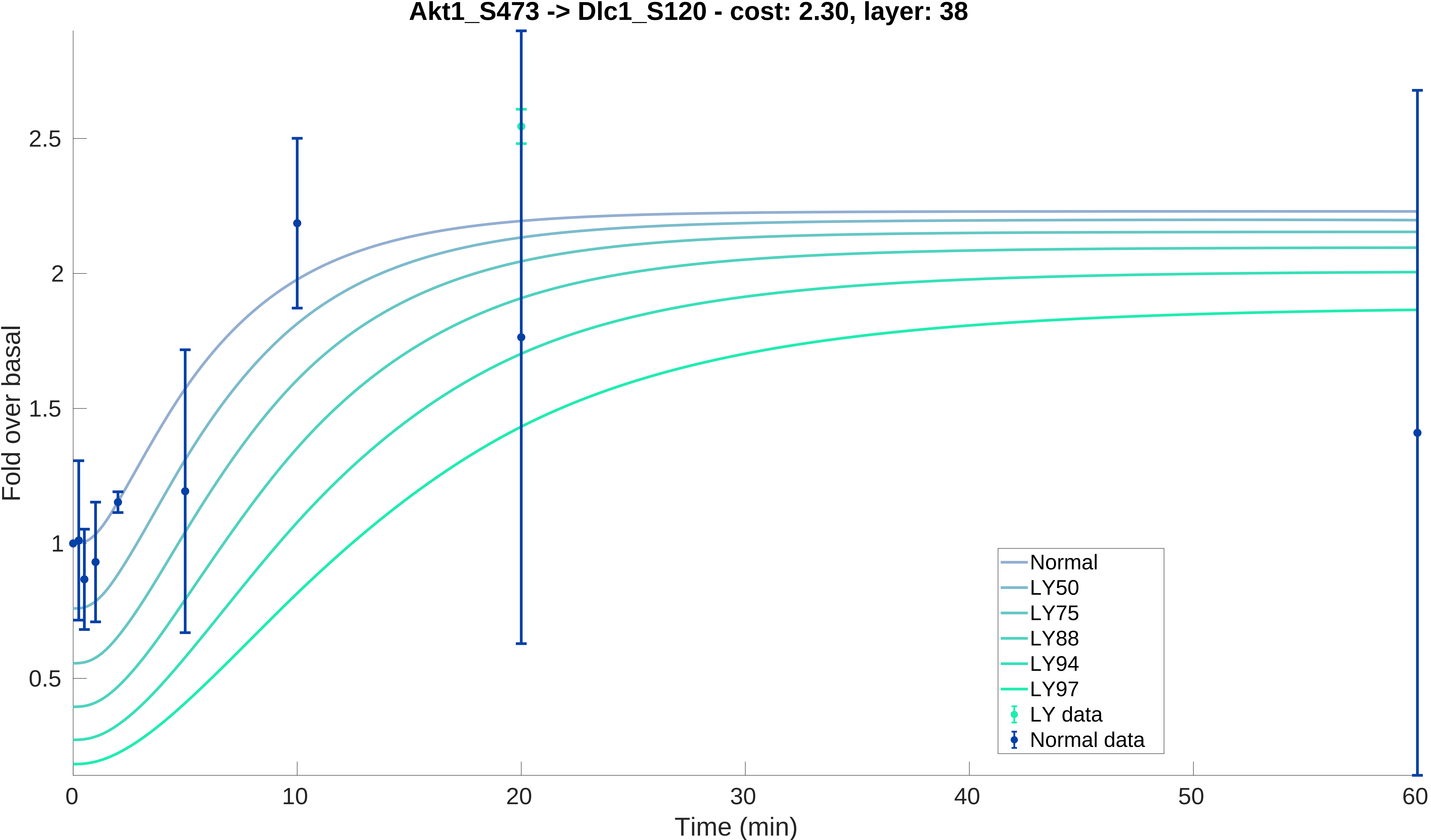

Akt1_S473 -> Dlc1_S120 - cost: 2.30, layer: 38
Fold over basal
Time (min)
0.5
1
1.5
2
2.5
0
10
20
30
40
50
60
Normal
LY50
LY75
LY88
LY94
LY97
LY data
Normal data

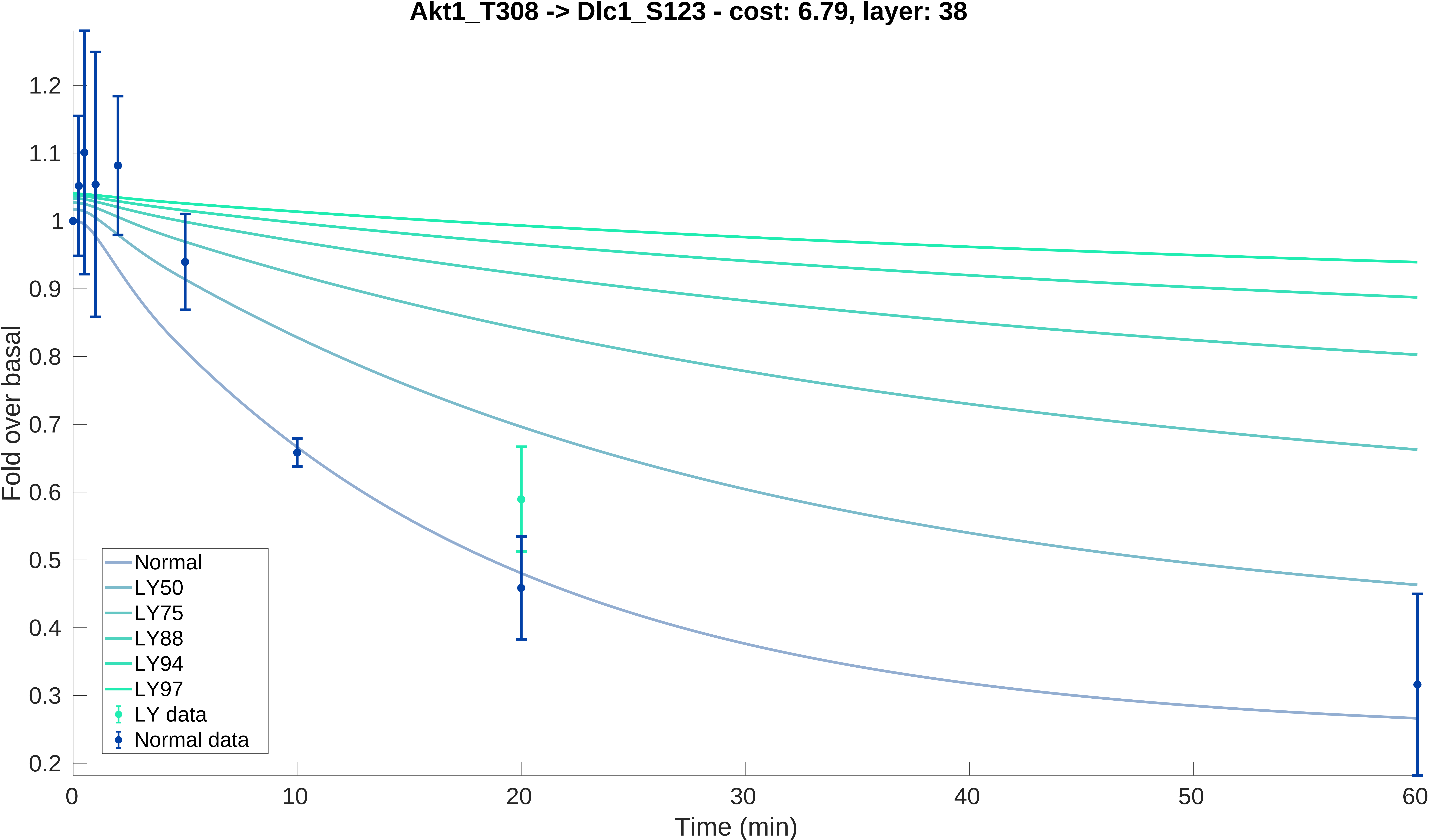
Akt1_T308 -> Dlc1_S123 - cost: 6.79, layer: 38
Fold over basal
Time (min)
0
10
20
30
40
50
60
0.2
0.3
0.4
0.5
0.6
0.7
0.8
0.9
1
1.1
1.2
Normal
LY50
LY75
LY88
LY94
LY97
LY data
Normal data

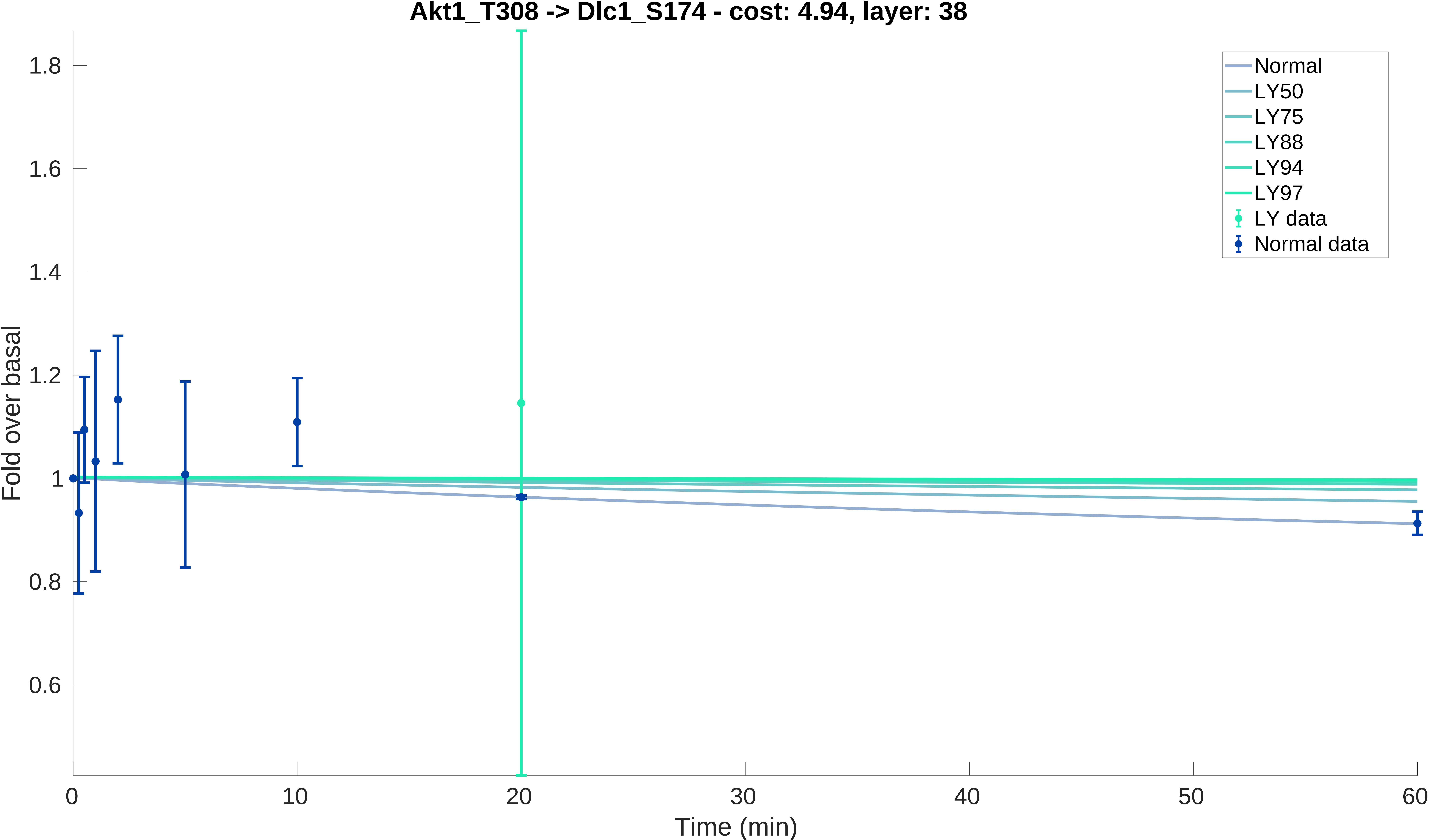

Akt1_T308 -> Dlc1_S174 - cost: 4.94, layer: 38
Normal
LY50
LY75
LY88
LY94
LY97
LY data
Normal data
Fold over basal
1.8
1.6
1.4
1.2
1
0.8
0.6
0
10
20
30
40
50
60
Time (min)

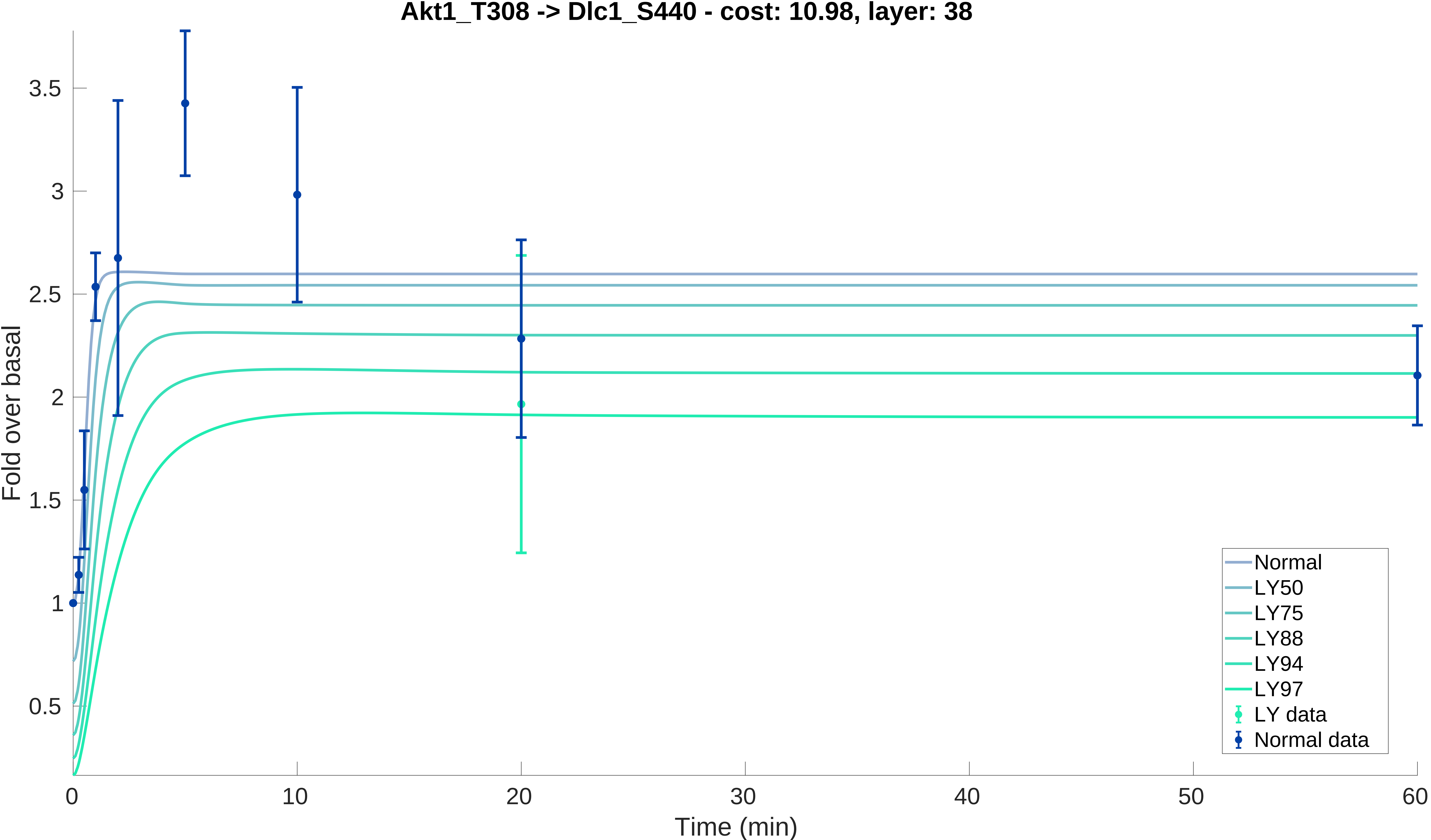

Akt1_T308 -> Dlc1_S440 - cost: 10.98, layer: 38
Fold over basal
Time (min)
0.5
1
1.5
2
2.5
3
3.5
0
10
20
30
40
50
60
Normal
LY50
LY75
LY88
LY94
LY97
LY data
Normal data

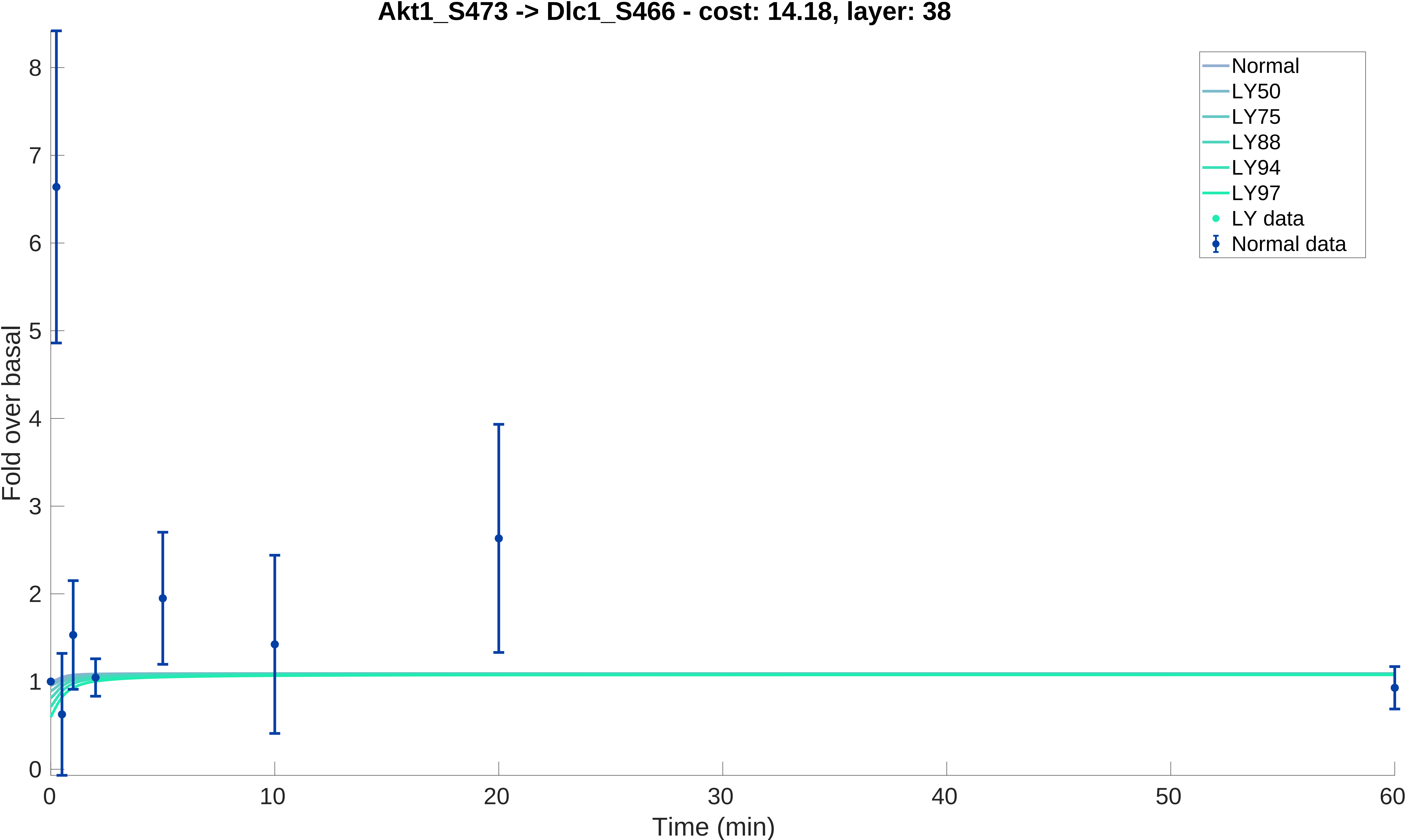

Akt1_S473 -> Dlc1_S466 - cost: 14.18, layer: 38
Fold over basal
Time (min)
0
1
2
3
4
5
6
7
8
0
10
20
30
40
50
60
Normal
LY50
LY75
LY88
LY94
LY97
LY data
Normal data

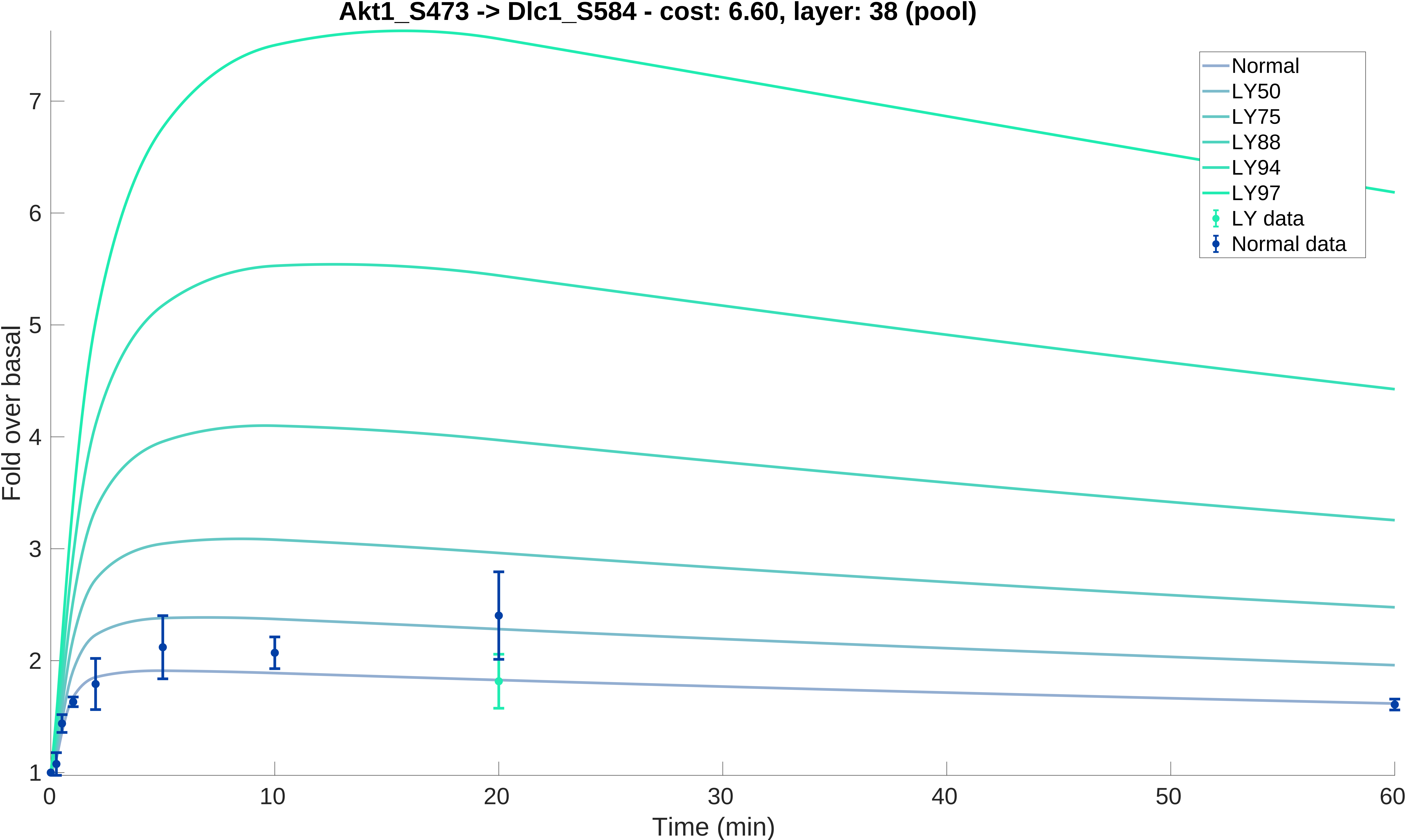

Akt1_S473 -> Dlc1_S584 - cost: 6.60, layer: 38 (pool)
Fold over basal
Time (min)
1
2
3
4
5
6
7
0
10
20
30
40
50
60
Normal
LY50
LY75
LY88
LY94
LY97
LY data
Normal data

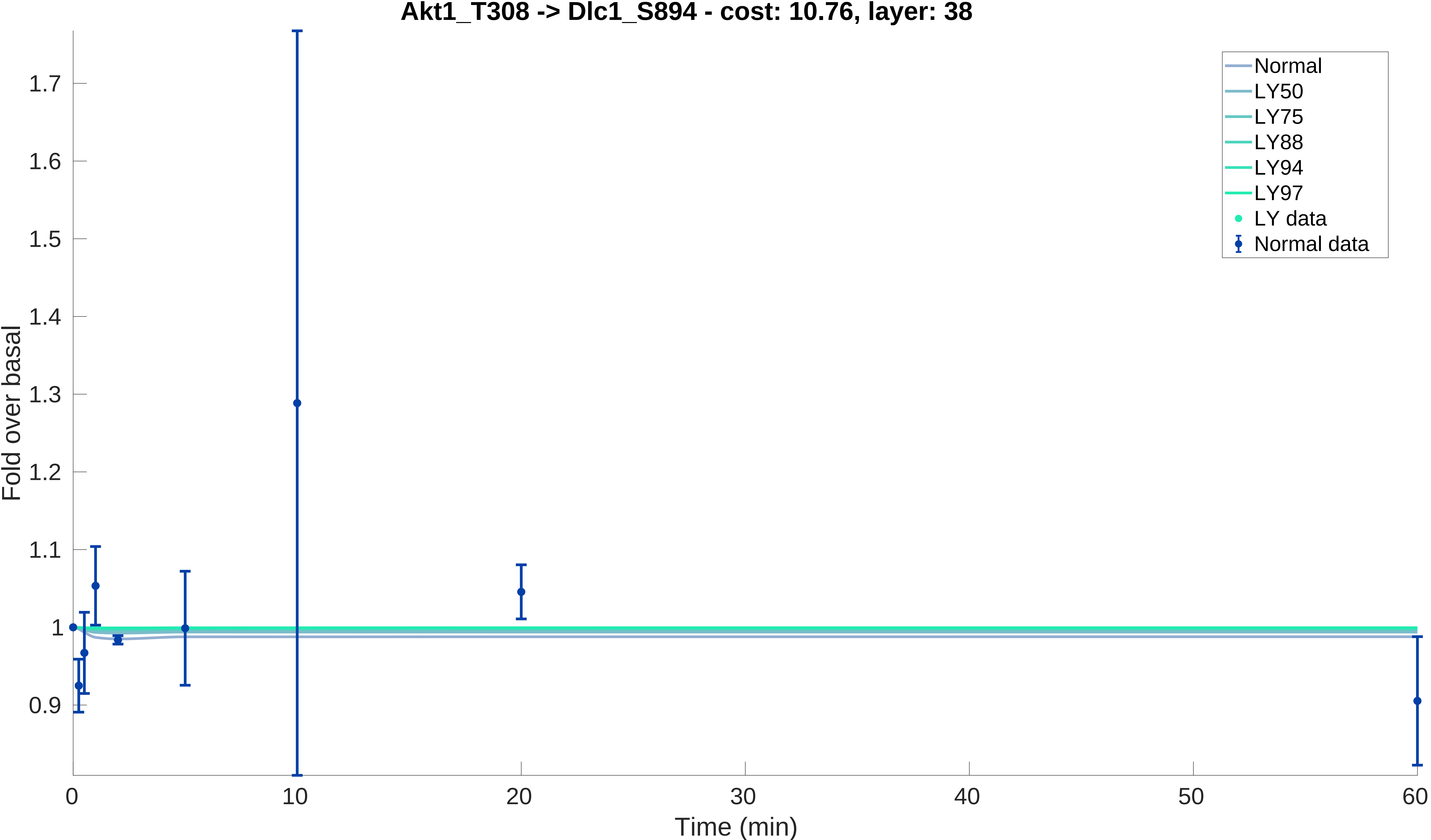

Akt1_T308 -> Dlc1_S894 - cost: 10.76, layer: 38
Fold over basal
Time (min)
1.7
1.6
1.5
1.4
1.3
1.2
1.1
1
0.9
0
10
20
30
40
50
60
Normal
LY50
LY75
LY88
LY94
LY97
LY data
Normal data

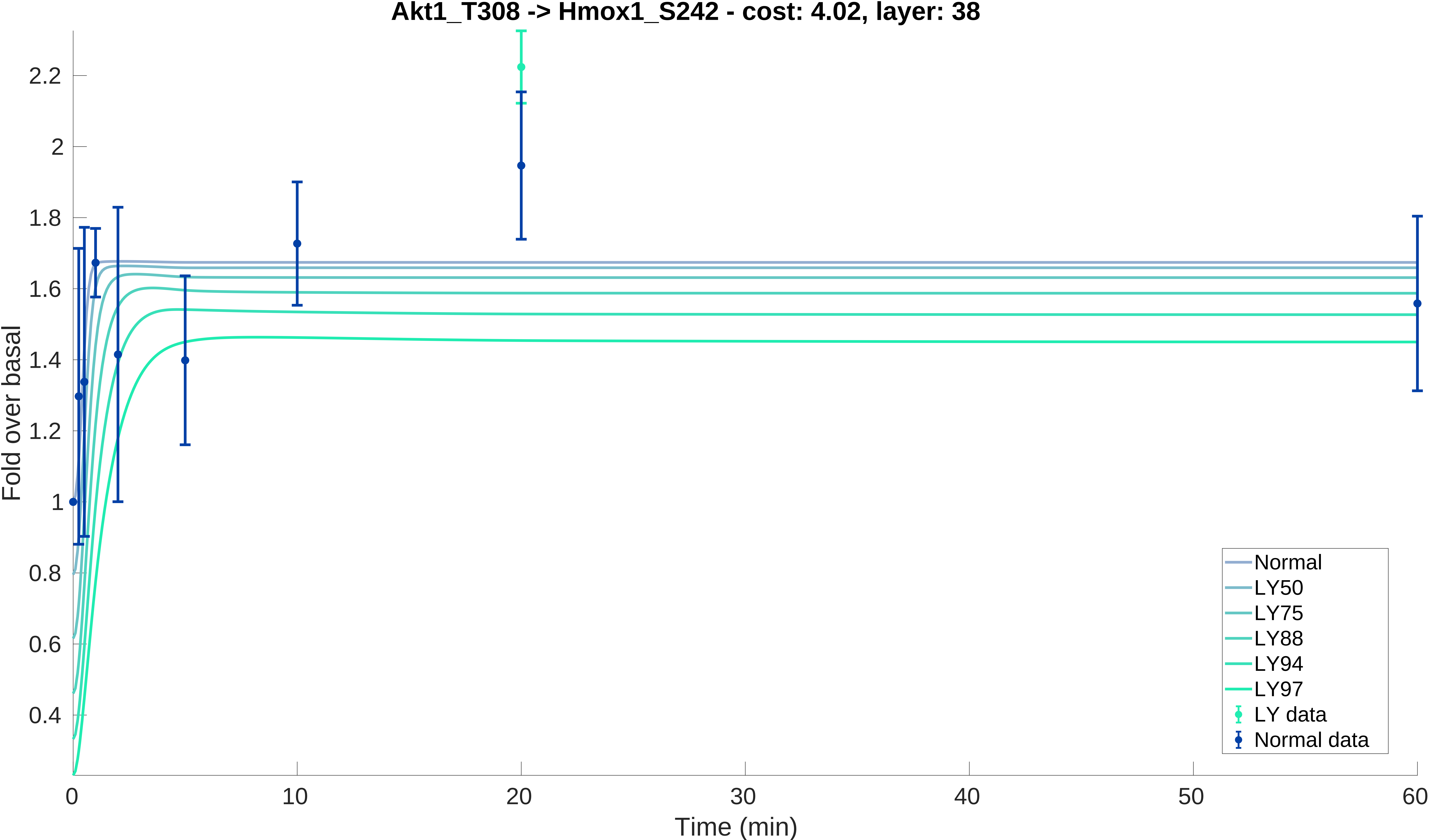
Akt1_T308 -> Hmox1_S242 - cost: 4.02, layer: 38
Fold over basal
2.2
2
1.8
1.6
1.4
1.2
1
0.8
0.6
0.4
0
10
20
30
40
50
60
Time (min)
Normal
LY50
LY75
LY88
LY94
LY97
LY data
Normal data

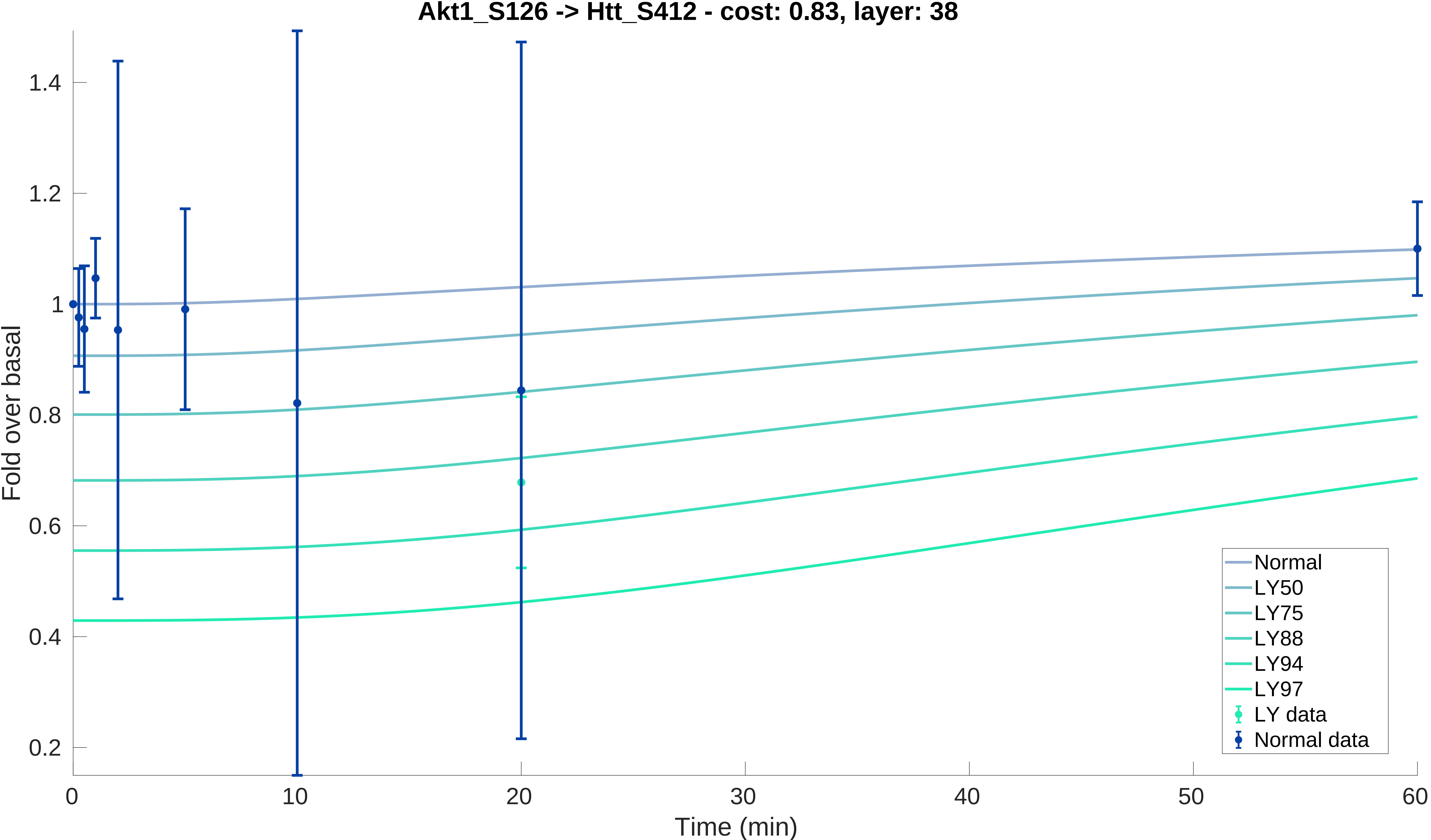
Akt1_S126 -> Htt_S412 - cost: 0.83, layer: 38
Fold over basal
Time (min)
0.2
0.4
0.6
0.8
1
1.2
1.4
0
10
20
30
40
50
60
Normal
LY50
LY75
LY88
LY94
LY97
LY data
Normal data

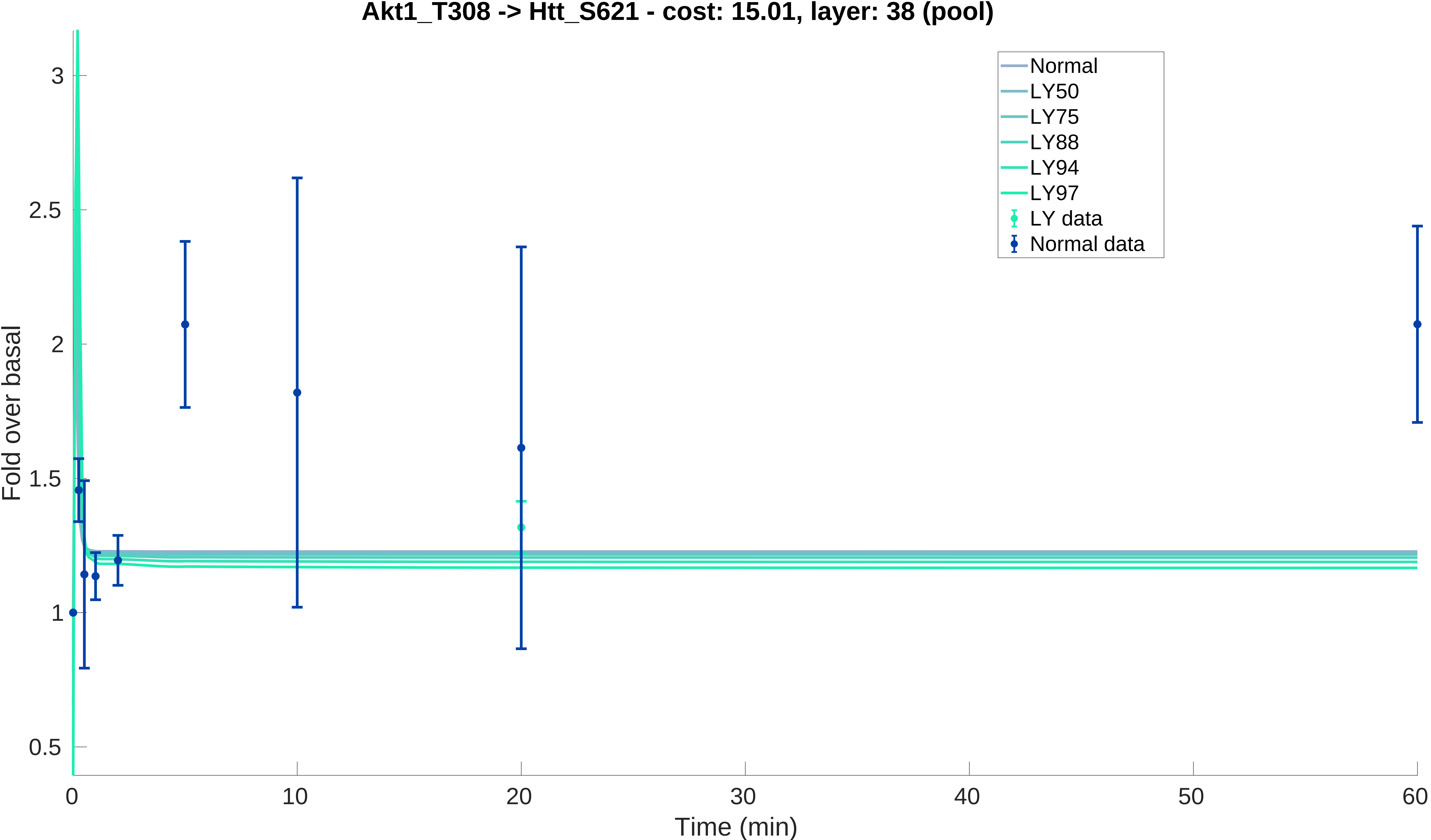
Akt1_T308 -> Htt_S621 - cost: 15.01, layer: 38 (pool)
Normal
LY50
LY75
LY88
LY94
LY97
LY data
Normal data
Fold over basal
0.5
1
1.5
2
2.5
3
0
10
20
30
40
50
60
Time (min)

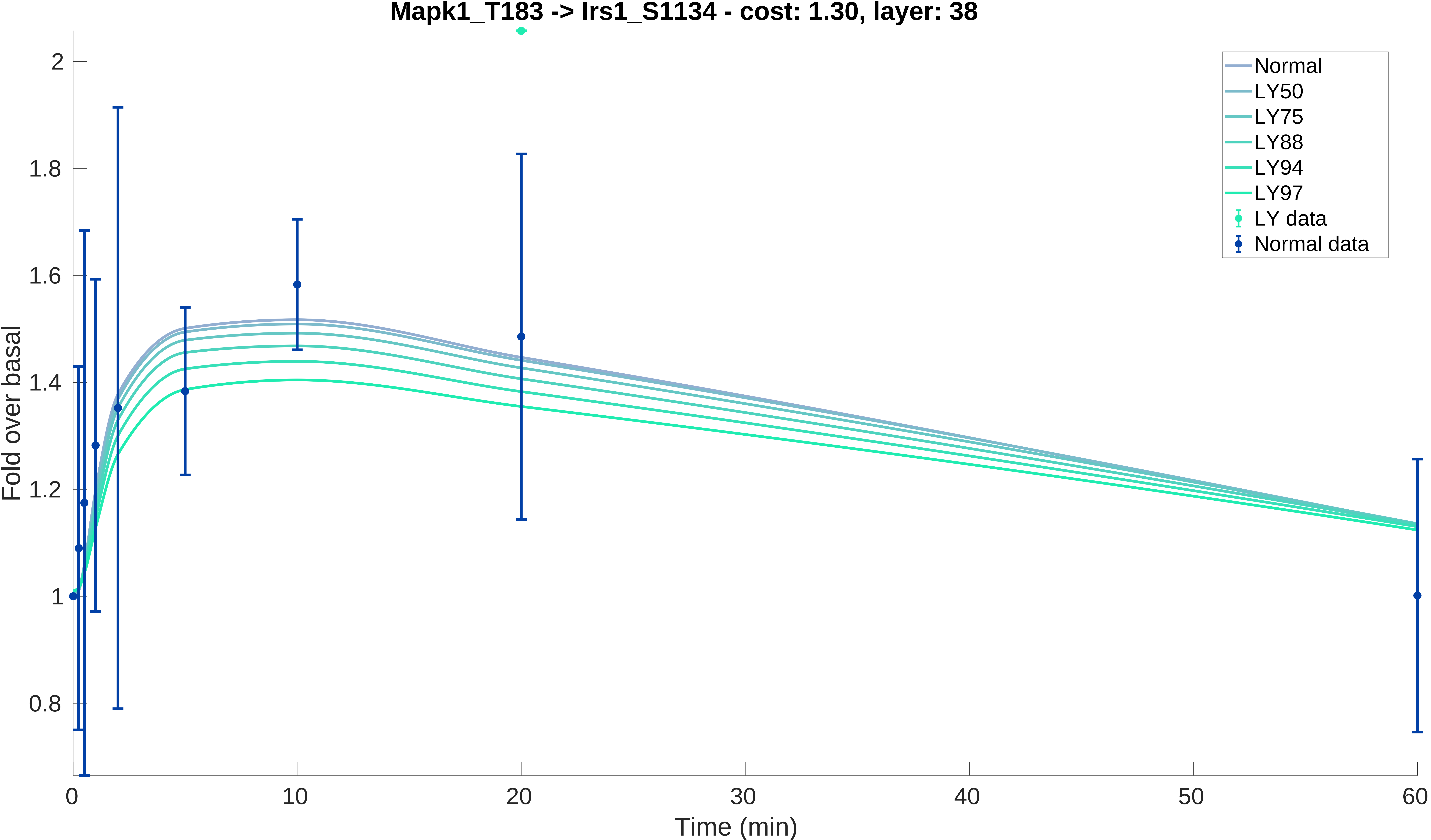

Mapk1_T183 -> Irs1_S1134 - cost: 1.30, layer: 38
Fold over basal
Time (min)
0.8
1
1.2
1.4
1.6
1.8
2
0
10
20
30
40
50
60
Normal
LY50
LY75
LY88
LY94
LY97
LY data
Normal data

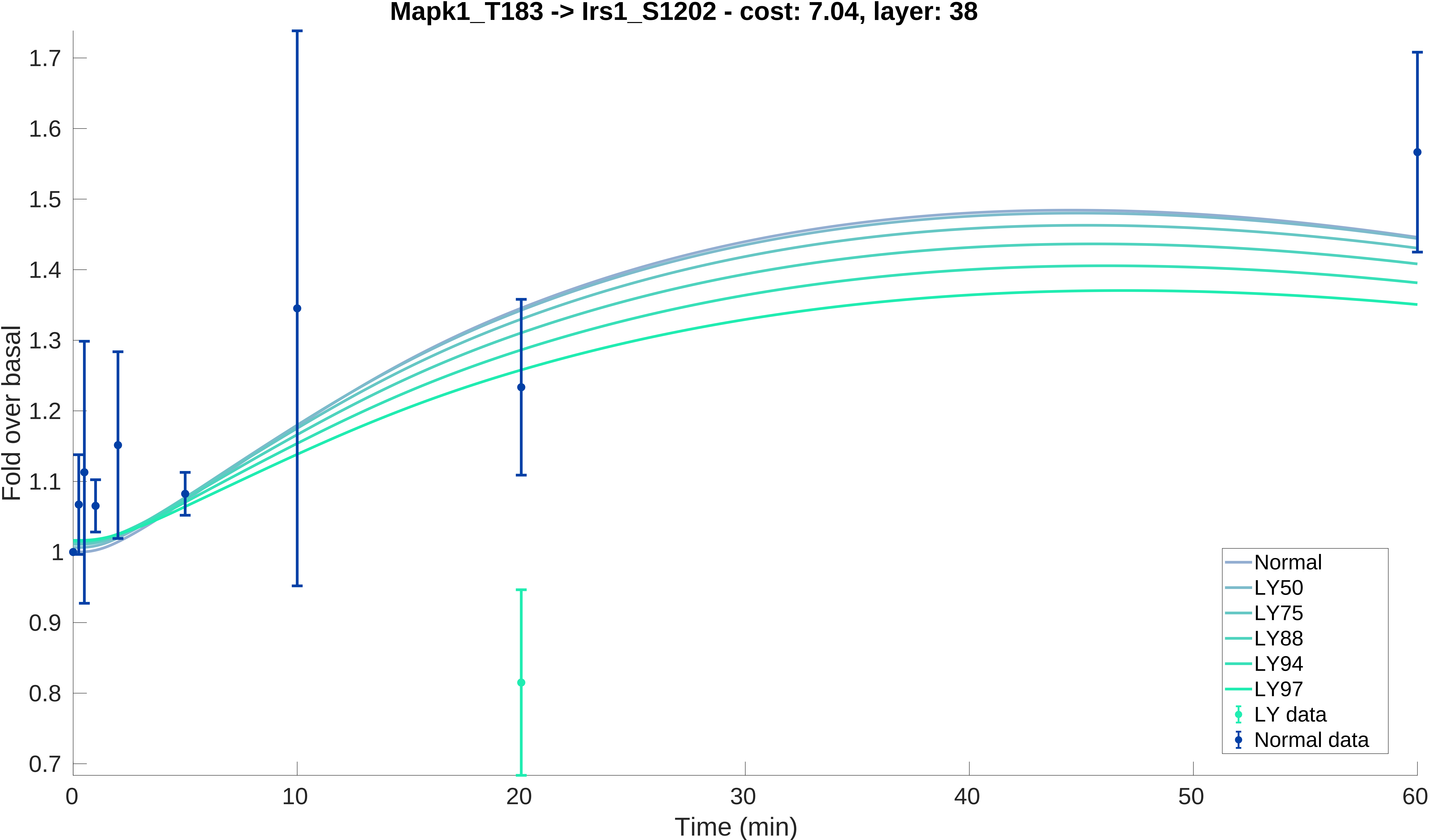

Mapk1_T183 -> Irs1_S1202 - cost: 7.04, layer: 38
Fold over basal
1.7
1.6
1.5
1.4
1.3
1.2
1.1
1
0.9
0.8
0.7
0
10
20
30
40
50
60
Time (min)
Normal
LY50
LY75
LY88
LY94
LY97
LY data
Normal data

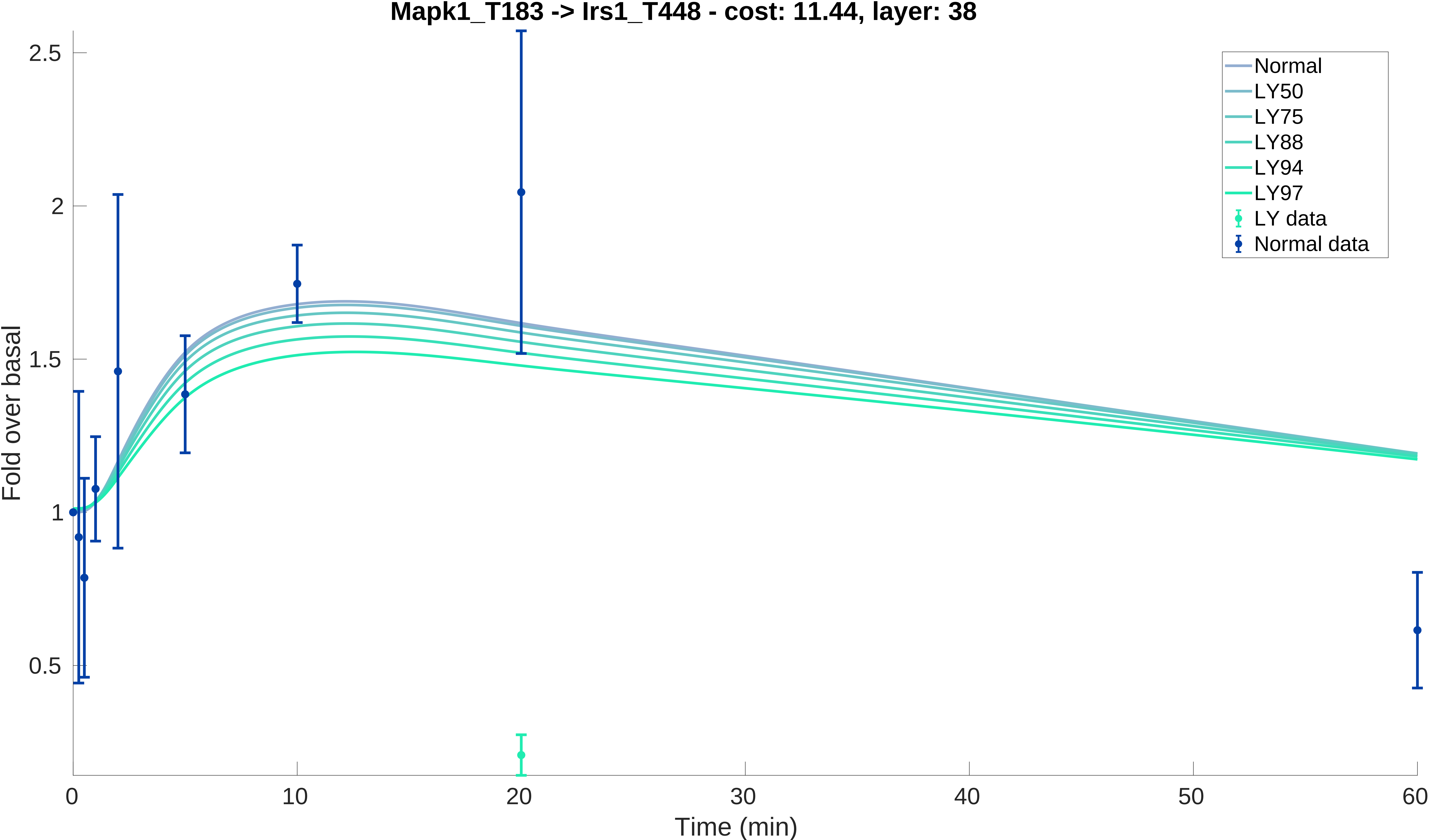
Mapk1_T183 -> Irs1_T448 - cost: 11.44, layer: 38
Fold over basal
Time (min)
0
10
20
30
40
50
60
0.5
1
1.5
2
2.5
Normal
LY50
LY75
LY88
LY94
LY97
LY data
Normal data

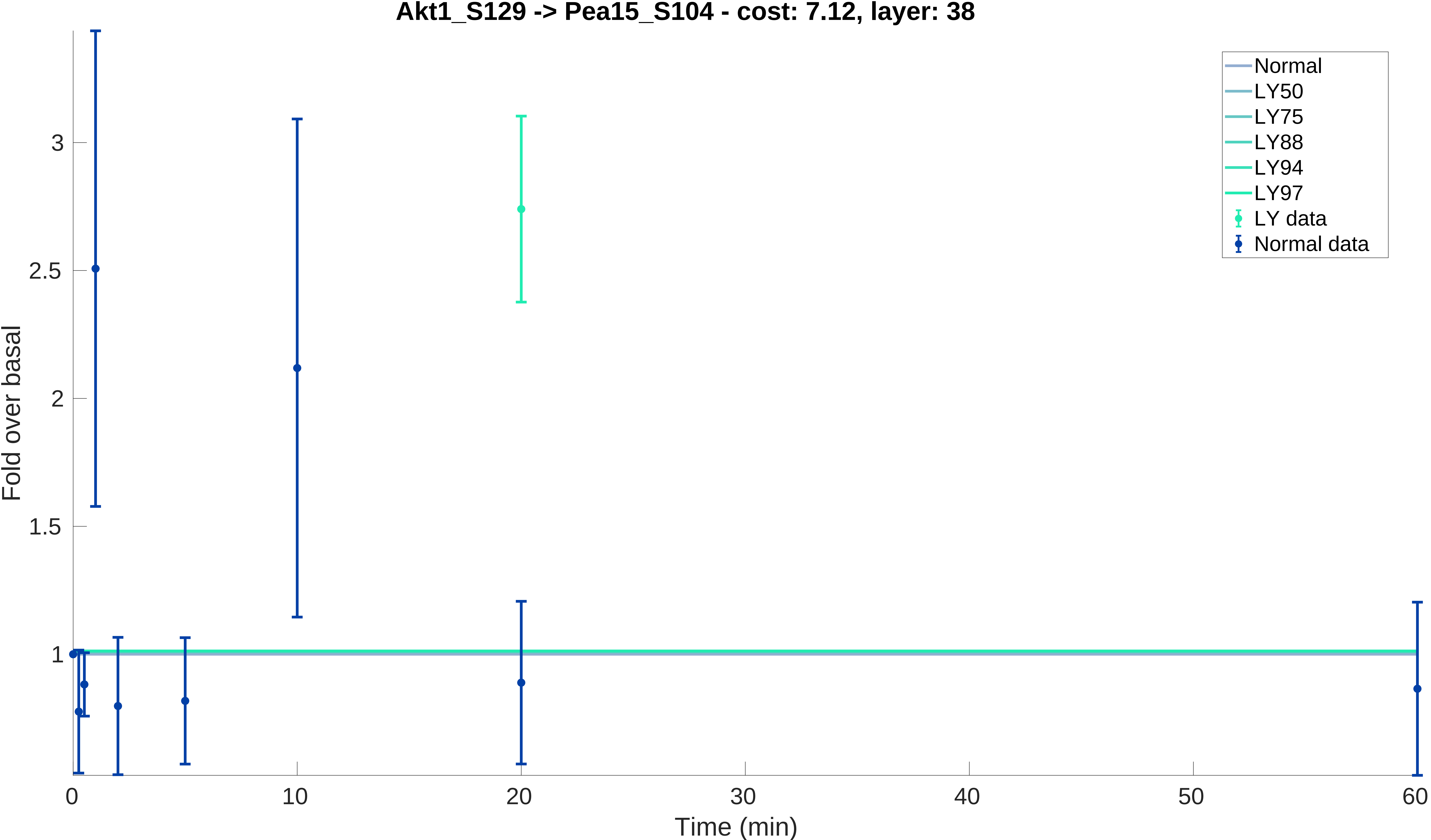

Akt1_S129 -> Pea15_S104 - cost: 7.12, layer: 38
Fold over basal
Time (min)
1
1.5
2
2.5
3
0
10
20
30
40
50
60
Normal
LY50
LY75
LY88
LY94
LY97
LY data
Normal data

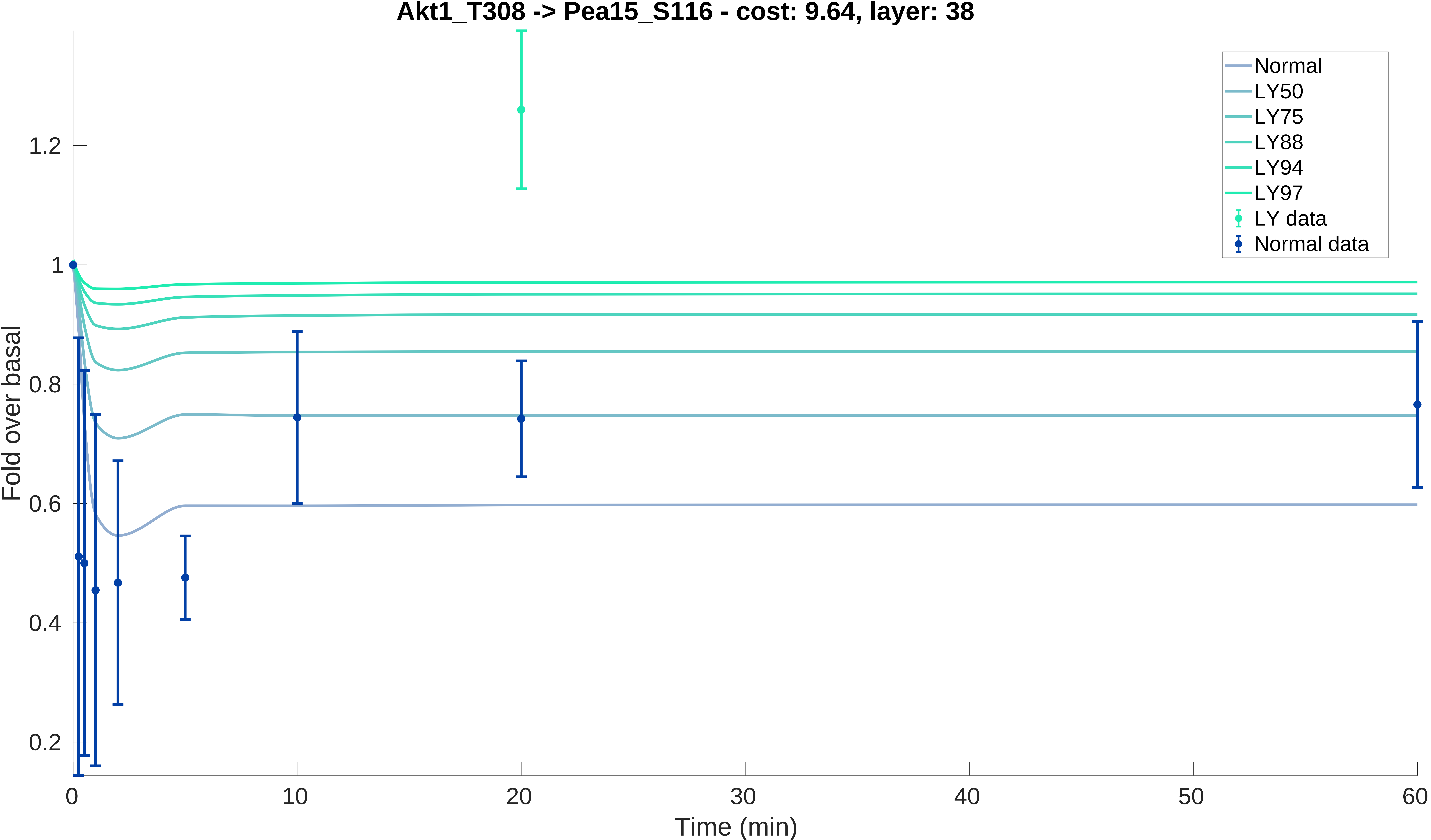
Akt1_T308 -> Pea15_S116 - cost: 9.64, layer: 38
Fold over basal
Time (min)
0
10
20
30
40
50
60
0.2
0.4
0.6
0.8
1
1.2
Normal
LY50
LY75
LY88
LY94
LY97
LY data
Normal data

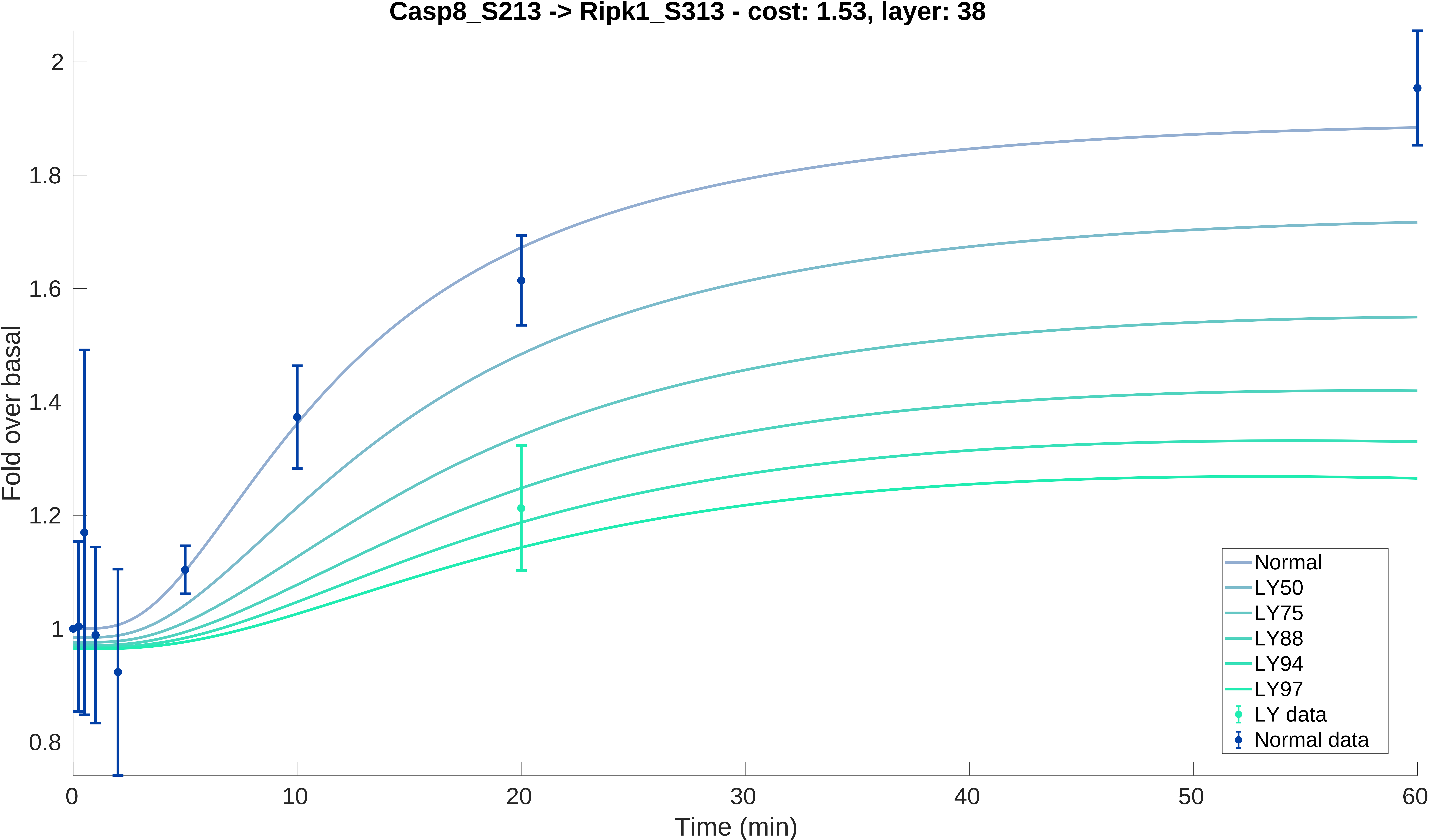

Casp8_S213 -> Ripk1_S313 - cost: 1.53, layer: 38
Fold over basal
Time (min)
0.8
1
1.2
1.4
1.6
1.8
2
0
10
20
30
40
50
60
Normal
LY50
LY75
LY88
LY94
LY97
LY data
Normal data

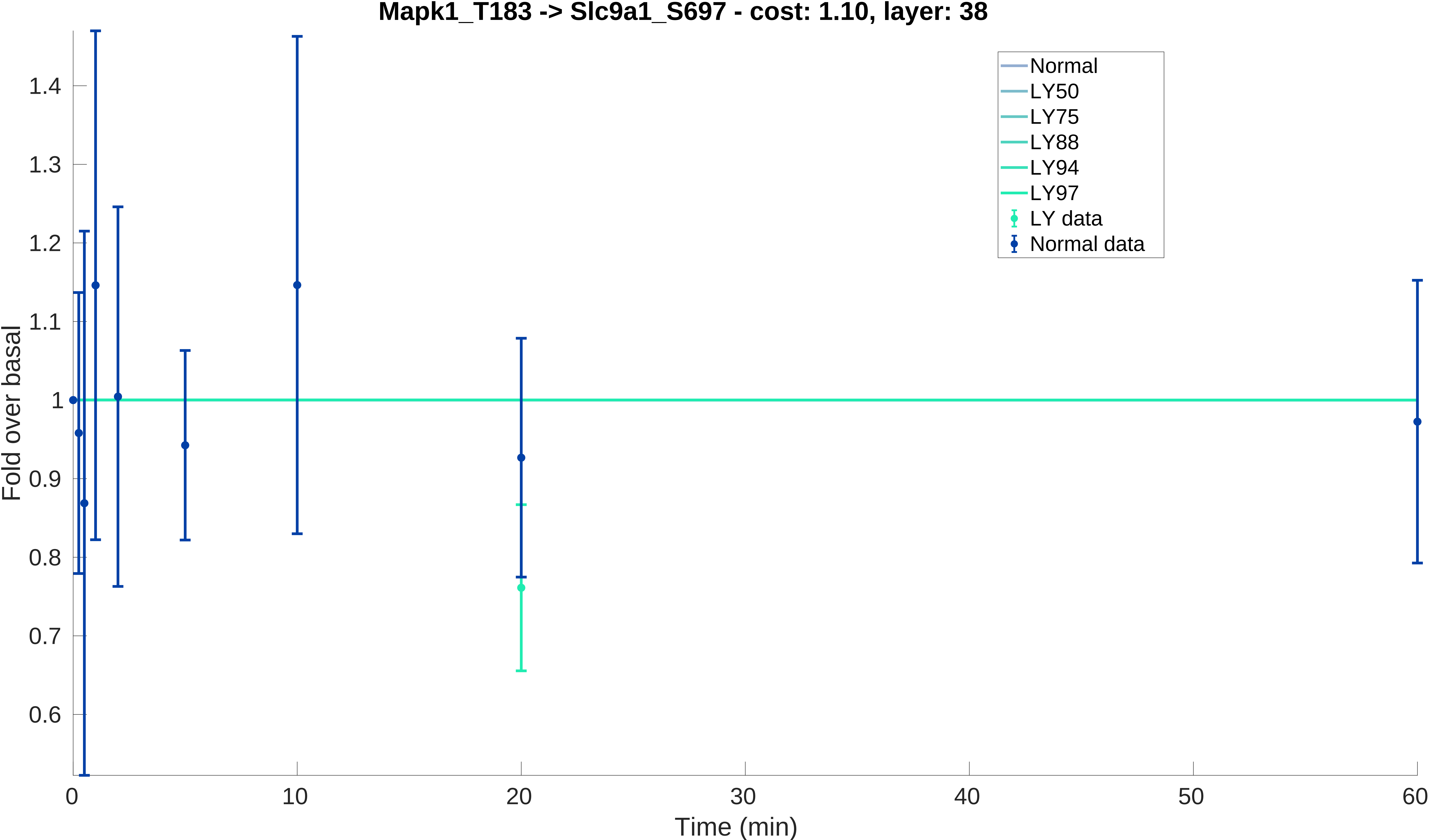

Mapk1_T183 -> Slc9a1_S697 - cost: 1.10, layer: 38
Normal
LY50
LY75
LY88
LY94
LY97
LY data
Normal data
Fold over basal
Time (min)
0.6
0.7
0.8
0.9
1
1.1
1.2
1.3
1.4
0
10
20
30
40
50
60

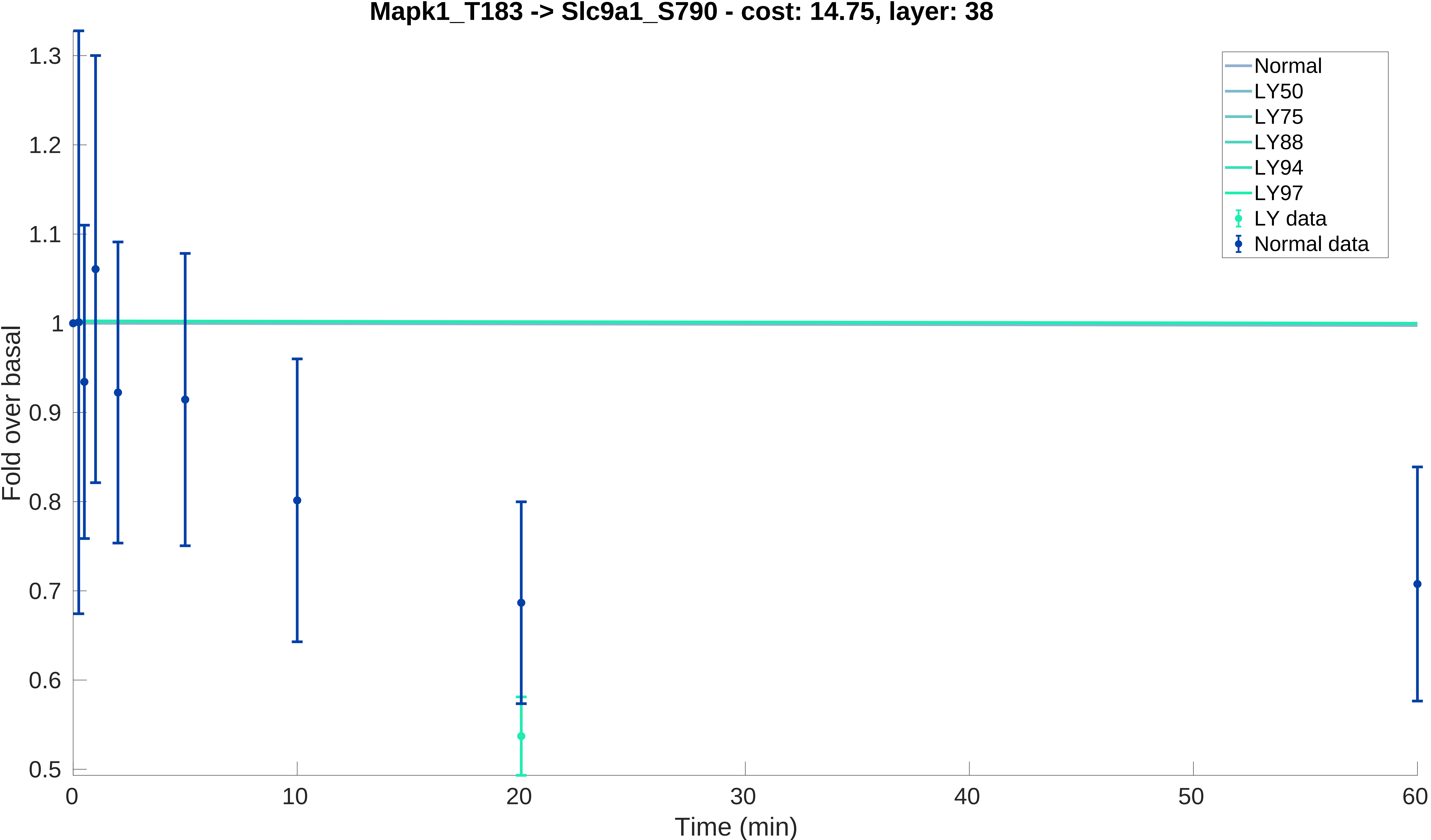

Mapk1_T183 -> Slc9a1_S790 - cost: 14.75, layer: 38
Normal
LY50
LY75
LY88
LY94
LY97
LY data
Normal data
Fold over basal
1.3
1.2
1.1
1
0.9
0.8
0.7
0.6
0.5
0
10
20
30
40
50
60
Time (min)

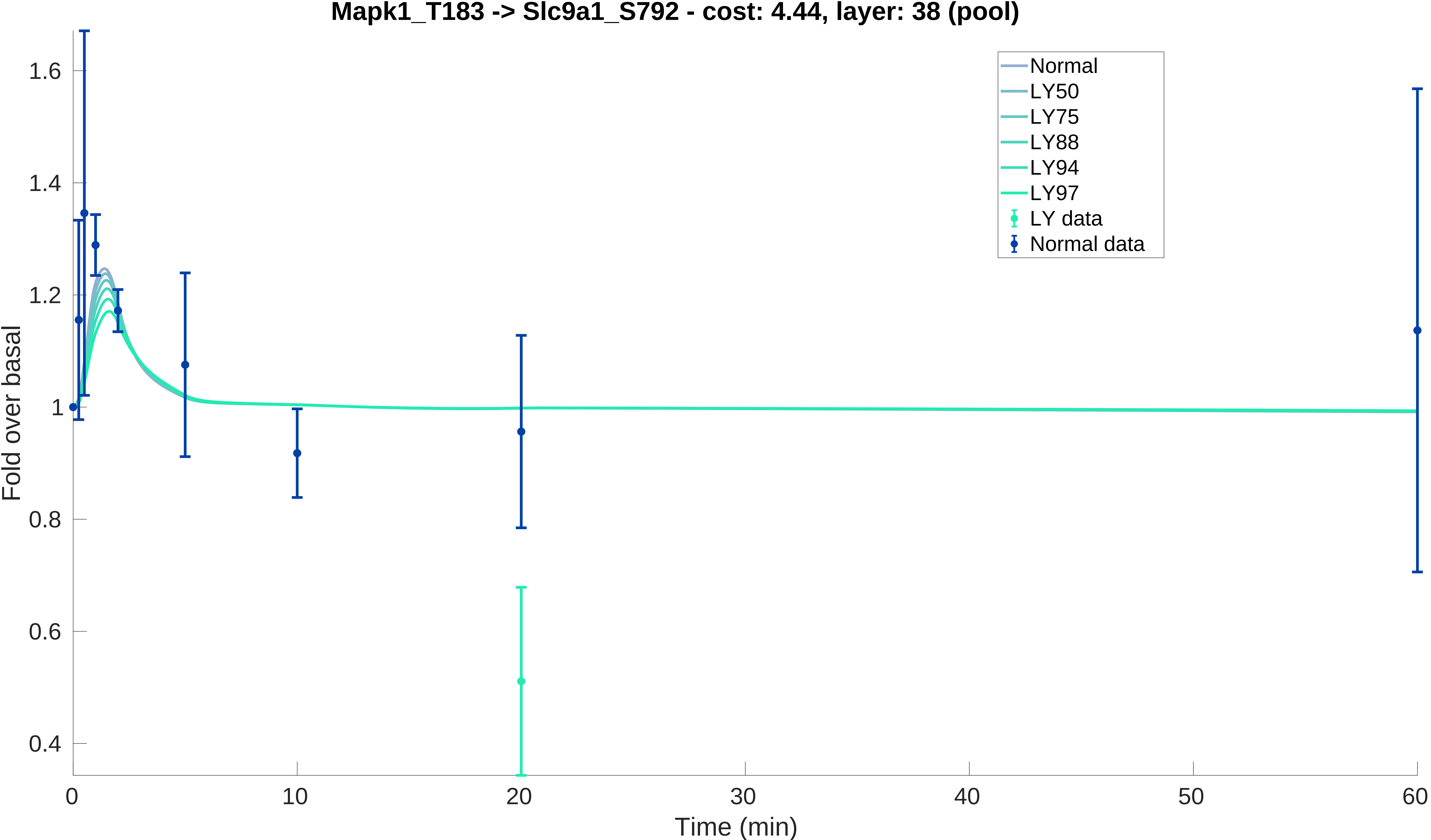
Mapk1_T183 -> Slc9a1_S792 - cost: 4.44, layer: 38 (pool)
Fold over basal
Time (min)
0.4
0.6
0.8
1
1.2
1.4
1.6
0
10
20
30
40
50
60
Normal
LY50
LY75
LY88
LY94
LY97
LY data
Normal data

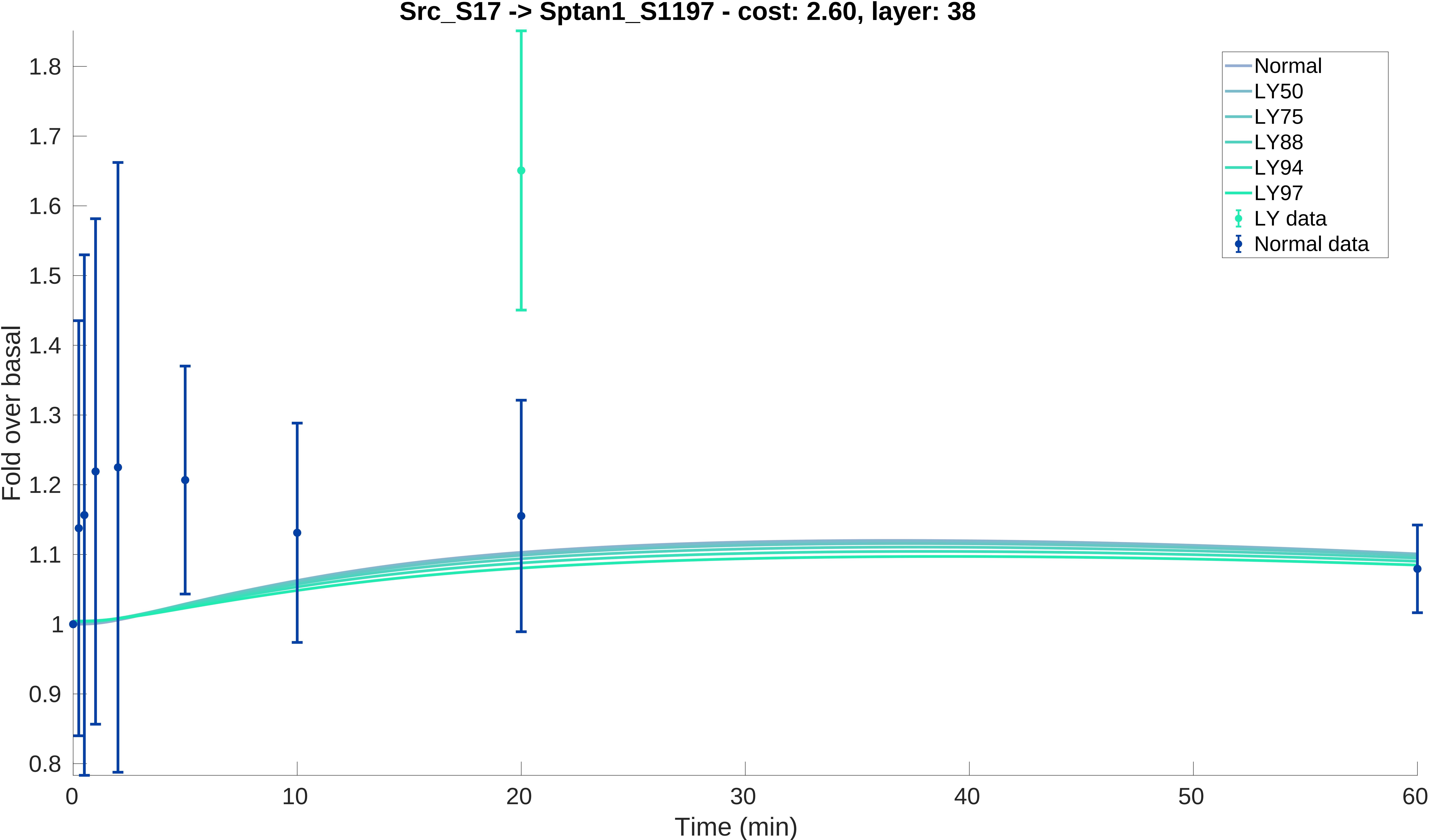

Src_S17 -> Sptan1_S1197 - cost: 2.60, layer: 38
Fold over basal
Time (min)
Normal
LY50
LY75
LY88
LY94
LY97
LY data
Normal data
0.8
0.9
1
1.1
1.2
1.3
1.4
1.5
1.6
1.7
1.8
0
10
20
30
40
50
60

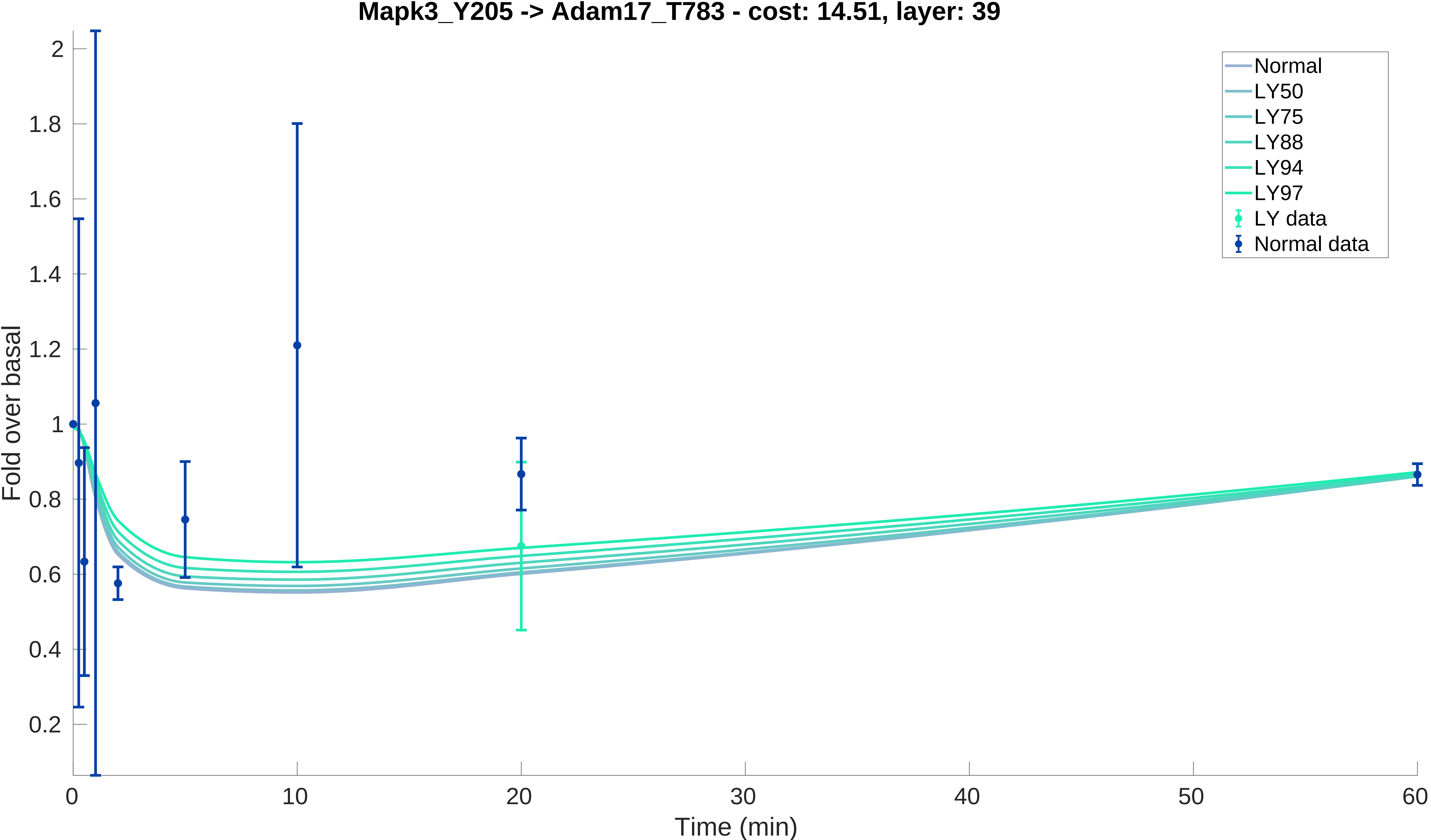

Mapk3_Y205 -> Adam17_T783 - cost: 14.51, layer: 39
Fold over basal
Time (min)
0
10
20
30
40
50
60
0.2
0.4
0.6
0.8
1
1.2
1.4
1.6
1.8
2
Normal
LY50
LY75
LY88
LY94
LY97
LY data
Normal data

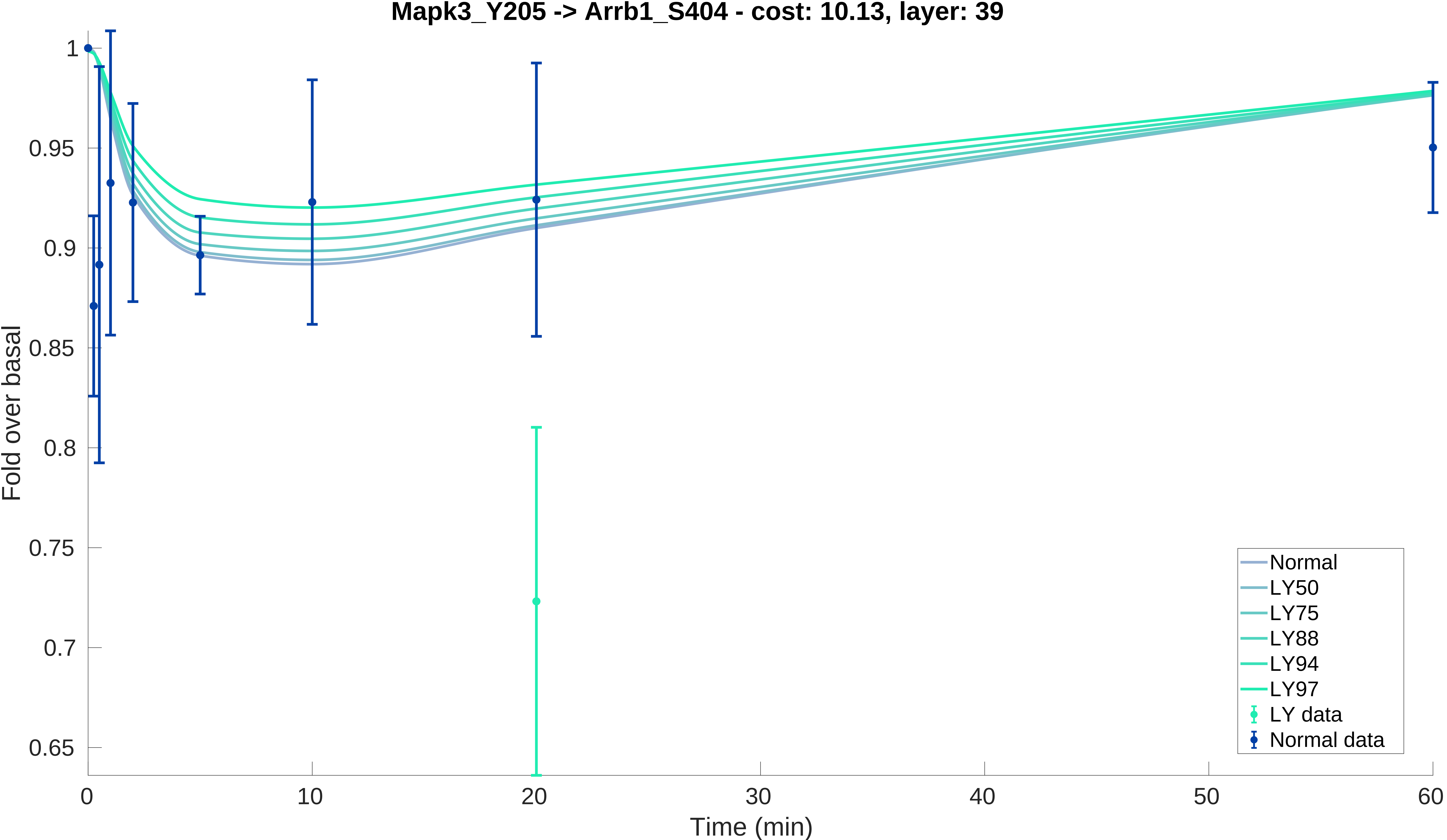

Mapk3_Y205 -> Arrb1_S404 - cost: 10.13, layer: 39
Fold over basal
1
0.95
0.9
0.85
0.8
0.75
0.7
0.65
0
10
20
30
40
50
60
Time (min)
Normal
LY50
LY75
LY88
LY94
LY97
LY data
Normal data

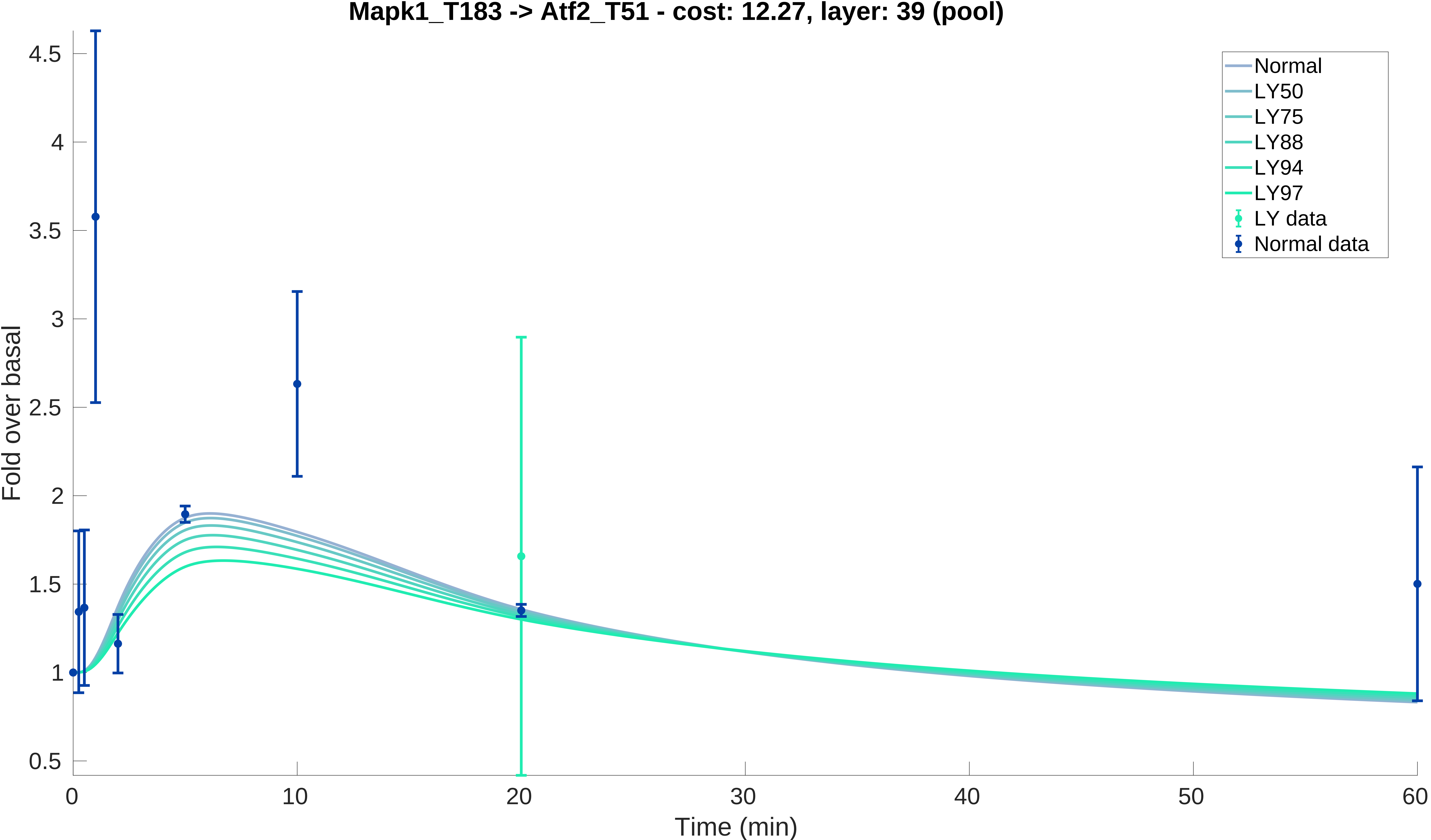
Mapk1_T183 -> Atf2_T51 - cost: 12.27, layer: 39 (pool)
Fold over basal
Time (min)
0.5
1
1.5
2
2.5
3
3.5
4
4.5
0
10
20
30
40
50
60
Normal
LY50
LY75
LY88
LY94
LY97
LY data
Normal data

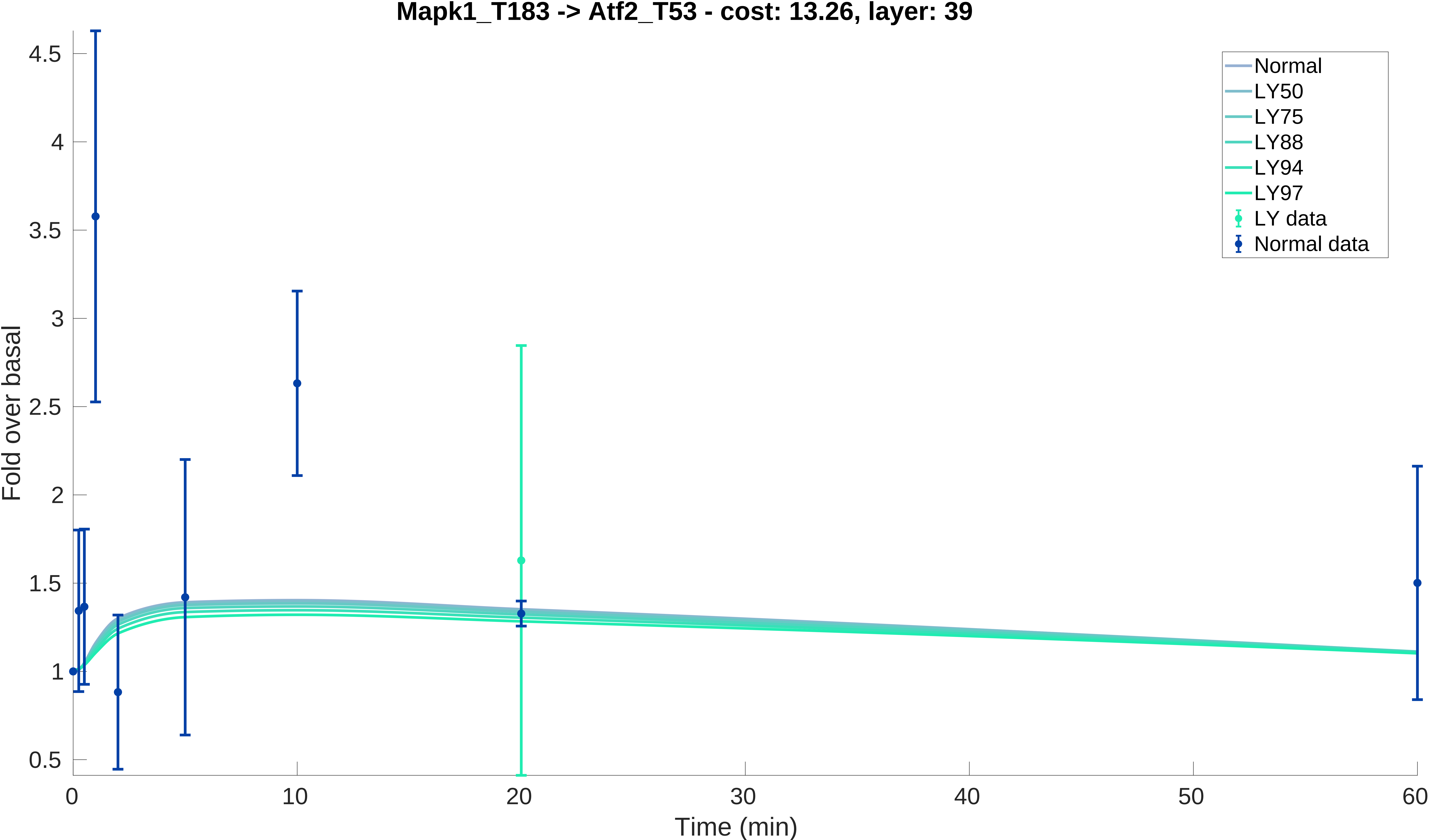

Mapk1_T183 -> Atf2_T53 - cost: 13.26, layer: 39
Fold over basal
Time (min)
Normal
LY50
LY75
LY88
LY94
LY97
LY data
Normal data
4.5
4
3.5
3
2.5
2
1.5
1
0.5
0
10
20
30
40
50
60

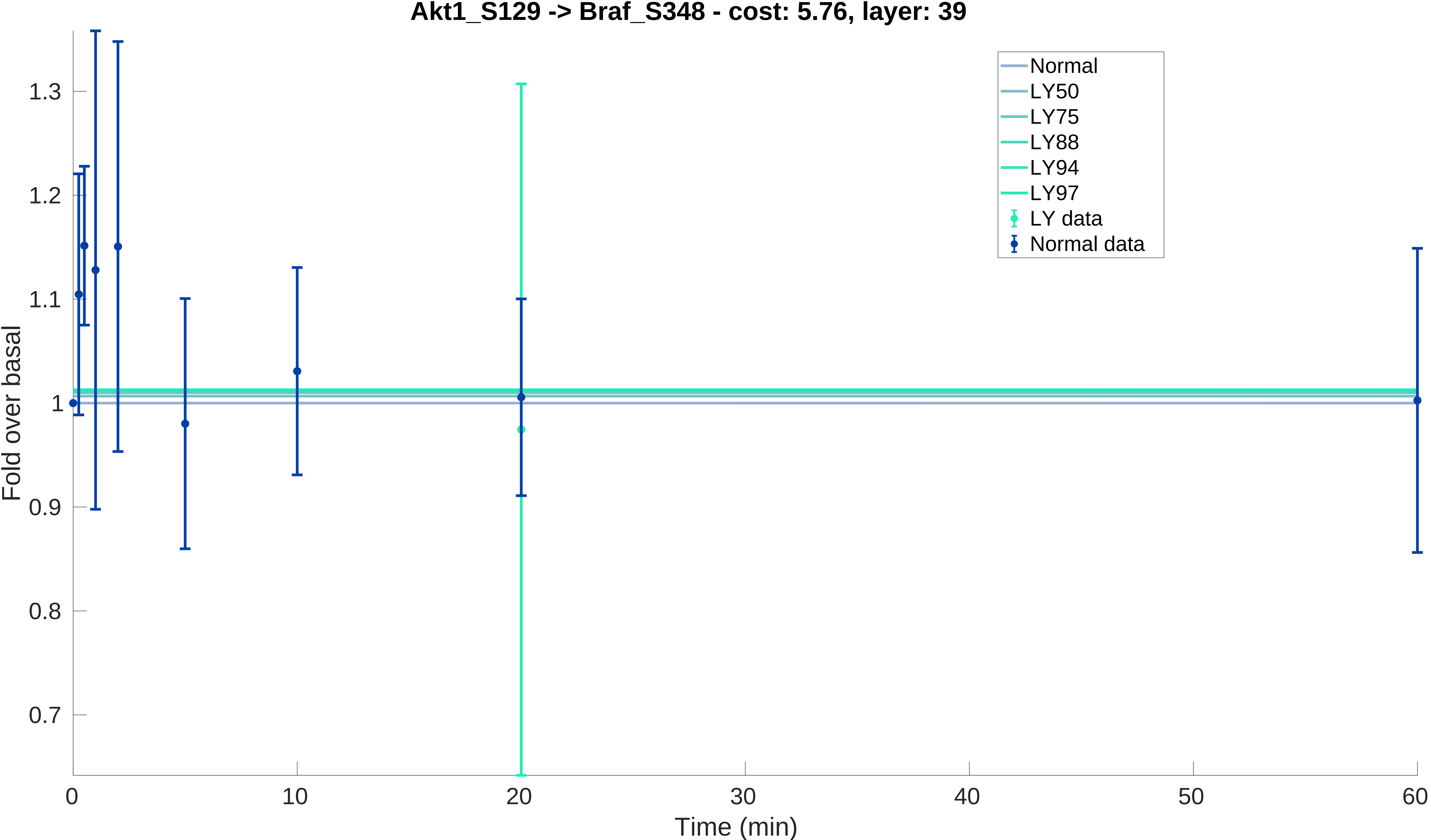

Akt1_S129 -> Braf_S348 - cost: 5.76, layer: 39
Normal
LY50
LY75
LY88
LY94
LY97
LY data
Normal data
Fold over basal
1.3
1.2
1.1
1
0.9
0.8
0.7
0
10
20
30
40
50
60
Time (min)

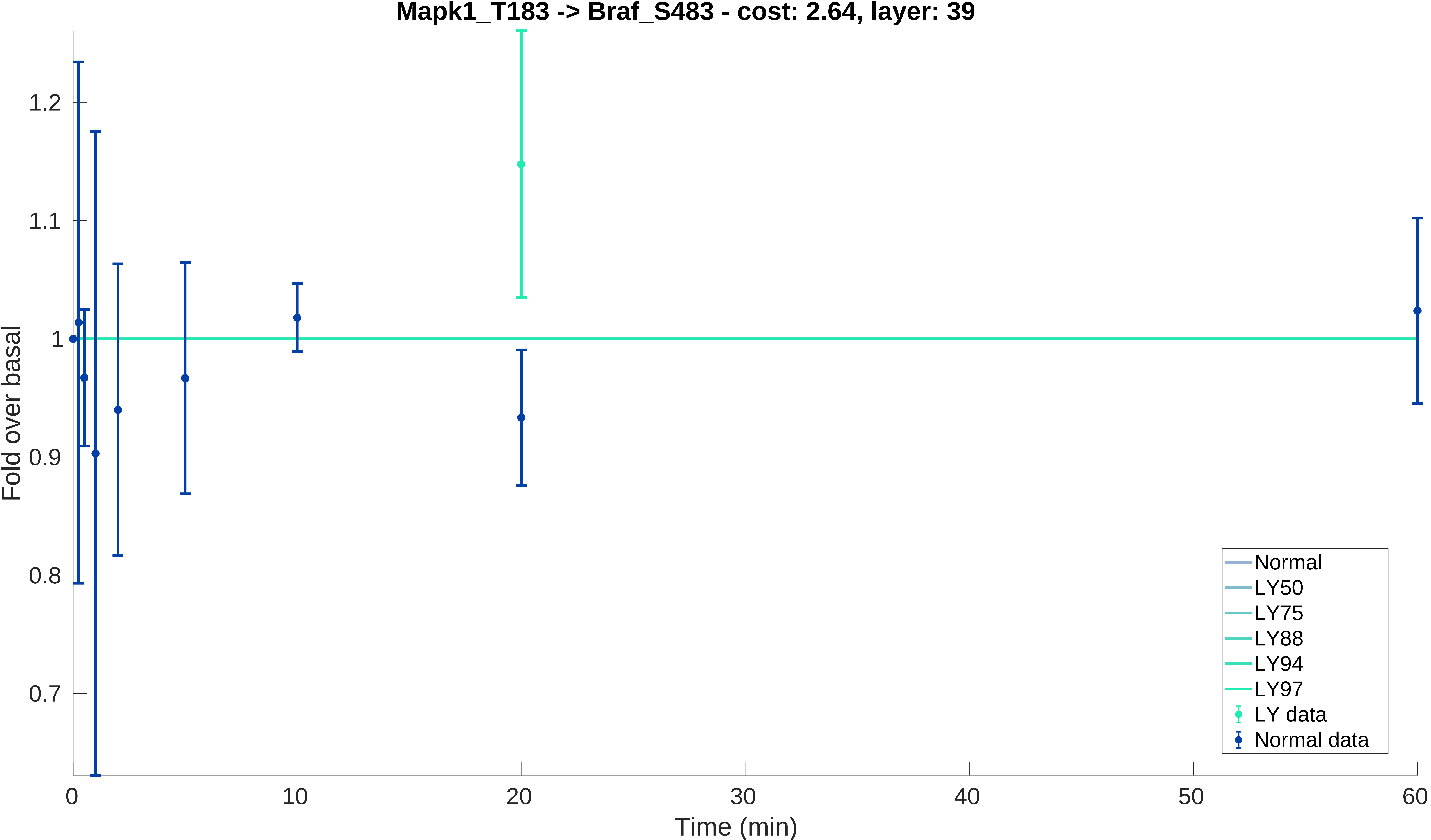

Mapk1_T183 -> Braf_S483 - cost: 2.64, layer: 39
Fold over basal
1.2
1.1
1
0.9
0.8
0.7
0
10
20
30
40
50
60
Time (min)
Normal
LY50
LY75
LY88
LY94
LY97
LY data
Normal data

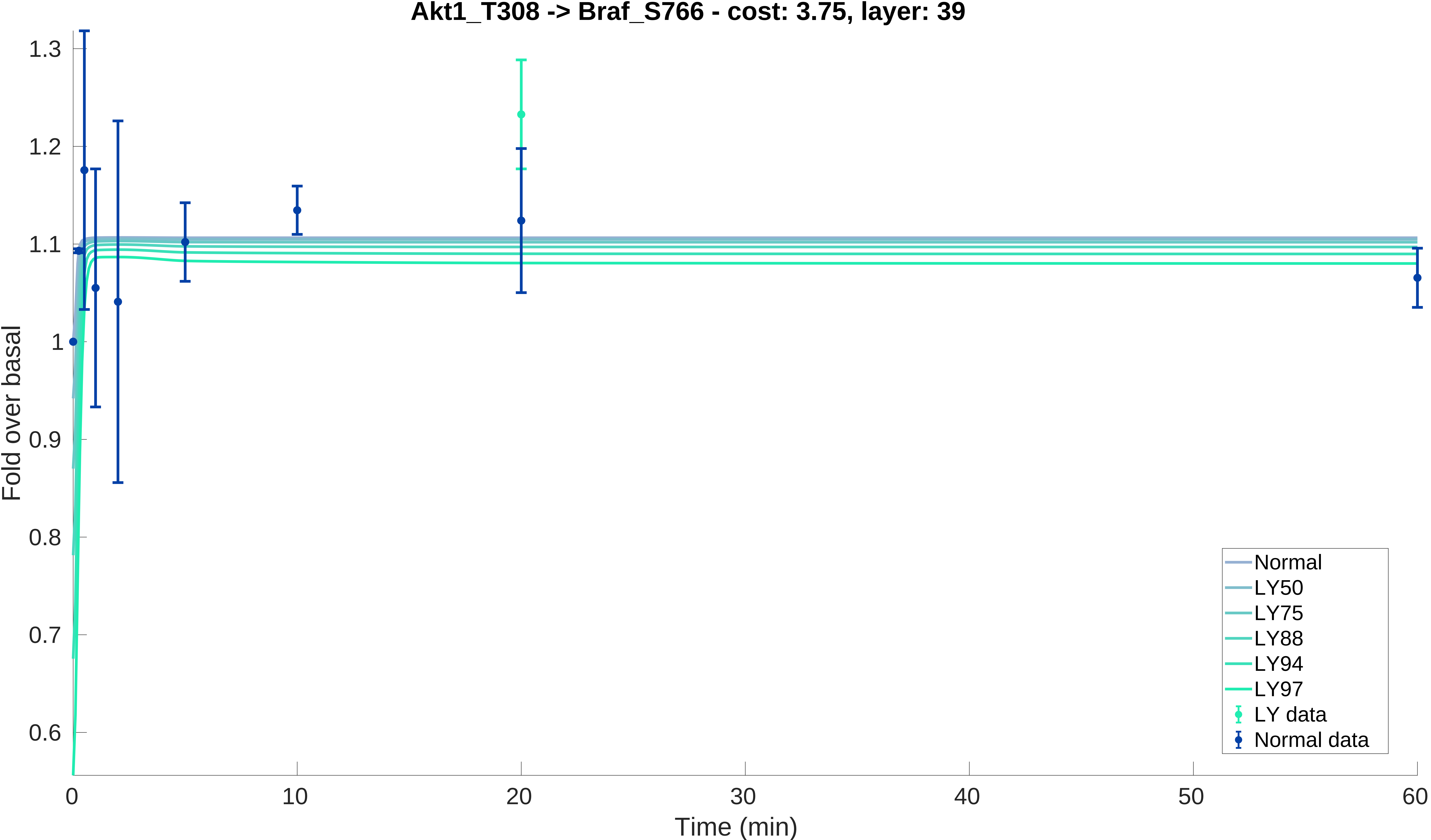

Akt1_T308 -> Braf_S766 - cost: 3.75, layer: 39
Fold over basal
Time (min)
1.3
1.2
1.1
1
0.9
0.8
0.7
0.6
0
10
20
30
40
50
60
Normal
LY50
LY75
LY88
LY94
LY97
LY data
Normal data

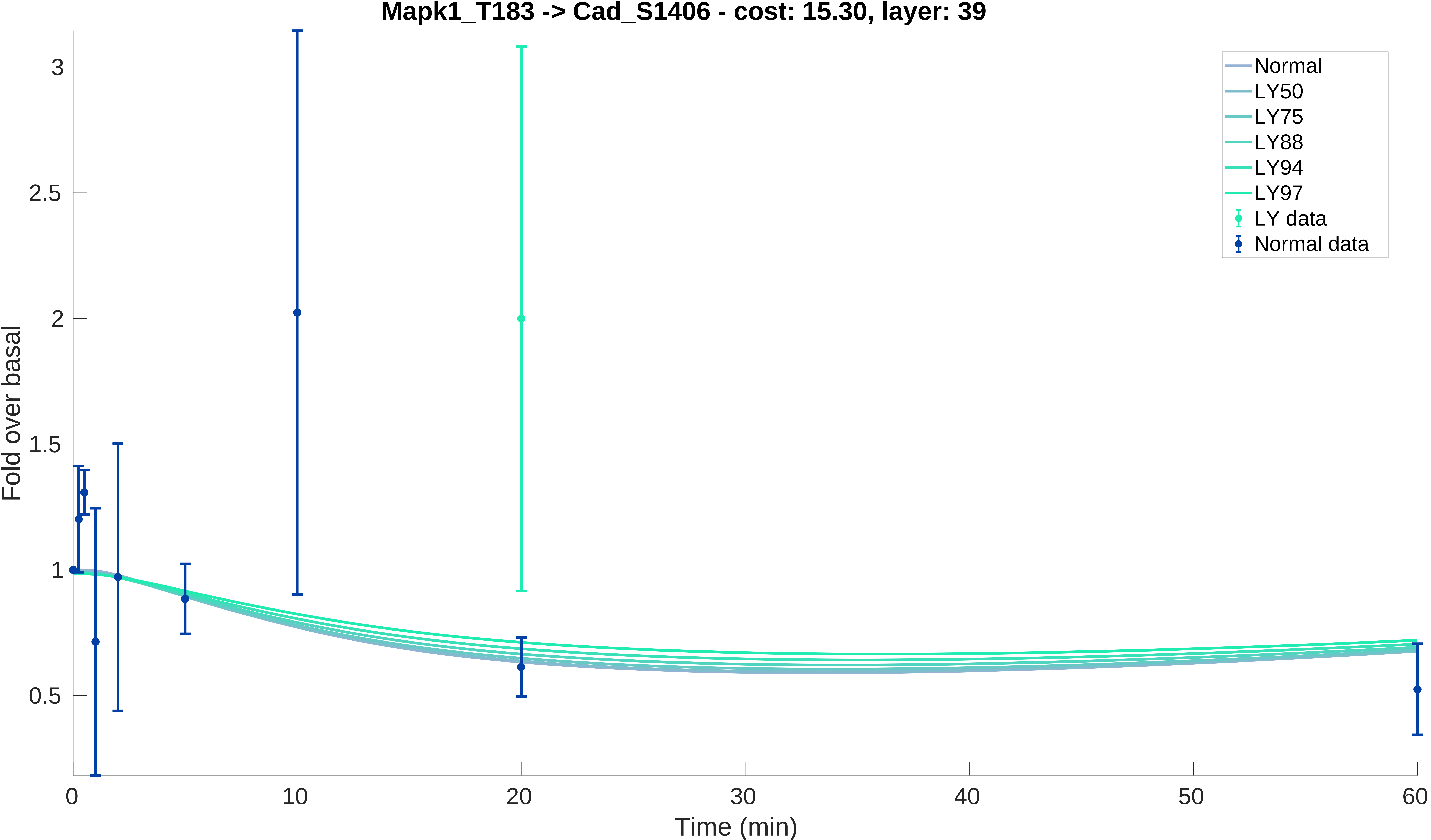

Mapk1_T183 -> Cad_S1406 - cost: 15.30, layer: 39
Normal
LY50
LY75
LY88
LY94
LY97
LY data
Normal data
Fold over basal
3
2.5
2
1.5
1
0.5
0
10
20
30
40
50
60
Time (min)

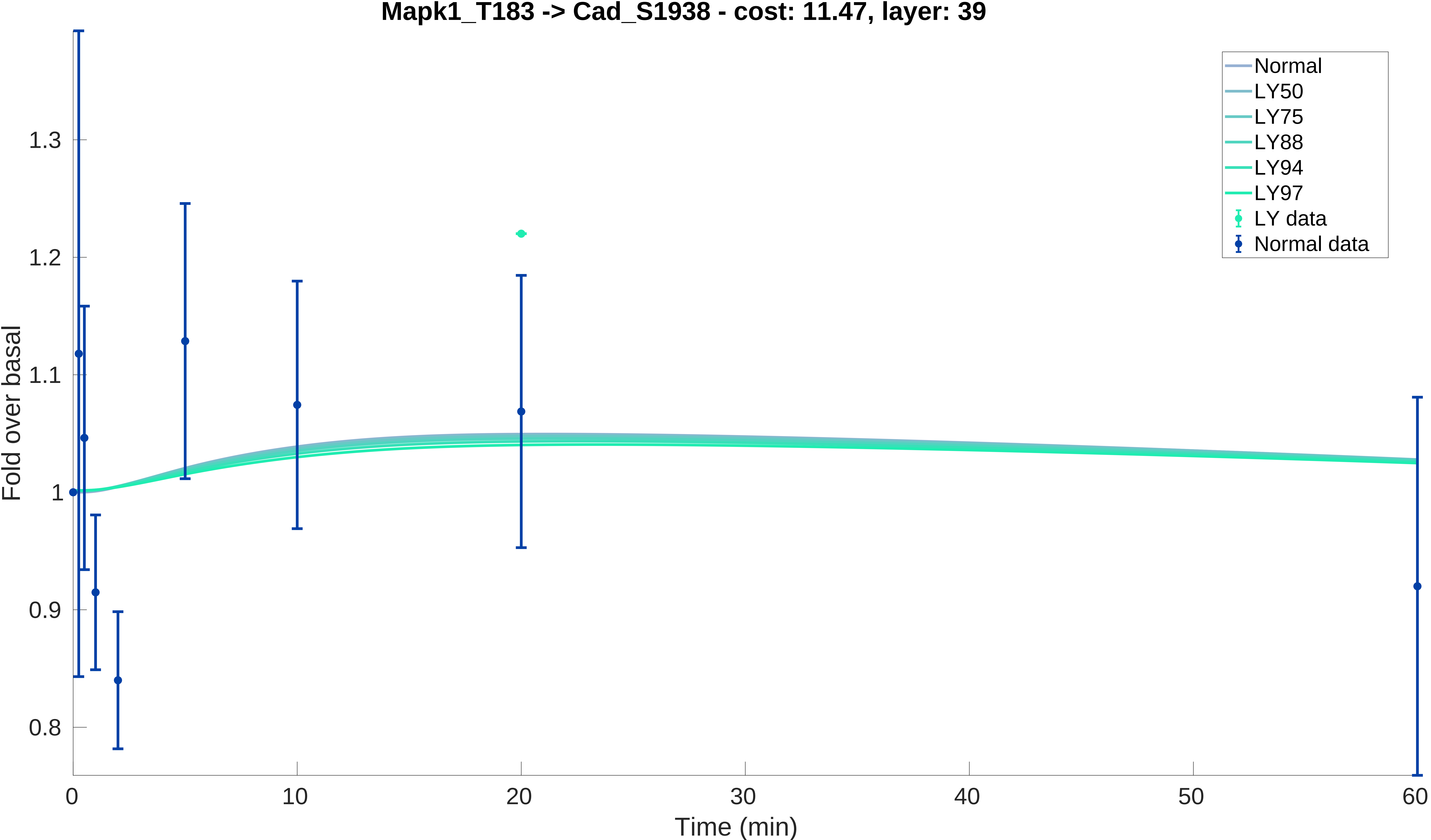

Mapk1_T183 -> Cad_S1938 - cost: 11.47, layer: 39
Normal
LY50
LY75
LY88
LY94
LY97
LY data
Normal data
1.3
1.2
1.1
1
0.9
0.8
Fold over basal
0
10
20
30
40
50
60
Time (min)

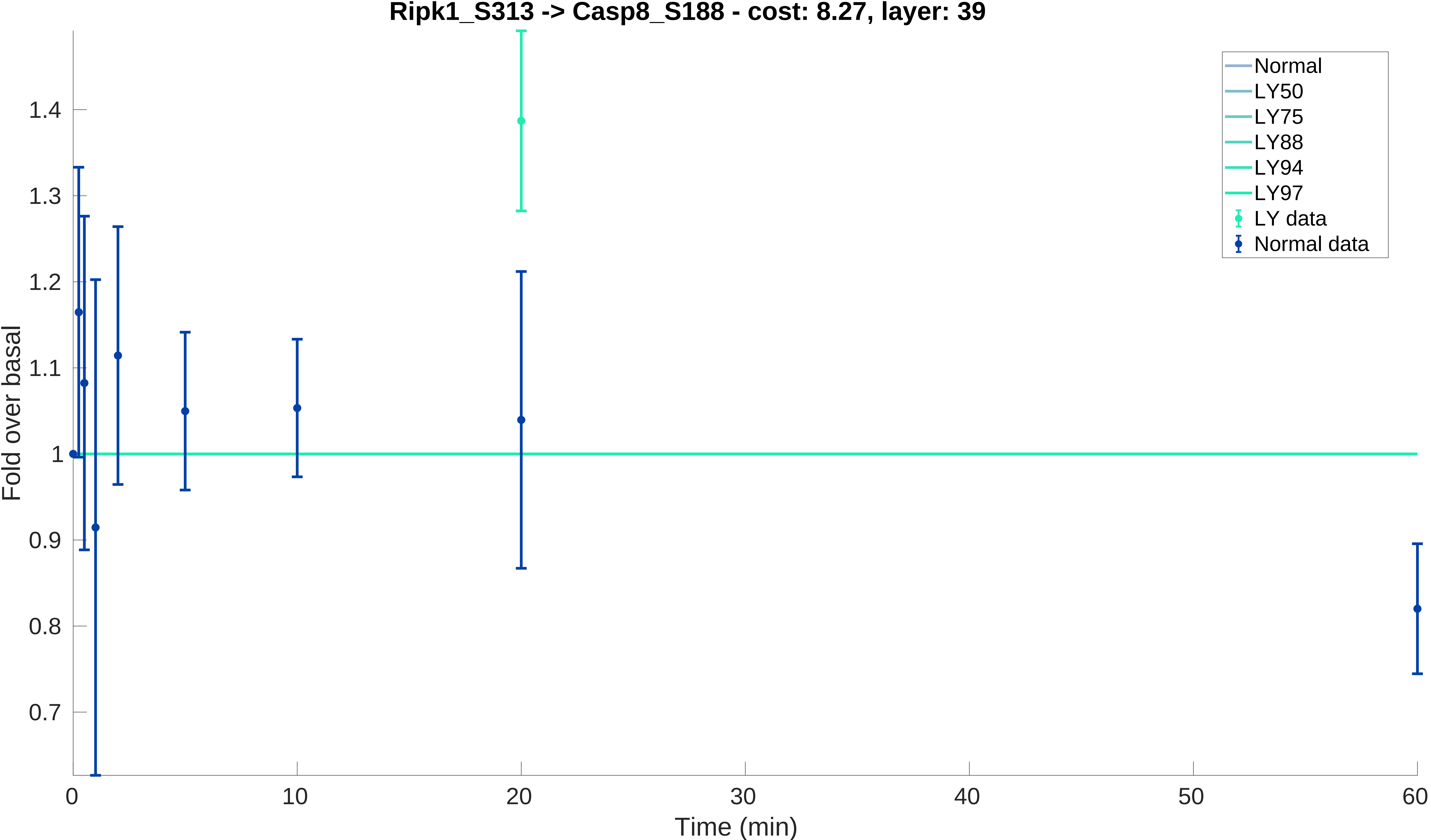

Ripk1_S313 -> Casp8_S188 - cost: 8.27, layer: 39
Fold over basal
1.4
1.3
1.2
1.1
1
0.9
0.8
0.7
0
10
20
30
40
50
60
Time (min)
Normal
LY50
LY75
LY88
LY94
LY97
LY data
Normal data

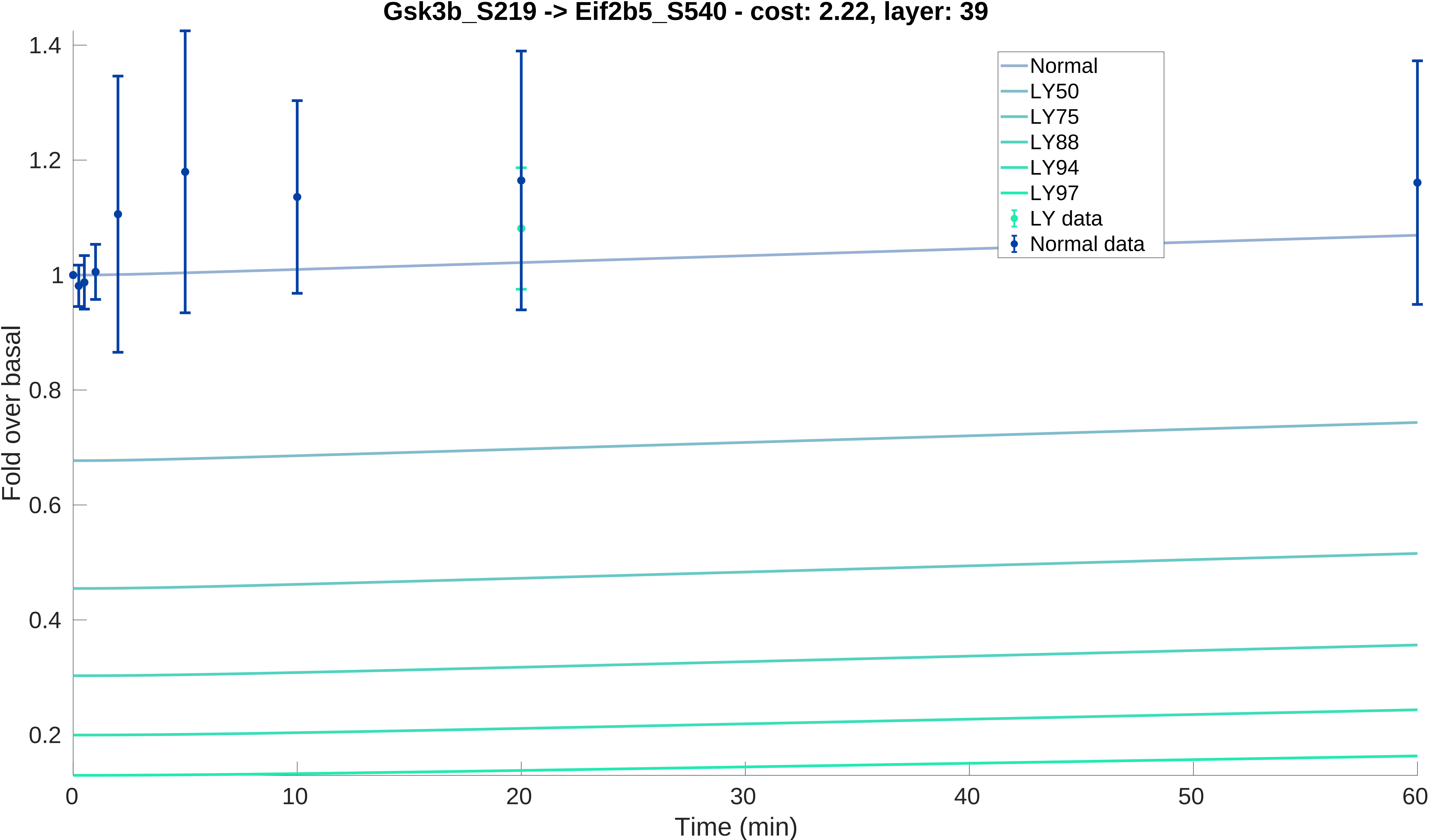

Gsk3b_S219 -> Eif2b5_S540 - cost: 2.22, layer: 39
Fold over basal
Time (min)
Normal
LY50
LY75
LY88
LY94
LY97
LY data
Normal data
1.4
1.2
1
0.8
0.6
0.4
0.2
0
10
20
30
40
50
60

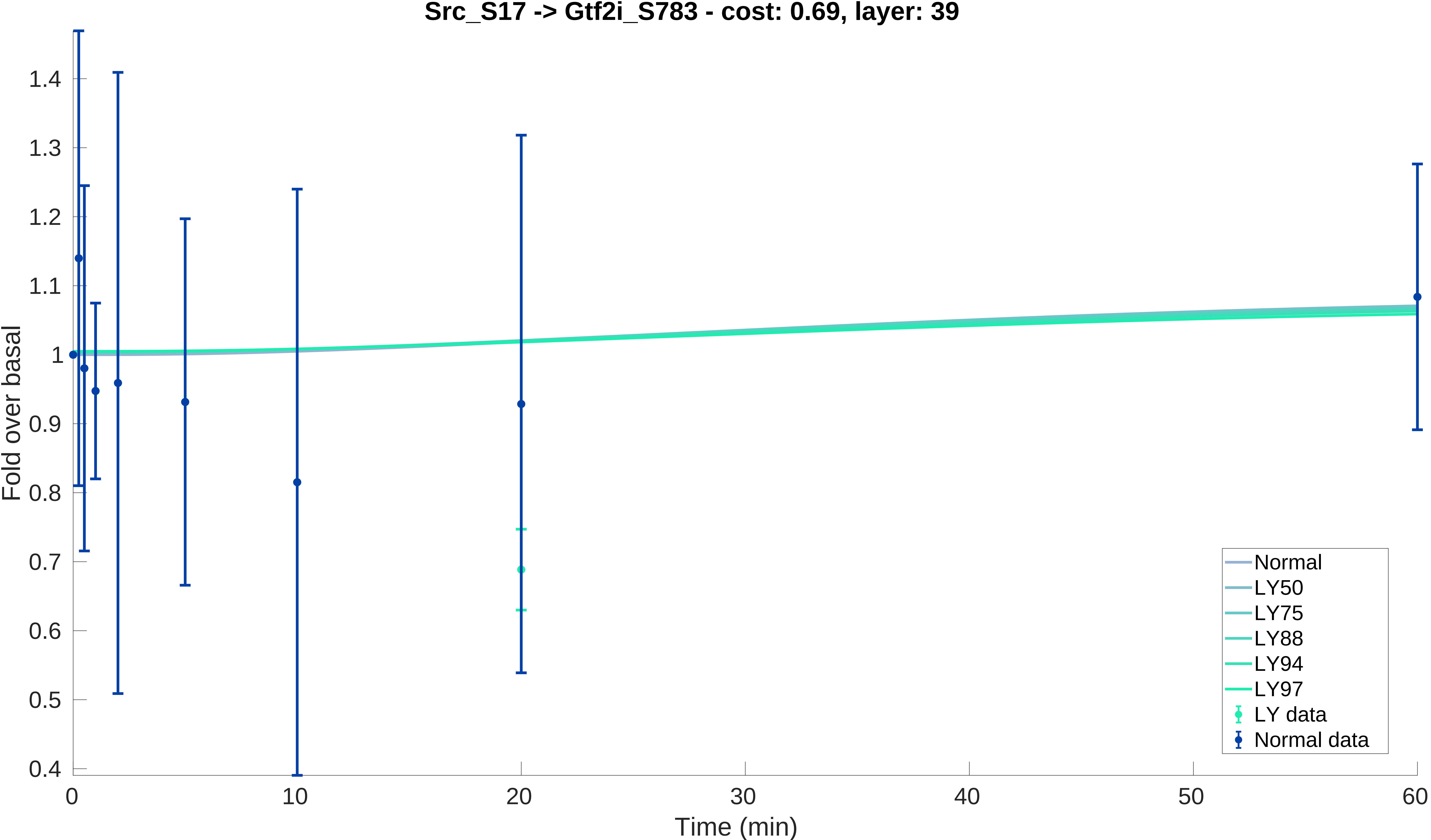

Src_S17 -> Gtf2i_S783 - cost: 0.69, layer: 39
Fold over basal
1.4
1.3
1.2
1.1
1
0.9
0.8
0.7
0.6
0.5
0.4
0
10
20
30
40
50
60
Time (min)
Normal
LY50
LY75
LY88
LY94
LY97
LY data
Normal data

Ripk1_S313 -> Ikbkg_S398 - cost: 3.06, layer: 39

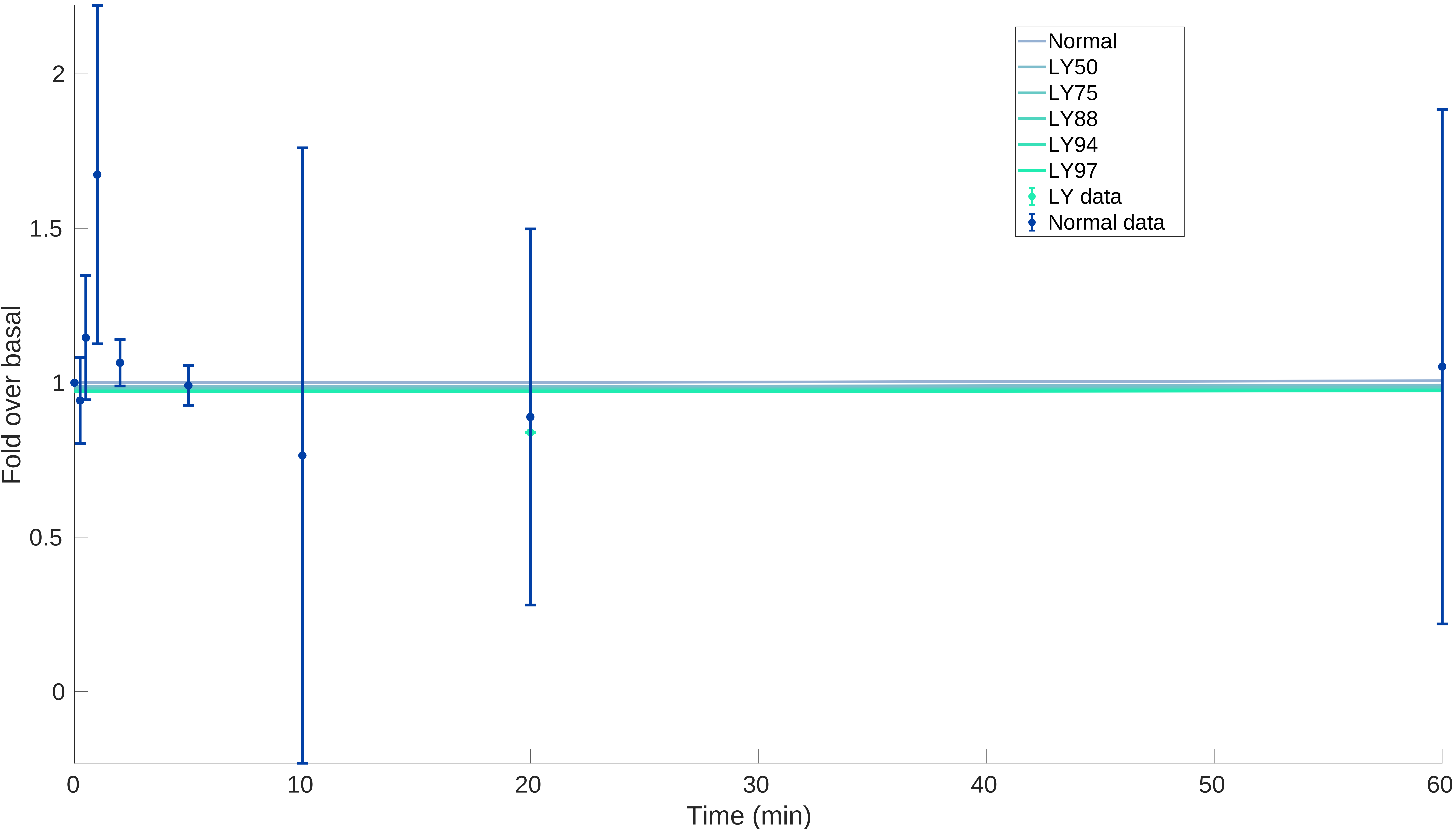

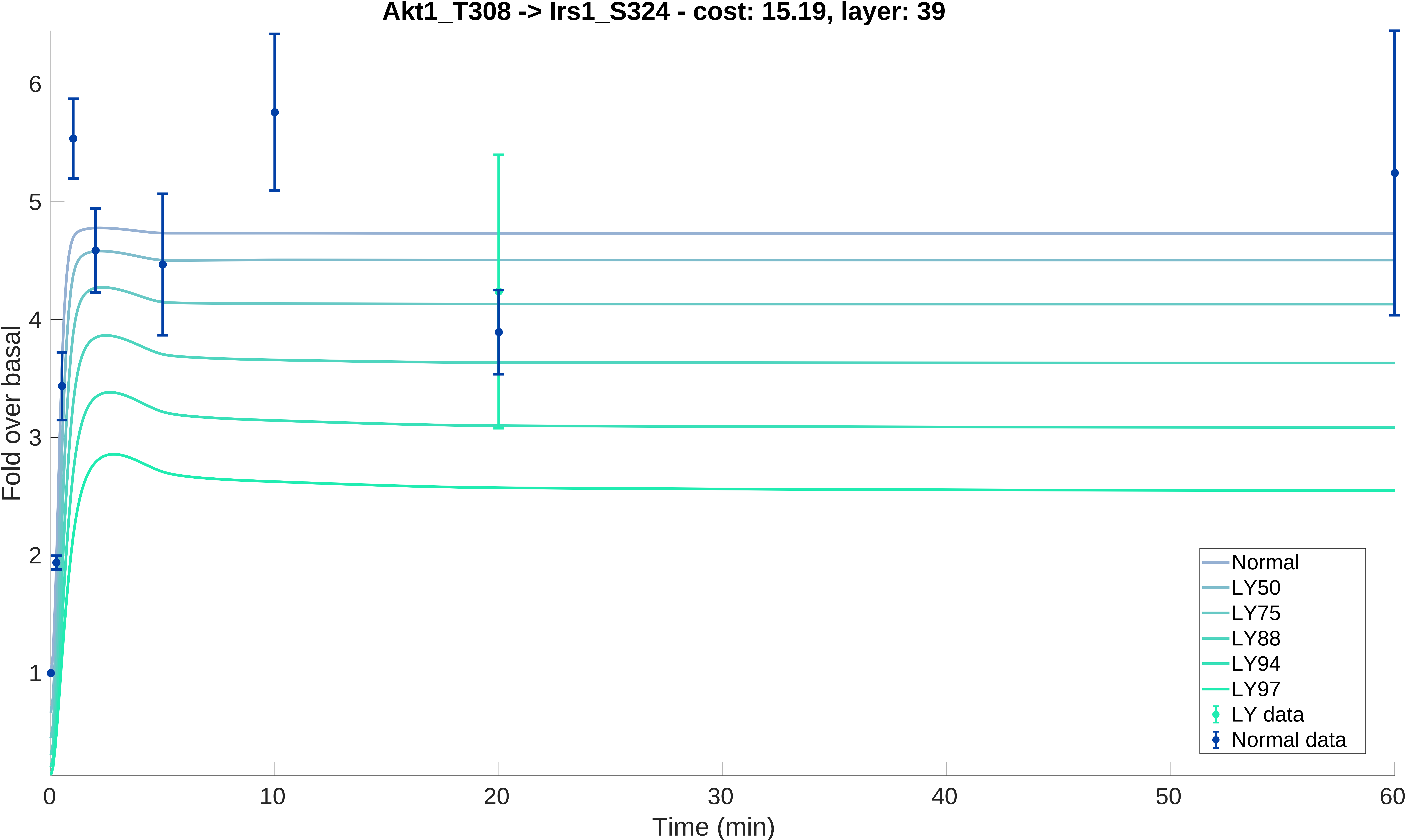

Akt1_T308 -> Irs1_S324 - cost: 15.19, layer: 39
Fold over basal
Time (min)
0
10
20
30
40
50
60
1
2
3
4
5
6
Normal
LY50
LY75
LY88
LY94
LY97
LY data
Normal data

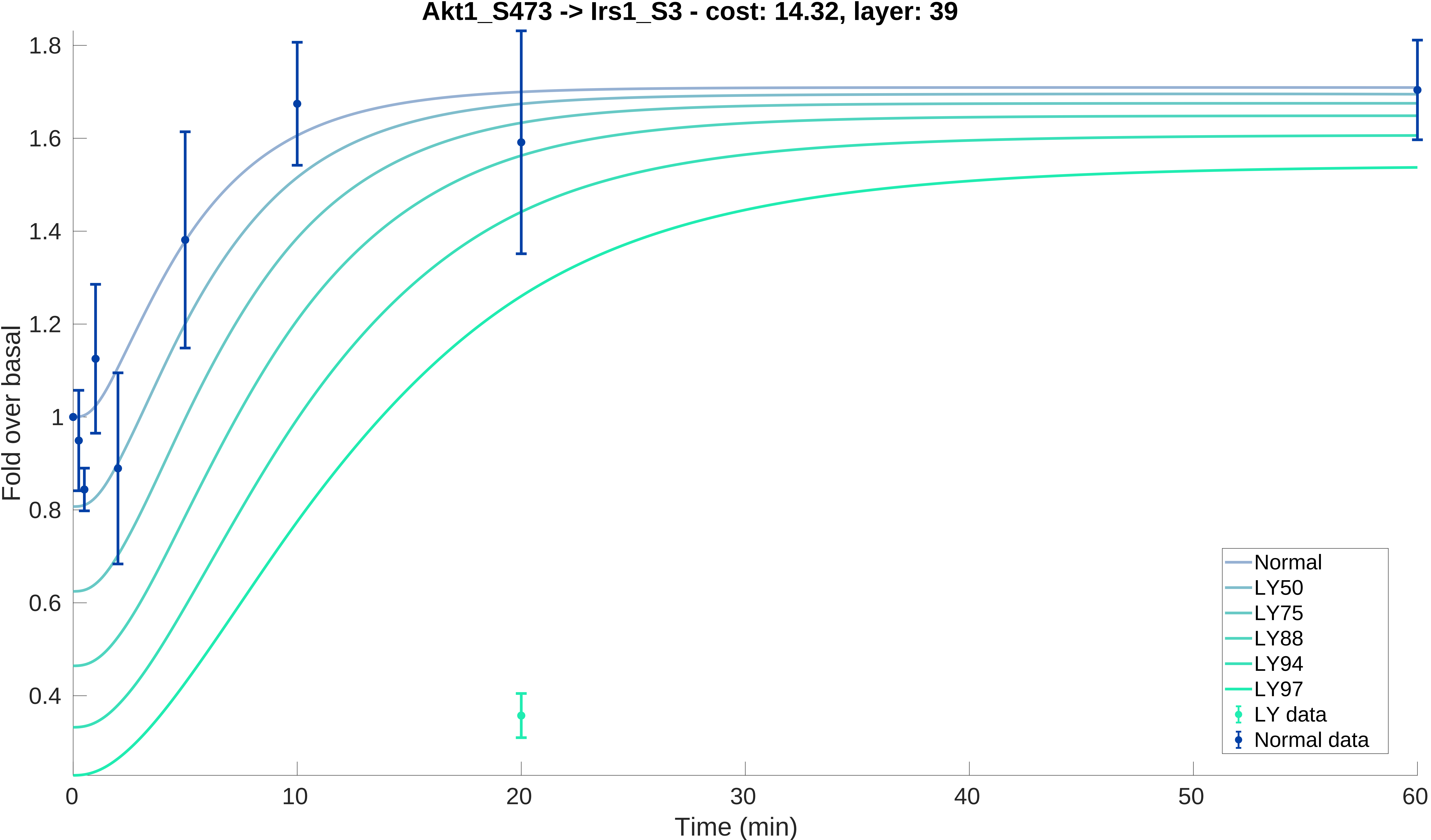

Akt1_S473 -> Irs1_S3 - cost: 14.32, layer: 39
Fold over basal
1.8
1.6
1.4
1.2
1
0.8
0.6
0.4
0
10
20
30
40
50
60
Time (min)
Normal
LY50
LY75
LY88
LY94
LY97
LY data
Normal data

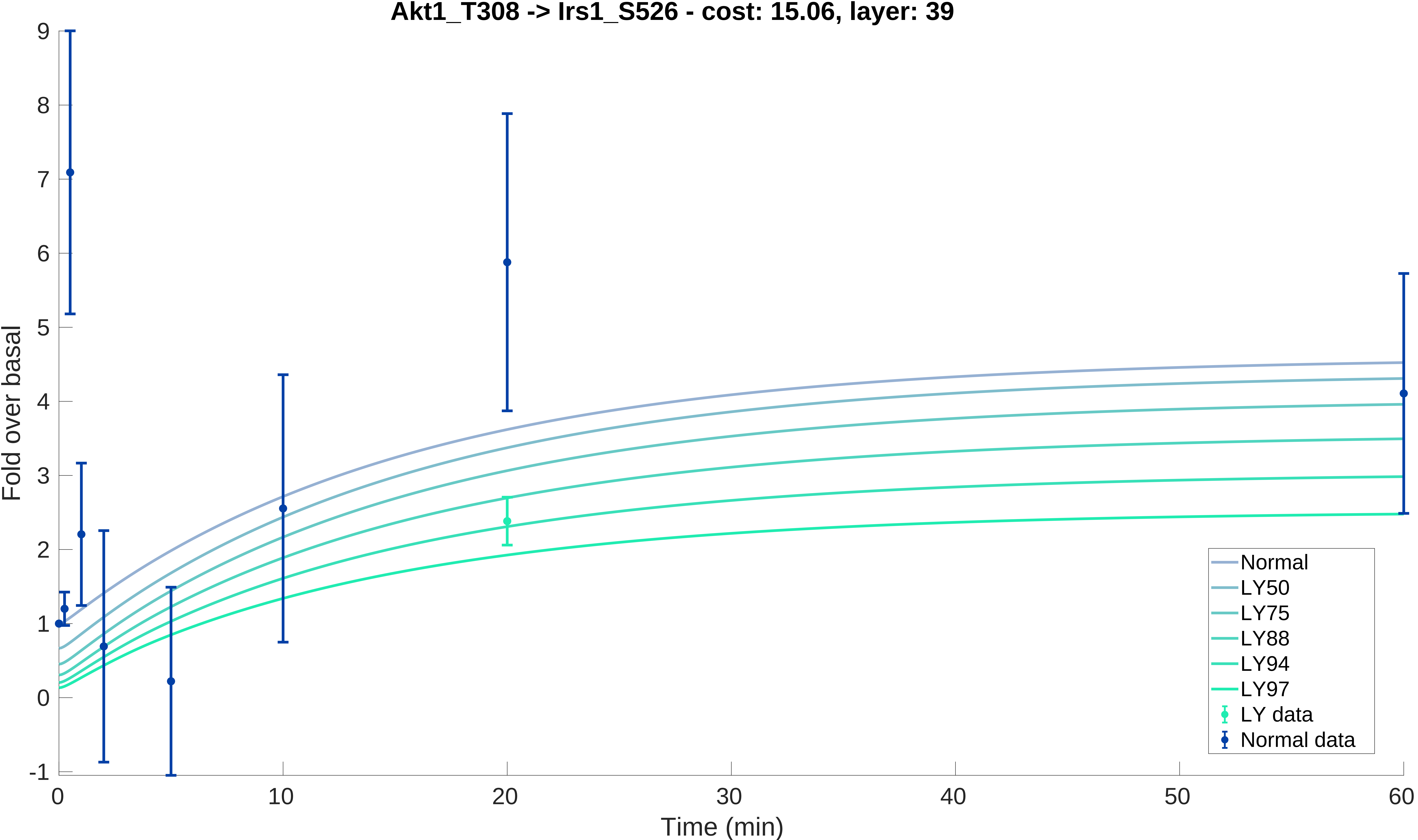
Akt1_T308 -> Irs1_S526 - cost: 15.06, layer: 39
Fold over basal
Time (min)
9
8
7
6
5
4
3
2
1
0
-1
0
10
20
30
40
50
60
Normal
LY50
LY75
LY88
LY94
LY97
LY data
Normal data

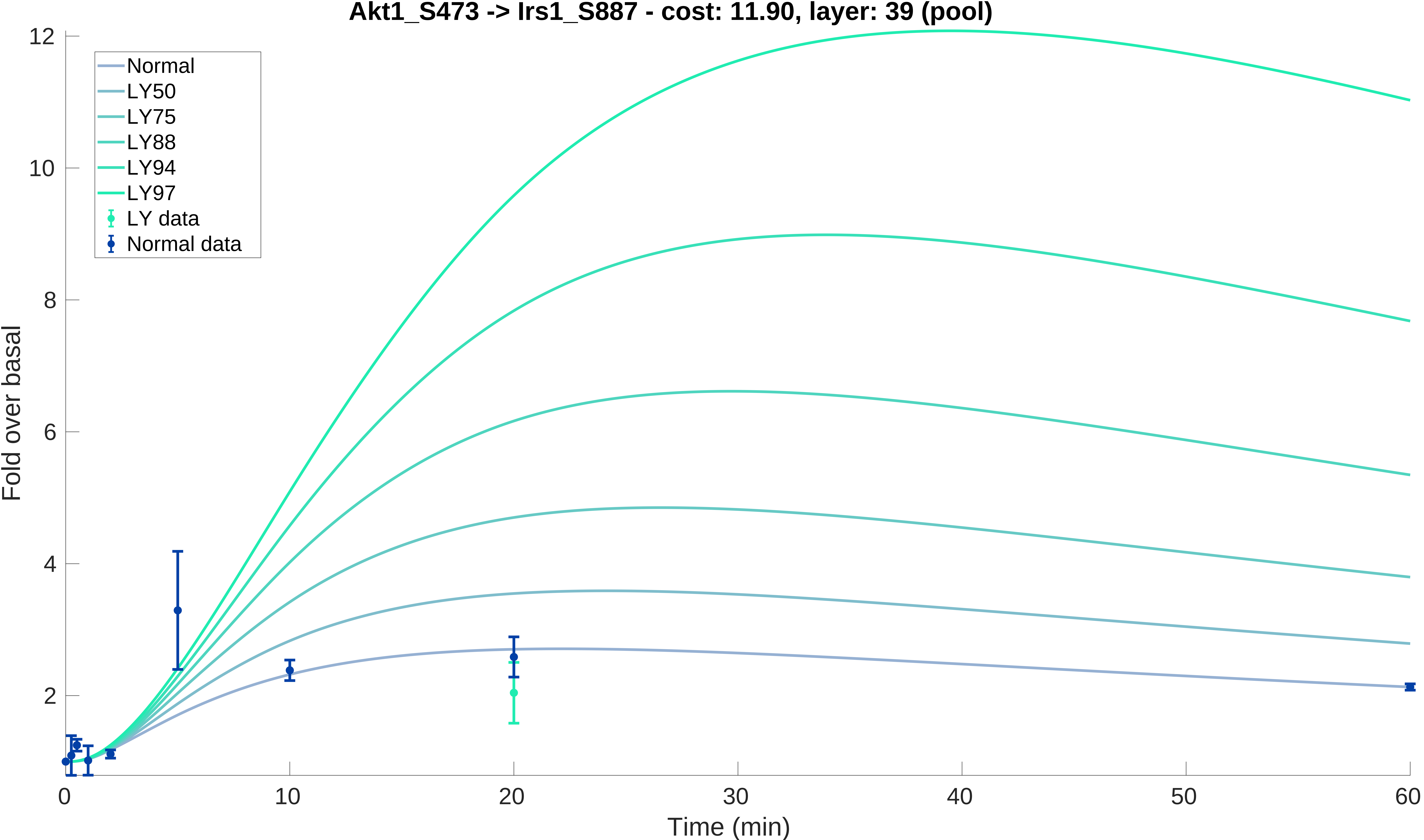

Akt1_S473 -> Irs1_S887 - cost: 11.90, layer: 39 (pool)
Normal
LY50
LY75
LY88
LY94
LY97
LY data
Normal data
Fold over basal
Time (min)
0
10
20
30
40
50
60
2
4
6
8
10
12

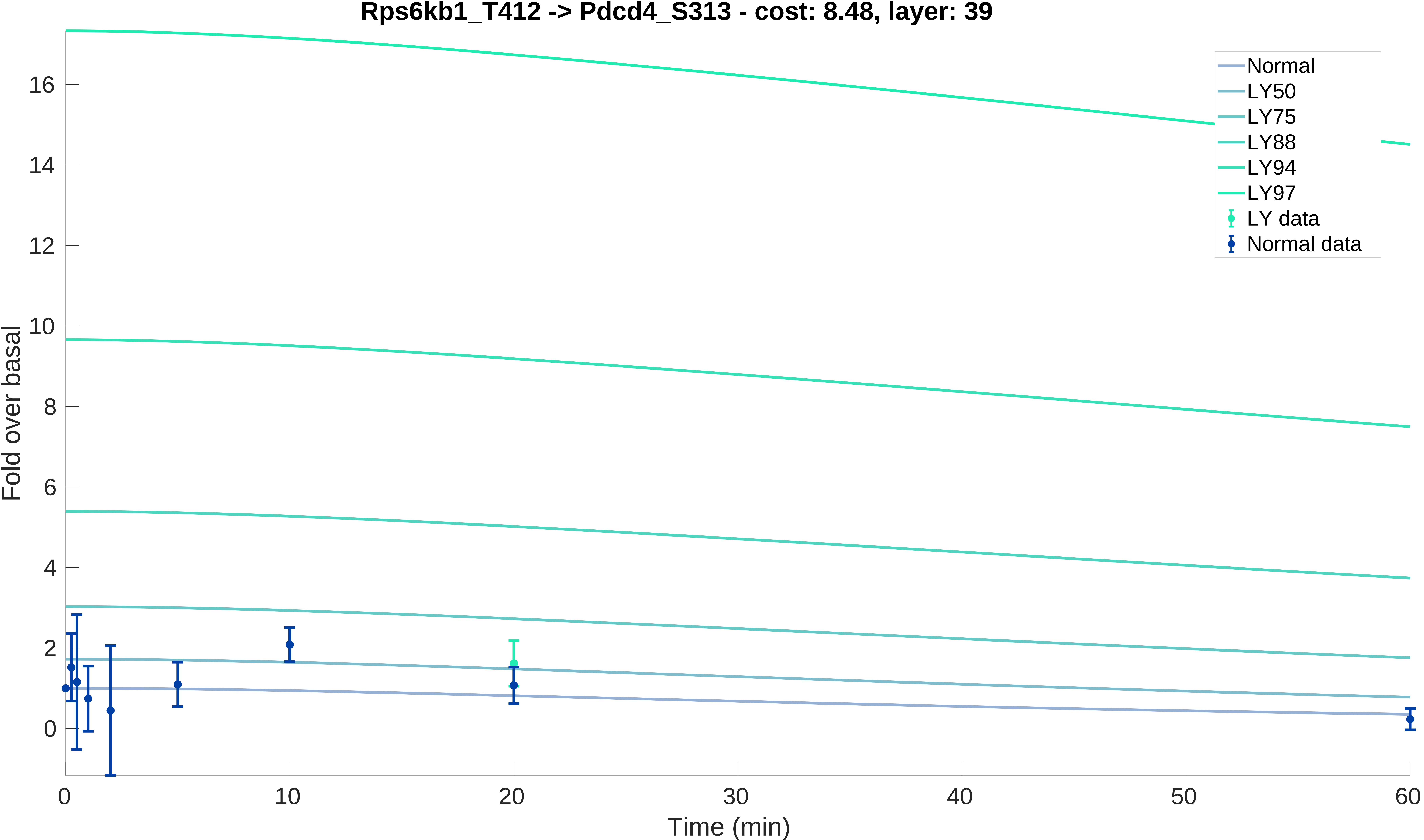

Rps6kb1_T412 -> Pdcd4_S313 - cost: 8.48, layer: 39
Fold over basal
Time (min)
0
2
4
6
8
10
12
14
16
0
10
20
30
40
50
60
Normal
LY50
LY75
LY88
LY94
LY97
LY data
Normal data

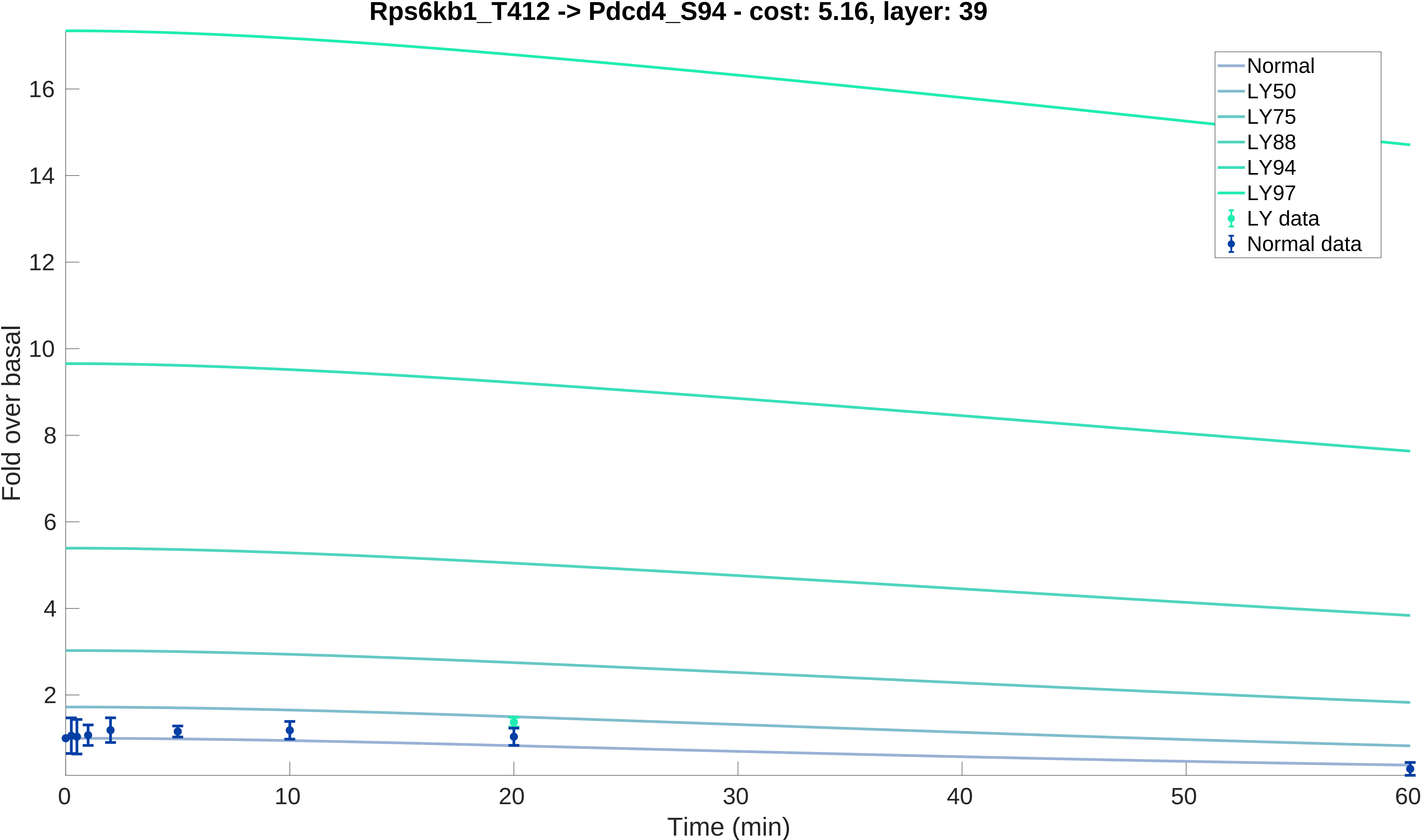

Rps6kb1_T412 -> Pdcd4_S94 - cost: 5.16, layer: 39
Fold over basal
16
14
12
10
8
6
4
2
0
10
20
30
40
50
60
Time (min)
Normal
LY50
LY75
LY88
LY94
LY97
LY data
Normal data

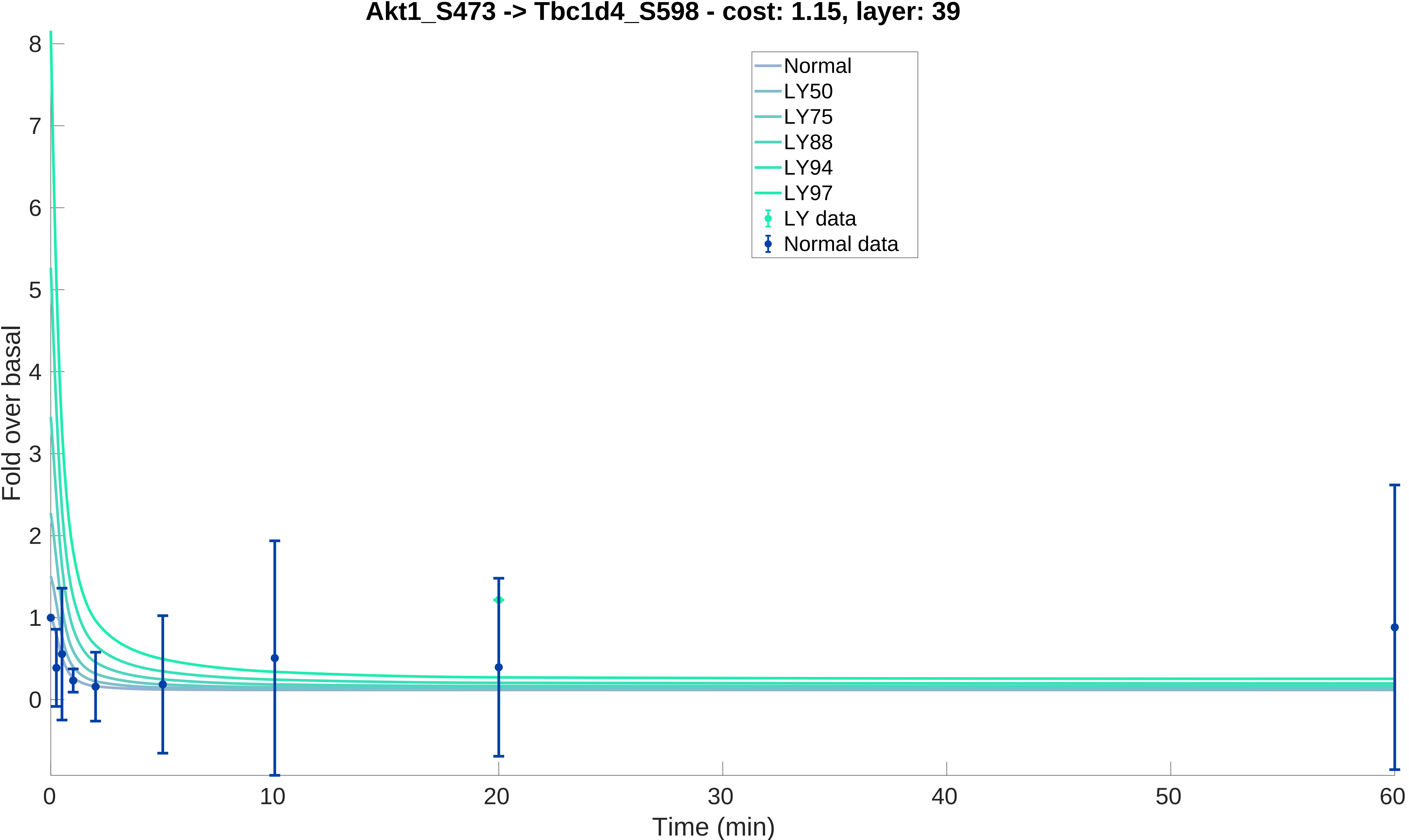

Akt1_S473 -> Tbc1d4_S598 - cost: 1.15, layer: 39
Normal
LY50
LY75
LY88
LY94
LY97
LY data
Normal data
Fold over basal
Time (min)
0
1
2
3
4
5
6
7
8
0
10
20
30
40
50
60

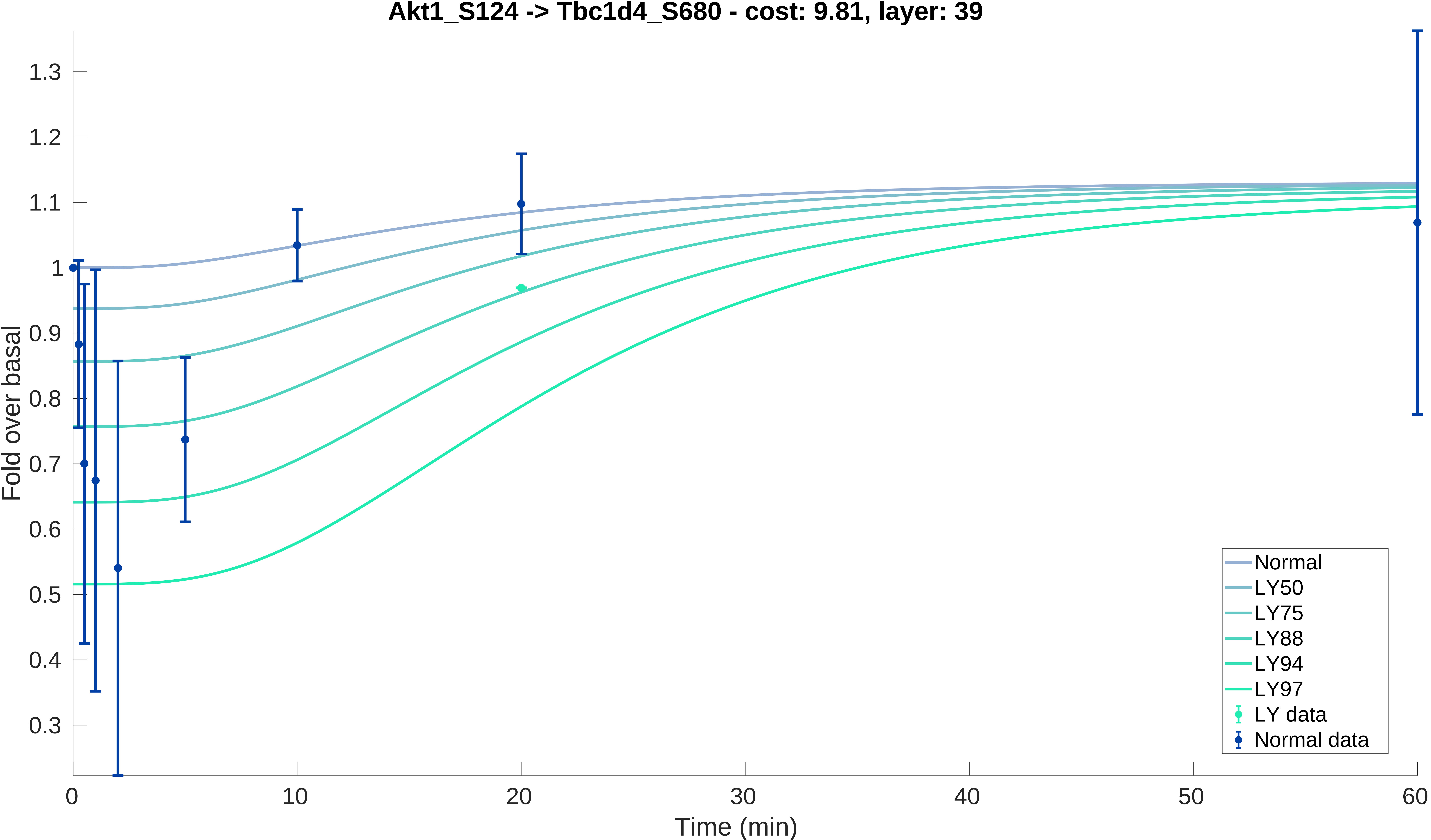

Akt1_S124 -> Tbc1d4_S680 - cost: 9.81, layer: 39
Fold over basal
Time (min)
1.3
1.2
1.1
1
0.9
0.8
0.7
0.6
0.5
0.4
0.3
0
10
20
30
40
50
60
Normal
LY50
LY75
LY88
LY94
LY97
LY data
Normal data

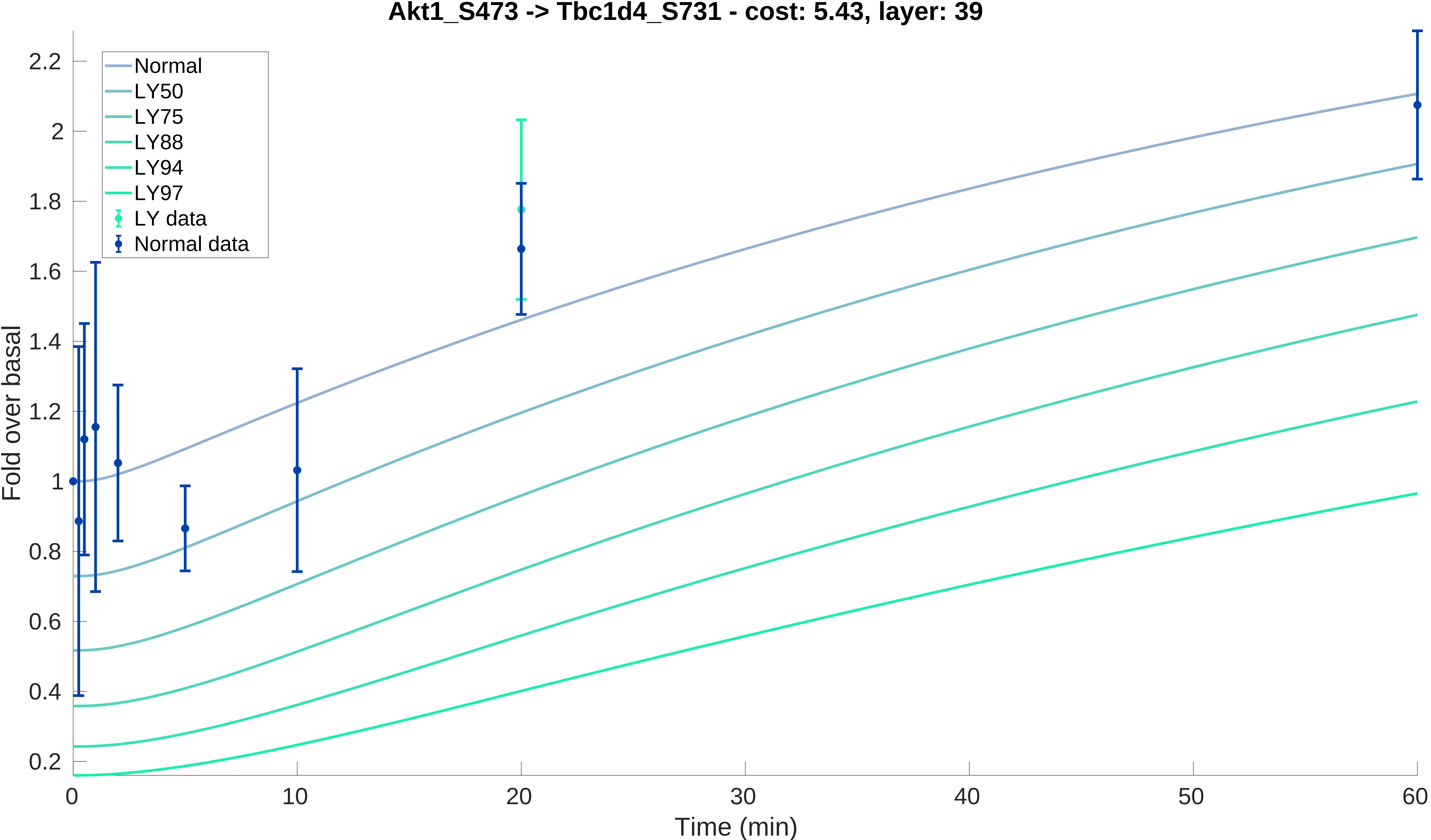

Akt1_S473 -> Tbc1d4_S731 - cost: 5.43, layer: 39
Normal
LY50
LY75
LY88
LY94
LY97
LY data
Normal data
Fold over basal
Time (min)
0.2
0.4
0.6
0.8
1
1.2
1.4
1.6
1.8
2
2.2
0
10
20
30
40
50
60

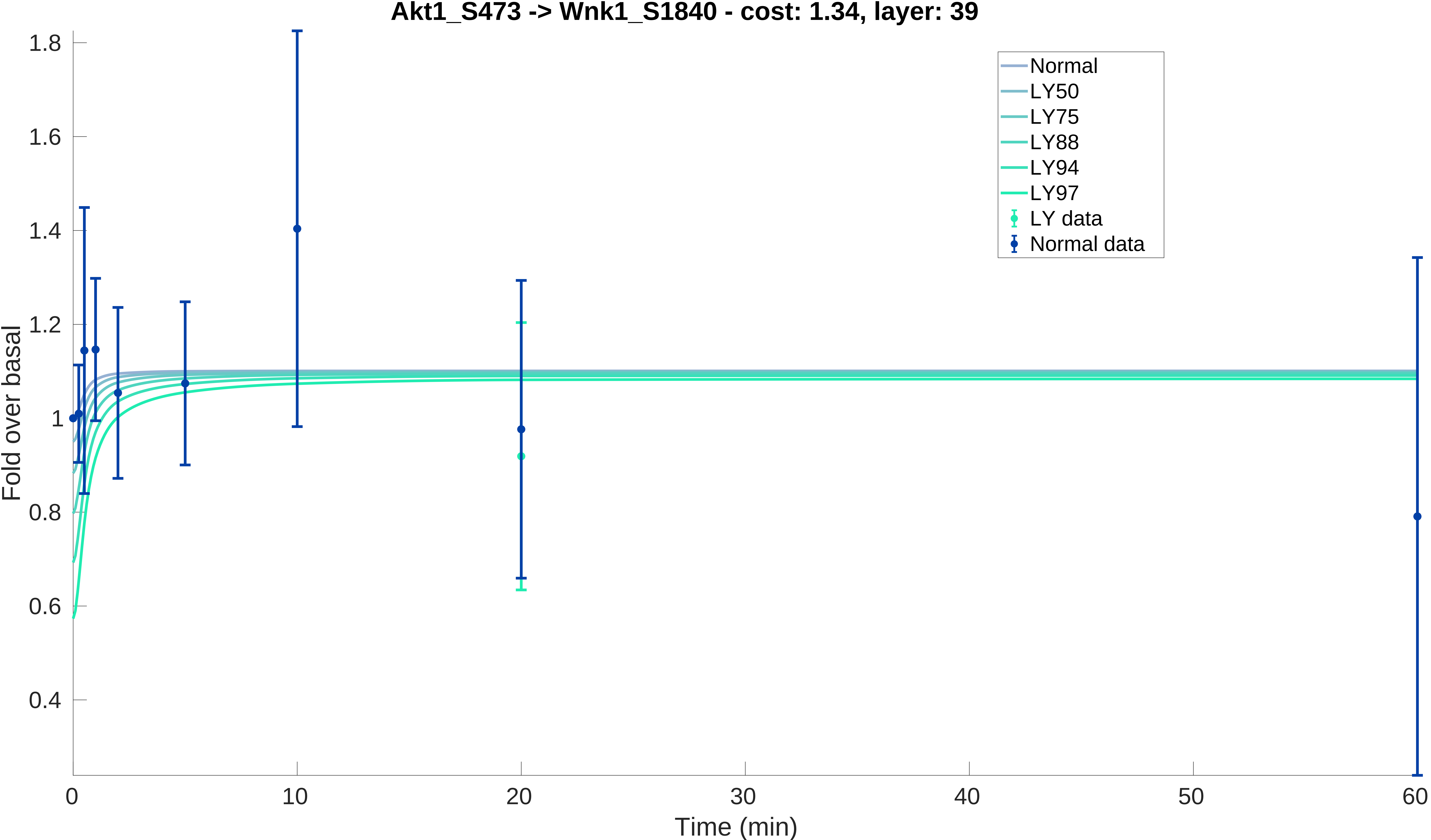

Akt1_S473 -> Wnk1_S1840 - cost: 1.34, layer: 39
Normal
LY50
LY75
LY88
LY94
LY97
LY data
Normal data
Fold over basal
Time (min)
1.8
1.6
1.4
1.2
1
0.8
0.6
0.4
0
10
20
30
40
50
60

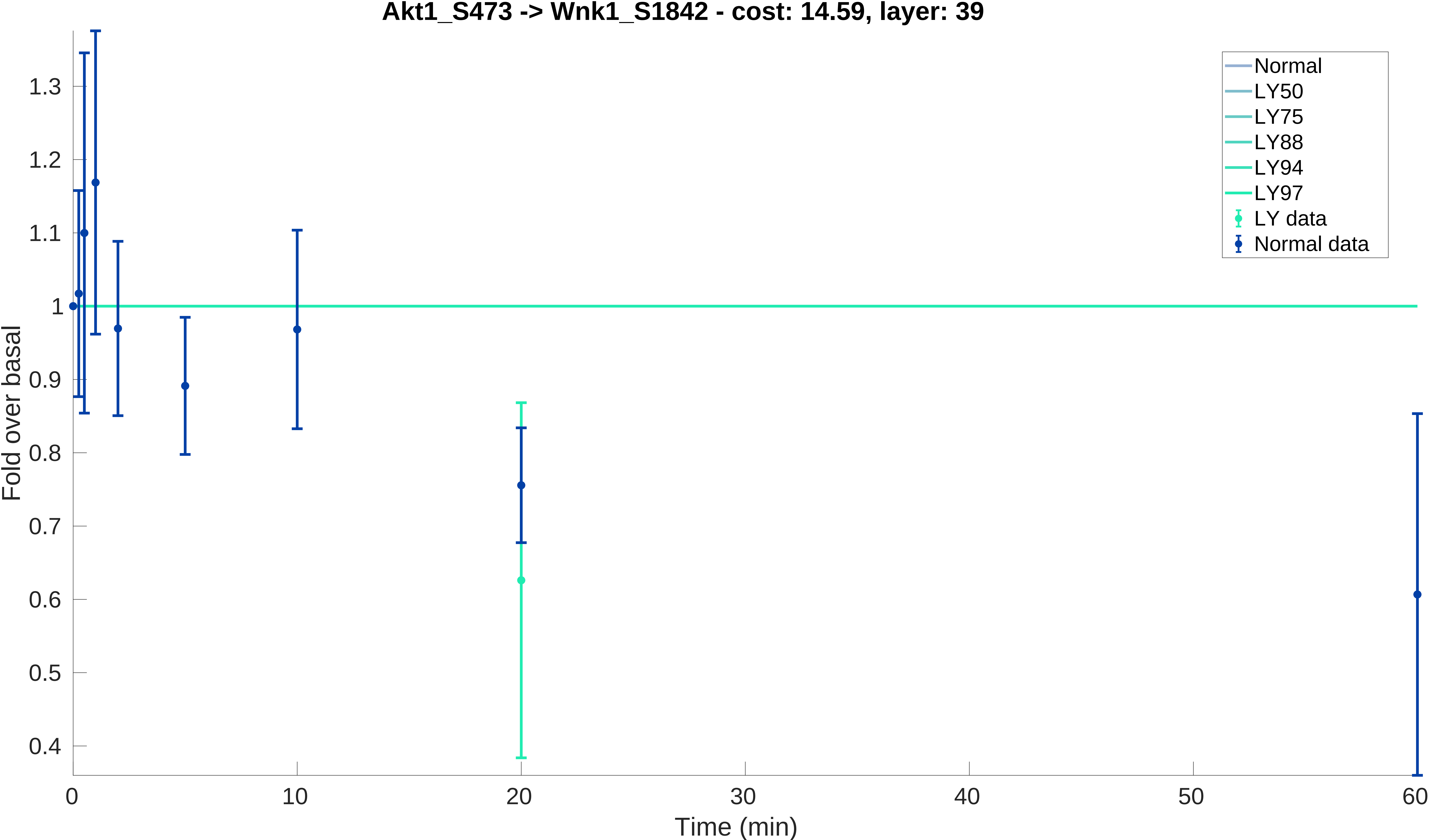
Akt1_S473 -> Wnk1_S1842 - cost: 14.59, layer: 39
Fold over basal
Time (min)
0
10
20
30
40
50
60
0.4
0.5
0.6
0.7
0.8
0.9
1
1.1
1.2
1.3
Normal
LY50
LY75
LY88
LY94
LY97
LY data
Normal data

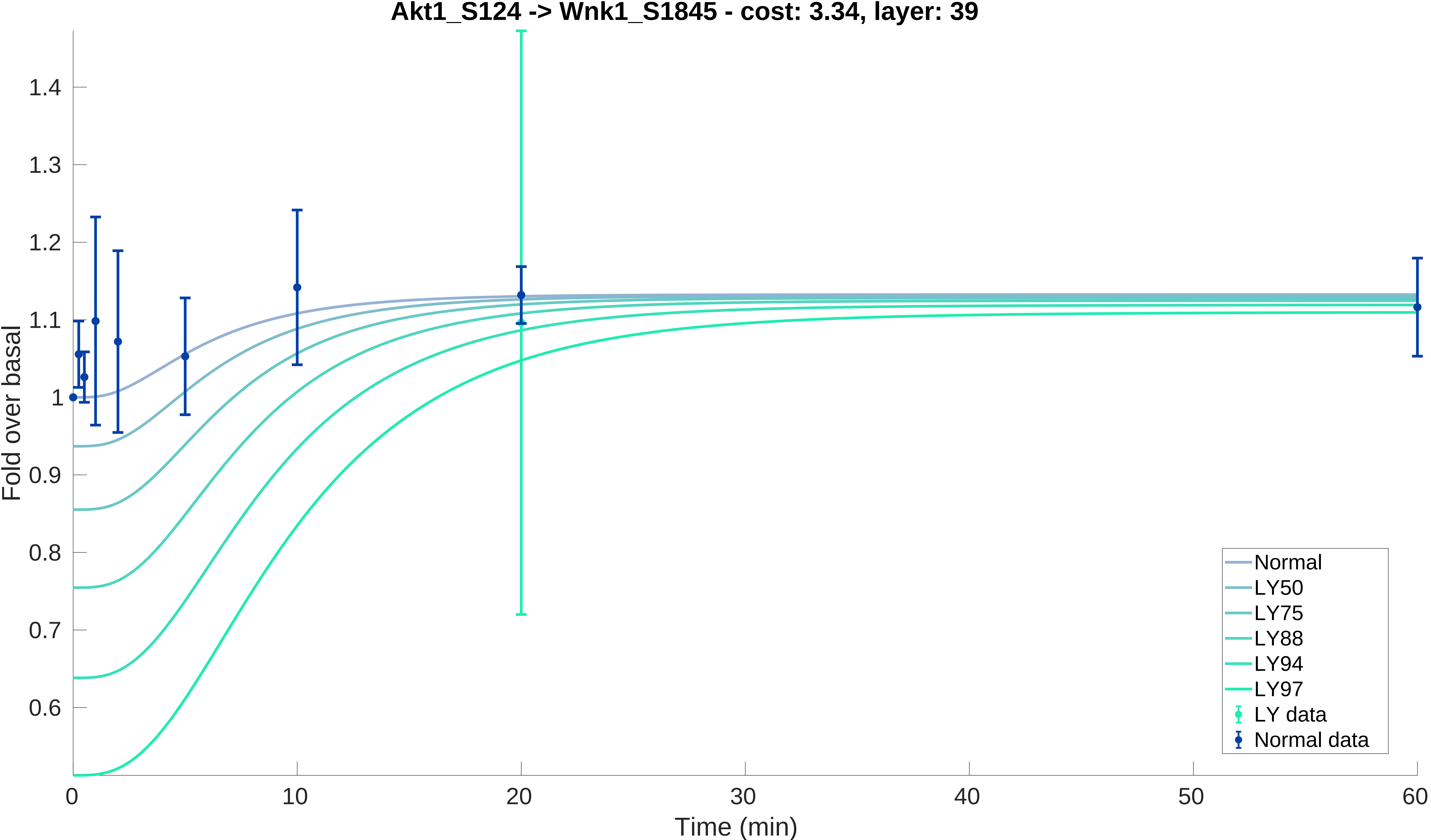

Akt1_S124 -> Wnk1_S1845 - cost: 3.34, layer: 39
Fold over basal
1.4
1.3
1.2
1.1
1
0.9
0.8
0.7
0.6
0
10
20
30
40
50
60
Time (min)
Normal
LY50
LY75
LY88
LY94
LY97
LY data
Normal data

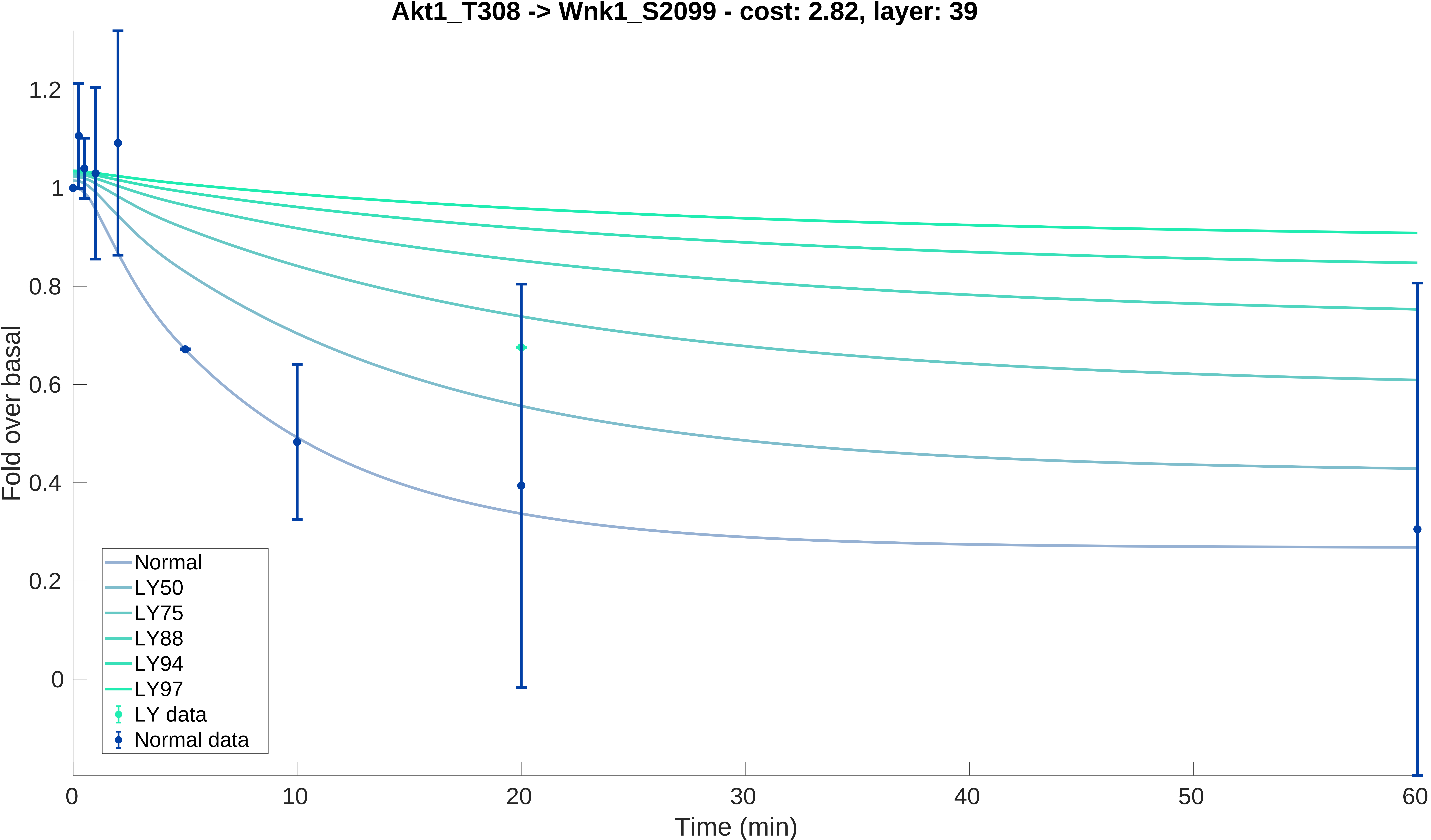

Akt1_T308 -> Wnk1_S2099 - cost: 2.82, layer: 39
Fold over basal
Time (min)
0
0.2
0.4
0.6
0.8
1
1.2
0
10
20
30
40
50
60
Normal
LY50
LY75
LY88
LY94
LY97
LY data
Normal data

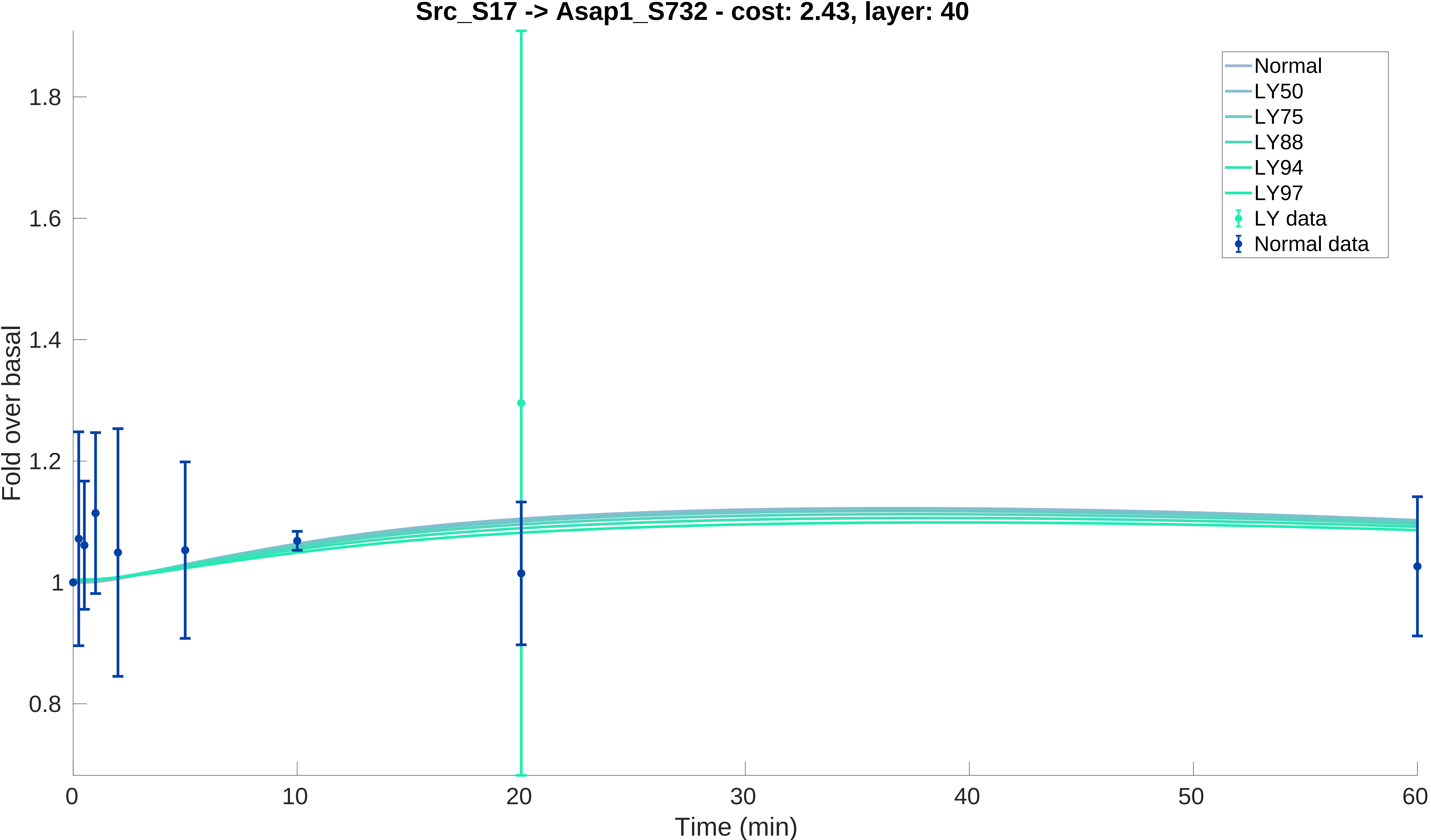

Src_S17 -> Asap1_S732 - cost: 2.43, layer: 40
Normal
LY50
LY75
LY88
LY94
LY97
LY data
Normal data
Fold over basal
1.8
1.6
1.4
1.2
1
0.8
0
10
20
30
40
50
60
Time (min)

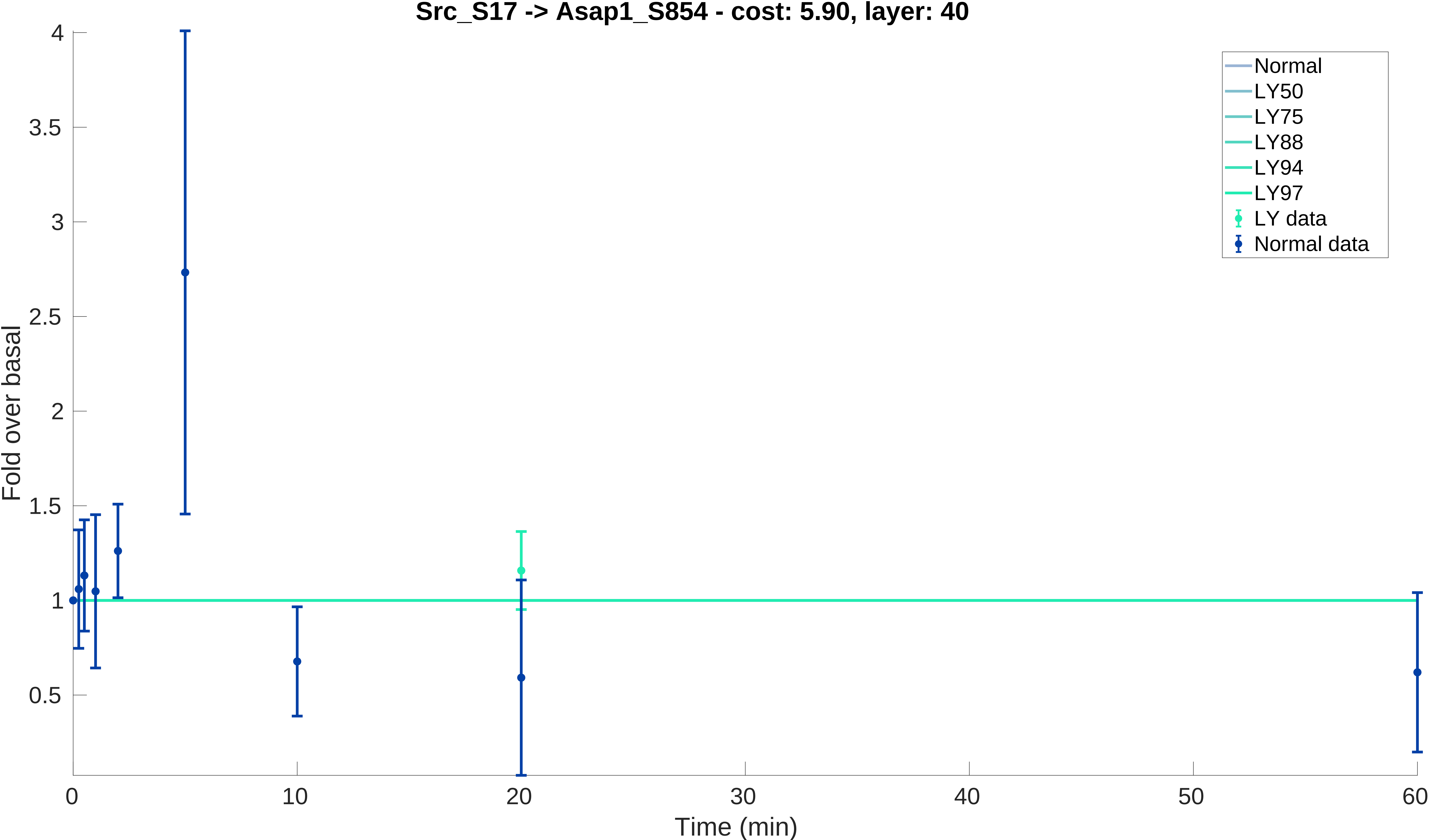
Src_S17 -> Asap1_S854 - cost: 5.90, layer: 40
Fold over basal
Time (min)
Normal
LY50
LY75
LY88
LY94
LY97
LY data
Normal data
0
10
20
30
40
50
60
0.5
1
1.5
2
2.5
3
3.5
4

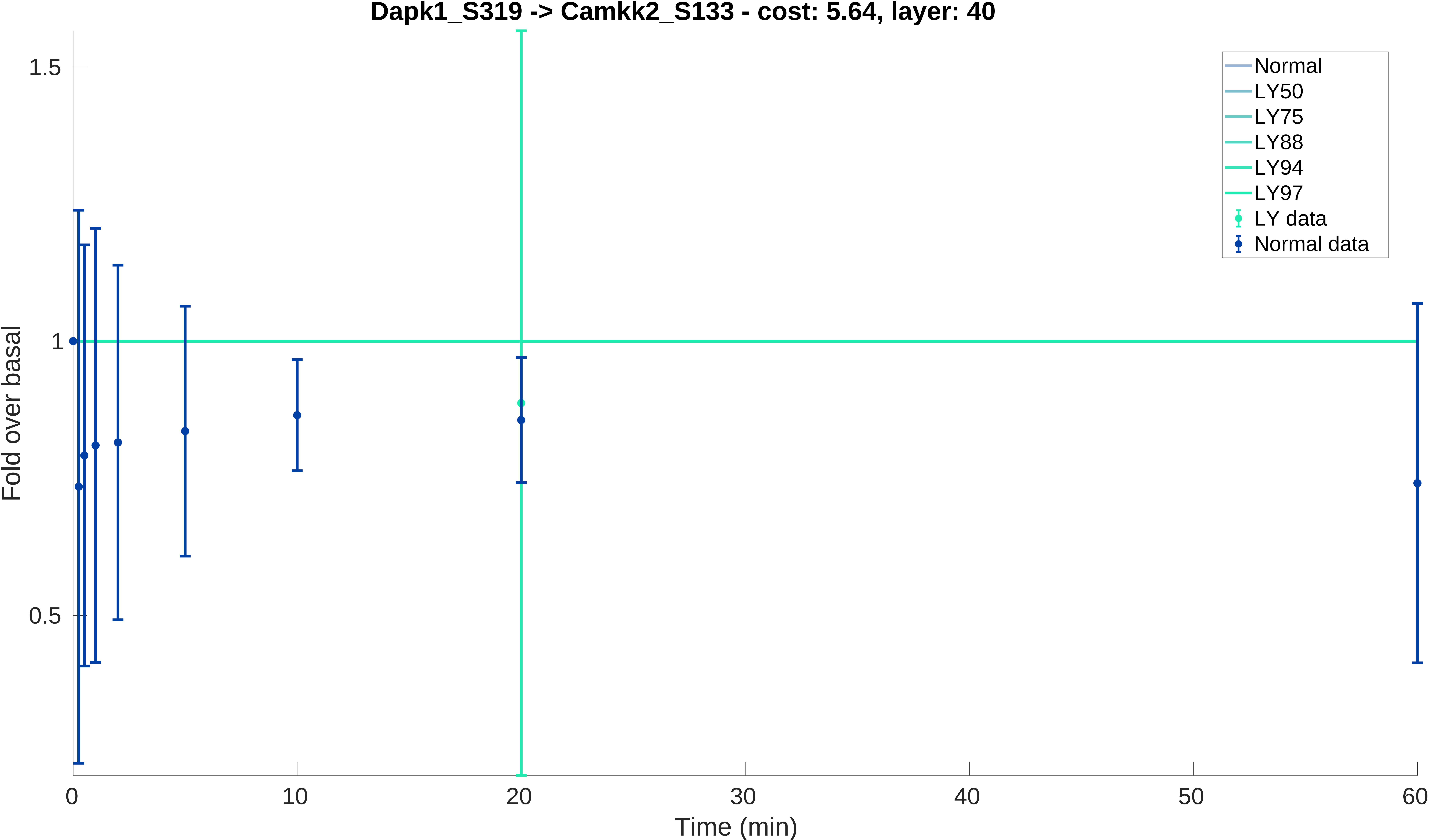

Dapk1_S319 -> Camkk2_S133 - cost: 5.64, layer: 40
Normal
LY50
LY75
LY88
LY94
LY97
LY data
Normal data
Fold over basal
1.5
1
0.5
0
10
20
30
40
50
60
Time (min)

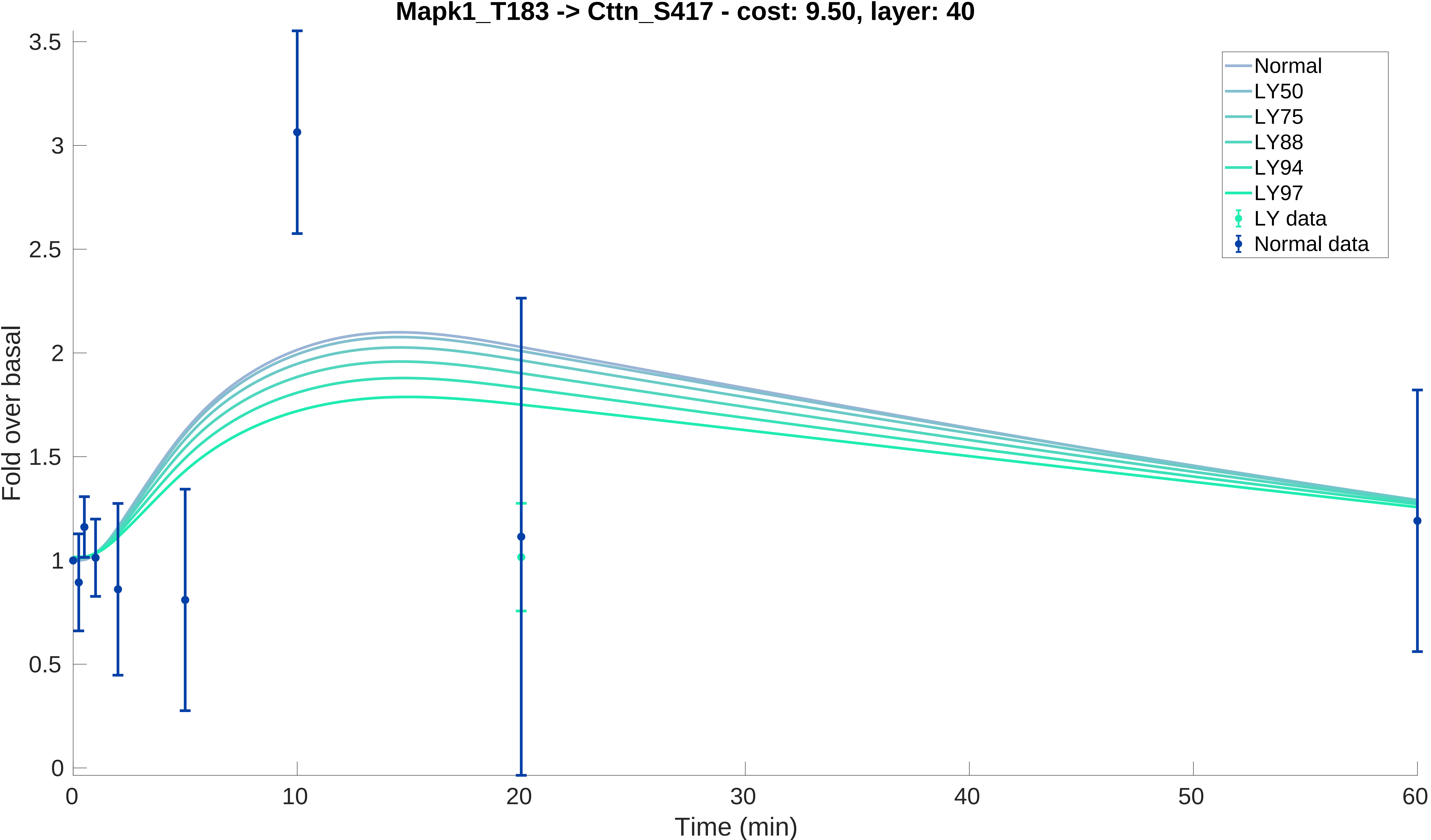

Mapk1_T183 -> Cttn_S417 - cost: 9.50, layer: 40
Fold over basal
Time (min)
0
0.5
1
1.5
2
2.5
3
3.5
0
10
20
30
40
50
60
Normal
LY50
LY75
LY88
LY94
LY97
LY data
Normal data

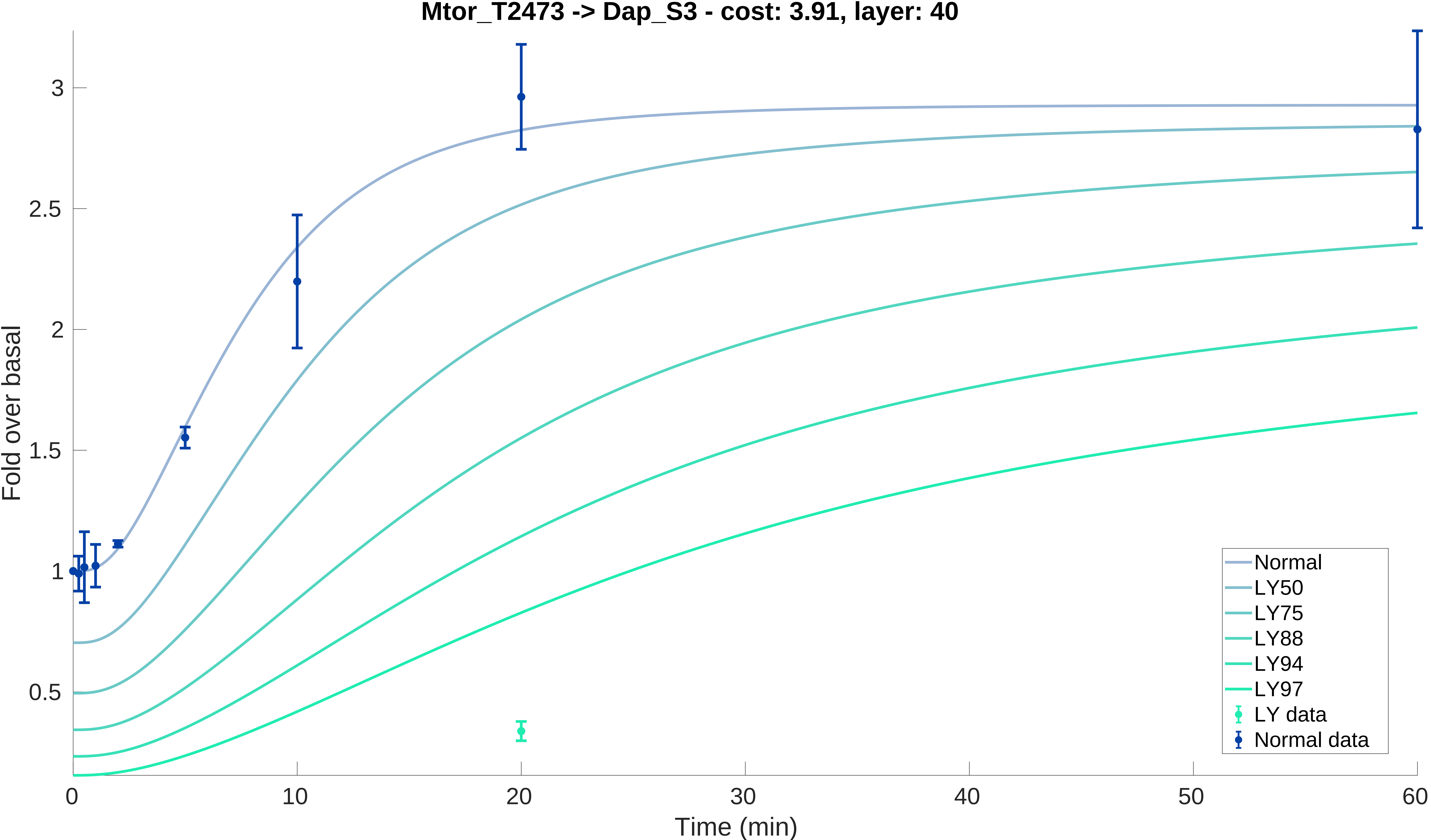
Mtor_T2473 -> Dap_S3 - cost: 3.91, layer: 40
Fold over basal
Time (min)
0.5
1
1.5
2
2.5
3
0
10
20
30
40
50
60
Normal
LY50
LY75
LY88
LY94
LY97
LY data
Normal data

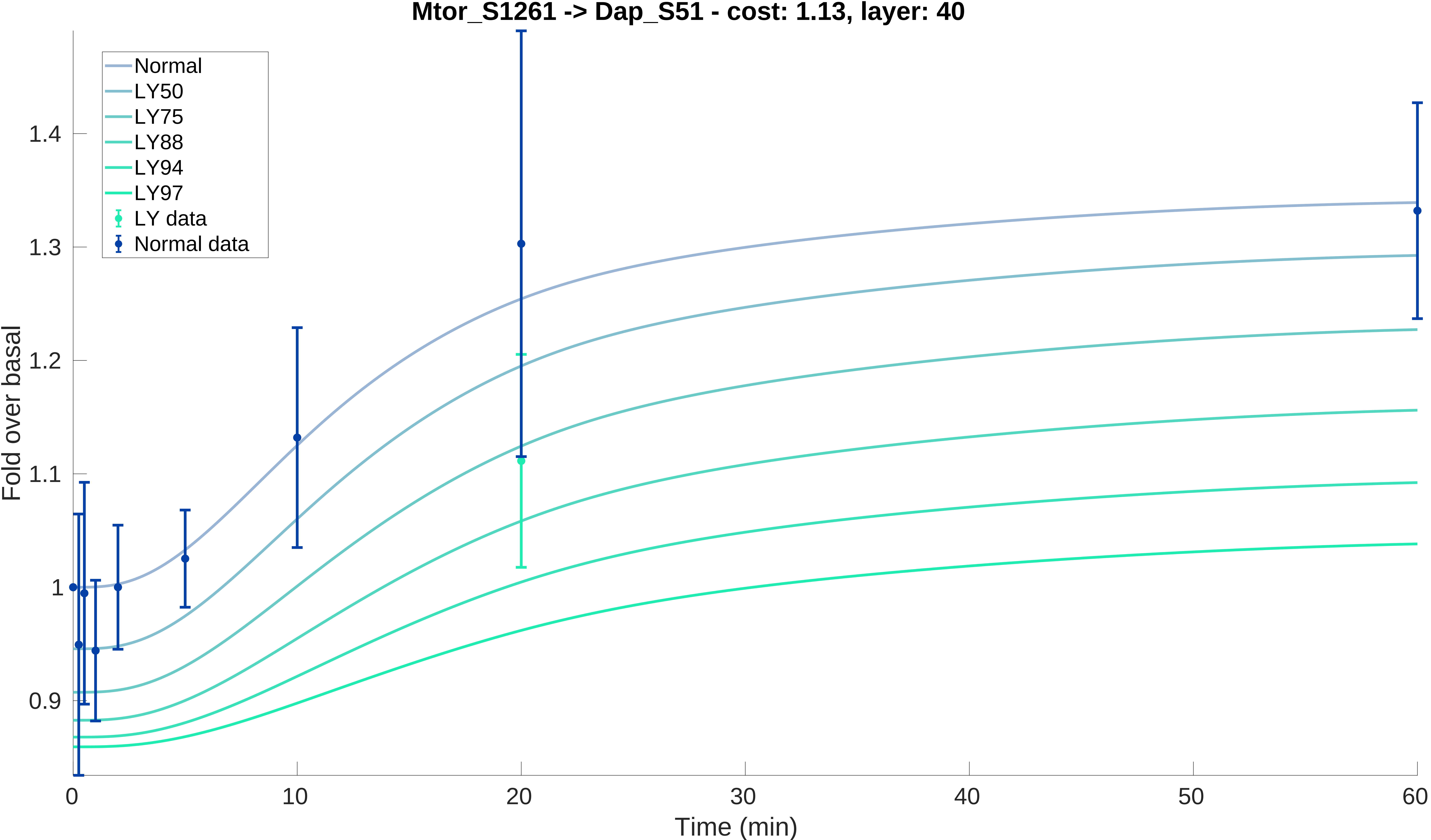

Mtor_S1261 -> Dap_S51 - cost: 1.13, layer: 40
Normal
LY50
LY75
LY88
LY94
LY97
LY data
Normal data
Fold over basal
0.9
1
1.1
1.2
1.3
1.4
0
10
20
30
40
50
60
Time (min)

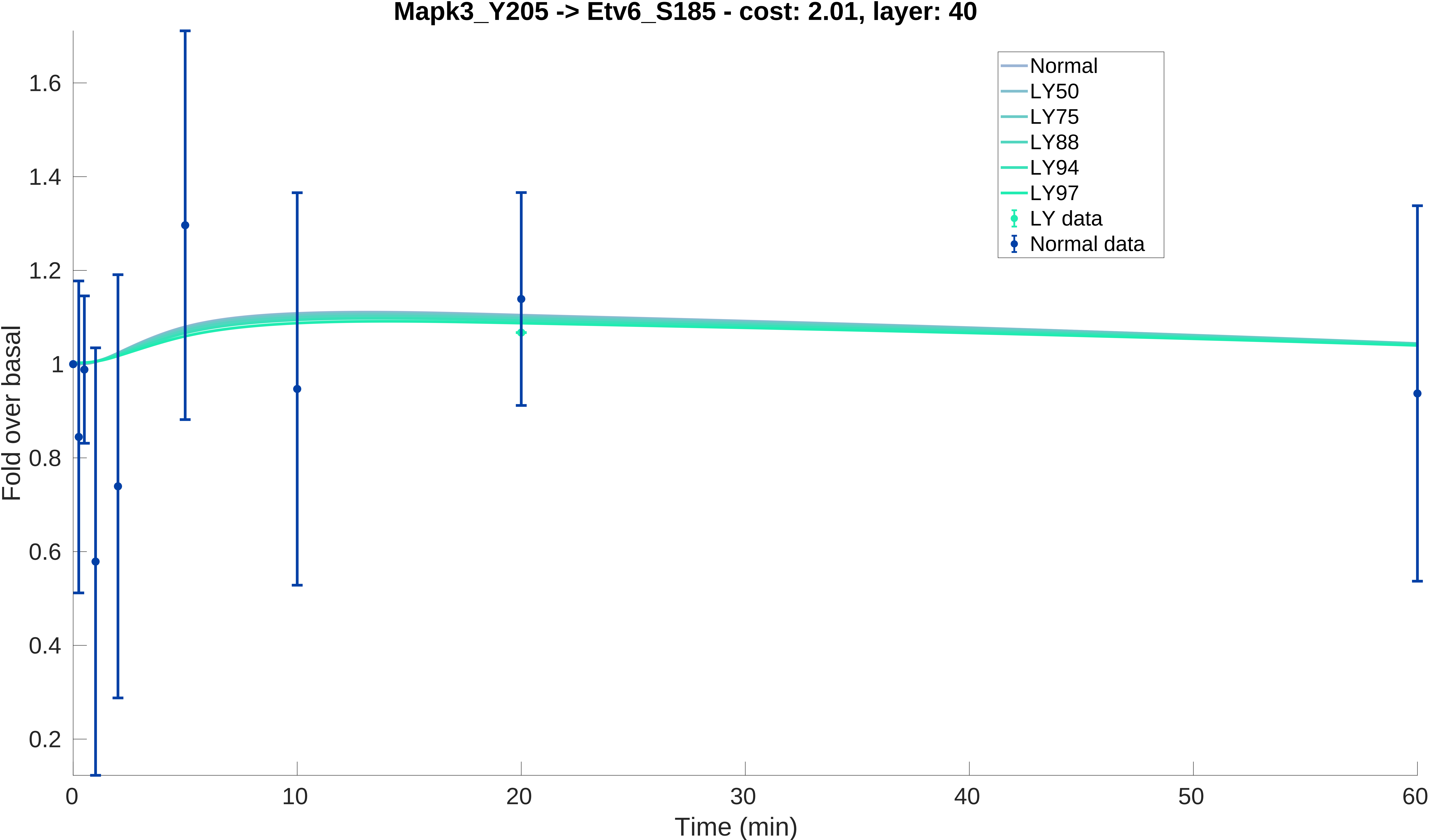

Mapk3_Y205 -> Etv6_S185 - cost: 2.01, layer: 40
Normal
LY50
LY75
LY88
LY94
LY97
LY data
Normal data
Fold over basal
1.6
1.4
1.2
1
0.8
0.6
0.4
0.2
0
10
20
30
40
50
60
Time (min)

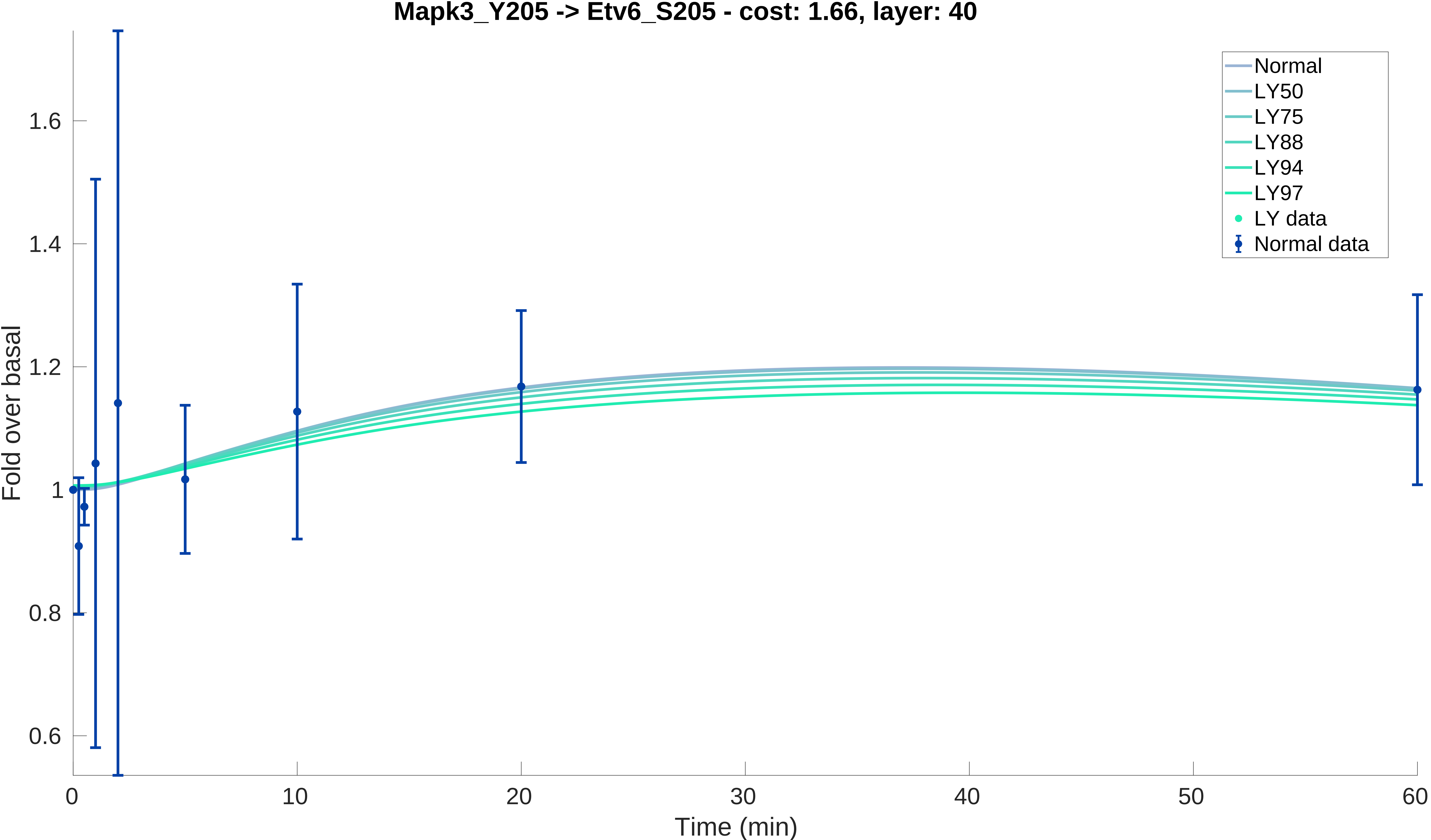

Mapk3_Y205 -> Etv6_S205 - cost: 1.66, layer: 40
Normal
LY50
LY75
LY88
LY94
LY97
LY data
Normal data
Fold over basal
1.6
1.4
1.2
1
0.8
0.6
0
10
20
30
40
50
60
Time (min)

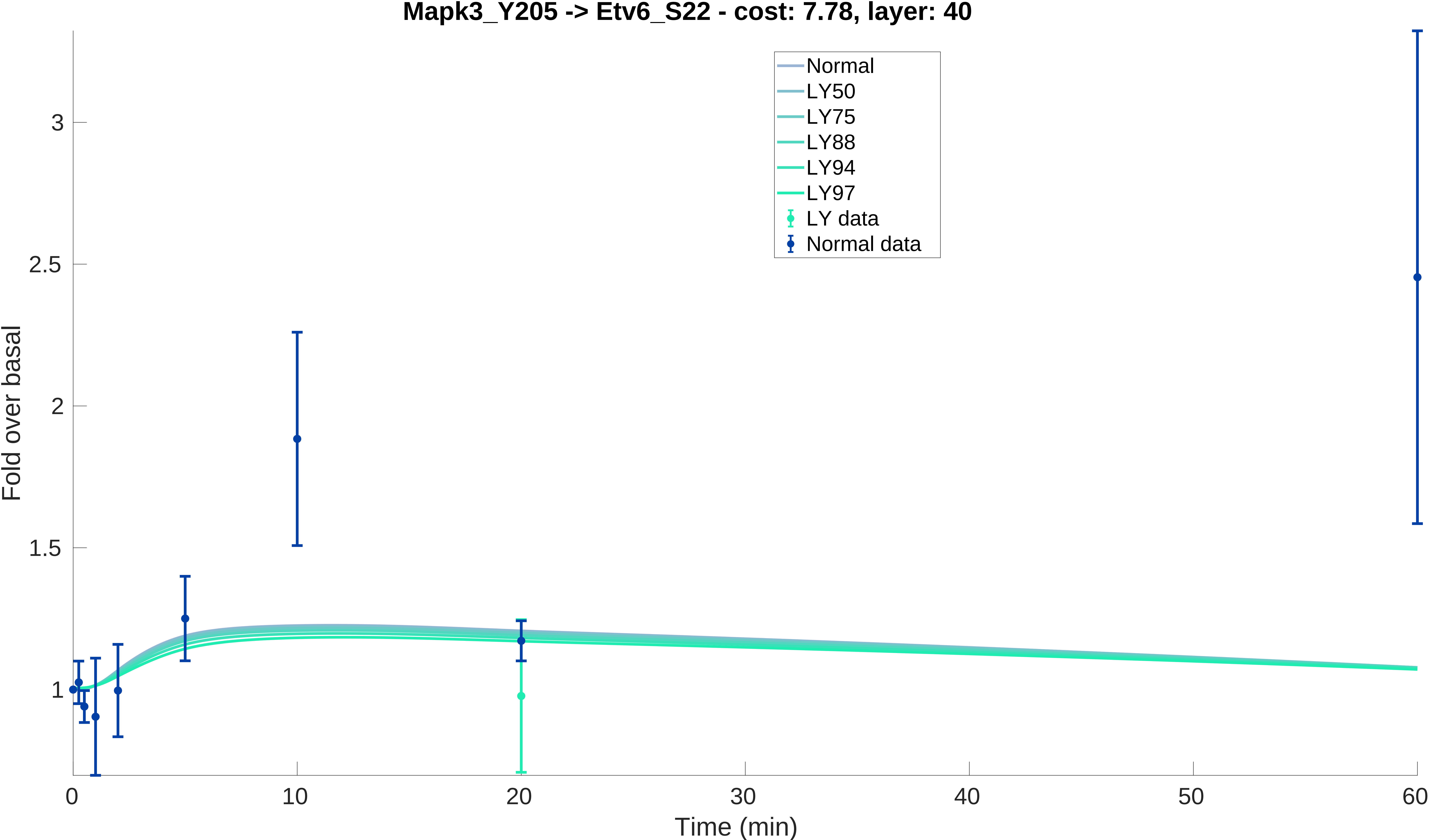

Mapk3_Y205 -> Etv6_S22 - cost: 7.78, layer: 40
Normal
LY50
LY75
LY88
LY94
LY97
LY data
Normal data
Fold over basal
3
2.5
2
1.5
1
0
10
20
30
40
50
60
Time (min)

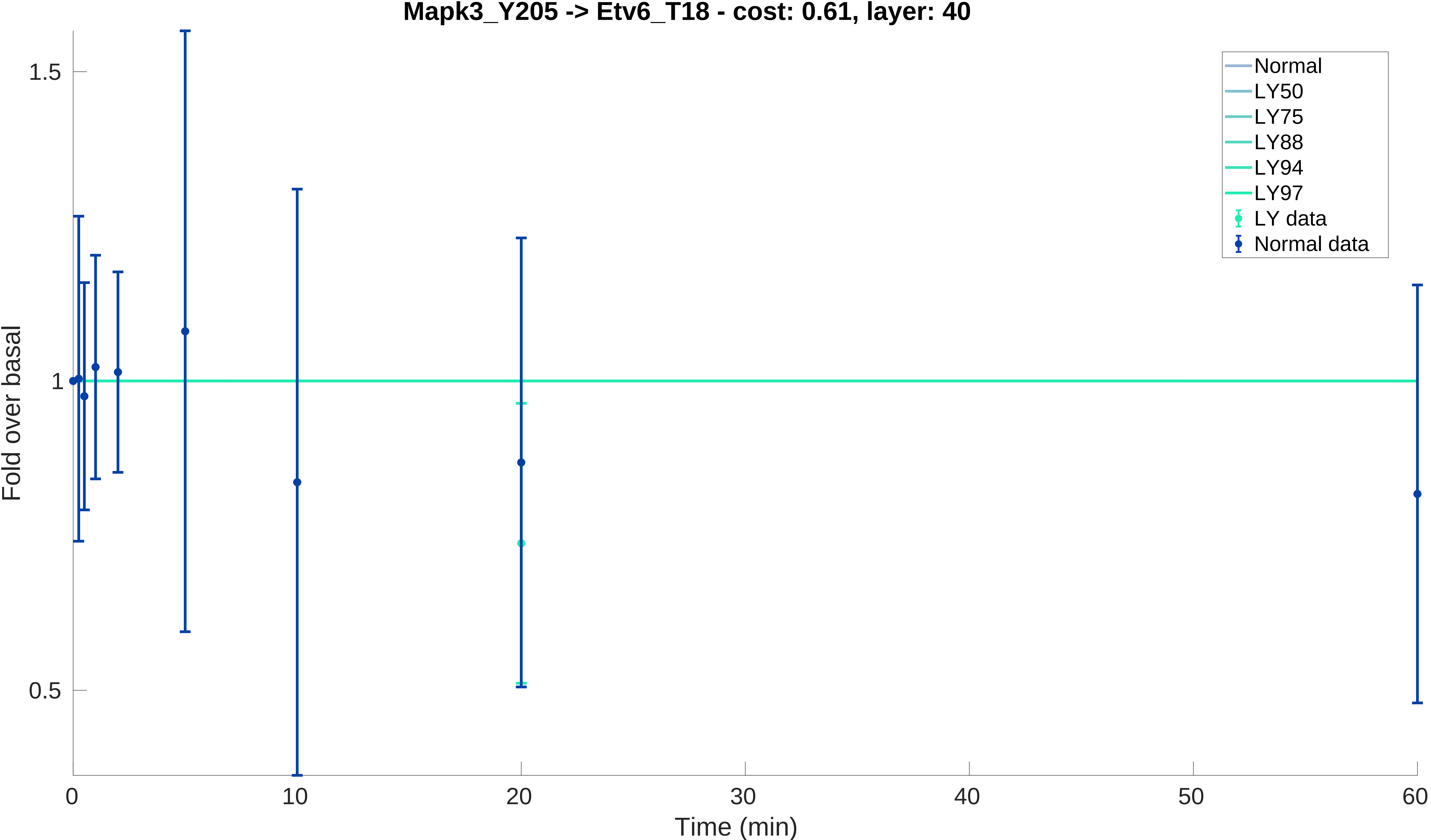
Mapk3_Y205 -> Etv6_T18 - cost: 0.61, layer: 40
Fold over basal
Time (min)
1.5
1
0.5
0
10
20
30
40
50
60
Normal
LY50
LY75
LY88
LY94
LY97
LY data
Normal data

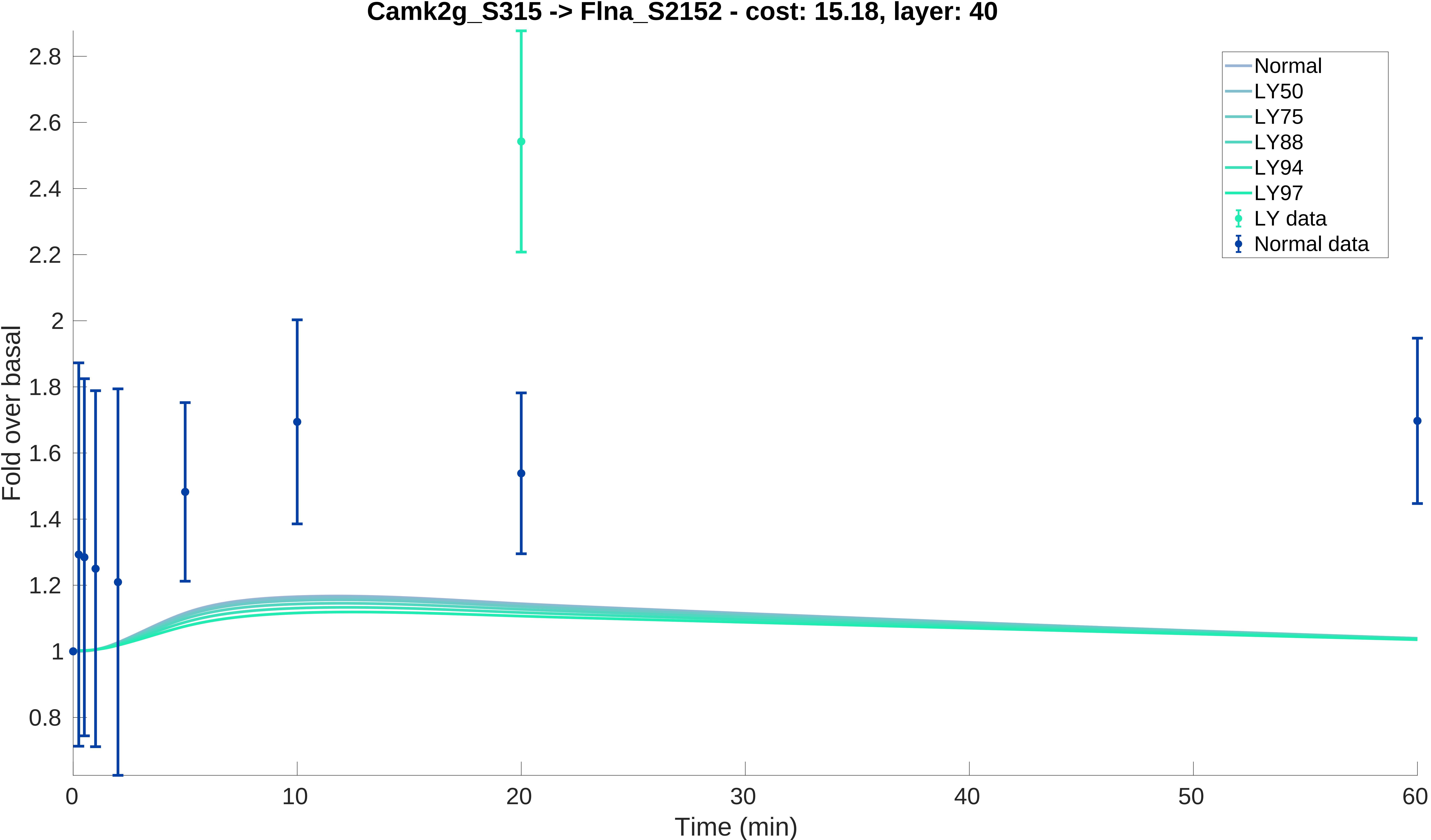

Camk2g_S315 -> Flna_S2152 - cost: 15.18, layer: 40
Fold over basal
Time (min)
0.8
1
1.2
1.4
1.6
1.8
2
2.2
2.4
2.6
2.8
0
10
20
30
40
50
60
Normal
LY50
LY75
LY88
LY94
LY97
LY data
Normal data

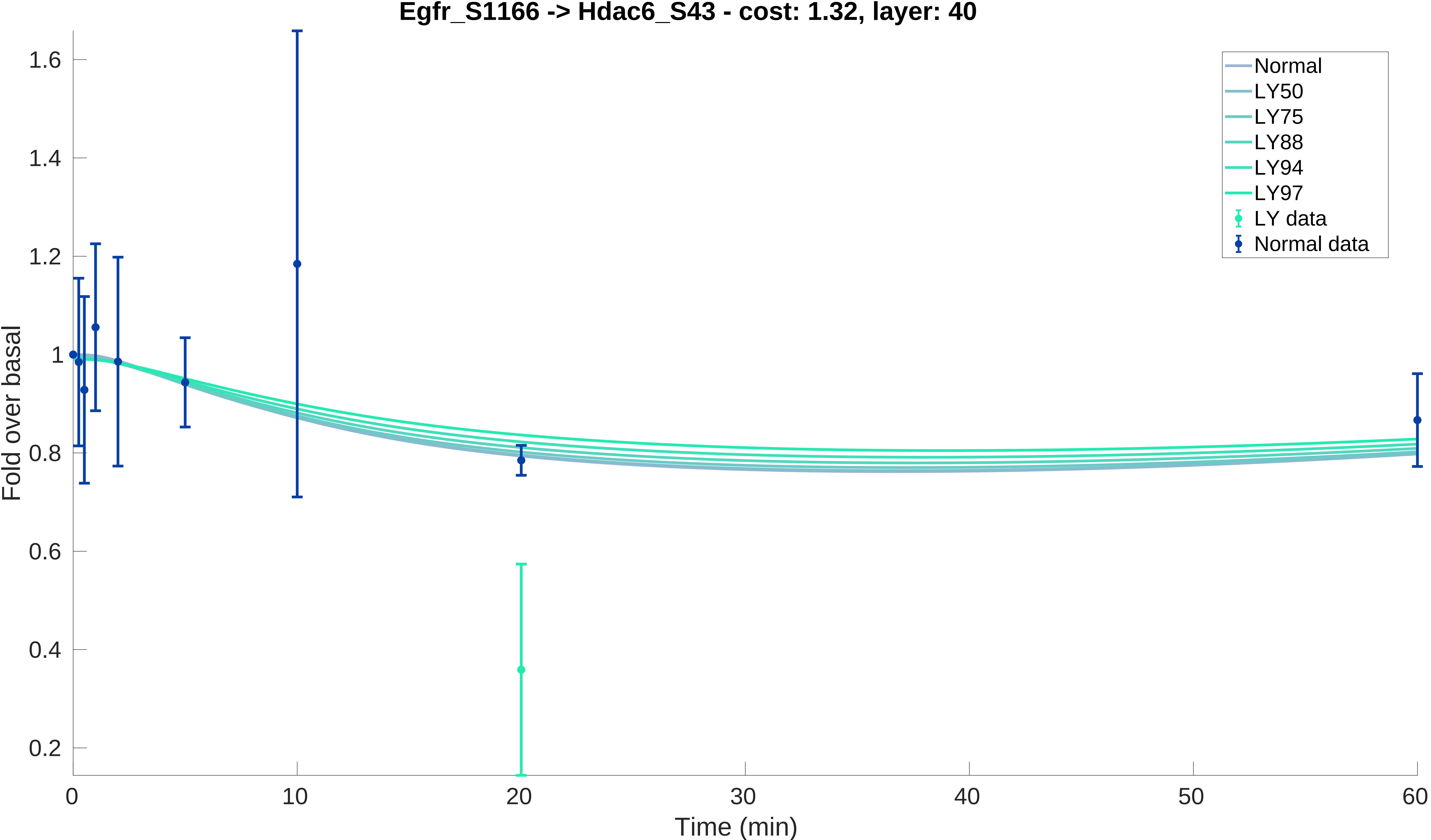
Egfr_S1166 -> Hdac6_S43 - cost: 1.32, layer: 40
Fold over basal
Time (min)
0.2
0.4
0.6
0.8
1
1.2
1.4
1.6
0
10
20
30
40
50
60
Normal
LY50
LY75
LY88
LY94
LY97
LY data
Normal data

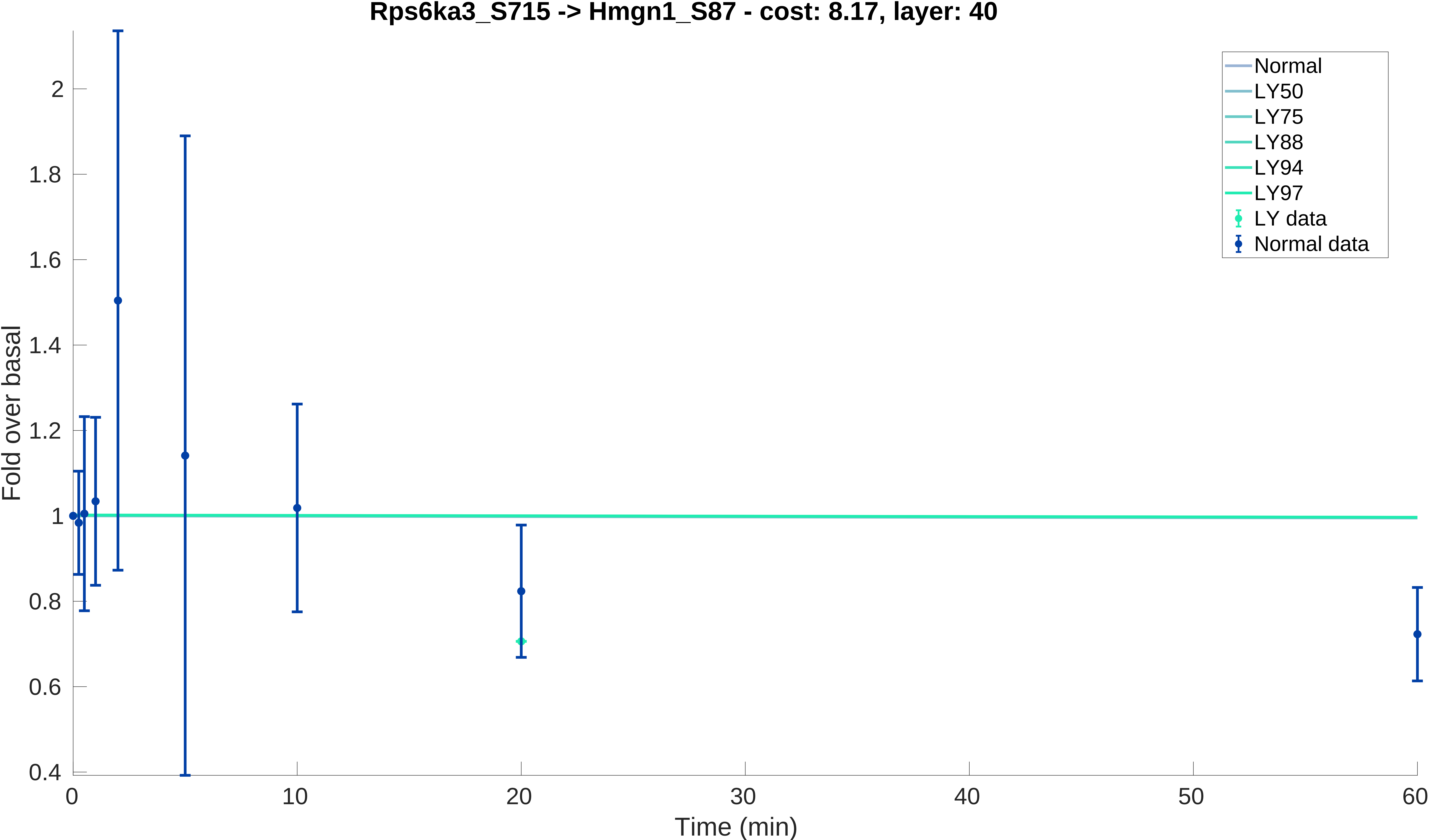

Rps6ka3_S715 -> Hmgn1_S87 - cost: 8.17, layer: 40
Fold over basal
Time (min)
0.4
0.6
0.8
1
1.2
1.4
1.6
1.8
2
0
10
20
30
40
50
60
Normal
LY50
LY75
LY88
LY94
LY97
LY data
Normal data

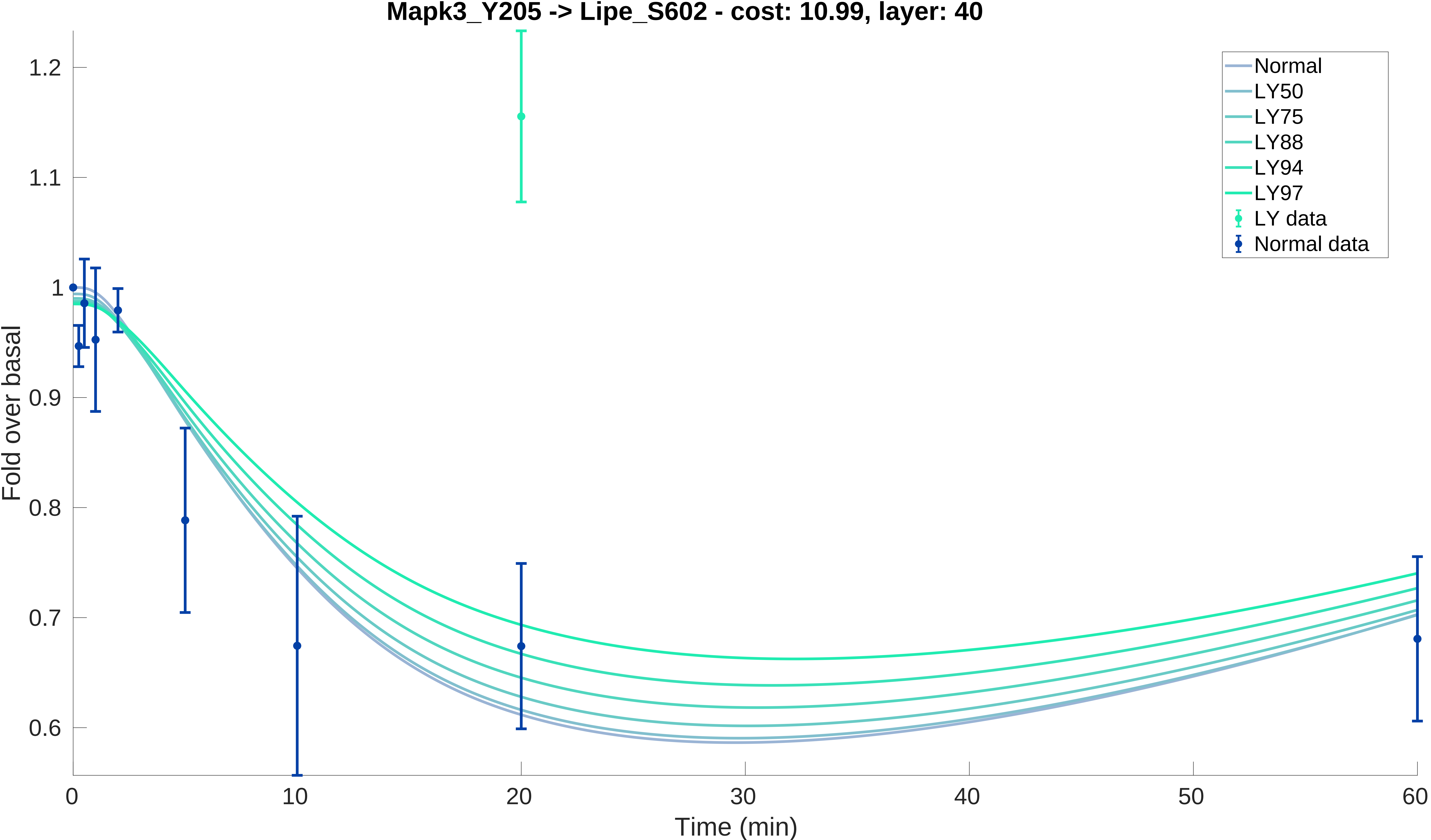

Mapk3_Y205 -> Lipe_S602 - cost: 10.99, layer: 40
Fold over basal
Time (min)
1.2
1.1
1
0.9
0.8
0.7
0.6
0
10
20
30
40
50
60
Normal
LY50
LY75
LY88
LY94
LY97
LY data
Normal data

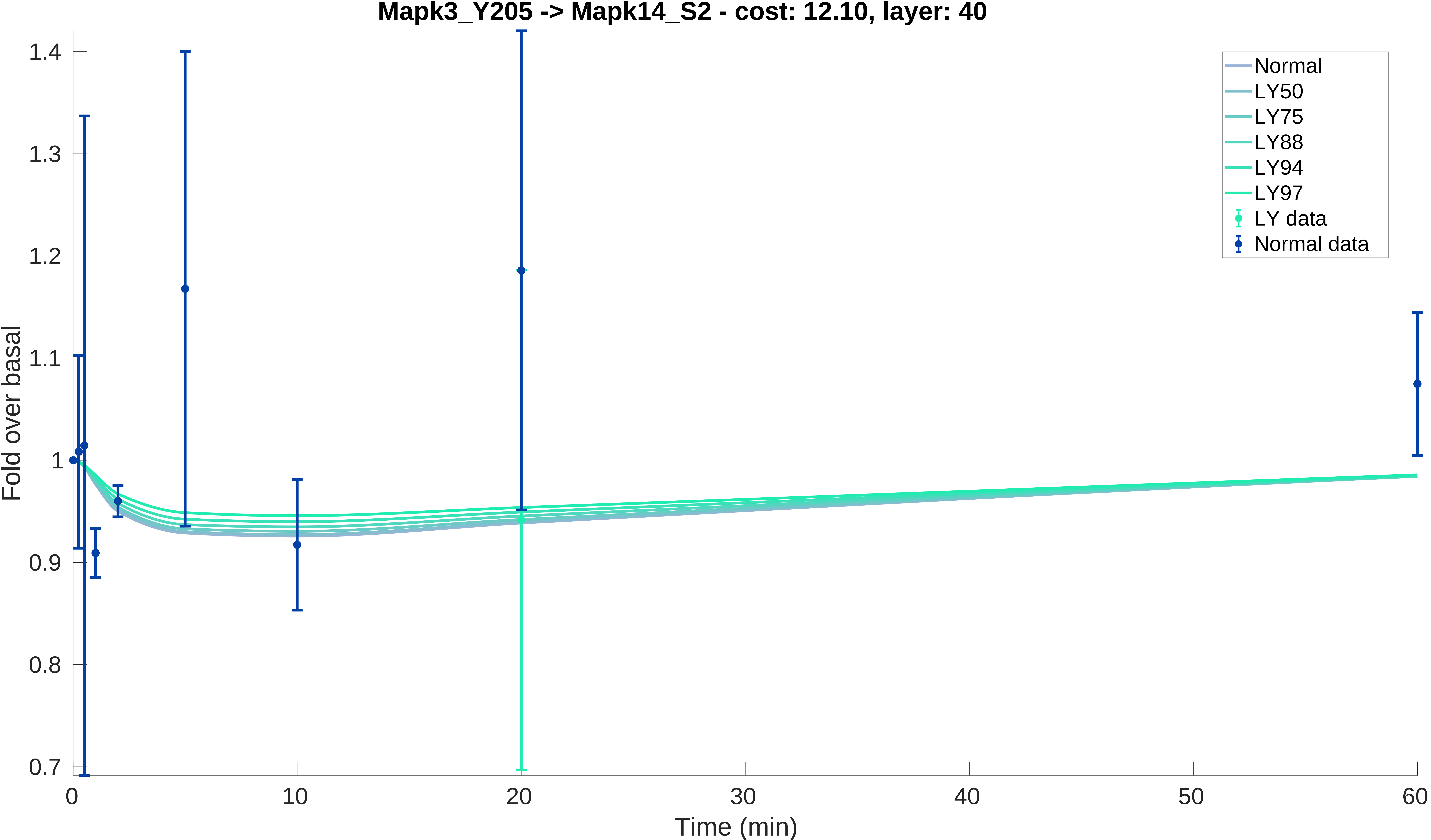
Mapk3_Y205 -> Mapk14_S2 - cost: 12.10, layer: 40
Fold over basal
Time (min)
1.4
1.3
1.2
1.1
1
0.9
0.8
0.7
0
10
20
30
40
50
60
Normal
LY50
LY75
LY88
LY94
LY97
LY data
Normal data

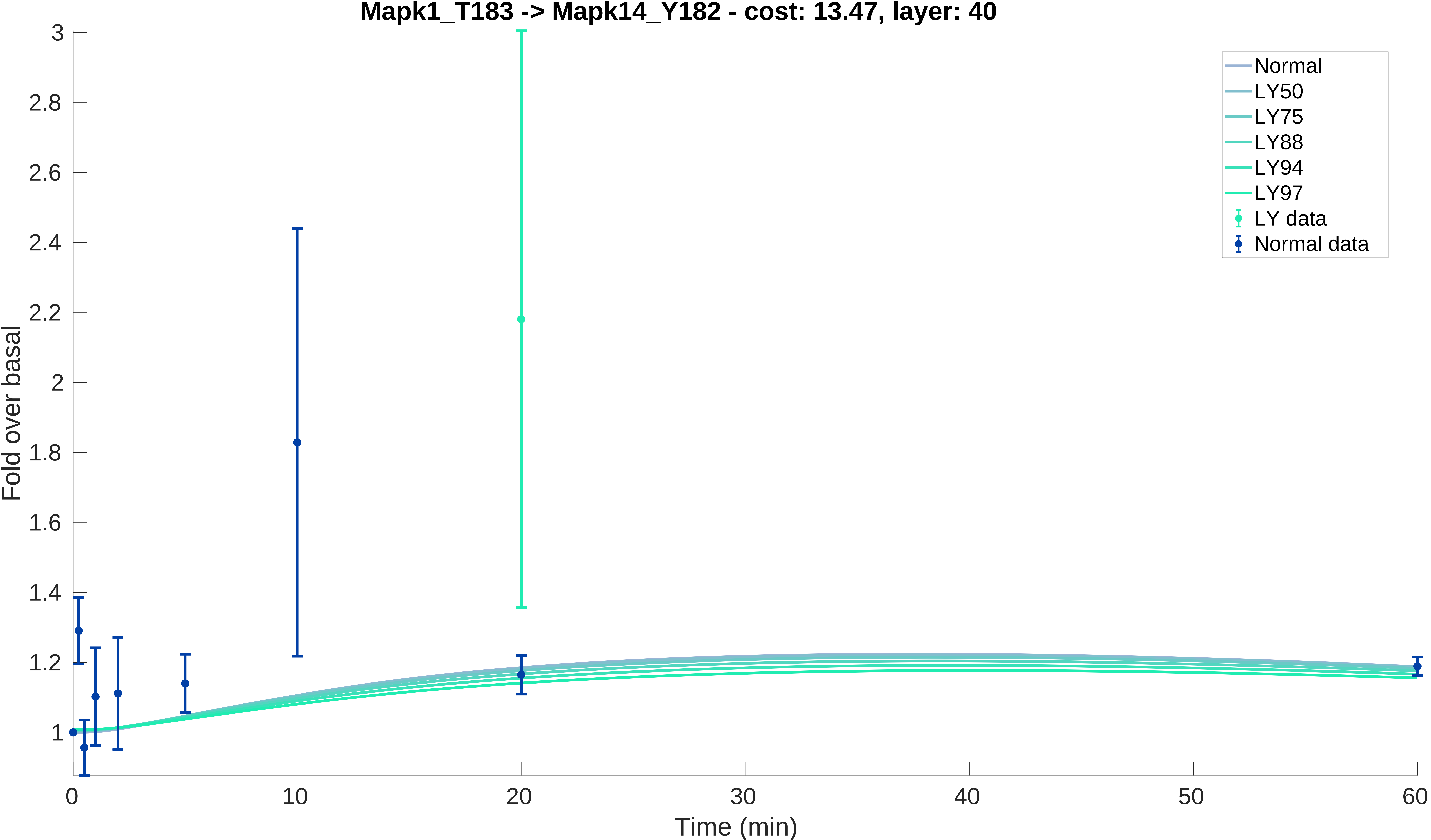

Mapk1_T183 -> Mapk14_Y182 - cost: 13.47, layer: 40
Fold over basal
Time (min)
3
2.8
2.6
2.4
2.2
2
1.8
1.6
1.4
1.2
1
0
10
20
30
40
50
60
Normal
LY50
LY75
LY88
LY94
LY97
LY data
Normal data

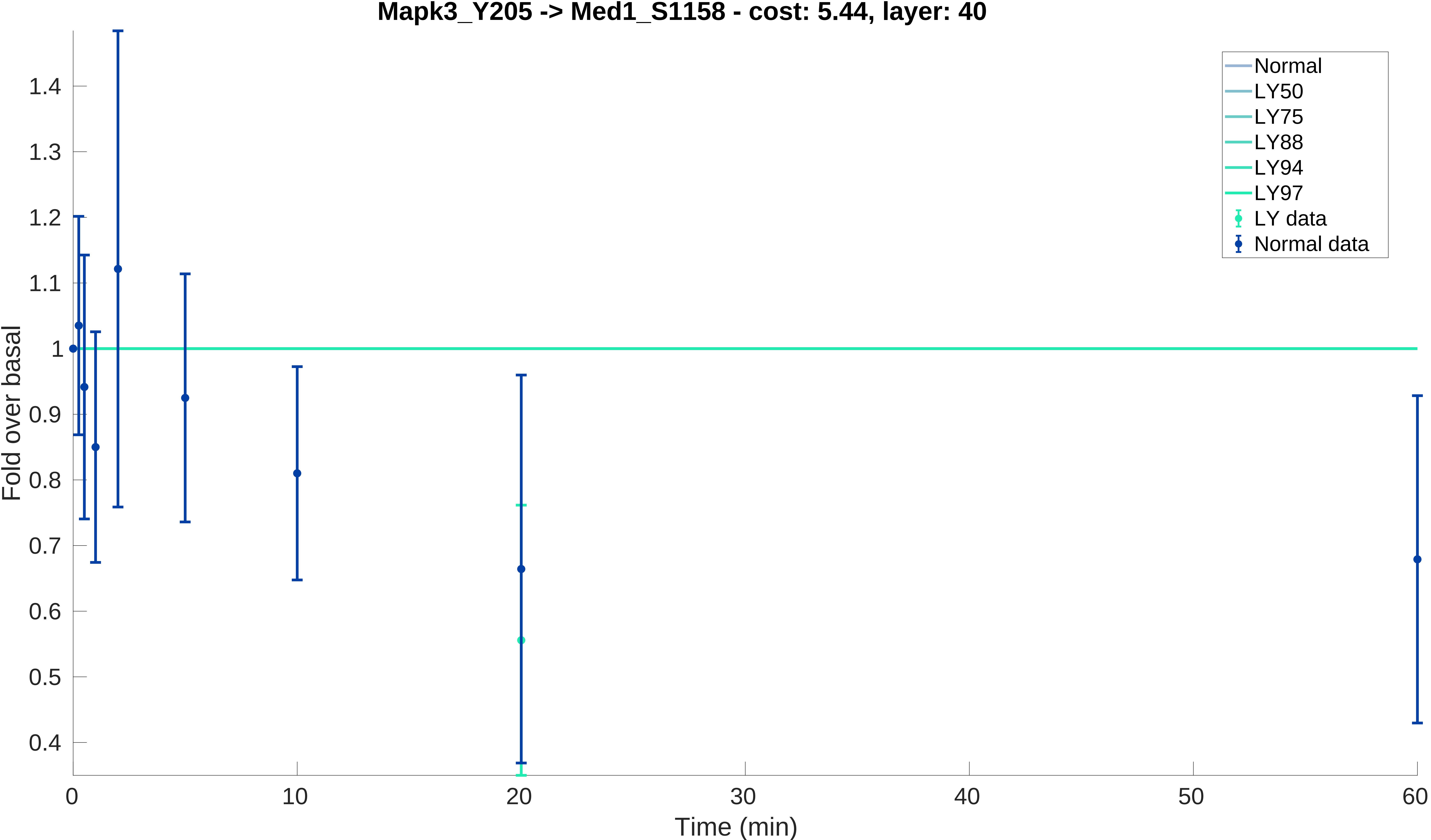

Mapk3_Y205 -> Med1_S1158 - cost: 5.44, layer: 40
Normal
LY50
LY75
LY88
LY94
LY97
LY data
Normal data
Fold over basal
0.4
0.5
0.6
0.7
0.8
0.9
1
1.1
1.2
1.3
1.4
0
10
20
30
40
50
60
Time (min)

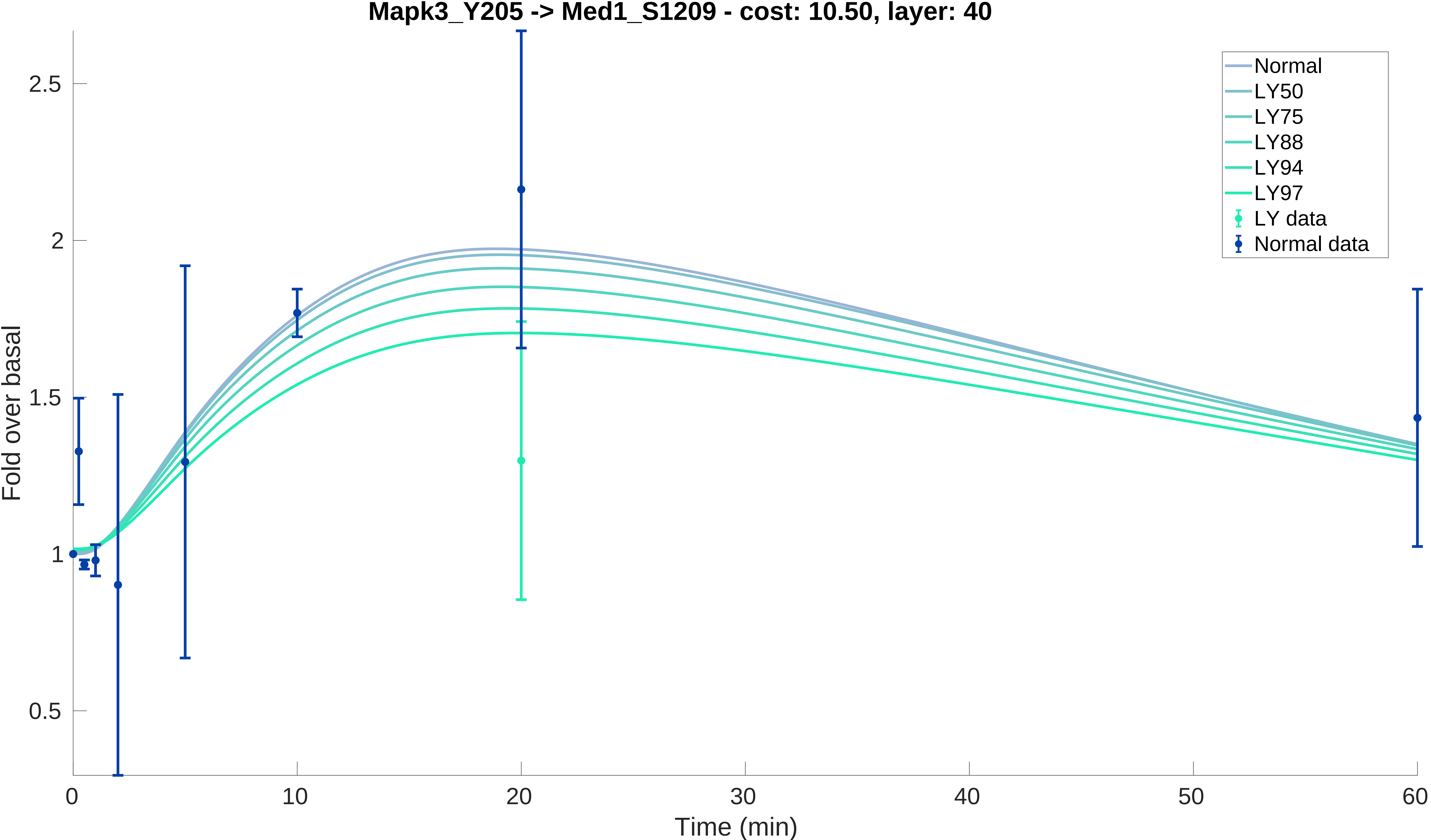

Mapk3_Y205 -> Med1_S1209 - cost: 10.50, layer: 40
Fold over basal
Time (min)
0.5
1
1.5
2
2.5
0
10
20
30
40
50
60
Normal
LY50
LY75
LY88
LY94
LY97
LY data
Normal data

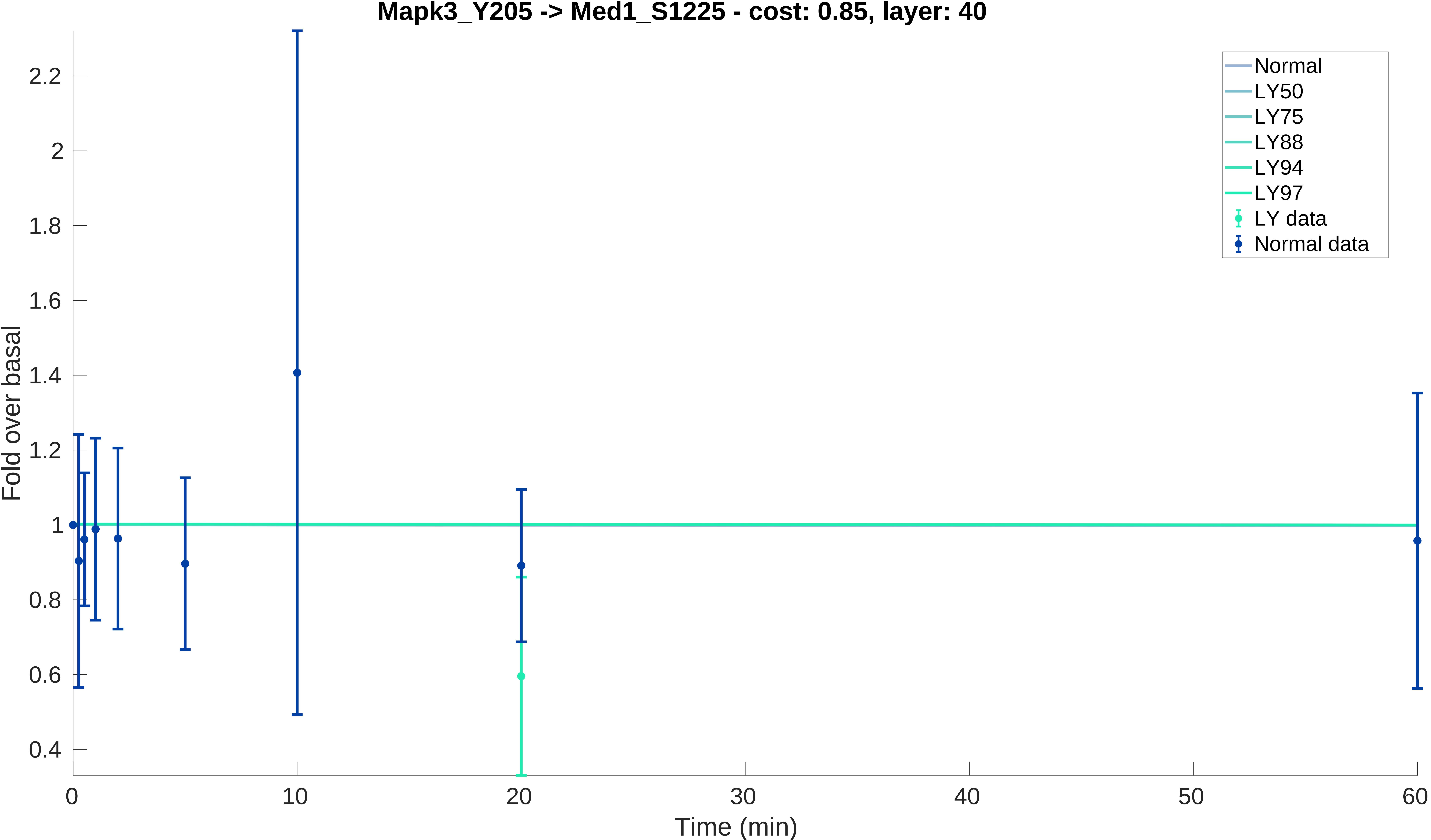
Mapk3_Y205 -> Med1_S1225 - cost: 0.85, layer: 40
Normal
LY50
LY75
LY88
LY94
LY97
LY data
Normal data
Fold over basal
2.2
2
1.8
1.6
1.4
1.2
1
0.8
0.6
0.4
0
10
20
30
40
50
60
Time (min)

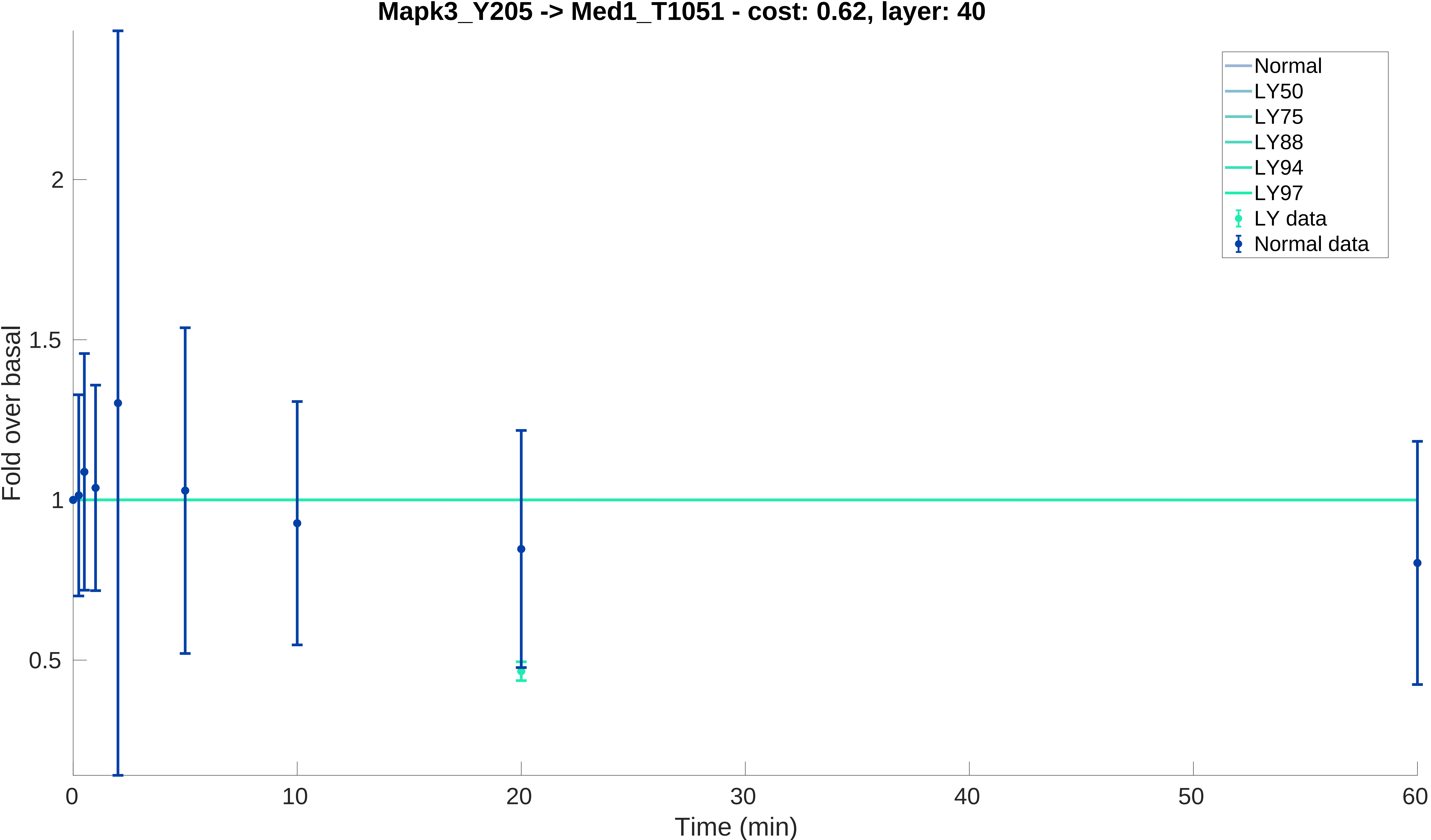

Mapk3_Y205 -> Med1_T1051 - cost: 0.62, layer: 40
Normal
LY50
LY75
LY88
LY94
LY97
LY data
Normal data
Fold over basal
2
1.5
1
0.5
0
10
20
30
40
50
60
Time (min)

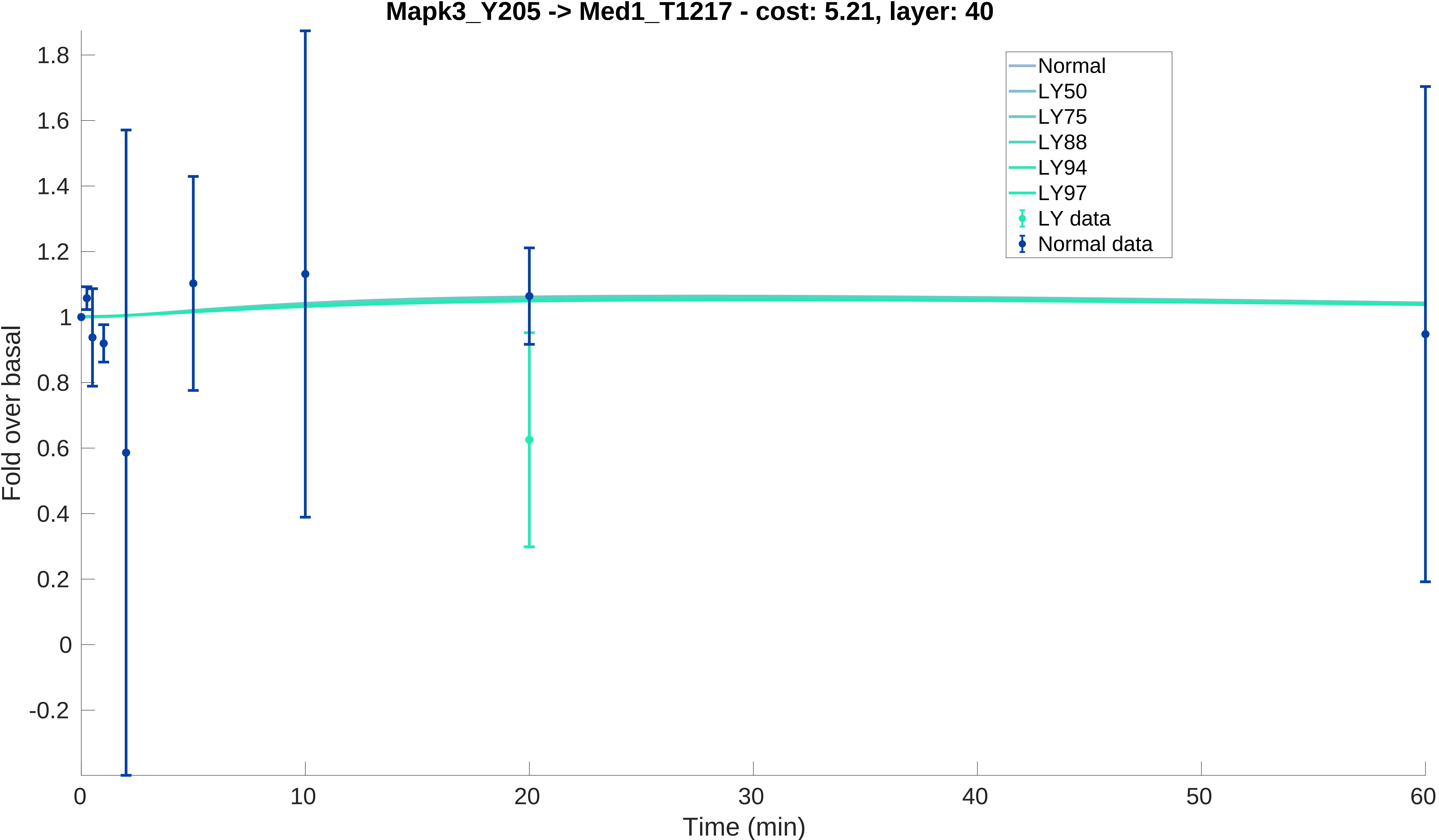

Mapk3_Y205 -> Med1_T1217 - cost: 5.21, layer: 40
Fold over basal
1.8
1.6
1.4
1.2
1
0.8
0.6
0.4
0.2
0
-0.2
0
10
20
30
40
50
60
Time (min)
Normal
LY50
LY75
LY88
LY94
LY97
LY data
Normal data

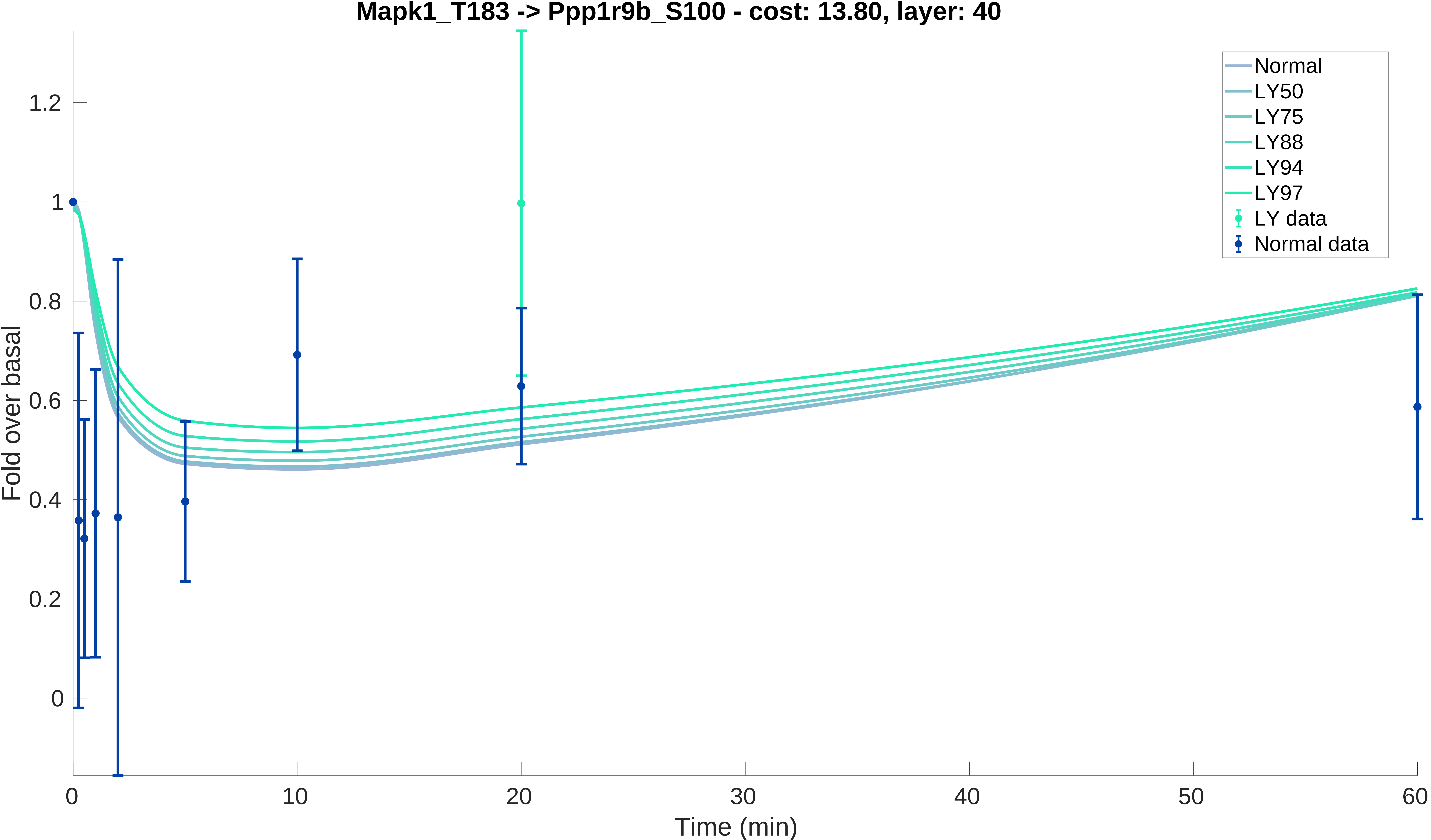

Mapk1_T183 -> Ppp1r9b_S100 - cost: 13.80, layer: 40
Normal
LY50
LY75
LY88
LY94
LY97
LY data
Normal data
Fold over basal
1.2
1
0.8
0.6
0.4
0.2
0
0
10
20
30
40
50
60
Time (min)

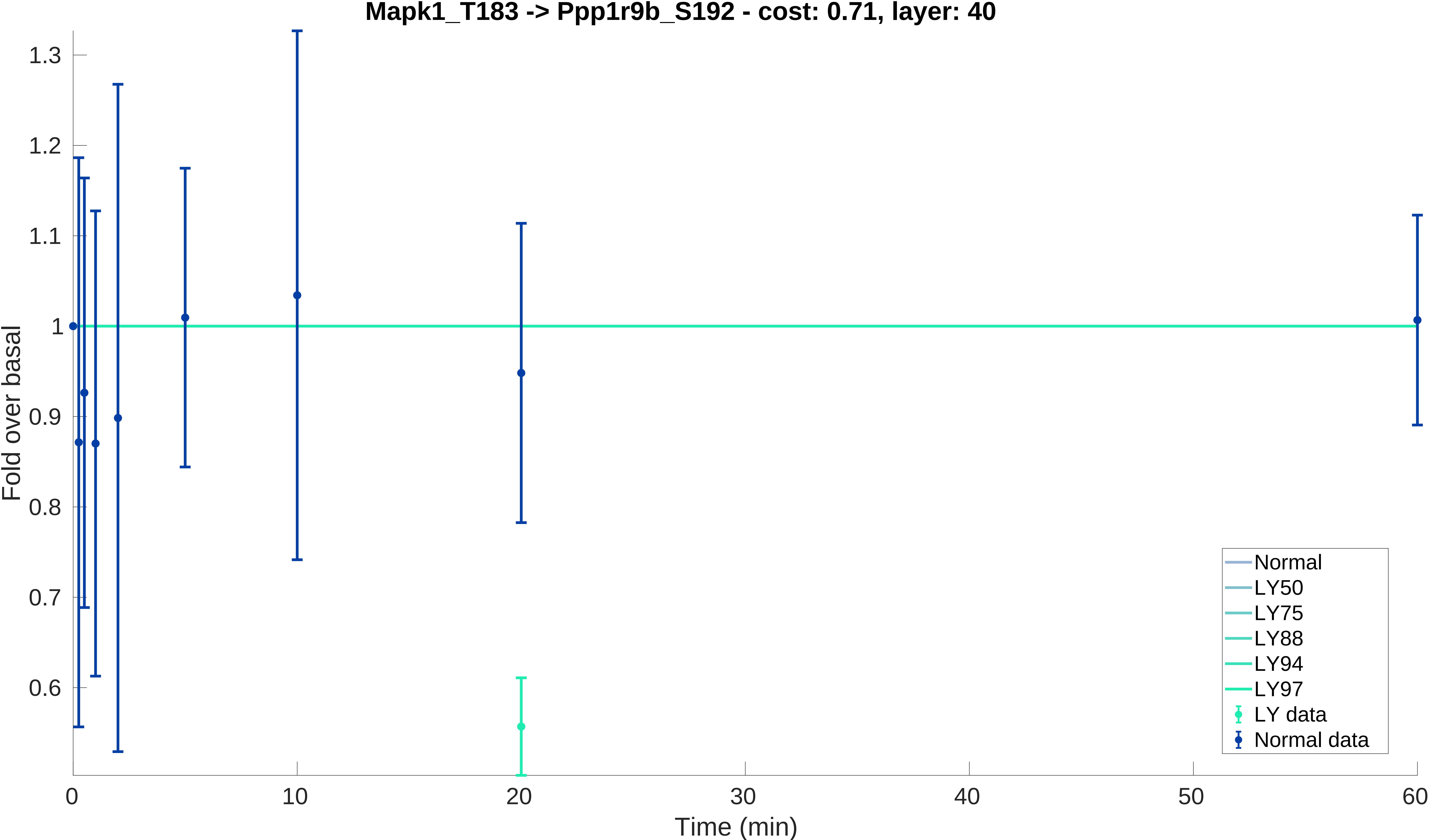

Mapk1_T183 -> Ppp1r9b_S192 - cost: 0.71, layer: 40
Fold over basal
Time (min)
1.3
1.2
1.1
1
0.9
0.8
0.7
0.6
0
10
20
30
40
50
60
Normal
LY50
LY75
LY88
LY94
LY97
LY data
Normal data

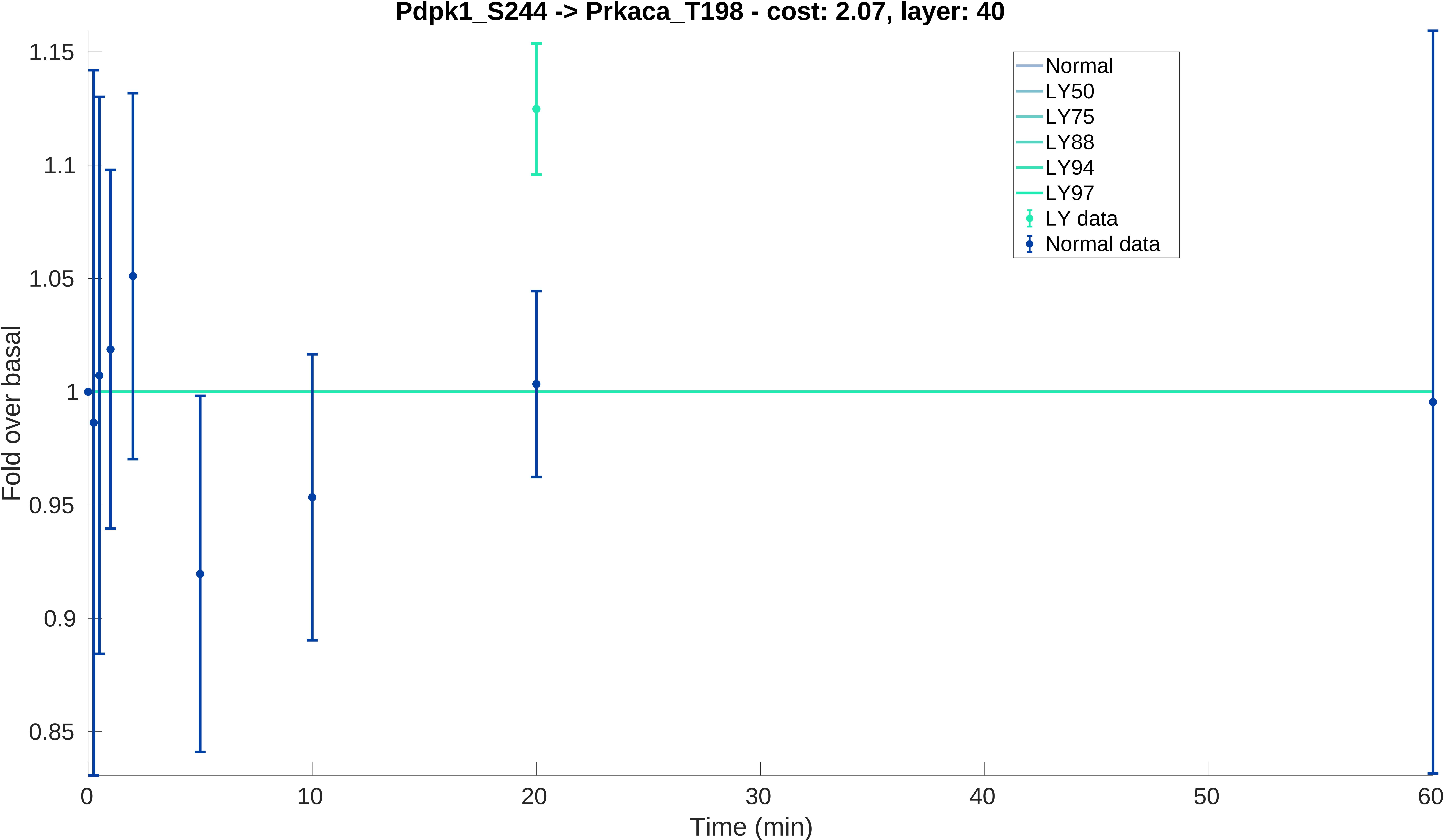

Pdpk1_S244 -> Prkaca_T198 - cost: 2.07, layer: 40
Fold over basal
1.15
1.1
1.05
1
0.95
0.9
0.85
0
10
20
30
40
50
60
Time (min)
Normal
LY50
LY75
LY88
LY94
LY97
LY data
Normal data

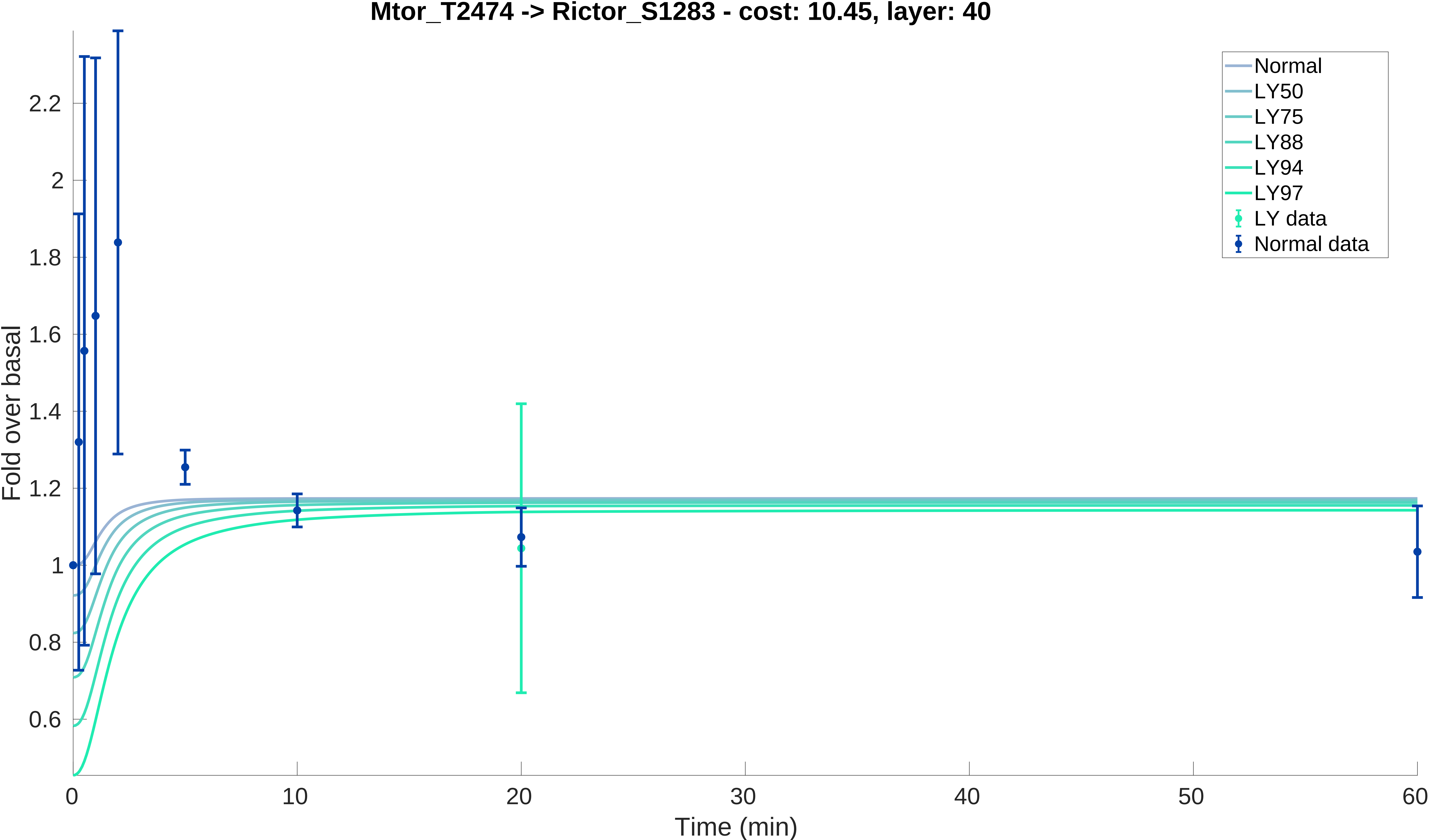

Mtor_T2474 -> Rictor_S1283 - cost: 10.45, layer: 40
Fold over basal
Time (min)
0.6
0.8
1
1.2
1.4
1.6
1.8
2
2.2
0
10
20
30
40
50
60
Normal
LY50
LY75
LY88
LY94
LY97
LY data
Normal data

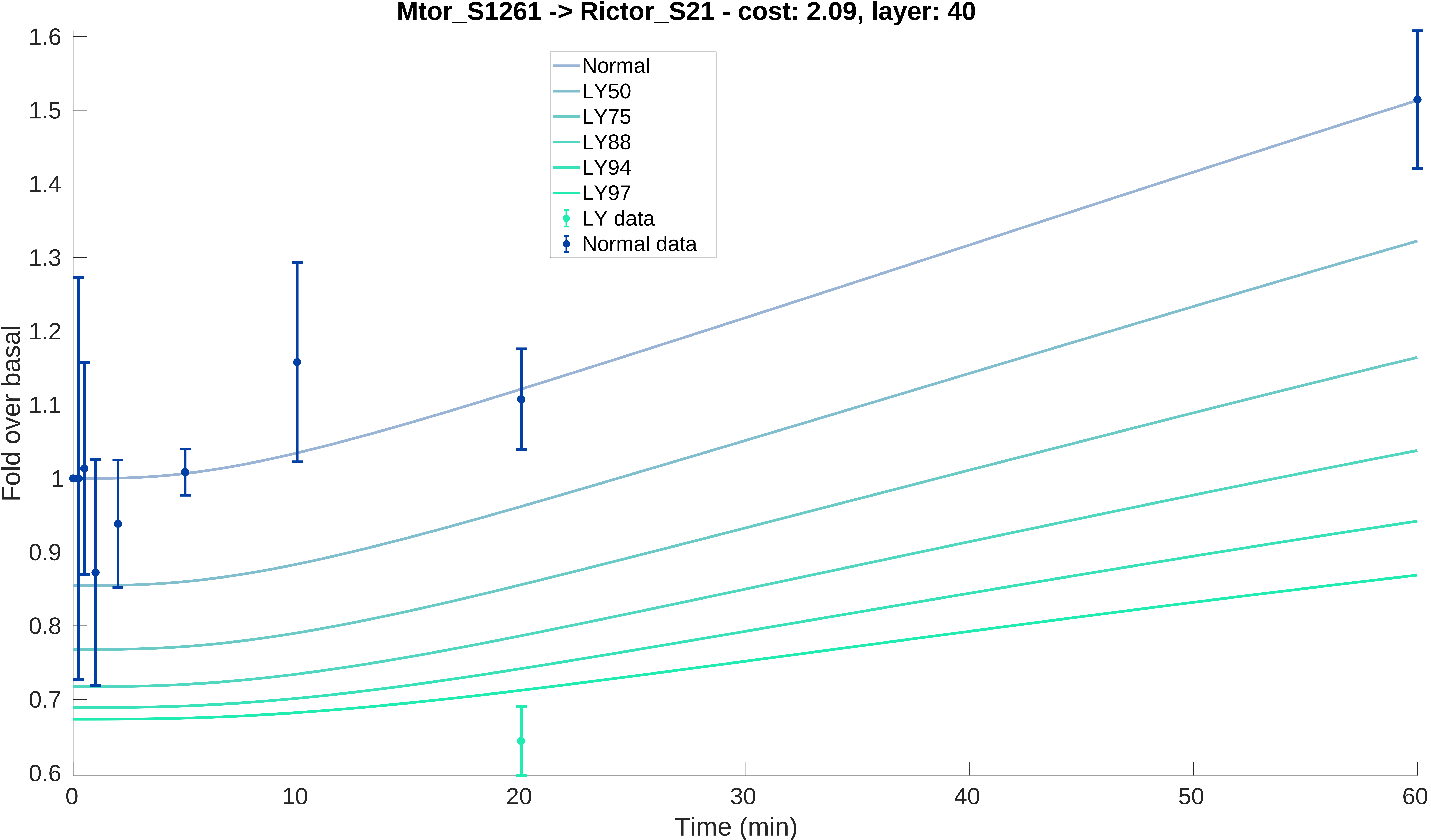

Mtor_S1261 -> Rictor_S21 - cost: 2.09, layer: 40
Normal
LY50
LY75
LY88
LY94
LY97
LY data
Normal data
Fold over basal
Time (min)
1.6
1.5
1.4
1.3
1.2
1.1
1
0.9
0.8
0.7
0.6
0
10
20
30
40
50
60

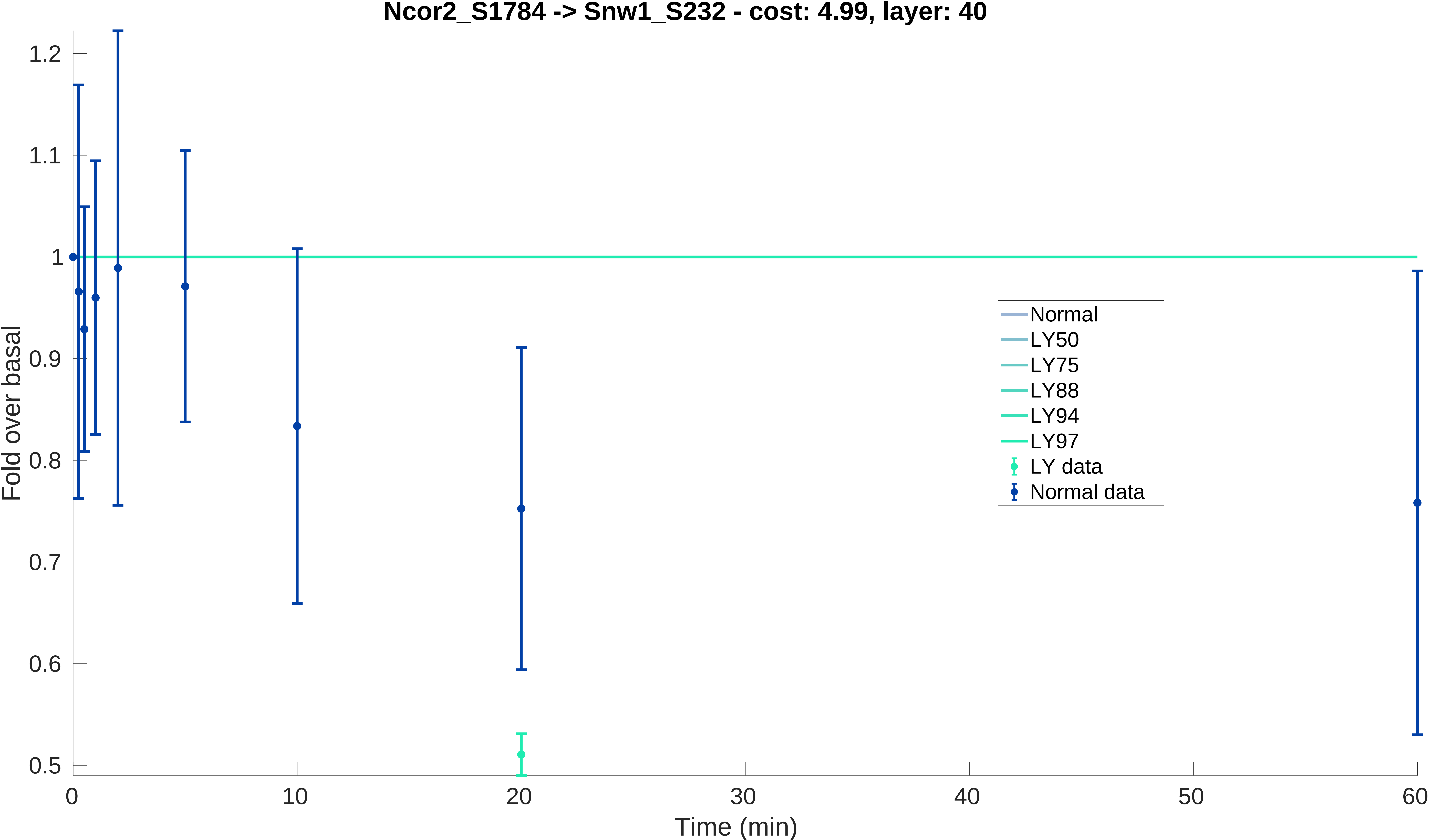
Ncor2_S1784 -> Snw1_S232 - cost: 4.99, layer: 40
Fold over basal
Time (min)
0.5
0.6
0.7
0.8
0.9
1
1.1
1.2
0
10
20
30
40
50
60
Normal
LY50
LY75
LY88
LY94
LY97
LY data
Normal data

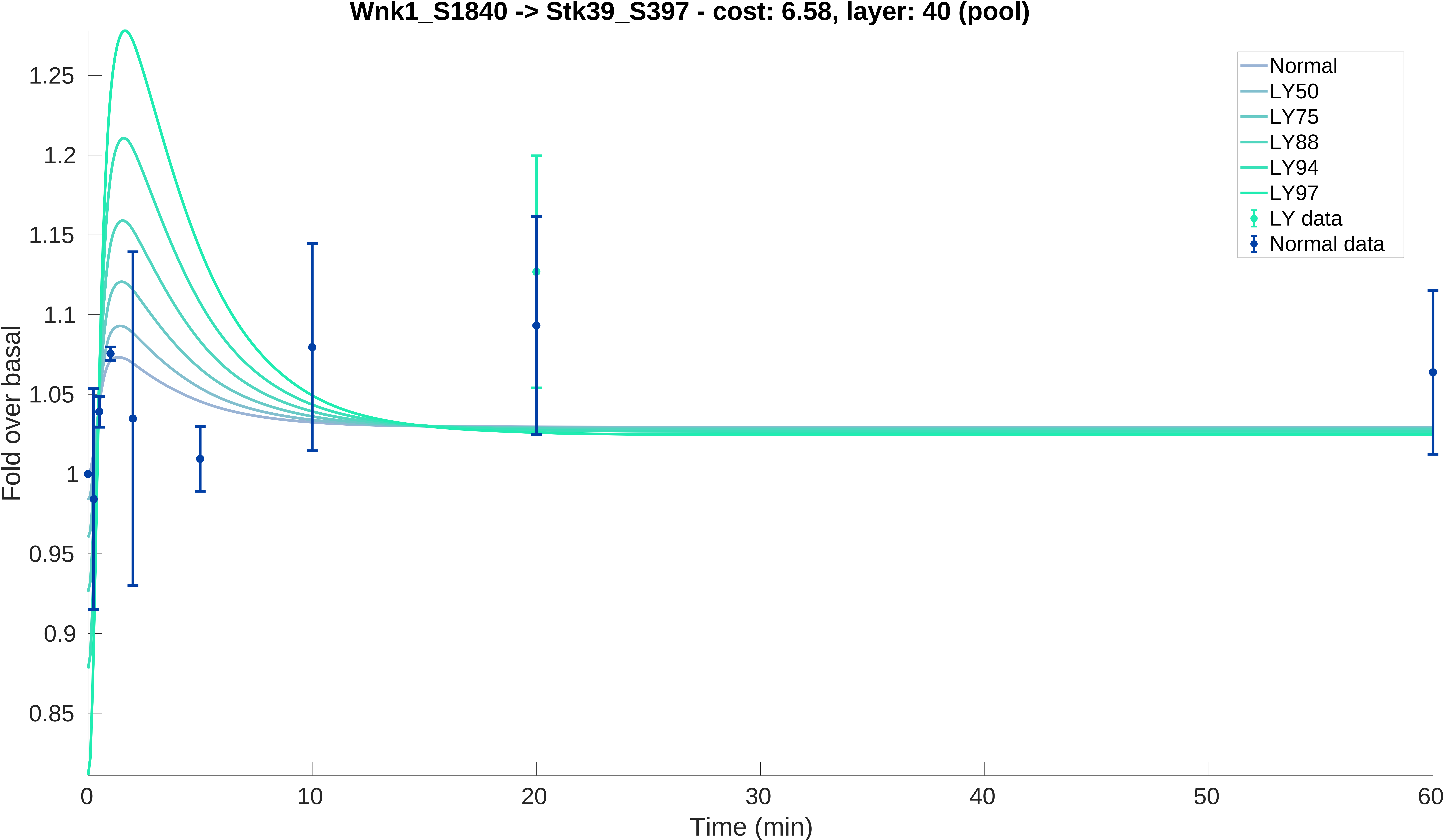

Wnk1_S1840 -> Stk39_S397 - cost: 6.58, layer: 40 (pool)
Fold over basal
Time (min)
1.25
1.2
1.15
1.1
1.05
1
0.95
0.9
0.85
0
10
20
30
40
50
60
Normal
LY50
LY75
LY88
LY94
LY97
LY data
Normal data

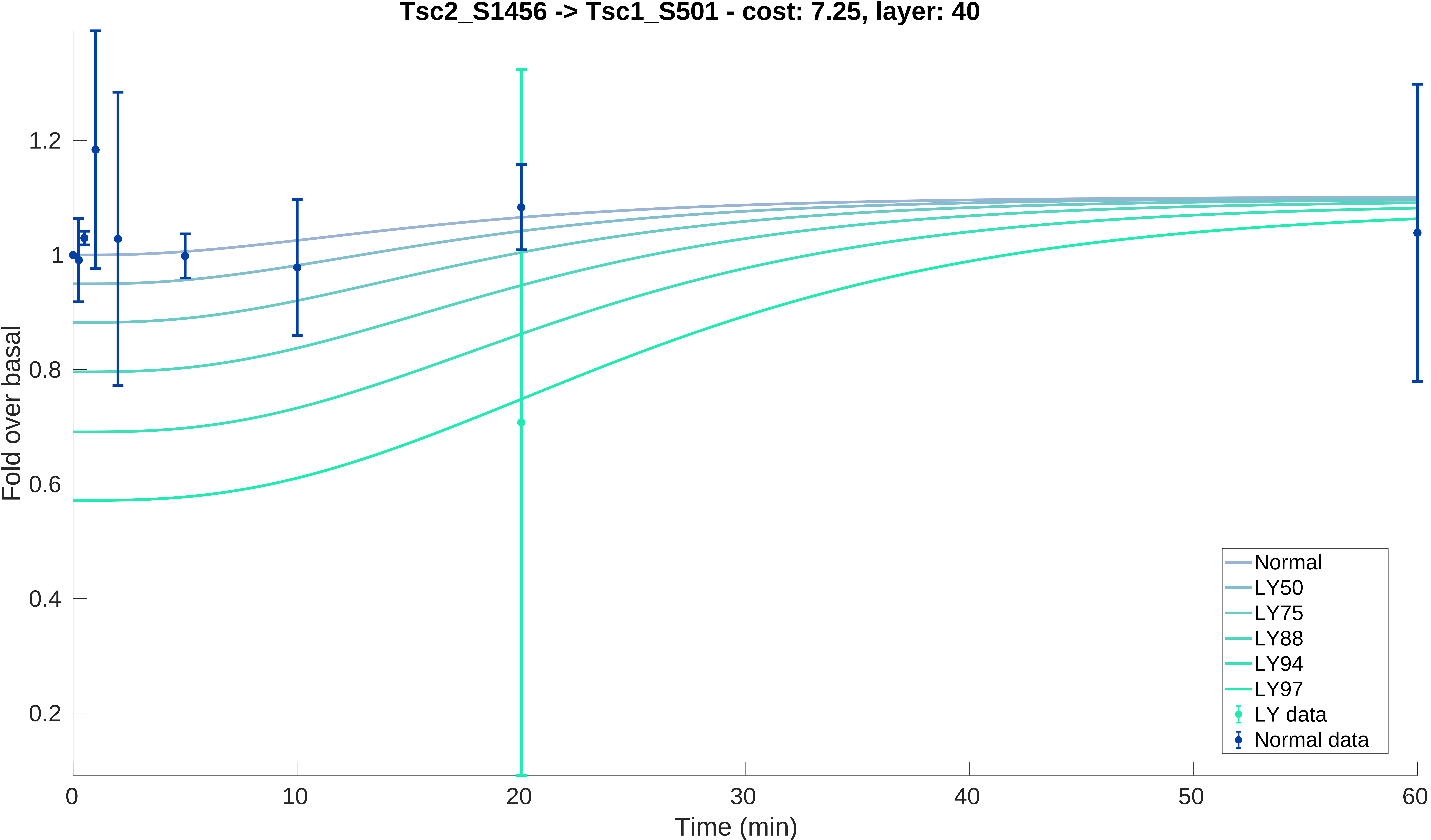
Tsc2_S1456 -> Tsc1_S501 - cost: 7.25, layer: 40
Fold over basal
Time (min)
0.2
0.4
0.6
0.8
1
1.2
0
10
20
30
40
50
60
Normal
LY50
LY75
LY88
LY94
LY97
LY data
Normal data

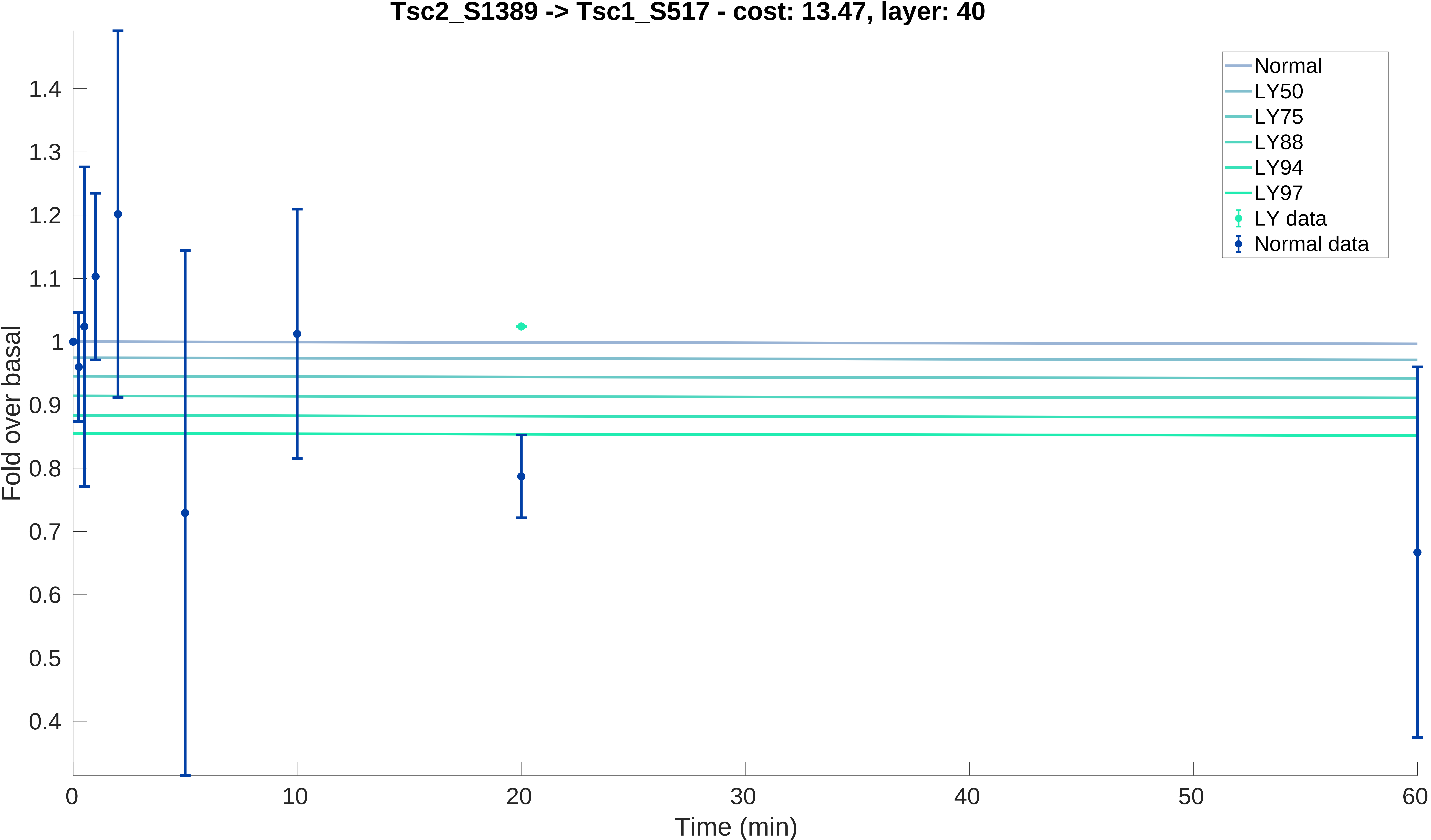

Tsc2_S1389 -> Tsc1_S517 - cost: 13.47, layer: 40
Fold over basal
Time (min)
0
10
20
30
40
50
60
1.4
1.3
1.2
1.1
1
0.9
0.8
0.7
0.6
0.5
0.4
Normal
LY50
LY75
LY88
LY94
LY97
LY data
Normal data

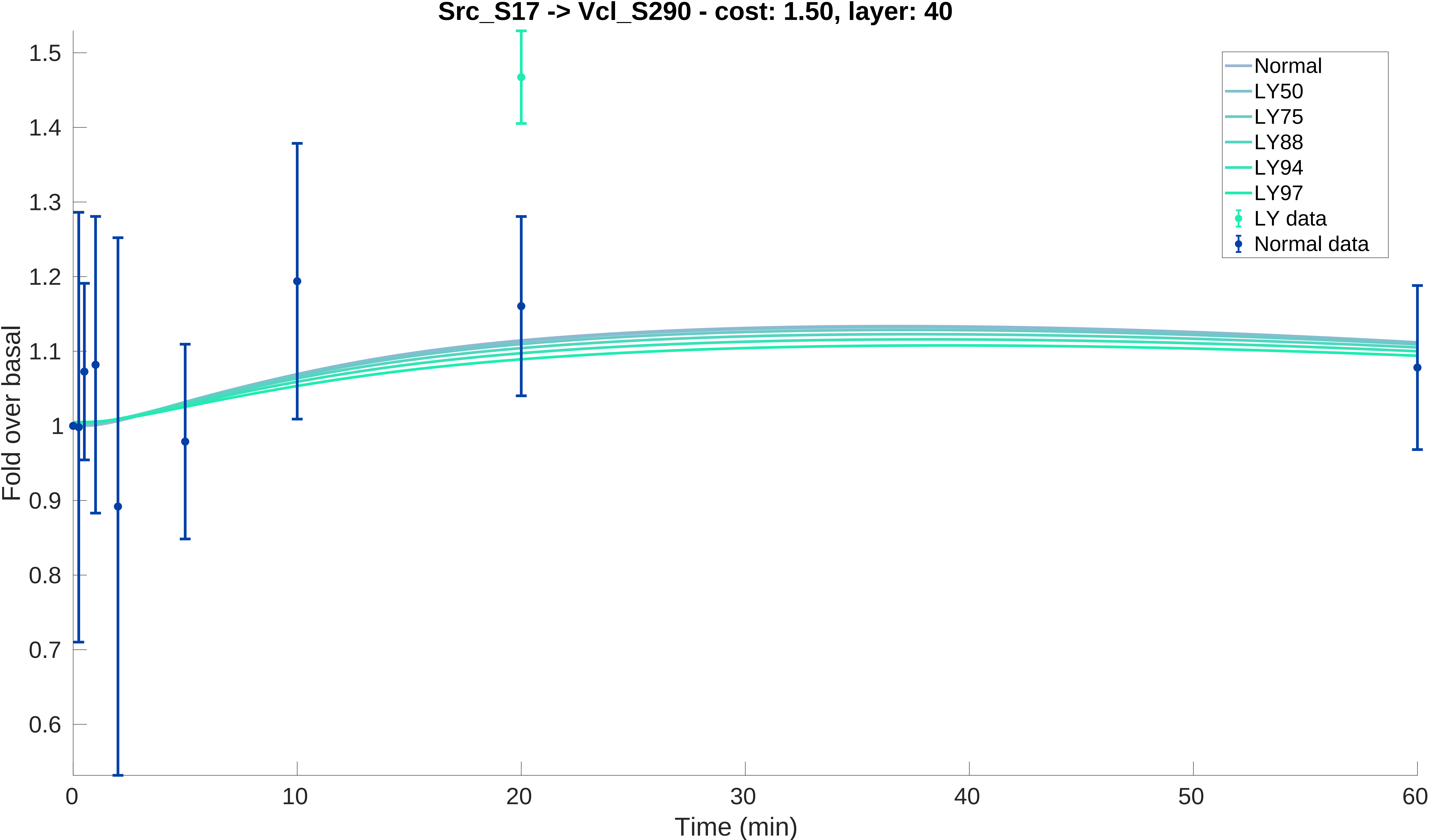
Src_S17 -> Vcl_S290 - cost: 1.50, layer: 40
Fold over basal
Time (min)
Normal
LY50
LY75
LY88
LY94
LY97
LY data
Normal data
1.5
1.4
1.3
1.2
1.1
1
0.9
0.8
0.7
0.6
0
10
20
30
40
50
60

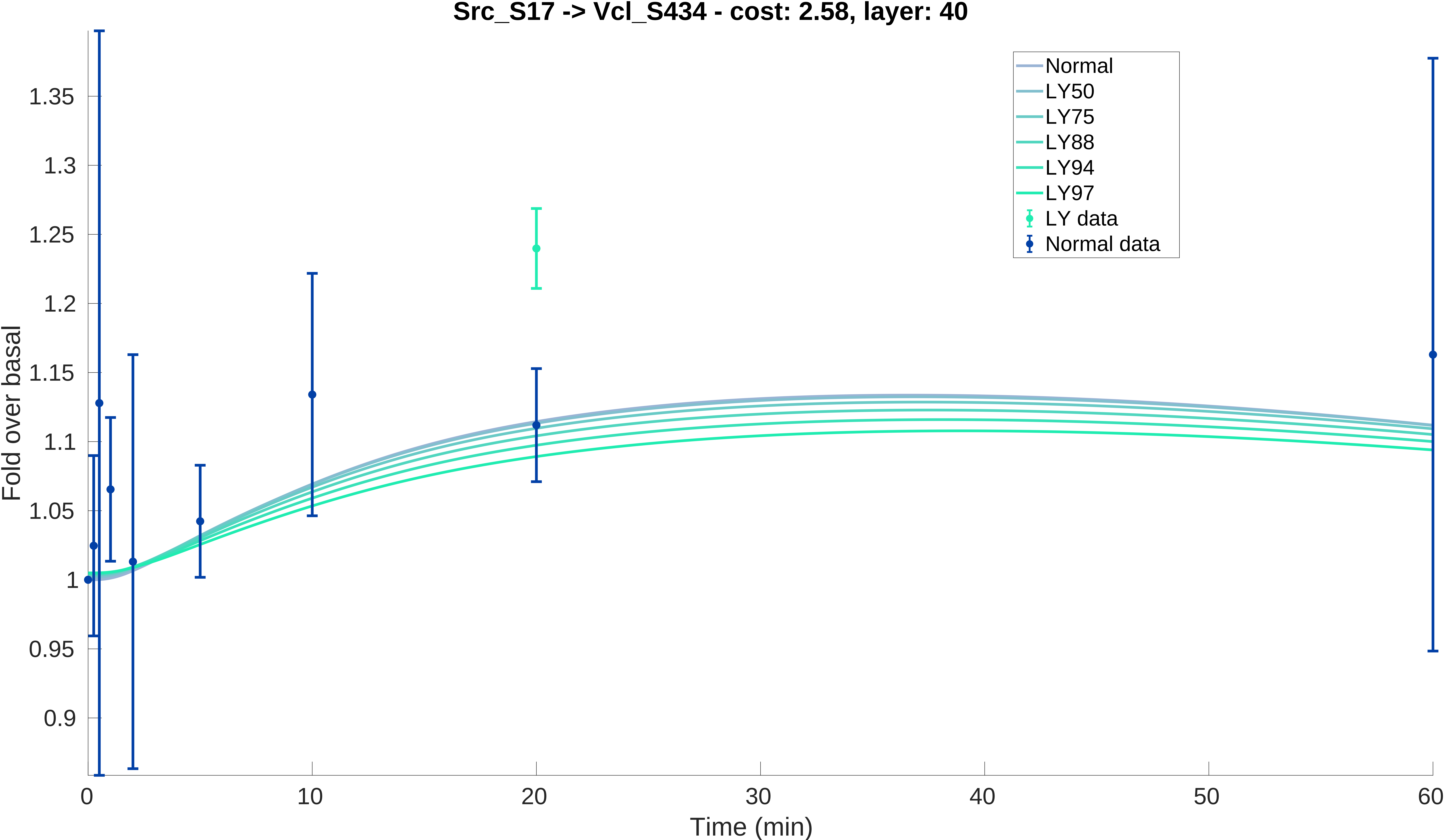

Src_S17 -> Vcl_S434 - cost: 2.58, layer: 40
Normal
LY50
LY75
LY88
LY94
LY97
LY data
Normal data
Fold over basal
1.35
1.3
1.25
1.2
1.15
1.1
1.05
1
0.95
0.9
0
10
20
30
40
50
60
Time (min)

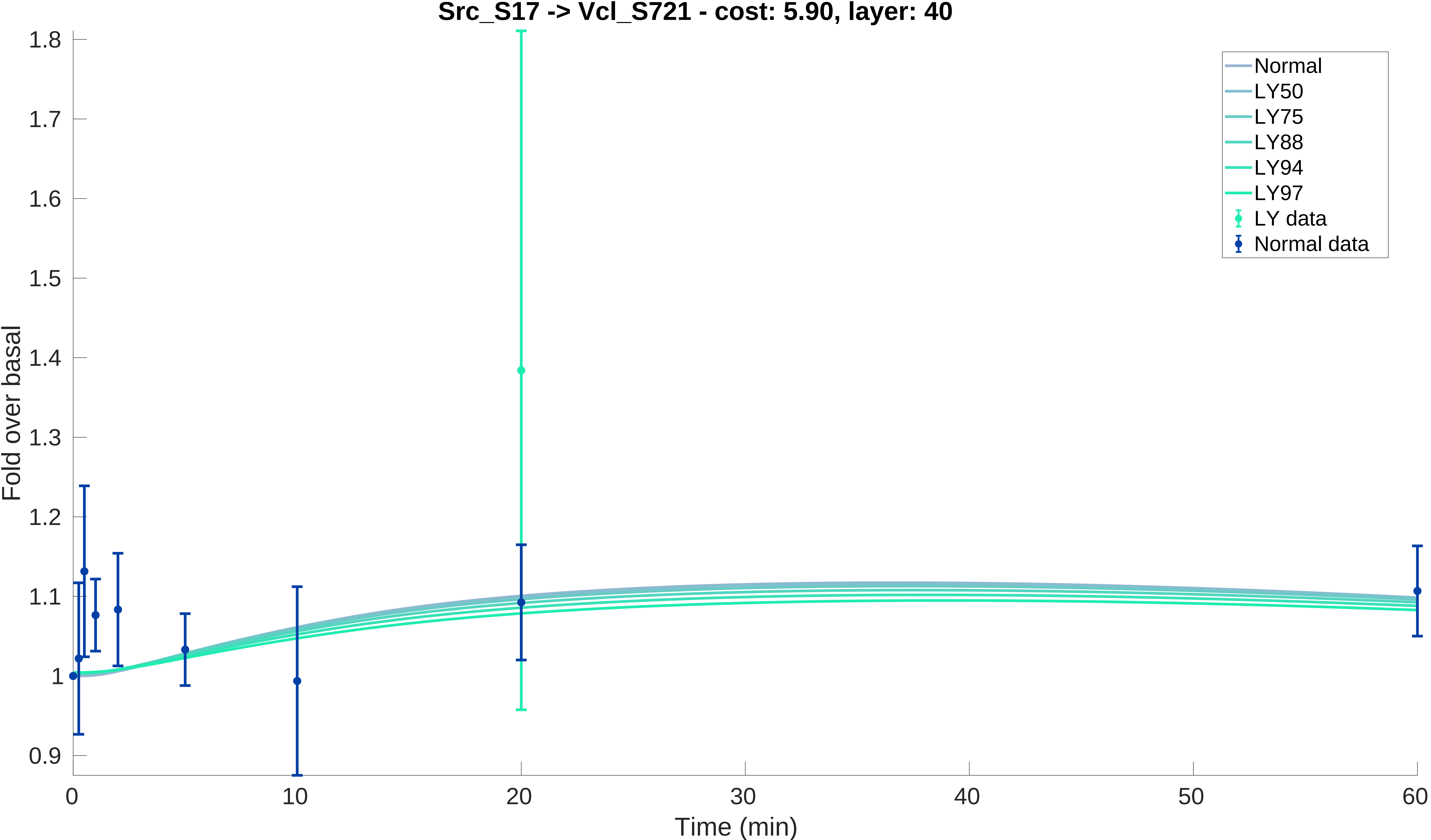

Src_S17 -> Vcl_S721 - cost: 5.90, layer: 40
Fold over basal
Time (min)
Normal
LY50
LY75
LY88
LY94
LY97
LY data
Normal data
0.9
1
1.1
1.2
1.3
1.4
1.5
1.6
1.7
1.8
0
10
20
30
40
50
60

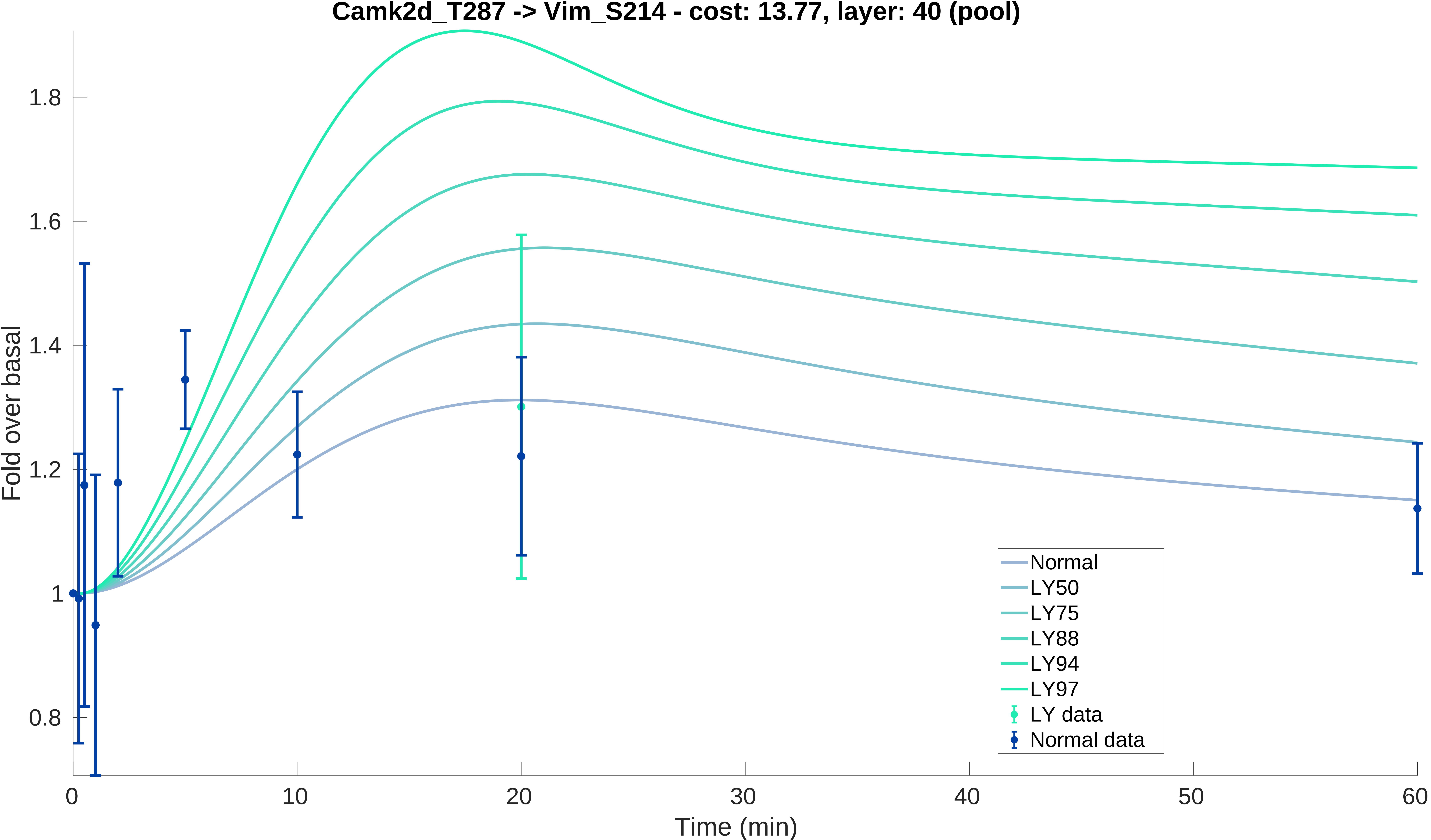

Camk2d_T287 -> Vim_S214 - cost: 13.77, layer: 40 (pool)
Fold over basal
0.8
1
1.2
1.4
1.6
1.8
0
10
20
30
40
50
60
Time (min)
Normal
LY50
LY75
LY88
LY94
LY97
LY data
Normal data

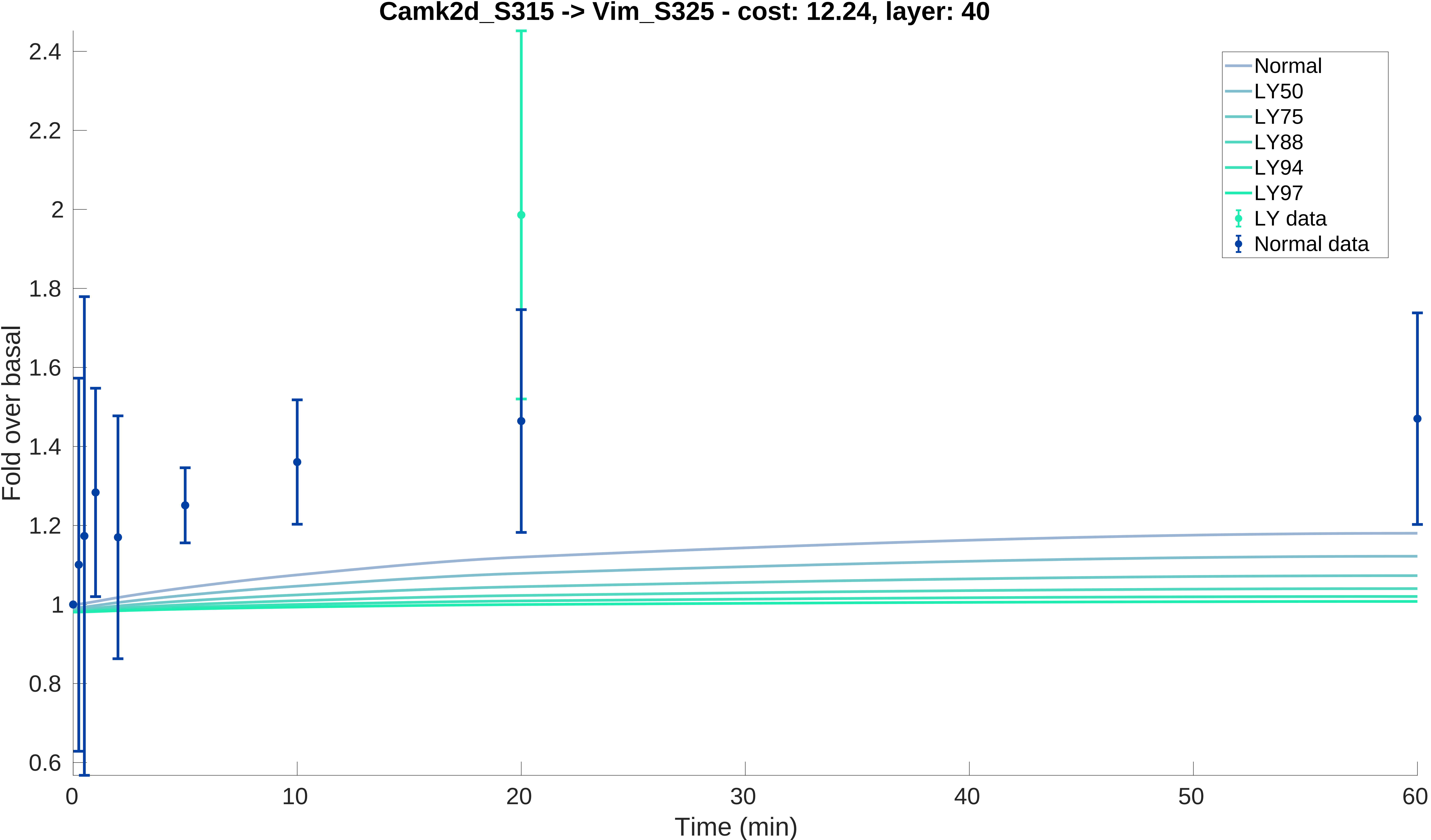

Camk2d_S315 -> Vim_S325 - cost: 12.24, layer: 40
Fold over basal
Time (min)
Normal
LY50
LY75
LY88
LY94
LY97
LY data
Normal data
0.6
0.8
1
1.2
1.4
1.6
1.8
2
2.2
2.4
0
10
20
30
40
50
60

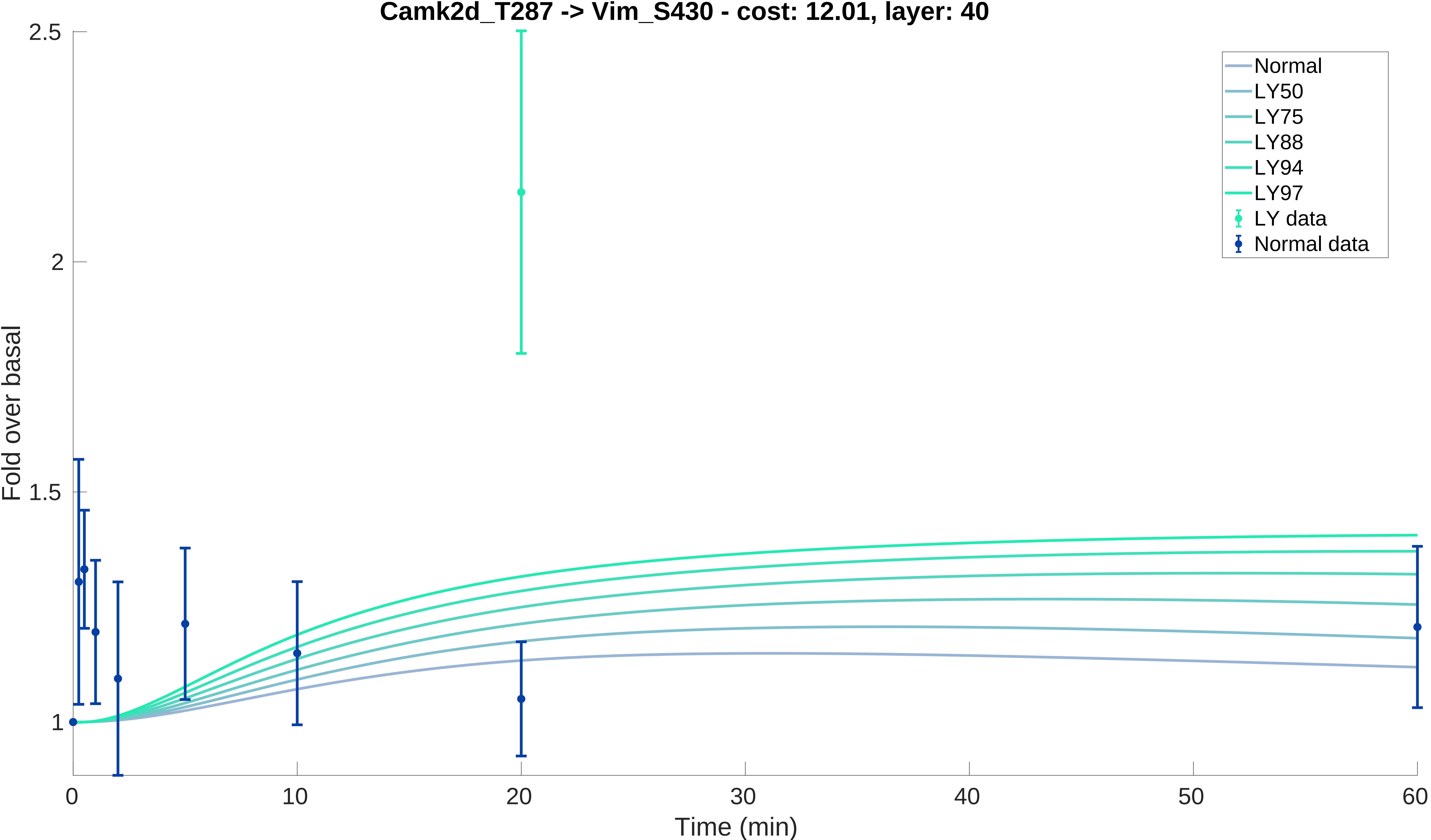

Camk2d_T287 -> Vim_S430 - cost: 12.01, layer: 40
Fold over basal
Time (min)
Normal
LY50
LY75
LY88
LY94
LY97
LY data
Normal data
2.5
2
1.5
1
0
10
20
30
40
50
60

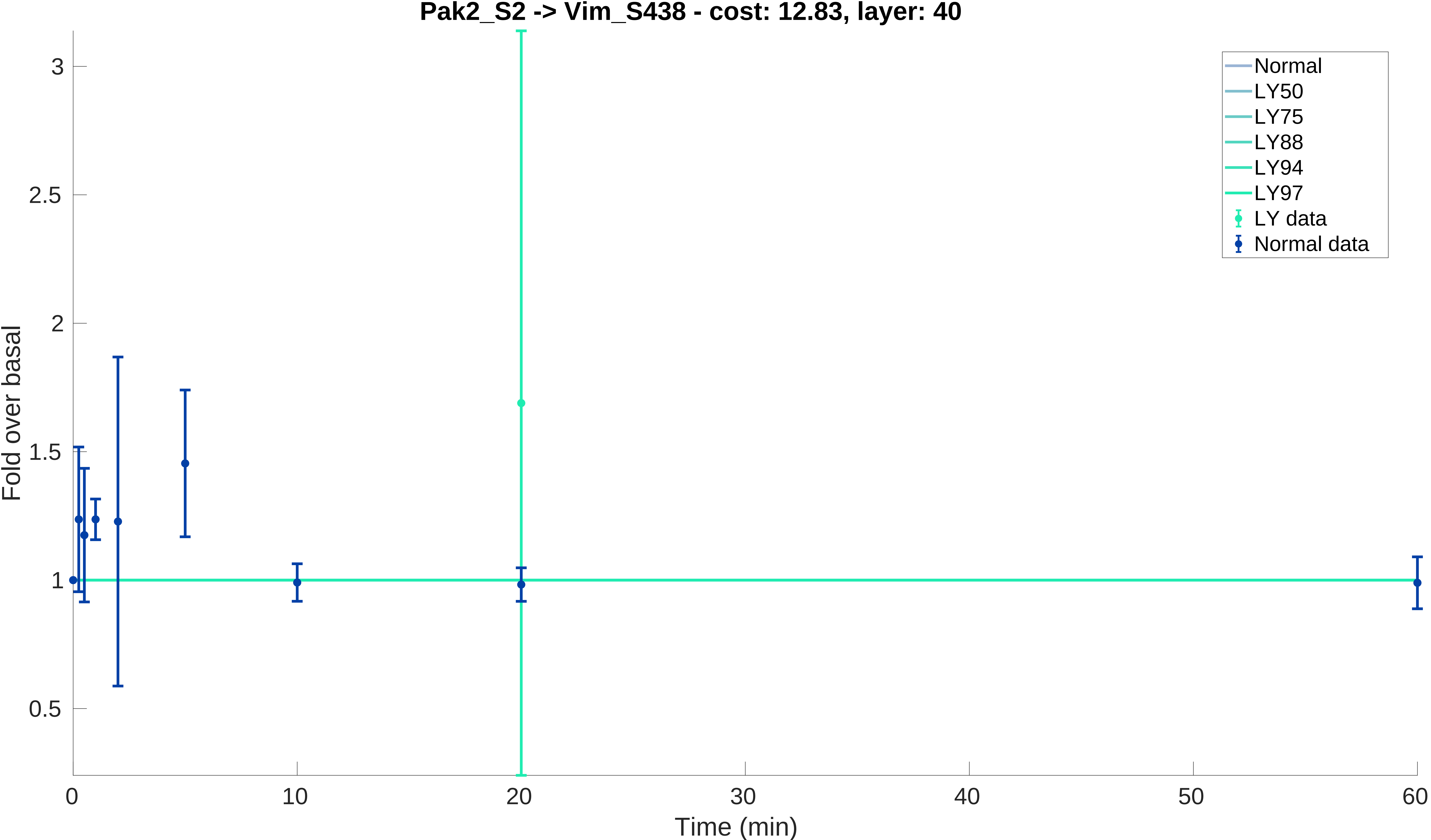

Pak2_S2 -> Vim_S438 - cost: 12.83, layer: 40
Normal
LY50
LY75
LY88
LY94
LY97
LY data
Normal data
Fold over basal
0.5
1
1.5
2
2.5
3
0
10
20
30
40
50
60
Time (min)

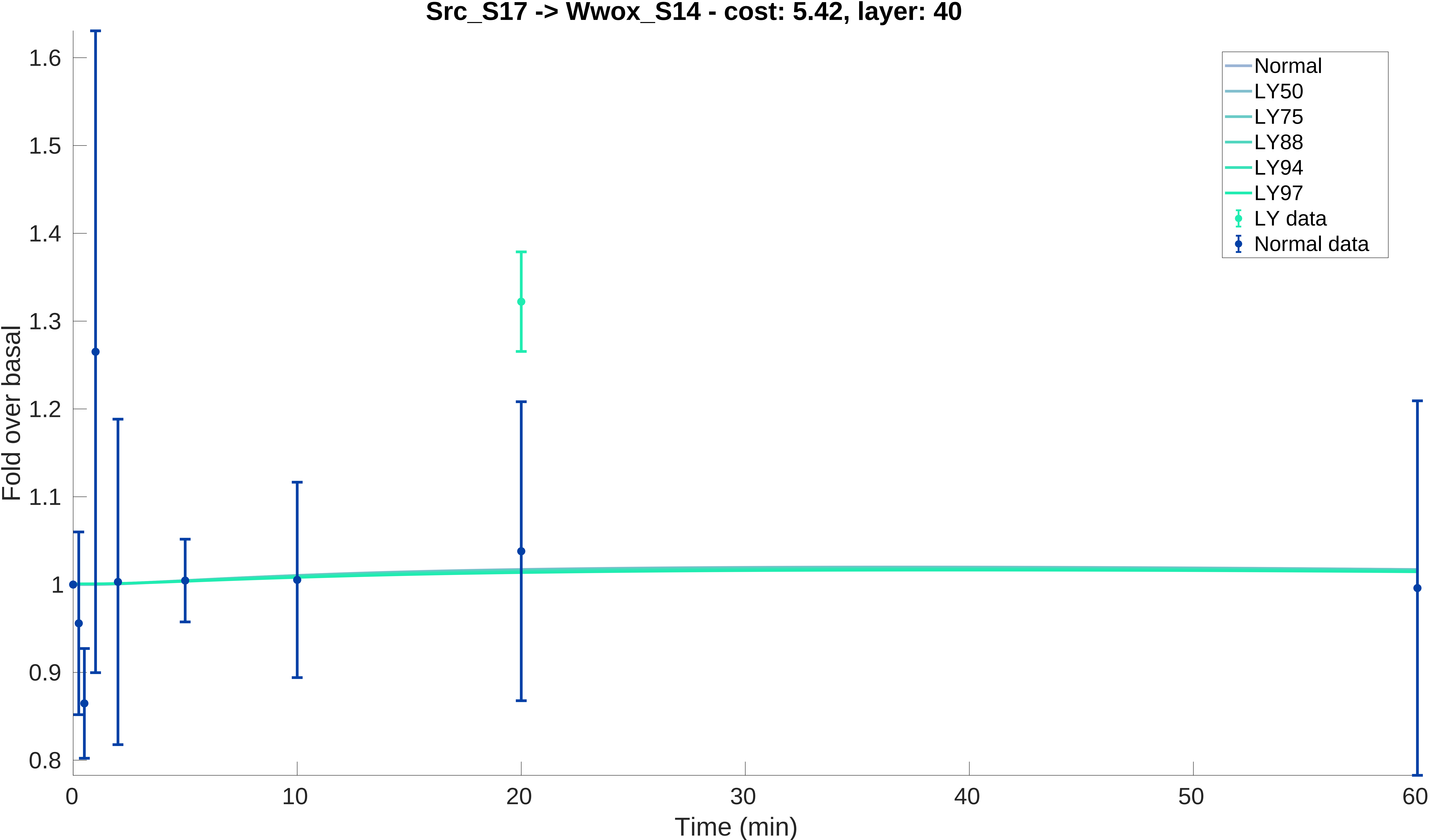

Src_S17 -> Wwox_S14 - cost: 5.42, layer: 40
Fold over basal
Time (min)
0.8
0.9
1
1.1
1.2
1.3
1.4
1.5
1.6
0
10
20
30
40
50
60
Normal
LY50
LY75
LY88
LY94
LY97
LY data
Normal data

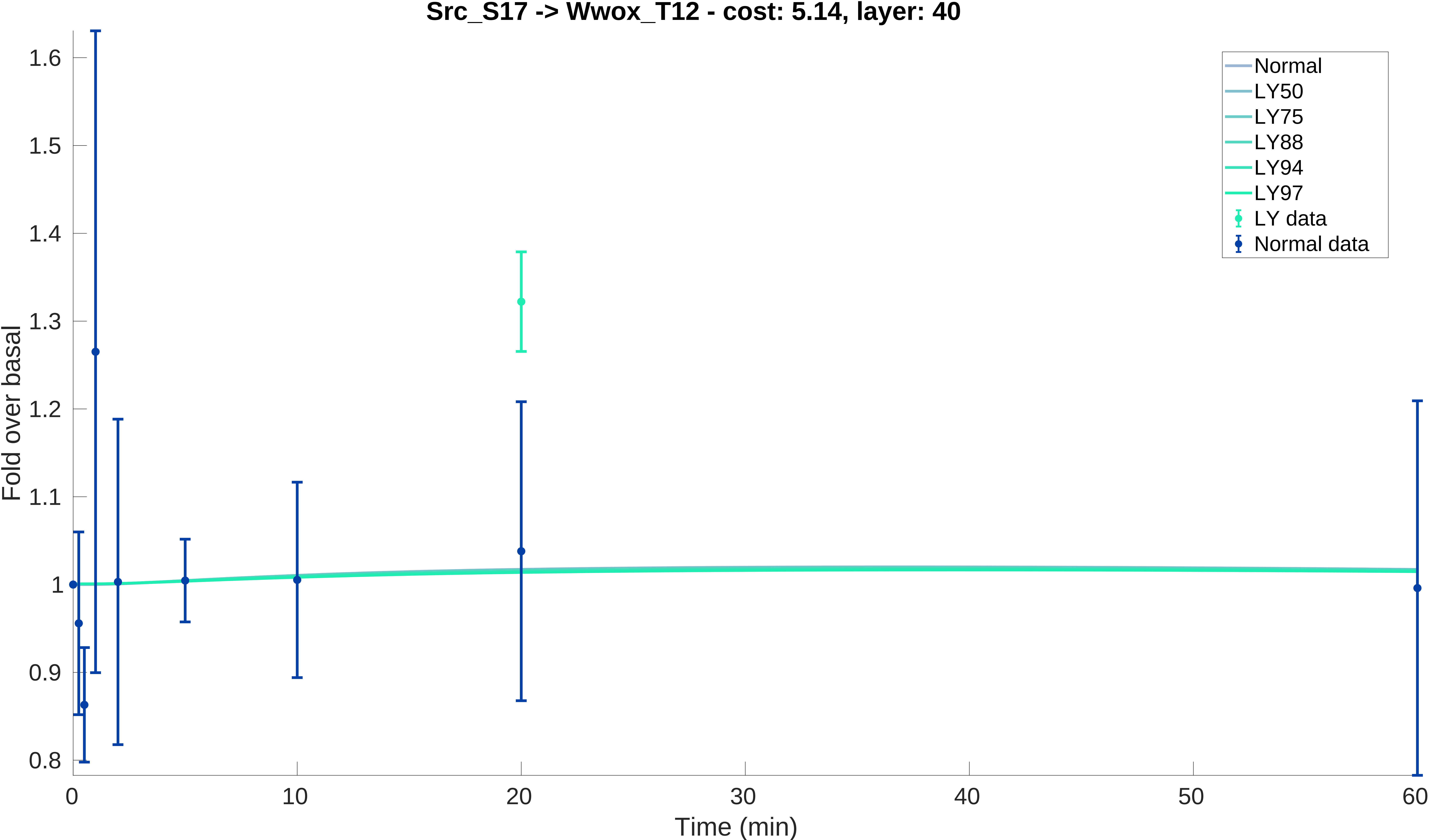

Src_S17 -> Wwox_T12 - cost: 5.14, layer: 40
Normal
LY50
LY75
LY88
LY94
LY97
LY data
Normal data
Fold over basal
1.6
1.5
1.4
1.3
1.2
1.1
1
0.9
0.8
0
10
20
30
40
50
60
Time (min)

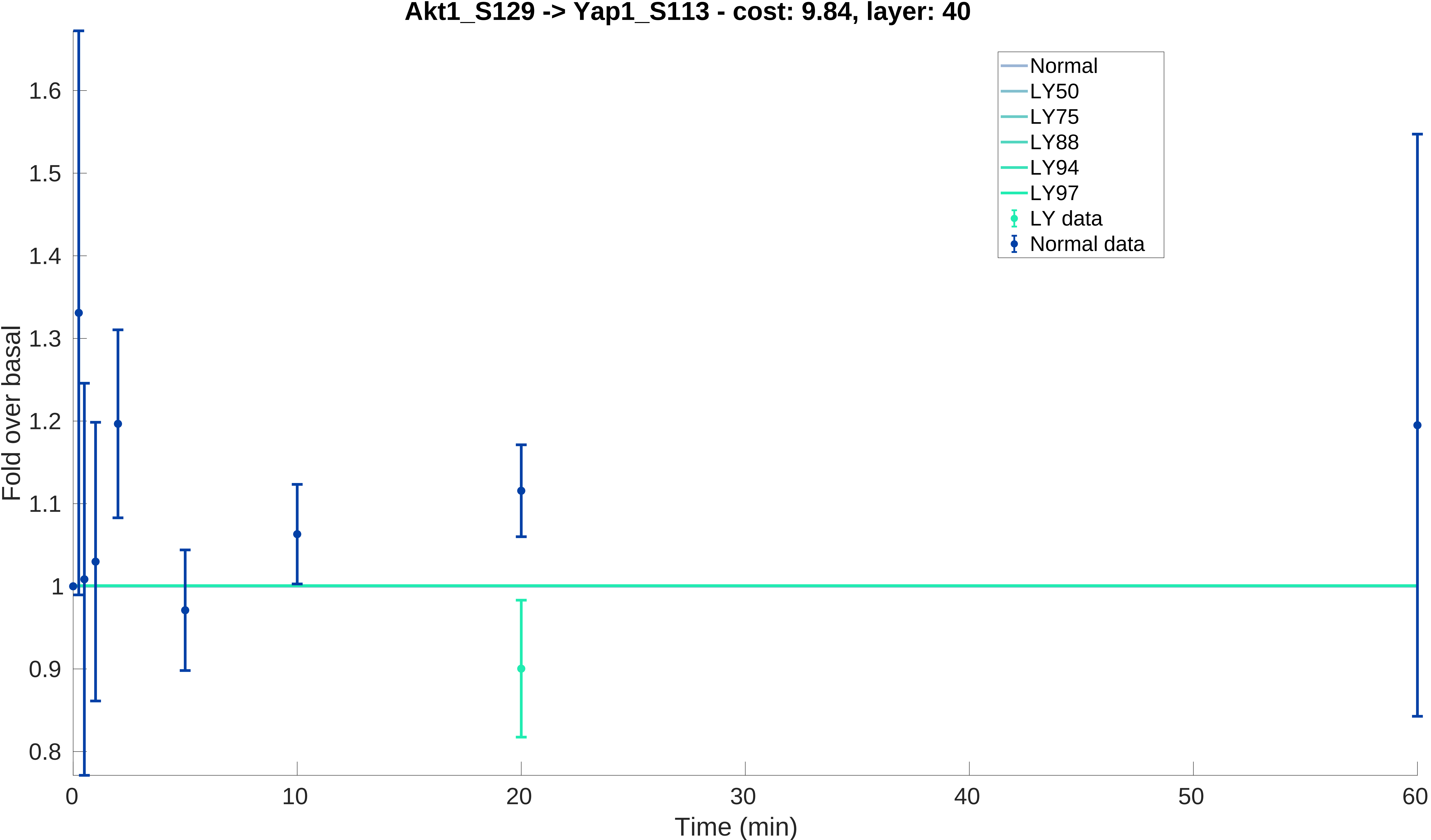
Akt1_S129 -> Yap1_S113 - cost: 9.84, layer: 40
Fold over basal
Time (min)
0.8
0.9
1
1.1
1.2
1.3
1.4
1.5
1.6
0
10
20
30
40
50
60
Normal
LY50
LY75
LY88
LY94
LY97
LY data
Normal data

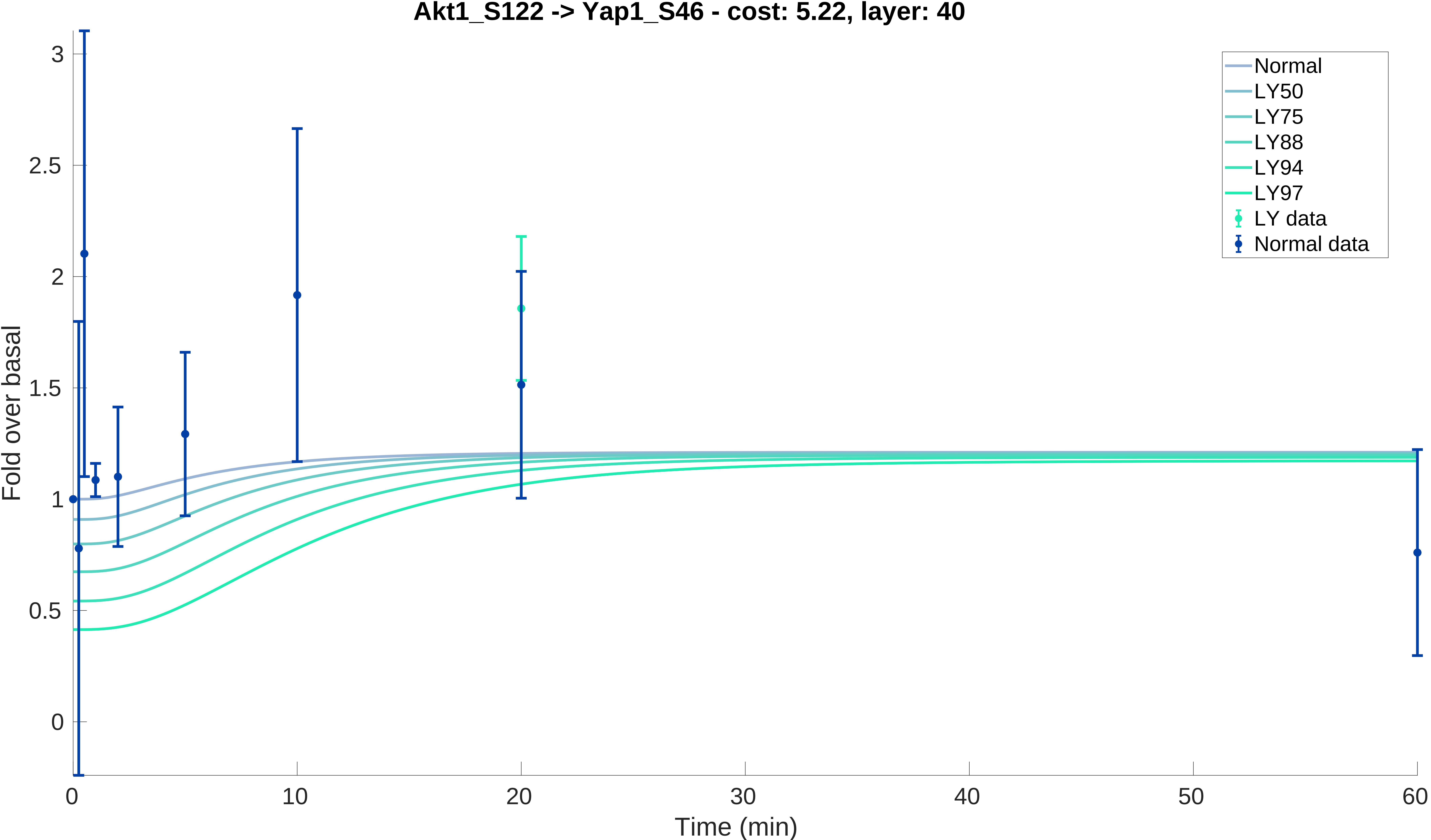

Akt1_S122 -> Yap1_S46 - cost: 5.22, layer: 40
Fold over basal
Time (min)
0
0.5
1
1.5
2
2.5
3
0
10
20
30
40
50
60
Normal
LY50
LY75
LY88
LY94
LY97
LY data
Normal data

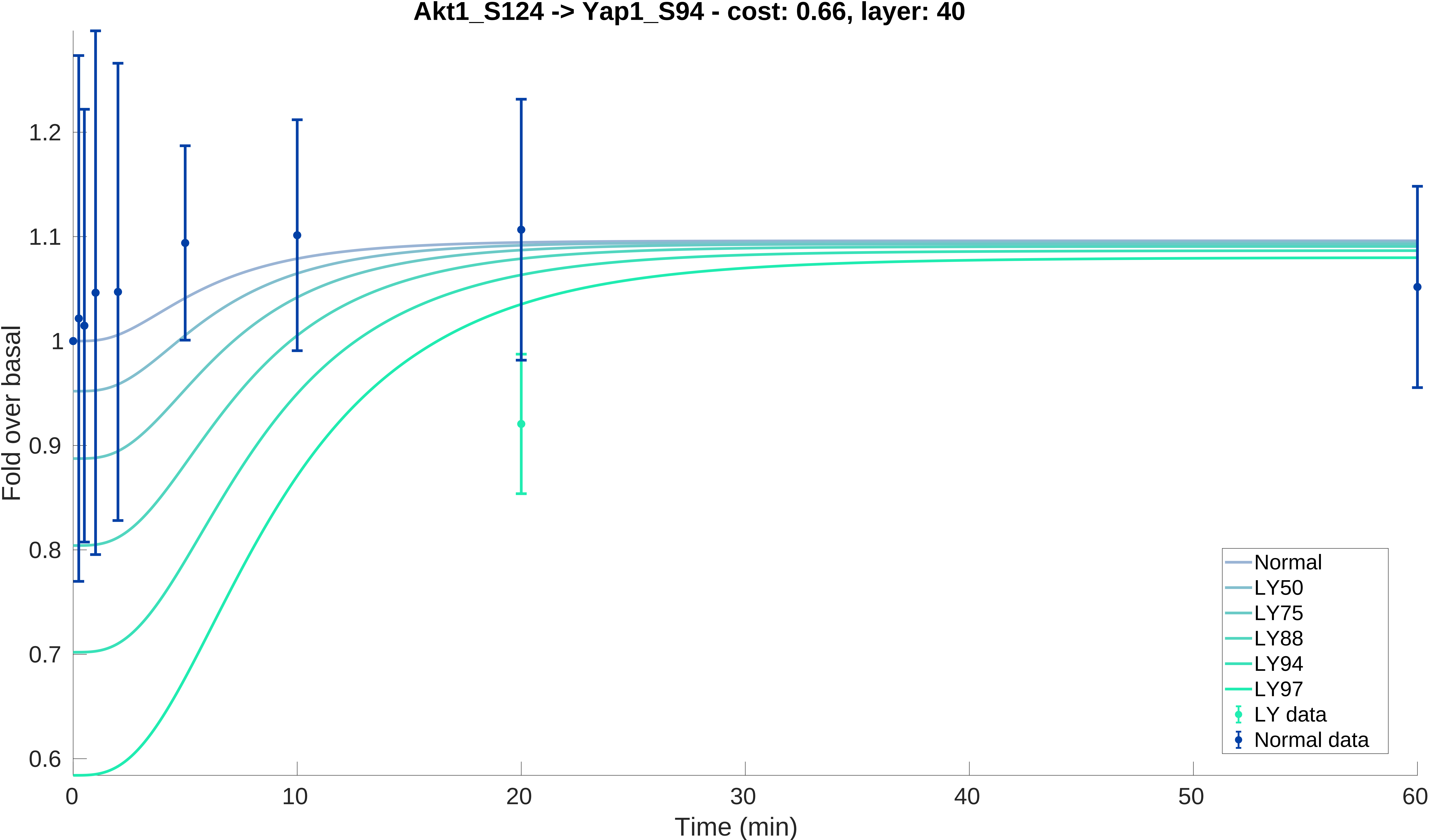
Akt1_S124 -> Yap1_S94 - cost: 0.66, layer: 40
Fold over basal
1.2
1.1
1
0.9
0.8
0.7
0.6
0
10
20
30
40
50
60
Time (min)
Normal
LY50
LY75
LY88
LY94
LY97
LY data
Normal data

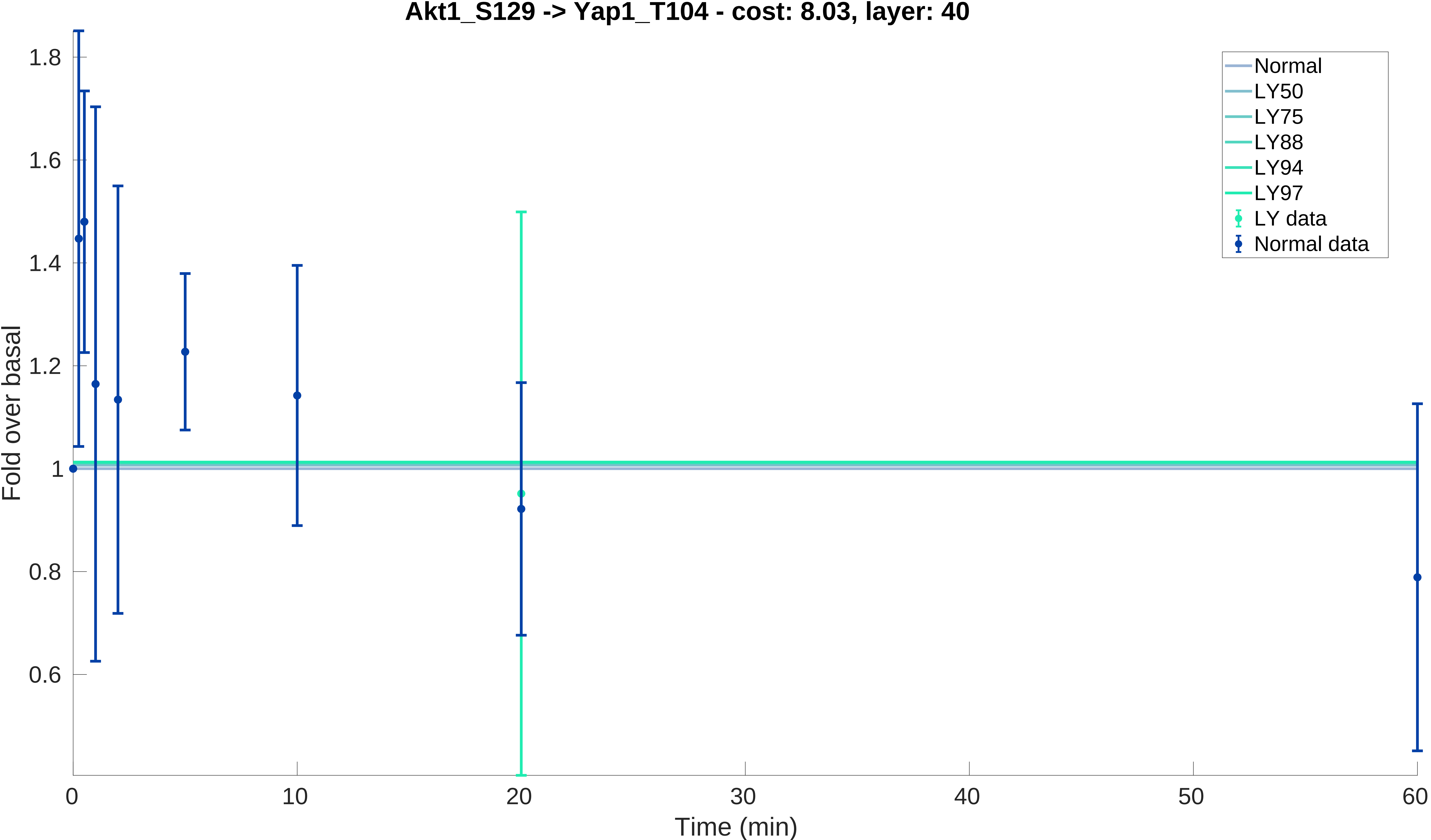
Akt1_S129 -> Yap1_T104 - cost: 8.03, layer: 40
Fold over basal
1.8
1.6
1.4
1.2
1
0.8
0.6
0
10
20
30
40
50
60
Time (min)
Normal
LY50
LY75
LY88
LY94
LY97
LY data
Normal data

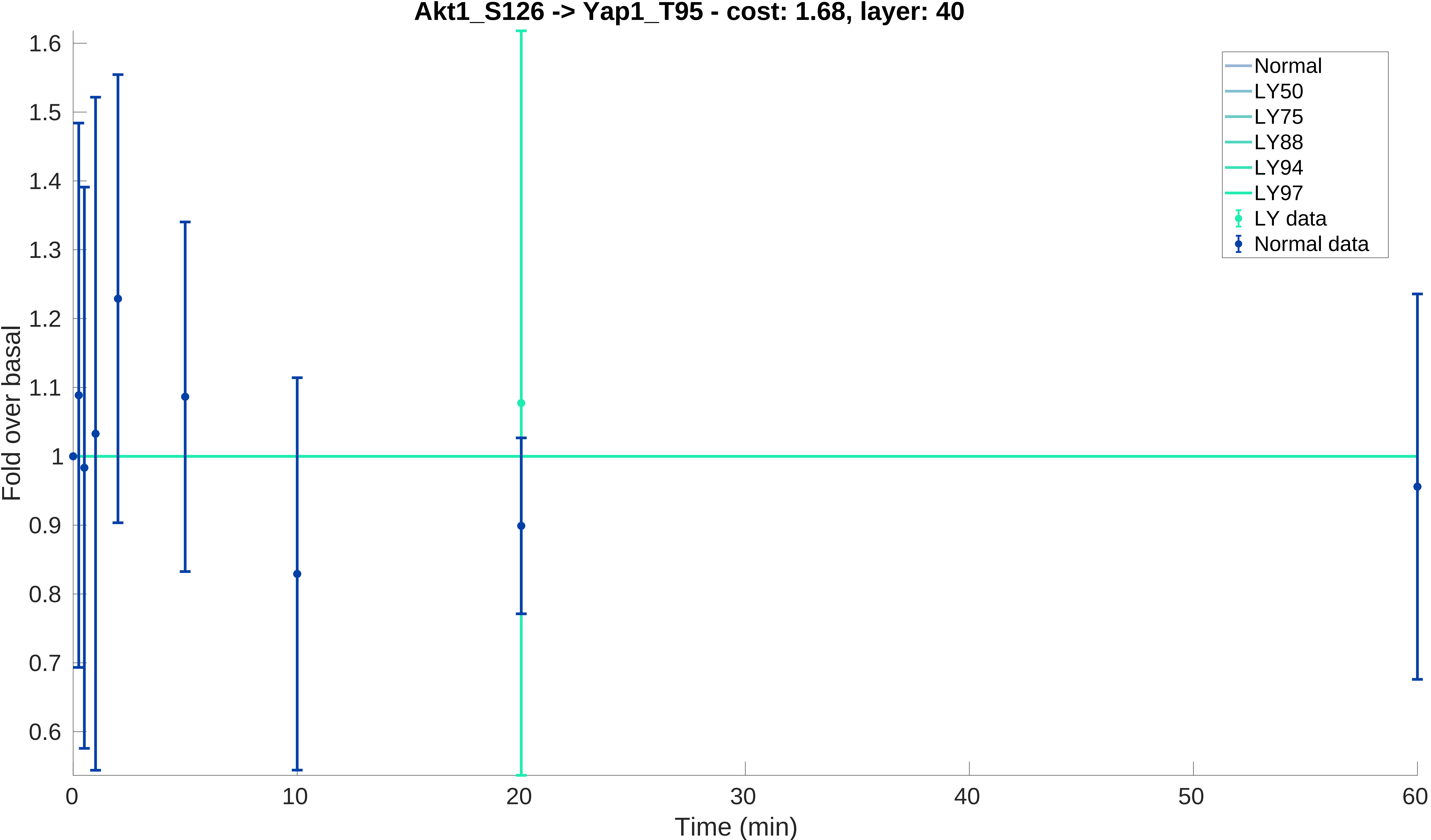

Akt1_S126 -> Yap1_T95 - cost: 1.68, layer: 40
Fold over basal
Time (min)
0
10
20
30
40
50
60
0.6
0.7
0.8
0.9
1
1.1
1.2
1.3
1.4
1.5
1.6
Normal
LY50
LY75
LY88
LY94
LY97
LY data
Normal data

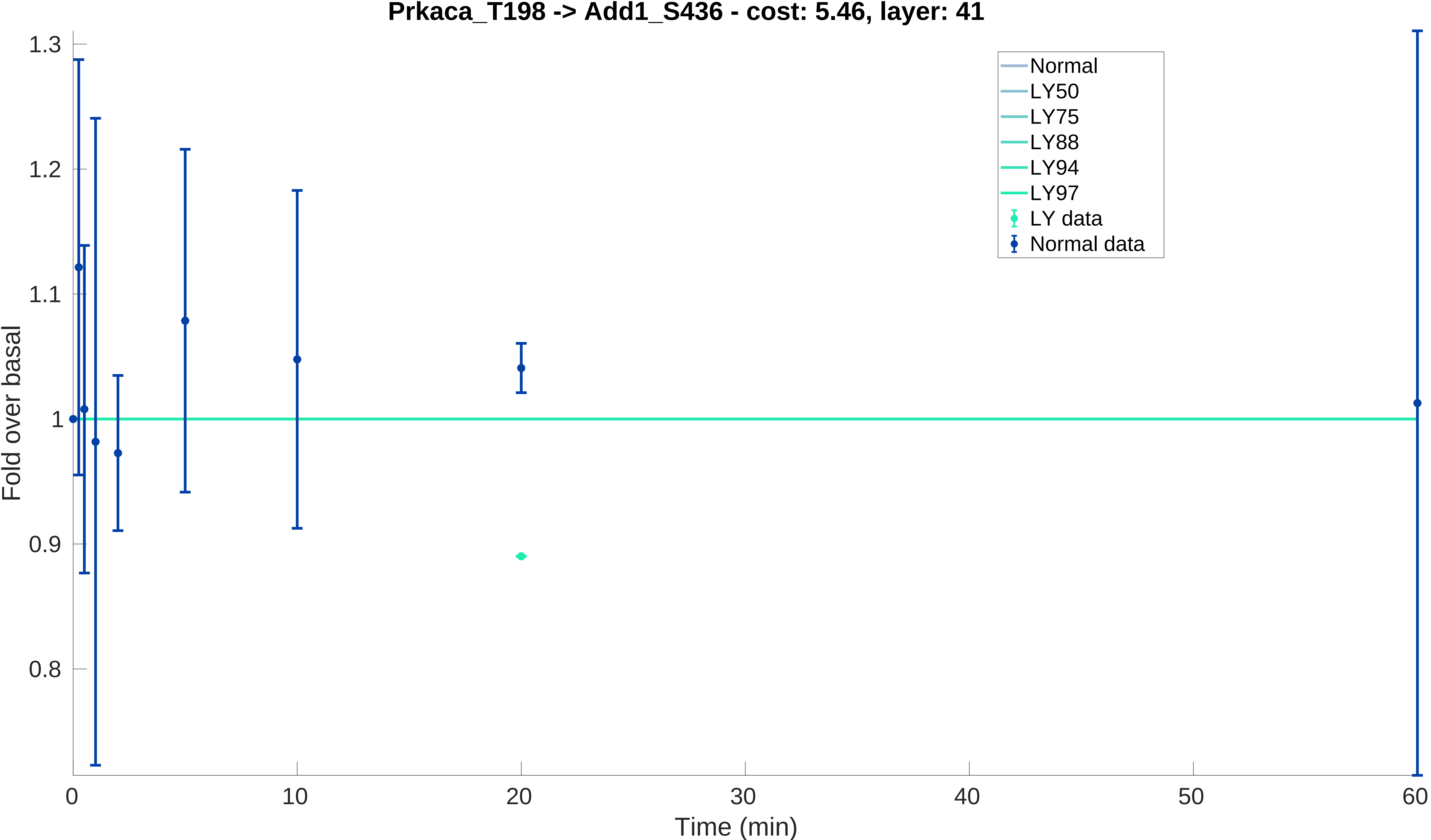

Prrkaca_T198 -> Add1_S436 - cost: 5.46, layer: 41
Fold over basal
1.3
1.2
1.1
1
0.9
0.8
0
10
20
30
40
50
60
Time (min)
Normal
LY50
LY75
LY88
LY94
LY97
LY data
Normal data

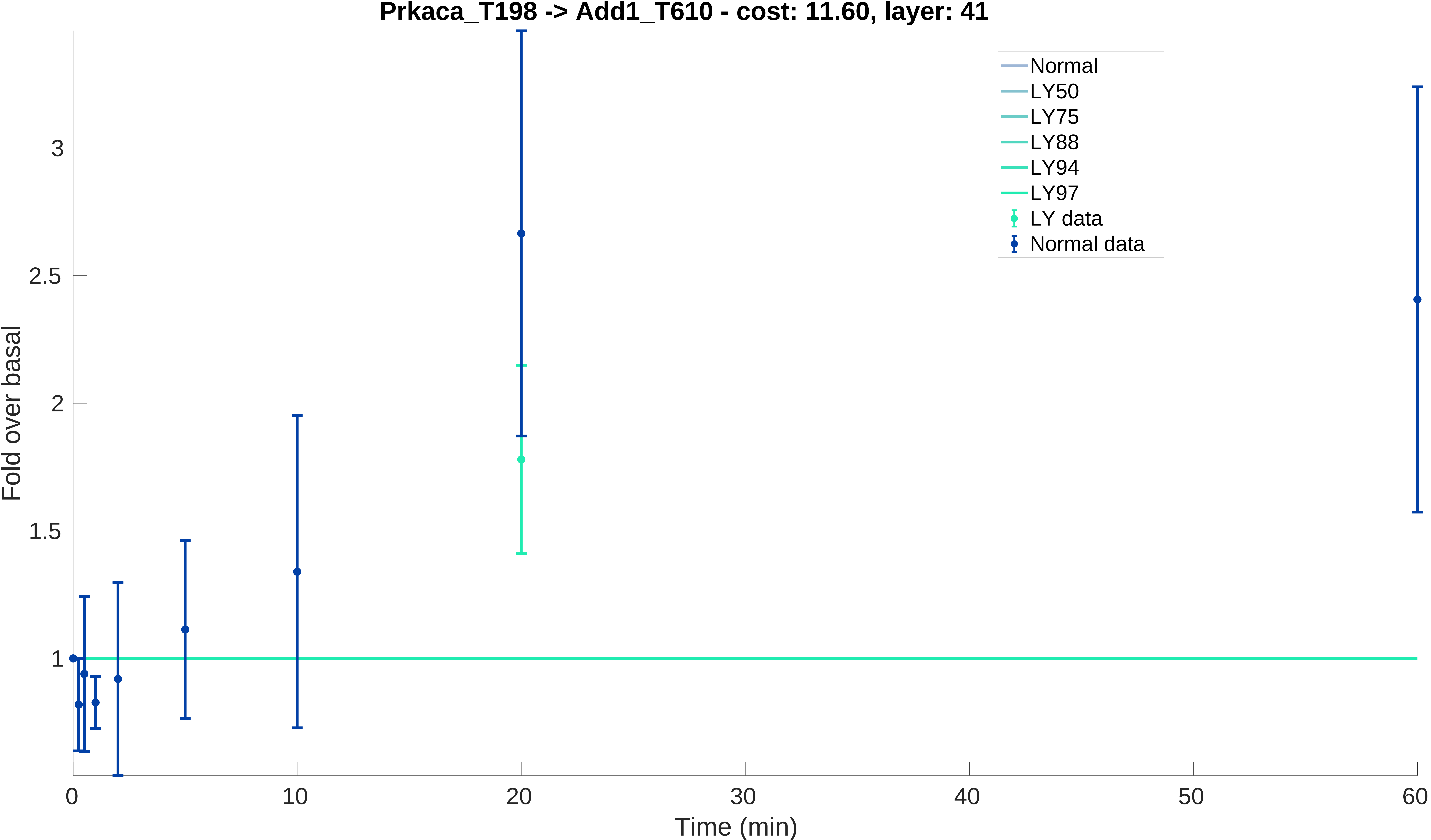
Prkaca_T198 -> Add1_T610 - cost: 11.60, layer: 41
Normal
LY50
LY75
LY88
LY94
LY97
LY data
Normal data
Fold over basal
Time (min)
0
10
20
30
40
50
60
1
1.5
2
2.5
3

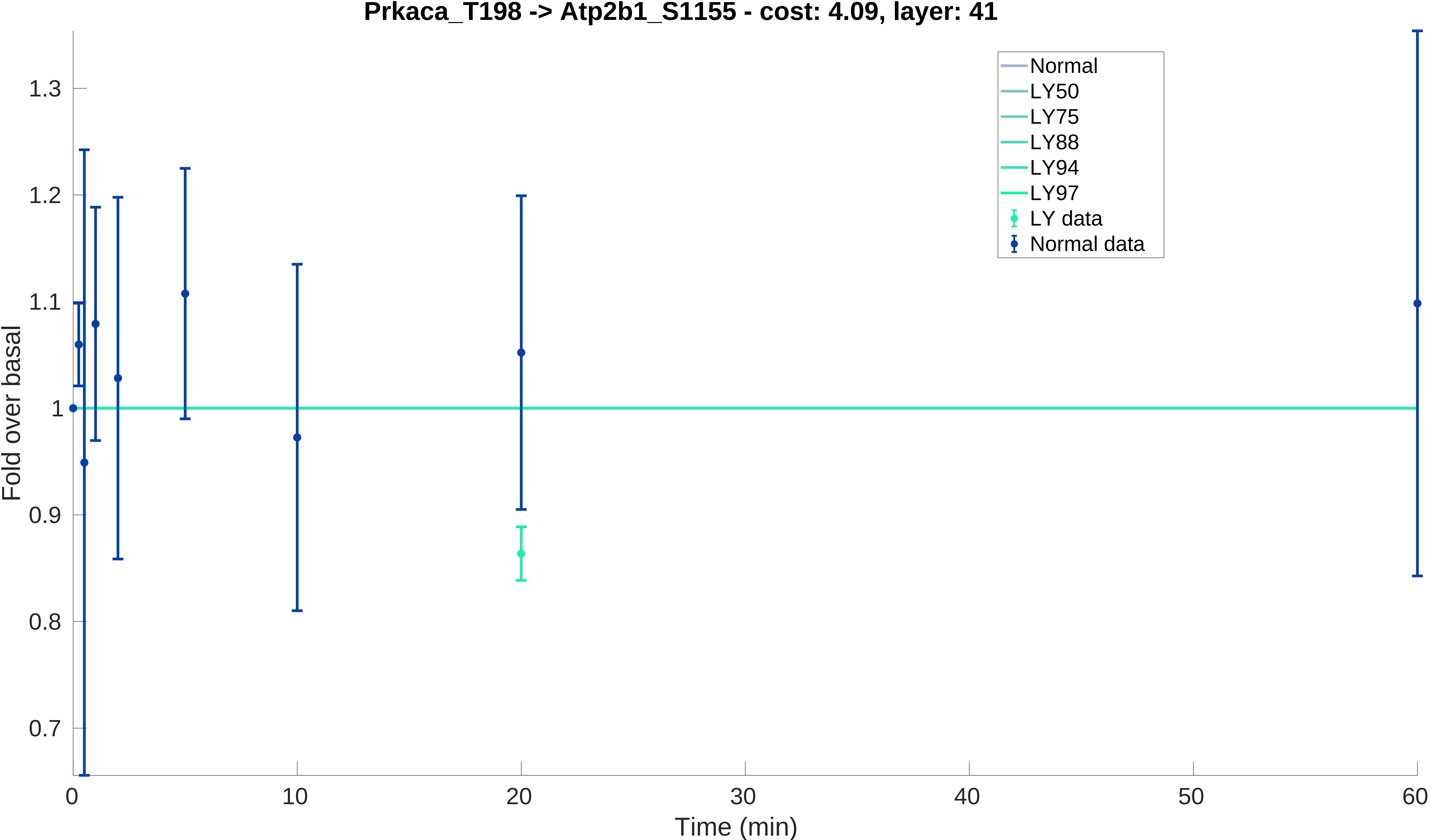
Prkaca_T198 -> Atp2b1_S1155 - cost: 4.09, layer: 41
Fold over basal
Time (min)
Normal
LY50
LY75
LY88
LY94
LY97
LY data
Normal data
1.3
1.2
1.1
1
0.9
0.8
0.7
0
10
20
30
40
50
60

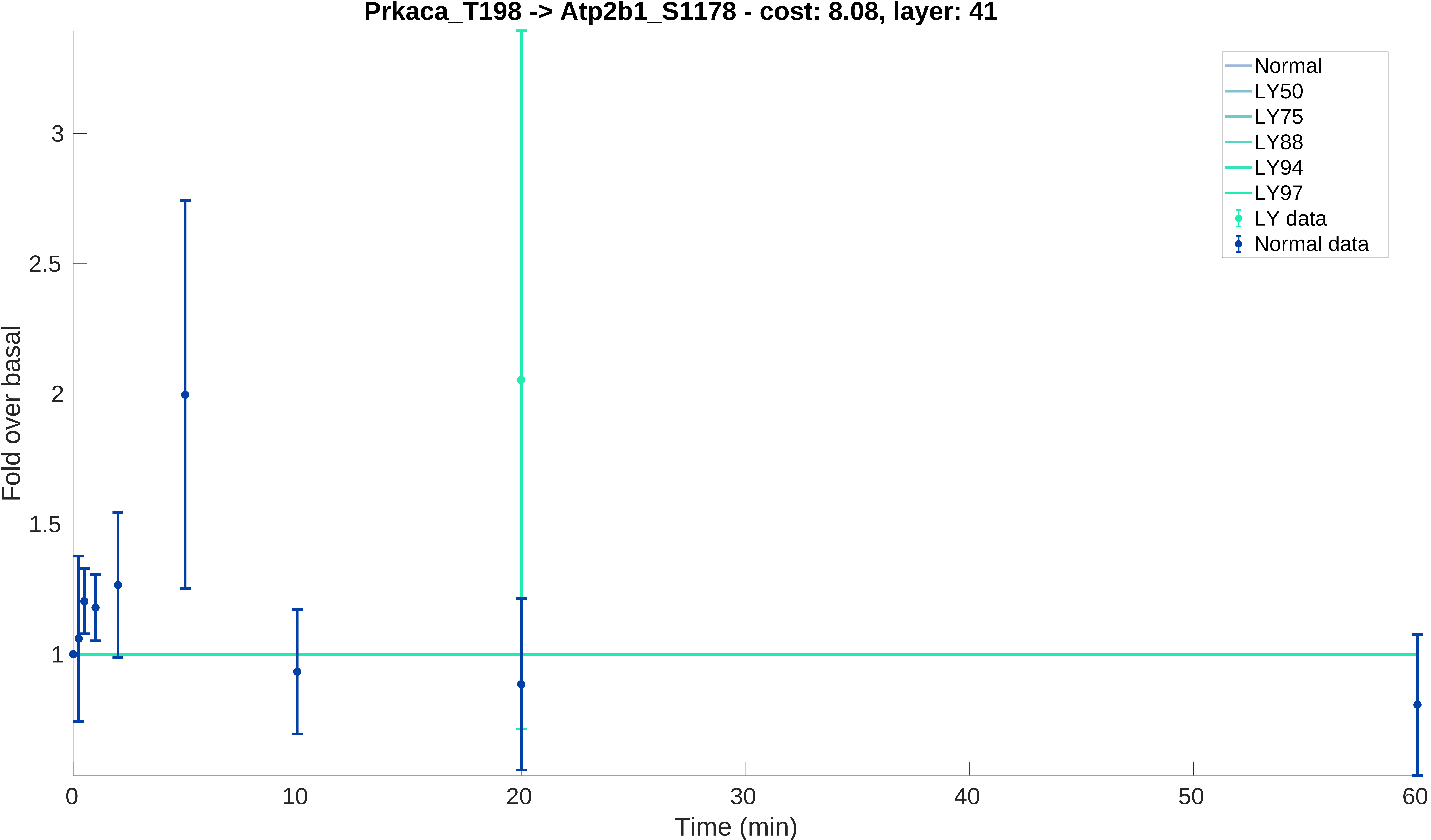
Prkaca_T198 -> Atp2b1_S1178 - cost: 8.08, layer: 41
Fold over basal
Time (min)
0
10
20
30
40
50
60
1
1.5
2
2.5
3
Normal
LY50
LY75
LY88
LY94
LY97
LY data
Normal data

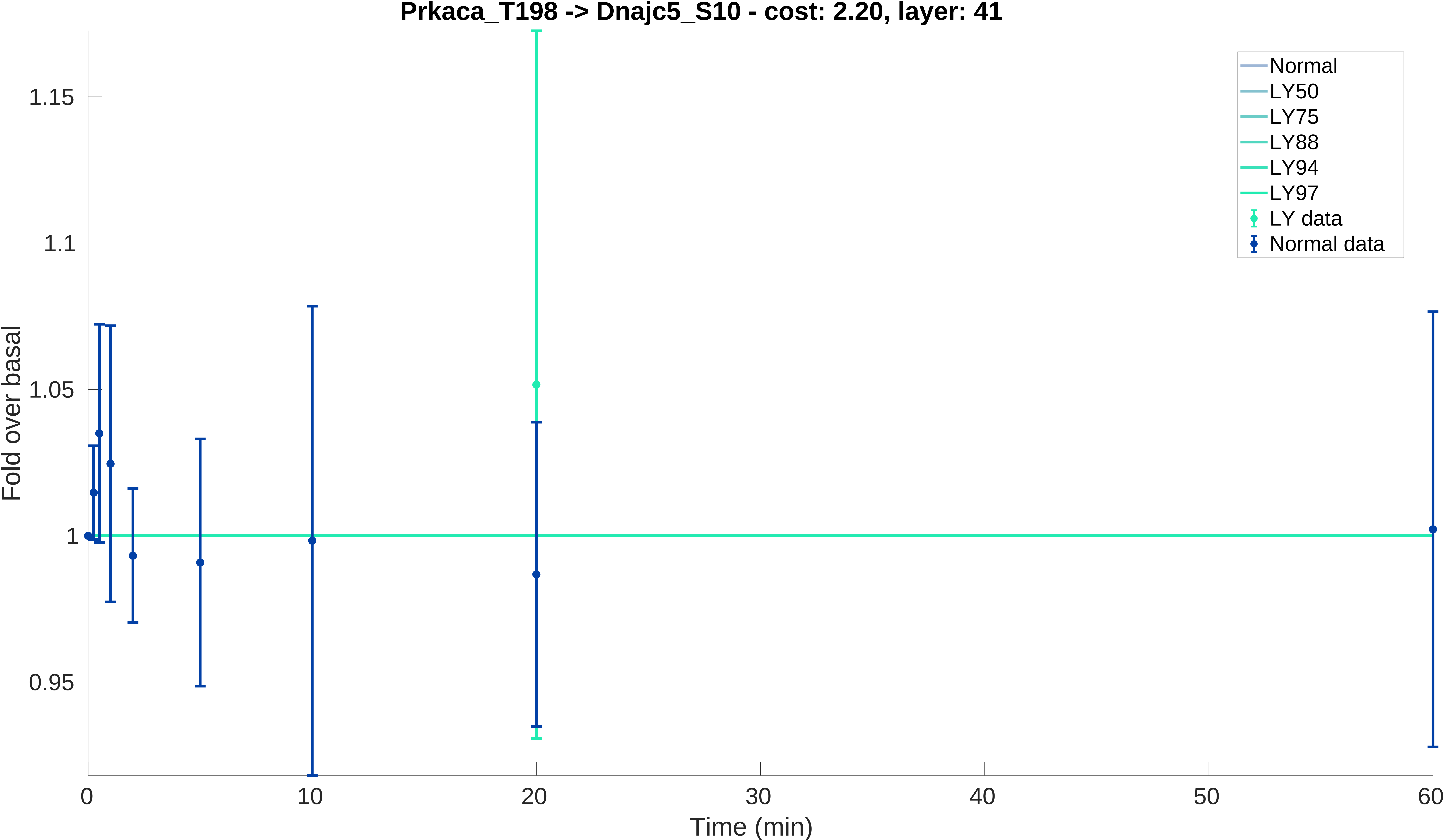

Prkaca_T198 -> Dnajc5_S10 - cost: 2.20, layer: 41
Fold over basal
1.15
1.1
1.05
1
0.95
0
10
20
30
40
50
60
Time (min)
Normal
LY50
LY75
LY88
LY94
LY97
LY data
Normal data

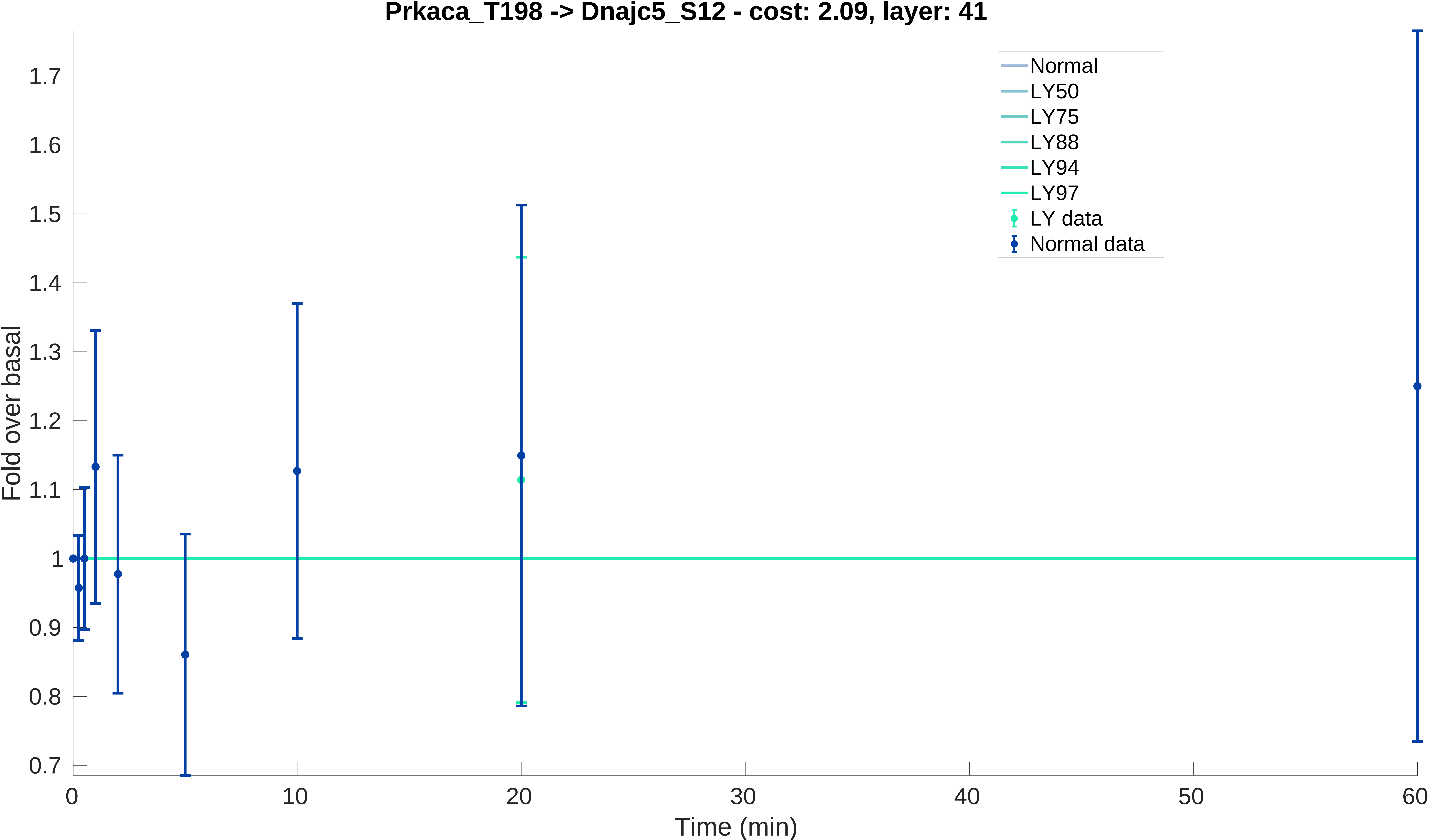

Prkaca_T198 -> Dnajc5_S12 - cost: 2.09, layer: 41
Normal
LY50
LY75
LY88
LY94
LY97
LY data
Normal data
Fold over basal
1.7
1.6
1.5
1.4
1.3
1.2
1.1
1
0.9
0.8
0.7
0
10
20
30
40
50
60
Time (min)

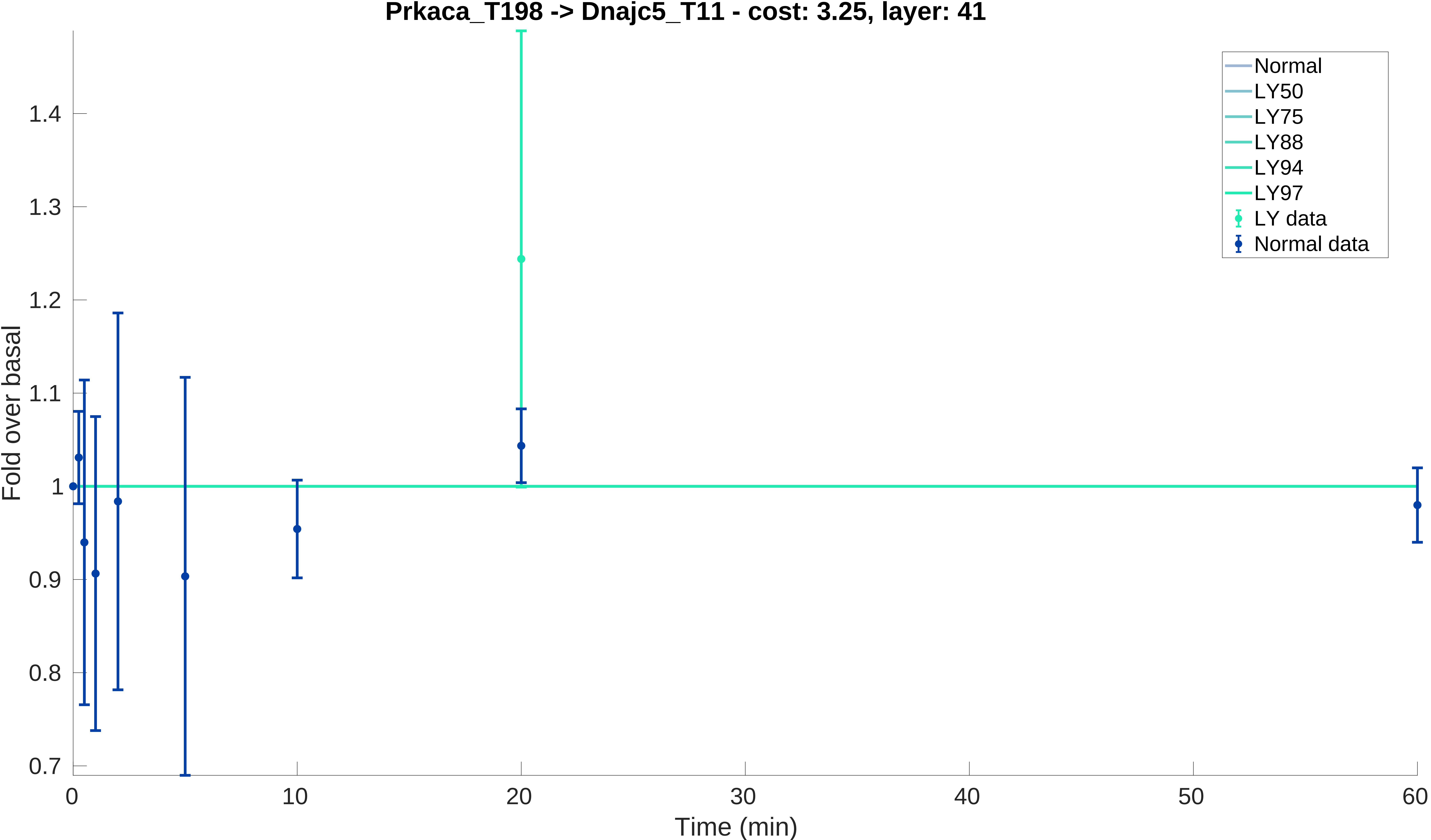
Prkaca_T198 -> Dnajc5_T11 - cost: 3.25, layer: 41
Normal
LY50
LY75
LY88
LY94
LY97
LY data
Normal data
Fold over basal
1.4
1.3
1.2
1.1
1
0.9
0.8
0.7
0
10
20
30
40
50
60
Time (min)

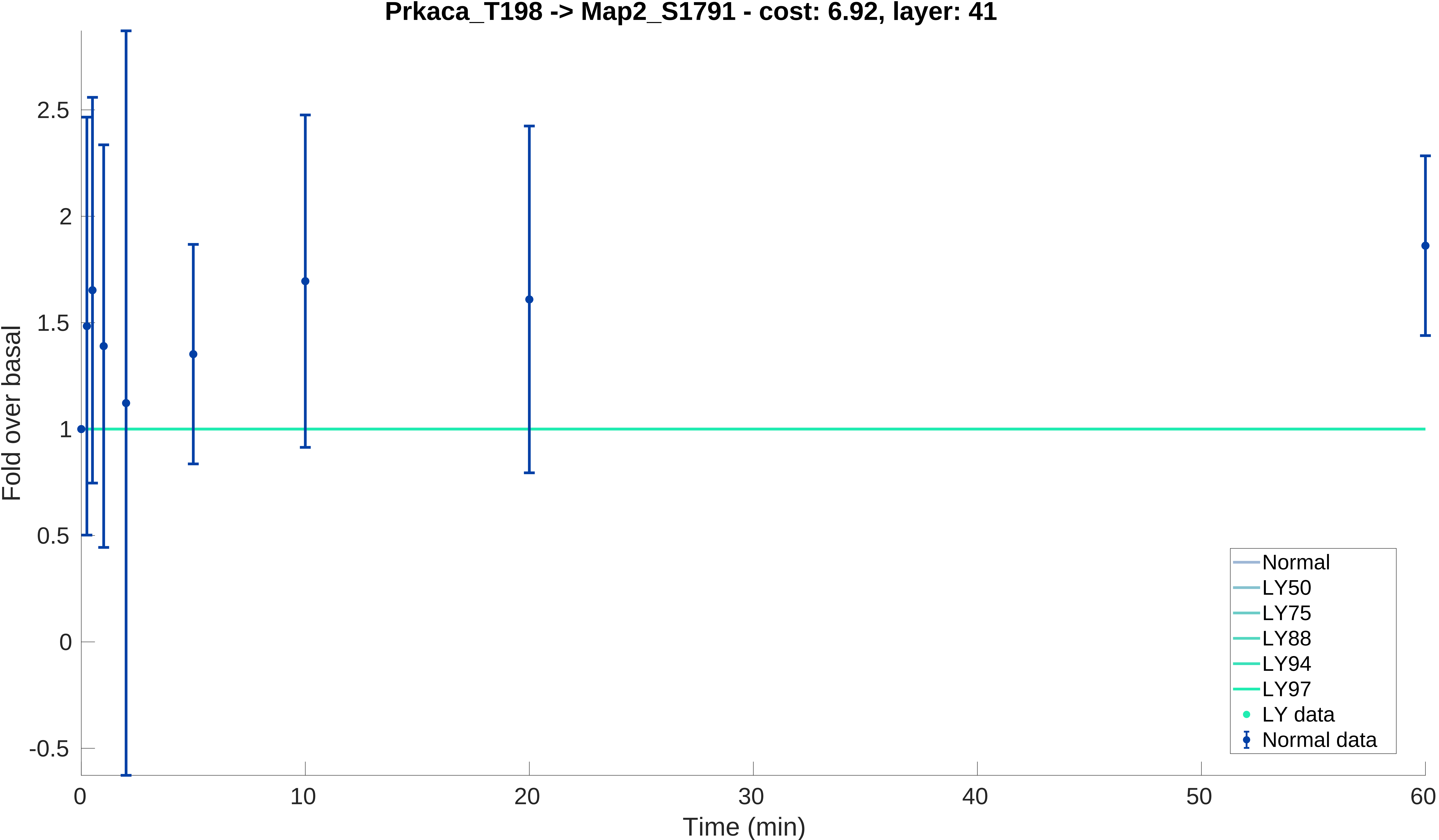

Prkaca_T198 -> Map2_S1791 - cost: 6.92, layer: 41
Fold over basal
Time (min)
2.5
2
1.5
1
0.5
0
-0.5
0
10
20
30
40
50
60
Normal
LY50
LY75
LY88
LY94
LY97
LY data
Normal data

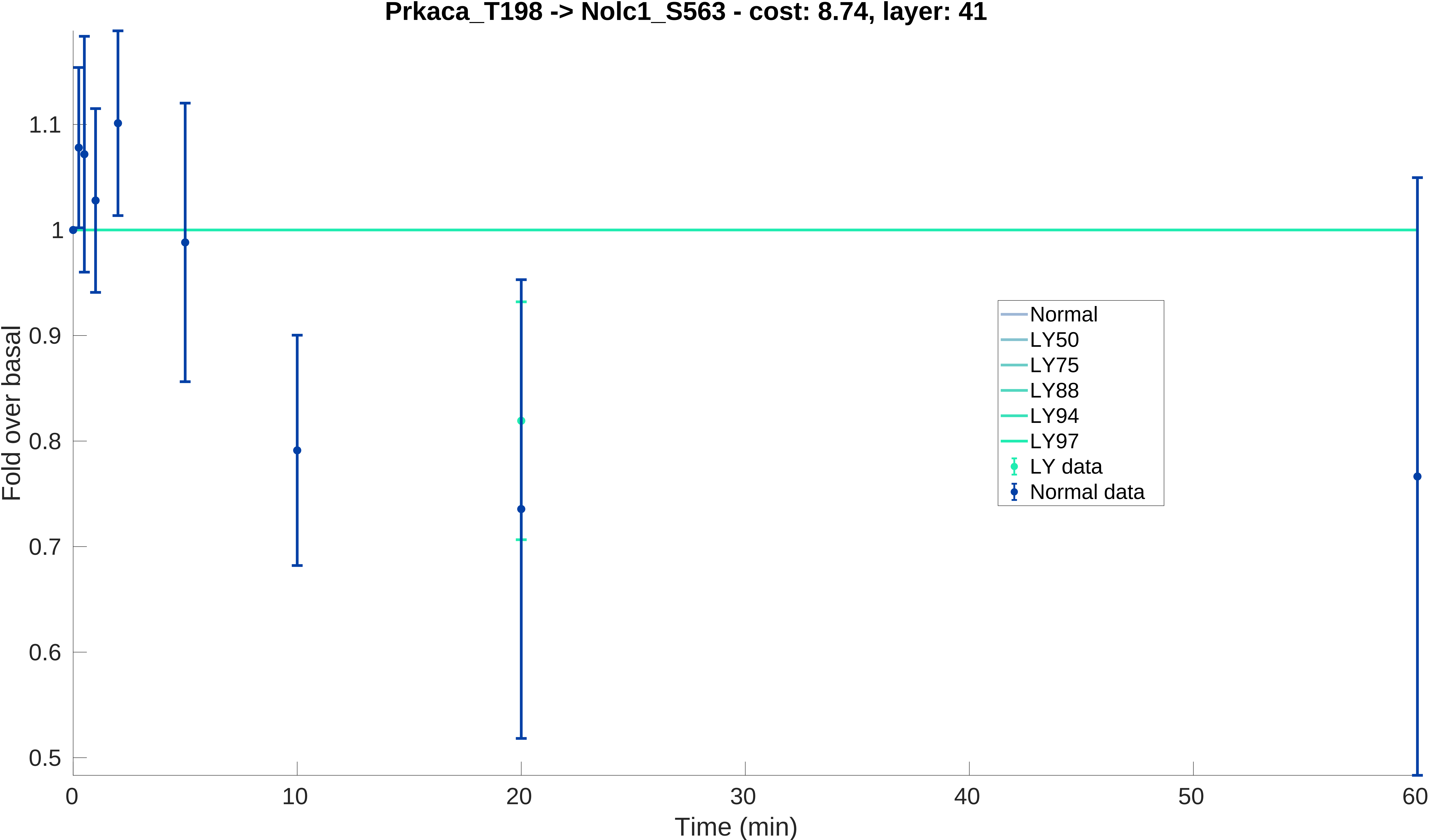

Prkaca_T198 -> Nolc1_S563 - cost: 8.74, layer: 41
Fold over basal
1.1
1
0.9
0.8
0.7
0.6
0.5
0
10
20
30
40
50
60
Time (min)
Normal
LY50
LY75
LY88
LY94
LY97
LY data
Normal data

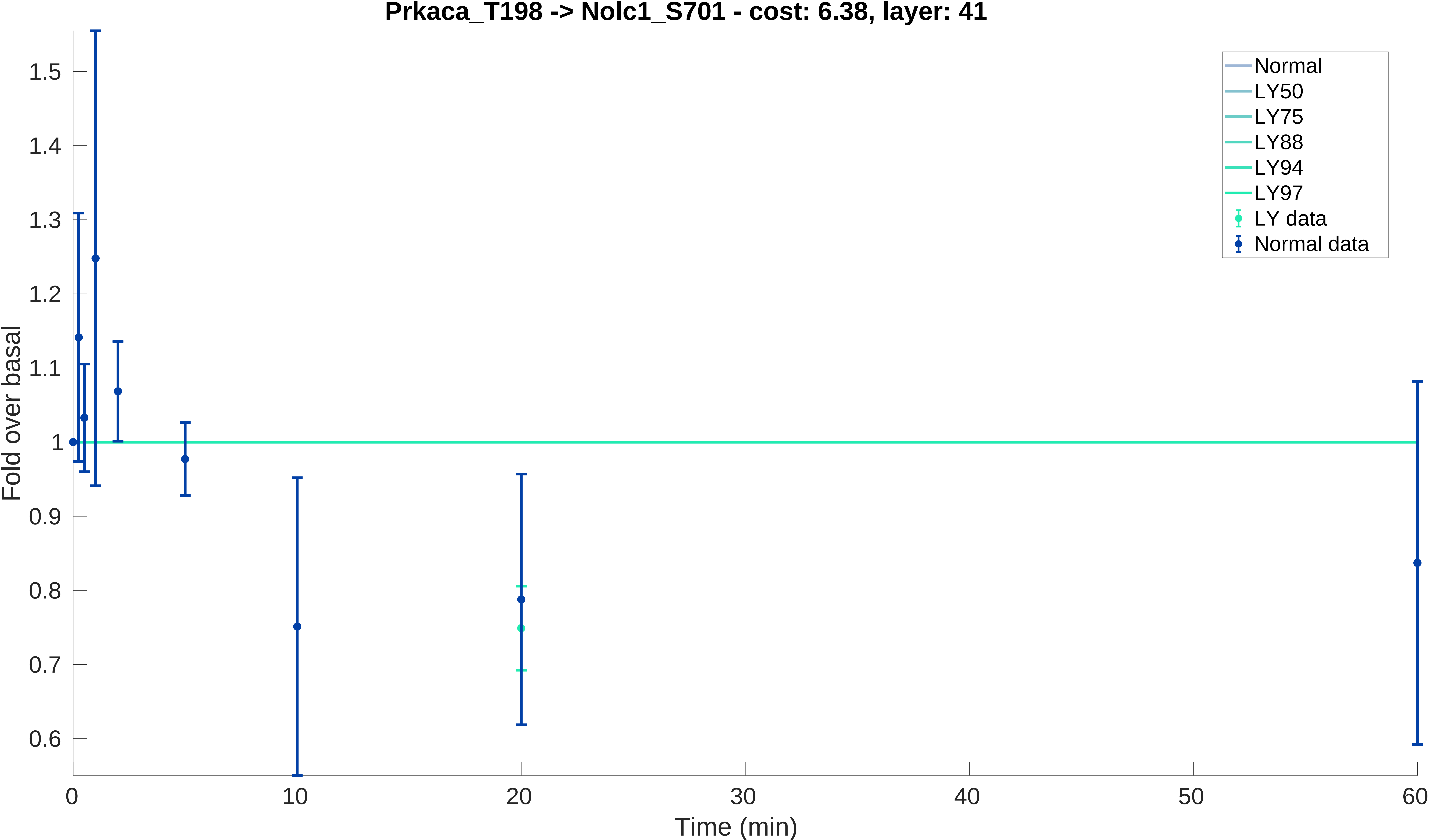

Prkaca_T198 -> Nolc1_S701 - cost: 6.38, layer: 41
Fold over basal
Time (min)
1.5
1.4
1.3
1.2
1.1
1
0.9
0.8
0.7
0.6
0
10
20
30
40
50
60
Normal
LY50
LY75
LY88
LY94
LY97
LY data
Normal data

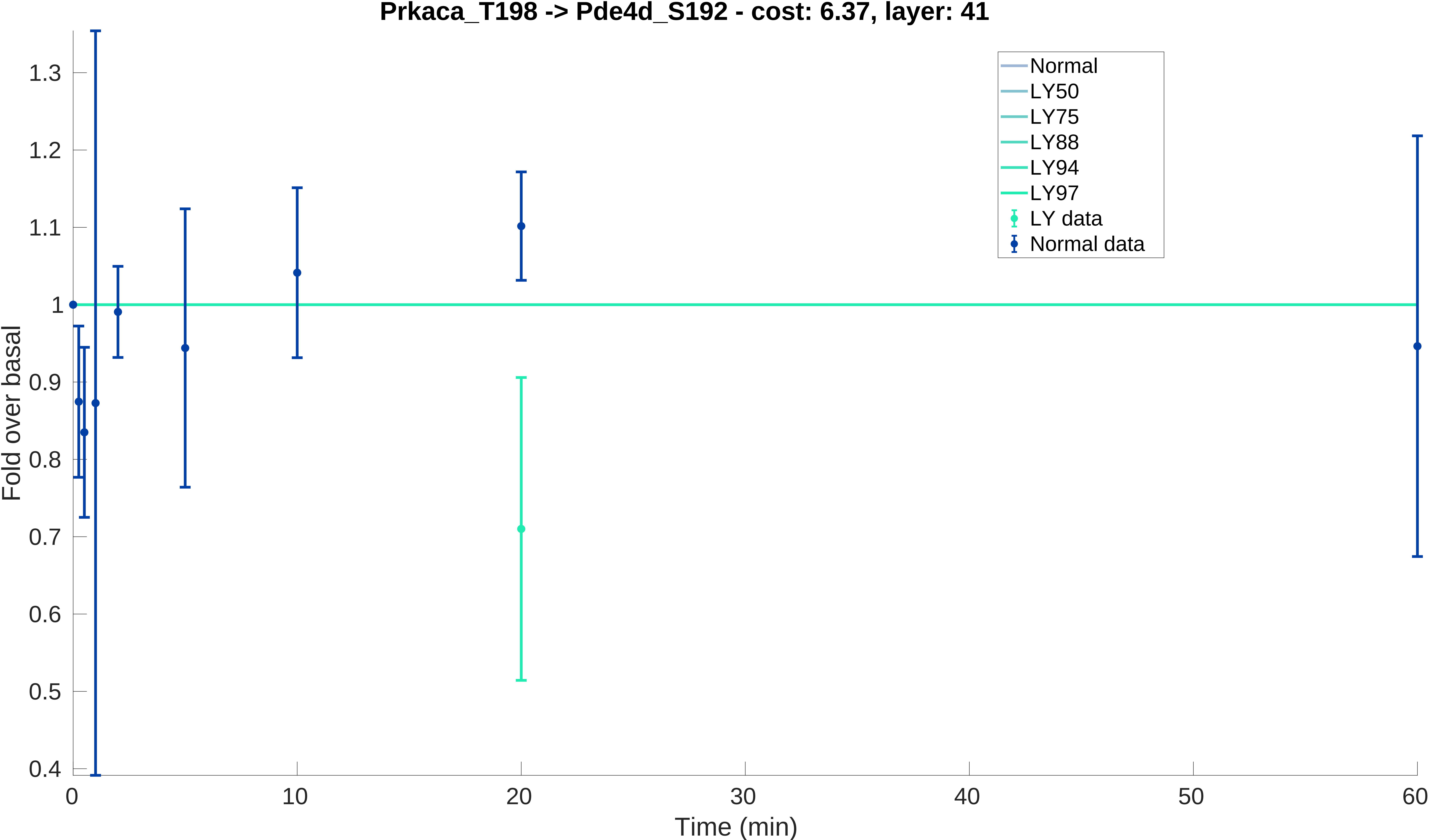
Prkaca_T198 -> Pde4d_S192 - cost: 6.37, layer: 41
Normal
LY50
LY75
LY88
LY94
LY97
LY data
Normal data
Fold over basal
1.3
1.2
1.1
1
0.9
0.8
0.7
0.6
0.5
0.4
0
10
20
30
40
50
60
Time (min)

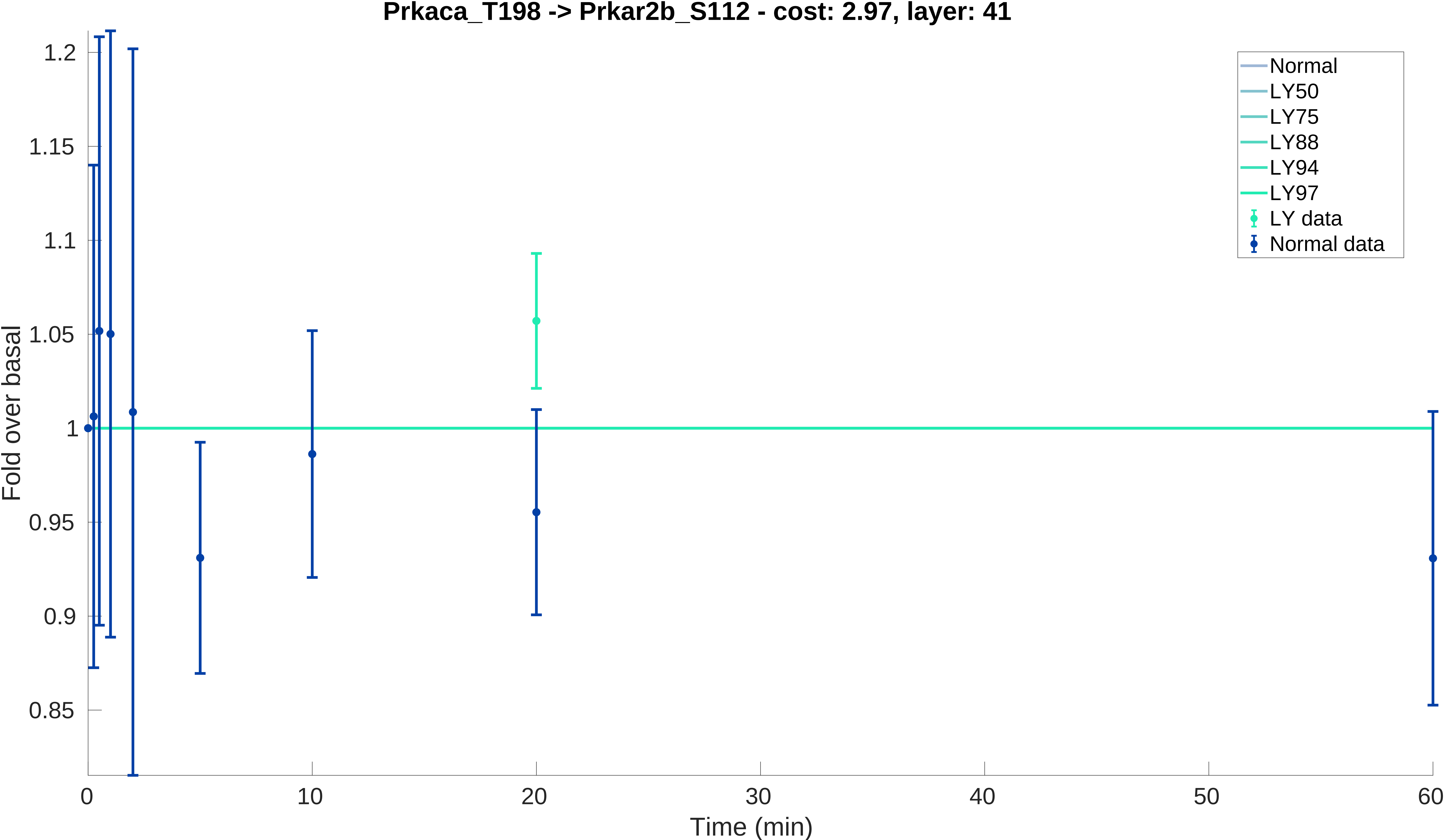

Prkaca_T198 -> Prkar2b_S112 - cost: 2.97, layer: 41
Fold over basal
Time (min)
1.2
1.15
1.1
1.05
1
0.95
0.9
0.85
0
10
20
30
40
50
60
Normal
LY50
LY75
LY88
LY94
LY97
LY data
Normal data

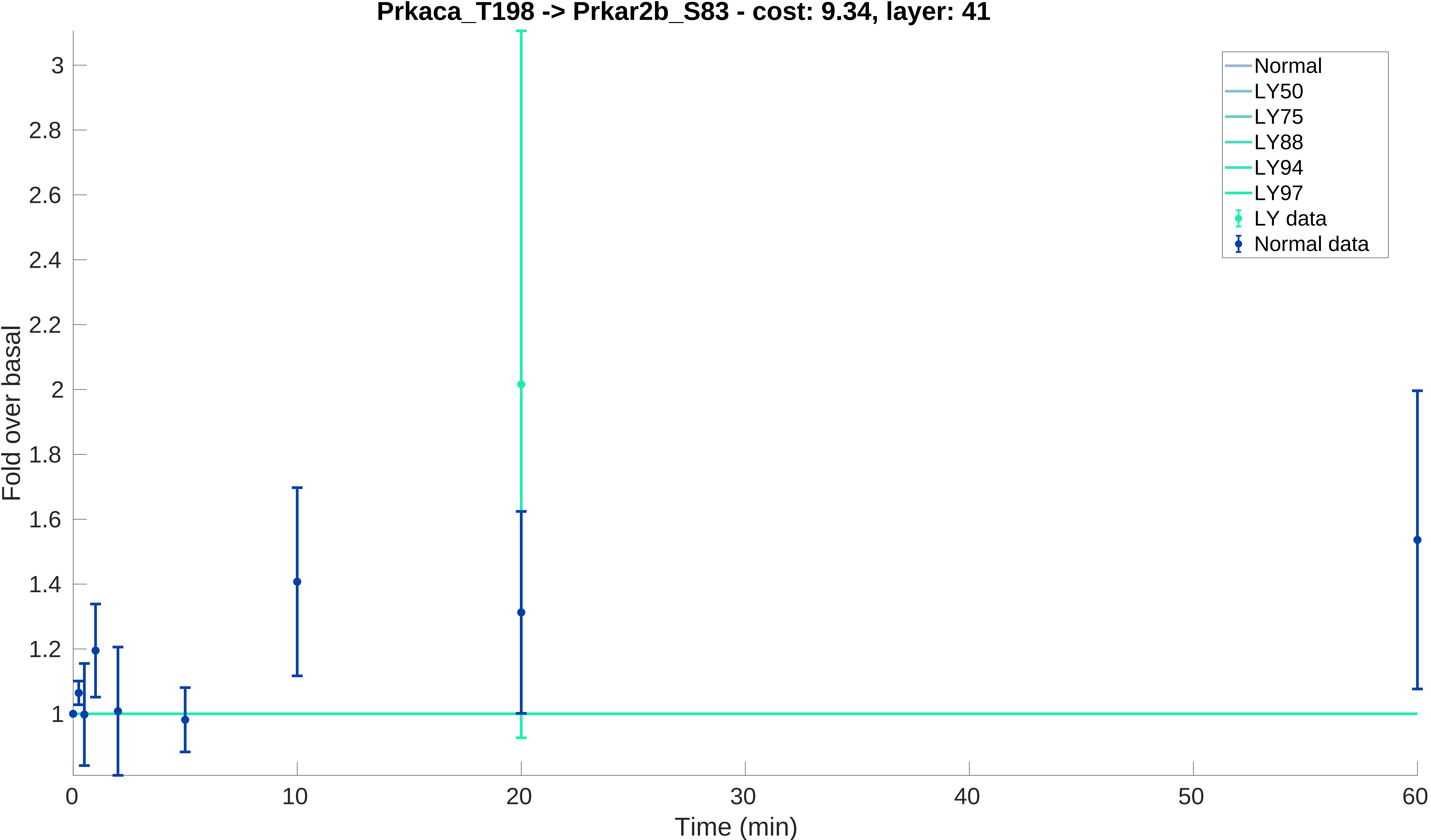

Prkaca_T198 -> Prkar2b_S83 - cost: 9.34, layer: 41
Fold over basal
Time (min)
Normal
LY50
LY75
LY88
LY94
LY97
LY data
Normal data
3
2.8
2.6
2.4
2.2
2
1.8
1.6
1.4
1.2
1
0
10
20
30
40
50
60

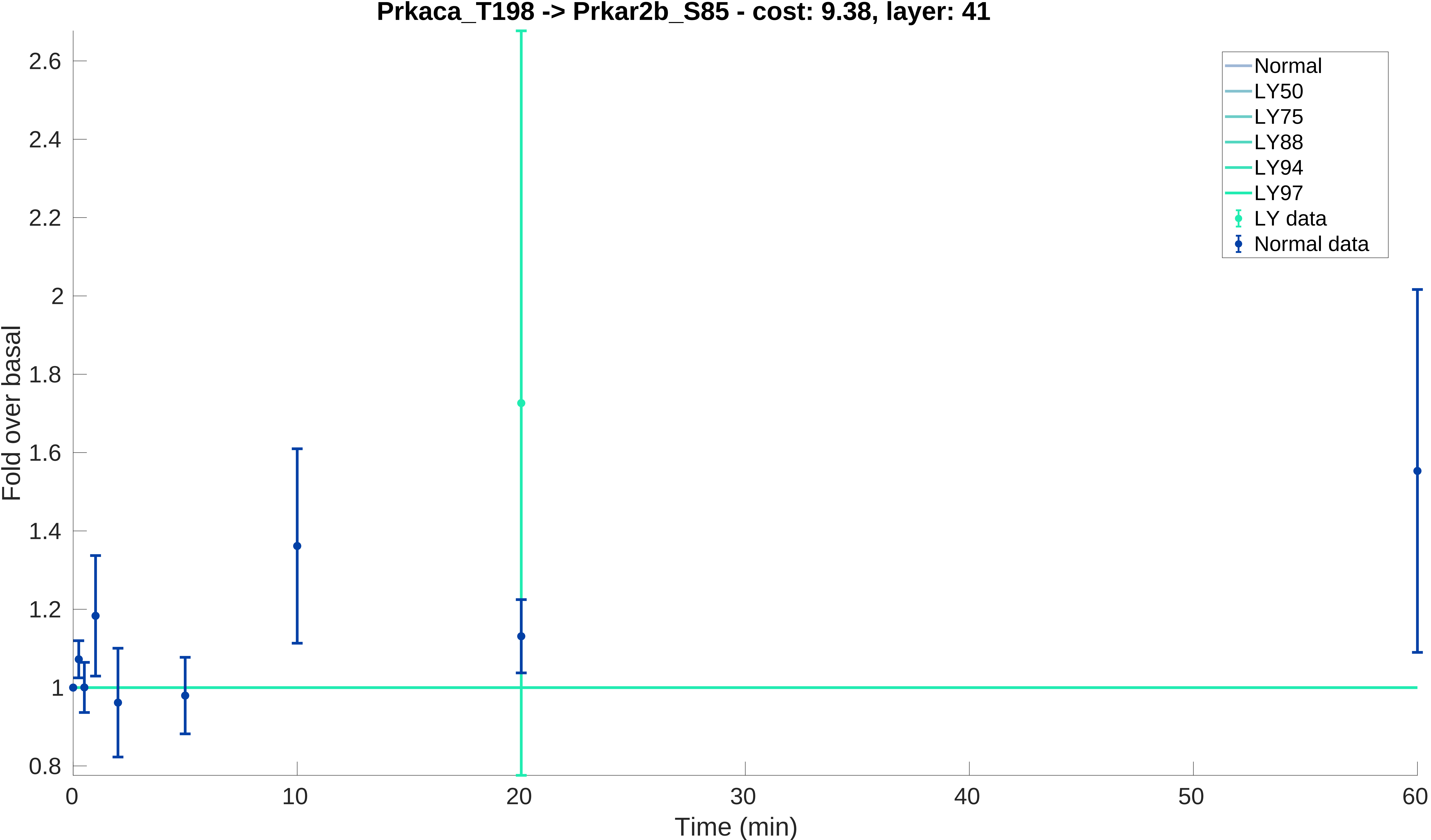
Prkaca_T198 -> Prkar2b_S85 - cost: 9.38, layer: 41
Fold over basal
Time (min)
2.6
2.4
2.2
2
1.8
1.6
1.4
1.2
1
0.8
0
10
20
30
40
50
60
Normal
LY50
LY75
LY88
LY94
LY97
LY data
Normal data

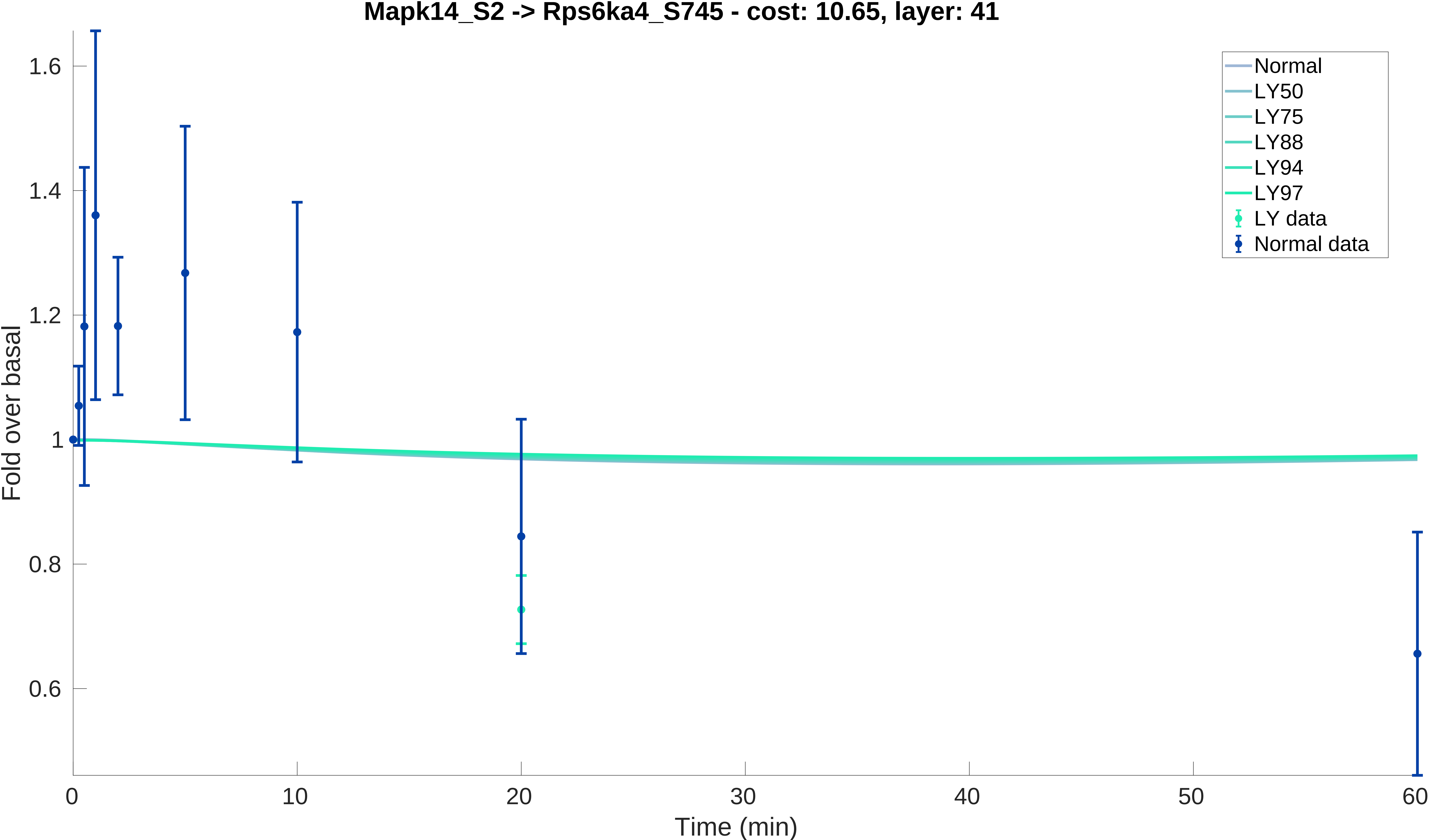
Mapk14_S2 -> Rps6ka4_S745 - cost: 10.65, layer: 41
Fold over basal
1.6
1.4
1.2
1
0.8
0.6
0
10
20
30
40
50
60
Time (min)
Normal
LY50
LY75
LY88
LY94
LY97
LY data
Normal data

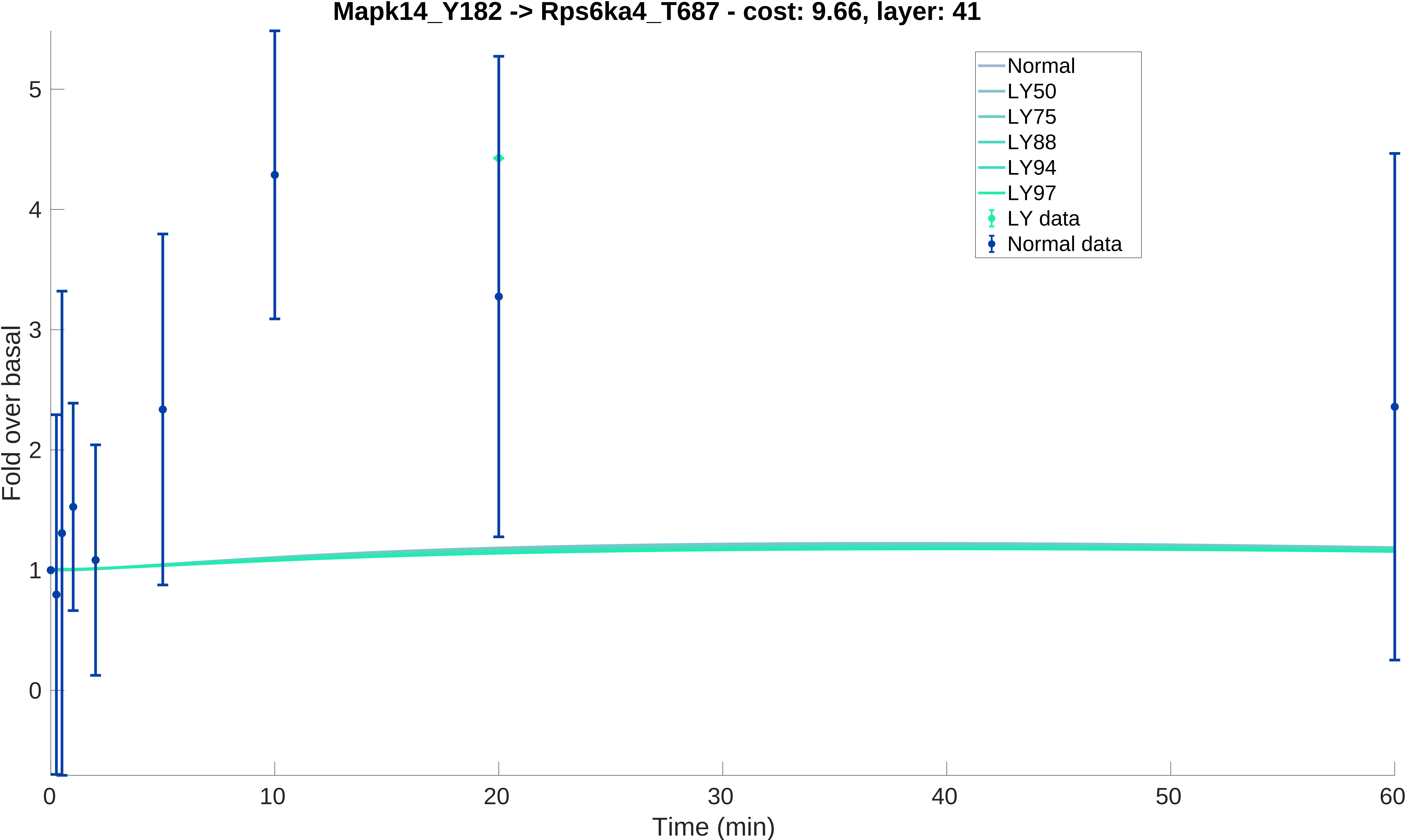

Mapk14_Y182 -> Rps6ka4_T687 - cost: 9.66, layer: 41
Fold over basal
Time (min)
0
1
2
3
4
5
0
10
20
30
40
50
60
Normal
LY50
LY75
LY88
LY94
LY97
LY data
Normal data

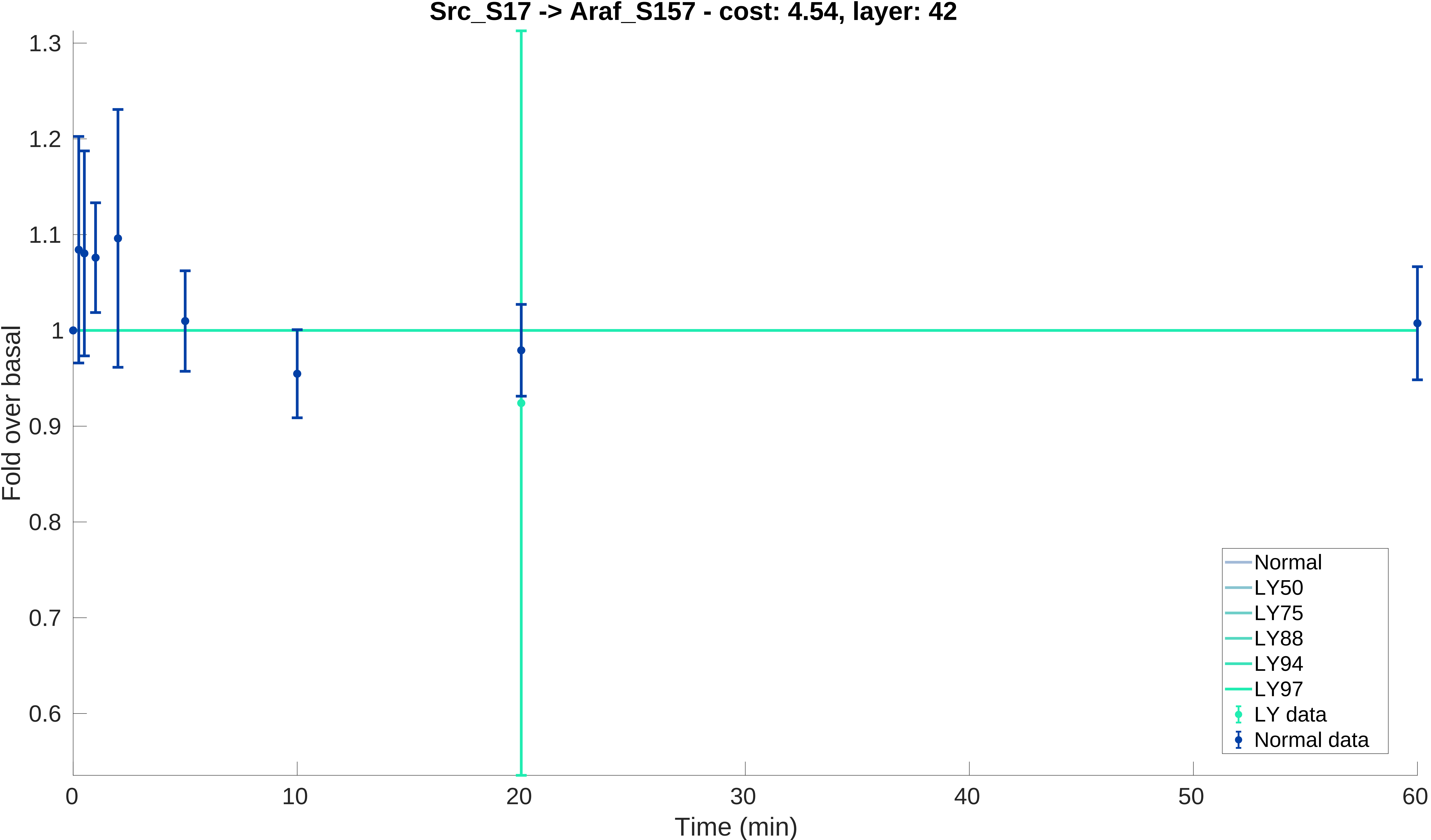

Src_S17 -> Araf_S157 - cost: 4.54, layer: 42
Fold over basal
Time (min)
1.3
1.2
1.1
1
0.9
0.8
0.7
0.6
0
10
20
30
40
50
60
Normal
LY50
LY75
LY88
LY94
LY97
LY data
Normal data

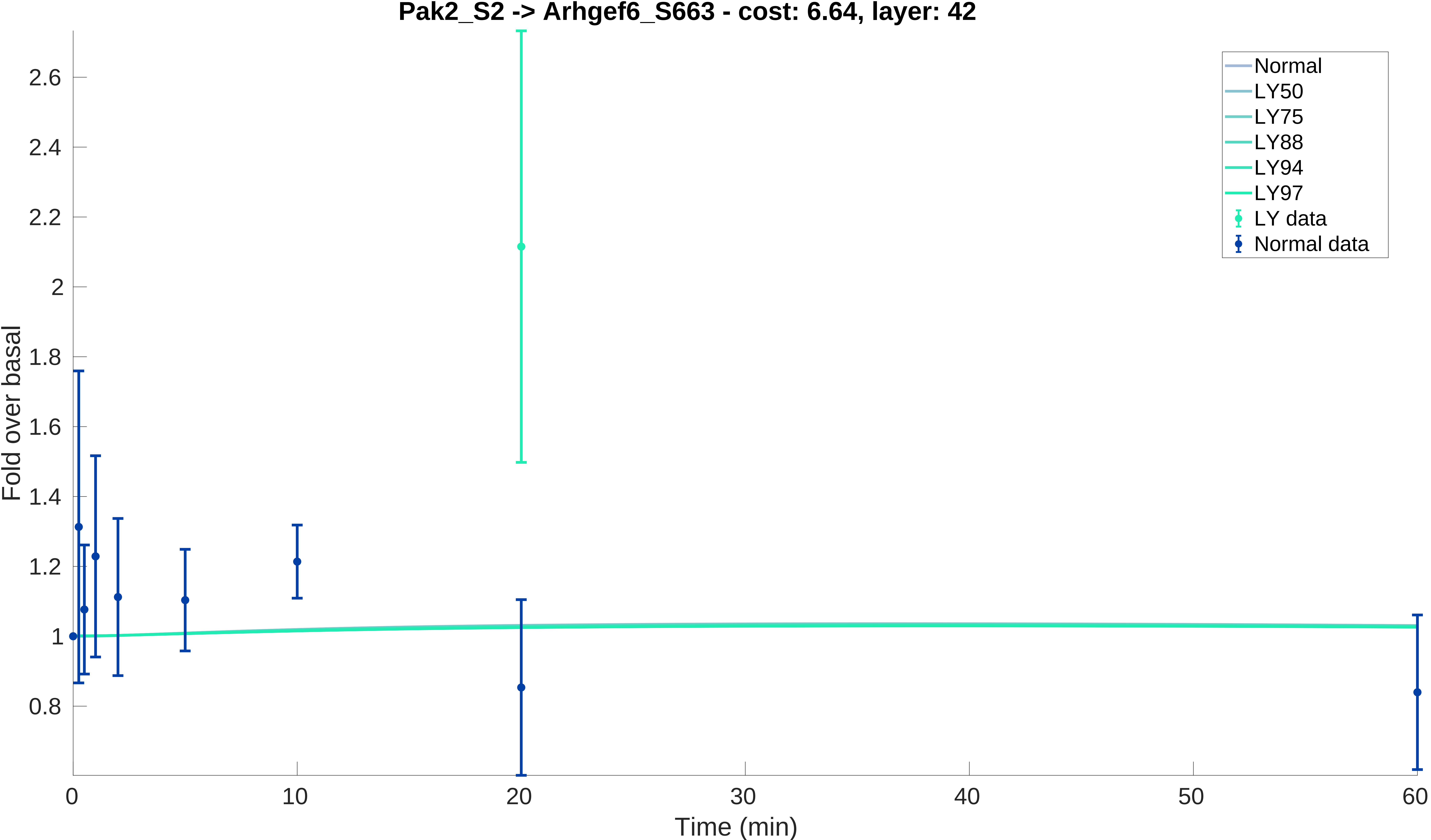

Pak2_S2 -> Arhgef6_S663 - cost: 6.64, layer: 42
Normal
LY50
LY75
LY88
LY94
LY97
LY data
Normal data
Fold over basal
2.6
2.4
2.2
2
1.8
1.6
1.4
1.2
1
0.8
0
10
20
30
40
50
60
Time (min)

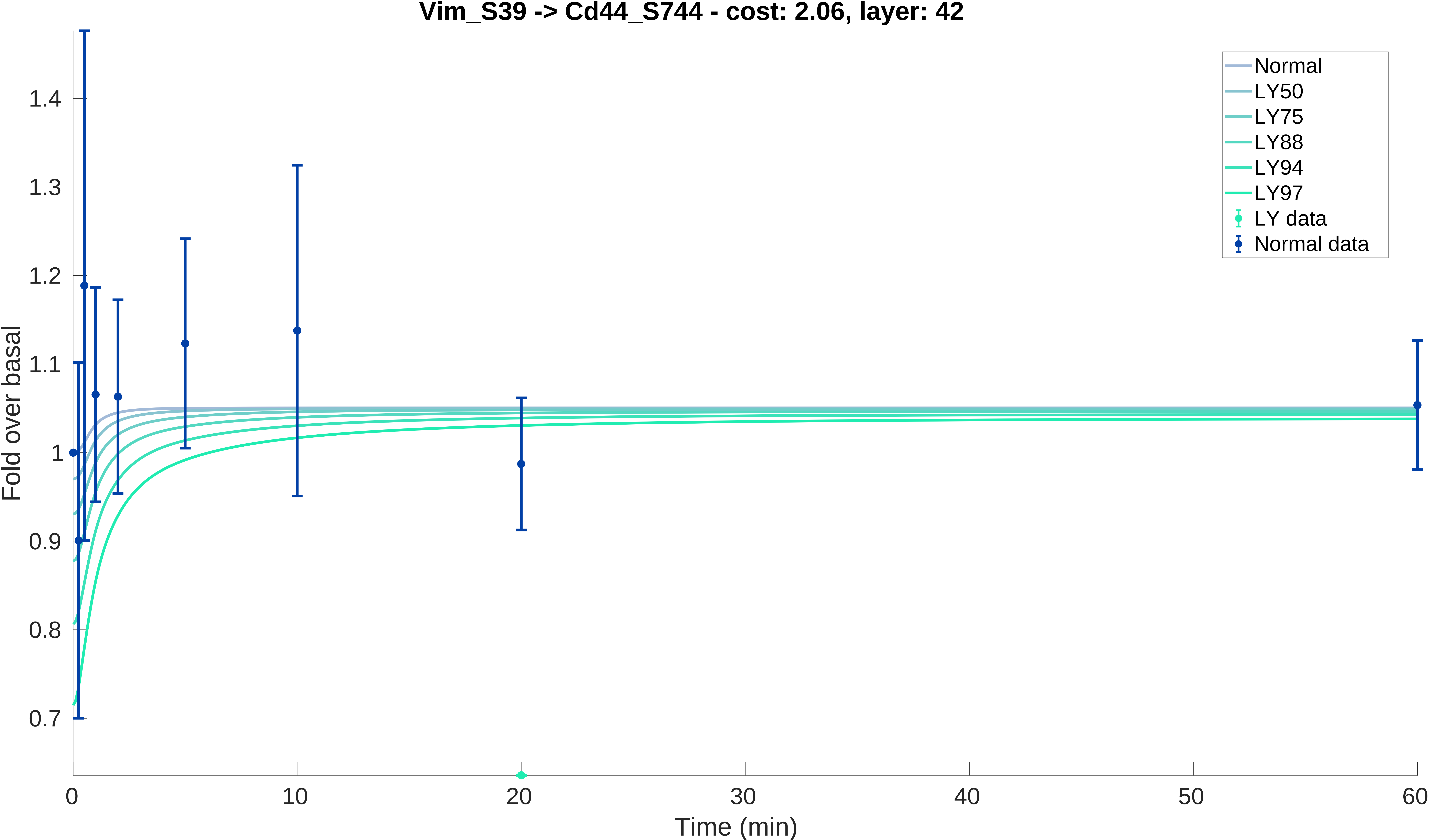

Vim_S39 -> Cd44_S744 - cost: 2.06, layer: 42
Normal
LY50
LY75
LY88
LY94
LY97
LY data
Normal data
Fold over basal
Time (min)
0.7
0.8
0.9
1
1.1
1.2
1.3
1.4
0
10
20
30
40
50
60

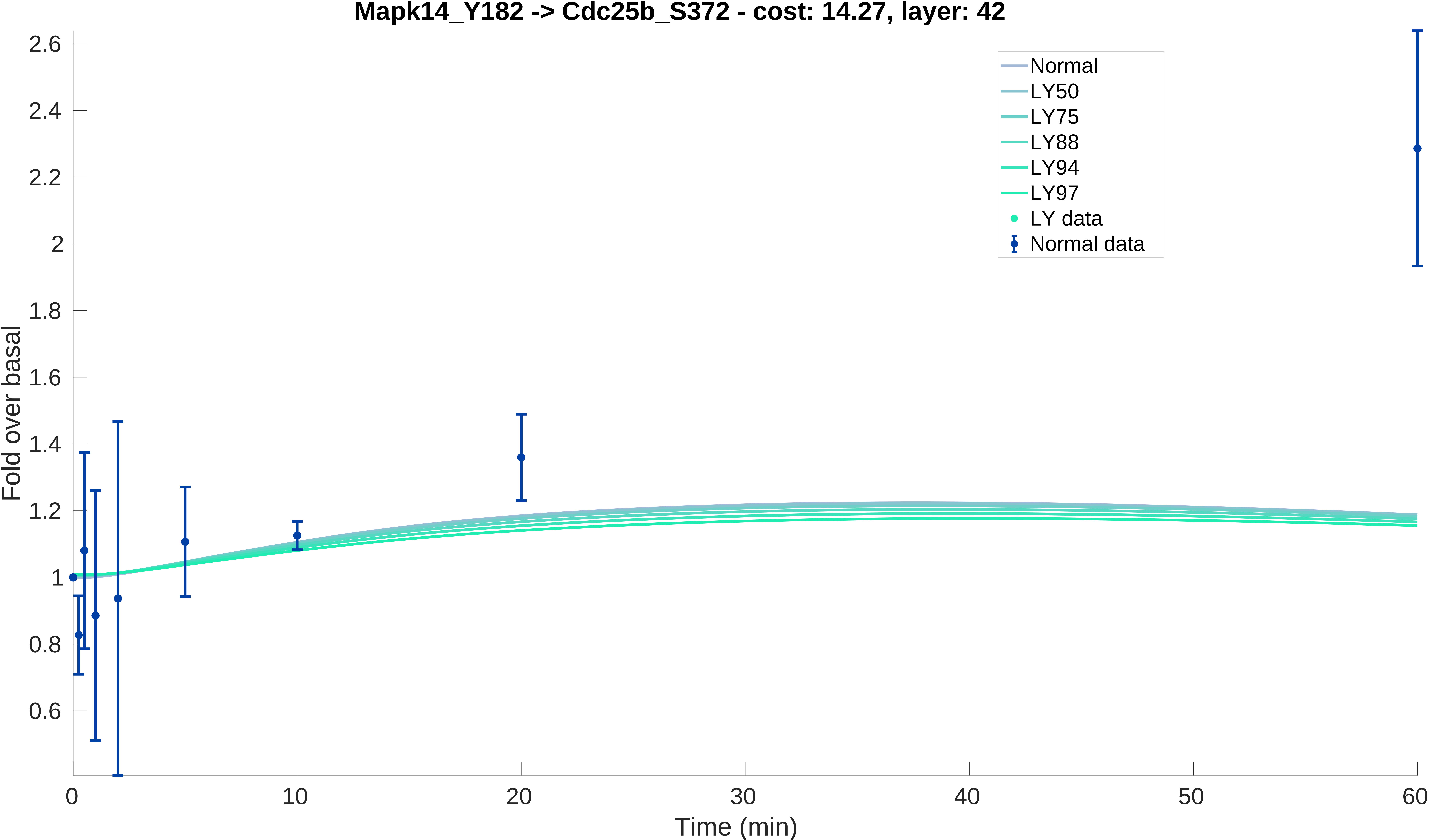

Mapk14_Y182 -> Cdc25b_S372 - cost: 14.27, layer: 42
Fold over basal
2.6
2.4
2.2
2
1.8
1.6
1.4
1.2
1
0.8
0.6
0
10
20
30
40
50
60
Time (min)
Normal
LY50
LY75
LY88
LY94
LY97
LY data
Normal data

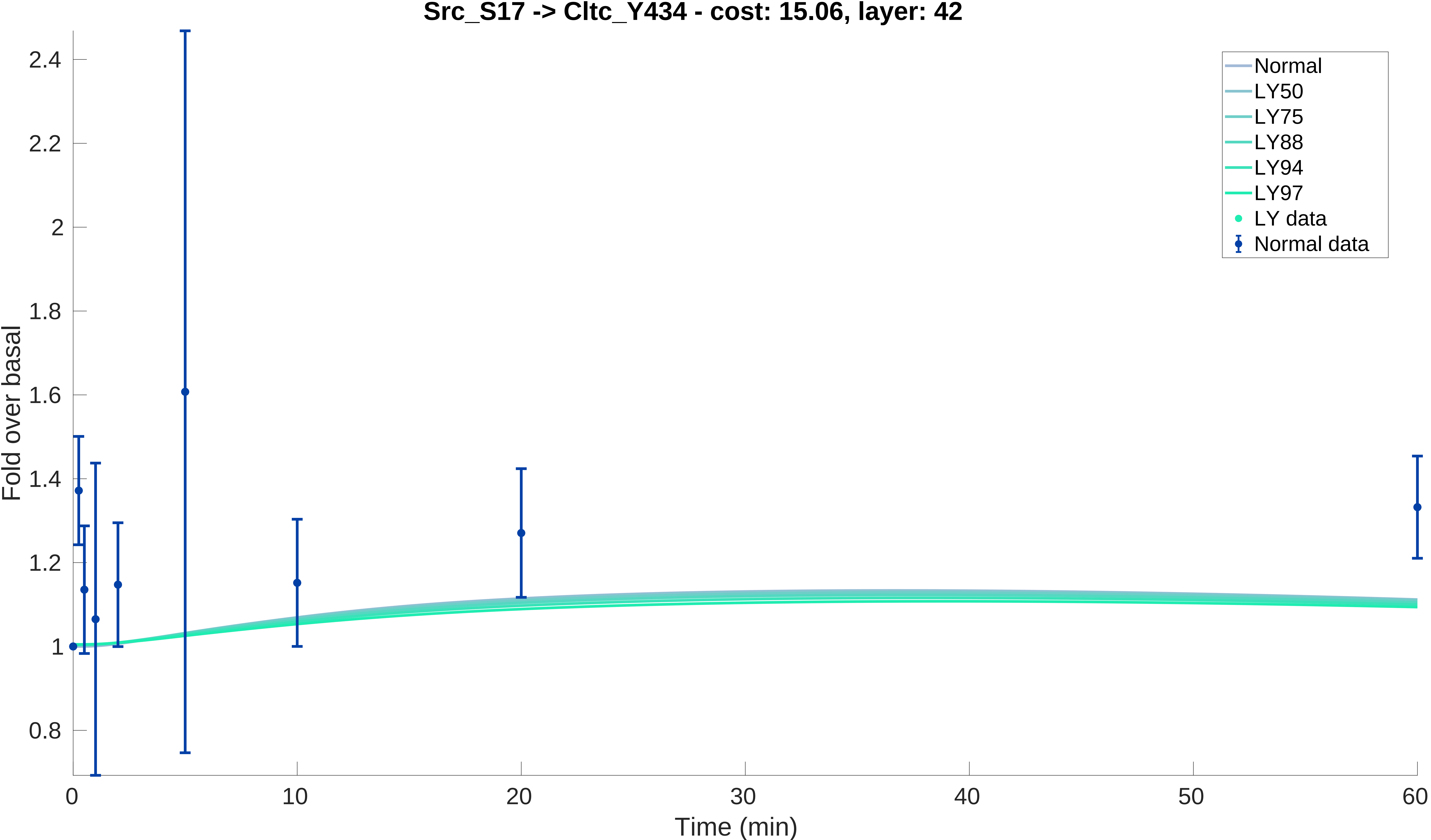

Src_S17 -> Cltc_Y434 - cost: 15.06, layer: 42
Fold over basal
Time (min)
Normal
LY50
LY75
LY88
LY94
LY97
LY data
Normal data
2.4
2.2
2
1.8
1.6
1.4
1.2
1
0.8
0
10
20
30
40
50
60

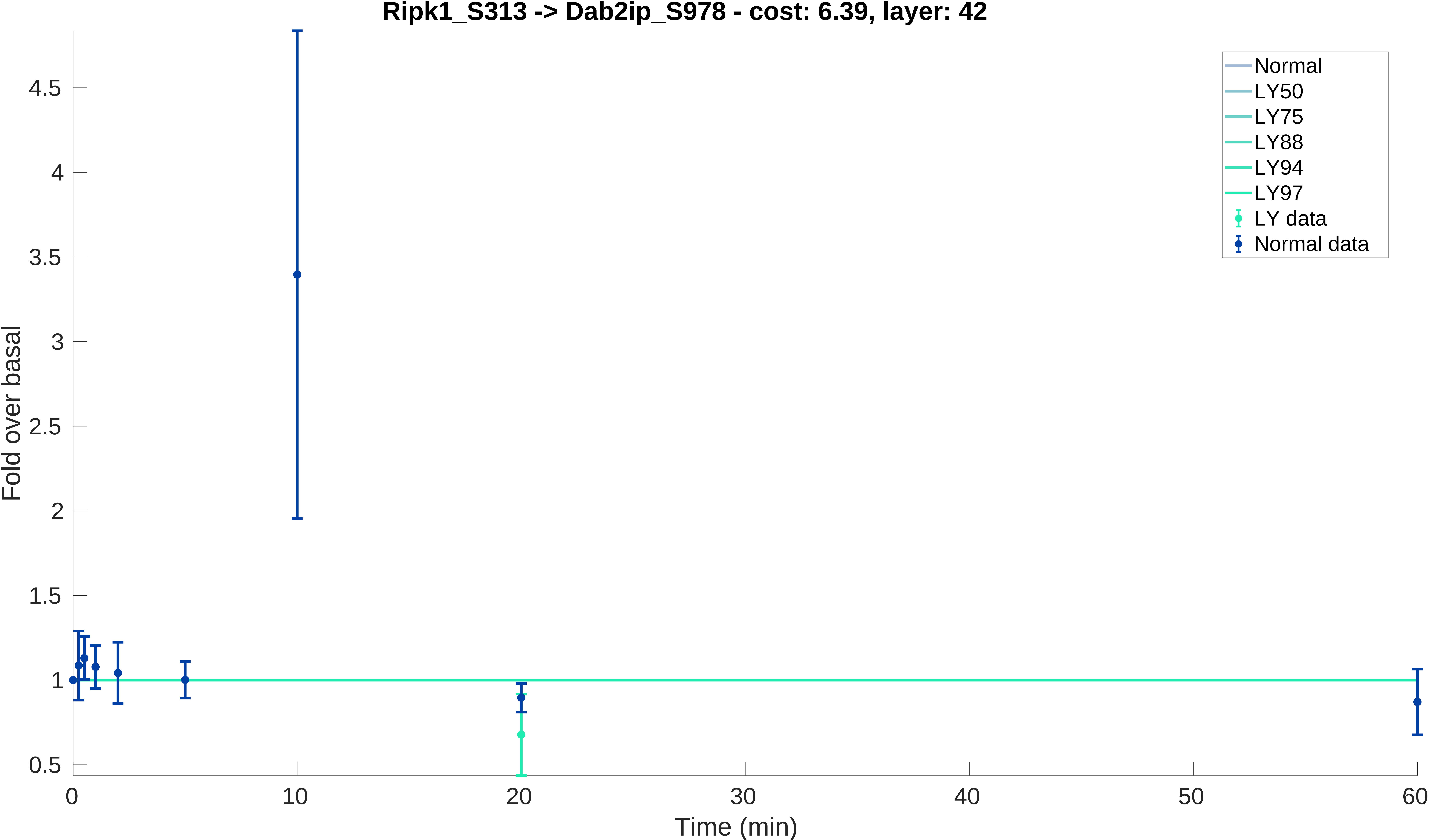

Ripk1_S313 -> Dab2ip_S978 - cost: 6.39, layer: 42
Fold over basal
4.5
4
3.5
3
2.5
2
1.5
1
0.5
0
10
20
30
40
50
60
Time (min)
Normal
LY50
LY75
LY88
LY94
LY97
LY data
Normal data

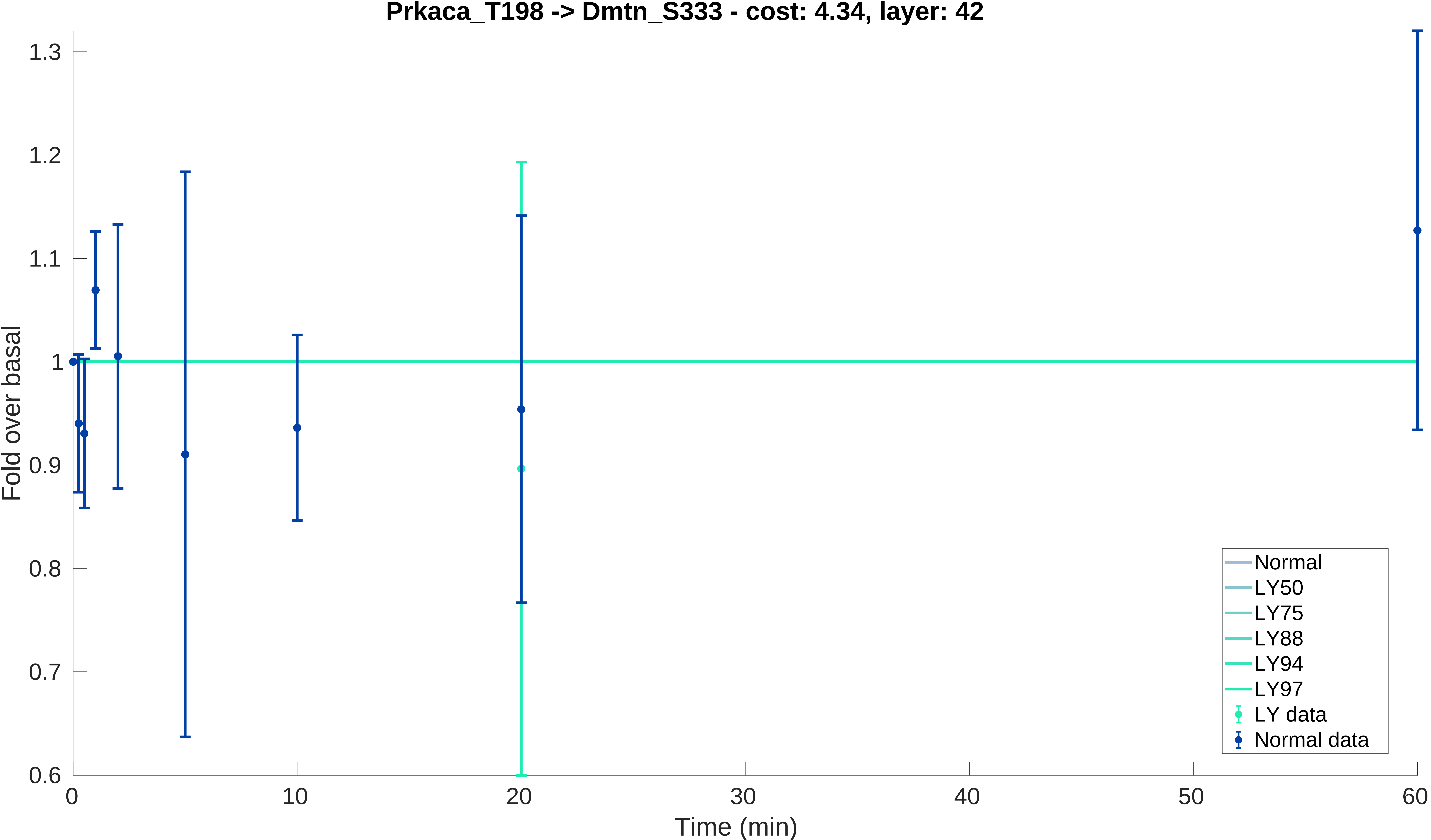

Prkaca_T198 -> Dmtn_S333 - cost: 4.34, layer: 42
Fold over basal
1.3
1.2
1.1
1
0.9
0.8
0.7
0.6
0
10
20
30
40
50
60
Time (min)
Normal
LY50
LY75
LY88
LY94
LY97
LY data
Normal data

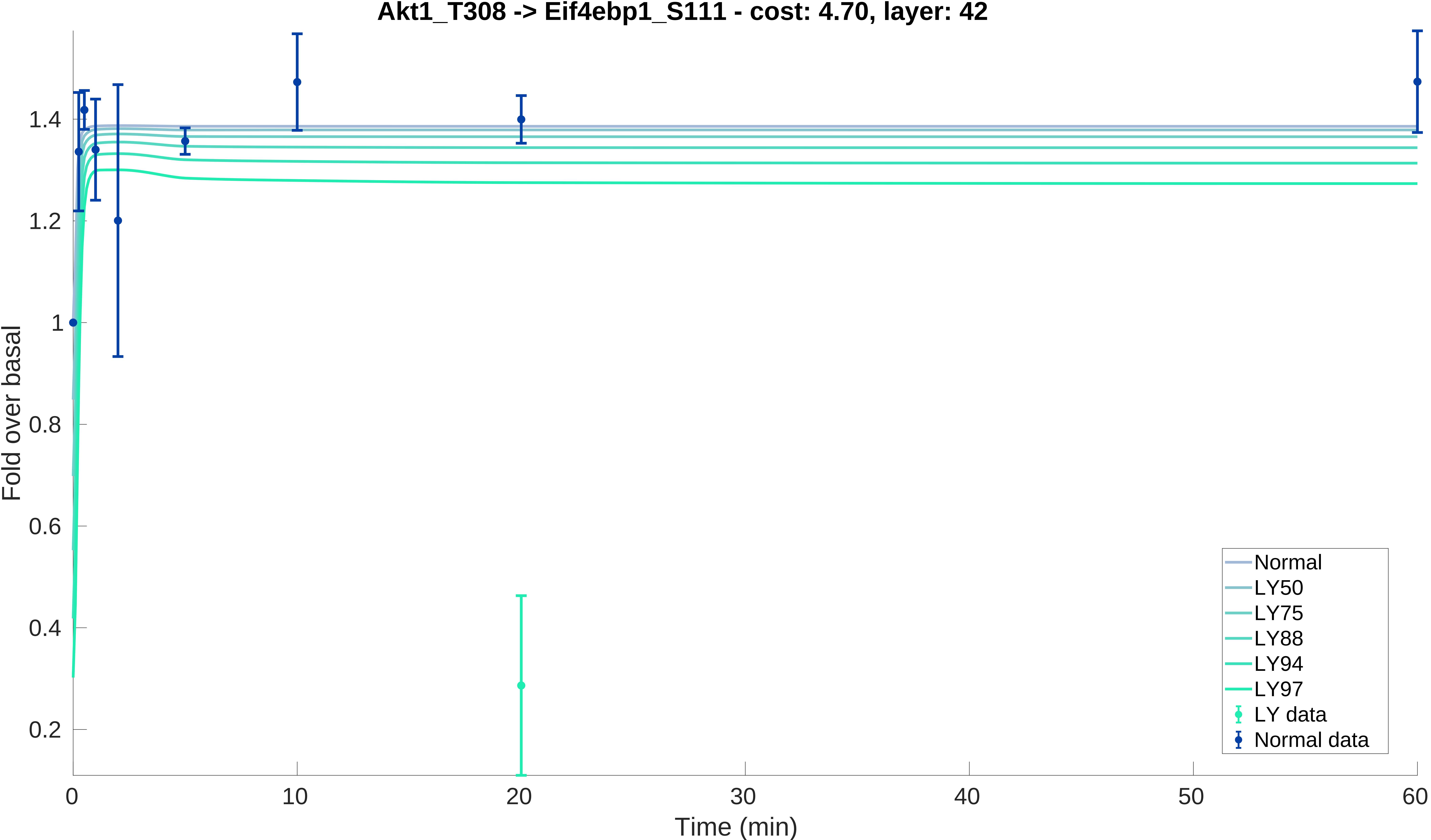

Akt1_T308 -> Eif4ebp1_S111 - cost: 4.70, layer: 42
Fold over basal
Time (min)
0.2
0.4
0.6
0.8
1
1.2
1.4
0
10
20
30
40
50
60
Normal
LY50
LY75
LY88
LY94
LY97
LY data
Normal data

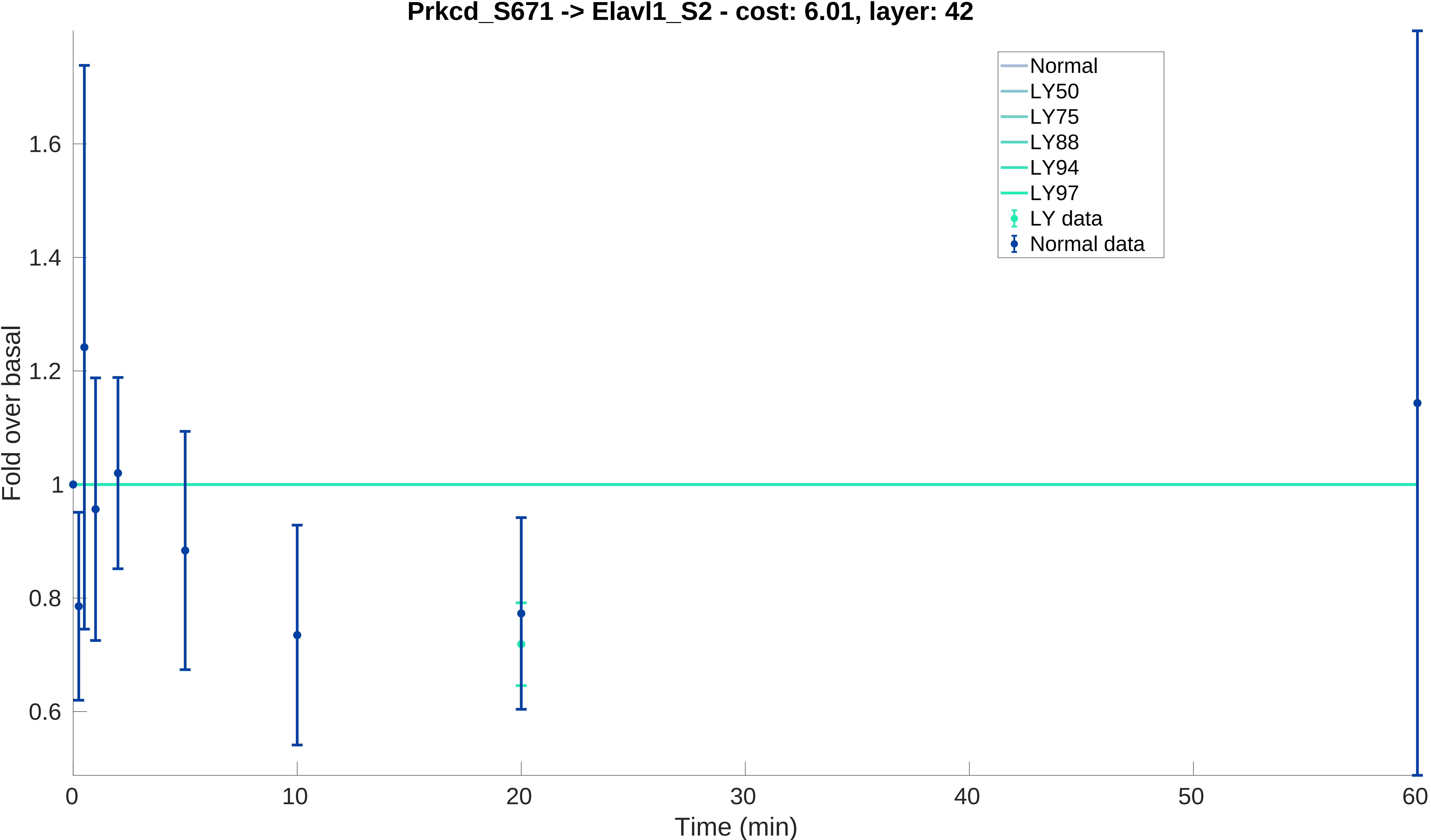

Prkcd_S671 -> Elavl1_S2 - cost: 6.01, layer: 42
Fold over basal
1.6
1.4
1.2
1
0.8
0.6
0
10
20
30
40
50
60
Time (min)
Normal
LY50
LY75
LY88
LY94
LY97
LY data
Normal data

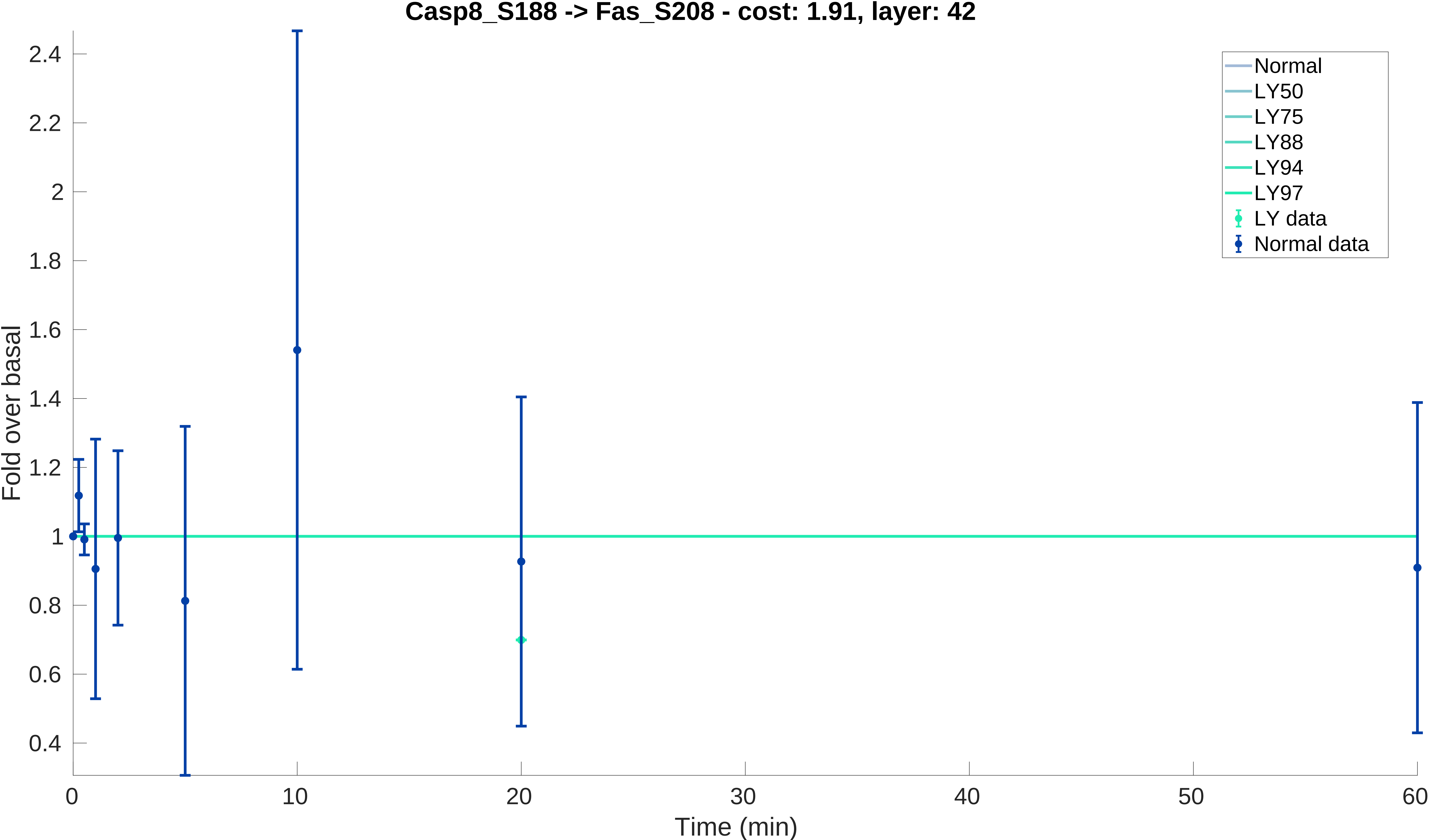

Casp8_S188 -> Fas_S208 - cost: 1.91, layer: 42
Fold over basal
2.4
2.2
2
1.8
1.6
1.4
1.2
1
0.8
0.6
0.4
0
10
20
30
40
50
60
Time (min)
Normal
LY50
LY75
LY88
LY94
LY97
LY data
Normal data

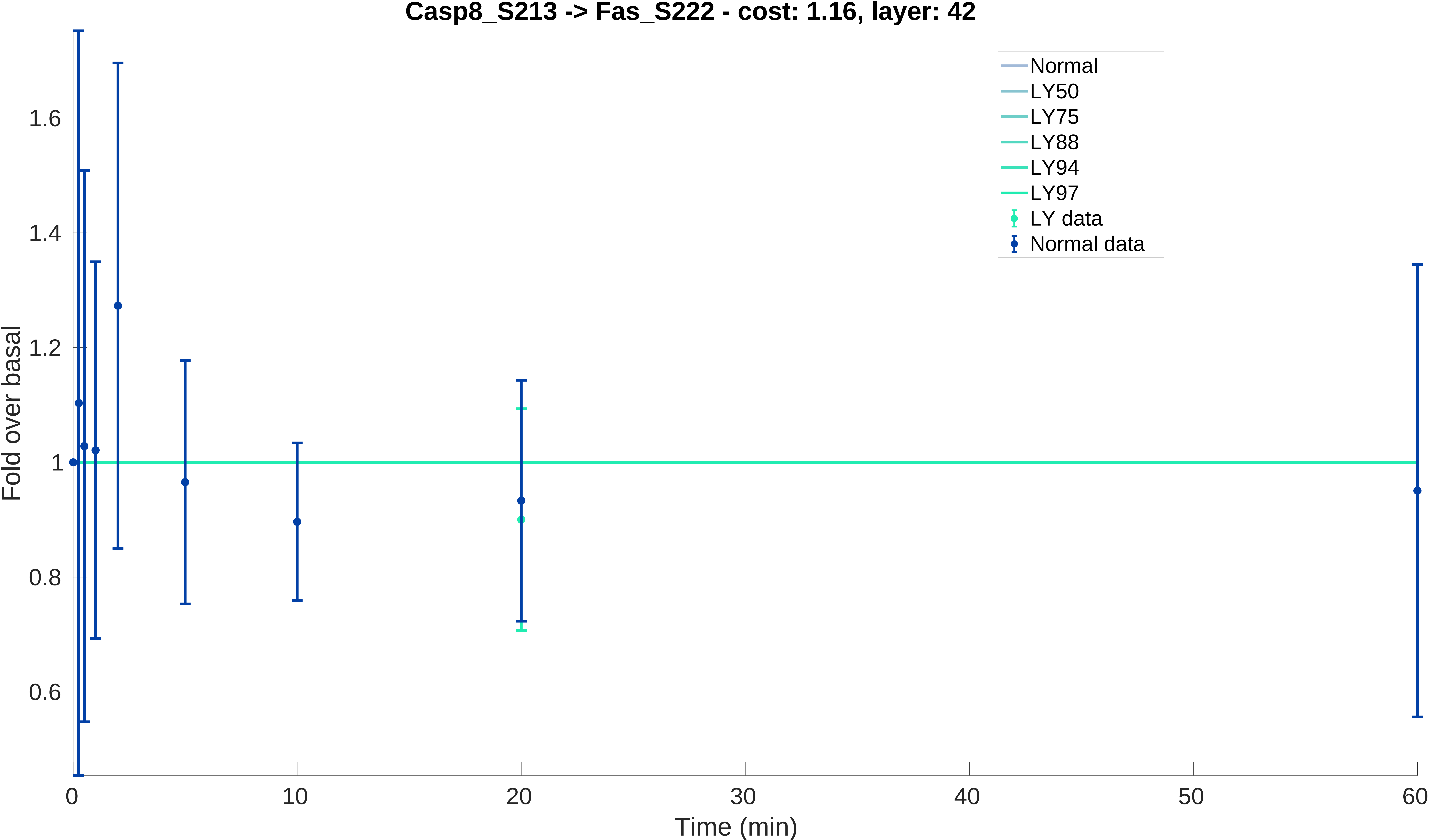

Casp8_S213 -> Fas_S222 - cost: 1.16, layer: 42
Normal
LY50
LY75
LY88
LY94
LY97
LY data
Normal data
Fold over basal
1.6
1.4
1.2
1
0.8
0.6
0
10
20
30
40
50
60
Time (min)

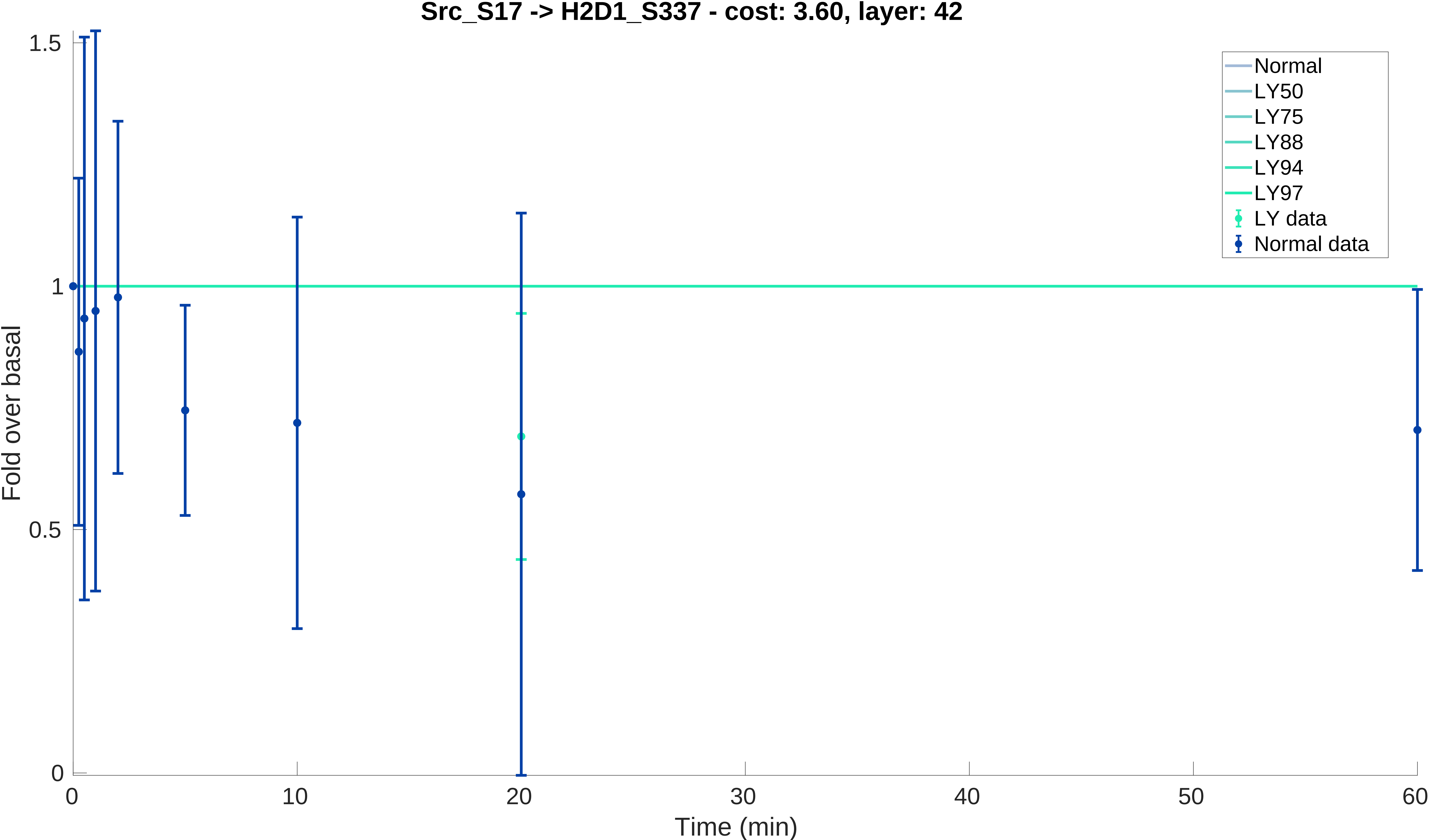

Src_S17 -> H2D1_S337 - cost: 3.60, layer: 42
Fold over basal
Time (min)
0
0.5
1
1.5
0
10
20
30
40
50
60
Normal
LY50
LY75
LY88
LY94
LY97
LY data
Normal data

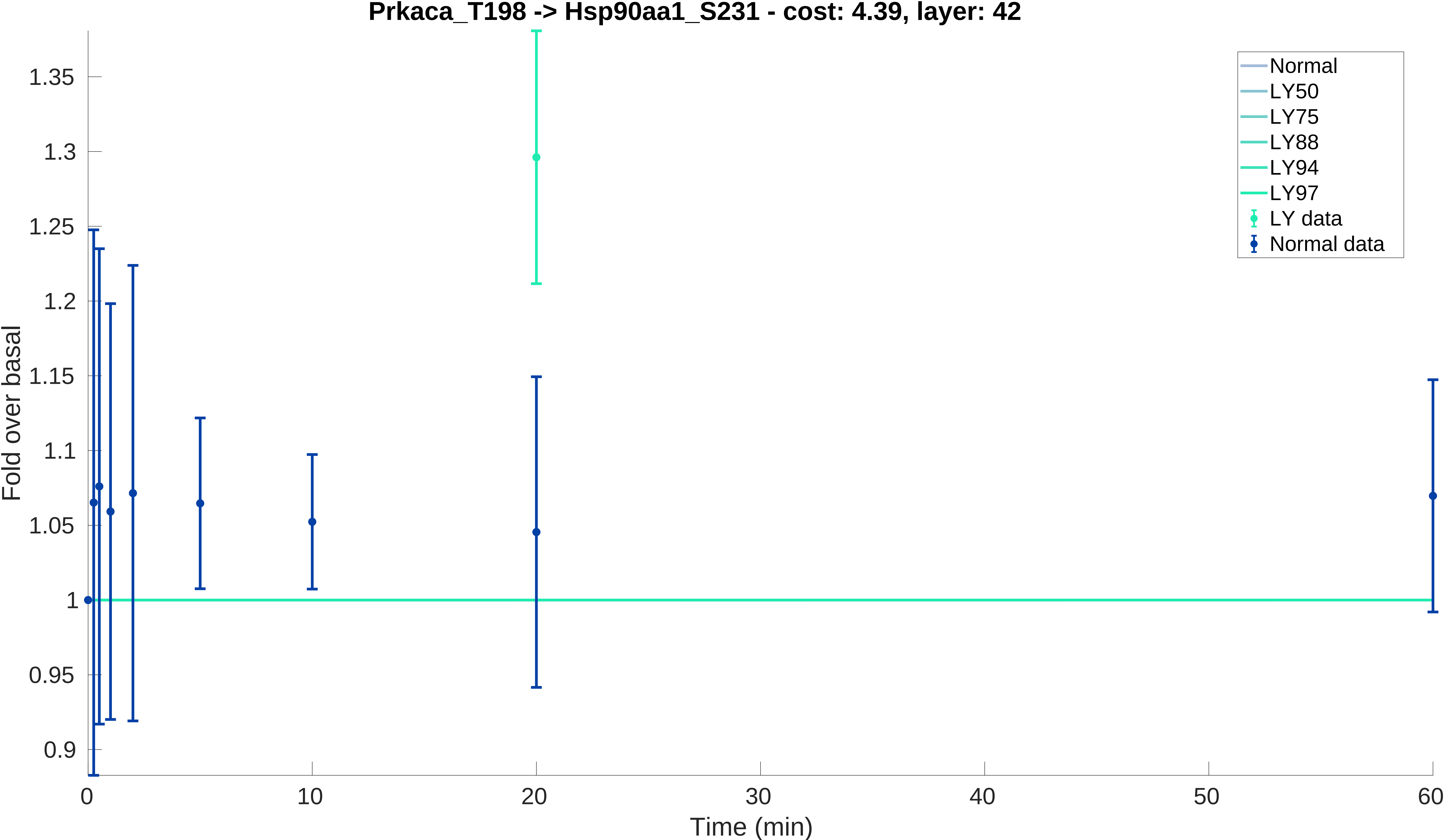
Prkaca_T198 -> Hsp90aa1_S231 - cost: 4.39, layer: 42
Fold over basal
Time (min)
1.35
1.3
1.25
1.2
1.15
1.1
1.05
1
0.95
0.9
0
10
20
30
40
50
60
Normal
LY50
LY75
LY88
LY94
LY97
LY data
Normal data

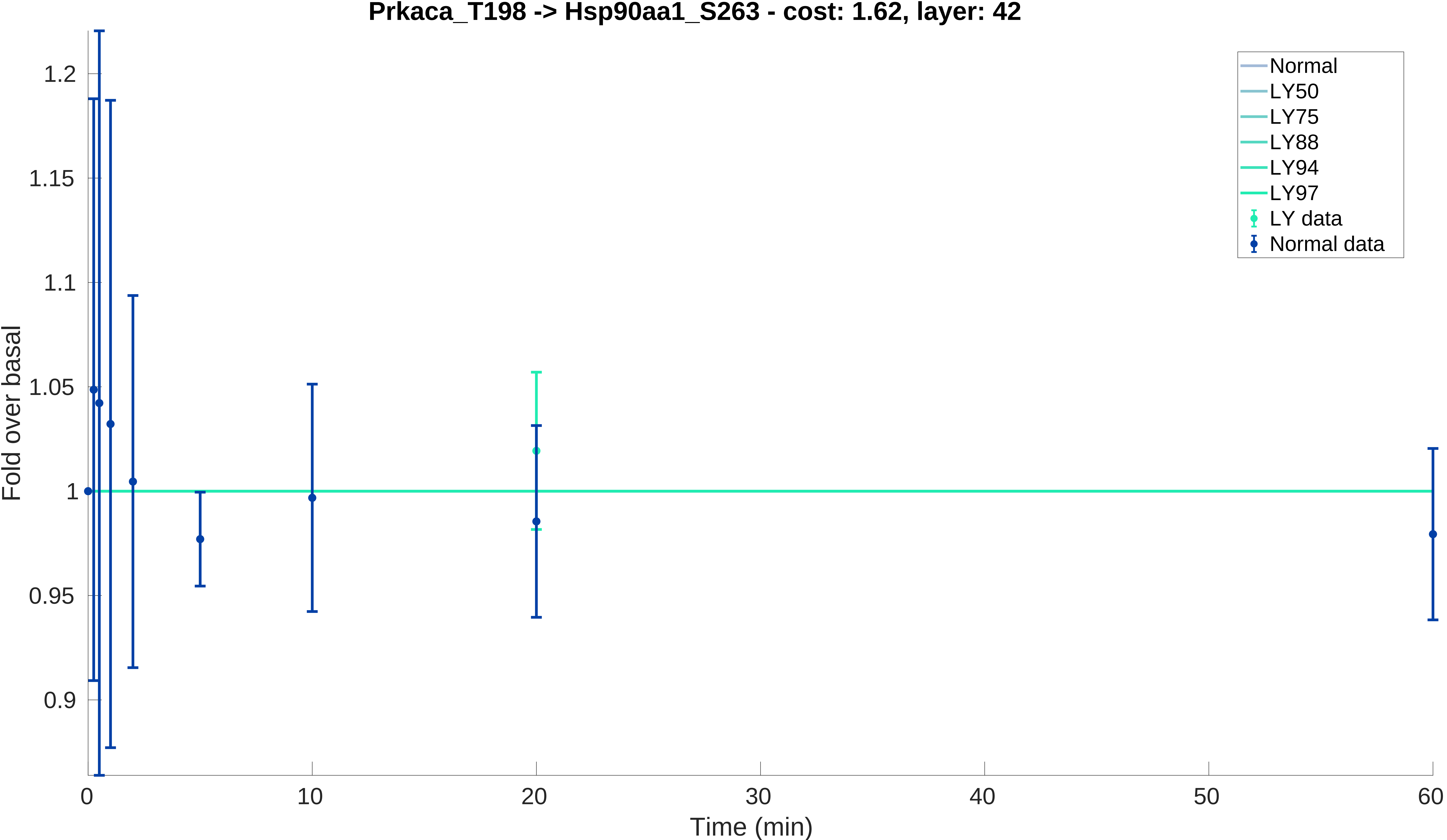

Prkaca_T198 -> Hsp90aa1_S263 - cost: 1.62, layer: 42
Fold over basal
1.2
1.15
1.1
1.05
1
0.95
0.9
0
10
20
30
40
50
60
Time (min)
Normal
LY50
LY75
LY88
LY94
LY97
LY data
Normal data

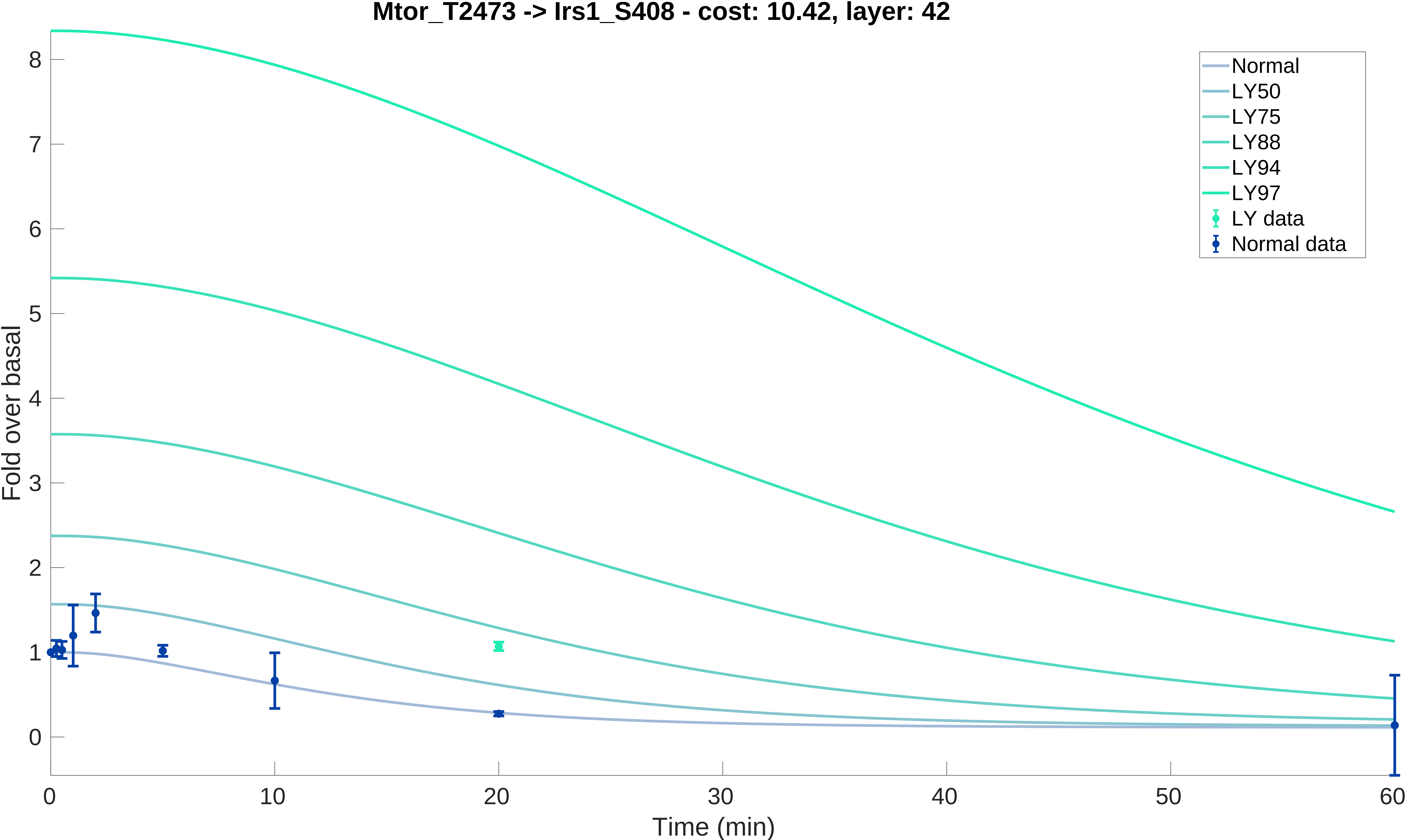

Mtor_T2473 -> Irs1_S408 - cost: 10.42, layer: 42
Fold over basal
Time (min)
0
1
2
3
4
5
6
7
8
0
10
20
30
40
50
60
Normal
LY50
LY75
LY88
LY94
LY97
LY data
Normal data

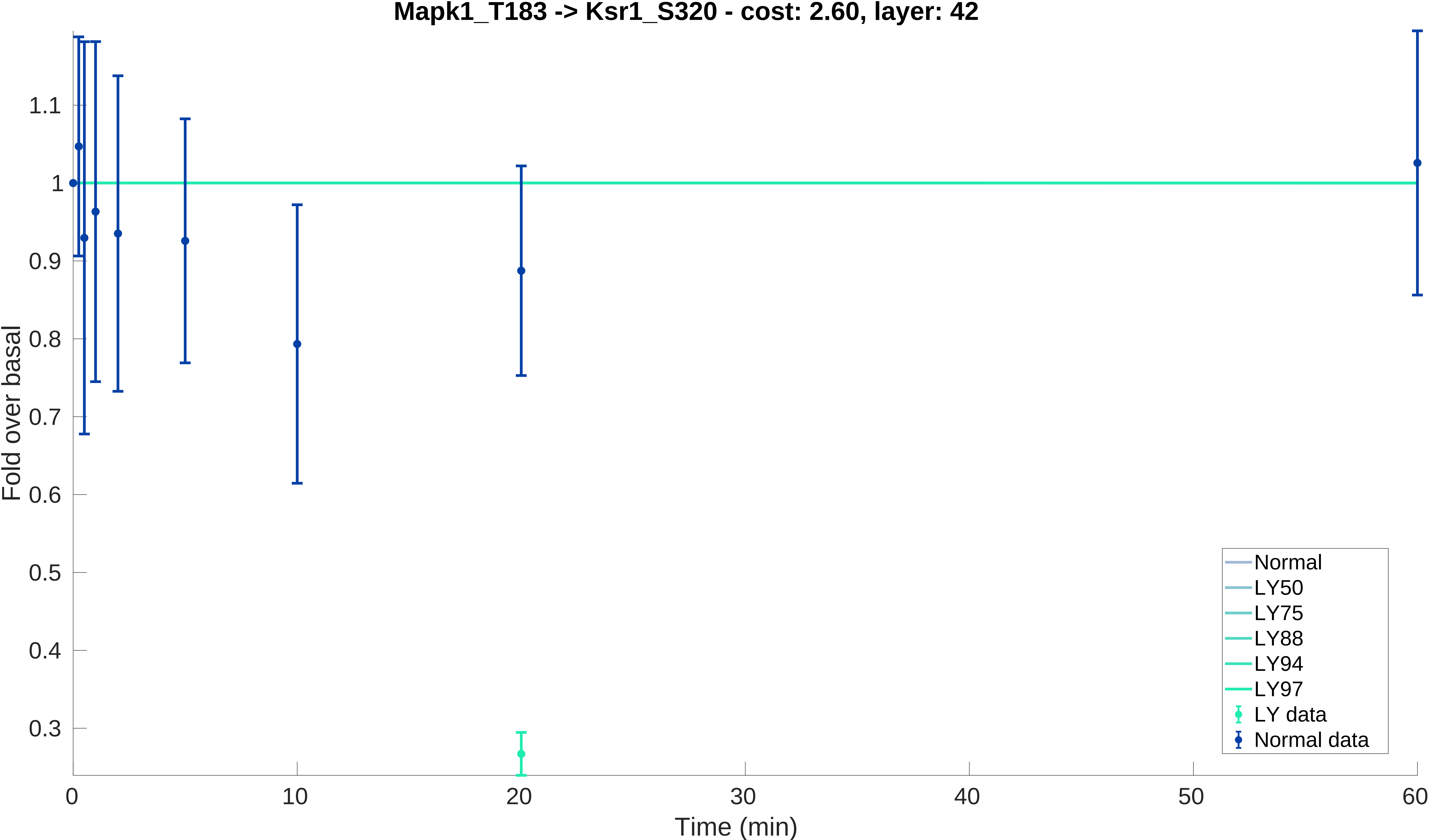
Mapk1_T183 -> Ksr1_S320 - cost: 2.60, layer: 42
Fold over basal
1.1
1
0.9
0.8
0.7
0.6
0.5
0.4
0.3
0
10
20
30
40
50
60
Time (min)
Normal
LY50
LY75
LY88
LY94
LY97
LY data
Normal data

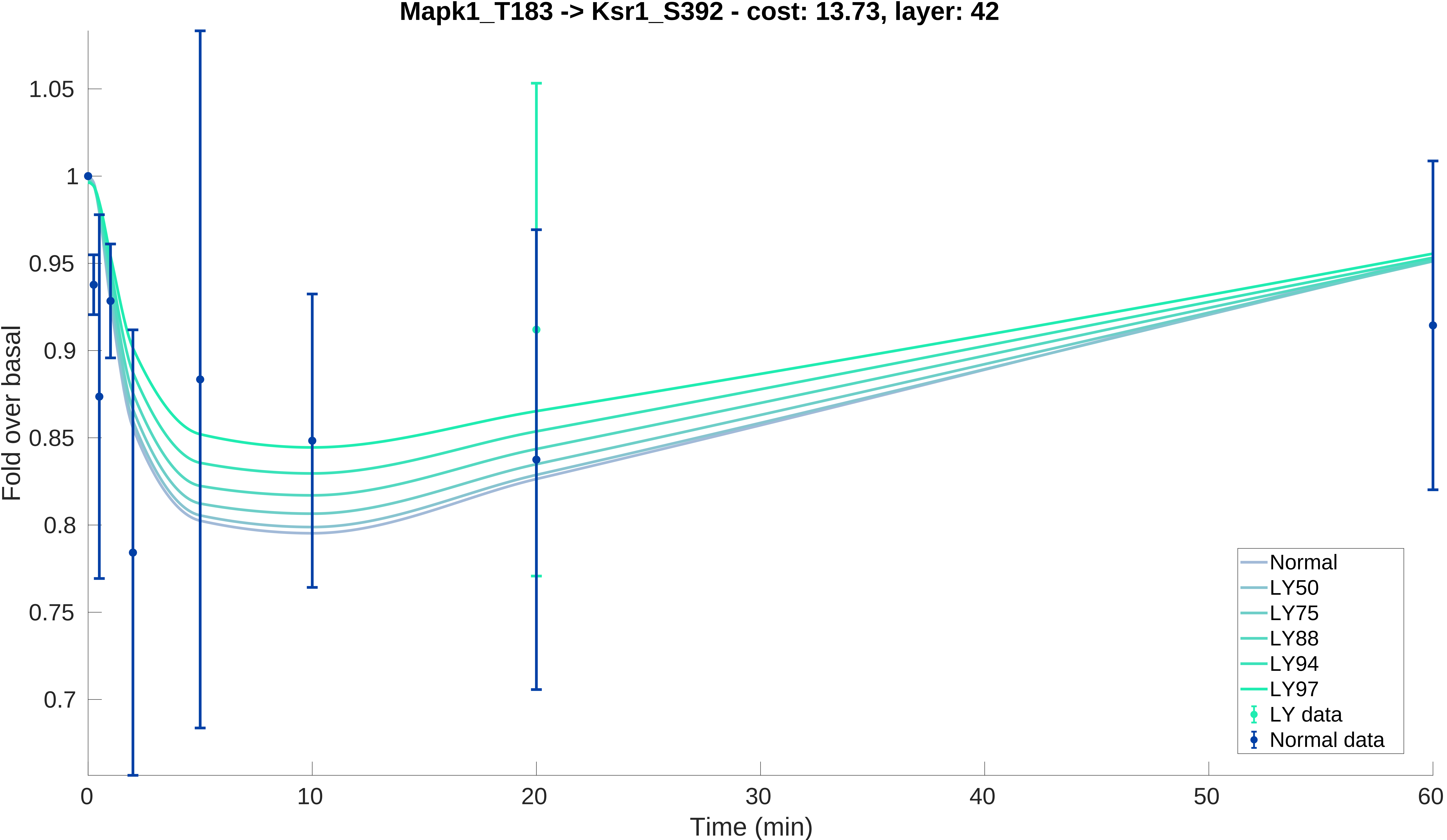

Mapk1_T183 -> Ksr1_S392 - cost: 13.73, layer: 42
Fold over basal
Time (min)
1.05
1
0.95
0.9
0.85
0.8
0.75
0.7
0
10
20
30
40
50
60
Normal
LY50
LY75
LY88
LY94
LY97
LY data
Normal data

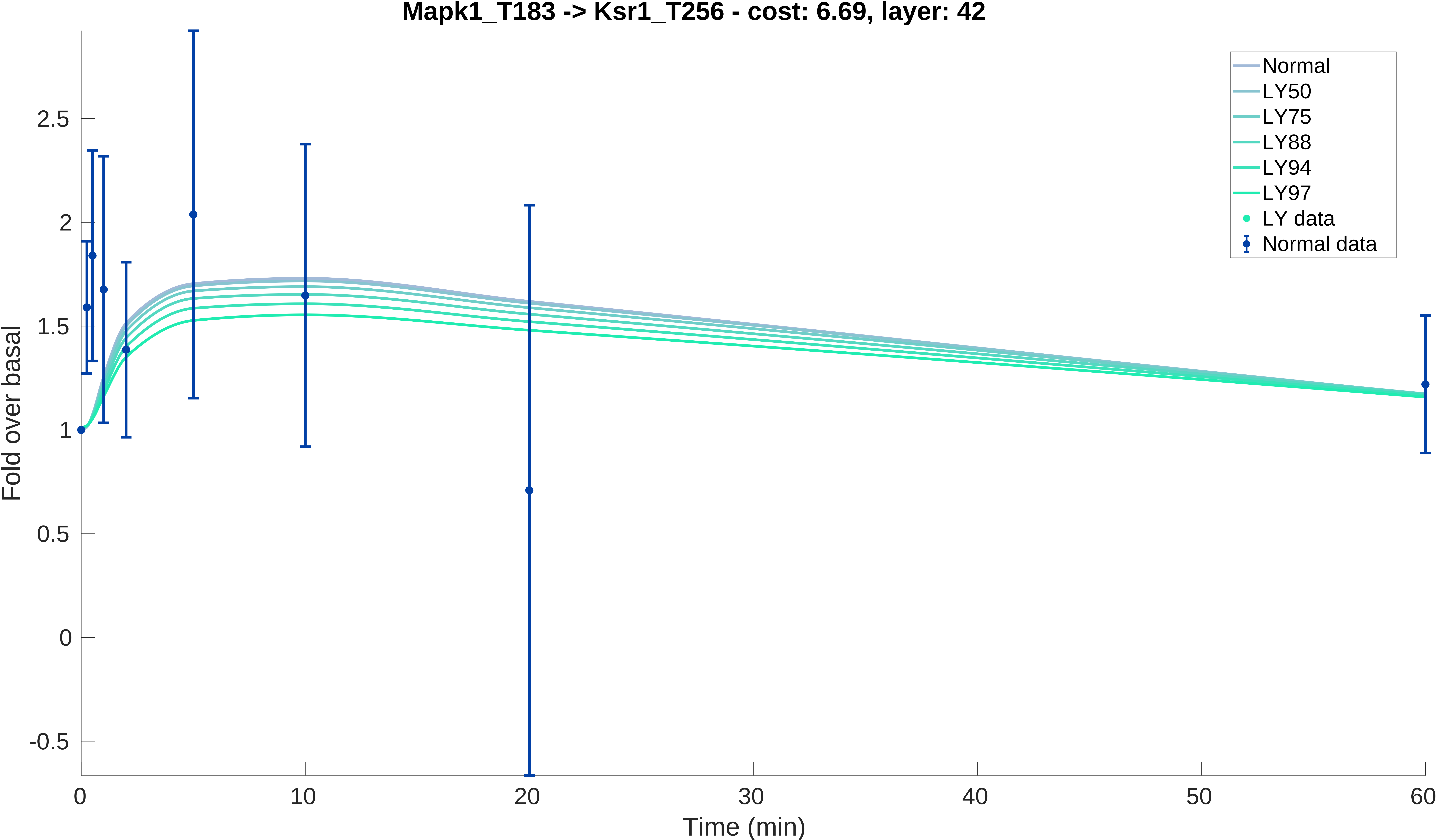

Mapk1_T183 -> Ksr1_T256 - cost: 6.69, layer: 42
Normal
LY50
LY75
LY88
LY94
LY97
LY data
Normal data
Fold over basal
2.5
2
1.5
1
0.5
0
-0.5
0
10
20
30
40
50
60
Time (min)

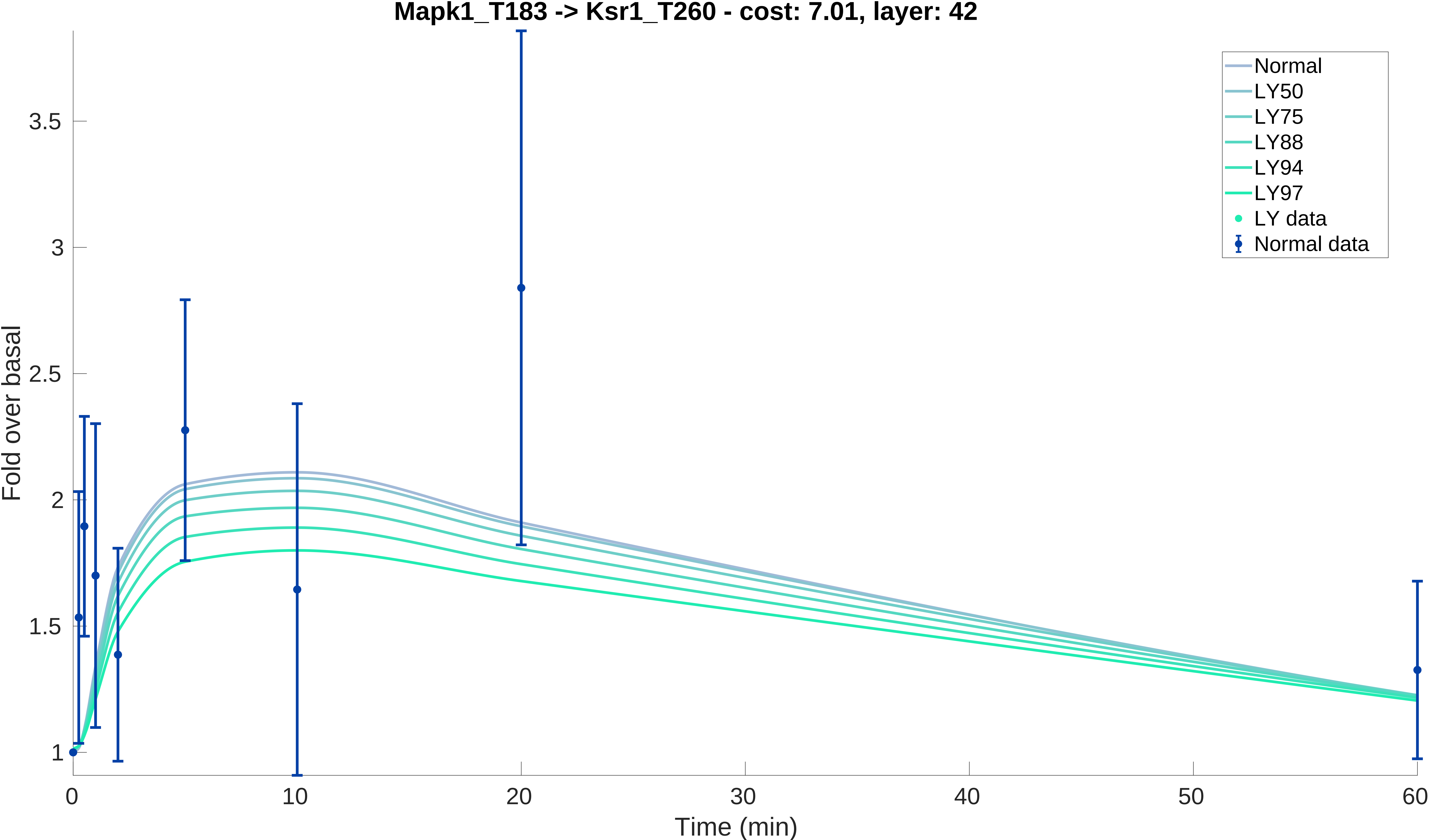

Mapk1_T183 -> Ksr1_T260 - cost: 7.01, layer: 42
Fold over basal
3.5
3
2.5
2
1.5
1
0
10
20
30
40
50
60
Time (min)
Normal
LY50
LY75
LY88
LY94
LY97
LY data
Normal data

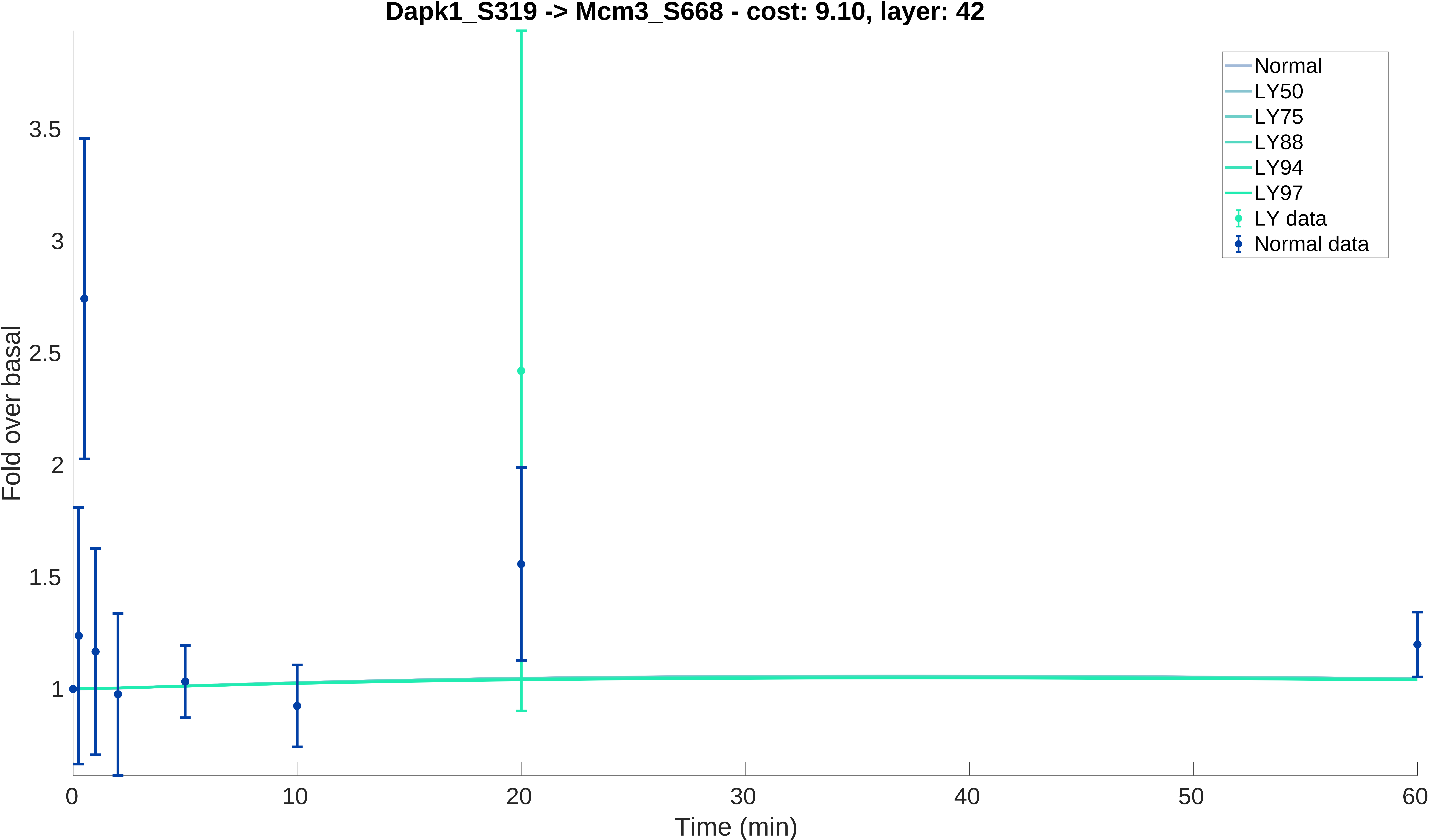
Dapk1_S319 -> Mcm3_S668 - cost: 9.10, layer: 42
Fold over basal
3.5
3
2.5
2
1.5
1
0
10
20
30
40
50
60
Time (min)
Normal
LY50
LY75
LY88
LY94
LY97
LY data
Normal data

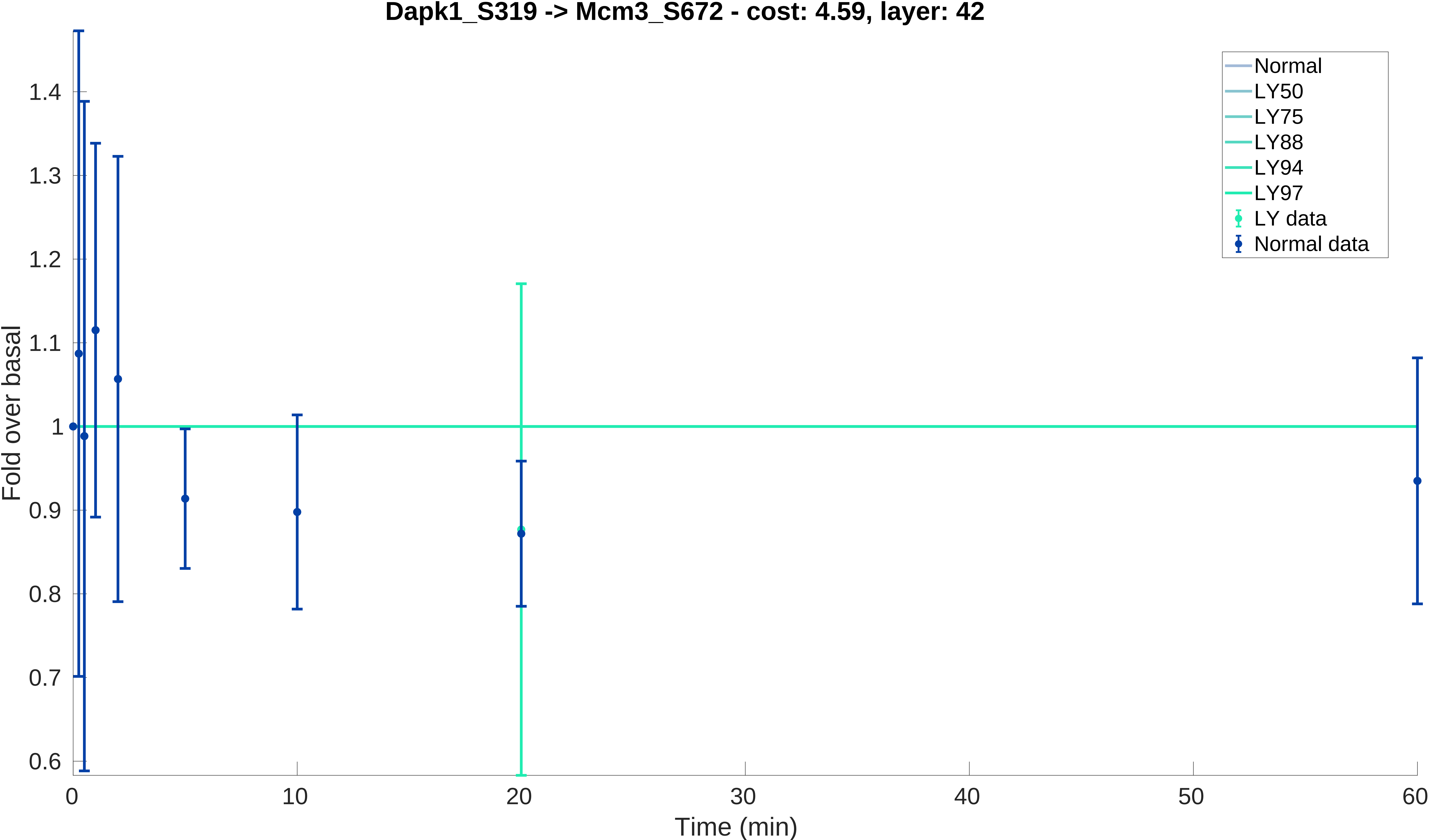

Dapk1_S319 -> Mcm3_S672 - cost: 4.59, layer: 42
Normal
LY50
LY75
LY88
LY94
LY97
LY data
Normal data
Fold over basal
1.4
1.3
1.2
1.1
1
0.9
0.8
0.7
0.6
0
10
20
30
40
50
60
Time (min)

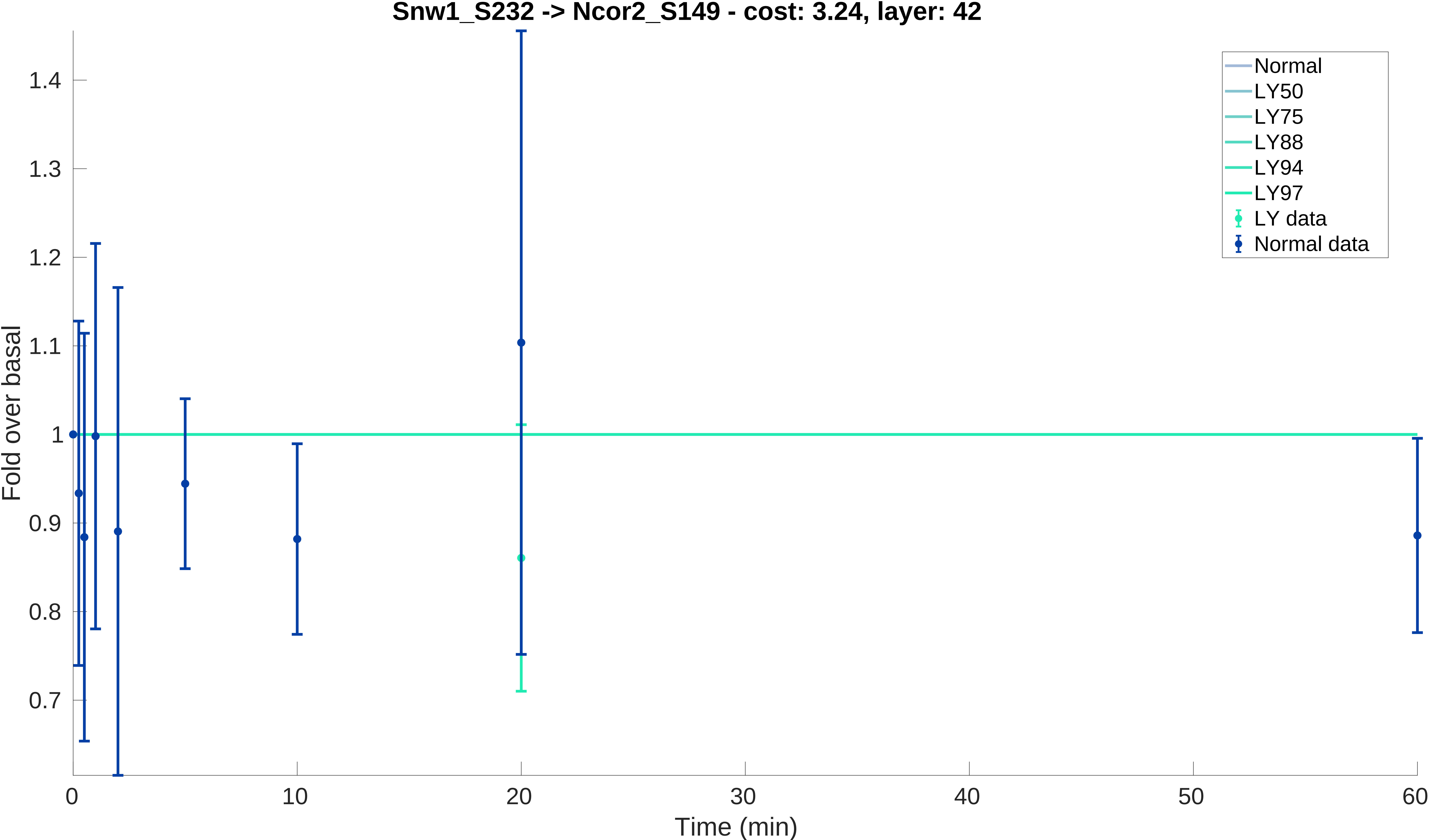

Snw1_S232 -> Ncor2_S149 - cost: 3.24, layer: 42
Normal
LY50
LY75
LY88
LY94
LY97
LY data
Normal data
Fold over basal
1.4
1.3
1.2
1.1
1
0.9
0.8
0.7
0
10
20
30
40
50
60
Time (min)

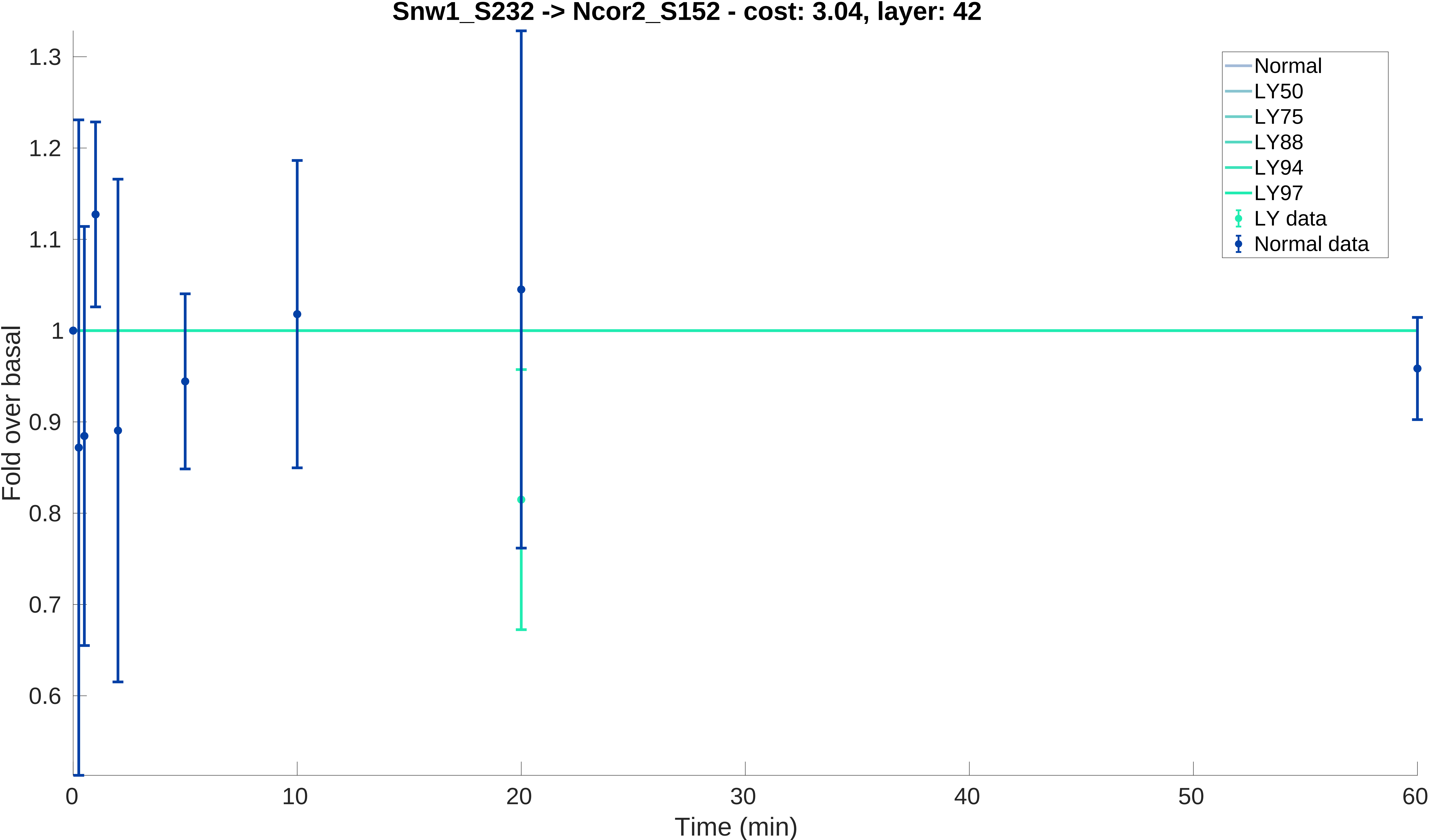

Snw1_S232 -> Ncor2_S152 - cost: 3.04, layer: 42
Fold over basal
1.3
1.2
1.1
1
0.9
0.8
0.7
0.6
0
10
20
30
40
50
60
Time (min)
Normal
LY50
LY75
LY88
LY94
LY97
LY data
Normal data

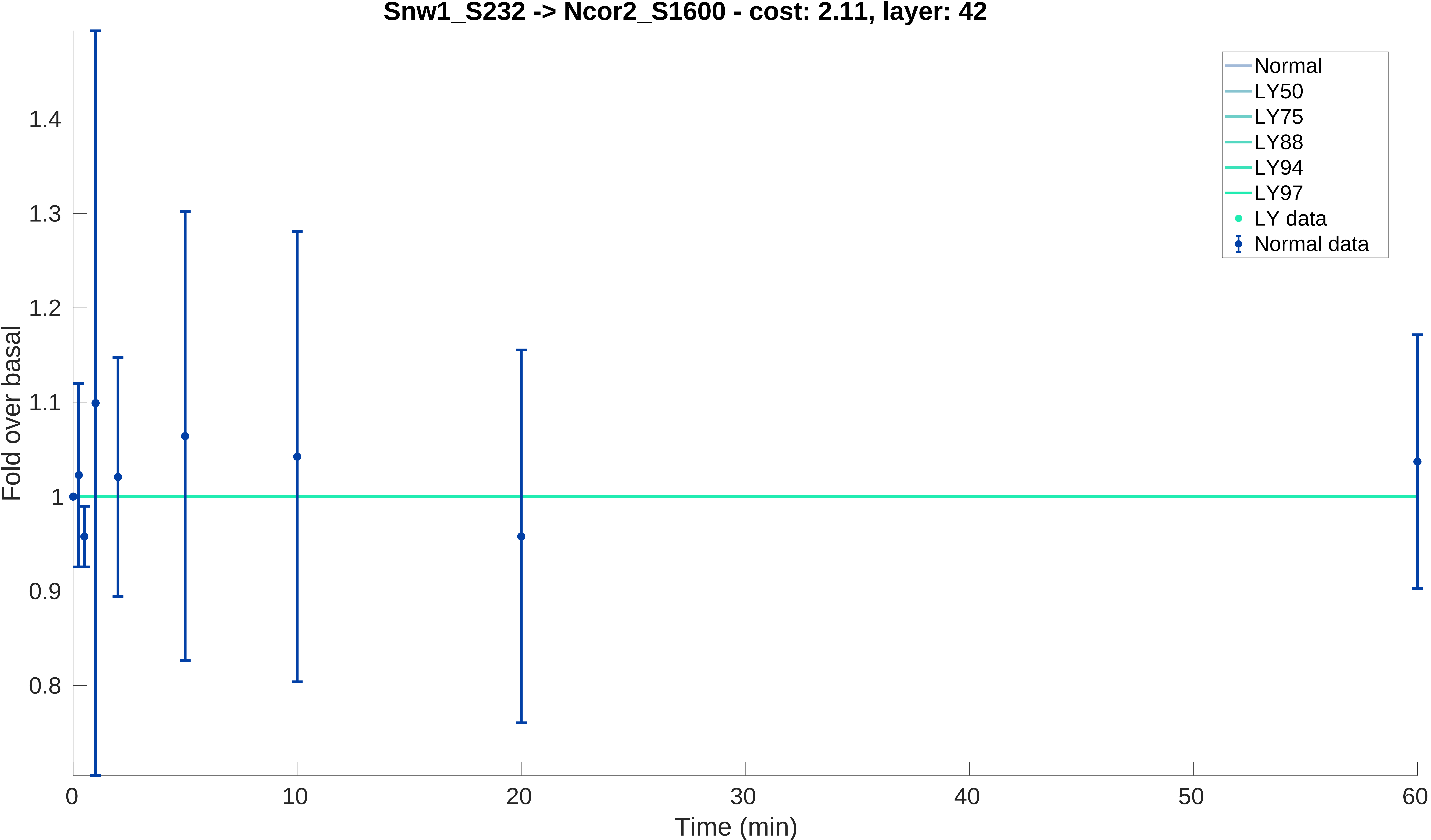
Snw1_S232 -> Ncor2_S1600 - cost: 2.11, layer: 42
Fold over basal
Time (min)
0.8
0.9
1
1.1
1.2
1.3
1.4
0
10
20
30
40
50
60
Normal
LY50
LY75
LY88
LY94
LY97
LY data
Normal data

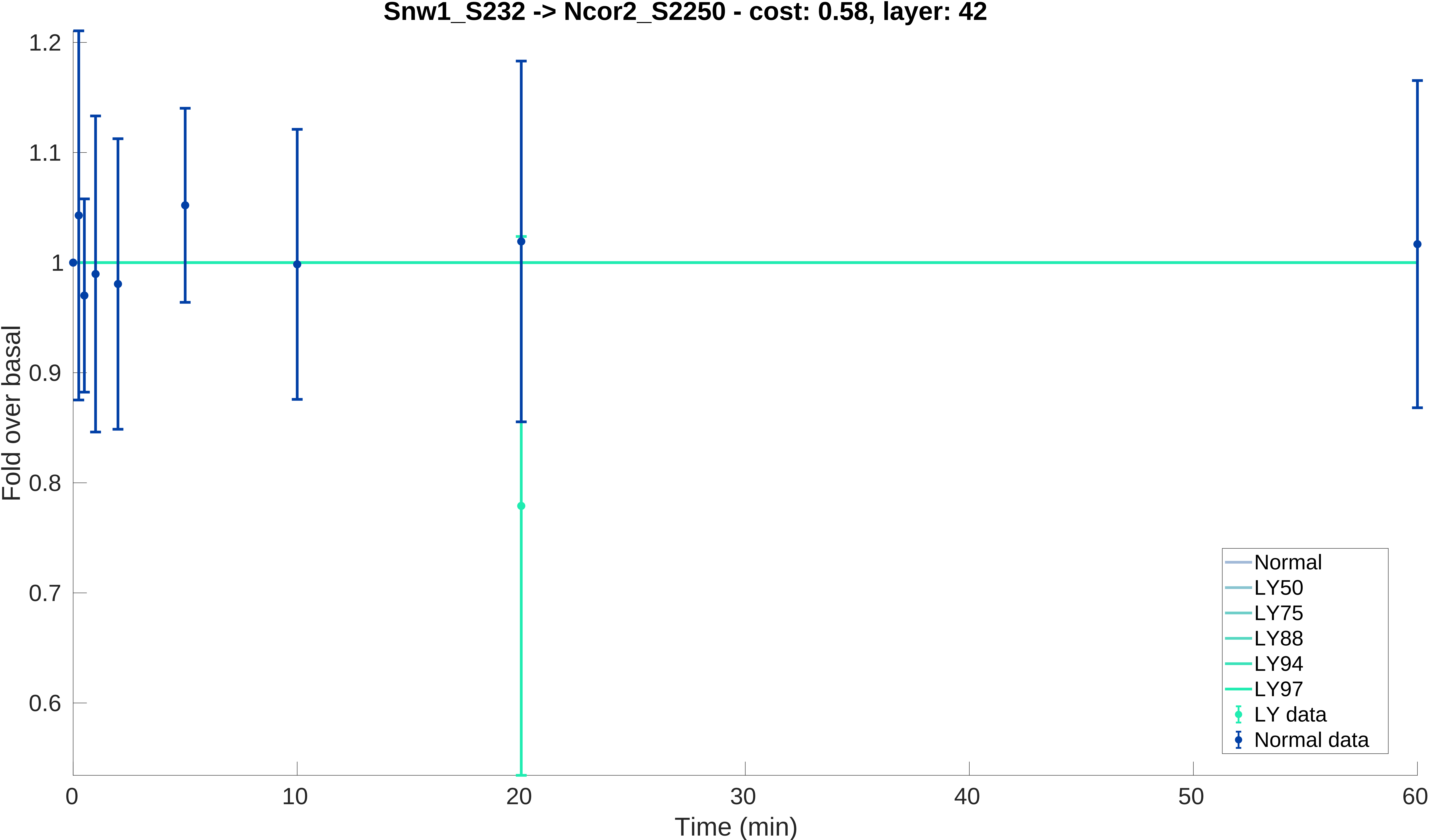
Snw1_S232 -> Ncor2_S2250 - cost: 0.58, layer: 42
Fold over basal
1.2
1.1
1
0.9
0.8
0.7
0.6
0
10
20
30
40
50
60
Time (min)
Normal
LY50
LY75
LY88
LY94
LY97
LY data
Normal data

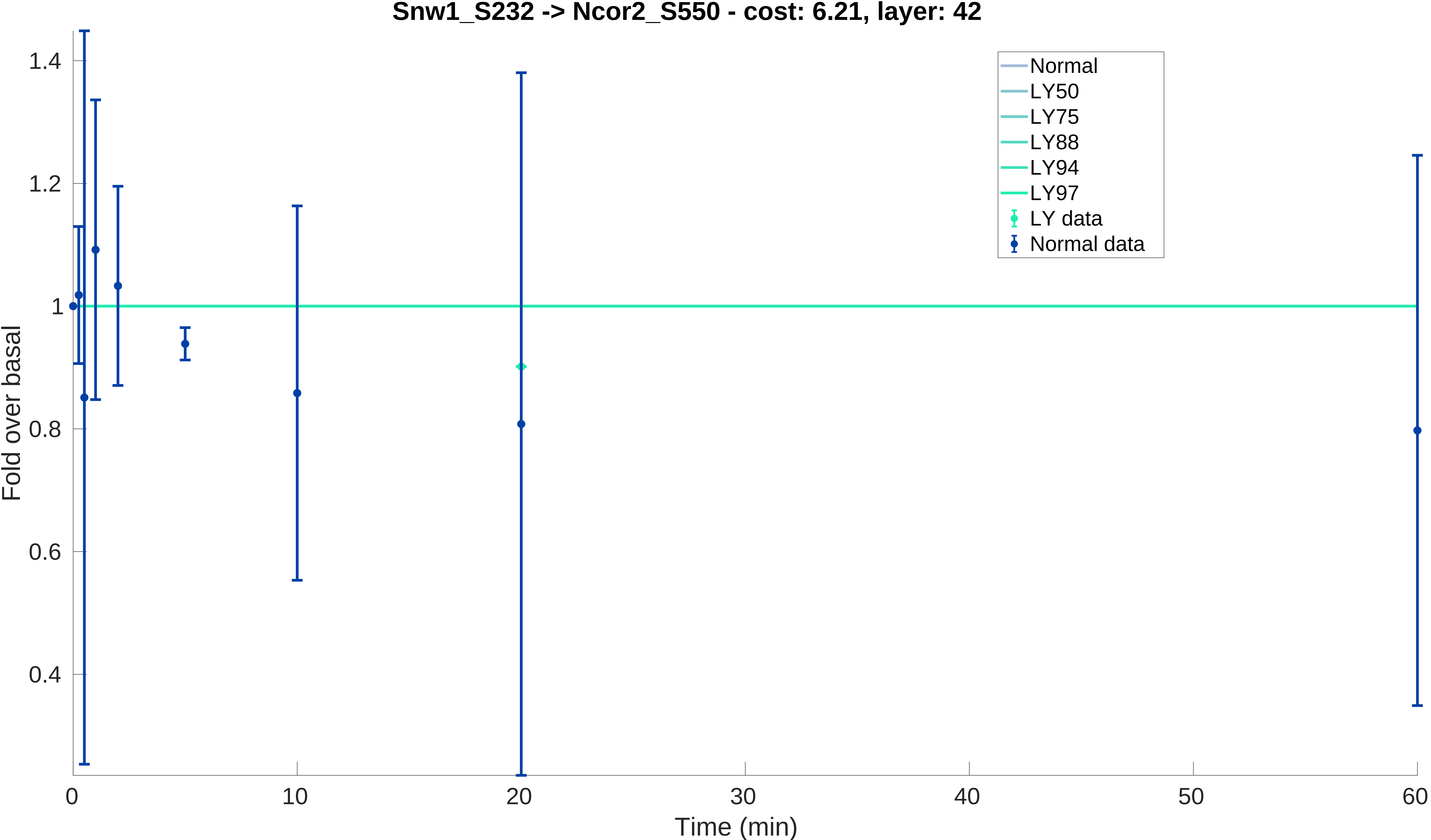

Snw1_S232 -> Ncor2_S550 - cost: 6.21, layer: 42
Normal
LY50
LY75
LY88
LY94
LY97
LY data
Normal data
Fold over basal
1.4
1.2
1
0.8
0.6
0.4
0
10
20
30
40
50
60
Time (min)

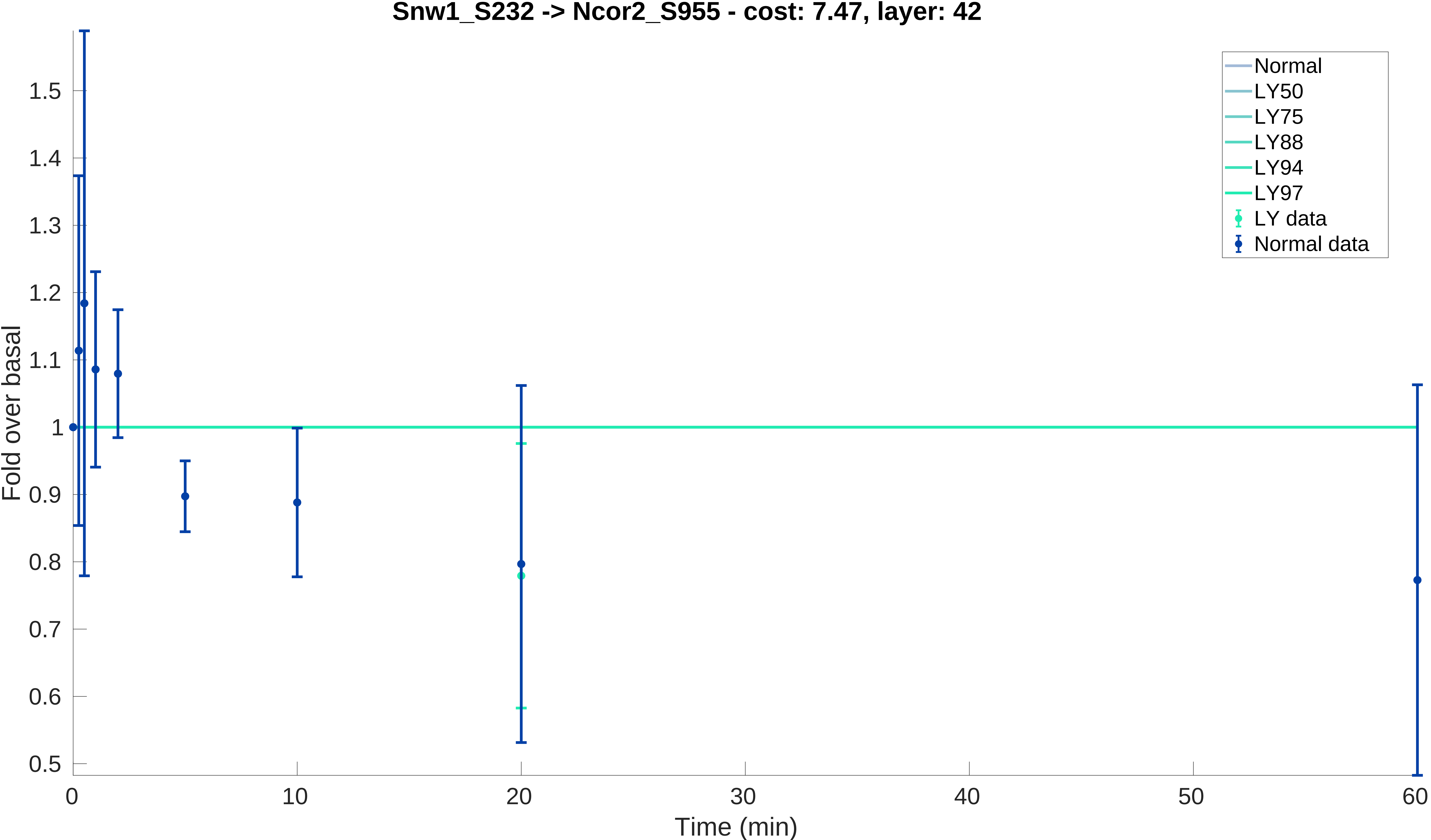

Snw1_S232 -> Ncor2_S955 - cost: 7.47, layer: 42
Fold over basal
Time (min)
Normal
LY50
LY75
LY88
LY94
LY97
LY data
Normal data
1.5
1.4
1.3
1.2
1.1
1
0.9
0.8
0.7
0.6
0.5
0
10
20
30
40
50
60

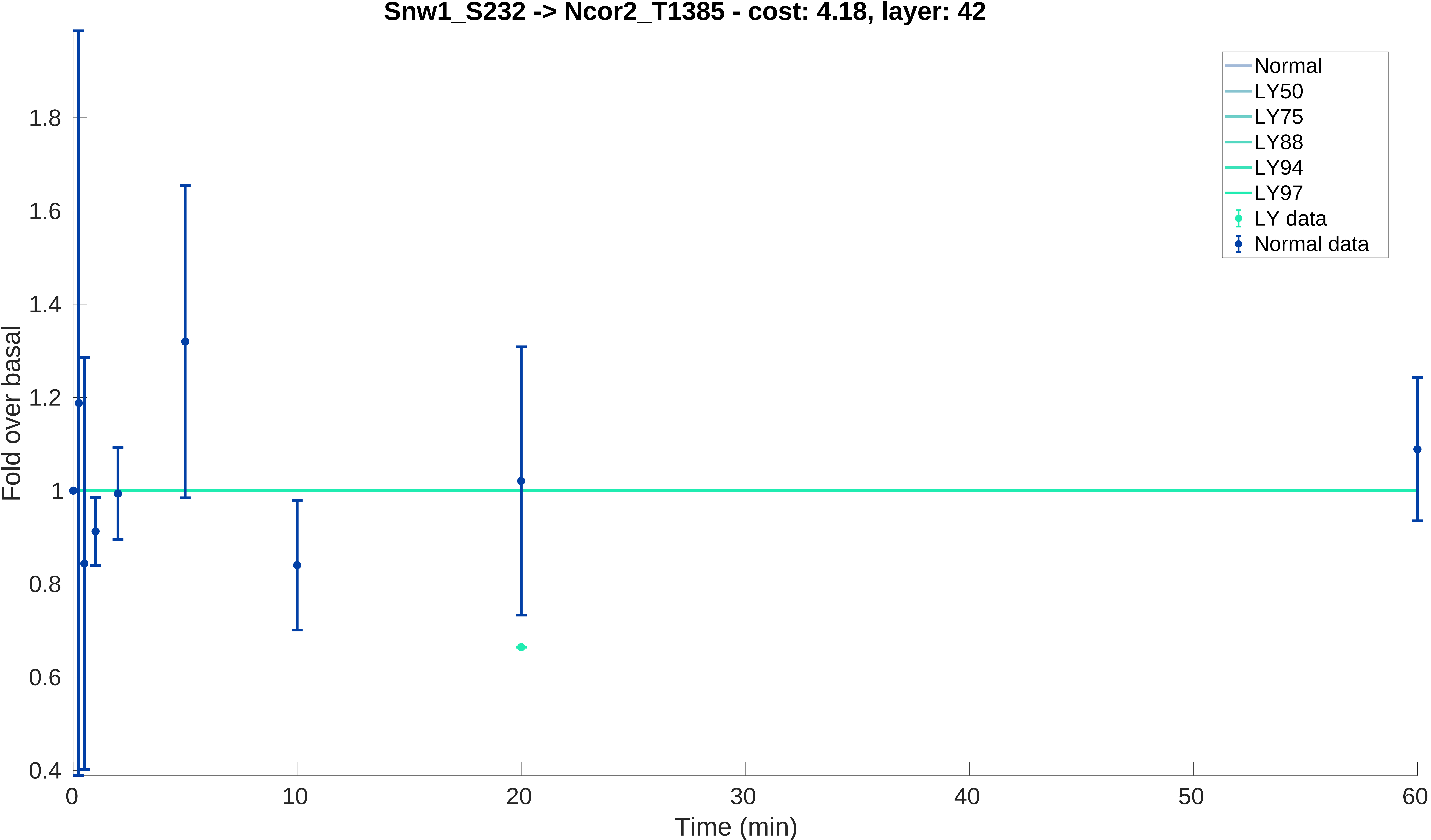

Snw1_S232 -> Ncor2_T1385 - cost: 4.18, layer: 42
Normal
LY50
LY75
LY88
LY94
LY97
LY data
Normal data
Fold over basal
1.8
1.6
1.4
1.2
1
0.8
0.6
0.4
0
10
20
30
40
50
60
Time (min)

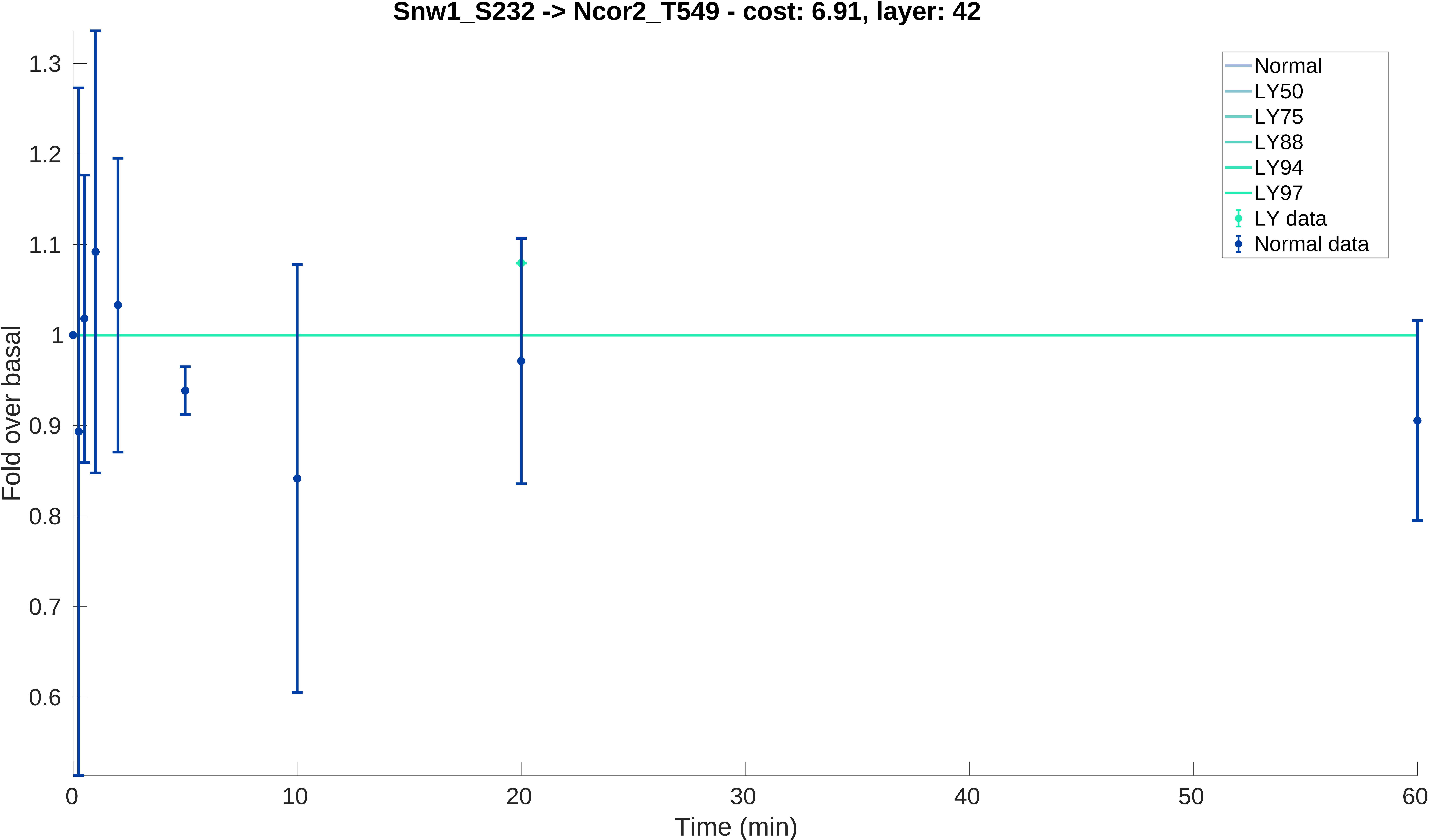

Snw1_S232 -> Ncor2_T549 - cost: 6.91, layer: 42
Fold over basal
Time (min)
1.3
1.2
1.1
1
0.9
0.8
0.7
0.6
0
10
20
30
40
50
60
Normal
LY50
LY75
LY88
LY94
LY97
LY data
Normal data

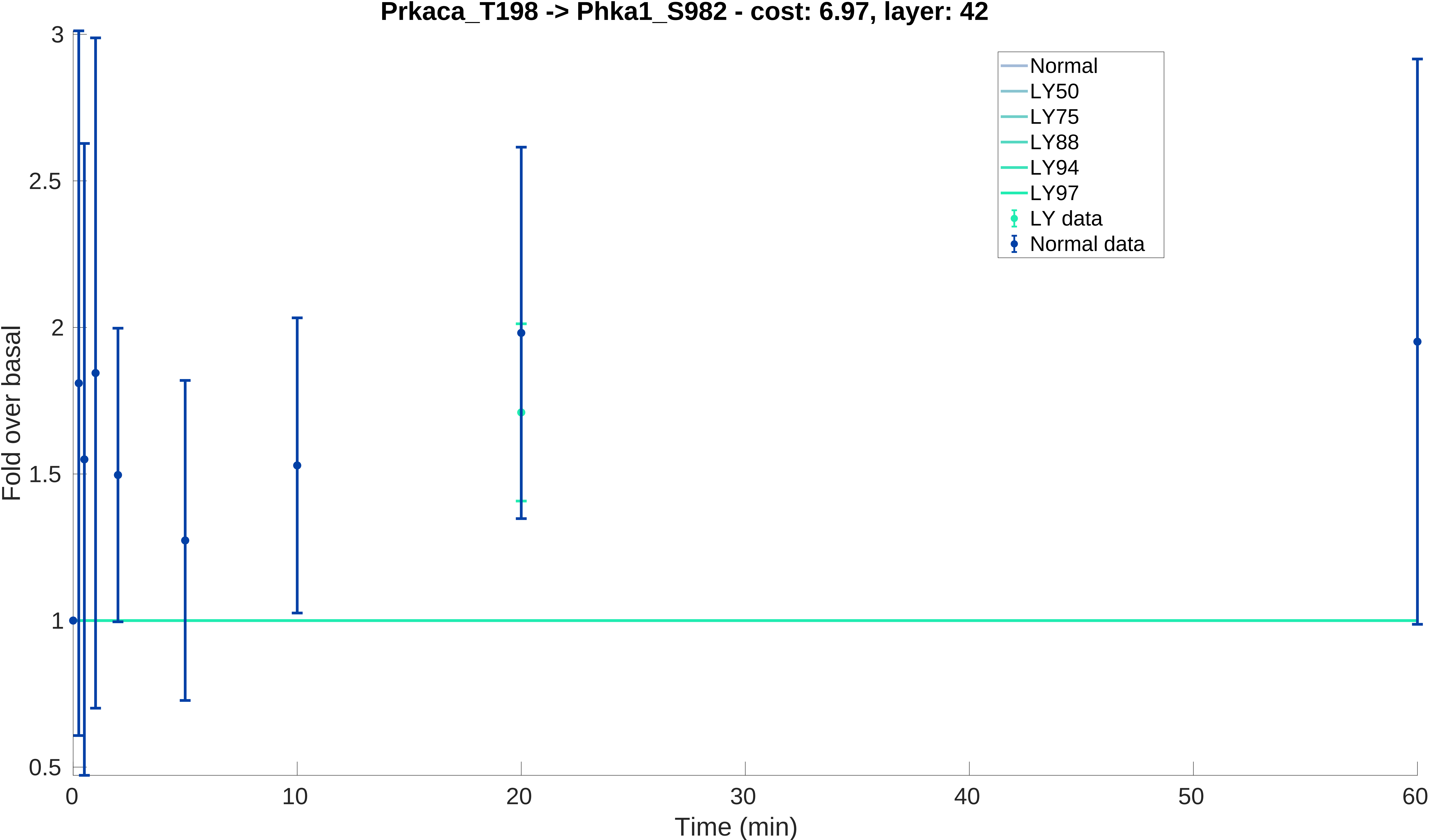
Prkaca_T198 -> Phka1_S982 - cost: 6.97, layer: 42
Normal
LY50
LY75
LY88
LY94
LY97
LY data
Normal data
Fold over basal
Time (min)
0.5
1
1.5
2
2.5
3
0
10
20
30
40
50
60

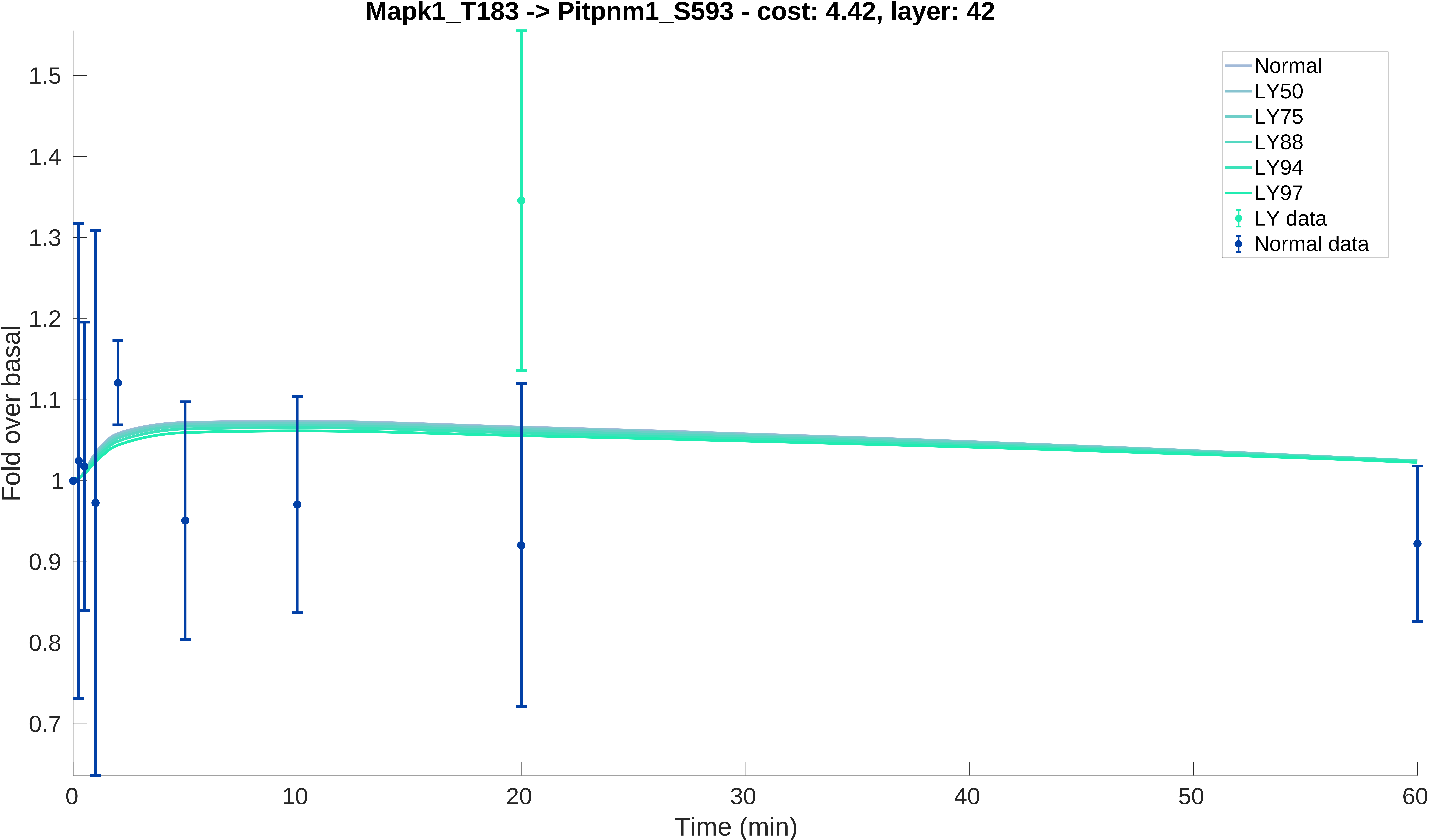

Mapk1_T183 -> Pitpnm1_S593 - cost: 4.42, layer: 42
Normal
LY50
LY75
LY88
LY94
LY97
LY data
Normal data
Fold over basal
1.5
1.4
1.3
1.2
1.1
1
0.9
0.8
0.7
0
10
20
30
40
50
60
Time (min)

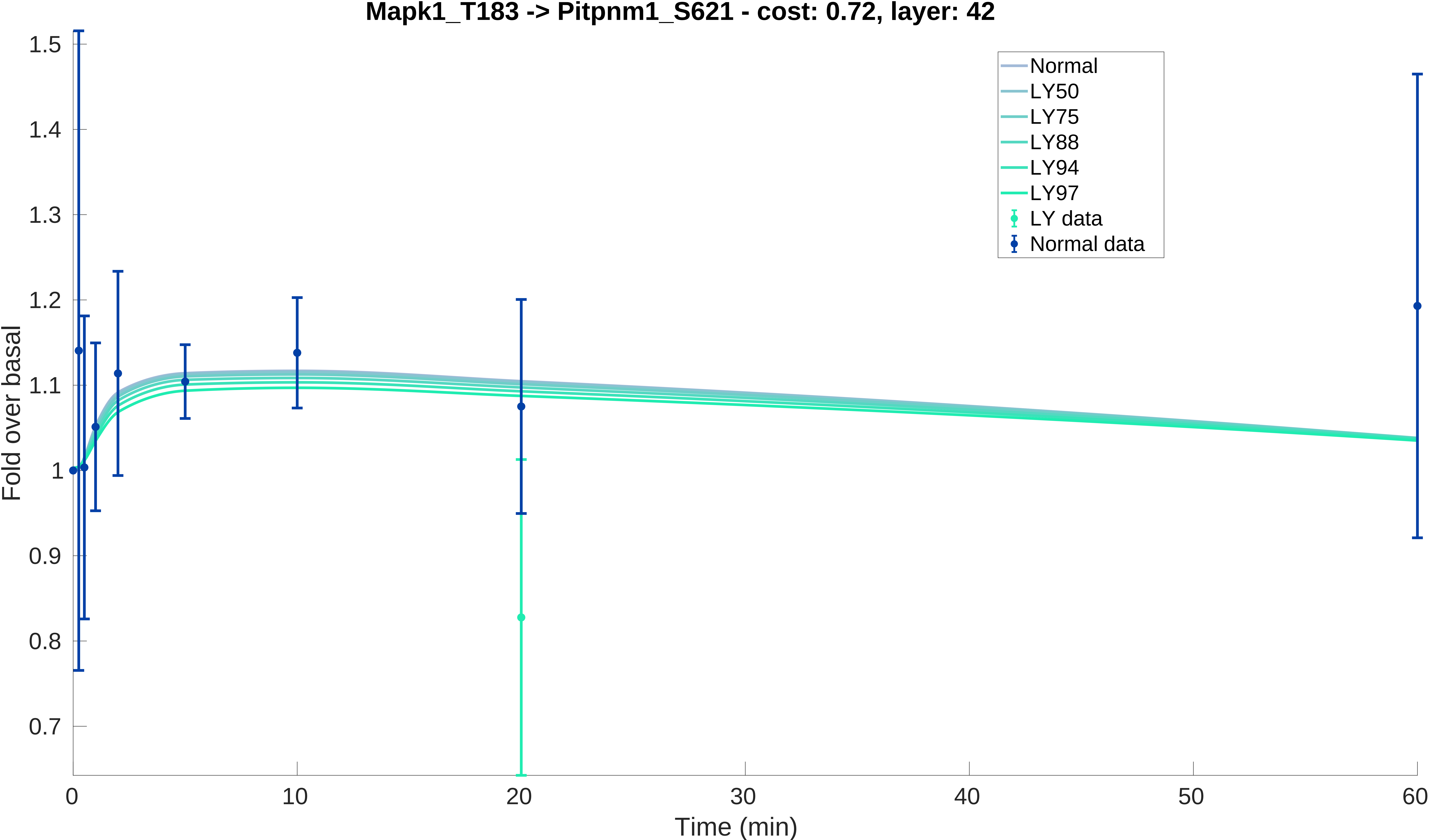
Mapk1_T183 -> Pitpnm1_S621 - cost: 0.72, layer: 42
Normal
LY50
LY75
LY88
LY94
LY97
LY data
Normal data
Fold over basal
1.5
1.4
1.3
1.2
1.1
1
0.9
0.8
0.7
0
10
20
30
40
50
60
Time (min)

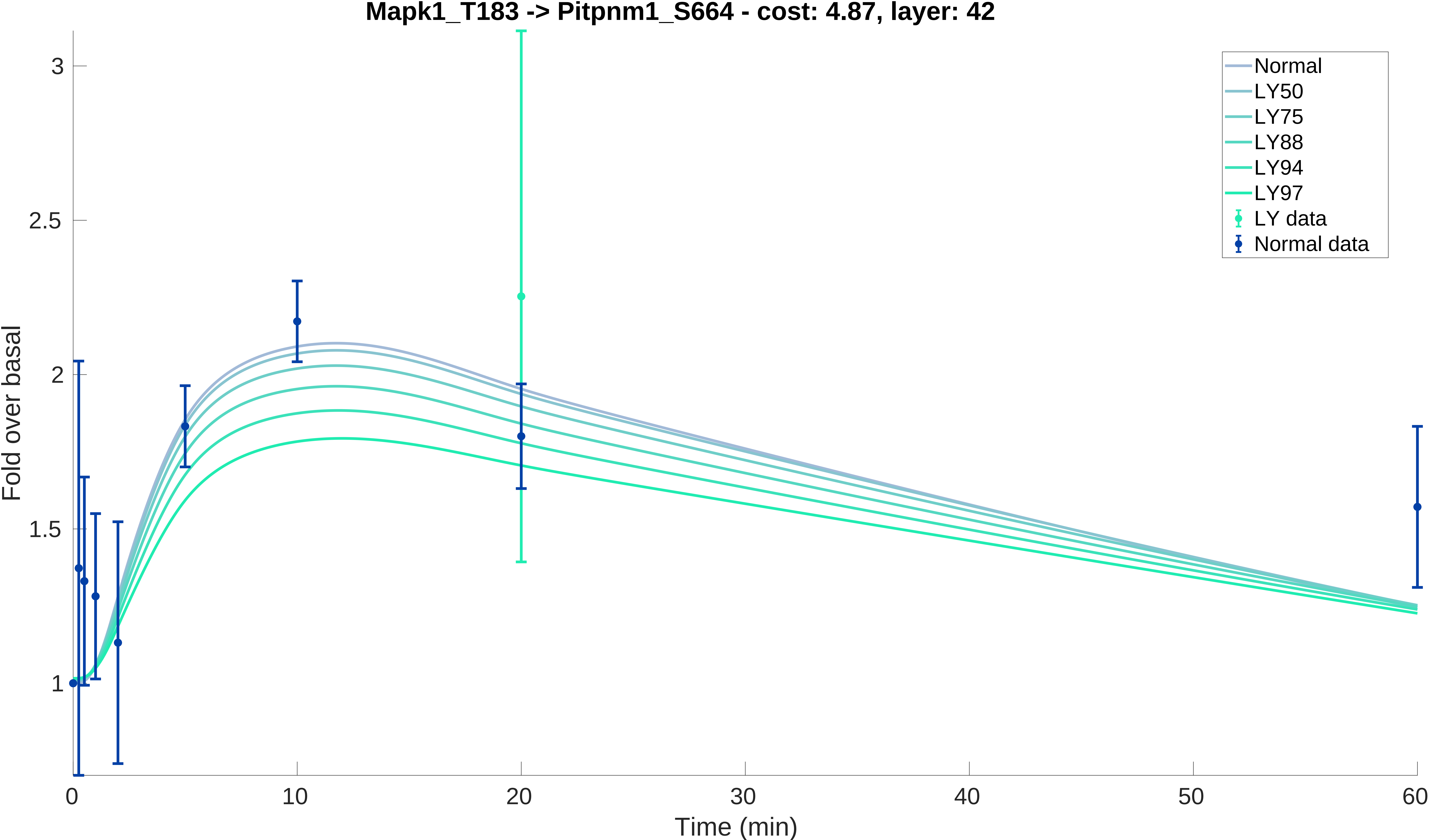

Mapk1_T183 -> Pitpnm1_S664 - cost: 4.87, layer: 42
Normal
LY50
LY75
LY88
LY94
LY97
LY data
Normal data
Fold over basal
3
2.5
2
1.5
1
0
10
20
30
40
50
60
Time (min)

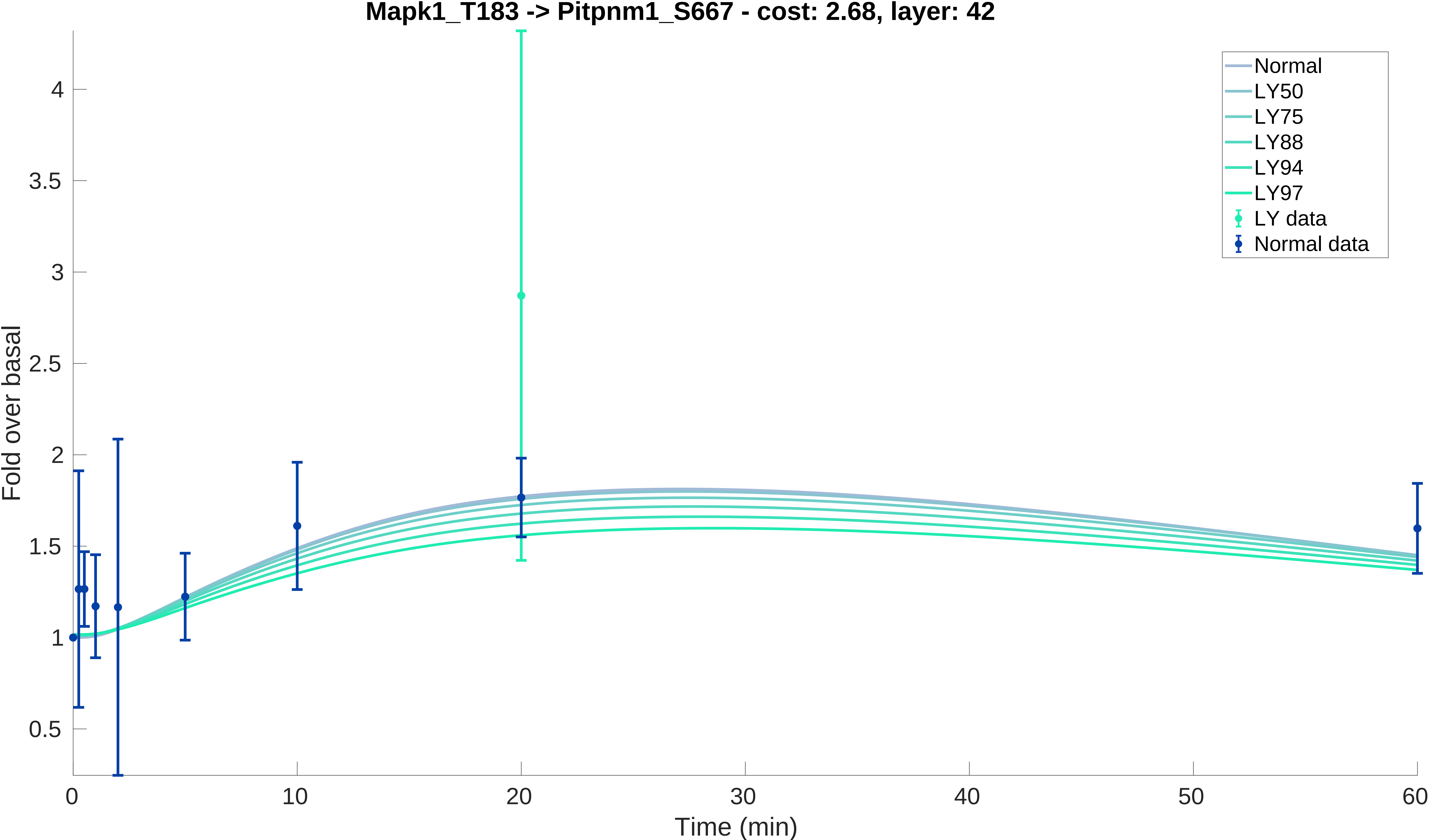

Mapk1_T183 -> Pitpnm1_S667 - cost: 2.68, layer: 42
Fold over basal
Time (min)
Normal
LY50
LY75
LY88
LY94
LY97
LY data
Normal data
0.5
1
1.5
2
2.5
3
3.5
4
0
10
20
30
40
50
60

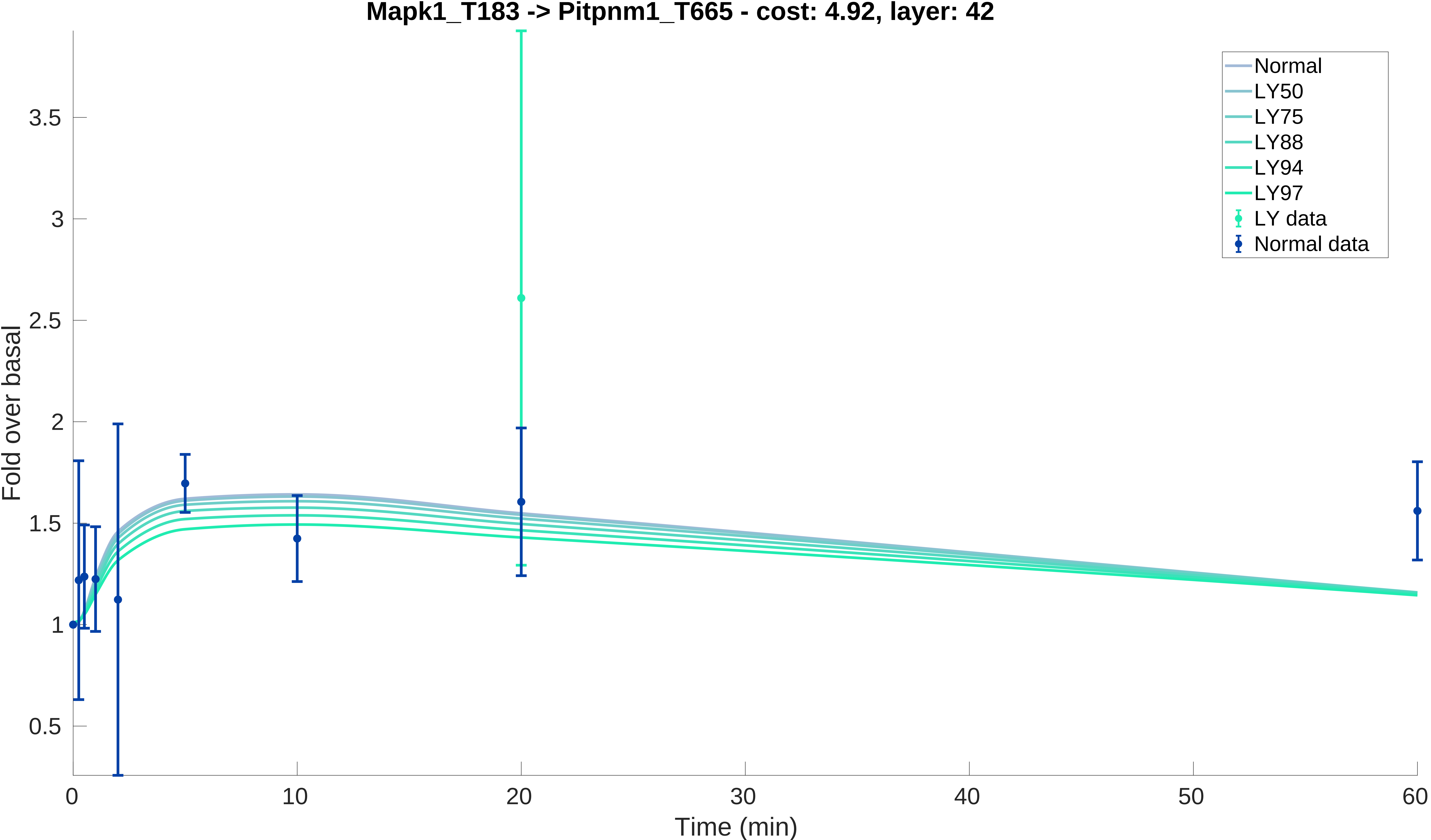

Mapk1_T183 -> Pitpnm1_T665 - cost: 4.92, layer: 42
Fold over basal
Time (min)
0.5
1
1.5
2
2.5
3
3.5
0
10
20
30
40
50
60
Normal
LY50
LY75
LY88
LY94
LY97
LY data
Normal data

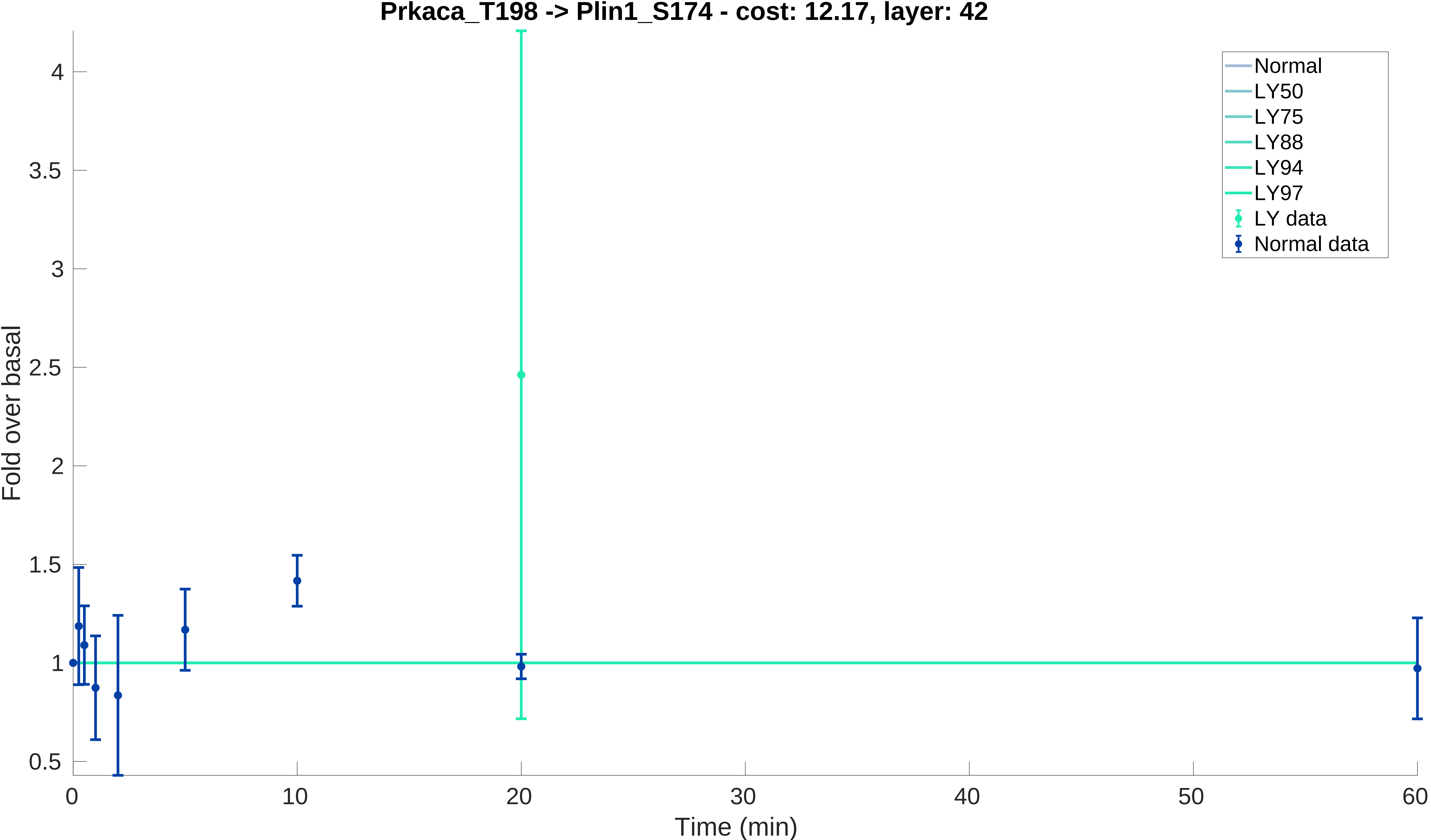

Prkaca_T198 -> Plin1_S174 - cost: 12.17, layer: 42
Fold over basal
Time (min)
0.5
1
1.5
2
2.5
3
3.5
4
0
10
20
30
40
50
60
Normal
LY50
LY75
LY88
LY94
LY97
LY data
Normal data

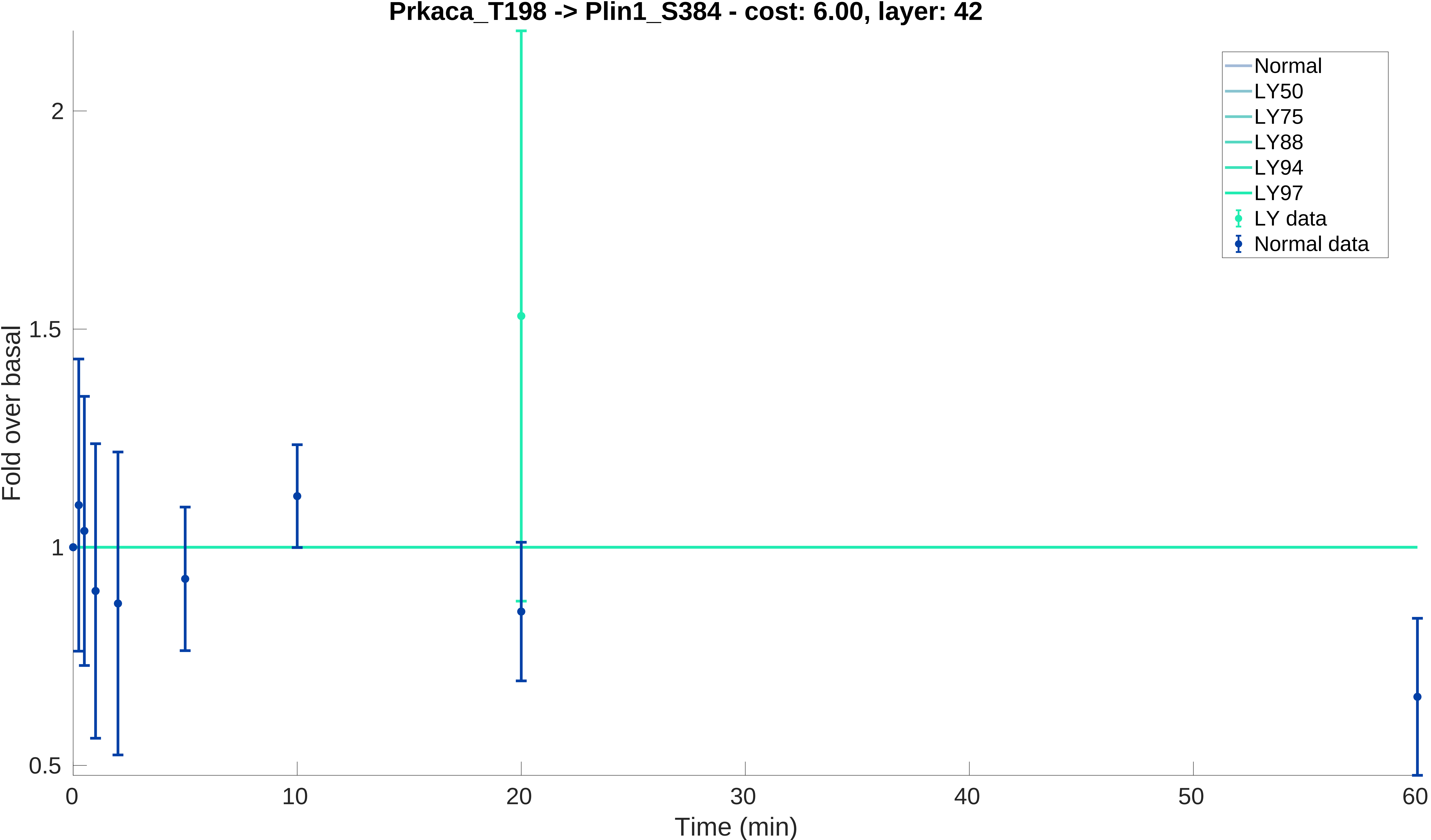

Prkaca_T198 -> Plin1_S384 - cost: 6.00, layer: 42
Normal
LY50
LY75
LY88
LY94
LY97
LY data
Normal data
Fold over basal
2
1.5
1
0.5
0
10
20
30
40
50
60
Time (min)

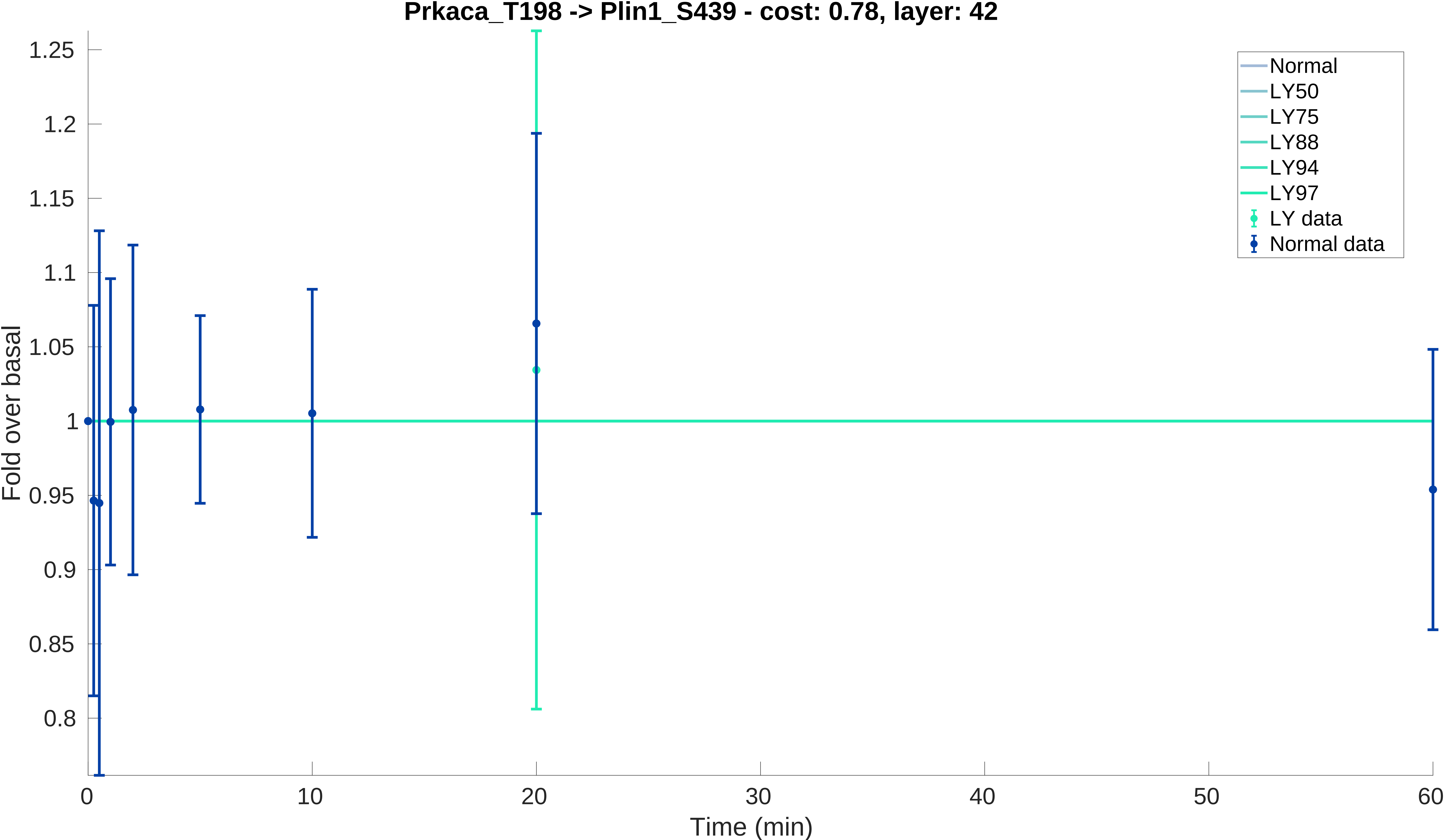

Prkaca_T198 -> Plin1_S439 - cost: 0.78, layer: 42
Fold over basal
1.25
1.2
1.15
1.1
1.05
1
0.95
0.9
0.85
0.8
0
10
20
30
40
50
60
Time (min)
Normal
LY50
LY75
LY88
LY94
LY97
LY data
Normal data

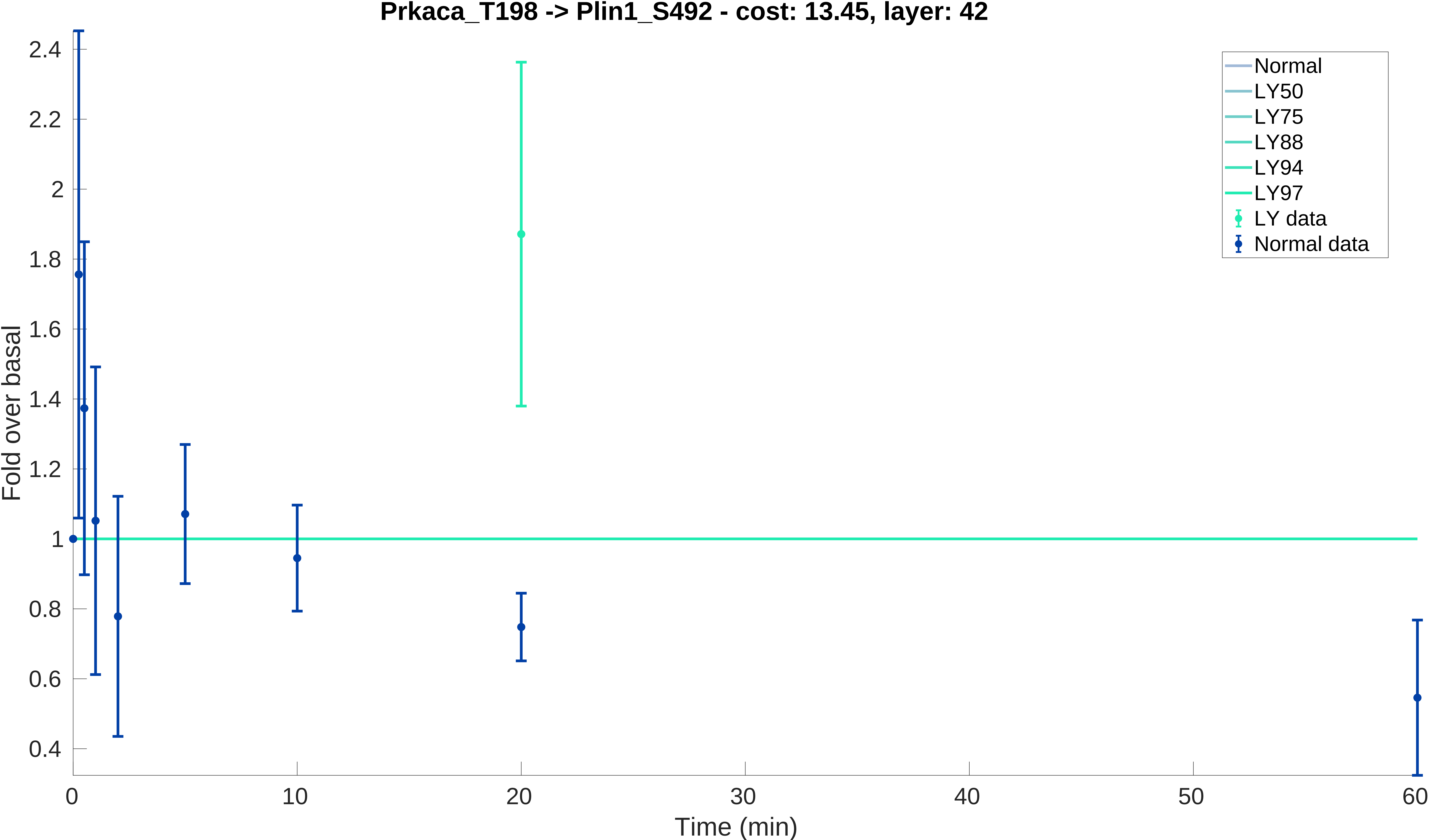

Prkaca_T198 -> Plin1_S492 - cost: 13.45, layer: 42
Fold over basal
2.4
2.2
2
1.8
1.6
1.4
1.2
1
0.8
0.6
0.4
0
10
20
30
40
50
60
Time (min)
Normal
LY50
LY75
LY88
LY94
LY97
LY data
Normal data

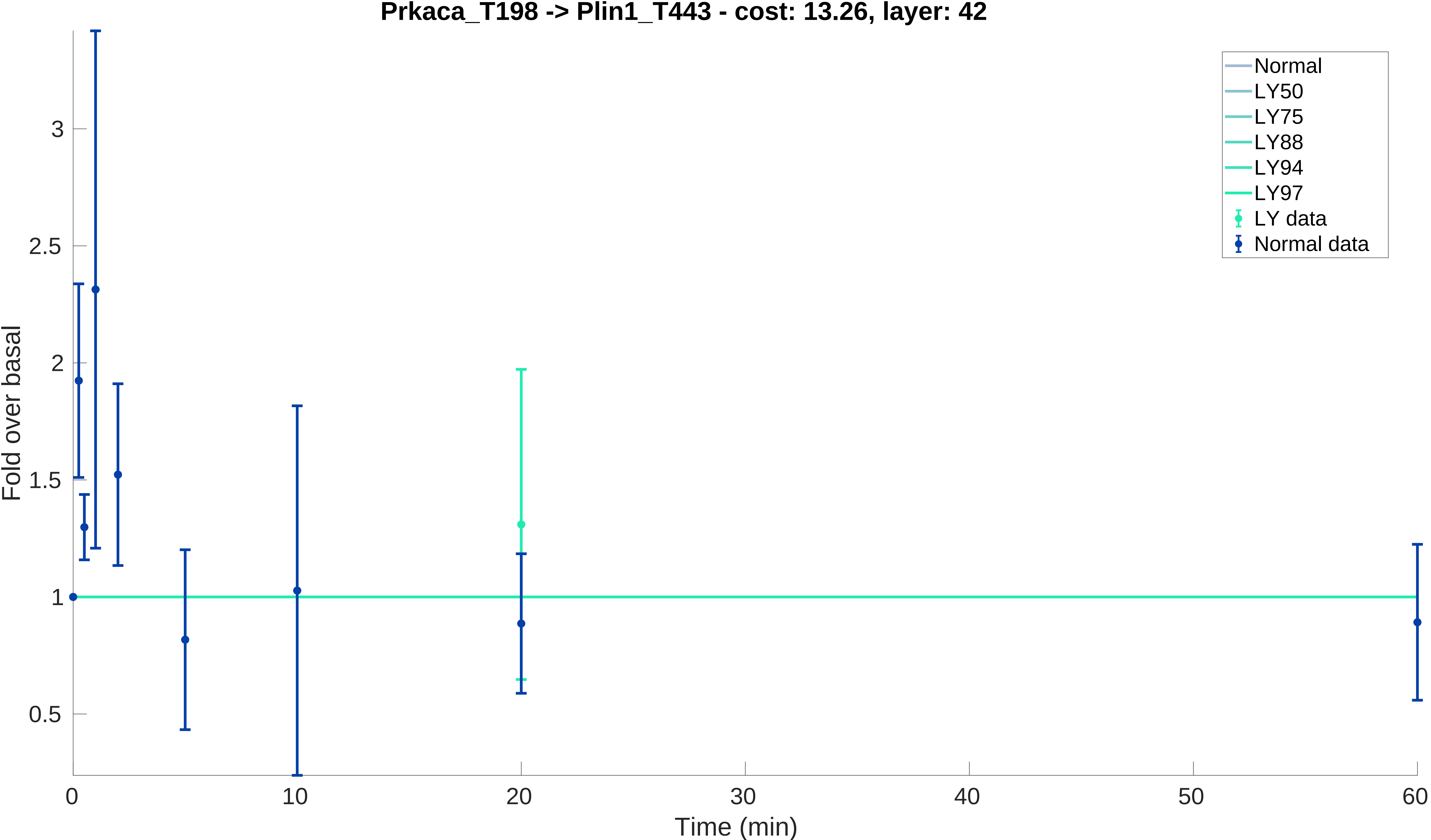
Prkaca_T198 -> Plin1_T443 - cost: 13.26, layer: 42
Normal
LY50
LY75
LY88
LY94
LY97
LY data
Normal data
Fold over basal
0.5
1
1.5
2
2.5
3
0
10
20
30
40
50
60
Time (min)

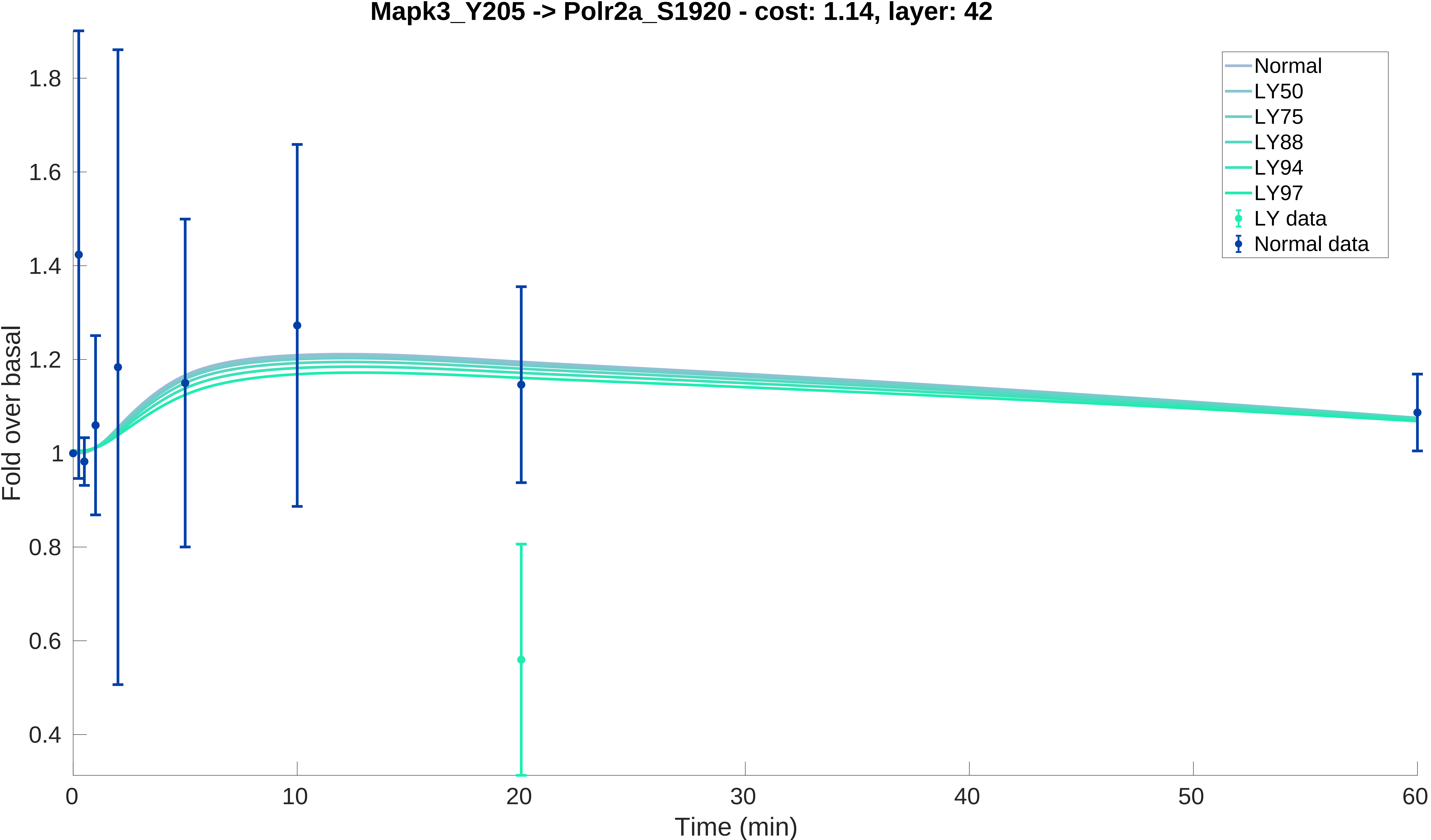

Mapk3_Y205 -> Polr2a_S1920 - cost: 1.14, layer: 42
Normal
LY50
LY75
LY88
LY94
LY97
LY data
Normal data
Fold over basal
0.4
0.6
0.8
1
1.2
1.4
1.6
1.8
0
10
20
30
40
50
60
Time (min)

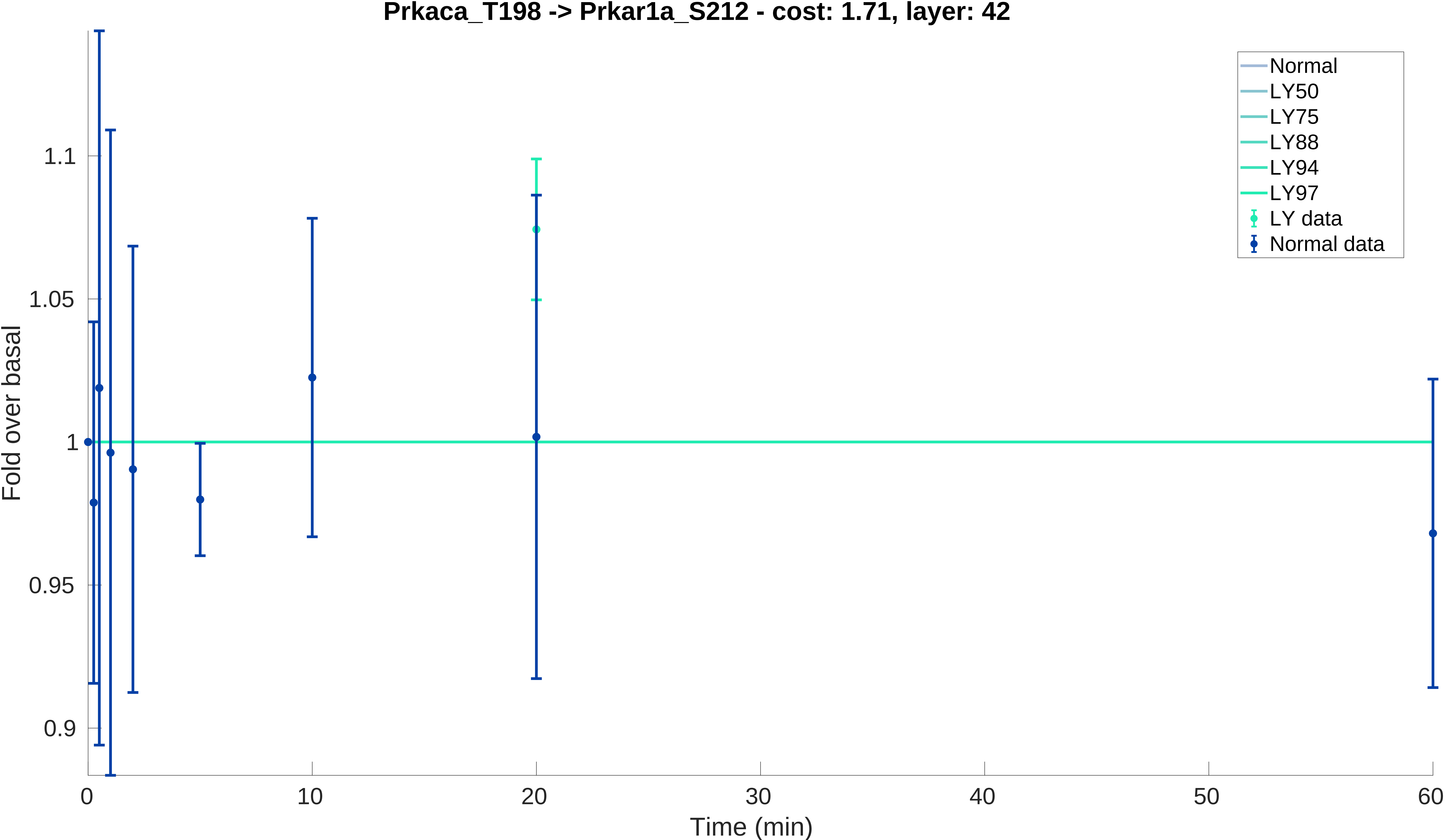
Prkaca_T198 -> Prkar1a_S212 - cost: 1.71, layer: 42
Fold over basal
1.1
1.05
1
0.95
0.9
0
10
20
30
40
50
60
Time (min)
Normal
LY50
LY75
LY88
LY94
LY97
LY data
Normal data

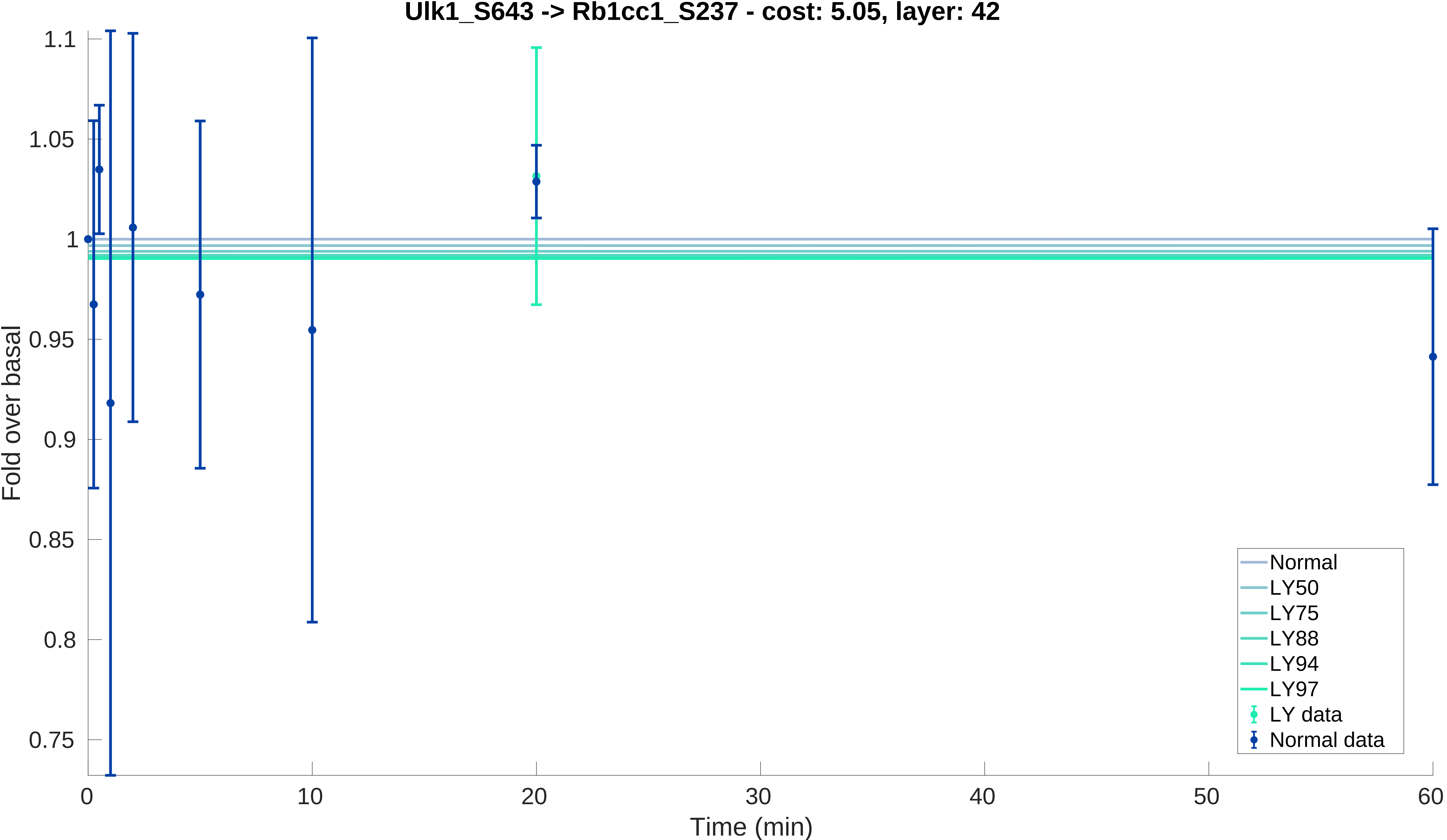

Ulk1_S643 -> Rb1cc1_S237 - cost: 5.05, layer: 42
Fold over basal
1.1
1.05
1
0.95
0.9
0.85
0.8
0.75
0
10
20
30
40
50
60
Time (min)
Normal
LY50
LY75
LY88
LY94
LY97
LY data
Normal data

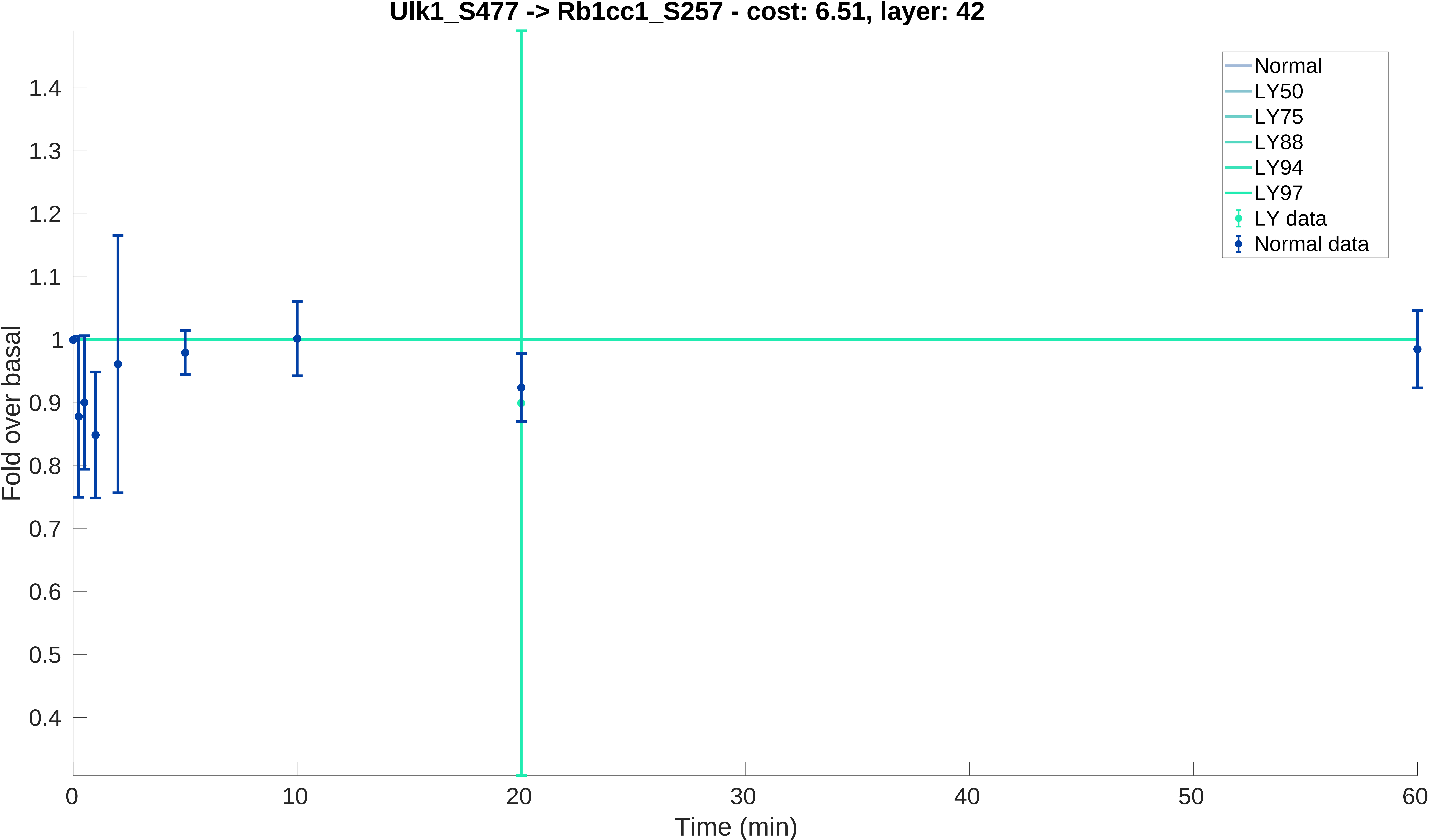

Ulk1_S477 -> Rb1cc1_S257 - cost: 6.51, layer: 42
Fold over basal
Time (min)
Normal
LY50
LY75
LY88
LY94
LY97
LY data
Normal data
1.4
1.3
1.2
1.1
1
0.9
0.8
0.7
0.6
0.5
0.4
0
10
20
30
40
50
60

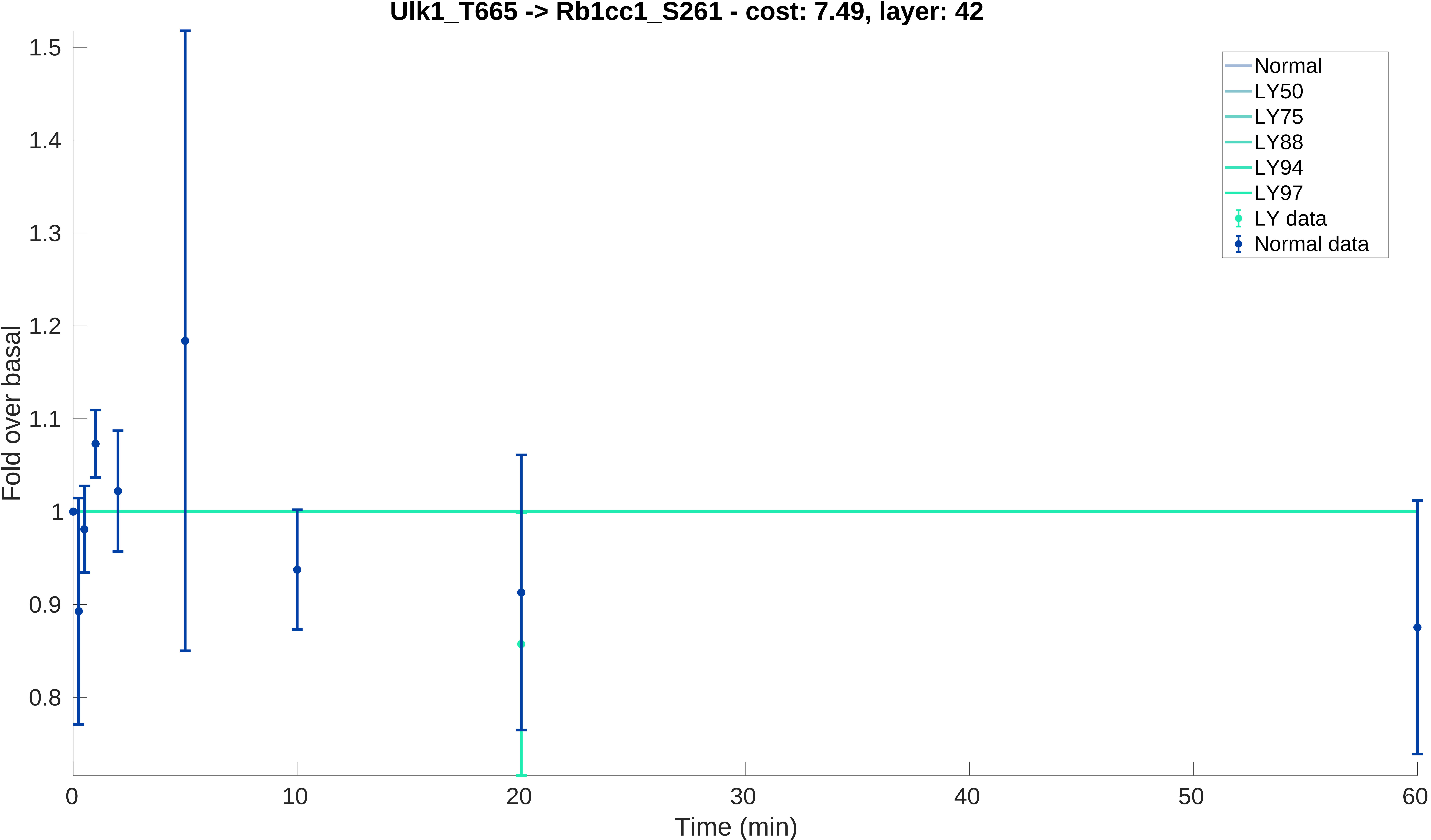
Ulk1_T665 -> Rb1cc1_S261 - cost: 7.49, layer: 42
Fold over basal
Time (min)
1.5
1.4
1.3
1.2
1.1
1
0.9
0.8
0
10
20
30
40
50
60
Normal
LY50
LY75
LY88
LY94
LY97
LY data
Normal data

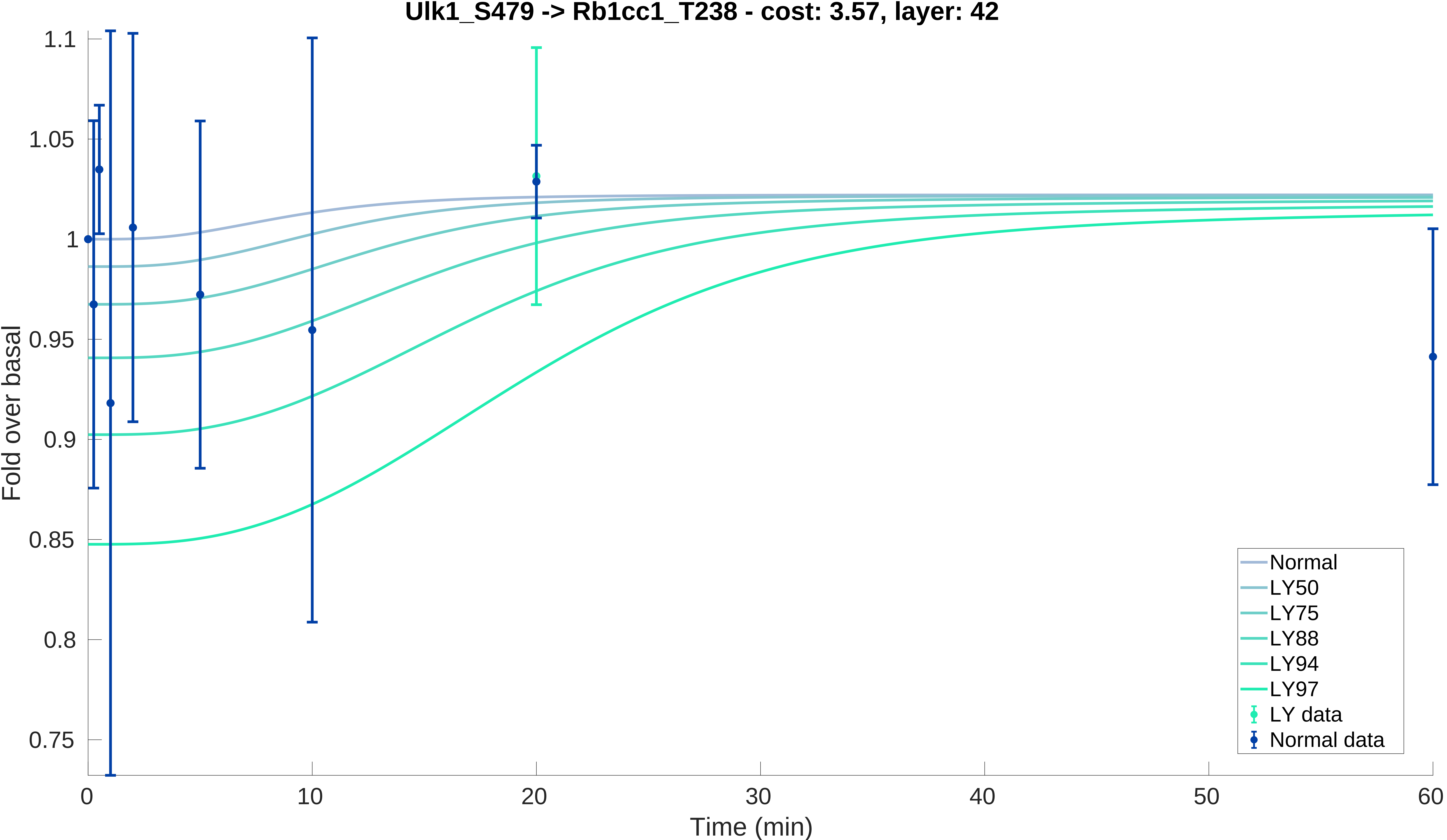
Ulk1_S479 -> Rb1cc1_T238 - cost: 3.57, layer: 42
Fold over basal
1.1
1.05
1
0.95
0.9
0.85
0.8
0.75
0
10
20
30
40
50
60
Time (min)
Normal
LY50
LY75
LY88
LY94
LY97
LY data
Normal data

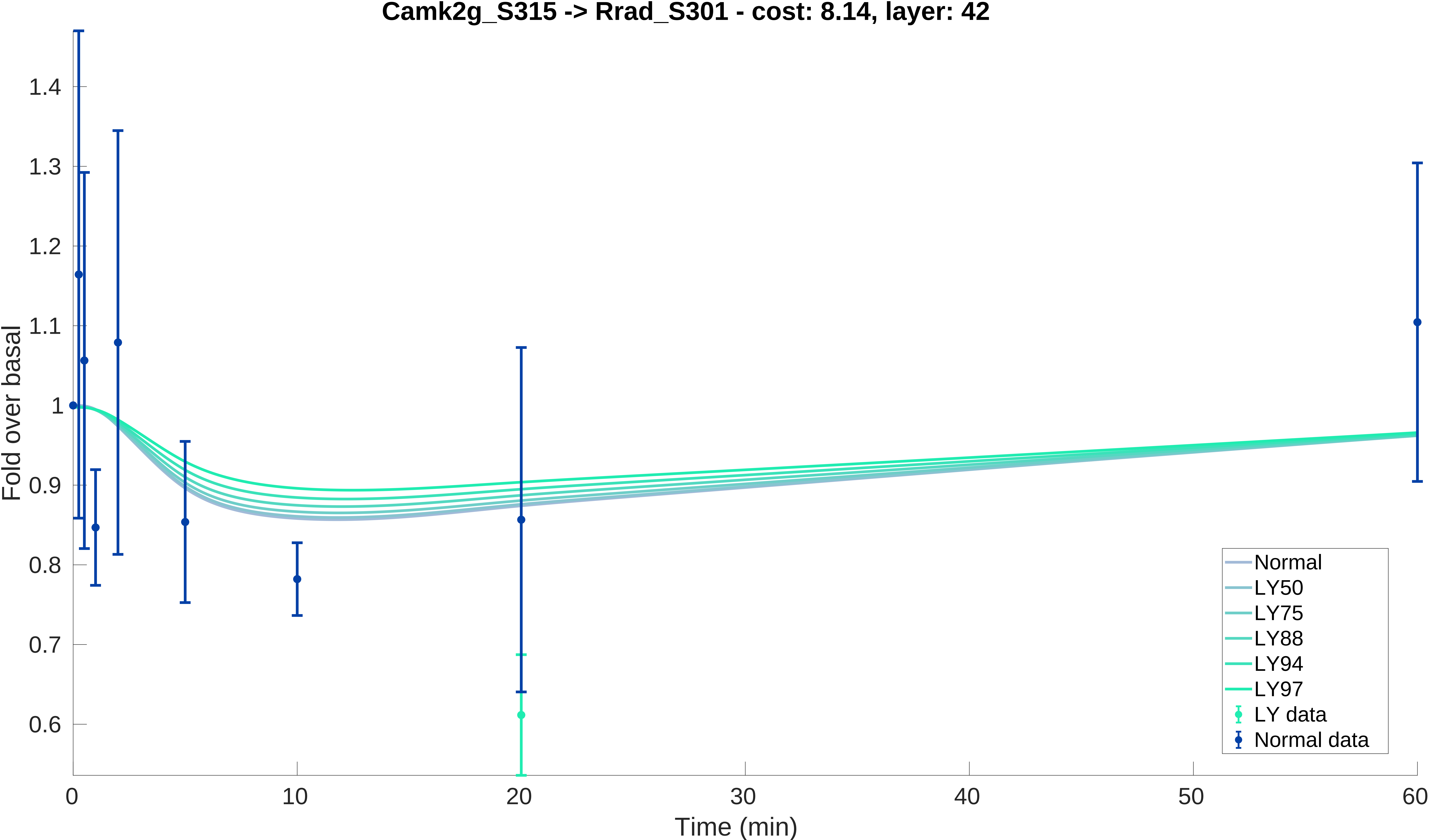
Camk2g_S315 -> Rrad_S301 - cost: 8.14, layer: 42
Fold over basal
1.4
1.3
1.2
1.1
1
0.9
0.8
0.7
0.6
0
10
20
30
40
50
60
Time (min)
Normal
LY50
LY75
LY88
LY94
LY97
LY data
Normal data

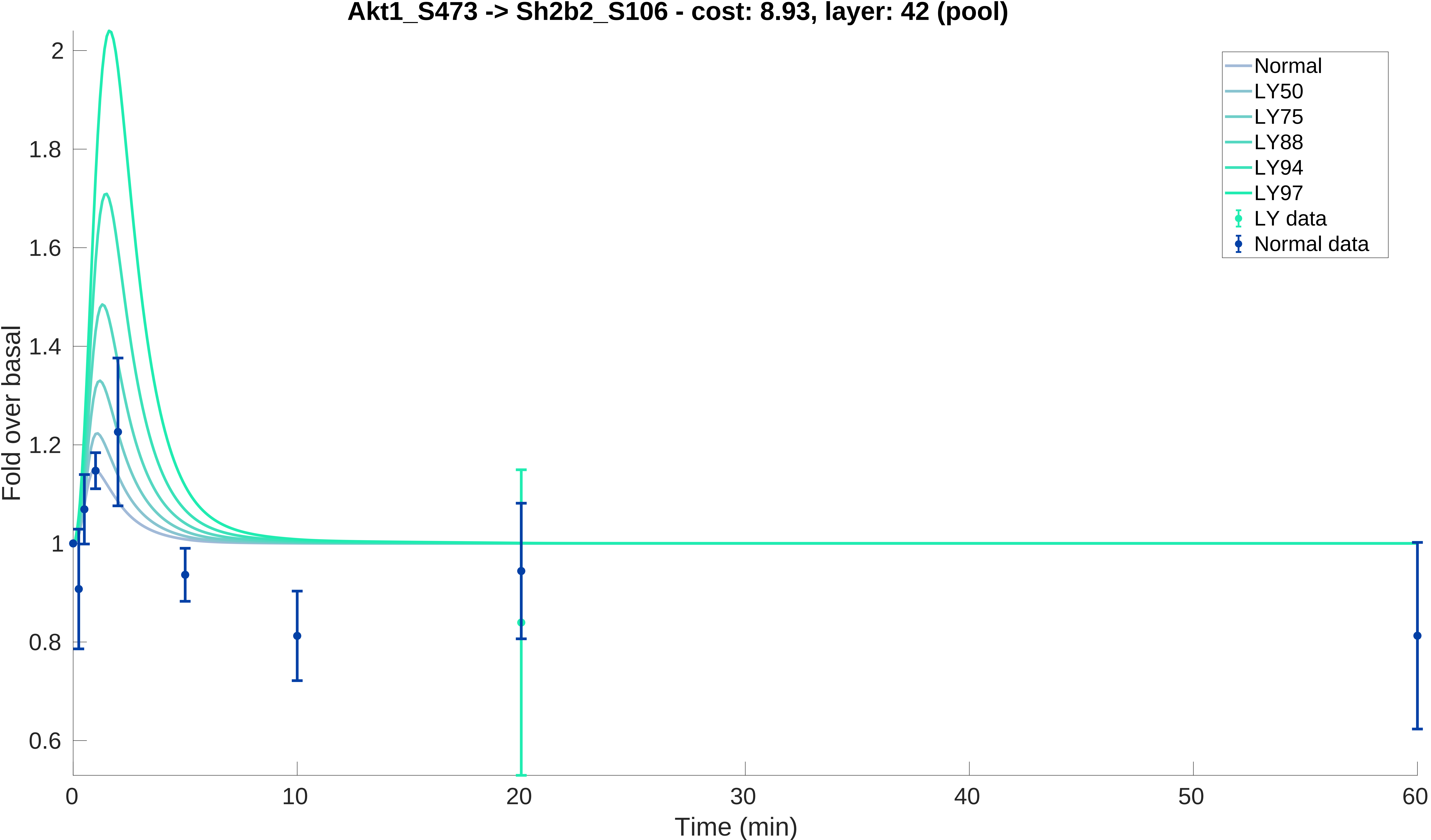

Akt1_S473 -> Sh2b2_S106 - cost: 8.93, layer: 42 (pool)
Fold over basal
Time (min)
Normal
LY50
LY75
LY88
LY94
LY97
LY data
Normal data
0.6
0.8
1
1.2
1.4
1.6
1.8
2
0
10
20
30
40
50
60

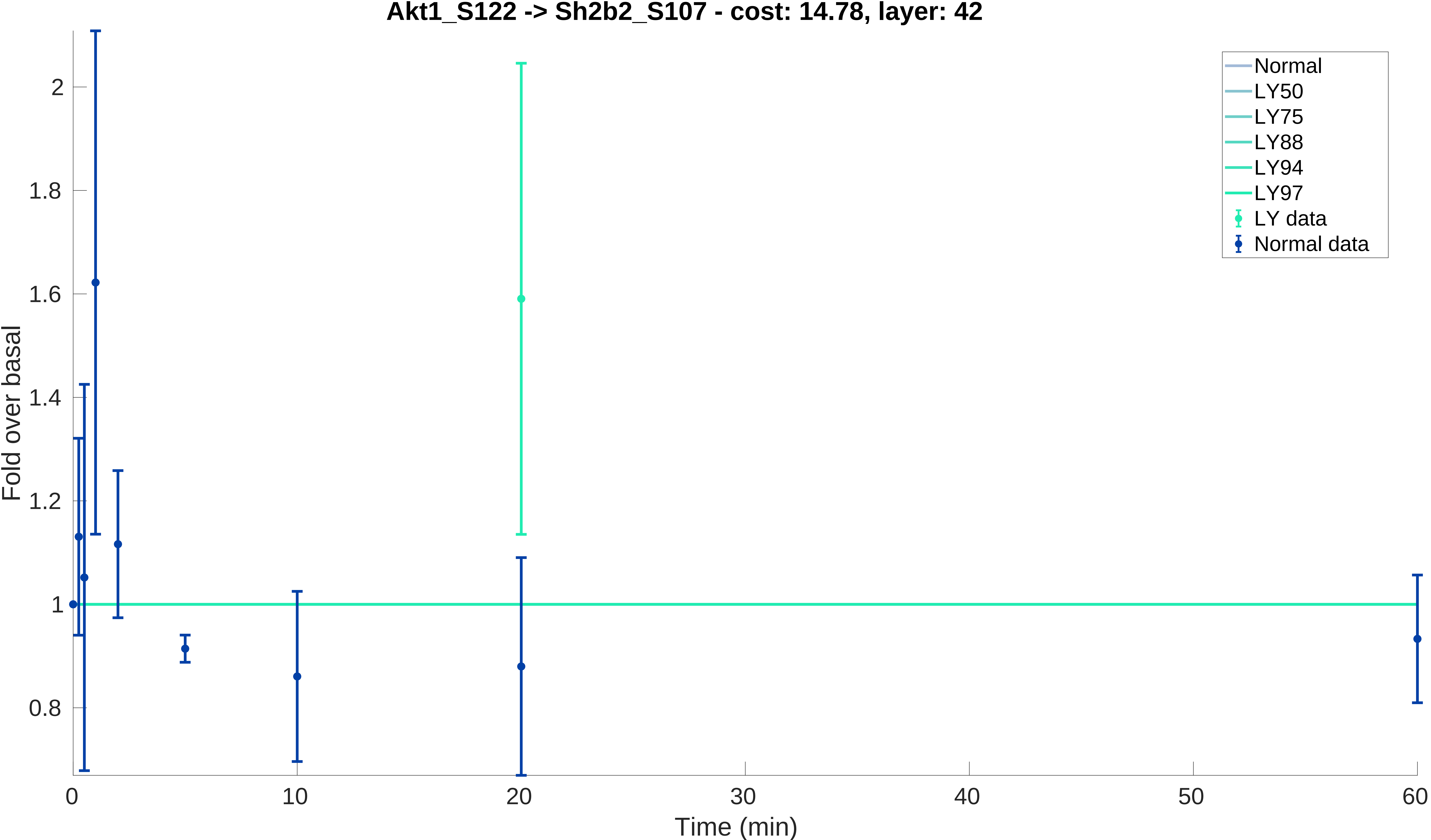

Akt1_S122 -> Sh2b2_S107 - cost: 14.78, layer: 42
Fold over basal
Time (min)
0.8
1
1.2
1.4
1.6
1.8
2
0
10
20
30
40
50
60
Normal
LY50
LY75
LY88
LY94
LY97
LY data
Normal data

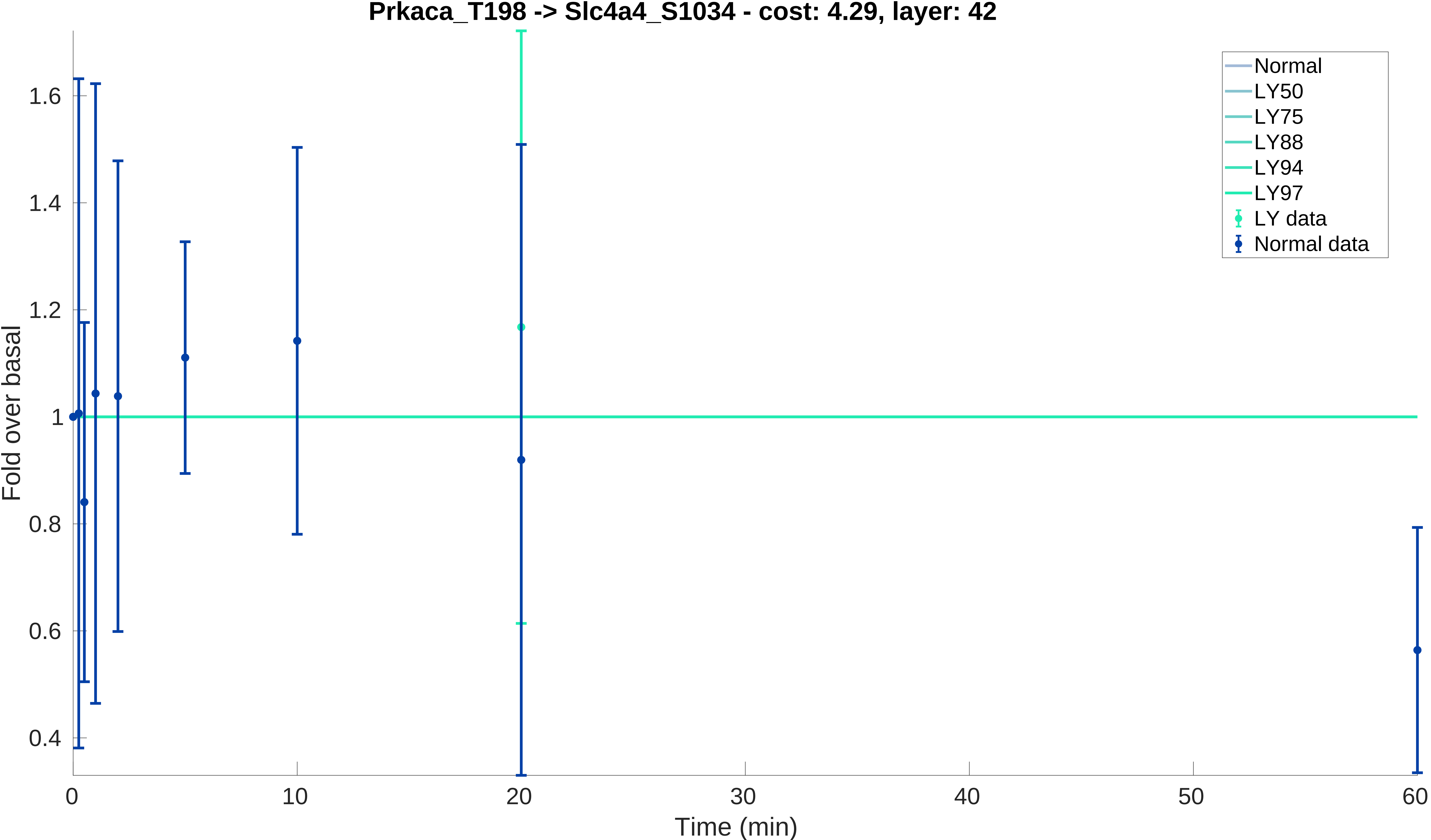

Prkaca_T198 -> Slc4a4_S1034 - cost: 4.29, layer: 42
Fold over basal
Time (min)
1.6
1.4
1.2
1
0.8
0.6
0.4
0
10
20
30
40
50
60
Normal
LY50
LY75
LY88
LY94
LY97
LY data
Normal data

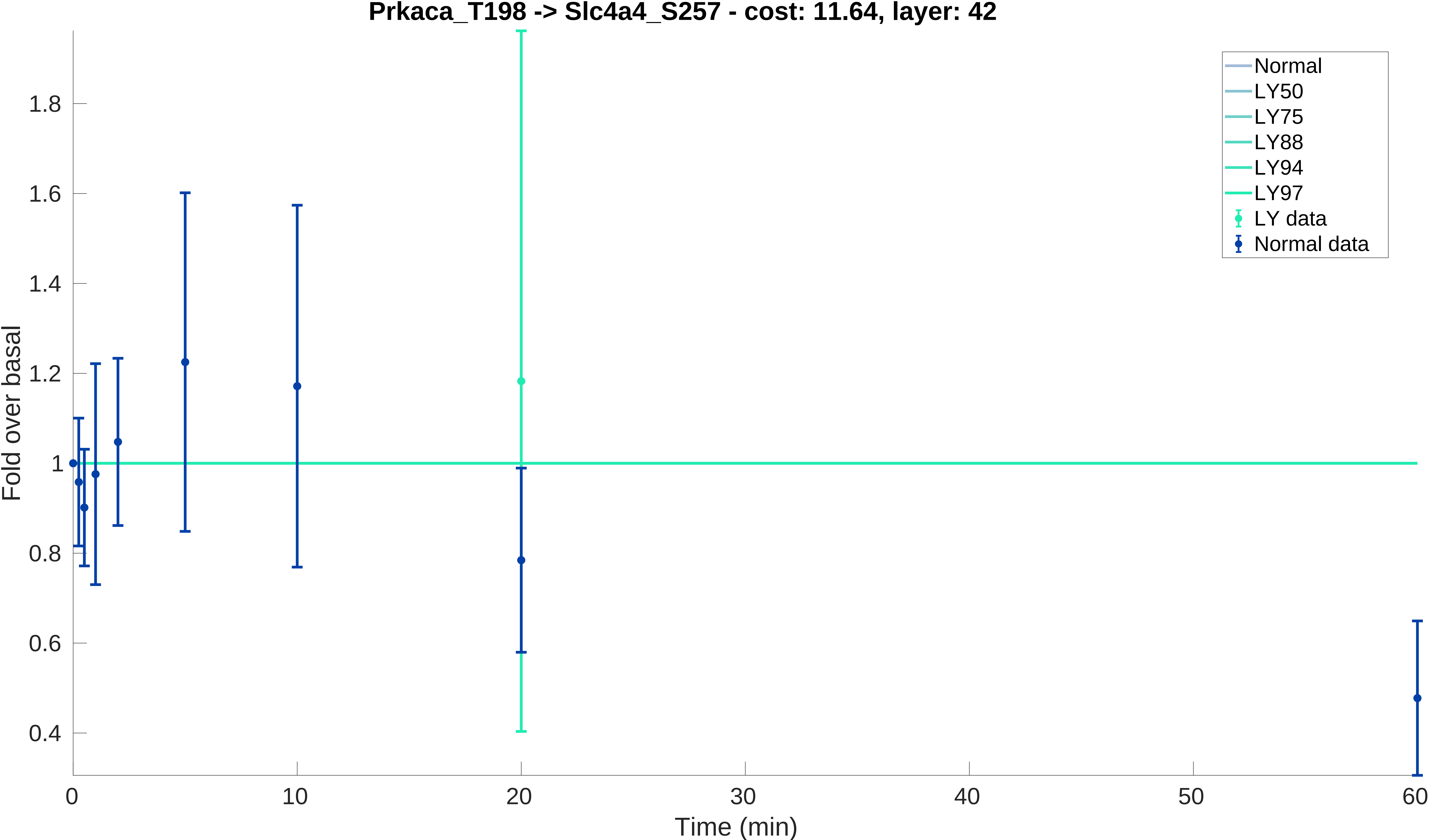

Prkaca_T198 -> Slc4a4_S257 - cost: 11.64, layer: 42
Fold over basal
Time (min)
0.4
0.6
0.8
1
1.2
1.4
1.6
1.8
0
10
20
30
40
50
60
Normal
LY50
LY75
LY88
LY94
LY97
LY data
Normal data

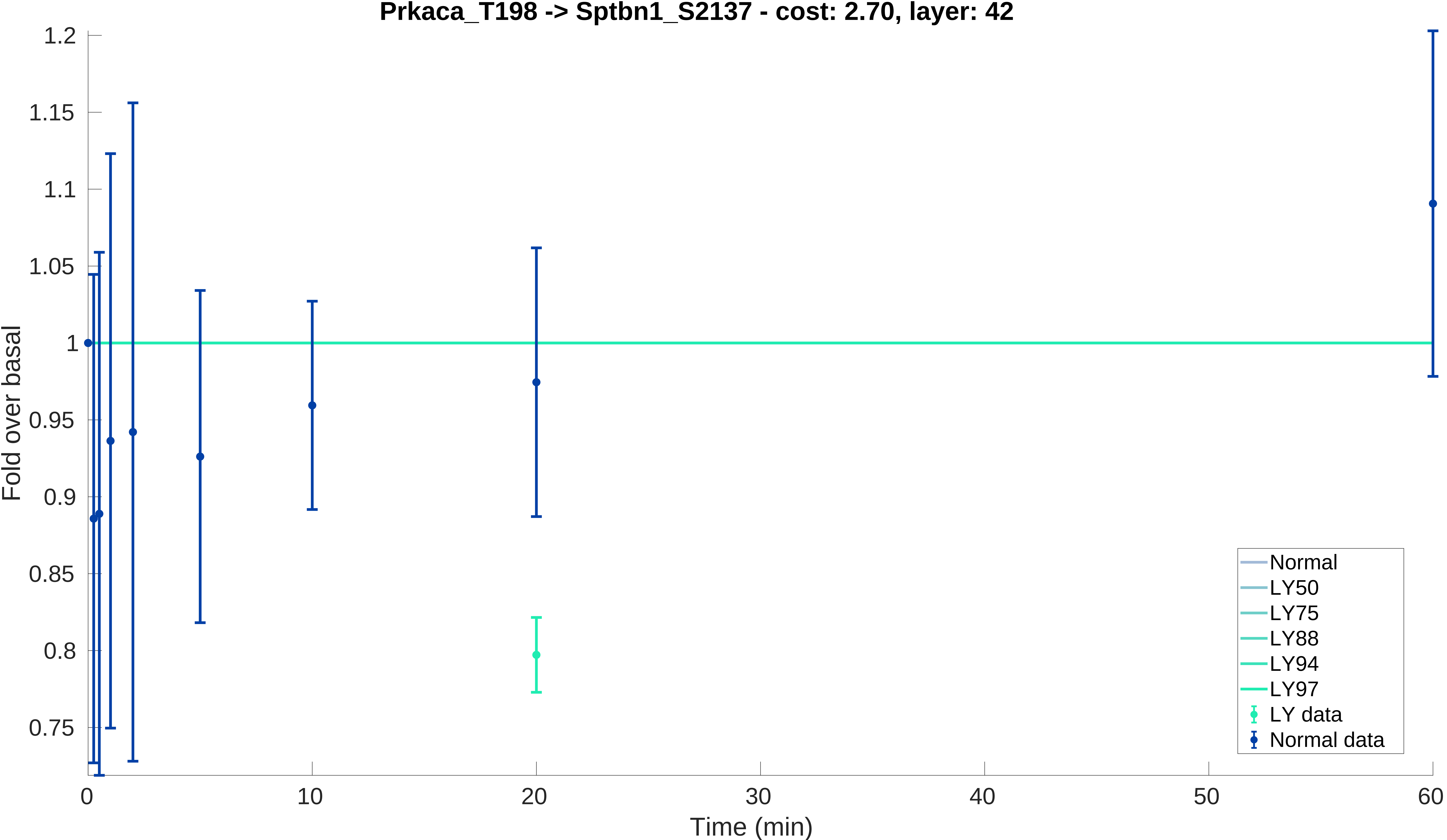
Prkaca_T198 -> Sptbn1_S2137 - cost: 2.70, layer: 42
Fold over basal
1.2
1.15
1.1
1.05
1
0.95
0.9
0.85
0.8
0.75
0
10
20
30
40
50
60
Time (min)
Normal
LY50
LY75
LY88
LY94
LY97
LY data
Normal data

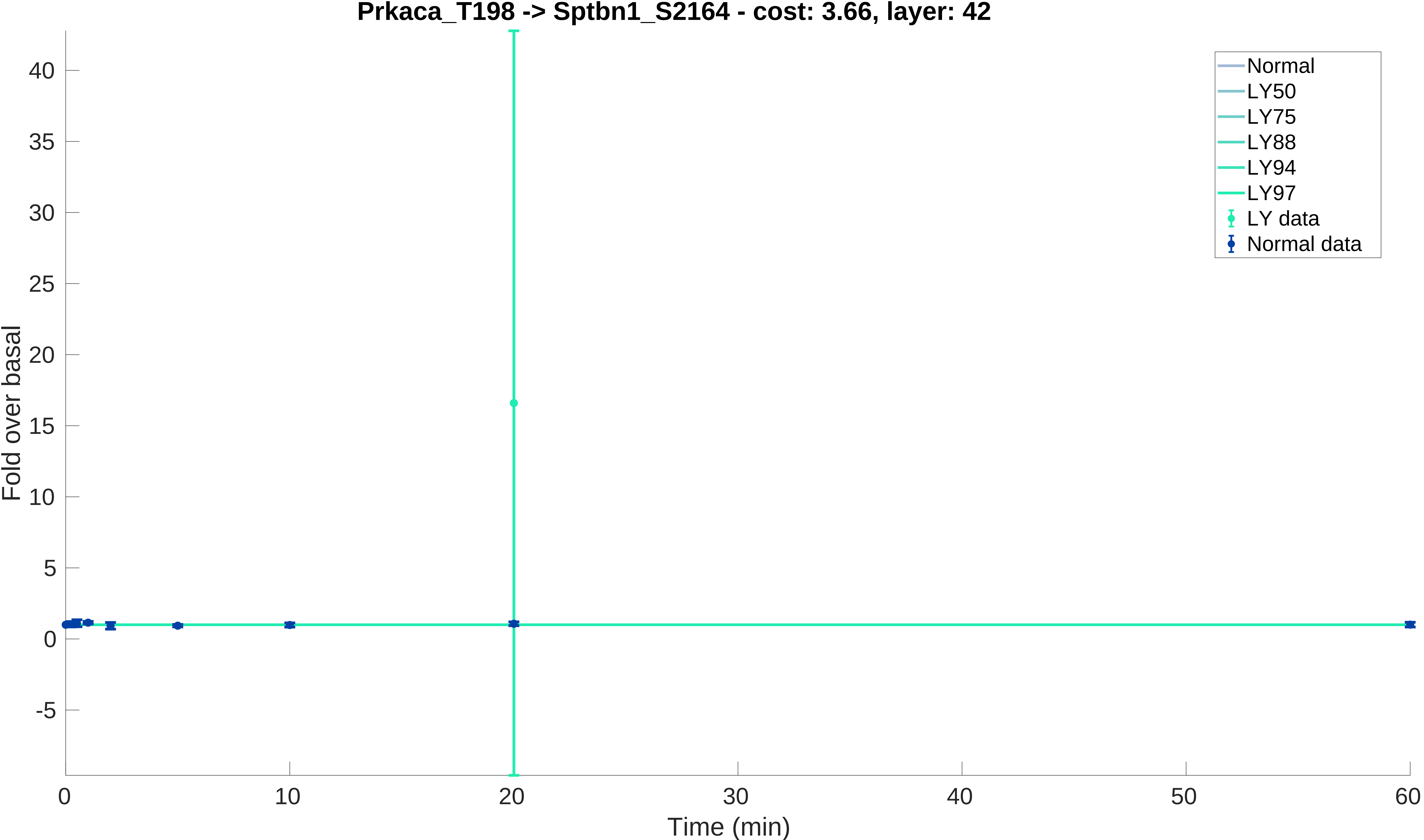

Prkaca_T198 -> Sptbn1_S2164 - cost: 3.66, layer: 42
Fold over basal
Time (min)
40
35
30
25
20
15
10
5
0
-5
0
10
20
30
40
50
60
Normal
LY50
LY75
LY88
LY94
LY97
LY data
Normal data

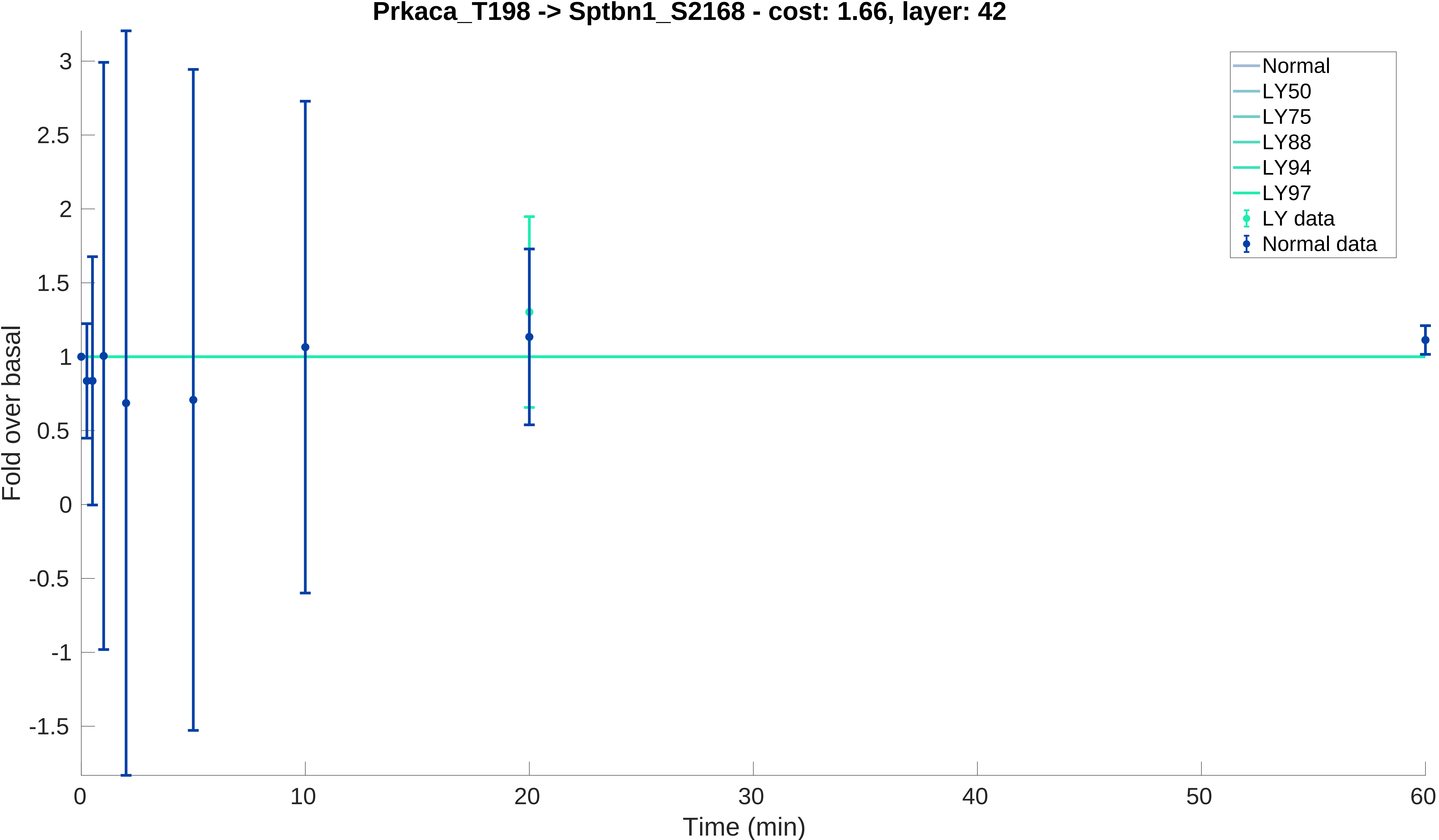

Prkaca_T198 -> Sptbn1_S2168 - cost: 1.66, layer: 42
Fold over basal
Time (min)
3
2.5
2
1.5
1
0.5
0
-0.5
-1
-1.5
0
10
20
30
40
50
60
Normal
LY50
LY75
LY88
LY94
LY97
LY data
Normal data

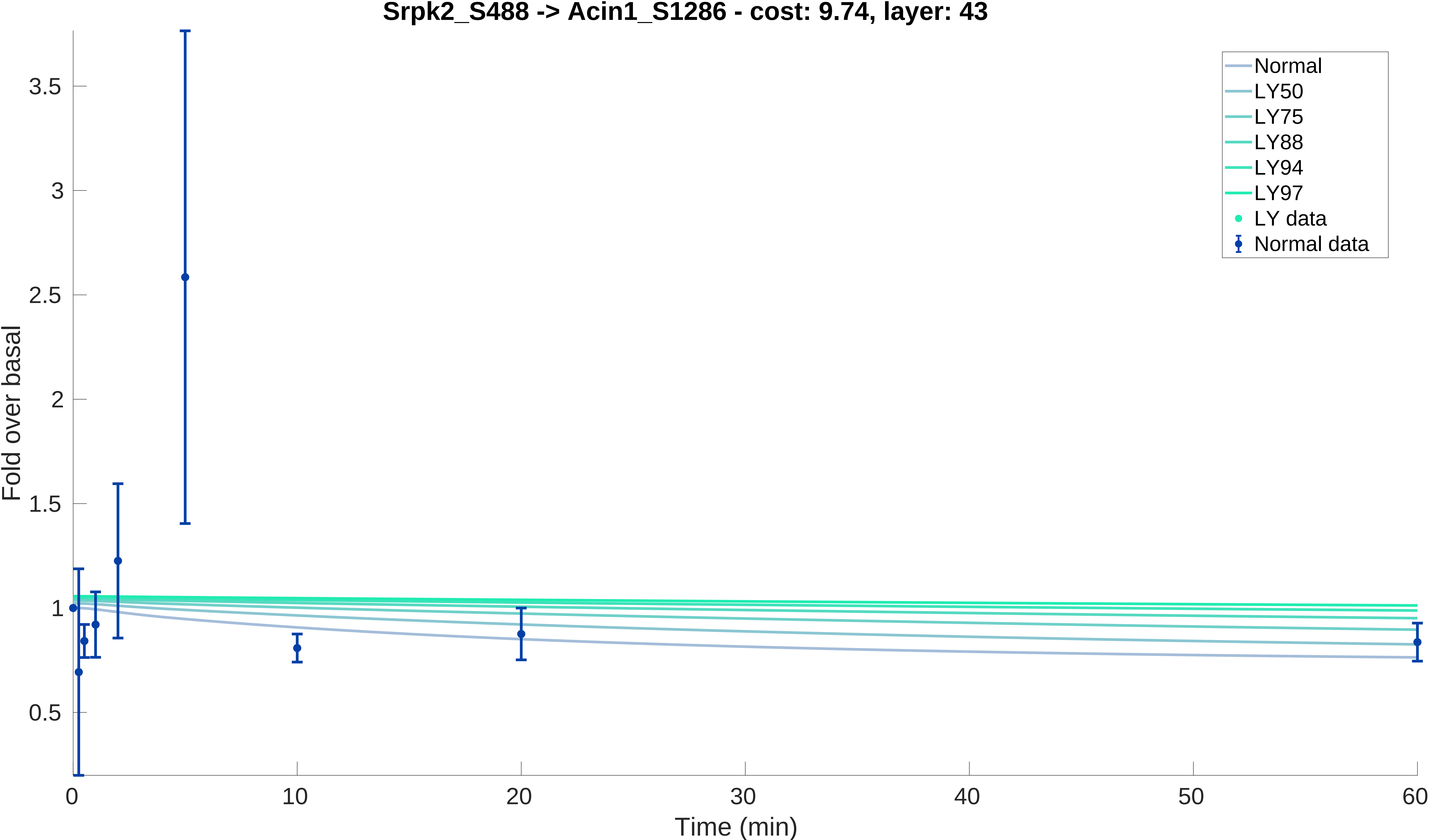

Srpk2_S488 -> Acin1_S1286 - cost: 9.74, layer: 43
Fold over basal
Time (min)
3.5
3
2.5
2
1.5
1
0.5
0
10
20
30
40
50
60
Normal
LY50
LY75
LY88
LY94
LY97
LY data
Normal data

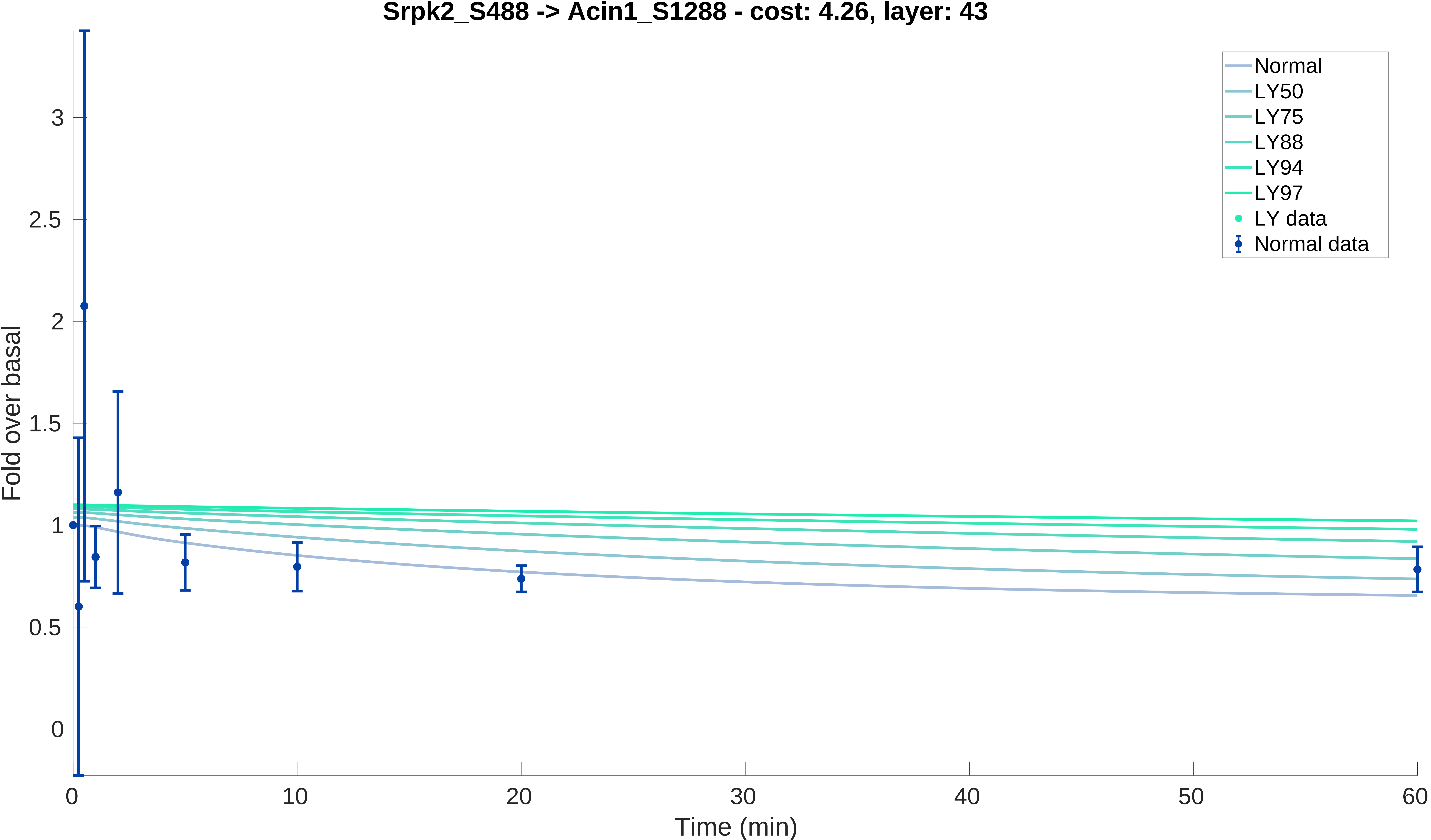

Srpk2_S488 -> Acin1_S1288 - cost: 4.26, layer: 43
Normal
LY50
LY75
LY88
LY94
LY97
LY data
Normal data
Fold over basal
Time (min)
0
0.5
1
1.5
2
2.5
3
0
10
20
30
40
50
60

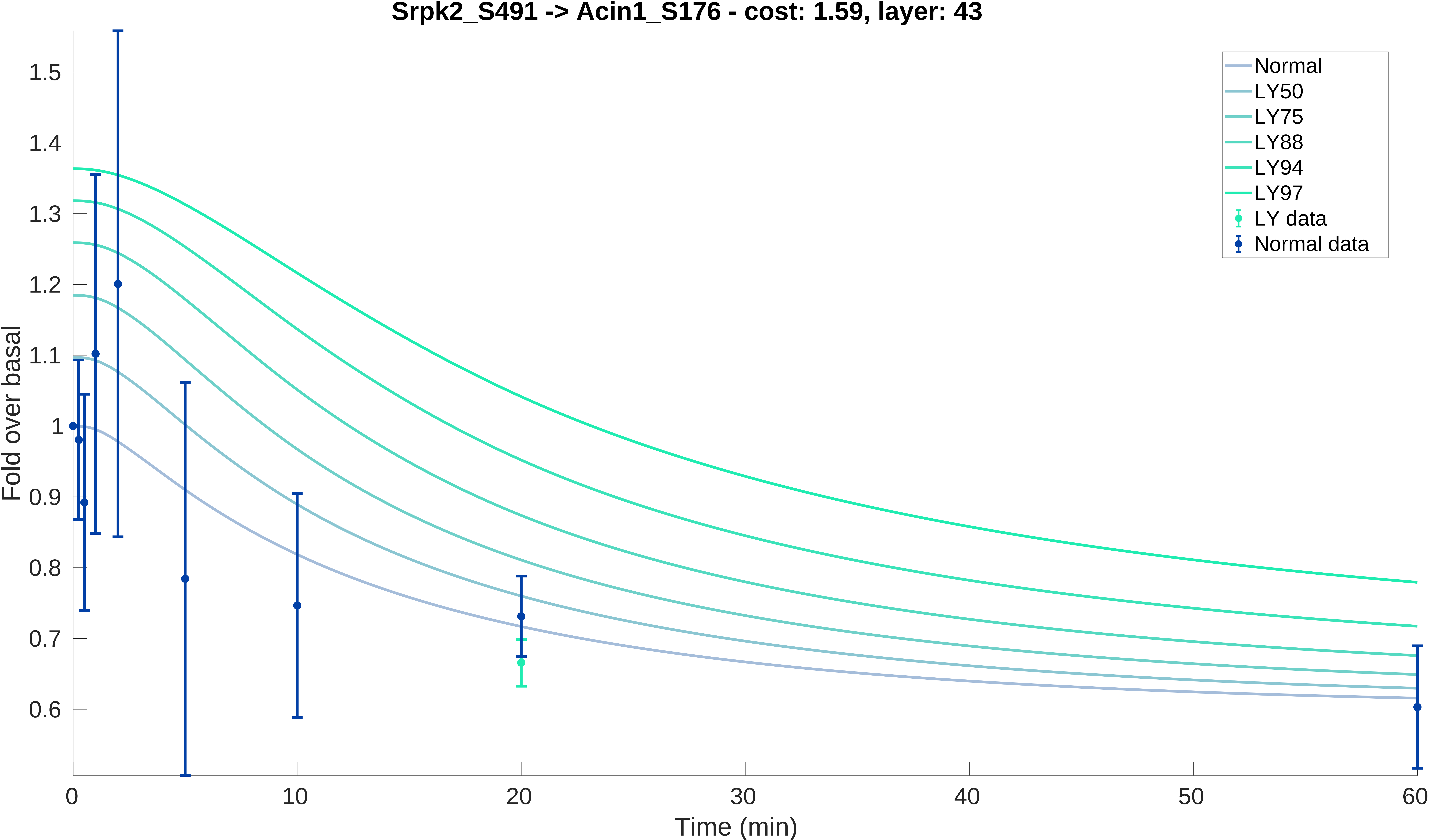

Srpk2_S491 -> Acin1_S176 - cost: 1.59, layer: 43
Normal
LY50
LY75
LY88
LY94
LY97
LY data
Normal data
Fold over basal
Time (min)
1.5
1.4
1.3
1.2
1.1
1
0.9
0.8
0.7
0.6
0
10
20
30
40
50
60

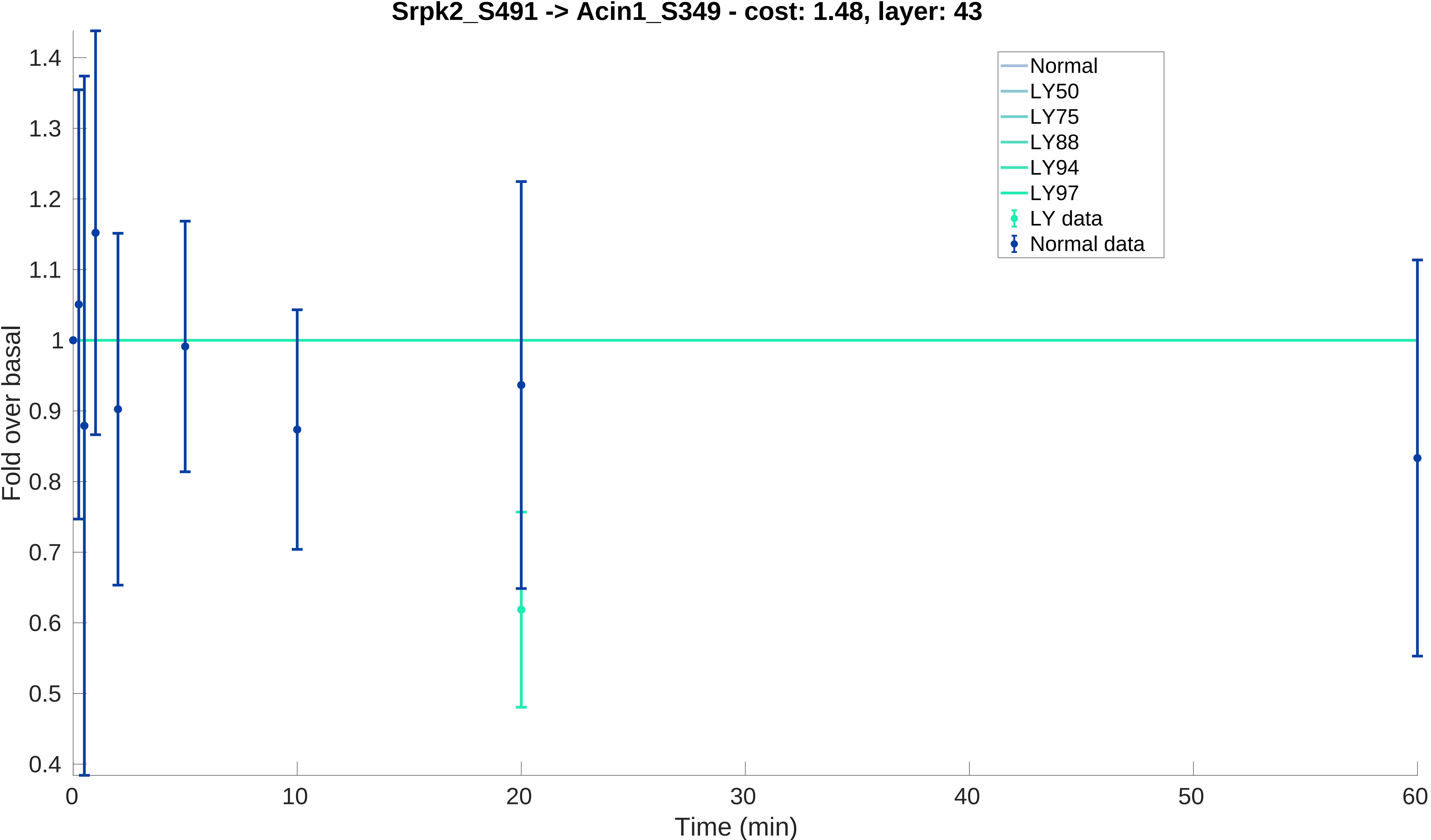

Srpk2_S491 -> Acin1_S349 - cost: 1.48, layer: 43
Normal
LY50
LY75
LY88
LY94
LY97
LY data
Normal data
Fold over basal
Time (min)
0.4
0.5
0.6
0.7
0.8
0.9
1
1.1
1.2
1.3
1.4
0
10
20
30
40
50
60

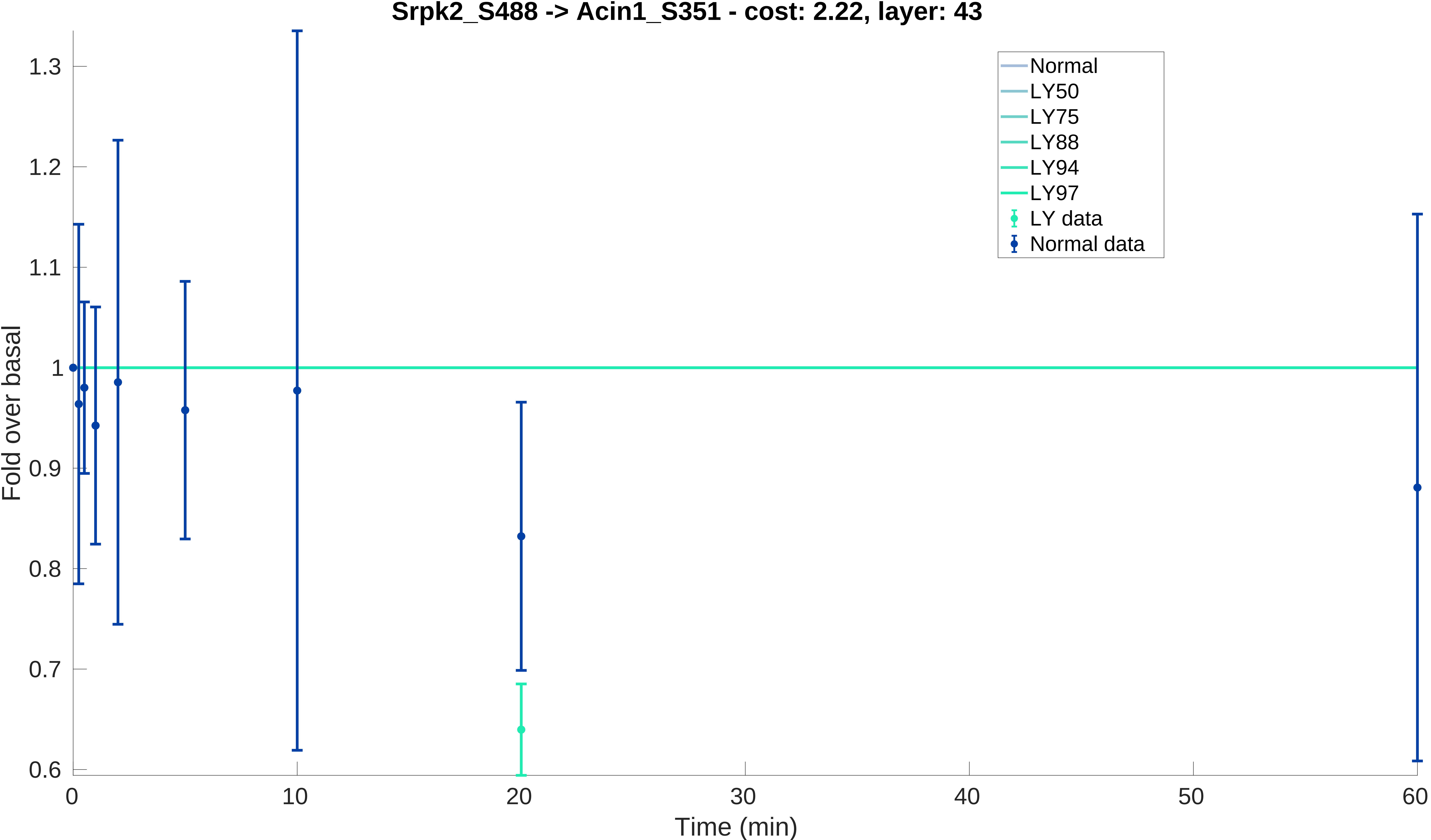

Srpk2_S488 -> Acin1_S351 - cost: 2.22, layer: 43
Fold over basal
Time (min)
1.3
1.2
1.1
1
0.9
0.8
0.7
0.6
0
10
20
30
40
50
60
Normal
LY50
LY75
LY88
LY94
LY97
LY data
Normal data

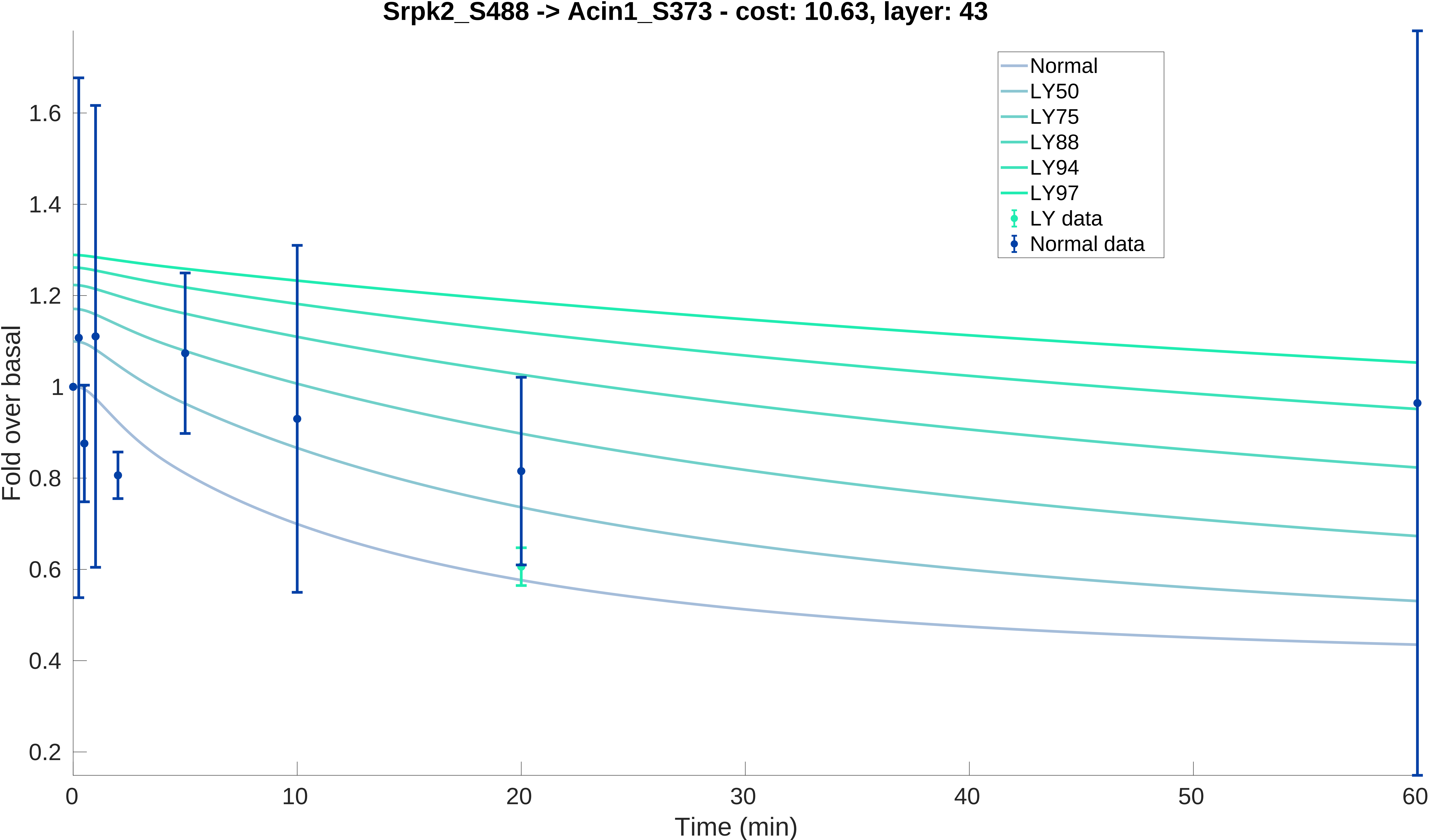

Srpk2_S488 -> Acin1_S373 - cost: 10.63, layer: 43
Normal
LY50
LY75
LY88
LY94
LY97
LY data
Normal data
Fold over basal
Time (min)
0.2
0.4
0.6
0.8
1
1.2
1.4
1.6
0
10
20
30
40
50
60

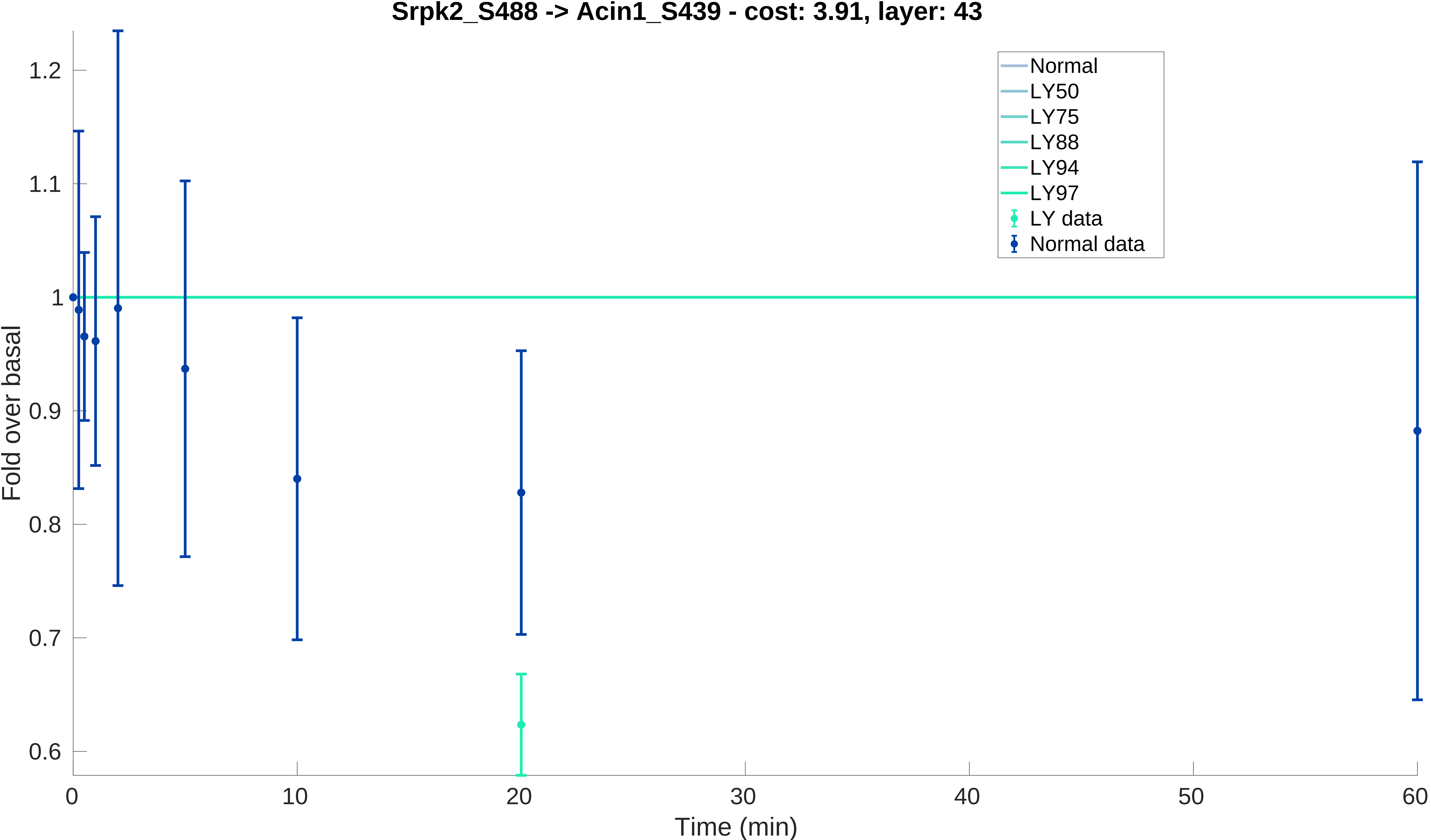
Srpk2_S488 -> Acin1_S439 - cost: 3.91, layer: 43
Normal
LY50
LY75
LY88
LY94
LY97
LY data
Normal data
Fold over basal
Time (min)
0.6
0.7
0.8
0.9
1
1.1
1.2
0
10
20
30
40
50
60

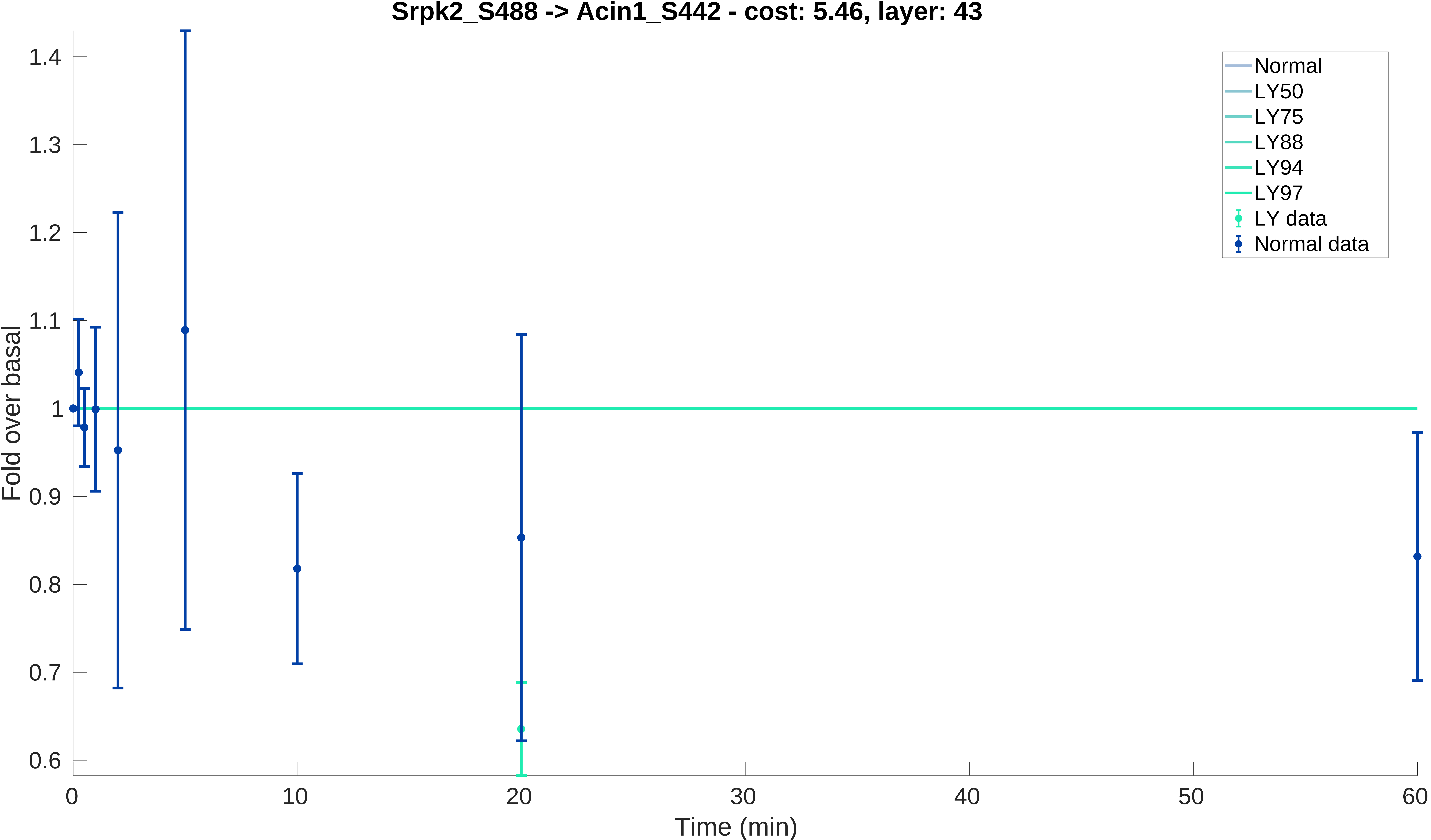

Srpk2_S488 -> Acin1_S442 - cost: 5.46, layer: 43
Fold over basal
Time (min)
1.4
1.3
1.2
1.1
1
0.9
0.8
0.7
0.6
0
10
20
30
40
50
60
Normal
LY50
LY75
LY88
LY94
LY97
LY data
Normal data

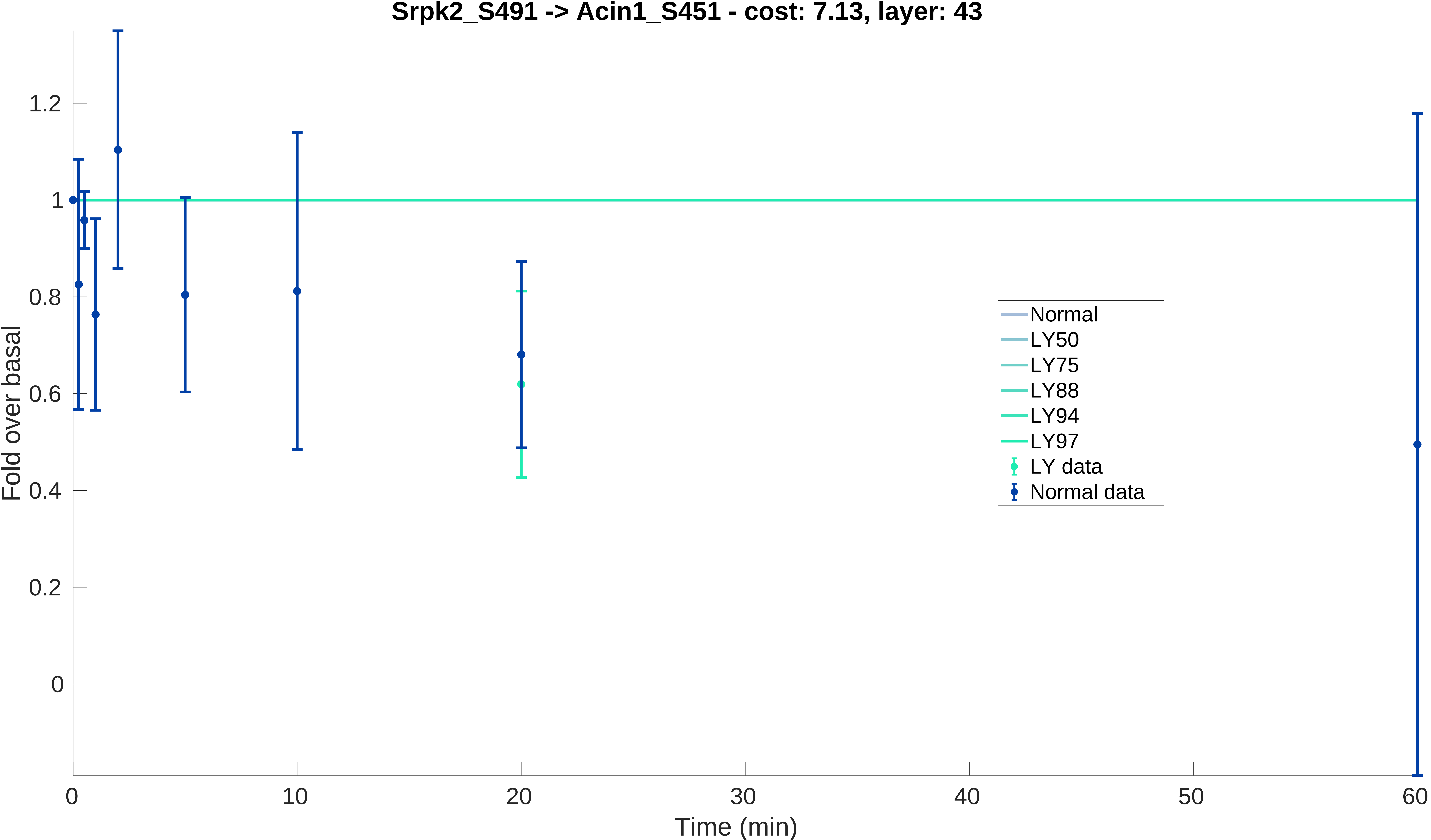
Srpk2_S491 -> Acin1_S451 - cost: 7.13, layer: 43
Fold over basal
Time (min)
1.2
1
0.8
0.6
0.4
0.2
0
0
10
20
30
40
50
60
Normal
LY50
LY75
LY88
LY94
LY97
LY data
Normal data

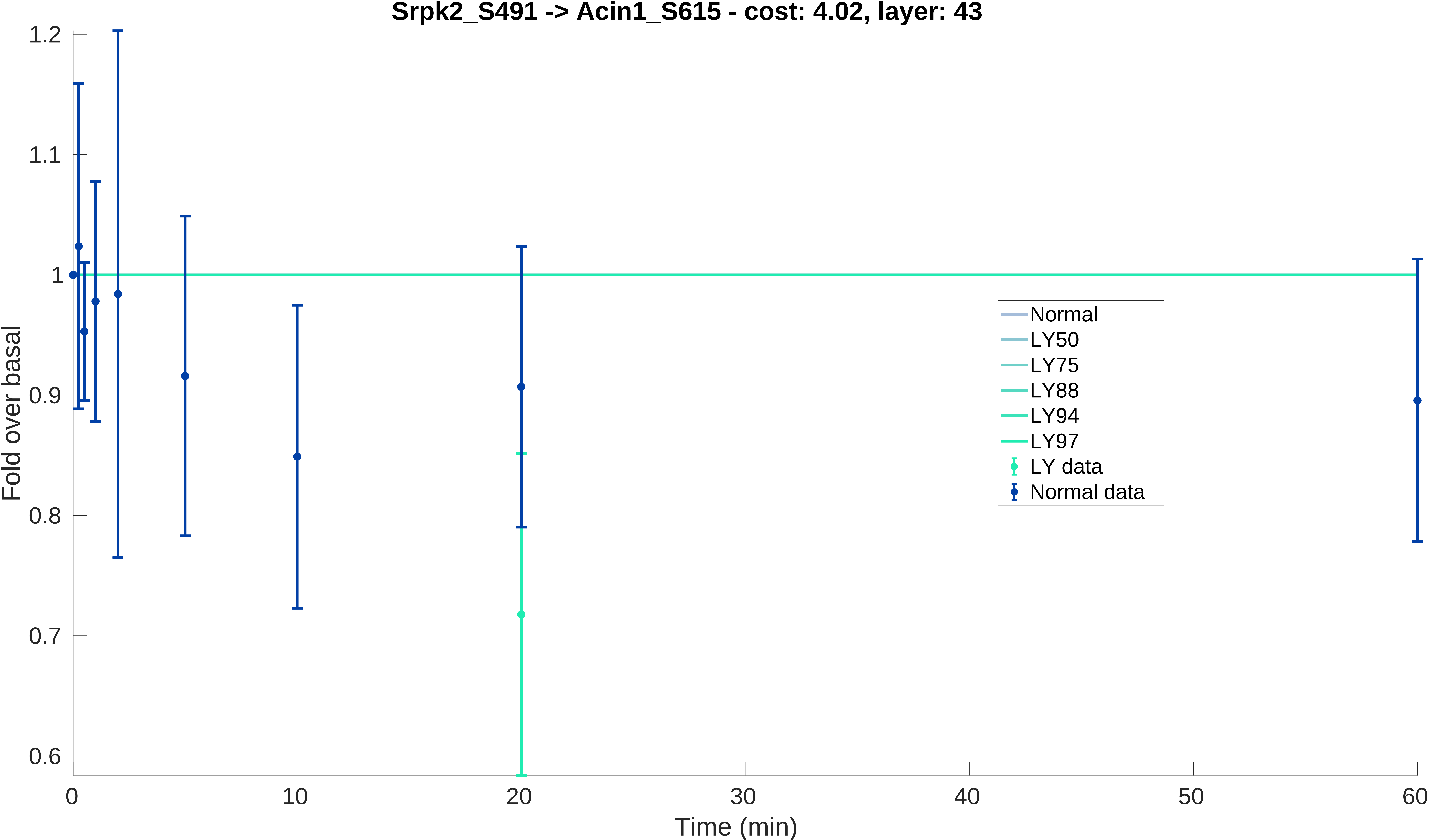

Srpk2_S491 -> Acin1_S615 - cost: 4.02, layer: 43
1.2
1.1
1
0.9
0.8
0.7
0.6
Fold over basal
0
10
20
30
40
50
60
Time (min)
Normal
LY50
LY75
LY88
LY94
LY97
LY data
Normal data

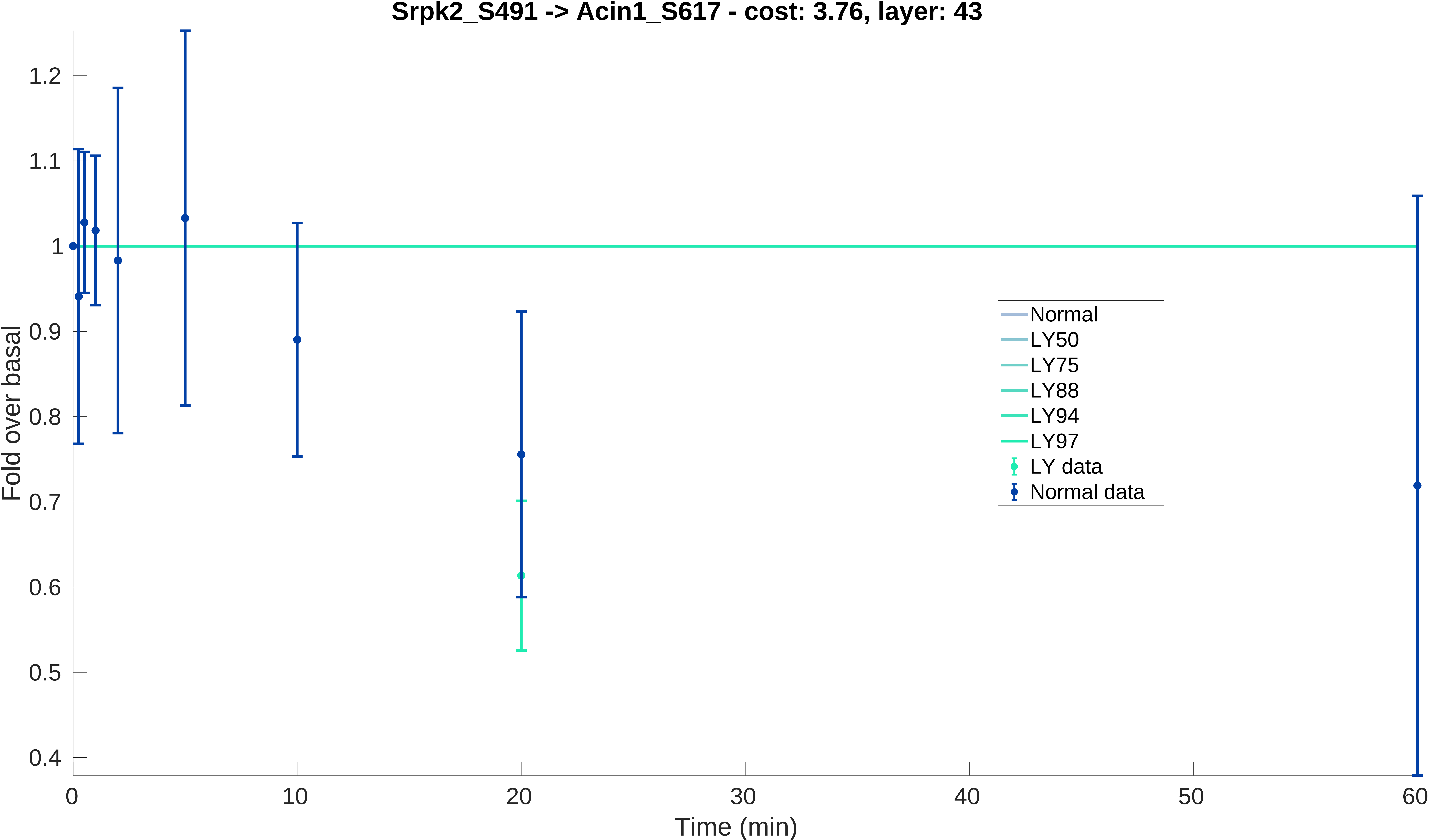
Srpk2_S491 -> Acin1_S617 - cost: 3.76, layer: 43
Fold over basal
1.2
1.1
1
0.9
0.8
0.7
0.6
0.5
0.4
0
10
20
30
40
50
60
Time (min)
Normal
LY50
LY75
LY88
LY94
LY97
LY data
Normal data

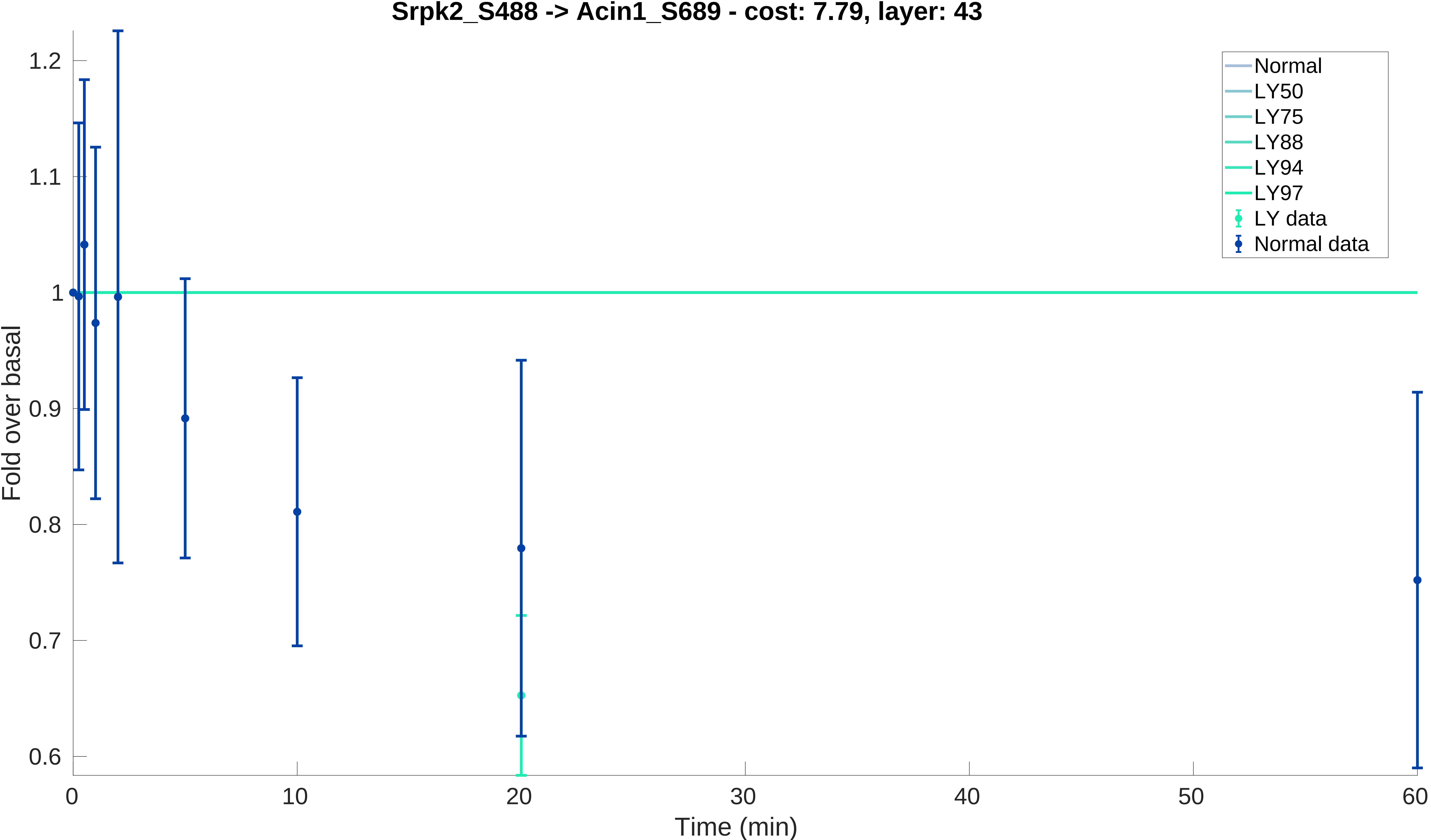

Srpk2_S488 -> Acin1_S689 - cost: 7.79, layer: 43
Fold over basal
Time (min)
Normal
LY50
LY75
LY88
LY94
LY97
LY data
Normal data
1.2
1.1
1
0.9
0.8
0.7
0.6
0
10
20
30
40
50
60

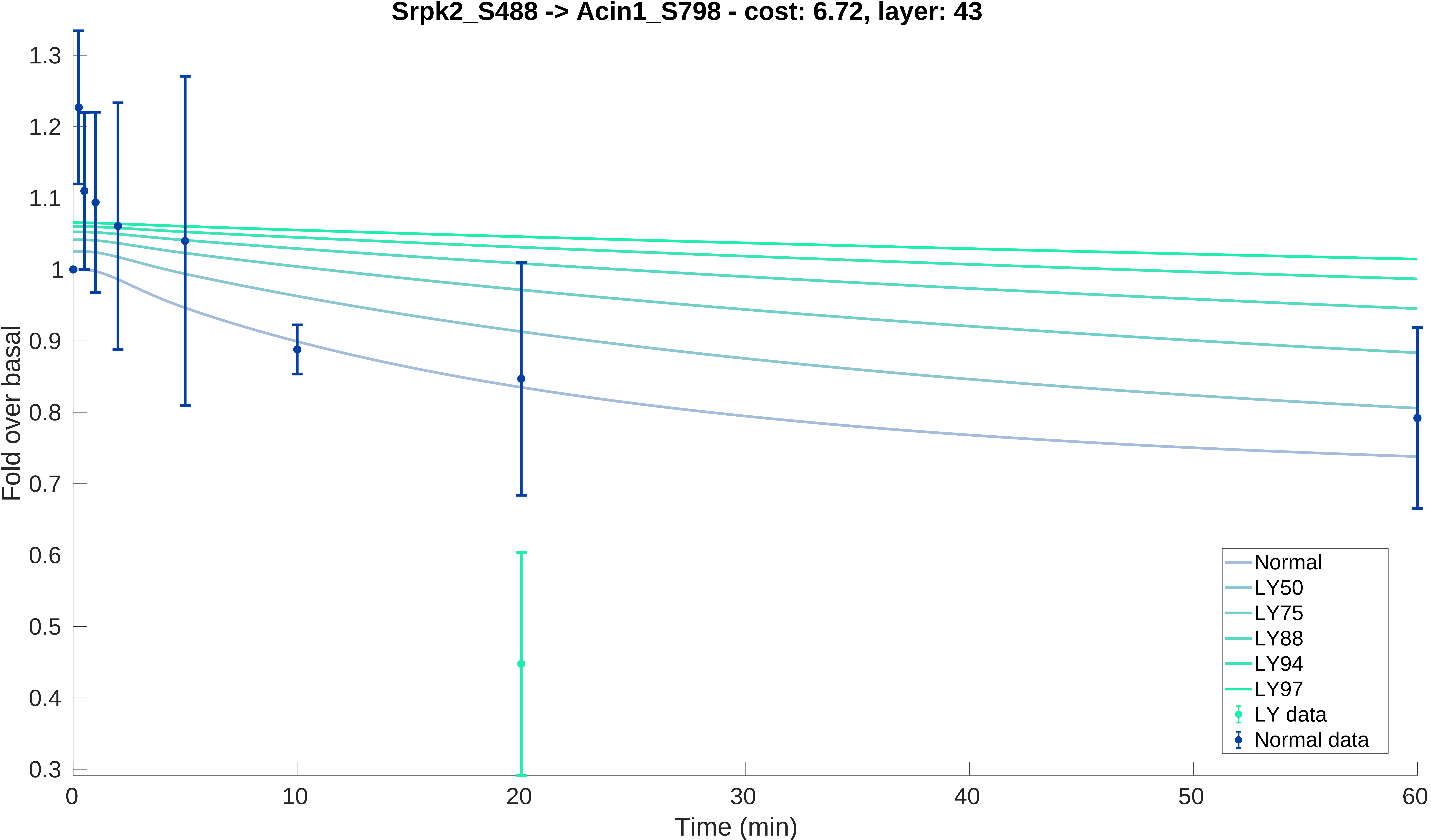

Srpk2_S488 -> Acin1_S798 - cost: 6.72, layer: 43
Fold over basal
1.3
1.2
1.1
1
0.9
0.8
0.7
0.6
0.5
0.4
0.3
0
10
20
30
40
50
60
Time (min)
Normal
LY50
LY75
LY88
LY94
LY97
LY data
Normal data

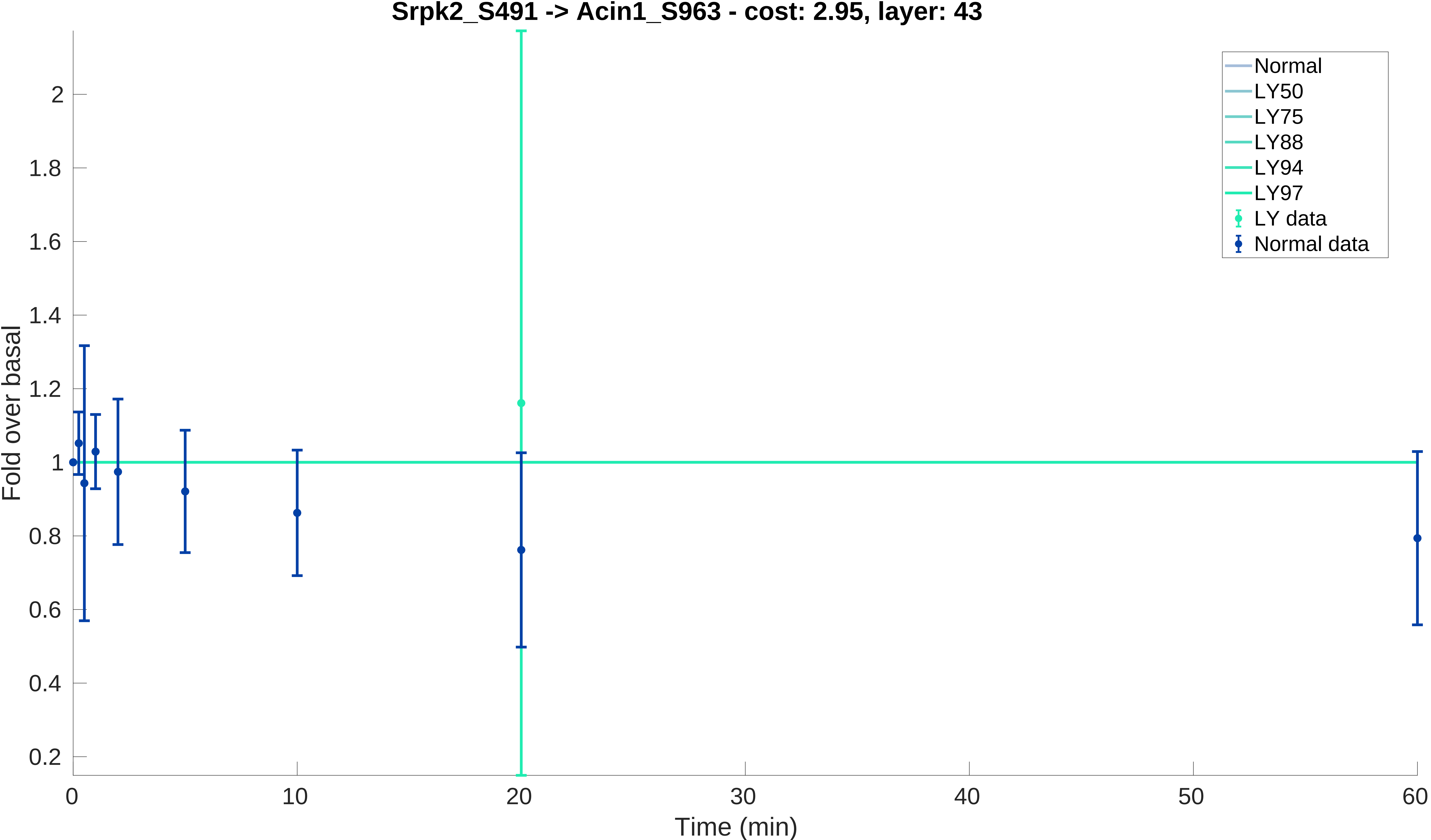

Srpk2_S491 -> Acin1_S963 - cost: 2.95, layer: 43
Normal
LY50
LY75
LY88
LY94
LY97
LY data
Normal data
Fold over basal
Time (min)
0.2
0.4
0.6
0.8
1
1.2
1.4
1.6
1.8
2
0
10
20
30
40
50
60

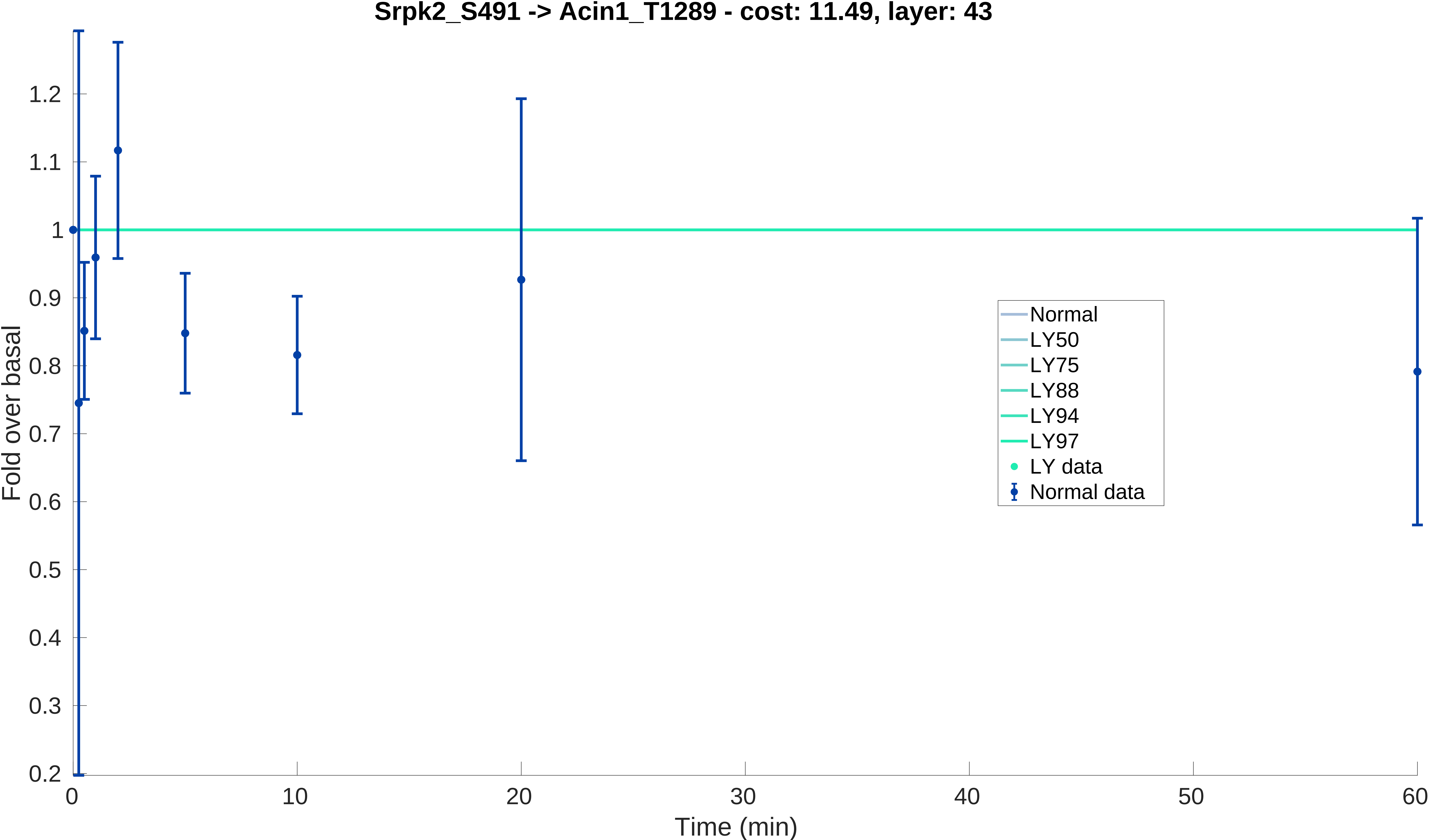
Srpk2_S491 -> Acin1_T1289 - cost: 11.49, layer: 43
Fold over basal
Time (min)
0
10
20
30
40
50
60
0.2
0.3
0.4
0.5
0.6
0.7
0.8
0.9
1
1.1
1.2
Normal
LY50
LY75
LY88
LY94
LY97
LY data
Normal data

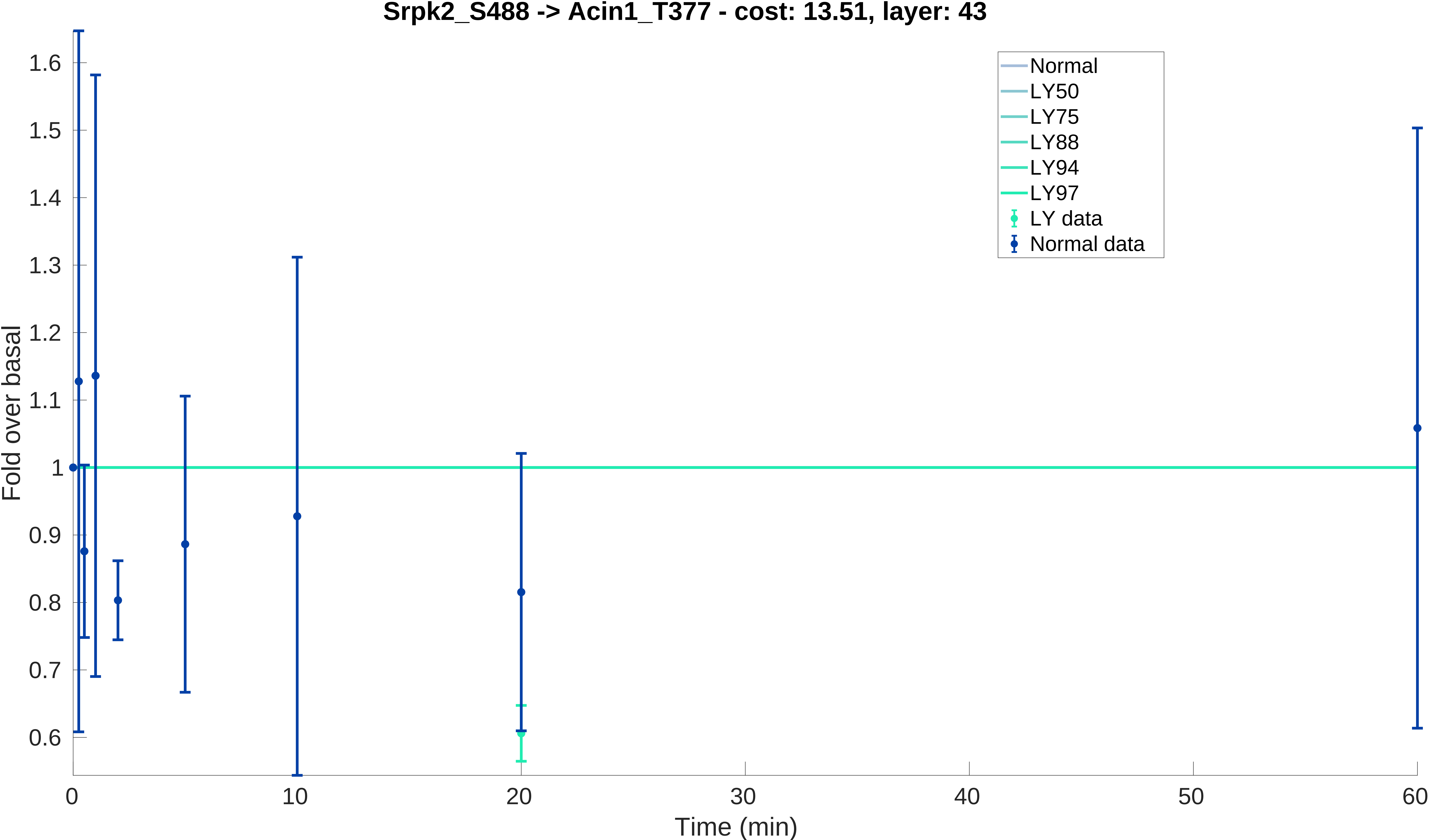

Srpk2_S488 -> Acin1_T377 - cost: 13.51, layer: 43
Fold over basal
Time (min)
Normal
LY50
LY75
LY88
LY94
LY97
LY data
Normal data
1.6
1.5
1.4
1.3
1.2
1.1
1
0.9
0.8
0.7
0.6
0
10
20
30
40
50
60

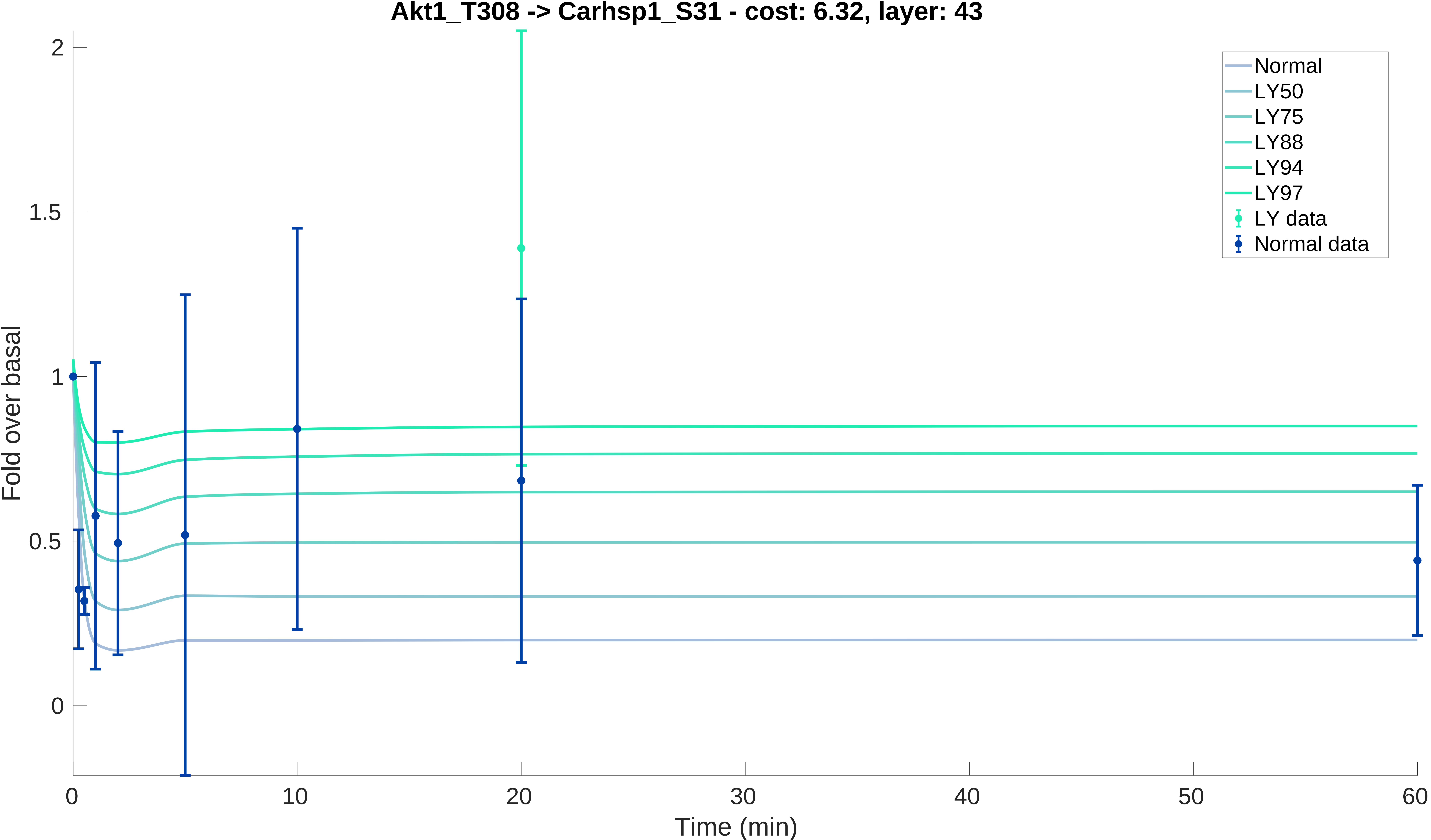

Akt1_T308 -> Carhsp1_S31 - cost: 6.32, layer: 43
Fold over basal
Time (min)
0
0.5
1
1.5
2
0
10
20
30
40
50
60
Normal
LY50
LY75
LY88
LY94
LY97
LY data
Normal data

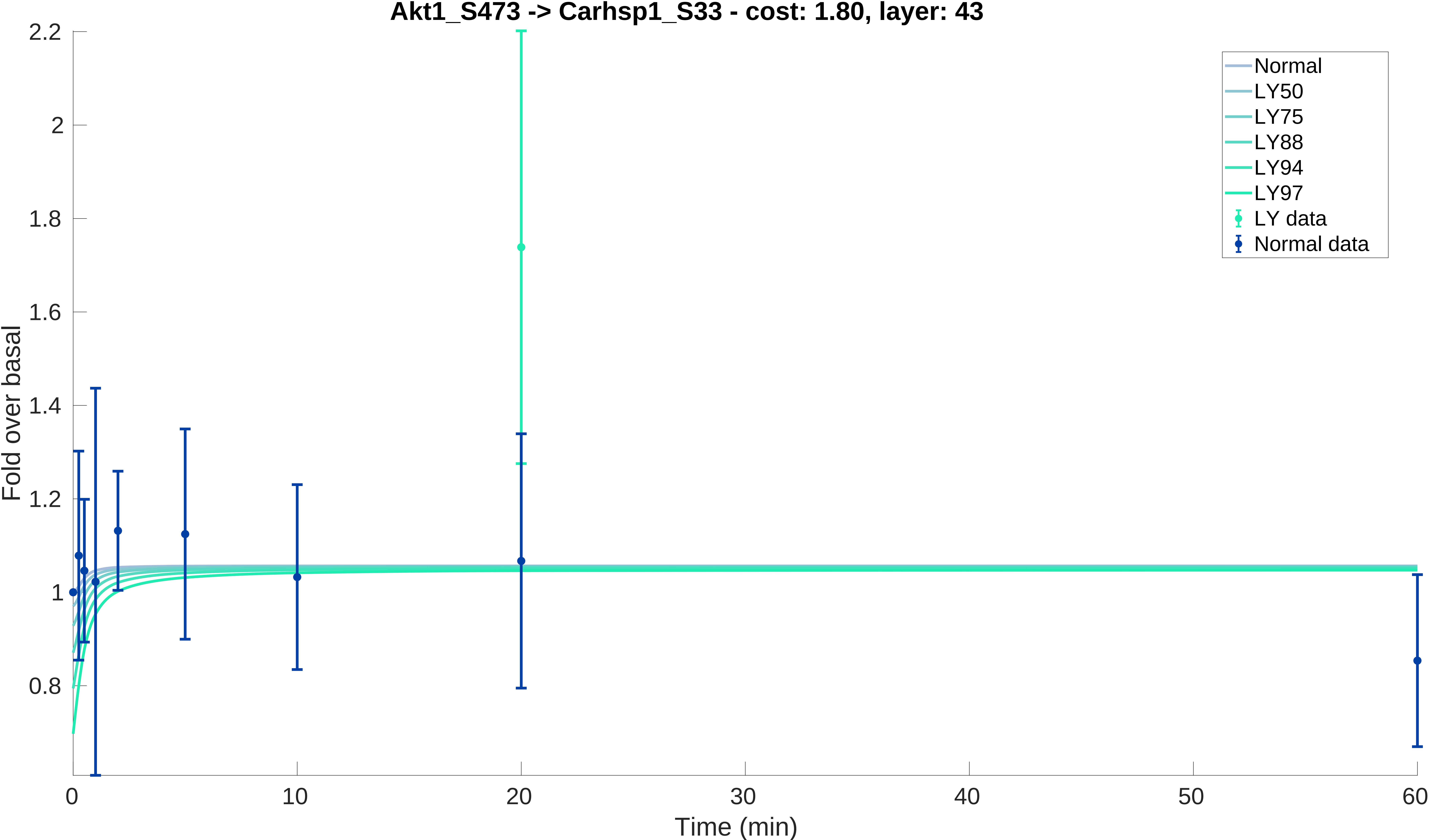
Akt1_S473 -> Carhsp1_S33 - cost: 1.80, layer: 43
Fold over basal
Time (min)
0.8
1
1.2
1.4
1.6
1.8
2
2.2
0
10
20
30
40
50
60
Normal
LY50
LY75
LY88
LY94
LY97
LY data
Normal data

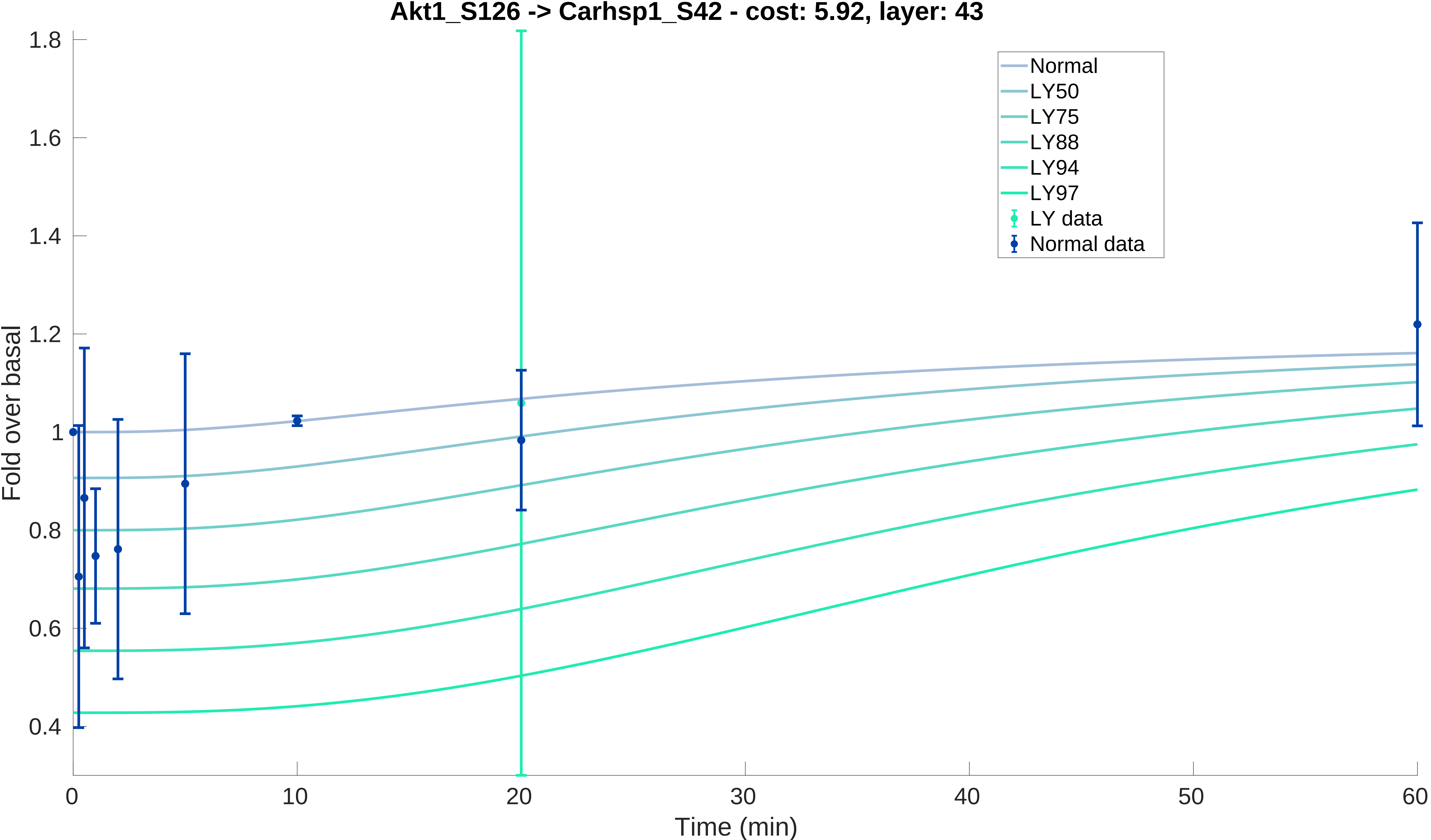

Akt1_S126 -> Carhsp1_S42 - cost: 5.92, layer: 43
Normal
LY50
LY75
LY88
LY94
LY97
LY data
Normal data
Fold over basal
Time (min)
0.4
0.6
0.8
1
1.2
1.4
1.6
1.8
0
10
20
30
40
50
60

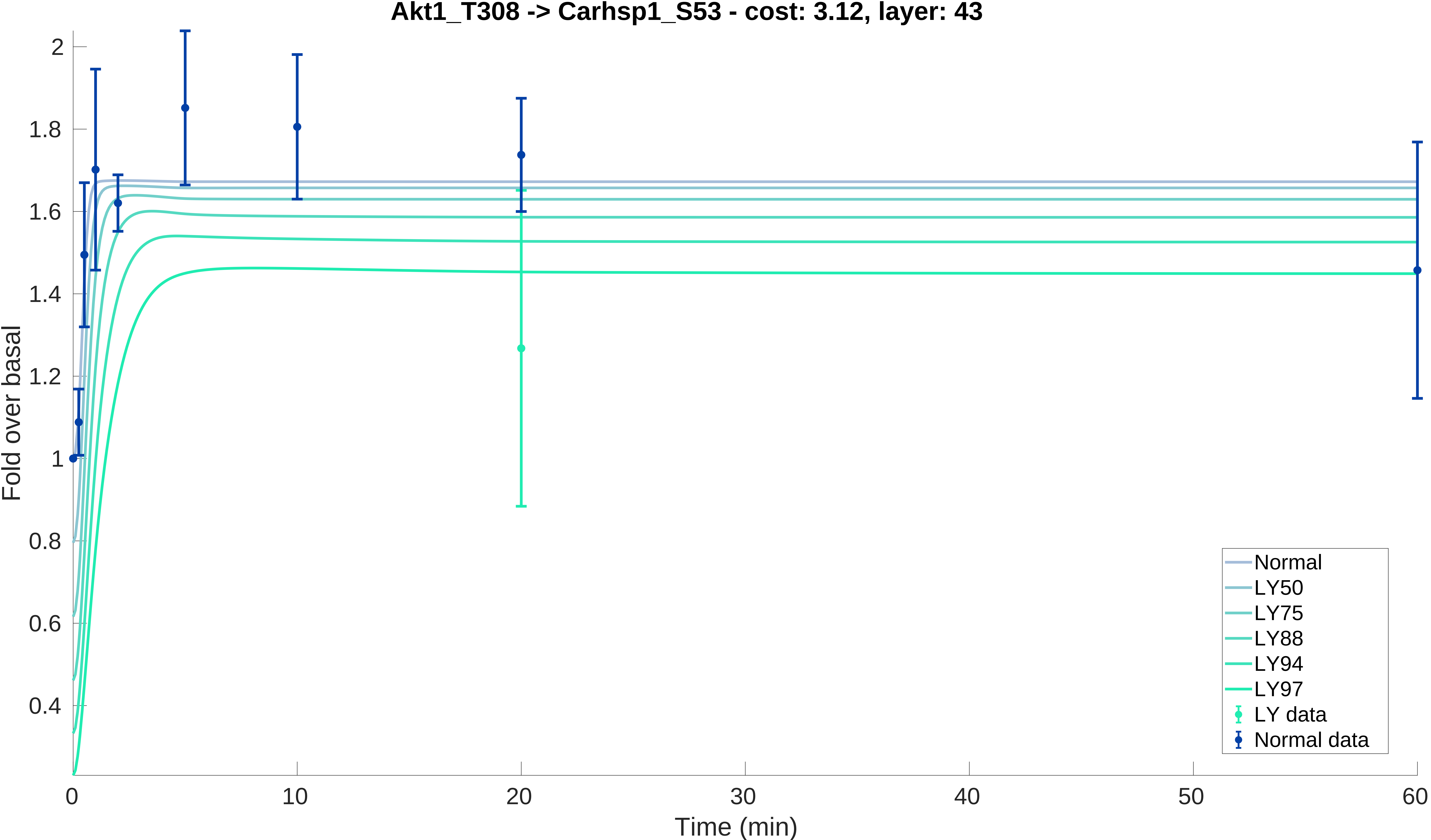

Akt1_T308 -> Carhsp1_S53 - cost: 3.12, layer: 43
Fold over basal
Time (min)
0
10
20
30
40
50
60
0.4
0.6
0.8
1
1.2
1.4
1.6
1.8
2
Normal
LY50
LY75
LY88
LY94
LY97
LY data
Normal data

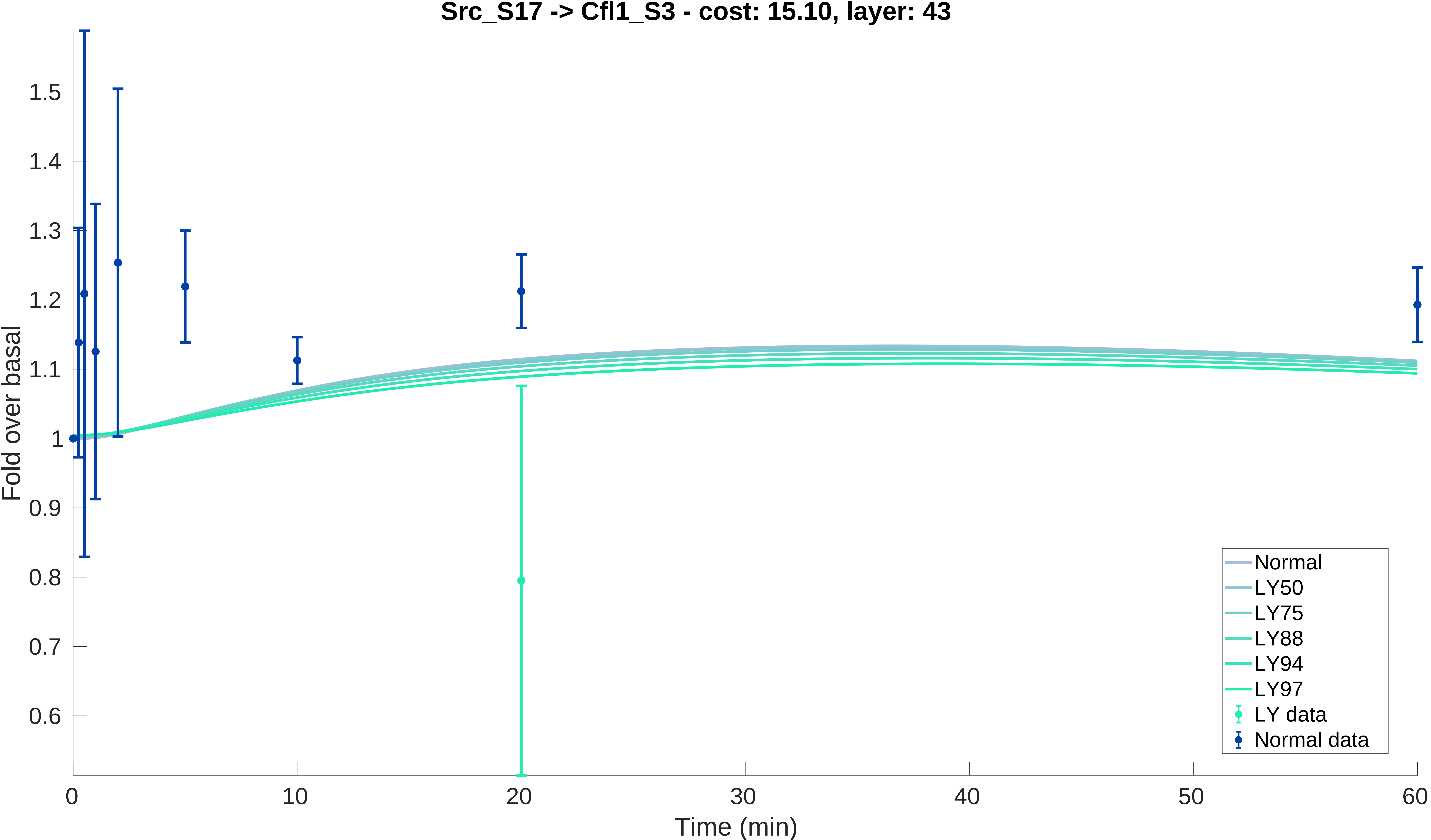

Src_S17 -> Cfl1_S3 - cost: 15.10, layer: 43
Fold over basal
1.5
1.4
1.3
1.2
1.1
1
0.9
0.8
0.7
0.6
0
10
20
30
40
50
60
Time (min)
Normal
LY50
LY75
LY88
LY94
LY97
LY data
Normal data

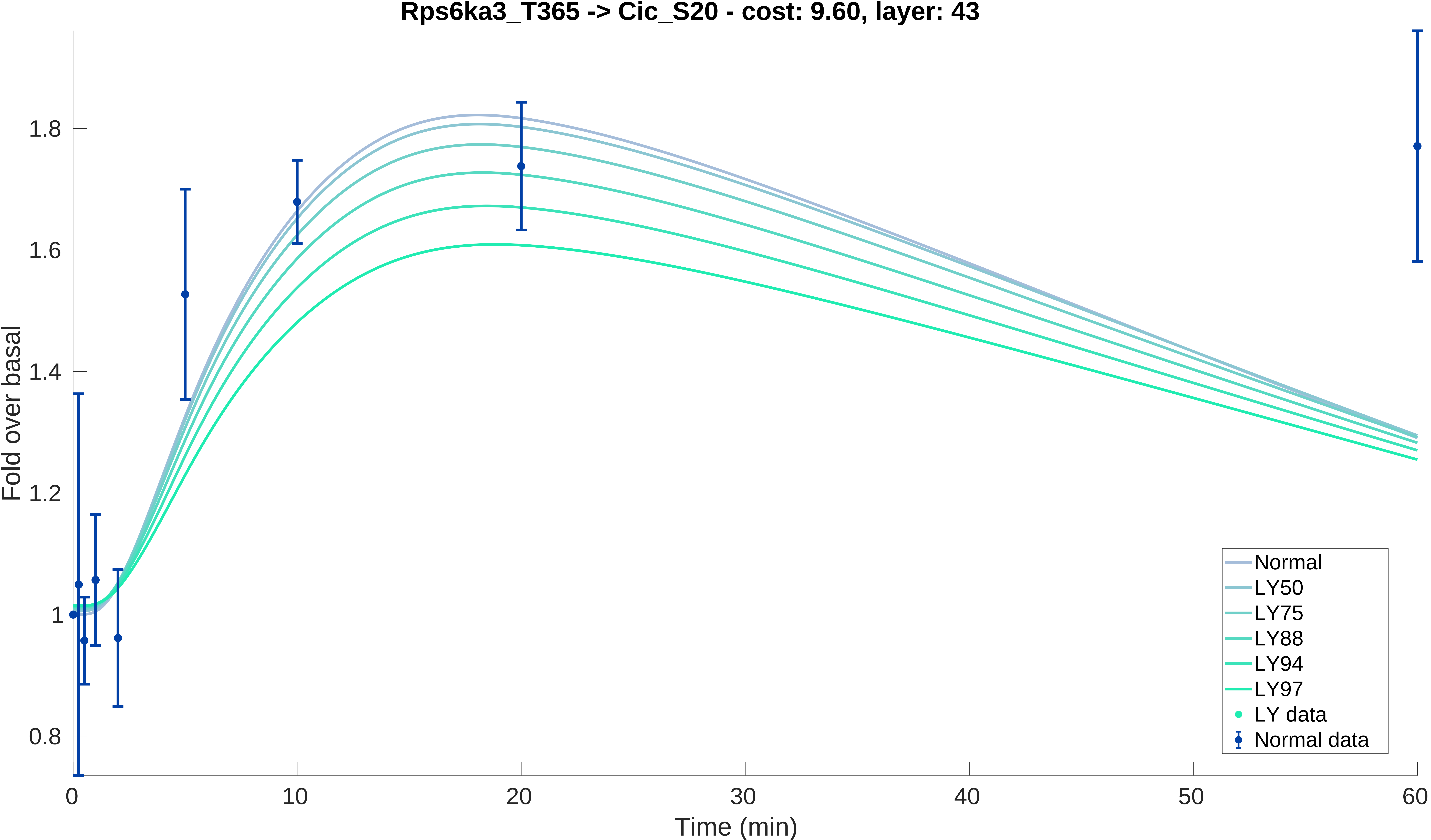

Rps6ka3_T365 -> Cic_S20 - cost: 9.60, layer: 43
Fold over basal
1.8
1.6
1.4
1.2
1
0.8
0
10
20
30
40
50
60
Time (min)
Normal
LY50
LY75
LY88
LY94
LY97
LY data
Normal data

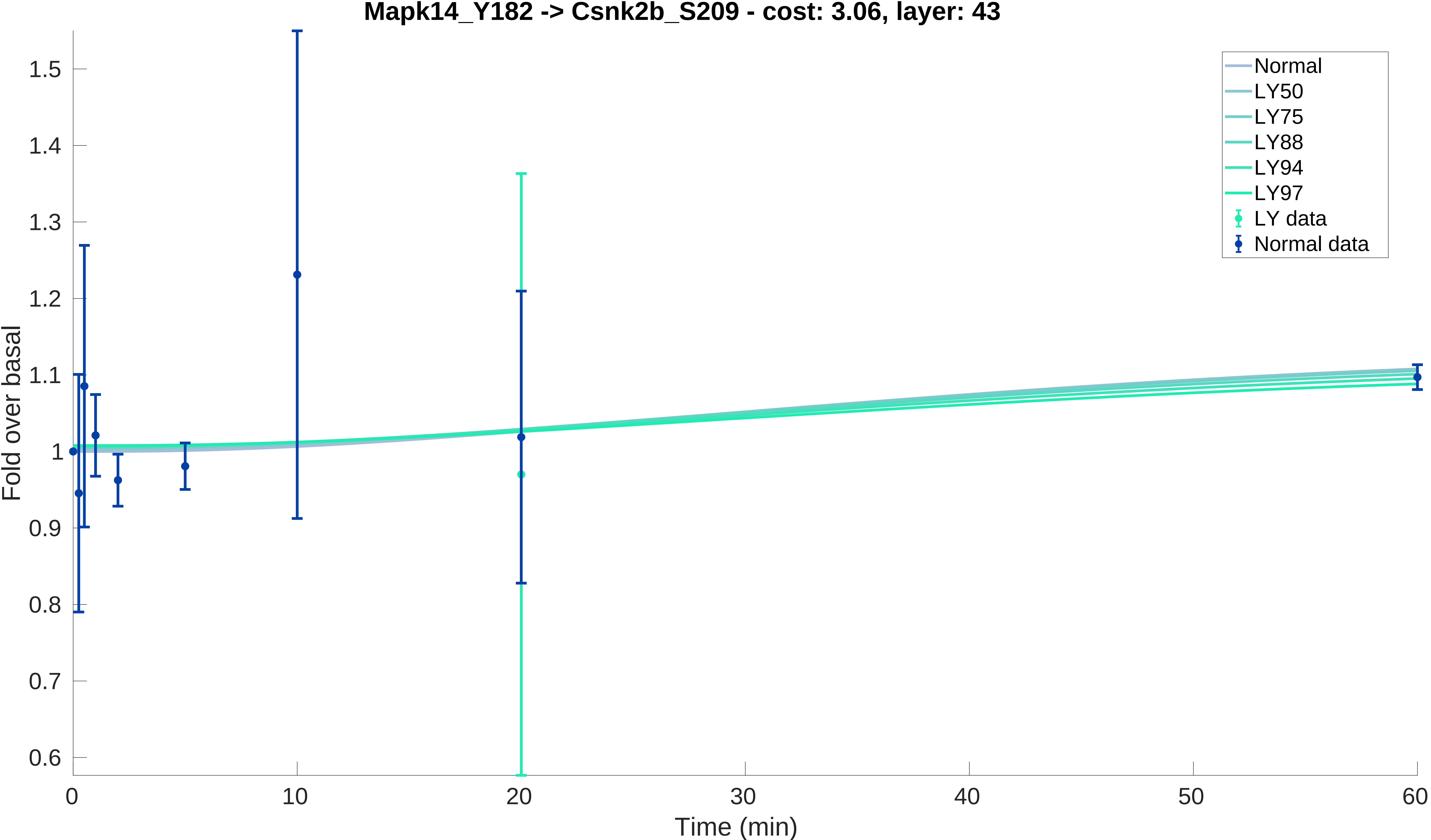
Mapk14_Y182 -> Csnk2b_S209 - cost: 3.06, layer: 43
Fold over basal
Time (min)
1.5
1.4
1.3
1.2
1.1
1
0.9
0.8
0.7
0.6
0
10
20
30
40
50
60
Normal
LY50
LY75
LY88
LY94
LY97
LY data
Normal data

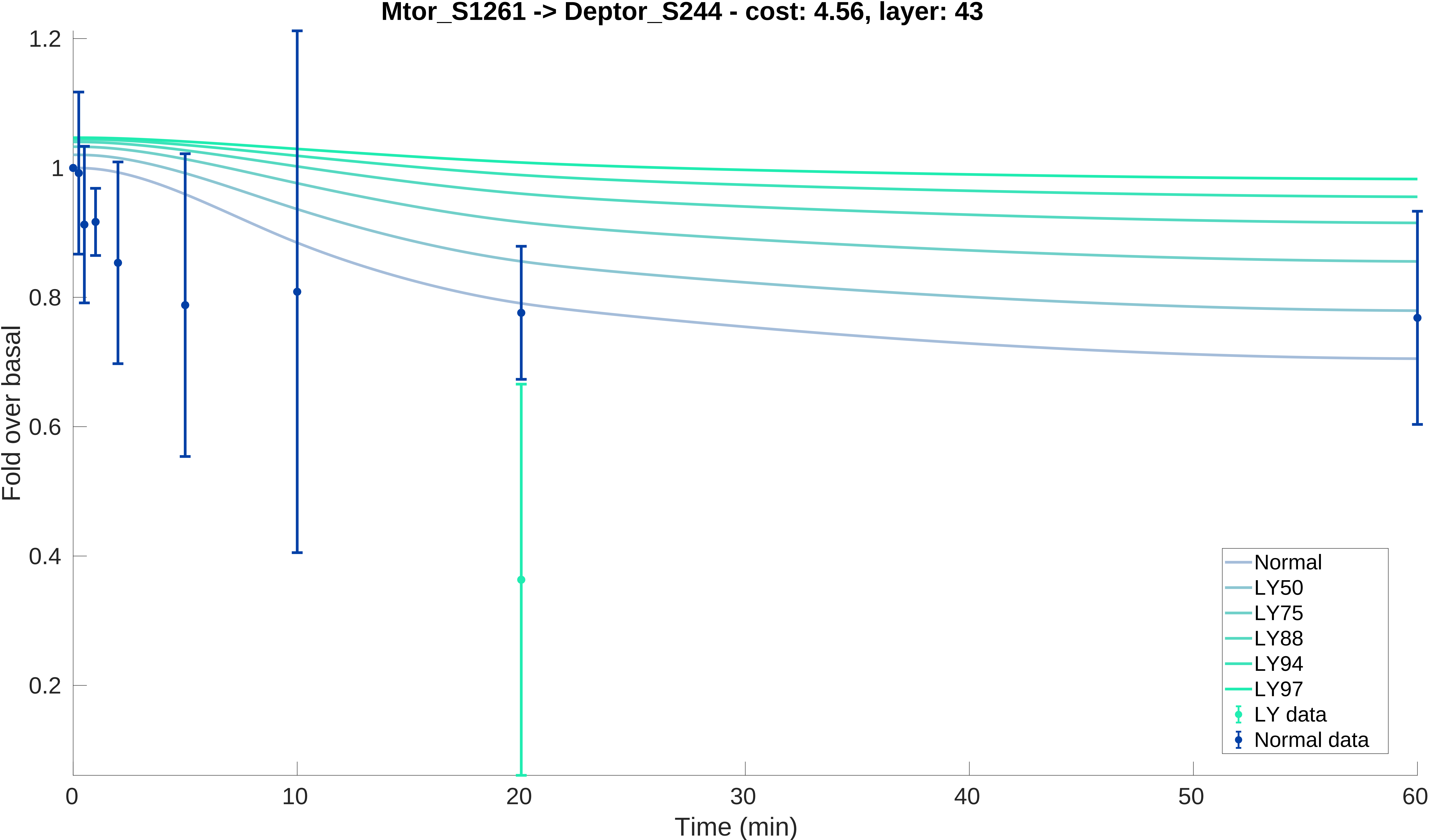

Mtor_S1261 -> Deptor_S244 - cost: 4.56, layer: 43
Fold over basal
Time (min)
0.2
0.4
0.6
0.8
1
1.2
0
10
20
30
40
50
60
Normal
LY50
LY75
LY88
LY94
LY97
LY data
Normal data

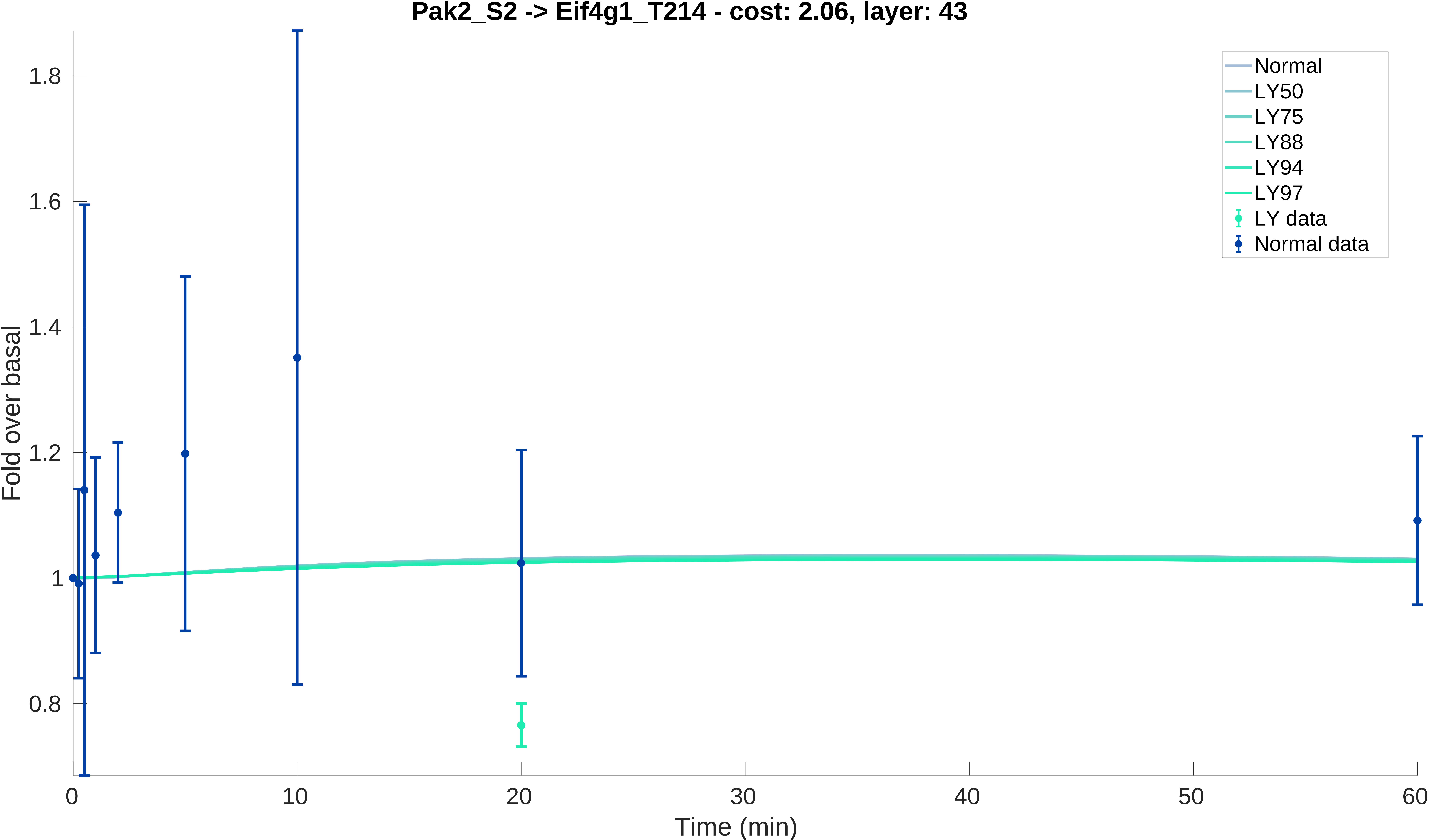
Pak2_S2 -> Eif4g1_T214 - cost: 2.06, layer: 43
Fold over basal
1.8
1.6
1.4
1.2
1
0.8
0
10
20
30
40
50
60
Time (min)
Normal
LY50
LY75
LY88
LY94
LY97
LY data
Normal data

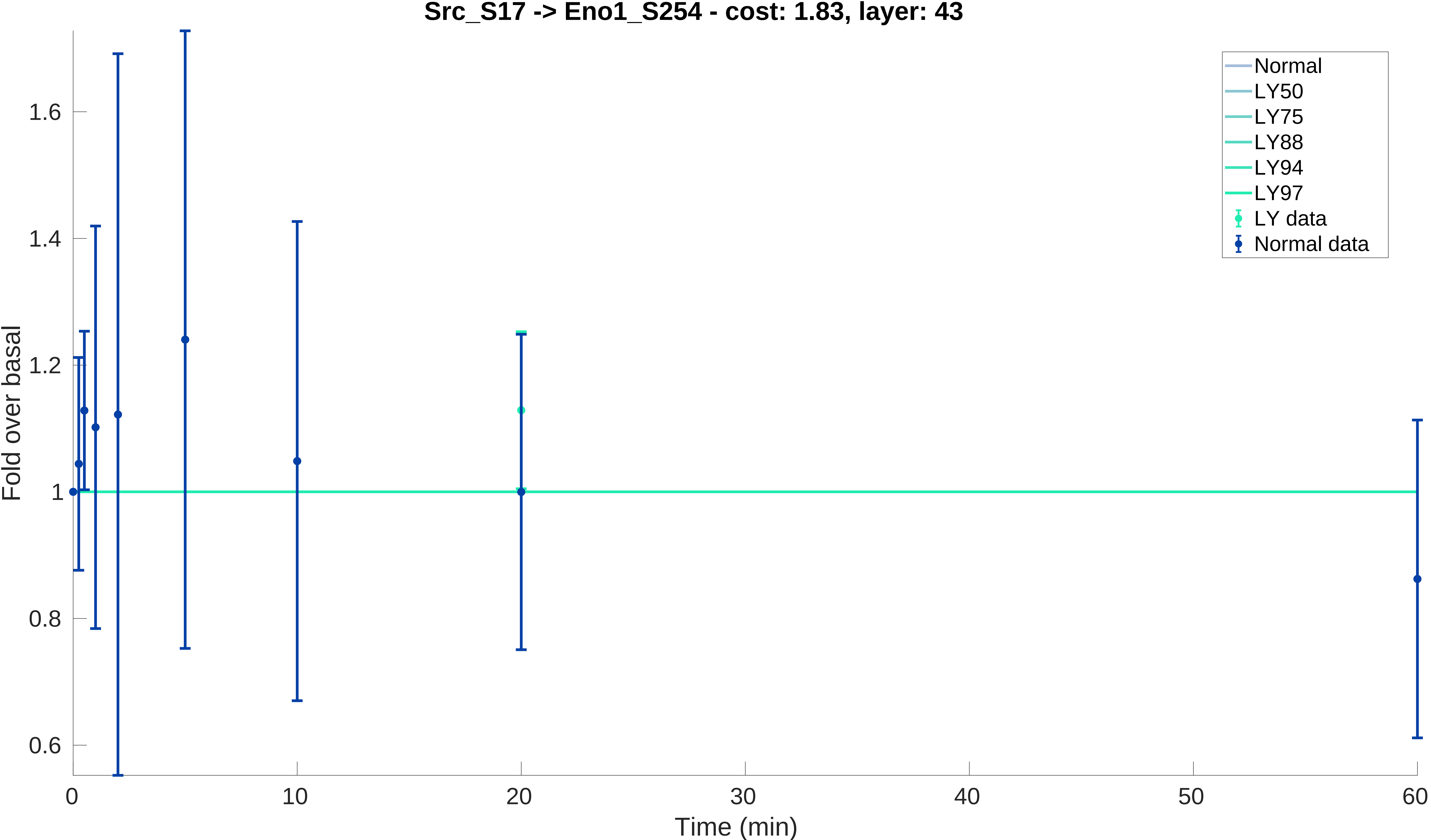
Src_S17 -> Eno1_S254 - cost: 1.83, layer: 43
Fold over basal
1.6
1.4
1.2
1
0.8
0.6
0
10
20
30
40
50
60
Time (min)
Normal
LY50
LY75
LY88
LY94
LY97
LY data
Normal data

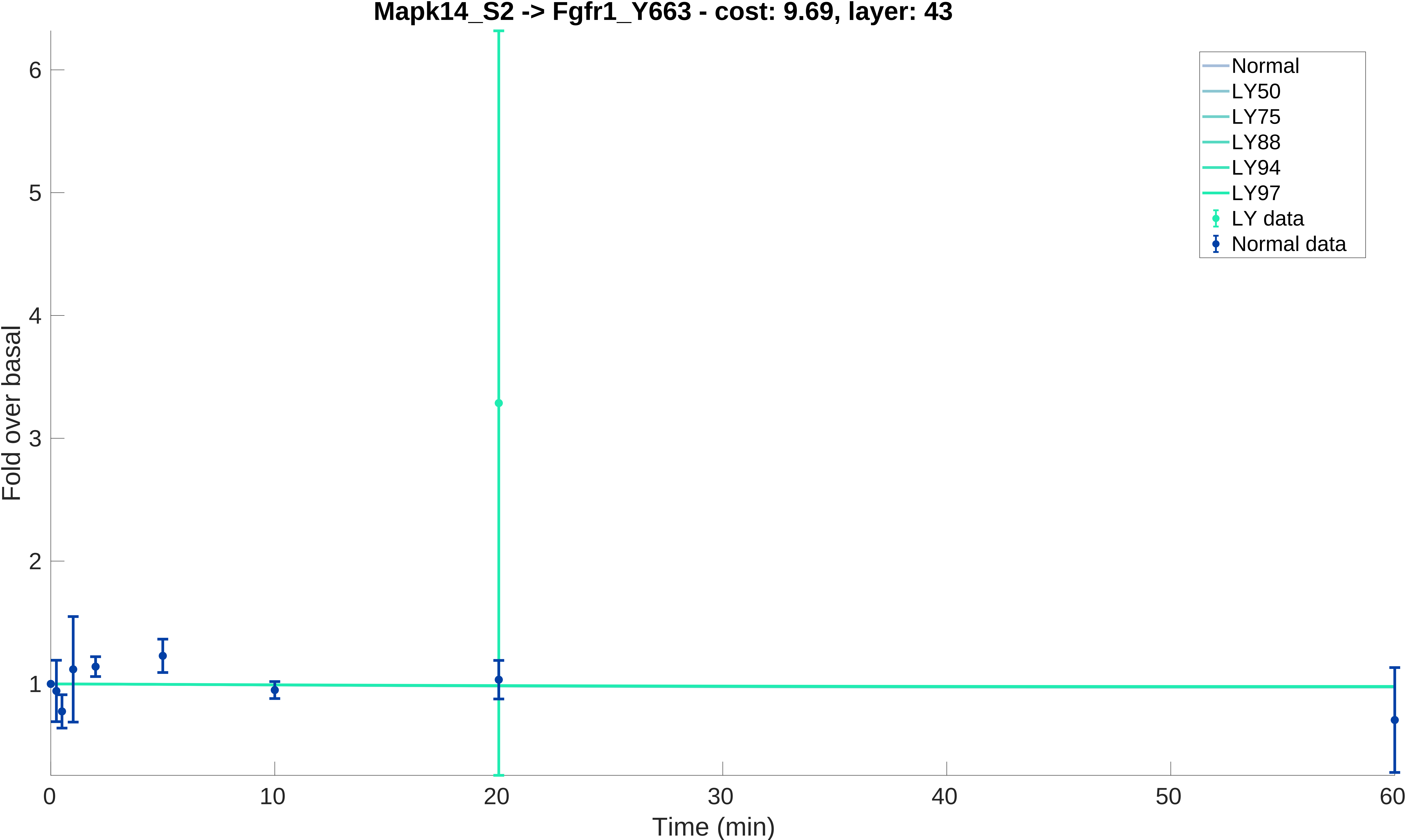

Mapk14_S2 -> Fgfr1_Y663 - cost: 9.69, layer: 43
Normal
LY50
LY75
LY88
LY94
LY97
LY data
Normal data
Fold over basal
Time (min)
0
10
20
30
40
50
60
1
2
3
4
5
6

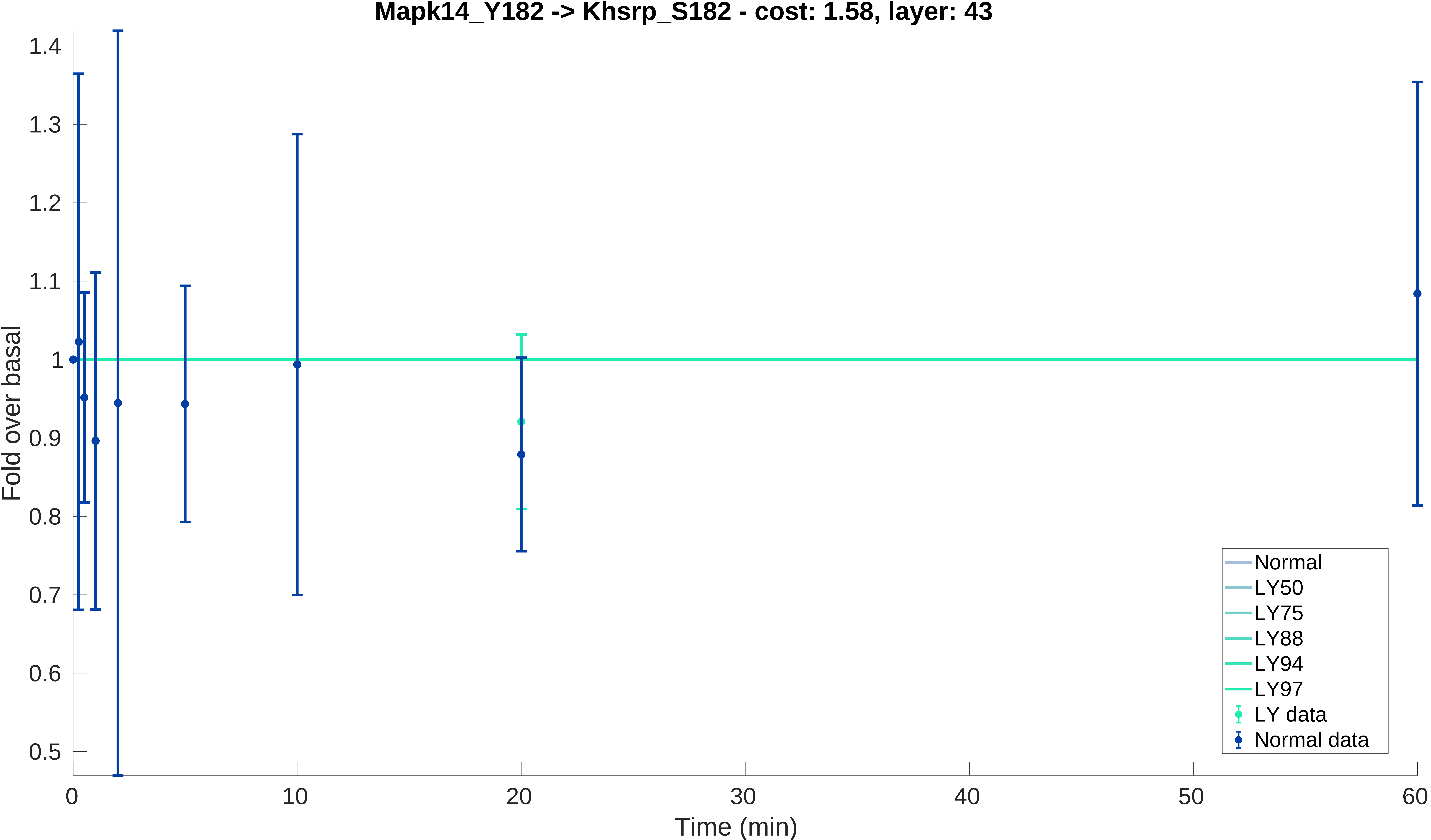

Mapk14_Y182 -> Khsrp_S182 - cost: 1.58, layer: 43
Fold over basal
Time (min)
1.4
1.3
1.2
1.1
1
0.9
0.8
0.7
0.6
0.5
0
10
20
30
40
50
60
Normal
LY50
LY75
LY88
LY94
LY97
LY data
Normal data

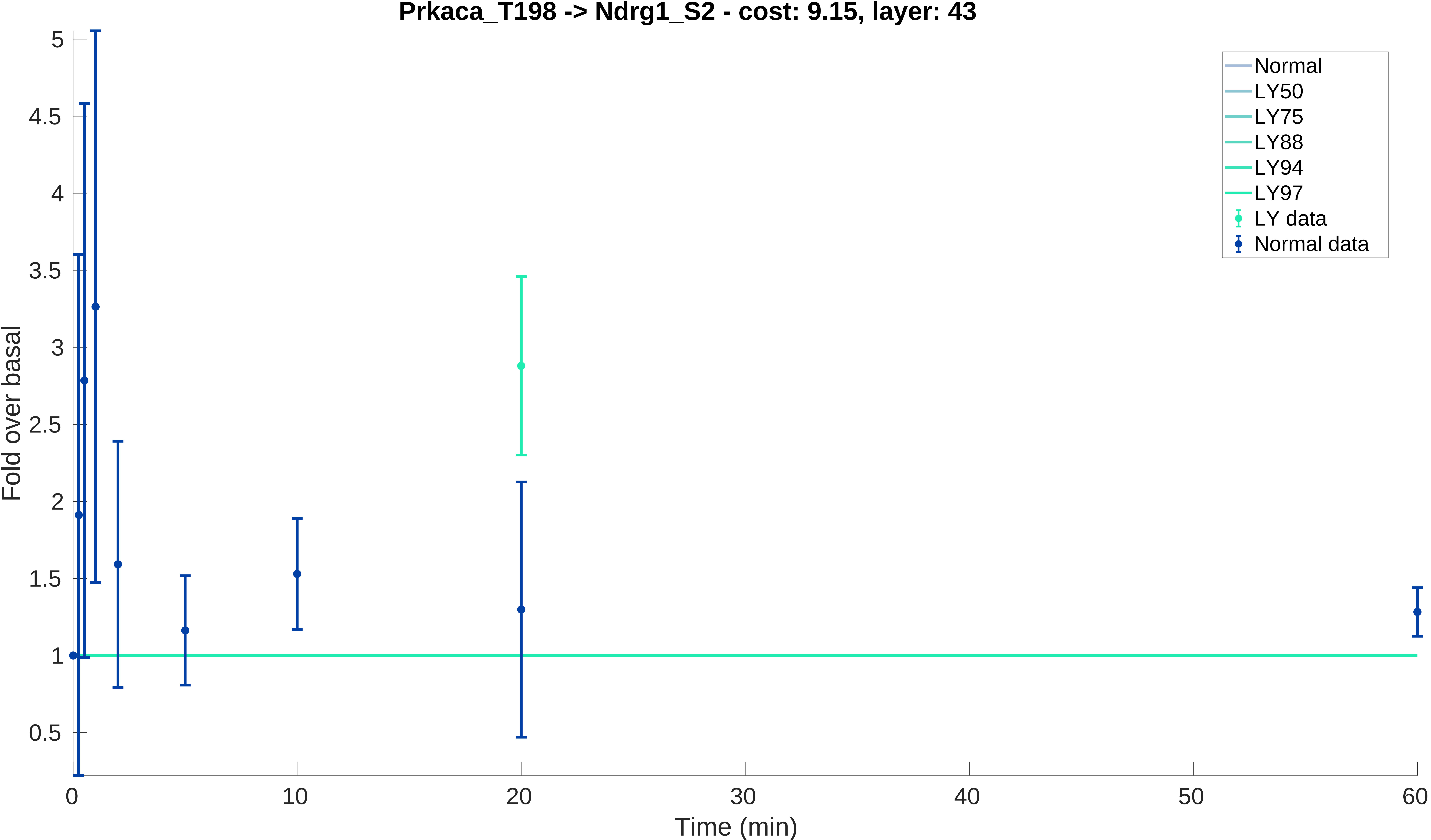

Prkaca_T198 -> Ndrg1_S2 - cost: 9.15, layer: 43
Fold over basal
Time (min)
0.5
1
1.5
2
2.5
3
3.5
4
4.5
5
0
10
20
30
40
50
60
Normal
LY50
LY75
LY88
LY94
LY97
LY data
Normal data

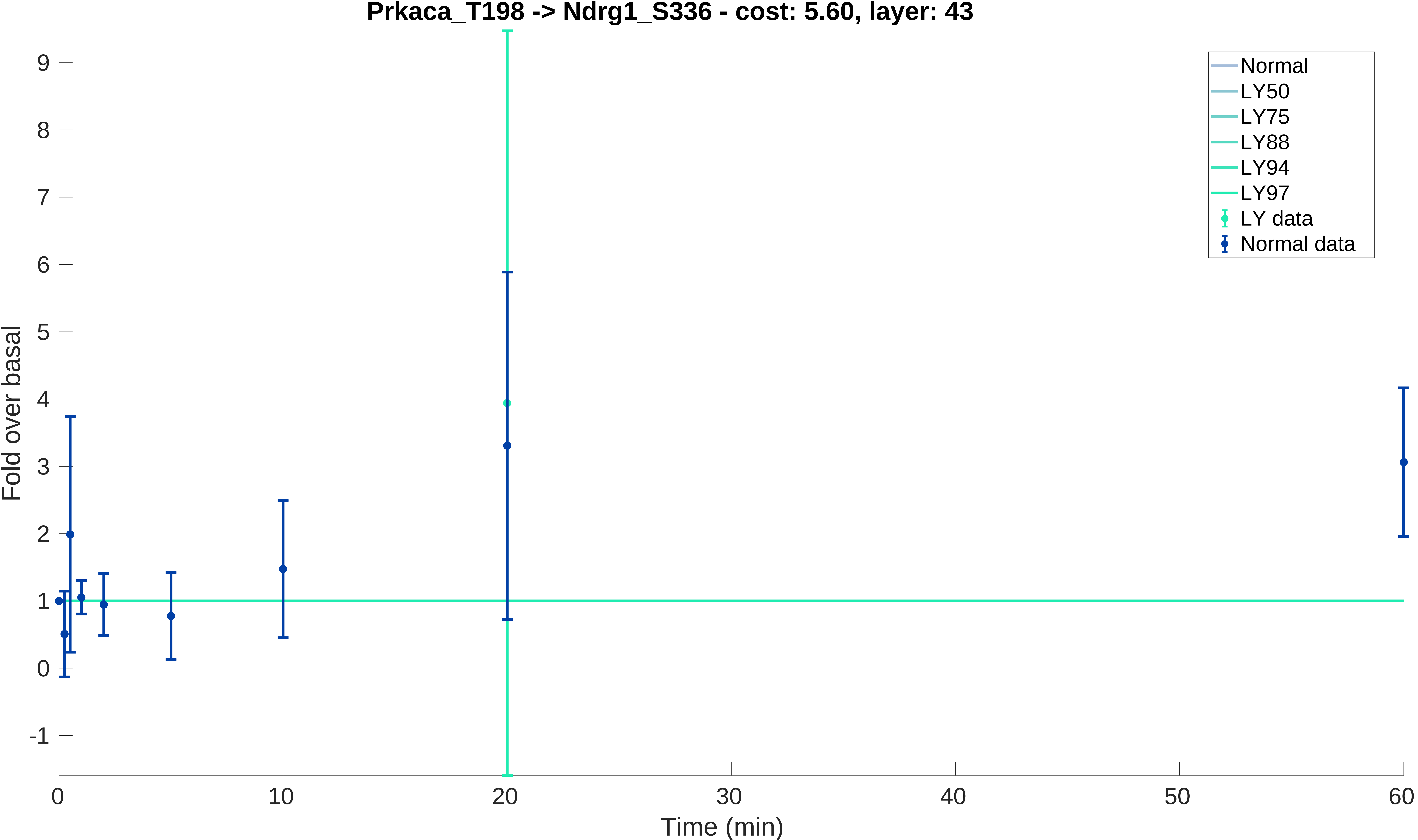

Prkaca_T198 -> Ndrg1_S336 - cost: 5.60, layer: 43
Normal
LY50
LY75
LY88
LY94
LY97
LY data
Normal data
Fold over basal
Time (min)
9
8
7
6
5
4
3
2
1
0
-1
0
10
20
30
40
50
60

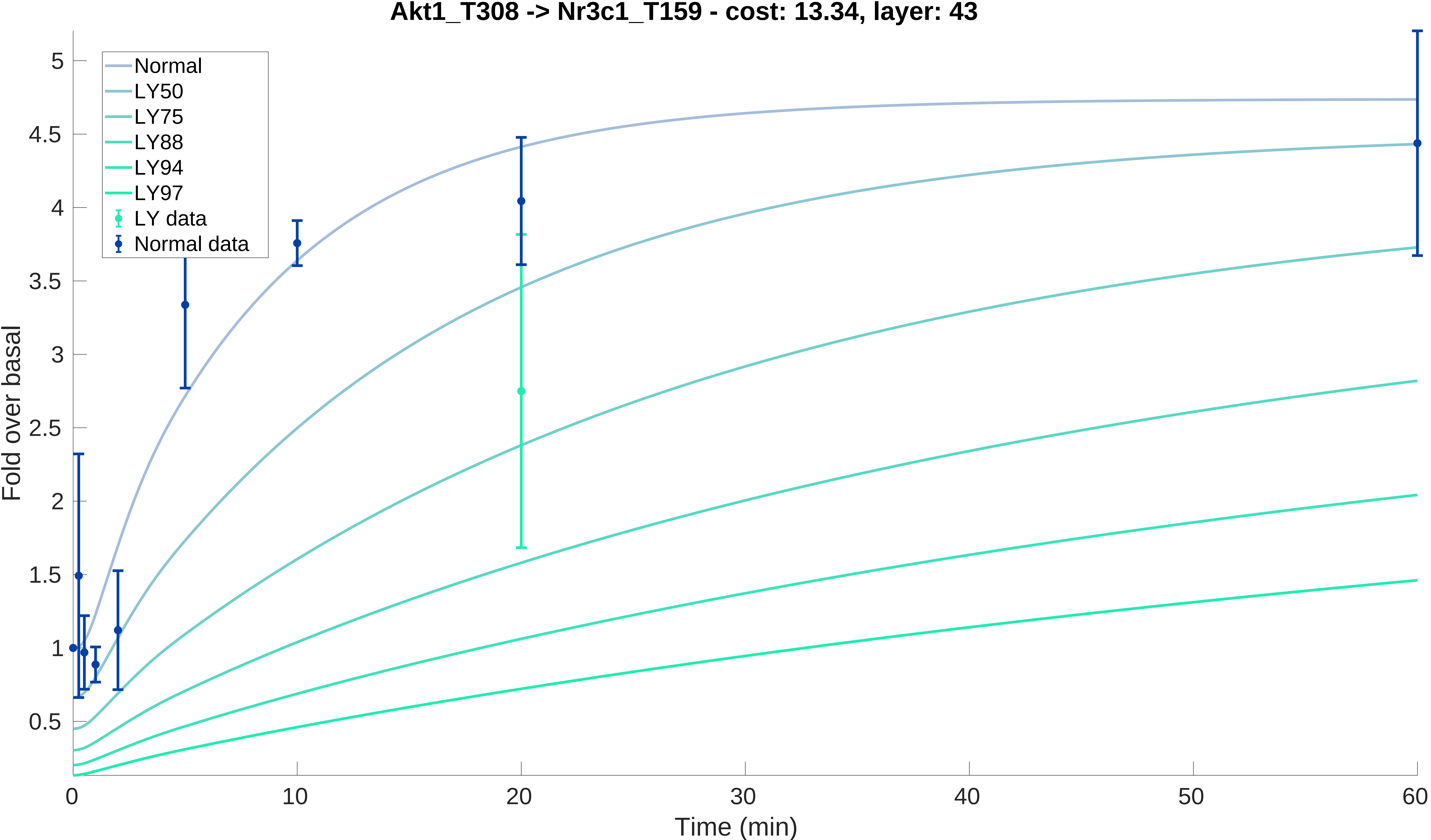
Akt1_T308 -> Nr3c1_T159 - cost: 13.34, layer: 43
Normal
LY50
LY75
LY88
LY94
LY97
LY data
Normal data
Fold over basal
Time (min)
0.5
1
1.5
2
2.5
3
3.5
4
4.5
5
0
10
20
30
40
50
60

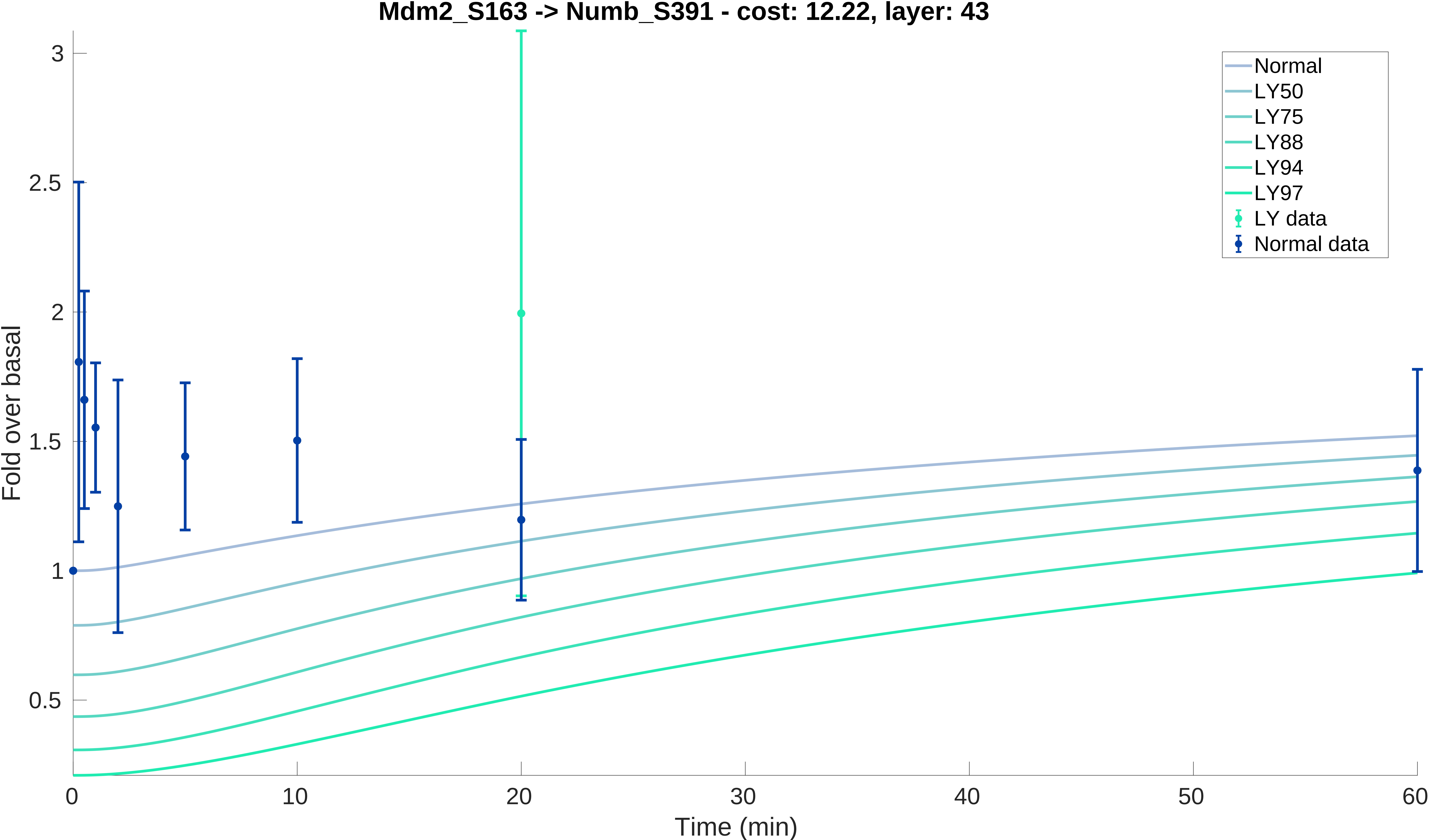

Mdm2_S163 -> Numb_S391 - cost: 12.22, layer: 43
Fold over basal
Time (min)
0
10
20
30
40
50
60
0.5
1
1.5
2
2.5
3
Normal
LY50
LY75
LY88
LY94
LY97
LY data
Normal data

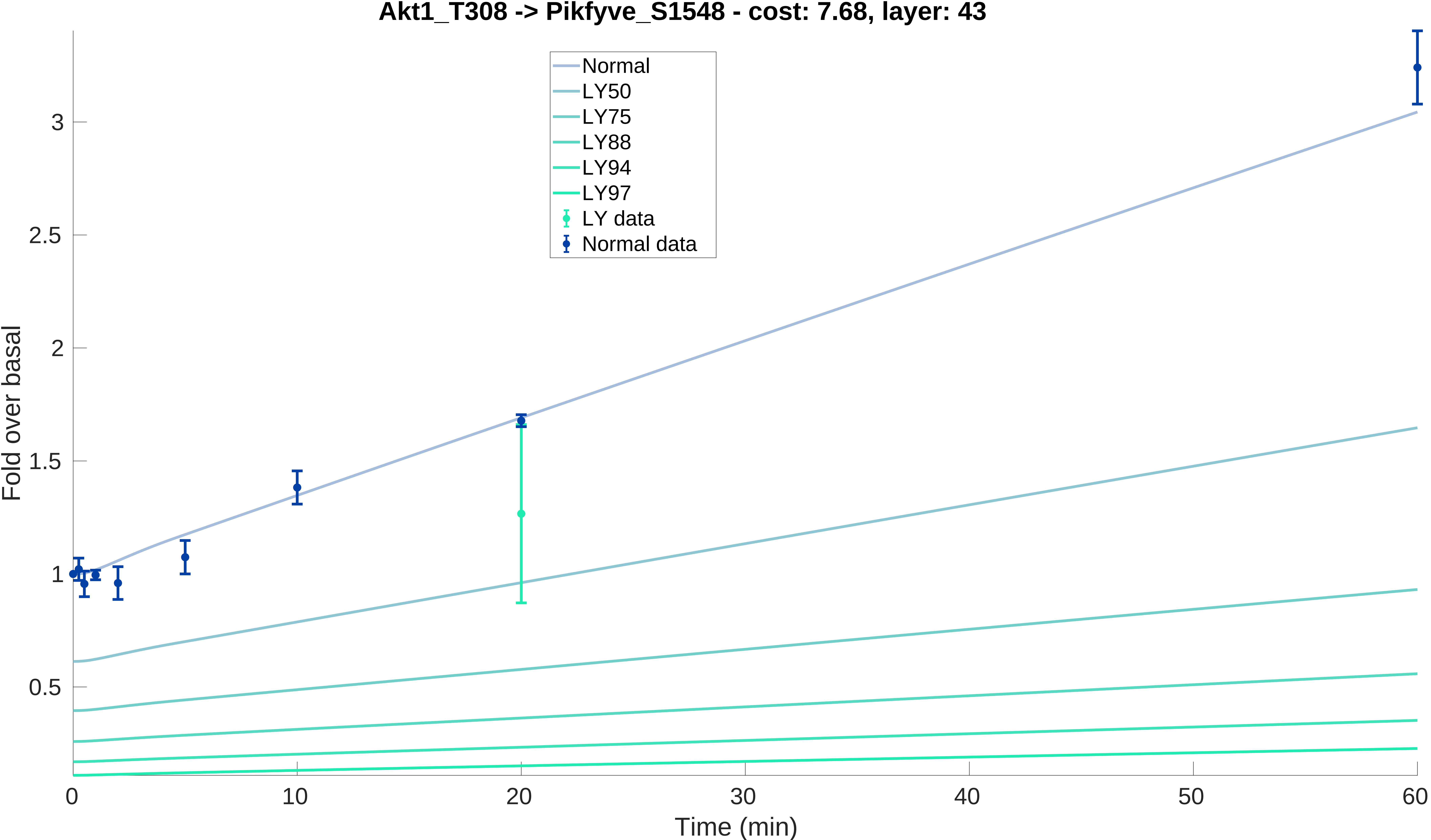

Akt1_T308 -> Pikfyve_S1548 - cost: 7.68, layer: 43
Normal
LY50
LY75
LY88
LY94
LY97
LY data
Normal data
Fold over basal
Time (min)
0
10
20
30
40
50
60
0.5
1
1.5
2
2.5
3

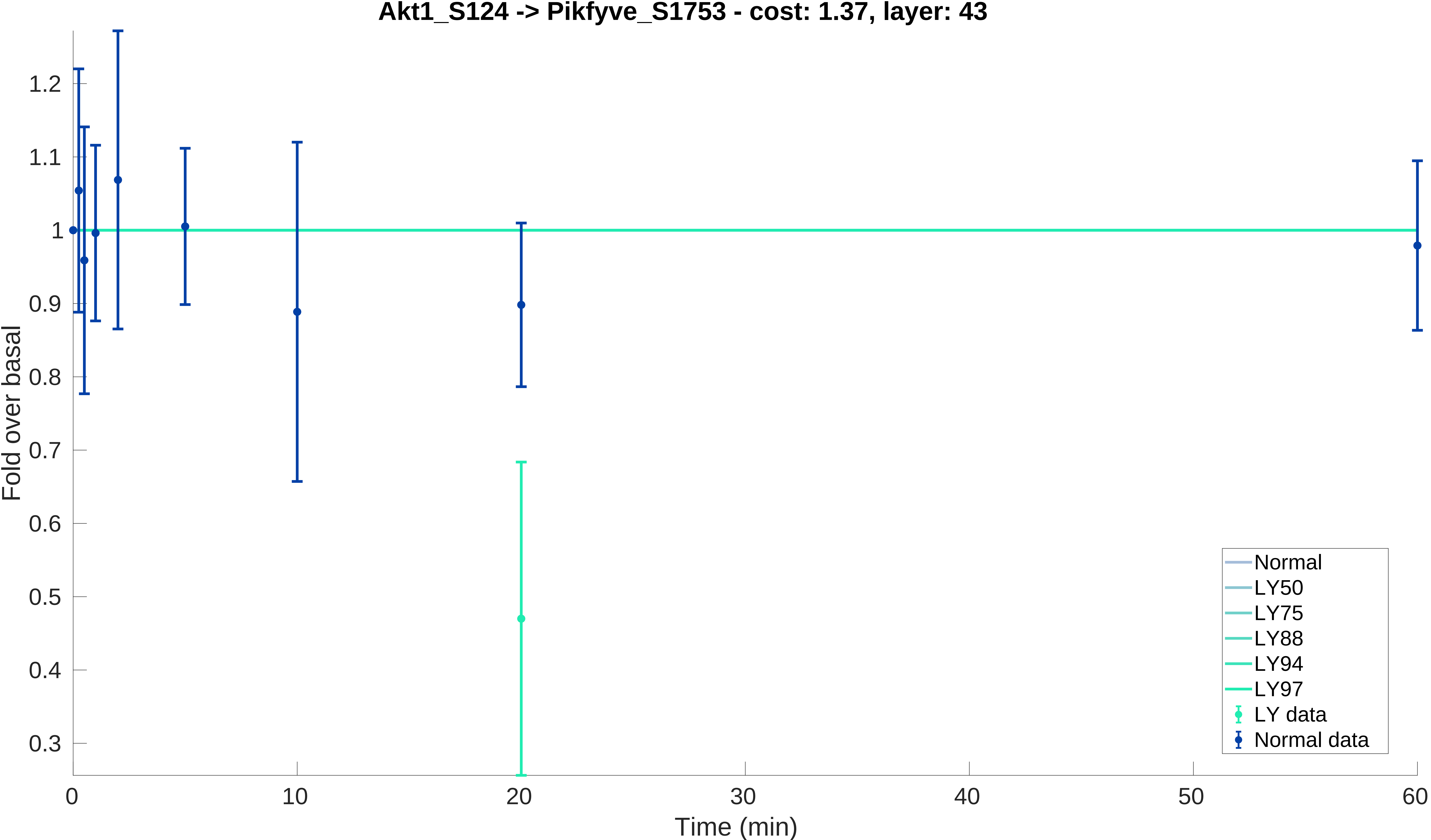

Akt1_S124 -> Pikfyve_S1753 - cost: 1.37, layer: 43
Fold over basal
1.2
1.1
1
0.9
0.8
0.7
0.6
0.5
0.4
0.3
0
10
20
30
40
50
60
Time (min)
Normal
LY50
LY75
LY88
LY94
LY97
LY data
Normal data

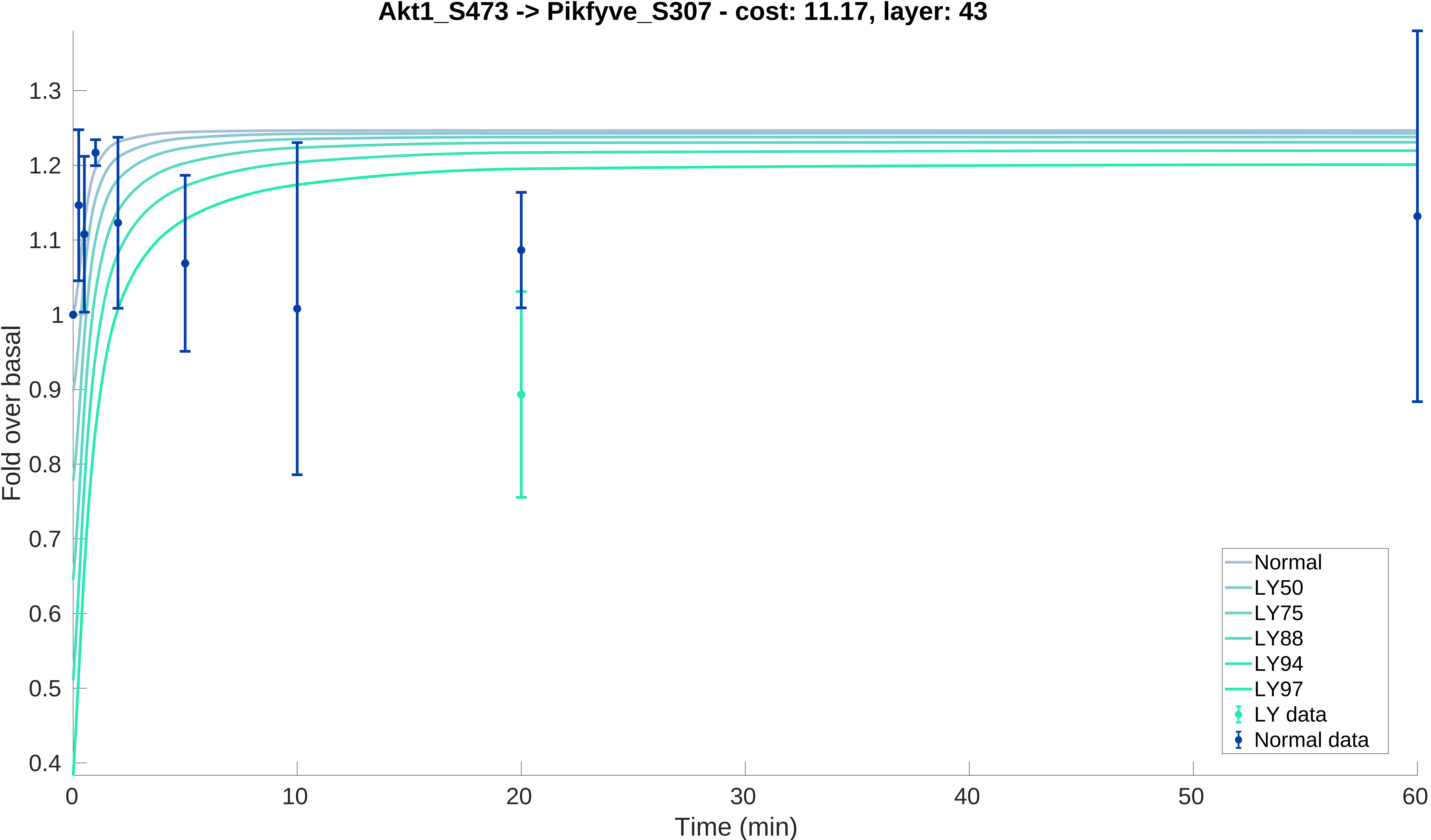

Akt1_S473 -> Pikfyve_S307 - cost: 11.17, layer: 43
Fold over basal
1.3
1.2
1.1
1
0.9
0.8
0.7
0.6
0.5
0.4
0
10
20
30
40
50
60
Time (min)
Normal
LY50
LY75
LY88
LY94
LY97
LY data
Normal data

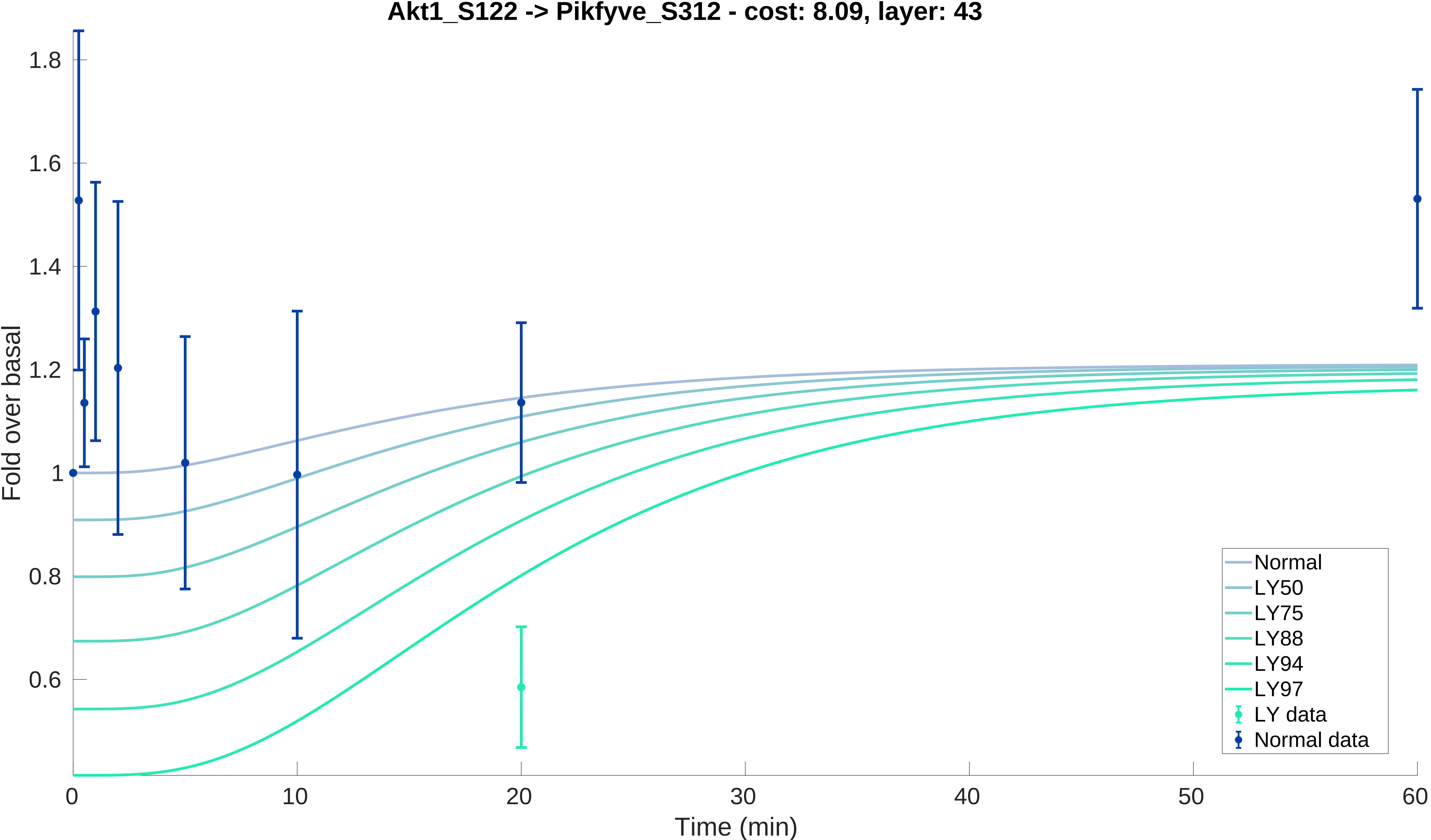

Akt1_S122 -> Pikfyve_S312 - cost: 8.09, layer: 43
1.8
1.6
1.4
1.2
1
0.8
0.6
Fold over basal
0
10
20
30
40
50
60
Time (min)
Normal
LY50
LY75
LY88
LY94
LY97
LY data
Normal data

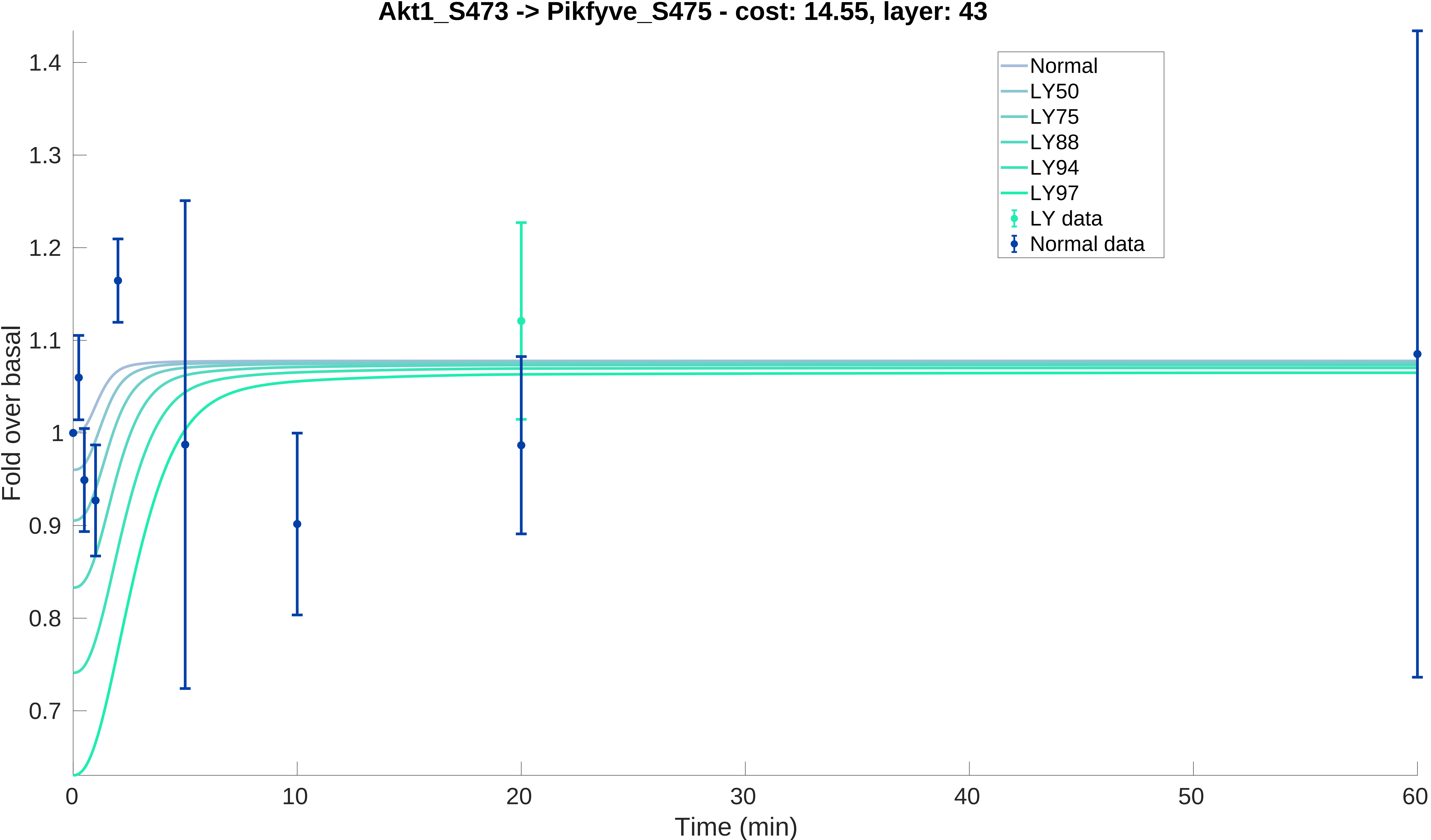

Akt1_S473 -> Pikfyve_S475 - cost: 14.55, layer: 43
Normal
LY50
LY75
LY88
LY94
LY97
LY data
Normal data
Fold over basal
1.4
1.3
1.2
1.1
1
0.9
0.8
0.7
0
10
20
30
40
50
60
Time (min)

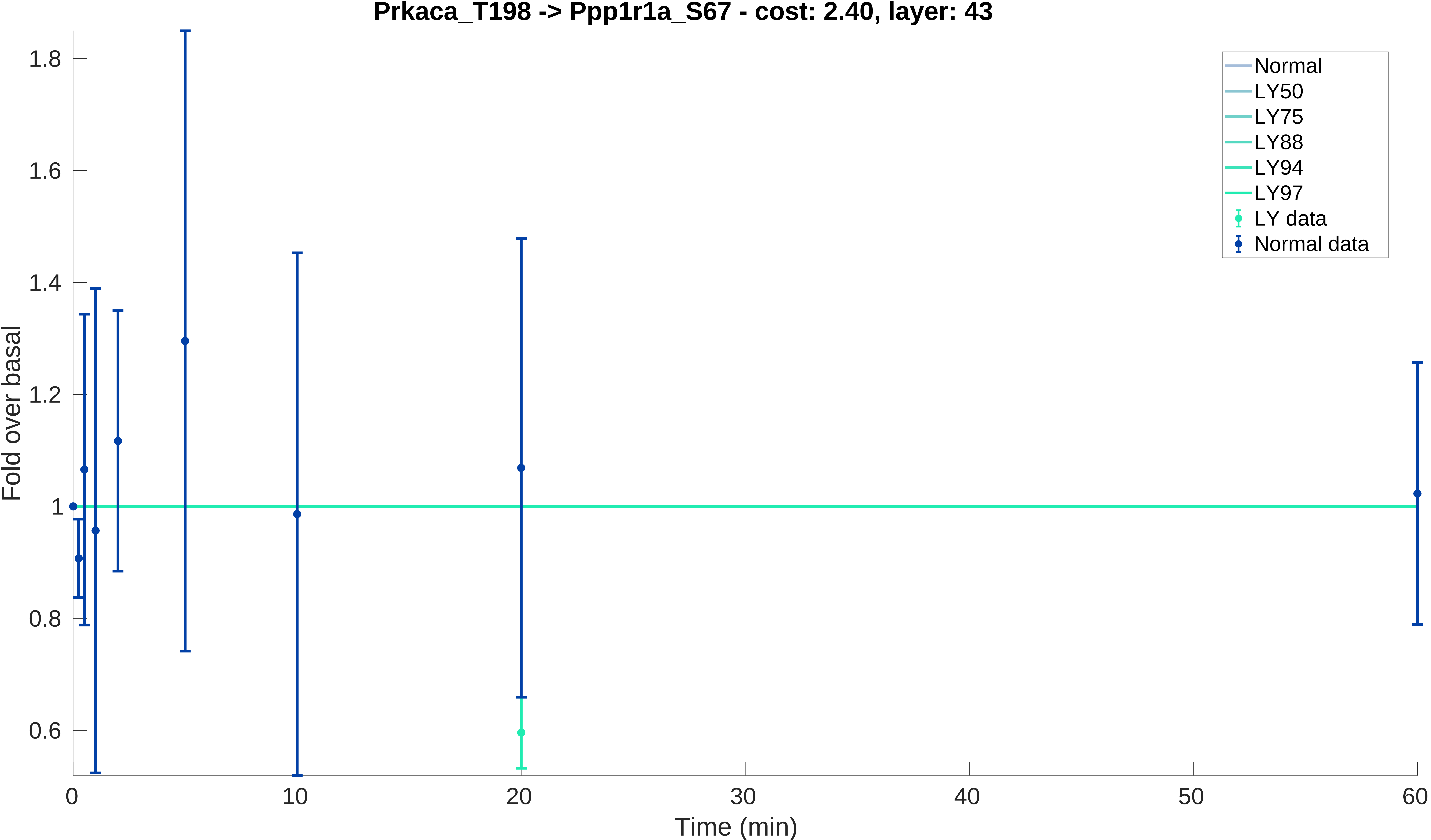

Prkaca_T198 -> Ppp1r1a_S67 - cost: 2.40, layer: 43
Fold over basal
Time (min)
1.8
1.6
1.4
1.2
1
0.8
0.6
0
10
20
30
40
50
60
Normal
LY50
LY75
LY88
LY94
LY97
LY data
Normal data

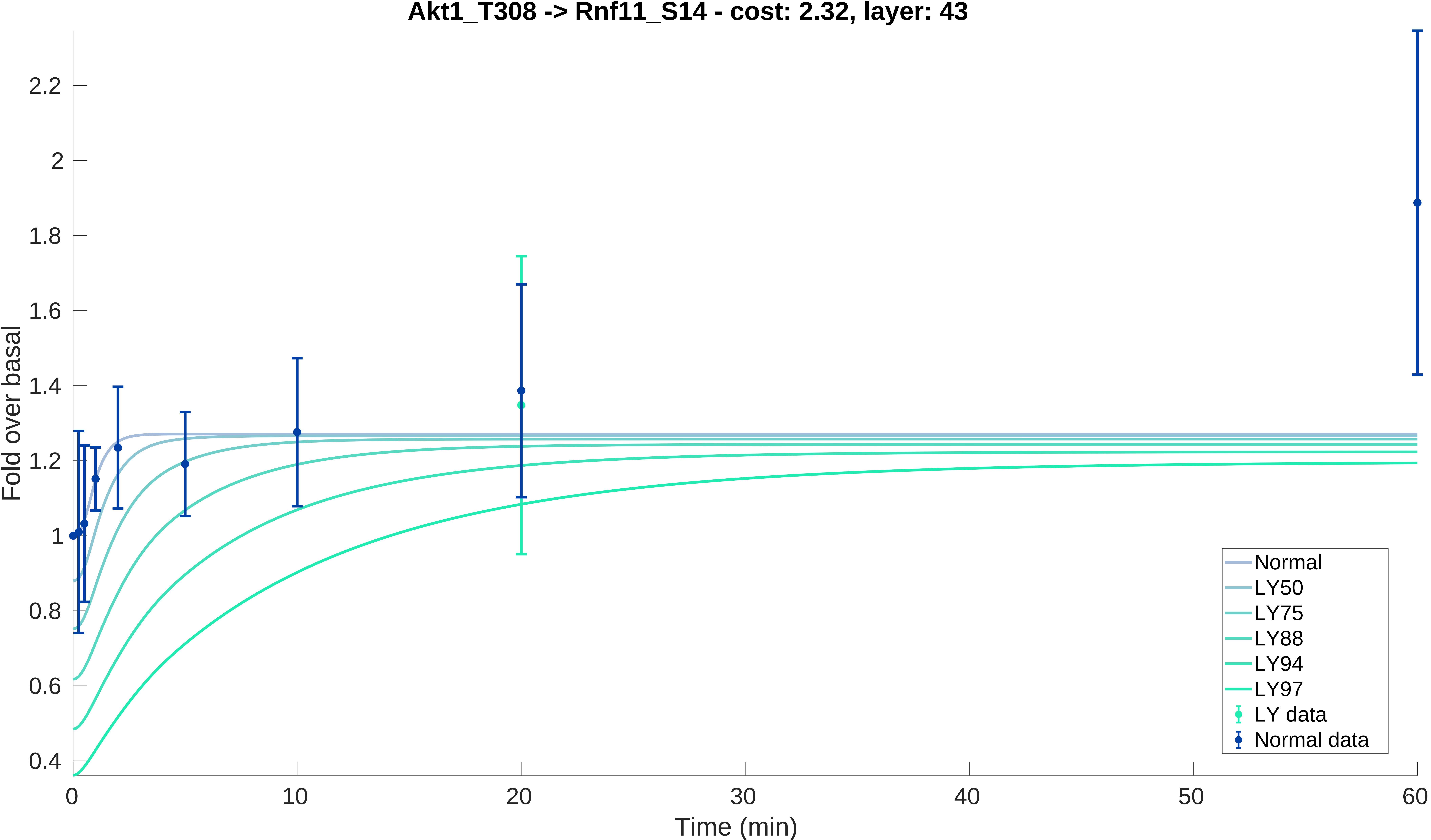

Akt1_T308 -> Rnf11_S14 - cost: 2.32, layer: 43
Fold over basal
Time (min)
0.4
0.6
0.8
1
1.2
1.4
1.6
1.8
2
2.2
0
10
20
30
40
50
60
Normal
LY50
LY75
LY88
LY94
LY97
LY data
Normal data

Src_S17 -> Tiam1_S231 - cost: 5.83, layer: 43

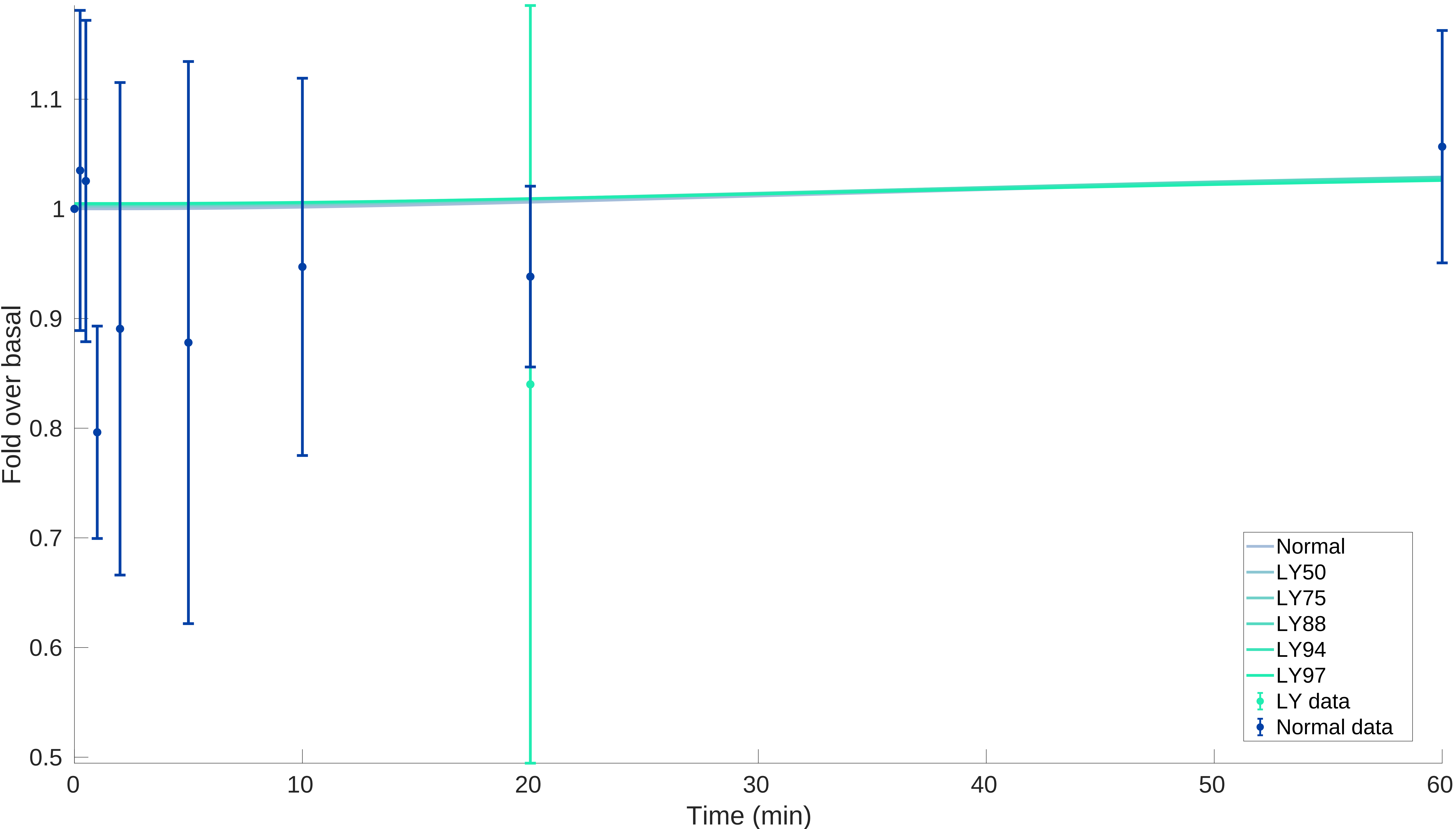

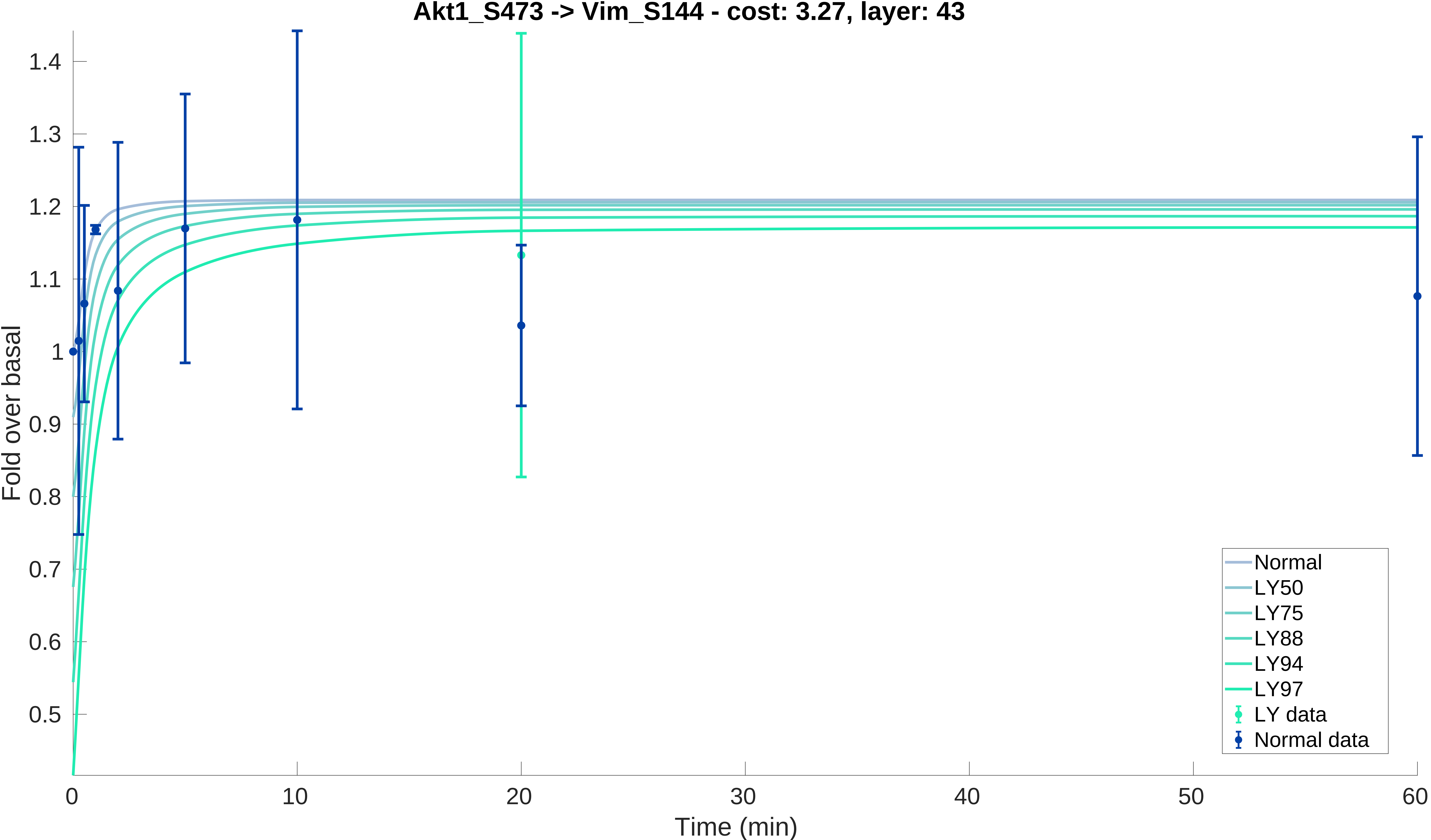

Akt1_S473 -> Vim_S144 - cost: 3.27, layer: 43
Fold over basal
Time (min)
0.5
0.6
0.7
0.8
0.9
1
1.1
1.2
1.3
1.4
0
10
20
30
40
50
60
Normal
LY50
LY75
LY88
LY94
LY97
LY data
Normal data

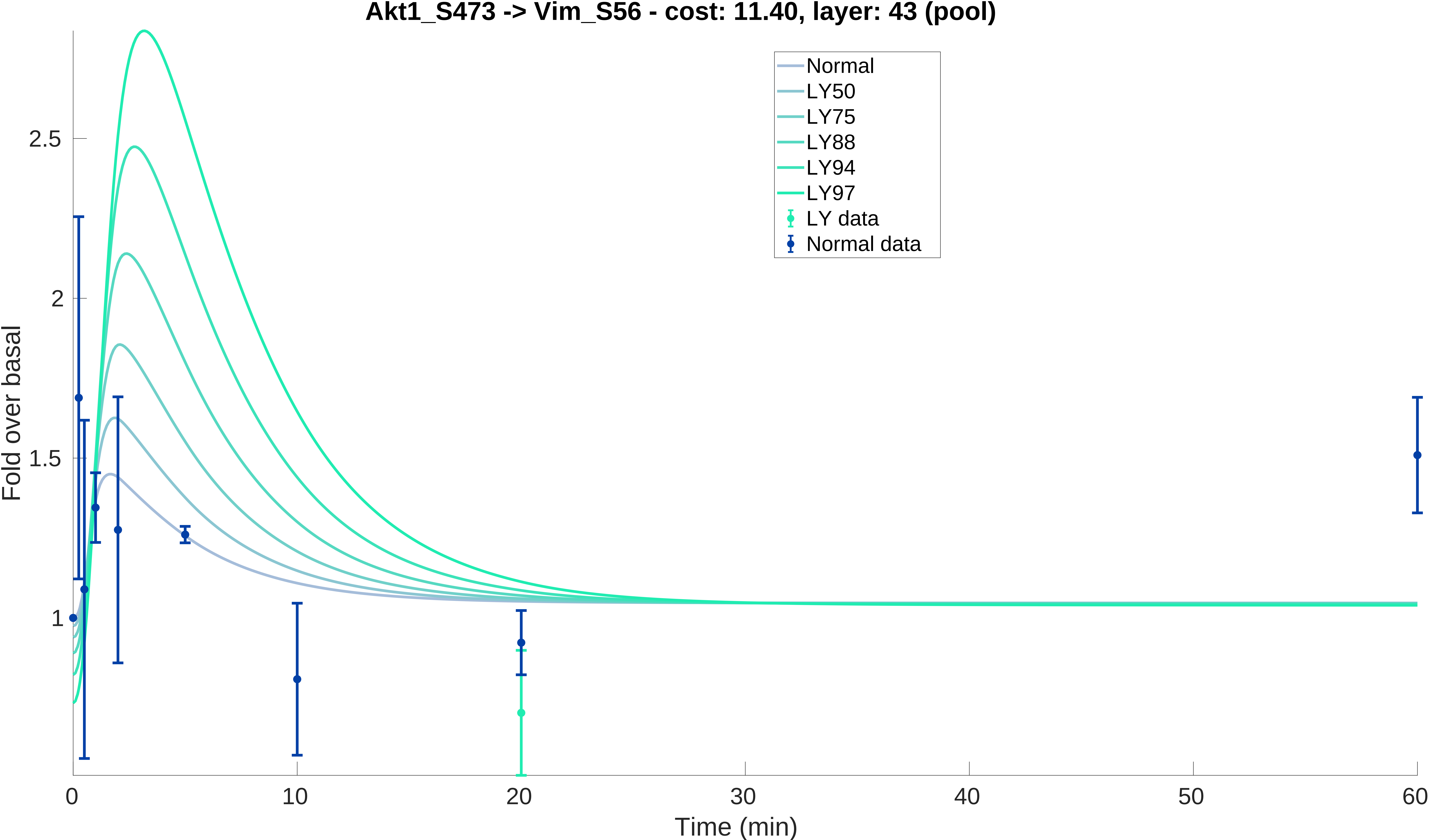

Akt1_S473 -> Vim_S56 - cost: 11.40, layer: 43 (pool)
Normal
LY50
LY75
LY88
LY94
LY97
LY data
Normal data
Fold over basal
2.5
2
1.5
1
0
10
20
30
40
50
60
Time (min)

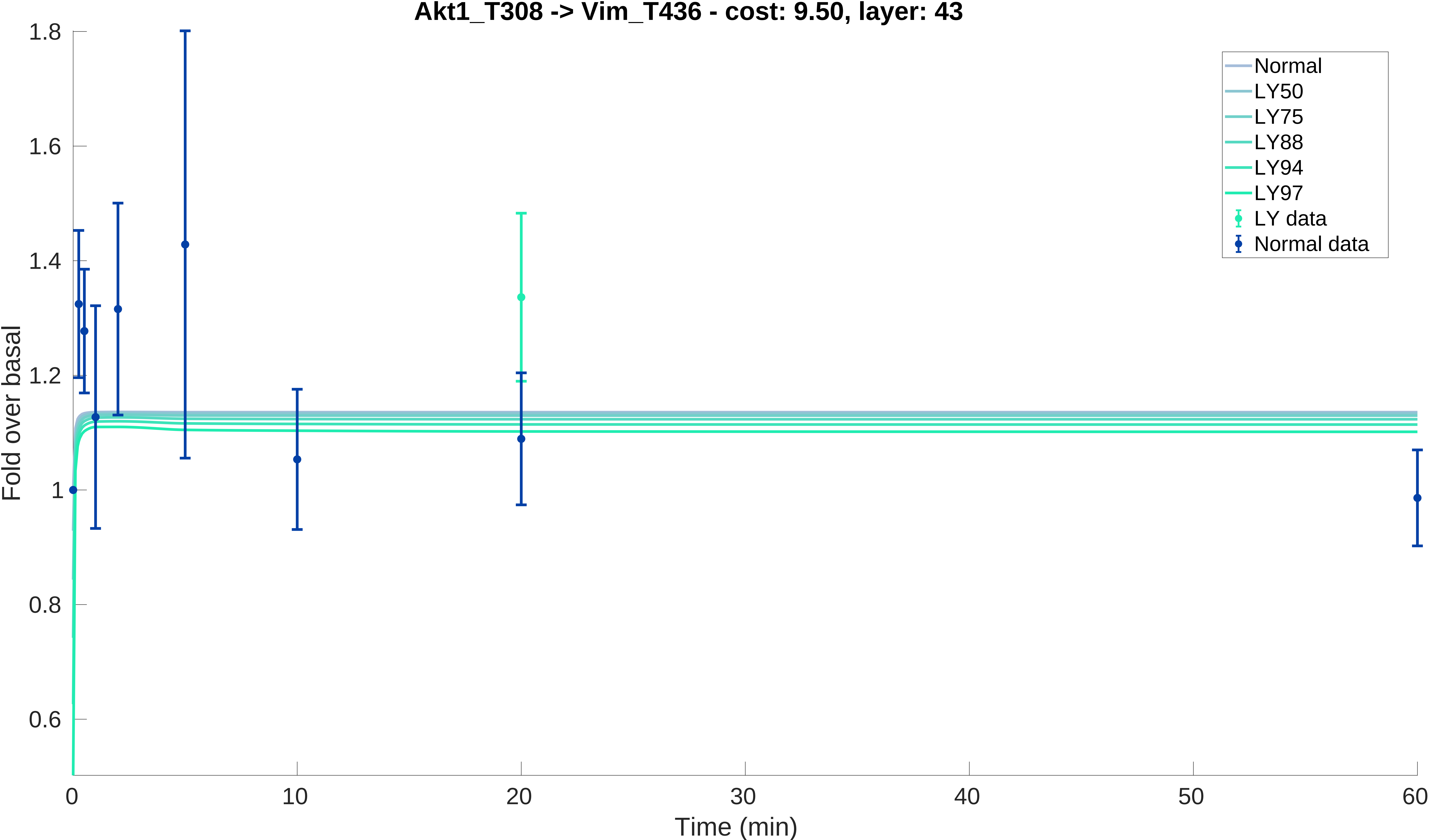

Akt1_T308 -> Vim_T436 - cost: 9.50, layer: 43
Fold over basal
Time (min)
1.8
1.6
1.4
1.2
1
0.8
0.6
0
10
20
30
40
50
60
Normal
LY50
LY75
LY88
LY94
LY97
LY data
Normal data

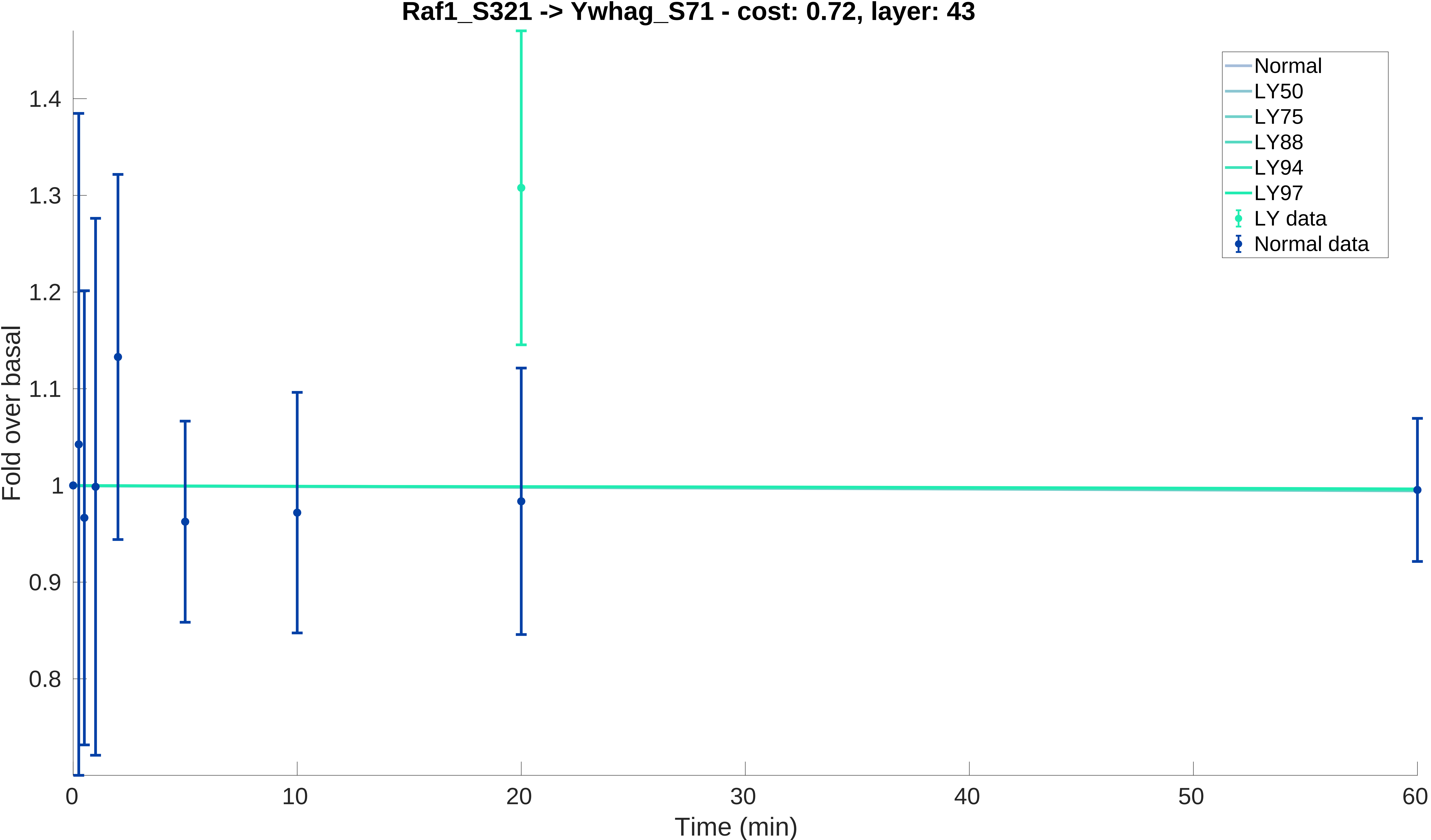

Raf1_S321 -> Ywhag_S71 - cost: 0.72, layer: 43
Normal
LY50
LY75
LY88
LY94
LY97
LY data
Normal data
Fold over basal
Time (min)
1.4
1.3
1.2
1.1
1
0.9
0.8
0
10
20
30
40
50
60

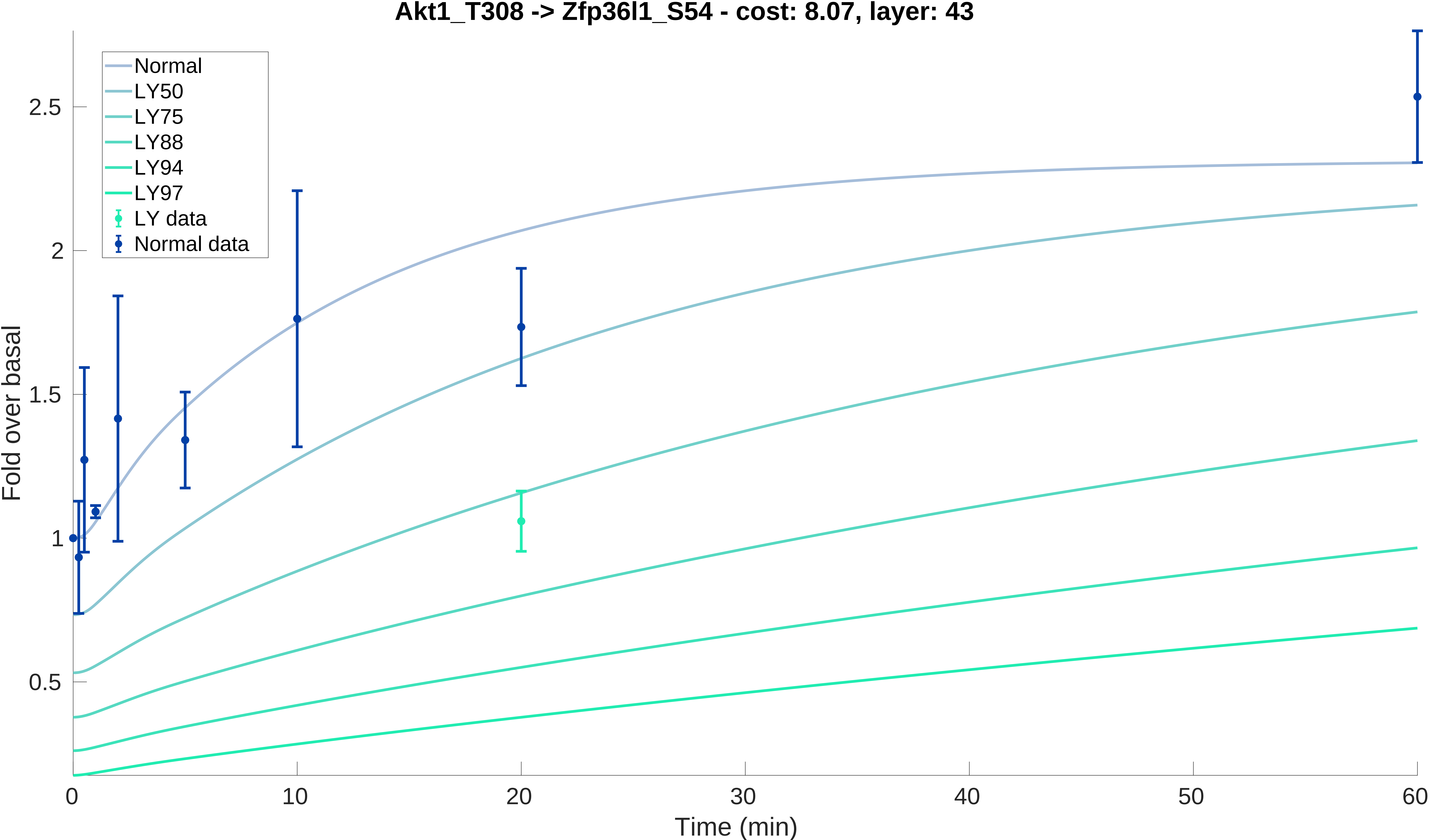
Akt1_T308 -> Zfp36l1_S54 - cost: 8.07, layer: 43
Normal
LY50
LY75
LY88
LY94
LY97
LY data
Normal data
Fold over basal
0.5
1
1.5
2
2.5
0
10
20
30
40
50
60
Time (min)

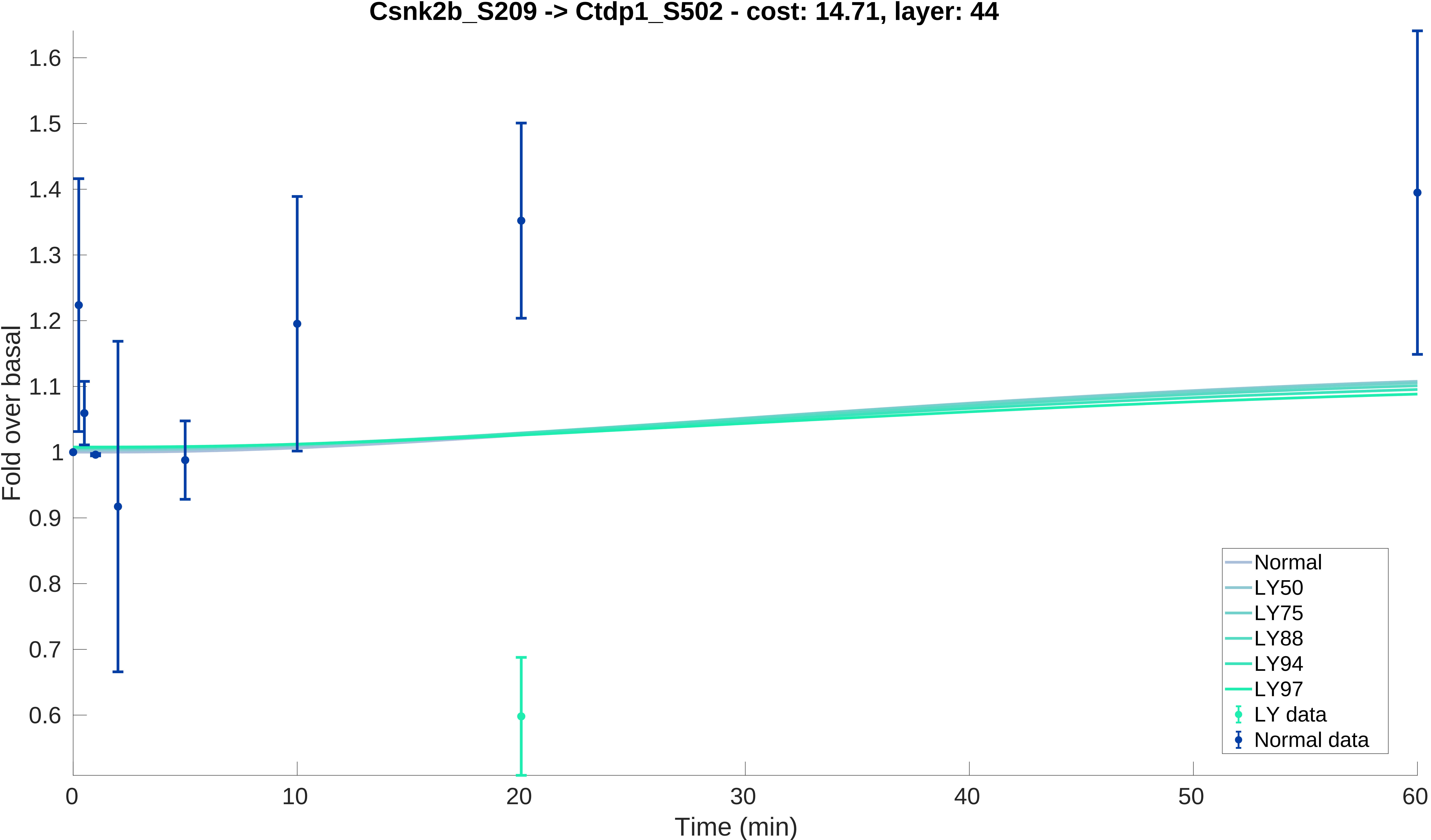

Csnk2b_S209 -> Ctdp1_S502 - cost: 14.71, layer: 44
Fold over basal
Time (min)
1.6
1.5
1.4
1.3
1.2
1.1
1
0.9
0.8
0.7
0.6
0
10
20
30
40
50
60
Normal
LY50
LY75
LY88
LY94
LY97
LY data
Normal data

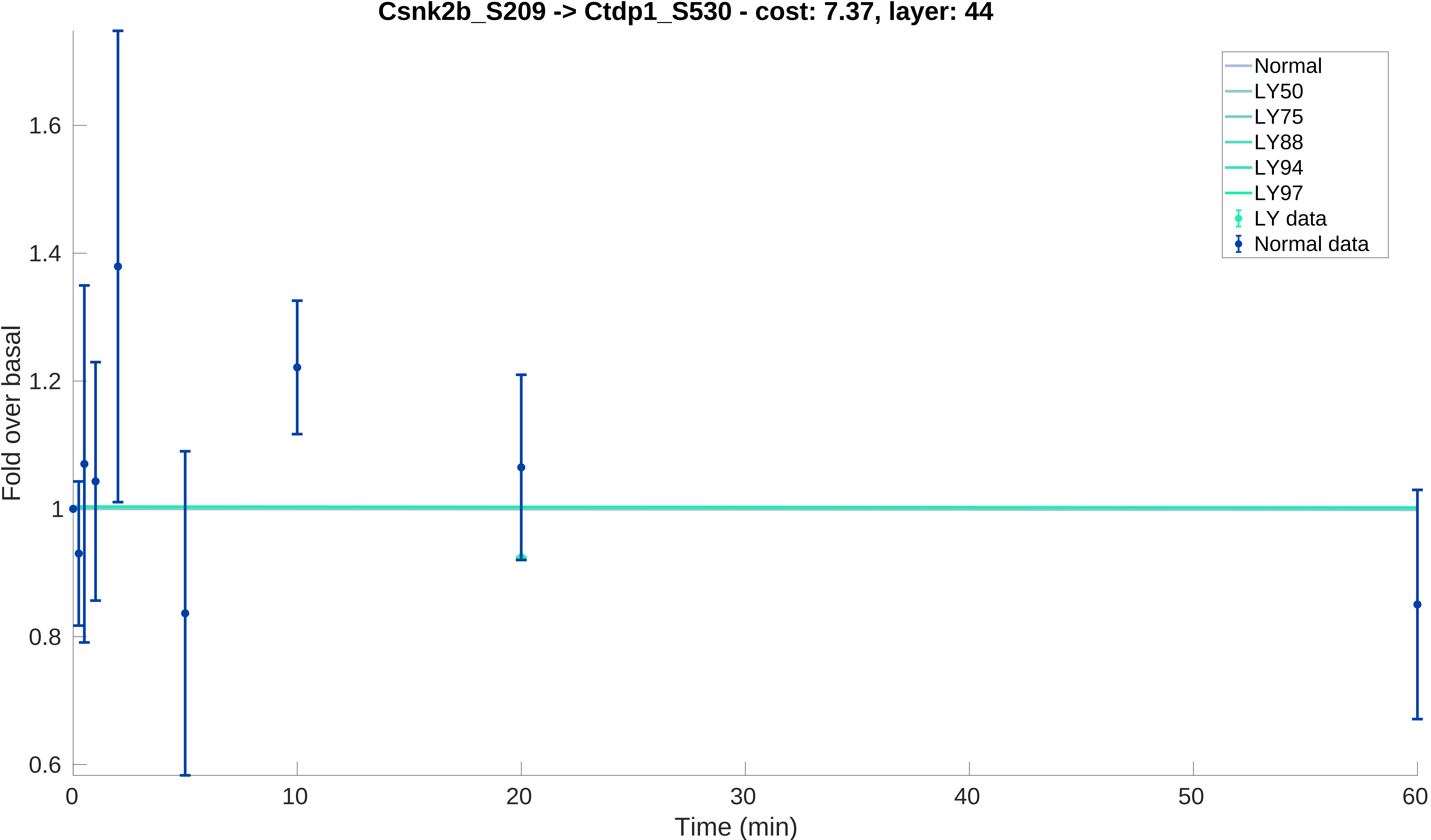

Csnk2b_S209 -> Ctdp1_S530 - cost: 7.37, layer: 44
Fold over basal
Time (min)
0.6
0.8
1
1.2
1.4
1.6
0
10
20
30
40
50
60
Normal
LY50
LY75
LY88
LY94
LY97
LY data
Normal data

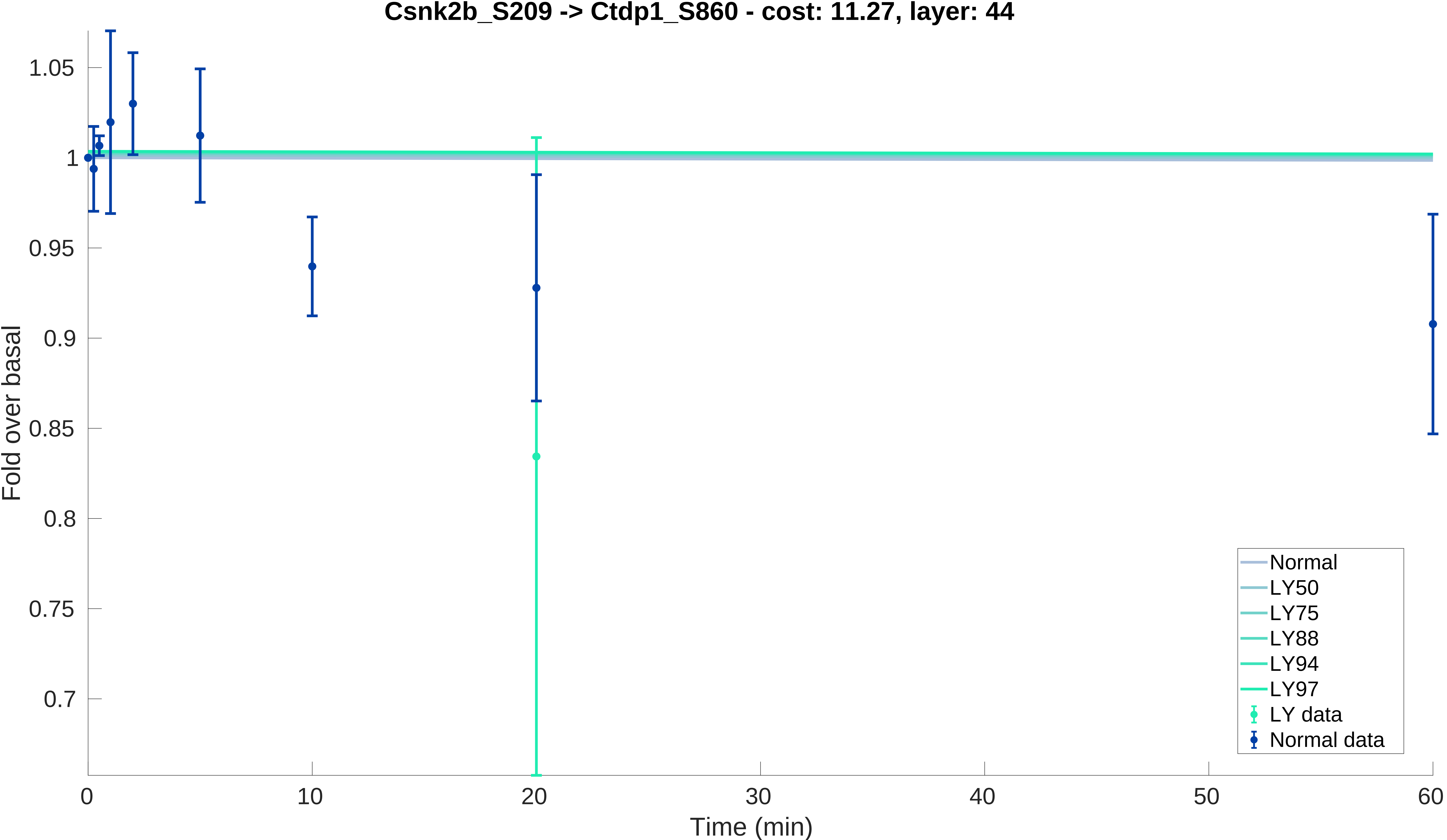

Csnk2b_S209 -> Ctdp1_S860 - cost: 11.27, layer: 44
Fold over basal
1.05
1
0.95
0.9
0.85
0.8
0.75
0.7
0
10
20
30
40
50
60
Time (min)
Normal
LY50
LY75
LY88
LY94
LY97
LY data
Normal data

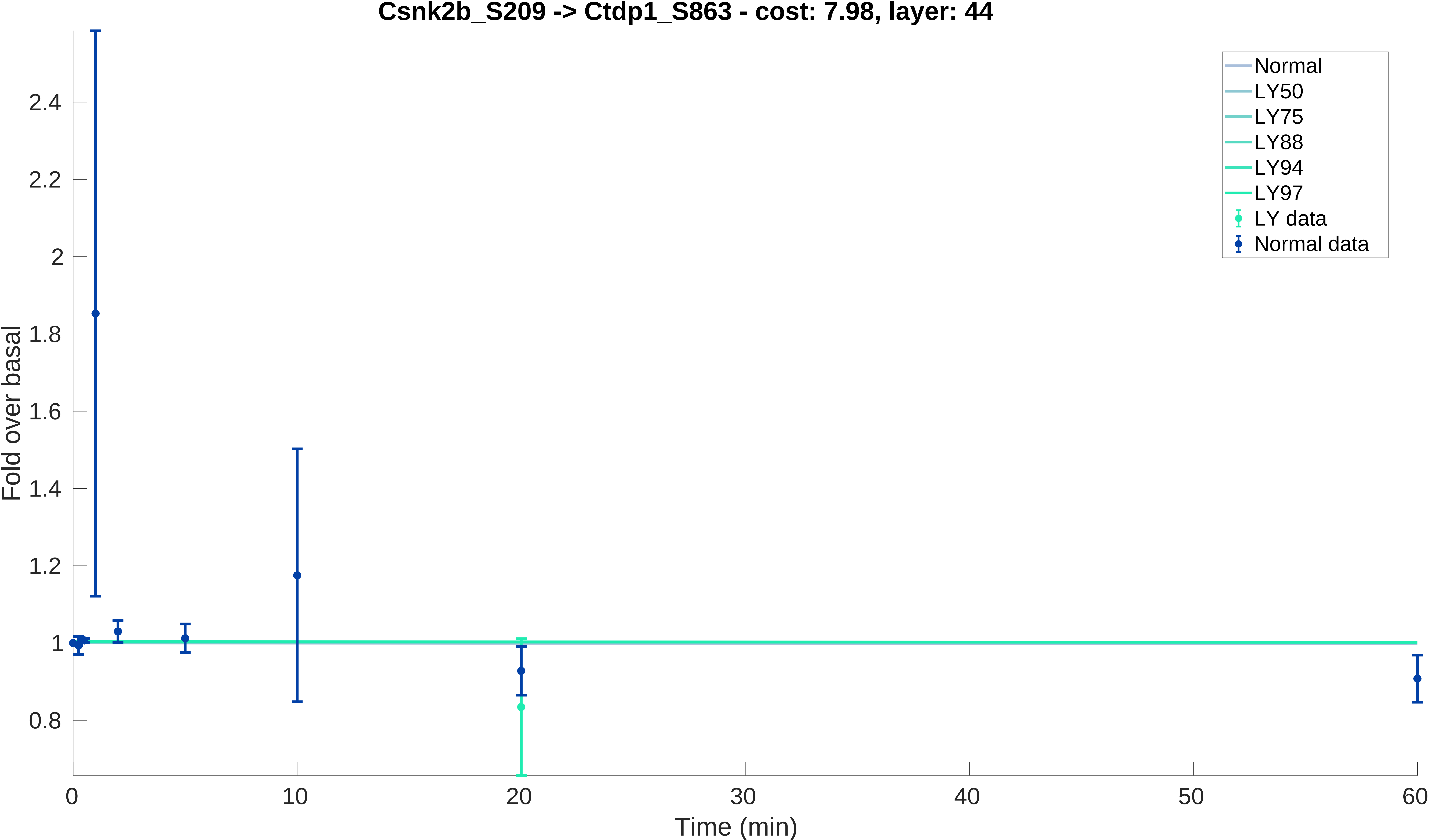

Csnk2b_S209 -> Ctdp1_S863 - cost: 7.98, layer: 44
Fold over basal
Time (min)
2.4
2.2
2
1.8
1.6
1.4
1.2
1
0.8
0
10
20
30
40
50
60
Normal
LY50
LY75
LY88
LY94
LY97
LY data
Normal data

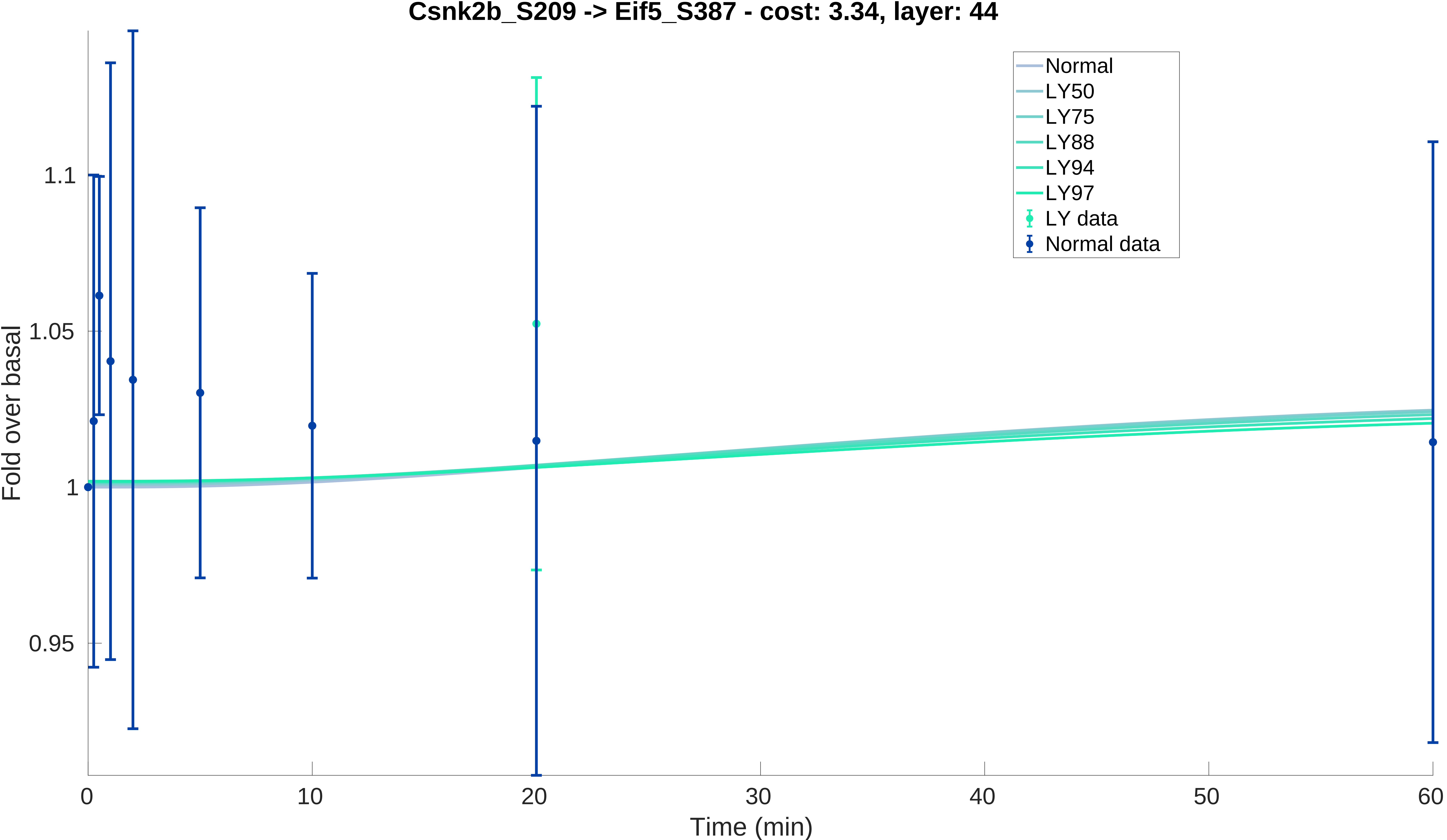
Csnk2b_S209 -> Eif5_S387 - cost: 3.34, layer: 44
Normal
LY50
LY75
LY88
LY94
LY97
LY data
Normal data
Fold over basal
1.1
1.05
1
0.95
0
10
20
30
40
50
60
Time (min)

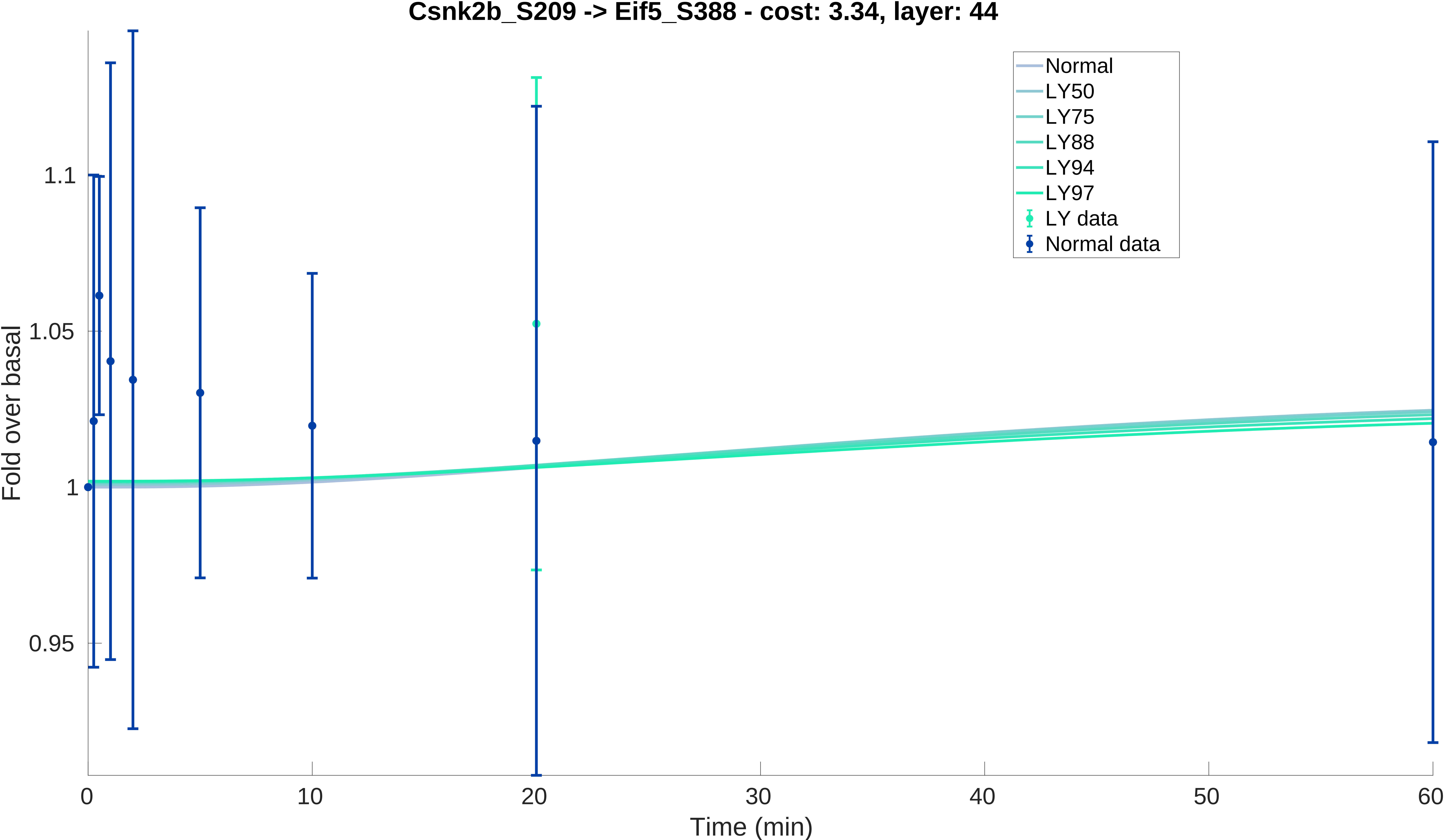
Csnk2b_S209 -> Eif5_S388 - cost: 3.34, layer: 44
Normal
LY50
LY75
LY88
LY94
LY97
LY data
Normal data
Fold over basal
1.1
1.05
1
0.95
0
10
20
30
40
50
60
Time (min)

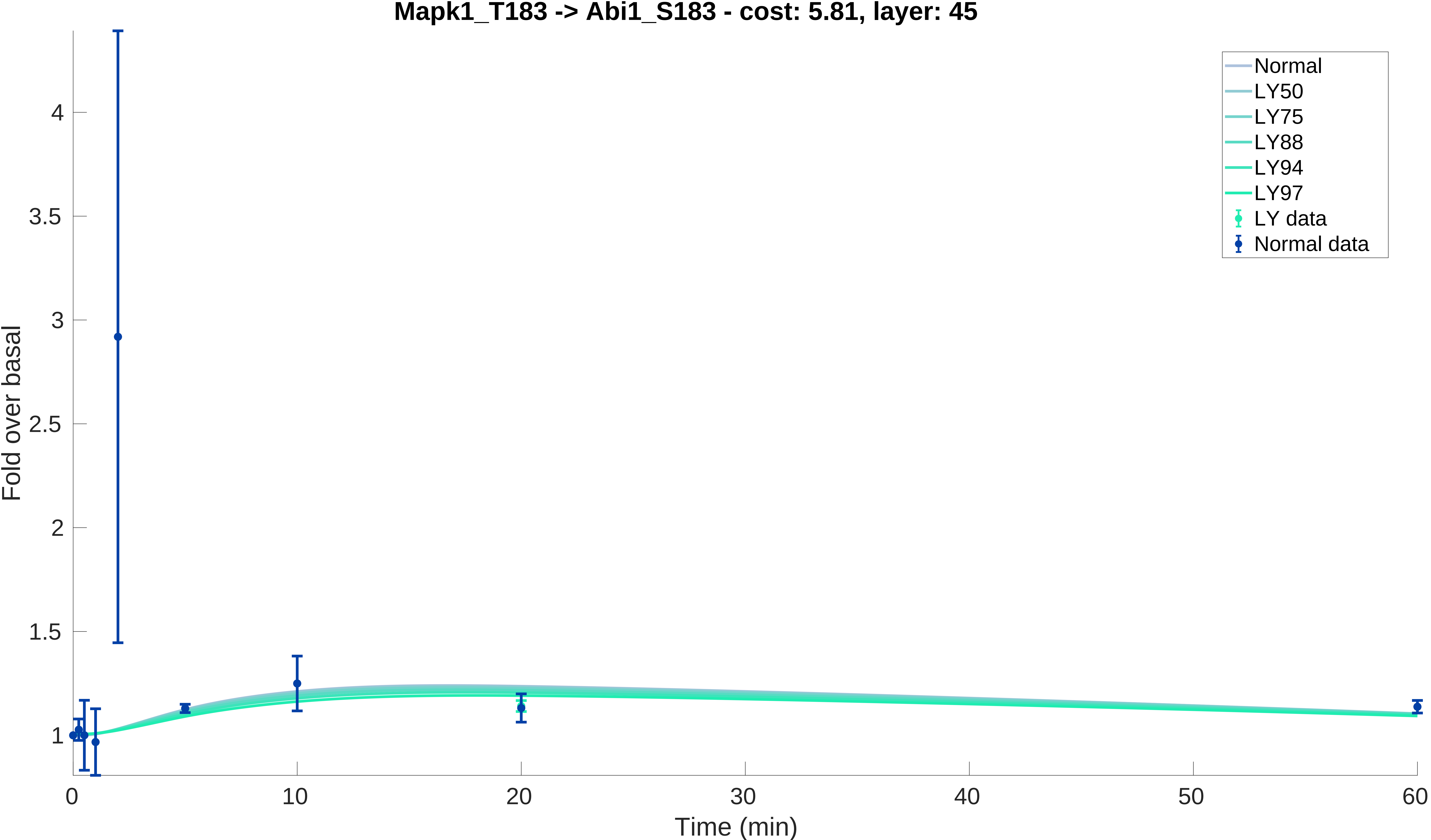

Mapk1_T183 -> Abi1_S183 - cost: 5.81, layer: 45
Fold over basal
Time (min)
4
3.5
3
2.5
2
1.5
1
0
10
20
30
40
50
60
Normal
LY50
LY75
LY88
LY94
LY97
LY data
Normal data

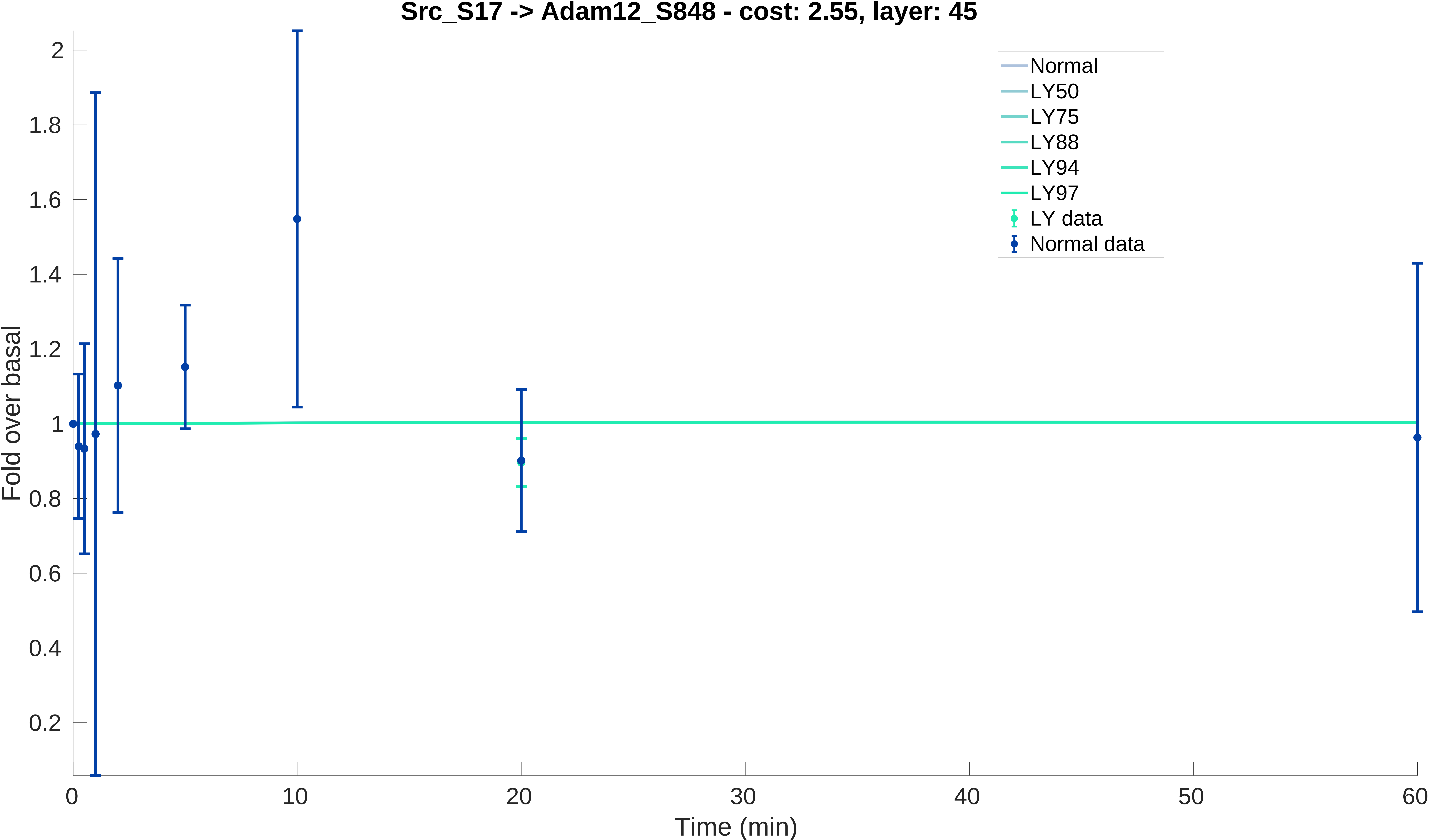
Src_S17 -> Adam12_S848 - cost: 2.55, layer: 45
Fold over basal
Time (min)
Normal
LY50
LY75
LY88
LY94
LY97
LY data
Normal data
0
10
20
30
40
50
60
0.2
0.4
0.6
0.8
1
1.2
1.4
1.6
1.8
2

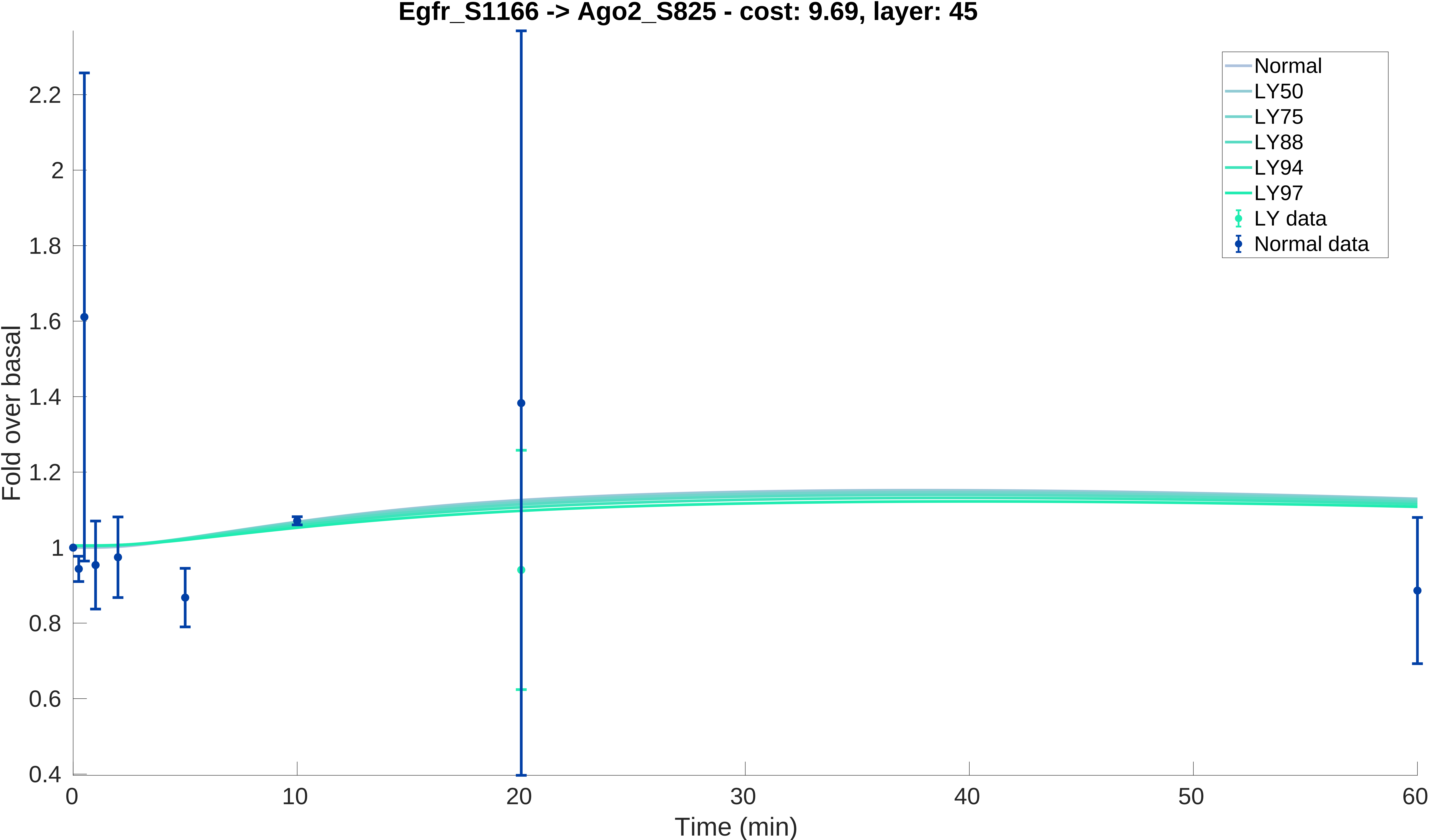

Egfr_S1166 -> Ago2_S825 - cost: 9.69, layer: 45
Fold over basal
Time (min)
Normal
LY50
LY75
LY88
LY94
LY97
LY data
Normal data
0.4
0.6
0.8
1
1.2
1.4
1.6
1.8
2
2.2
0
10
20
30
40
50
60

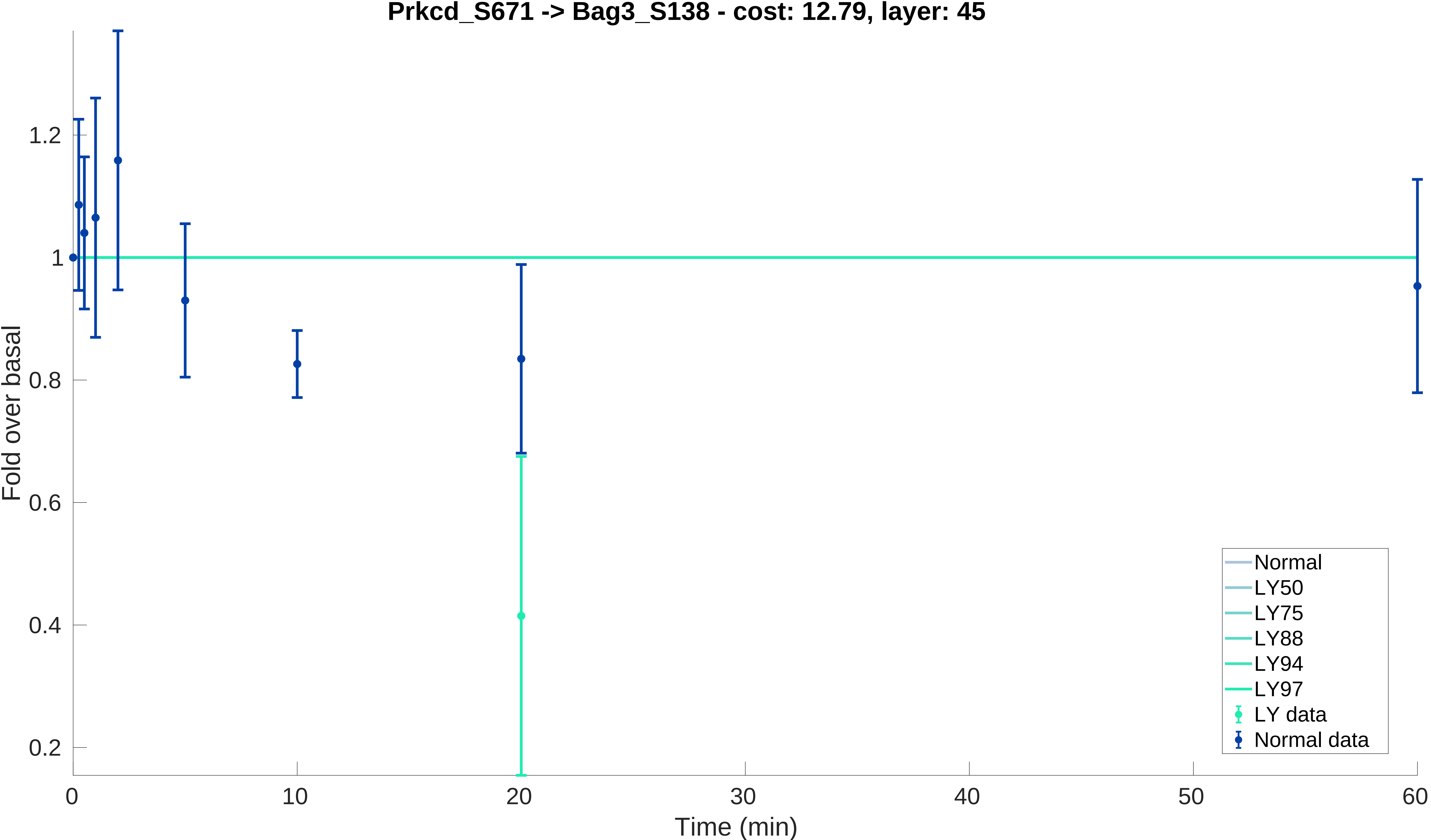

Prkcd_S671 -> Bag3_S138 - cost: 12.79, layer: 45
Fold over basal
Time (min)
0.2
0.4
0.6
0.8
1
1.2
0
10
20
30
40
50
60
Normal
LY50
LY75
LY88
LY94
LY97
LY data
Normal data

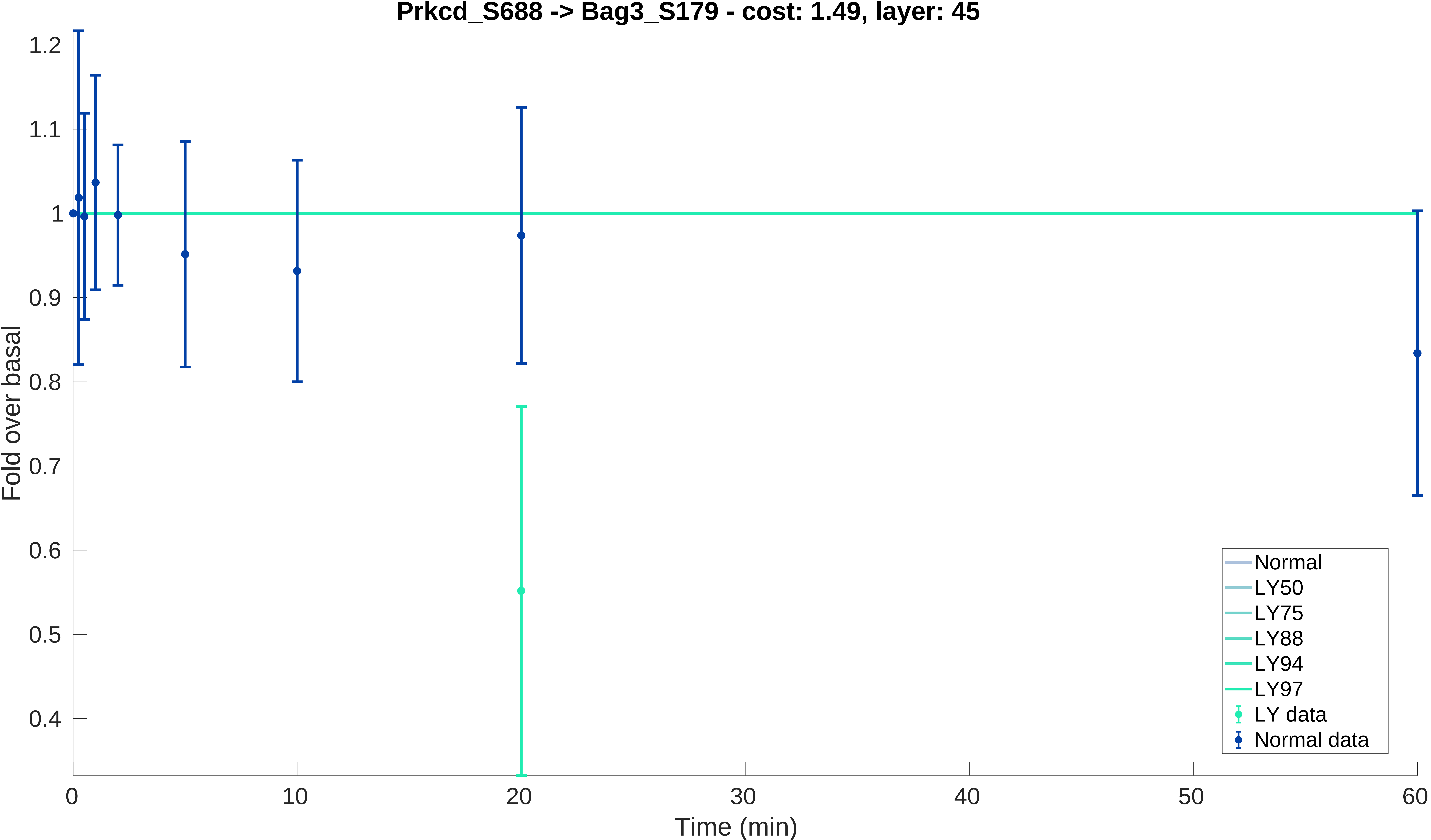

Prkcd_S688 -> Bag3_S179 - cost: 1.49, layer: 45
Fold over basal
Time (min)
1.2
1.1
1
0.9
0.8
0.7
0.6
0.5
0.4
0
10
20
30
40
50
60
Normal
LY50
LY75
LY88
LY94
LY97
LY data
Normal data

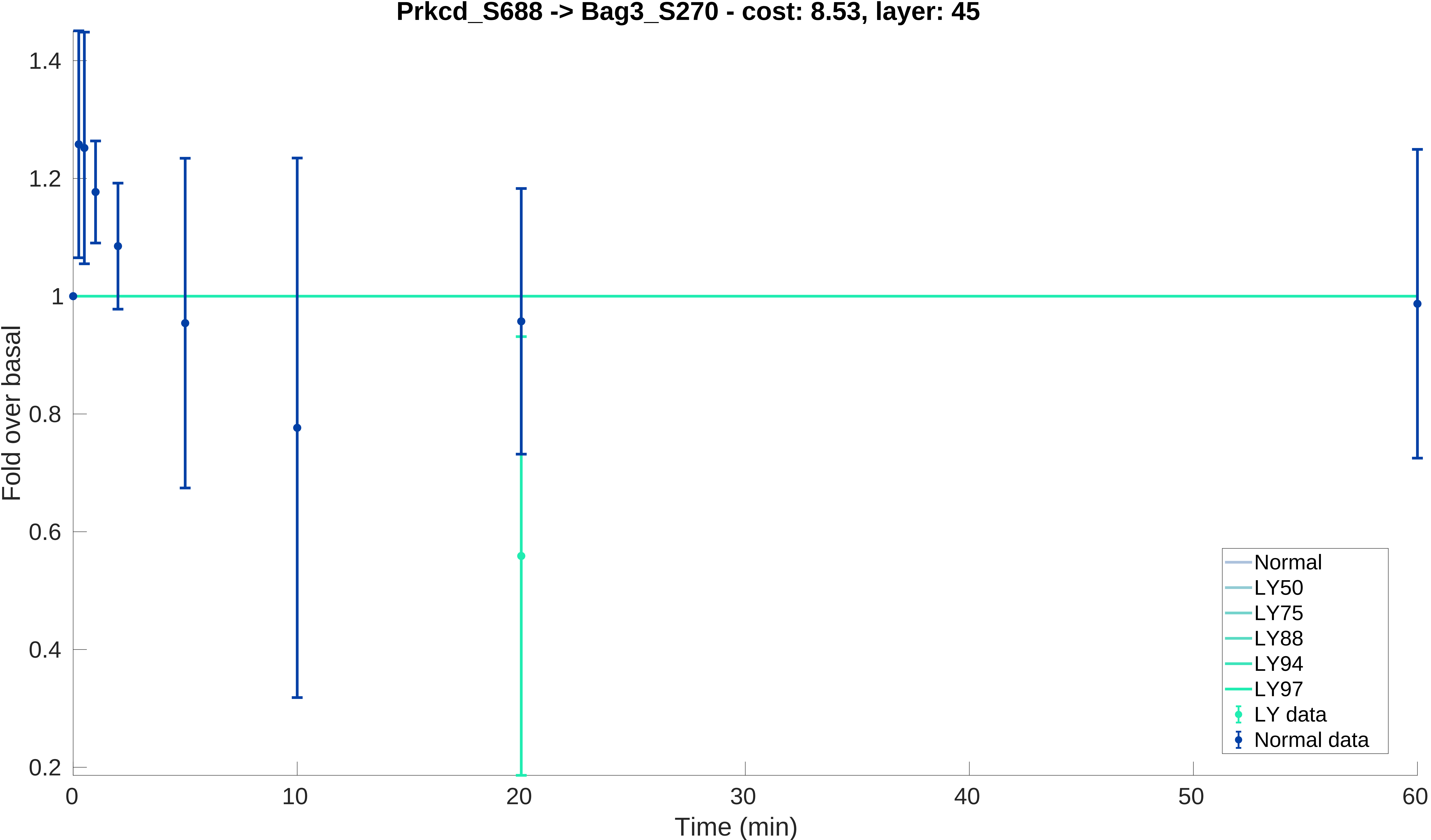

Prkcd_S688 -> Bag3_S270 - cost: 8.53, layer: 45
Fold over basal
1.4
1.2
1
0.8
0.6
0.4
0.2
0
10
20
30
40
50
60
Time (min)
Normal
LY50
LY75
LY88
LY94
LY97
LY data
Normal data

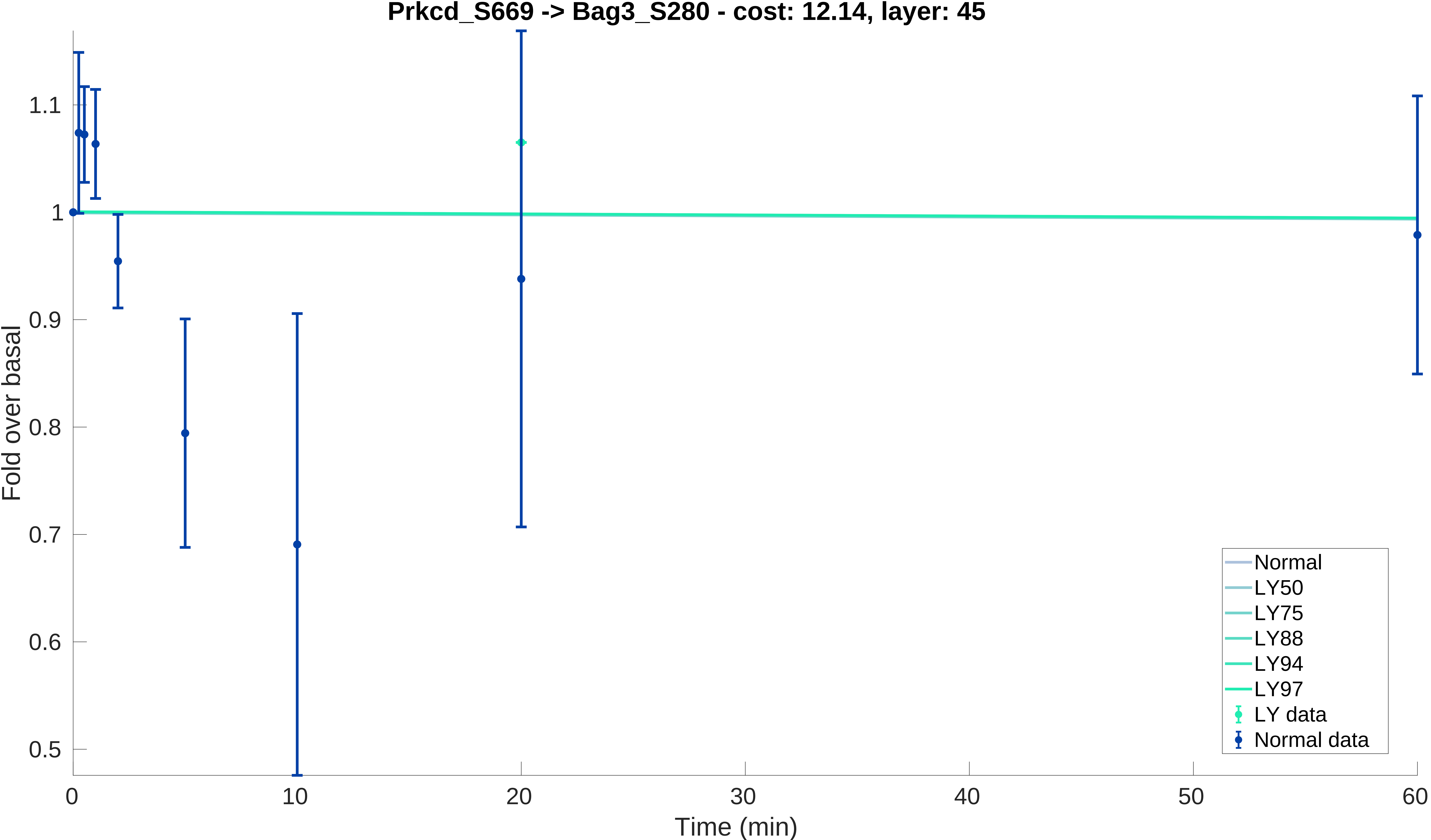
Prkcd_S669 -> Bag3_S280 - cost: 12.14, layer: 45
Fold over basal
1.1
1
0.9
0.8
0.7
0.6
0.5
0
10
20
30
40
50
60
Time (min)
Normal
LY50
LY75
LY88
LY94
LY97
LY data
Normal data

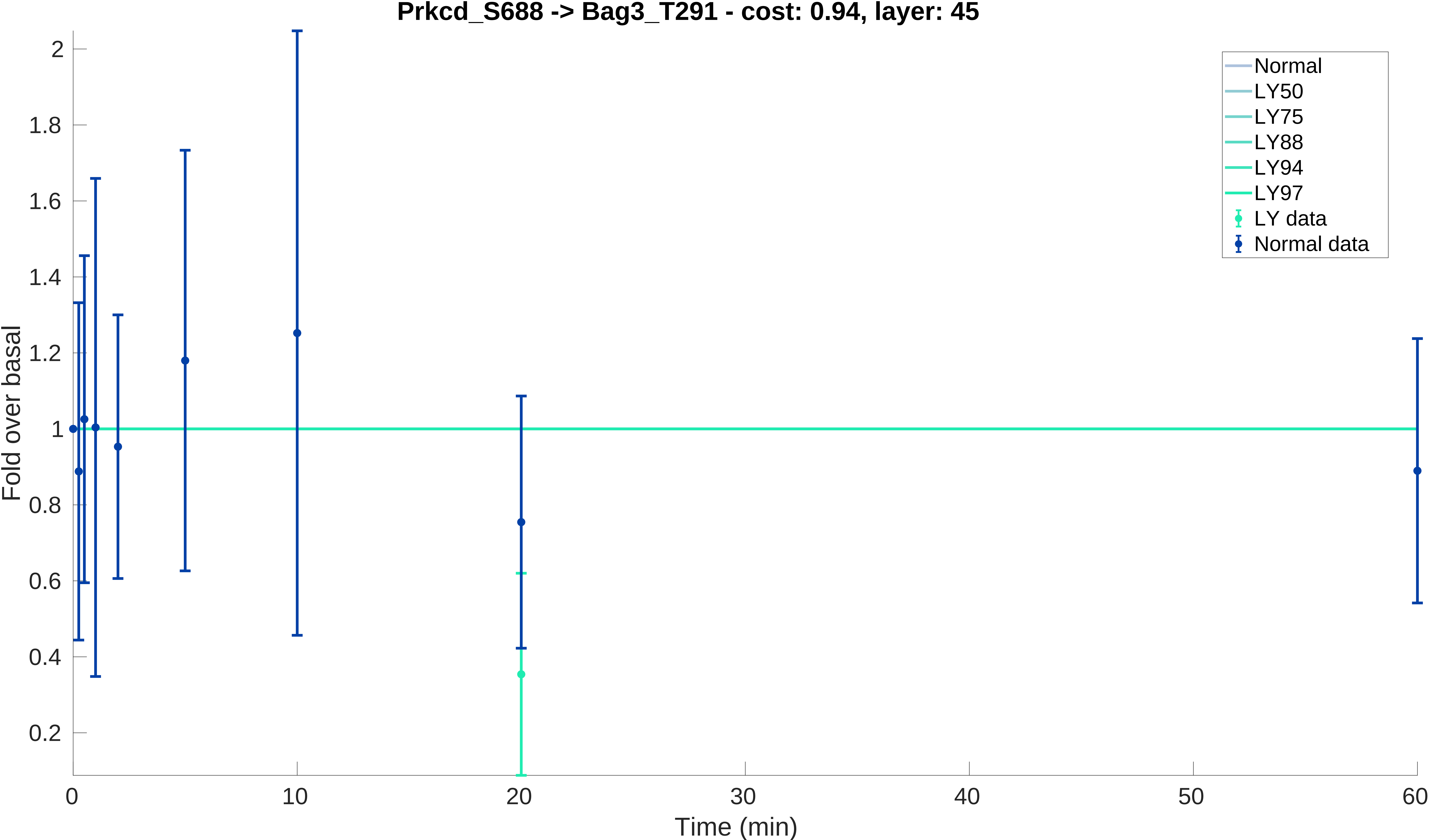

Prkcd_S688 -> Bag3_T291 - cost: 0.94, layer: 45
Fold over basal
Time (min)
Normal
LY50
LY75
LY88
LY94
LY97
LY data
Normal data
0
10
20
30
40
50
60
0.2
0.4
0.6
0.8
1
1.2
1.4
1.6
1.8
2

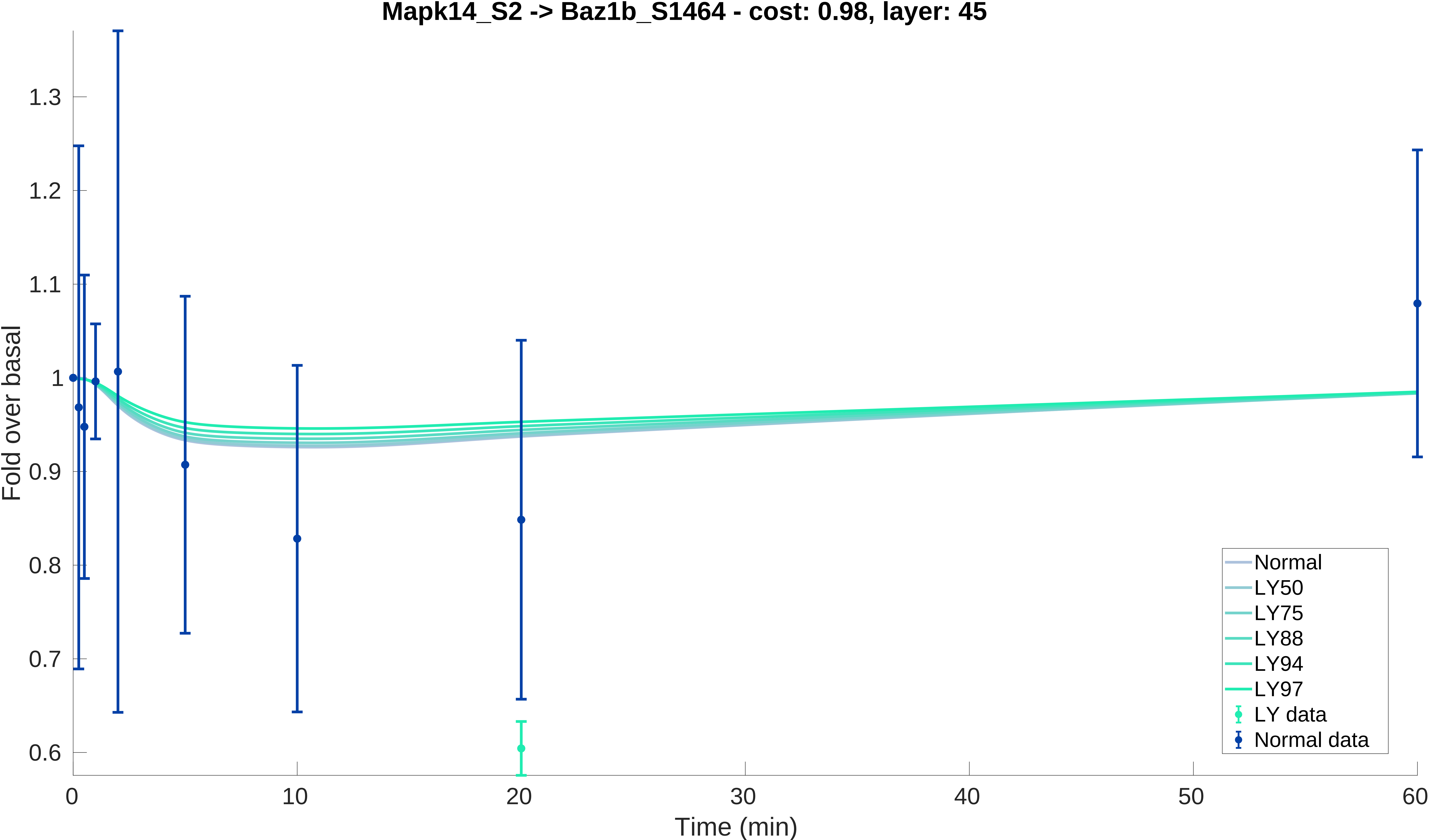

Mapk14_S2 -> Baz1b_S1464 - cost: 0.98, layer: 45
Fold over basal
1.3
1.2
1.1
1
0.9
0.8
0.7
0.6
0
10
20
30
40
50
60
Time (min)
Normal
LY50
LY75
LY88
LY94
LY97
LY data
Normal data

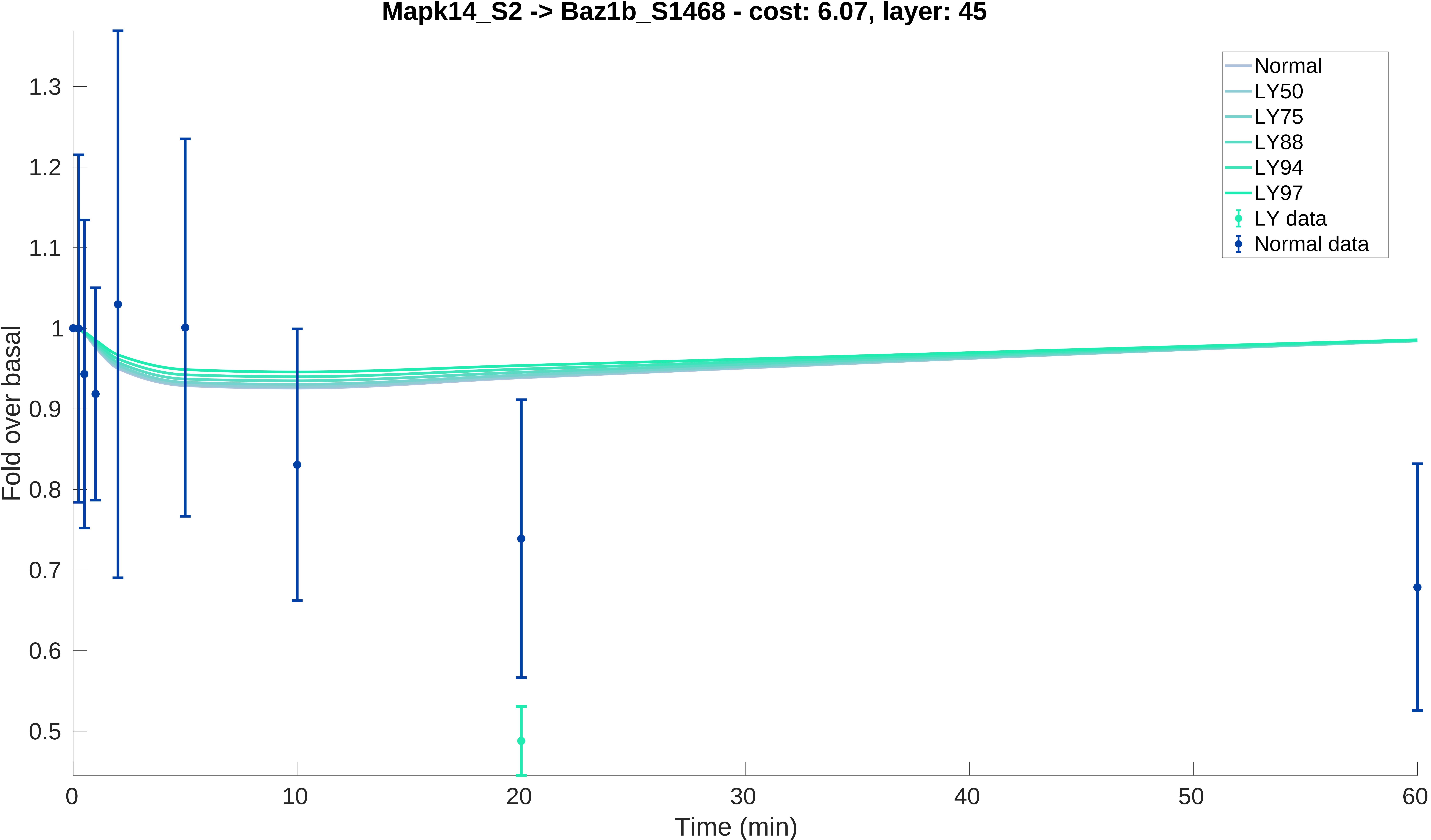

Mapk14_S2 -> Baz1b_S1468 - cost: 6.07, layer: 45
Fold over basal
Time (min)
0
10
20
30
40
50
60
0.5
0.6
0.7
0.8
0.9
1
1.1
1.2
1.3
Normal
LY50
LY75
LY88
LY94
LY97
LY data
Normal data

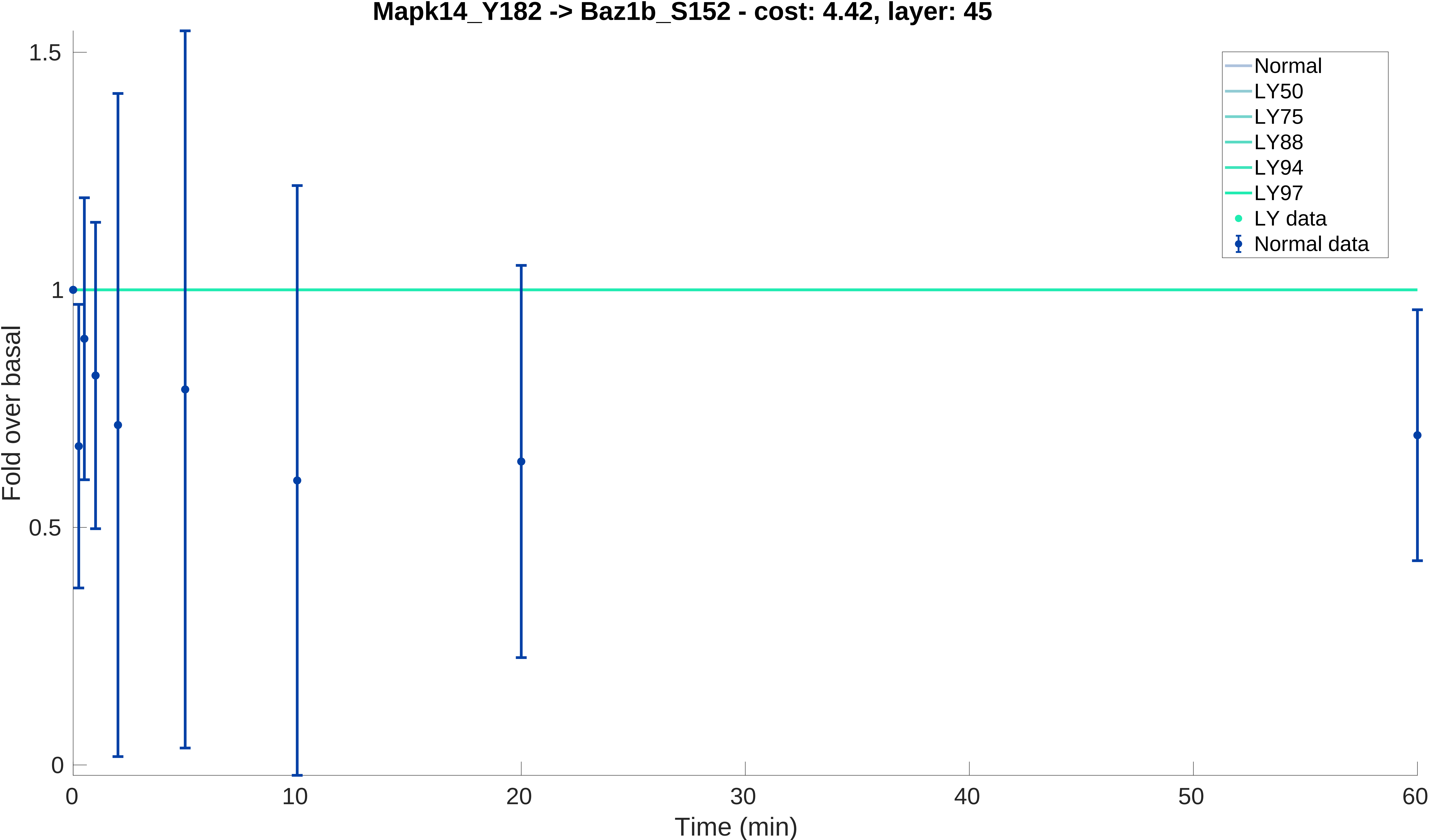

Mapk14_Y182 -> Baz1b_S152 - cost: 4.42, layer: 45
Fold over basal
1.5
1
0.5
0
0
10
20
30
40
50
60
Time (min)
Normal
LY50
LY75
LY88
LY94
LY97
LY data
Normal data

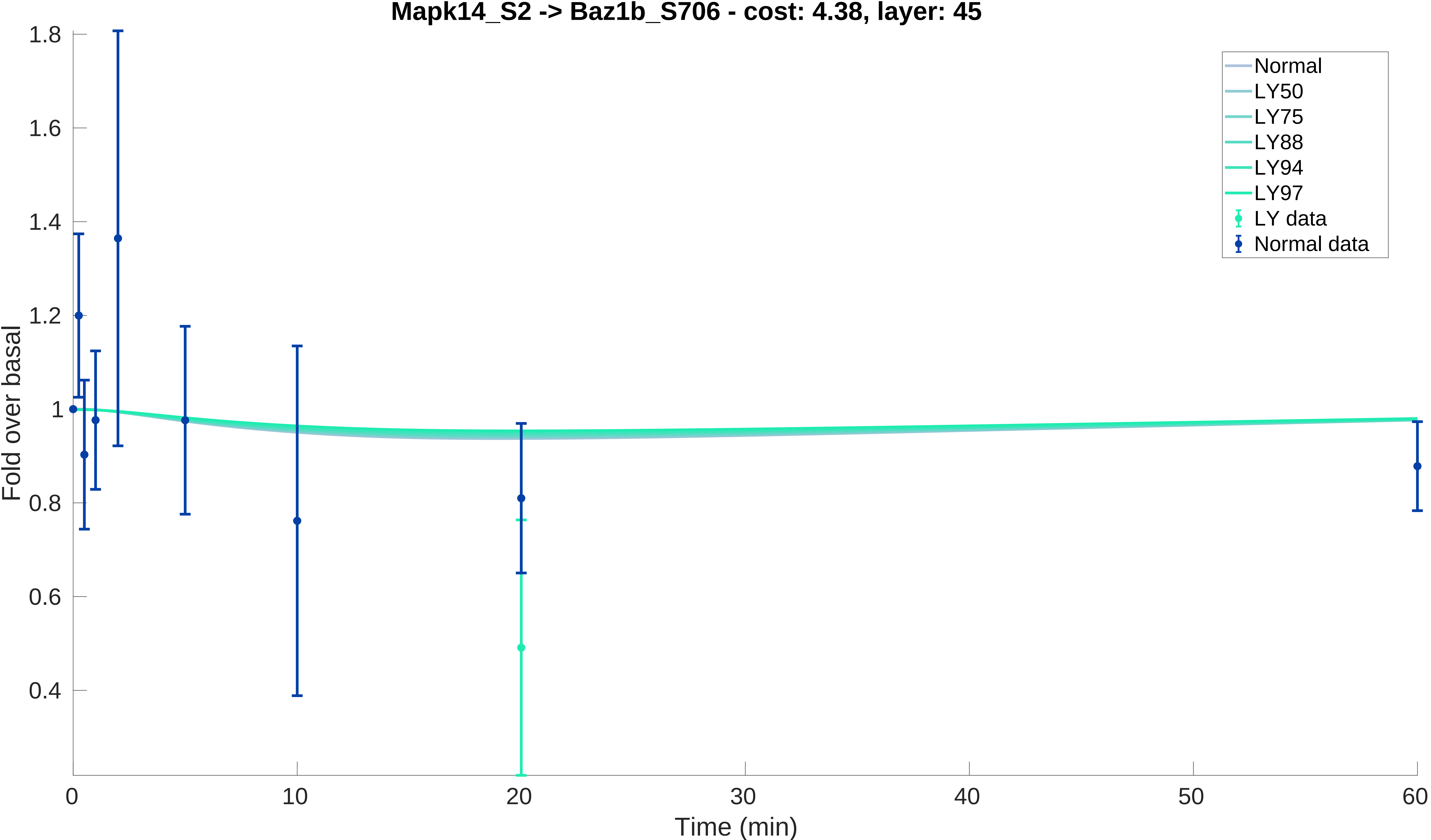

Mapk14_S2 -> Baz1b_S706 - cost: 4.38, layer: 45
Fold over basal
Time (min)
Normal
LY50
LY75
LY88
LY94
LY97
LY data
Normal data
0.4
0.6
0.8
1
1.2
1.4
1.6
1.8
0
10
20
30
40
50
60

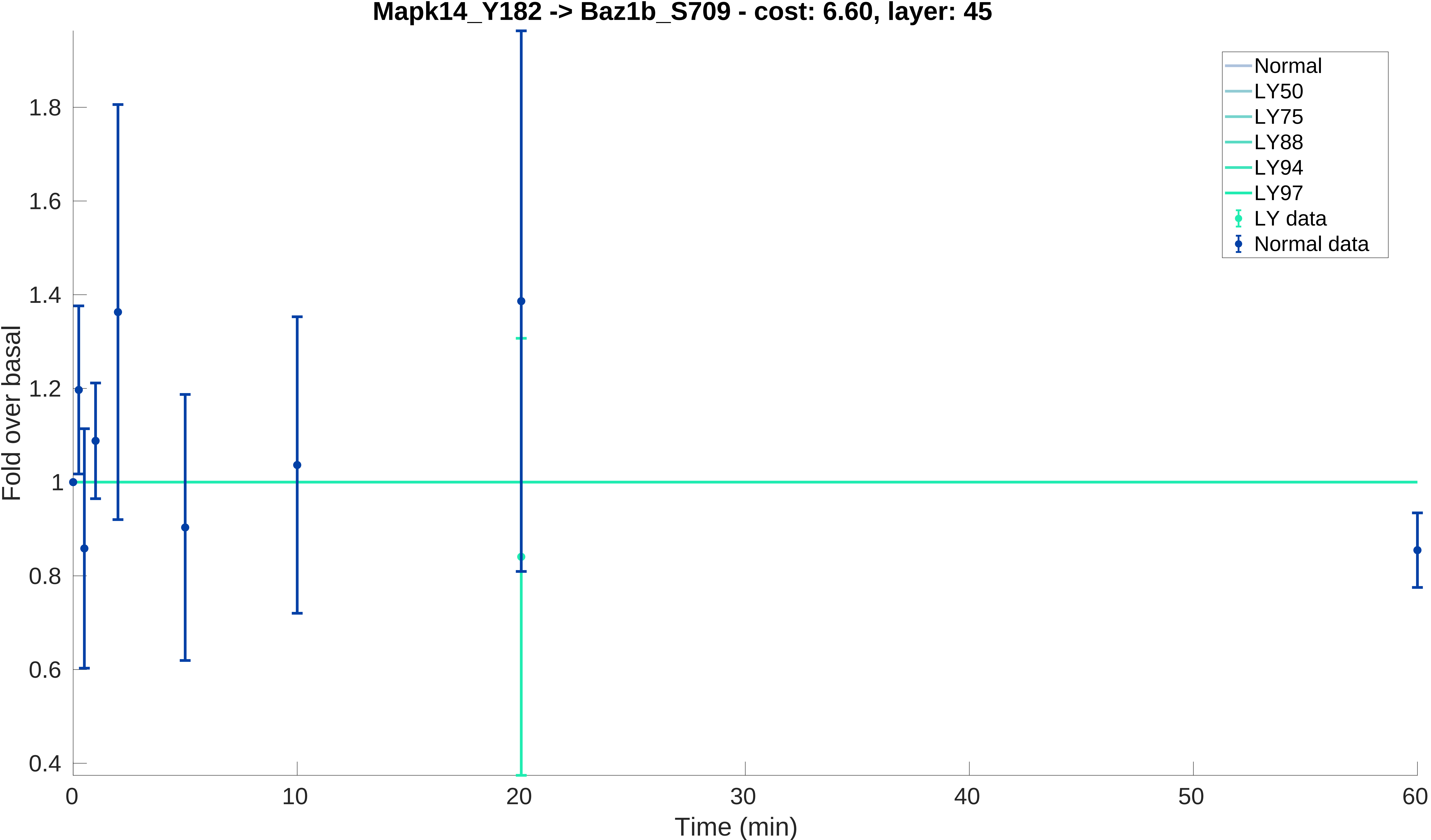

Mapk14_Y182 -> Baz1b_S709 - cost: 6.60, layer: 45
Fold over basal
Time (min)
0.4
0.6
0.8
1
1.2
1.4
1.6
1.8
0
10
20
30
40
50
60
Normal
LY50
LY75
LY88
LY94
LY97
LY data
Normal data

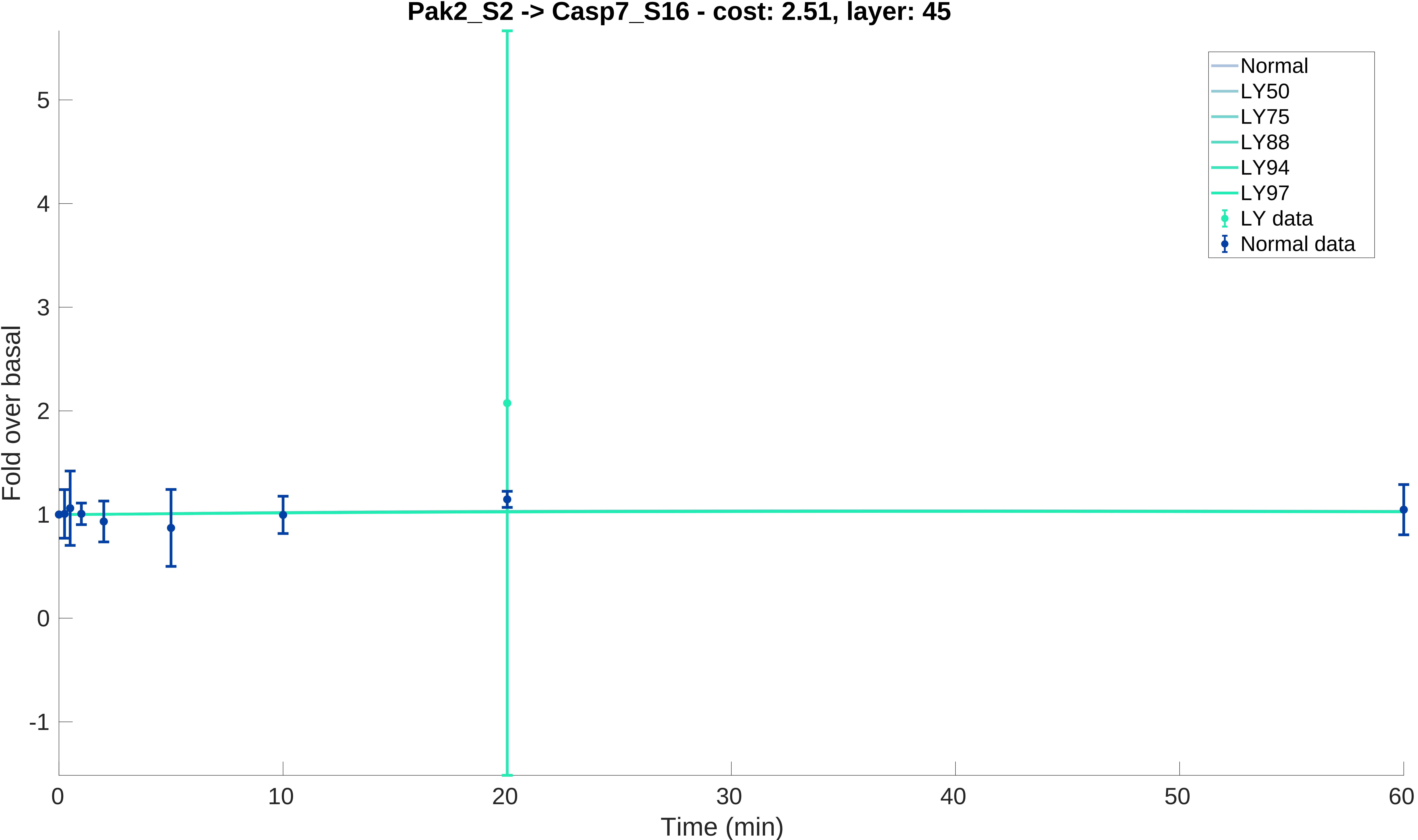

Pak2_S2 -> Casp7_S16 - cost: 2.51, layer: 45
Fold over basal
Time (min)
Normal
LY50
LY75
LY88
LY94
LY97
LY data
Normal data
5
4
3
2
1
0
-1
0
10
20
30
40
50
60

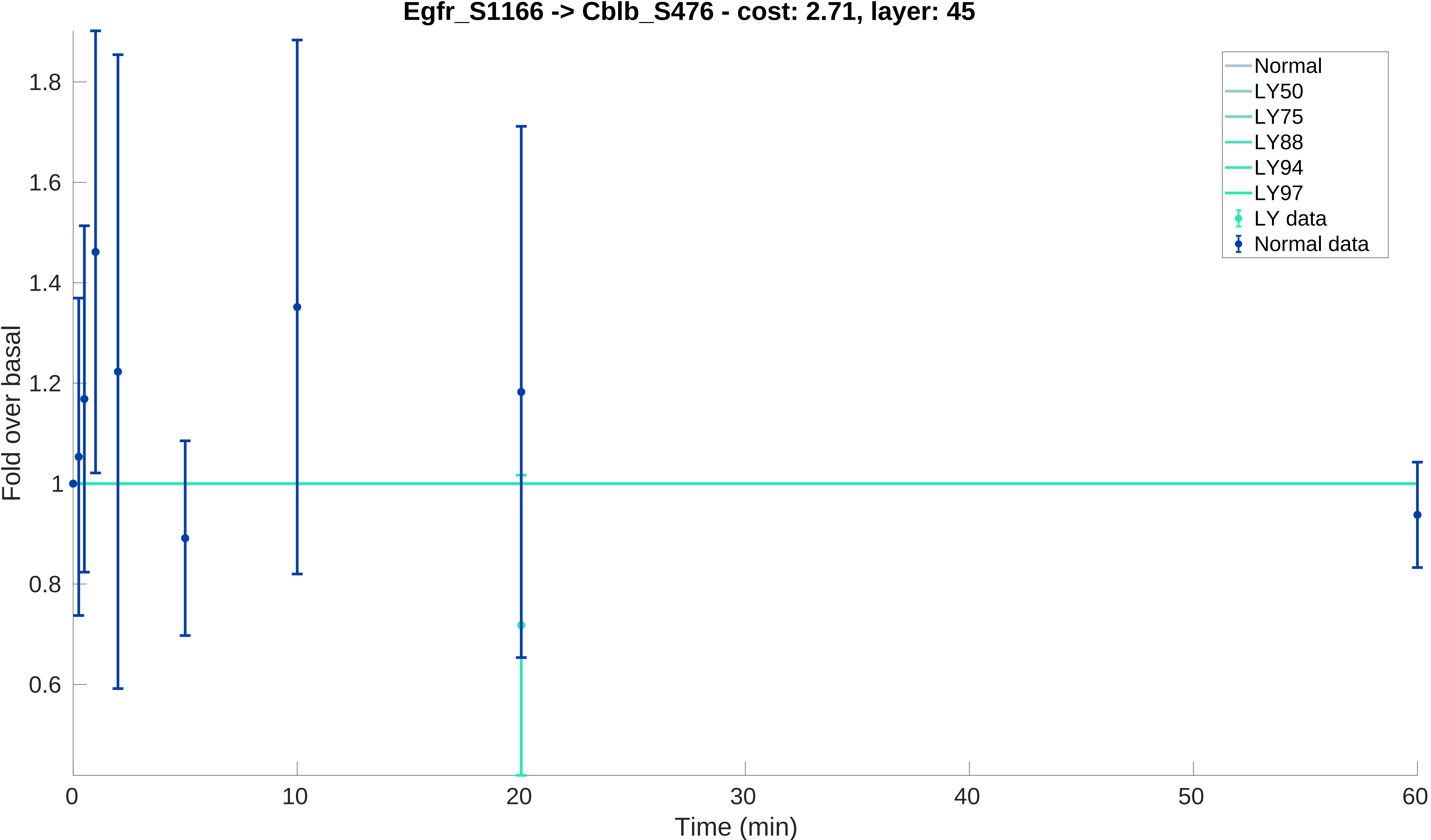

Egfr_S1166 -> Cblb_S476 - cost: 2.71, layer: 45
Fold over basal
Time (min)
0.6
0.8
1
1.2
1.4
1.6
1.8
0
10
20
30
40
50
60
Normal
LY50
LY75
LY88
LY94
LY97
LY data
Normal data

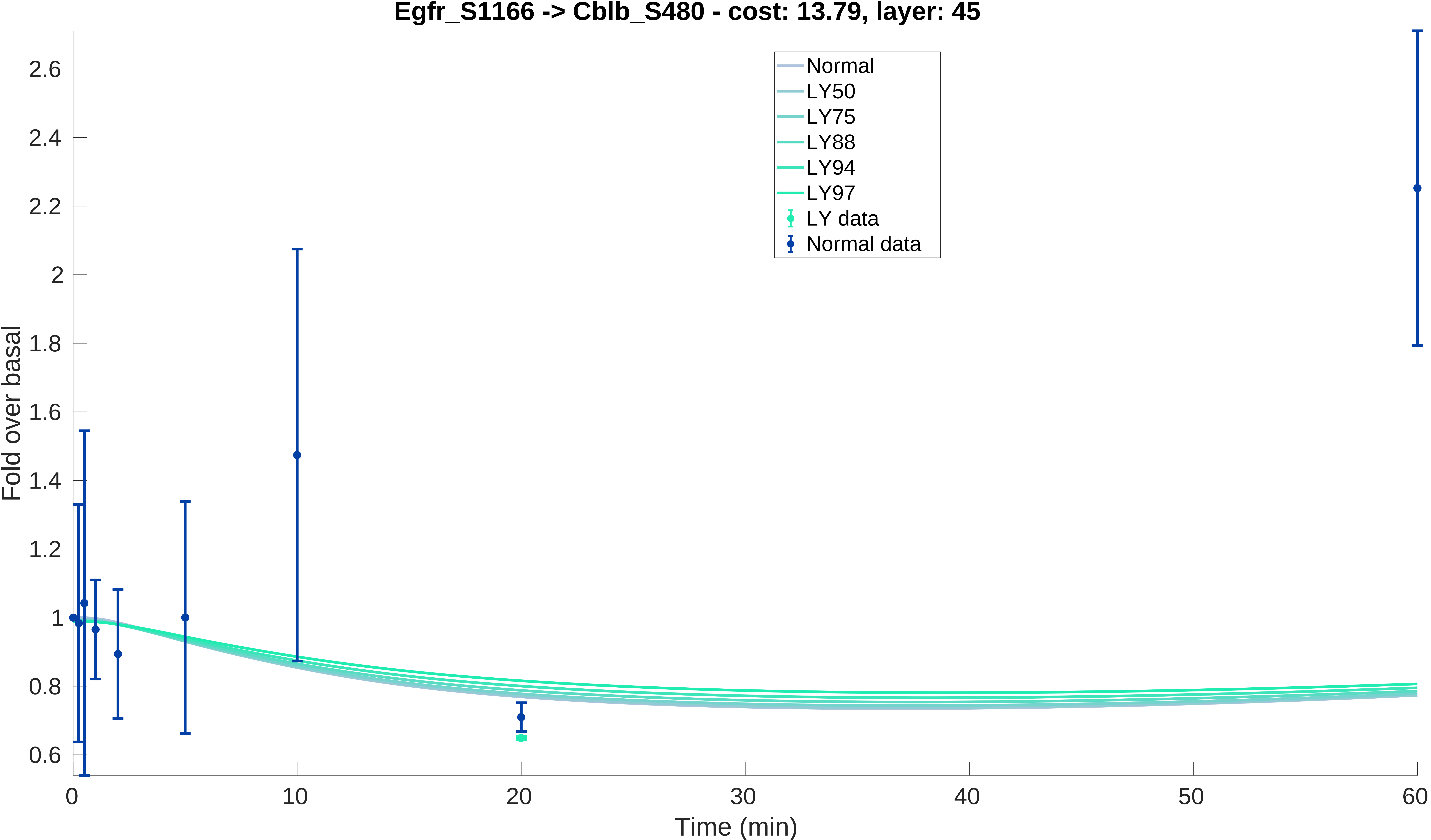

Egfr_S1166 -> Cblb_S480 - cost: 13.79, layer: 45
Normal
LY50
LY75
LY88
LY94
LY97
LY data
Normal data
Fold over basal
0.6
0.8
1
1.2
1.4
1.6
1.8
2
2.2
2.4
2.6
0
10
20
30
40
50
60
Time (min)

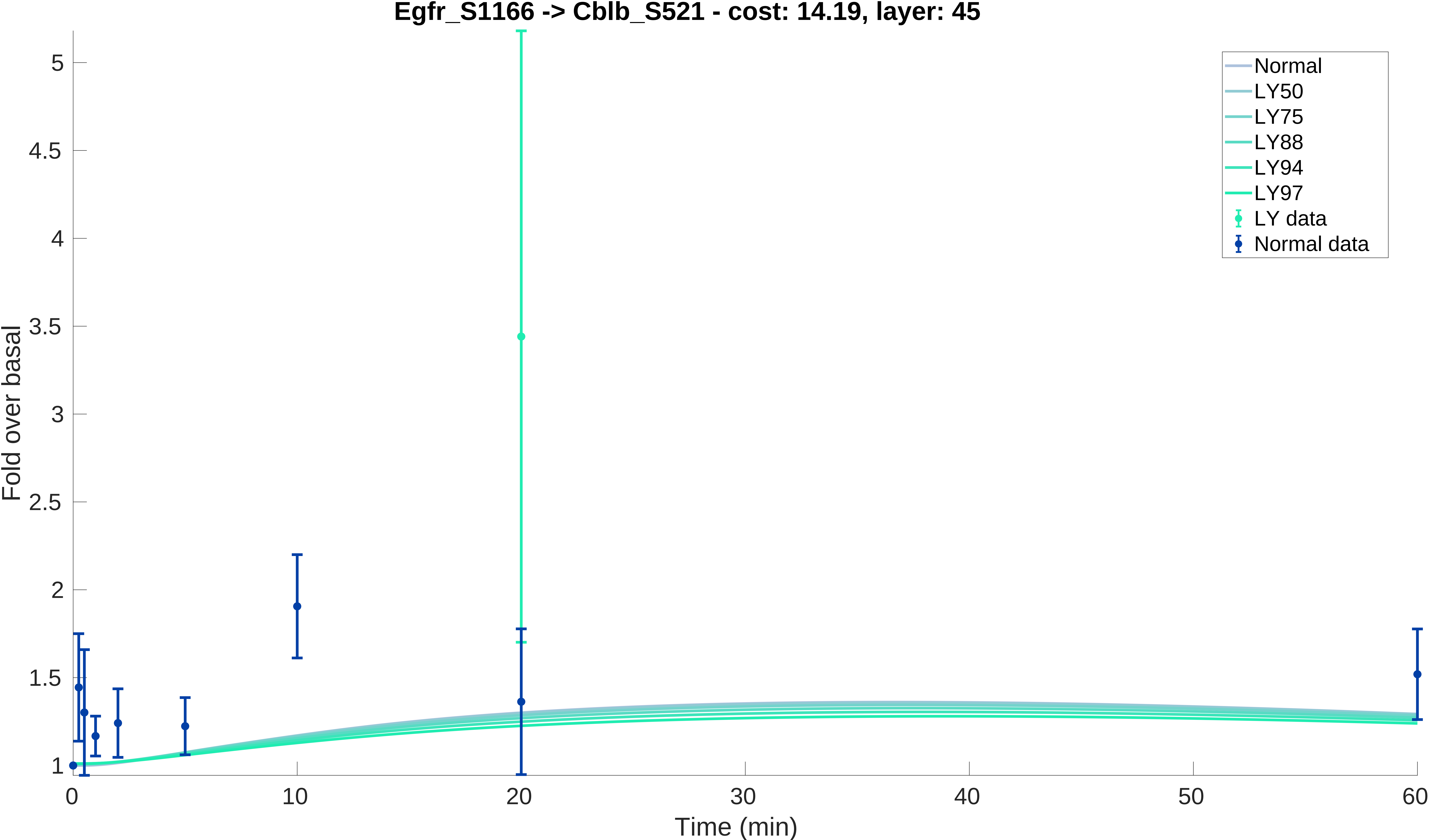

Egfr_S1166 -> Cblb_S521 - cost: 14.19, layer: 45
Fold over basal
Time (min)
0
10
20
30
40
50
60
1
1.5
2
2.5
3
3.5
4
4.5
5
Normal
LY50
LY75
LY88
LY94
LY97
LY data
Normal data

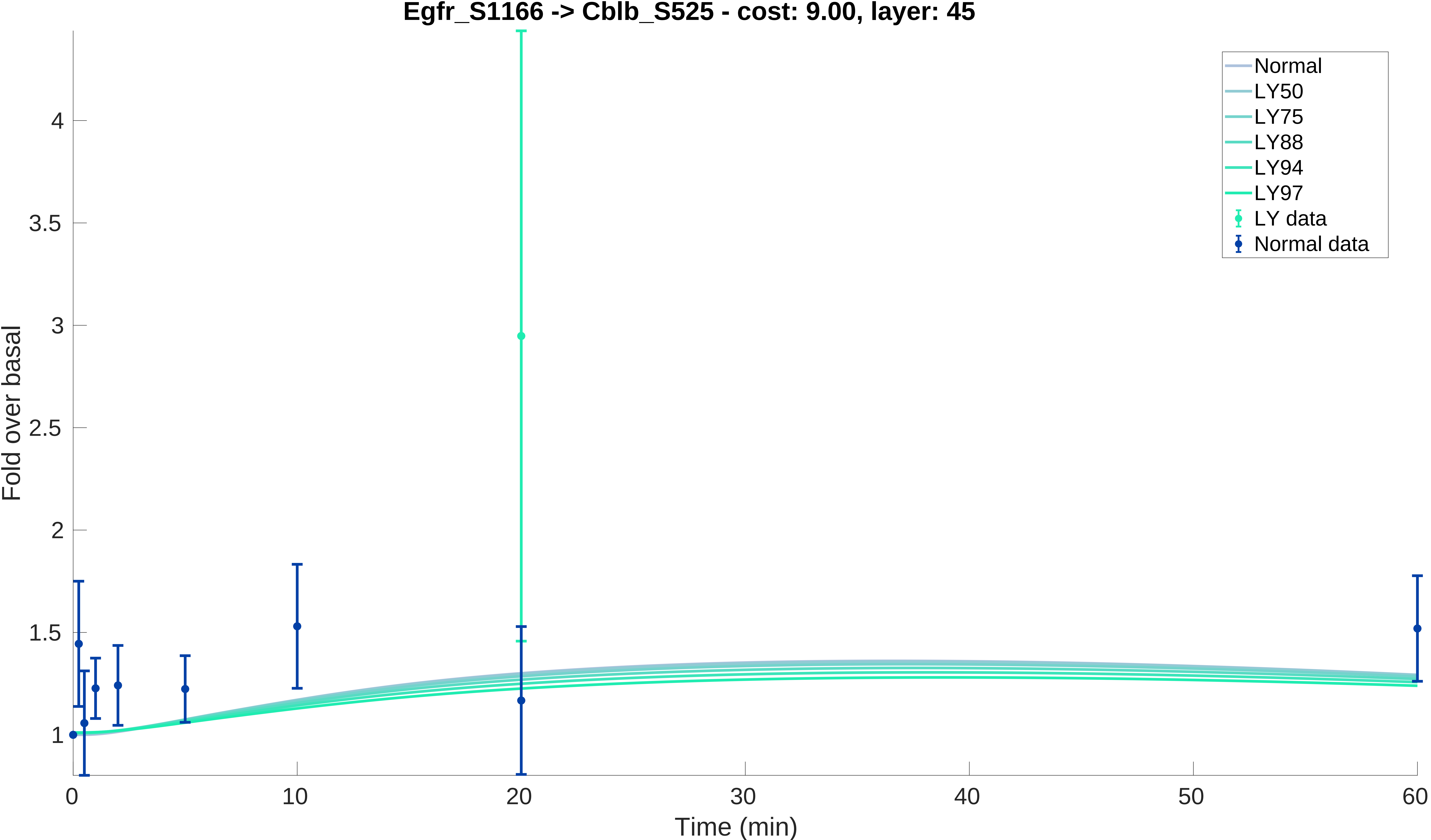

Egfr_S1166 -> Cblb_S525 - cost: 9.00, layer: 45
Fold over basal
Time (min)
Normal
LY50
LY75
LY88
LY94
LY97
LY data
Normal data
0
10
20
30
40
50
60
1
1.5
2
2.5
3
3.5
4

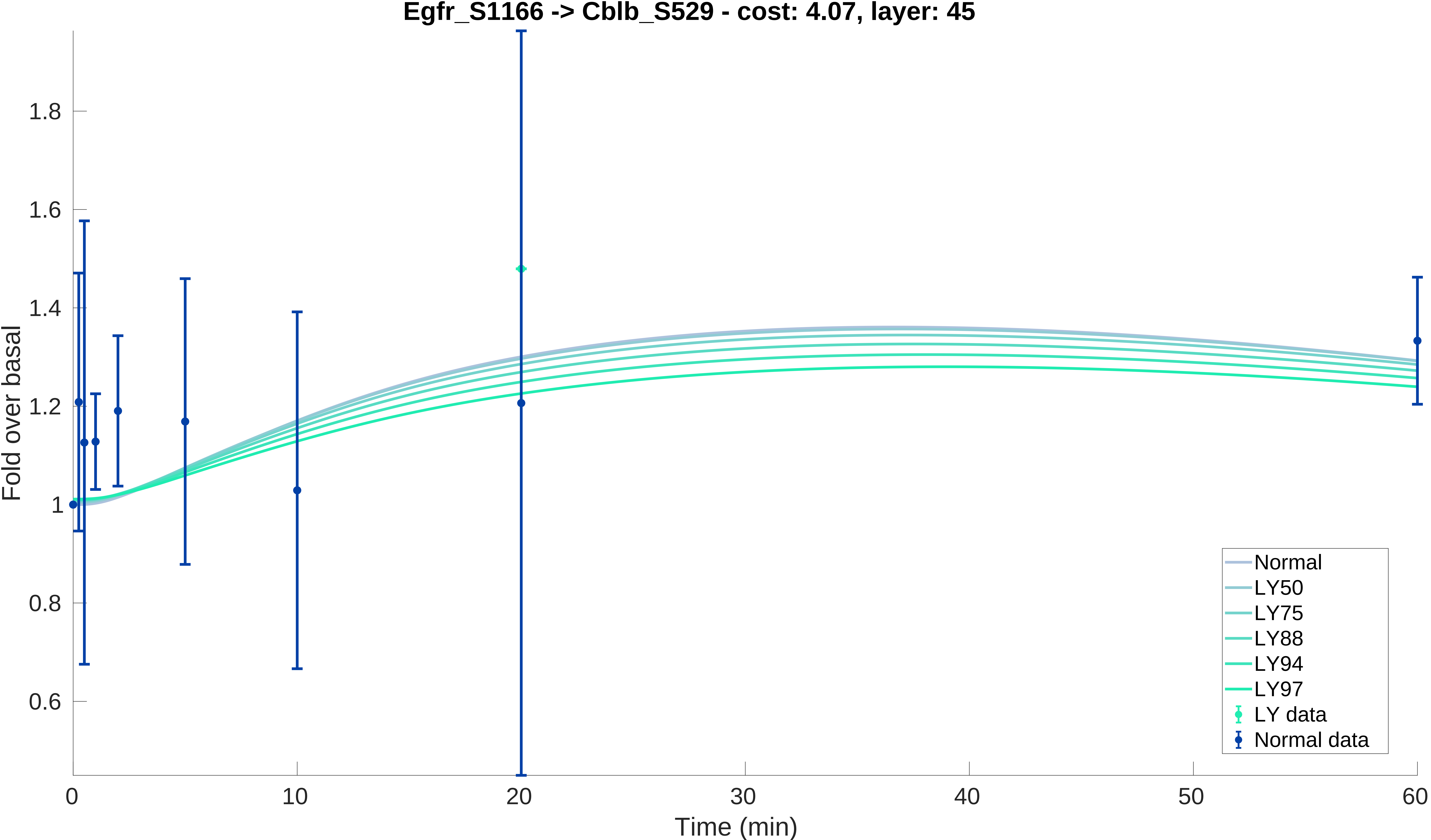
Egfr_S1166 -> Cblb_S529 - cost: 4.07, layer: 45
Fold over basal
Time (min)
0
10
20
30
40
50
60
0.6
0.8
1
1.2
1.4
1.6
1.8
Normal
LY50
LY75
LY88
LY94
LY97
LY data
Normal data

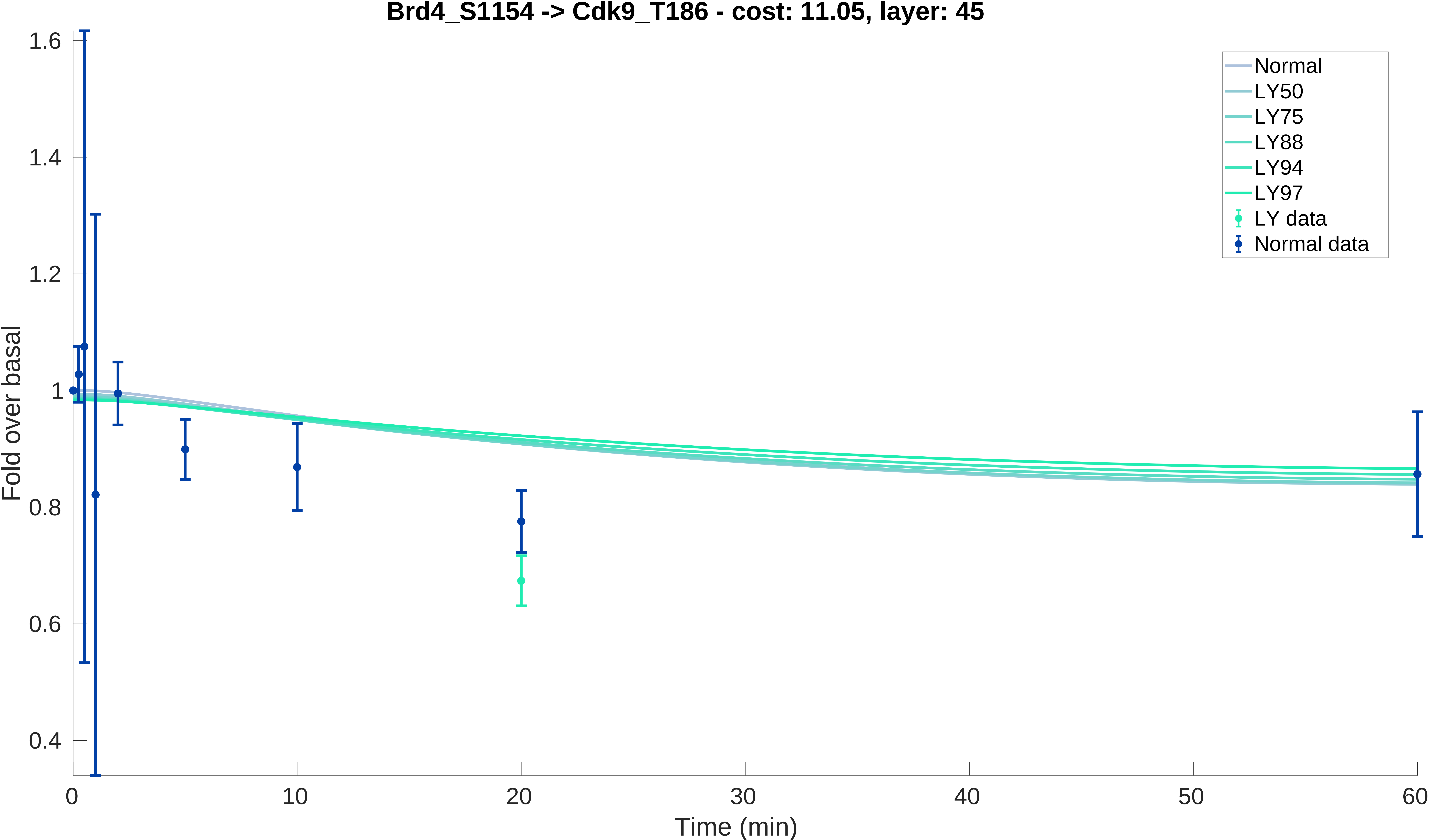
Brd4_S1154 -> Cdk9_T186 - cost: 11.05, layer: 45
Fold over basal
Time (min)
0
10
20
30
40
50
60
0.4
0.6
0.8
1
1.2
1.4
1.6
Normal
LY50
LY75
LY88
LY94
LY97
LY data
Normal data

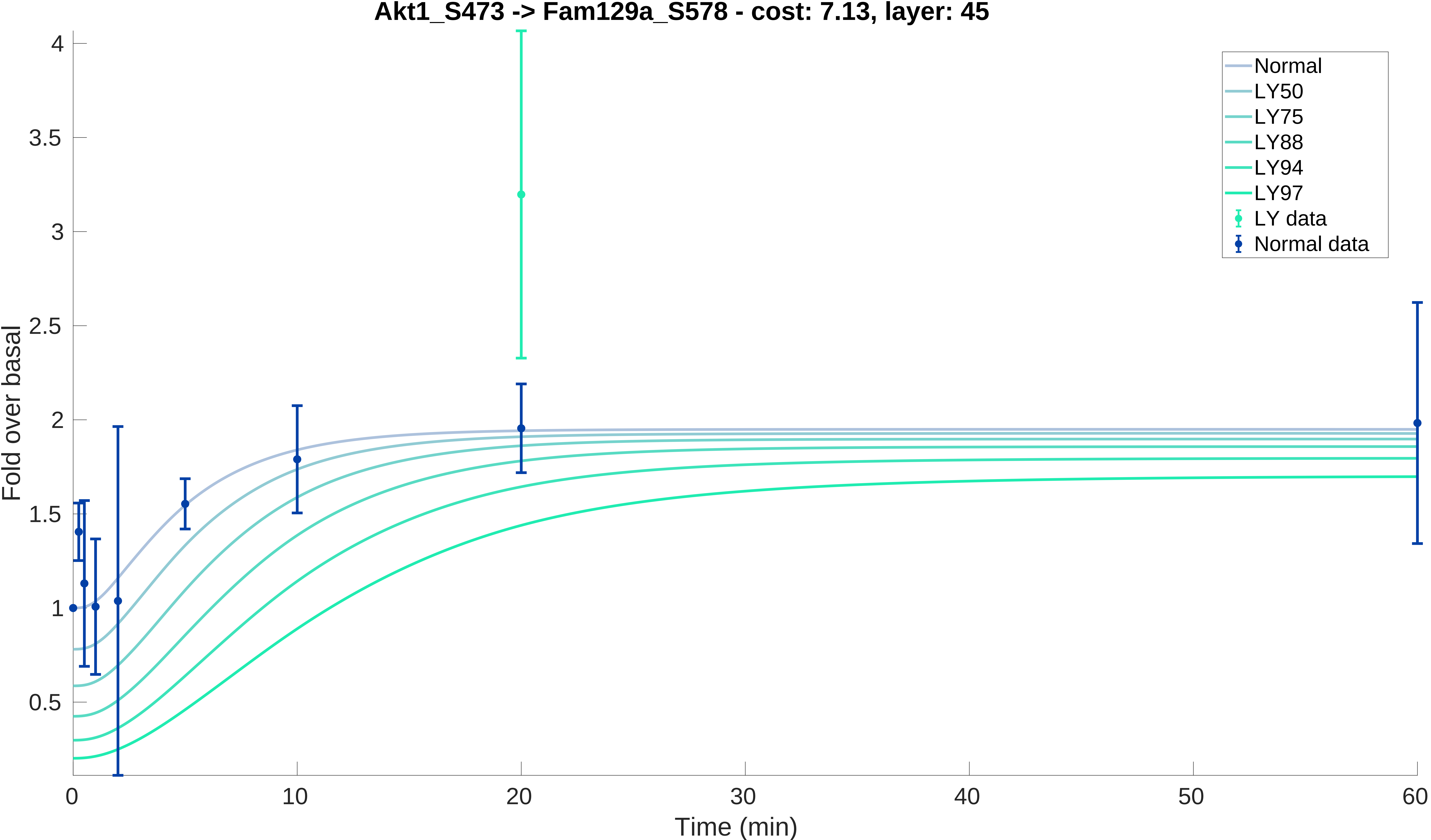
Akt1_S473 -> Fam129a_S578 - cost: 7.13, layer: 45
Fold over basal
Time (min)
0
10
20
30
40
50
60
0.5
1
1.5
2
2.5
3
3.5
4
Normal
LY50
LY75
LY88
LY94
LY97
LY data
Normal data

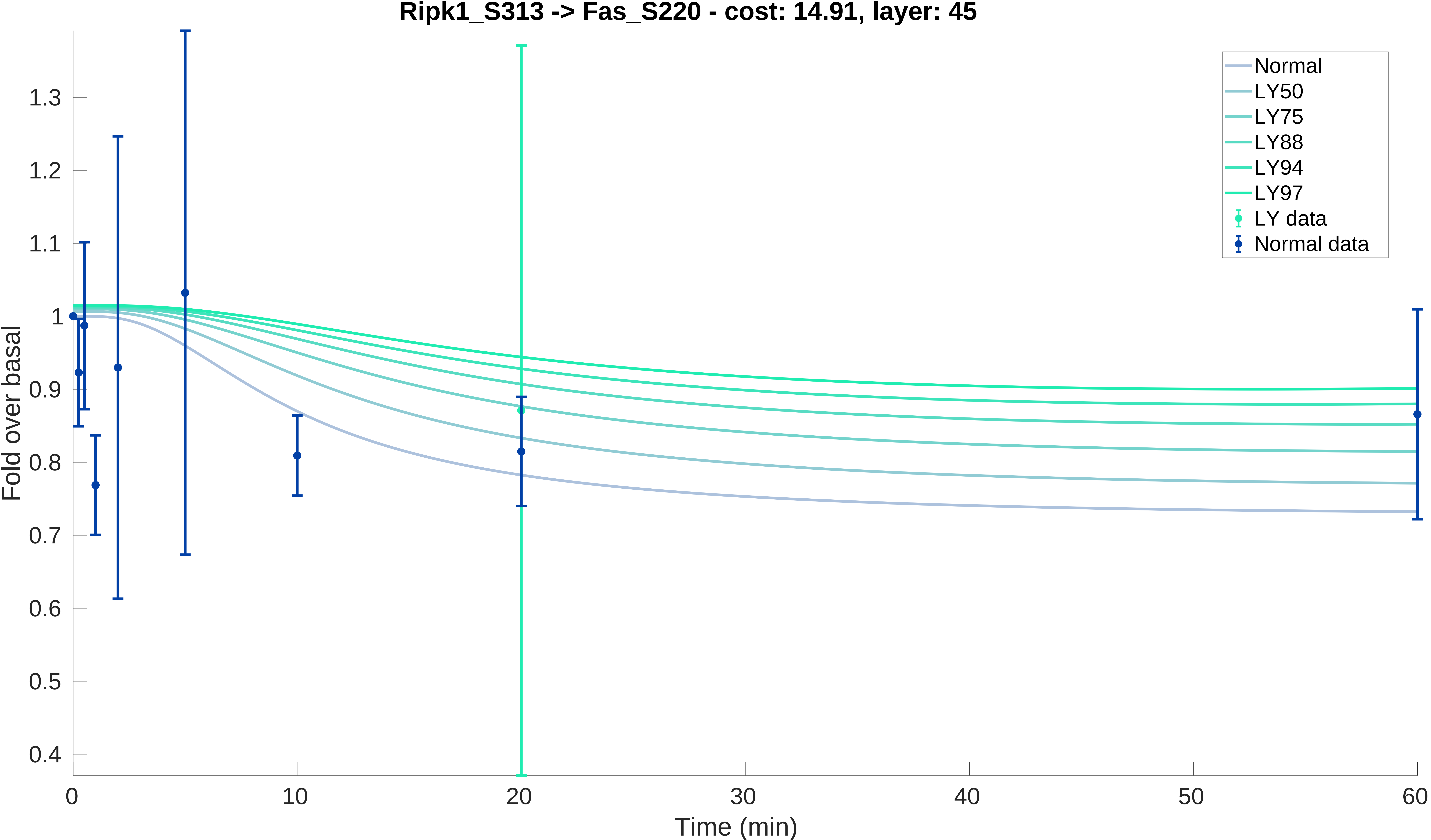
Ripk1_S313 -> Fas_S220 - cost: 14.91, layer: 45
Fold over basal
Time (min)
0
10
20
30
40
50
60
0.4
0.5
0.6
0.7
0.8
0.9
1
1.1
1.2
1.3
Normal
LY50
LY75
LY88
LY94
LY97
LY data
Normal data

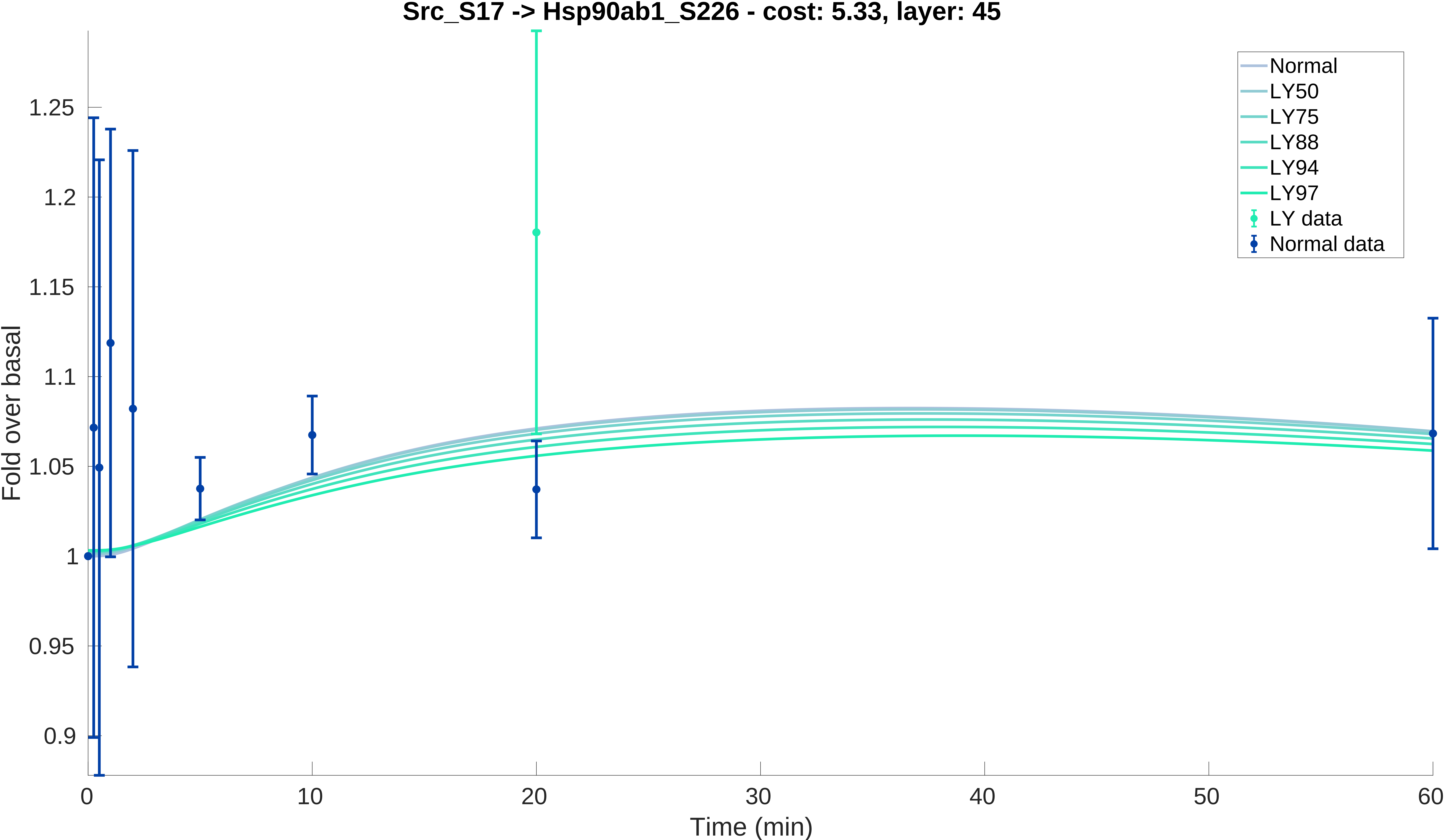
Src_S17 -> Hsp90ab1_S226 - cost: 5.33, layer: 45
Normal
LY50
LY75
LY88
LY94
LY97
LY data
Normal data
Fold over basal
Time (min)
1.25
1.2
1.15
1.1
1.05
1
0.95
0.9
0
10
20
30
40
50
60

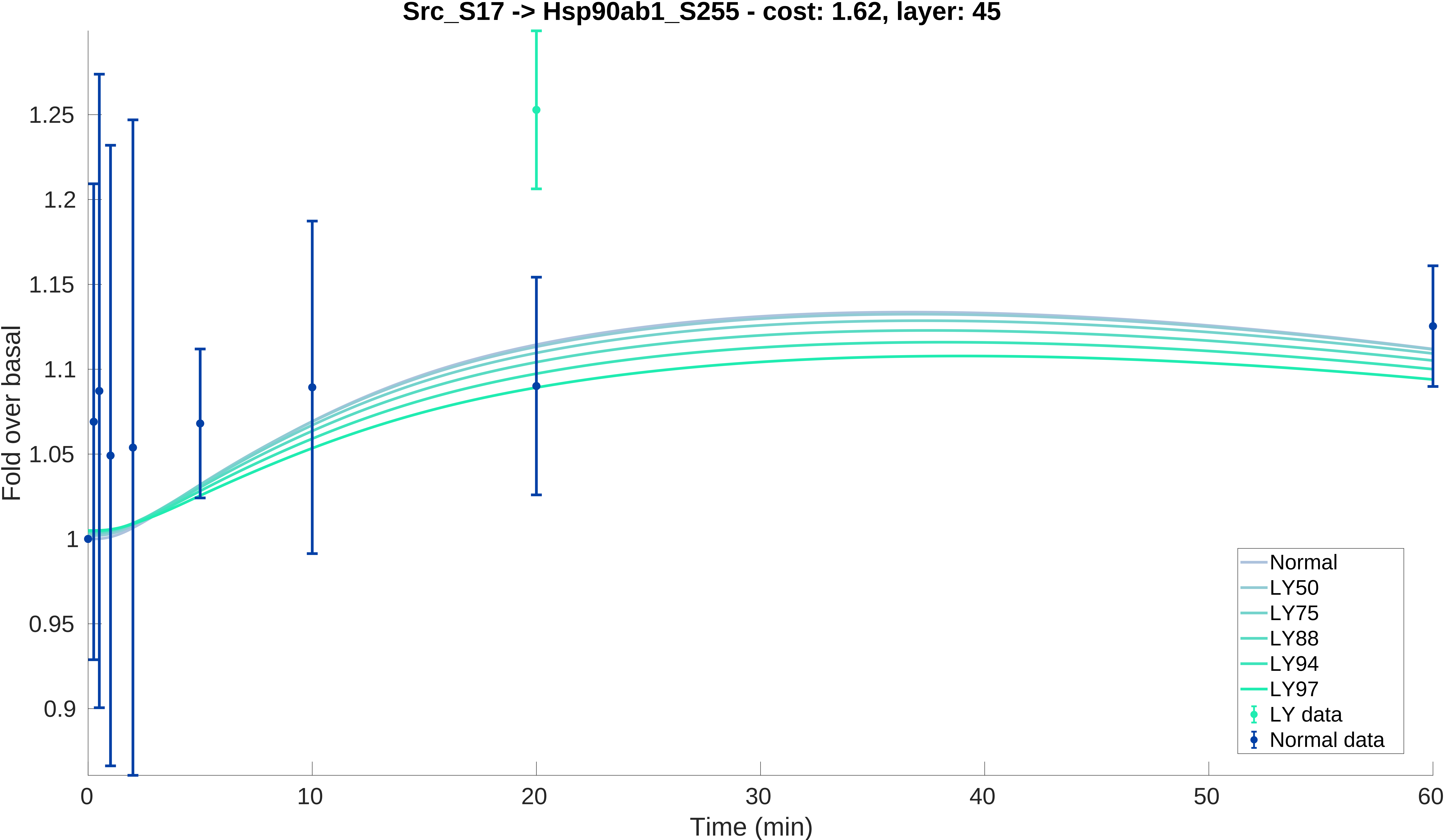

Src_S17 -> Hsp90ab1_S255 - cost: 1.62, layer: 45
Fold over basal
Time (min)
0.9
0.95
1
1.05
1.1
1.15
1.2
1.25
0
10
20
30
40
50
60
Normal
LY50
LY75
LY88
LY94
LY97
LY data
Normal data

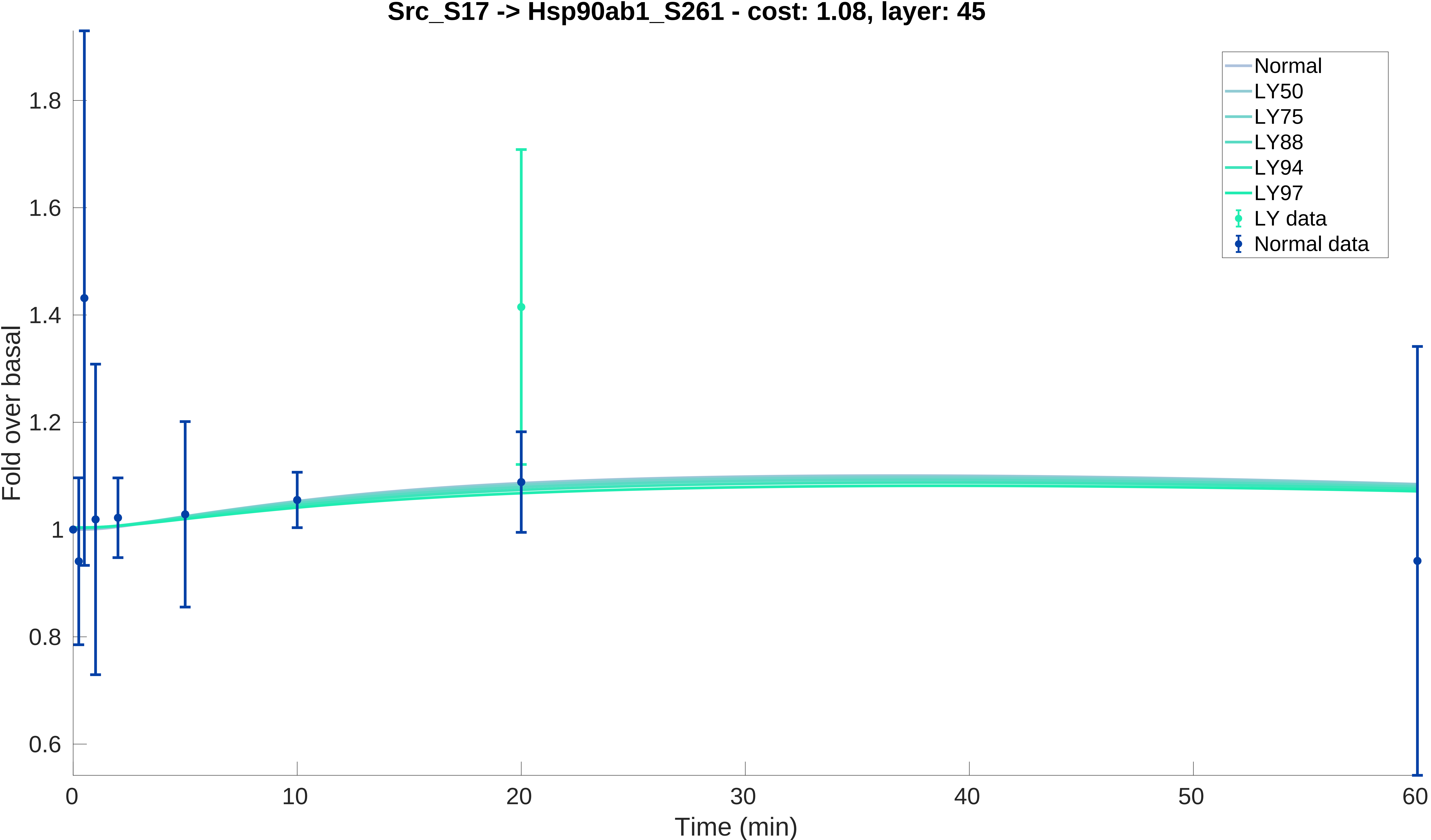

Src_S17 -> Hsp90ab1_S261 - cost: 1.08, layer: 45
Normal
LY50
LY75
LY88
LY94
LY97
LY data
Normal data
Fold over basal
1.8
1.6
1.4
1.2
1
0.8
0.6
0
10
20
30
40
50
60
Time (min)

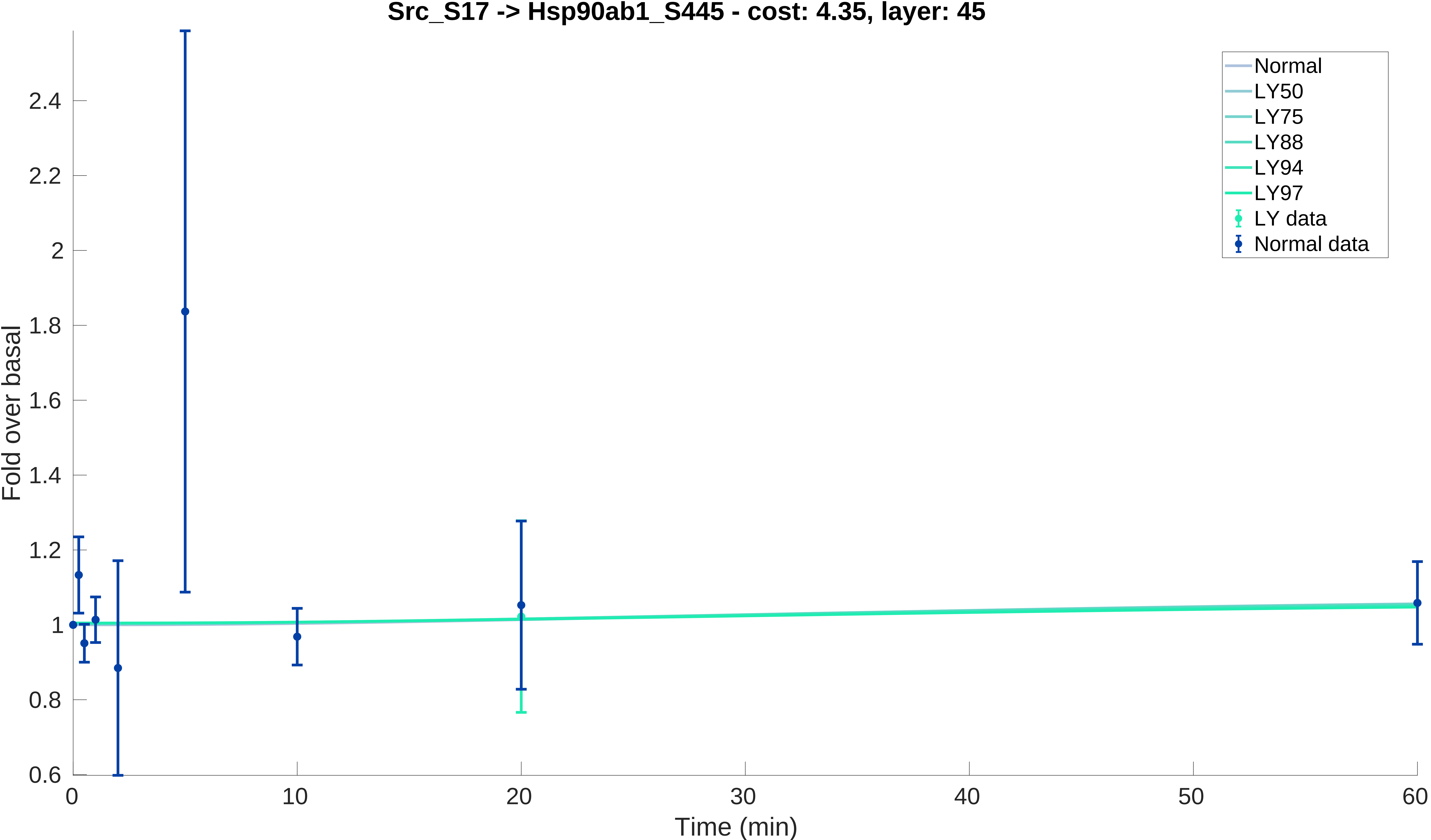
Src_S17 -> Hsp90ab1_S445 - cost: 4.35, layer: 45
Fold over basal
2.4
2.2
2
1.8
1.6
1.4
1.2
1
0.8
0.6
0
10
20
30
40
50
60
Time (min)
Normal
LY50
LY75
LY88
LY94
LY97
LY data
Normal data

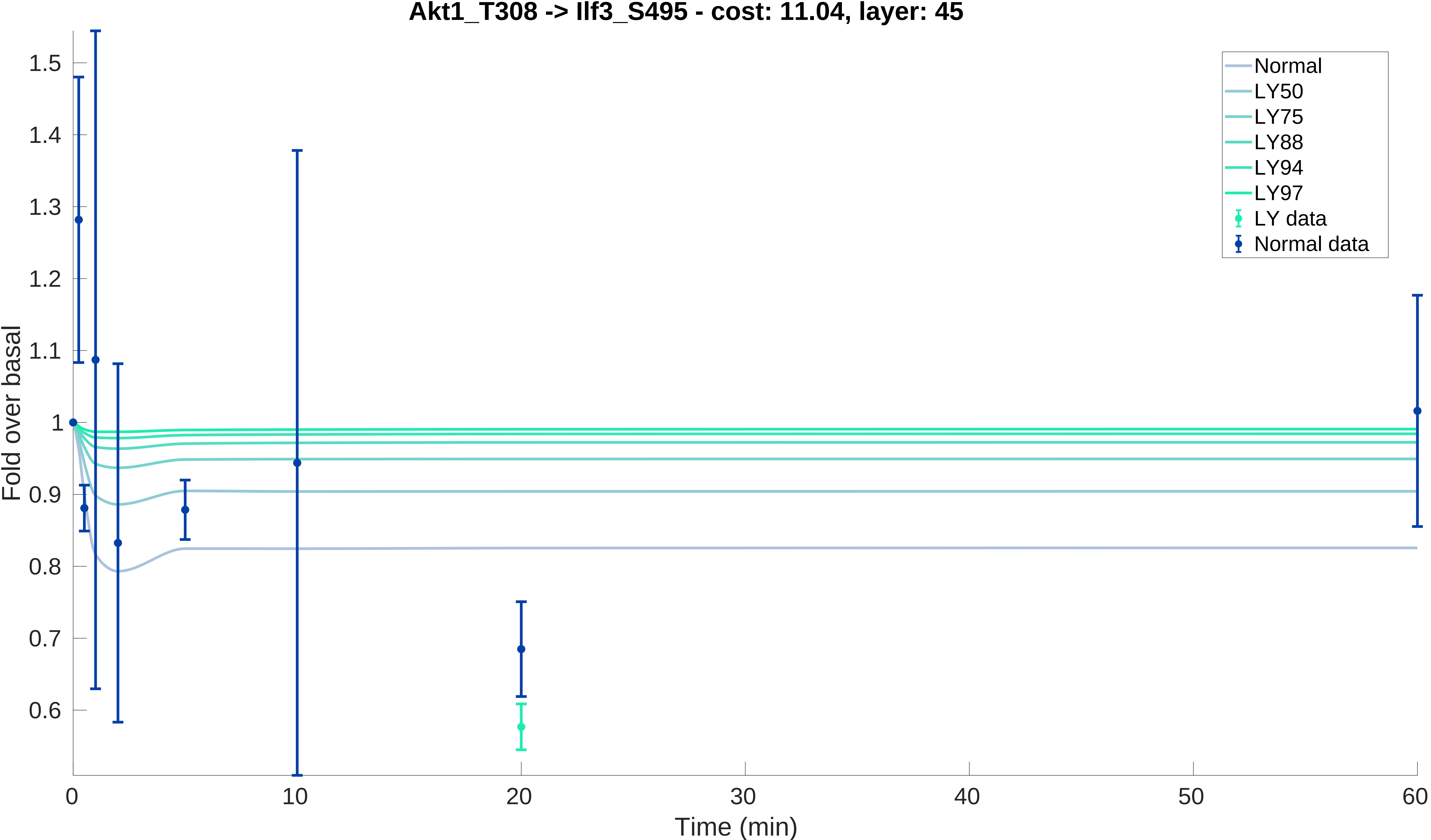

Akt1_T308 -> Ilf3_S495 - cost: 11.04, layer: 45
Fold over basal
Time (min)
0.6
0.7
0.8
0.9
1
1.1
1.2
1.3
1.4
1.5
0
10
20
30
40
50
60
Normal
LY50
LY75
LY88
LY94
LY97
LY data
Normal data

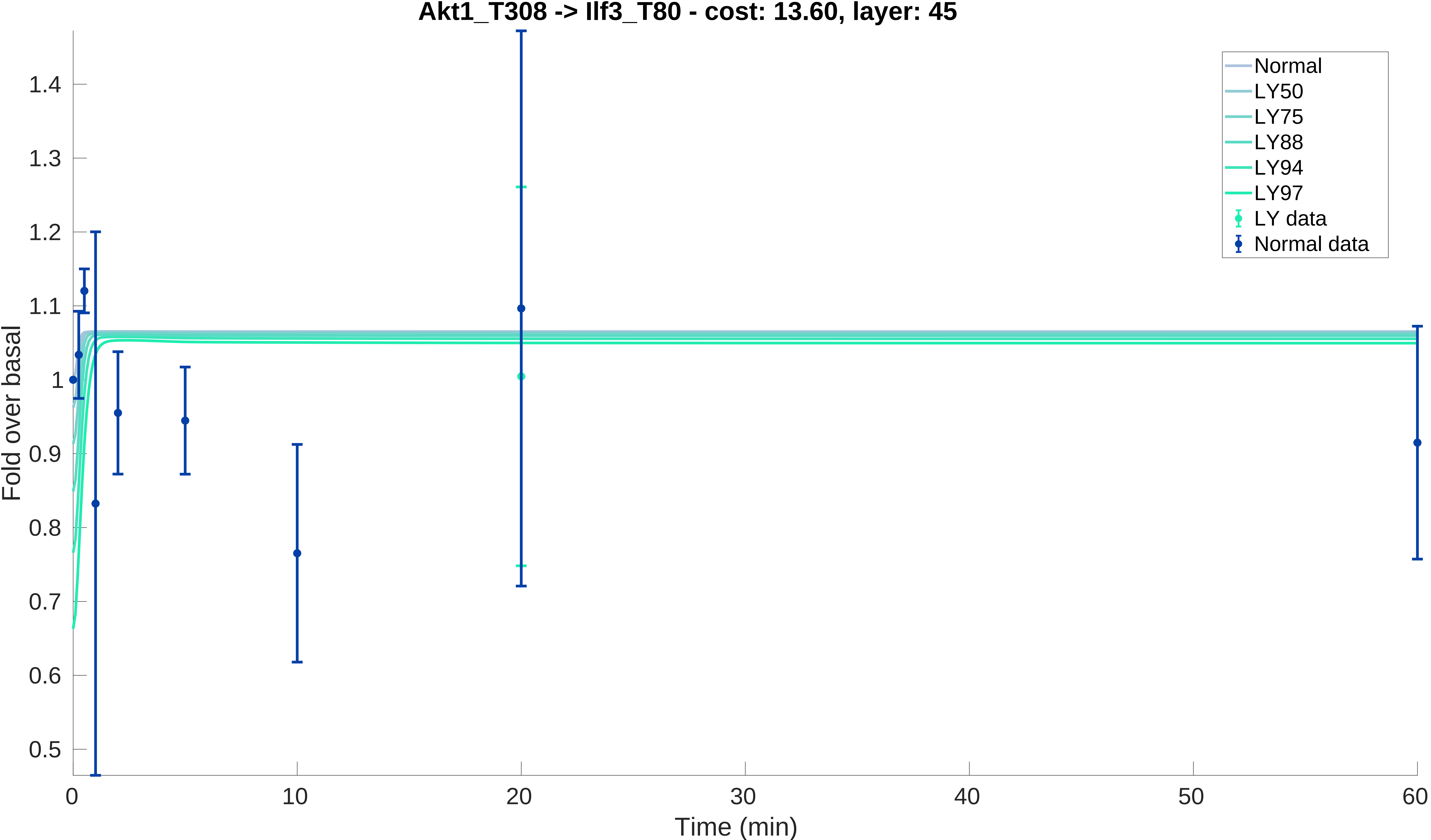

Akt1_T308 -> Ilf3_T80 - cost: 13.60, layer: 45
Fold over basal
Time (min)
Normal
LY50
LY75
LY88
LY94
LY97
LY data
Normal data
0.5
0.6
0.7
0.8
0.9
1
1.1
1.2
1.3
1.4
0
10
20
30
40
50
60

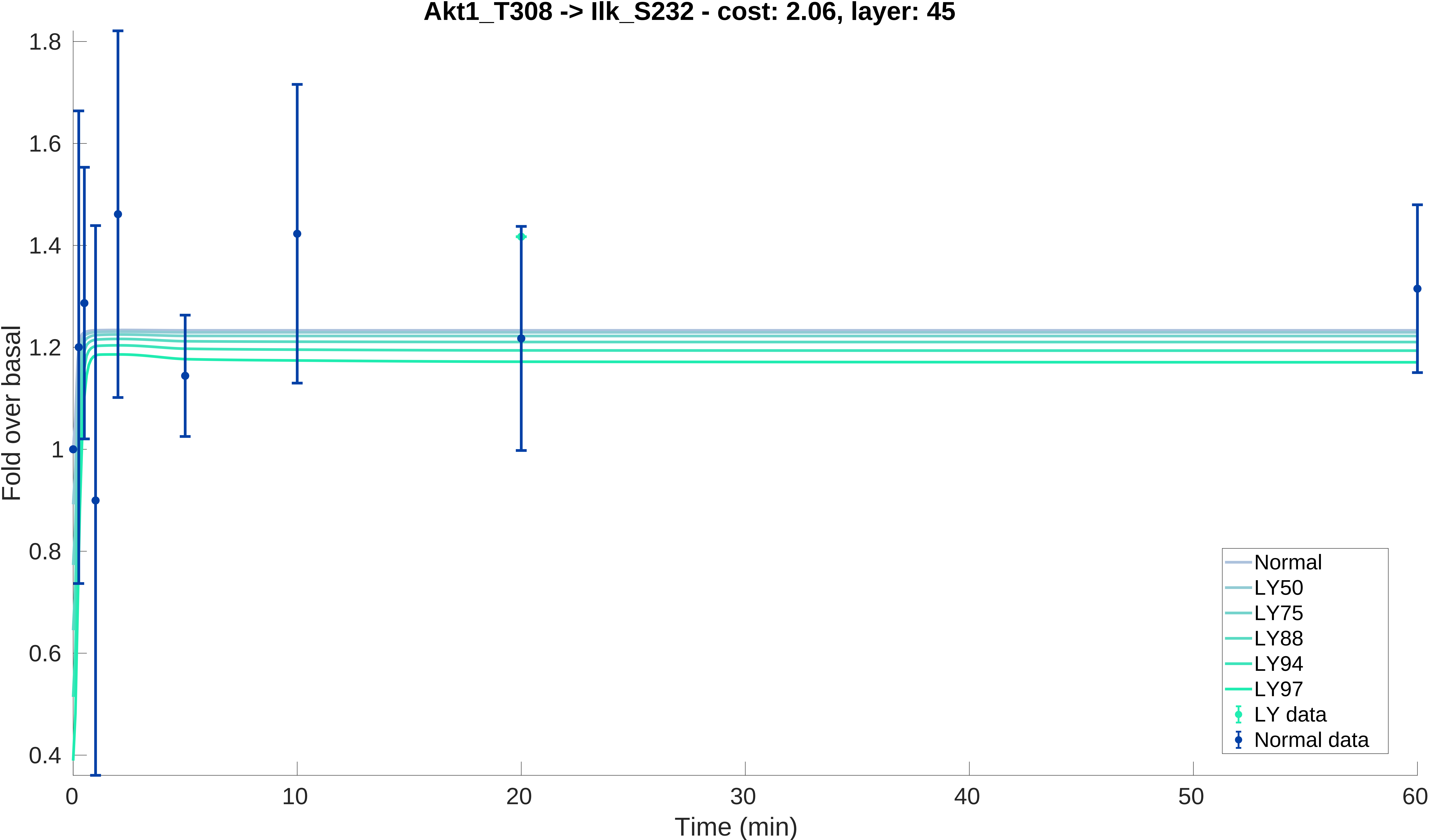
Akt1_T308 -> Ilk_S232 - cost: 2.06, layer: 45
Fold over basal
1.8
1.6
1.4
1.2
1
0.8
0.6
0.4
0
10
20
30
40
50
60
Time (min)
Normal
LY50
LY75
LY88
LY94
LY97
LY data
Normal data

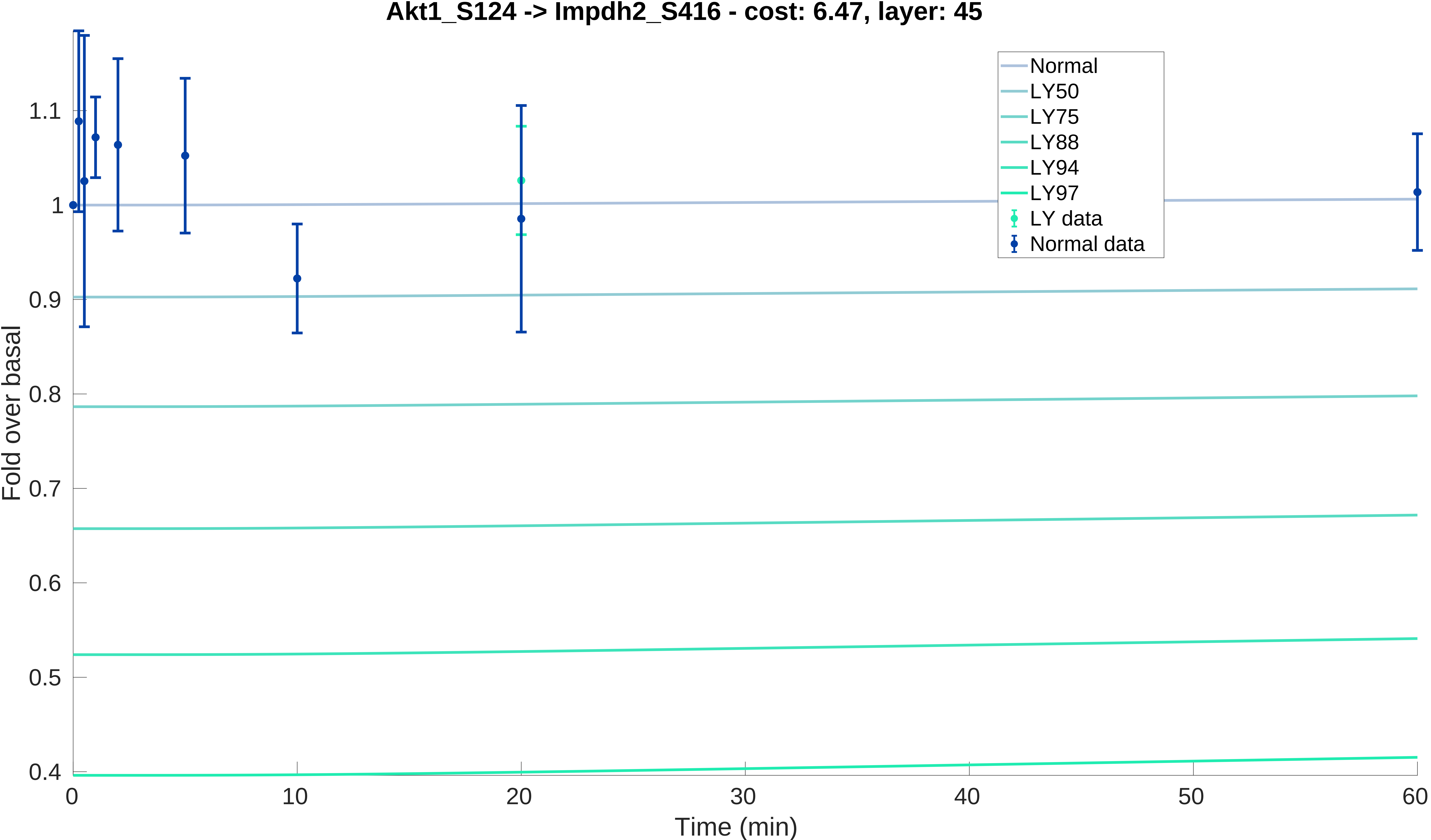

Akt1_S124 -> Impdh2_S416 - cost: 6.47, layer: 45
Normal
LY50
LY75
LY88
LY94
LY97
LY data
Normal data
Fold over basal
1.1
1
0.9
0.8
0.7
0.6
0.5
0.4
0
10
20
30
40
50
60
Time (min)

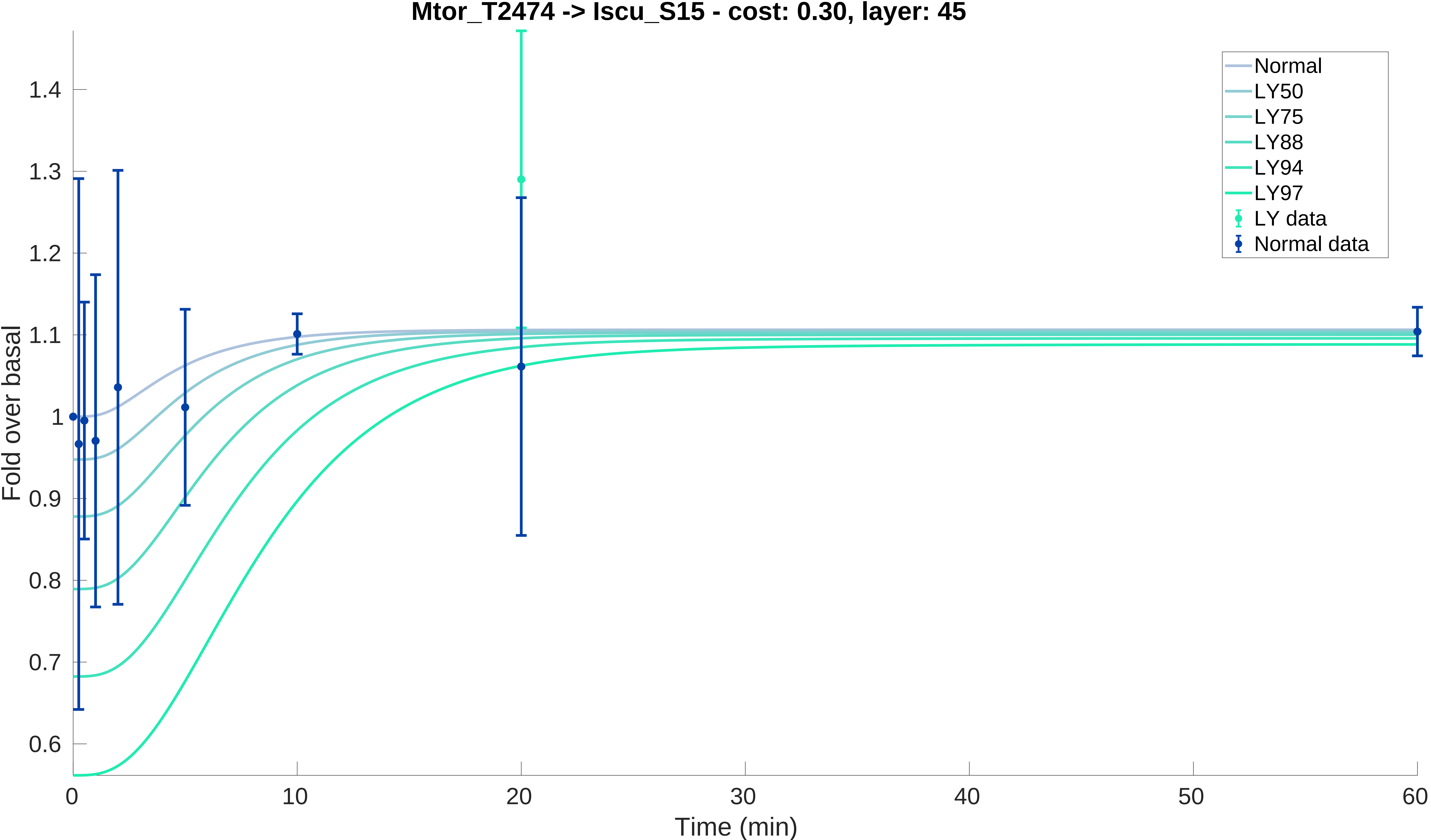

Mtor_T2474 -> Iscu_S15 - cost: 0.30, layer: 45
Fold over basal
Time (min)
1.4
1.3
1.2
1.1
1
0.9
0.8
0.7
0.6
0
10
20
30
40
50
60
Normal
LY50
LY75
LY88
LY94
LY97
LY data
Normal data

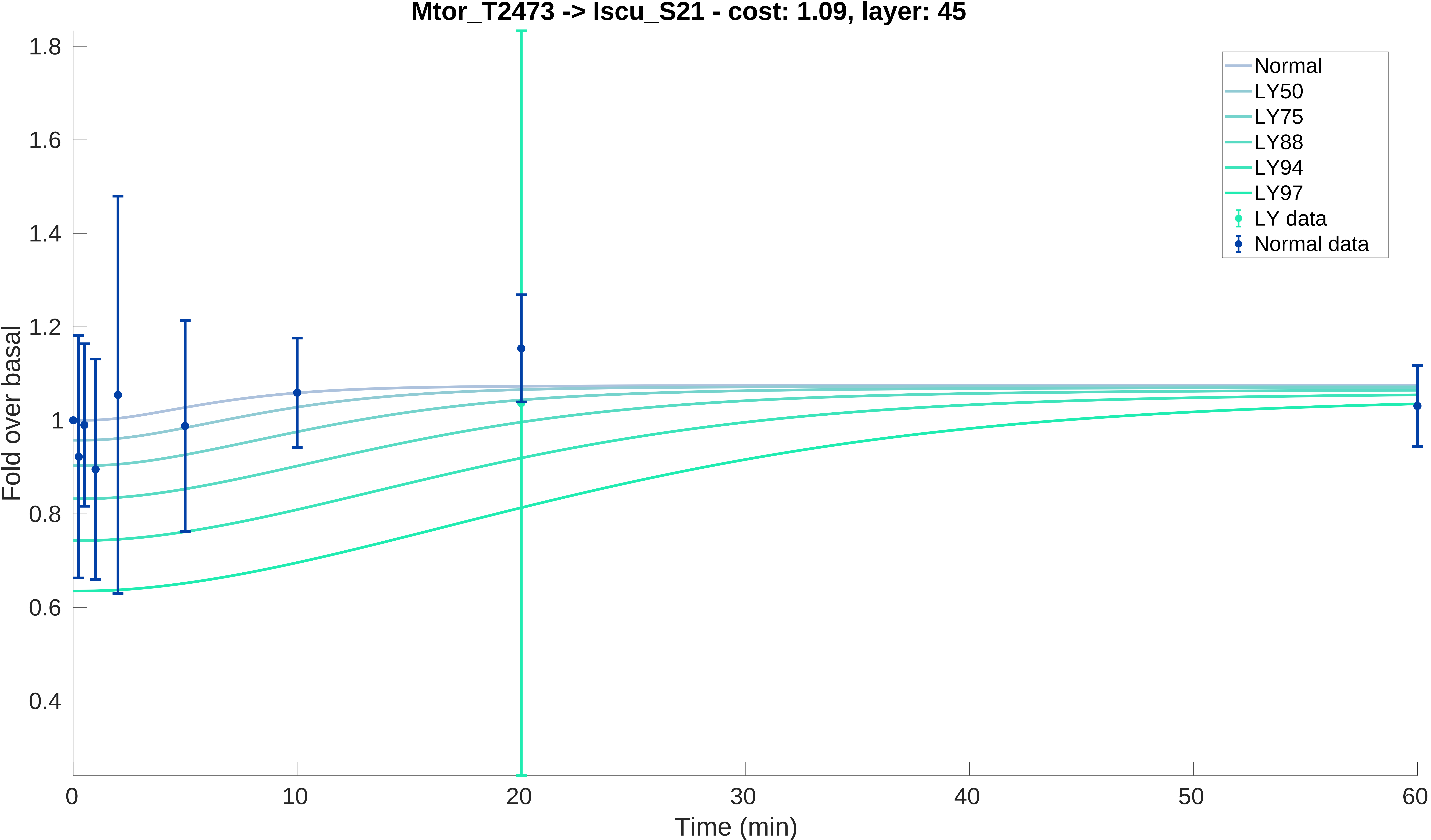

Mtor_T2473 -> Iscu_S21 - cost: 1.09, layer: 45
Fold over basal
Time (min)
1.8
1.6
1.4
1.2
1
0.8
0.6
0.4
0
10
20
30
40
50
60
Normal
LY50
LY75
LY88
LY94
LY97
LY data
Normal data

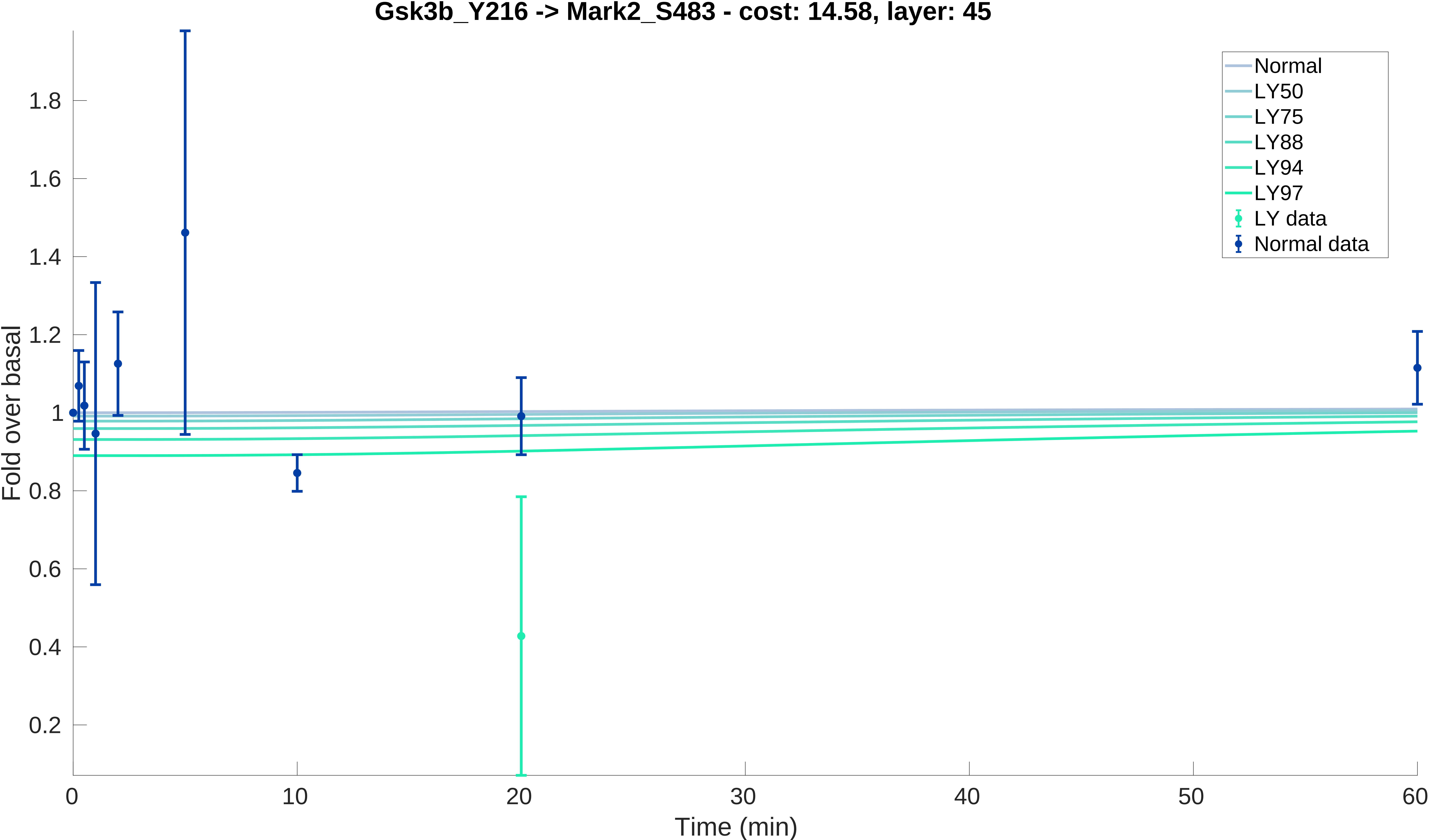

Gsk3b_Y216 -> Mark2_S483 - cost: 14.58, layer: 45
Fold over basal
Time (min)
1.8
1.6
1.4
1.2
1
0.8
0.6
0.4
0.2
0
10
20
30
40
50
60
Normal
LY50
LY75
LY88
LY94
LY97
LY data
Normal data

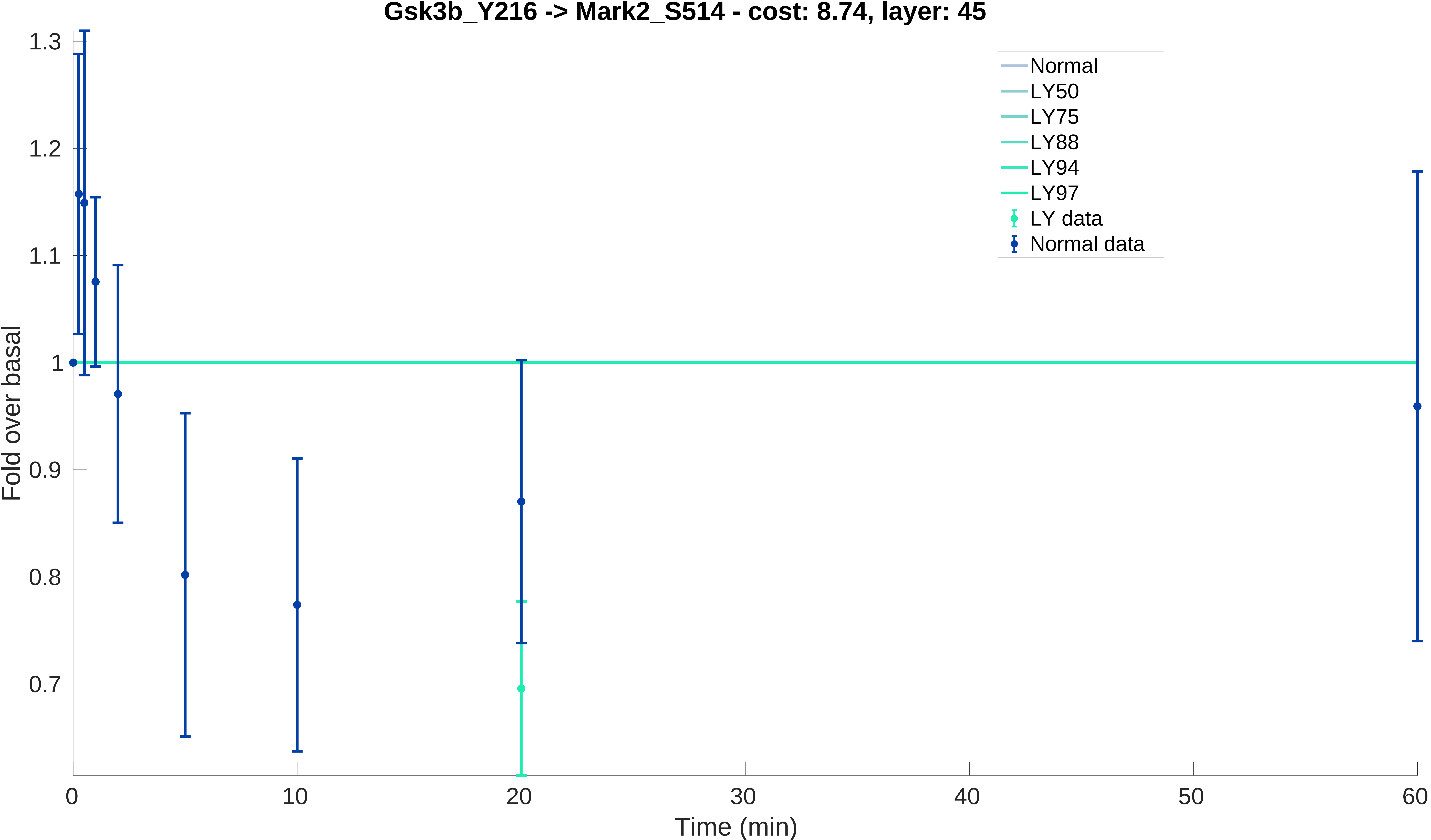

Gsk3b_Y216 -> Mark2_S514 - cost: 8.74, layer: 45
Fold over basal
Time (min)
Normal
LY50
LY75
LY88
LY94
LY97
LY data
Normal data
1.3
1.2
1.1
1
0.9
0.8
0.7
0
10
20
30
40
50
60

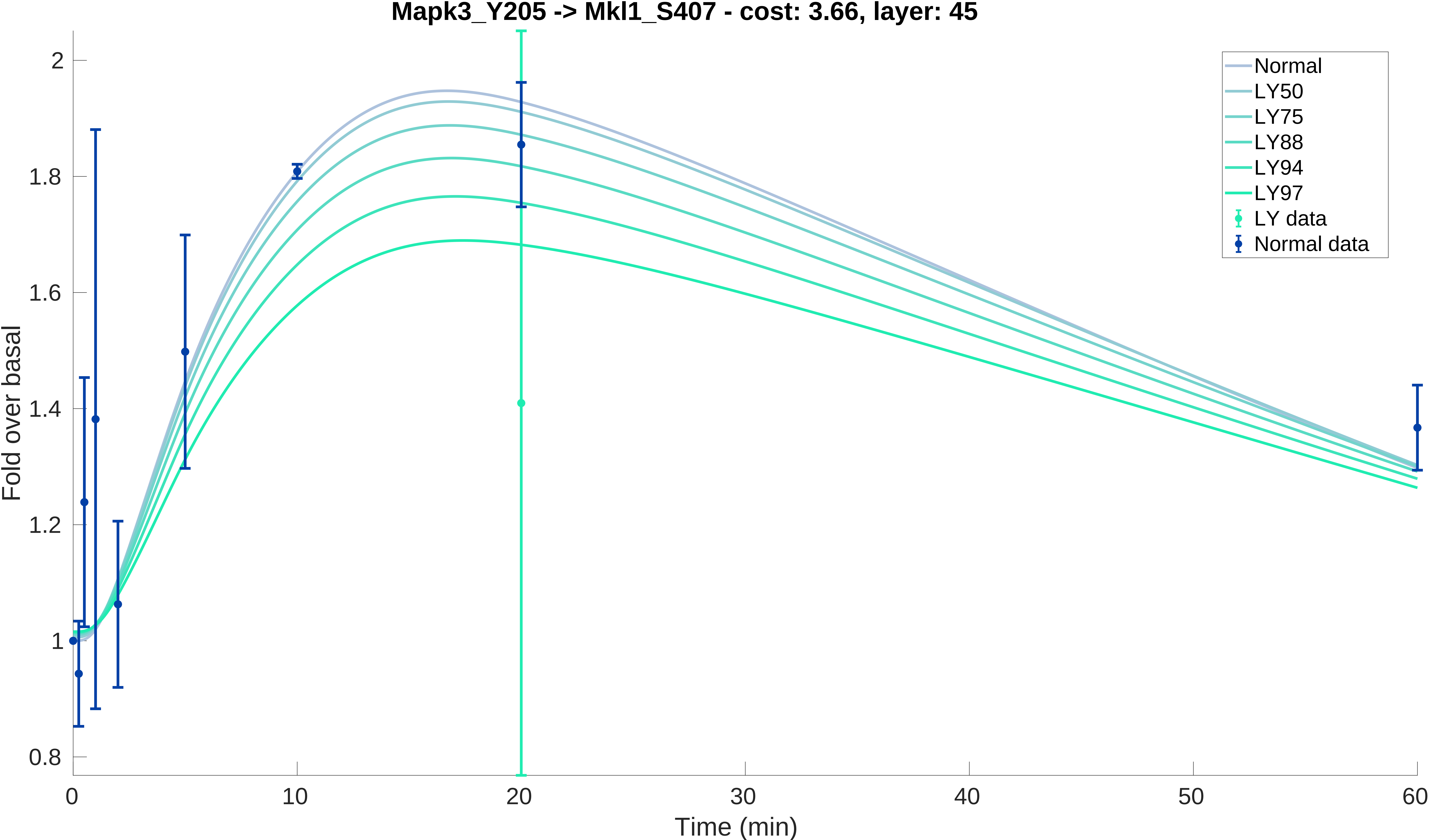

Mapk3_Y205 -> Mkl1_S407 - cost: 3.66, layer: 45
Fold over basal
Time (min)
0.8
1
1.2
1.4
1.6
1.8
2
0
10
20
30
40
50
60
Normal
LY50
LY75
LY88
LY94
LY97
LY data
Normal data

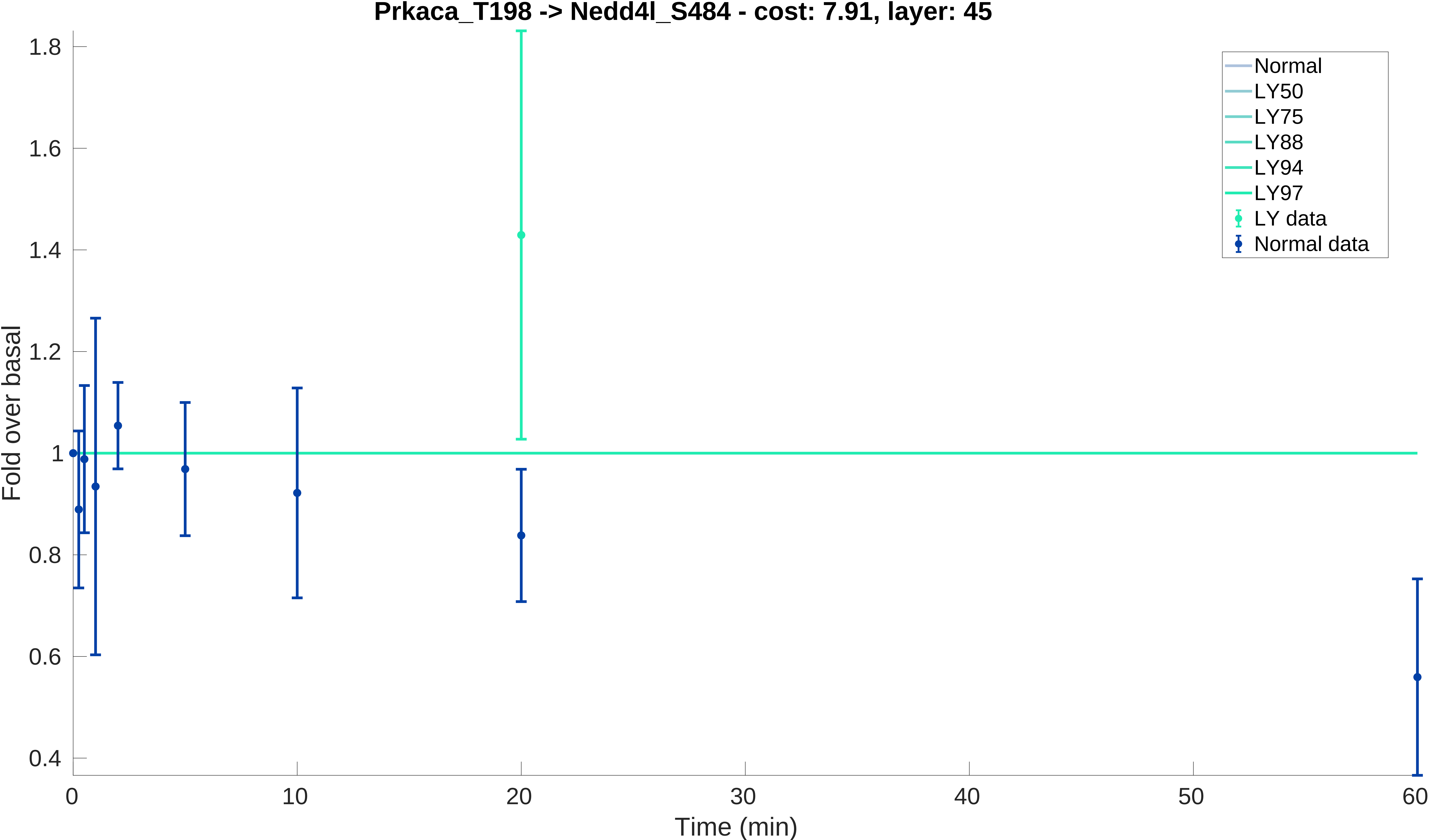
Prkaca_T198 -> Nedd4l_S484 - cost: 7.91, layer: 45
Fold over basal
Time (min)
0.4
0.6
0.8
1
1.2
1.4
1.6
1.8
0
10
20
30
40
50
60
Normal
LY50
LY75
LY88
LY94
LY97
LY data
Normal data

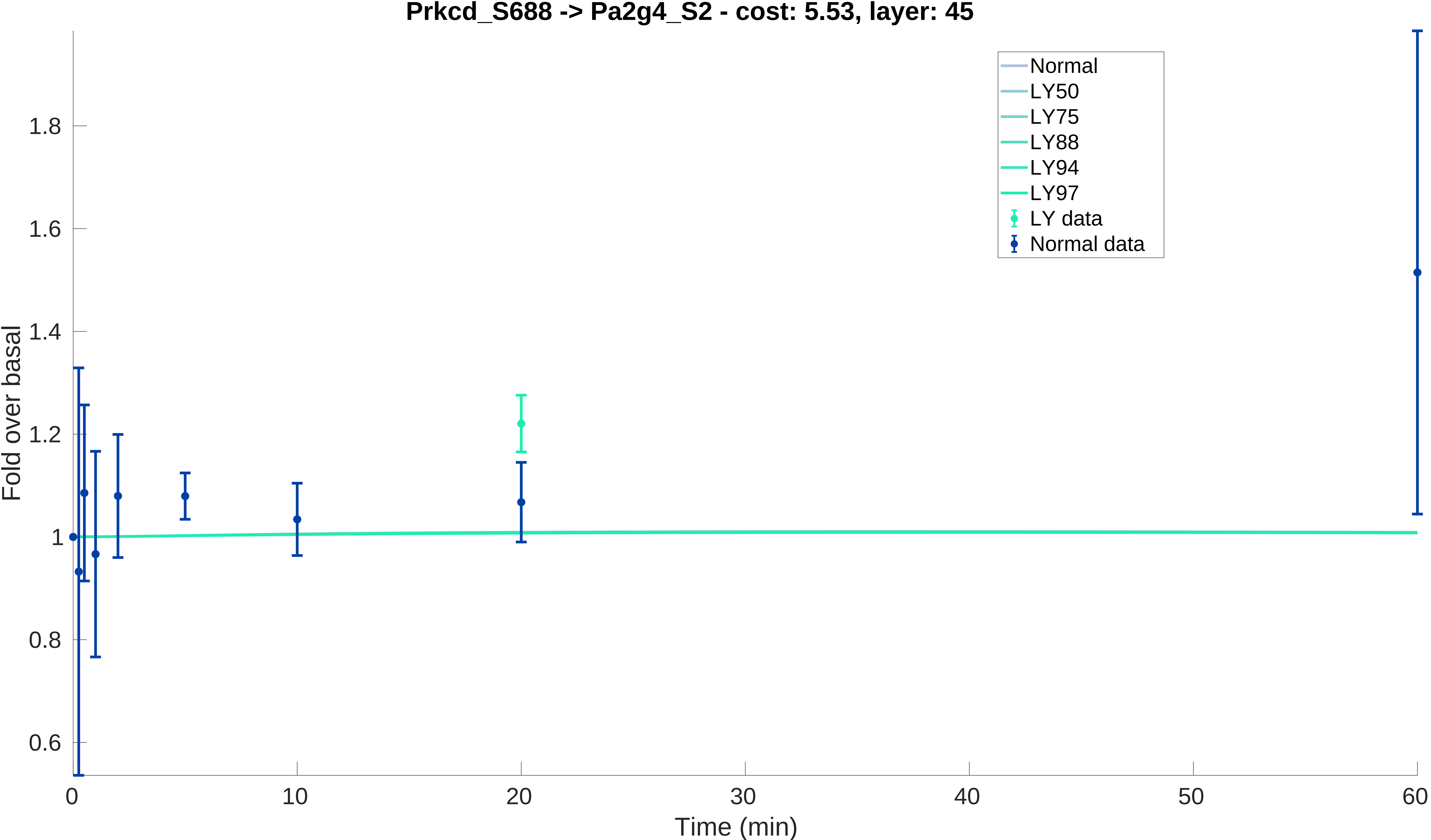

Prkcd_S688 -> Pa2g4_S2 - cost: 5.53, layer: 45
Normal
LY50
LY75
LY88
LY94
LY97
LY data
Normal data
Fold over basal
1.8
1.6
1.4
1.2
1
0.8
0.6
0
10
20
30
40
50
60
Time (min)

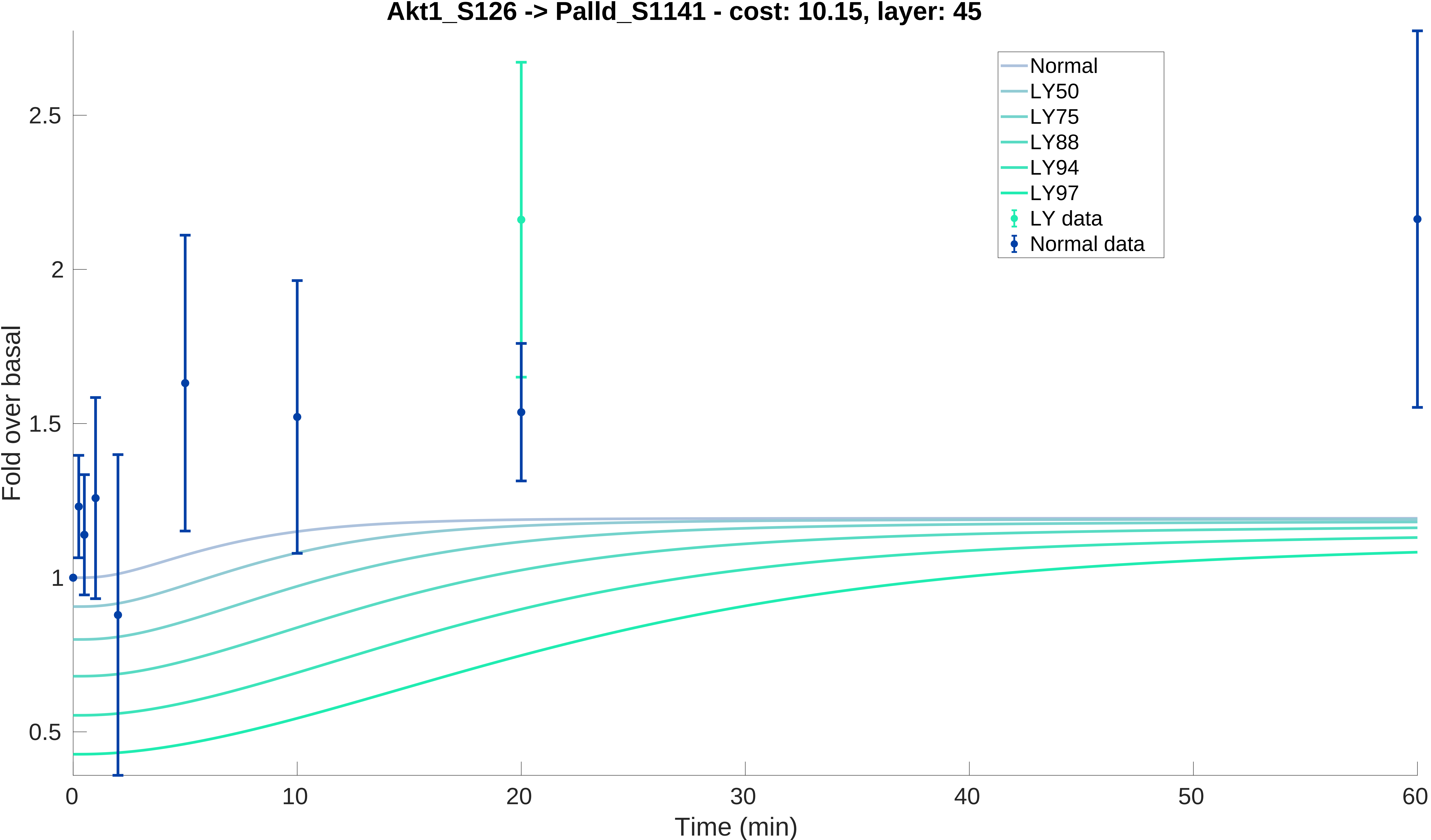
Akt1_S126 -> Palld_S1141 - cost: 10.15, layer: 45
Normal
LY50
LY75
LY88
LY94
LY97
LY data
Normal data
Fold over basal
Time (min)
0.5
1
1.5
2
2.5
0
10
20
30
40
50
60

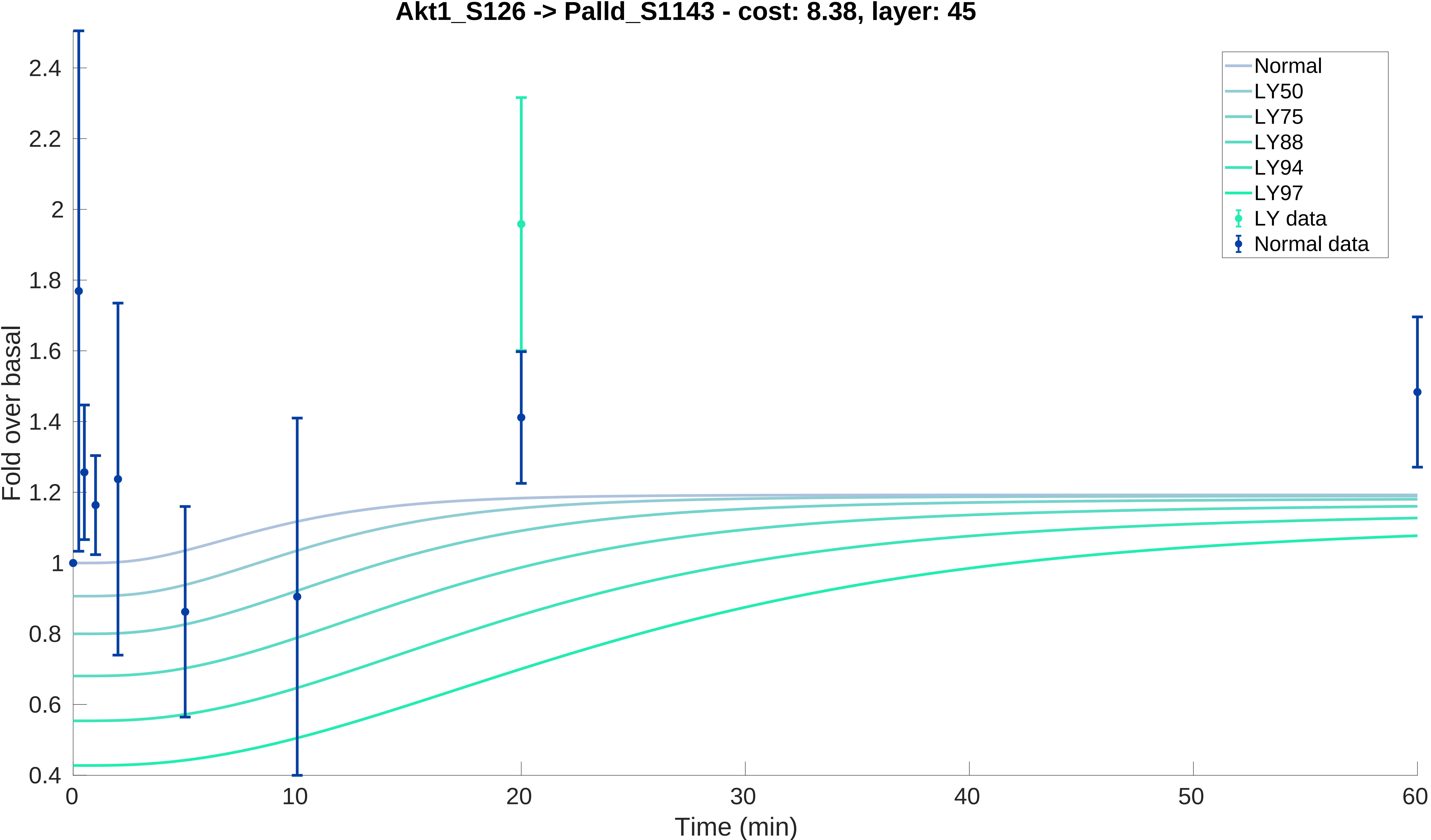

Akt1_S126 -> Palld_S1143 - cost: 8.38, layer: 45
Normal
LY50
LY75
LY88
LY94
LY97
LY data
Normal data
Fold over basal
Time (min)
0.4
0.6
0.8
1
1.2
1.4
1.6
1.8
2
2.2
2.4
0
10
20
30
40
50
60

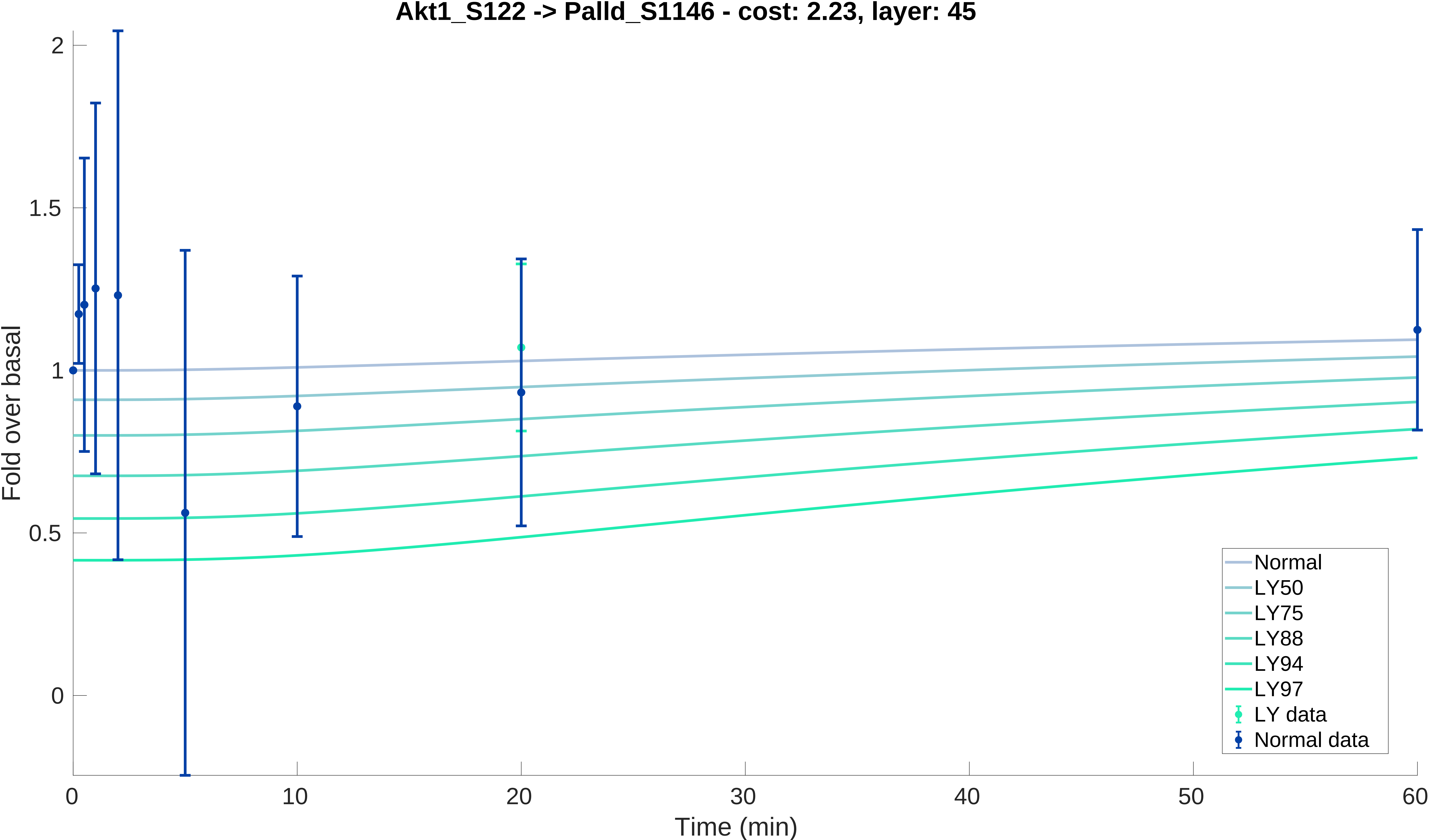

Akt1_S122 -> Palld_S1146 - cost: 2.23, layer: 45
Fold over basal
Time (min)
0
0.5
1
1.5
2
0
10
20
30
40
50
60
Normal
LY50
LY75
LY88
LY94
LY97
LY data
Normal data

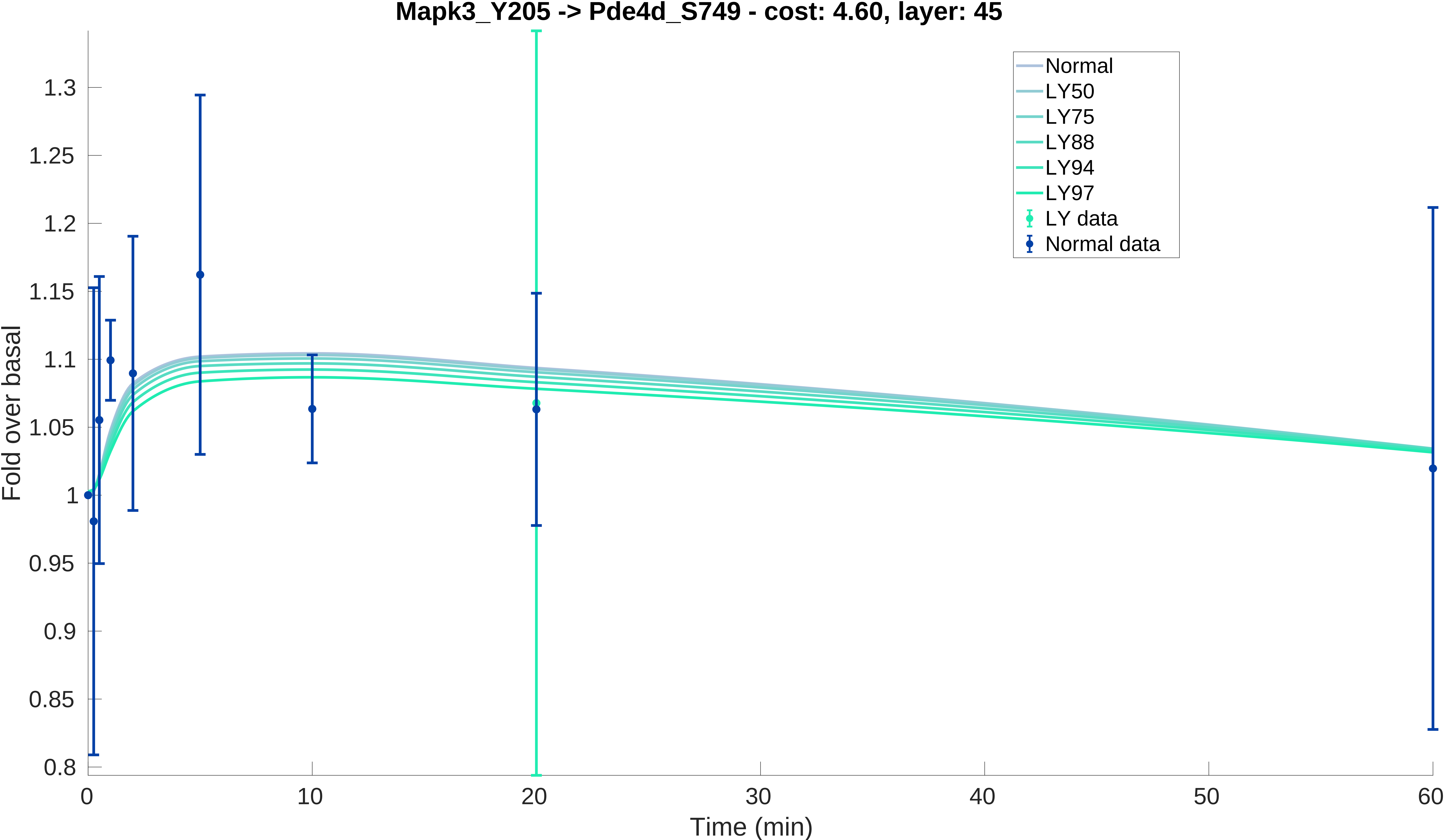

Mapk3_Y205 -> Pde4d_S749 - cost: 4.60, layer: 45
Fold over basal
1.3
1.25
1.2
1.15
1.1
1.05
1
0.95
0.9
0.85
0.8
0
10
20
30
40
50
60
Time (min)
Normal
LY50
LY75
LY88
LY94
LY97
LY data
Normal data

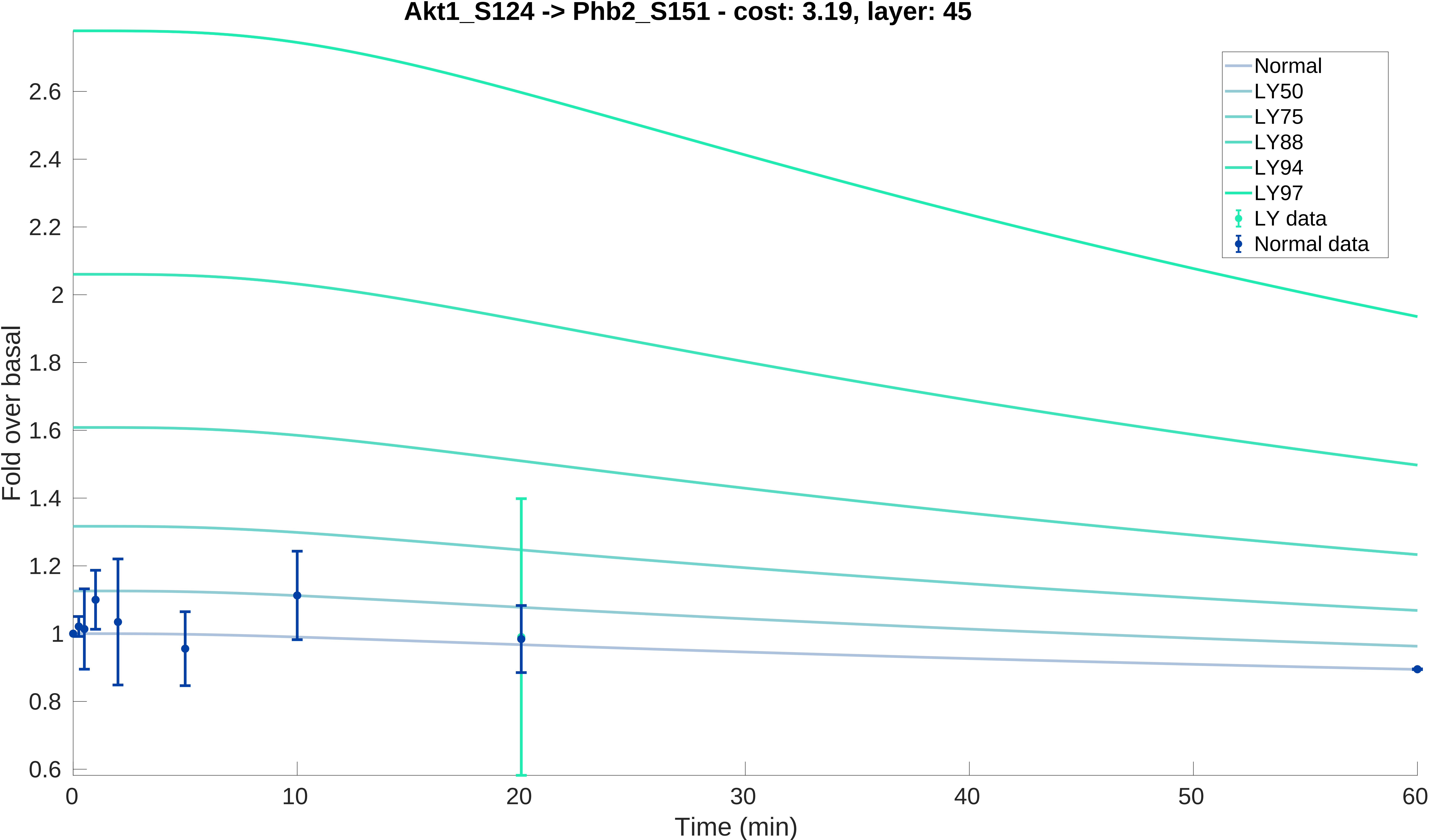
Akt1_S124 -> Phb2_S151 - cost: 3.19, layer: 45
Fold over basal
Time (min)
0
10
20
30
40
50
60
0.6
0.8
1
1.2
1.4
1.6
1.8
2
2.2
2.4
2.6
Normal
LY50
LY75
LY88
LY94
LY97
LY data
Normal data

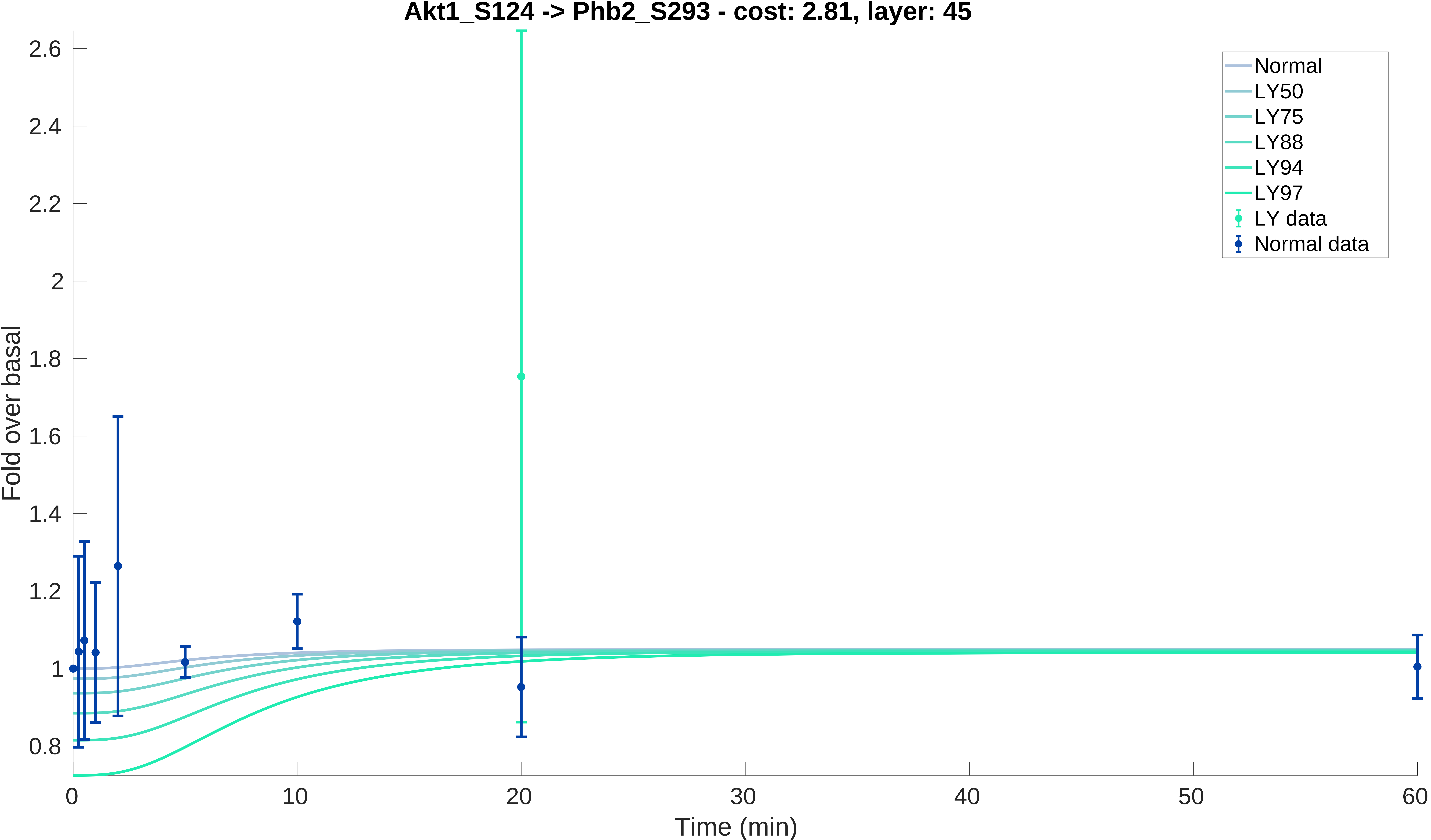

Akt1_S124 -> Phb2_S293 - cost: 2.81, layer: 45
Fold over basal
2.6
2.4
2.2
2
1.8
1.6
1.4
1.2
1
0.8
0
10
20
30
40
50
60
Time (min)
Normal
LY50
LY75
LY88
LY94
LY97
LY data
Normal data

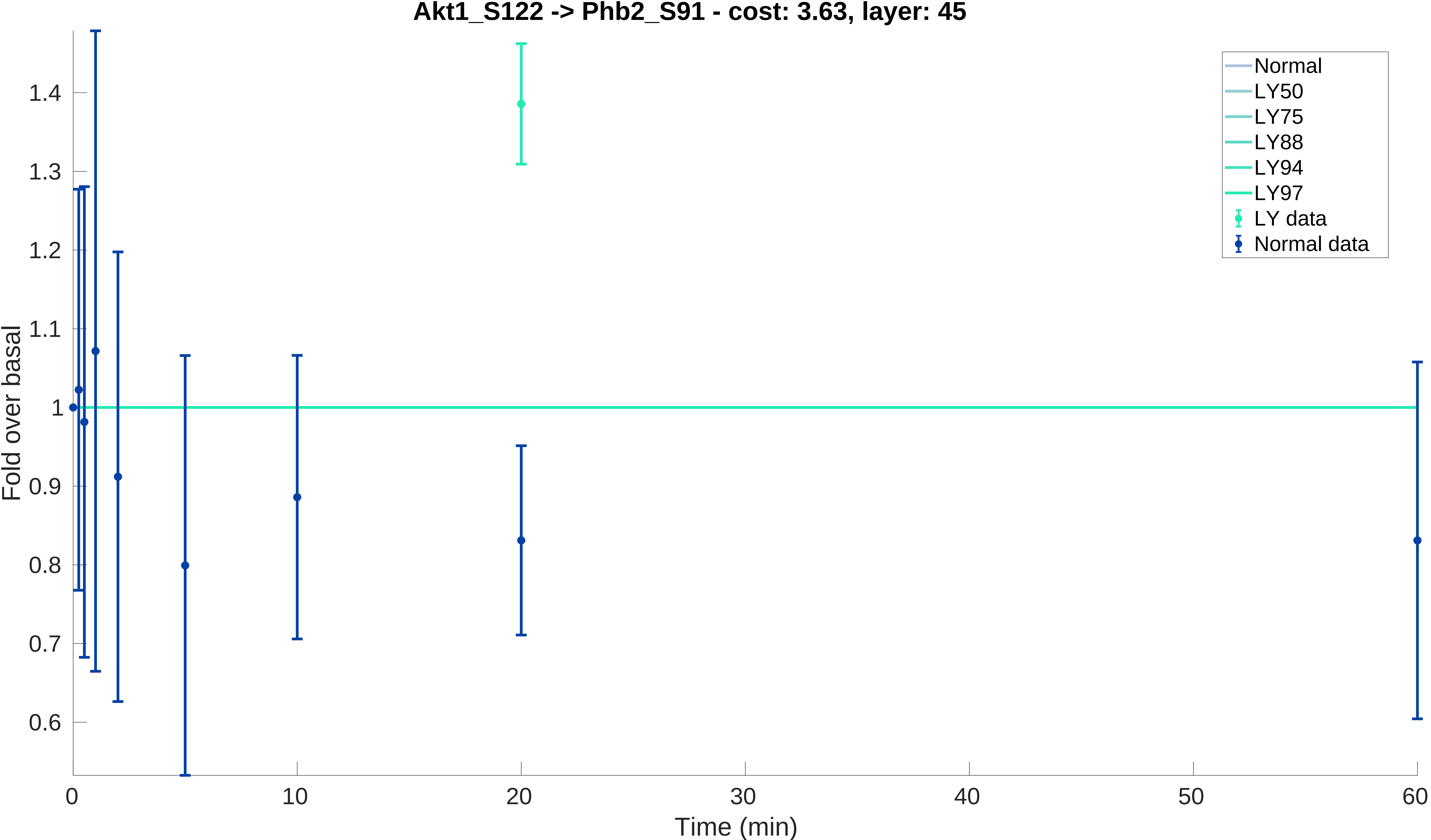

Akt1_S122 -> Phb2_S91 - cost: 3.63, layer: 45
Fold over basal
Time (min)
Normal
LY50
LY75
LY88
LY94
LY97
LY data
Normal data
1.4
1.3
1.2
1.1
1
0.9
0.8
0.7
0.6
0
10
20
30
40
50
60

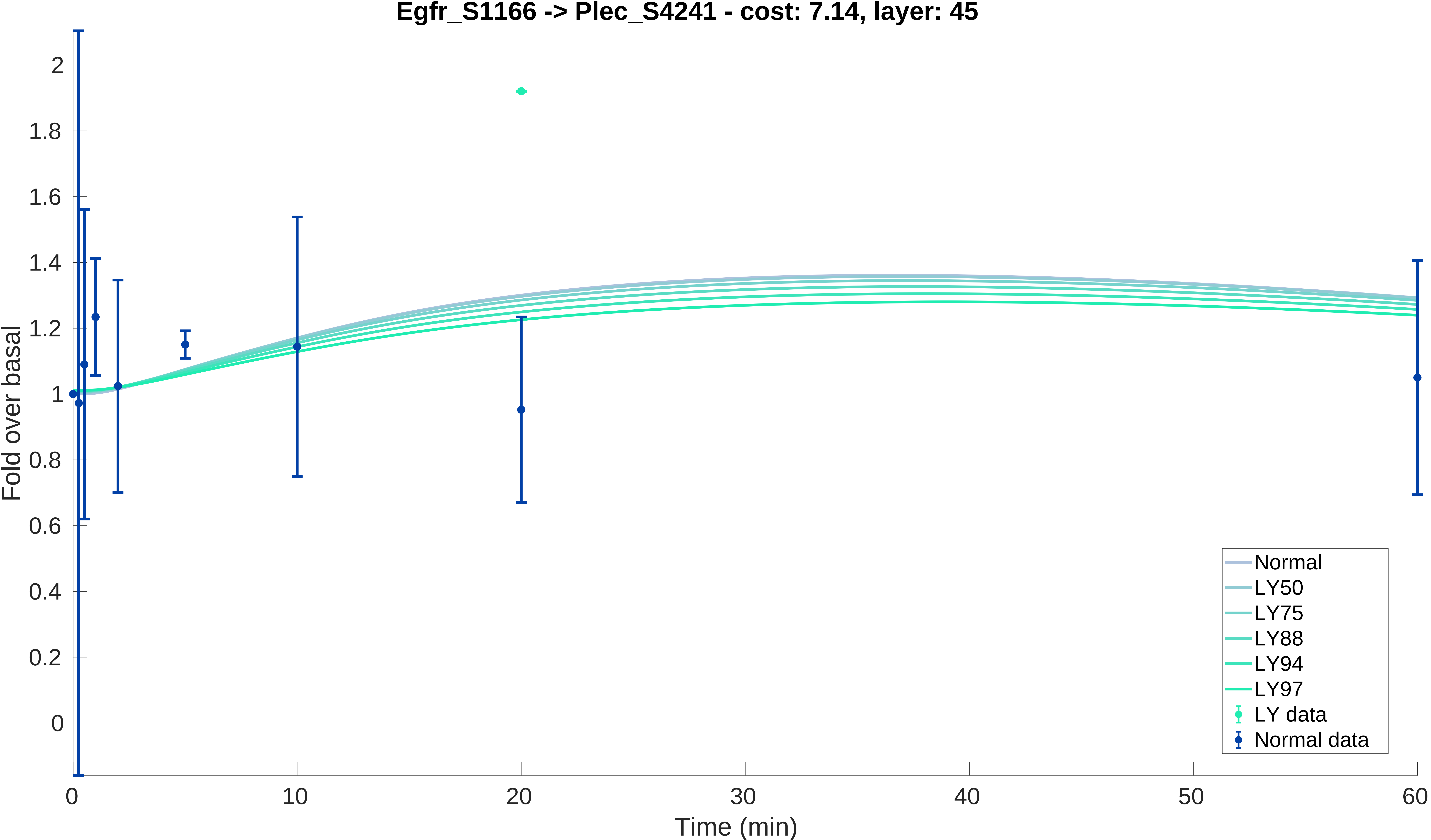

Egfr_S1166 -> Plec_S4241 - cost: 7.14, layer: 45
Fold over basal
Time (min)
0
10
20
30
40
50
60
0
0.2
0.4
0.6
0.8
1
1.2
1.4
1.6
1.8
2
Normal
LY50
LY75
LY88
LY94
LY97
LY data
Normal data

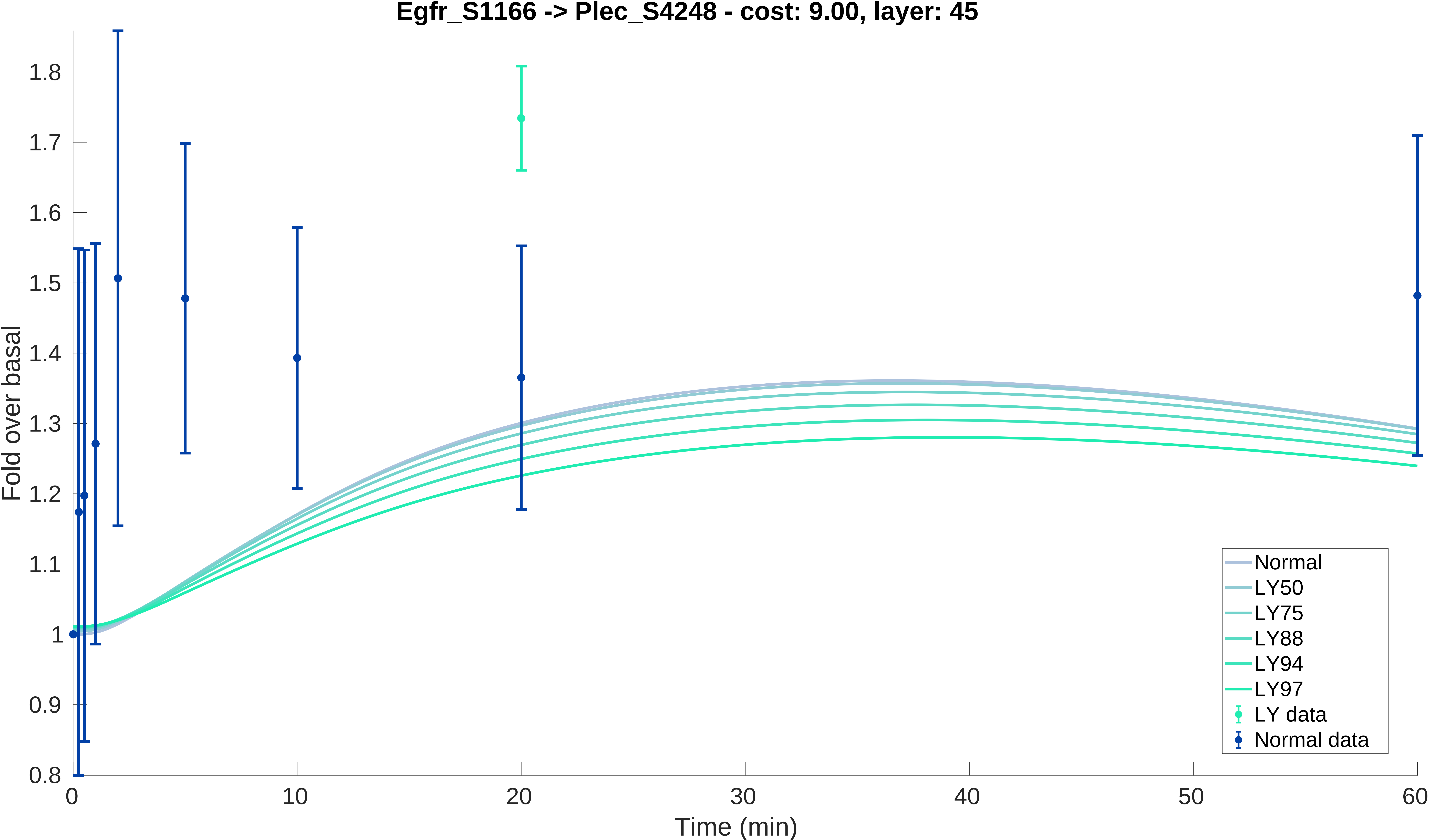

Egfr_S1166 -> Plec_S4248 - cost: 9.00, layer: 45
Fold over basal
1.8
1.7
1.6
1.5
1.4
1.3
1.2
1.1
1
0.9
0.8
0
10
20
30
40
50
60
Time (min)
Normal
LY50
LY75
LY88
LY94
LY97
LY data
Normal data

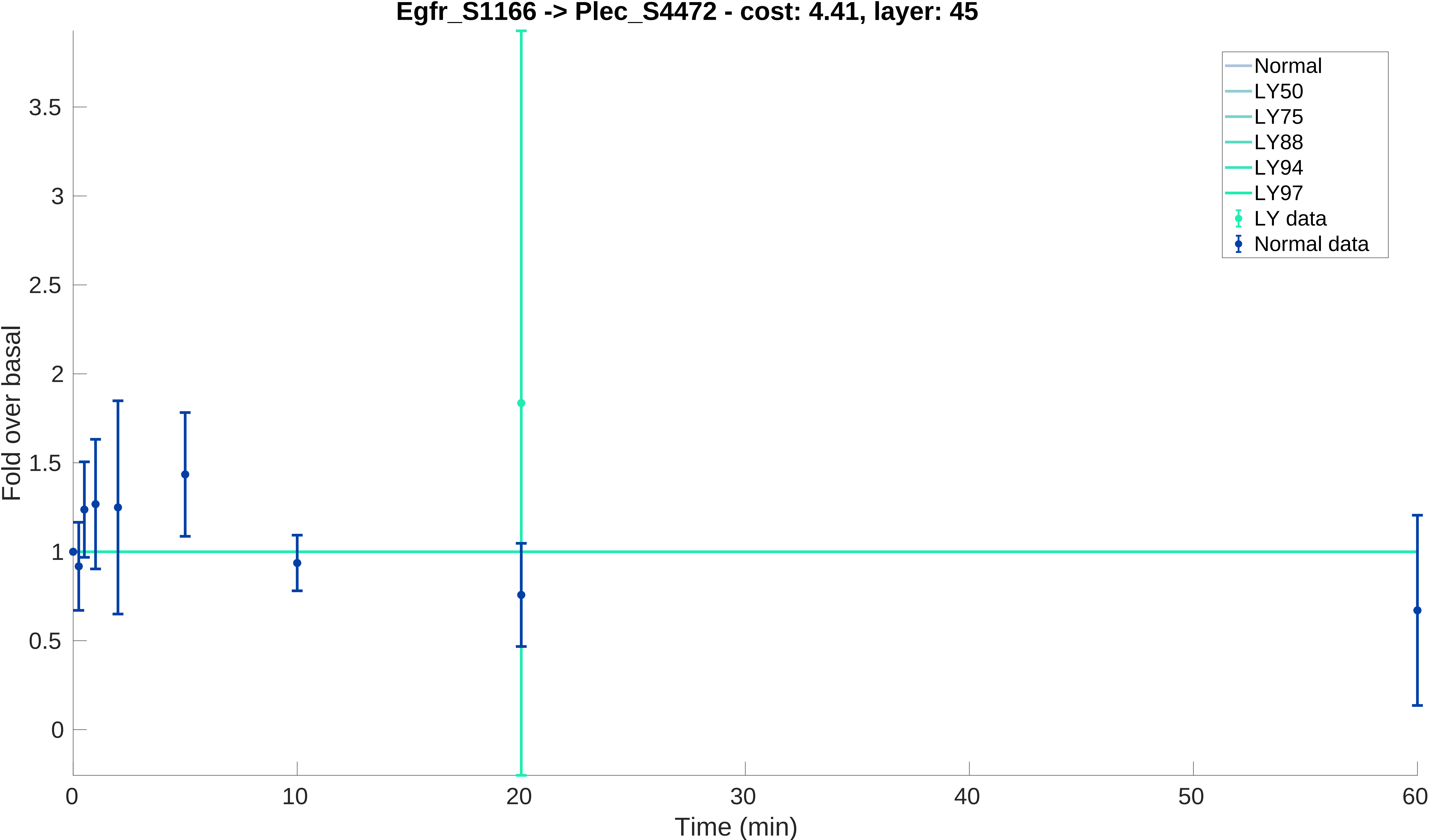

Egfr_S1166 -> Plec_S4472 - cost: 4.41, layer: 45
Normal
LY50
LY75
LY88
LY94
LY97
LY data
Normal data
Fold over basal
3.5
3
2.5
2
1.5
1
0.5
0
0
10
20
30
40
50
60
Time (min)

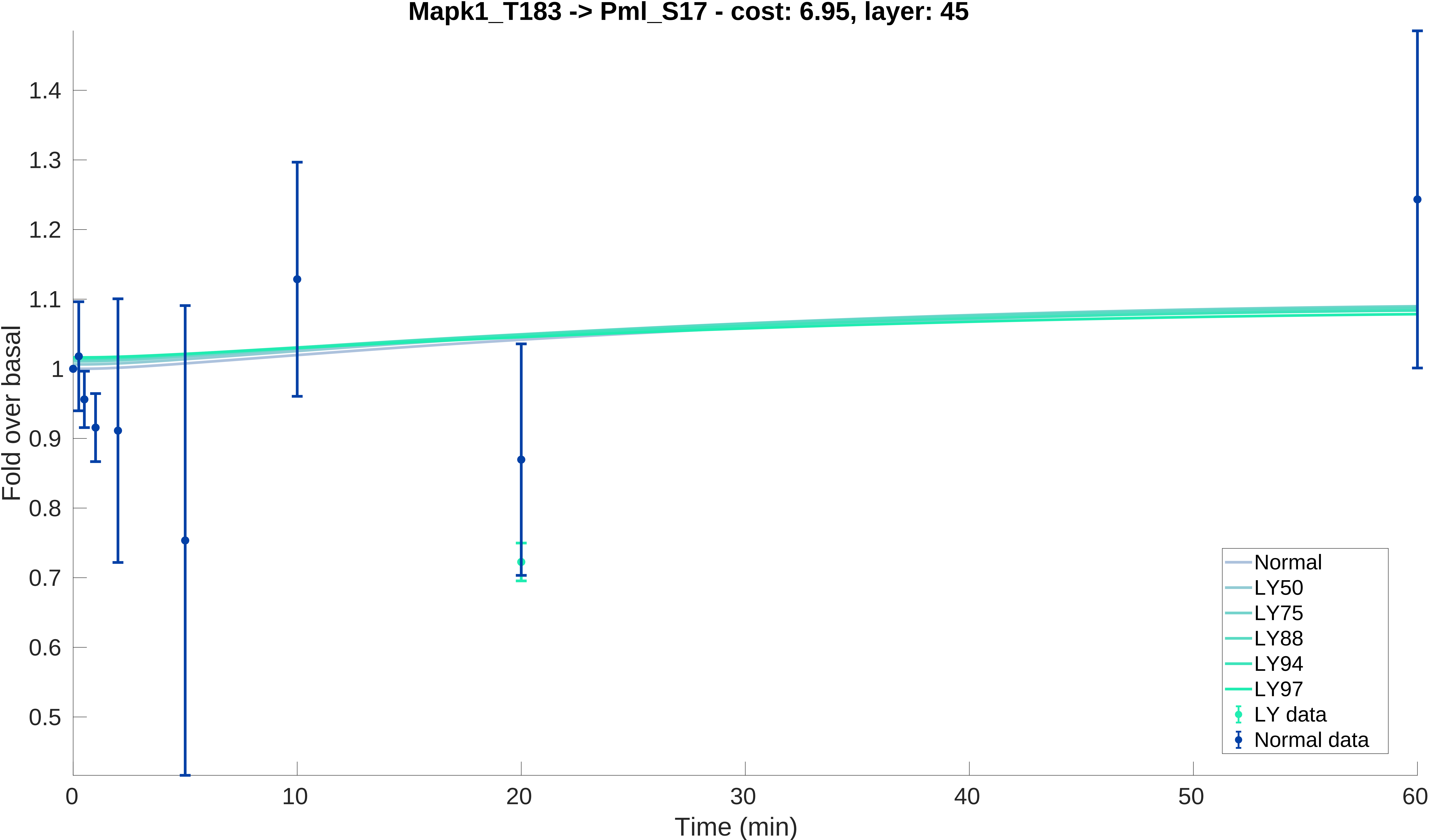

Mapk1_T183 -> Pml_S17 - cost: 6.95, layer: 45
Fold over basal
Time (min)
0.5
0.6
0.7
0.8
0.9
1
1.1
1.2
1.3
1.4
0
10
20
30
40
50
60
Normal
LY50
LY75
LY88
LY94
LY97
LY data
Normal data

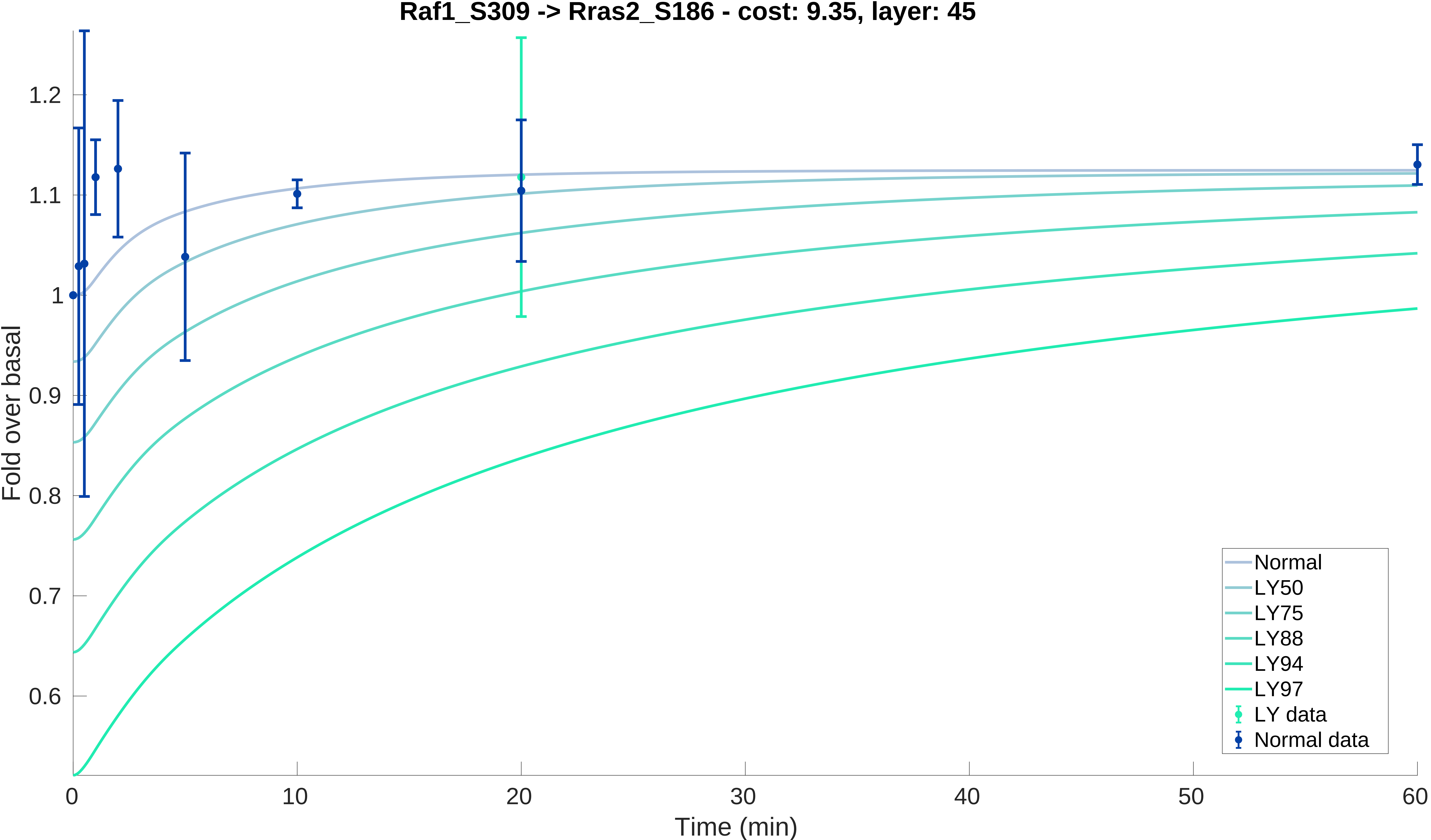
Raf1_S309 -> Rras2_S186 - cost: 9.35, layer: 45
Fold over basal
Time (min)
1.2
1.1
1
0.9
0.8
0.7
0.6
0
10
20
30
40
50
60
Normal
LY50
LY75
LY88
LY94
LY97
LY data
Normal data

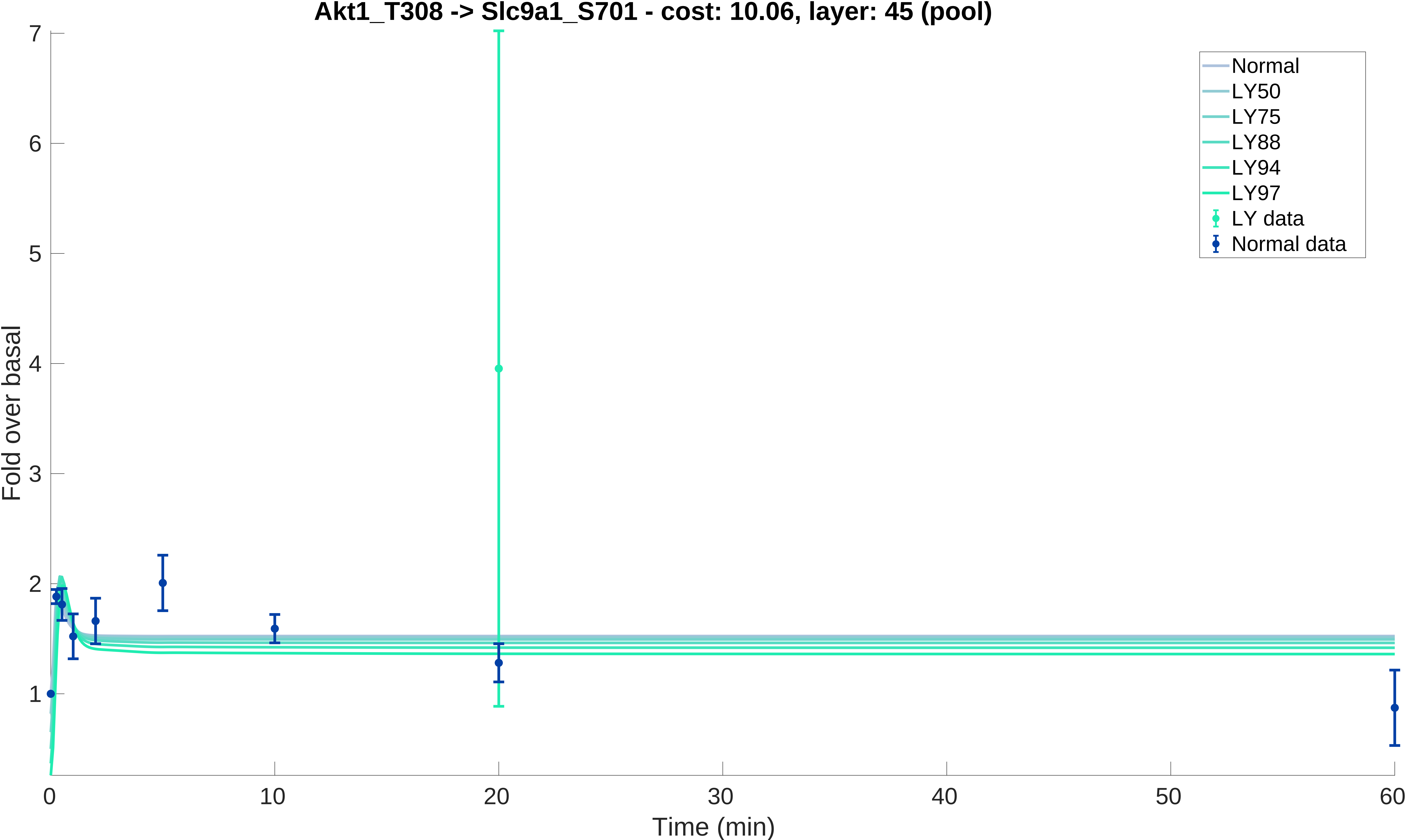

Akt1_T308 -> Slc9a1_S701 - cost: 10.06, layer: 45 (pool)
Fold over basal
Time (min)
0
10
20
30
40
50
60
1
2
3
4
5
6
7
Normal
LY50
LY75
LY88
LY94
LY97
LY data
Normal data

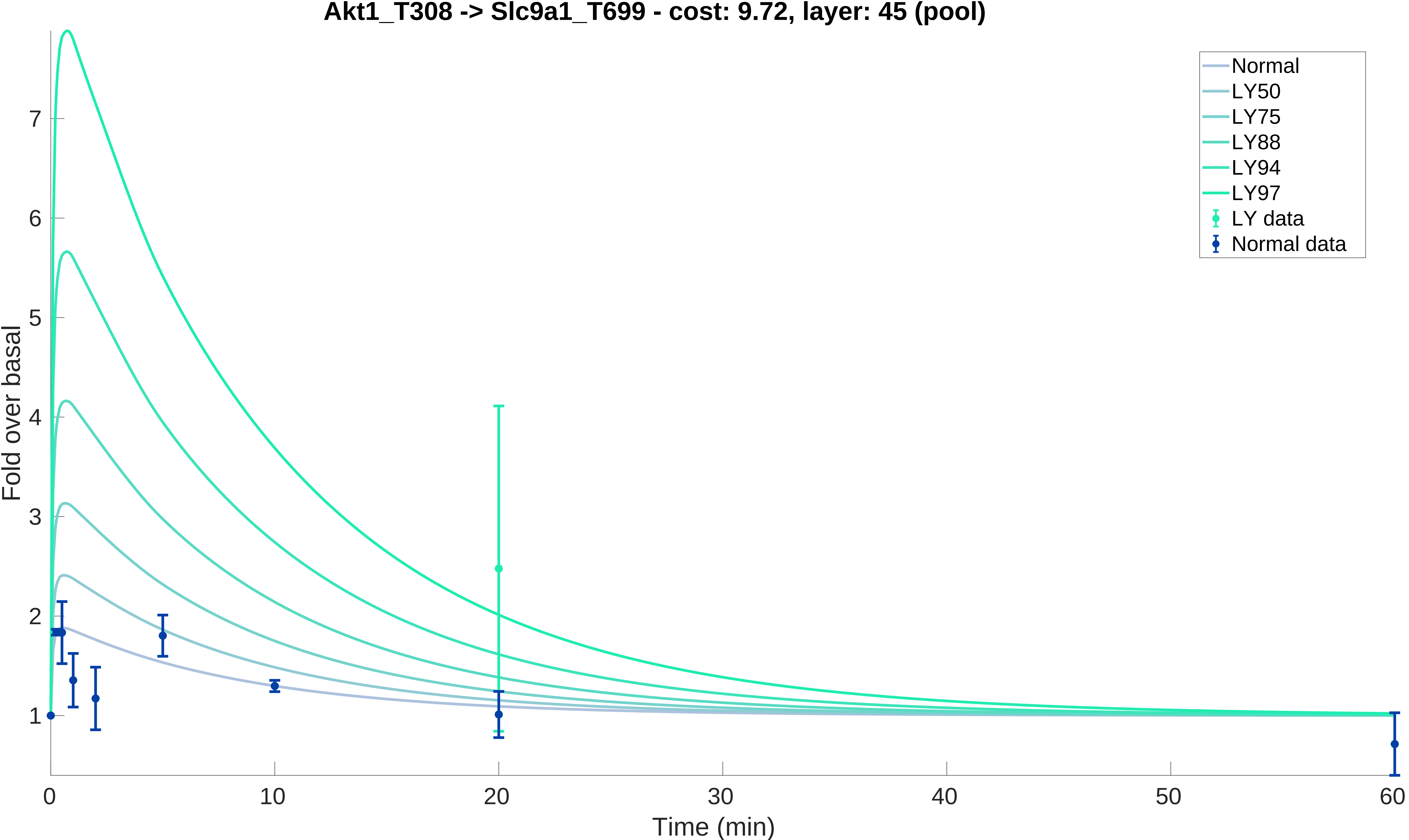

Akt1_T308 -> Slc9a1_T699 - cost: 9.72, layer: 45 (pool)
Fold over basal
Time (min)
Normal
LY50
LY75
LY88
LY94
LY97
LY data
Normal data
0
10
20
30
40
50
60
1
2
3
4
5
6
7

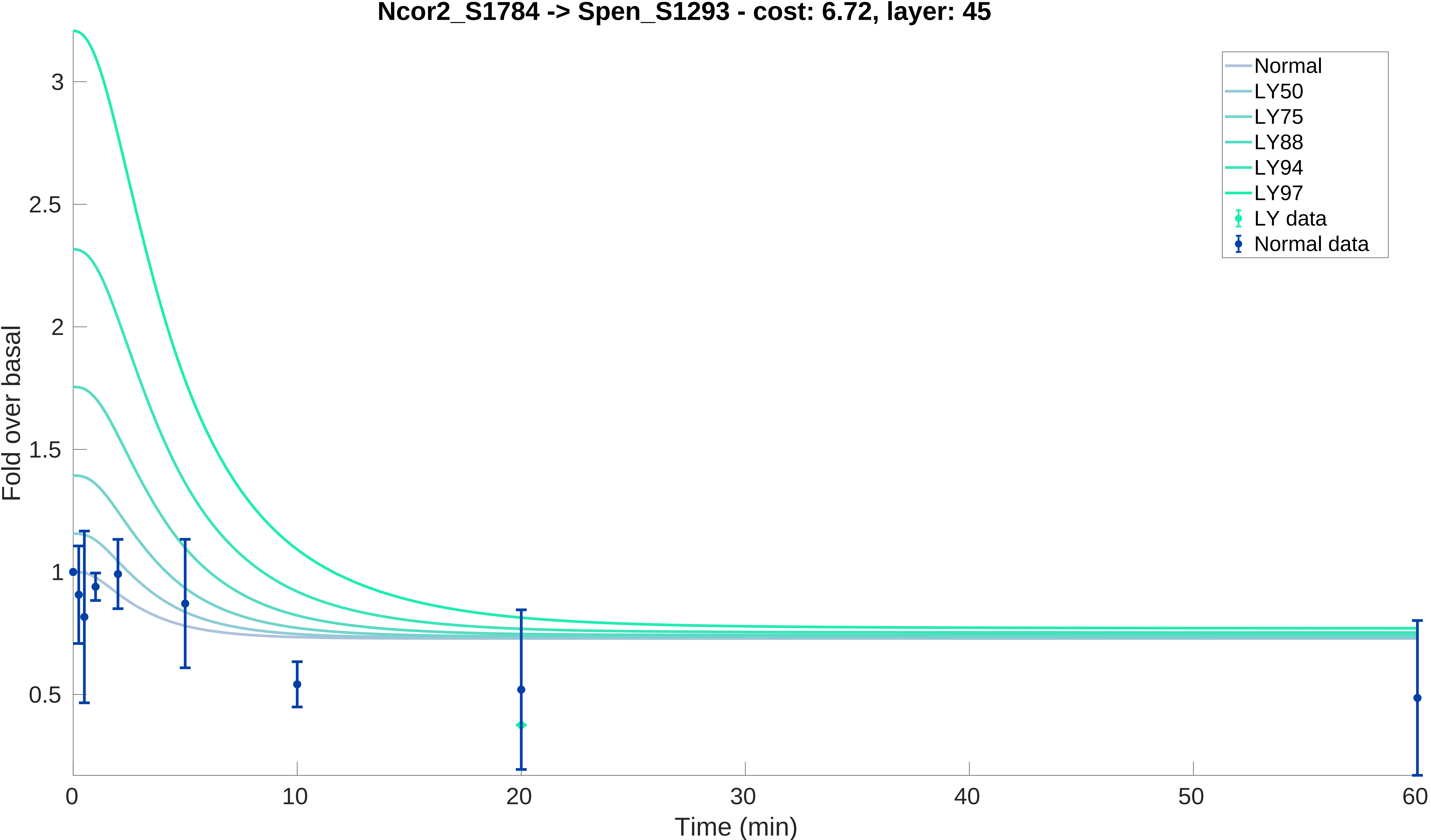

Ncor2_S1784 -> Spen_S1293 - cost: 6.72, layer: 45
Normal
LY50
LY75
LY88
LY94
LY97
LY data
Normal data
Fold over basal
3
2.5
2
1.5
1
0.5
0
10
20
30
40
50
60
Time (min)

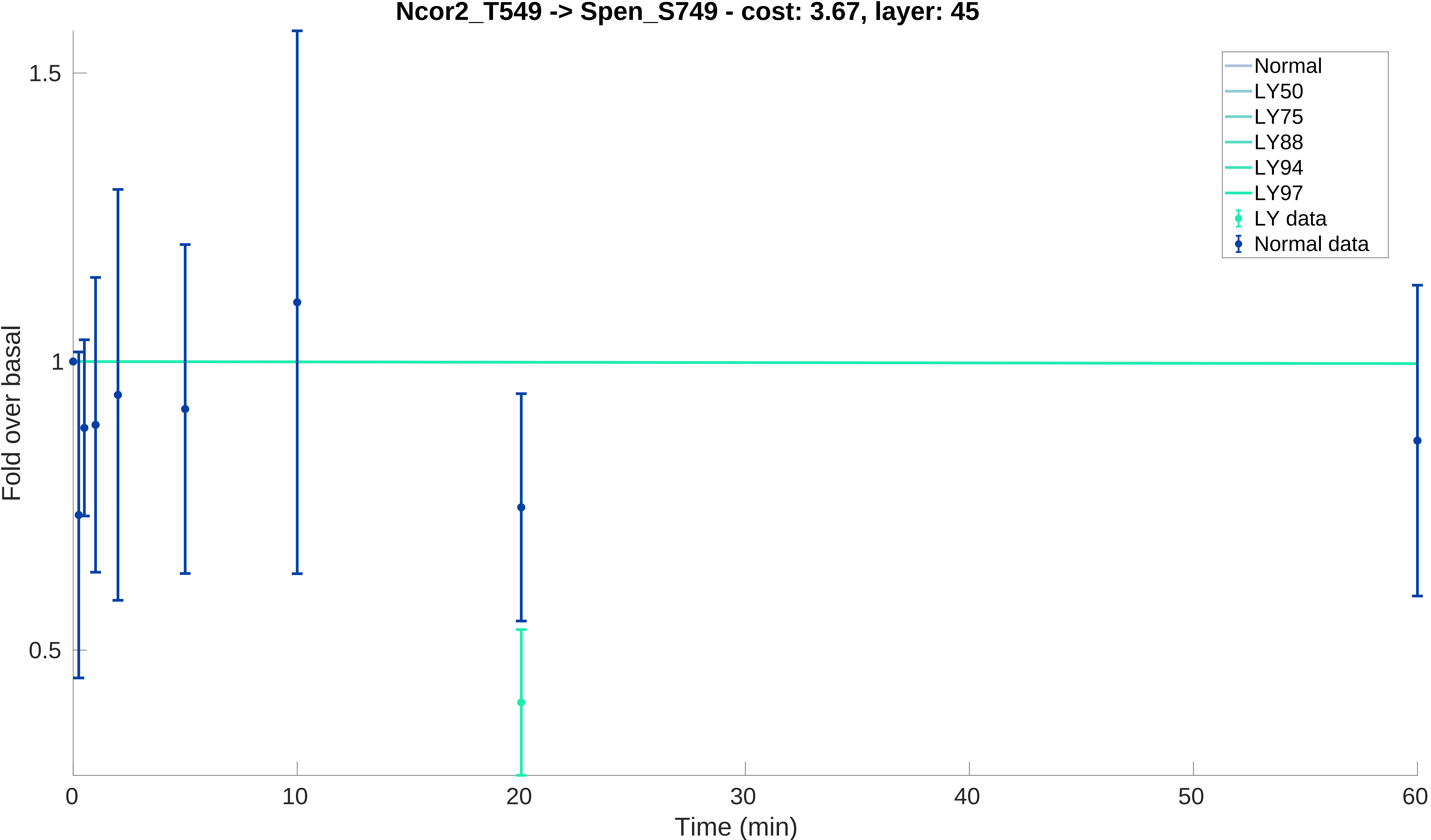

Ncor2_T549 -> Spen_S749 - cost: 3.67, layer: 45
Fold over basal
1.5
1
0.5
0
10
20
30
40
50
60
Time (min)
Normal
LY50
LY75
LY88
LY94
LY97
LY data
Normal data

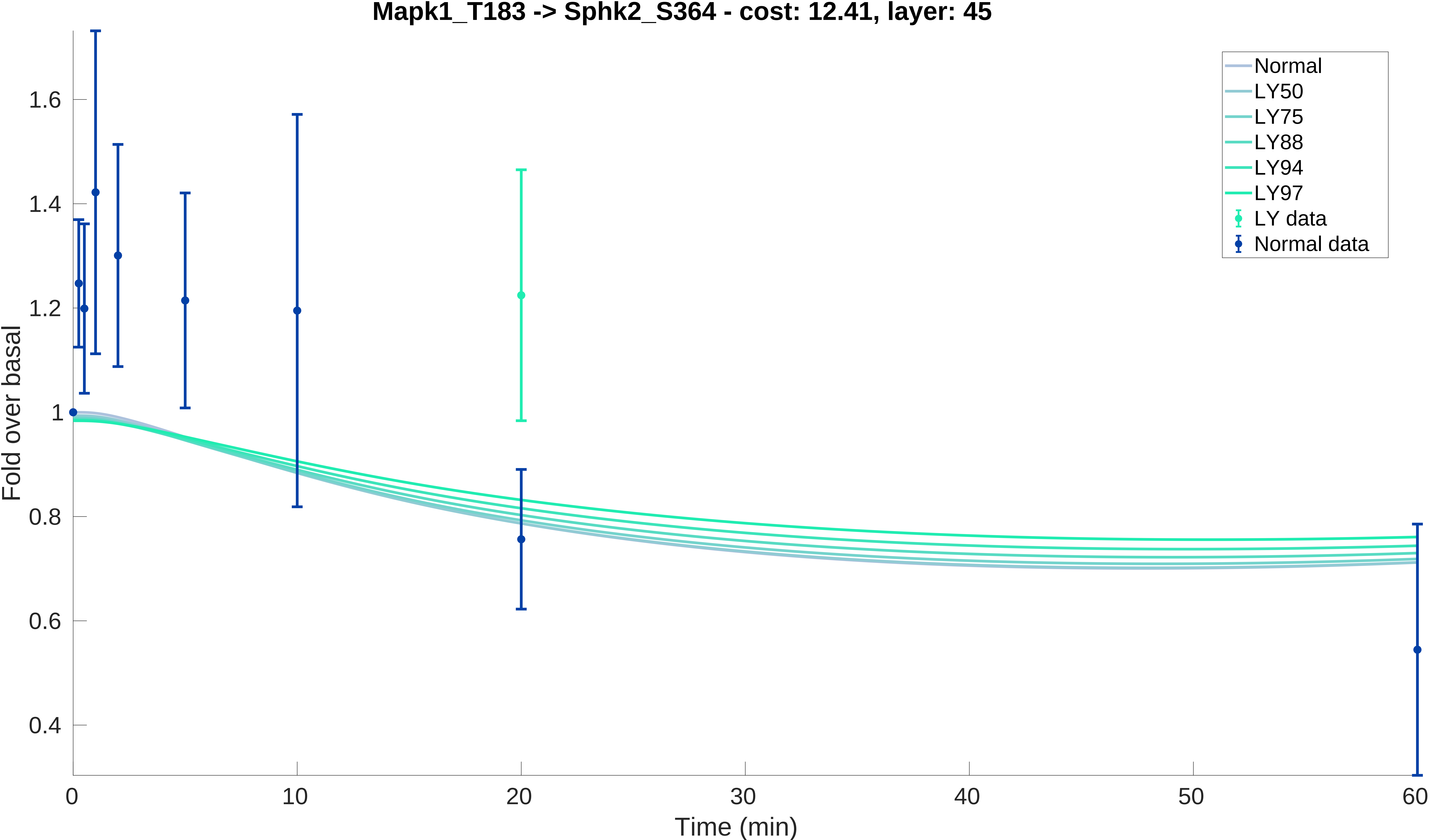

Mapk1_T183 -> Sphk2_S364 - cost: 12.41, layer: 45
Fold over basal
Time (min)
0.4
0.6
0.8
1
1.2
1.4
1.6
0
10
20
30
40
50
60
Normal
LY50
LY75
LY88
LY94
LY97
LY data
Normal data

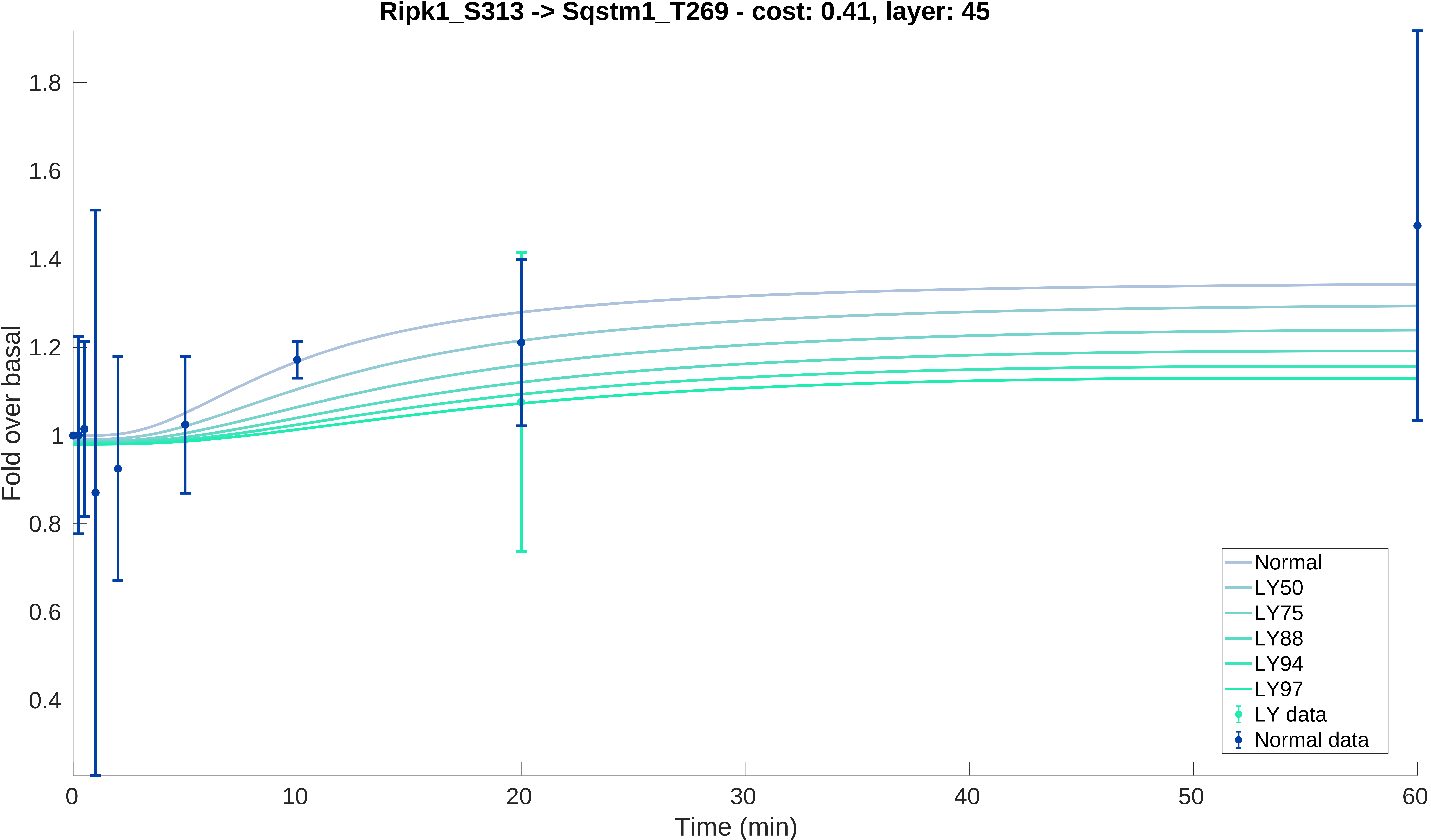

Ripk1_S313 -> Sqstm1_T269 - cost: 0.41, layer: 45
Fold over basal
1.8
1.6
1.4
1.2
1
0.8
0.6
0.4
0
10
20
30
40
50
60
Time (min)
Normal
LY50
LY75
LY88
LY94
LY97
LY data
Normal data

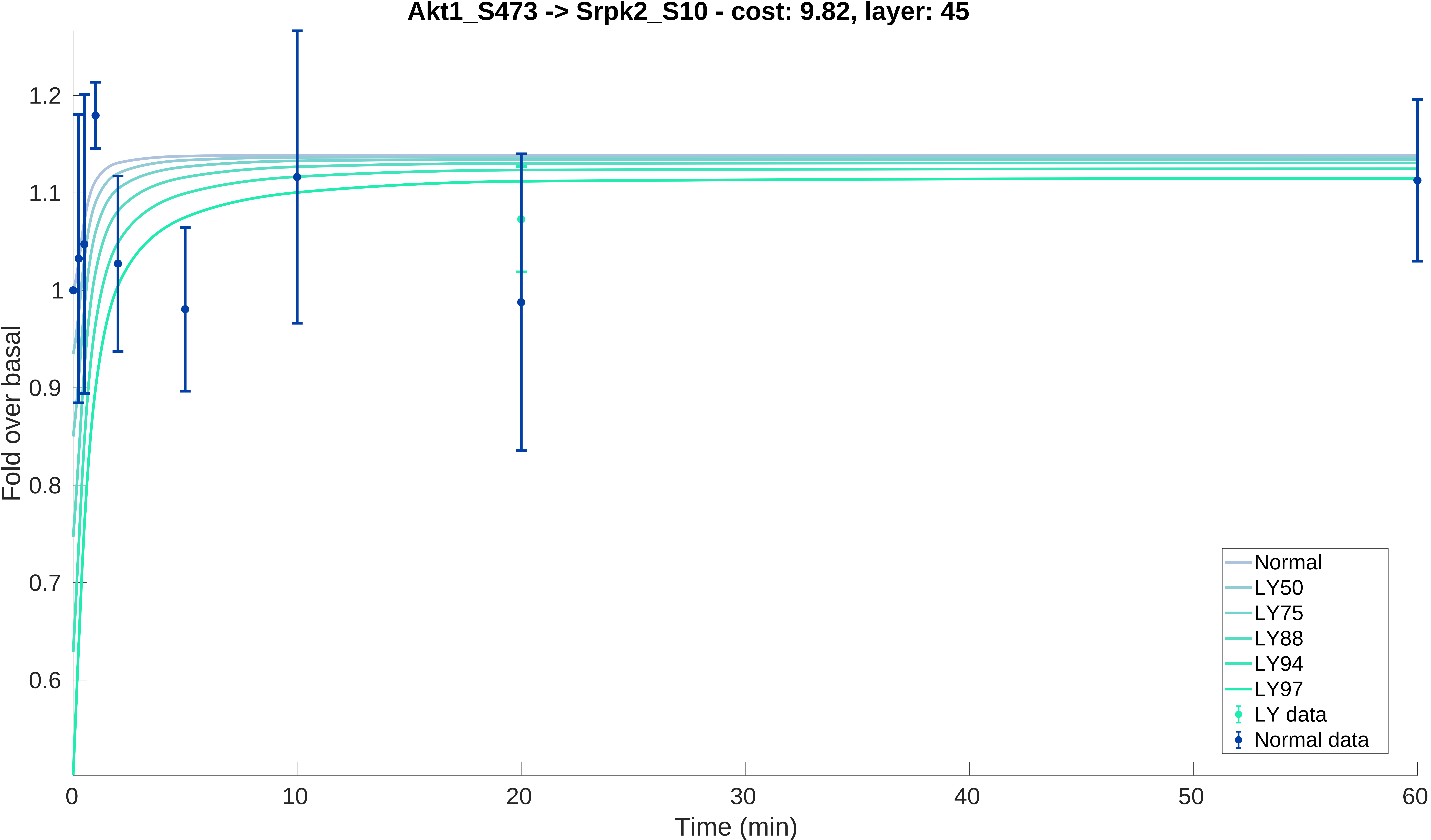

Akt1_S473 -> Srpk2_S10 - cost: 9.82, layer: 45
Fold over basal
Time (min)
0.6
0.7
0.8
0.9
1
1.1
1.2
0
10
20
30
40
50
60
Normal
LY50
LY75
LY88
LY94
LY97
LY data
Normal data

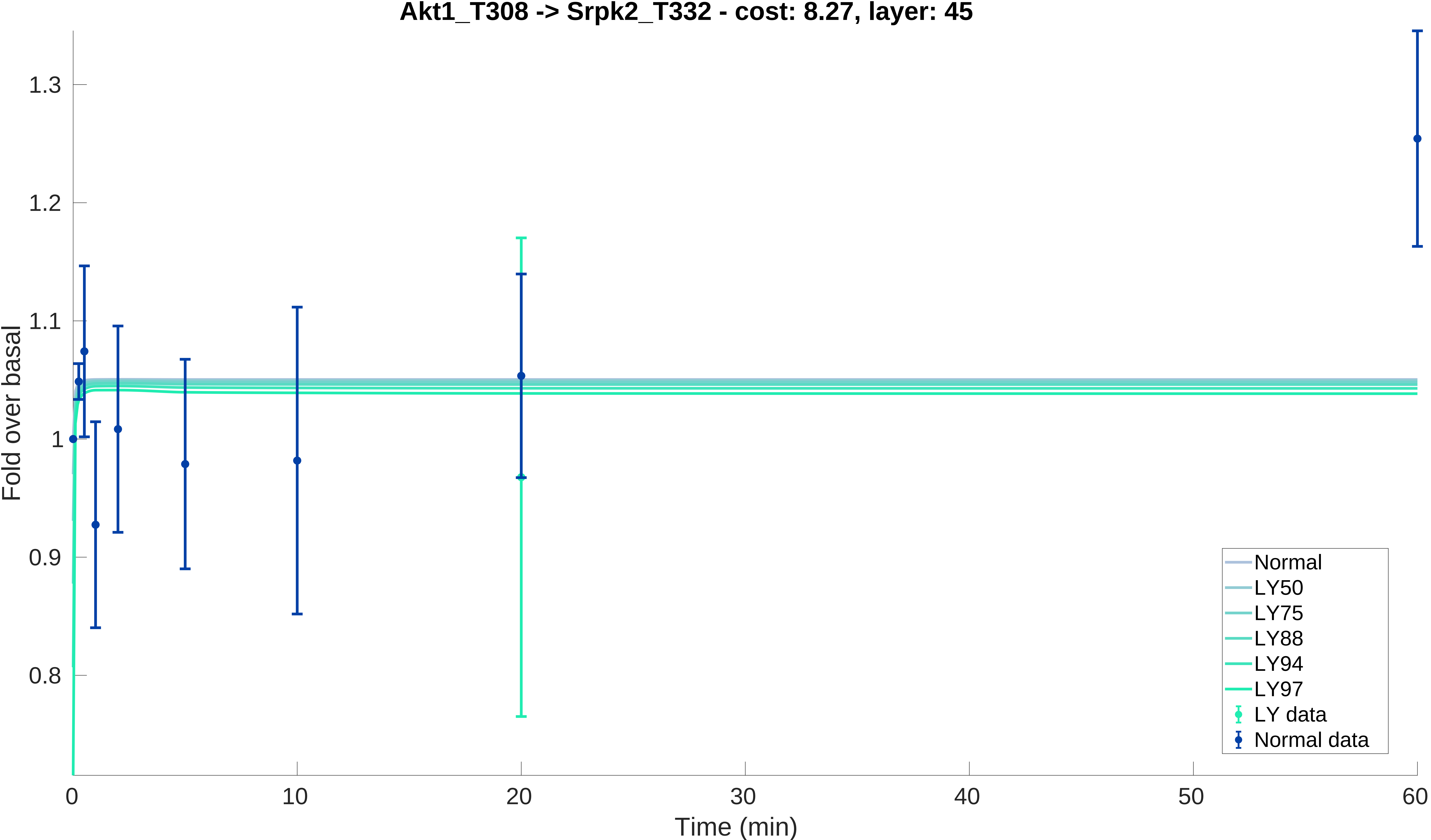
Akt1_T308 -> Srpk2_T332 - cost: 8.27, layer: 45
Fold over basal
Time (min)
1.3
1.2
1.1
1
0.9
0.8
0
10
20
30
40
50
60
Normal
LY50
LY75
LY88
LY94
LY97
LY data
Normal data

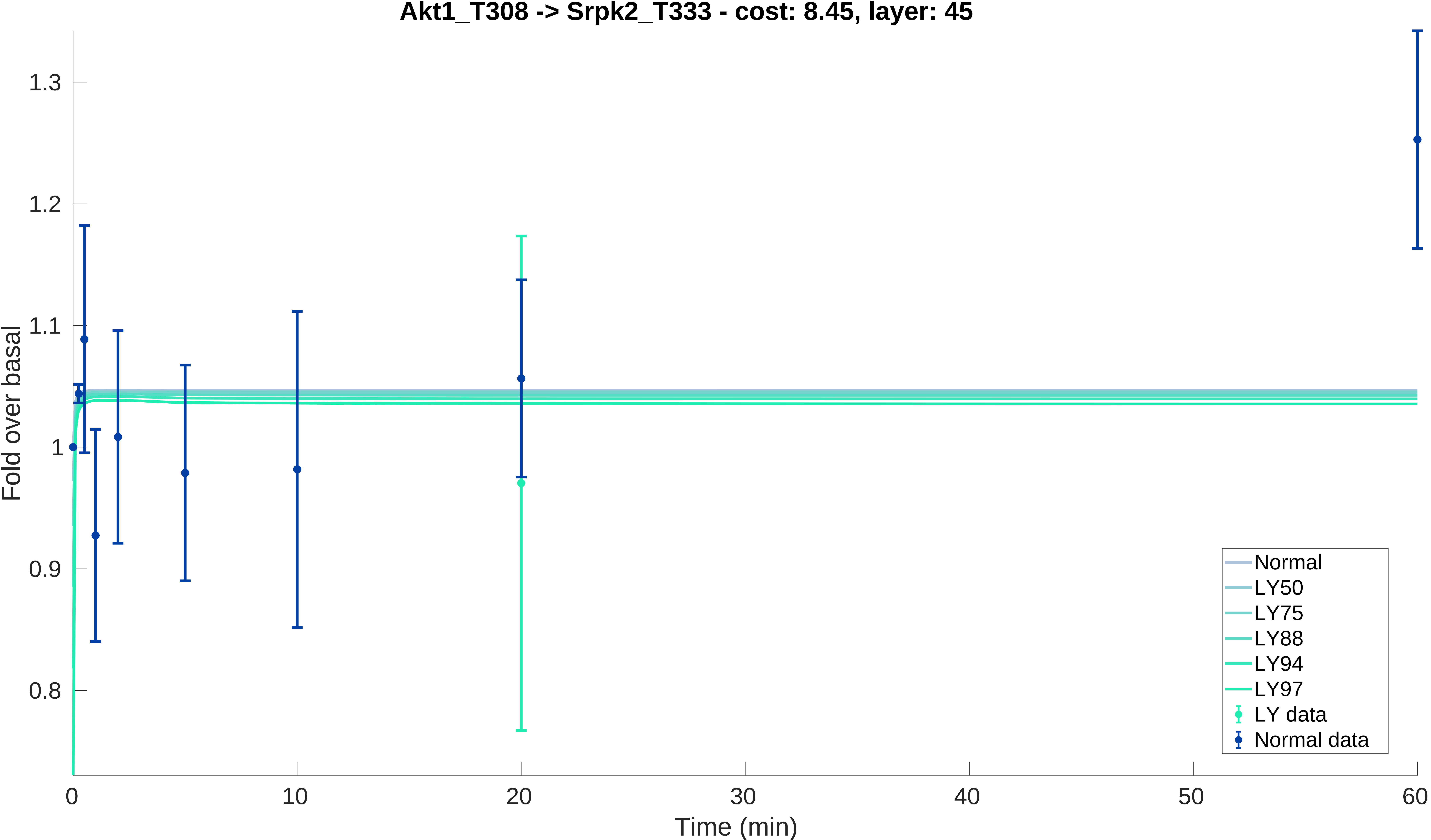

Akt1_T308 -> Srpk2_T333 - cost: 8.45, layer: 45
1.3
1.2
1.1
1
0.9
0.8
Fold over basal
0
10
20
30
40
50
60
Time (min)
Normal
LY50
LY75
LY88
LY94
LY97
LY data
Normal data

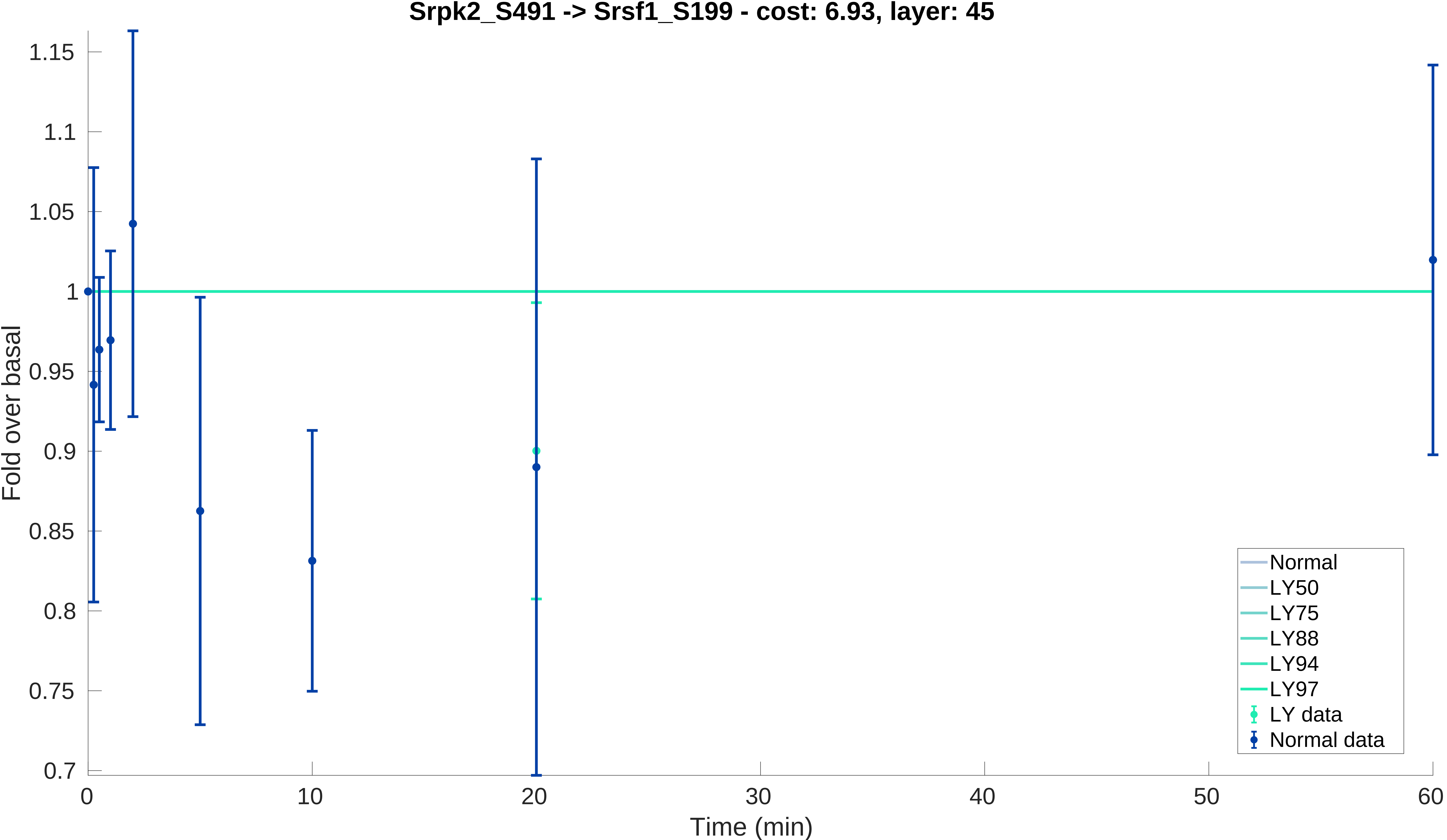

Srpk2_S491 -> Srsf1_S199 - cost: 6.93, layer: 45
Fold over basal
Time (min)
1.15
1.1
1.05
1
0.95
0.9
0.85
0.8
0.75
0.7
0
10
20
30
40
50
60
Normal
LY50
LY75
LY88
LY94
LY97
LY data
Normal data

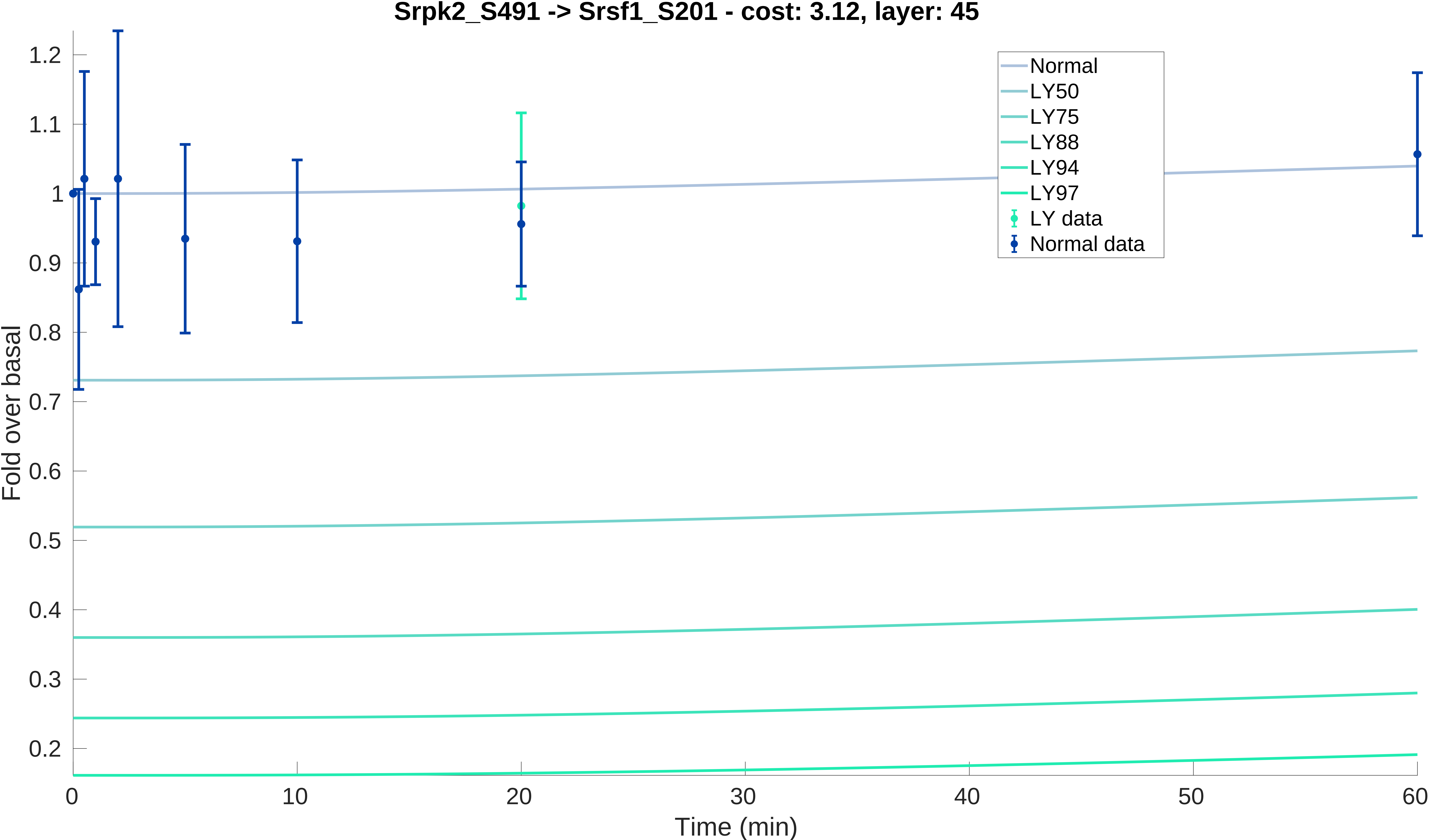
Srpk2_S491 -> Srsf1_S201 - cost: 3.12, layer: 45
Fold over basal
Time (min)
0
10
20
30
40
50
60
0.2
0.3
0.4
0.5
0.6
0.7
0.8
0.9
1
1.1
1.2
Normal
LY50
LY75
LY88
LY94
LY97
LY data
Normal data

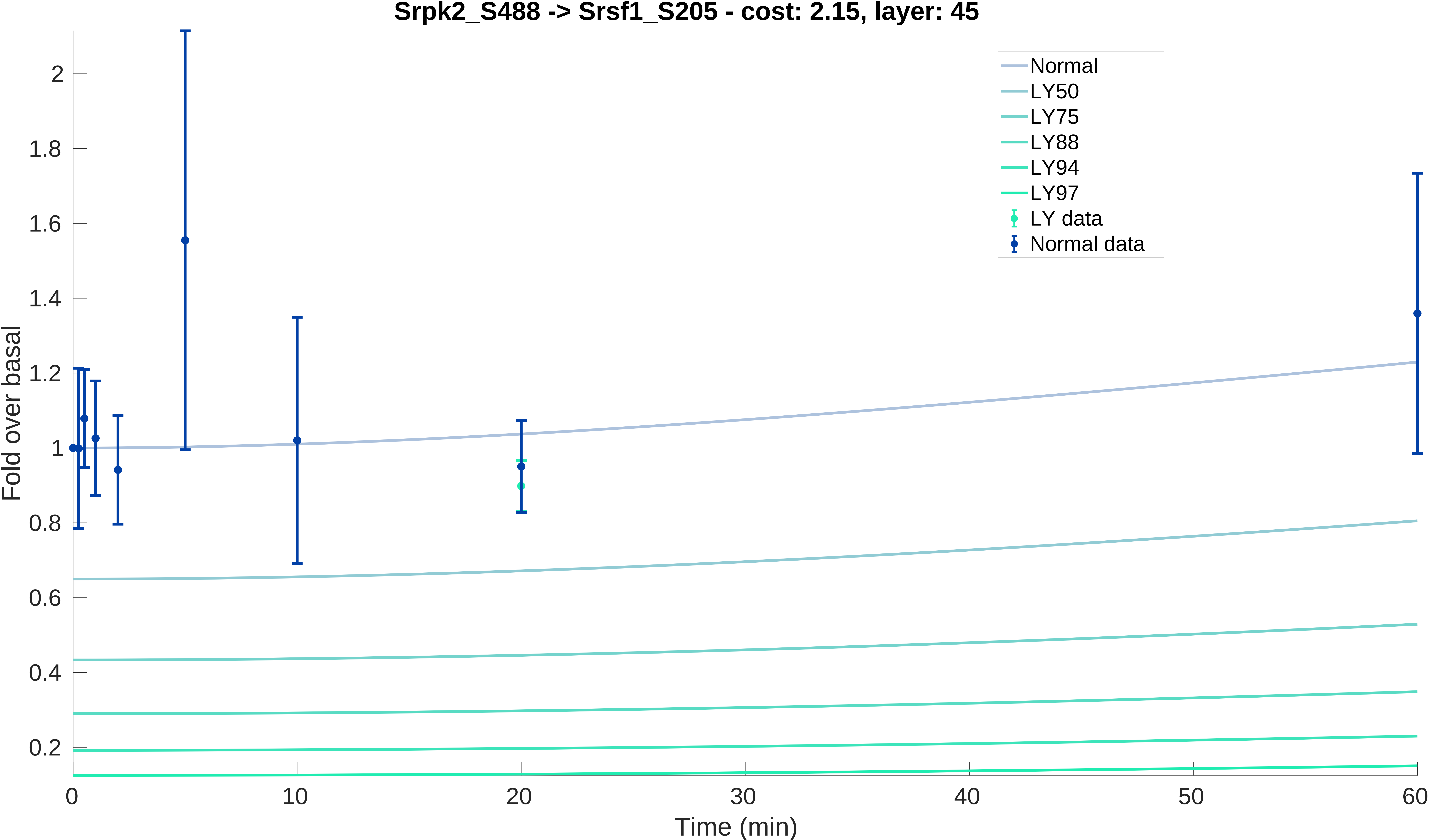

Srpk2_S488 -> Srsf1_S205 - cost: 2.15, layer: 45
Fold over basal
Time (min)
0
10
20
30
40
50
60
0.2
0.4
0.6
0.8
1
1.2
1.4
1.6
1.8
2
Normal
LY50
LY75
LY88
LY94
LY97
LY data
Normal data

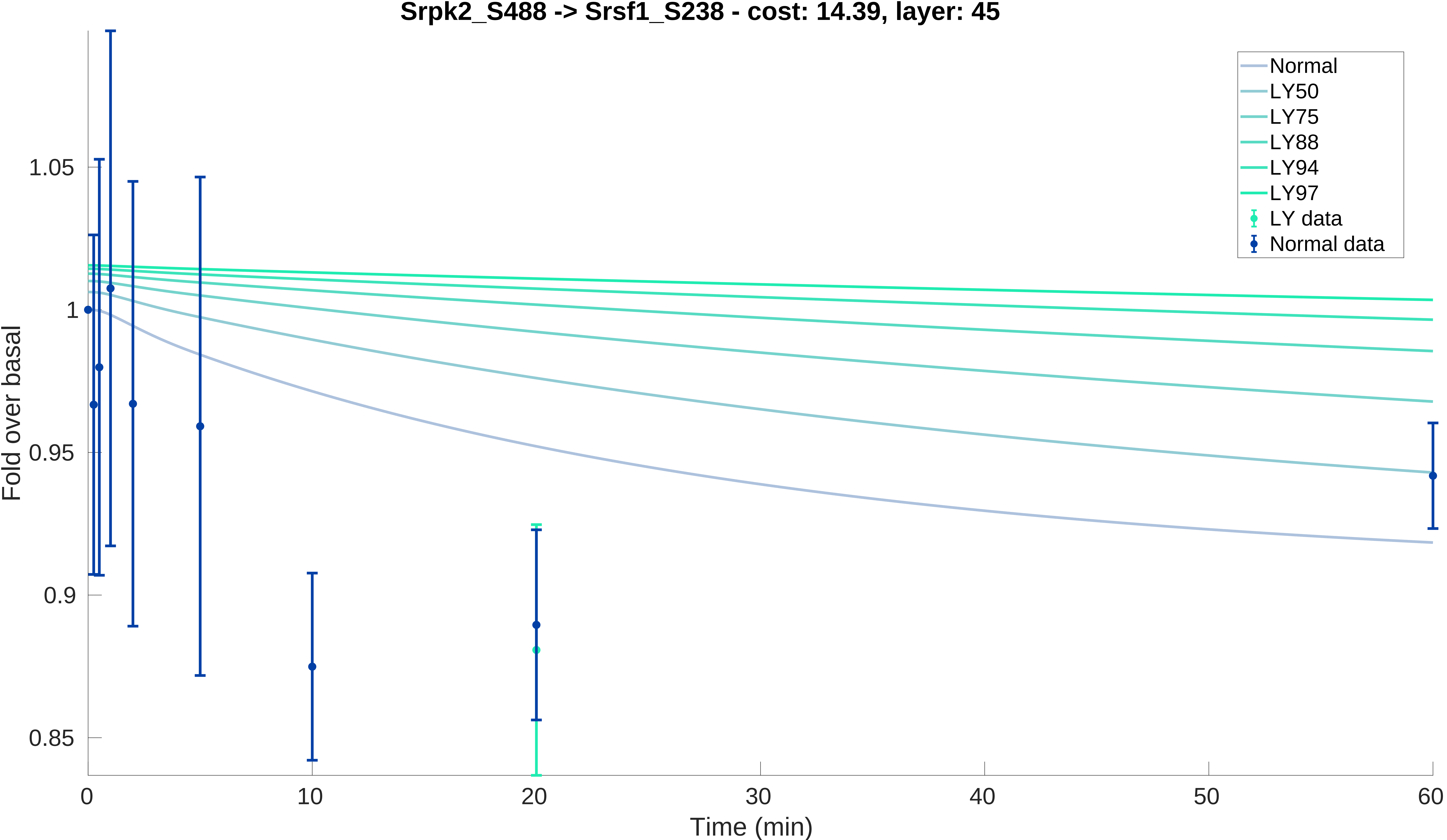

Srpk2_S488 -> Srsf1_S238 - cost: 14.39, layer: 45
Normal
LY50
LY75
LY88
LY94
LY97
LY data
Normal data
1.05
1
0.95
0.9
0.85
Fold over basal
0
10
20
30
40
50
60
Time (min)

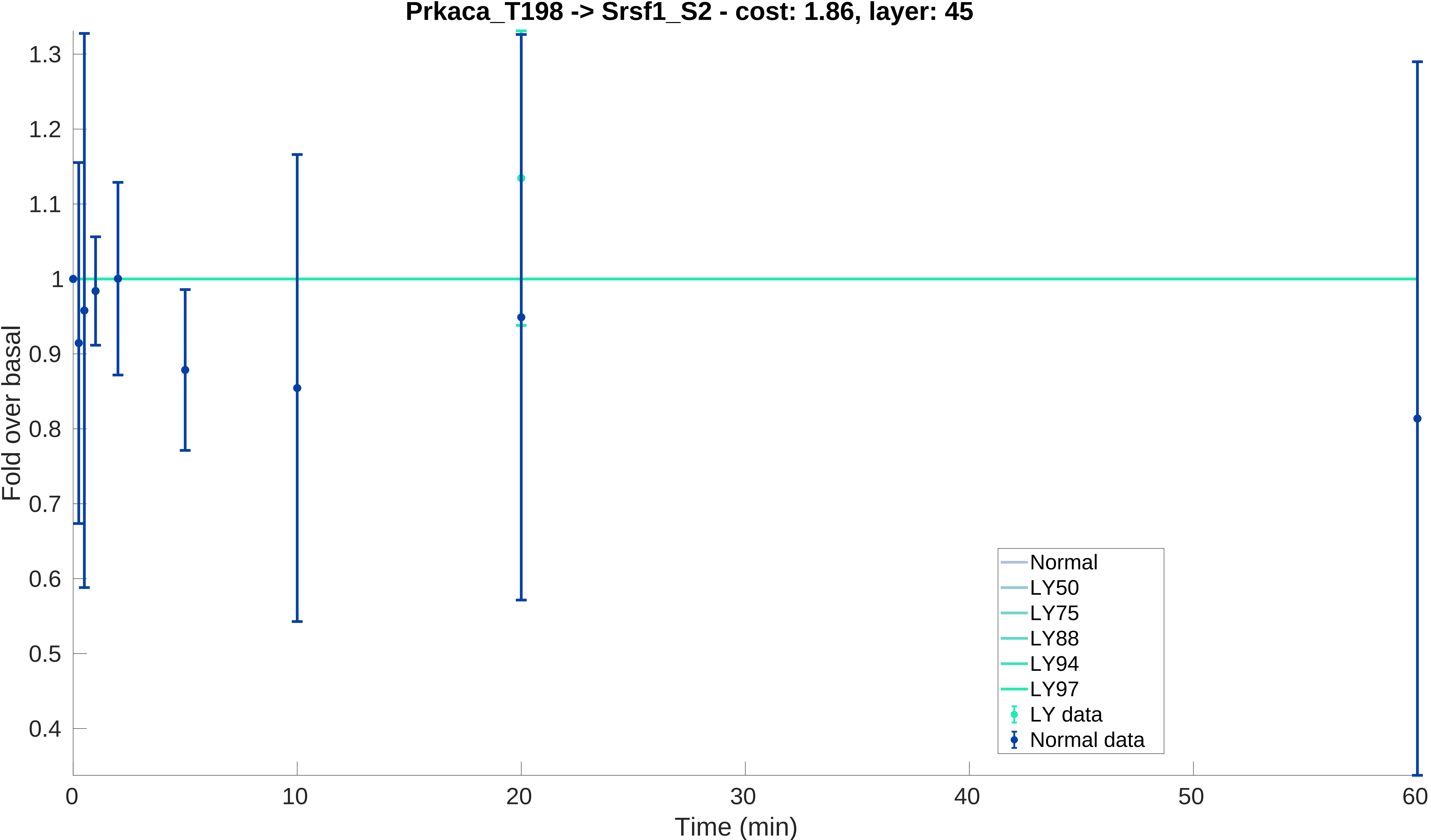
Prkaca_T198 -> Srsf1_S2 - cost: 1.86, layer: 45
Fold over basal
Time (min)
0
10
20
30
40
50
60
0.4
0.5
0.6
0.7
0.8
0.9
1
1.1
1.2
1.3
Normal
LY50
LY75
LY88
LY94
LY97
LY data
Normal data

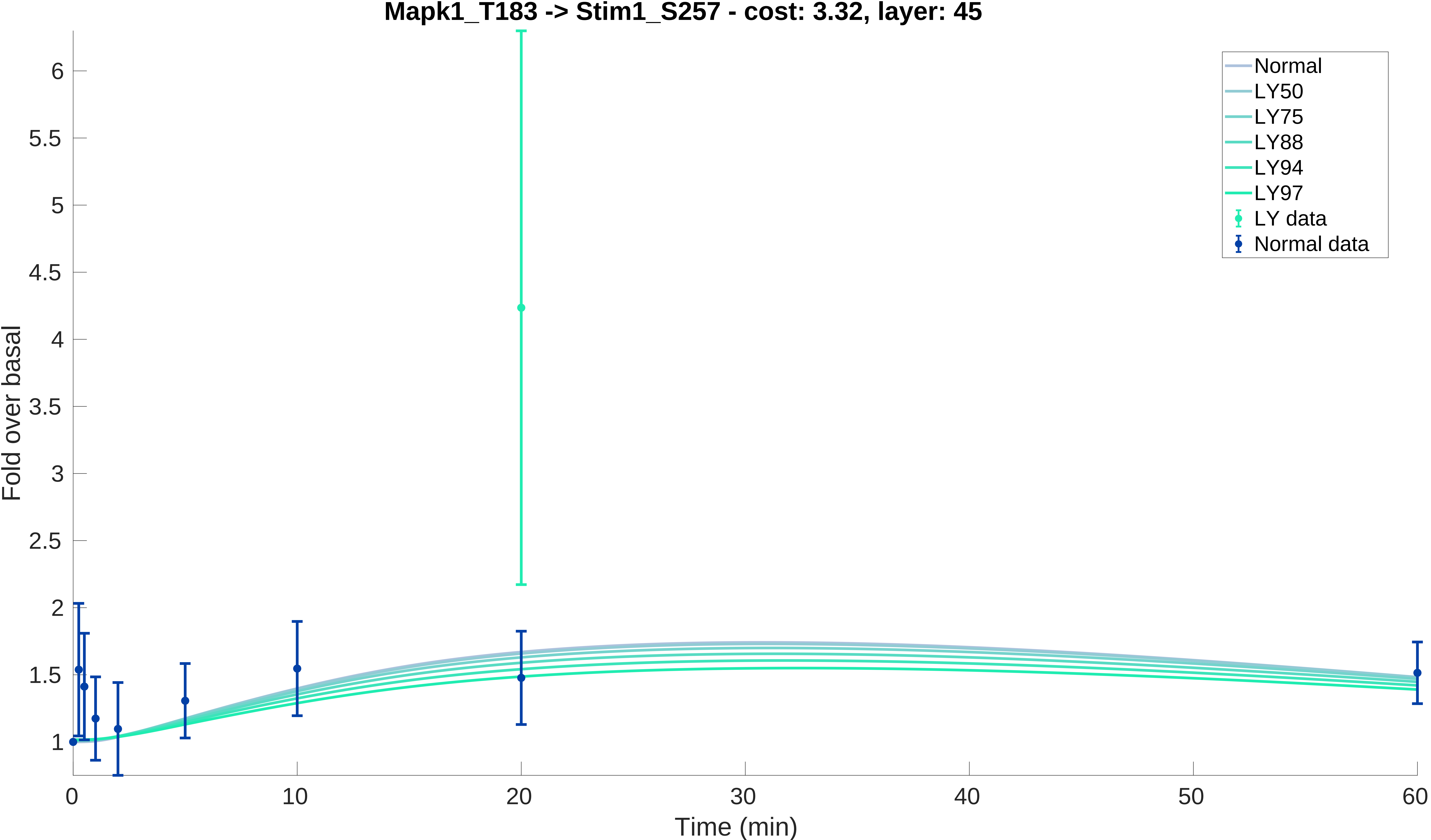

Mapk1_T183 -> Stim1_S257 - cost: 3.32, layer: 45
Fold over basal
Time (min)
0
10
20
30
40
50
60
1
1.5
2
2.5
3
3.5
4
4.5
5
5.5
6
Normal
LY50
LY75
LY88
LY94
LY97
LY data
Normal data

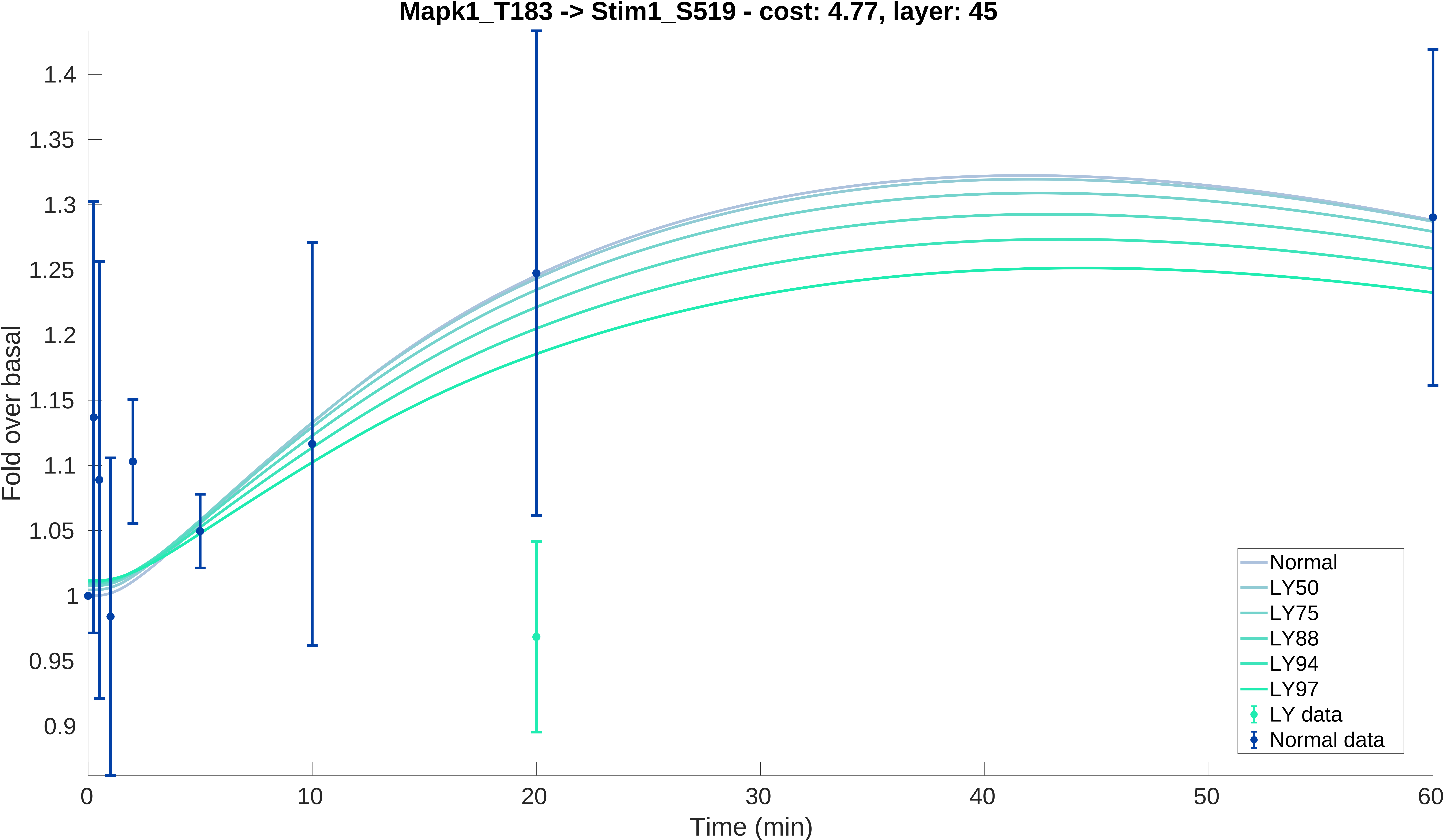
Mapk1_T183 -> Stim1_S519 - cost: 4.77, layer: 45
Fold over basal
1.4
1.35
1.3
1.25
1.2
1.15
1.1
1.05
1
0.95
0.9
0
10
20
30
40
50
60
Time (min)
Normal
LY50
LY75
LY88
LY94
LY97
LY data
Normal data

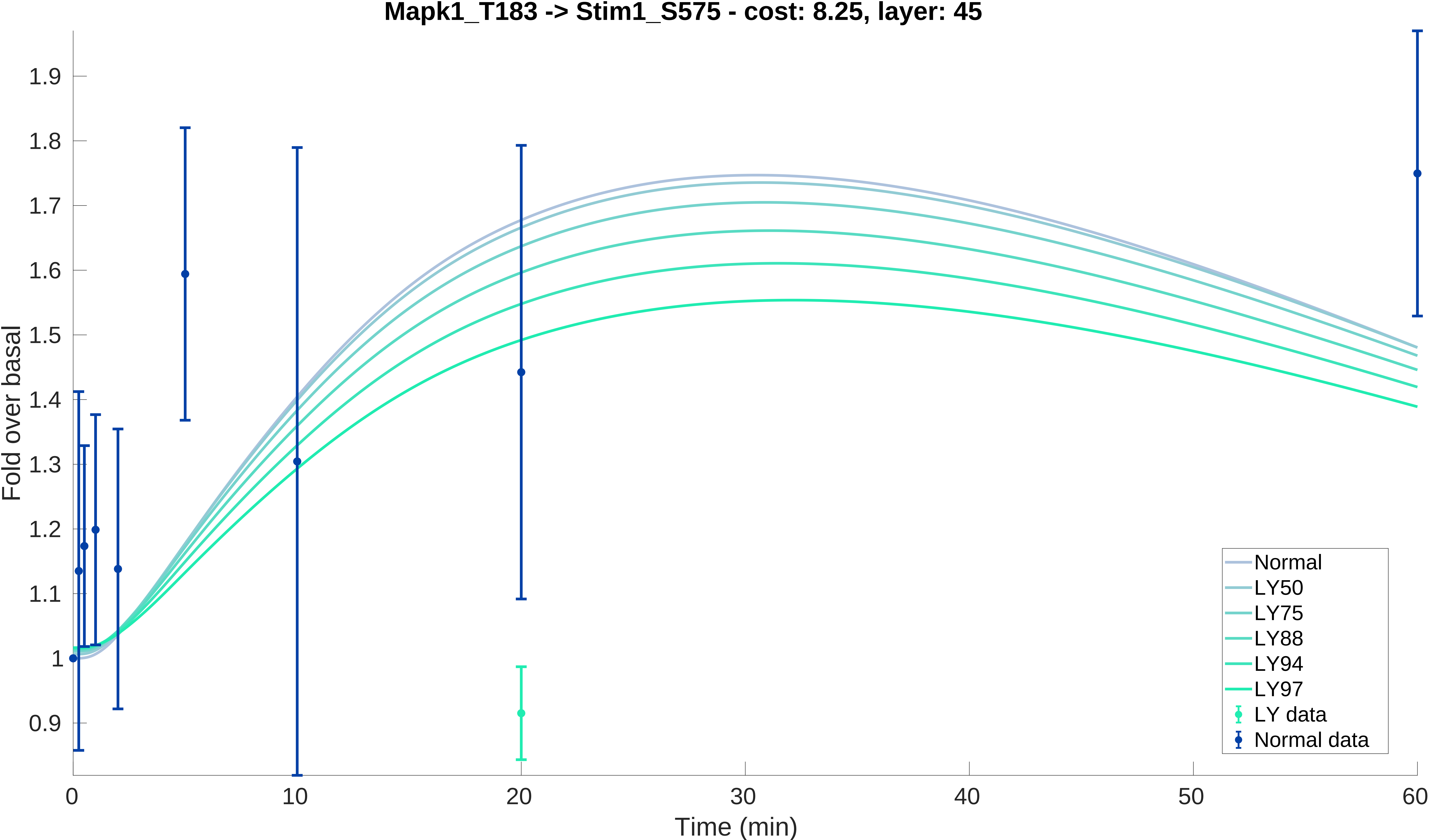

Mapk1_T183 -> Stim1_S575 - cost: 8.25, layer: 45
Fold over basal
Time (min)
0.9
1
1.1
1.2
1.3
1.4
1.5
1.6
1.7
1.8
1.9
0
10
20
30
40
50
60
Normal
LY50
LY75
LY88
LY94
LY97
LY data
Normal data

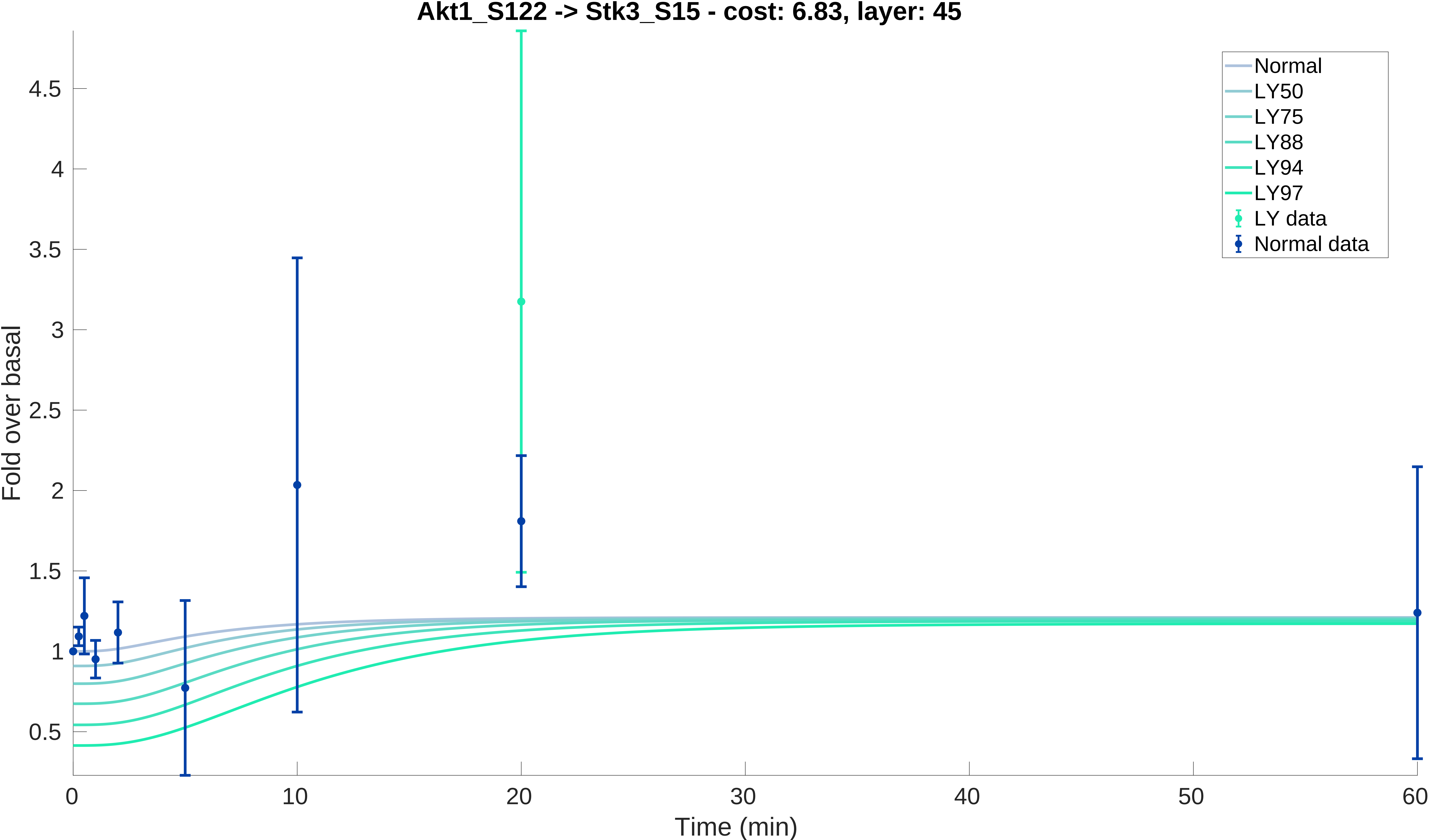
Akt1_S122 -> Stk3_S15 - cost: 6.83, layer: 45
Normal
LY50
LY75
LY88
LY94
LY97
LY data
Normal data
Fold over basal
Time (min)
0.5
1
1.5
2
2.5
3
3.5
4
4.5
0
10
20
30
40
50
60

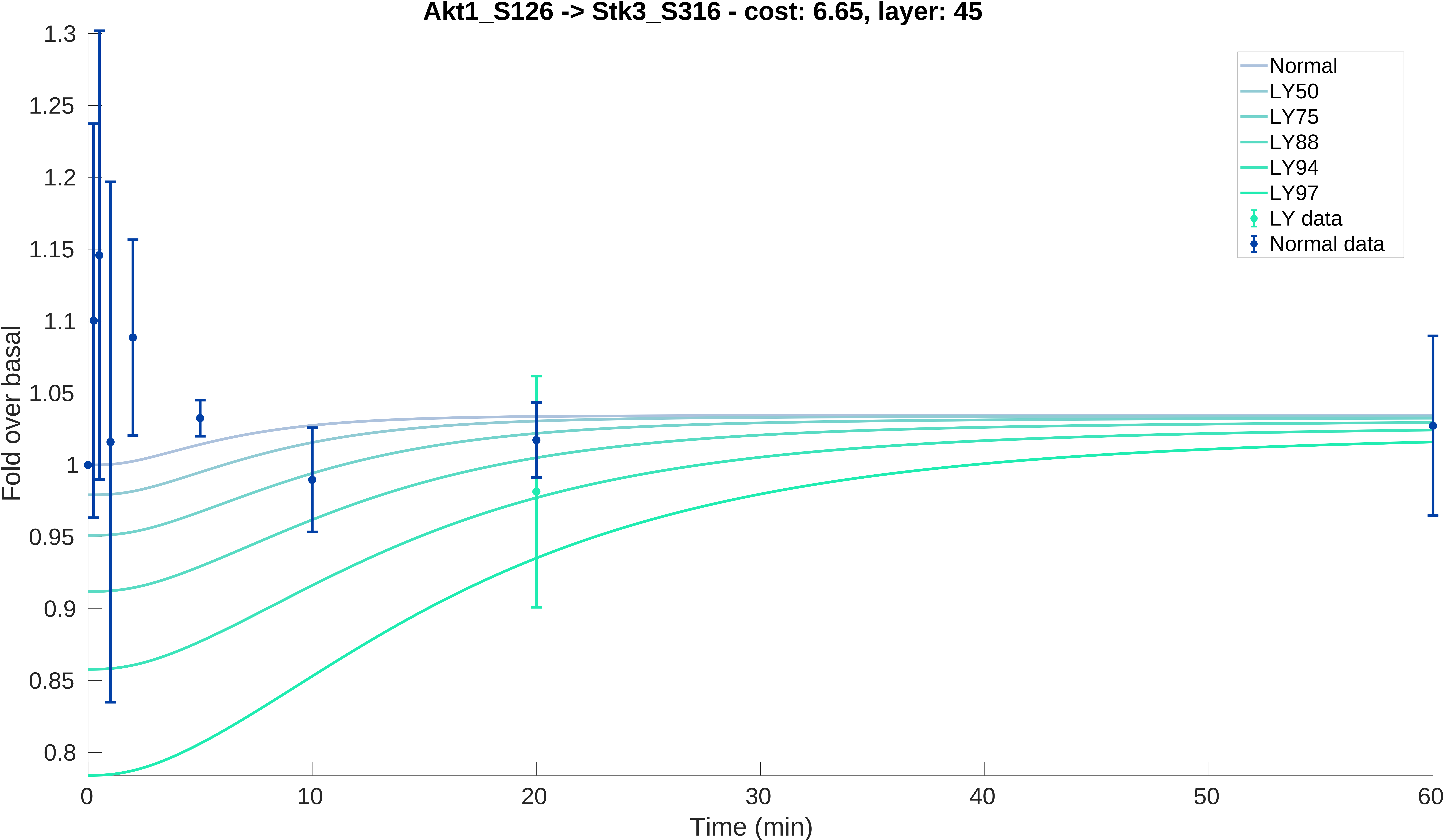

Akt1_S126 -> Stk3_S316 - cost: 6.65, layer: 45
Fold over basal
Time (min)
1.3
1.25
1.2
1.15
1.1
1.05
1
0.95
0.9
0.85
0.8
0
10
20
30
40
50
60
Normal
LY50
LY75
LY88
LY94
LY97
LY data
Normal data

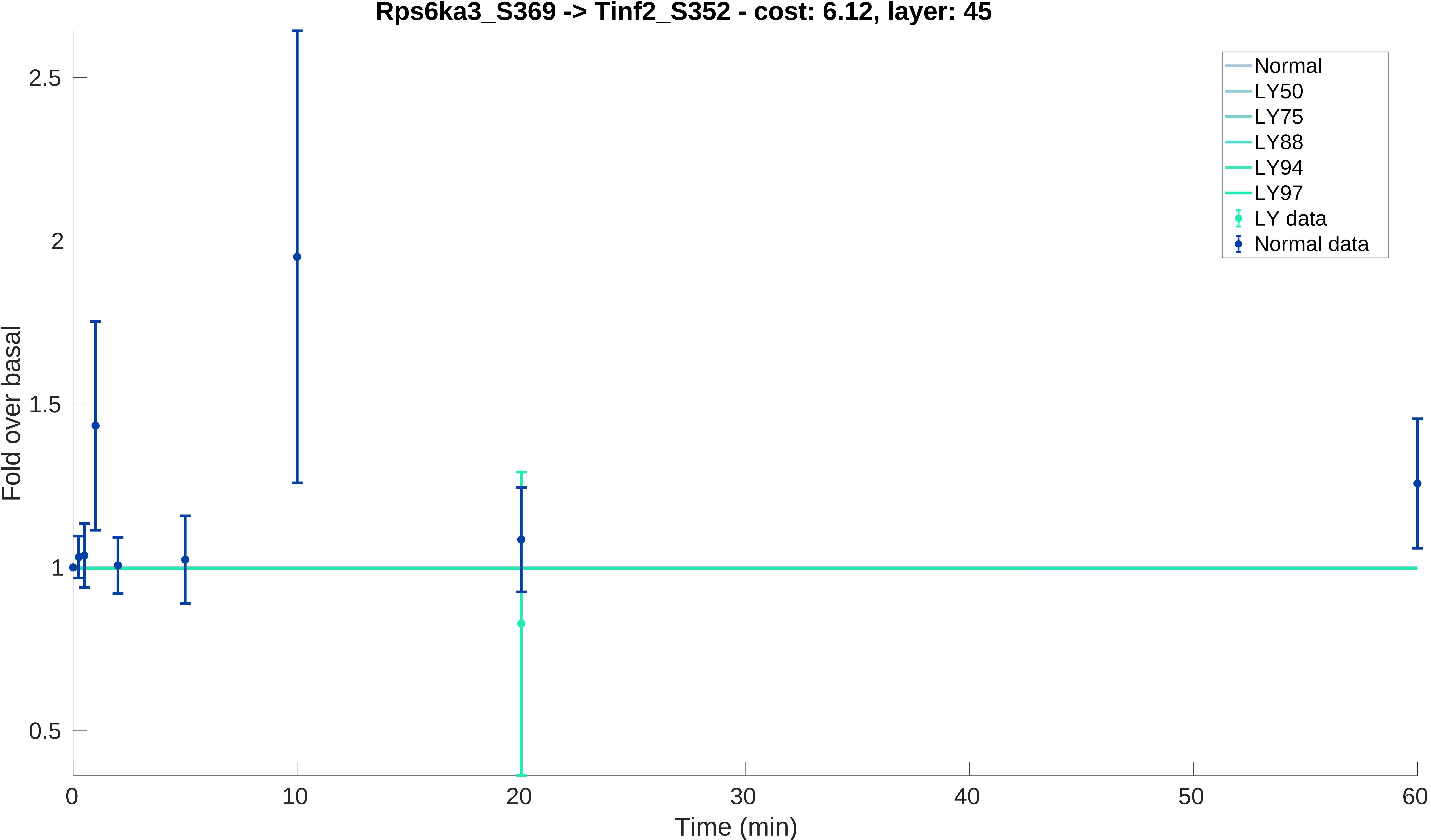

Rps6ka3_S369 -> Tinf2_S352 - cost: 6.12, layer: 45
Fold over basal
2.5
2
1.5
1
0.5
0
10
20
30
40
50
60
Time (min)
Normal
LY50
LY75
LY88
LY94
LY97
LY data
Normal data

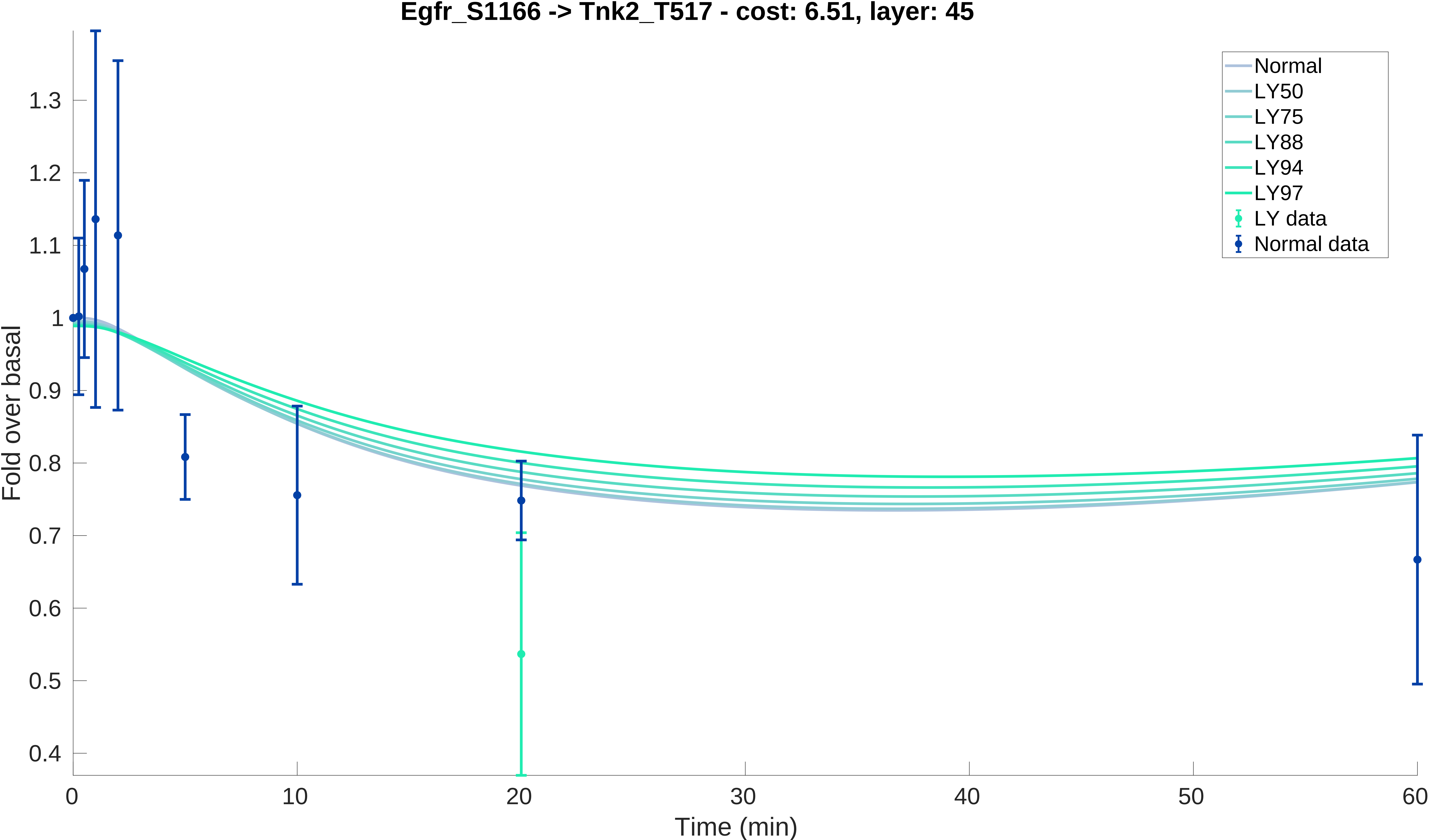

Egfr_S1166 -> Tnk2_T517 - cost: 6.51, layer: 45
Fold over basal
Time (min)
Normal
LY50
LY75
LY88
LY94
LY97
LY data
Normal data
1.3
1.2
1.1
1
0.9
0.8
0.7
0.6
0.5
0.4
0
10
20
30
40
50
60

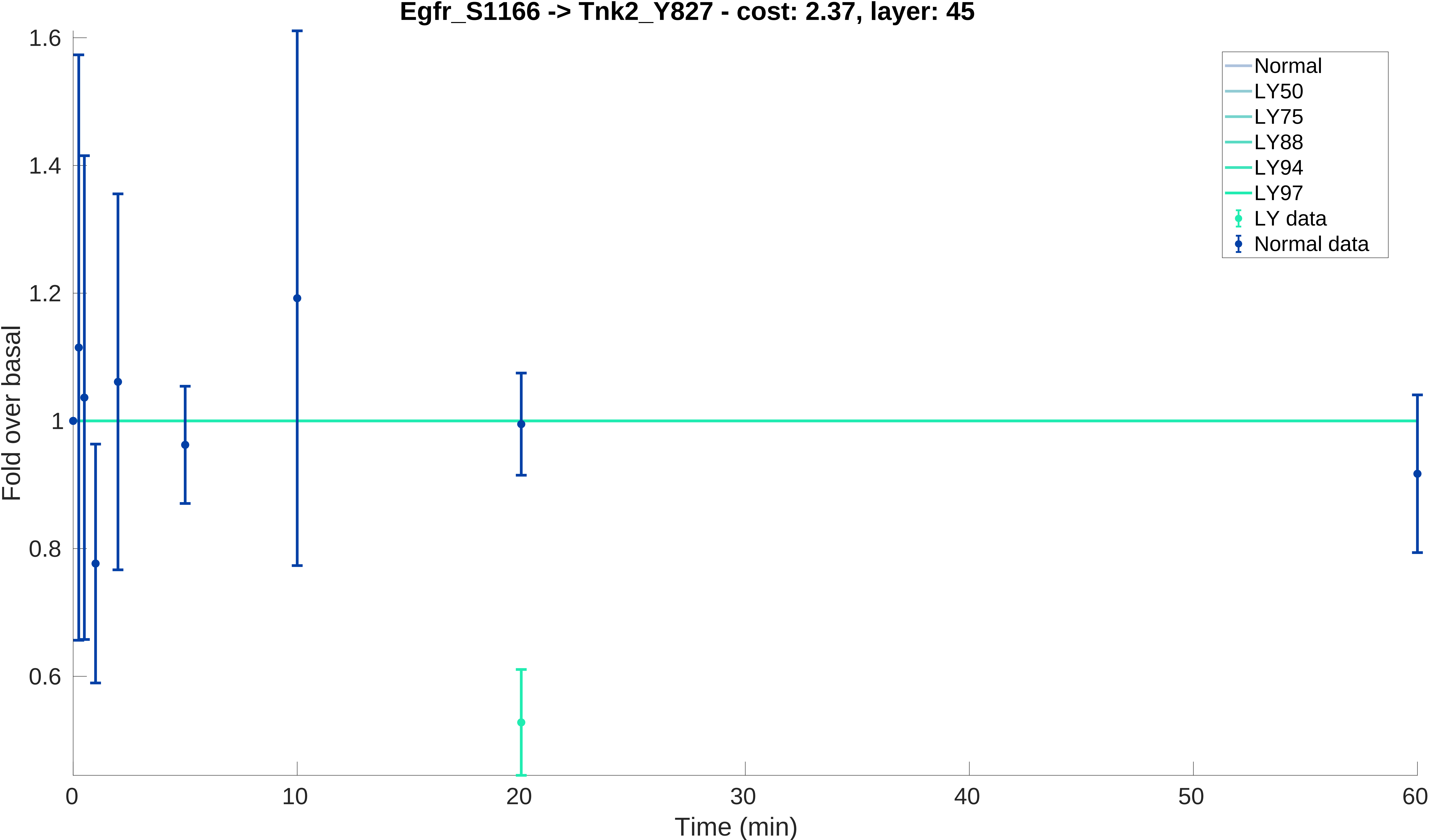

Egfr_S1166 -> Tnk2_Y827 - cost: 2.37, layer: 45
Fold over basal
Time (min)
1.6
1.4
1.2
1
0.8
0.6
0
10
20
30
40
50
60
Normal
LY50
LY75
LY88
LY94
LY97
LY data
Normal data

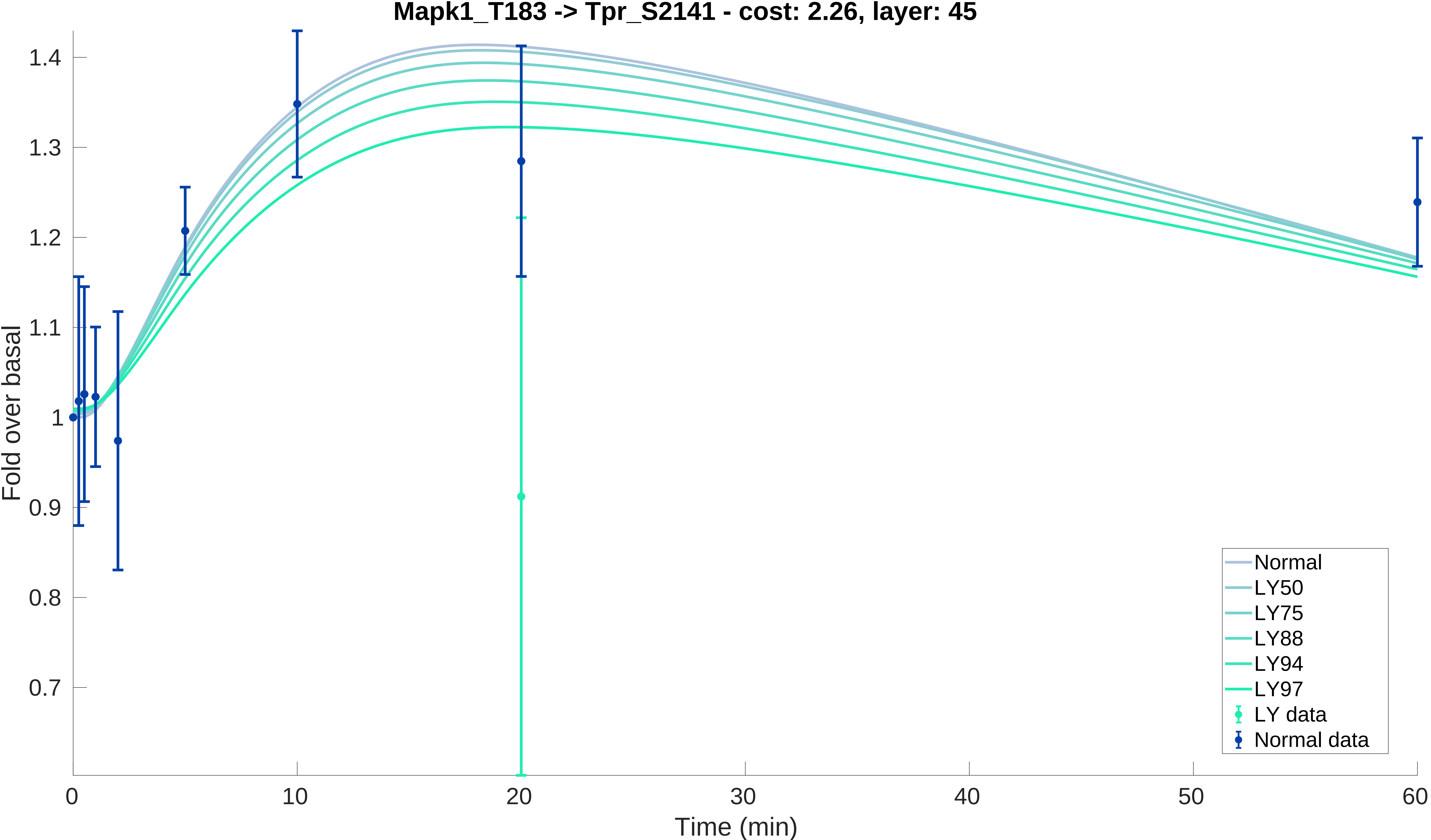

Mapk1_T183 -> Tpr_S2141 - cost: 2.26, layer: 45
Fold over basal
1.4
1.3
1.2
1.1
1
0.9
0.8
0.7
0
10
20
30
40
50
60
Time (min)
Normal
LY50
LY75
LY88
LY94
LY97
LY data
Normal data

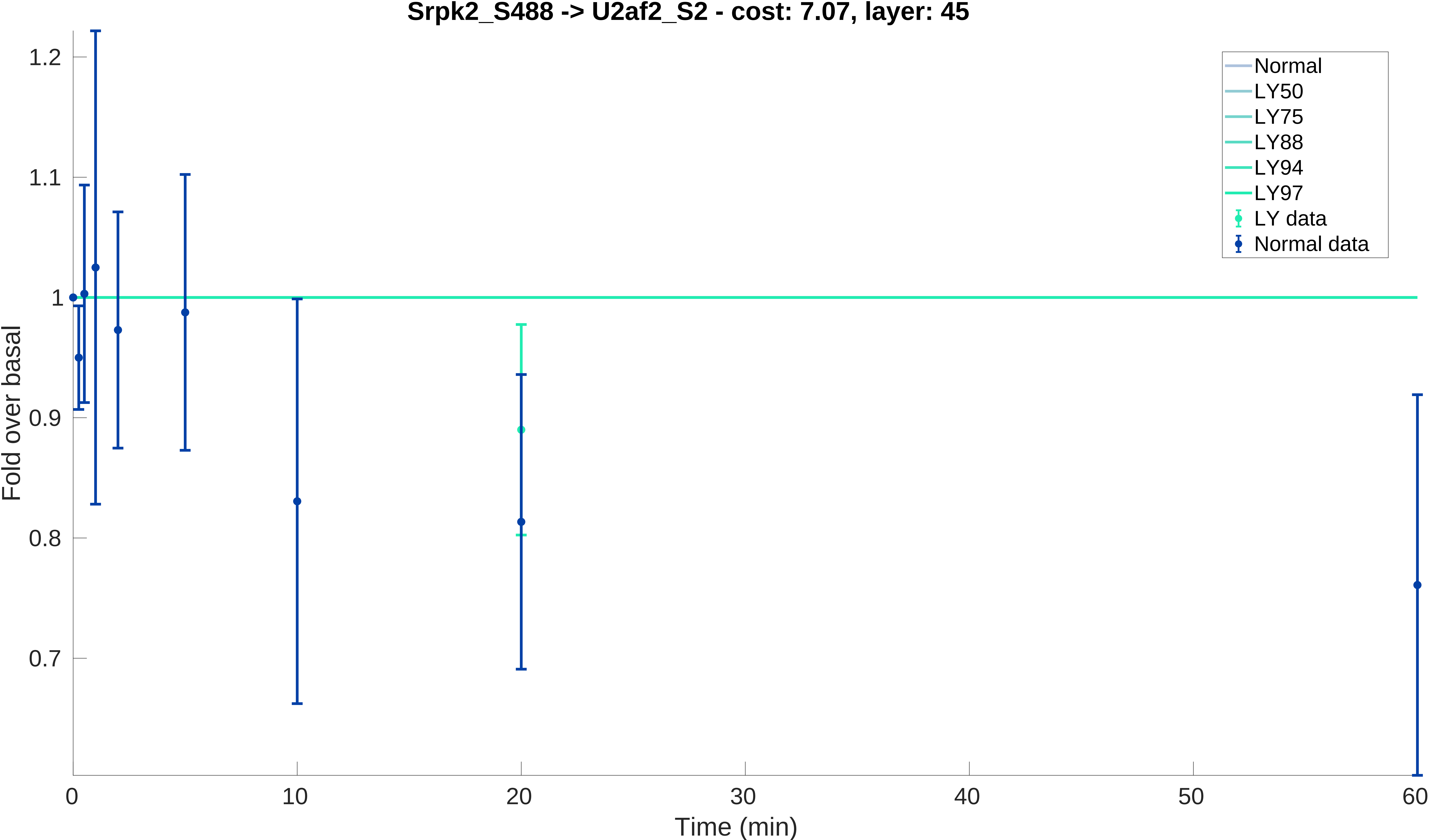
Srpk2_S488 -> U2af2_S2 - cost: 7.07, layer: 45
Fold over basal
Time (min)
1.2
1.1
1
0.9
0.8
0.7
0
10
20
30
40
50
60
Normal
LY50
LY75
LY88
LY94
LY97
LY data
Normal data

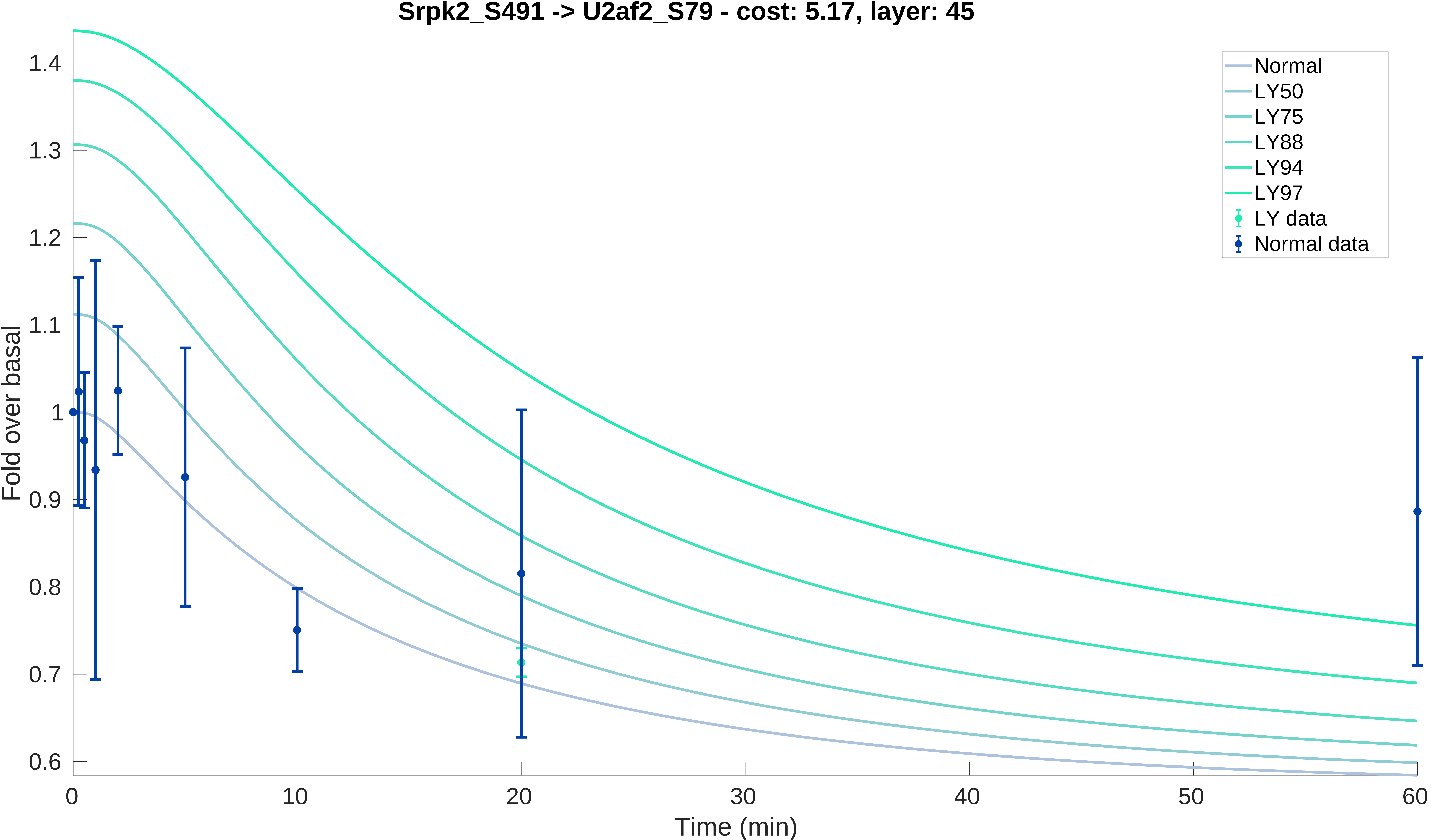

Srpk2_S491 -> U2af2_S79 - cost: 5.17, layer: 45
Fold over basal
1.4
1.3
1.2
1.1
1
0.9
0.8
0.7
0.6
0
10
20
30
40
50
60
Time (min)
Normal
LY50
LY75
LY88
LY94
LY97
LY data
Normal data

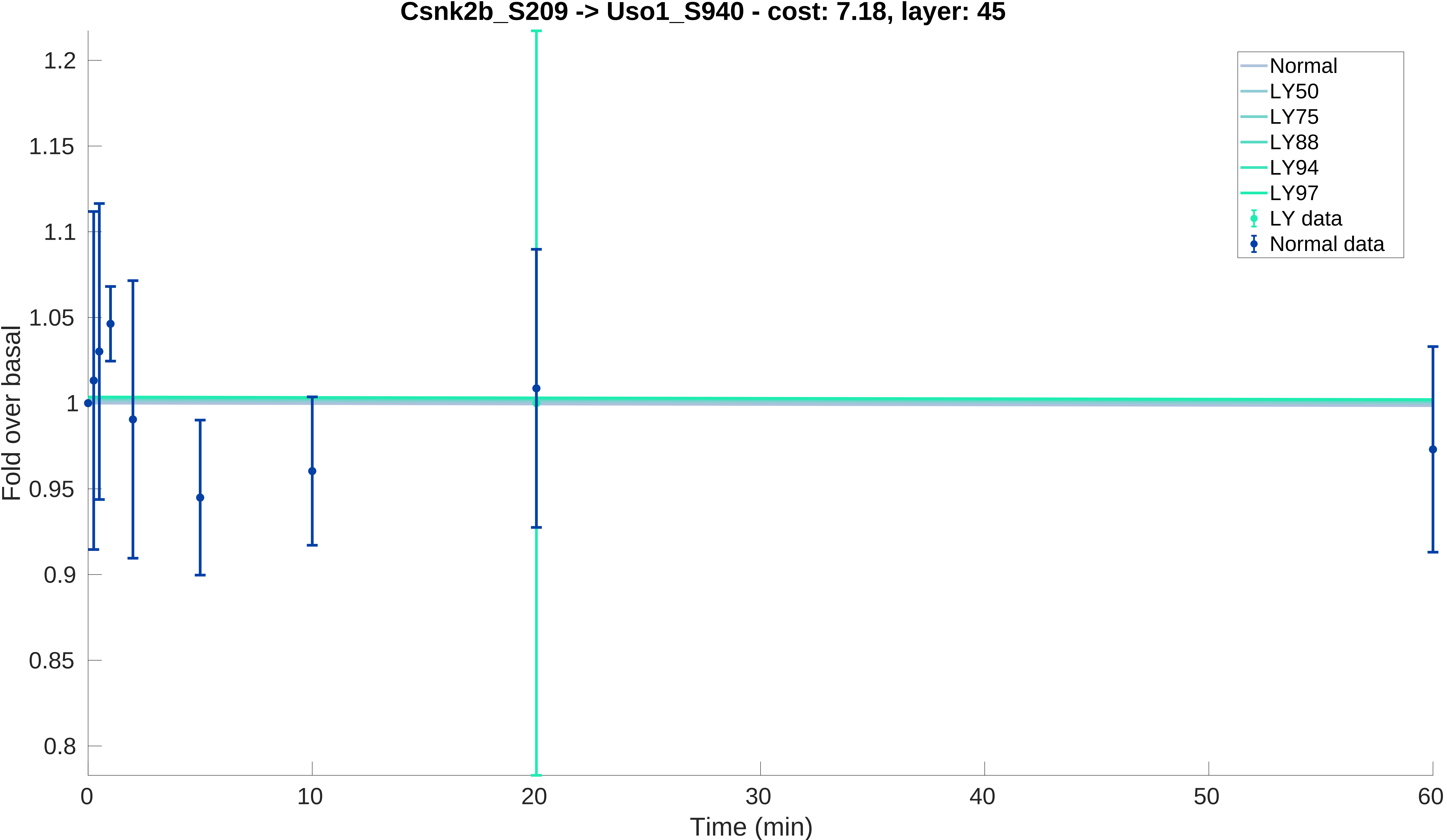
Csnk2b_S209 -> Uso1_S940 - cost: 7.18, layer: 45
Fold over basal
Time (min)
1.2
1.15
1.1
1.05
1
0.95
0.9
0.85
0.8
0
10
20
30
40
50
60
Normal
LY50
LY75
LY88
LY94
LY97
LY data
Normal data

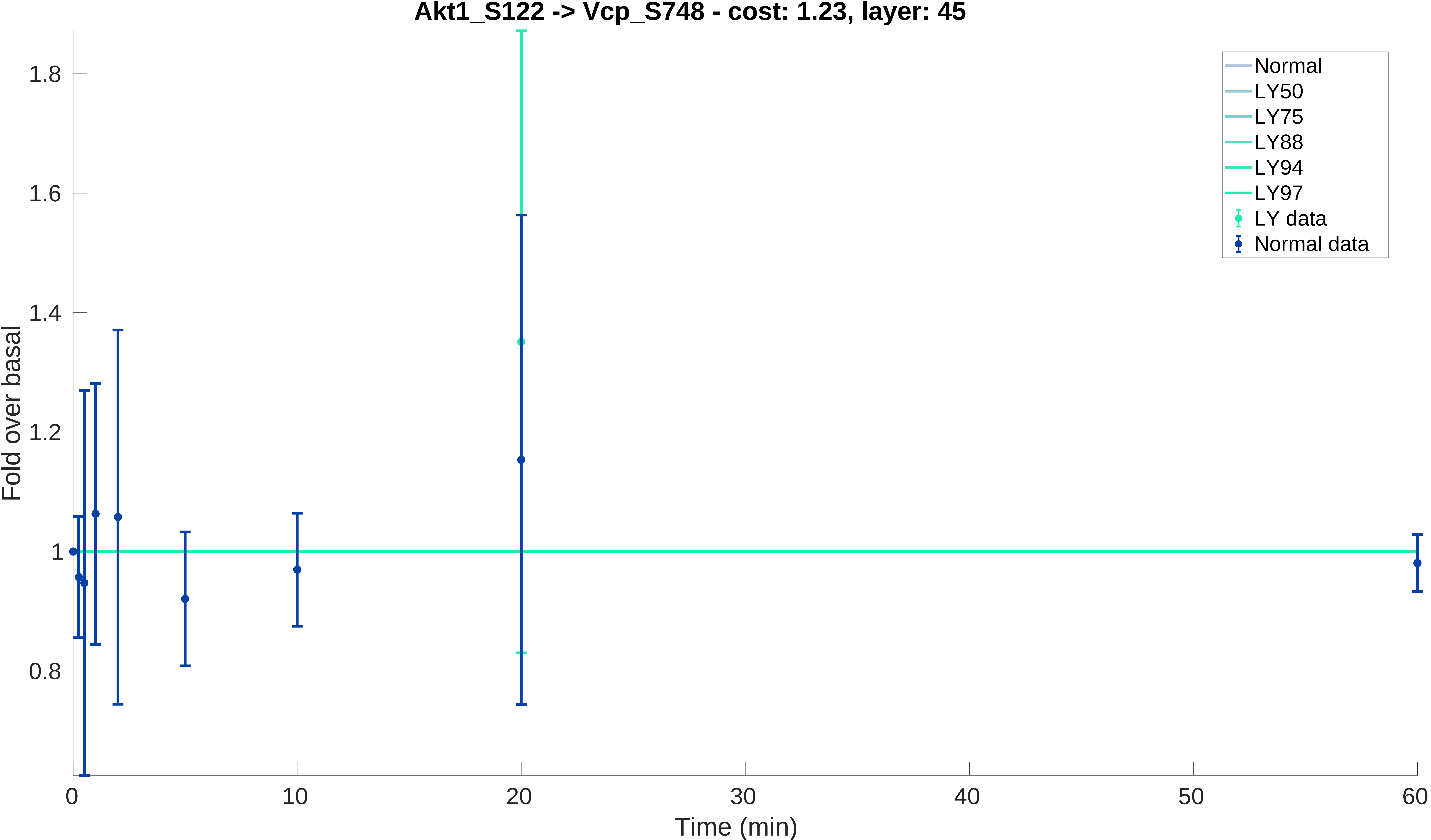

Akt1_S122 -> Vcp_S748 - cost: 1.23, layer: 45
Normal
LY50
LY75
LY88
LY94
LY97
LY data
Normal data
Fold over basal
1.8
1.6
1.4
1.2
1
0.8
0
10
20
30
40
50
60
Time (min)

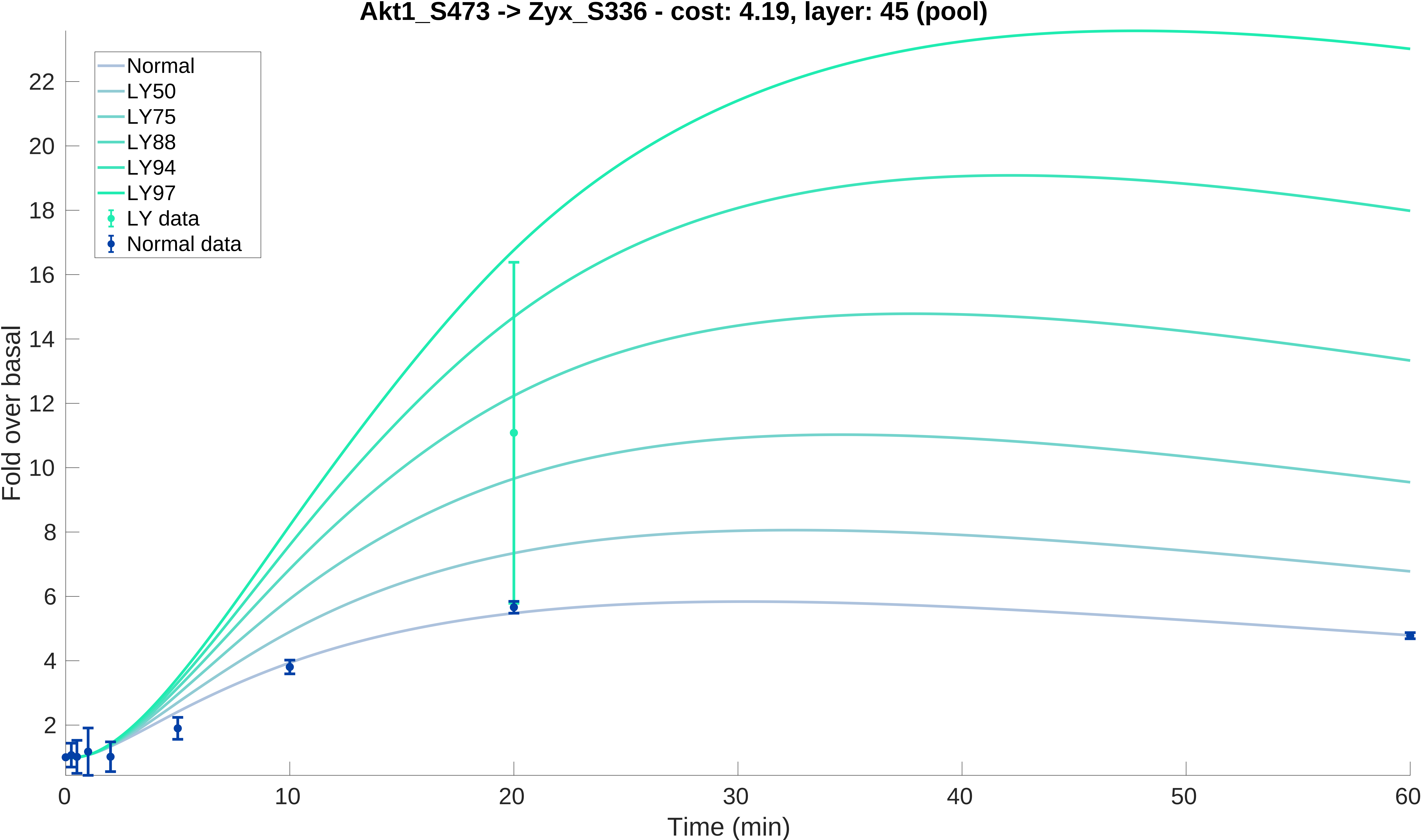

Akt1_S473 -> Zyx_S336 - cost: 4.19, layer: 45 (pool)
Normal
LY50
LY75
LY88
LY94
LY97
LY data
Normal data
Fold over basal
Time (min)
2
4
6
8
10
12
14
16
18
20
22
0
10
20
30
40
50
60

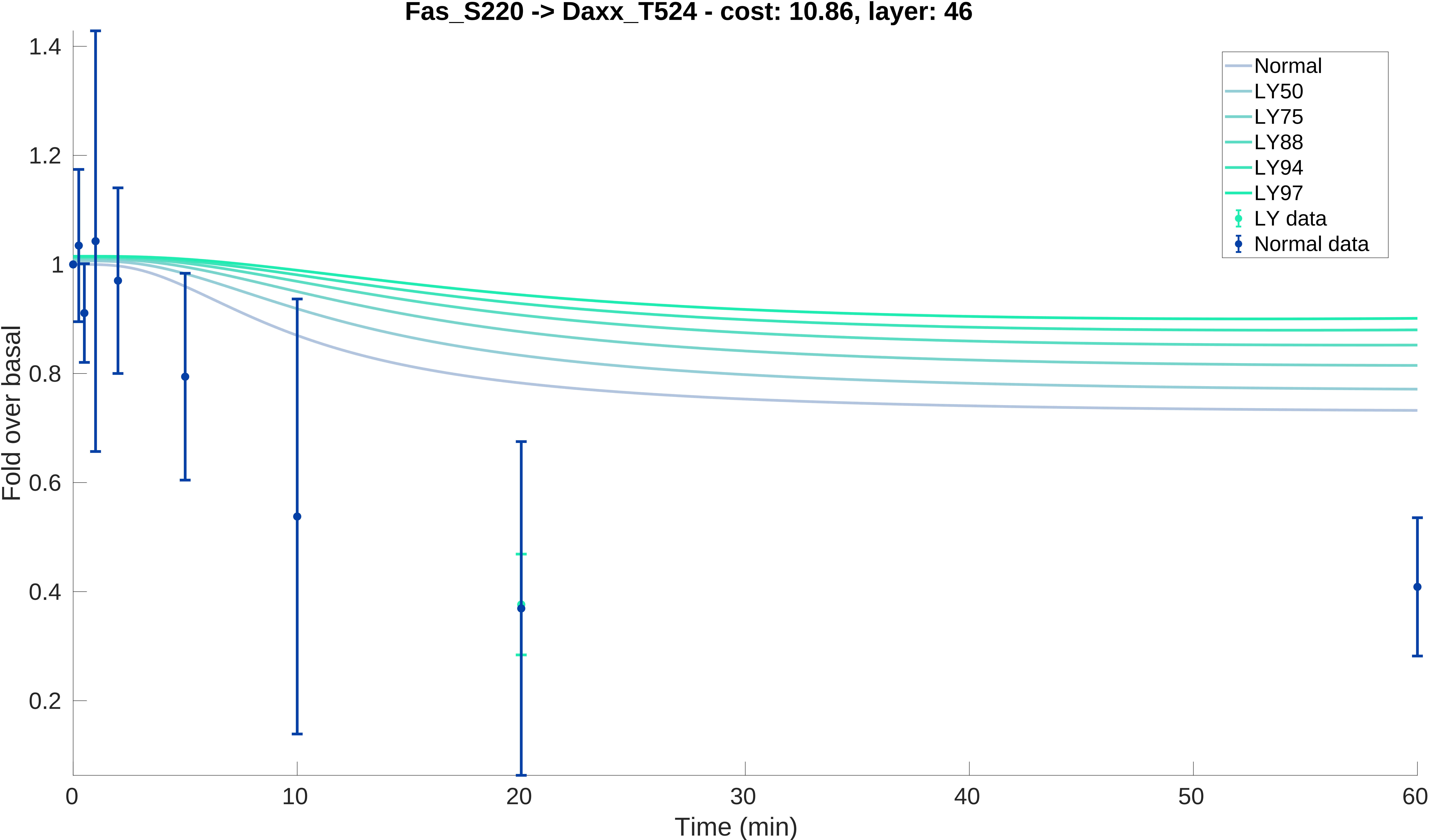

Fas_S220 -> Daxx_T524 - cost: 10.86, layer: 46
Fold over basal
Time (min)
0
10
20
30
40
50
60
0.2
0.4
0.6
0.8
1
1.2
1.4
Normal
LY50
LY75
LY88
LY94
LY97
LY data
Normal data

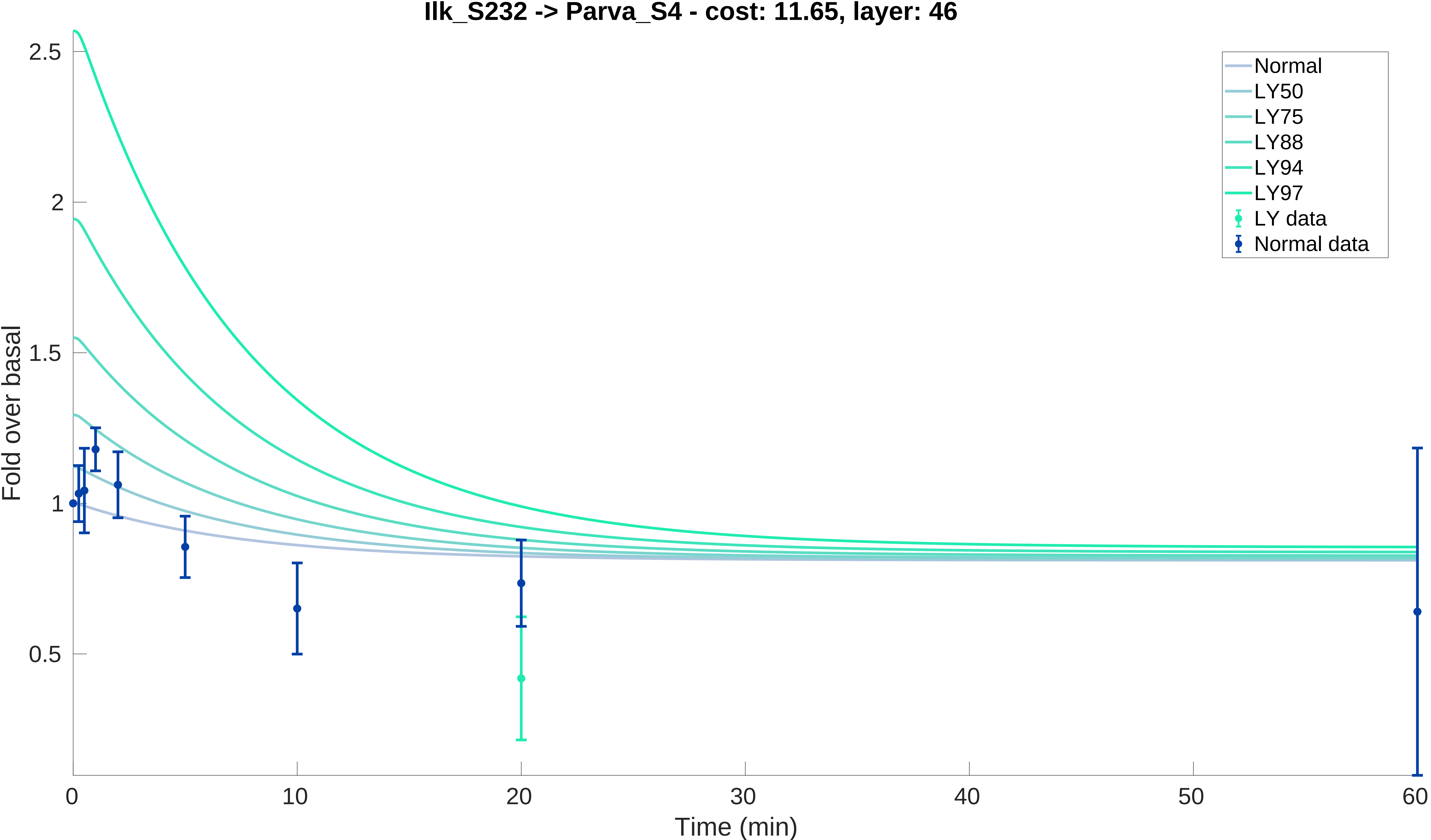

Ilk_S232 -> Parva_S4 - cost: 11.65, layer: 46
Fold over basal
Time (min)
0
10
20
30
40
50
60
0.5
1
1.5
2
2.5
Normal
LY50
LY75
LY88
LY94
LY97
LY data
Normal data

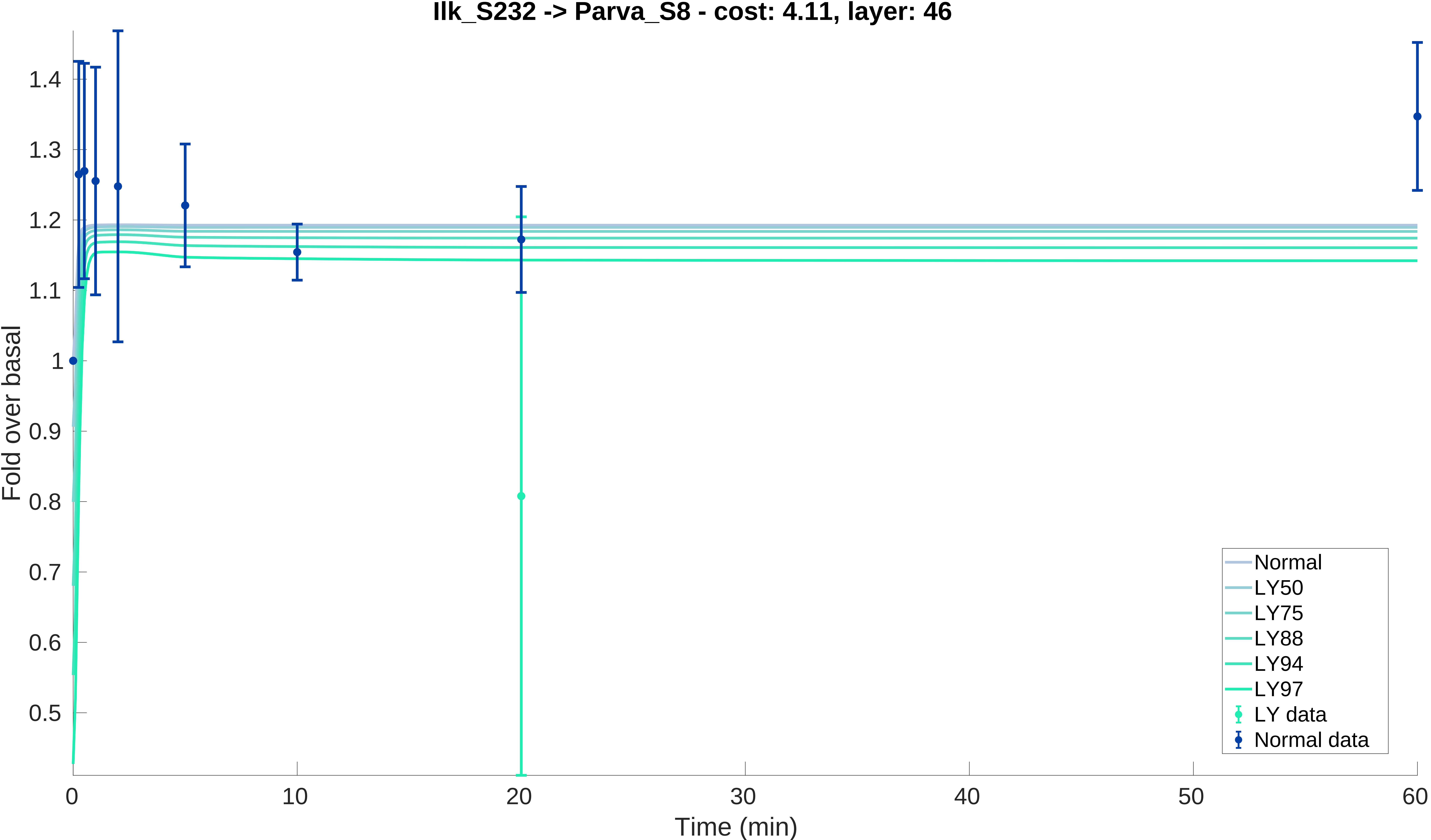
Ilk_S232 -> Parva_S8 - cost: 4.11, layer: 46
Fold over basal
1.4
1.3
1.2
1.1
1
0.9
0.8
0.7
0.6
0.5
0
10
20
30
40
50
60
Time (min)
Normal
LY50
LY75
LY88
LY94
LY97
LY data
Normal data

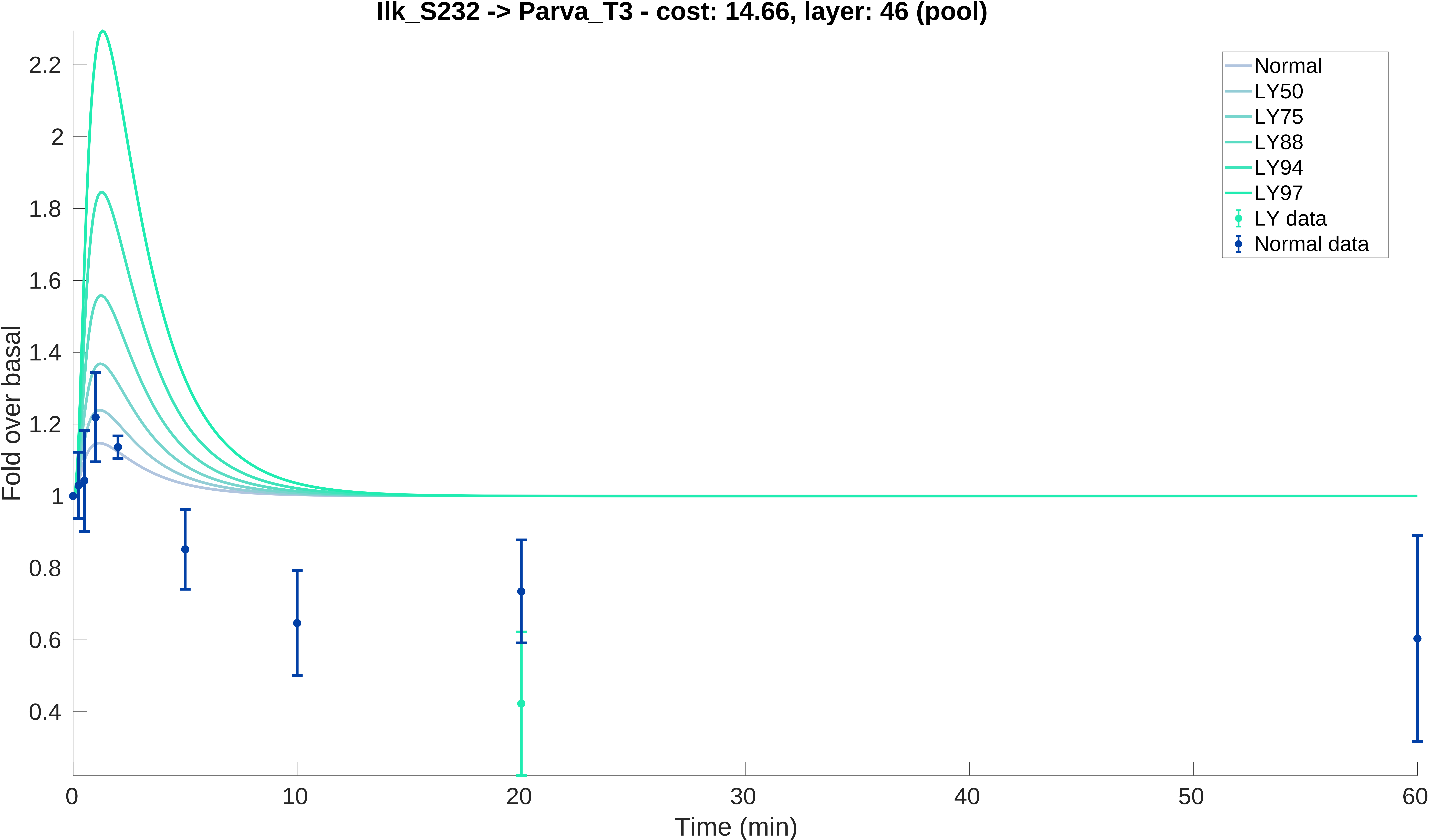

Ilk_S232 -> Parva_T3 - cost: 14.66, layer: 46 (pool)
Fold over basal
Time (min)
0
10
20
30
40
50
60
0.4
0.6
0.8
1
1.2
1.4
1.6
1.8
2
2.2
Normal
LY50
LY75
LY88
LY94
LY97
LY data
Normal data

### Ilk_S232 -> Ppp1r12a_S870 - cost: 4.96, layer: 46

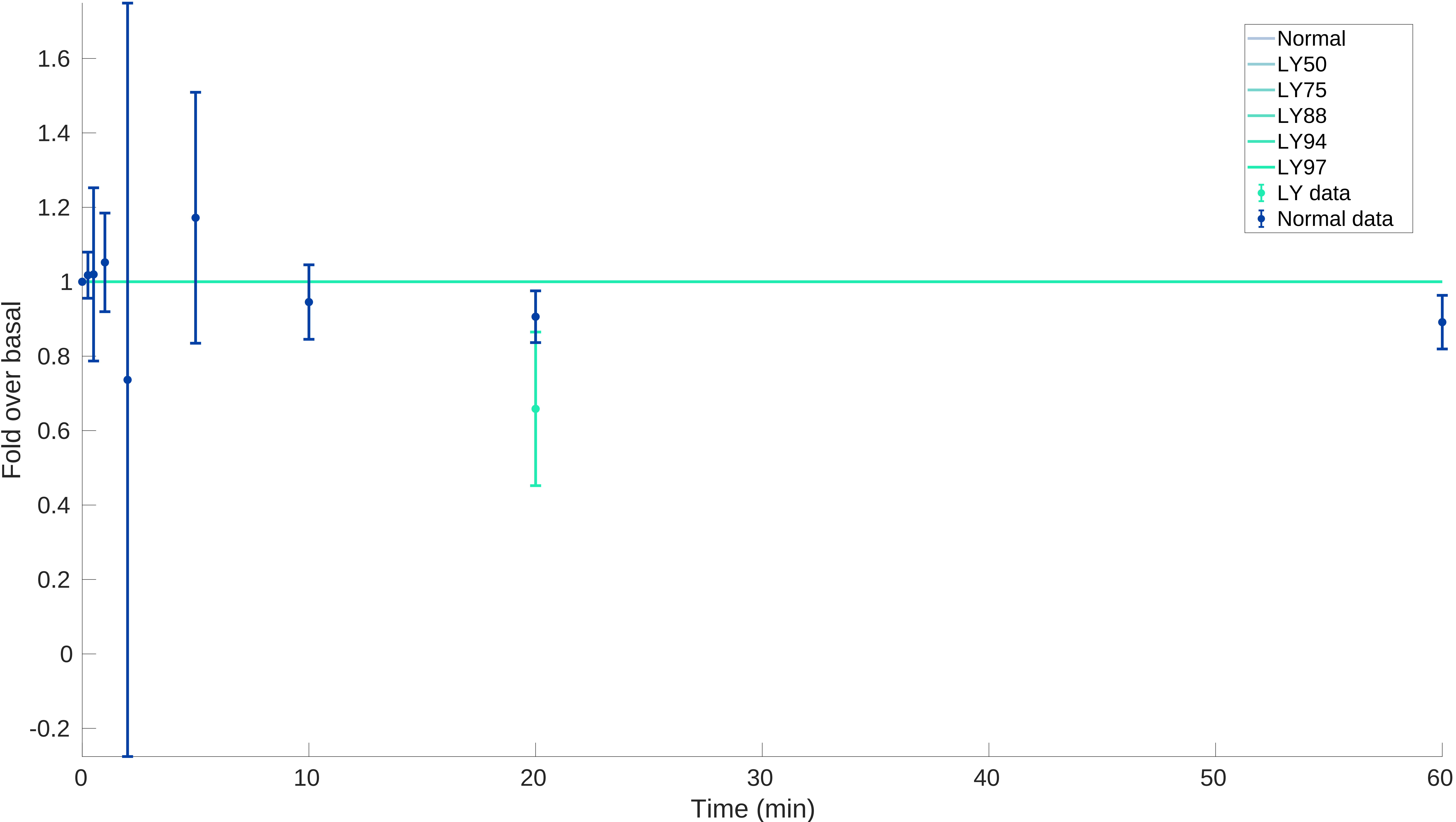

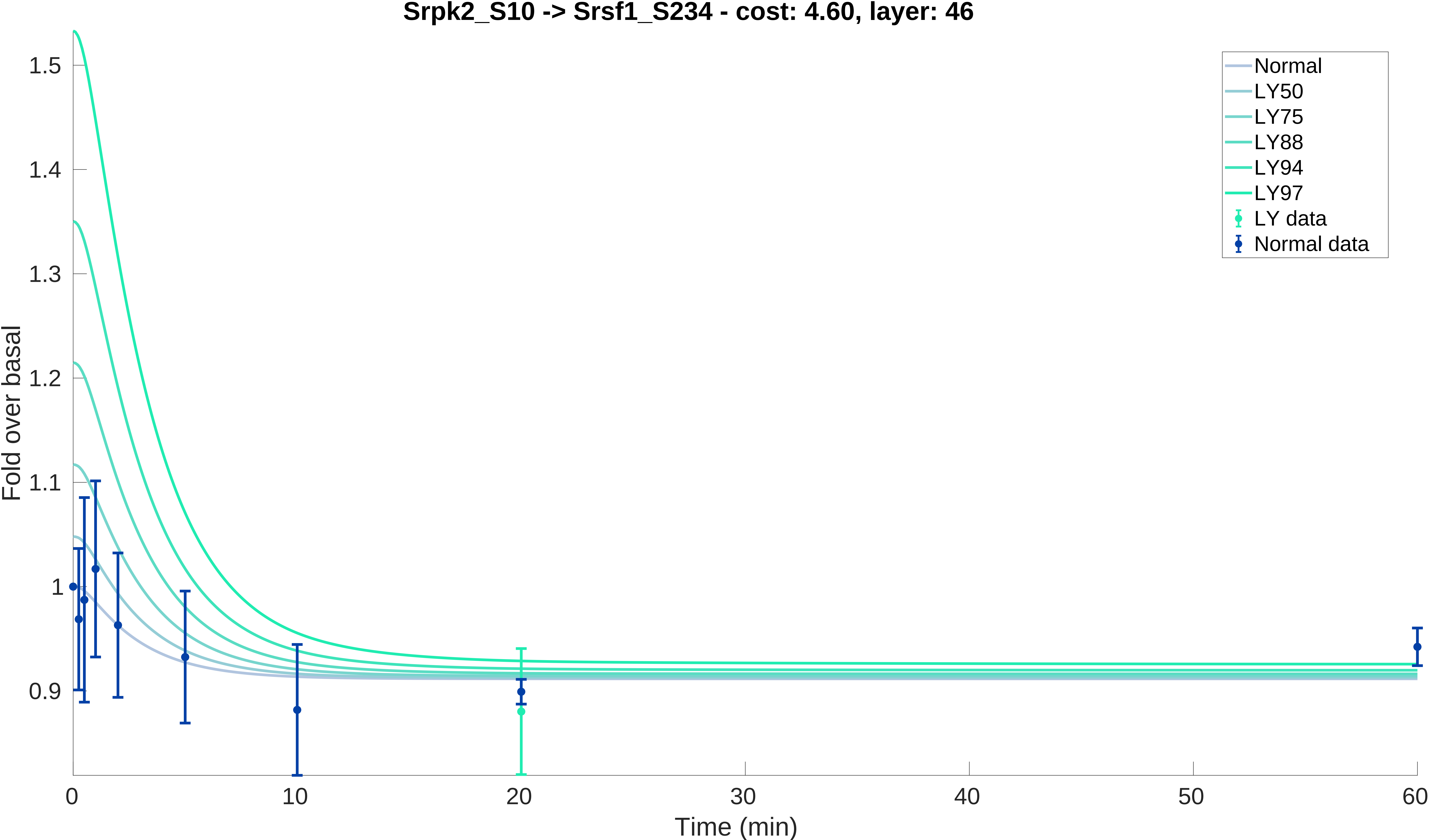

Srpk2_S10 -> Srsf1_S234 - cost: 4.60, layer: 46
Fold over basal
Time (min)
0.9
1
1.1
1.2
1.3
1.4
1.5
0
10
20
30
40
50
60
Normal
LY50
LY75
LY88
LY94
LY97
LY data
Normal data

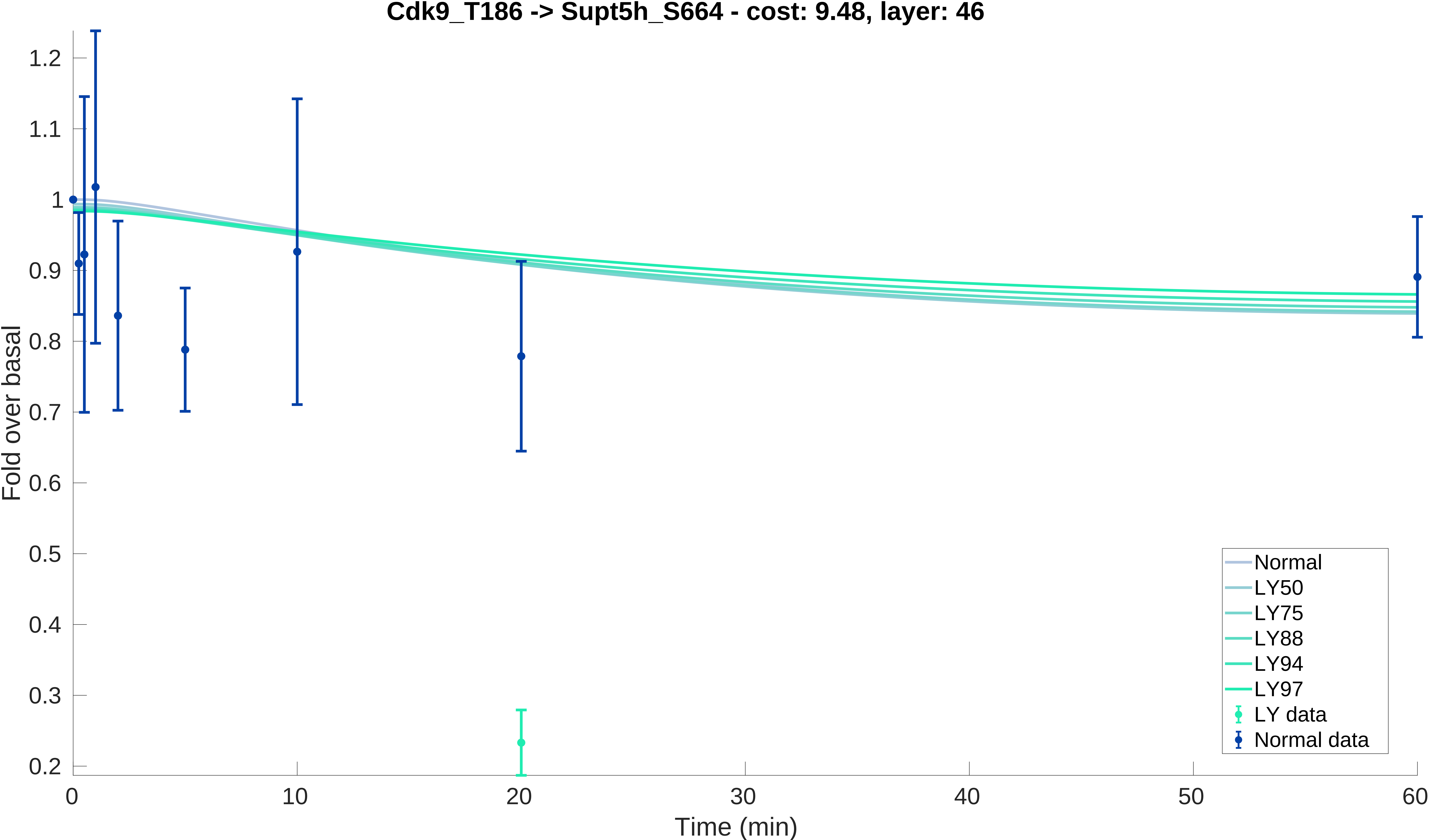

Cdk9_T186 -> Supt5h_S664 - cost: 9.48, layer: 46
Fold over basal
Time (min)
1.2
1.1
1
0.9
0.8
0.7
0.6
0.5
0.4
0.3
0.2
0
10
20
30
40
50
60
Normal
LY50
LY75
LY88
LY94
LY97
LY data
Normal data

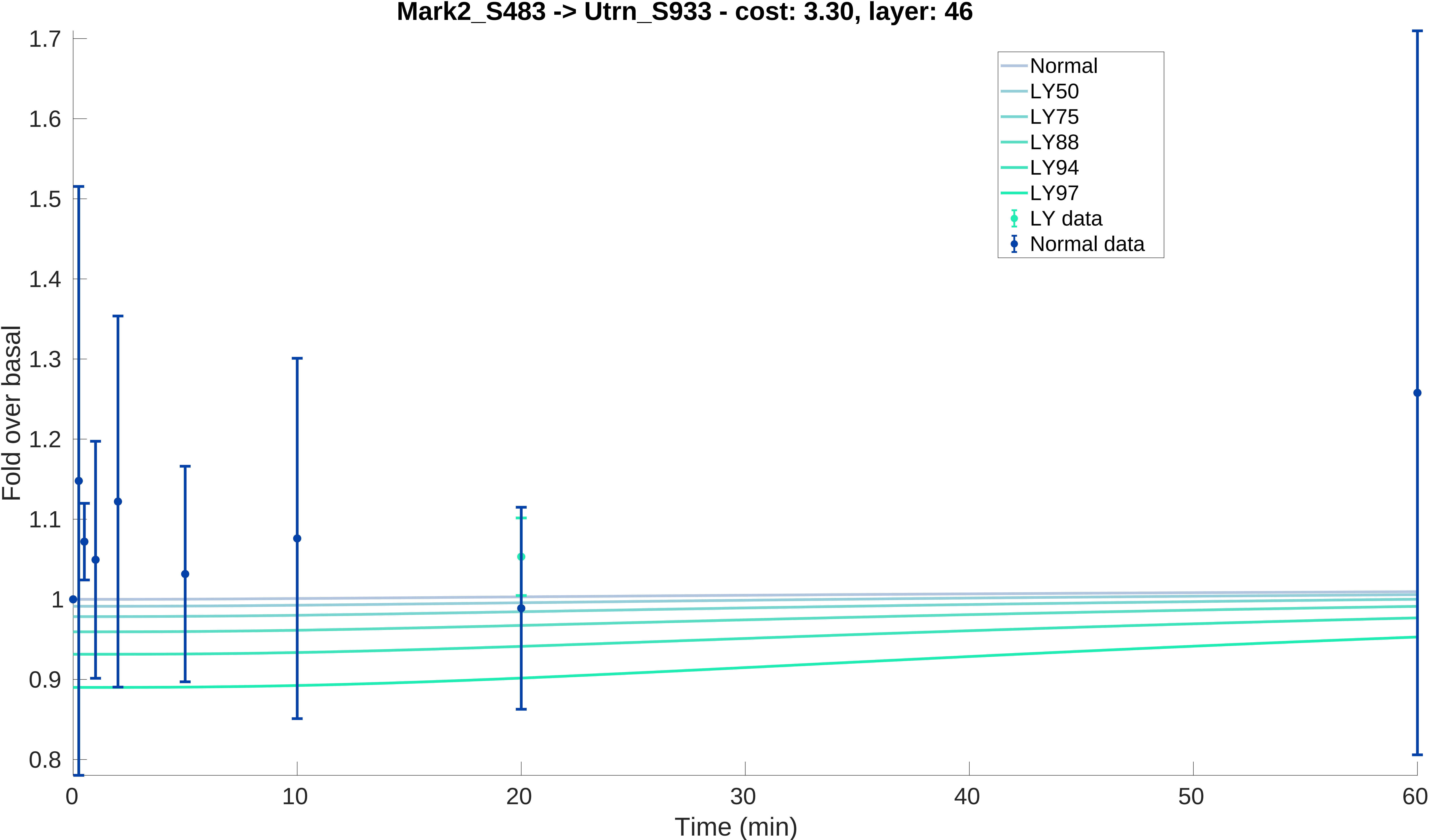

Mark2_S483 -> Utrn_S933 - cost: 3.30, layer: 46
Fold over basal
Time (min)
1.7
1.6
1.5
1.4
1.3
1.2
1.1
1
0.9
0.8
0
10
20
30
40
50
60
Normal
LY50
LY75
LY88
LY94
LY97
LY data
Normal data

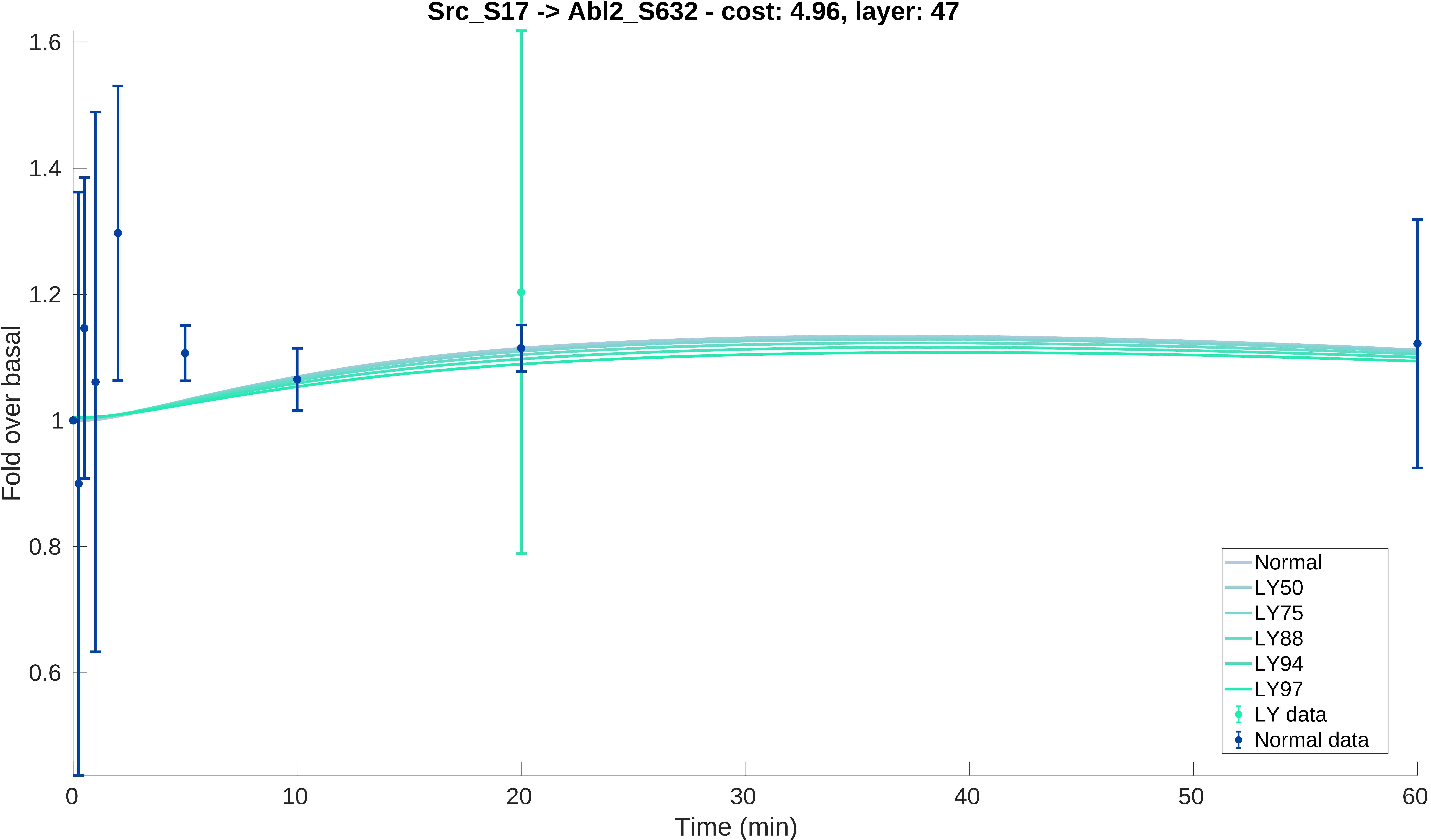

Src_S17 -> Abl2_S632 - cost: 4.96, layer: 47
Fold over basal
1.6
1.4
1.2
1
0.8
0.6
0
10
20
30
40
50
60
Time (min)
Normal
LY50
LY75
LY88
LY94
LY97
LY data
Normal data

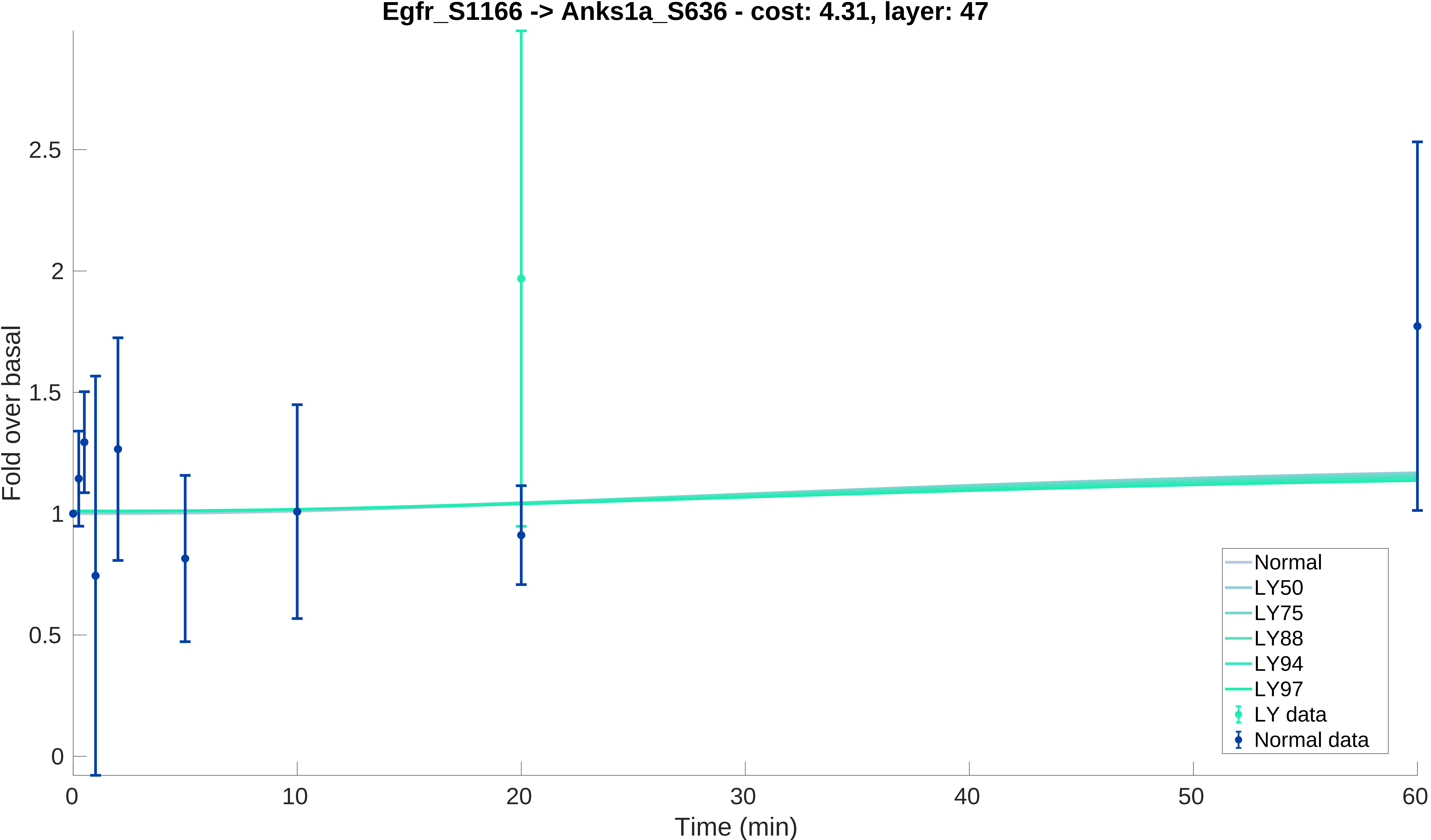

Egfr_S1166 -> Anks1a_S636 - cost: 4.31, layer: 47
Fold over basal
Time (min)
0
0.5
1
1.5
2
2.5
0
10
20
30
40
50
60
Normal
LY50
LY75
LY88
LY94
LY97
LY data
Normal data

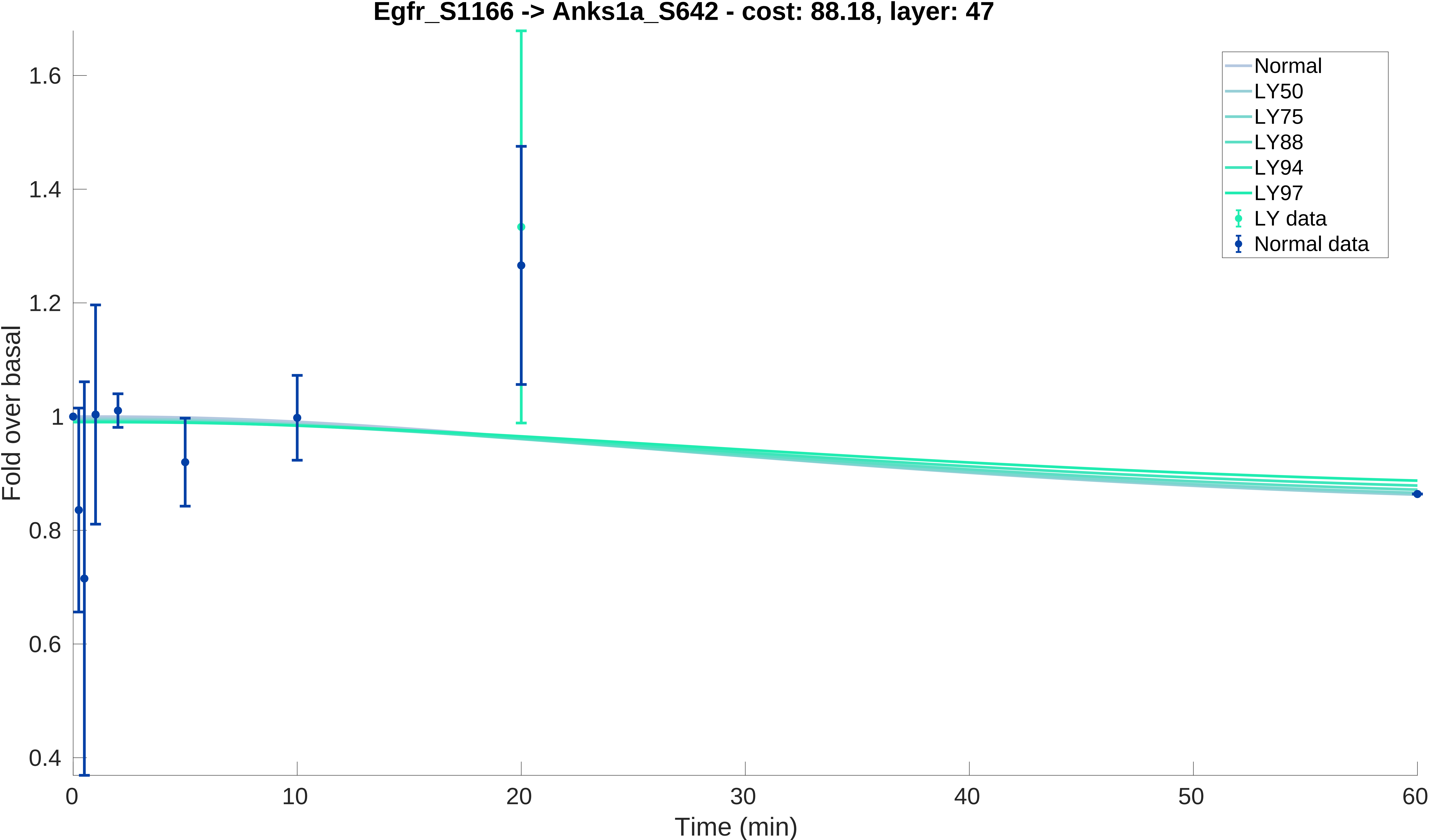

Egfr_S1166 -> Anks1a_S642 - cost: 88.18, layer: 47
Fold over basal
1.6
1.4
1.2
1
0.8
0.6
0.4
0
10
20
30
40
50
60
Time (min)
Normal
LY50
LY75
LY88
LY94
LY97
LY data
Normal data

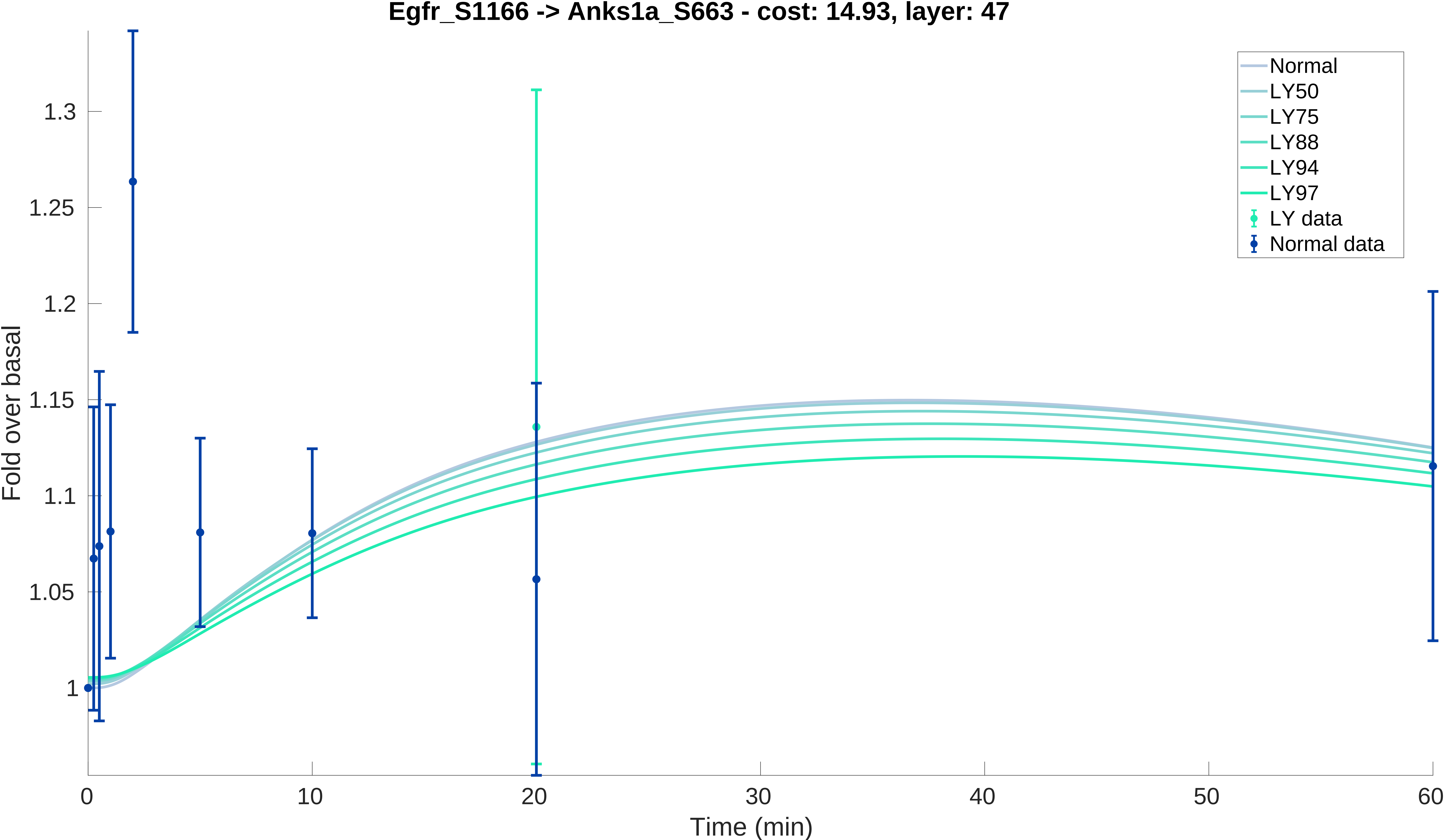
Egfr_S1166 -> Anks1a_S663 - cost: 14.93, layer: 47
Fold over basal
Time (min)
1
1.05
1.1
1.15
1.2
1.25
1.3
0
10
20
30
40
50
60
Normal
LY50
LY75
LY88
LY94
LY97
LY data
Normal data

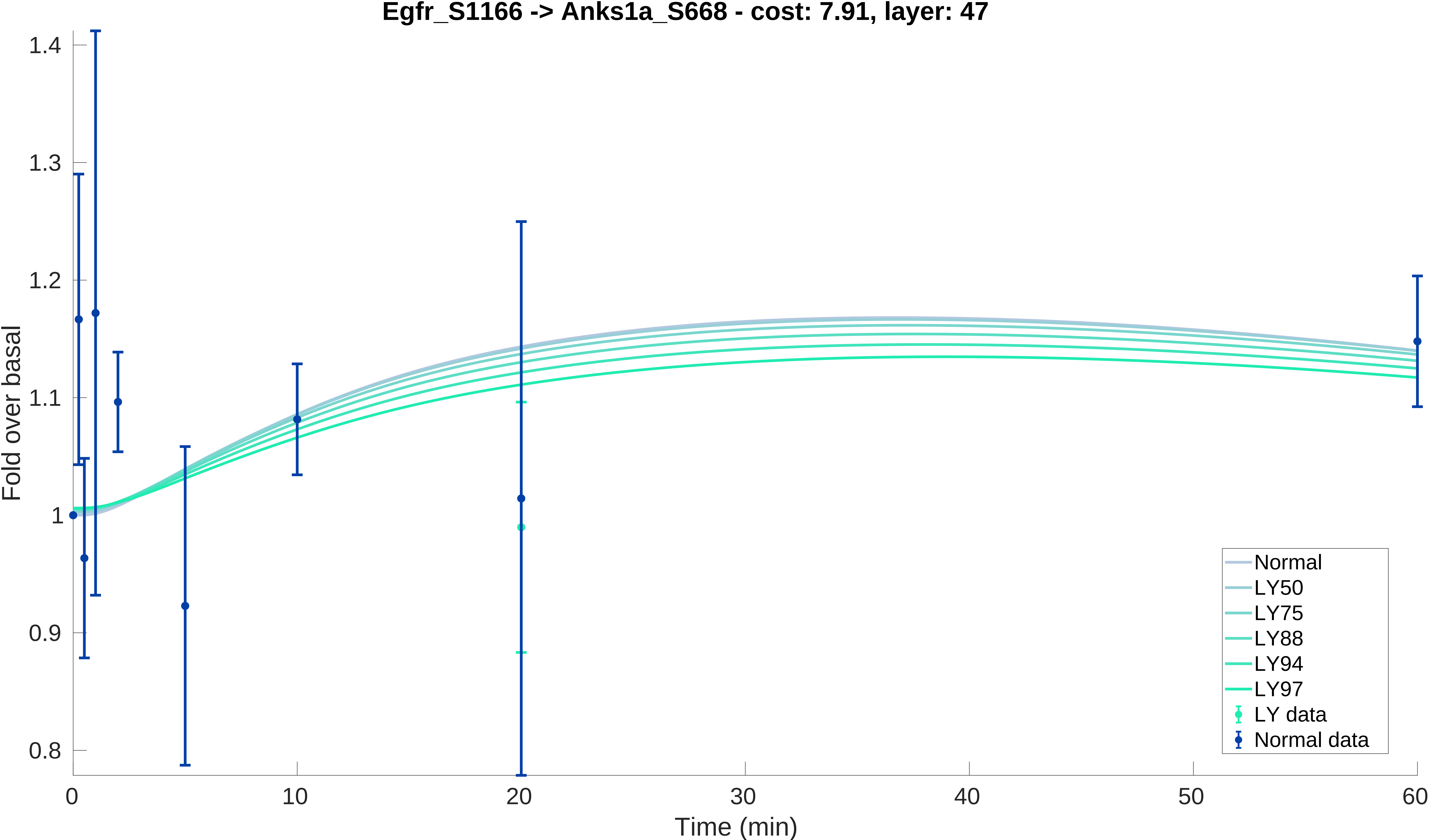

Egfr_S1166 -> Anks1a_S668 - cost: 7.91, layer: 47
Fold over basal
Time (min)
1.4
1.3
1.2
1.1
1
0.9
0.8
0
10
20
30
40
50
60
Normal
LY50
LY75
LY88
LY94
LY97
LY data
Normal data

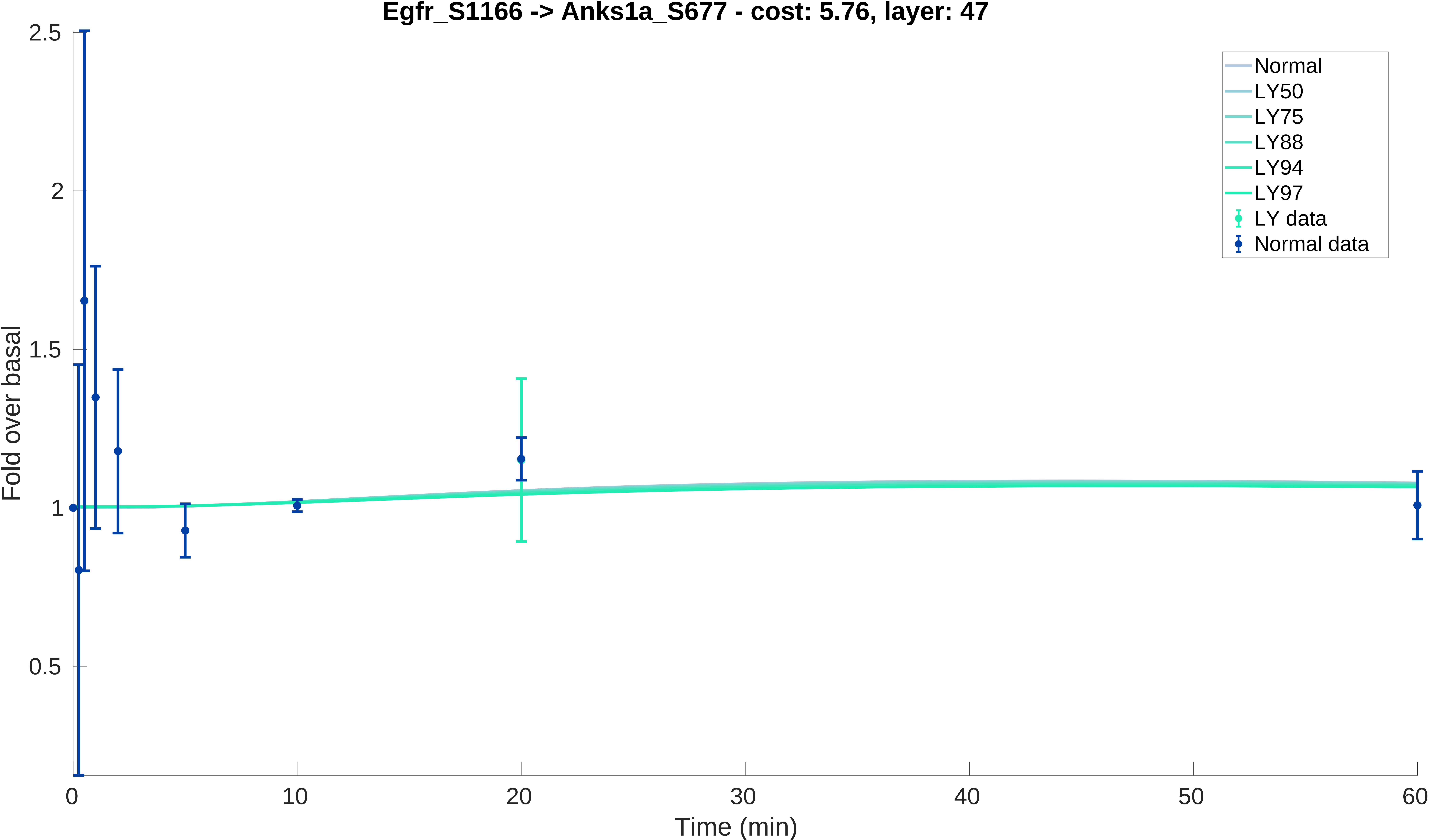

Egfr_S1166 -> Anks1a_S677 - cost: 5.76, layer: 47
Normal
LY50
LY75
LY88
LY94
LY97
LY data
Normal data
Fold over basal
2.5
2
1.5
1
0.5
0
10
20
30
40
50
60
Time (min)

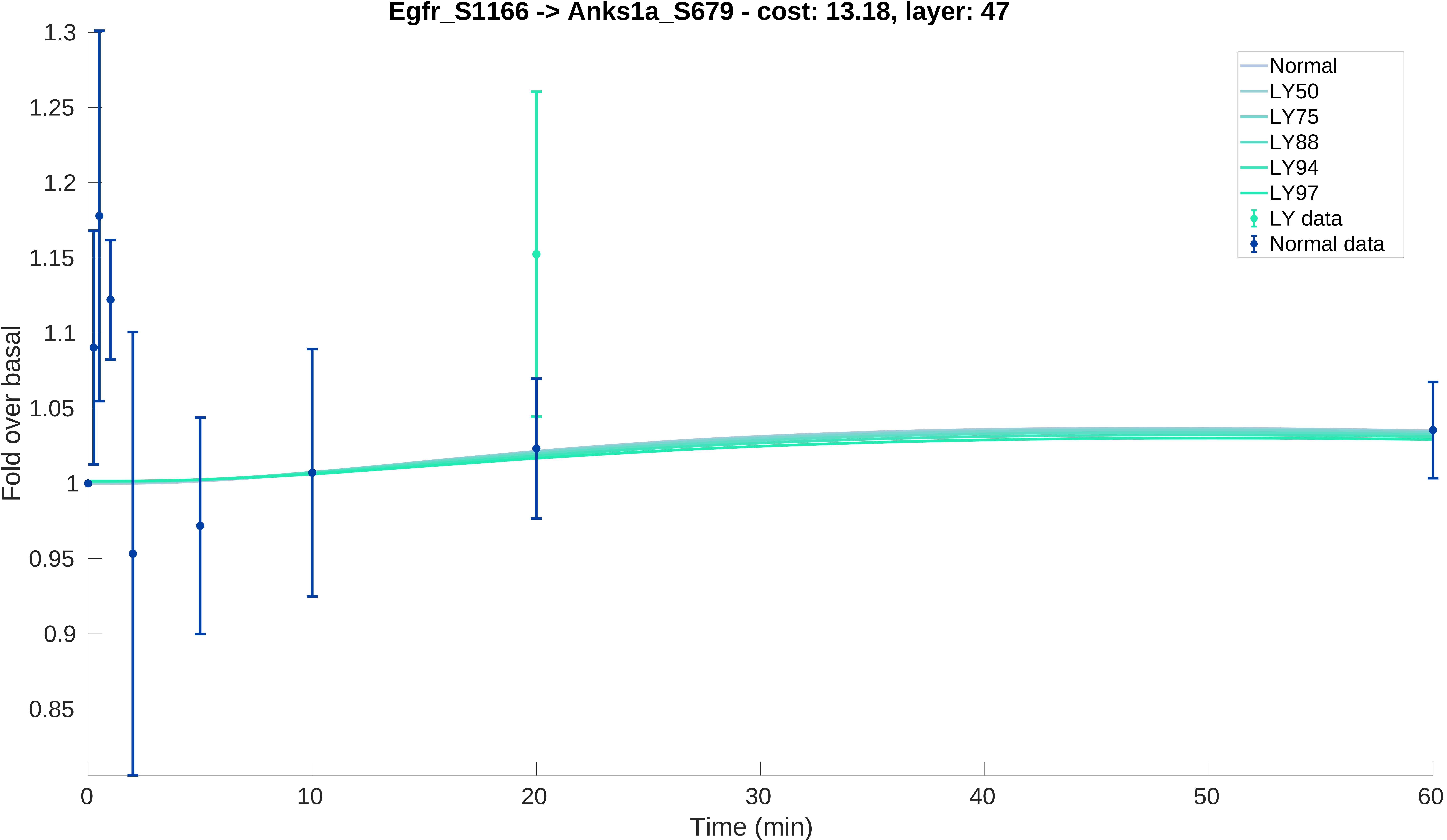

Egfr_S1166 -> Anks1a_S679 - cost: 13.18, layer: 47
Fold over basal
Time (min)
1.3
1.25
1.2
1.15
1.1
1.05
1
0.95
0.9
0.85
0
10
20
30
40
50
60
Normal
LY50
LY75
LY88
LY94
LY97
LY data
Normal data

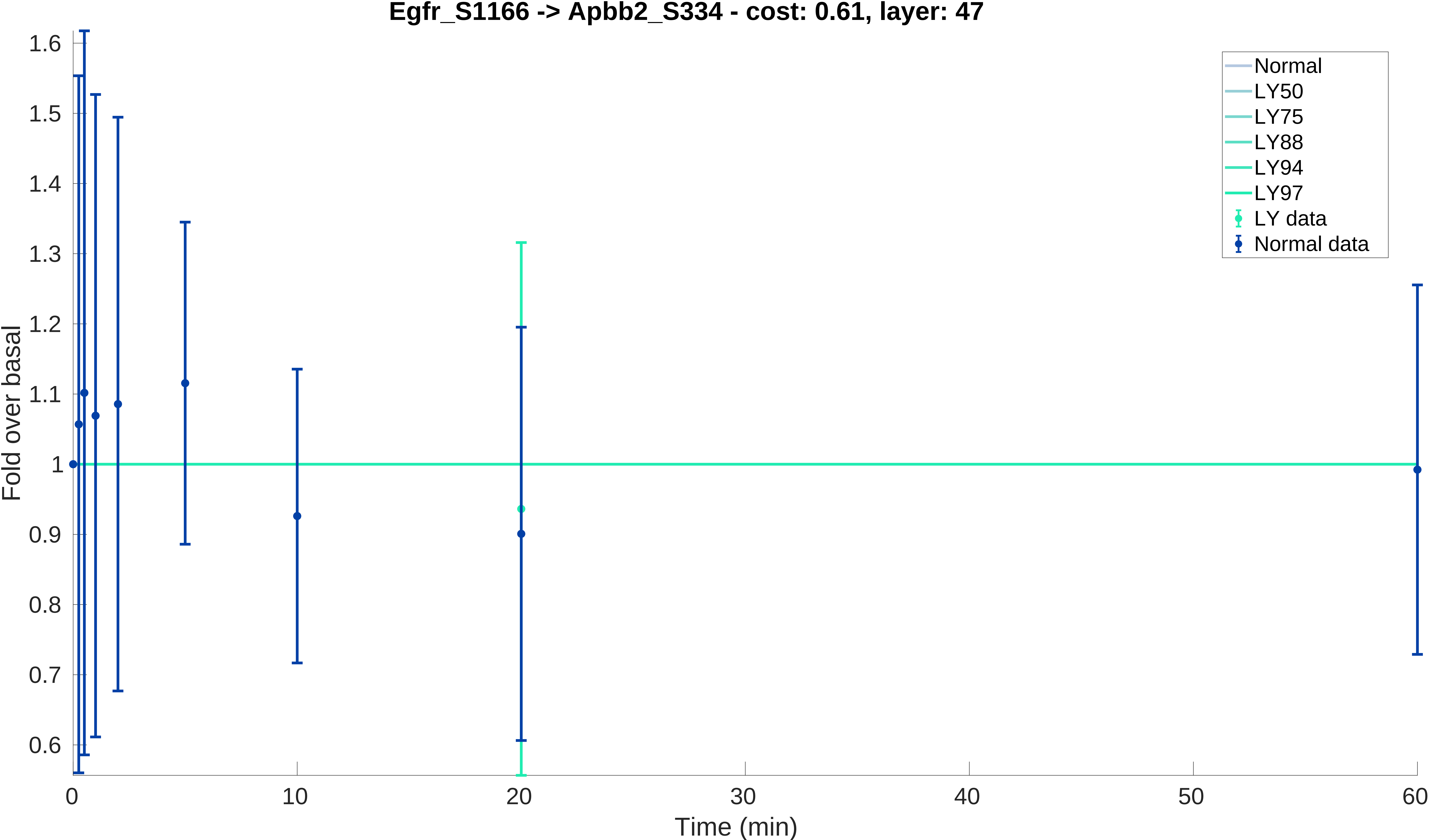

Egfr_S1166 -> Apbb2_S334 - cost: 0.61, layer: 47
Fold over basal
Time (min)
Normal
LY50
LY75
LY88
LY94
LY97
LY data
Normal data
1.6
1.5
1.4
1.3
1.2
1.1
1
0.9
0.8
0.7
0.6
0
10
20
30
40
50
60

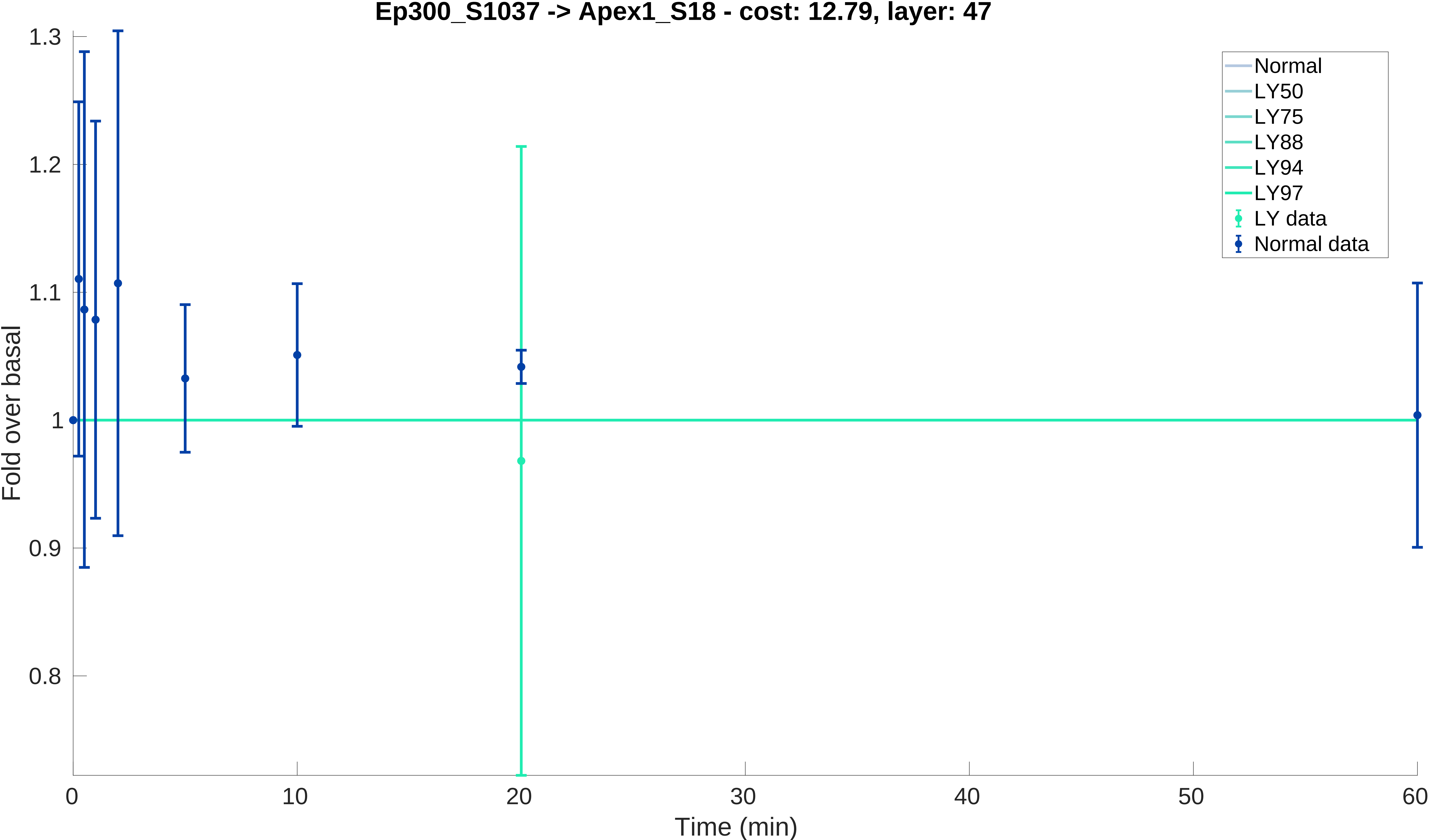

Ep300_S1037 -> Apex1_S18 - cost: 12.79, layer: 47
Fold over basal
1.3
1.2
1.1
1
0.9
0.8
0
10
20
30
40
50
60
Time (min)
Normal
LY50
LY75
LY88
LY94
LY97
LY data
Normal data

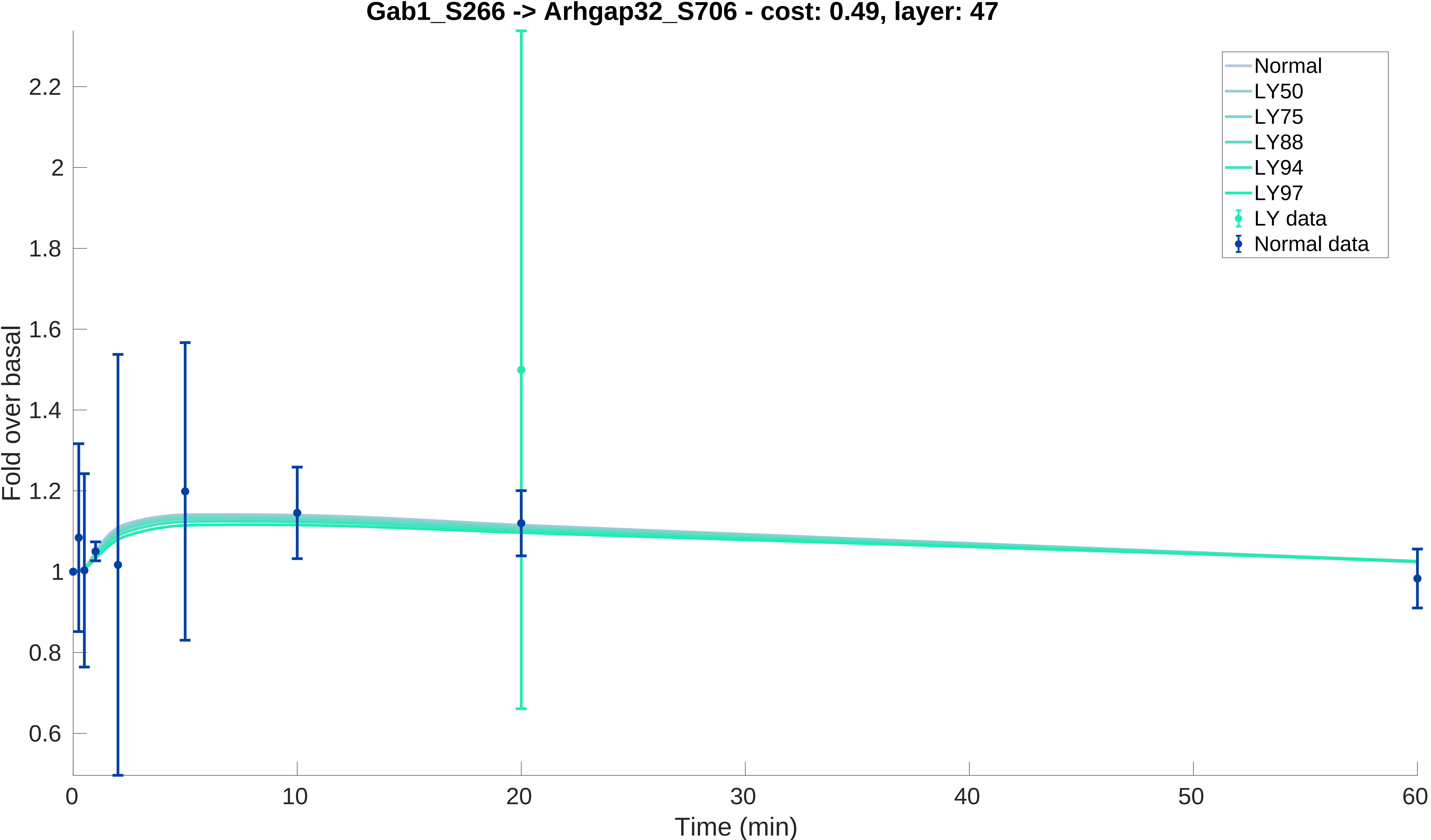

Gab1_S266 -> Arhgap32_S706 - cost: 0.49, layer: 47
Normal
LY50
LY75
LY88
LY94
LY97
LY data
Normal data
Fold over basal
0.6
0.8
1
1.2
1.4
1.6
1.8
2
2.2
0
10
20
30
40
50
60
Time (min)

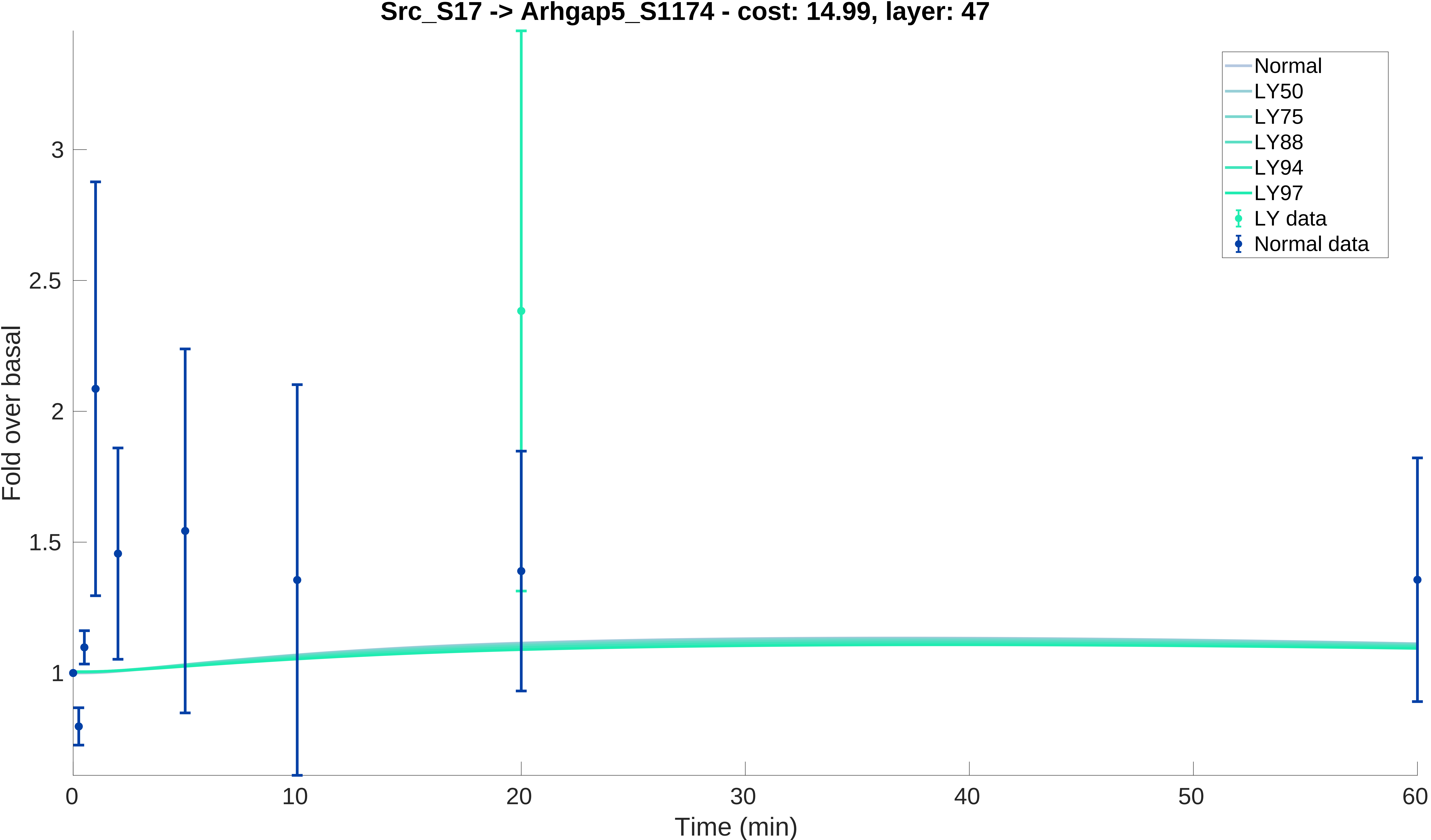

Src_S17 -> Arhgap5_S1174 - cost: 14.99, layer: 47
Normal
LY50
LY75
LY88
LY94
LY97
LY data
Normal data
Fold over basal
3
2.5
2
1.5
1
0
10
20
30
40
50
60
Time (min)

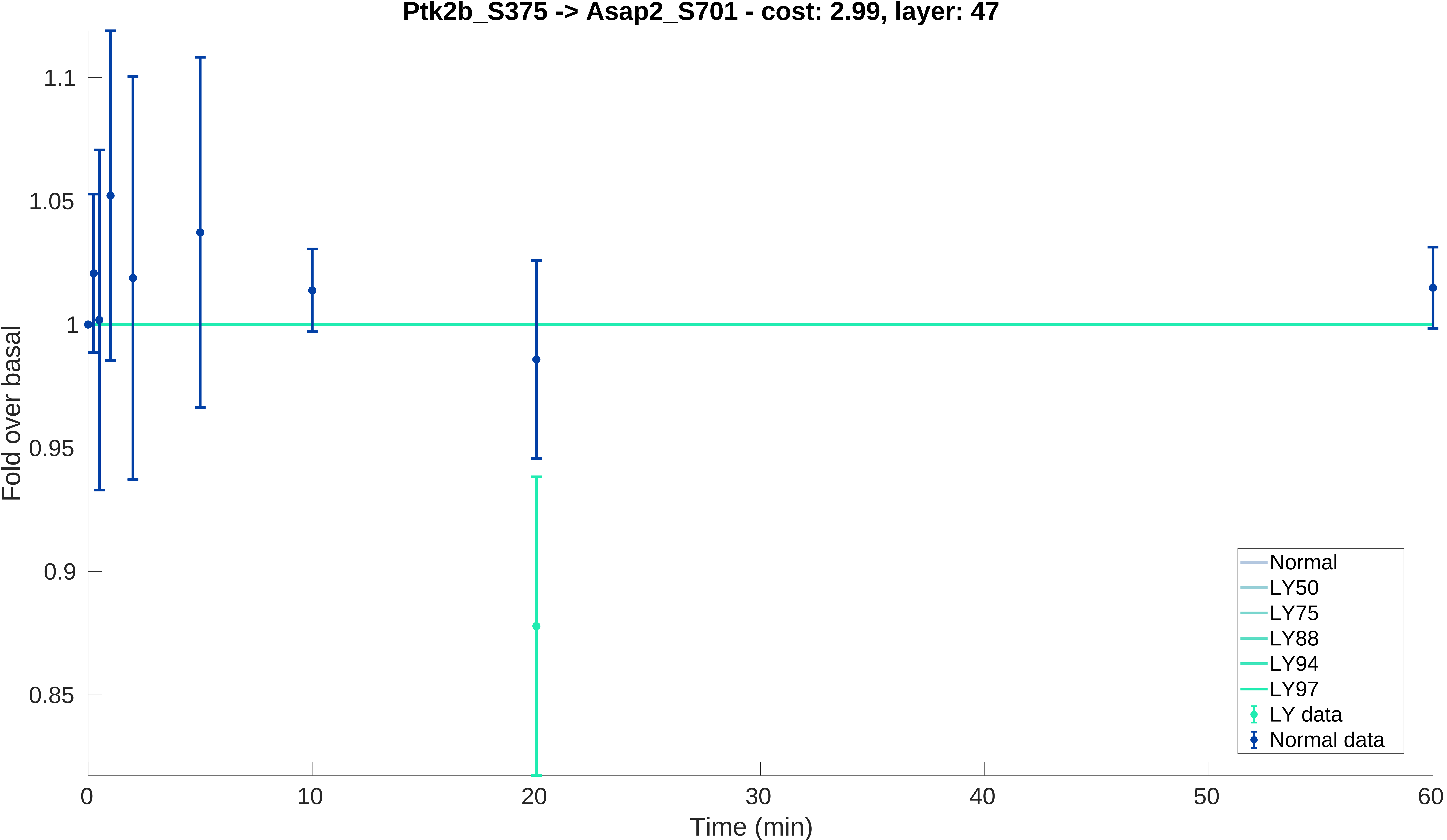

Ptk2b_S375 -> Asap2_S701 - cost: 2.99, layer: 47
Fold over basal
Time (min)
1.1
1.05
1
0.95
0.9
0.85
0
10
20
30
40
50
60
Normal
LY50
LY75
LY88
LY94
LY97
LY data
Normal data

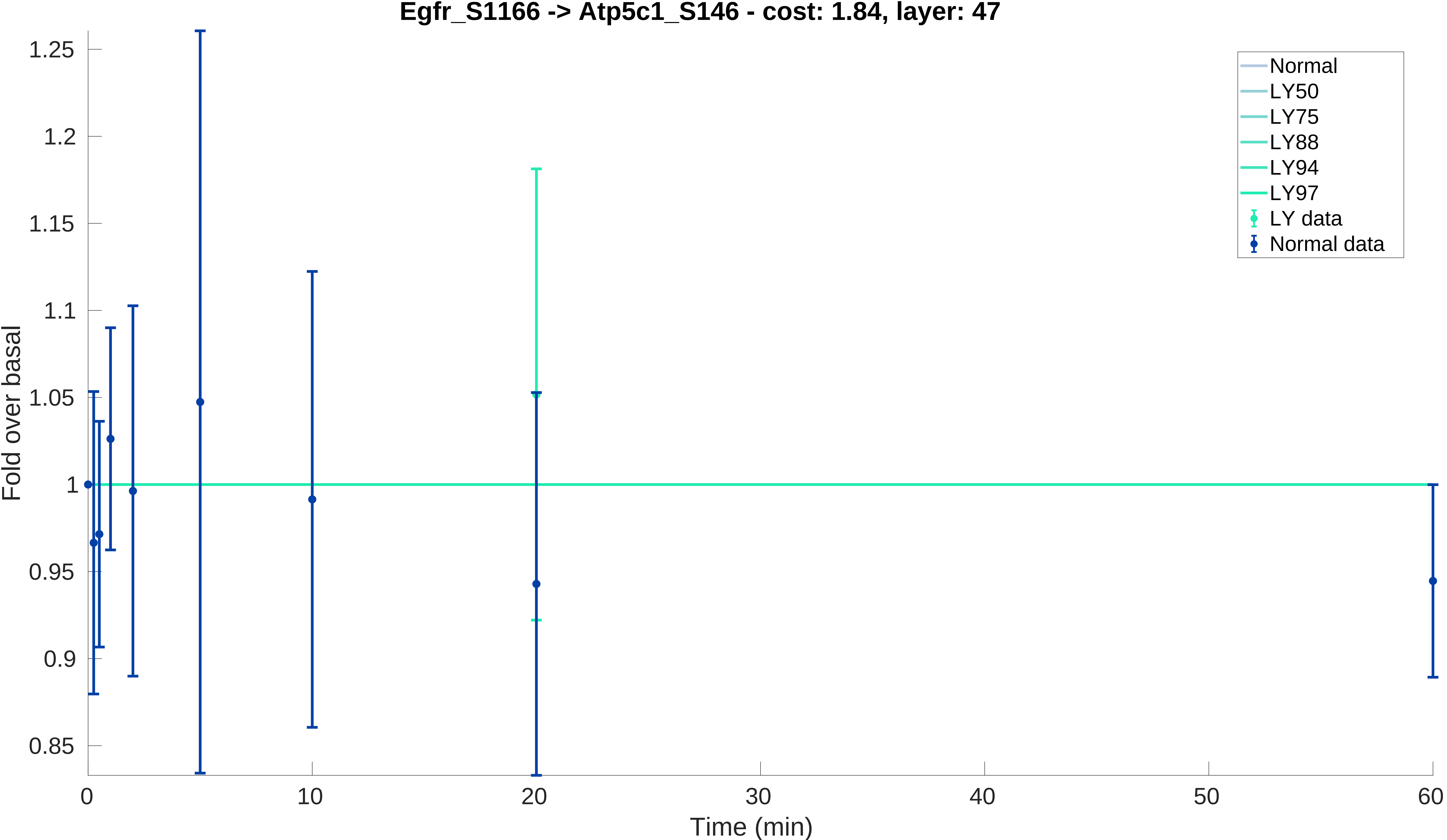
Egfr_S1166 -> Atp5c1_S146 - cost: 1.84, layer: 47
Fold over basal
Time (min)
1.25
1.2
1.15
1.1
1.05
1
0.95
0.9
0.85
0
10
20
30
40
50
60
Normal
LY50
LY75
LY88
LY94
LY97
LY data
Normal data

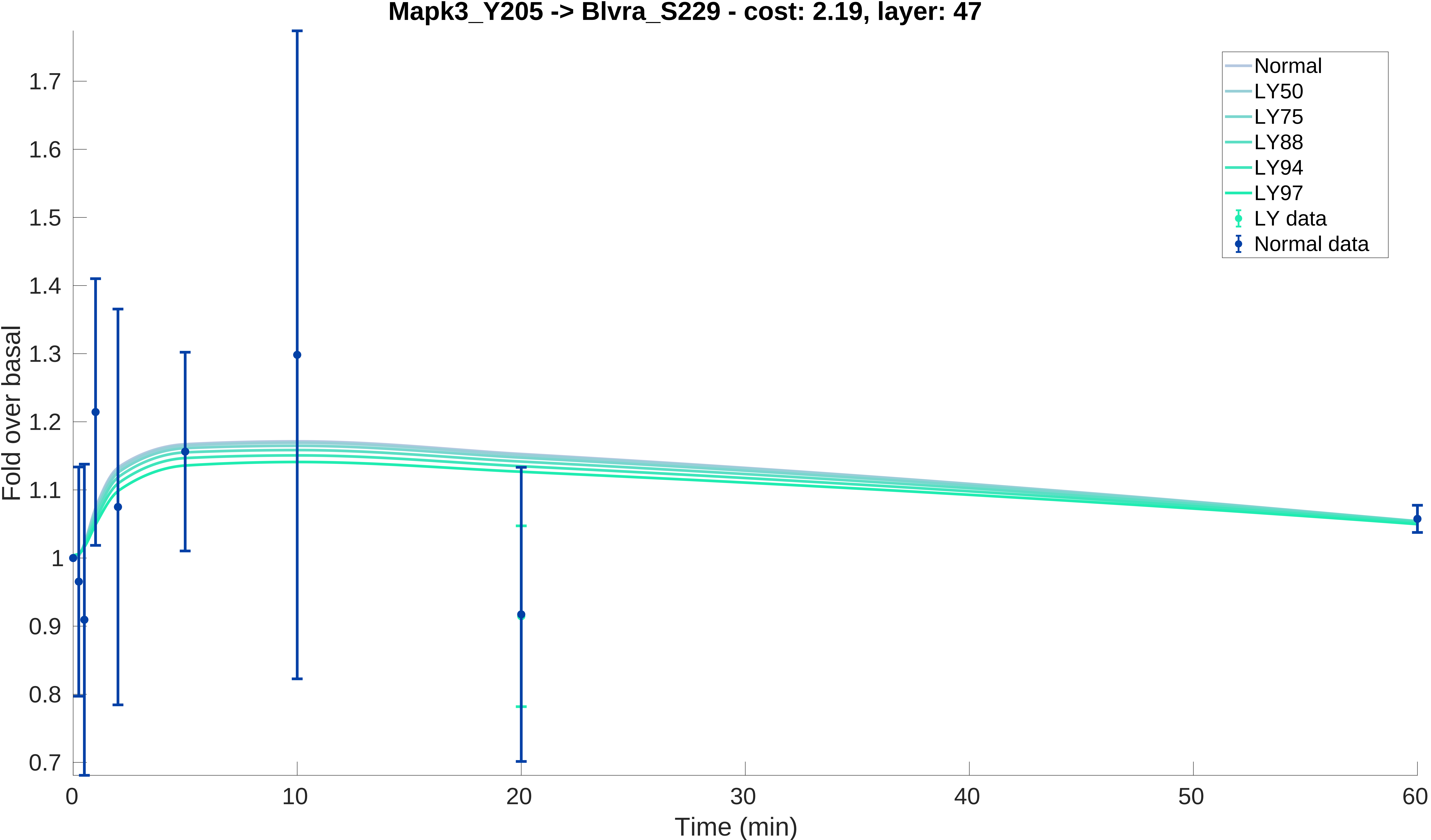

Mapk3_Y205 -> Blvra_S229 - cost: 2.19, layer: 47
Fold over basal
Time (min)
0.7
0.8
0.9
1
1.1
1.2
1.3
1.4
1.5
1.6
1.7
0
10
20
30
40
50
60
Normal
LY50
LY75
LY88
LY94
LY97
LY data
Normal data

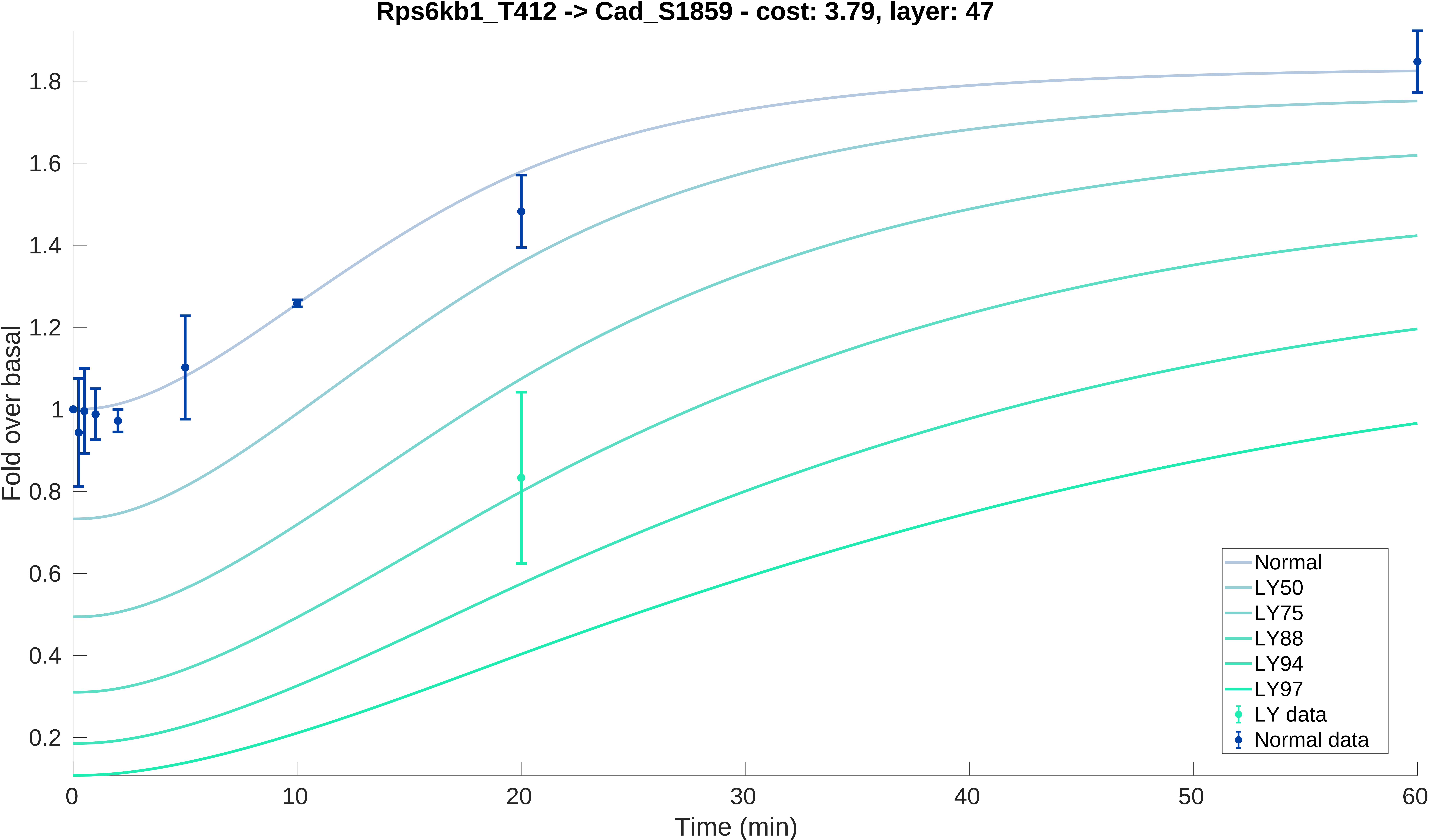

Rps6kb1_T412 -> Cad_S1859 - cost: 3.79, layer: 47
Fold over basal
0.2
0.4
0.6
0.8
1
1.2
1.4
1.6
1.8
Time (min)
0
10
20
30
40
50
60
Normal
LY50
LY75
LY88
LY94
LY97
LY data
Normal data

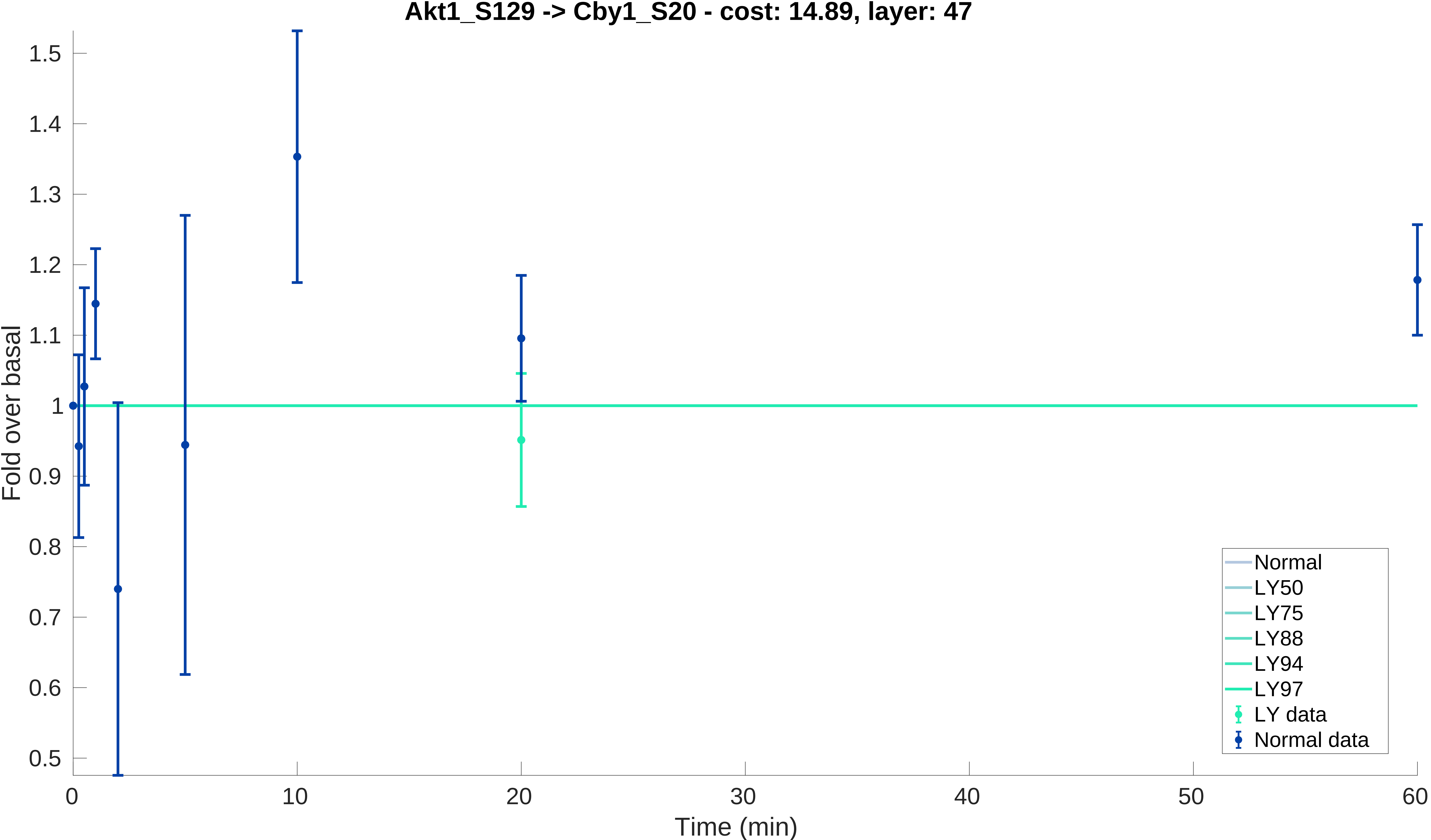
Akt1_S129 -> Cby1_S20 - cost: 14.89, layer: 47
Fold over basal
1.5
1.4
1.3
1.2
1.1
1
0.9
0.8
0.7
0.6
0.5
0
10
20
30
40
50
60
Time (min)
Normal
LY50
LY75
LY88
LY94
LY97
LY data
Normal data

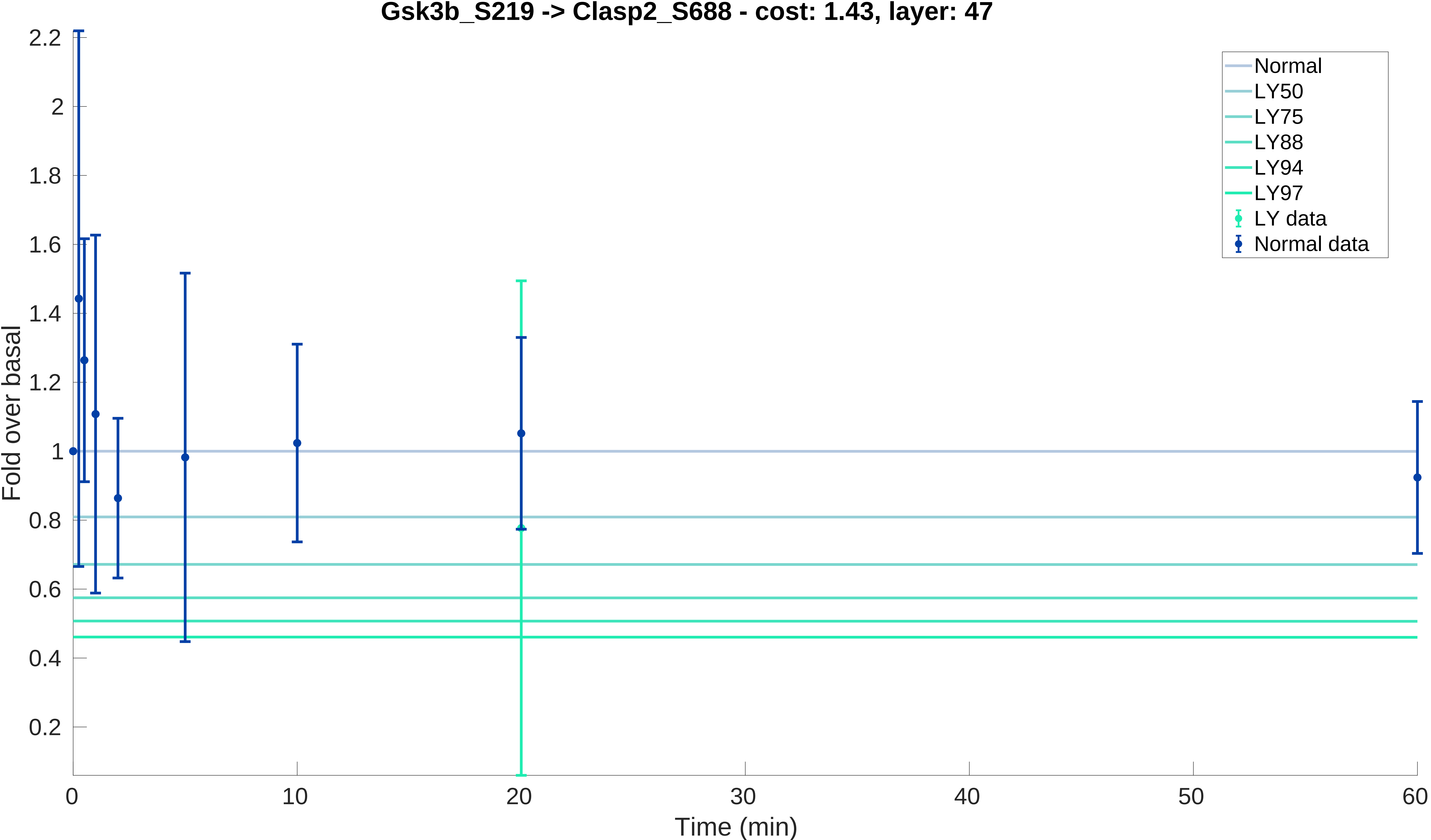
Gsk3b_S219 -> Clasp2_S688 - cost: 1.43, layer: 47
Fold over basal
Time (min)
2.2
2
1.8
1.6
1.4
1.2
1
0.8
0.6
0.4
0.2
0
10
20
30
40
50
60
Normal
LY50
LY75
LY88
LY94
LY97
LY data
Normal data

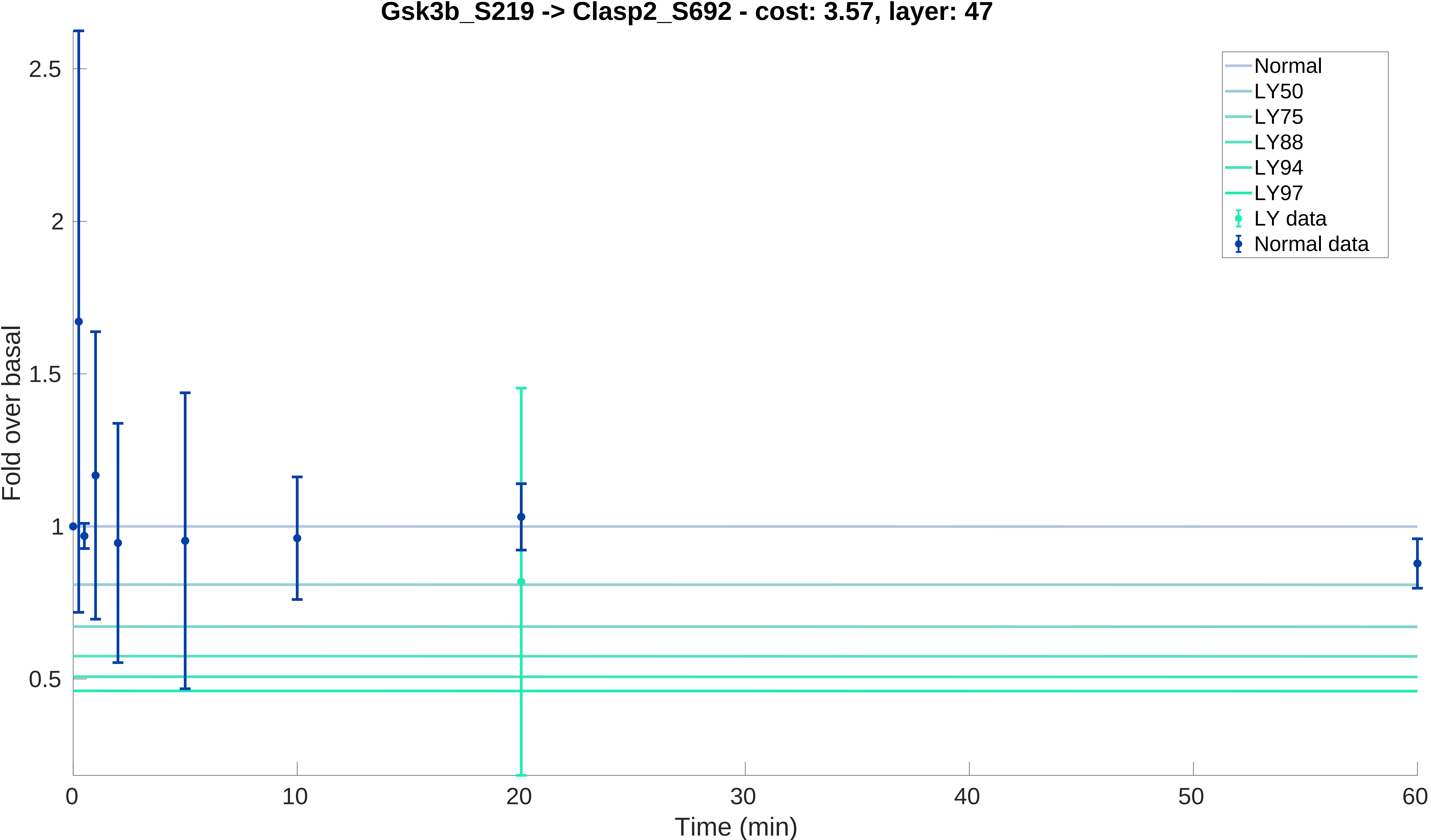

Gsk3b_S219 -> Clasp2_S692 - cost: 3.57, layer: 47
Fold over basal
Time (min)
2.5
2
1.5
1
0.5
0
10
20
30
40
50
60
Normal
LY50
LY75
LY88
LY94
LY97
LY data
Normal data

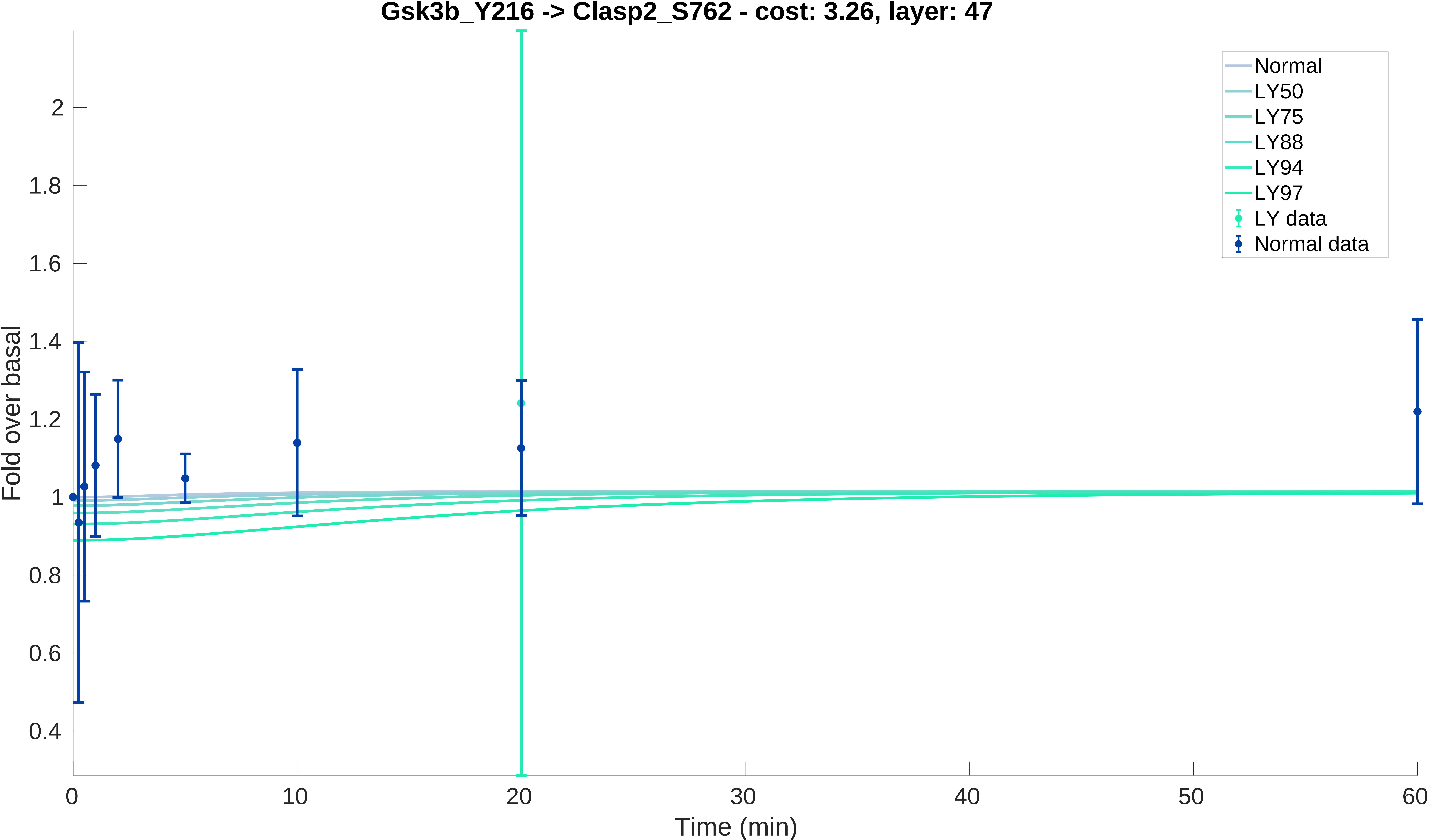
Gsk3b_Y216 -> Clasp2_S762 - cost: 3.26, layer: 47
Fold over basal
Time (min)
0
10
20
30
40
50
60
0.4
0.6
0.8
1
1.2
1.4
1.6
1.8
2
Normal
LY50
LY75
LY88
LY94
LY97
LY data
Normal data

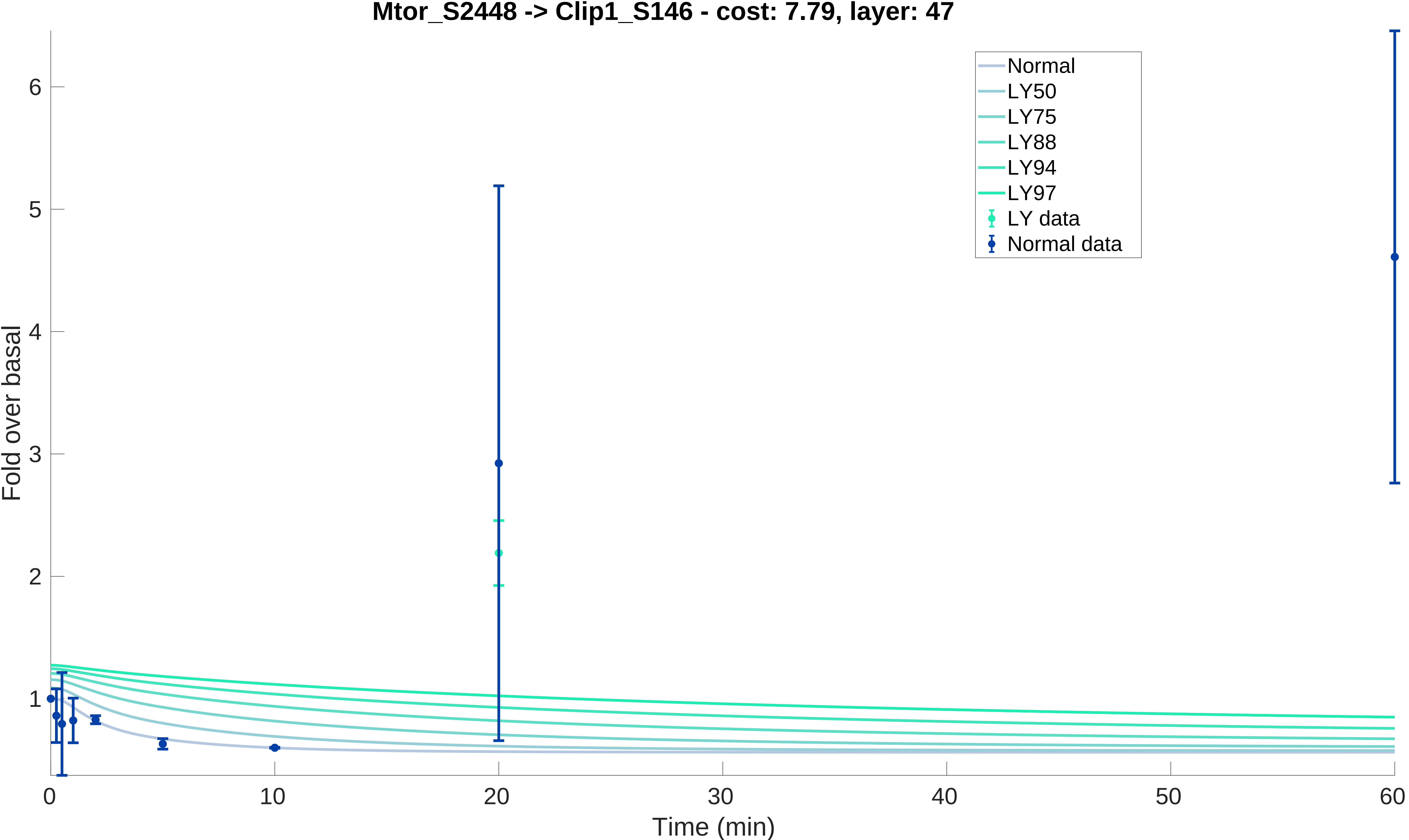
Mtor_S2448 -> Clip1_S146 - cost: 7.79, layer: 47
Normal
LY50
LY75
LY88
LY94
LY97
LY data
Normal data
Fold over basal
Time (min)
0
10
20
30
40
50
60
1
2
3
4
5
6

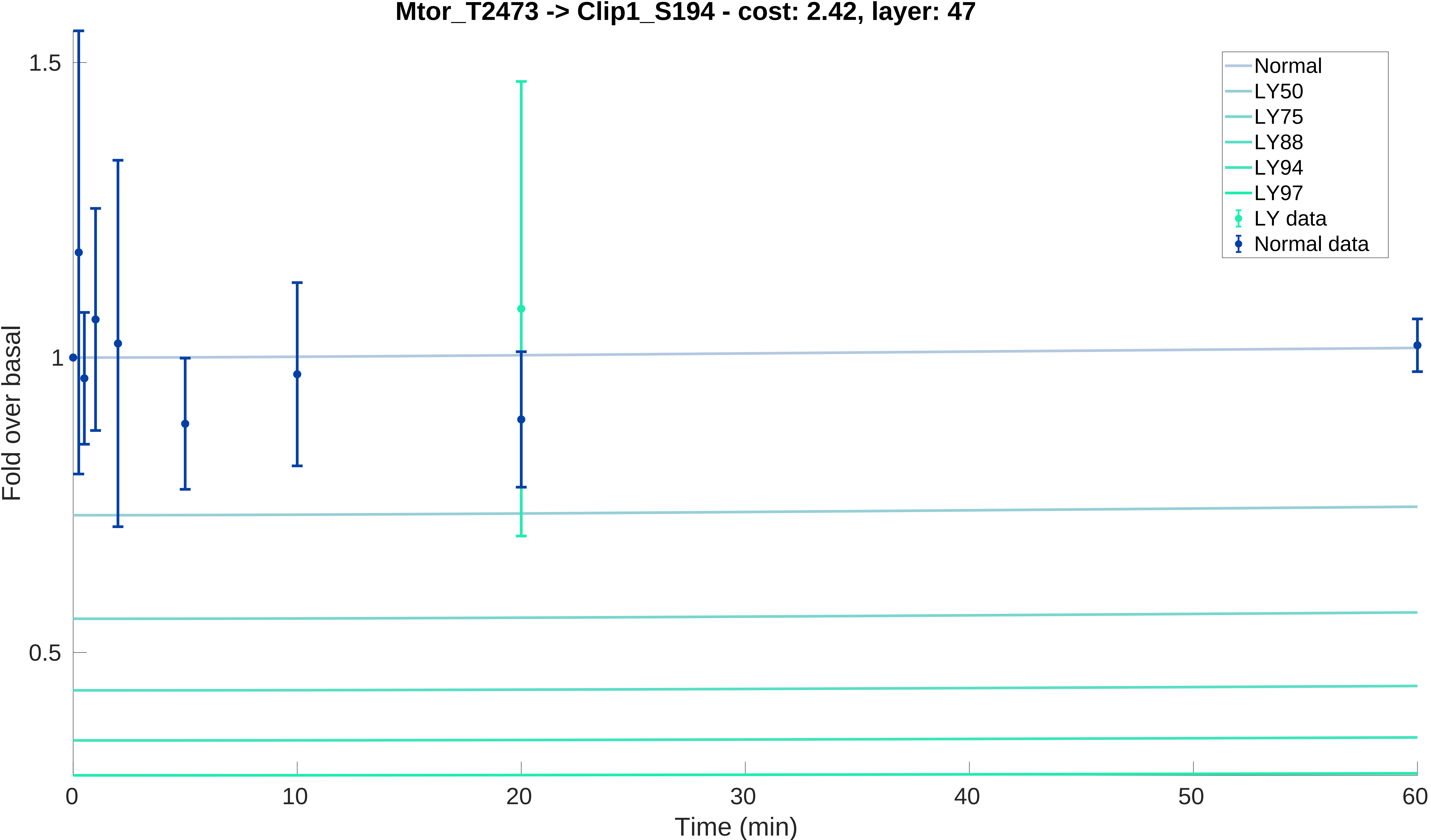

Mtor_T2473 -> Clip1_S194 - cost: 2.42, layer: 47
Fold over basal
1.5
1
0.5
0
10
20
30
40
50
60
Time (min)
Normal
LY50
LY75
LY88
LY94
LY97
LY data
Normal data

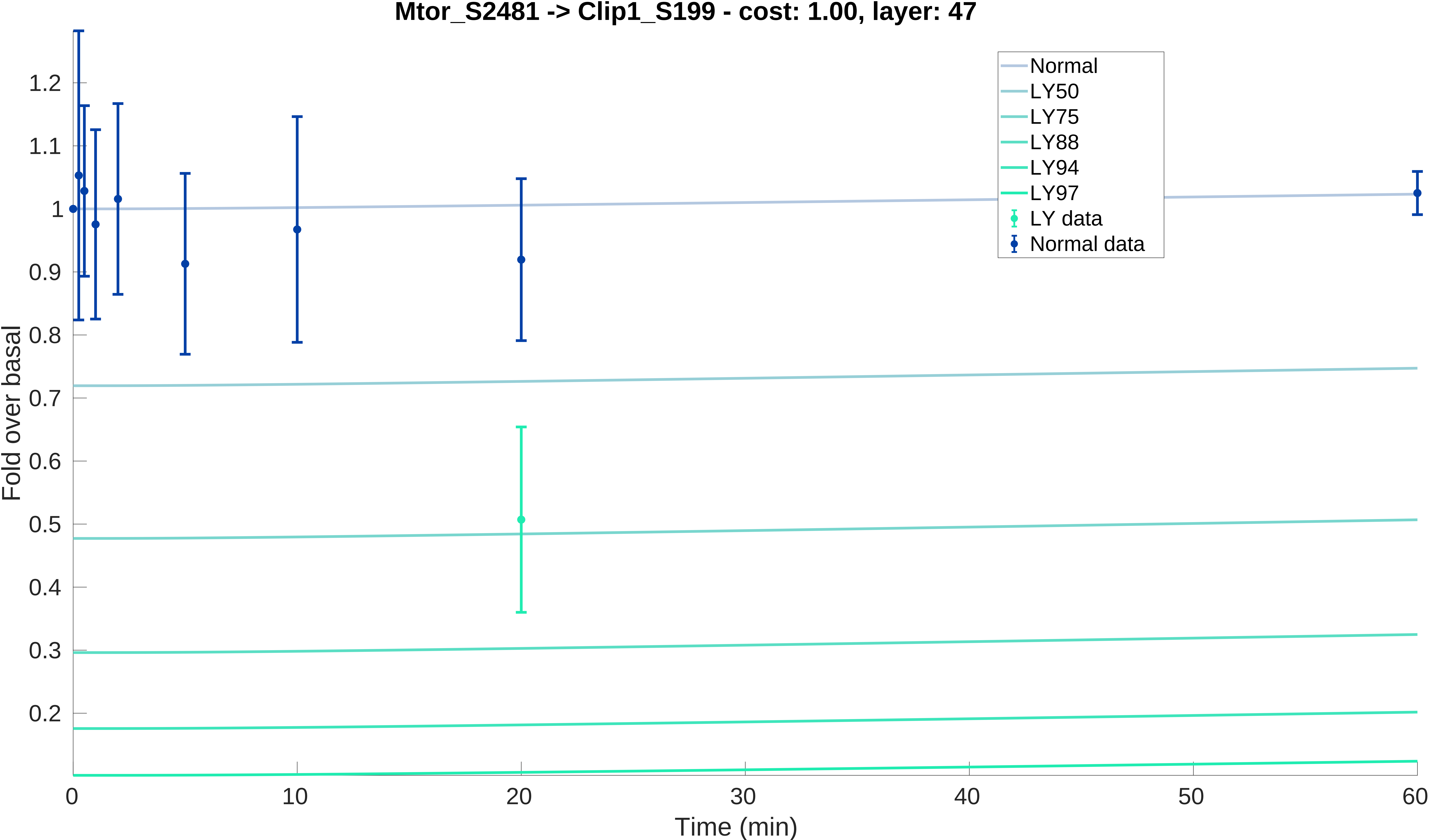

Mtor_S2481 -> Clip1_S199 - cost: 1.00, layer: 47
Normal
LY50
LY75
LY88
LY94
LY97
LY data
Normal data
Fold over basal
Time (min)
0
10
20
30
40
50
60
1.2
1.1
1
0.9
0.8
0.7
0.6
0.5
0.4
0.3
0.2

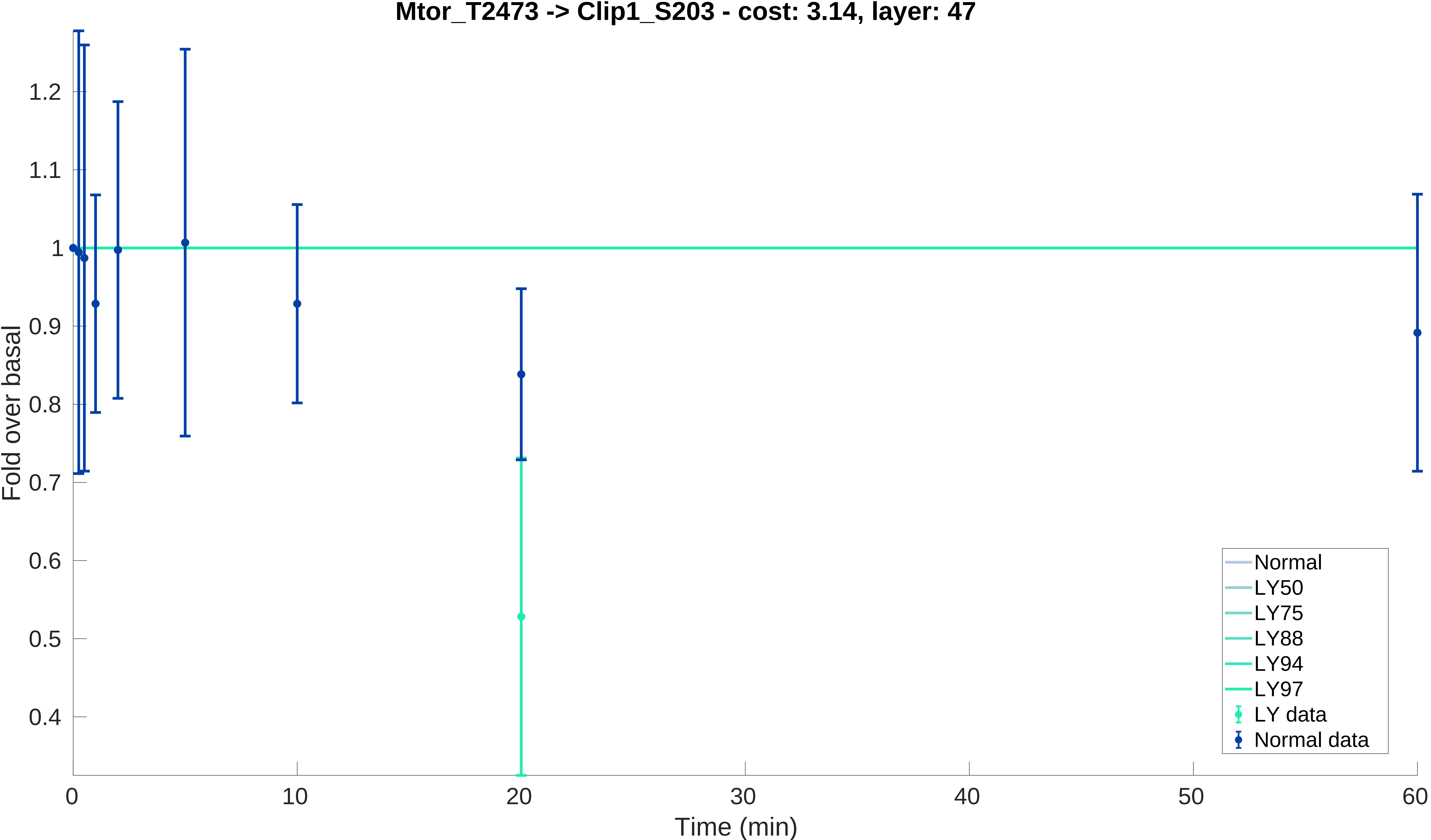

Mtor_T2473 -> Clip1_S203 - cost: 3.14, layer: 47
Fold over basal
1.2
1.1
1
0.9
0.8
0.7
0.6
0.5
0.4
0
10
20
30
40
50
60
Time (min)
Normal
LY50
LY75
LY88
LY94
LY97
LY data
Normal data

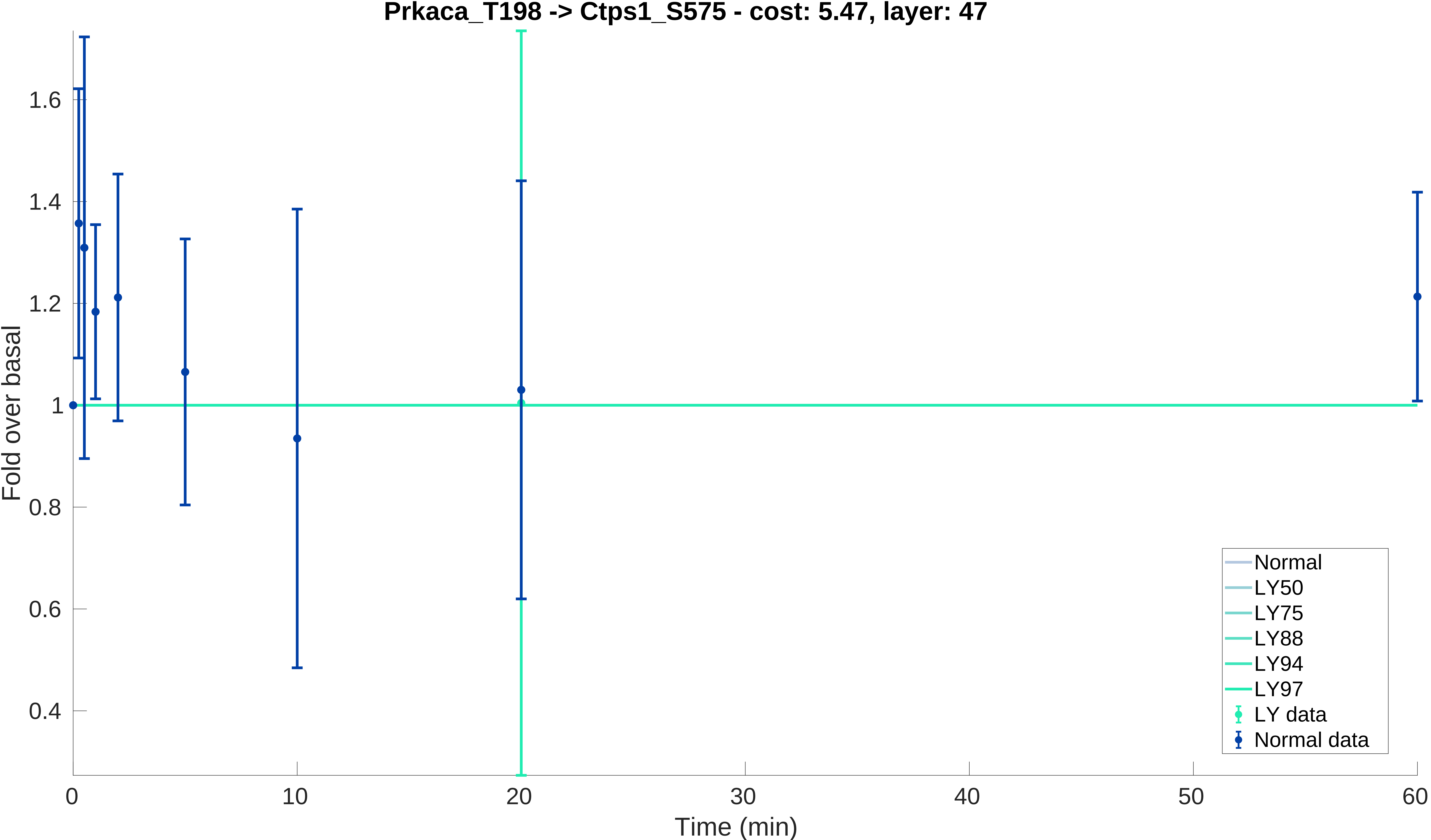

Prrkaca_T198 -> Ctps1_S575 - cost: 5.47, layer: 47
Fold over basal
Time (min)
0
10
20
30
40
50
60
0.4
0.6
0.8
1
1.2
1.4
1.6
Normal
LY50
LY75
LY88
LY94
LY97
LY data
Normal data

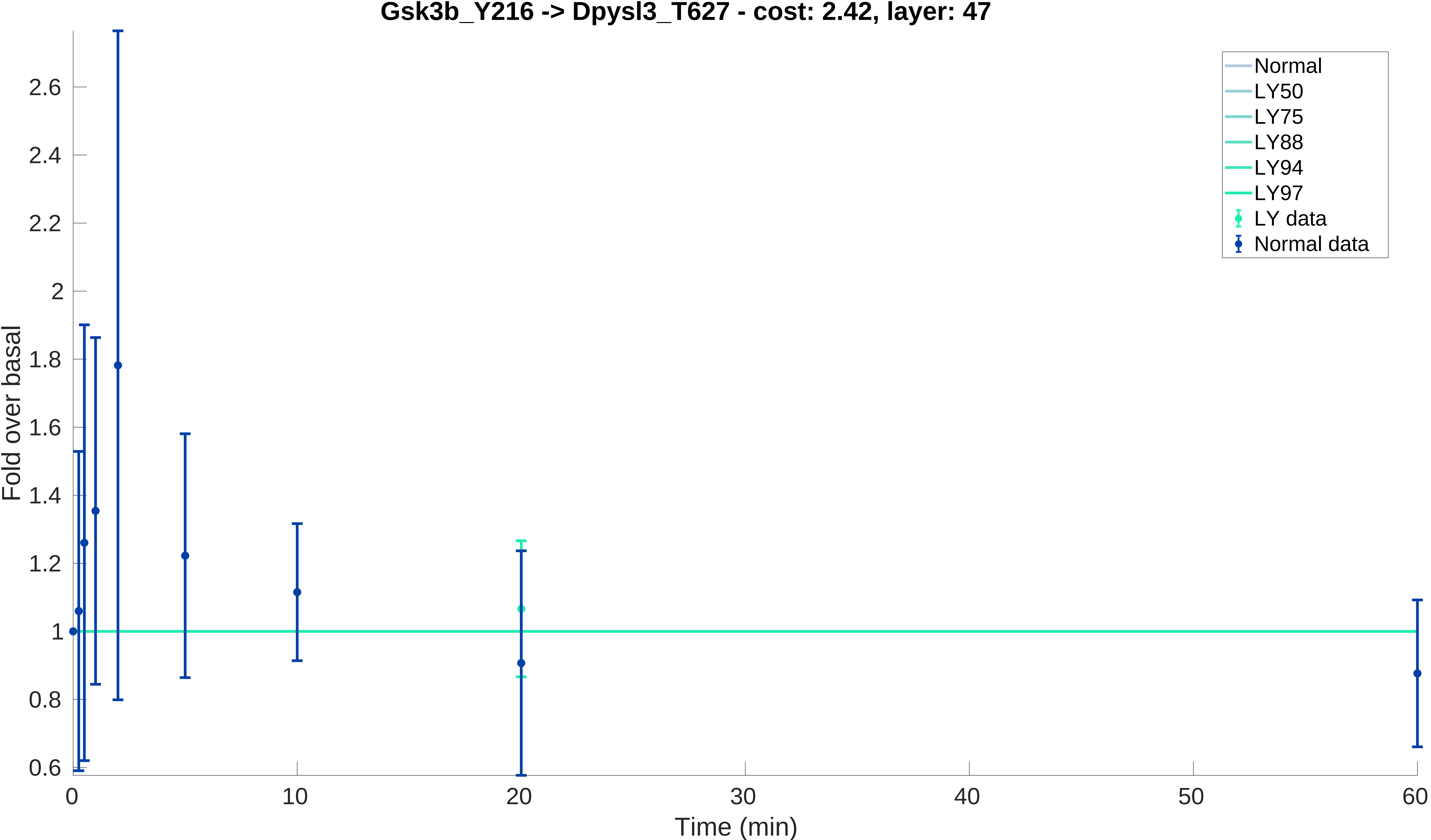

Gsk3b_Y216 -> Dpysl3_T627 - cost: 2.42, layer: 47
Fold over basal
Time (min)
0
10
20
30
40
50
60
0.6
0.8
1
1.2
1.4
1.6
1.8
2
2.2
2.4
2.6
Normal
LY50
LY75
LY88
LY94
LY97
LY data
Normal data

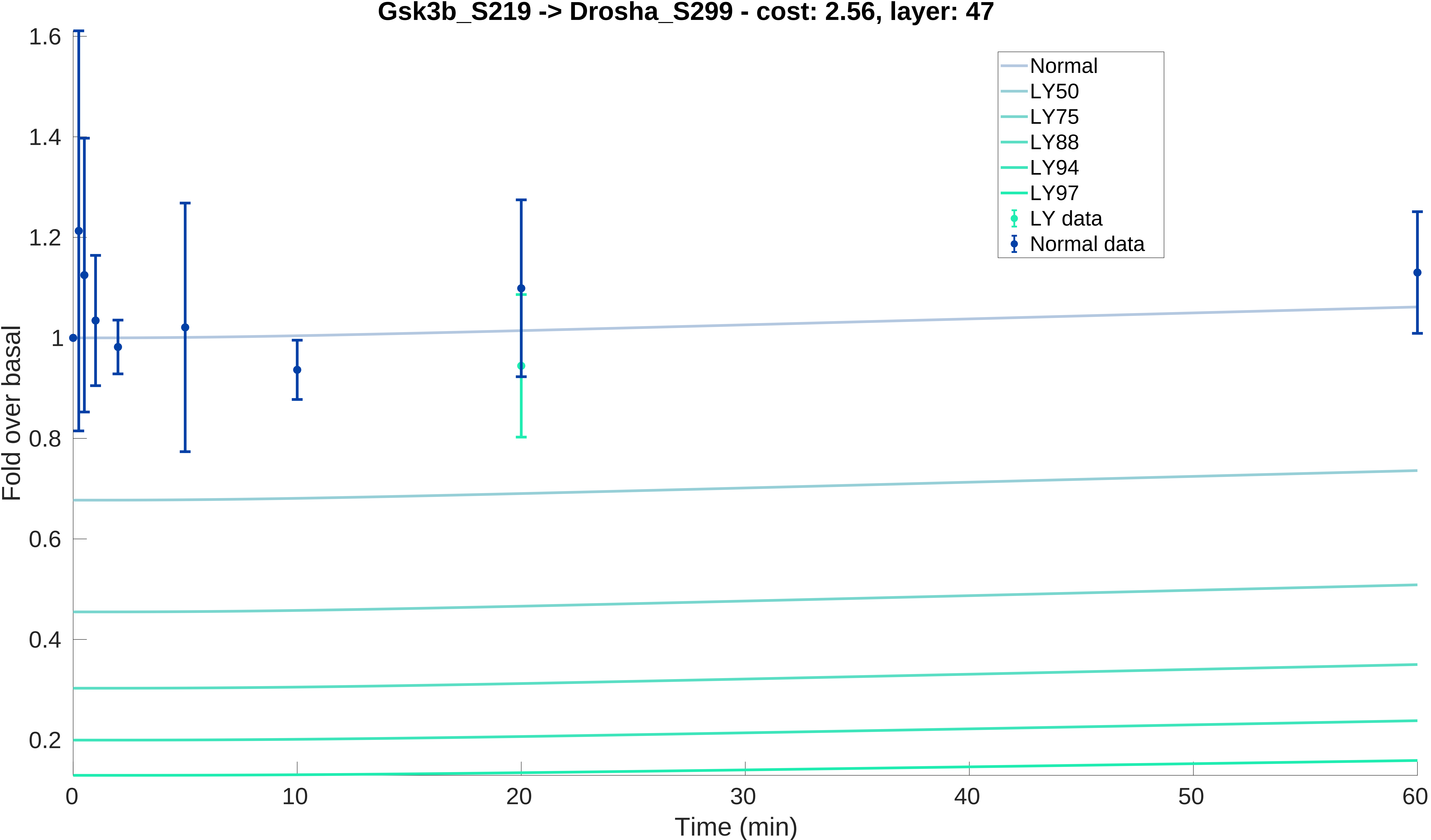

Gsk3b_S219 -> Drosha_S299 - cost: 2.56, layer: 47
Fold over basal
Time (min)
1.6
1.4
1.2
1
0.8
0.6
0.4
0.2
0
10
20
30
40
50
60
Normal
LY50
LY75
LY88
LY94
LY97
LY data
Normal data

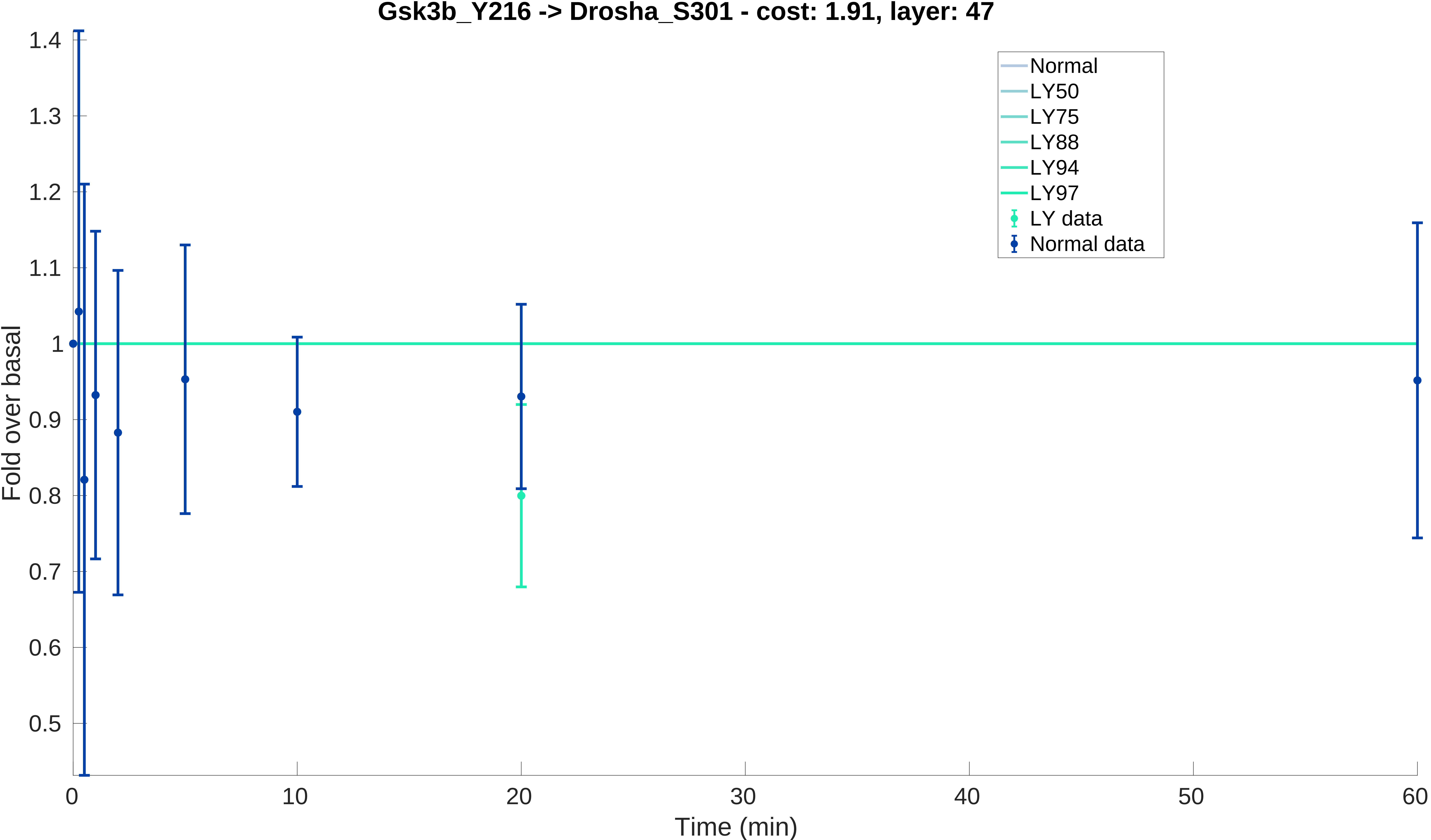

Gsk3b_Y216 -> Drosha_S301 - cost: 1.91, layer: 47
Fold over basal
Time (min)
1.4
1.3
1.2
1.1
1
0.9
0.8
0.7
0.6
0.5
0
10
20
30
40
50
60
Normal
LY50
LY75
LY88
LY94
LY97
LY data
Normal data

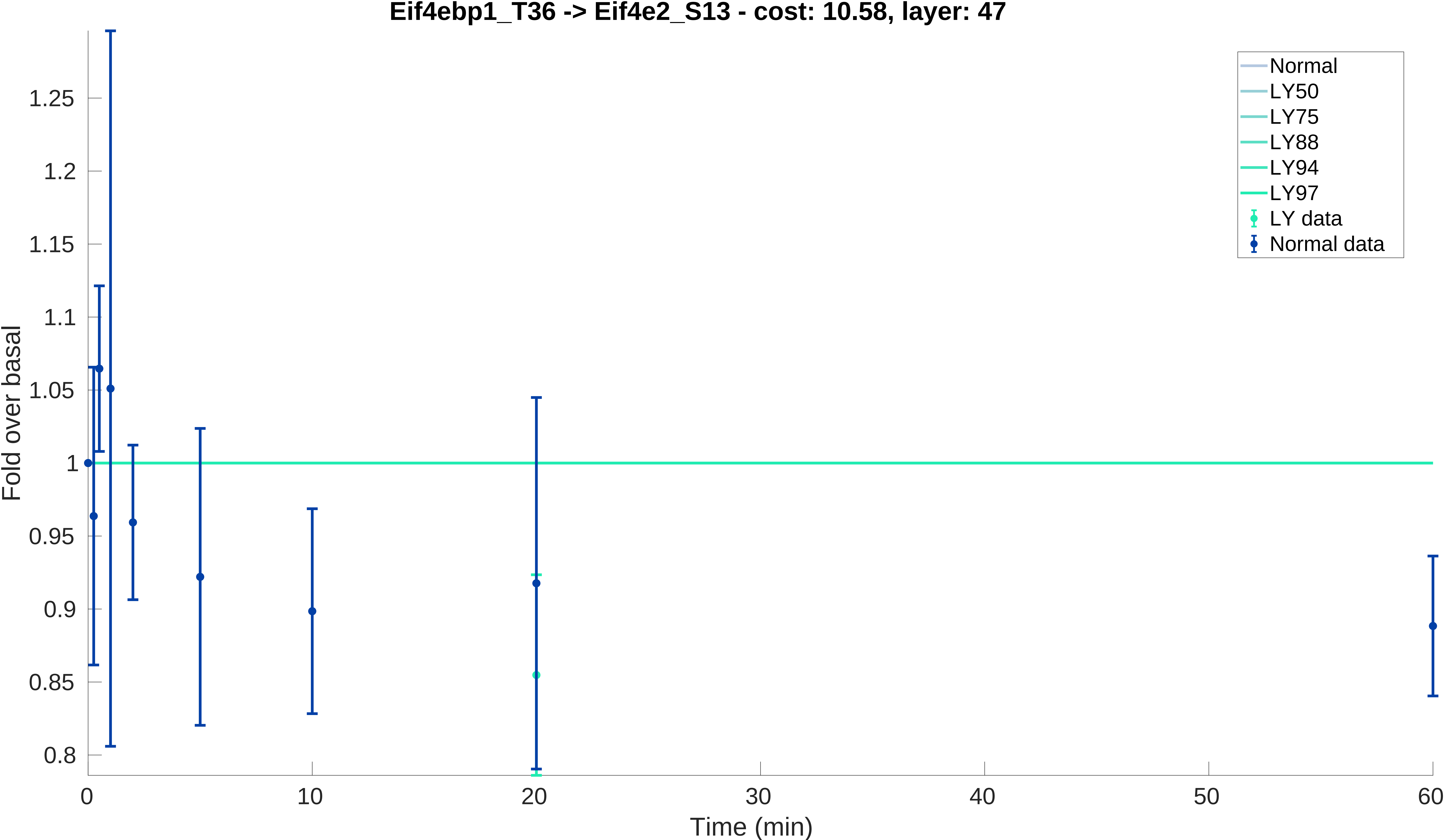

Eif4ebp1_T36 -> Eif4e2_S13 - cost: 10.58, layer: 47
Fold over basal
1.25
1.2
1.15
1.1
1.05
1
0.95
0.9
0.85
0.8
0
10
20
30
40
50
60
Time (min)
Normal
LY50
LY75
LY88
LY94
LY97
LY data
Normal data

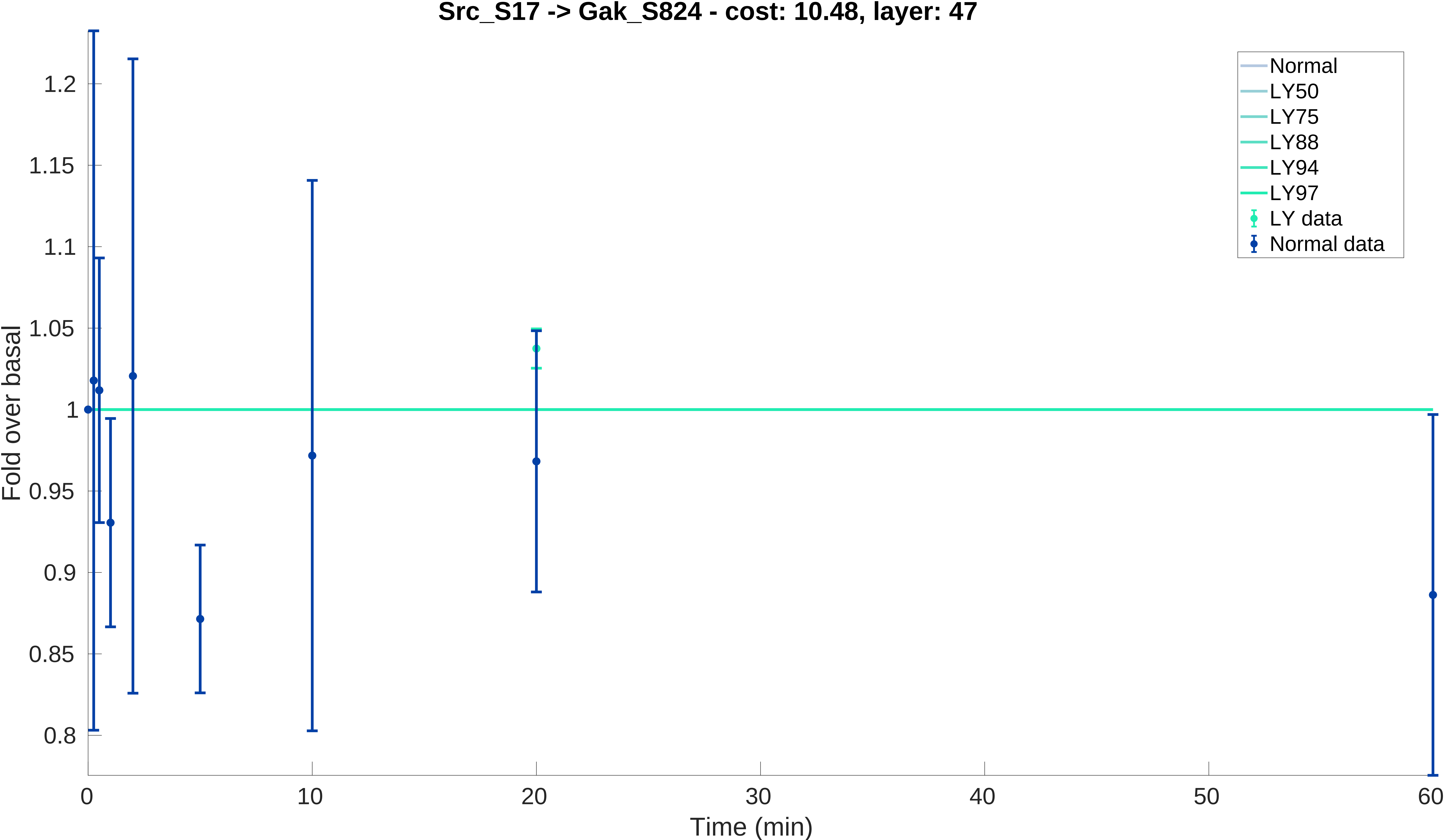

Src_S17 -> Gak_S824 - cost: 10.48, layer: 47
Normal
LY50
LY75
LY88
LY94
LY97
LY data
Normal data
Fold over basal
1.2
1.15
1.1
1.05
1
0.95
0.9
0.85
0.8
0
10
20
30
40
50
60
Time (min)

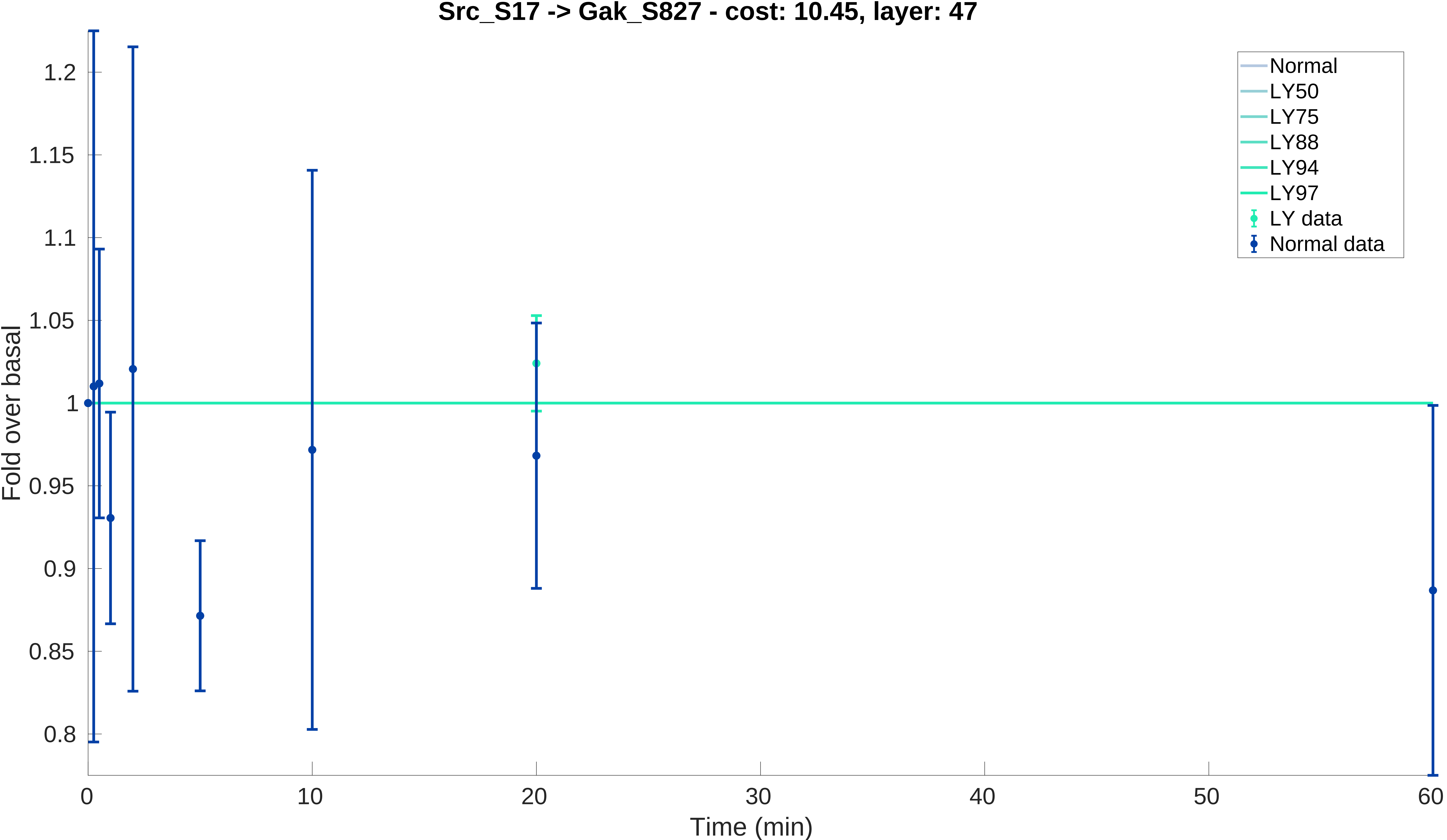

Src_S17 -> Gak_S827 - cost: 10.45, layer: 47
Normal
LY50
LY75
LY88
LY94
LY97
LY data
Normal data
Fold over basal
1.2
1.15
1.1
1.05
1
0.95
0.9
0.85
0.8
0
10
20
30
40
50
60
Time (min)

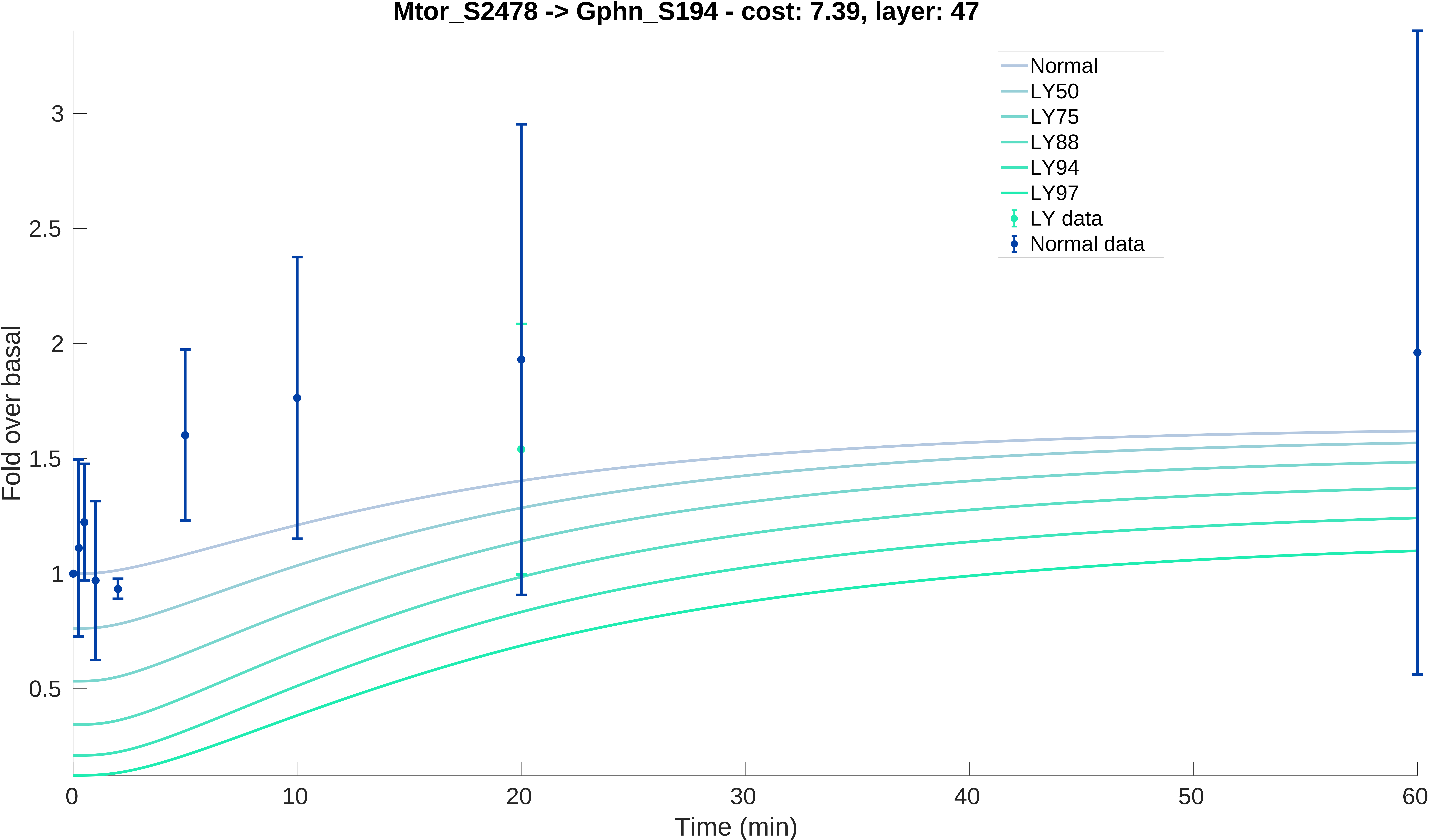

Mtor_S2478 -> Gphn_S194 - cost: 7.39, layer: 47
Normal
LY50
LY75
LY88
LY94
LY97
LY data
Normal data
Fold over basal
0.5
1
1.5
2
2.5
3
0
10
20
30
40
50
60
Time (min)

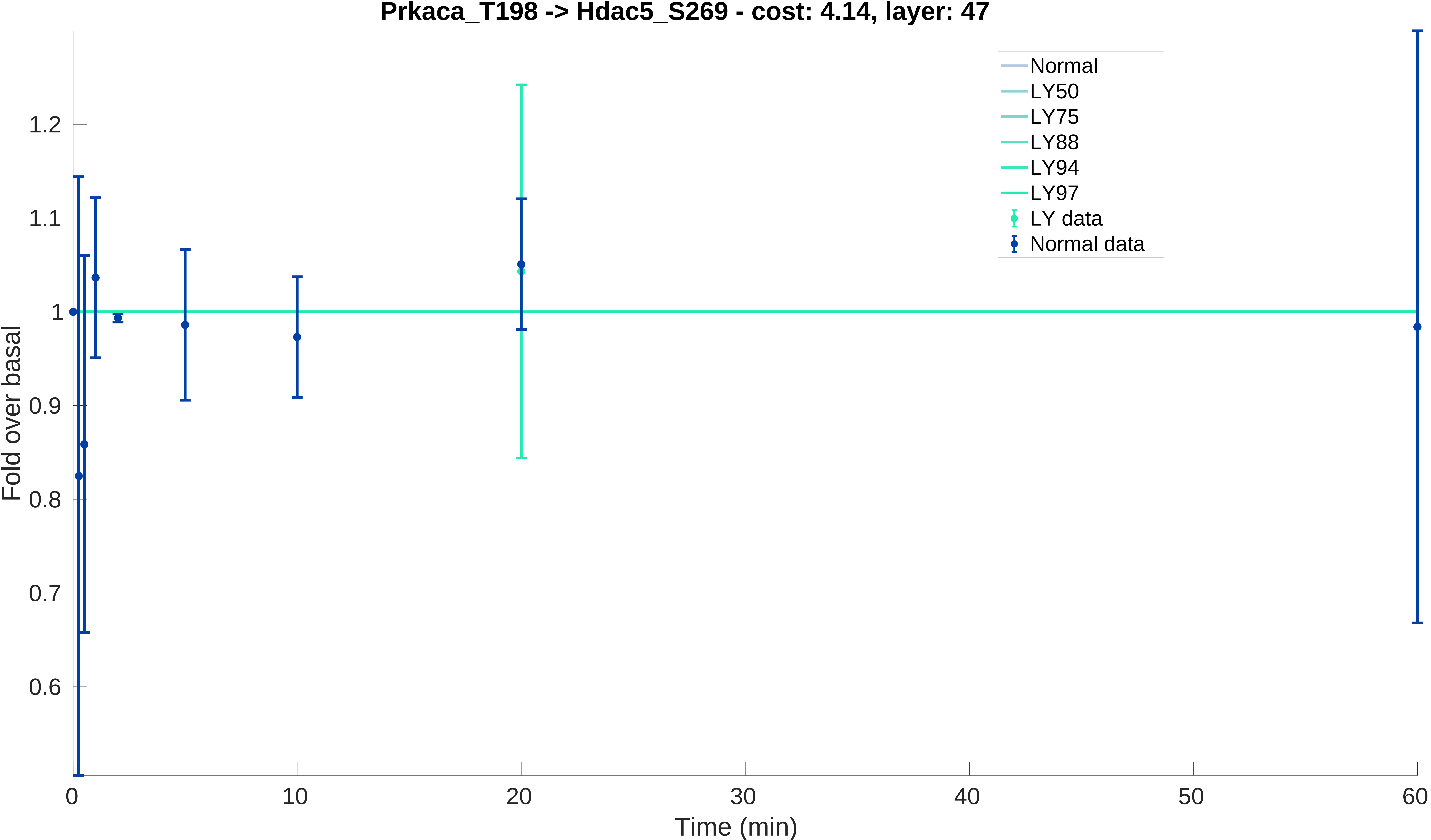

Prkaca_T198 -> Hdac5_S269 - cost: 4.14, layer: 47
Normal
LY50
LY75
LY88
LY94
LY97
LY data
Normal data
Fold over basal
1.2
1.1
1
0.9
0.8
0.7
0.6
0
10
20
30
40
50
60
Time (min)

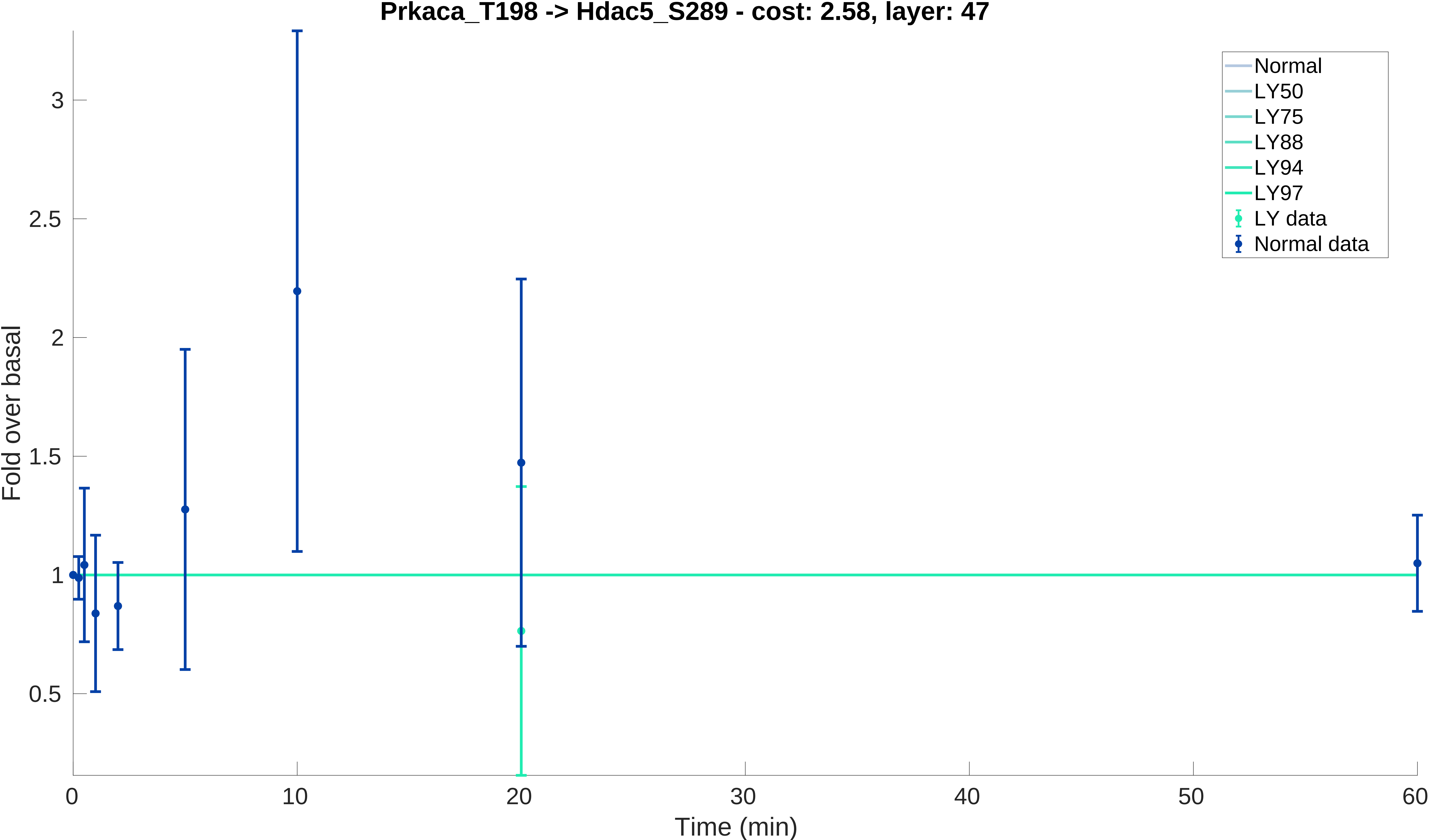

Prkaca_T198 -> Hdac5_S289 - cost: 2.58, layer: 47
Normal
LY50
LY75
LY88
LY94
LY97
LY data
Normal data
Fold over basal
0.5
1
1.5
2
2.5
3
0
10
20
30
40
50
60
Time (min)

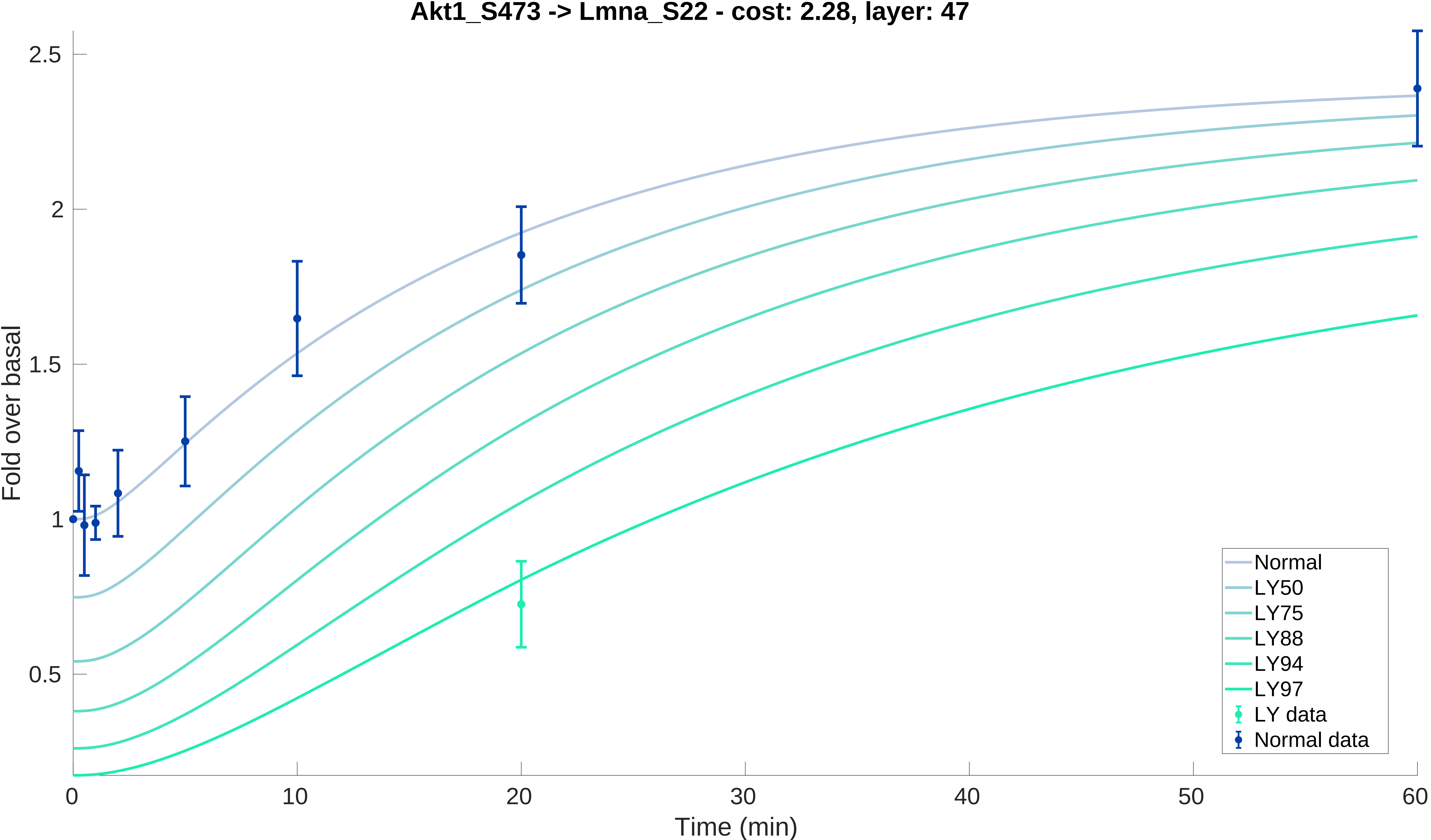

Akt1_S473 -> Lmna_S22 - cost: 2.28, layer: 47
Fold over basal
Time (min)
0.5
1
1.5
2
2.5
0
10
20
30
40
50
60
Normal
LY50
LY75
LY88
LY94
LY97
LY data
Normal data

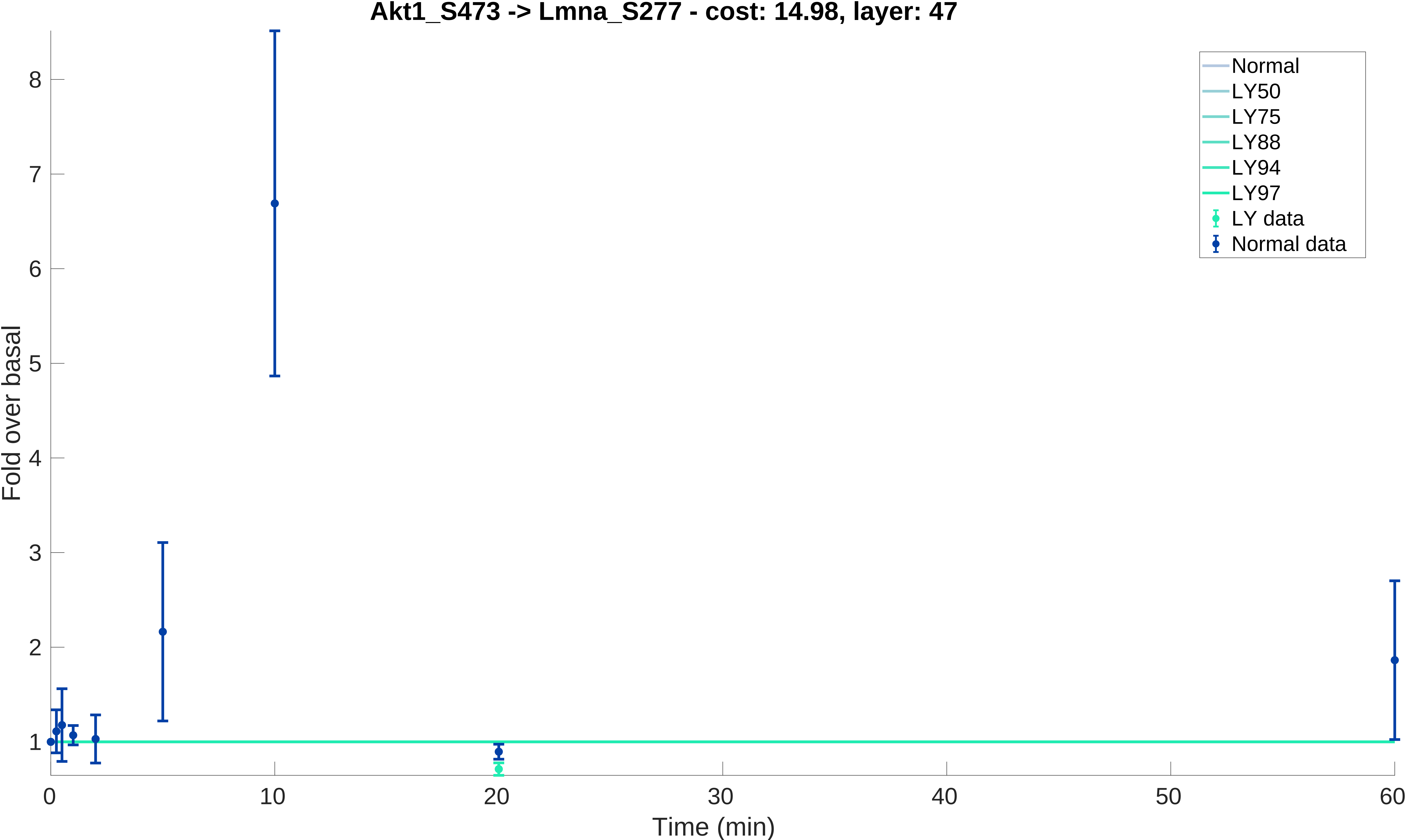
Akt1_S473 -> Lmna_S277 - cost: 14.98, layer: 47
Normal
LY50
LY75
LY88
LY94
LY97
LY data
Normal data
Fold over basal
Time (min)
1
2
3
4
5
6
7
8
0
10
20
30
40
50
60

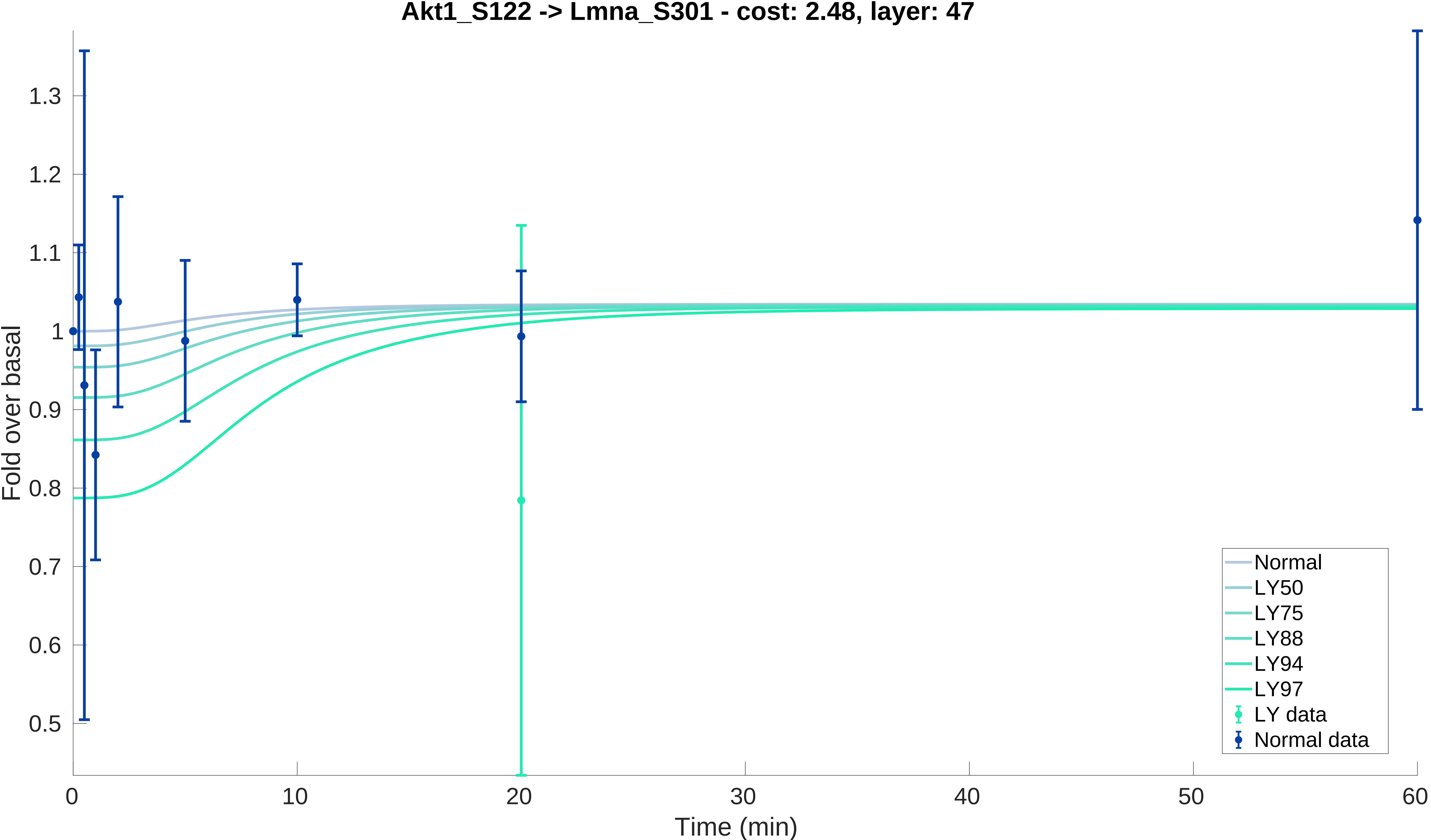

Akt1_S122 -> Lmna_S301 - cost: 2.48, layer: 47
Fold over basal
Time (min)
1.3
1.2
1.1
1
0.9
0.8
0.7
0.6
0.5
0
10
20
30
40
50
60
Normal
LY50
LY75
LY88
LY94
LY97
LY data
Normal data

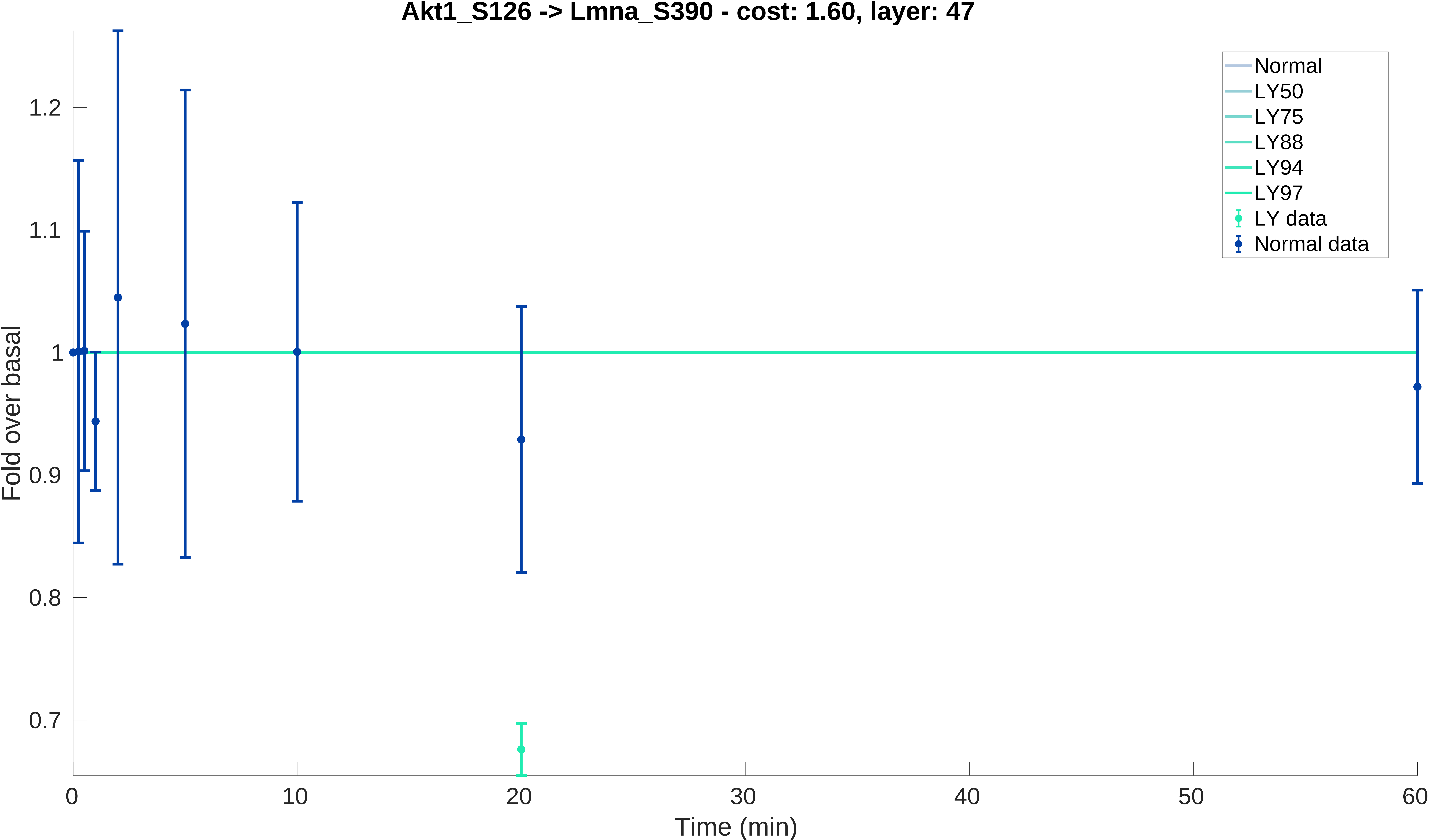
Akt1_S126 -> Lmna_S390 - cost: 1.60, layer: 47
Normal
LY50
LY75
LY88
LY94
LY97
LY data
Normal data
Fold over basal
1.2
1.1
1
0.9
0.8
0.7
0
10
20
30
40
50
60
Time (min)

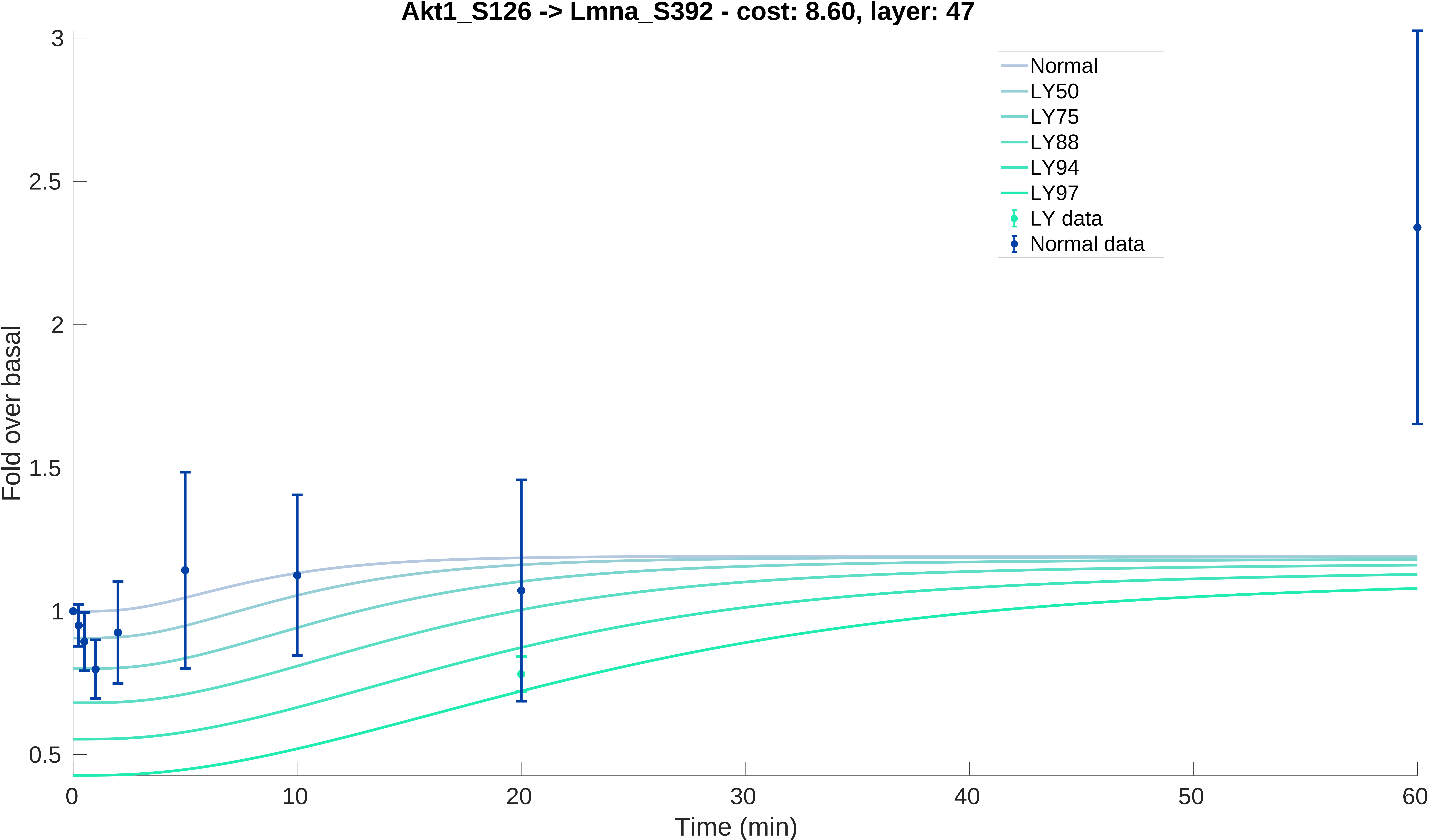

Akt1_S126 -> Lmna_S392 - cost: 8.60, layer: 47
Normal
LY50
LY75
LY88
LY94
LY97
LY data
Normal data
Fold over basal
Time (min)
0.5
1
1.5
2
2.5
3
0
10
20
30
40
50
60

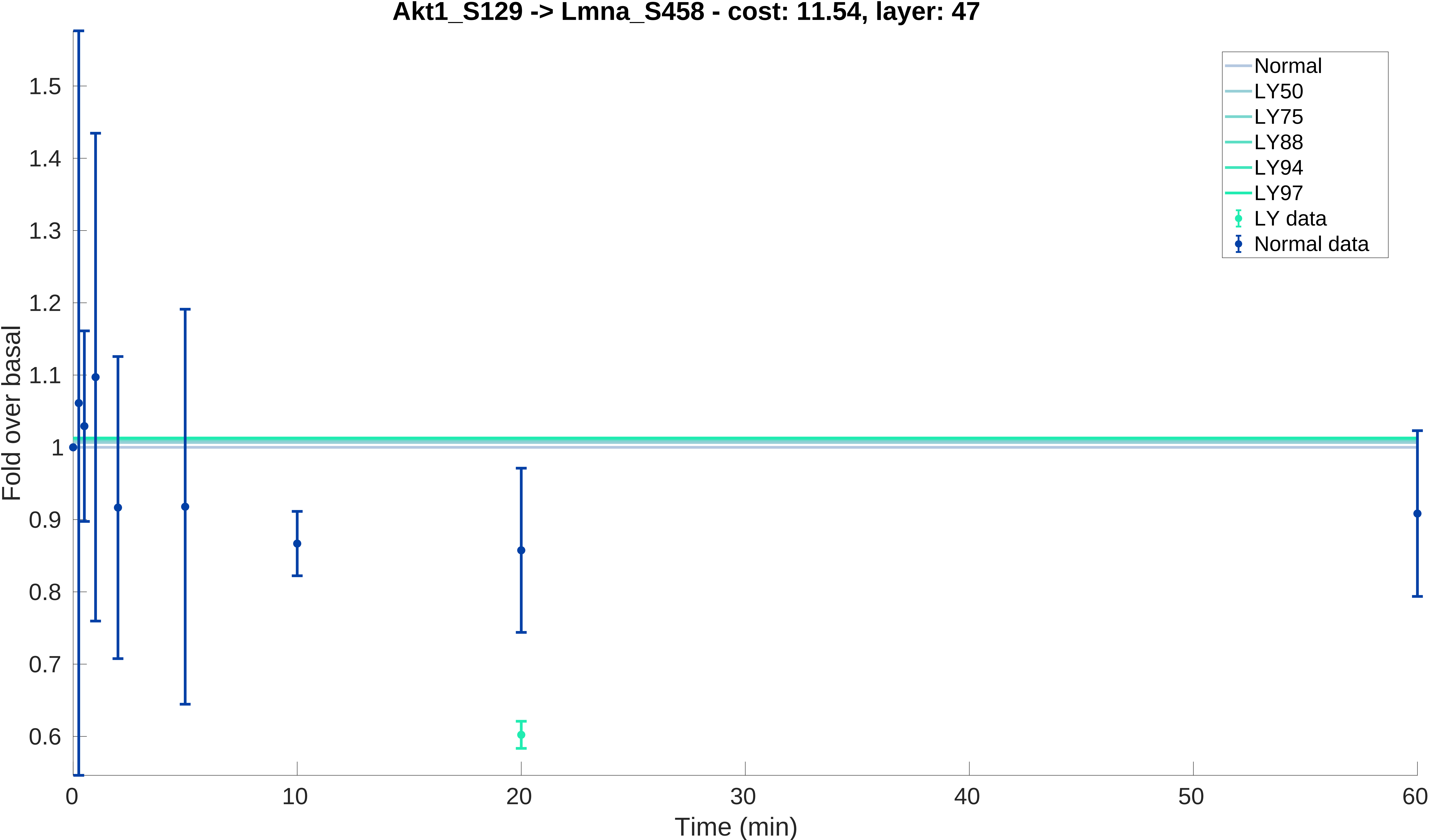

Akt1_S129 -> Lmna_S458 - cost: 11.54, layer: 47
Fold over basal
Time (min)
Normal
LY50
LY75
LY88
LY94
LY97
LY data
Normal data
1.5
1.4
1.3
1.2
1.1
1
0.9
0.8
0.7
0.6
0
10
20
30
40
50
60

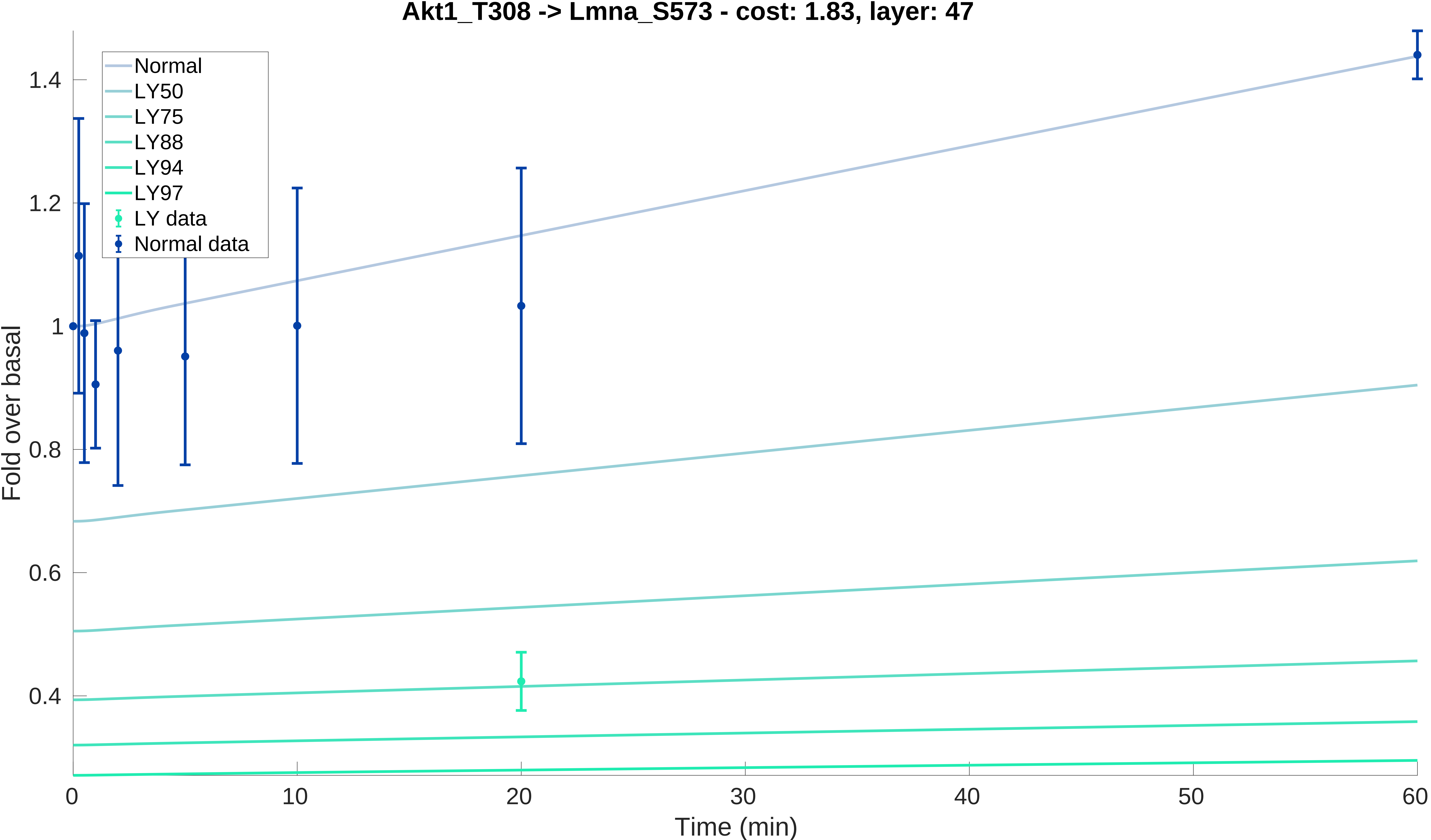

Akt1_T308 -> Lmna_S573 - cost: 1.83, layer: 47
Normal
LY50
LY75
LY88
LY94
LY97
LY data
Normal data
Fold over basal
1.4
1.2
1
0.8
0.6
0.4
0
10
20
30
40
50
60
Time (min)

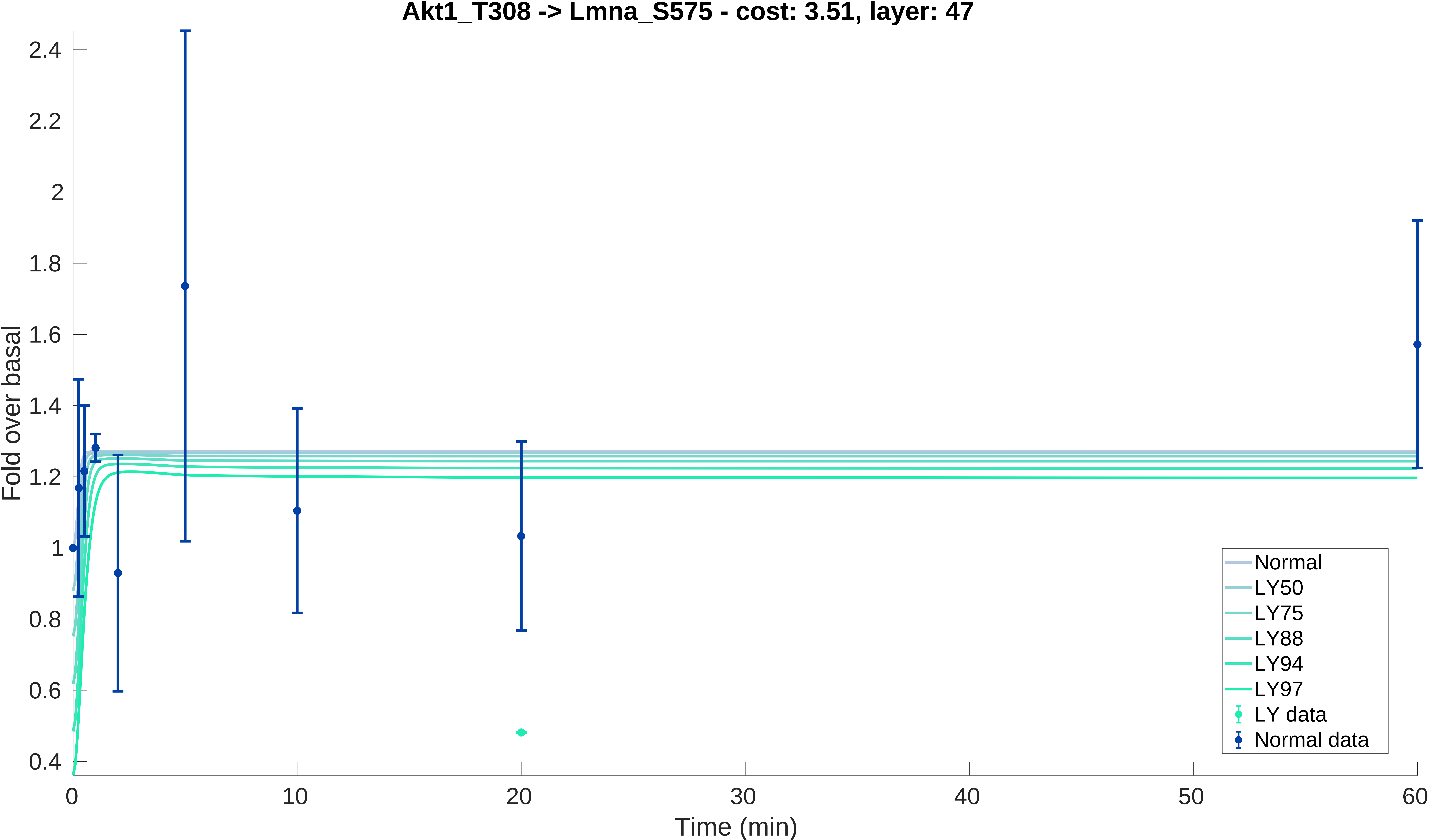

Akt1_T308 -> Lmna_S575 - cost: 3.51, layer: 47
Fold over basal
Time (min)
2.4
2.2
2
1.8
1.6
1.4
1.2
1
0.8
0.6
0.4
0
10
20
30
40
50
60
Normal
LY50
LY75
LY88
LY94
LY97
LY data
Normal data

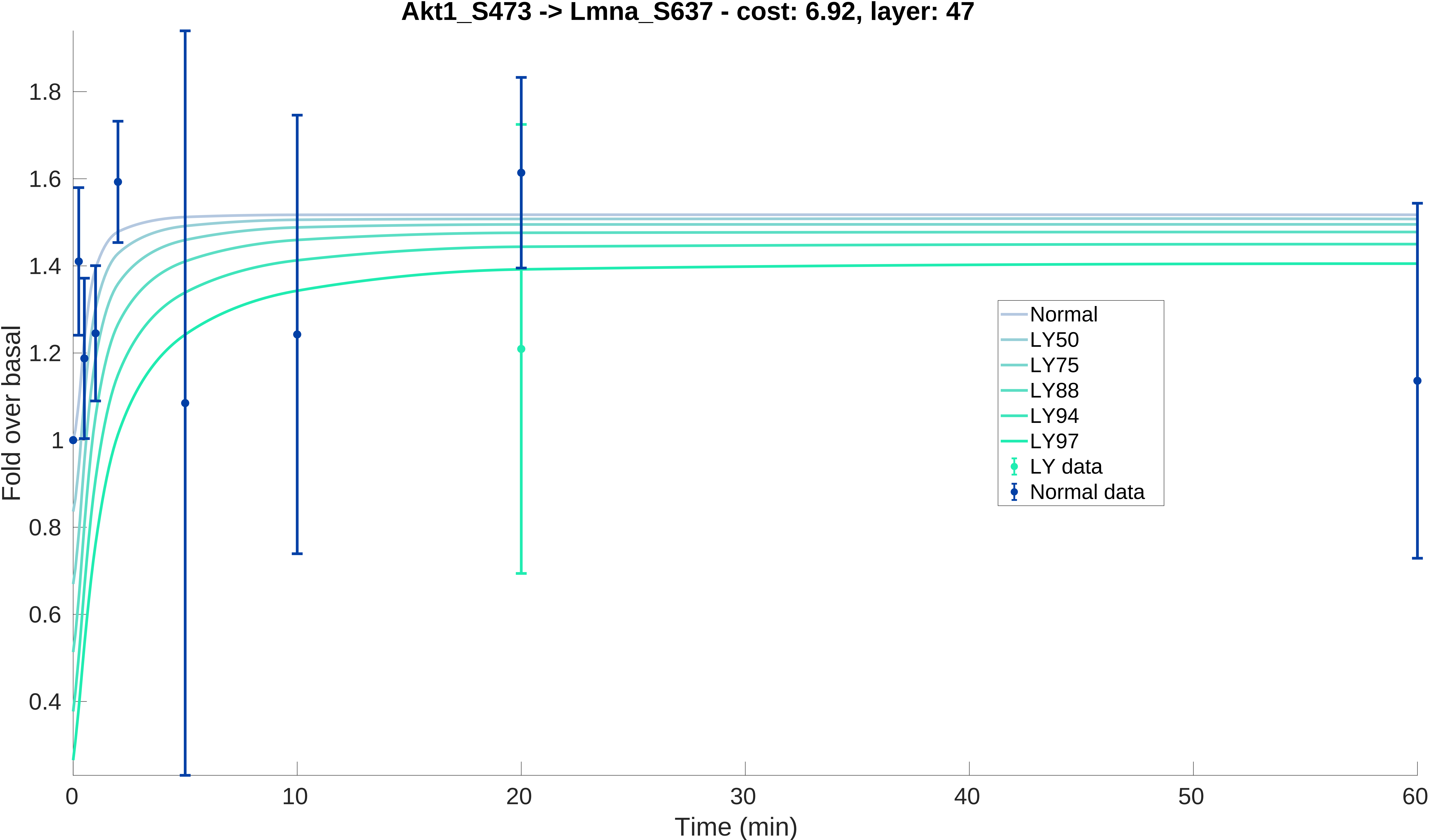

Akt1_S473 -> Lmna_S637 - cost: 6.92, layer: 47
Fold over basal
1.8
1.6
1.4
1.2
1
0.8
0.6
0.4
0
10
20
30
40
50
60
Time (min)
Normal
LY50
LY75
LY88
LY94
LY97
LY data
Normal data

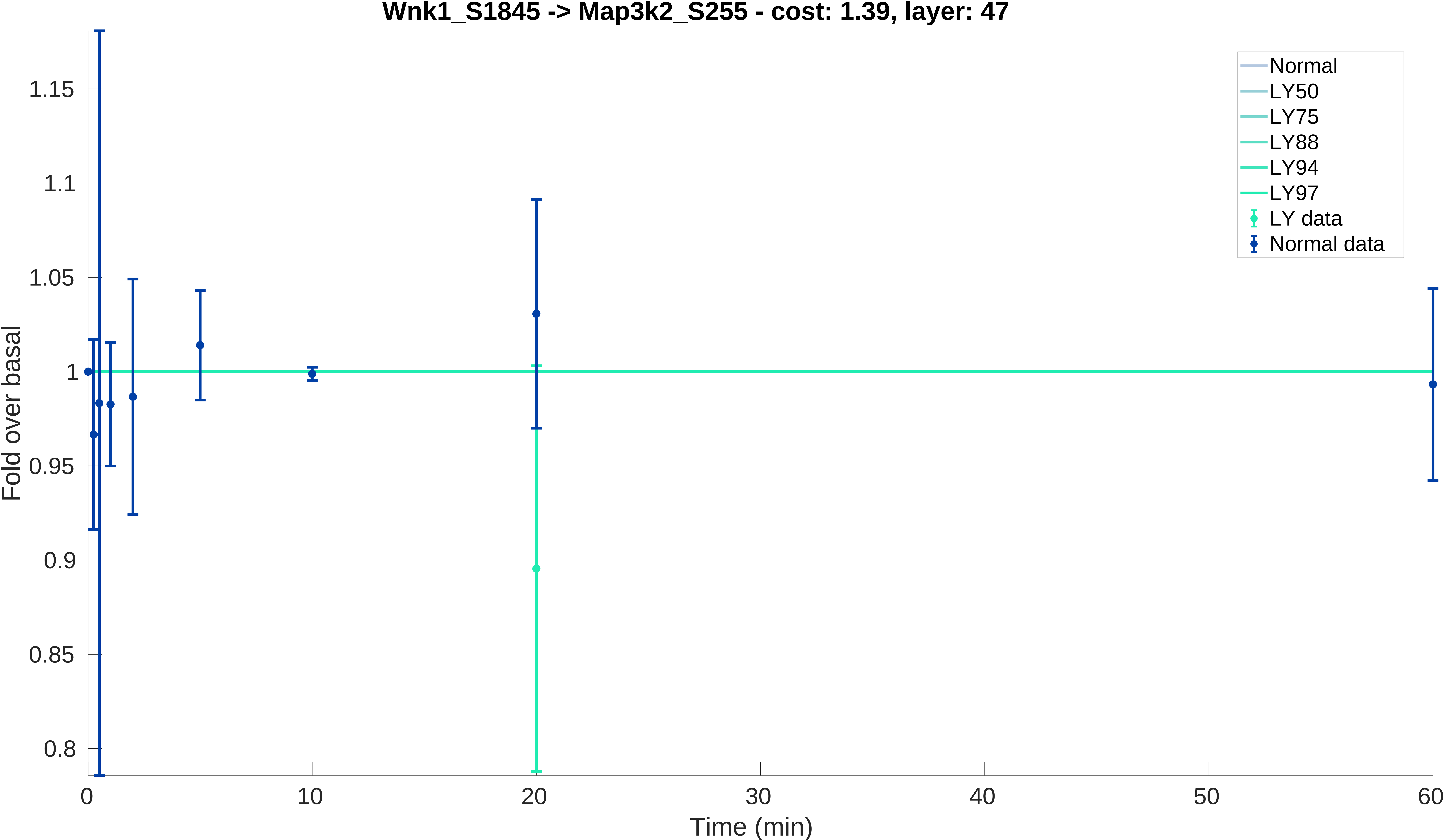

Wnk1_S1845 -> Map3k2_S255 - cost: 1.39, layer: 47
Normal
LY50
LY75
LY88
LY94
LY97
LY data
Normal data
Fold over basal
1.15
1.1
1.05
1
0.95
0.9
0.85
0.8
0
10
20
30
40
50
60
Time (min)

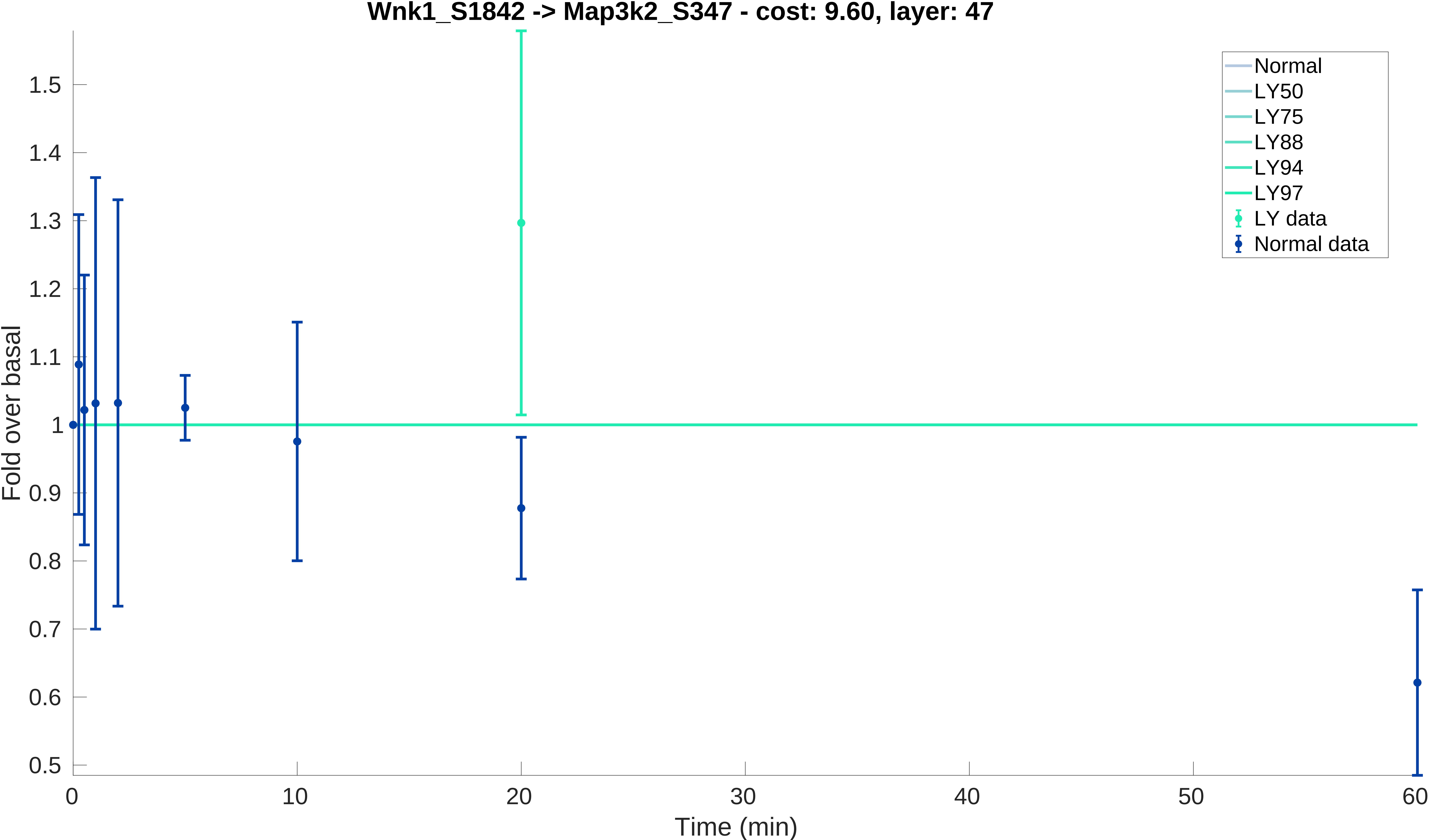

Wnk1_S1842 -> Map3k2_S347 - cost: 9.60, layer: 47
Fold over basal
Time (min)
Normal
LY50
LY75
LY88
LY94
LY97
LY data
Normal data
0.5
0.6
0.7
0.8
0.9
1
1.1
1.2
1.3
1.4
1.5
0
10
20
30
40
50
60

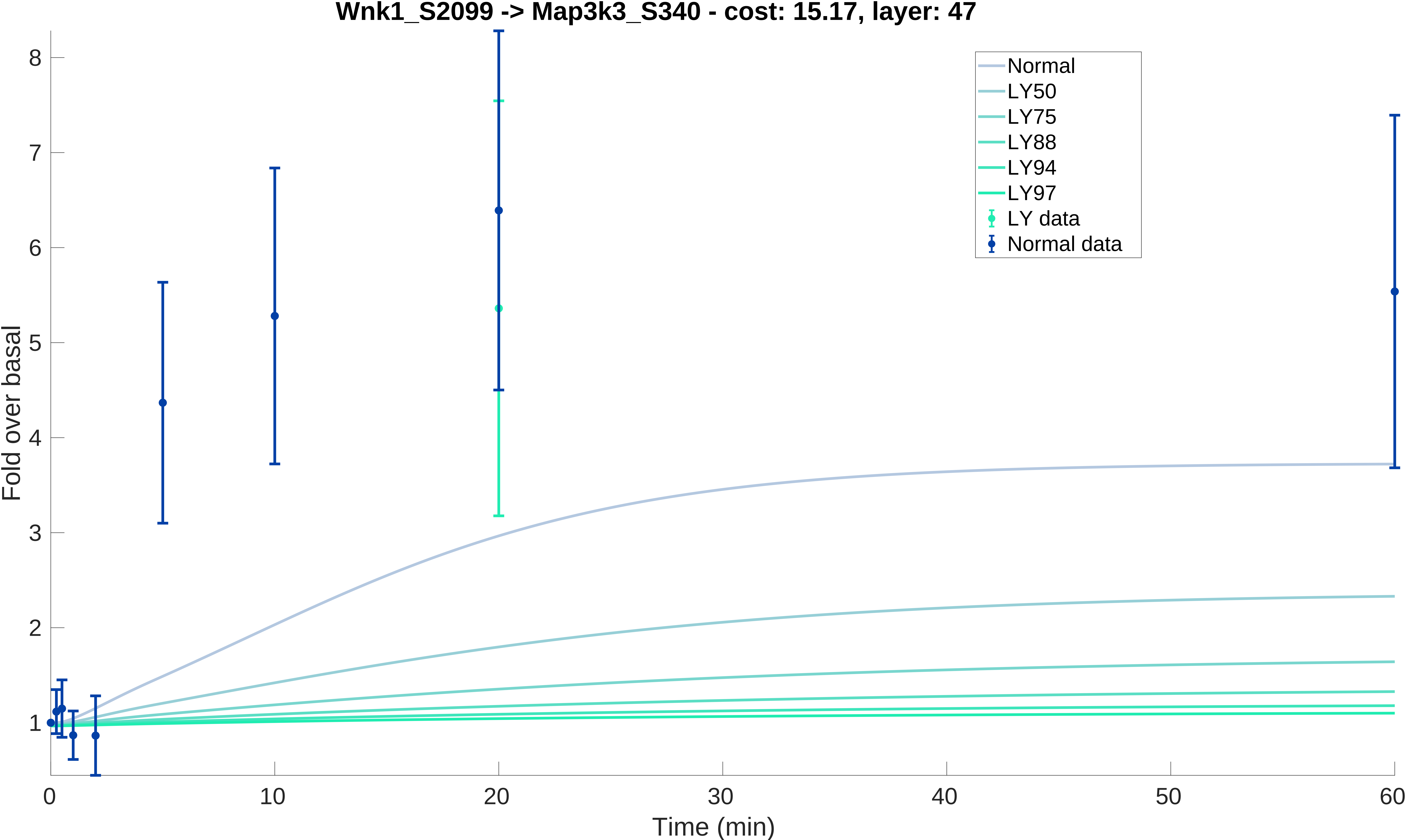

Wnk1_S2099 -> Map3k3_S340 - cost: 15.17, layer: 47
Fold over basal
Time (min)
Normal
LY50
LY75
LY88
LY94
LY97
LY data
Normal data
1
2
3
4
5
6
7
8
0
10
20
30
40
50
60

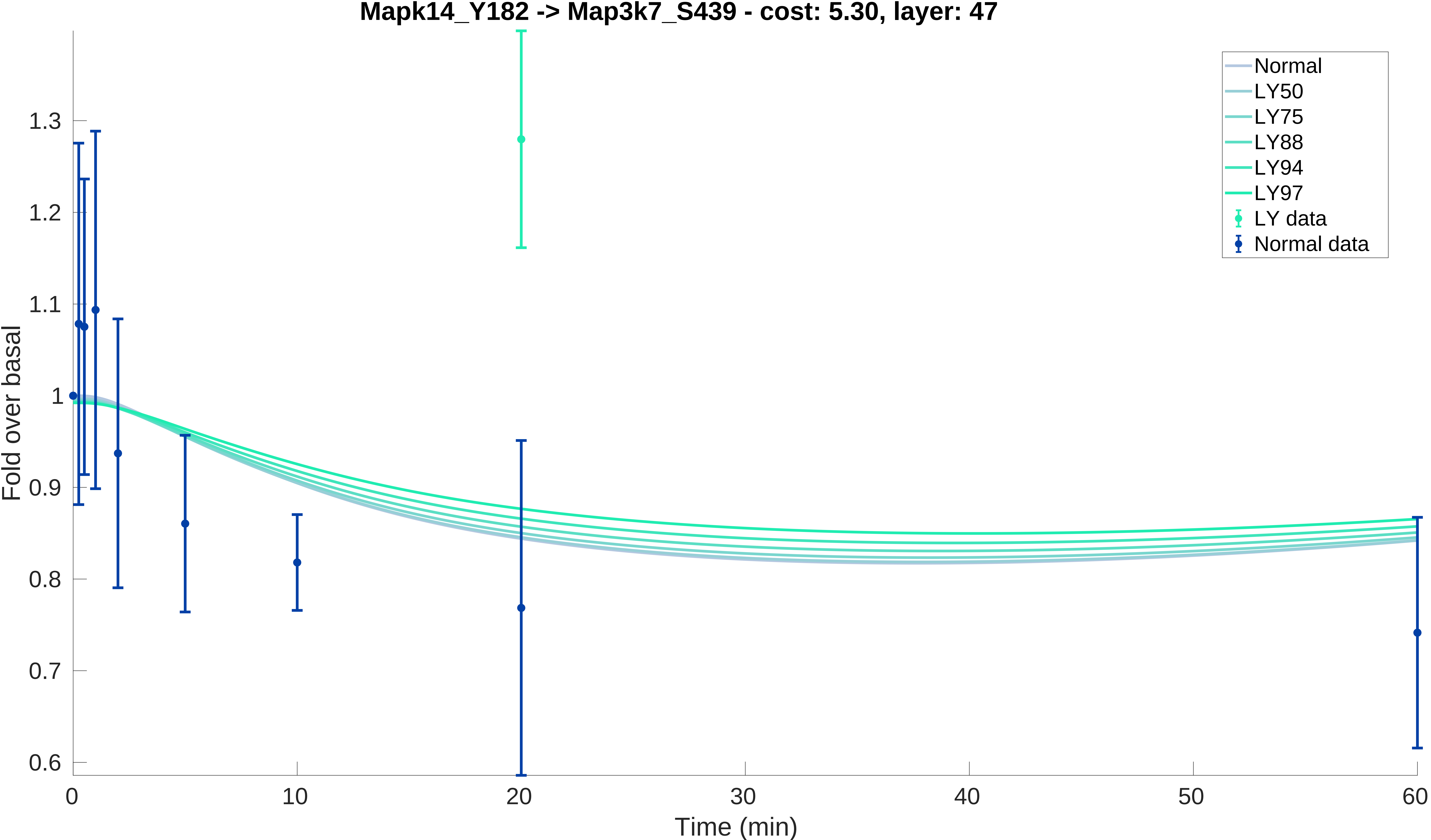
Mapk14_Y182 -> Map3k7_S439 - cost: 5.30, layer: 47
Fold over basal
Time (min)
0.6
0.7
0.8
0.9
1
1.1
1.2
1.3
0
10
20
30
40
50
60
Normal
LY50
LY75
LY88
LY94
LY97
LY data
Normal data

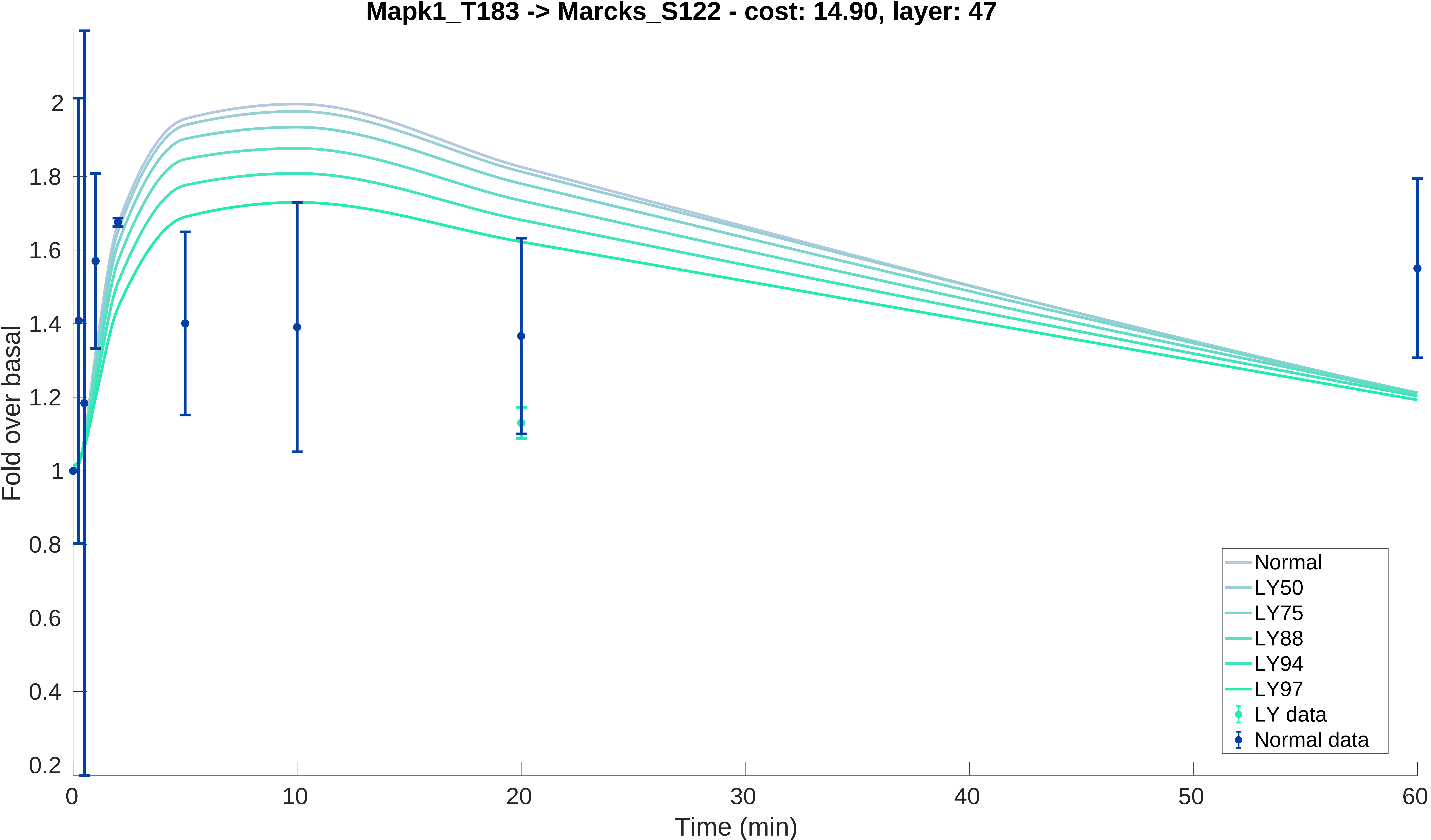

Mapk1_T183 -> Marcks_S122 - cost: 14.90, layer: 47
Fold over basal
Time (min)
0.2
0.4
0.6
0.8
1
1.2
1.4
1.6
1.8
2
0
10
20
30
40
50
60
Normal
LY50
LY75
LY88
LY94
LY97
LY data
Normal data

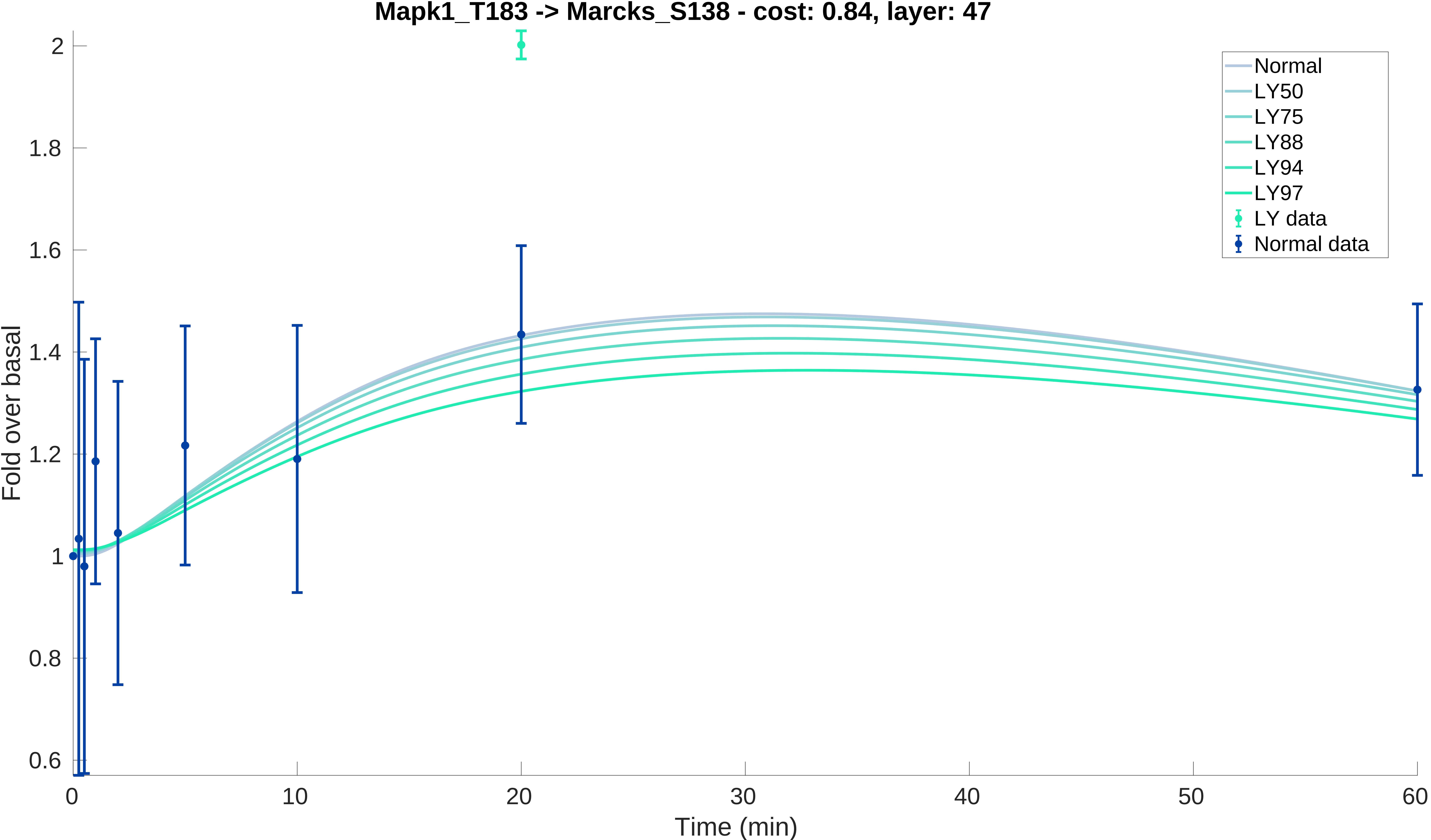

Mapk1_T183 -> Marcks_S138 - cost: 0.84, layer: 47
Fold over basal
Time (min)
Normal
LY50
LY75
LY88
LY94
LY97
LY data
Normal data
0.6
0.8
1
1.2
1.4
1.6
1.8
2
0
10
20
30
40
50
60

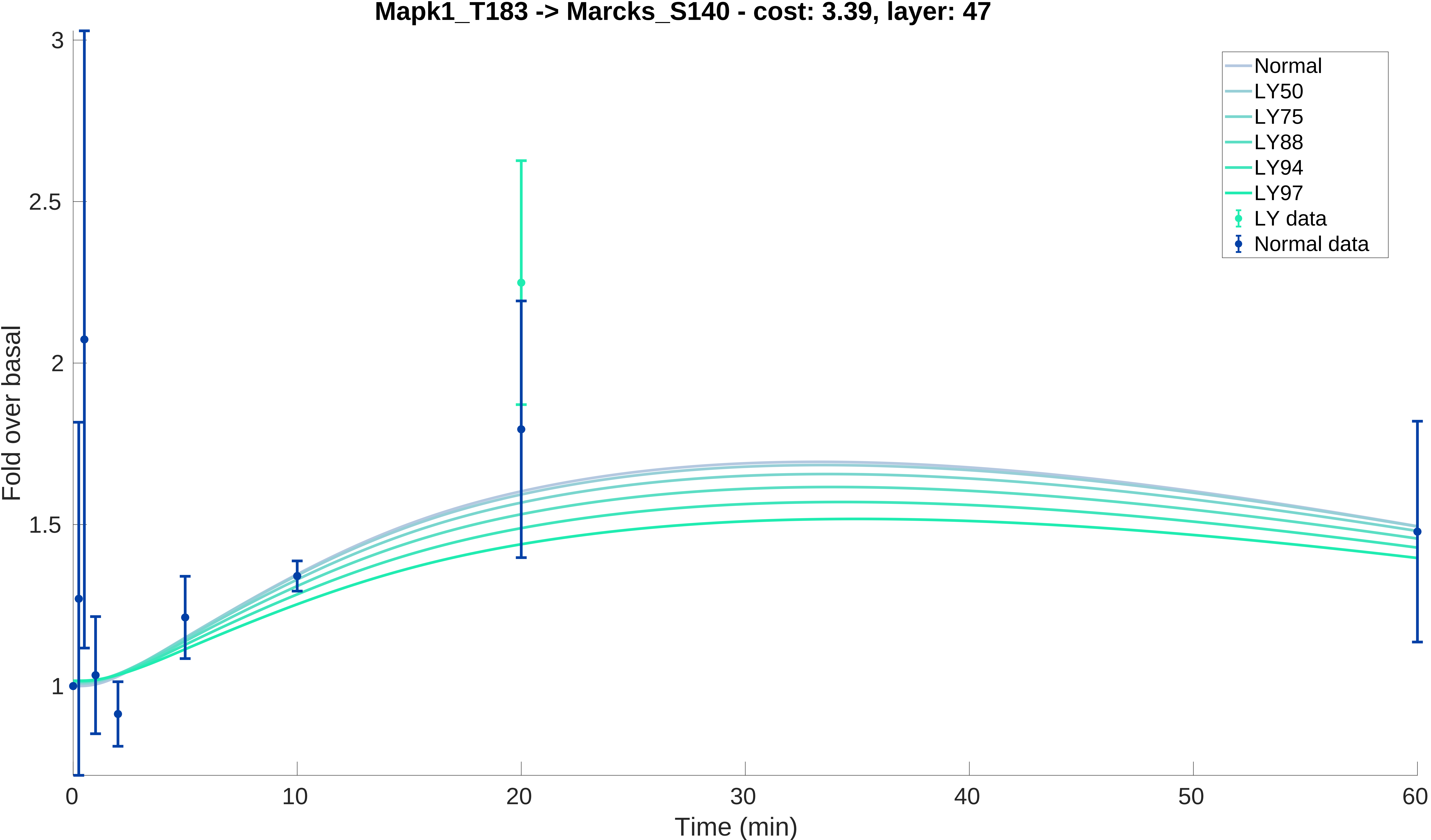

Mapk1_T183 -> Marcks_S140 - cost: 3.39, layer: 47
Fold over basal
Time (min)
Normal
LY50
LY75
LY88
LY94
LY97
LY data
Normal data
0
10
20
30
40
50
60
1
1.5
2
2.5
3

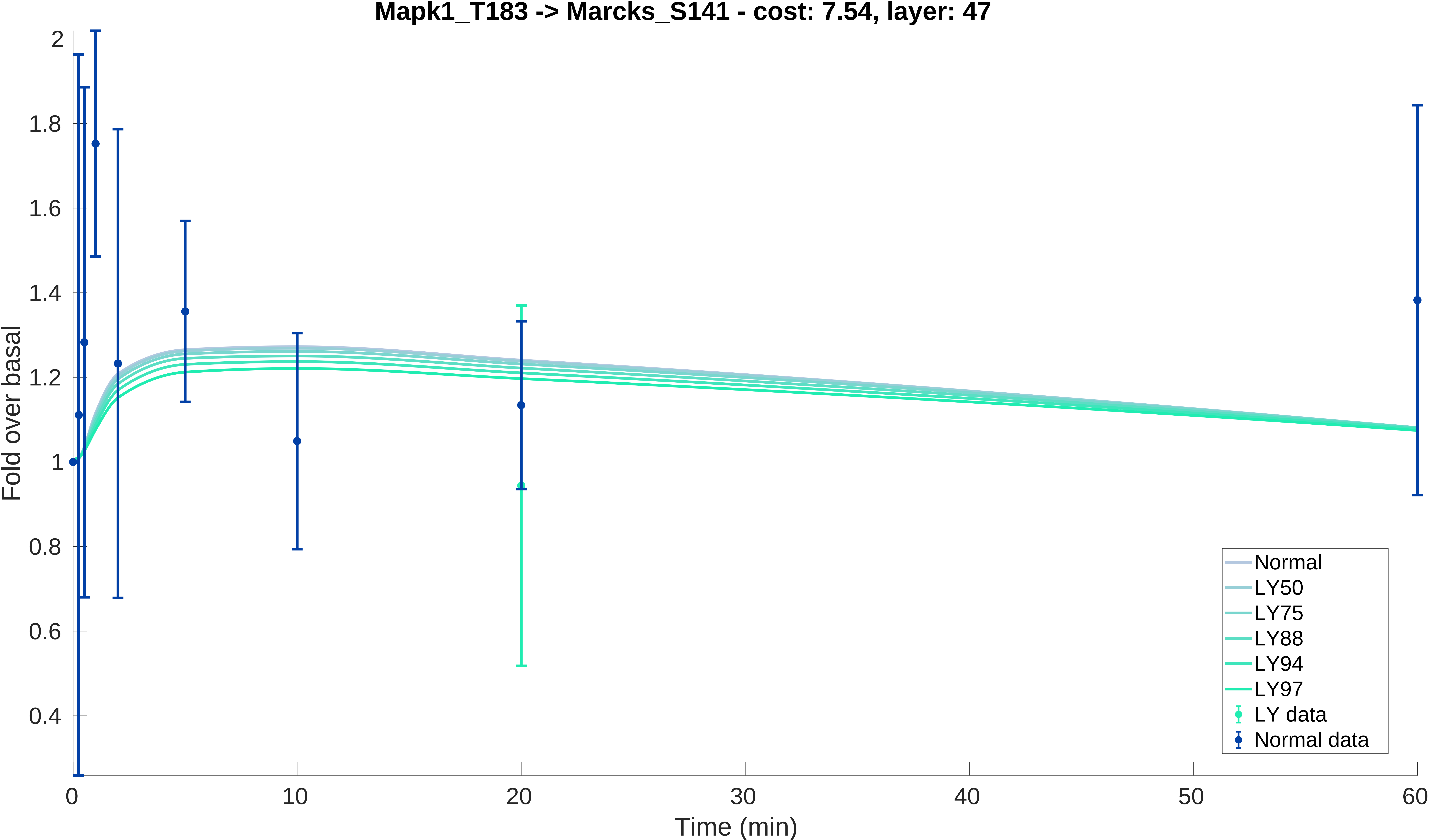
Mapk1_T183 -> Marcks_S141 - cost: 7.54, layer: 47
Fold over basal
Time (min)
0
10
20
30
40
50
60
0.4
0.6
0.8
1
1.2
1.4
1.6
1.8
2
Normal
LY50
LY75
LY88
LY94
LY97
LY data
Normal data

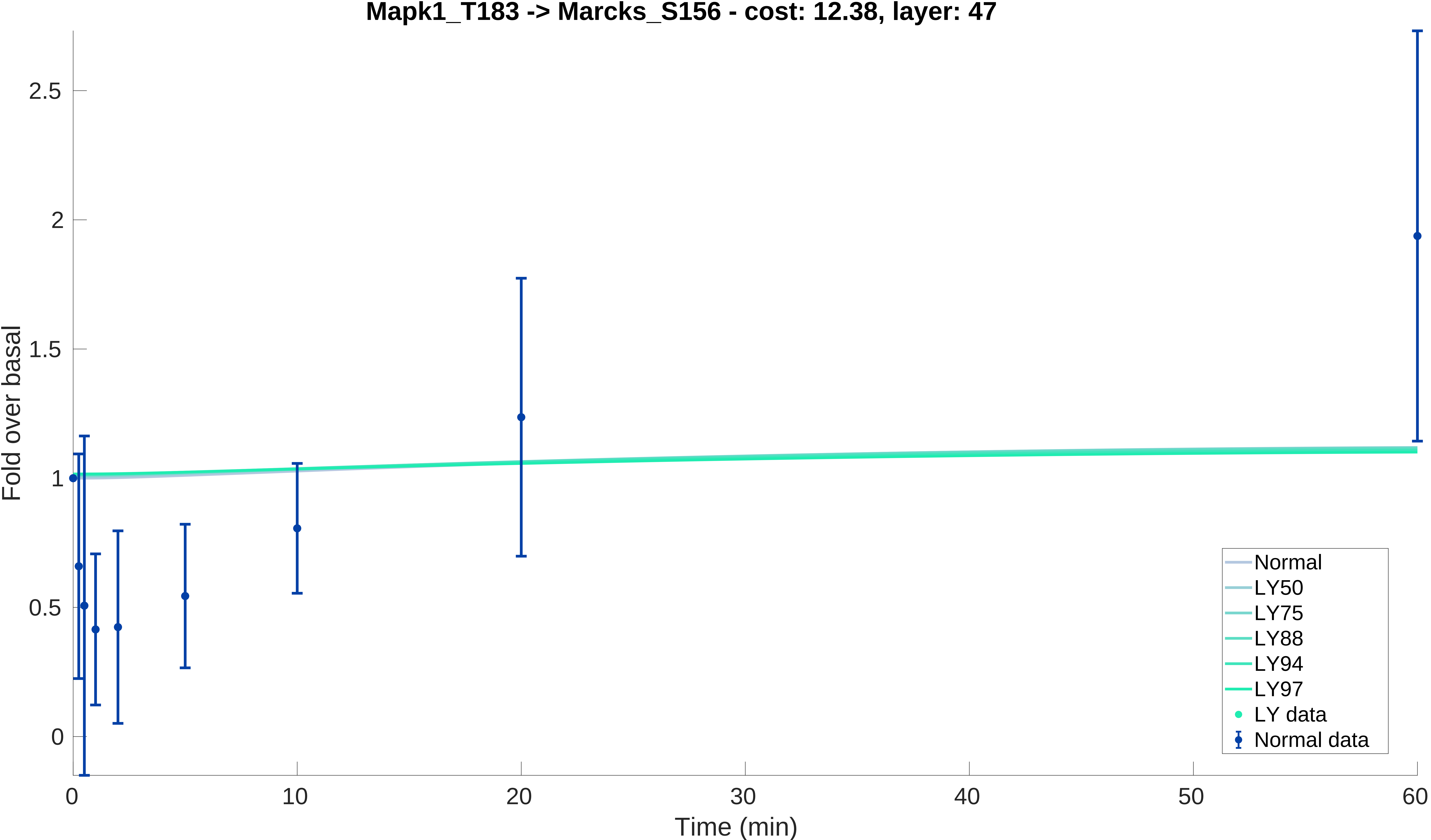

Mapk1_T183 -> Marcks_S156 - cost: 12.38, layer: 47
Fold over basal
2.5
2
1.5
1
0.5
0
0
10
20
30
40
50
60
Time (min)
Normal
LY50
LY75
LY88
LY94
LY97
LY data
Normal data

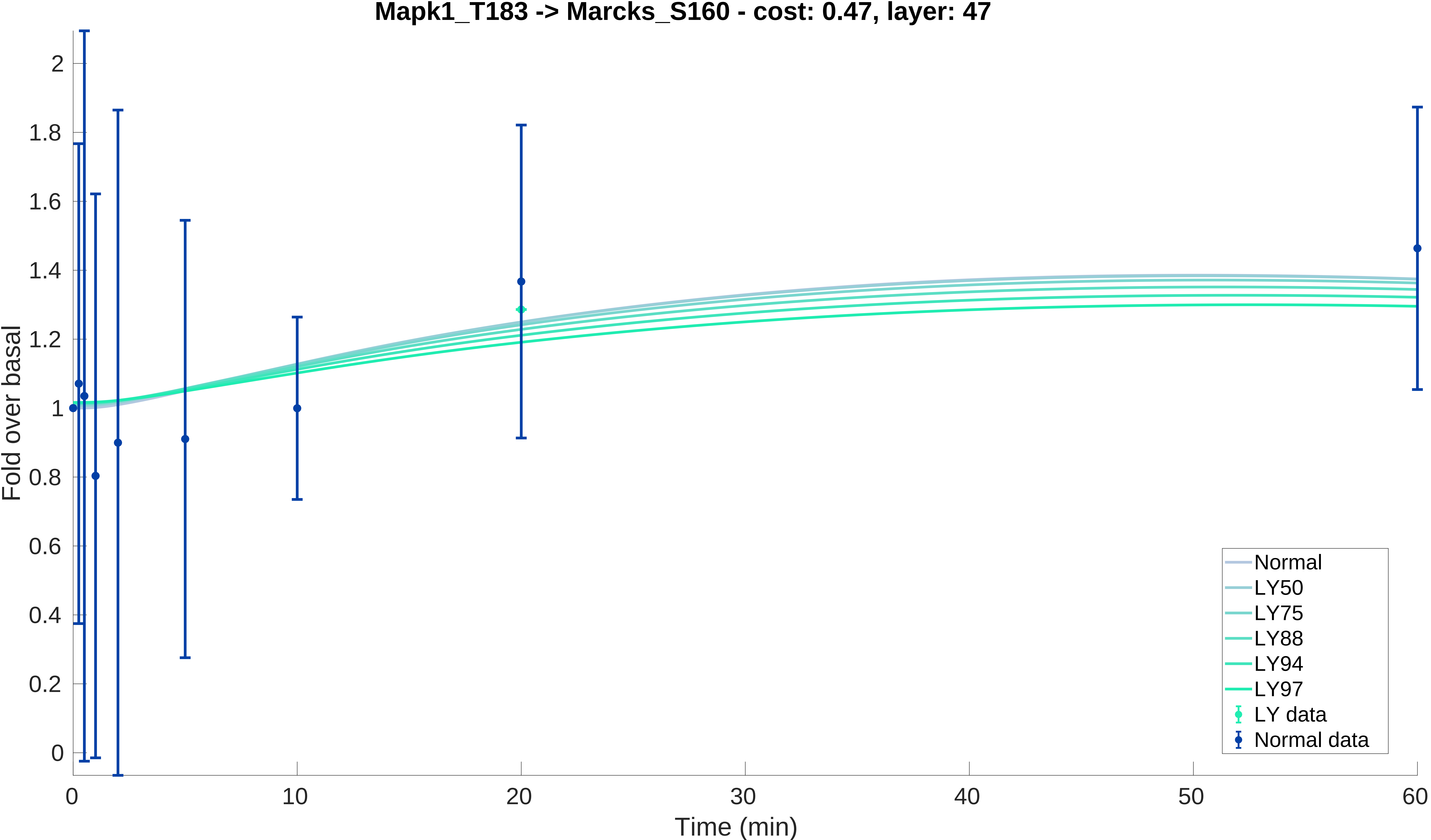

Mapk1_T183 -> Marcks_S160 - cost: 0.47, layer: 47
Fold over basal
Time (min)
0
0.2
0.4
0.6
0.8
1
1.2
1.4
1.6
1.8
2
0
10
20
30
40
50
60
Normal
LY50
LY75
LY88
LY94
LY97
LY data
Normal data

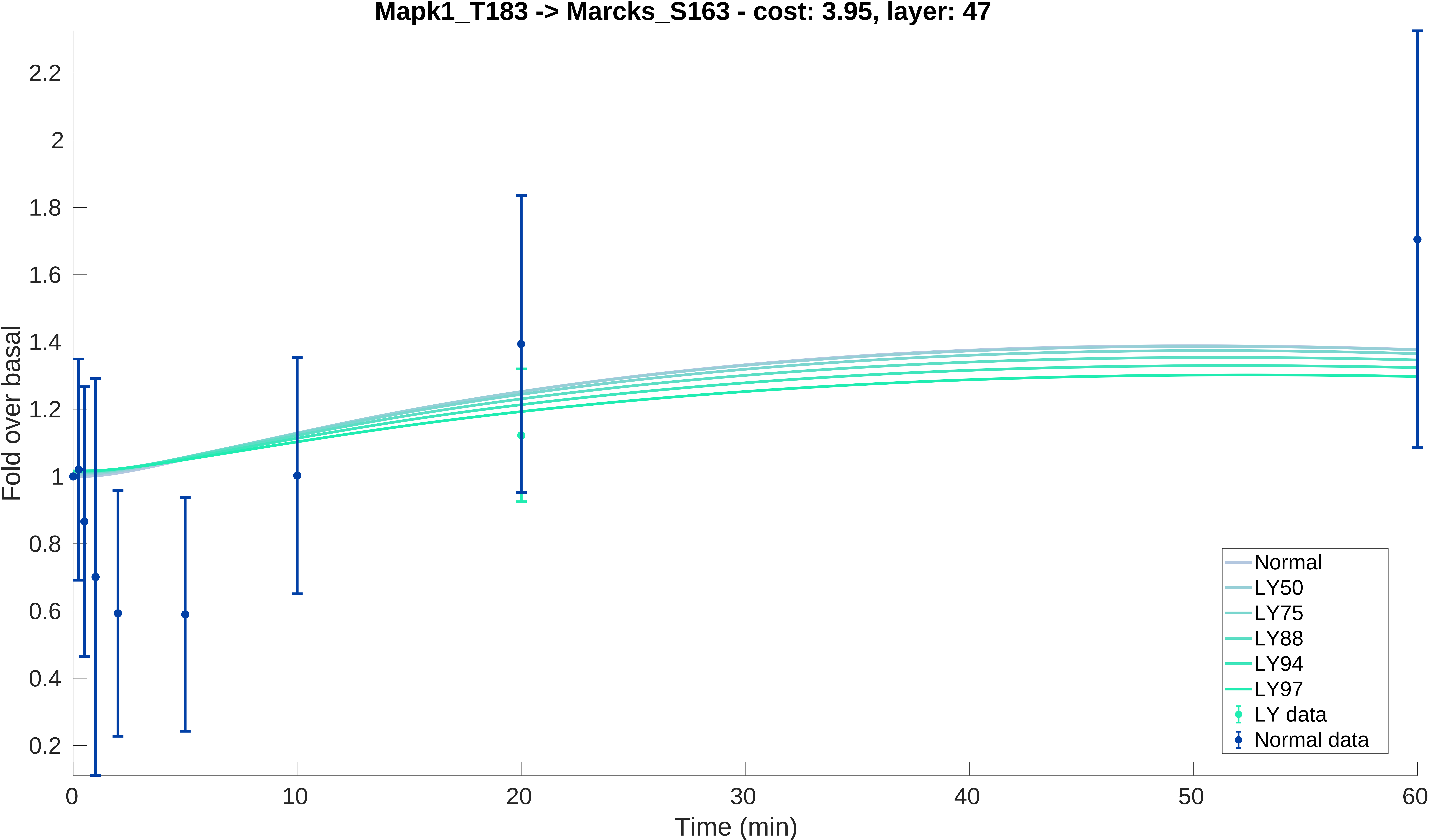

Mapk1_T183 -> Marcks_S163 - cost: 3.95, layer: 47
Fold over basal
Time (min)
0.2
0.4
0.6
0.8
1
1.2
1.4
1.6
1.8
2
2.2
0
10
20
30
40
50
60
Normal
LY50
LY75
LY88
LY94
LY97
LY data
Normal data

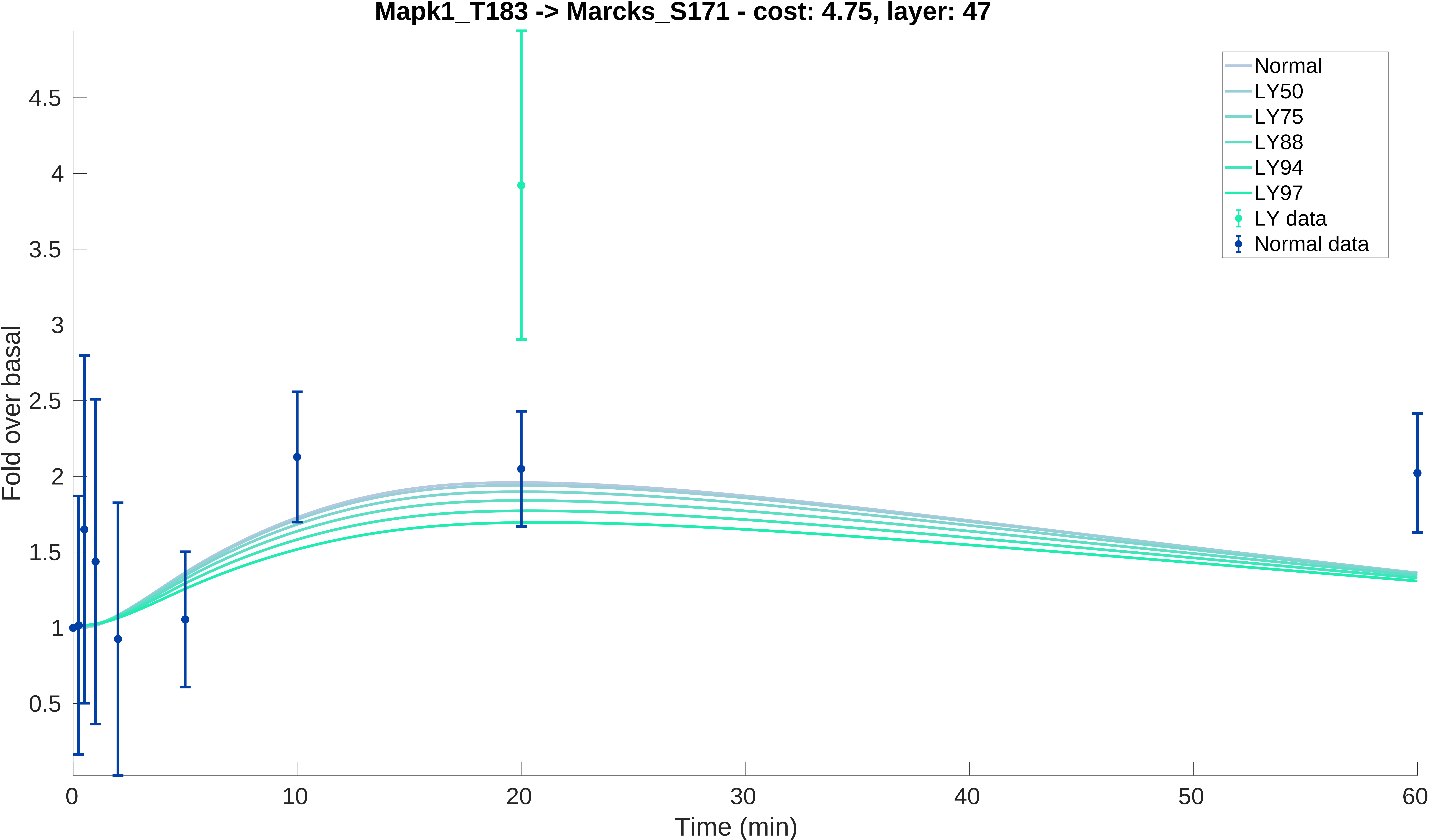
Mapk1_T183 -> Marcks_S171 - cost: 4.75, layer: 47
Fold over basal
Time (min)
0.5
1
1.5
2
2.5
3
3.5
4
4.5
0
10
20
30
40
50
60
Normal
LY50
LY75
LY88
LY94
LY97
LY data
Normal data

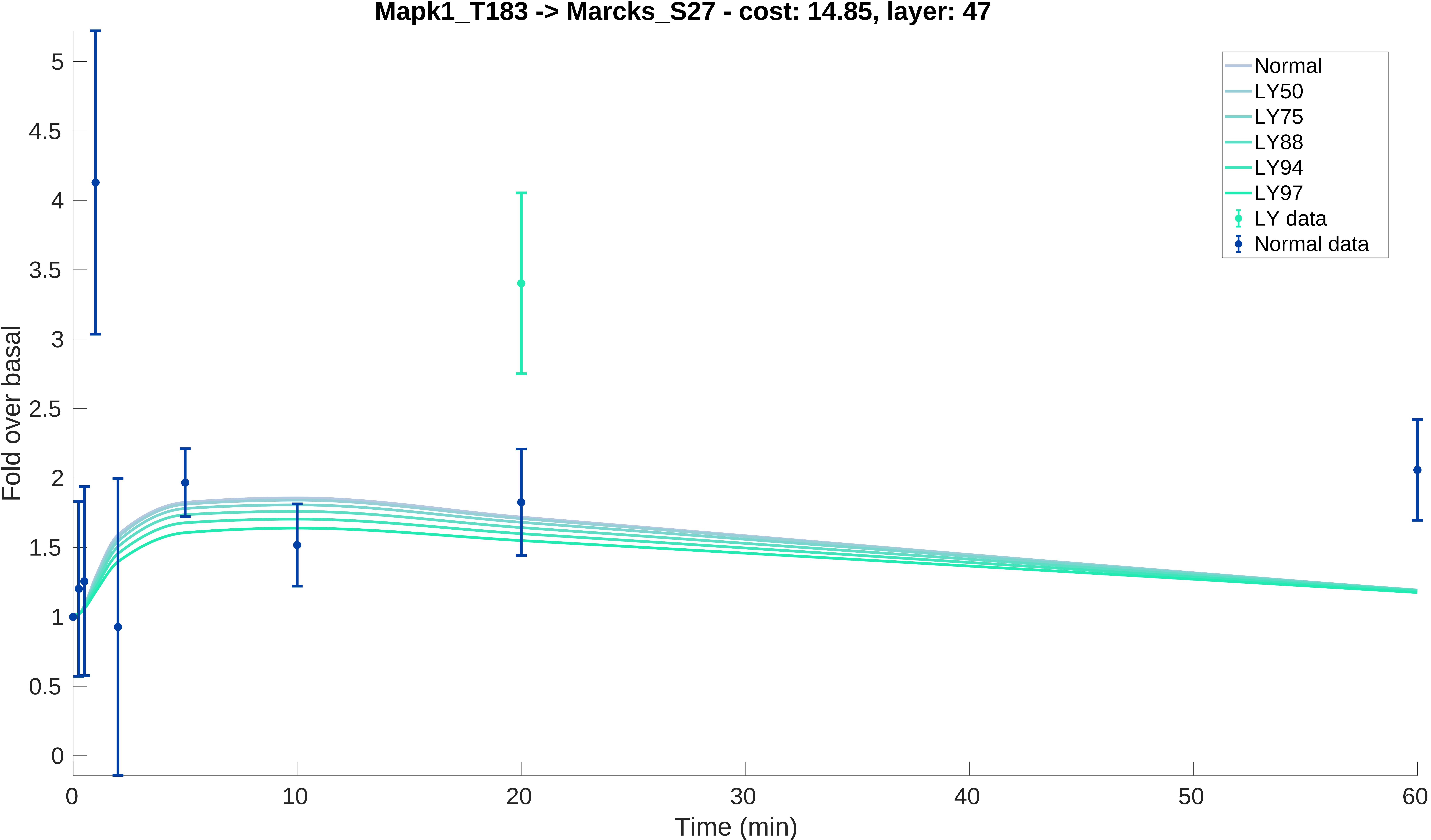

Mapk1_T183 -> Marcks_S27 - cost: 14.85, layer: 47
Normal
LY50
LY75
LY88
LY94
LY97
LY data
Normal data
Fold over basal
Time (min)
0
0.5
1
1.5
2
2.5
3
3.5
4
4.5
5
0
10
20
30
40
50
60

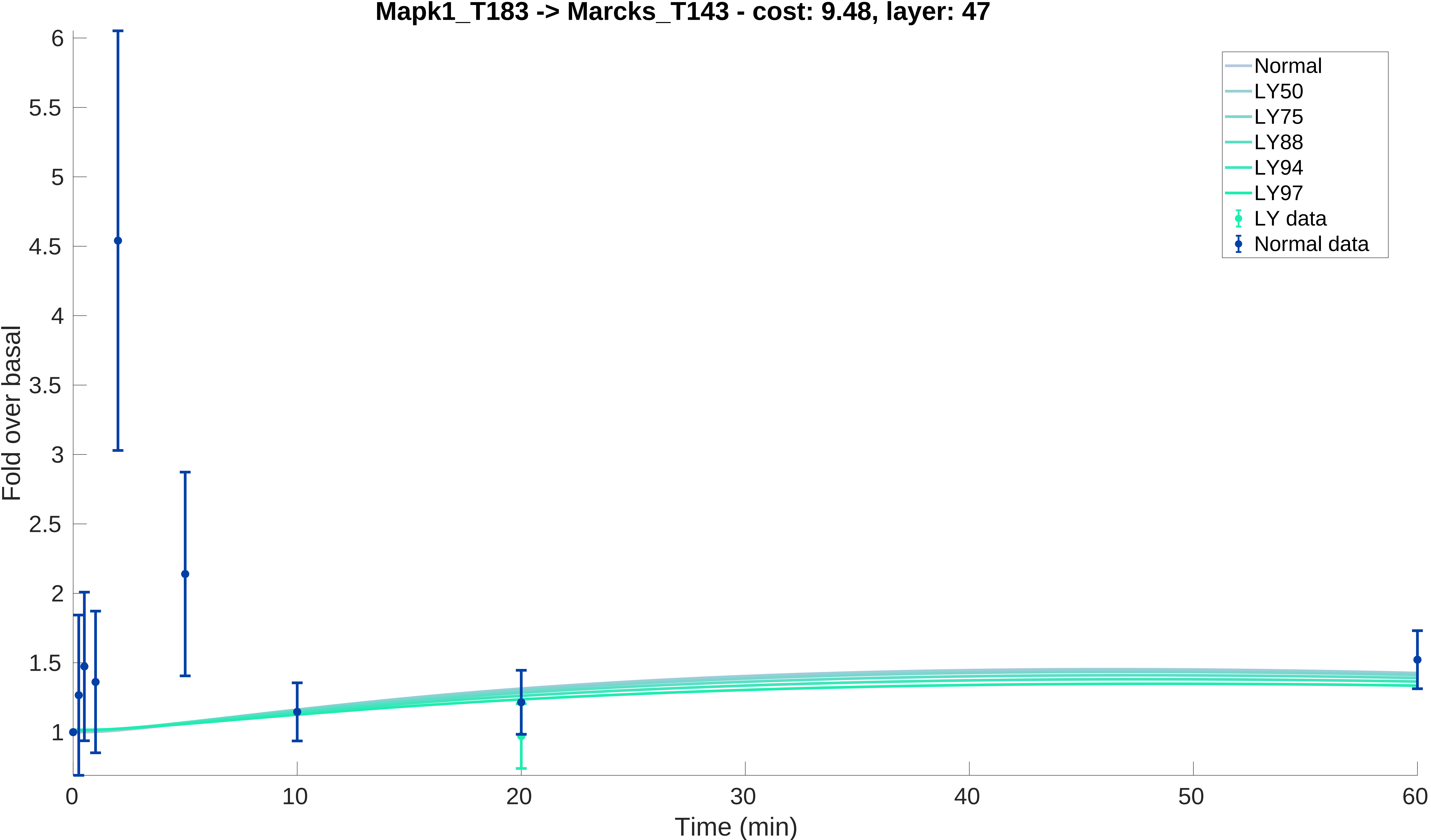

Mapk1_T183 -> Marcks_T143 - cost: 9.48, layer: 47
Fold over basal
6
5.5
5
4.5
4
3.5
3
2.5
2
1.5
1
0
10
20
30
40
50
60
Time (min)
Normal
LY50
LY75
LY88
LY94
LY97
LY data
Normal data

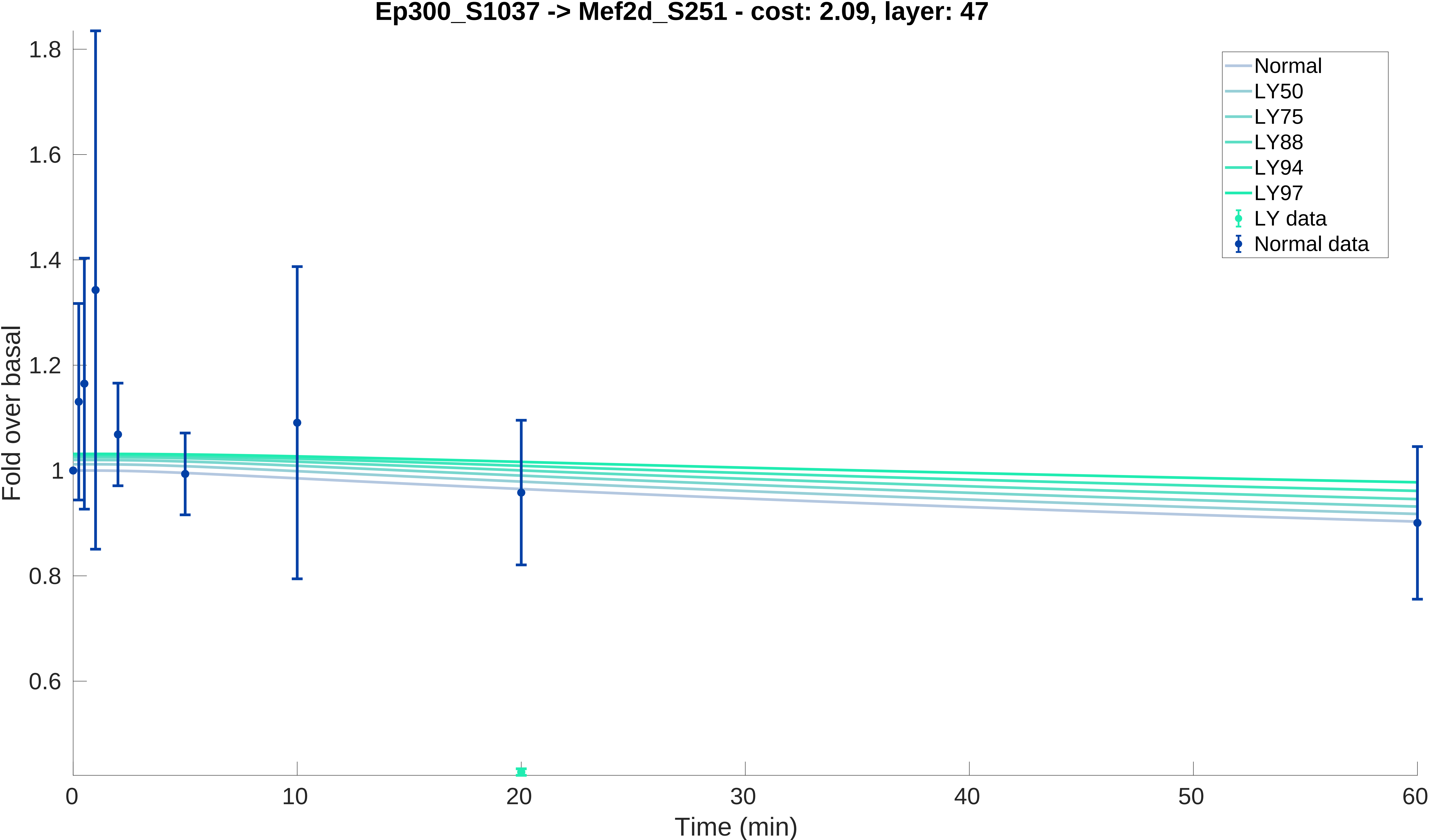
Ep300_S1037 -> Mef2d_S251 - cost: 2.09, layer: 47
Fold over basal
1.8
1.6
1.4
1.2
1
0.8
0.6
0
10
20
30
40
50
60
Time (min)
Normal
LY50
LY75
LY88
LY94
LY97
LY data
Normal data

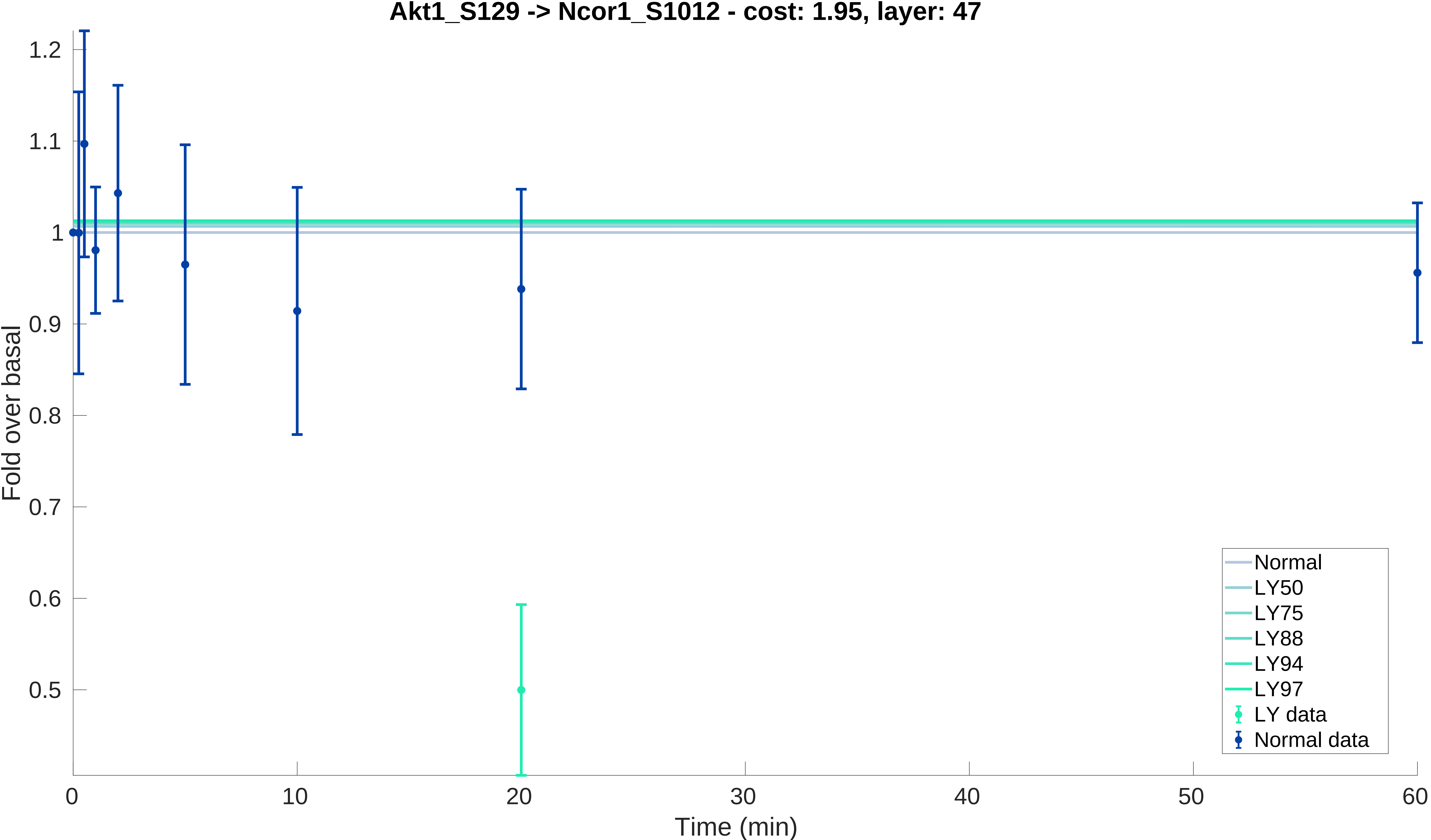

Akt1_S129 -> Ncor1_S1012 - cost: 1.95, layer: 47
Fold over basal
1.2
1.1
1
0.9
0.8
0.7
0.6
0.5
0
10
20
30
40
50
60
Time (min)
Normal
LY50
LY75
LY88
LY94
LY97
LY data
Normal data

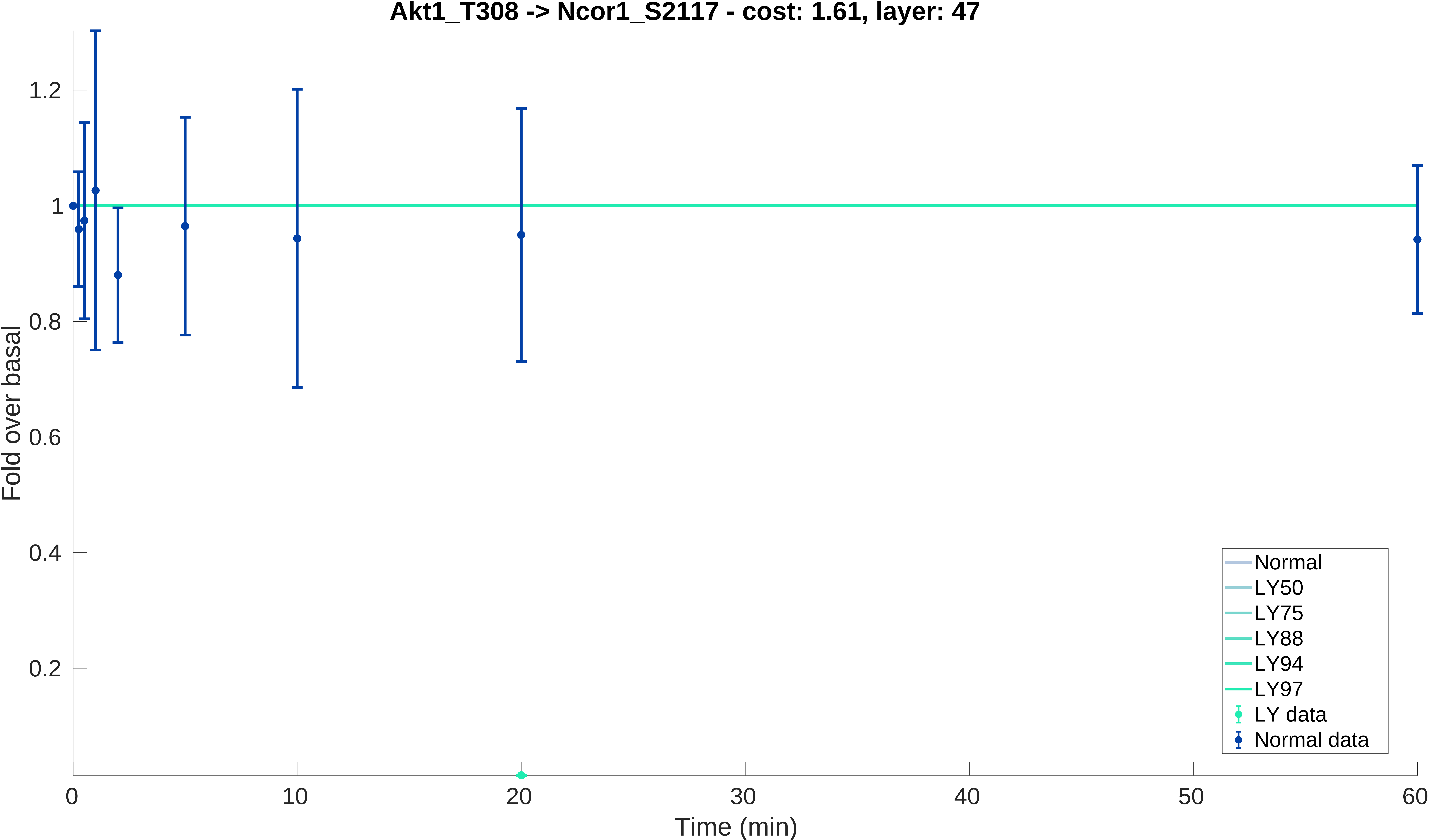
Akt1_T308 -> Ncor1_S2117 - cost: 1.61, layer: 47
Fold over basal
1.2
1
0.8
0.6
0.4
0.2
0
10
20
30
40
50
60
Time (min)
Normal
LY50
LY75
LY88
LY94
LY97
LY data
Normal data

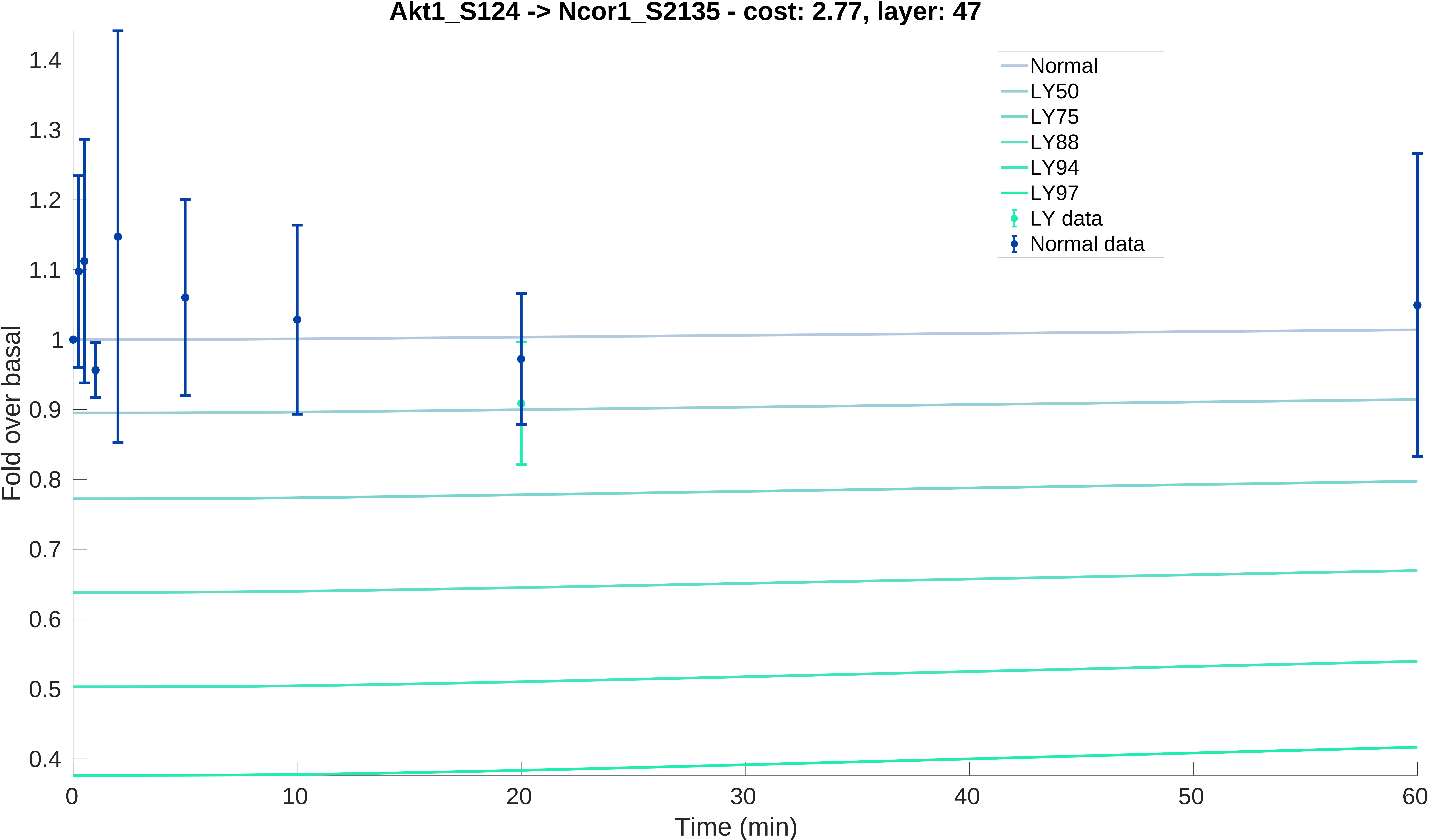
Akt1_S124 -> Ncor1_S2135 - cost: 2.77, layer: 47
Normal
LY50
LY75
LY88
LY94
LY97
LY data
Normal data
Fold over basal
1.4
1.3
1.2
1.1
1
0.9
0.8
0.7
0.6
0.5
0.4
0
10
20
30
40
50
60
Time (min)

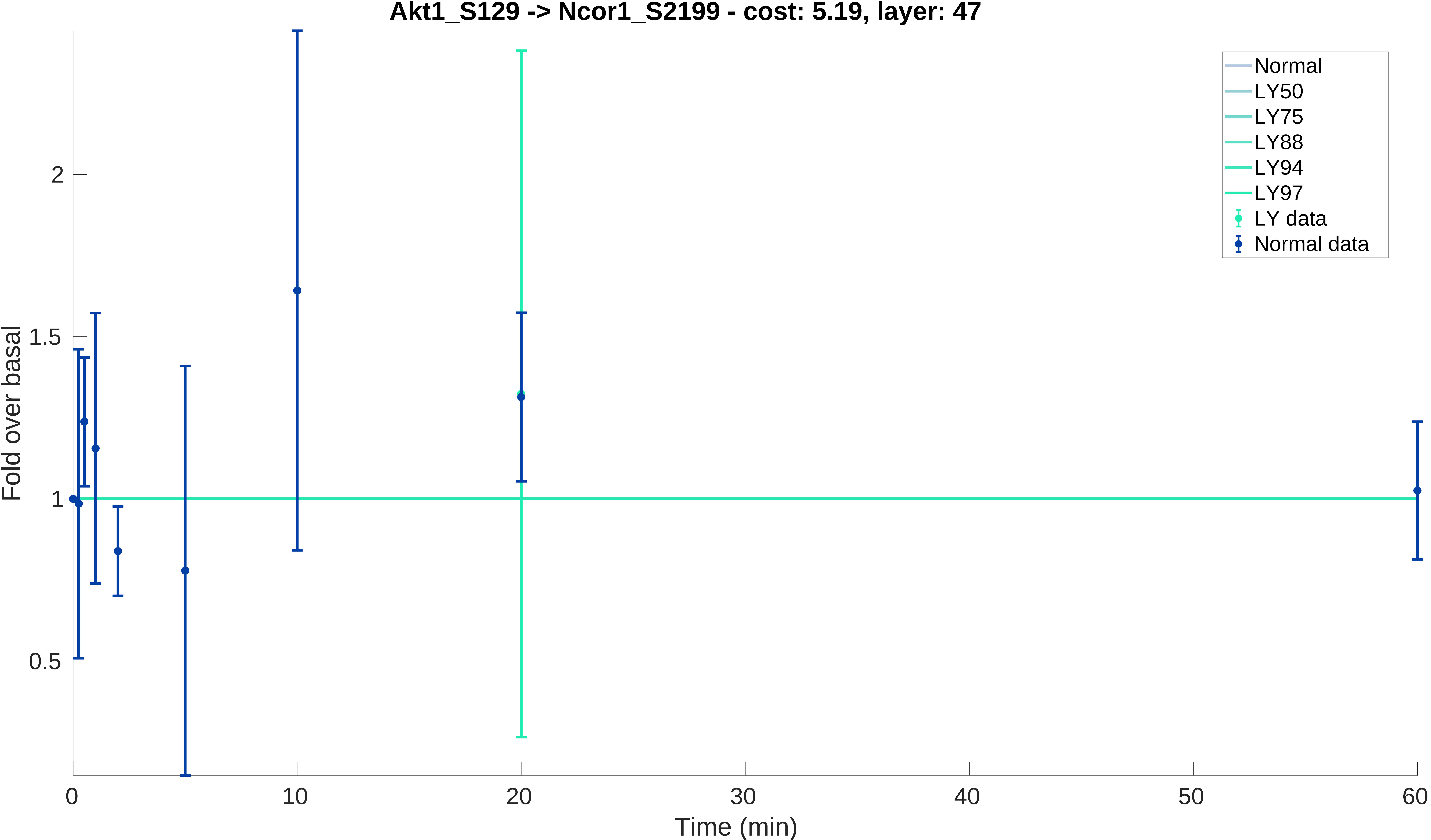
Akt1_S129 -> Ncor1_S2199 - cost: 5.19, layer: 47
Fold over basal
Time (min)
Normal
LY50
LY75
LY88
LY94
LY97
LY data
Normal data
0.5
1
1.5
2
0
10
20
30
40
50
60

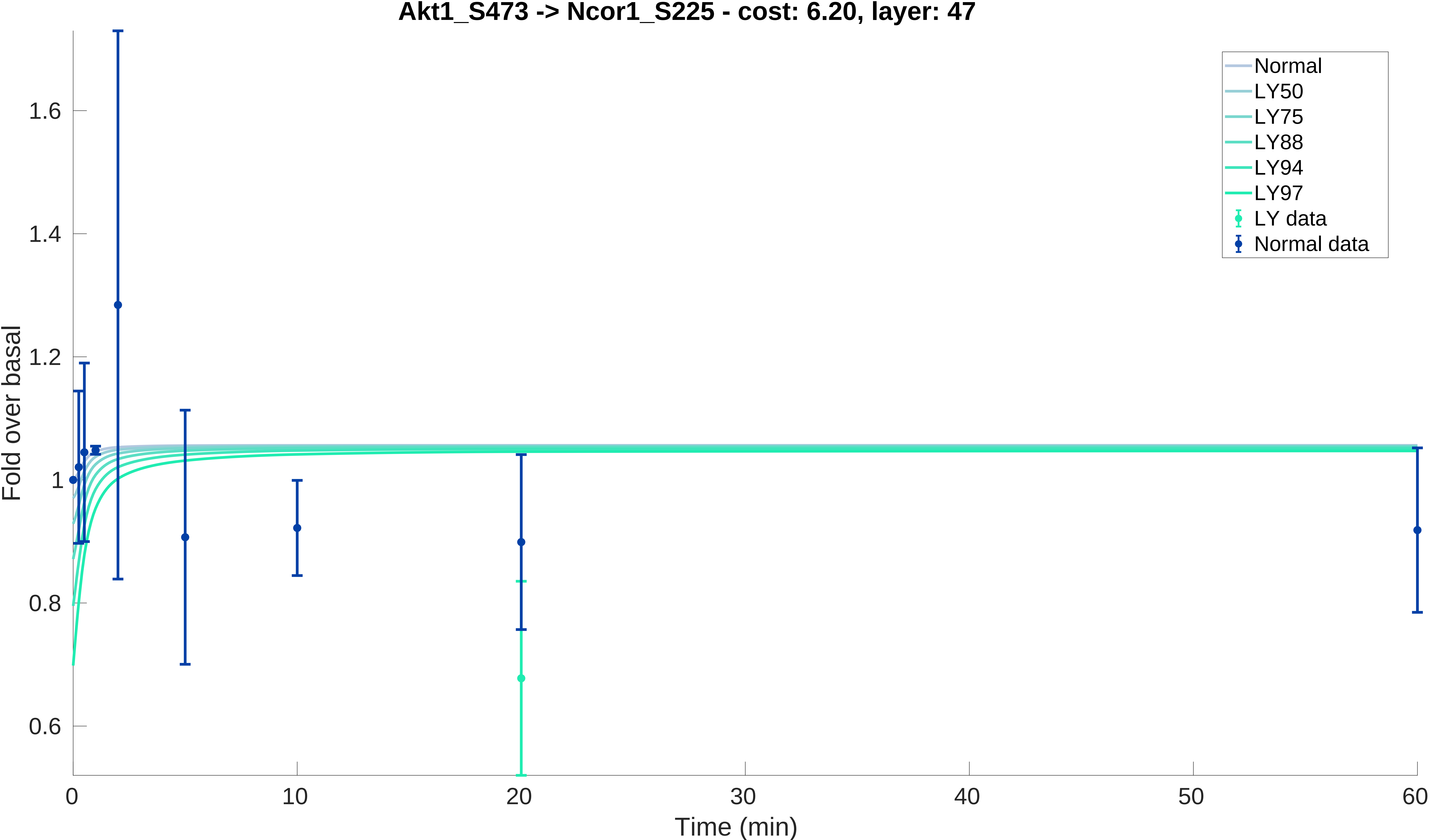

Akt1_S473 -> Ncor1_S225 - cost: 6.20, layer: 47
Normal
LY50
LY75
LY88
LY94
LY97
LY data
Normal data
Fold over basal
1.6
1.4
1.2
1
0.8
0.6
0
10
20
30
40
50
60
Time (min)

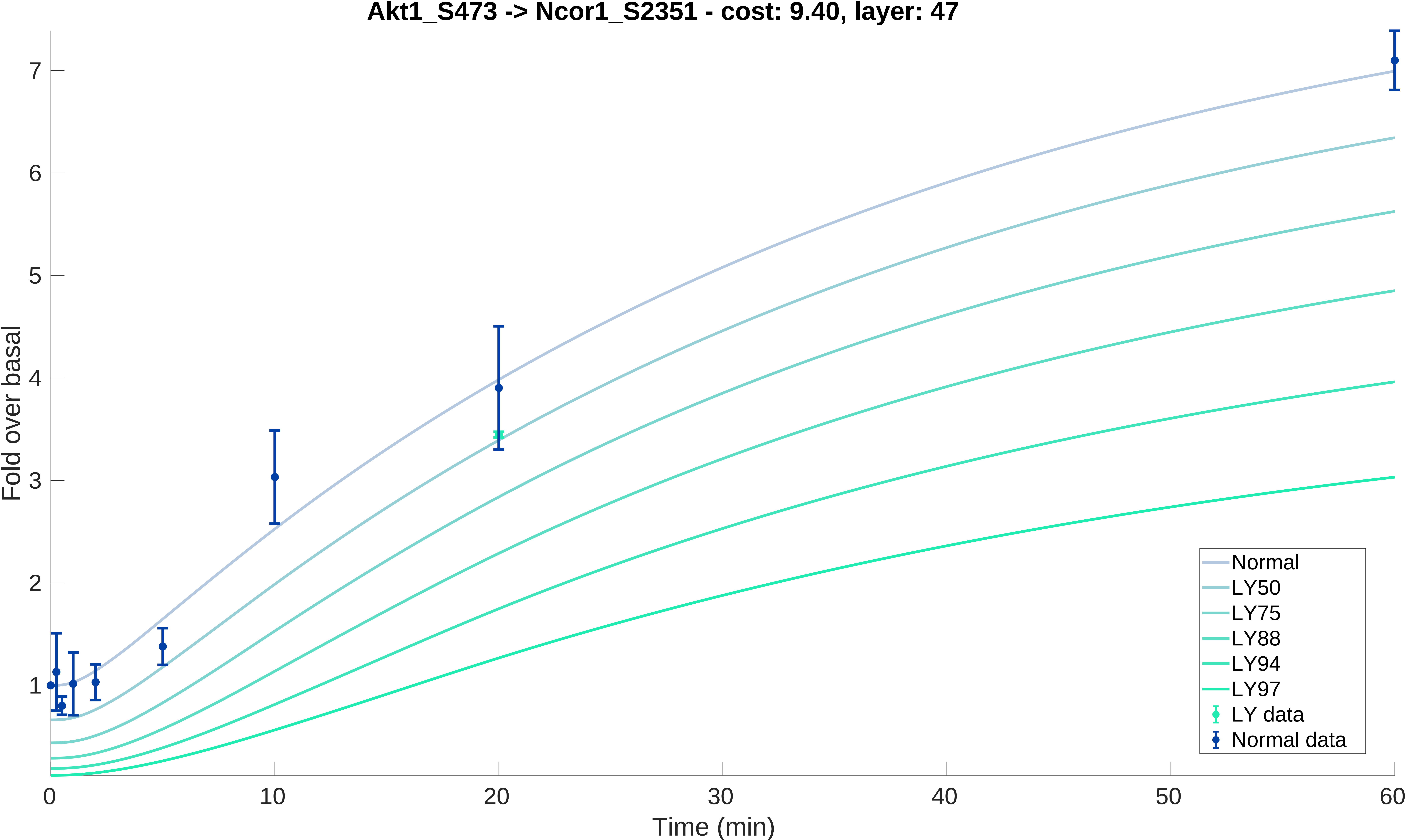

Akt1_S473 -> Ncor1_S2351 - cost: 9.40, layer: 47
Fold over basal
Time (min)
0
10
20
30
40
50
60
1
2
3
4
5
6
7
Normal
LY50
LY75
LY88
LY94
LY97
LY data
Normal data

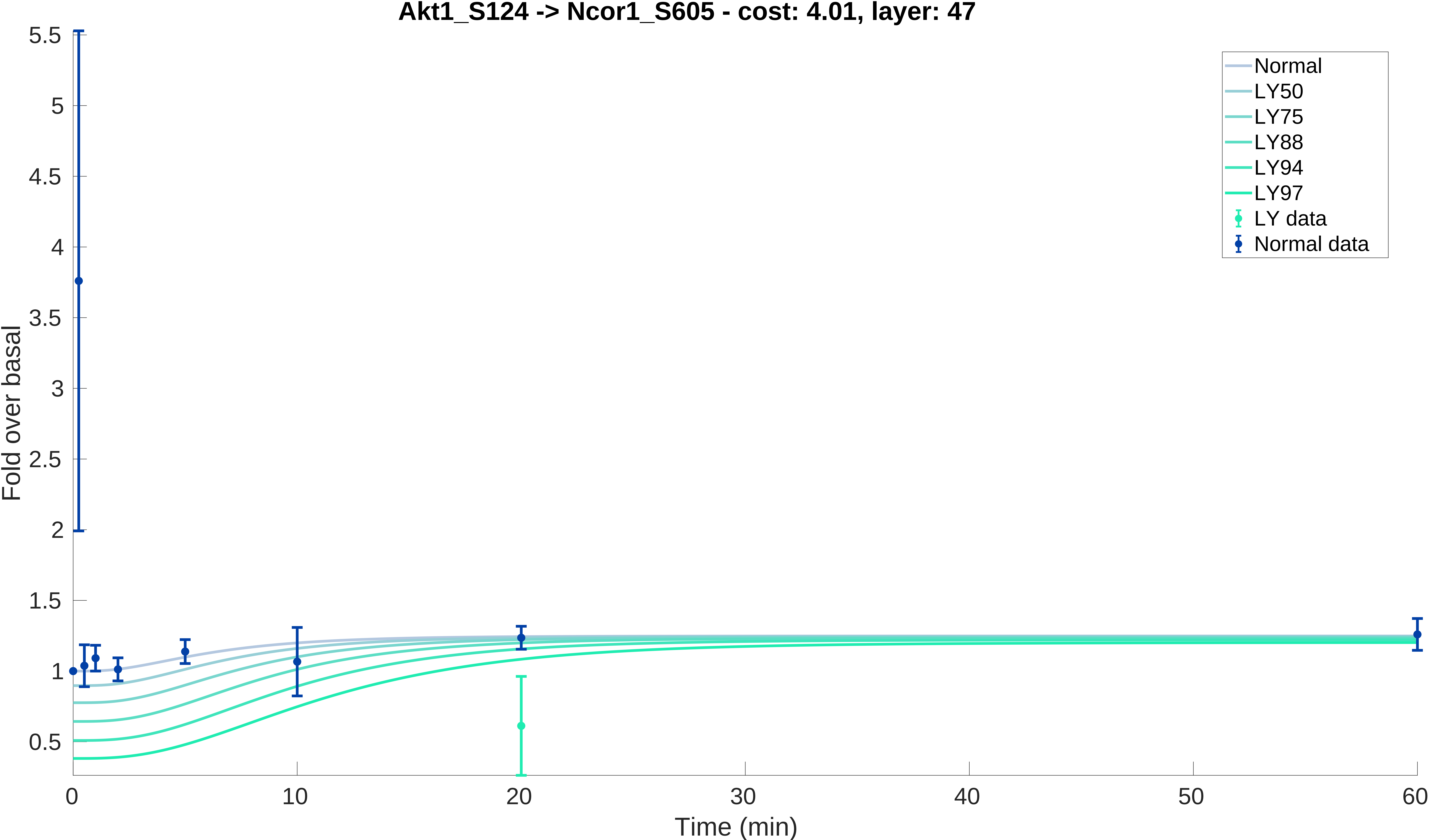

Akt1_S124 -> Ncor1_S605 - cost: 4.01, layer: 47
Fold over basal
Time (min)
0
10
20
30
40
50
60
0.5
1
1.5
2
2.5
3
3.5
4
4.5
5
5.5
Normal
LY50
LY75
LY88
LY94
LY97
LY data
Normal data

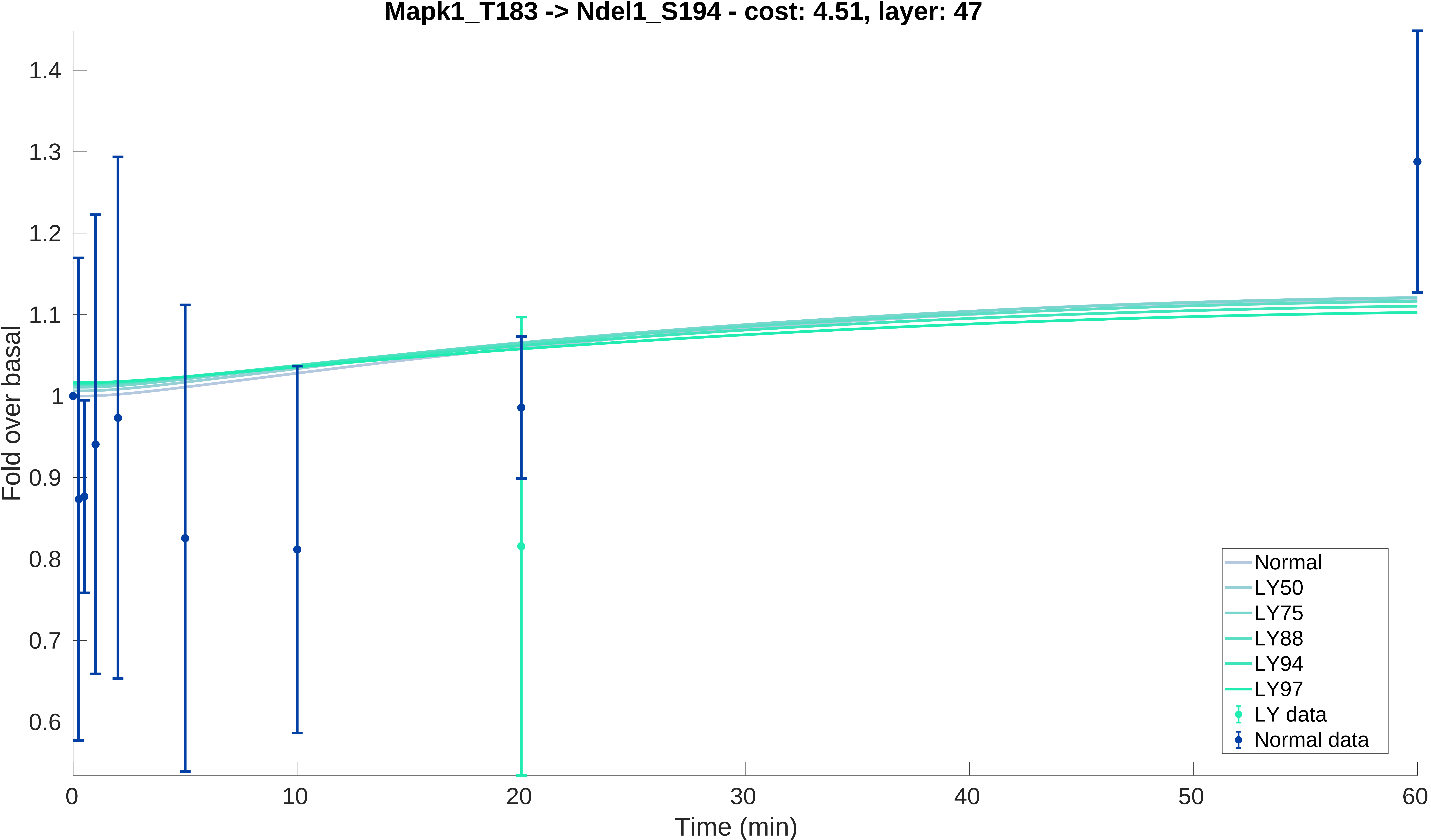

Mapk1_T183 -> Ndel1_S194 - cost: 4.51, layer: 47
Fold over basal
1.4
1.3
1.2
1.1
1
0.9
0.8
0.7
0.6
0
10
20
30
40
50
60
Time (min)
Normal
LY50
LY75
LY88
LY94
LY97
LY data
Normal data

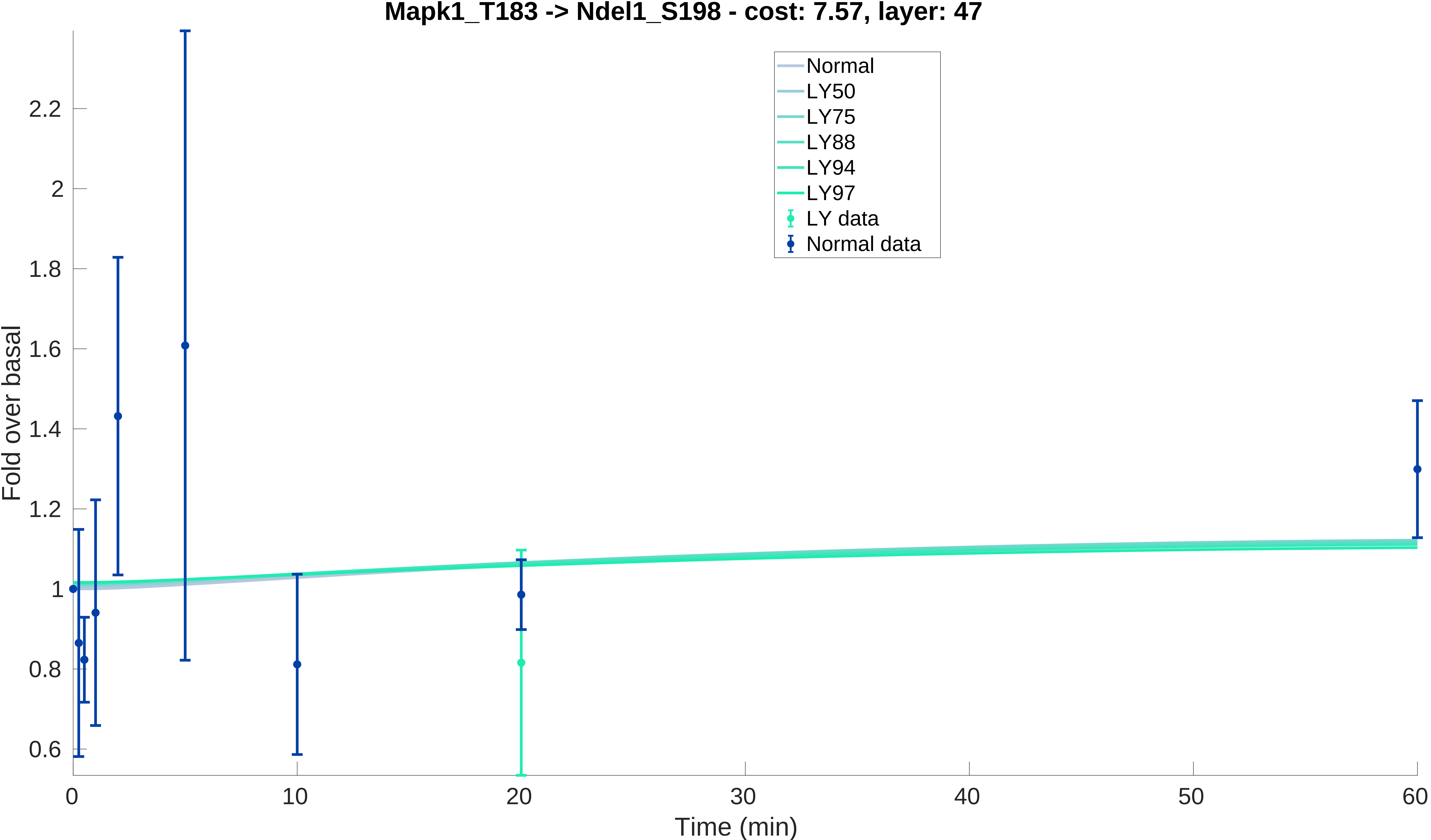

Mapk1_T183 -> Ndel1_S198 - cost: 7.57, layer: 47
Normal
LY50
LY75
LY88
LY94
LY97
LY data
Normal data
Fold over basal
2.2
2
1.8
1.6
1.4
1.2
1
0.8
0.6
0
10
20
30
40
50
60
Time (min)

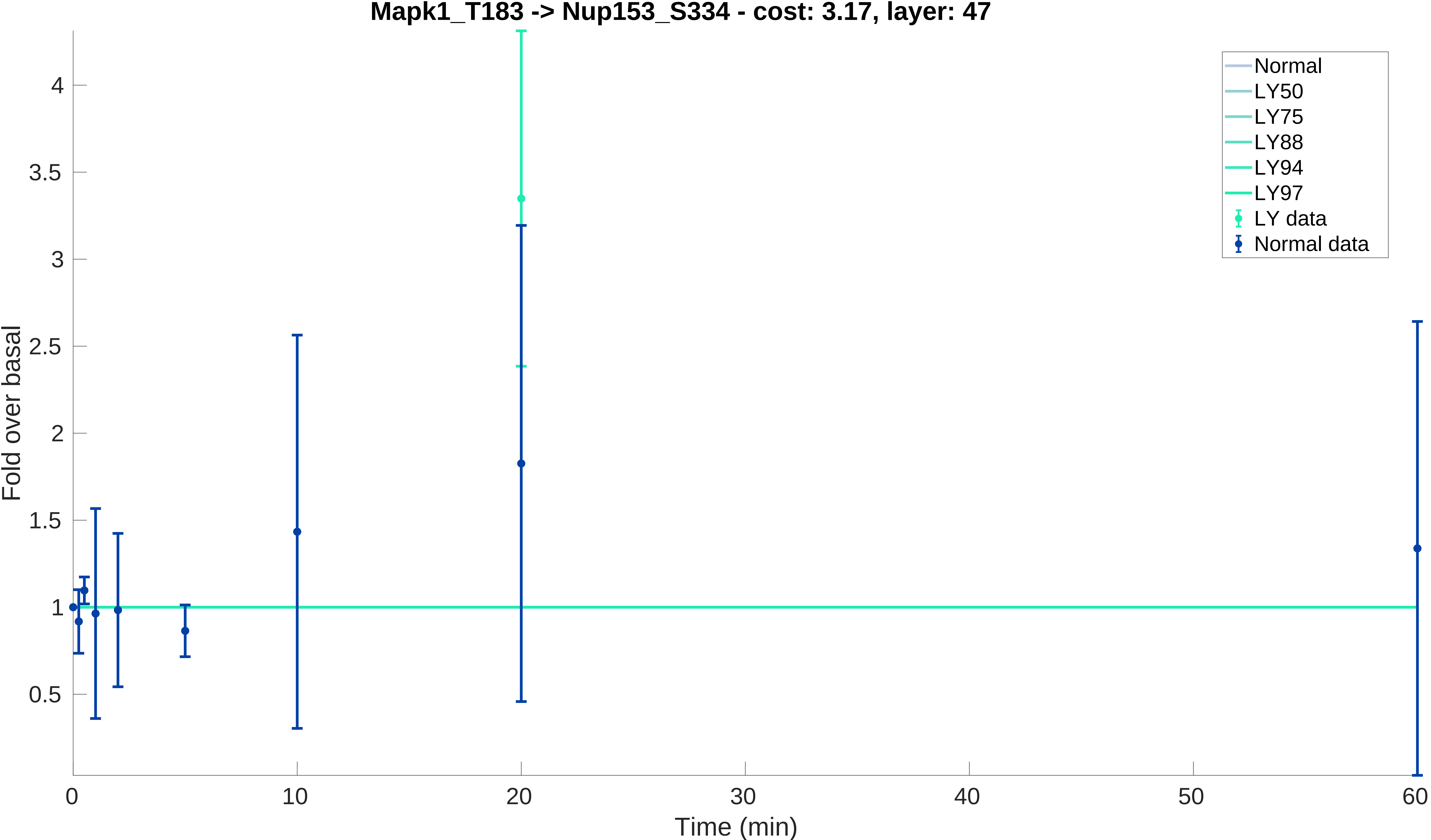

Mapk1_T183 -> Nup153_S334 - cost: 3.17, layer: 47
Normal
LY50
LY75
LY88
LY94
LY97
LY data
Normal data
Fold over basal
Time (min)
0.5
1
1.5
2
2.5
3
3.5
4
0
10
20
30
40
50
60

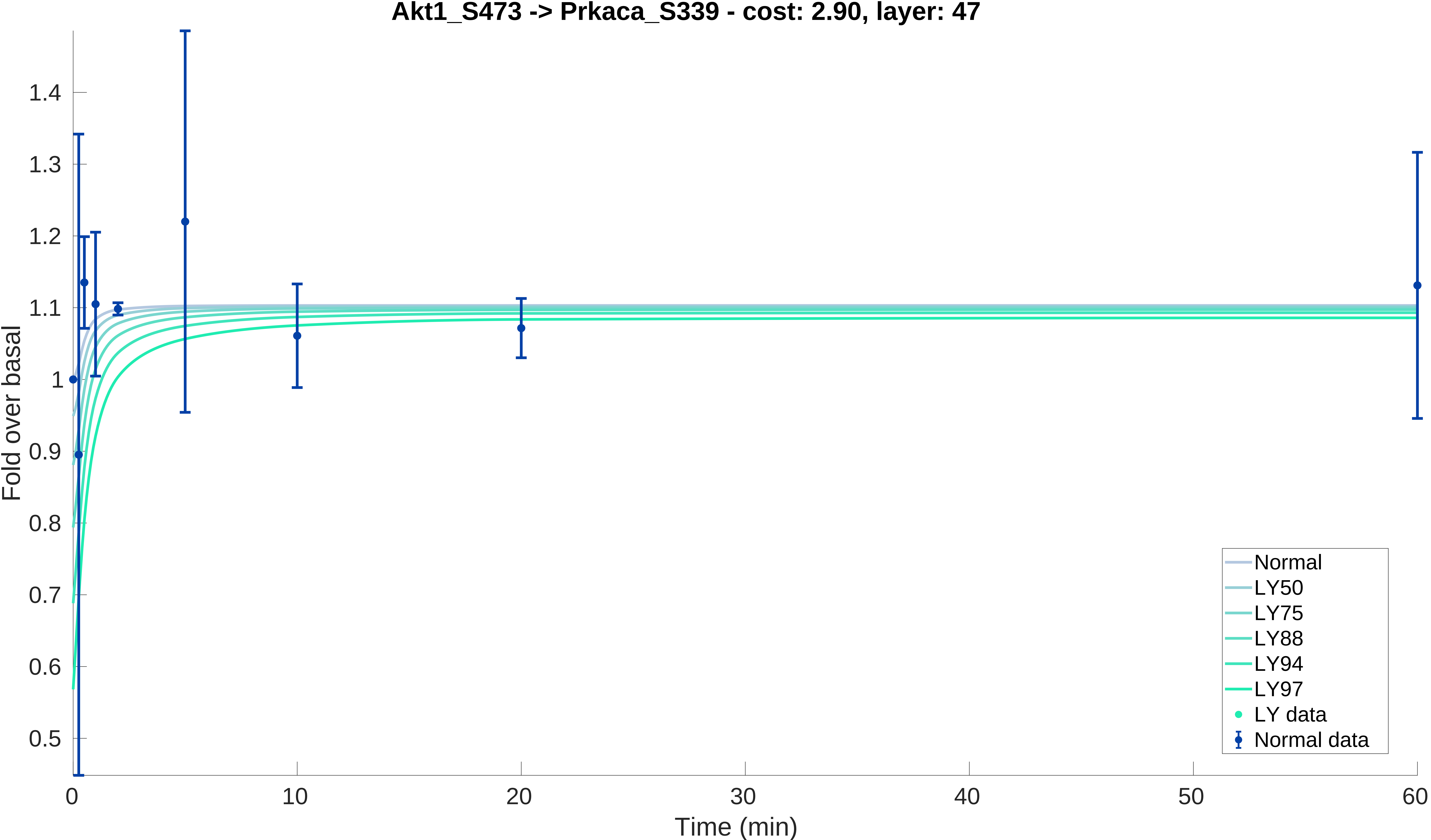

Akt1_S473 -> Prkaca_S339 - cost: 2.90, layer: 47
Fold over basal
1.4
1.3
1.2
1.1
1
0.9
0.8
0.7
0.6
0.5
0
10
20
30
40
50
60
Time (min)
Normal
LY50
LY75
LY88
LY94
LY97
LY data
Normal data

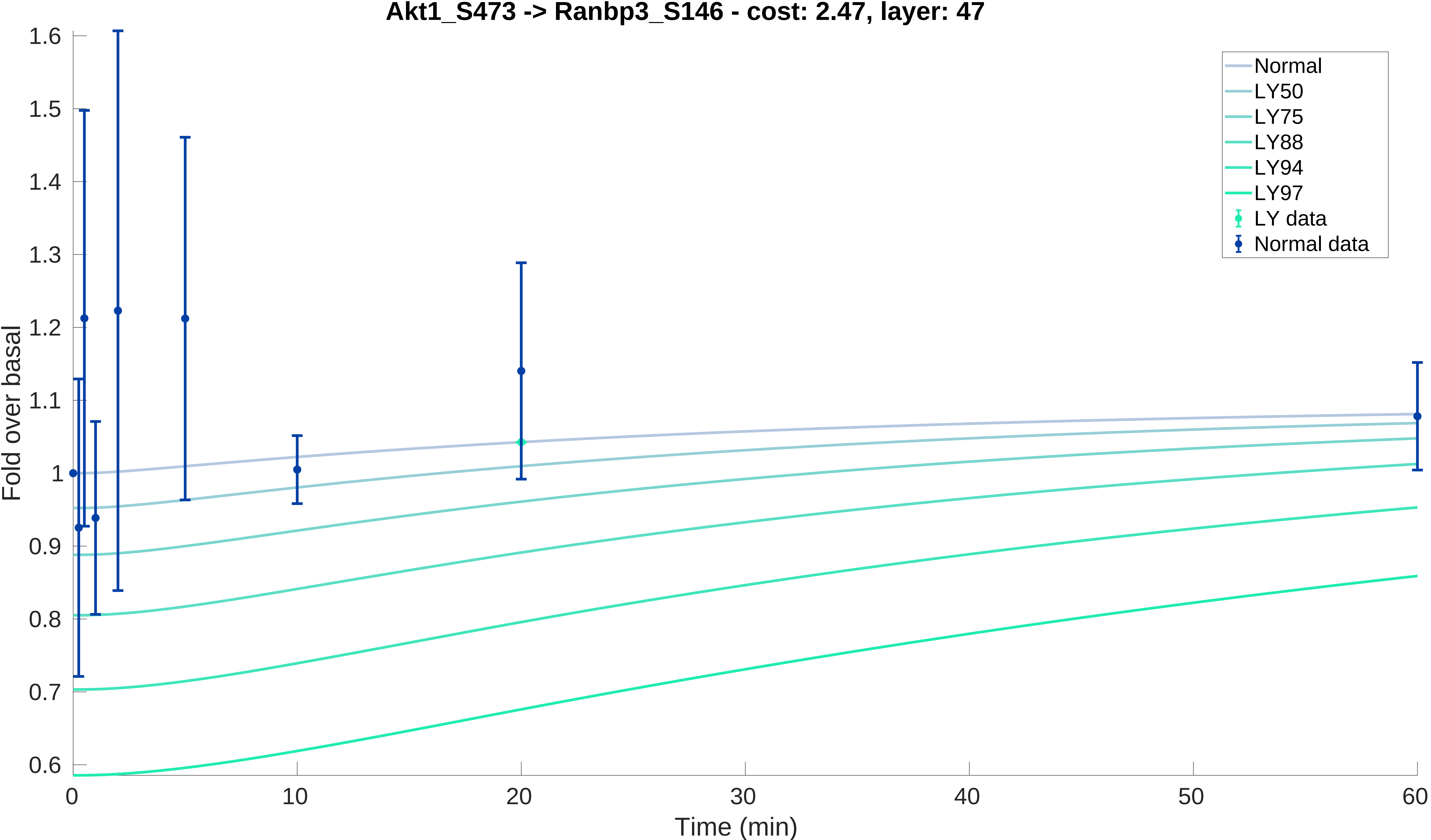

Akt1_S473 -> Ranbp3_S146 - cost: 2.47, layer: 47
Fold over basal
Time (min)
Normal
LY50
LY75
LY88
LY94
LY97
LY data
Normal data
1.6
1.5
1.4
1.3
1.2
1.1
1
0.9
0.8
0.7
0.6
0
10
20
30
40
50
60

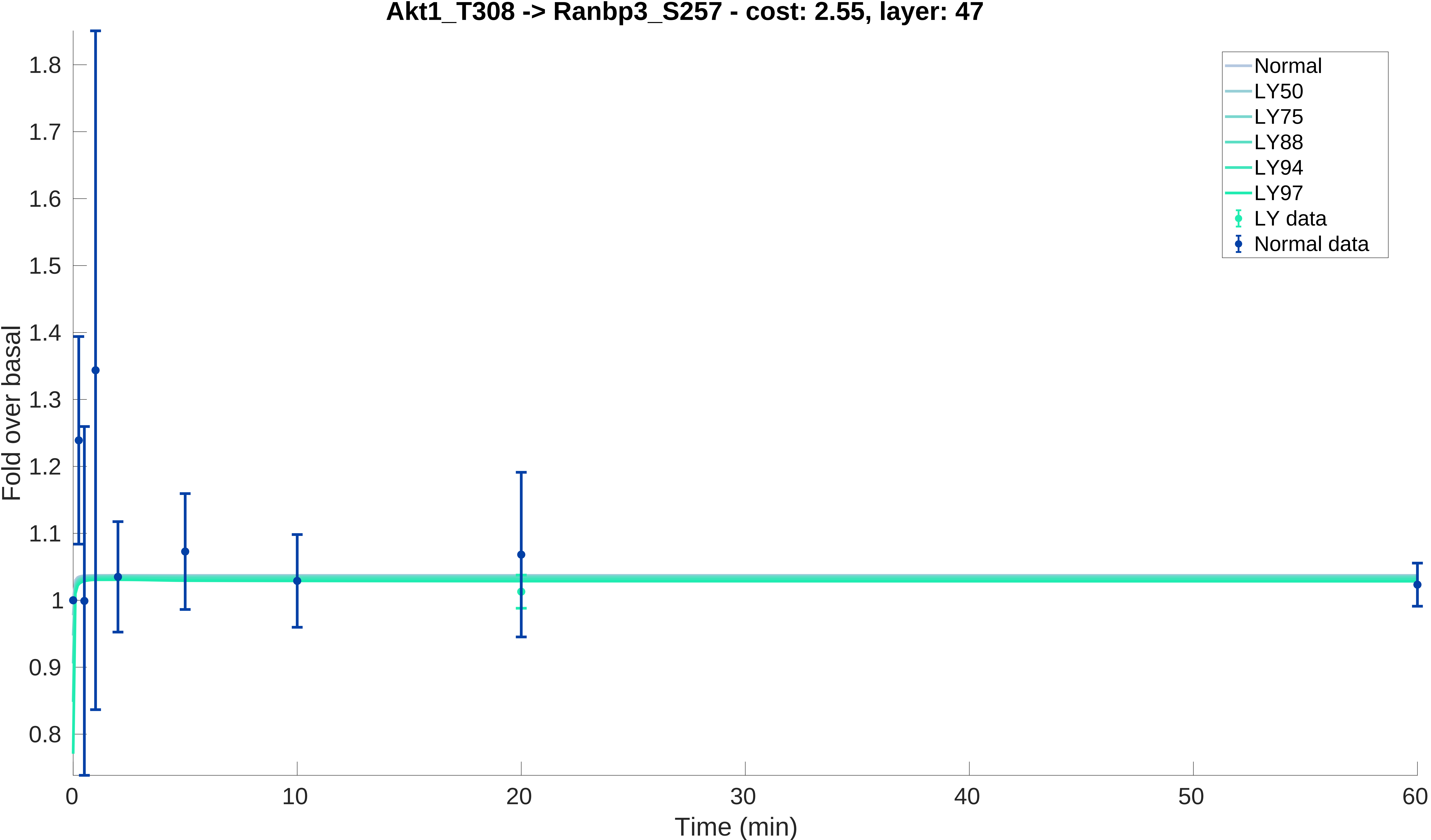
Akt1_T308 -> Ranbp3_S257 - cost: 2.55, layer: 47
Fold over basal
Time (min)
Normal
LY50
LY75
LY88
LY94
LY97
LY data
Normal data
0.8
0.9
1
1.1
1.2
1.3
1.4
1.5
1.6
1.7
1.8
0
10
20
30
40
50
60

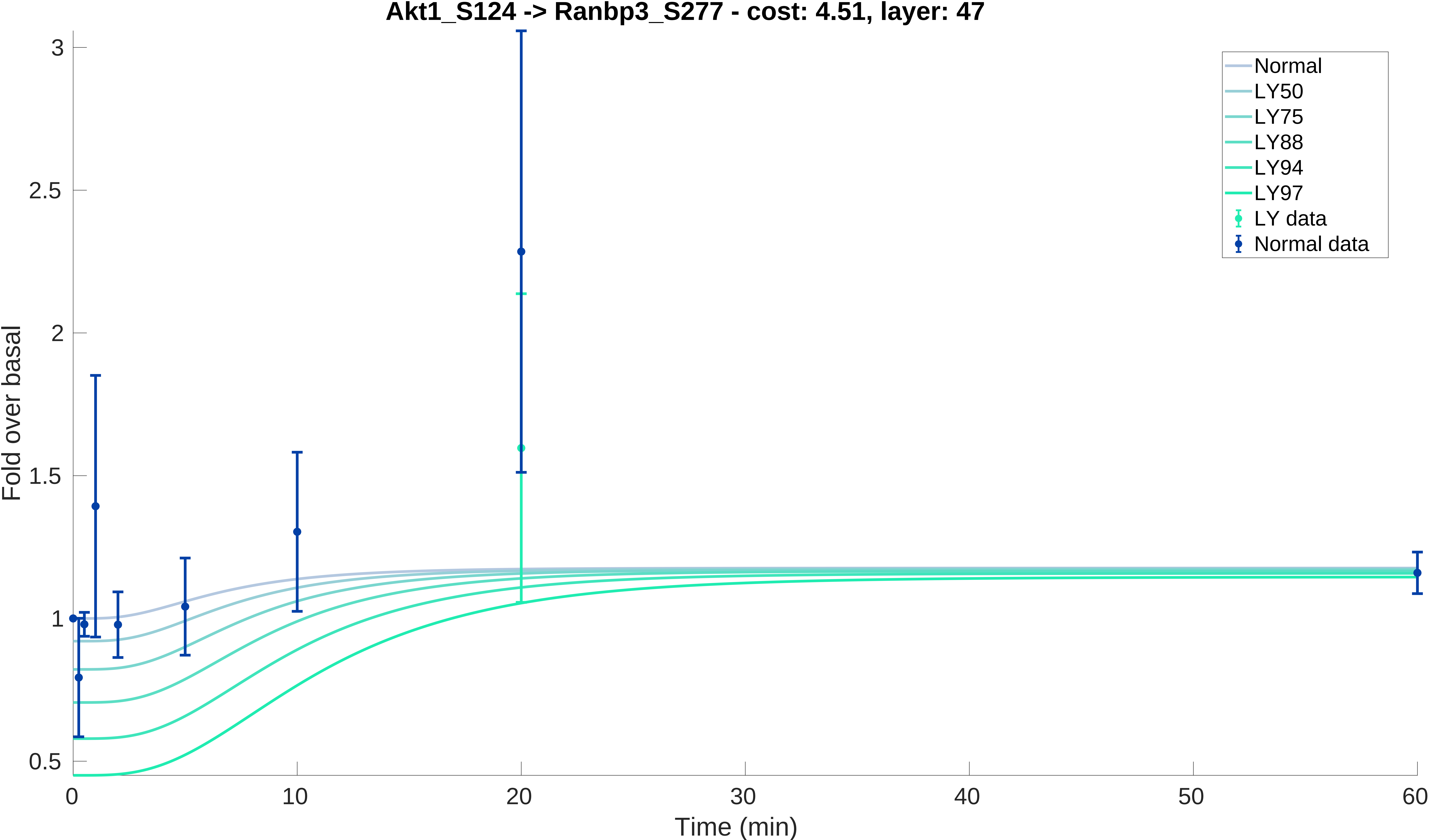

Akt1_S124 -> Ranbp3_S277 - cost: 4.51, layer: 47
Fold over basal
Time (min)
0.5
1
1.5
2
2.5
3
0
10
20
30
40
50
60
Normal
LY50
LY75
LY88
LY94
LY97
LY data
Normal data

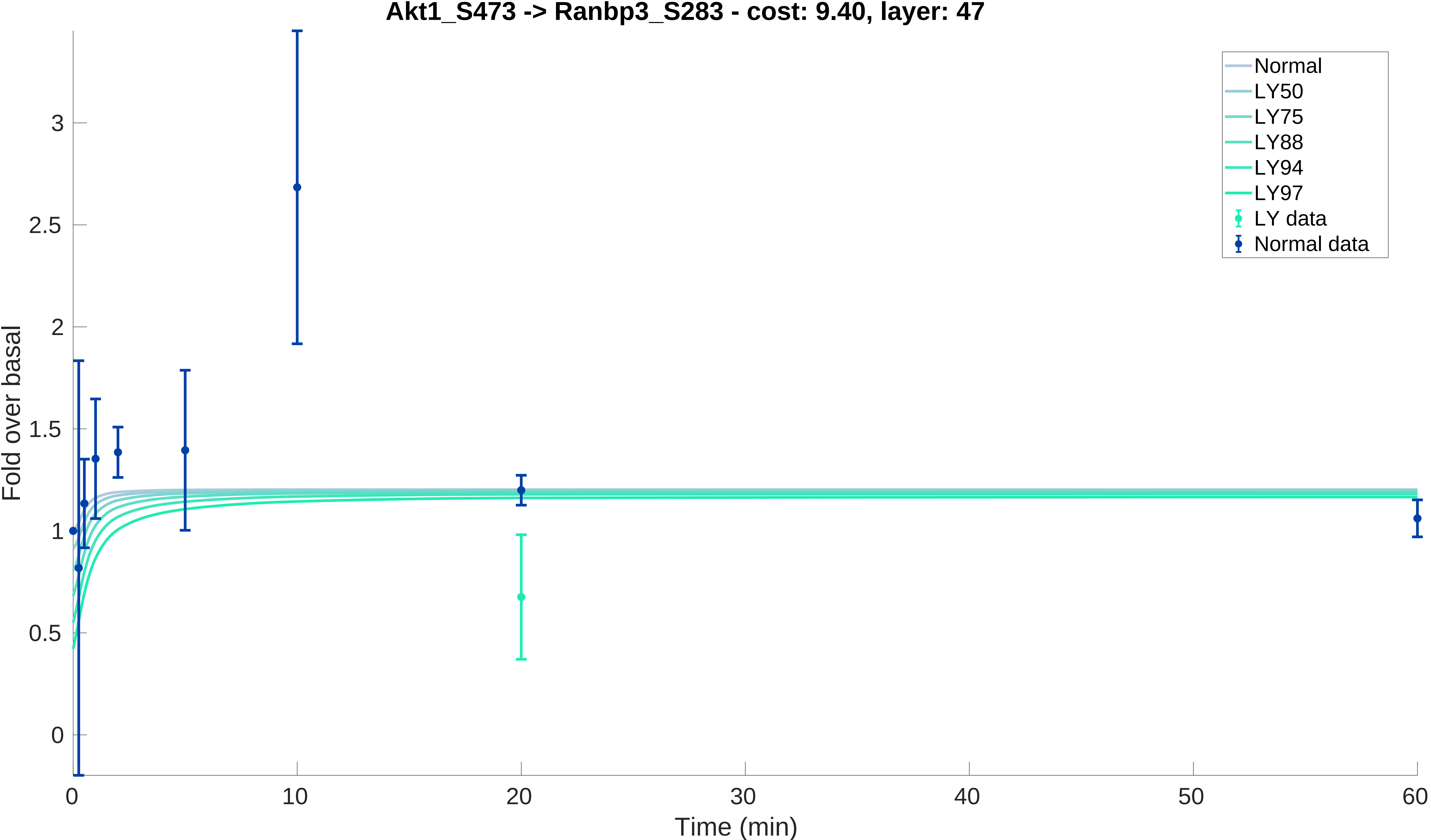

Akt1_S473 -> Ranbp3_S283 - cost: 9.40, layer: 47
Normal
LY50
LY75
LY88
LY94
LY97
LY data
Normal data
Fold over basal
Time (min)
0
0.5
1
1.5
2
2.5
3
0
10
20
30
40
50
60

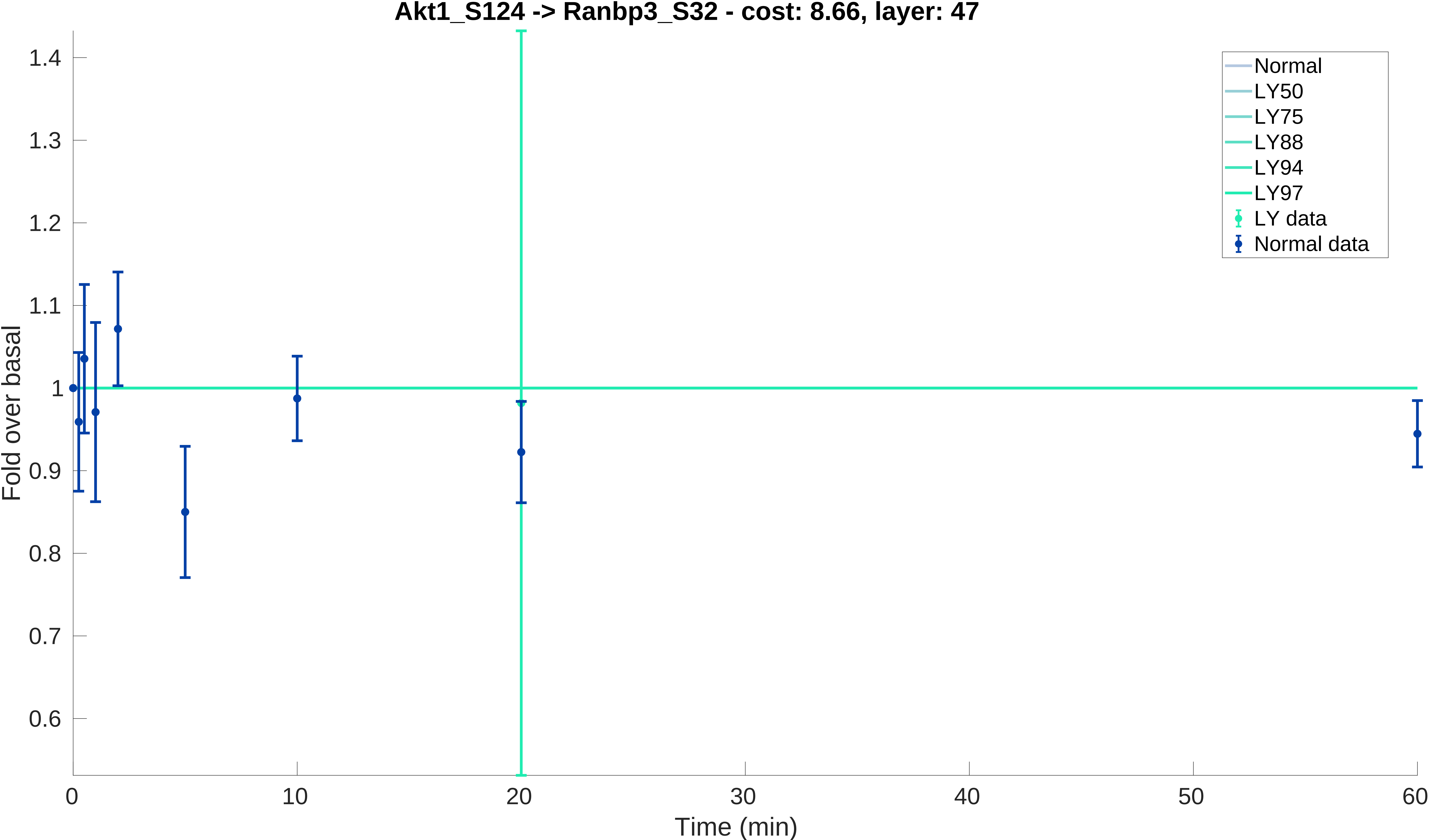

Akt1_S124 -> Ranbp3_S32 - cost: 8.66, layer: 47
Fold over basal
Time (min)
1.4
1.3
1.2
1.1
1
0.9
0.8
0.7
0.6
0
10
20
30
40
50
60
Normal
LY50
LY75
LY88
LY94
LY97
LY data
Normal data

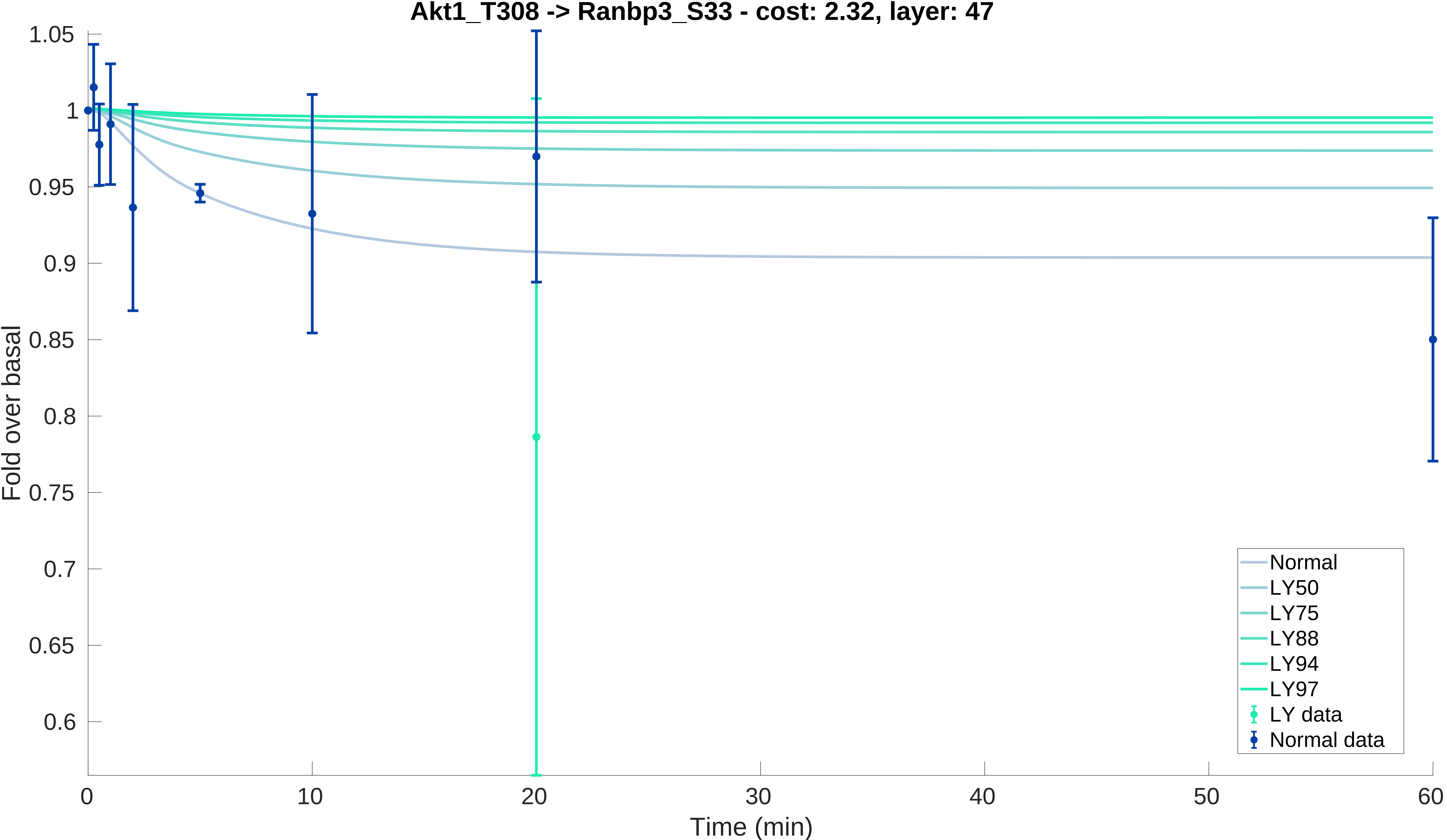

Akt1_T308 -> Ranbp3_S33 - cost: 2.32, layer: 47
1.05
1
0.95
0.9
0.85
0.8
0.75
0.7
0.65
0.6
Fold over basal
0
10
20
30
40
50
60
Time (min)
Normal
LY50
LY75
LY88
LY94
LY97
LY data
Normal data

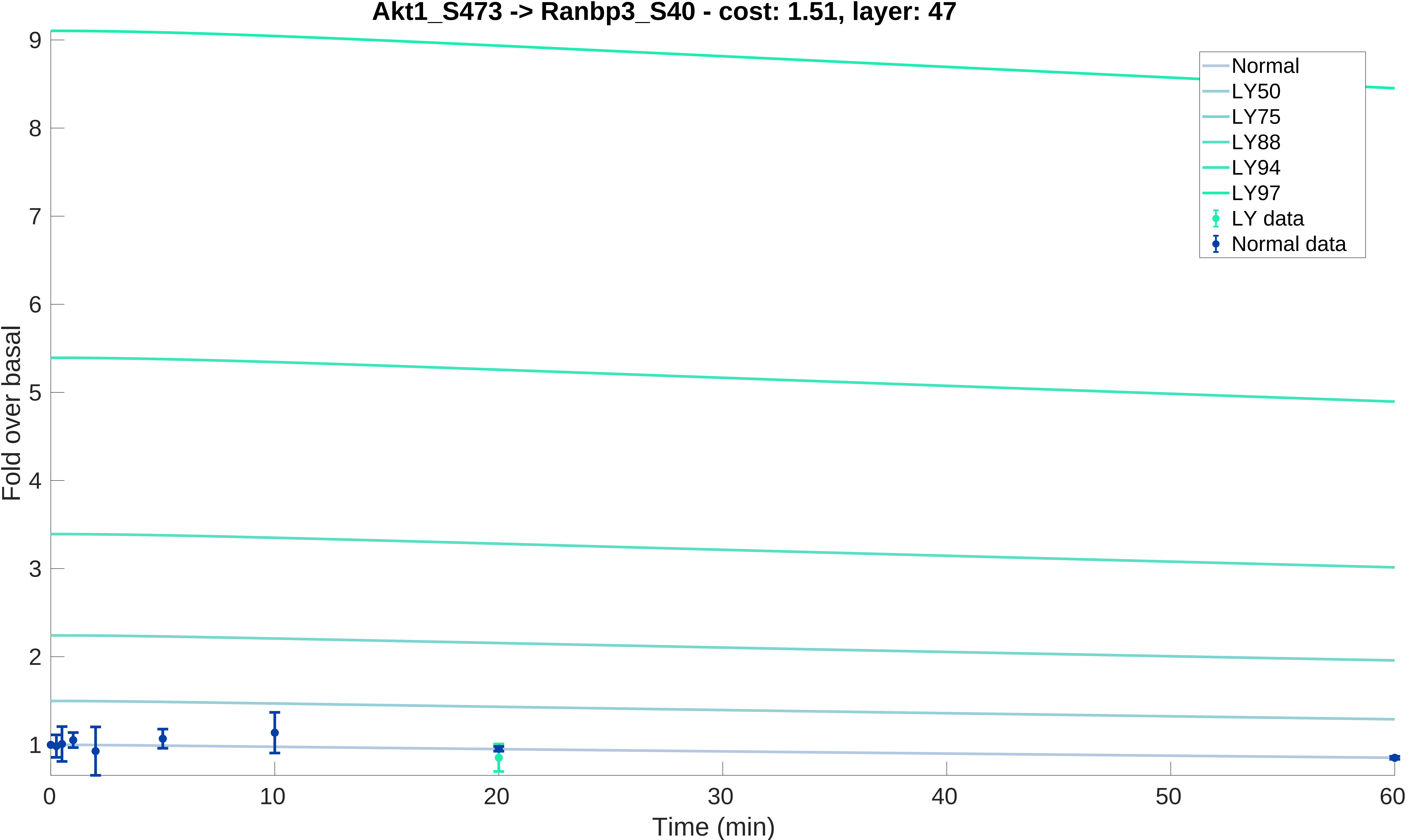

Akt1_S473 -> Ranbp3_S40 - cost: 1.51, layer: 47
Fold over basal
Time (min)
0
10
20
30
40
50
60
1
2
3
4
5
6
7
8
9
Normal
LY50
LY75
LY88
LY94
LY97
LY data
Normal data

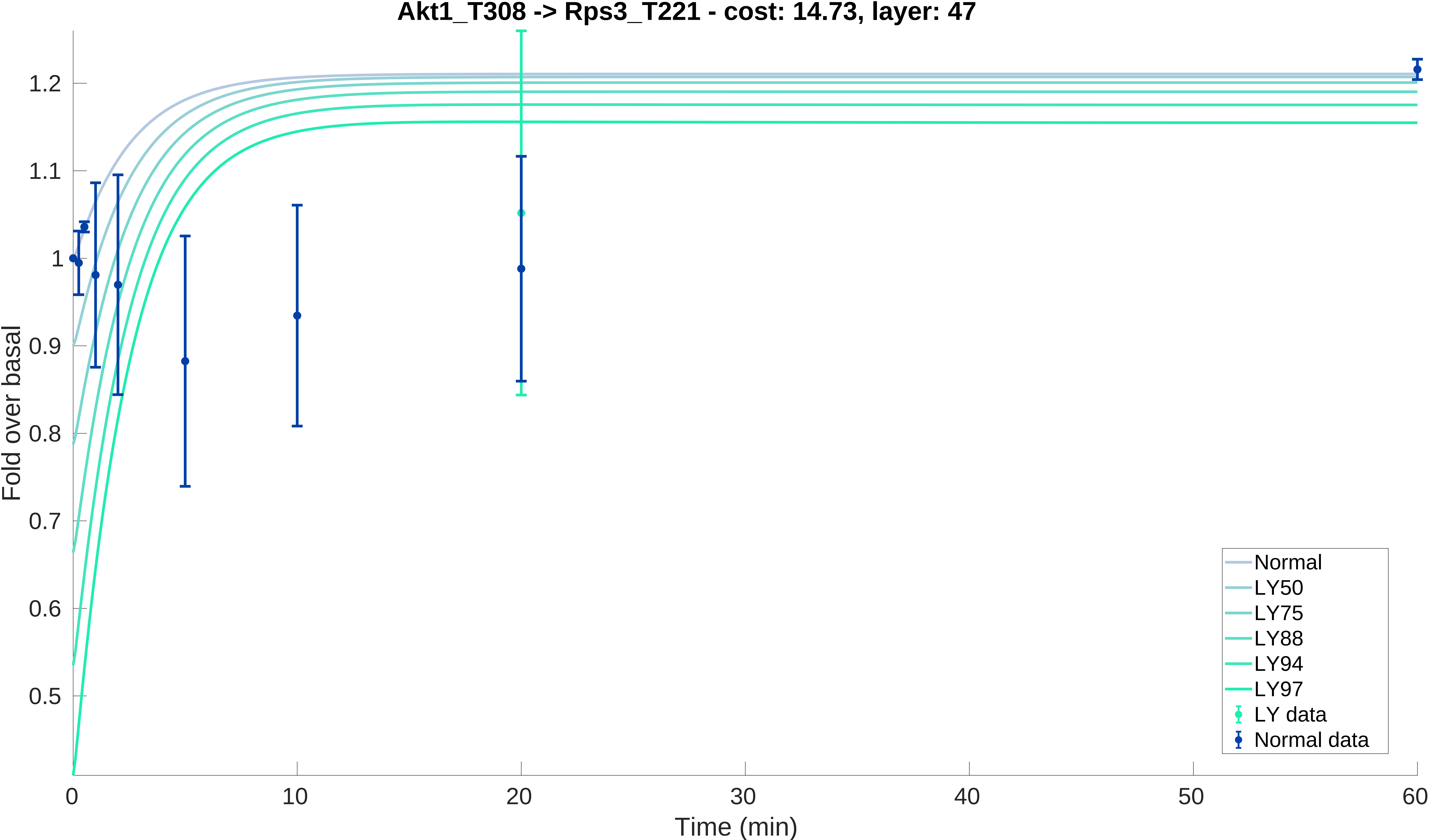

Akt1_T308 -> Rps3_T221 - cost: 14.73, layer: 47
Fold over basal
Time (min)
0
10
20
30
40
50
60
0.5
0.6
0.7
0.8
0.9
1
1.1
1.2
Normal
LY50
LY75
LY88
LY94
LY97
LY data
Normal data

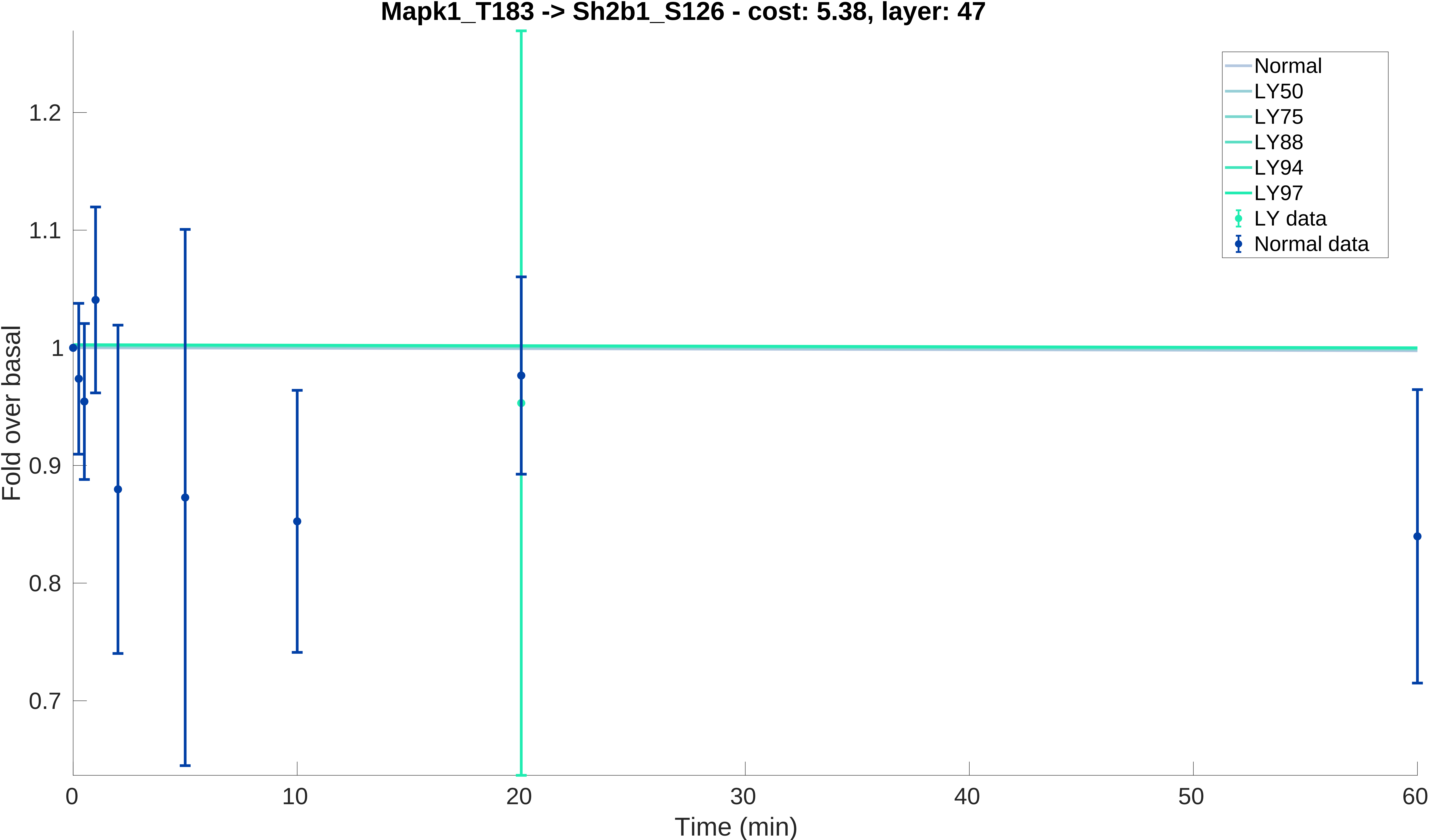

Mapk1_T183 -> Sh2b1_S126 - cost: 5.38, layer: 47
Normal
LY50
LY75
LY88
LY94
LY97
LY data
Normal data
Fold over basal
1.2
1.1
1
0.9
0.8
0.7
0
10
20
30
40
50
60
Time (min)

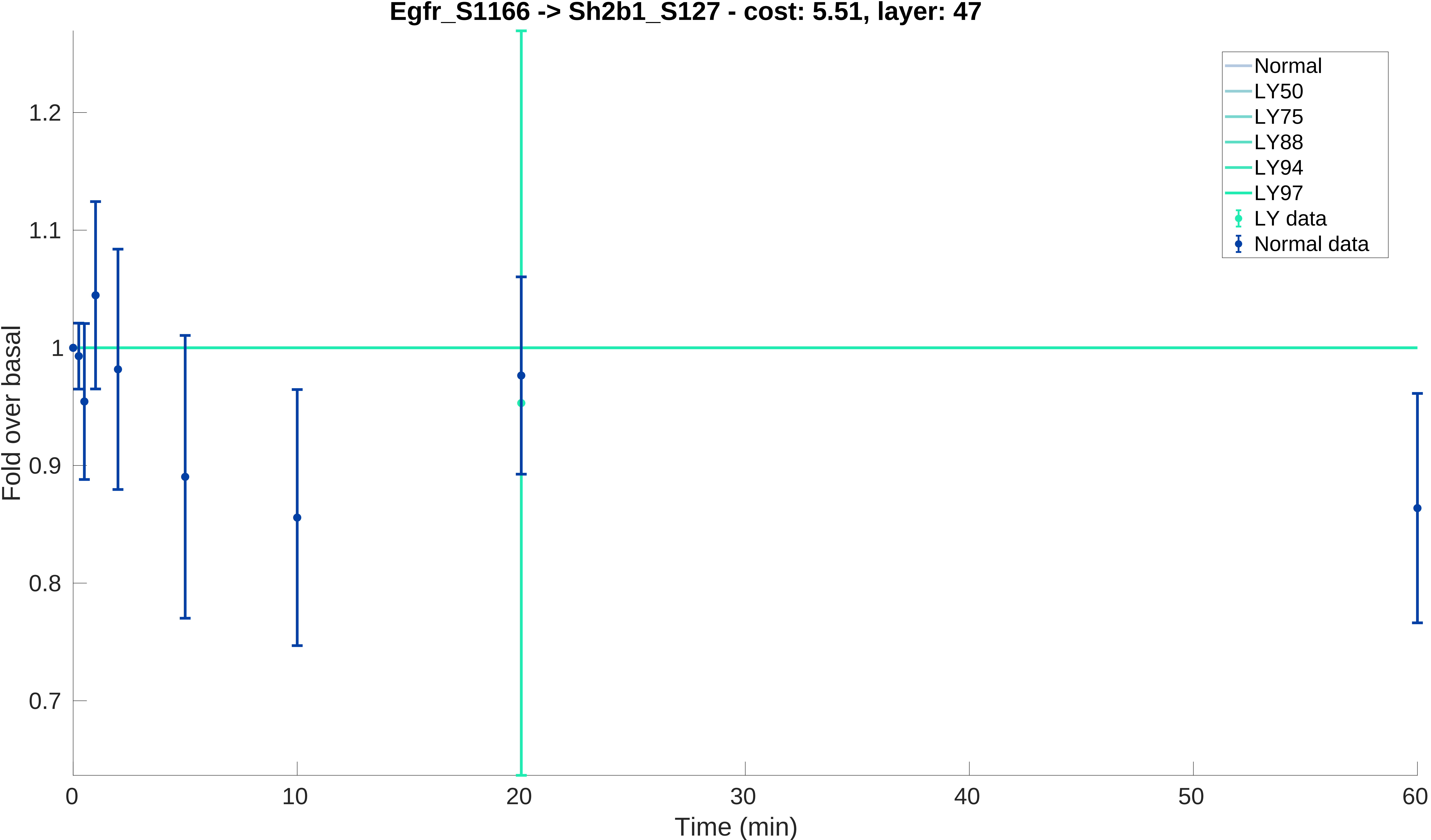
Egfr_S1166 -> Sh2b1_S127 - cost: 5.51, layer: 47
Fold over basal
Time (min)
1.2
1.1
1
0.9
0.8
0.7
0
10
20
30
40
50
60
Normal
LY50
LY75
LY88
LY94
LY97
LY data
Normal data

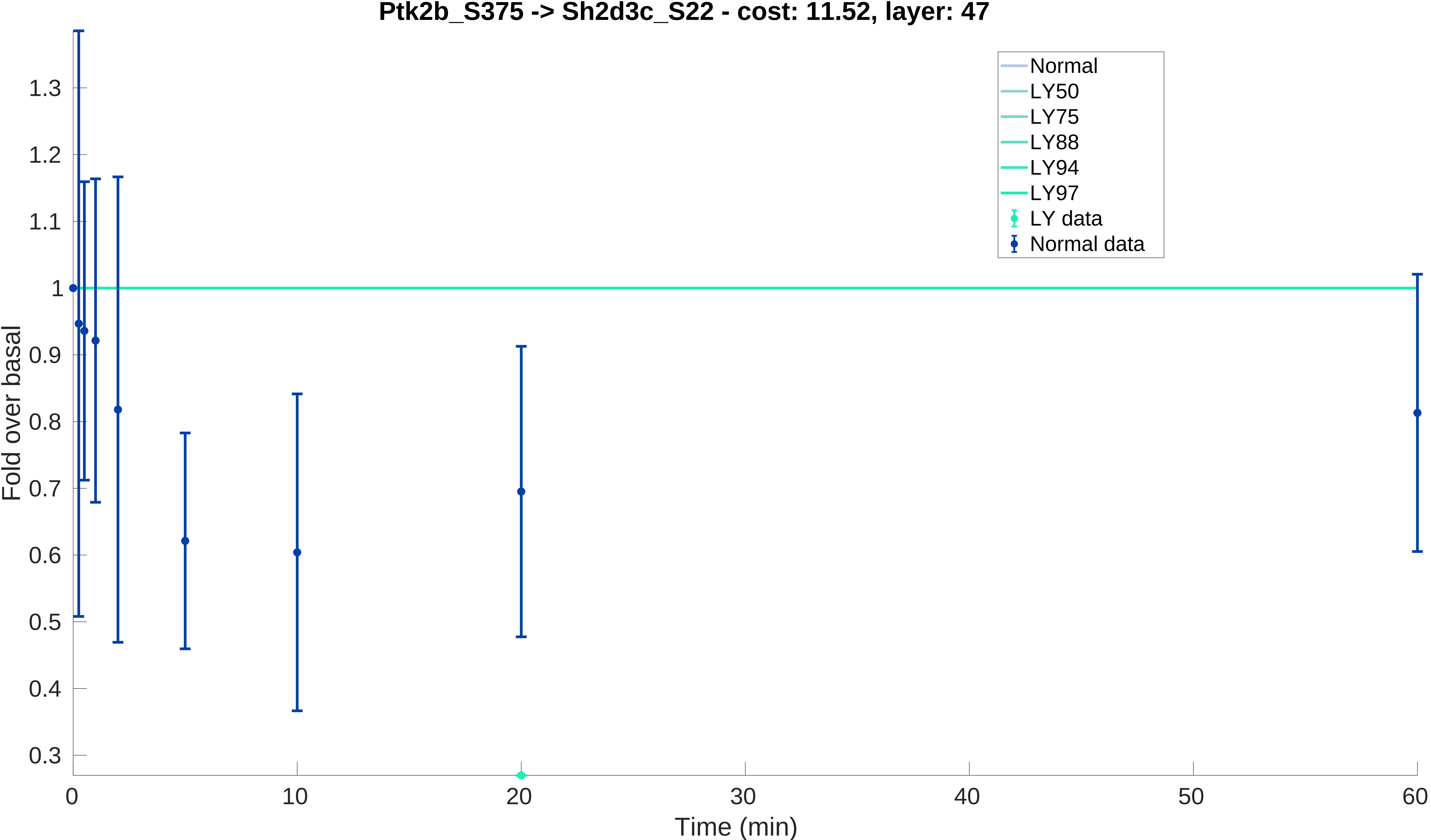

Ptk2b_S375 -> Sh2d3c_S22 - cost: 11.52, layer: 47
Normal
LY50
LY75
LY88
LY94
LY97
LY data
Normal data
Fold over basal
1.3
1.2
1.1
1
0.9
0.8
0.7
0.6
0.5
0.4
0.3
0
10
20
30
40
50
60
Time (min)

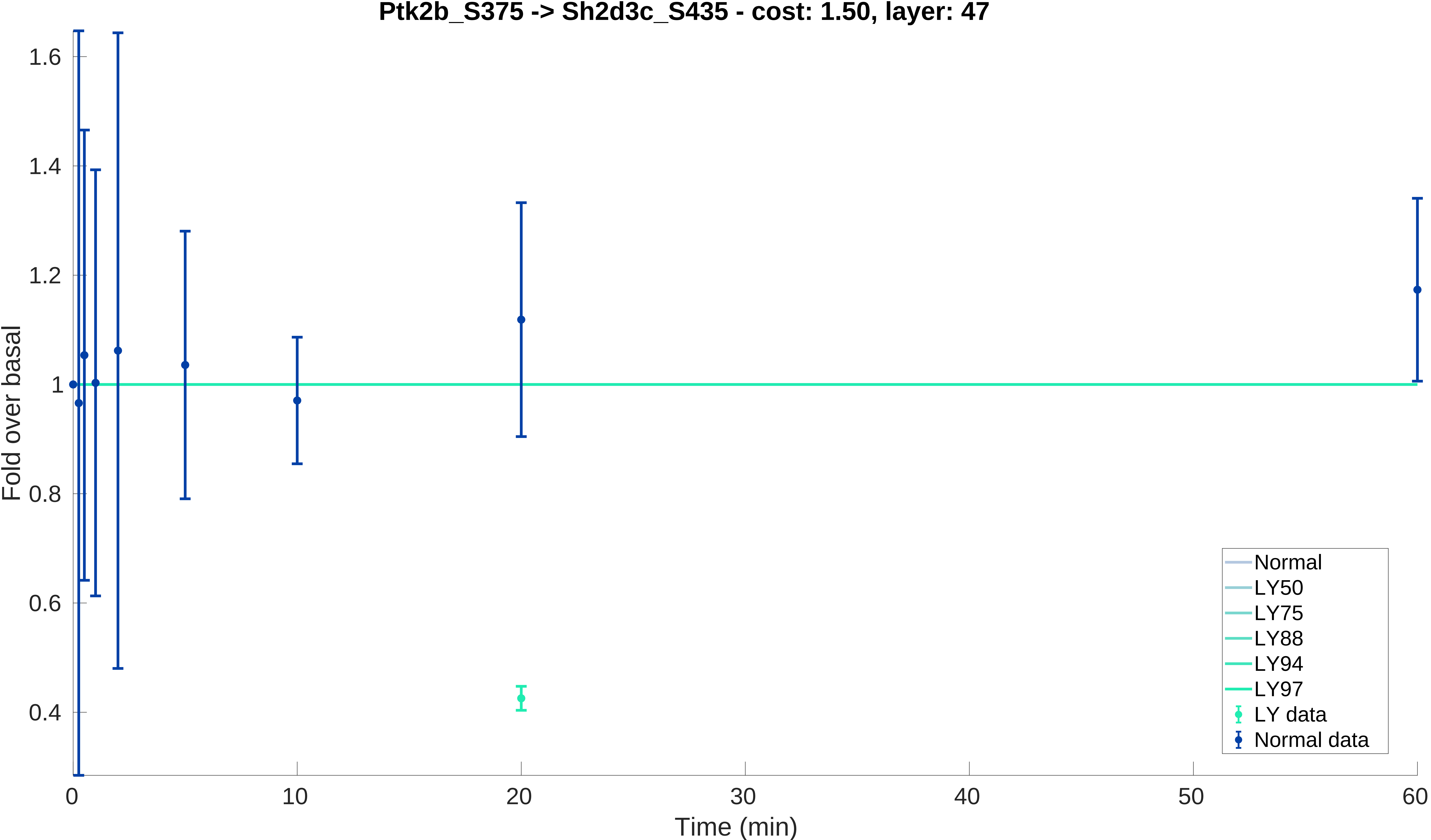

Ptk2b_S375 -> Sh2d3c_S435 - cost: 1.50, layer: 47
Fold over basal
1.6
1.4
1.2
1
0.8
0.6
0.4
0
10
20
30
40
50
60
Time (min)
Normal
LY50
LY75
LY88
LY94
LY97
LY data
Normal data

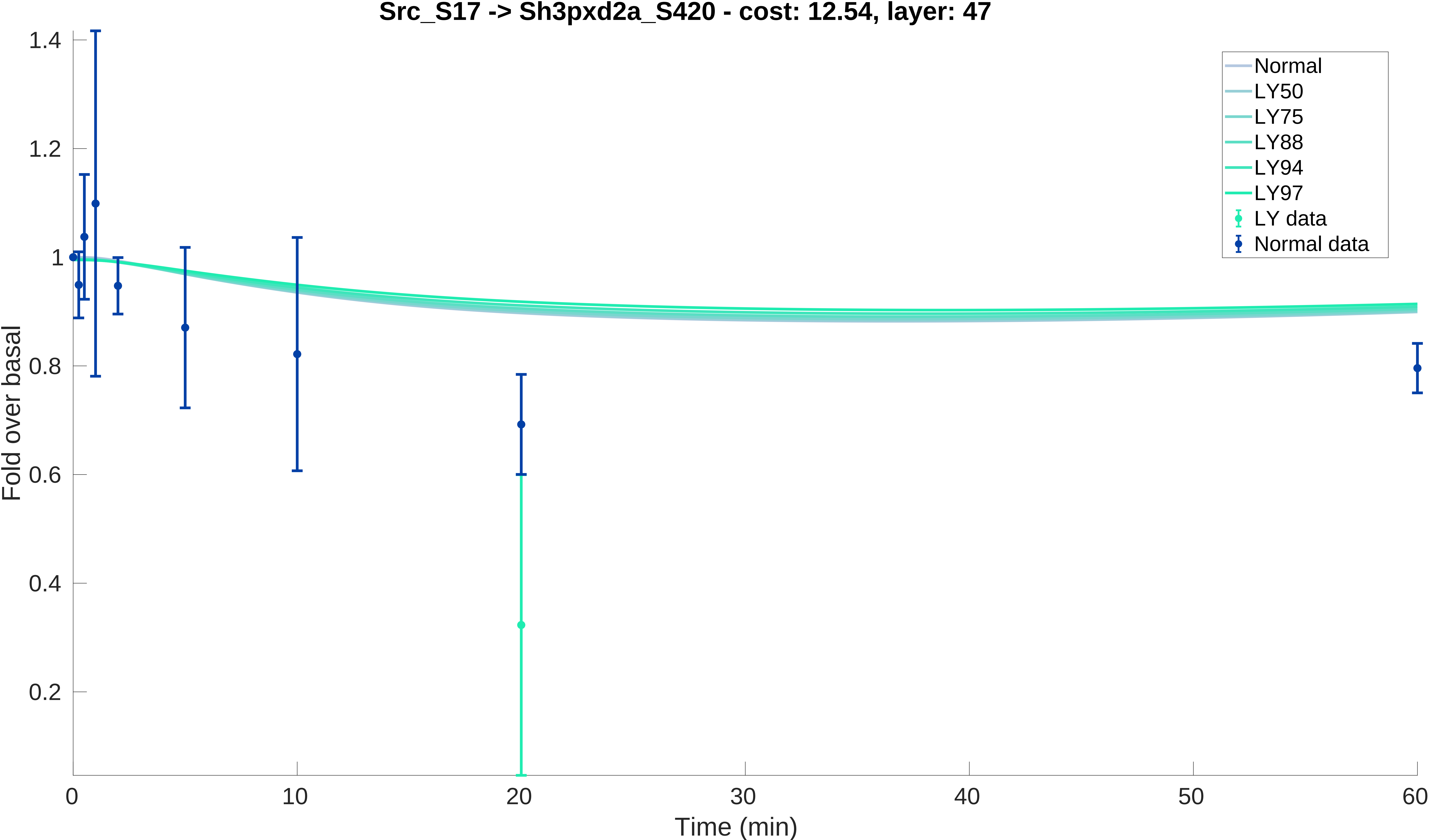

Src_S17 -> Sh3pxd2a_S420 - cost: 12.54, layer: 47
Fold over basal
Time (min)
0
10
20
30
40
50
60
0.2
0.4
0.6
0.8
1
1.2
1.4
Normal
LY50
LY75
LY88
LY94
LY97
LY data
Normal data

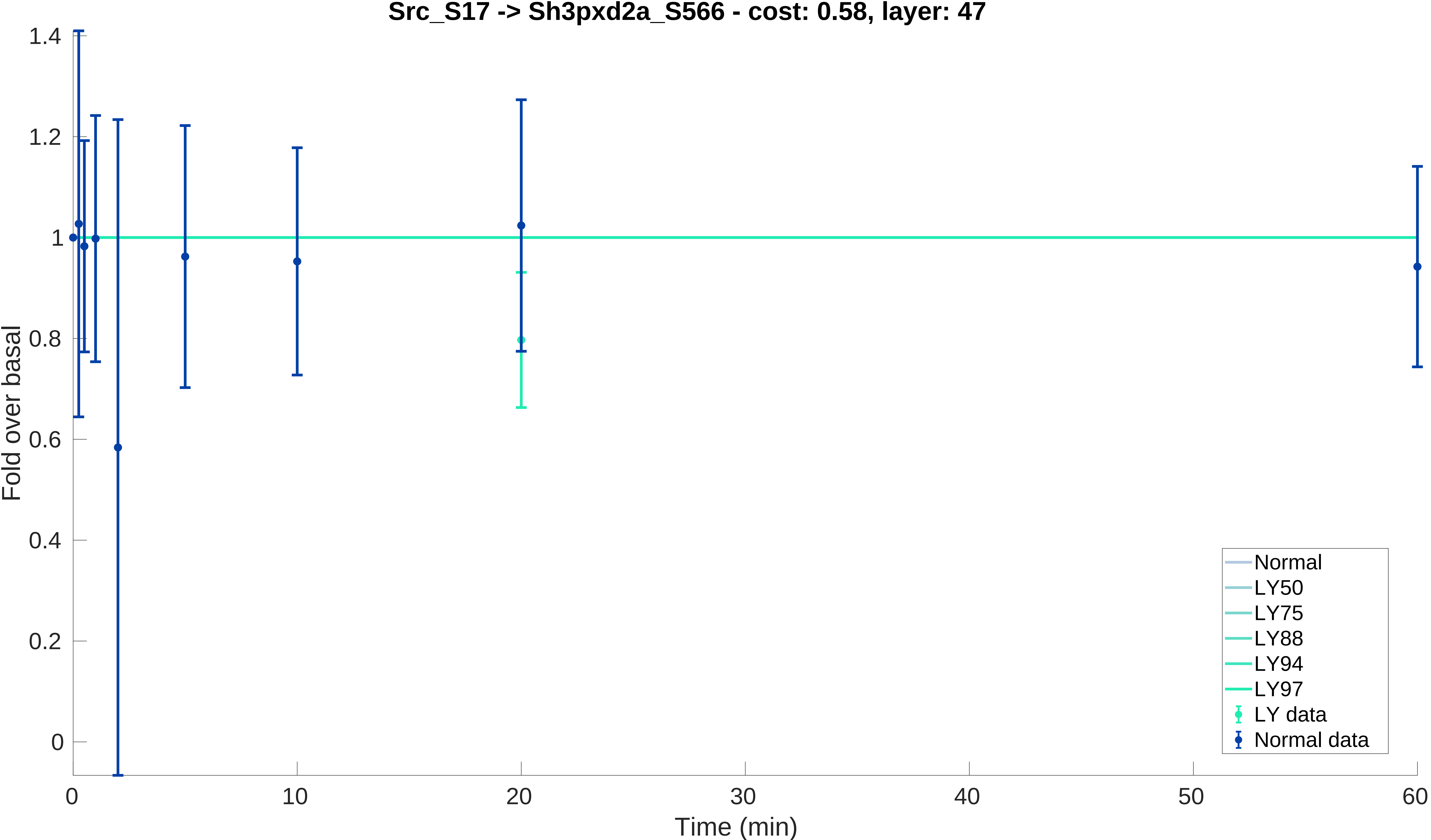

Src_S17 -> Sh3pxd2a_S566 - cost: 0.58, layer: 47
Fold over basal
1.4
1.2
1
0.8
0.6
0.4
0.2
0
0
10
20
30
40
50
60
Time (min)
Normal
LY50
LY75
LY88
LY94
LY97
LY data
Normal data

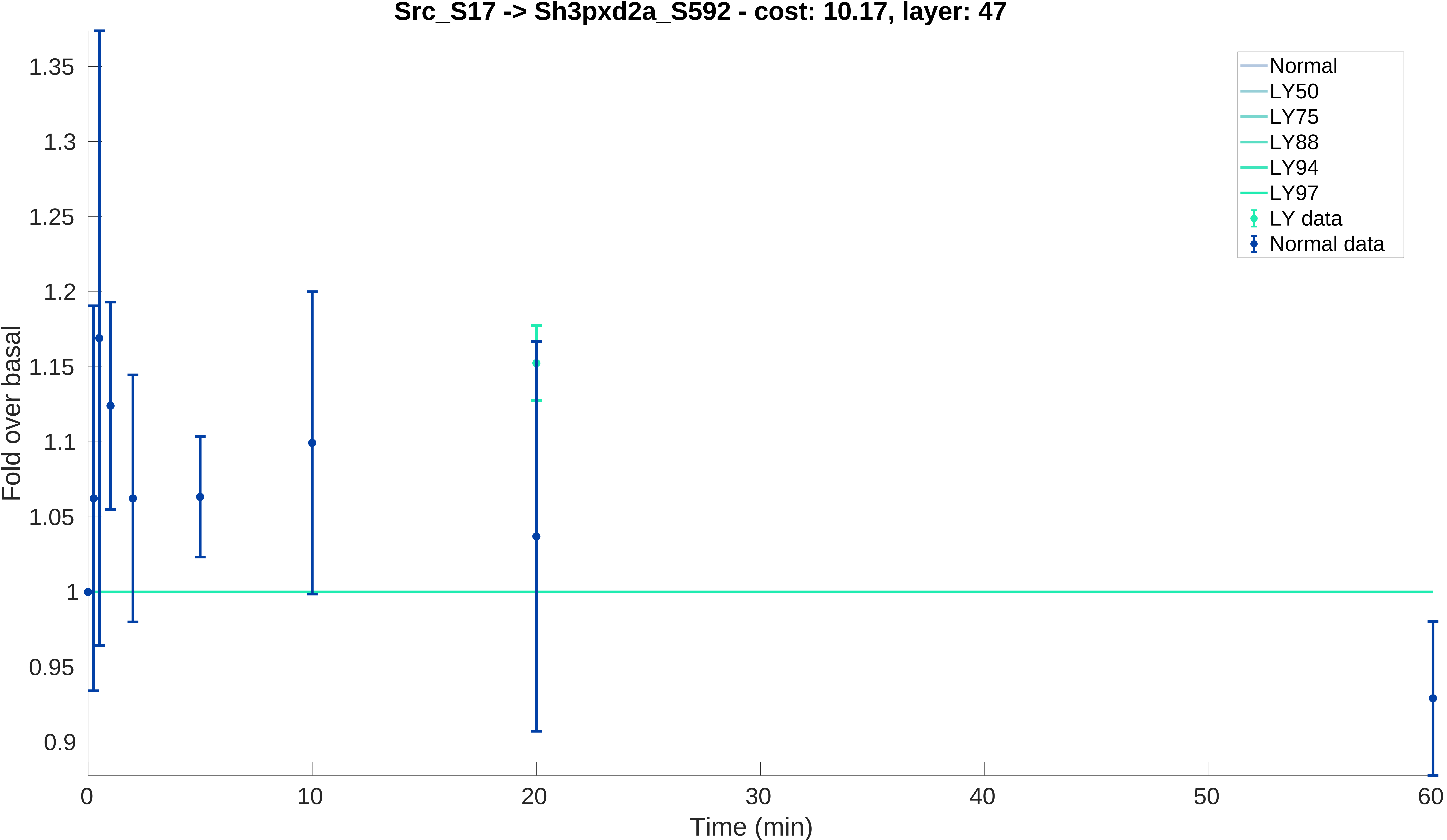

Src_S17 -> Sh3pxd2a_S592 - cost: 10.17, layer: 47
Fold over basal
Time (min)
1.35
1.3
1.25
1.2
1.15
1.1
1.05
1
0.95
0.9
0
10
20
30
40
50
60
Normal
LY50
LY75
LY88
LY94
LY97
LY data
Normal data

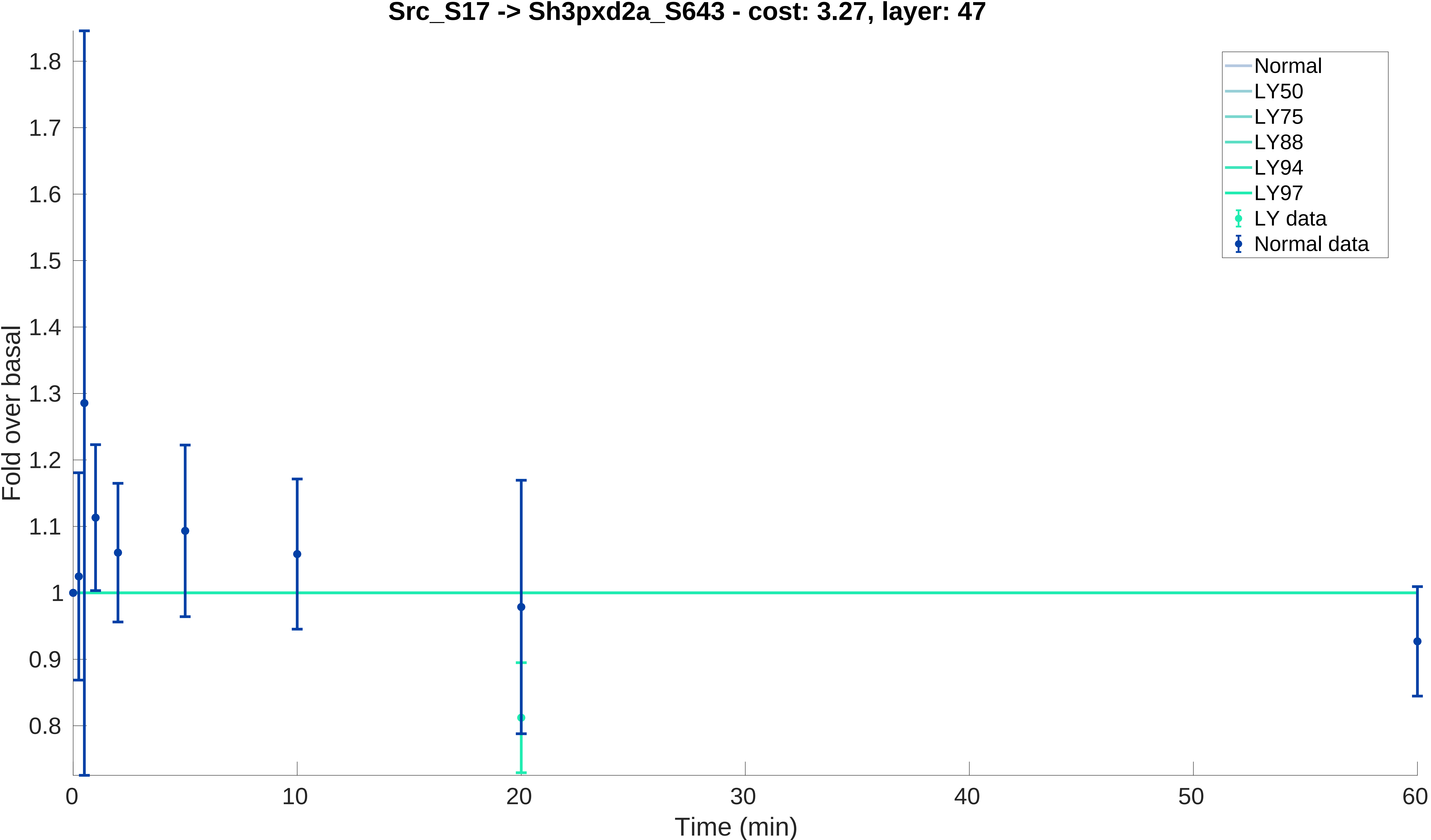

Src_S17 -> Sh3pxd2a_S643 - cost: 3.27, layer: 47
Fold over basal
Time (min)
Normal
LY50
LY75
LY88
LY94
LY97
LY data
Normal data
1.8
1.7
1.6
1.5
1.4
1.3
1.2
1.1
1
0.9
0.8
0
10
20
30
40
50
60

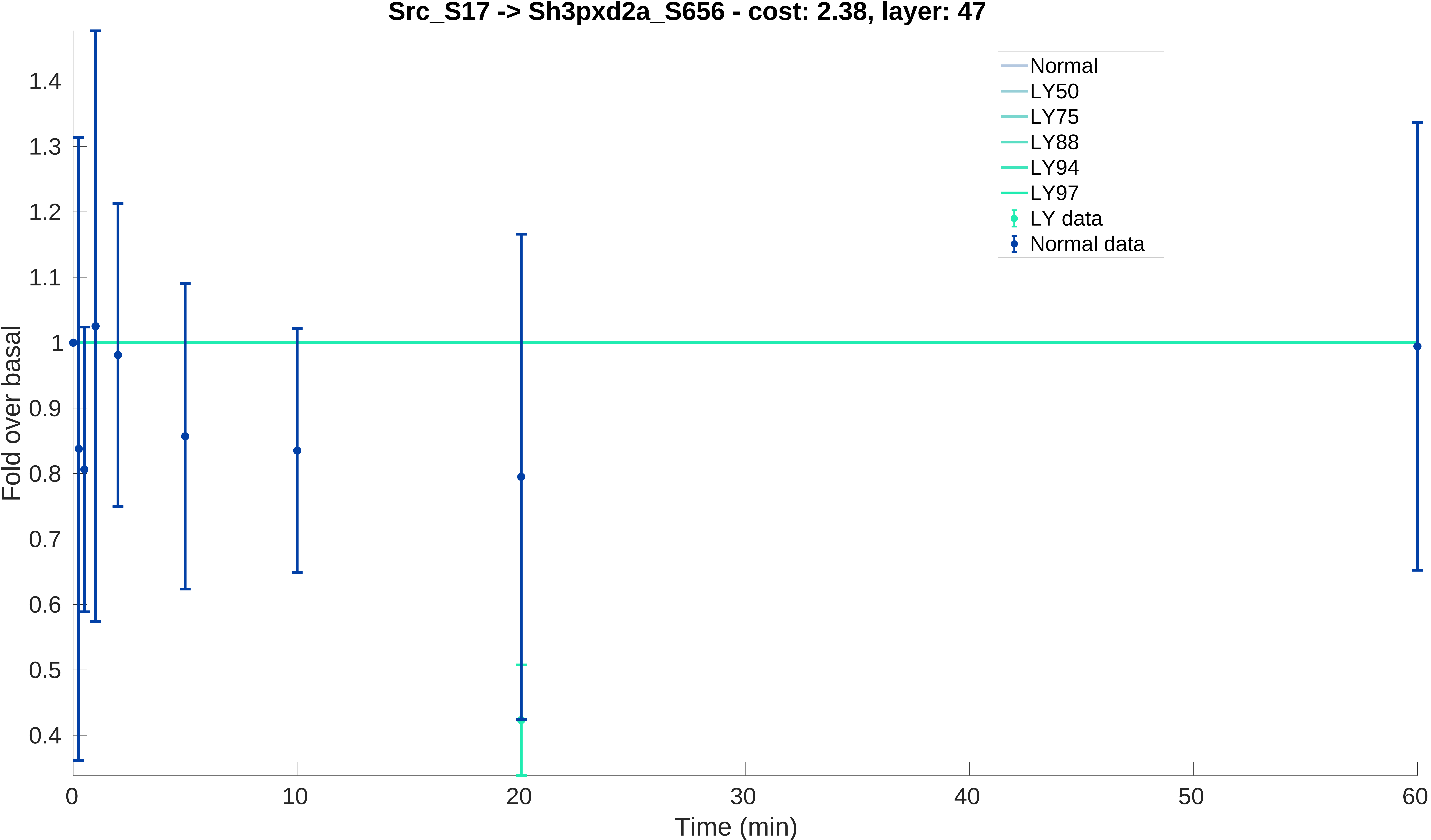

Src_S17 -> Sh3pxd2a_S656 - cost: 2.38, layer: 47
Normal
LY50
LY75
LY88
LY94
LY97
LY data
Normal data
Fold over basal
1.4
1.3
1.2
1.1
1
0.9
0.8
0.7
0.6
0.5
0.4
0
10
20
30
40
50
60
Time (min)

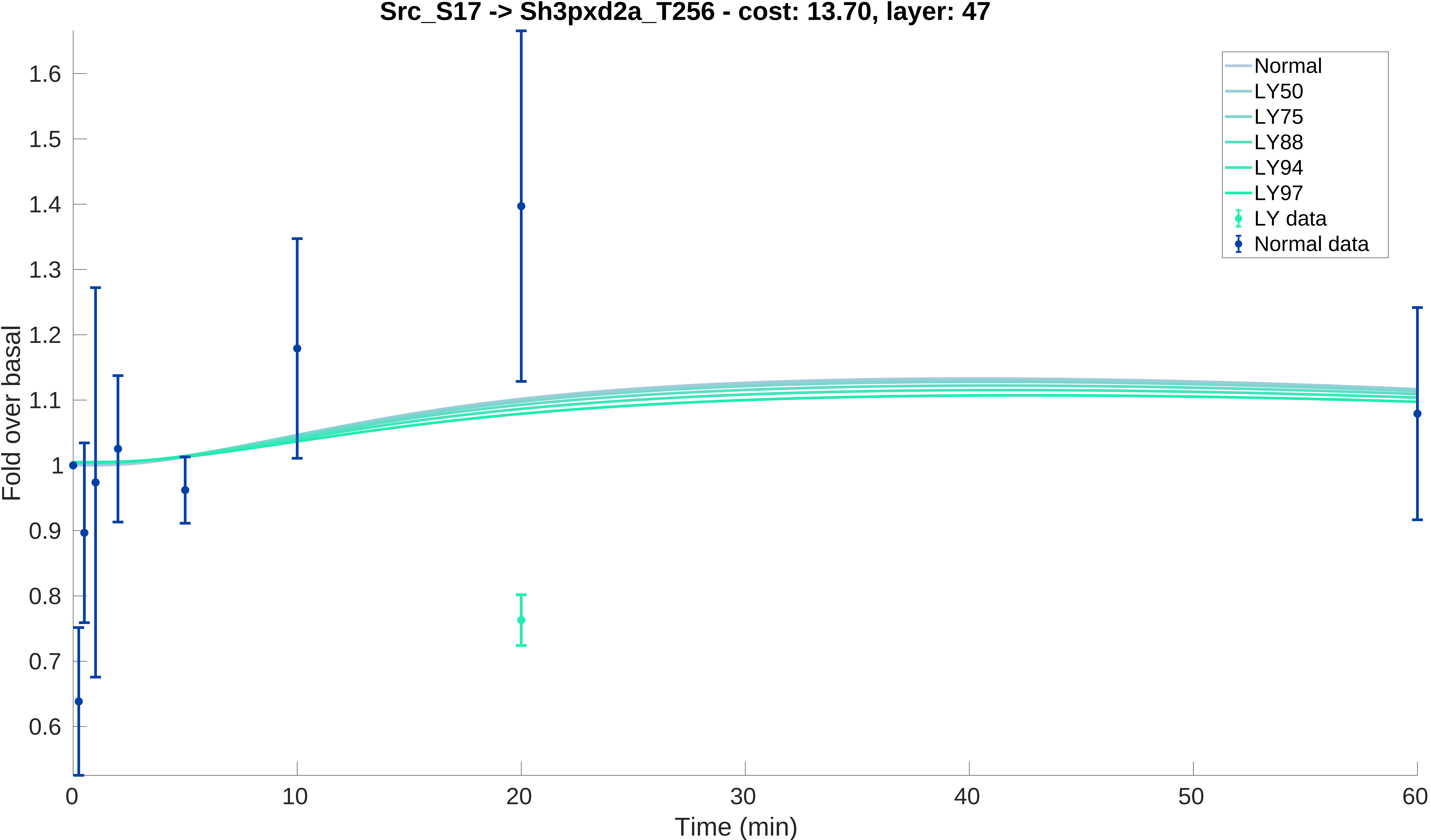

Src_S17 -> Sh3pxd2a_T256 - cost: 13.70, layer: 47
Normal
LY50
LY75
LY88
LY94
LY97
LY data
Normal data
Fold over basal
Time (min)
0
10
20
30
40
50
60
0.6
0.7
0.8
0.9
1
1.1
1.2
1.3
1.4
1.5
1.6

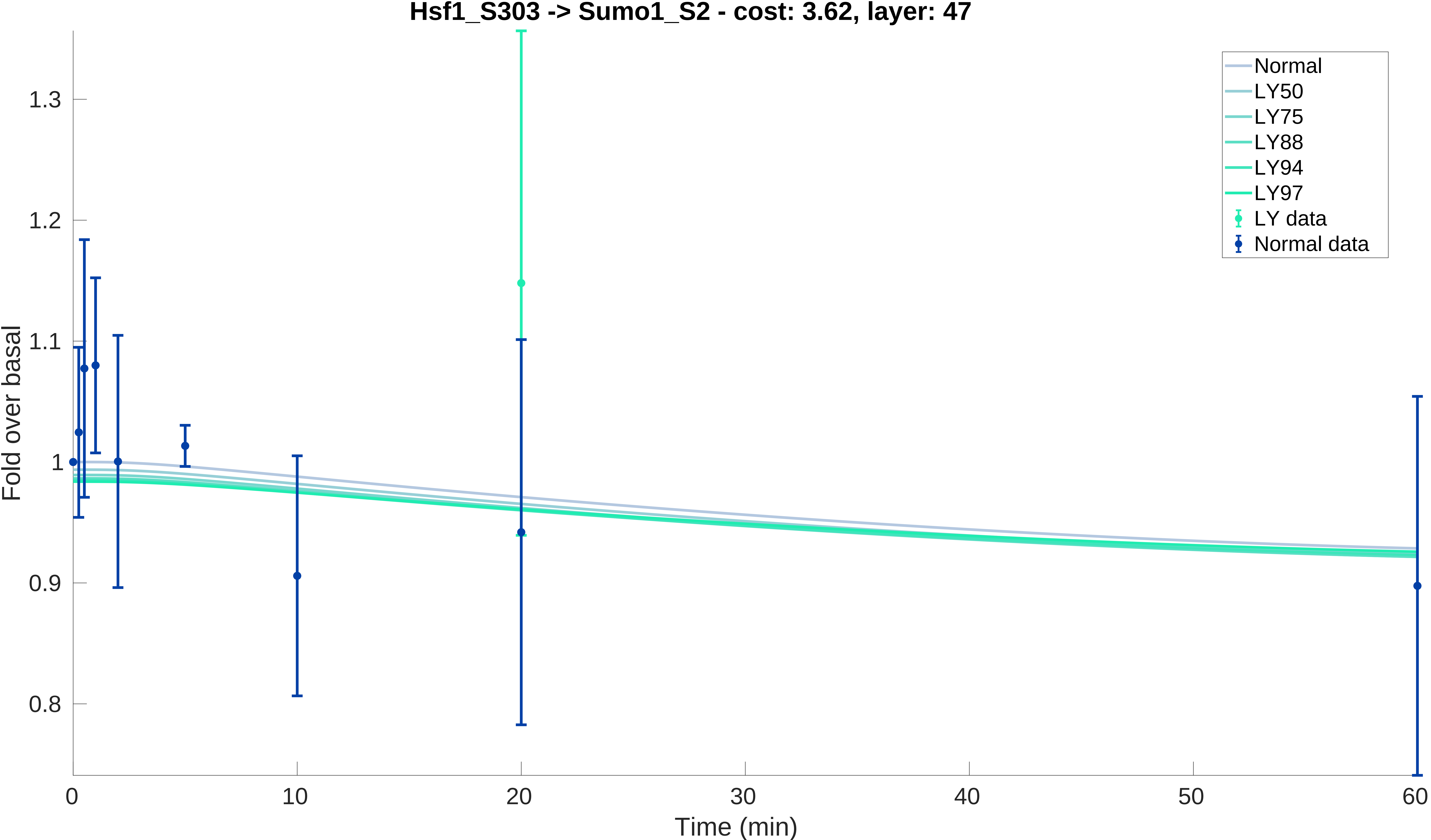

Hsf1_S303 -> Sumo1_S2 - cost: 3.62, layer: 47
Normal
LY50
LY75
LY88
LY94
LY97
LY data
Normal data
Fold over basal
1.3
1.2
1.1
1
0.9
0.8
0
10
20
30
40
50
60
Time (min)

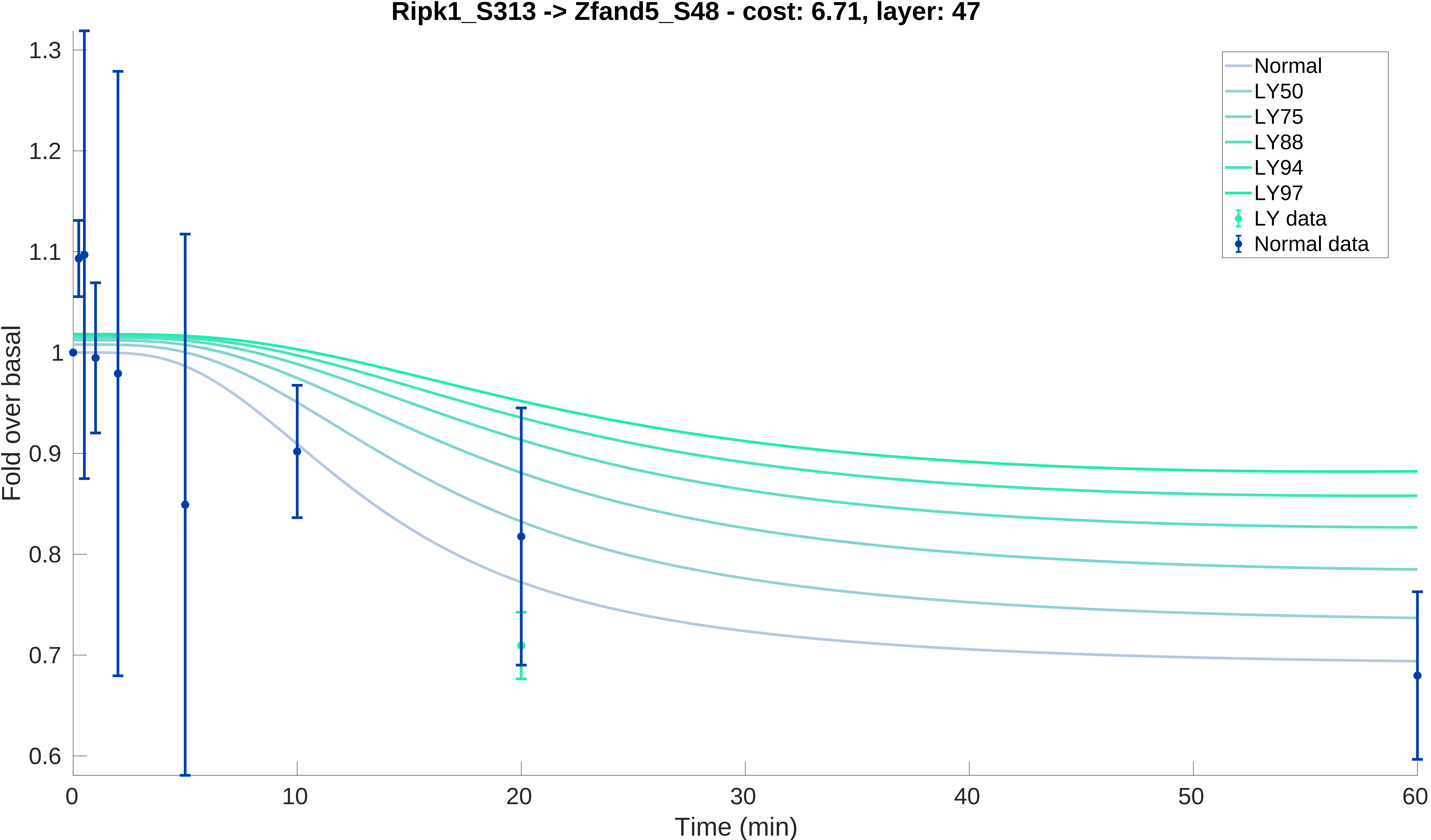
Ripk1_S313 -> Zfand5_S48 - cost: 6.71, layer: 47
Fold over basal
Time (min)
0
10
20
30
40
50
60
0.6
0.7
0.8
0.9
1
1.1
1.2
1.3
Normal
LY50
LY75
LY88
LY94
LY97
LY data
Normal data

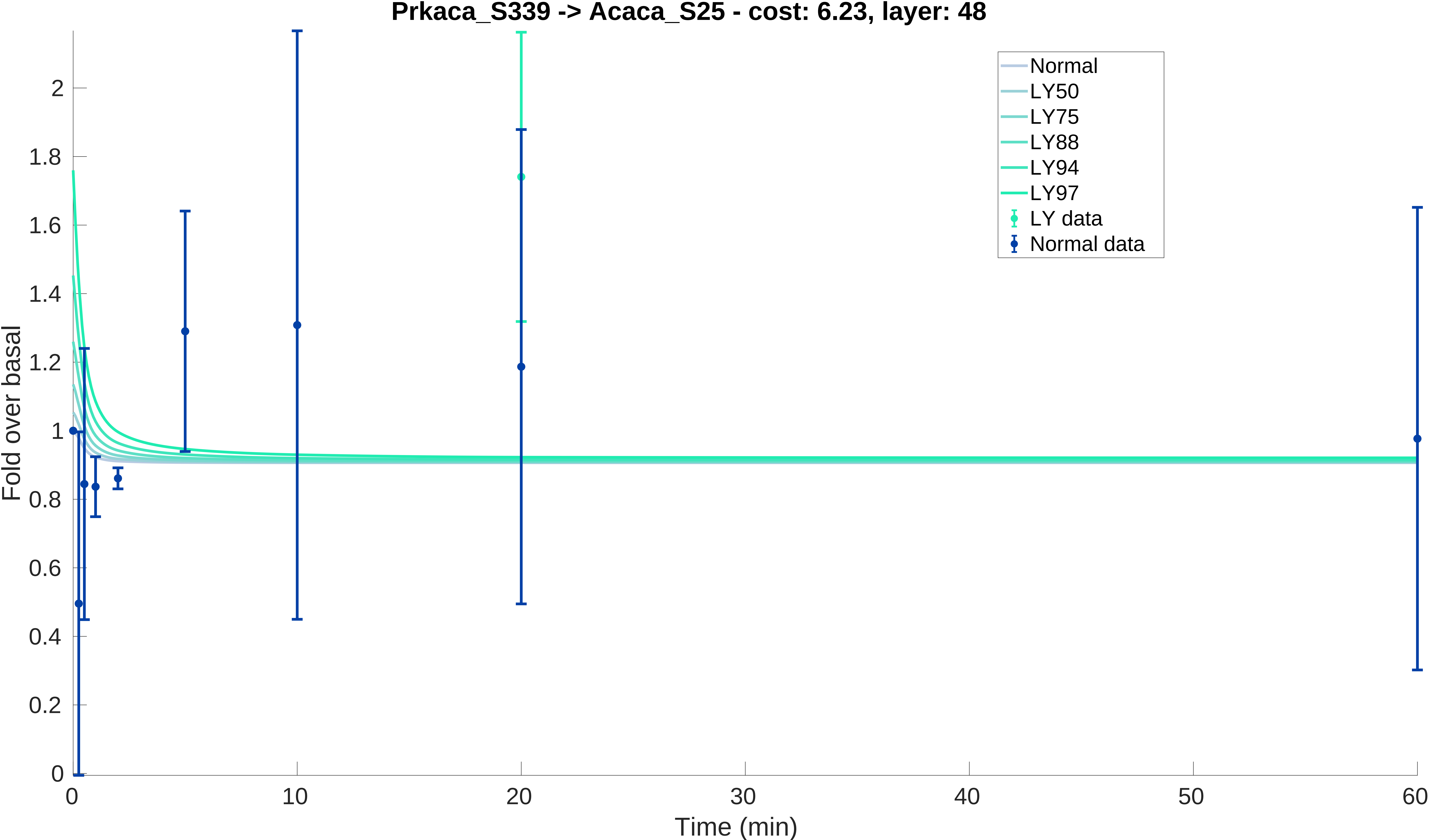

Prkaca_S339 -> Acaca_S25 - cost: 6.23, layer: 48
Fold over basal
Time (min)
0
0.2
0.4
0.6
0.8
1
1.2
1.4
1.6
1.8
2
0
10
20
30
40
50
60
Normal
LY50
LY75
LY88
LY94
LY97
LY data
Normal data

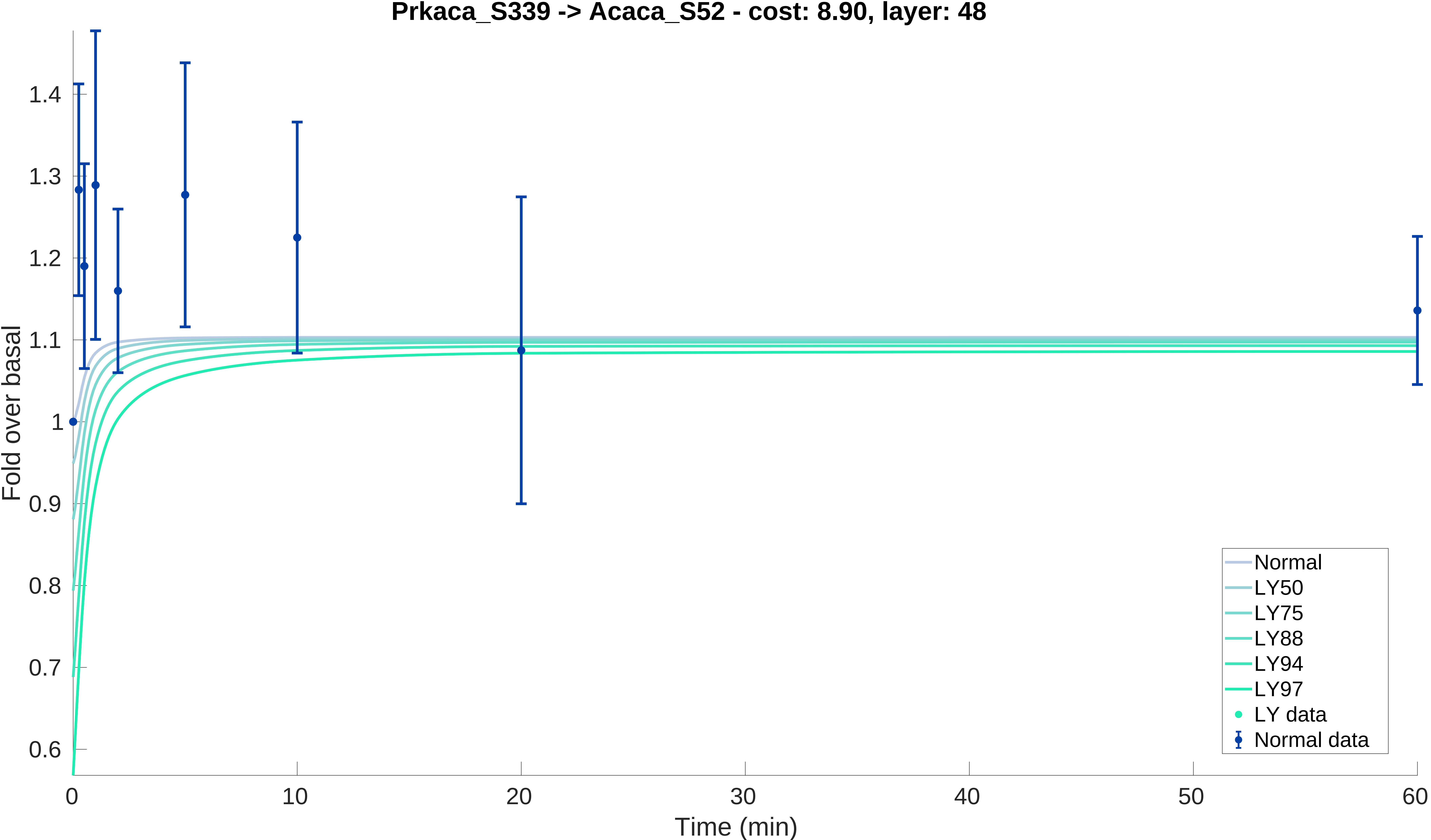

Prkaca_S339 -> Acaca_S52 - cost: 8.90, layer: 48
Fold over basal
Time (min)
1.4
1.3
1.2
1.1
1
0.9
0.8
0.7
0.6
0
10
20
30
40
50
60
Normal
LY50
LY75
LY88
LY94
LY97
LY data
Normal data

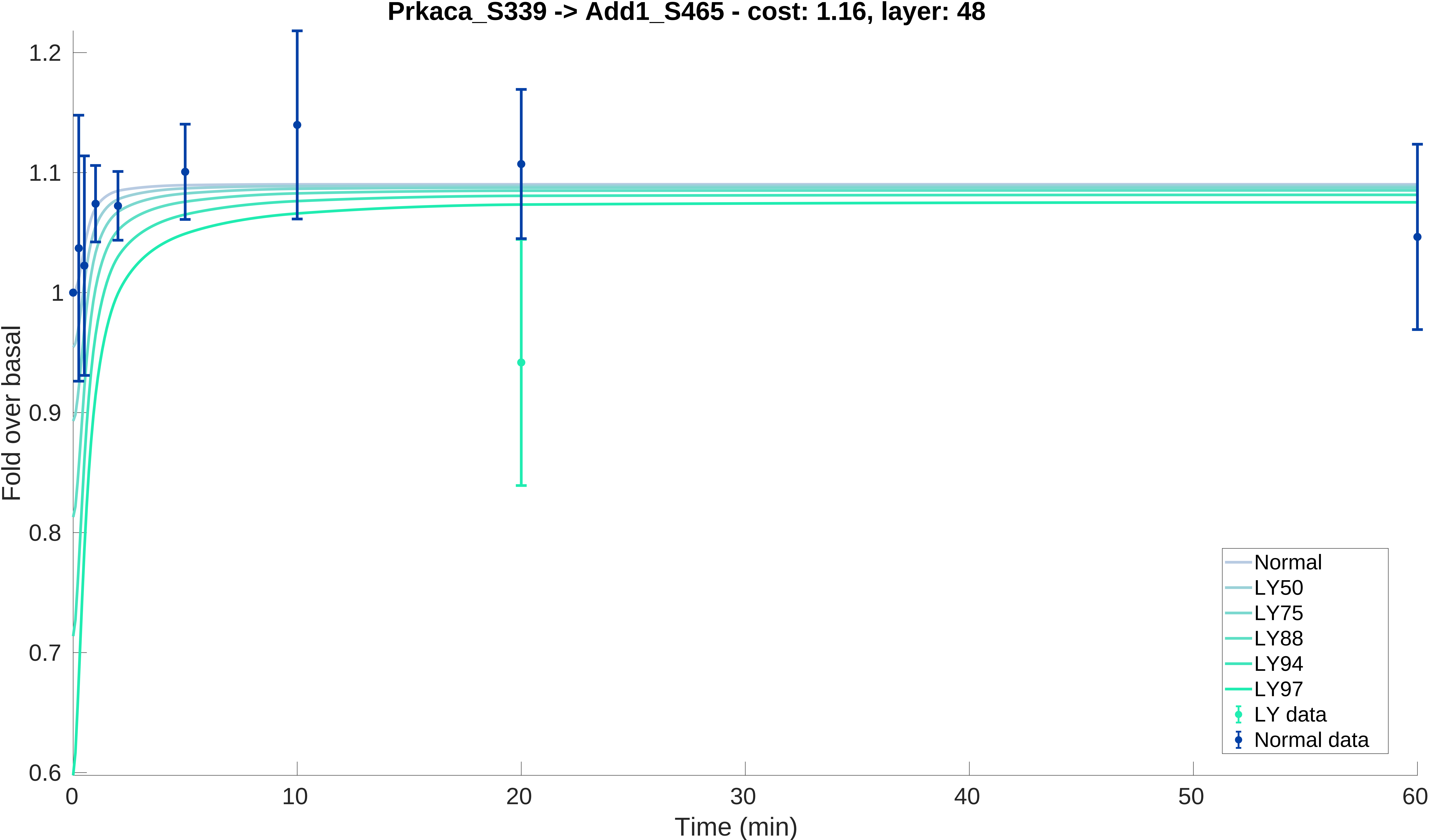
Prrkaca_S339 -> Add1_S465 - cost: 1.16, layer: 48
Fold over basal
1.2
1.1
1
0.9
0.8
0.7
0.6
0
10
20
30
40
50
60
Time (min)
Normal
LY50
LY75
LY88
LY94
LY97
LY data
Normal data

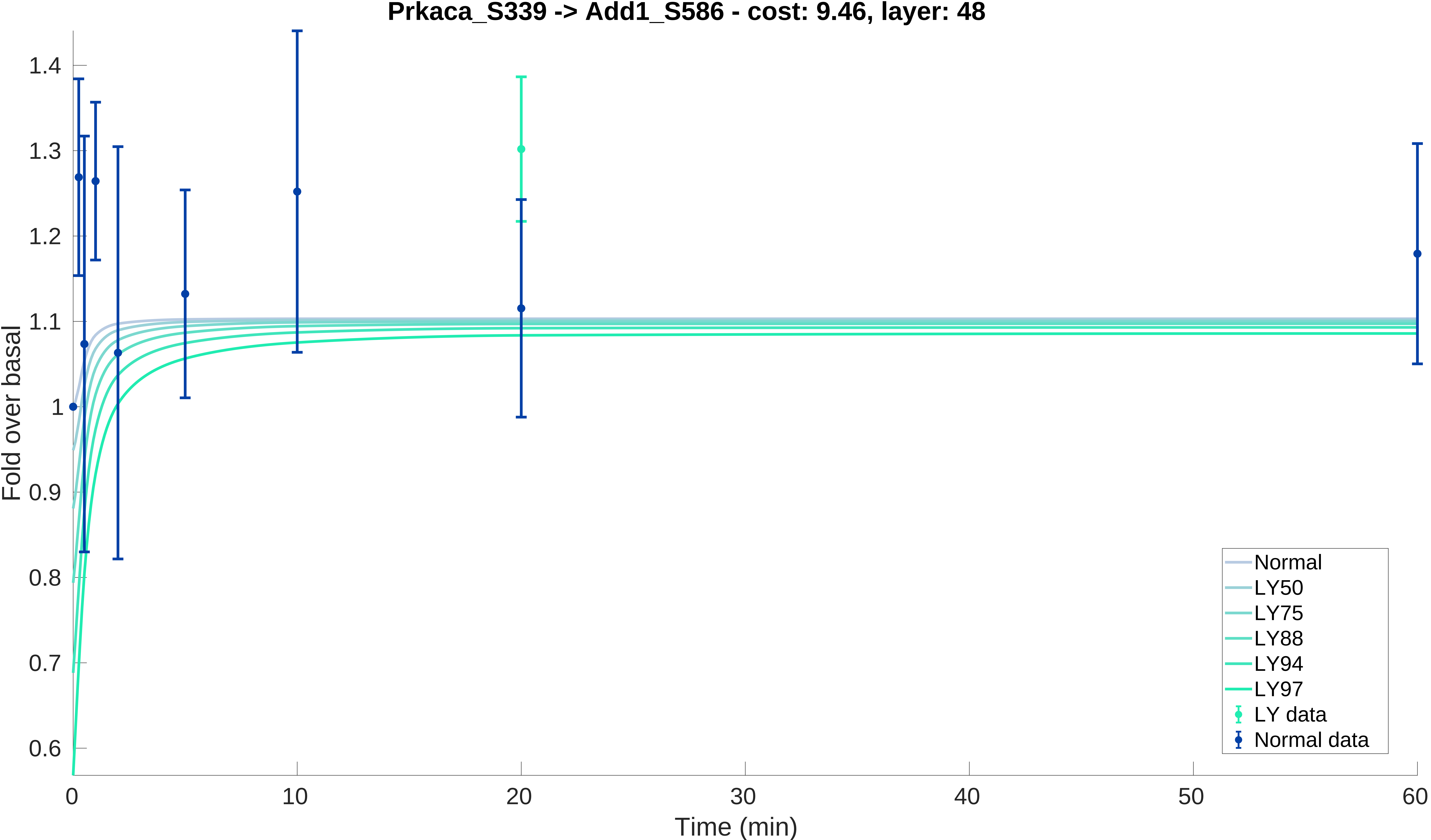

Prkaca_S339 -> Add1_S586 - cost: 9.46, layer: 48
Fold over basal
Time (min)
0.6
0.7
0.8
0.9
1
1.1
1.2
1.3
1.4
0
10
20
30
40
50
60
Normal
LY50
LY75
LY88
LY94
LY97
LY data
Normal data

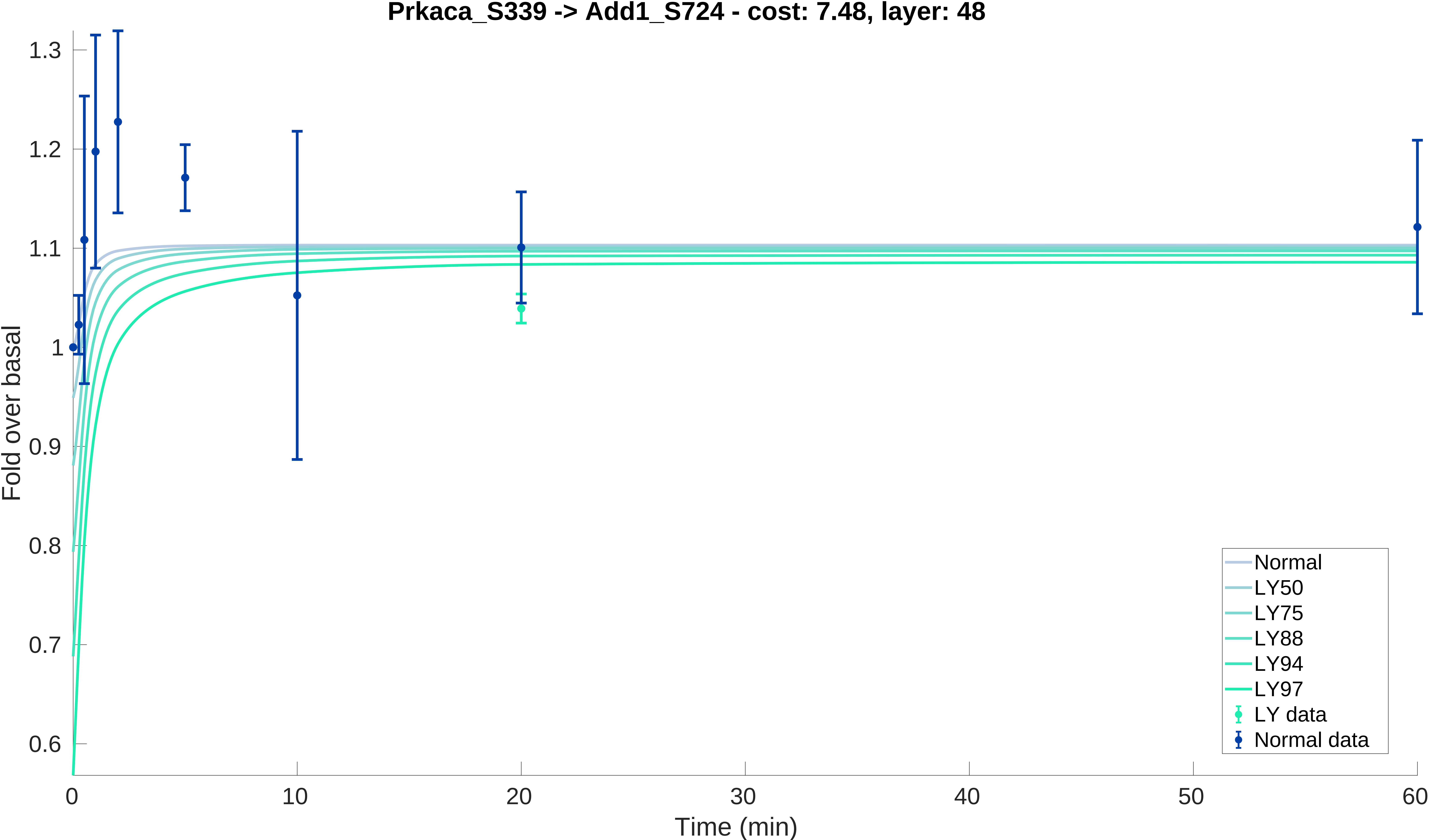

Prkaca_S339 -> Add1_S724 - cost: 7.48, layer: 48
Fold over basal
1.3
1.2
1.1
1
0.9
0.8
0.7
0.6
0
10
20
30
40
50
60
Time (min)
Normal
LY50
LY75
LY88
LY94
LY97
LY data
Normal data

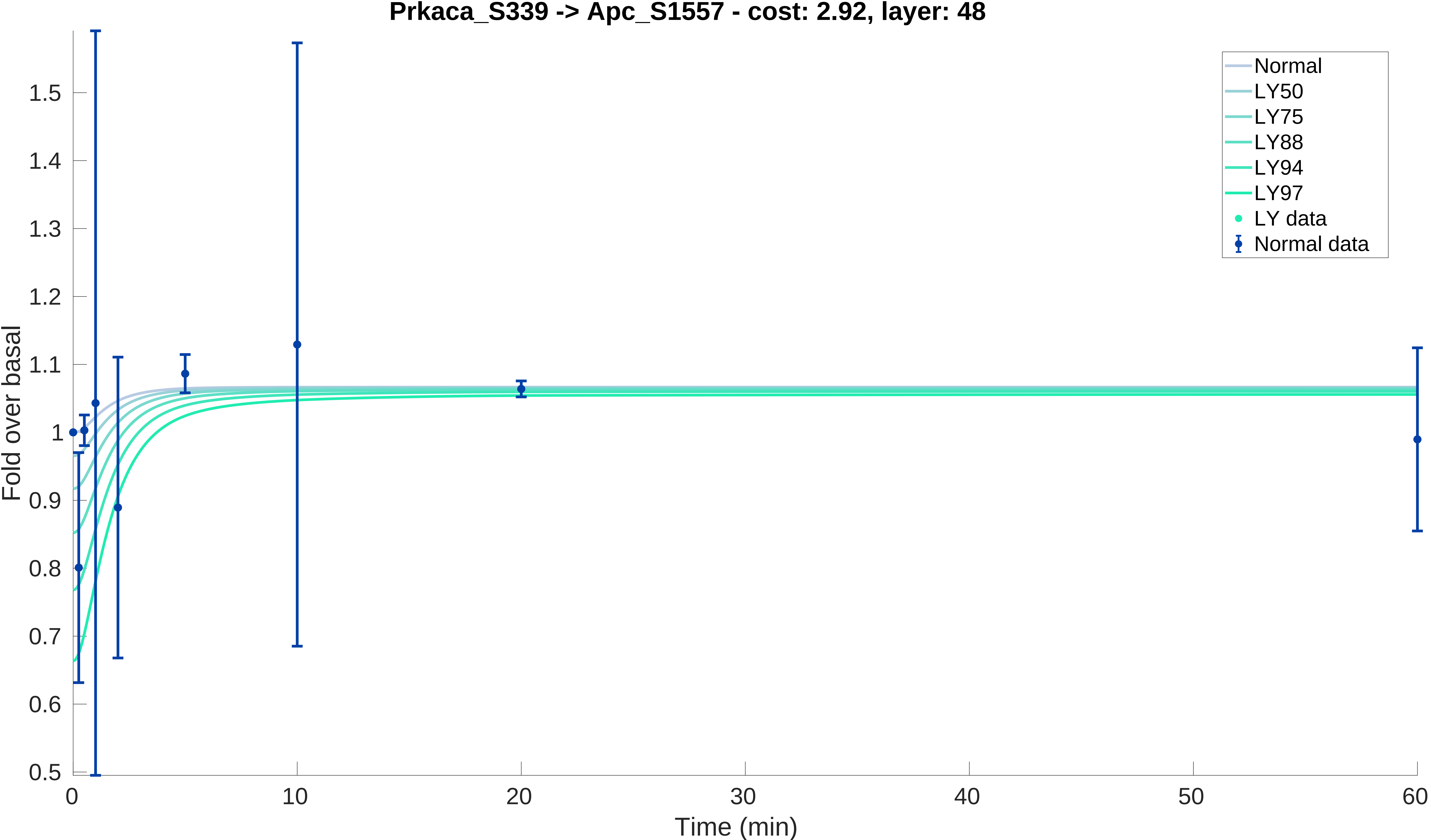

Prkaca_S339 -> Apc_S1557 - cost: 2.92, layer: 48
Normal
LY50
LY75
LY88
LY94
LY97
LY data
Normal data
Fold over basal
Time (min)
0.5
0.6
0.7
0.8
0.9
1
1.1
1.2
1.3
1.4
1.5
0
10
20
30
40
50
60

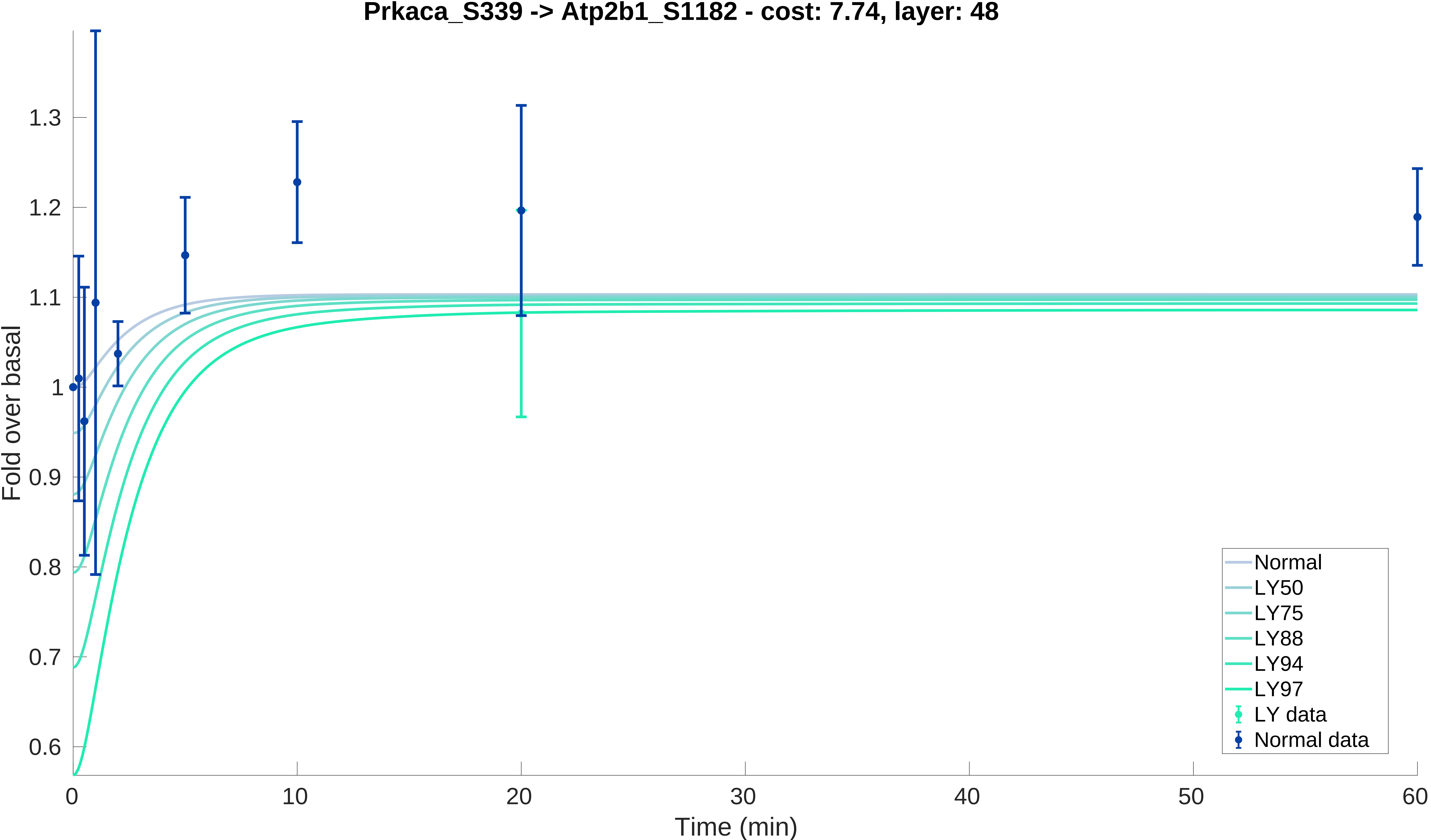

Prkaca_S339 -> Atp2b1_S1182 - cost: 7.74, layer: 48
Fold over basal
Time (min)
0.6
0.7
0.8
0.9
1
1.1
1.2
1.3
0
10
20
30
40
50
60
Normal
LY50
LY75
LY88
LY94
LY97
LY data
Normal data

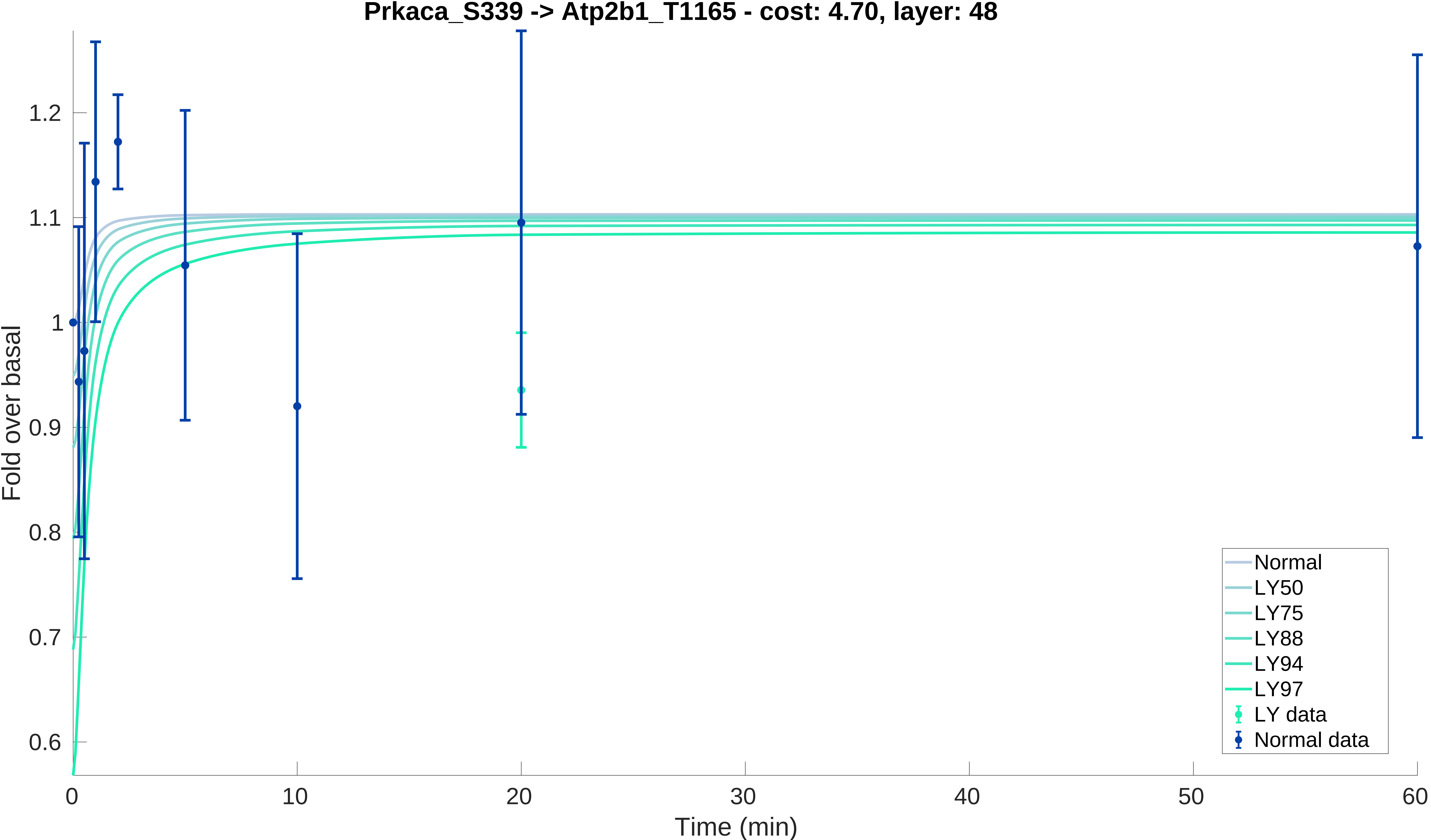
Prkaca_S339 -> Atp2b1_T1165 - cost: 4.70, layer: 48
Fold over basal
Time (min)
0.6
0.7
0.8
0.9
1
1.1
1.2
0
10
20
30
40
50
60
Normal
LY50
LY75
LY88
LY94
LY97
LY data
Normal data

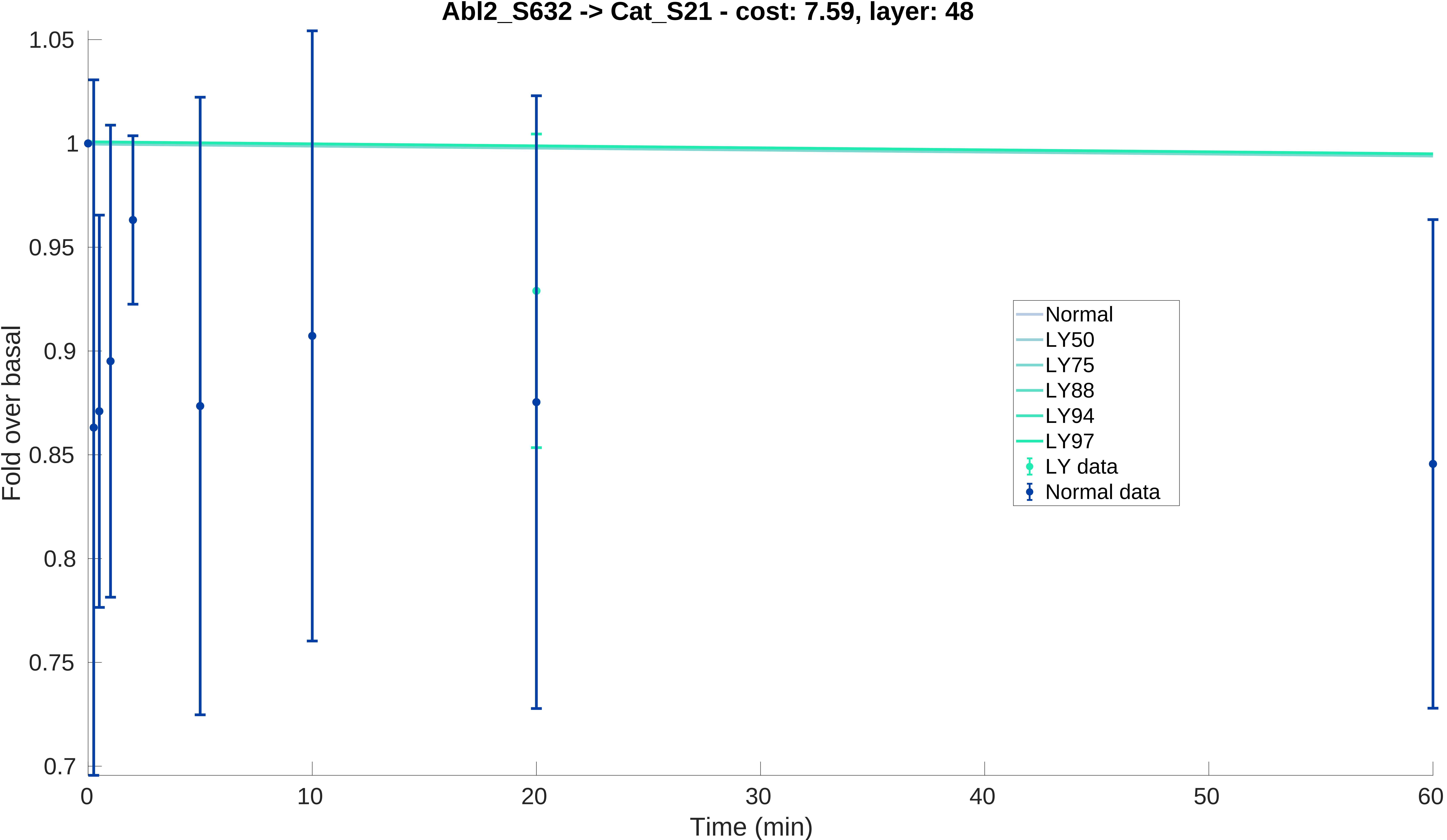

Abl2_S632 -> Cat_S21 - cost: 7.59, layer: 48
1.05
1
0.95
0.9
0.85
0.8
0.75
0.7
Fold over basal
0
10
20
30
40
50
60
Time (min)
Normal
LY50
LY75
LY88
LY94
LY97
LY data
Normal data

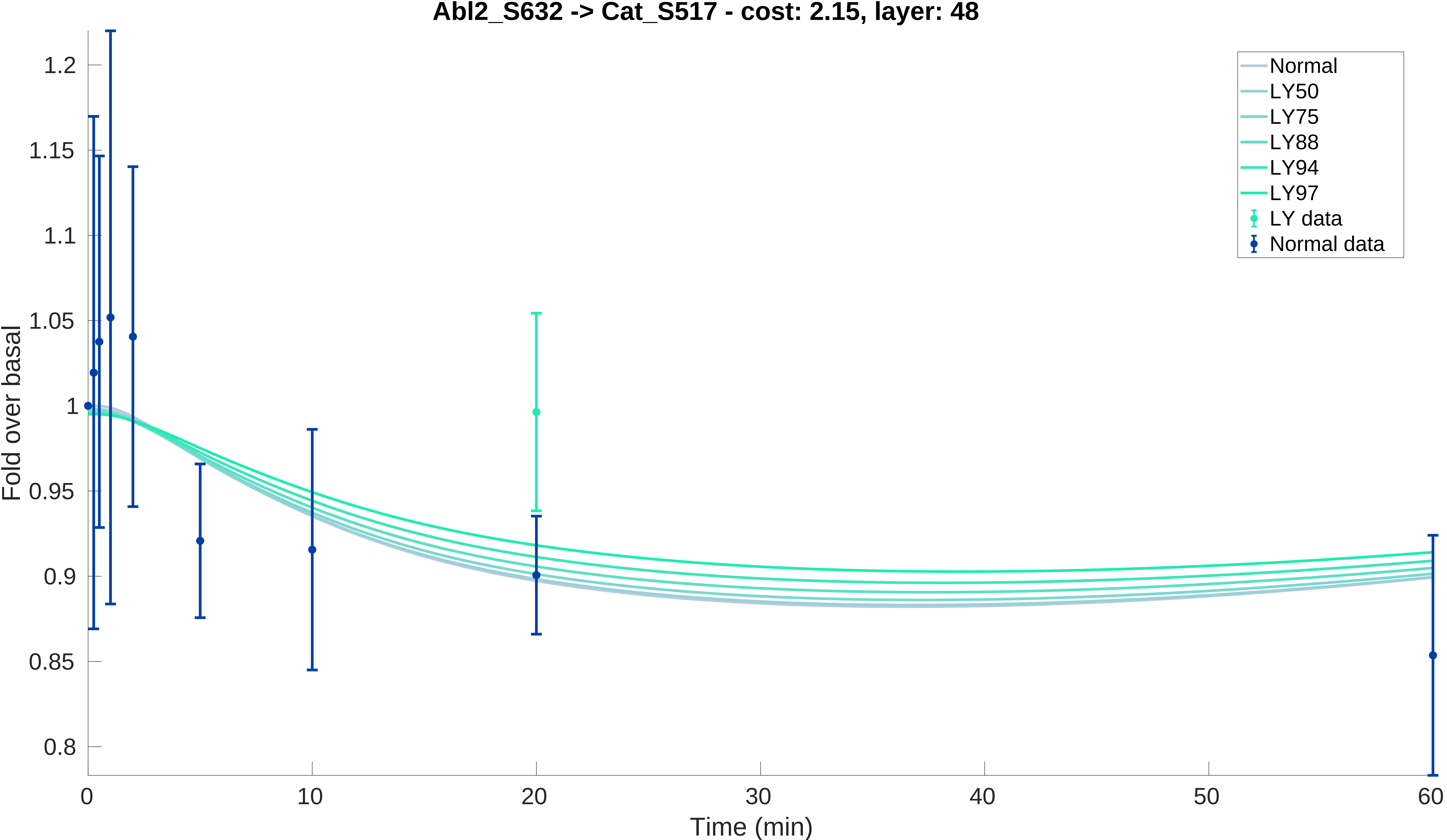
Abl2_S632 -> Cat_S517 - cost: 2.15, layer: 48
Fold over basal
Time (min)
1.2
1.15
1.1
1.05
1
0.95
0.9
0.85
0.8
0
10
20
30
40
50
60
Normal
LY50
LY75
LY88
LY94
LY97
LY data
Normal data

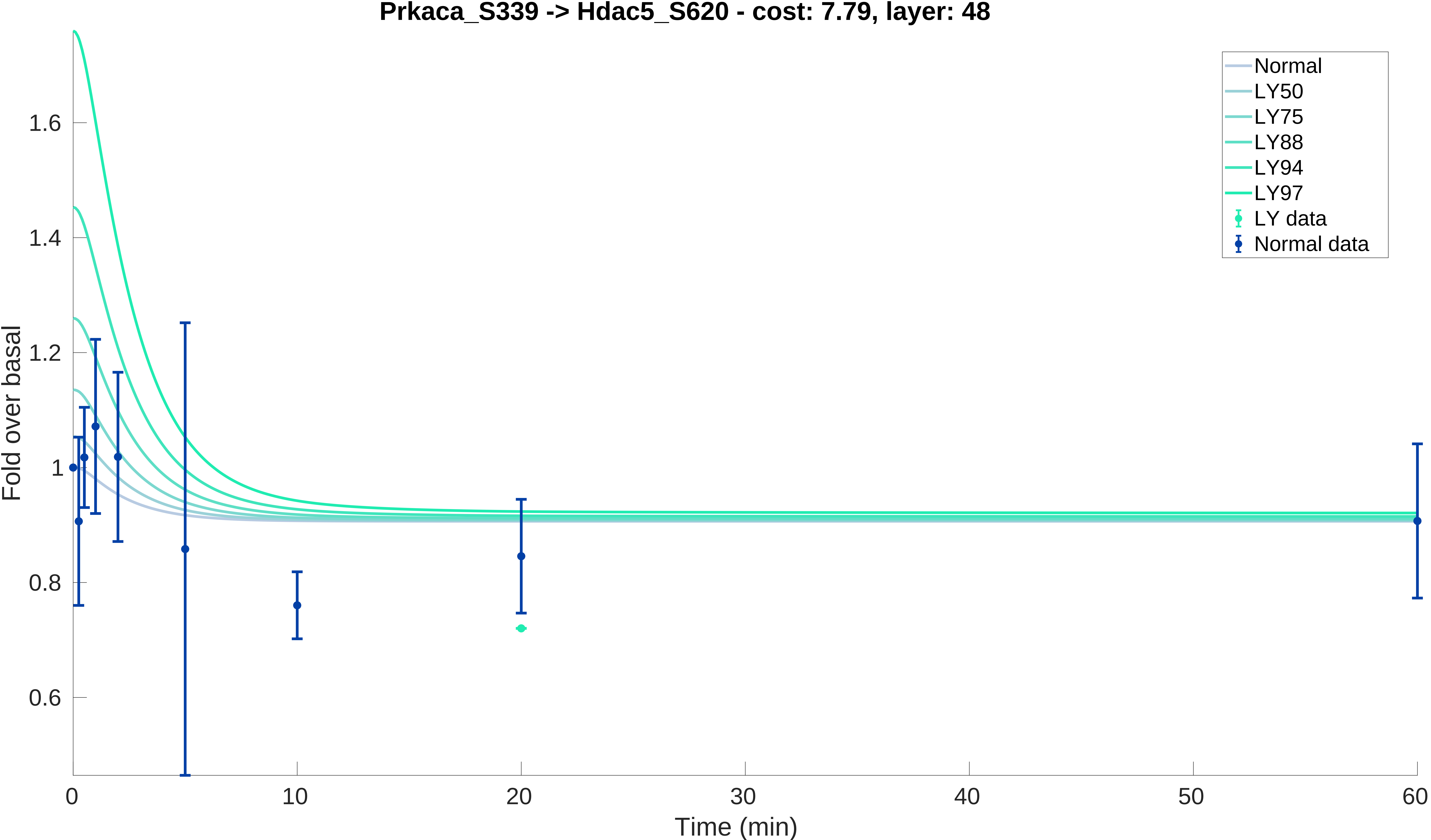

Prkaca_S339 -> Hdac5_S620 - cost: 7.79, layer: 48
Normal
LY50
LY75
LY88
LY94
LY97
LY data
Normal data
Fold over basal
1.6
1.4
1.2
1
0.8
0.6
0
10
20
30
40
50
60
Time (min)

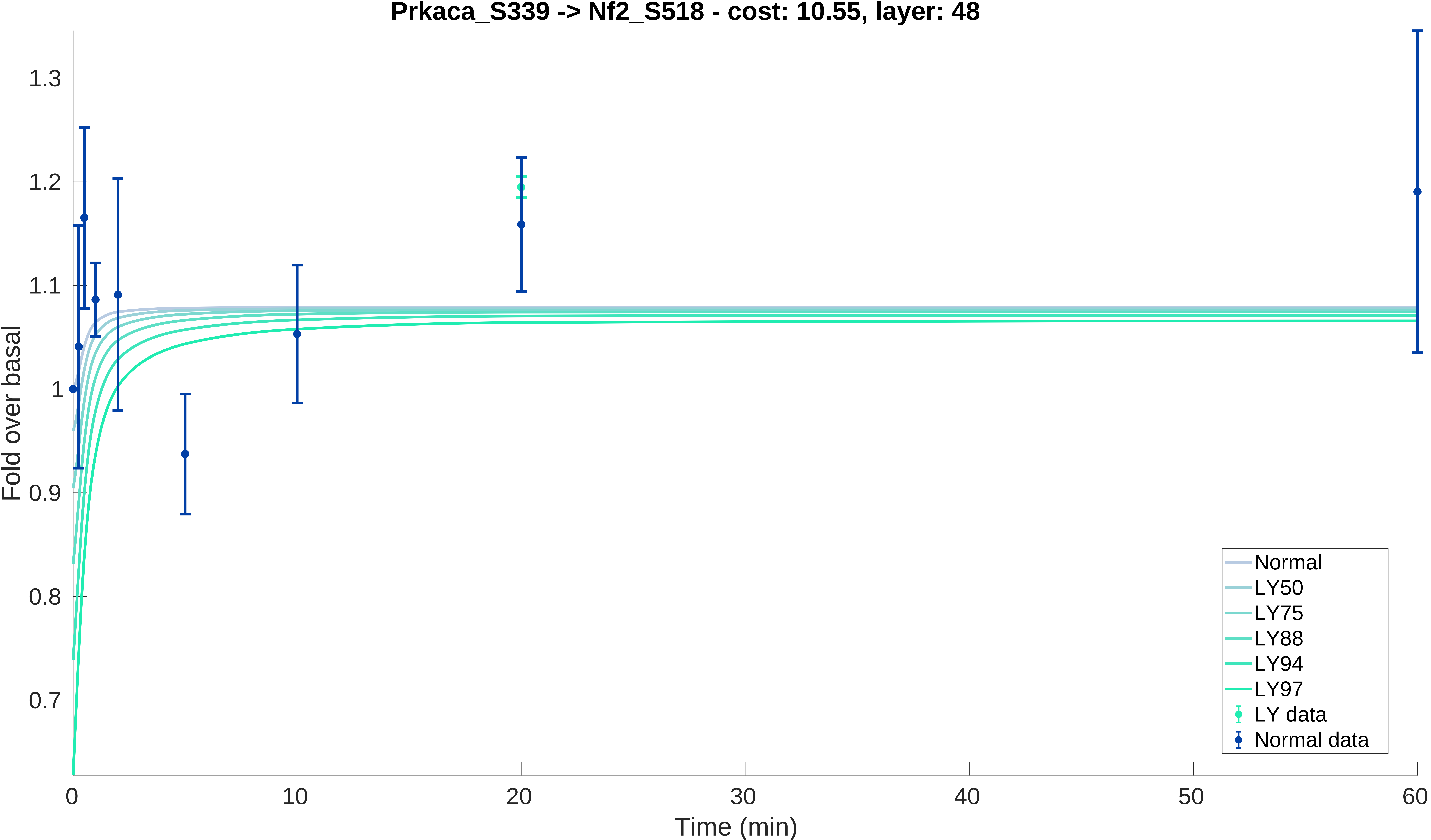

Prkaca_S339 -> Nf2_S518 - cost: 10.55, layer: 48
Fold over basal
Time (min)
1.3
1.2
1.1
1
0.9
0.8
0.7
0
10
20
30
40
50
60
Normal
LY50
LY75
LY88
LY94
LY97
LY data
Normal data

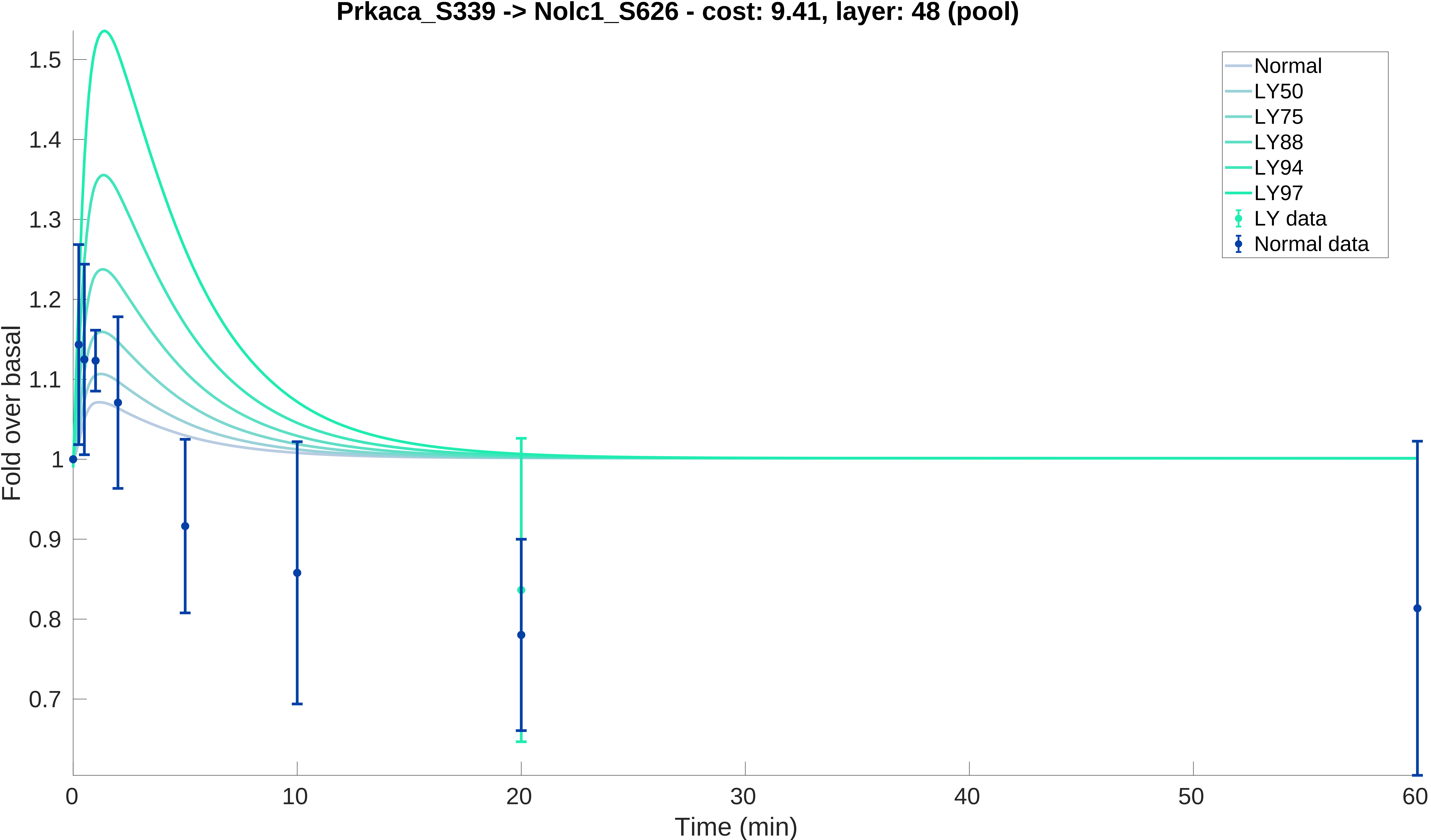
Prkaca_S339 -> Nolc1_S626 - cost: 9.41, layer: 48 (pool)
Normal
LY50
LY75
LY88
LY94
LY97
LY data
Normal data
Fold over basal
Time (min)
1.5
1.4
1.3
1.2
1.1
1
0.9
0.8
0.7
0
10
20
30
40
50
60

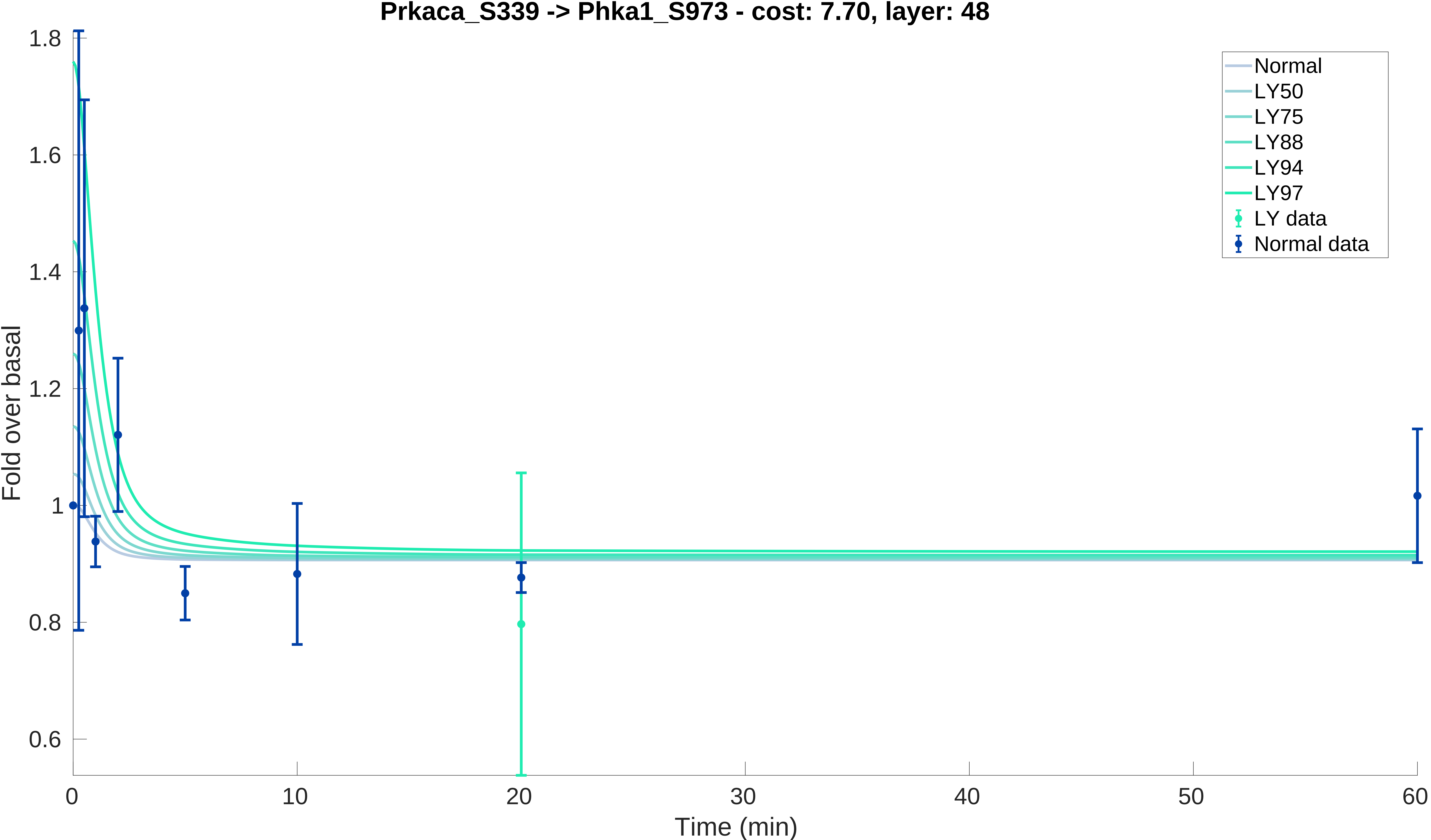

Prkaca_S339 -> Phka1_S973 - cost: 7.70, layer: 48
Fold over basal
Time (min)
Normal
LY50
LY75
LY88
LY94
LY97
LY data
Normal data
1.8
1.6
1.4
1.2
1
0.8
0.6
0
10
20
30
40
50
60

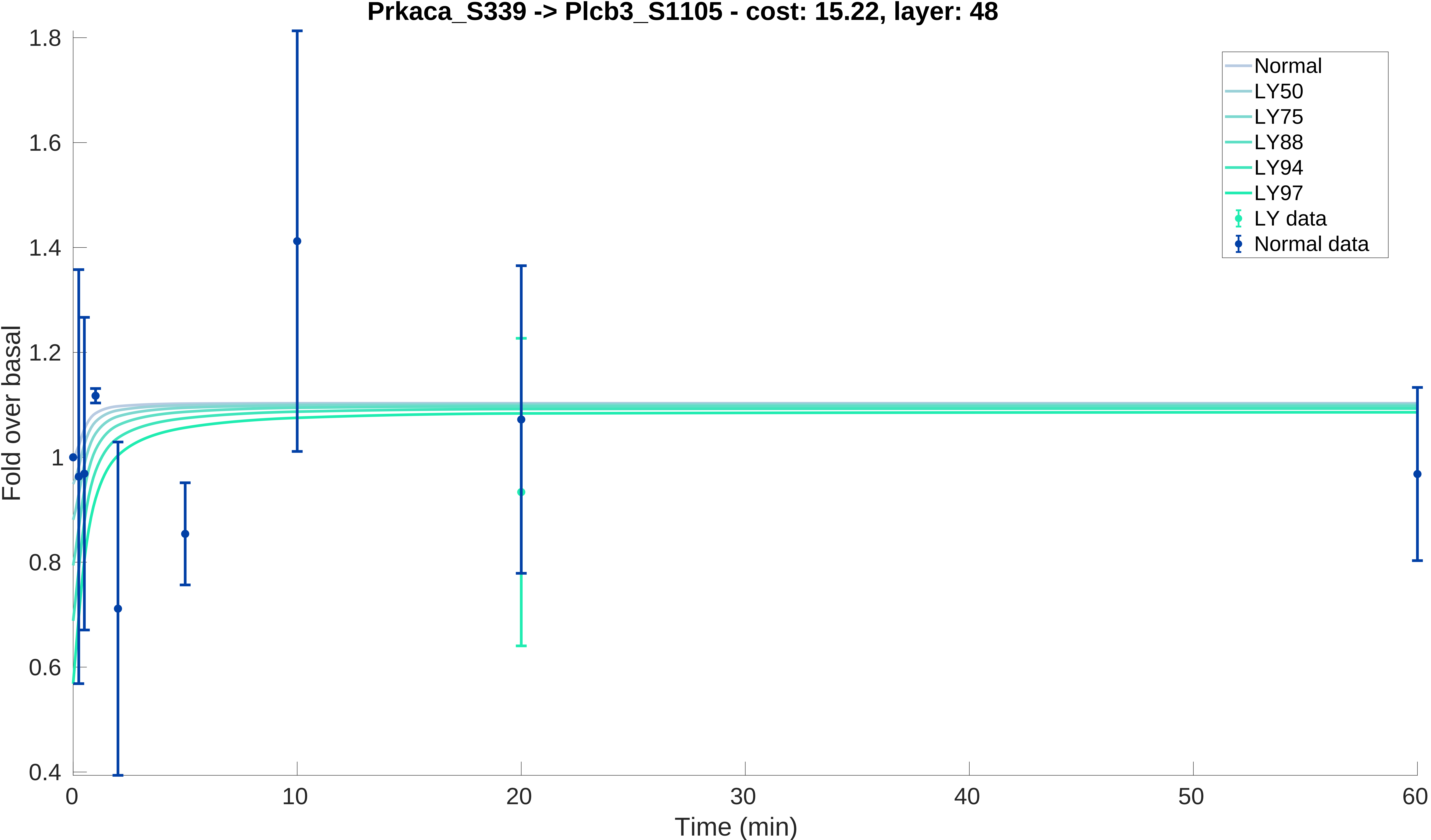

Prrkaca_S339 -> Plcb3_S1105 - cost: 15.22, layer: 48
Normal
LY50
LY75
LY88
LY94
LY97
LY data
Normal data
Fold over basal
Time (min)
0
10
20
30
40
50
60
0.4
0.6
0.8
1
1.2
1.4
1.6
1.8

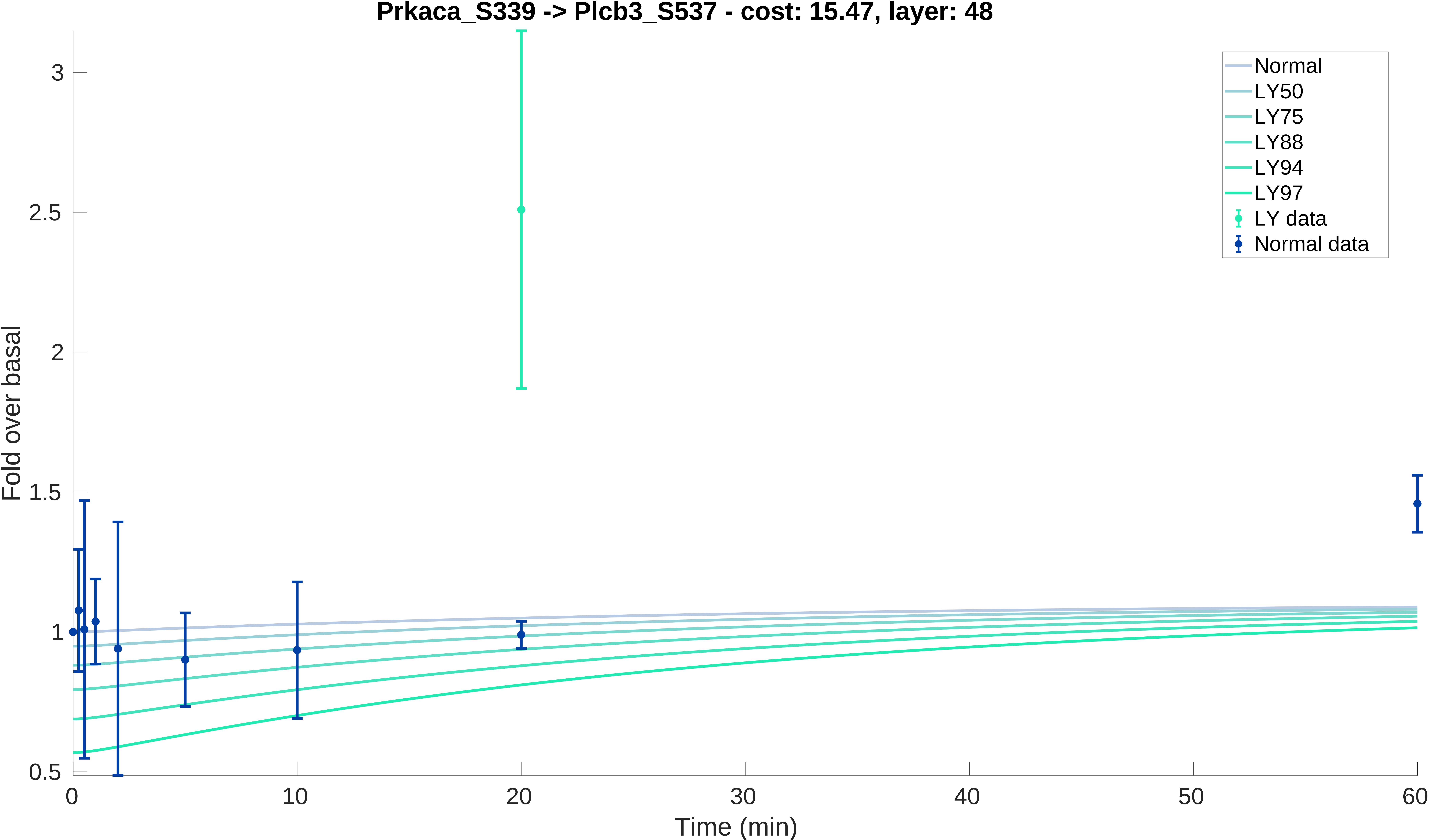

Prkaca_S339 -> Plcb3_S537 - cost: 15.47, layer: 48
Fold over basal
Time (min)
0.5
1
1.5
2
2.5
3
0
10
20
30
40
50
60
Normal
LY50
LY75
LY88
LY94
LY97
LY data
Normal data

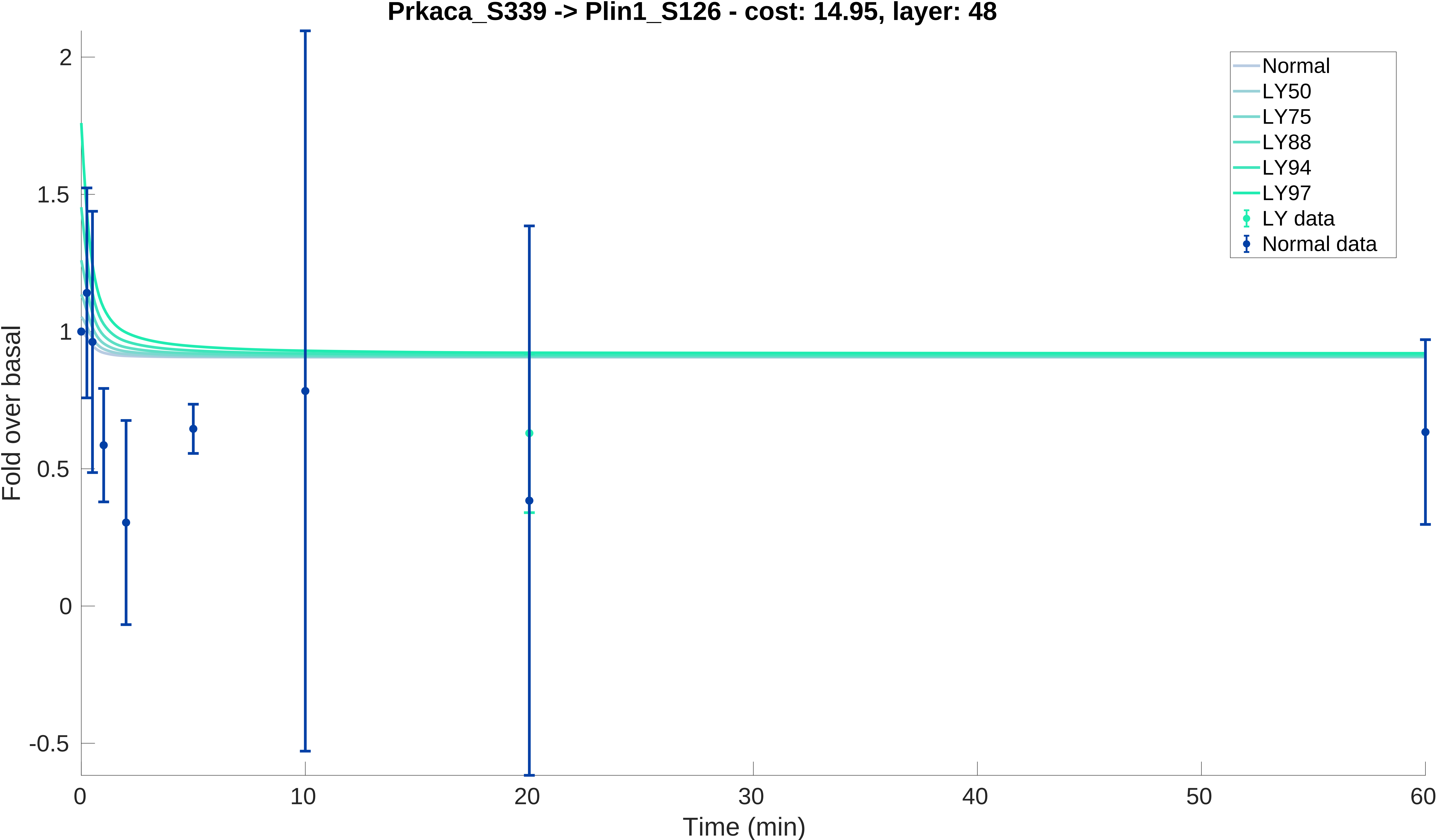
Prkaca_S339 -> Plin1_S126 - cost: 14.95, layer: 48
Fold over basal
Time (min)
2
1.5
1
0.5
0
-0.5
0
10
20
30
40
50
60
Normal
LY50
LY75
LY88
LY94
LY97
LY data
Normal data

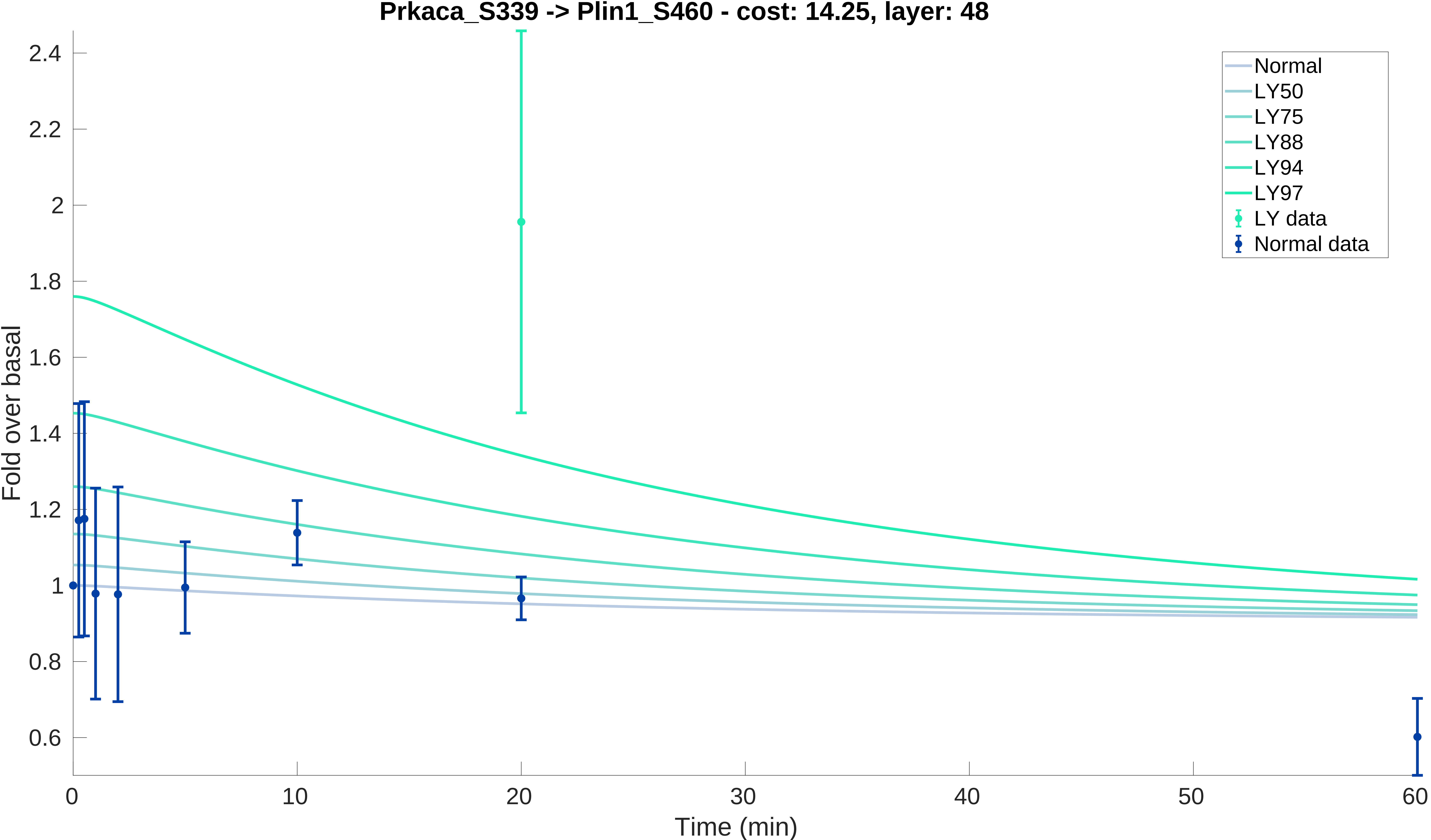

Prkaca_S339 -> Plin1_S460 - cost: 14.25, layer: 48
Fold over basal
Time (min)
0.6
0.8
1
1.2
1.4
1.6
1.8
2
2.2
2.4
0
10
20
30
40
50
60
Normal
LY50
LY75
LY88
LY94
LY97
LY data
Normal data

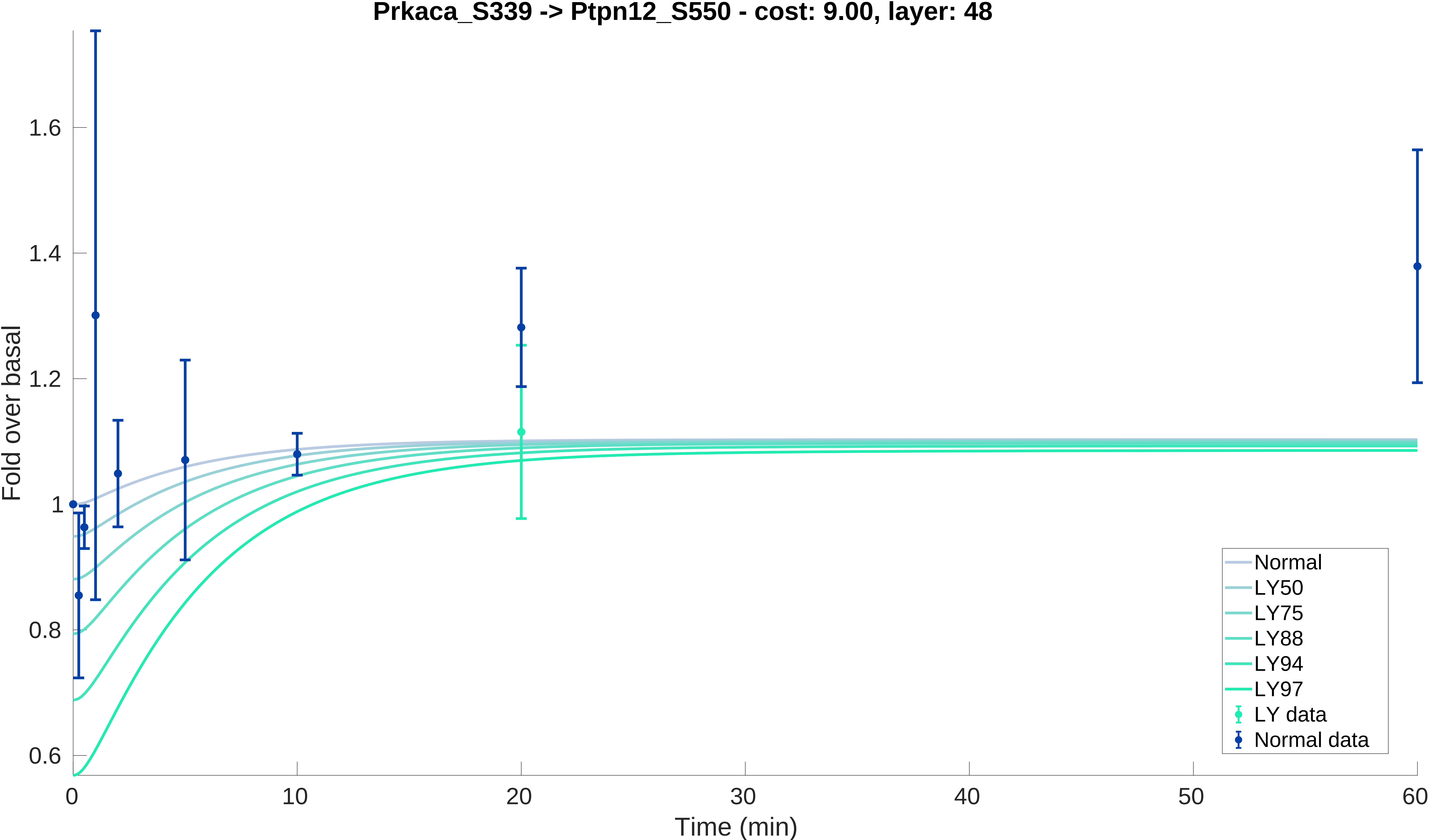
Prkaca_S339 -> Ptpn12_S550 - cost: 9.00, layer: 48
Fold over basal
1.6
1.4
1.2
1
0.8
0.6
0
10
20
30
40
50
60
Time (min)
Normal
LY50
LY75
LY88
LY94
LY97
LY data
Normal data

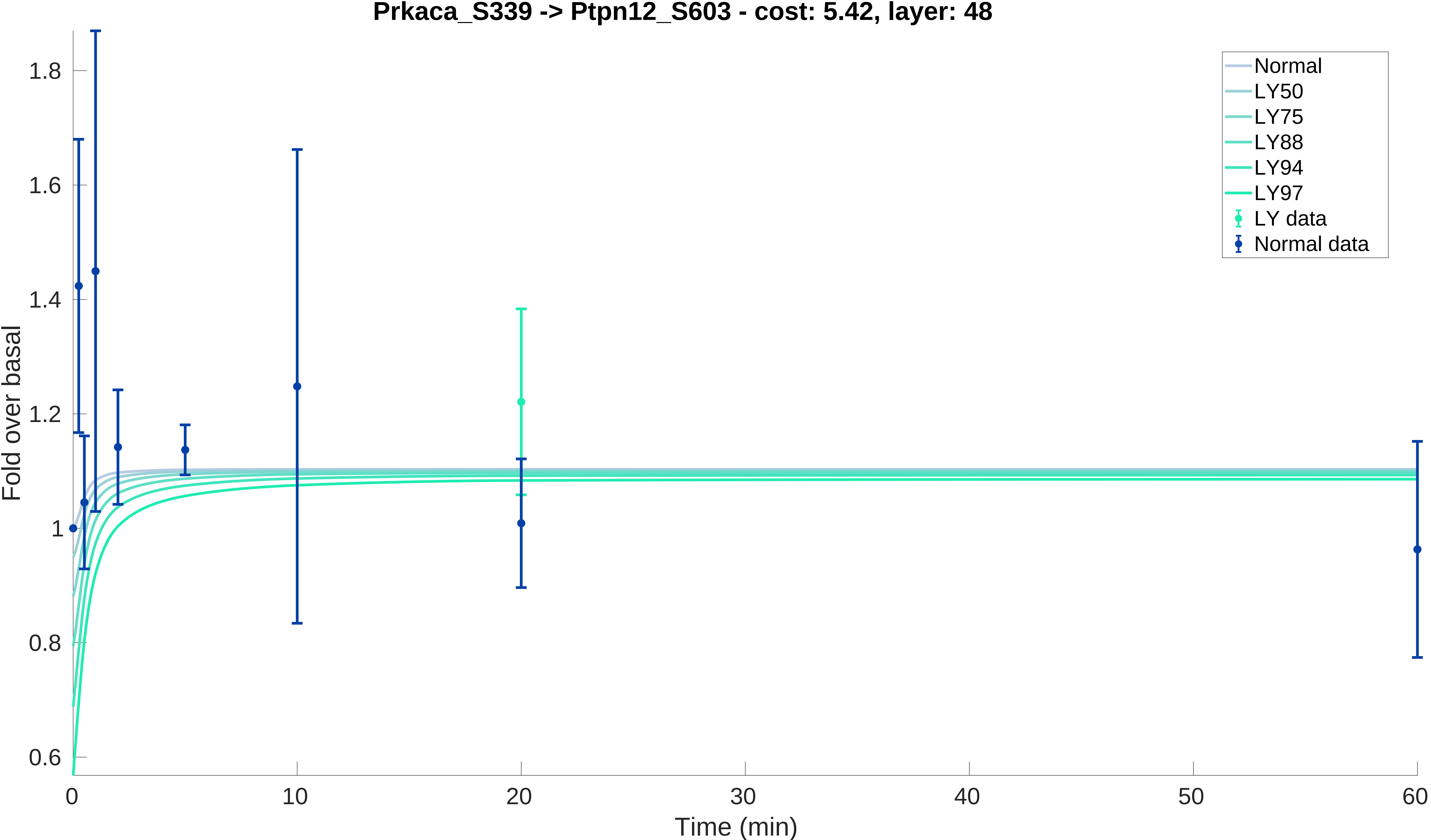
Prkaca_S339 -> Ptpn12_S603 - cost: 5.42, layer: 48
Fold over basal
Time (min)
0.6
0.8
1
1.2
1.4
1.6
1.8
0
10
20
30
40
50
60
Normal
LY50
LY75
LY88
LY94
LY97
LY data
Normal data

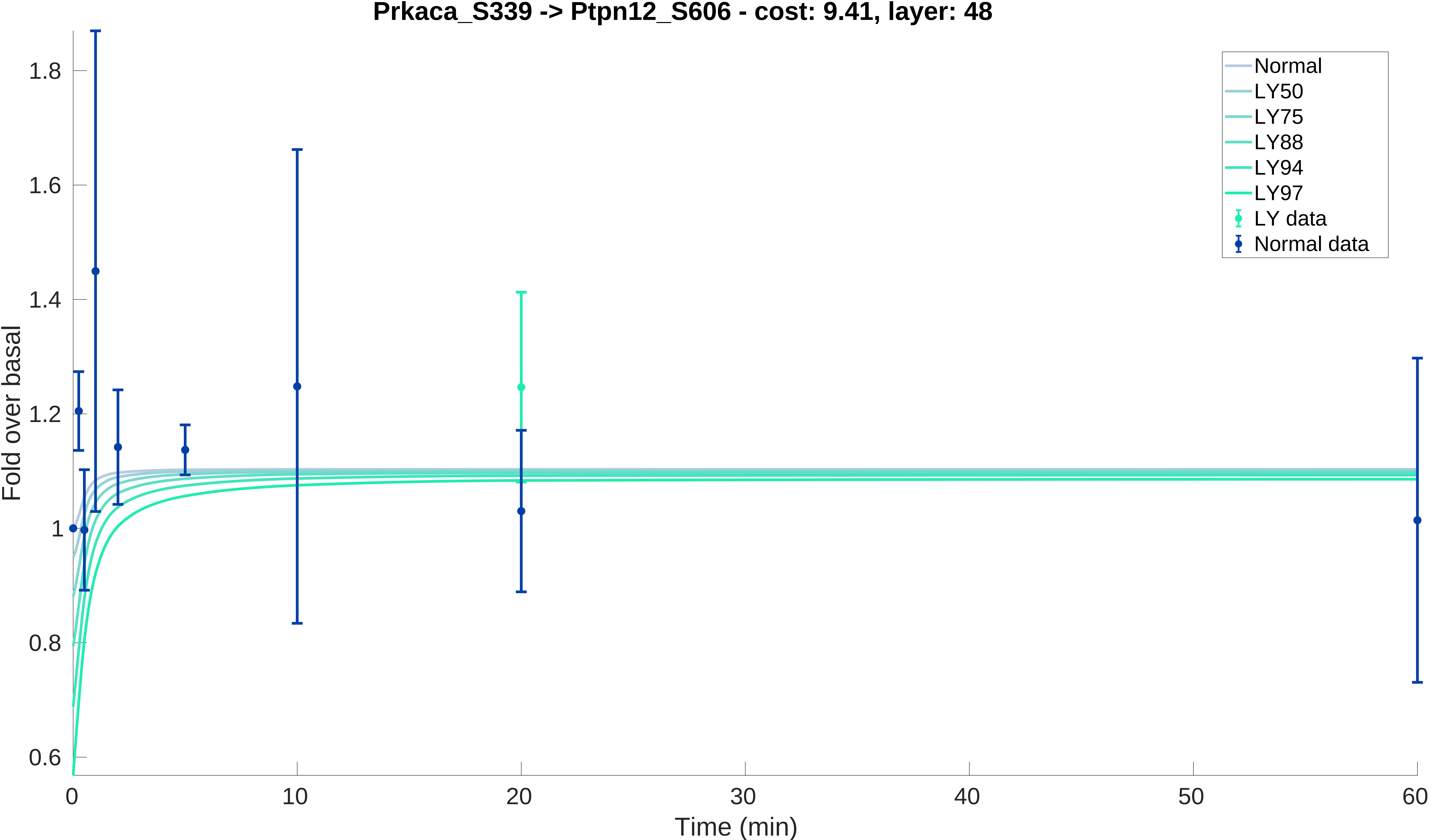

Prkaca_S339 -> Ptpn12_S606 - cost: 9.41, layer: 48
Fold over basal
Time (min)
1.8
1.6
1.4
1.2
1
0.8
0.6
0
10
20
30
40
50
60
Normal
LY50
LY75
LY88
LY94
LY97
LY data
Normal data

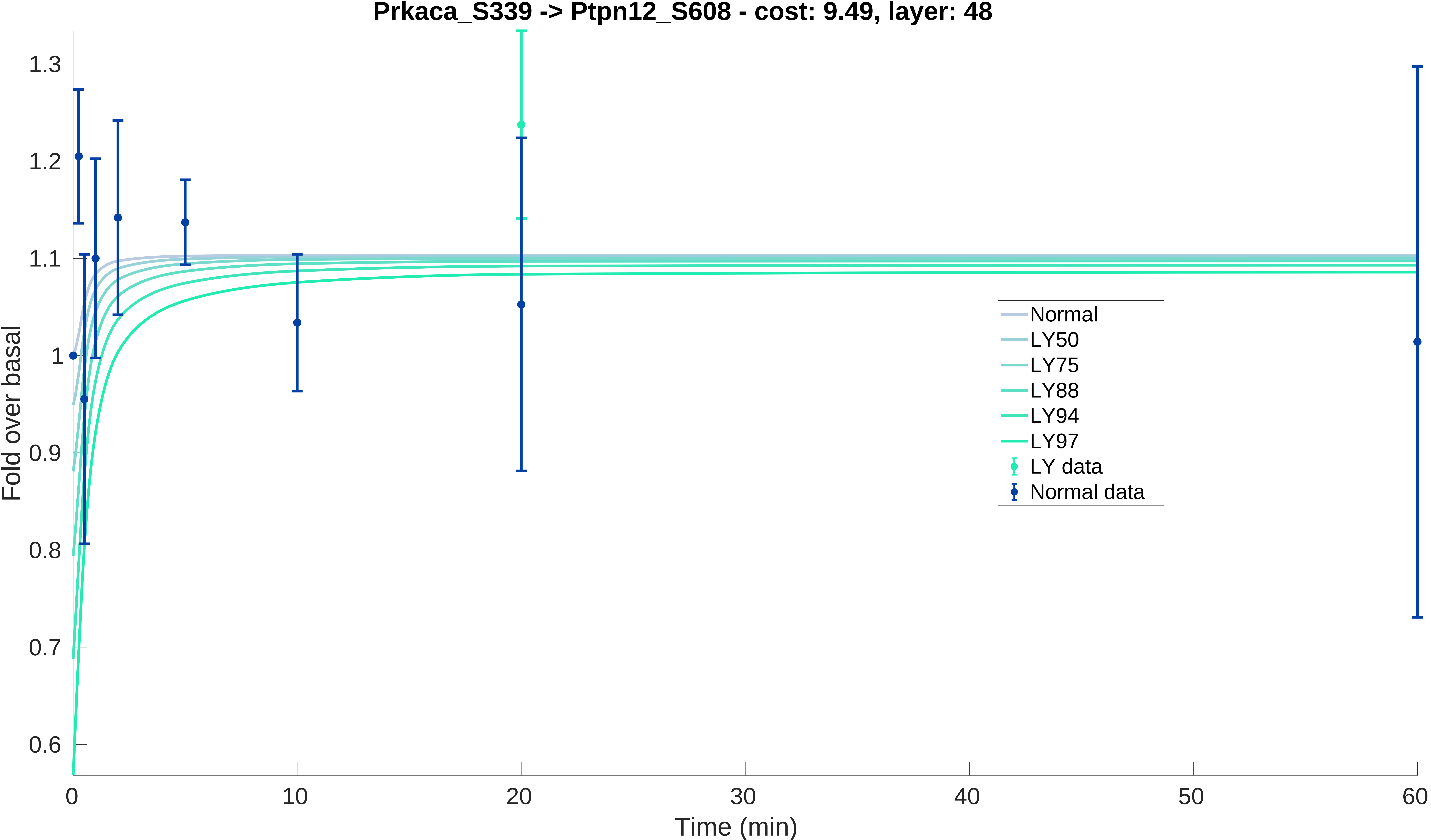

Prkaca_S339 -> Ptpn12_S608 - cost: 9.49, layer: 48
Fold over basal
1.3
1.2
1.1
1
0.9
0.8
0.7
0.6
0
10
20
30
40
50
60
Time (min)
Normal
LY50
LY75
LY88
LY94
LY97
LY data
Normal data

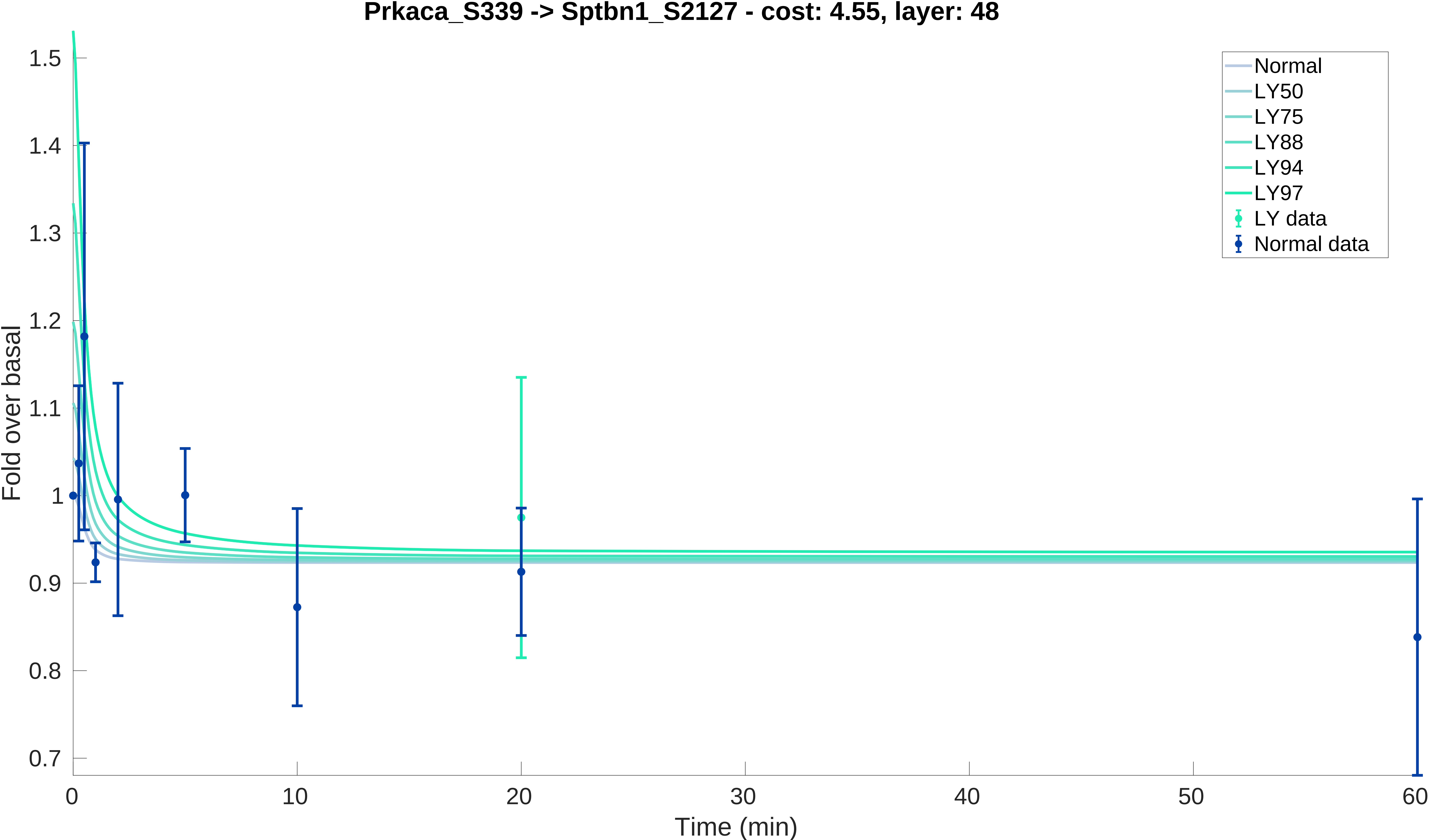

Prrkaca_S339 -> Sptbn1_S2127 - cost: 4.55, layer: 48
Fold over basal
Time (min)
1.5
1.4
1.3
1.2
1.1
1
0.9
0.8
0.7
0
10
20
30
40
50
60
Normal
LY50
LY75
LY88
LY94
LY97
LY data
Normal data

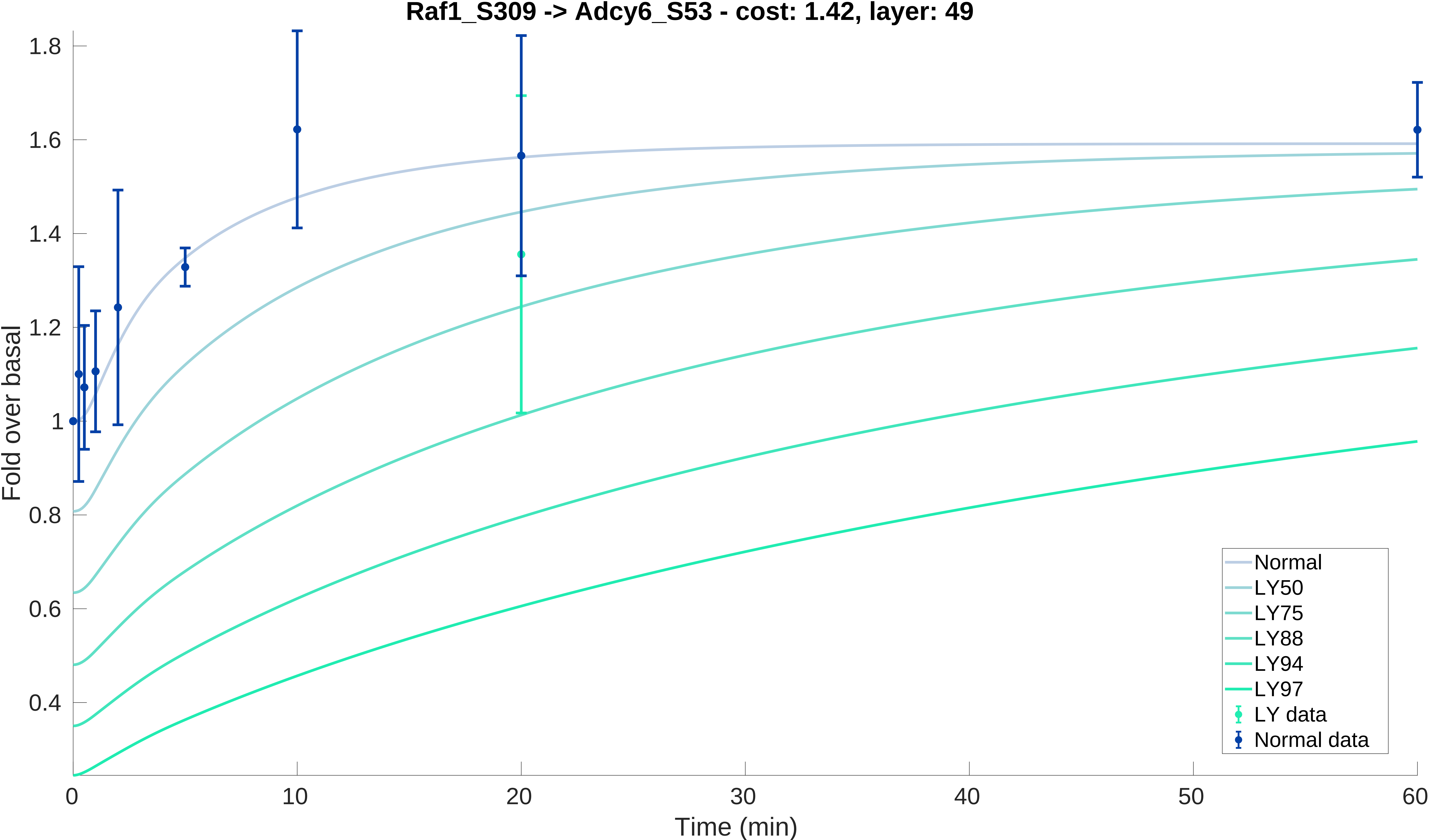

Raf1_S309 -> Adcy6_S53 - cost: 1.42, layer: 49
Fold over basal
1.8
1.6
1.4
1.2
1
0.8
0.6
0.4
0
10
20
30
40
50
60
Time (min)
Normal
LY50
LY75
LY88
LY94
LY97
LY data
Normal data

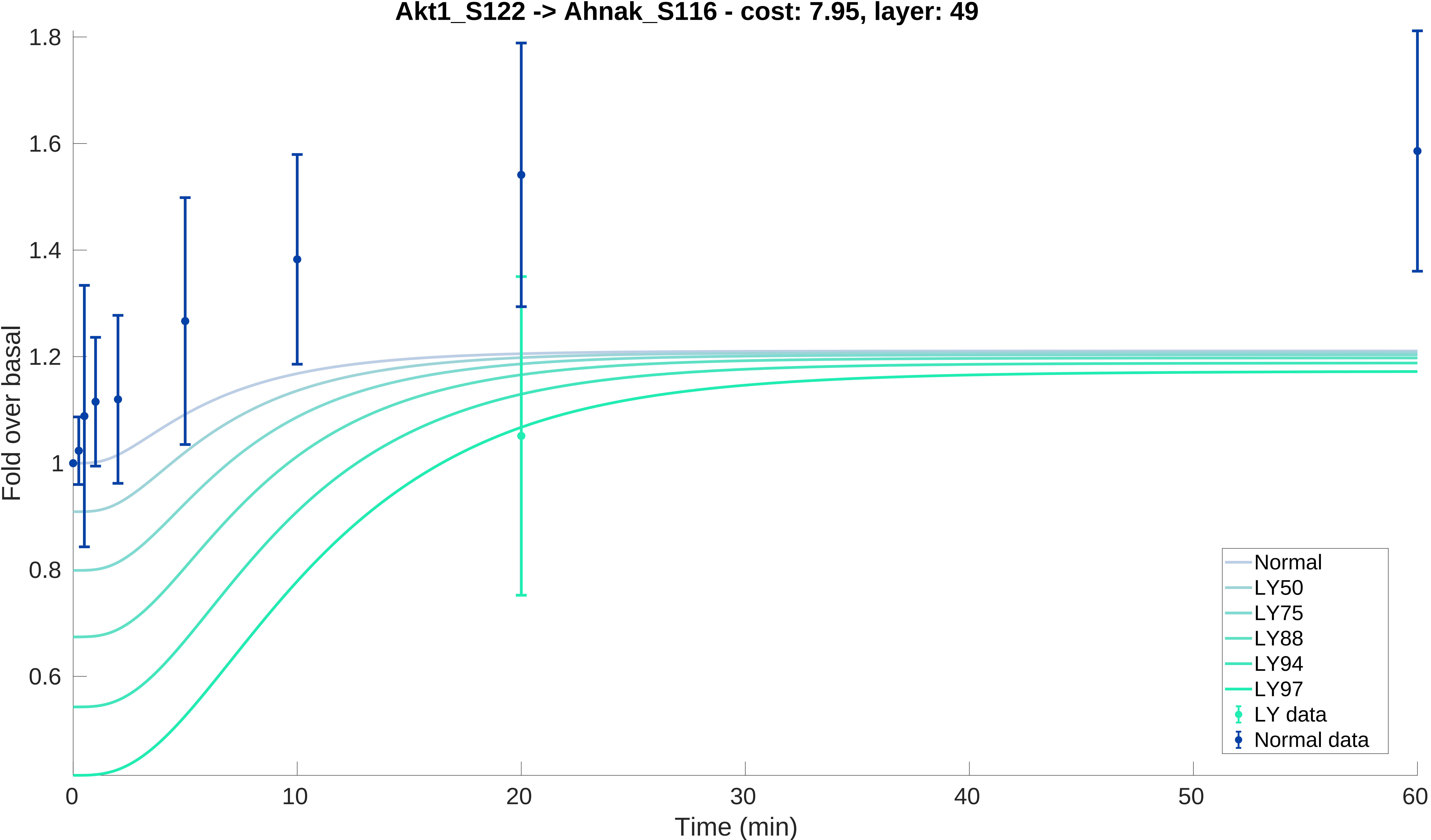

Akt1_S122 -> Ahnak_S116 - cost: 7.95, layer: 49
Fold over basal
1.8
1.6
1.4
1.2
1
0.8
0.6
0
10
20
30
40
50
60
Time (min)
Normal
LY50
LY75
LY88
LY94
LY97
LY data
Normal data

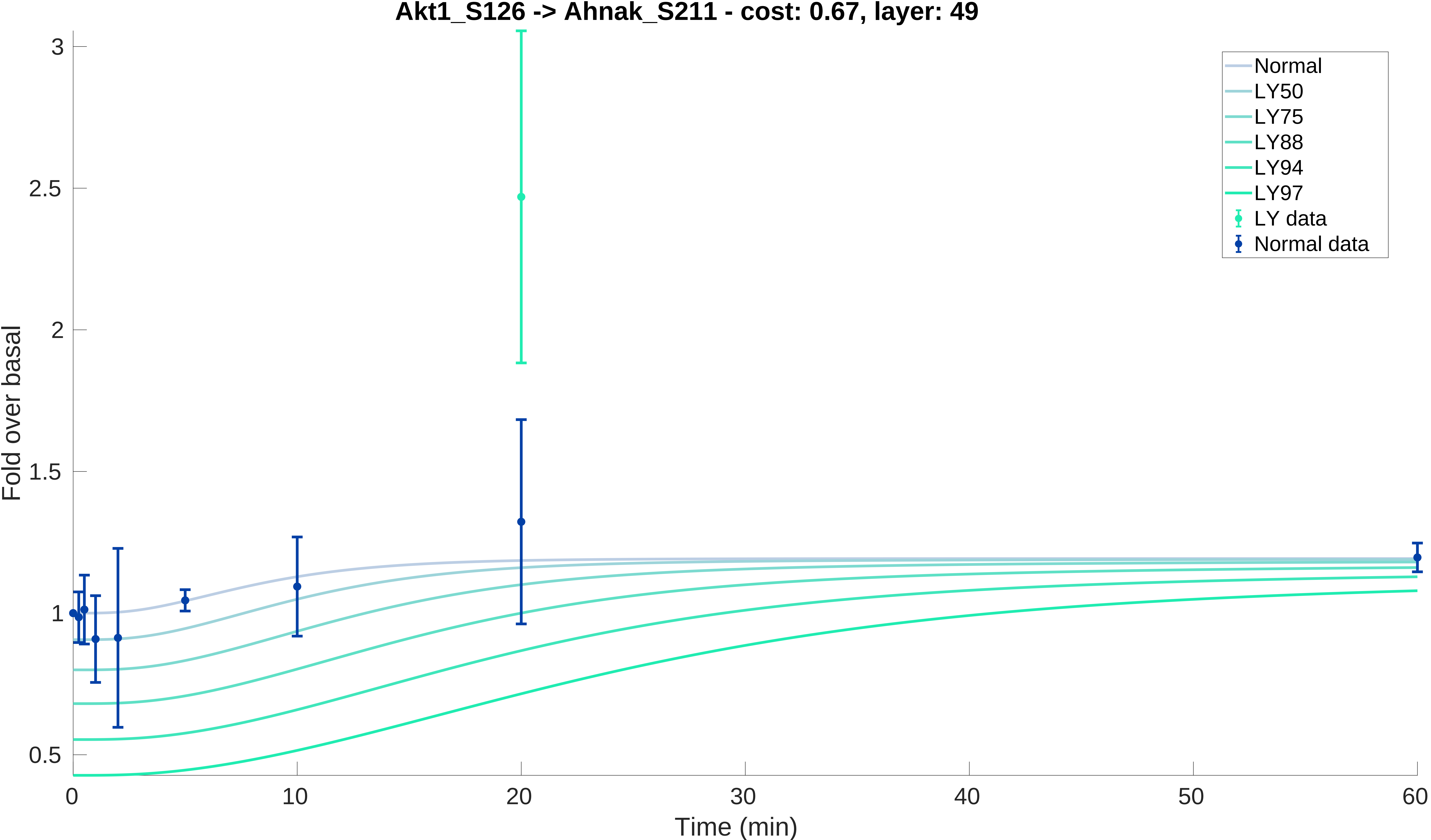

Akt1_S126 -> Ahnak_S211 - cost: 0.67, layer: 49
Fold over basal
Time (min)
0.5
1
1.5
2
2.5
3
0
10
20
30
40
50
60
Normal
LY50
LY75
LY88
LY94
LY97
LY data
Normal data

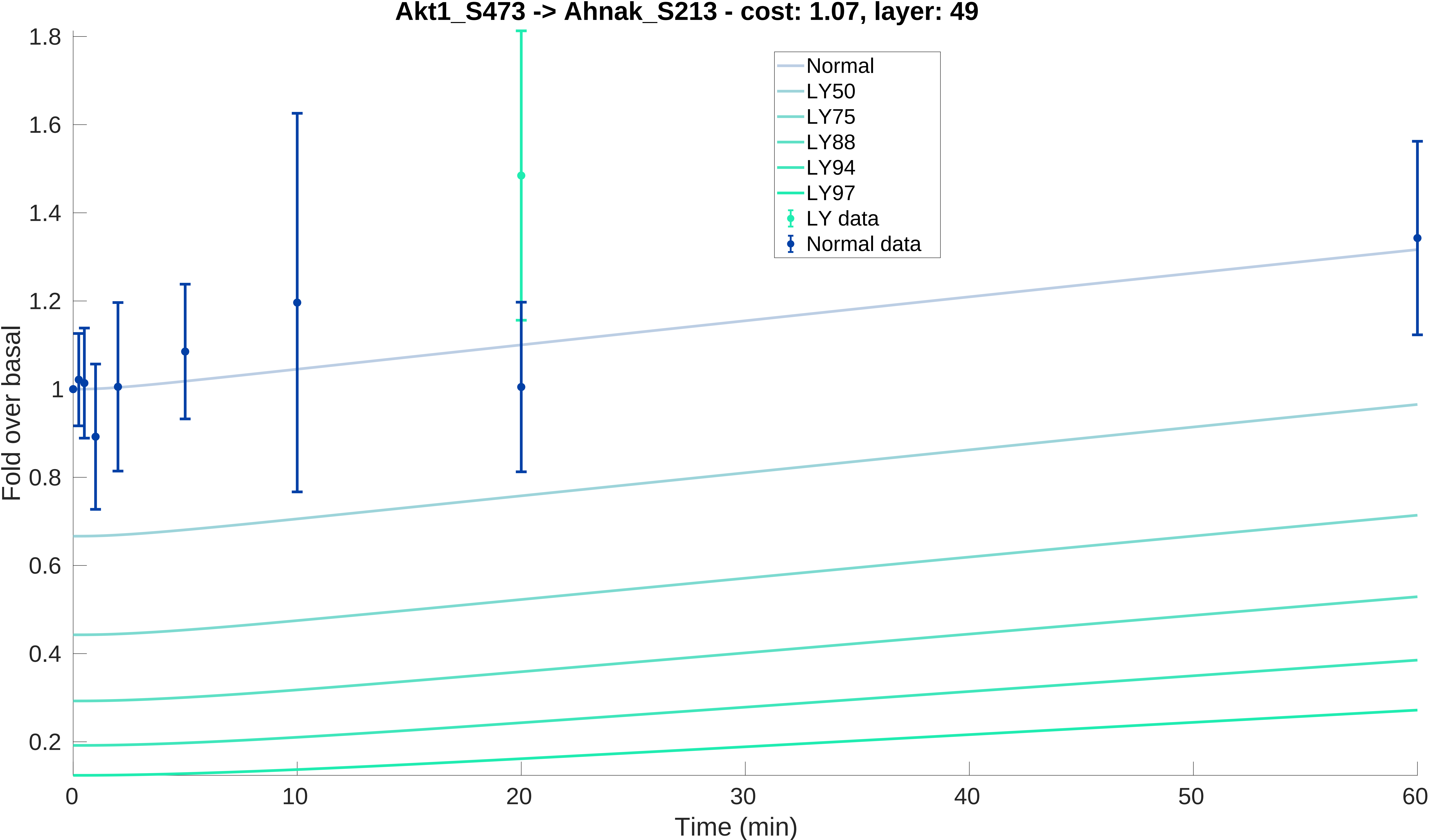
Akt1_S473 -> Ahnak_S213 - cost: 1.07, layer: 49
Normal
LY50
LY75
LY88
LY94
LY97
LY data
Normal data
Fold over basal
Time (min)
0
10
20
30
40
50
60
0.2
0.4
0.6
0.8
1
1.2
1.4
1.6
1.8

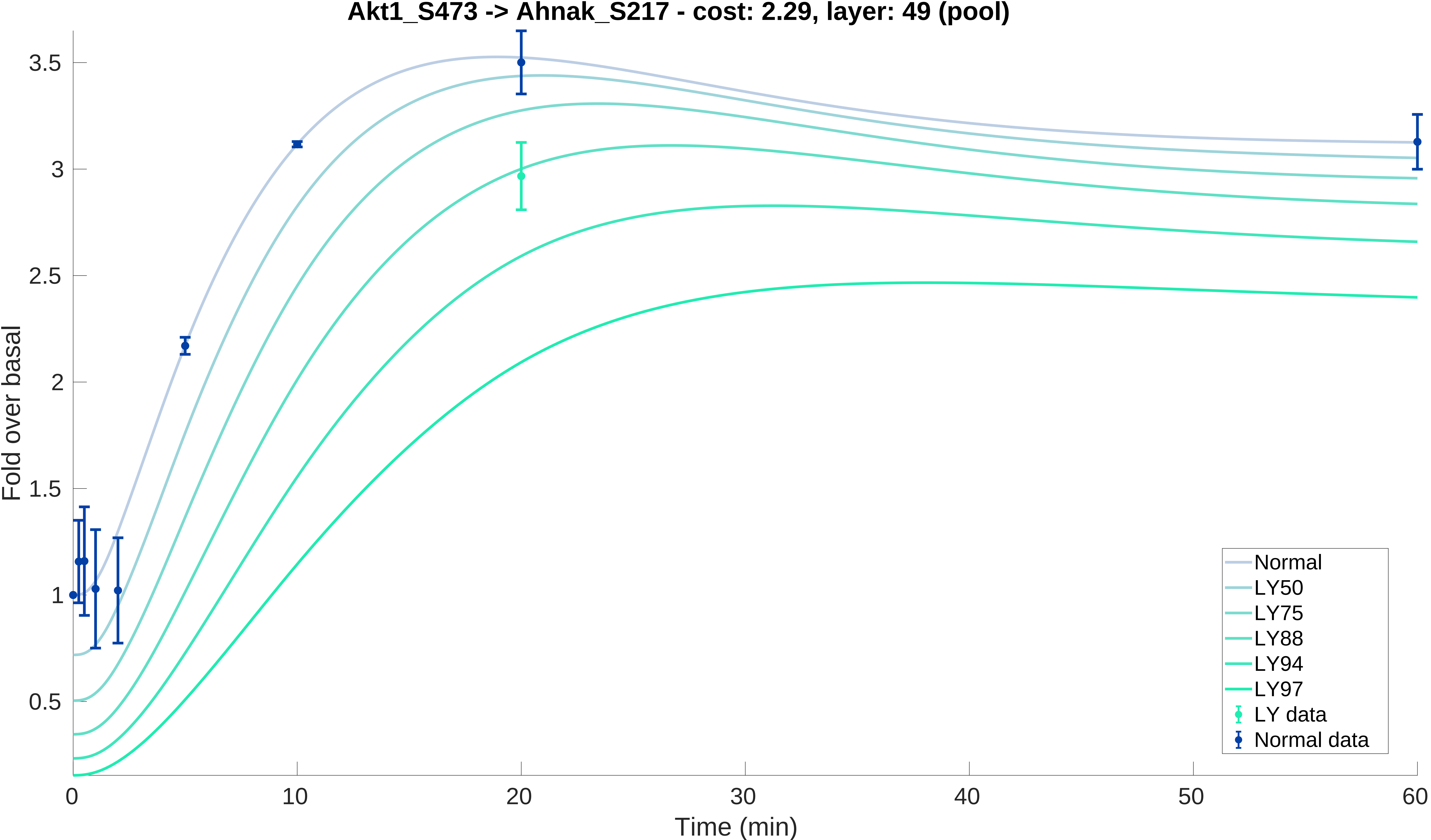

Akt1_S473 -> Ahnak_S217 - cost: 2.29, layer: 49 (pool)
Fold over basal
3.5
3
2.5
2
1.5
1
0.5
0
10
20
30
40
50
60
Time (min)
Normal
LY50
LY75
LY88
LY94
LY97
LY data
Normal data

Akt1_T308 -> Ahnak_S2948 - cost: 13.78, layer: 49

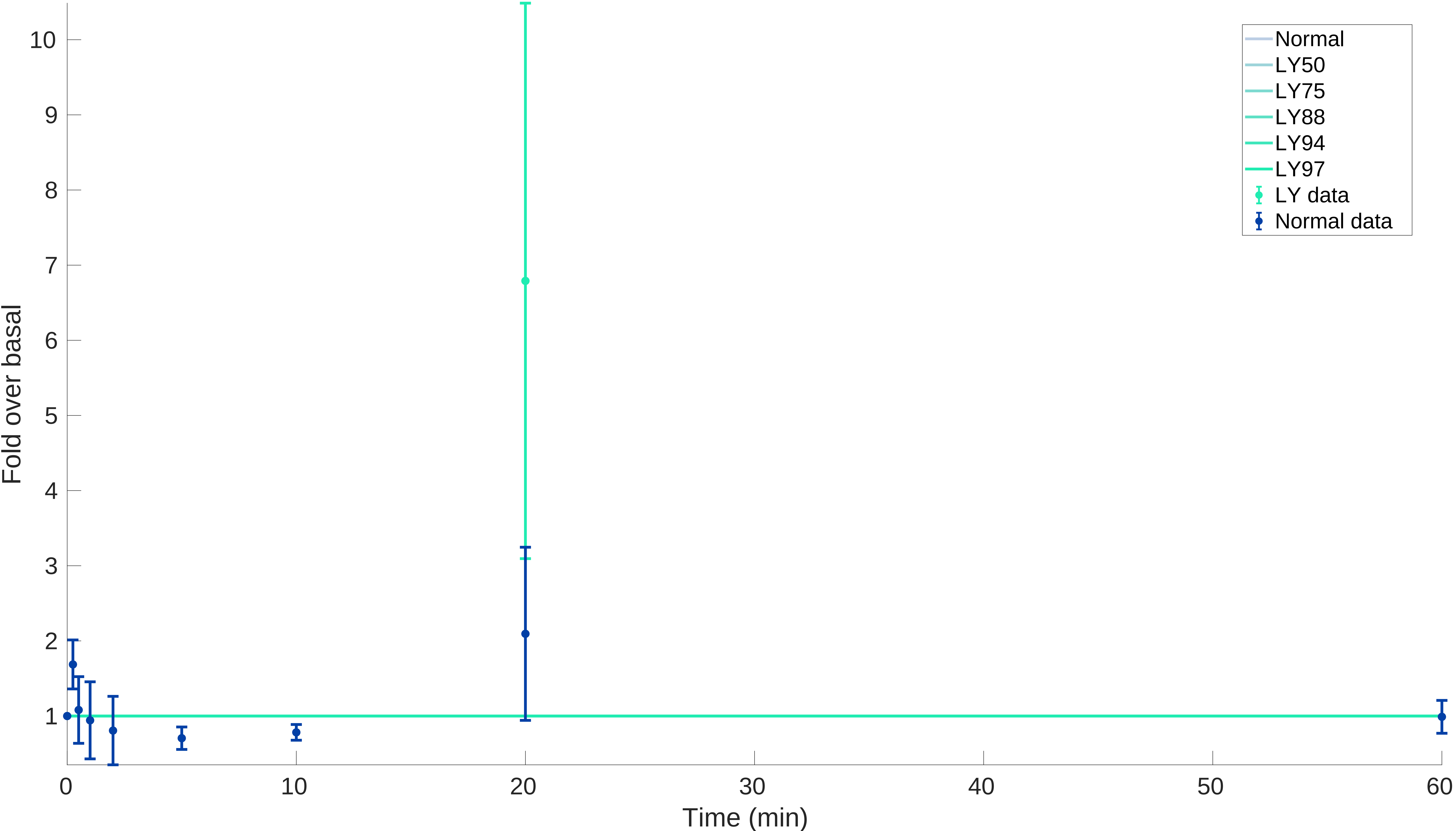

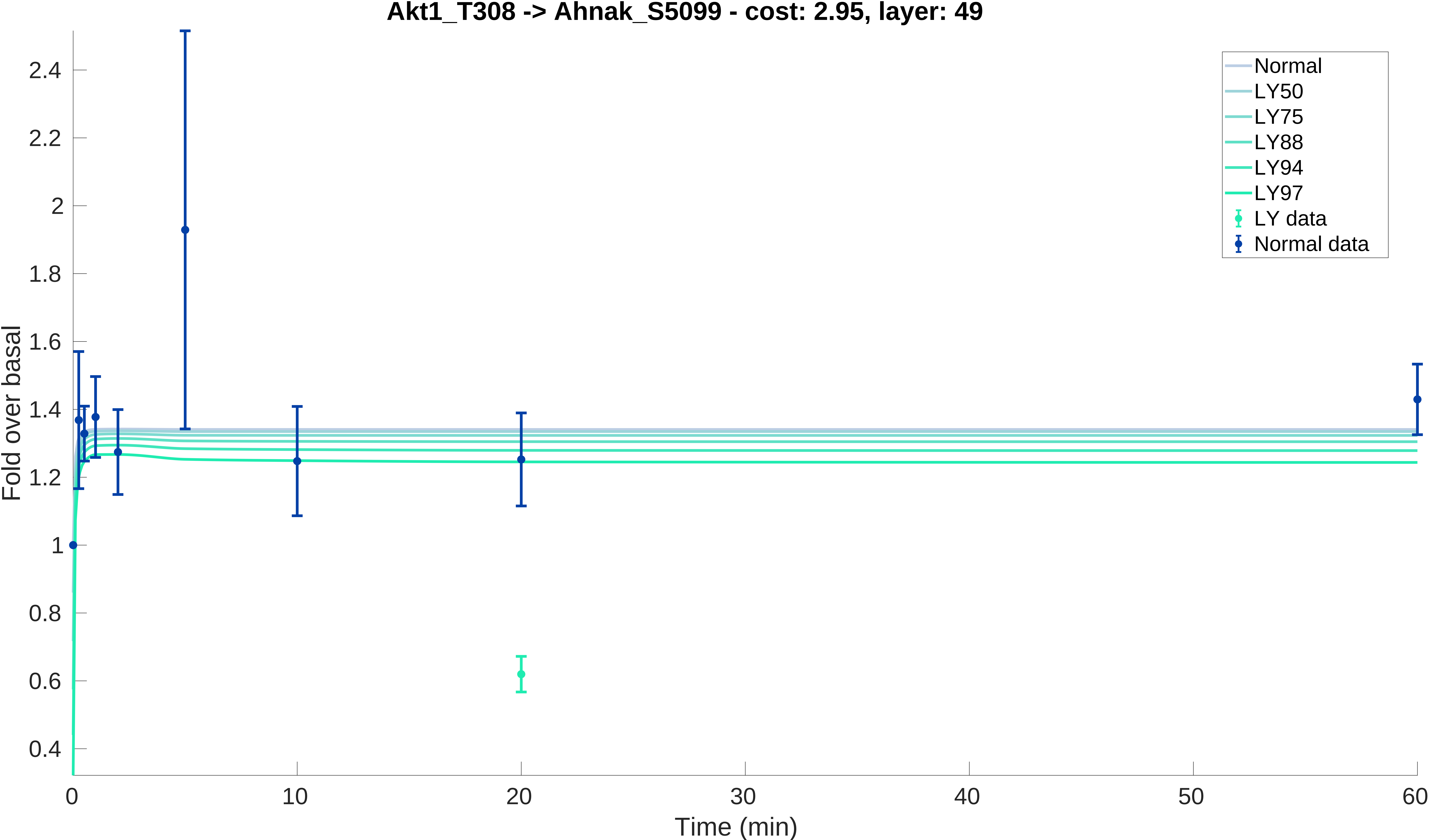

Akt1_T308 -> Ahnak_S5099 - cost: 2.95, layer: 49
Fold over basal
Time (min)
Normal
LY50
LY75
LY88
LY94
LY97
LY data
Normal data
2.4
2.2
2
1.8
1.6
1.4
1.2
1
0.8
0.6
0.4
0
10
20
30
40
50
60

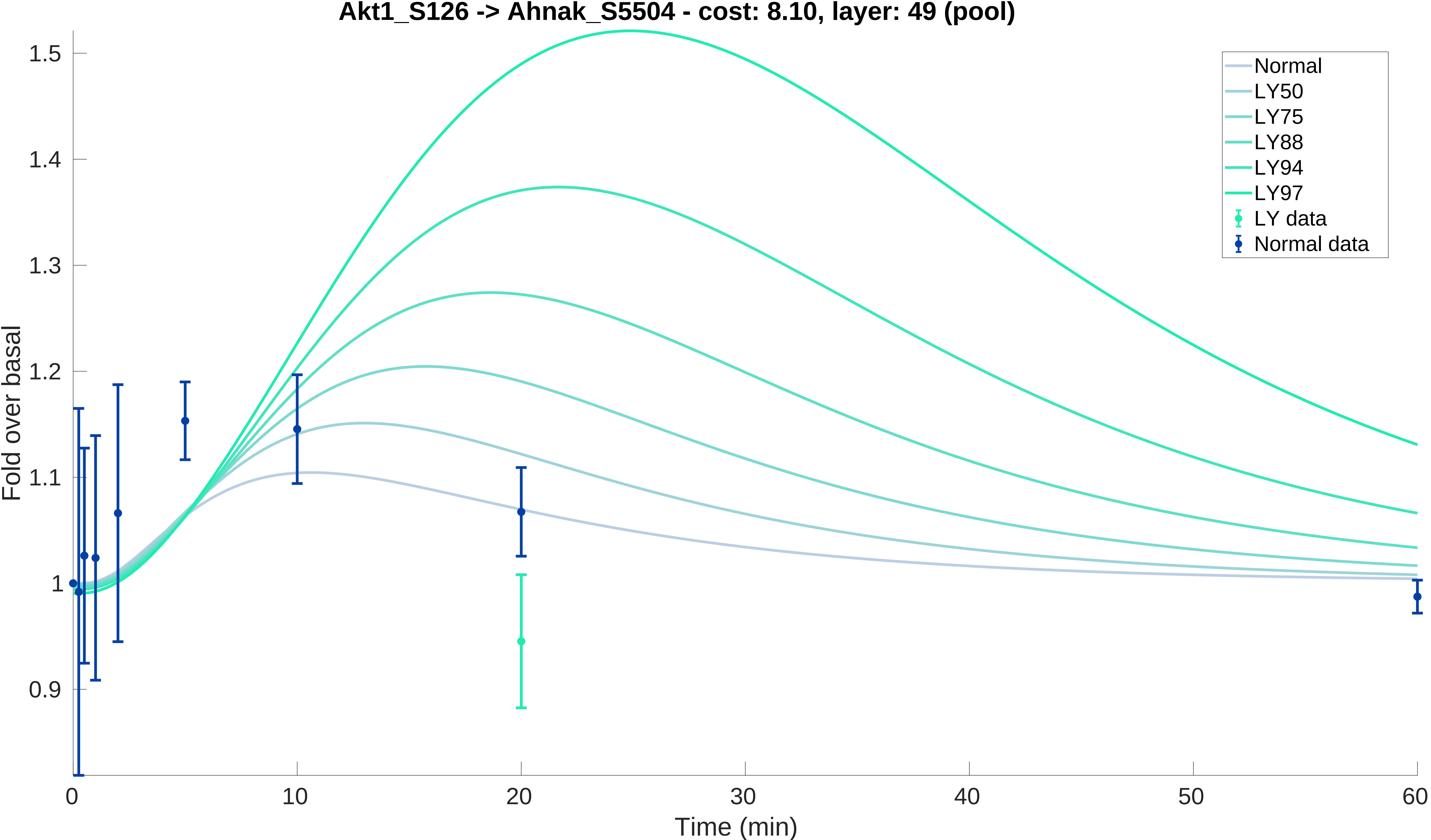
Akt1_S126 -> Ahnak_S5504 - cost: 8.10, layer: 49 (pool)
Fold over basal
1.5
1.4
1.3
1.2
1.1
1
0.9
0
10
20
30
40
50
60
Time (min)
Normal
LY50
LY75
LY88
LY94
LY97
LY data
Normal data

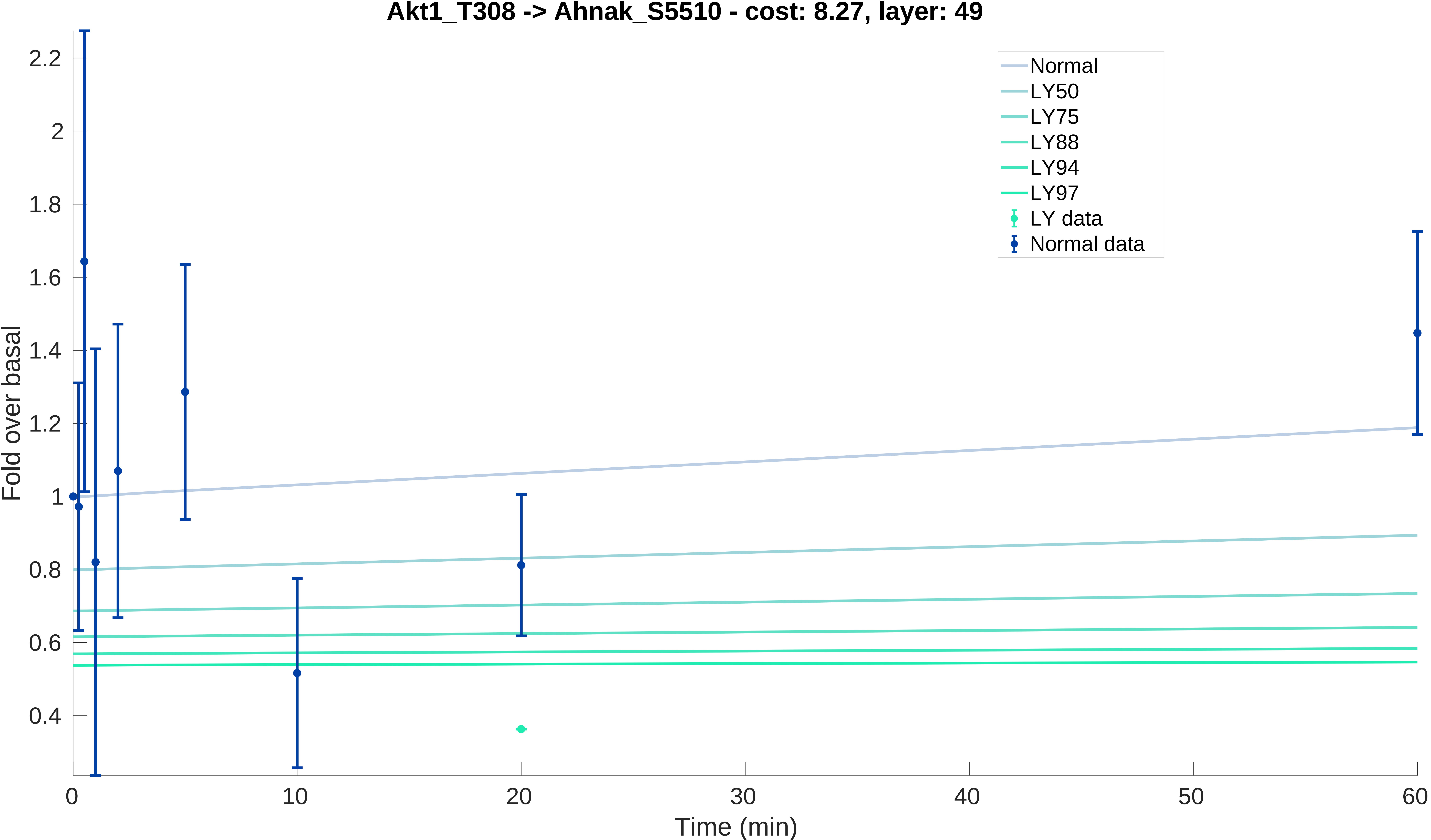

Akt1_T308 -> Ahnak_S5510 - cost: 8.27, layer: 49
Fold over basal
Time (min)
0.4
0.6
0.8
1
1.2
1.4
1.6
1.8
2
2.2
0
10
20
30
40
50
60
Normal
LY50
LY75
LY88
LY94
LY97
LY data
Normal data

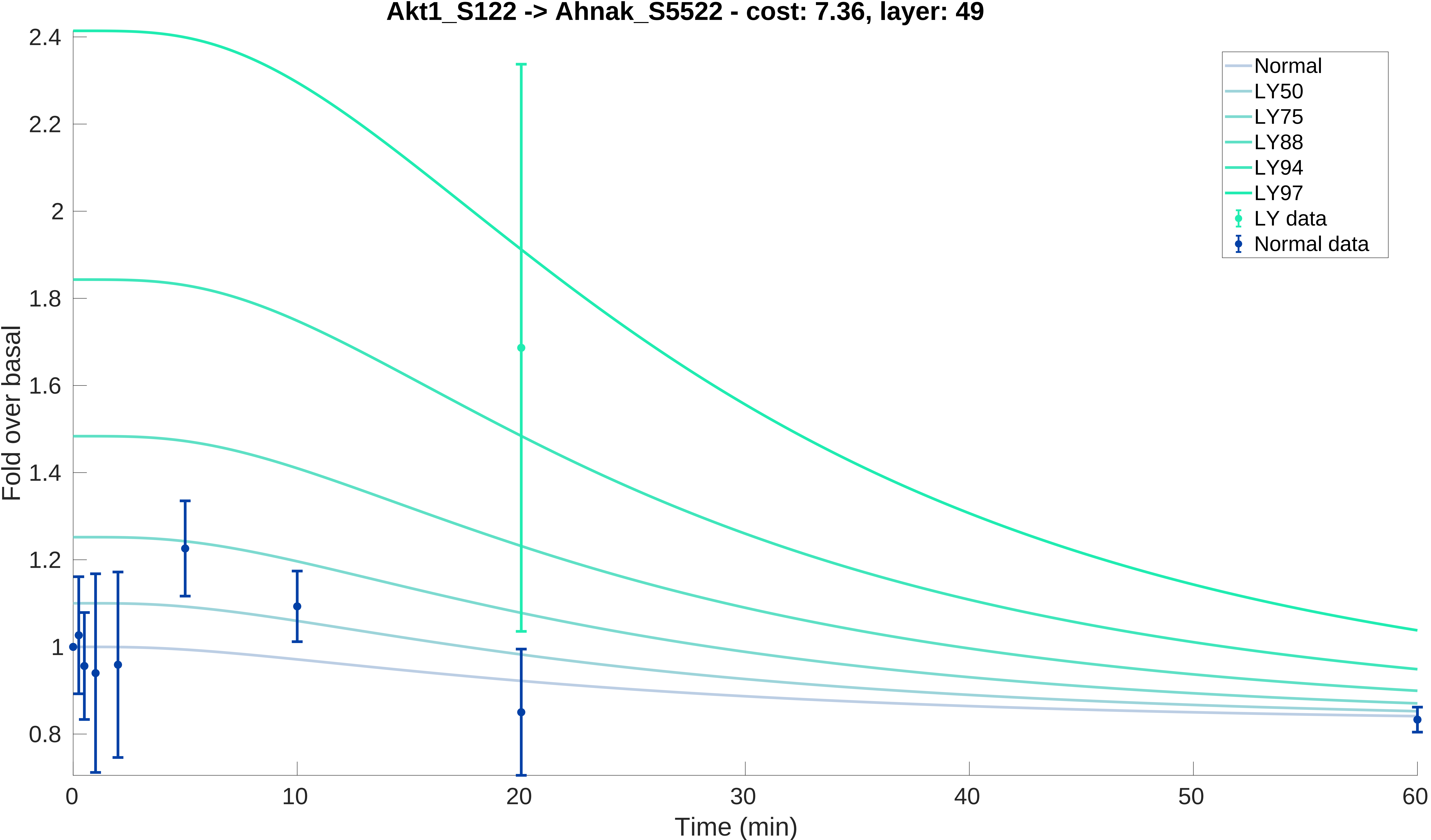

Akt1_S122 -> Ahnak_S5522 - cost: 7.36, layer: 49
Fold over basal
Time (min)
2.4
2.2
2
1.8
1.6
1.4
1.2
1
0.8
0
10
20
30
40
50
60
Normal
LY50
LY75
LY88
LY94
LY97
LY data
Normal data

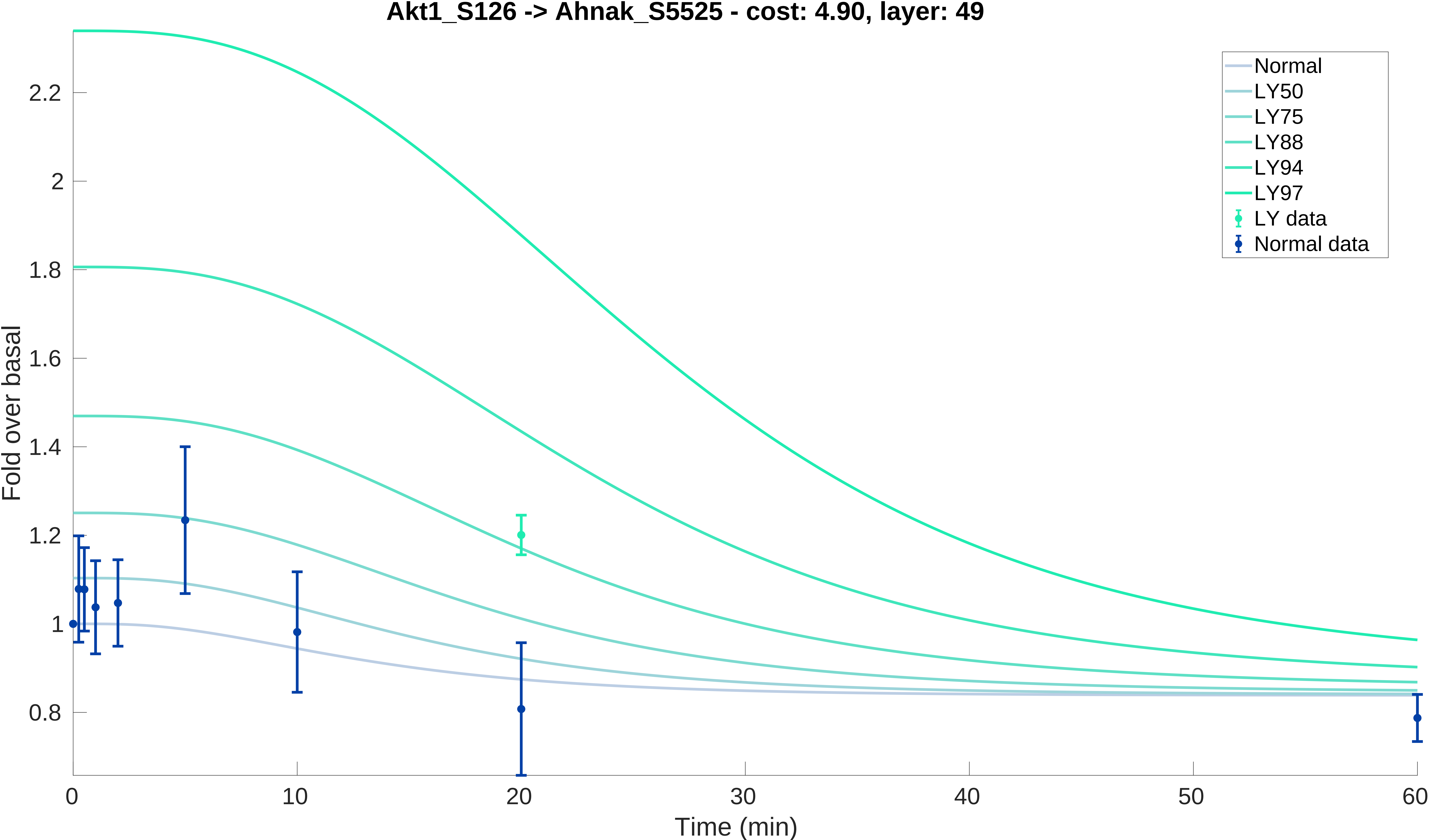

Akt1_S126 -> Ahnak_S5525 - cost: 4.90, layer: 49
Fold over basal
Time (min)
Normal
LY50
LY75
LY88
LY94
LY97
LY data
Normal data
2.2
2
1.8
1.6
1.4
1.2
1
0.8
0
10
20
30
40
50
60

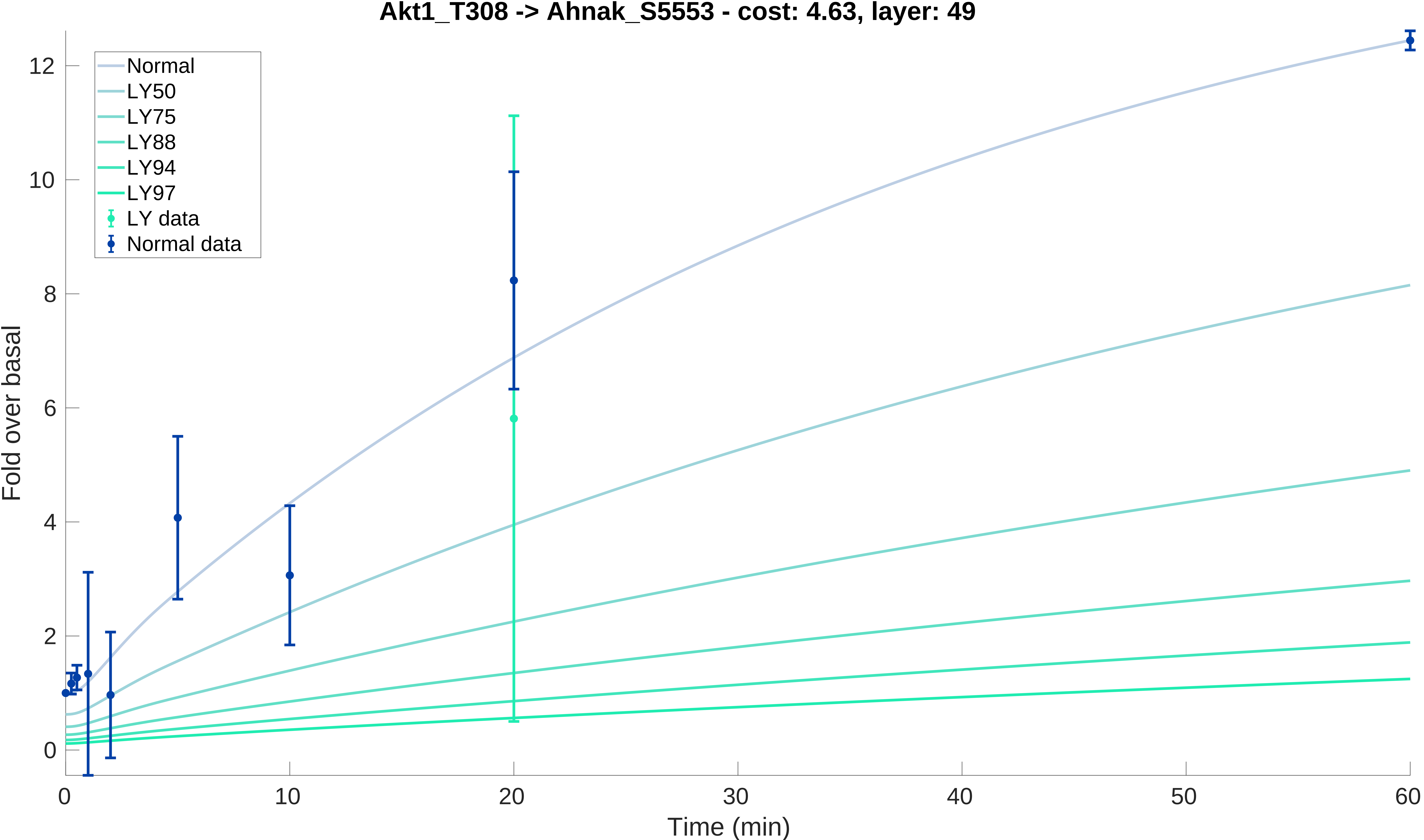

Akt1_T308 -> Ahnak_S5553 - cost: 4.63, layer: 49
Normal
LY50
LY75
LY88
LY94
LY97
LY data
Normal data
Fold over basal
Time (min)
0
2
4
6
8
10
12
0
10
20
30
40
50
60

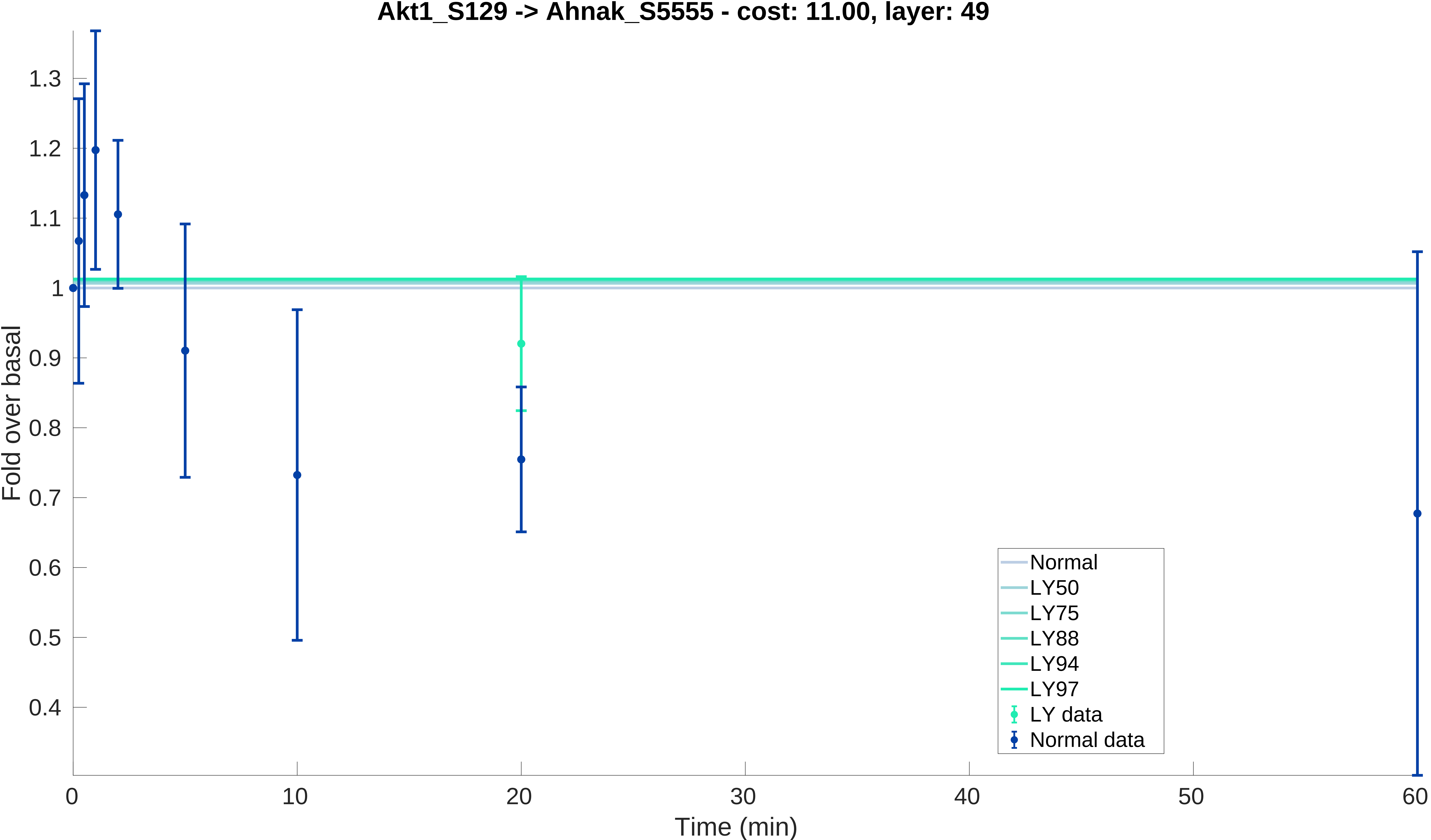
Akt1_S129 -> Ahnak_S5555 - cost: 11.00, layer: 49
Fold over basal
Time (min)
1.3
1.2
1.1
1
0.9
0.8
0.7
0.6
0.5
0.4
0
10
20
30
40
50
60
Normal
LY50
LY75
LY88
LY94
LY97
LY data
Normal data

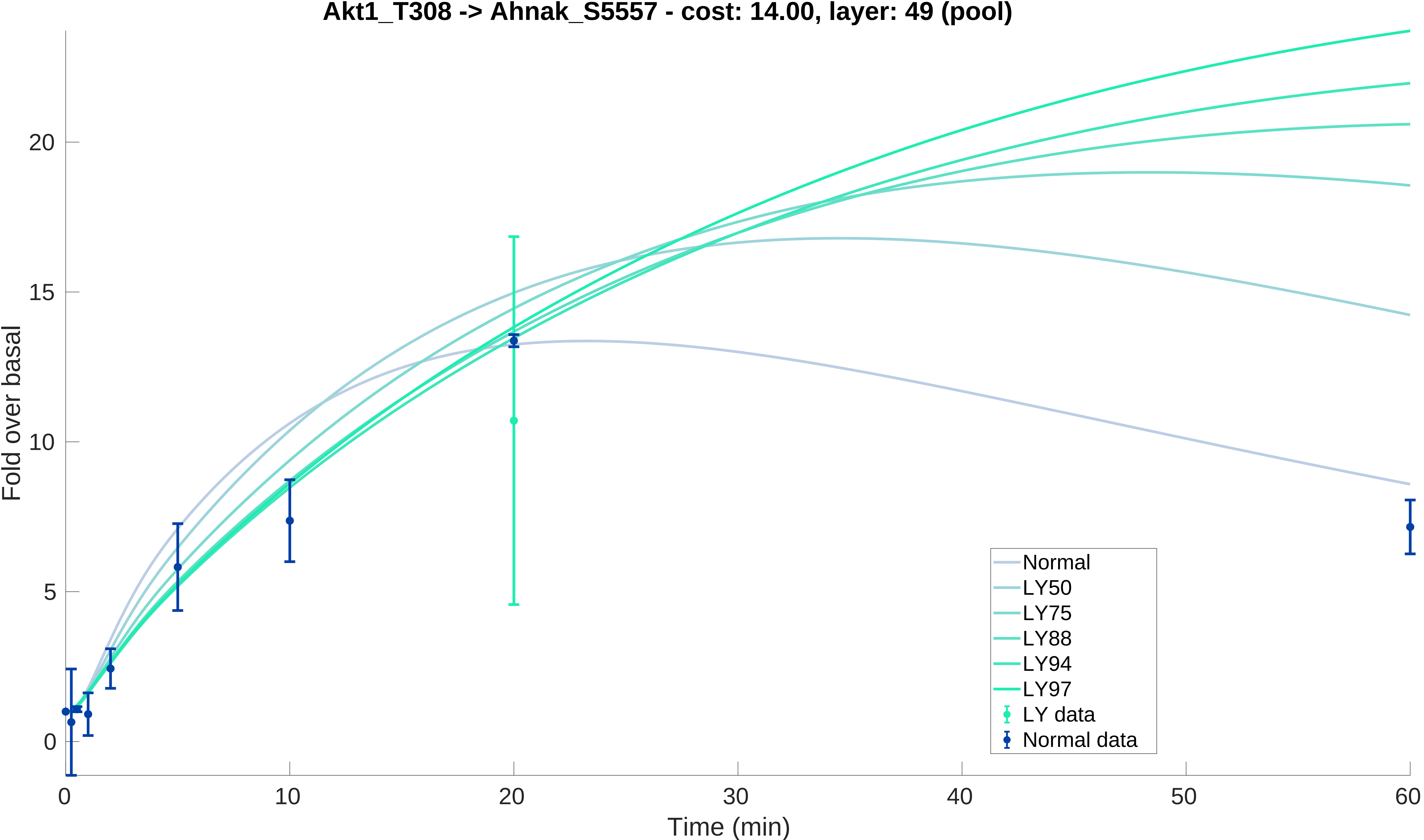

Akt1_T308 -> Ahnak_S5557 - cost: 14.00, layer: 49 (pool)
Fold over basal
Time (min)
0
5
10
15
20
0
10
20
30
40
50
60
Normal
LY50
LY75
LY88
LY94
LY97
LY data
Normal data

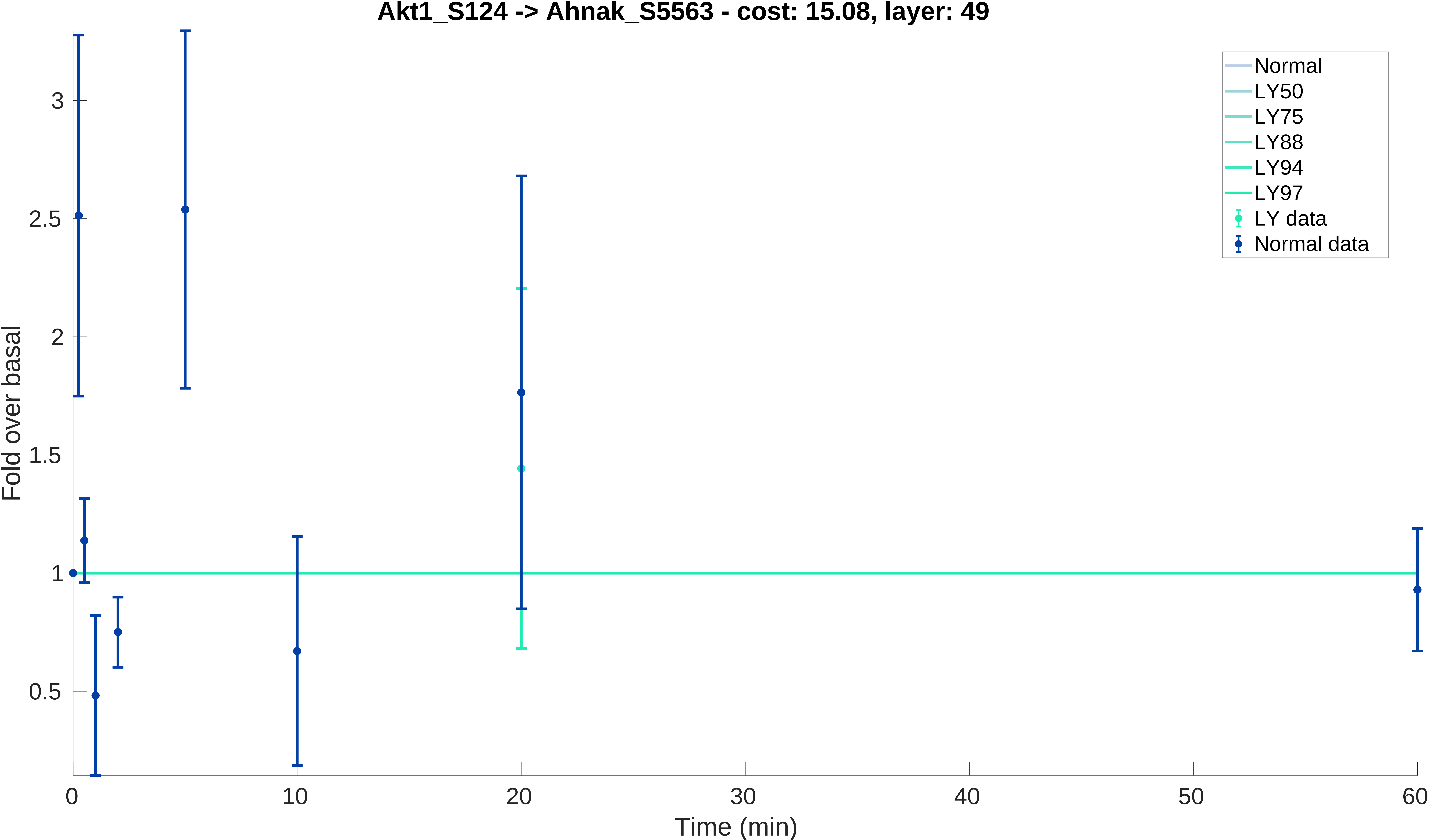

Akt1_S124 -> Ahnak_S5563 - cost: 15.08, layer: 49
Fold over basal
Time (min)
Normal
LY50
LY75
LY88
LY94
LY97
LY data
Normal data
0.5
1
1.5
2
2.5
3
0
10
20
30
40
50
60

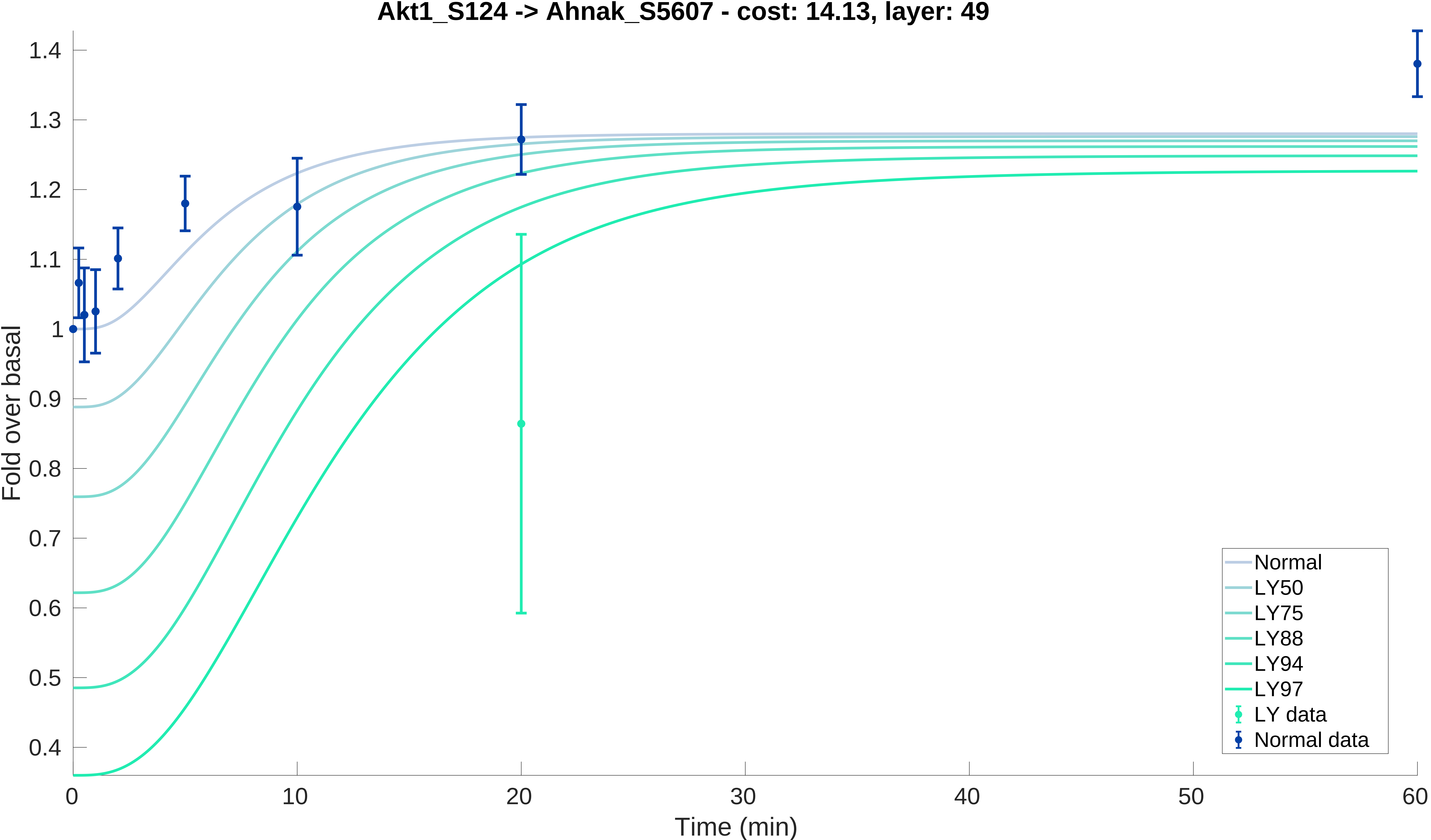

Akt1_S124 -> Ahnak_S5607 - cost: 14.13, layer: 49
Fold over basal
Time (min)
0.4
0.5
0.6
0.7
0.8
0.9
1
1.1
1.2
1.3
1.4
0
10
20
30
40
50
60
Normal
LY50
LY75
LY88
LY94
LY97
LY data
Normal data

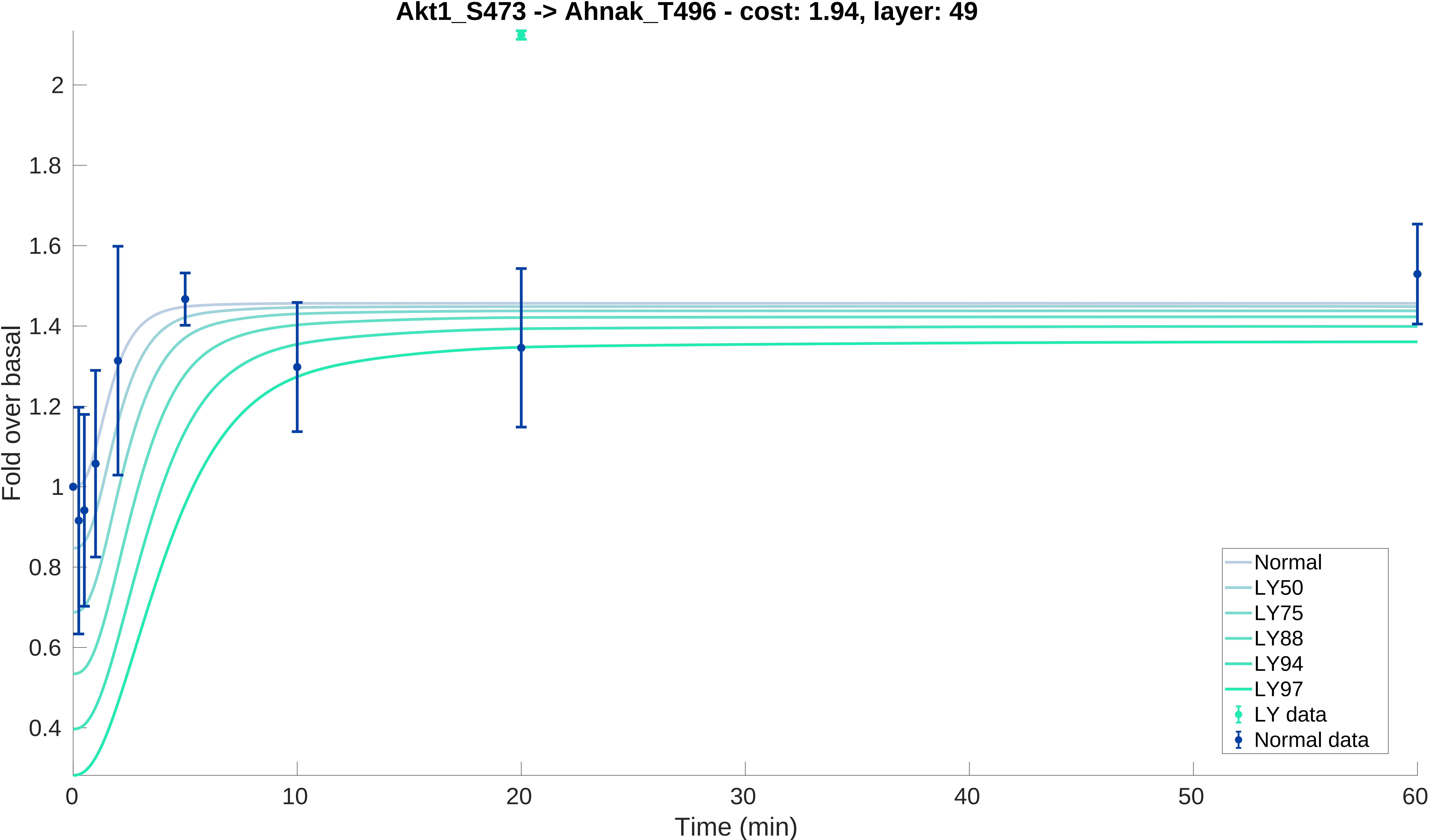
Akt1_S473 -> Ahnak_T496 - cost: 1.94, layer: 49
Fold over basal
2
1.8
1.6
1.4
1.2
1
0.8
0.6
0.4
0
10
20
30
40
50
60
Time (min)
Normal
LY50
LY75
LY88
LY94
LY97
LY data
Normal data

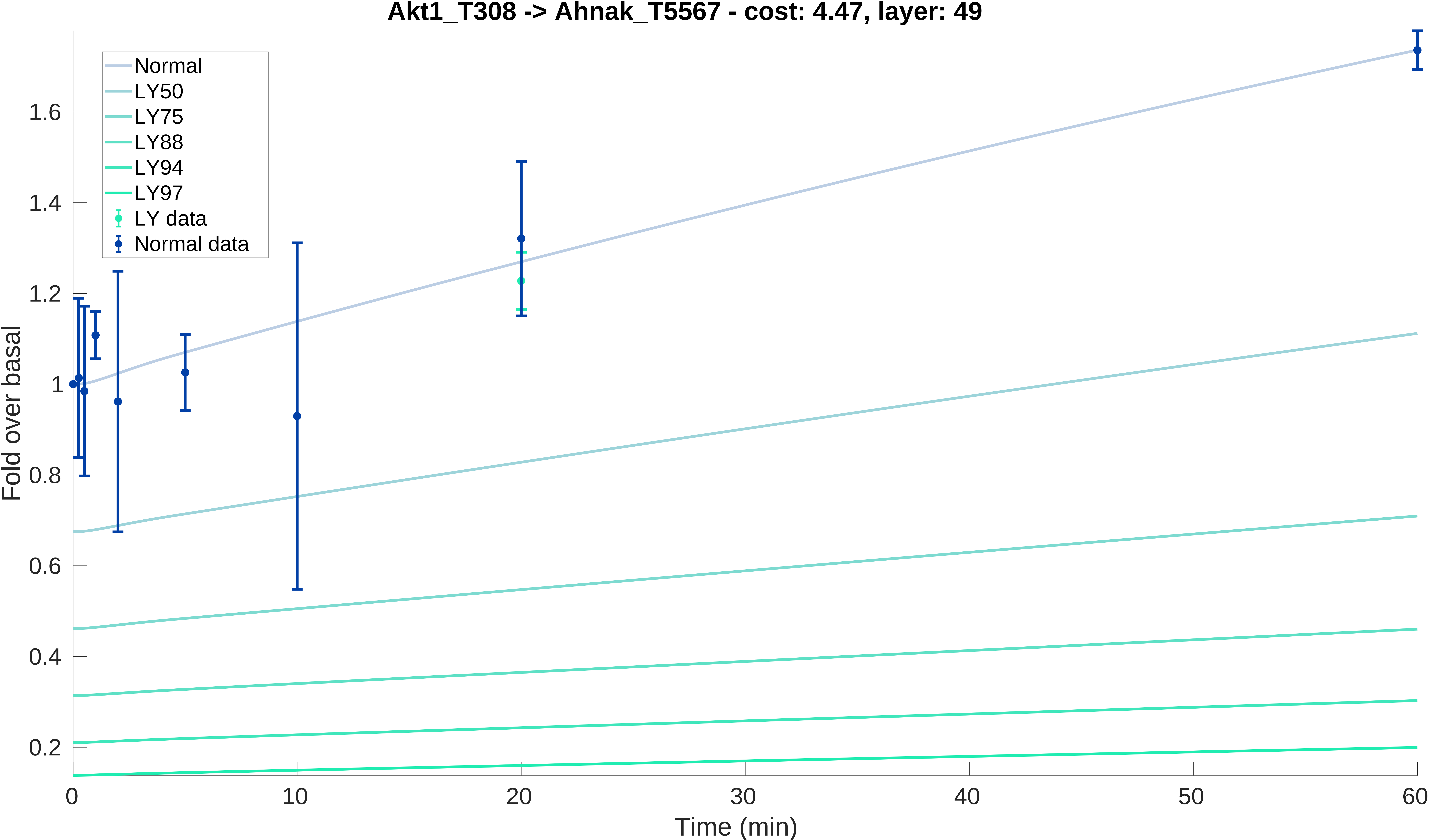

Akt1_T308 -> Ahnak_T5567 - cost: 4.47, layer: 49
Normal
LY50
LY75
LY88
LY94
LY97
LY data
Normal data
Fold over basal
Time (min)
0
10
20
30
40
50
60
0.2
0.4
0.6
0.8
1
1.2
1.4
1.6

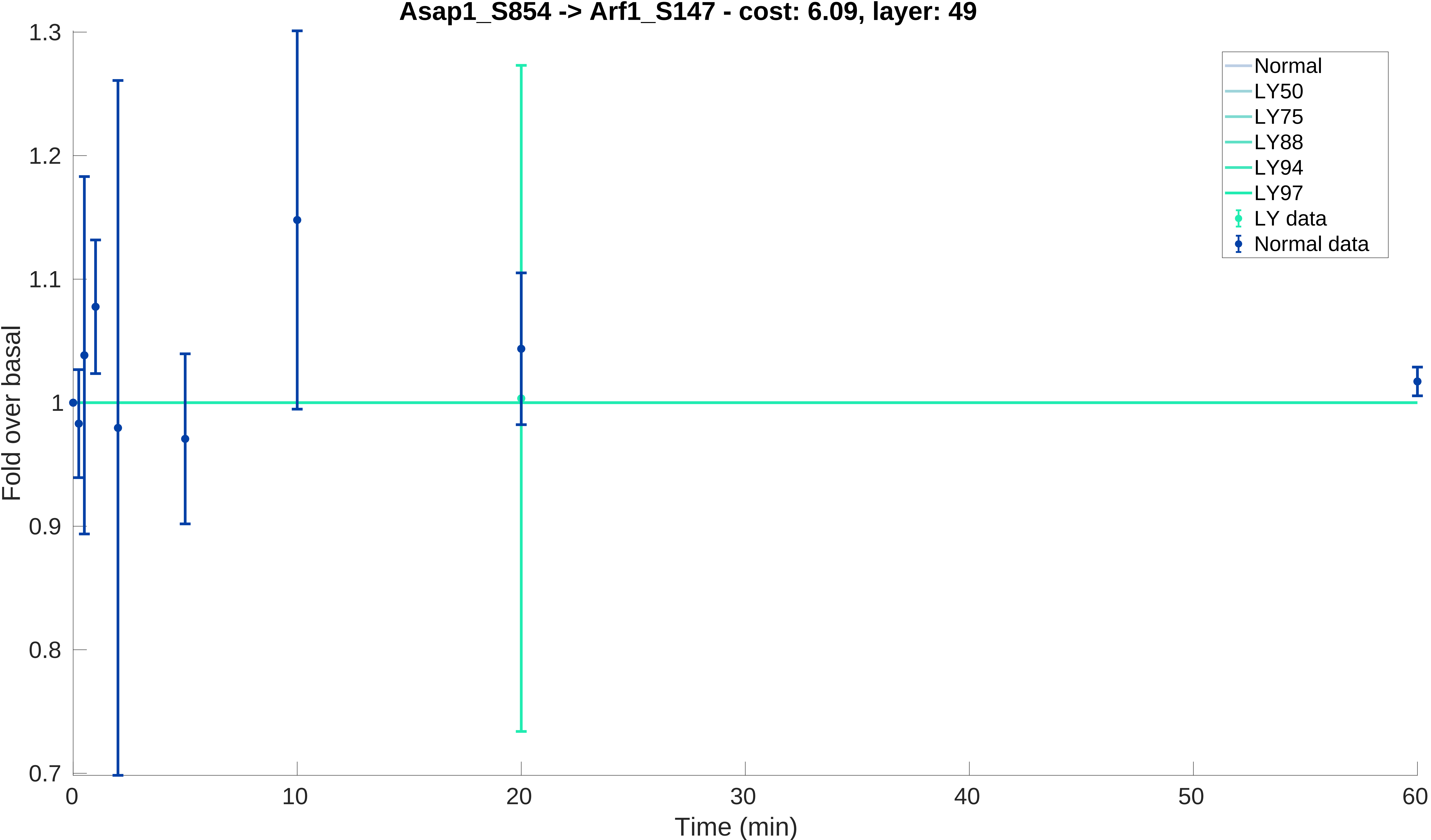
Asap1_S854 -> Arf1_S147 - cost: 6.09, layer: 49
Fold over basal
Time (min)
1.3
1.2
1.1
1
0.9
0.8
0.7
0
10
20
30
40
50
60
Normal
LY50
LY75
LY88
LY94
LY97
LY data
Normal data

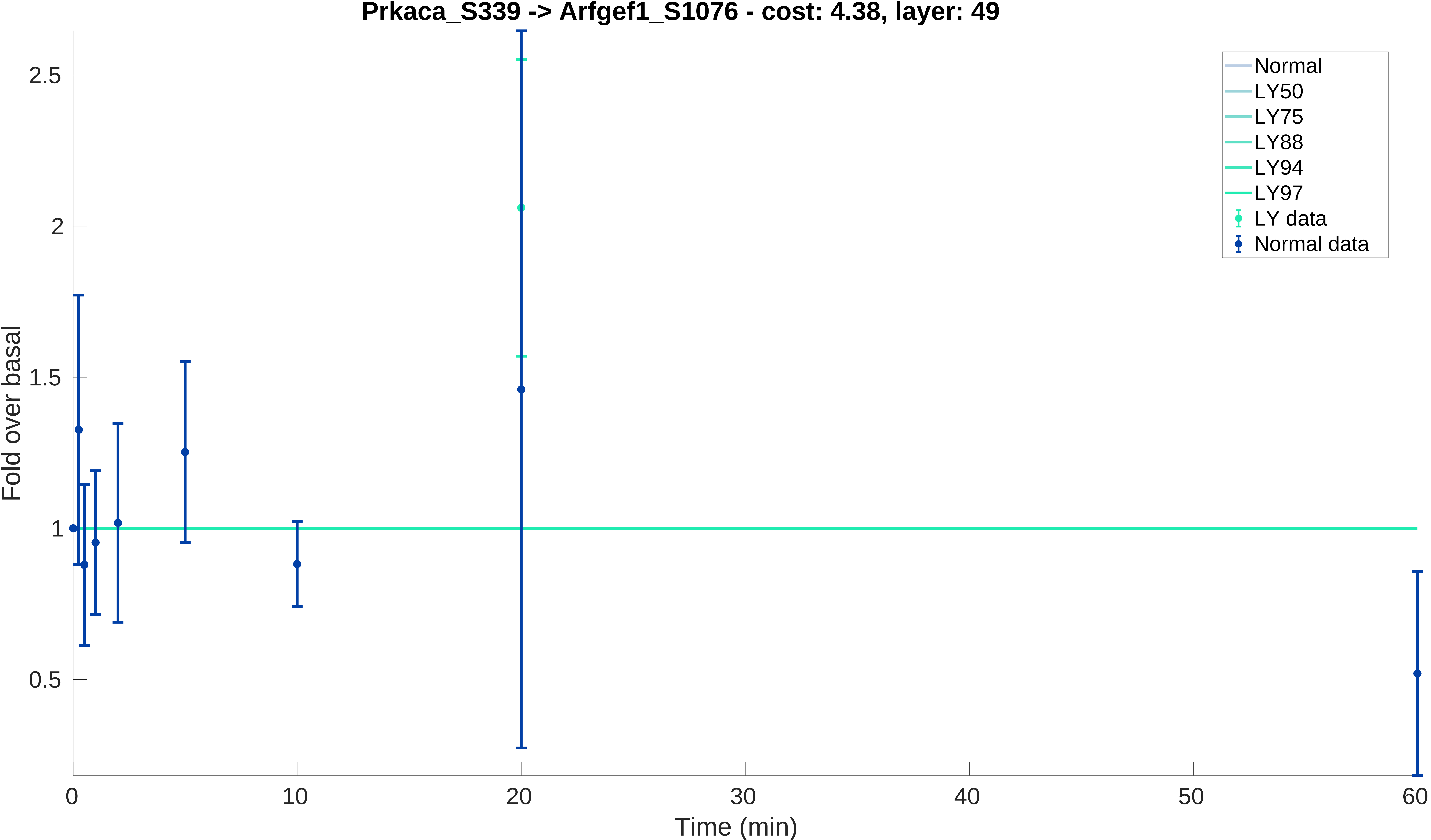
Prkaca_S339 -> Arfgef1_S1076 - cost: 4.38, layer: 49
Fold over basal
Time (min)
0.5
1
1.5
2
2.5
0
10
20
30
40
50
60
Normal
LY50
LY75
LY88
LY94
LY97
LY data
Normal data

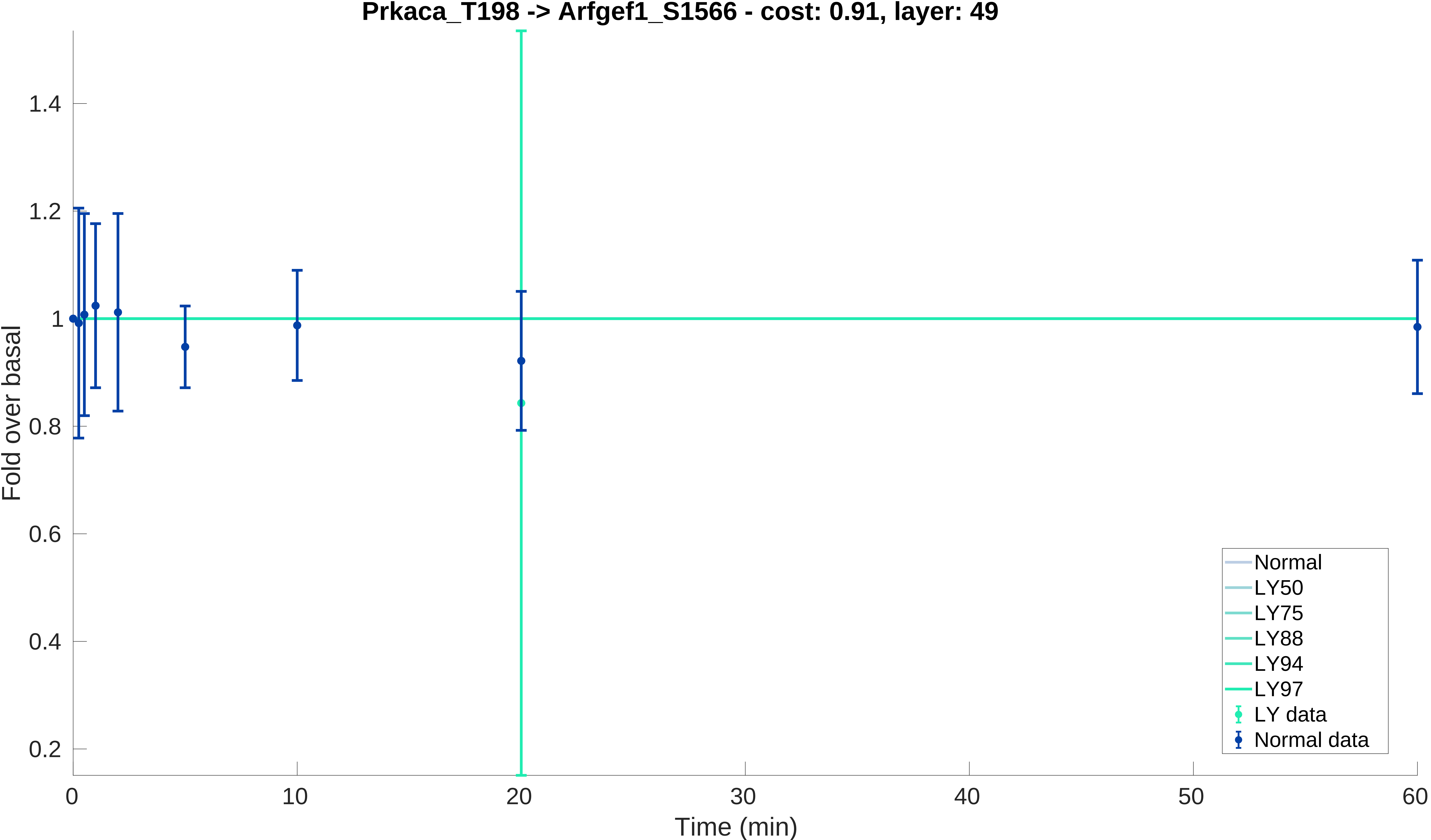

Prkaca_T198 -> Arfgef1_S1566 - cost: 0.91, layer: 49
Fold over basal
1.4
1.2
1
0.8
0.6
0.4
0.2
0
10
20
30
40
50
60
Time (min)
Normal
LY50
LY75
LY88
LY94
LY97
LY data
Normal data

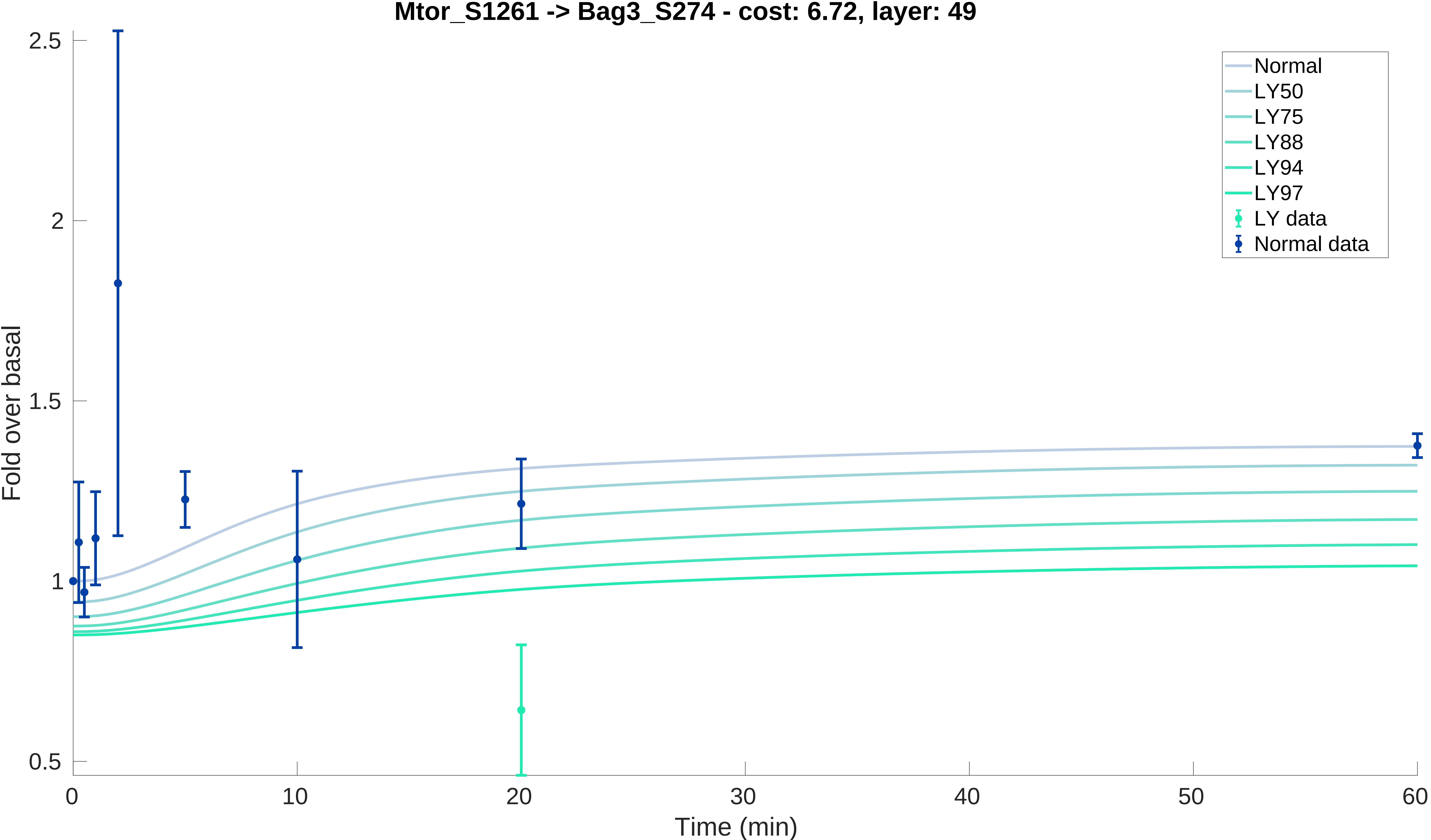

Mtor_S1261 -> Bag3_S274 - cost: 6.72, layer: 49
Fold over basal
Time (min)
2.5
2
1.5
1
0.5
0
10
20
30
40
50
60
Normal
LY50
LY75
LY88
LY94
LY97
LY data
Normal data

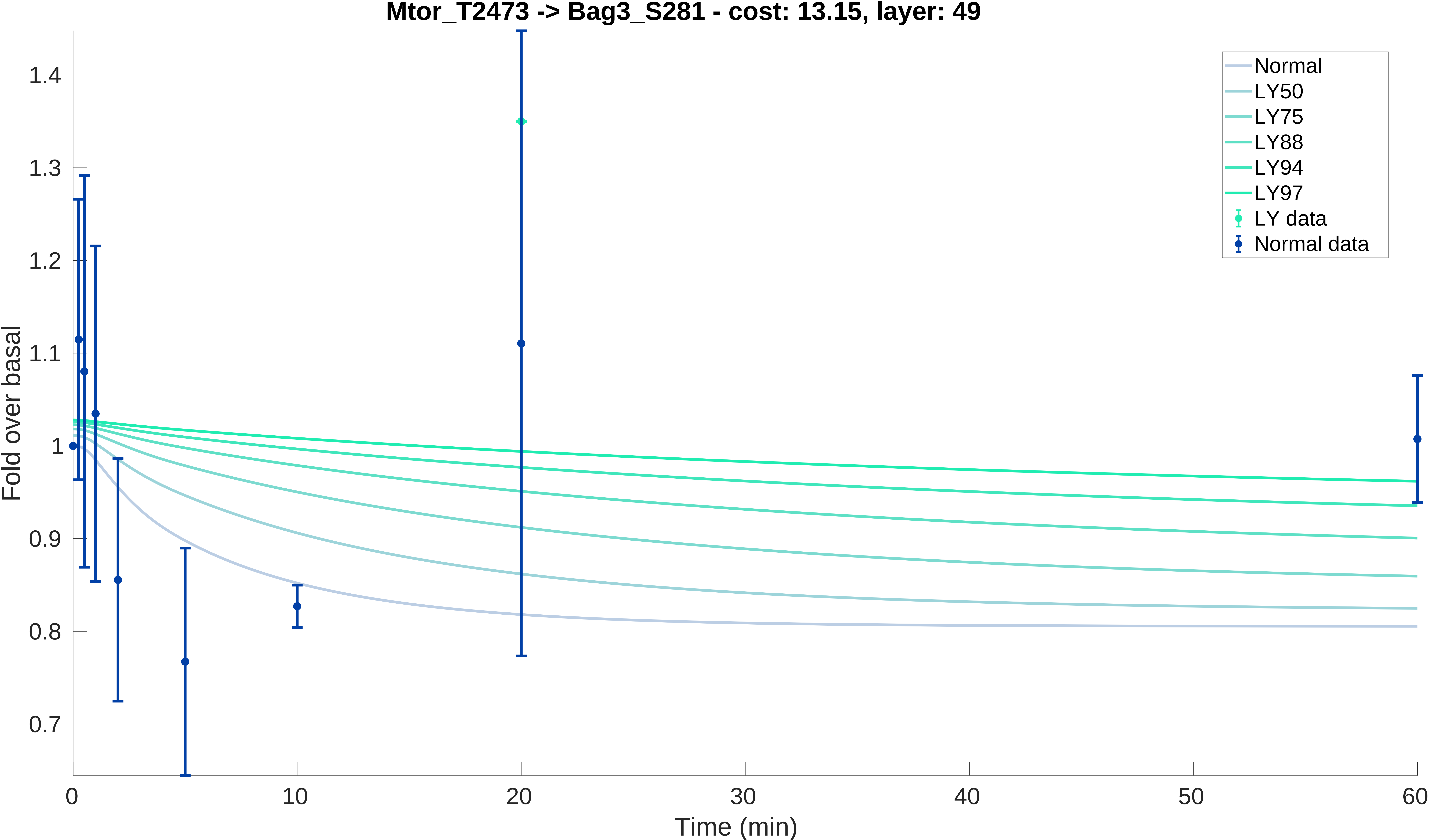

Mtor_T2473 -> Bag3_S281 - cost: 13.15, layer: 49
Fold over basal
1.4
1.3
1.2
1.1
1
0.9
0.8
0.7
0
10
20
30
40
50
60
Time (min)
Normal
LY50
LY75
LY88
LY94
LY97
LY data
Normal data

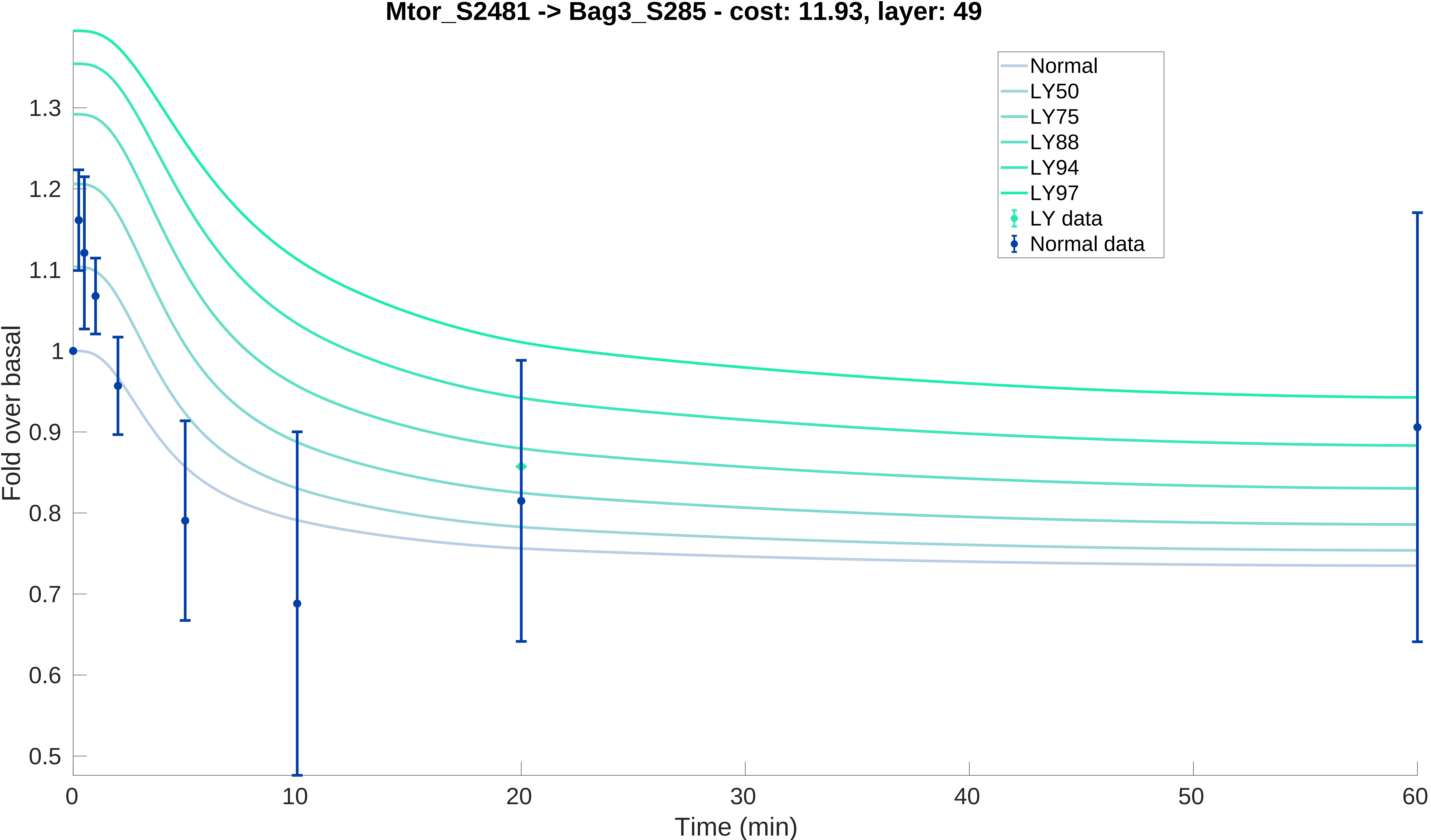

Mtor_S2481 -> Bag3_S285 - cost: 11.93, layer: 49
Normal
LY50
LY75
LY88
LY94
LY97
LY data
Normal data
Fold over basal
1.3
1.2
1.1
1
0.9
0.8
0.7
0.6
0.5
0
10
20
30
40
50
60
Time (min)

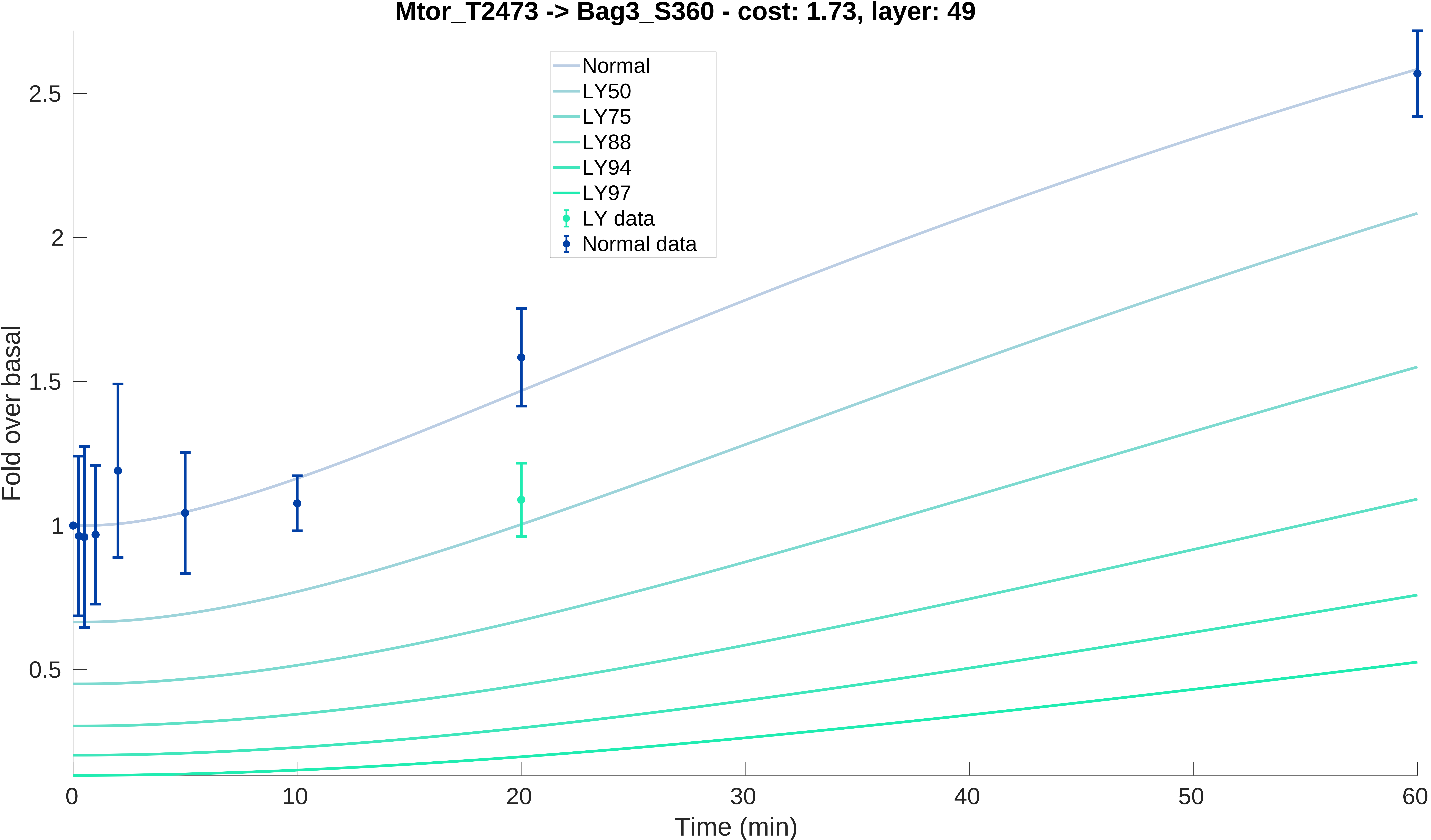

Mtor_T2473 -> Bag3_S360 - cost: 1.73, layer: 49
Normal
LY50
LY75
LY88
LY94
LY97
LY data
Normal data
2.5
2
1.5
1
0.5
Fold over basal
0
10
20
30
40
50
60
Time (min)

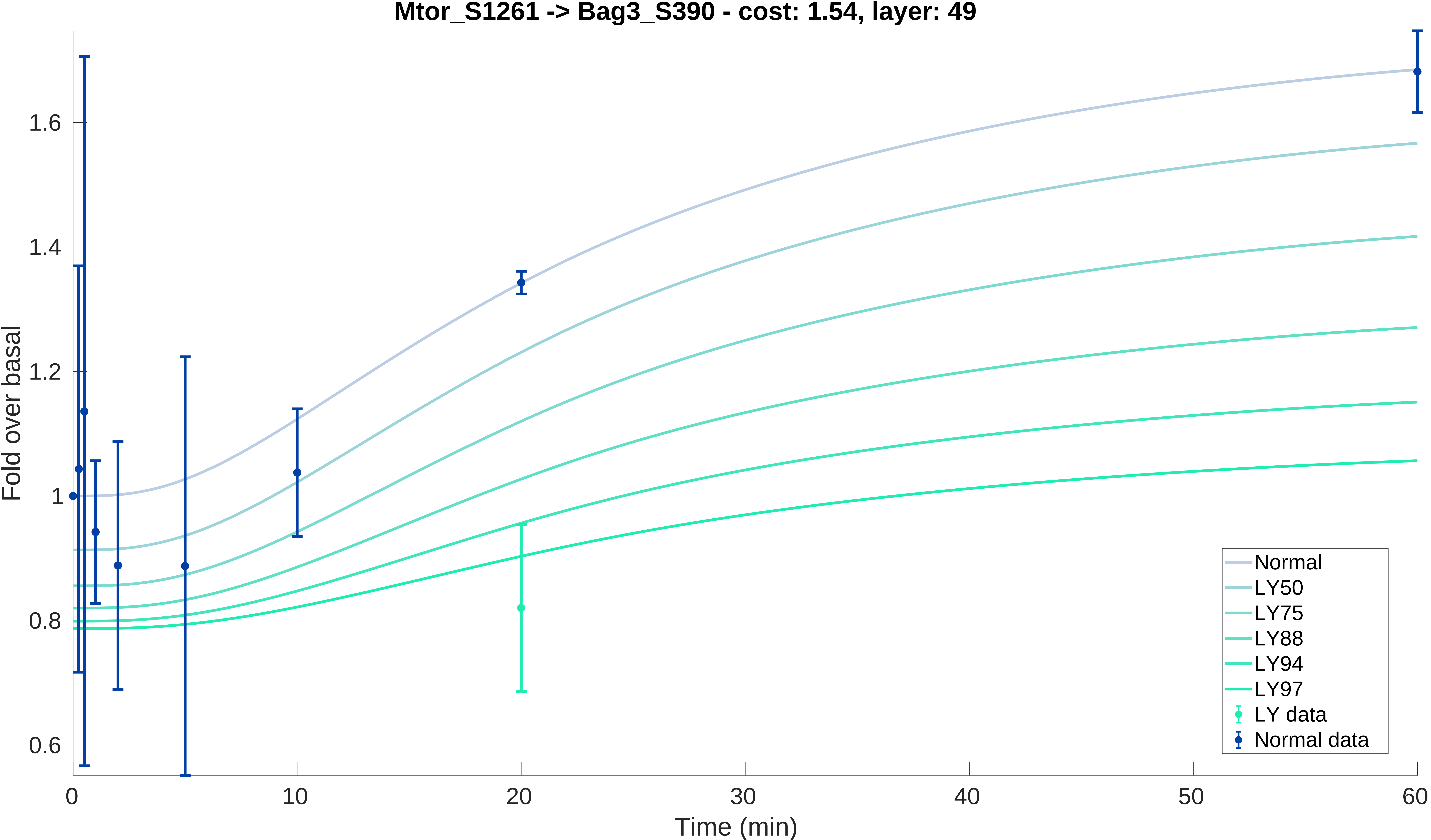

Mtor_S1261 -> Bag3_S390 - cost: 1.54, layer: 49
Fold over basal
1.6
1.4
1.2
1
0.8
0.6
0
10
20
30
40
50
60
Time (min)
Normal
LY50
LY75
LY88
LY94
LY97
LY data
Normal data

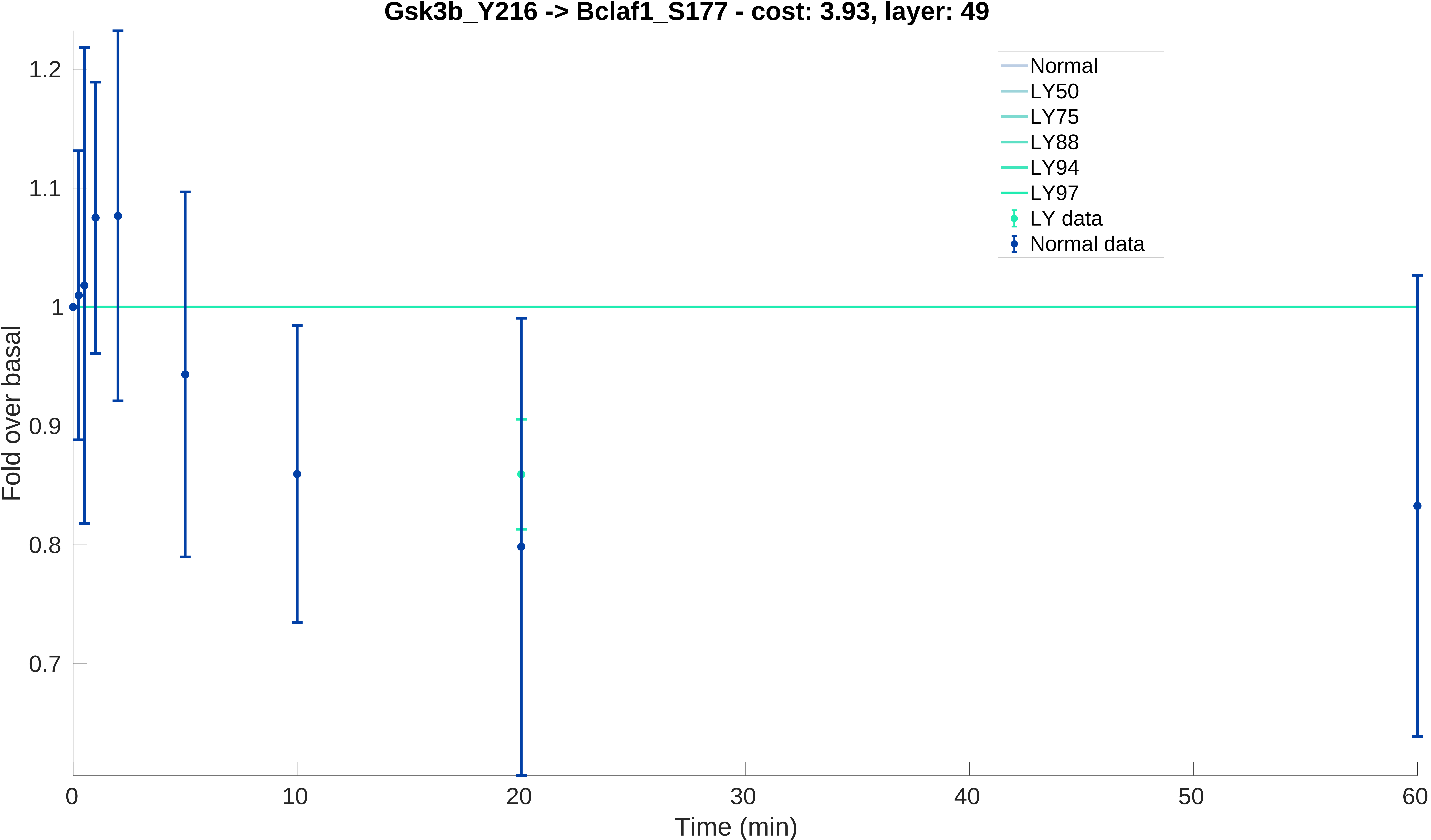
Gsk3b_Y216 -> Bclaf1_S177 - cost: 3.93, layer: 49
Normal
LY50
LY75
LY88
LY94
LY97
LY data
Normal data
Fold over basal
1.2
1.1
1
0.9
0.8
0.7
0
10
20
30
40
50
60
Time (min)

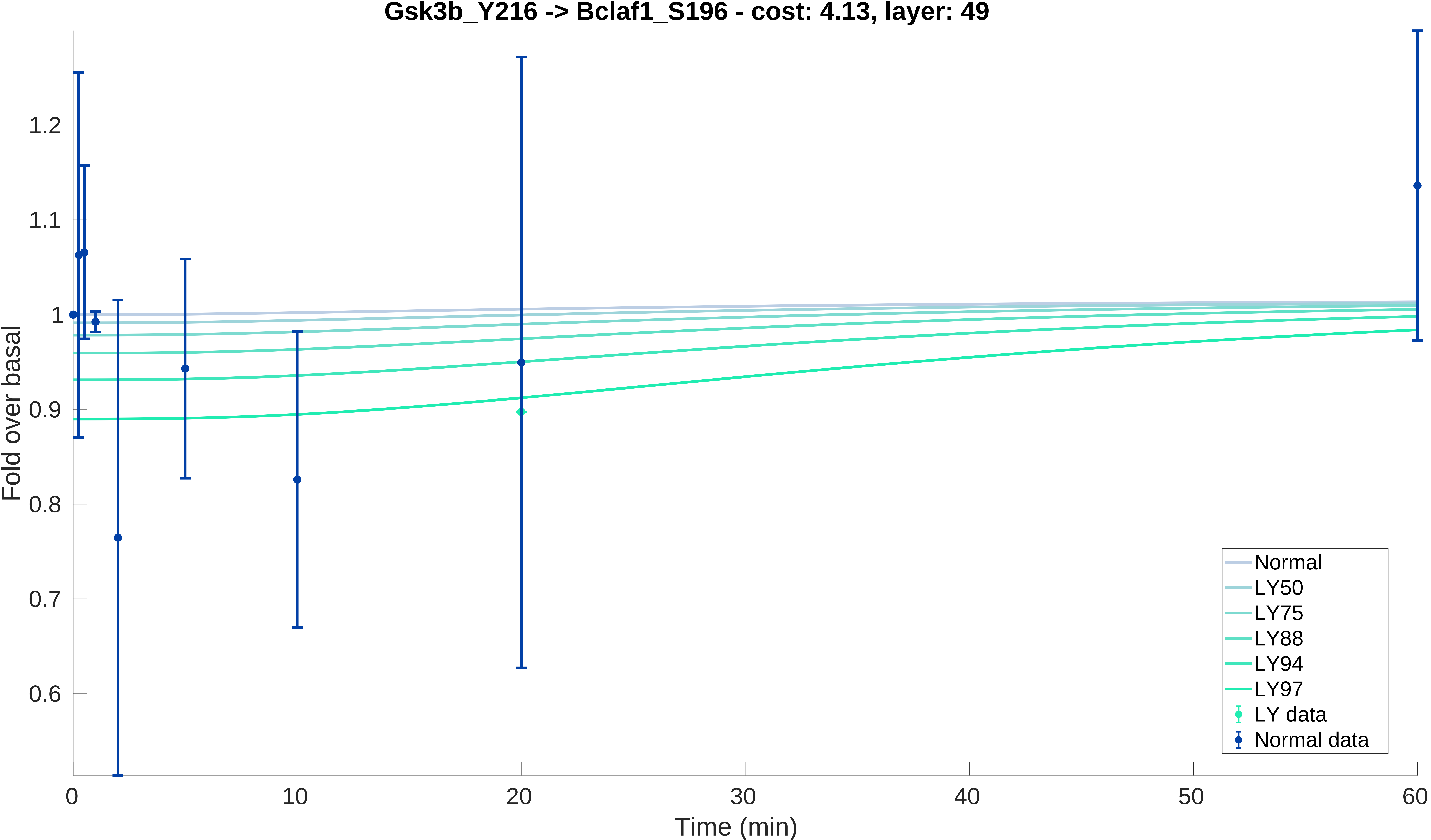

Gsk3b_Y216 -> Bclaf1_S196 - cost: 4.13, layer: 49
Fold over basal
1.2
1.1
1
0.9
0.8
0.7
0.6
0
10
20
30
40
50
60
Time (min)
Normal
LY50
LY75
LY88
LY94
LY97
LY data
Normal data

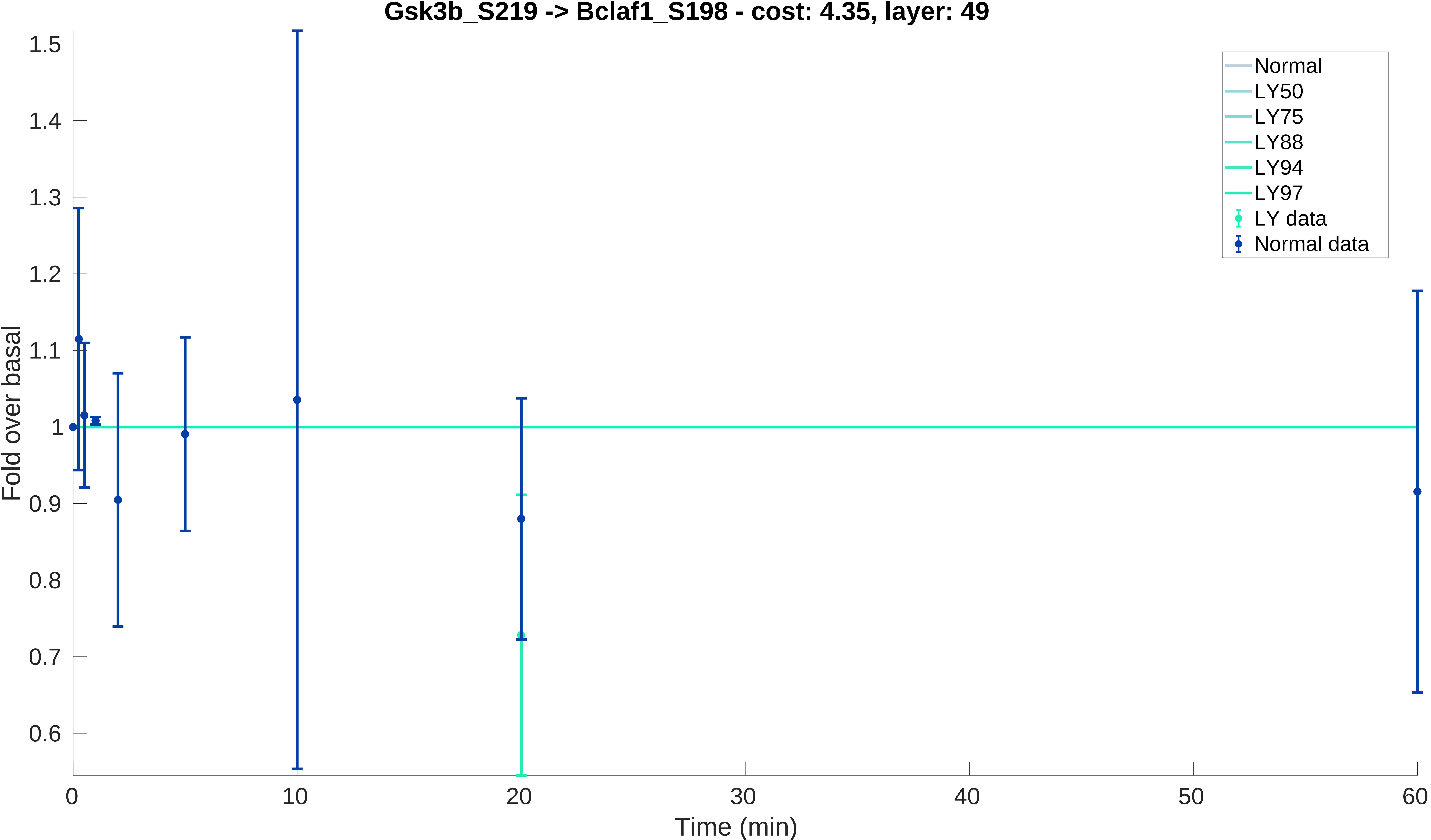

Gsk3b_S219 -> Bclaf1_S198 - cost: 4.35, layer: 49
Fold over basal
Time (min)
1.5
1.4
1.3
1.2
1.1
1
0.9
0.8
0.7
0.6
0
10
20
30
40
50
60
Normal
LY50
LY75
LY88
LY94
LY97
LY data
Normal data

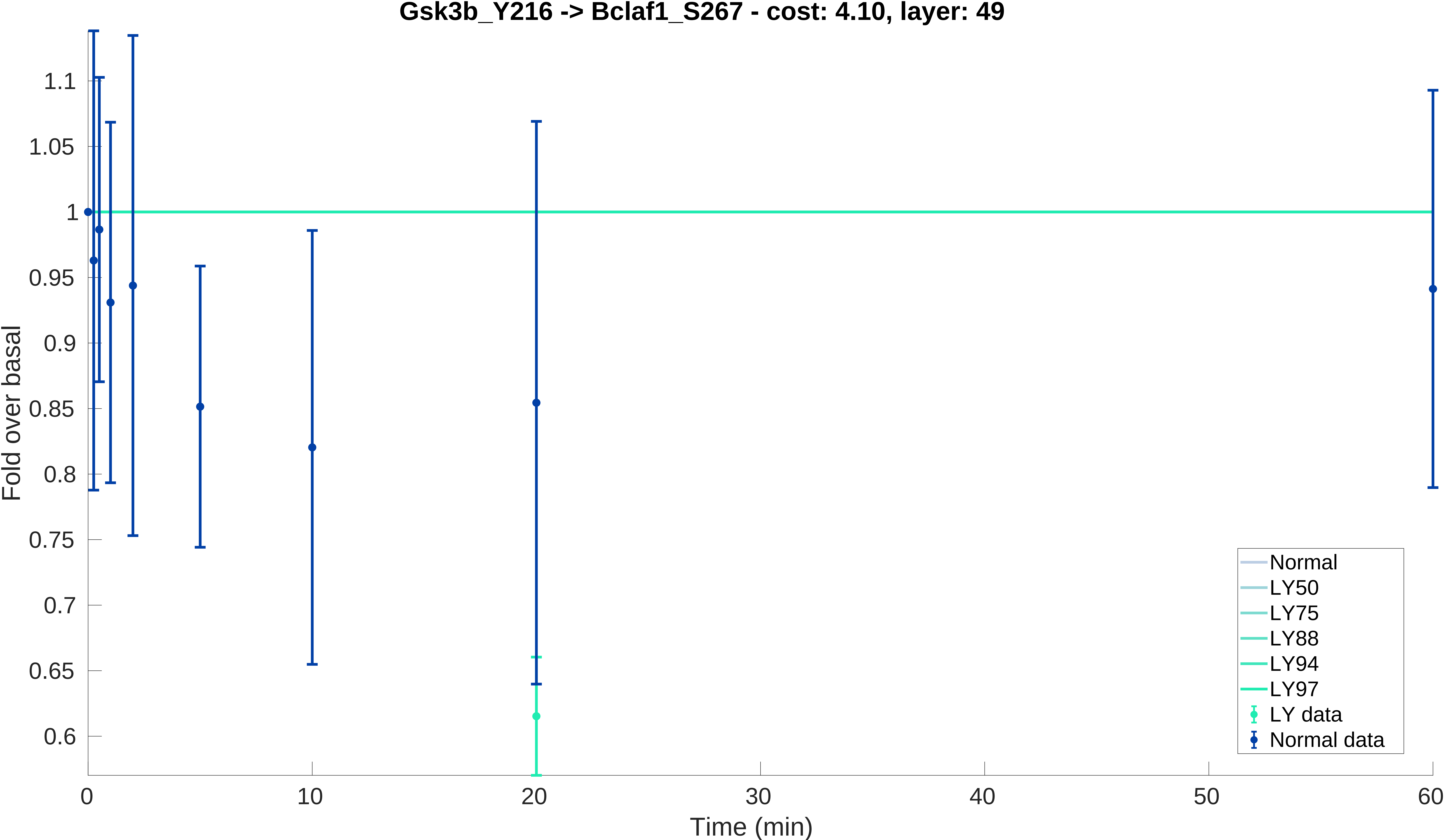

Gsk3b_Y216 -> Bclaf1_S267 - cost: 4.10, layer: 49
Fold over basal
Time (min)
1.1
1.05
1
0.95
0.9
0.85
0.8
0.75
0.7
0.65
0.6
0
10
20
30
40
50
60
Normal
LY50
LY75
LY88
LY94
LY97
LY data
Normal data

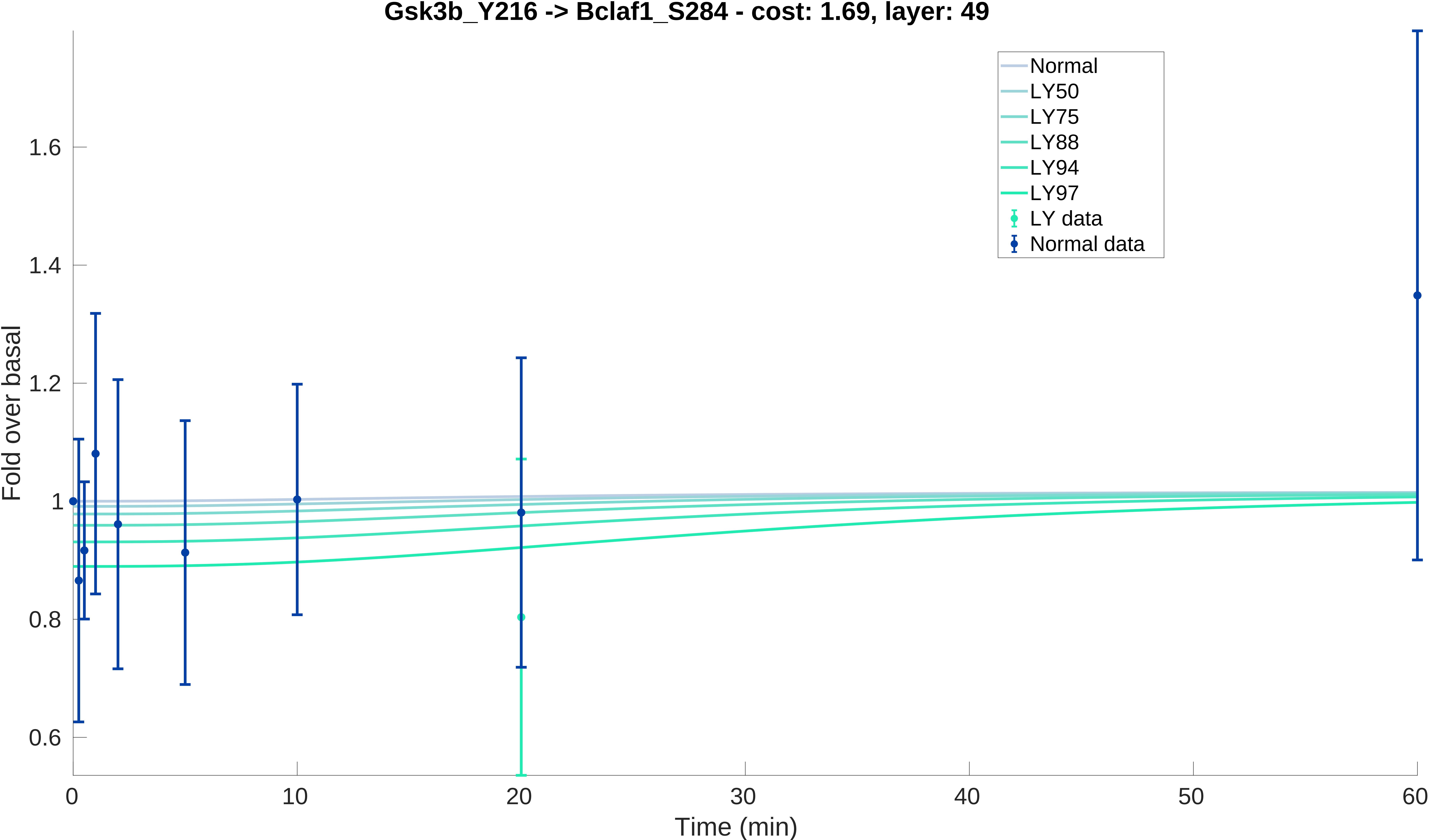

Gsk3b_Y216 -> Bclaf1_S284 - cost: 1.69, layer: 49
Normal
LY50
LY75
LY88
LY94
LY97
LY data
Normal data
Fold over basal
1.6
1.4
1.2
1
0.8
0.6
0
10
20
30
40
50
60
Time (min)

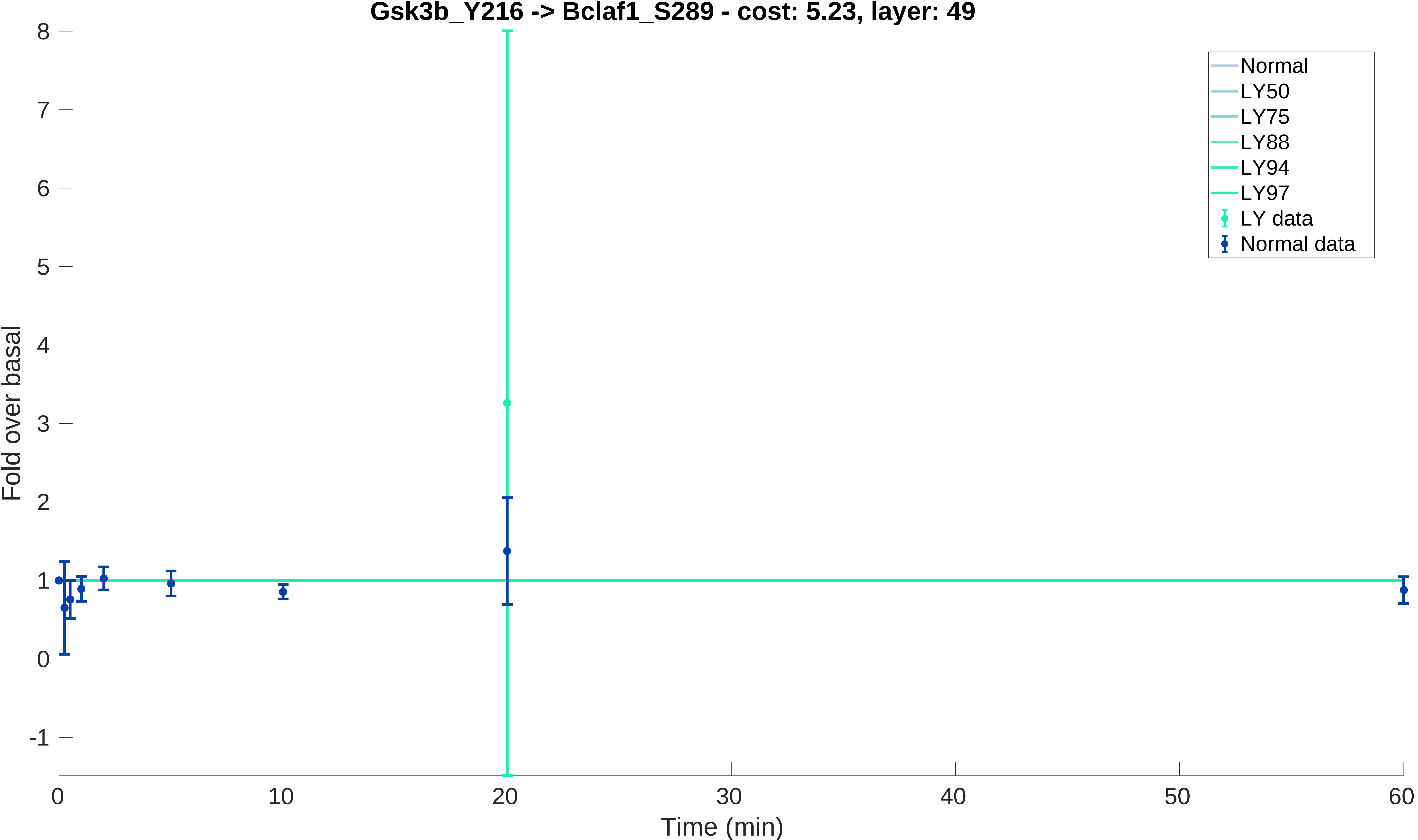

Gsk3b_Y216 -> Bclaf1_S289 - cost: 5.23, layer: 49
Fold over basal
Time (min)
8
7
6
5
4
3
2
1
0
-1
0
10
20
30
40
50
60
Normal
LY50
LY75
LY88
LY94
LY97
LY data
Normal data

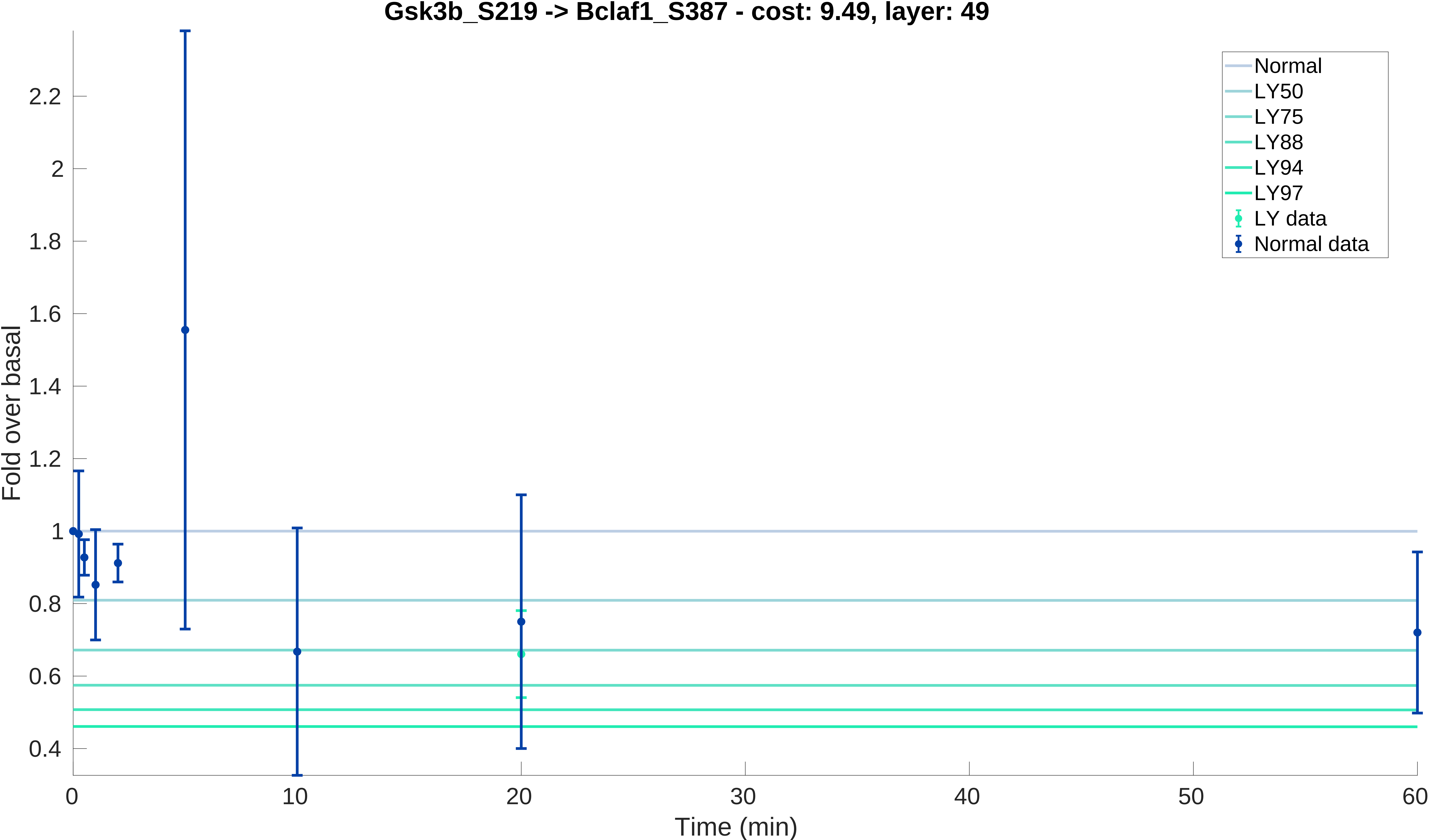

Gsk3b_S219 -> Bclaf1_S387 - cost: 9.49, layer: 49
Fold over basal
Time (min)
Normal
LY50
LY75
LY88
LY94
LY97
LY data
Normal data
2.2
2
1.8
1.6
1.4
1.2
1
0.8
0.6
0.4
0
10
20
30
40
50
60

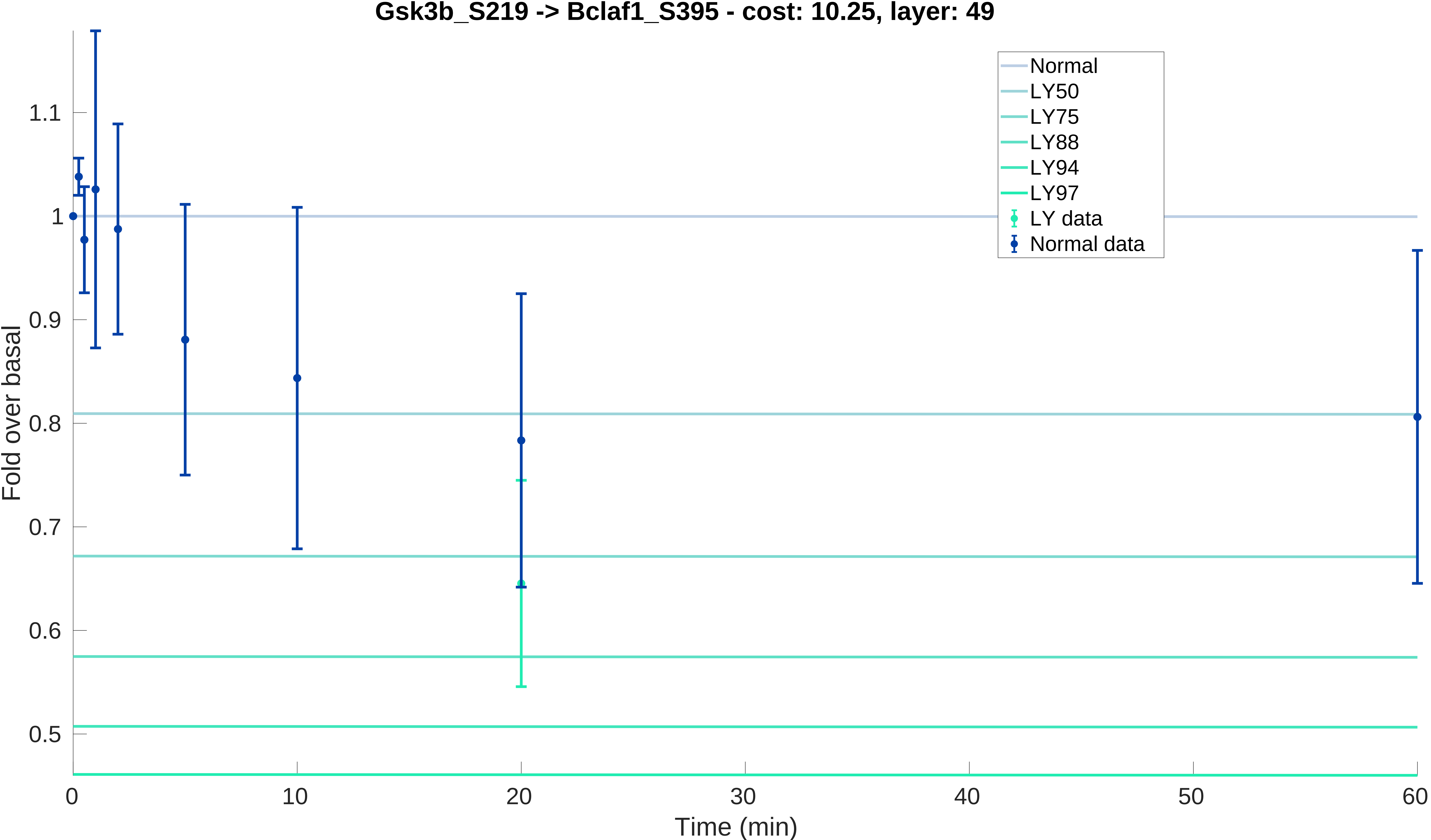
Gsk3b_S219 -> Bclaf1_S395 - cost: 10.25, layer: 49
Fold over basal
Time (min)
0.5
0.6
0.7
0.8
0.9
1
1.1
0
10
20
30
40
50
60
Normal
LY50
LY75
LY88
LY94
LY97
LY data
Normal data

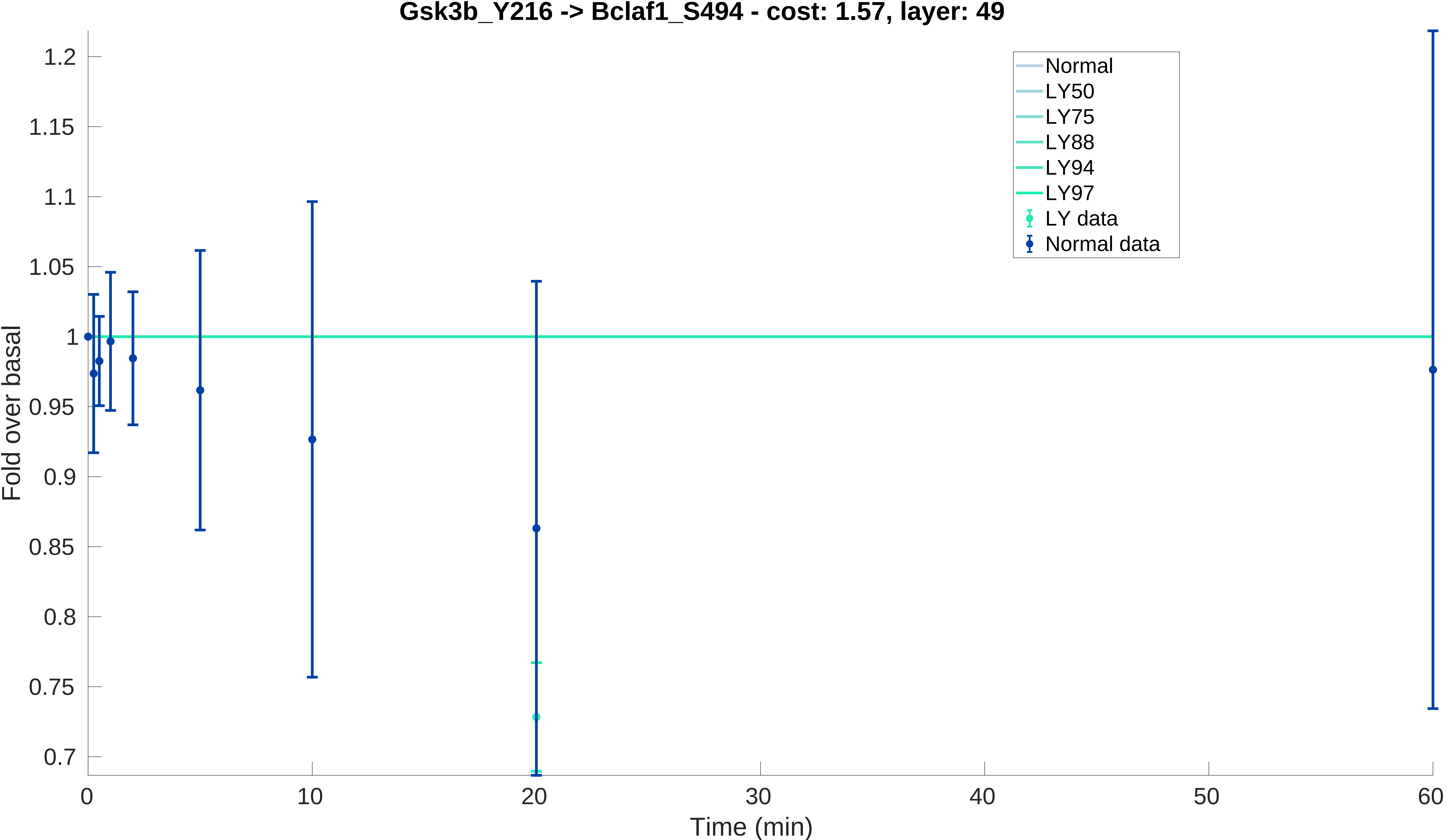

Gsk3b_Y216 -> Bclaf1_S494 - cost: 1.57, layer: 49
Fold over basal
Time (min)
1.2
1.15
1.1
1.05
1
0.95
0.9
0.85
0.8
0.75
0.7
0
10
20
30
40
50
60
Normal
LY50
LY75
LY88
LY94
LY97
LY data
Normal data

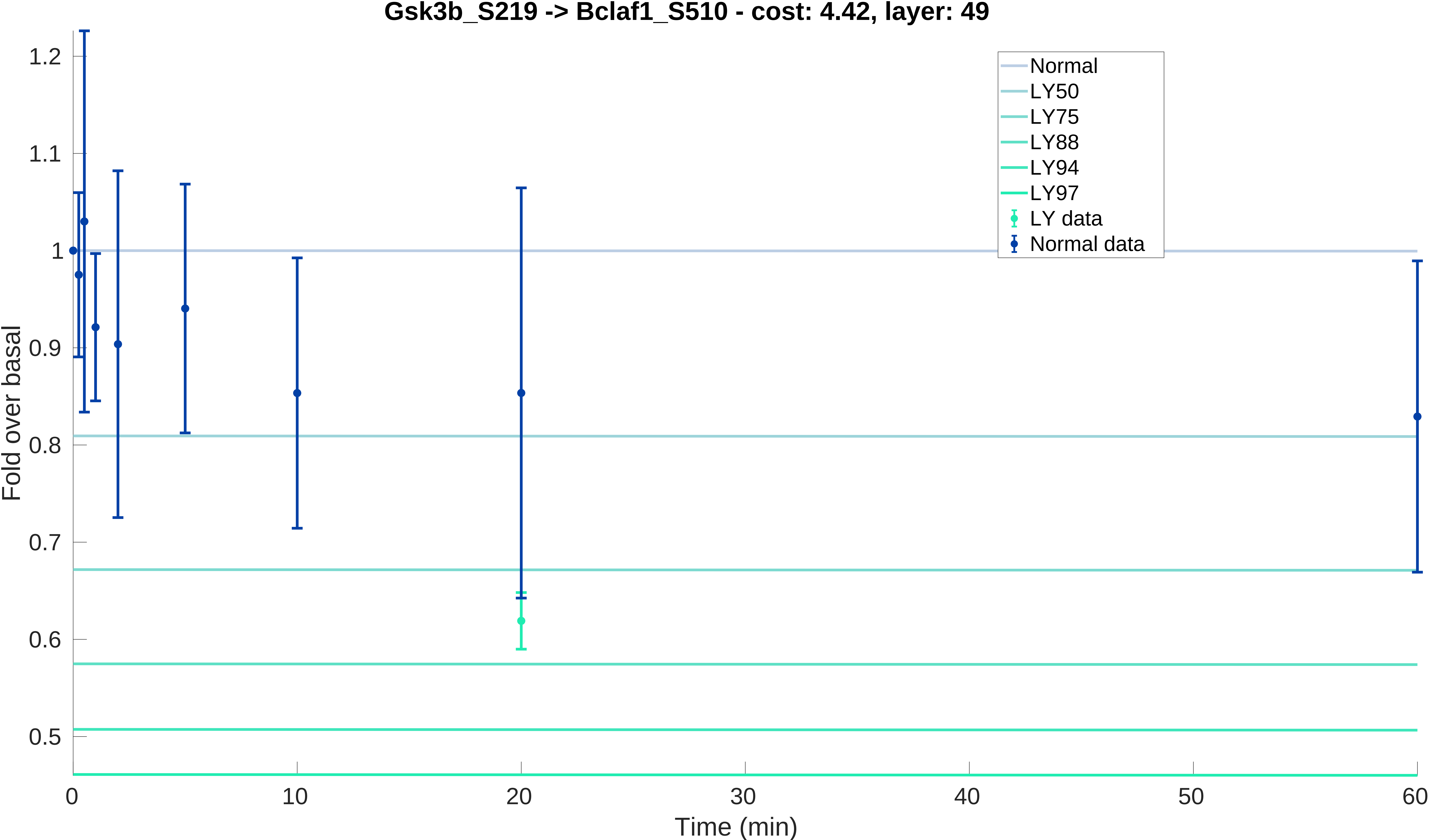

Gsk3b_S219 -> Bclaf1_S510 - cost: 4.42, layer: 49
Fold over basal
1.2
1.1
1
0.9
0.8
0.7
0.6
0.5
0
10
20
30
40
50
60
Time (min)
Normal
LY50
LY75
LY88
LY94
LY97
LY data
Normal data

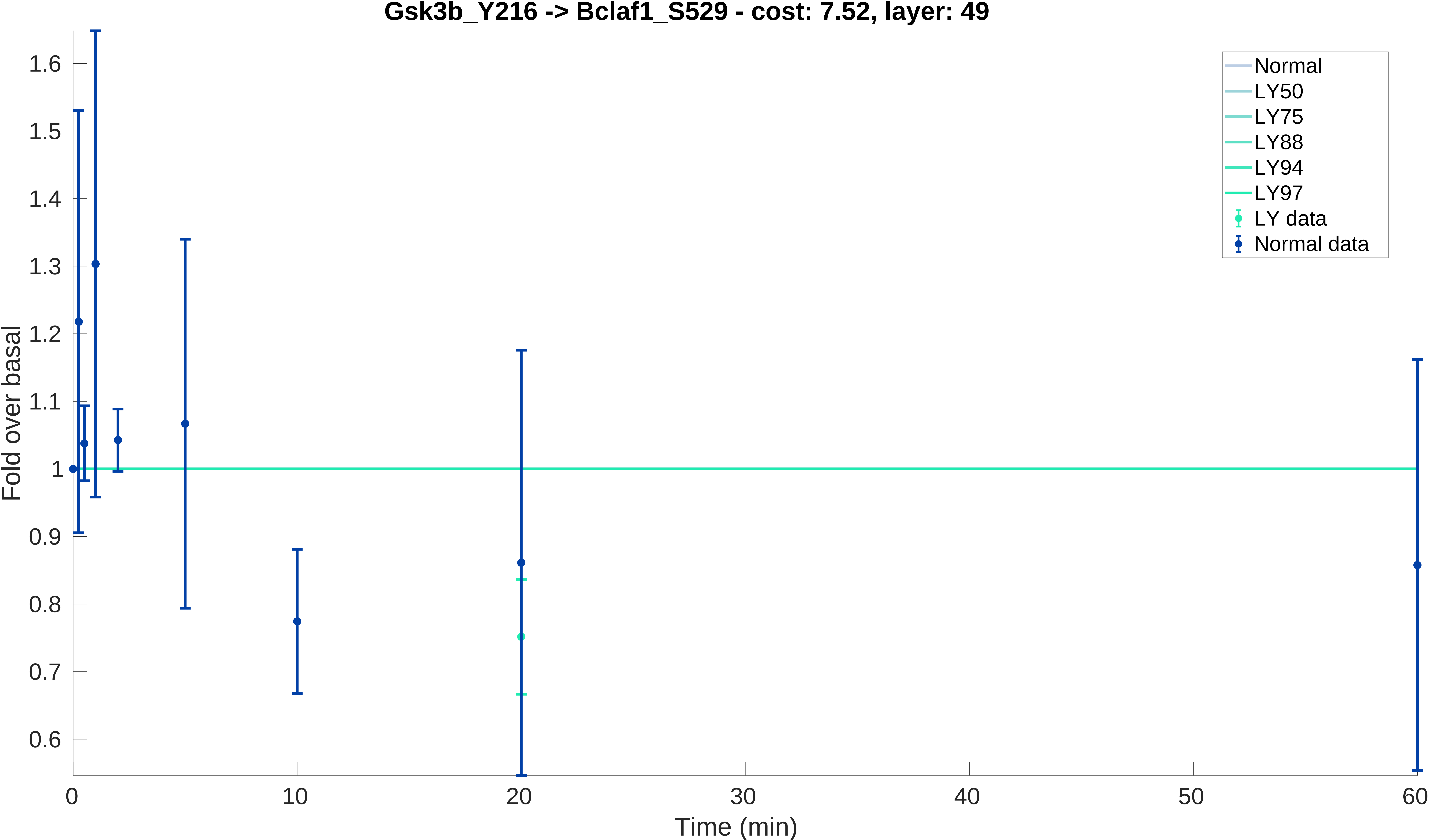

Gsk3b_Y216 -> Bclaf1_S529 - cost: 7.52, layer: 49
Normal
LY50
LY75
LY88
LY94
LY97
LY data
Normal data
Fold over basal
Time (min)
0
10
20
30
40
50
60
0.6
0.7
0.8
0.9
1
1.1
1.2
1.3
1.4
1.5
1.6

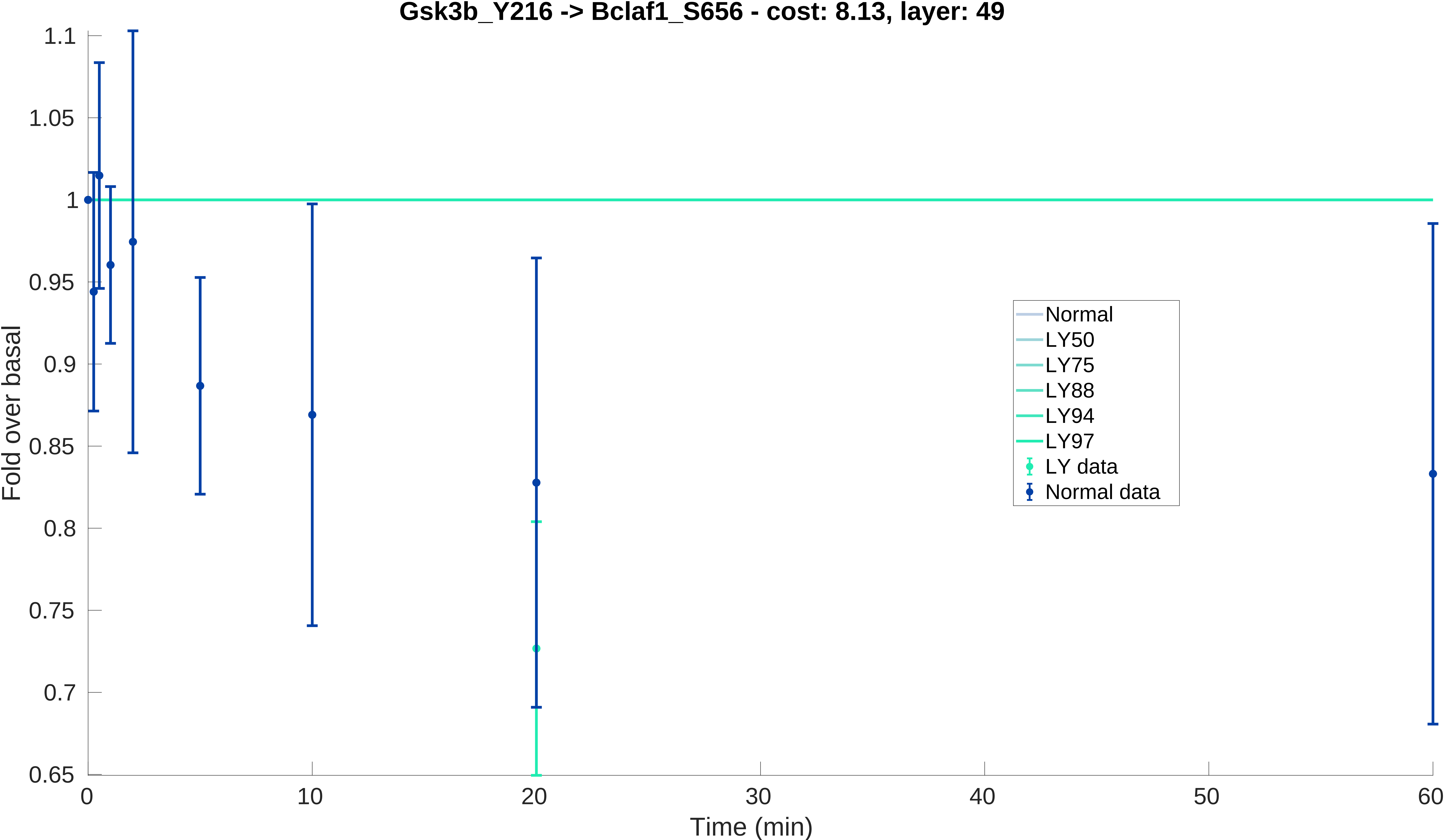

Gsk3b_Y216 -> Bclaf1_S656 - cost: 8.13, layer: 49
Fold over basal
1.1
1.05
1
0.95
0.9
0.85
0.8
0.75
0.7
0.65
0
10
20
30
40
50
60
Time (min)
Normal
LY50
LY75
LY88
LY94
LY97
LY data
Normal data

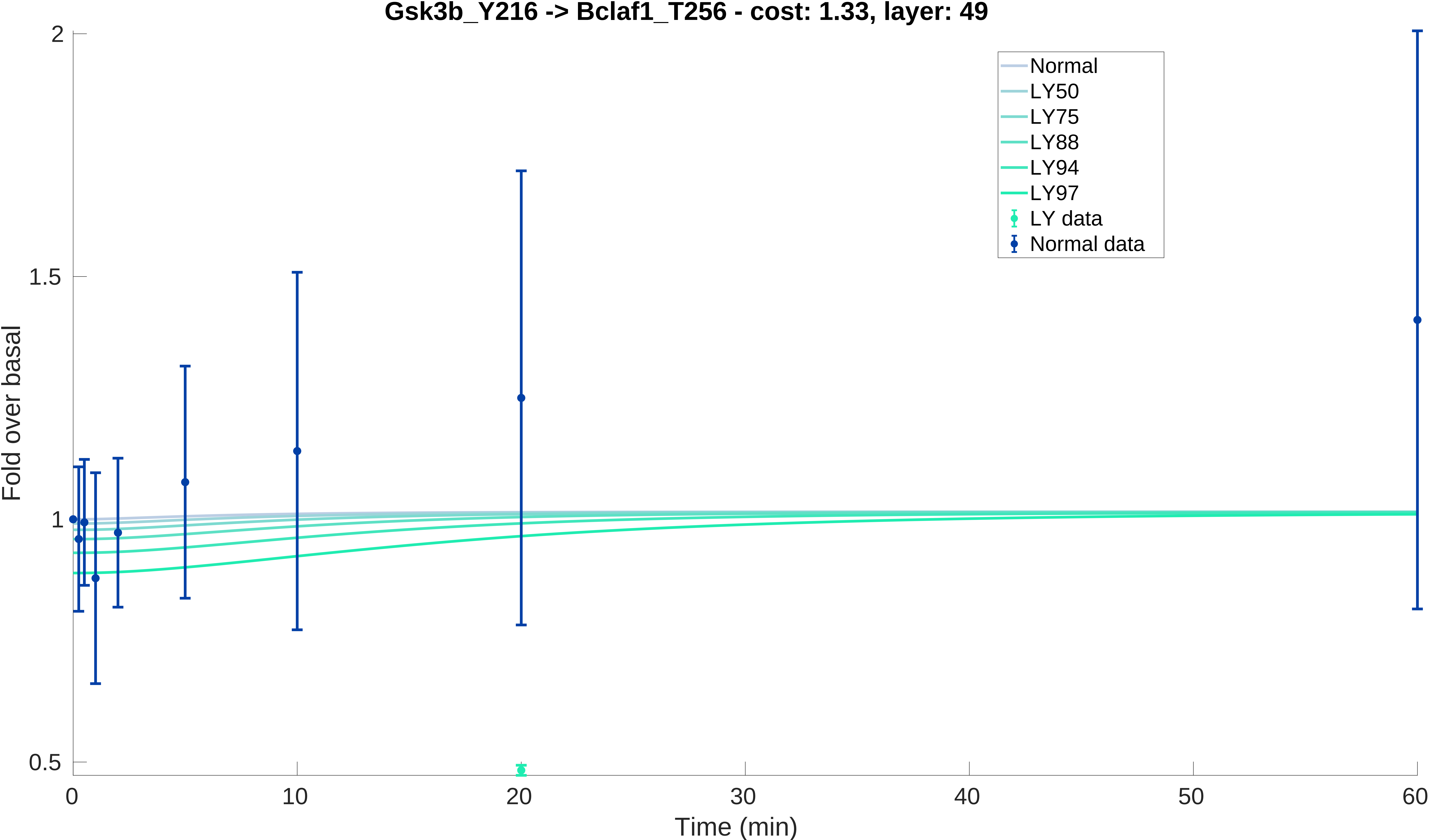

Gsk3b_Y216 -> Bclaf1_T256 - cost: 1.33, layer: 49
Normal
LY50
LY75
LY88
LY94
LY97
LY data
Normal data
Fold over basal
Time (min)
2
1.5
1
0.5
0
10
20
30
40
50
60

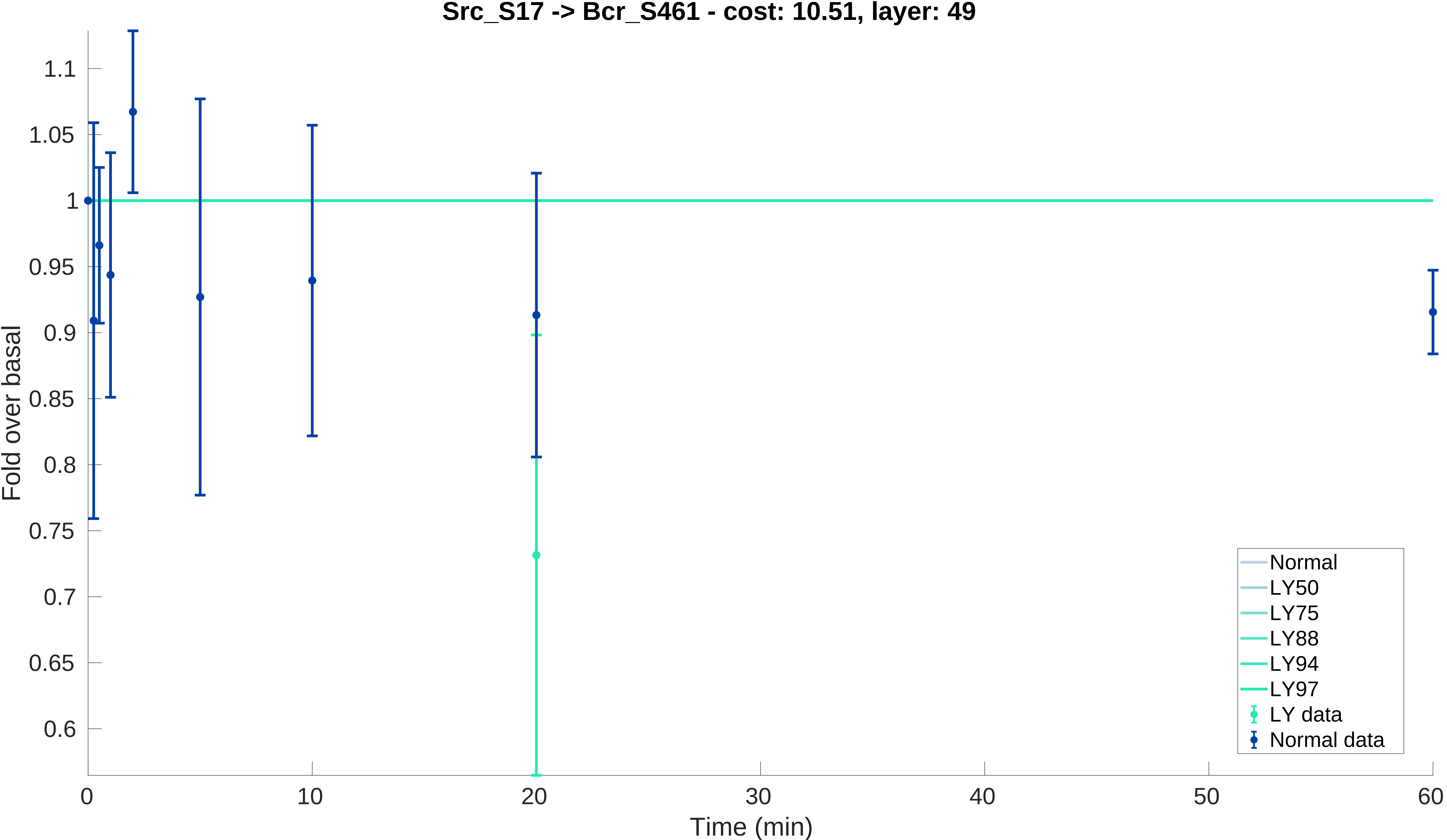
Src_S17 -> Bcr_S461 - cost: 10.51, layer: 49
Fold over basal
1.1
1.05
1
0.95
0.9
0.85
0.8
0.75
0.7
0.65
0.6
0
10
20
30
40
50
60
Time (min)
Normal
LY50
LY75
LY88
LY94
LY97
LY data
Normal data

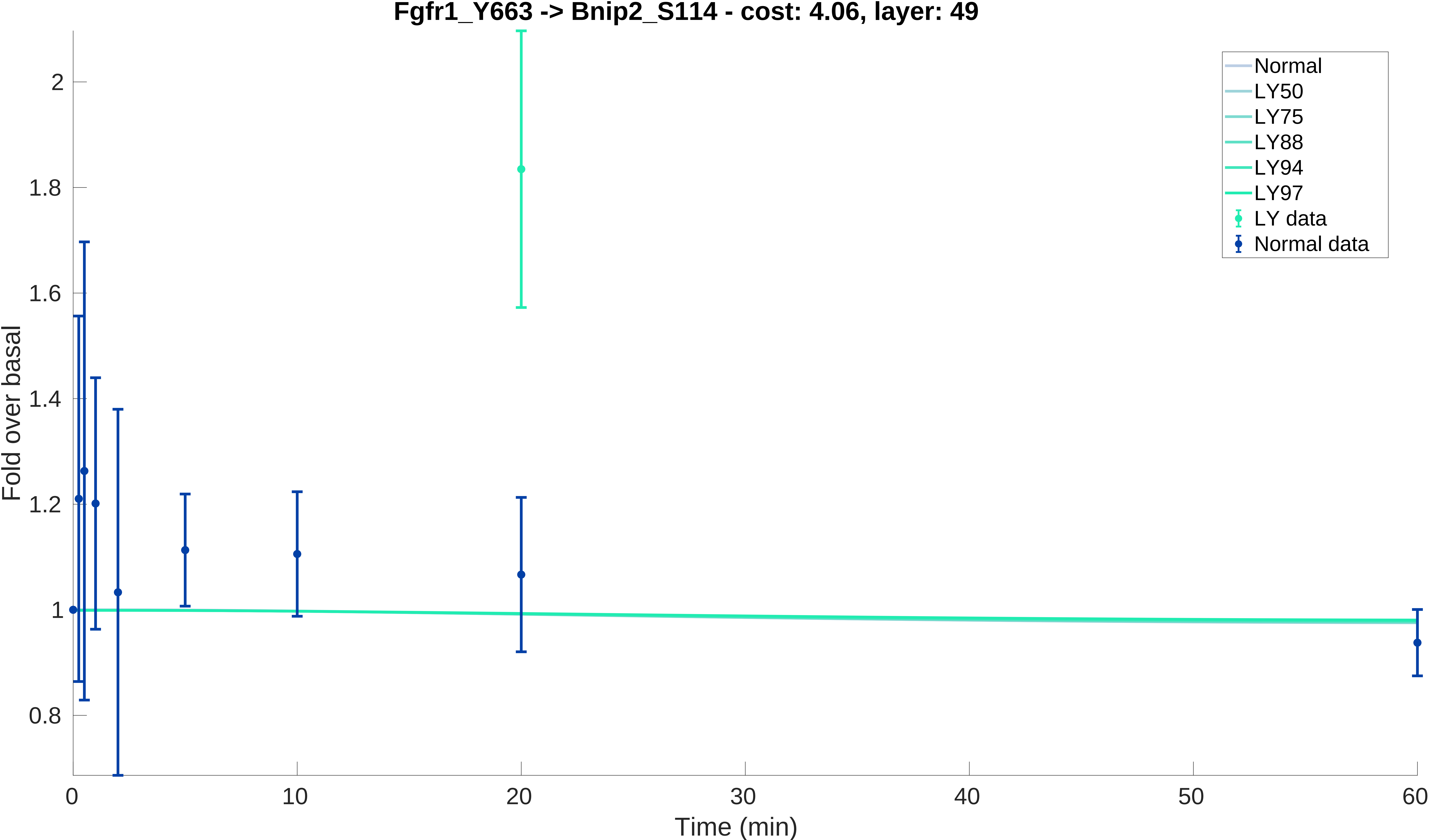

Fgfr1_Y663 -> Bnip2_S114 - cost: 4.06, layer: 49
Fold over basal
Time (min)
0.8
1
1.2
1.4
1.6
1.8
2
0
10
20
30
40
50
60
Normal
LY50
LY75
LY88
LY94
LY97
LY data
Normal data

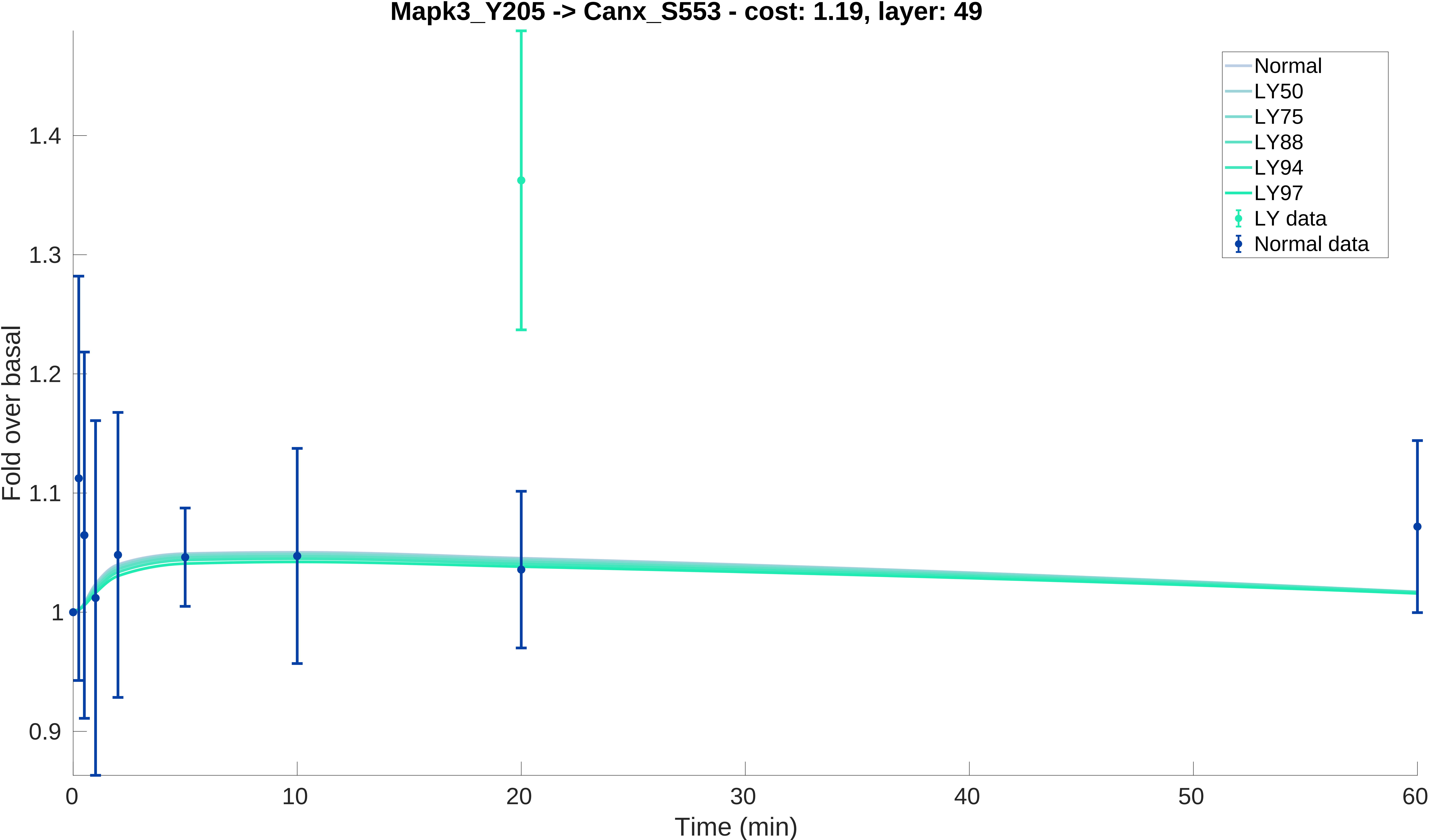

Mapk3_Y205 -> Canx_S553 - cost: 1.19, layer: 49
Fold over basal
Time (min)
0.9
1
1.1
1.2
1.3
1.4
0
10
20
30
40
50
60
Normal
LY50
LY75
LY88
LY94
LY97
LY data
Normal data

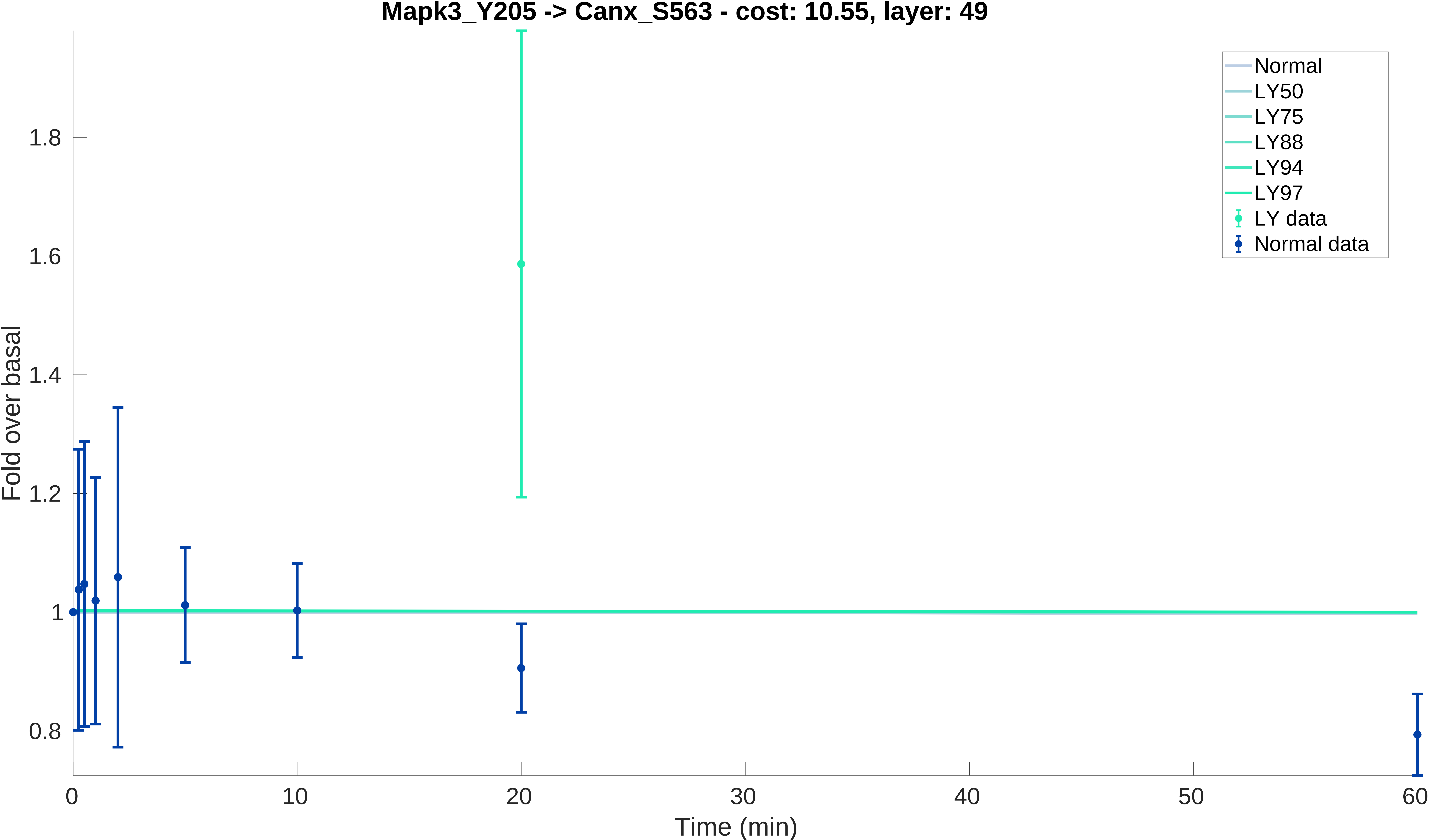
Mapk3_Y205 -> Canx_S563 - cost: 10.55, layer: 49
Fold over basal
Time (min)
0.8
1
1.2
1.4
1.6
1.8
0
10
20
30
40
50
60
Normal
LY50
LY75
LY88
LY94
LY97
LY data
Normal data

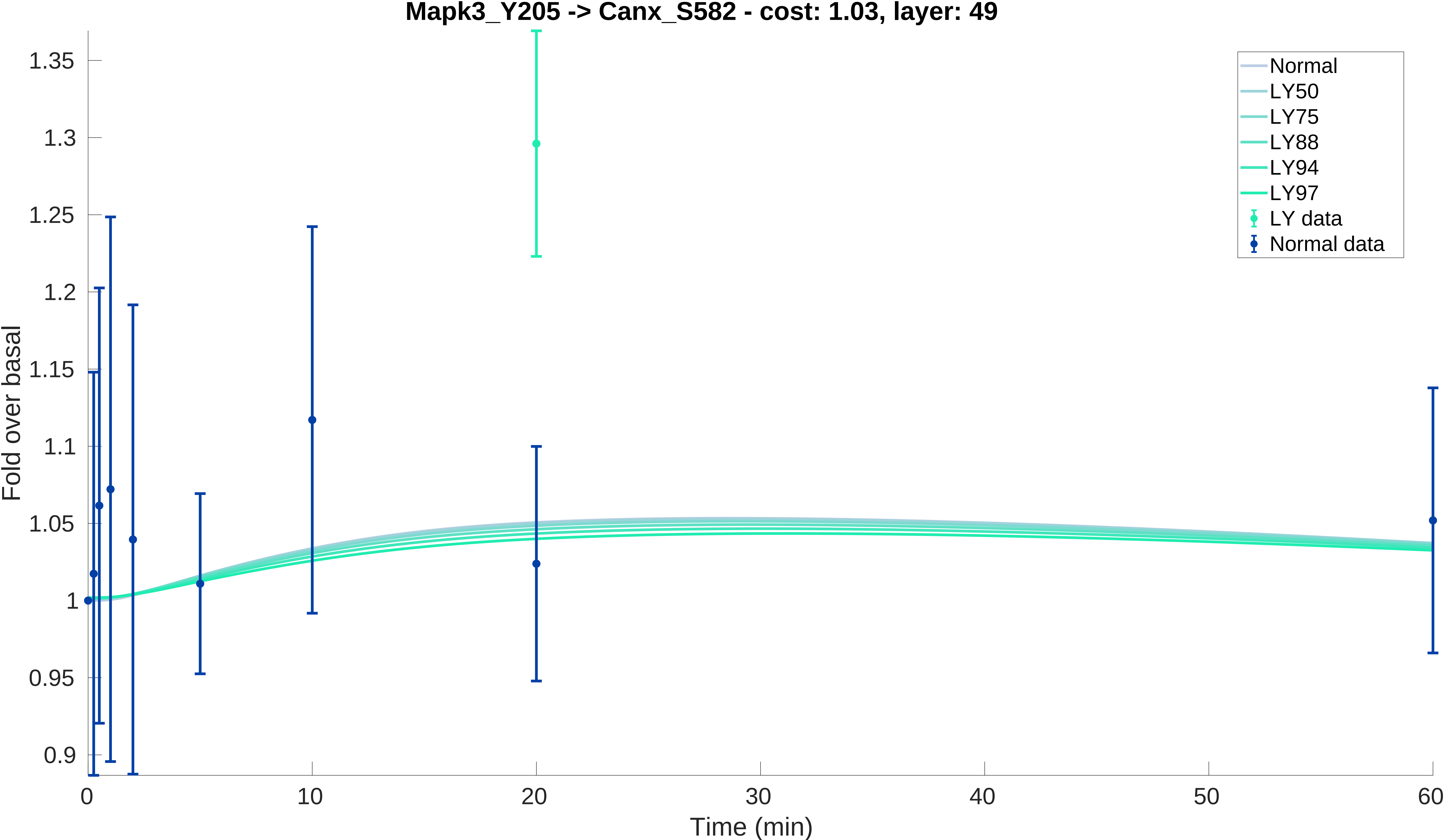

Mapk3_Y205 -> Canx_S582 - cost: 1.03, layer: 49
Normal
LY50
LY75
LY88
LY94
LY97
LY data
Normal data
Fold over basal
Time (min)
1.35
1.3
1.25
1.2
1.15
1.1
1.05
1
0.95
0.9
0
10
20
30
40
50
60

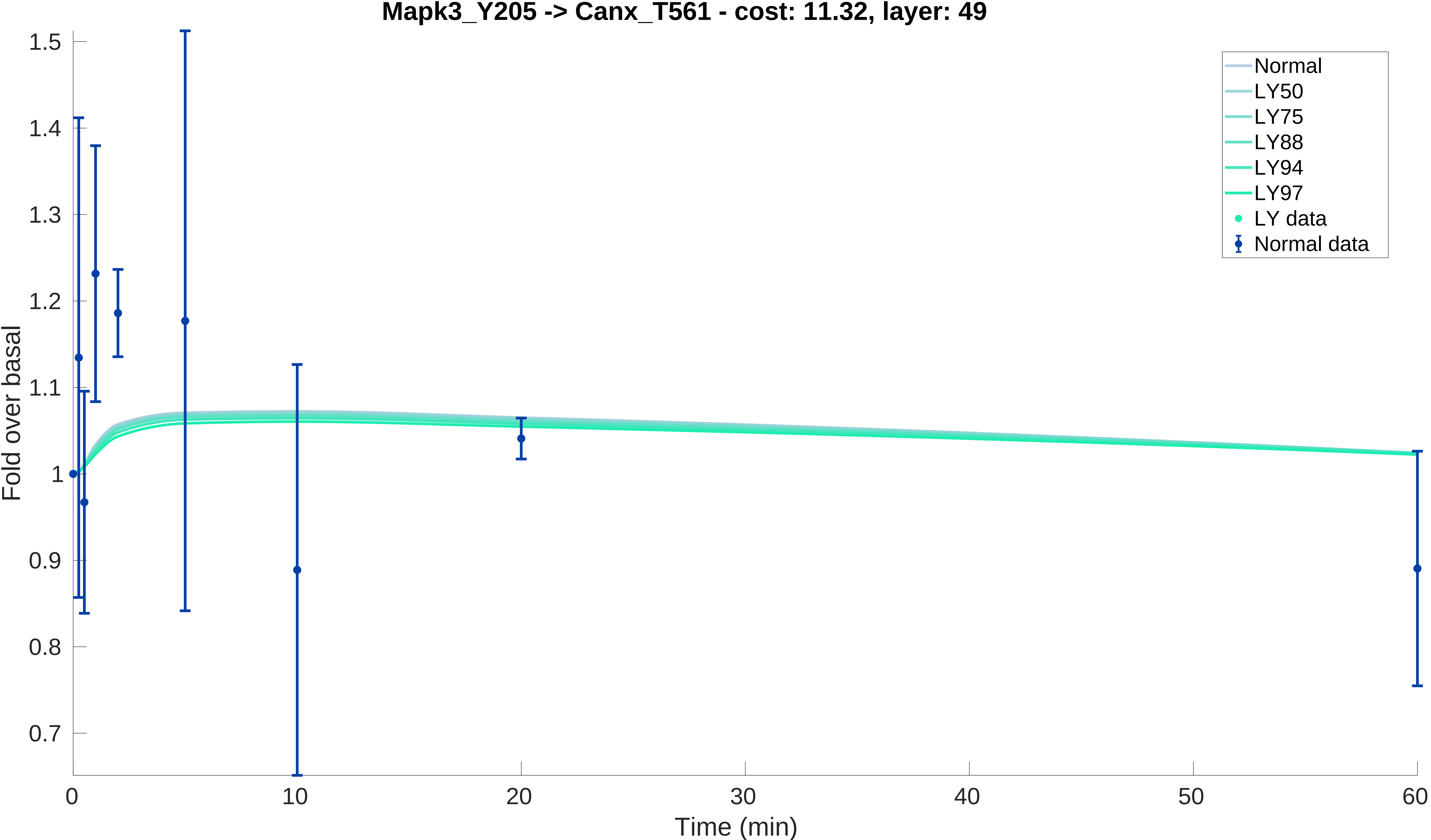

Mapk3_Y205 -> Canx_T561 - cost: 11.32, layer: 49
Fold over basal
Time (min)
0
10
20
30
40
50
60
0.7
0.8
0.9
1
1.1
1.2
1.3
1.4
1.5
Normal
LY50
LY75
LY88
LY94
LY97
LY data
Normal data

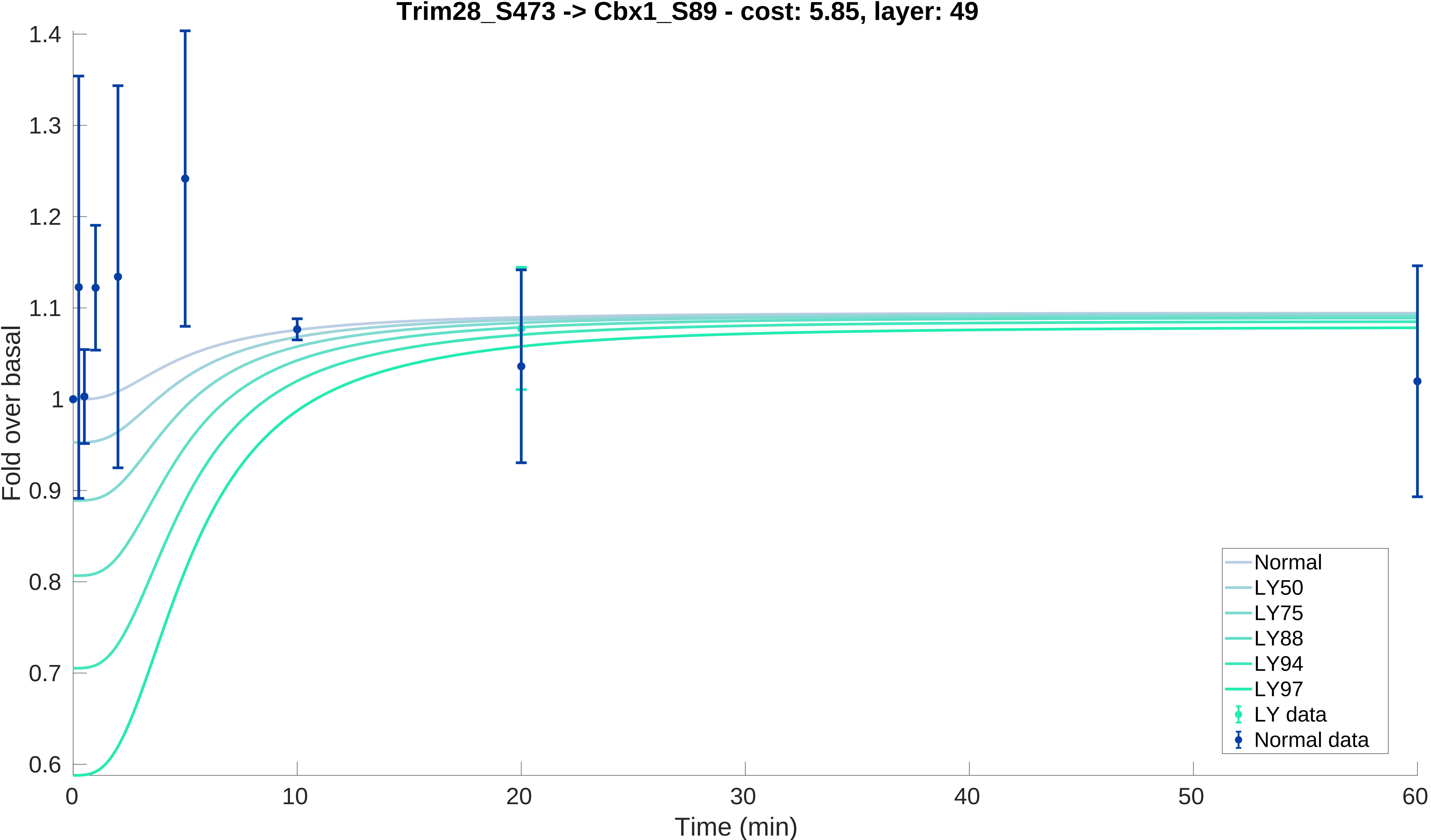

Trim28_S473 -> Cbx1_S89 - cost: 5.85, layer: 49
Fold over basal
1.4
1.3
1.2
1.1
1
0.9
0.8
0.7
0.6
0
10
20
30
40
50
60
Time (min)
Normal
LY50
LY75
LY88
LY94
LY97
LY data
Normal data

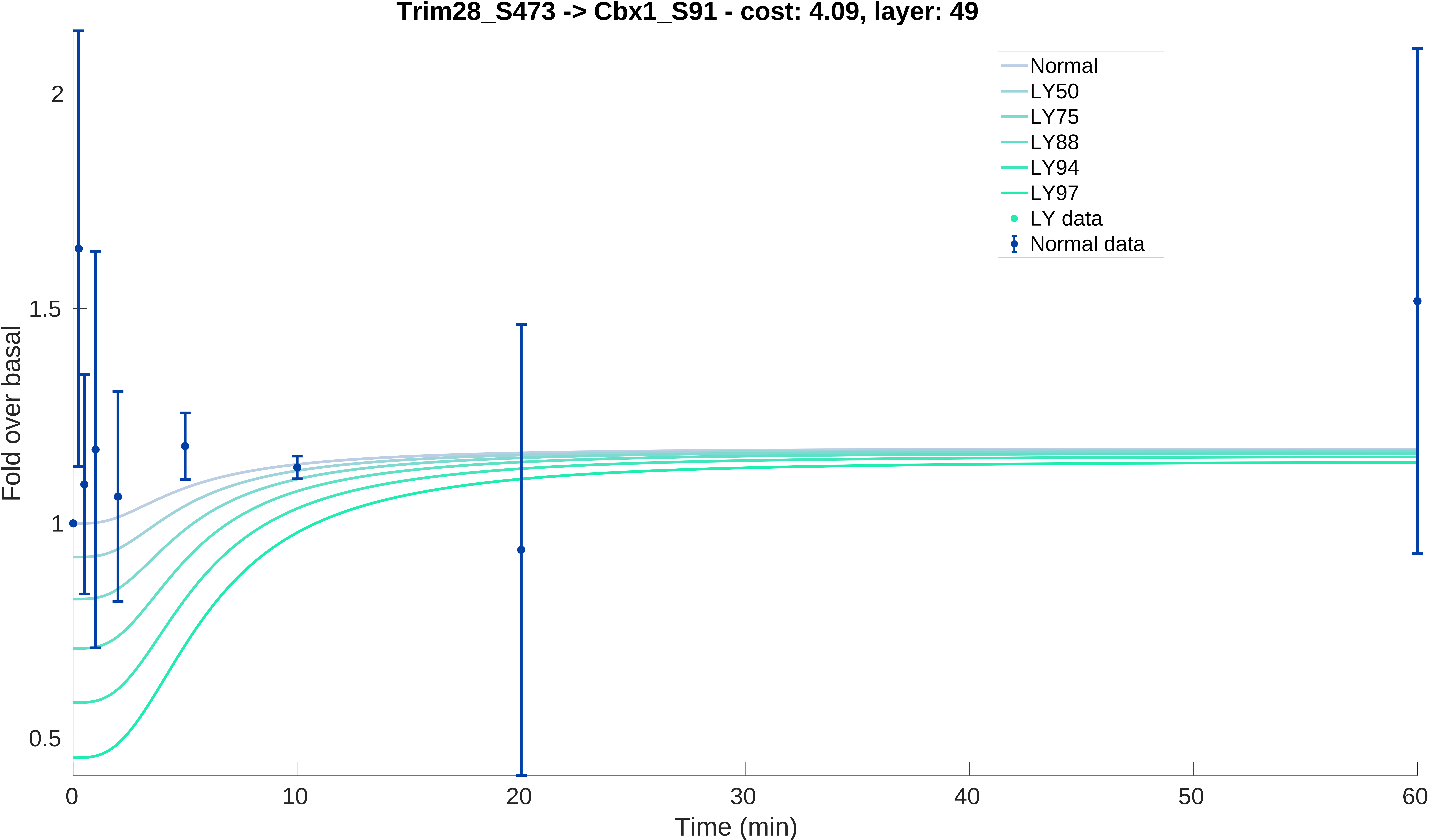

Trim28_S473 -> Cbx1_S91 - cost: 4.09, layer: 49
Normal
LY50
LY75
LY88
LY94
LY97
LY data
Normal data
Fold over basal
2
1.5
1
0.5
0
10
20
30
40
50
60
Time (min)

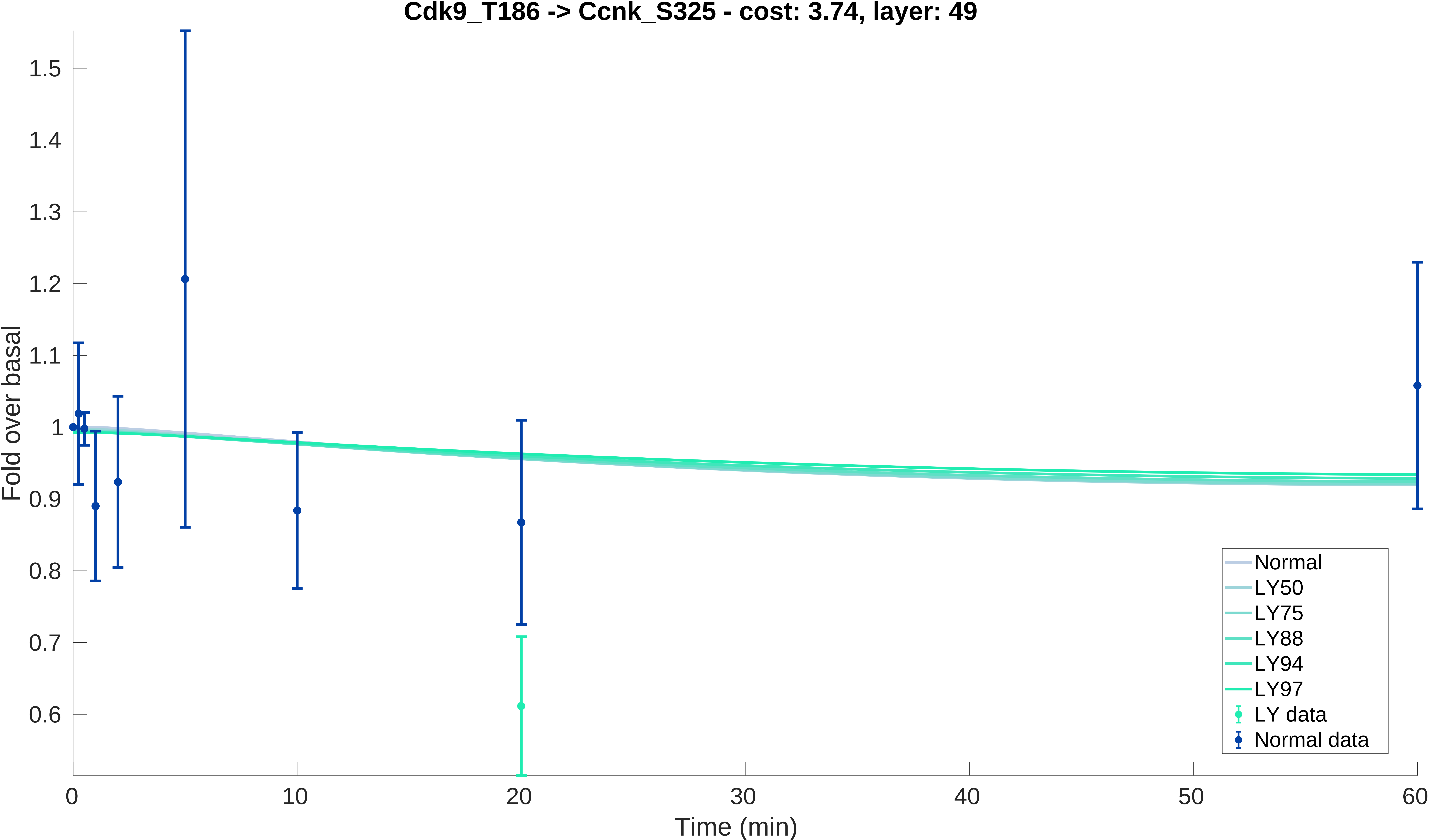

Cdk9_T186 -> Ccnk_S325 - cost: 3.74, layer: 49
Fold over basal
Time (min)
1.5
1.4
1.3
1.2
1.1
1
0.9
0.8
0.7
0.6
0
10
20
30
40
50
60
Normal
LY50
LY75
LY88
LY94
LY97
LY data
Normal data

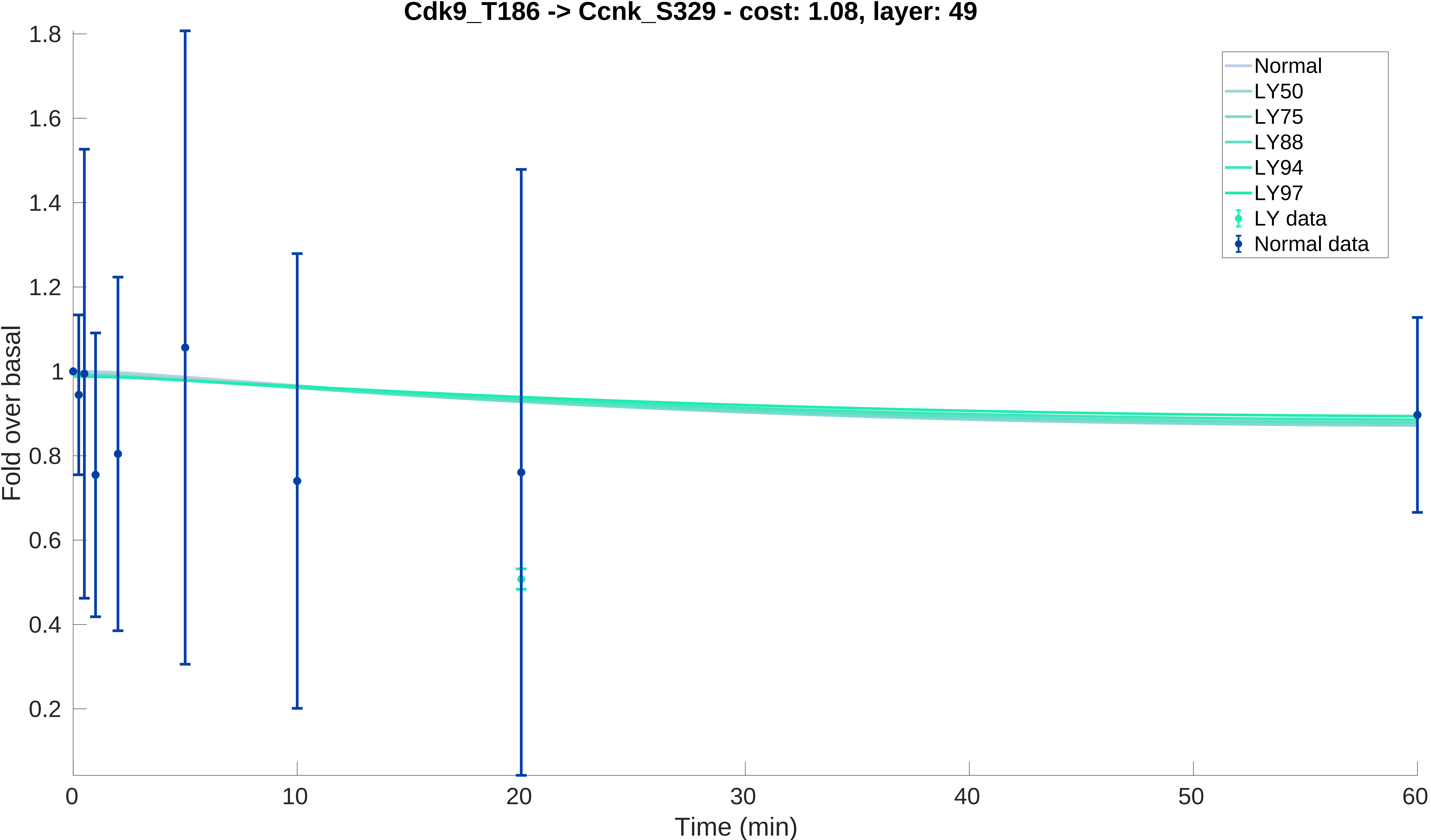

Cdk9_T186 -> Ccnk_S329 - cost: 1.08, layer: 49
Fold over basal
Time (min)
1.8
1.6
1.4
1.2
1
0.8
0.6
0.4
0.2
0
10
20
30
40
50
60
Normal
LY50
LY75
LY88
LY94
LY97
LY data
Normal data

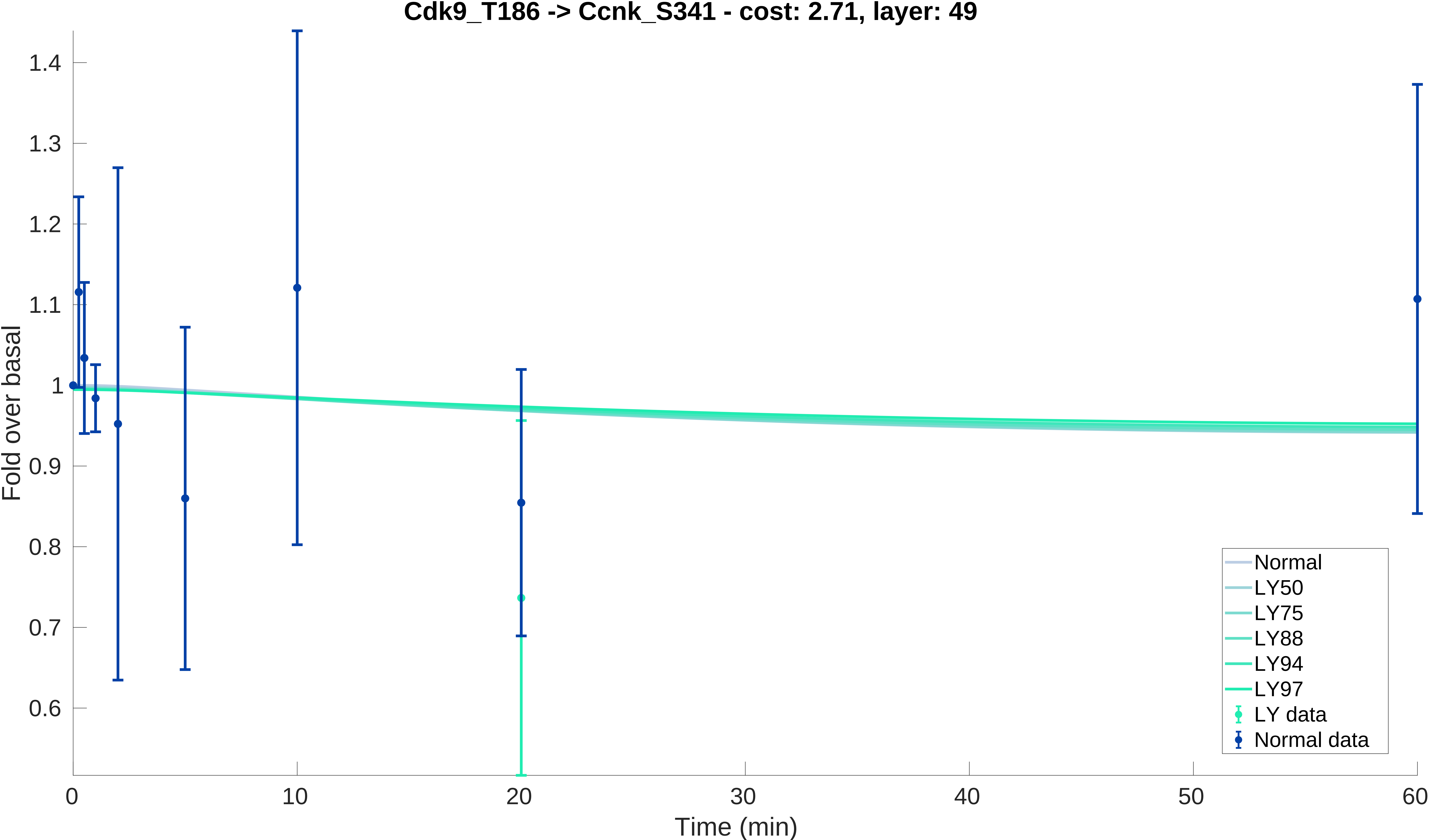

Cdk9_T186 -> Ccnk_S341 - cost: 2.71, layer: 49
Fold over basal
Time (min)
1.4
1.3
1.2
1.1
1
0.9
0.8
0.7
0.6
0
10
20
30
40
50
60
Normal
LY50
LY75
LY88
LY94
LY97
LY data
Normal data

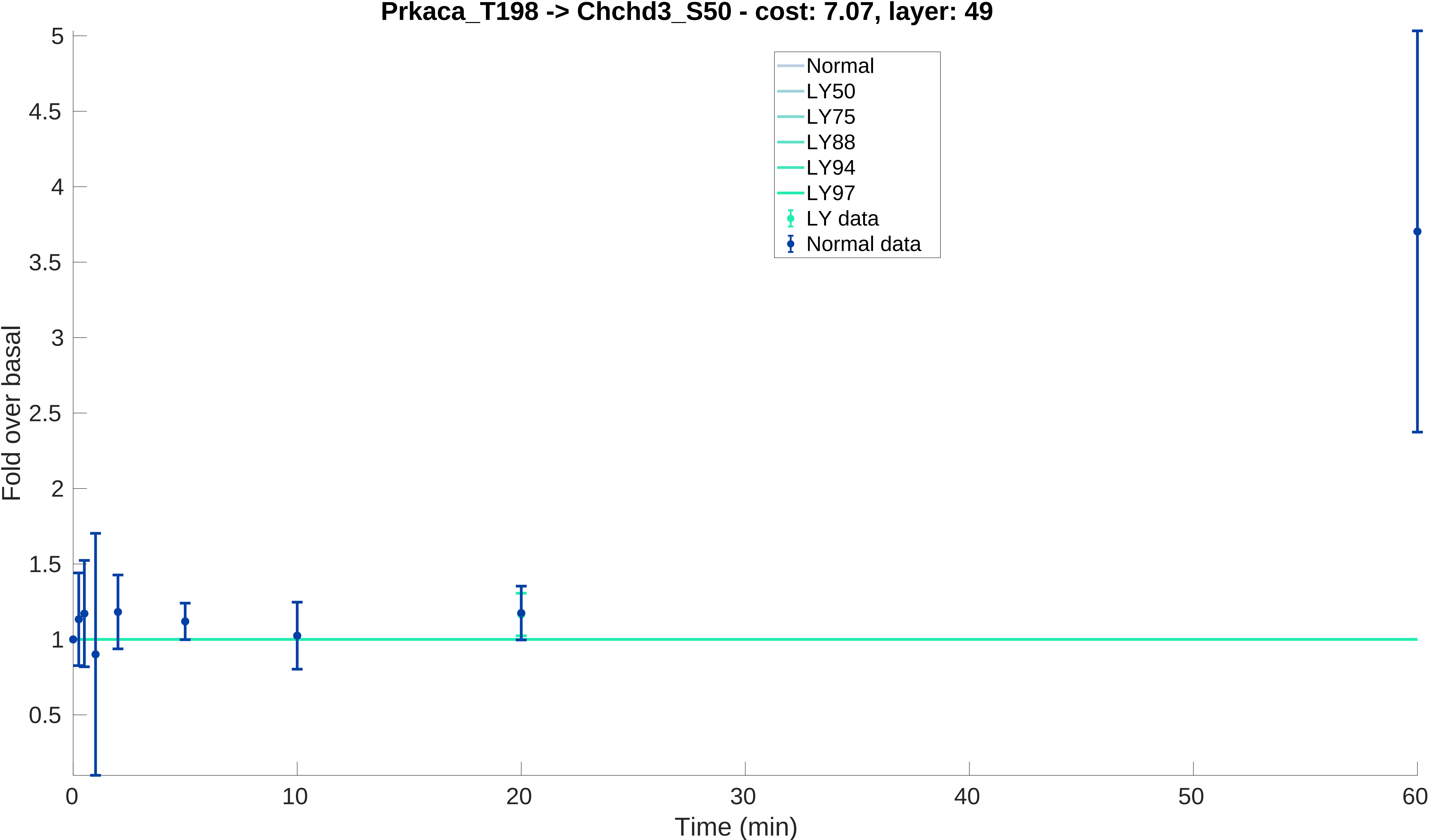

Prkaca_T198 -> Chchd3_S50 - cost: 7.07, layer: 49
Fold over basal
Time (min)
0.5
1
1.5
2
2.5
3
3.5
4
4.5
5
0
10
20
30
40
50
60
Normal
LY50
LY75
LY88
LY94
LY97
LY data
Normal data

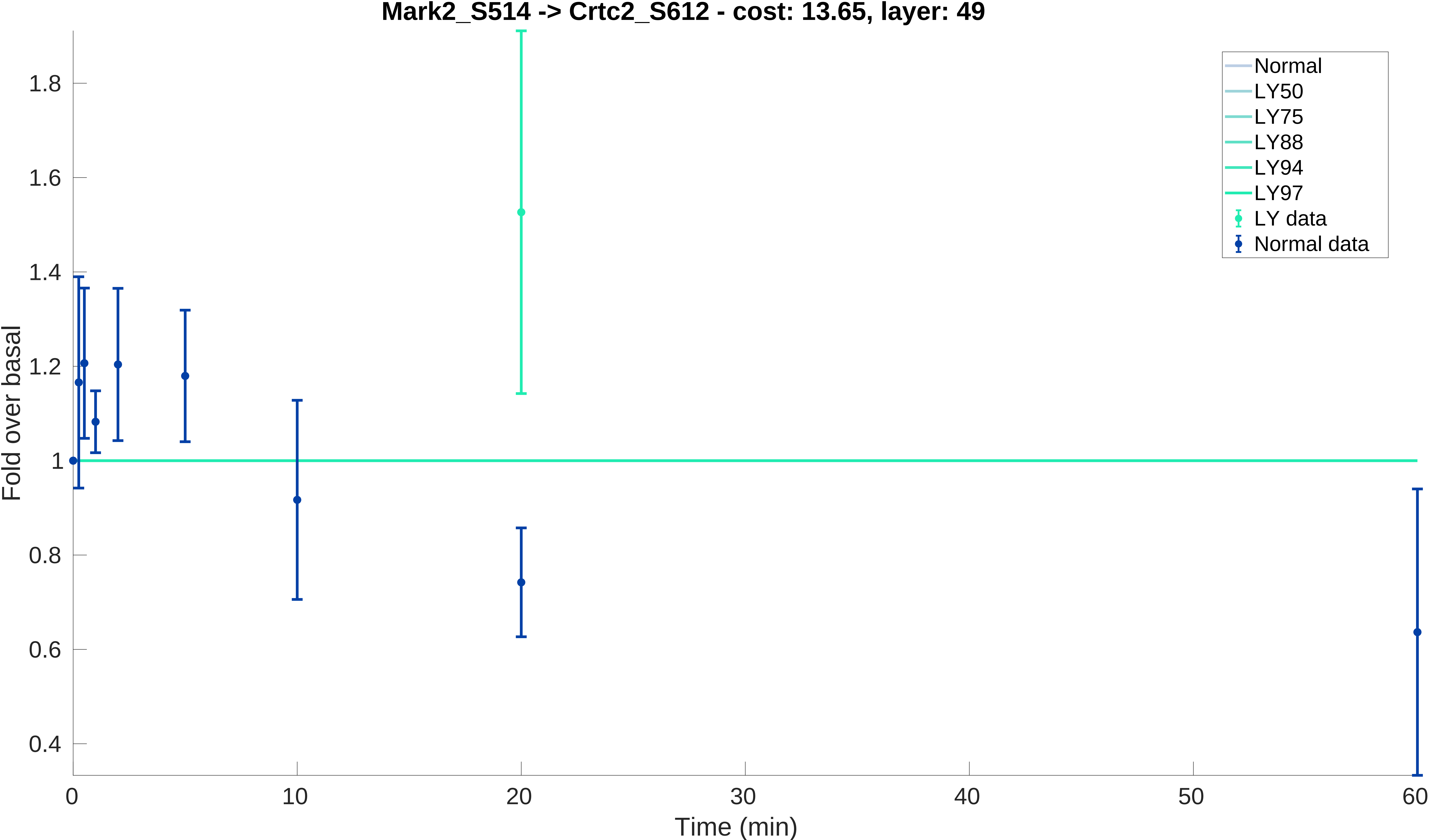

Mark2_S514 -> Crtc2_S612 - cost: 13.65, layer: 49
Normal
LY50
LY75
LY88
LY94
LY97
LY data
Normal data
Fold over basal
1.8
1.6
1.4
1.2
1
0.8
0.6
0.4
0
10
20
30
40
50
60
Time (min)

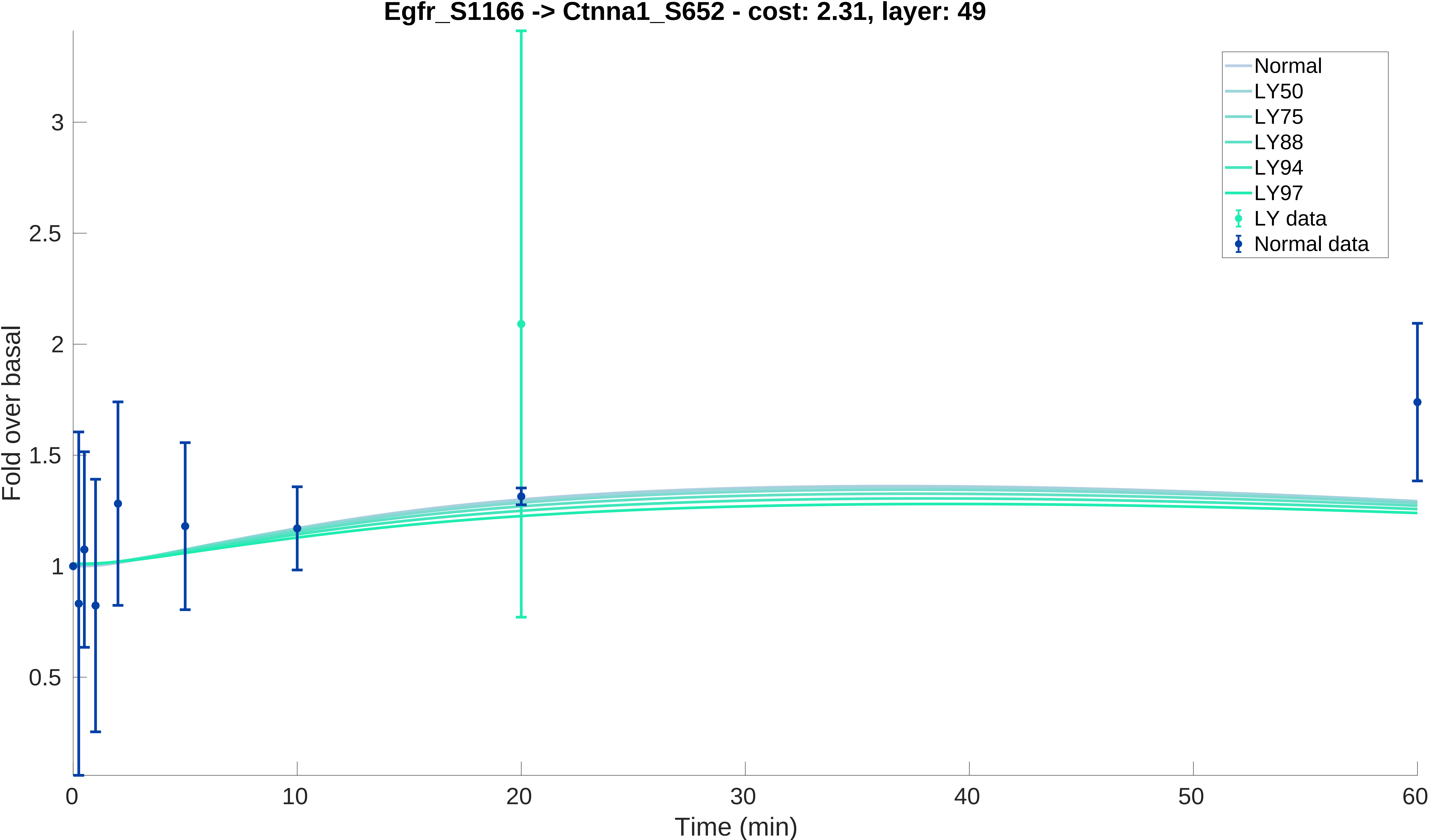
Egfr_S1166 -> Ctnna1_S652 - cost: 2.31, layer: 49
Normal
LY50
LY75
LY88
LY94
LY97
LY data
Normal data
Fold over basal
Time (min)
0.5
1
1.5
2
2.5
3
0
10
20
30
40
50
60

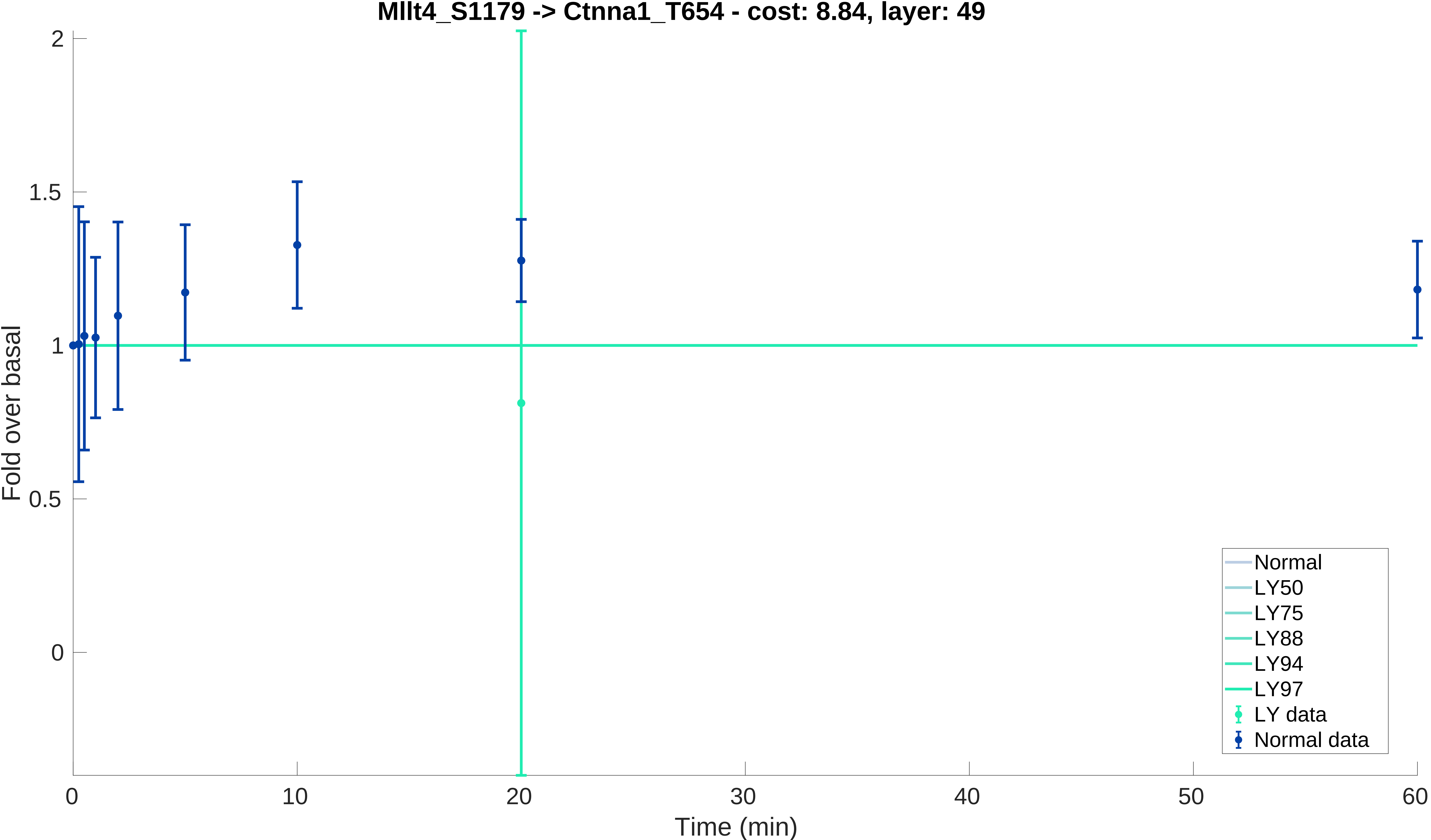

Mllt4_S1179 -> Ctnna1_T654 - cost: 8.84, layer: 49
Fold over basal
Time (min)
0
0.5
1
1.5
2
0
10
20
30
40
50
60
Normal
LY50
LY75
LY88
LY94
LY97
LY data
Normal data

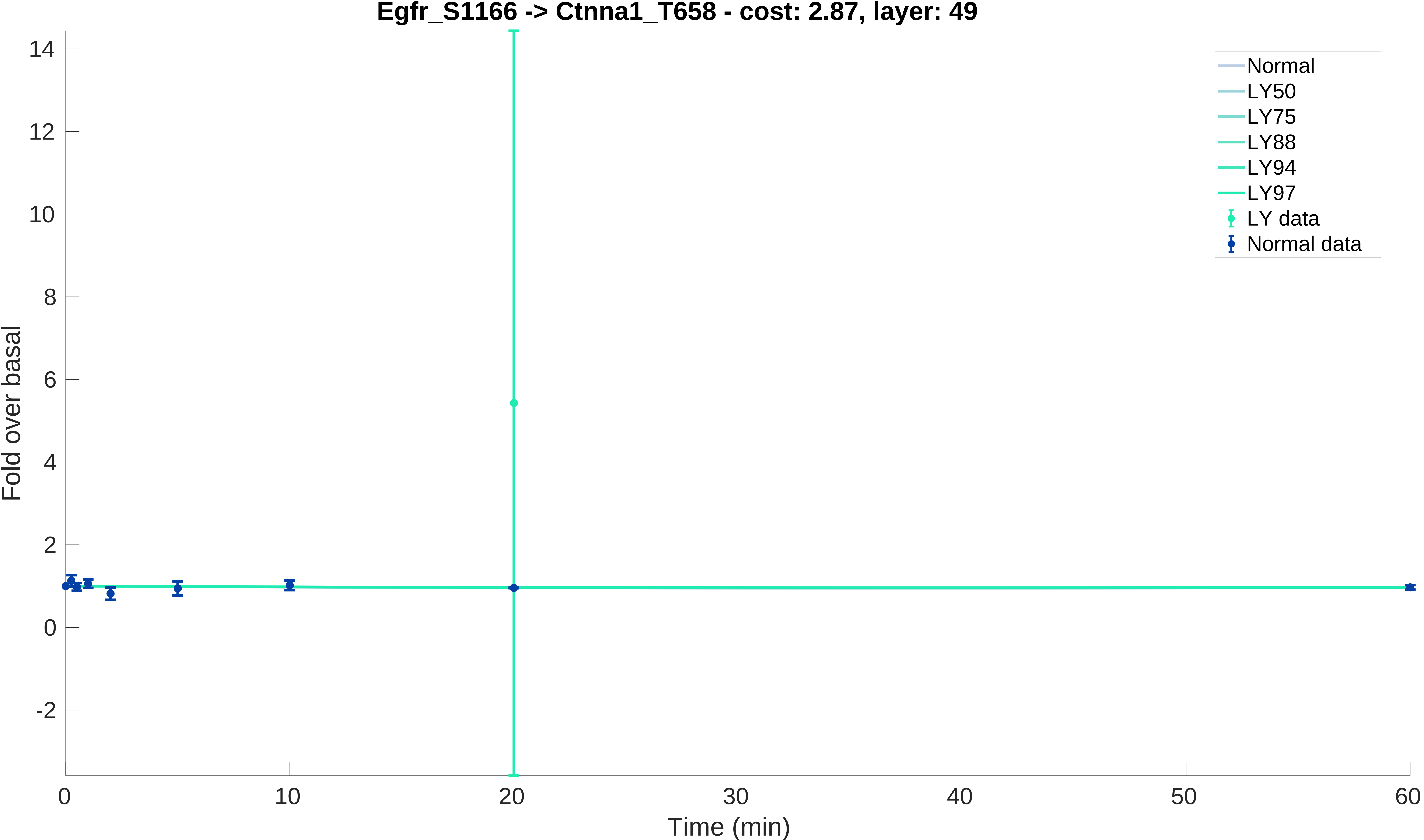

Egfr_S1166 -> Ctnna1_T658 - cost: 2.87, layer: 49
Fold over basal
Time (min)
14
12
10
8
6
4
2
0
-2
0
10
20
30
40
50
60
Normal
LY50
LY75
LY88
LY94
LY97
LY data
Normal data

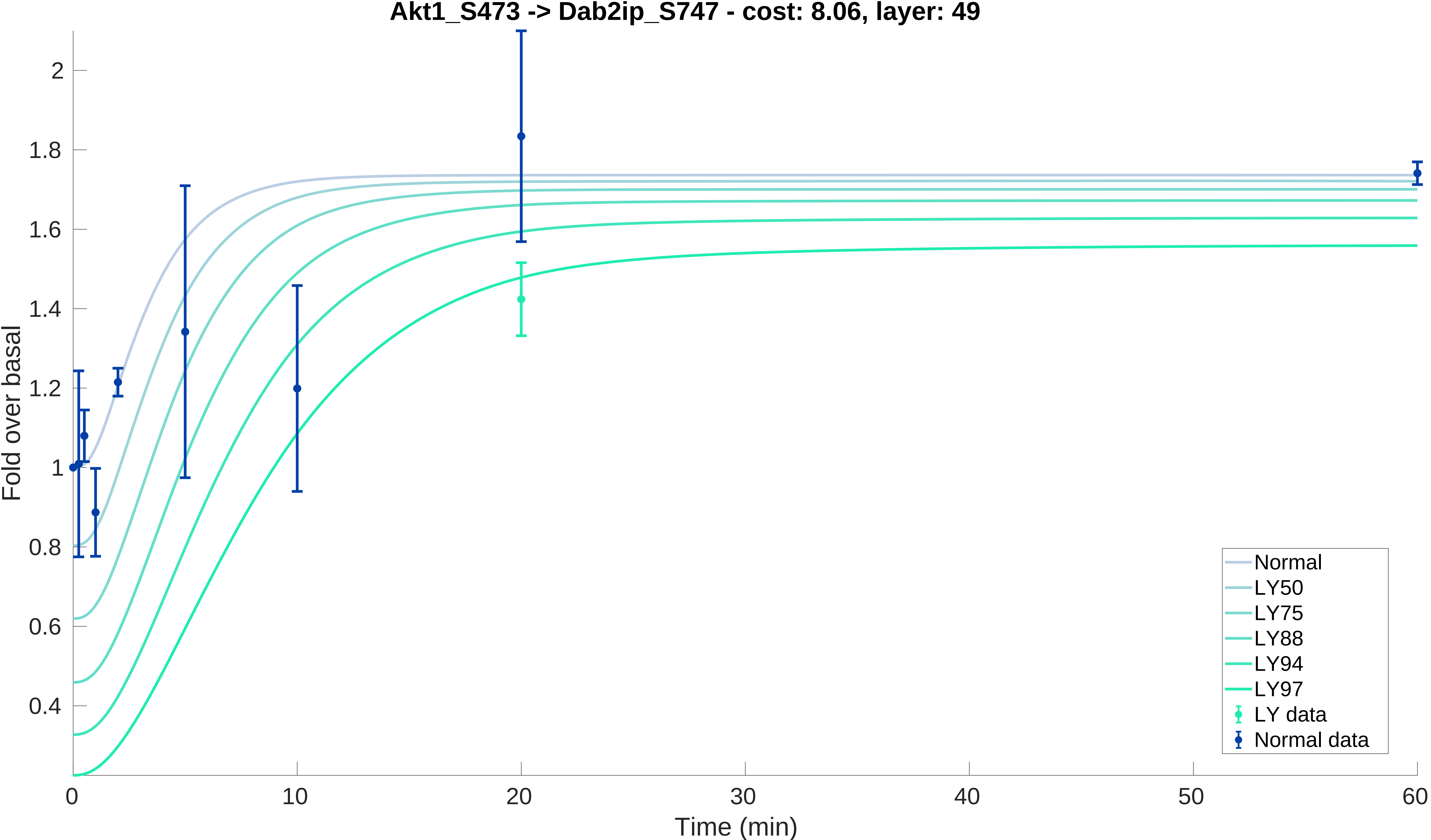

Akt1_S473 -> Dab2ip_S747 - cost: 8.06, layer: 49
Fold over basal
Time (min)
0
10
20
30
40
50
60
0.4
0.6
0.8
1
1.2
1.4
1.6
1.8
2
Normal
LY50
LY75
LY88
LY94
LY97
LY data
Normal data

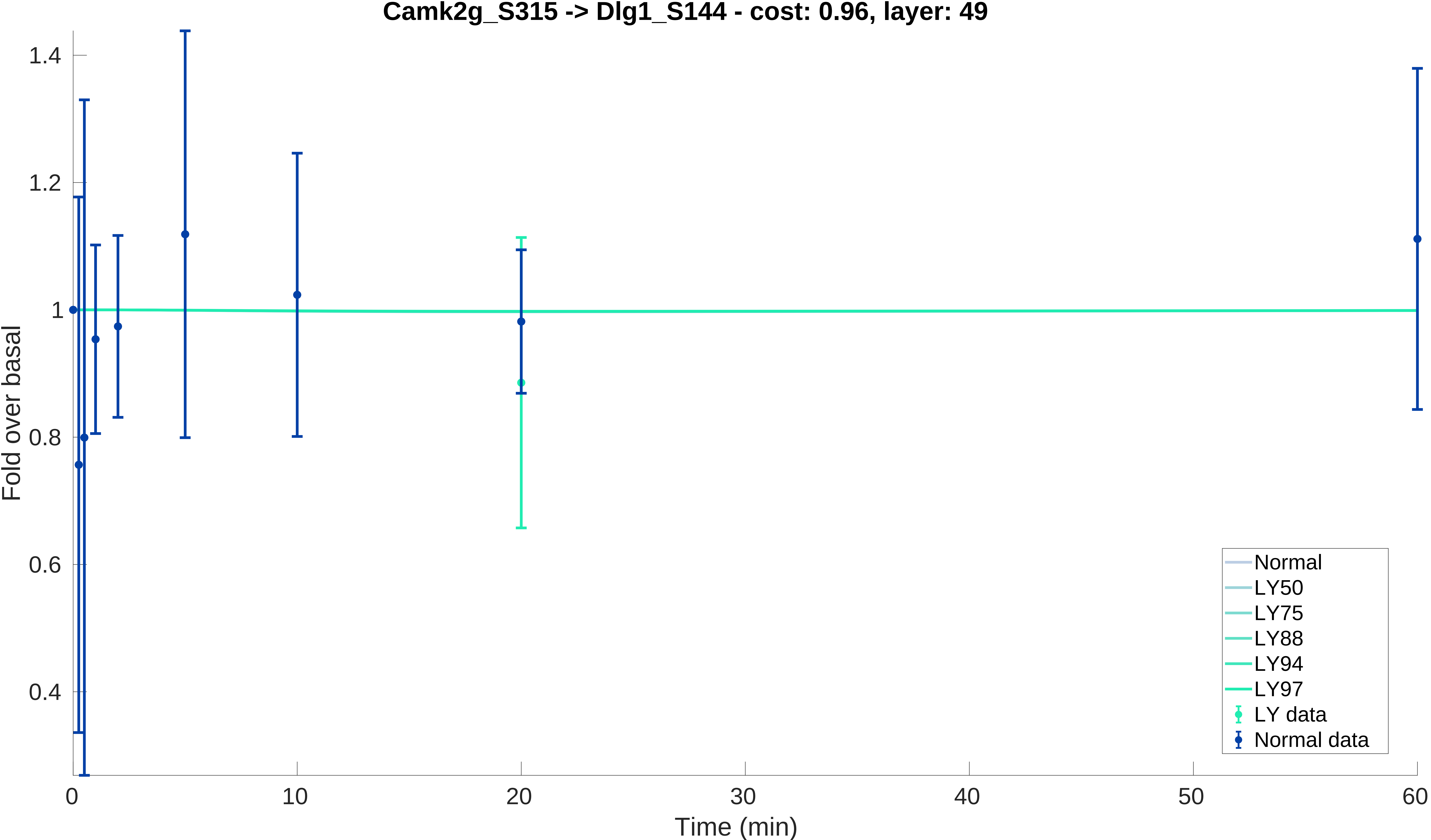

Camk2g_S315 -> Dlg1_S144 - cost: 0.96, layer: 49
Fold over basal
Time (min)
1.4
1.2
1
0.8
0.6
0.4
0
10
20
30
40
50
60
Normal
LY50
LY75
LY88
LY94
LY97
LY data
Normal data

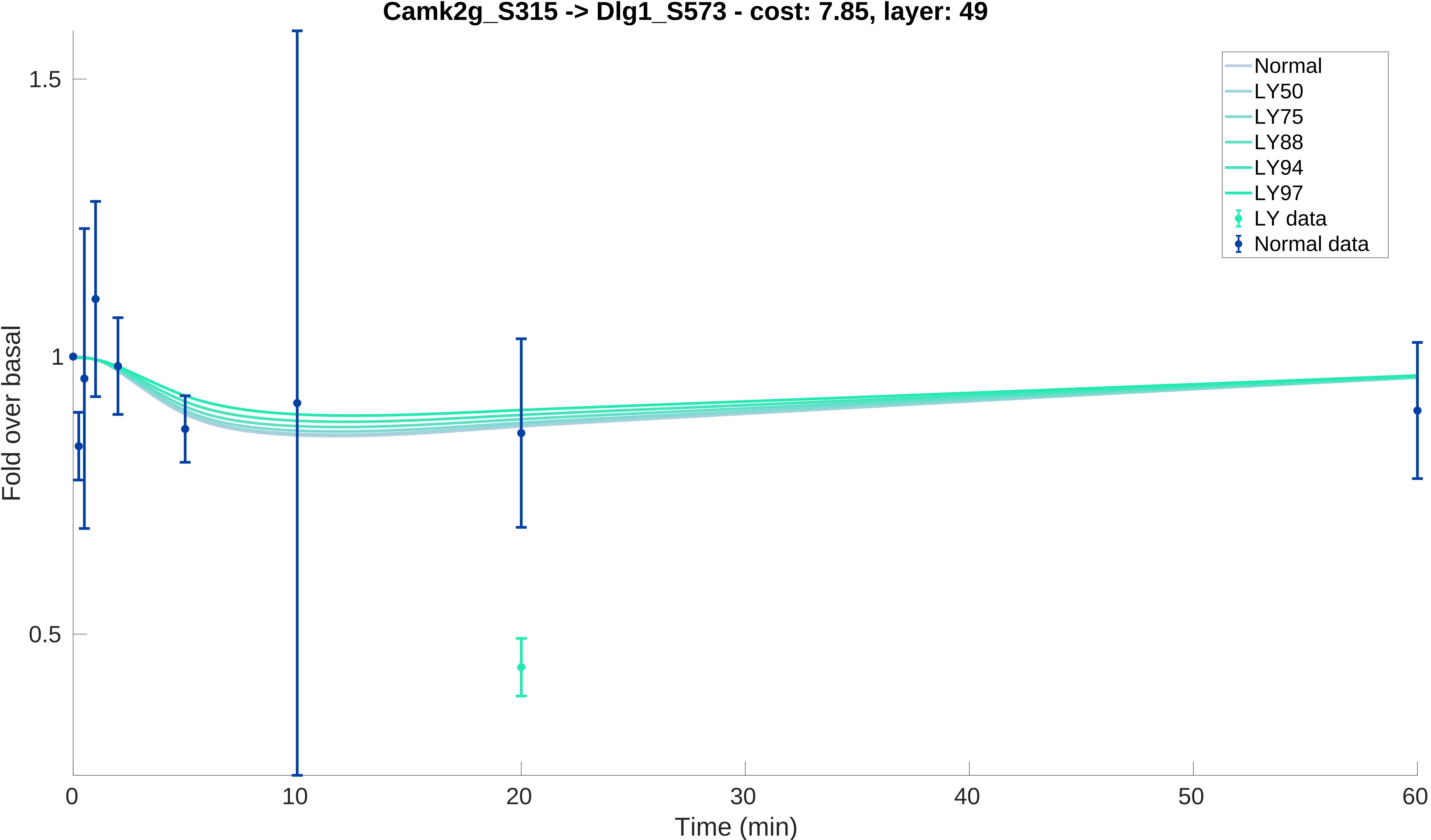
Camk2g_S315 -> Dlg1_S573 - cost: 7.85, layer: 49
Fold over basal
1.5
1
0.5
0
10
20
30
40
50
60
Time (min)
Normal
LY50
LY75
LY88
LY94
LY97
LY data
Normal data

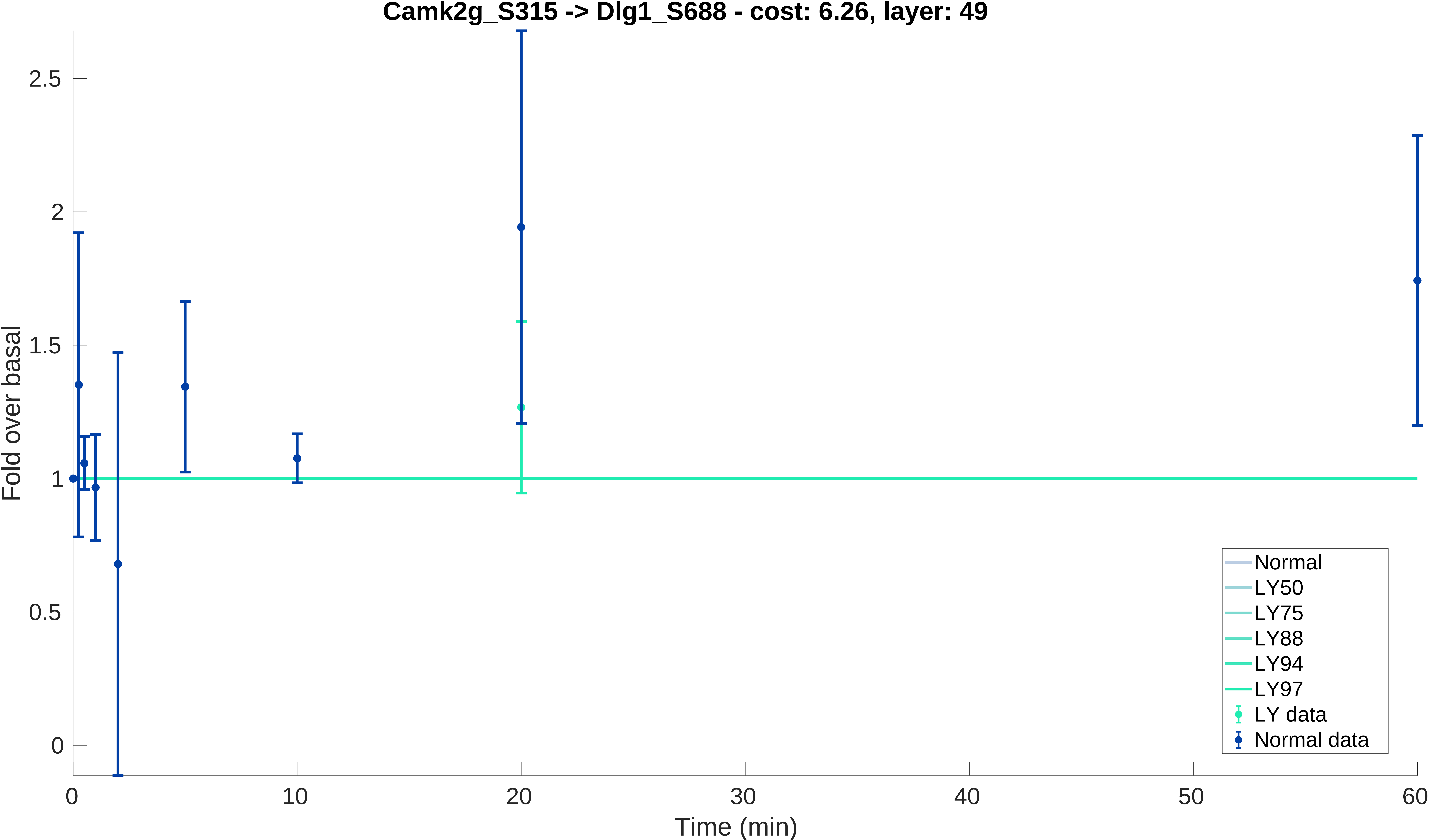
Camk2g_S315 -> Dlg1_S688 - cost: 6.26, layer: 49
Fold over basal
Time (min)
0
0.5
1
1.5
2
2.5
0
10
20
30
40
50
60
Normal
LY50
LY75
LY88
LY94
LY97
LY data
Normal data

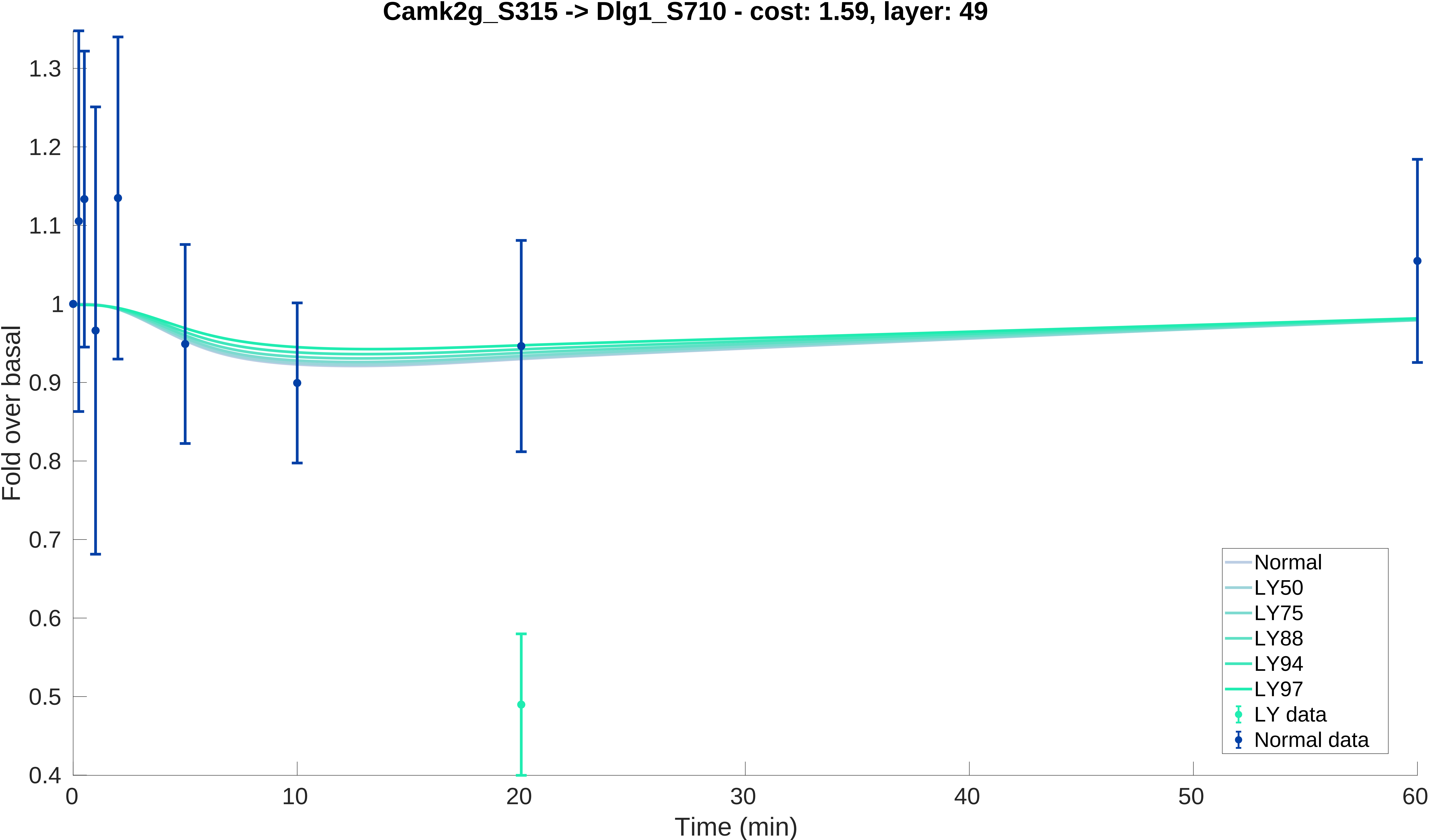

Camk2g_S315 -> Dlg1_S710 - cost: 1.59, layer: 49
1.3
1.2
1.1
1
0.9
0.8
0.7
0.6
0.5
0.4
Fold over basal
0
10
20
30
40
50
60
Time (min)
Normal
LY50
LY75
LY88
LY94
LY97
LY data
Normal data

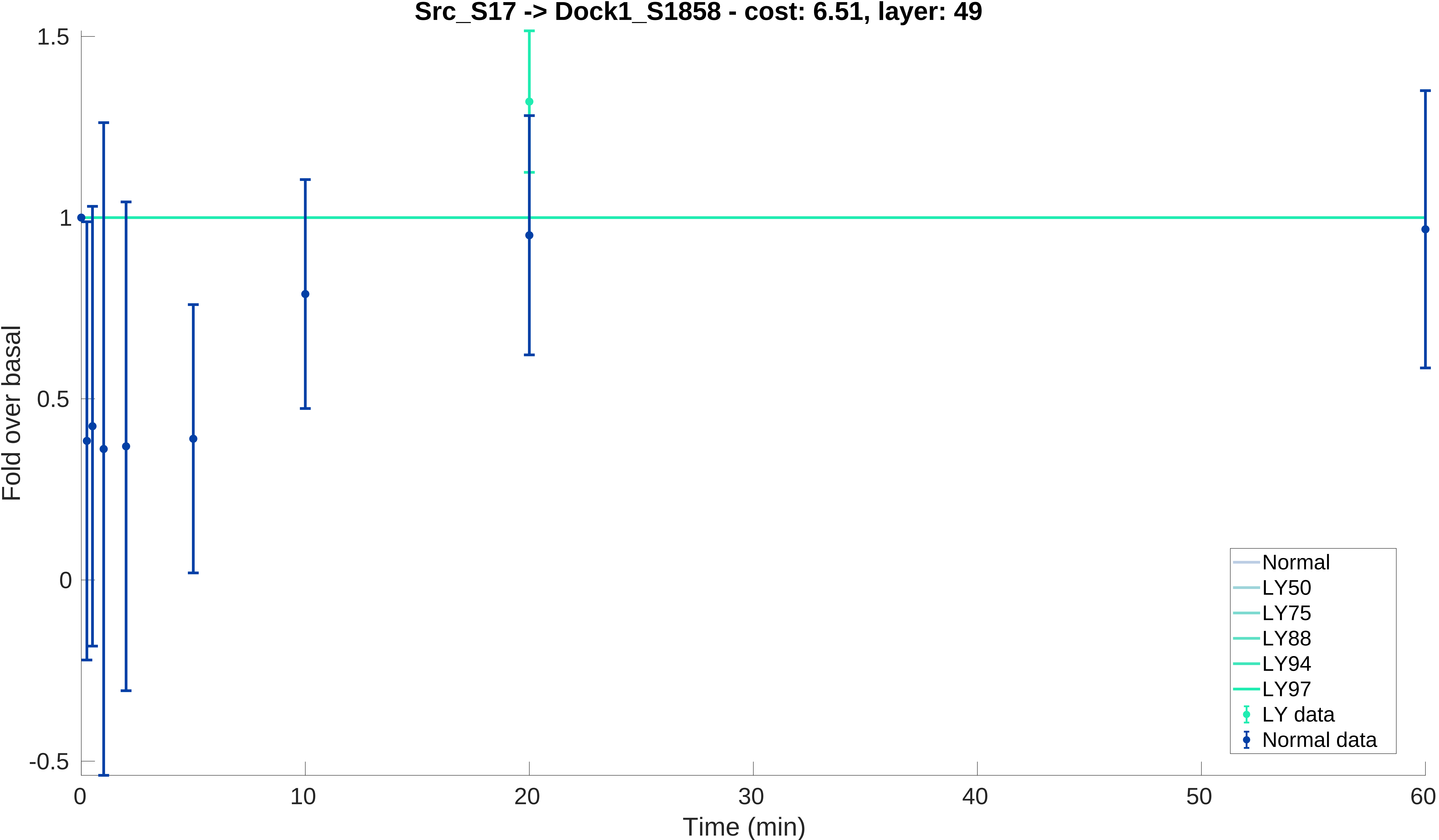

Src_S17 -> Dock1_S1858 - cost: 6.51, layer: 49
1.5
1
0.5
0
-0.5
Fold over basal
0
10
20
30
40
50
60
Time (min)
Normal
LY50
LY75
LY88
LY94
LY97
LY data
Normal data

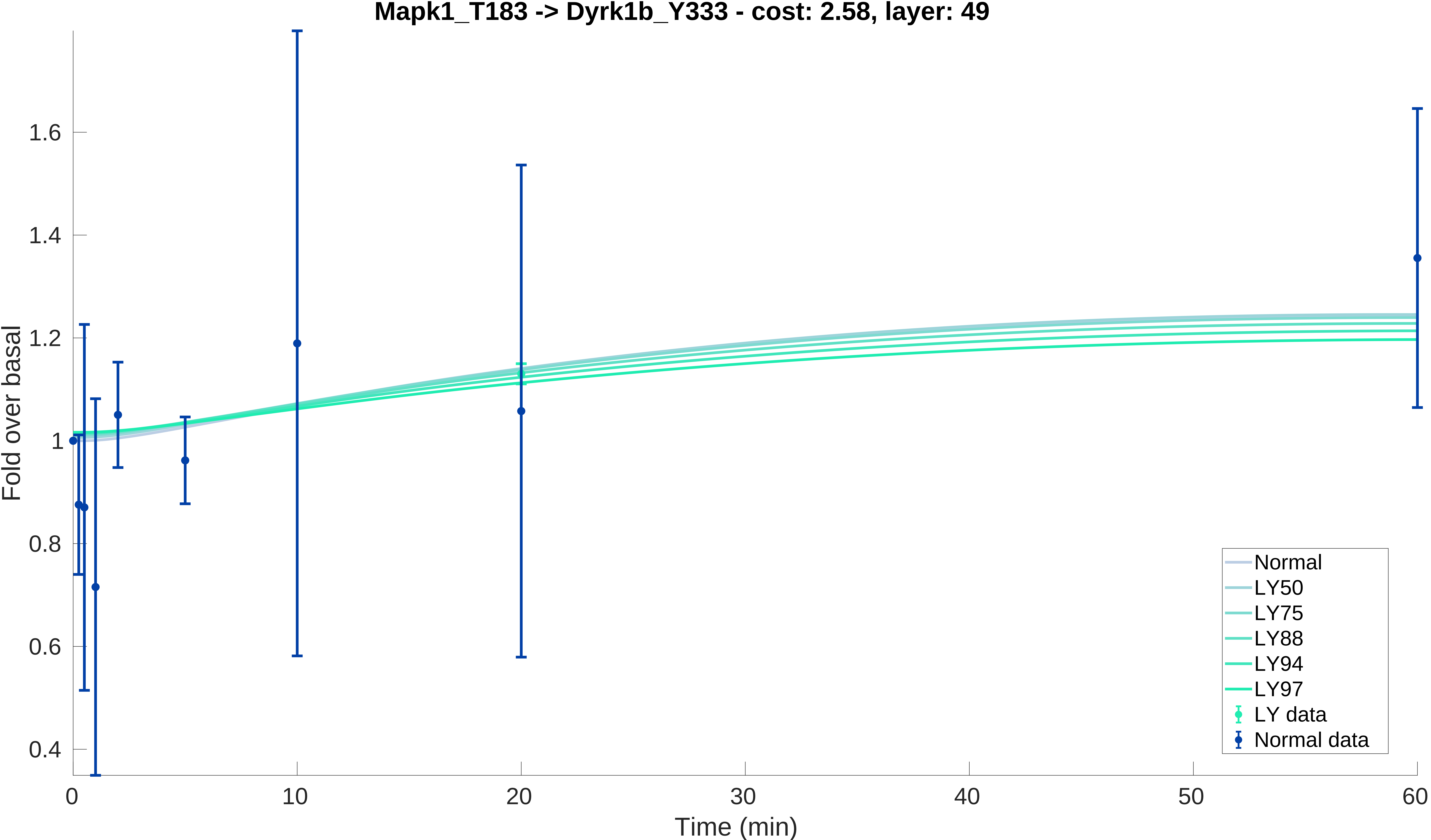

Mapk1_T183 -> Dyrk1b_Y333 - cost: 2.58, layer: 49
Fold over basal
Time (min)
0.4
0.6
0.8
1
1.2
1.4
1.6
0
10
20
30
40
50
60
Normal
LY50
LY75
LY88
LY94
LY97
LY data
Normal data

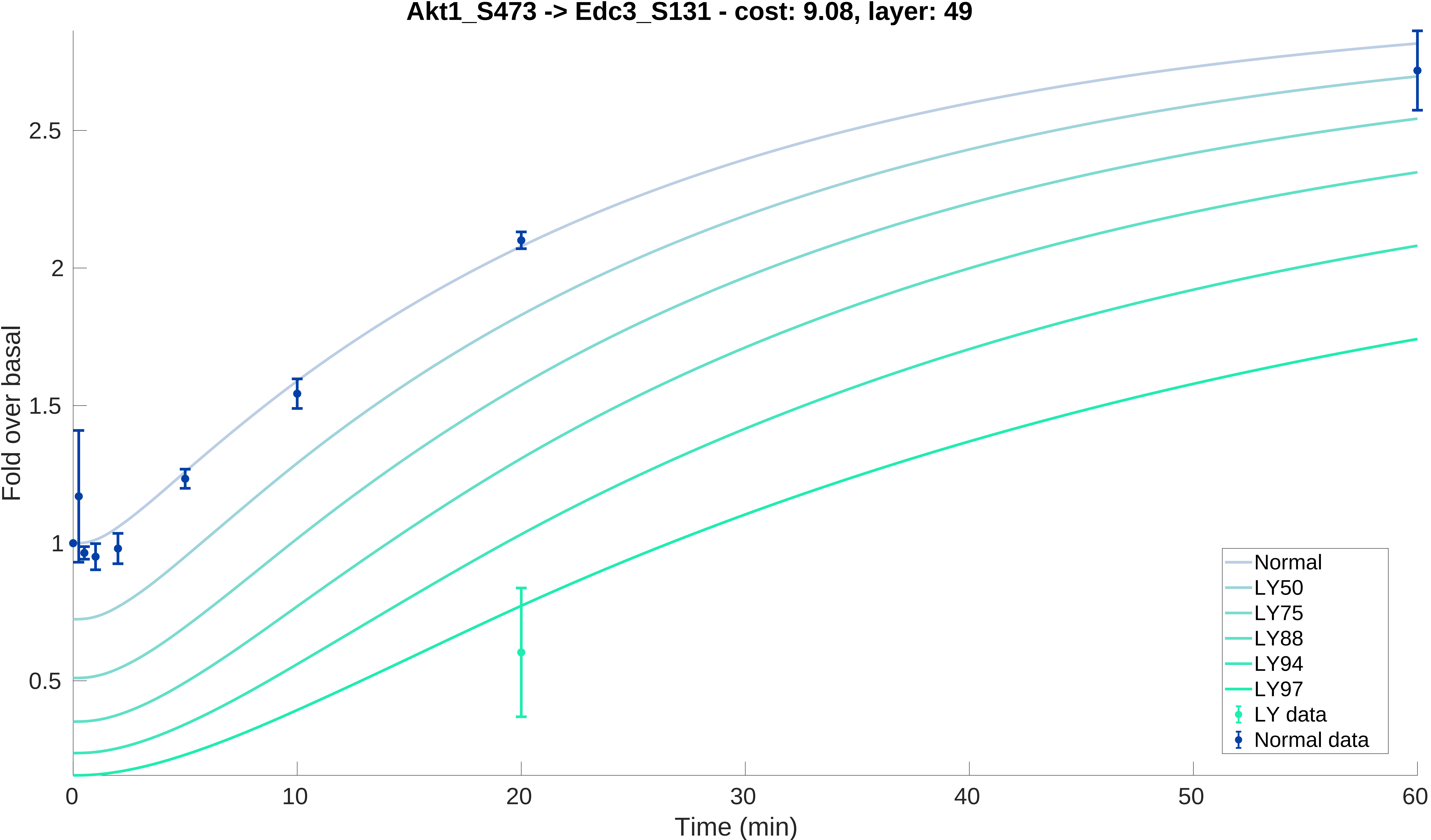
Akt1_S473 -> Edc3_S131 - cost: 9.08, layer: 49
Fold over basal
2.5
2
1.5
1
0.5
0
10
20
30
40
50
60
Time (min)
Normal
LY50
LY75
LY88
LY94
LY97
LY data
Normal data

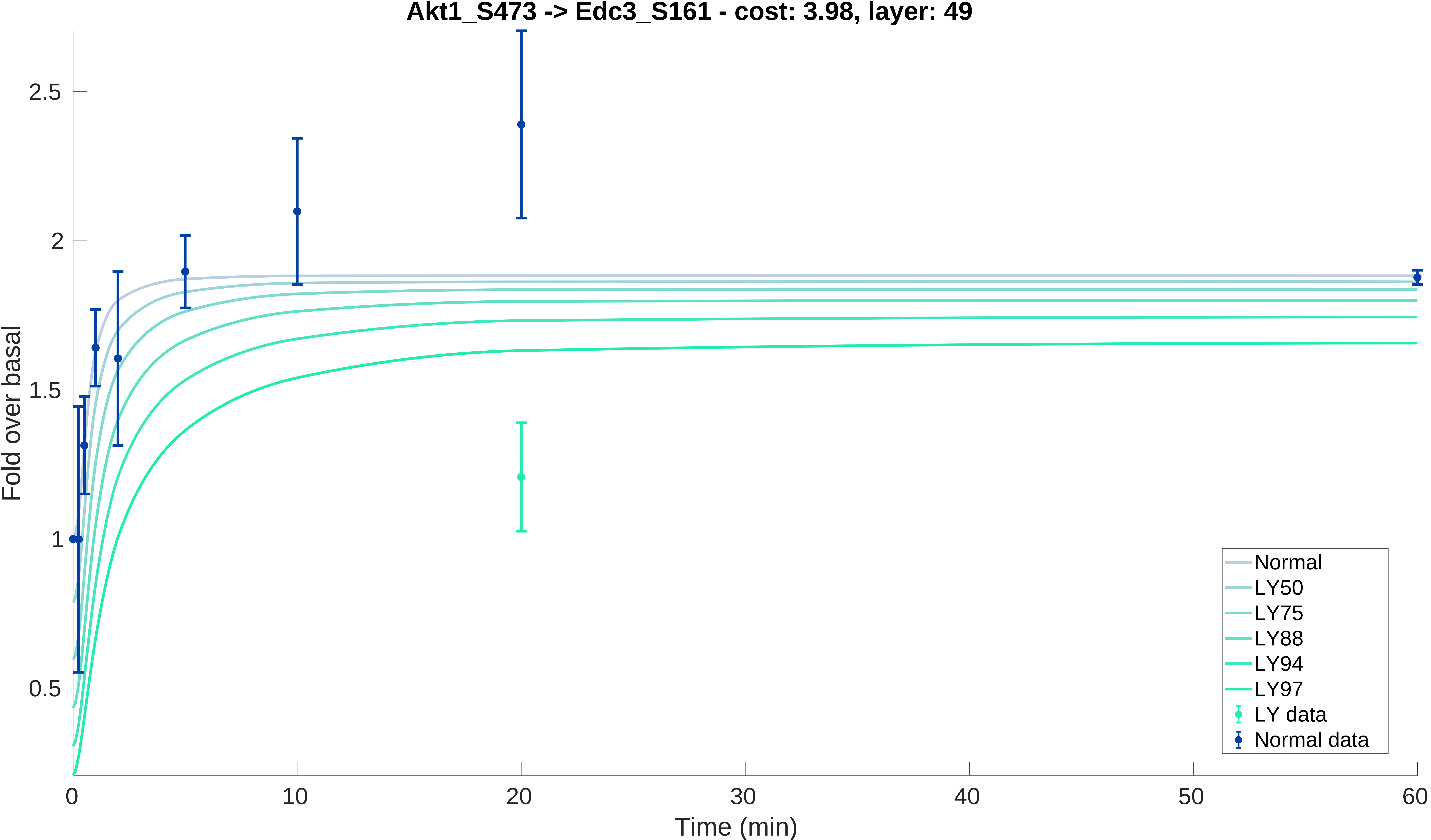

Akt1_S473 -> Edc3_S161 - cost: 3.98, layer: 49
Fold over basal
Time (min)
0.5
1
1.5
2
2.5
0
10
20
30
40
50
60
Normal
LY50
LY75
LY88
LY94
LY97
LY data
Normal data

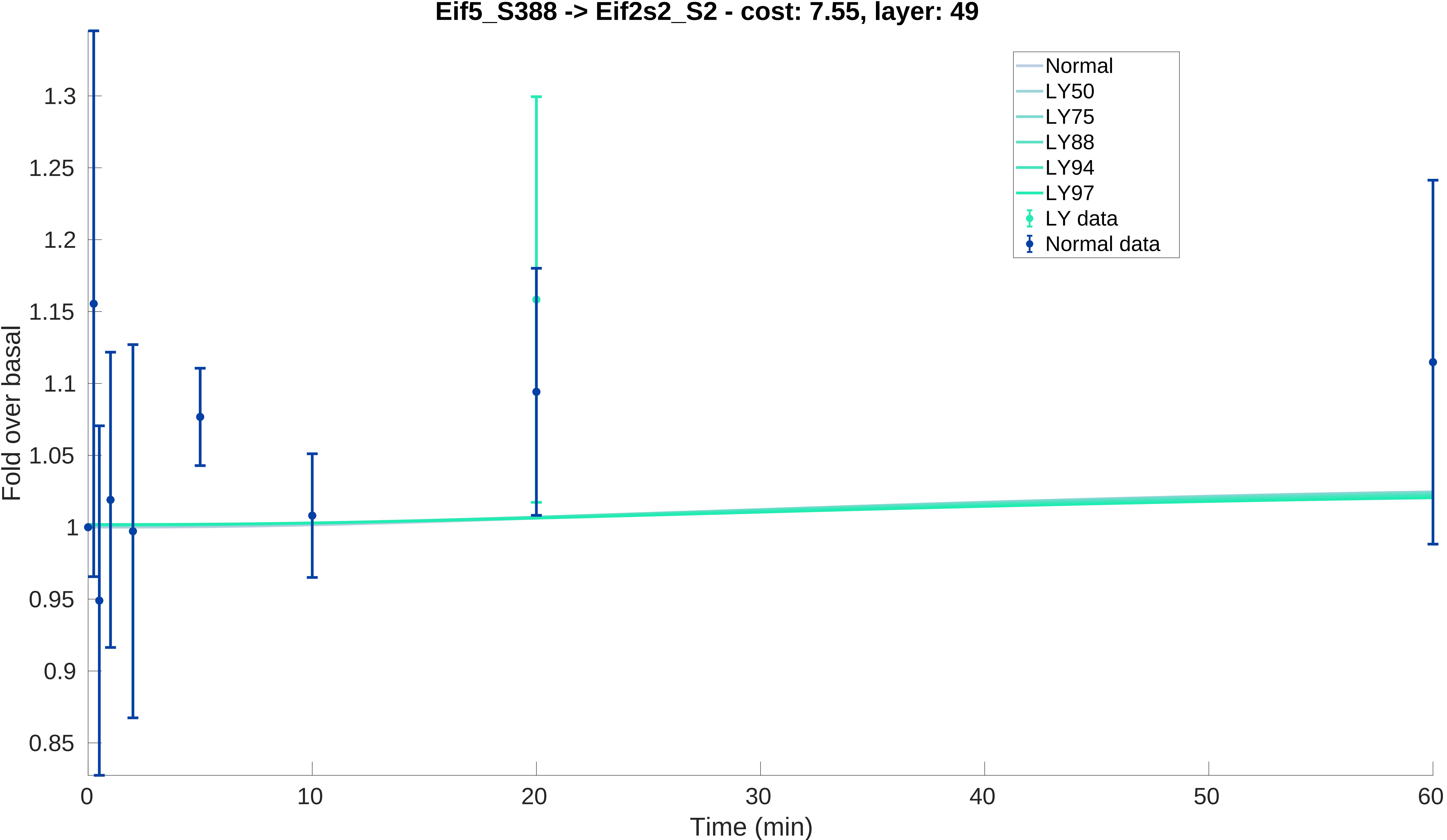

Eif5_S388 -> Eif2s2_S2 - cost: 7.55, layer: 49
Normal
LY50
LY75
LY88
LY94
LY97
LY data
Normal data
Fold over basal
1.3
1.25
1.2
1.15
1.1
1.05
1
0.95
0.9
0.85
0
10
20
30
40
50
60
Time (min)

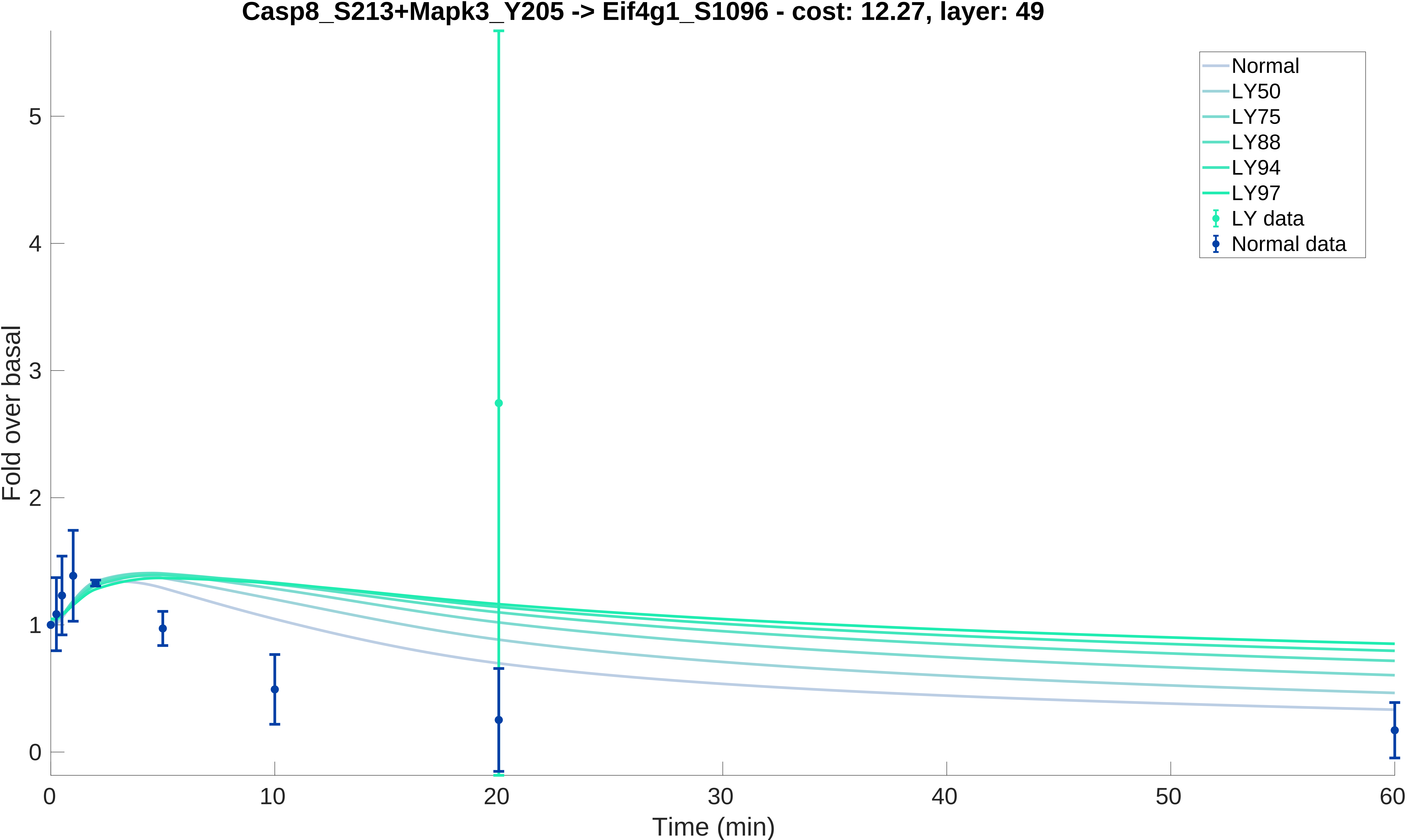

Casp8_S213+Mapk3_Y205 -> Eif4g1_S1096 - cost: 12.27, layer: 49
Fold over basal
Time (min)
0
1
2
3
4
5
0
10
20
30
40
50
60
Normal
LY50
LY75
LY88
LY94
LY97
LY data
Normal data

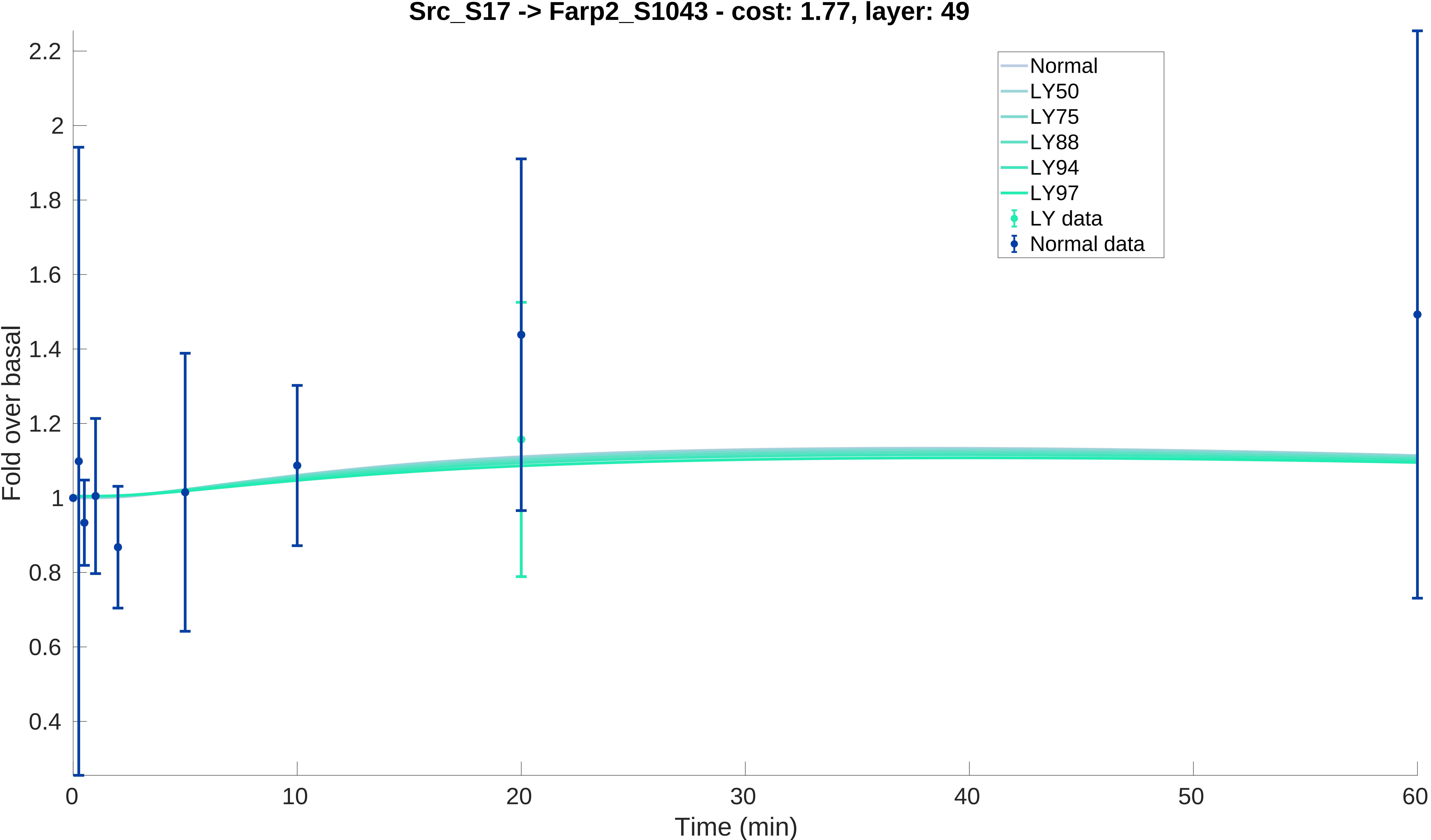

Src_S17 -> Farp2_S1043 - cost: 1.77, layer: 49
Normal
LY50
LY75
LY88
LY94
LY97
LY data
Normal data
Fold over basal
2.2
2
1.8
1.6
1.4
1.2
1
0.8
0.6
0.4
0
10
20
30
40
50
60
Time (min)

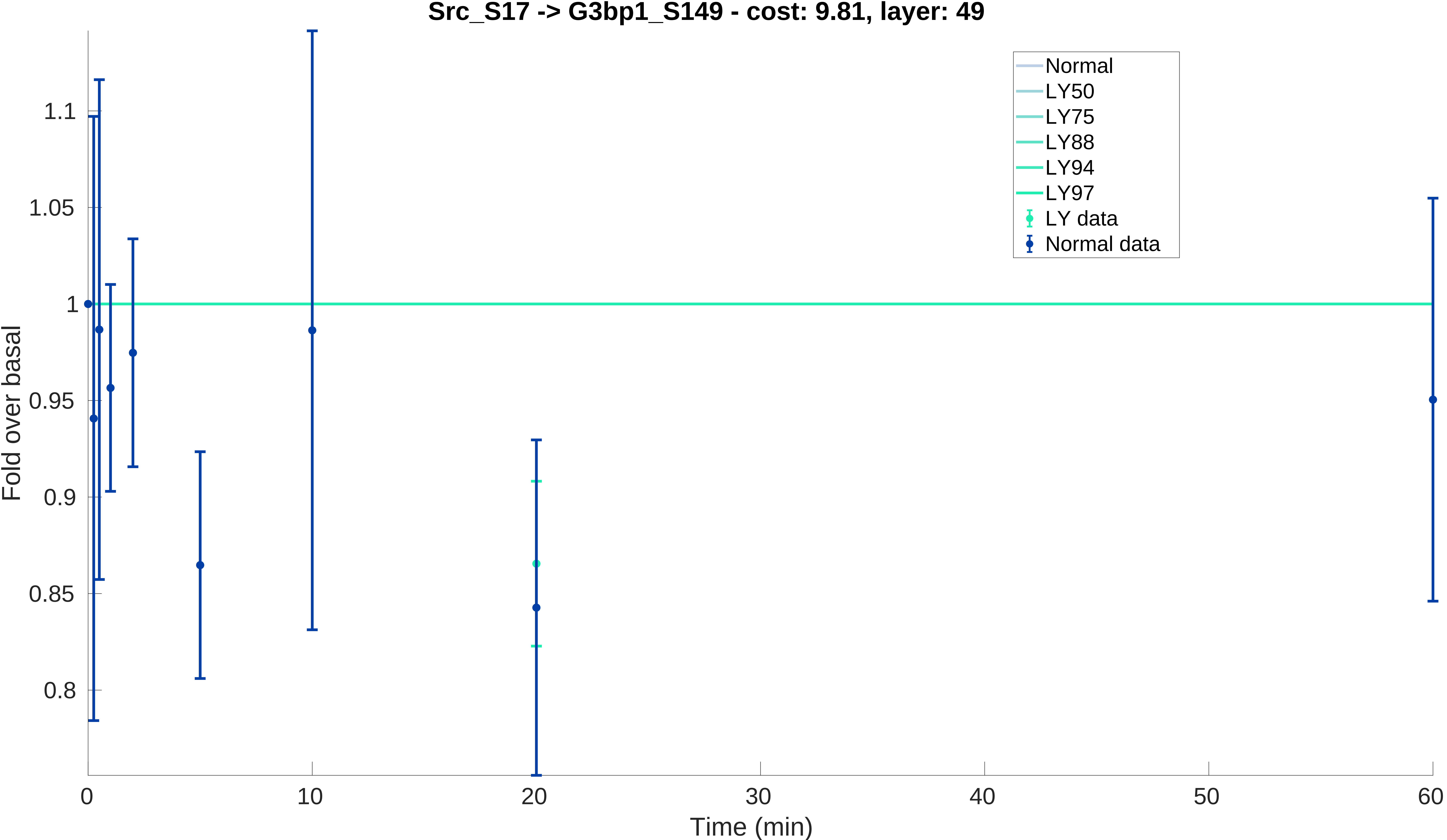

Src_S17 -> G3bp1_S149 - cost: 9.81, layer: 49
Normal
LY50
LY75
LY88
LY94
LY97
LY data
Normal data
Fold over basal
1.1
1.05
1
0.95
0.9
0.85
0.8
0
10
20
30
40
50
60
Time (min)

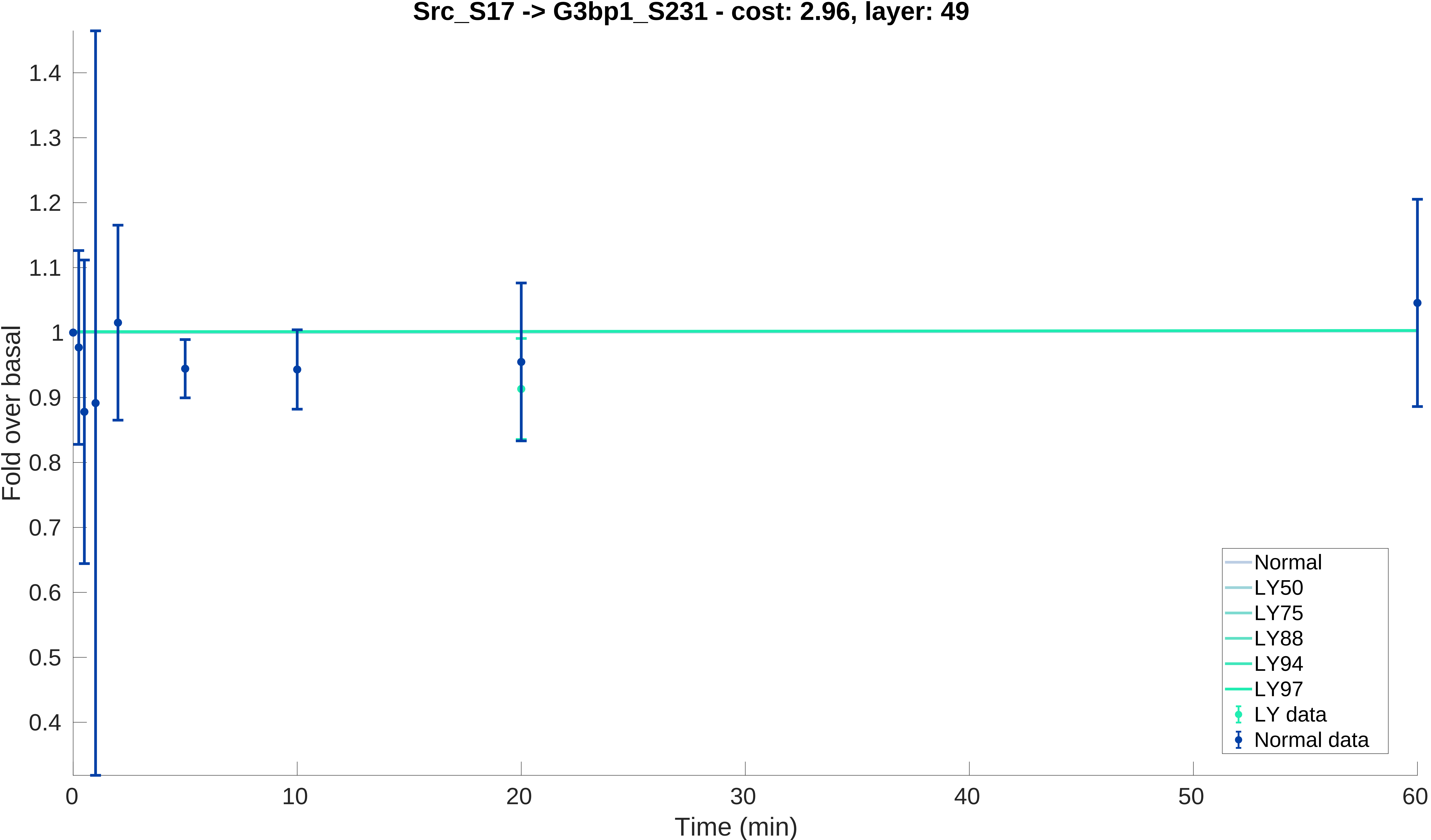

Src_S17 -> G3bp1_S231 - cost: 2.96, layer: 49
Fold over basal
Time (min)
1.4
1.3
1.2
1.1
1
0.9
0.8
0.7
0.6
0.5
0.4
0
10
20
30
40
50
60
Normal
LY50
LY75
LY88
LY94
LY97
LY data
Normal data

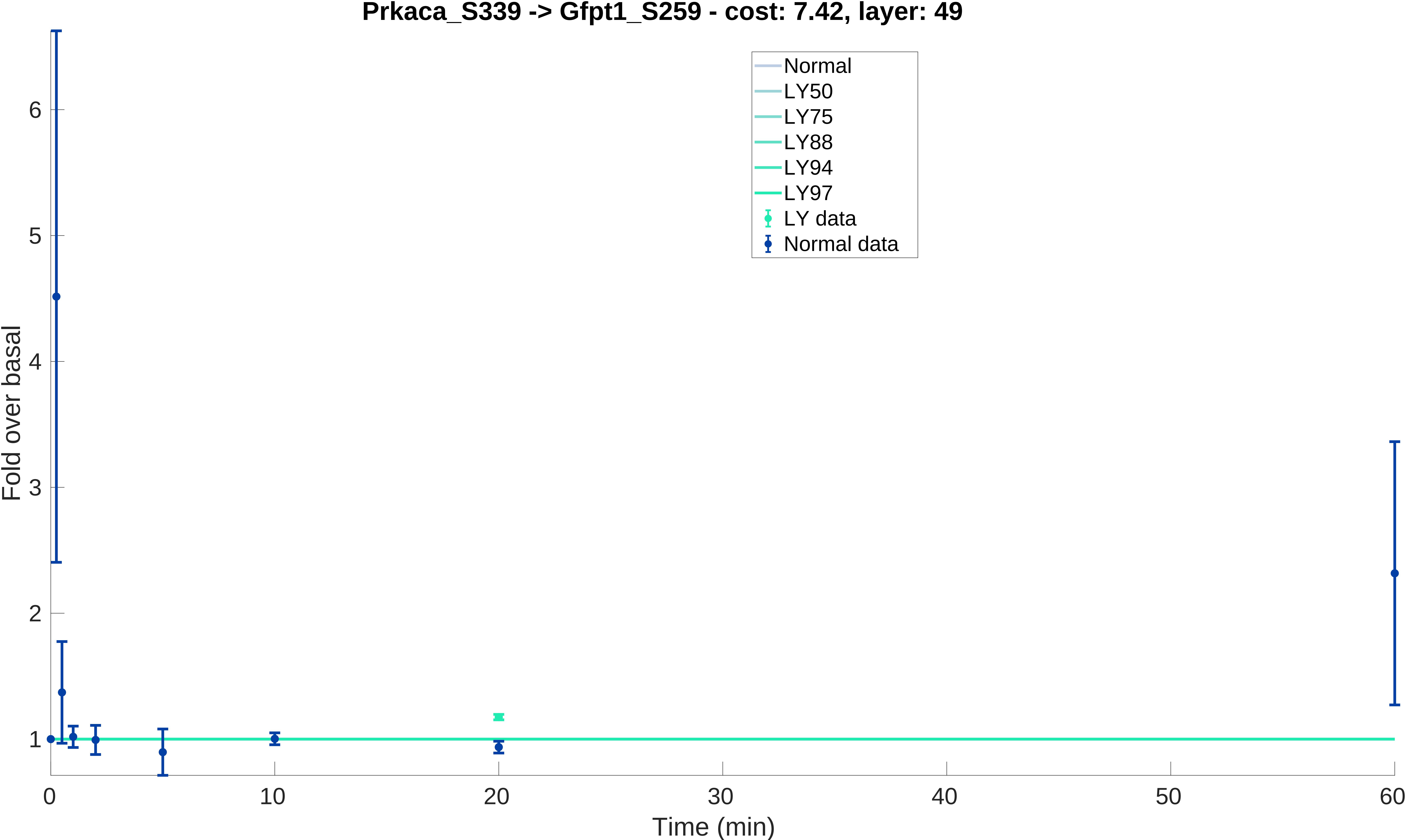
Prkaca_S339 -> Gfpt1_S259 - cost: 7.42, layer: 49
Normal
LY50
LY75
LY88
LY94
LY97
LY data
Normal data
Fold over basal
Time (min)
0
10
20
30
40
50
60
1
2
3
4
5
6

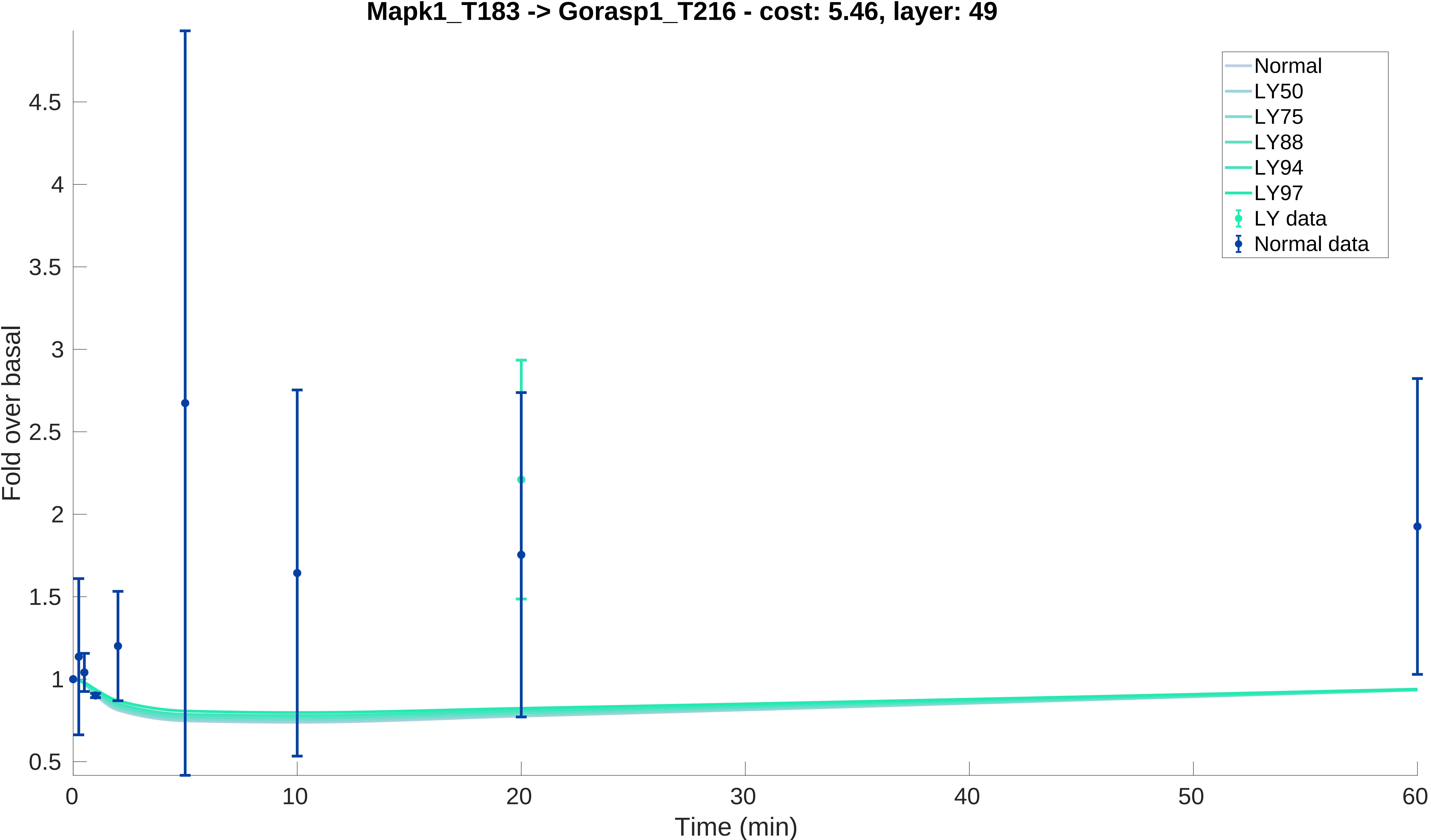
Mapk1_T183 -> Gorasp1_T216 - cost: 5.46, layer: 49
Fold over basal
Time (min)
0.5
1
1.5
2
2.5
3
3.5
4
4.5
0
10
20
30
40
50
60
Normal
LY50
LY75
LY88
LY94
LY97
LY data
Normal data

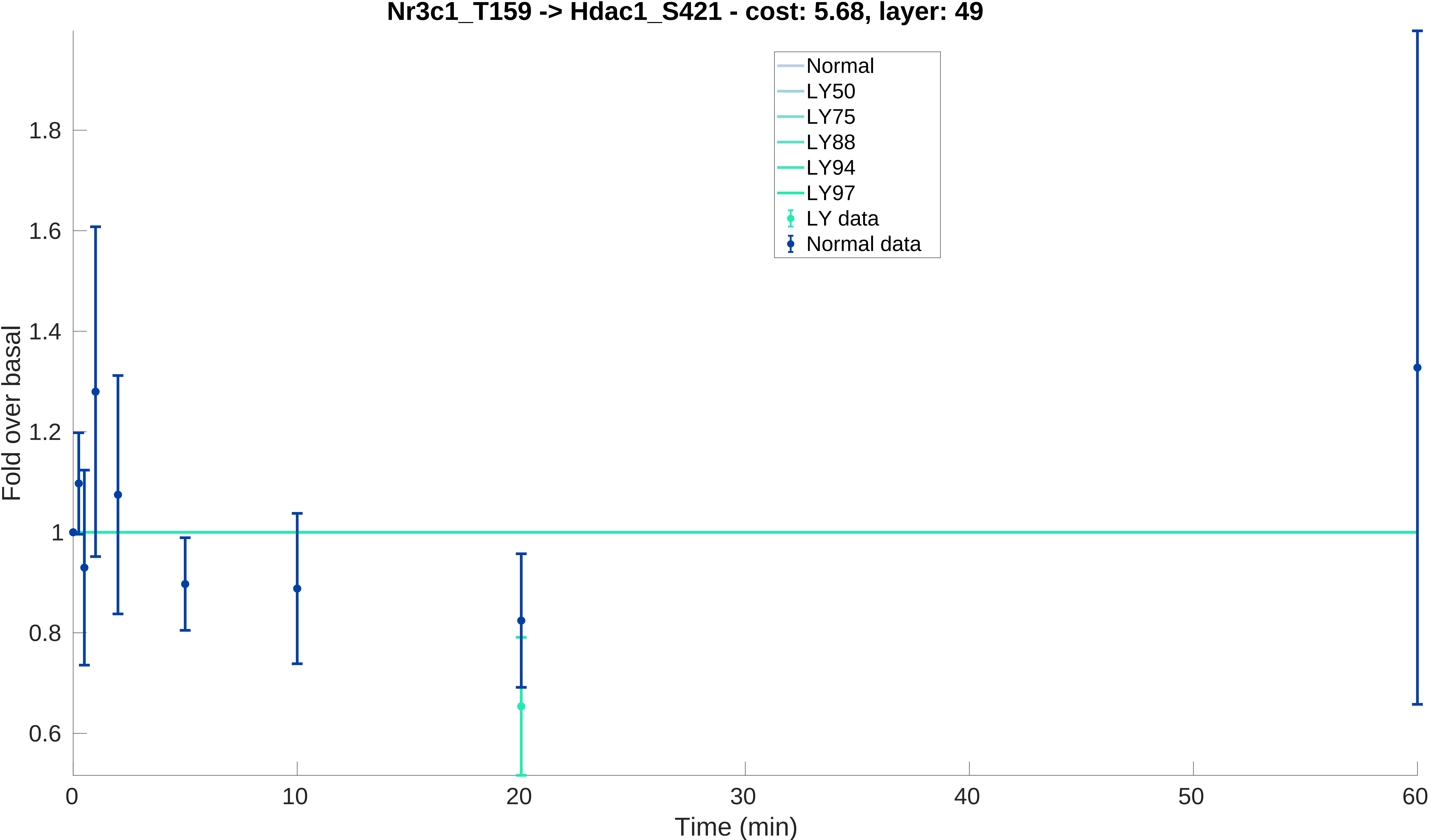
Nr3c1_T159 -> Hdac1_S421 - cost: 5.68, layer: 49
Normal
LY50
LY75
LY88
LY94
LY97
LY data
Normal data
Fold over basal
1.8
1.6
1.4
1.2
1
0.8
0.6
0
10
20
30
40
50
60
Time (min)

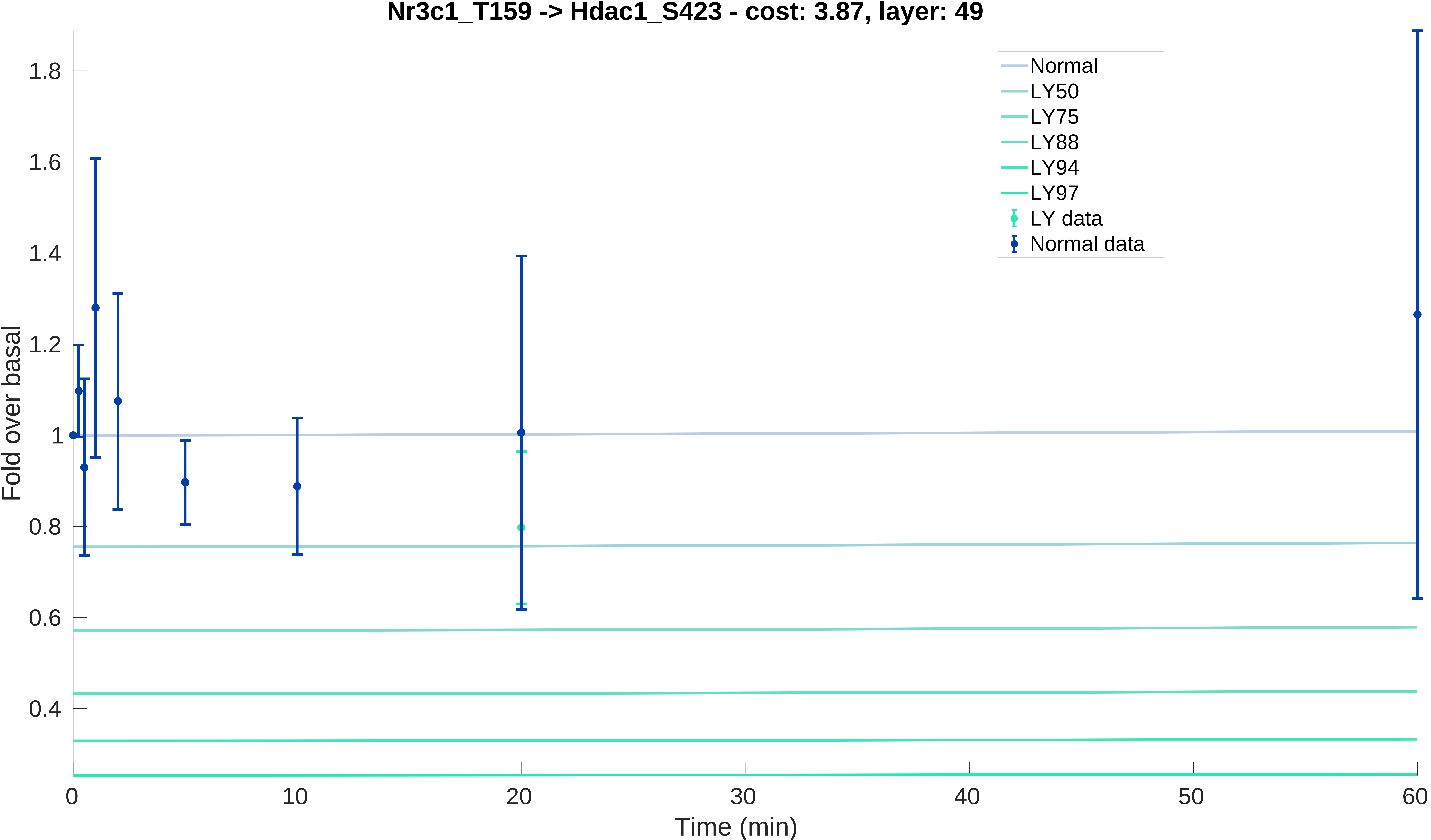

Nr3c1_T159 -> Hdac1_S423 - cost: 3.87, layer: 49
Fold over basal
Time (min)
Normal
LY50
LY75
LY88
LY94
LY97
LY data
Normal data
0.4
0.6
0.8
1
1.2
1.4
1.6
1.8
0
10
20
30
40
50
60

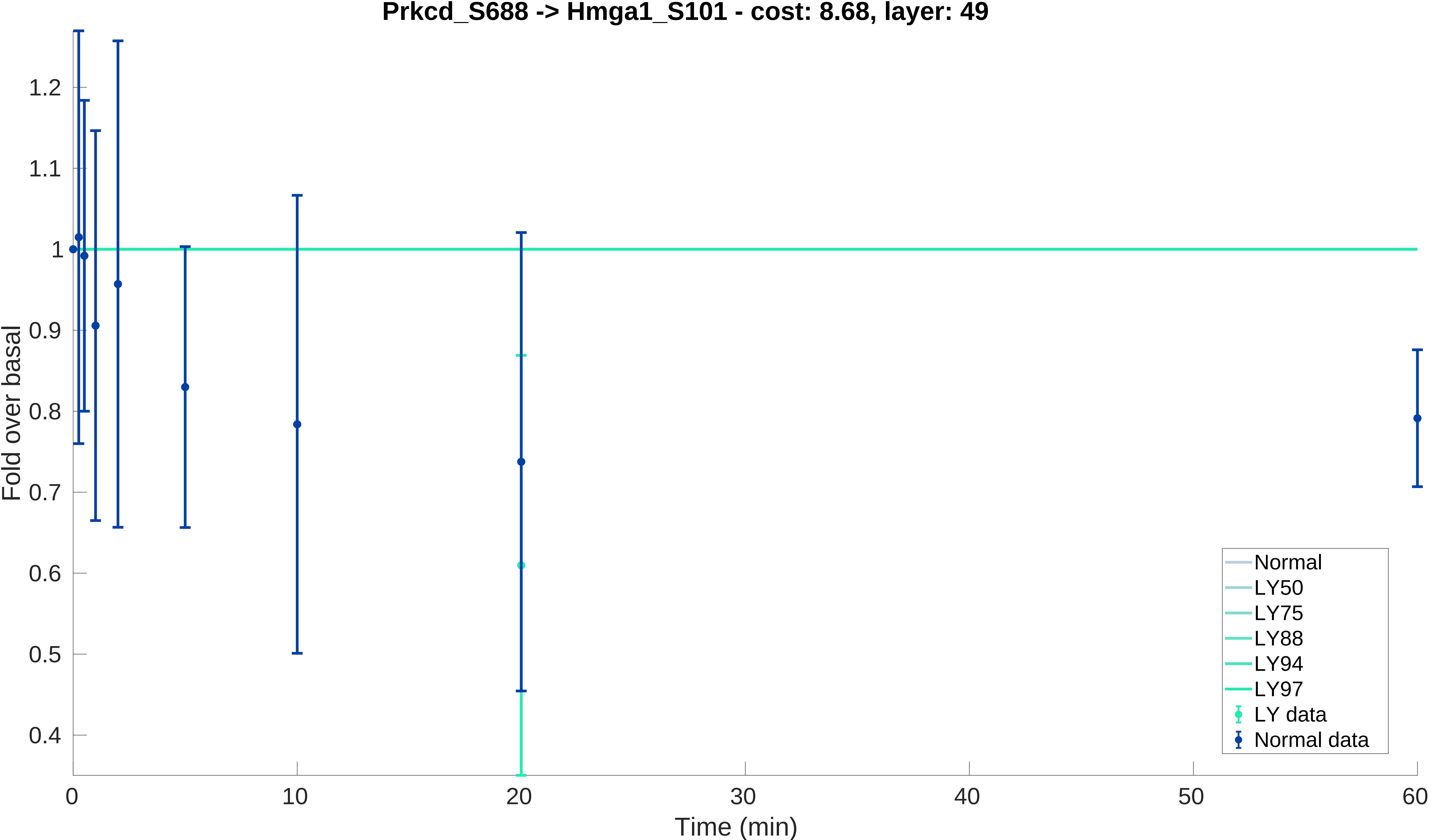

Prkcd_S688 -> Hmga1_S101 - cost: 8.68, layer: 49
Fold over basal
1.2
1.1
1
0.9
0.8
0.7
0.6
0.5
0.4
0
10
20
30
40
50
60
Time (min)
Normal
LY50
LY75
LY88
LY94
LY97
LY data
Normal data

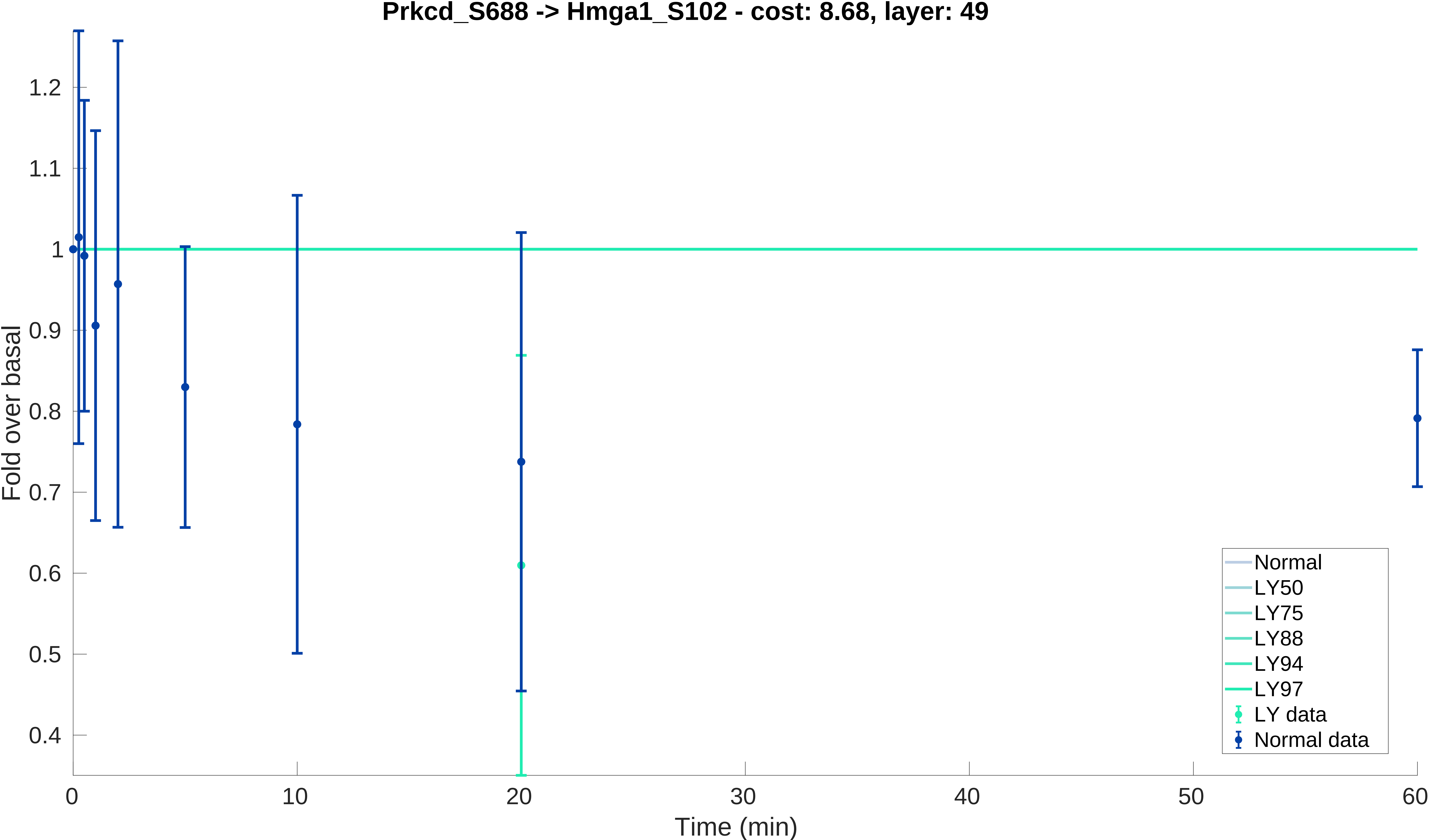

Prkcd_S688 -> Hmga1_S102 - cost: 8.68, layer: 49
Fold over basal
1.2
1.1
1
0.9
0.8
0.7
0.6
0.5
0.4
0
10
20
30
40
50
60
Time (min)
Normal
LY50
LY75
LY88
LY94
LY97
LY data
Normal data

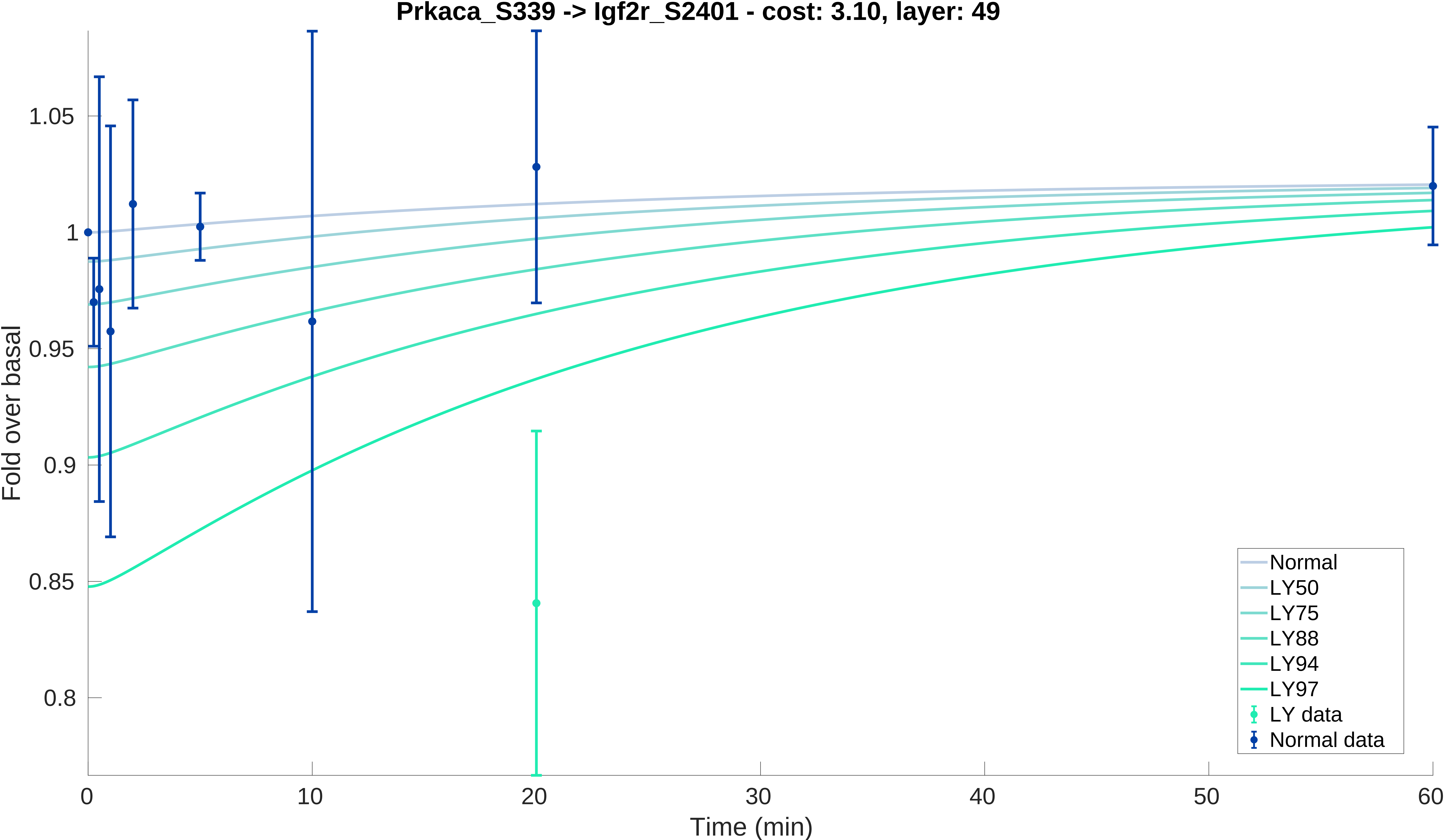

Prkaca_S339 -> Igf2r_S2401 - cost: 3.10, layer: 49
Fold over basal
1.05
1
0.95
0.9
0.85
0.8
0
10
20
30
40
50
60
Time (min)
Normal
LY50
LY75
LY88
LY94
LY97
LY data
Normal data

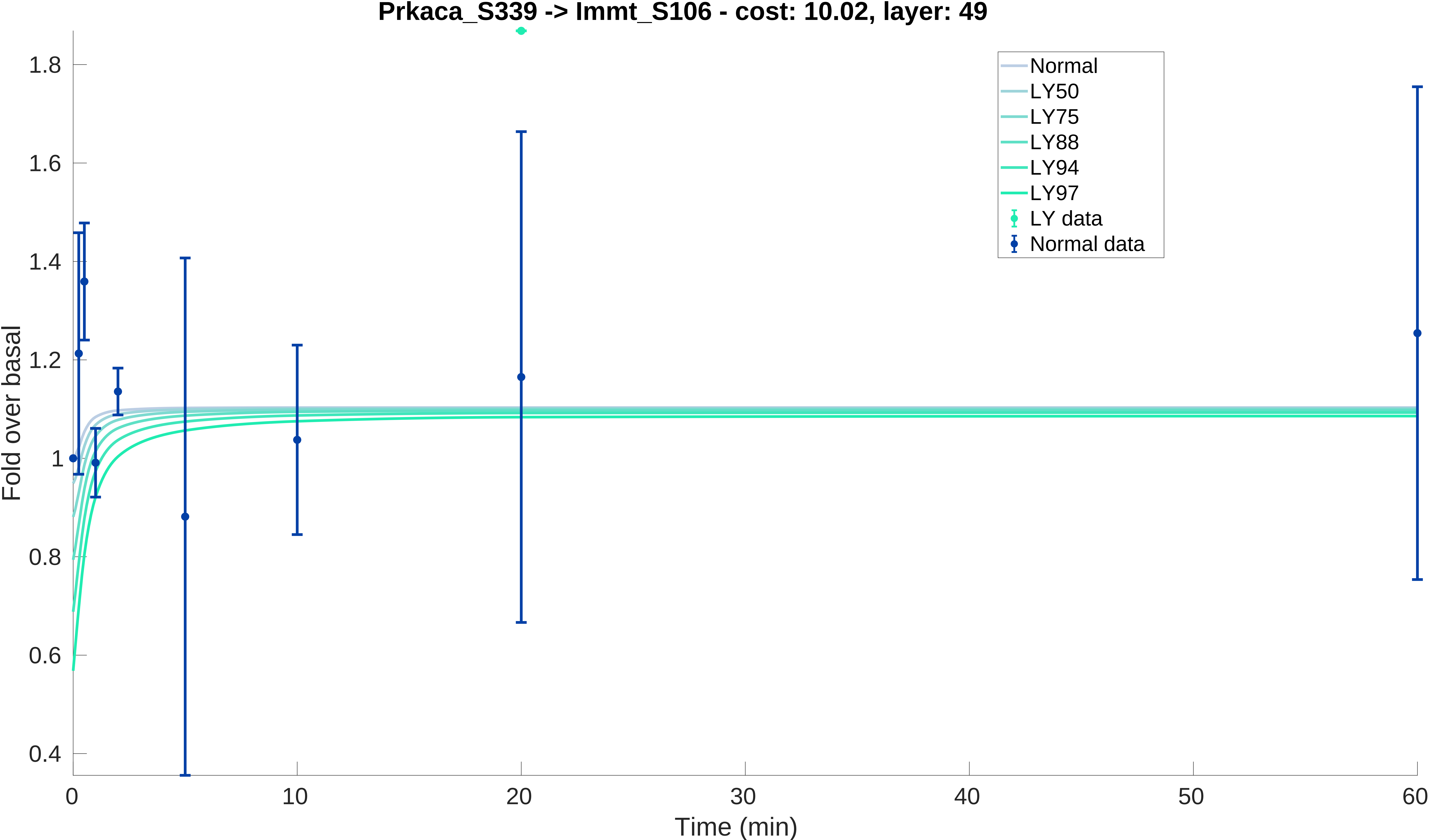

Prkaca_S339 -> Immt_S106 - cost: 10.02, layer: 49
Fold over basal
Time (min)
0
10
20
30
40
50
60
0.4
0.6
0.8
1
1.2
1.4
1.6
1.8
Normal
LY50
LY75
LY88
LY94
LY97
LY data
Normal data

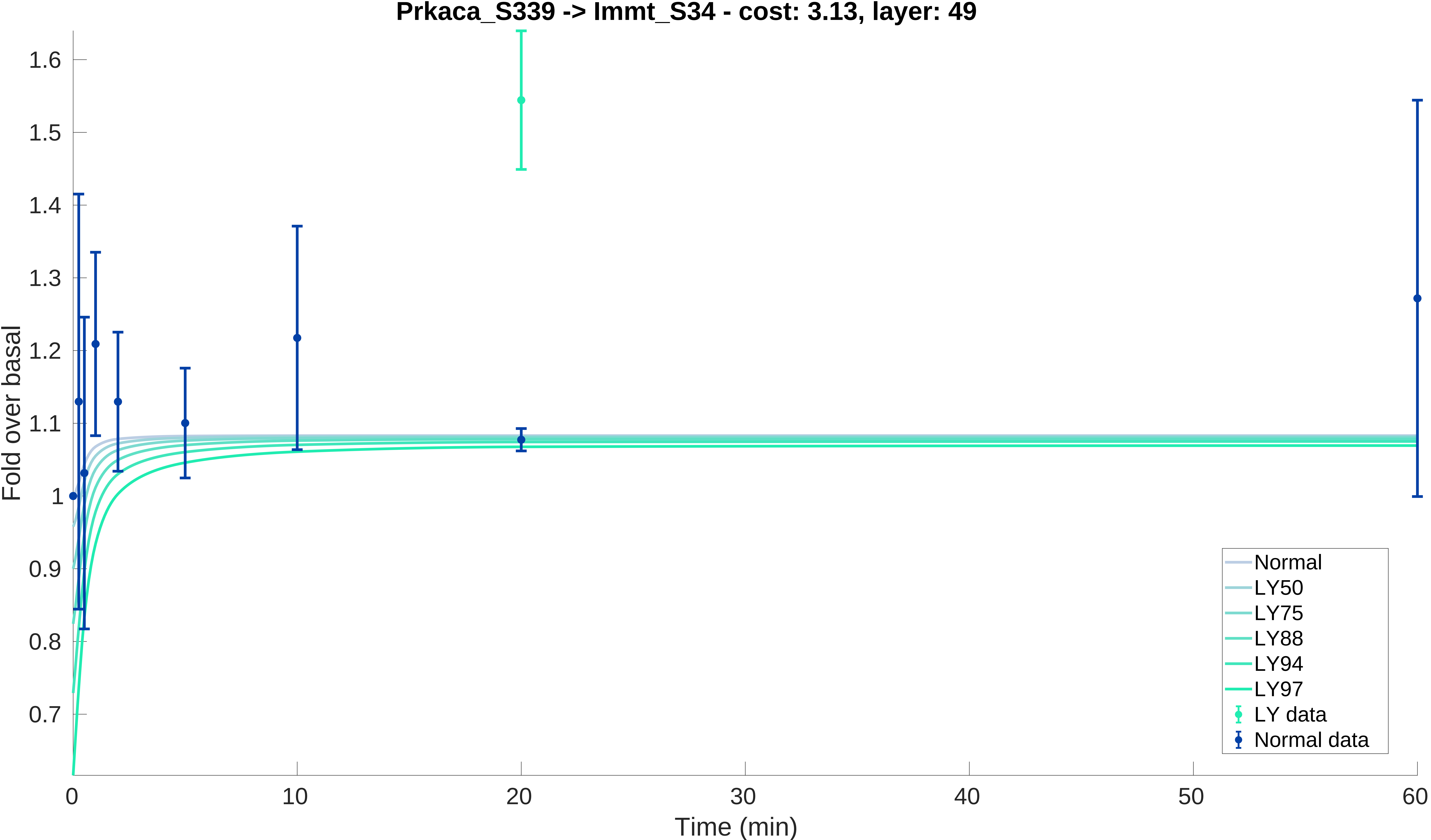

Prkaca_S339 -> Immt_S34 - cost: 3.13, layer: 49
Fold over basal
1.6
1.5
1.4
1.3
1.2
1.1
1
0.9
0.8
0.7
0
10
20
30
40
50
60
Time (min)
Normal
LY50
LY75
LY88
LY94
LY97
LY data
Normal data

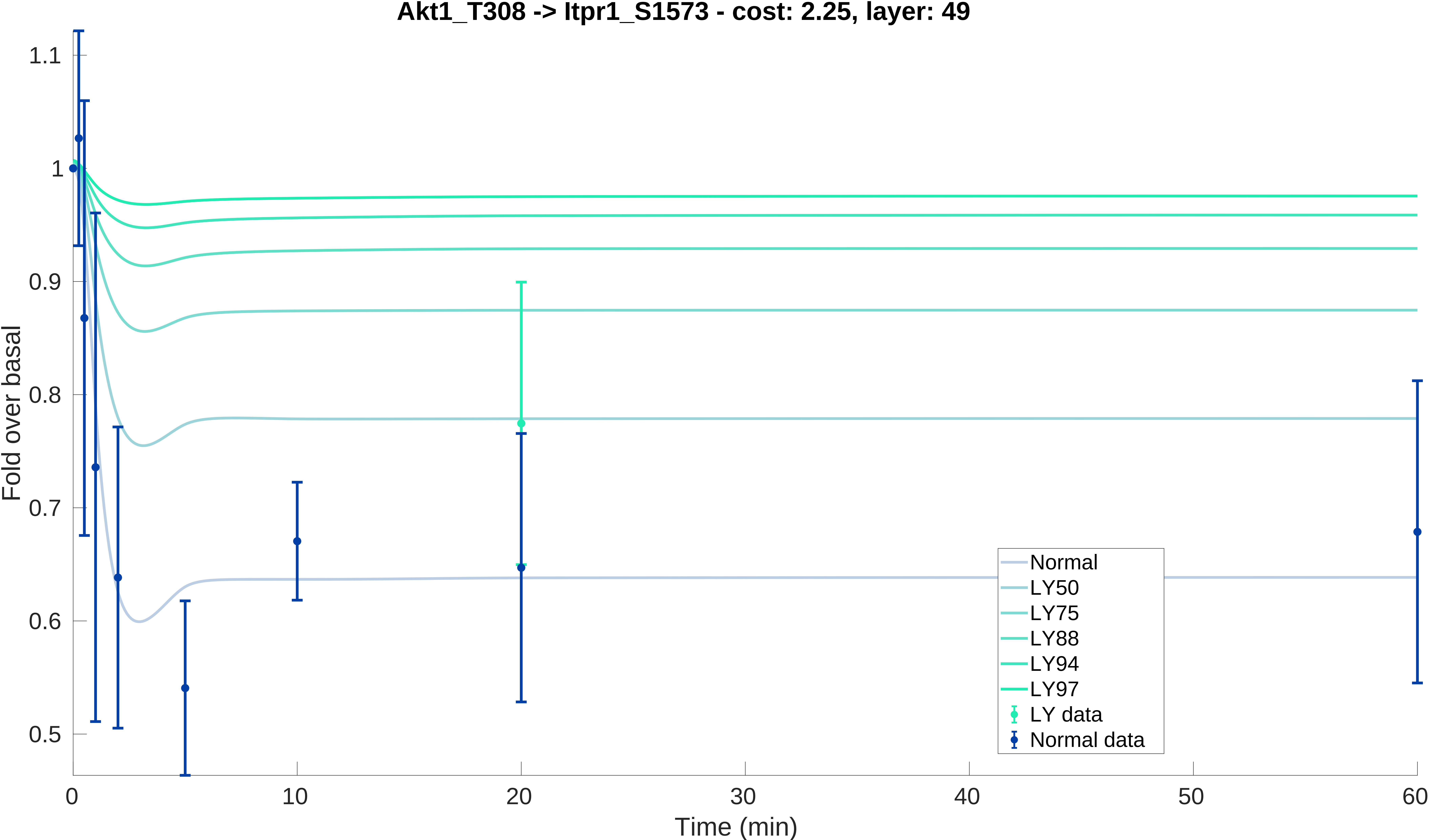

Akt1_T308 -> Itpr1_S1573 - cost: 2.25, layer: 49
Fold over basal
1.1
1
0.9
0.8
0.7
0.6
0.5
0
10
20
30
40
50
60
Time (min)
Normal
LY50
LY75
LY88
LY94
LY97
LY data
Normal data

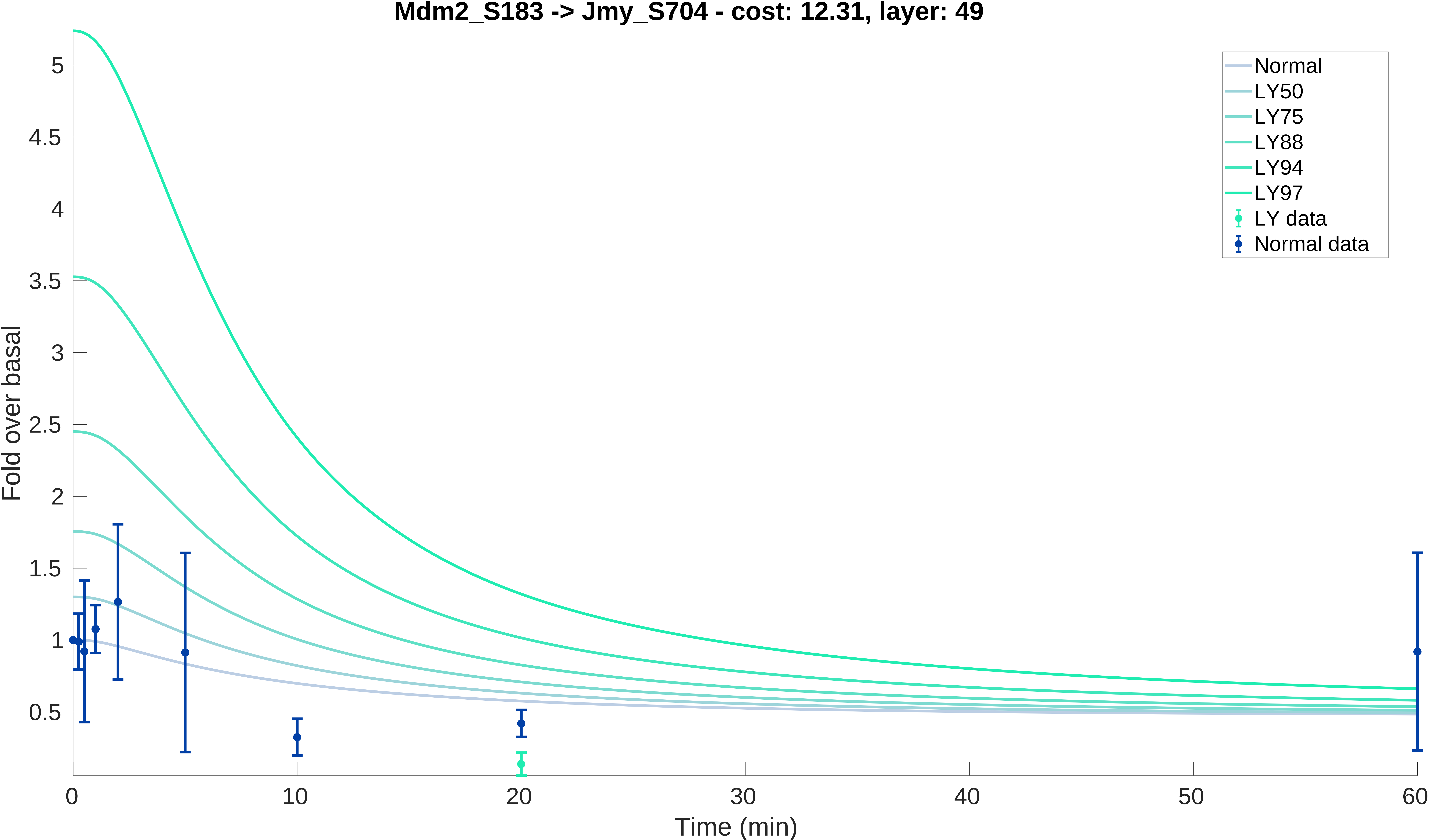

Mdm2_S183 -> Jmy_S704 - cost: 12.31, layer: 49
Fold over basal
Time (min)
Normal
LY50
LY75
LY88
LY94
LY97
LY data
Normal data
5
4.5
4
3.5
3
2.5
2
1.5
1
0.5
0
10
20
30
40
50
60

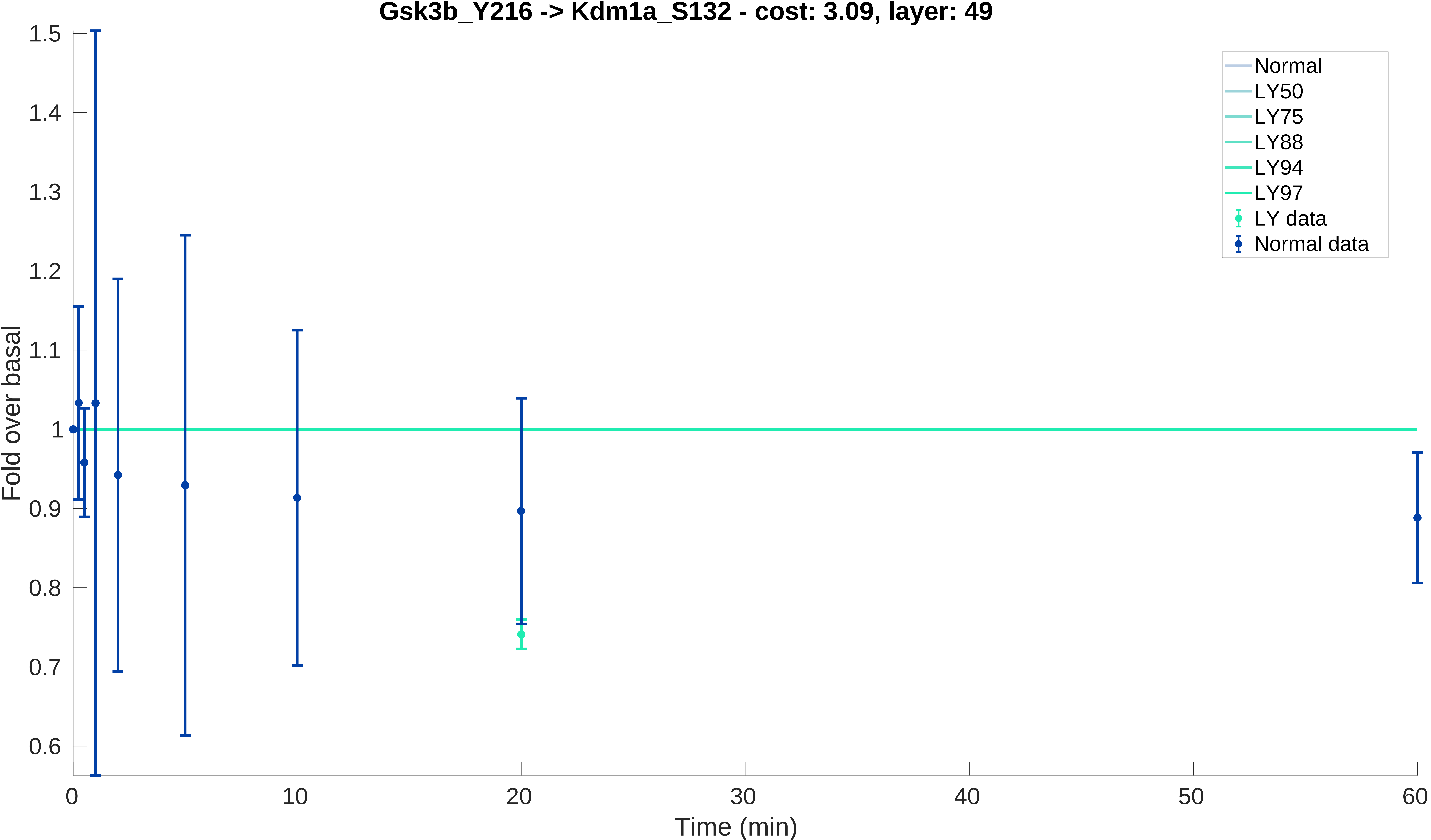

Gsk3b_Y216 -> Kdm1a_S132 - cost: 3.09, layer: 49
Fold over basal
Time (min)
0.6
0.7
0.8
0.9
1
1.1
1.2
1.3
1.4
1.5
0
10
20
30
40
50
60
Normal
LY50
LY75
LY88
LY94
LY97
LY data
Normal data

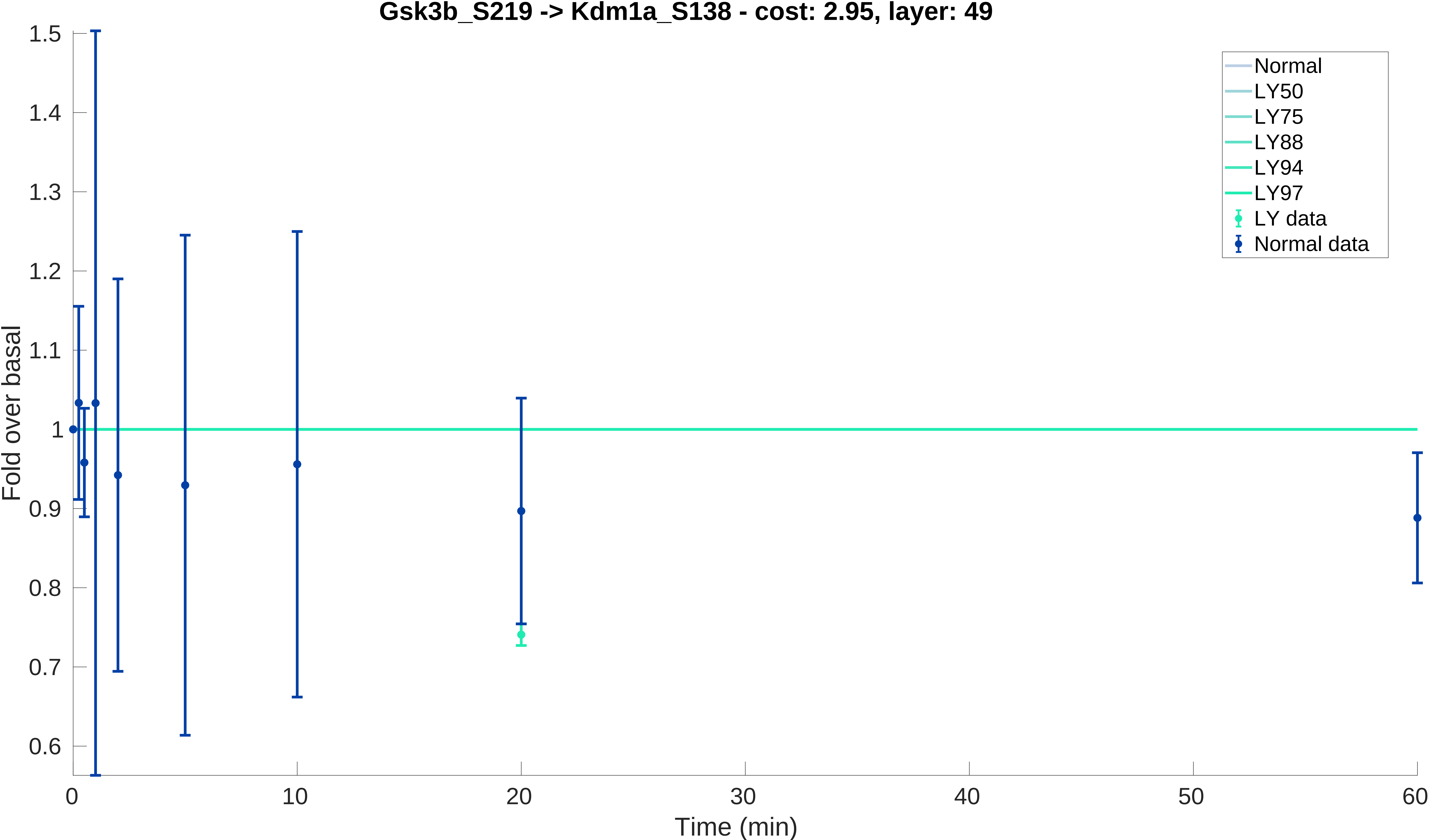

Gsk3b_S219 -> Kdm1a_S138 - cost: 2.95, layer: 49
Fold over basal
Time (min)
1.5
1.4
1.3
1.2
1.1
1
0.9
0.8
0.7
0.6
0
10
20
30
40
50
60
Normal
LY50
LY75
LY88
LY94
LY97
LY data
Normal data

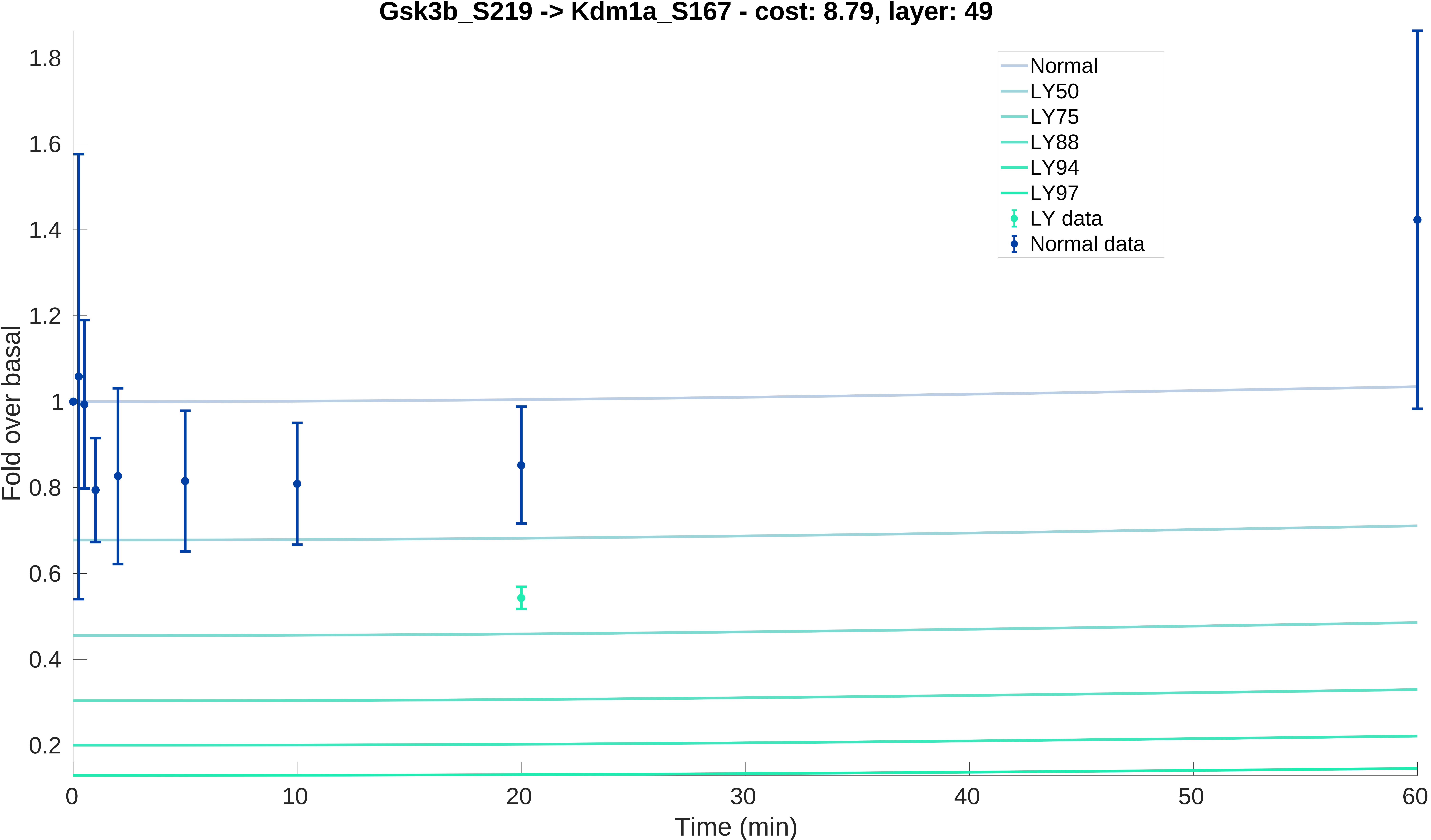

Gsk3b_S219 -> Kdm1a_S167 - cost: 8.79, layer: 49
Fold over basal
Time (min)
0.2
0.4
0.6
0.8
1
1.2
1.4
1.6
1.8
0
10
20
30
40
50
60
Normal
LY50
LY75
LY88
LY94
LY97
LY data
Normal data

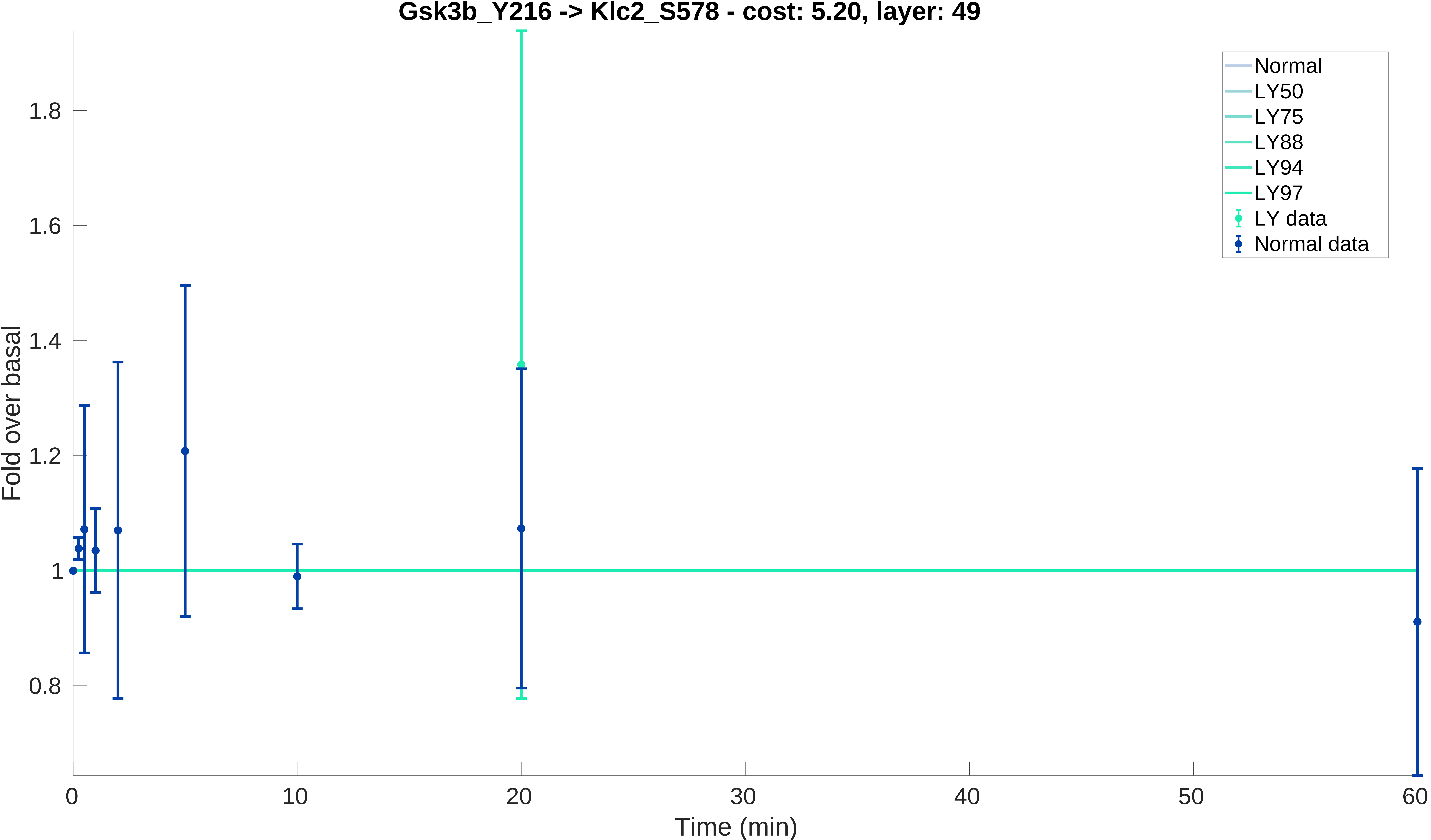
Gsk3b_Y216 -> Klc2_S578 - cost: 5.20, layer: 49
Fold over basal
1.8
1.6
1.4
1.2
1
0.8
0
10
20
30
40
50
60
Time (min)
Normal
LY50
LY75
LY88
LY94
LY97
LY data
Normal data

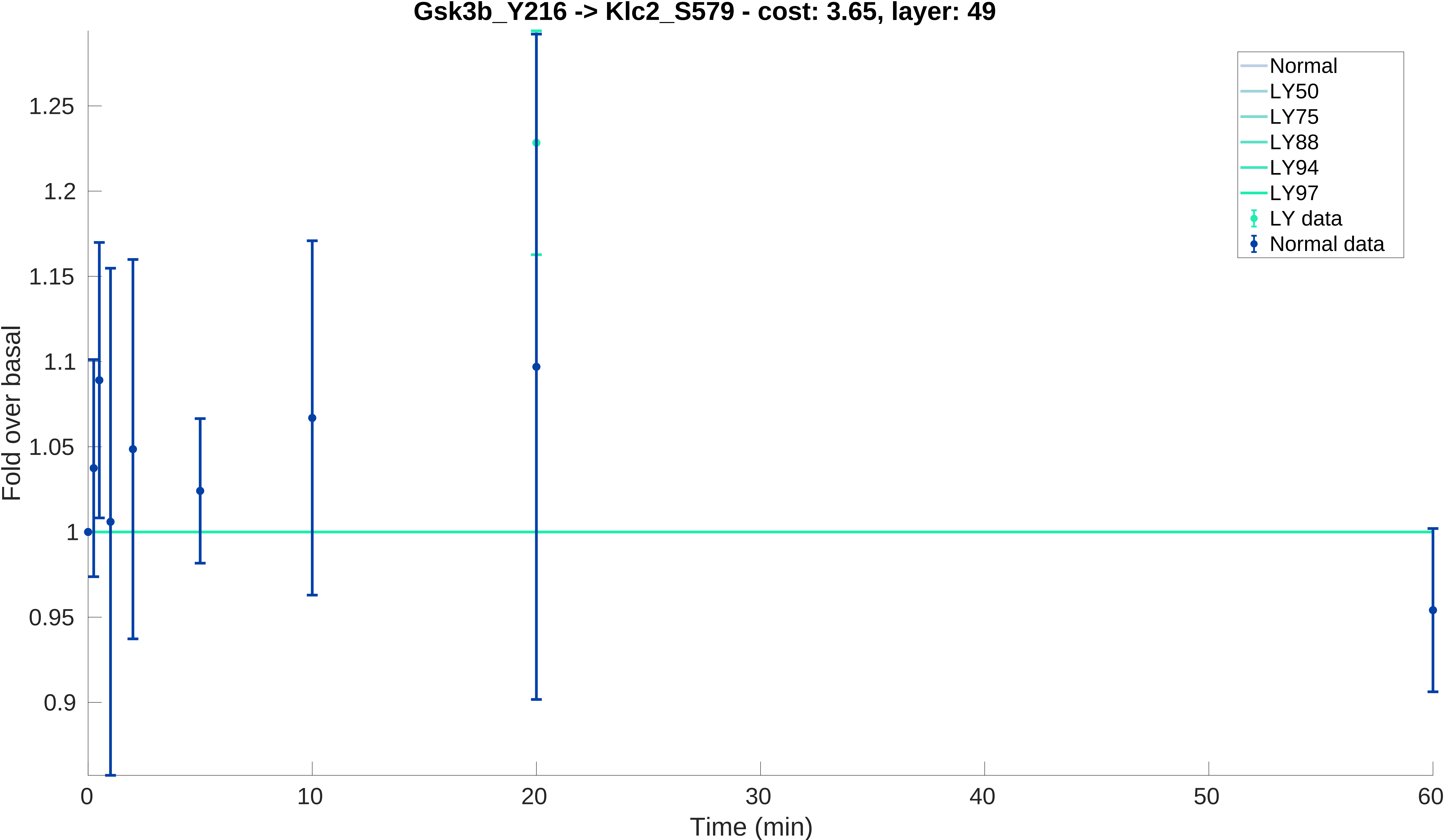

Gsk3b_Y216 -> Klc2_S579 - cost: 3.65, layer: 49
Normal
LY50
LY75
LY88
LY94
LY97
LY data
Normal data
1.25
1.2
1.15
1.1
1.05
1
0.95
0.9
Fold over basal
0
10
20
30
40
50
60
Time (min)

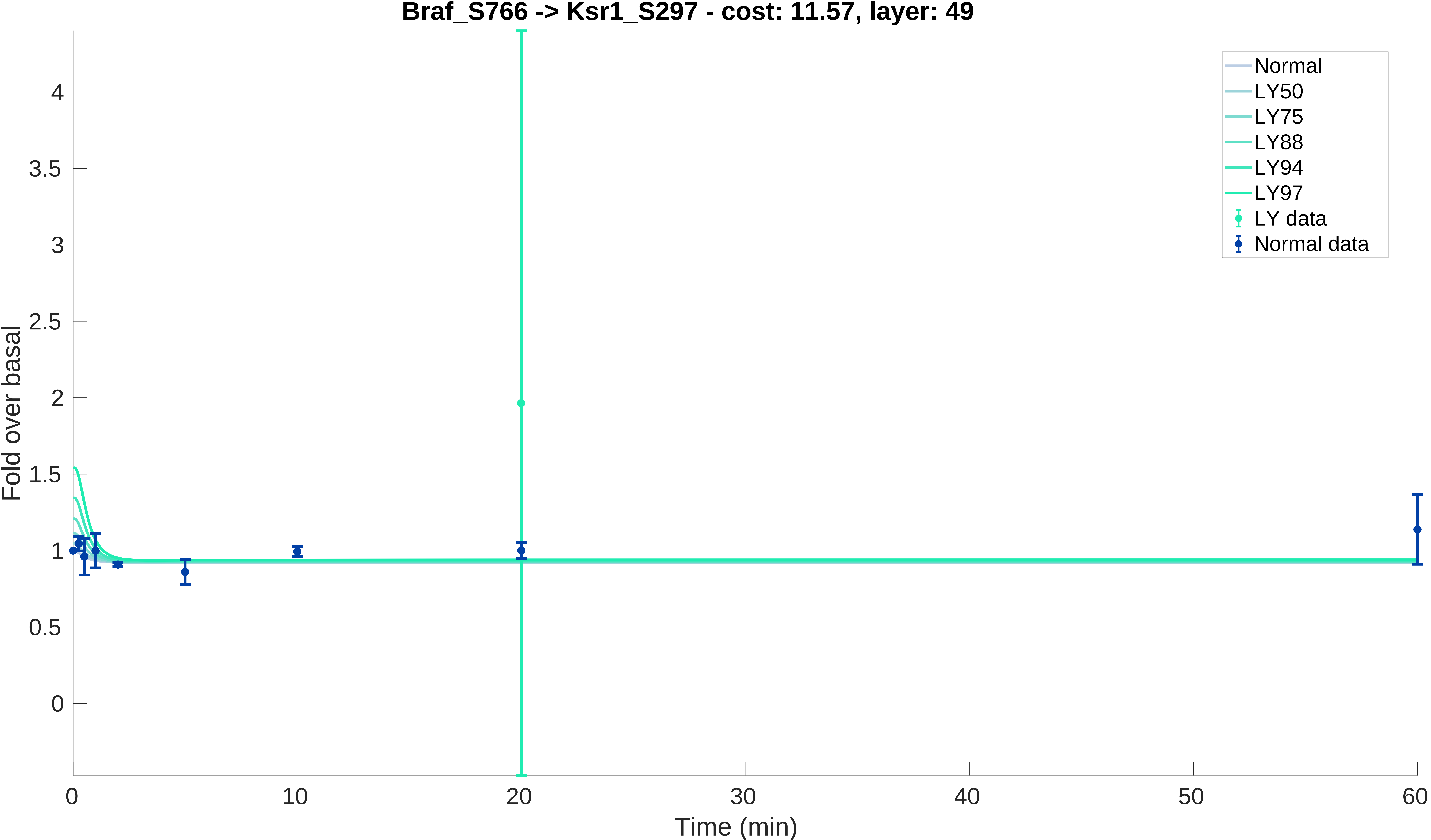

Braf_S766 -> Ksr1_S297 - cost: 11.57, layer: 49
Normal
LY50
LY75
LY88
LY94
LY97
LY data
Normal data
Fold over basal
Time (min)
0
0.5
1
1.5
2
2.5
3
3.5
4
0
10
20
30
40
50
60

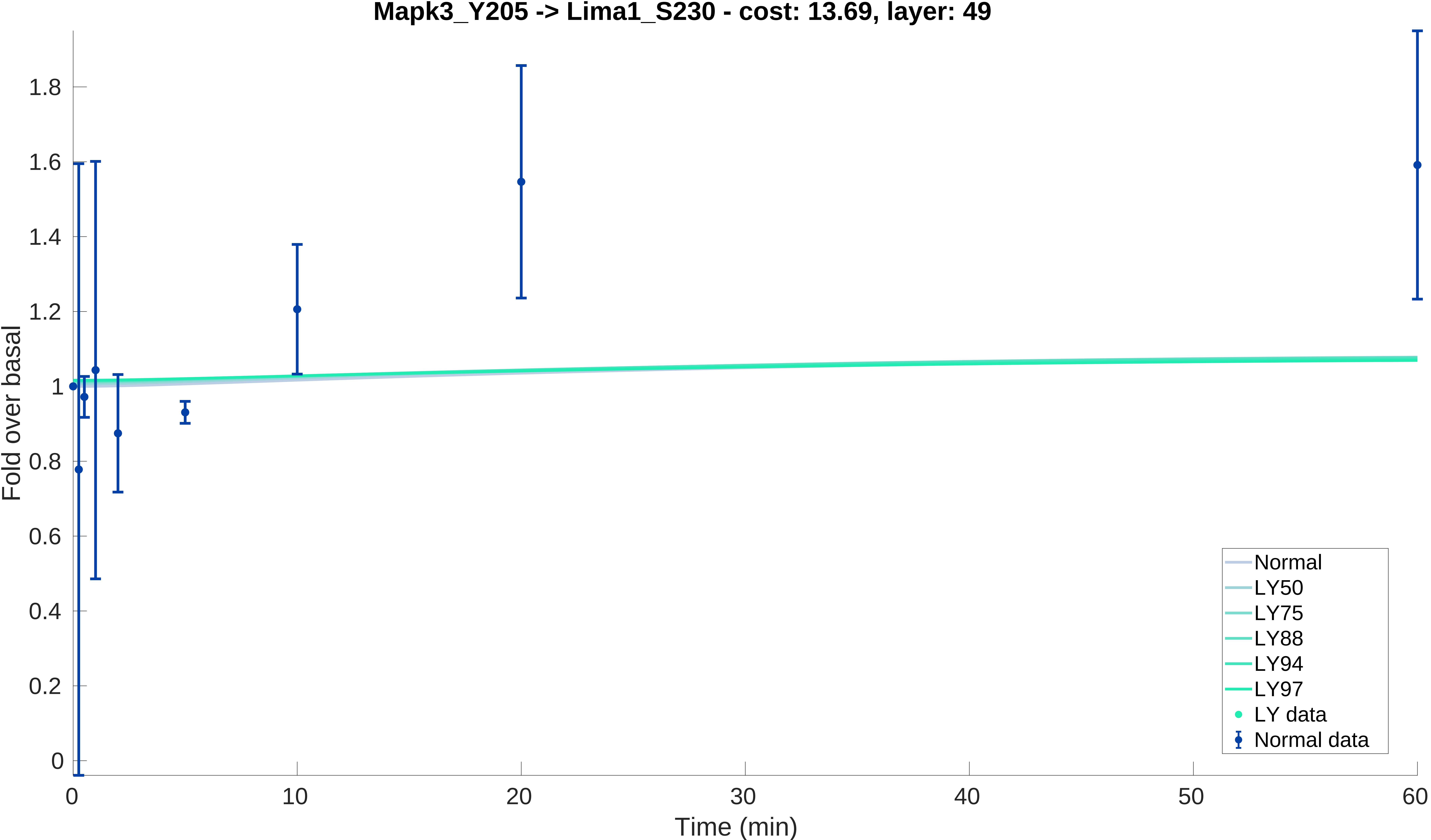

Mapk3_Y205 -> Lima1_S230 - cost: 13.69, layer: 49
Fold over basal
Time (min)
0
0.2
0.4
0.6
0.8
1
1.2
1.4
1.6
1.8
0
10
20
30
40
50
60
Normal
LY50
LY75
LY88
LY94
LY97
LY data
Normal data

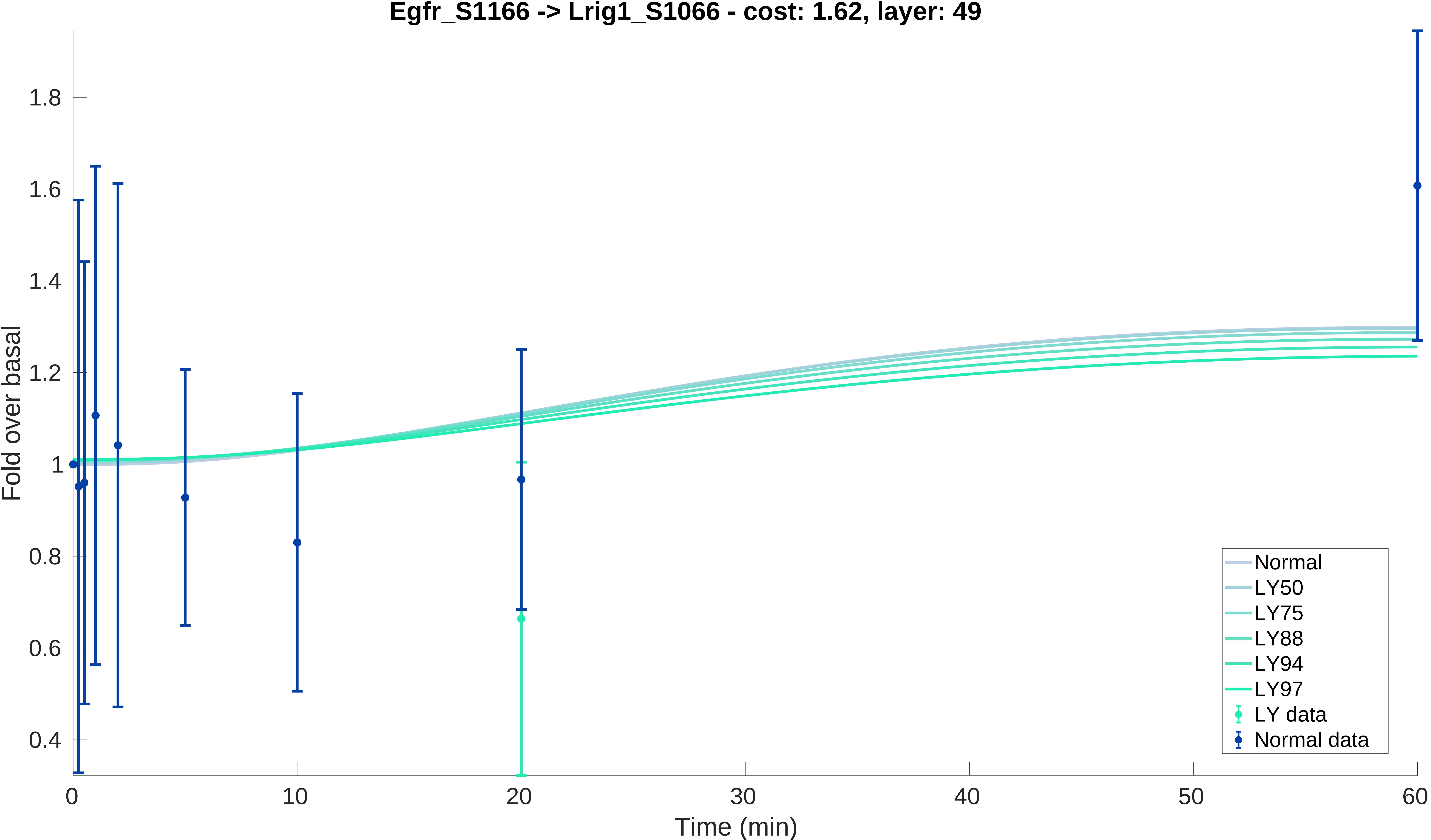

Egfr_S1166 -> Lrig1_S1066 - cost: 1.62, layer: 49
Fold over basal
1.8
1.6
1.4
1.2
1
0.8
0.6
0.4
0
10
20
30
40
50
60
Time (min)
Normal
LY50
LY75
LY88
LY94
LY97
LY data
Normal data

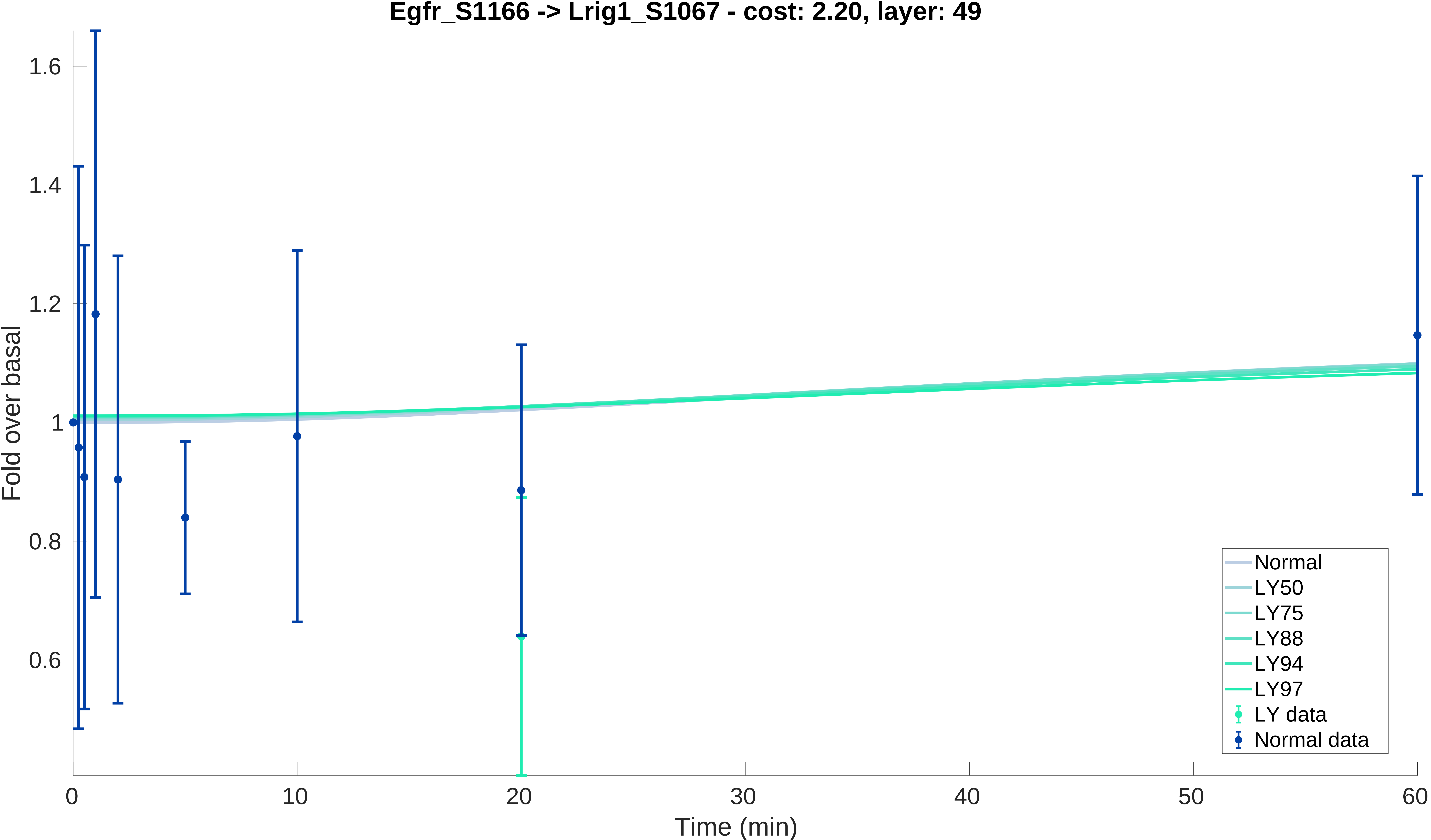

Egfr_S1166 -> Lrig1_S1067 - cost: 2.20, layer: 49
Fold over basal
1.6
1.4
1.2
1
0.8
0.6
0
10
20
30
40
50
60
Time (min)
Normal
LY50
LY75
LY88
LY94
LY97
LY data
Normal data

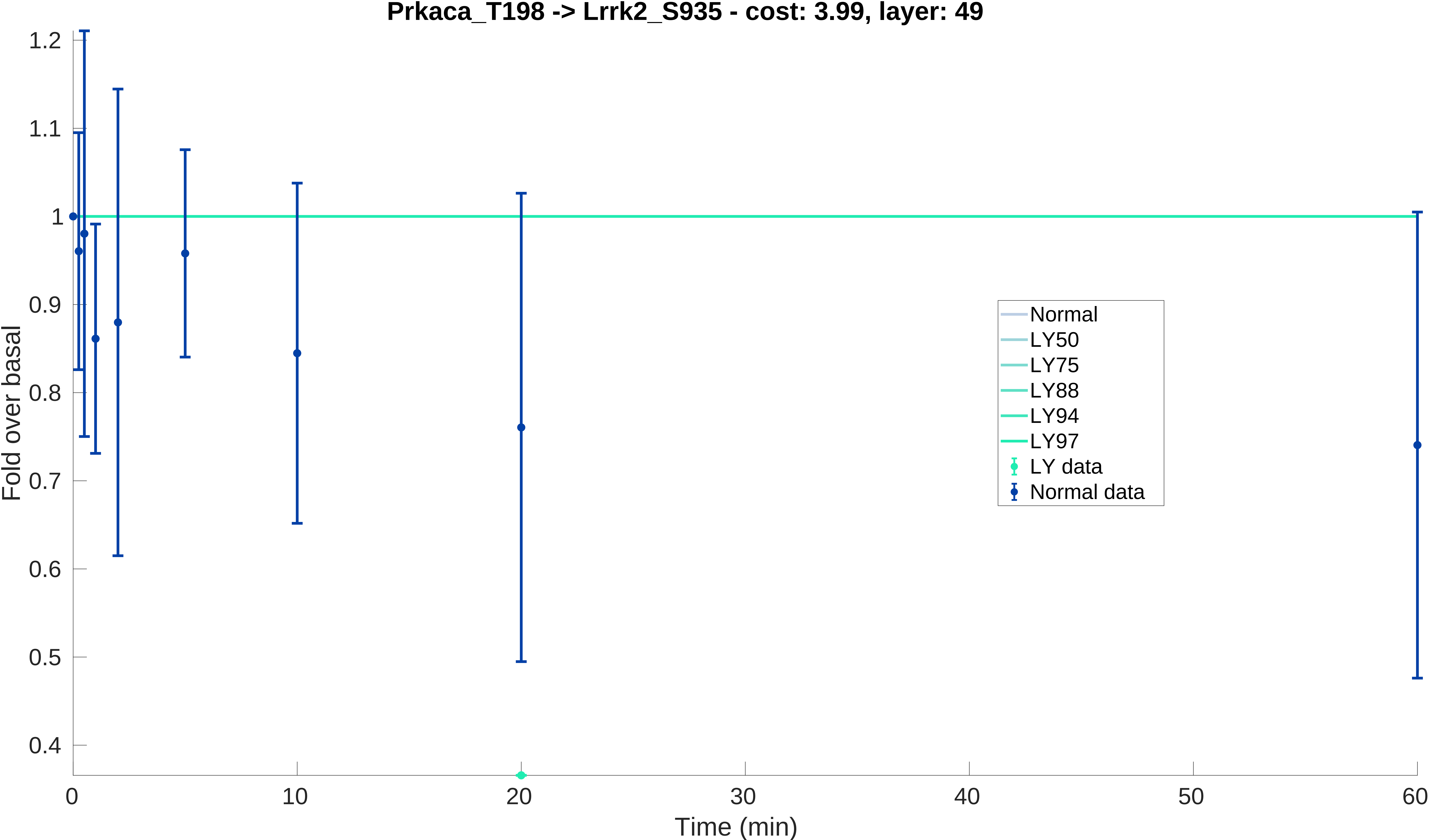
Prkaca_T198 -> Lrrk2_S935 - cost: 3.99, layer: 49
Fold over basal
Time (min)
1.2
1.1
1
0.9
0.8
0.7
0.6
0.5
0.4
0
10
20
30
40
50
60
Normal
LY50
LY75
LY88
LY94
LY97
LY data
Normal data

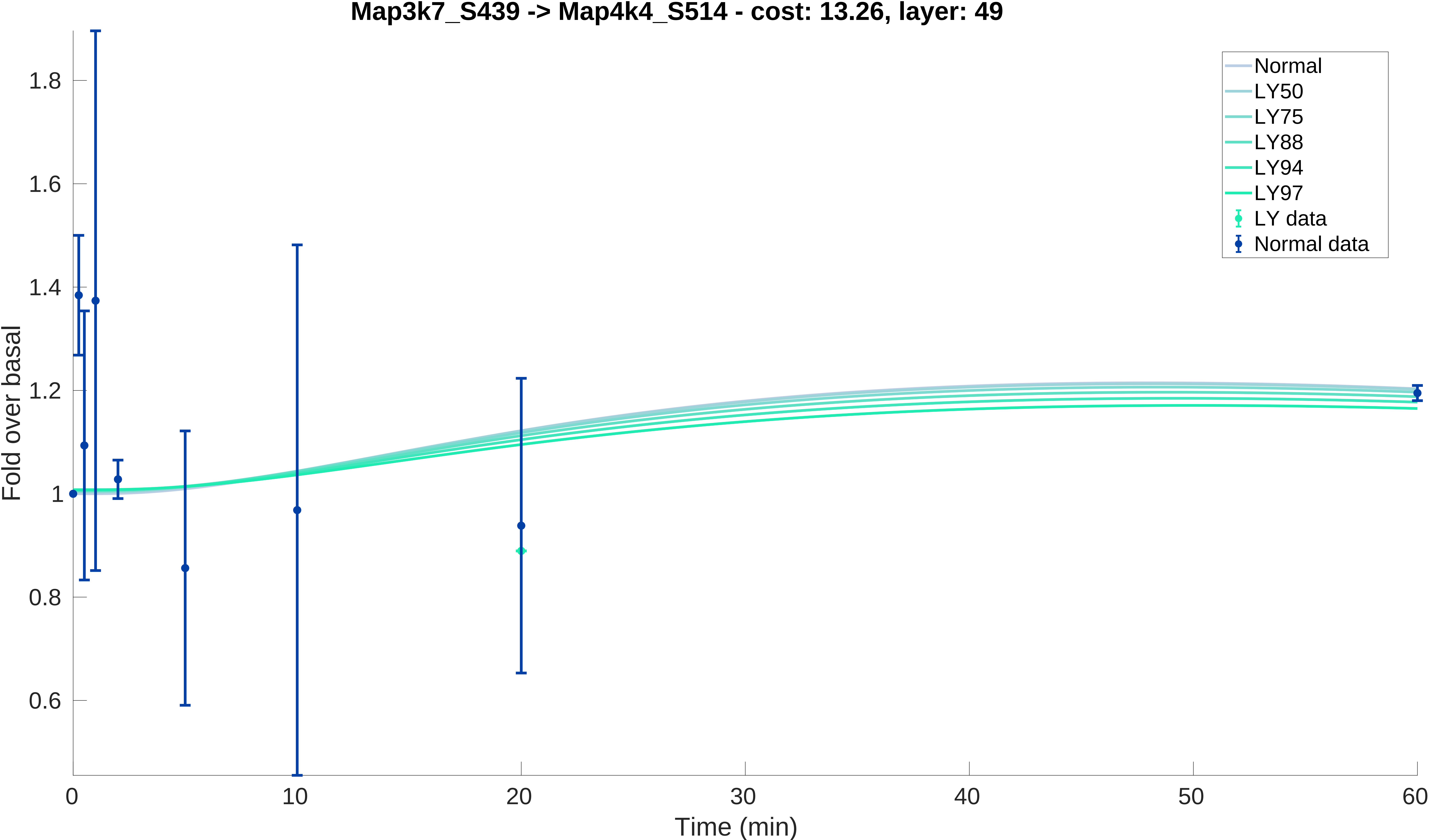

Map3k7_S439 -> Map4k4_S514 - cost: 13.26, layer: 49
Normal
LY50
LY75
LY88
LY94
LY97
LY data
Normal data
Fold over basal
1.8
1.6
1.4
1.2
1
0.8
0.6
0
10
20
30
40
50
60
Time (min)

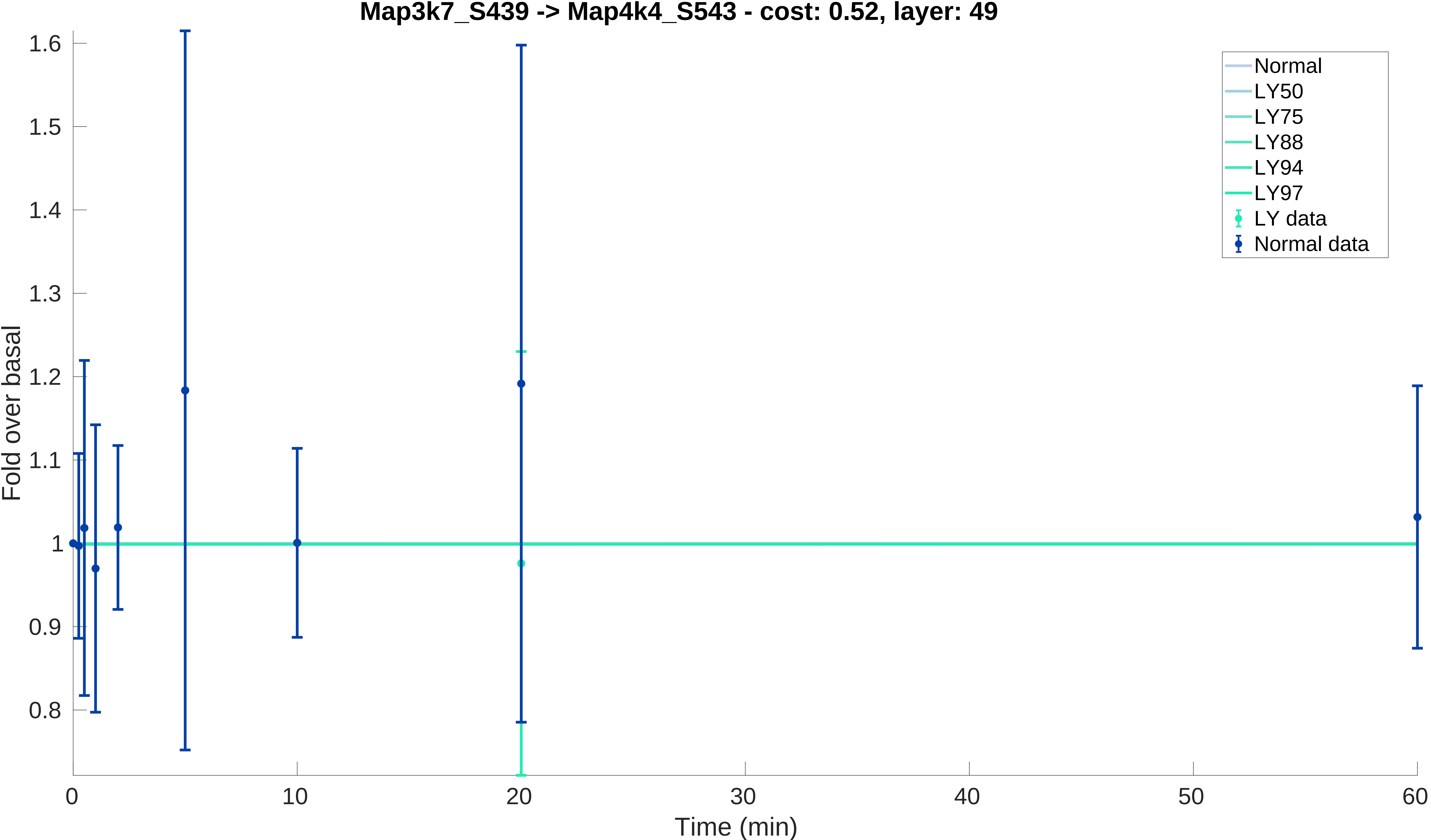

Map3k7_S439 -> Map4k4_S543 - cost: 0.52, layer: 49
Fold over basal
Time (min)
0.8
0.9
1
1.1
1.2
1.3
1.4
1.5
1.6
0
10
20
30
40
50
60
Normal
LY50
LY75
LY88
LY94
LY97
LY data
Normal data

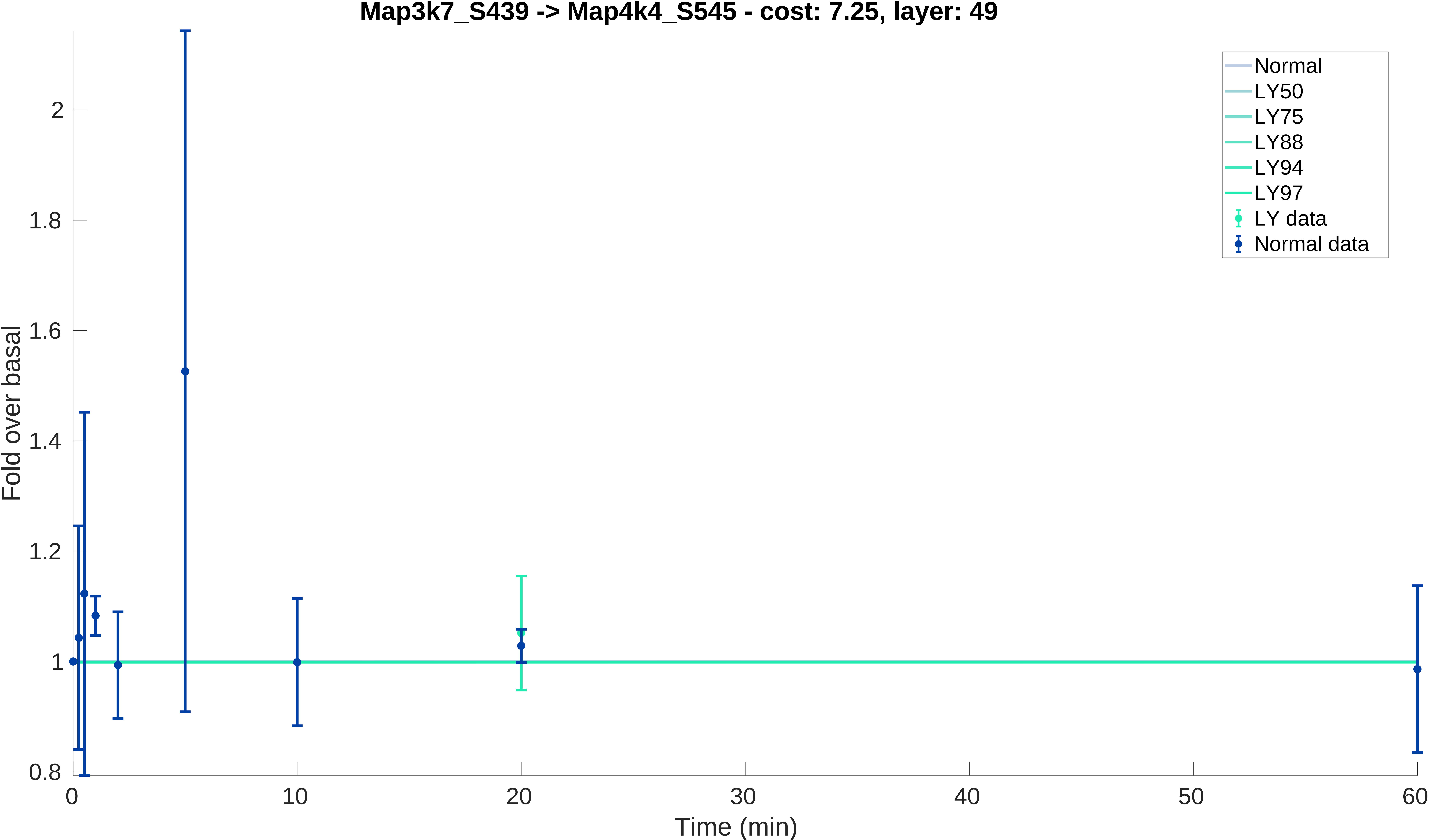

Map3k7_S439 -> Map4k4_S545 - cost: 7.25, layer: 49
Fold over basal
Time (min)
Normal
LY50
LY75
LY88
LY94
LY97
LY data
Normal data
0.8
1
1.2
1.4
1.6
1.8
2
0
10
20
30
40
50
60

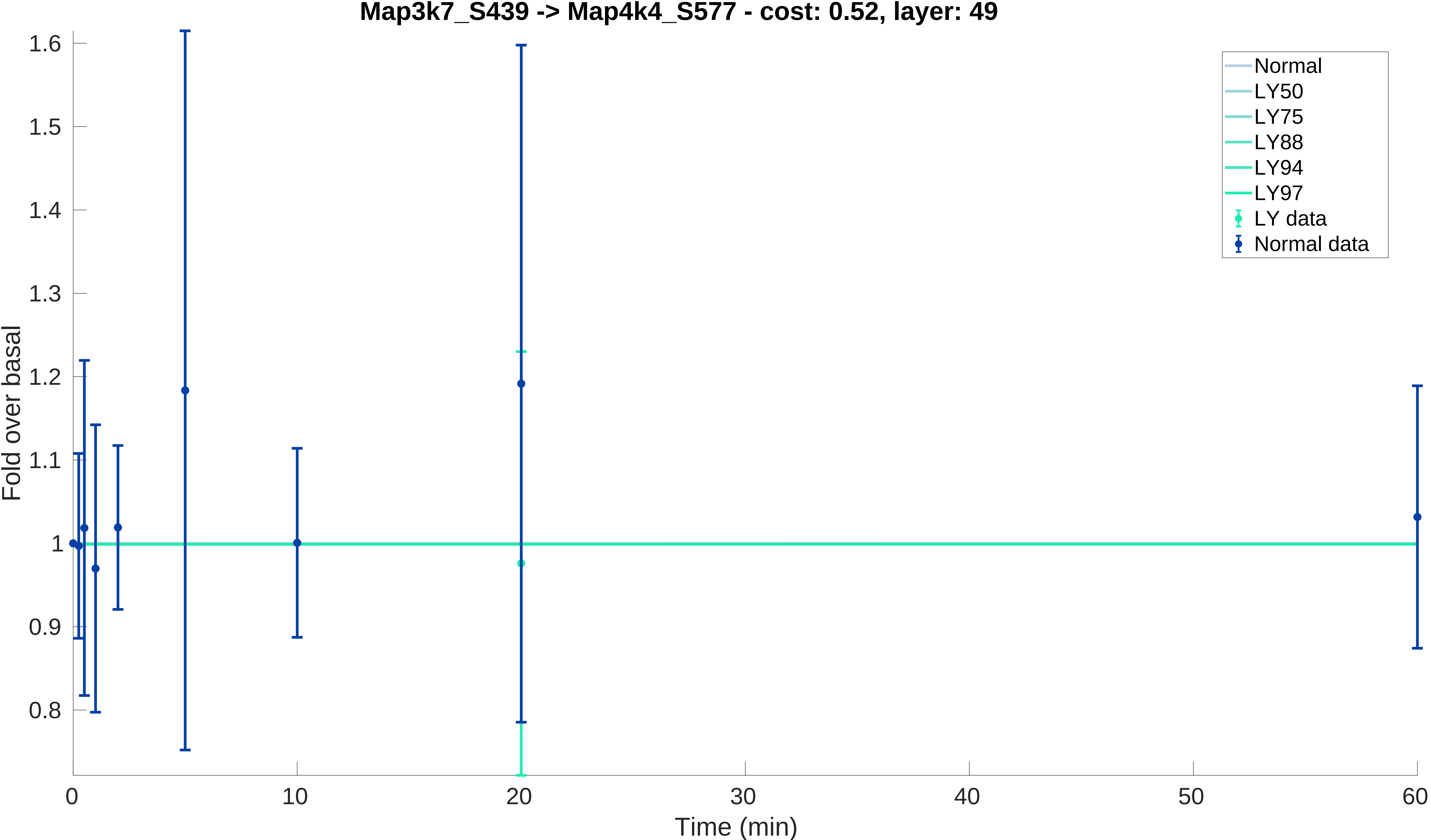

Map3k7_S439 -> Map4k4_S577 - cost: 0.52, layer: 49
Fold over basal
Time (min)
1.6
1.5
1.4
1.3
1.2
1.1
1
0.9
0.8
0
10
20
30
40
50
60
Normal
LY50
LY75
LY88
LY94
LY97
LY data
Normal data

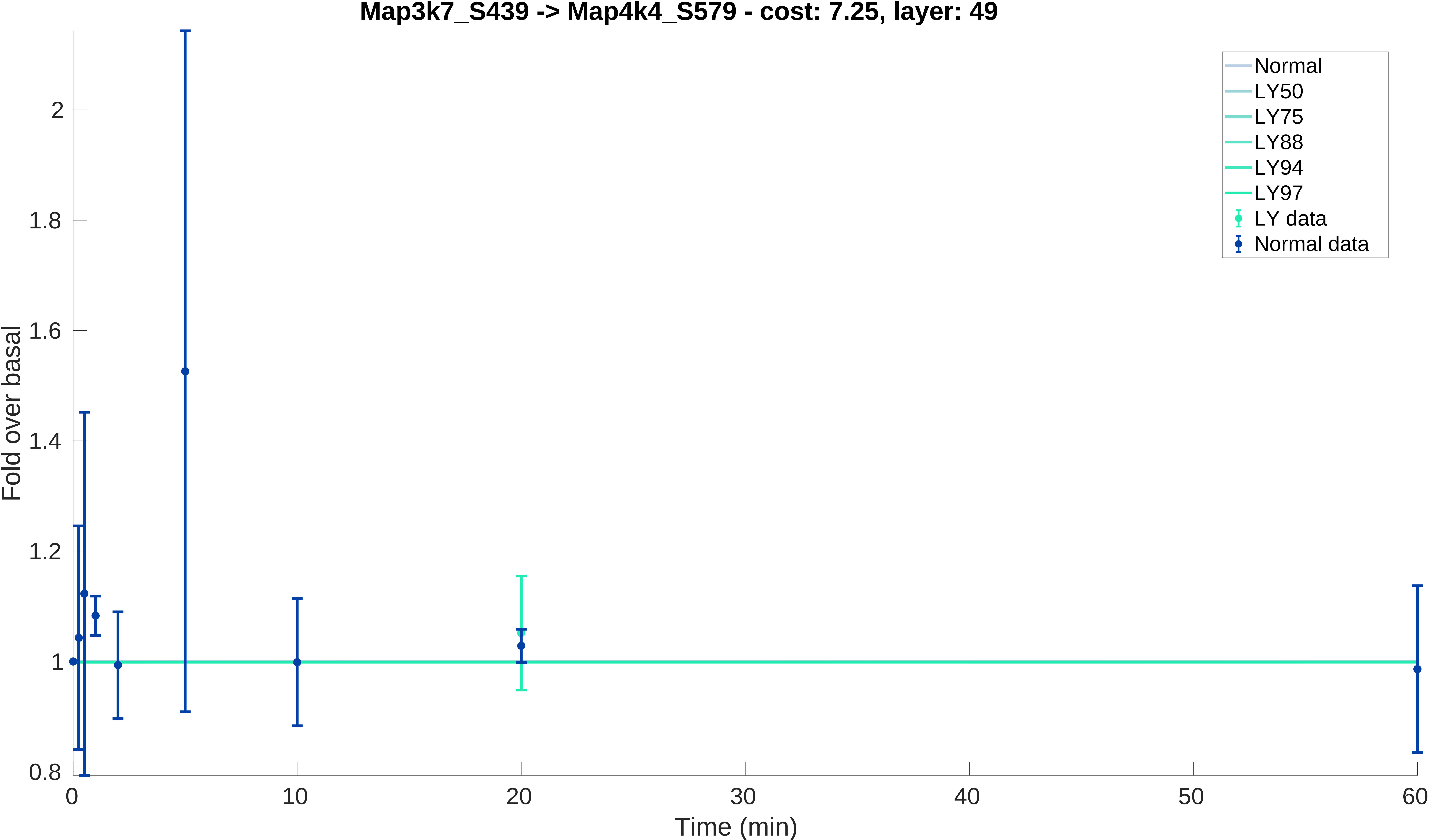

Map3k7_S439 -> Map4k4_S579 - cost: 7.25, layer: 49
Normal
LY50
LY75
LY88
LY94
LY97
LY data
Normal data
Fold over basal
2
1.8
1.6
1.4
1.2
1
0.8
0
10
20
30
40
50
60
Time (min)

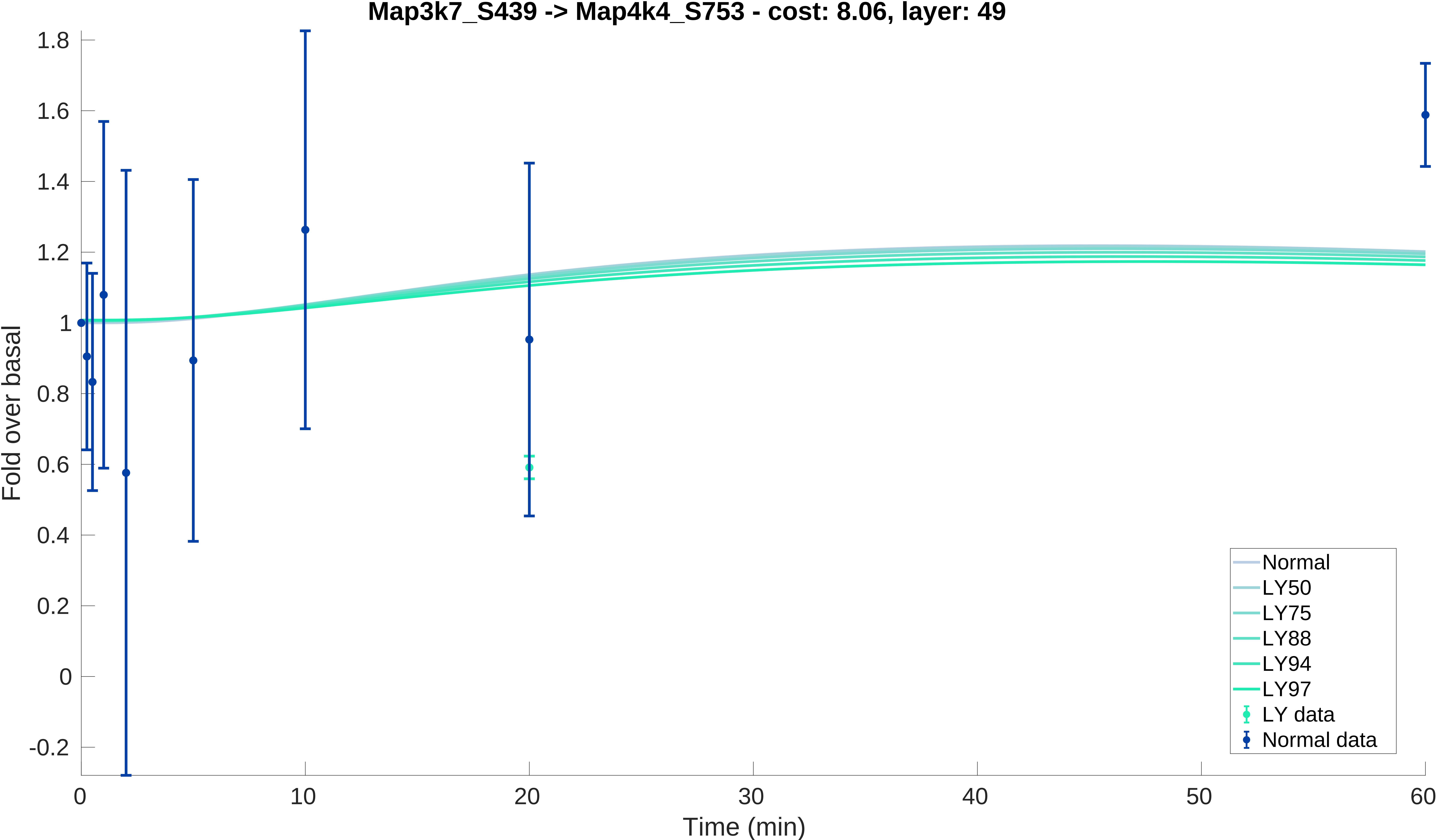

Map3k7_S439 -> Map4k4_S753 - cost: 8.06, layer: 49
Fold over basal
Time (min)
1.8
1.6
1.4
1.2
1
0.8
0.6
0.4
0.2
0
-0.2
0
10
20
30
40
50
60
Normal
LY50
LY75
LY88
LY94
LY97
LY data
Normal data

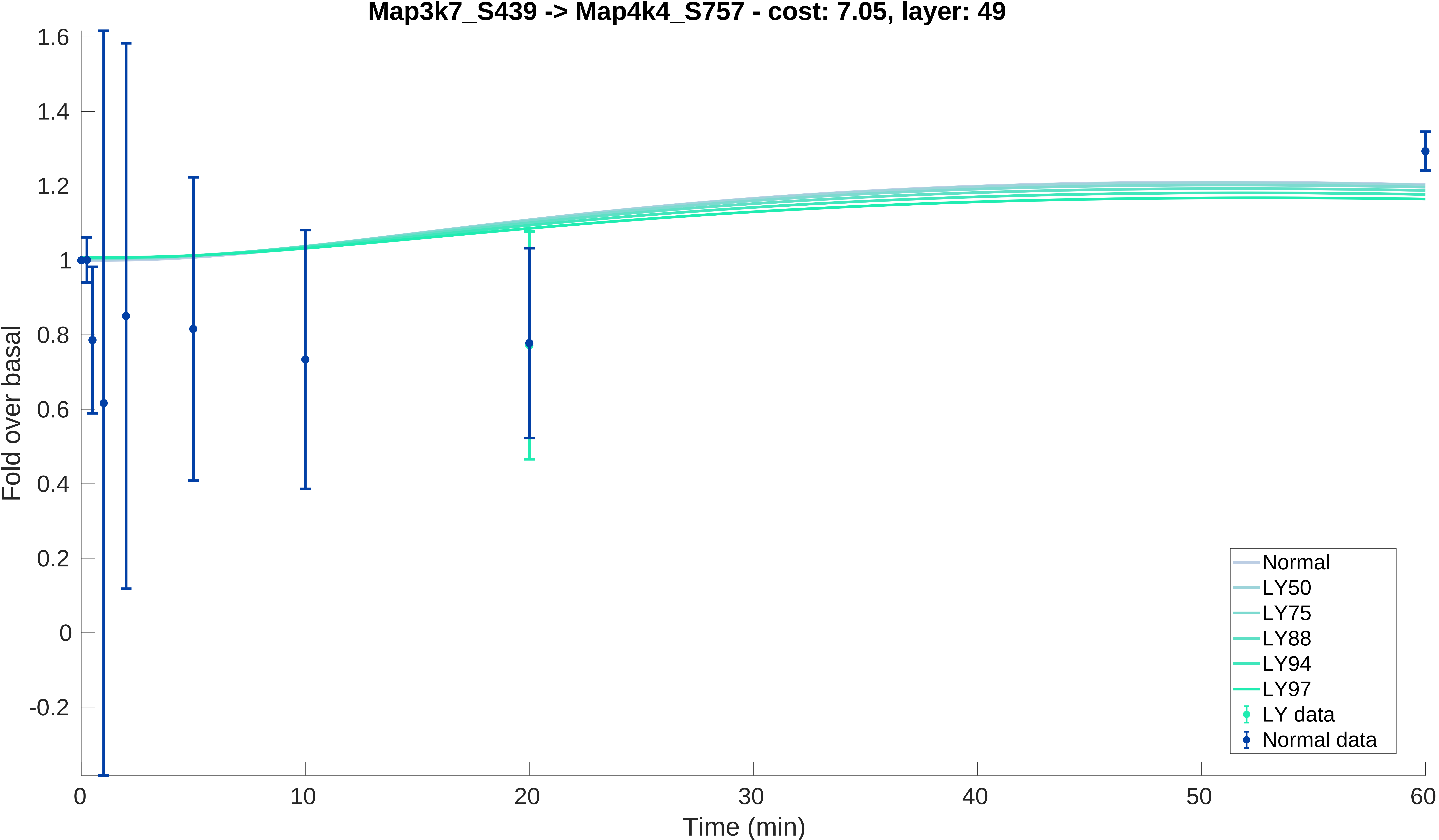

Map3k7_S439 -> Map4k4_S757 - cost: 7.05, layer: 49
Fold over basal
1.6
1.4
1.2
1
0.8
0.6
0.4
0.2
0
-0.2
0
10
20
30
40
50
60
Time (min)
Normal
LY50
LY75
LY88
LY94
LY97
LY data
Normal data

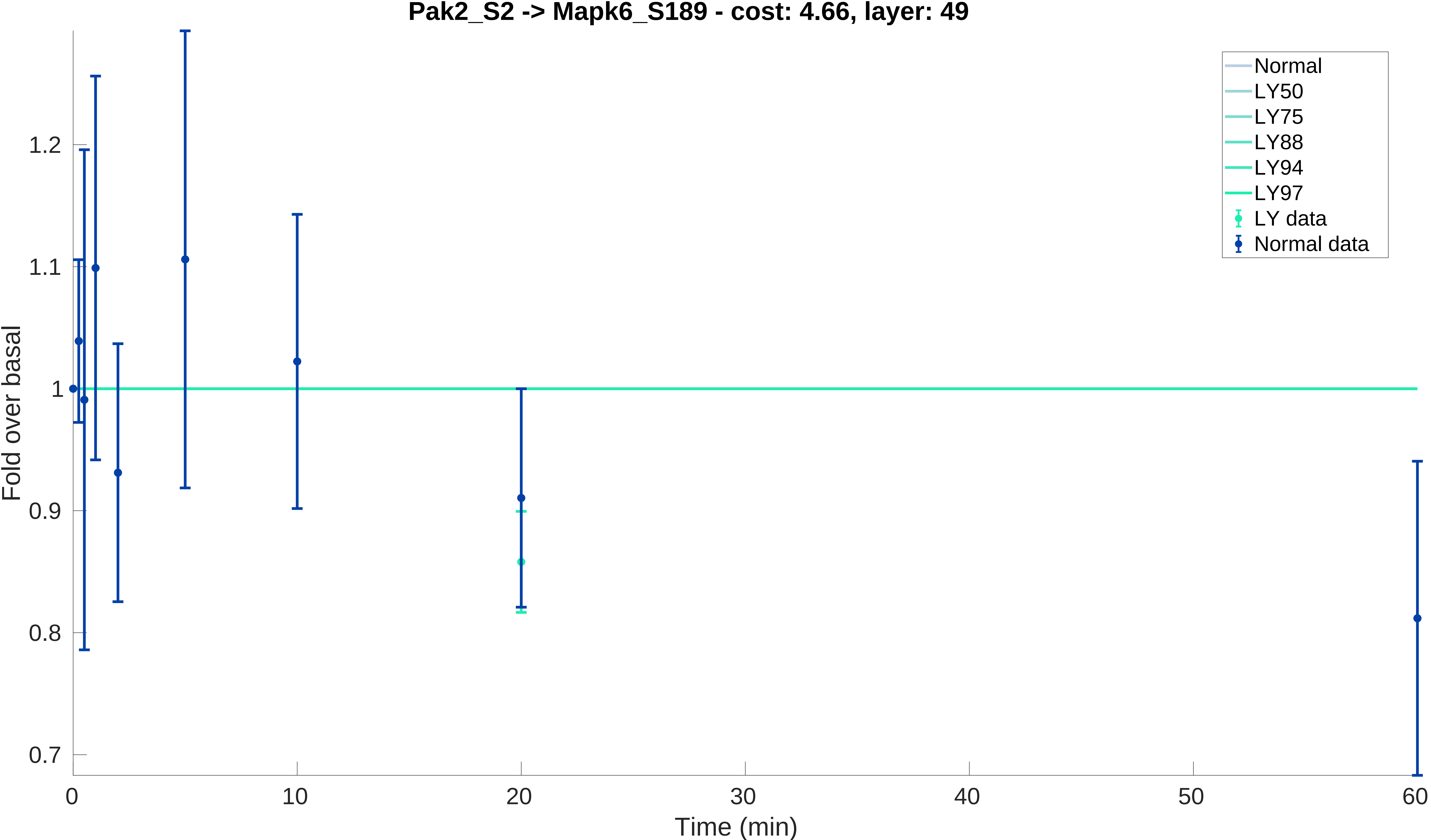

Pak2_S2 -> Mapk6_S189 - cost: 4.66, layer: 49
Fold over basal
1.2
1.1
1
0.9
0.8
0.7
0
10
20
30
40
50
60
Time (min)
Normal
LY50
LY75
LY88
LY94
LY97
LY data
Normal data

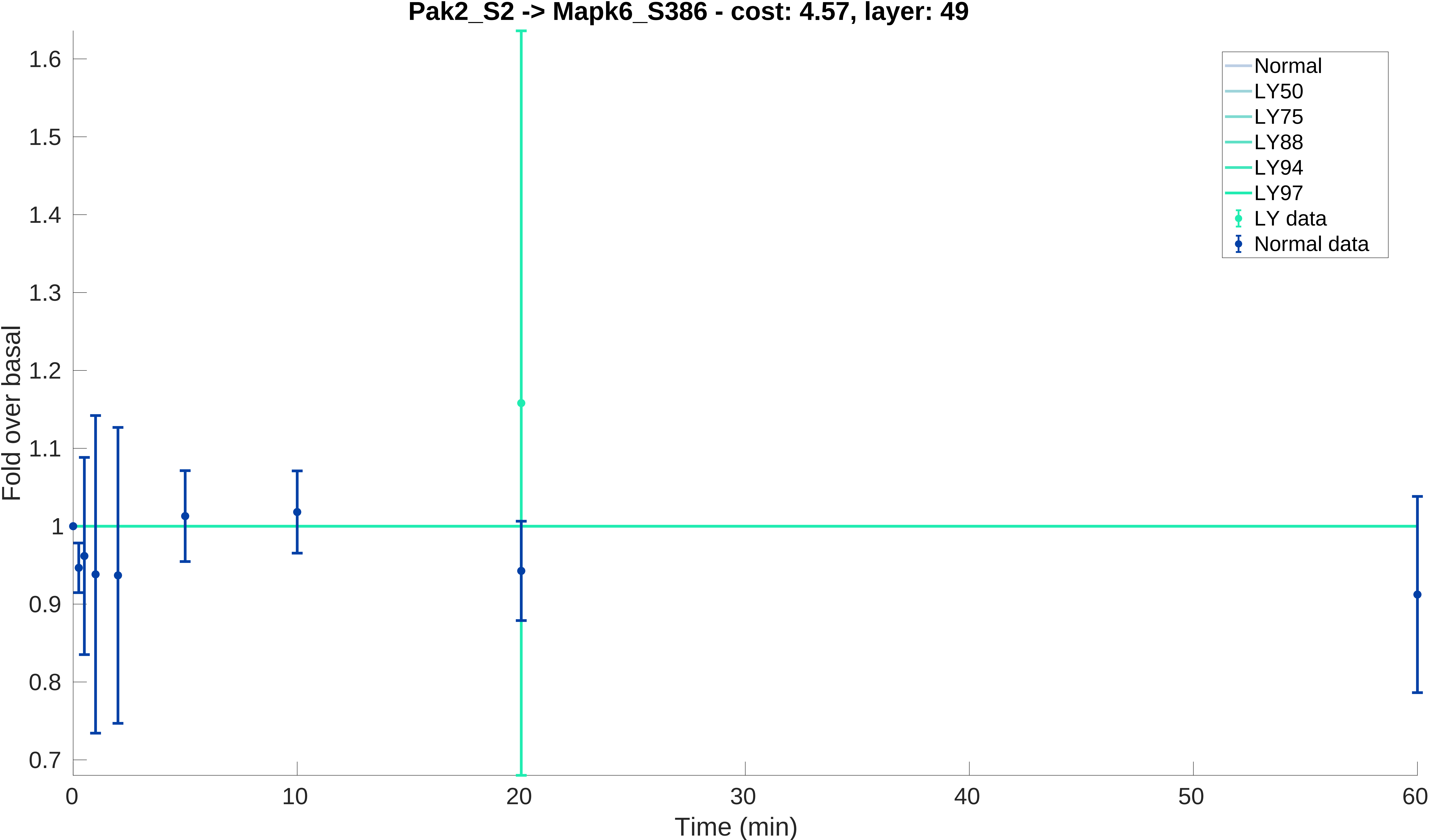

Pak2_S2 -> Mapk6_S386 - cost: 4.57, layer: 49
Normal
LY50
LY75
LY88
LY94
LY97
LY data
Normal data
Fold over basal
1.6
1.5
1.4
1.3
1.2
1.1
1
0.9
0.8
0.7
0
10
20
30
40
50
60
Time (min)

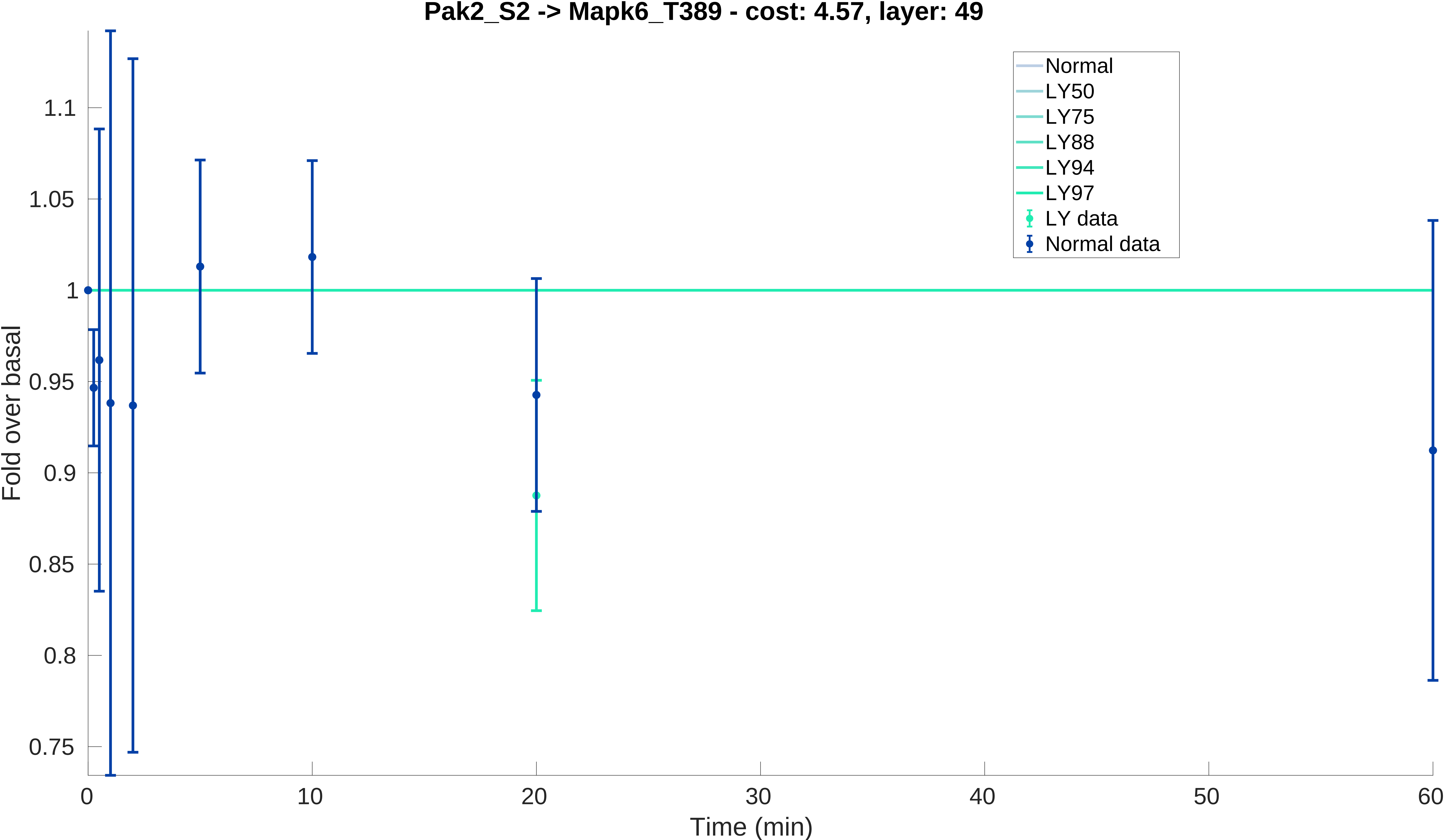

Pak2_S2 -> Mapk6_T389 - cost: 4.57, layer: 49
Normal
LY50
LY75
LY88
LY94
LY97
LY data
Normal data
Fold over basal
1.1
1.05
1
0.95
0.9
0.85
0.8
0.75
0
10
20
30
40
50
60
Time (min)

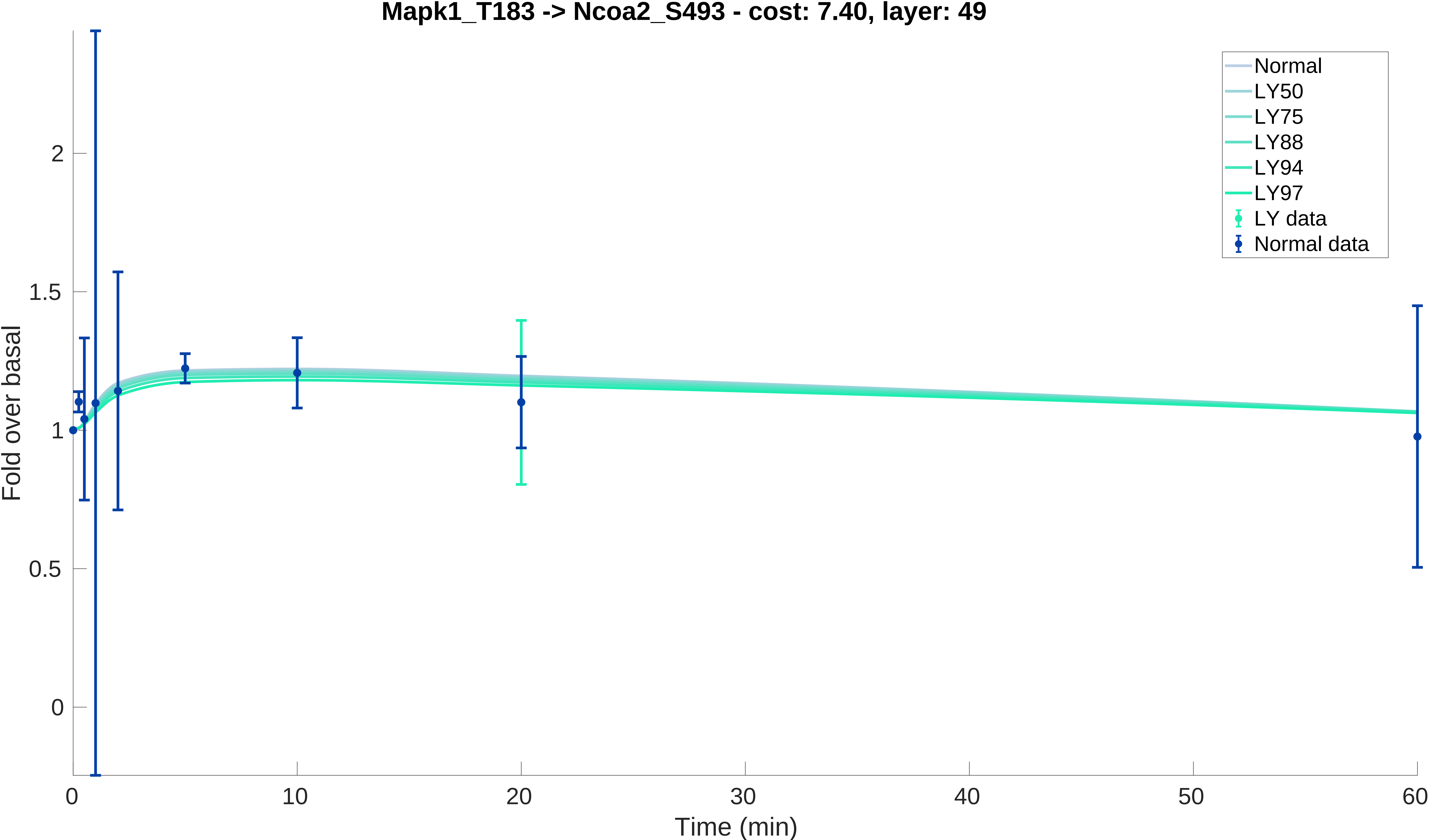

Mapk1_T183 -> Ncoa2_S493 - cost: 7.40, layer: 49
Normal
LY50
LY75
LY88
LY94
LY97
LY data
Normal data
Fold over basal
Time (min)
0
0.5
1
1.5
2
0
10
20
30
40
50
60

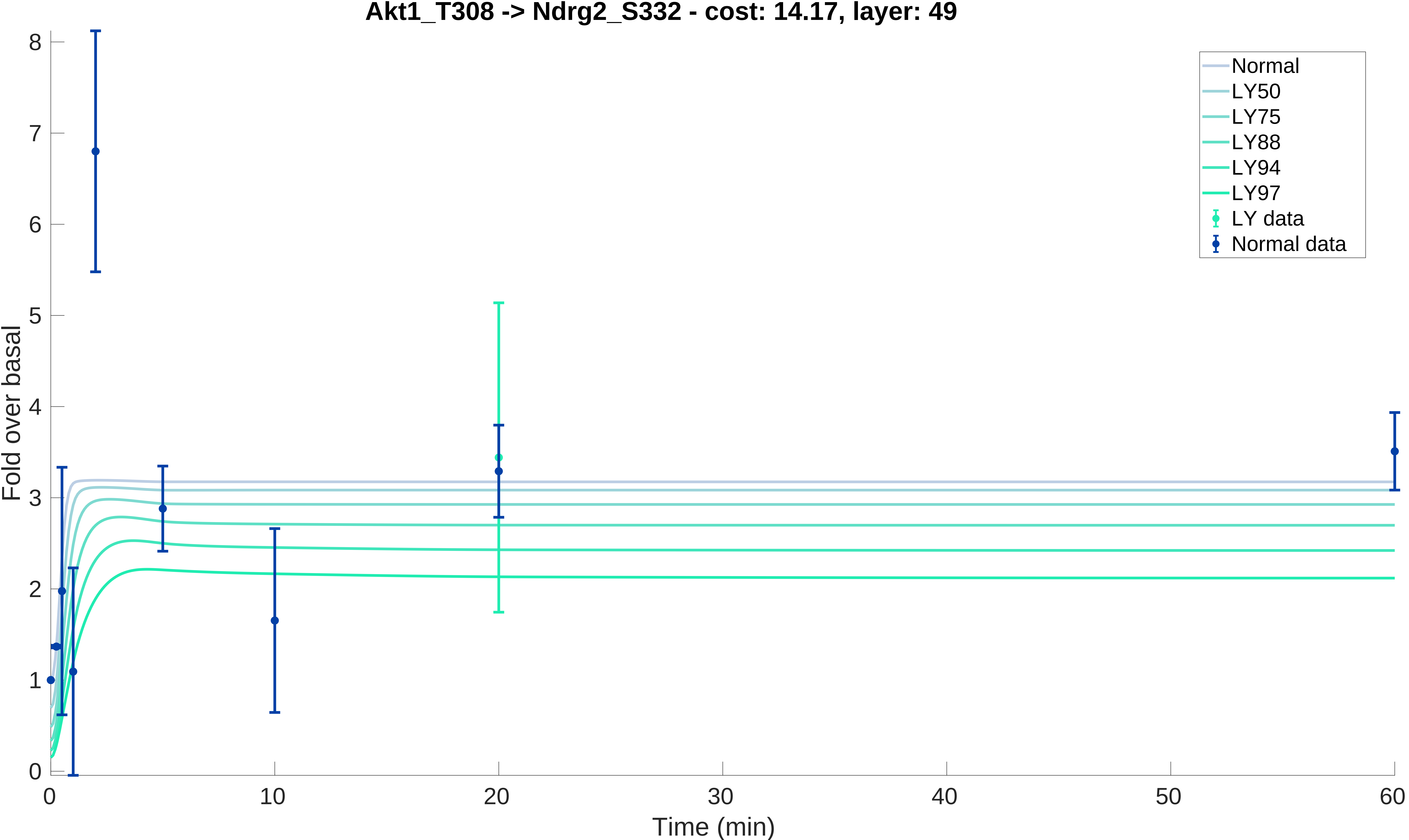
Akt1_T308 -> Ndrg2_S332 - cost: 14.17, layer: 49
Fold over basal
Time (min)
0
1
2
3
4
5
6
7
8
0
10
20
30
40
50
60
Normal
LY50
LY75
LY88
LY94
LY97
LY data
Normal data

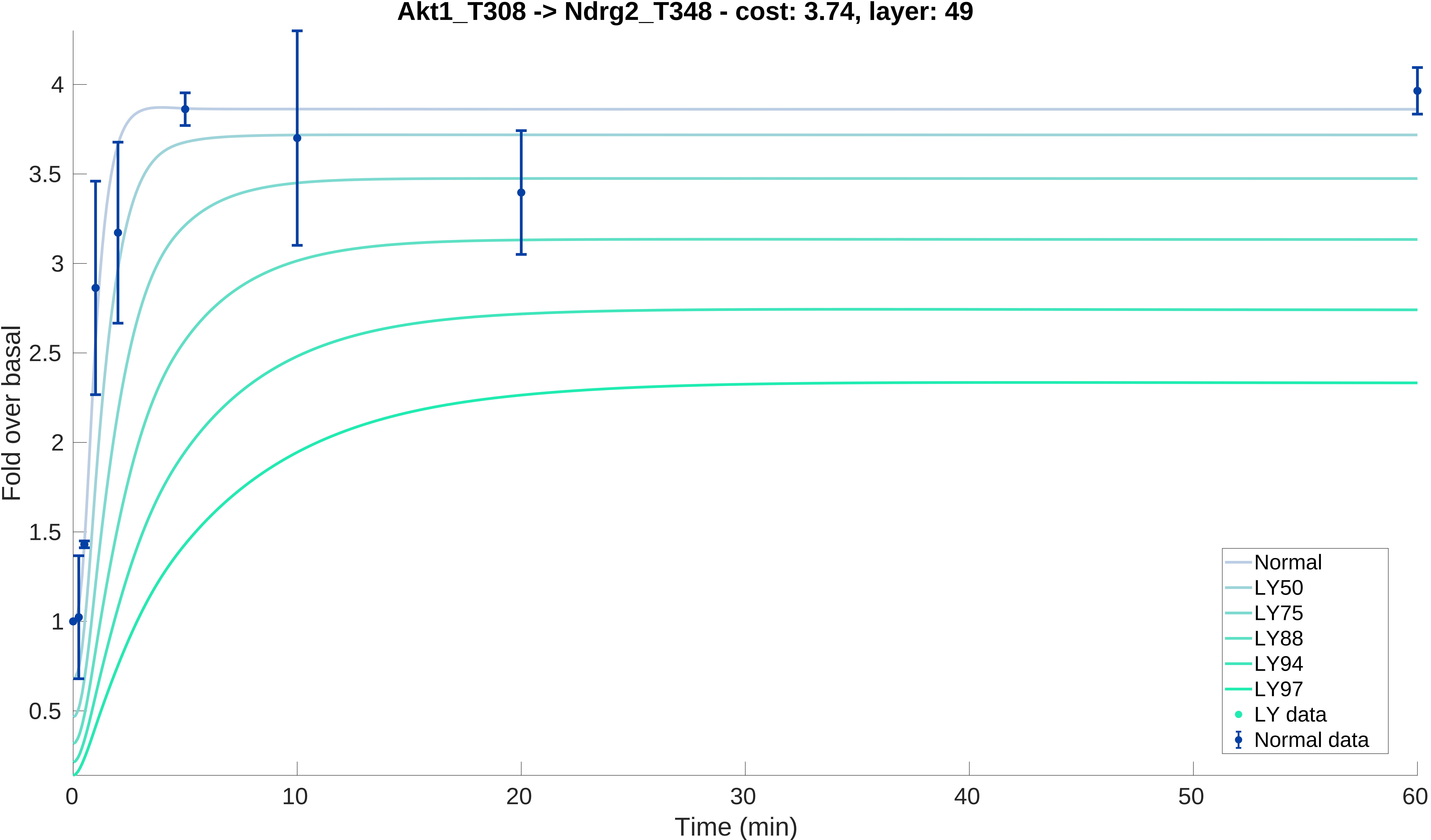

Akt1_T308 -> Ndrg2_T348 - cost: 3.74, layer: 49
Fold over basal
0.5
1
1.5
2
2.5
3
3.5
4
0
10
20
30
40
50
60
Time (min)
Normal
LY50
LY75
LY88
LY94
LY97
LY data
Normal data

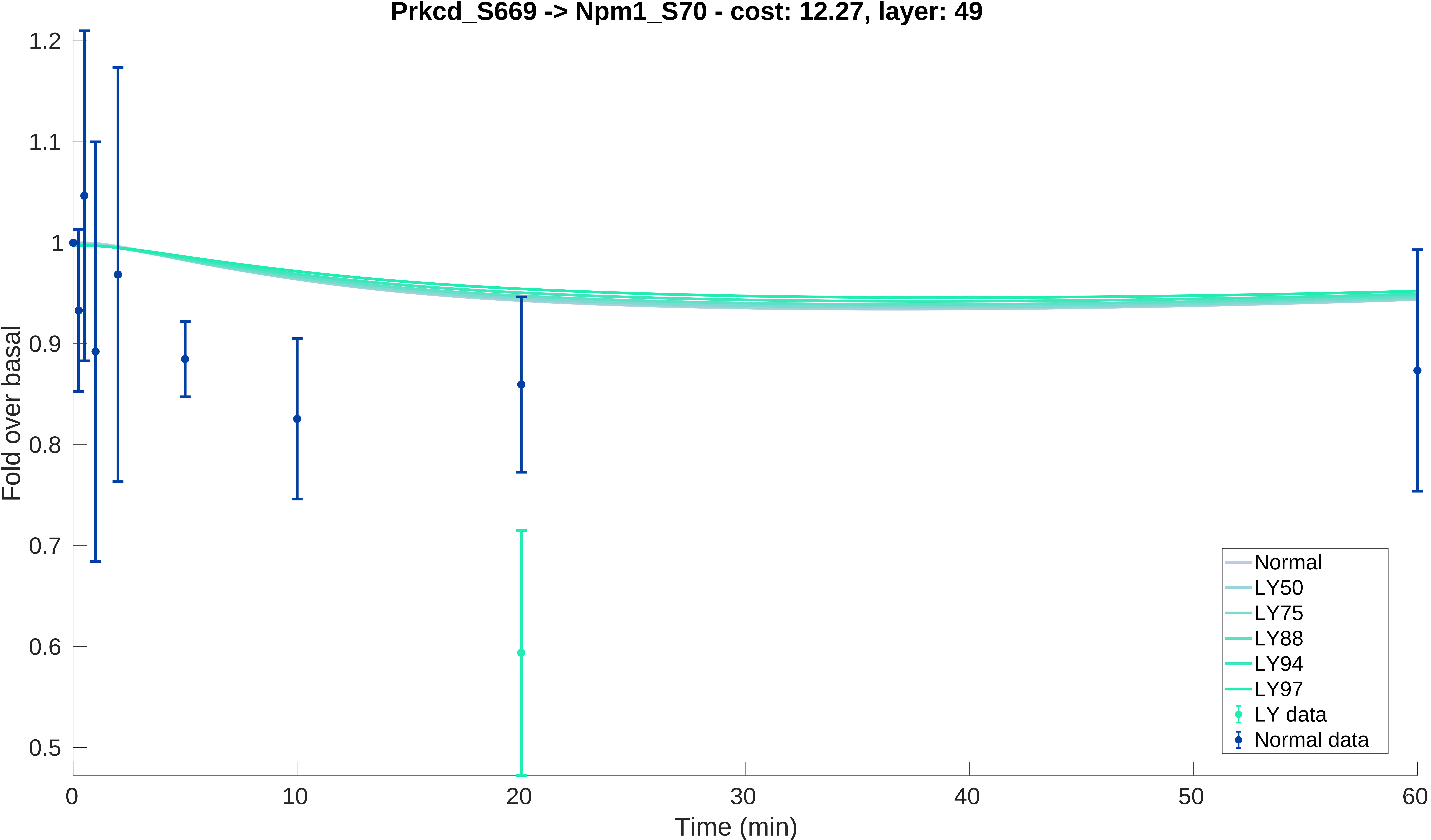
Prkcd_S669 -> Npm1_S70 - cost: 12.27, layer: 49
Fold over basal
Time (min)
1.2
1.1
1
0.9
0.8
0.7
0.6
0.5
0
10
20
30
40
50
60
Normal
LY50
LY75
LY88
LY94
LY97
LY data
Normal data

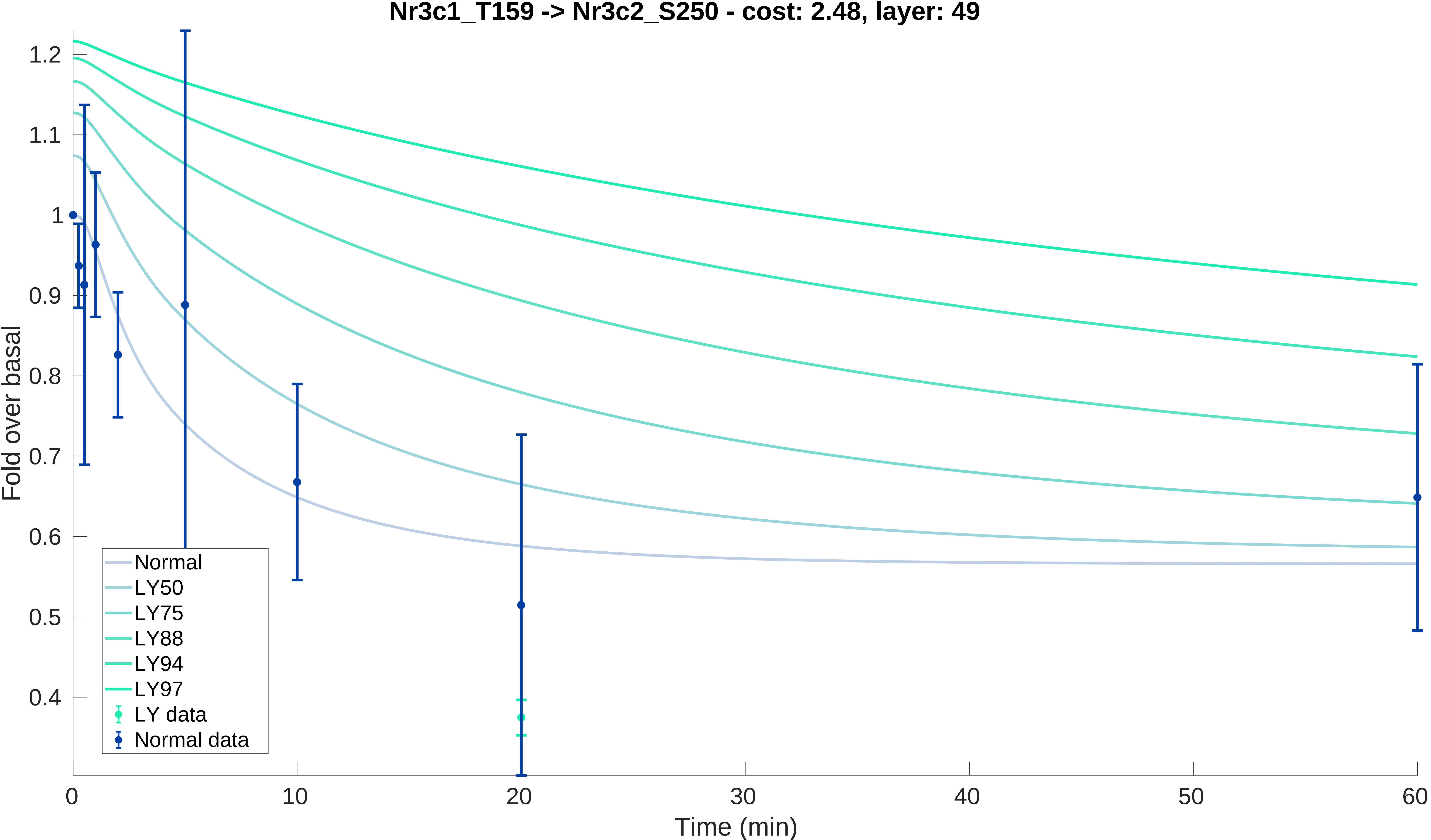

Nr3c1_T159 -> Nr3c2_S250 - cost: 2.48, layer: 49
Fold over basal
Time (min)
1.2
1.1
1
0.9
0.8
0.7
0.6
0.5
0.4
0
10
20
30
40
50
60
Normal
LY50
LY75
LY88
LY94
LY97
LY data
Normal data

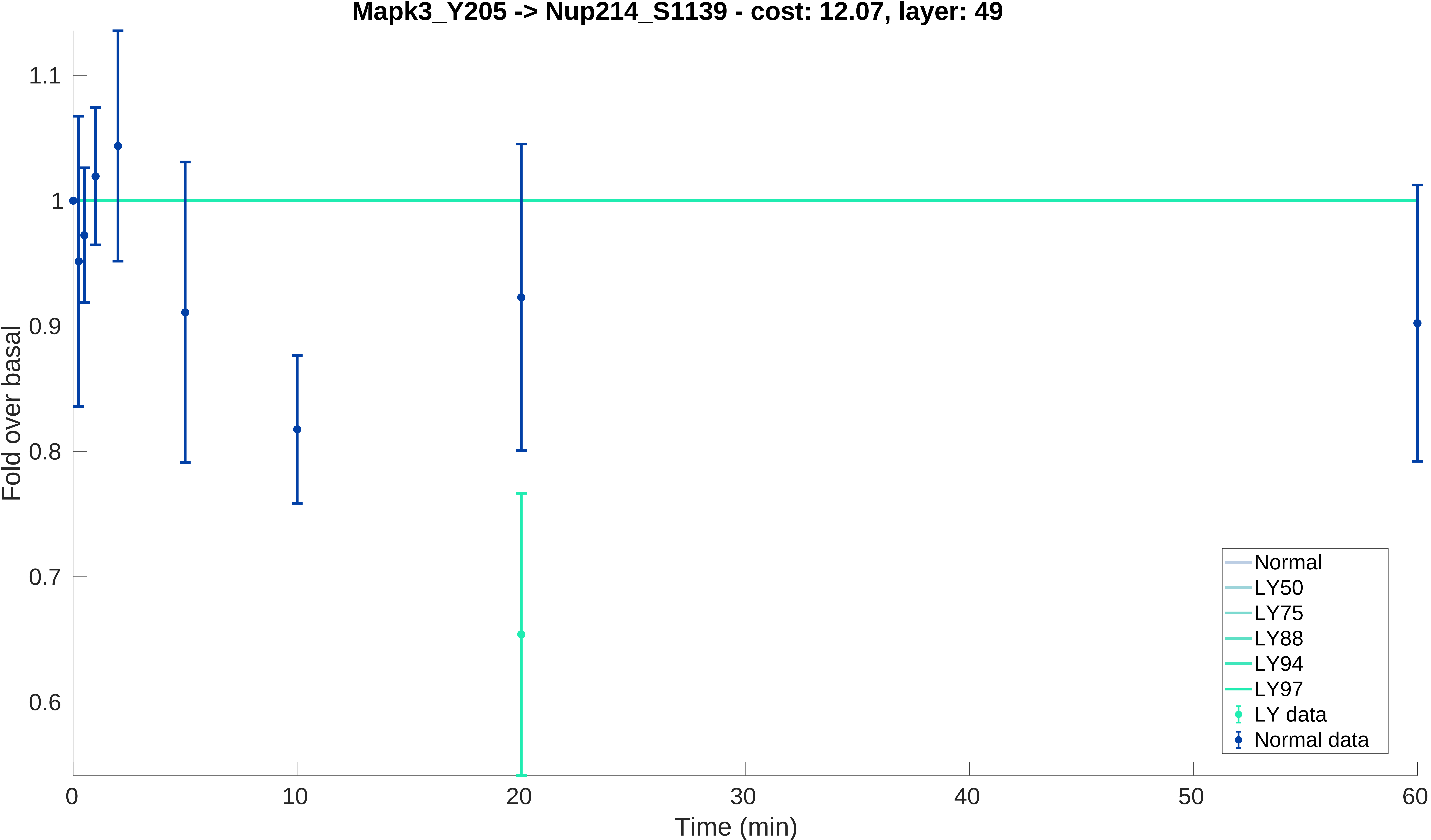
Mapk3_Y205 -> Nup214_S1139 - cost: 12.07, layer: 49
Fold over basal
1.1
1
0.9
0.8
0.7
0.6
0
10
20
30
40
50
60
Time (min)
Normal
LY50
LY75
LY88
LY94
LY97
LY data
Normal data

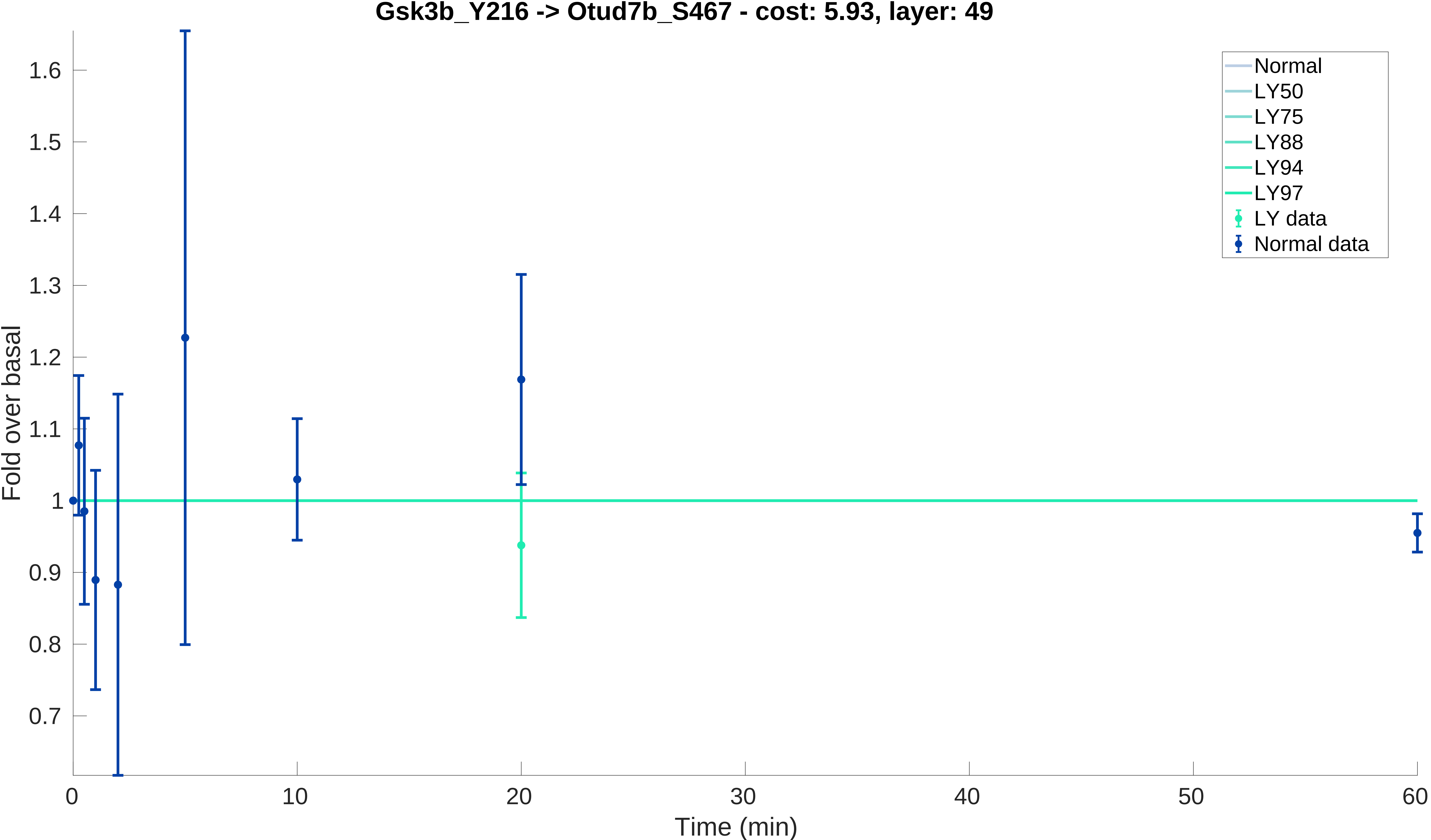

Gsk3b_Y216 -> Otud7b_S467 - cost: 5.93, layer: 49
Fold over basal
1.6
1.5
1.4
1.3
1.2
1.1
1
0.9
0.8
0.7
0
10
20
30
40
50
60
Time (min)
Normal
LY50
LY75
LY88
LY94
LY97
LY data
Normal data

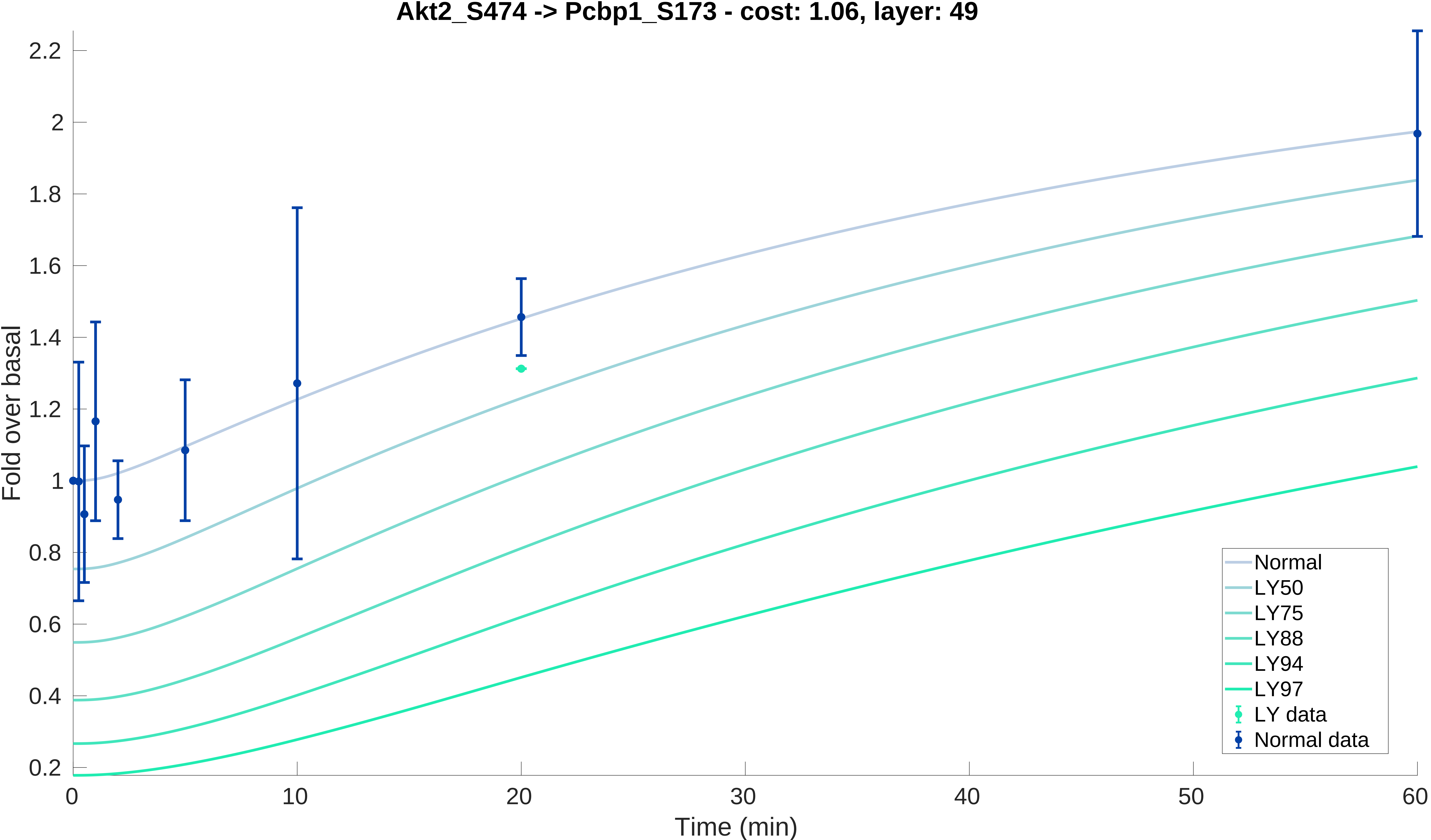

Akt2_S474 -> Pcbp1_S173 - cost: 1.06, layer: 49
Fold over basal
Time (min)
0
10
20
30
40
50
60
0.2
0.4
0.6
0.8
1
1.2
1.4
1.6
1.8
2
2.2
Normal
LY50
LY75
LY88
LY94
LY97
LY data
Normal data

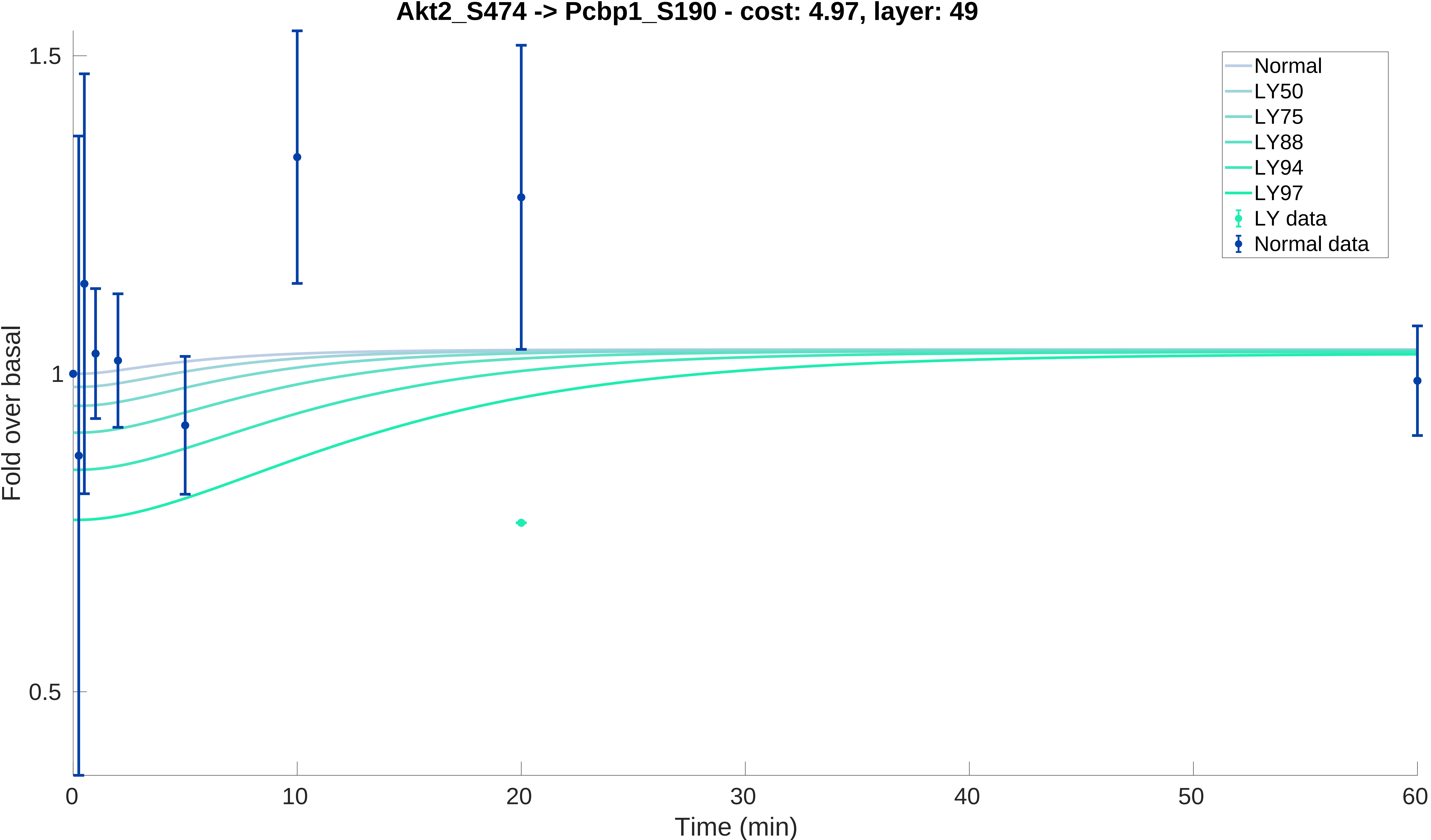

Akt2_S474 -> Pcbp1_S190 - cost: 4.97, layer: 49
Fold over basal
Time (min)
1.5
1
0.5
0
10
20
30
40
50
60
Normal
LY50
LY75
LY88
LY94
LY97
LY data
Normal data

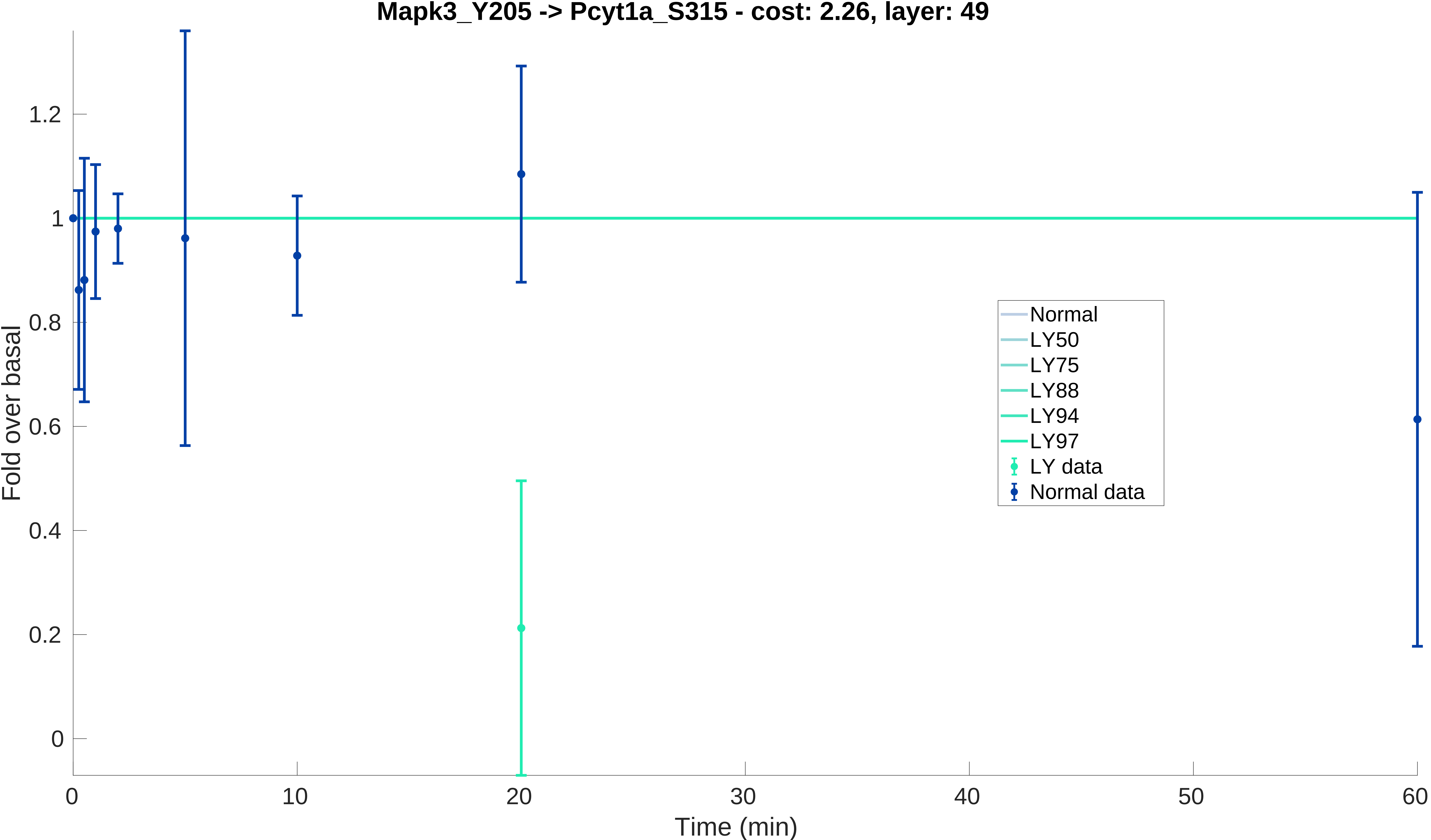
Mapk3_Y205 -> Pcyt1a_S315 - cost: 2.26, layer: 49
Fold over basal
1.2
1
0.8
0.6
0.4
0.2
0
0
10
20
30
40
50
60
Time (min)
Normal
LY50
LY75
LY88
LY94
LY97
LY data
Normal data

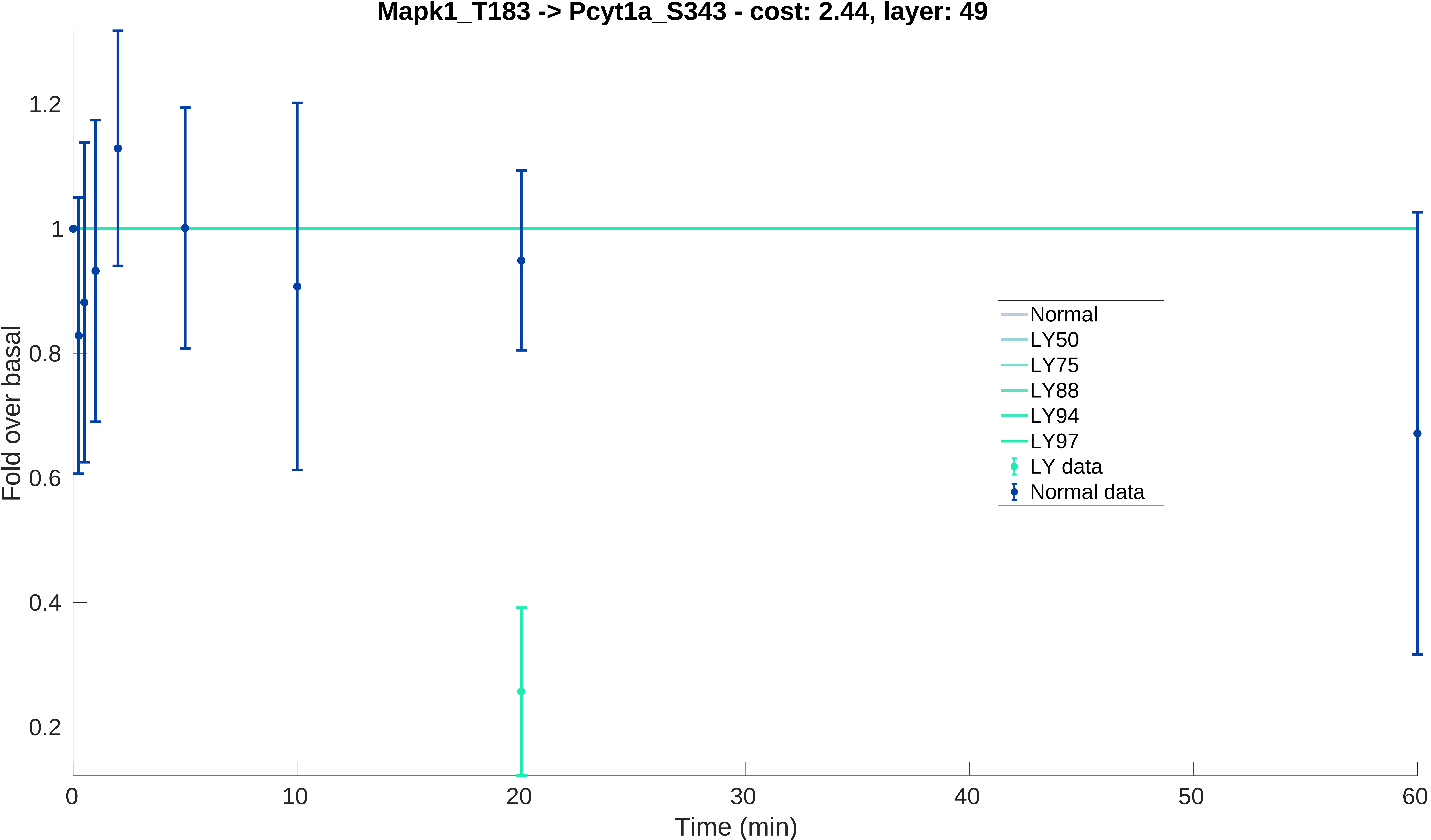

Mapk1_T183 -> Pcyt1a_S343 - cost: 2.44, layer: 49
Fold over basal
1.2
1
0.8
0.6
0.4
0.2
0
10
20
30
40
50
60
Time (min)
Normal
LY50
LY75
LY88
LY94
LY97
LY data
Normal data

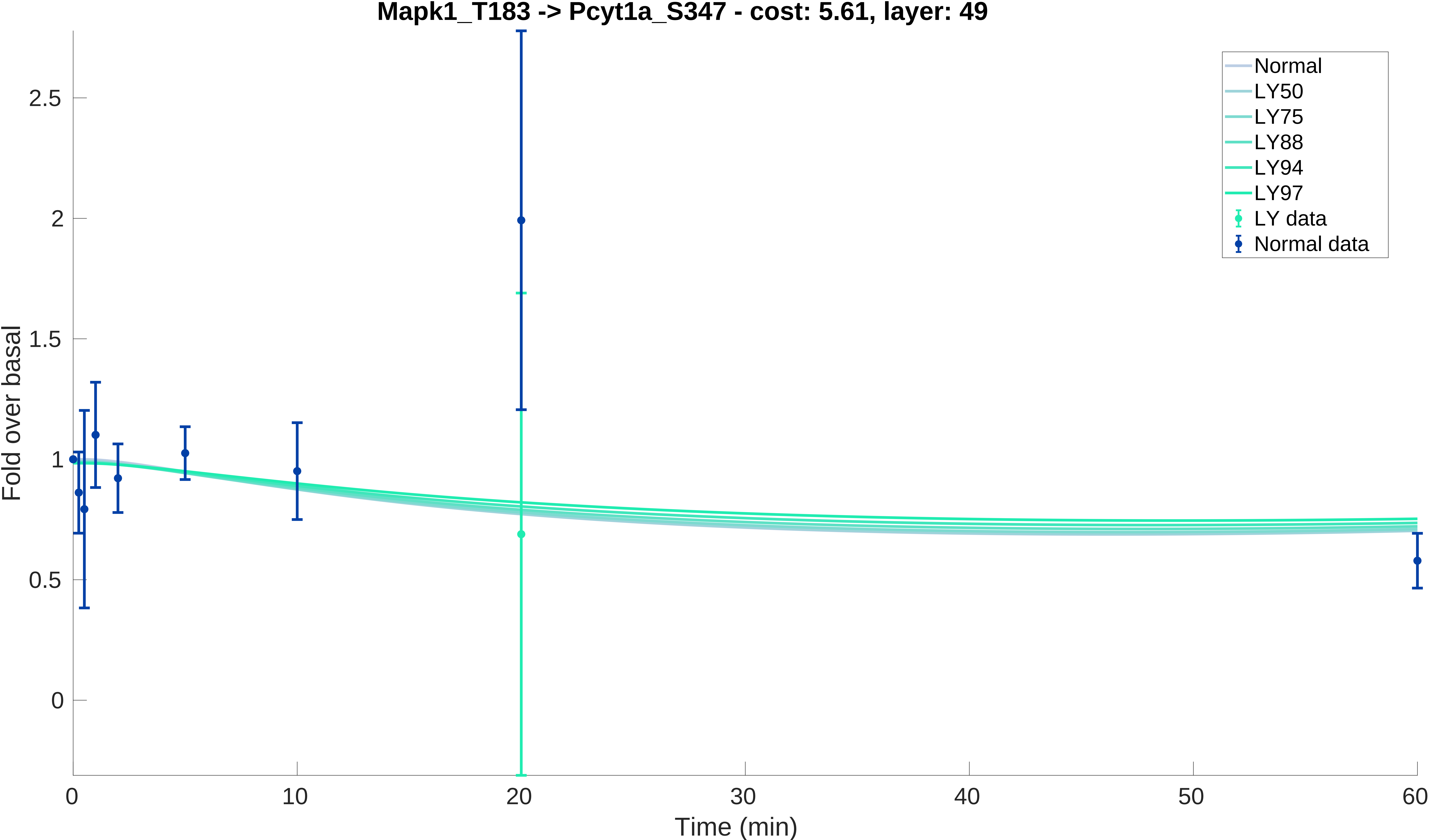

Mapk1_T183 -> Pcyt1a_S347 - cost: 5.61, layer: 49
Fold over basal
Time (min)
0
10
20
30
40
50
60
0
0.5
1
1.5
2
2.5
Normal
LY50
LY75
LY88
LY94
LY97
LY data
Normal data

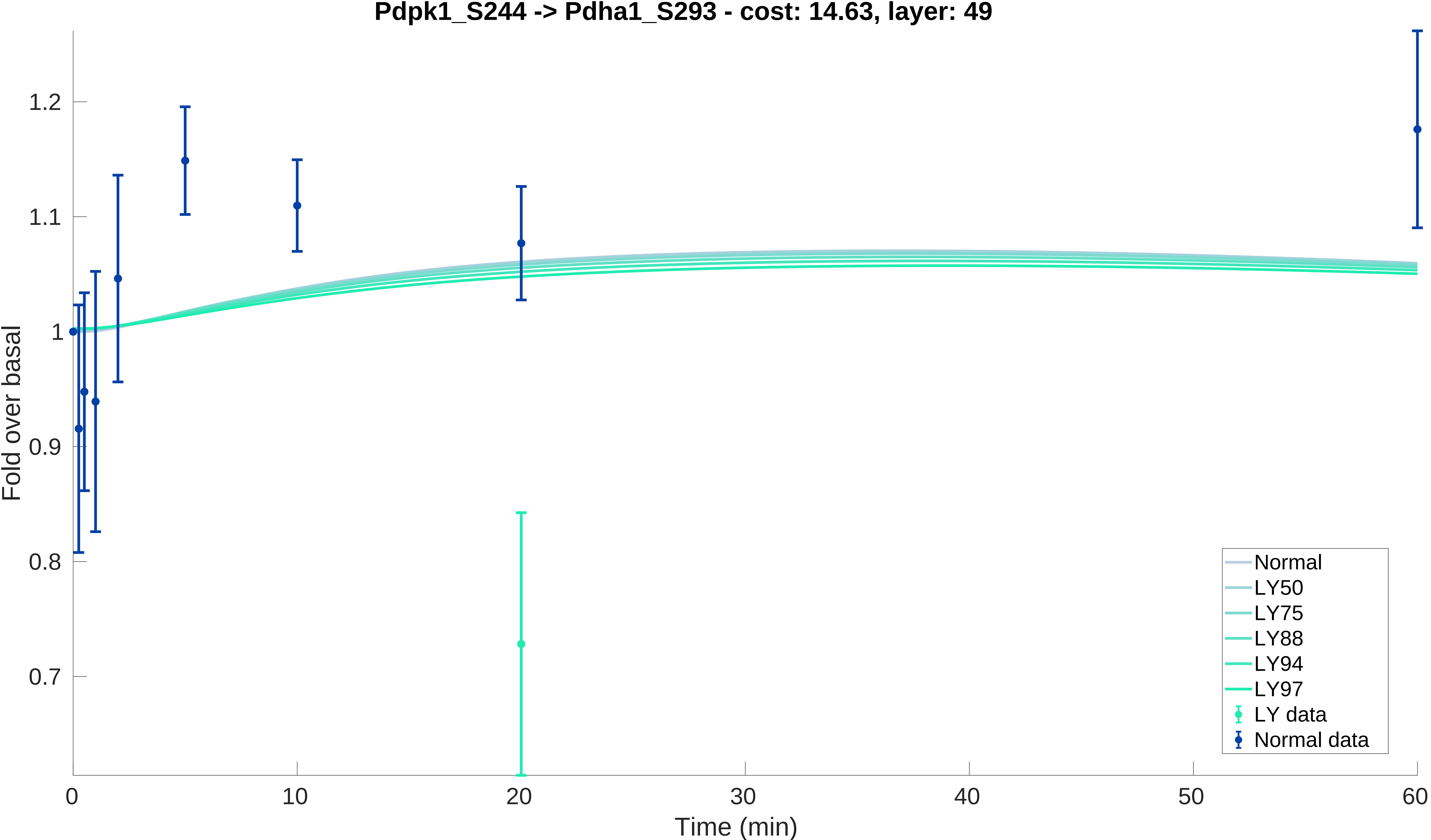

Pdpk1_S244 -> Pdha1_S293 - cost: 14.63, layer: 49
Fold over basal
Time (min)
1.2
1.1
1
0.9
0.8
0.7
0
10
20
30
40
50
60
Normal
LY50
LY75
LY88
LY94
LY97
LY data
Normal data

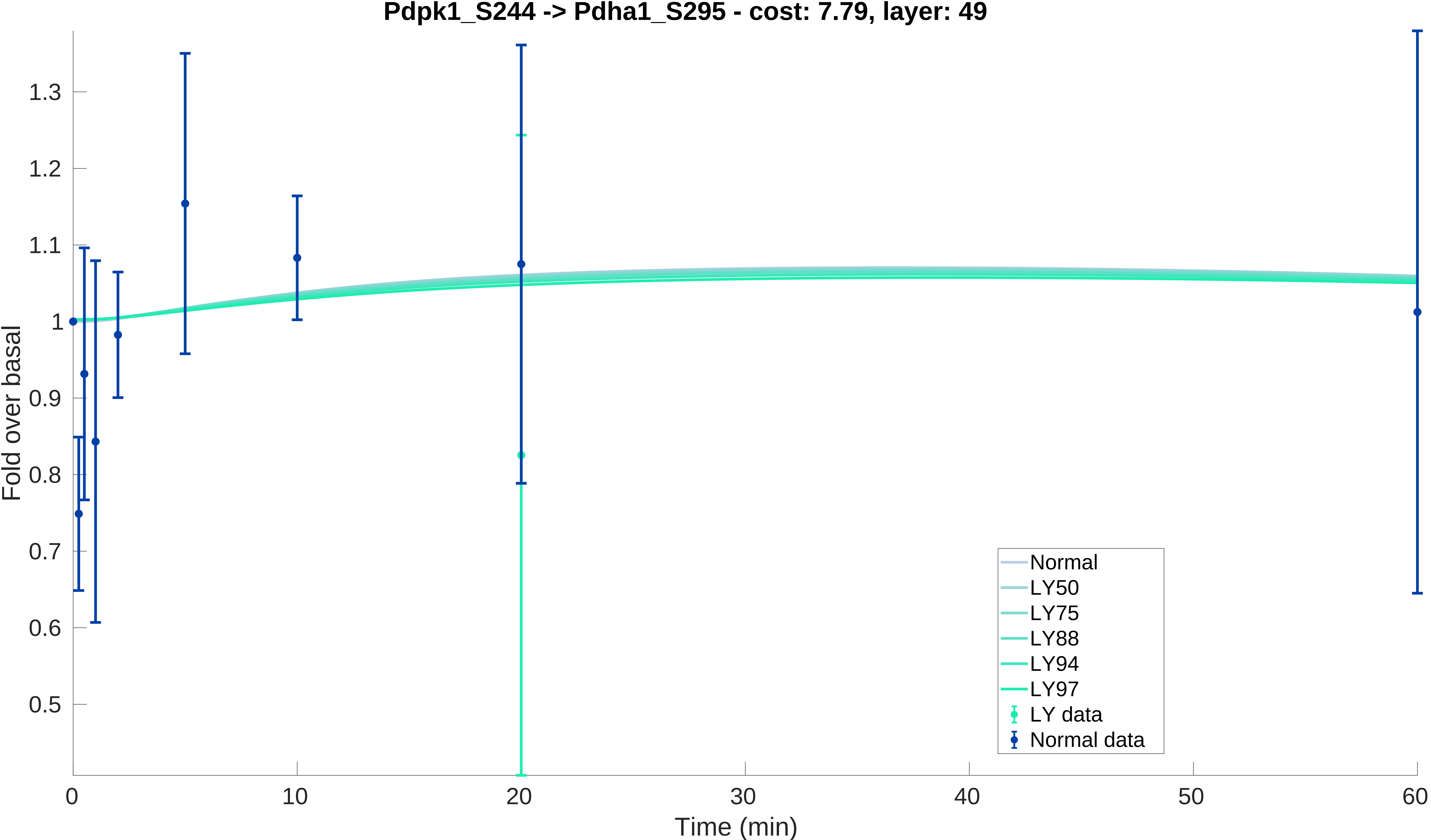
Pdpk1_S244 -> Pdha1_S295 - cost: 7.79, layer: 49
Fold over basal
Time (min)
0
10
20
30
40
50
60
0.5
0.6
0.7
0.8
0.9
1
1.1
1.2
1.3
Normal
LY50
LY75
LY88
LY94
LY97
LY data
Normal data

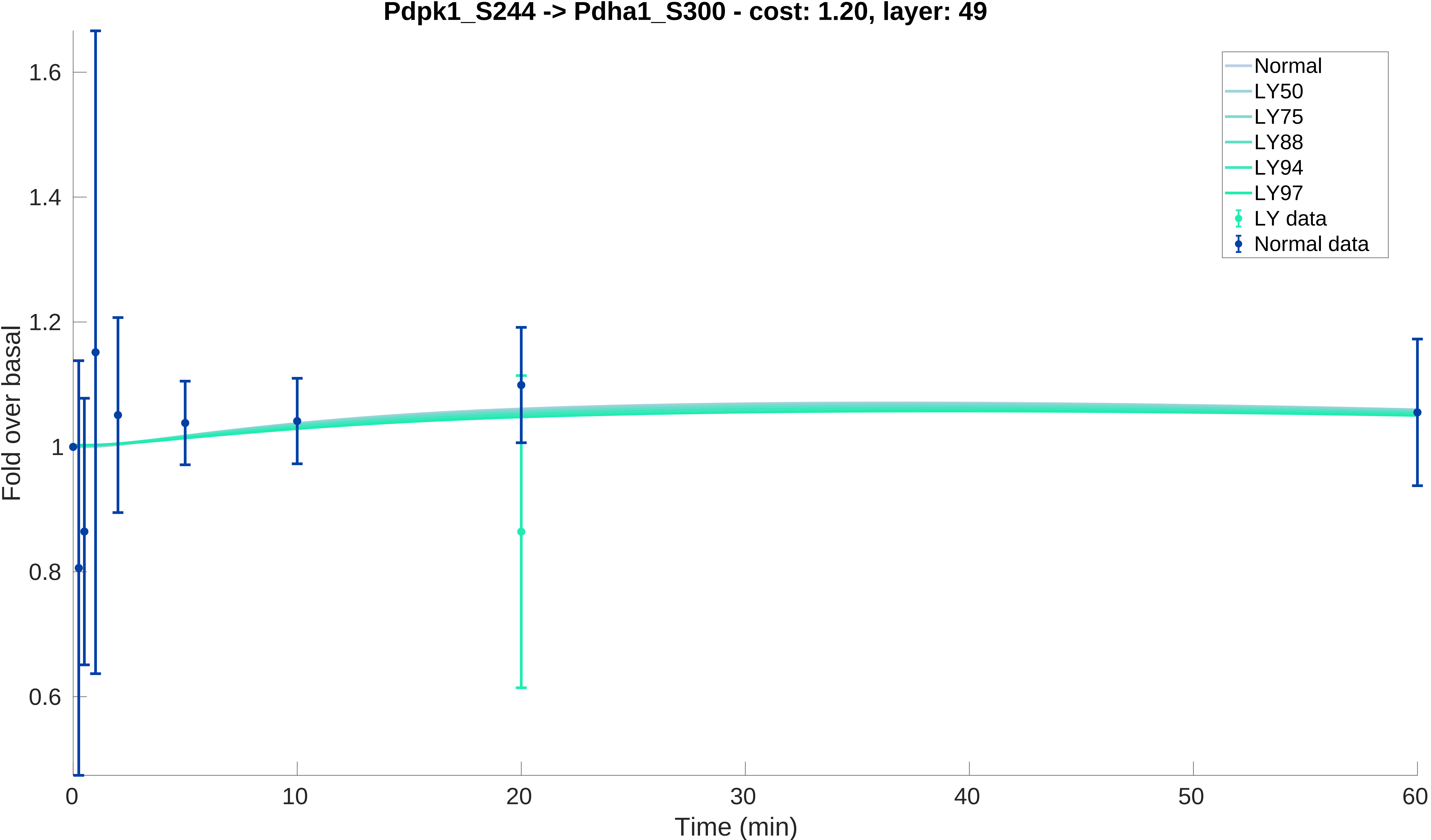

Pdpk1_S244 -> Pdha1_S300 - cost: 1.20, layer: 49
Fold over basal
1.6
1.4
1.2
1
0.8
0.6
0
10
20
30
40
50
60
Time (min)
Normal
LY50
LY75
LY88
LY94
LY97
LY data
Normal data

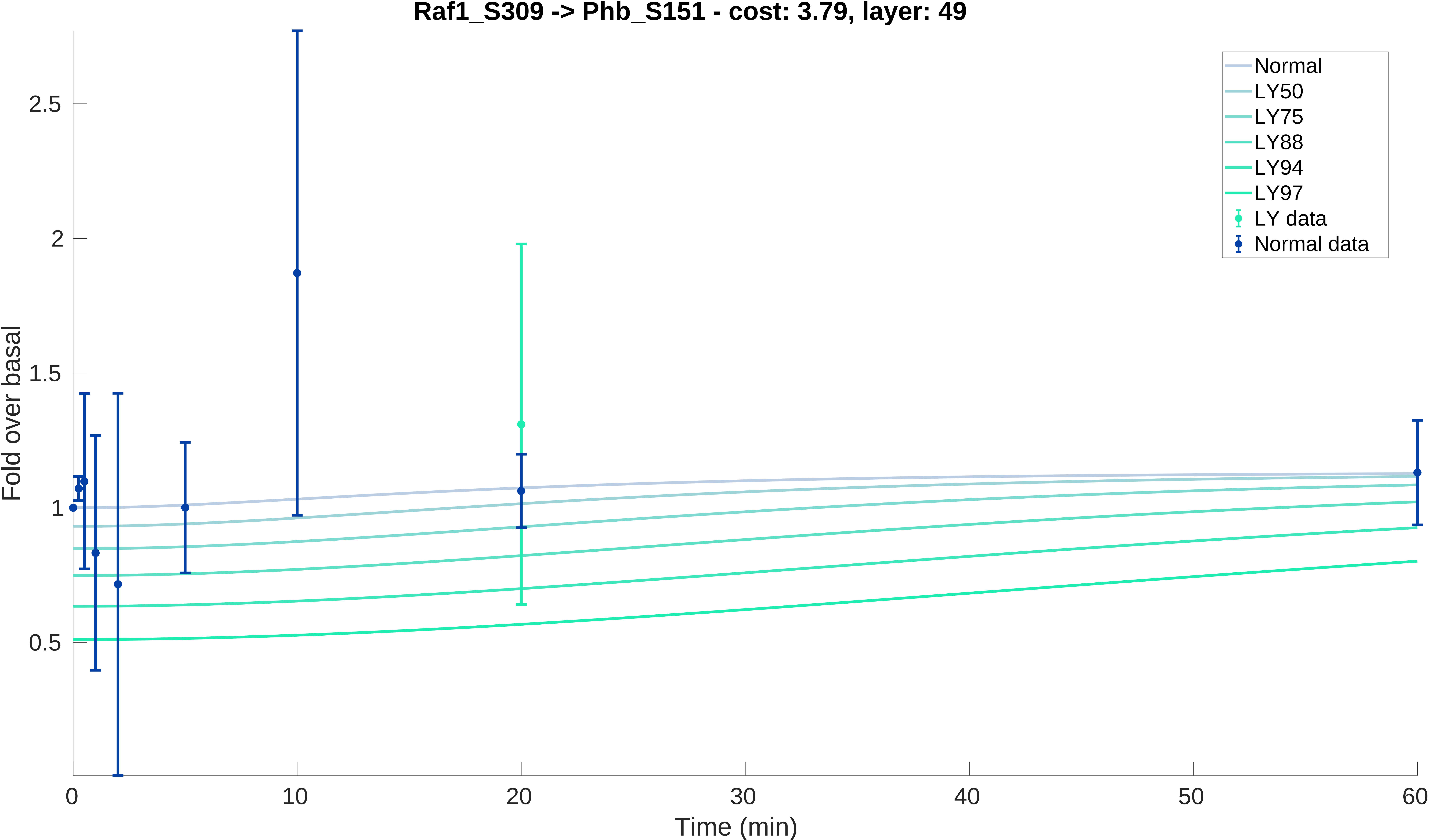

Raf1_S309 -> Phb_S151 - cost: 3.79, layer: 49
Normal
LY50
LY75
LY88
LY94
LY97
LY data
Normal data
Fold over basal
2.5
2
1.5
1
0.5
0
10
20
30
40
50
60
Time (min)

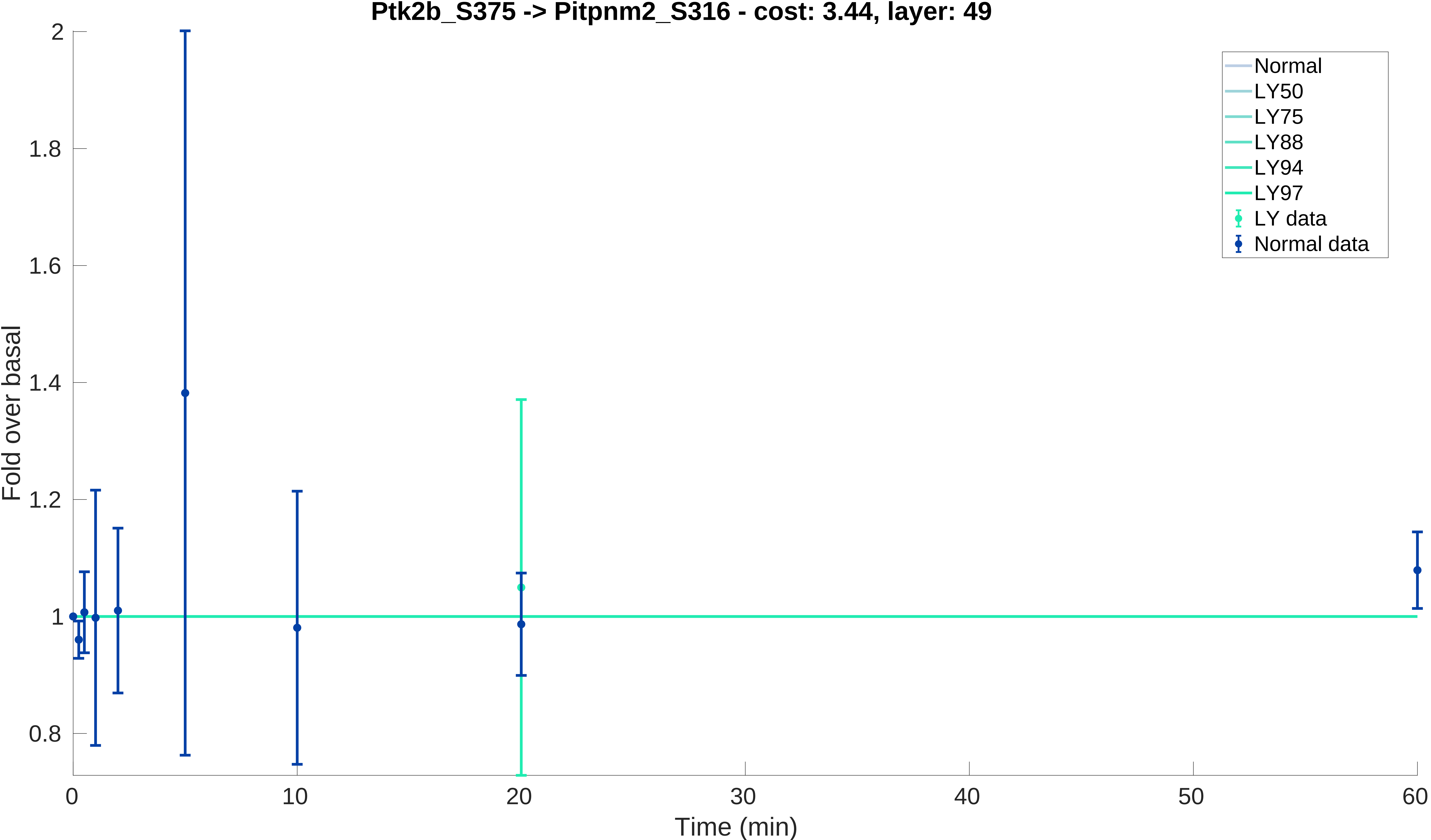
Ptk2b_S375 -> Pitpnm2_S316 - cost: 3.44, layer: 49
Fold over basal
Time (min)
2
1.8
1.6
1.4
1.2
1
0.8
0
10
20
30
40
50
60
Normal
LY50
LY75
LY88
LY94
LY97
LY data
Normal data

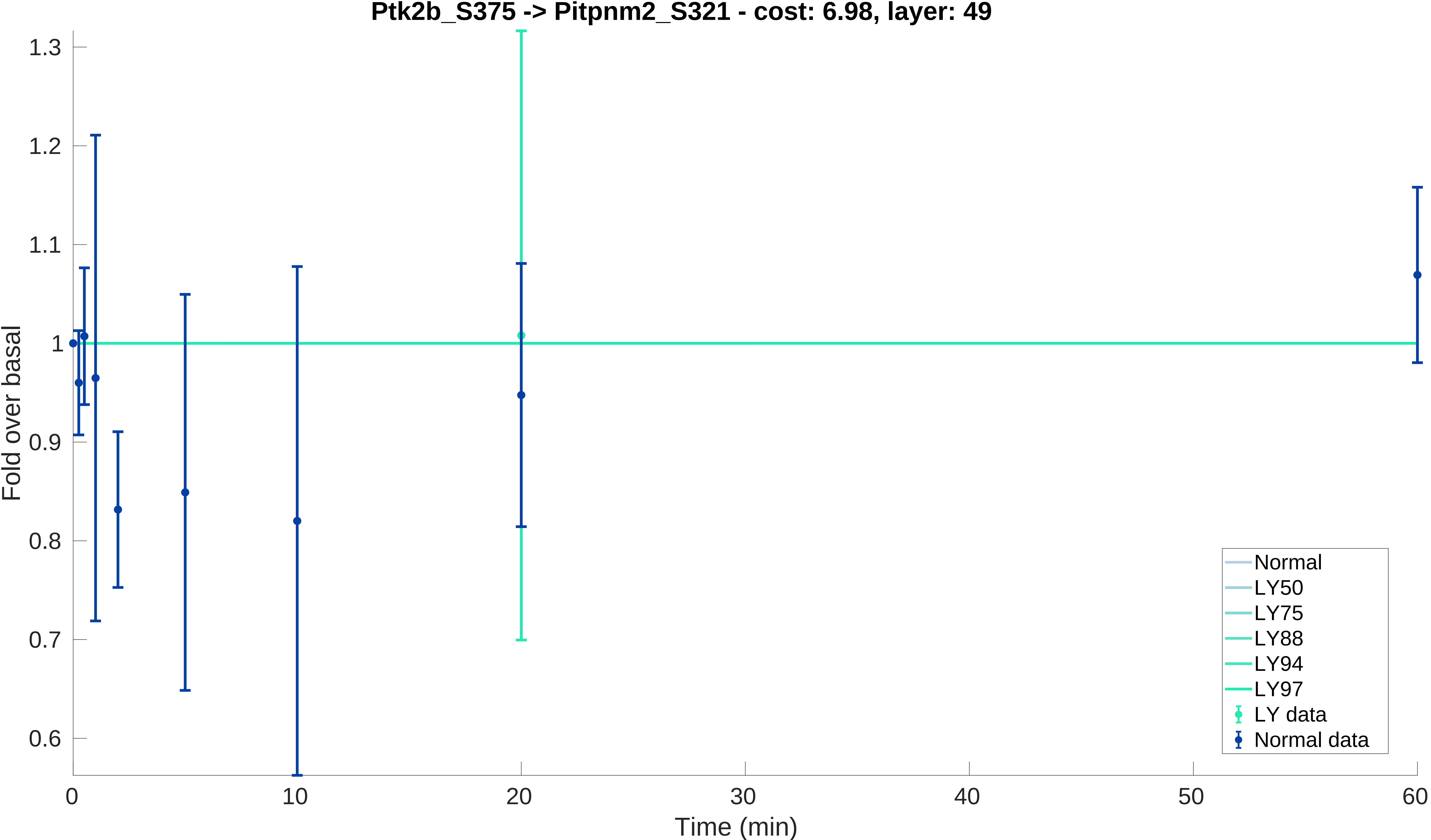

Ptk2b_S375 -> Pitpnm2_S321 - cost: 6.98, layer: 49
Fold over basal
Time (min)
1.3
1.2
1.1
1
0.9
0.8
0.7
0.6
0
10
20
30
40
50
60
Normal
LY50
LY75
LY88
LY94
LY97
LY data
Normal data

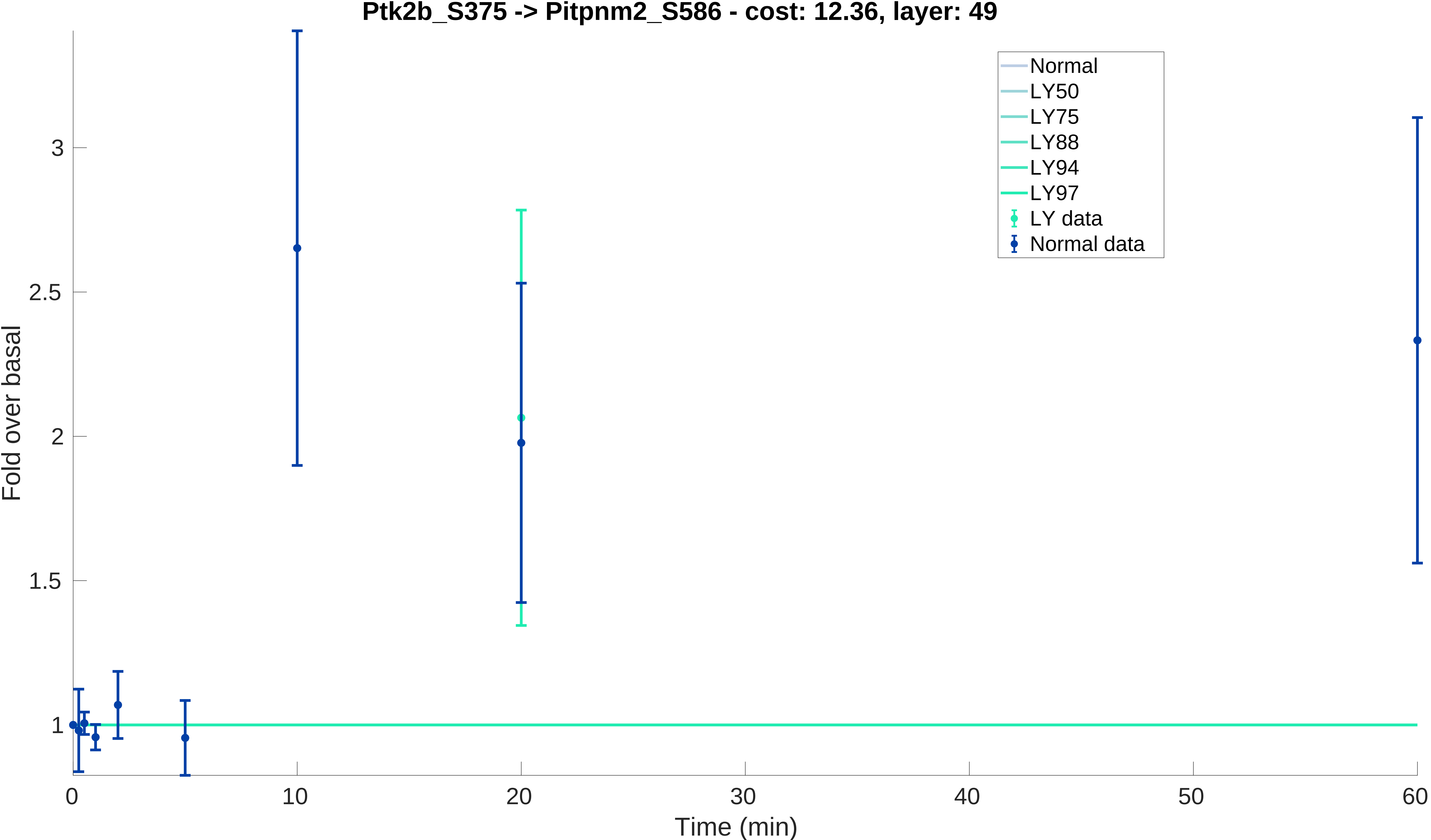

Ptk2b_S375 -> Pitpnm2_S586 - cost: 12.36, layer: 49
Normal
LY50
LY75
LY88
LY94
LY97
LY data
Normal data
Fold over basal
Time (min)
1
1.5
2
2.5
3
0
10
20
30
40
50
60

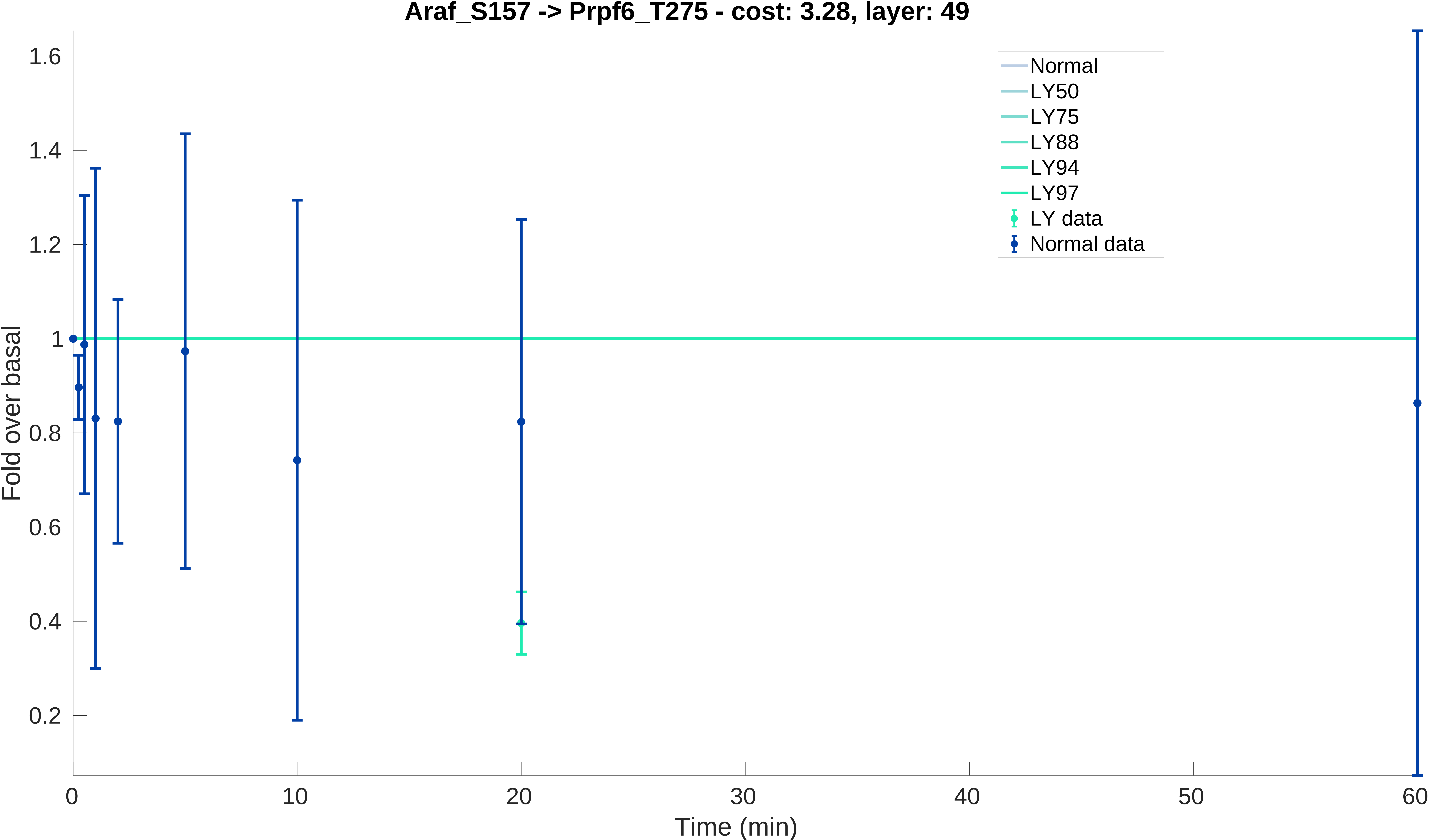

Araf_S157 -> Prpf6_T275 - cost: 3.28, layer: 49
Fold over basal
Time (min)
1.6
1.4
1.2
1
0.8
0.6
0.4
0.2
0
10
20
30
40
50
60
Normal
LY50
LY75
LY88
LY94
LY97
LY data
Normal data

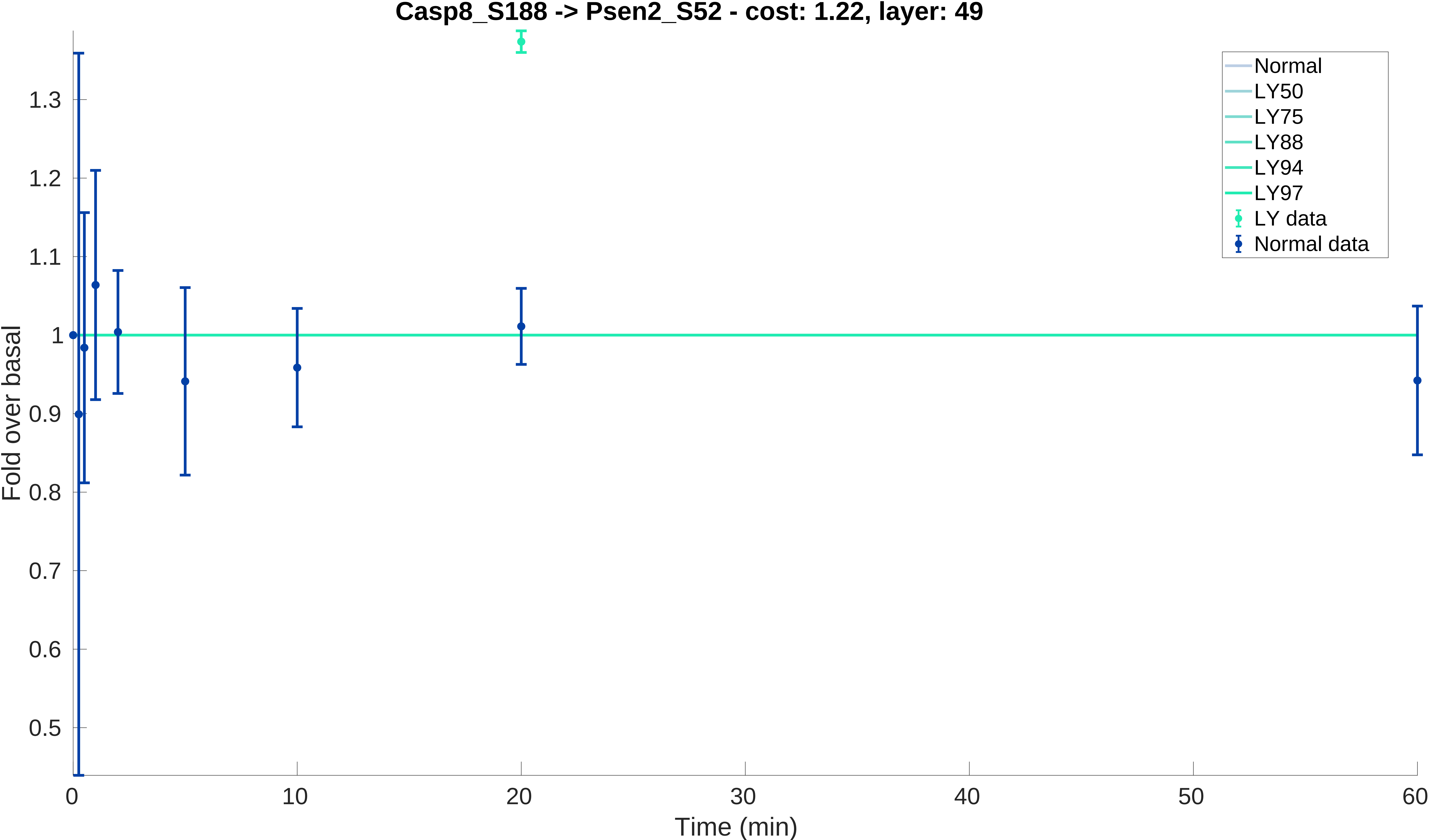

Casp8_S188 -> Psen2_S52 - cost: 1.22, layer: 49
Fold over basal
Time (min)
0.5
0.6
0.7
0.8
0.9
1
1.1
1.2
1.3
0
10
20
30
40
50
60
Normal
LY50
LY75
LY88
LY94
LY97
LY data
Normal data

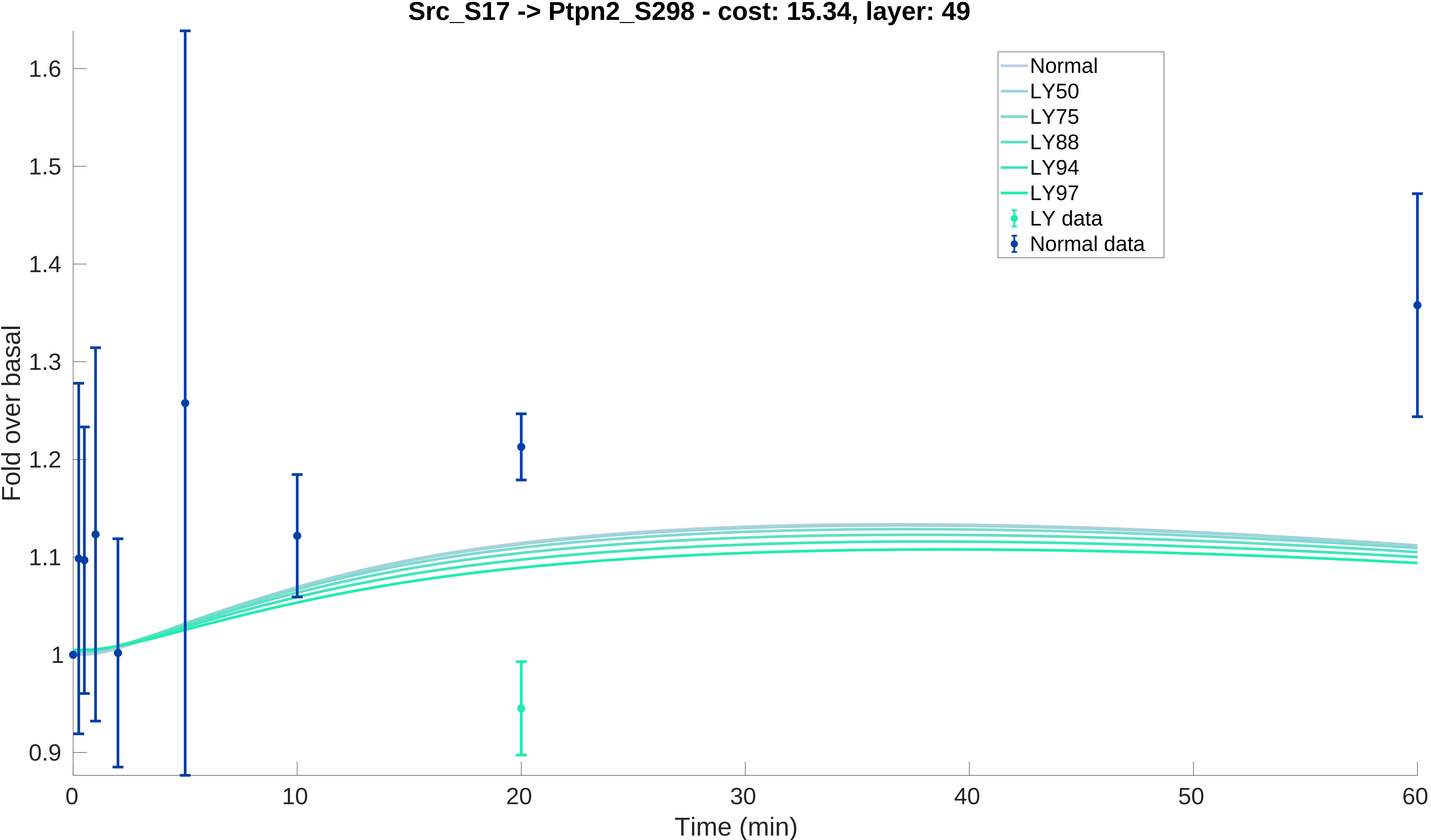

Src_S17 -> Ptpn2_S298 - cost: 15.34, layer: 49
Normal
LY50
LY75
LY88
LY94
LY97
LY data
Normal data
Fold over basal
1.6
1.5
1.4
1.3
1.2
1.1
1
0.9
0
10
20
30
40
50
60
Time (min)

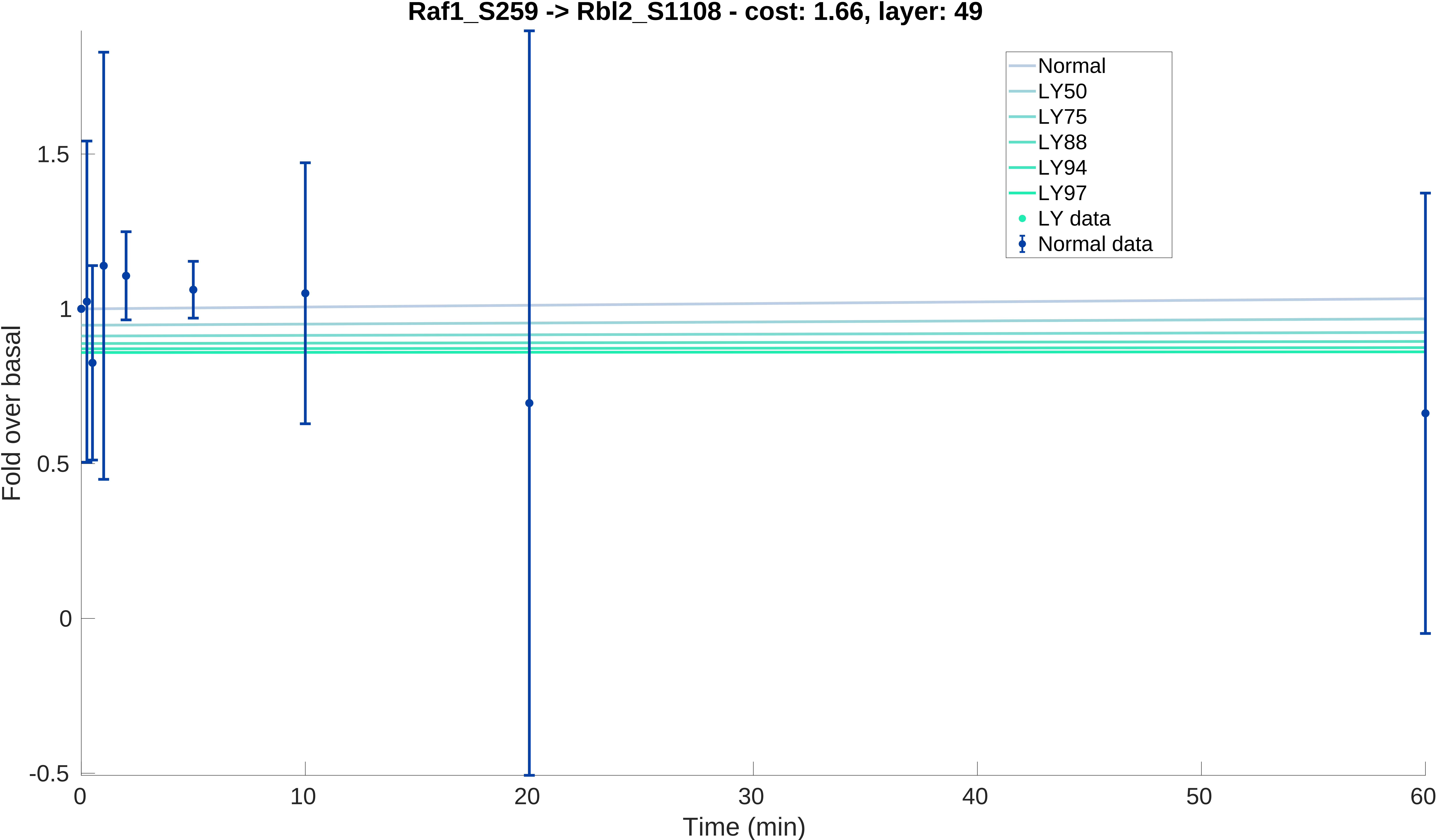

Raf1_S259 -> Rbl2_S1108 - cost: 1.66, layer: 49
Normal
LY50
LY75
LY88
LY94
LY97
LY data
Normal data
Fold over basal
1.5
1
0.5
0
-0.5
0
10
20
30
40
50
60
Time (min)

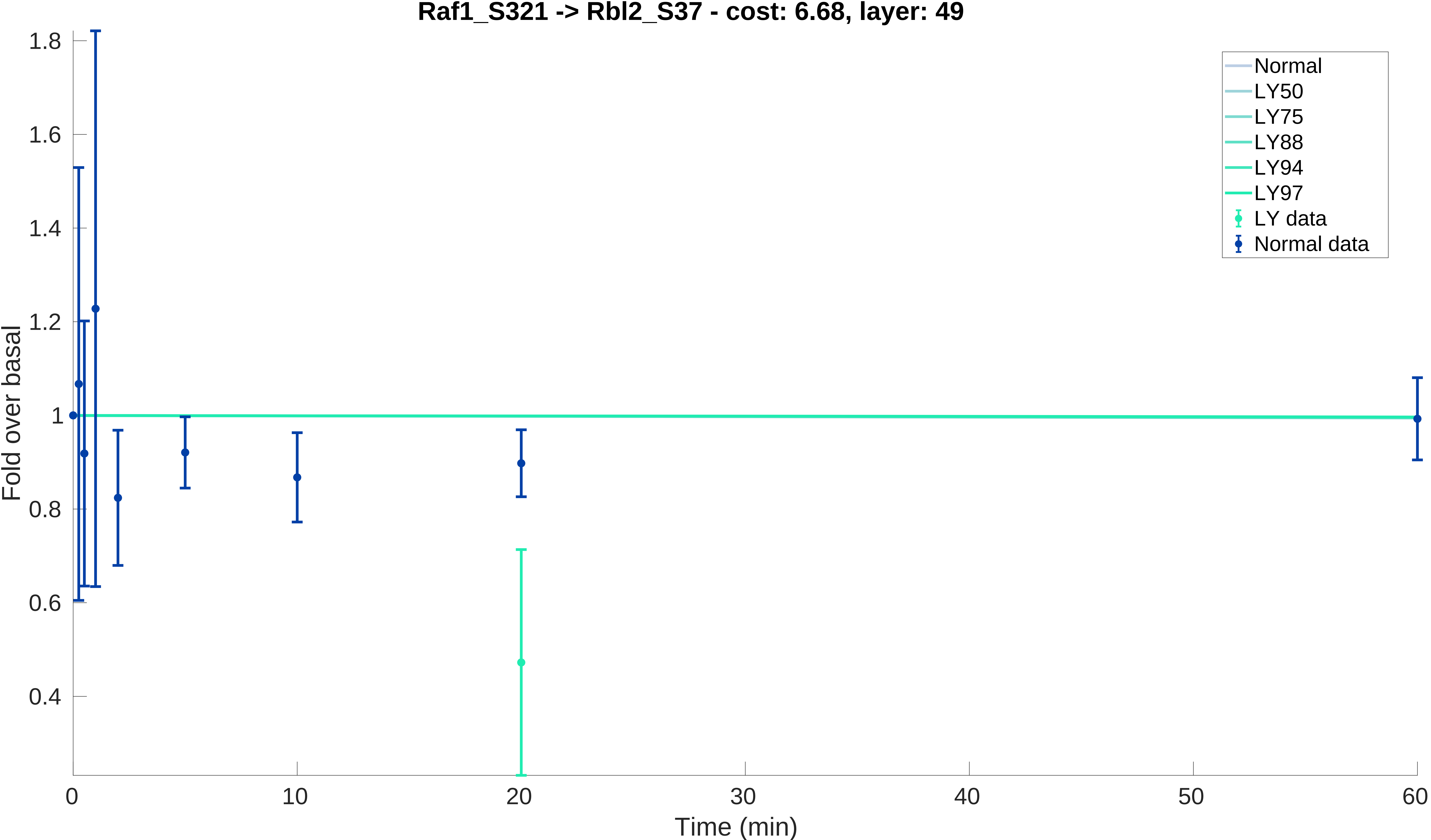
Raf1_S321 -> Rbl2_S37 - cost: 6.68, layer: 49
Fold over basal
Time (min)
1.8
1.6
1.4
1.2
1
0.8
0.6
0.4
0
10
20
30
40
50
60
Normal
LY50
LY75
LY88
LY94
LY97
LY data
Normal data

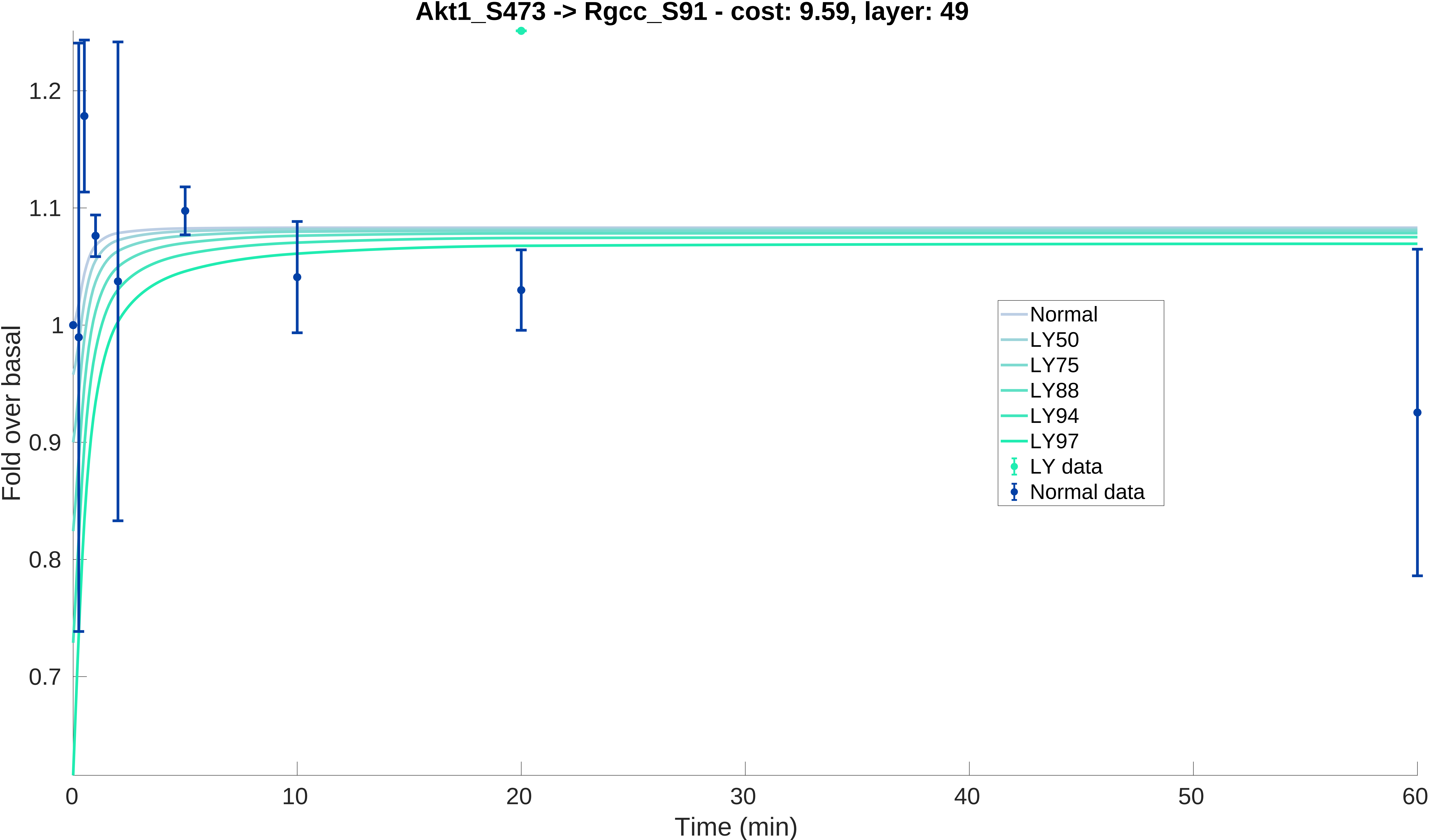

Akt1_S473 -> Rgcc_S91 - cost: 9.59, layer: 49
Fold over basal
1.2
1.1
1
0.9
0.8
0.7
0
10
20
30
40
50
60
Time (min)
Normal
LY50
LY75
LY88
LY94
LY97
LY data
Normal data

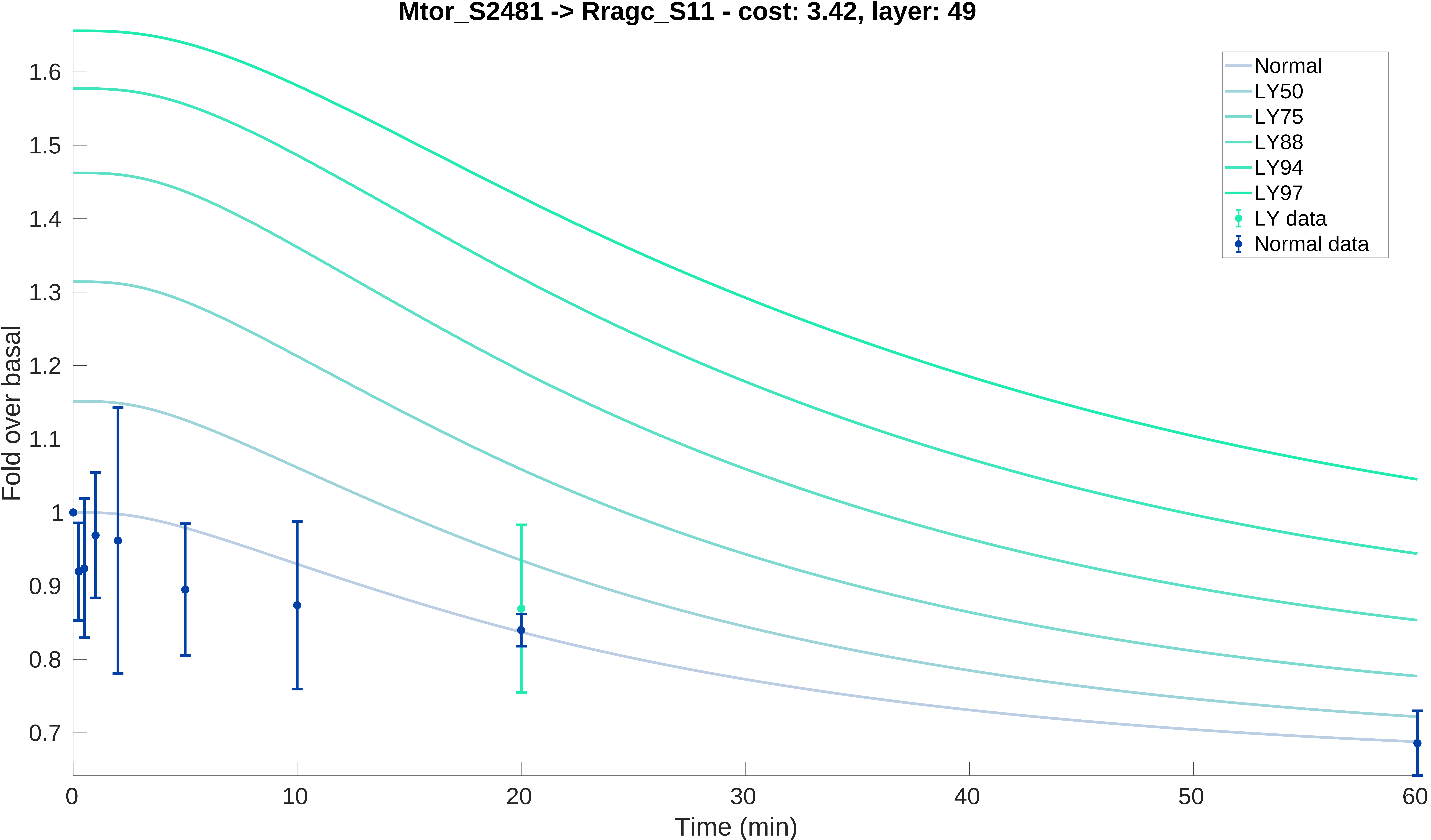

Mtor_S2481 -> Rragc_S11 - cost: 3.42, layer: 49
Fold over basal
Time (min)
Normal
LY50
LY75
LY88
LY94
LY97
LY data
Normal data
0.7
0.8
0.9
1
1.1
1.2
1.3
1.4
1.5
1.6
0
10
20
30
40
50
60

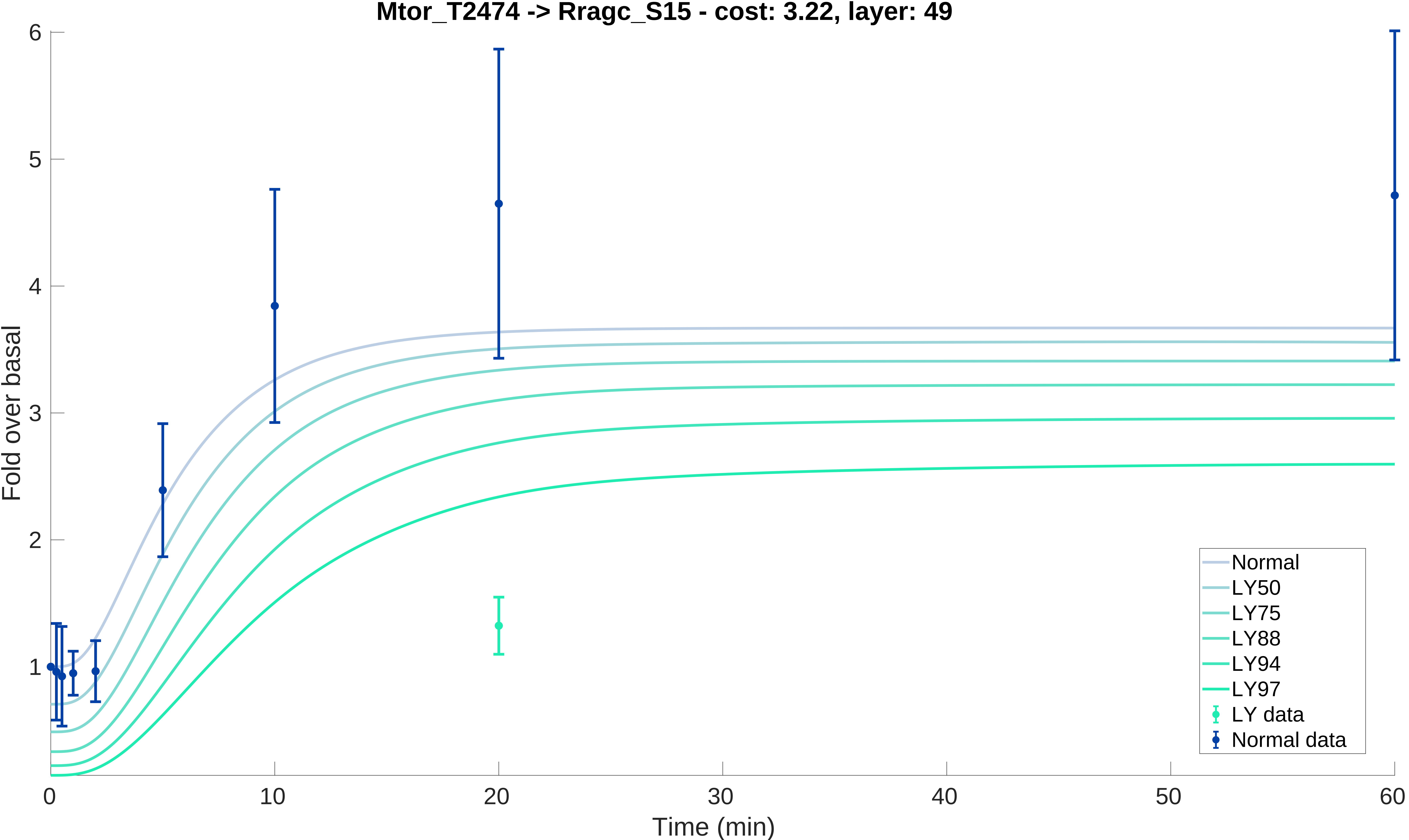

Mtor_T2474 -> Rragc_S15 - cost: 3.22, layer: 49
6
5
4
3
2
1
Fold over basal
0
10
20
30
40
50
60
Time (min)
Normal
LY50
LY75
LY88
LY94
LY97
LY data
Normal data

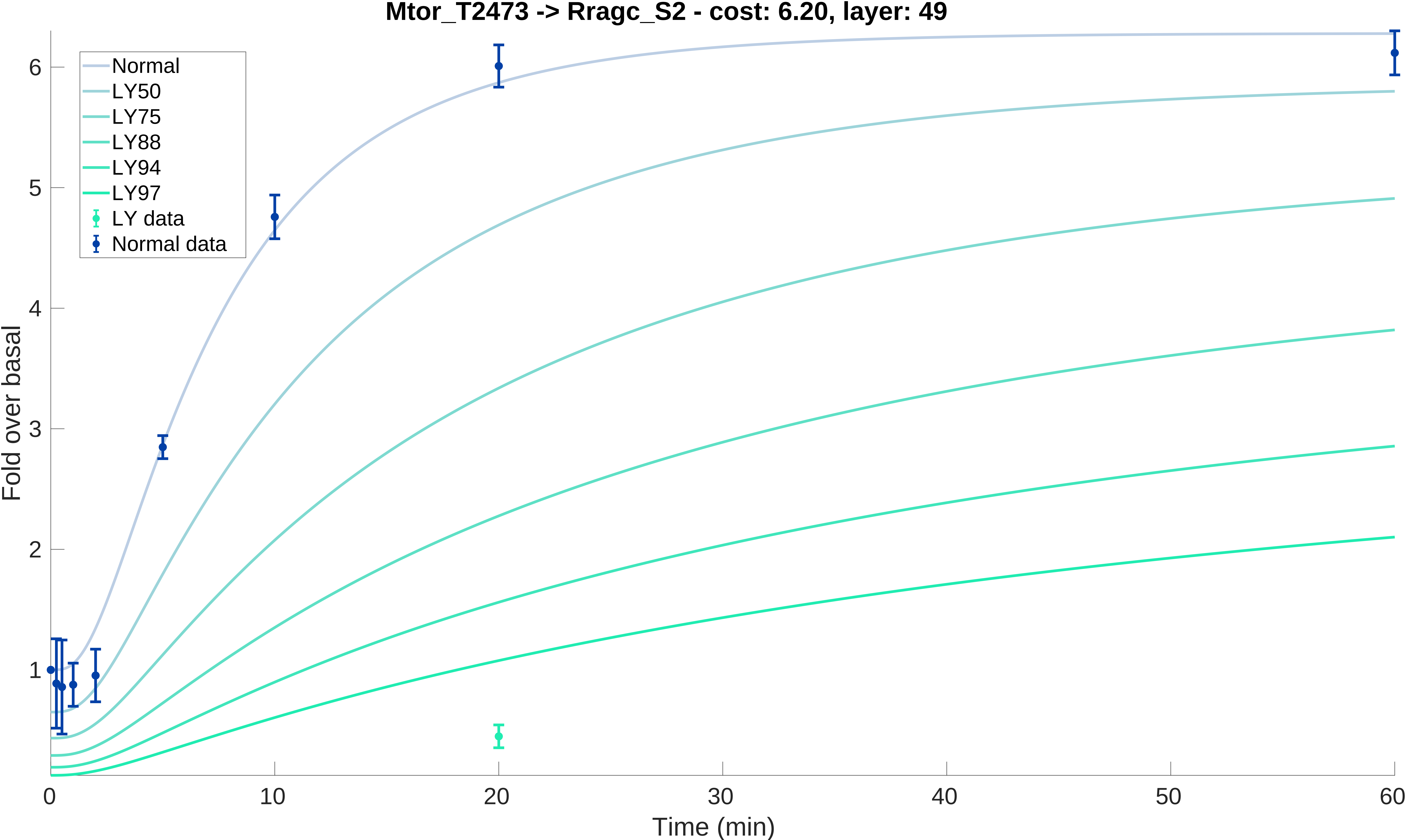

Mtor_T2473 -> Rragc_S2 - cost: 6.20, layer: 49
Normal
LY50
LY75
LY88
LY94
LY97
LY data
Normal data
Fold over basal
Time (min)
0
10
20
30
40
50
60
1
2
3
4
5
6

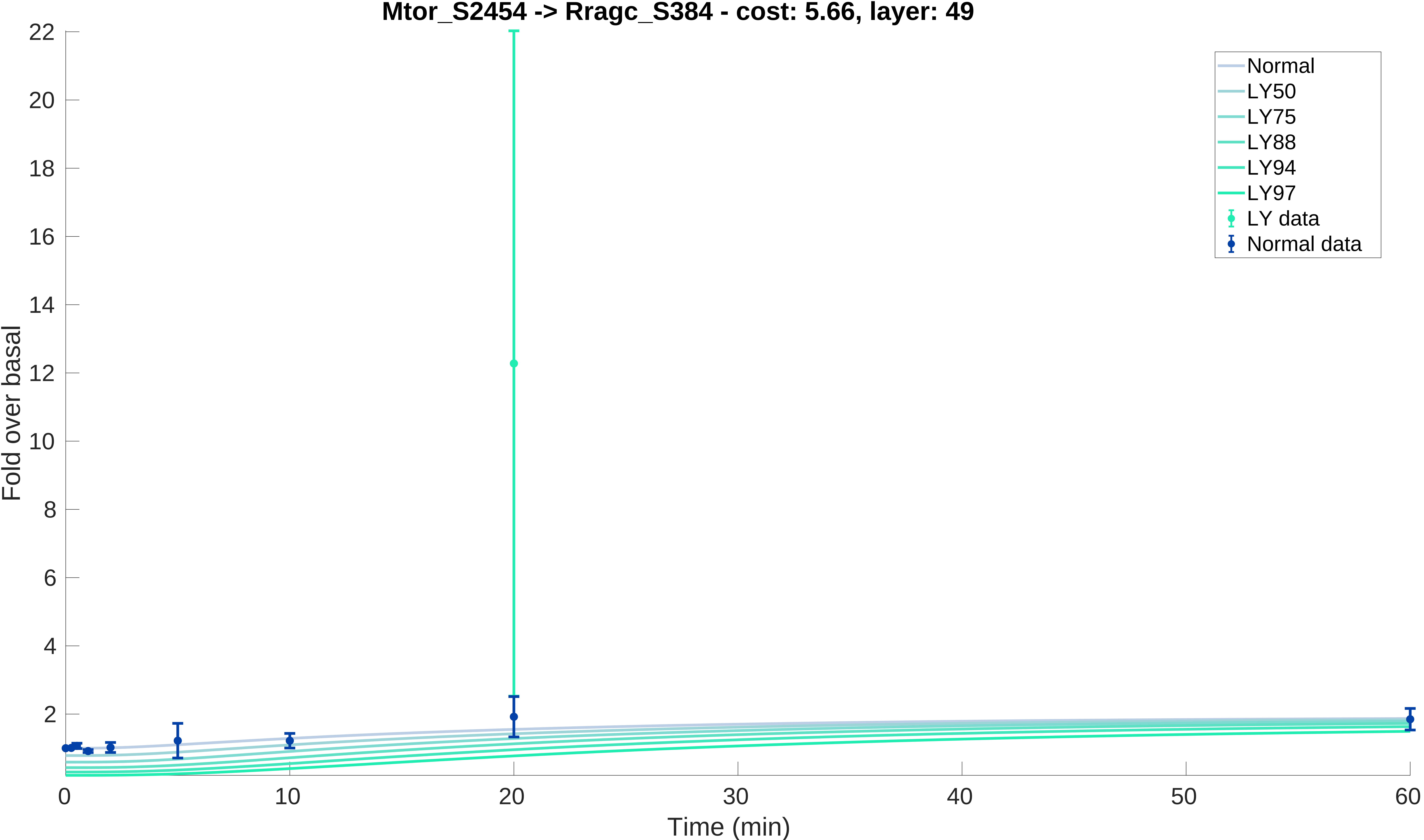

Mtor_S2454 -> Rragc_S384 - cost: 5.66, layer: 49
Fold over basal
Time (min)
0
10
20
30
40
50
60
2
4
6
8
10
12
14
16
18
20
22
Normal
LY50
LY75
LY88
LY94
LY97
LY data
Normal data

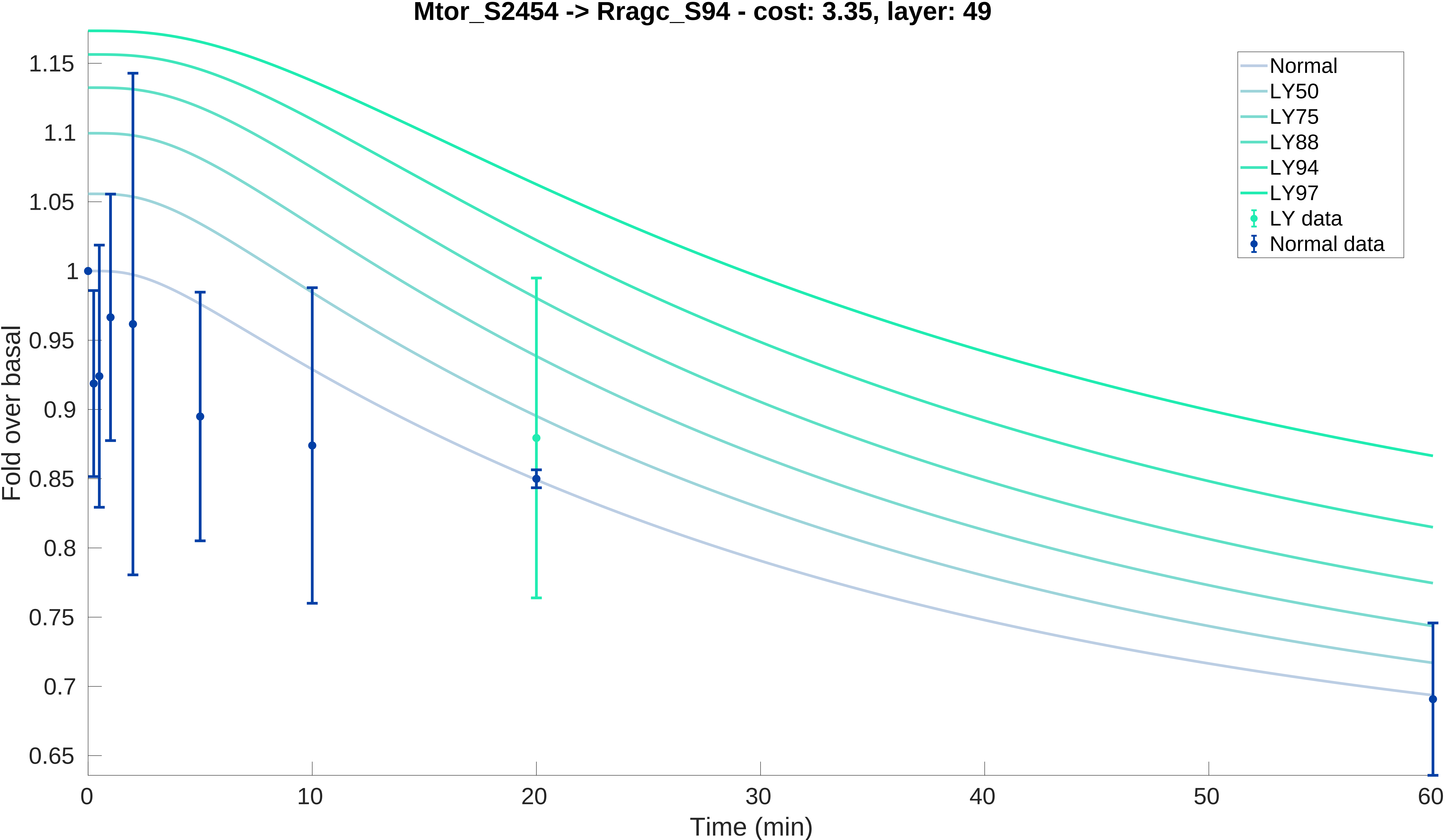

Mtor_S2454 -> Rragc_S94 - cost: 3.35, layer: 49
Normal
LY50
LY75
LY88
LY94
LY97
LY data
Normal data
Fold over basal
1.15
1.1
1.05
1
0.95
0.9
0.85
0.8
0.75
0.7
0.65
0
10
20
30
40
50
60
Time (min)

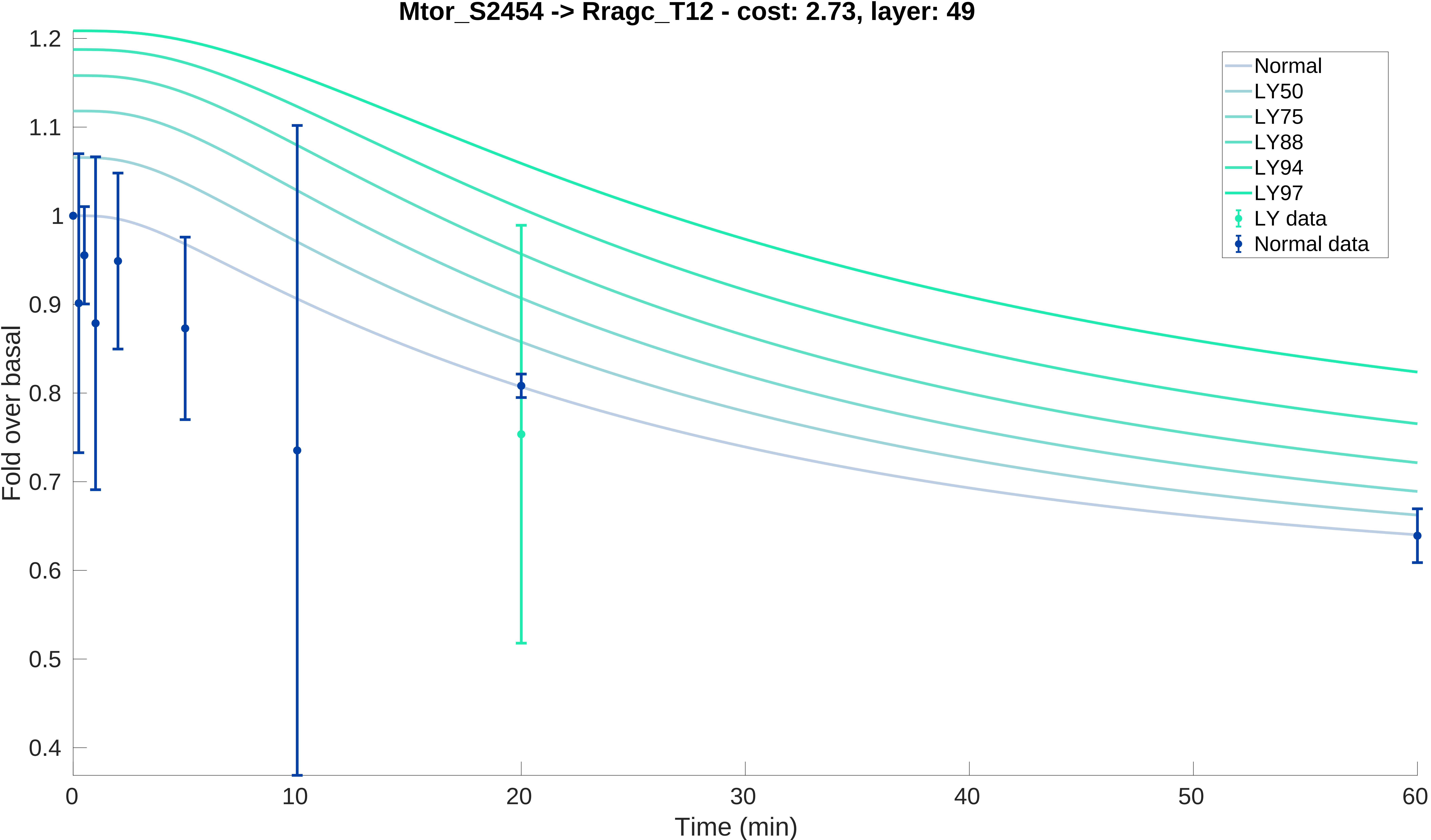

Mtor_S2454 -> Rragc_T12 - cost: 2.73, layer: 49
Fold over basal
Time (min)
0
10
20
30
40
50
60
0.4
0.5
0.6
0.7
0.8
0.9
1
1.1
1.2
Normal
LY50
LY75
LY88
LY94
LY97
LY data
Normal data

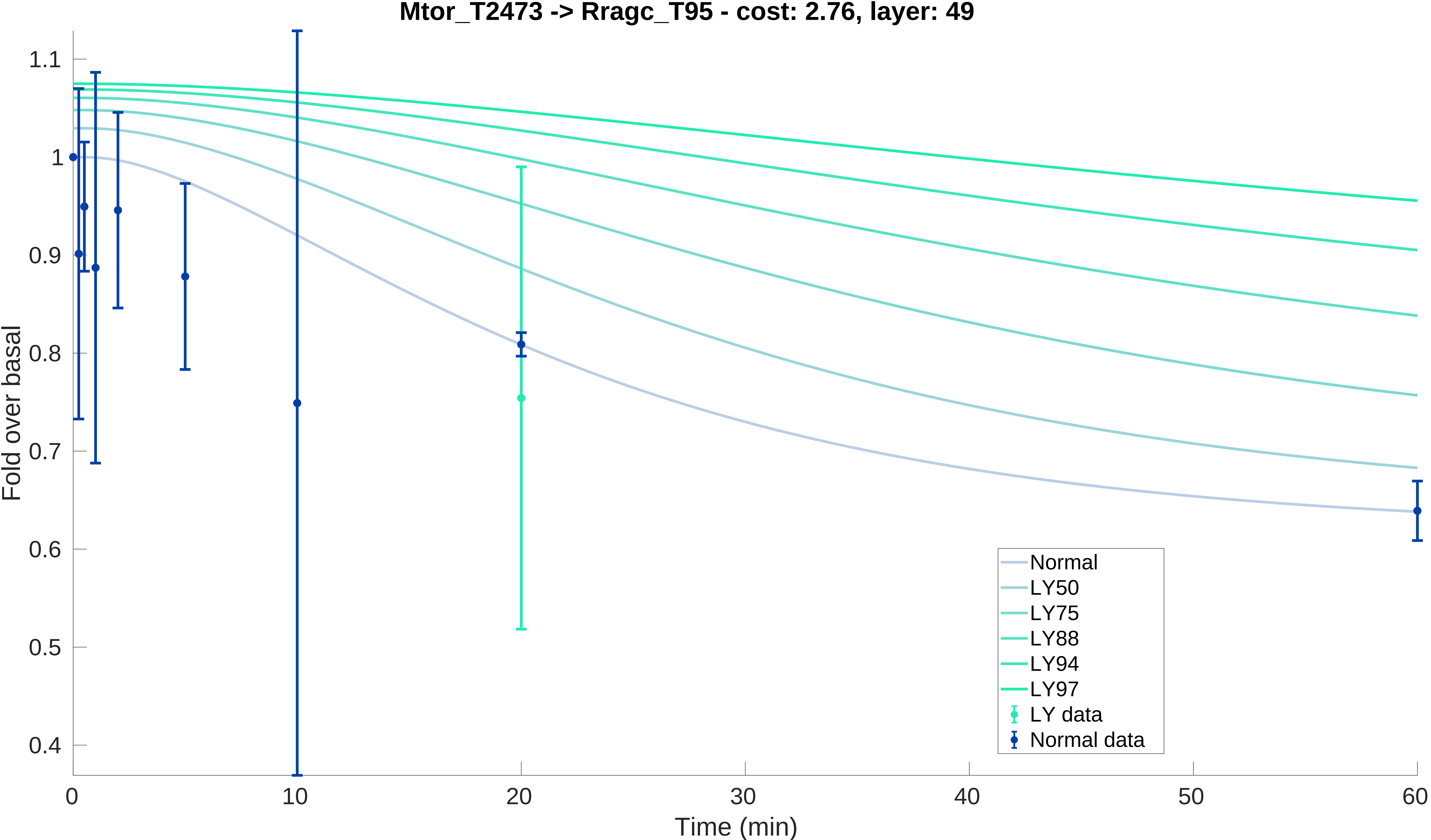

Mtor_T2473 -> Rragc_T95 - cost: 2.76, layer: 49
Fold over basal
1.1
1
0.9
0.8
0.7
0.6
0.5
0.4
0
10
20
30
40
50
60
Time (min)
Normal
LY50
LY75
LY88
LY94
LY97
LY data
Normal data

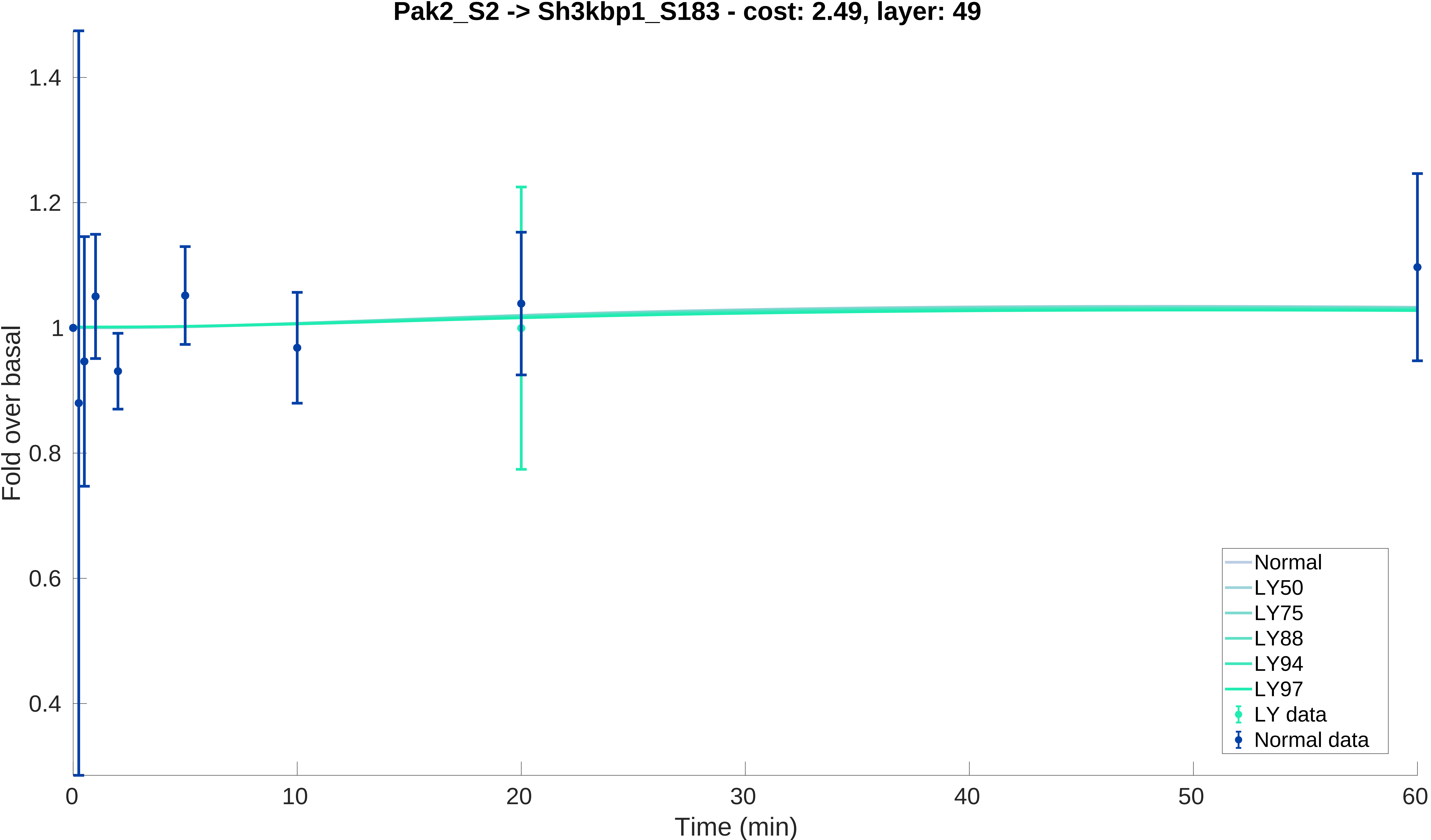
Pak2_S2 -> Sh3kbp1_S183 - cost: 2.49, layer: 49
Fold over basal
1.4
1.2
1
0.8
0.6
0.4
0
10
20
30
40
50
60
Time (min)
Normal
LY50
LY75
LY88
LY94
LY97
LY data
Normal data

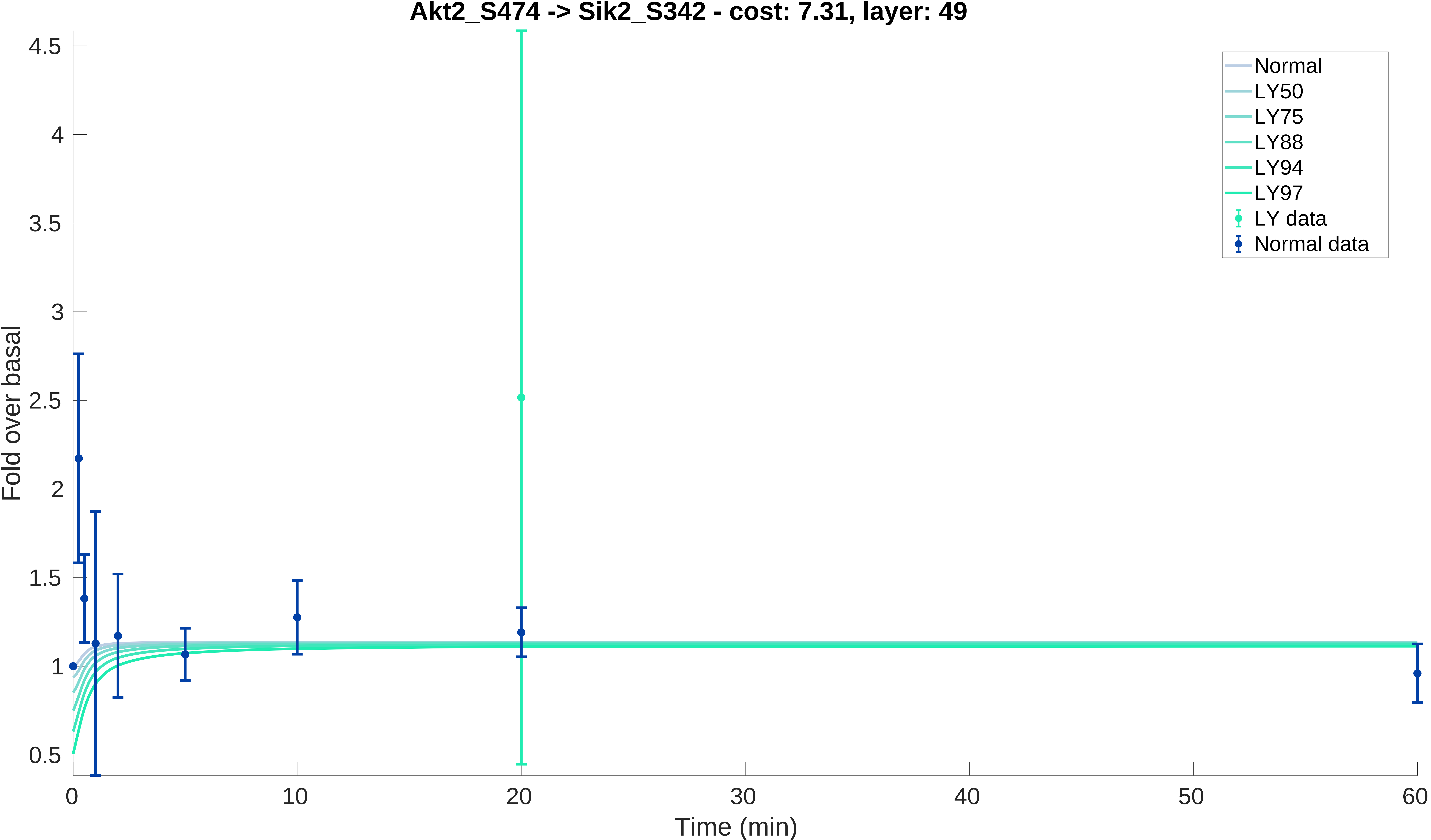

Akt2_S474 -> Sik2_S342 - cost: 7.31, layer: 49
Fold over basal
4.5
4
3.5
3
2.5
2
1.5
1
0.5
0
10
20
30
40
50
60
Time (min)
Normal
LY50
LY75
LY88
LY94
LY97
LY data
Normal data

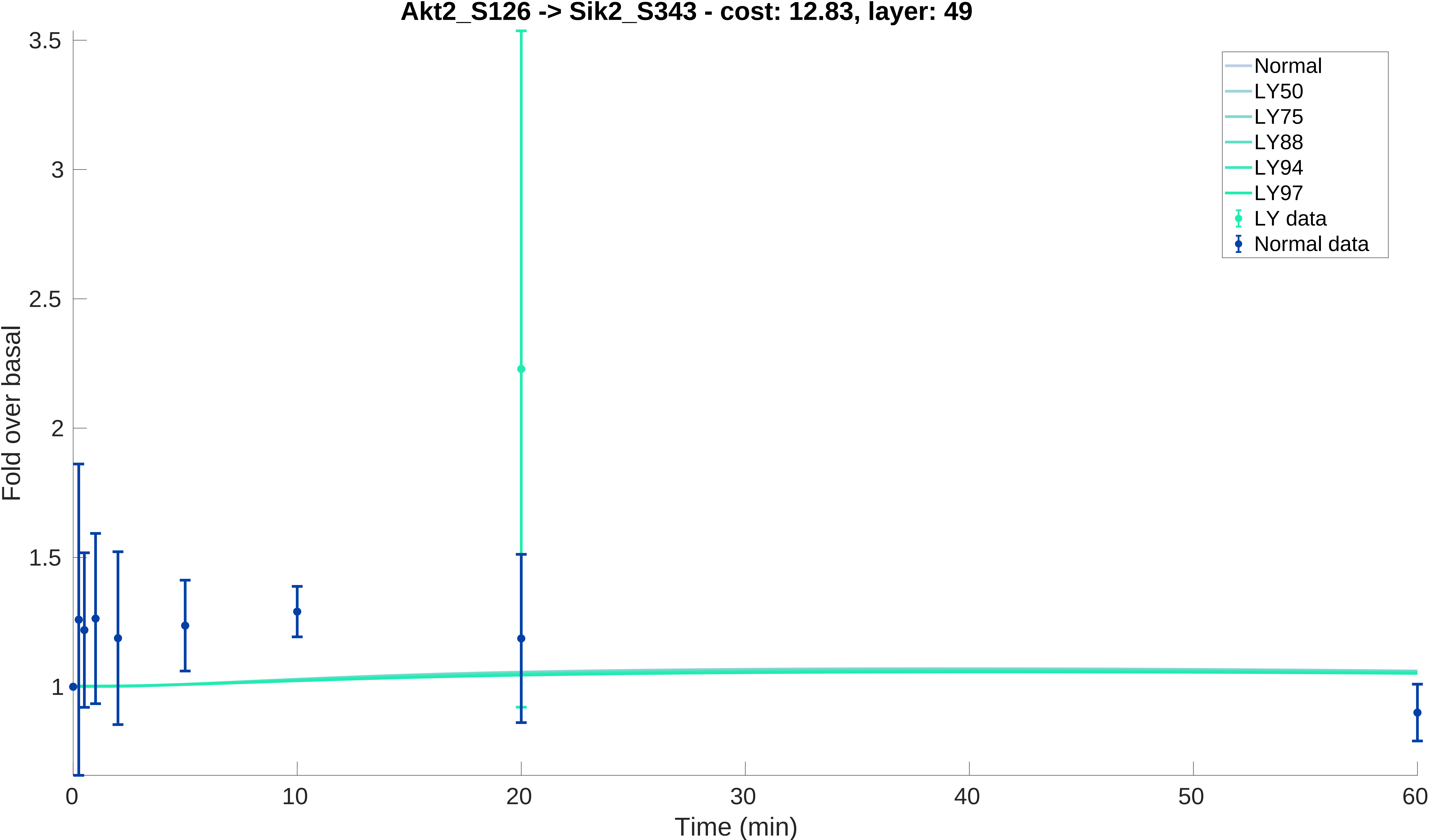
Akt2_S126 -> Sik2_S343 - cost: 12.83, layer: 49
Fold over basal
3.5
3
2.5
2
1.5
1
Time (min)
0
10
20
30
40
50
60
Normal
LY50
LY75
LY88
LY94
LY97
LY data
Normal data

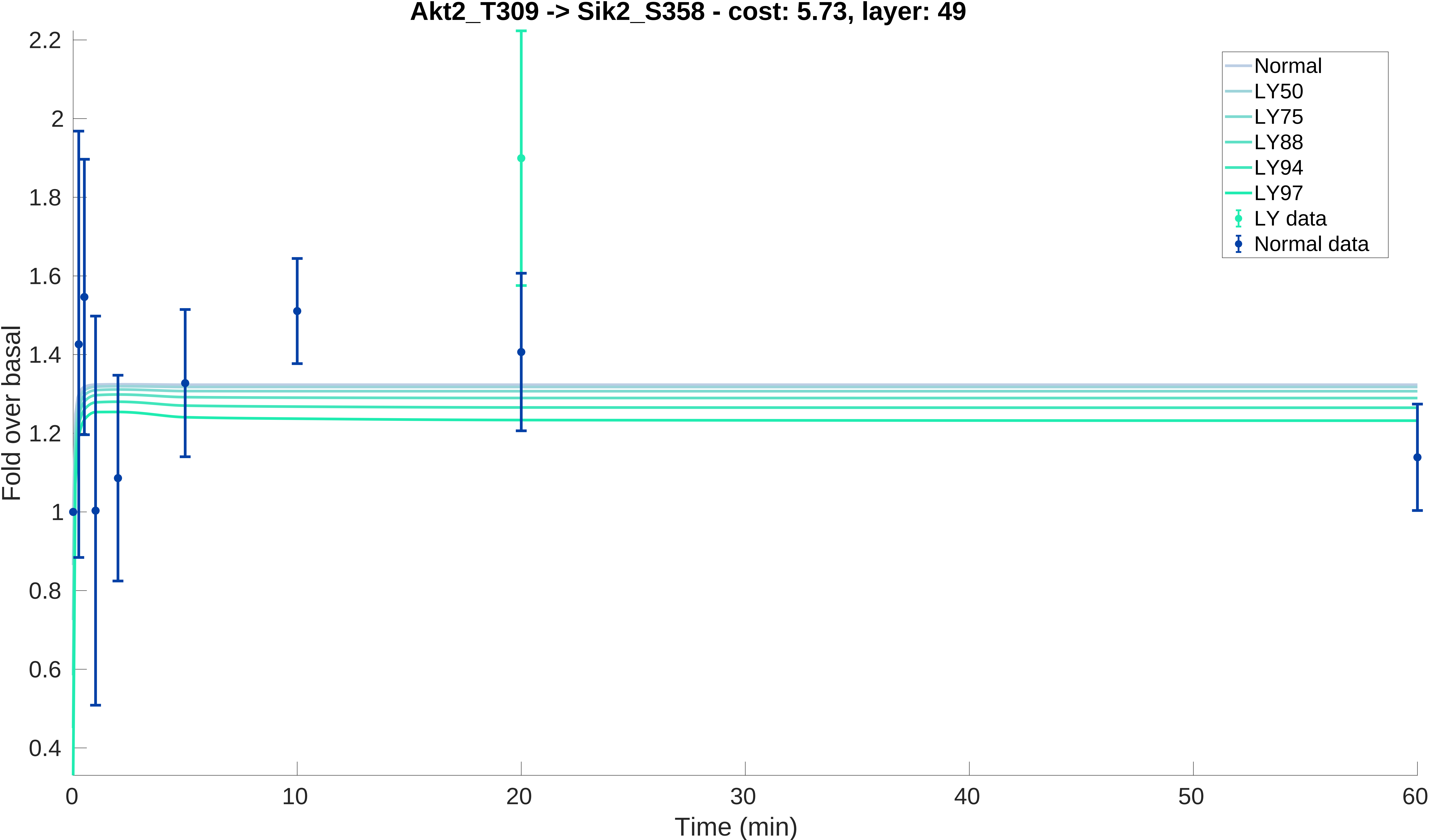

Akt2_T309 -> Sik2_S358 - cost: 5.73, layer: 49
Fold over basal
Time (min)
0.4
0.6
0.8
1
1.2
1.4
1.6
1.8
2
2.2
0
10
20
30
40
50
60
Normal
LY50
LY75
LY88
LY94
LY97
LY data
Normal data

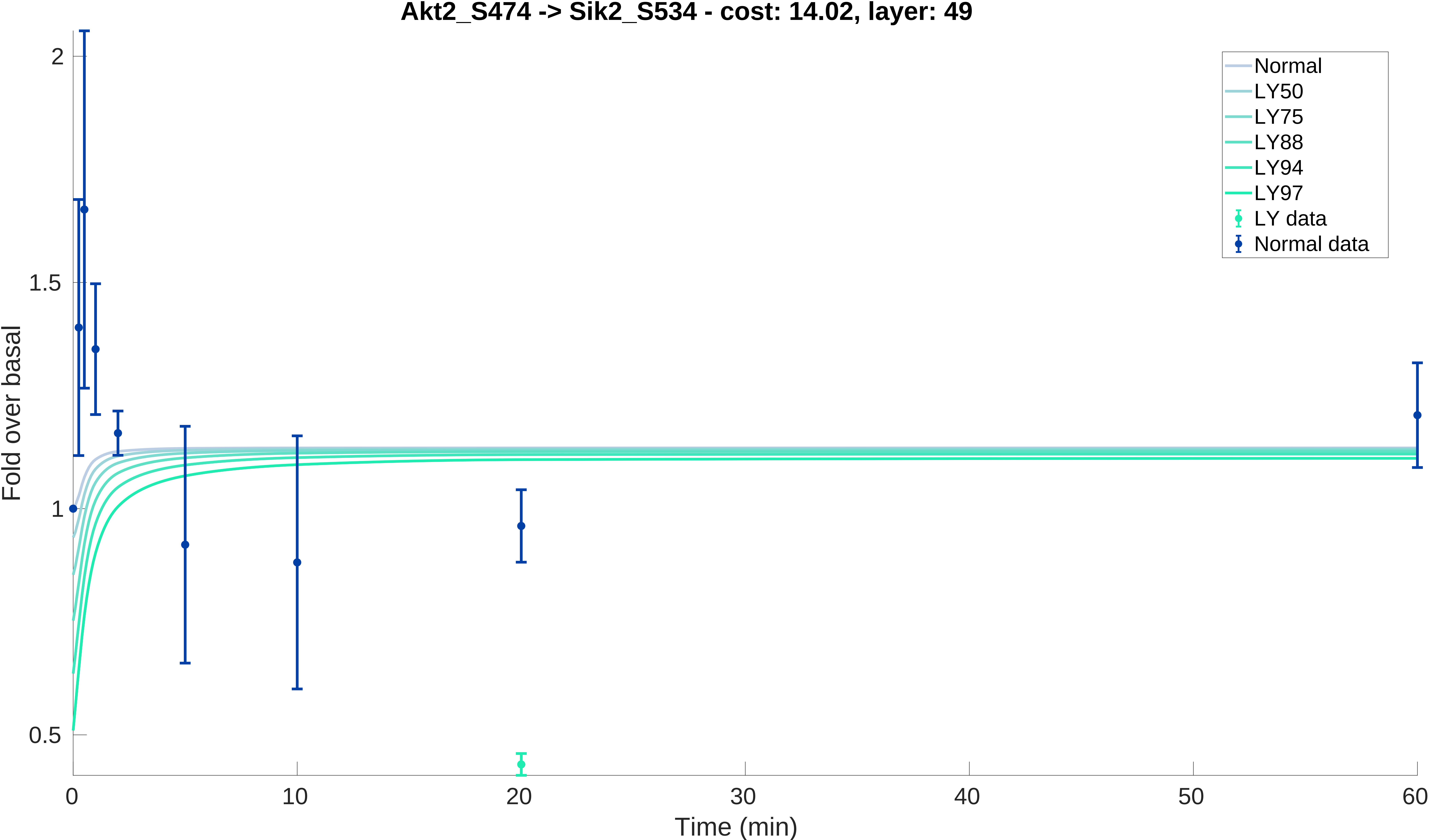
Akt2_S474 -> Sik2_S534 - cost: 14.02, layer: 49
Fold over basal
Time (min)
0.5
1
1.5
2
0
10
20
30
40
50
60
Normal
LY50
LY75
LY88
LY94
LY97
LY data
Normal data

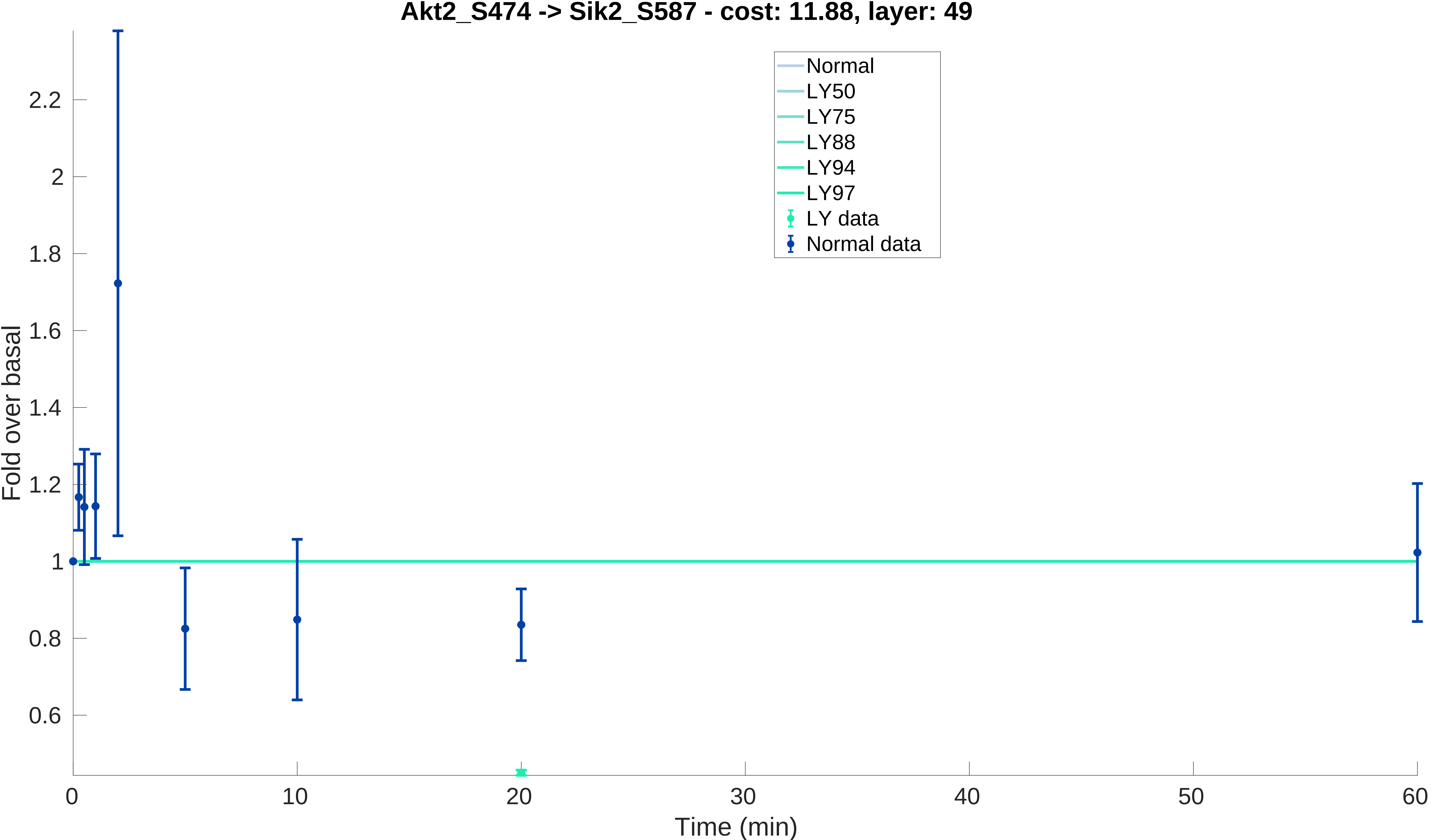
Akt2_S474 -> Sik2_S587 - cost: 11.88, layer: 49
Normal
LY50
LY75
LY88
LY94
LY97
LY data
Normal data
Fold over basal
2.2
2
1.8
1.6
1.4
1.2
1
0.8
0.6
0
10
20
30
40
50
60
Time (min)

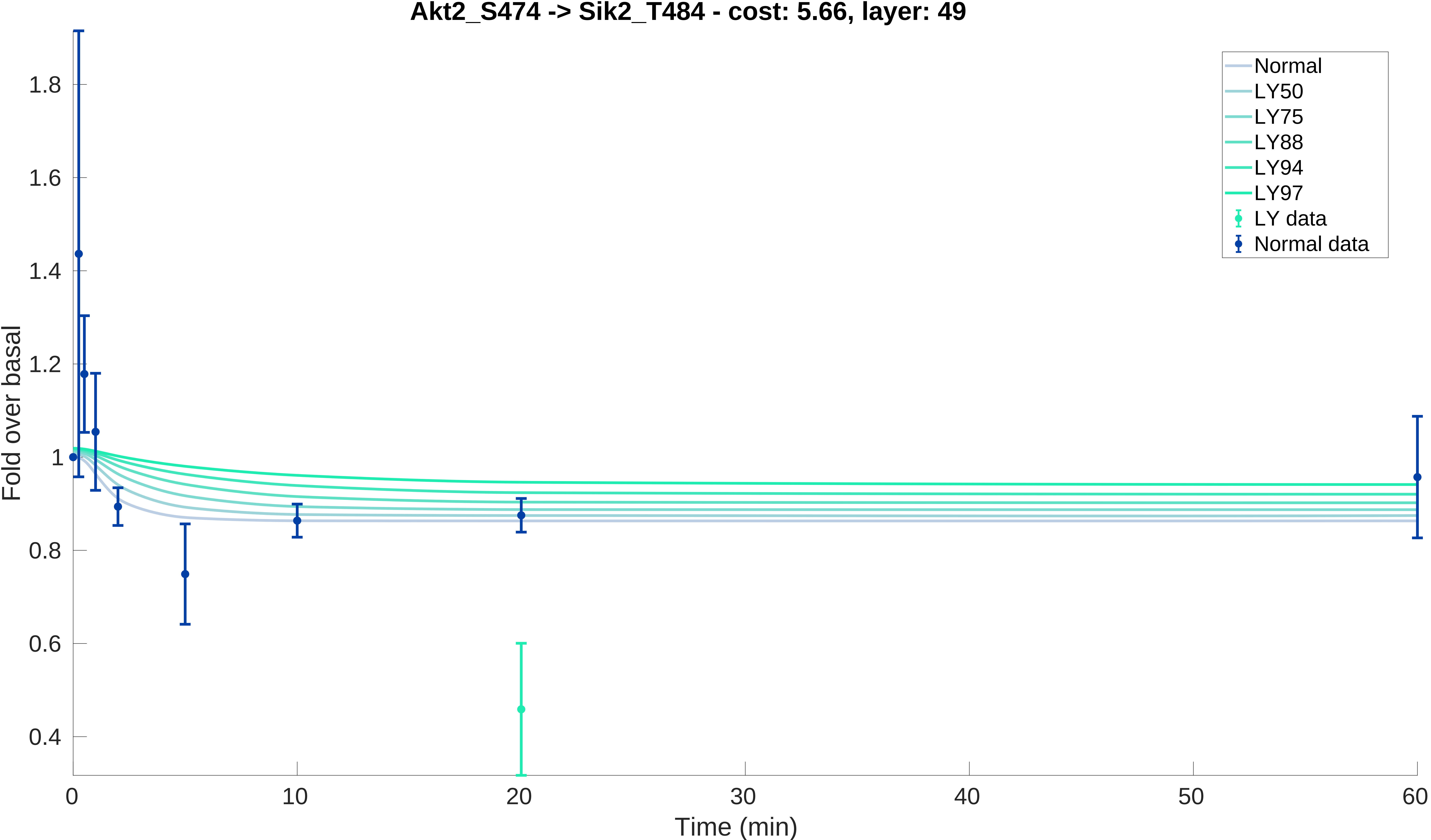

Akt2_S474 -> Sik2_T484 - cost: 5.66, layer: 49
Fold over basal
Time (min)
0.4
0.6
0.8
1
1.2
1.4
1.6
1.8
0
10
20
30
40
50
60
Normal
LY50
LY75
LY88
LY94
LY97
LY data
Normal data

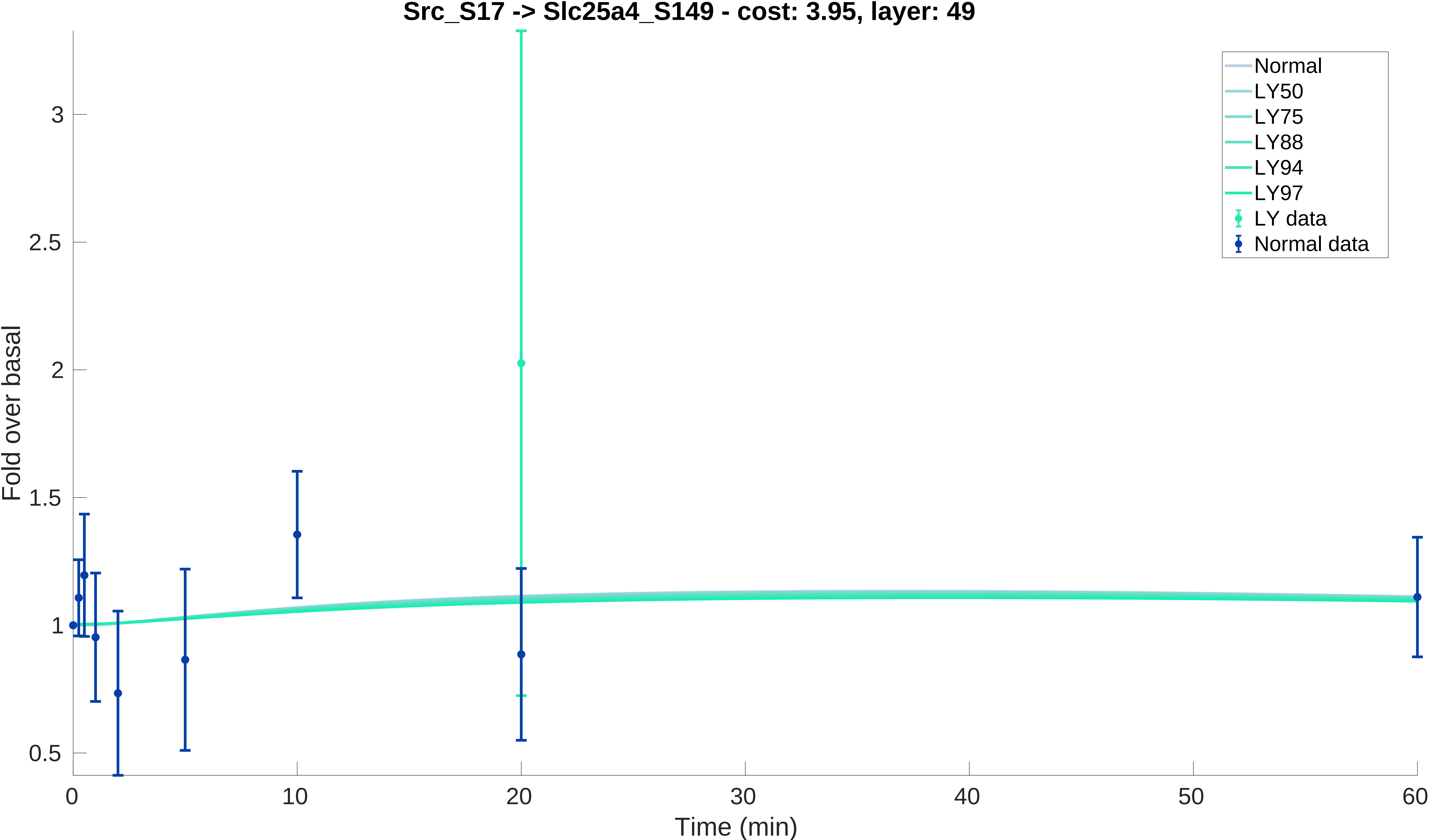

Src_S17 -> Slc25a4_S149 - cost: 3.95, layer: 49
Normal
LY50
LY75
LY88
LY94
LY97
LY data
Normal data
Fold over basal
3
2.5
2
1.5
1
0.5
0
10
20
30
40
50
60
Time (min)

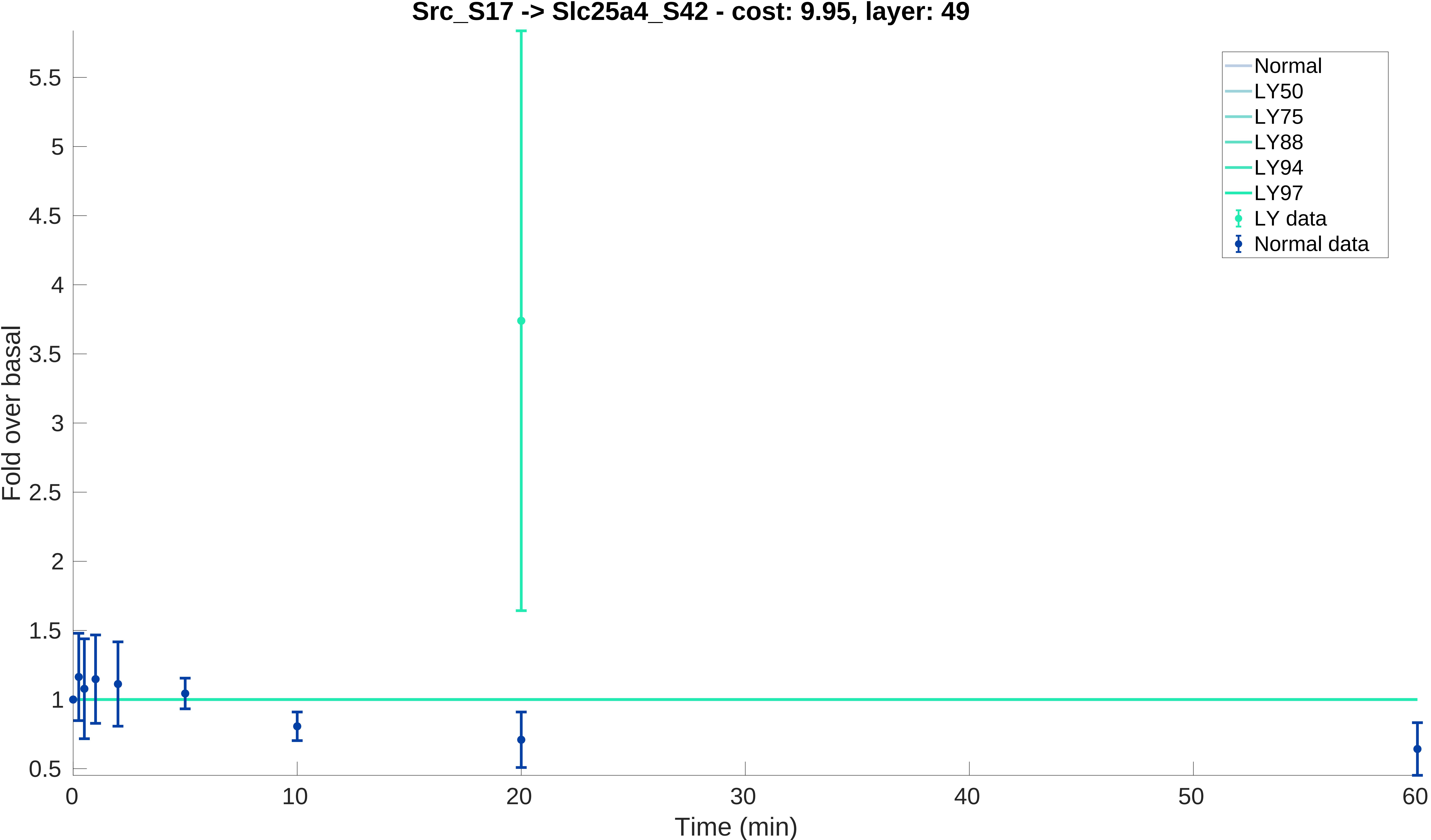
Src_S17 -> Slc25a4_S42 - cost: 9.95, layer: 49
Fold over basal
Time (min)
0.5
1
1.5
2
2.5
3
3.5
4
4.5
5
5.5
0
10
20
30
40
50
60
Normal
LY50
LY75
LY88
LY94
LY97
LY data
Normal data

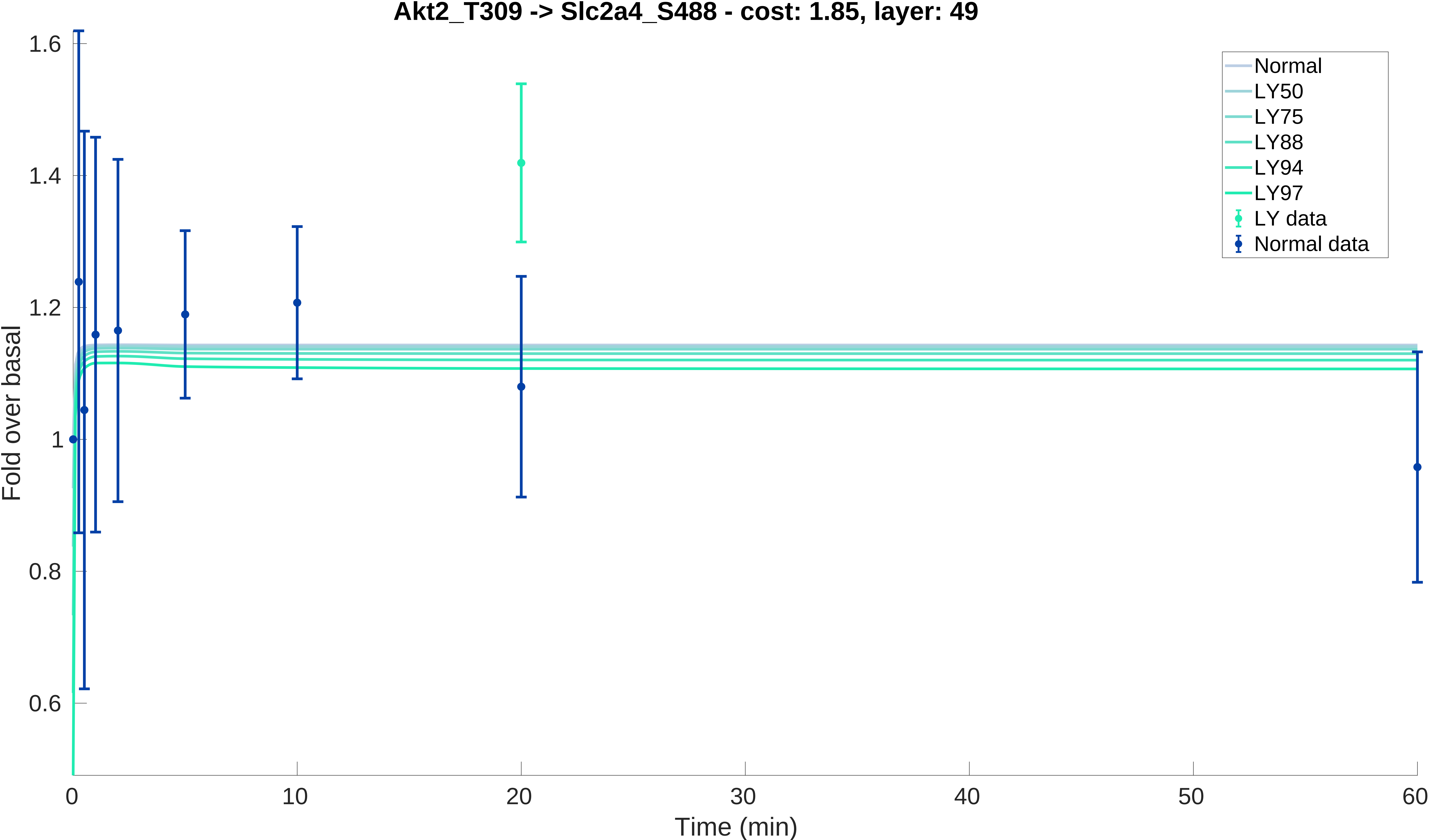
Akt2_T309 -> Slc2a4_S488 - cost: 1.85, layer: 49
Fold over basal
Time (min)
0
10
20
30
40
50
60
0.6
0.8
1
1.2
1.4
1.6
Normal
LY50
LY75
LY88
LY94
LY97
LY data
Normal data

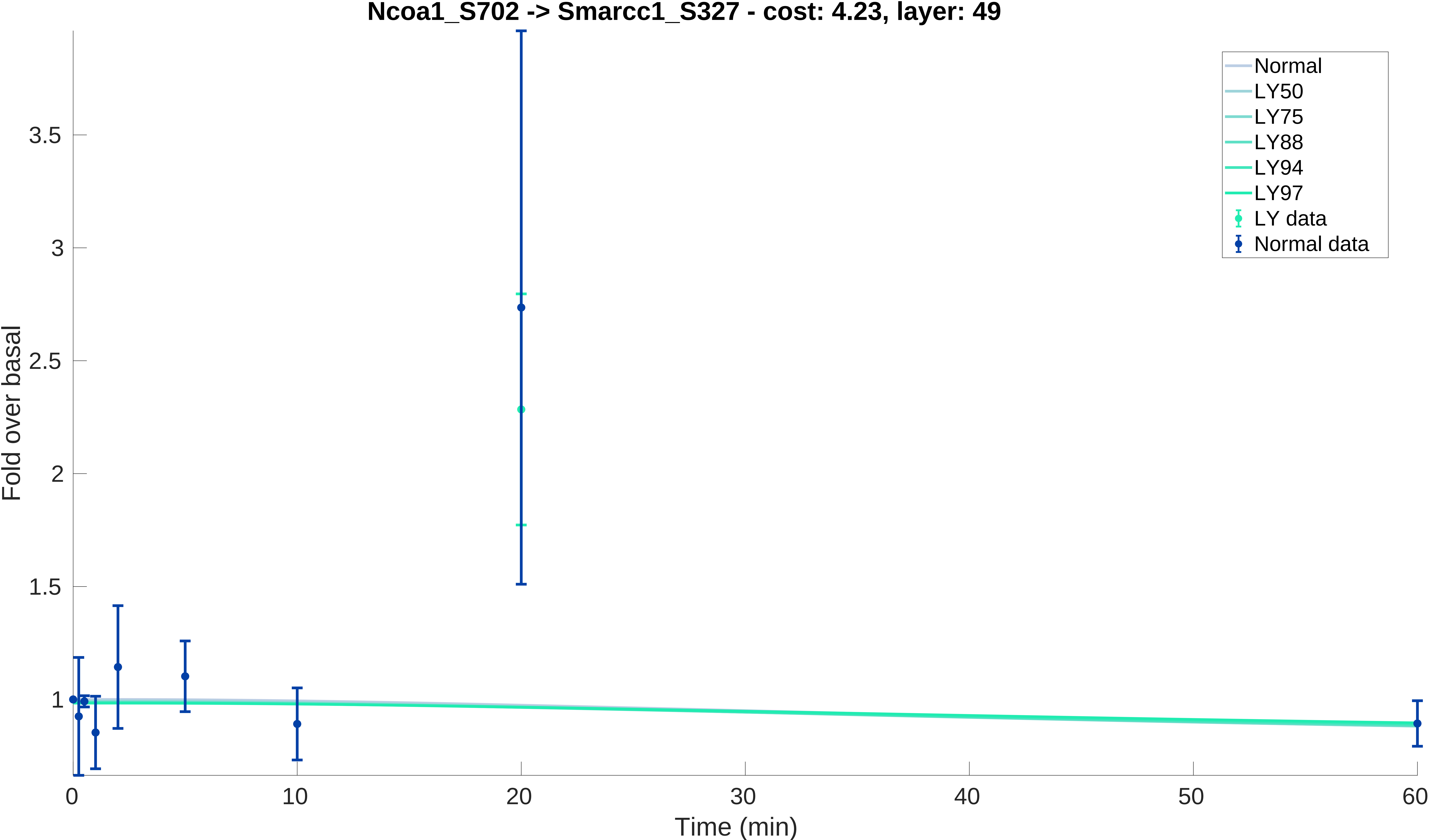

Ncoa1_S702 -> Smarcc1_S327 - cost: 4.23, layer: 49
Fold over basal
Time (min)
3.5
3
2.5
2
1.5
1
0
10
20
30
40
50
60
Normal
LY50
LY75
LY88
LY94
LY97
LY data
Normal data

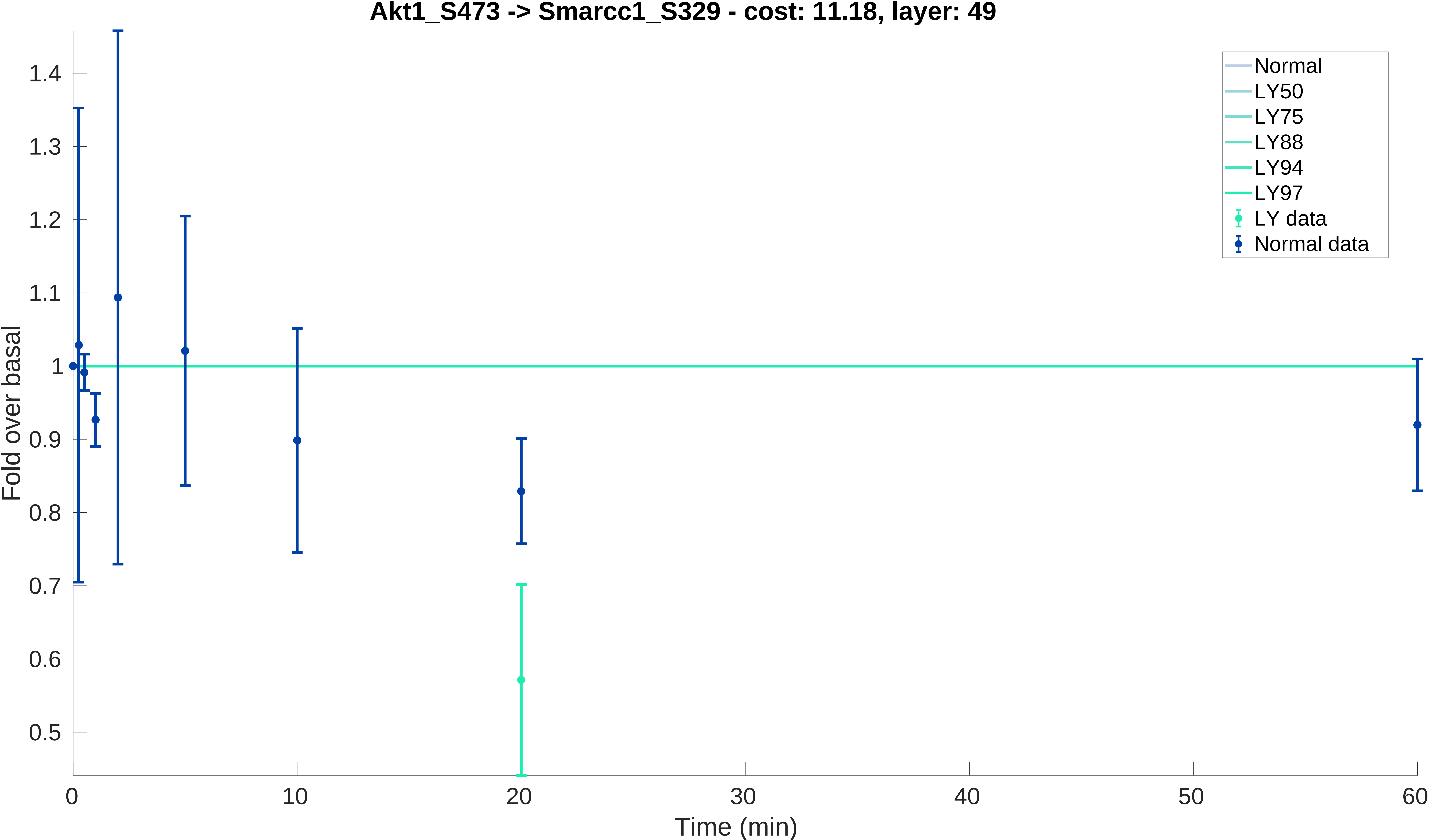

Akt1_S473 -> Smarcc1_S329 - cost: 11.18, layer: 49
Fold over basal
Time (min)
0
10
20
30
40
50
60
0.5
0.6
0.7
0.8
0.9
1
1.1
1.2
1.3
1.4
Normal
LY50
LY75
LY88
LY94
LY97
LY data
Normal data

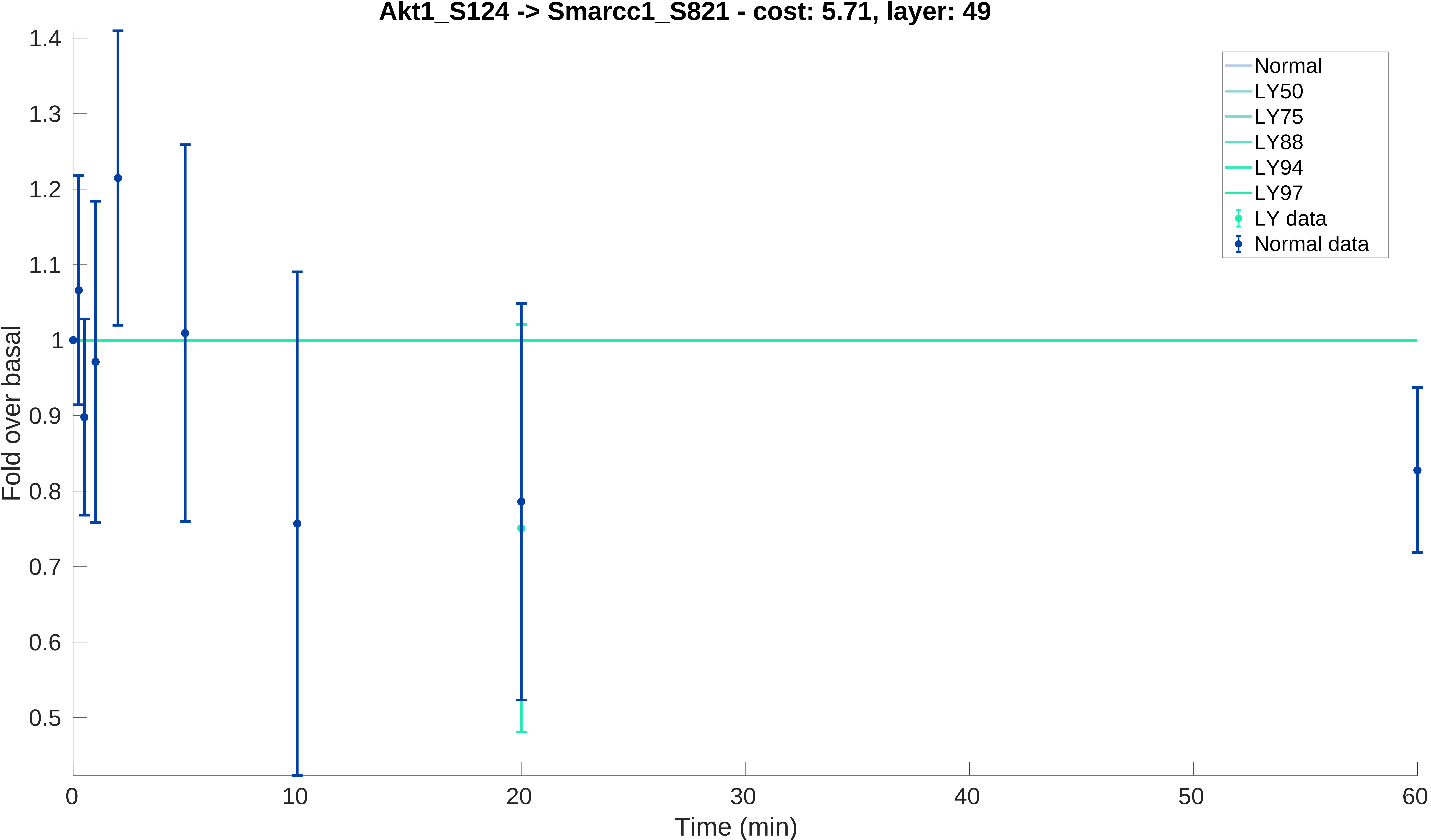

Akt1_S124 -> Smarcc1_S821 - cost: 5.71, layer: 49
Fold over basal
Time (min)
1.4
1.3
1.2
1.1
1
0.9
0.8
0.7
0.6
0.5
0
10
20
30
40
50
60
Normal
LY50
LY75
LY88
LY94
LY97
LY data
Normal data

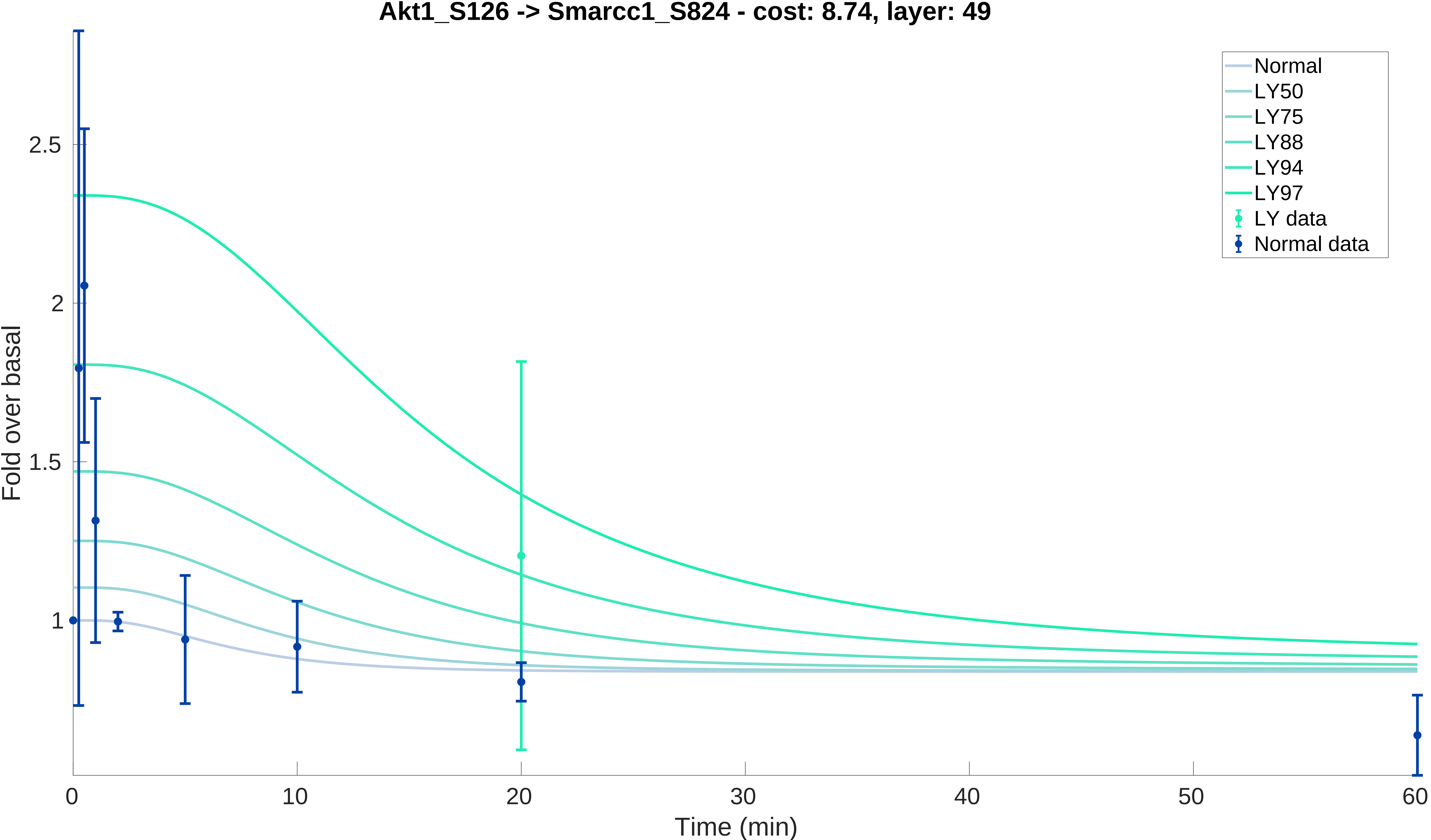
Akt1_S126 -> Smarcc1_S824 - cost: 8.74, layer: 49
Fold over basal
Time (min)
0
10
20
30
40
50
60
1
1.5
2
2.5
Normal
LY50
LY75
LY88
LY94
LY97
LY data
Normal data

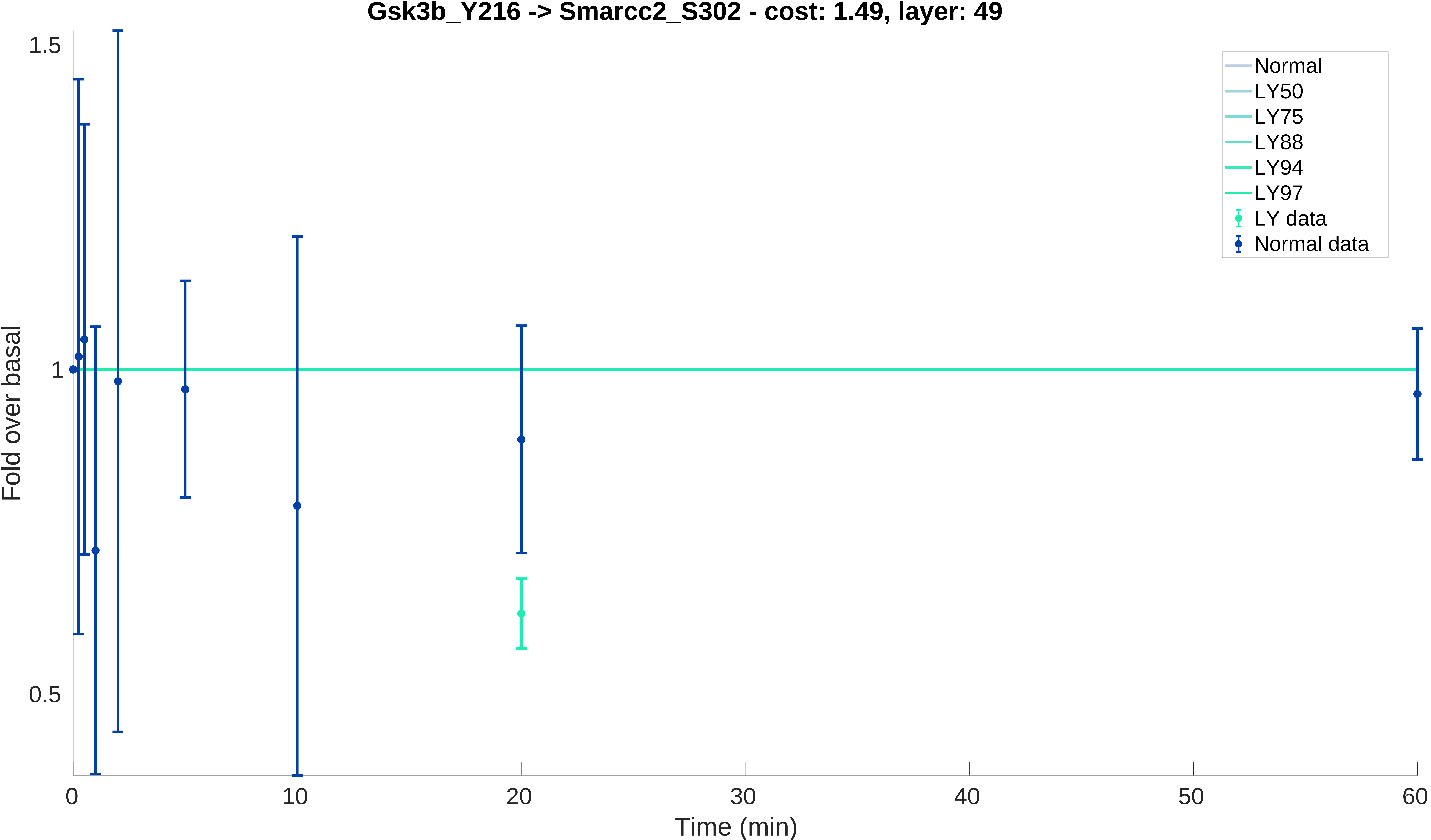
Gsk3b_Y216 -> Smarcc2_S302 - cost: 1.49, layer: 49
Fold over basal
1.5
1
0.5
0
10
20
30
40
50
60
Time (min)
Normal
LY50
LY75
LY88
LY94
LY97
LY data
Normal data

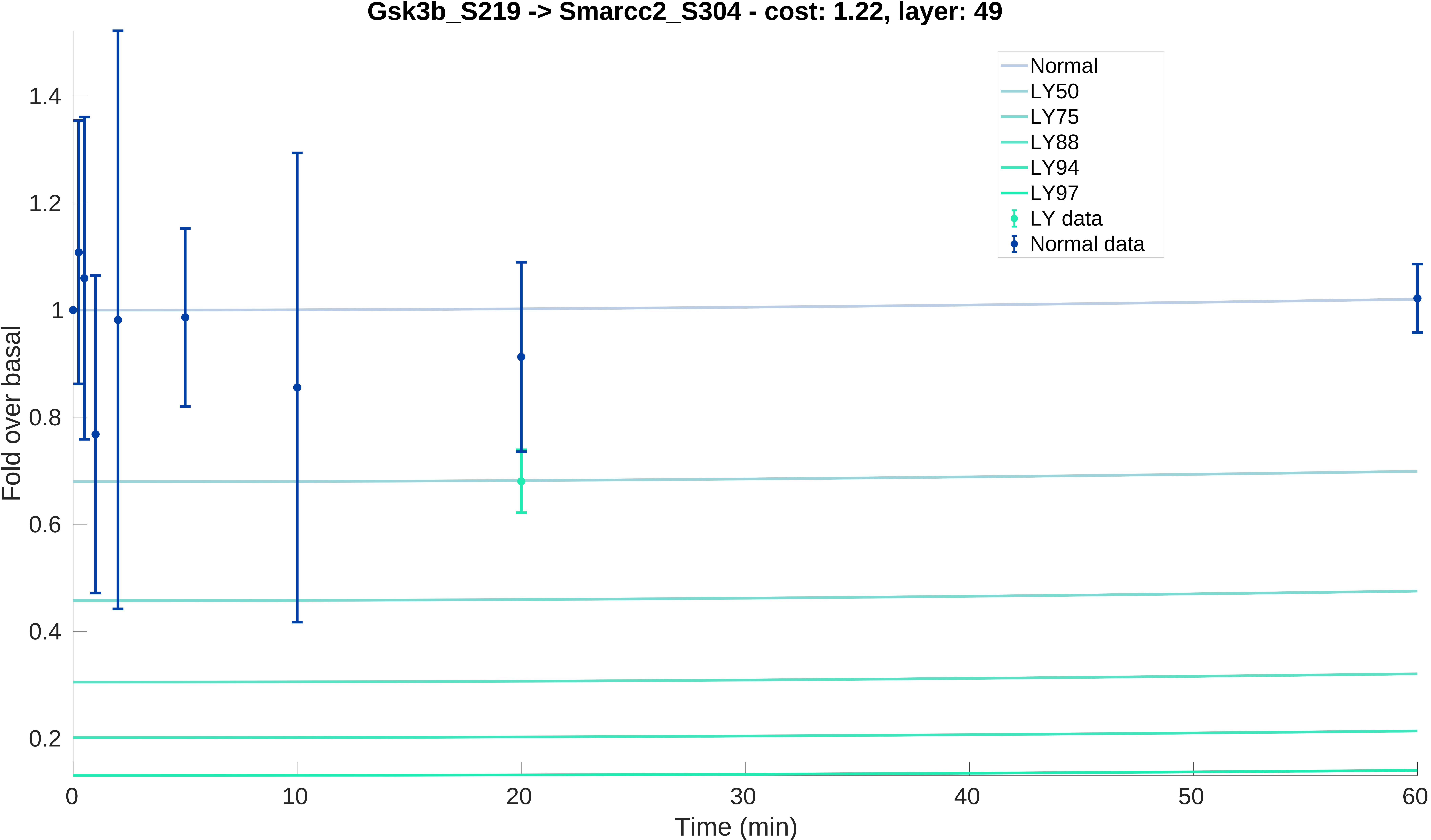

Gsk3b_S219 -> Smarcc2_S304 - cost: 1.22, layer: 49
Fold over basal
Time (min)
Normal
LY50
LY75
LY88
LY94
LY97
LY data
Normal data
1.4
1.2
1
0.8
0.6
0.4
0.2
0
10
20
30
40
50
60

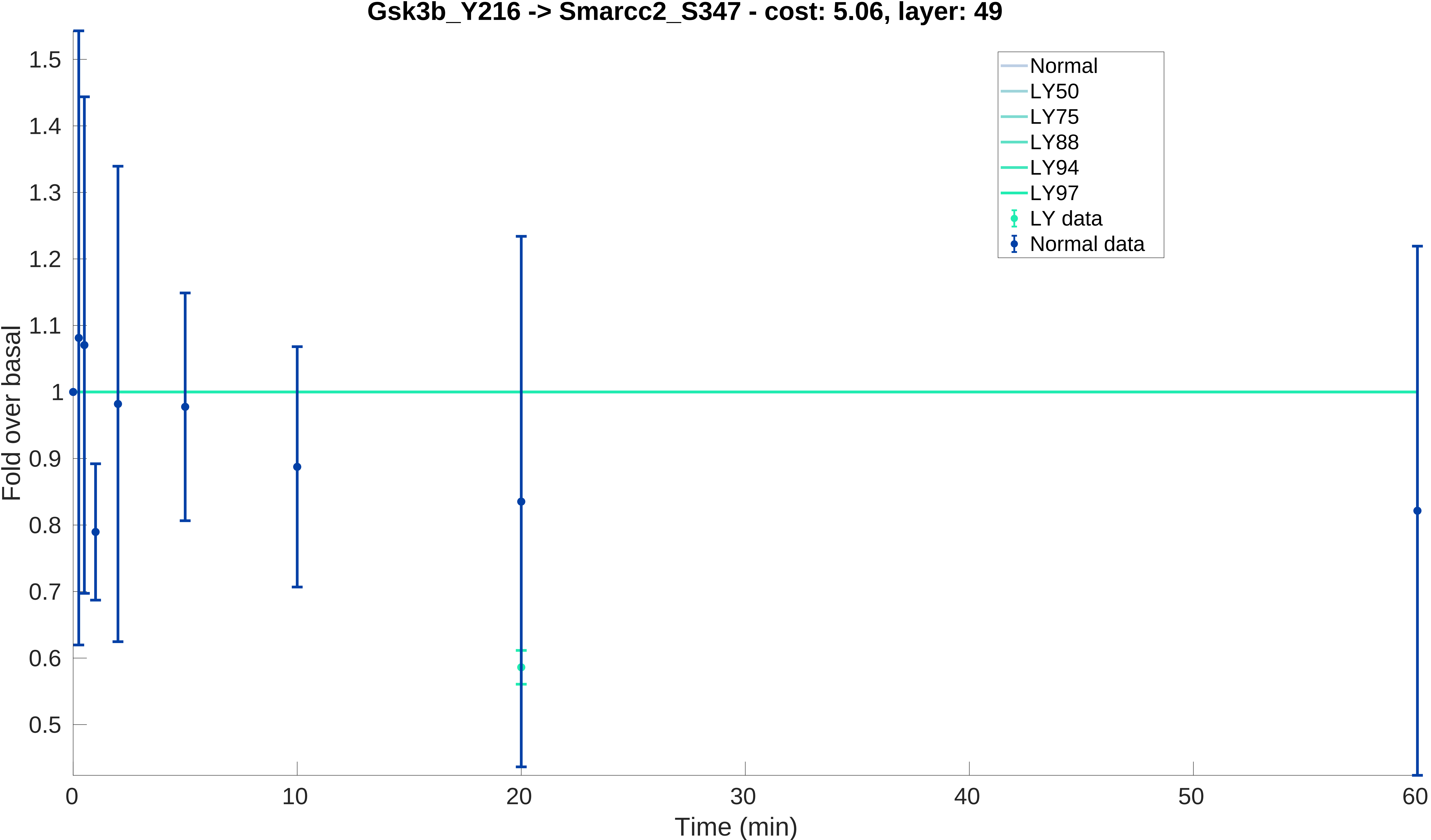

Gsk3b_Y216 -> Smarcc2_S347 - cost: 5.06, layer: 49
Normal
LY50
LY75
LY88
LY94
LY97
LY data
Normal data
Fold over basal
1.5
1.4
1.3
1.2
1.1
1
0.9
0.8
0.7
0.6
0.5
0
10
20
30
40
50
60
Time (min)

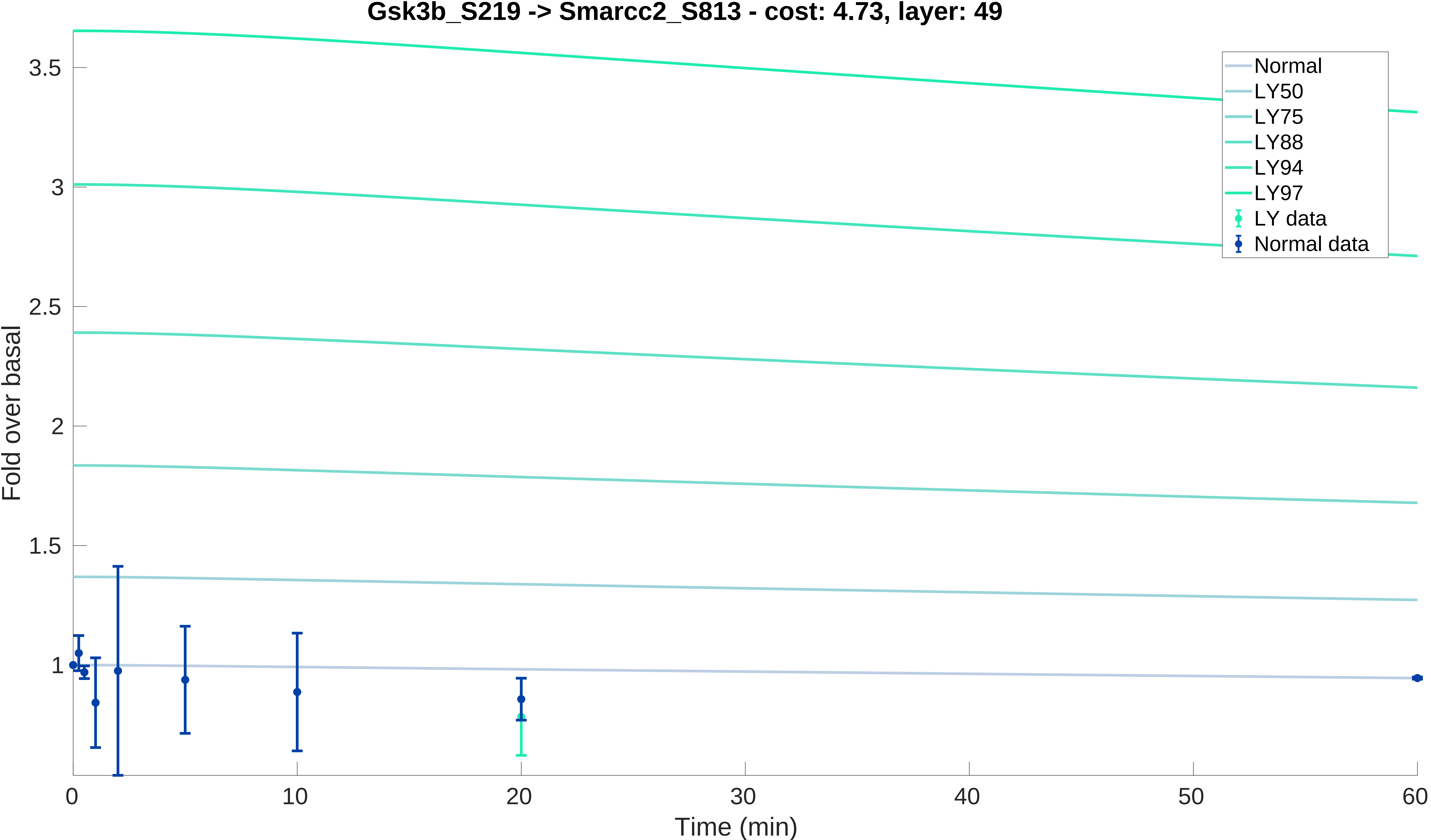
Gsk3b_S219 -> Smarcc2_S813 - cost: 4.73, layer: 49
Fold over basal
Time (min)
3.5
3
2.5
2
1.5
1
0
10
20
30
40
50
60
Normal
LY50
LY75
LY88
LY94
LY97
LY data
Normal data

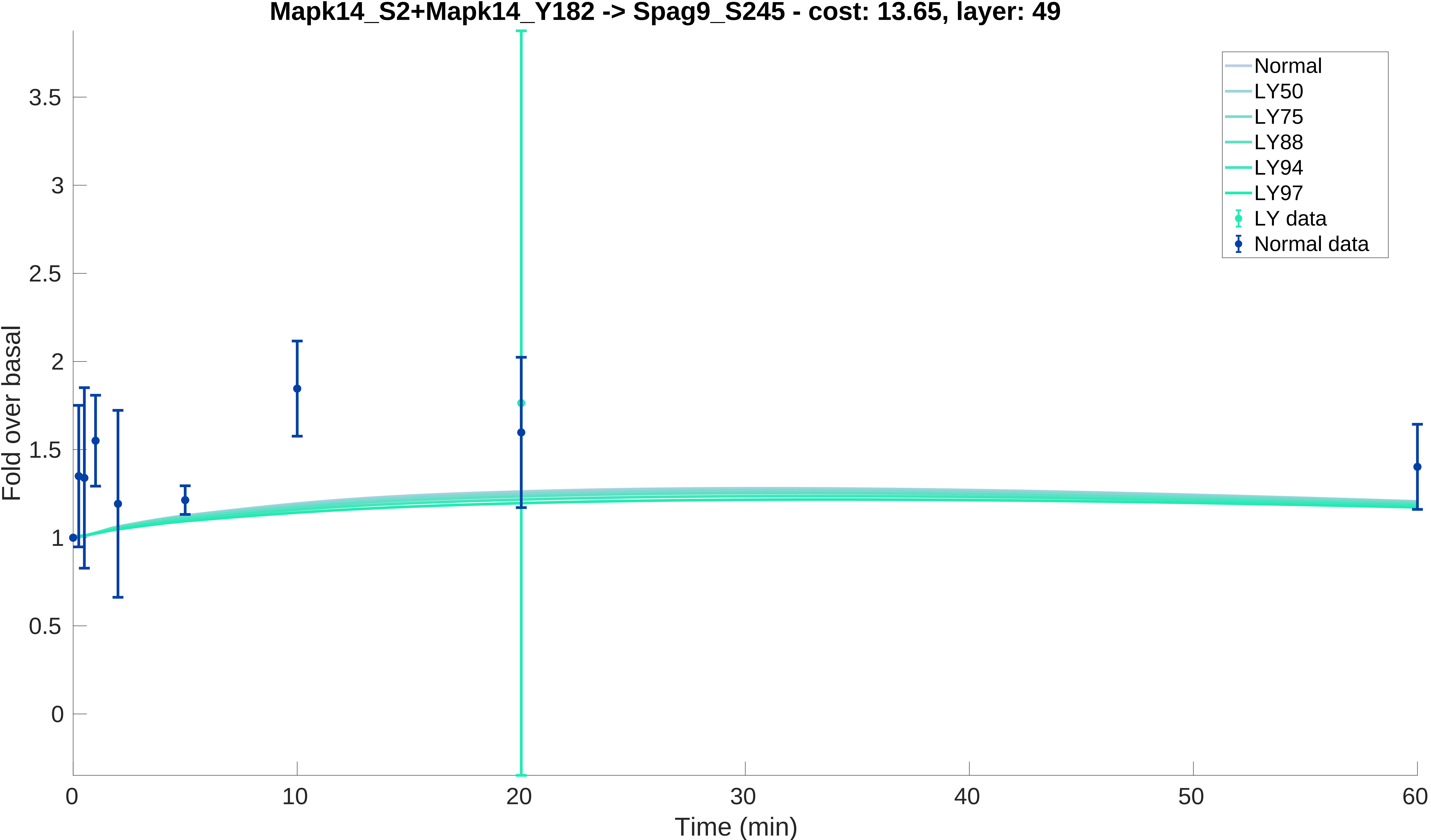

Mapk14_S2+Mapk14_Y182 -> Spag9_S245 - cost: 13.65, layer: 49
Fold over basal
Time (min)
0
0.5
1
1.5
2
2.5
3
3.5
0
10
20
30
40
50
60
Normal
LY50
LY75
LY88
LY94
LY97
LY data
Normal data

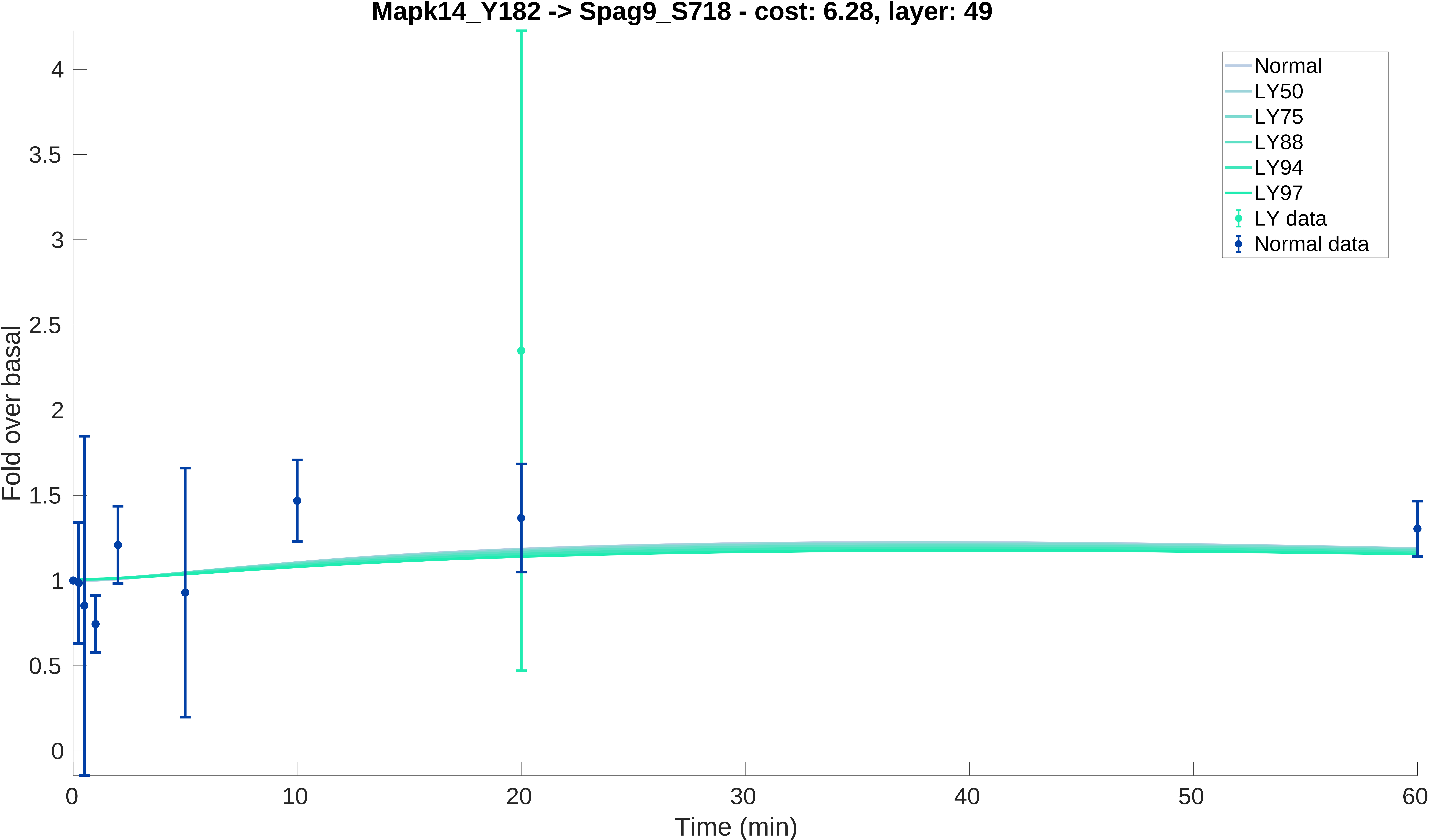

Mapk14_Y182 -> Spag9_S718 - cost: 6.28, layer: 49
Fold over basal
Time (min)
0
0.5
1
1.5
2
2.5
3
3.5
4
0
10
20
30
40
50
60
Normal
LY50
LY75
LY88
LY94
LY97
LY data
Normal data

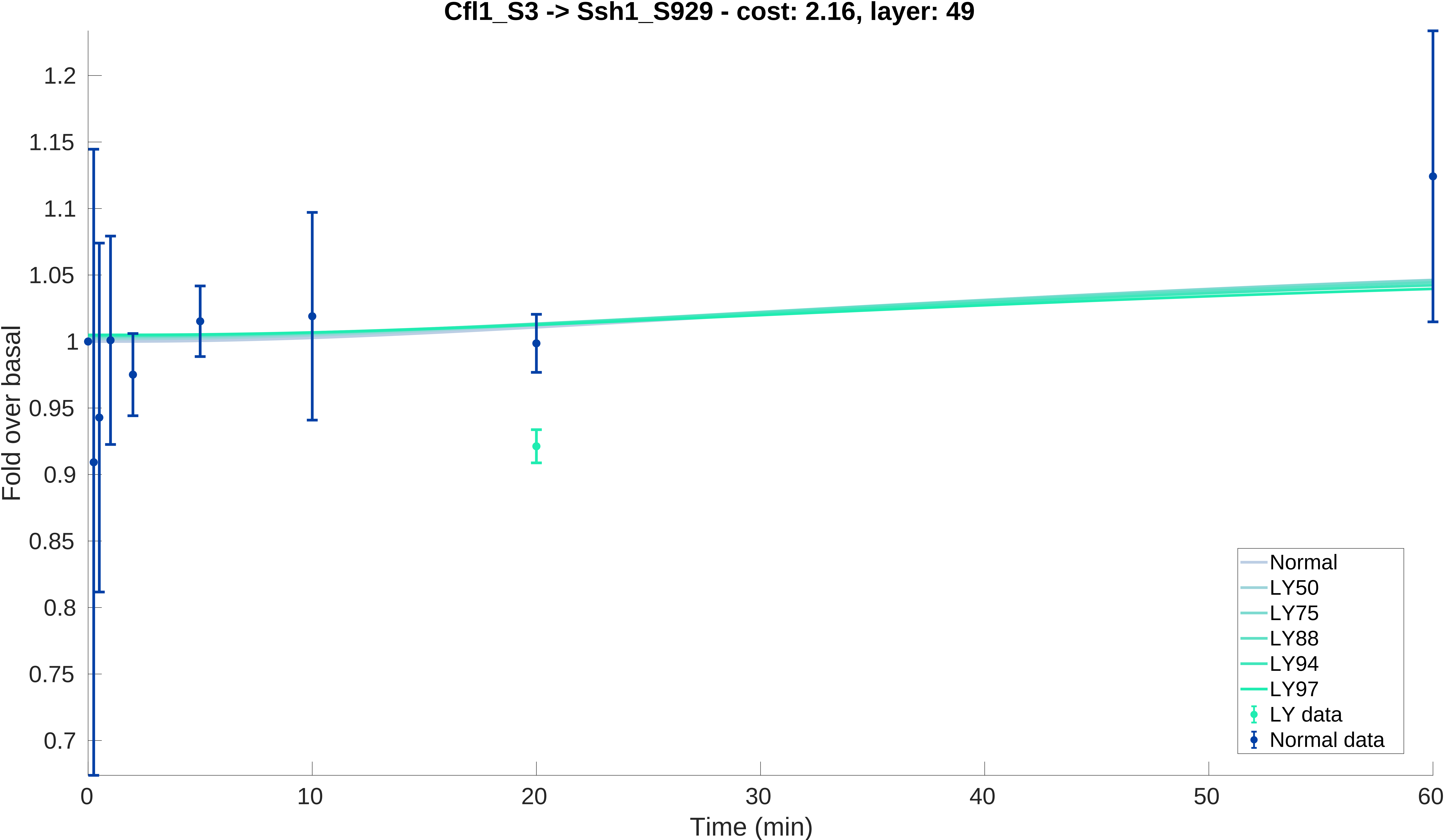

Cfl1_S3 -> Ssh1_S929 - cost: 2.16, layer: 49
Fold over basal
Time (min)
1.2
1.15
1.1
1.05
1
0.95
0.9
0.85
0.8
0.75
0.7
0
10
20
30
40
50
60
Normal
LY50
LY75
LY88
LY94
LY97
LY data
Normal data

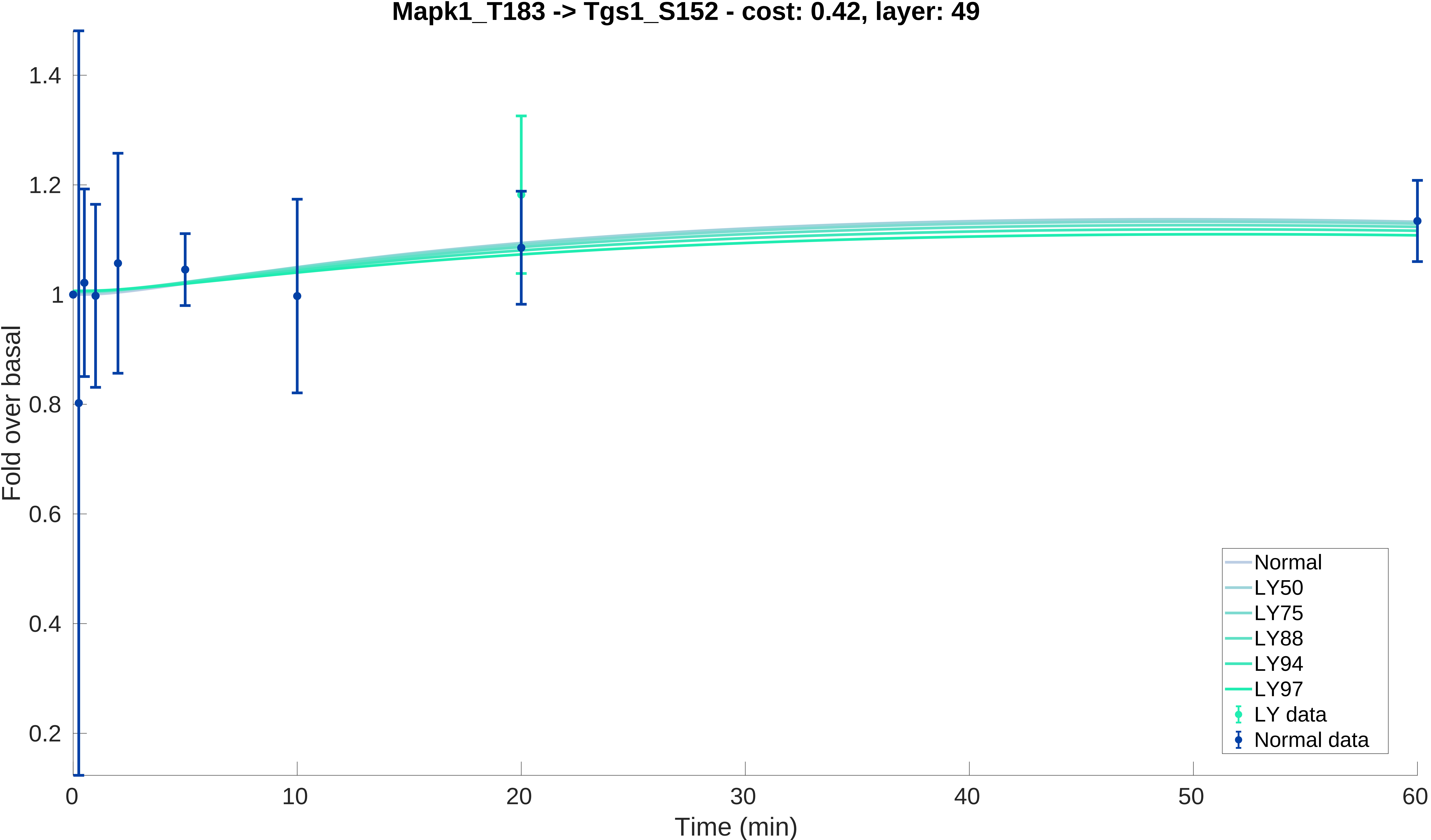
Mapk1_T183 -> Tgs1_S152 - cost: 0.42, layer: 49
Fold over basal
Time (min)
0
10
20
30
40
50
60
0.2
0.4
0.6
0.8
1
1.2
1.4
Normal
LY50
LY75
LY88
LY94
LY97
LY data
Normal data

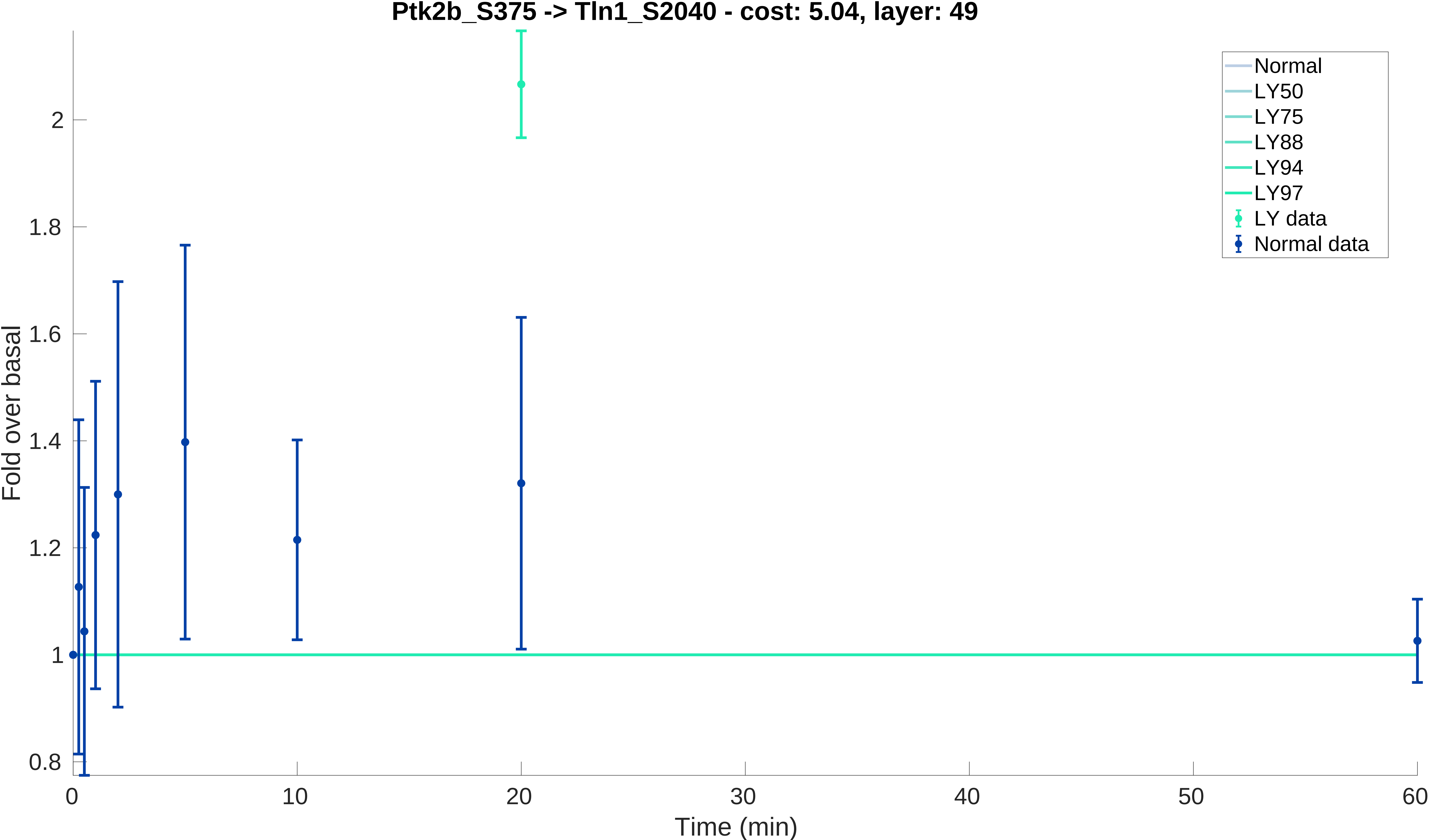

Ptk2b_S375 -> Tln1_S2040 - cost: 5.04, layer: 49
Fold over basal
Time (min)
0.8
1
1.2
1.4
1.6
1.8
2
0
10
20
30
40
50
60
Normal
LY50
LY75
LY88
LY94
LY97
LY data
Normal data

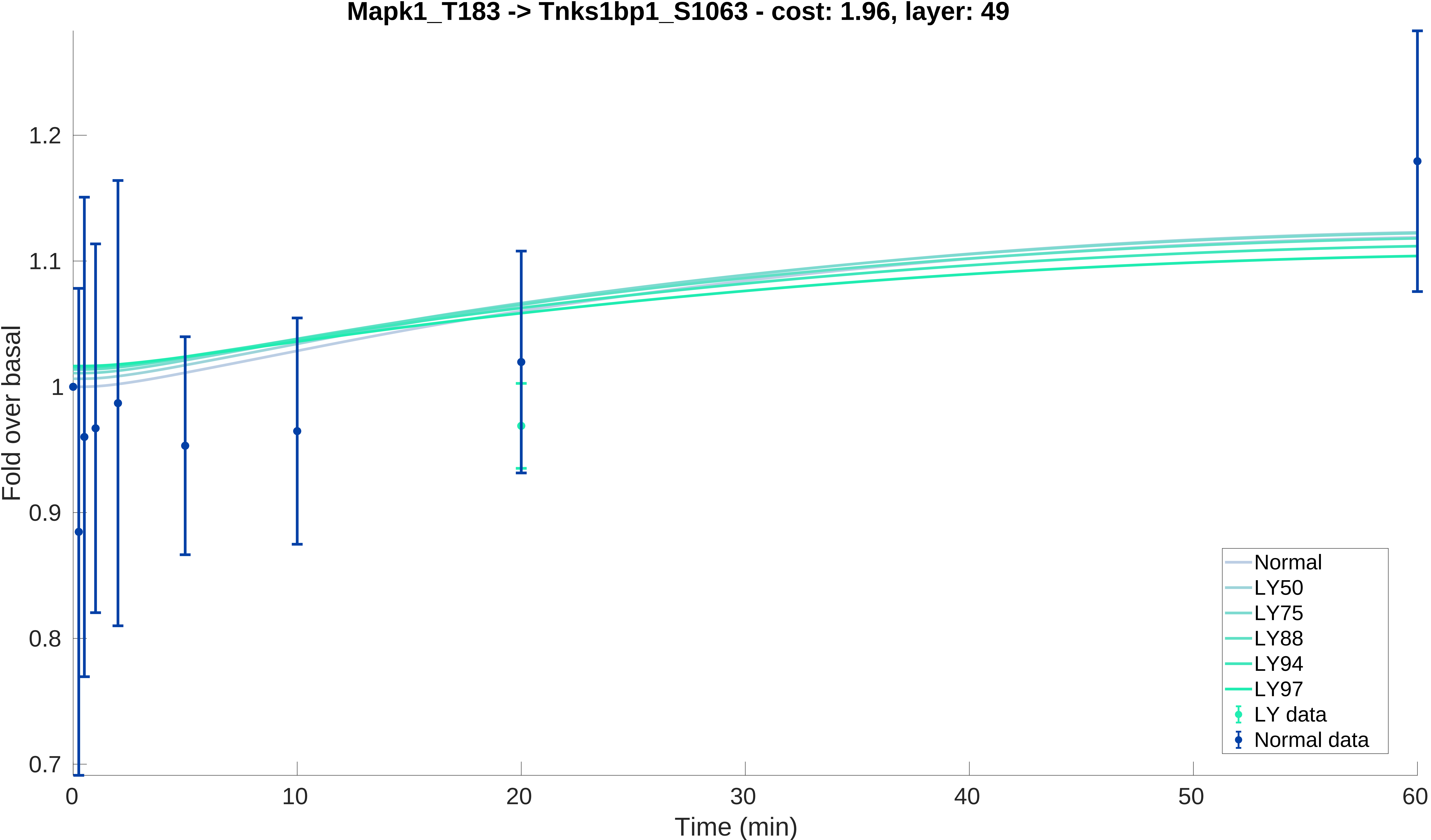

Mapk1_T183 -> Tnks1bp1_S1063 - cost: 1.96, layer: 49
Fold over basal
1.2
1.1
1
0.9
0.8
0.7
0
10
20
30
40
50
60
Time (min)
Normal
LY50
LY75
LY88
LY94
LY97
LY data
Normal data

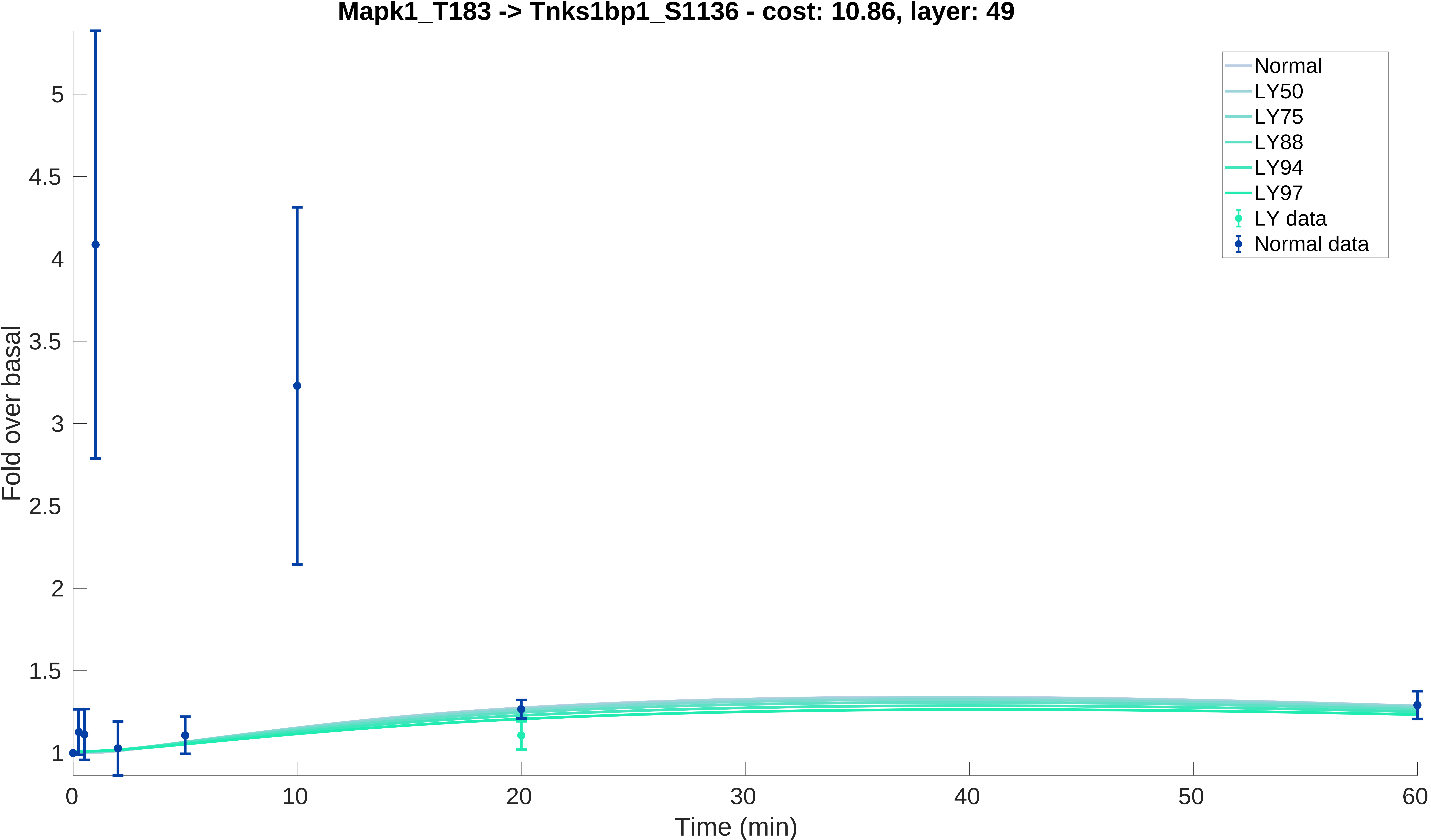

Mapk1_T183 -> Tnks1bp1_S1136 - cost: 10.86, layer: 49
Fold over basal
1
1.5
2
2.5
3
3.5
4
4.5
5
0
10
20
30
40
50
60
Time (min)
Normal
LY50
LY75
LY88
LY94
LY97
LY data
Normal data

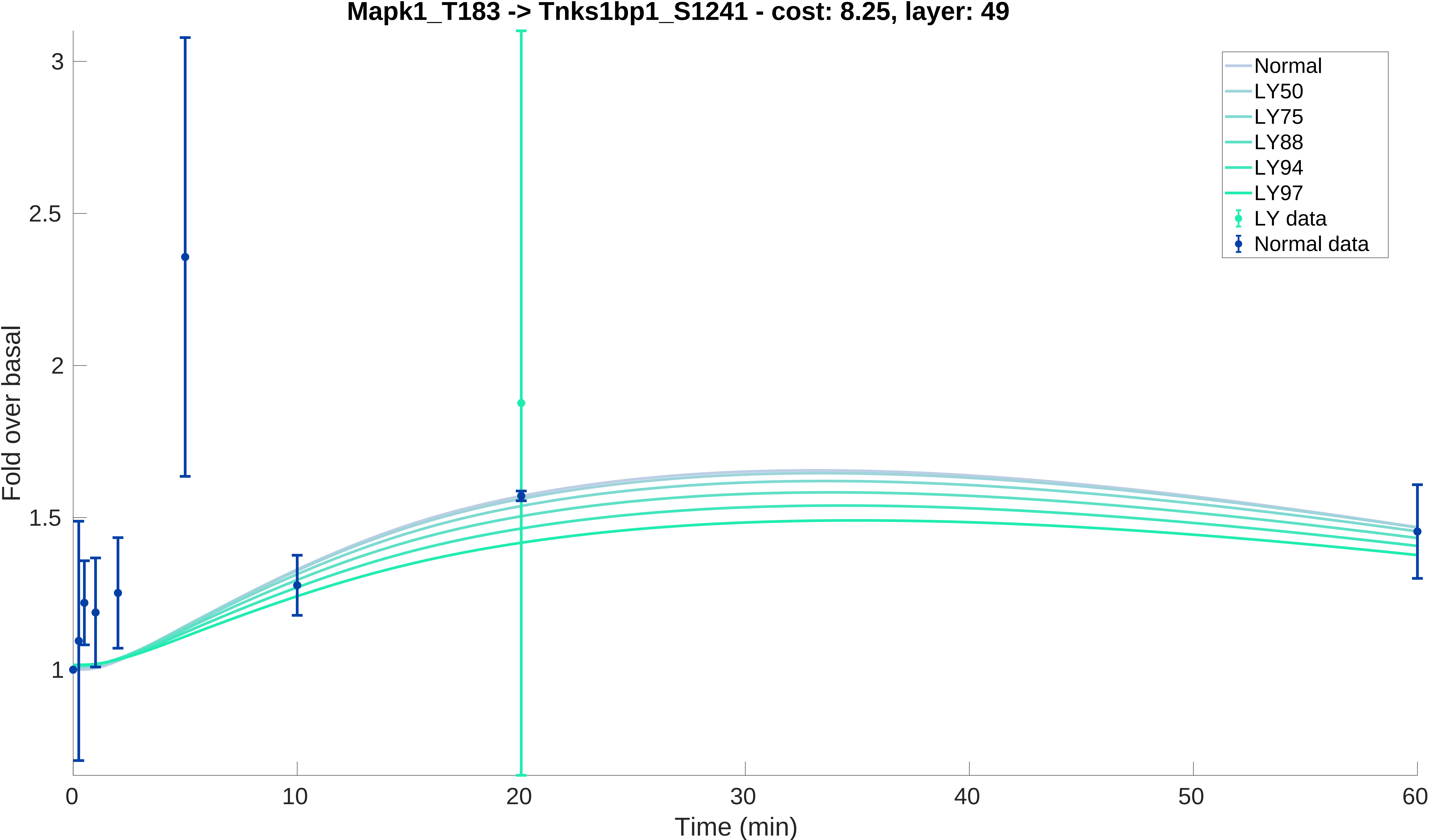

Mapk1_T183 -> Tnks1bp1_S1241 - cost: 8.25, layer: 49
Fold over basal
Time (min)
Normal
LY50
LY75
LY88
LY94
LY97
LY data
Normal data
3
2.5
2
1.5
1
0
10
20
30
40
50
60

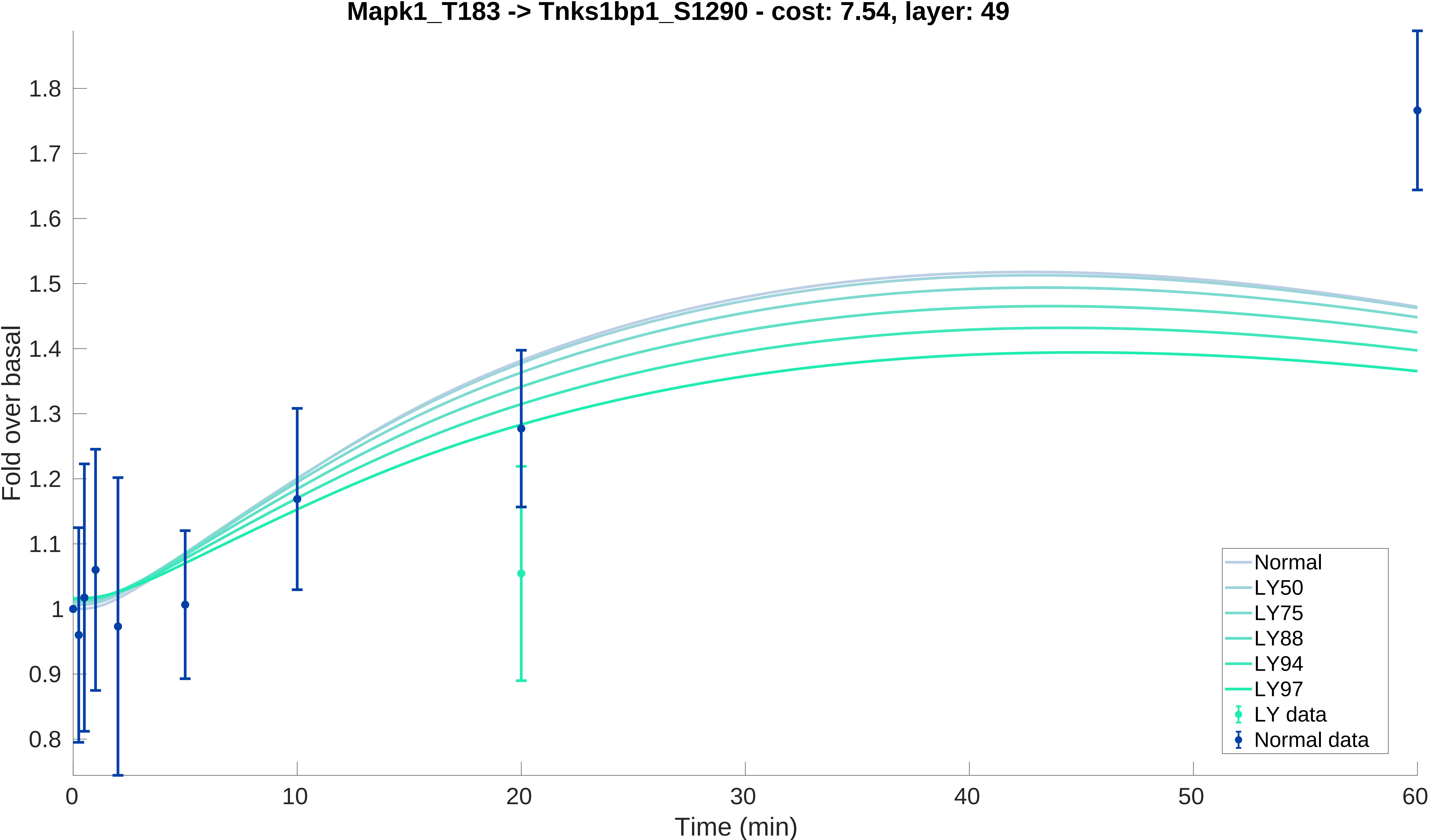

Mapk1_T183 -> Tnks1bp1_S1290 - cost: 7.54, layer: 49
Fold over basal
Time (min)
0.8
0.9
1
1.1
1.2
1.3
1.4
1.5
1.6
1.7
1.8
0
10
20
30
40
50
60
Normal
LY50
LY75
LY88
LY94
LY97
LY data
Normal data

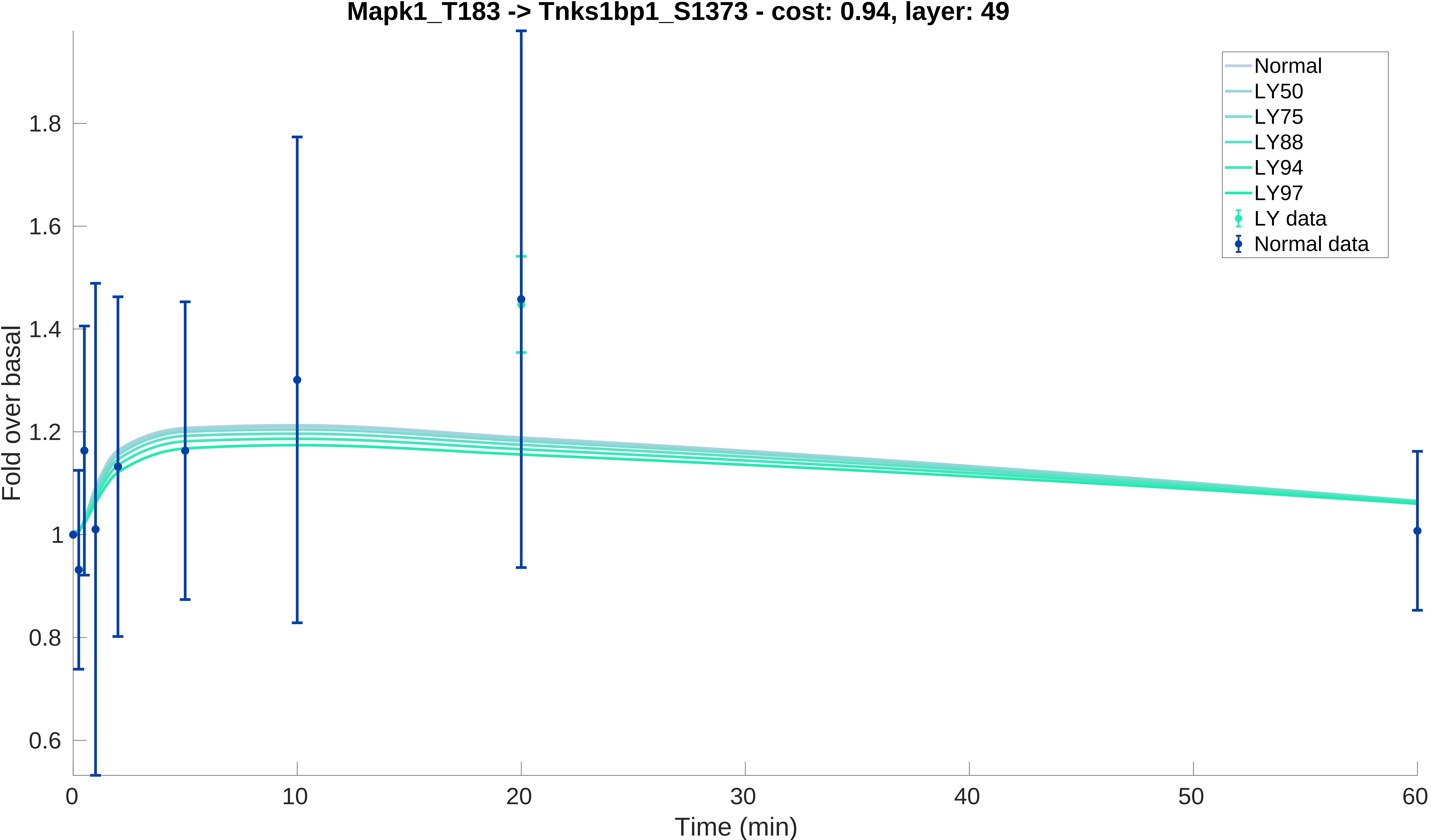
Mapk1_T183 -> Tnks1bp1_S1373 - cost: 0.94, layer: 49
Fold over basal
Time (min)
0.6
0.8
1
1.2
1.4
1.6
1.8
0
10
20
30
40
50
60
Normal
LY50
LY75
LY88
LY94
LY97
LY data
Normal data

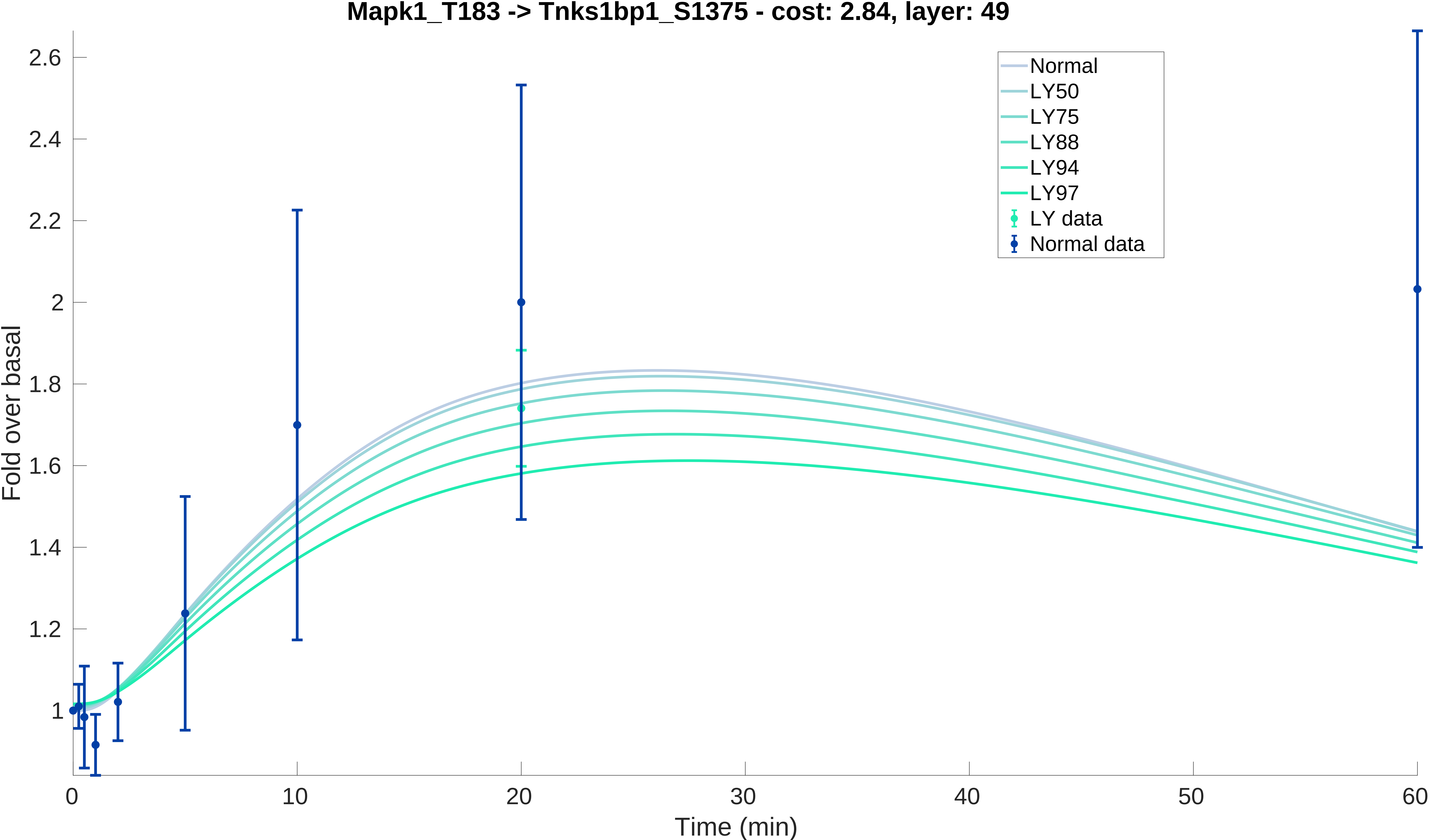

Mapk1_T183 -> Tnks1bp1_S1375 - cost: 2.84, layer: 49
Fold over basal
Time (min)
0
10
20
30
40
50
60
1
1.2
1.4
1.6
1.8
2
2.2
2.4
2.6
Normal
LY50
LY75
LY88
LY94
LY97
LY data
Normal data

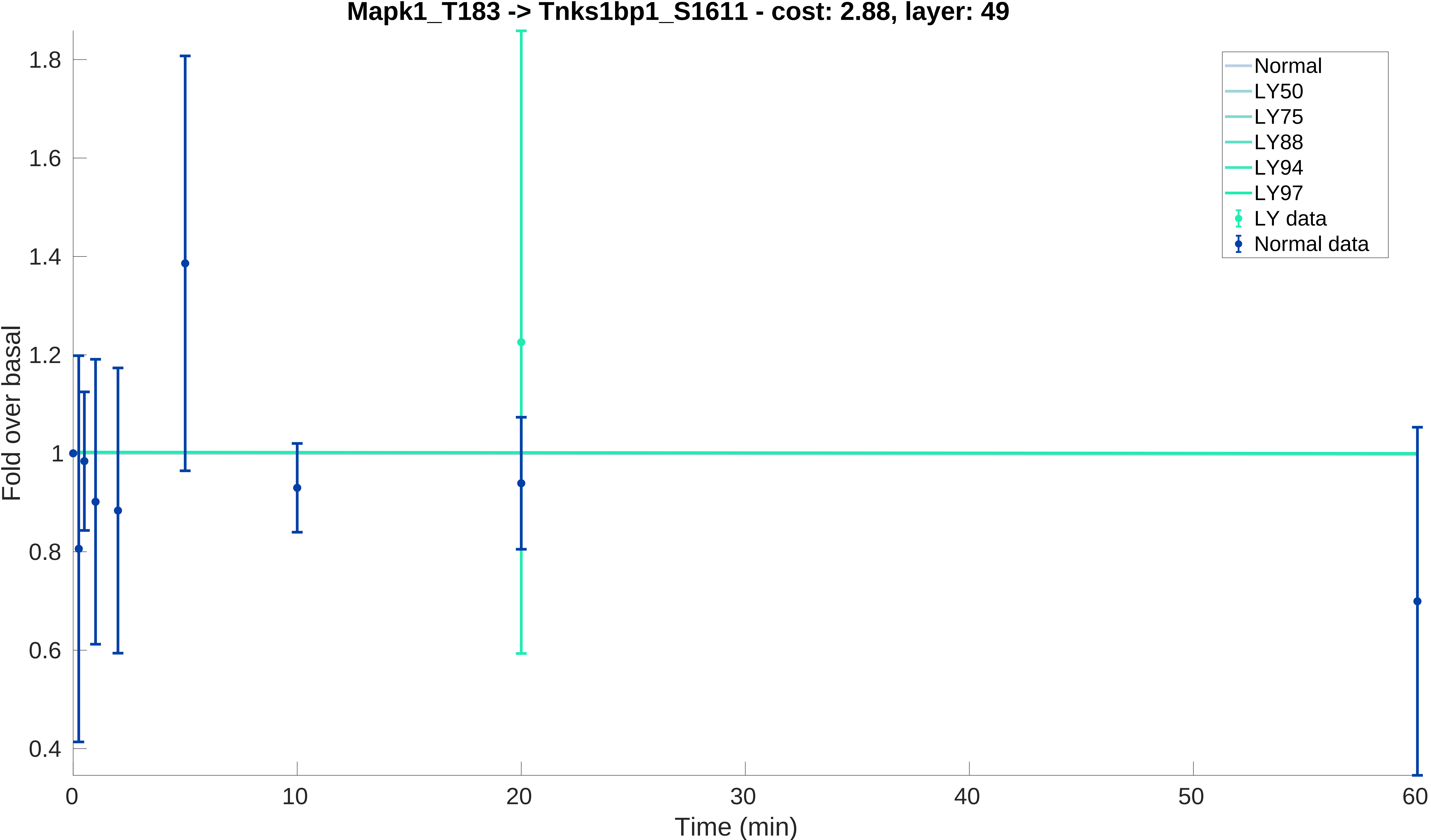
Mapk1_T183 -> Tnks1bp1_S1611 - cost: 2.88, layer: 49
Fold over basal
Time (min)
0.4
0.6
0.8
1
1.2
1.4
1.6
1.8
0
10
20
30
40
50
60
Normal
LY50
LY75
LY88
LY94
LY97
LY data
Normal data

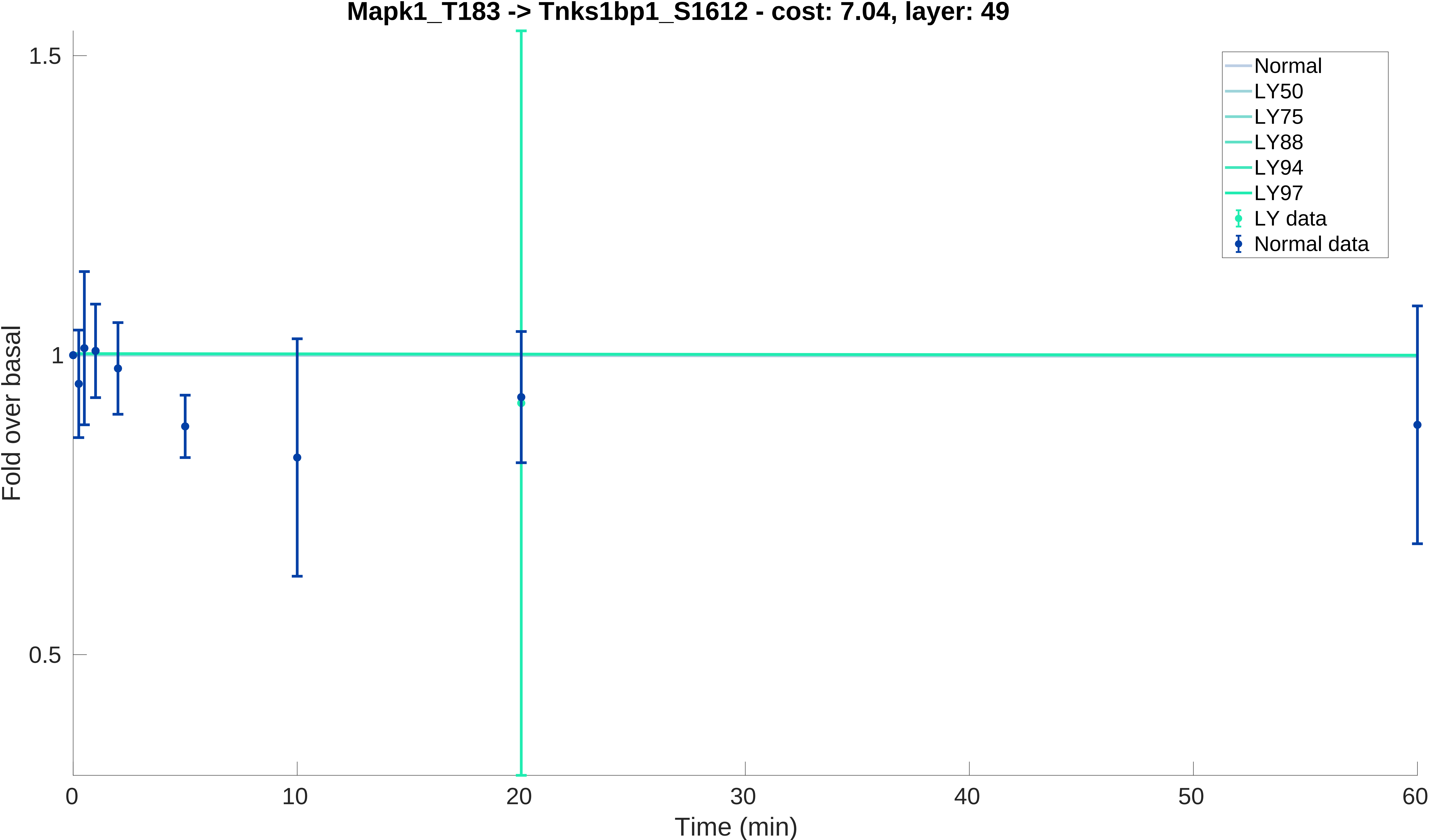

Mapk1_T183 -> Tnks1bp1_S1612 - cost: 7.04, layer: 49
Fold over basal
1.5
1
0.5
0
10
20
30
40
50
60
Time (min)
Normal
LY50
LY75
LY88
LY94
LY97
LY data
Normal data

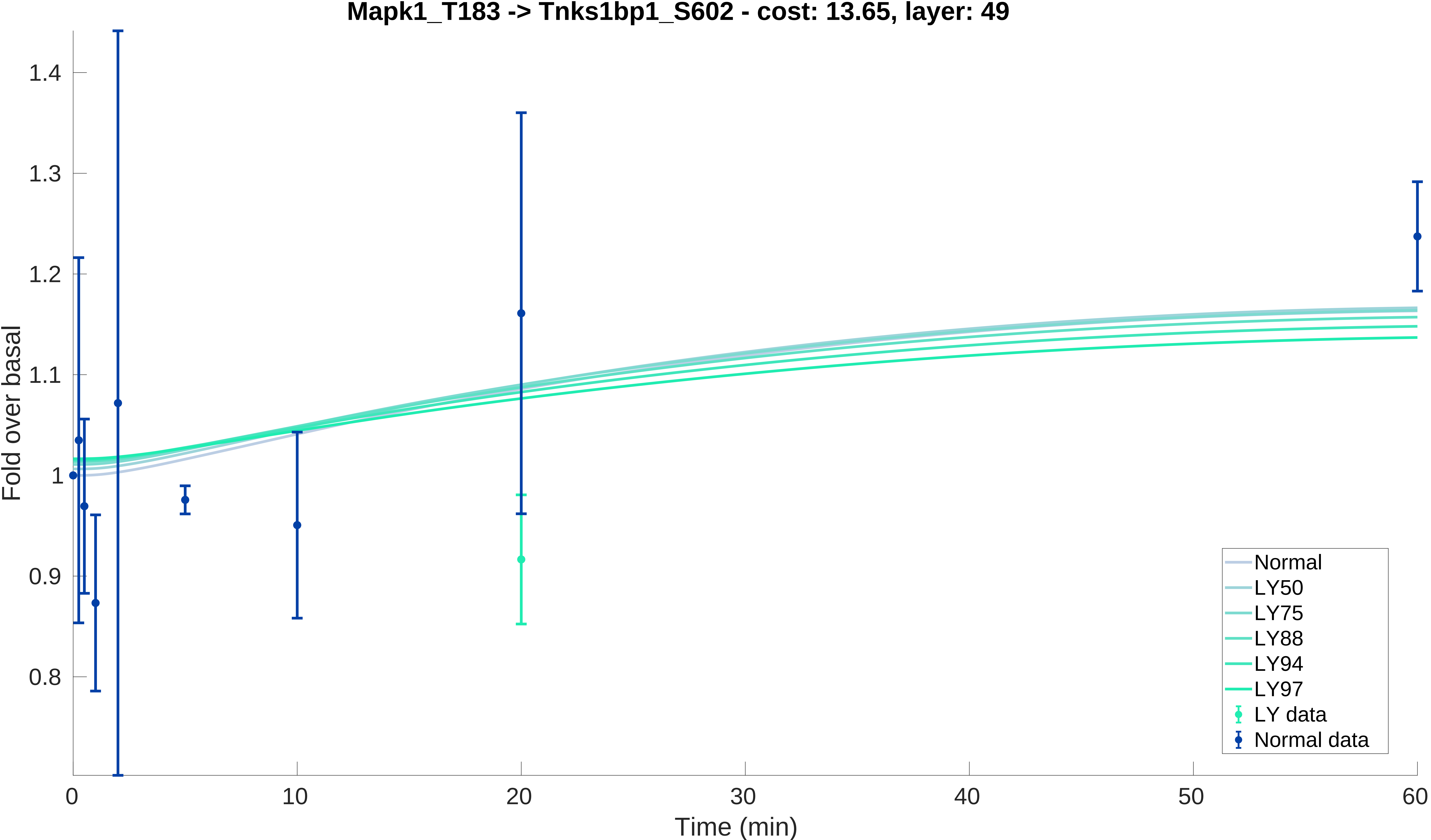

Mapk1_T183 -> Tnks1bp1_S602 - cost: 13.65, layer: 49
Fold over basal
1.4
1.3
1.2
1.1
1
0.9
0.8
0
10
20
30
40
50
60
Time (min)
Normal
LY50
LY75
LY88
LY94
LY97
LY data
Normal data

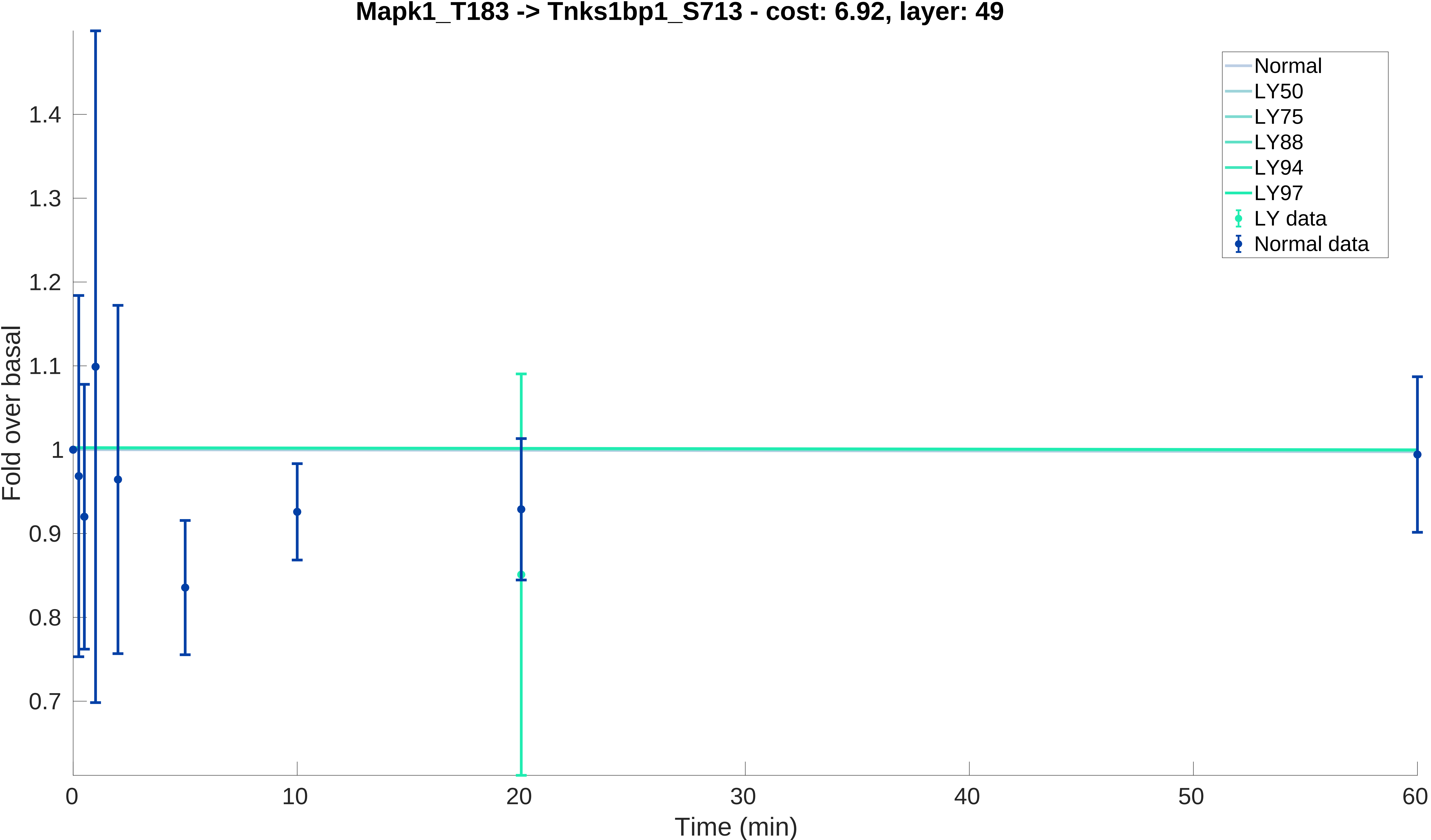

Mapk1_T183 -> Tnks1bp1_S713 - cost: 6.92, layer: 49
Normal
LY50
LY75
LY88
LY94
LY97
LY data
Normal data
Fold over basal
1.4
1.3
1.2
1.1
1
0.9
0.8
0.7
0
10
20
30
40
50
60
Time (min)

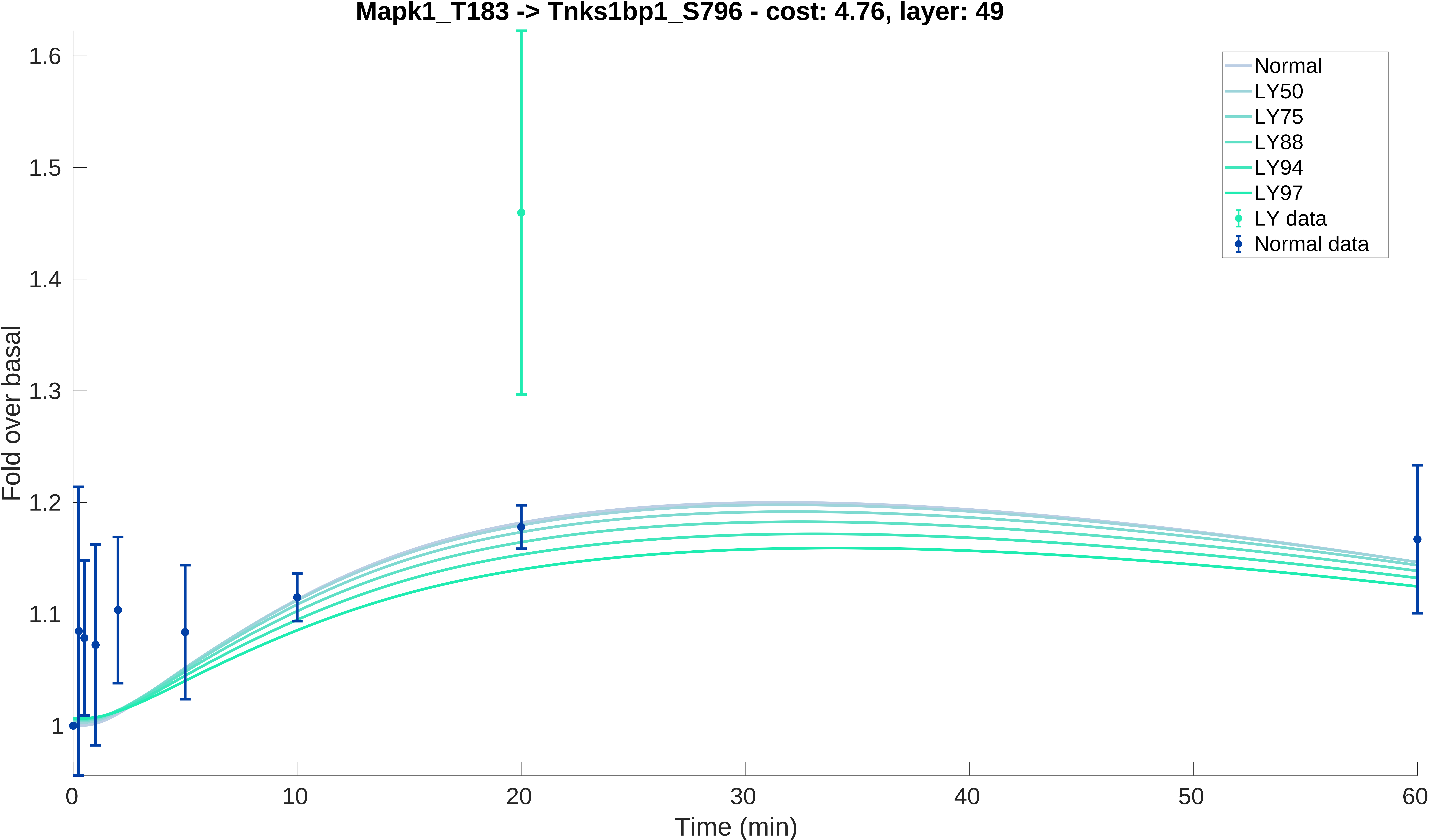
Mapk1_T183 -> Tnks1bp1_S796 - cost: 4.76, layer: 49
Fold over basal
Time (min)
1.6
1.5
1.4
1.3
1.2
1.1
1
0
10
20
30
40
50
60
Normal
LY50
LY75
LY88
LY94
LY97
LY data
Normal data

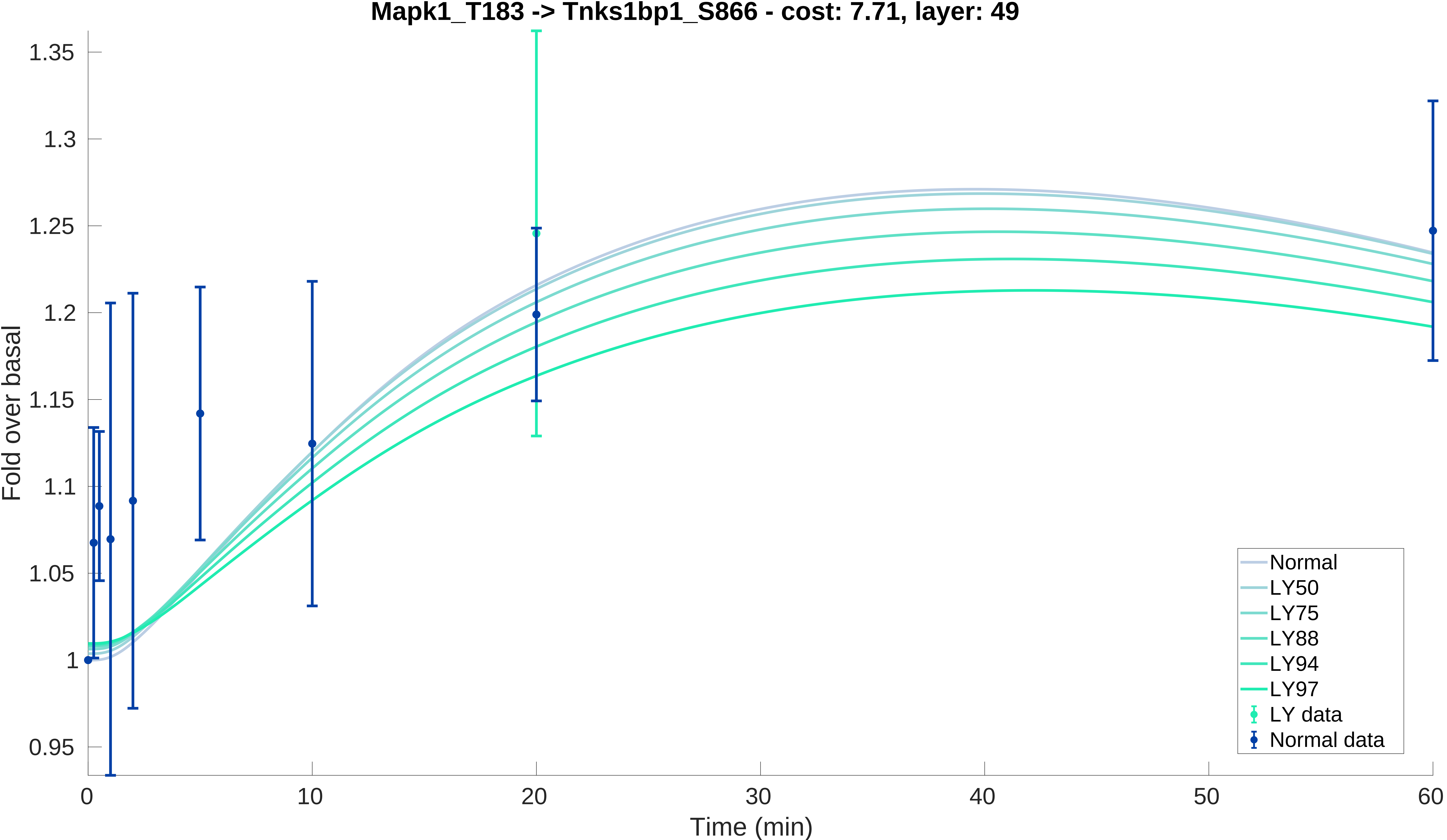

Mapk1_T183 -> Tnks1bp1_S866 - cost: 7.71, layer: 49
Fold over basal
1.35
1.3
1.25
1.2
1.15
1.1
1.05
1
0.95
0
10
20
30
40
50
60
Time (min)
Normal
LY50
LY75
LY88
LY94
LY97
LY data
Normal data

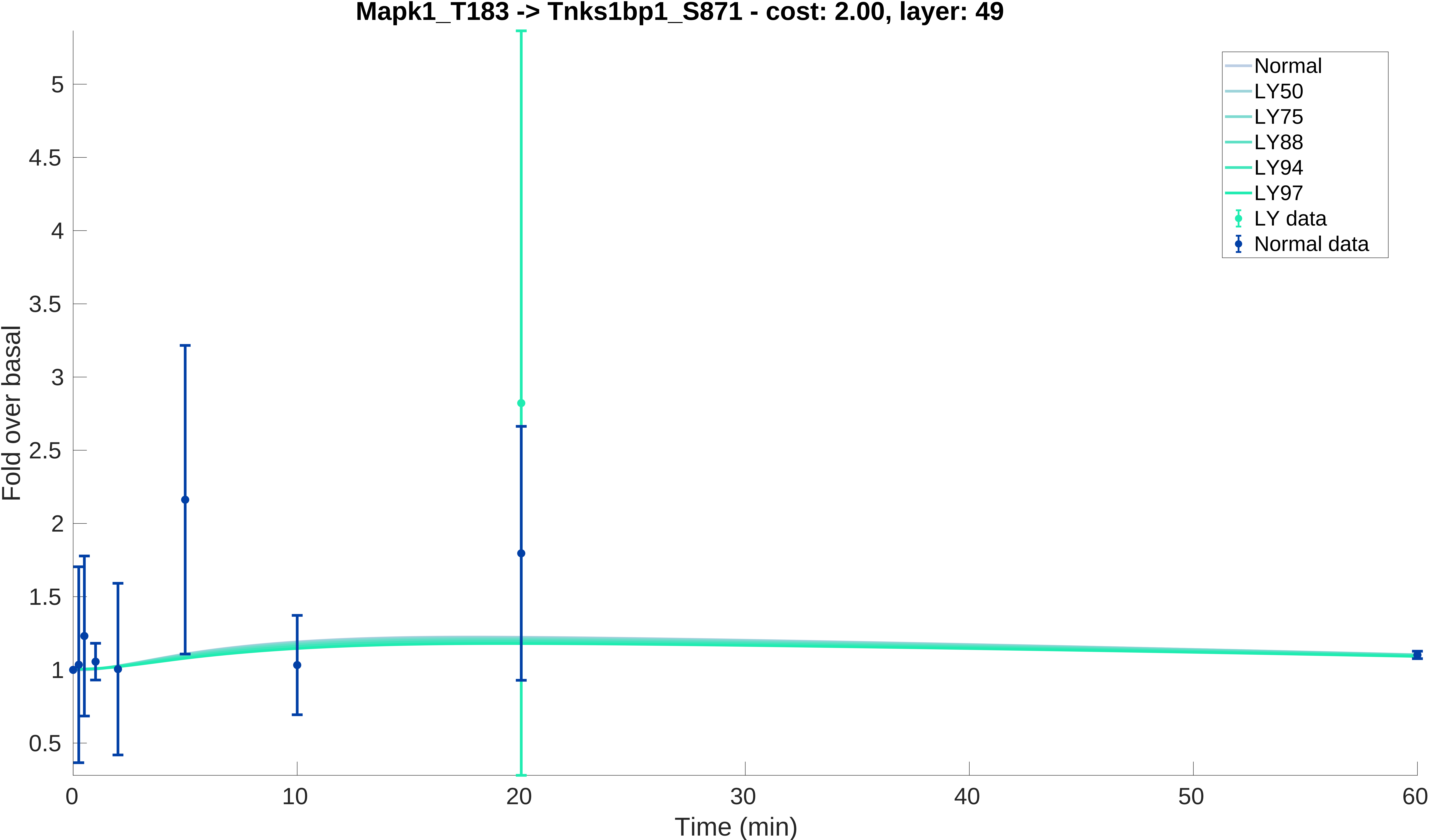

Mapk1_T183 -> Tnks1bp1_S871 - cost: 2.00, layer: 49
Fold over basal
Time (min)
0.5
1
1.5
2
2.5
3
3.5
4
4.5
5
0
10
20
30
40
50
60
Normal
LY50
LY75
LY88
LY94
LY97
LY data
Normal data

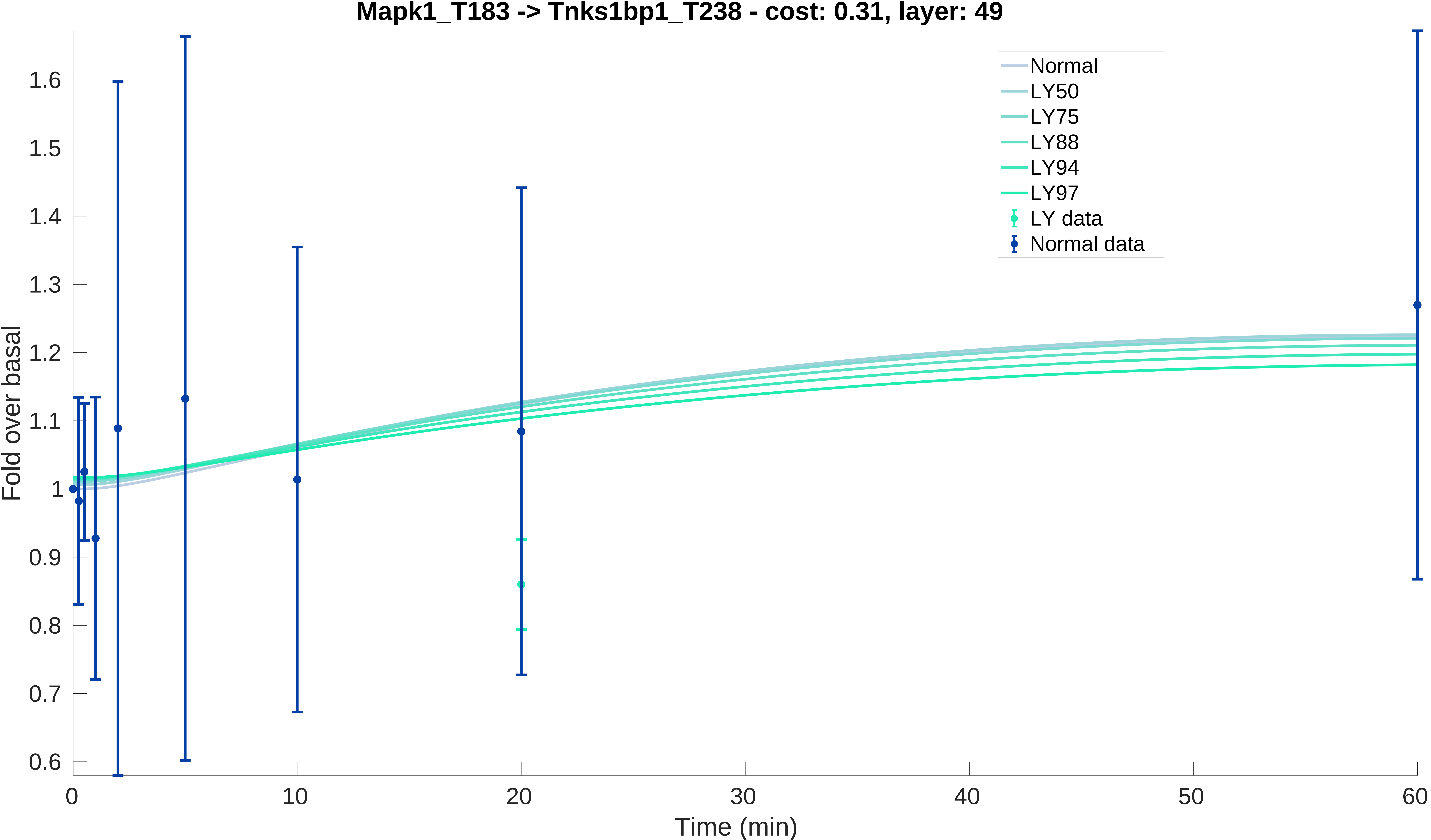

Mapk1_T183 -> Tnks1bp1_T238 - cost: 0.31, layer: 49
Fold over basal
Time (min)
Normal
LY50
LY75
LY88
LY94
LY97
LY data
Normal data
0.6
0.7
0.8
0.9
1
1.1
1.2
1.3
1.4
1.5
1.6
0
10
20
30
40
50
60

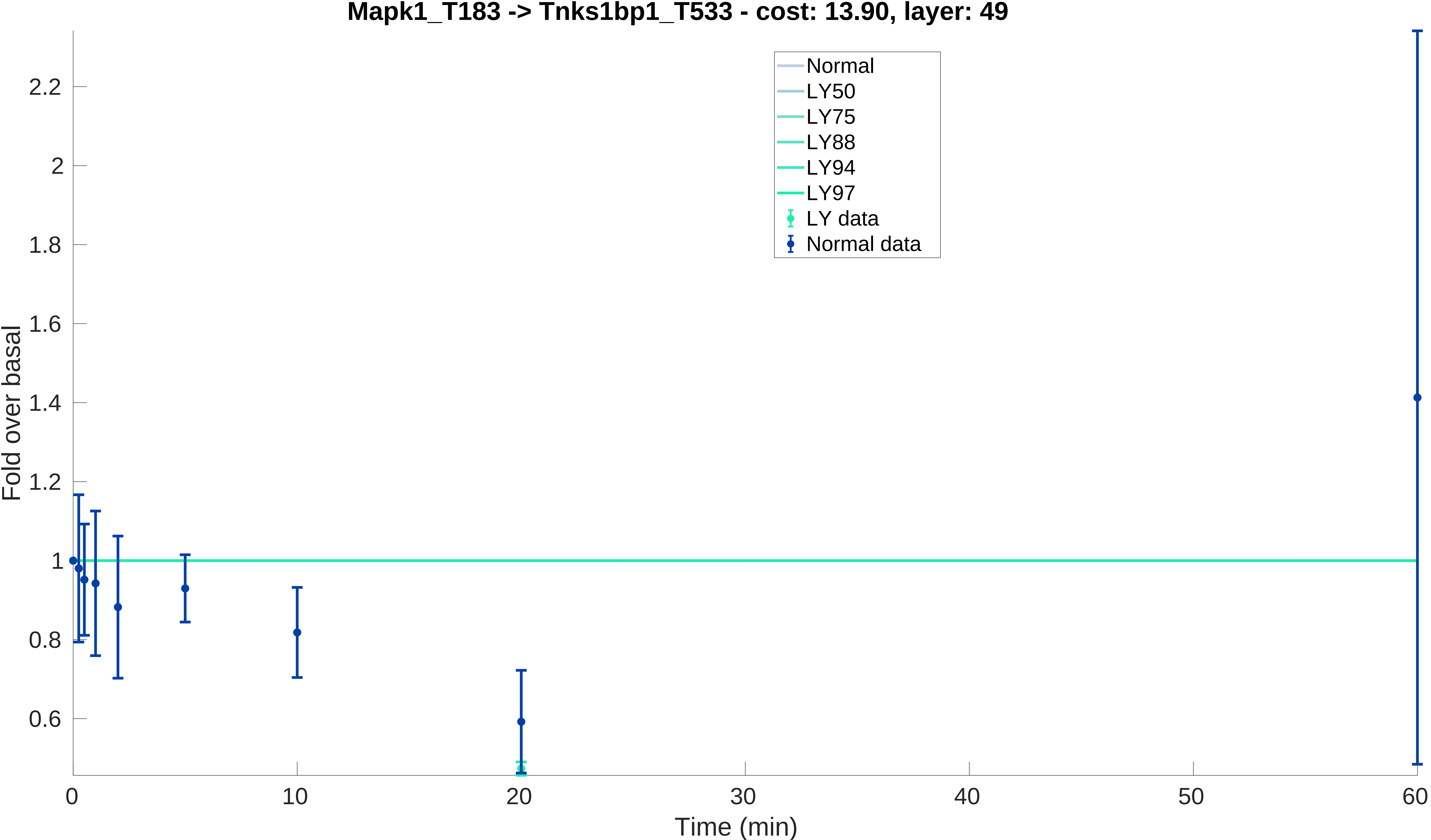

Mapk1_T183 -> Tnks1bp1_T533 - cost: 13.90, layer: 49
Normal
LY50
LY75
LY88
LY94
LY97
LY data
Normal data
Fold over basal
Time (min)
0.6
0.8
1
1.2
1.4
1.6
1.8
2
2.2
0
10
20
30
40
50
60

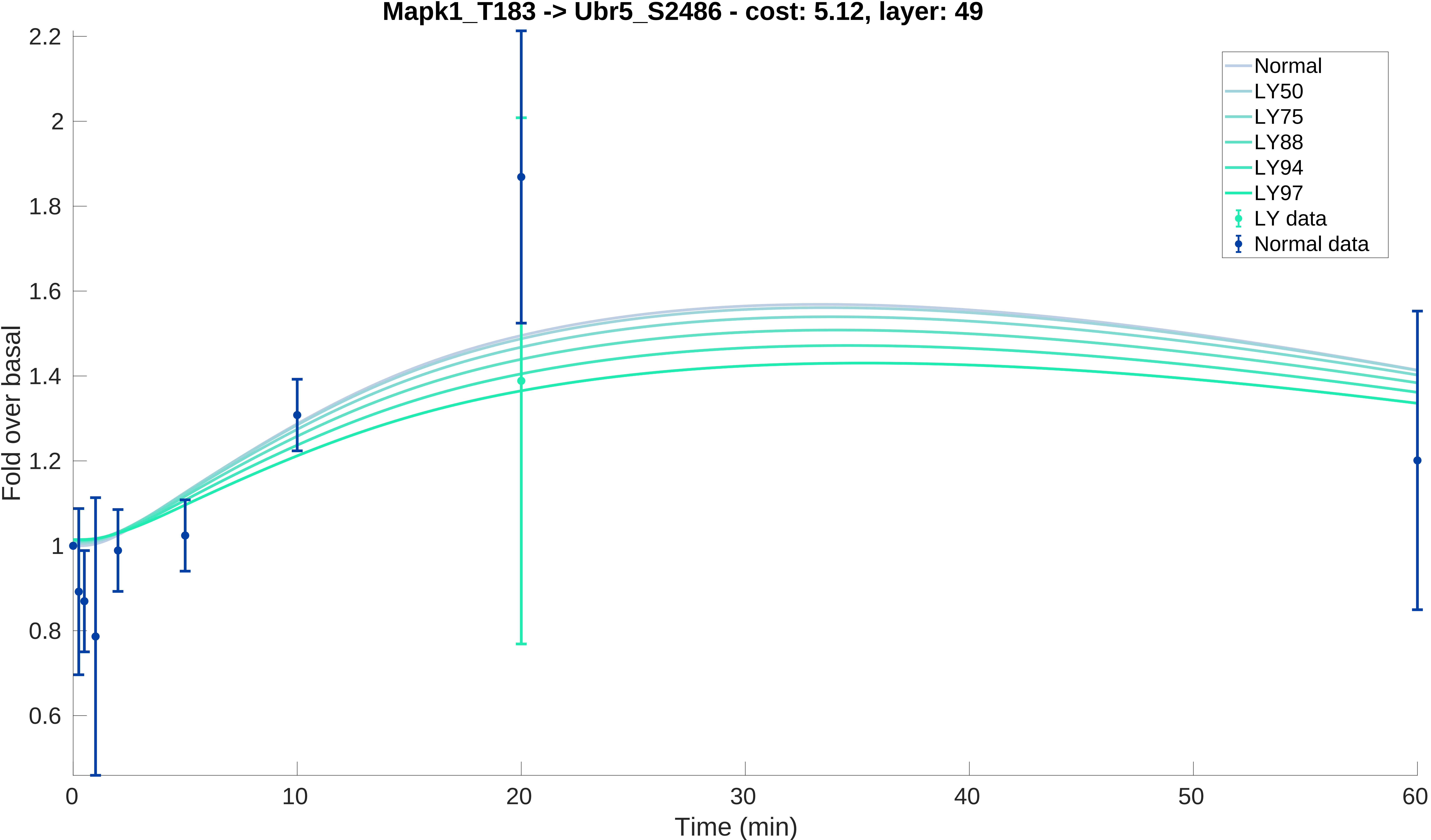

Mapk1_T183 -> Ubr5_S2486 - cost: 5.12, layer: 49
Normal
LY50
LY75
LY88
LY94
LY97
LY data
Normal data
Fold over basal
Time (min)
0
10
20
30
40
50
60
0.6
0.8
1
1.2
1.4
1.6
1.8
2
2.2

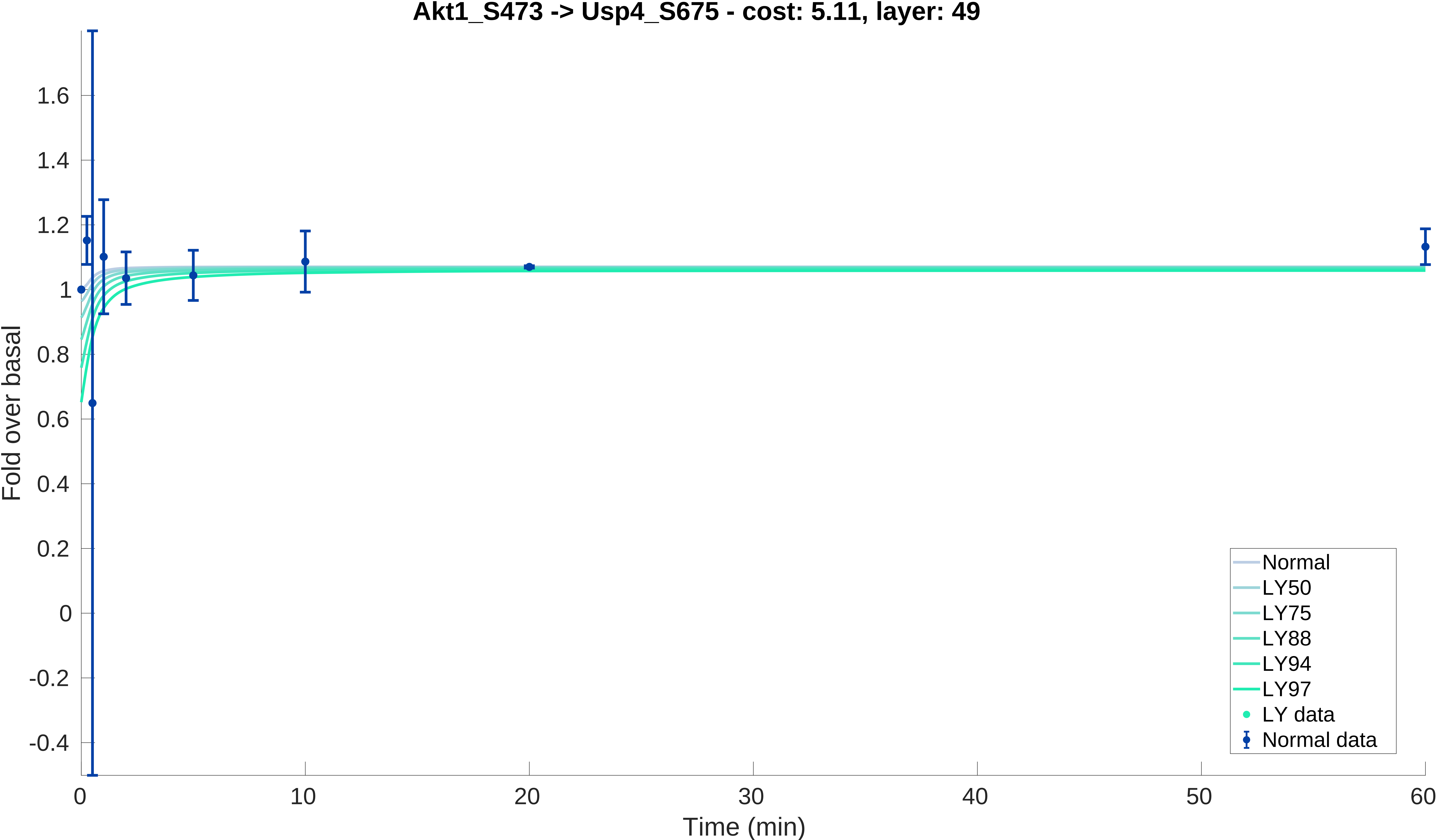

Akt1_S473 -> Usp4_S675 - cost: 5.11, layer: 49
Fold over basal
1.6
1.4
1.2
1
0.8
0.6
0.4
0.2
0
-0.2
-0.4
0
10
20
30
40
50
60
Time (min)
Normal
LY50
LY75
LY88
LY94
LY97
LY data
Normal data

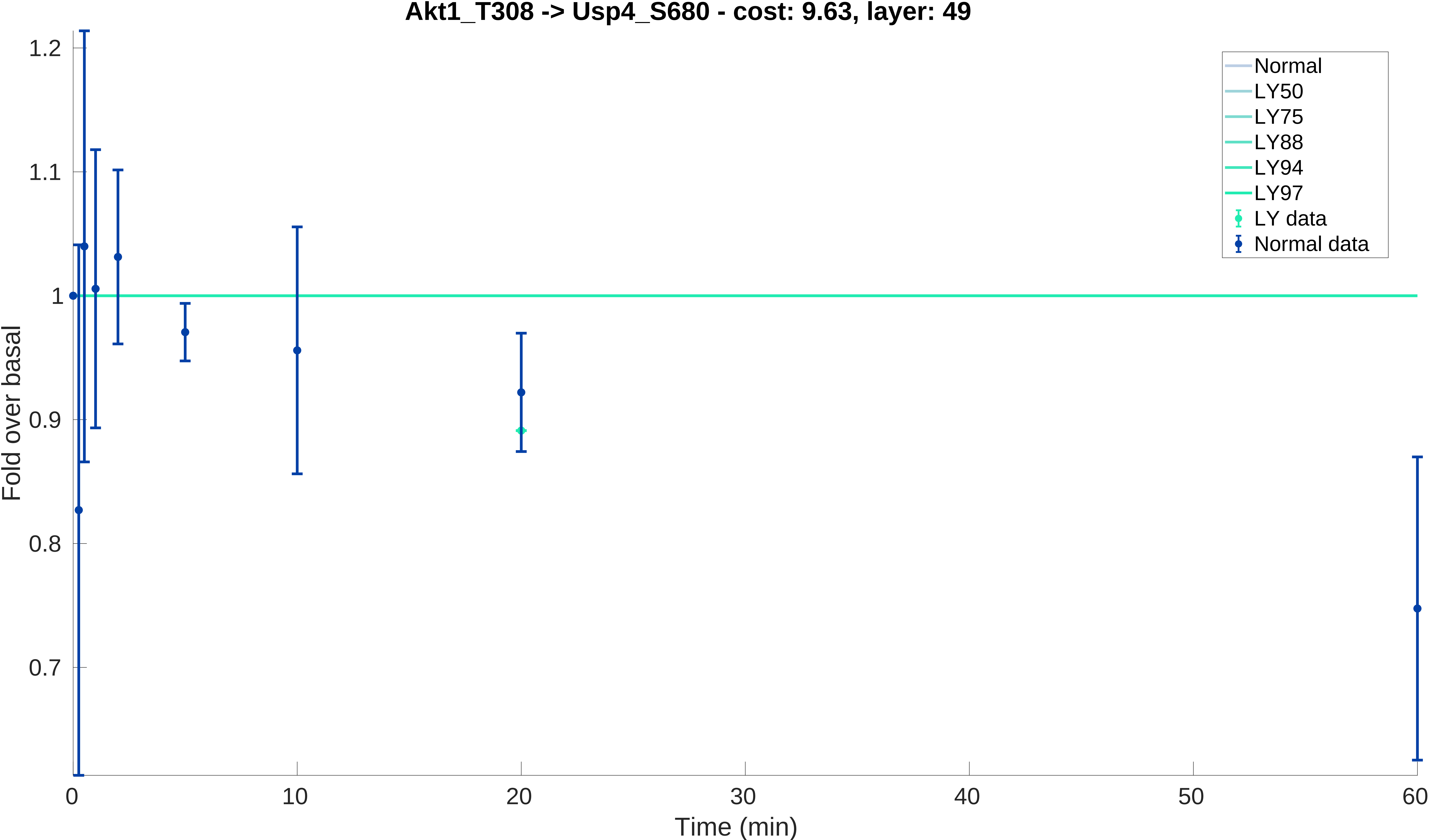

Akt1_T308 -> Usp4_S680 - cost: 9.63, layer: 49
Normal
LY50
LY75
LY88
LY94
LY97
LY data
Normal data
Fold over basal
1.2
1.1
1
0.9
0.8
0.7
0
10
20
30
40
50
60
Time (min)

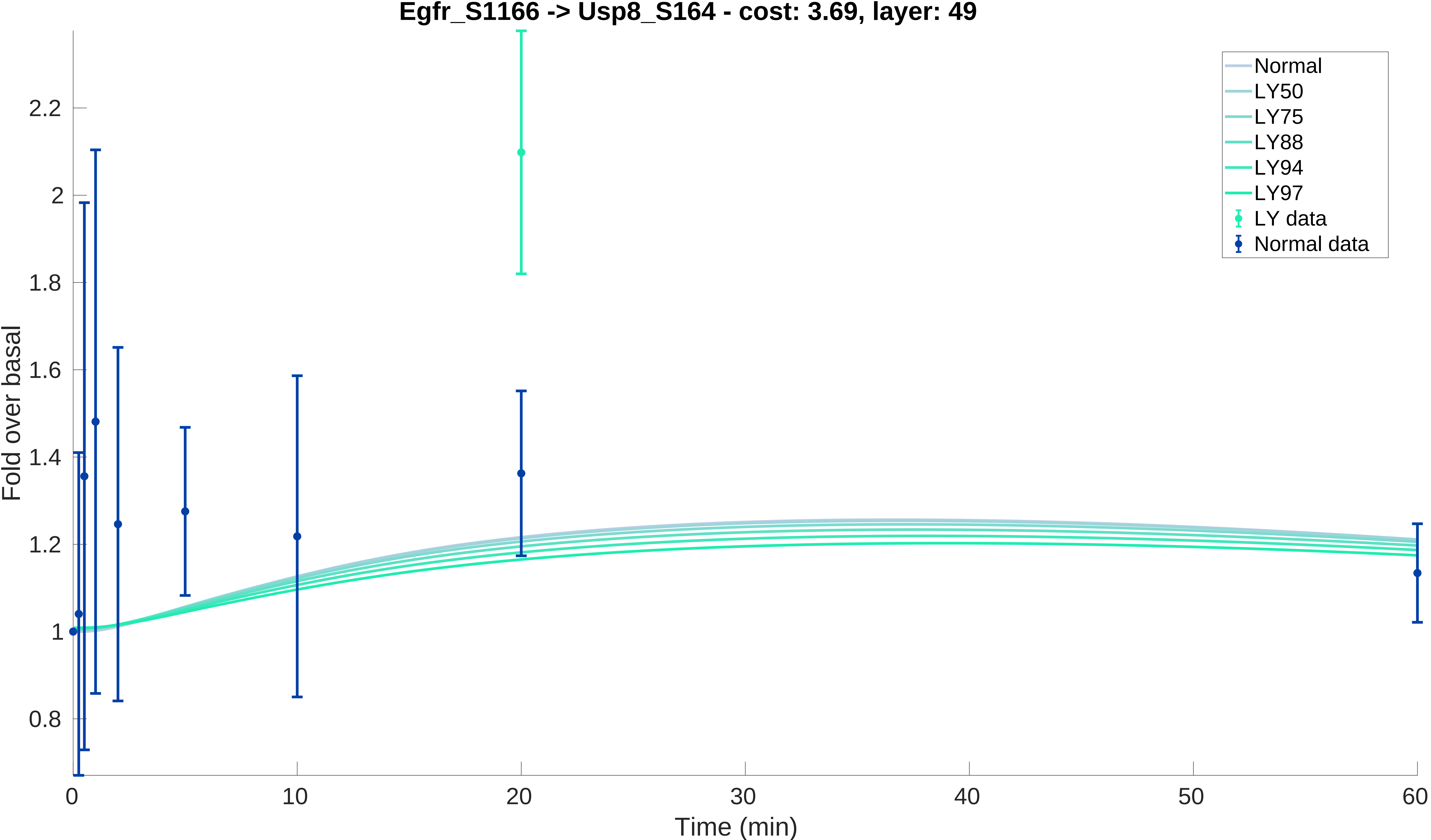

Egfr_S1166 -> Usp8_S164 - cost: 3.69, layer: 49
Fold over basal
Time (min)
0.8
1
1.2
1.4
1.6
1.8
2
2.2
0
10
20
30
40
50
60
Normal
LY50
LY75
LY88
LY94
LY97
LY data
Normal data

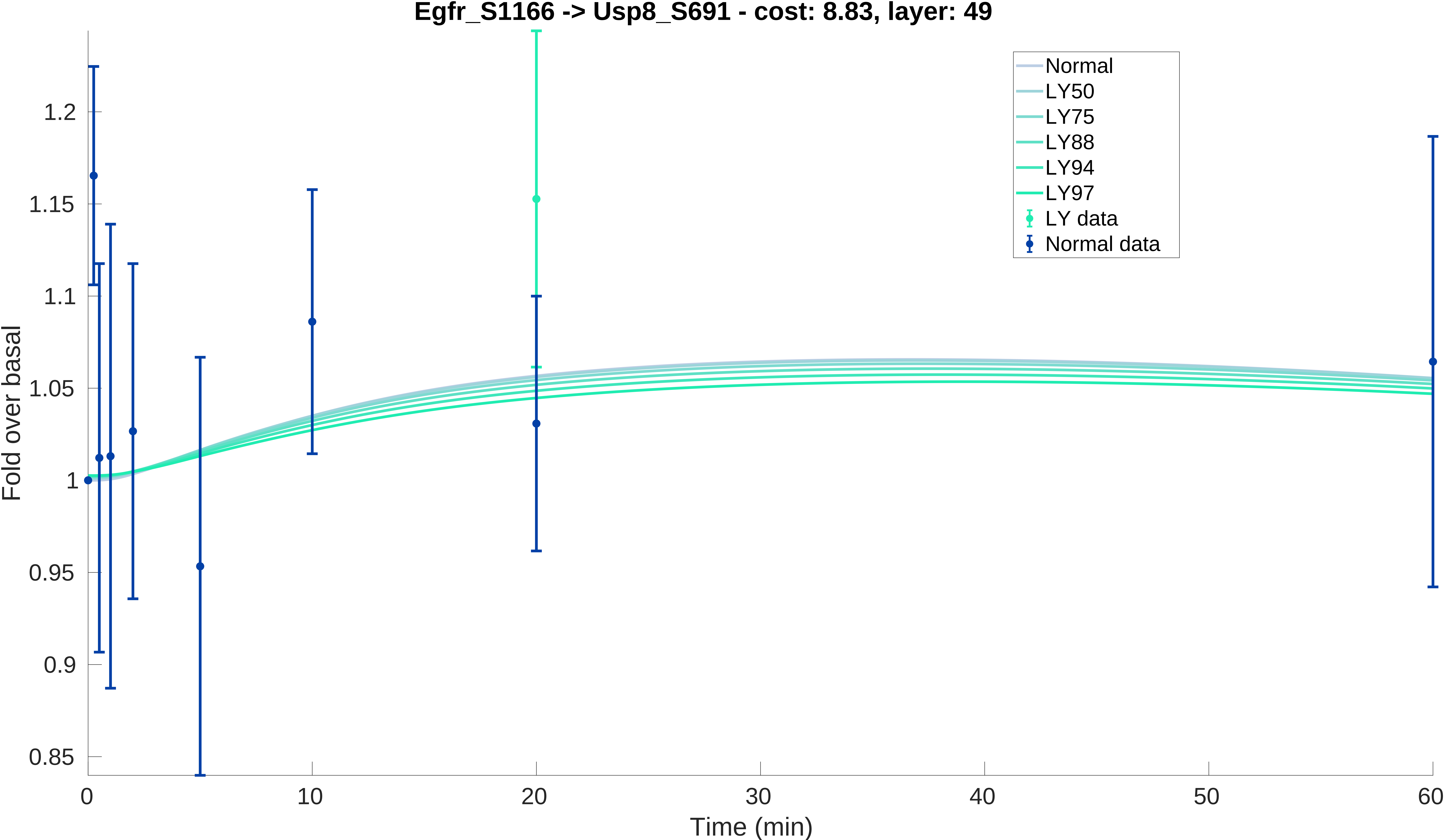
Egfr_S1166 -> Usp8_S691 - cost: 8.83, layer: 49
Normal
LY50
LY75
LY88
LY94
LY97
LY data
Normal data
Fold over basal
1.2
1.15
1.1
1.05
1
0.95
0.9
0.85
0
10
20
30
40
50
60
Time (min)

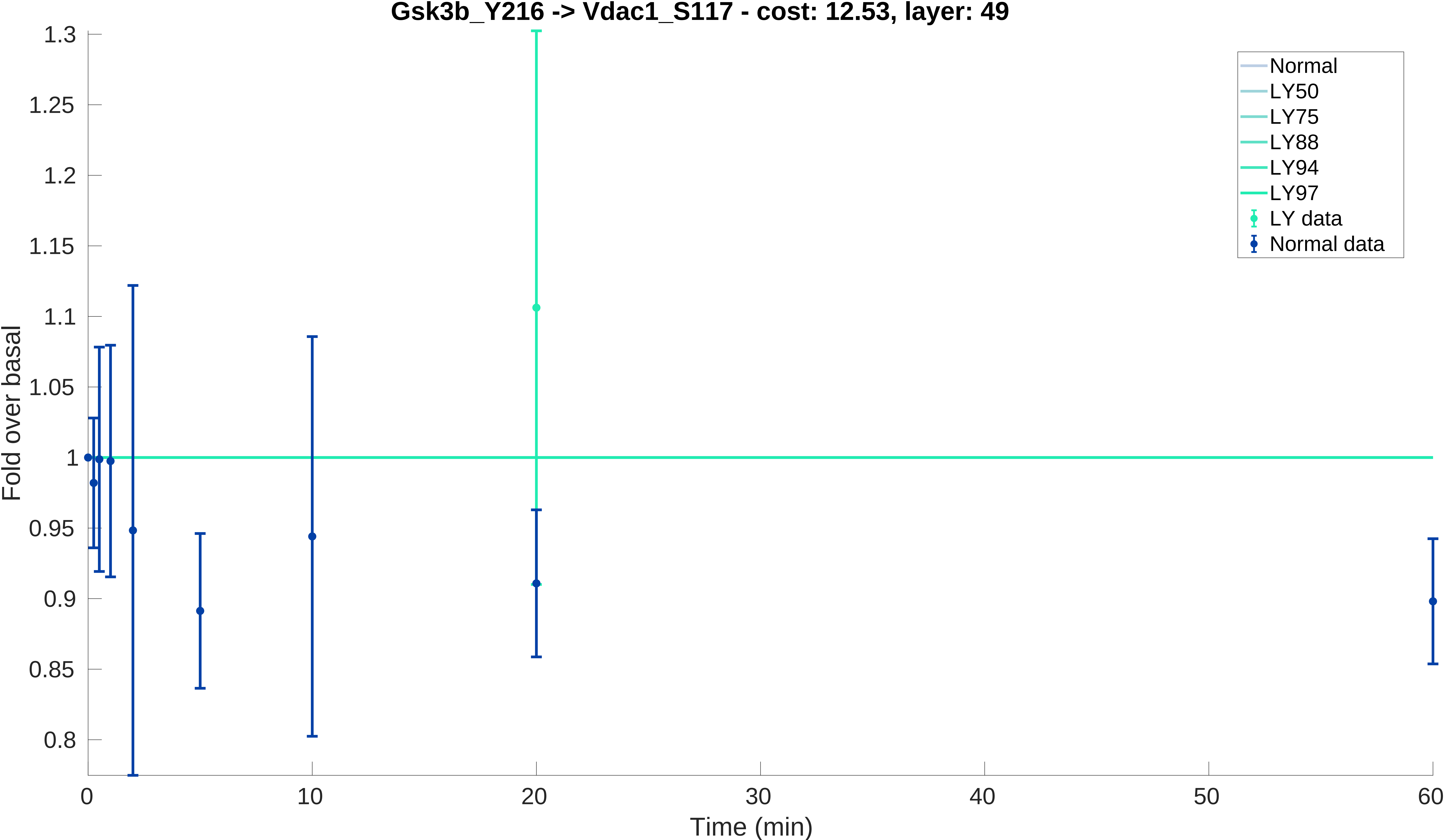

Gsk3b_Y216 -> Vdac1_S117 - cost: 12.53, layer: 49
Fold over basal
Time (min)
Normal
LY50
LY75
LY88
LY94
LY97
LY data
Normal data
1.3
1.25
1.2
1.15
1.1
1.05
1
0.95
0.9
0.85
0.8
0
10
20
30
40
50
60

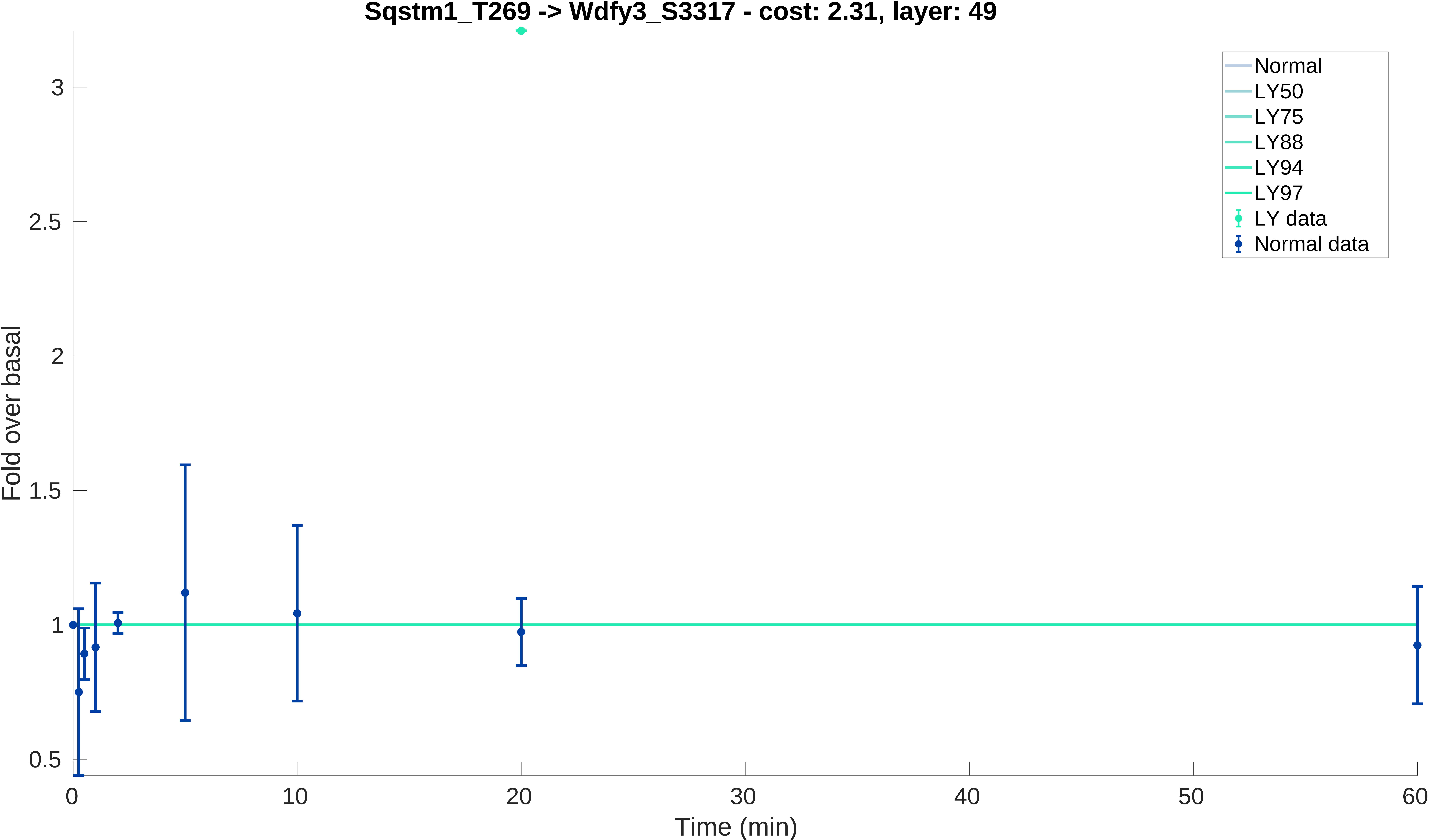

Sqstm1_T269 -> Wdfy3_S3317 - cost: 2.31, layer: 49
Fold over basal
Time (min)
0.5
1
1.5
2
2.5
3
0
10
20
30
40
50
60
Normal
LY50
LY75
LY88
LY94
LY97
LY data
Normal data

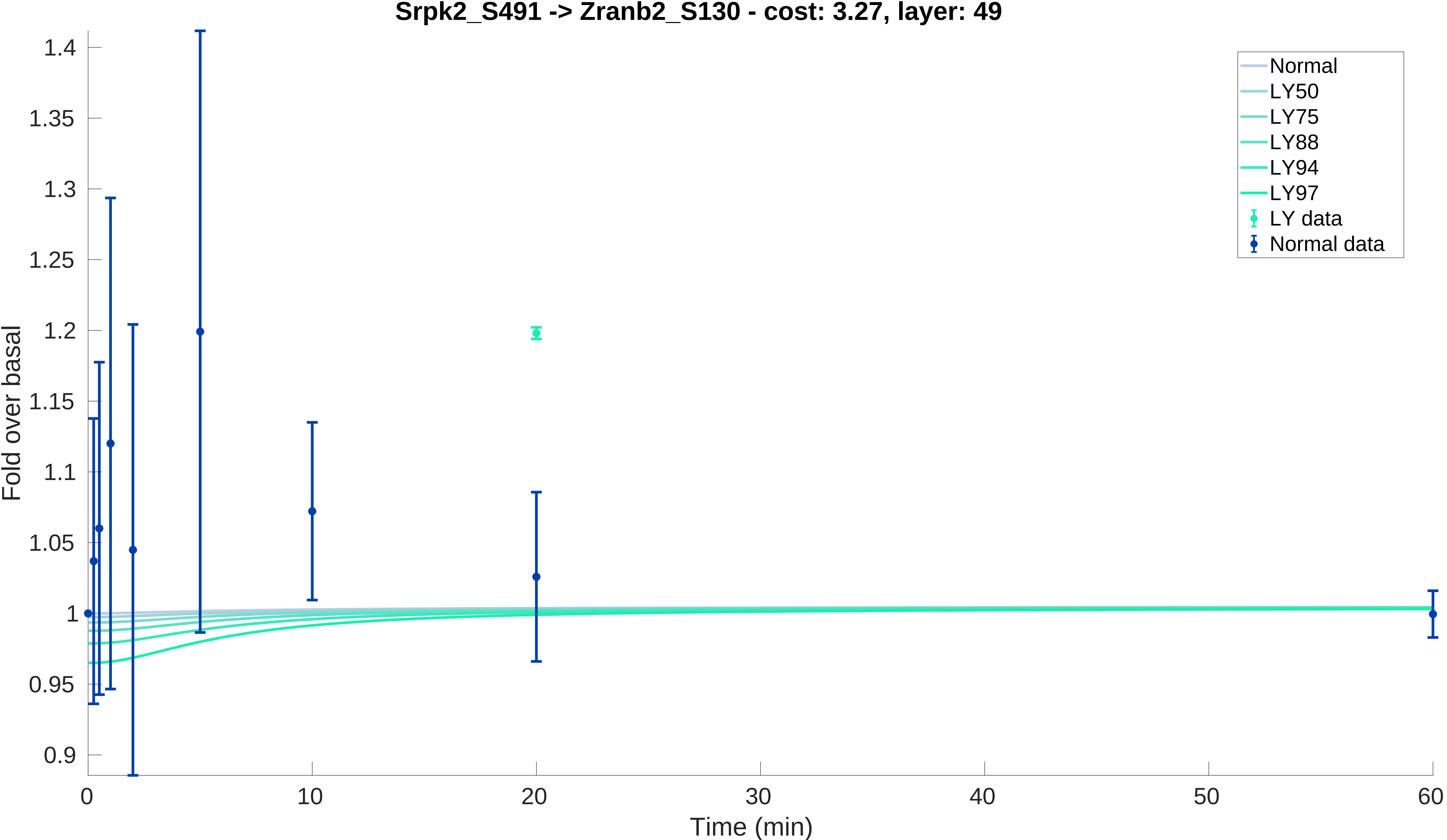

Srpk2_S491 -> Zranb2_S130 - cost: 3.27, layer: 49
Fold over basal
Time (min)
Normal
LY50
LY75
LY88
LY94
LY97
LY data
Normal data
1.4
1.35
1.3
1.25
1.2
1.15
1.1
1.05
1
0.95
0.9
0
10
20
30
40
50
60

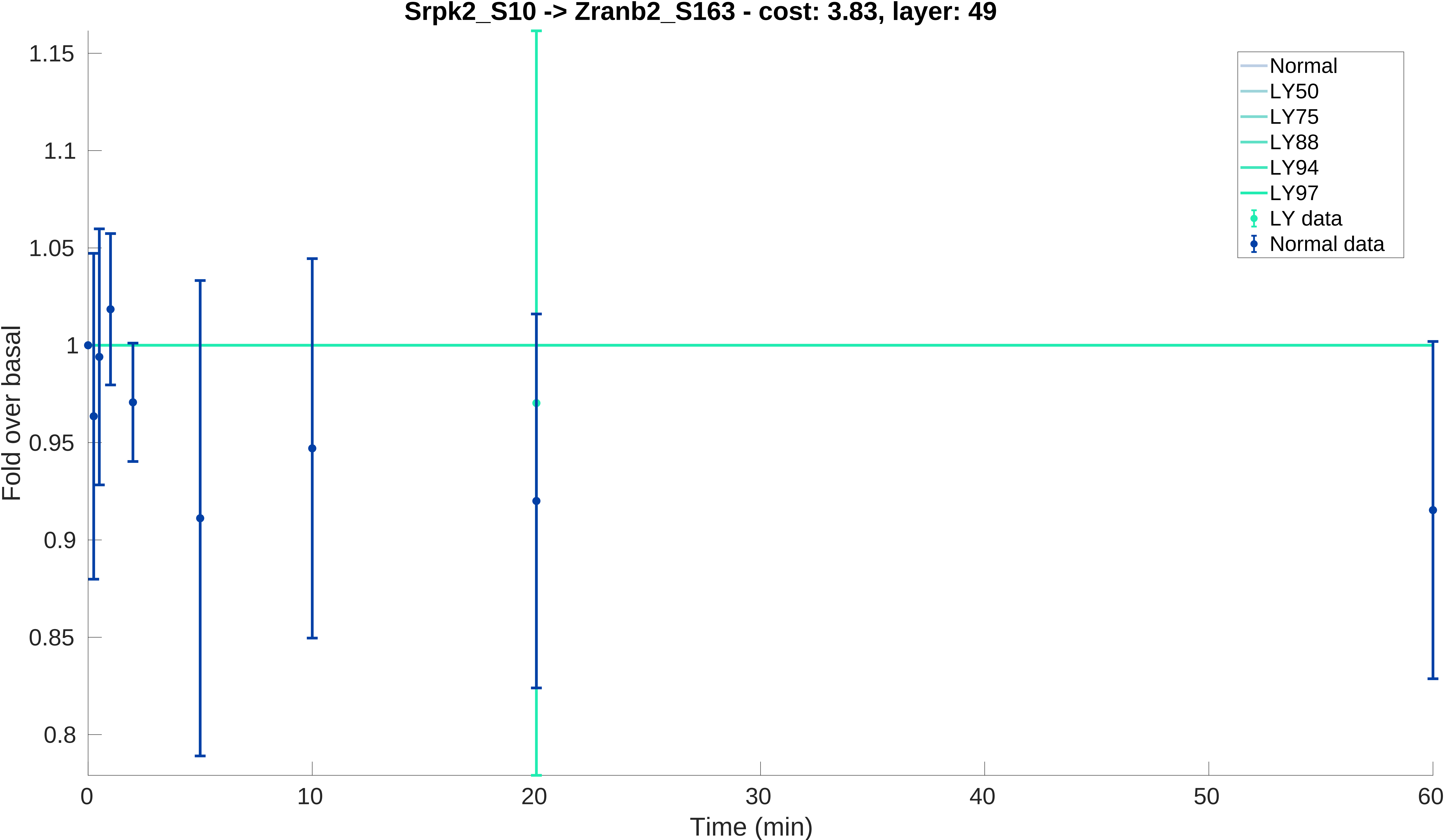

Srpk2_S10 -> Zranb2_S163 - cost: 3.83, layer: 49
Fold over basal
Time (min)
1.15
1.1
1.05
1
0.95
0.9
0.85
0.8
0
10
20
30
40
50
60
Normal
LY50
LY75
LY88
LY94
LY97
LY data
Normal data

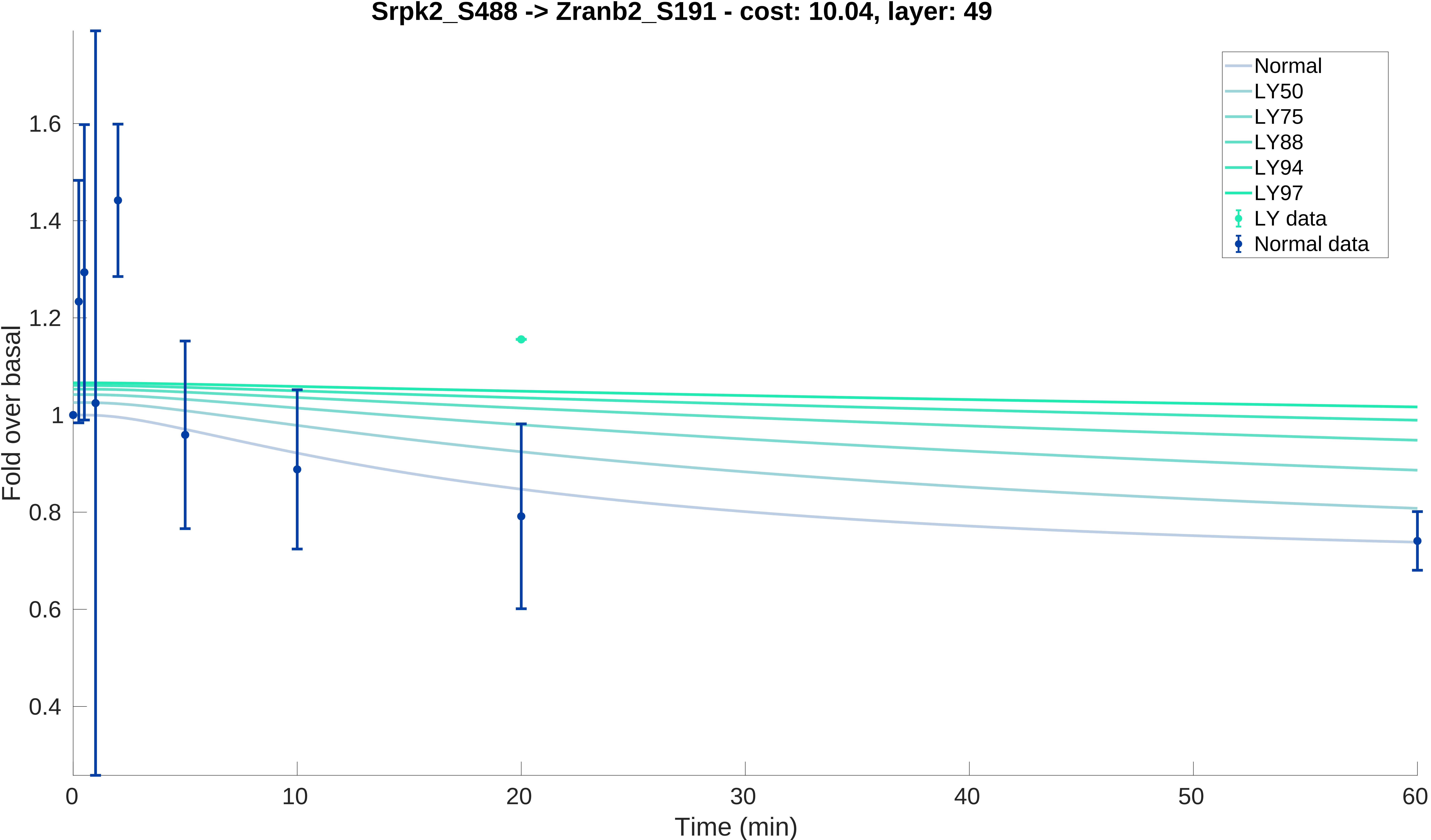

Srpk2_S488 -> Zranb2_S191 - cost: 10.04, layer: 49
Normal
LY50
LY75
LY88
LY94
LY97
LY data
Normal data
Fold over basal
Time (min)
0.4
0.6
0.8
1
1.2
1.4
1.6
0
10
20
30
40
50
60

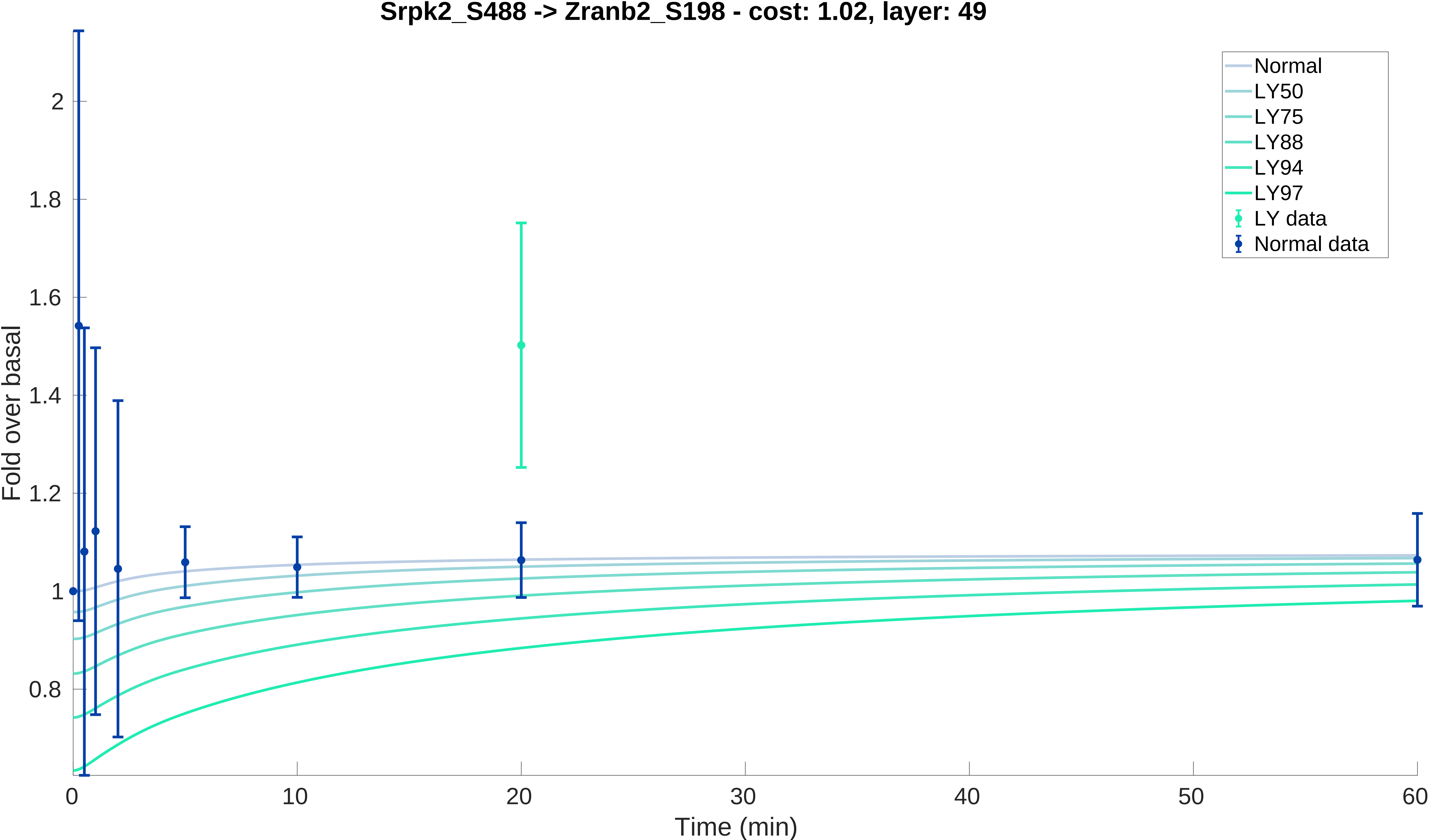
Srpk2_S488 -> Zranb2_S198 - cost: 1.02, layer: 49
Normal
LY50
LY75
LY88
LY94
LY97
LY data
Normal data
Fold over basal
2
1.8
1.6
1.4
1.2
1
0.8
0
10
20
30
40
50
60
Time (min)

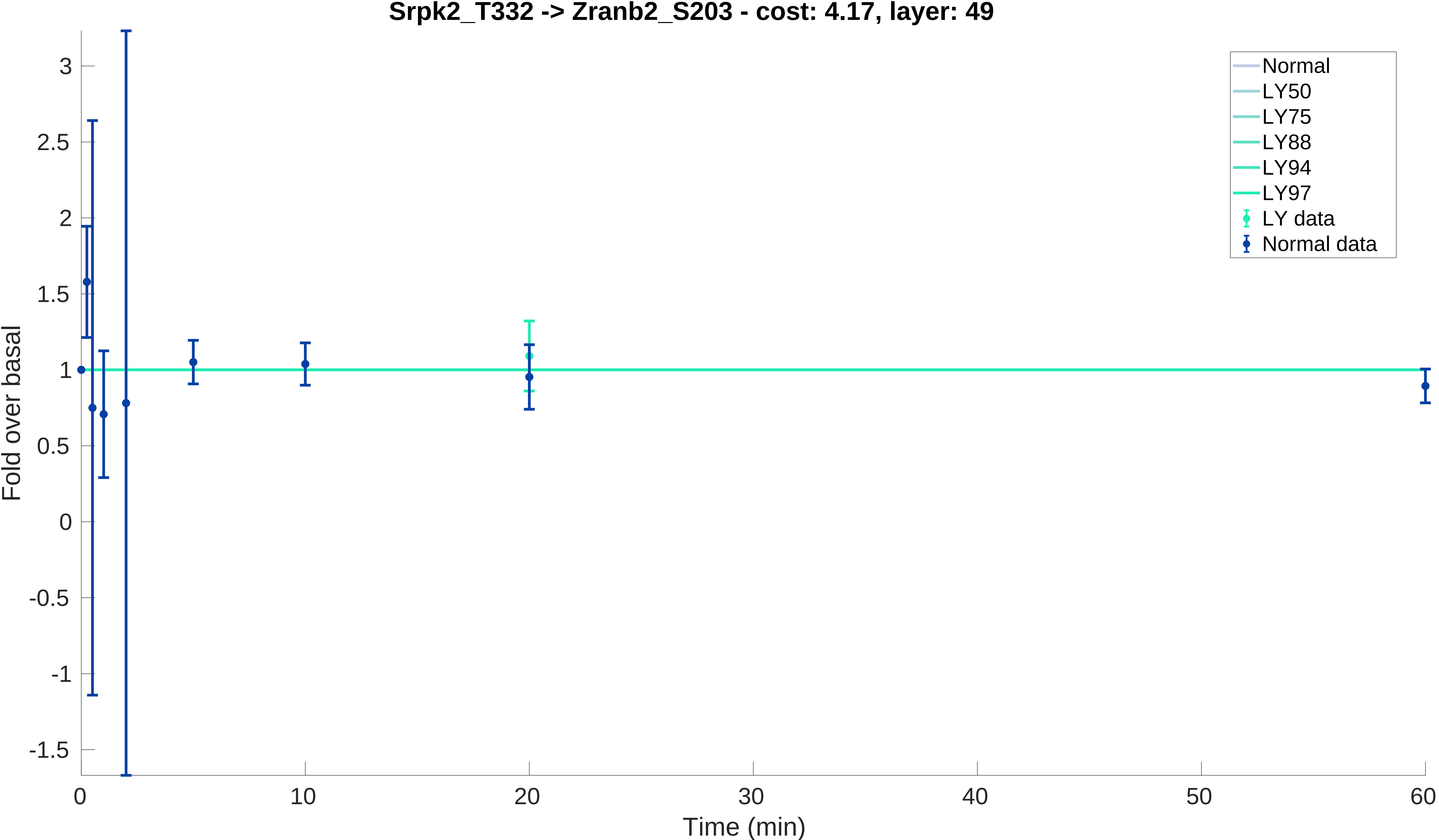

Srpk2_T332 -> Zranb2_S203 - cost: 4.17, layer: 49
Fold over basal
Time (min)
Normal
LY50
LY75
LY88
LY94
LY97
LY data
Normal data
3
2.5
2
1.5
1
0.5
0
-0.5
-1
-1.5
0
10
20
30
40
50
60

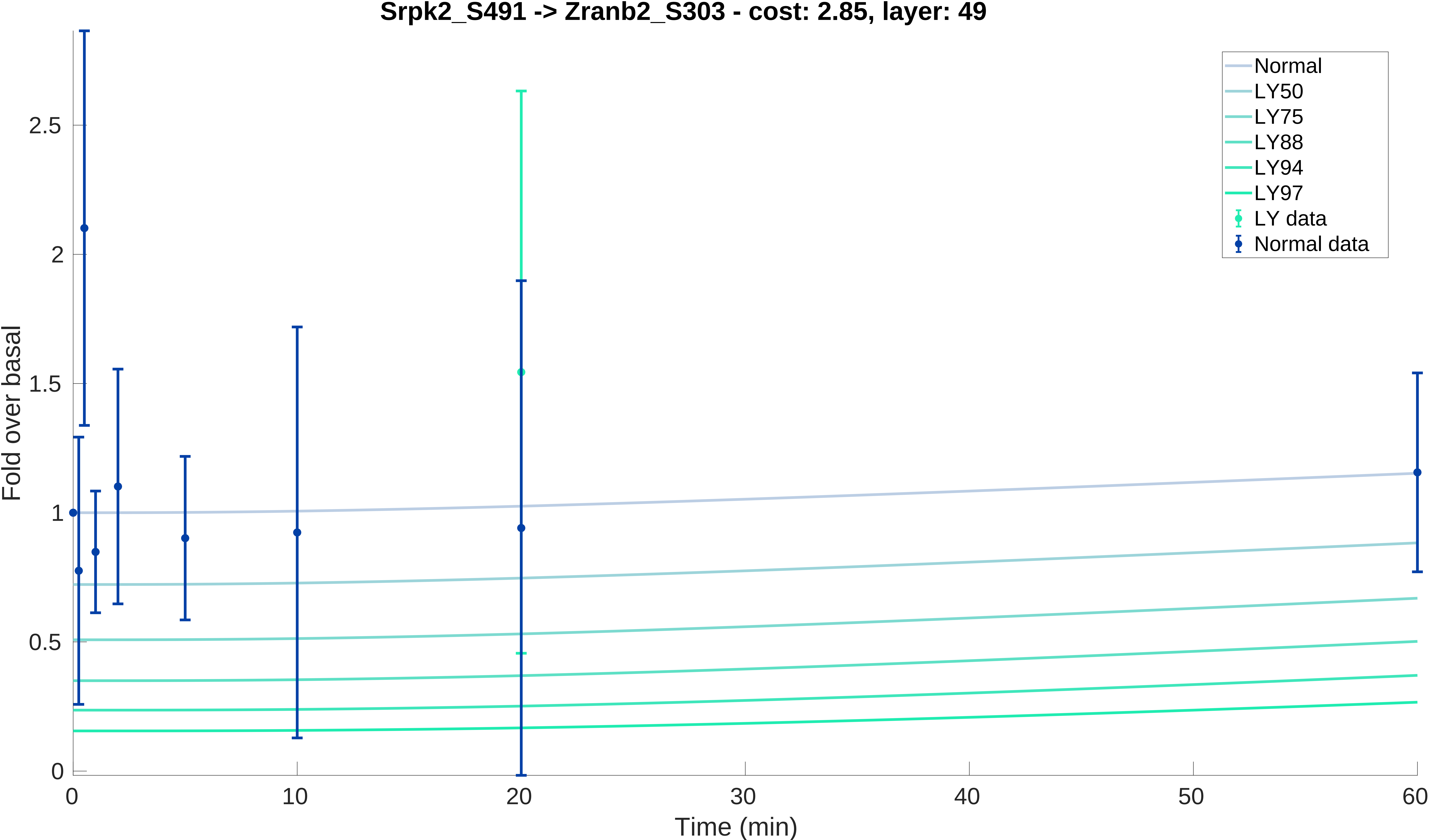

Srpk2_S491 -> Zranb2_S303 - cost: 2.85, layer: 49
Fold over basal
Time (min)
0
0.5
1
1.5
2
2.5
0
10
20
30
40
50
60
Normal
LY50
LY75
LY88
LY94
LY97
LY data
Normal data

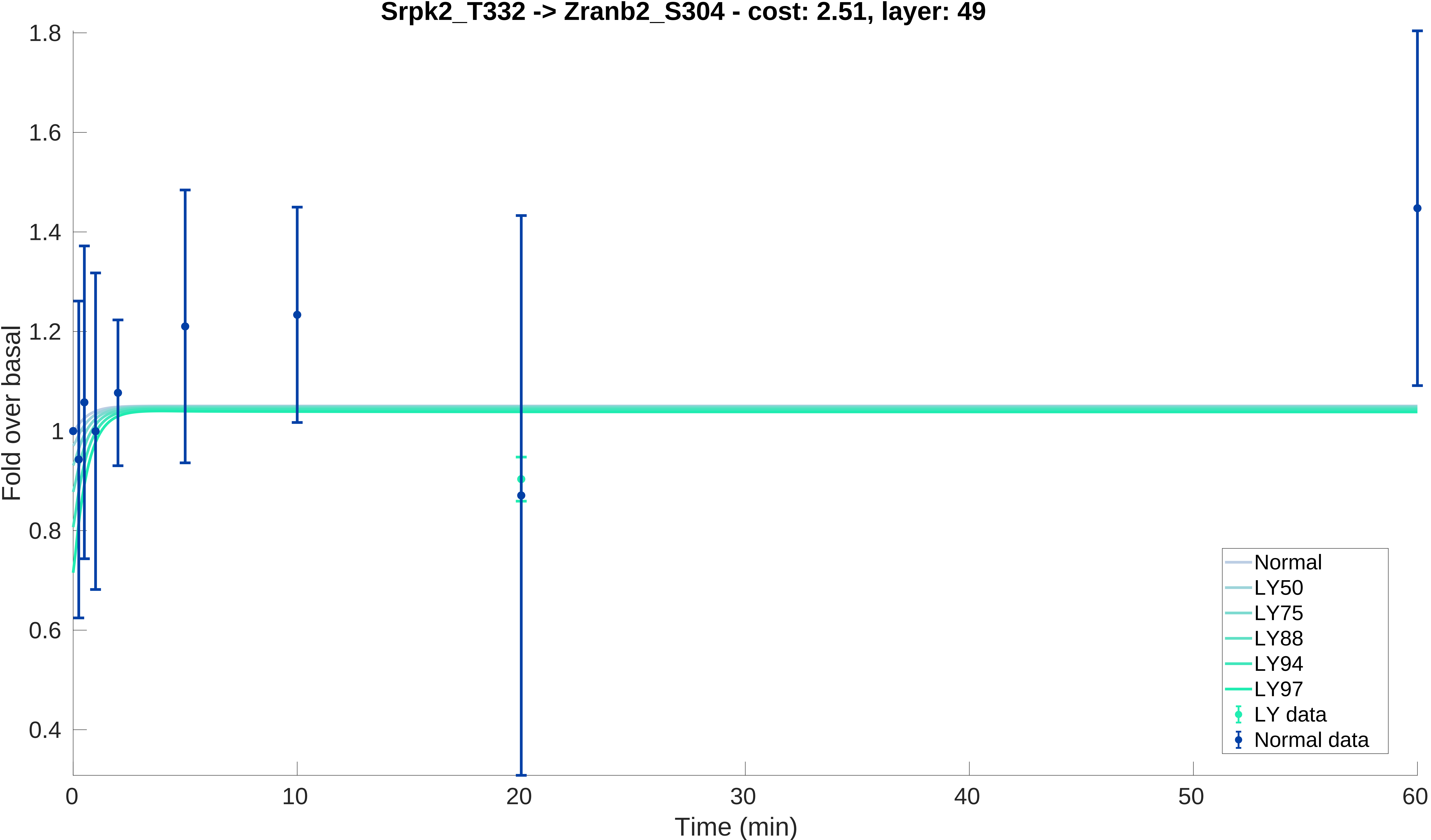

Srpk2_T332 -> Zranb2_S304 - cost: 2.51, layer: 49
Fold over basal
Time (min)
1.8
1.6
1.4
1.2
1
0.8
0.6
0.4
0
10
20
30
40
50
60
Normal
LY50
LY75
LY88
LY94
LY97
LY data
Normal data

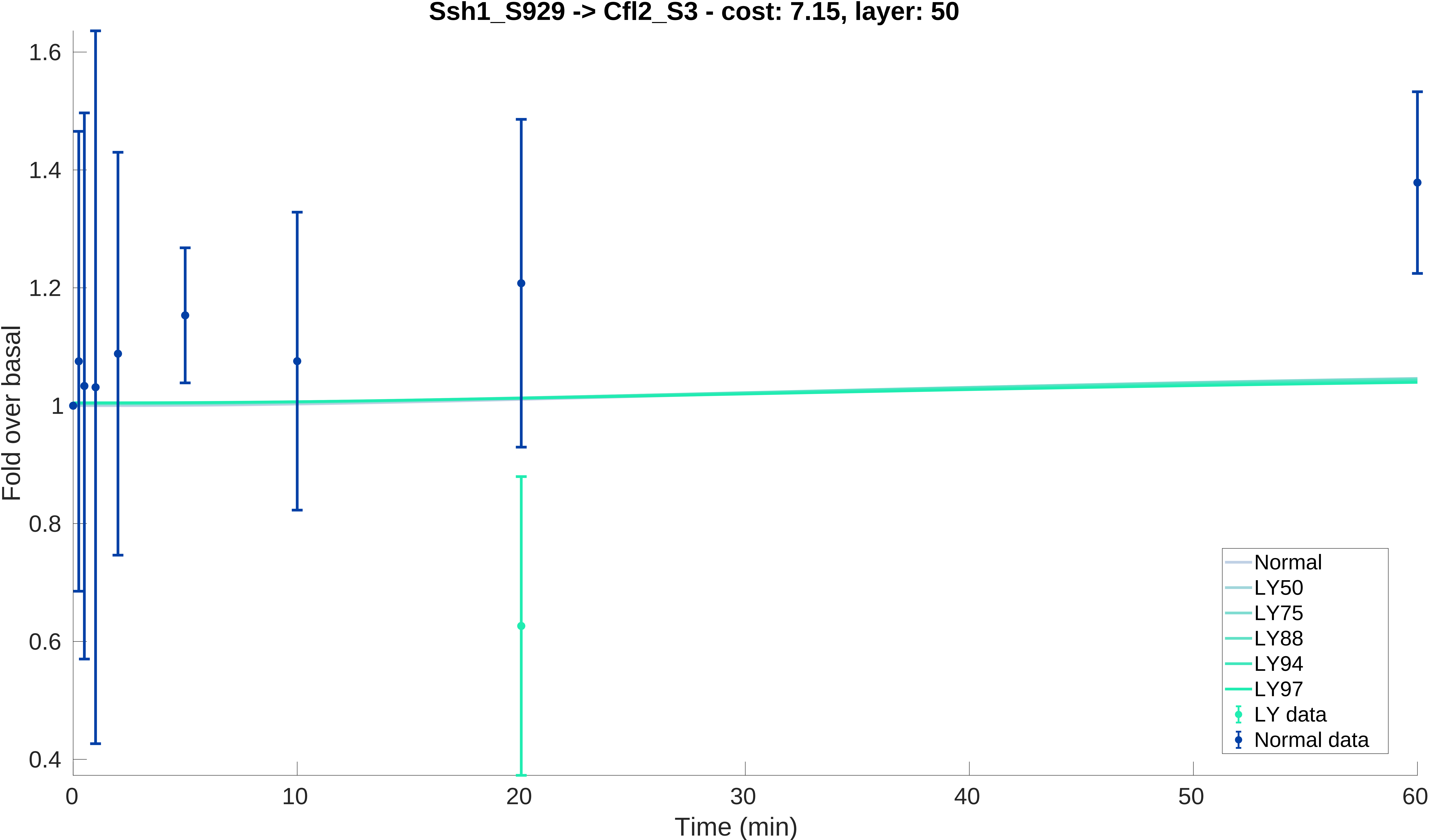

Ssh1_S929 -> Cfl2_S3 - cost: 7.15, layer: 50
Fold over basal
Time (min)
1.6
1.4
1.2
1
0.8
0.6
0.4
0
10
20
30
40
50
60
Normal
LY50
LY75
LY88
LY94
LY97
LY data
Normal data

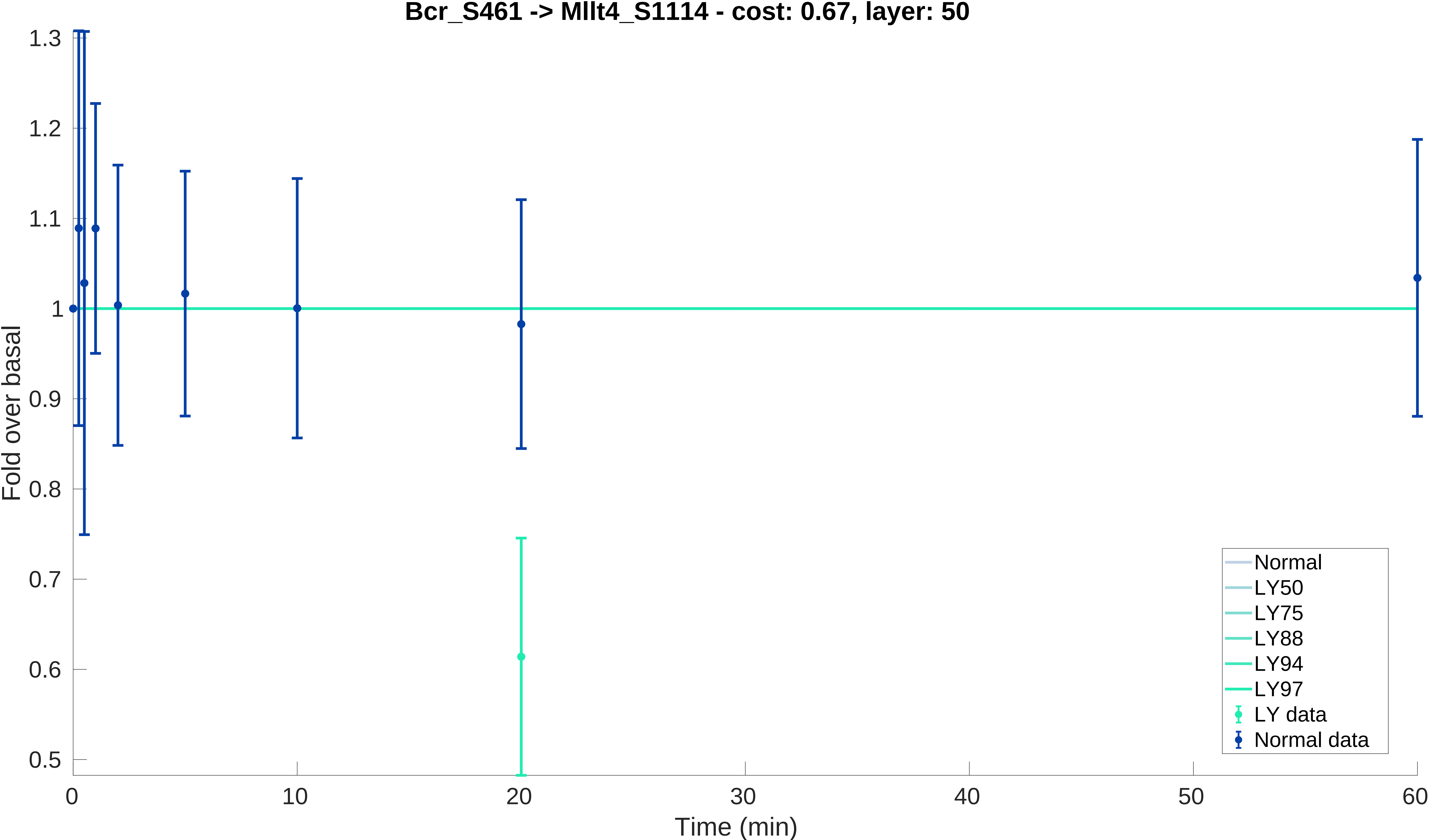
Bcr_S461 -> Mllt4_S1114 - cost: 0.67, layer: 50
Fold over basal
1.3
1.2
1.1
1
0.9
0.8
0.7
0.6
0.5
0
10
20
30
40
50
60
Time (min)
Normal
LY50
LY75
LY88
LY94
LY97
LY data
Normal data

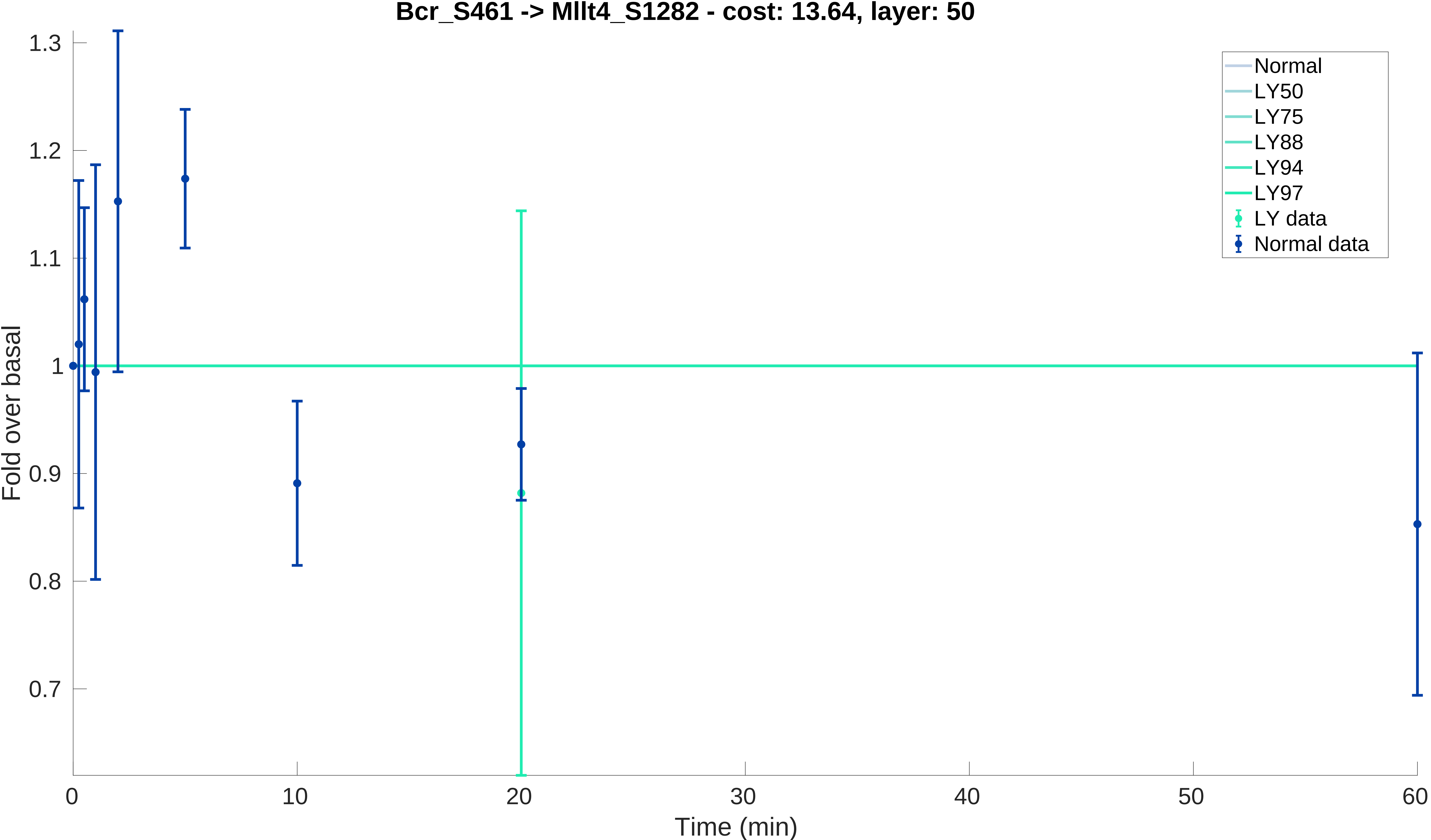

Bcr_S461 -> Mllt4_S1282 - cost: 13.64, layer: 50
Fold over basal
Time (min)
0
10
20
30
40
50
60
0.7
0.8
0.9
1
1.1
1.2
1.3
Normal
LY50
LY75
LY88
LY94
LY97
LY data
Normal data

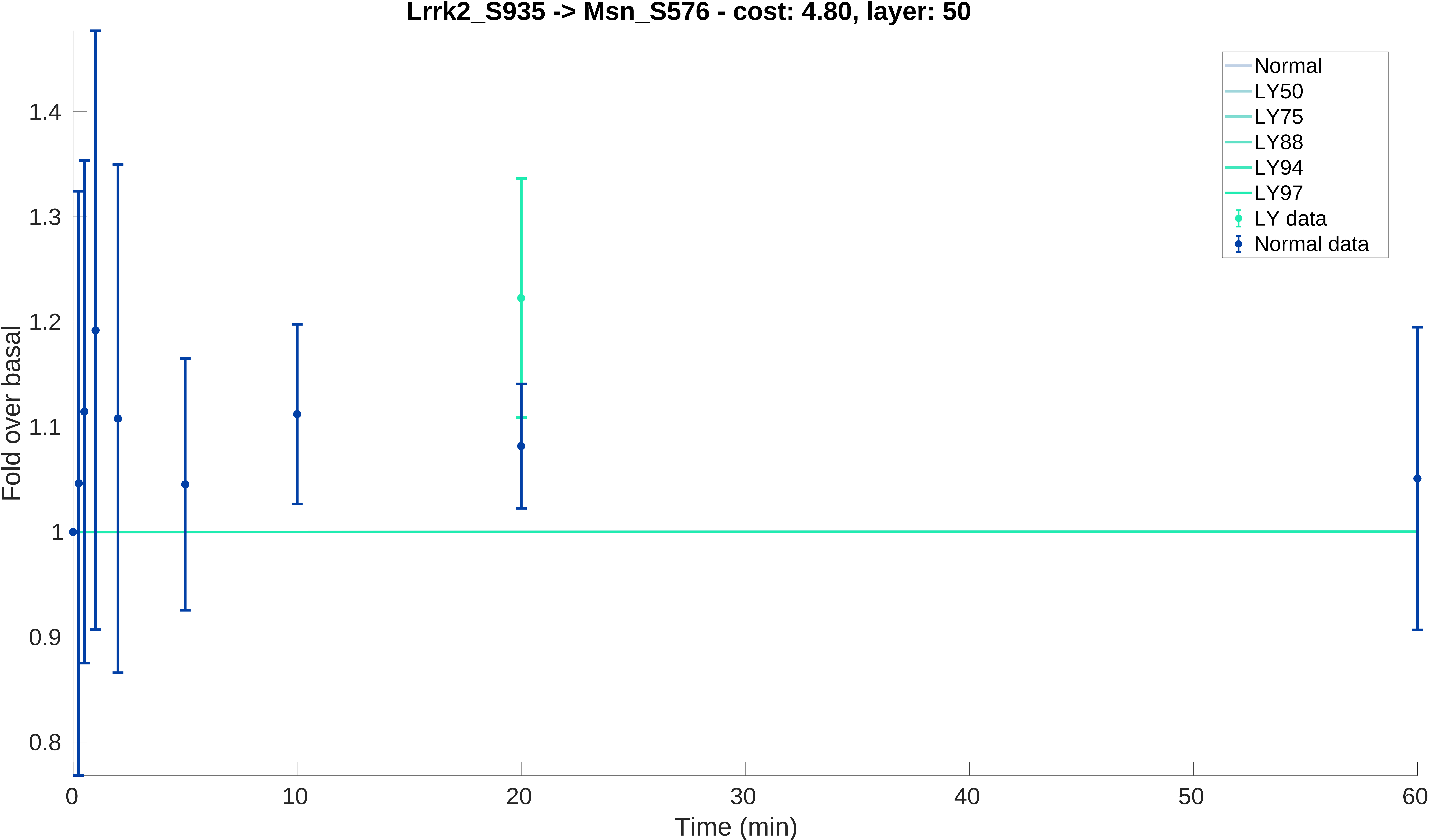

Lrrk2_S935 -> Msn_S576 - cost: 4.80, layer: 50
Normal
LY50
LY75
LY88
LY94
LY97
LY data
Normal data
Fold over basal
1.4
1.3
1.2
1.1
1
0.9
0.8
0
10
20
30
40
50
60
Time (min)

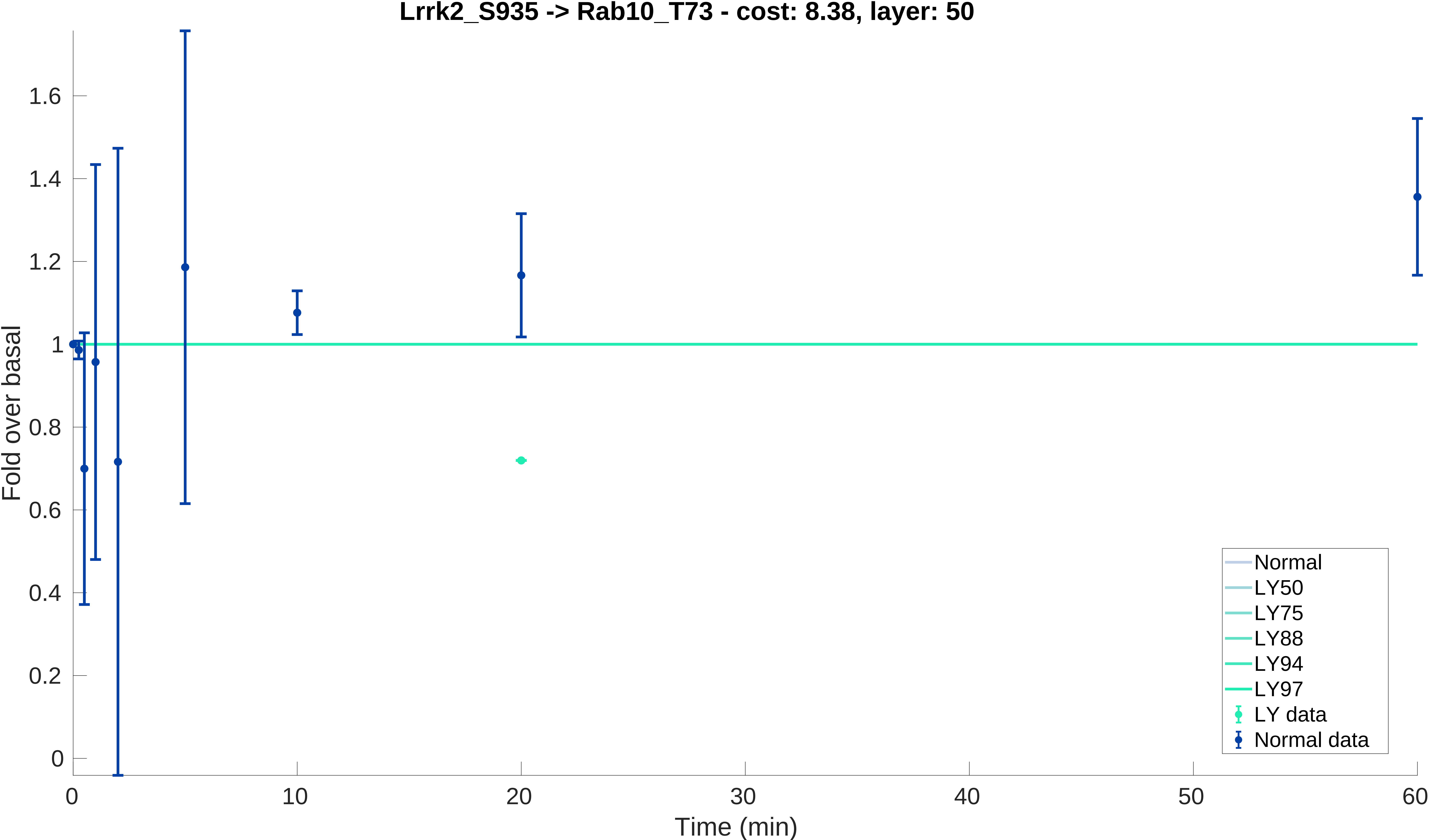
Lrrk2_S935 -> Rab10_T73 - cost: 8.38, layer: 50
Fold over basal
Time (min)
0
0.2
0.4
0.6
0.8
1
1.2
1.4
1.6
0
10
20
30
40
50
60
Normal
LY50
LY75
LY88
LY94
LY97
LY data
Normal data

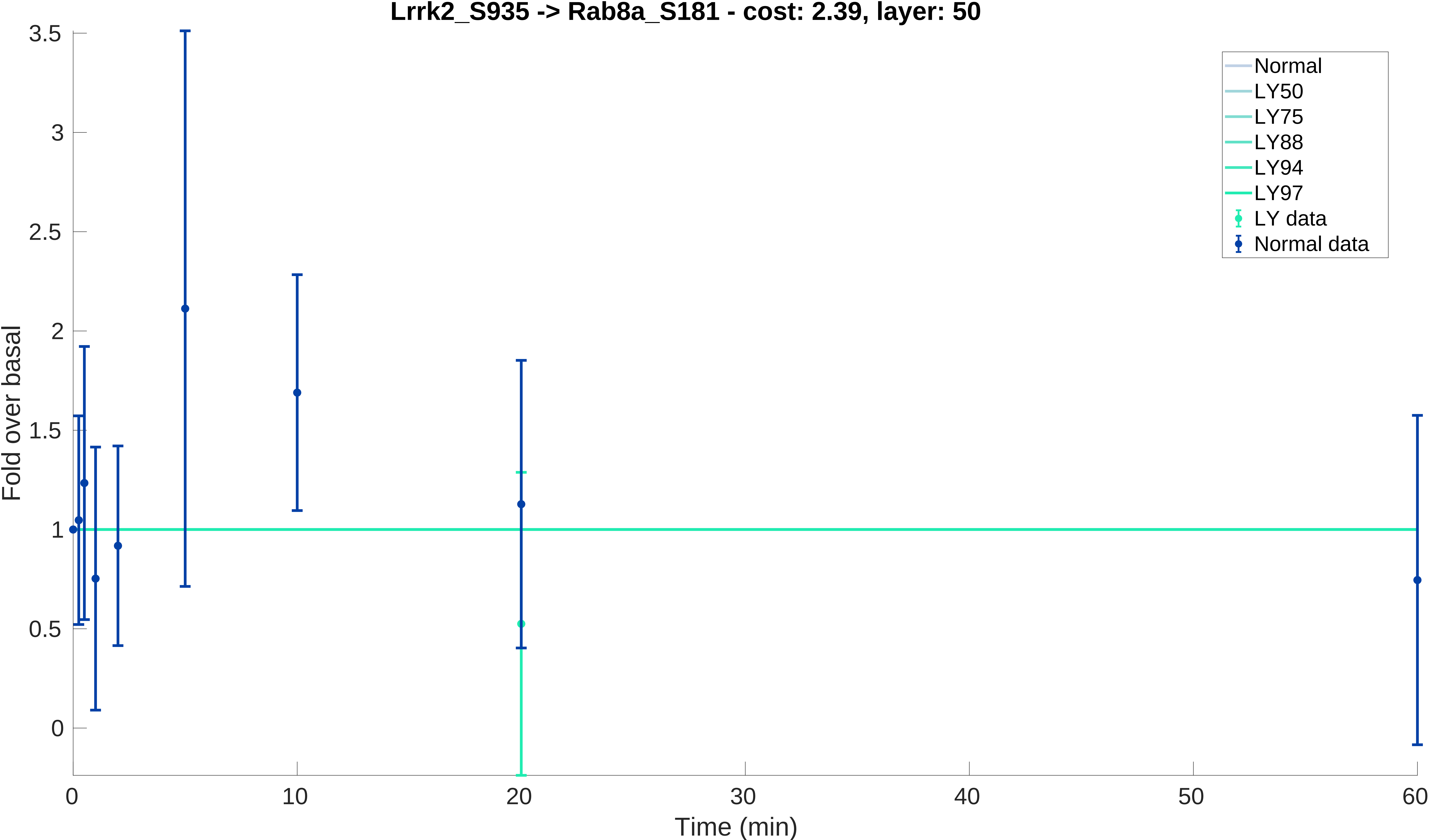
Lrrk2_S935 -> Rab8a_S181 - cost: 2.39, layer: 50
Fold over basal
Time (min)
0
0.5
1
1.5
2
2.5
3
3.5
0
10
20
30
40
50
60
Normal
LY50
LY75
LY88
LY94
LY97
LY data
Normal data

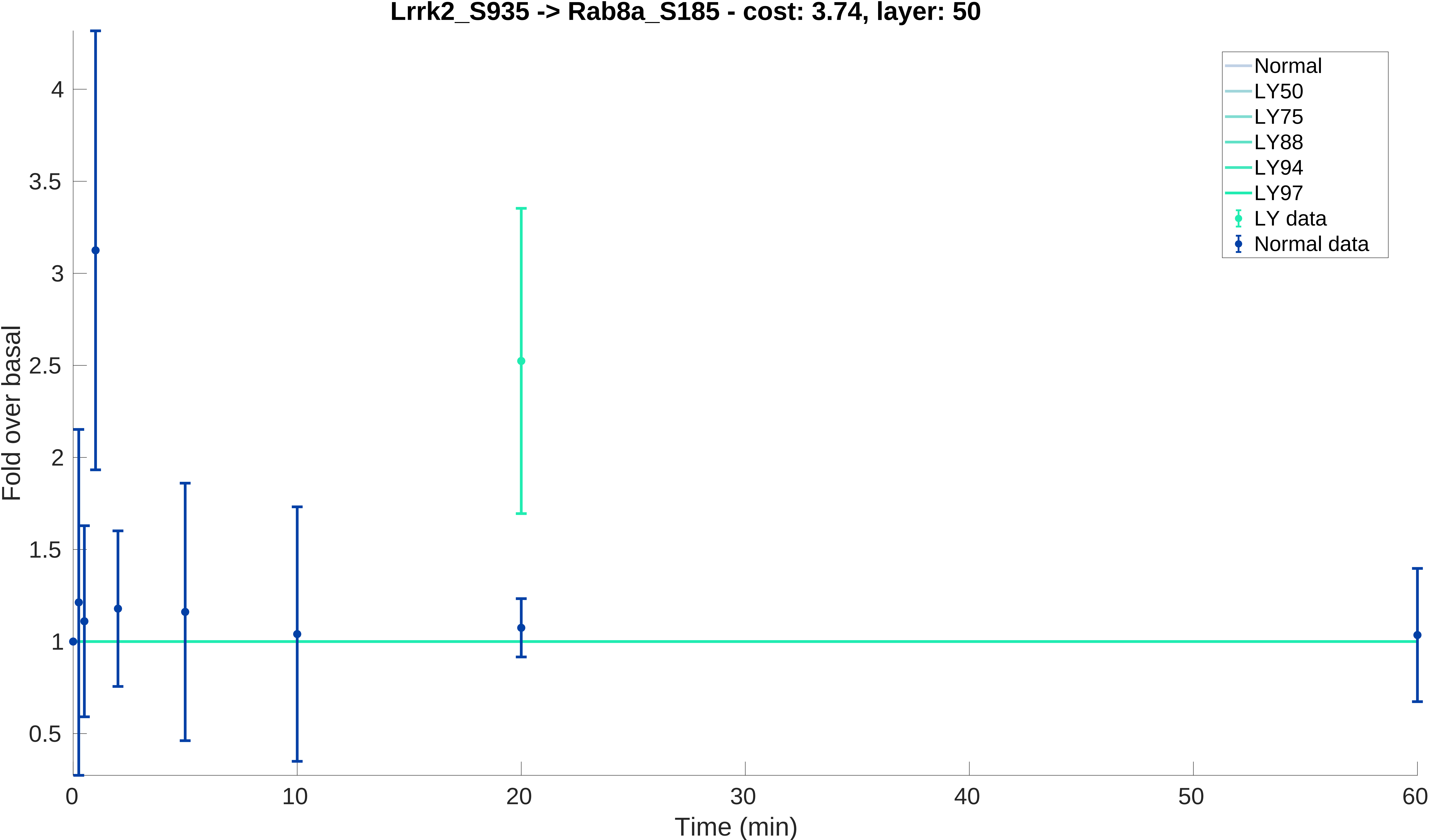

Lrrk2_S935 -> Rab8a_S185 - cost: 3.74, layer: 50
Fold over basal
Time (min)
0.5
1
1.5
2
2.5
3
3.5
4
0
10
20
30
40
50
60
Normal
LY50
LY75
LY88
LY94
LY97
LY data
Normal data

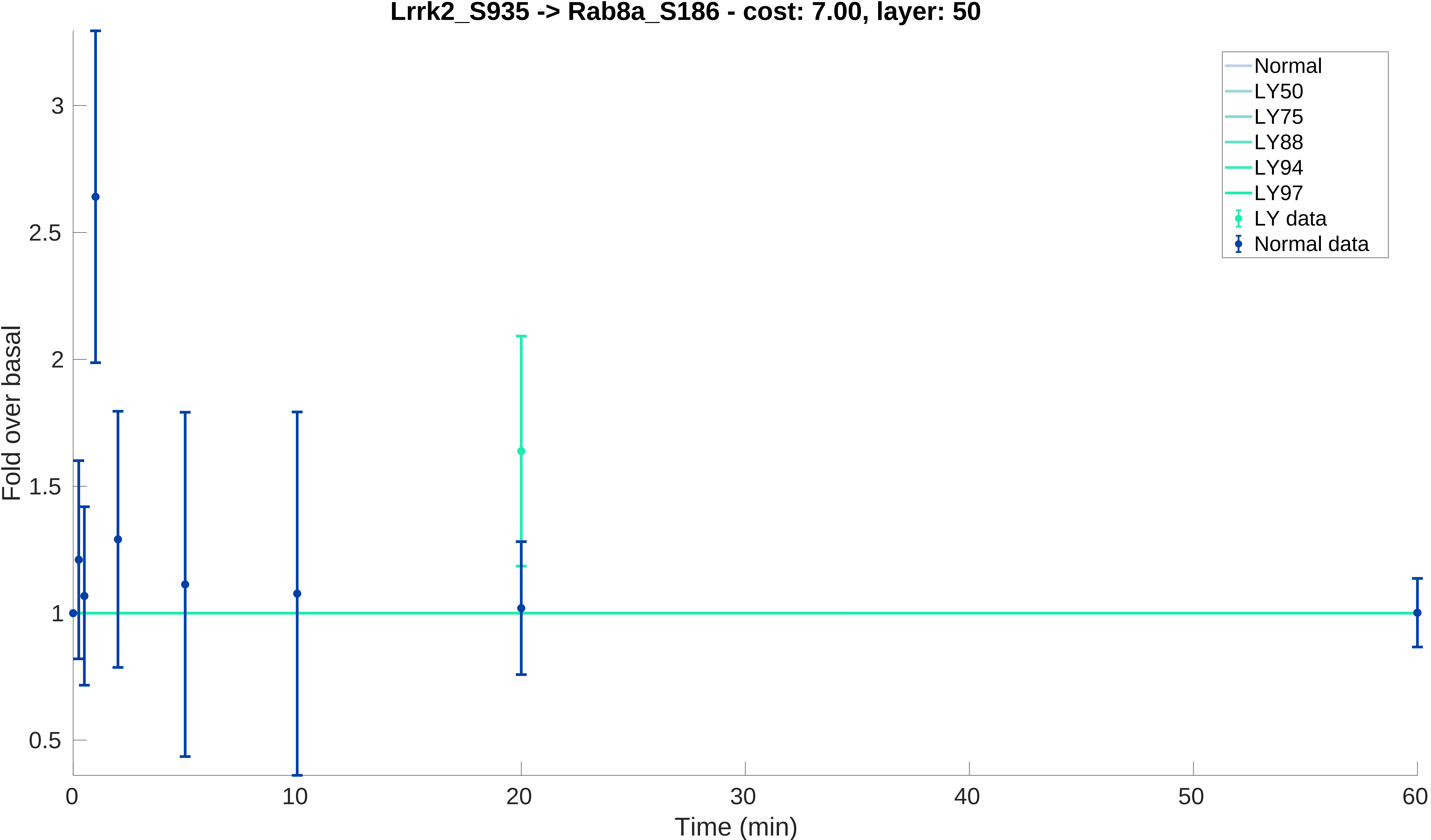

Lrrk2_S935 -> Rab8a_S186 - cost: 7.00, layer: 50
Normal
LY50
LY75
LY88
LY94
LY97
LY data
Normal data
Fold over basal
Time (min)
0.5
1
1.5
2
2.5
3
0
10
20
30
40
50
60

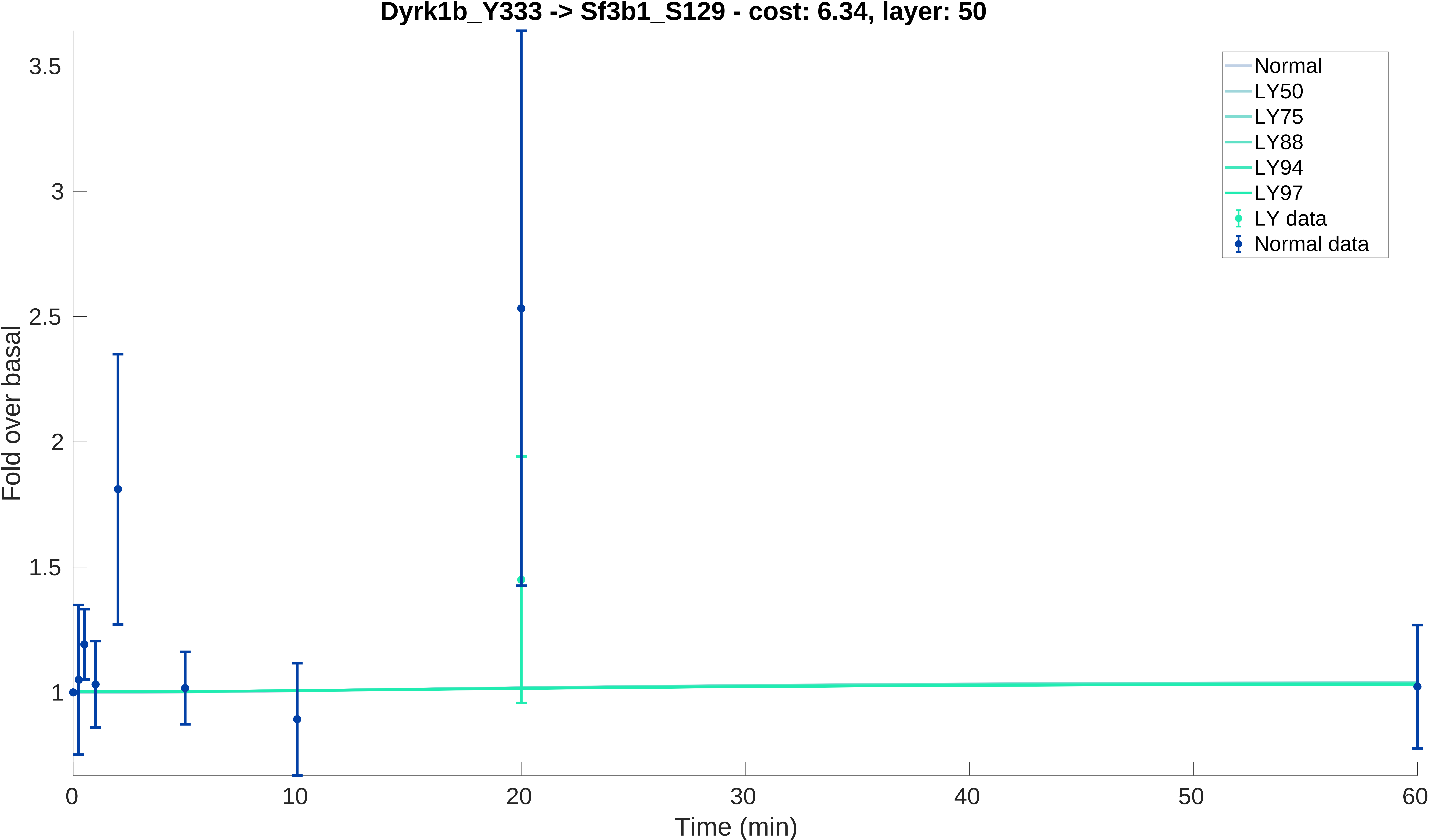
Dyrk1b_Y333 -> Sf3b1_S129 - cost: 6.34, layer: 50
Fold over basal
Time (min)
3.5
3
2.5
2
1.5
1
0
10
20
30
40
50
60
Normal
LY50
LY75
LY88
LY94
LY97
LY data
Normal data

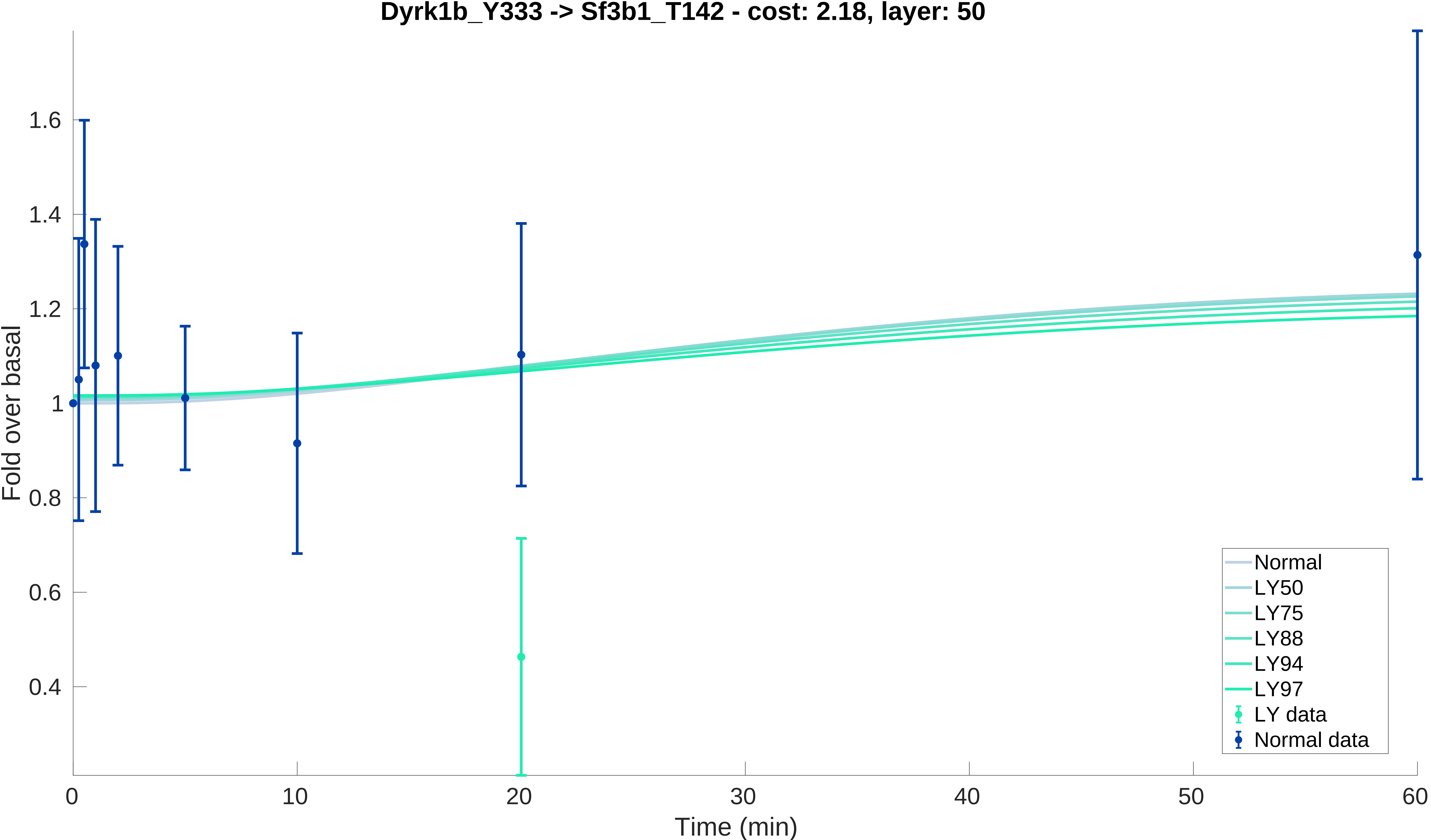

Dyrk1b_Y333 -> Sf3b1_T142 - cost: 2.18, layer: 50
Fold over basal
1.6
1.4
1.2
1
0.8
0.6
0.4
0
10
20
30
40
50
60
Time (min)
Normal
LY50
LY75
LY88
LY94
LY97
LY data
Normal data

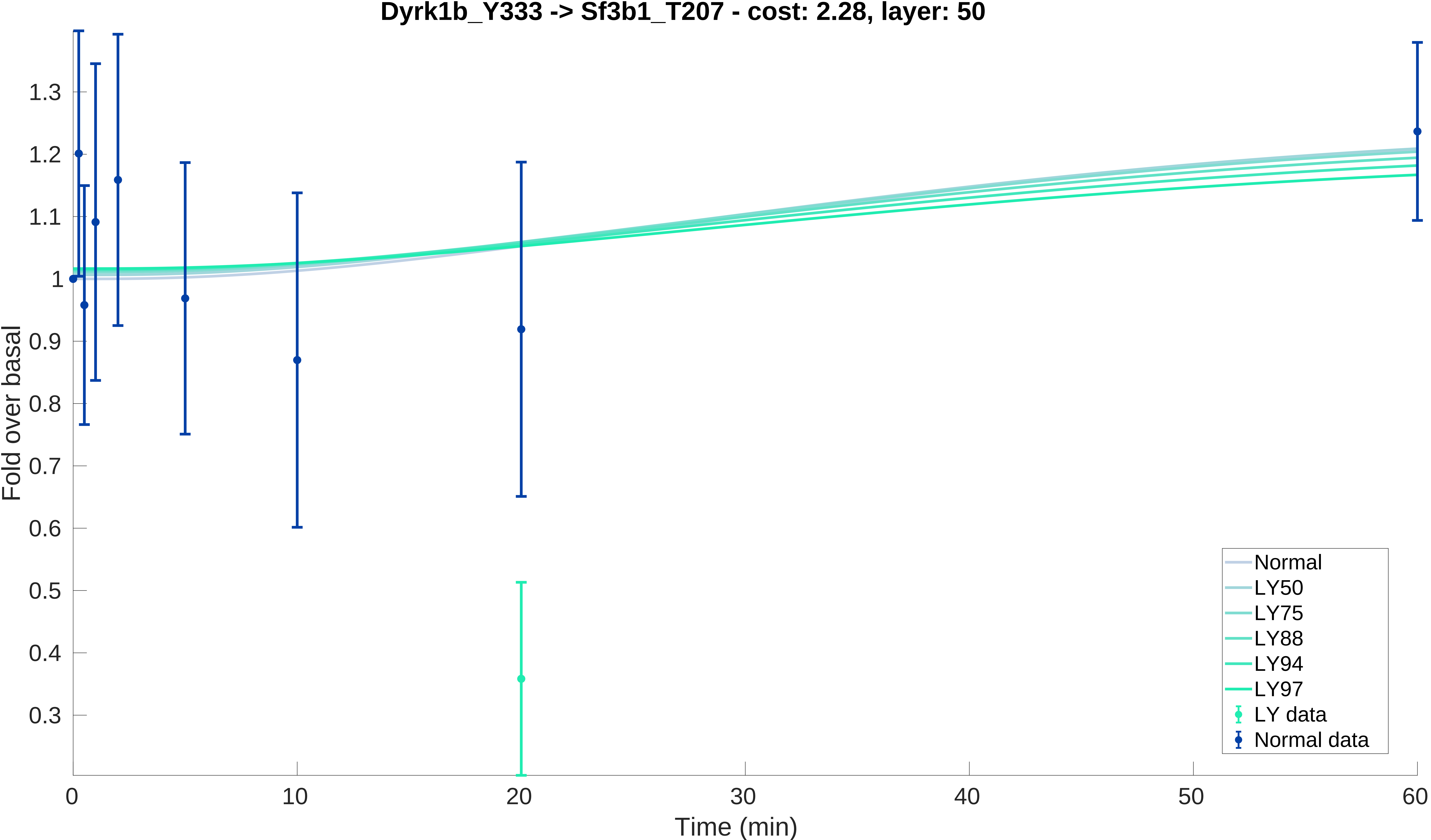

Dyrk1b_Y333 -> Sf3b1_T207 - cost: 2.28, layer: 50
Fold over basal
Time (min)
1.3
1.2
1.1
1
0.9
0.8
0.7
0.6
0.5
0.4
0.3
0
10
20
30
40
50
60
Normal
LY50
LY75
LY88
LY94
LY97
LY data
Normal data

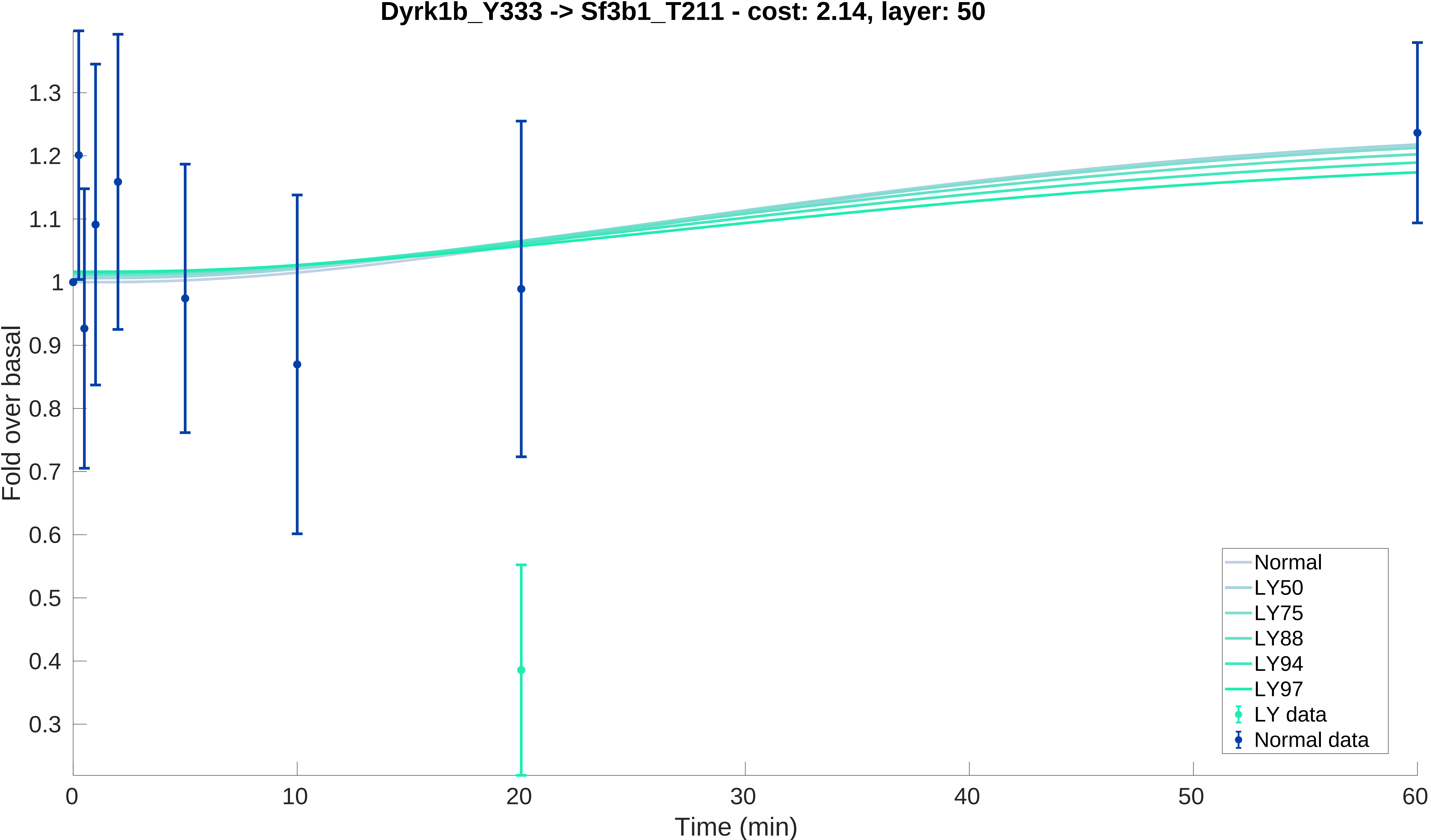
Dyrk1b_Y333 -> Sf3b1_T211 - cost: 2.14, layer: 50
Fold over basal
Time (min)
0.3
0.4
0.5
0.6
0.7
0.8
0.9
1
1.1
1.2
1.3
0
10
20
30
40
50
60
Normal
LY50
LY75
LY88
LY94
LY97
LY data
Normal data

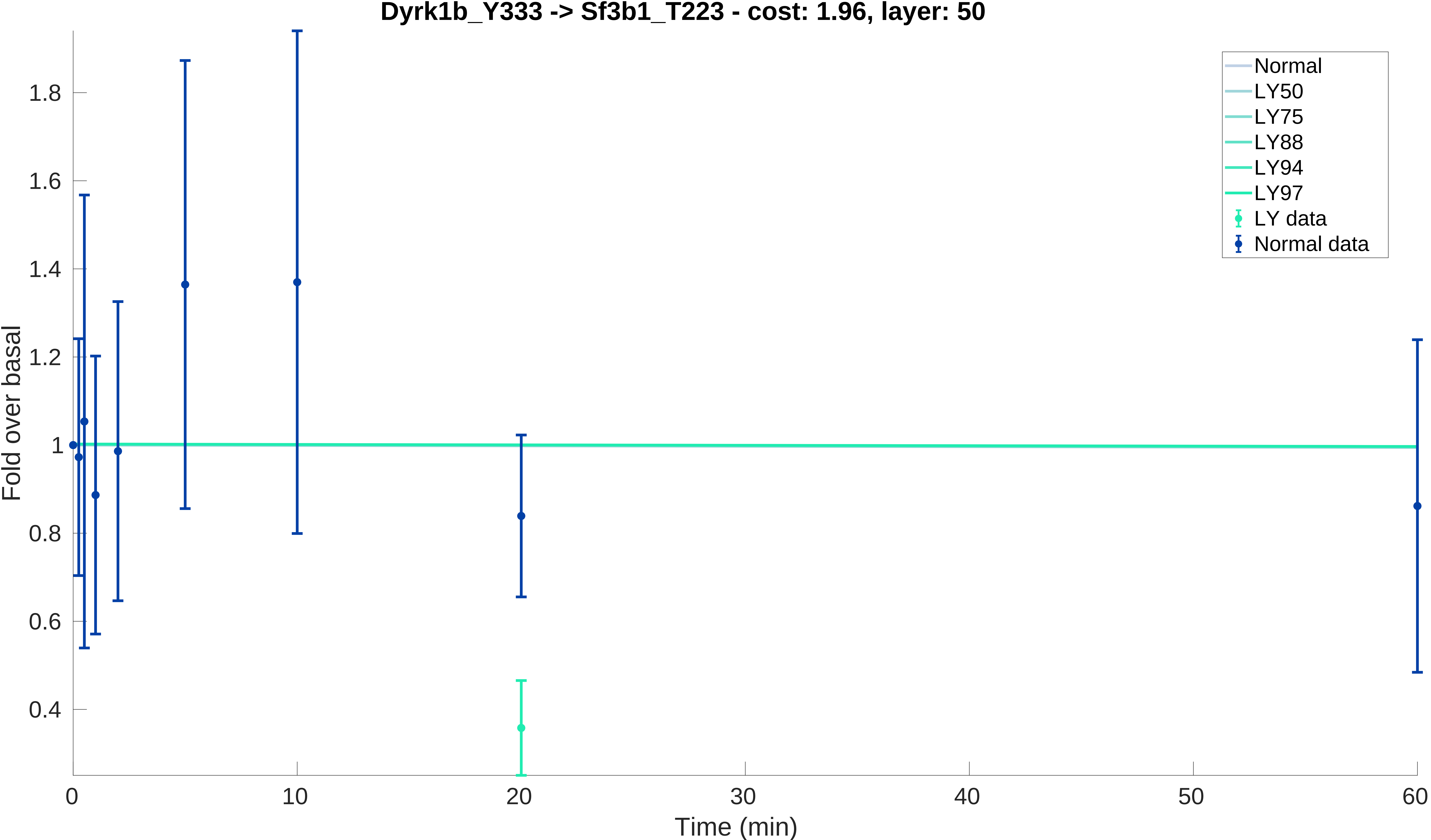

Dyrk1b_Y333 -> Sf3b1_T223 - cost: 1.96, layer: 50
Fold over basal
Time (min)
Normal
LY50
LY75
LY88
LY94
LY97
LY data
Normal data
0.4
0.6
0.8
1
1.2
1.4
1.6
1.8
0
10
20
30
40
50
60

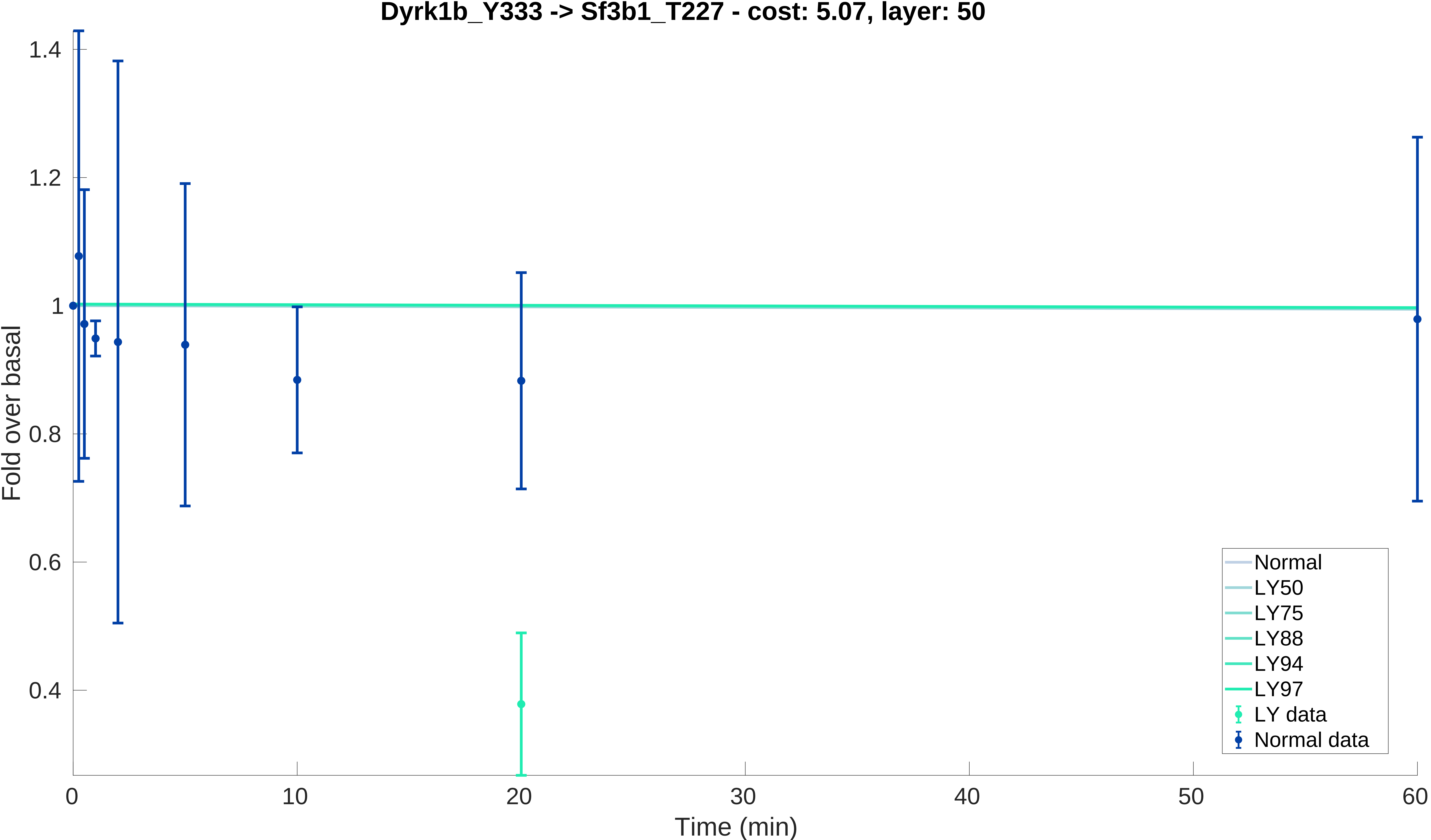

Dyrk1b_Y333 -> Sf3b1_T227 - cost: 5.07, layer: 50
Fold over basal
1.4
1.2
1
0.8
0.6
0.4
0
10
20
30
40
50
60
Time (min)
Normal
LY50
LY75
LY88
LY94
LY97
LY data
Normal data

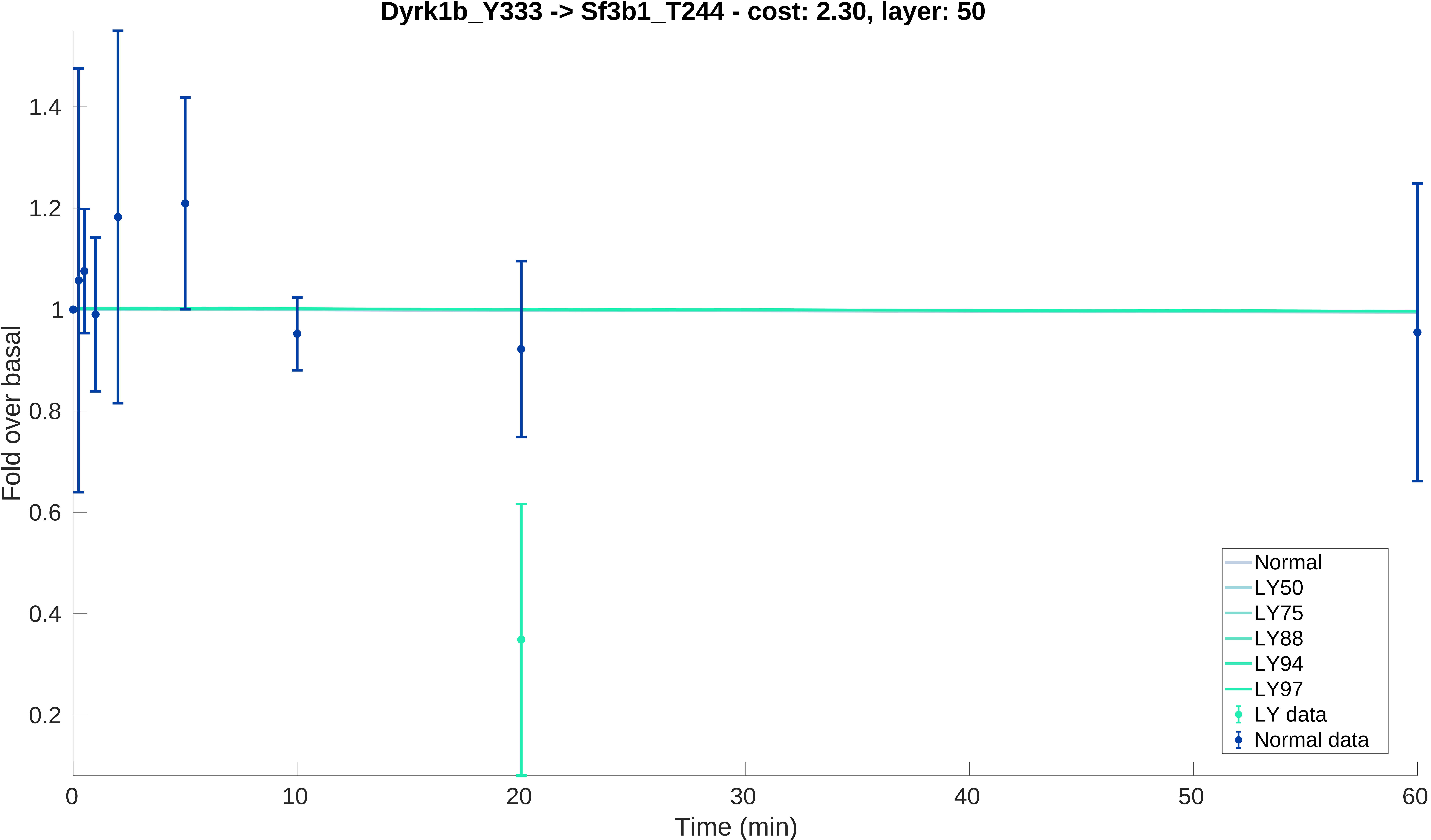

Dyrk1b_Y333 -> Sf3b1_T244 - cost: 2.30, layer: 50
Fold over basal
1.4
1.2
1
0.8
0.6
0.4
0.2
0
10
20
30
40
50
60
Time (min)
Normal
LY50
LY75
LY88
LY94
LY97
LY data
Normal data

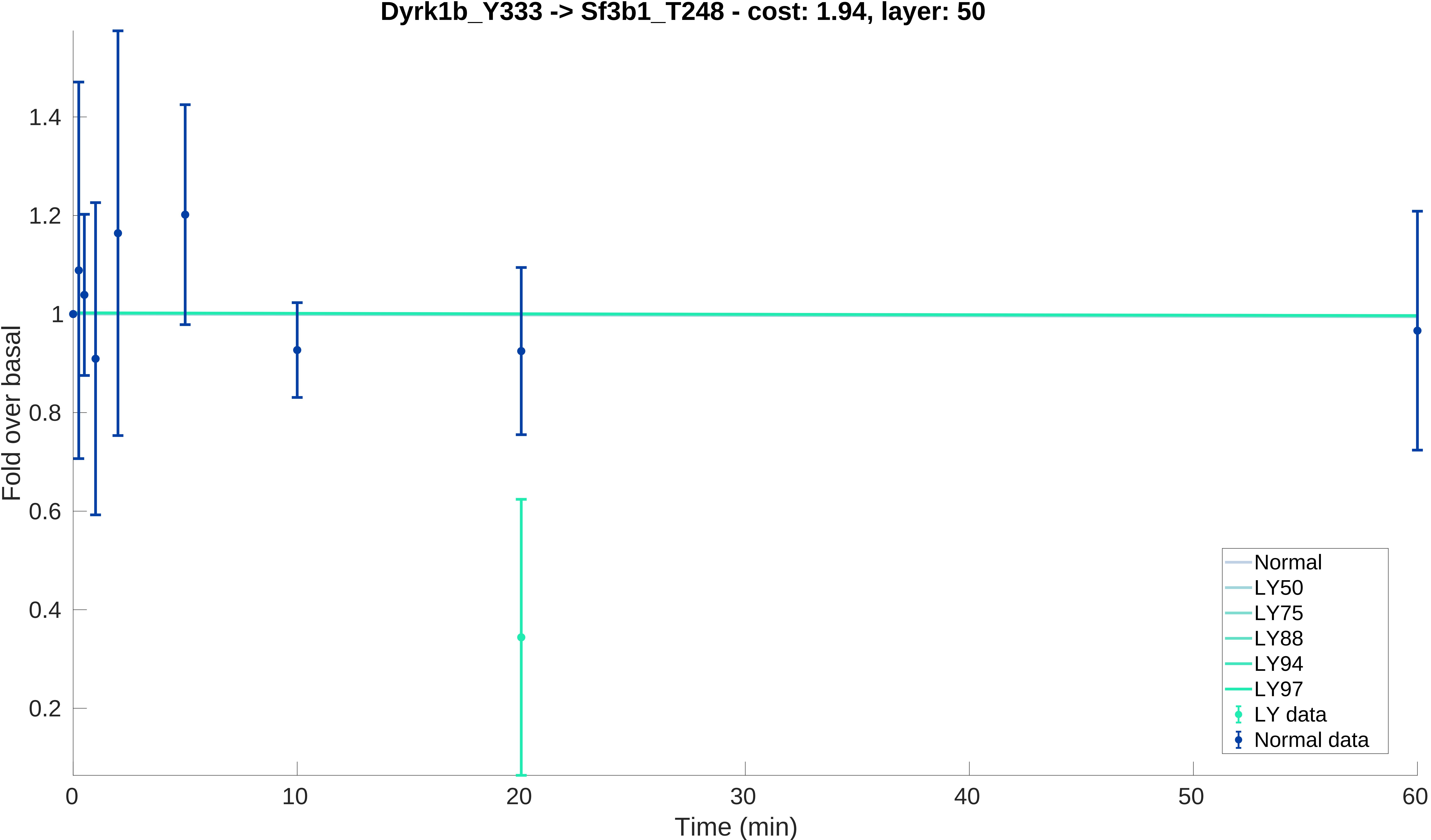

Dyrk1b_Y333 -> Sf3b1_T248 - cost: 1.94, layer: 50
Fold over basal
1.4
1.2
1
0.8
0.6
0.4
0.2
0
10
20
30
40
50
60
Time (min)
Normal
LY50
LY75
LY88
LY94
LY97
LY data
Normal data

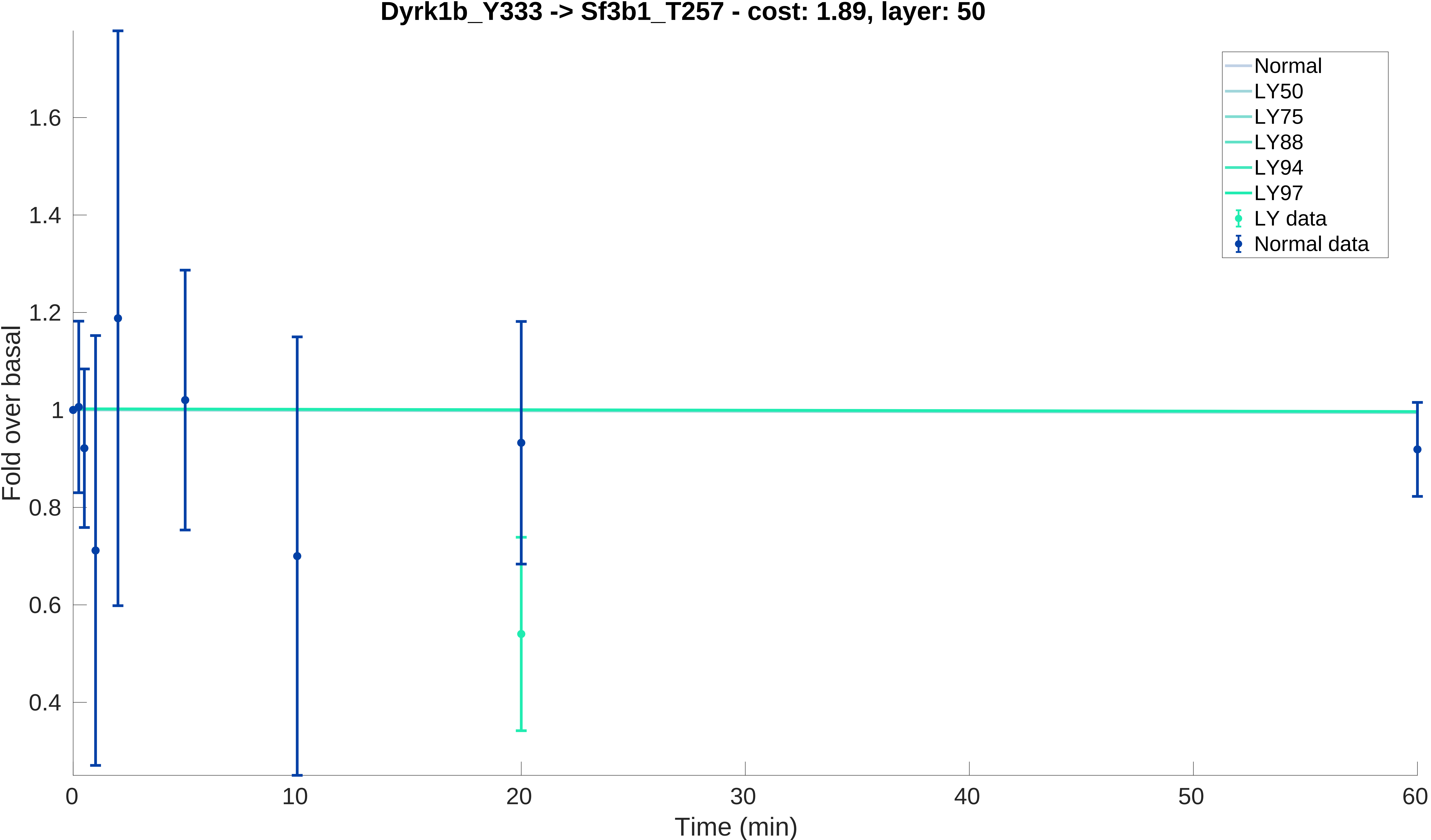
Dyrk1b_Y333 -> Sf3b1_T257 - cost: 1.89, layer: 50
Normal
LY50
LY75
LY88
LY94
LY97
LY data
Normal data
Fold over basal
1.6
1.4
1.2
1
0.8
0.6
0.4
0
10
20
30
40
50
60
Time (min)

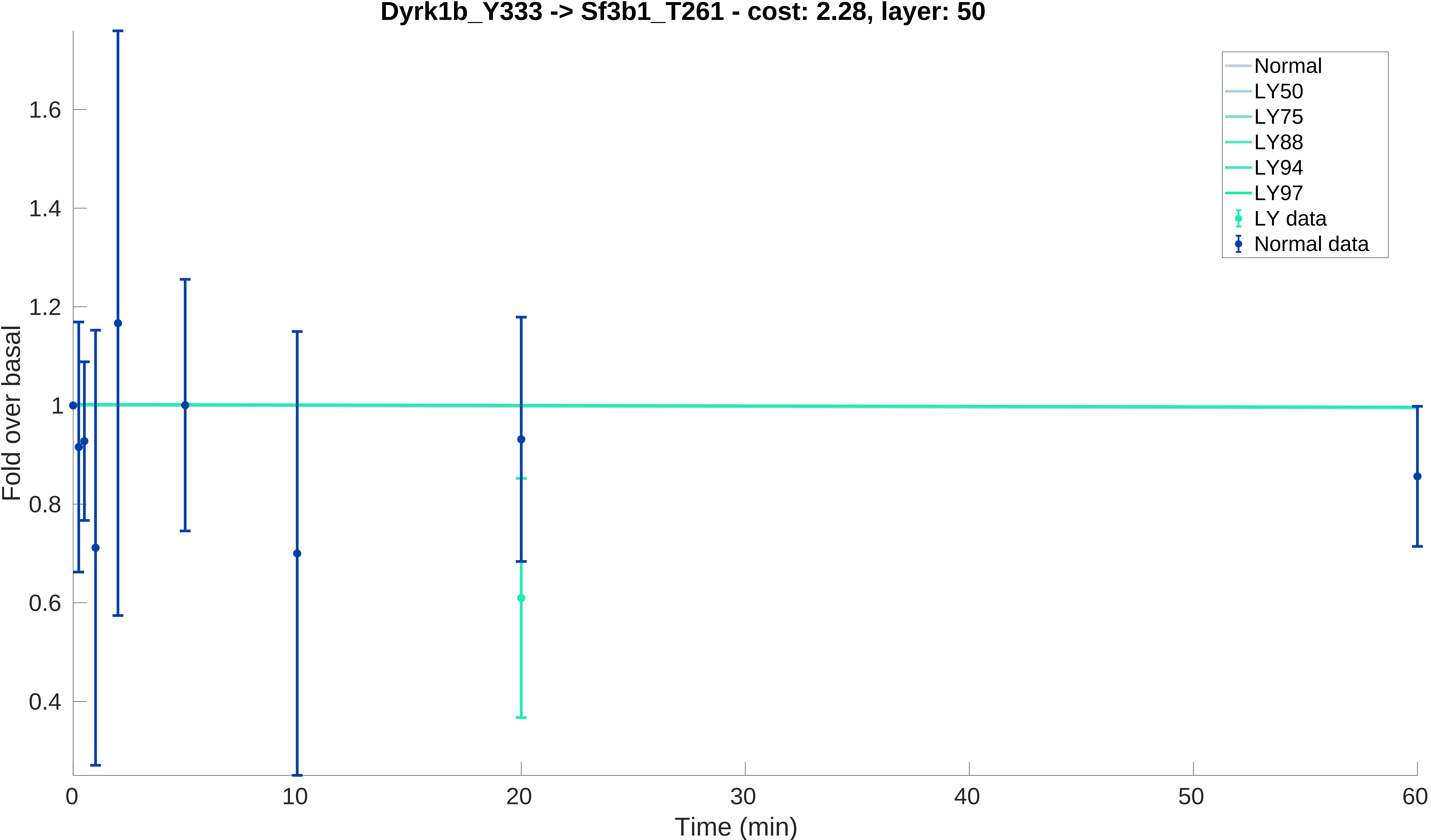

Dyrk1b_Y333 -> Sf3b1_T261 - cost: 2.28, layer: 50
Fold over basal
Time (min)
0.4
0.6
0.8
1
1.2
1.4
1.6
0
10
20
30
40
50
60
Normal
LY50
LY75
LY88
LY94
LY97
LY data
Normal data

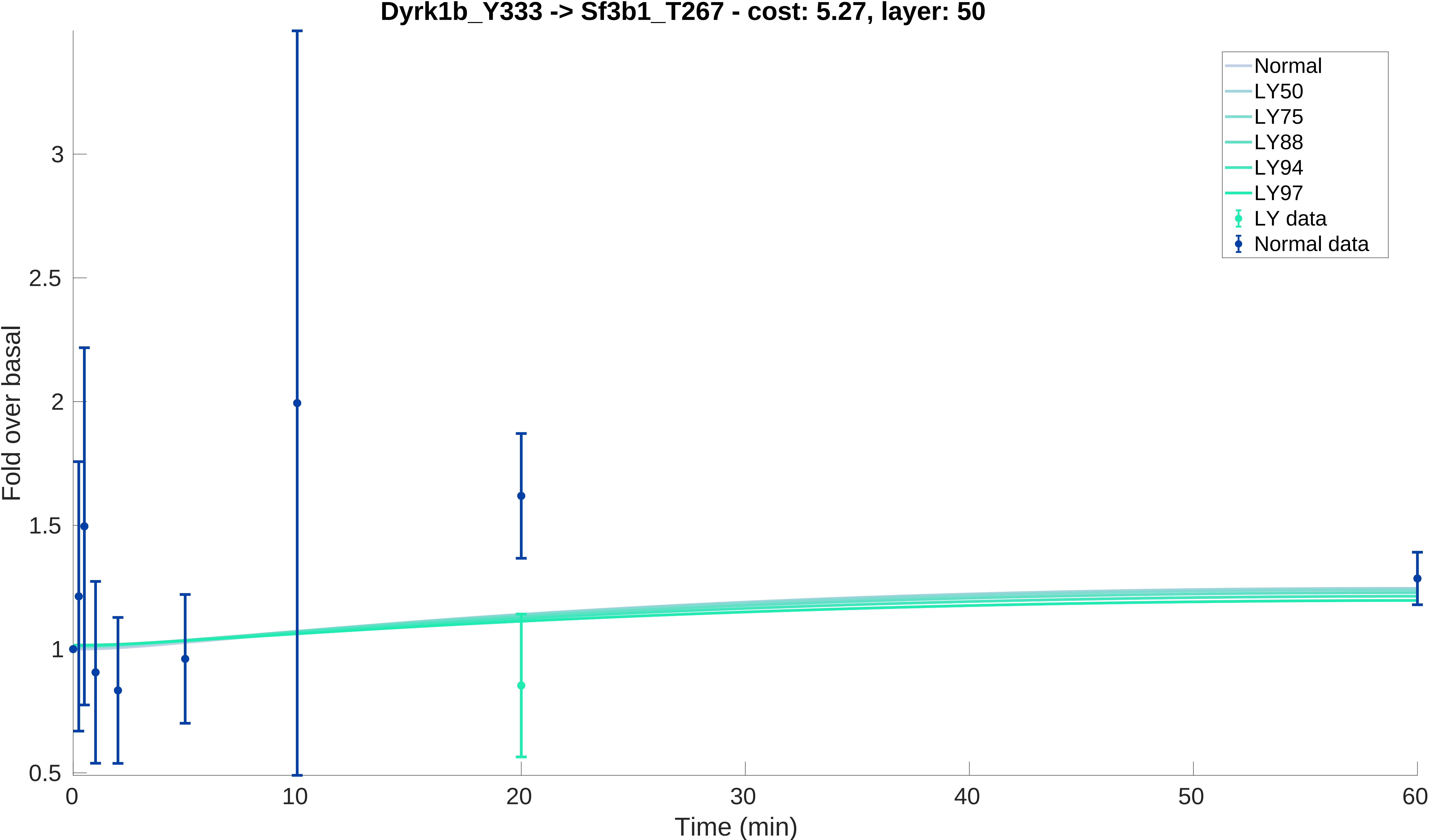
Dyrk1b_Y333 -> Sf3b1_T267 - cost: 5.27, layer: 50
Fold over basal
Time (min)
0.5
1
1.5
2
2.5
3
0
10
20
30
40
50
60
Normal
LY50
LY75
LY88
LY94
LY97
LY data
Normal data

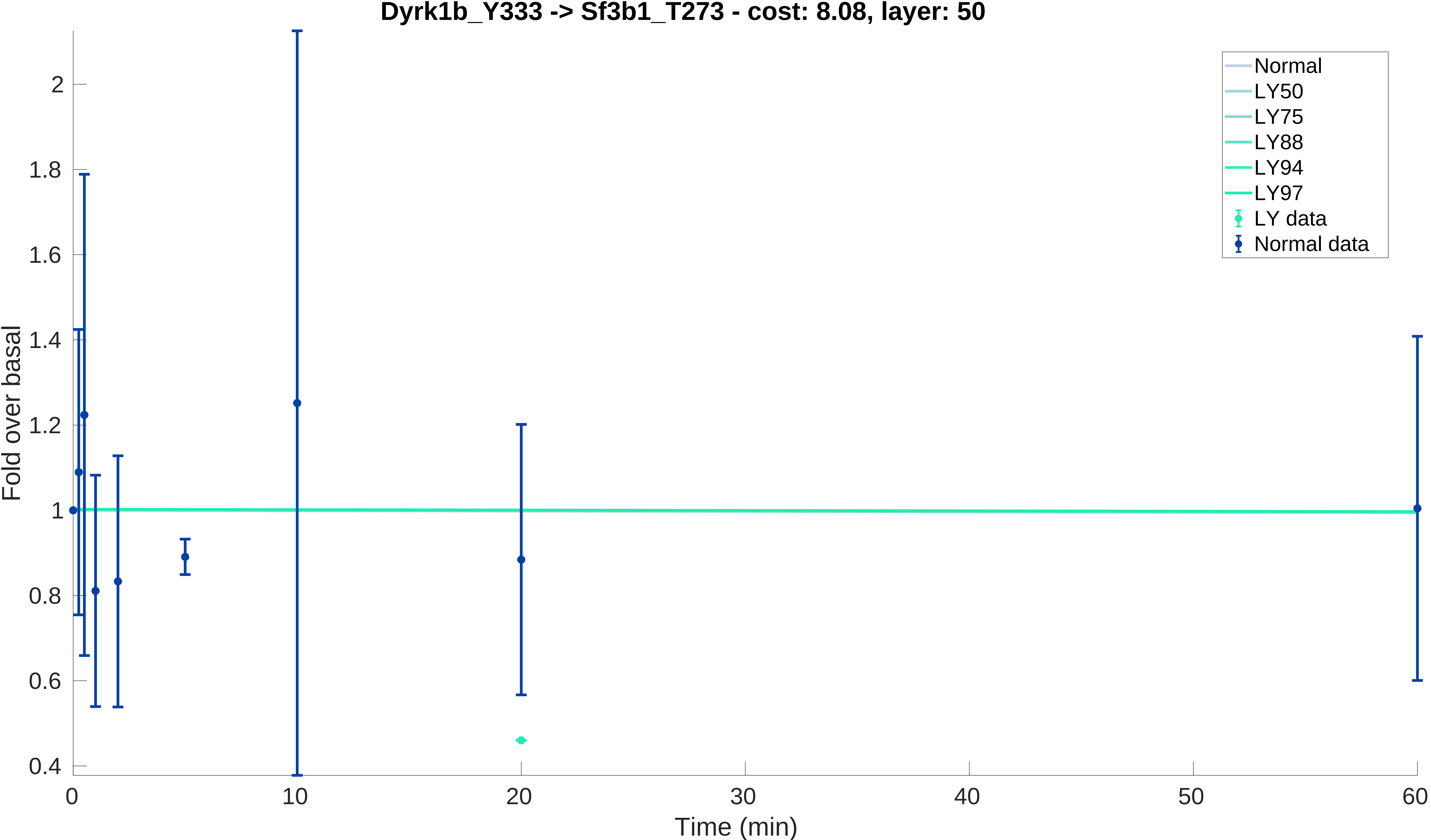
Dyrk1b_Y333 -> Sf3b1_T273 - cost: 8.08, layer: 50
Fold over basal
Time (min)
0.4
0.6
0.8
1
1.2
1.4
1.6
1.8
2
0
10
20
30
40
50
60
Normal
LY50
LY75
LY88
LY94
LY97
LY data
Normal data

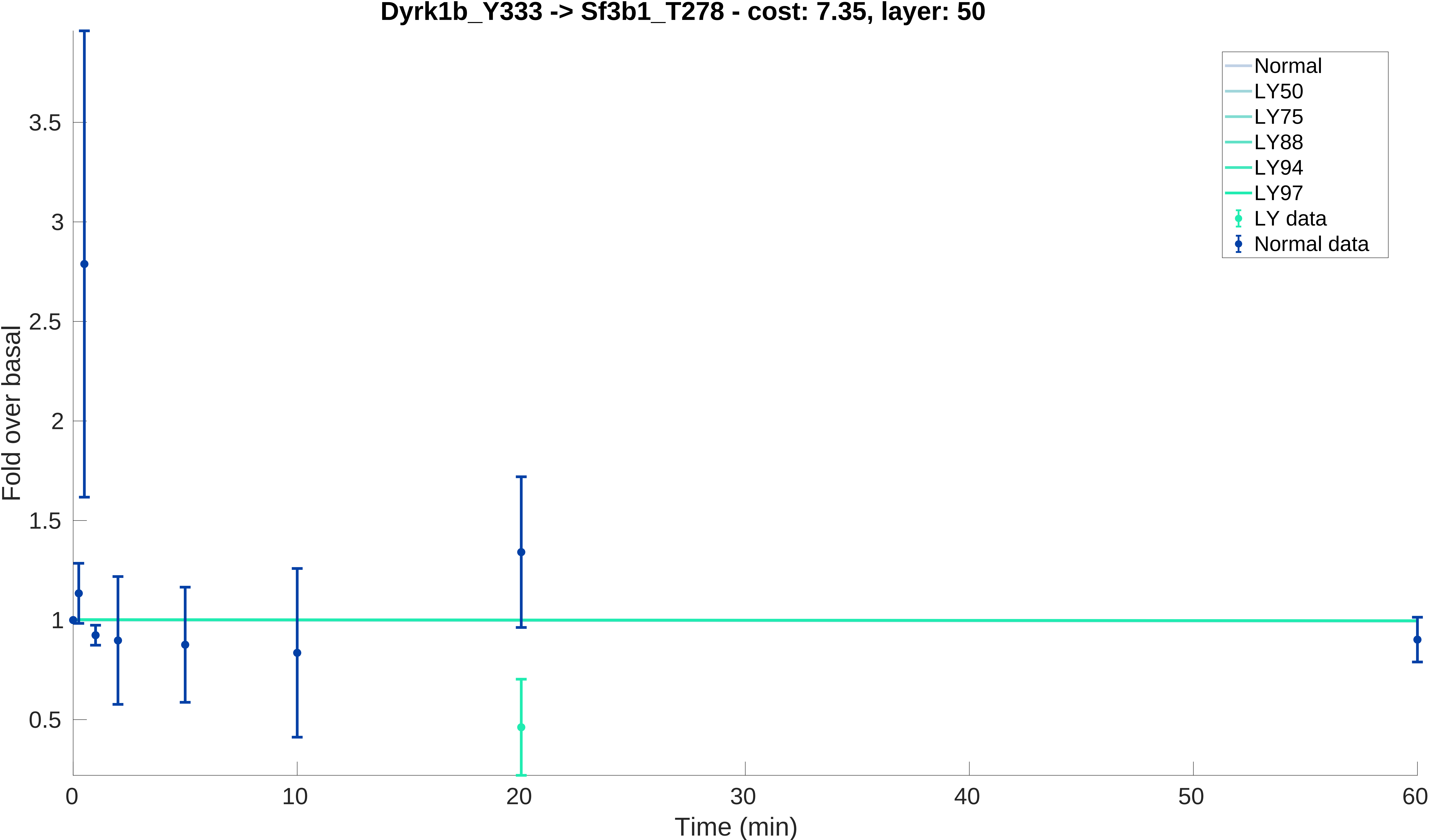

Dyrk1b_Y333 -> Sf3b1_T278 - cost: 7.35, layer: 50
Fold over basal
Time (min)
0.5
1
1.5
2
2.5
3
3.5
0
10
20
30
40
50
60
Normal
LY50
LY75
LY88
LY94
LY97
LY data
Normal data

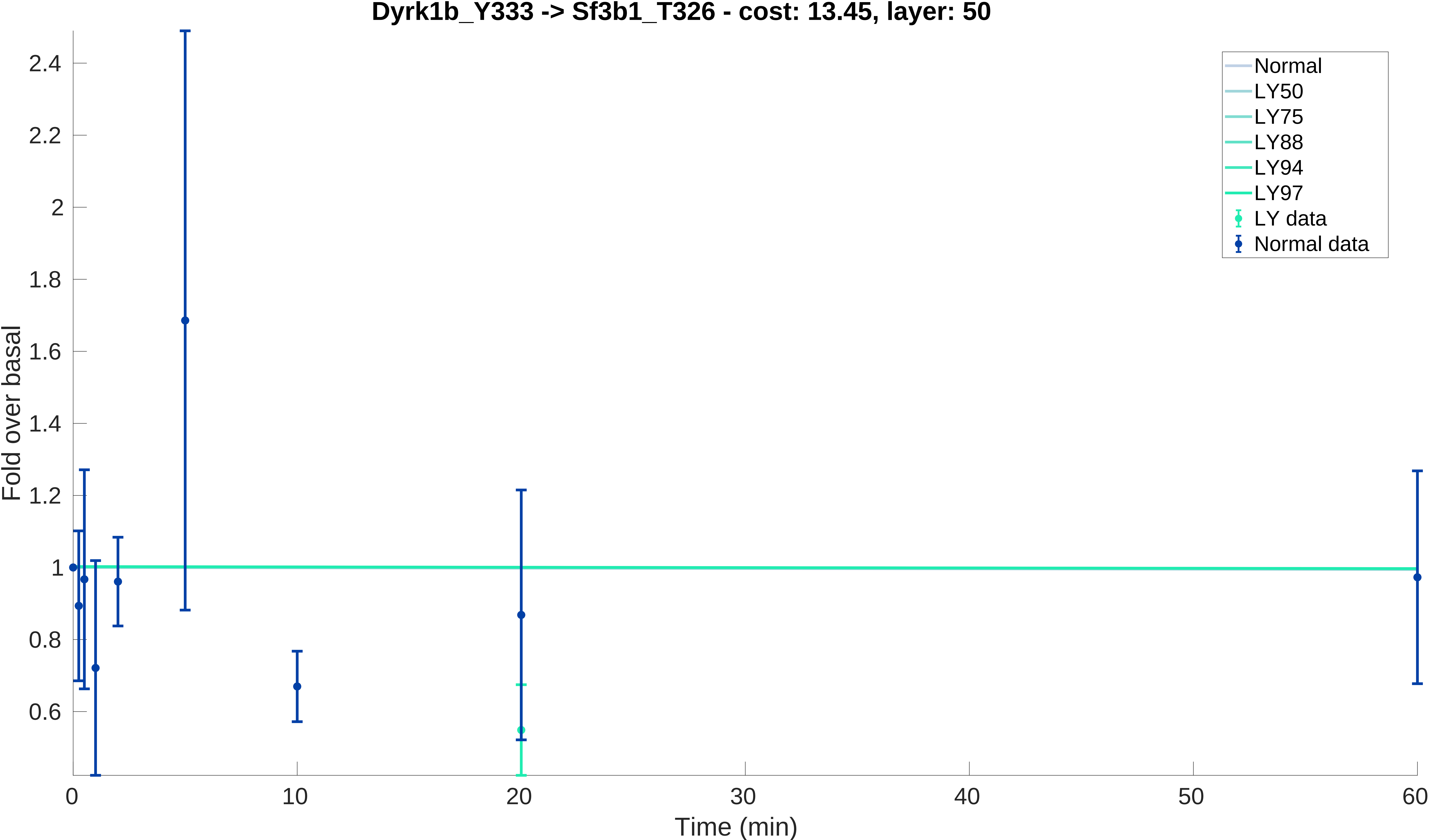
Dyrk1b_Y333 -> Sf3b1_T326 - cost: 13.45, layer: 50
Fold over basal
Time (min)
Normal
LY50
LY75
LY88
LY94
LY97
LY data
Normal data
2.4
2.2
2
1.8
1.6
1.4
1.2
1
0.8
0.6
0
10
20
30
40
50
60

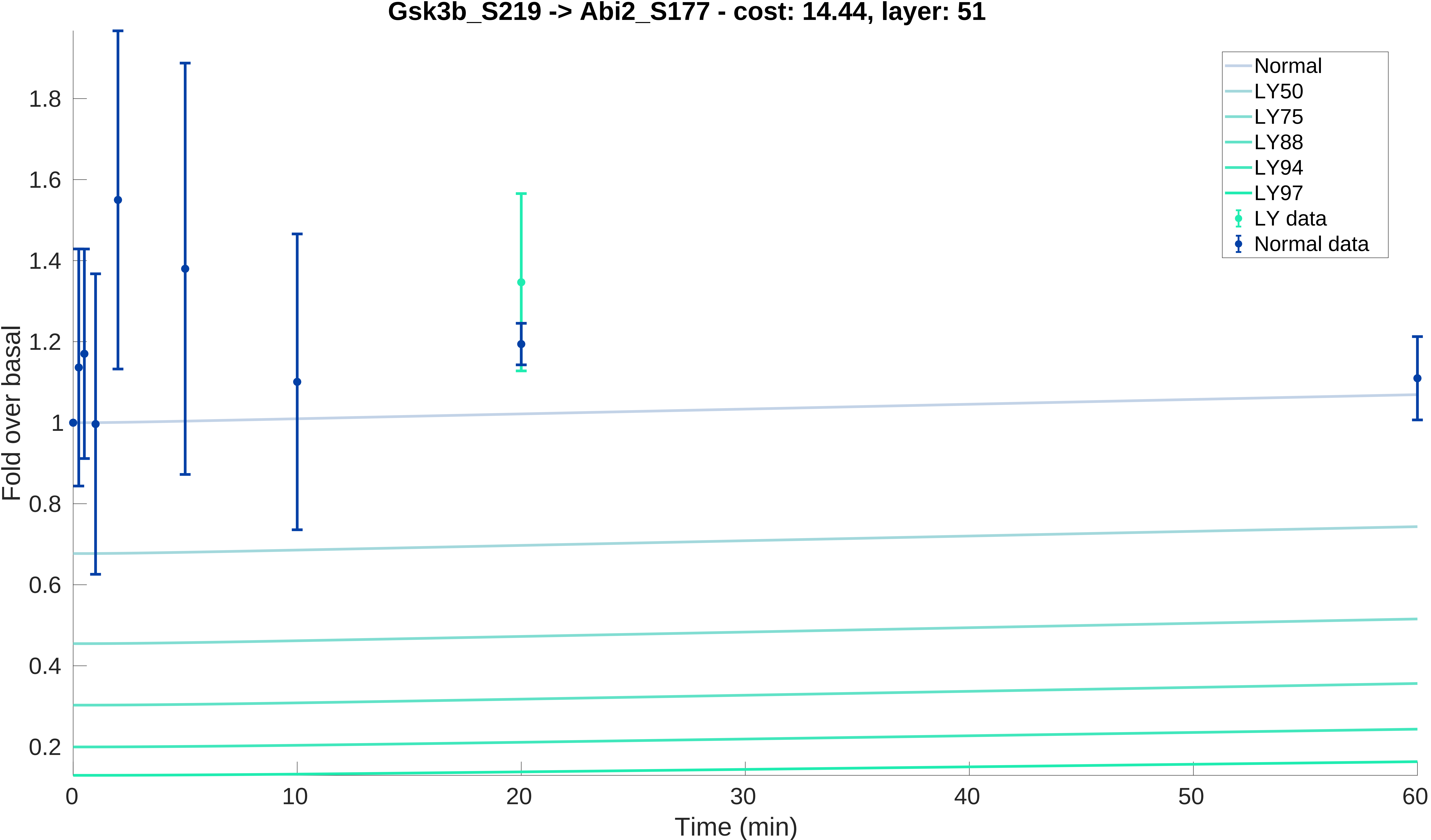
Gsk3b_S219 -> Abi2_S177 - cost: 14.44, layer: 51
Fold over basal
Time (min)
0.2
0.4
0.6
0.8
1
1.2
1.4
1.6
1.8
0
10
20
30
40
50
60
Normal
LY50
LY75
LY88
LY94
LY97
LY data
Normal data

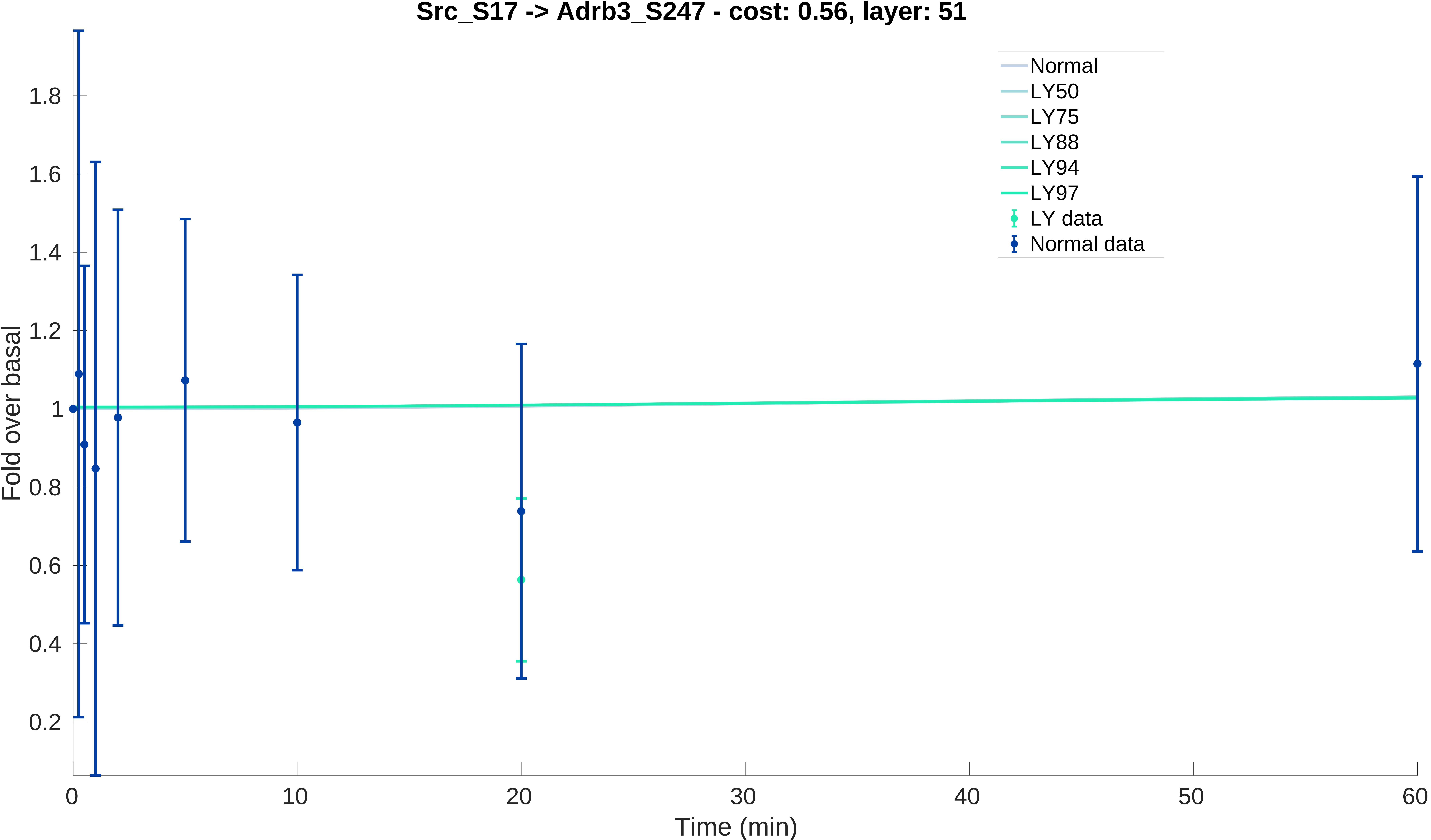

Src_S17 -> Adrb3_S247 - cost: 0.56, layer: 51
Normal
LY50
LY75
LY88
LY94
LY97
LY data
Normal data
Fold over basal
1.8
1.6
1.4
1.2
1
0.8
0.6
0.4
0.2
0
10
20
30
40
50
60
Time (min)

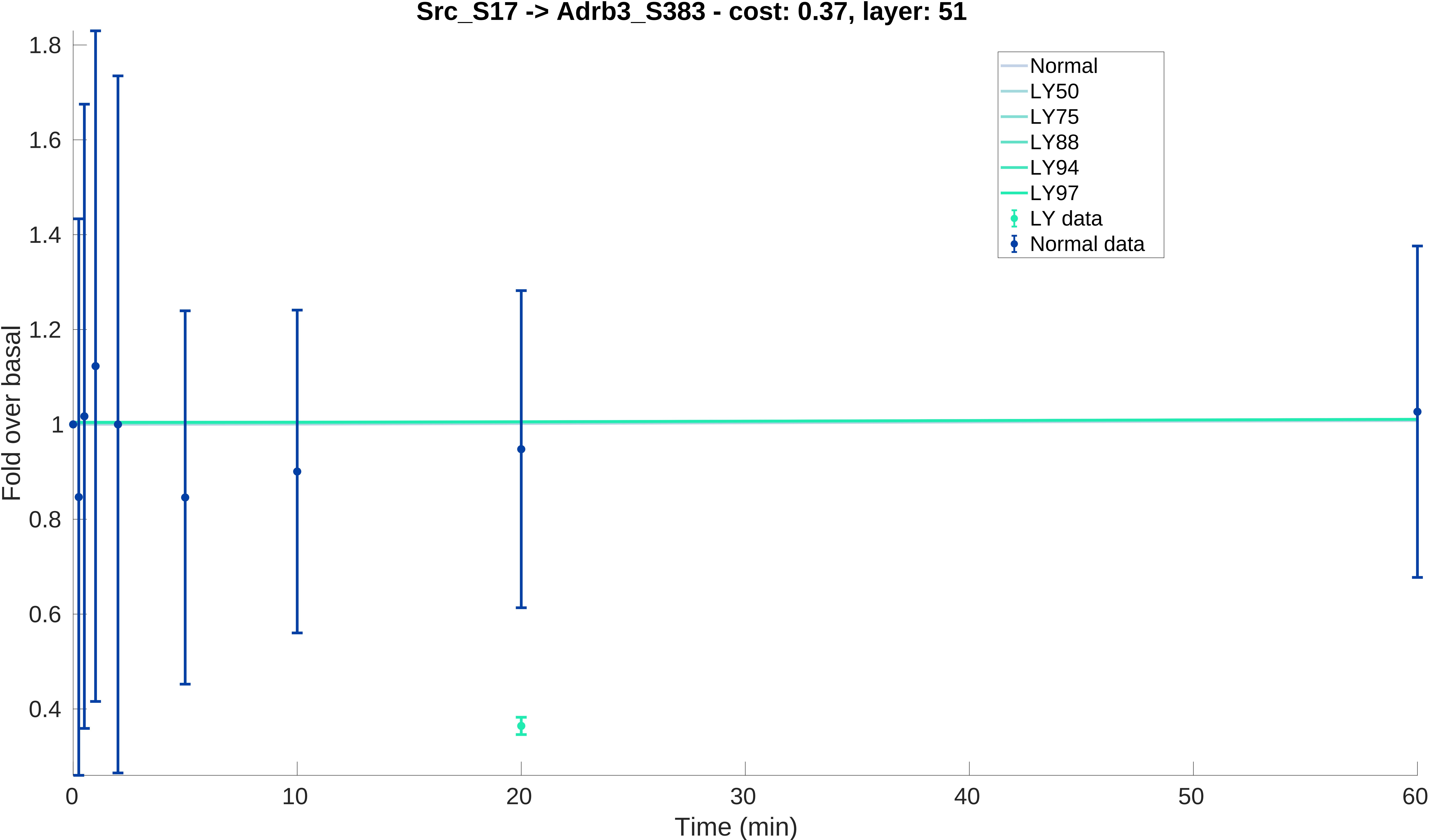

Src_S17 -> Adrb3_S383 - cost: 0.37, layer: 51
Fold over basal
Time (min)
Normal
LY50
LY75
LY88
LY94
LY97
LY data
Normal data
1.8
1.6
1.4
1.2
1
0.8
0.6
0.4
0
10
20
30
40
50
60

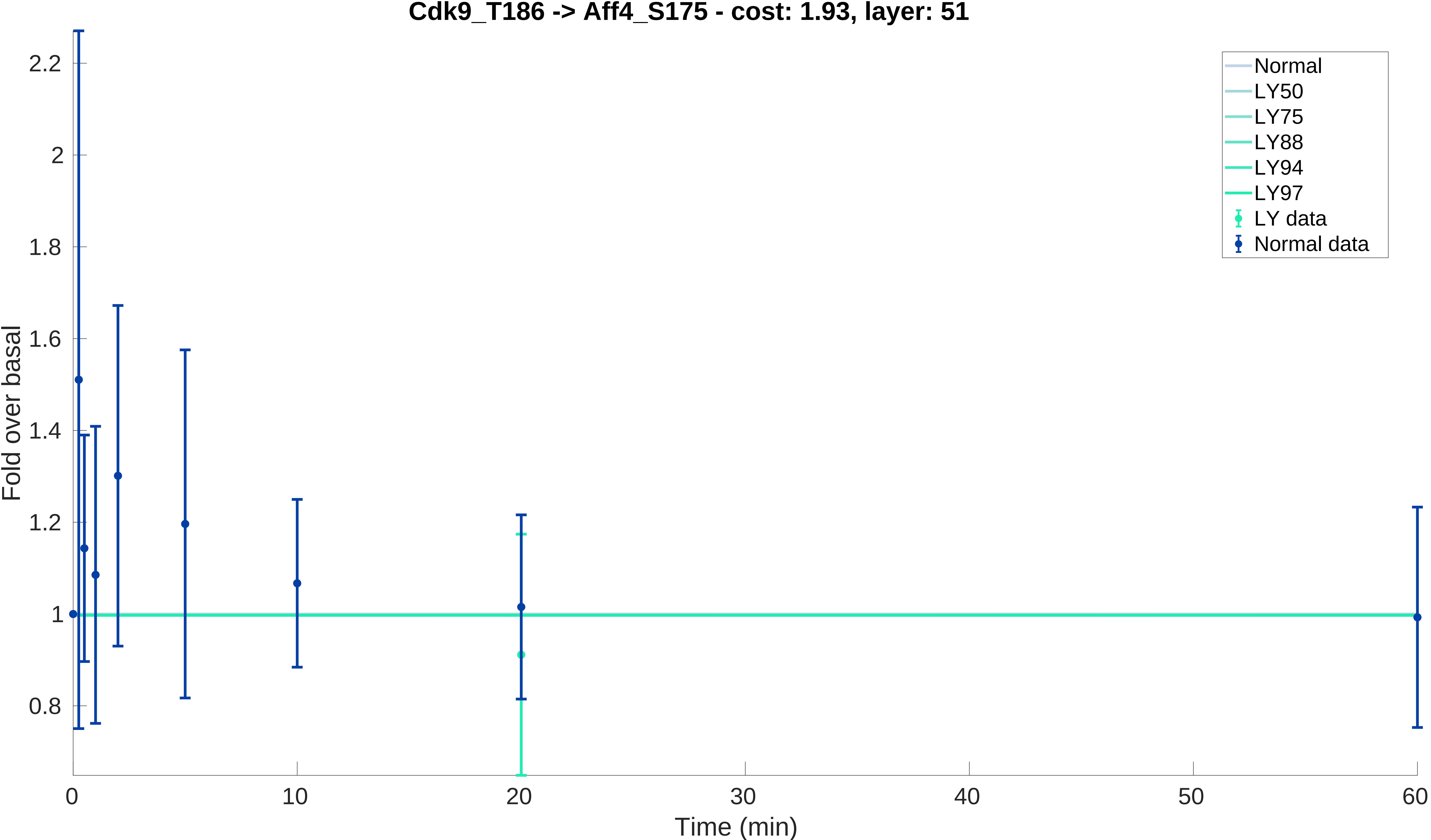
Cdk9_T186 -> Aff4_S175 - cost: 1.93, layer: 51
Fold over basal
Time (min)
Normal
LY50
LY75
LY88
LY94
LY97
LY data
Normal data
2.2
2
1.8
1.6
1.4
1.2
1
0.8
0
10
20
30
40
50
60

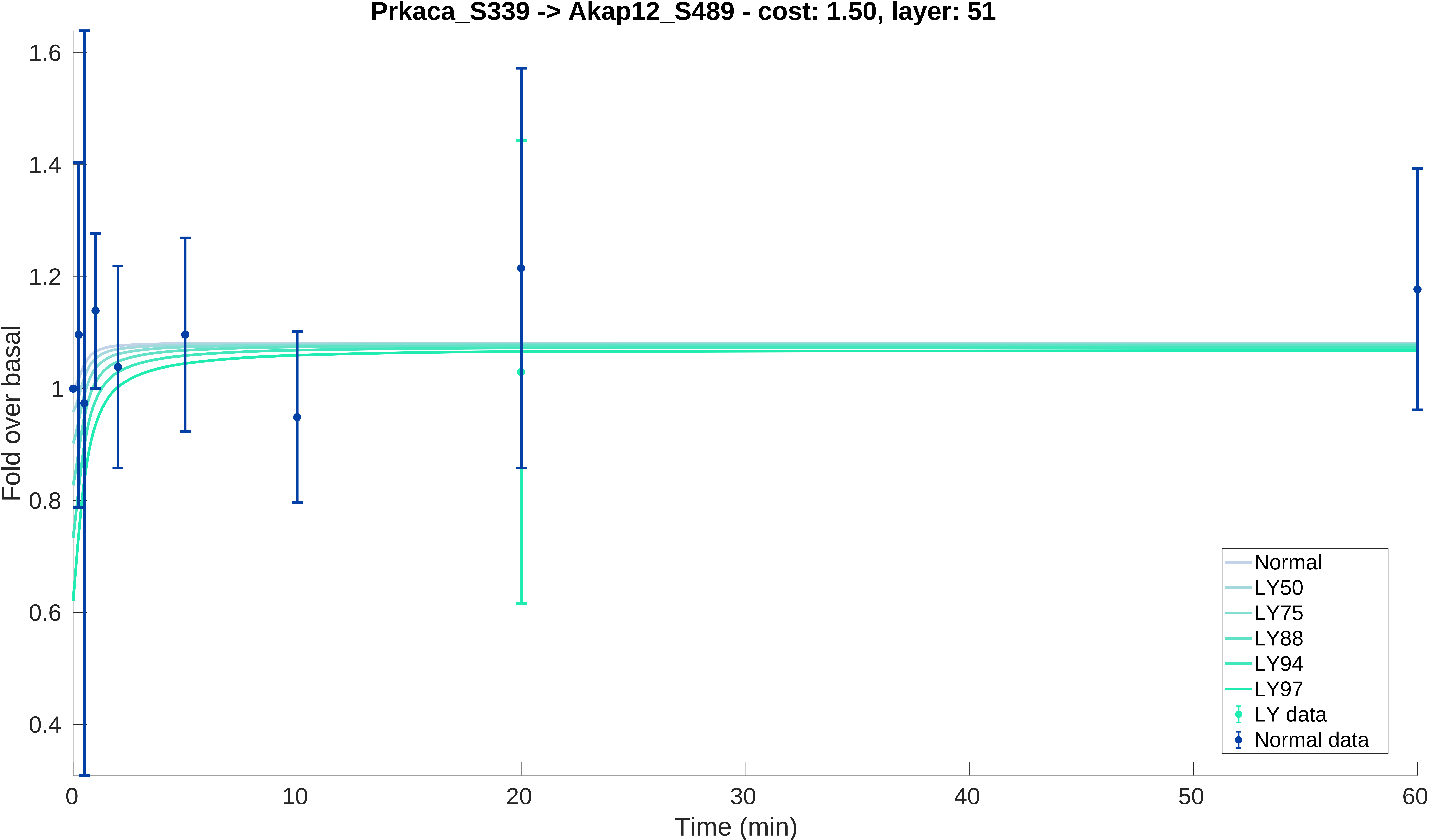

Prkaca_S339 -> Akap12_S489 - cost: 1.50, layer: 51
Fold over basal
1.6
1.4
1.2
1
0.8
0.6
0.4
0
10
20
30
40
50
60
Time (min)
Normal
LY50
LY75
LY88
LY94
LY97
LY data
Normal data

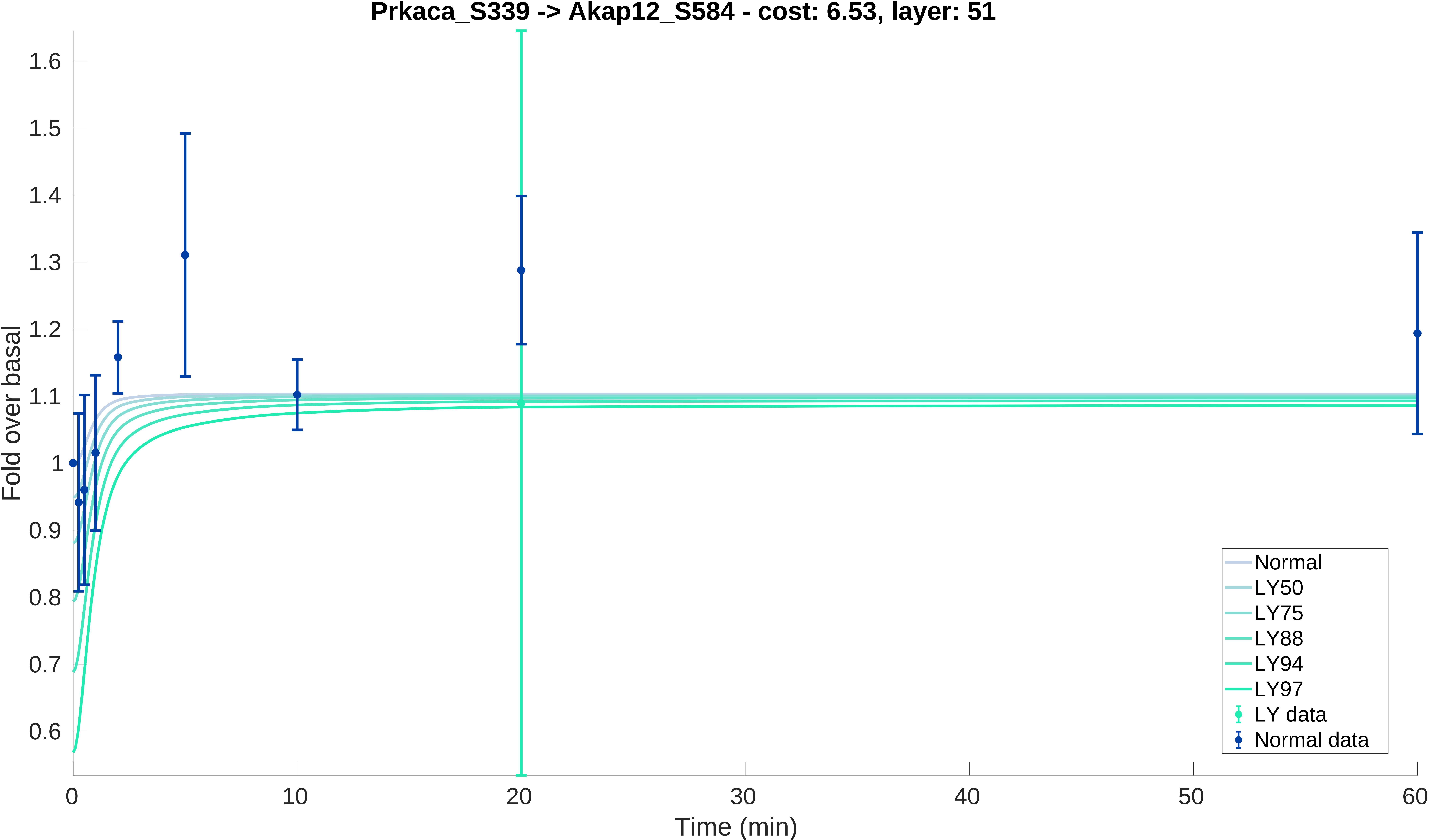

Prkaca_S339 -> Akap12_S584 - cost: 6.53, layer: 51
Fold over basal
1.6
1.5
1.4
1.3
1.2
1.1
1
0.9
0.8
0.7
0.6
0
10
20
30
40
50
60
Time (min)
Normal
LY50
LY75
LY88
LY94
LY97
LY data
Normal data

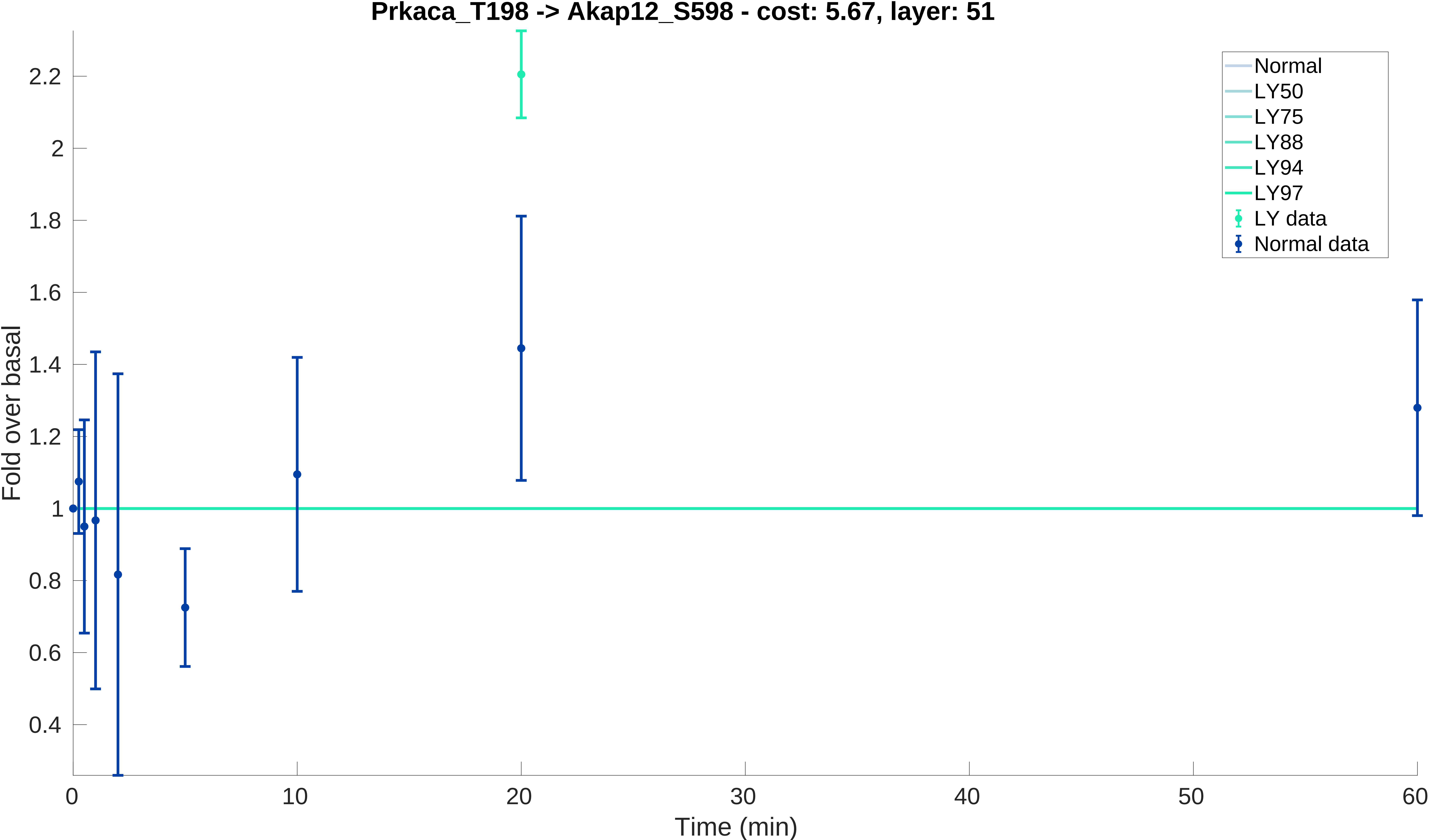

Prkaca_T198 -> Akap12_S598 - cost: 5.67, layer: 51
Fold over basal
Time (min)
Normal
LY50
LY75
LY88
LY94
LY97
LY data
Normal data
0.4
0.6
0.8
1
1.2
1.4
1.6
1.8
2
2.2
0
10
20
30
40
50
60

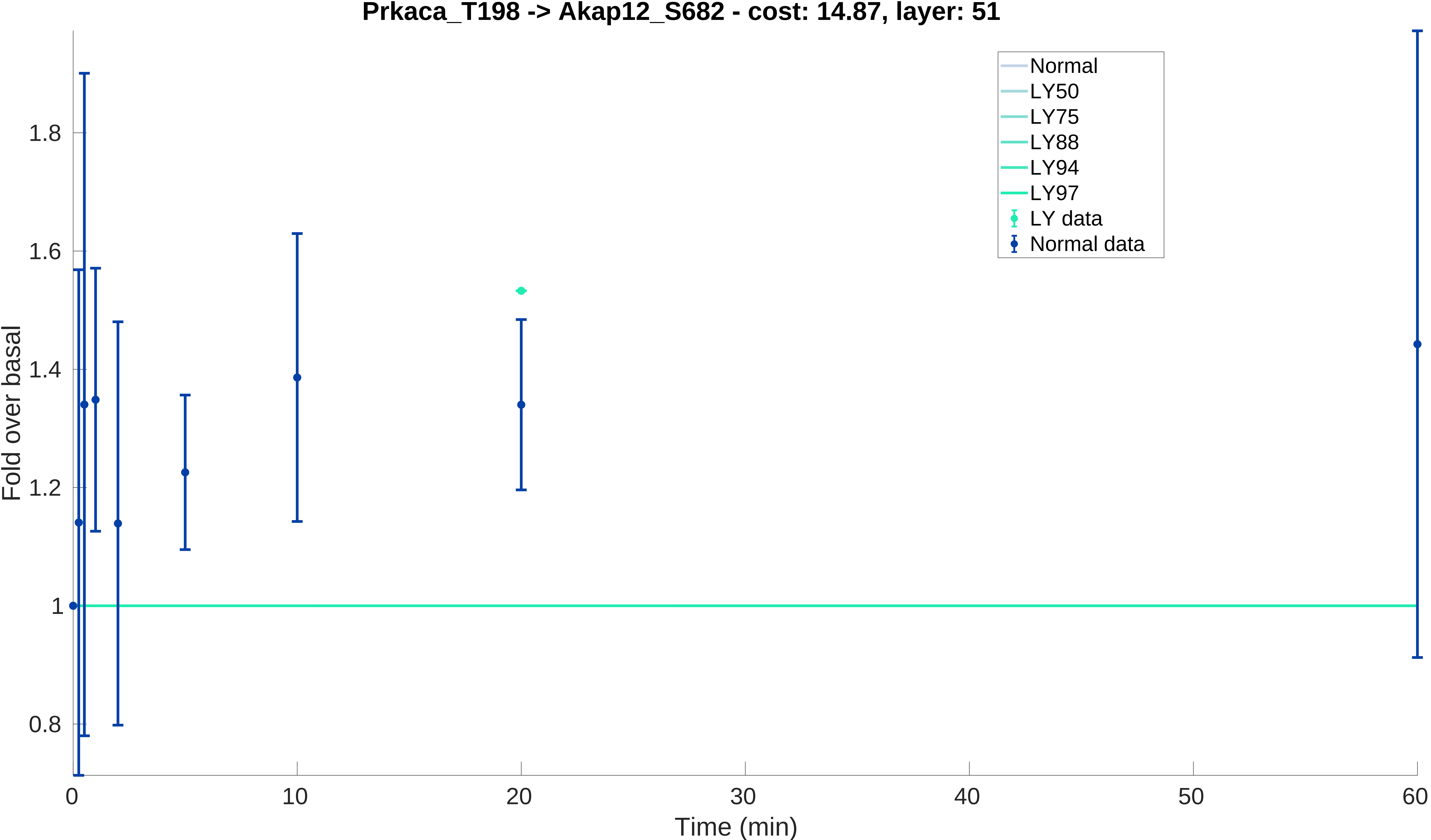

Prrkaca_T198 -> Akap12_S682 - cost: 14.87, layer: 51
Fold over basal
Time (min)
0.8
1
1.2
1.4
1.6
1.8
0
10
20
30
40
50
60
Normal
LY50
LY75
LY88
LY94
LY97
LY data
Normal data

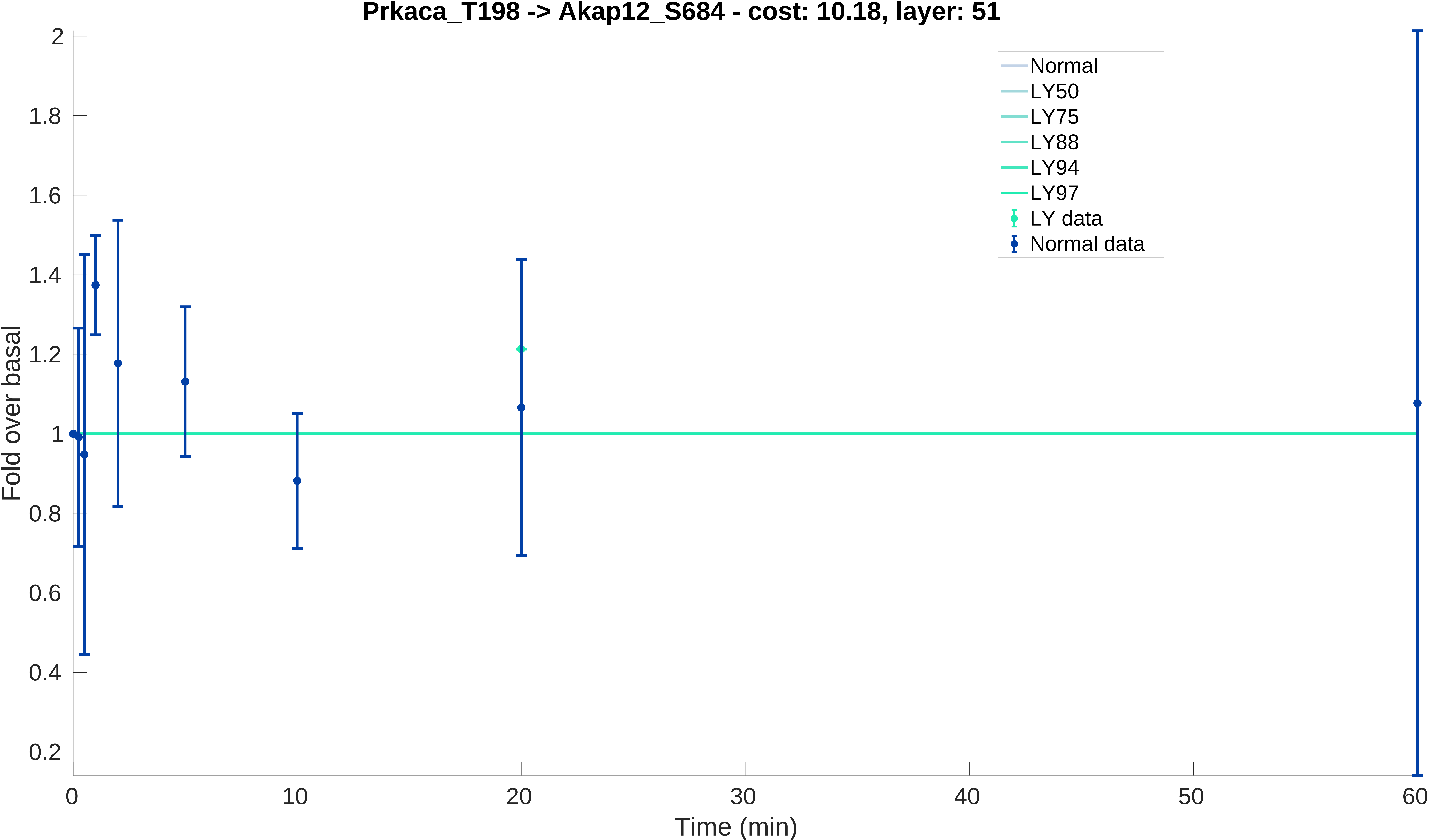
Prkaca_T198 -> Akap12_S684 - cost: 10.18, layer: 51
Fold over basal
Time (min)
0
10
20
30
40
50
60
0.2
0.4
0.6
0.8
1
1.2
1.4
1.6
1.8
2
Normal
LY50
LY75
LY88
LY94
LY97
LY data
Normal data

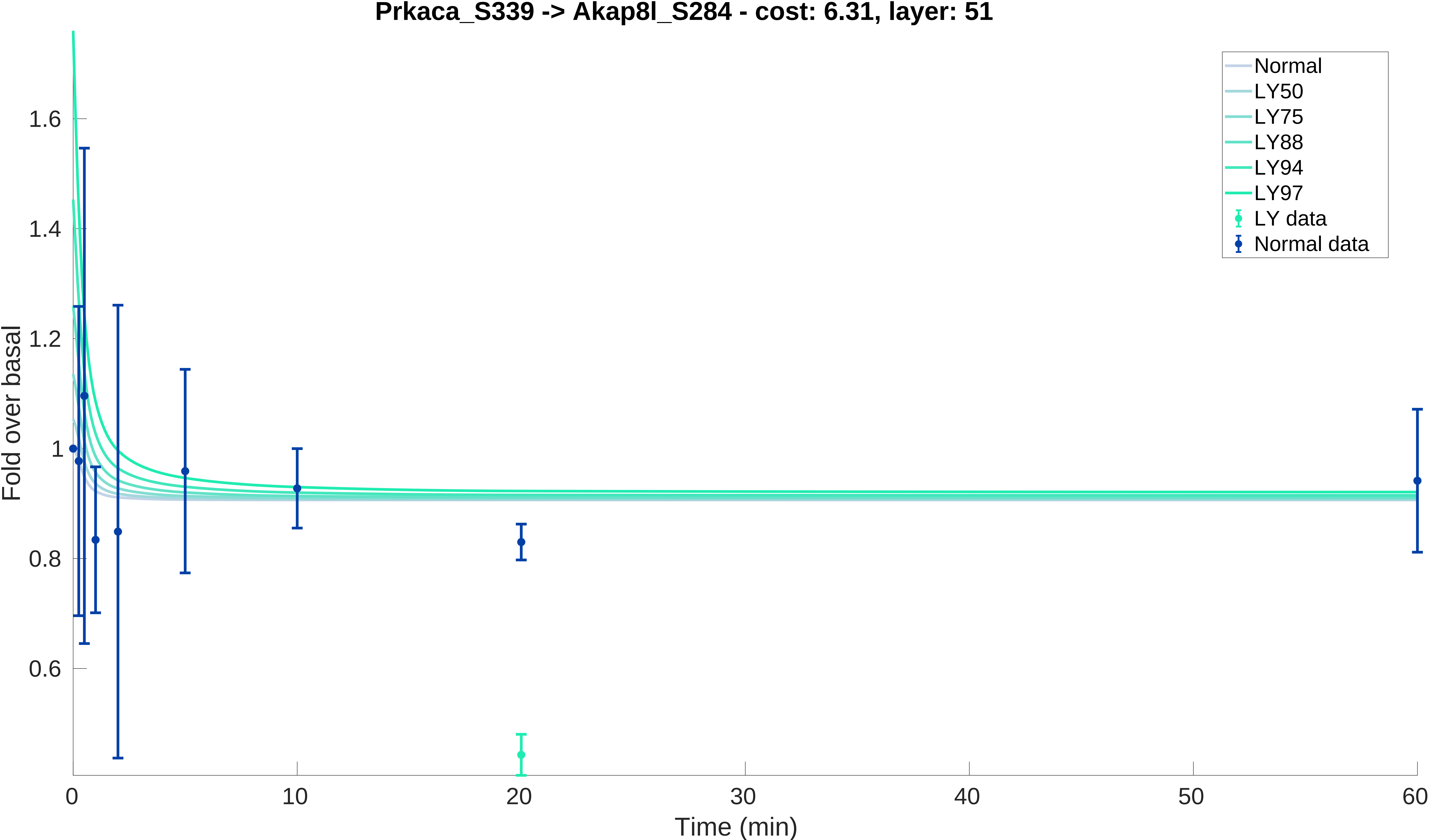
Prkaca_S339 -> Akap8l_S284 - cost: 6.31, layer: 51
Fold over basal
Time (min)
0.6
0.8
1
1.2
1.4
1.6
0
10
20
30
40
50
60
Normal
LY50
LY75
LY88
LY94
LY97
LY data
Normal data

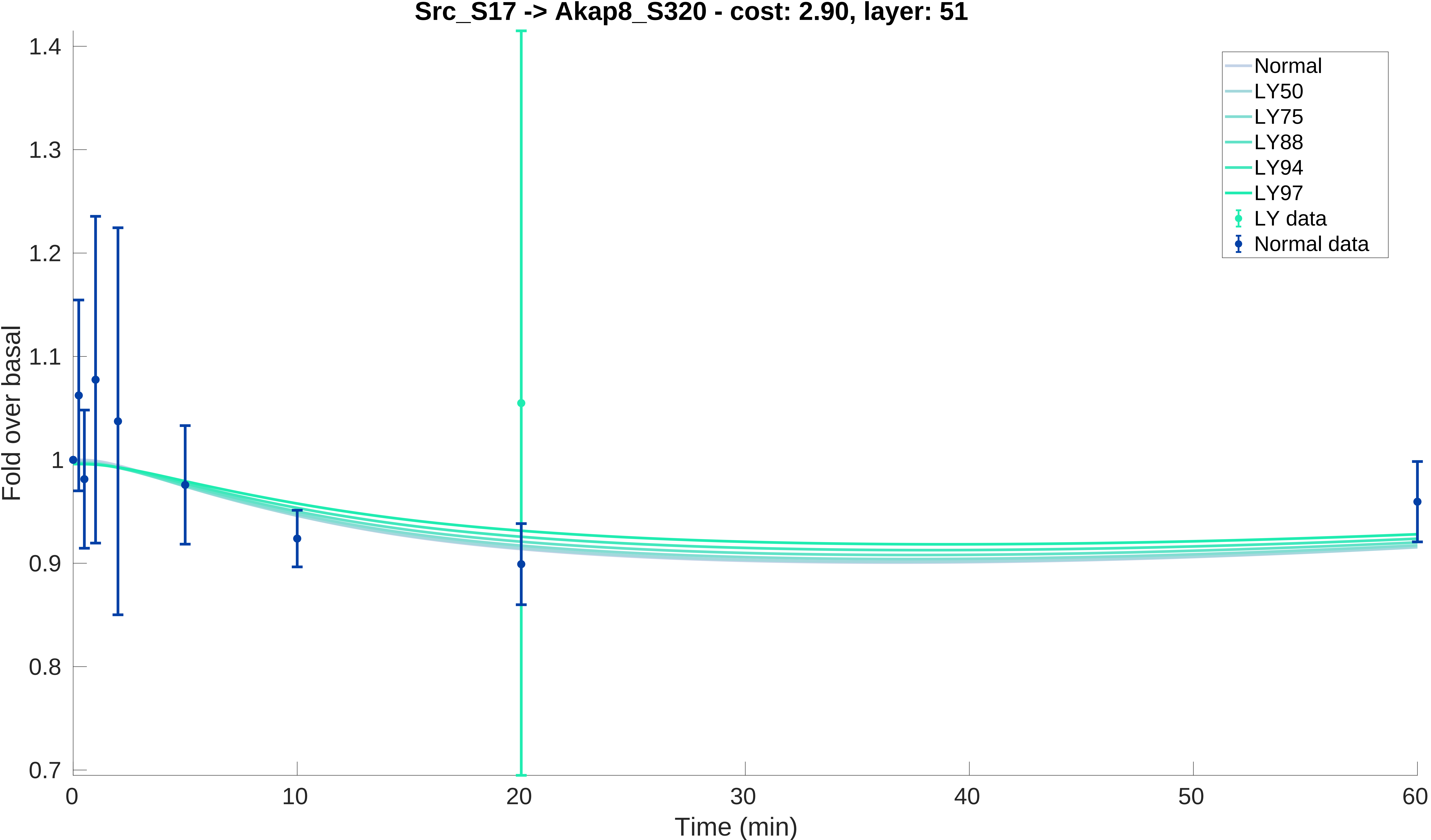
Src_S17 -> Akap8_S320 - cost: 2.90, layer: 51
Fold over basal
Time (min)
1.4
1.3
1.2
1.1
1
0.9
0.8
0.7
0
10
20
30
40
50
60
Normal
LY50
LY75
LY88
LY94
LY97
LY data
Normal data

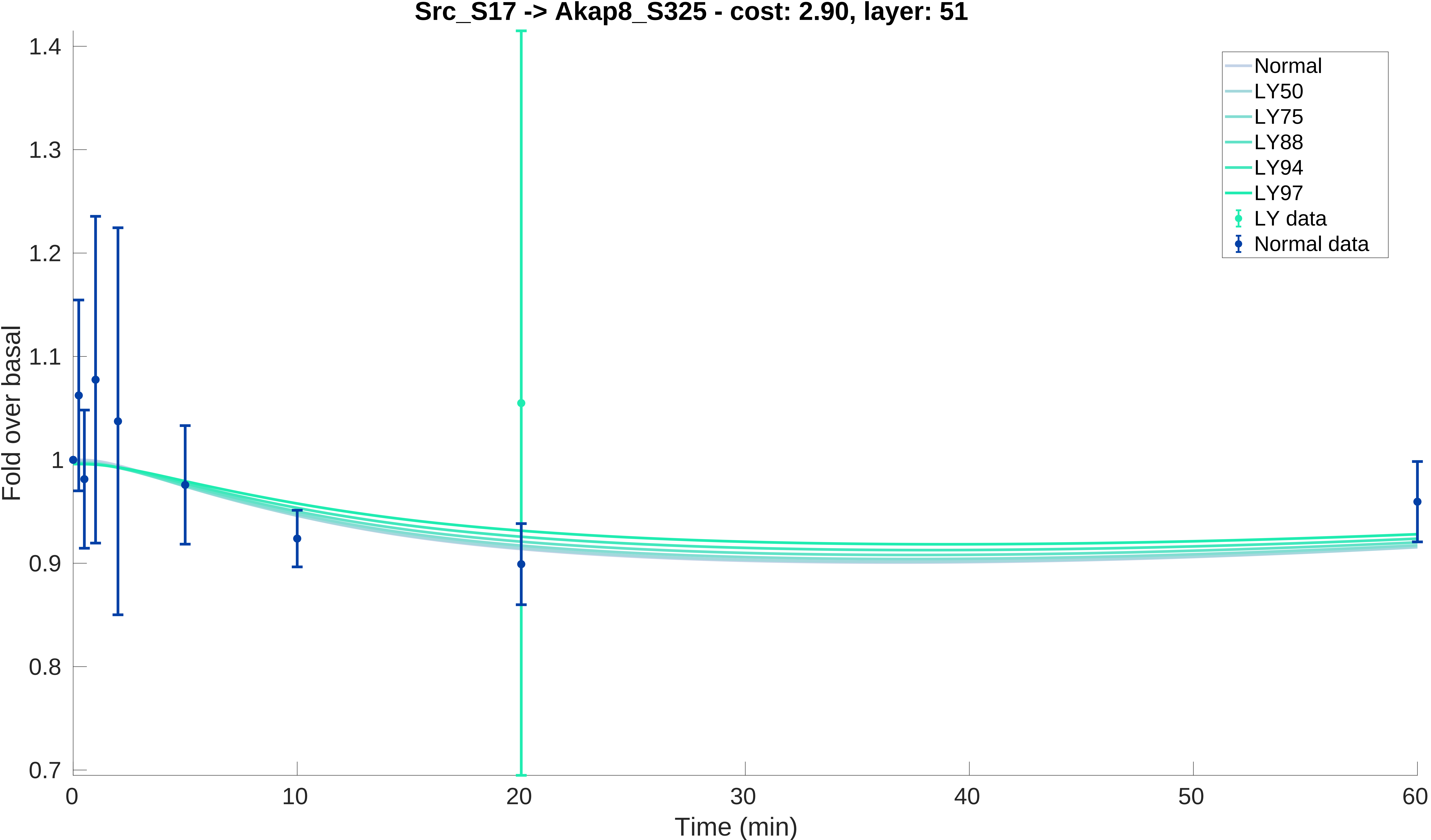
Src_S17 -> Akap8_S325 - cost: 2.90, layer: 51
Fold over basal
Time (min)
0.7
0.8
0.9
1
1.1
1.2
1.3
1.4
0
10
20
30
40
50
60
Normal
LY50
LY75
LY88
LY94
LY97
LY data
Normal data

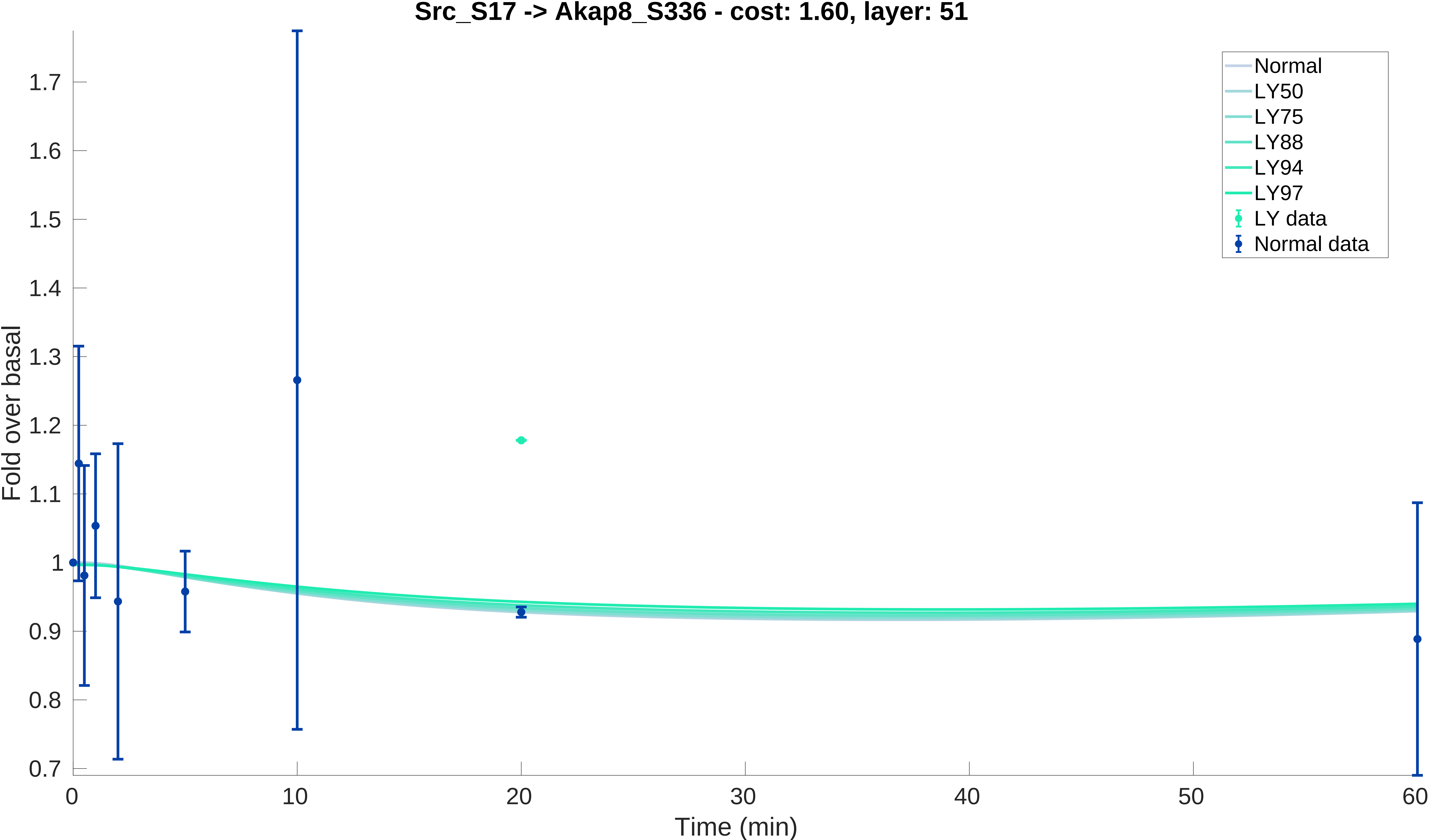

Src_S17 -> Akap8_S336 - cost: 1.60, layer: 51
Fold over basal
1.7
1.6
1.5
1.4
1.3
1.2
1.1
1
0.9
0.8
0.7
0
10
20
30
40
50
60
Time (min)
Normal
LY50
LY75
LY88
LY94
LY97
LY data
Normal data

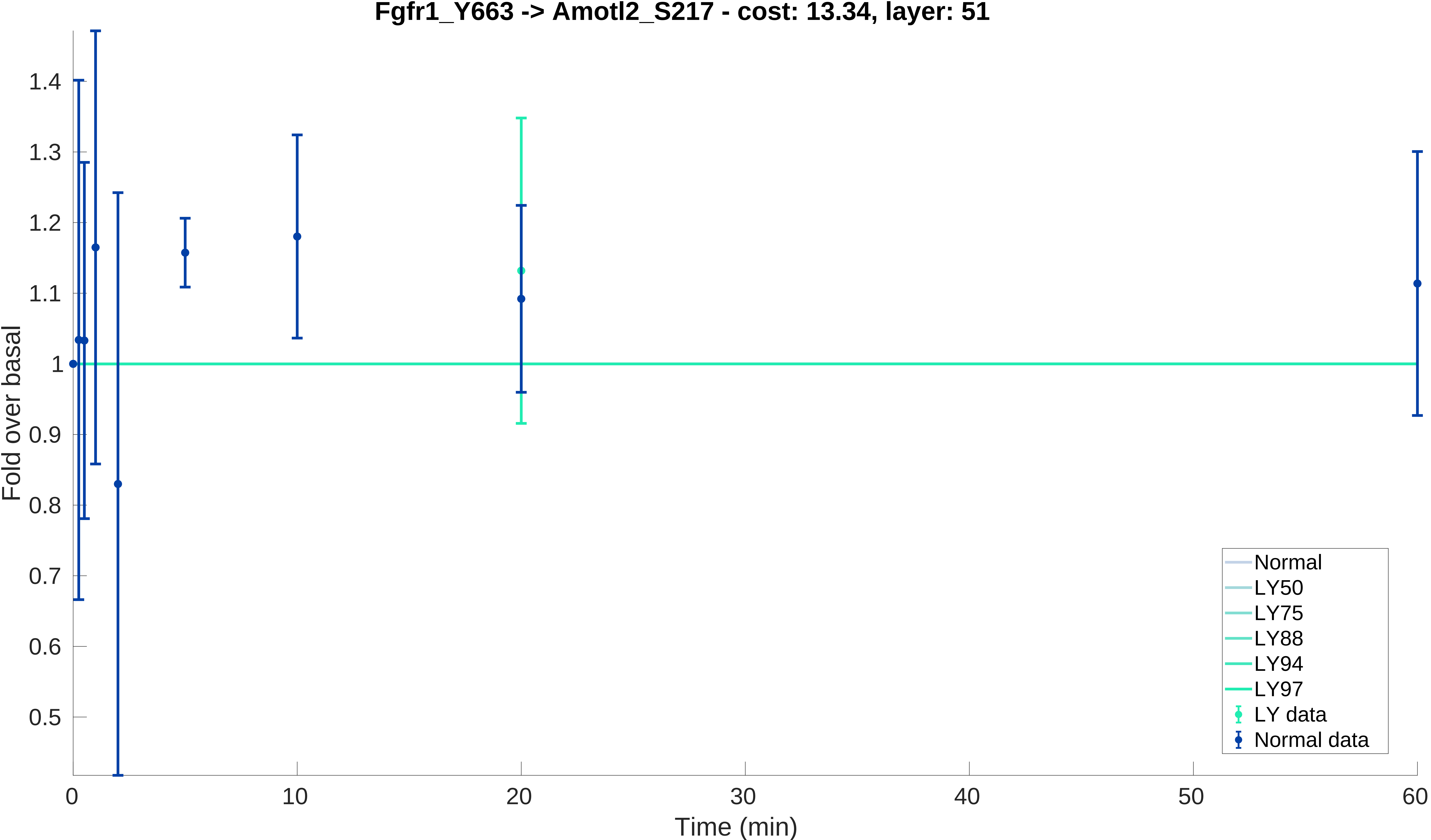

Fgfr1_Y663 -> Amotl2_S217 - cost: 13.34, layer: 51
Fold over basal
1.4
1.3
1.2
1.1
1
0.9
0.8
0.7
0.6
0.5
0
10
20
30
40
50
60
Time (min)
Normal
LY50
LY75
LY88
LY94
LY97
LY data
Normal data

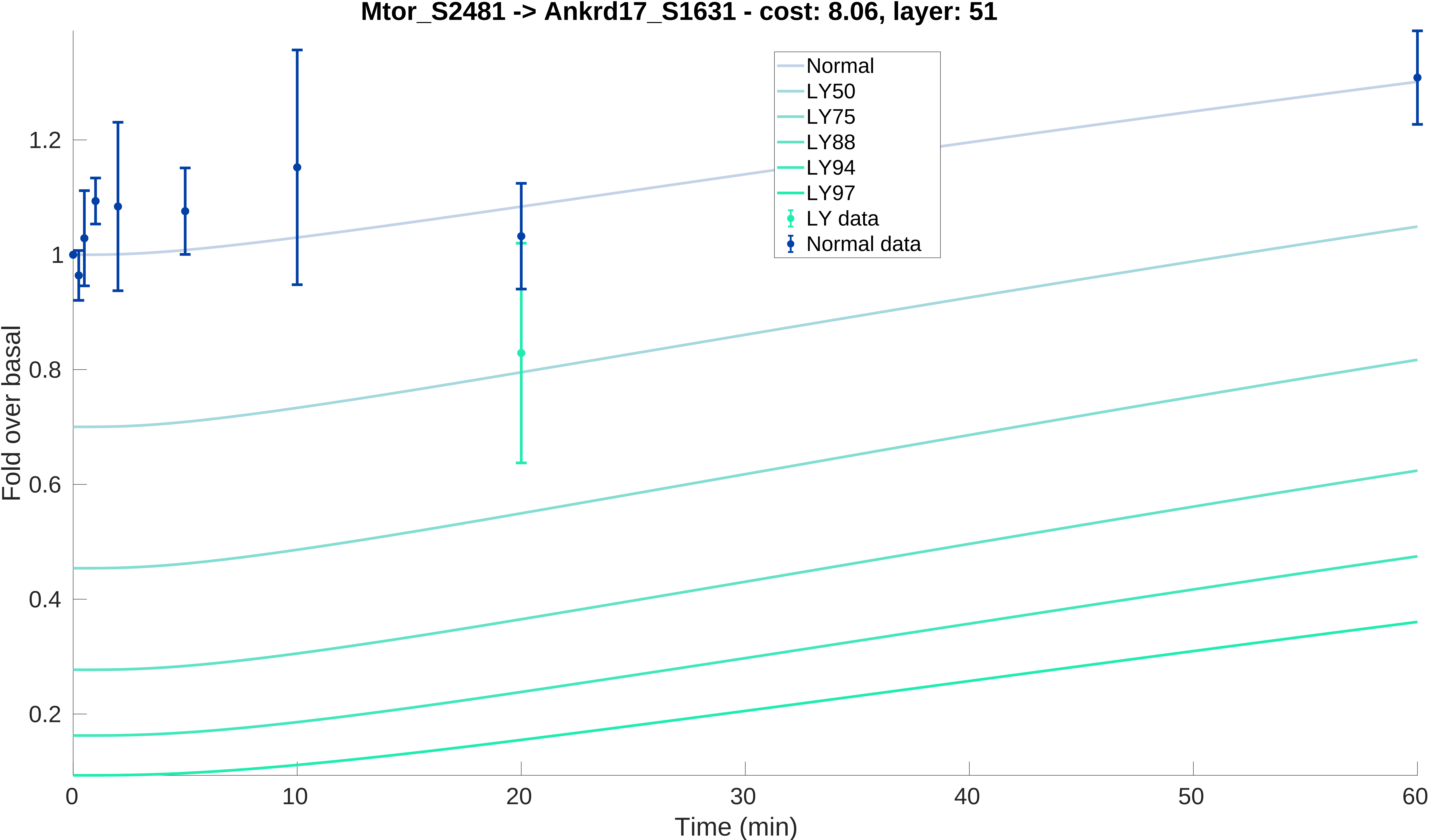

Mtor_S2481 -> Ankrd17_S1631 - cost: 8.06, layer: 51
Normal
LY50
LY75
LY88
LY94
LY97
LY data
Normal data
Fold over basal
0.2
0.4
0.6
0.8
1
1.2
0
10
20
30
40
50
60
Time (min)

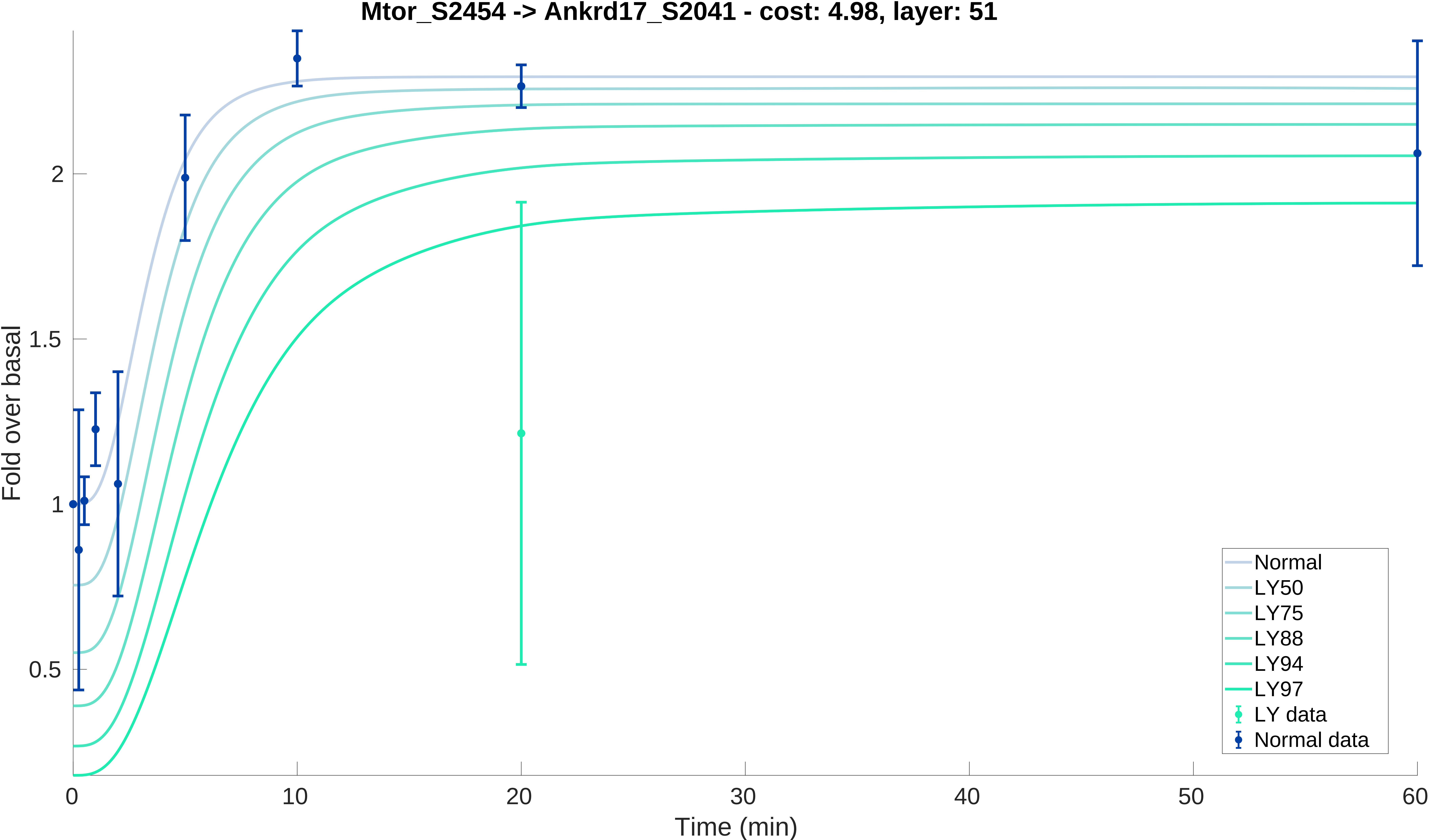

Mtor_S2454 -> Ankrd17_S2041 - cost: 4.98, layer: 51
Fold over basal
2
1.5
1
0.5
0
10
20
30
40
50
60
Time (min)
Normal
LY50
LY75
LY88
LY94
LY97
LY data
Normal data

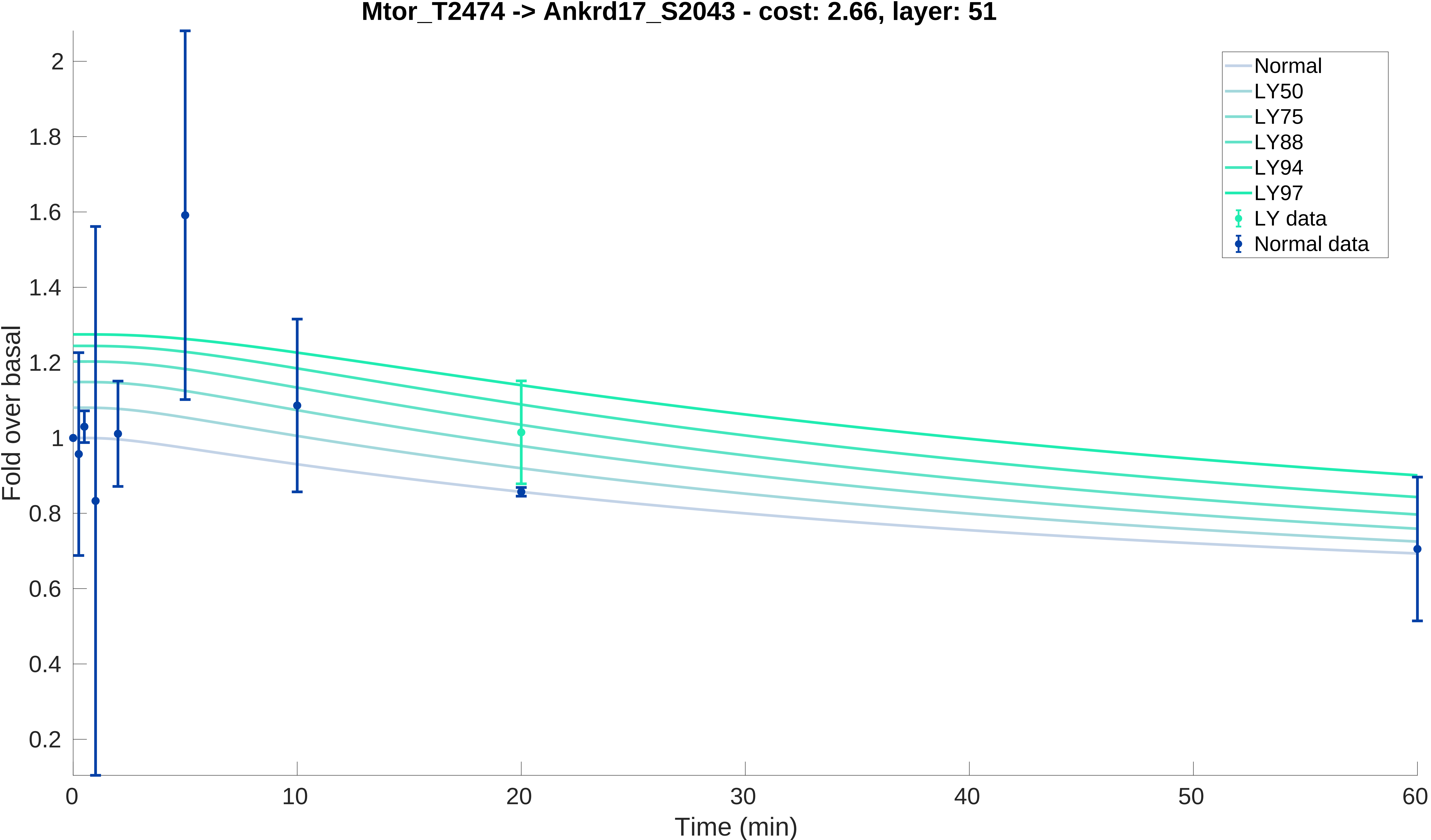

Mtor_T2474 -> Ankrd17_S2043 - cost: 2.66, layer: 51
Fold over basal
Time (min)
0
10
20
30
40
50
60
0.2
0.4
0.6
0.8
1
1.2
1.4
1.6
1.8
2
Normal
LY50
LY75
LY88
LY94
LY97
LY data
Normal data

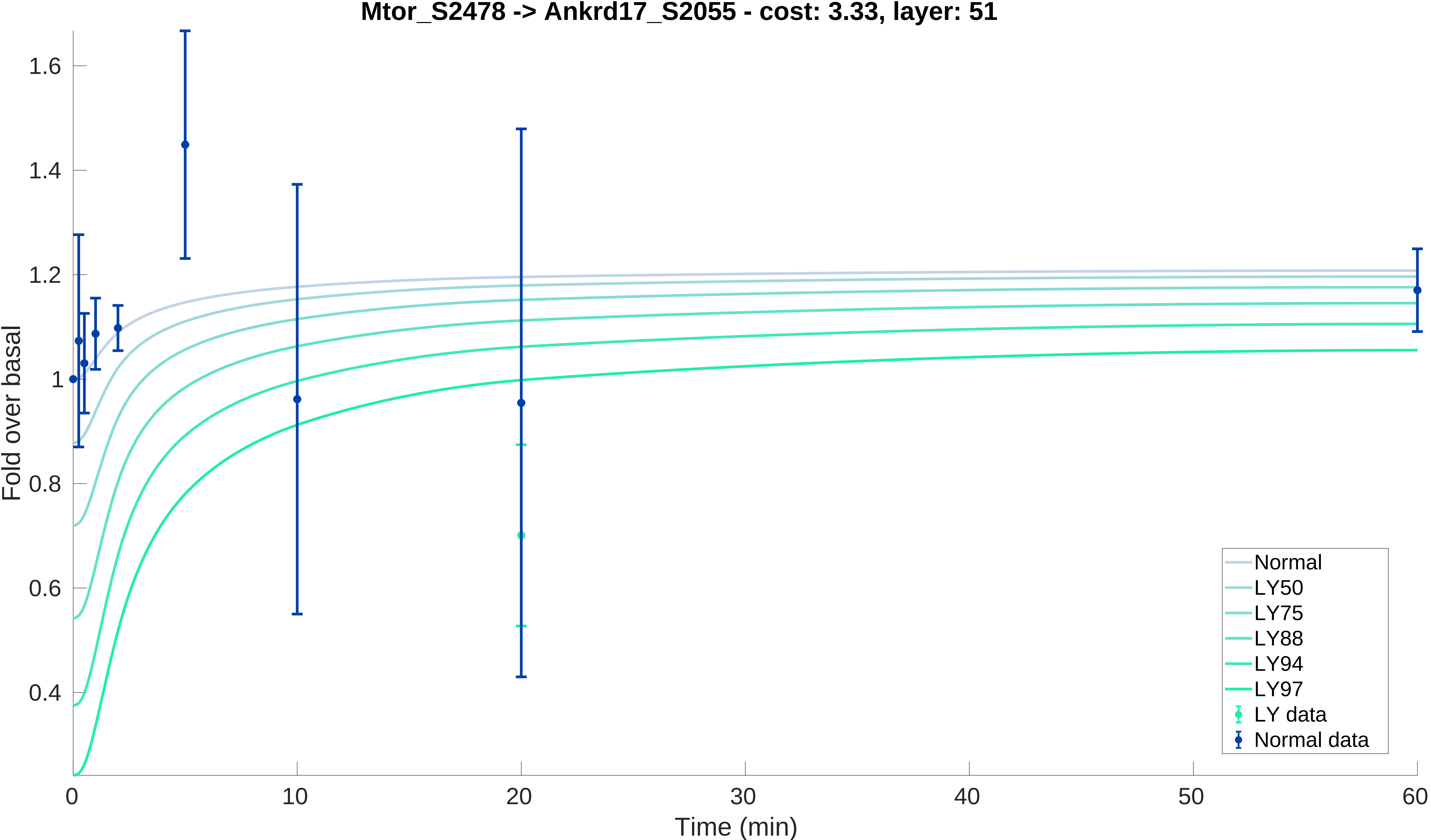
Mtor_S2478 -> Ankrd17_S2055 - cost: 3.33, layer: 51
Fold over basal
1.6
1.4
1.2
1
0.8
0.6
0.4
0
10
20
30
40
50
60
Time (min)
Normal
LY50
LY75
LY88
LY94
LY97
LY data
Normal data

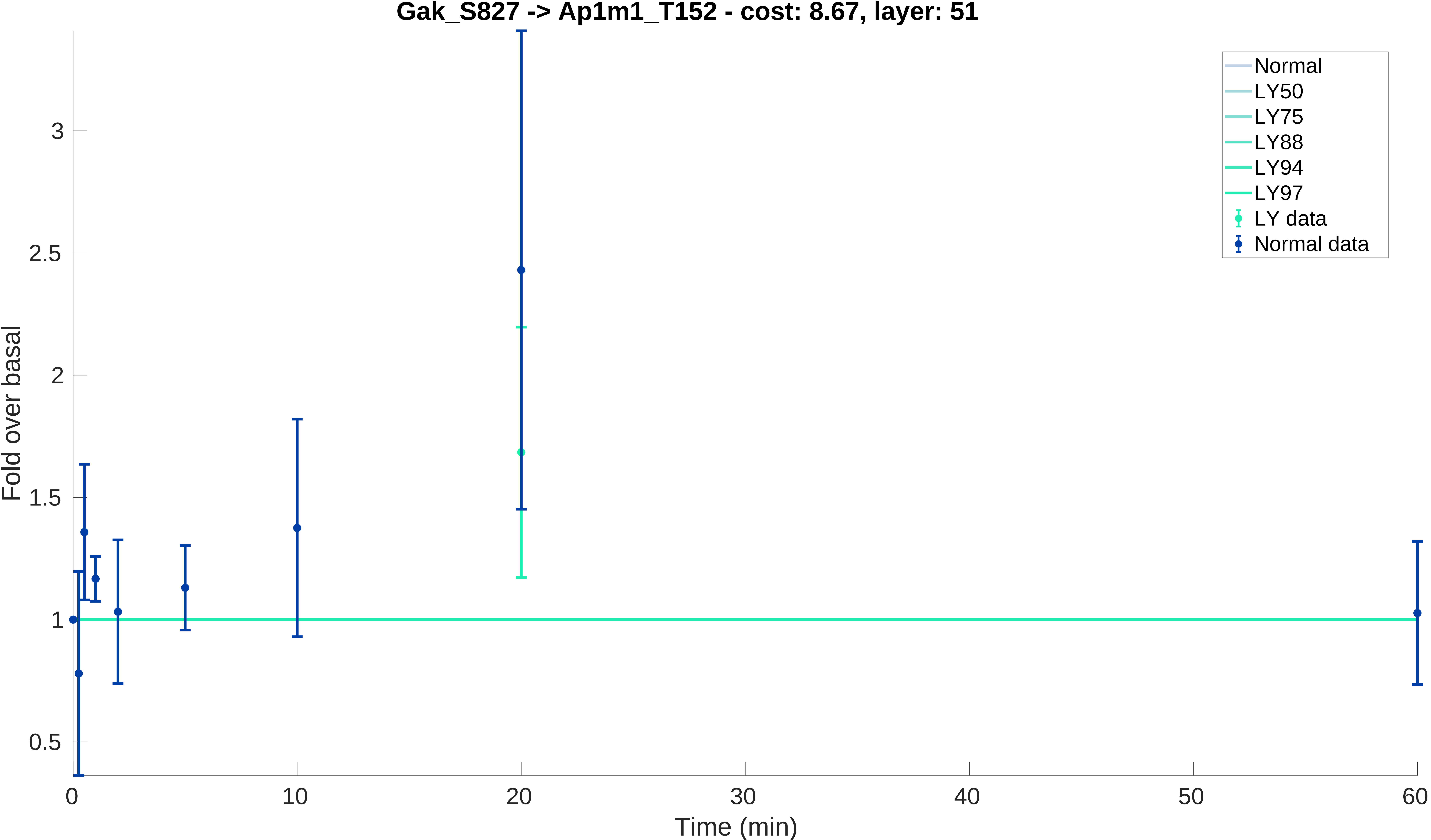

Gak_S827 -> Ap1m1_T152 - cost: 8.67, layer: 51
Normal
LY50
LY75
LY88
LY94
LY97
LY data
Normal data
Fold over basal
Time (min)
0.5
1
1.5
2
2.5
3
0
10
20
30
40
50
60

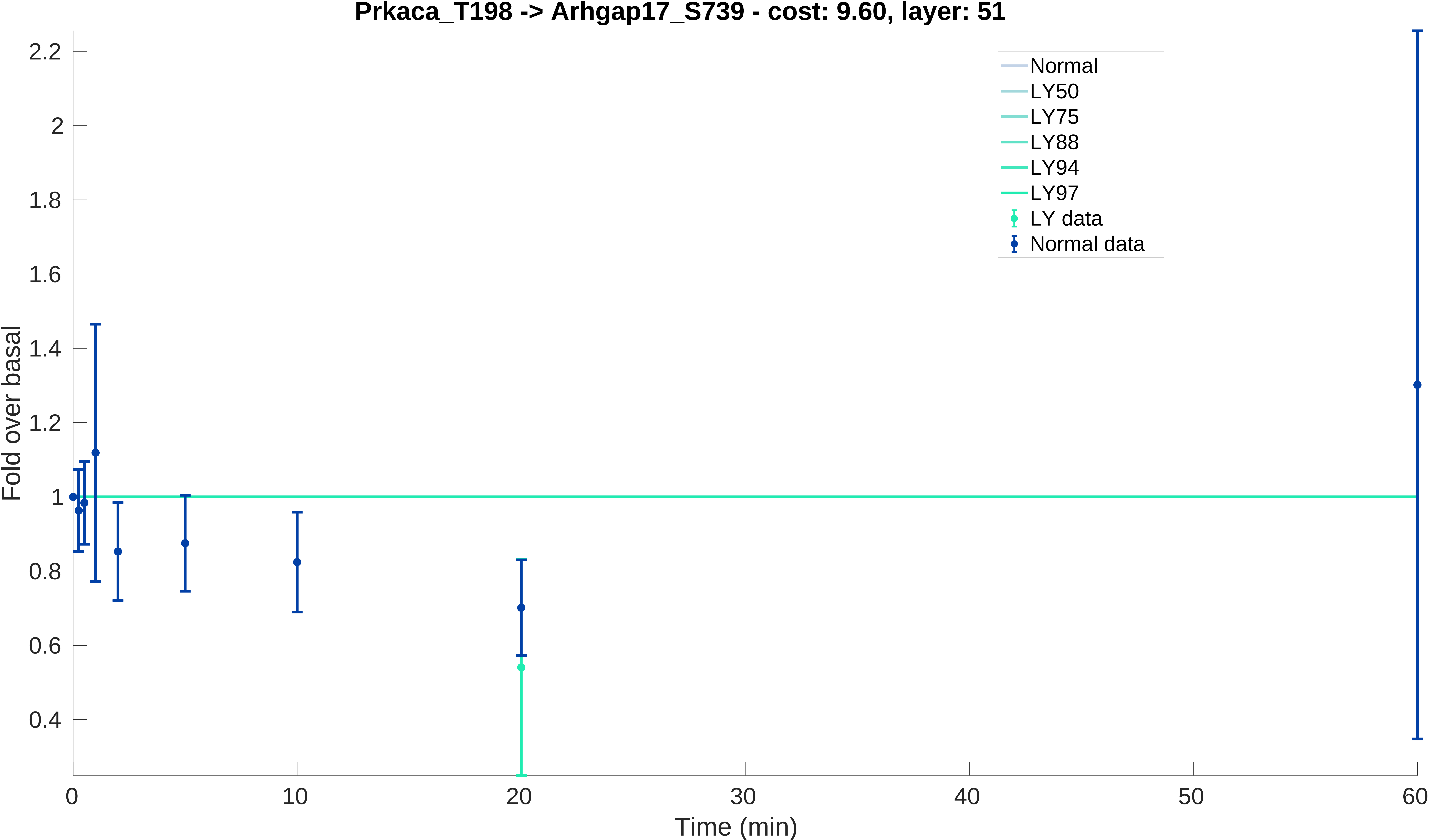
Prkaca_T198 -> Arhgap17_S739 - cost: 9.60, layer: 51
Fold over basal
2.2
2
1.8
1.6
1.4
1.2
1
0.8
0.6
0.4
0
10
20
30
40
50
60
Time (min)
Normal
LY50
LY75
LY88
LY94
LY97
LY data
Normal data

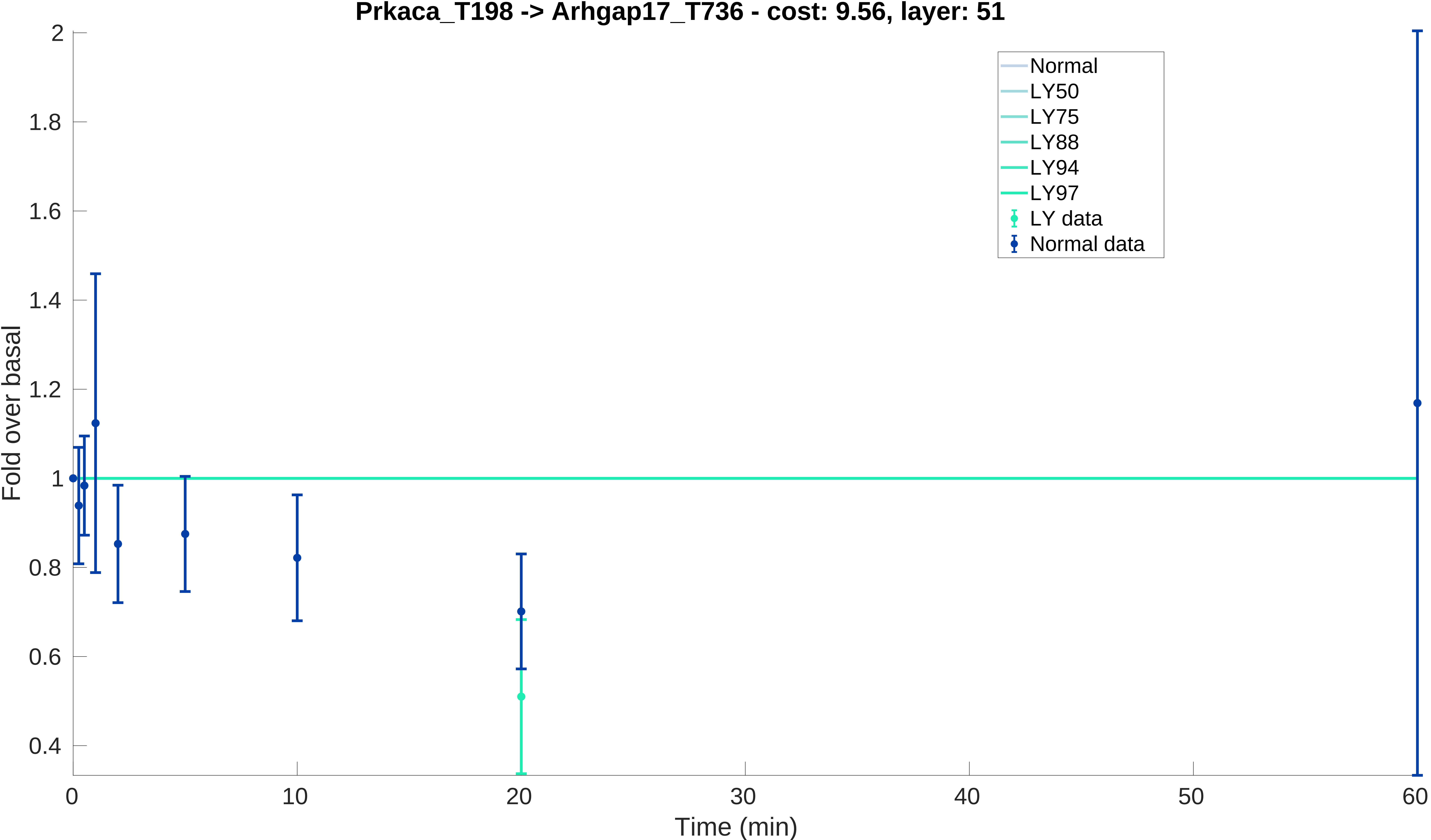

Prkaca_T198 -> Arhgap17_T736 - cost: 9.56, layer: 51
Fold over basal
Time (min)
2
1.8
1.6
1.4
1.2
1
0.8
0.6
0.4
0
10
20
30
40
50
60
Normal
LY50
LY75
LY88
LY94
LY97
LY data
Normal data

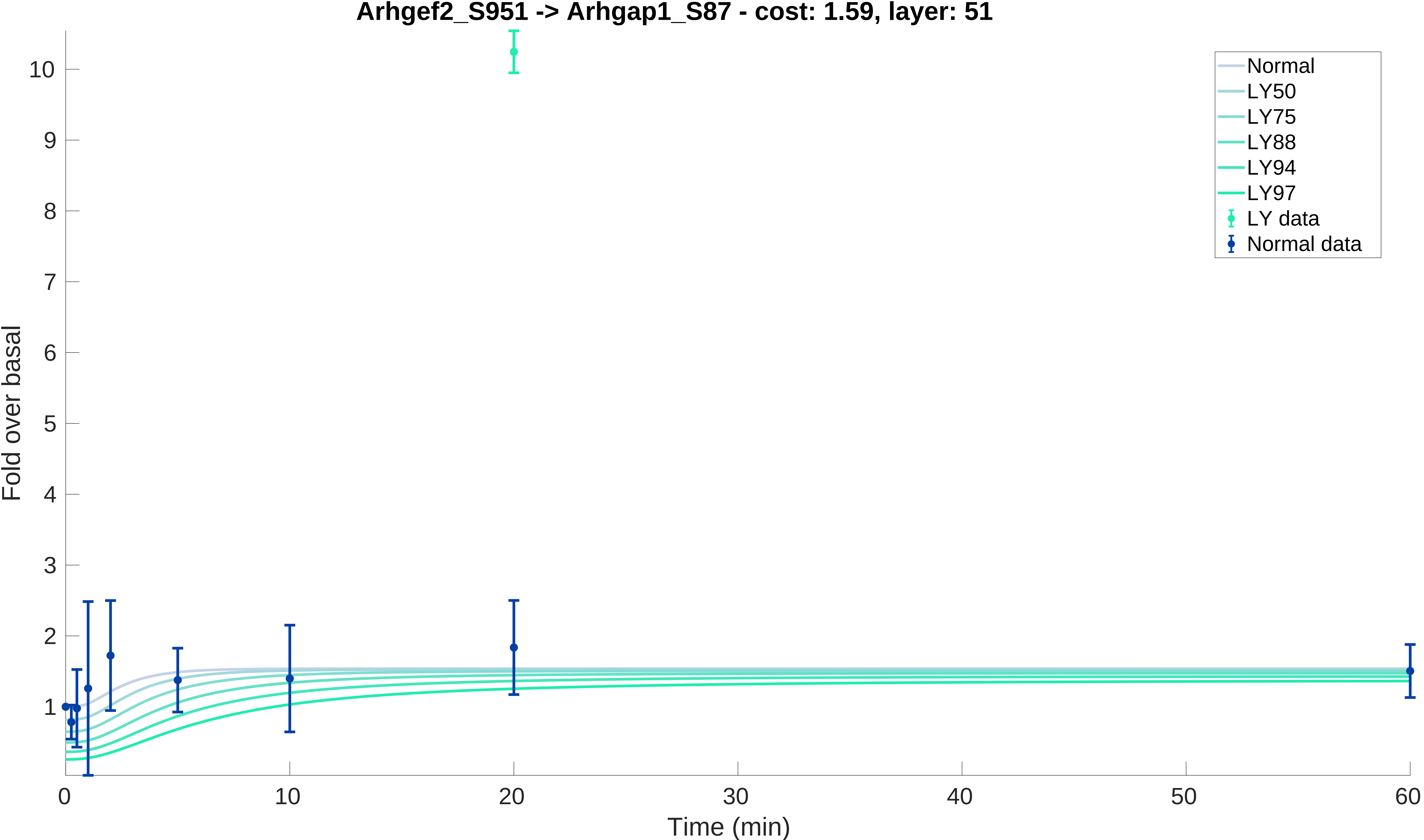

Arhgef2_S951 -> Arhgap1_S87 - cost: 1.59, layer: 51
Fold over basal
Time (min)
0
10
20
30
40
50
60
1
2
3
4
5
6
7
8
9
10
Normal
LY50
LY75
LY88
LY94
LY97
LY data
Normal data

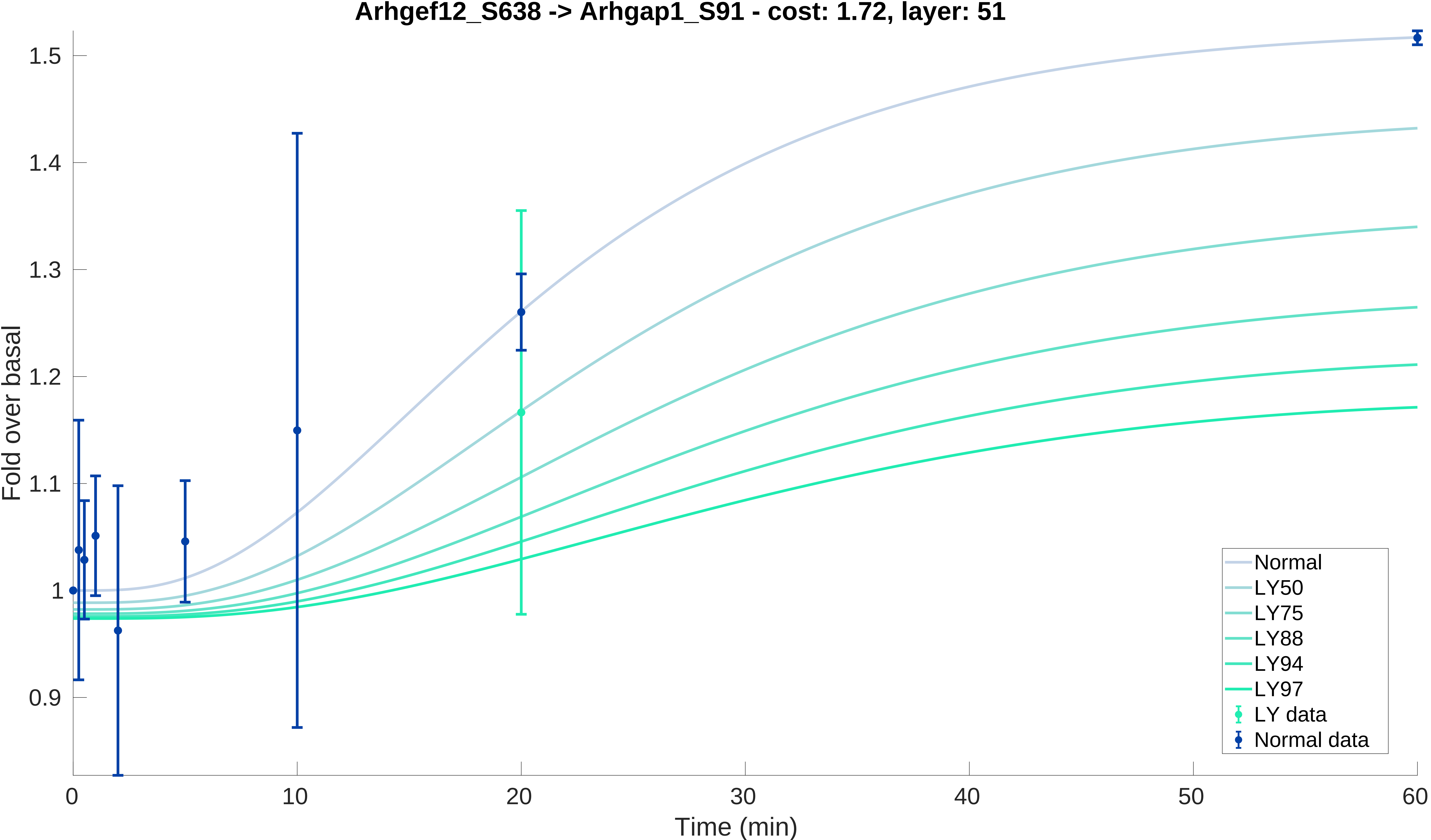

Arhgef12_S638 -> Arhgap1_S91 - cost: 1.72, layer: 51
Fold over basal
Time (min)
1.5
1.4
1.3
1.2
1.1
1
0.9
0
10
20
30
40
50
60
Normal
LY50
LY75
LY88
LY94
LY97
LY data
Normal data

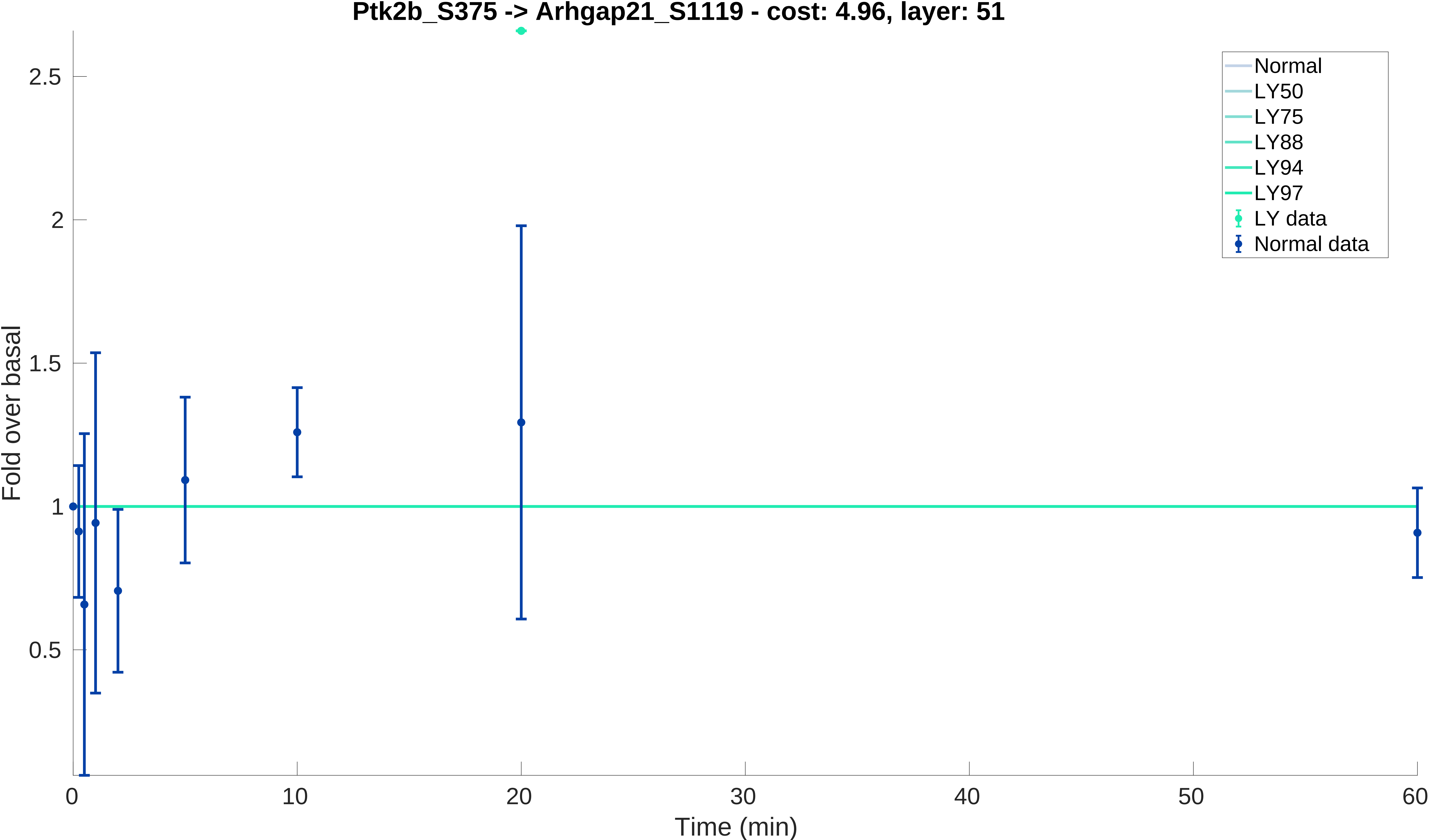

Ptk2b_S375 -> Arhgap21_S1119 - cost: 4.96, layer: 51
Fold over basal
Time (min)
0
10
20
30
40
50
60
0.5
1
1.5
2
2.5
Normal
LY50
LY75
LY88
LY94
LY97
LY data
Normal data

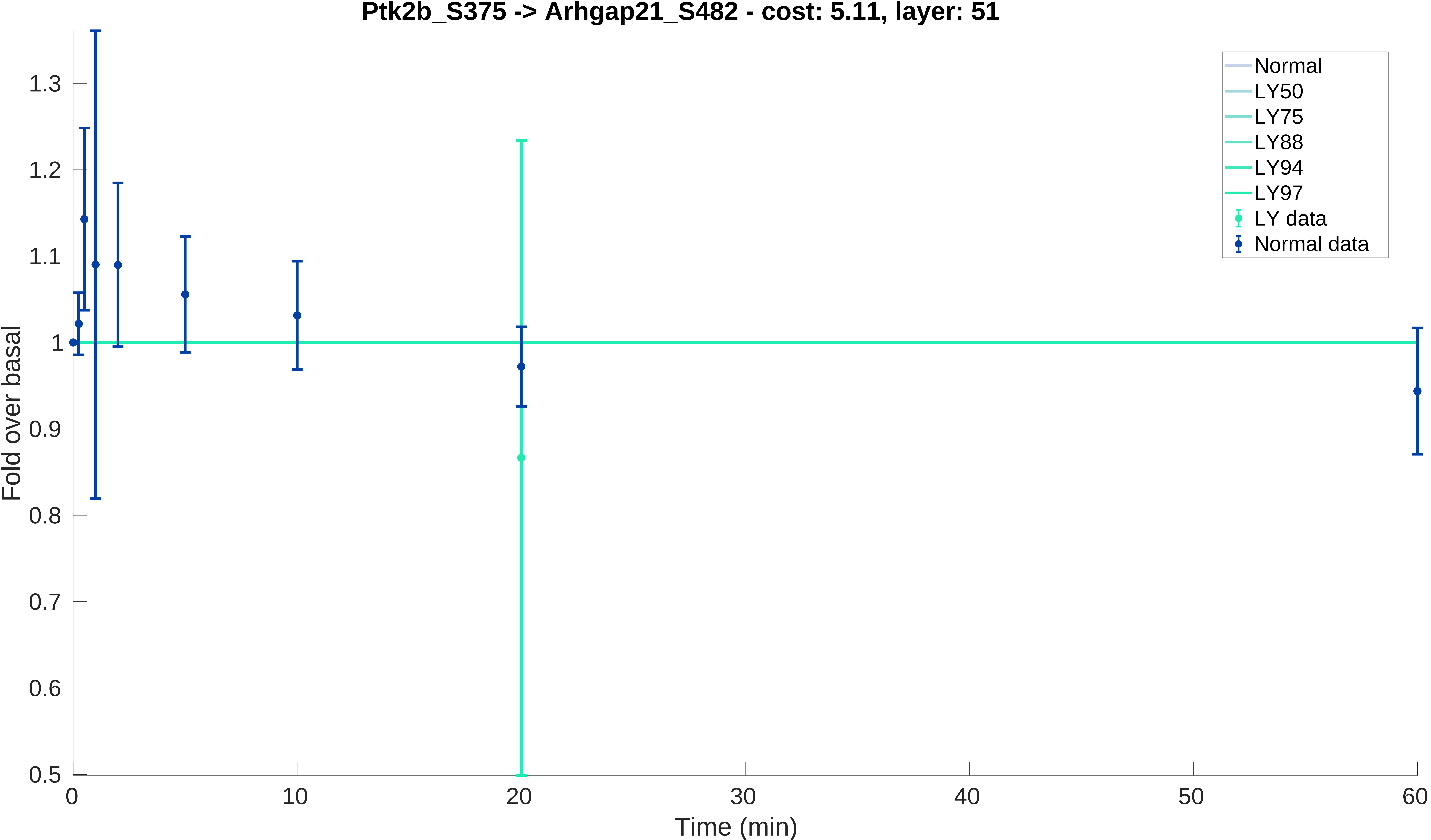

Ptk2b_S375 -> Arhgap21_S482 - cost: 5.11, layer: 51
Normal
LY50
LY75
LY88
LY94
LY97
LY data
Normal data
Fold over basal
Time (min)
1.3
1.2
1.1
1
0.9
0.8
0.7
0.6
0.5
0
10
20
30
40
50
60

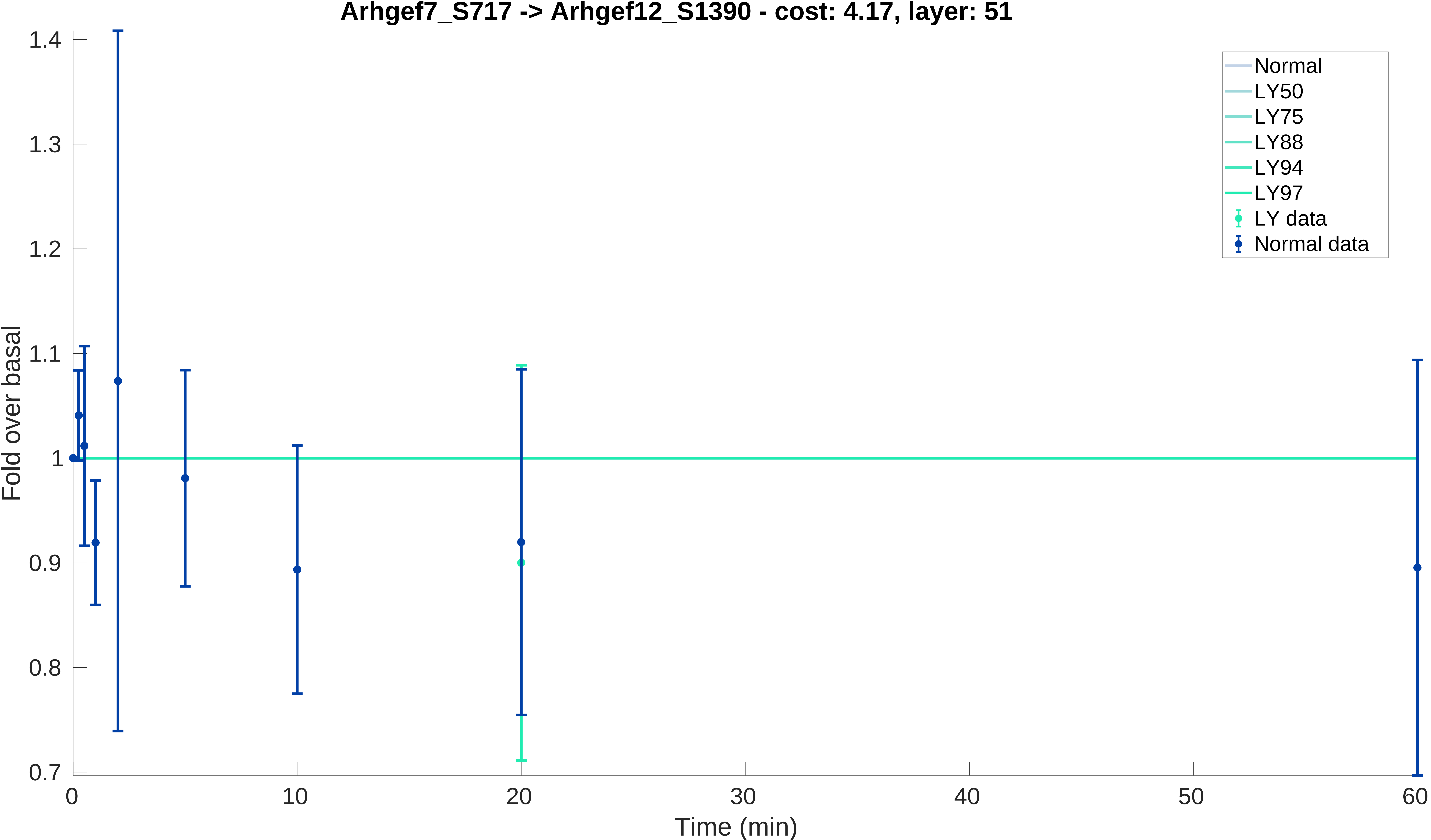

Arhgef7_S717 -> Arhgef12_S1390 - cost: 4.17, layer: 51
Fold over basal
Time (min)
0.7
0.8
0.9
1
1.1
1.2
1.3
1.4
0
10
20
30
40
50
60
Normal
LY50
LY75
LY88
LY94
LY97
LY data
Normal data

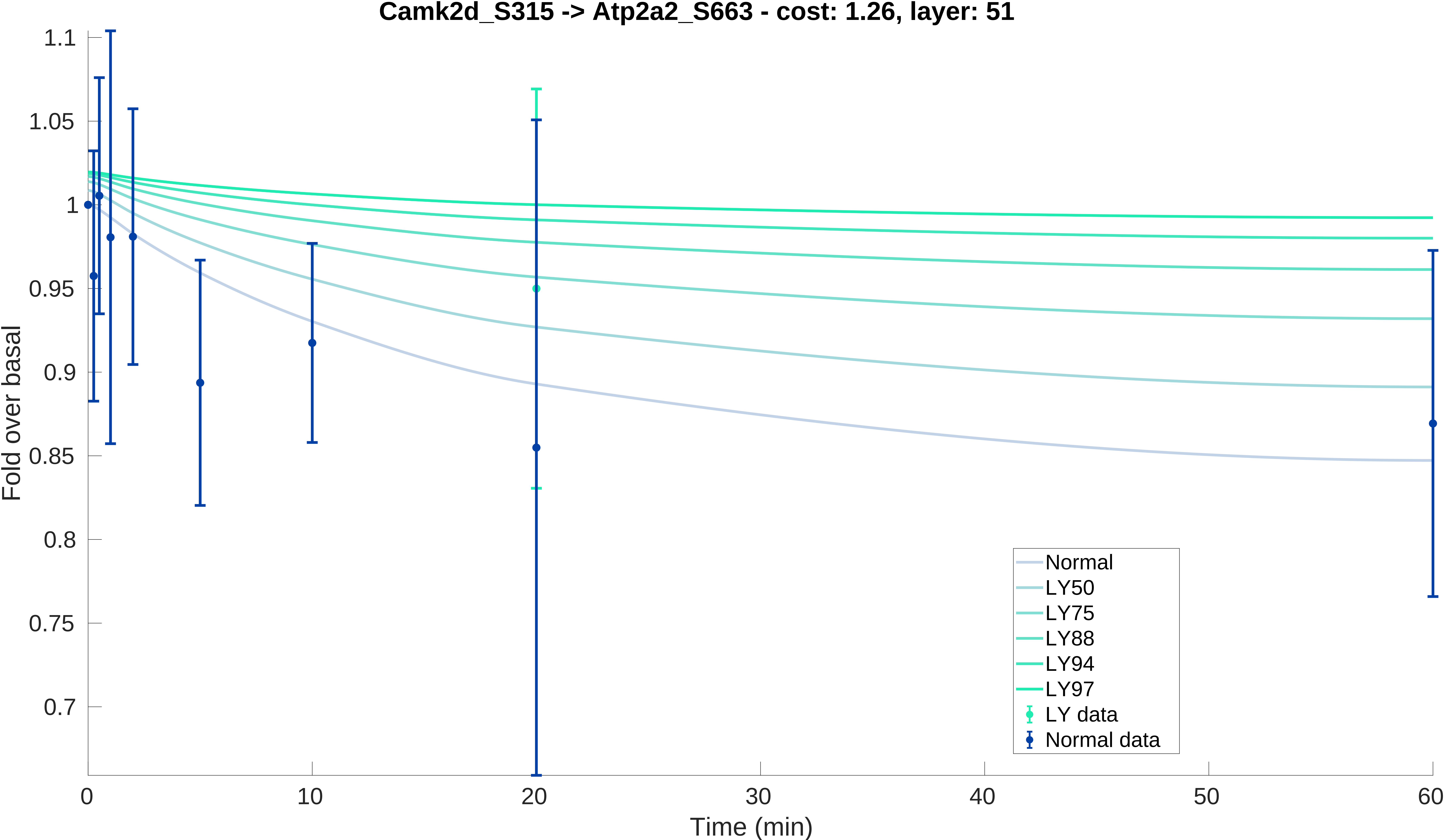

Camk2d_S315 -> Atp2a2_S663 - cost: 1.26, layer: 51
Fold over basal
Time (min)
1.1
1.05
1
0.95
0.9
0.85
0.8
0.75
0.7
0
10
20
30
40
50
60
Normal
LY50
LY75
LY88
LY94
LY97
LY data
Normal data

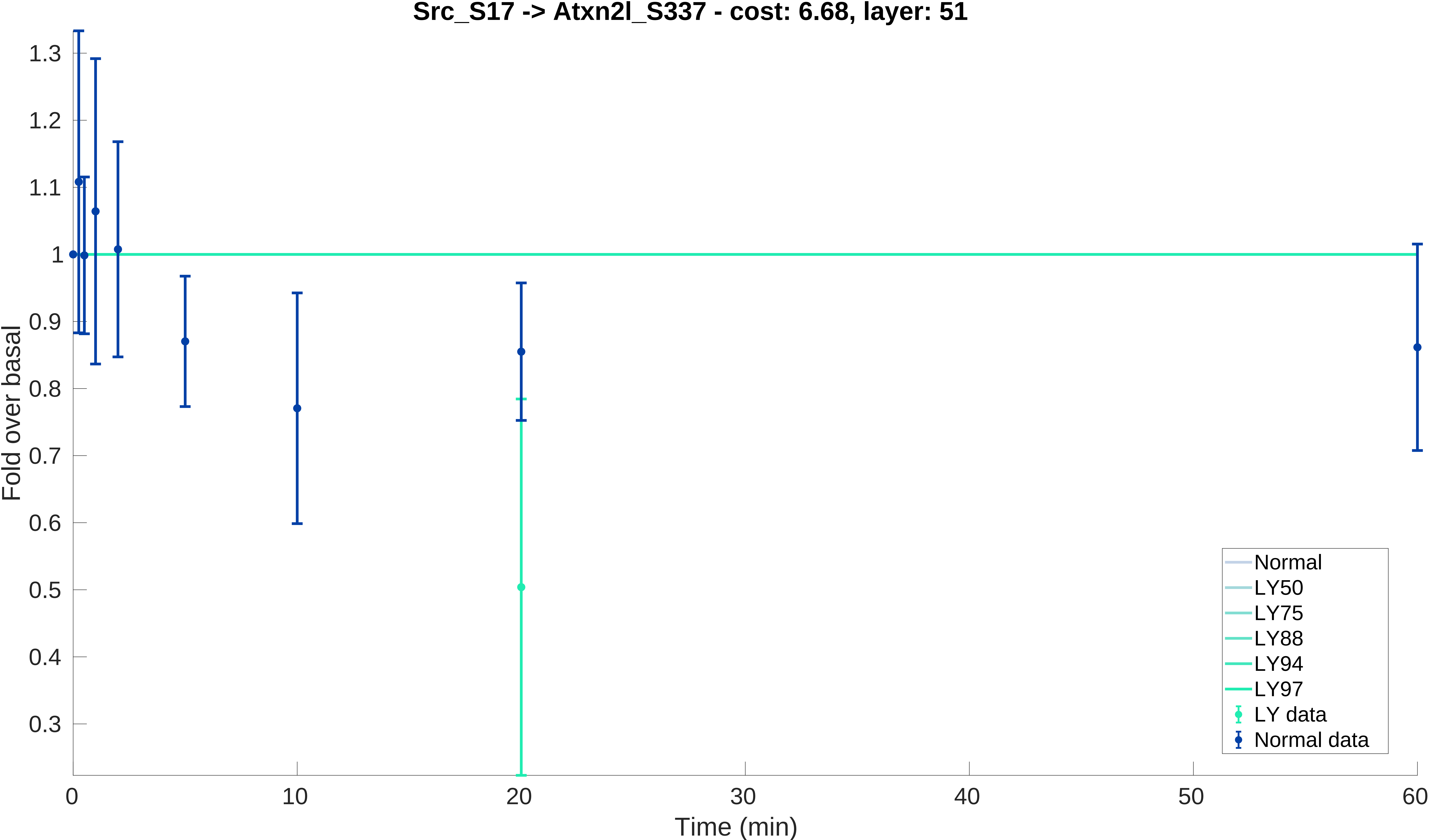

Src_S17 -> Atxn2l_S337 - cost: 6.68, layer: 51
Fold over basal
1.3
1.2
1.1
1
0.9
0.8
0.7
0.6
0.5
0.4
0.3
0
10
20
30
40
50
60
Time (min)
Normal
LY50
LY75
LY88
LY94
LY97
LY data
Normal data

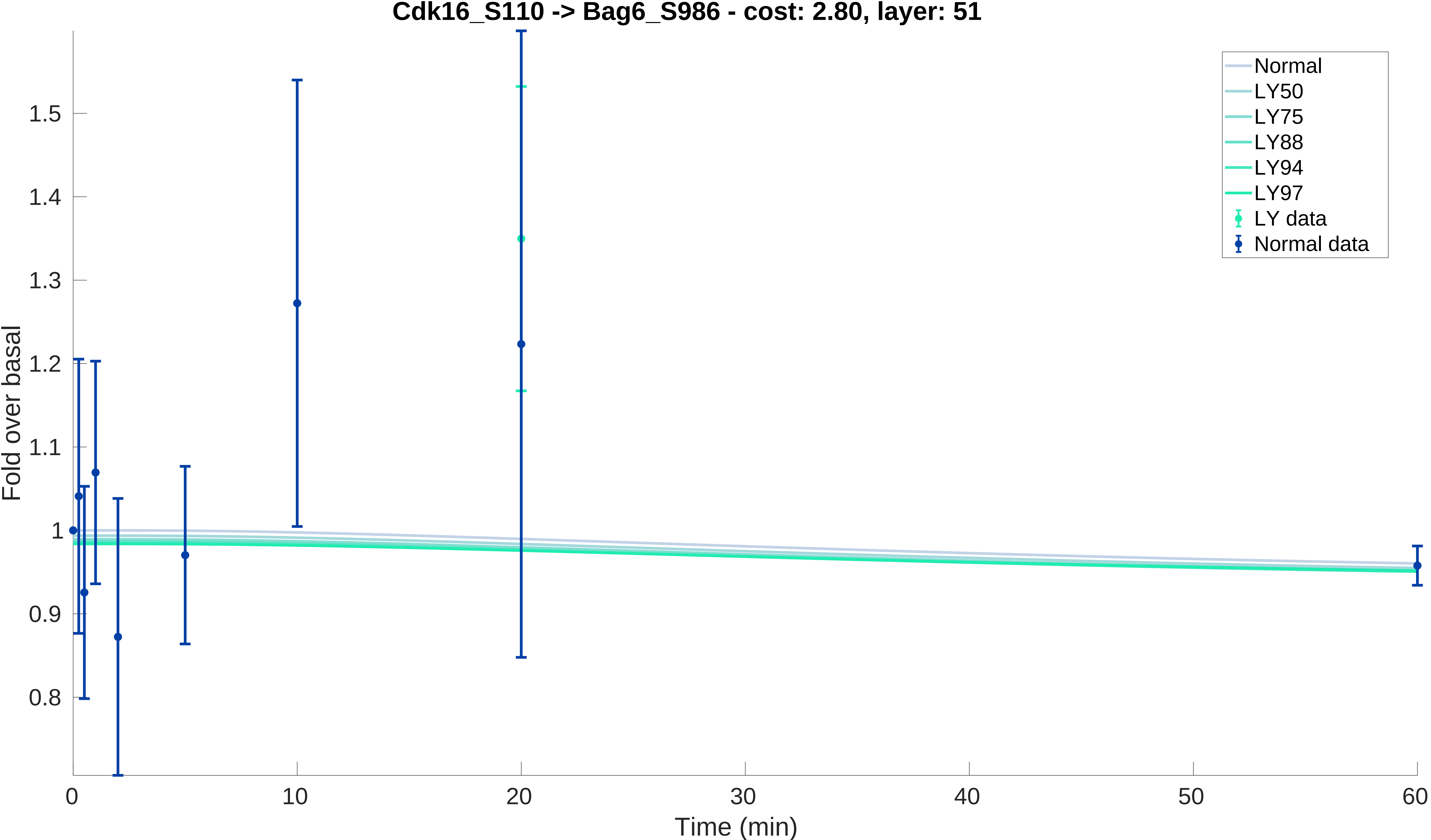

Cdk16_S110 -> Bag6_S986 - cost: 2.80, layer: 51
Fold over basal
Time (min)
1.5
1.4
1.3
1.2
1.1
1
0.9
0.8
0
10
20
30
40
50
60
Normal
LY50
LY75
LY88
LY94
LY97
LY data
Normal data

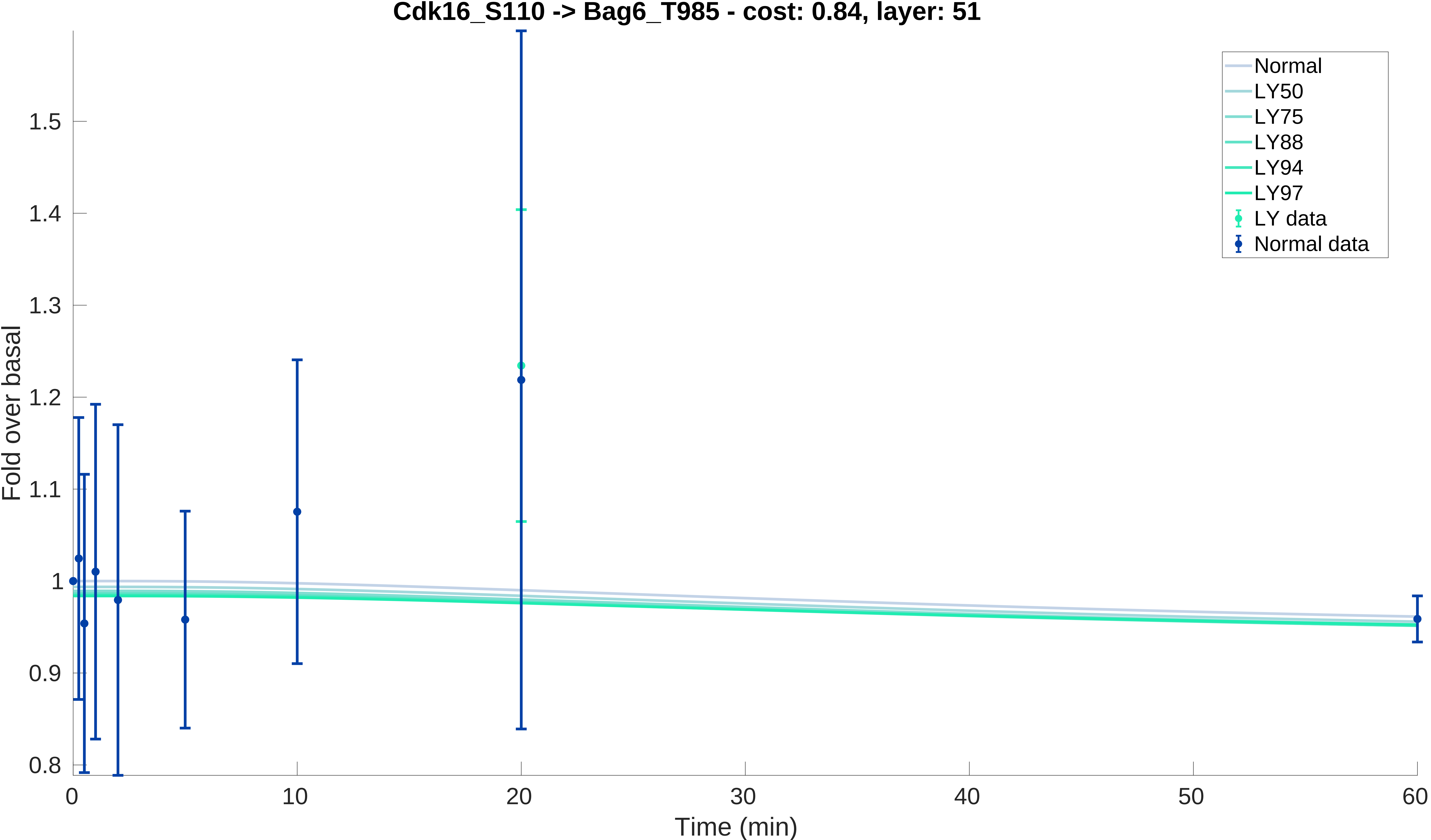

Cdk16_S110 -> Bag6_T985 - cost: 0.84, layer: 51
Normal
LY50
LY75
LY88
LY94
LY97
LY data
Normal data
Fold over basal
1.5
1.4
1.3
1.2
1.1
1
0.9
0.8
0
10
20
30
40
50
60
Time (min)

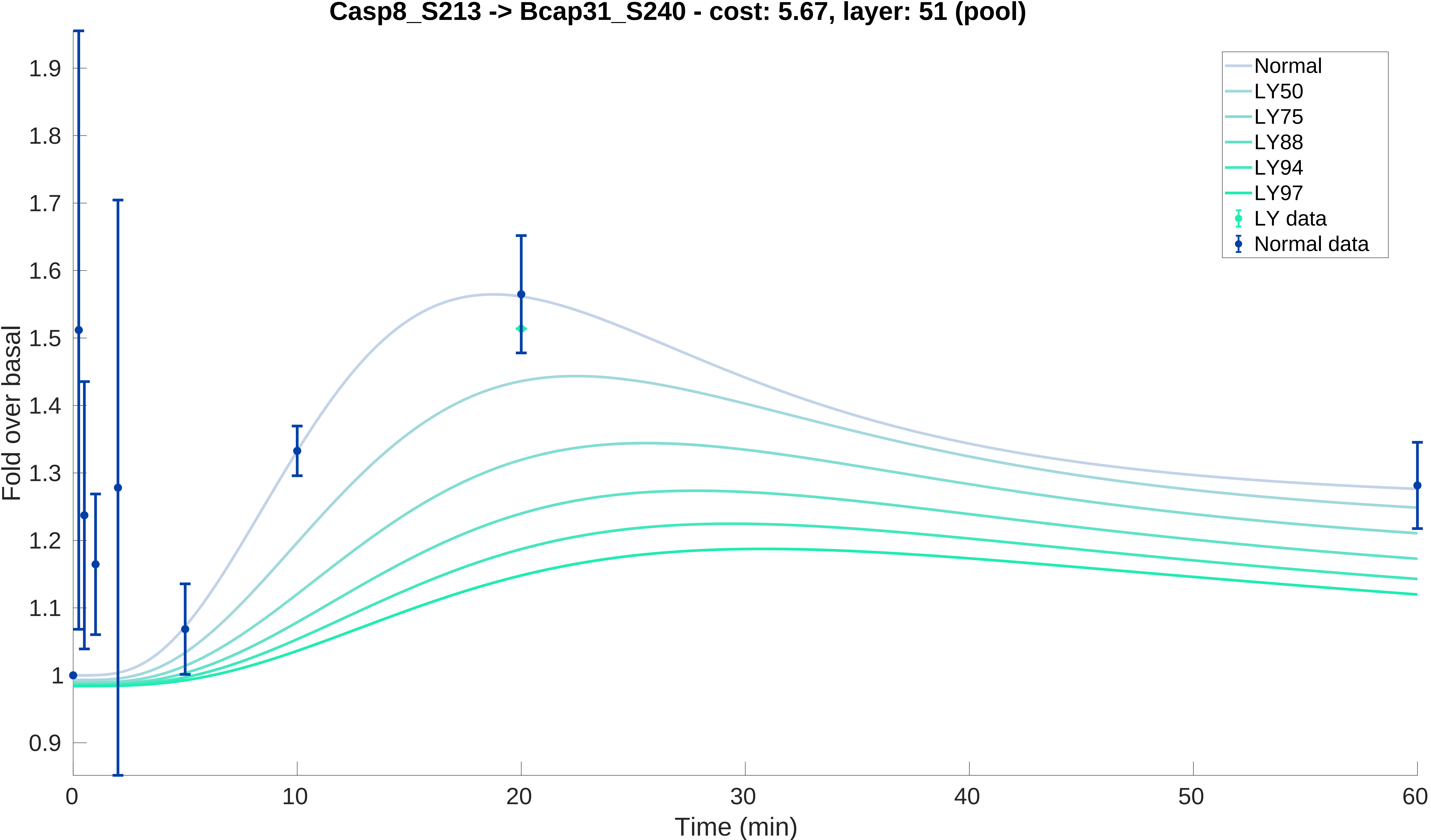

Casp8_S213 -> Bcap31_S240 - cost: 5.67, layer: 51 (pool)
Normal
LY50
LY75
LY88
LY94
LY97
LY data
Normal data
Fold over basal
1.9
1.8
1.7
1.6
1.5
1.4
1.3
1.2
1.1
1
0.9
0
10
20
30
40
50
60
Time (min)

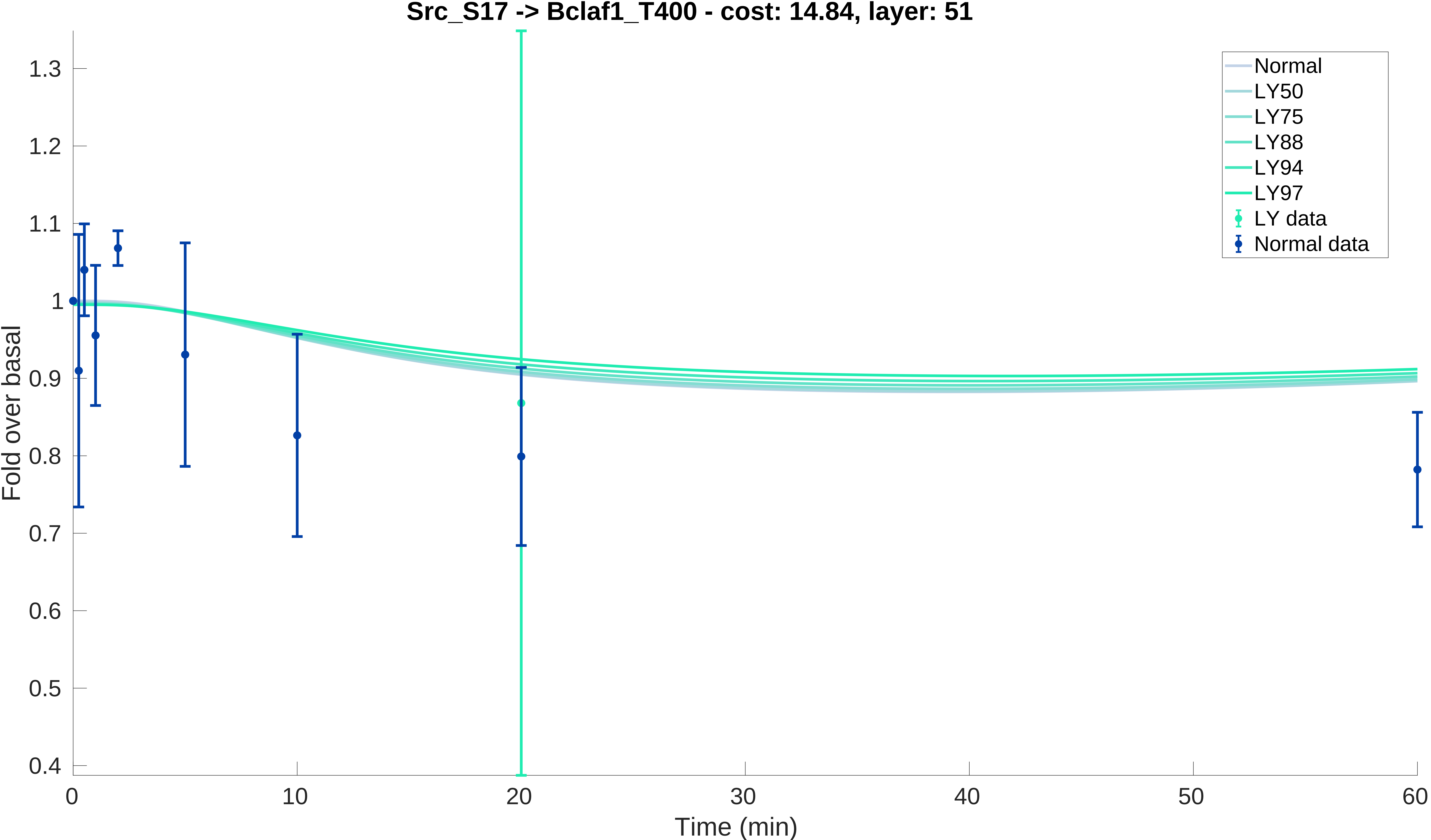
Src_S17 -> Bclaf1_T400 - cost: 14.84, layer: 51
Normal
LY50
LY75
LY88
LY94
LY97
LY data
Normal data
Fold over basal
Time (min)
1.3
1.2
1.1
1
0.9
0.8
0.7
0.6
0.5
0.4
0
10
20
30
40
50
60

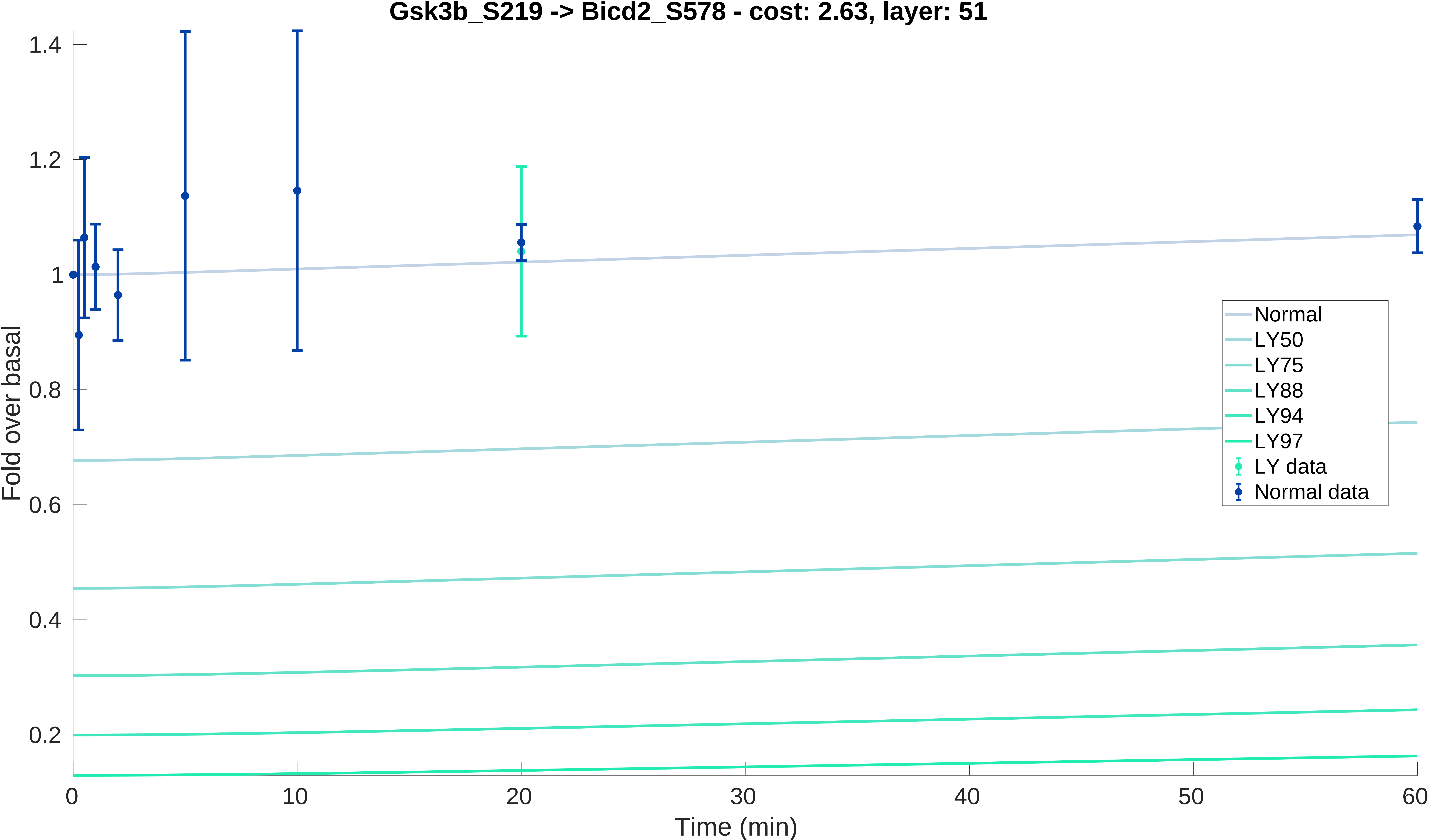

Gsk3b_S219 -> Bicd2_S578 - cost: 2.63, layer: 51
1.4
1.2
1
0.8
0.6
0.4
0.2
Fold over basal
0
10
20
30
40
50
60
Time (min)
Normal
LY50
LY75
LY88
LY94
LY97
LY data
Normal data

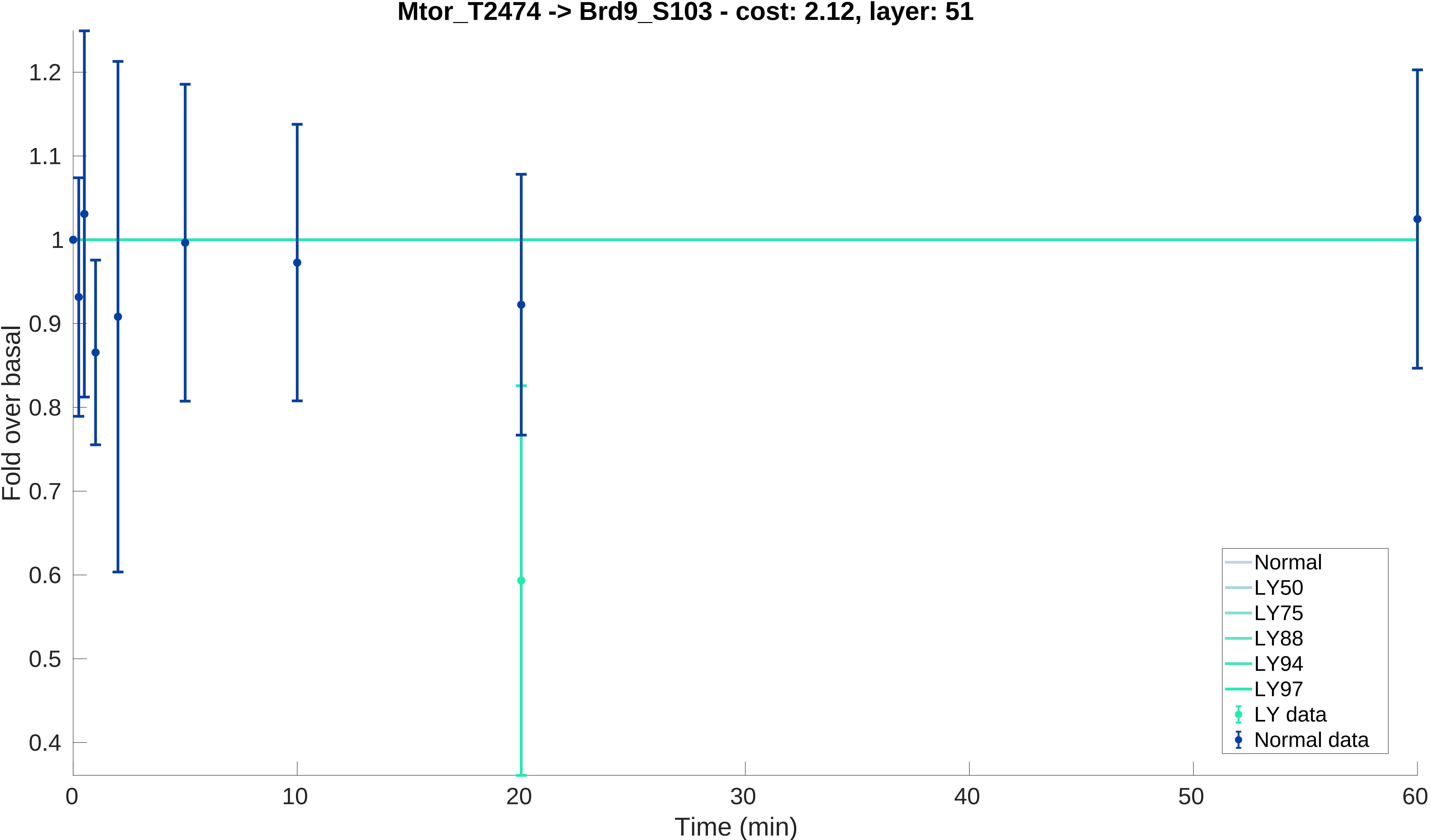

Mtor_T2474 -> Brd9_S103 - cost: 2.12, layer: 51
Fold over basal
1.2
1.1
1
0.9
0.8
0.7
0.6
0.5
0.4
0
10
20
30
40
50
60
Time (min)
Normal
LY50
LY75
LY88
LY94
LY97
LY data
Normal data

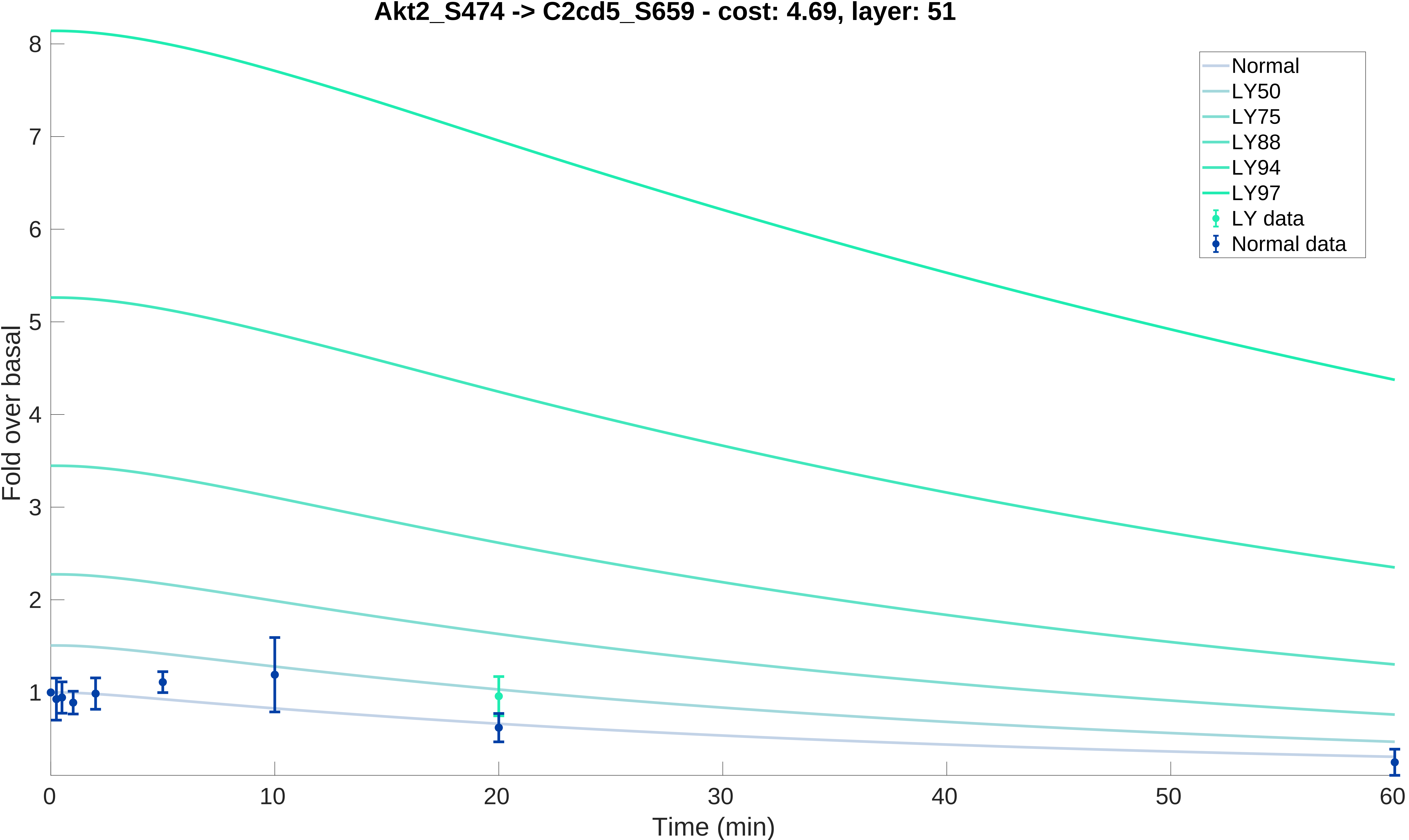

Akt2_S474 -> C2cd5_S659 - cost: 4.69, layer: 51
Fold over basal
Time (min)
Normal
LY50
LY75
LY88
LY94
LY97
LY data
Normal data
0
10
20
30
40
50
60
1
2
3
4
5
6
7
8

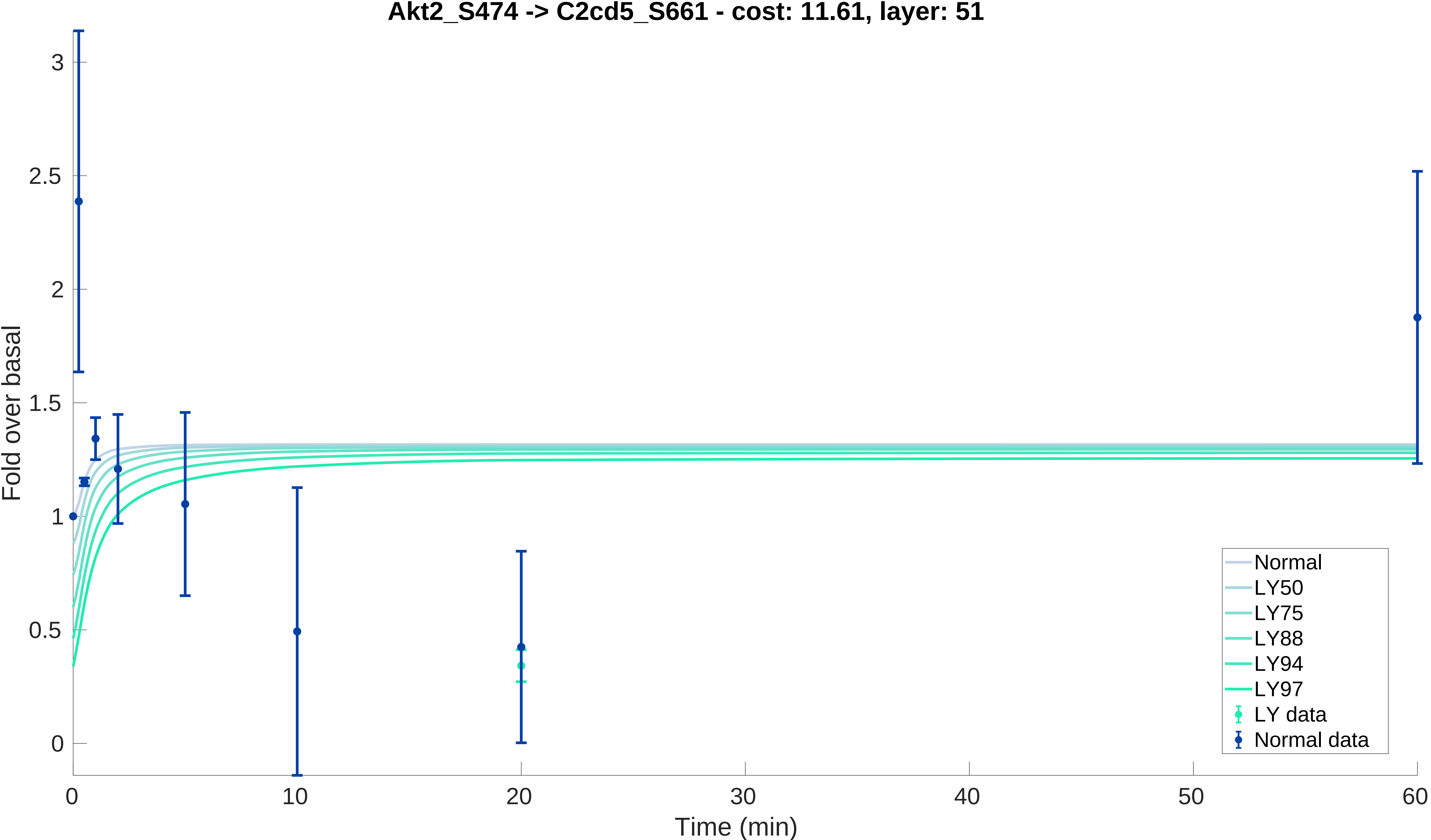

Akt2_S474 -> C2cd5_S661 - cost: 11.61, layer: 51
Fold over basal
Time (min)
0
0.5
1
1.5
2
2.5
3
0
10
20
30
40
50
60
Normal
LY50
LY75
LY88
LY94
LY97
LY data
Normal data

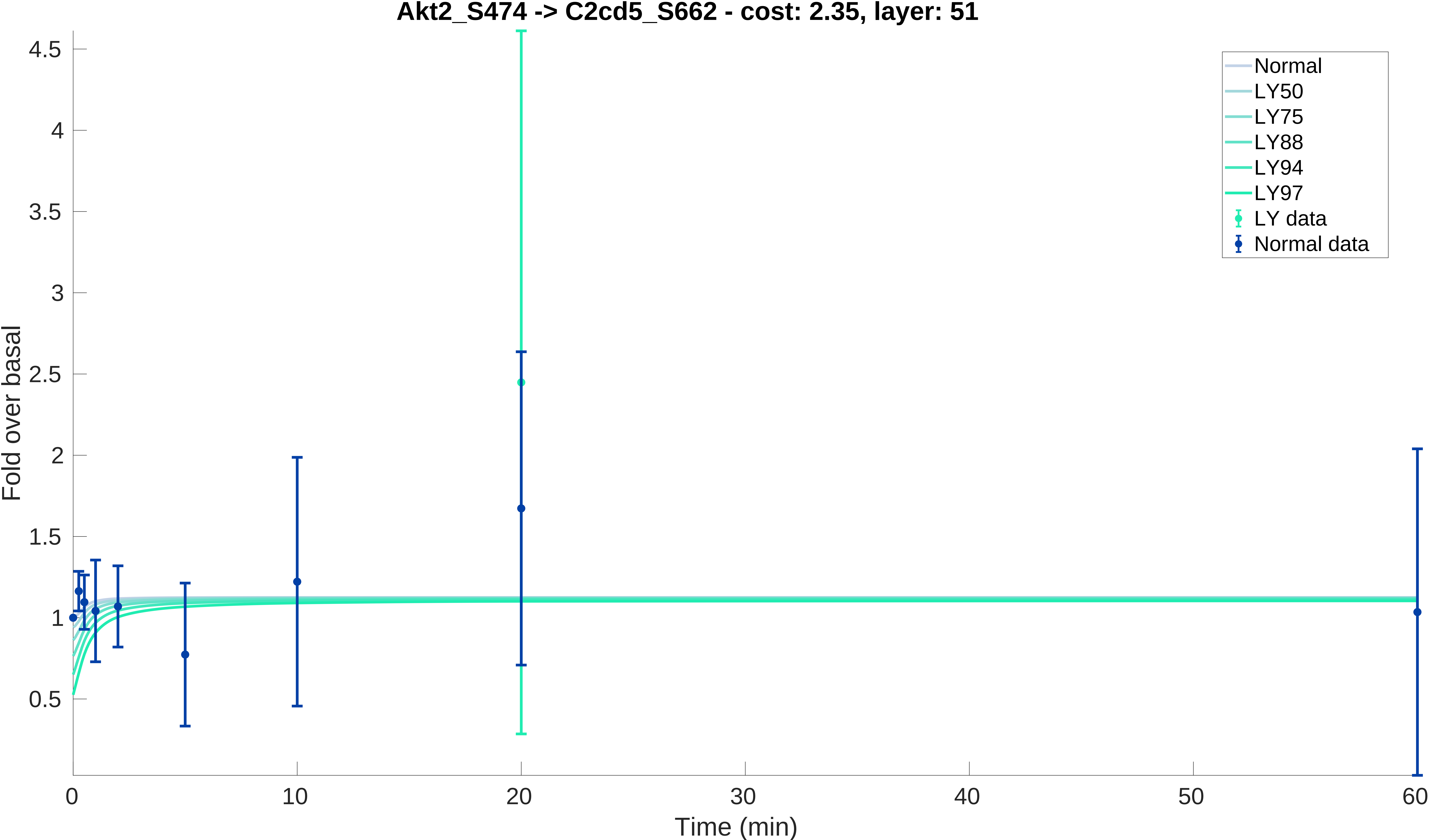
Akt2_S474 -> C2cd5_S662 - cost: 2.35, layer: 51
Fold over basal
4.5
4
3.5
3
2.5
2
1.5
1
0.5
0
10
20
30
40
50
60
Time (min)
Normal
LY50
LY75
LY88
LY94
LY97
LY data
Normal data

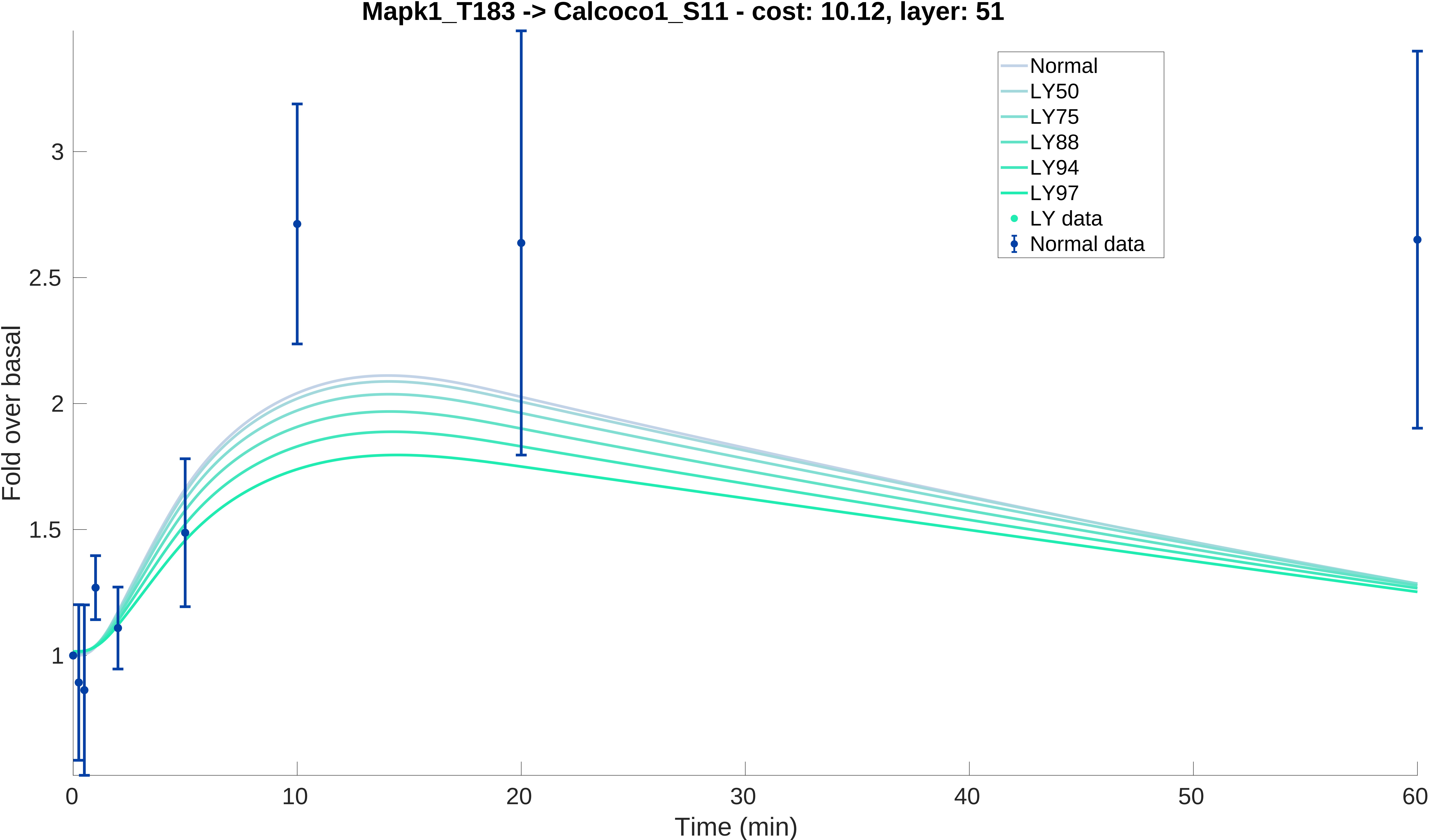

Mapk1_T183 -> Calcoco1_S11 - cost: 10.12, layer: 51
Normal
LY50
LY75
LY88
LY94
LY97
LY data
Normal data
Fold over basal
Time (min)
1
1.5
2
2.5
3
0
10
20
30
40
50
60

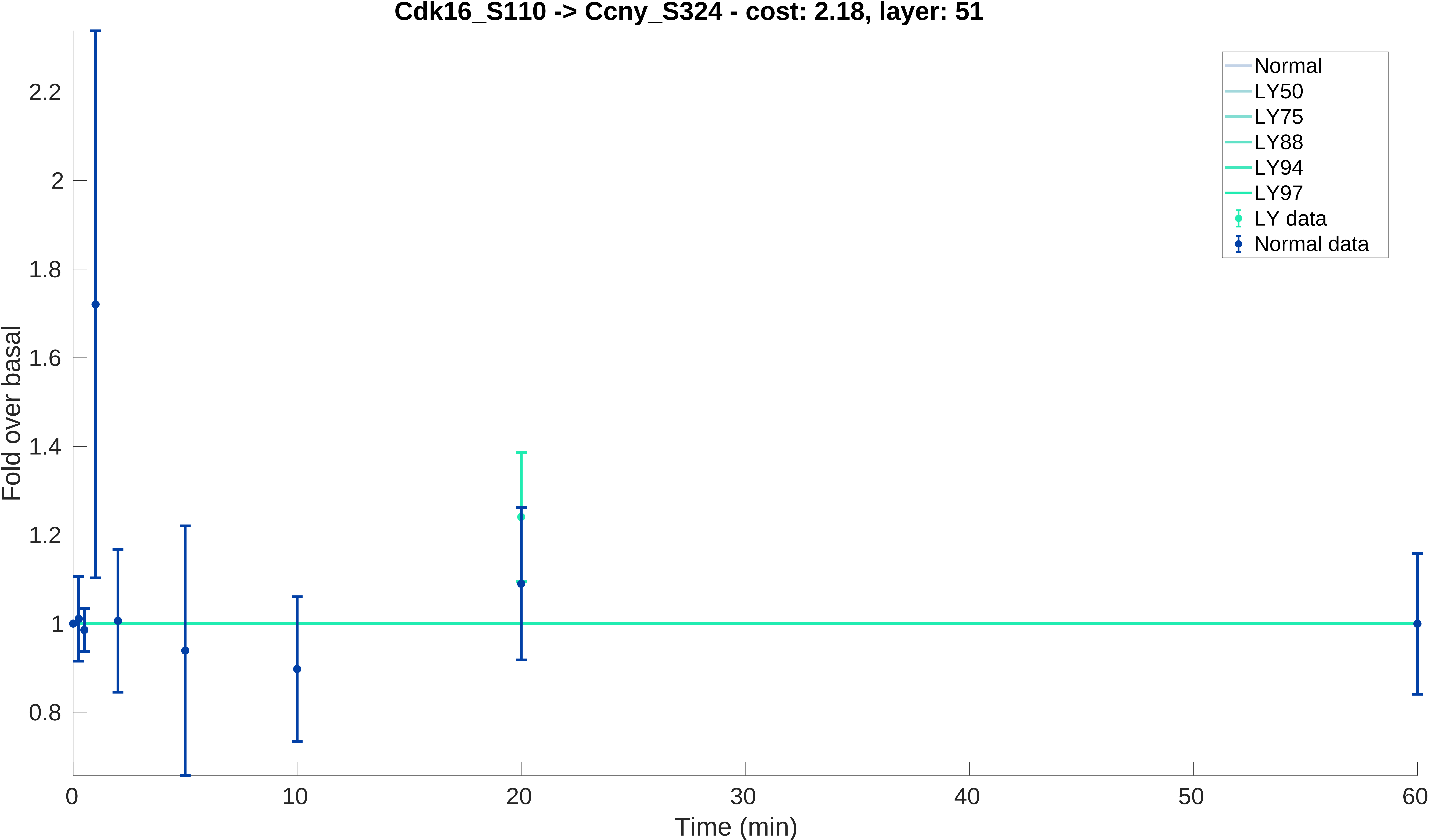
Cdk16_S110 -> Ccny_S324 - cost: 2.18, layer: 51
Fold over basal
Time (min)
0
10
20
30
40
50
60
0.8
1
1.2
1.4
1.6
1.8
2
2.2
Normal
LY50
LY75
LY88
LY94
LY97
LY data
Normal data

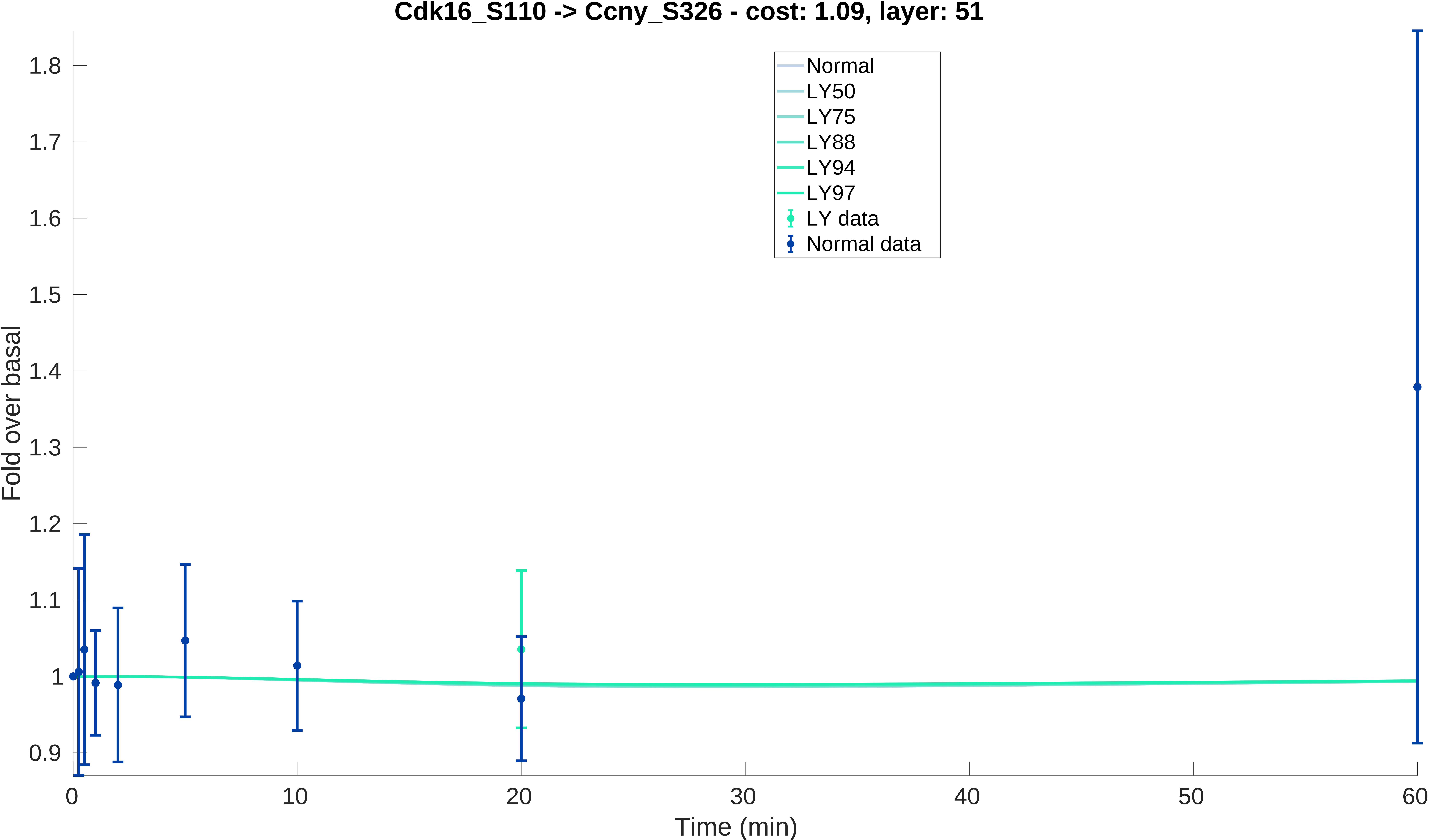
Cdk16_S110 -> Ccny_S326 - cost: 1.09, layer: 51
Normal
LY50
LY75
LY88
LY94
LY97
LY data
Normal data
Fold over basal
1.8
1.7
1.6
1.5
1.4
1.3
1.2
1.1
1
0.9
0
10
20
30
40
50
60
Time (min)

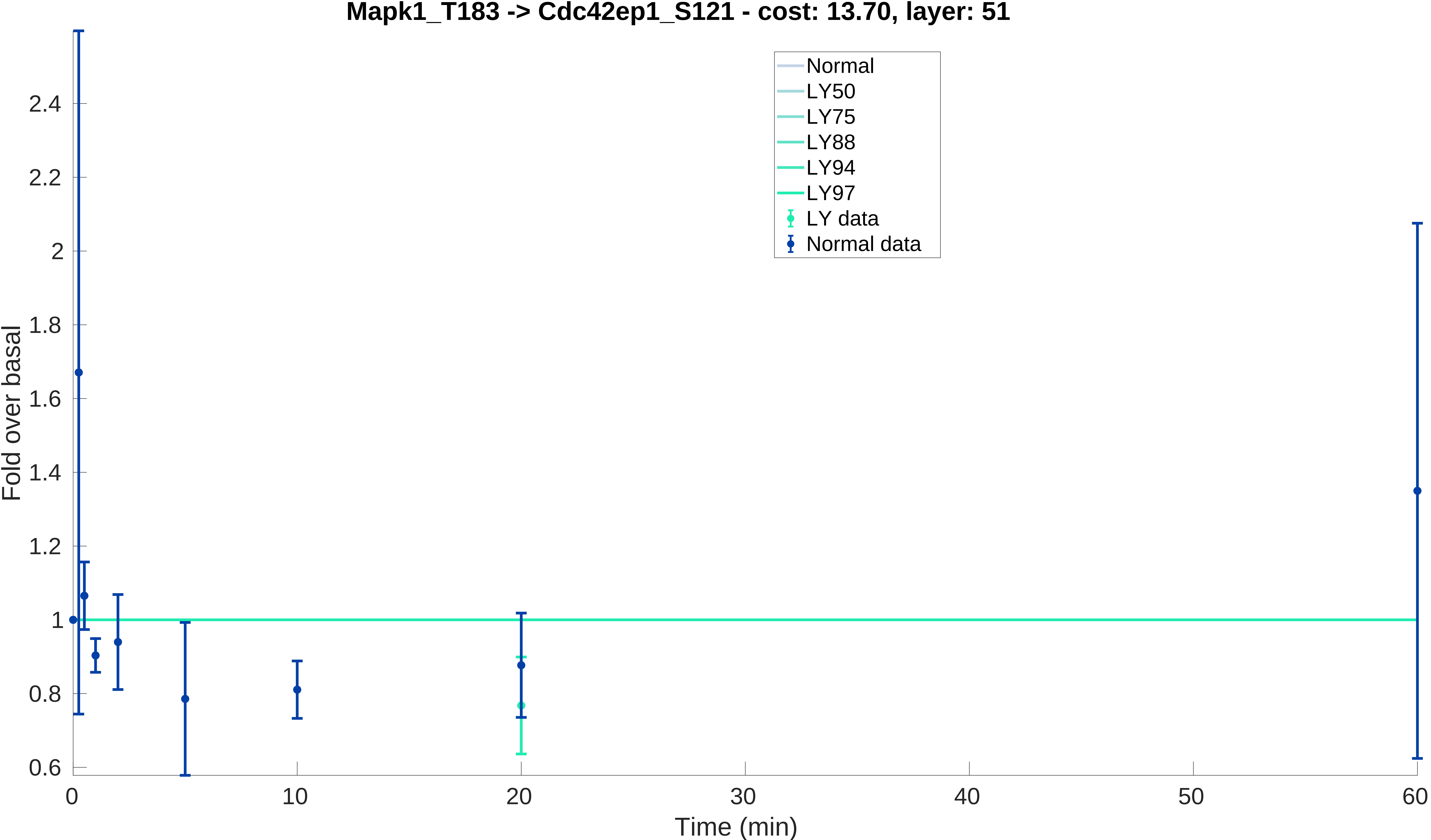

Mapk1_T183 -> Cdc42ep1_S121 - cost: 13.70, layer: 51
Normal
LY50
LY75
LY88
LY94
LY97
LY data
Normal data
Fold over basal
2.4
2.2
2
1.8
1.6
1.4
1.2
1
0.8
0.6
0
10
20
30
40
50
60
Time (min)

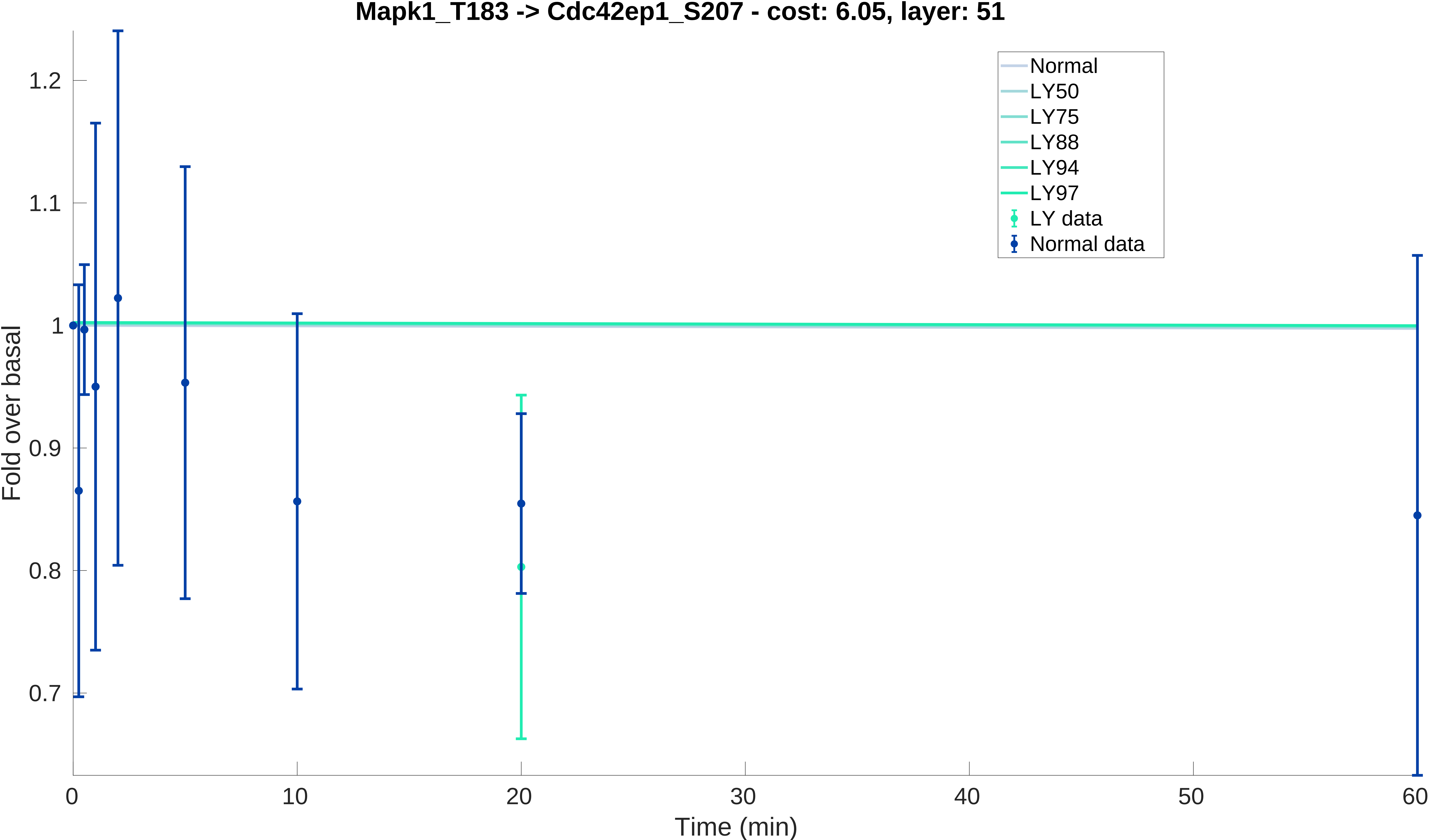

Mapk1_T183 -> Cdc42ep1_S207 - cost: 6.05, layer: 51
Normal
LY50
LY75
LY88
LY94
LY97
LY data
Normal data
Fold over basal
Time (min)
1.2
1.1
1
0.9
0.8
0.7
0
10
20
30
40
50
60

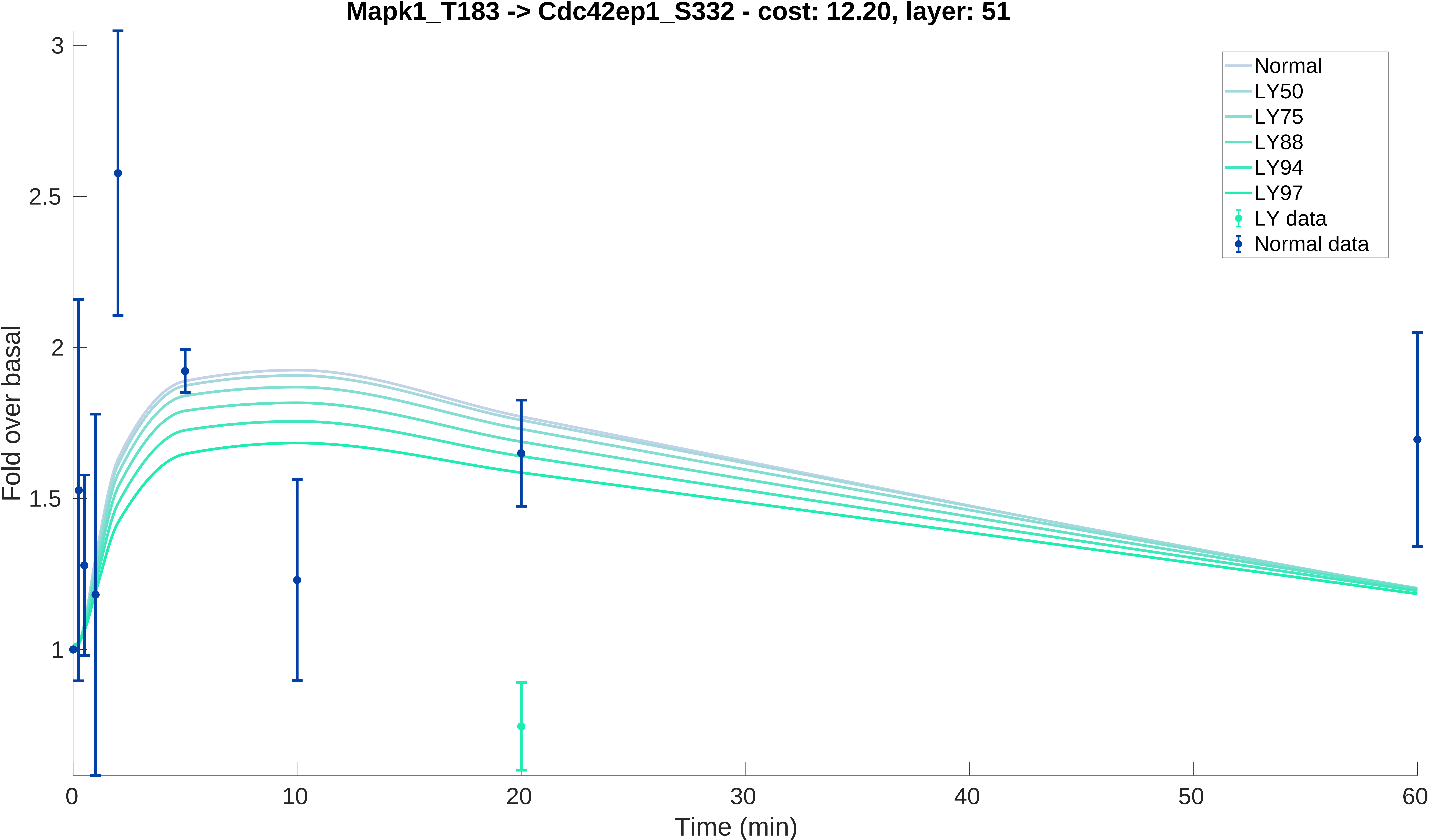

Mapk1_T183 -> Cdc42ep1_S332 - cost: 12.20, layer: 51
Fold over basal
Time (min)
0
10
20
30
40
50
60
1
1.5
2
2.5
3
Normal
LY50
LY75
LY88
LY94
LY97
LY data
Normal data

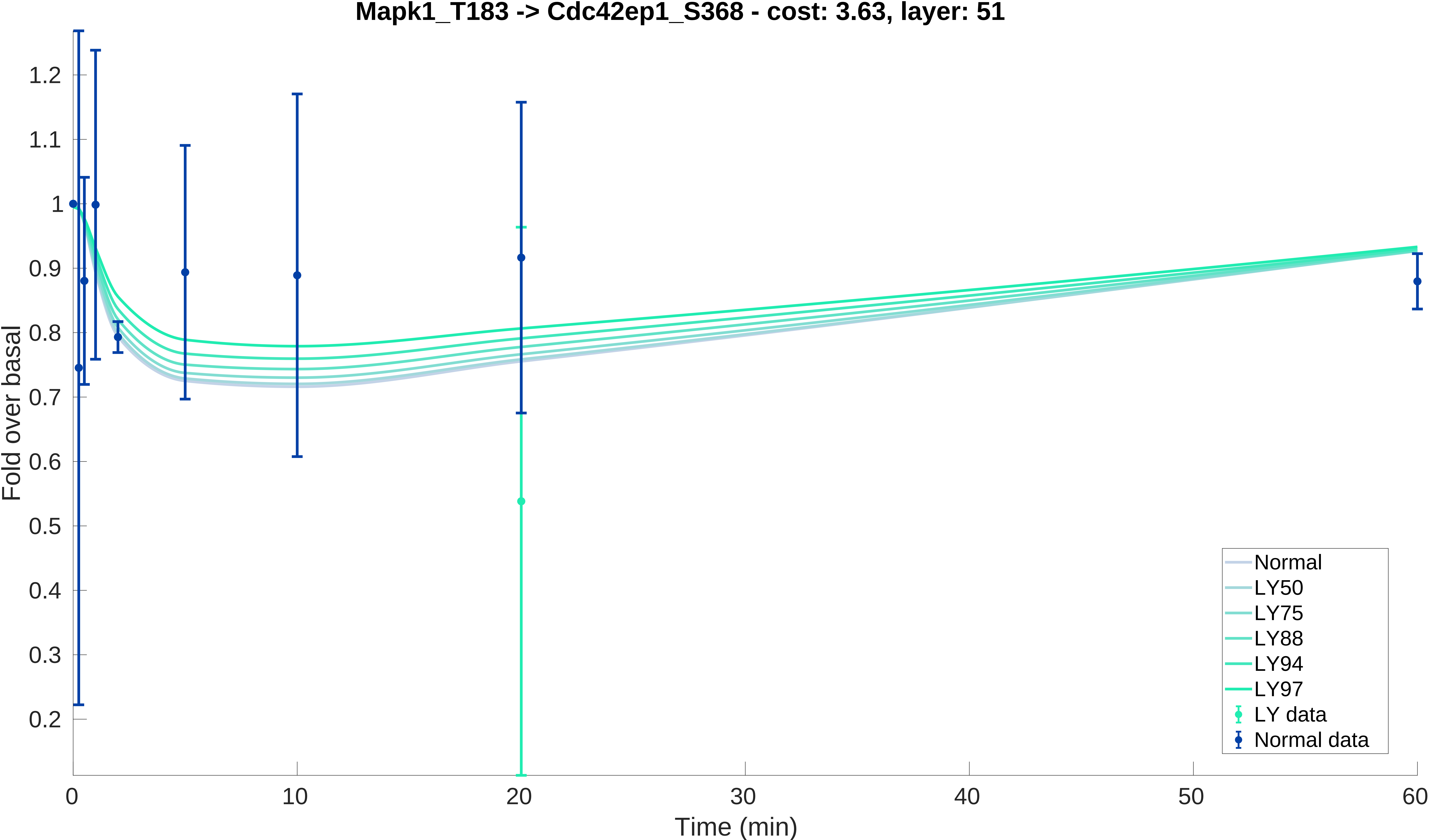
Mapk1_T183 -> Cdc42ep1_S368 - cost: 3.63, layer: 51
Fold over basal
Time (min)
1.2
1.1
1
0.9
0.8
0.7
0.6
0.5
0.4
0.3
0.2
0
10
20
30
40
50
60
Normal
LY50
LY75
LY88
LY94
LY97
LY data
Normal data

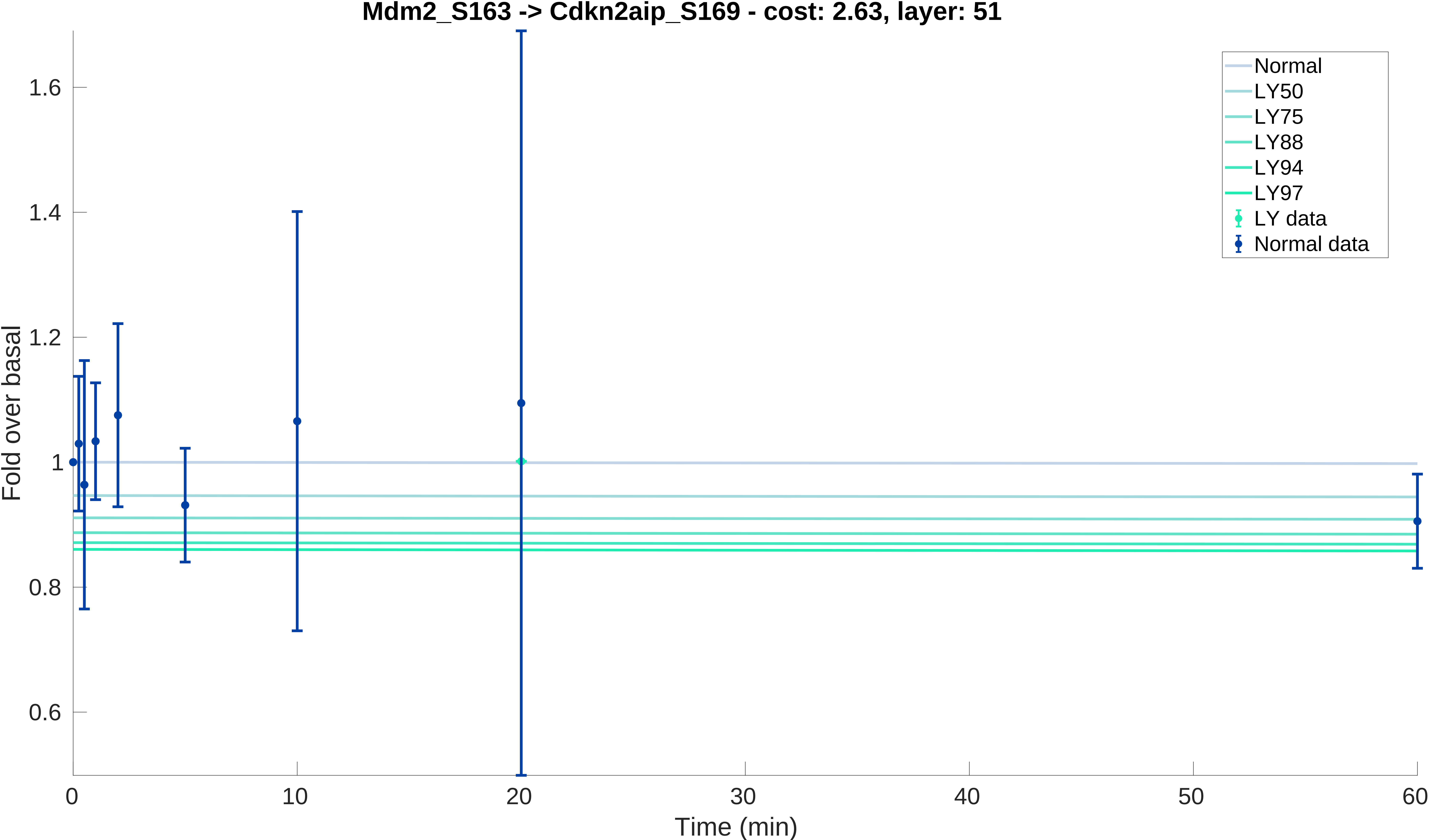

Mdm2_S163 -> Cdkn2aip_S169 - cost: 2.63, layer: 51
Fold over basal
Time (min)
1.6
1.4
1.2
1
0.8
0.6
0
10
20
30
40
50
60
Normal
LY50
LY75
LY88
LY94
LY97
LY data
Normal data

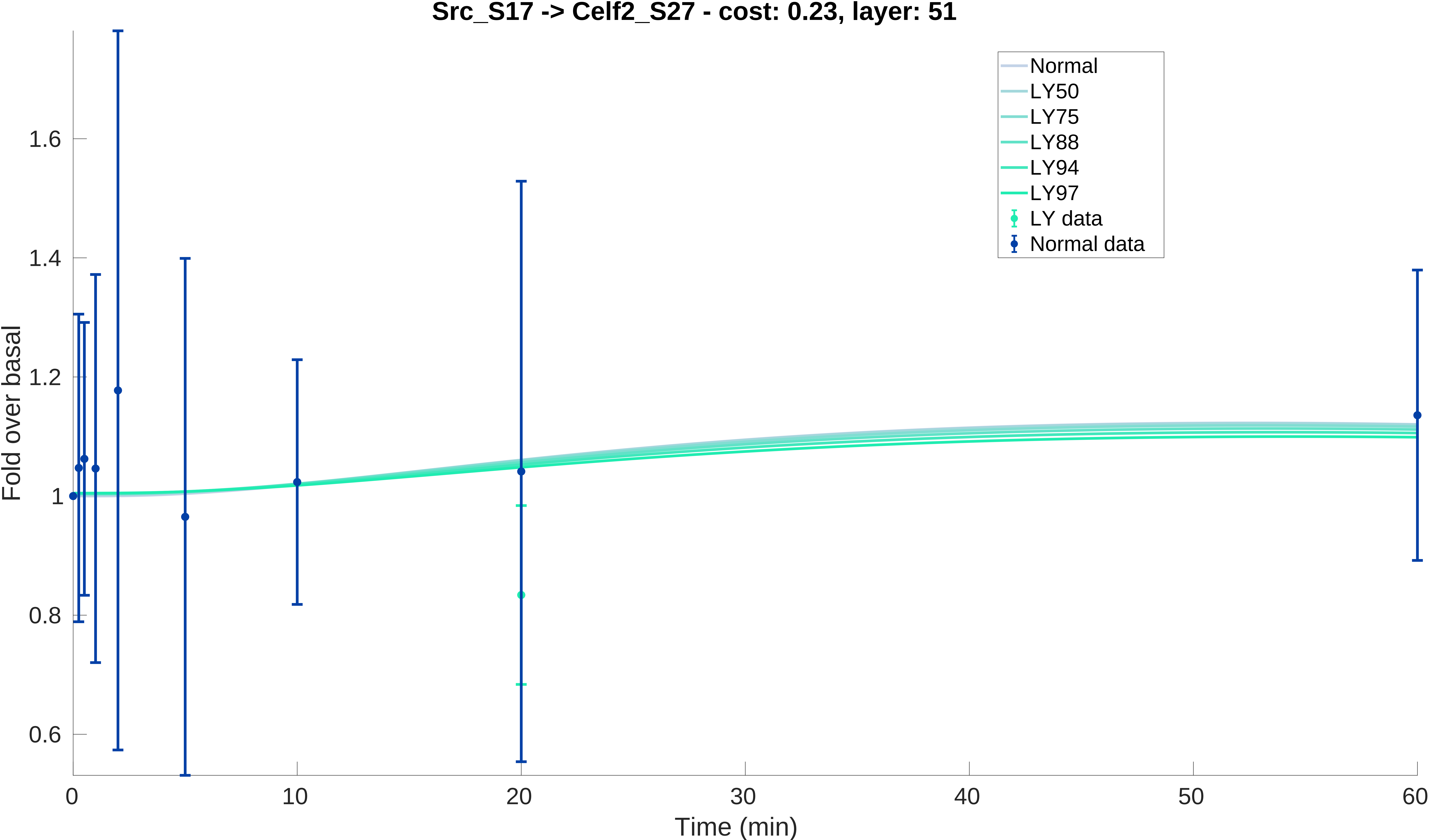

Src_S17 -> Celf2_S27 - cost: 0.23, layer: 51
Normal
LY50
LY75
LY88
LY94
LY97
LY data
Normal data
1.6
1.4
1.2
1
0.8
0.6
Fold over basal
0
10
20
30
40
50
60
Time (min)

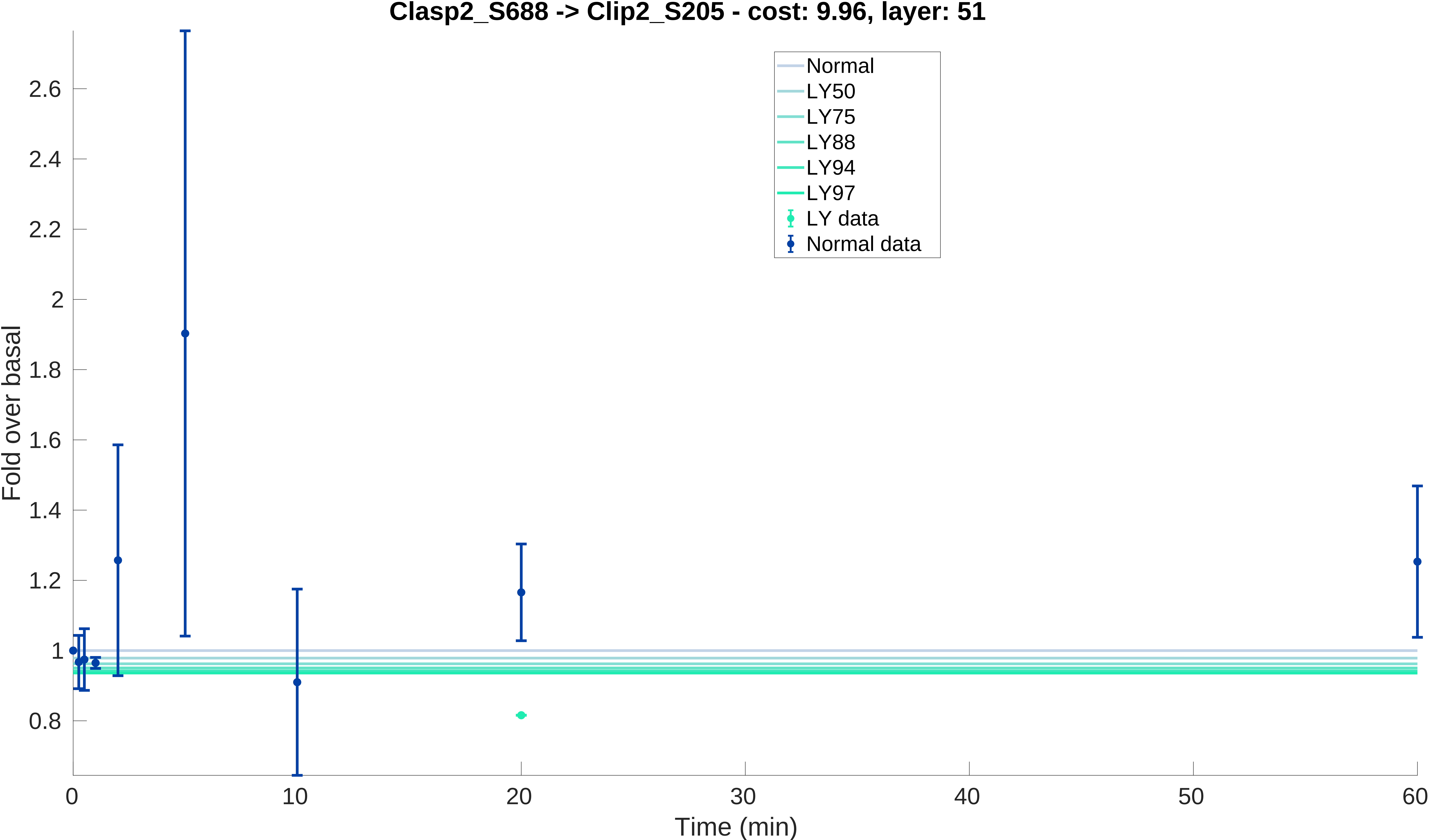

Clasp2_S688 -> Clip2_S205 - cost: 9.96, layer: 51
Normal
LY50
LY75
LY88
LY94
LY97
LY data
Normal data
Fold over basal
Time (min)
0
10
20
30
40
50
60
0.8
1
1.2
1.4
1.6
1.8
2
2.2
2.4
2.6

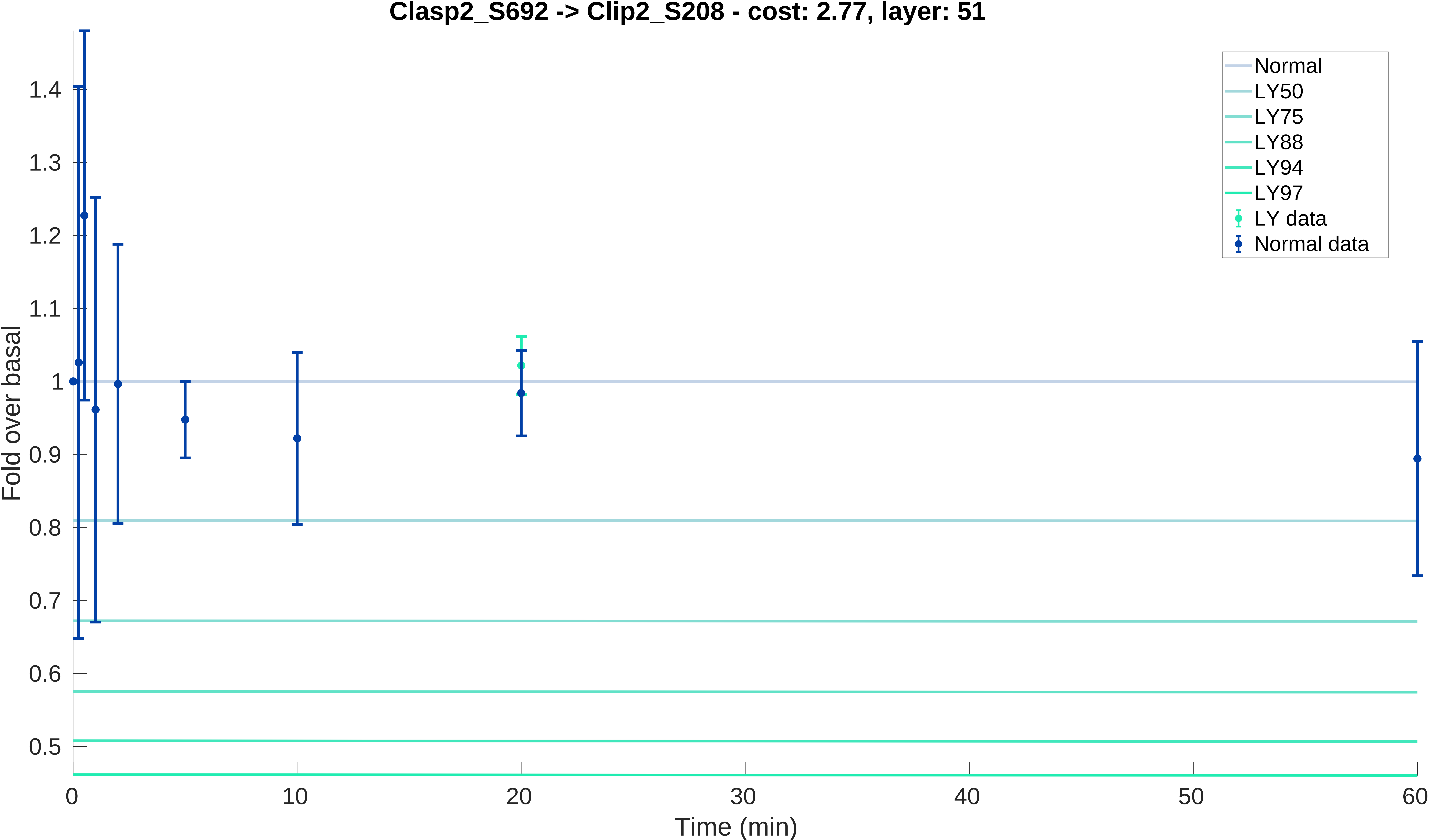
Clasp2_S692 -> Clip2_S208 - cost: 2.77, layer: 51
Fold over basal
Time (min)
1.4
1.3
1.2
1.1
1
0.9
0.8
0.7
0.6
0.5
0
10
20
30
40
50
60
Normal
LY50
LY75
LY88
LY94
LY97
LY data
Normal data

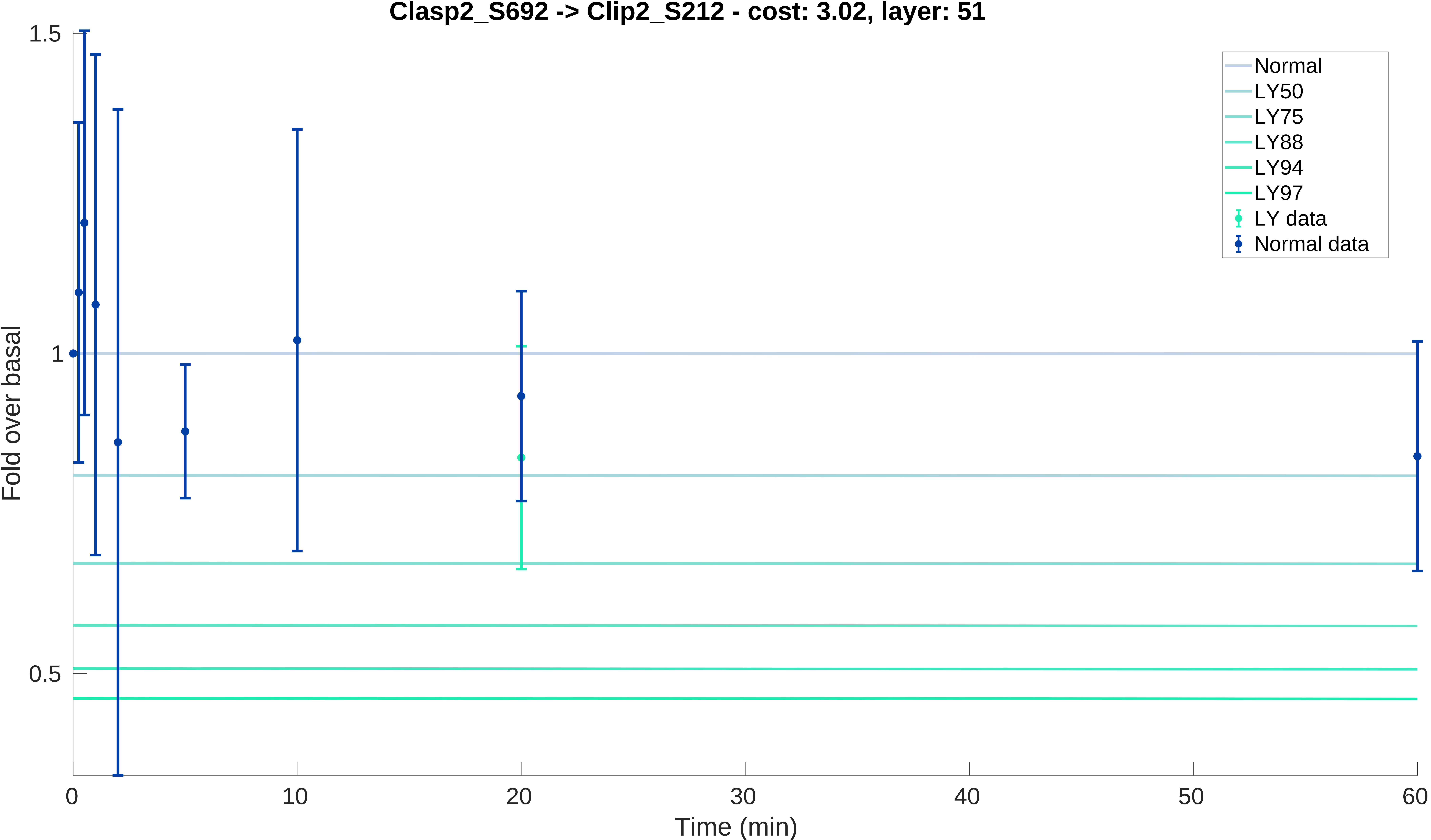

Clasp2_S692 -> Clip2_S212 - cost: 3.02, layer: 51
Fold over basal
1.5
1
0.5
Time (min)
0
10
20
30
40
50
60
Normal
LY50
LY75
LY88
LY94
LY97
LY data
Normal data

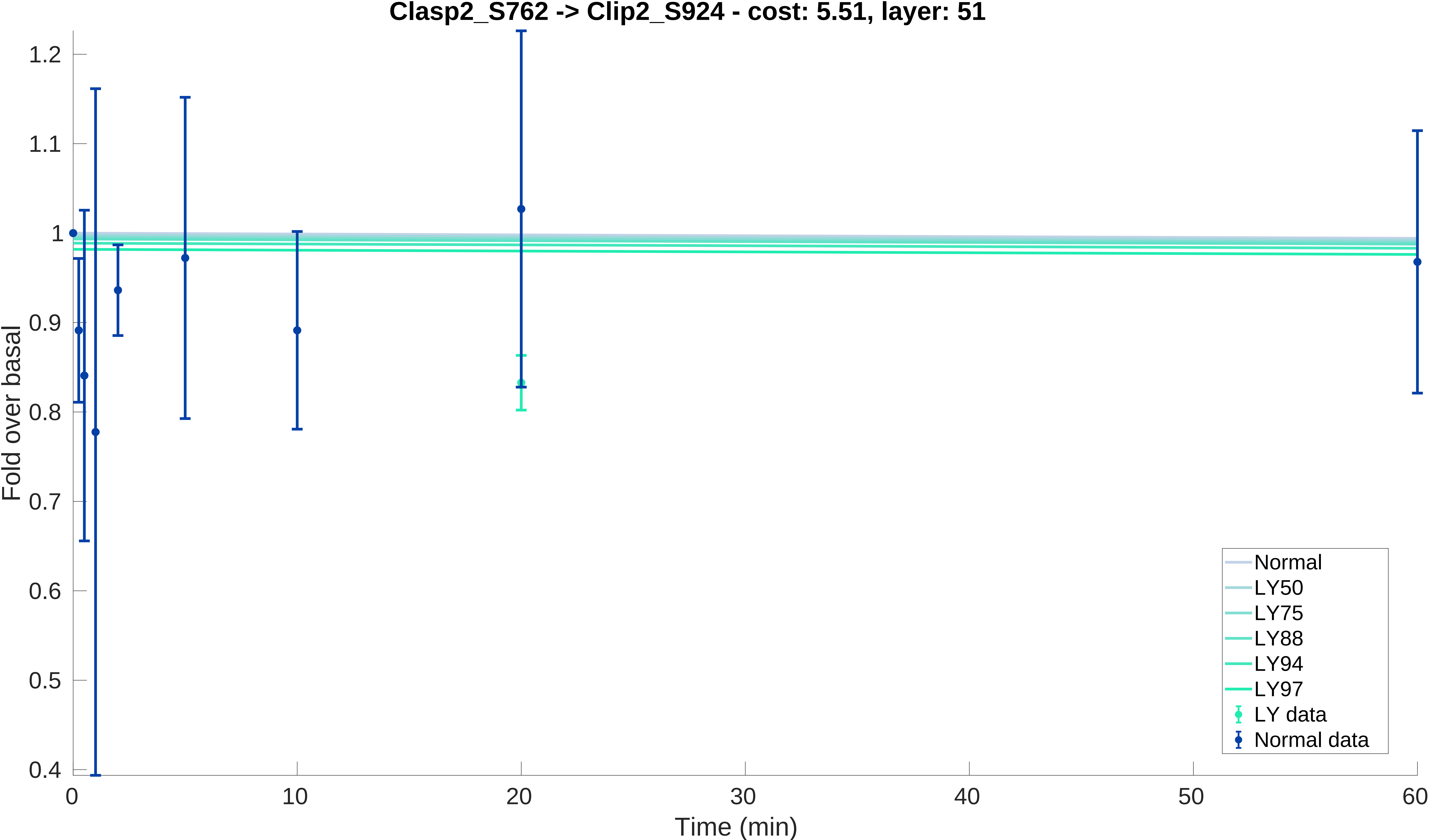

Clasp2_S762 -> Clip2_S924 - cost: 5.51, layer: 51
Fold over basal
Time (min)
1.2
1.1
1
0.9
0.8
0.7
0.6
0.5
0.4
0
10
20
30
40
50
60
Normal
LY50
LY75
LY88
LY94
LY97
LY data
Normal data

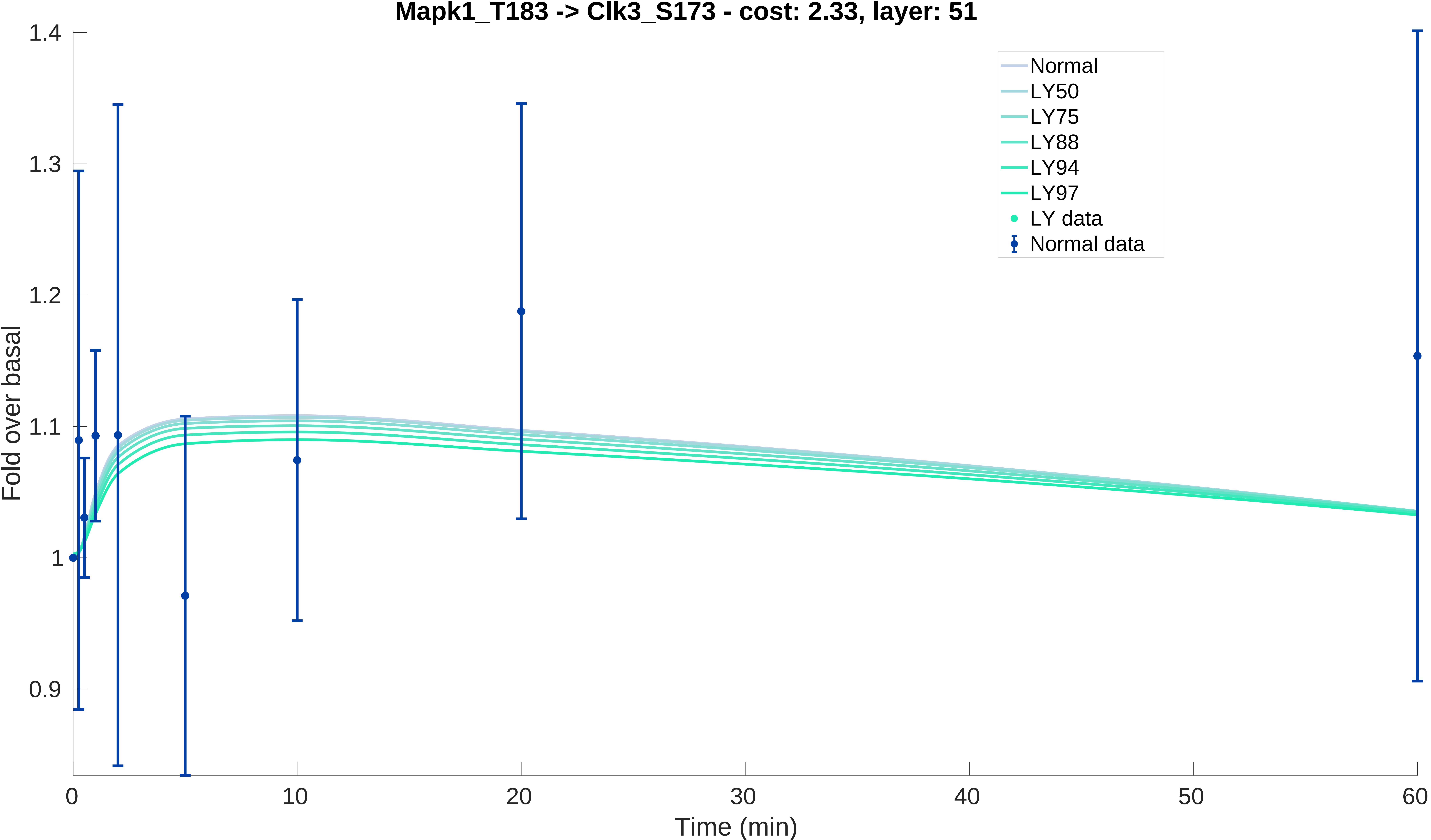

Mapk1_T183 -> Clk3_S173 - cost: 2.33, layer: 51
Fold over basal
1.4
1.3
1.2
1.1
1
0.9
0
10
20
30
40
50
60
Time (min)
Normal
LY50
LY75
LY88
LY94
LY97
LY data
Normal data

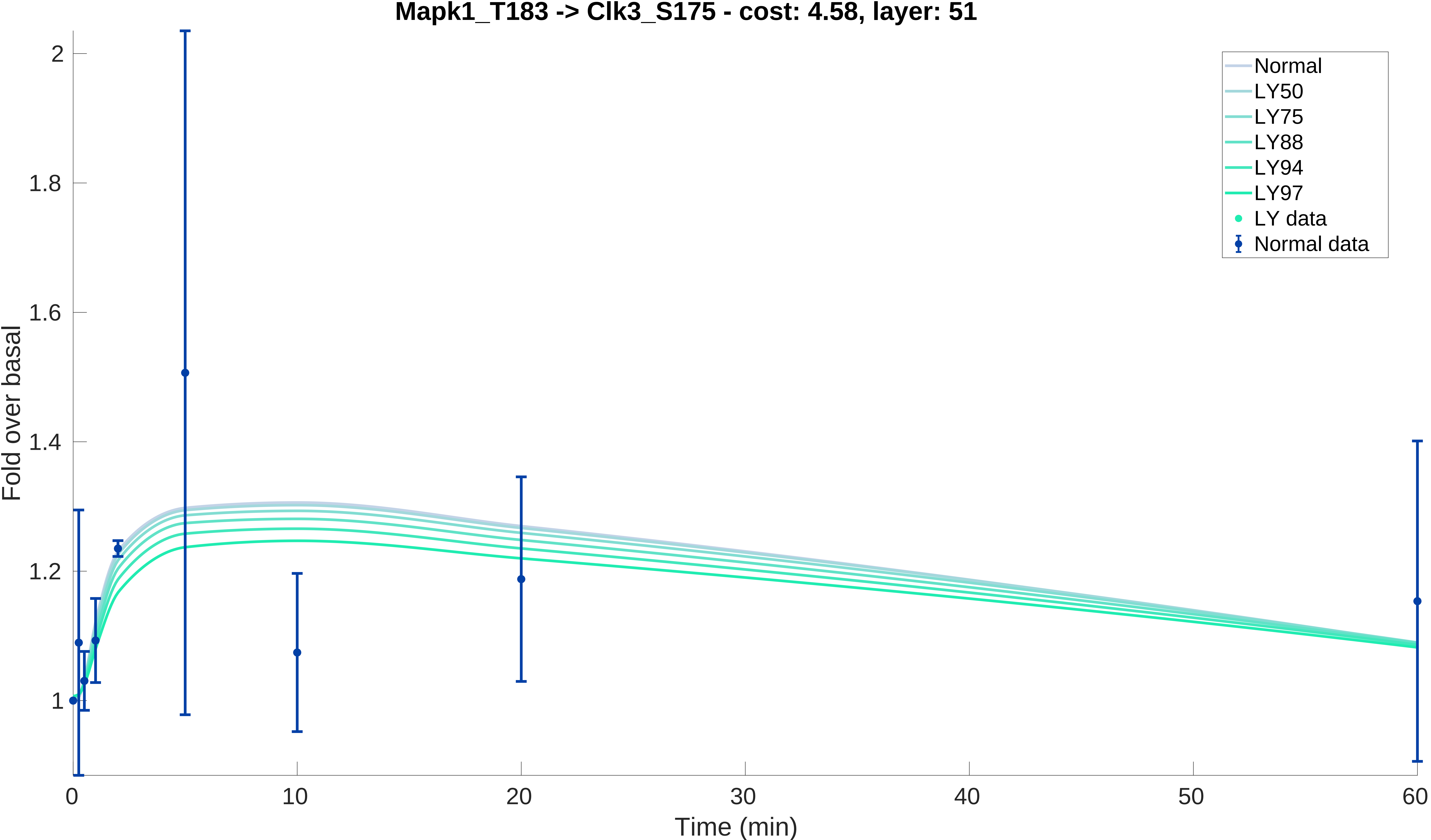

Mapk1_T183 -> Clk3_S175 - cost: 4.58, layer: 51
Normal
LY50
LY75
LY88
LY94
LY97
LY data
Normal data
Fold over basal
2
1.8
1.6
1.4
1.2
1
0
10
20
30
40
50
60
Time (min)

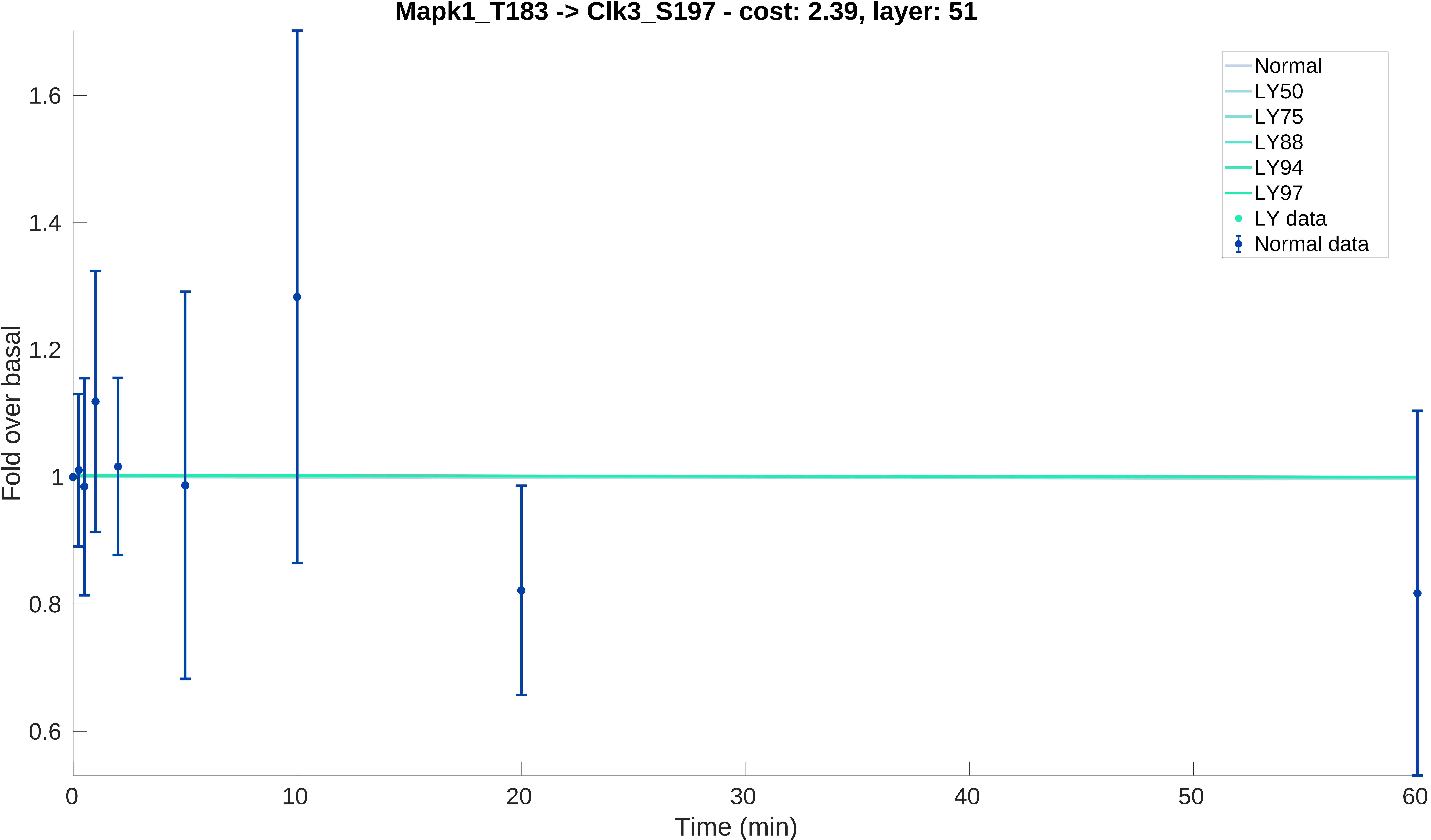

Mapk1_T183 -> Clk3_S197 - cost: 2.39, layer: 51
Normal
LY50
LY75
LY88
LY94
LY97
LY data
Normal data
Fold over basal
1.6
1.4
1.2
1
0.8
0.6
0
10
20
30
40
50
60
Time (min)

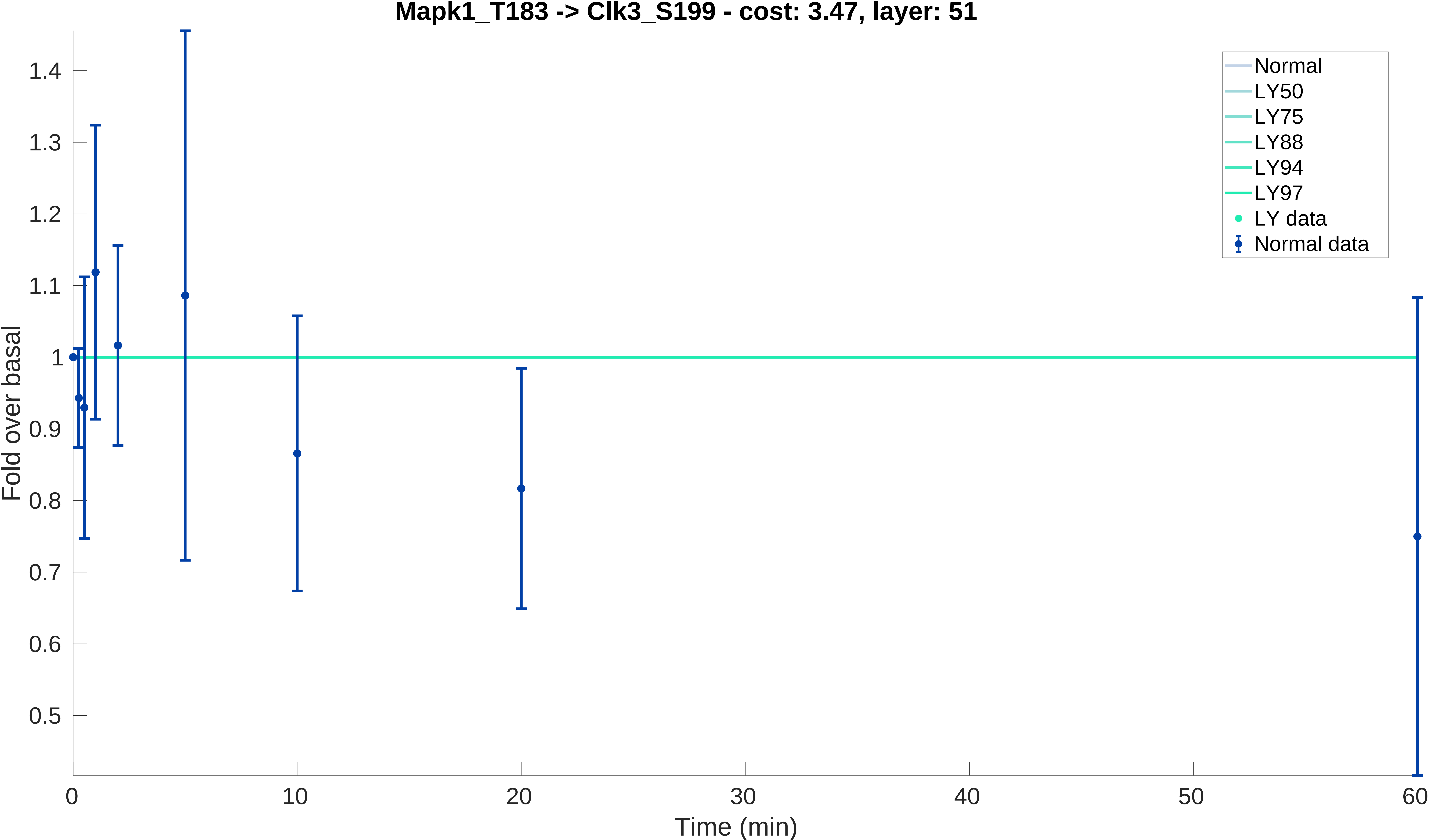
Mapk1_T183 -> Clk3_S199 - cost: 3.47, layer: 51
Fold over basal
Time (min)
1.4
1.3
1.2
1.1
1
0.9
0.8
0.7
0.6
0.5
0
10
20
30
40
50
60
Normal
LY50
LY75
LY88
LY94
LY97
LY data
Normal data

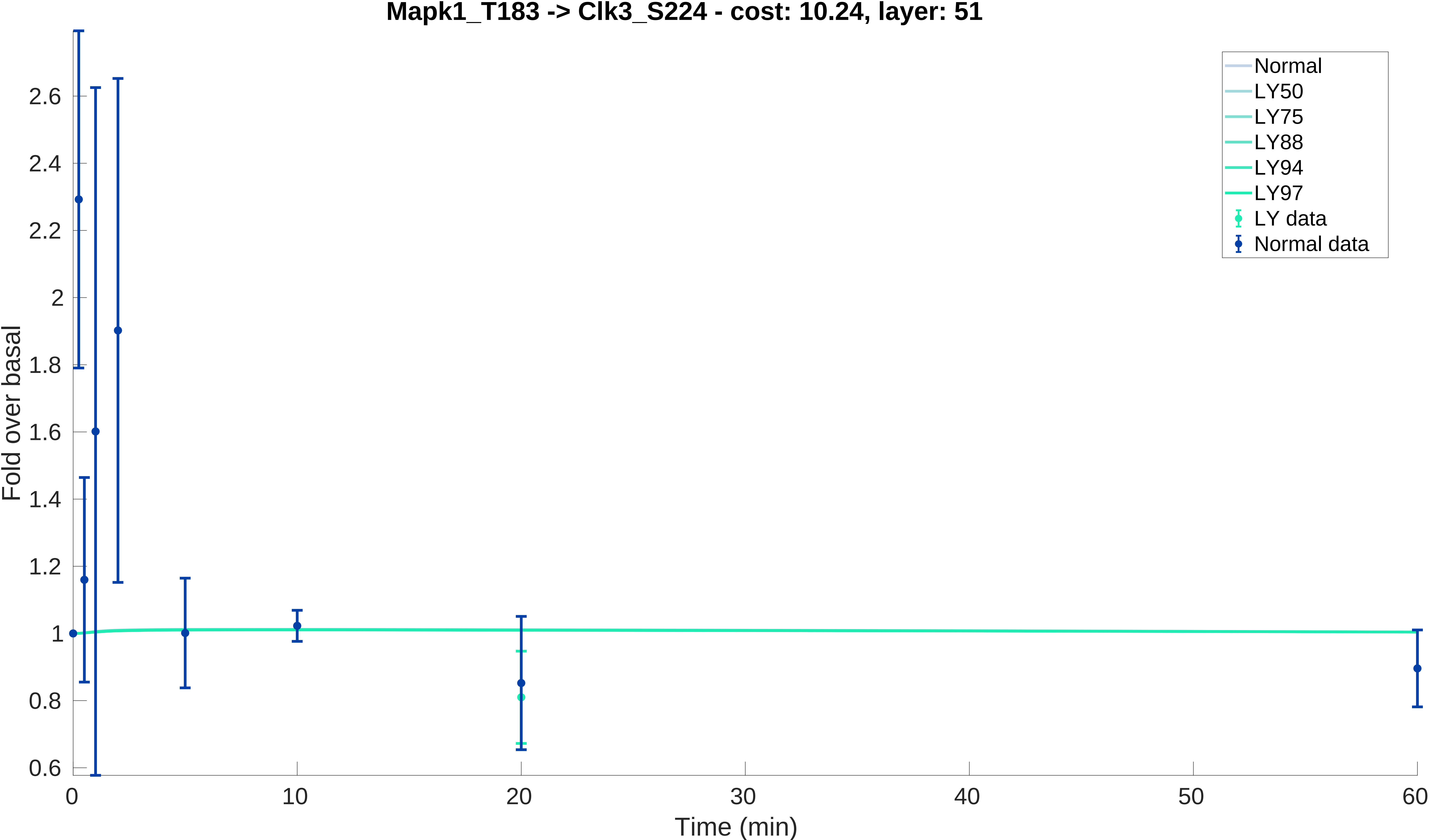

Mapk1_T183 -> Clk3_S224 - cost: 10.24, layer: 51
Normal
LY50
LY75
LY88
LY94
LY97
LY data
Normal data
Fold over basal
2.6
2.4
2.2
2
1.8
1.6
1.4
1.2
1
0.8
0.6
0
10
20
30
40
50
60
Time (min)

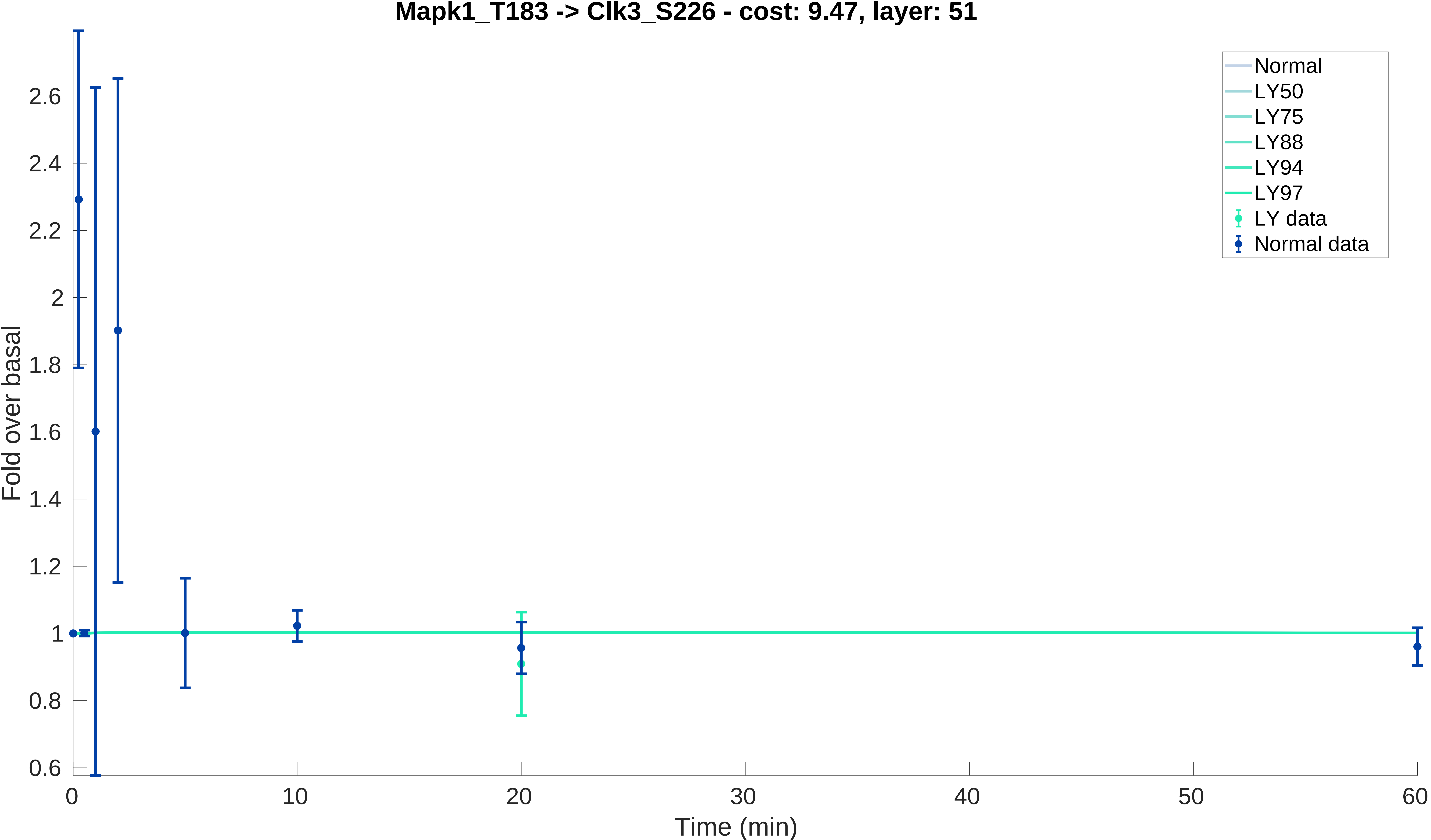

Mapk1_T183 -> Clk3_S226 - cost: 9.47, layer: 51
Normal
LY50
LY75
LY88
LY94
LY97
LY data
Normal data
Fold over basal
Time (min)
2.6
2.4
2.2
2
1.8
1.6
1.4
1.2
1
0.8
0.6
0
10
20
30
40
50
60

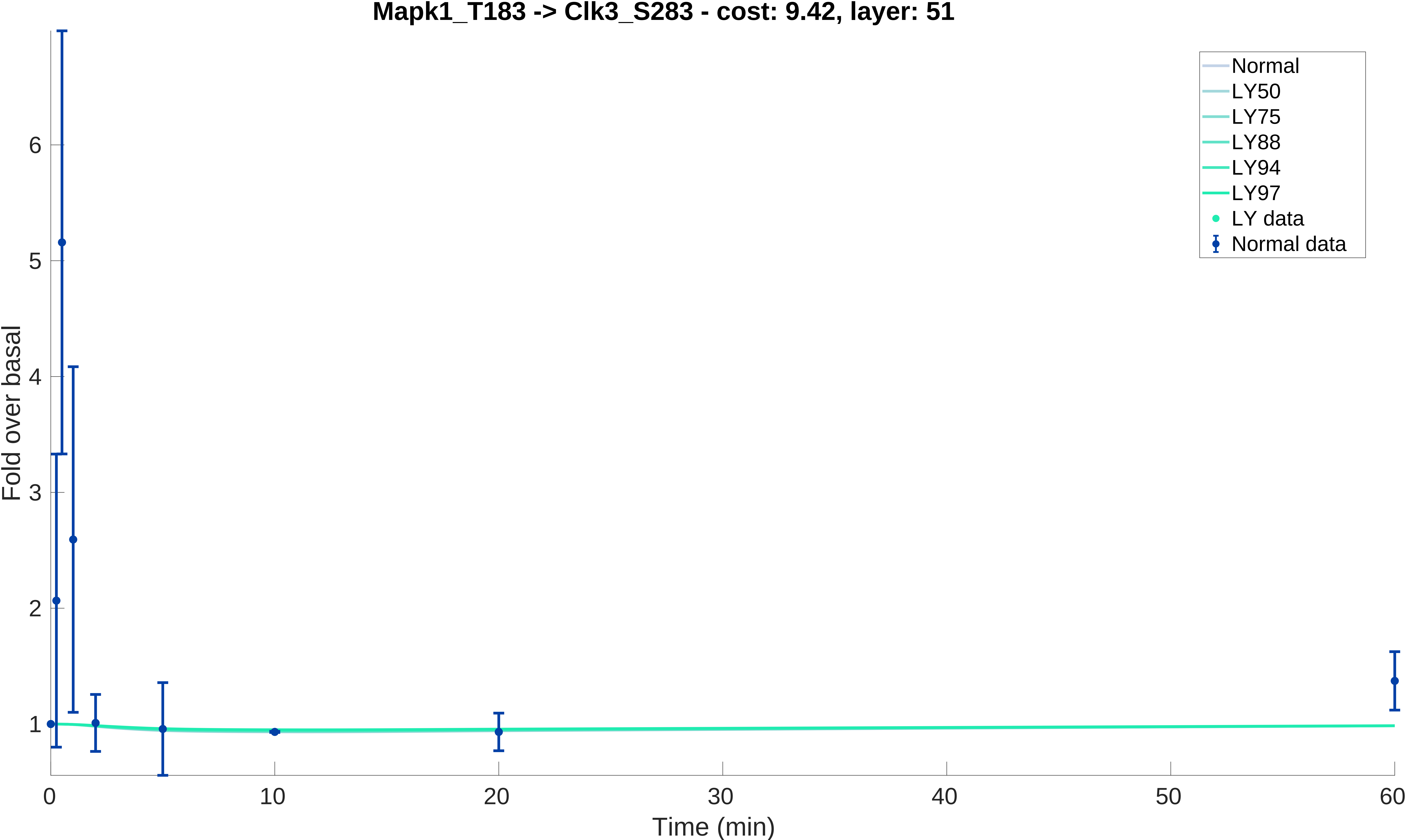

Mapk1_T183 -> Clk3_S283 - cost: 9.42, layer: 51
Fold over basal
Time (min)
1
2
3
4
5
6
0
10
20
30
40
50
60
Normal
LY50
LY75
LY88
LY94
LY97
LY data
Normal data

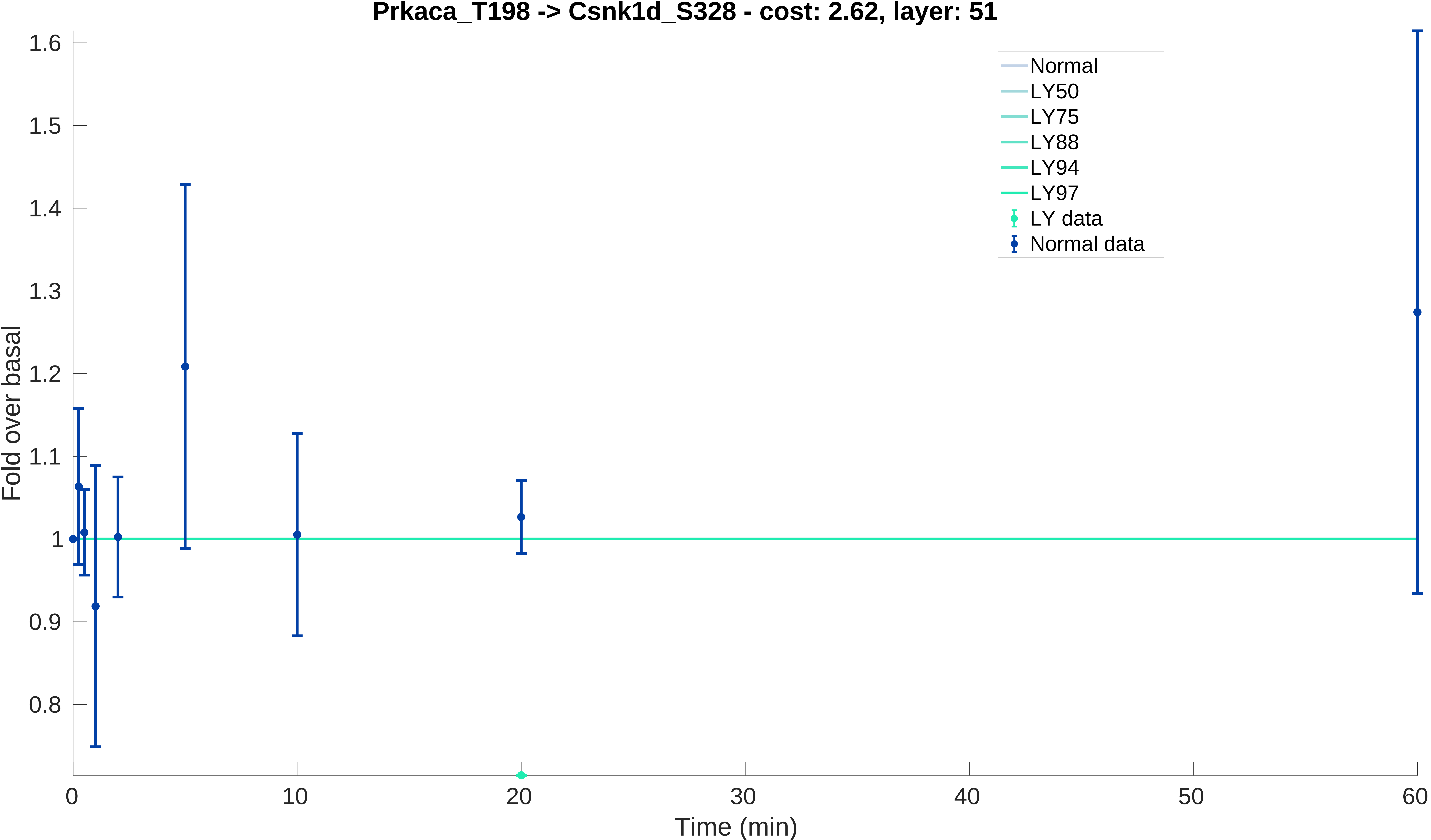
Prkaca_T198 -> Csnk1d_S328 - cost: 2.62, layer: 51
Fold over basal
Time (min)
1.6
1.5
1.4
1.3
1.2
1.1
1
0.9
0.8
0
10
20
30
40
50
60
Normal
LY50
LY75
LY88
LY94
LY97
LY data
Normal data

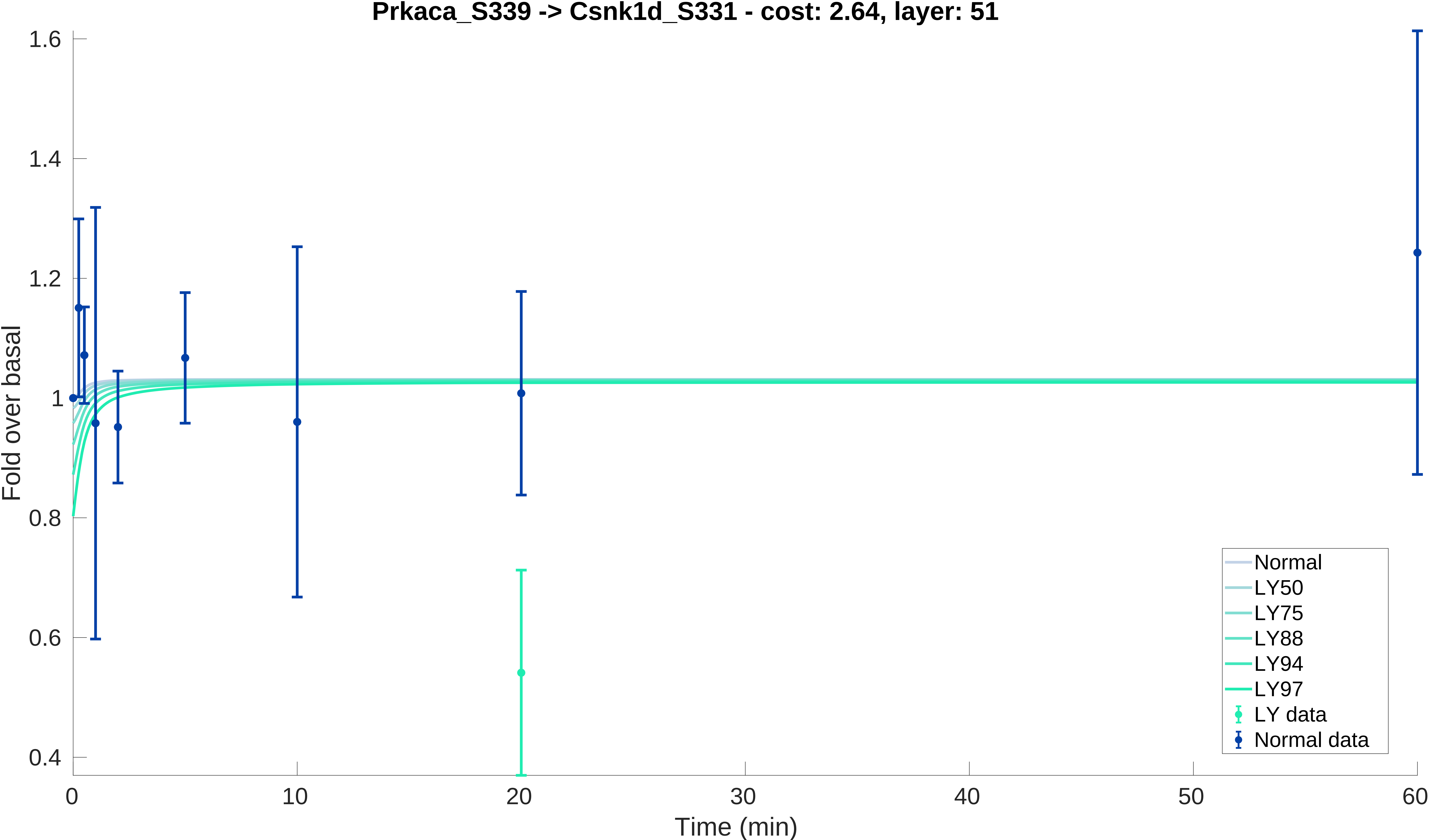

Prkaca_S339 -> Csnk1d_S331 - cost: 2.64, layer: 51
Fold over basal
1.6
1.4
1.2
1
0.8
0.6
0.4
0
10
20
30
40
50
60
Time (min)
Normal
LY50
LY75
LY88
LY94
LY97
LY data
Normal data

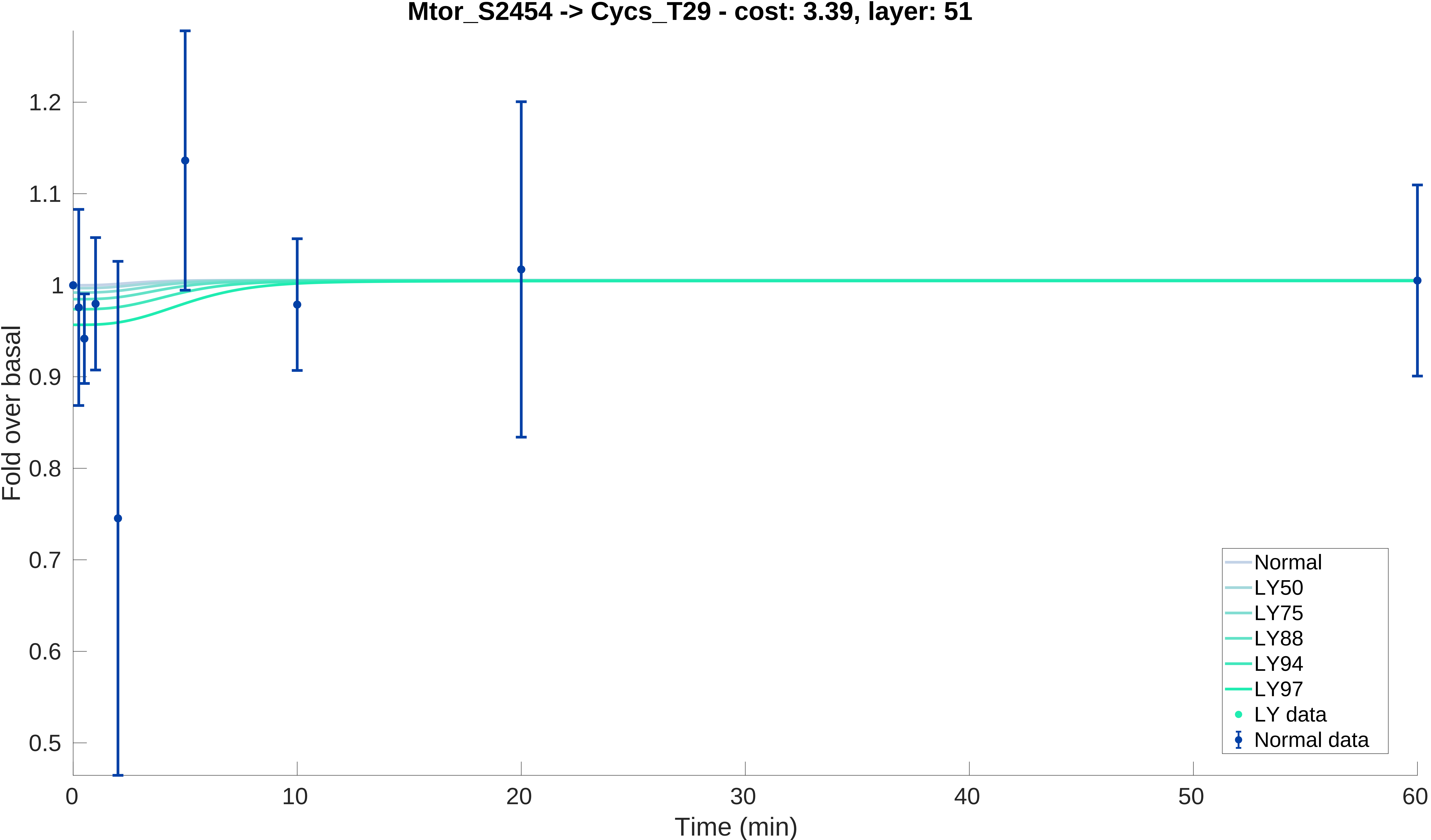

Mtor_S2454 -> Cycs_T29 - cost: 3.39, layer: 51
Fold over basal
1.2
1.1
1
0.9
0.8
0.7
0.6
0.5
0
10
20
30
40
50
60
Time (min)
Normal
LY50
LY75
LY88
LY94
LY97
LY data
Normal data

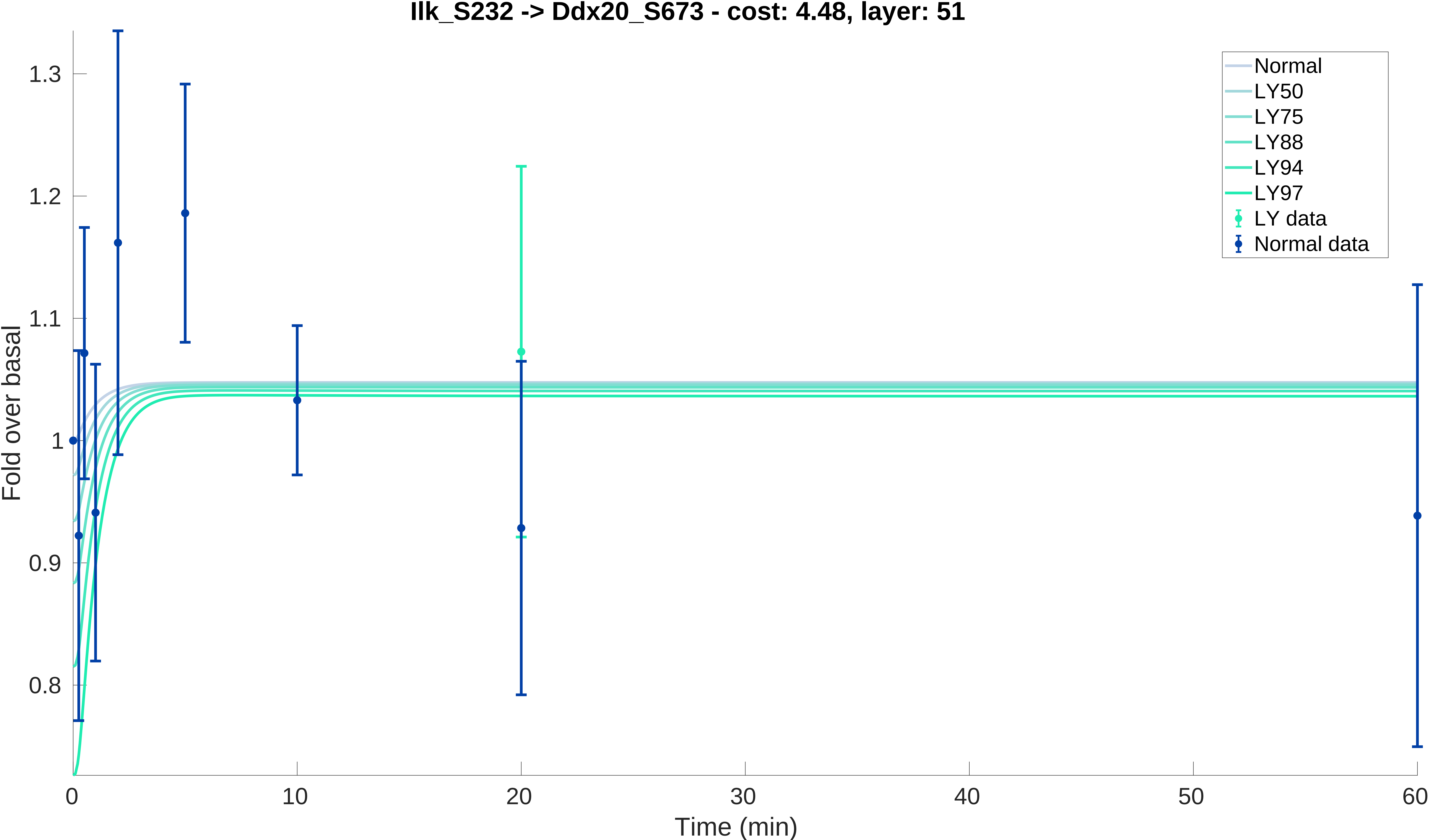

Ilk_S232 -> Ddx20_S673 - cost: 4.48, layer: 51
Normal
LY50
LY75
LY88
LY94
LY97
LY data
Normal data
Fold over basal
1.3
1.2
1.1
1
0.9
0.8
0
10
20
30
40
50
60
Time (min)

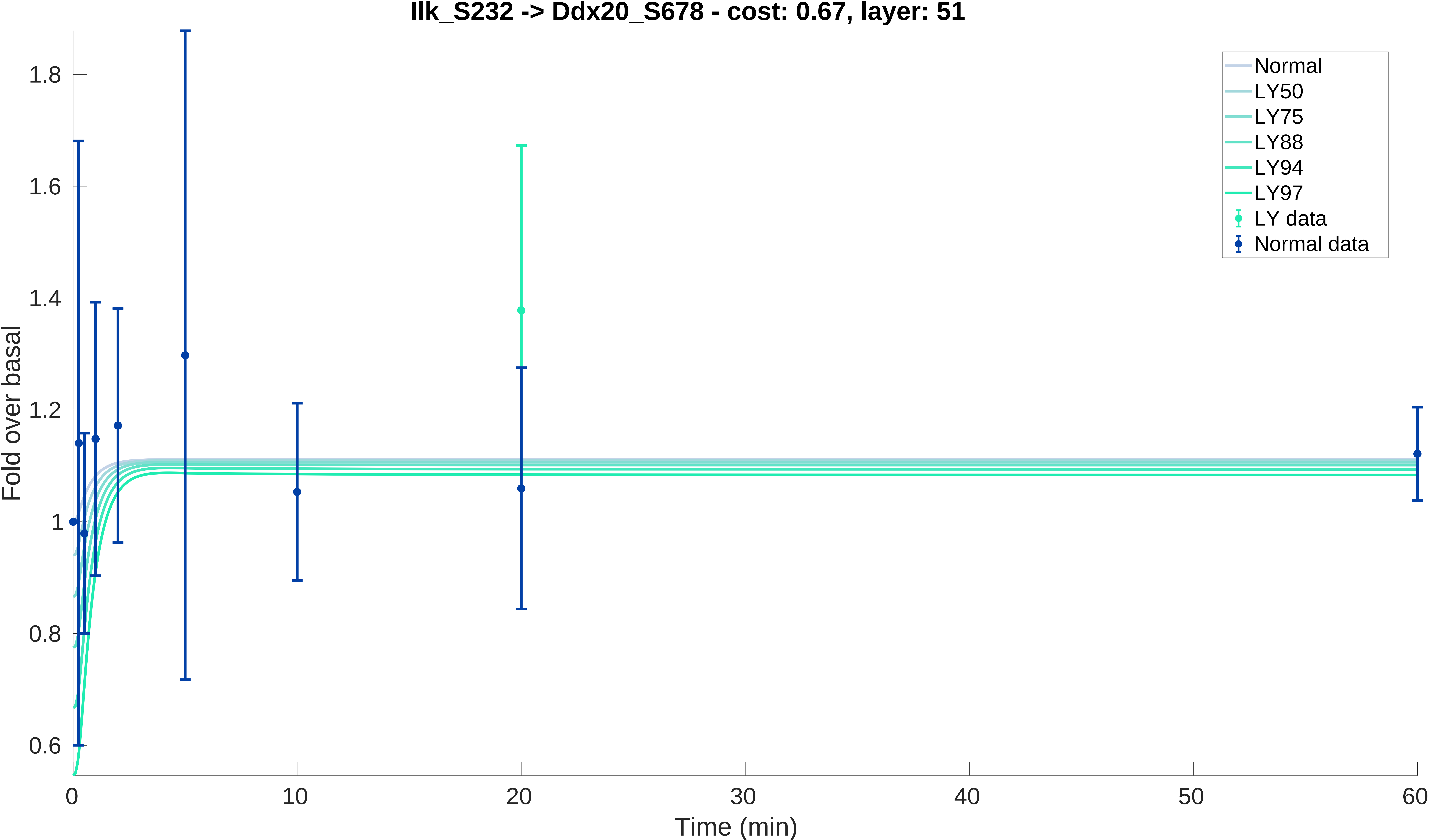

Ilk_S232 -> Ddx20_S678 - cost: 0.67, layer: 51
Fold over basal
Time (min)
0.6
0.8
1
1.2
1.4
1.6
1.8
0
10
20
30
40
50
60
Normal
LY50
LY75
LY88
LY94
LY97
LY data
Normal data

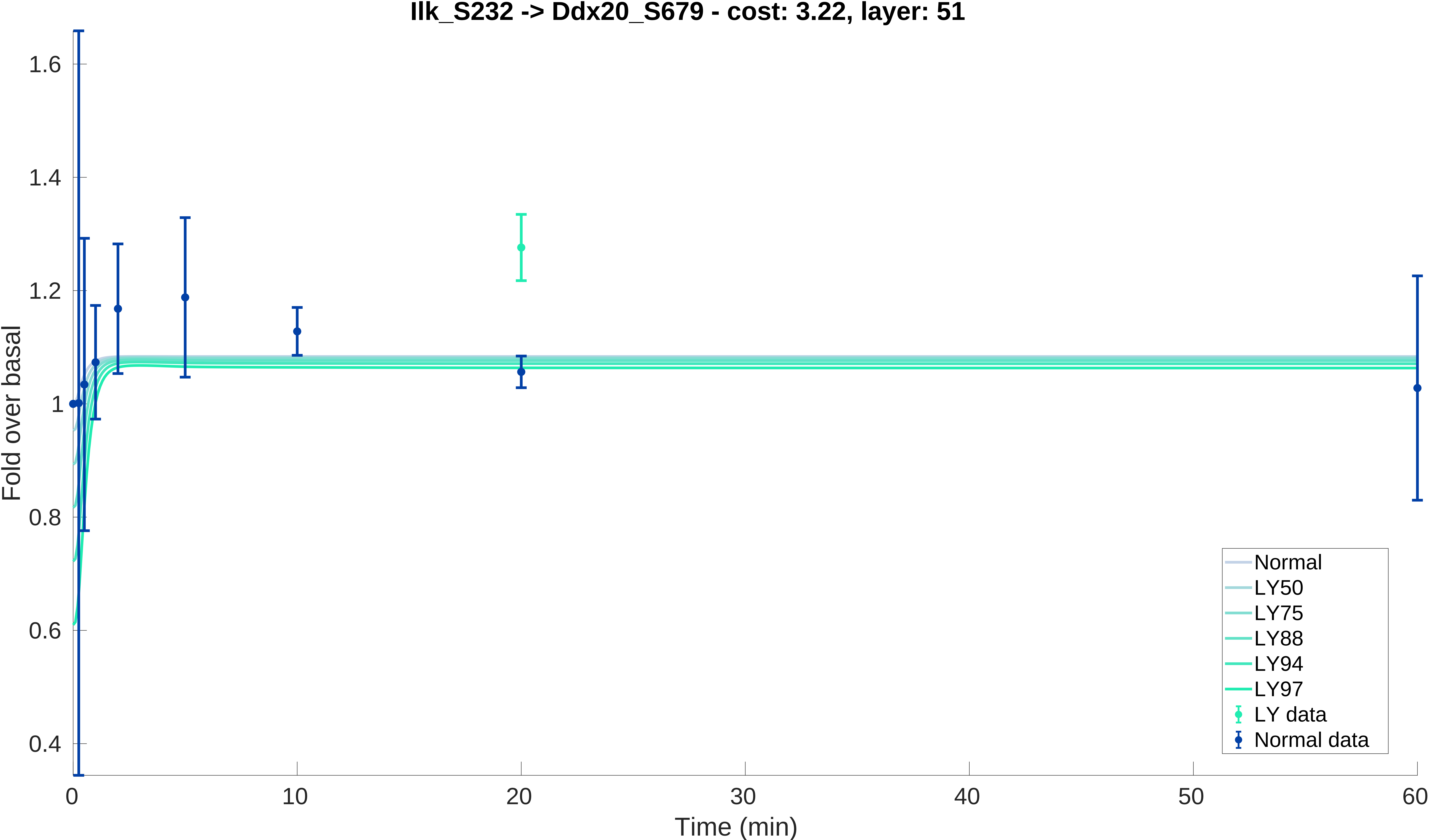

Ilk_S232 -> Ddx20_S679 - cost: 3.22, layer: 51
1.6
1.4
1.2
1
0.8
0.6
0.4
Fold over basal
0
10
20
30
40
50
60
Time (min)
Normal
LY50
LY75
LY88
LY94
LY97
LY data
Normal data

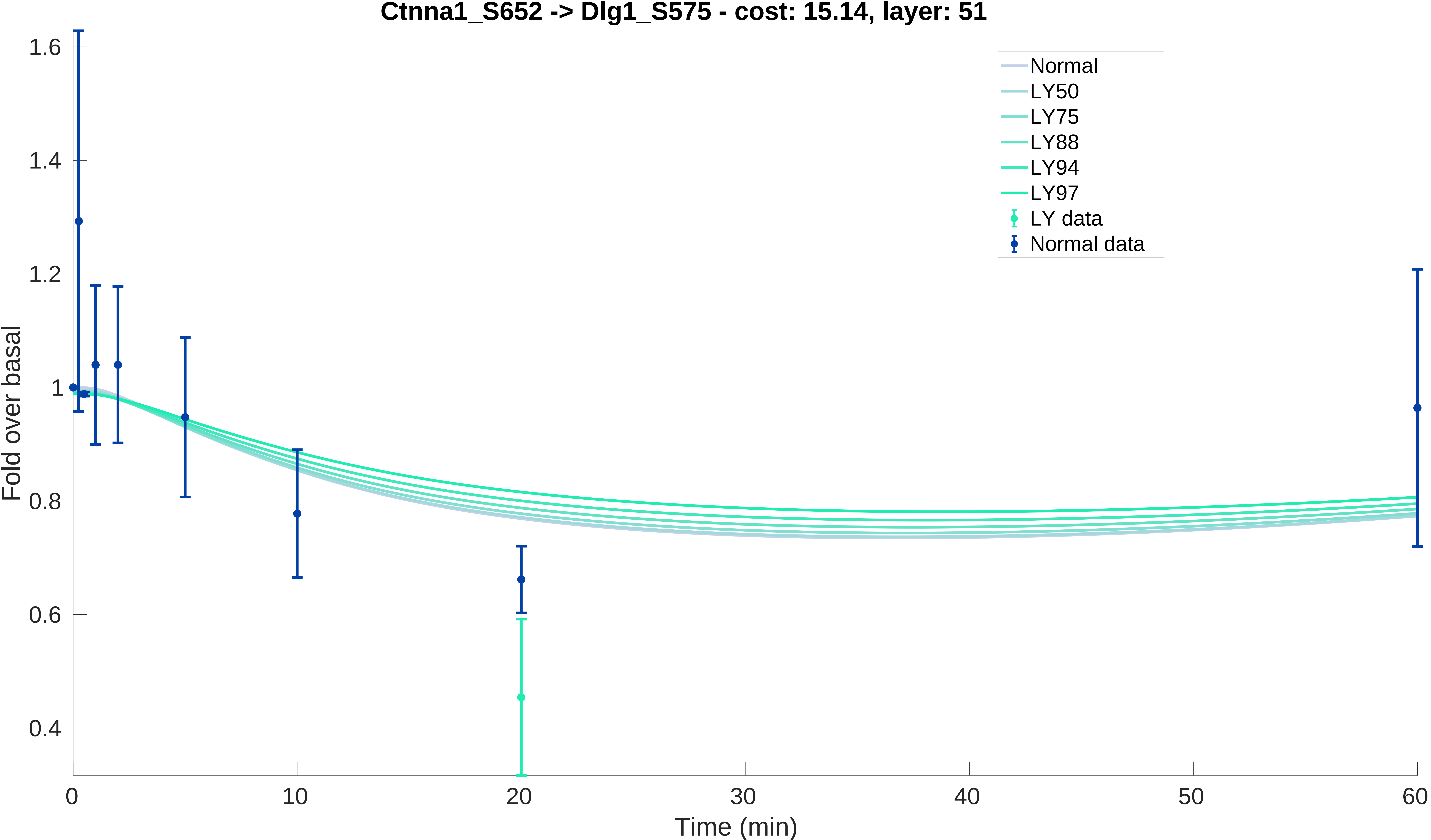

Ctnna1_S652 -> Dlg1_S575 - cost: 15.14, layer: 51
Fold over basal
Time (min)
1.6
1.4
1.2
1
0.8
0.6
0.4
0
10
20
30
40
50
60
Normal
LY50
LY75
LY88
LY94
LY97
LY data
Normal data

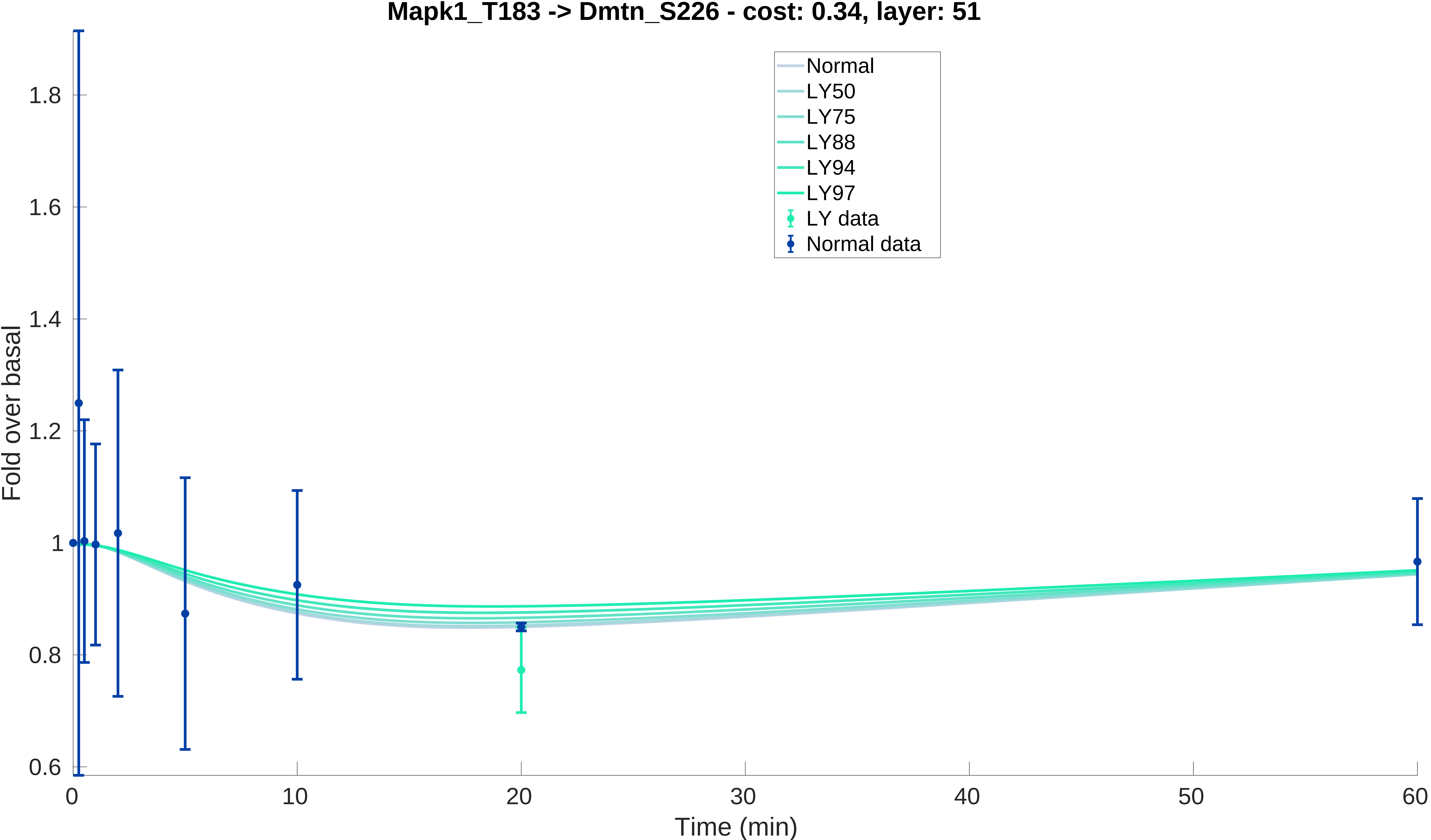
Mapk1_T183 -> Dmtn_S226 - cost: 0.34, layer: 51
Normal
LY50
LY75
LY88
LY94
LY97
LY data
Normal data
Fold over basal
Time (min)
0.6
0.8
1
1.2
1.4
1.6
1.8
0
10
20
30
40
50
60

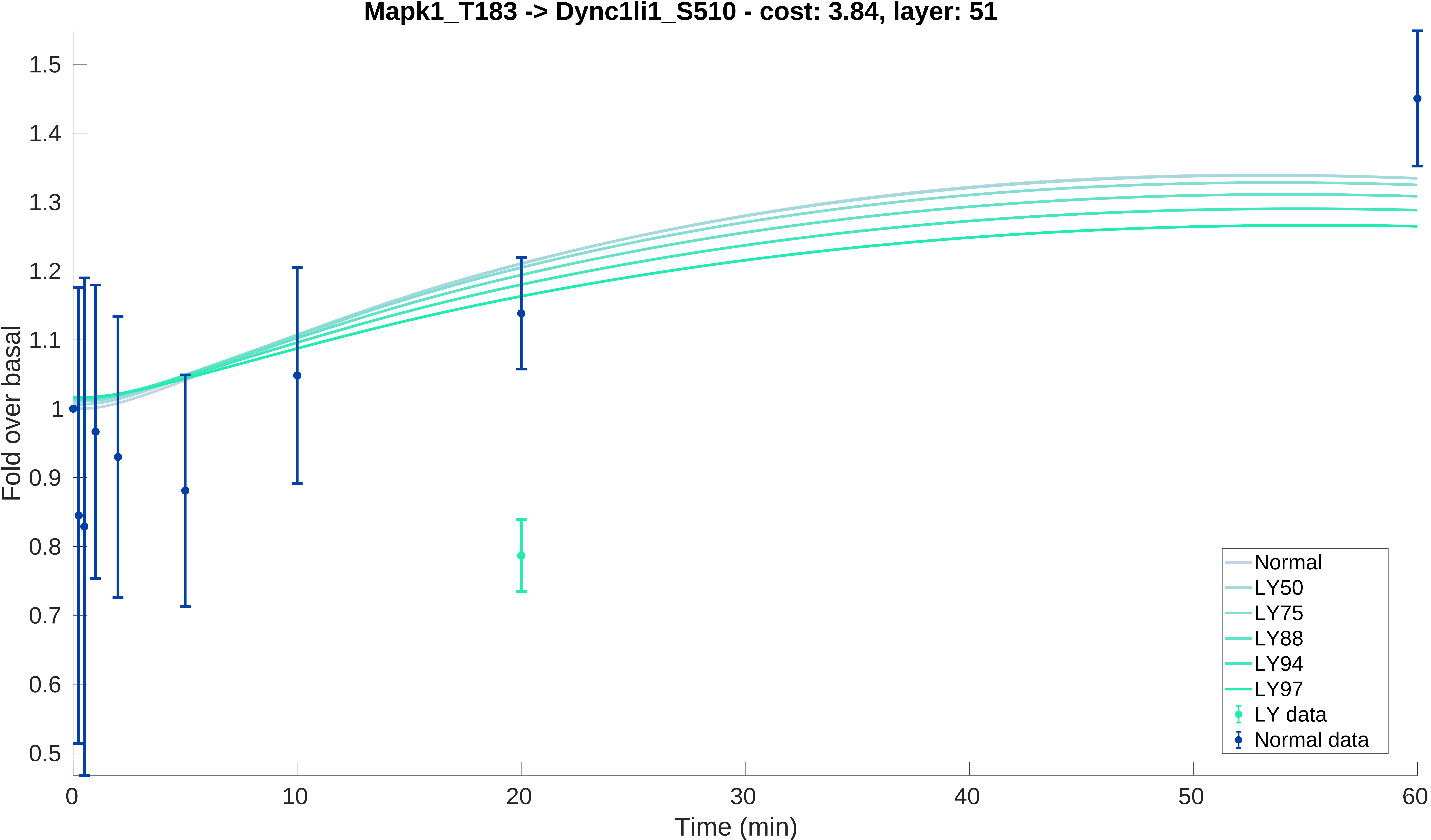
Mapk1_T183 -> Dync1li1_S510 - cost: 3.84, layer: 51
Fold over basal
1.5
1.4
1.3
1.2
1.1
1
0.9
0.8
0.7
0.6
0.5
0
10
20
30
40
50
60
Time (min)
Normal
LY50
LY75
LY88
LY94
LY97
LY data
Normal data

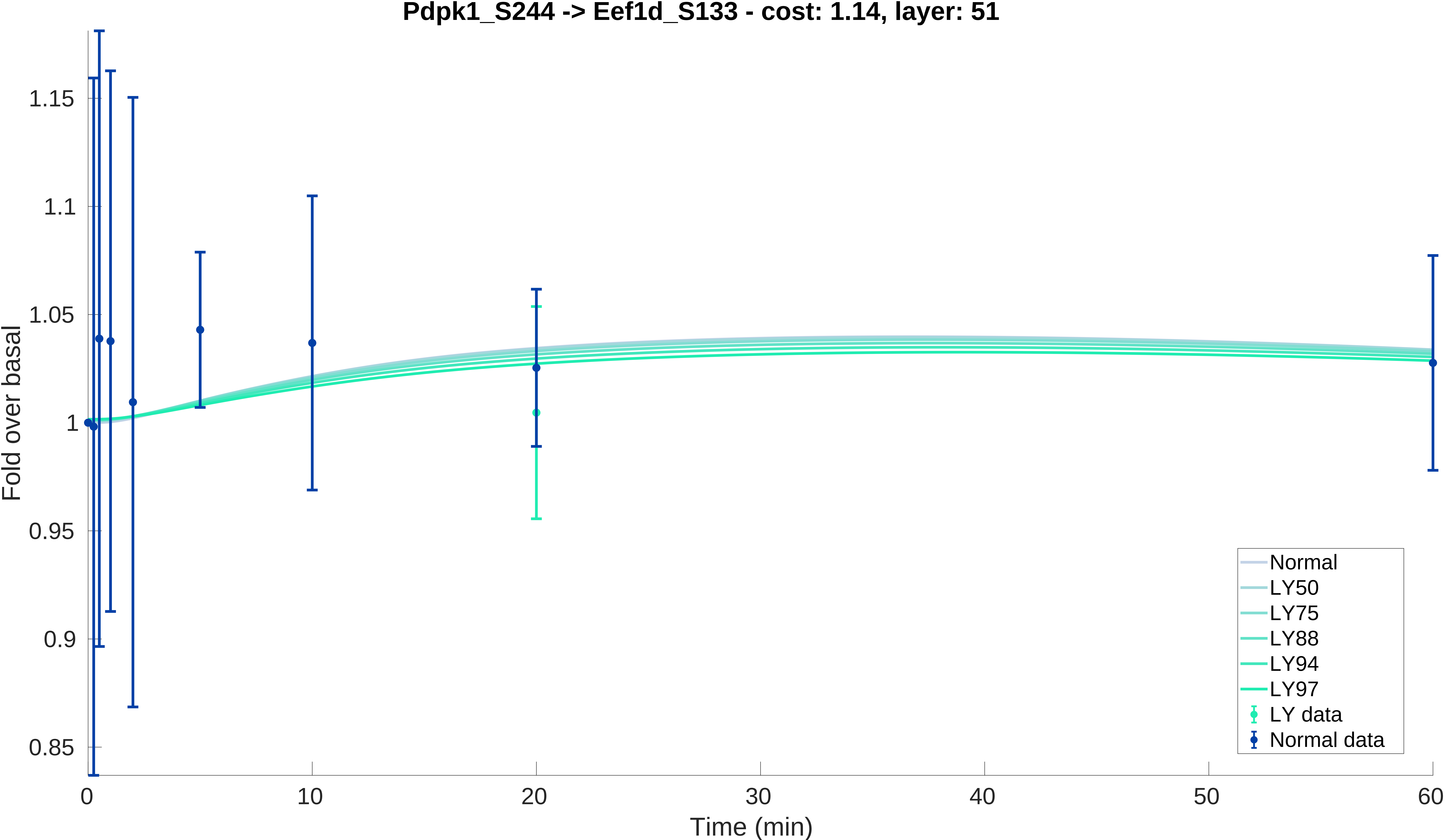

Pdpk1_S244 -> Eef1d_S133 - cost: 1.14, layer: 51
Fold over basal
1.15
1.1
1.05
1
0.95
0.9
0.85
0
10
20
30
40
50
60
Time (min)
Normal
LY50
LY75
LY88
LY94
LY97
LY data
Normal data

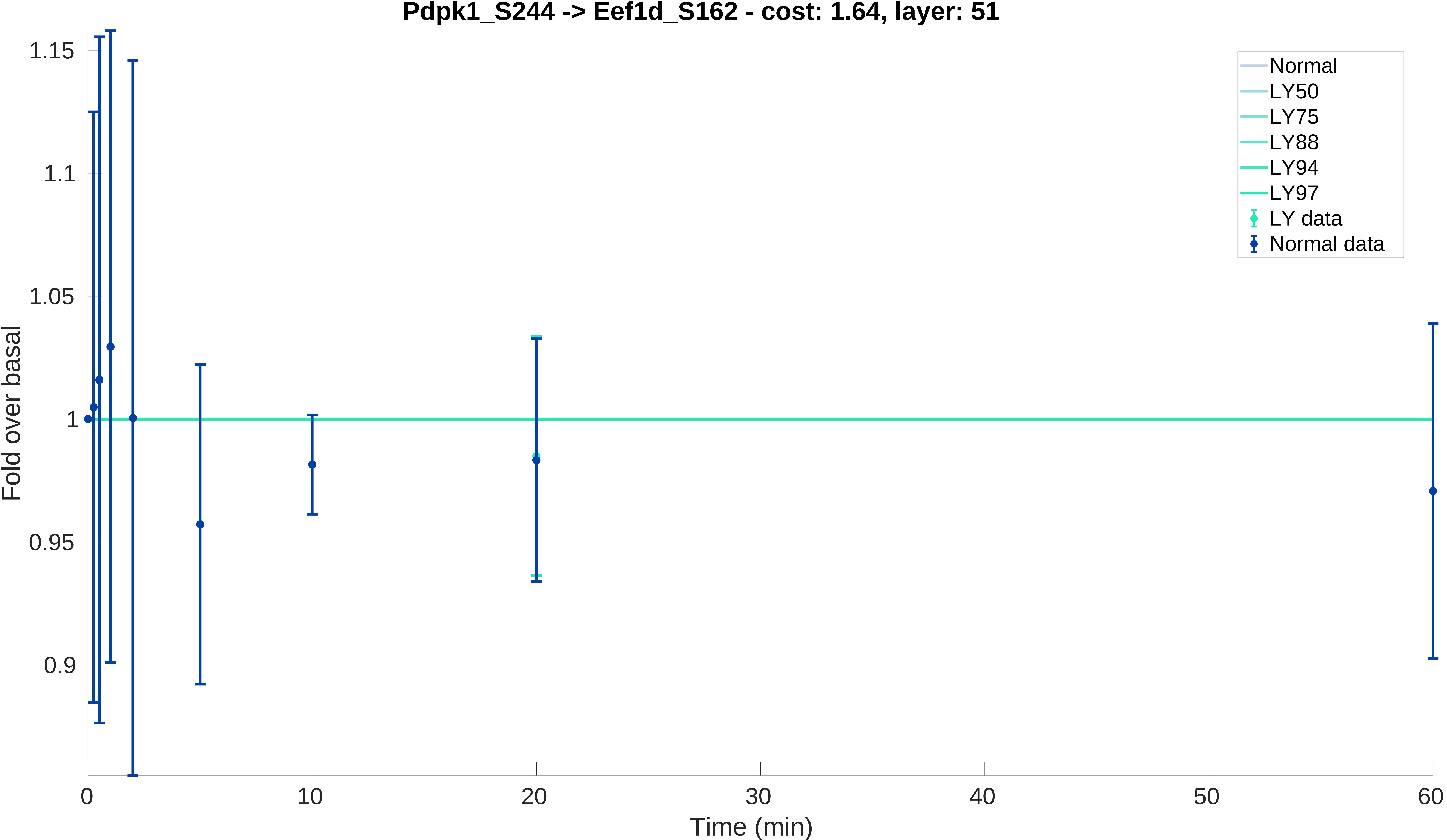
Pdpk1_S244 -> Eef1d_S162 - cost: 1.64, layer: 51
Fold over basal
1.15
1.1
1.05
1
0.95
0.9
0
10
20
30
40
50
60
Time (min)
Normal
LY50
LY75
LY88
LY94
LY97
LY data
Normal data

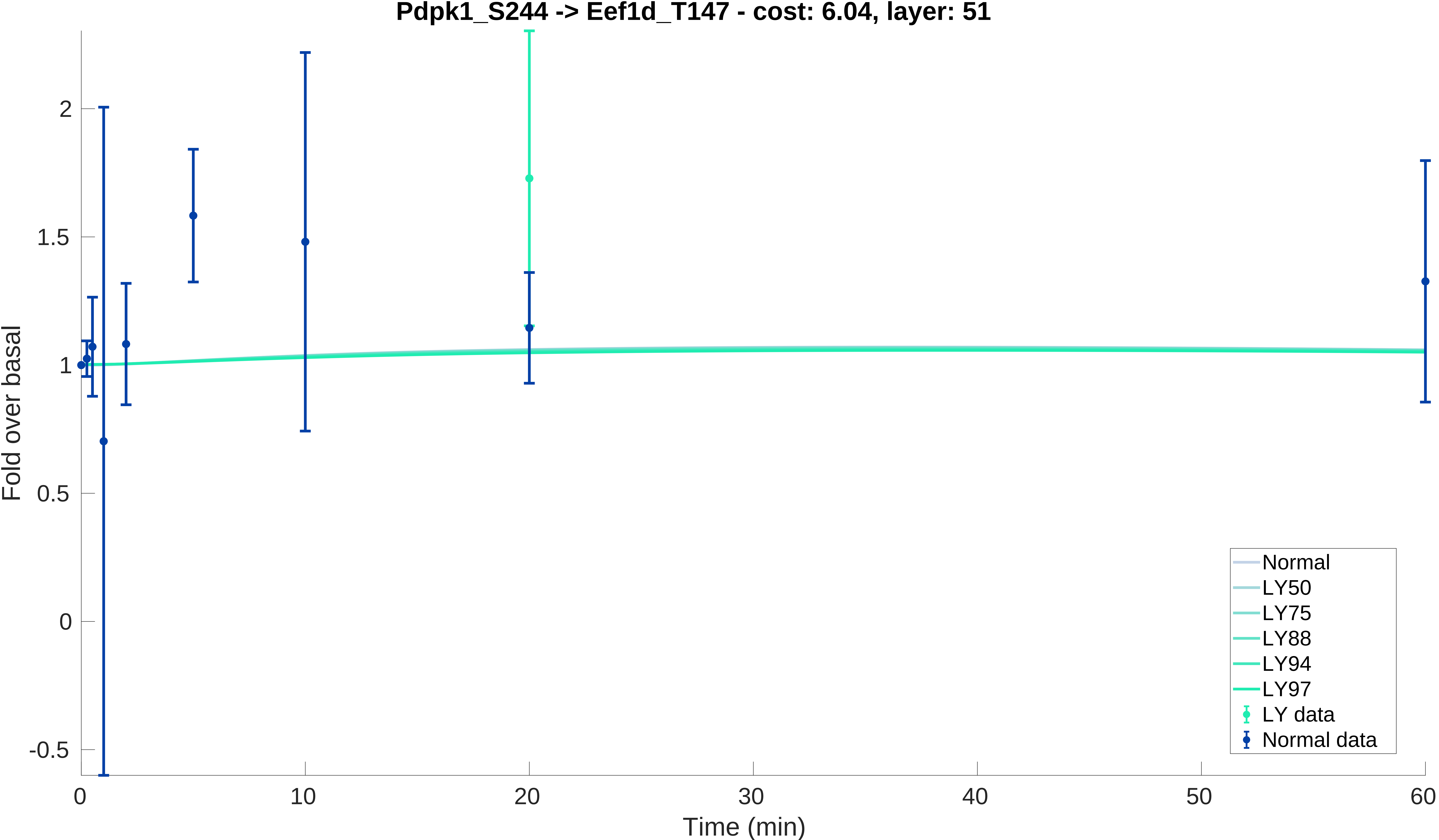

Pdpk1_S244 -> Eef1d_T147 - cost: 6.04, layer: 51
Fold over basal
Time (min)
2
1.5
1
0.5
0
-0.5
0
10
20
30
40
50
60
Normal
LY50
LY75
LY88
LY94
LY97
LY data
Normal data

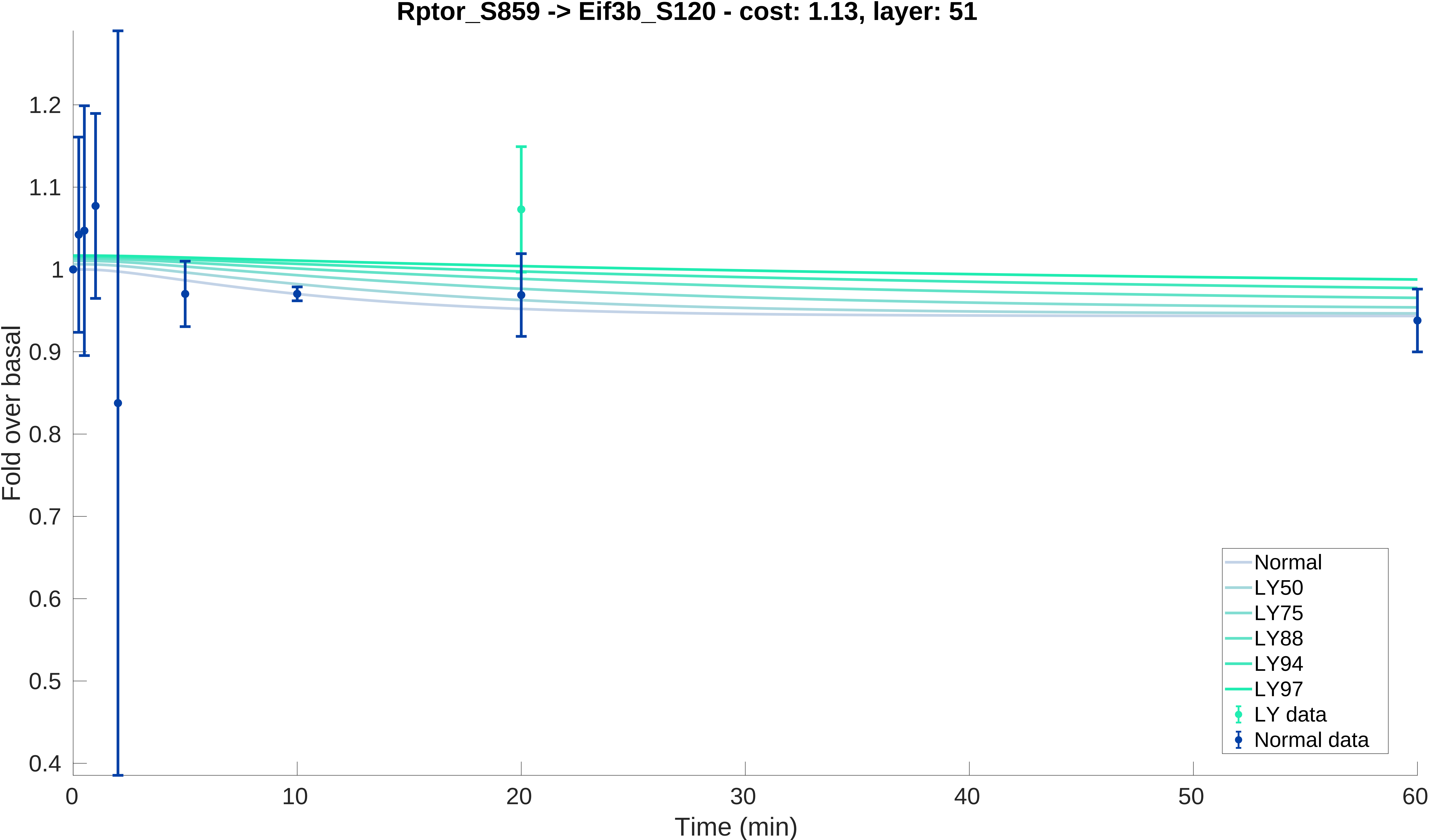

Rptor_S859 -> Eif3b_S120 - cost: 1.13, layer: 51
Fold over basal
1.2
1.1
1
0.9
0.8
0.7
0.6
0.5
0.4
0
10
20
30
40
50
60
Time (min)
Normal
LY50
LY75
LY88
LY94
LY97
LY data
Normal data

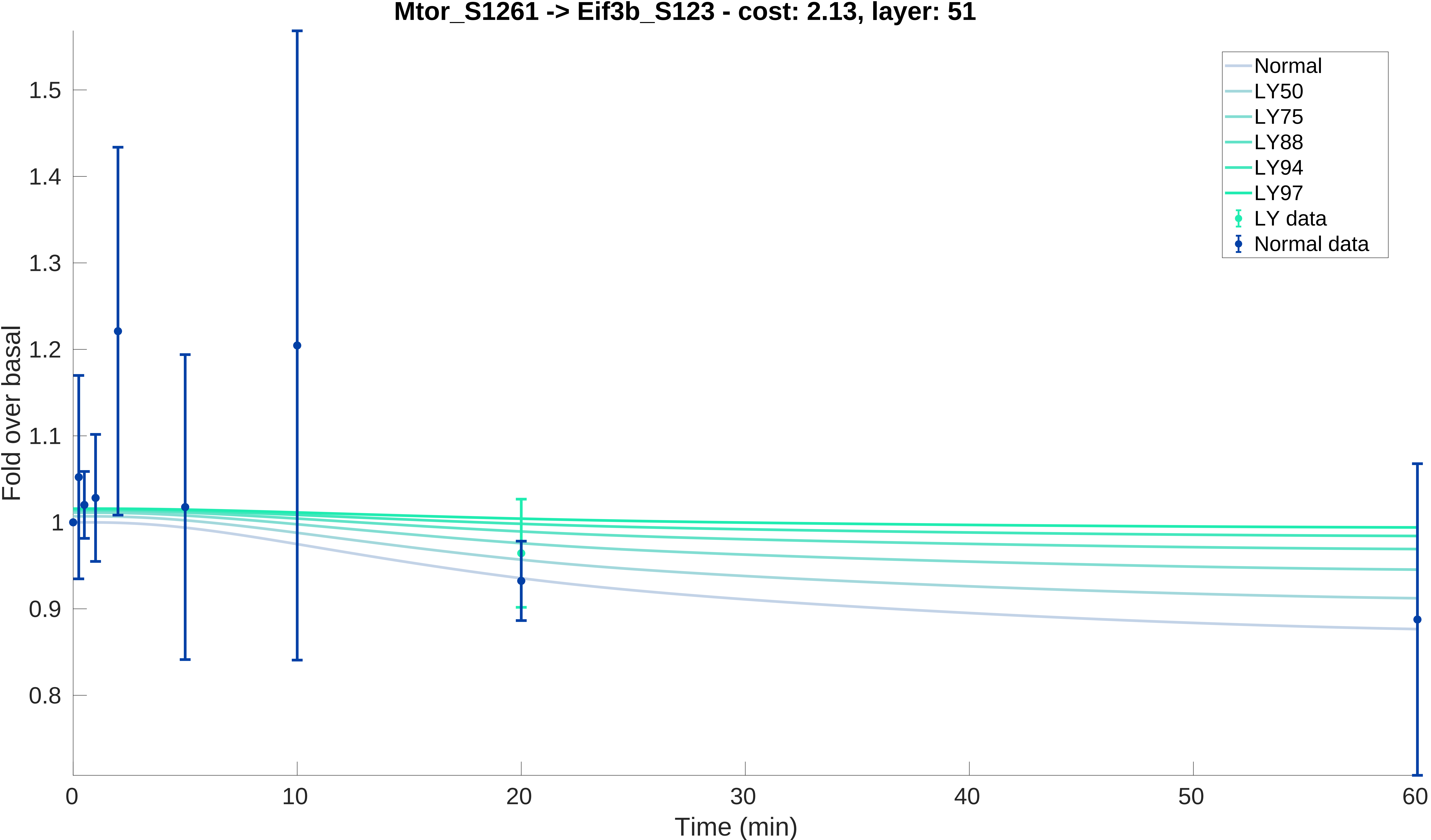
Mtor_S1261 -> Eif3b_S123 - cost: 2.13, layer: 51
Fold over basal
Time (min)
0.8
0.9
1
1.1
1.2
1.3
1.4
1.5
0
10
20
30
40
50
60
Normal
LY50
LY75
LY88
LY94
LY97
LY data
Normal data

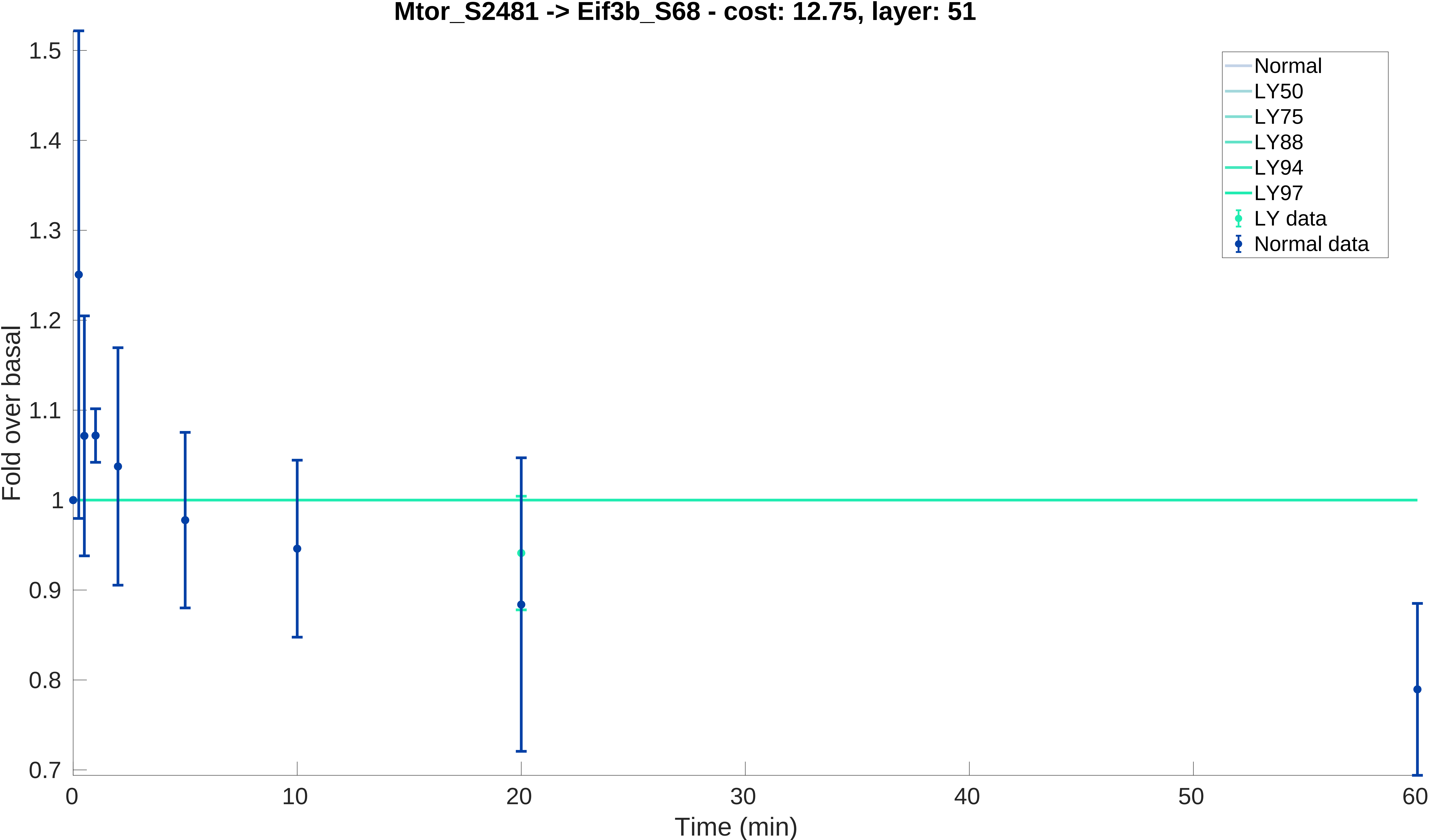

Mtor_S2481 -> Eif3b_S68 - cost: 12.75, layer: 51
Fold over basal
1.5
1.4
1.3
1.2
1.1
1
0.9
0.8
0.7
0
10
20
30
40
50
60
Time (min)
Normal
LY50
LY75
LY88
LY94
LY97
LY data
Normal data

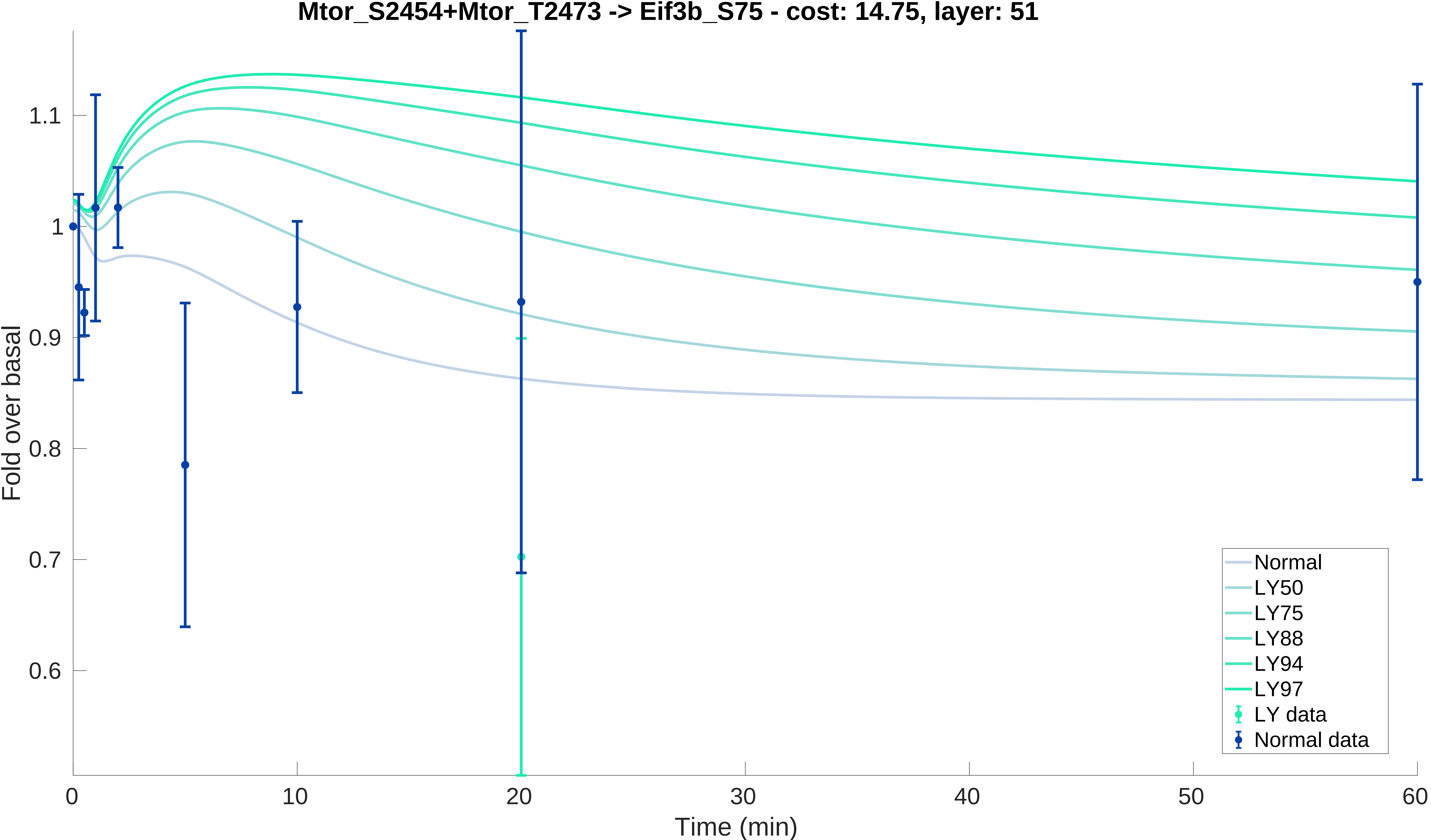

Mtor_S2454+Mtor_T2473 -> Eif3b_S75 - cost: 14.75, layer: 51
Fold over basal
Time (min)
0.6
0.7
0.8
0.9
1
1.1
0
10
20
30
40
50
60
Normal
LY50
LY75
LY88
LY94
LY97
LY data
Normal data

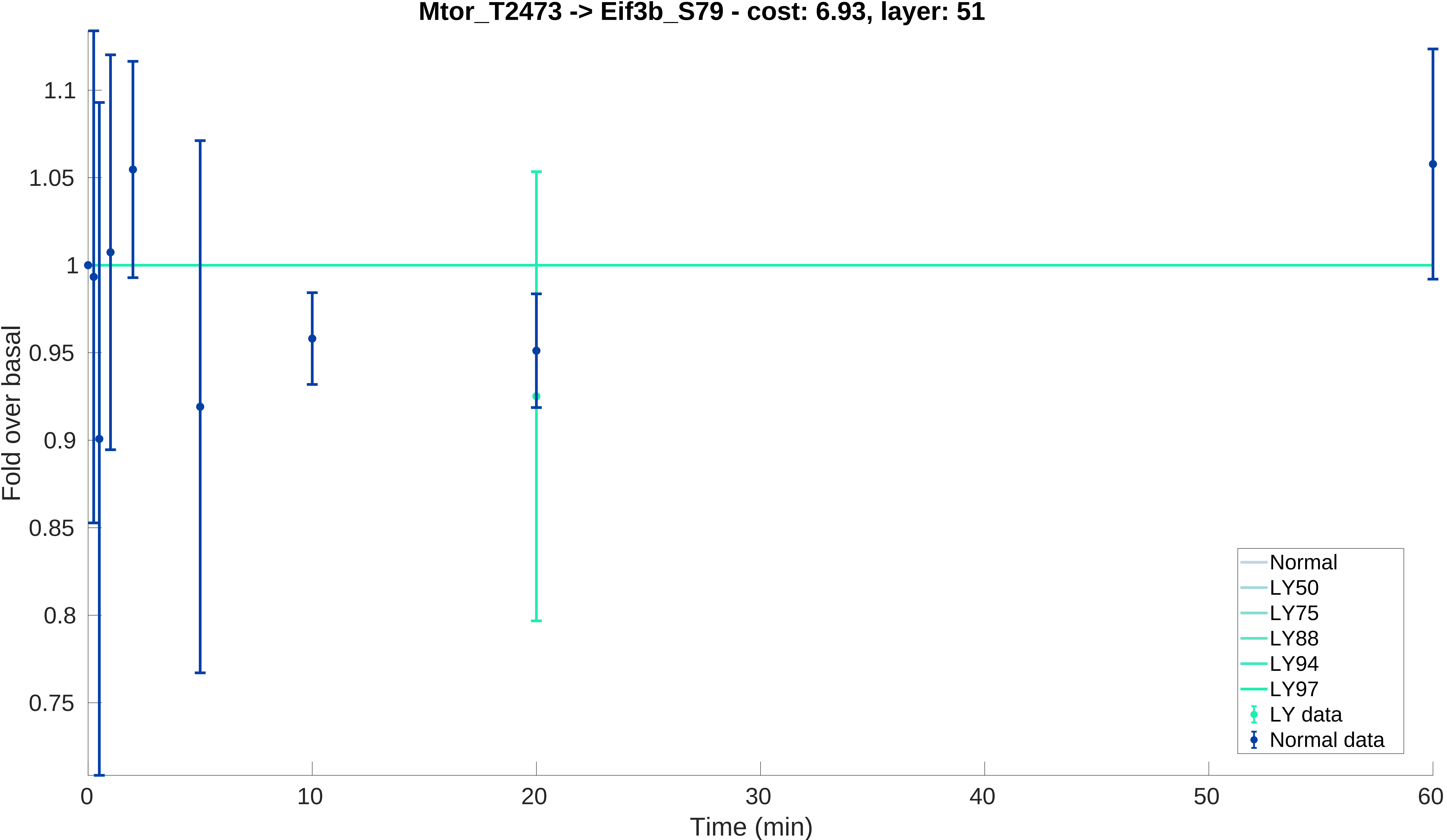

Mtor_T2473 -> Eif3b_S79 - cost: 6.93, layer: 51
Fold over basal
1.1
1.05
1
0.95
0.9
0.85
0.8
0.75
0
10
20
30
40
50
60
Time (min)
Normal
LY50
LY75
LY88
LY94
LY97
LY data
Normal data

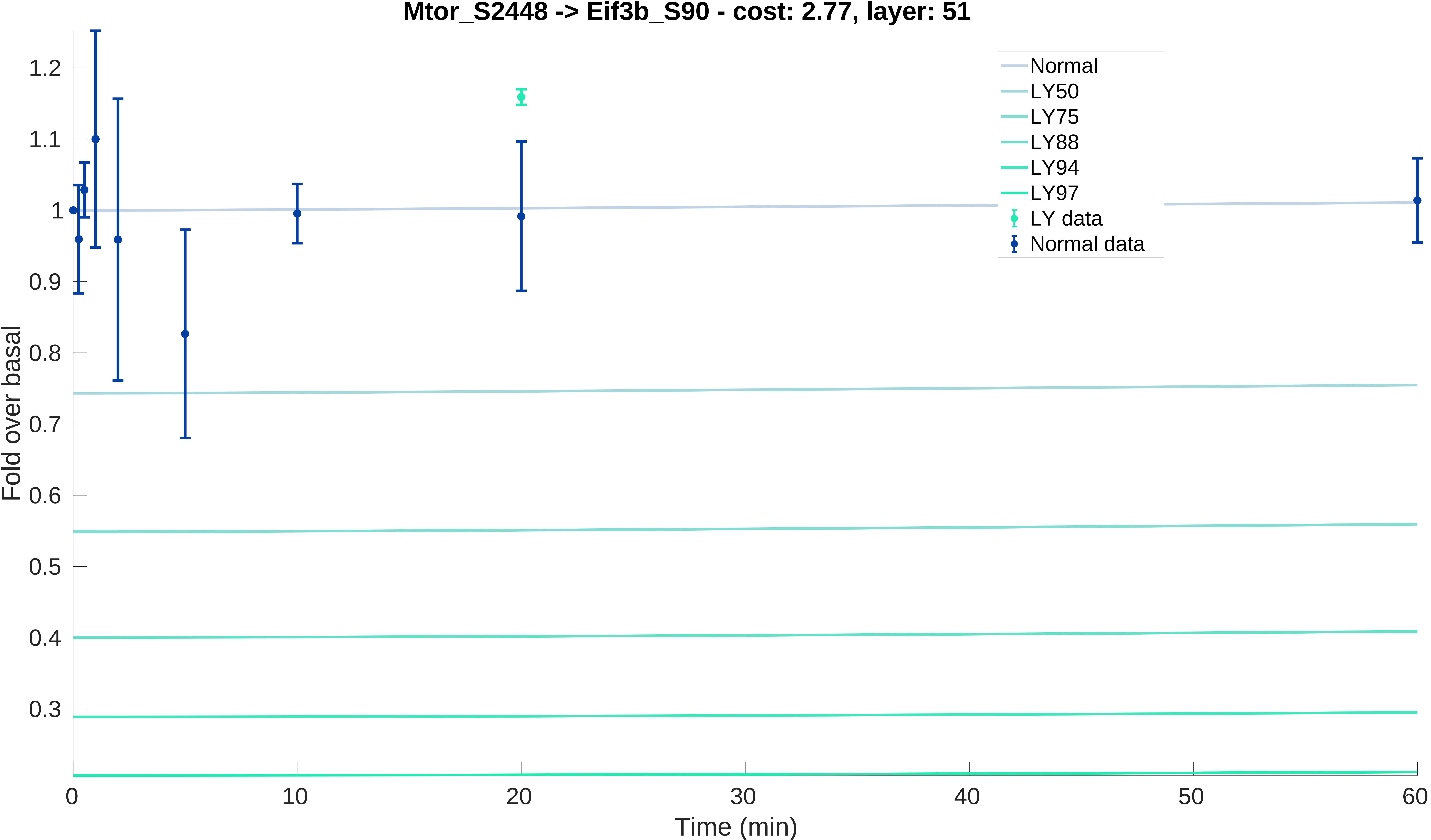

Mtor_S2448 -> Eif3b_S90 - cost: 2.77, layer: 51
Normal
LY50
LY75
LY88
LY94
LY97
LY data
Normal data
Fold over basal
Time (min)
1.2
1.1
1
0.9
0.8
0.7
0.6
0.5
0.4
0.3
0
10
20
30
40
50
60

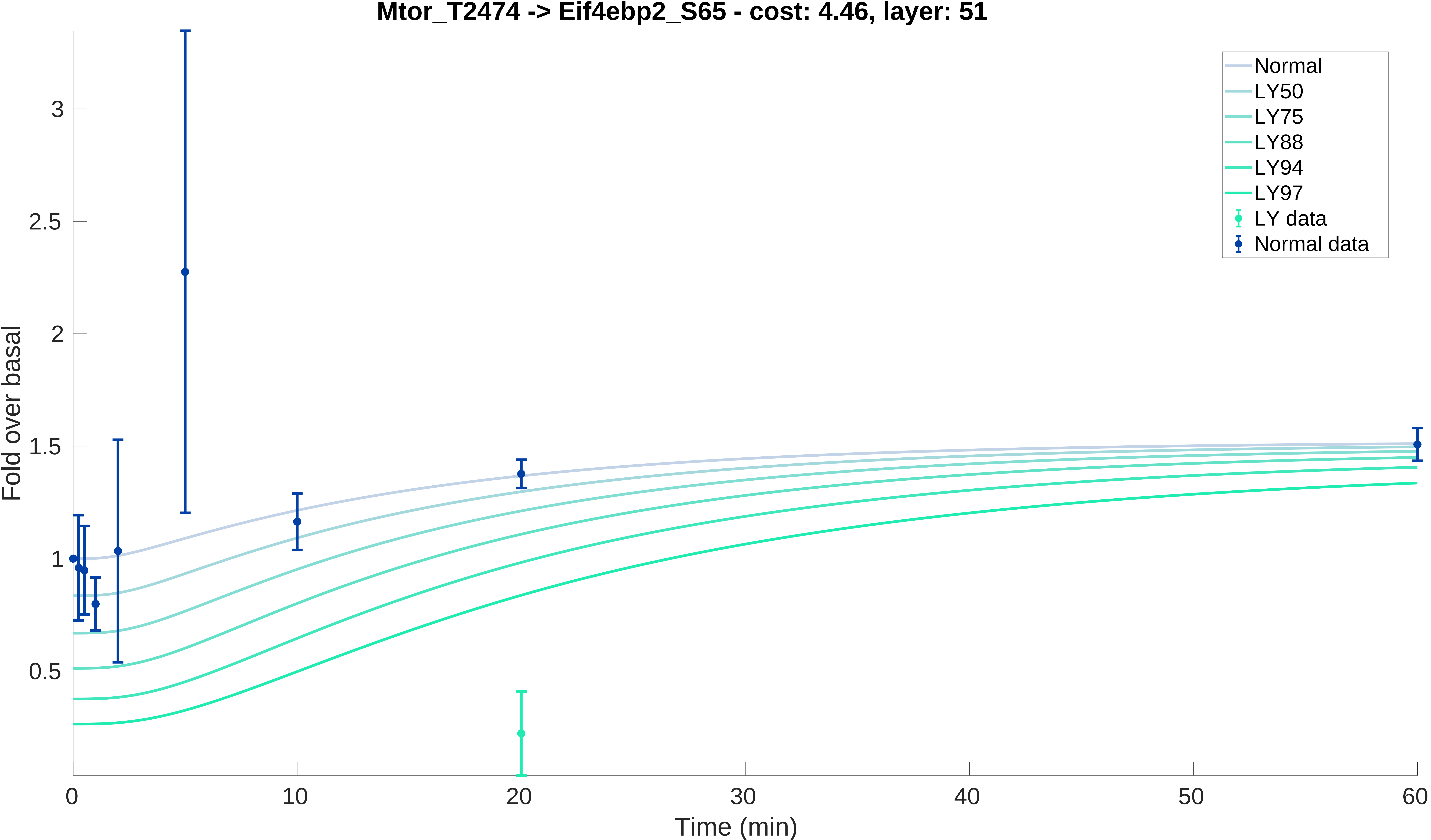
Mtor_T2474 -> Eif4ebp2_S65 - cost: 4.46, layer: 51
Fold over basal
Time (min)
0.5
1
1.5
2
2.5
3
0
10
20
30
40
50
60
Normal
LY50
LY75
LY88
LY94
LY97
LY data
Normal data

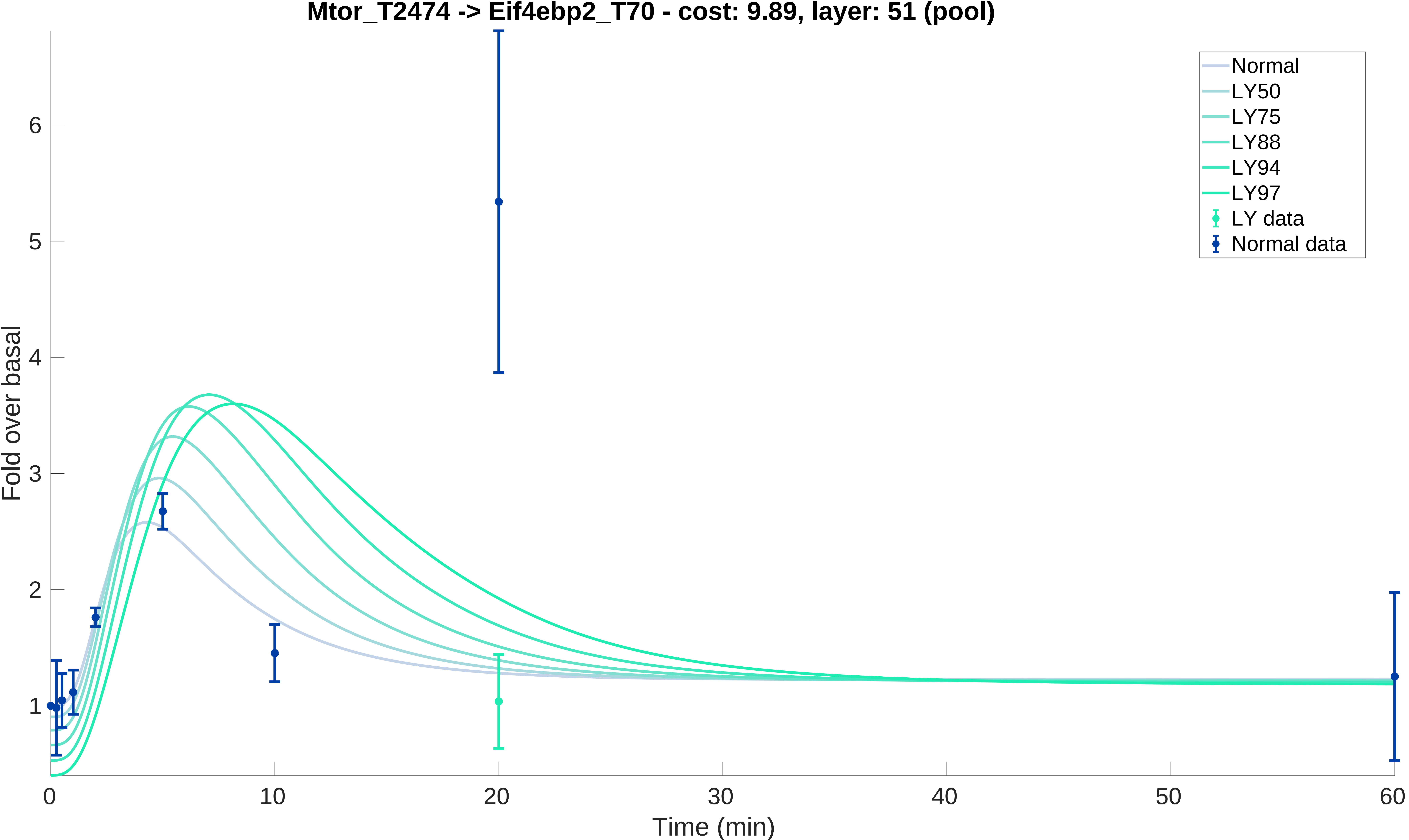

Mtor_T2474 -> Eif4ebp2_T70 - cost: 9.89, layer: 51 (pool)
Fold over basal
Time (min)
0
10
20
30
40
50
60
1
2
3
4
5
6
Normal
LY50
LY75
LY88
LY94
LY97
LY data
Normal data

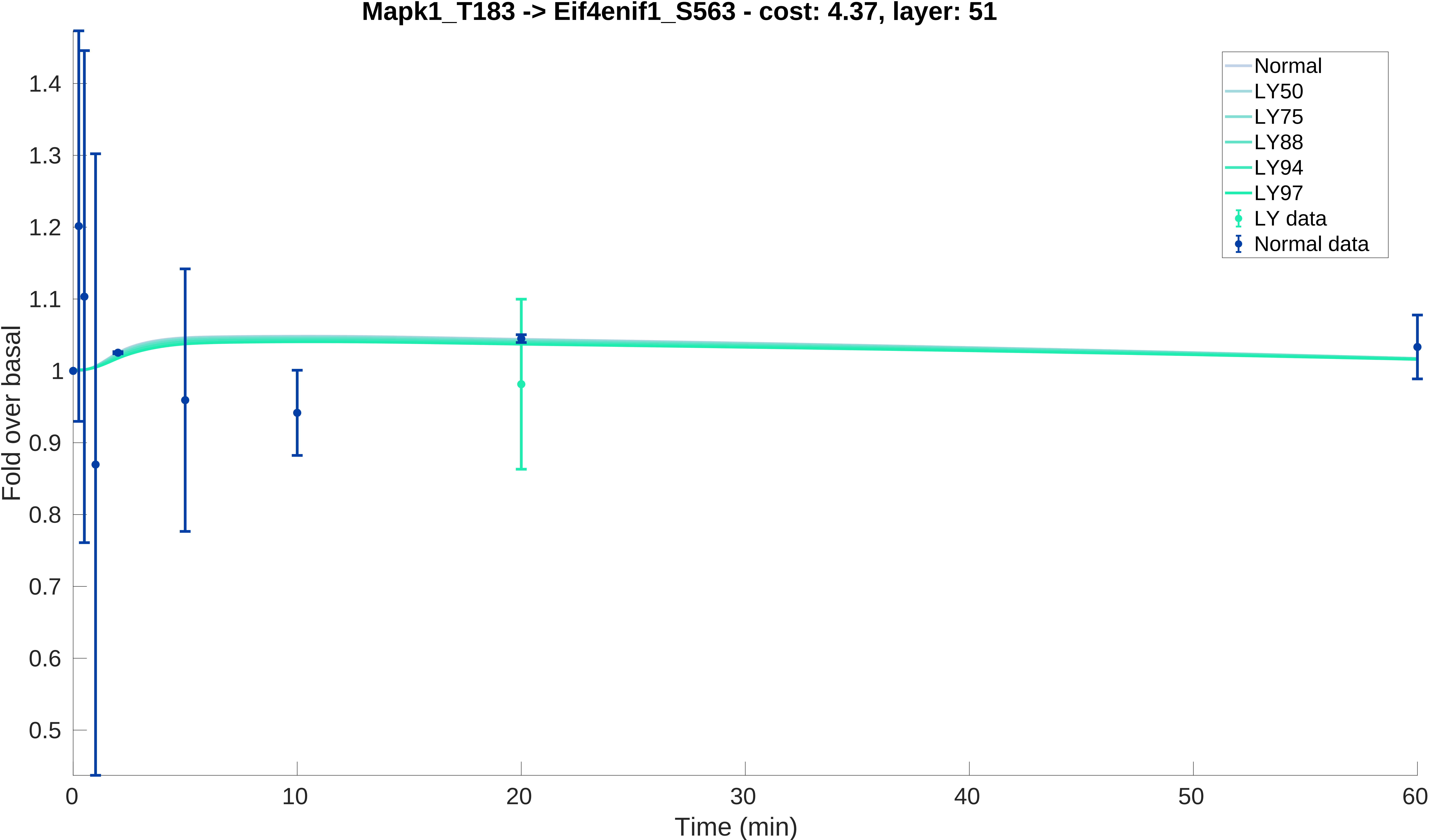

Mapk1_T183 -> Eif4enif1_S563 - cost: 4.37, layer: 51
Normal
LY50
LY75
LY88
LY94
LY97
LY data
Normal data
Fold over basal
1.4
1.3
1.2
1.1
1
0.9
0.8
0.7
0.6
0.5
0
10
20
30
40
50
60
Time (min)

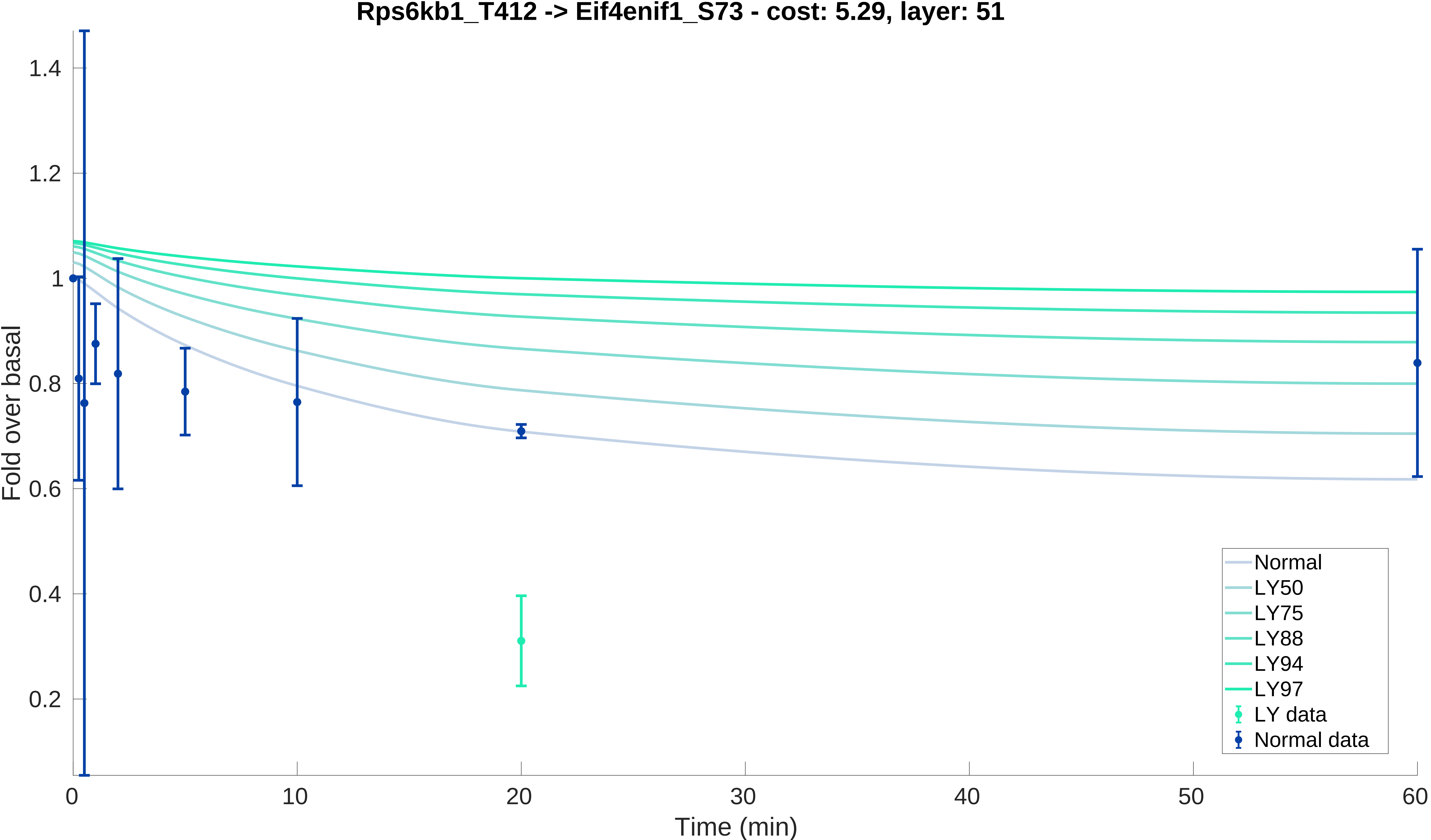
Rps6kb1_T412 -> Eif4enif1_S73 - cost: 5.29, layer: 51
Fold over basal
1.4
1.2
1
0.8
0.6
0.4
0.2
0
10
20
30
40
50
60
Time (min)
Normal
LY50
LY75
LY88
LY94
LY97
LY data
Normal data

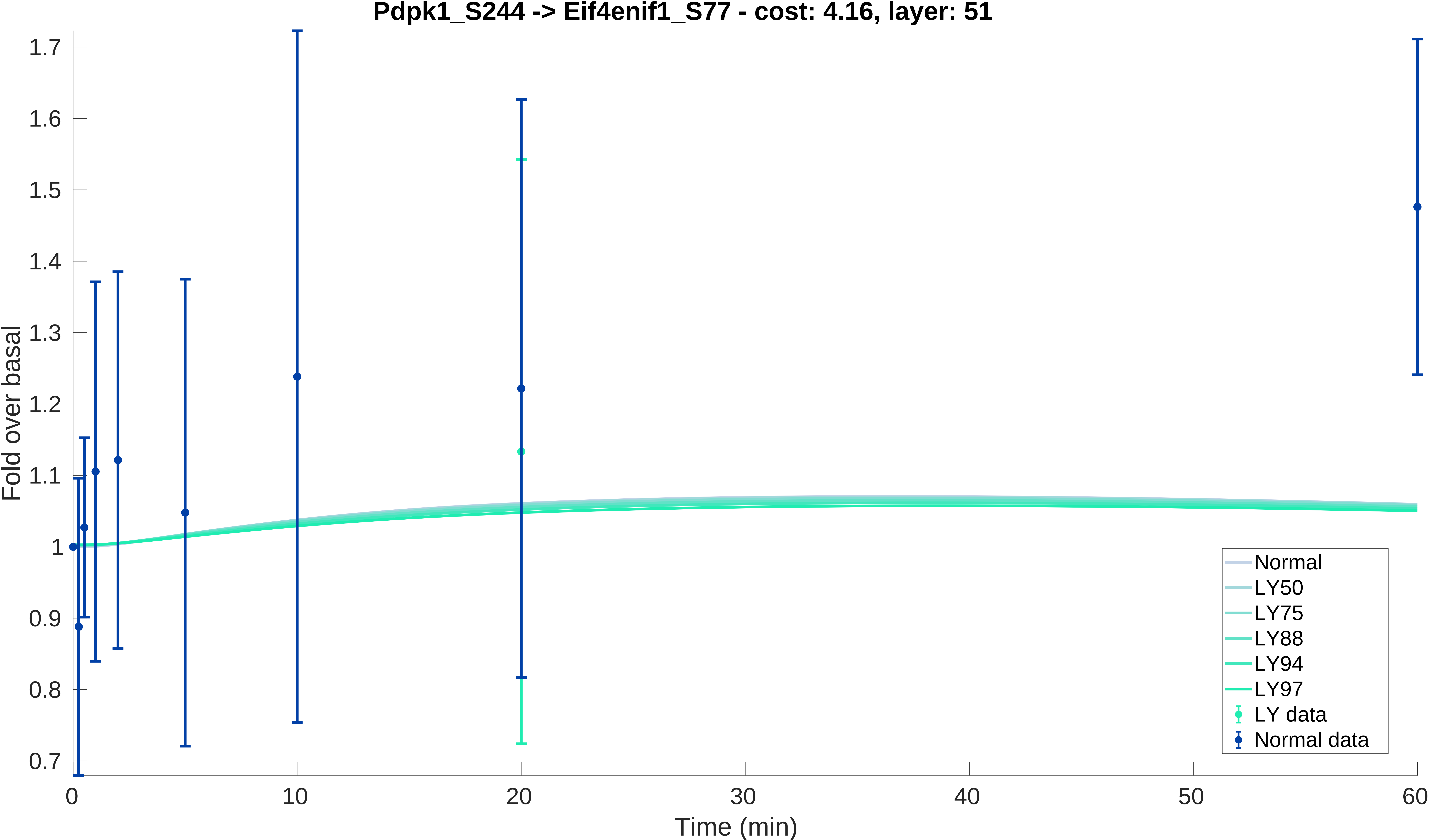

Pdpk1_S244 -> Eif4enif1_S77 - cost: 4.16, layer: 51
Fold over basal
Time (min)
0.7
0.8
0.9
1
1.1
1.2
1.3
1.4
1.5
1.6
1.7
0
10
20
30
40
50
60
Normal
LY50
LY75
LY88
LY94
LY97
LY data
Normal data

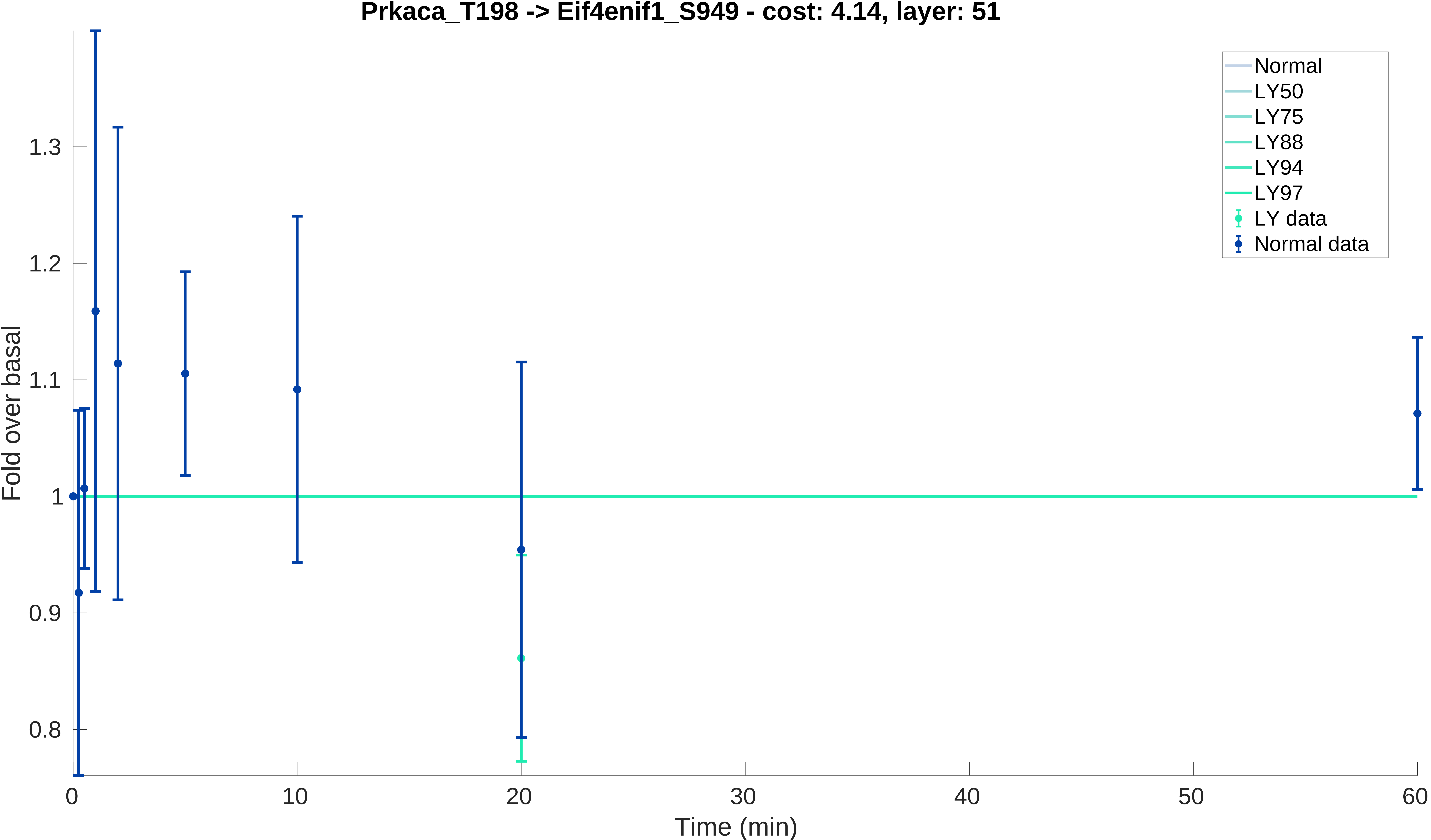
Prkaca_T198 -> Eif4enif1_S949 - cost: 4.14, layer: 51
Fold over basal
Time (min)
1.3
1.2
1.1
1
0.9
0.8
0
10
20
30
40
50
60
Normal
LY50
LY75
LY88
LY94
LY97
LY data
Normal data

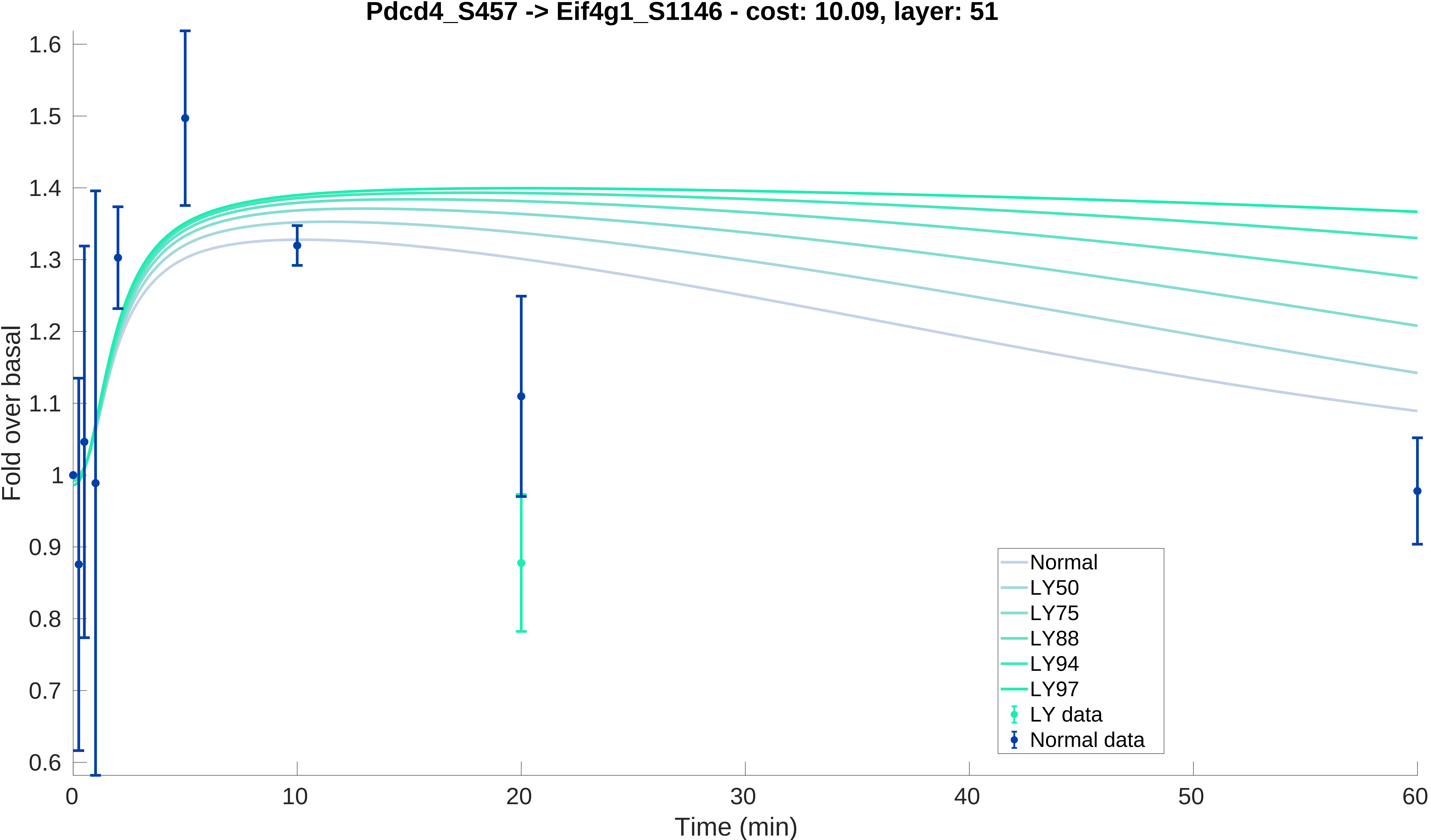
Pdcd4_S457 -> Eif4g1_S1146 - cost: 10.09, layer: 51
Fold over basal
1.6
1.5
1.4
1.3
1.2
1.1
1
0.9
0.8
0.7
0.6
0
10
20
30
40
50
60
Time (min)
Normal
LY50
LY75
LY88
LY94
LY97
LY data
Normal data

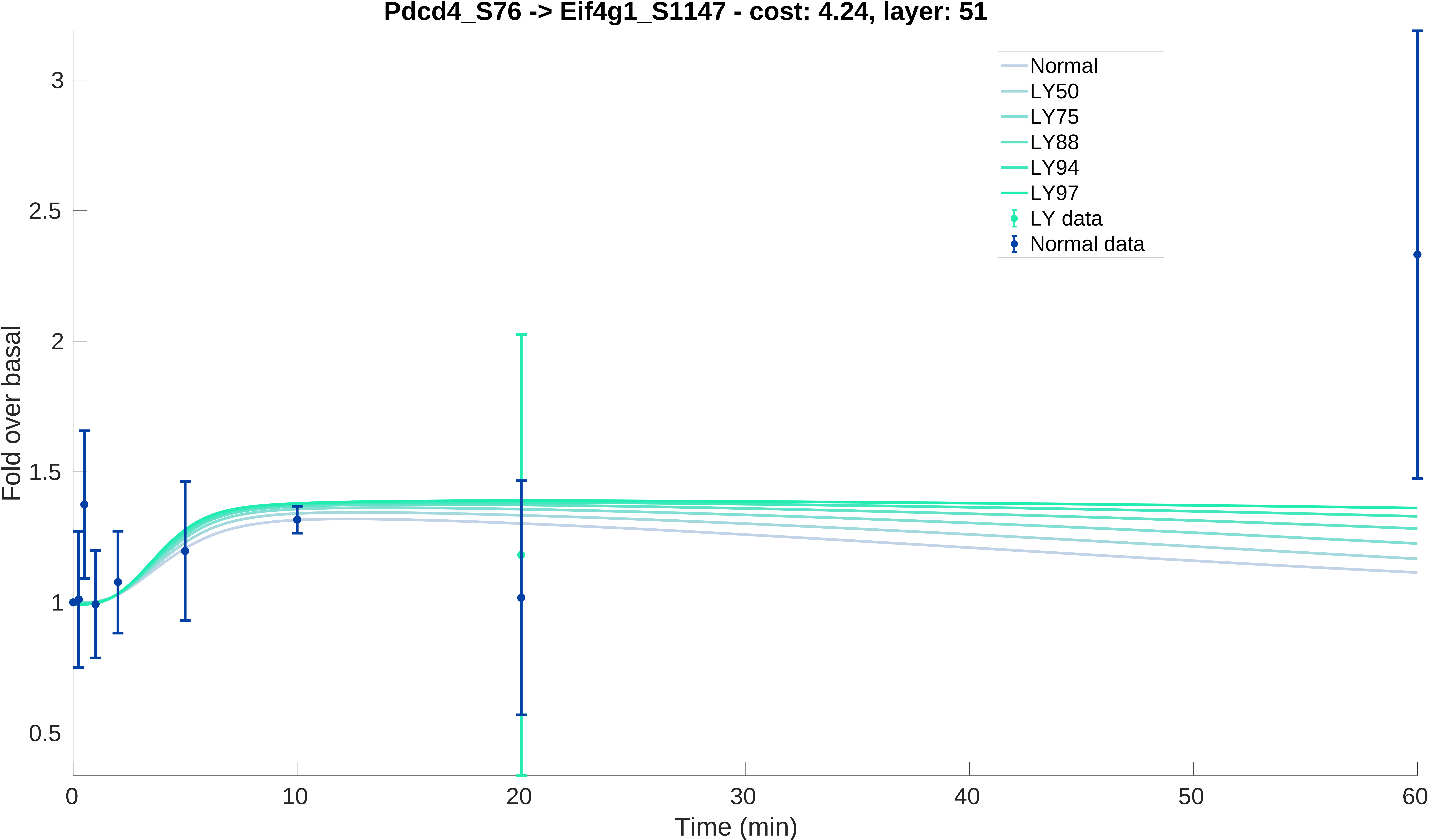

Pdcd4_S76 -> Eif4g1_S1147 - cost: 4.24, layer: 51
Normal
LY50
LY75
LY88
LY94
LY97
LY data
Normal data
Fold over basal
3
2.5
2
1.5
1
0.5
0
10
20
30
40
50
60
Time (min)

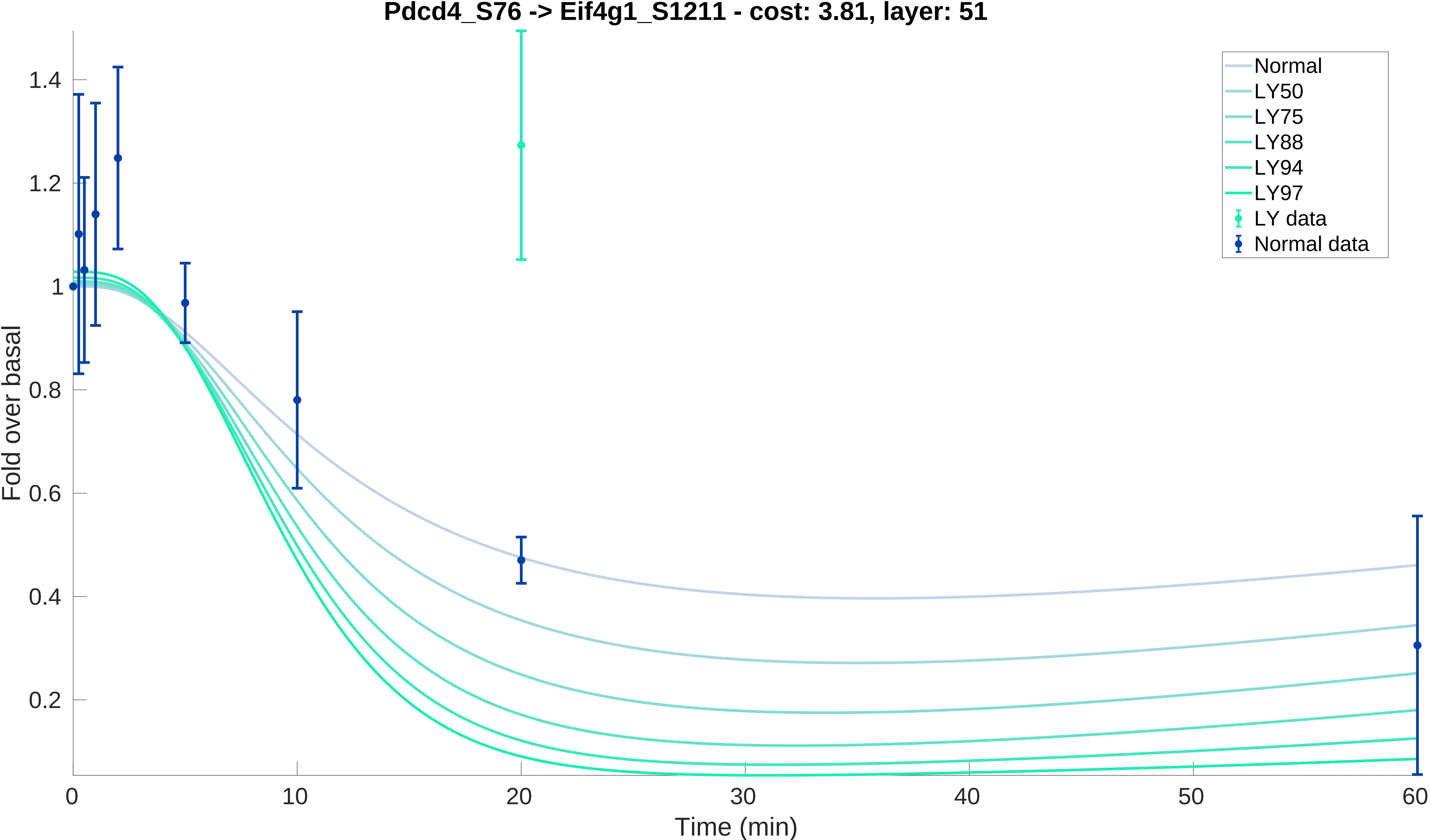

Pdcd4_S76 -> Eif4g1_S1211 - cost: 3.81, layer: 51
Fold over basal
Time (min)
1.4
1.2
1
0.8
0.6
0.4
0.2
0
10
20
30
40
50
60
Normal
LY50
LY75
LY88
LY94
LY97
LY data
Normal data

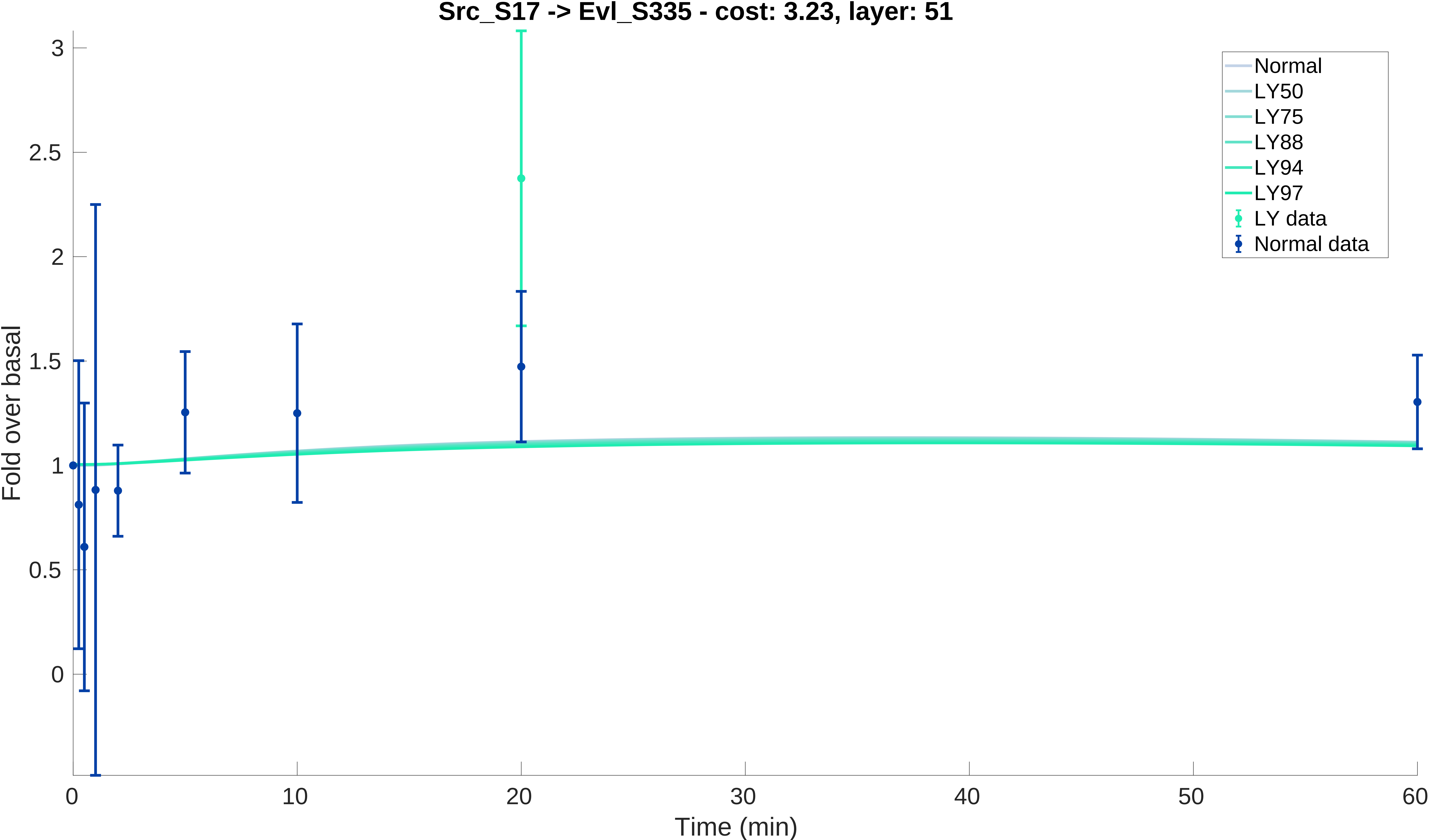

Src_S17 -> Evl_S335 - cost: 3.23, layer: 51
Fold over basal
Time (min)
0
0.5
1
1.5
2
2.5
3
0
10
20
30
40
50
60
Normal
LY50
LY75
LY88
LY94
LY97
LY data
Normal data

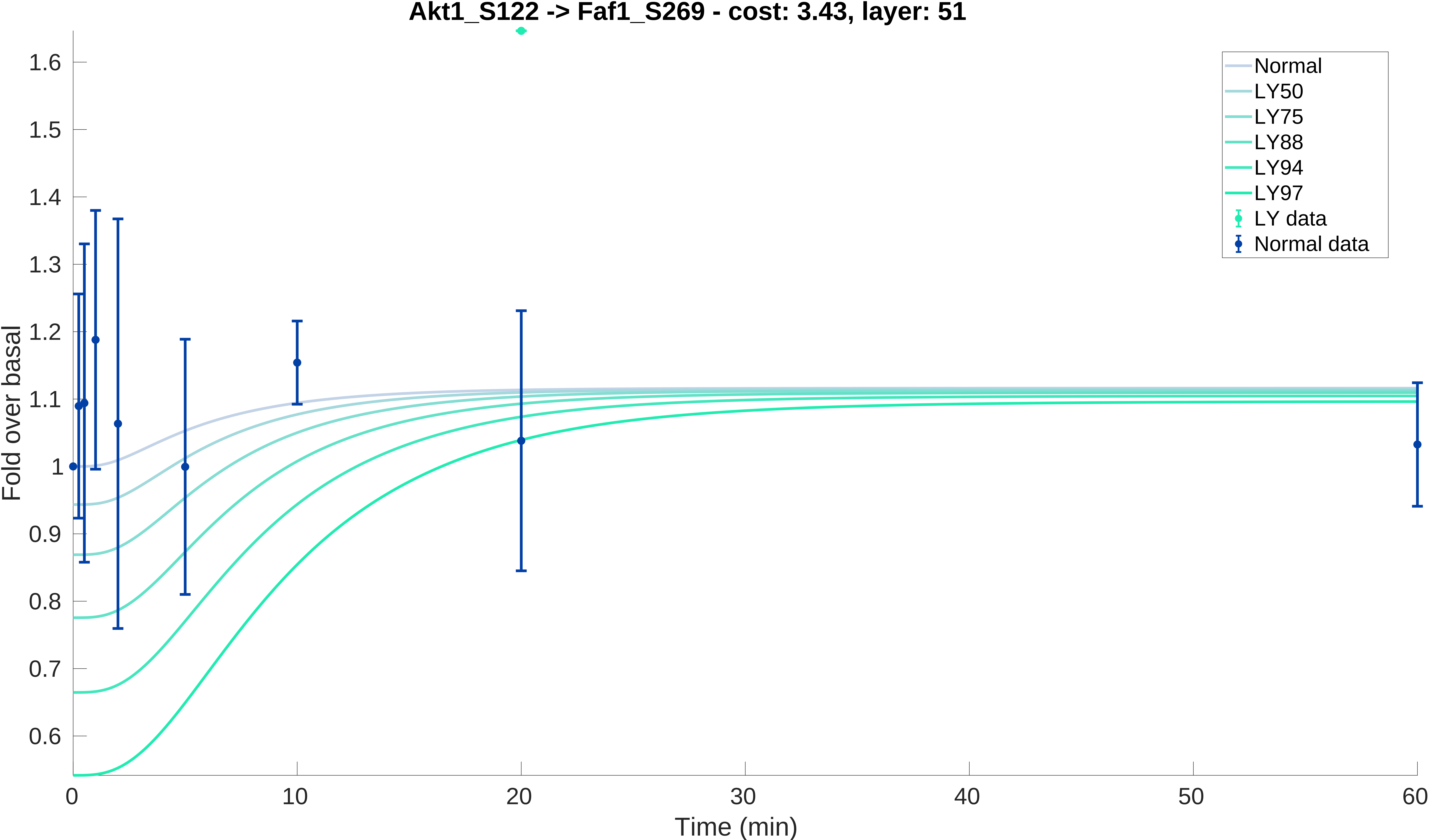

Akt1_S122 -> Faf1_S269 - cost: 3.43, layer: 51
Fold over basal
Time (min)
1.6
1.5
1.4
1.3
1.2
1.1
1
0.9
0.8
0.7
0.6
0
10
20
30
40
50
60
Normal
LY50
LY75
LY88
LY94
LY97
LY data
Normal data

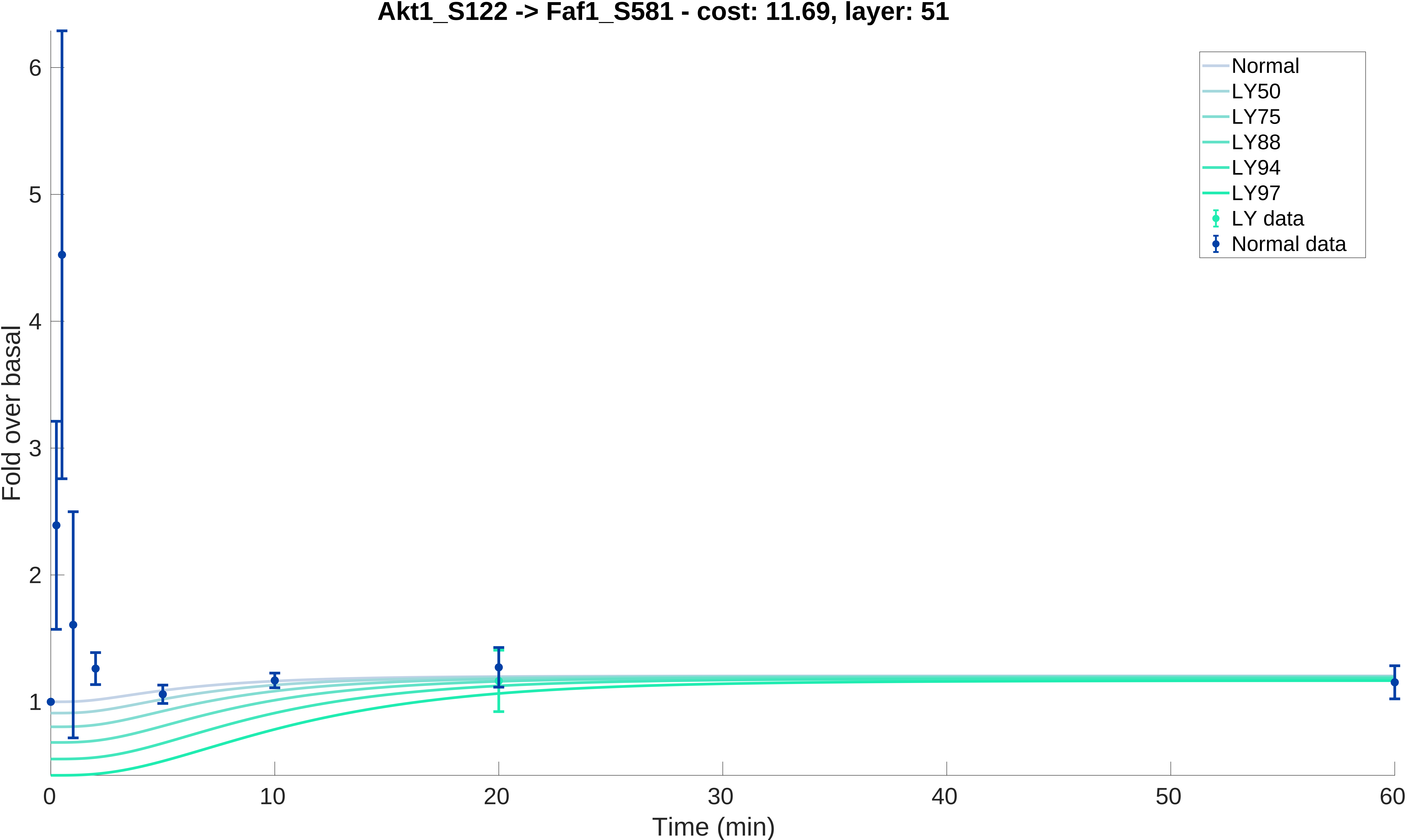

Akt1_S122 -> Faf1_S581 - cost: 11.69, layer: 51
Fold over basal
Time (min)
0
10
20
30
40
50
60
1
2
3
4
5
6
Normal
LY50
LY75
LY88
LY94
LY97
LY data
Normal data

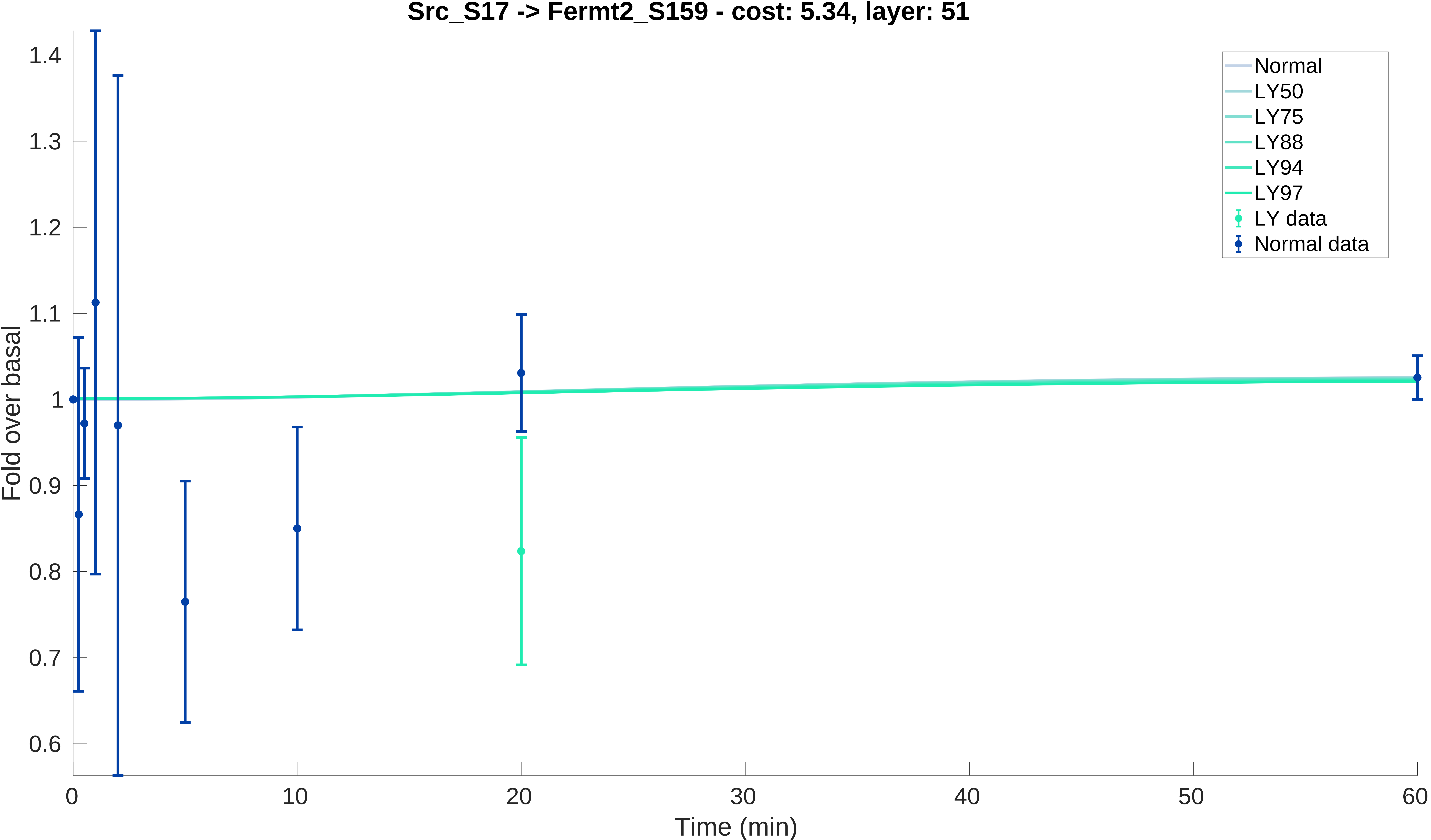

Src_S17 -> Fermt2_S159 - cost: 5.34, layer: 51
Fold over basal
Time (min)
1.4
1.3
1.2
1.1
1
0.9
0.8
0.7
0.6
0
10
20
30
40
50
60
Normal
LY50
LY75
LY88
LY94
LY97
LY data
Normal data

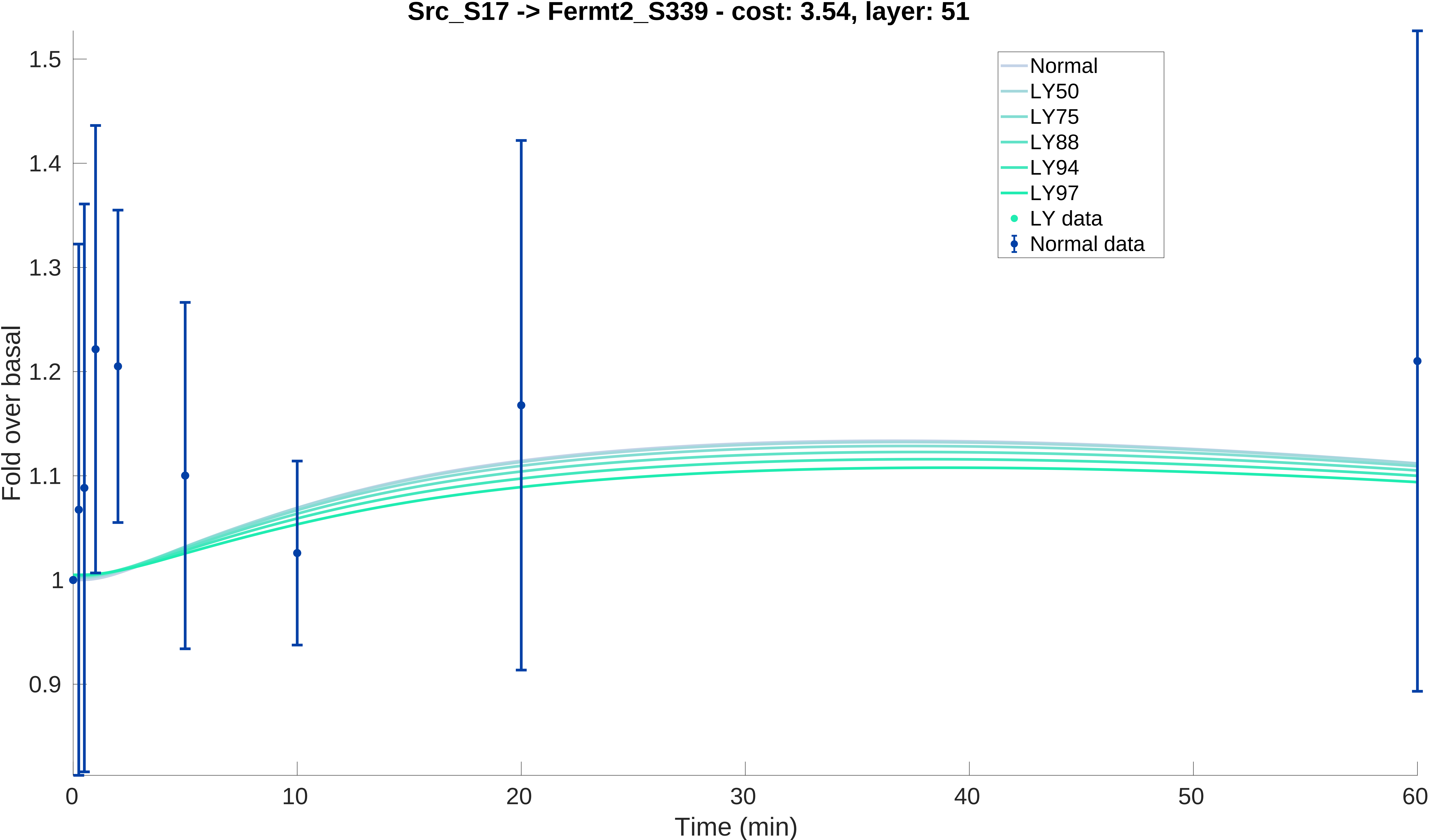

Src_S17 -> Fermt2_S339 - cost: 3.54, layer: 51
Normal
LY50
LY75
LY88
LY94
LY97
LY data
Normal data
Fold over basal
Time (min)
0.9
1
1.1
1.2
1.3
1.4
1.5
0
10
20
30
40
50
60

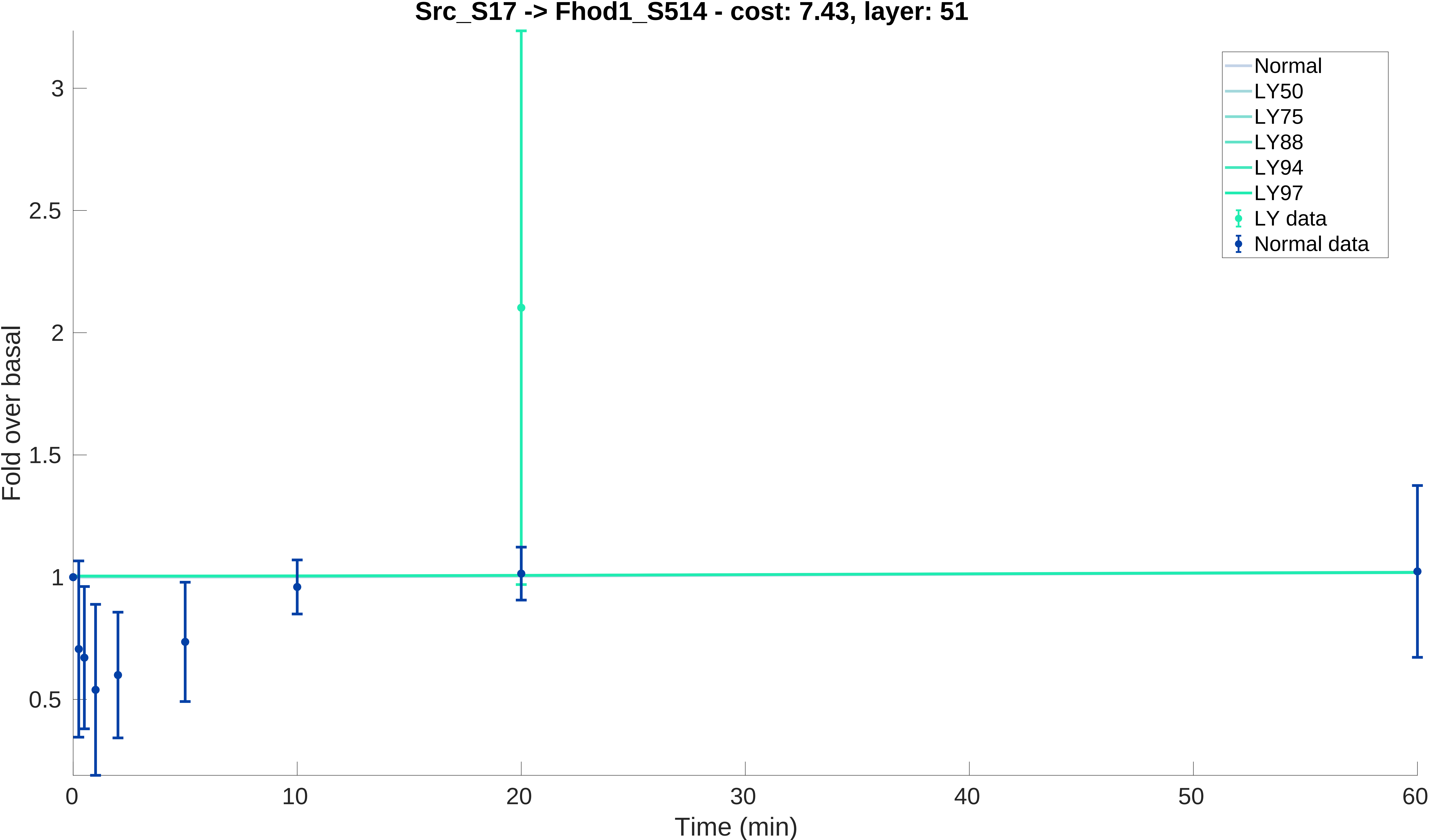

Src_S17 -> Fhod1_S514 - cost: 7.43, layer: 51
Fold over basal
Time (min)
0.5
1
1.5
2
2.5
3
0
10
20
30
40
50
60
Normal
LY50
LY75
LY88
LY94
LY97
LY data
Normal data

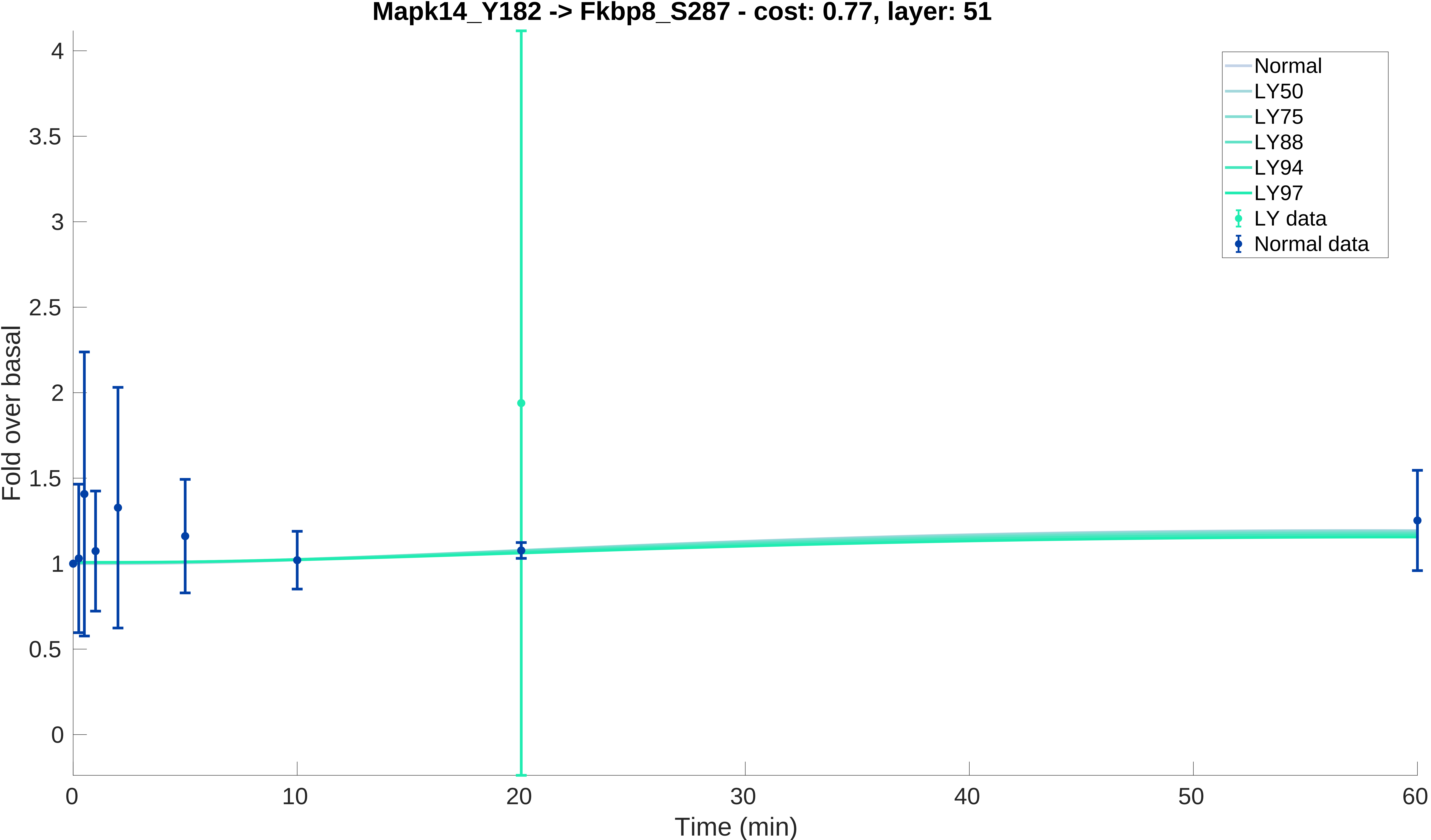

Mapk14_Y182 -> Fkbp8_S287 - cost: 0.77, layer: 51
Fold over basal
Time (min)
0
0.5
1
1.5
2
2.5
3
3.5
4
0
10
20
30
40
50
60
Normal
LY50
LY75
LY88
LY94
LY97
LY data
Normal data

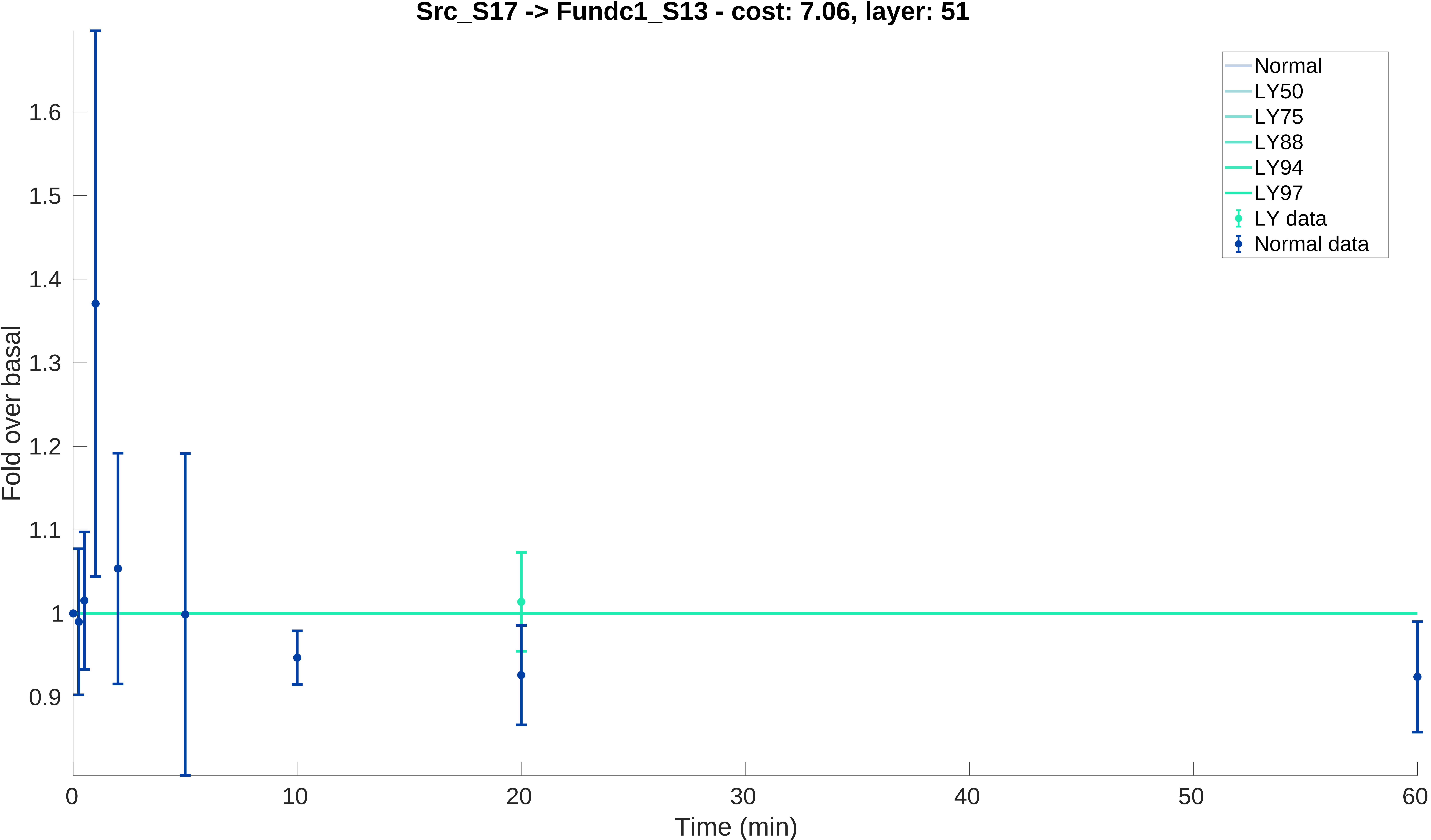
Src_S17 -> Fundc1_S13 - cost: 7.06, layer: 51
Fold over basal
1.6
1.5
1.4
1.3
1.2
1.1
1
0.9
0
10
20
30
40
50
60
Time (min)
Normal
LY50
LY75
LY88
LY94
LY97
LY data
Normal data

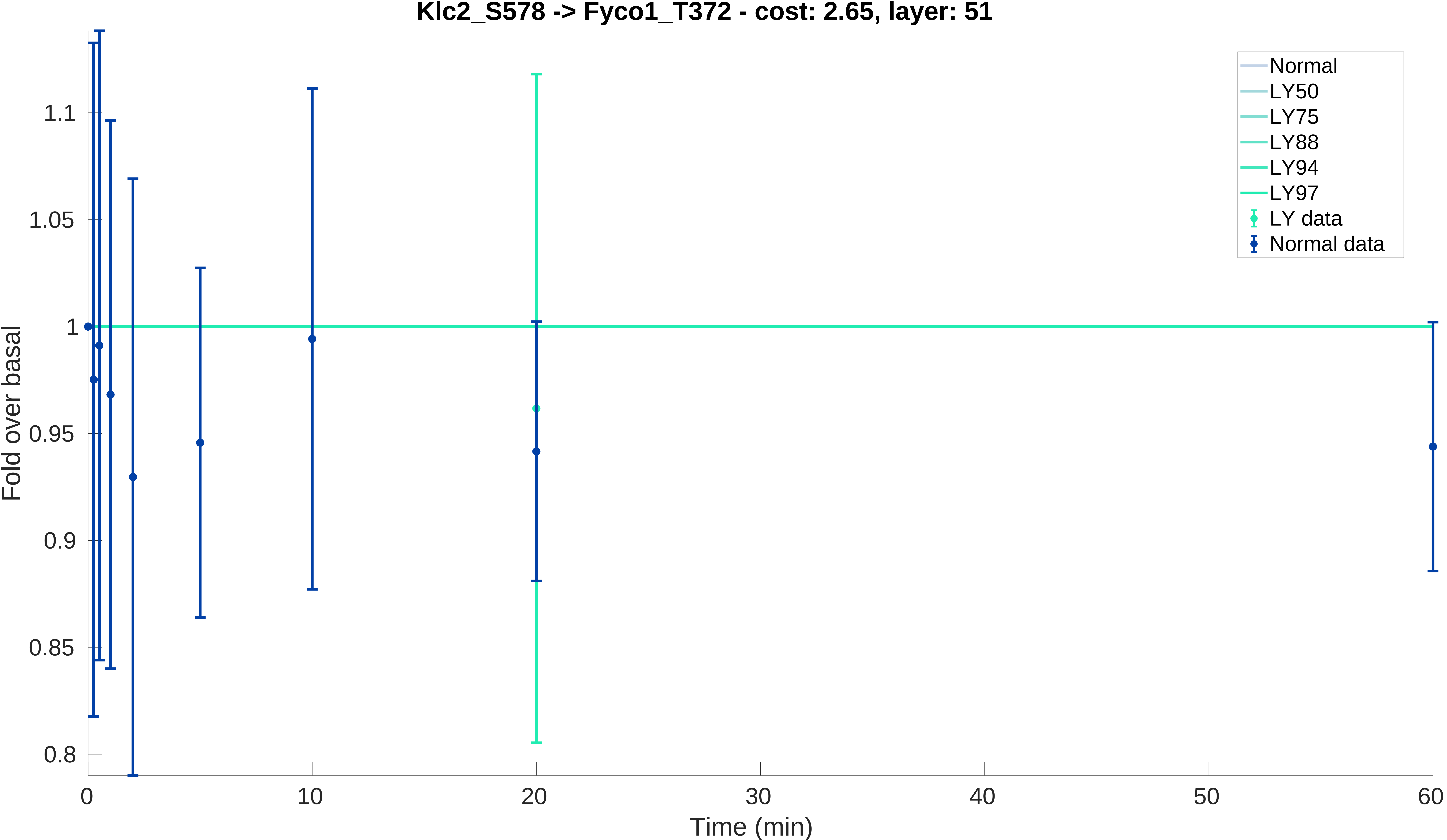

Klc2_S578 -> Fyco1_T372 - cost: 2.65, layer: 51
Normal
LY50
LY75
LY88
LY94
LY97
LY data
Normal data
Fold over basal
1.1
1.05
1
0.95
0.9
0.85
0.8
0
10
20
30
40
50
60
Time (min)

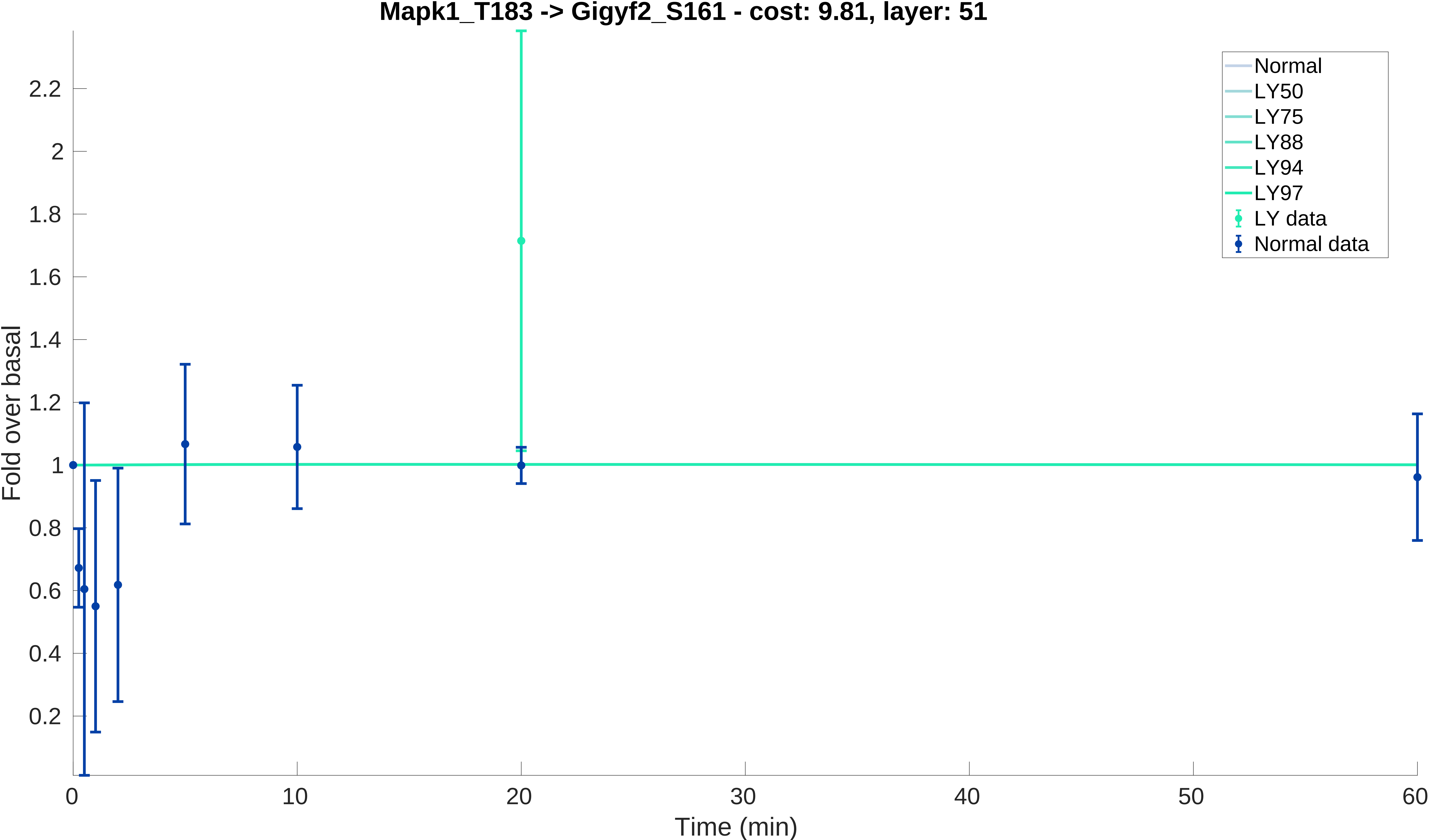
Mapk1_T183 -> Gigyf2_S161 - cost: 9.81, layer: 51
Fold over basal
Time (min)
Normal
LY50
LY75
LY88
LY94
LY97
LY data
Normal data
0.2
0.4
0.6
0.8
1
1.2
1.4
1.6
1.8
2
2.2
0
10
20
30
40
50
60

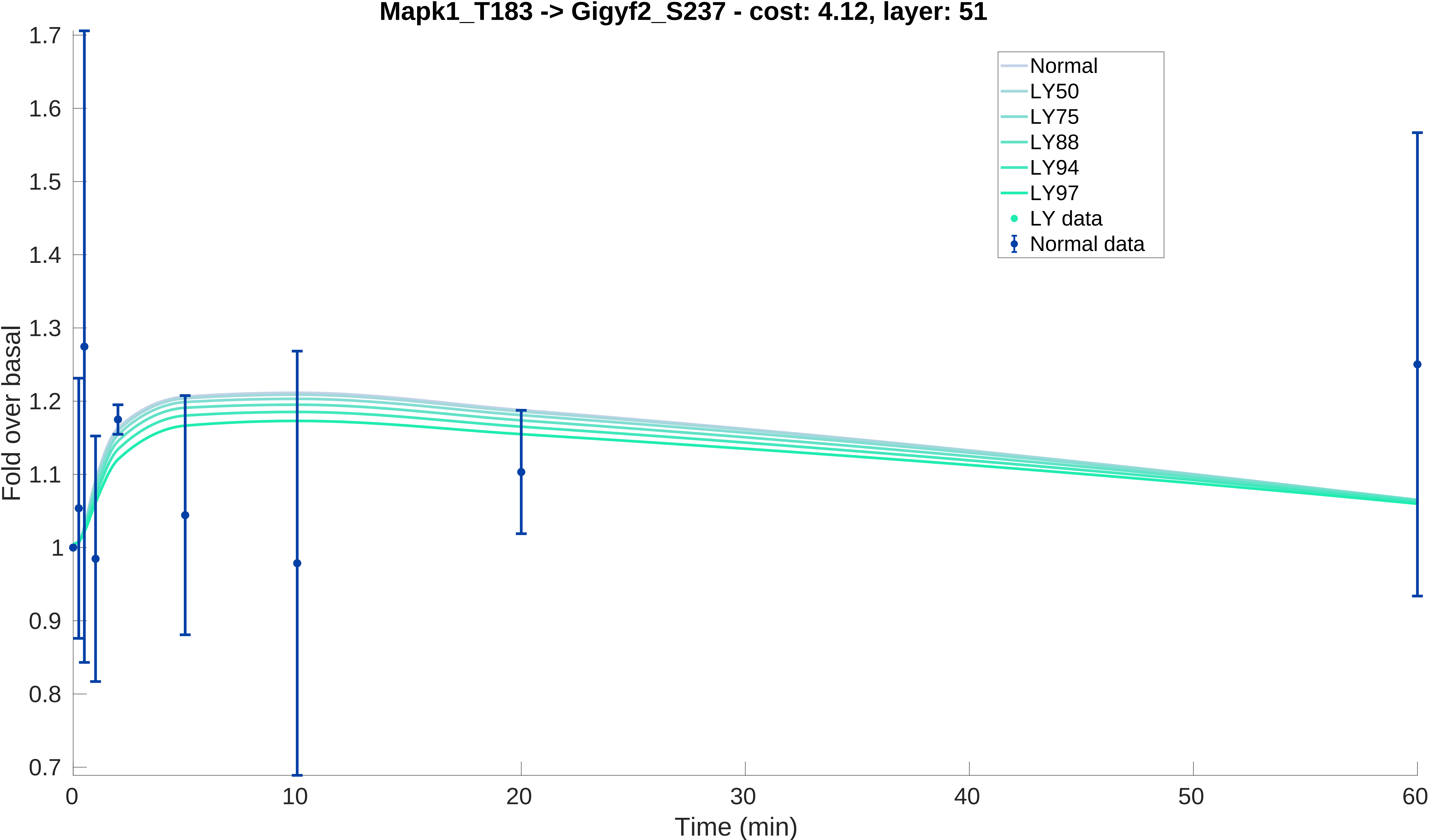

Mapk1_T183 -> Gigyf2_S237 - cost: 4.12, layer: 51
Fold over basal
Time (min)
Normal
LY50
LY75
LY88
LY94
LY97
LY data
Normal data
1.7
1.6
1.5
1.4
1.3
1.2
1.1
1
0.9
0.8
0.7
0
10
20
30
40
50
60

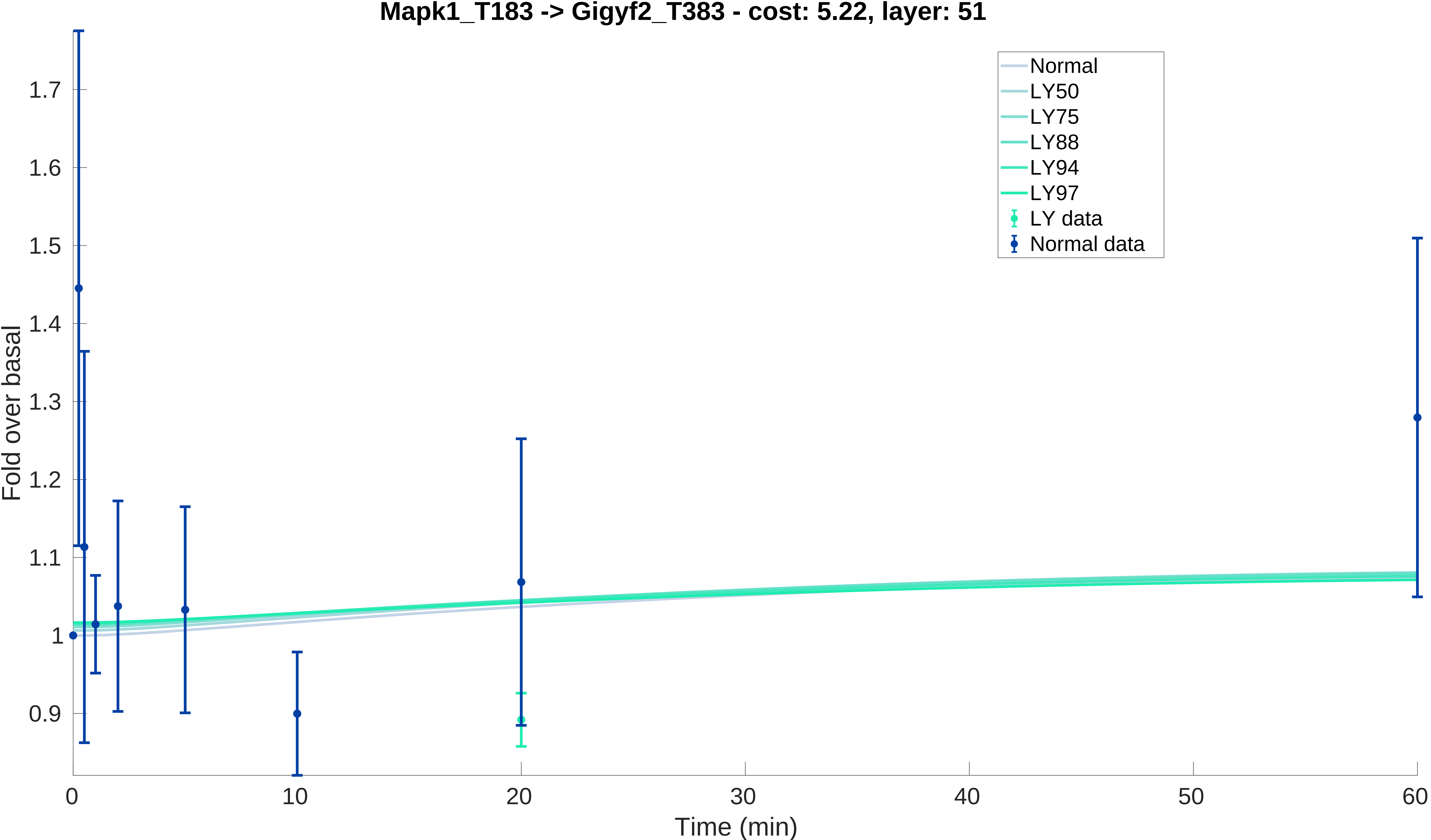

Mapk1_T183 -> Gigyf2_T383 - cost: 5.22, layer: 51
Fold over basal
Time (min)
0.9
1
1.1
1.2
1.3
1.4
1.5
1.6
1.7
0
10
20
30
40
50
60
Normal
LY50
LY75
LY88
LY94
LY97
LY data
Normal data

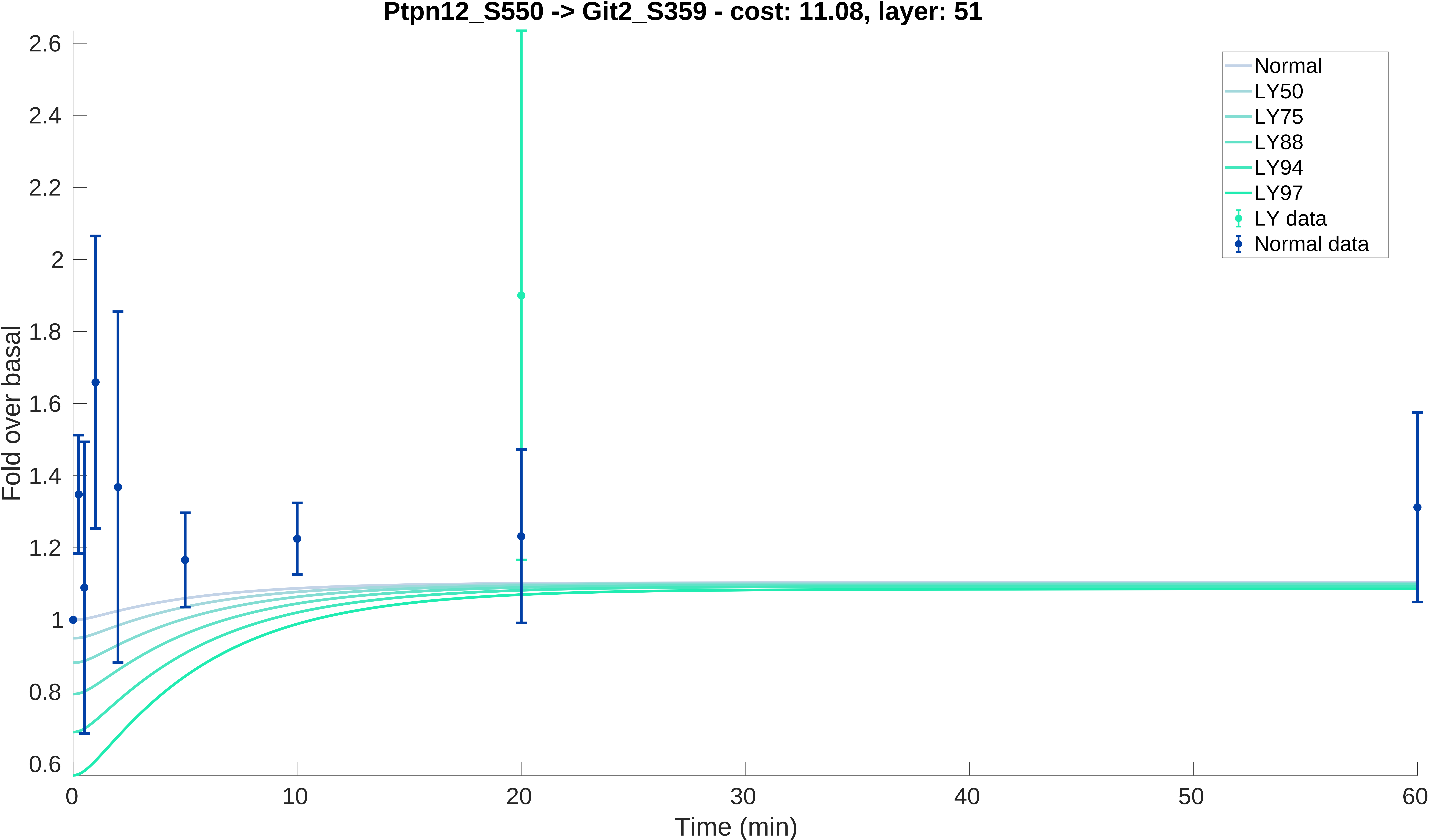
Ptpn12_S550 -> Git2_S359 - cost: 11.08, layer: 51
Normal
LY50
LY75
LY88
LY94
LY97
LY data
Normal data
Fold over basal
Time (min)
0.6
0.8
1
1.2
1.4
1.6
1.8
2
2.2
2.4
2.6
0
10
20
30
40
50
60

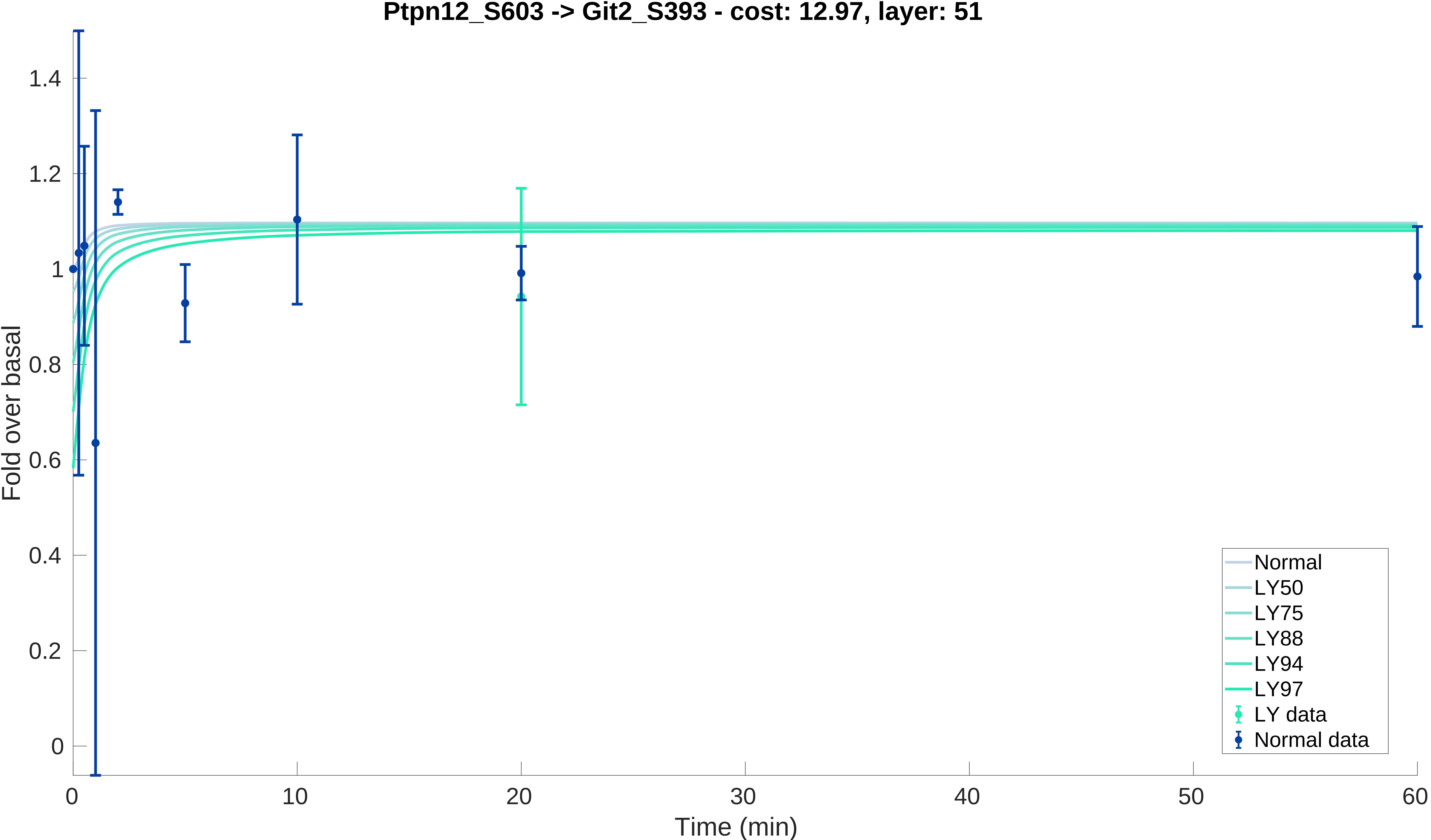

Ptpn12_S603 -> Git2_S393 - cost: 12.97, layer: 51
Fold over basal
Time (min)
0
0.2
0.4
0.6
0.8
1
1.2
1.4
0
10
20
30
40
50
60
Normal
LY50
LY75
LY88
LY94
LY97
LY data
Normal data

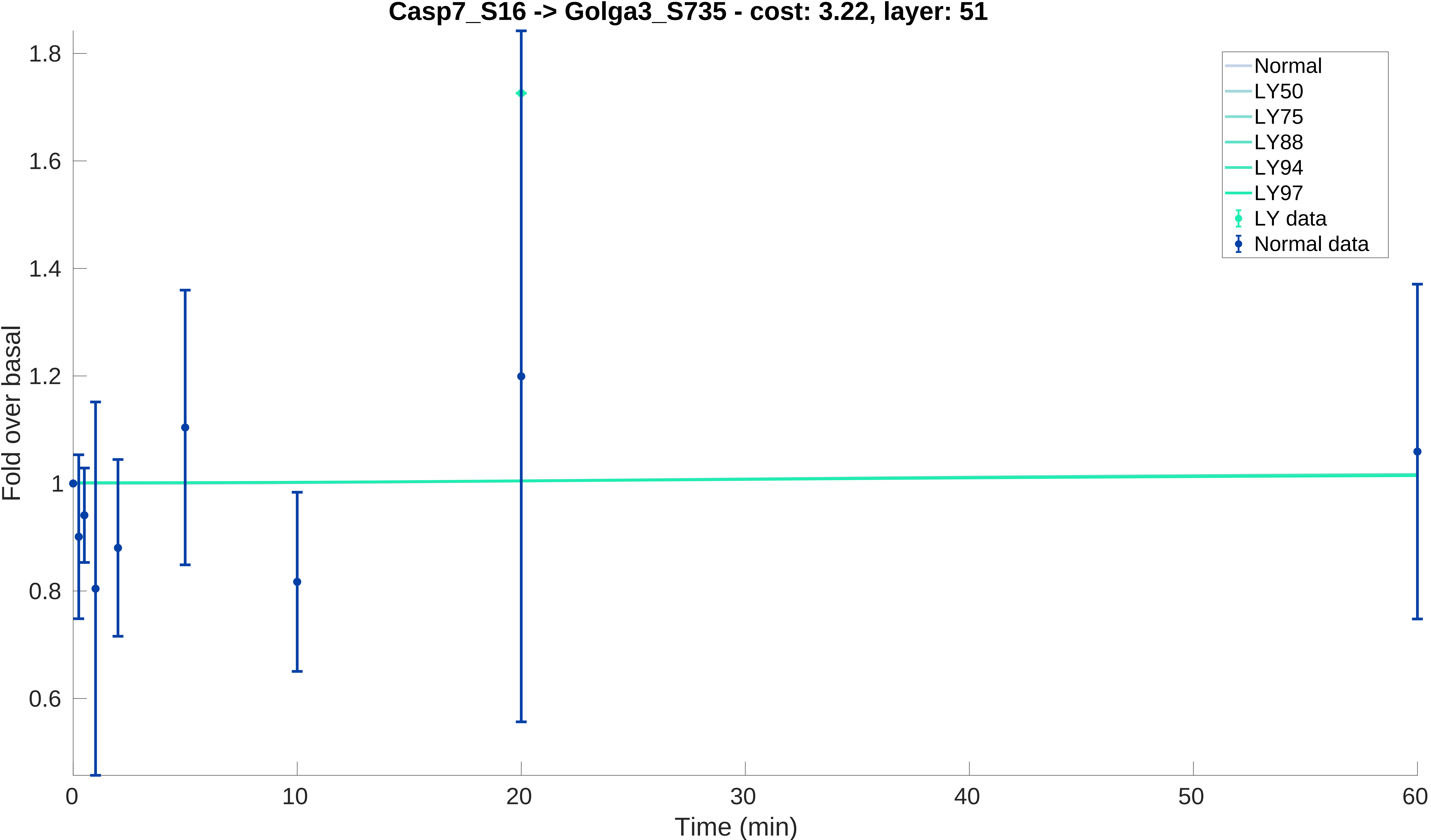

Casp7_S16 -> Golga3_S735 - cost: 3.22, layer: 51
Fold over basal
1.8
1.6
1.4
1.2
1
0.8
0.6
0
10
20
30
40
50
60
Time (min)
Normal
LY50
LY75
LY88
LY94
LY97
LY data
Normal data

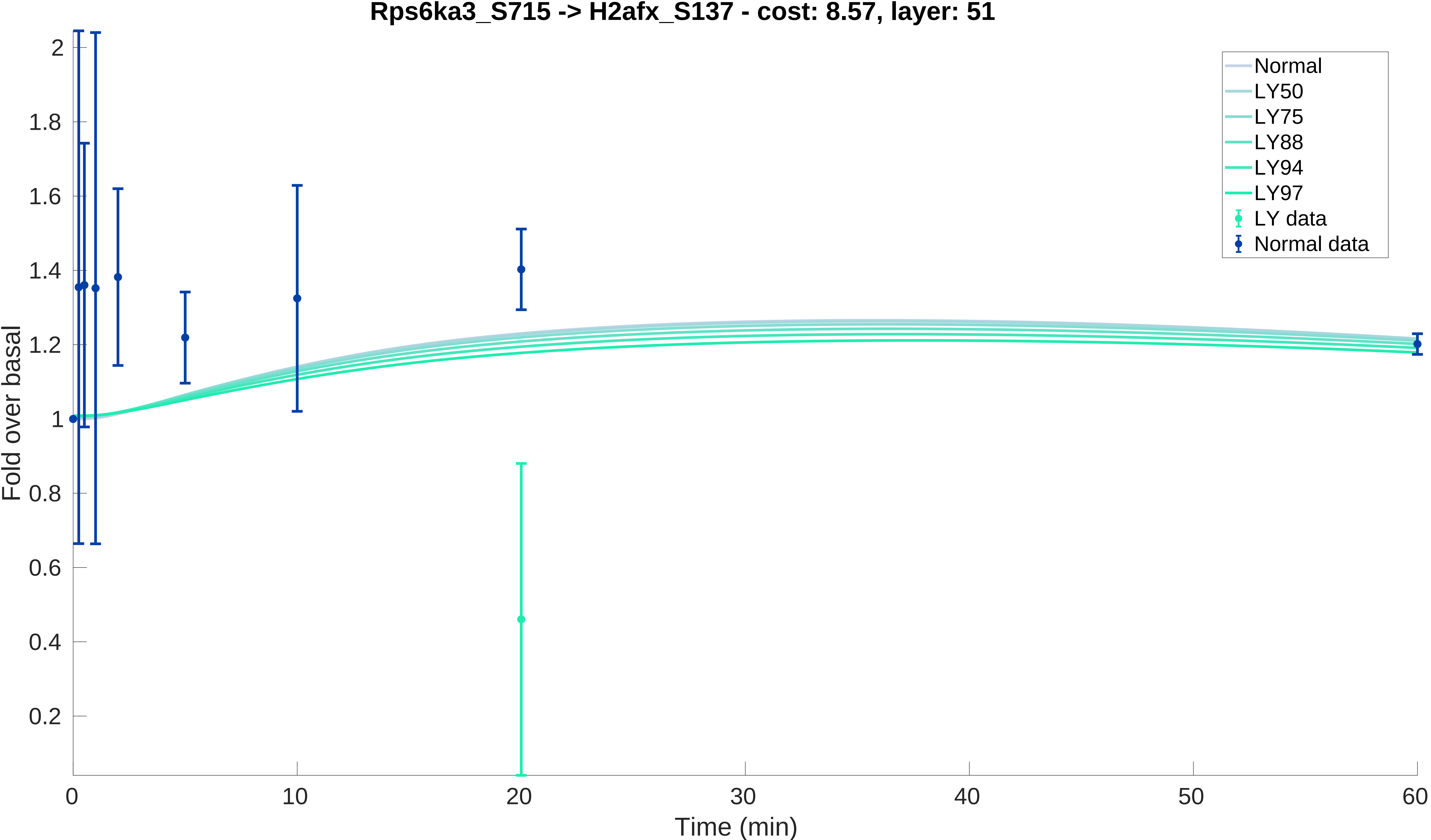

Rps6ka3_S715 -> H2afx_S137 - cost: 8.57, layer: 51
Fold over basal
Time (min)
2
1.8
1.6
1.4
1.2
1
0.8
0.6
0.4
0.2
0
10
20
30
40
50
60
Normal
LY50
LY75
LY88
LY94
LY97
LY data
Normal data

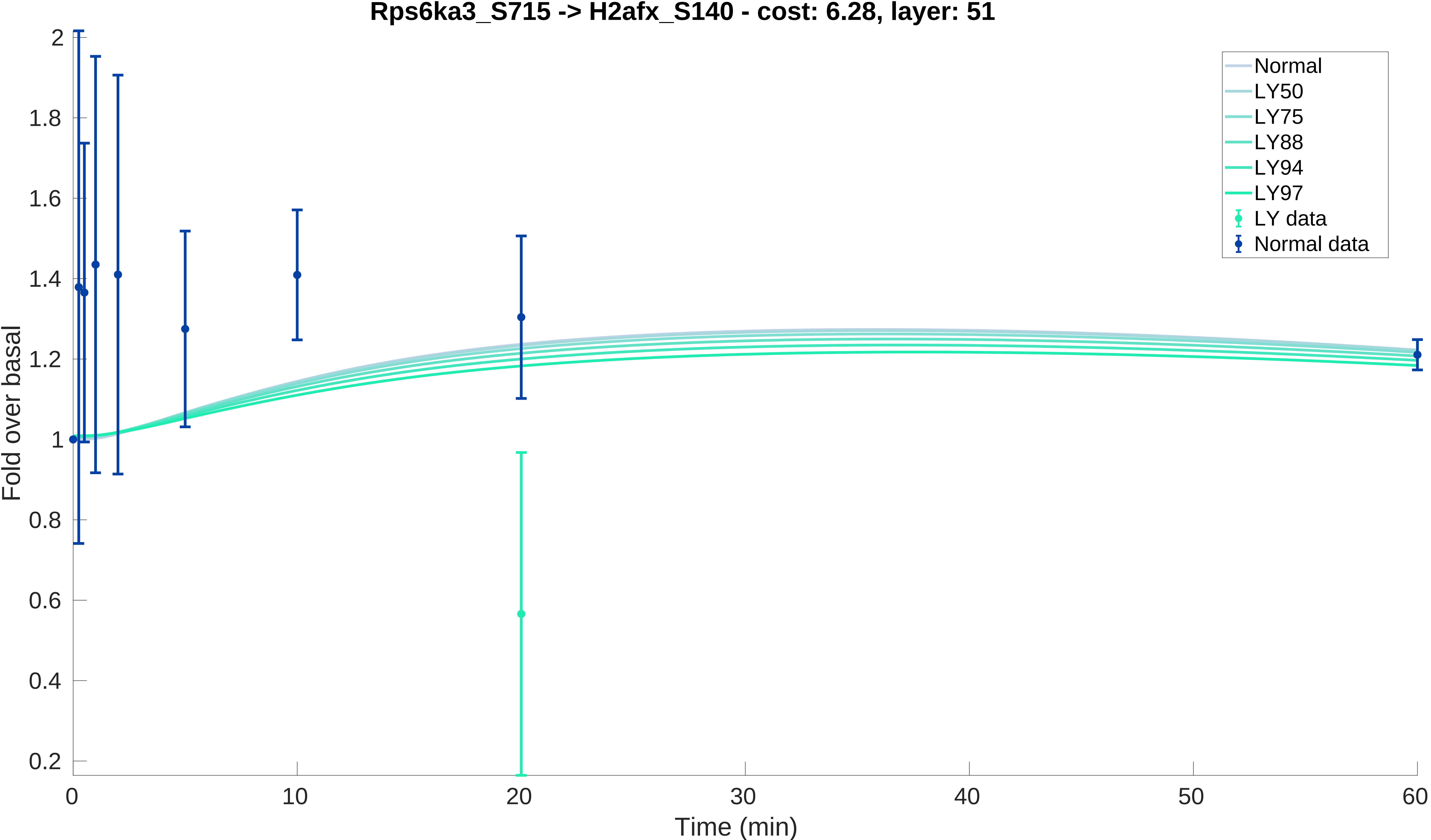

Rps6ka3_S715 -> H2afx_S140 - cost: 6.28, layer: 51
Fold over basal
Time (min)
0
10
20
30
40
50
60
0.2
0.4
0.6
0.8
1
1.2
1.4
1.6
1.8
2
Normal
LY50
LY75
LY88
LY94
LY97
LY data
Normal data

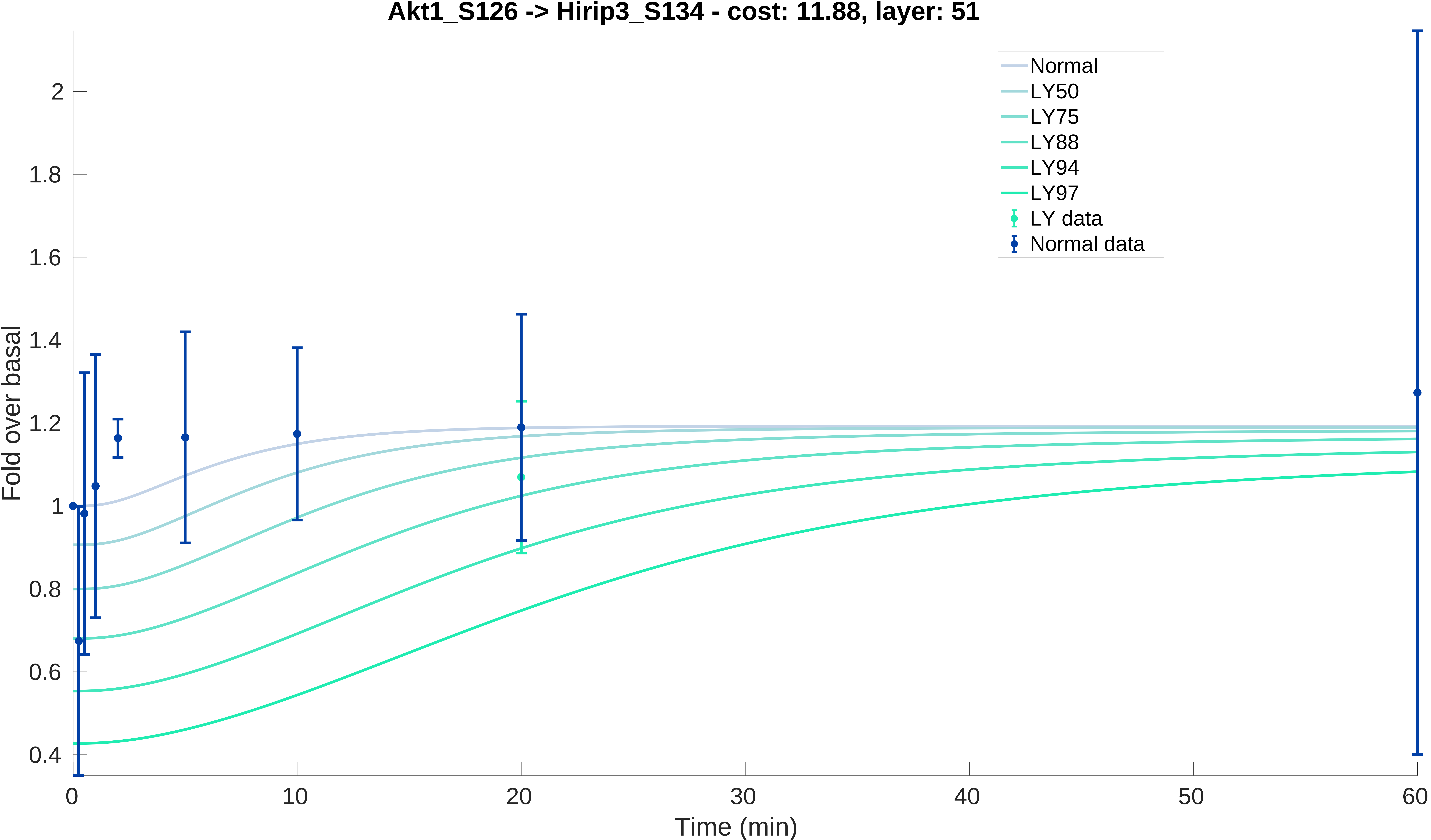

Akt1_S126 -> Hirip3_S134 - cost: 11.88, layer: 51
Normal
LY50
LY75
LY88
LY94
LY97
LY data
Normal data
Fold over basal
Time (min)
0.4
0.6
0.8
1
1.2
1.4
1.6
1.8
2
0
10
20
30
40
50
60

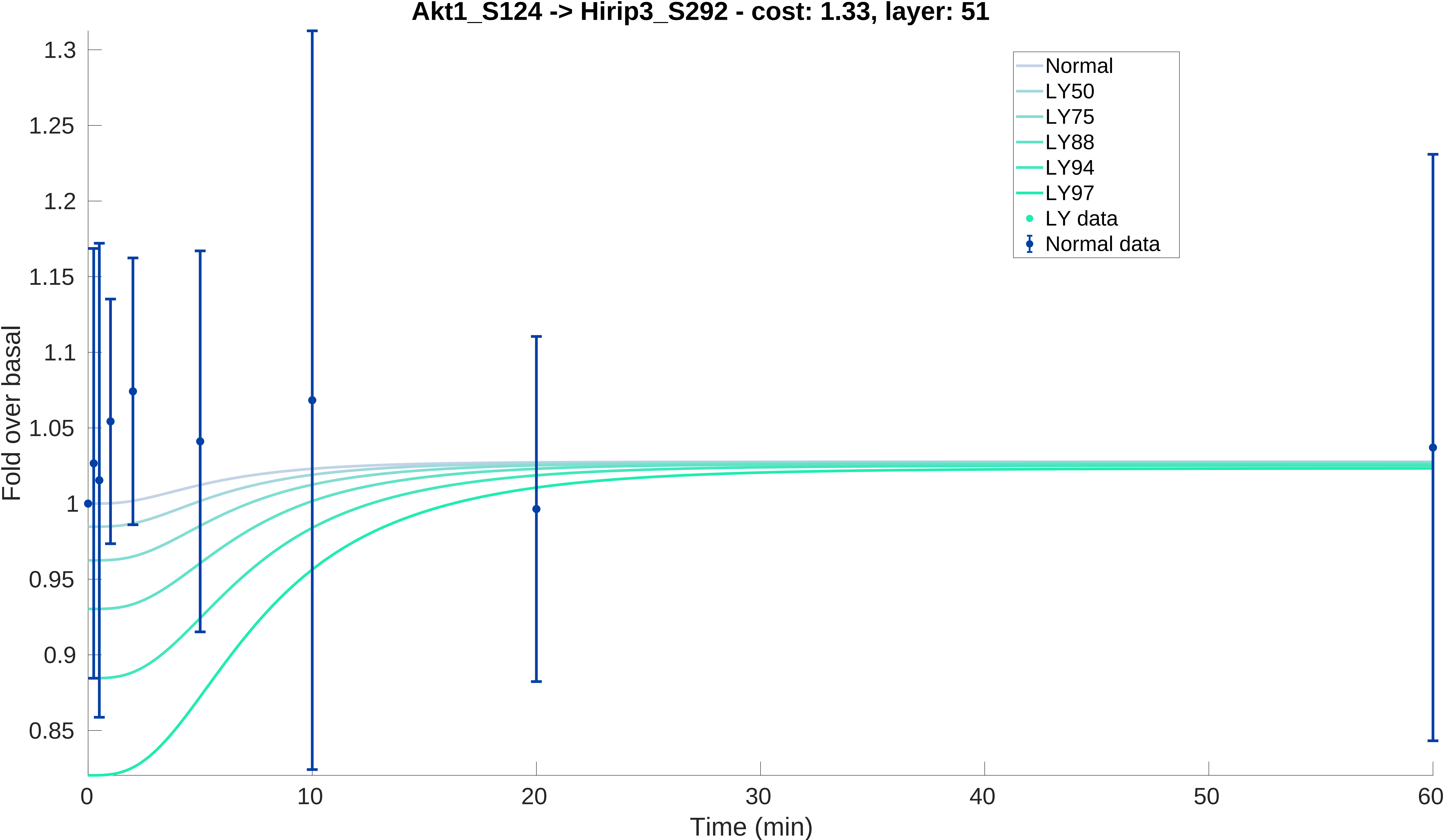

Akt1_S124 -> Hirip3_S292 - cost: 1.33, layer: 51
Fold over basal
Time (min)
1.3
1.25
1.2
1.15
1.1
1.05
1
0.95
0.9
0.85
0
10
20
30
40
50
60
Normal
LY50
LY75
LY88
LY94
LY97
LY data
Normal data

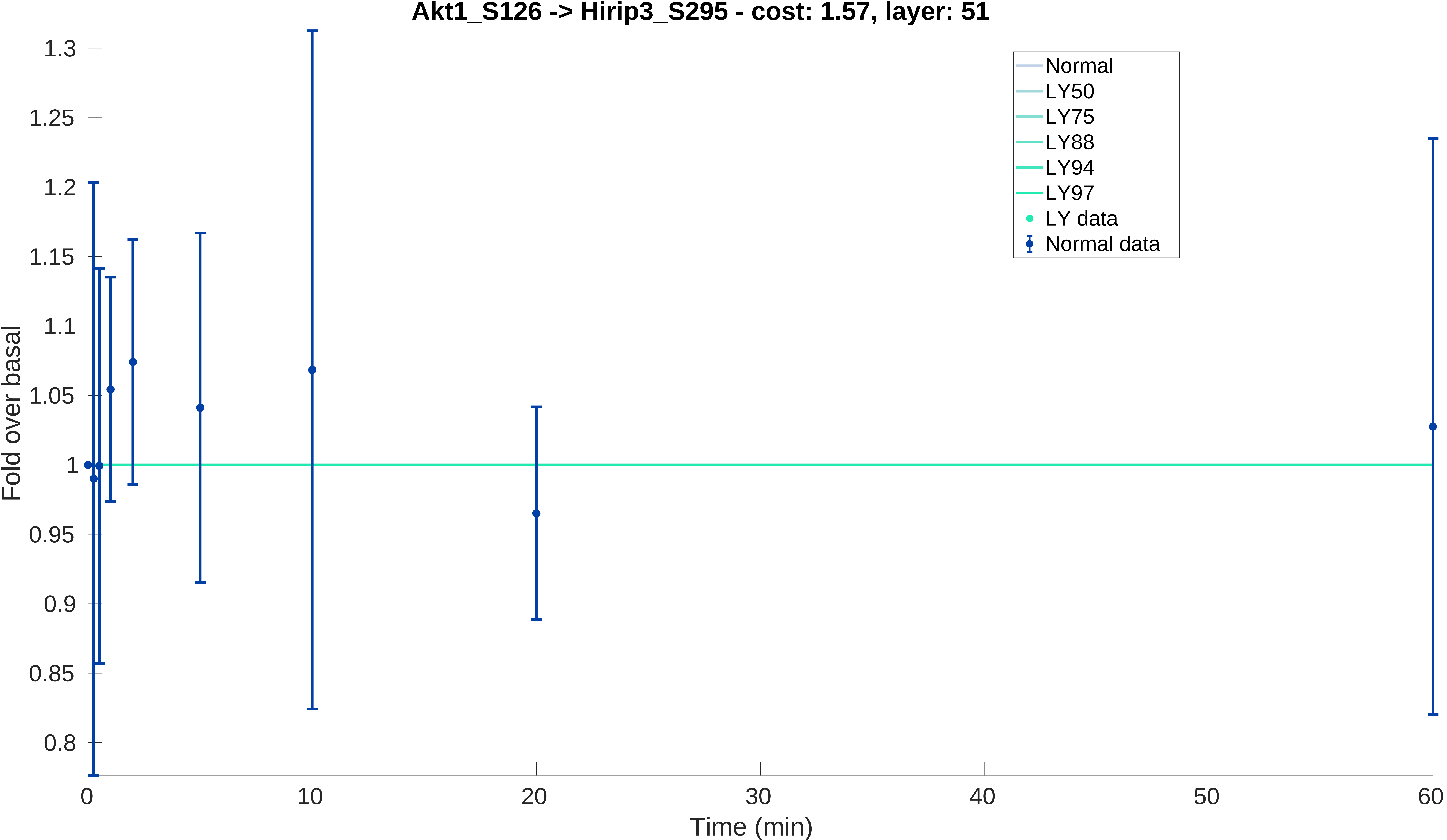

Akt1_S126 -> Hirip3_S295 - cost: 1.57, layer: 51
Normal
LY50
LY75
LY88
LY94
LY97
LY data
Normal data
Fold over basal
1.3
1.25
1.2
1.15
1.1
1.05
1
0.95
0.9
0.85
0.8
0
10
20
30
40
50
60
Time (min)

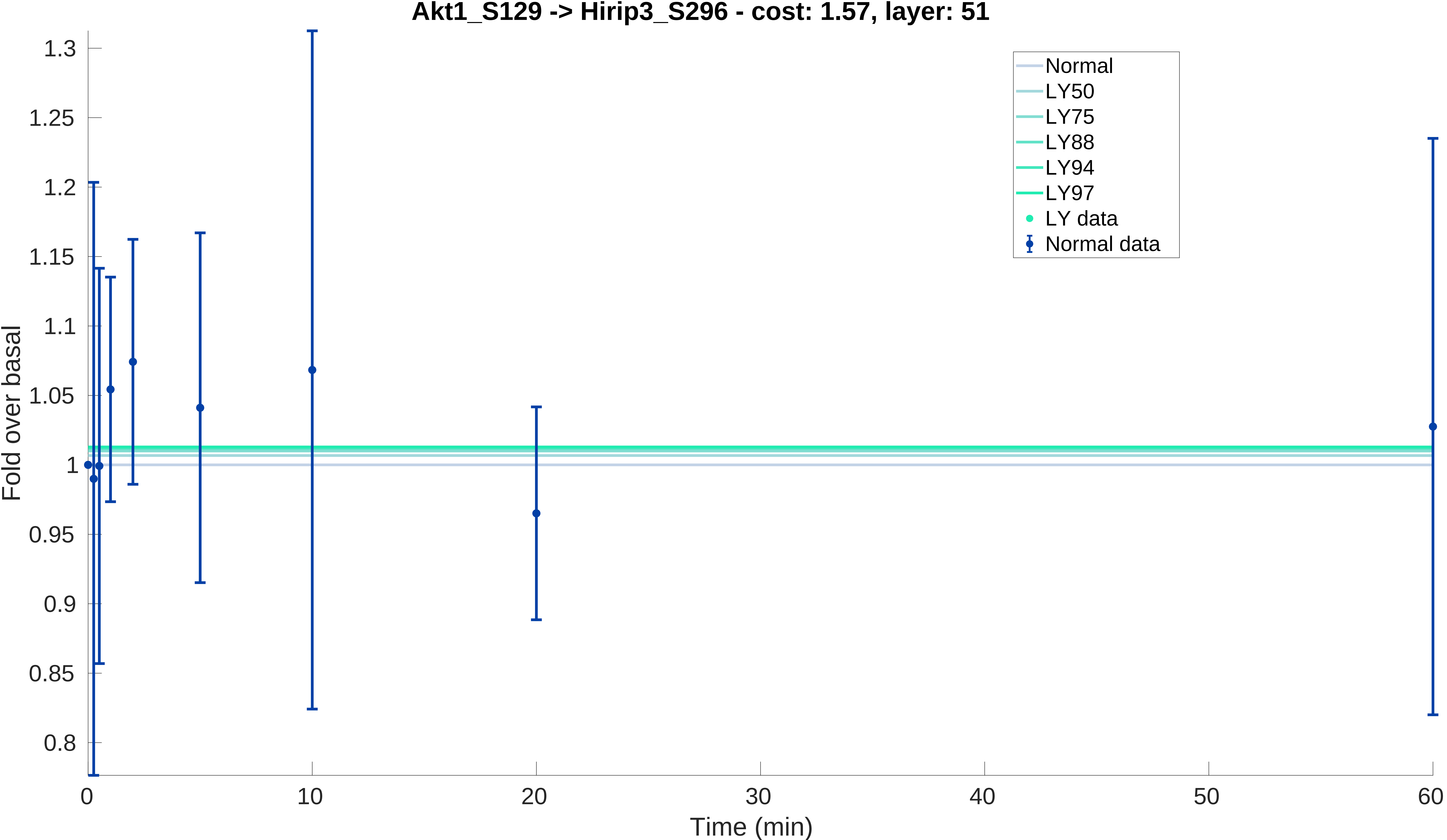

Akt1_S129 -> Hirip3_S296 - cost: 1.57, layer: 51
Fold over basal
Time (min)
1.3
1.25
1.2
1.15
1.1
1.05
1
0.95
0.9
0.85
0.8
0
10
20
30
40
50
60
Normal
LY50
LY75
LY88
LY94
LY97
LY data
Normal data

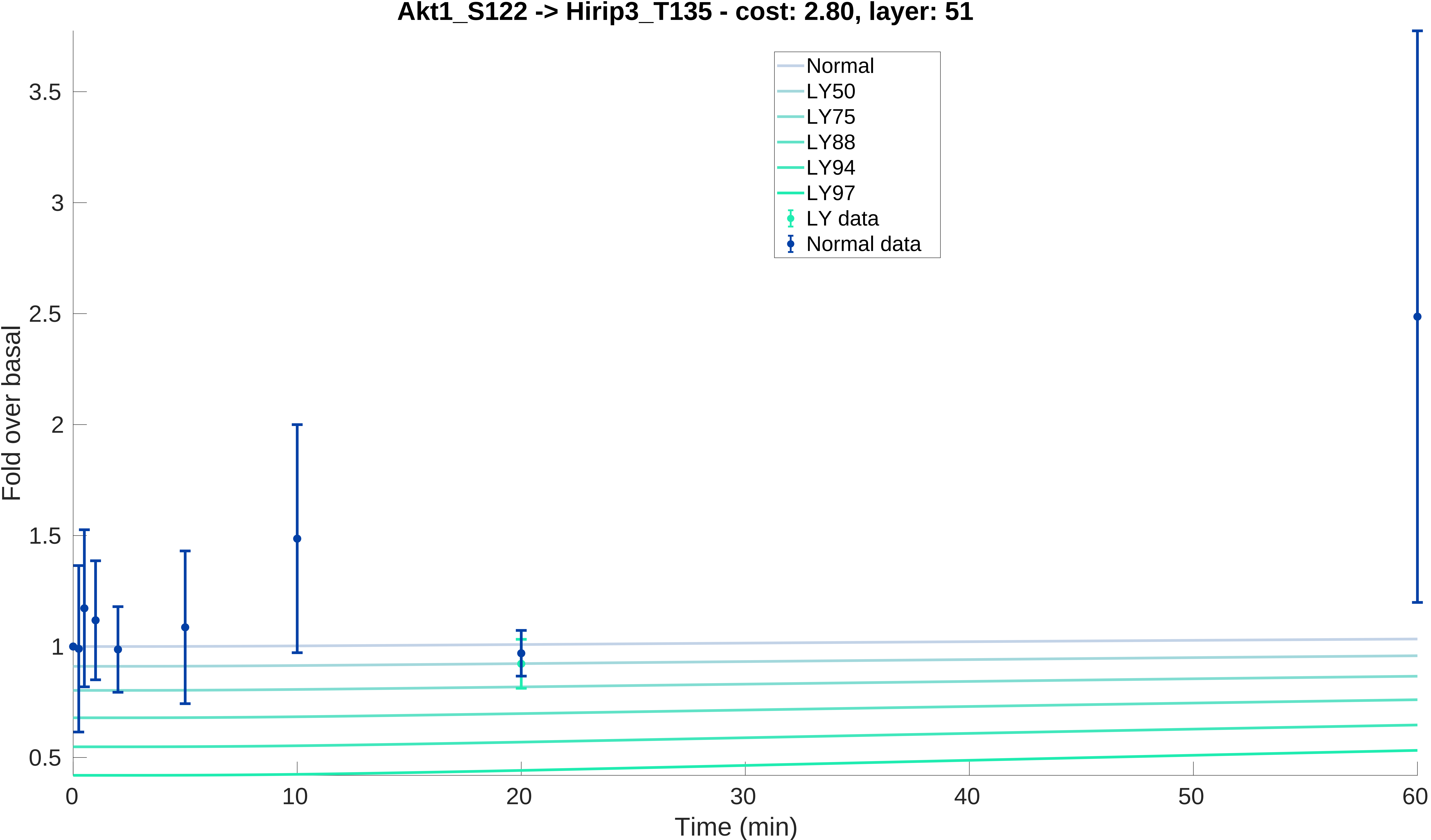
Akt1_S122 -> Hirip3_T135 - cost: 2.80, layer: 51
Normal
LY50
LY75
LY88
LY94
LY97
LY data
Normal data
Fold over basal
Time (min)
0
10
20
30
40
50
60
0.5
1
1.5
2
2.5
3
3.5

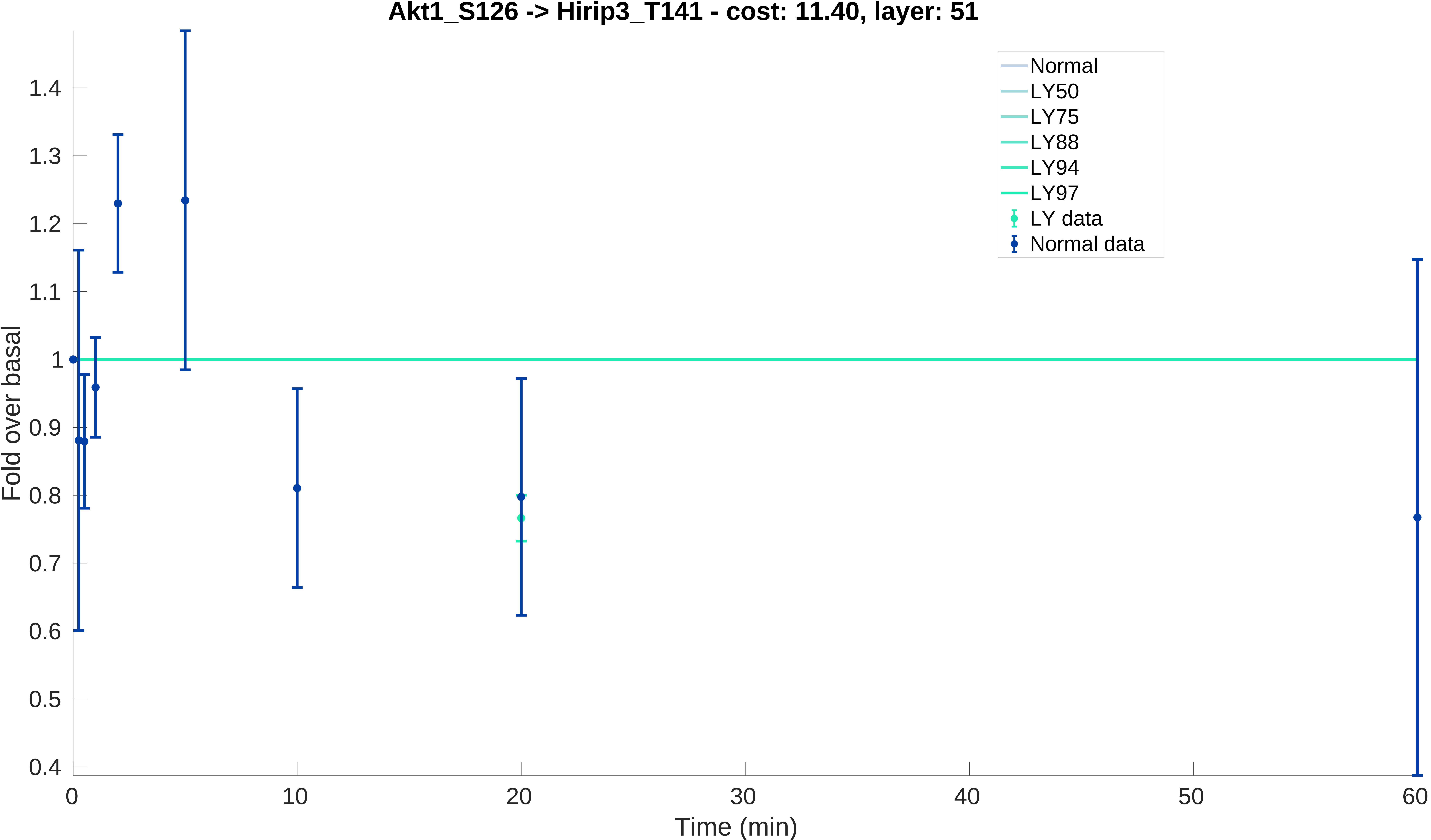

Akt1_S126 -> Hirip3_T141 - cost: 11.40, layer: 51
Normal
LY50
LY75
LY88
LY94
LY97
LY data
Normal data
Fold over basal
1.4
1.3
1.2
1.1
1
0.9
0.8
0.7
0.6
0.5
0.4
0
10
20
30
40
50
60
Time (min)

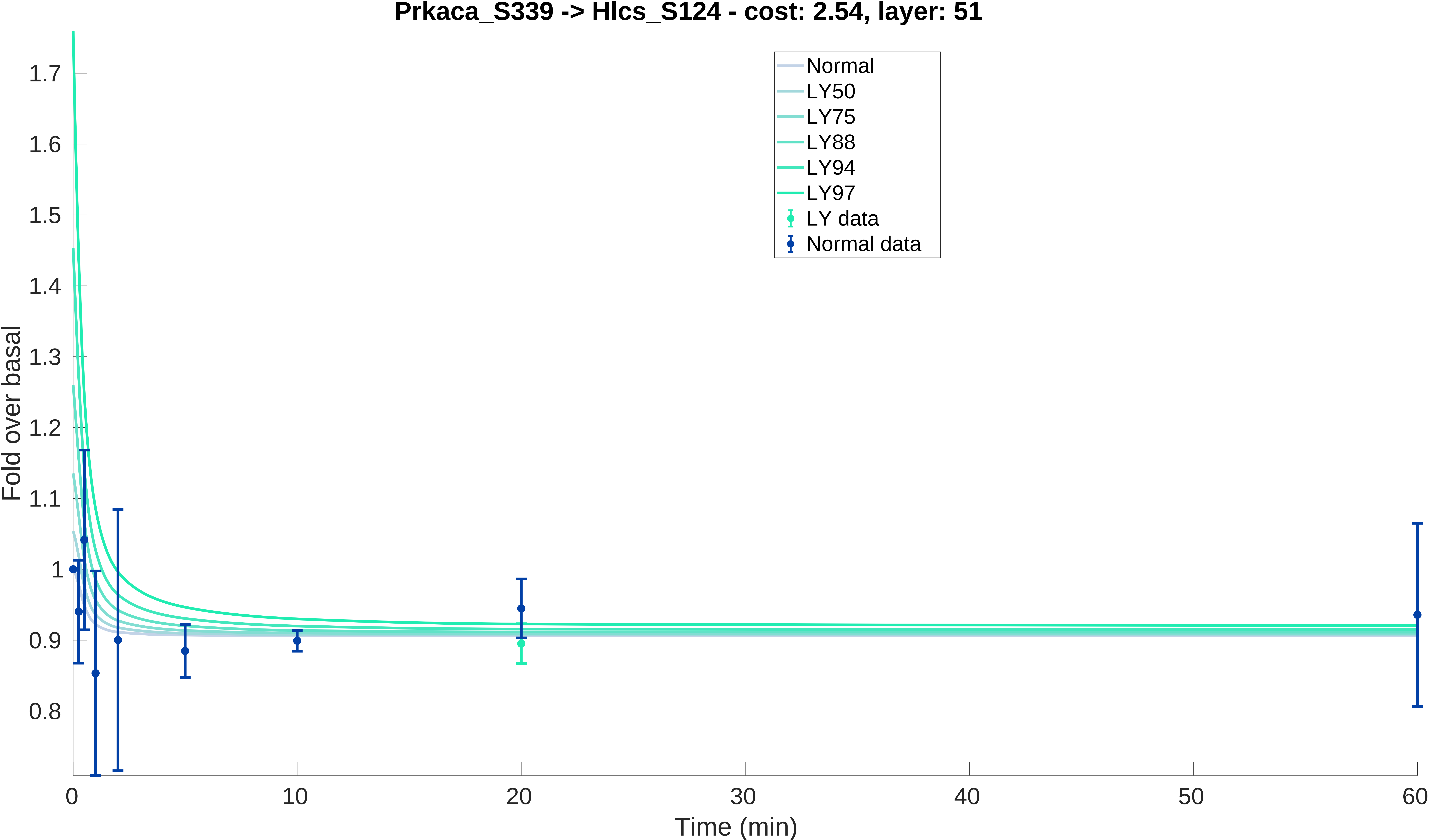

Prkaca_S339 -> Hlcs_S124 - cost: 2.54, layer: 51
Normal
LY50
LY75
LY88
LY94
LY97
LY data
Normal data
Fold over basal
1.7
1.6
1.5
1.4
1.3
1.2
1.1
1
0.9
0.8
0
10
20
30
40
50
60
Time (min)

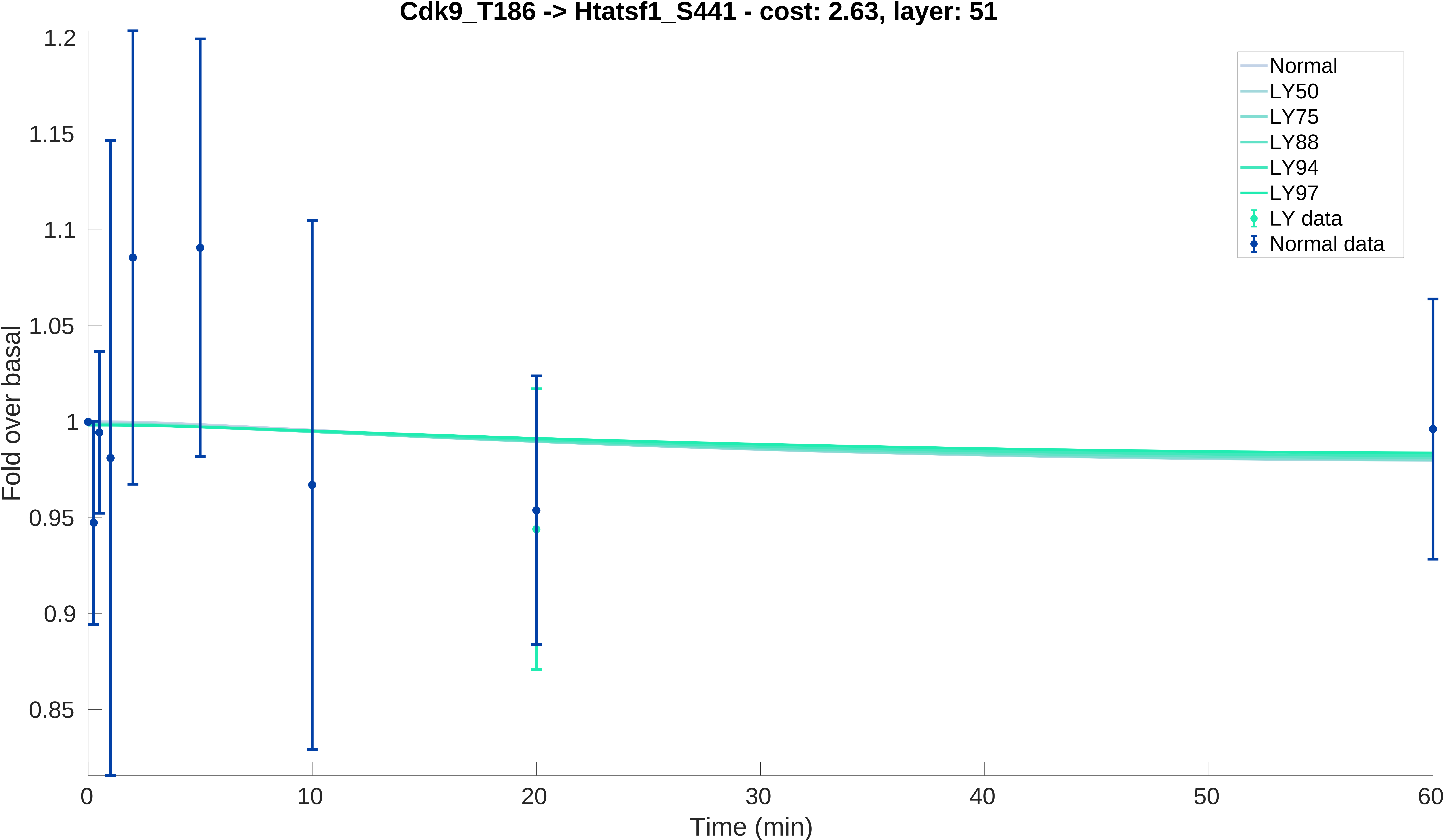

Cdk9_T186 -> Htatsf1_S441 - cost: 2.63, layer: 51
Fold over basal
Time (min)
0.85
0.9
0.95
1
1.05
1.1
1.15
1.2
0
10
20
30
40
50
60
Normal
LY50
LY75
LY88
LY94
LY97
LY data
Normal data

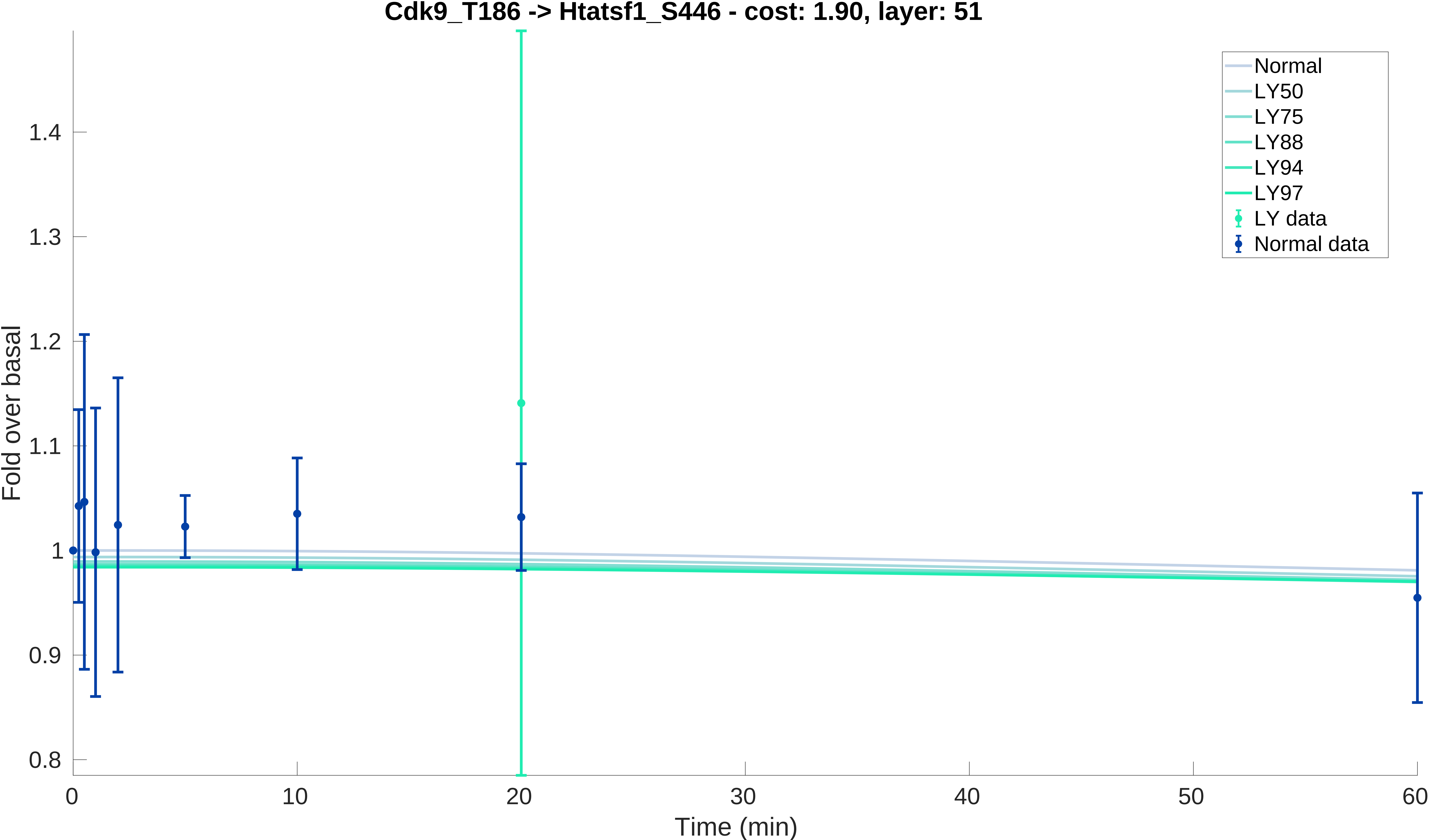

Cdk9_T186 -> Htatsf1_S446 - cost: 1.90, layer: 51
Normal
LY50
LY75
LY88
LY94
LY97
LY data
Normal data
Fold over basal
1.4
1.3
1.2
1.1
1
0.9
0.8
0
10
20
30
40
50
60
Time (min)

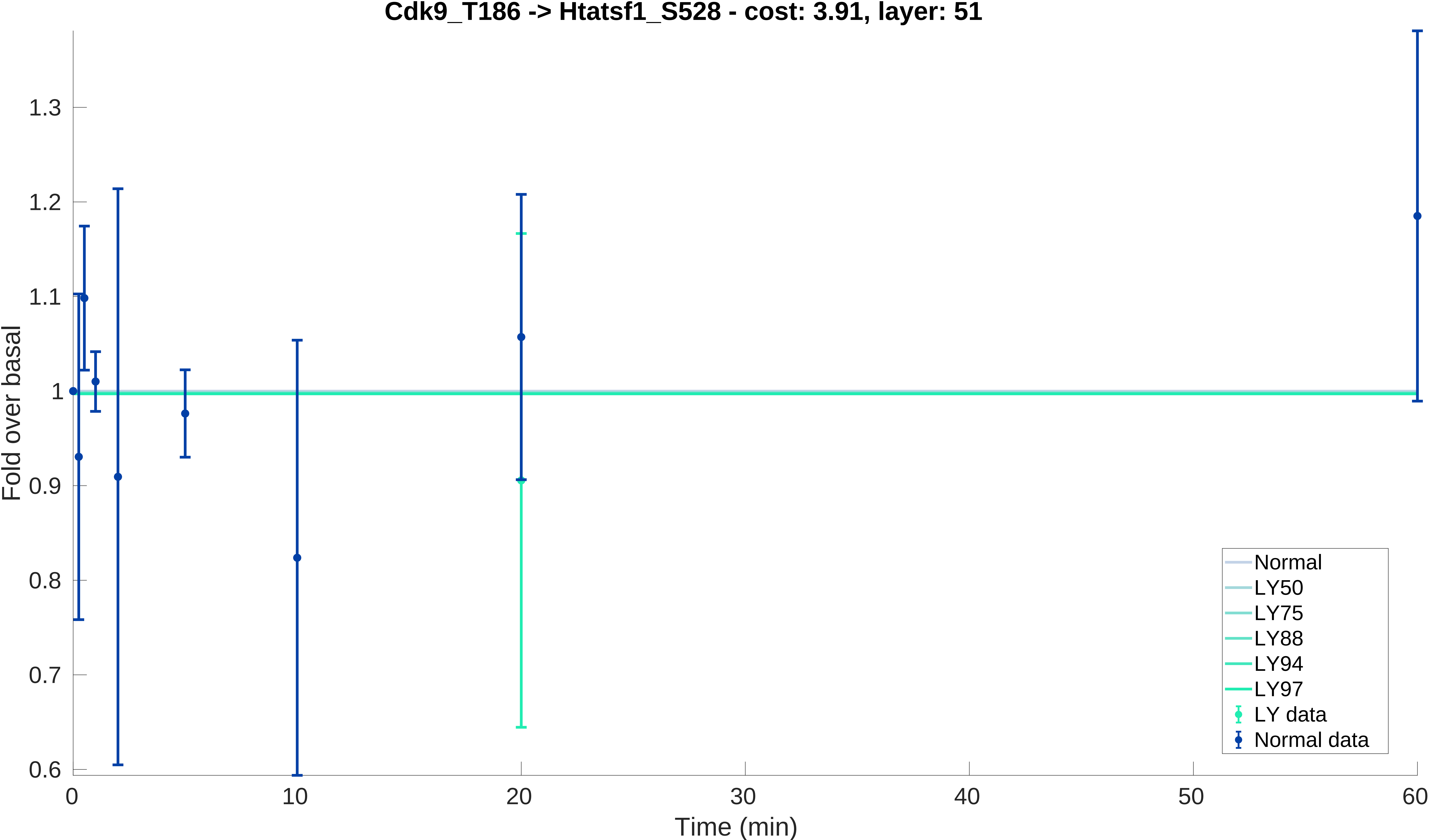

Cdk9_T186 -> Htatsf1_S528 - cost: 3.91, layer: 51
Fold over basal
Time (min)
1.3
1.2
1.1
1
0.9
0.8
0.7
0.6
0
10
20
30
40
50
60
Normal
LY50
LY75
LY88
LY94
LY97
LY data
Normal data

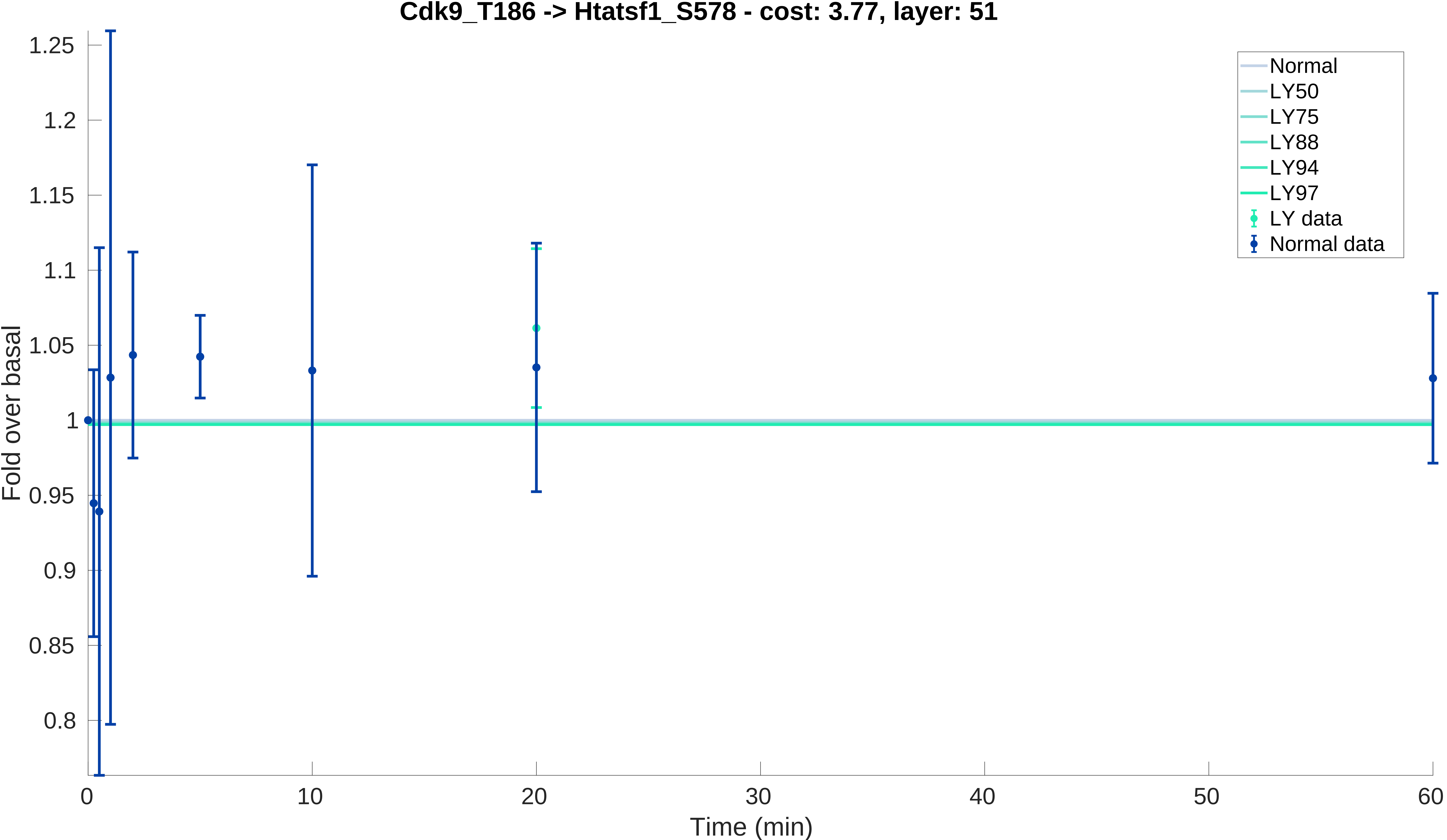

Cdk9_T186 -> Htatsf1_S578 - cost: 3.77, layer: 51
Normal
LY50
LY75
LY88
LY94
LY97
LY data
Normal data
Fold over basal
Time (min)
1.25
1.2
1.15
1.1
1.05
1
0.95
0.9
0.85
0.8
0
10
20
30
40
50
60

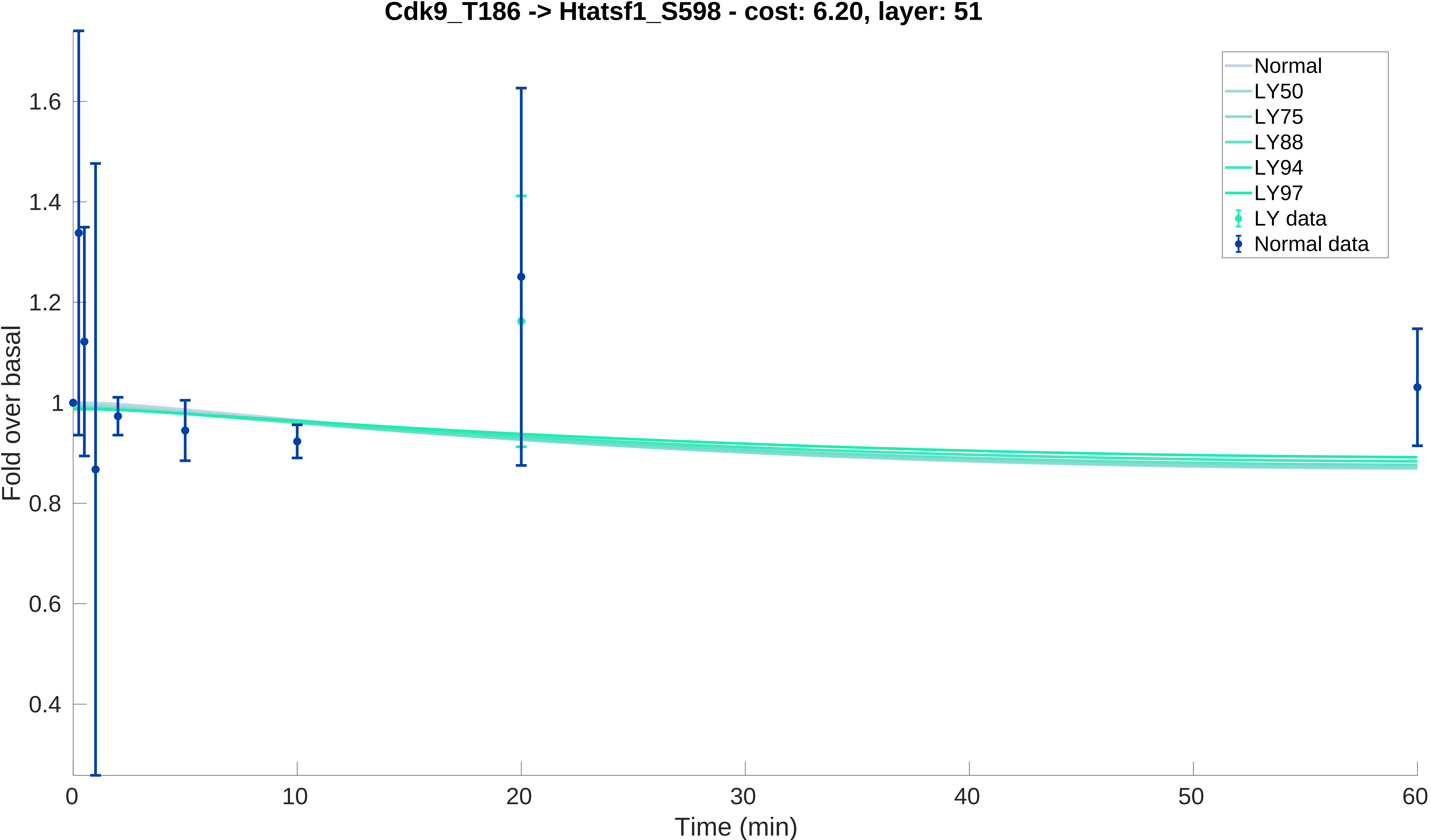

Cdk9_T186 -> Htatsf1_S598 - cost: 6.20, layer: 51
Normal
LY50
LY75
LY88
LY94
LY97
LY data
Normal data
Fold over basal
1.6
1.4
1.2
1
0.8
0.6
0.4
0
10
20
30
40
50
60
Time (min)

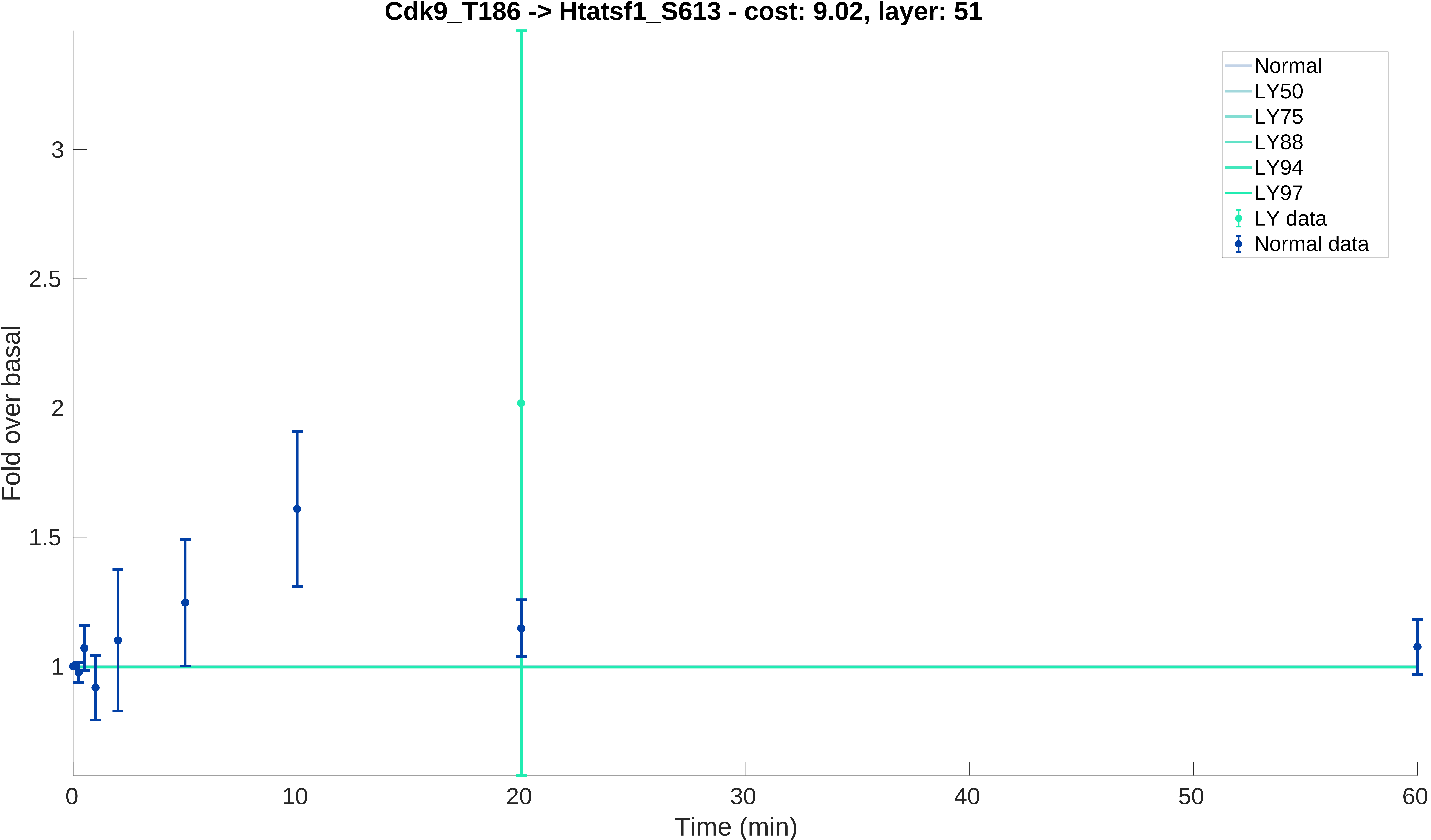

Cdk9_T186 -> Htatsf1_S613 - cost: 9.02, layer: 51
Fold over basal
Time (min)
0
10
20
30
40
50
60
1
1.5
2
2.5
3
Normal
LY50
LY75
LY88
LY94
LY97
LY data
Normal data

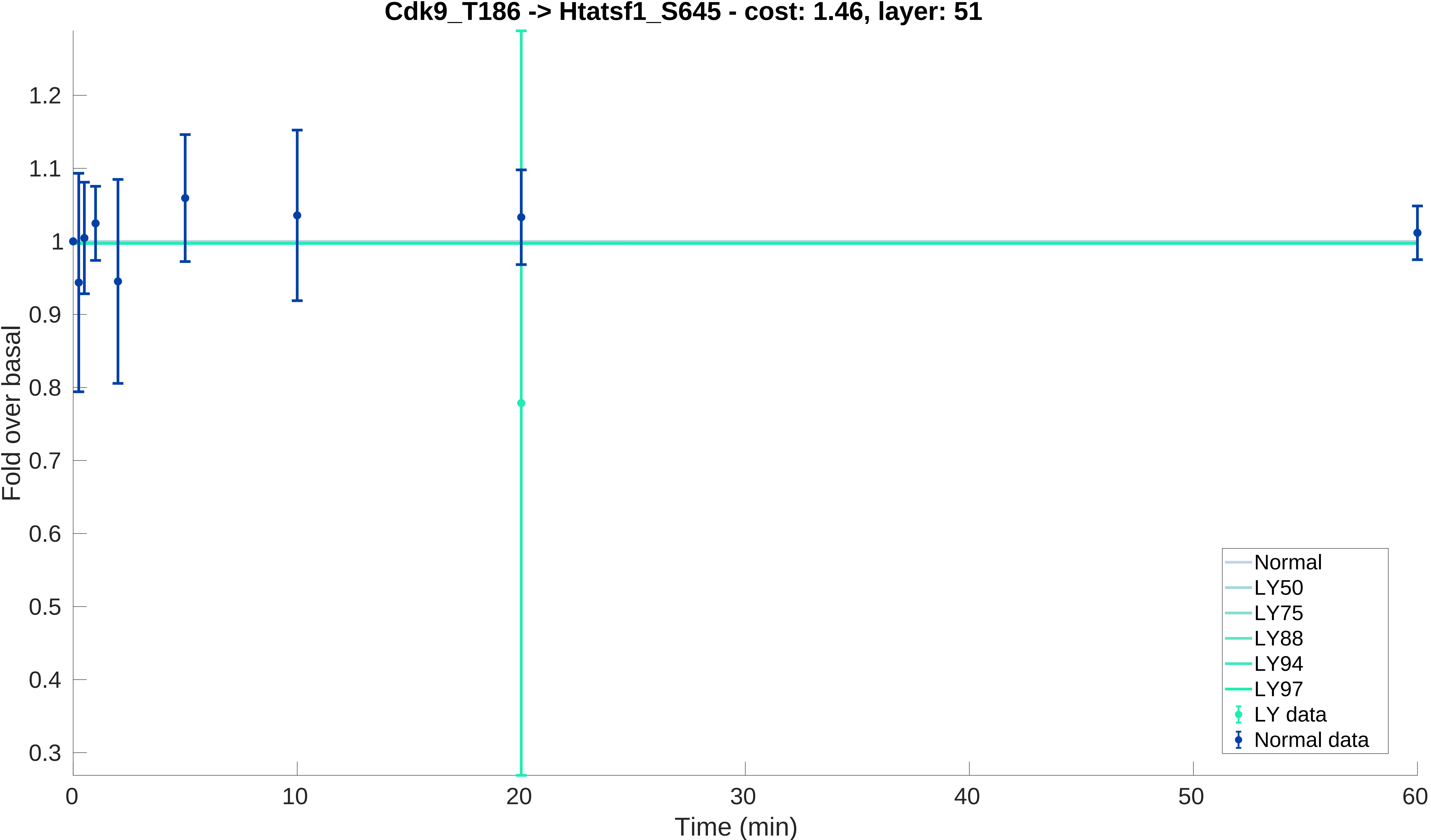

Cdk9_T186 -> Htatsf1_S645 - cost: 1.46, layer: 51
Fold over basal
1.2
1.1
1
0.9
0.8
0.7
0.6
0.5
0.4
0.3
0
10
20
30
40
50
60
Time (min)
Normal
LY50
LY75
LY88
LY94
LY97
LY data
Normal data

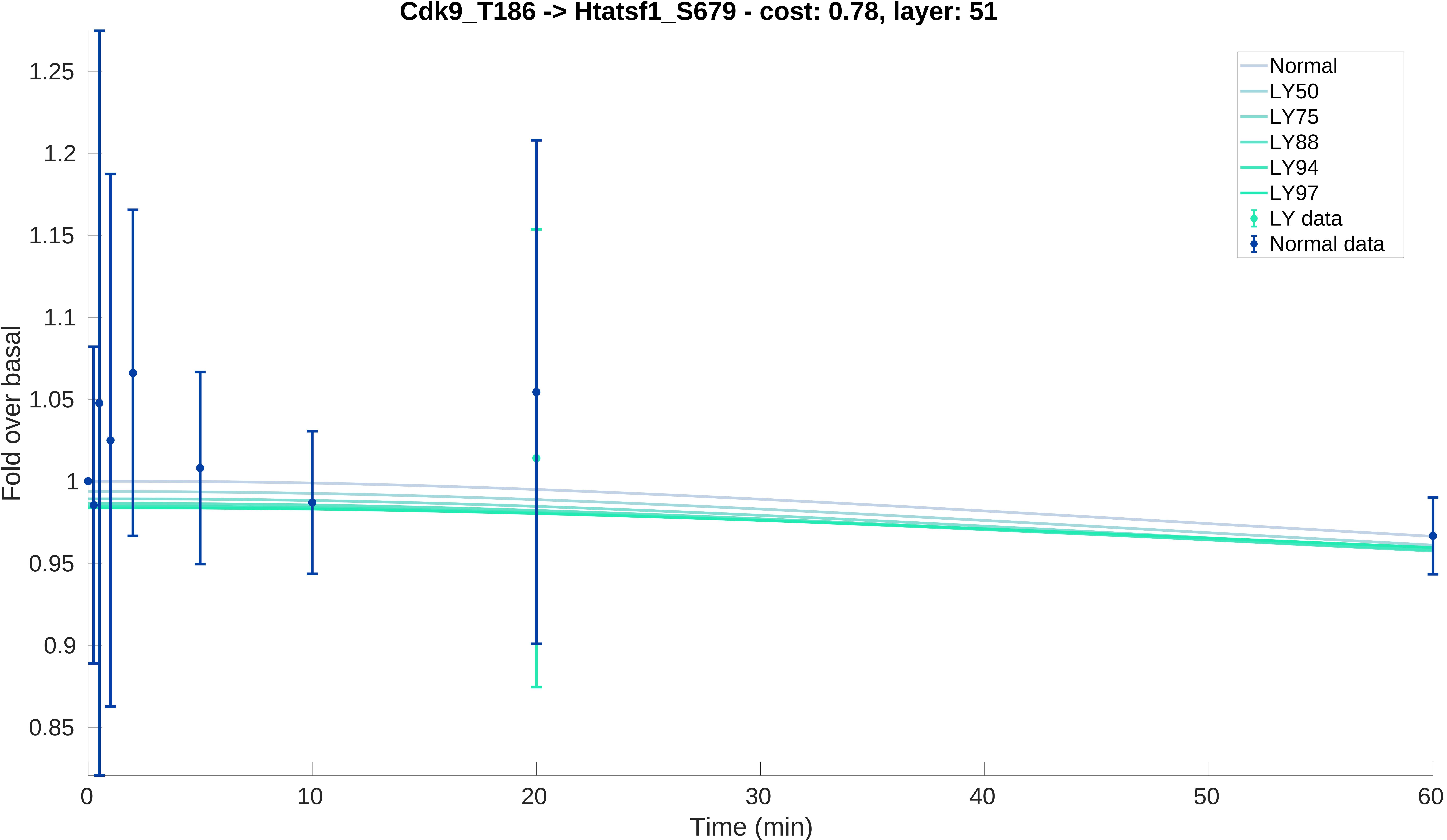

Cdk9_T186 -> Htatsf1_S679 - cost: 0.78, layer: 51
Fold over basal
Time (min)
1.25
1.2
1.15
1.1
1.05
1
0.95
0.9
0.85
0
10
20
30
40
50
60
Normal
LY50
LY75
LY88
LY94
LY97
LY data
Normal data

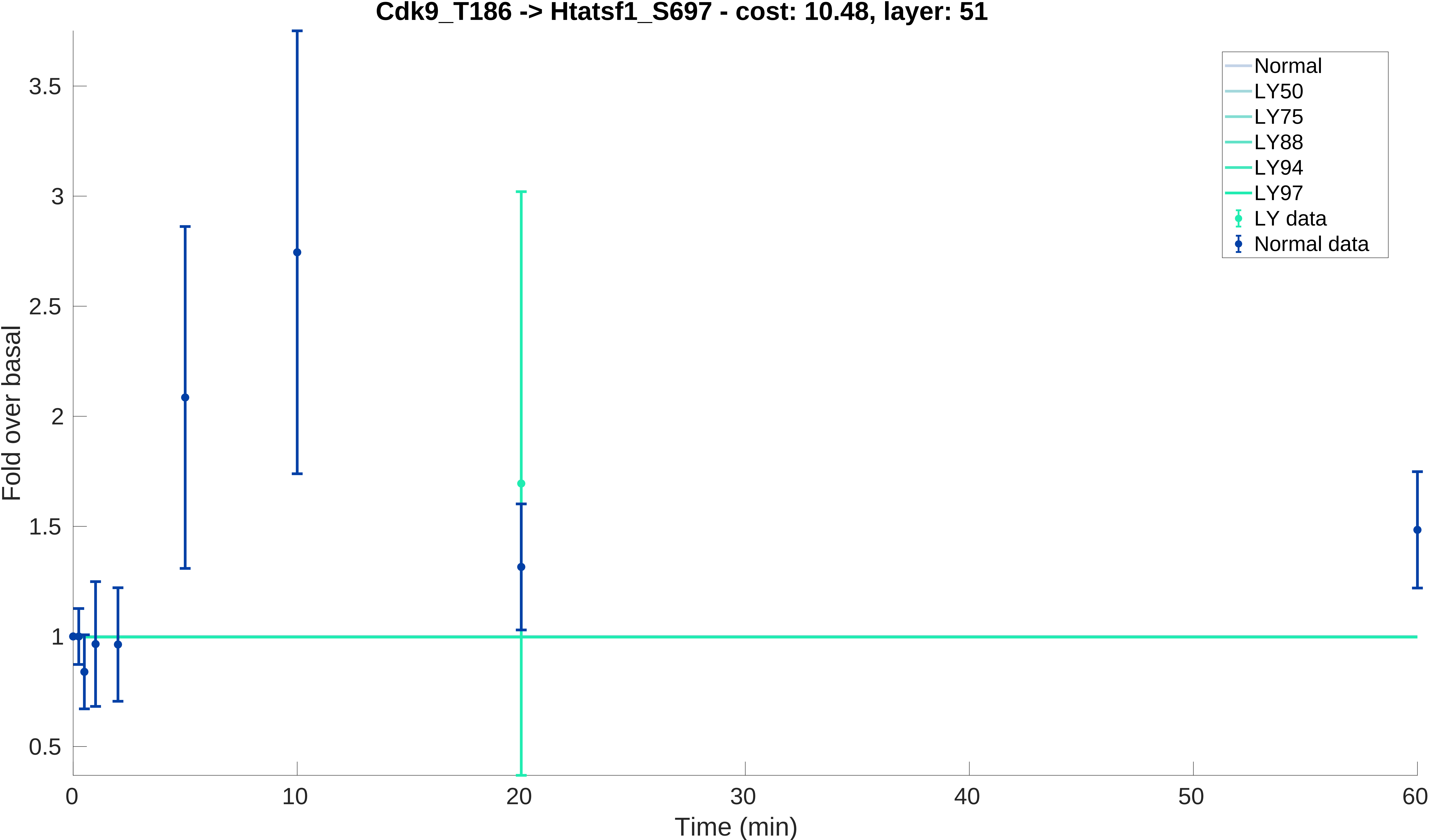

Cdk9_T186 -> Htatsf1_S697 - cost: 10.48, layer: 51
Fold over basal
Time (min)
0.5
1
1.5
2
2.5
3
3.5
0
10
20
30
40
50
60
Normal
LY50
LY75
LY88
LY94
LY97
LY data
Normal data

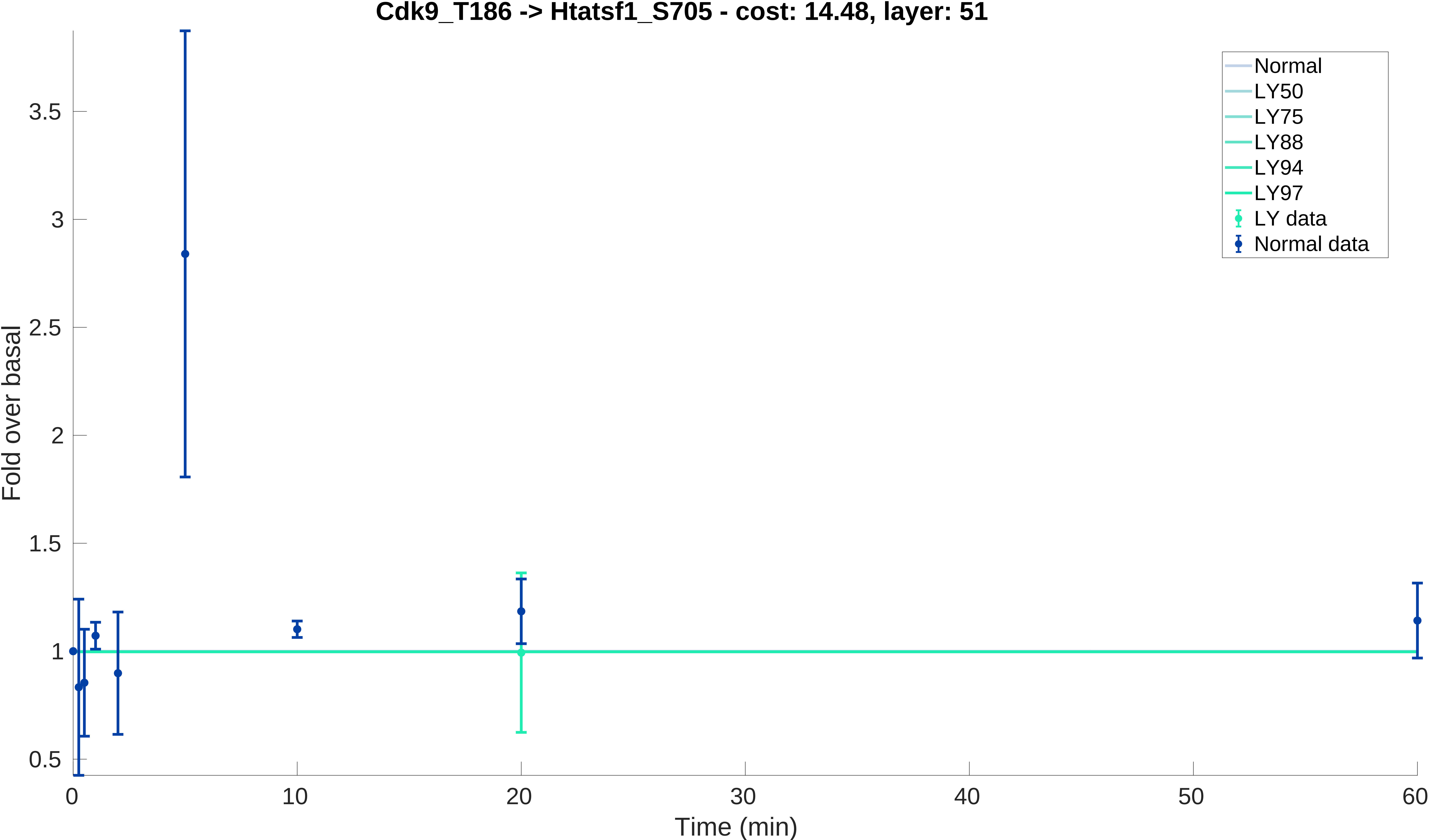

Cdk9_T186 -> Htatsf1_S705 - cost: 14.48, layer: 51
Normal
LY50
LY75
LY88
LY94
LY97
LY data
Normal data
Fold over basal
3.5
3
2.5
2
1.5
1
0.5
0
10
20
30
40
50
60
Time (min)

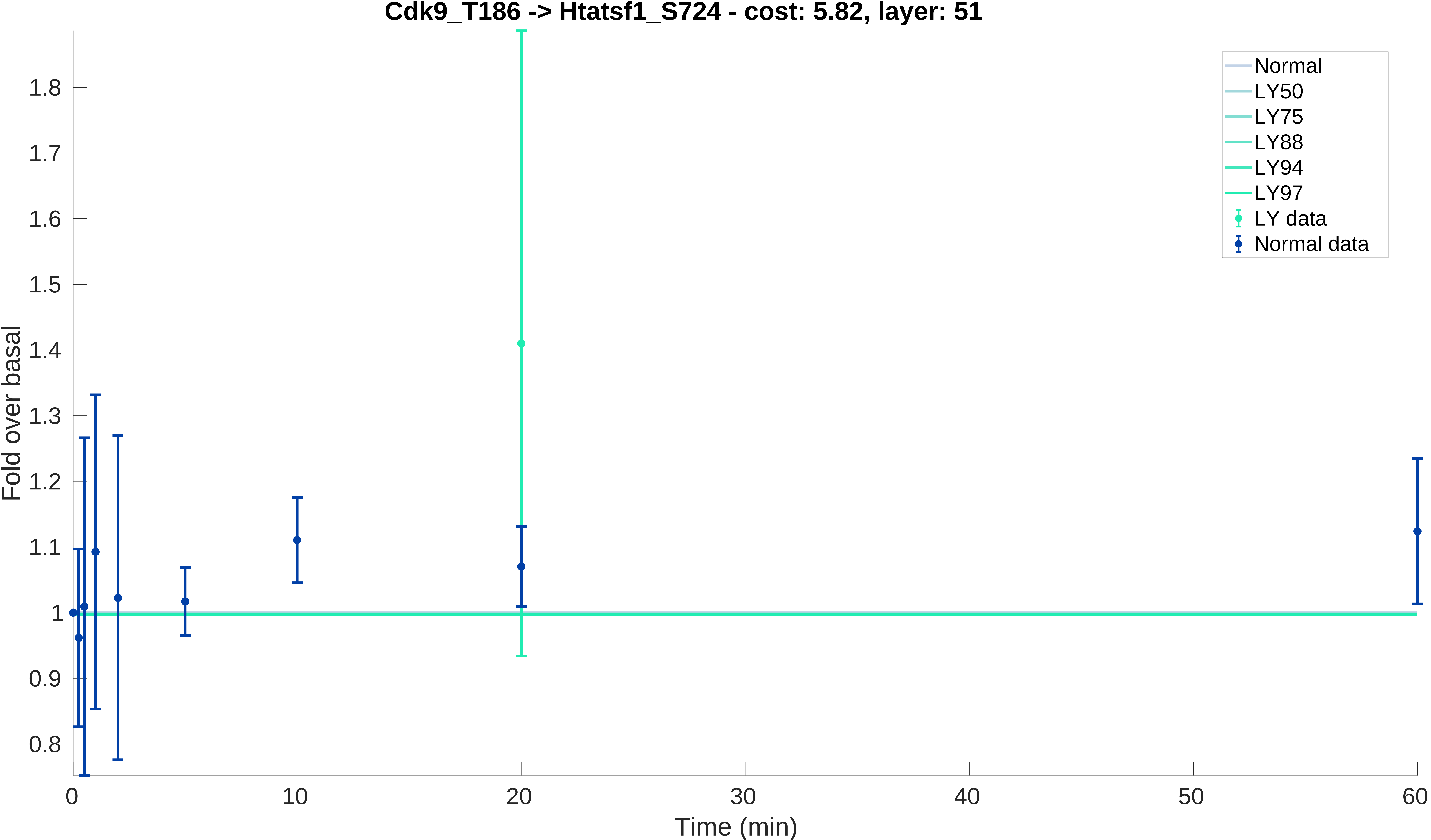

Cdk9_T186 -> Htatsf1_S724 - cost: 5.82, layer: 51
Fold over basal
Time (min)
1.8
1.7
1.6
1.5
1.4
1.3
1.2
1.1
1
0.9
0.8
0
10
20
30
40
50
60
Normal
LY50
LY75
LY88
LY94
LY97
LY data
Normal data

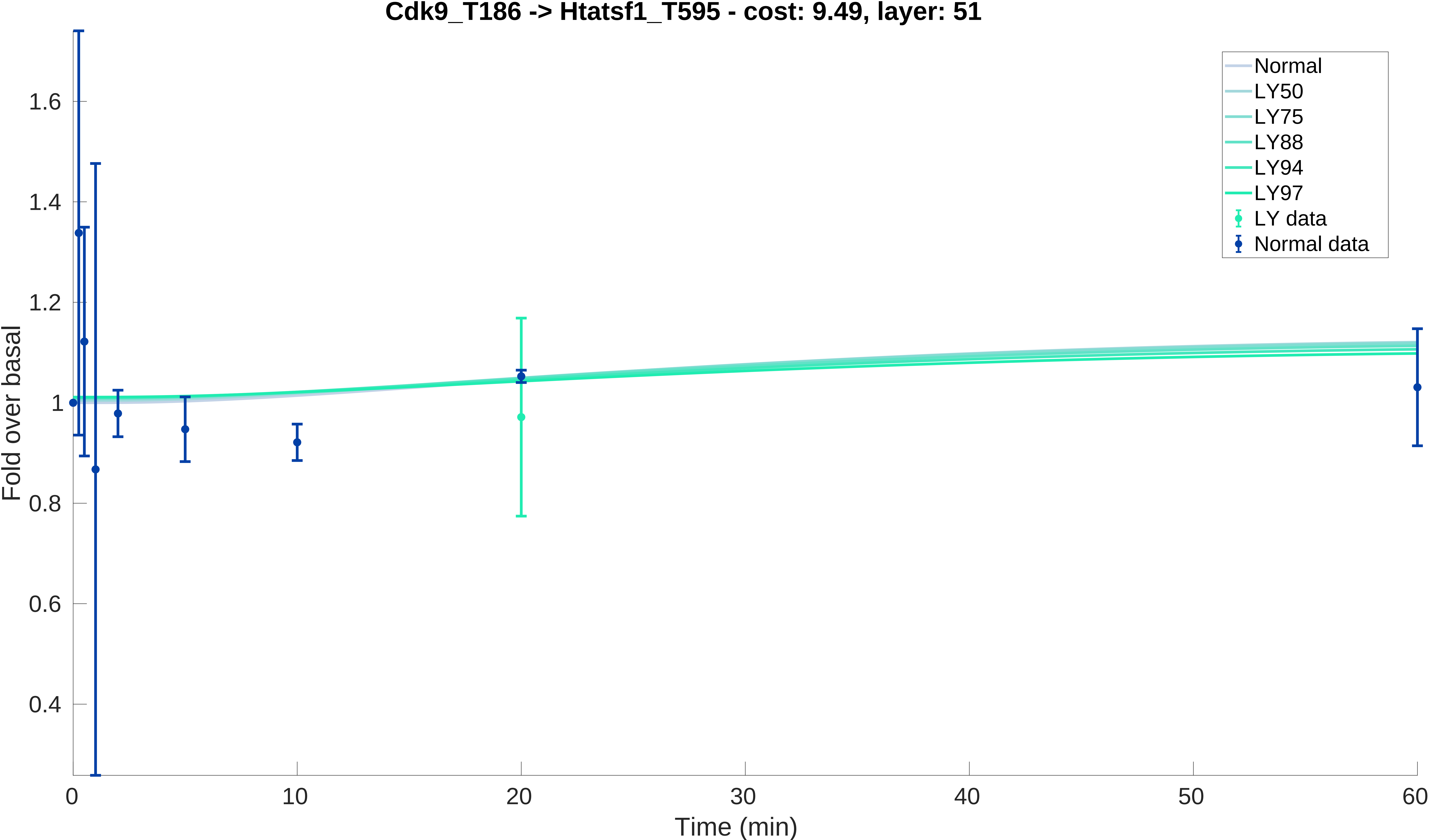

Cdk9_T186 -> Htatsf1_T595 - cost: 9.49, layer: 51
Normal
LY50
LY75
LY88
LY94
LY97
LY data
Normal data
Fold over basal
1.6
1.4
1.2
1
0.8
0.6
0.4
0
10
20
30
40
50
60
Time (min)

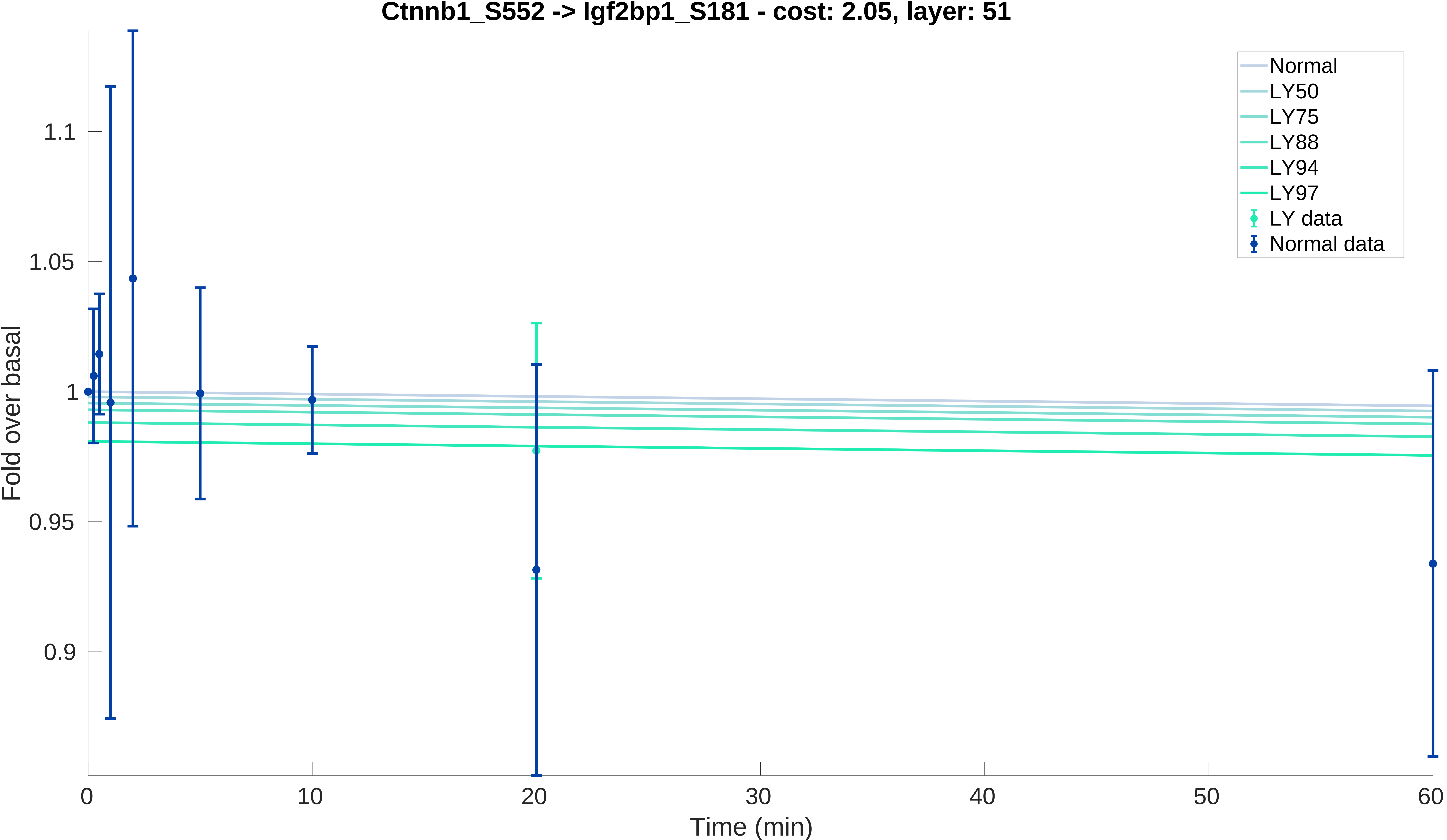
Ctnnb1_S552 -> Igf2bp1_S181 - cost: 2.05, layer: 51
Normal
LY50
LY75
LY88
LY94
LY97
LY data
Normal data
Fold over basal
1.1
1.05
1
0.95
0.9
0
10
20
30
40
50
60
Time (min)

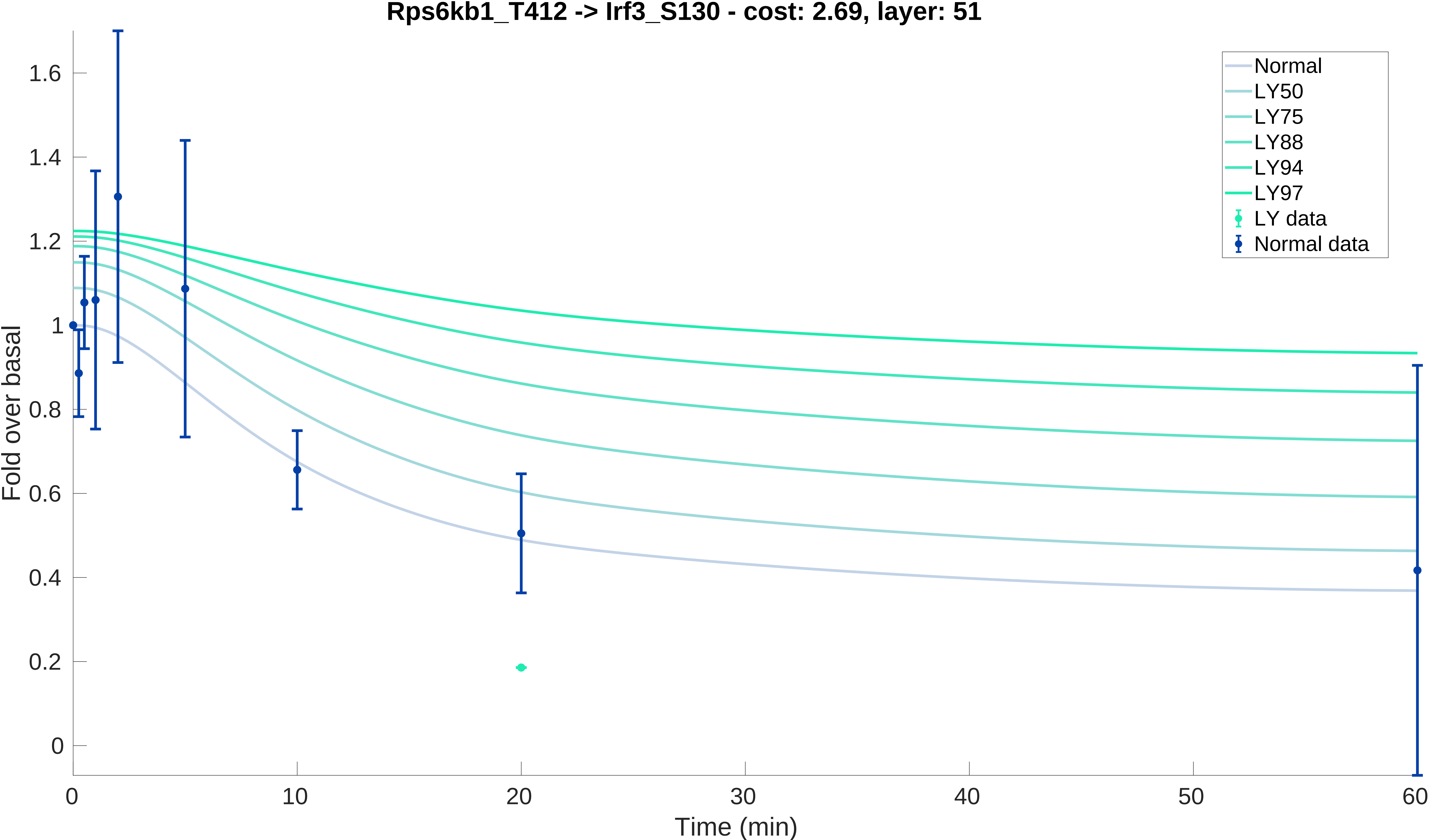
Rps6kb1_T412 -> Irf3_S130 - cost: 2.69, layer: 51
Fold over basal
Time (min)
0
0.2
0.4
0.6
0.8
1
1.2
1.4
1.6
0
10
20
30
40
50
60
Normal
LY50
LY75
LY88
LY94
LY97
LY data
Normal data

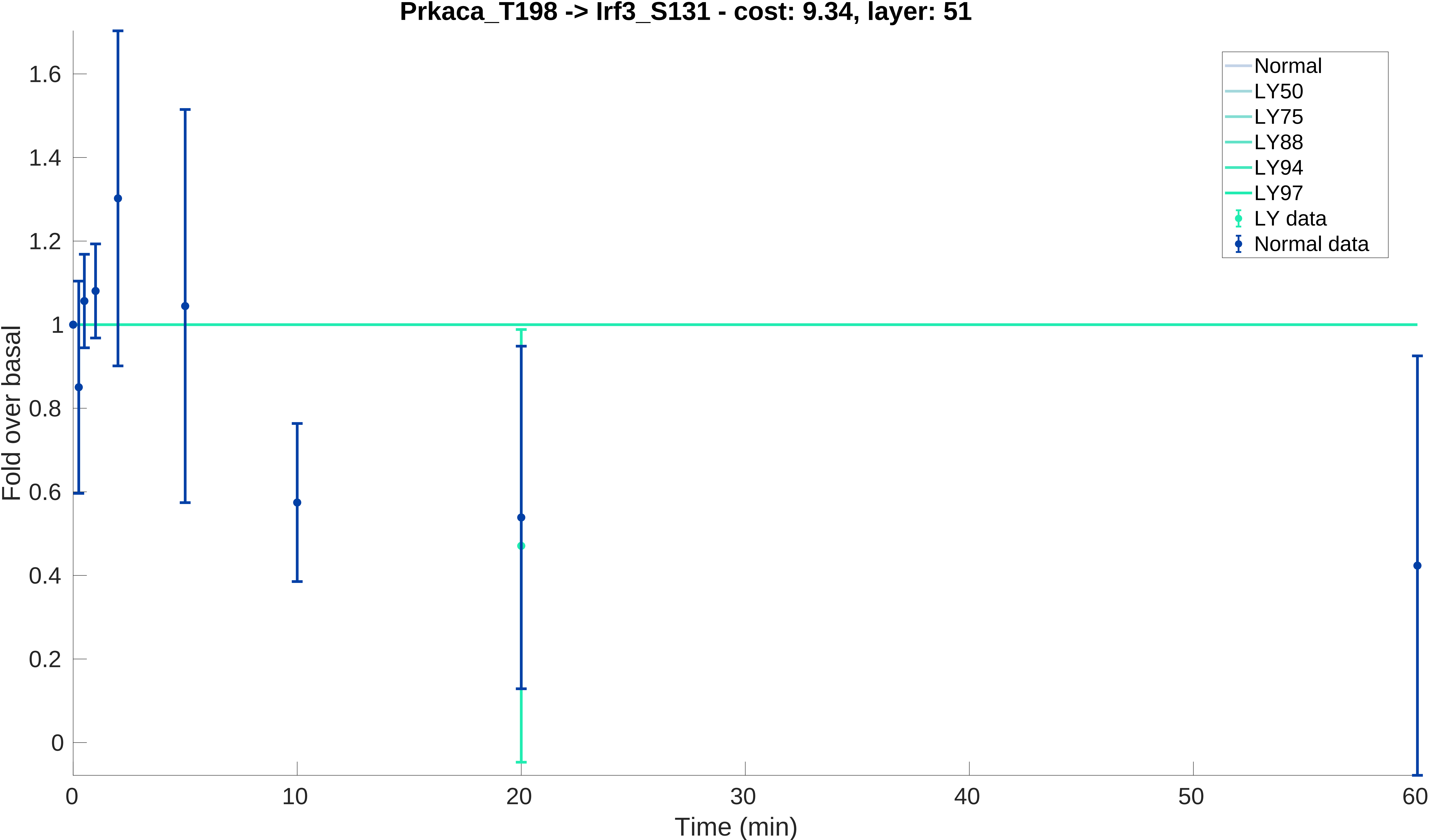

Prkaca_T198 -> Irf3_S131 - cost: 9.34, layer: 51
Fold over basal
Time (min)
0
0.2
0.4
0.6
0.8
1
1.2
1.4
1.6
0
10
20
30
40
50
60
Normal
LY50
LY75
LY88
LY94
LY97
LY data
Normal data

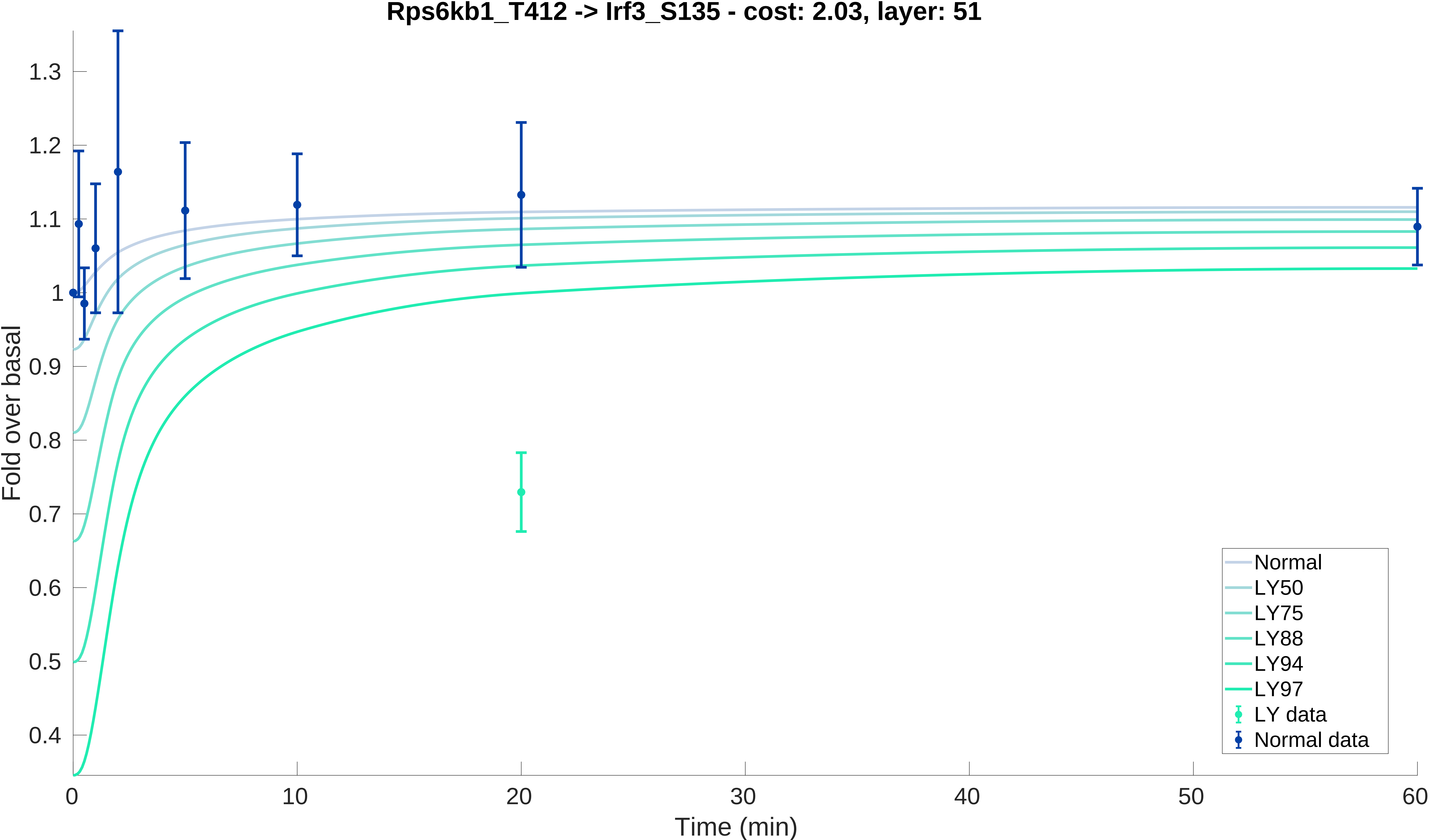

Rps6kb1_T412 -> Irf3_S135 - cost: 2.03, layer: 51
Fold over basal
Time (min)
0
10
20
30
40
50
60
0.4
0.5
0.6
0.7
0.8
0.9
1
1.1
1.2
1.3
Normal
LY50
LY75
LY88
LY94
LY97
LY data
Normal data

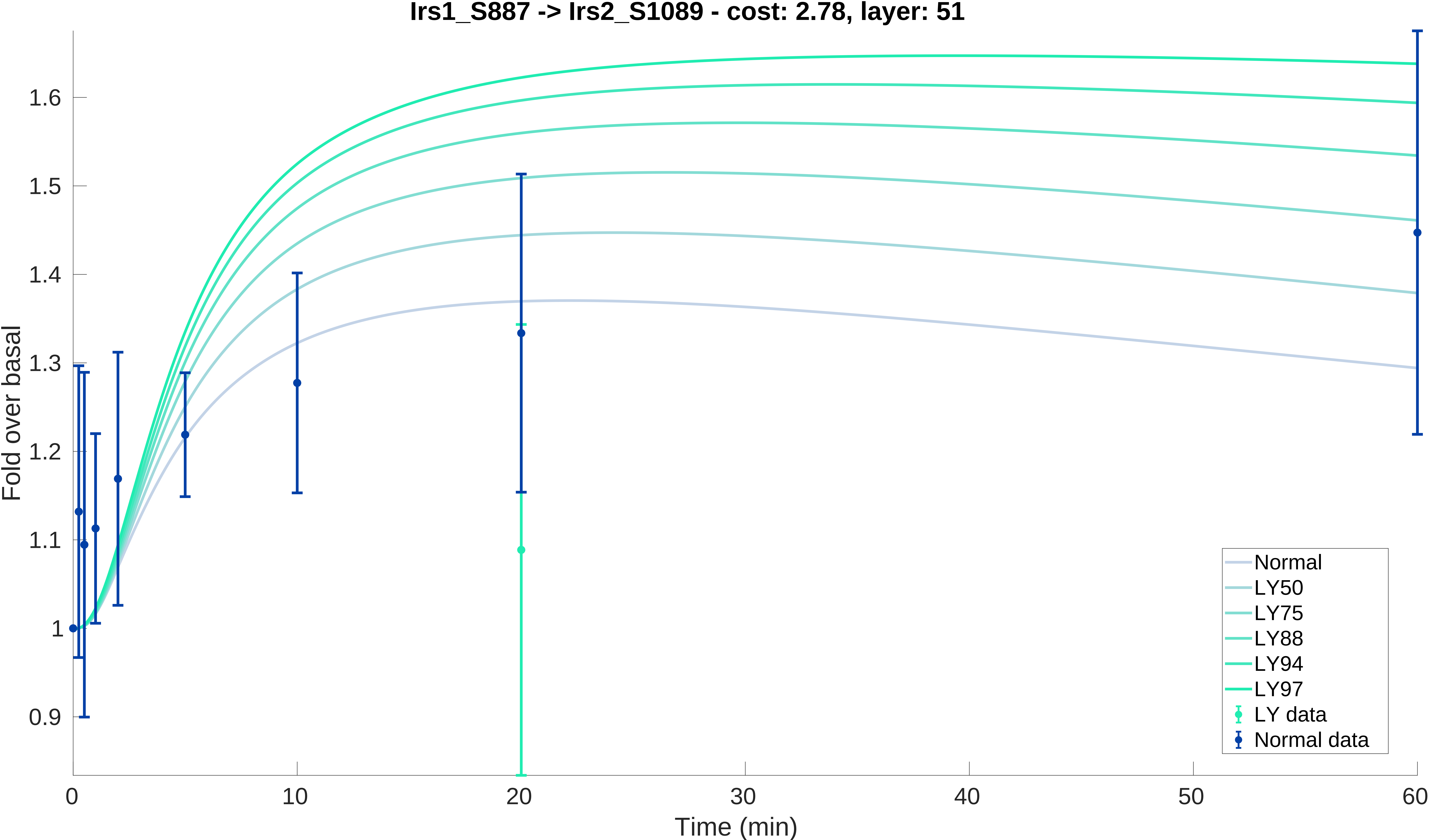
Irs1_S887 -> Irs2_S1089 - cost: 2.78, layer: 51
Fold over basal
Time (min)
0.9
1
1.1
1.2
1.3
1.4
1.5
1.6
0
10
20
30
40
50
60
Normal
LY50
LY75
LY88
LY94
LY97
LY data
Normal data

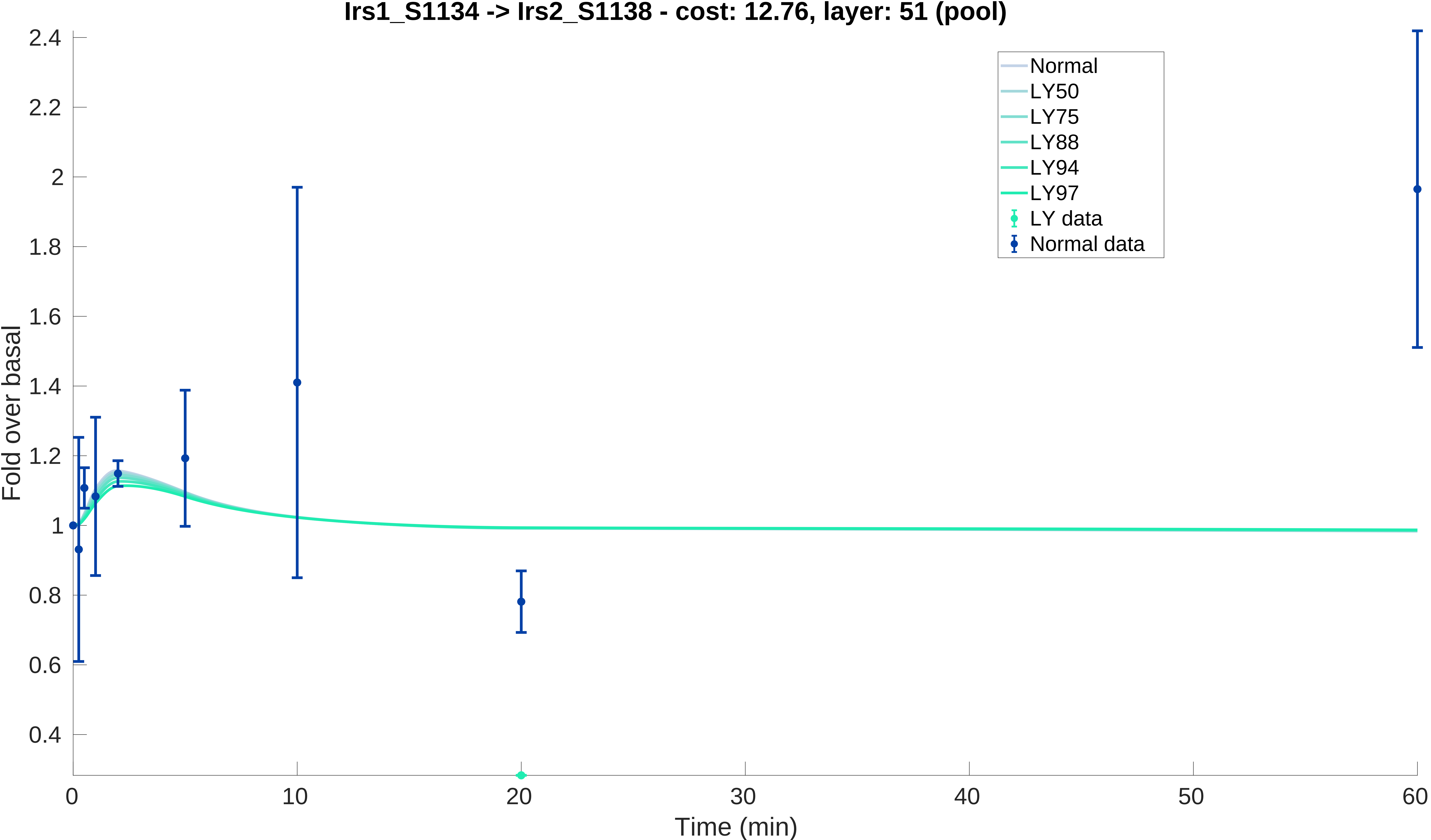

Irs1_S1134 -> Irs2_S1138 - cost: 12.76, layer: 51 (pool)
Fold over basal
Time (min)
2.4
2.2
2
1.8
1.6
1.4
1.2
1
0.8
0.6
0.4
0
10
20
30
40
50
60
Normal
LY50
LY75
LY88
LY94
LY97
LY data
Normal data

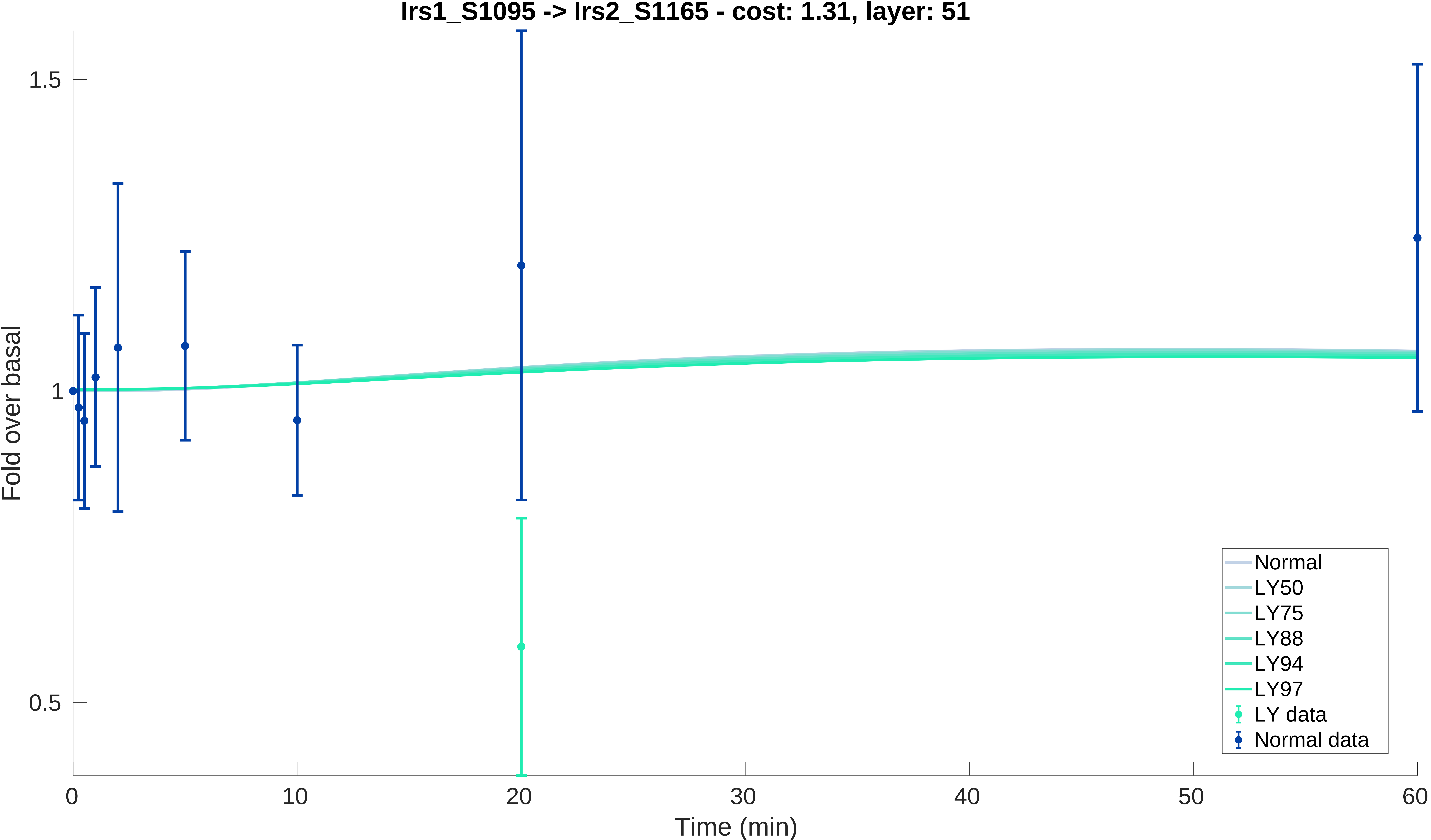

Irs1_S1095 -> Irs2_S1165 - cost: 1.31, layer: 51
Fold over basal
1.5
1
0.5
0
10
20
30
40
50
60
Time (min)
Normal
LY50
LY75
LY88
LY94
LY97
LY data
Normal data

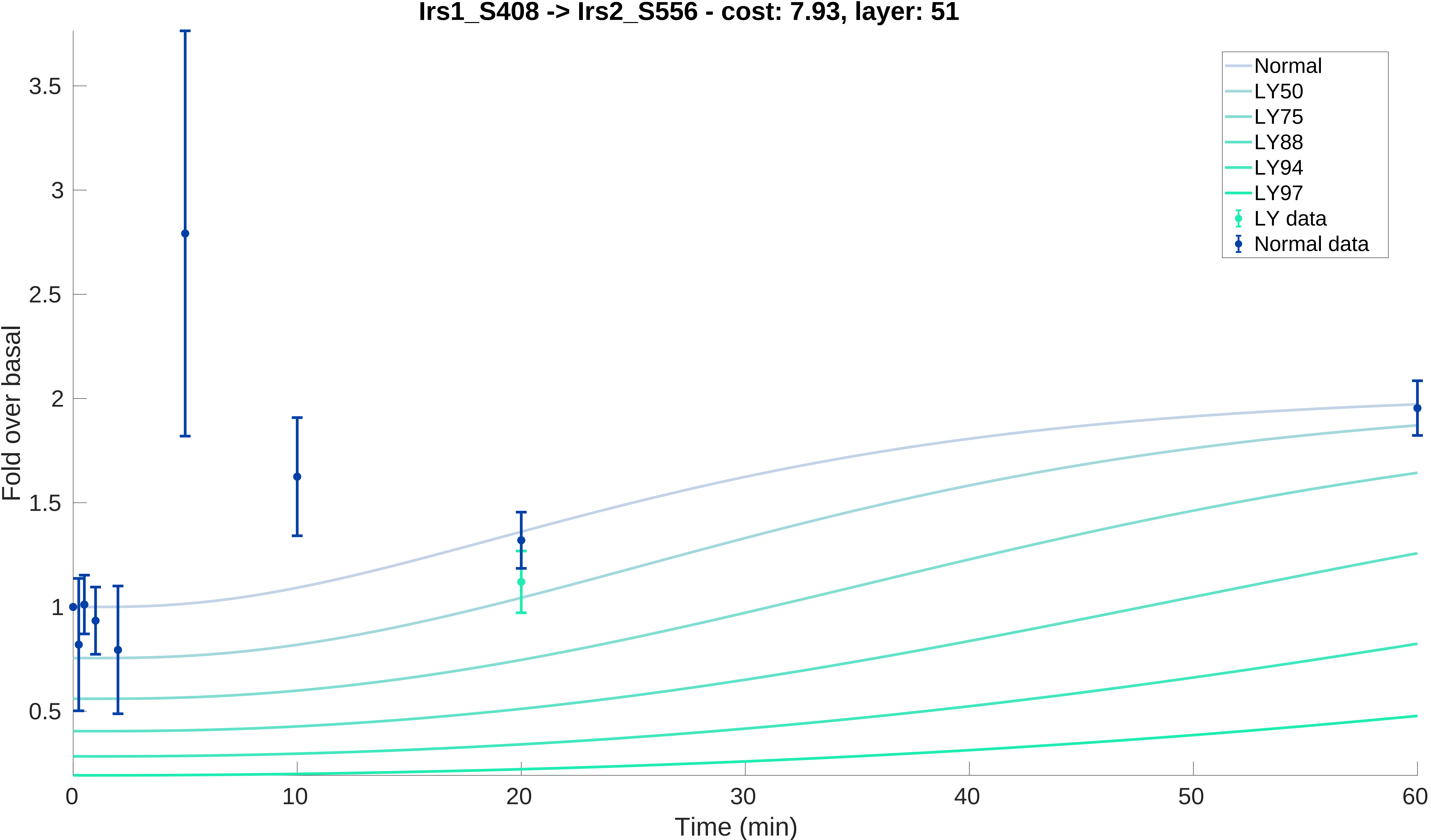

Irs1_S408 -> Irs2_S556 - cost: 7.93, layer: 51
Fold over basal
Time (min)
0.5
1
1.5
2
2.5
3
3.5
0
10
20
30
40
50
60
Normal
LY50
LY75
LY88
LY94
LY97
LY data
Normal data

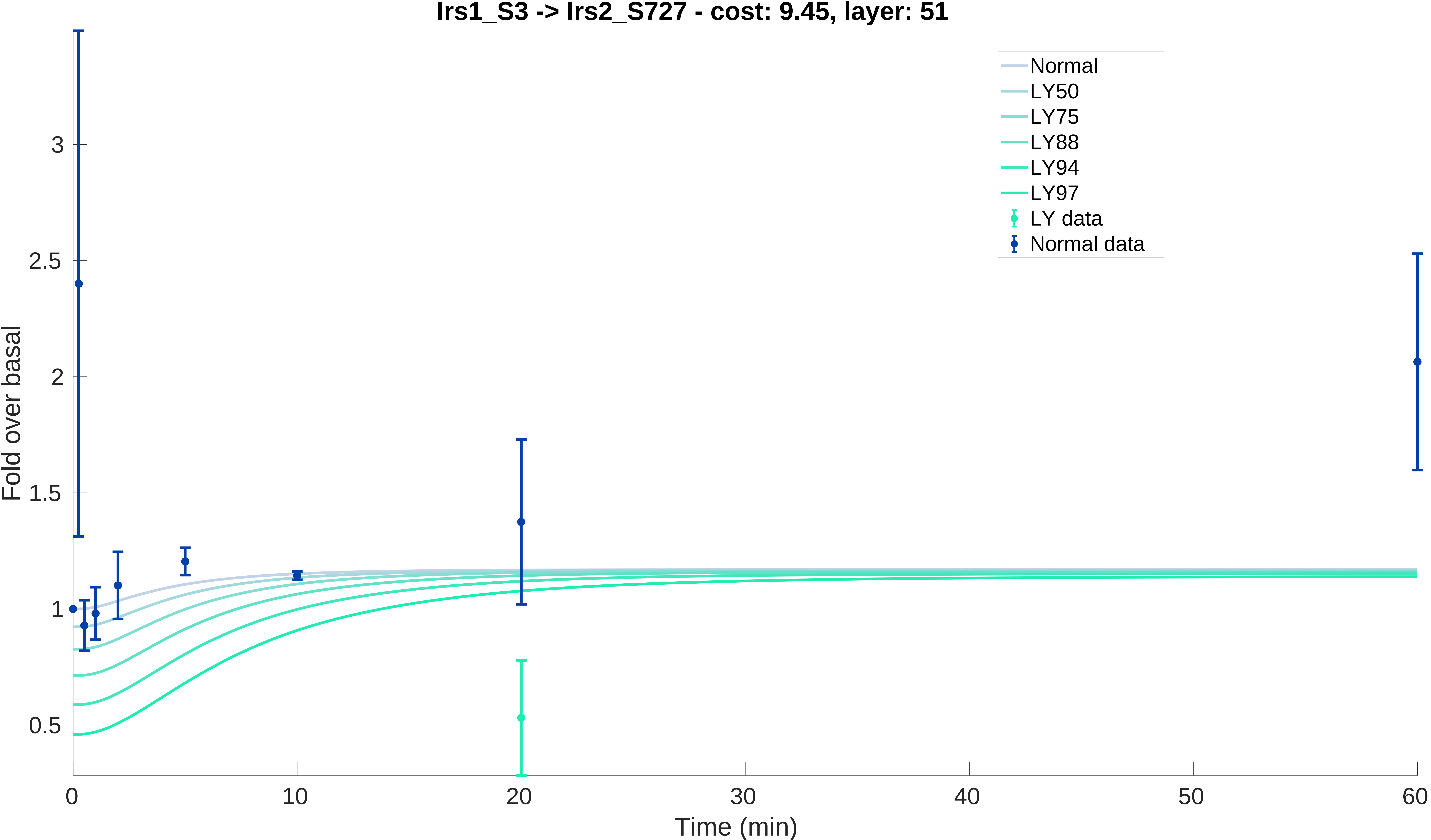

Irs1_S3 -> Irs2_S727 - cost: 9.45, layer: 51
Normal
LY50
LY75
LY88
LY94
LY97
LY data
Normal data
Fold over basal
3
2.5
2
1.5
1
0.5
0
10
20
30
40
50
60
Time (min)

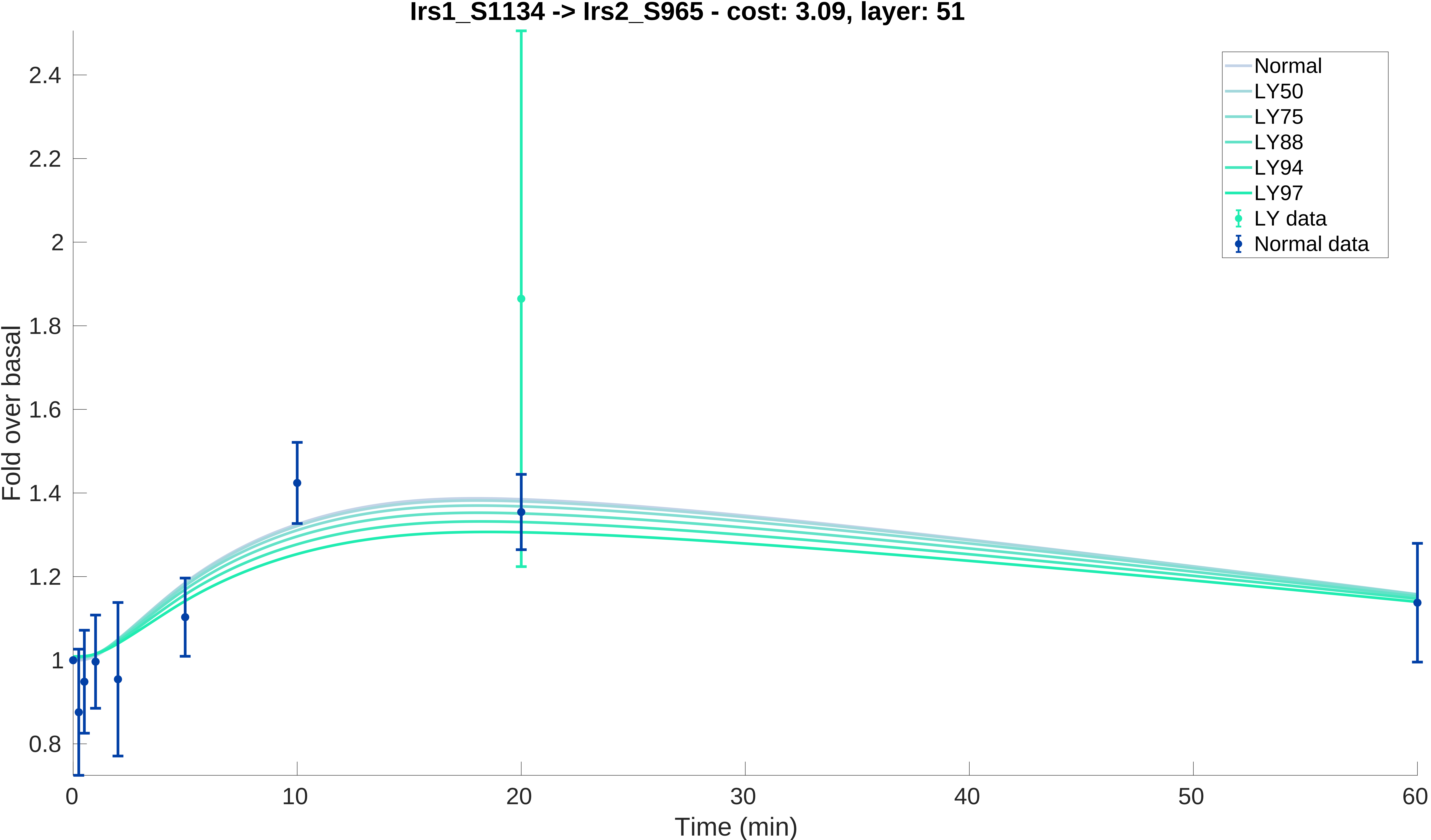

Irs1_S1134 -> Irs2_S965 - cost: 3.09, layer: 51
Fold over basal
Time (min)
0
10
20
30
40
50
60
0.8
1
1.2
1.4
1.6
1.8
2
2.2
2.4
Normal
LY50
LY75
LY88
LY94
LY97
LY data
Normal data

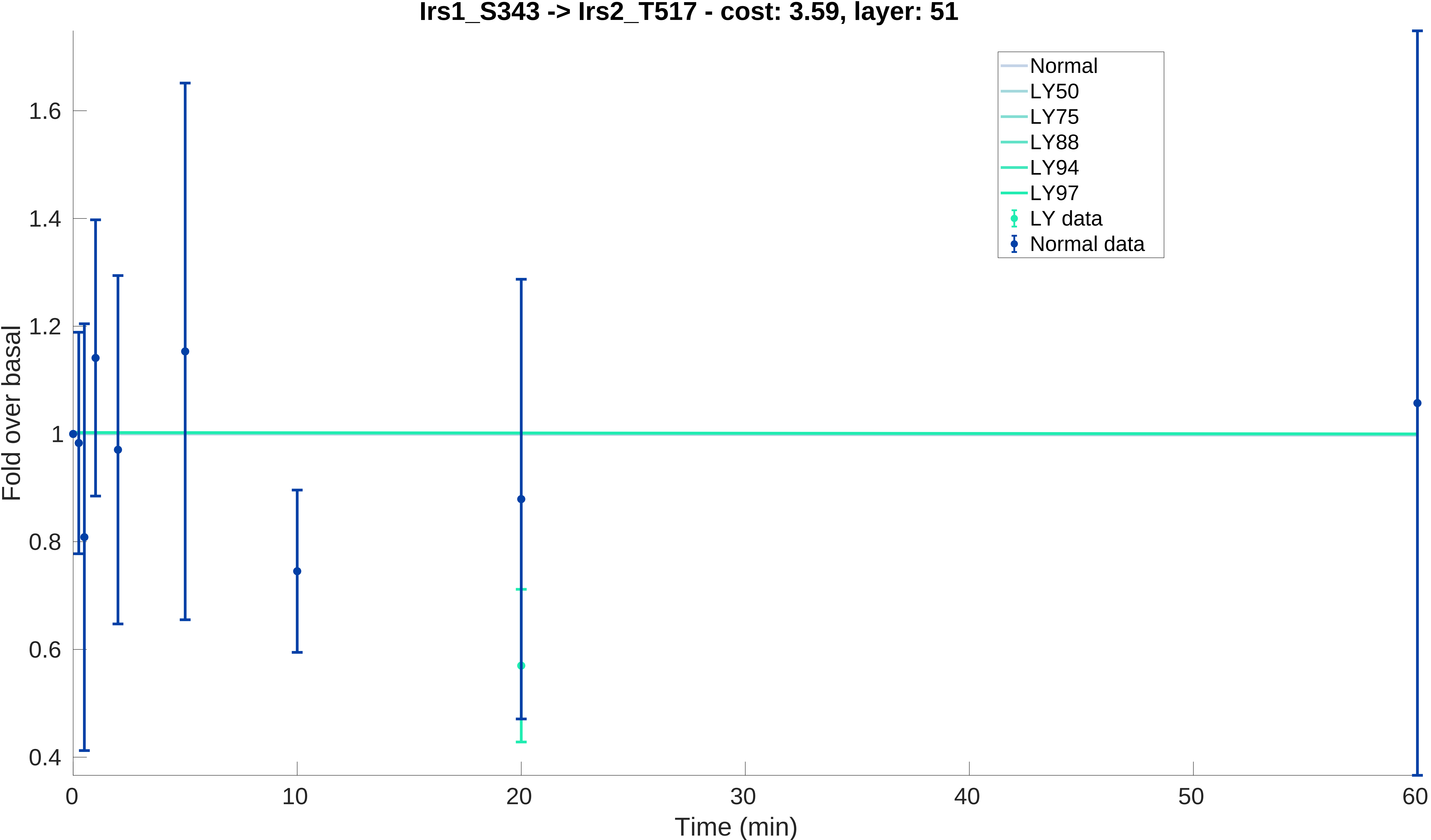

Irs1_S343 -> Irs2_T517 - cost: 3.59, layer: 51
Normal
LY50
LY75
LY88
LY94
LY97
LY data
Normal data
Fold over basal
1.6
1.4
1.2
1
0.8
0.6
0.4
0
10
20
30
40
50
60
Time (min)

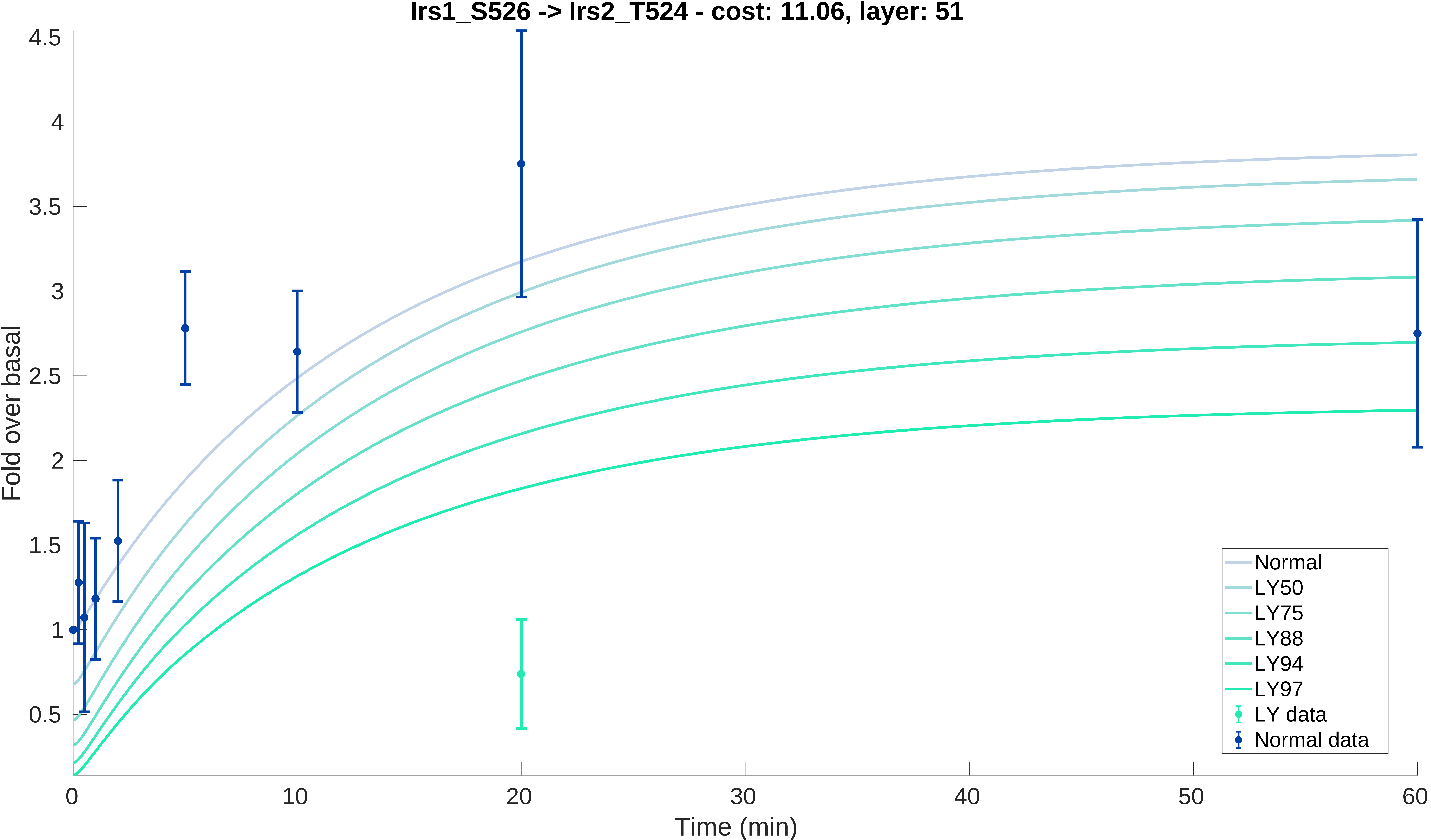

Irs1_S526 -> Irs2_T524 - cost: 11.06, layer: 51
Fold over basal
4.5
4
3.5
3
2.5
2
1.5
1
0.5
0
10
20
30
40
50
60
Time (min)
Normal
LY50
LY75
LY88
LY94
LY97
LY data
Normal data

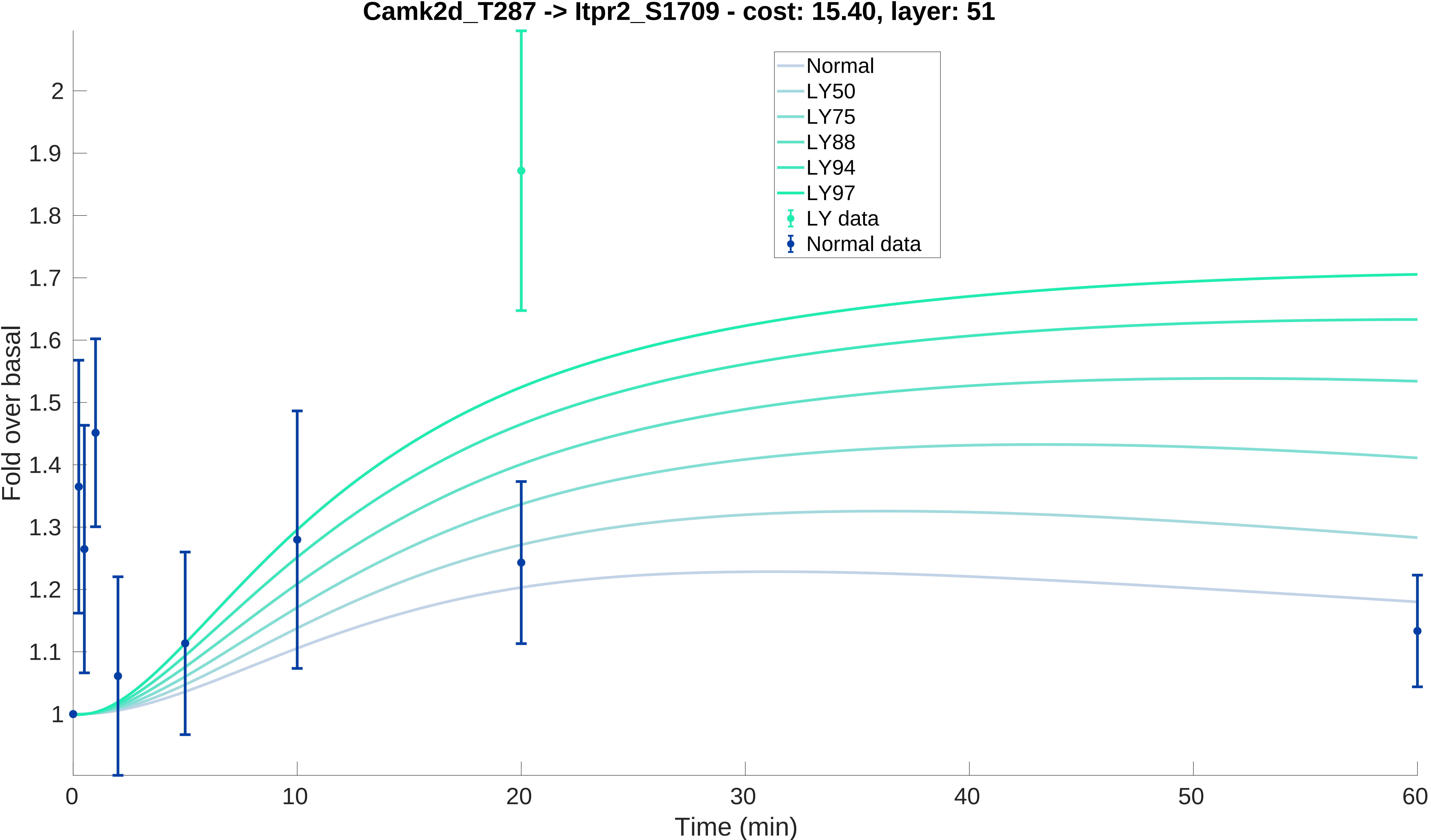

Camk2d_T287 -> Itpr2_S1709 - cost: 15.40, layer: 51
Normal
LY50
LY75
LY88
LY94
LY97
LY data
Normal data
Fold over basal
2
1.9
1.8
1.7
1.6
1.5
1.4
1.3
1.2
1.1
1
0
10
20
30
40
50
60
Time (min)

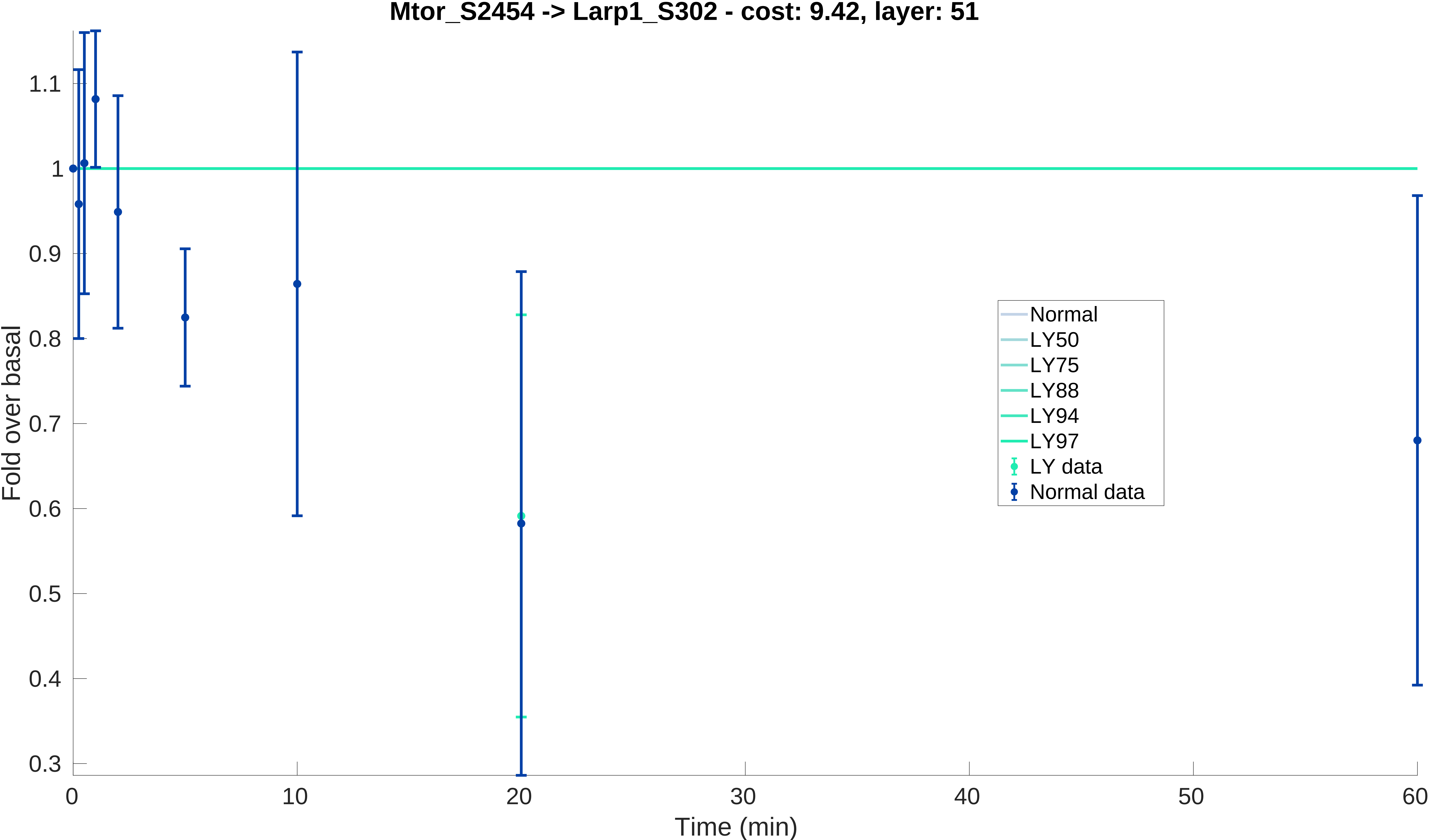

Mtor_S2454 -> Larp1_S302 - cost: 9.42, layer: 51
Fold over basal
1.1
1
0.9
0.8
0.7
0.6
0.5
0.4
0.3
0
10
20
30
40
50
60
Time (min)
Normal
LY50
LY75
LY88
LY94
LY97
LY data
Normal data

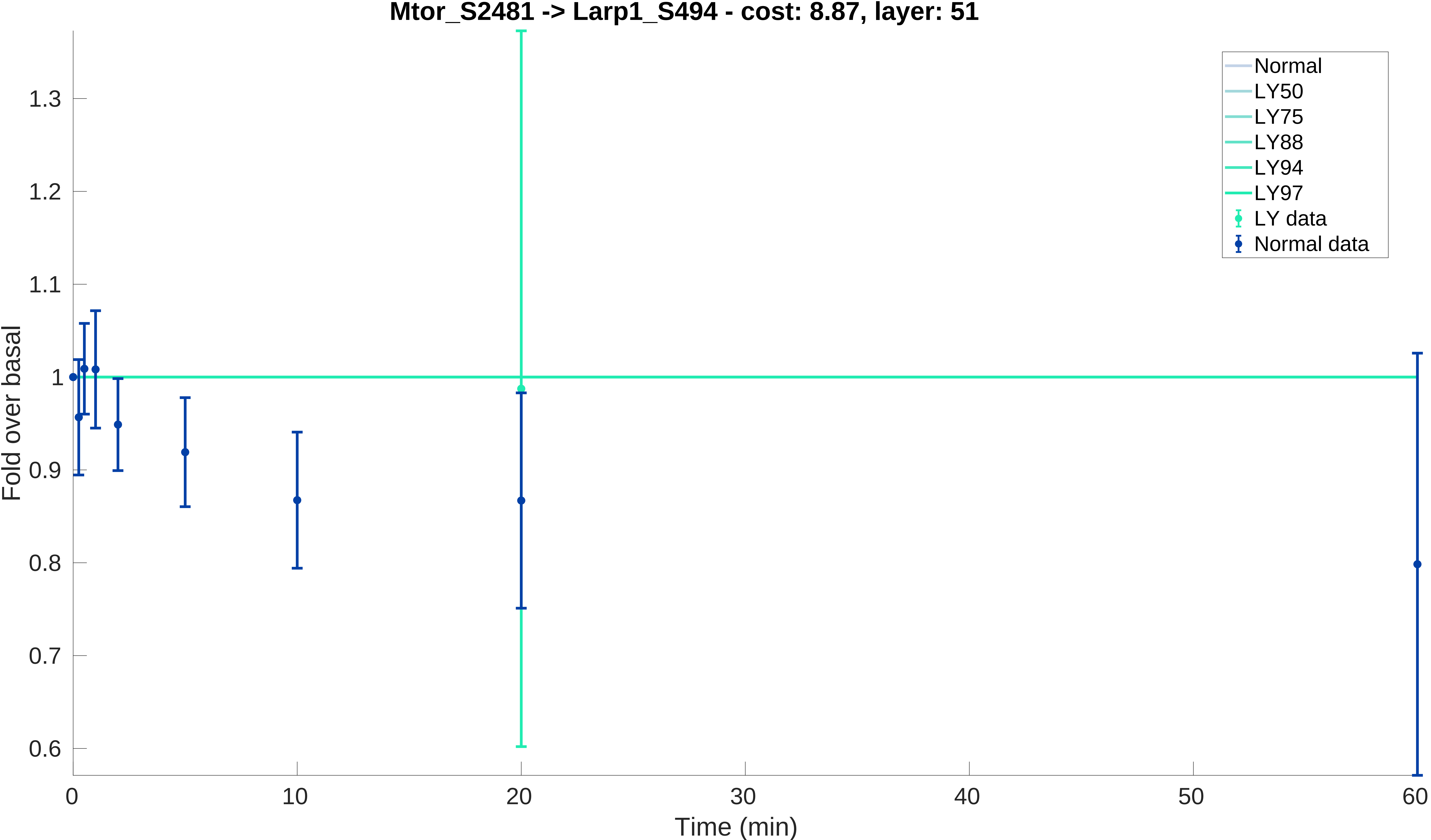

Mtor_S2481 -> Larp1_S494 - cost: 8.87, layer: 51
Fold over basal
Time (min)
0.6
0.7
0.8
0.9
1
1.1
1.2
1.3
0
10
20
30
40
50
60
Normal
LY50
LY75
LY88
LY94
LY97
LY data
Normal data

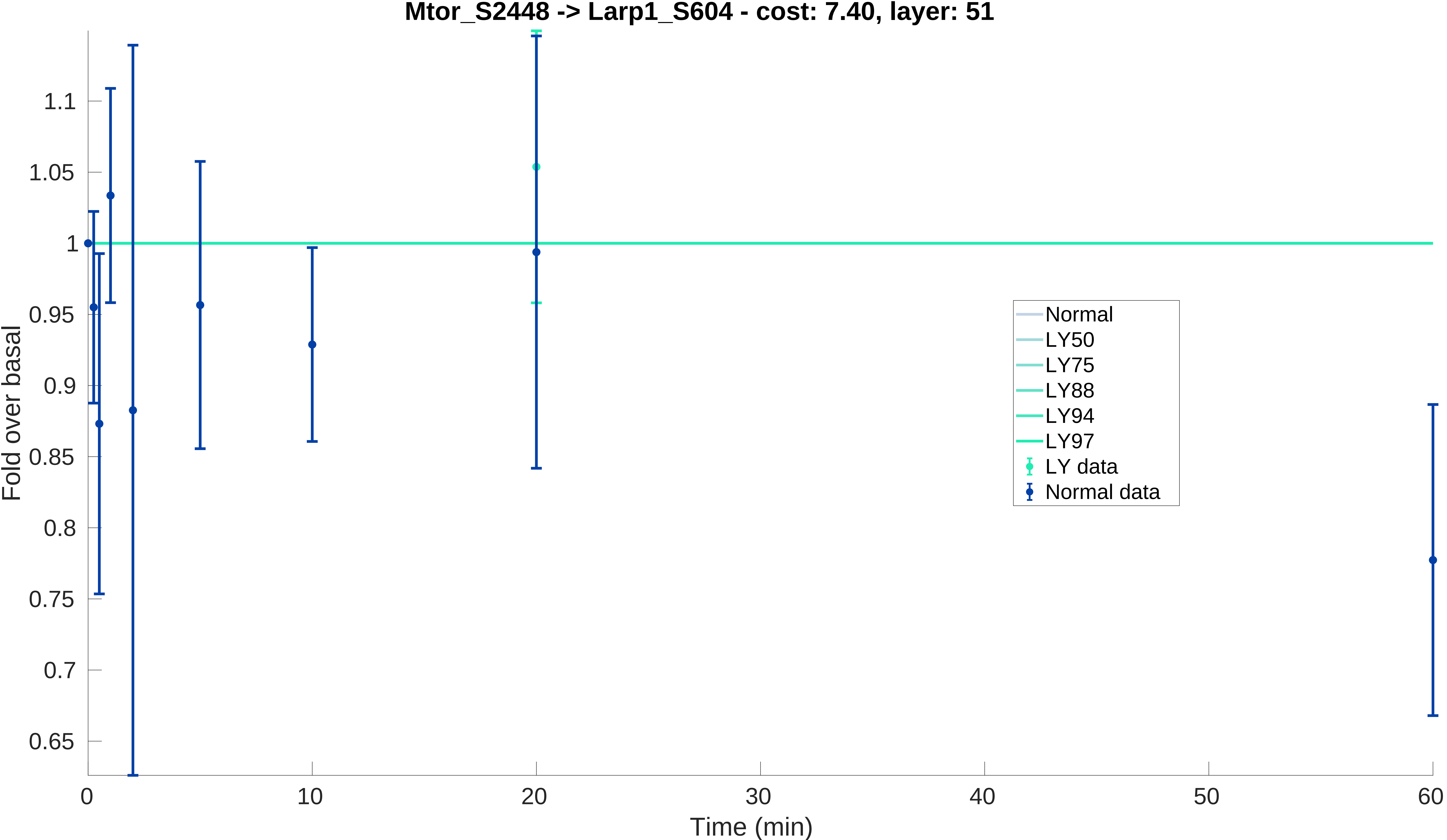

Mtor_S2448 -> Larp1_S604 - cost: 7.40, layer: 51
Fold over basal
1.1
1.05
1
0.95
0.9
0.85
0.8
0.75
0.7
0.65
0
10
20
30
40
50
60
Time (min)
Normal
LY50
LY75
LY88
LY94
LY97
LY data
Normal data

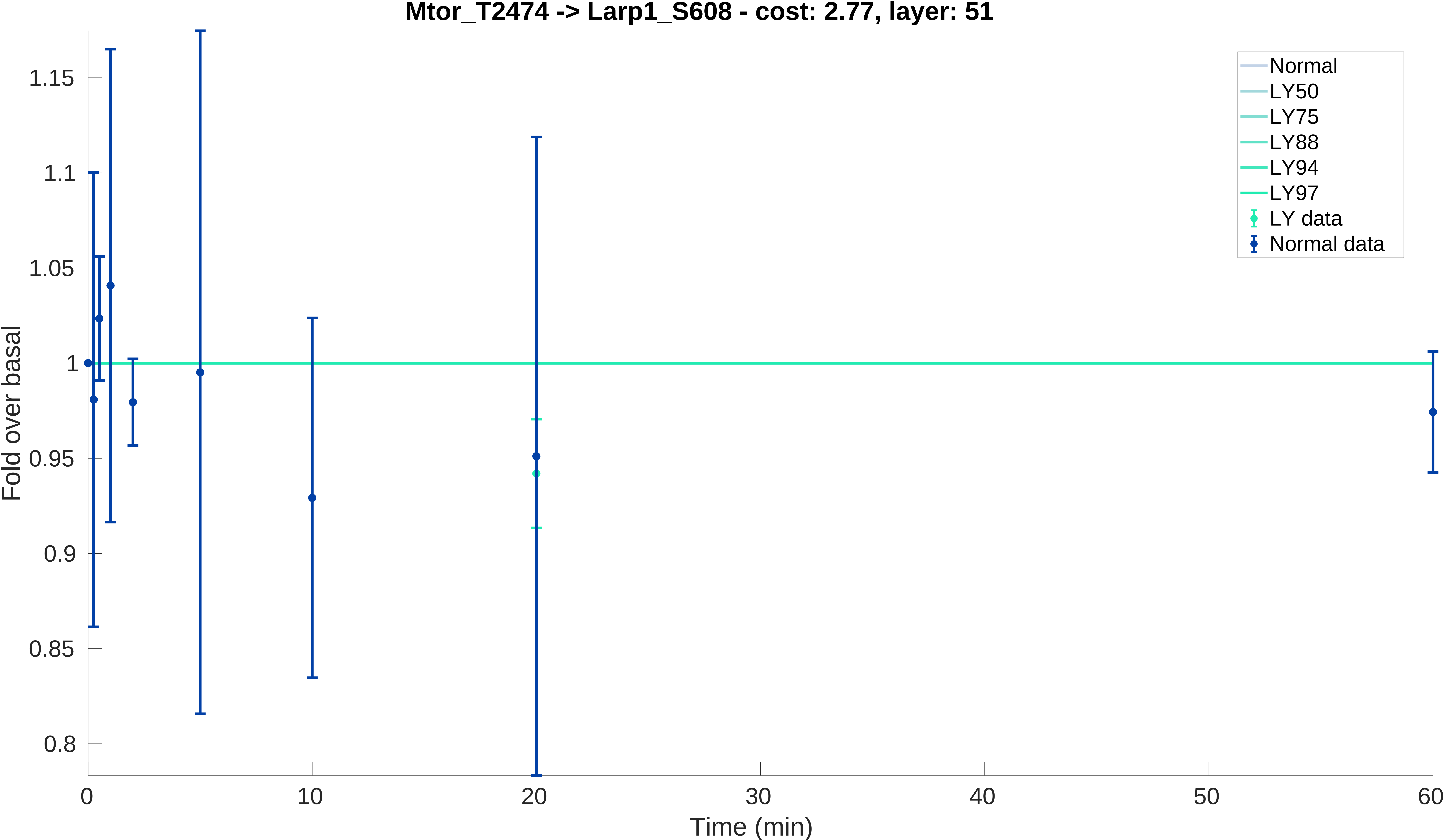

Mtor_T2474 -> Larp1_S608 - cost: 2.77, layer: 51
Normal
LY50
LY75
LY88
LY94
LY97
LY data
Normal data
Fold over basal
1.15
1.1
1.05
1
0.95
0.9
0.85
0.8
0
10
20
30
40
50
60
Time (min)

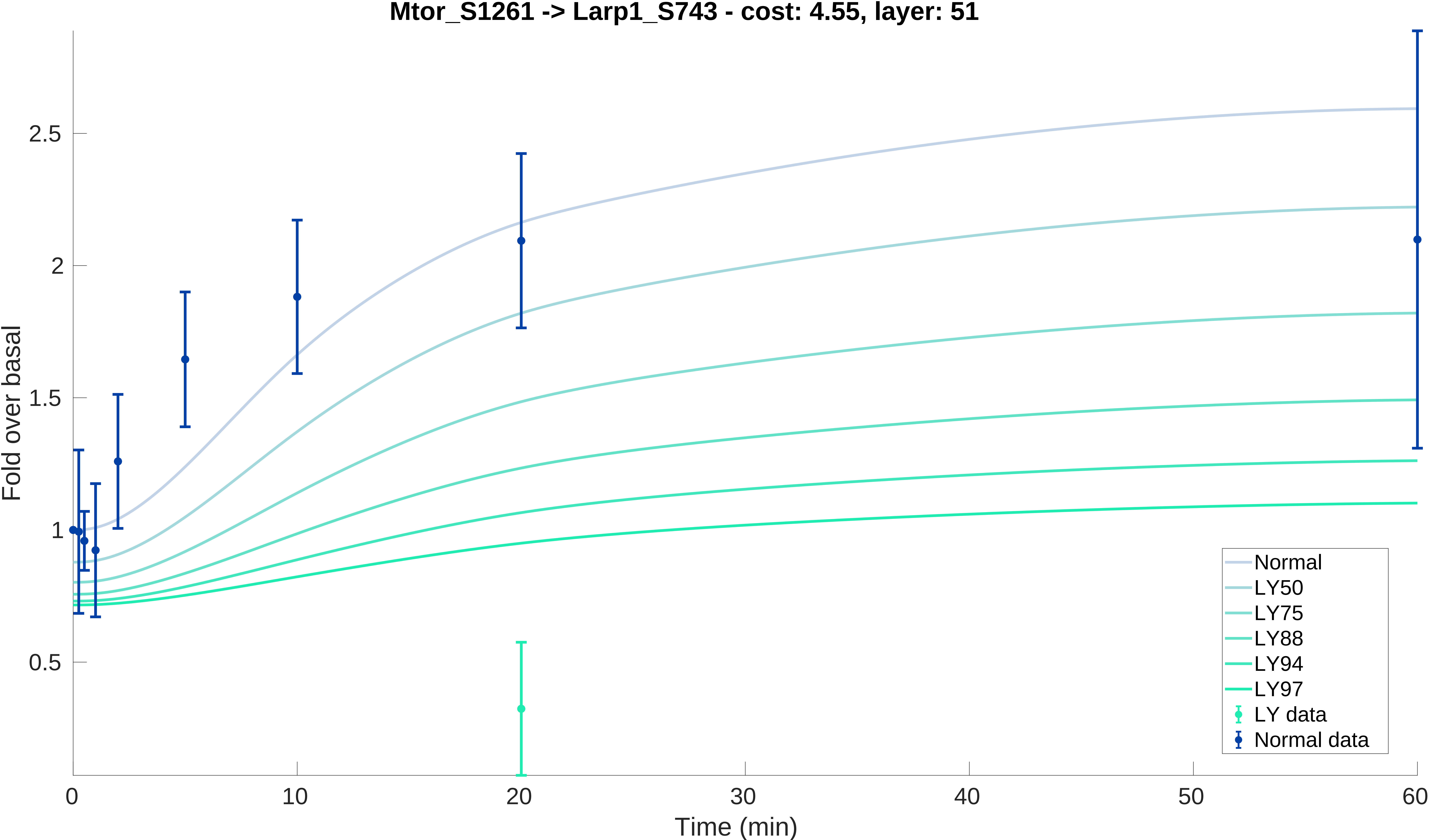

Mtor_S1261 -> Larp1_S743 - cost: 4.55, layer: 51
Fold over basal
Time (min)
2.5
2
1.5
1
0.5
0
10
20
30
40
50
60
Normal
LY50
LY75
LY88
LY94
LY97
LY data
Normal data

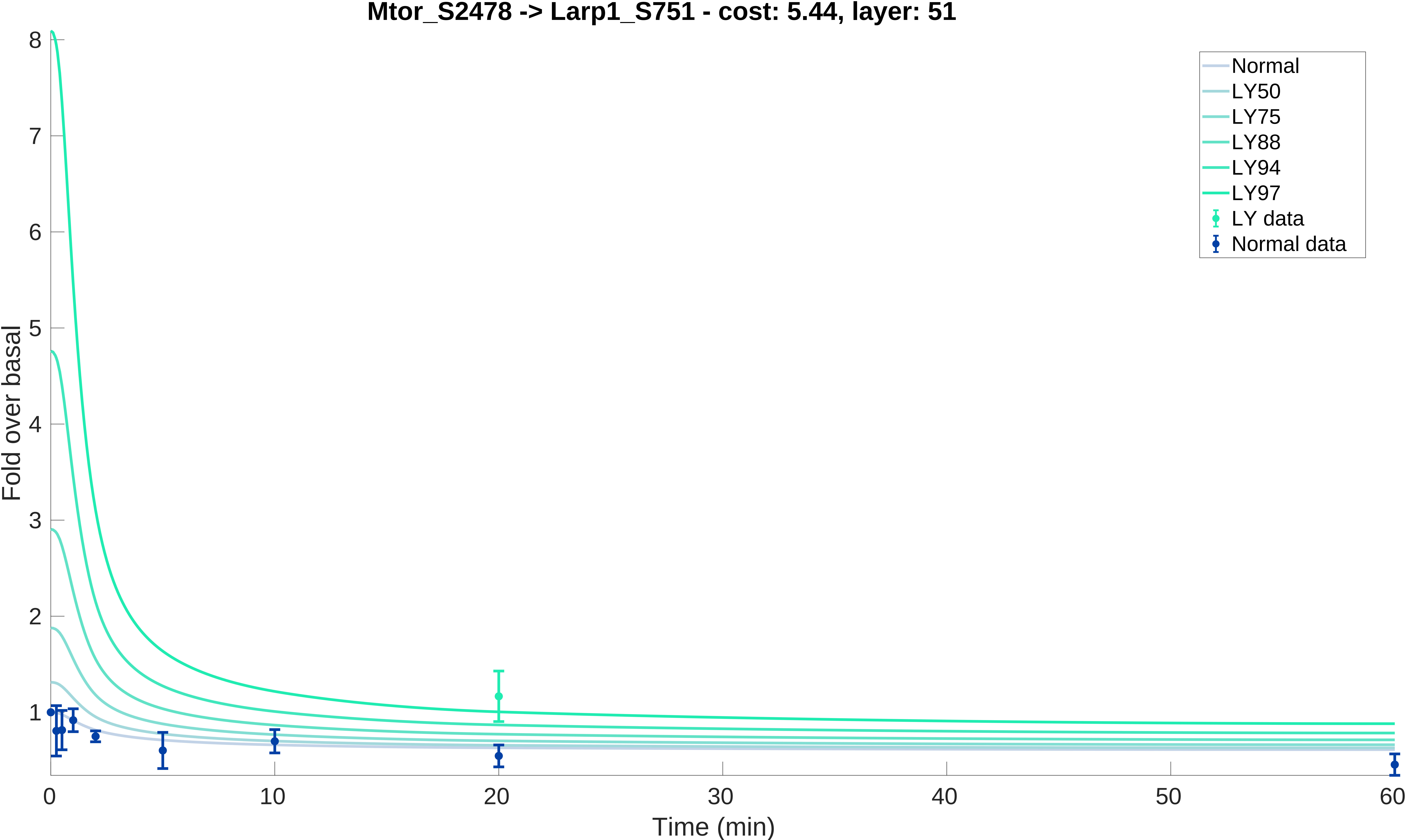
Mtor_S2478 -> Larp1_S751 - cost: 5.44, layer: 51
Fold over basal
Time (min)
Normal
LY50
LY75
LY88
LY94
LY97
LY data
Normal data
0
10
20
30
40
50
60
1
2
3
4
5
6
7
8

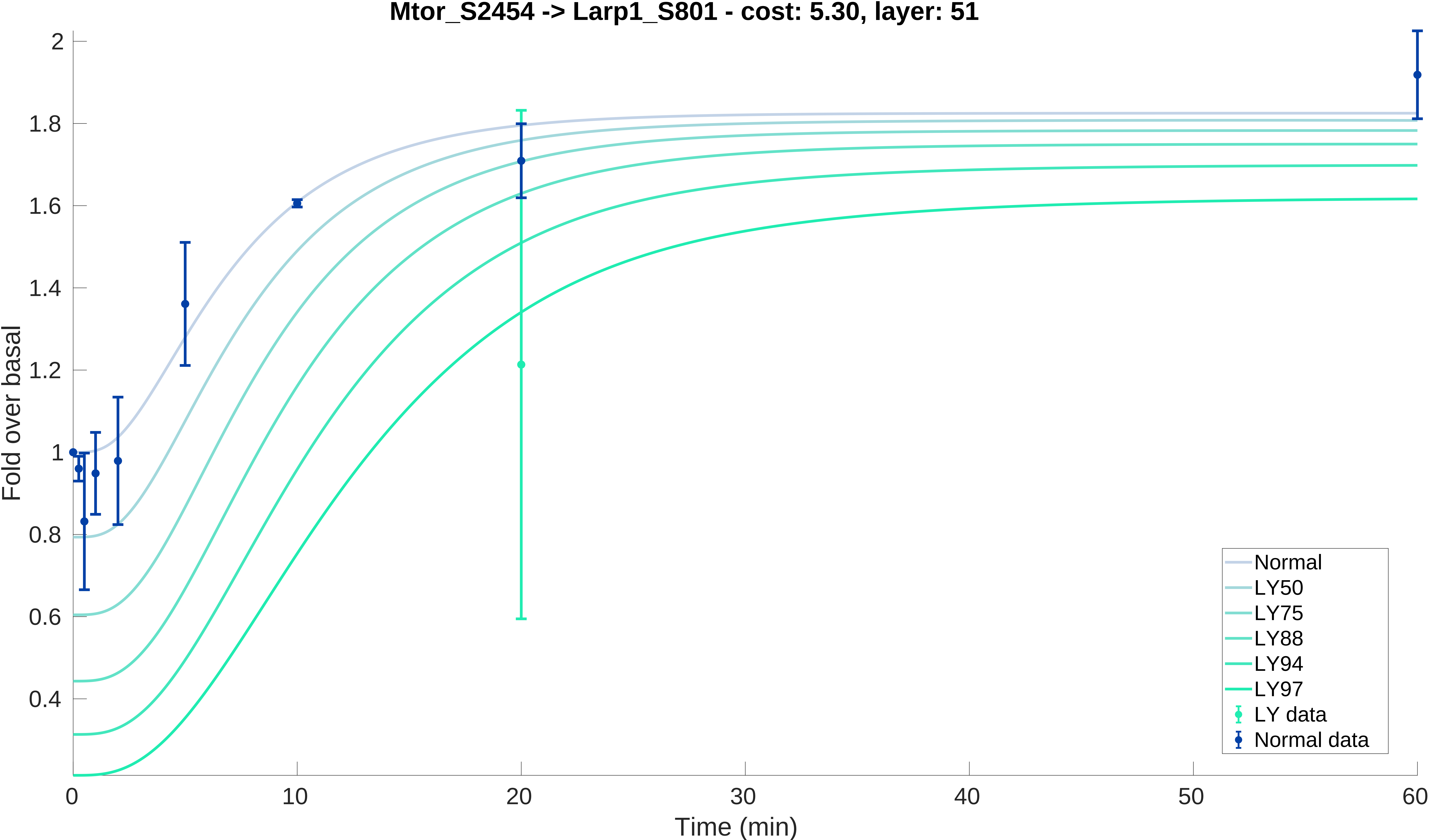
Mtor_S2454 -> Larp1_S801 - cost: 5.30, layer: 51
Fold over basal
2
1.8
1.6
1.4
1.2
1
0.8
0.6
0.4
0
10
20
30
40
50
60
Time (min)
Normal
LY50
LY75
LY88
LY94
LY97
LY data
Normal data

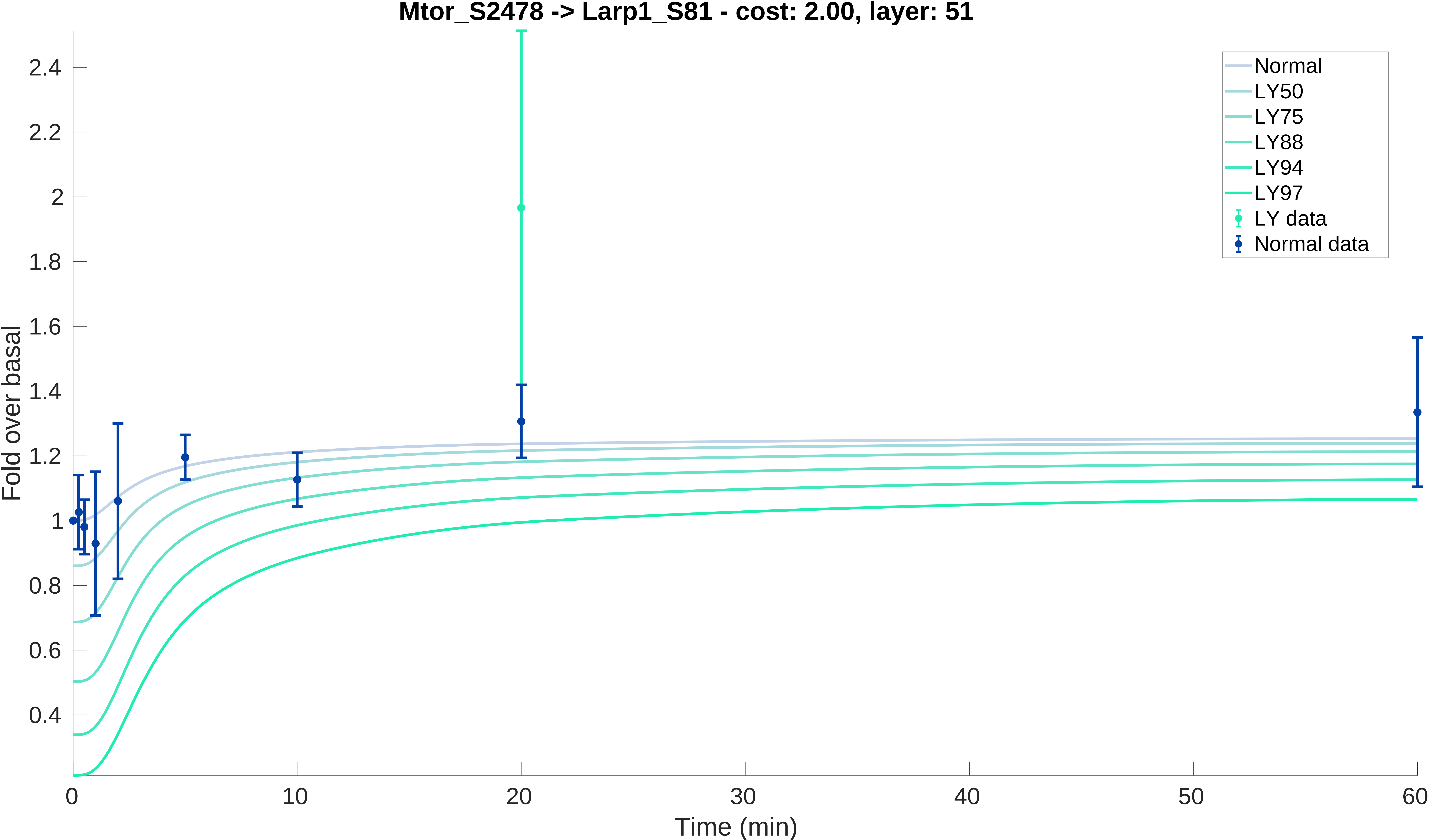
Mtor_S2478 -> Larp1_S81 - cost: 2.00, layer: 51
Fold over basal
Time (min)
0.4
0.6
0.8
1
1.2
1.4
1.6
1.8
2
2.2
2.4
0
10
20
30
40
50
60
Normal
LY50
LY75
LY88
LY94
LY97
LY data
Normal data

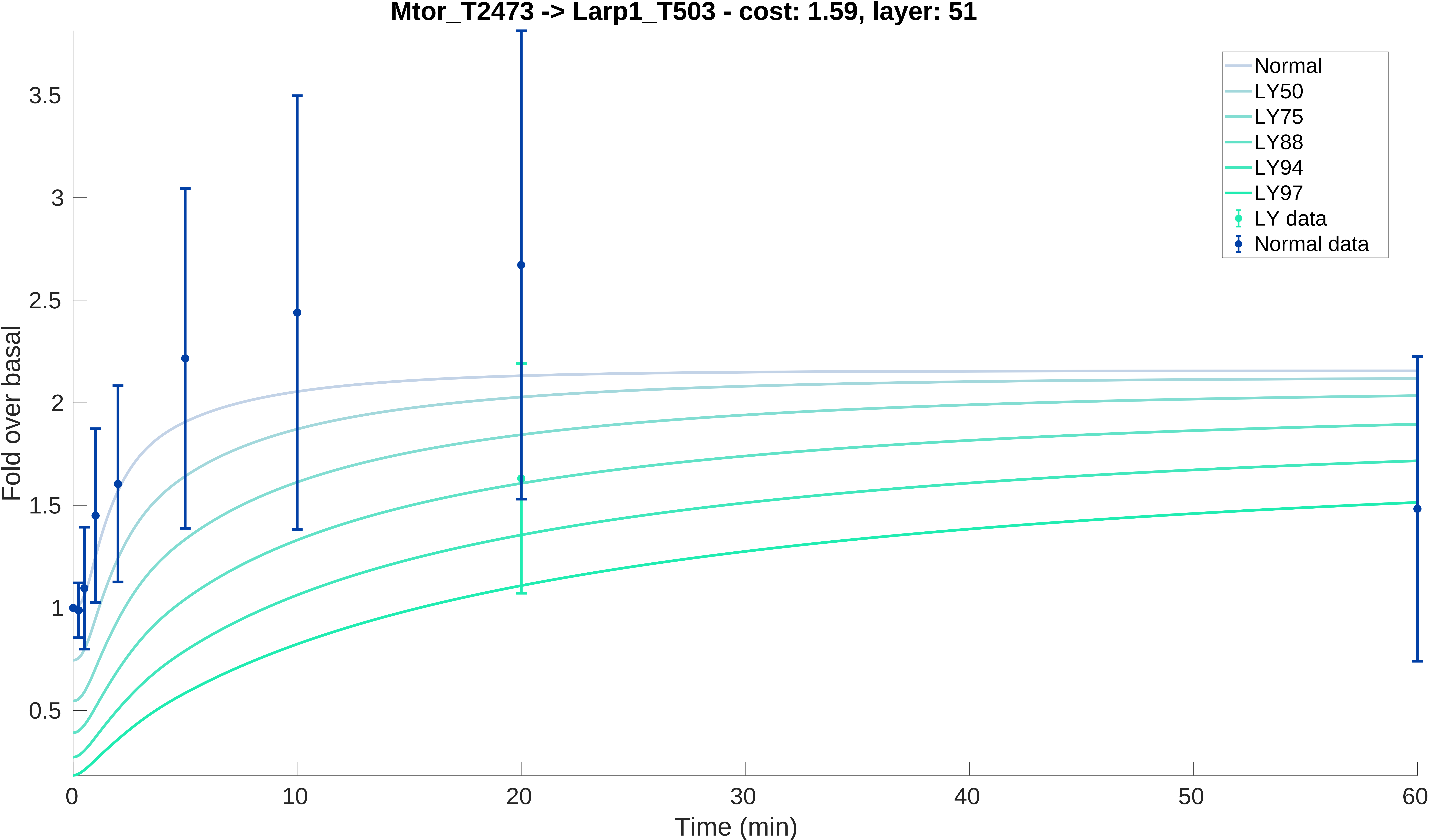
Mtor_T2473 -> Larp1_T503 - cost: 1.59, layer: 51
Fold over basal
Time (min)
0.5
1
1.5
2
2.5
3
3.5
0
10
20
30
40
50
60
Normal
LY50
LY75
LY88
LY94
LY97
LY data
Normal data

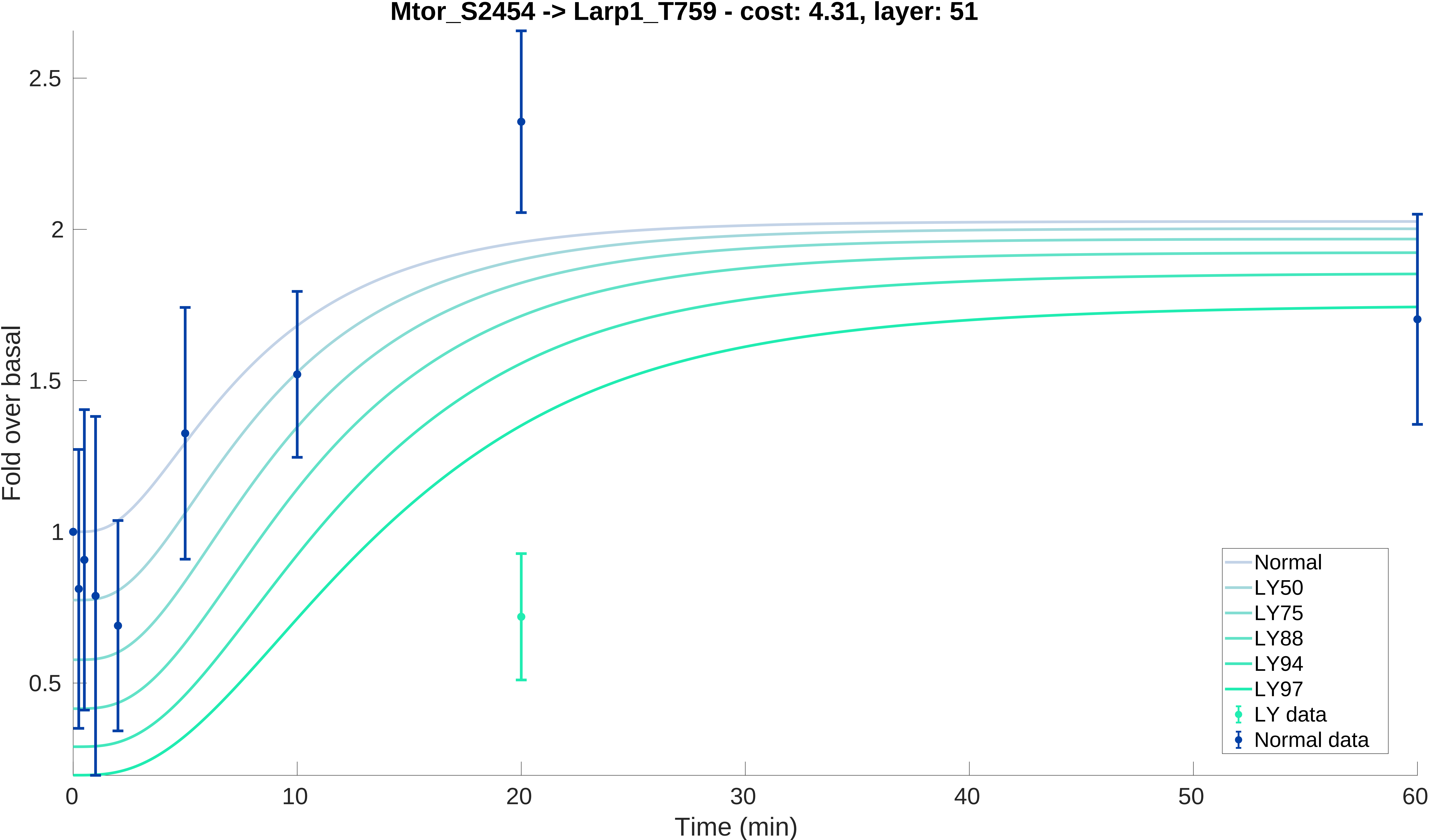

Mtor_S2454 -> Larp1_T759 - cost: 4.31, layer: 51
Fold over basal
Time (min)
0.5
1
1.5
2
2.5
0
10
20
30
40
50
60
Normal
LY50
LY75
LY88
LY94
LY97
LY data
Normal data

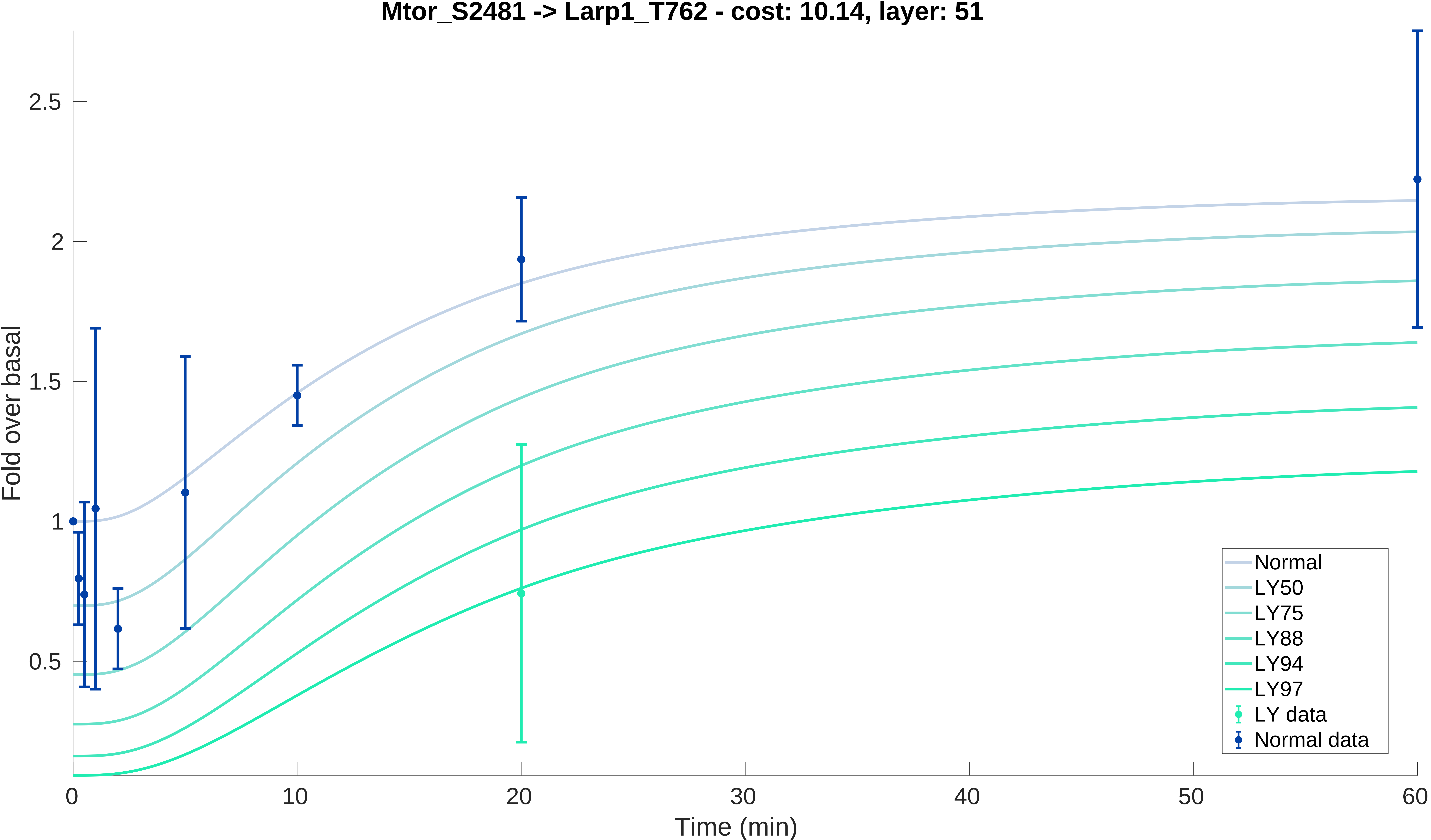

Mtor_S2481 -> Larp1_T762 - cost: 10.14, layer: 51
Fold over basal
Time (min)
2.5
2
1.5
1
0.5
0
10
20
30
40
50
60
Normal
LY50
LY75
LY88
LY94
LY97
LY data
Normal data

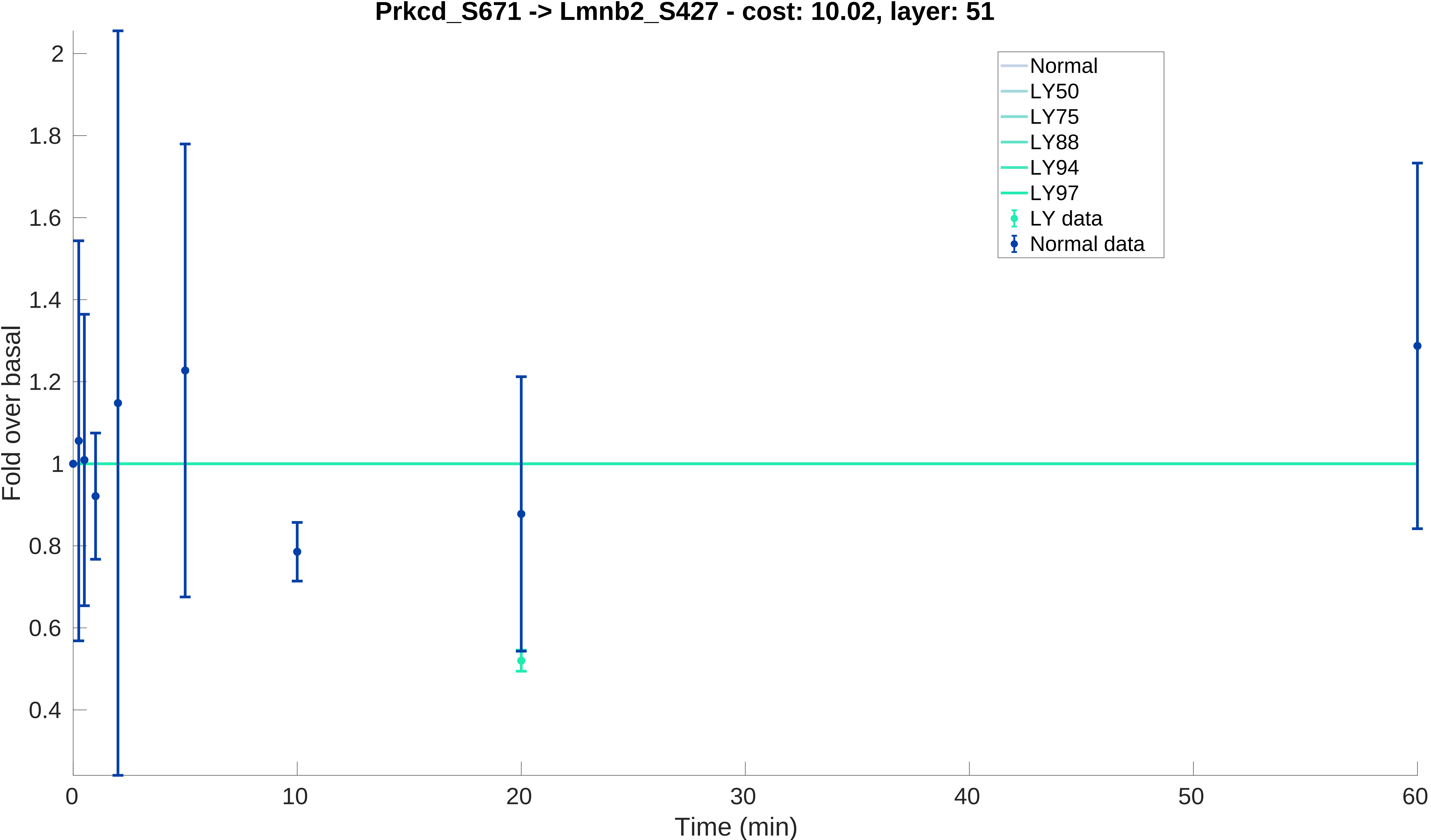

Prkcd_S671 -> Lmnb2_S427 - cost: 10.02, layer: 51
Normal
LY50
LY75
LY88
LY94
LY97
LY data
Normal data
Fold over basal
Time (min)
0
10
20
30
40
50
60
0.4
0.6
0.8
1
1.2
1.4
1.6
1.8
2

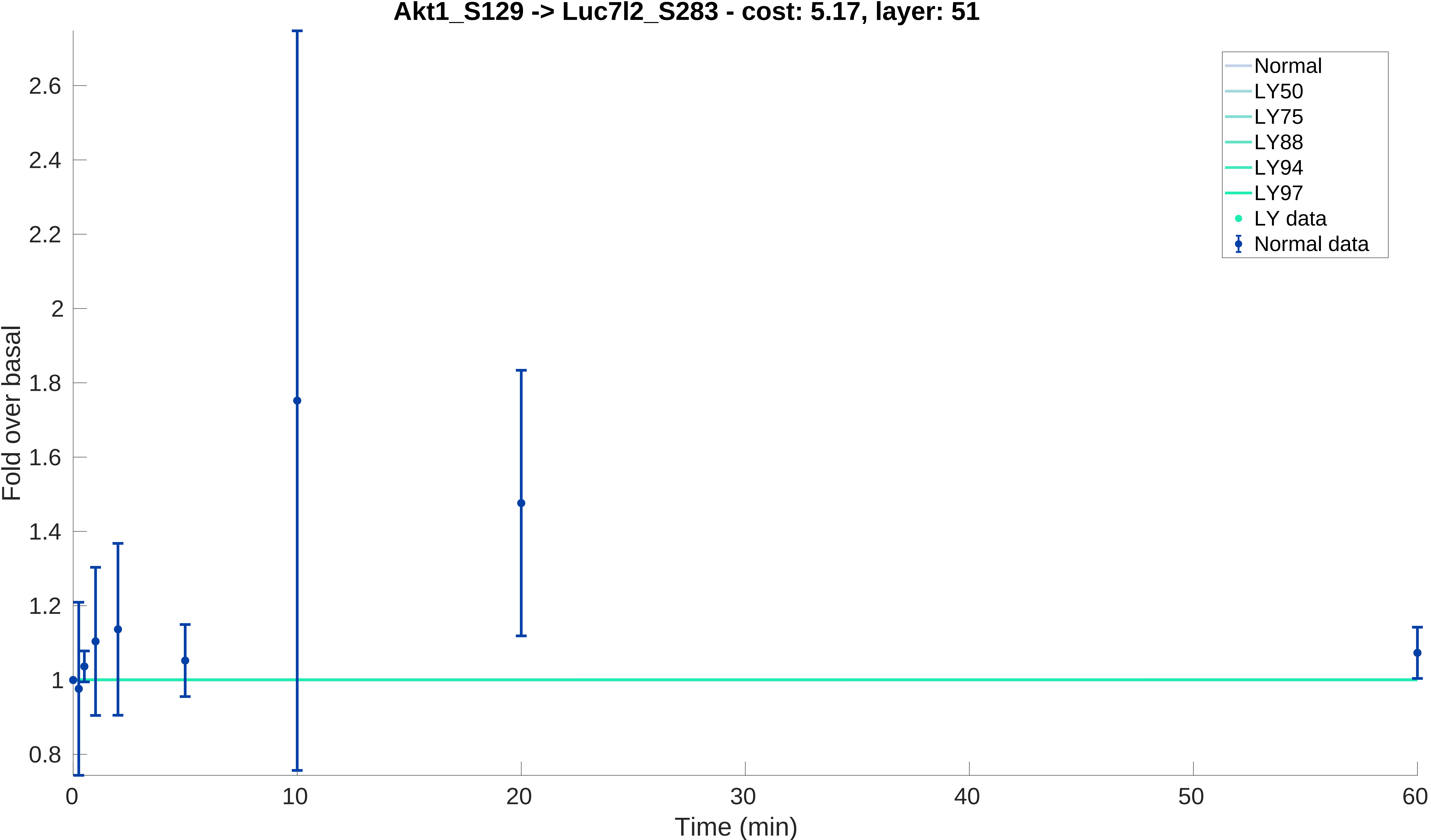

Akt1_S129 -> Luc7l2_S283 - cost: 5.17, layer: 51
Fold over basal
Time (min)
2.6
2.4
2.2
2
1.8
1.6
1.4
1.2
1
0.8
0
10
20
30
40
50
60
Normal
LY50
LY75
LY88
LY94
LY97
LY data
Normal data

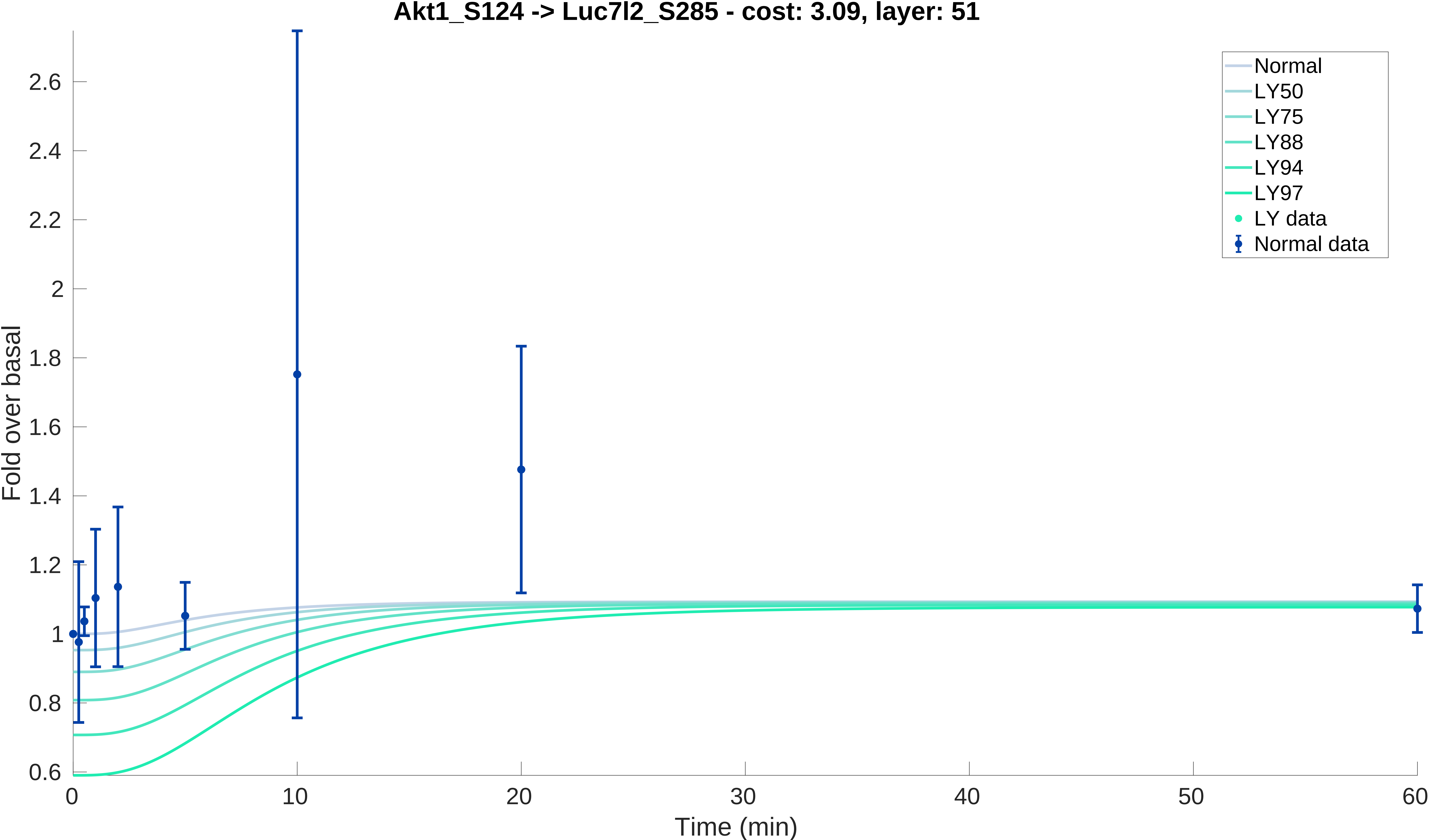

Akt1_S124 -> Luc7l2_S285 - cost: 3.09, layer: 51
Fold over basal
2.6
2.4
2.2
2
1.8
1.6
1.4
1.2
1
0.8
0.6
0
10
20
30
40
50
60
Time (min)
Normal
LY50
LY75
LY88
LY94
LY97
LY data
Normal data

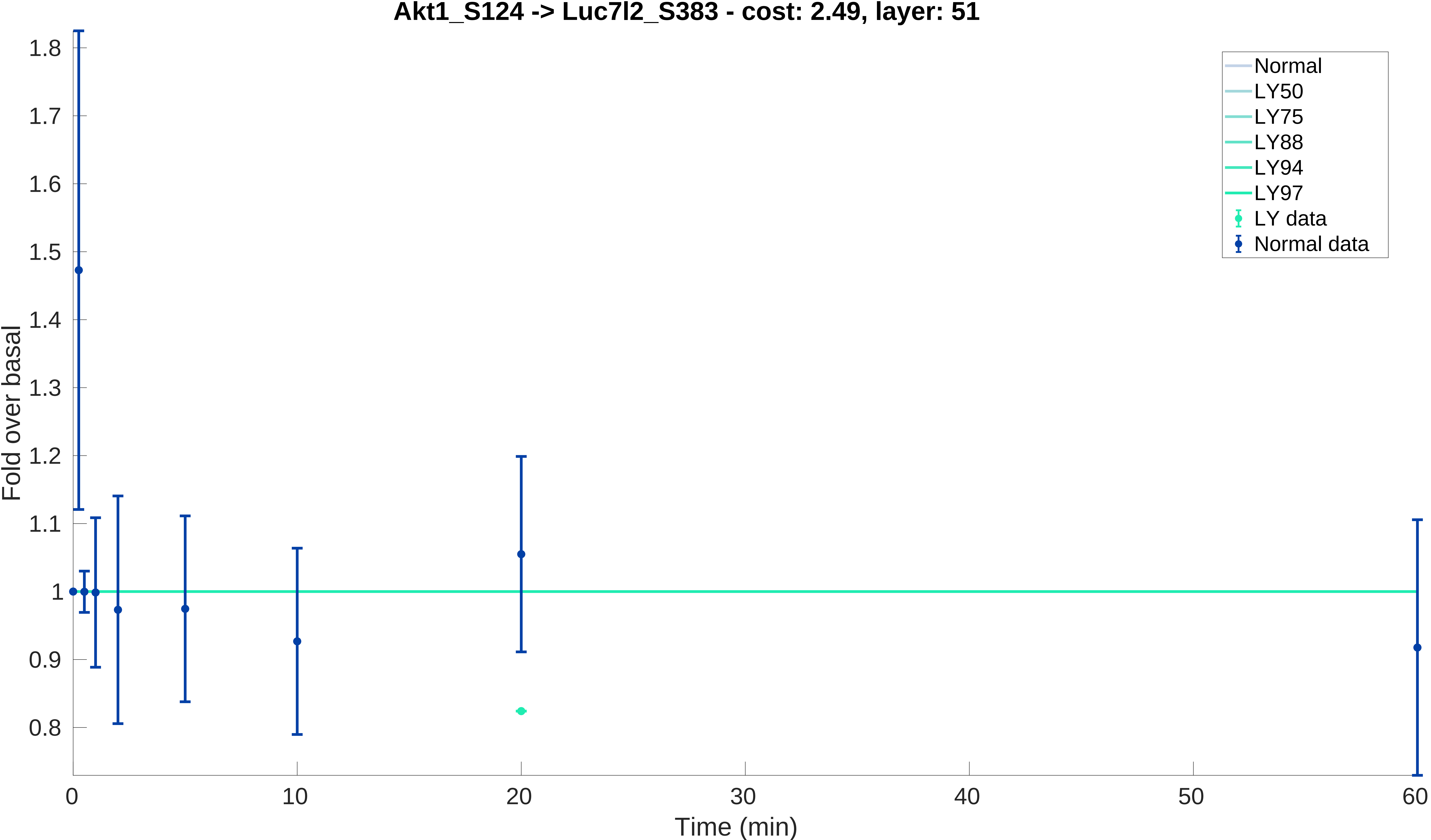
Akt1_S124 -> Luc7l2_S383 - cost: 2.49, layer: 51
Fold over basal
Time (min)
1.8
1.7
1.6
1.5
1.4
1.3
1.2
1.1
1
0.9
0.8
0
10
20
30
40
50
60
Normal
LY50
LY75
LY88
LY94
LY97
LY data
Normal data

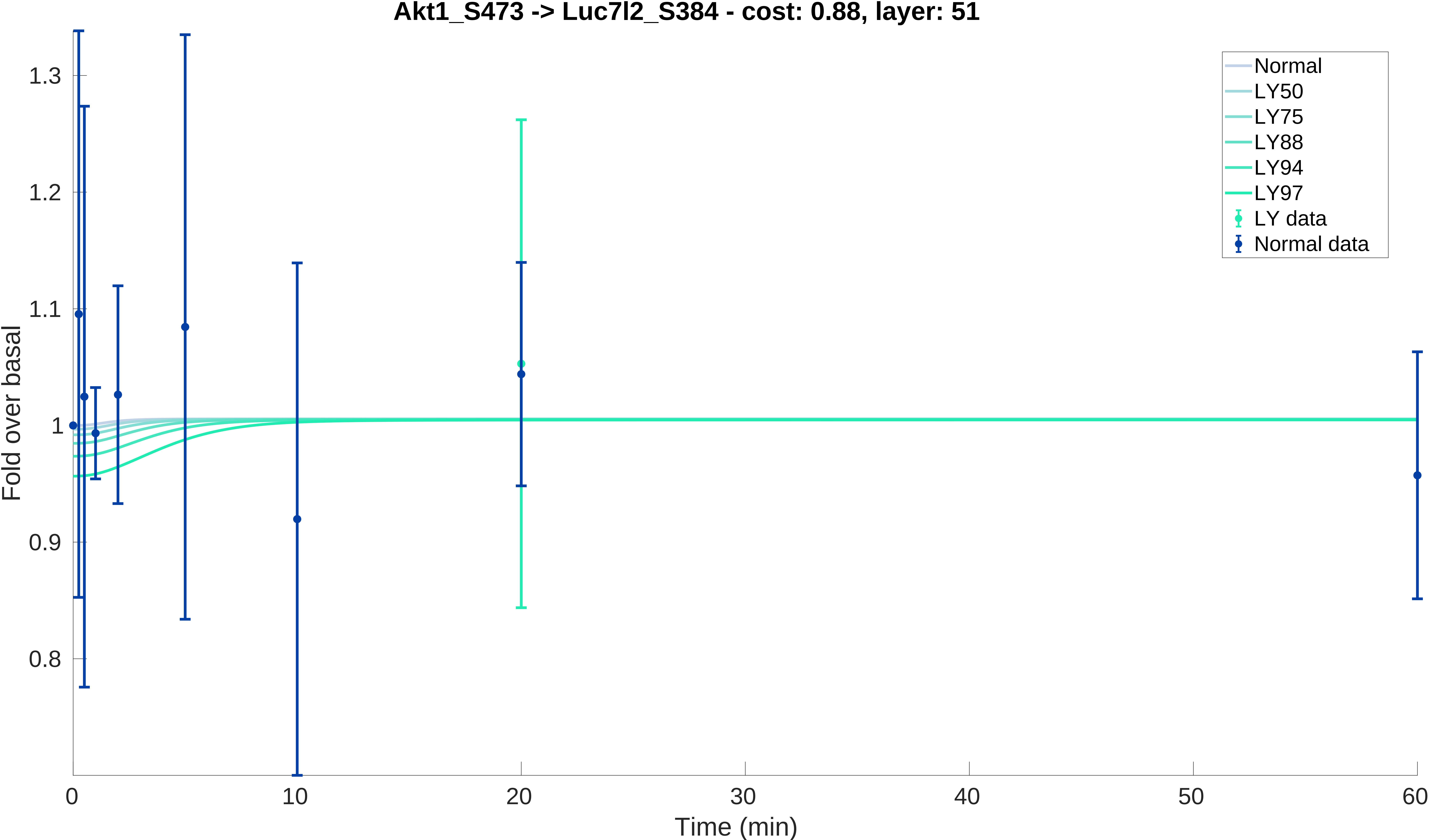

Akt1_S473 -> Luc7l2_S384 - cost: 0.88, layer: 51
Normal
LY50
LY75
LY88
LY94
LY97
LY data
Normal data
1.3
1.2
1.1
1
0.9
0.8
Fold over basal
0
10
20
30
40
50
60
Time (min)

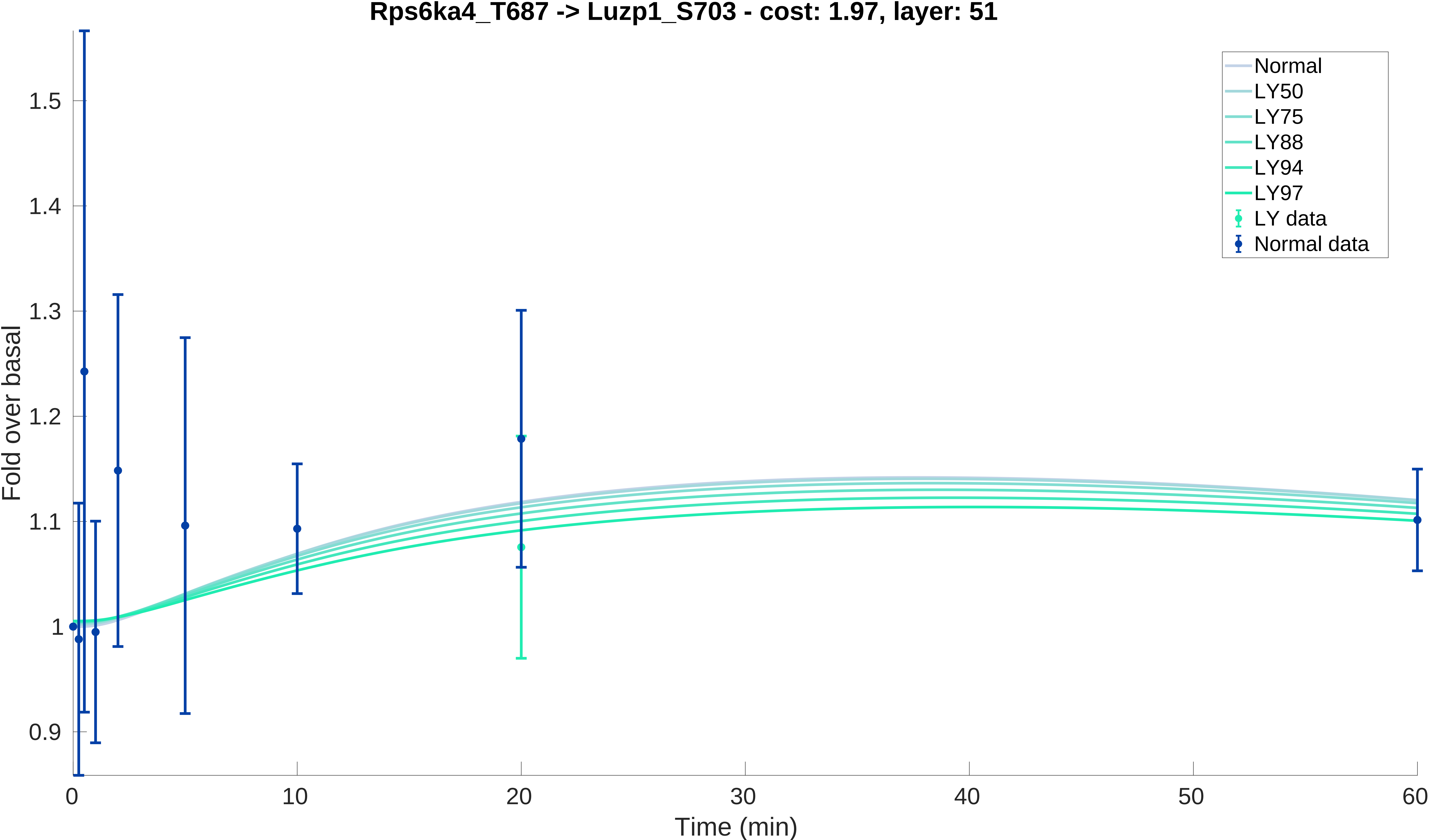

Rps6ka4_T687 -> Luzp1_S703 - cost: 1.97, layer: 51
Fold over basal
Time (min)
1.5
1.4
1.3
1.2
1.1
1
0.9
0
10
20
30
40
50
60
Normal
LY50
LY75
LY88
LY94
LY97
LY data
Normal data

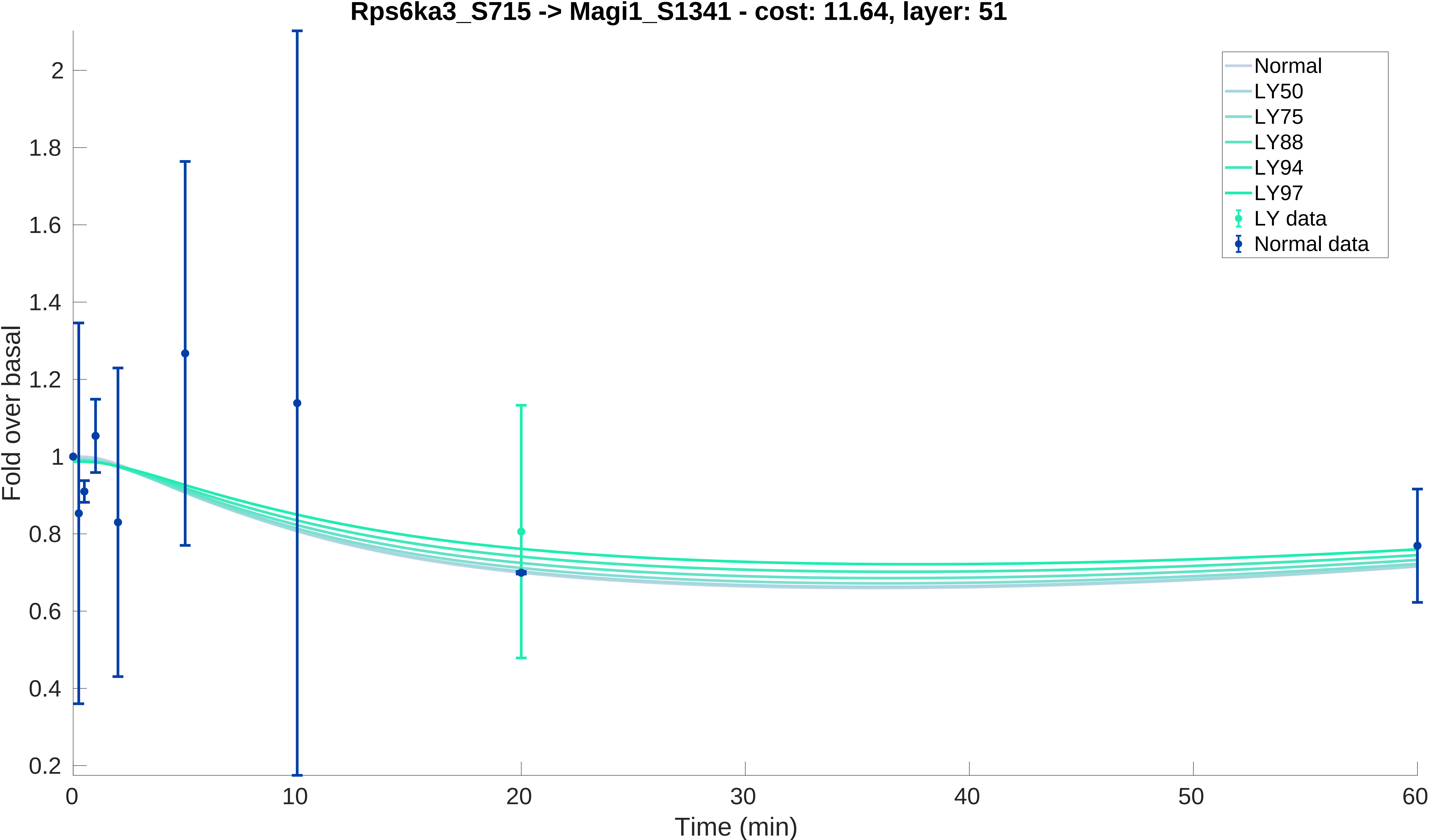
Rps6ka3_S715 -> Magi1_S1341 - cost: 11.64, layer: 51
Fold over basal
Time (min)
Normal
LY50
LY75
LY88
LY94
LY97
LY data
Normal data
0.2
0.4
0.6
0.8
1
1.2
1.4
1.6
1.8
2
0
10
20
30
40
50
60

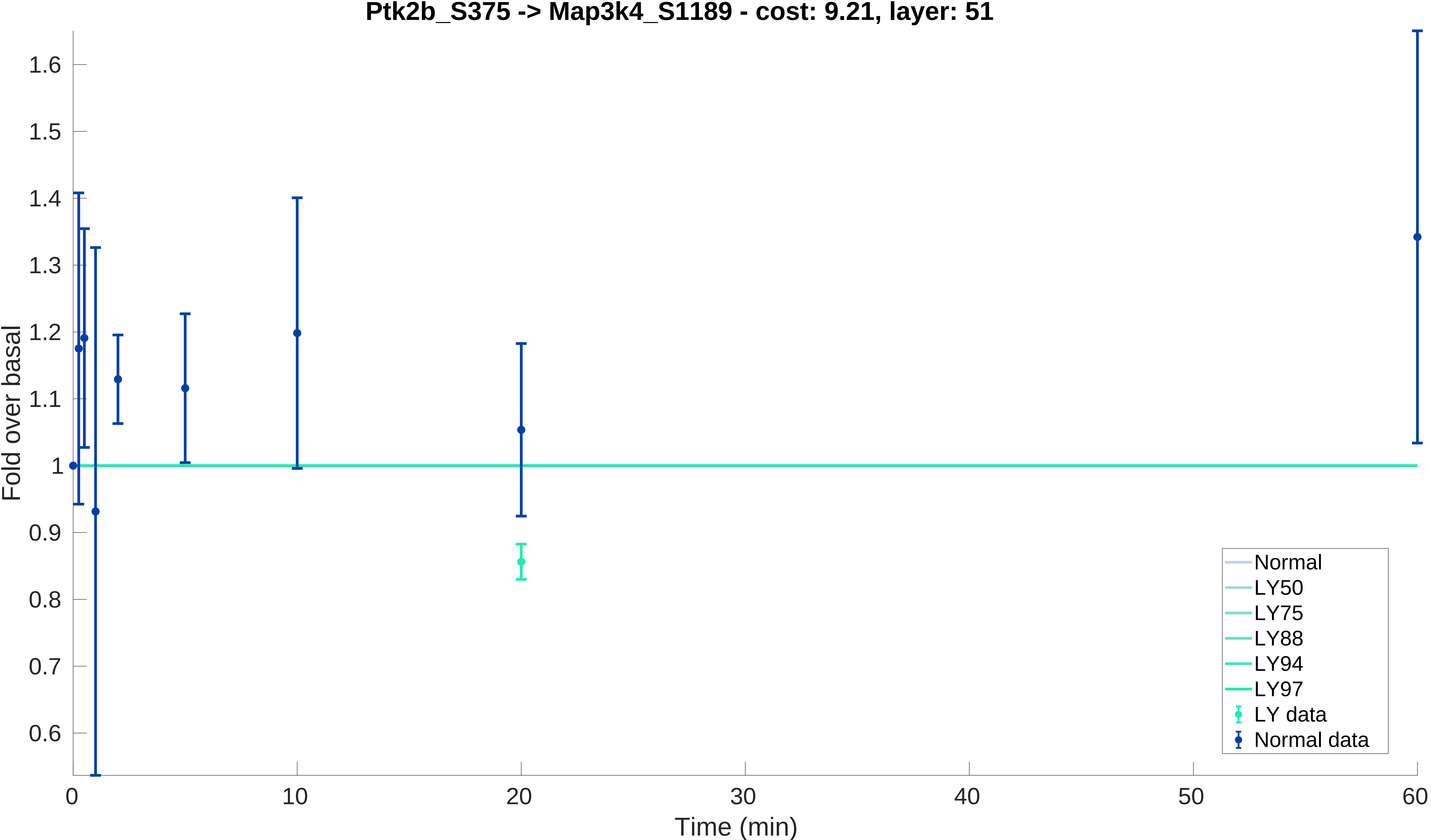

Ptk2b_S375 -> Map3k4_S1189 - cost: 9.21, layer: 51
Fold over basal
Time (min)
1.6
1.5
1.4
1.3
1.2
1.1
1
0.9
0.8
0.7
0.6
0
10
20
30
40
50
60
Normal
LY50
LY75
LY88
LY94
LY97
LY data
Normal data

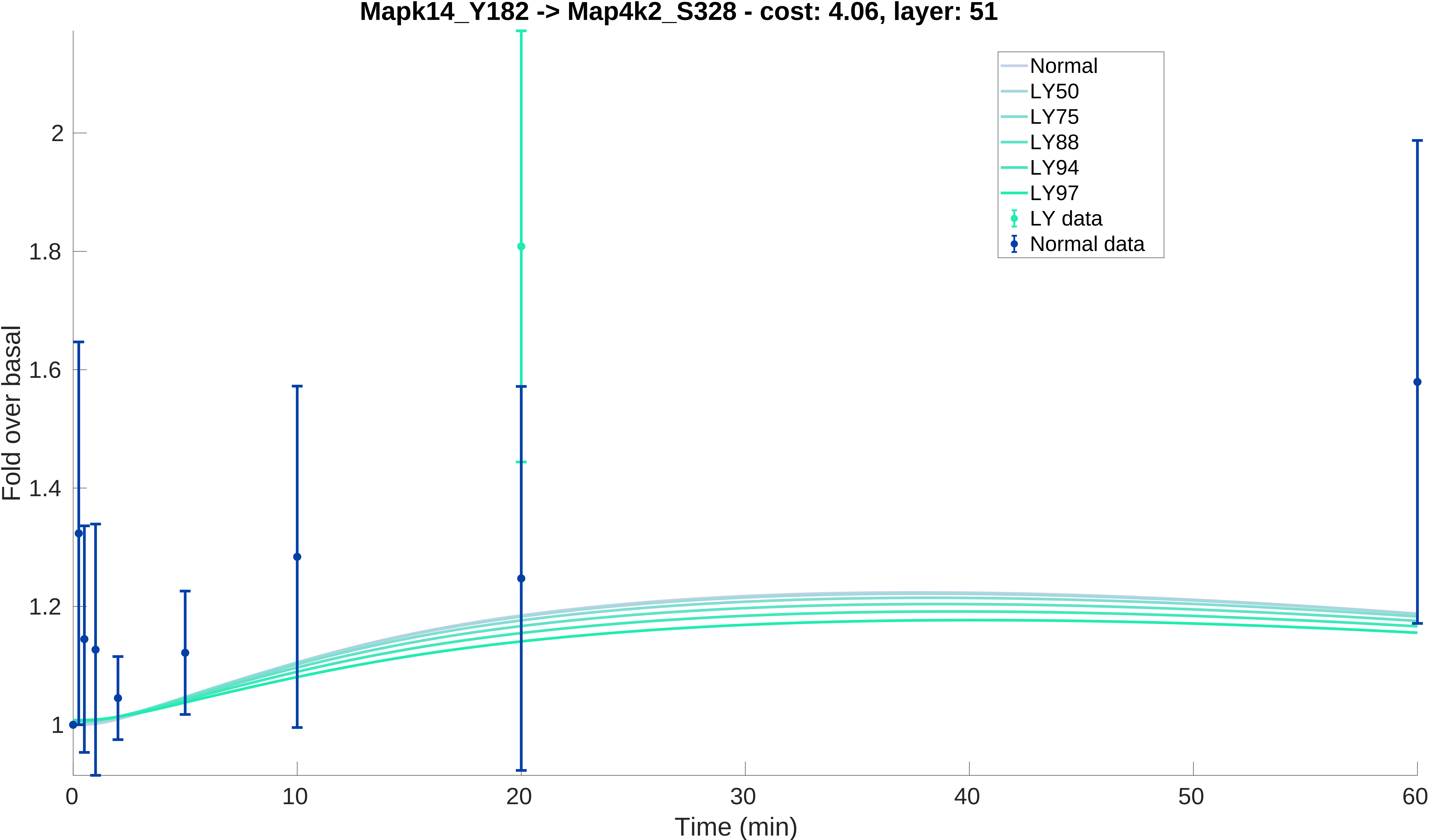

Mapk14_Y182 -> Map4k2_S328 - cost: 4.06, layer: 51
Fold over basal
Time (min)
0
10
20
30
40
50
60
1
1.2
1.4
1.6
1.8
2
Normal
LY50
LY75
LY88
LY94
LY97
LY data
Normal data

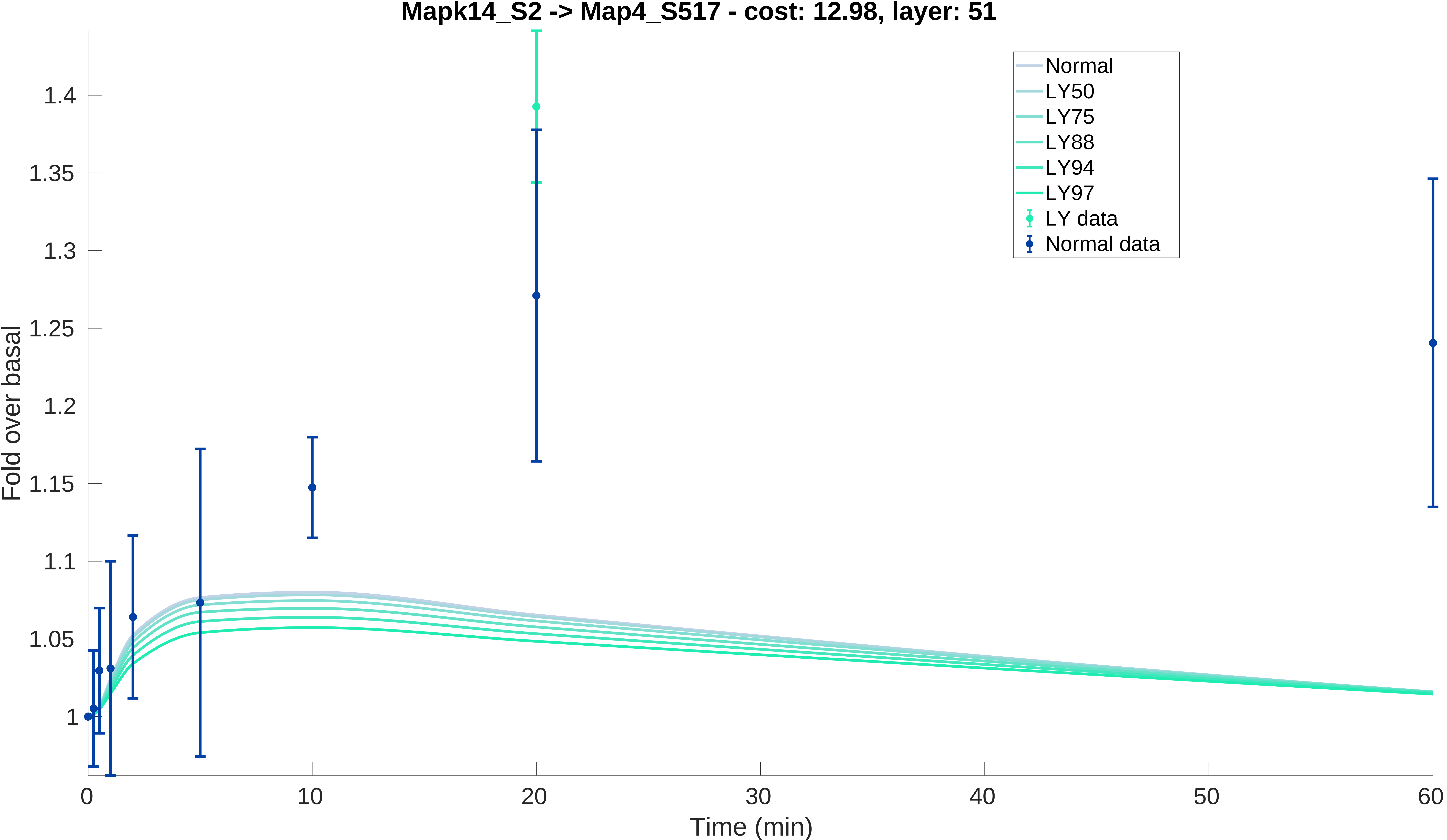
Mapk14_S2 -> Map4_S517 - cost: 12.98, layer: 51
Fold over basal
Time (min)
Normal
LY50
LY75
LY88
LY94
LY97
LY data
Normal data
1
1.05
1.1
1.15
1.2
1.25
1.3
1.35
1.4
0
10
20
30
40
50
60

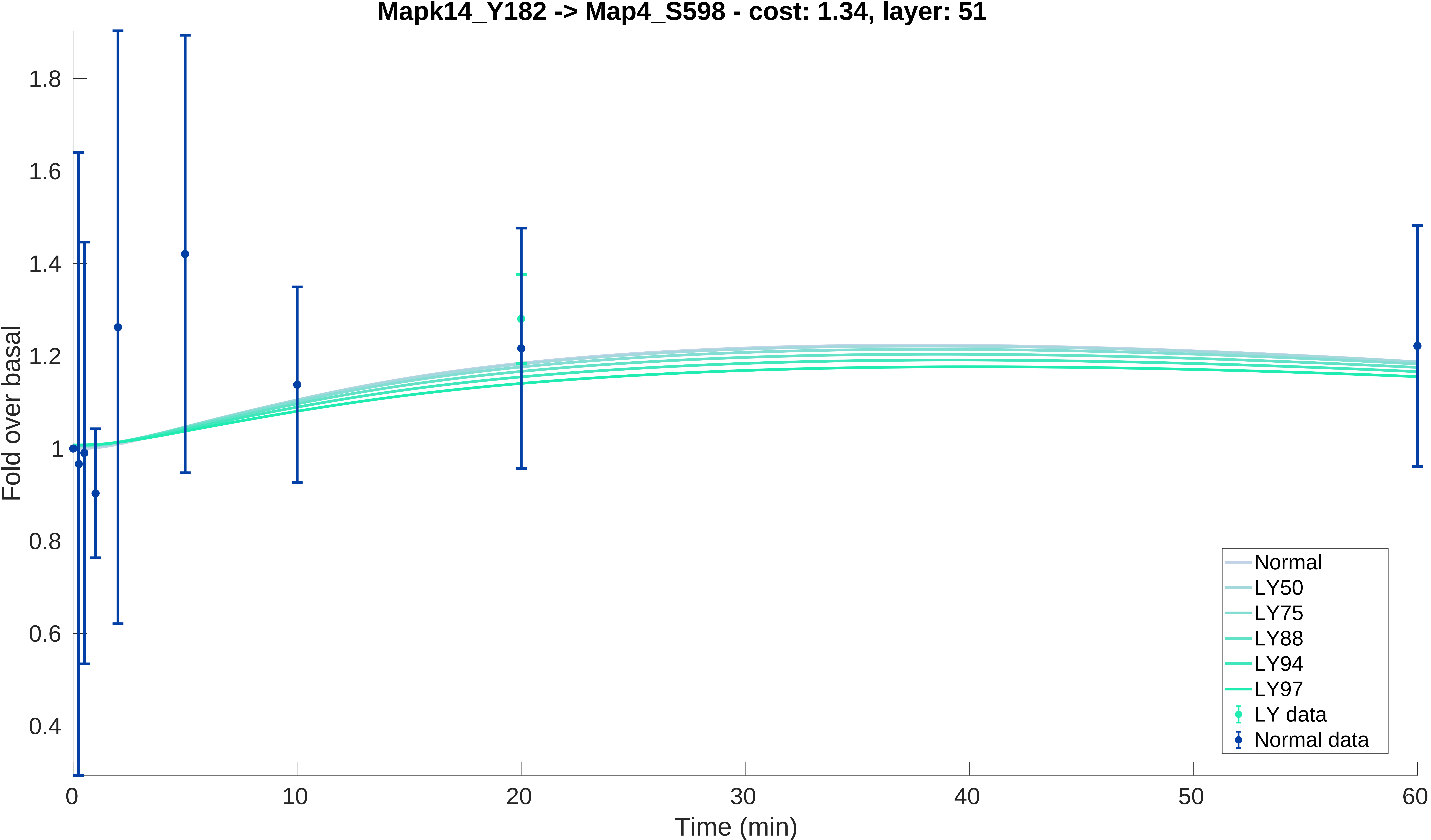

Mapk14_Y182 -> Map4_S598 - cost: 1.34, layer: 51
Fold over basal
1.8
1.6
1.4
1.2
1
0.8
0.6
0.4
0
10
20
30
40
50
60
Time (min)
Normal
LY50
LY75
LY88
LY94
LY97
LY data
Normal data

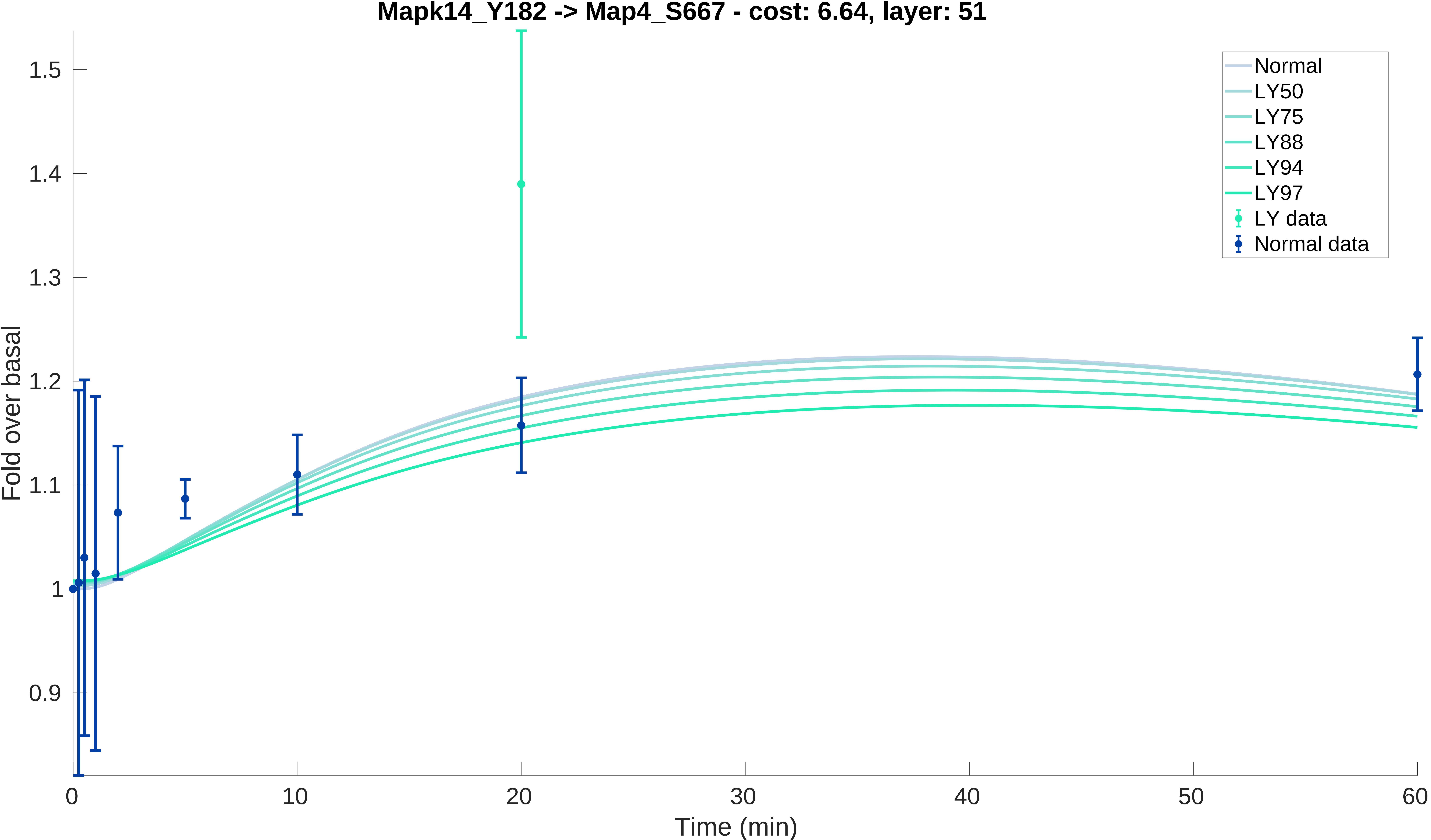

Mapk14_Y182 -> Map4_S667 - cost: 6.64, layer: 51
Fold over basal
1.5
1.4
1.3
1.2
1.1
1
0.9
0
10
20
30
40
50
60
Time (min)
Normal
LY50
LY75
LY88
LY94
LY97
LY data
Normal data

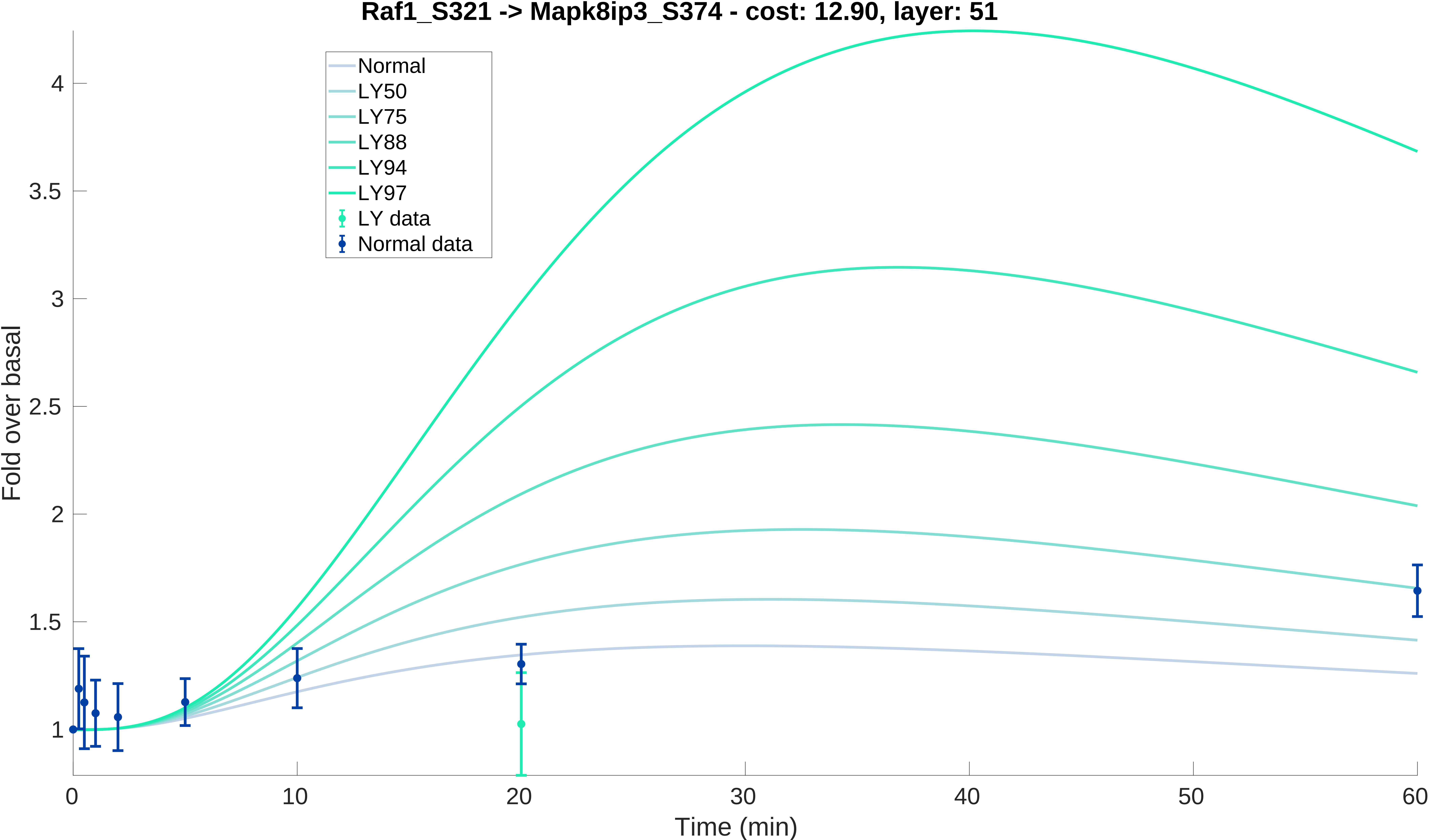

Raf1_S321 -> Mapk8ip3_S374 - cost: 12.90, layer: 51
Normal
LY50
LY75
LY88
LY94
LY97
LY data
Normal data
Fold over basal
Time (min)
1
1.5
2
2.5
3
3.5
4
0
10
20
30
40
50
60

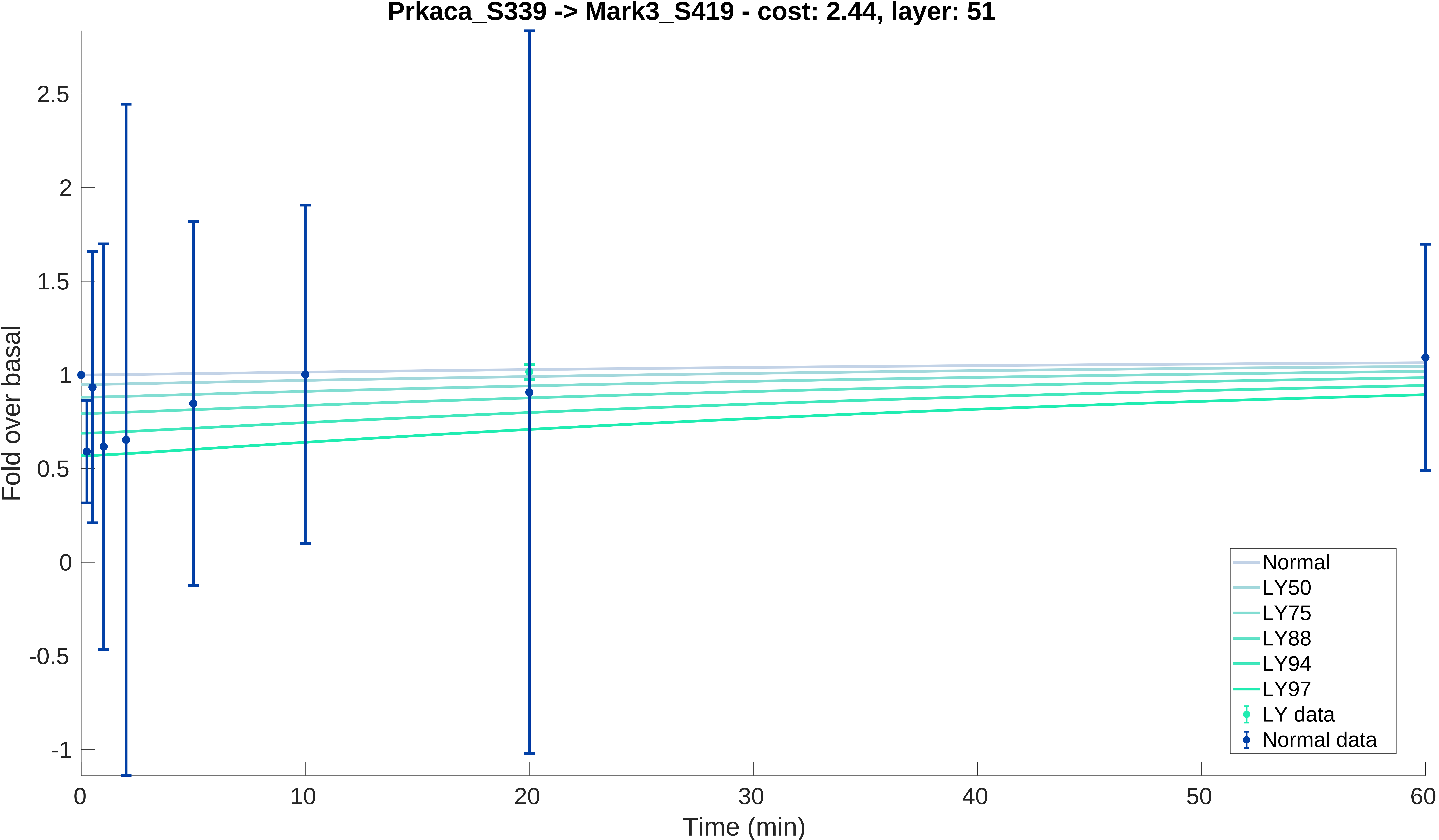

Prkaca_S339 -> Mark3_S419 - cost: 2.44, layer: 51
Fold over basal
Time (min)
2.5
2
1.5
1
0.5
0
-0.5
-1
0
10
20
30
40
50
60
Normal
LY50
LY75
LY88
LY94
LY97
LY data
Normal data

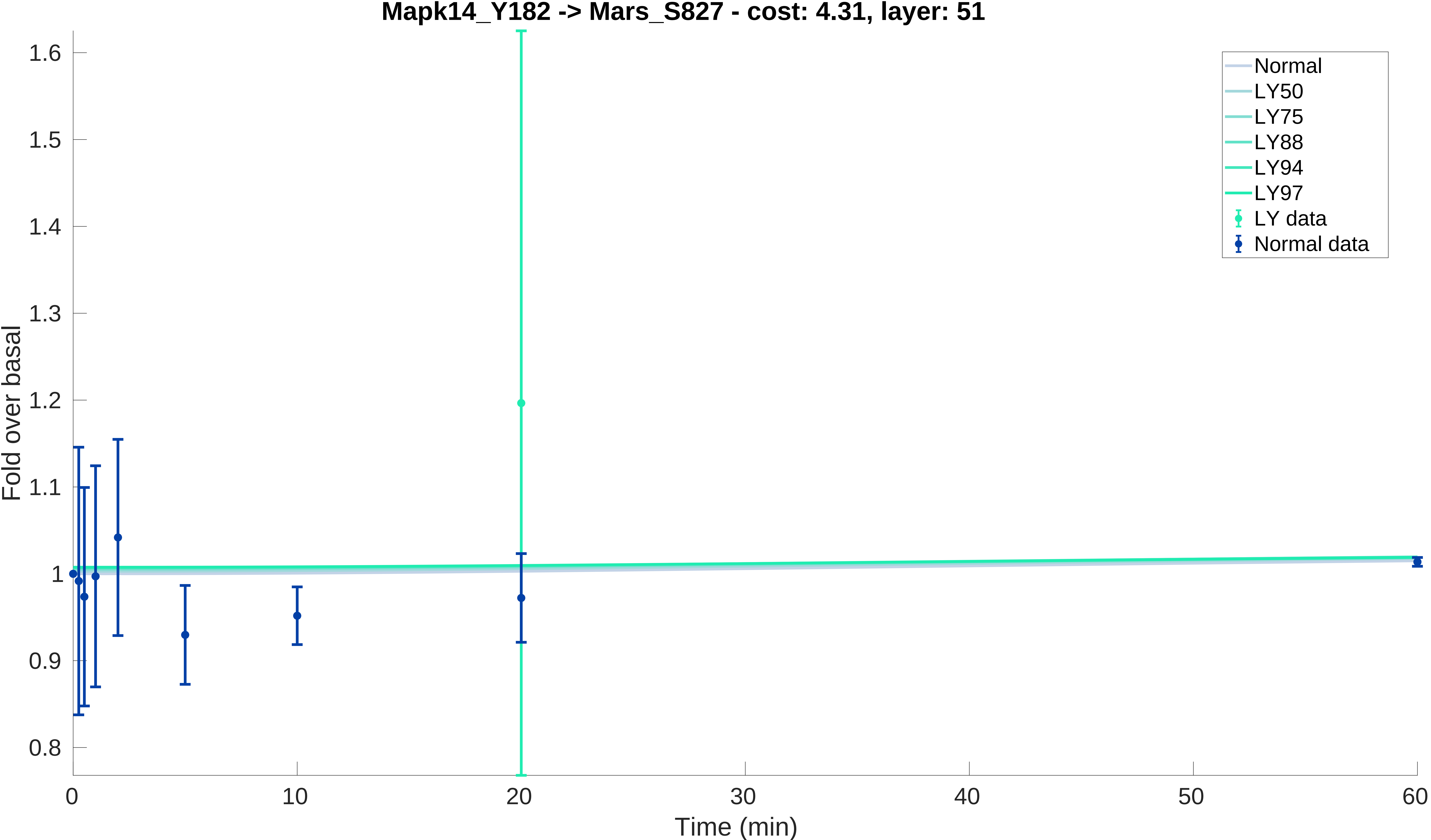

Mapk14_Y182 -> Mars_S827 - cost: 4.31, layer: 51
Fold over basal
1.6
1.5
1.4
1.3
1.2
1.1
1
0.9
0.8
0
10
20
30
40
50
60
Time (min)
Normal
LY50
LY75
LY88
LY94
LY97
LY data
Normal data

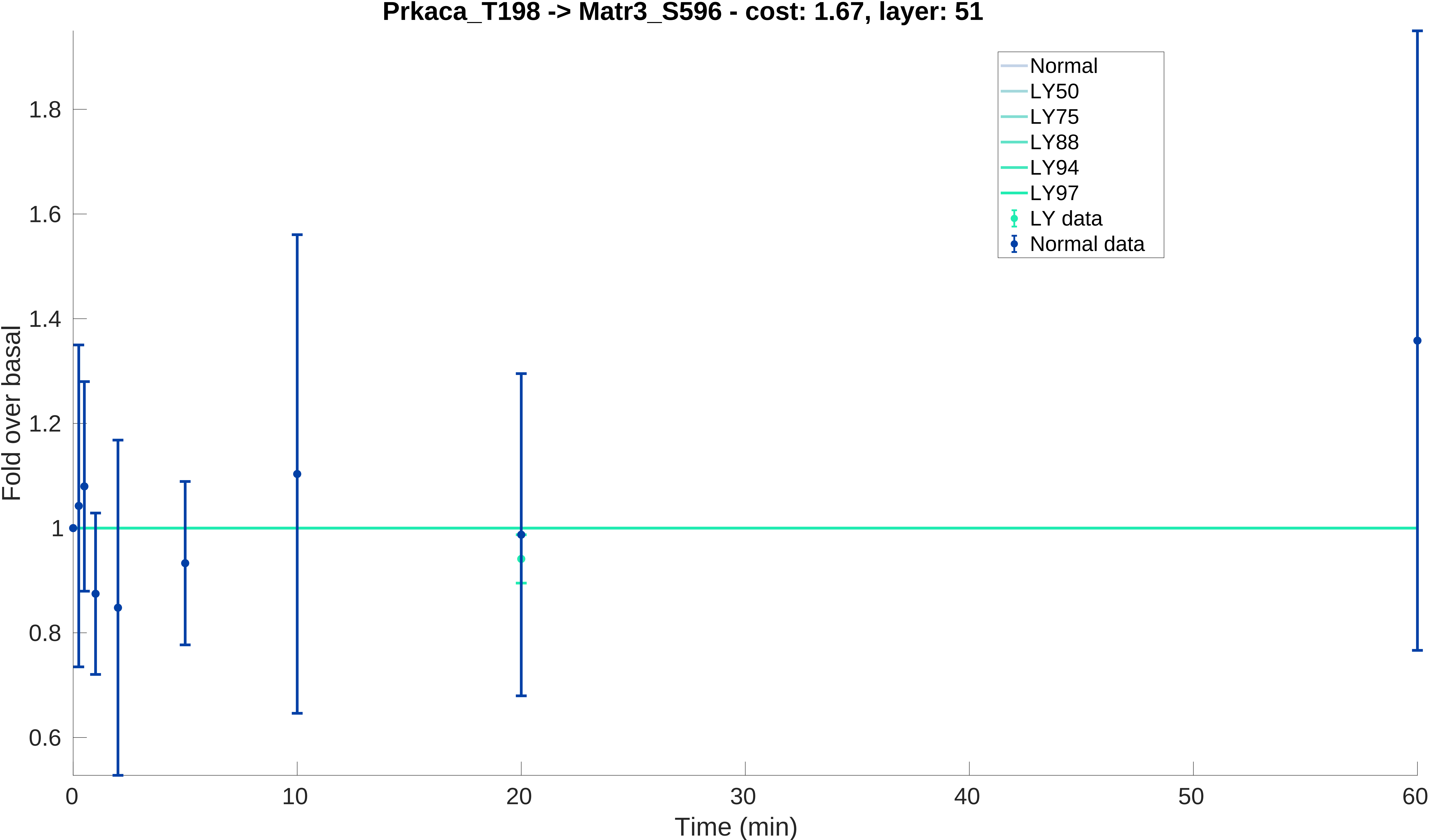

Prrkaca_T198 -> Matr3_S596 - cost: 1.67, layer: 51
Normal
LY50
LY75
LY88
LY94
LY97
LY data
Normal data
Fold over basal
1.8
1.6
1.4
1.2
1
0.8
0.6
0
10
20
30
40
50
60
Time (min)

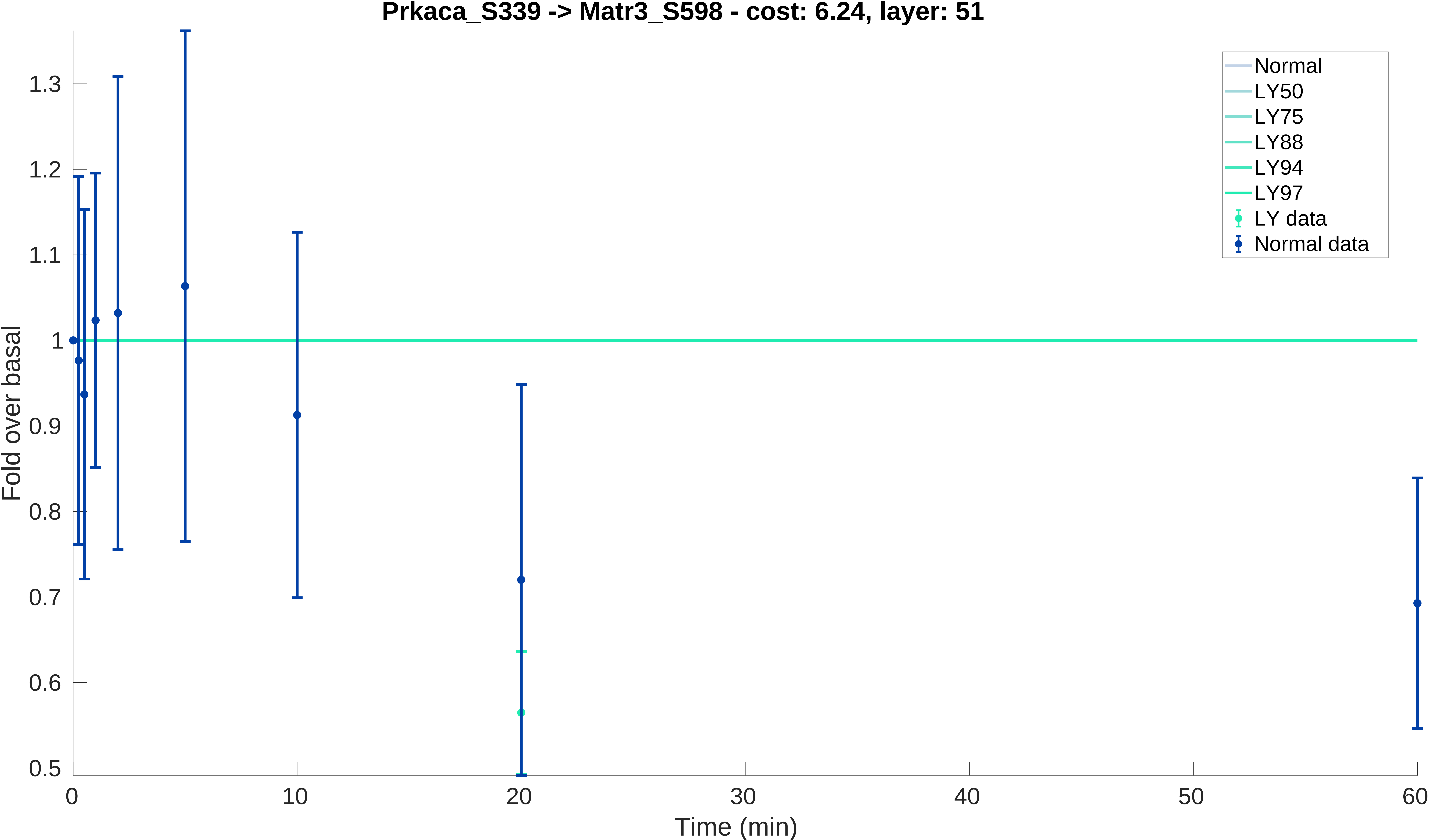

Prkaca_S339 -> Matr3_S598 - cost: 6.24, layer: 51
Normal
LY50
LY75
LY88
LY94
LY97
LY data
Normal data
Fold over basal
1.3
1.2
1.1
1
0.9
0.8
0.7
0.6
0.5
0
10
20
30
40
50
60
Time (min)

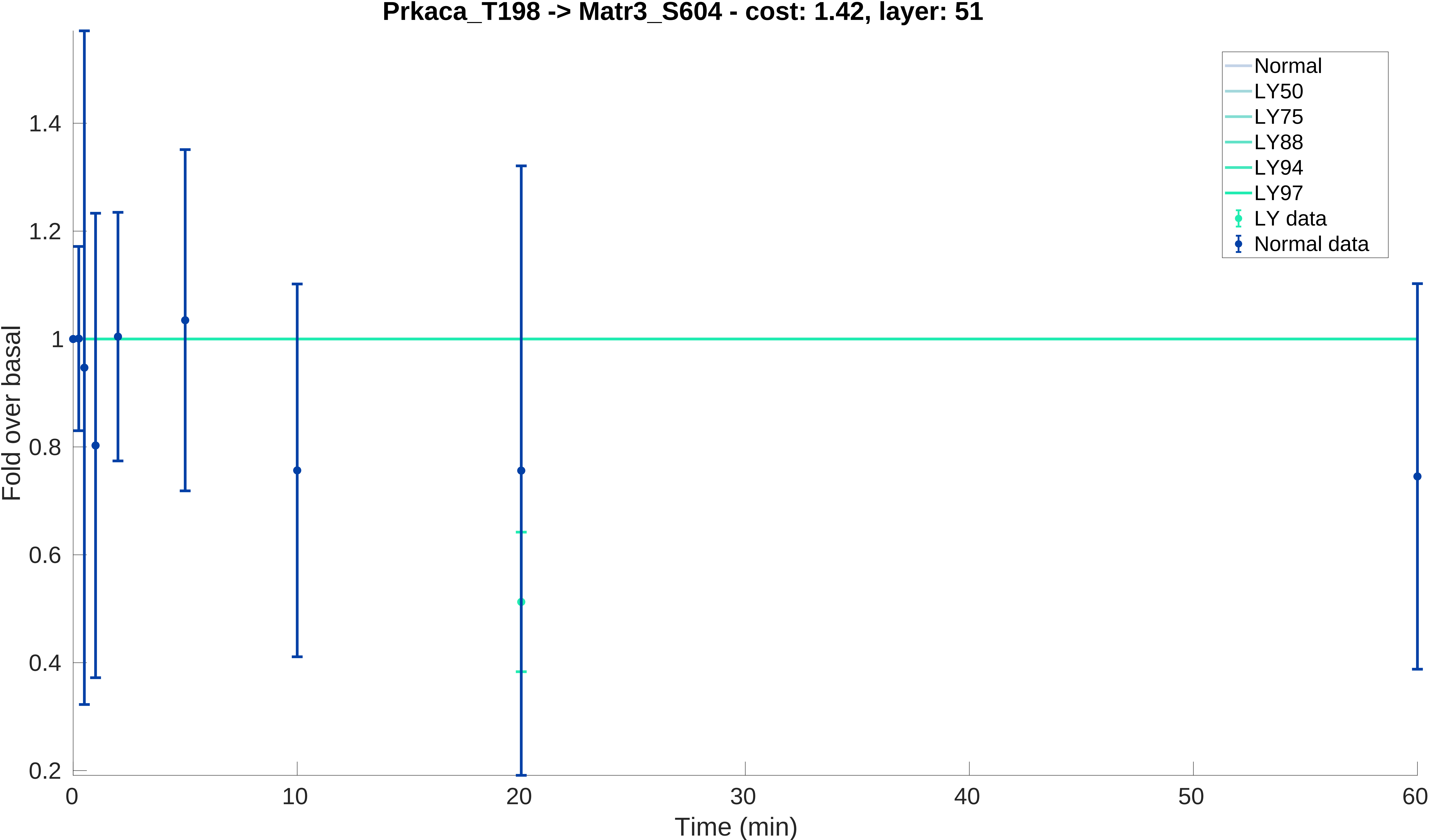

Prkaca_T198 -> Matr3_S604 - cost: 1.42, layer: 51
Normal
LY50
LY75
LY88
LY94
LY97
LY data
Normal data
Fold over basal
1.4
1.2
1
0.8
0.6
0.4
0.2
0
10
20
30
40
50
60
Time (min)

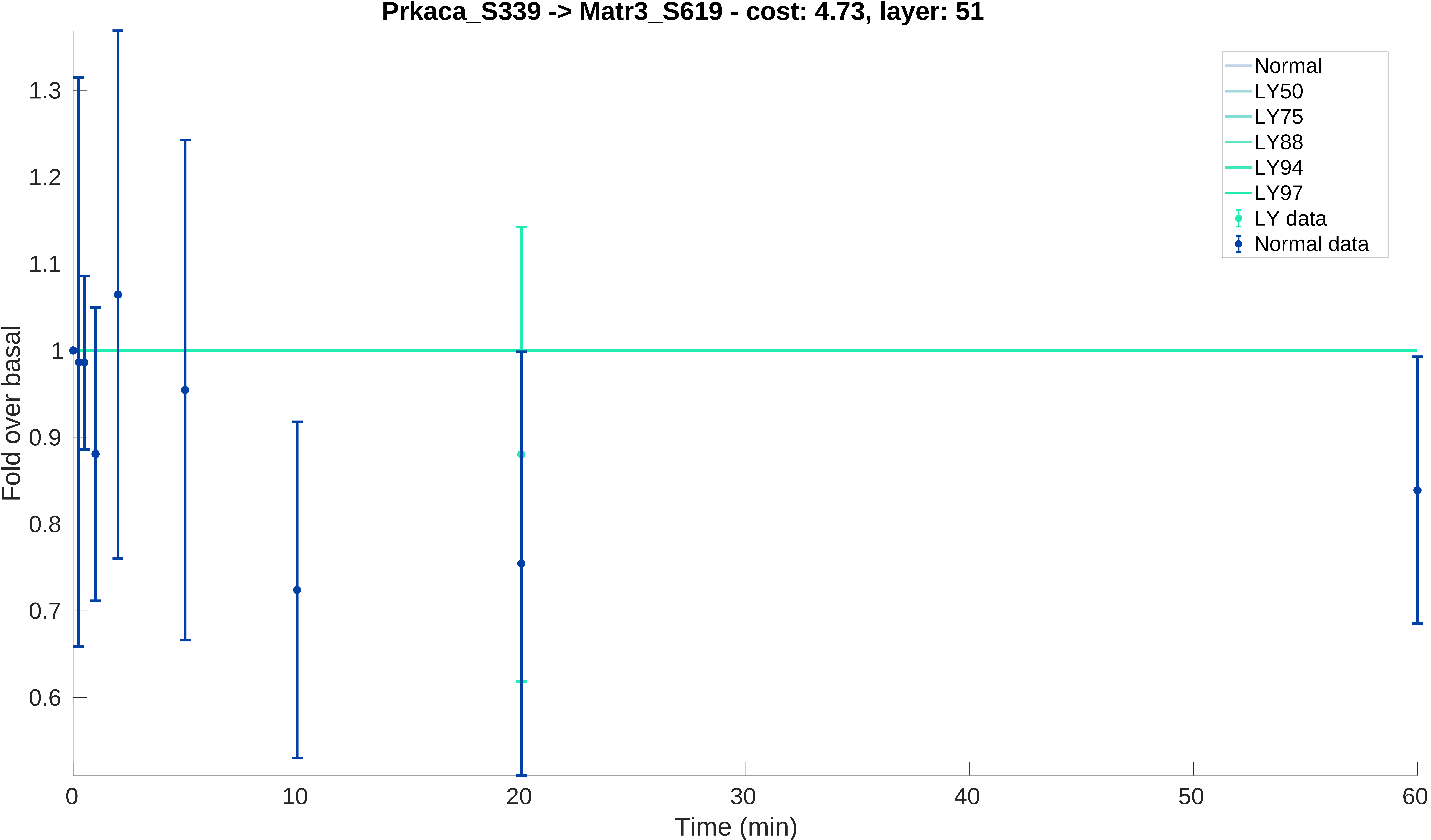
Prkaca_S339 -> Matr3_S619 - cost: 4.73, layer: 51
Fold over basal
Time (min)
Normal
LY50
LY75
LY88
LY94
LY97
LY data
Normal data
1.3
1.2
1.1
1
0.9
0.8
0.7
0.6
0
10
20
30
40
50
60

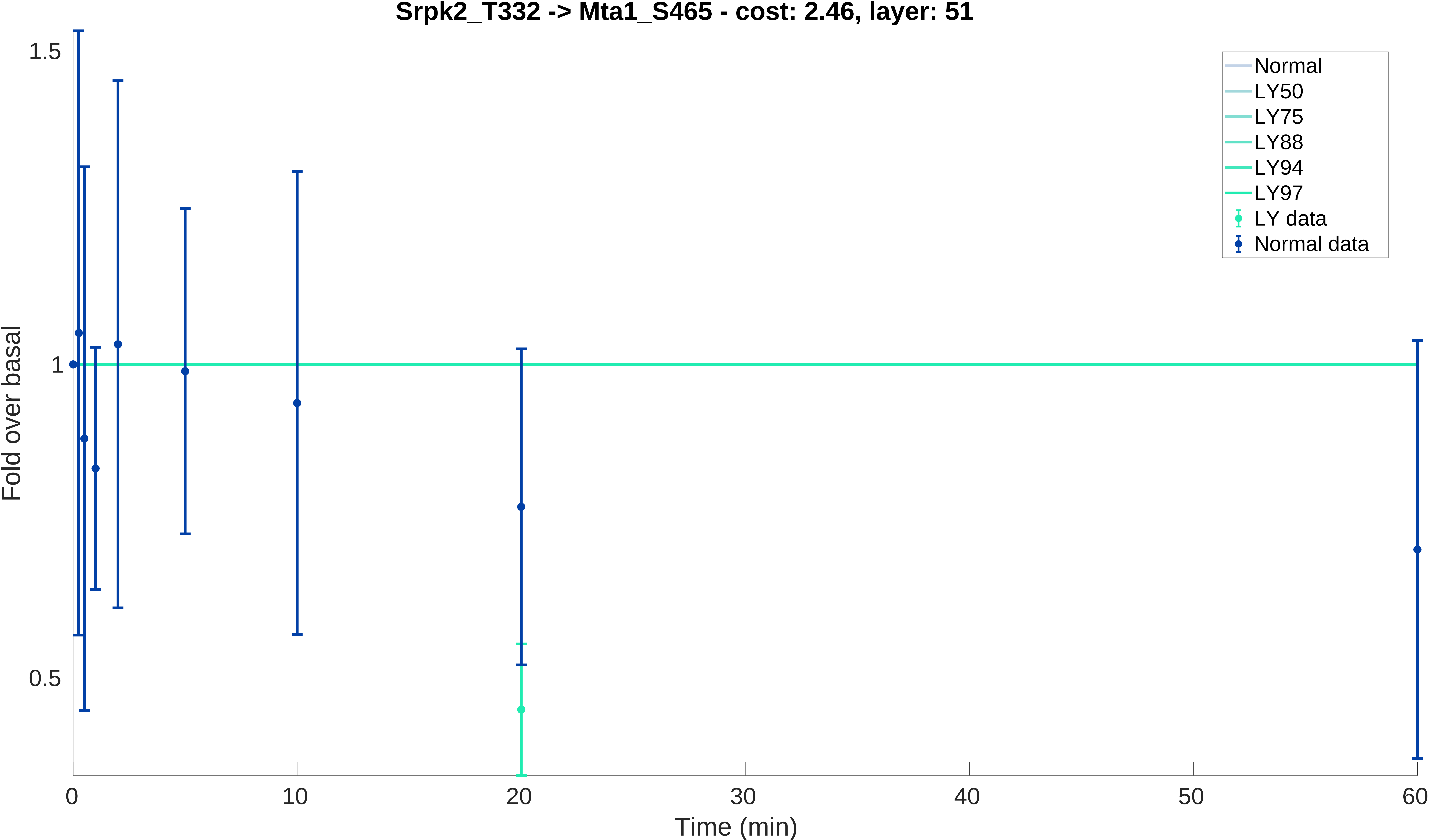

Srpk2_T332 -> Mta1_S465 - cost: 2.46, layer: 51
Fold over basal
1.5
1
0.5
0
10
20
30
40
50
60
Time (min)
Normal
LY50
LY75
LY88
LY94
LY97
LY data
Normal data

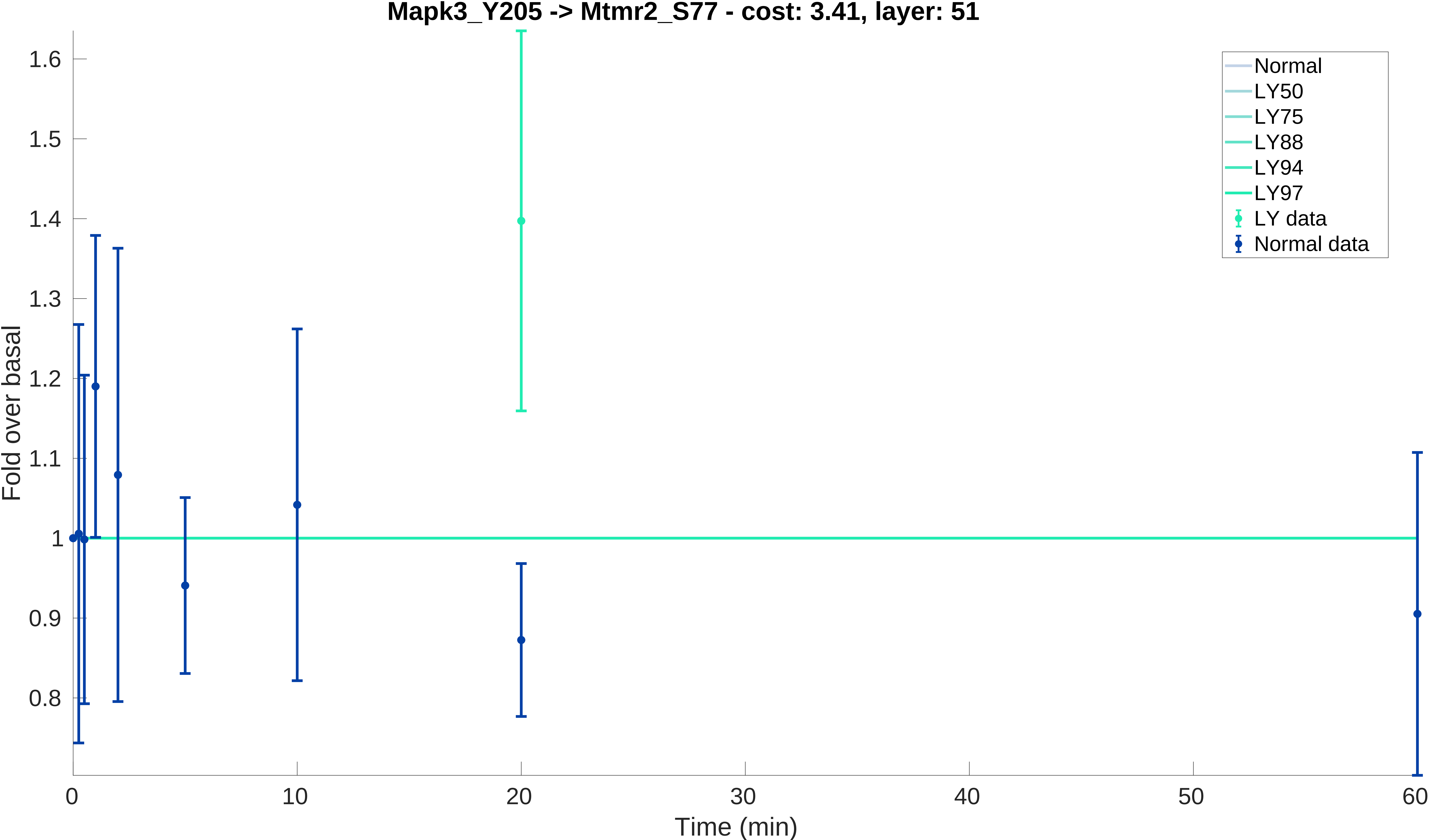
Mapk3_Y205 -> Mtmr2_S77 - cost: 3.41, layer: 51
Fold over basal
Time (min)
1.6
1.5
1.4
1.3
1.2
1.1
1
0.9
0.8
0
10
20
30
40
50
60
Normal
LY50
LY75
LY88
LY94
LY97
LY data
Normal data

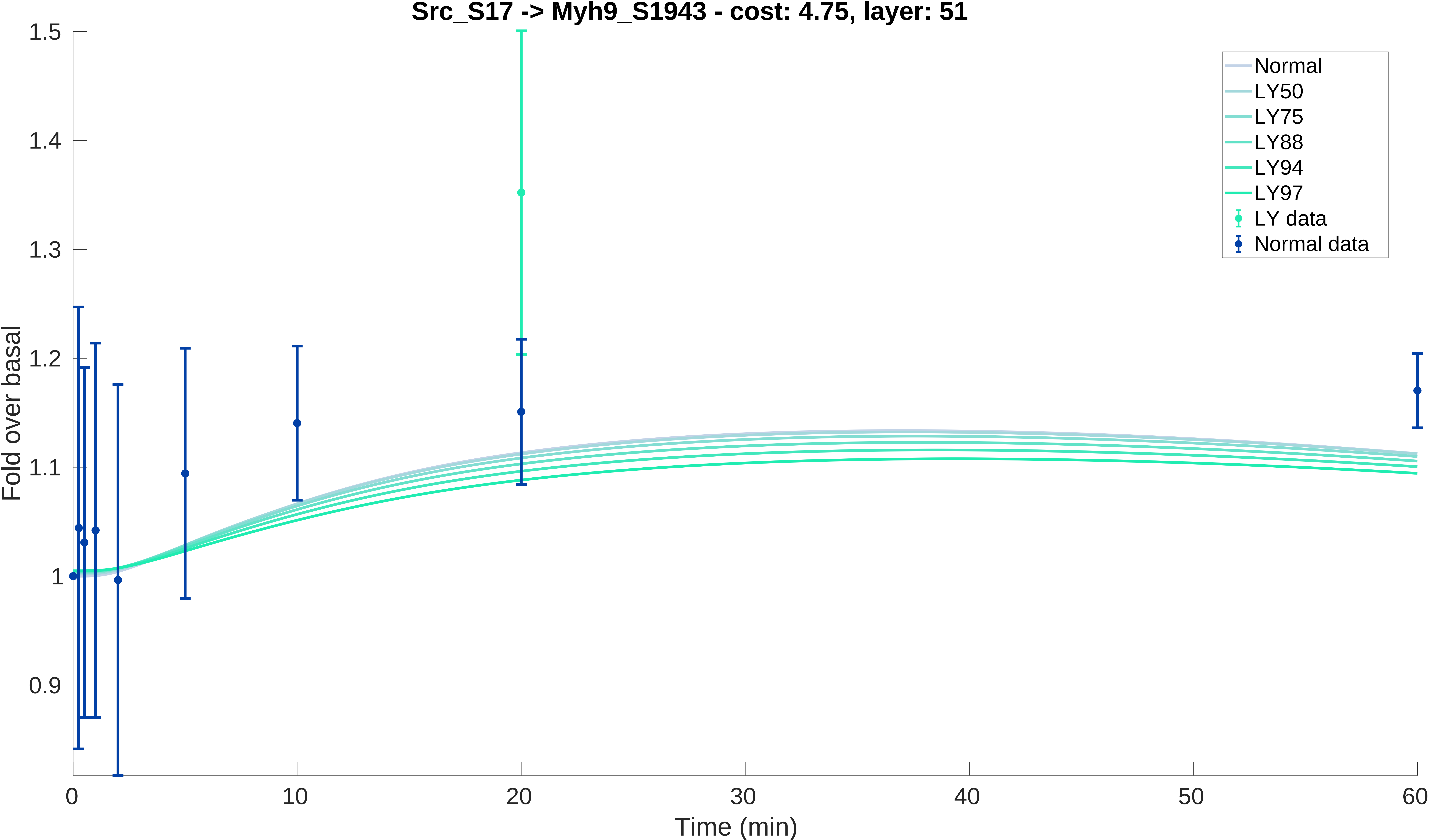
Src_S17 -> Myh9_S1943 - cost: 4.75, layer: 51
Fold over basal
Time (min)
1.5
1.4
1.3
1.2
1.1
1
0.9
0
10
20
30
40
50
60
Normal
LY50
LY75
LY88
LY94
LY97
LY data
Normal data

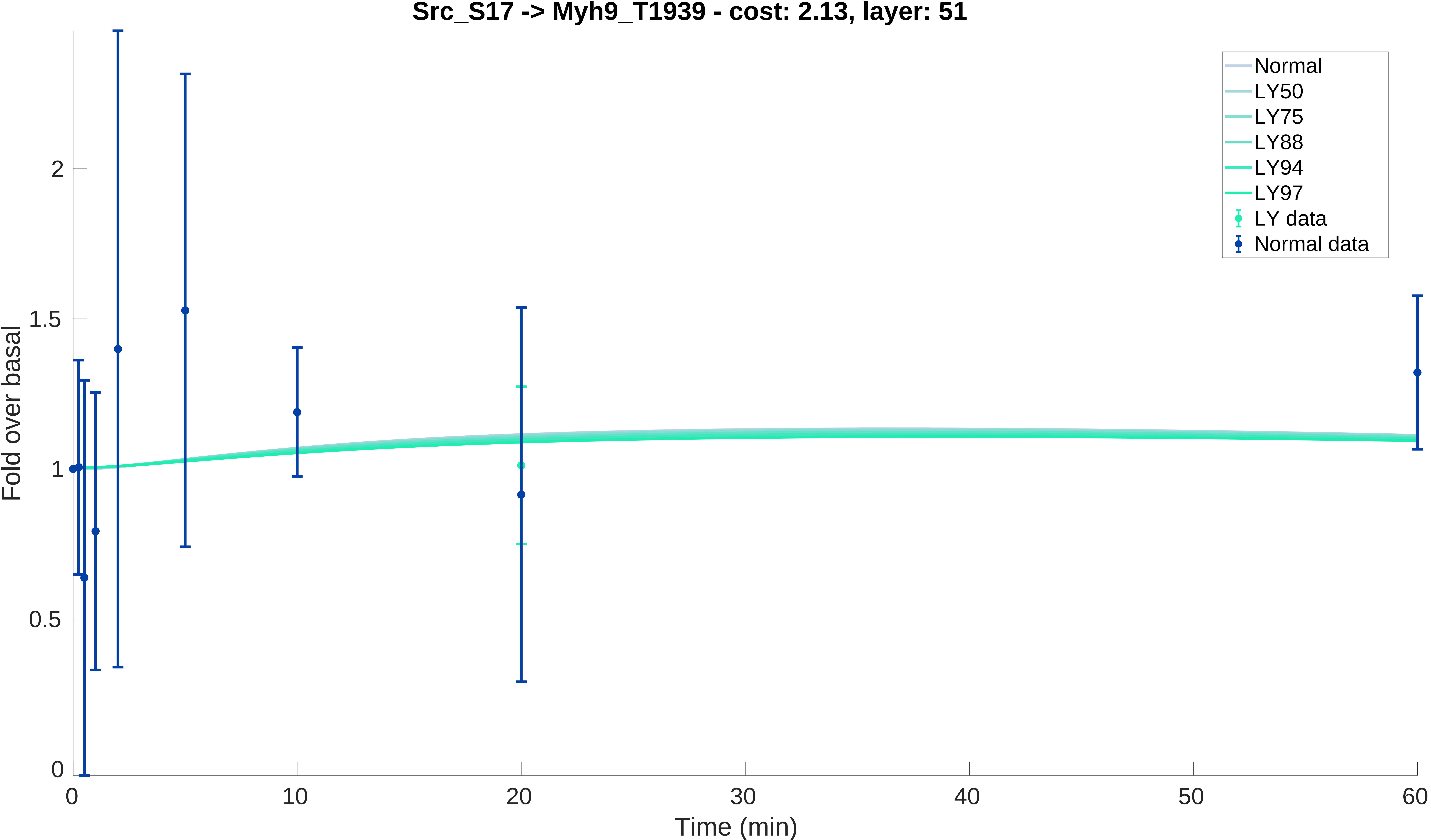

Src_S17 -> Myh9_T1939 - cost: 2.13, layer: 51
Normal
LY50
LY75
LY88
LY94
LY97
LY data
Normal data
Fold over basal
0
0.5
1
1.5
2
0
10
20
30
40
50
60
Time (min)

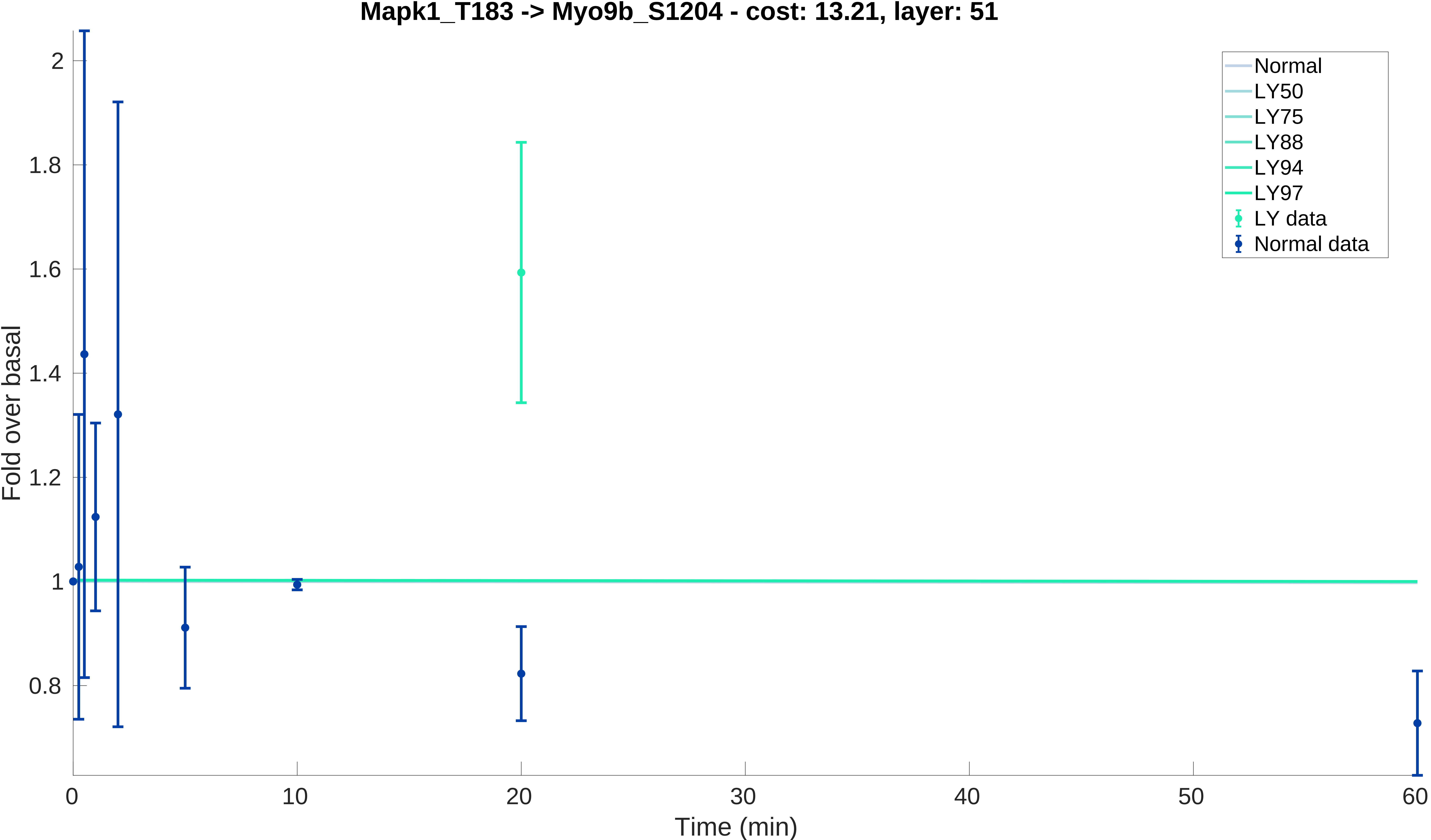

Mapk1_T183 -> Myo9b_S1204 - cost: 13.21, layer: 51
Fold over basal
2
1.8
1.6
1.4
1.2
1
0.8
0
10
20
30
40
50
60
Time (min)
Normal
LY50
LY75
LY88
LY94
LY97
LY data
Normal data

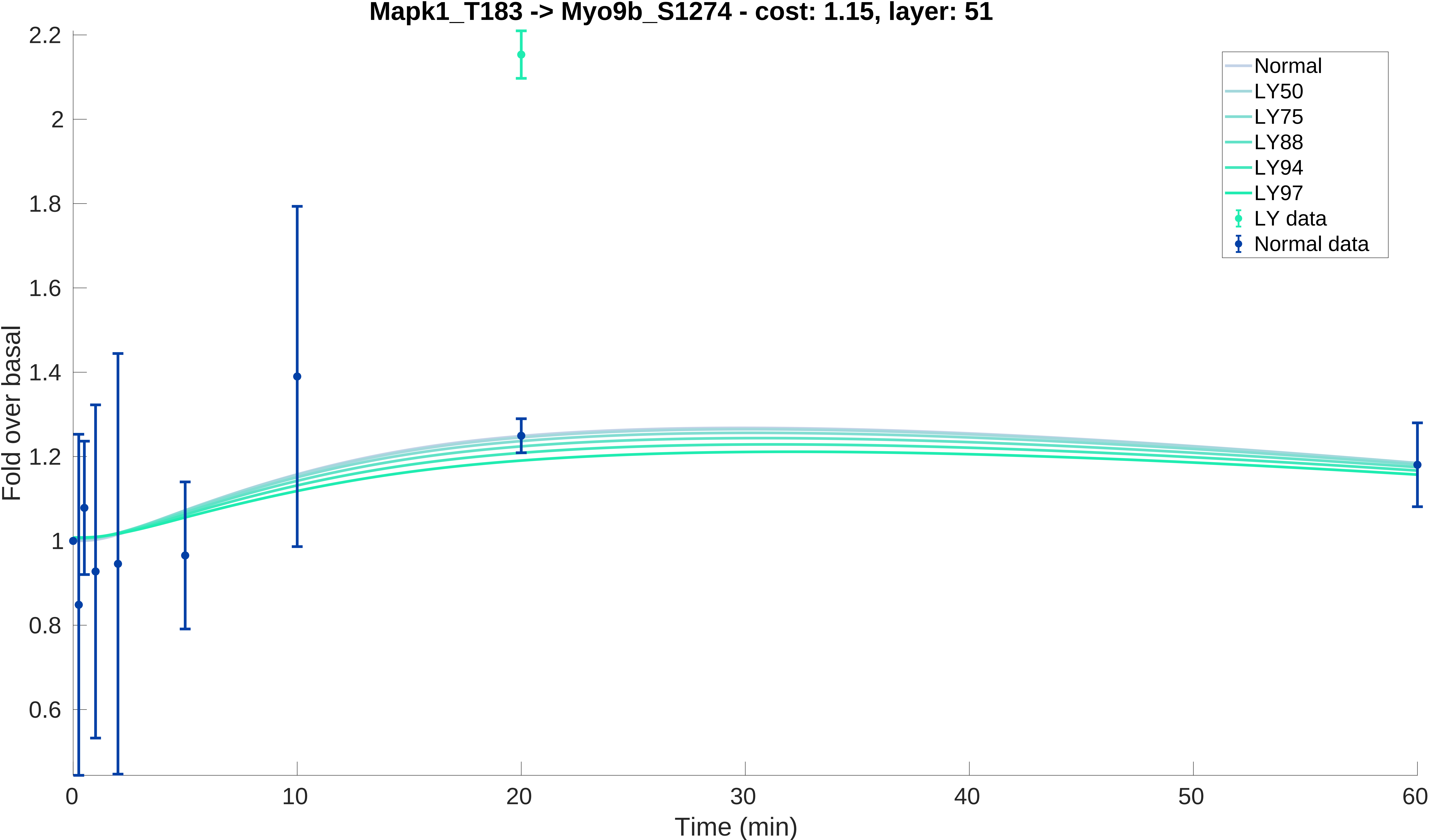

Mapk1_T183 -> Myo9b_S1274 - cost: 1.15, layer: 51
Fold over basal
Time (min)
2.2
2
1.8
1.6
1.4
1.2
1
0.8
0.6
0
10
20
30
40
50
60
Normal
LY50
LY75
LY88
LY94
LY97
LY data
Normal data

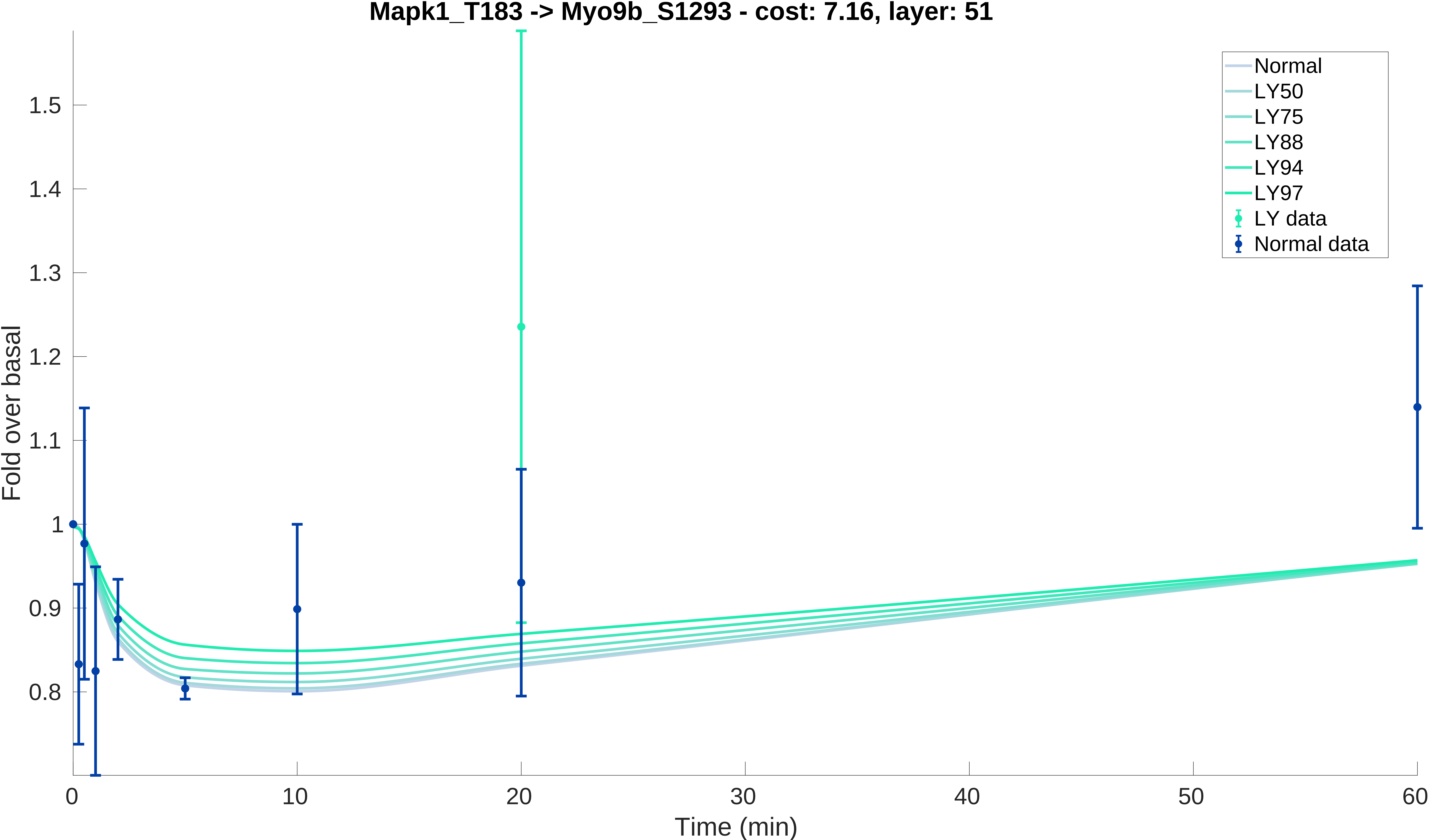

Mapk1_T183 -> Myo9b_S1293 - cost: 7.16, layer: 51
Fold over basal
Time (min)
0.8
0.9
1
1.1
1.2
1.3
1.4
1.5
0
10
20
30
40
50
60
Normal
LY50
LY75
LY88
LY94
LY97
LY data
Normal data

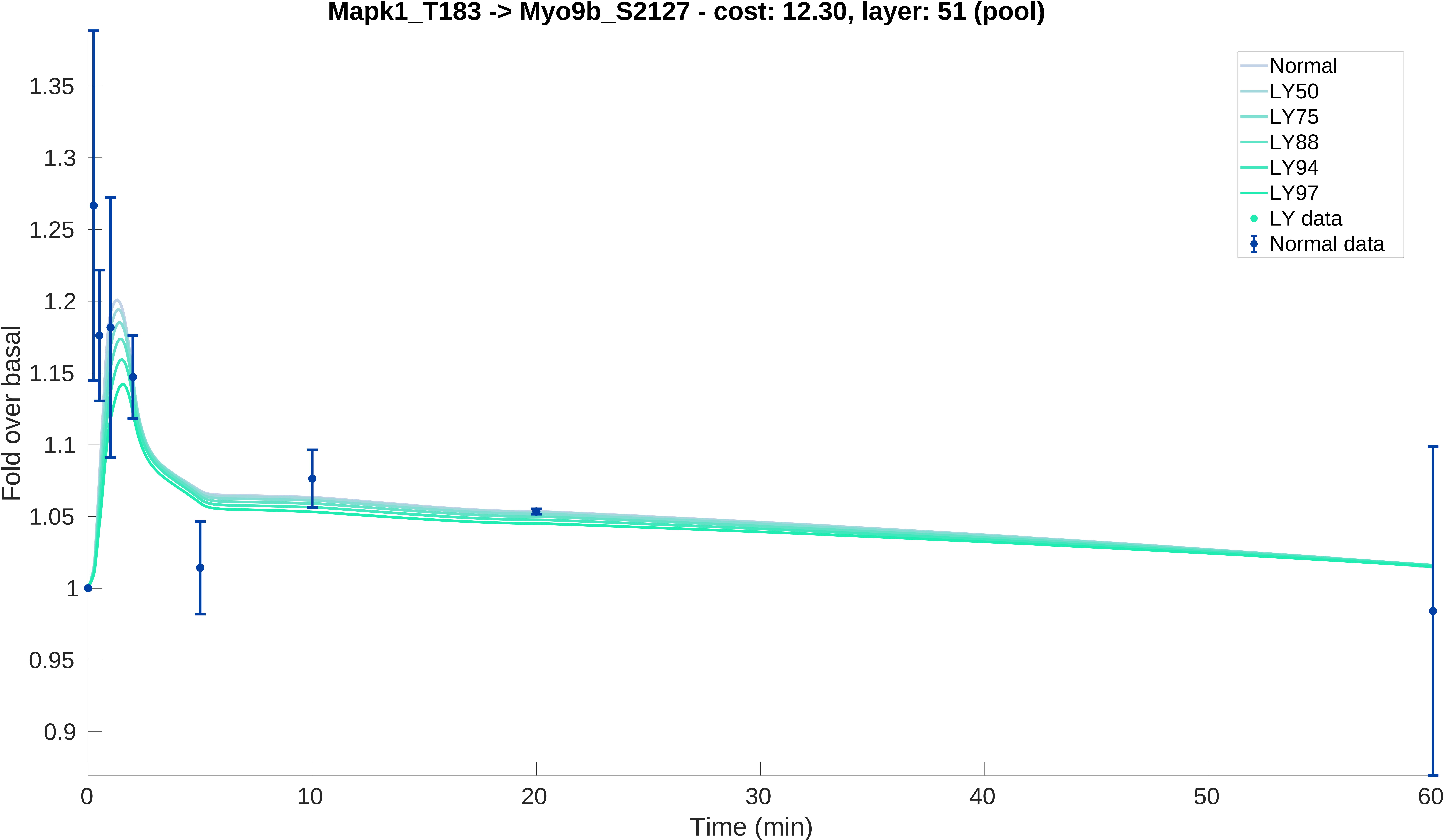

Mapk1_T183 -> Myo9b_S2127 - cost: 12.30, layer: 51 (pool)
Normal
LY50
LY75
LY88
LY94
LY97
LY data
Normal data
Fold over basal
1.35
1.3
1.25
1.2
1.15
1.1
1.05
1
0.95
0.9
0
10
20
30
40
50
60
Time (min)

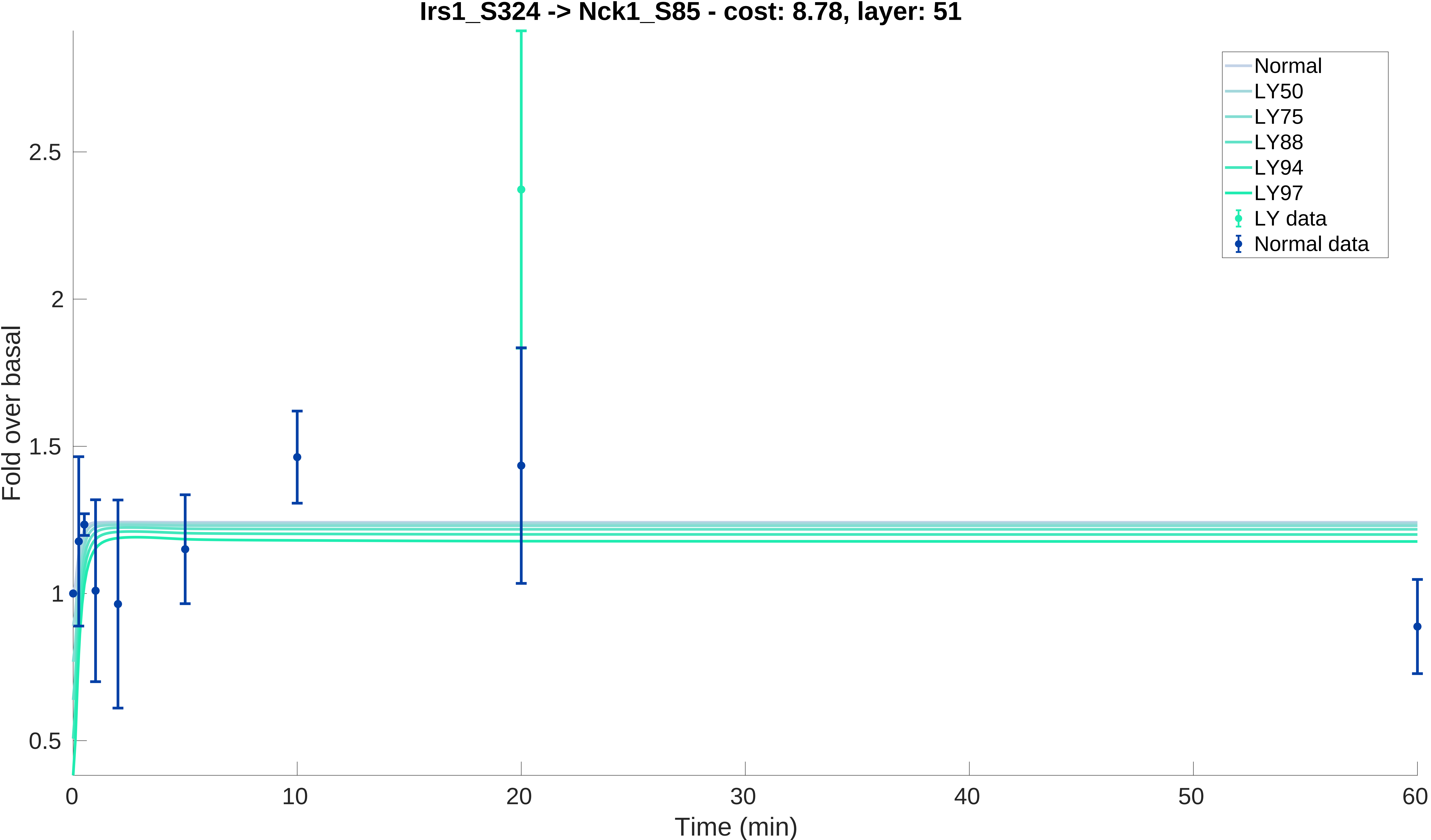

Irs1_S324 -> Nck1_S85 - cost: 8.78, layer: 51
Fold over basal
Time (min)
0.5
1
1.5
2
2.5
0
10
20
30
40
50
60
Normal
LY50
LY75
LY88
LY94
LY97
LY data
Normal data

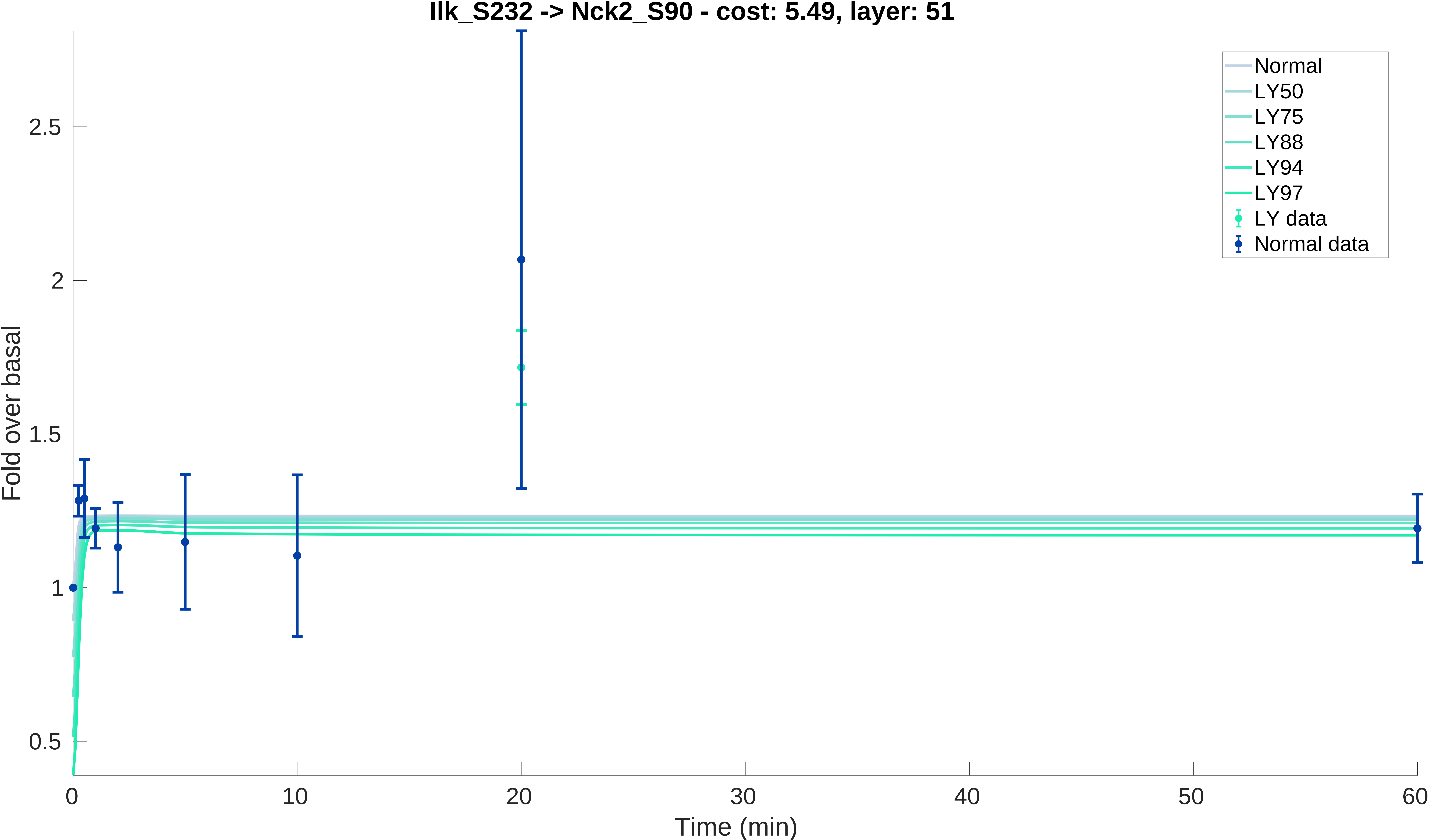

Ilk_S232 -> Nck2_S90 - cost: 5.49, layer: 51
Fold over basal
Time (min)
0.5
1
1.5
2
2.5
0
10
20
30
40
50
60
Normal
LY50
LY75
LY88
LY94
LY97
LY data
Normal data

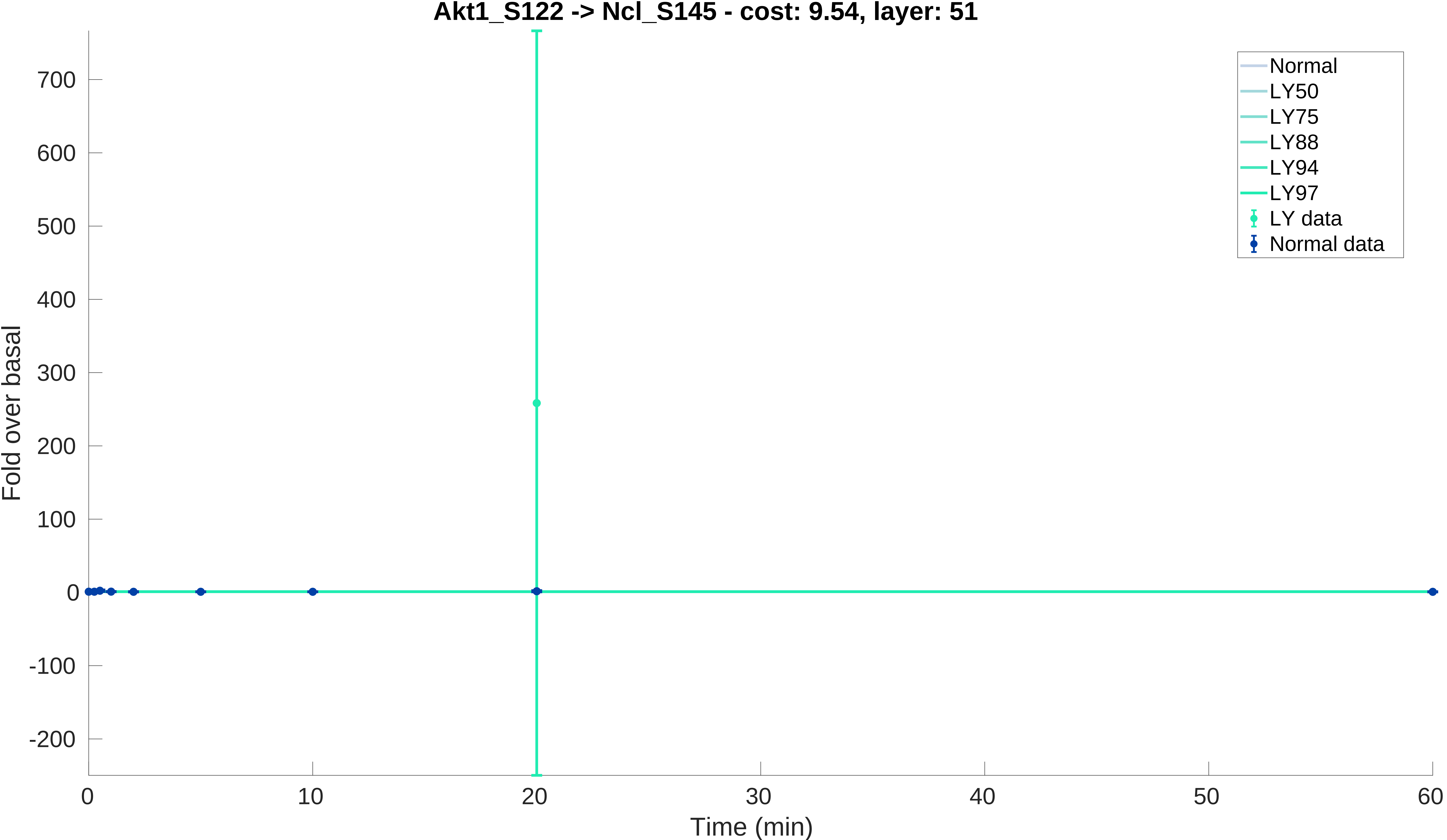

Akt1_S122 -> Ncl_S145 - cost: 9.54, layer: 51
Fold over basal
700
600
500
400
300
200
100
0
-100
-200
0
10
20
30
40
50
60
Time (min)
Normal
LY50
LY75
LY88
LY94
LY97
LY data
Normal data

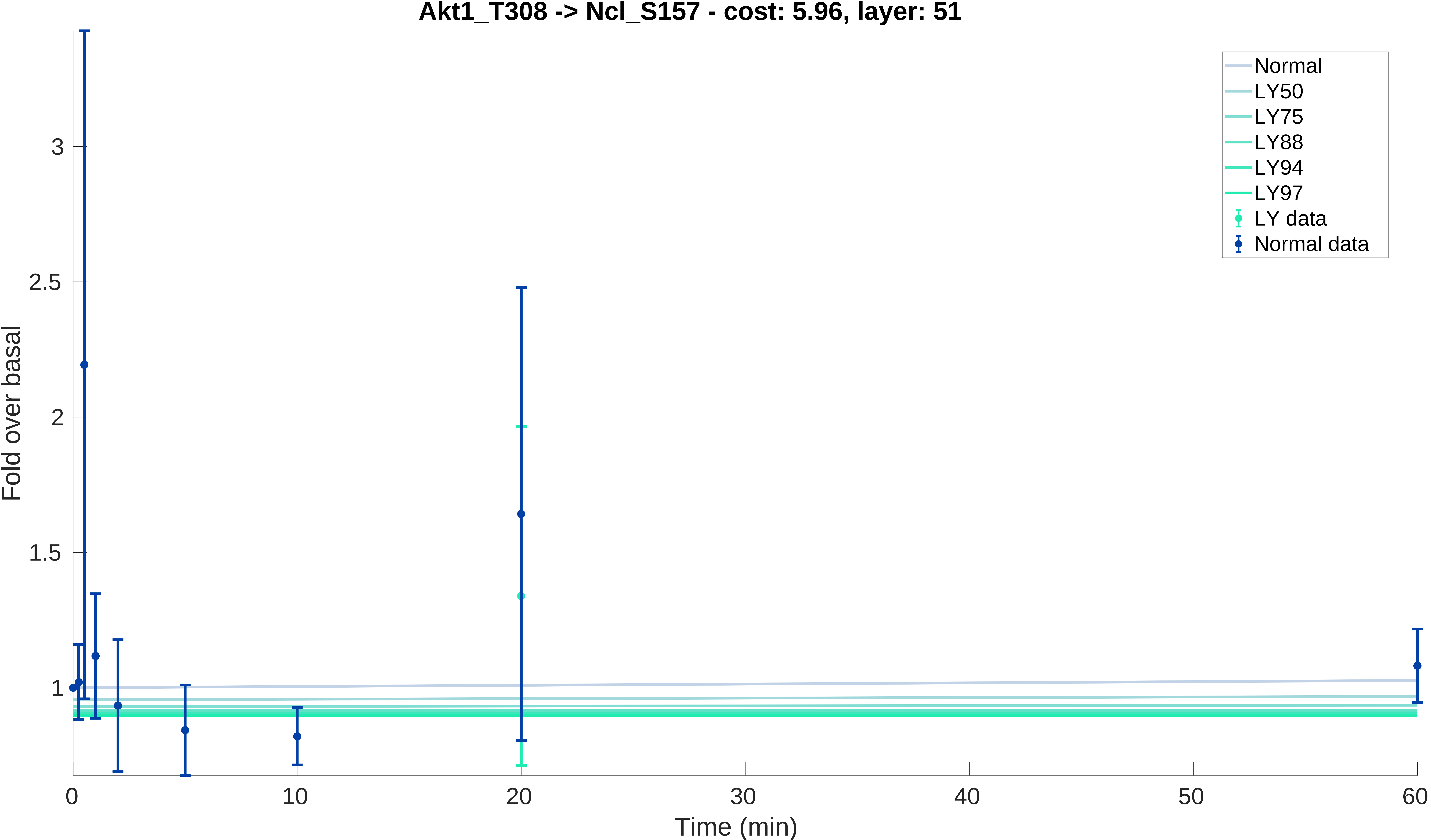

Akt1_T308 -> Ncl_S157 - cost: 5.96, layer: 51
Fold over basal
Time (min)
3
2.5
2
1.5
1
0
10
20
30
40
50
60
Normal
LY50
LY75
LY88
LY94
LY97
LY data
Normal data

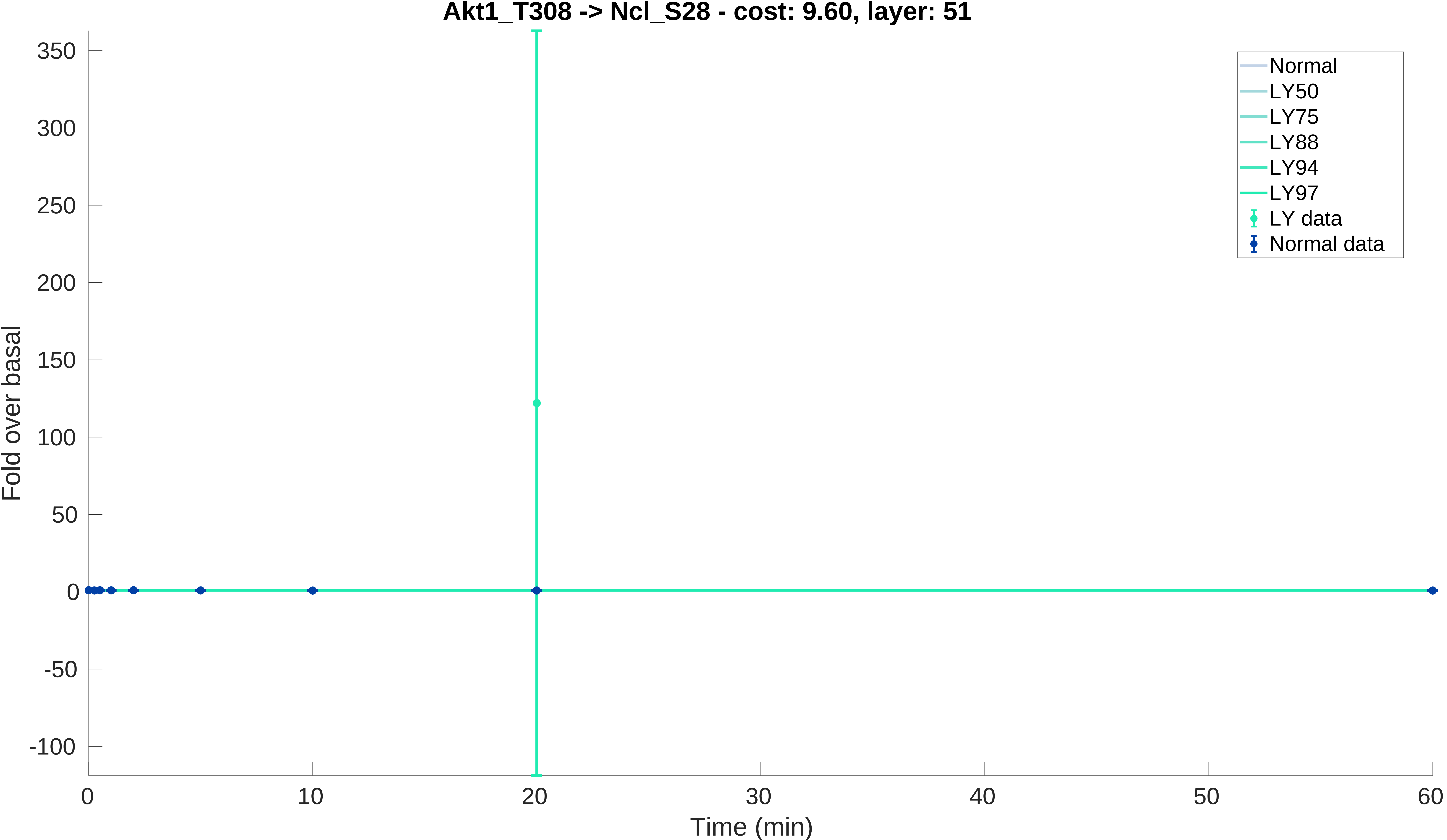

Akt1_T308 -> Ncl_S28 - cost: 9.60, layer: 51
Fold over basal
350
300
250
200
150
100
50
0
-50
-100
0
10
20
30
40
50
60
Time (min)
Normal
LY50
LY75
LY88
LY94
LY97
LY data
Normal data

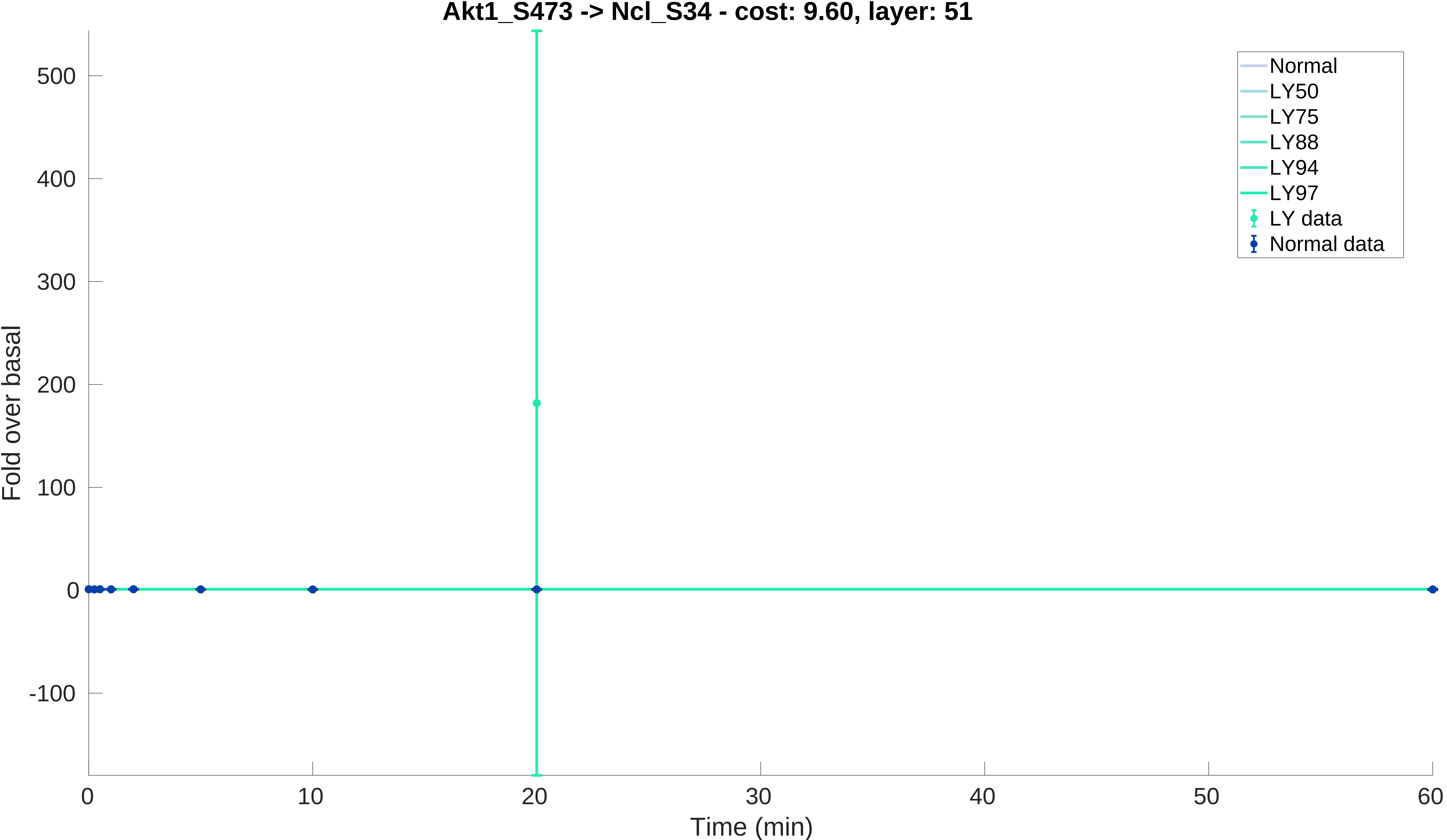

Akt1_S473 -> Ncl_S34 - cost: 9.60, layer: 51
Fold over basal
500
400
300
200
100
0
-100
0
10
20
30
40
50
60
Time (min)
Normal
LY50
LY75
LY88
LY94
LY97
LY data
Normal data

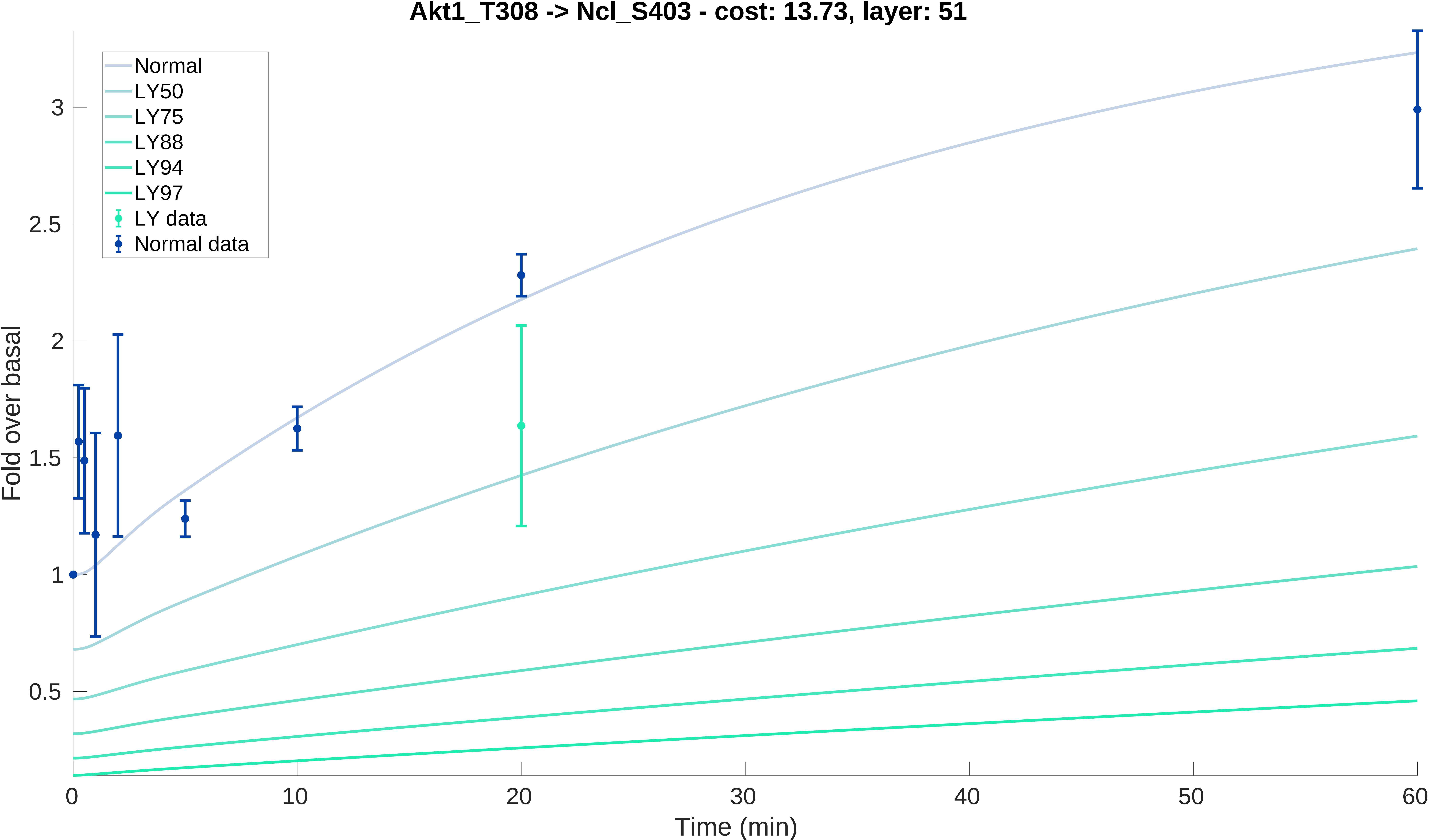

Akt1_T308 -> Ncl_S403 - cost: 13.73, layer: 51
Normal
LY50
LY75
LY88
LY94
LY97
LY data
Normal data
Fold over basal
Time (min)
0.5
1
1.5
2
2.5
3
0
10
20
30
40
50
60

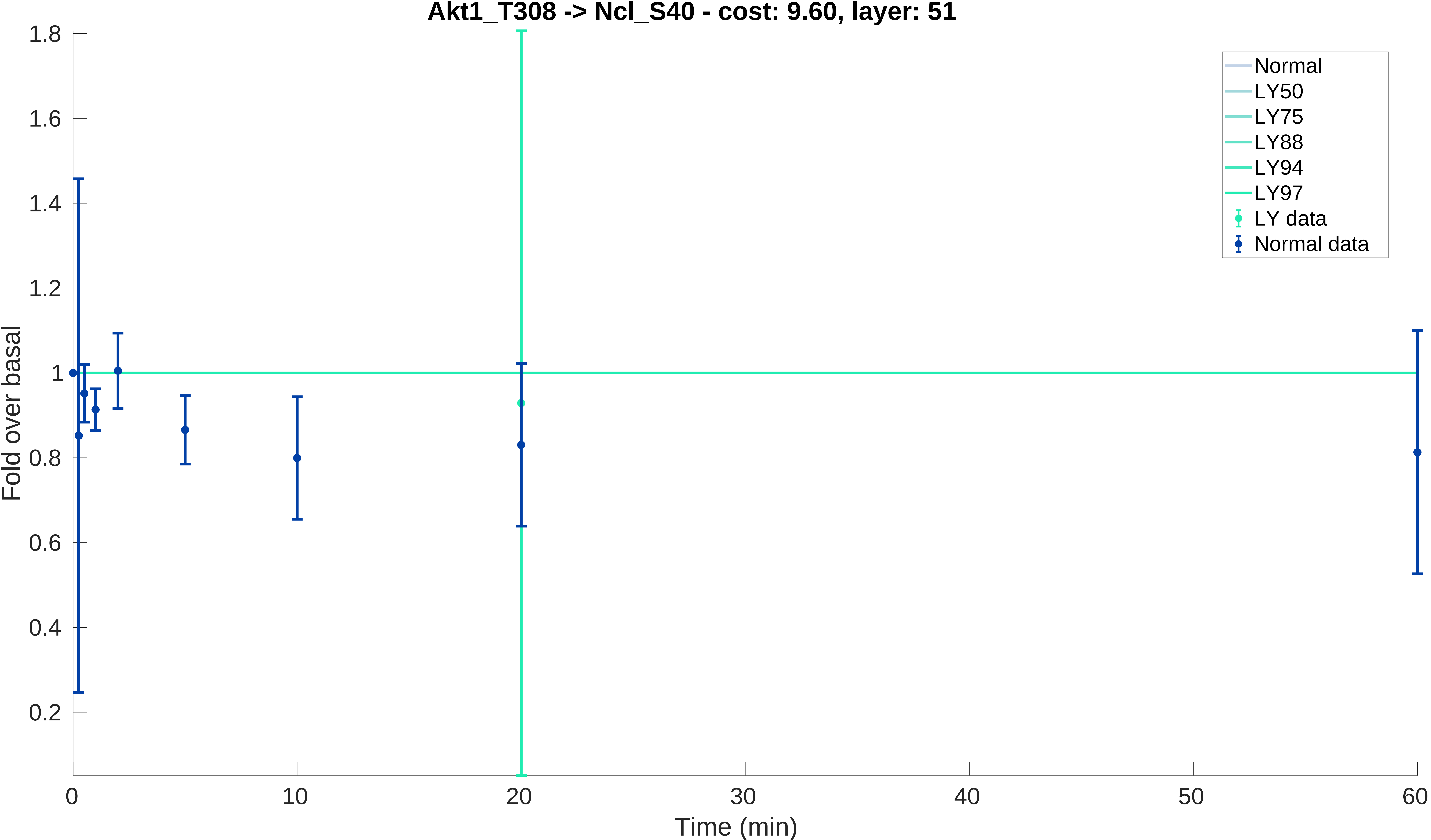

Akt1_T308 -> Ncl_S40 - cost: 9.60, layer: 51
Fold over basal
1.8
1.6
1.4
1.2
1
0.8
0.6
0.4
0.2
0
10
20
30
40
50
60
Time (min)
Normal
LY50
LY75
LY88
LY94
LY97
LY data
Normal data

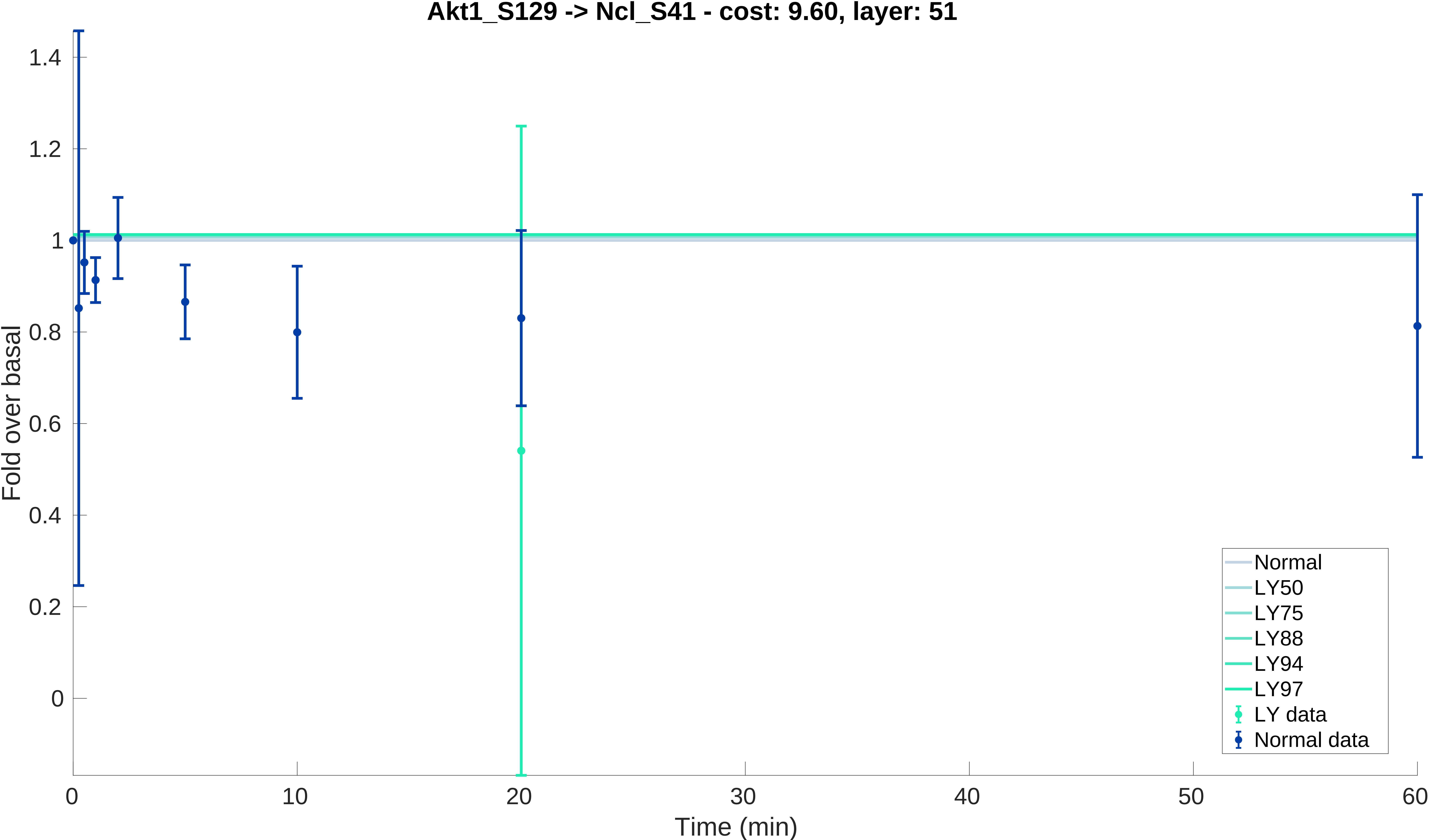

Akt1_S129 -> Ncl_S41 - cost: 9.60, layer: 51
Fold over basal
Time (min)
1.4
1.2
1
0.8
0.6
0.4
0.2
0
0
10
20
30
40
50
60
Normal
LY50
LY75
LY88
LY94
LY97
LY data
Normal data

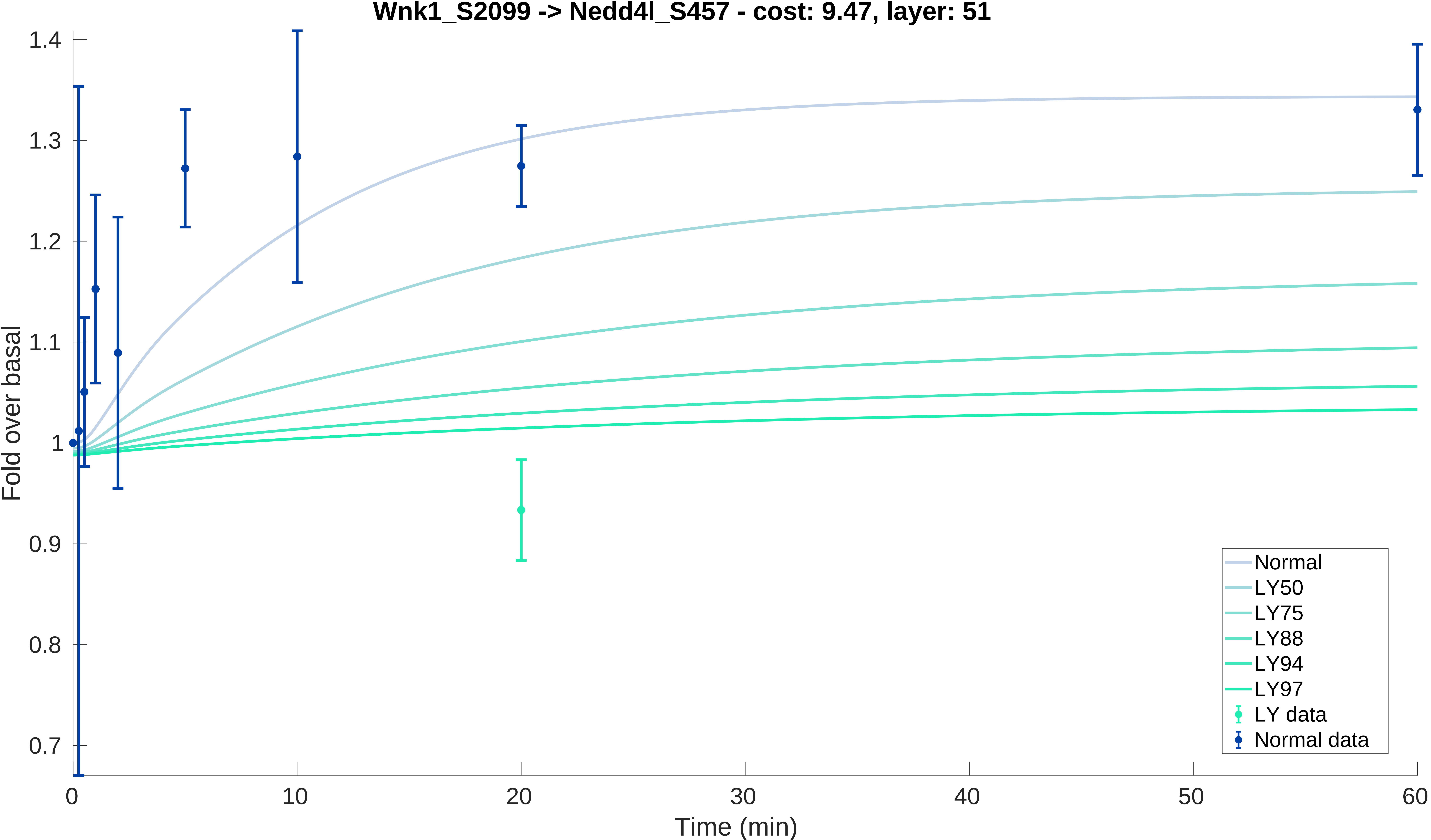

Wnk1_S2099 -> Nedd4l_S457 - cost: 9.47, layer: 51
Fold over basal
1.4
1.3
1.2
1.1
1
0.9
0.8
0.7
0
10
20
30
40
50
60
Time (min)
Normal
LY50
LY75
LY88
LY94
LY97
LY data
Normal data

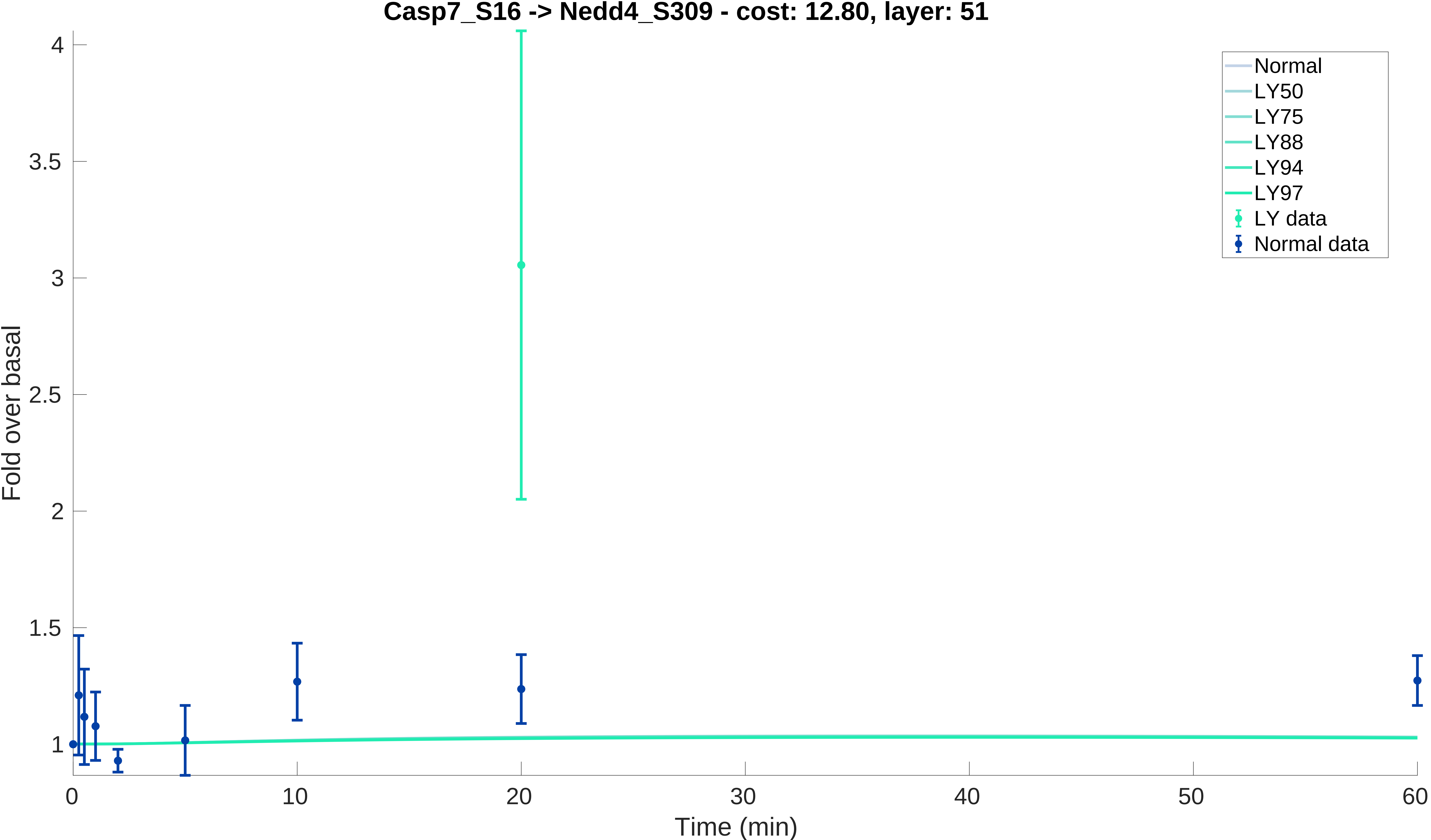

Casp7_S16 -> Nedd4_S309 - cost: 12.80, layer: 51
Fold over basal
Time (min)
4
3.5
3
2.5
2
1.5
1
0
10
20
30
40
50
60
Normal
LY50
LY75
LY88
LY94
LY97
LY data
Normal data

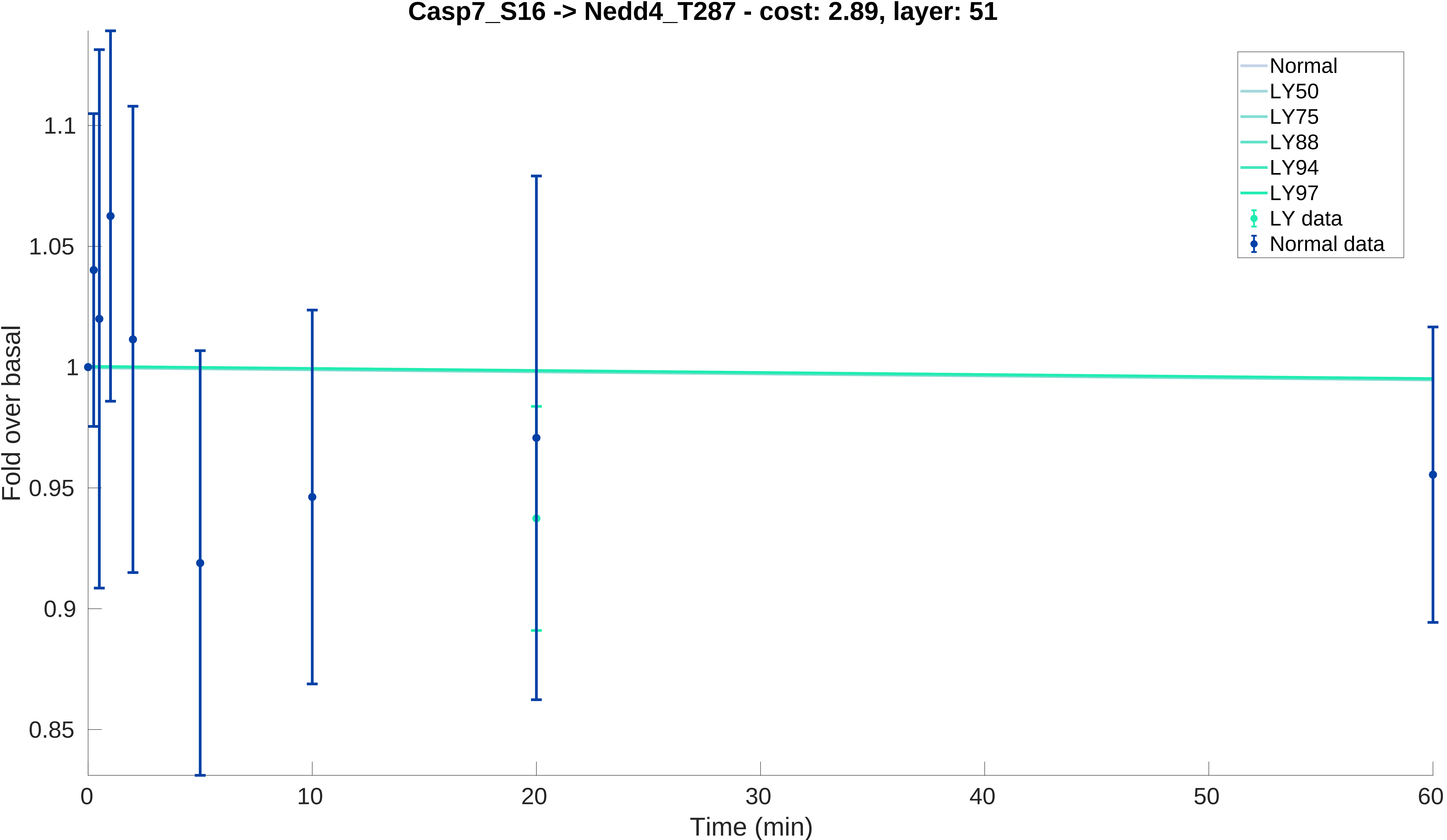

Casp7_S16 -> Nedd4_T287 - cost: 2.89, layer: 51
Fold over basal
1.1
1.05
1
0.95
0.9
0.85
0
10
20
30
40
50
60
Time (min)
Normal
LY50
LY75
LY88
LY94
LY97
LY data
Normal data

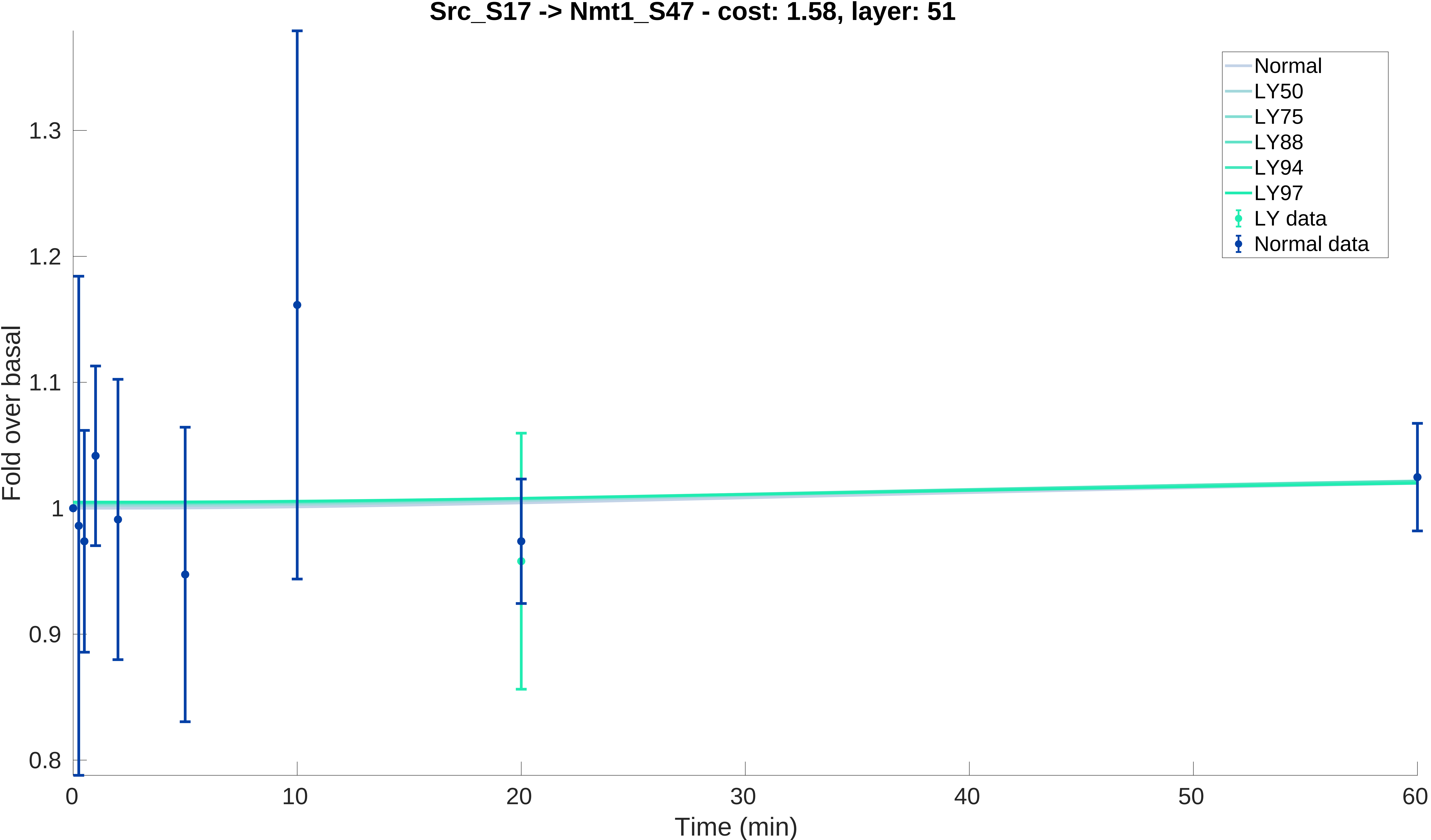
Src_S17 -> Nmt1_S47 - cost: 1.58, layer: 51
Normal
LY50
LY75
LY88
LY94
LY97
LY data
Normal data
Fold over basal
1.3
1.2
1.1
1
0.9
0.8
0
10
20
30
40
50
60
Time (min)

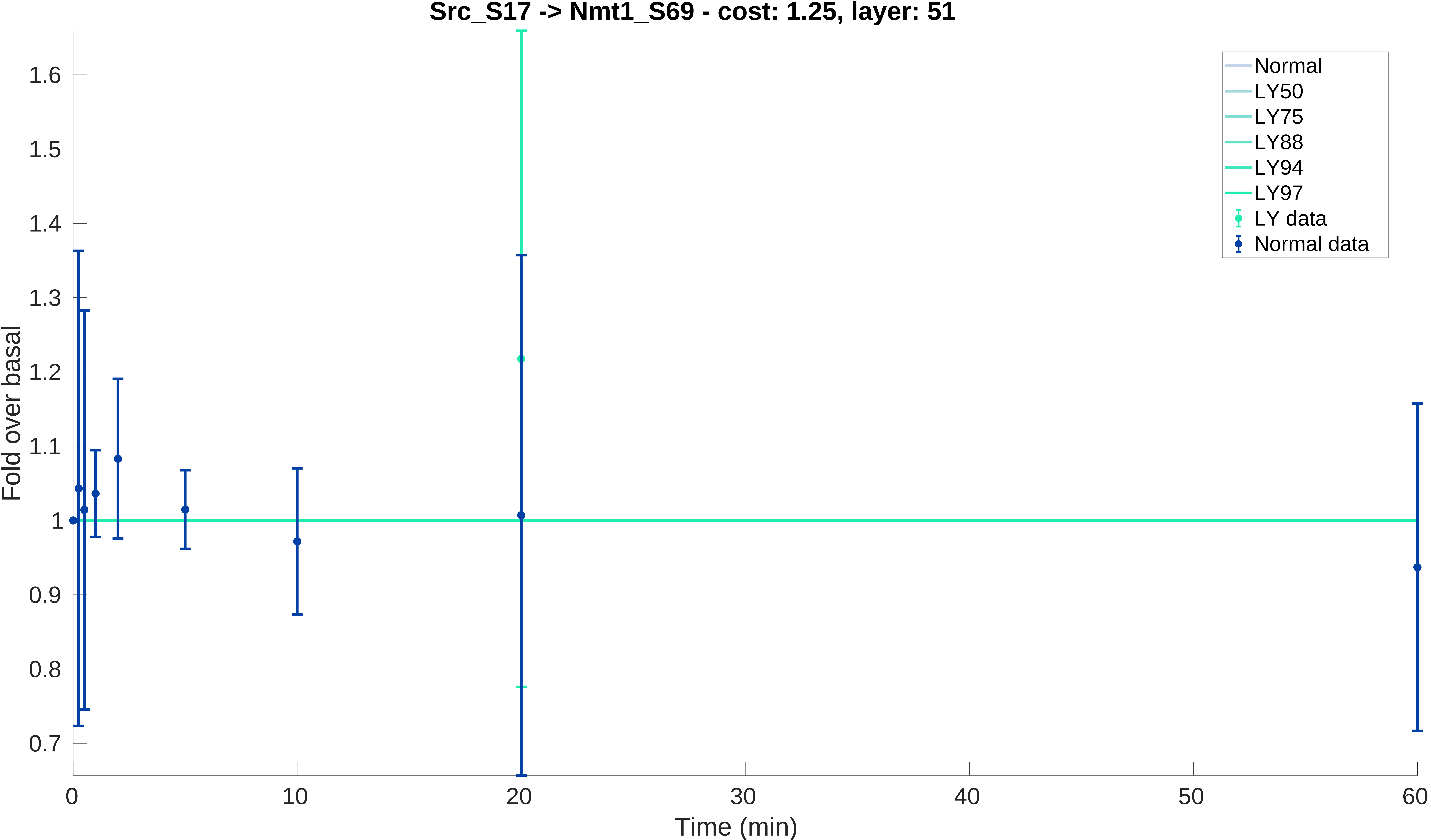

Src_S17 -> Nmt1_S69 - cost: 1.25, layer: 51
Normal
LY50
LY75
LY88
LY94
LY97
LY data
Normal data
Fold over basal
1.6
1.5
1.4
1.3
1.2
1.1
1
0.9
0.8
0.7
0
10
20
30
40
50
60
Time (min)

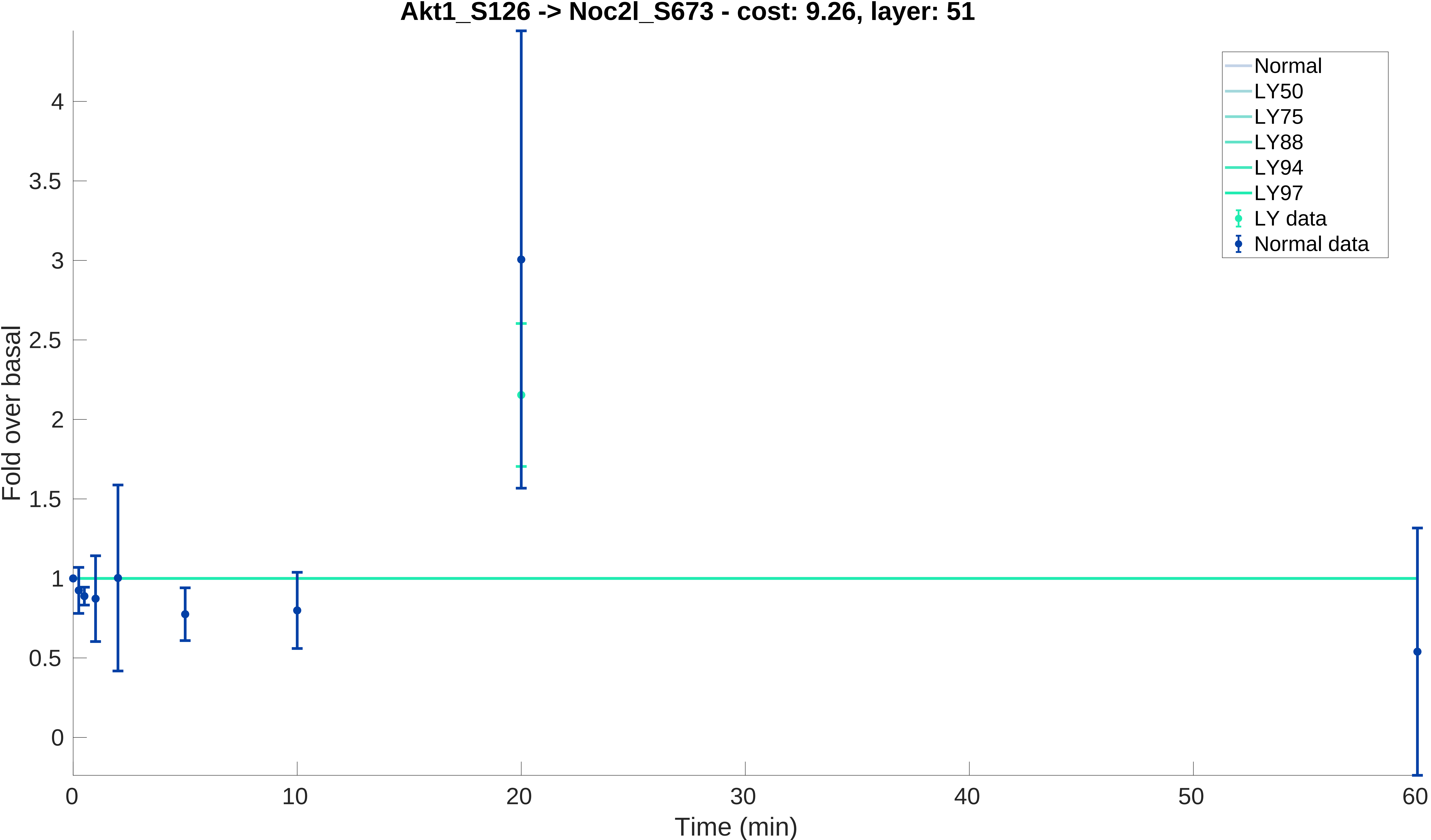

Akt1_S126 -> Noc2l_S673 - cost: 9.26, layer: 51
Fold over basal
Time (min)
0
0.5
1
1.5
2
2.5
3
3.5
4
0
10
20
30
40
50
60
Normal
LY50
LY75
LY88
LY94
LY97
LY data
Normal data

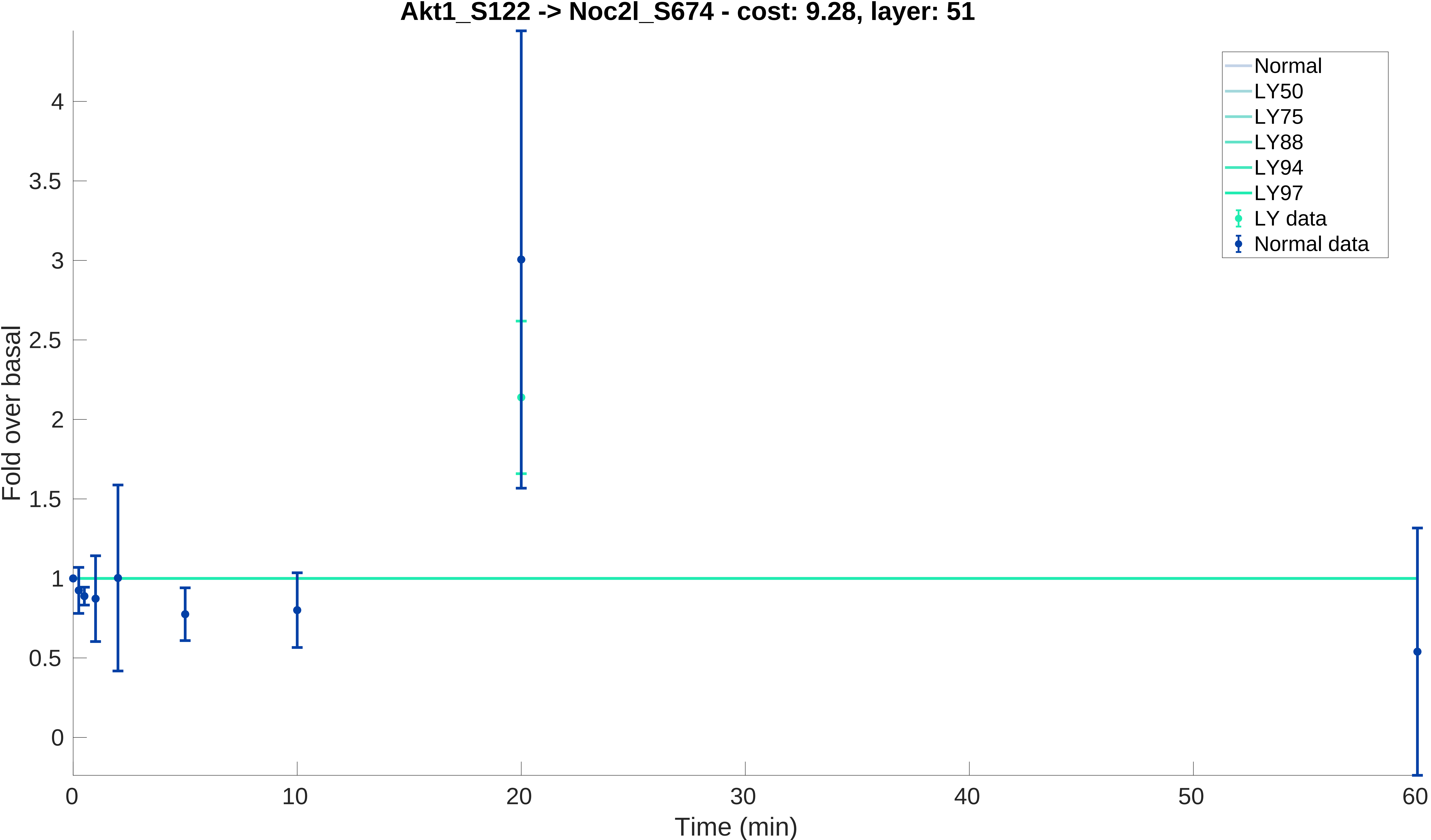

Akt1_S122 -> Noc2l_S674 - cost: 9.28, layer: 51
Fold over basal
Time (min)
0
0.5
1
1.5
2
2.5
3
3.5
4
0
10
20
30
40
50
60
Normal
LY50
LY75
LY88
LY94
LY97
LY data
Normal data

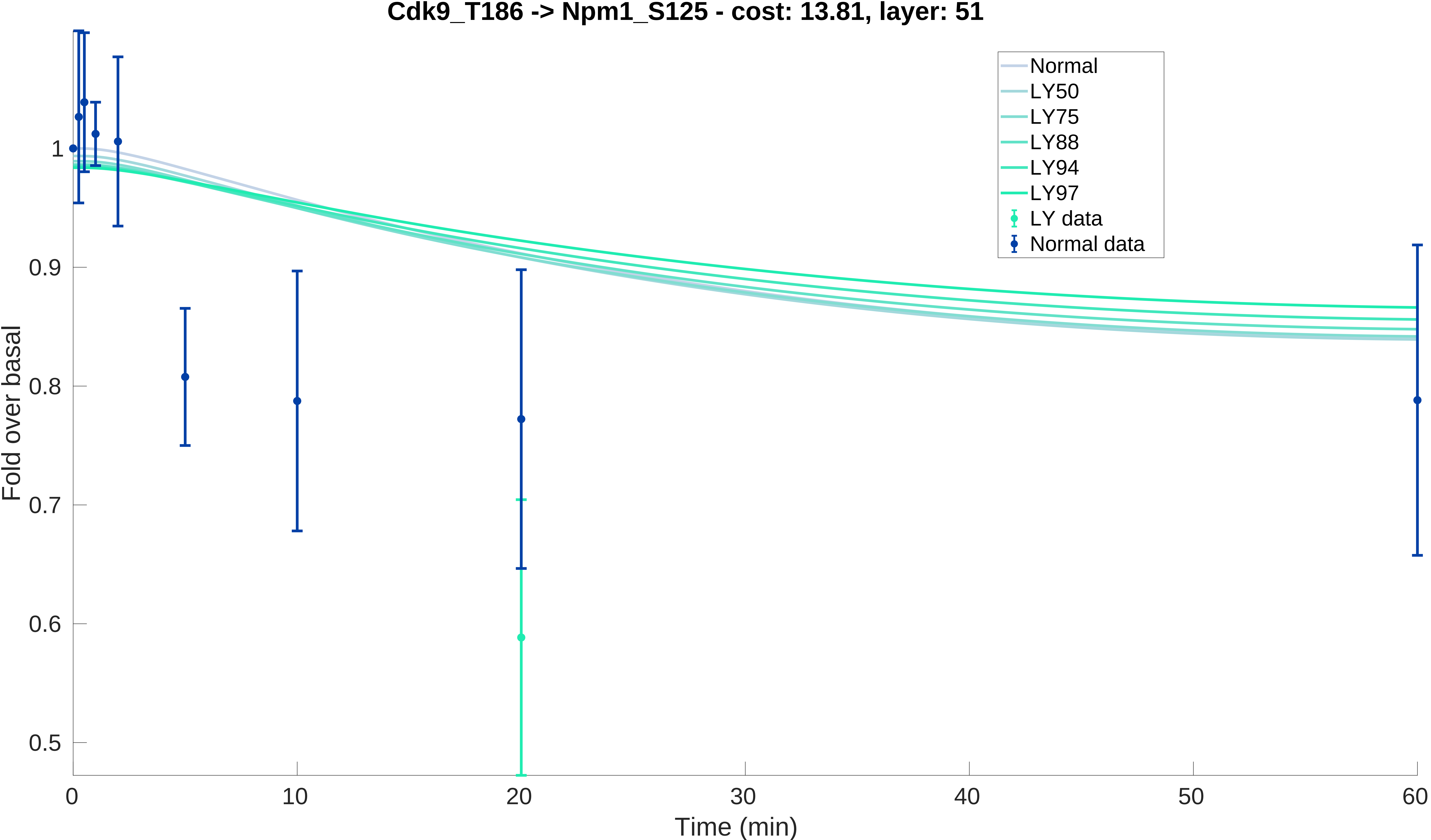

Cdk9_T186 -> Npm1_S125 - cost: 13.81, layer: 51
Normal
LY50
LY75
LY88
LY94
LY97
LY data
Normal data
Fold over basal
1
0.9
0.8
0.7
0.6
0.5
0
10
20
30
40
50
60
Time (min)

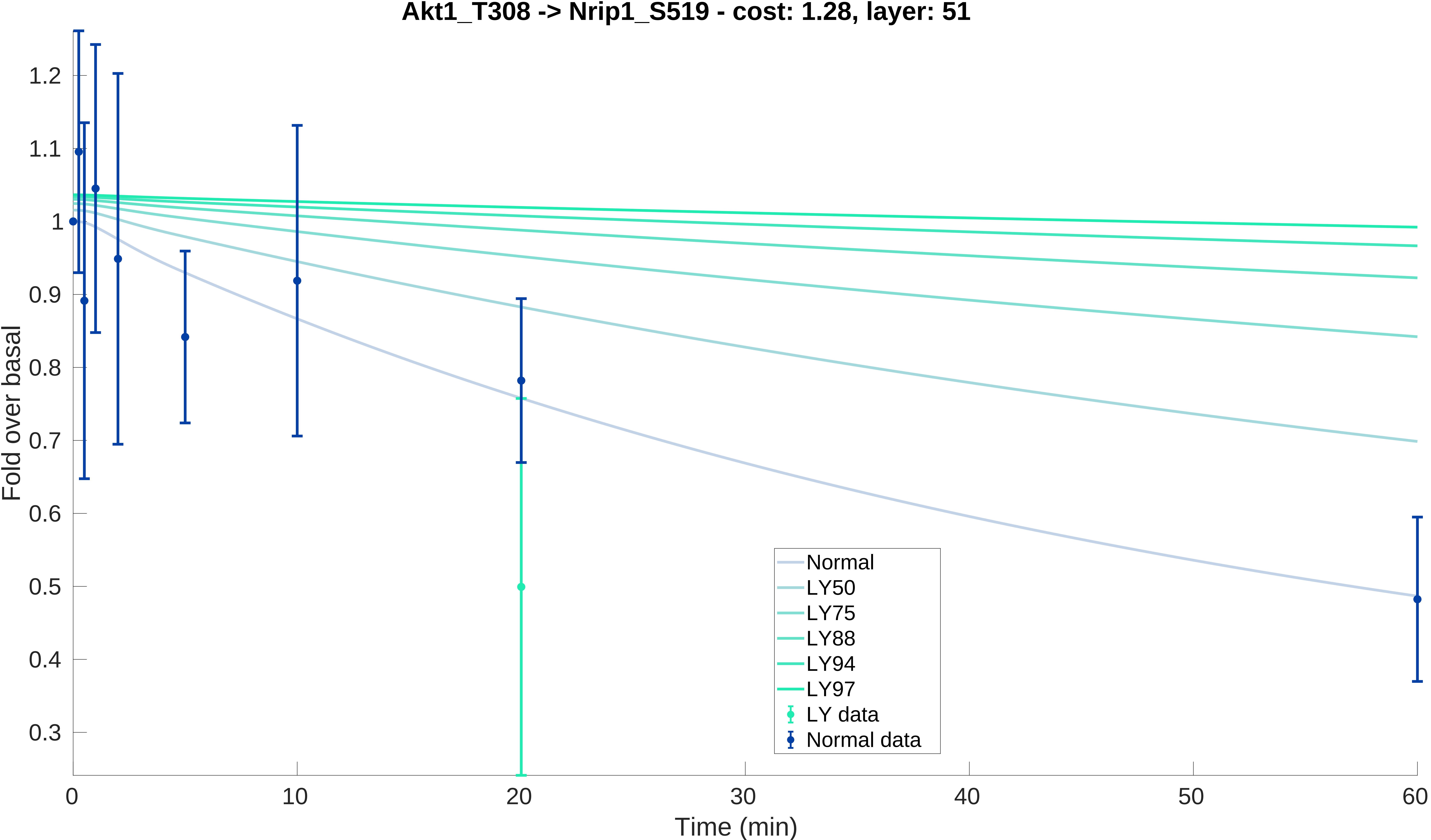

Akt1_T308 -> Nrip1_S519 - cost: 1.28, layer: 51
Fold over basal
Time (min)
1.2
1.1
1
0.9
0.8
0.7
0.6
0.5
0.4
0.3
0
10
20
30
40
50
60
Normal
LY50
LY75
LY88
LY94
LY97
LY data
Normal data

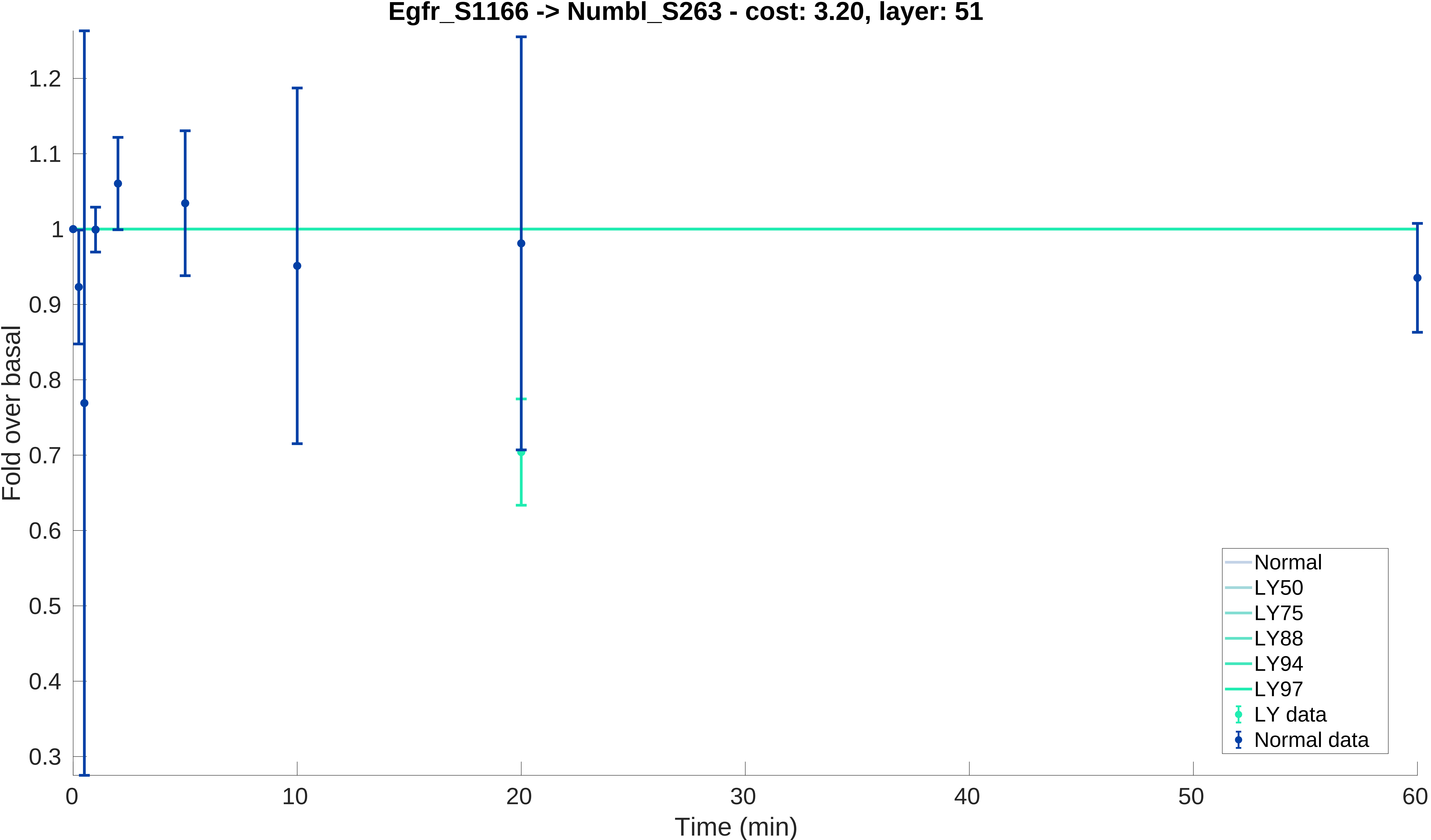
Egfr_S1166 -> Numbl_S263 - cost: 3.20, layer: 51
Fold over basal
1.2
1.1
1
0.9
0.8
0.7
0.6
0.5
0.4
0.3
0
10
20
30
40
50
60
Time (min)
Normal
LY50
LY75
LY88
LY94
LY97
LY data
Normal data

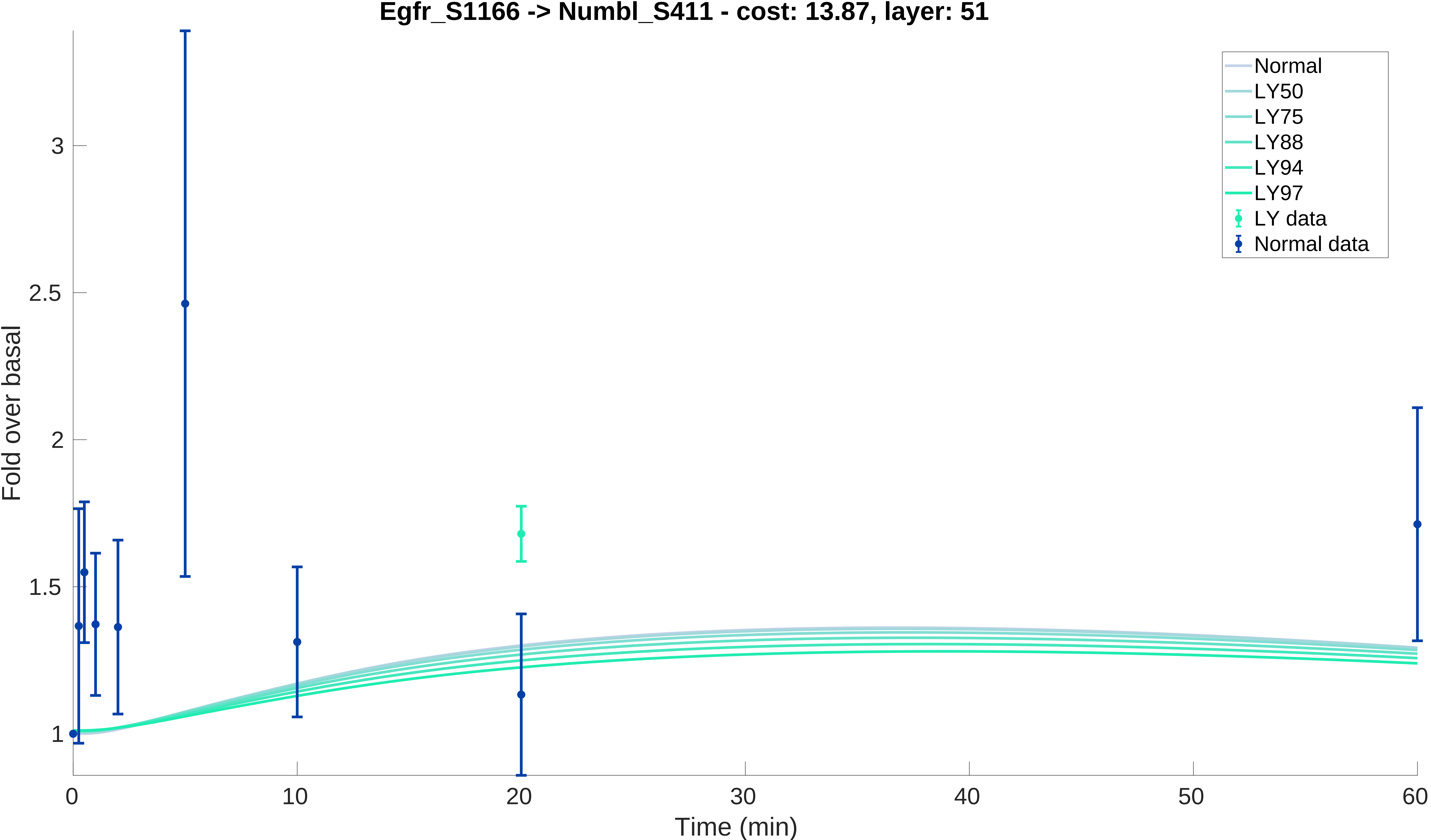

Egfr_S1166 -> Numbl_S411 - cost: 13.87, layer: 51
Normal
LY50
LY75
LY88
LY94
LY97
LY data
Normal data
Fold over basal
Time (min)
3
2.5
2
1.5
1
0
10
20
30
40
50
60

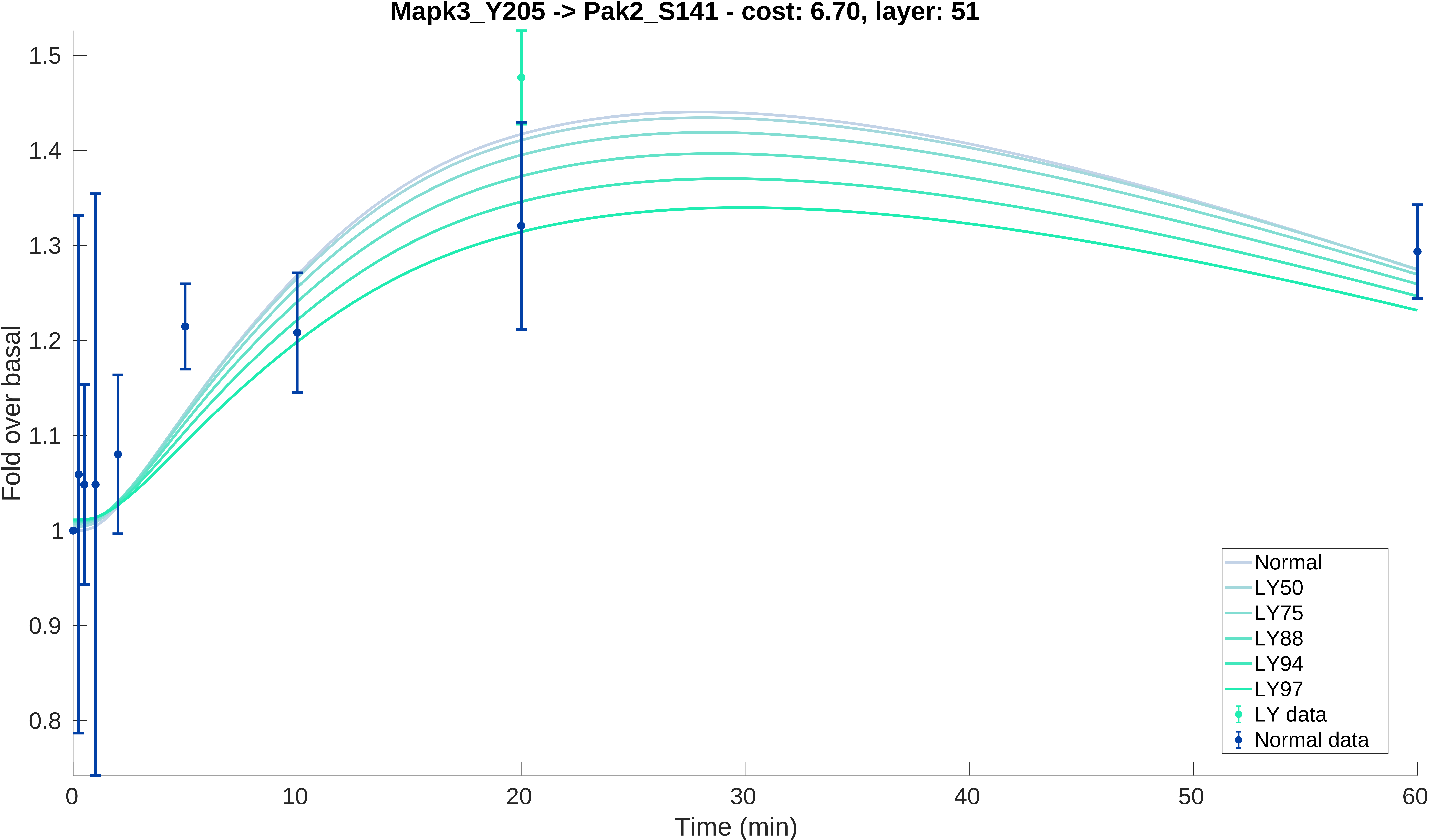

Mapk3_Y205 -> Pak2_S141 - cost: 6.70, layer: 51
Fold over basal
1.5
1.4
1.3
1.2
1.1
1
0.9
0.8
0
10
20
30
40
50
60
Time (min)
Normal
LY50
LY75
LY88
LY94
LY97
LY data
Normal data

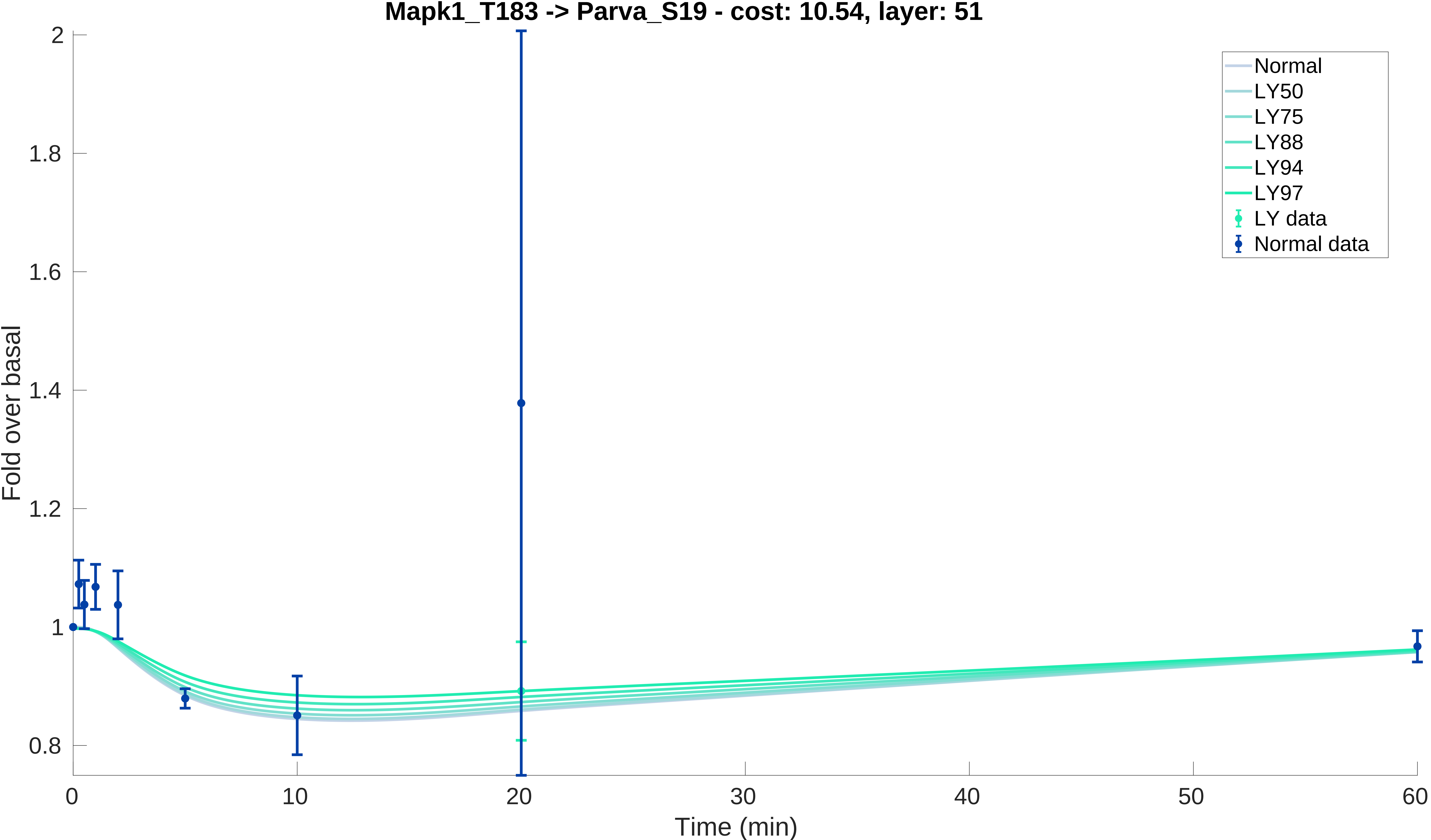

Mapk1_T183 -> Parva_S19 - cost: 10.54, layer: 51
Fold over basal
Time (min)
2
1.8
1.6
1.4
1.2
1
0.8
0
10
20
30
40
50
60
Normal
LY50
LY75
LY88
LY94
LY97
LY data
Normal data

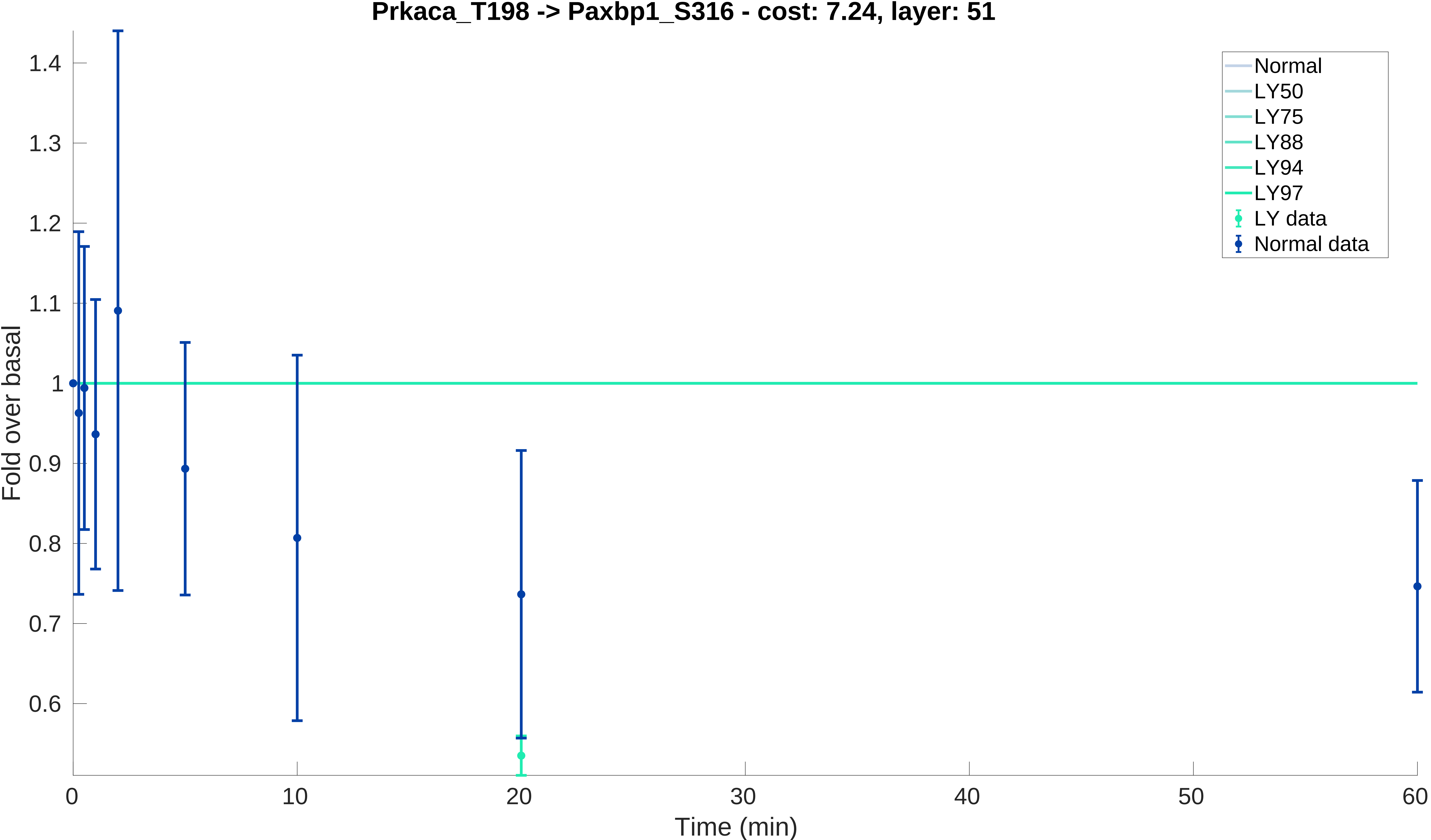

Prkaca_T198 -> Paxbp1_S316 - cost: 7.24, layer: 51
Fold over basal
Time (min)
1.4
1.3
1.2
1.1
1
0.9
0.8
0.7
0.6
0
10
20
30
40
50
60
Normal
LY50
LY75
LY88
LY94
LY97
LY data
Normal data

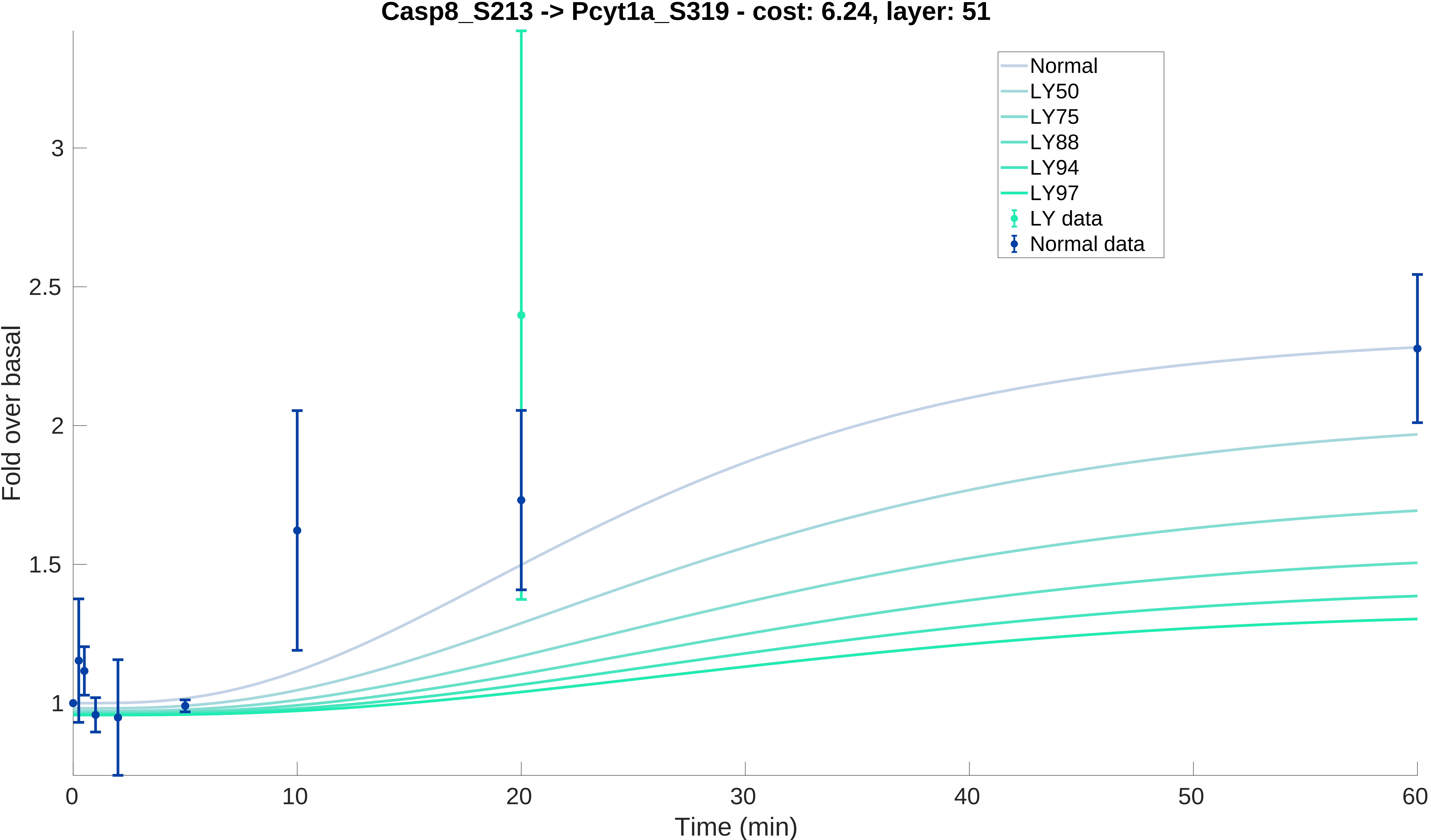

Casp8_S213 -> Pcyt1a_S319 - cost: 6.24, layer: 51
Fold over basal
Time (min)
0
10
20
30
40
50
60
1
1.5
2
2.5
3
Normal
LY50
LY75
LY88
LY94
LY97
LY data
Normal data

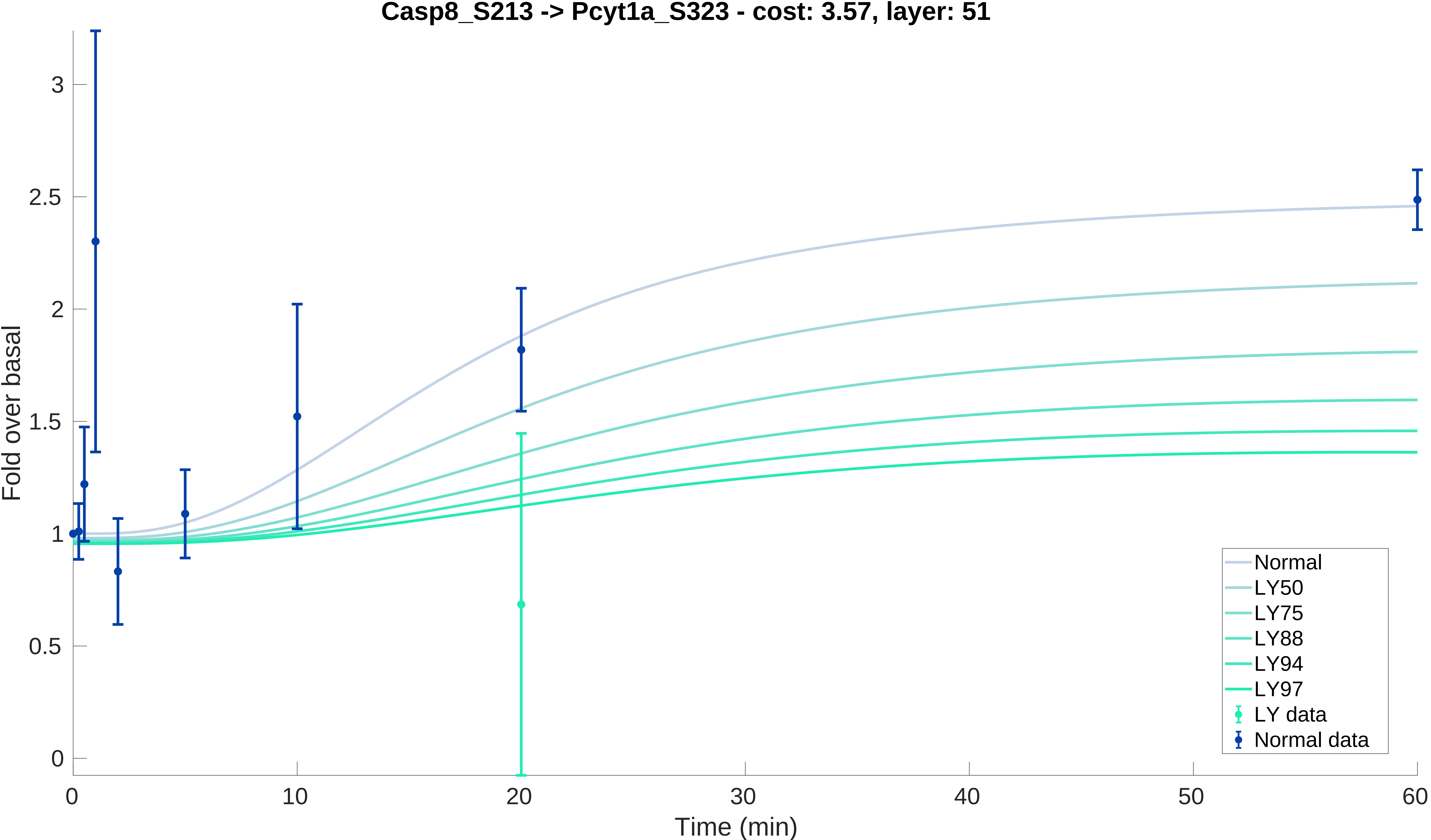

Casp8_S213 -> Pcyt1a_S323 - cost: 3.57, layer: 51
Fold over basal
Time (min)
0
0.5
1
1.5
2
2.5
3
0
10
20
30
40
50
60
Normal
LY50
LY75
LY88
LY94
LY97
LY data
Normal data

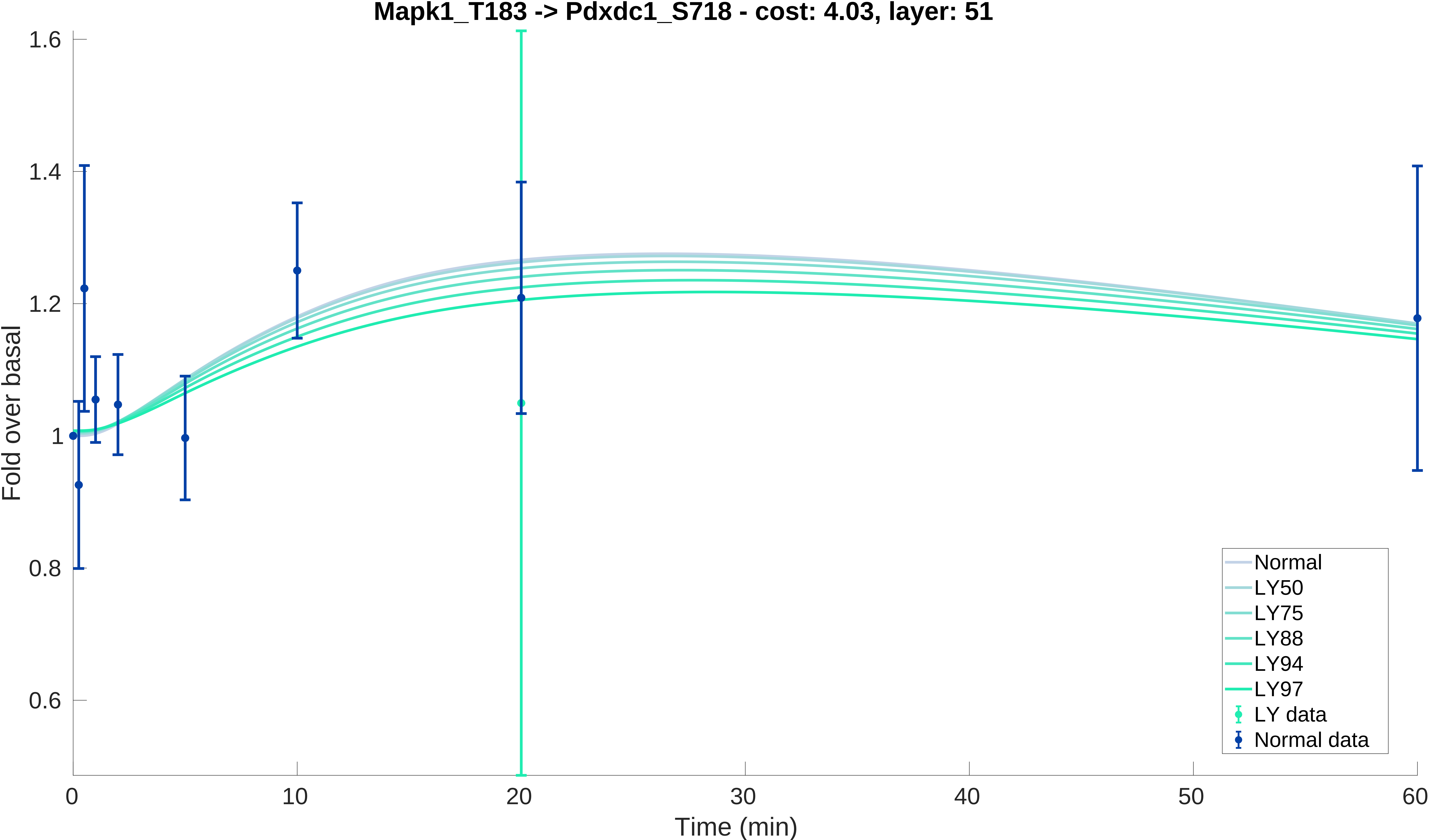

Mapk1_T183 -> Pdxdc1_S718 - cost: 4.03, layer: 51
Fold over basal
Time (min)
1.6
1.4
1.2
1
0.8
0.6
0
10
20
30
40
50
60
Normal
LY50
LY75
LY88
LY94
LY97
LY data
Normal data

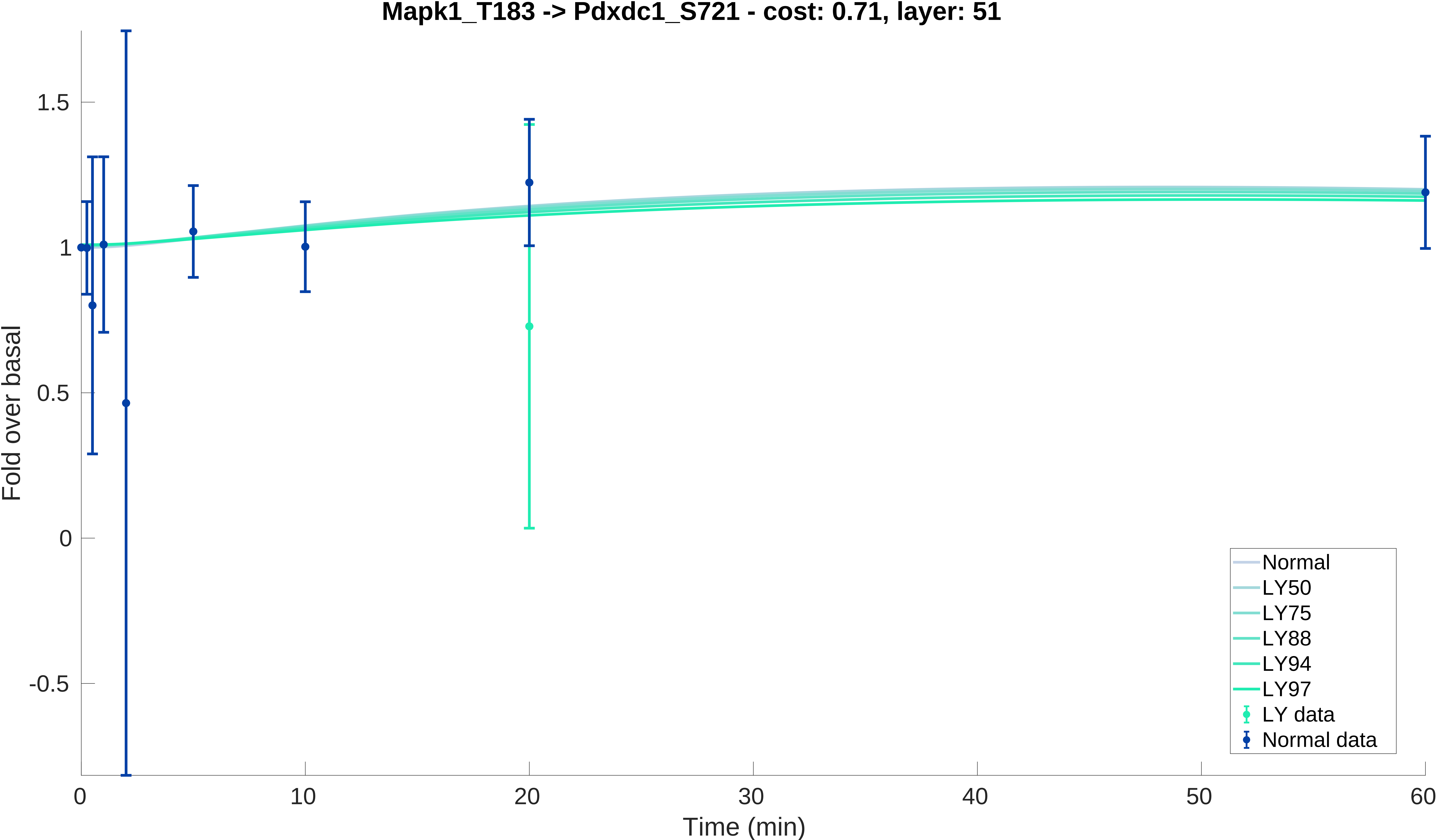

Mapk1_T183 -> Pdxdc1_S721 - cost: 0.71, layer: 51
Fold over basal
1.5
1
0.5
0
-0.5
0
10
20
30
40
50
60
Time (min)
Normal
LY50
LY75
LY88
LY94
LY97
LY data
Normal data

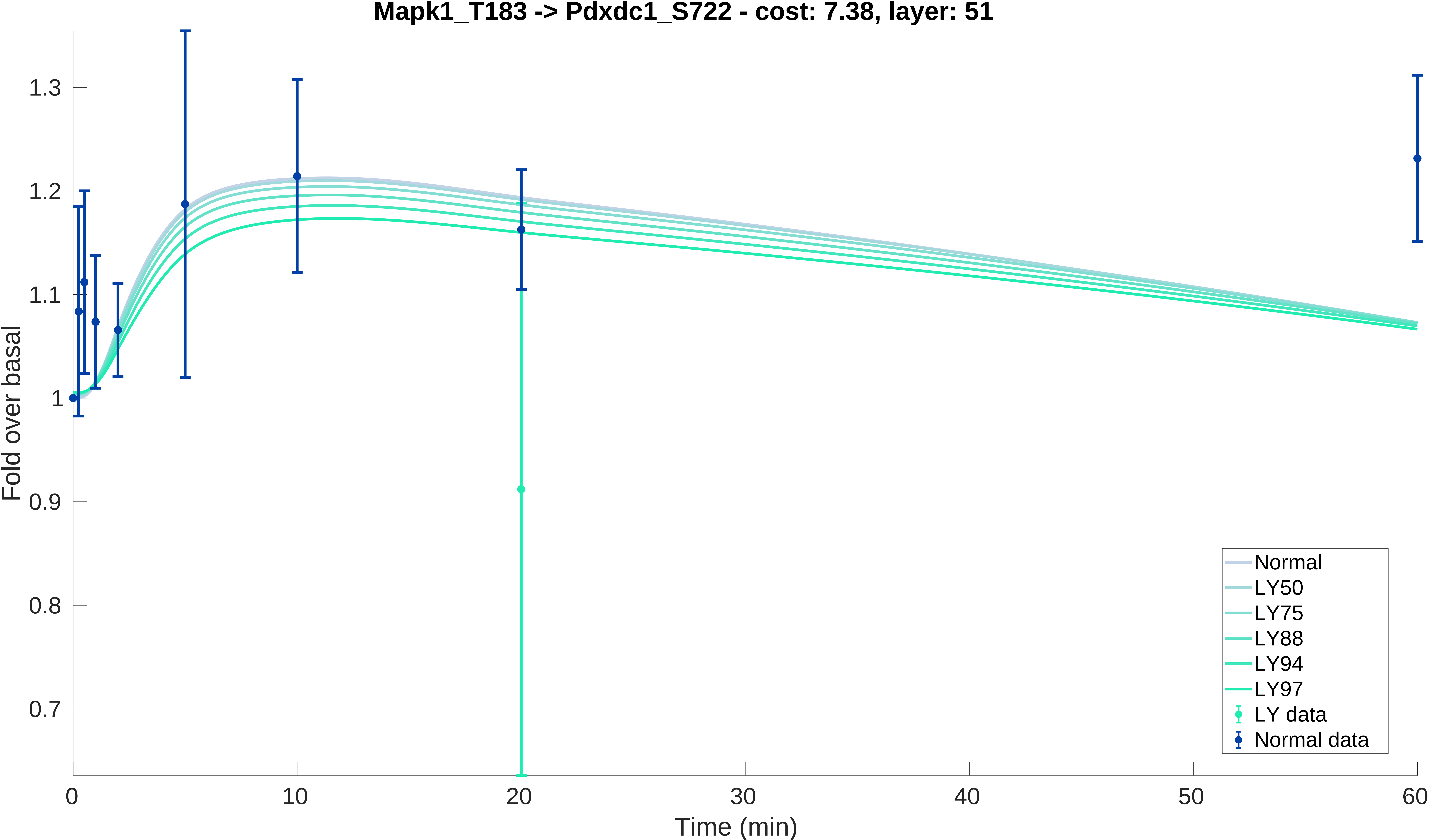
Mapk1_T183 -> Pdxdc1_S722 - cost: 7.38, layer: 51
Fold over basal
1.3
1.2
1.1
1
0.9
0.8
0.7
0
10
20
30
40
50
60
Time (min)
Normal
LY50
LY75
LY88
LY94
LY97
LY data
Normal data

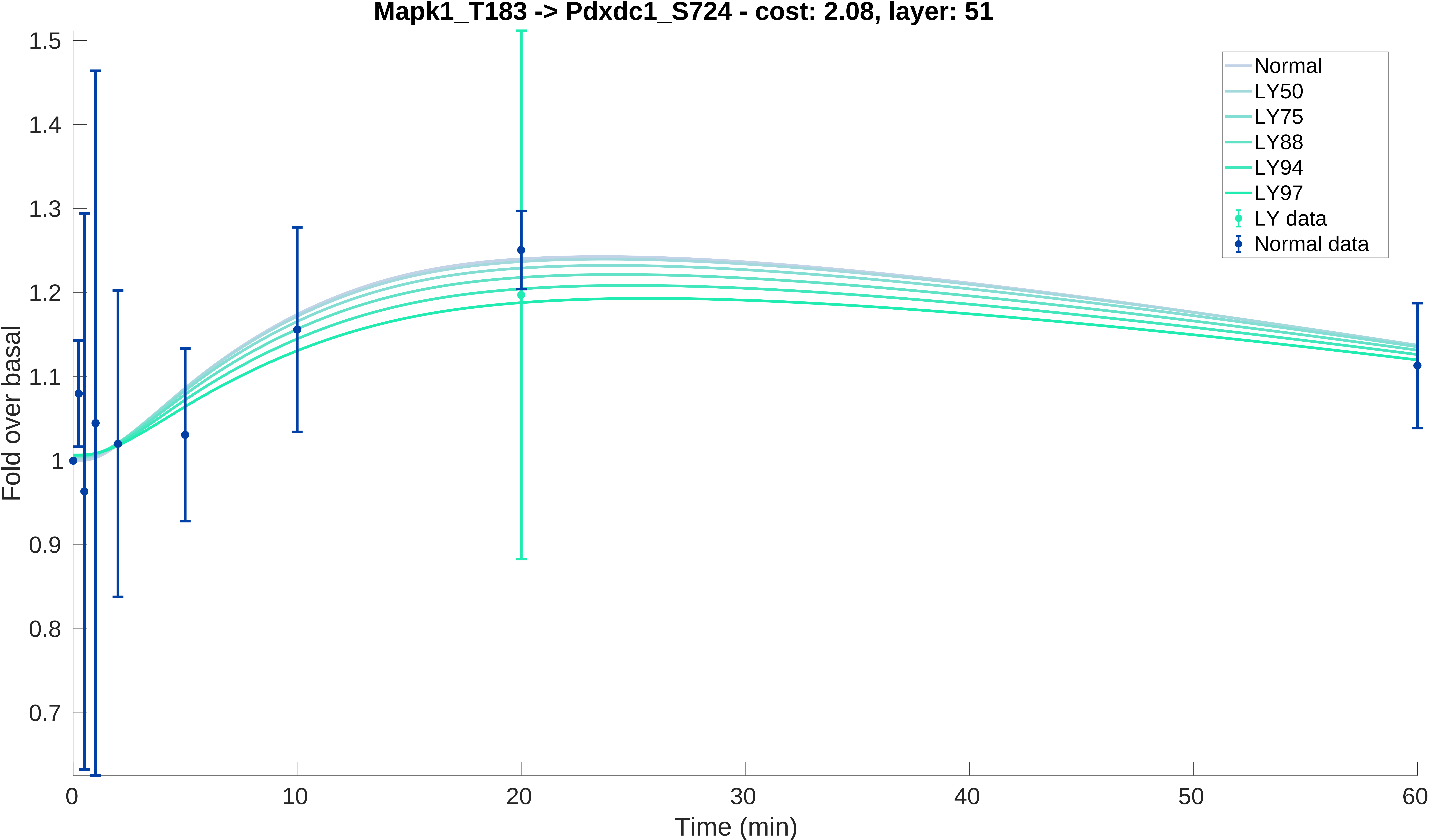

Mapk1_T183 -> Pdxdc1_S724 - cost: 2.08, layer: 51
Fold over basal
Time (min)
1.5
1.4
1.3
1.2
1.1
1
0.9
0.8
0.7
0
10
20
30
40
50
60
Normal
LY50
LY75
LY88
LY94
LY97
LY data
Normal data

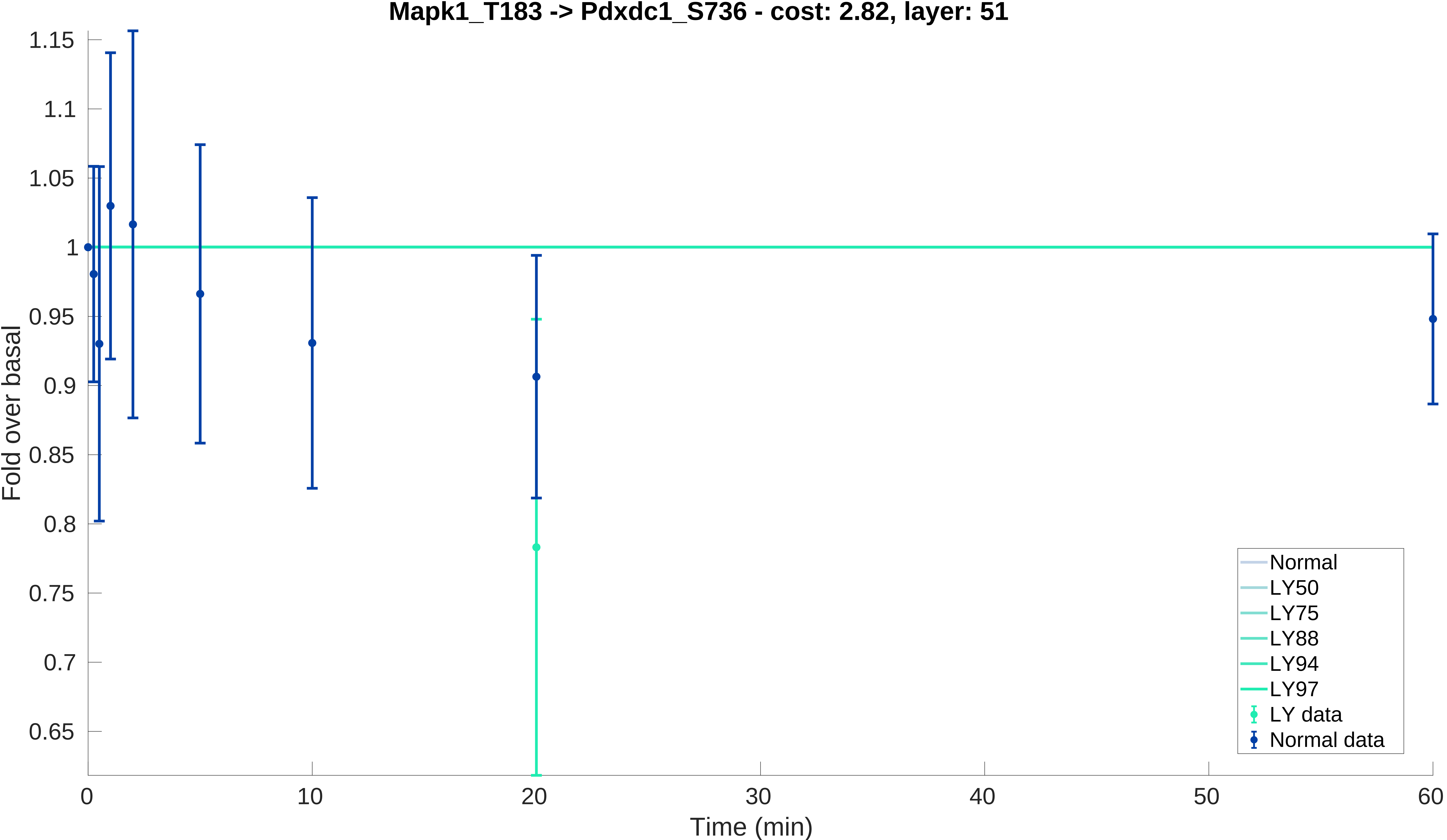

Mapk1_T183 -> Pdxdc1_S736 - cost: 2.82, layer: 51
Fold over basal
1.15
1.1
1.05
1
0.95
0.9
0.85
0.8
0.75
0.7
0.65
0
10
20
30
40
50
60
Time (min)
Normal
LY50
LY75
LY88
LY94
LY97
LY data
Normal data

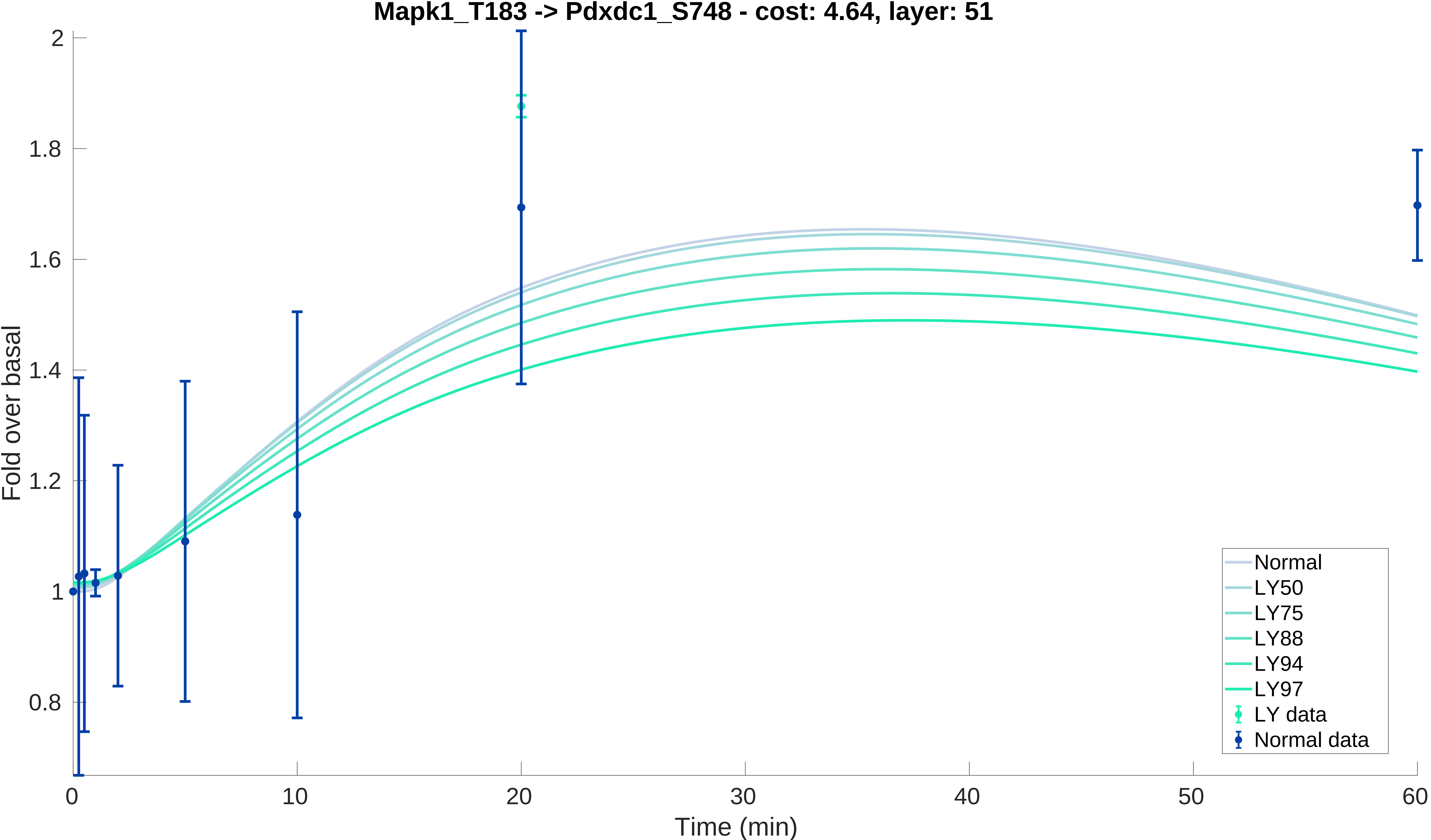

Mapk1_T183 -> Pdxdc1_S748 - cost: 4.64, layer: 51
Fold over basal
Time (min)
0.8
1
1.2
1.4
1.6
1.8
2
0
10
20
30
40
50
60
Normal
LY50
LY75
LY88
LY94
LY97
LY data
Normal data

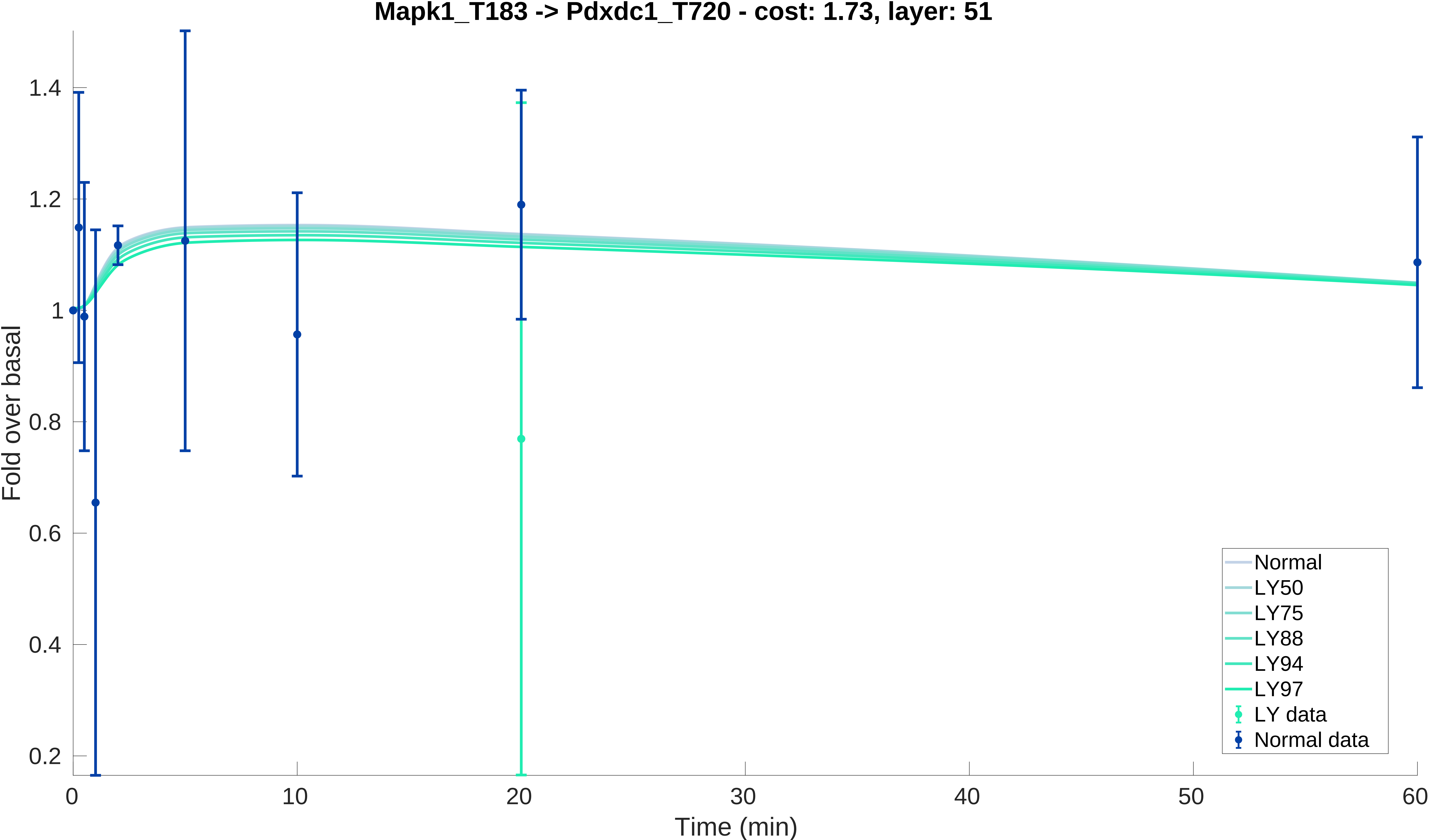

Mapk1_T183 -> Pdxdc1_T720 - cost: 1.73, layer: 51
Fold over basal
Time (min)
0
10
20
30
40
50
60
0.2
0.4
0.6
0.8
1
1.2
1.4
Normal
LY50
LY75
LY88
LY94
LY97
LY data
Normal data

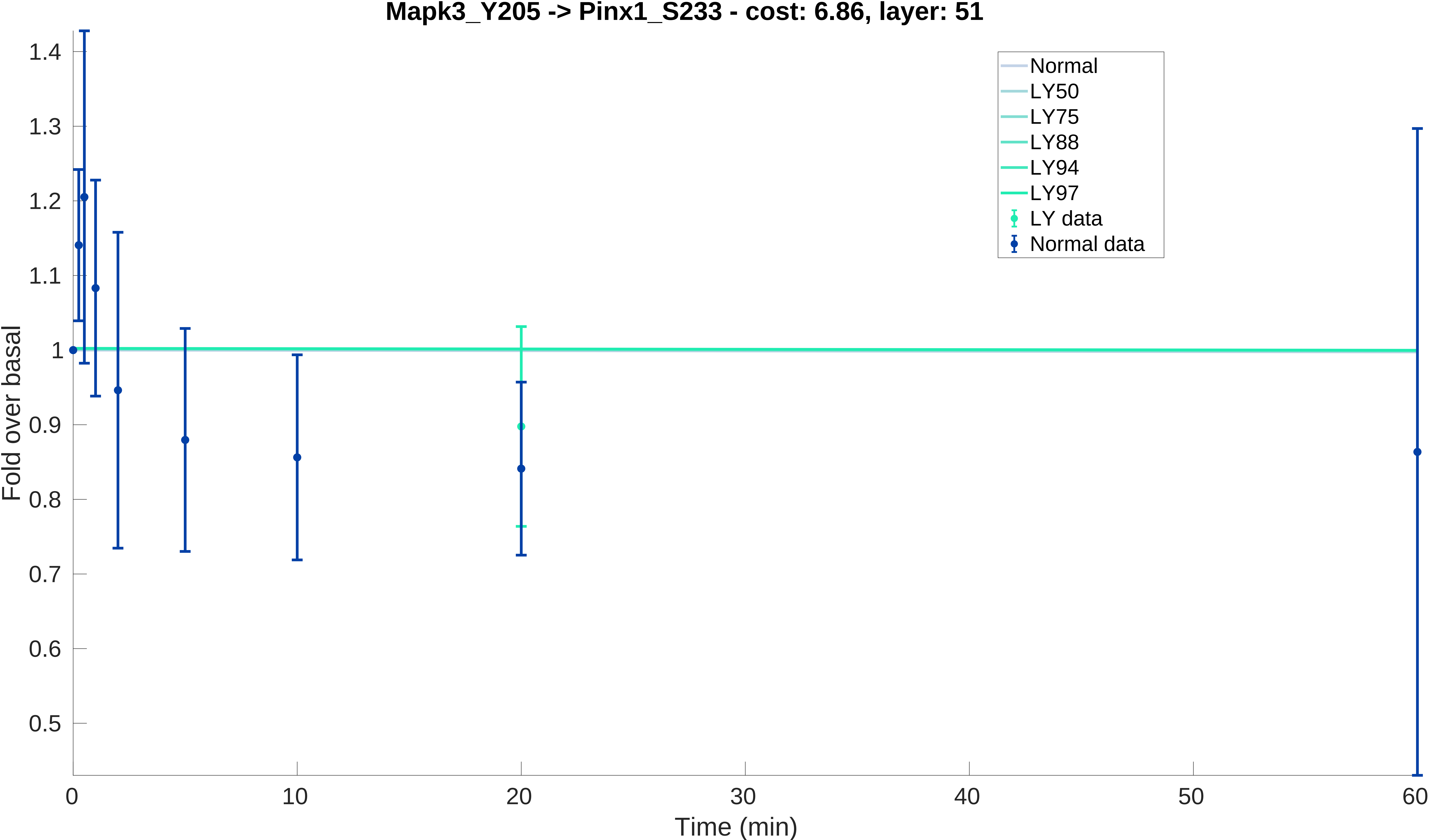
Mapk3_Y205 -> Pinx1_S233 - cost: 6.86, layer: 51
Normal
LY50
LY75
LY88
LY94
LY97
LY data
Normal data
Fold over basal
Time (min)
1.4
1.3
1.2
1.1
1
0.9
0.8
0.7
0.6
0.5
0
10
20
30
40
50
60

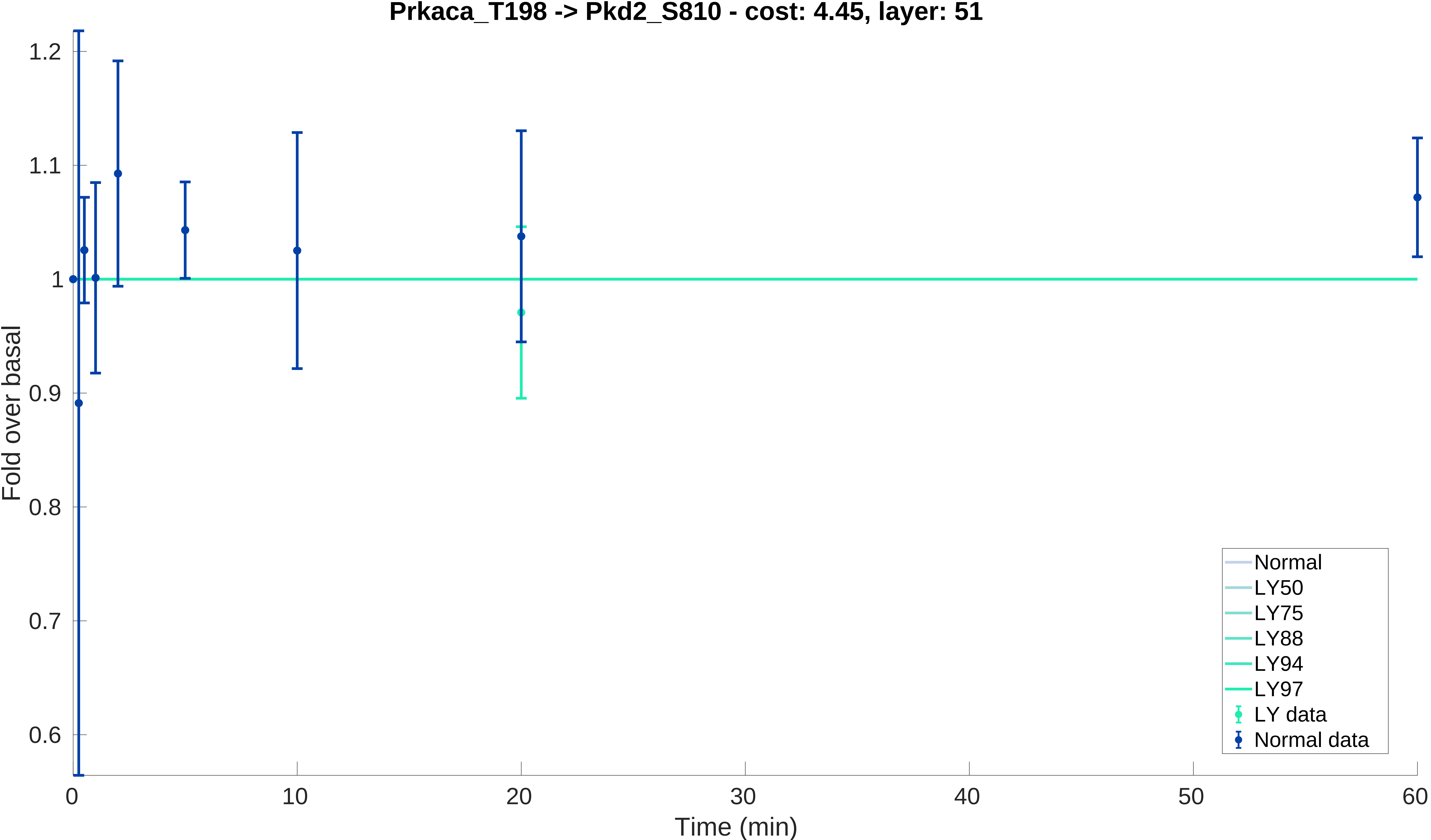

Prkaca_T198 -> Pkd2_S810 - cost: 4.45, layer: 51
Fold over basal
1.2
1.1
1
0.9
0.8
0.7
0.6
0
10
20
30
40
50
60
Time (min)
Normal
LY50
LY75
LY88
LY94
LY97
LY data
Normal data

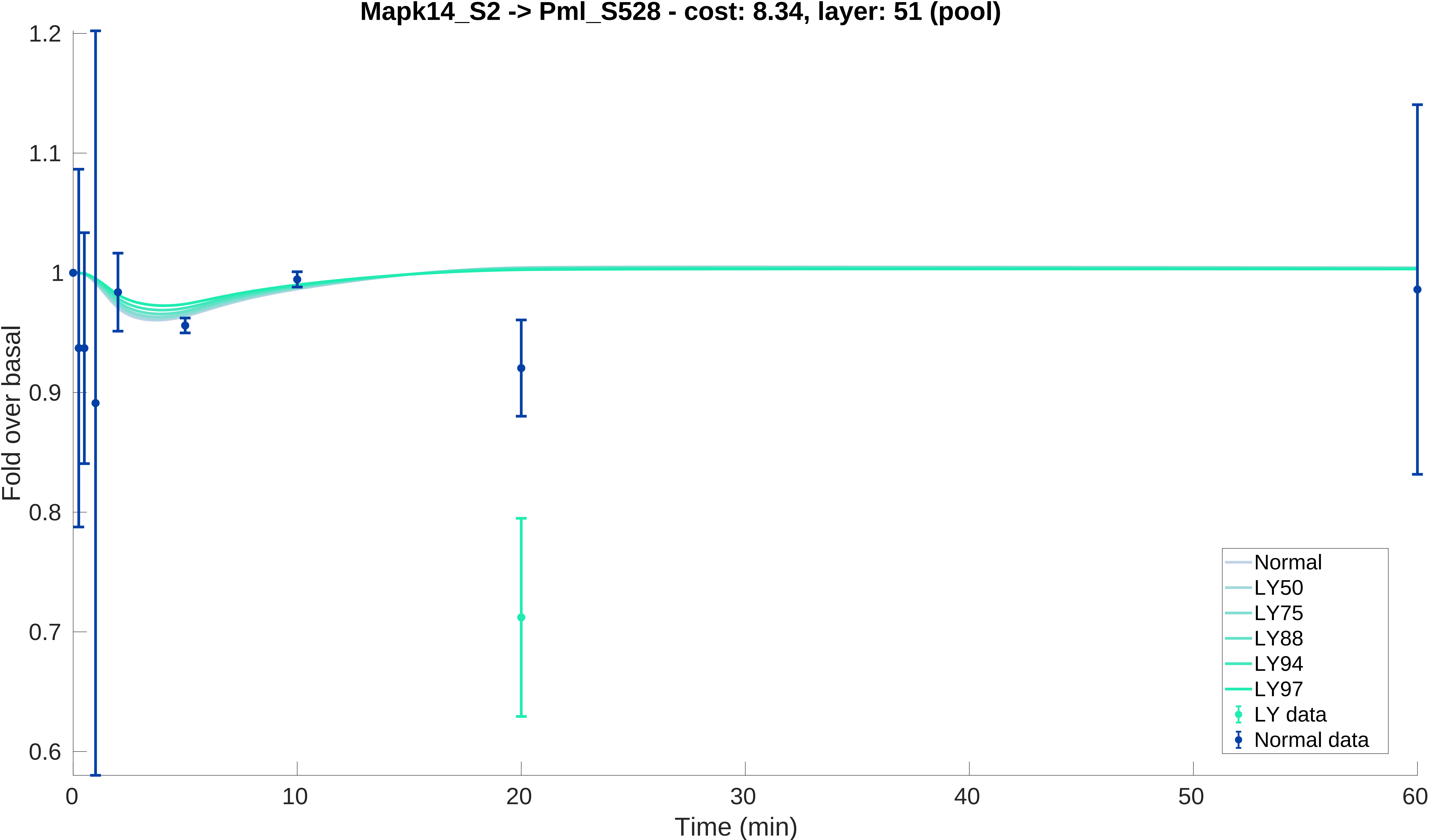

Mapk14_S2 -> Pml_S528 - cost: 8.34, layer: 51 (pool)
Fold over basal
Time (min)
1.2
1.1
1
0.9
0.8
0.7
0.6
0
10
20
30
40
50
60
Normal
LY50
LY75
LY88
LY94
LY97
LY data
Normal data

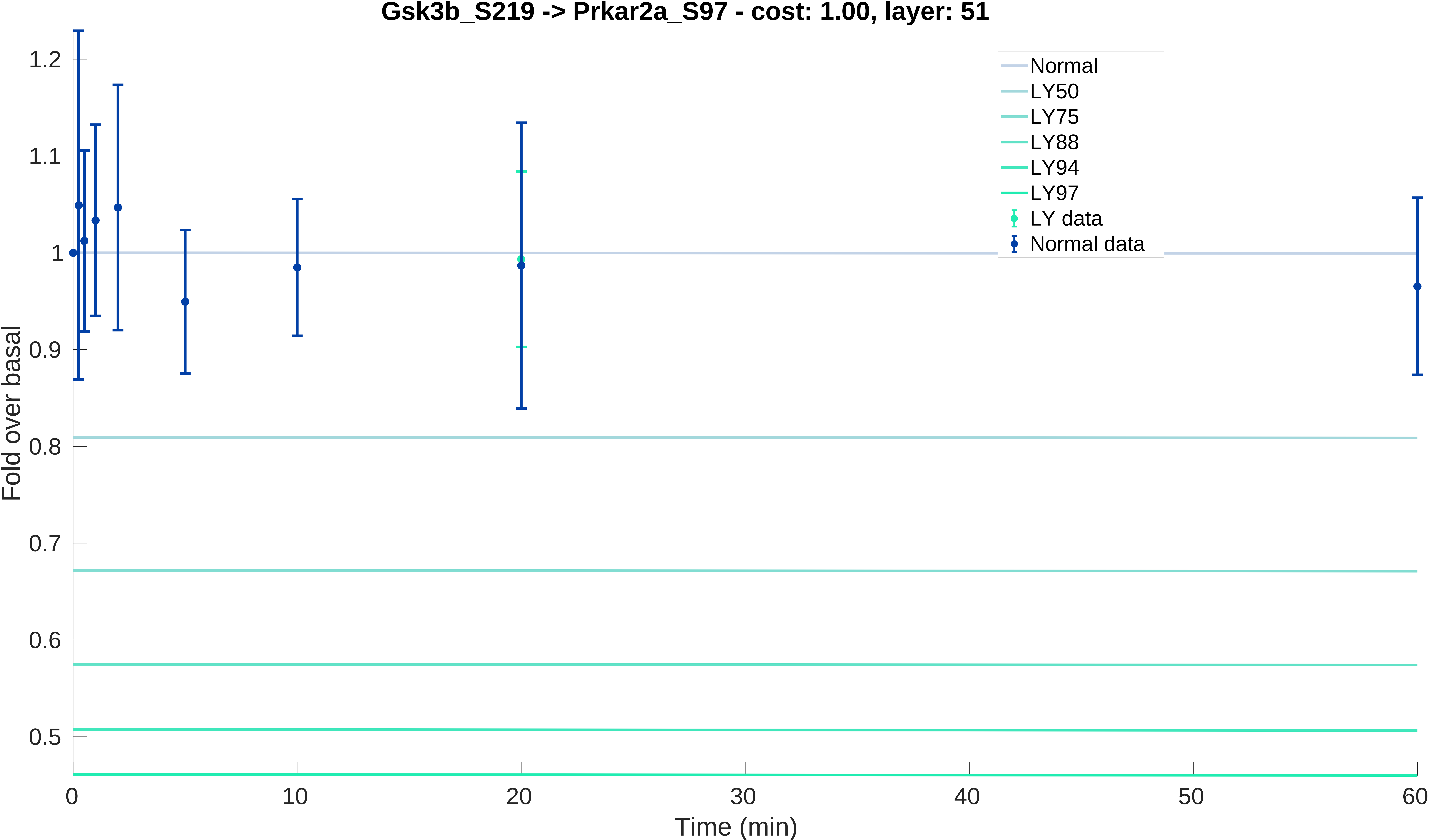

Gsk3b_S219 -> Prkar2a_S97 - cost: 1.00, layer: 51
Fold over basal
Time (min)
1.2
1.1
1
0.9
0.8
0.7
0.6
0.5
0
10
20
30
40
50
60
Normal
LY50
LY75
LY88
LY94
LY97
LY data
Normal data

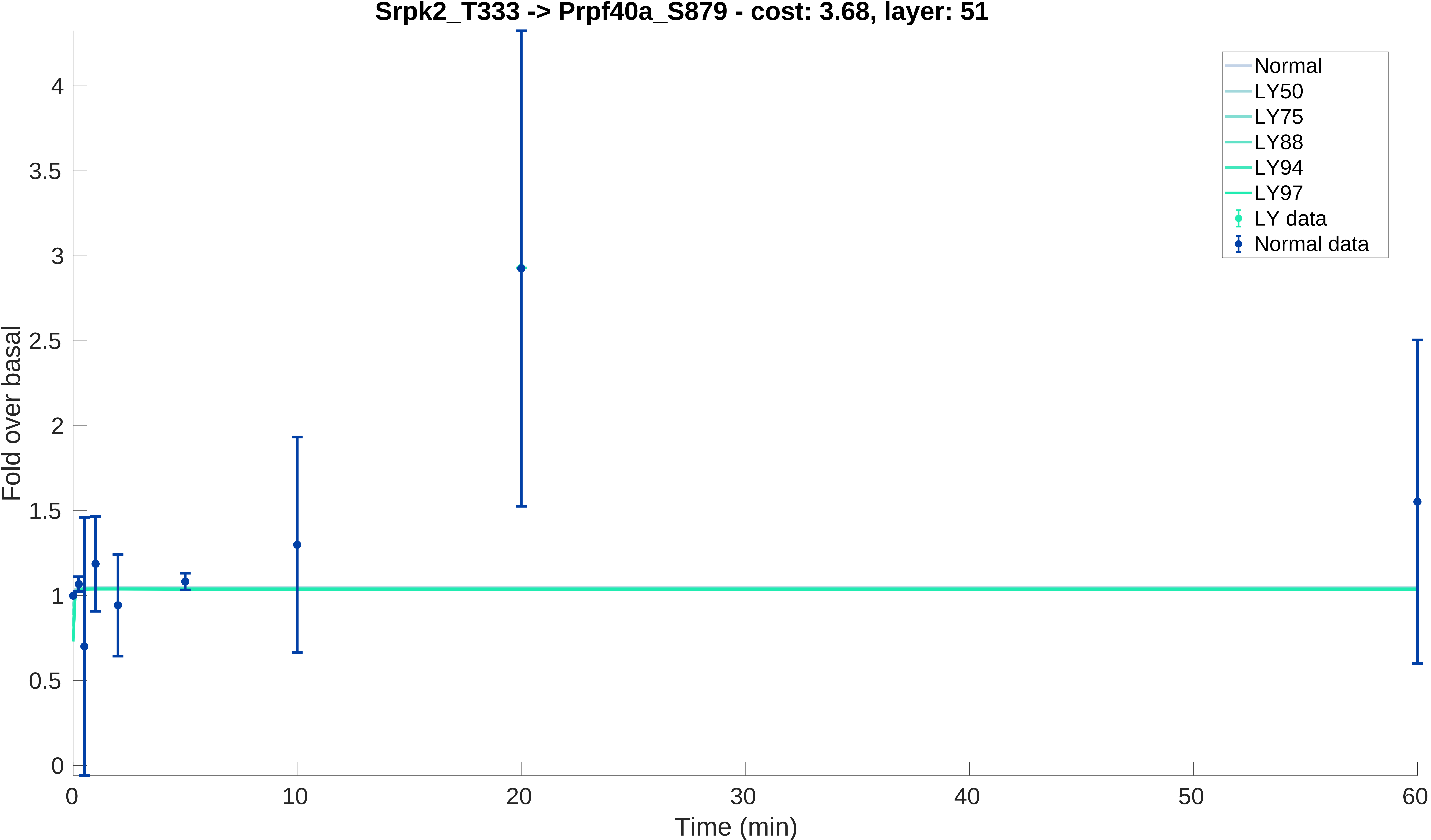

Srpk2_T333 -> Prpf40a_S879 - cost: 3.68, layer: 51
Fold over basal
Time (min)
0
0.5
1
1.5
2
2.5
3
3.5
4
0
10
20
30
40
50
60
Normal
LY50
LY75
LY88
LY94
LY97
LY data
Normal data

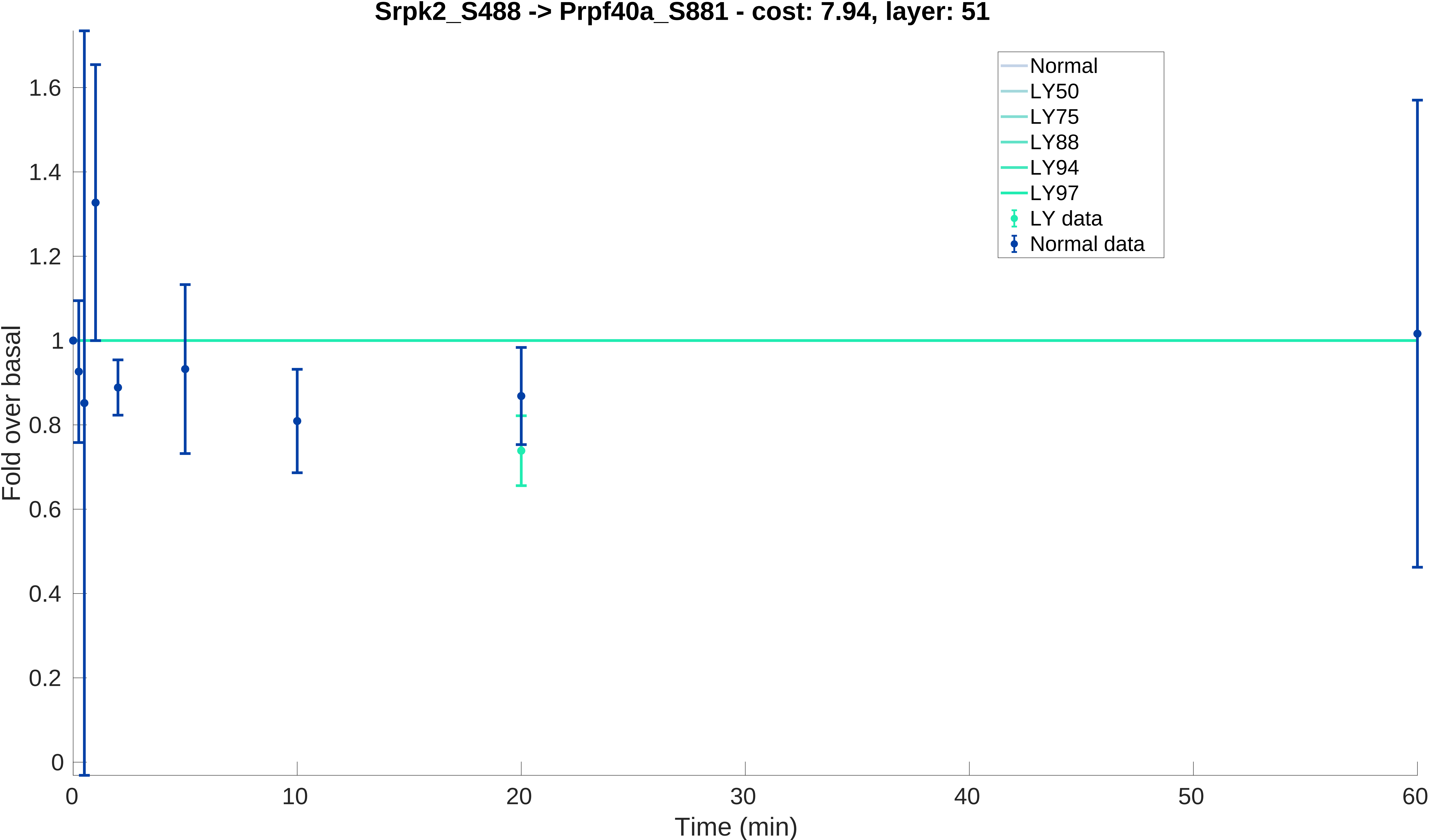

Srpk2_S488 -> Prpf40a_S881 - cost: 7.94, layer: 51
Normal
LY50
LY75
LY88
LY94
LY97
LY data
Normal data
Fold over basal
Time (min)
0
0.2
0.4
0.6
0.8
1
1.2
1.4
1.6
0
10
20
30
40
50
60

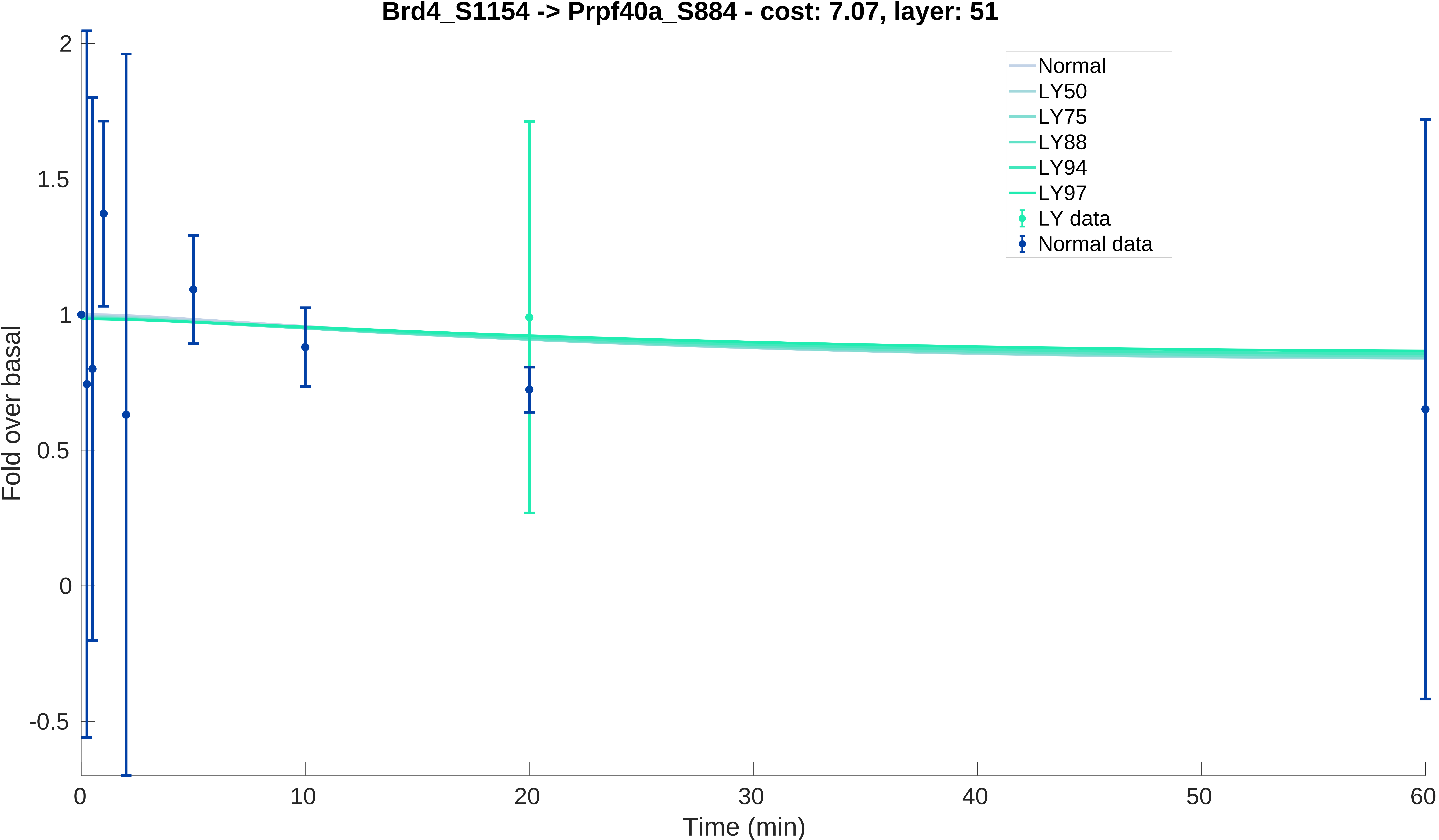

Brd4_S1154 -> Prpf40a_S884 - cost: 7.07, layer: 51
Normal
LY50
LY75
LY88
LY94
LY97
LY data
Normal data
Fold over basal
Time (min)
2
1.5
1
0.5
0
-0.5
0
10
20
30
40
50
60

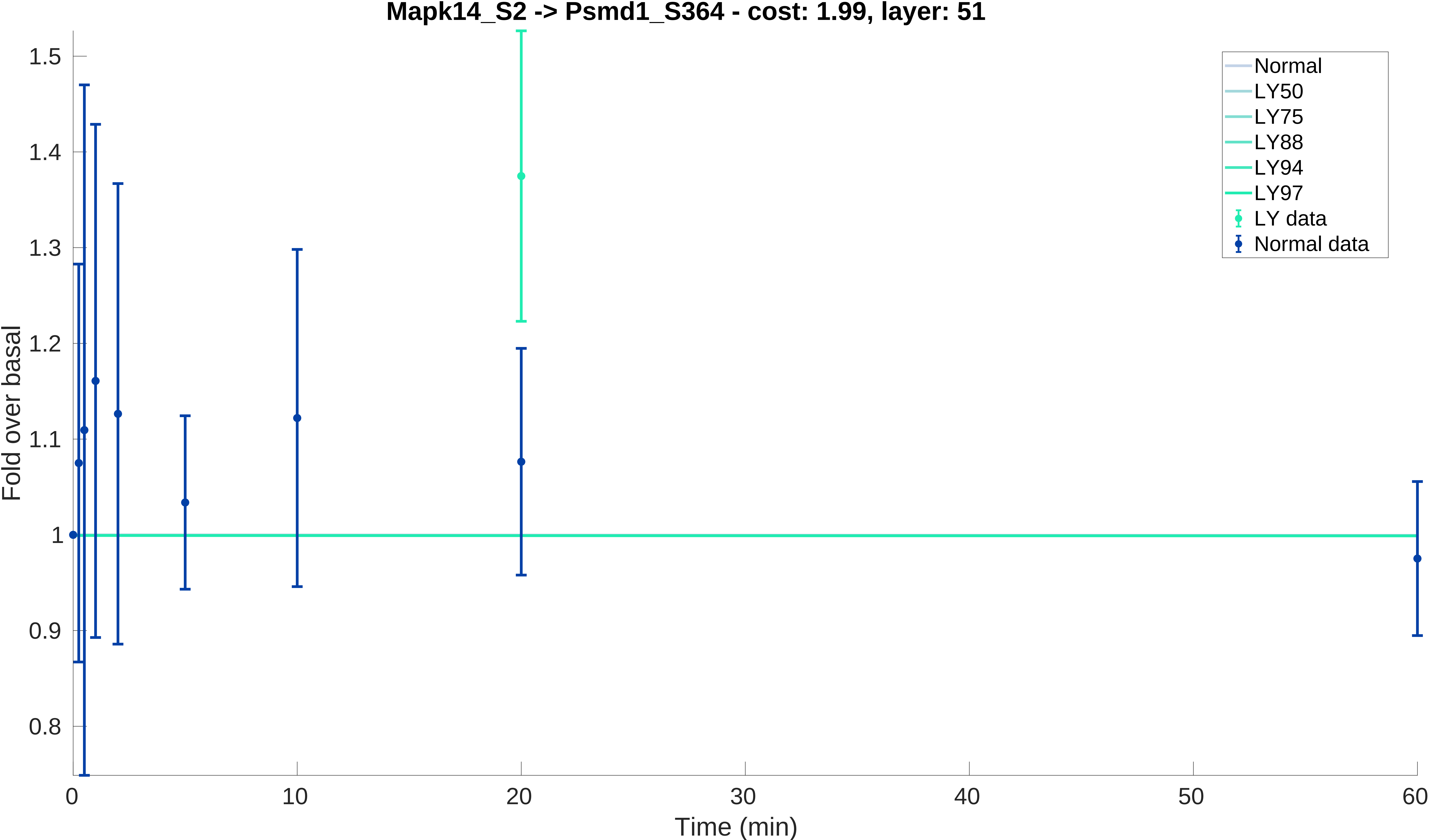
Mapk14_S2 -> Psmd1_S364 - cost: 1.99, layer: 51
Fold over basal
1.5
1.4
1.3
1.2
1.1
1
0.9
0.8
0
10
20
30
40
50
60
Time (min)
Normal
LY50
LY75
LY88
LY94
LY97
LY data
Normal data

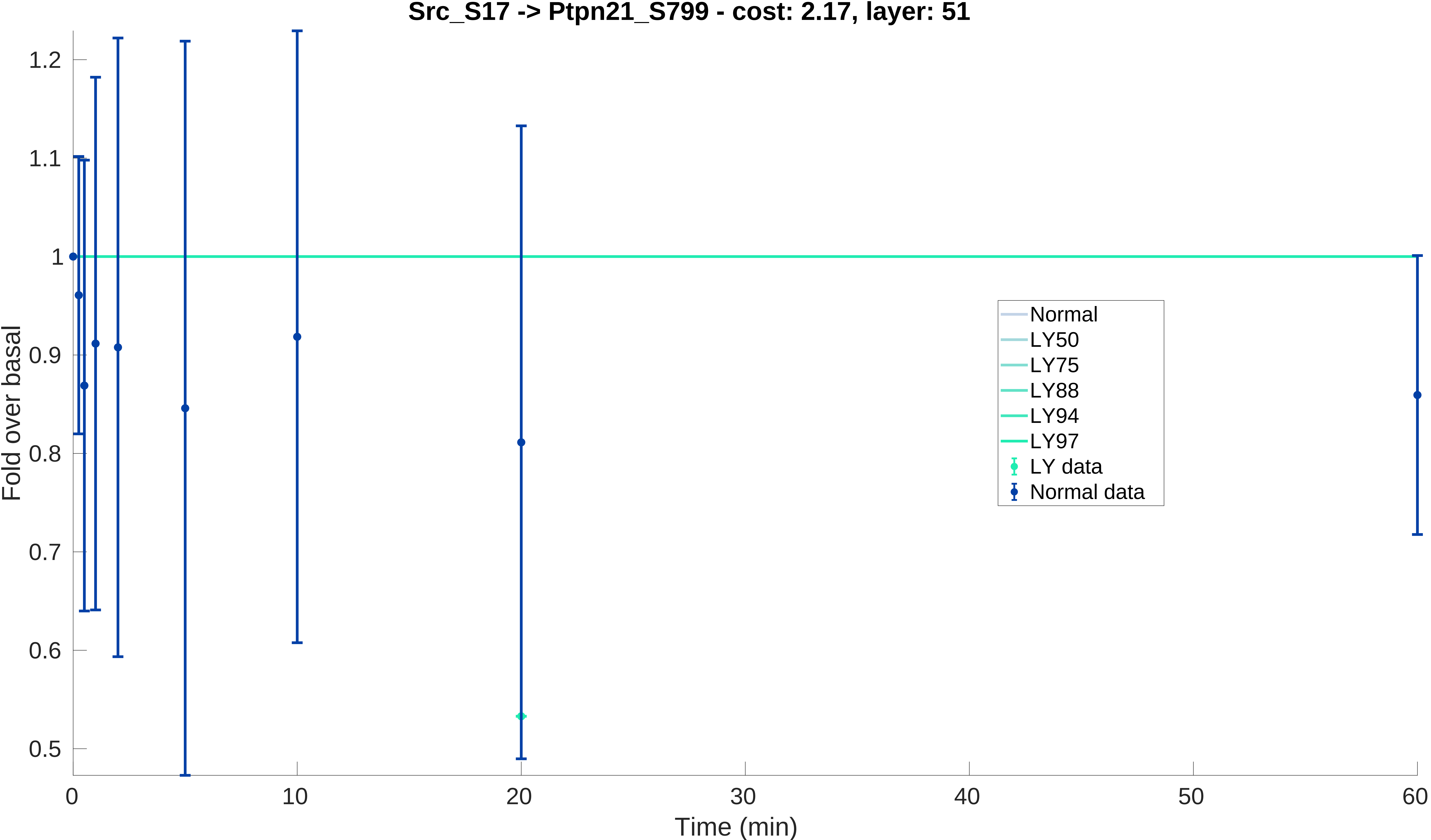

Src_S17 -> Ptpn21_S799 - cost: 2.17, layer: 51
Fold over basal
1.2
1.1
1
0.9
0.8
0.7
0.6
0.5
0
10
20
30
40
50
60
Time (min)
Normal
LY50
LY75
LY88
LY94
LY97
LY data
Normal data

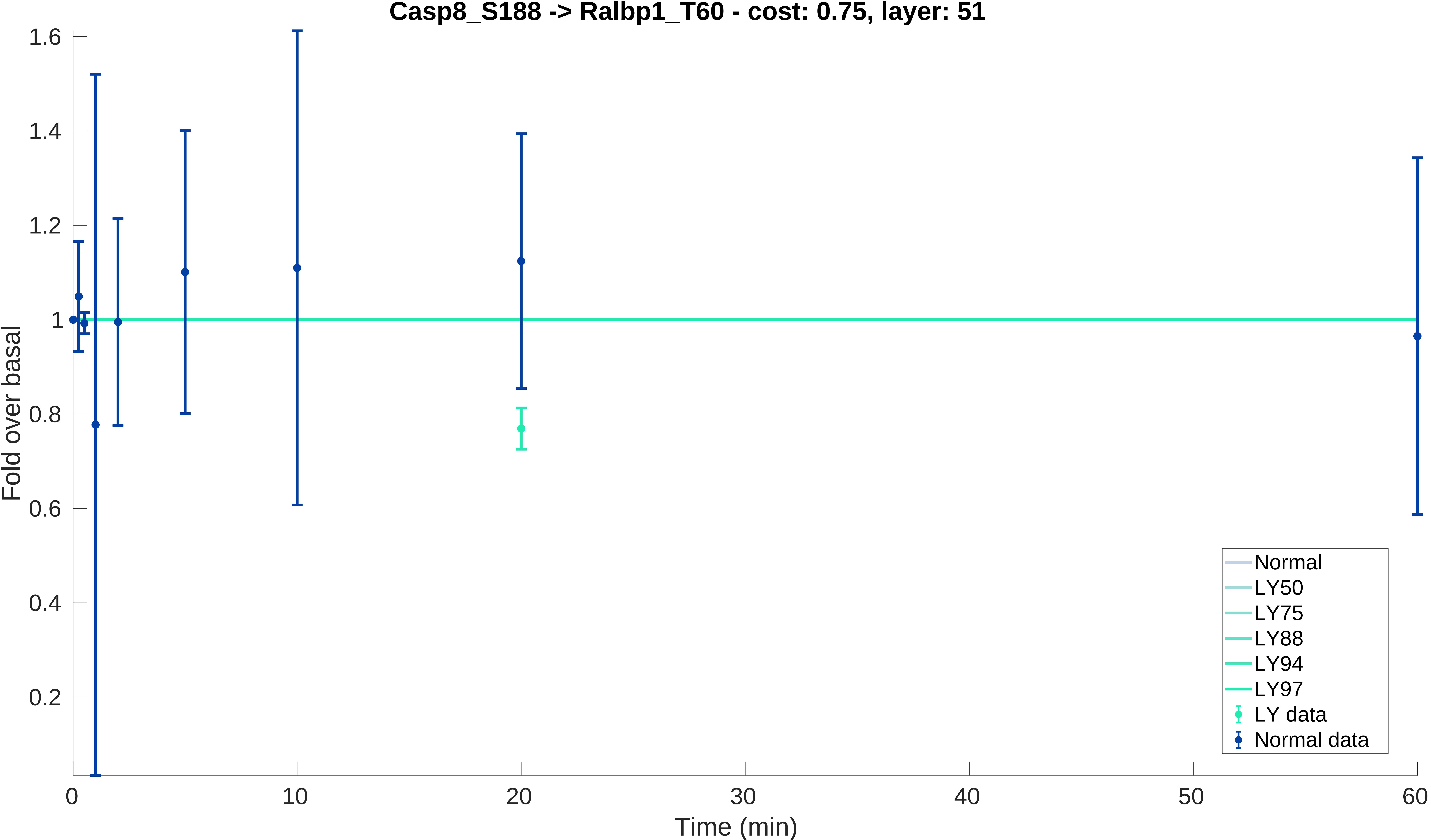

Casp8_S188 -> Ralbp1_T60 - cost: 0.75, layer: 51
Fold over basal
Time (min)
0
10
20
30
40
50
60
0.2
0.4
0.6
0.8
1
1.2
1.4
1.6
Normal
LY50
LY75
LY88
LY94
LY97
LY data
Normal data

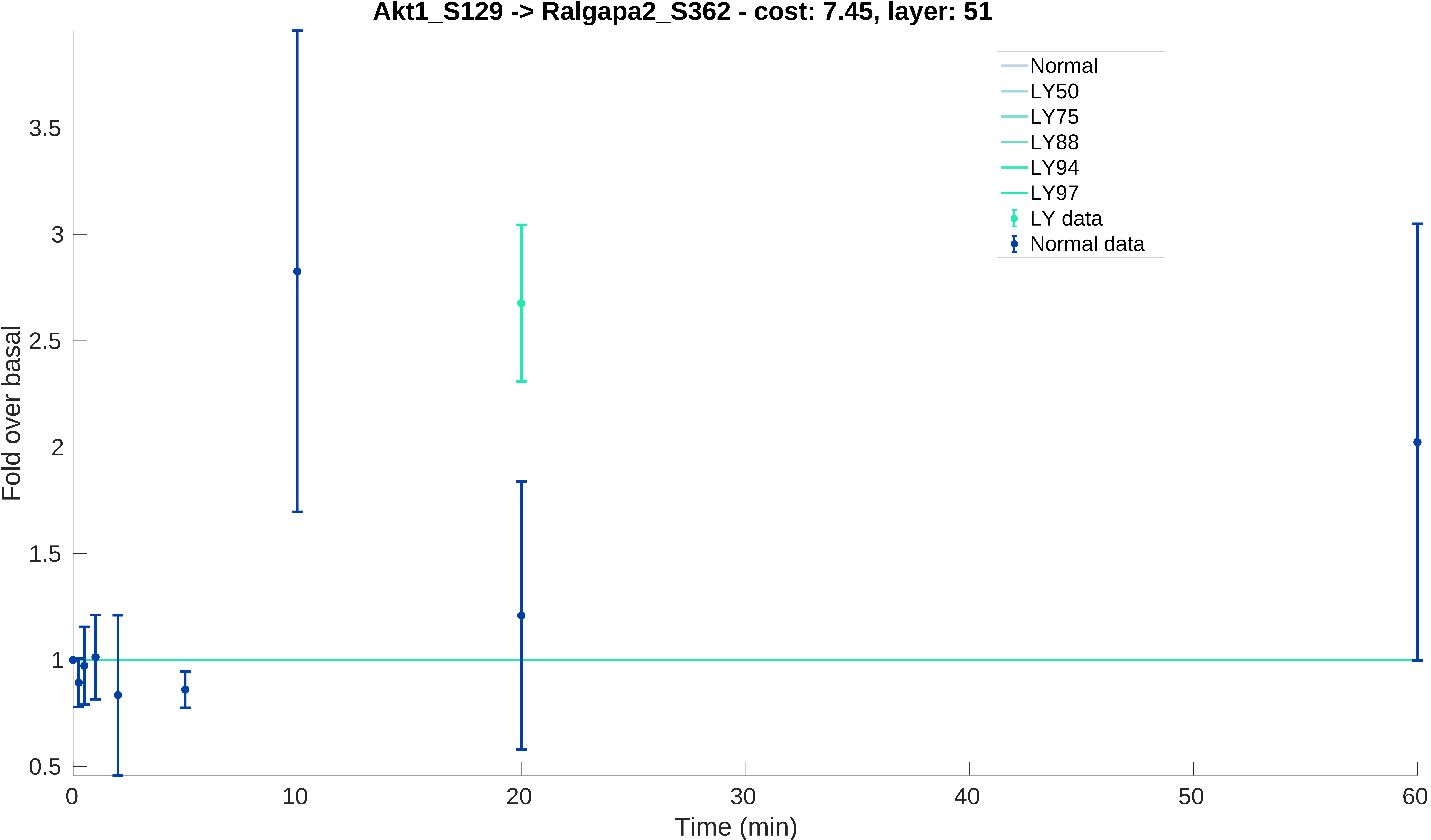
Akt1_S129 -> Ralgapa2_S362 - cost: 7.45, layer: 51
Normal
LY50
LY75
LY88
LY94
LY97
LY data
Normal data
Fold over basal
0.5
1
1.5
2
2.5
3
3.5
0
10
20
30
40
50
60
Time (min)

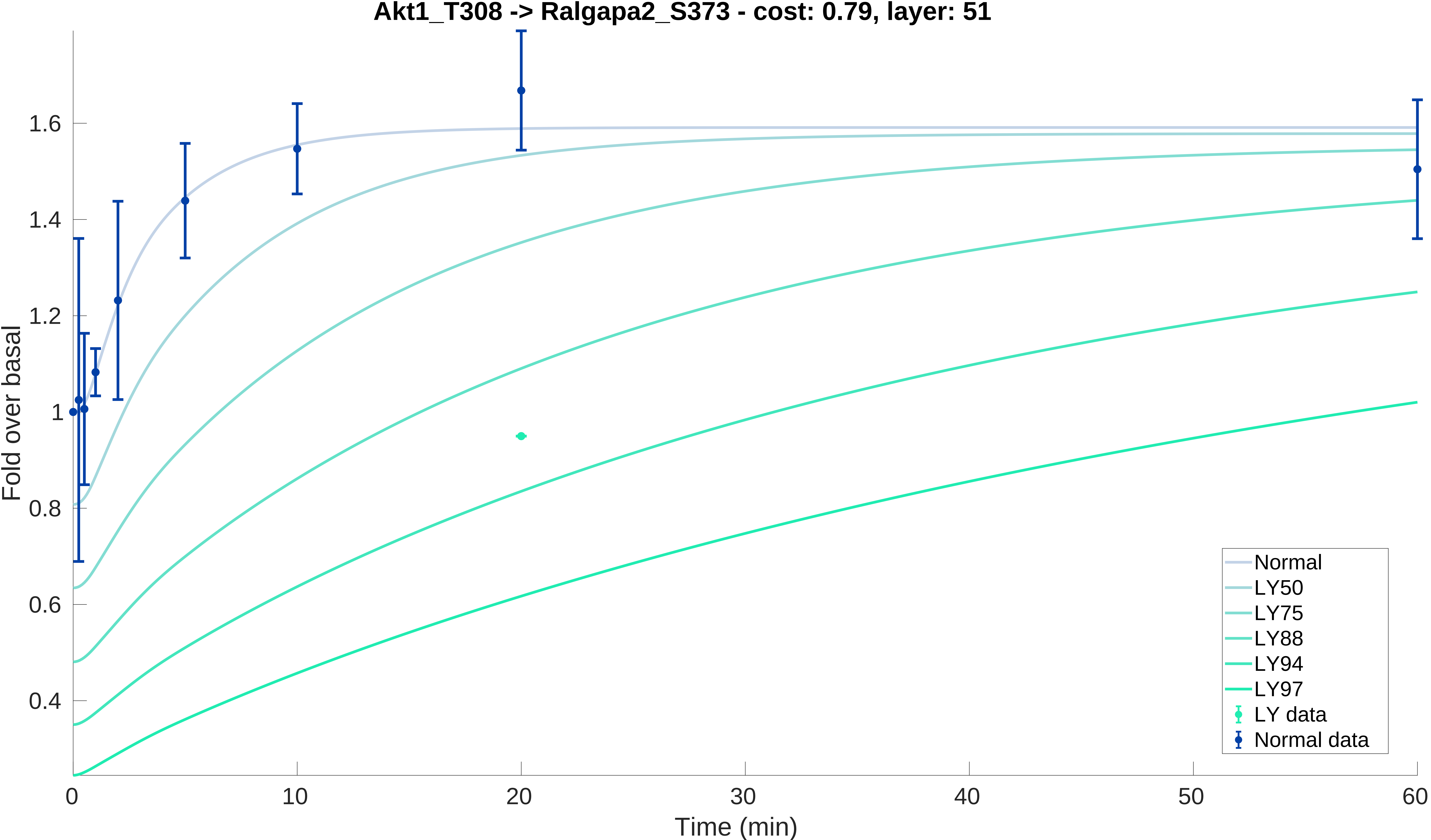
Akt1_T308 -> Ralgapa2_S373 - cost: 0.79, layer: 51
Fold over basal
Time (min)
0.4
0.6
0.8
1
1.2
1.4
1.6
0
10
20
30
40
50
60
Normal
LY50
LY75
LY88
LY94
LY97
LY data
Normal data

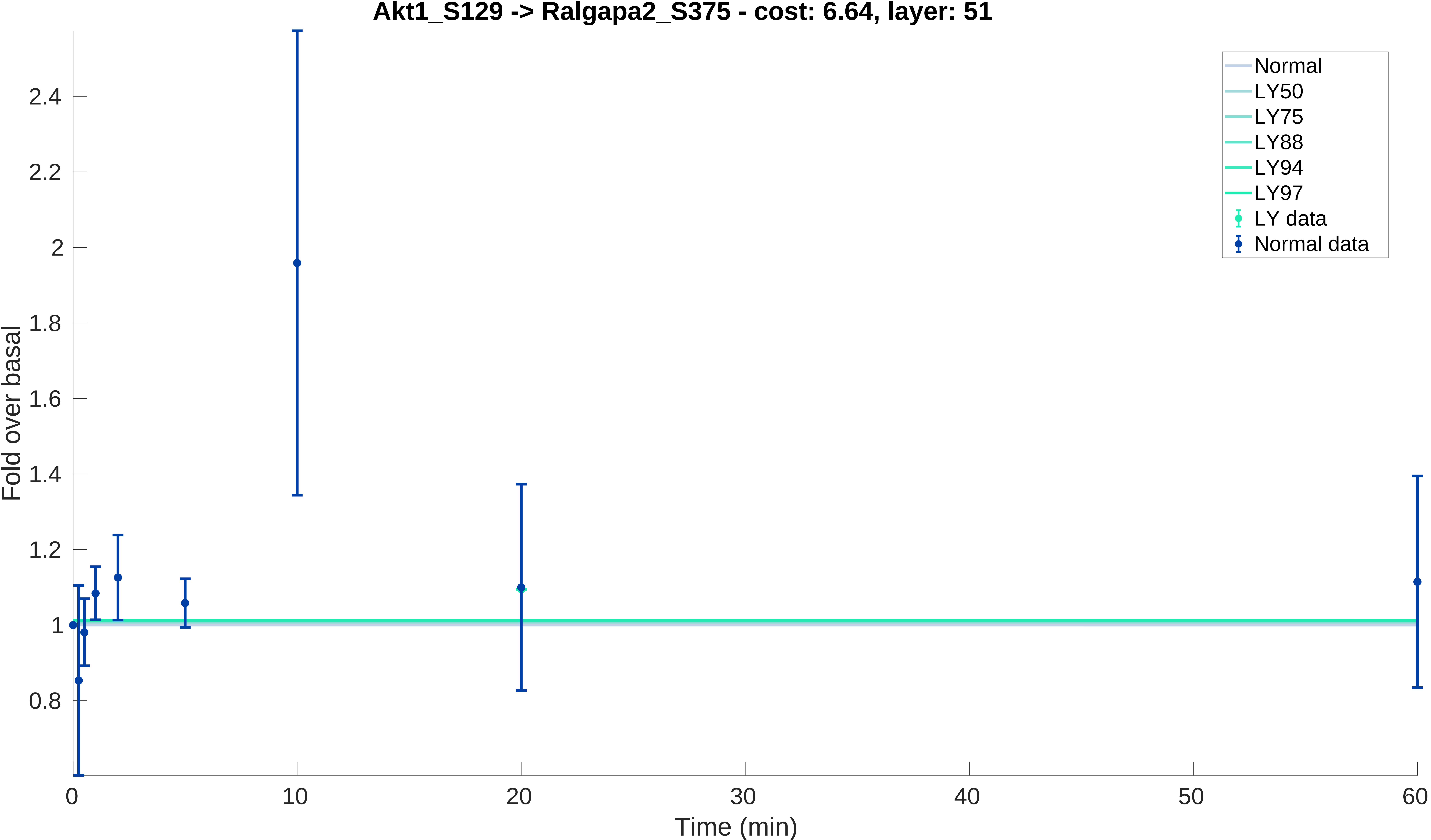

Akt1_S129 -> Ralgapa2_S375 - cost: 6.64, layer: 51
Fold over basal
Time (min)
0.8
1
1.2
1.4
1.6
1.8
2
2.2
2.4
0
10
20
30
40
50
60
Normal
LY50
LY75
LY88
LY94
LY97
LY data
Normal data

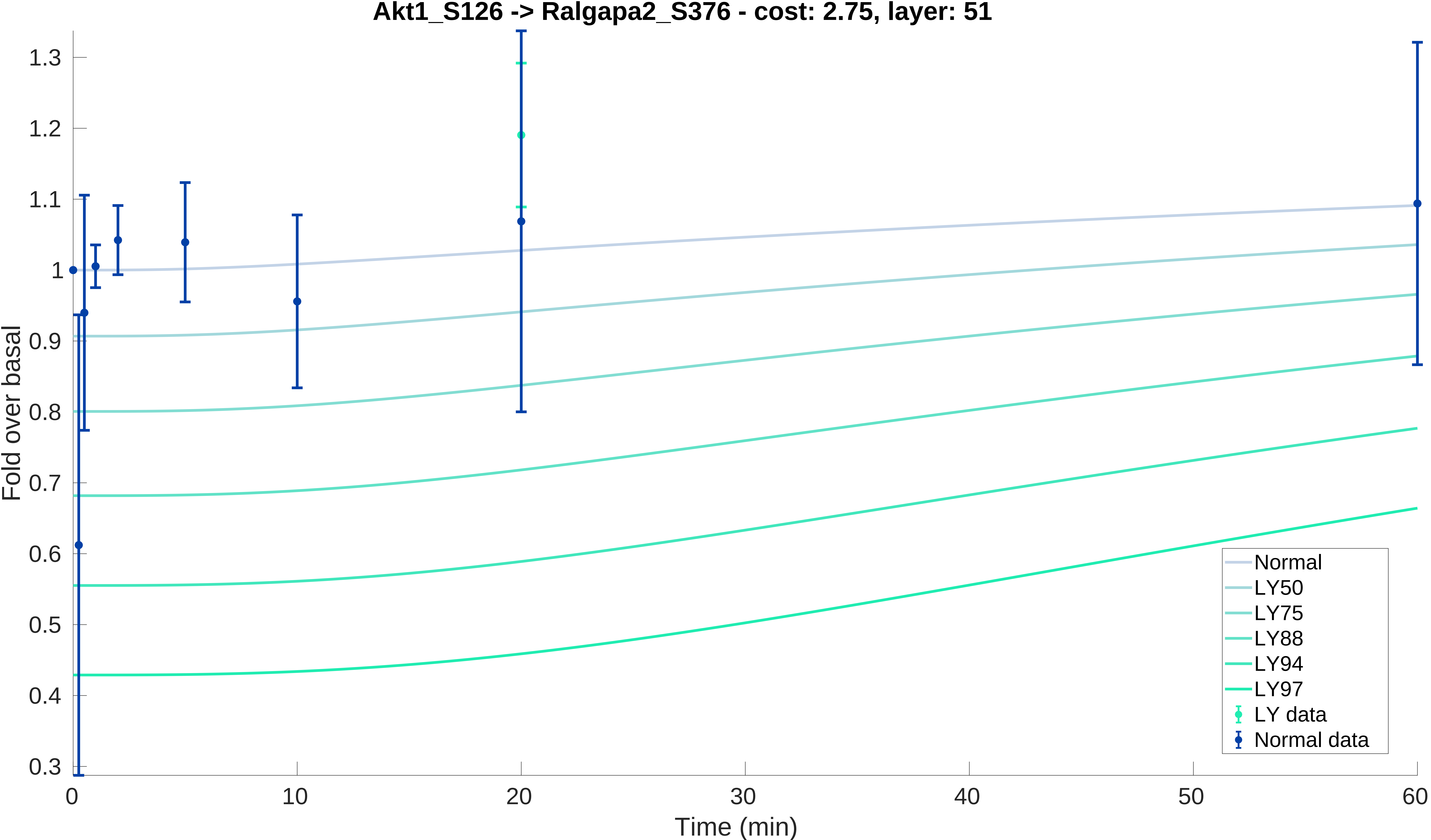

Akt1_S126 -> Ralgapa2_S376 - cost: 2.75, layer: 51
Fold over basal
Time (min)
0.3
0.4
0.5
0.6
0.7
0.8
0.9
1
1.1
1.2
1.3
0
10
20
30
40
50
60
Normal
LY50
LY75
LY88
LY94
LY97
LY data
Normal data

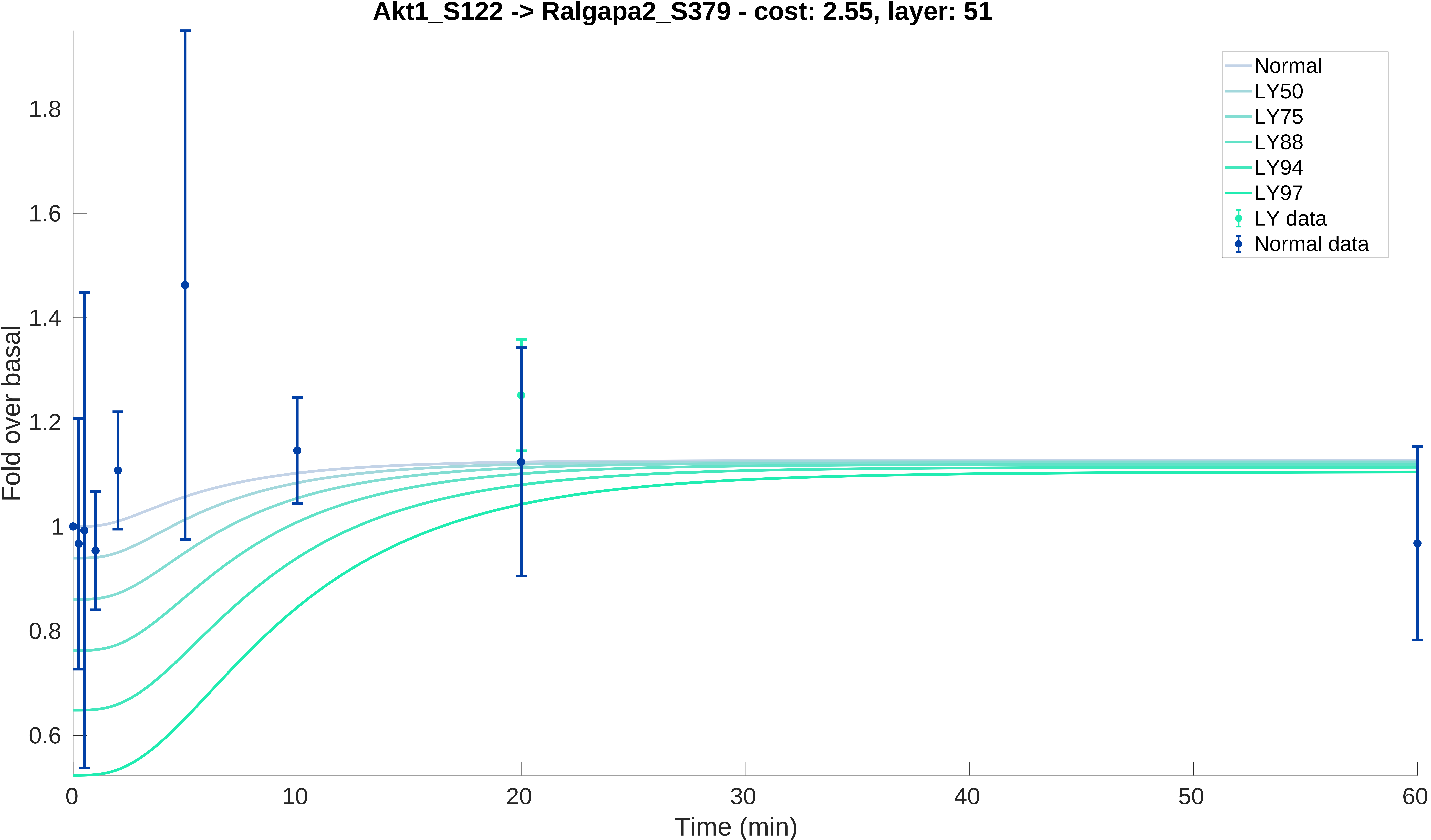

Akt1_S122 -> Ralgapa2_S379 - cost: 2.55, layer: 51
Fold over basal
Time (min)
0.6
0.8
1
1.2
1.4
1.6
1.8
0
10
20
30
40
50
60
Normal
LY50
LY75
LY88
LY94
LY97
LY data
Normal data

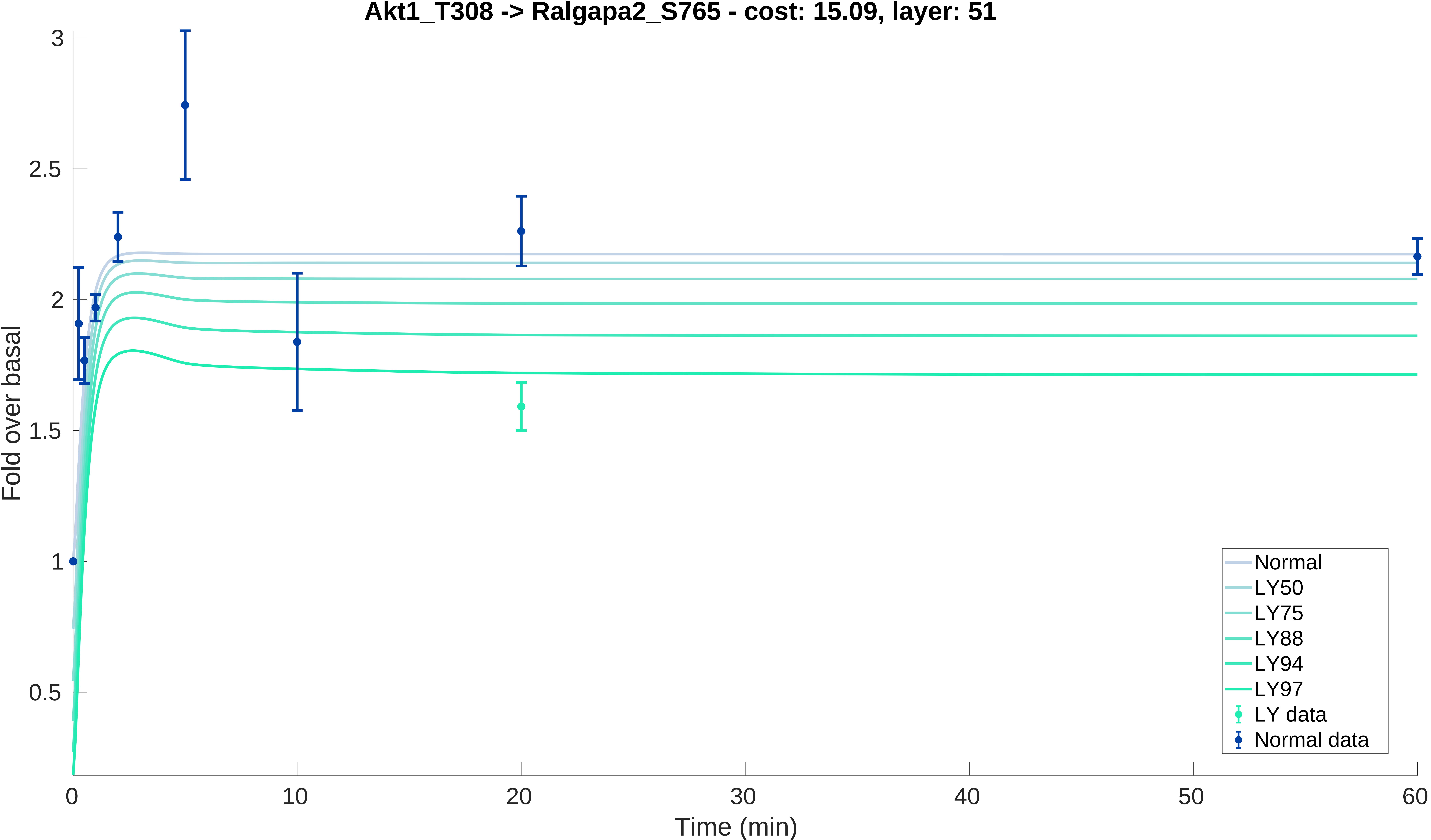

Akt1_T308 -> Ralgapa2_S765 - cost: 15.09, layer: 51
Fold over basal
Time (min)
0
10
20
30
40
50
60
0.5
1
1.5
2
2.5
3
Normal
LY50
LY75
LY88
LY94
LY97
LY data
Normal data

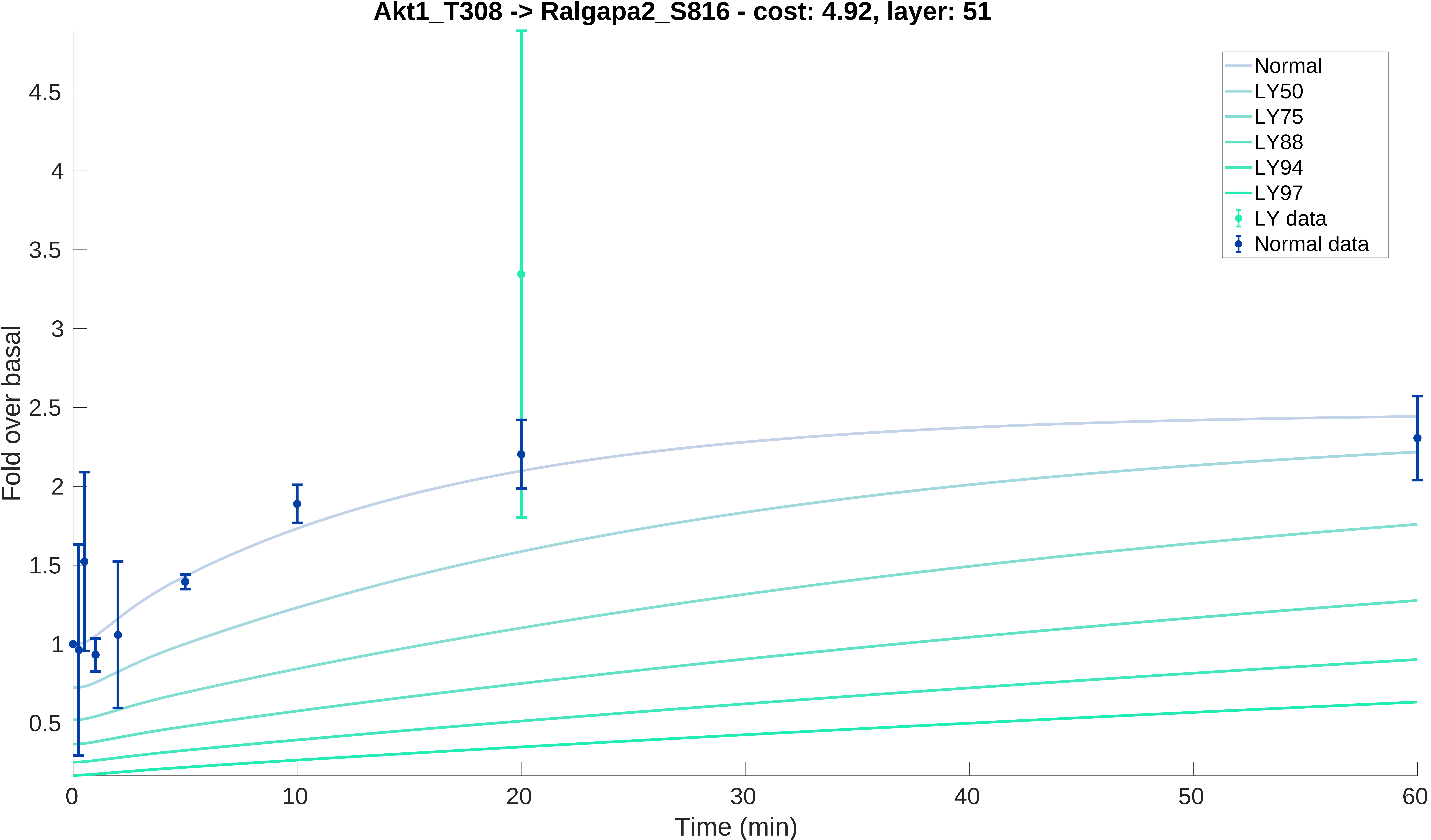

Akt1_T308 -> Ralgapa2_S816 - cost: 4.92, layer: 51
Fold over basal
Time (min)
0.5
1
1.5
2
2.5
3
3.5
4
4.5
0
10
20
30
40
50
60
Normal
LY50
LY75
LY88
LY94
LY97
LY data
Normal data

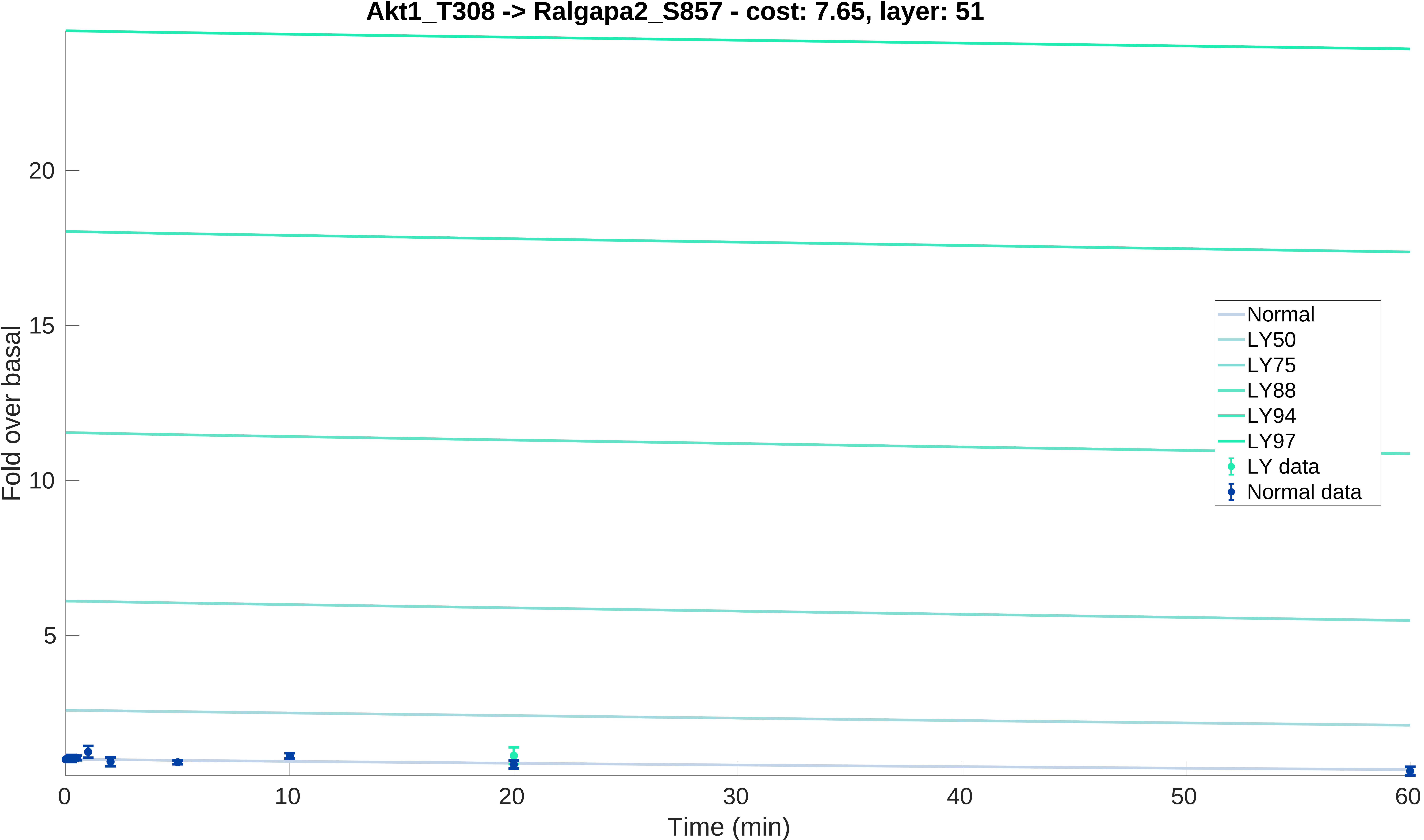

Akt1_T308 -> Ralgapa2_S857 - cost: 7.65, layer: 51
Fold over basal
20
15
10
5
0
10
20
30
40
50
60
Time (min)
Normal
LY50
LY75
LY88
LY94
LY97
LY data
Normal data

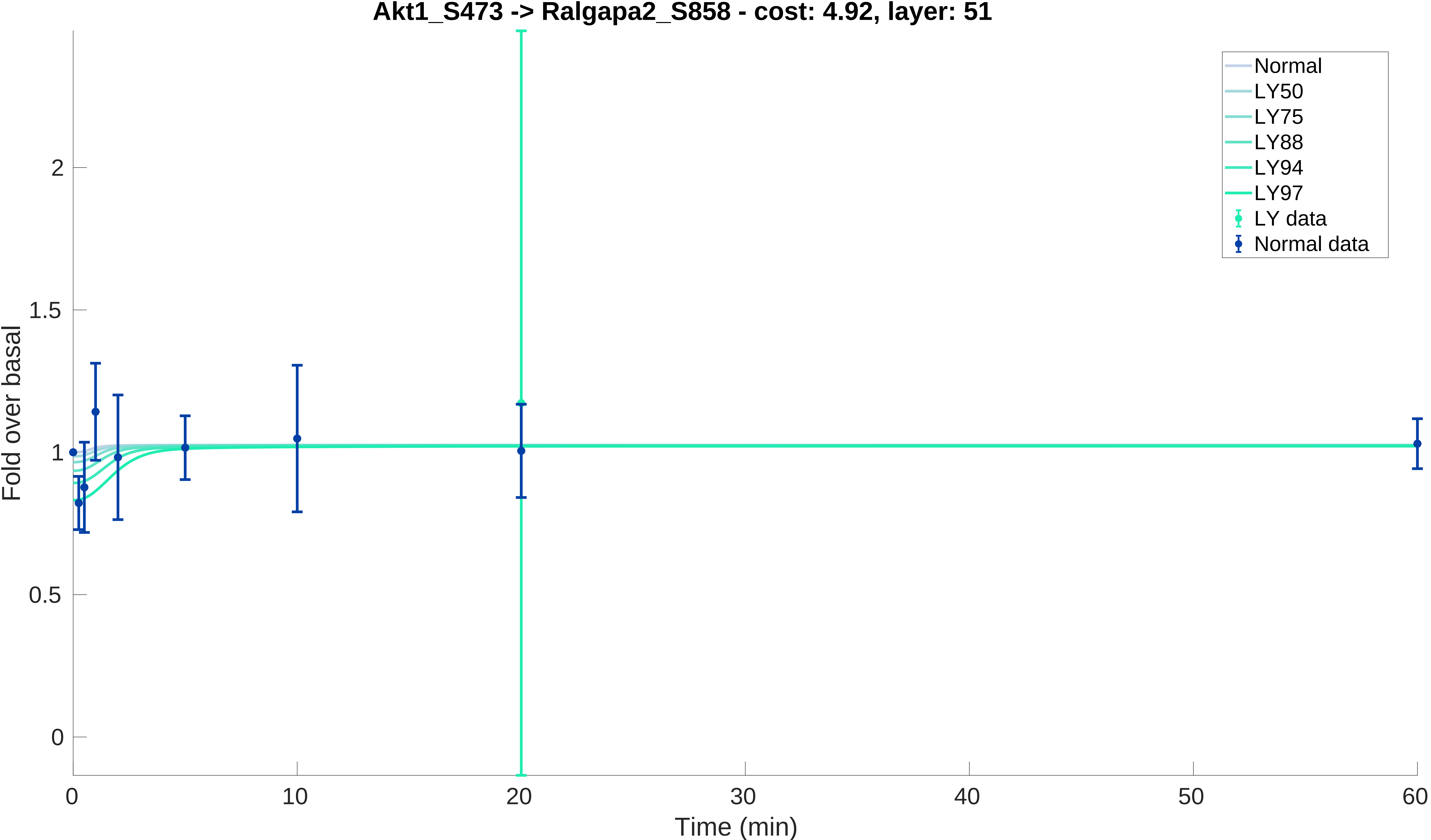

Akt1_S473 -> Ralgapa2_S858 - cost: 4.92, layer: 51
Fold over basal
Time (min)
0
0.5
1
1.5
2
0
10
20
30
40
50
60
Normal
LY50
LY75
LY88
LY94
LY97
LY data
Normal data

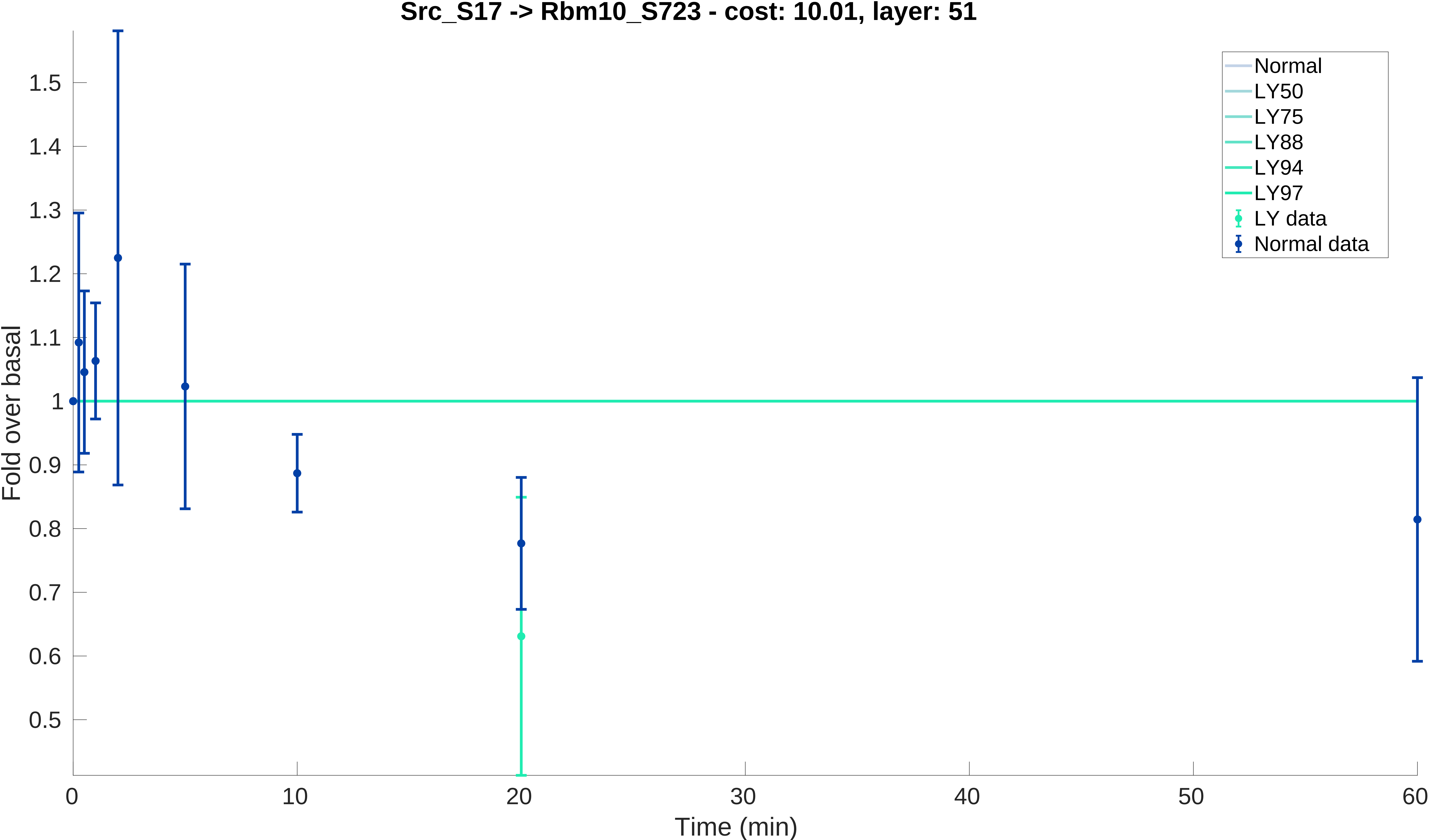

Src_S17 -> Rbm10_S723 - cost: 10.01, layer: 51
Fold over basal
Time (min)
0
10
20
30
40
50
60
0.5
0.6
0.7
0.8
0.9
1
1.1
1.2
1.3
1.4
1.5
Normal
LY50
LY75
LY88
LY94
LY97
LY data
Normal data

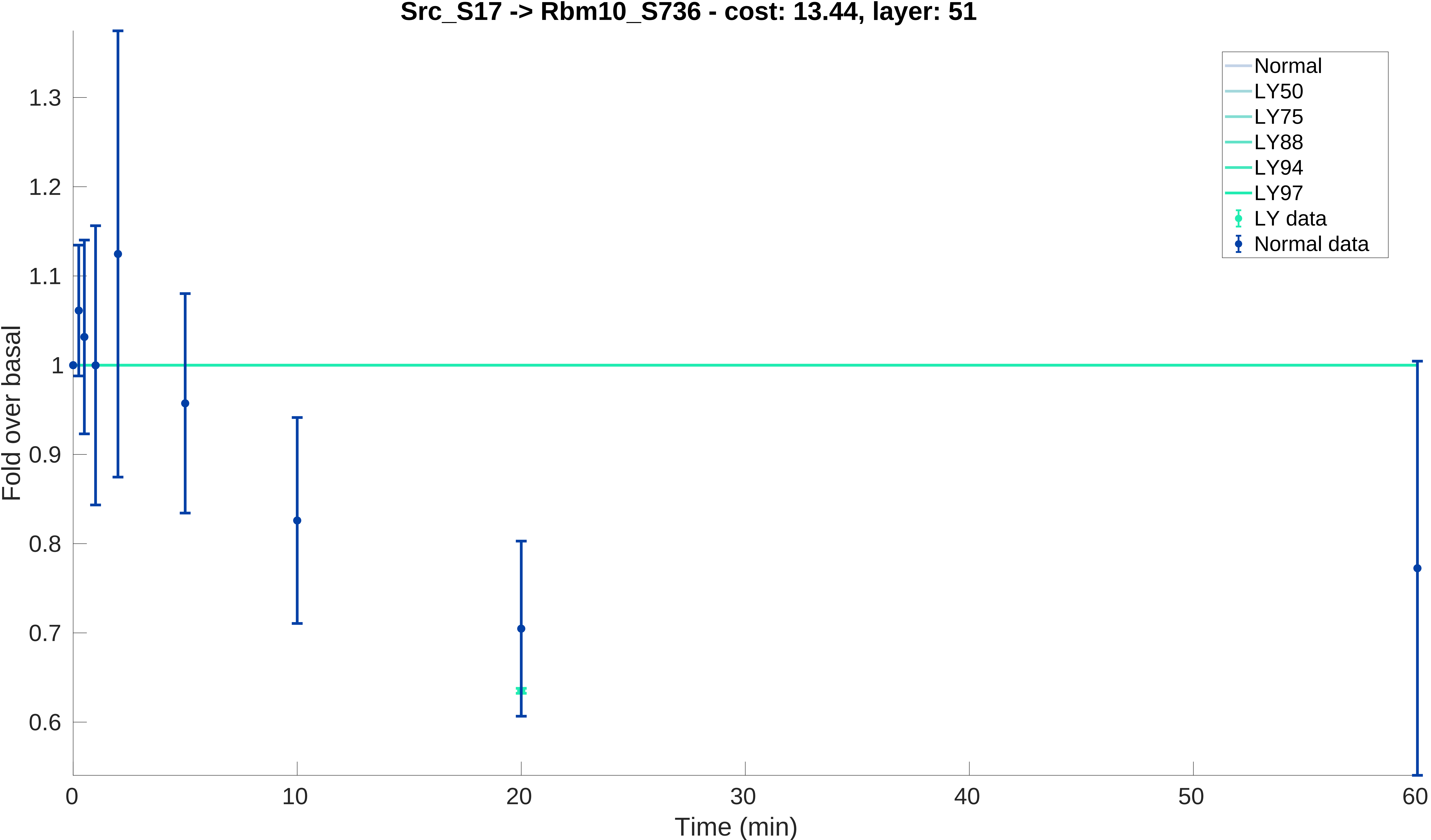

Src_S17 -> Rbm10_S736 - cost: 13.44, layer: 51
Fold over basal
Time (min)
0
10
20
30
40
50
60
0.6
0.7
0.8
0.9
1
1.1
1.2
1.3
Normal
LY50
LY75
LY88
LY94
LY97
LY data
Normal data

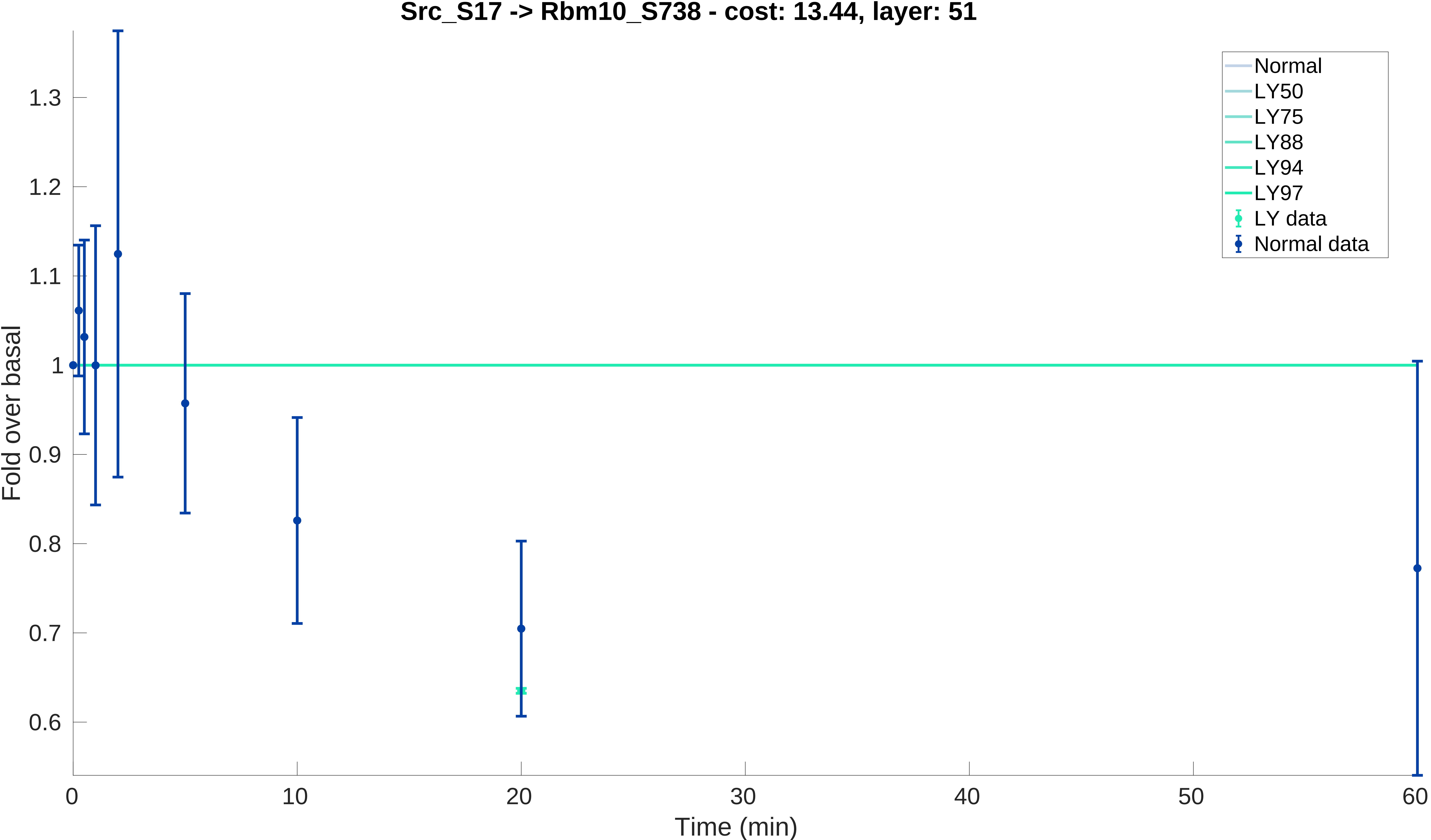

Src_S17 -> Rbm10_S738 - cost: 13.44, layer: 51
Normal
LY50
LY75
LY88
LY94
LY97
LY data
Normal data
Fold over basal
1.3
1.2
1.1
1
0.9
0.8
0.7
0.6
0
10
20
30
40
50
60
Time (min)

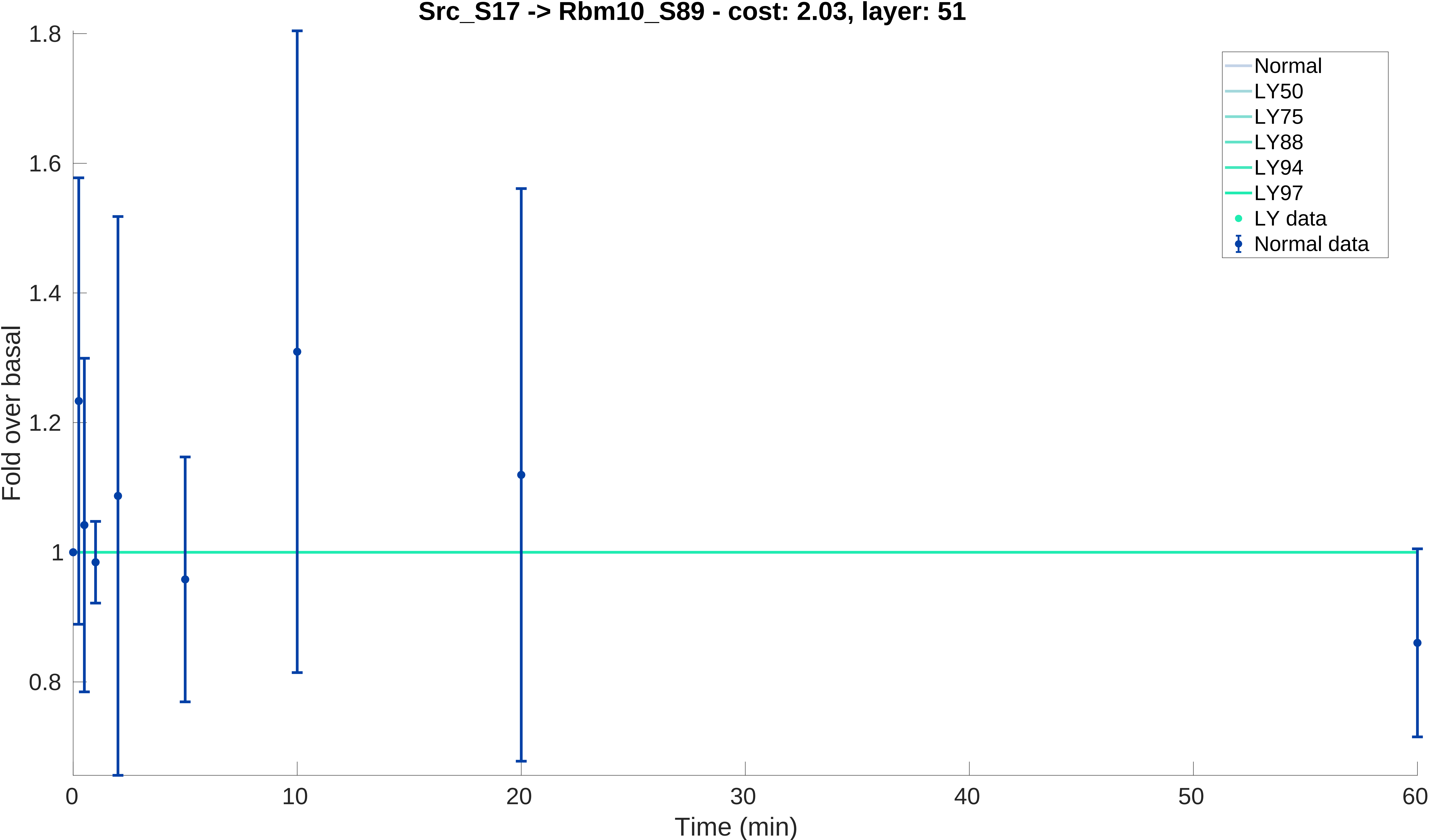

Src_S17 -> Rbm10_S89 - cost: 2.03, layer: 51
Fold over basal
Time (min)
0.8
1
1.2
1.4
1.6
1.8
0
10
20
30
40
50
60
Normal
LY50
LY75
LY88
LY94
LY97
LY data
Normal data

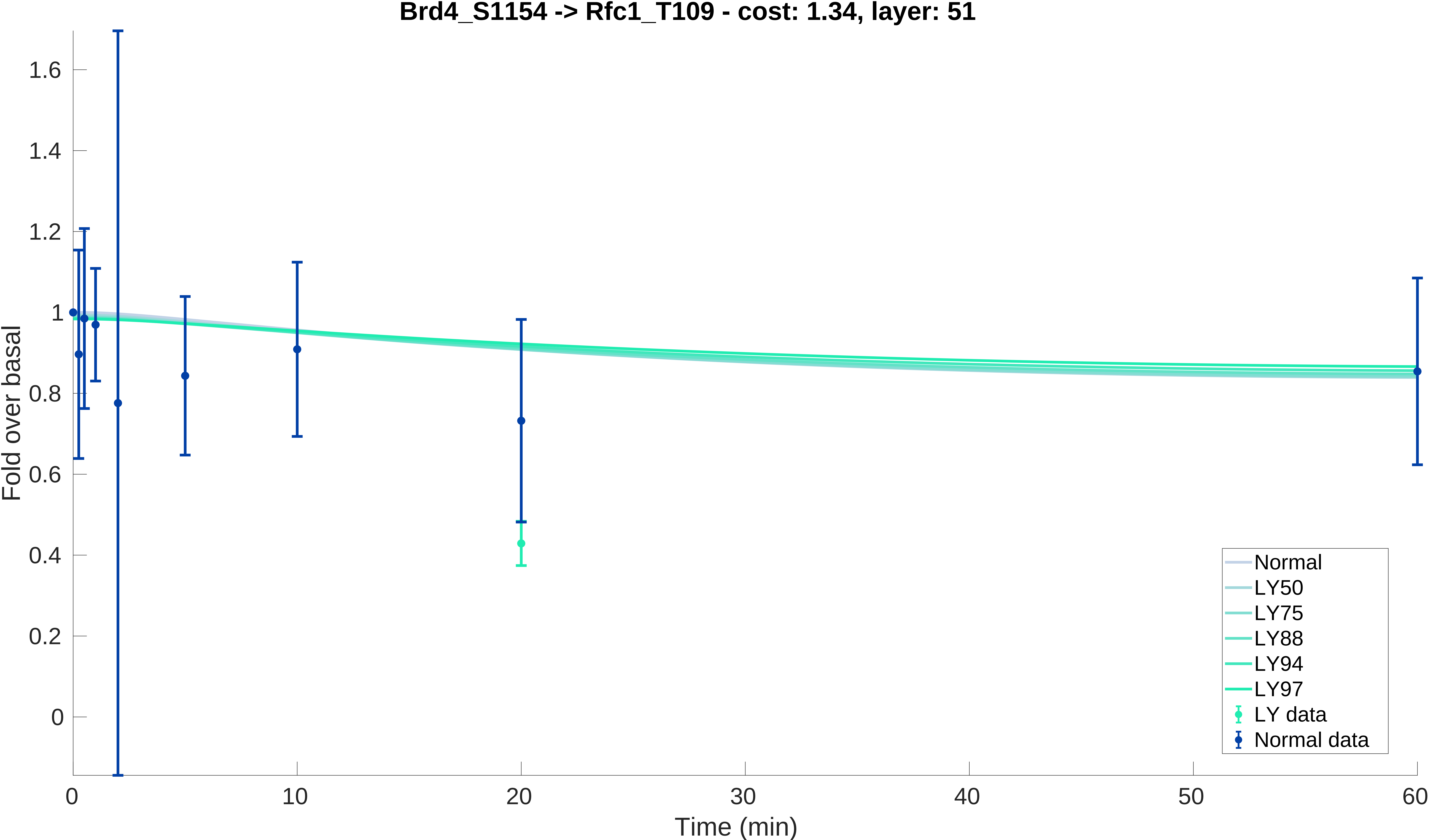

Brd4_S1154 -> Rfc1_T109 - cost: 1.34, layer: 51
Fold over basal
Time (min)
0
0.2
0.4
0.6
0.8
1
1.2
1.4
1.6
0
10
20
30
40
50
60
Normal
LY50
LY75
LY88
LY94
LY97
LY data
Normal data

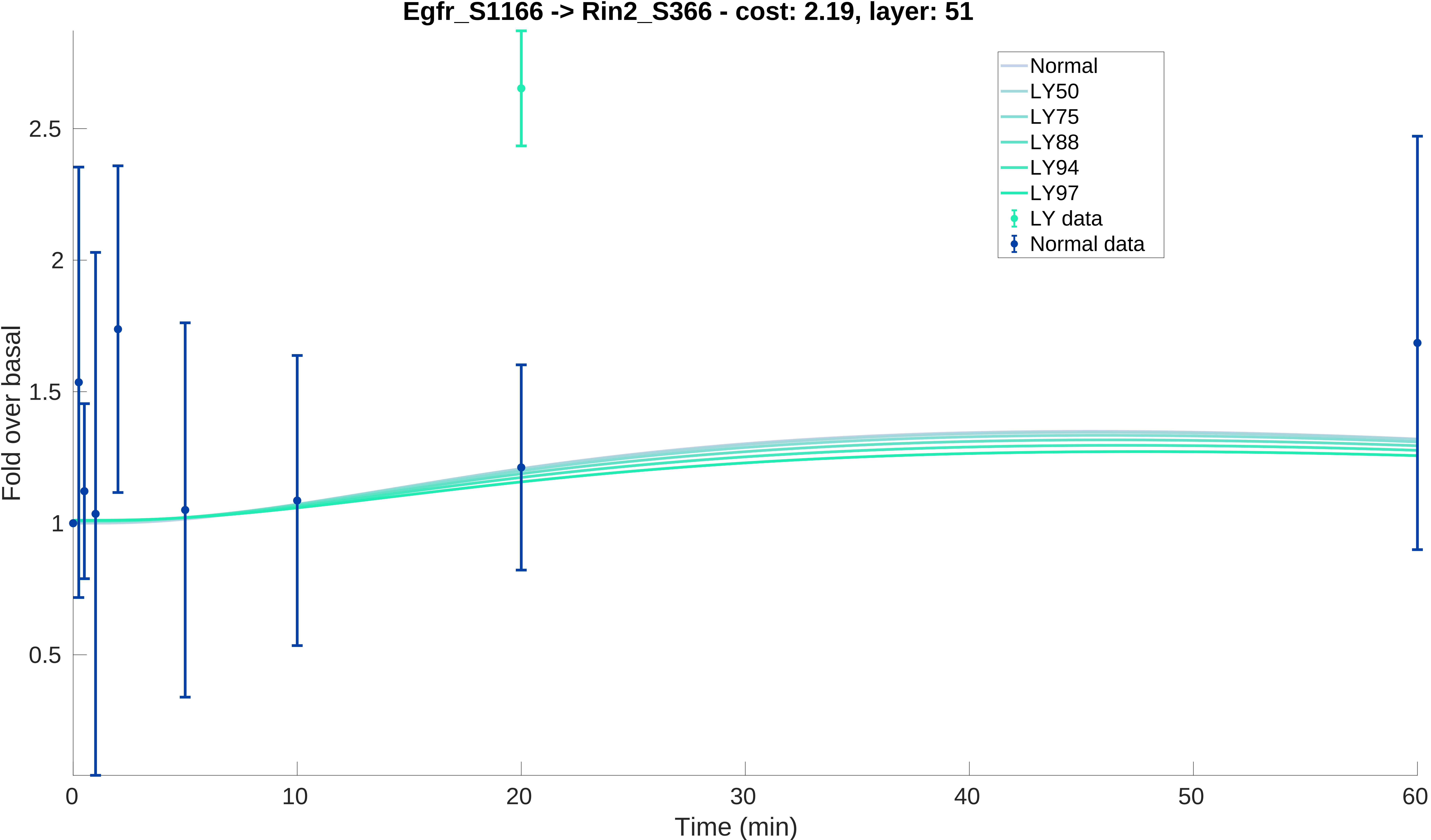

Egfr_S1166 -> Rin2_S366 - cost: 2.19, layer: 51
Normal
LY50
LY75
LY88
LY94
LY97
LY data
Normal data
Fold over basal
Time (min)
2.5
2
1.5
1
0.5
0
10
20
30
40
50
60

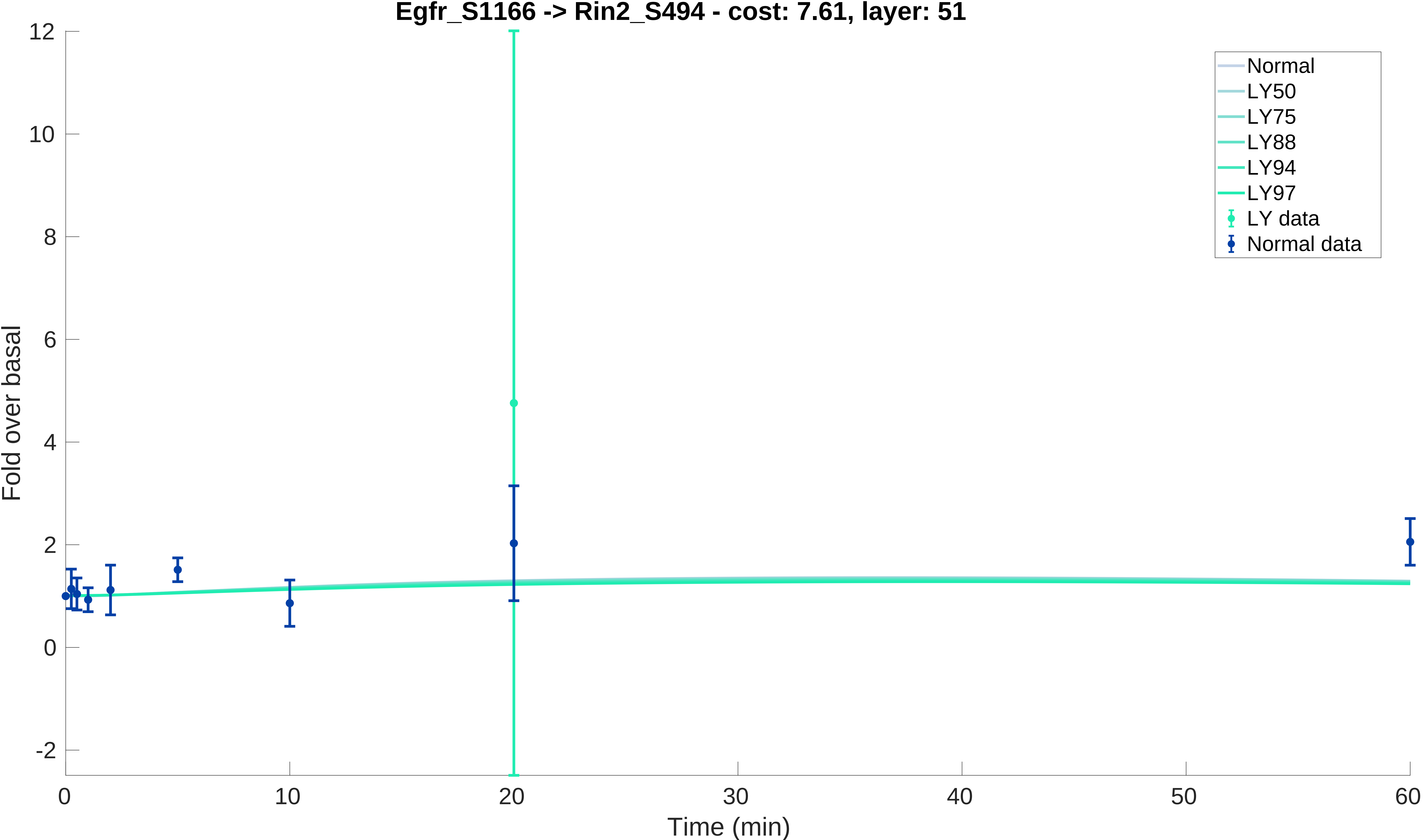

Egfr_S1166 -> Rin2_S494 - cost: 7.61, layer: 51
Fold over basal
Time (min)
Normal
LY50
LY75
LY88
LY94
LY97
LY data
Normal data
12
10
8
6
4
2
0
-2
0
10
20
30
40
50
60

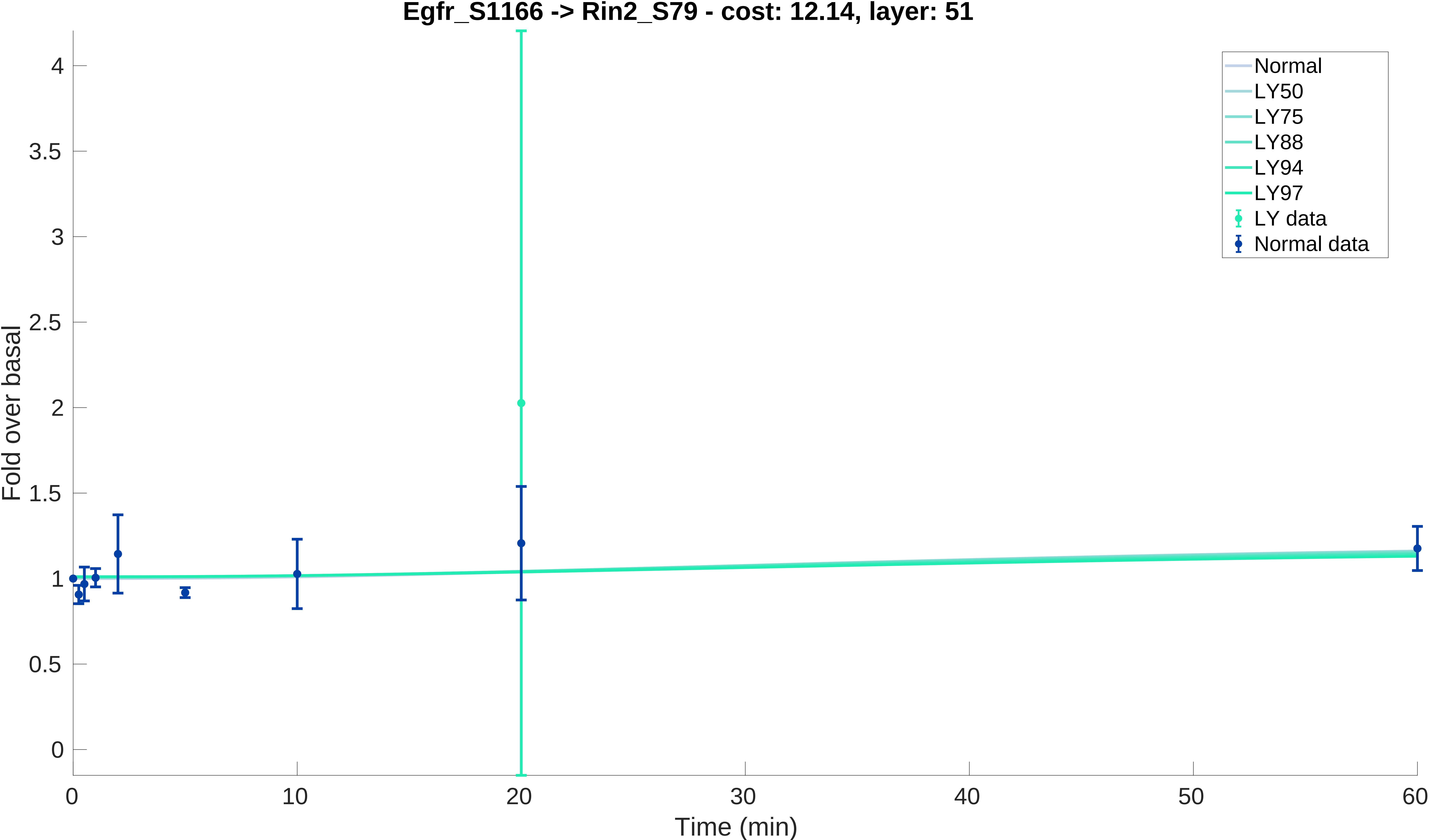

Egfr_S1166 -> Rin2_S79 - cost: 12.14, layer: 51
Fold over basal
Time (min)
0
0.5
1
1.5
2
2.5
3
3.5
4
0
10
20
30
40
50
60
Normal
LY50
LY75
LY88
LY94
LY97
LY data
Normal data

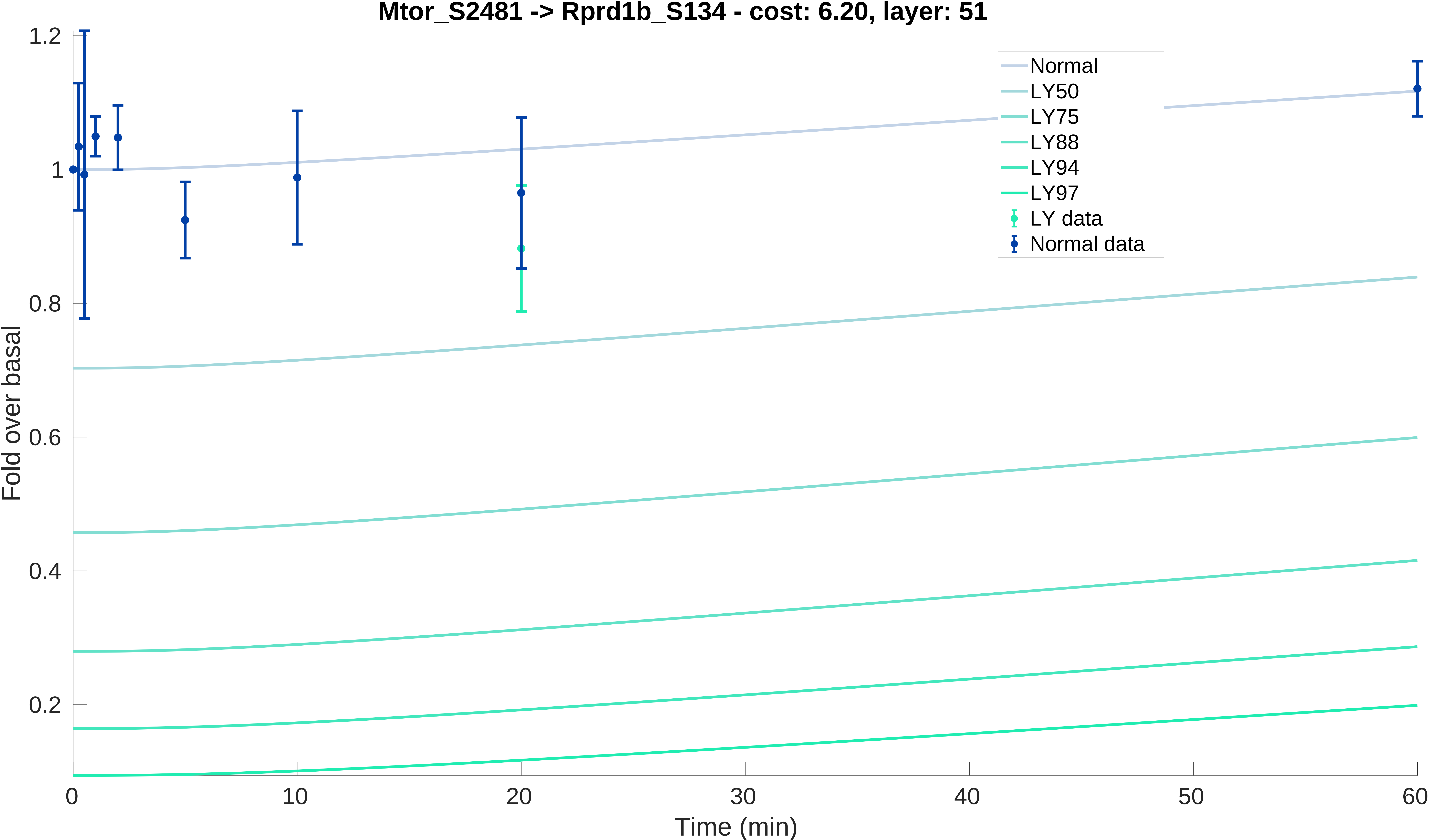

Mtor_S2481 -> Rprd1b_S134 - cost: 6.20, layer: 51
Normal
LY50
LY75
LY88
LY94
LY97
LY data
Normal data
Fold over basal
Time (min)
1.2
1
0.8
0.6
0.4
0.2
0
10
20
30
40
50
60

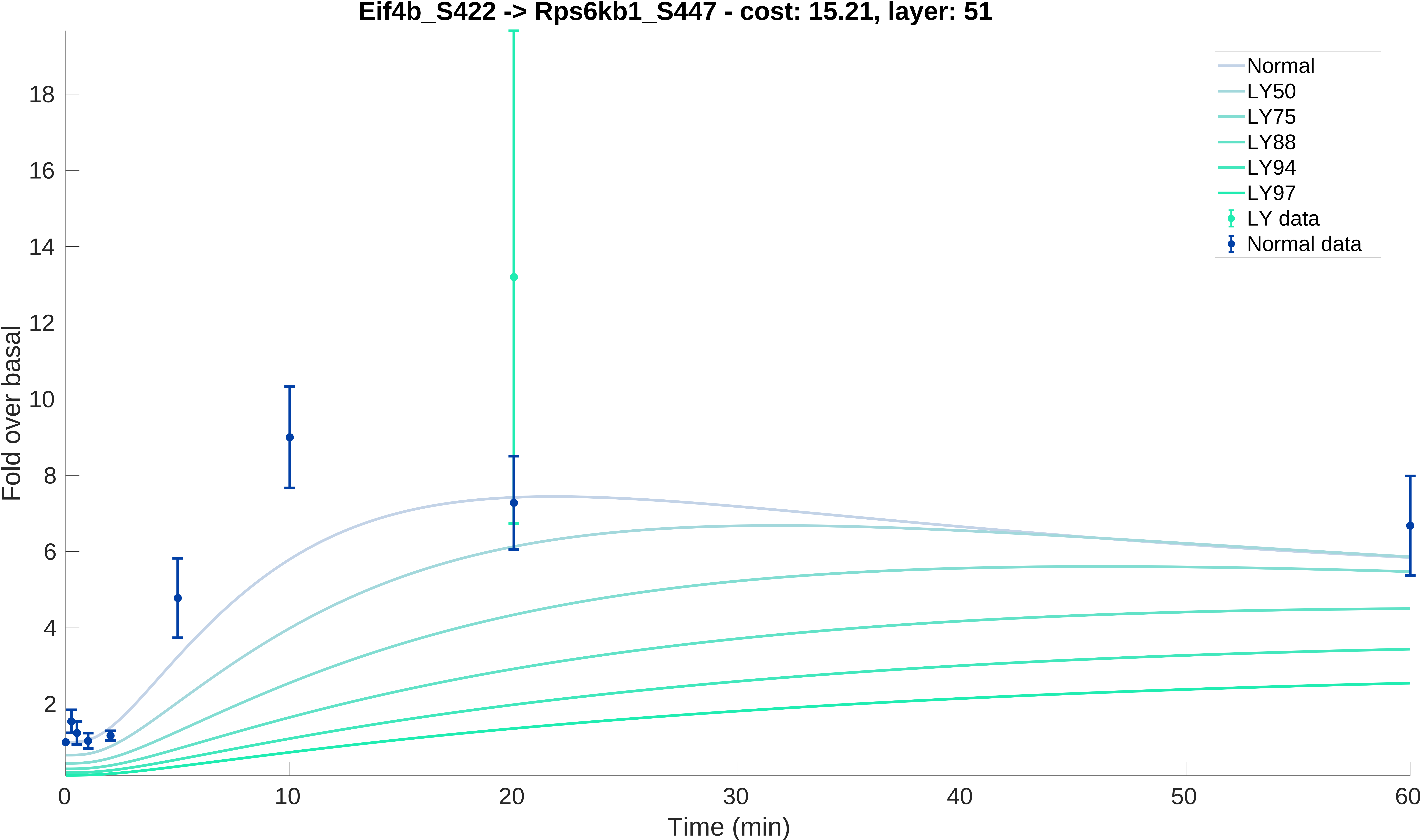

Eif4b_S422 -> Rps6kb1_S447 - cost: 15.21, layer: 51
Fold over basal
Time (min)
Normal
LY50
LY75
LY88
LY94
LY97
LY data
Normal data
0
10
20
30
40
50
60
2
4
6
8
10
12
14
16
18

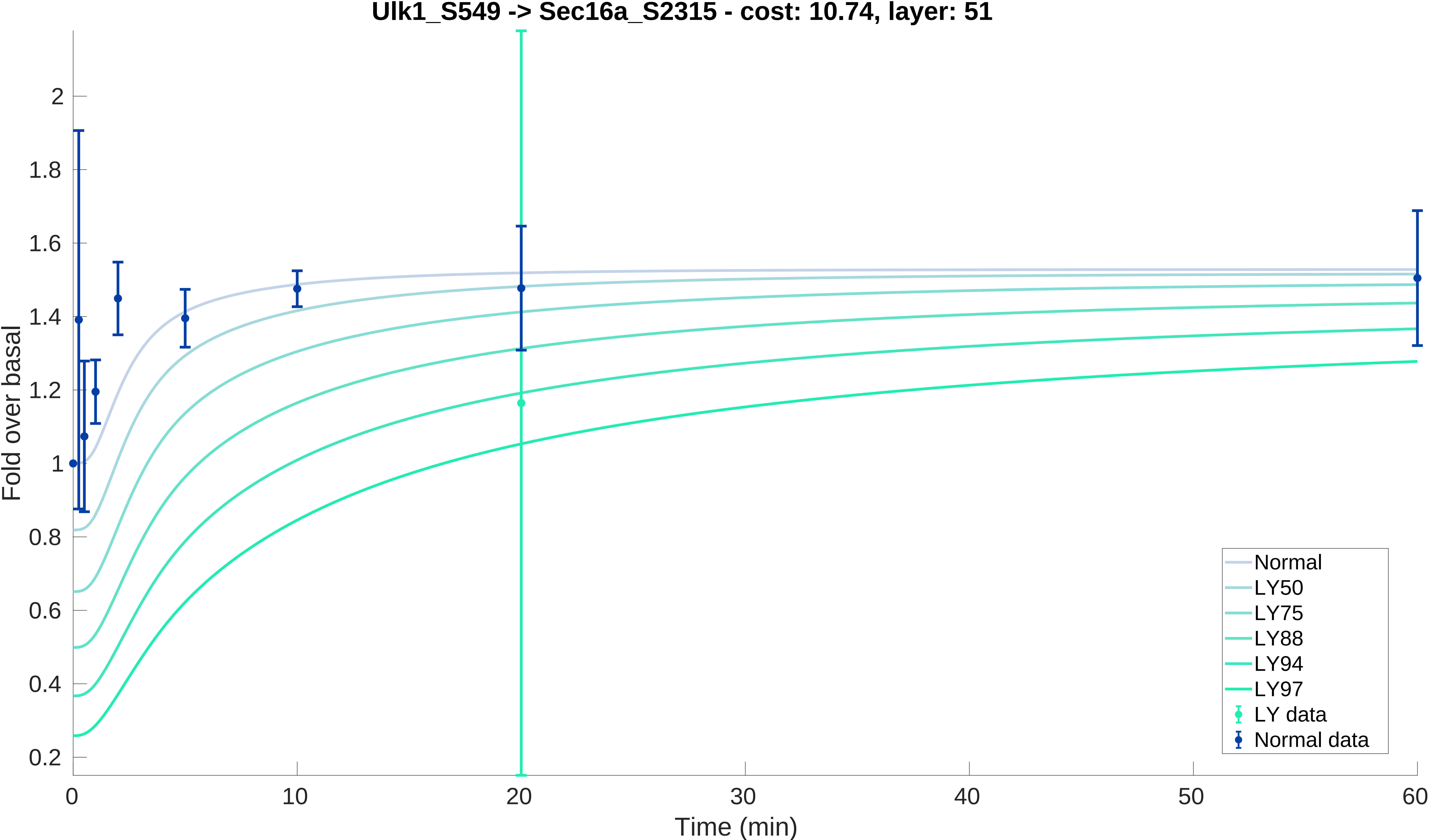
Ulk1_S549 -> Sec16a_S2315 - cost: 10.74, layer: 51
Fold over basal
2
1.8
1.6
1.4
1.2
1
0.8
0.6
0.4
0.2
0
10
20
30
40
50
60
Time (min)
Normal
LY50
LY75
LY88
LY94
LY97
LY data
Normal data

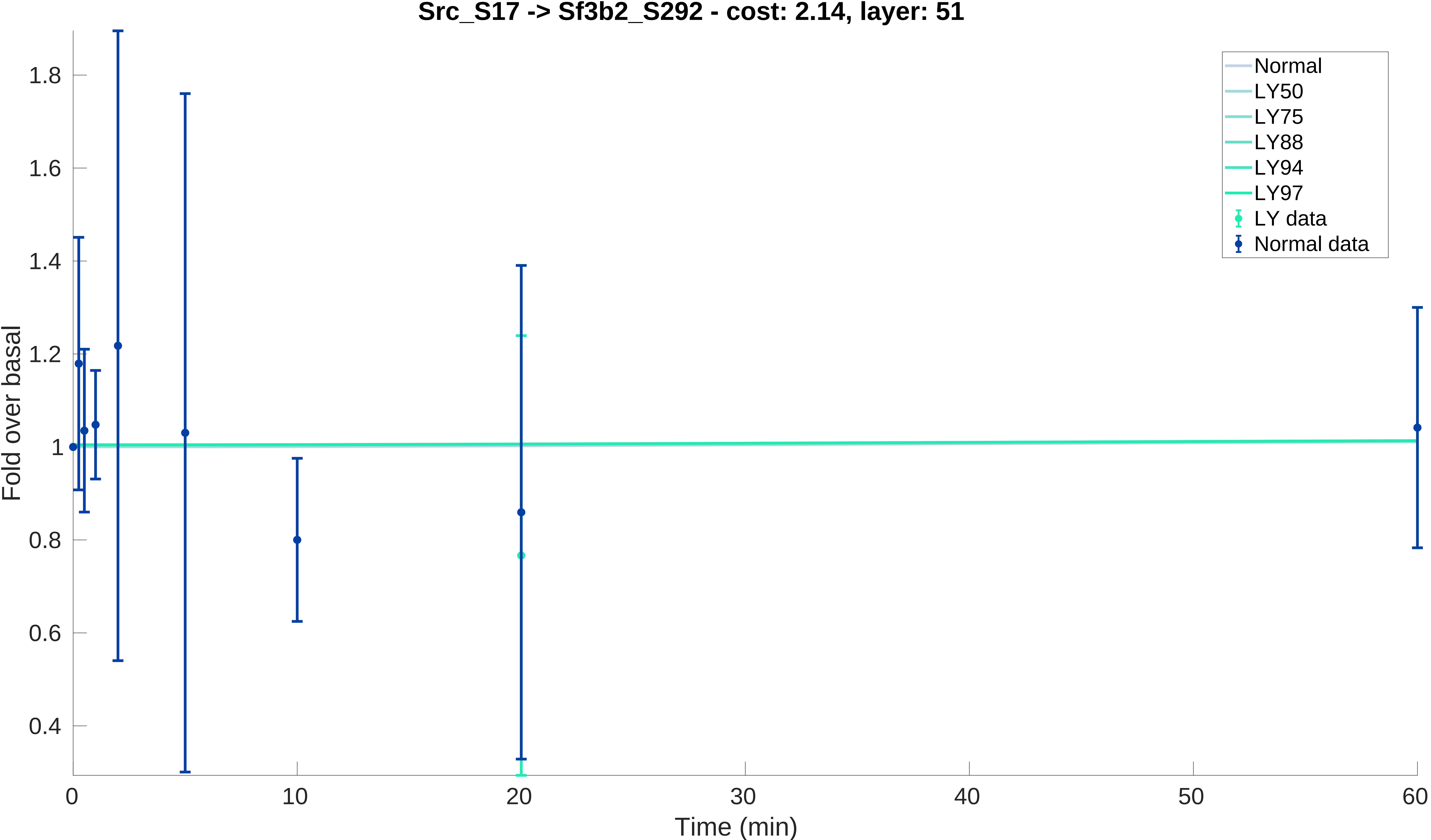

Src_S17 -> Sf3b2_S292 - cost: 2.14, layer: 51
Normal
LY50
LY75
LY88
LY94
LY97
LY data
Normal data
Fold over basal
Time (min)
0
10
20
30
40
50
60
0.4
0.6
0.8
1
1.2
1.4
1.6
1.8

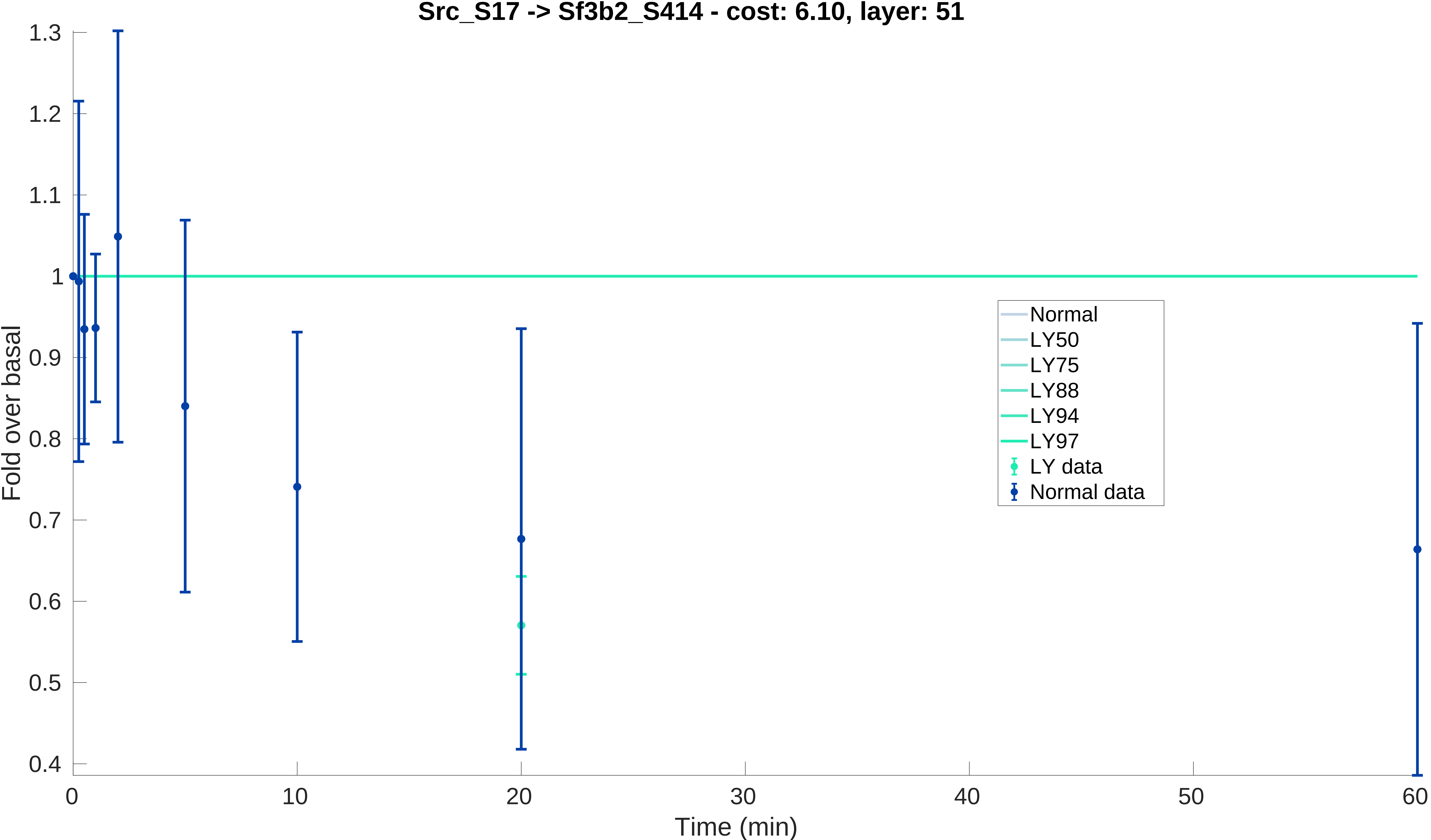

Src_S17 -> Sf3b2_S414 - cost: 6.10, layer: 51
Fold over basal
1.3
1.2
1.1
1
0.9
0.8
0.7
0.6
0.5
0.4
0
10
20
30
40
50
60
Time (min)
Normal
LY50
LY75
LY88
LY94
LY97
LY data
Normal data

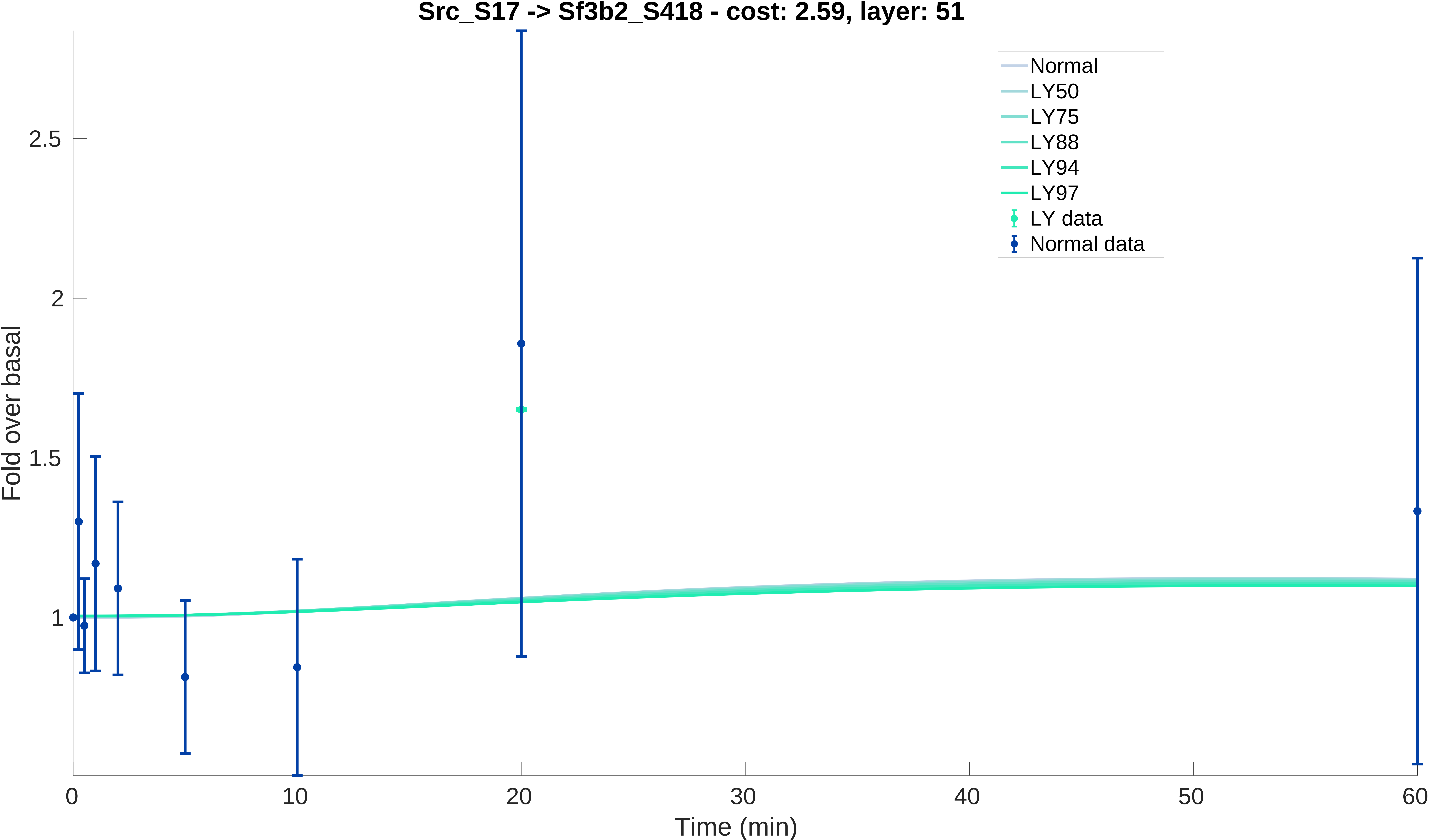

Src_S17 -> Sf3b2_S418 - cost: 2.59, layer: 51
Normal
LY50
LY75
LY88
LY94
LY97
LY data
Normal data
Fold over basal
Time (min)
2.5
2
1.5
1
0
10
20
30
40
50
60

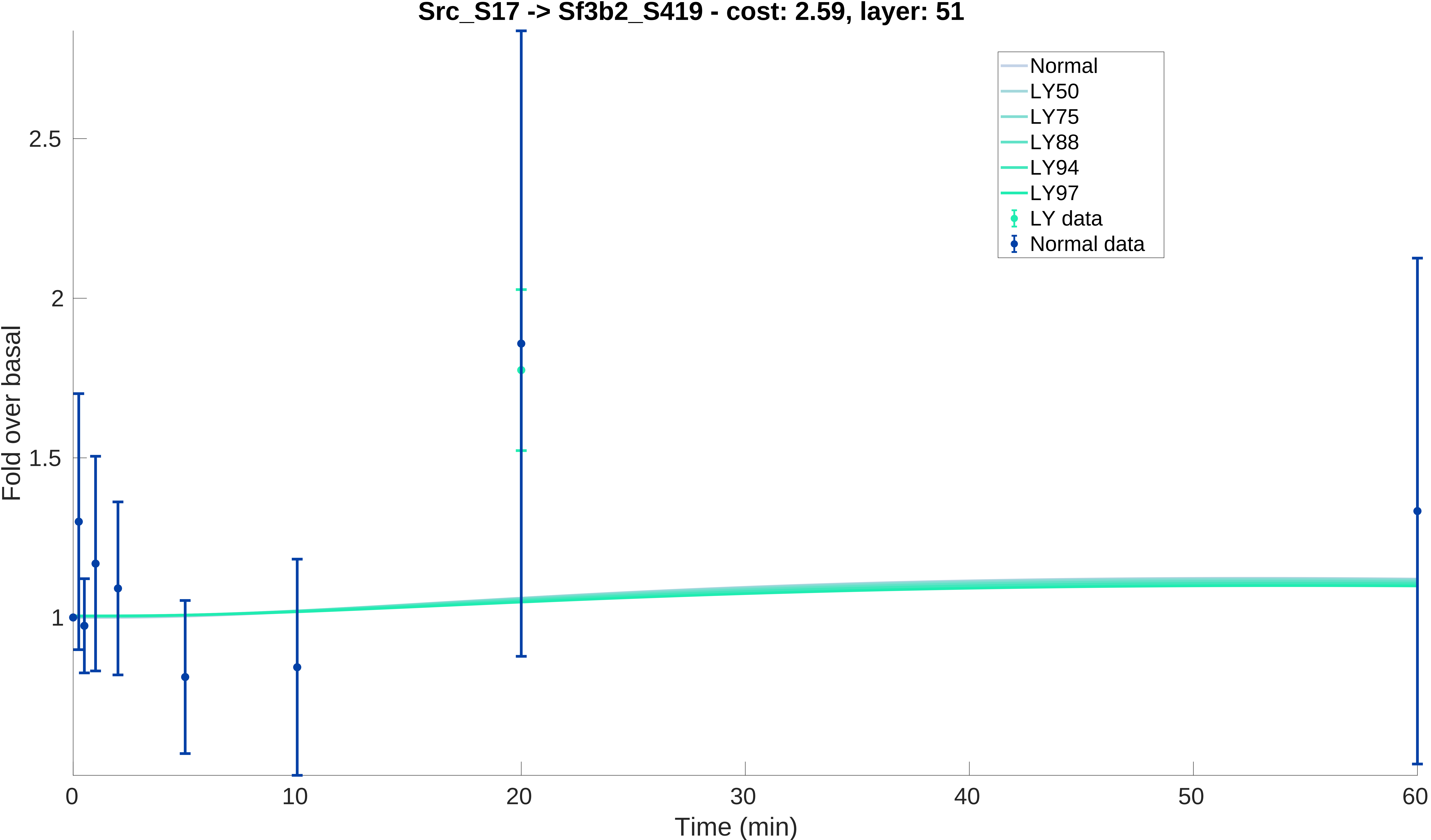

Src_S17 -> Sf3b2_S419 - cost: 2.59, layer: 51
Normal
LY50
LY75
LY88
LY94
LY97
LY data
Normal data
Fold over basal
Time (min)
1
1.5
2
2.5
0
10
20
30
40
50
60

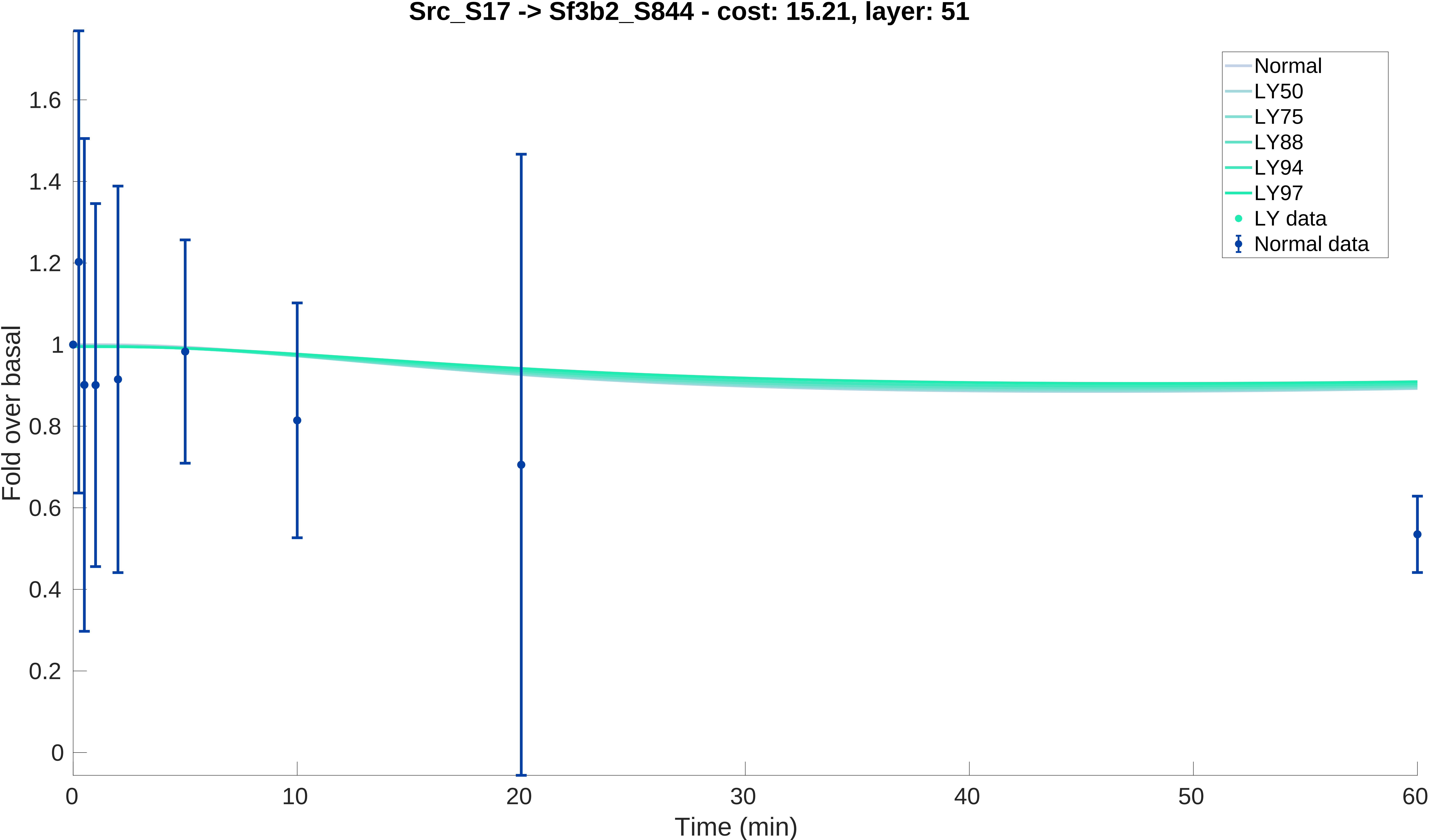

Src_S17 -> Sf3b2_S844 - cost: 15.21, layer: 51
Normal
LY50
LY75
LY88
LY94
LY97
LY data
Normal data
Fold over basal
Time (min)
0
0.2
0.4
0.6
0.8
1
1.2
1.4
1.6
0
10
20
30
40
50
60

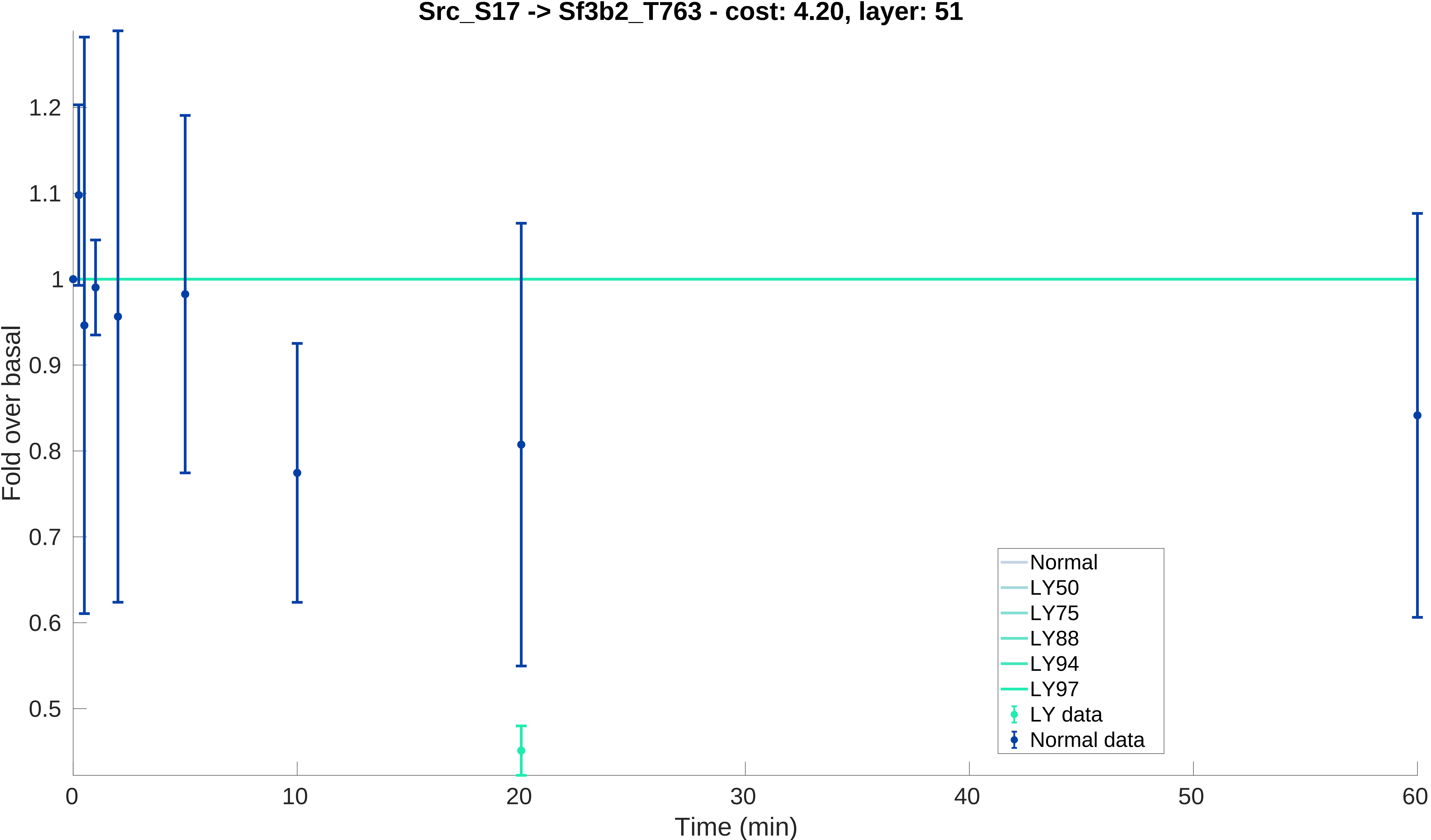

Src_S17 -> Sf3b2_T763 - cost: 4.20, layer: 51
Fold over basal
Time (min)
1.2
1.1
1
0.9
0.8
0.7
0.6
0.5
0
10
20
30
40
50
60
Normal
LY50
LY75
LY88
LY94
LY97
LY data
Normal data

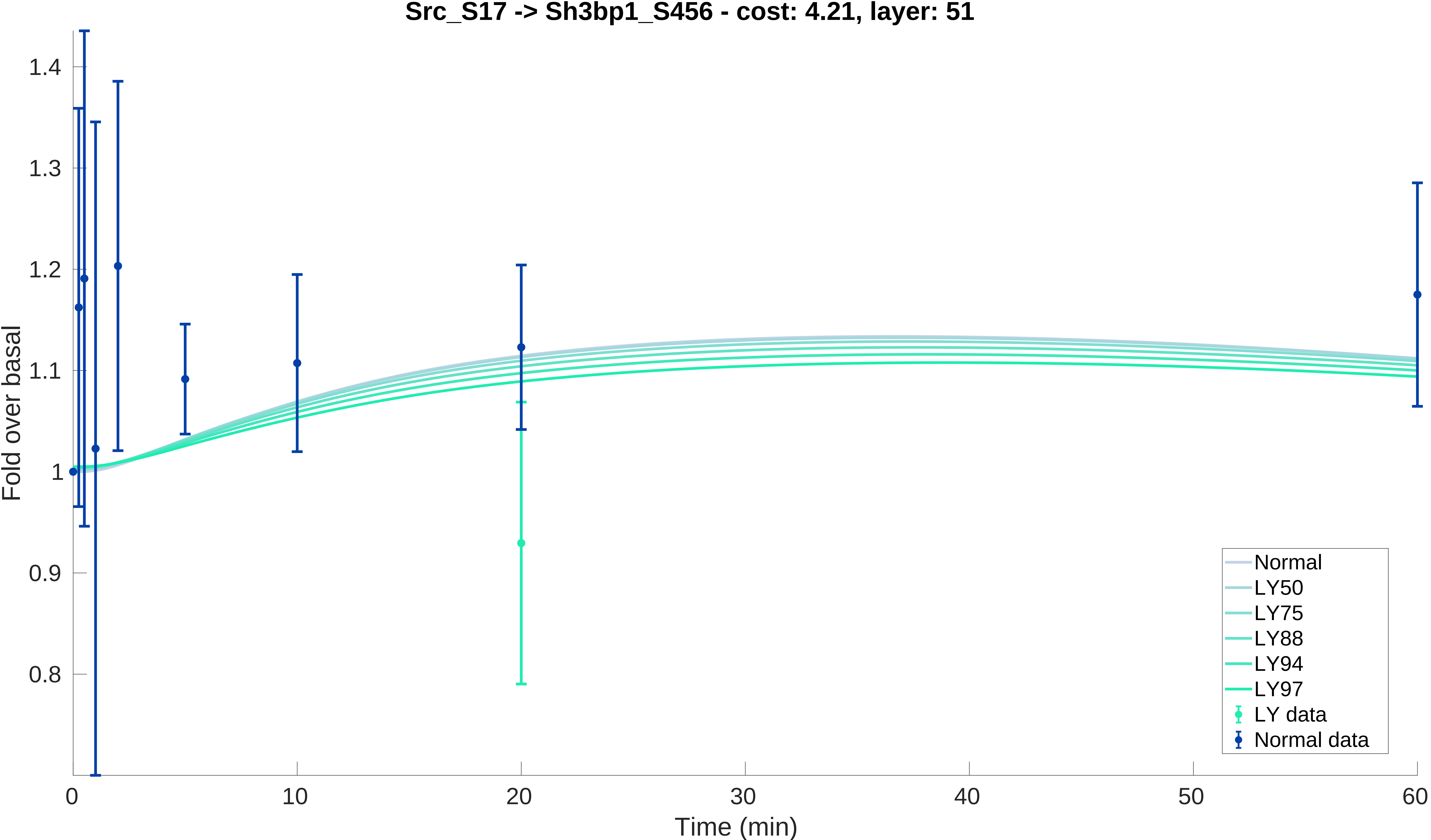
Src_S17 -> Sh3bp1_S456 - cost: 4.21, layer: 51
Fold over basal
Time (min)
0.8
0.9
1
1.1
1.2
1.3
1.4
0
10
20
30
40
50
60
Normal
LY50
LY75
LY88
LY94
LY97
LY data
Normal data

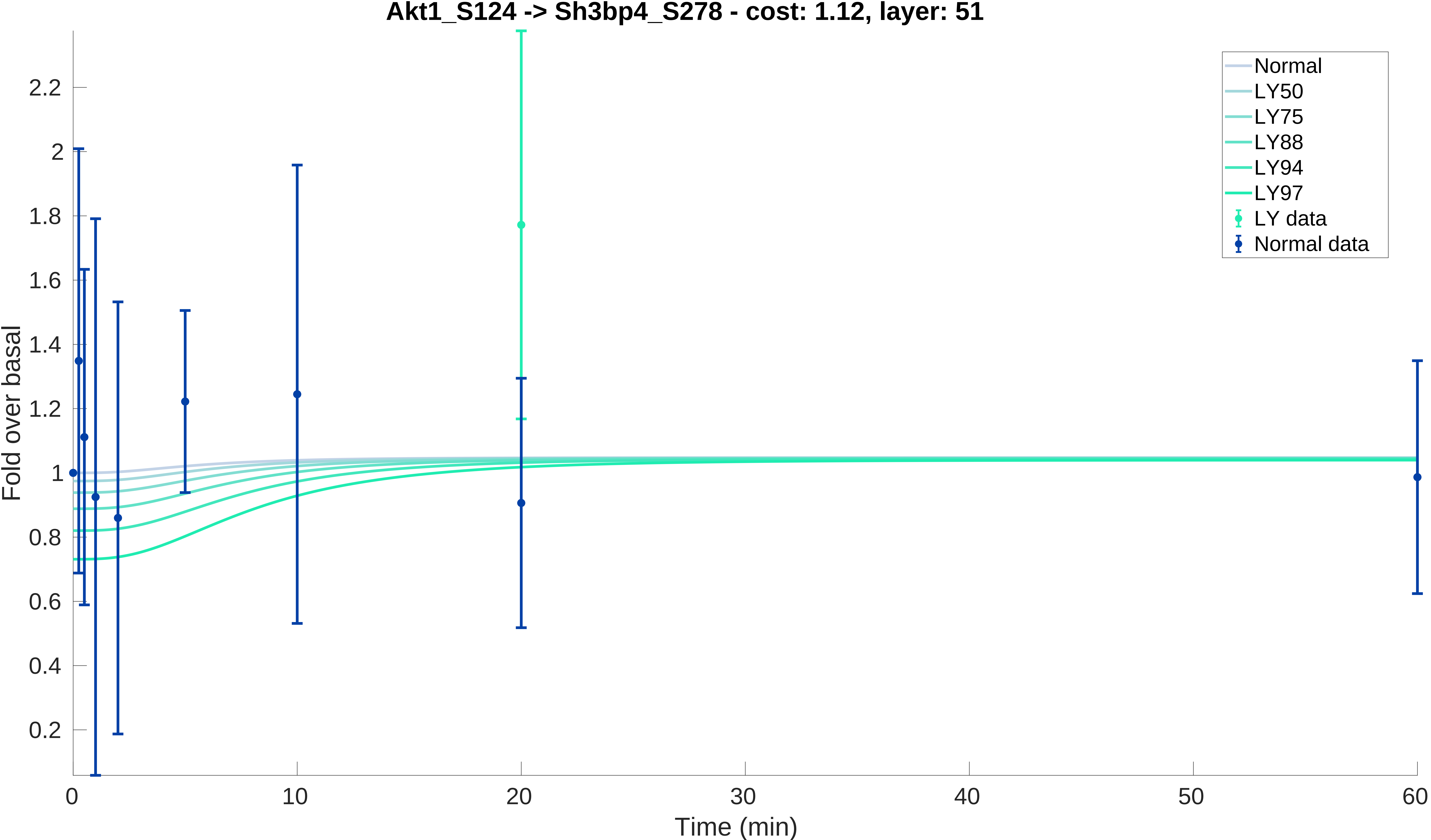

Akt1_S124 -> Sh3bp4_S278 - cost: 1.12, layer: 51
Fold over basal
Time (min)
0.2
0.4
0.6
0.8
1
1.2
1.4
1.6
1.8
2
2.2
0
10
20
30
40
50
60
Normal
LY50
LY75
LY88
LY94
LY97
LY data
Normal data

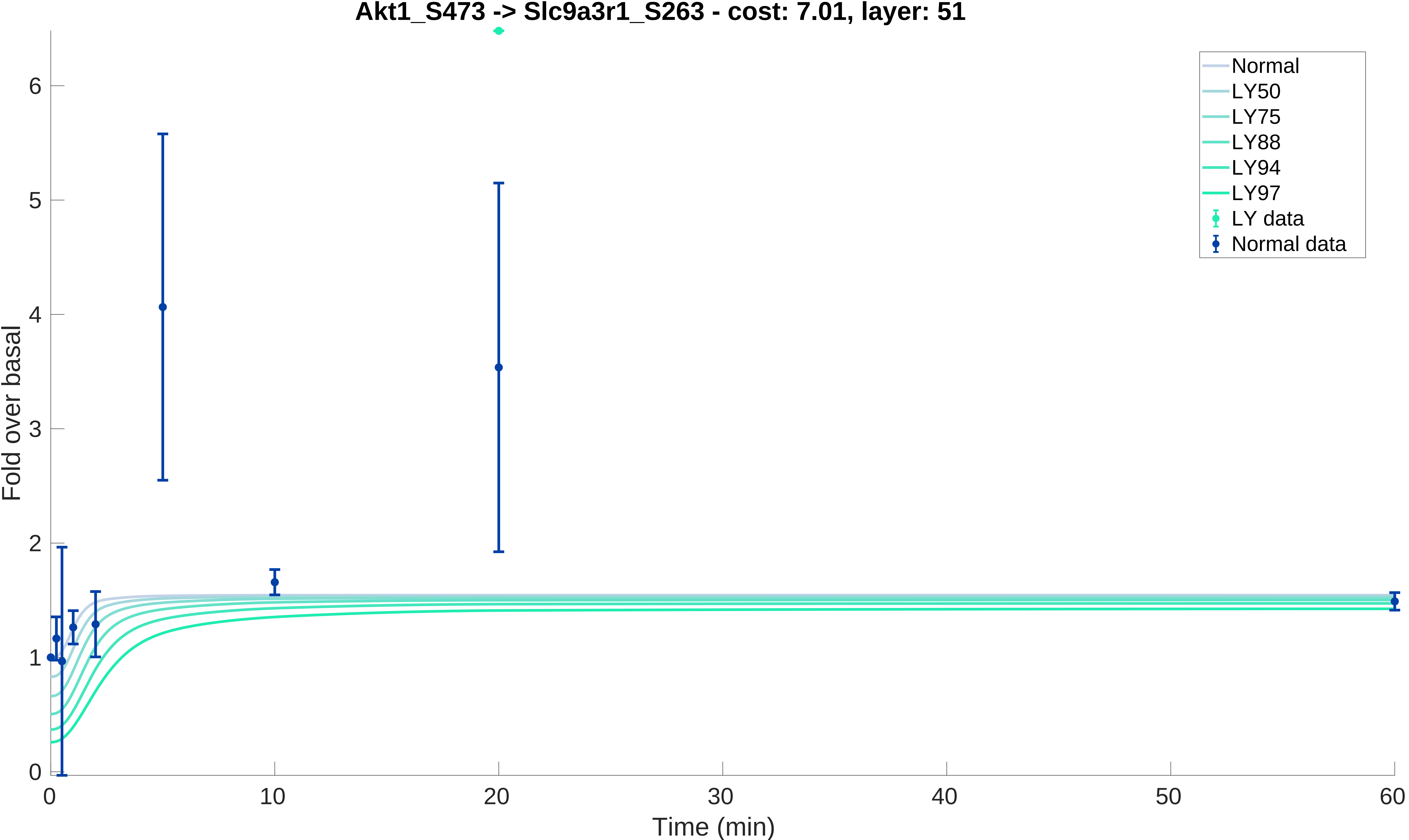

Akt1_S473 -> Slc9a3r1_S263 - cost: 7.01, layer: 51
Fold over basal
Time (min)
0
1
2
3
4
5
6
0
10
20
30
40
50
60
Normal
LY50
LY75
LY88
LY94
LY97
LY data
Normal data

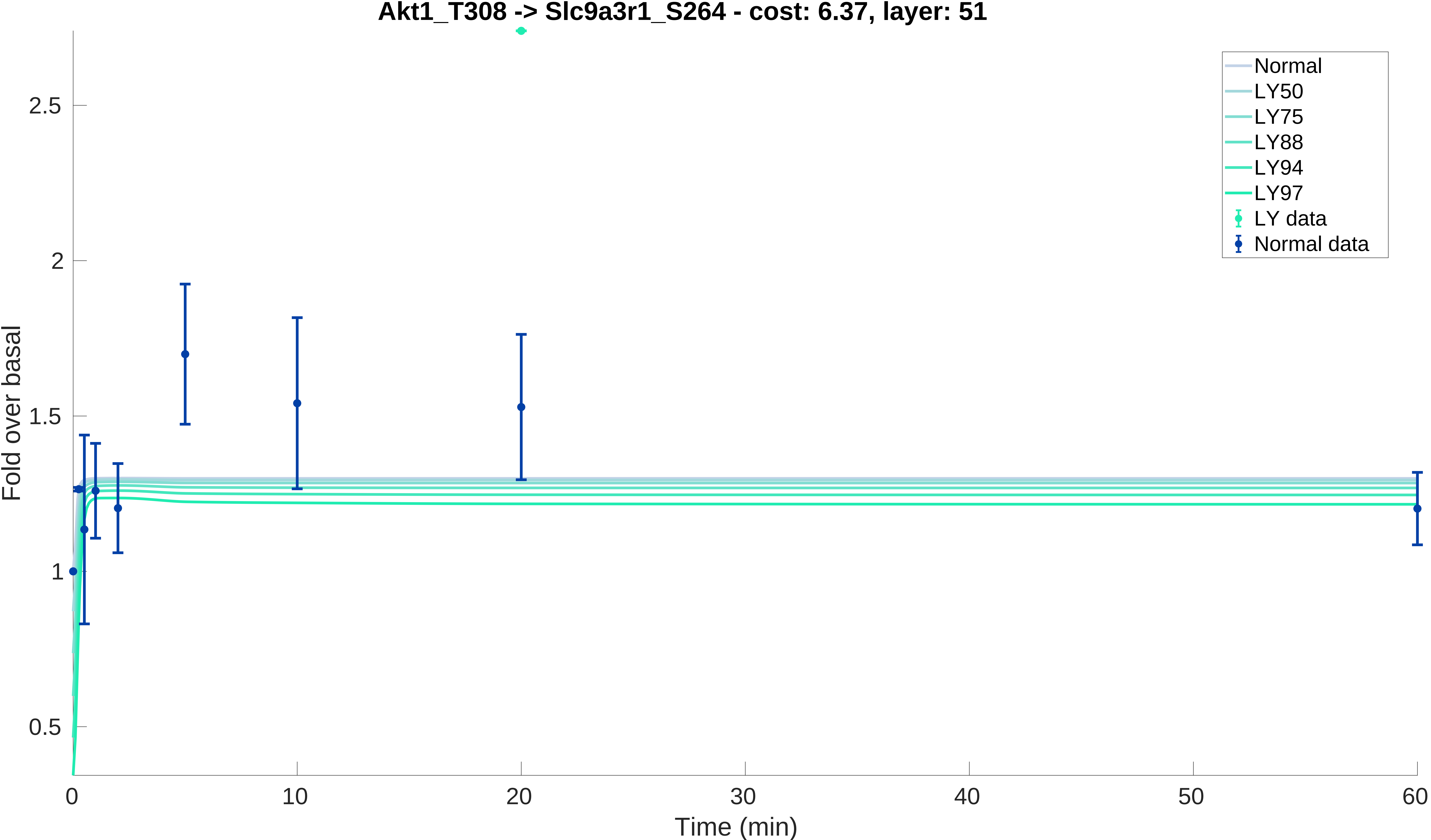
Akt1_T308 -> Slc9a3r1_S264 - cost: 6.37, layer: 51
Normal
LY50
LY75
LY88
LY94
LY97
LY data
Normal data
Fold over basal
Time (min)
0.5
1
1.5
2
2.5
0
10
20
30
40
50
60

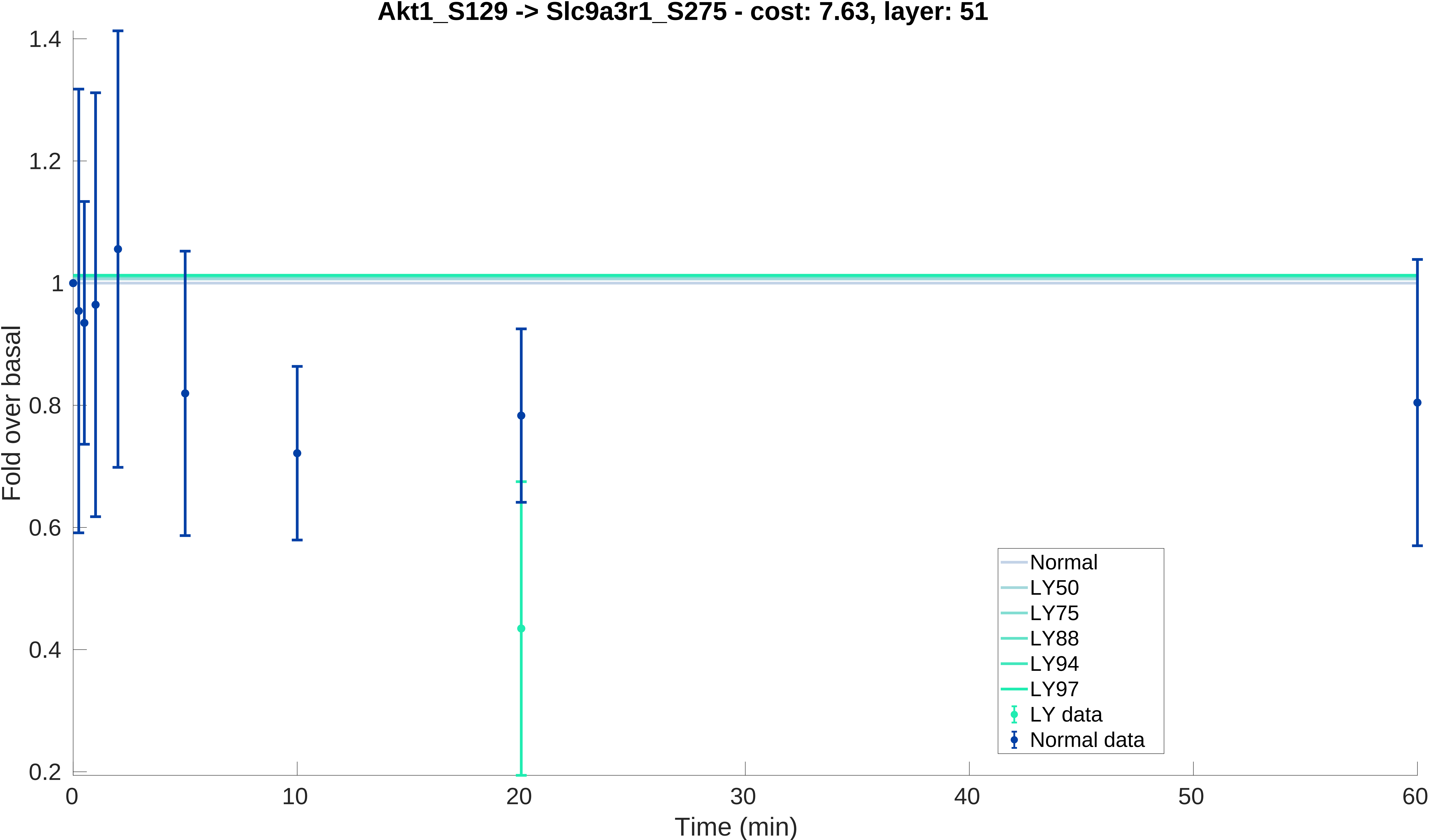
Akt1_S129 -> Slc9a3r1_S275 - cost: 7.63, layer: 51
Fold over basal
Time (min)
0
10
20
30
40
50
60
0.2
0.4
0.6
0.8
1
1.2
1.4
Normal
LY50
LY75
LY88
LY94
LY97
LY data
Normal data

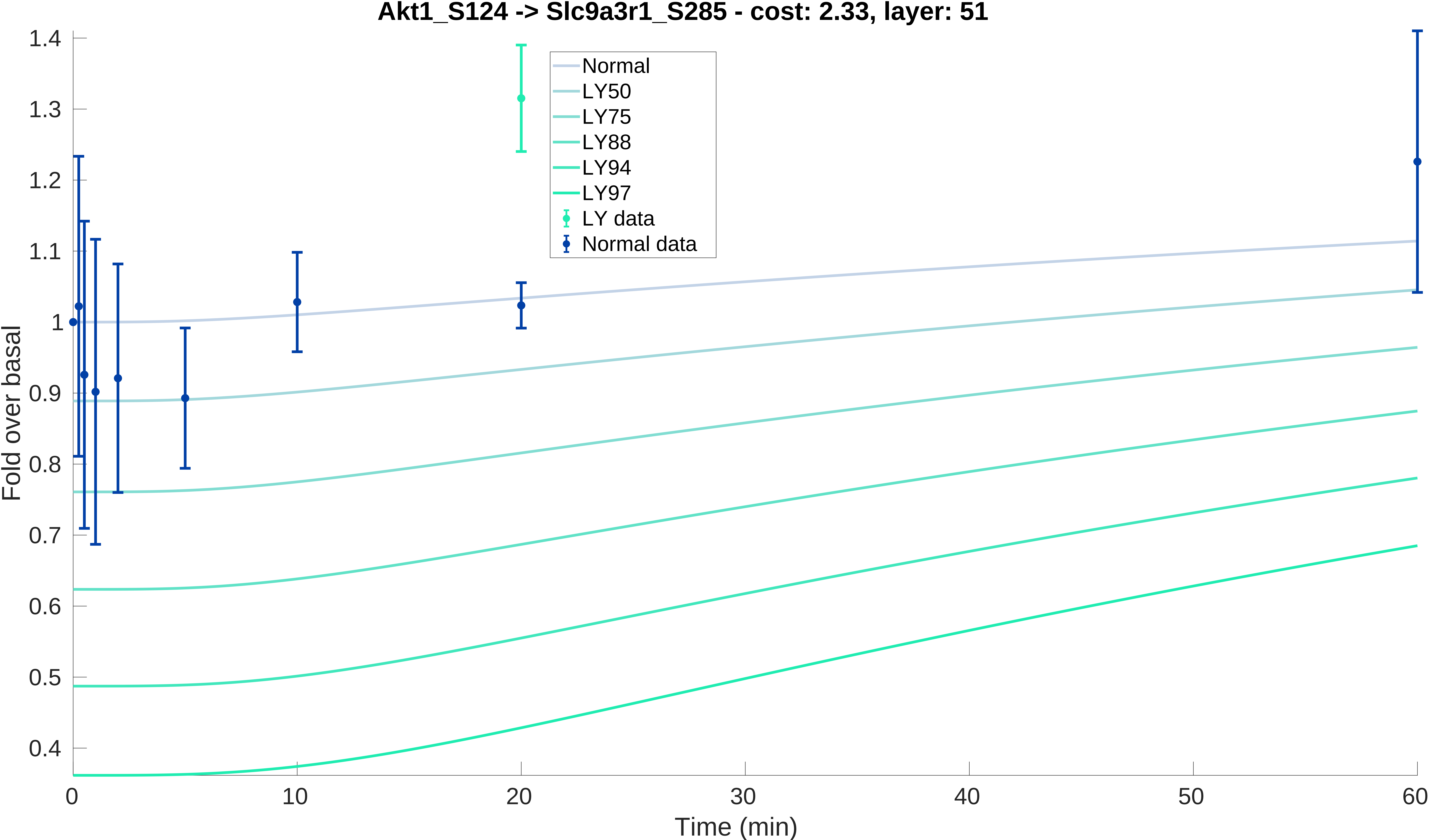

Akt1_S124 -> Slc9a3r1_S285 - cost: 2.33, layer: 51
Normal
LY50
LY75
LY88
LY94
LY97
LY data
Normal data
Fold over basal
Time (min)
0
10
20
30
40
50
60
0.4
0.5
0.6
0.7
0.8
0.9
1
1.1
1.2
1.3
1.4

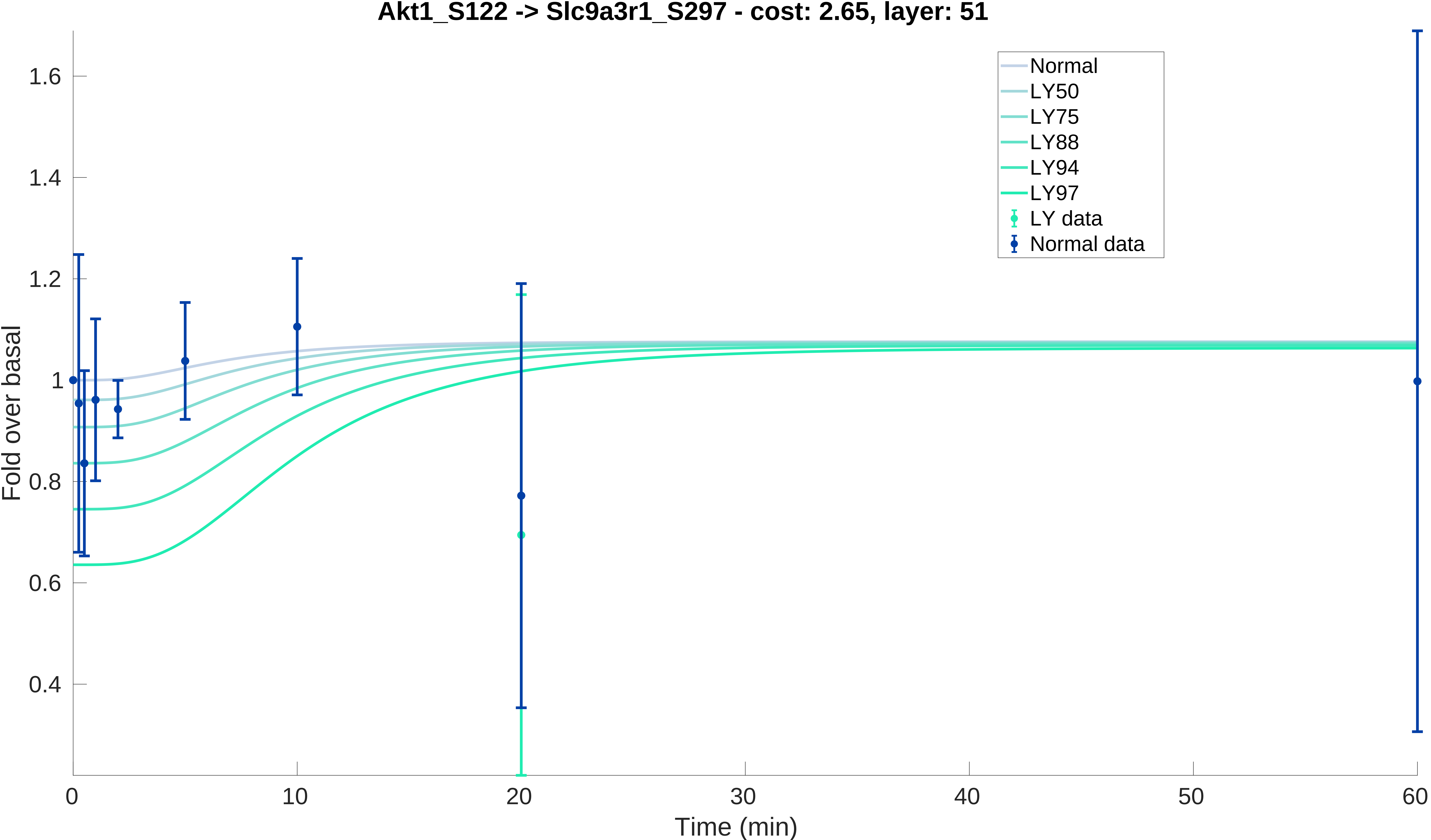

Akt1_S122 -> Slc9a3r1_S297 - cost: 2.65, layer: 51
Normal
LY50
LY75
LY88
LY94
LY97
LY data
Normal data
Fold over basal
1.6
1.4
1.2
1
0.8
0.6
0.4
0
10
20
30
40
50
60
Time (min)

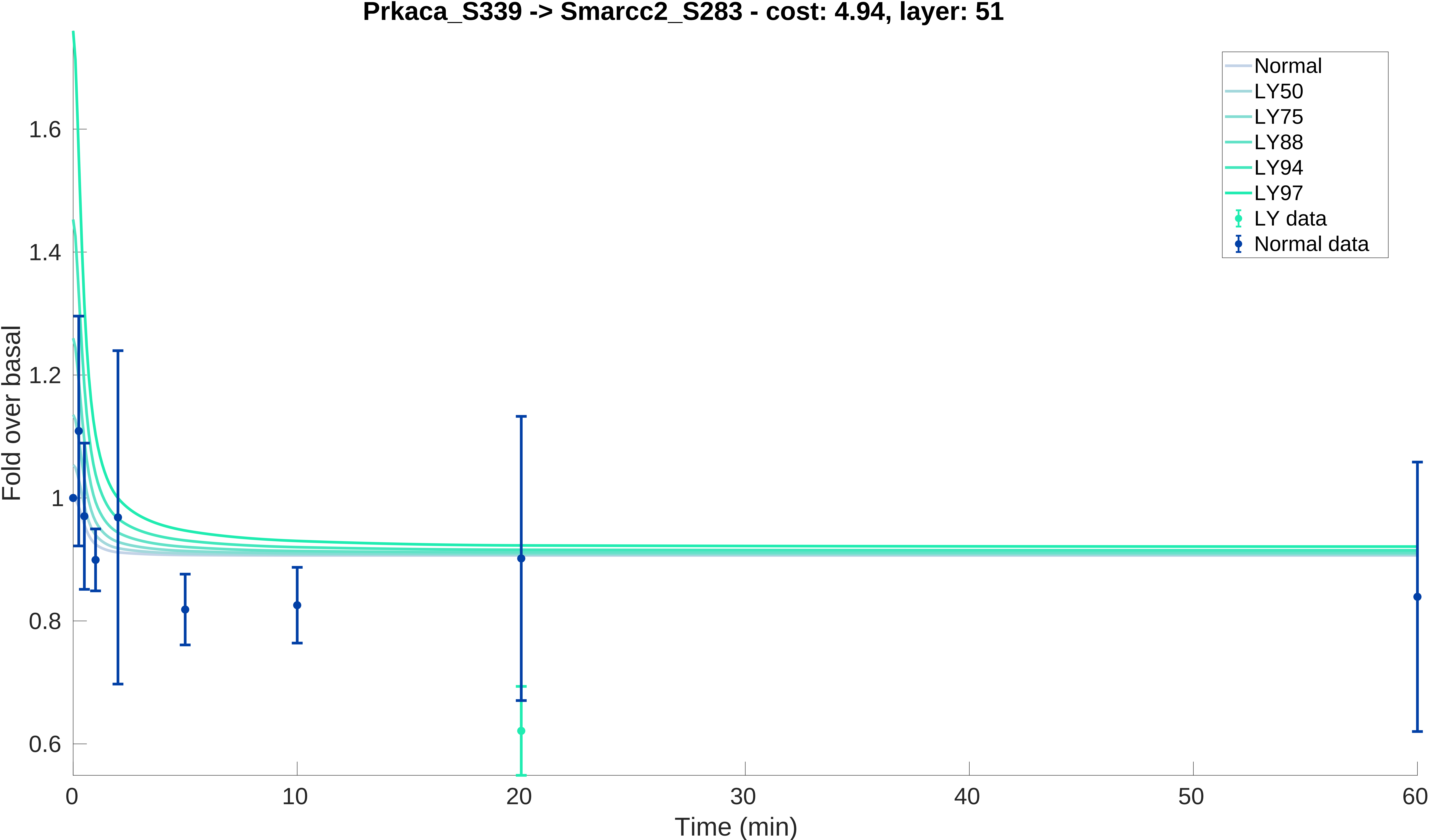

Prkaca_S339 -> Smarcc2_S283 - cost: 4.94, layer: 51
Fold over basal
1.6
1.4
1.2
1
0.8
0.6
0
10
20
30
40
50
60
Time (min)
Normal
LY50
LY75
LY88
LY94
LY97
LY data
Normal data

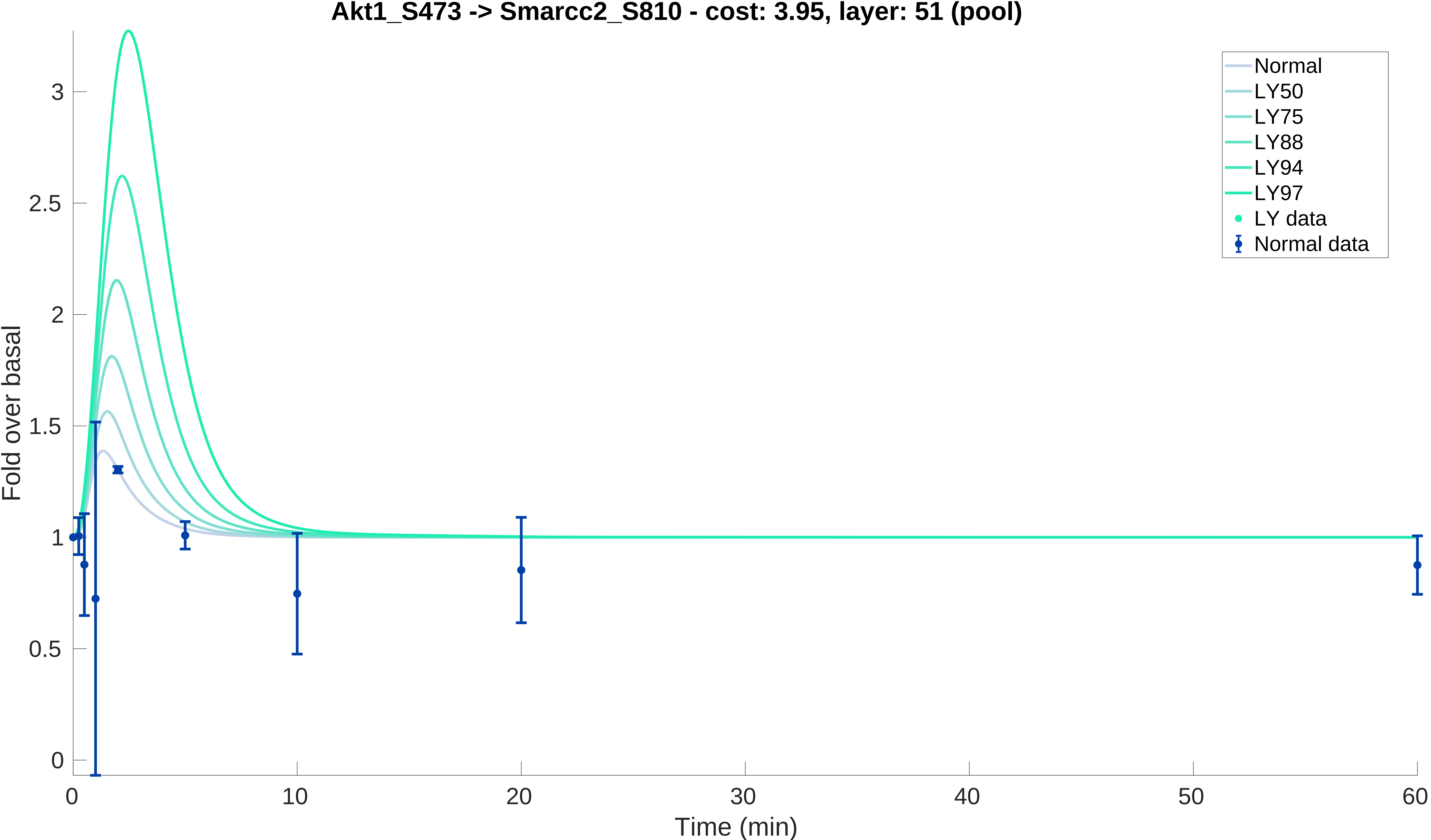

Akt1_S473 -> Smarcc2_S810 - cost: 3.95, layer: 51 (pool)
Fold over basal
Time (min)
Normal
LY50
LY75
LY88
LY94
LY97
LY data
Normal data
0
0.5
1
1.5
2
2.5
3
0
10
20
30
40
50
60

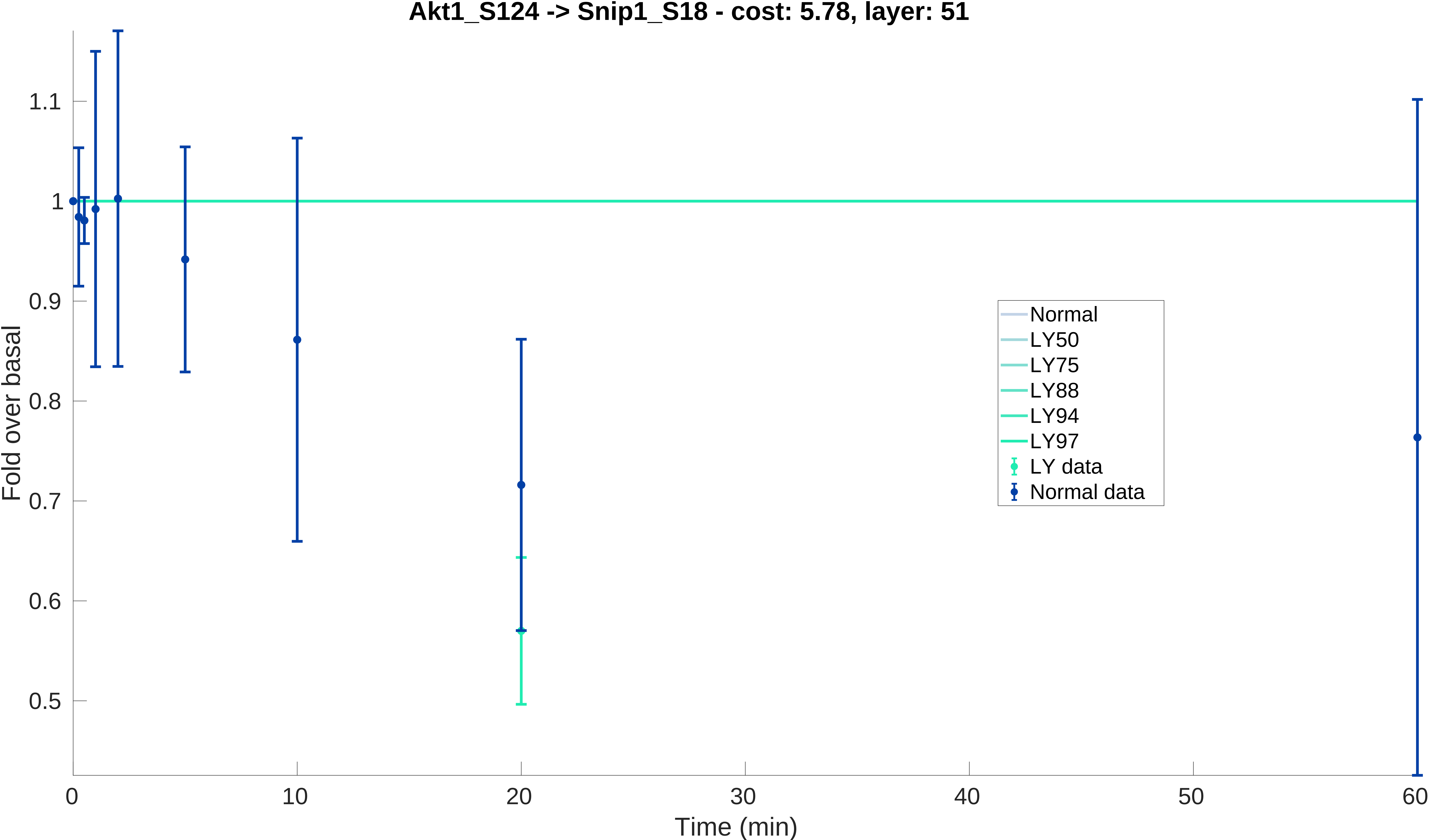
Akt1_S124 -> Snip1_S18 - cost: 5.78, layer: 51
Fold over basal
1.1
1
0.9
0.8
0.7
0.6
0.5
0
10
20
30
40
50
60
Time (min)
Normal
LY50
LY75
LY88
LY94
LY97
LY data
Normal data

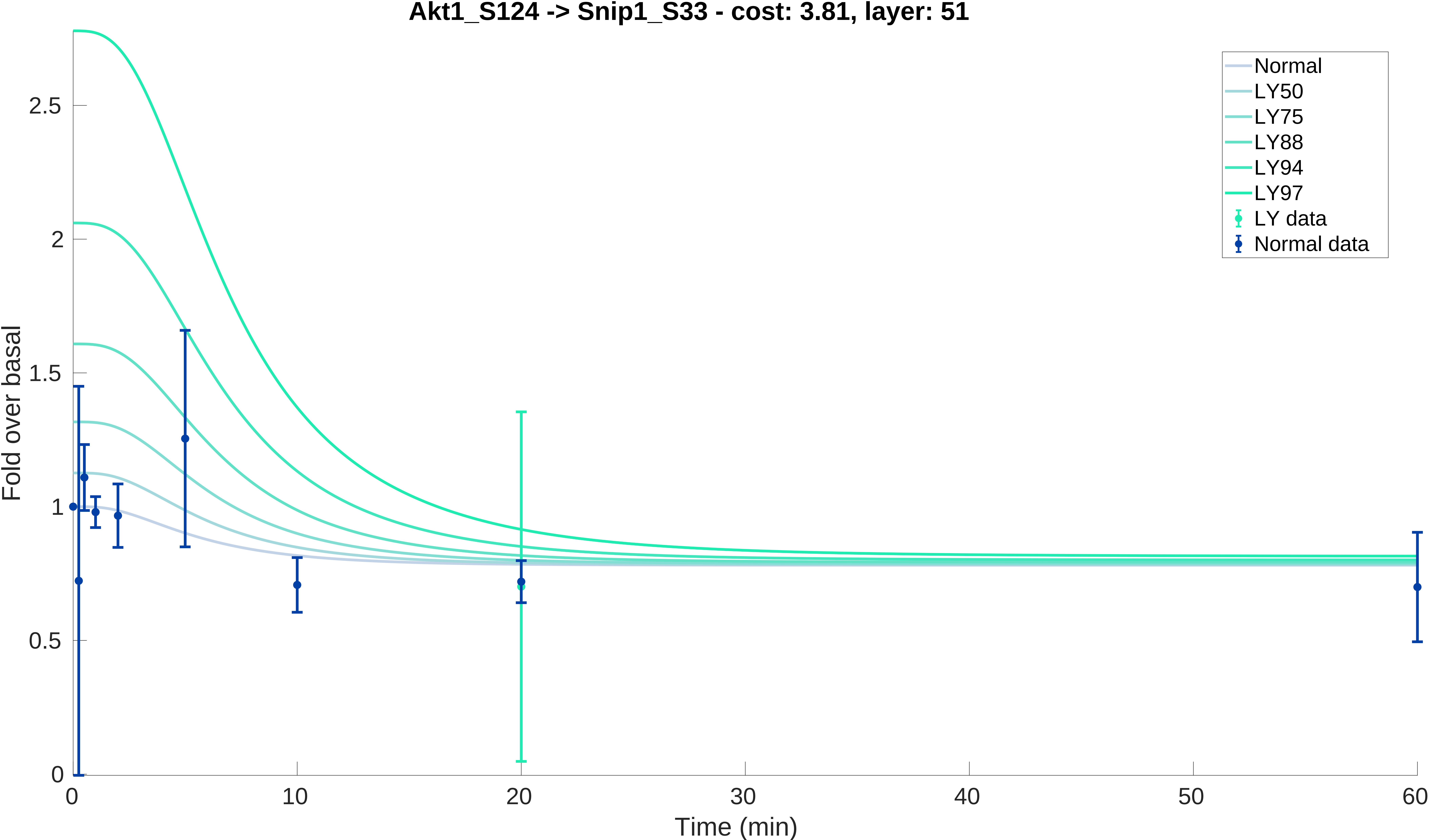

Akt1_S124 -> Snip1_S33 - cost: 3.81, layer: 51
Normal
LY50
LY75
LY88
LY94
LY97
LY data
Normal data
Fold over basal
Time (min)
0
0.5
1
1.5
2
2.5
0
10
20
30
40
50
60

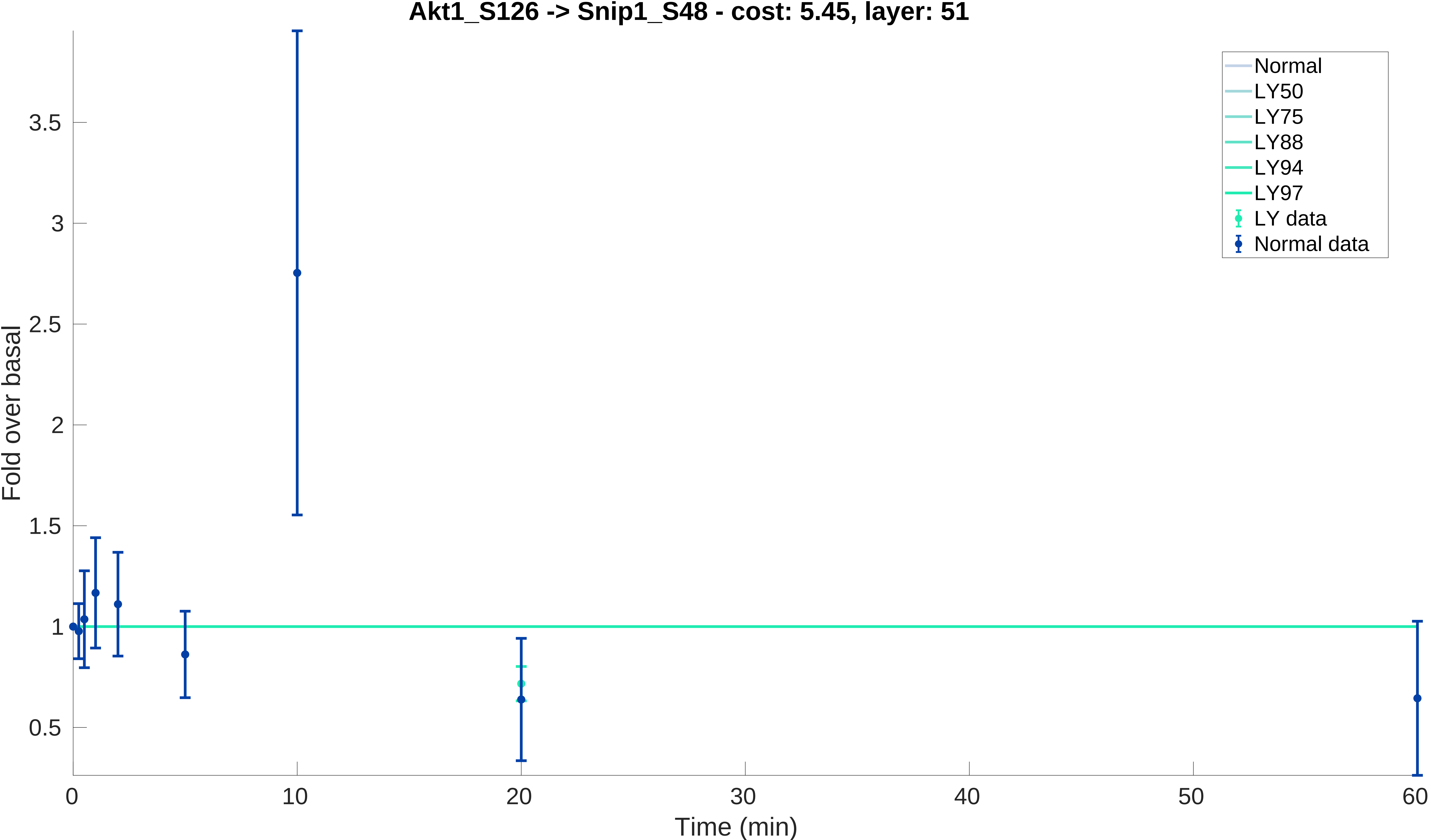

Akt1_S126 -> Snip1_S48 - cost: 5.45, layer: 51
Normal
LY50
LY75
LY88
LY94
LY97
LY data
Normal data
Fold over basal
3.5
3
2.5
2
1.5
1
0.5
0
10
20
30
40
50
60
Time (min)

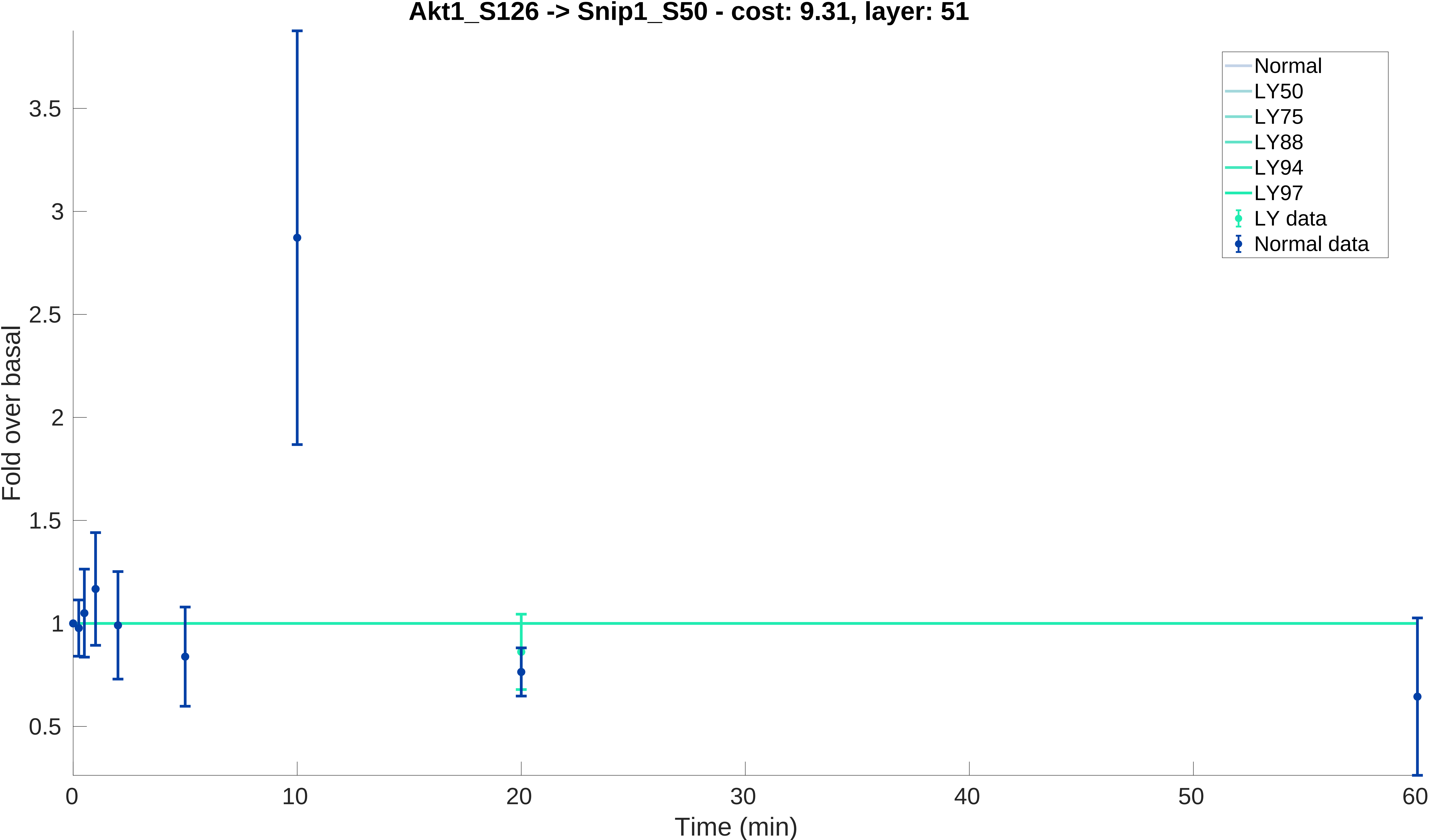
Akt1_S126 -> Snip1_S50 - cost: 9.31, layer: 51
Fold over basal
Time (min)
0.5
1
1.5
2
2.5
3
3.5
0
10
20
30
40
50
60
Normal
LY50
LY75
LY88
LY94
LY97
LY data
Normal data

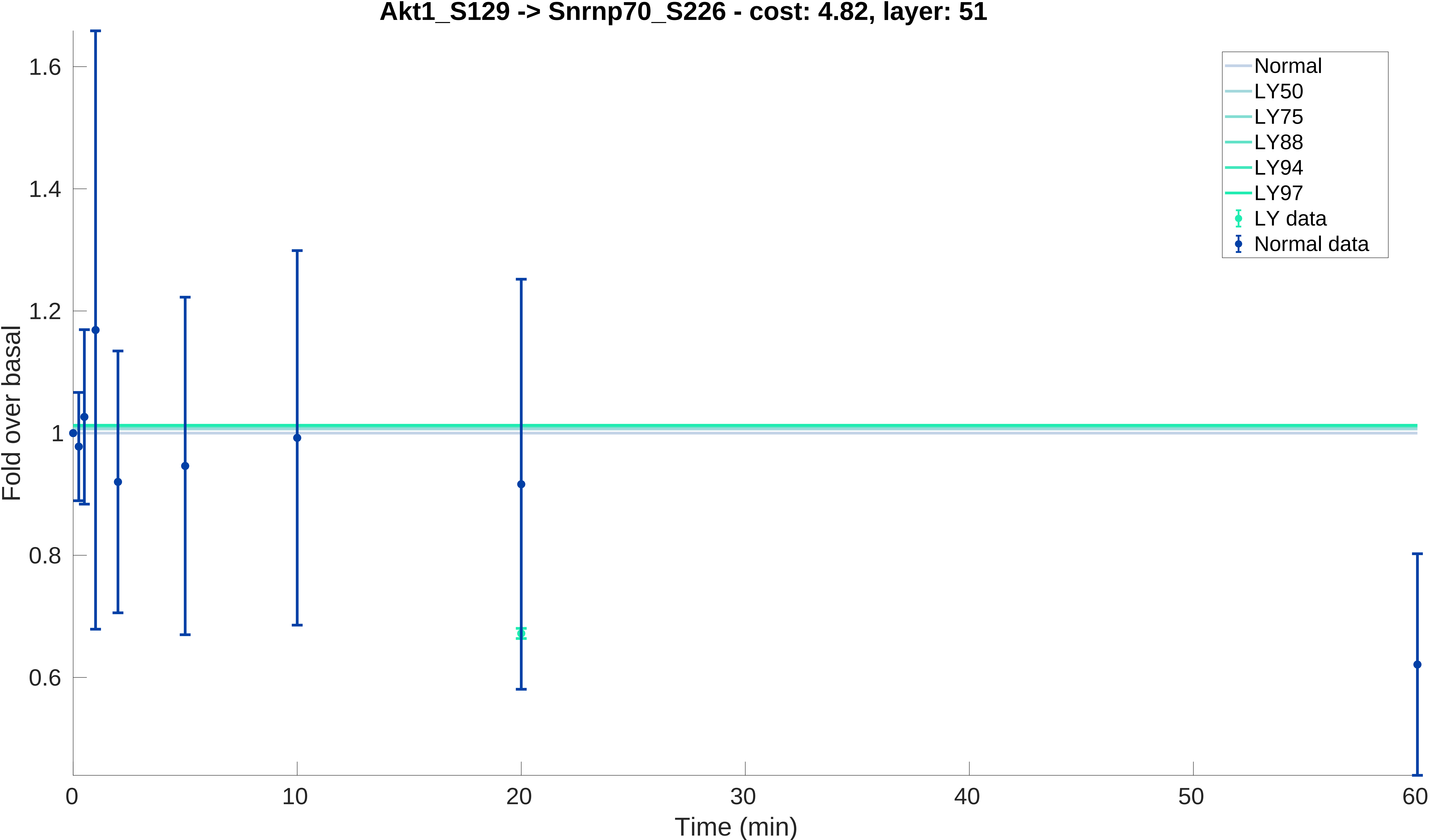

Akt1_S129 -> Snrnp70_S226 - cost: 4.82, layer: 51
Fold over basal
Time (min)
1.6
1.4
1.2
1
0.8
0.6
0
10
20
30
40
50
60
Normal
LY50
LY75
LY88
LY94
LY97
LY data
Normal data

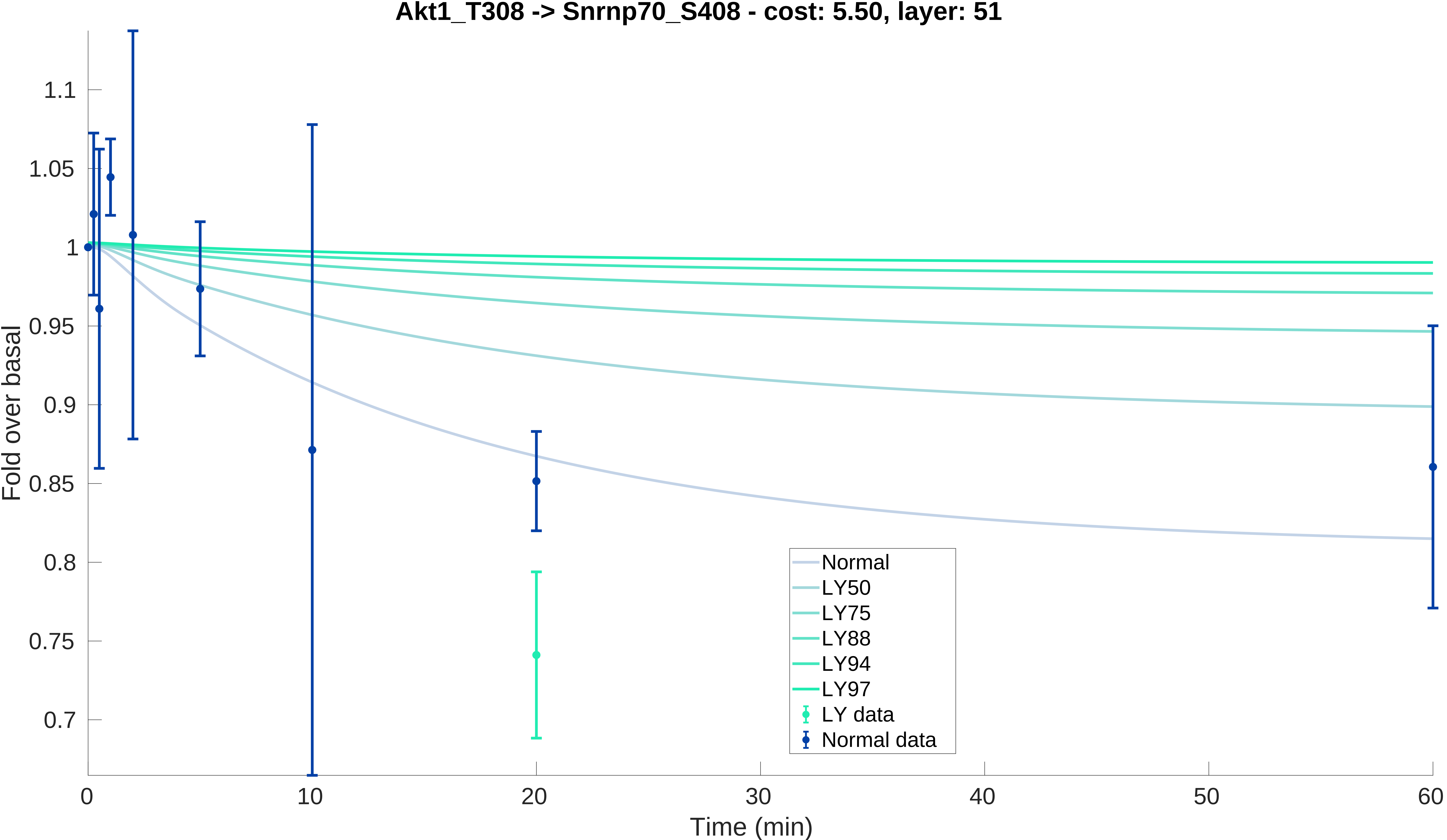

Akt1_T308 -> Snrnp70_S408 - cost: 5.50, layer: 51
Fold over basal
1.1
1.05
1
0.95
0.9
0.85
0.8
0.75
0.7
0
10
20
30
40
50
60
Time (min)
Normal
LY50
LY75
LY88
LY94
LY97
LY data
Normal data

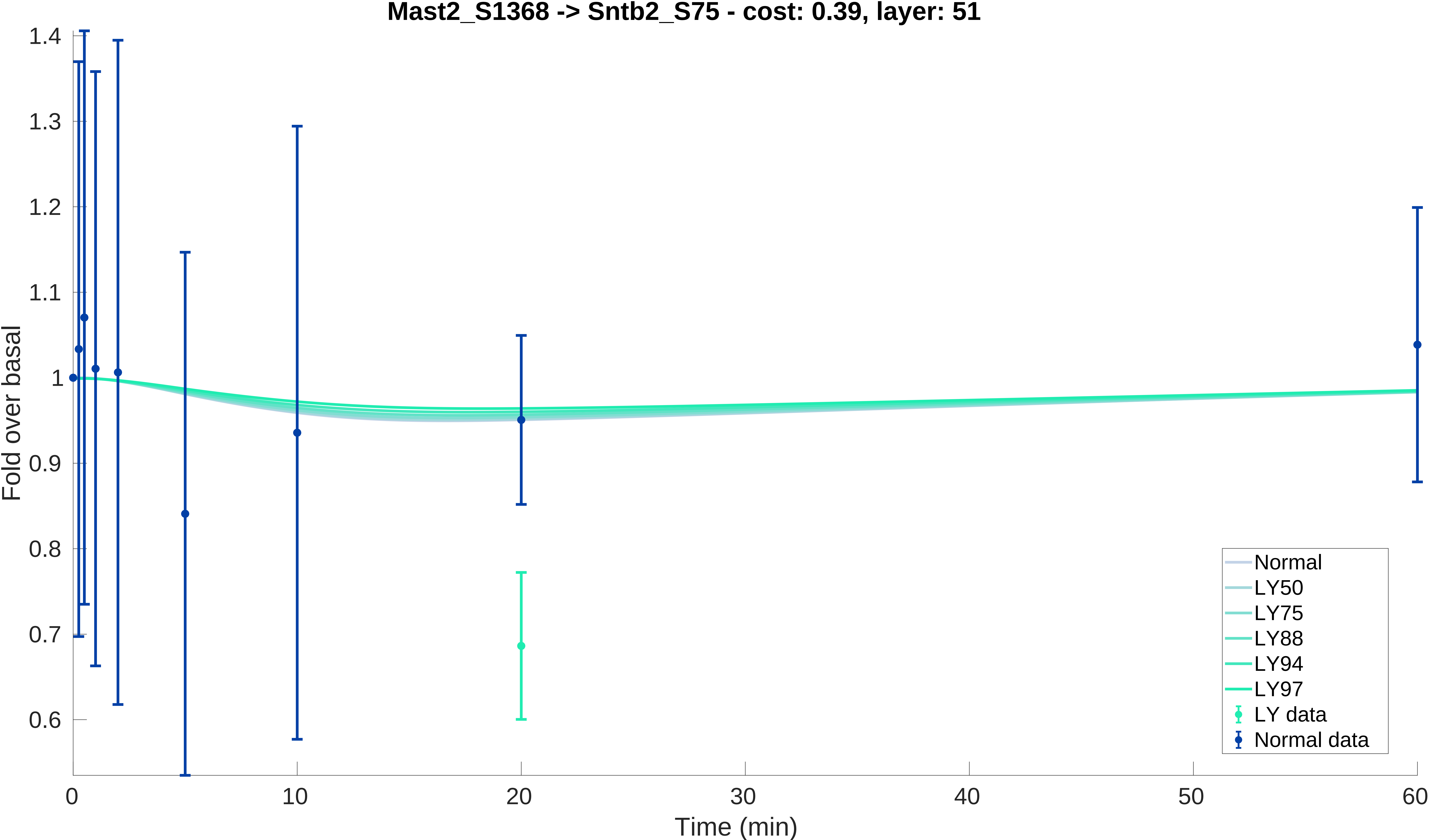

Mast2_S1368 -> Sntb2_S75 - cost: 0.39, layer: 51
Fold over basal
1.4
1.3
1.2
1.1
1
0.9
0.8
0.7
0.6
0
10
20
30
40
50
60
Time (min)
Normal
LY50
LY75
LY88
LY94
LY97
LY data
Normal data

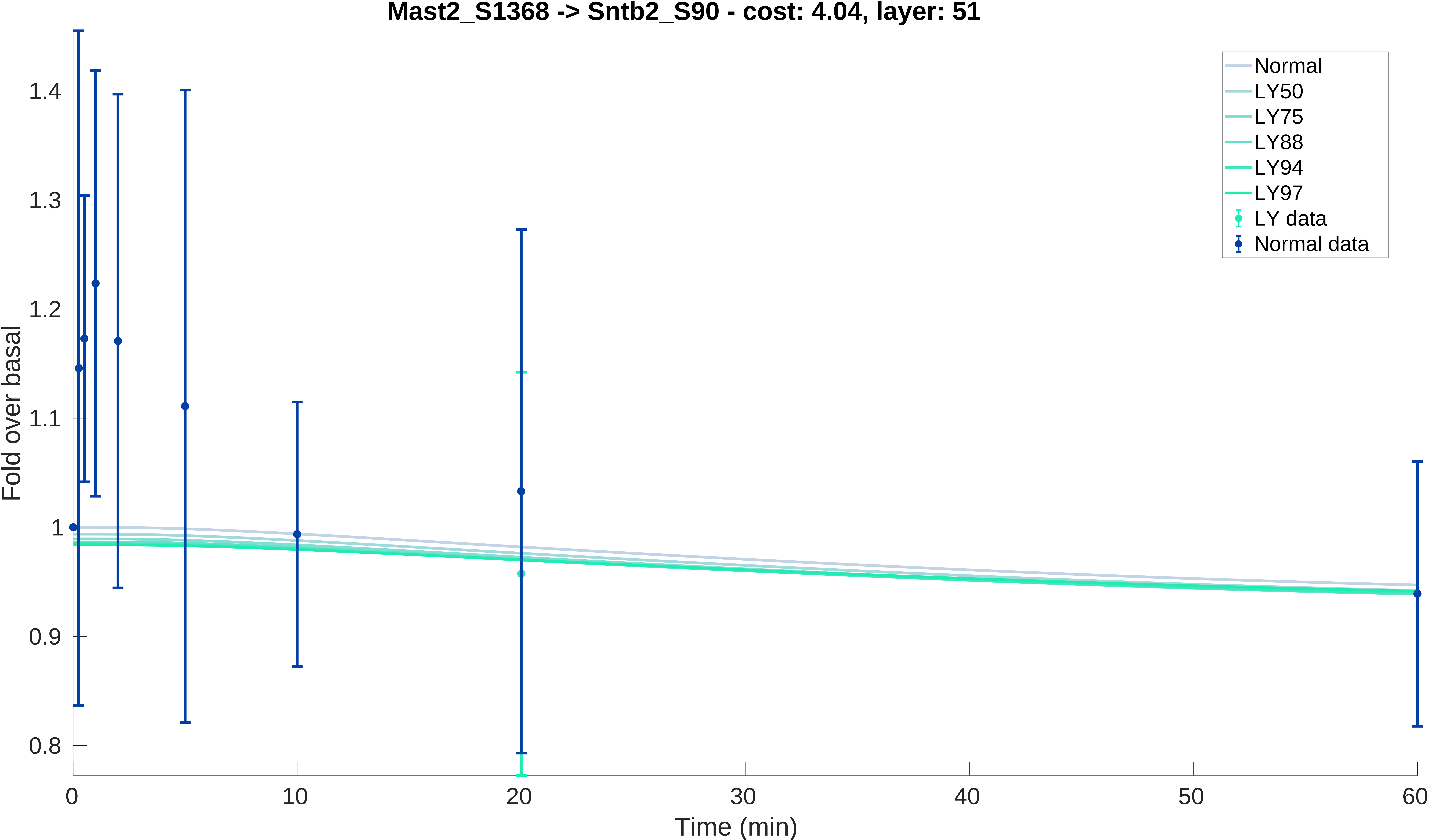

Mast2_S1368 -> Sntb2_S90 - cost: 4.04, layer: 51
Fold over basal
Time (min)
0.8
0.9
1
1.1
1.2
1.3
1.4
0
10
20
30
40
50
60
Normal
LY50
LY75
LY88
LY94
LY97
LY data
Normal data

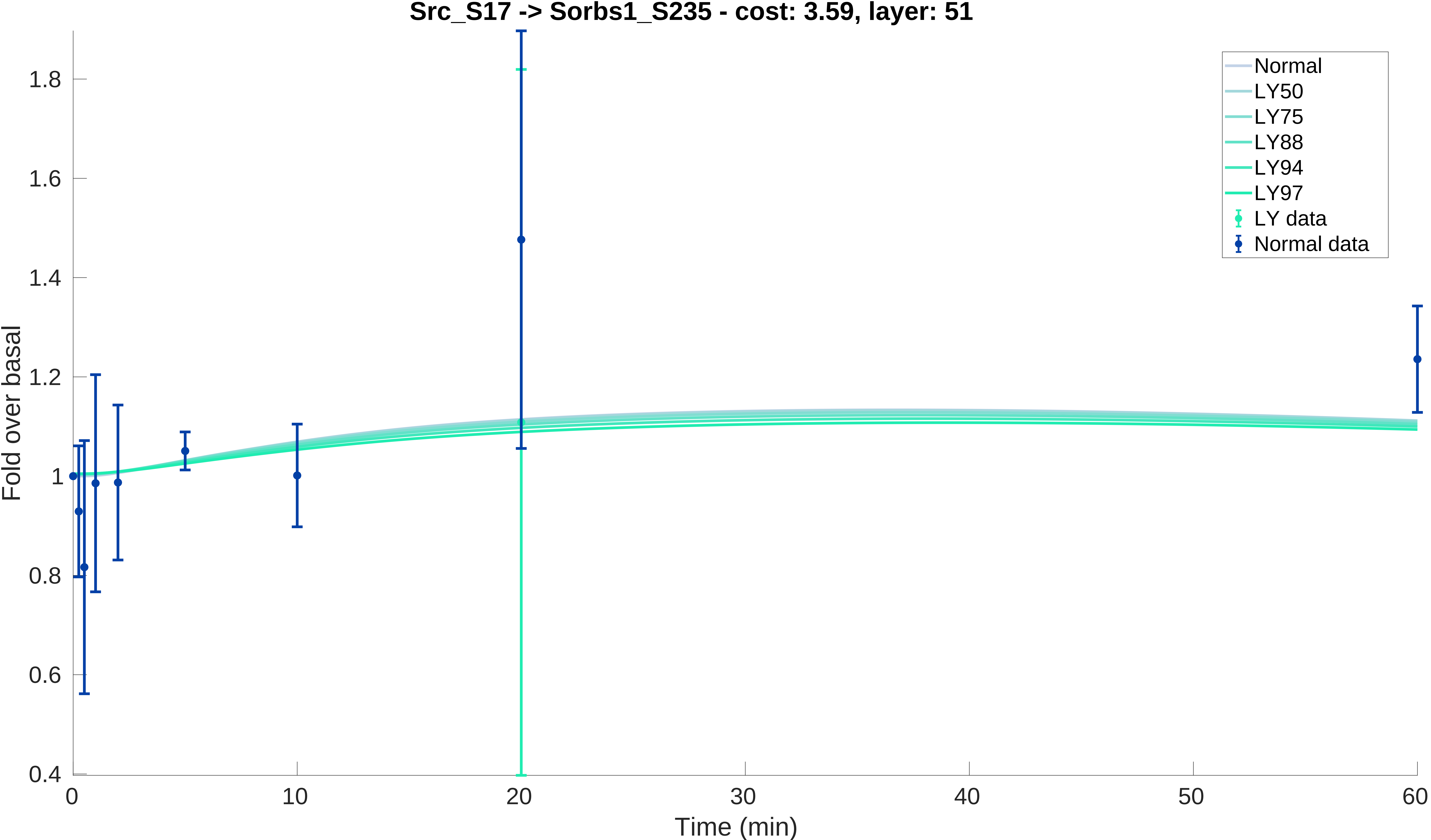
Src_S17 -> Sorbs1_S235 - cost: 3.59, layer: 51
Fold over basal
Time (min)
0.4
0.6
0.8
1
1.2
1.4
1.6
1.8
0
10
20
30
40
50
60
Normal
LY50
LY75
LY88
LY94
LY97
LY data
Normal data

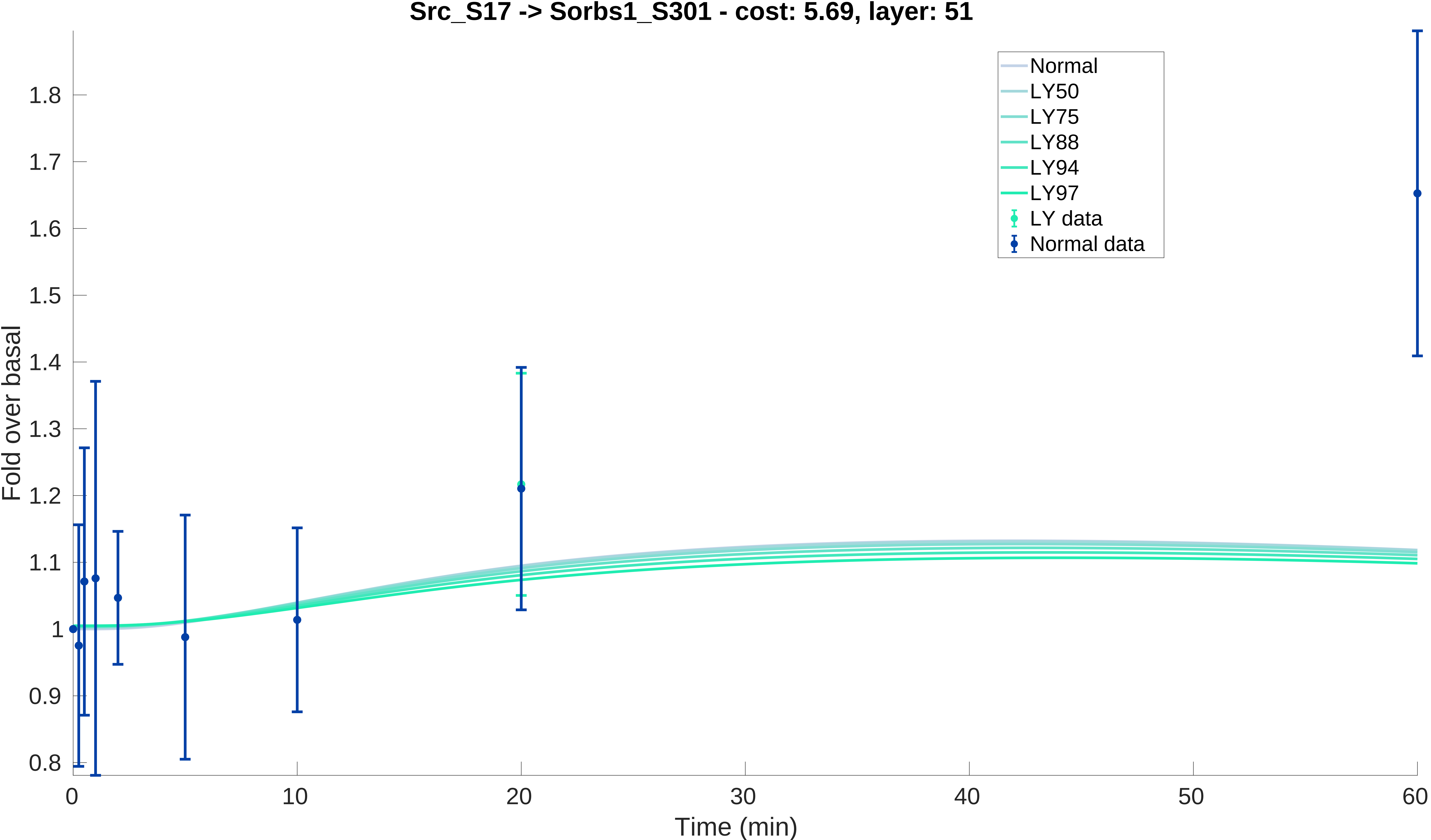
Src_S17 -> Sorbs1_S301 - cost: 5.69, layer: 51
Normal
LY50
LY75
LY88
LY94
LY97
LY data
Normal data
Fold over basal
Time (min)
0.8
0.9
1
1.1
1.2
1.3
1.4
1.5
1.6
1.7
1.8
0
10
20
30
40
50
60

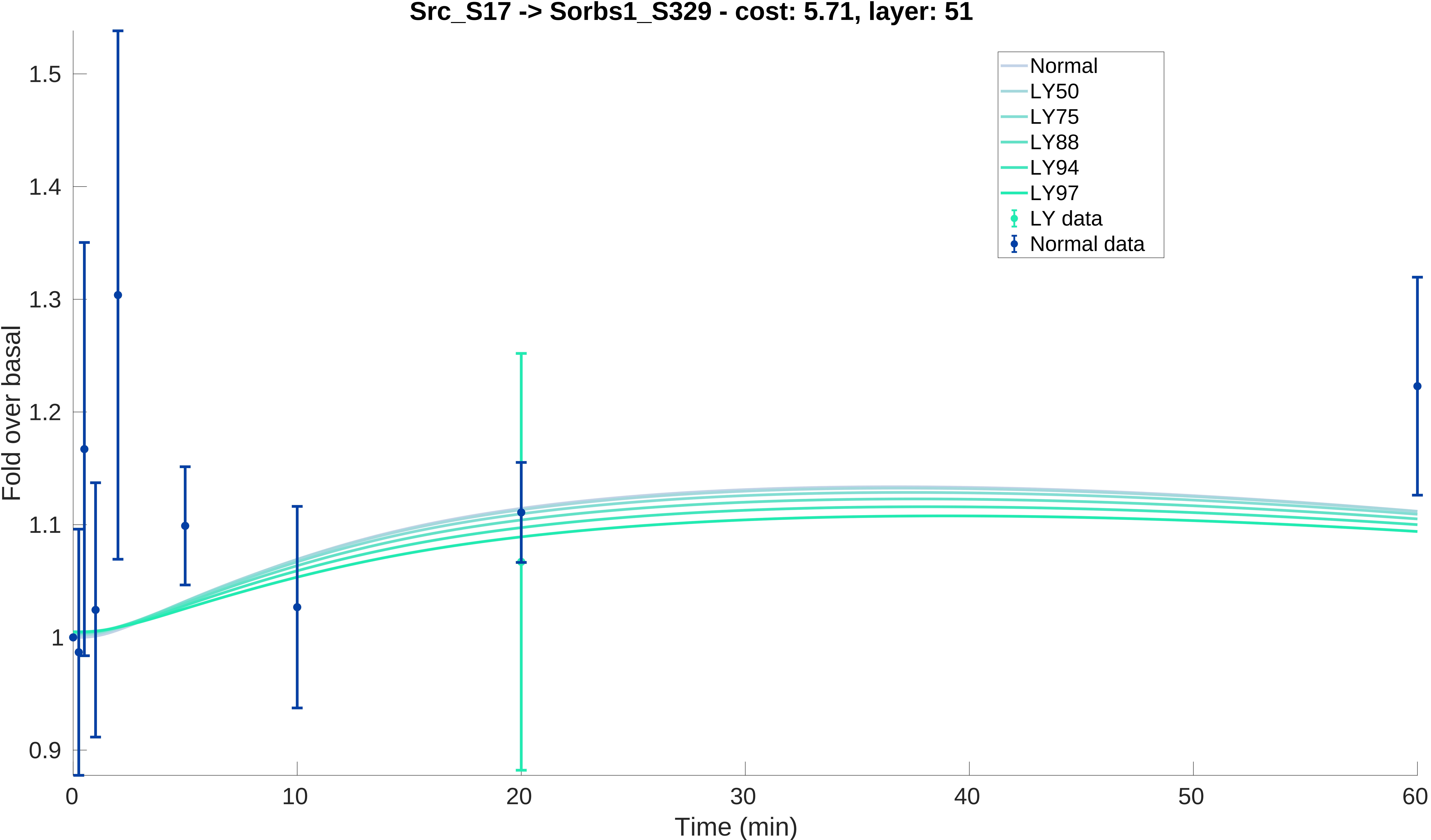
Src_S17 -> Sorbs1_S329 - cost: 5.71, layer: 51
Normal
LY50
LY75
LY88
LY94
LY97
LY data
Normal data
Fold over basal
1.5
1.4
1.3
1.2
1.1
1
0.9
0
10
20
30
40
50
60
Time (min)

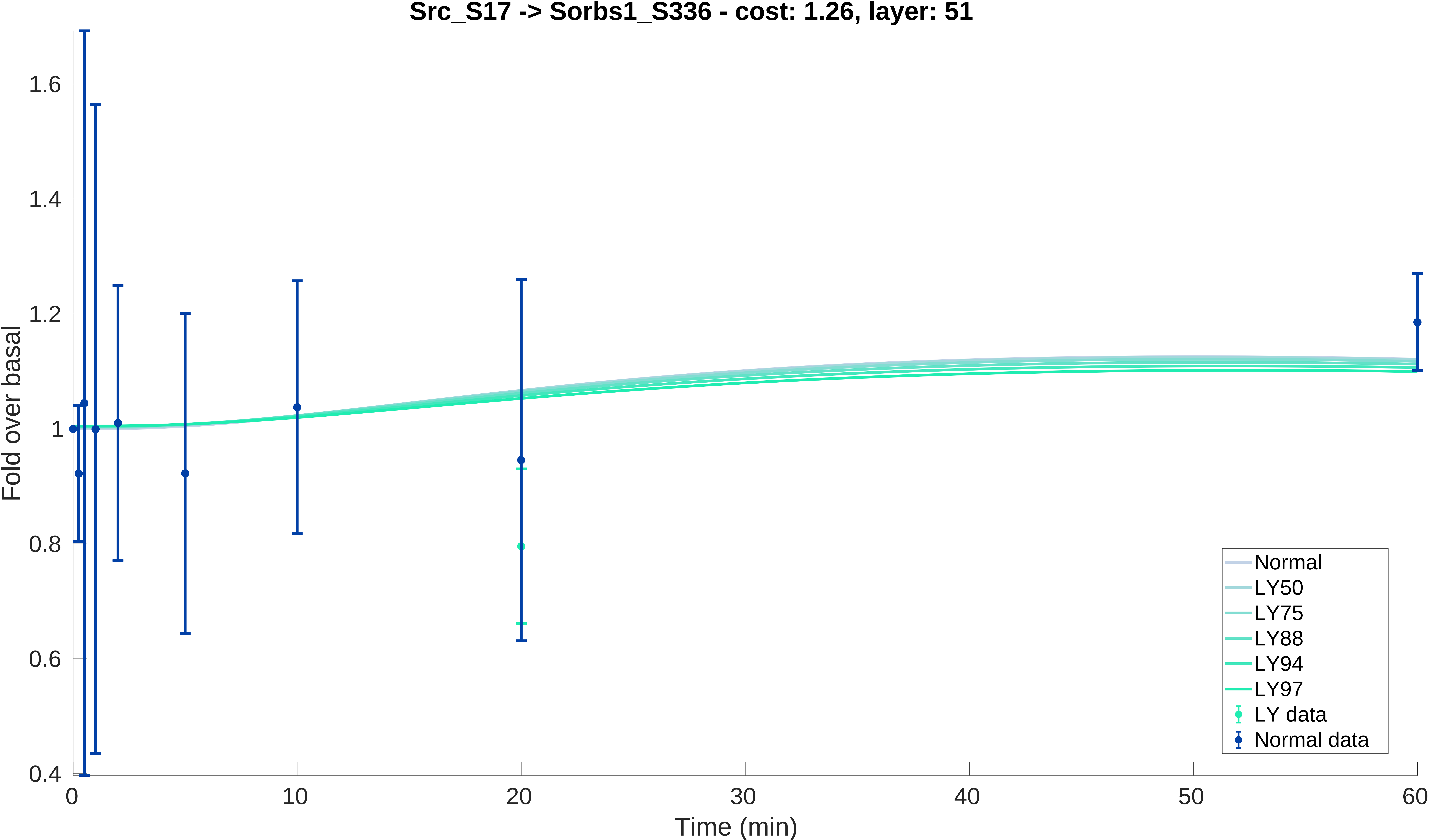
Src_S17 -> Sorbs1_S336 - cost: 1.26, layer: 51
Fold over basal
1.6
1.4
1.2
1
0.8
0.6
0.4
0
10
20
30
40
50
60
Time (min)
Normal
LY50
LY75
LY88
LY94
LY97
LY data
Normal data

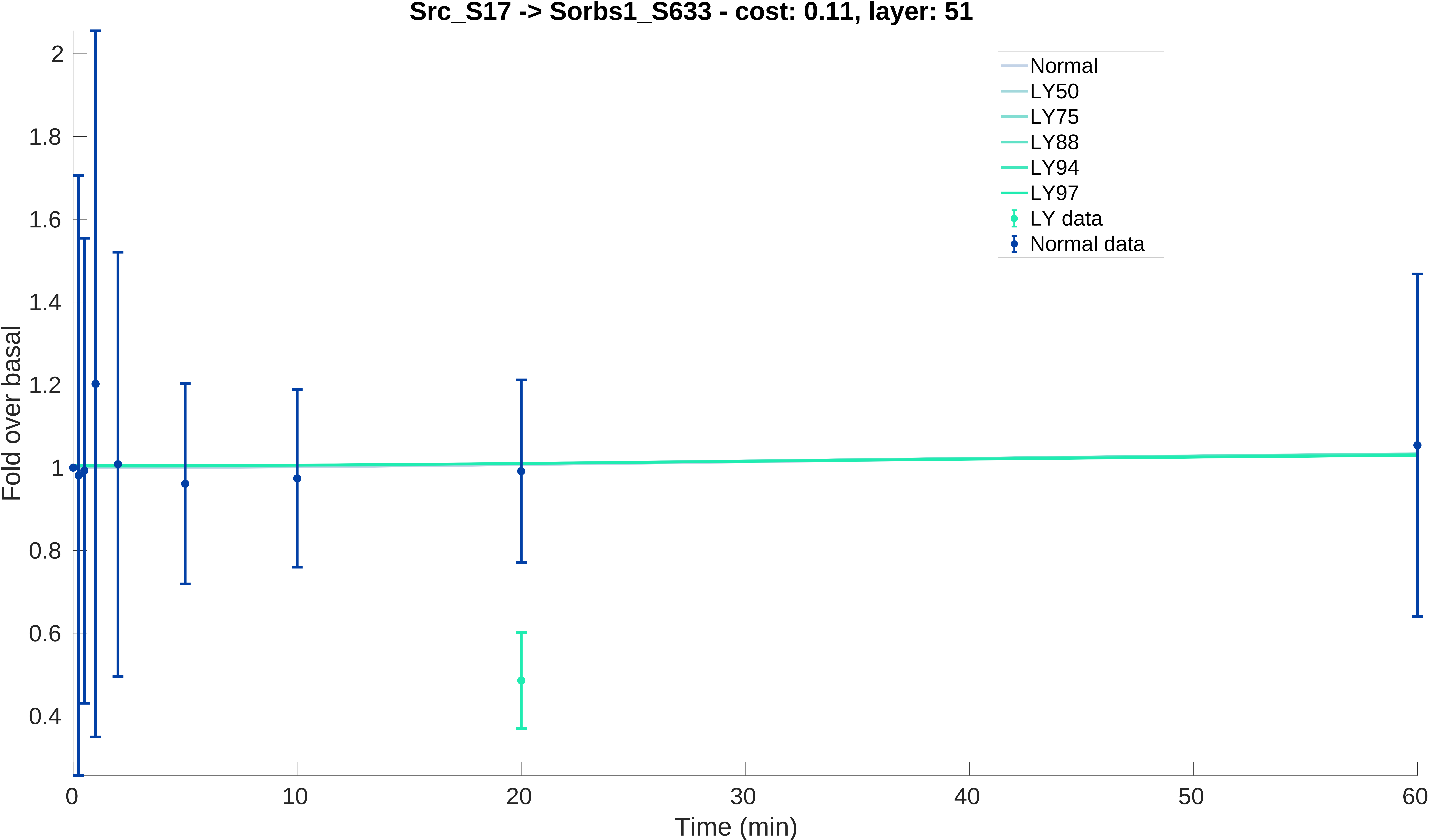
Src_S17 -> Sorbs1_S633 - cost: 0.11, layer: 51
Fold over basal
Time (min)
0
10
20
30
40
50
60
0.4
0.6
0.8
1
1.2
1.4
1.6
1.8
2
Normal
LY50
LY75
LY88
LY94
LY97
LY data
Normal data

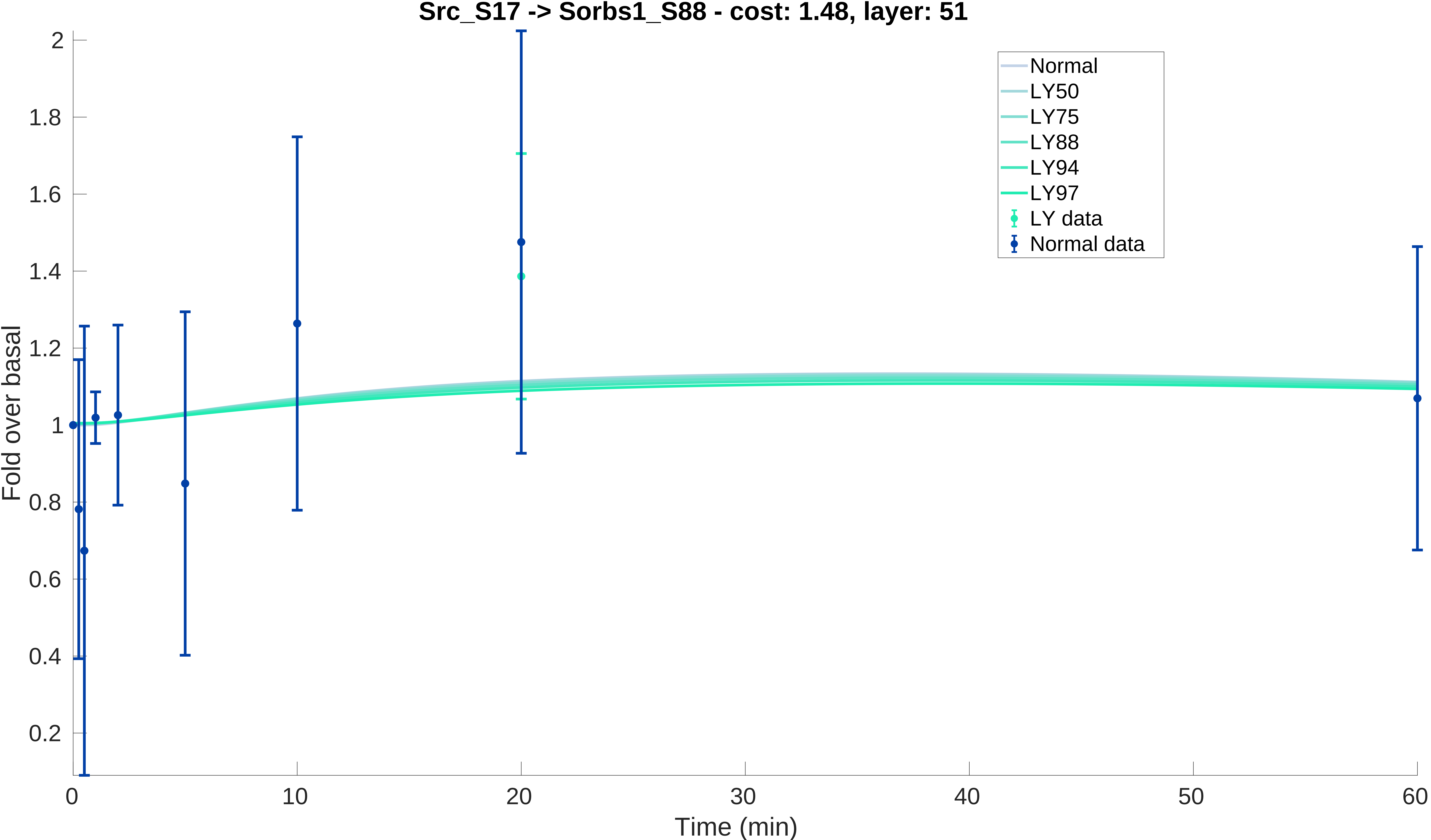

Src_S17 -> Sorbs1_S88 - cost: 1.48, layer: 51
Normal
LY50
LY75
LY88
LY94
LY97
LY data
Normal data
Fold over basal
Time (min)
0
10
20
30
40
50
60
0.2
0.4
0.6
0.8
1
1.2
1.4
1.6
1.8
2

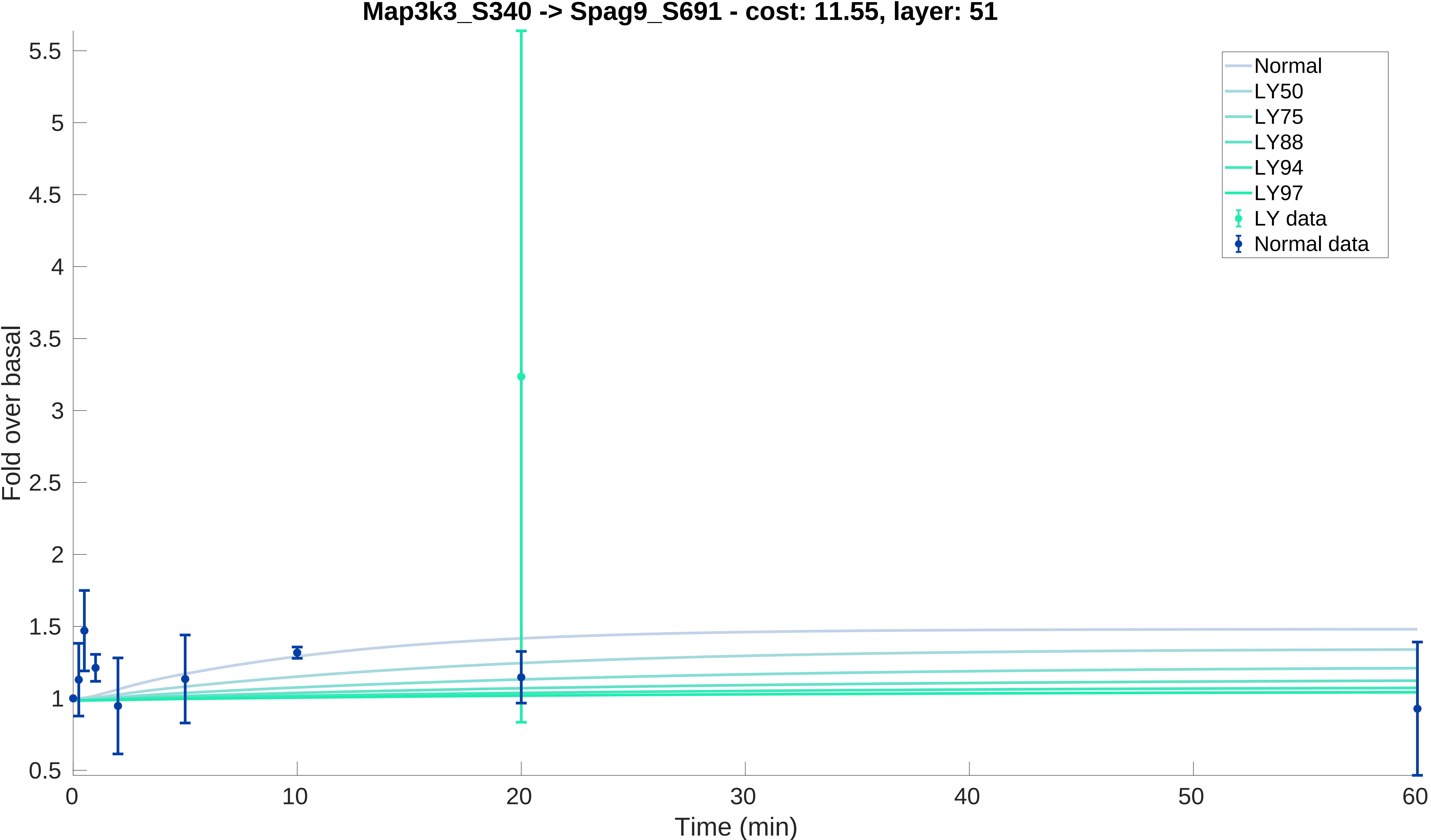

Map3k3_S340 -> Spag9_S691 - cost: 11.55, layer: 51
Fold over basal
Time (min)
0.5
1
1.5
2
2.5
3
3.5
4
4.5
5
5.5
0
10
20
30
40
50
60
Normal
LY50
LY75
LY88
LY94
LY97
LY data
Normal data

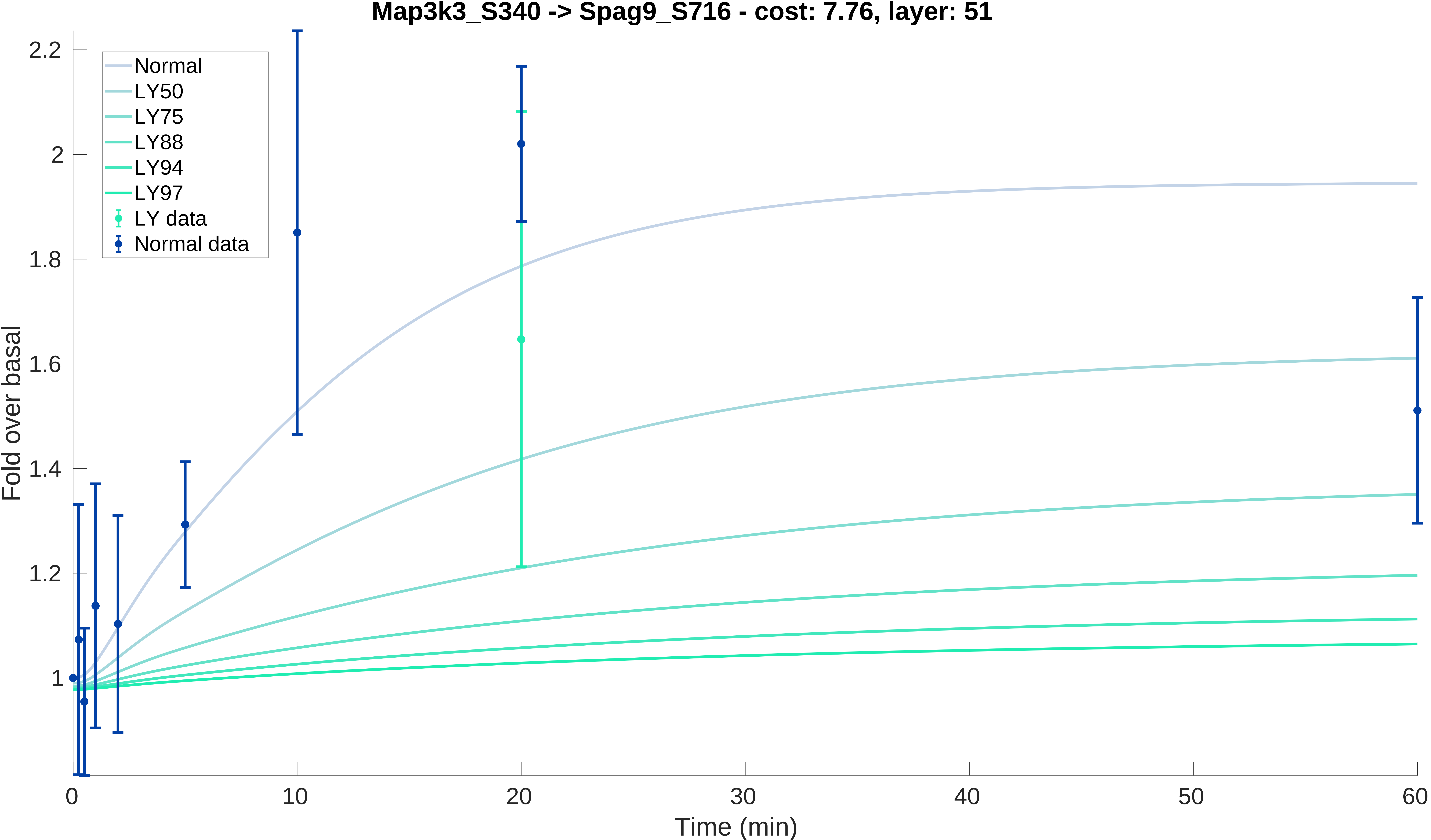

Map3k3_S340 -> Spag9_S716 - cost: 7.76, layer: 51
Normal
LY50
LY75
LY88
LY94
LY97
LY data
Normal data
Fold over basal
Time (min)
2.2
2
1.8
1.6
1.4
1.2
1
0
10
20
30
40
50
60

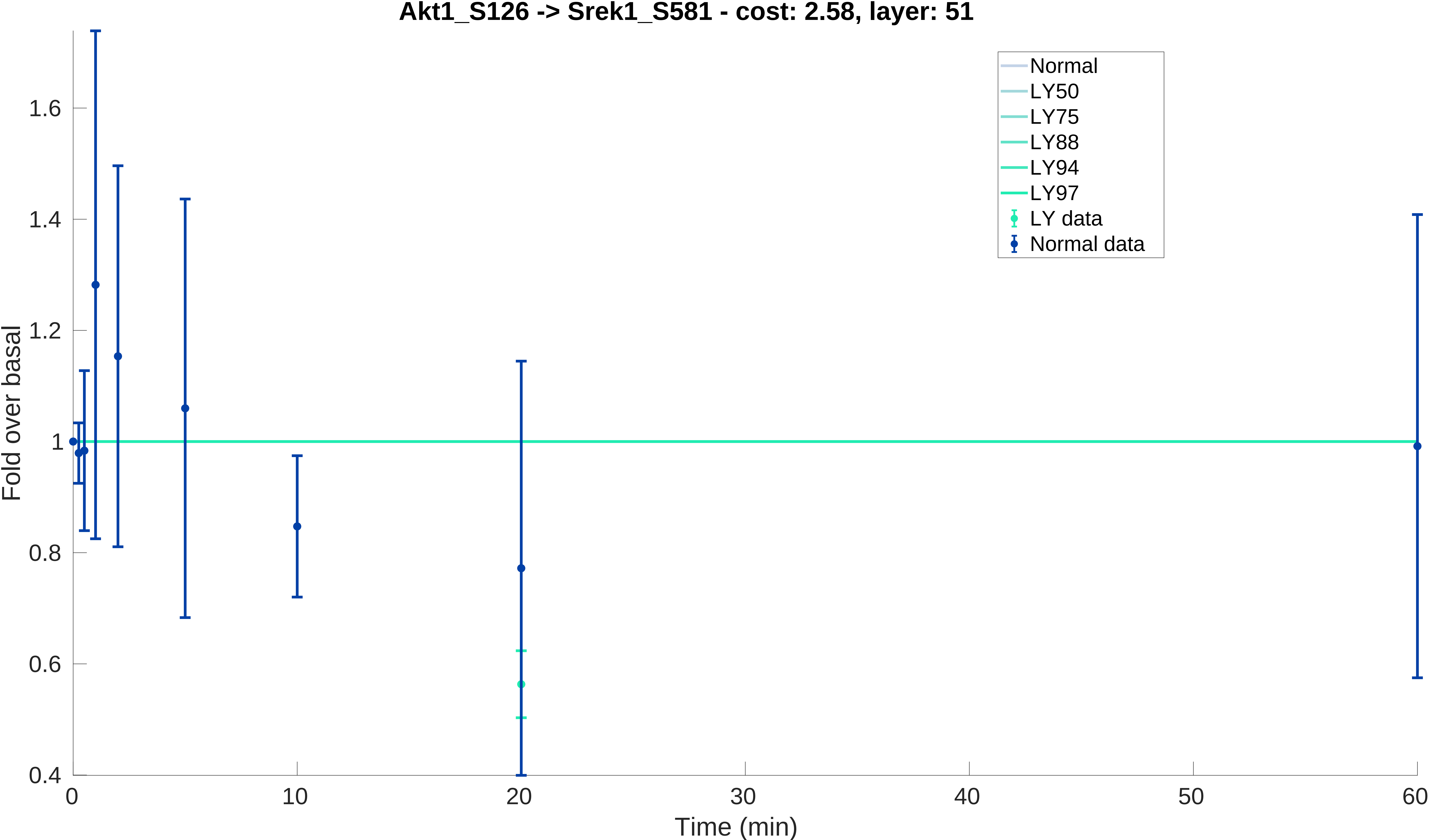
Akt1_S126 -> Srek1_S581 - cost: 2.58, layer: 51
Fold over basal
Time (min)
1.6
1.4
1.2
1
0.8
0.6
0.4
0
10
20
30
40
50
60
Normal
LY50
LY75
LY88
LY94
LY97
LY data
Normal data

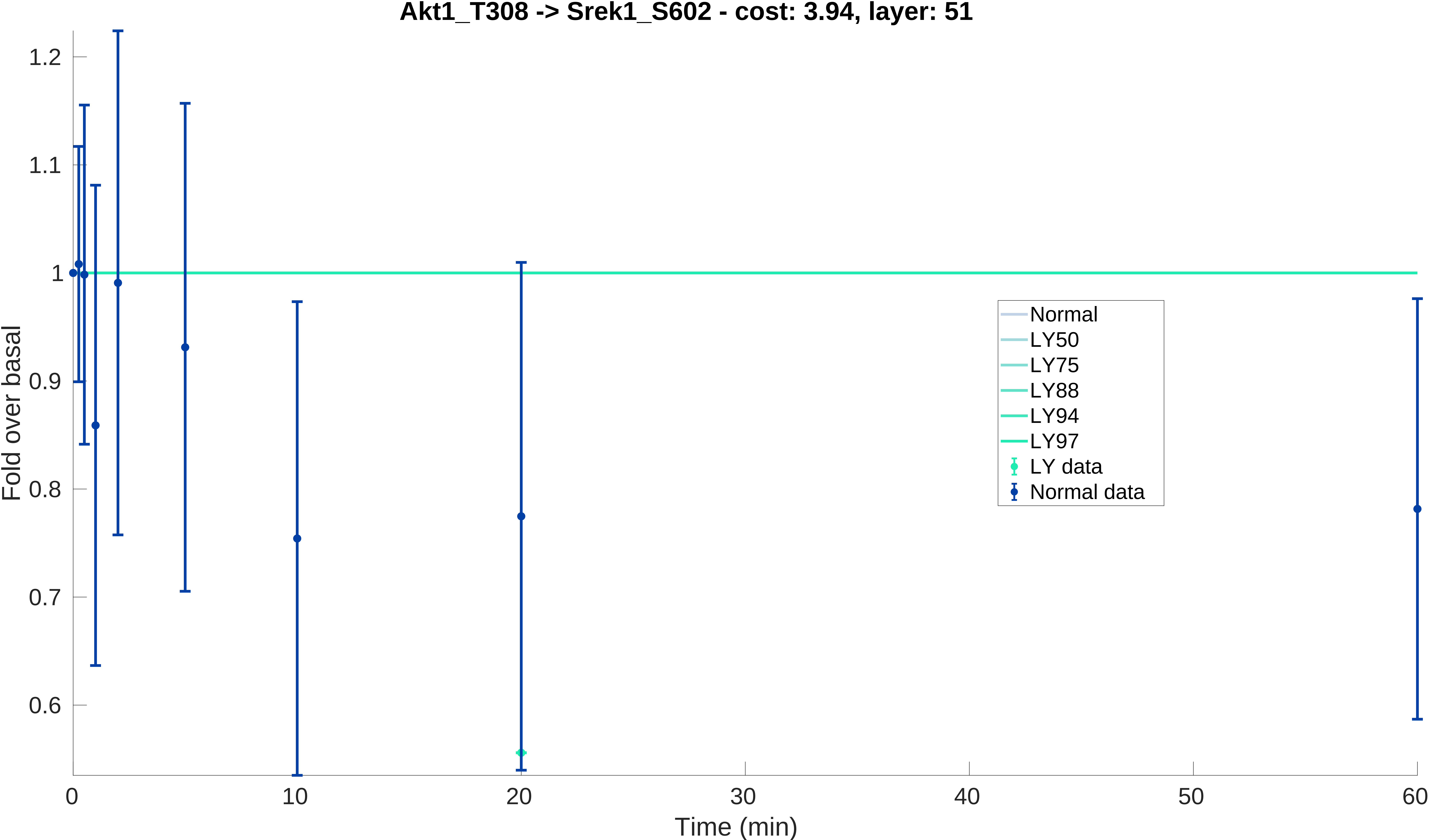
Akt1_T308 -> Srek1_S602 - cost: 3.94, layer: 51
Fold over basal
Time (min)
0
10
20
30
40
50
60
0.6
0.7
0.8
0.9
1
1.1
1.2
Normal
LY50
LY75
LY88
LY94
LY97
LY data
Normal data

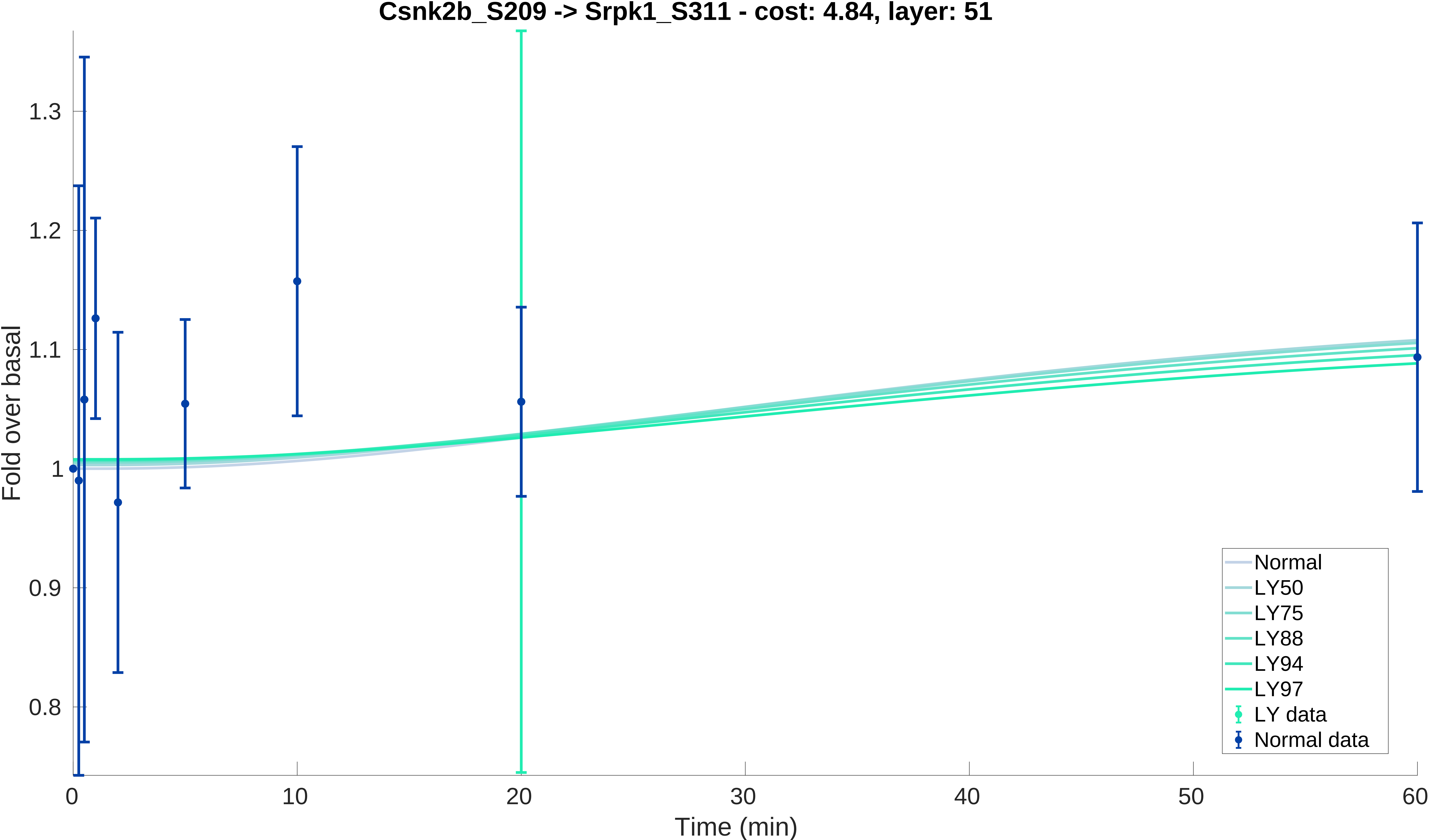
Csnk2b_S209 -> Srpk1_S311 - cost: 4.84, layer: 51
Fold over basal
1.3
1.2
1.1
1
0.9
0.8
0
10
20
30
40
50
60
Time (min)
Normal
LY50
LY75
LY88
LY94
LY97
LY data
Normal data

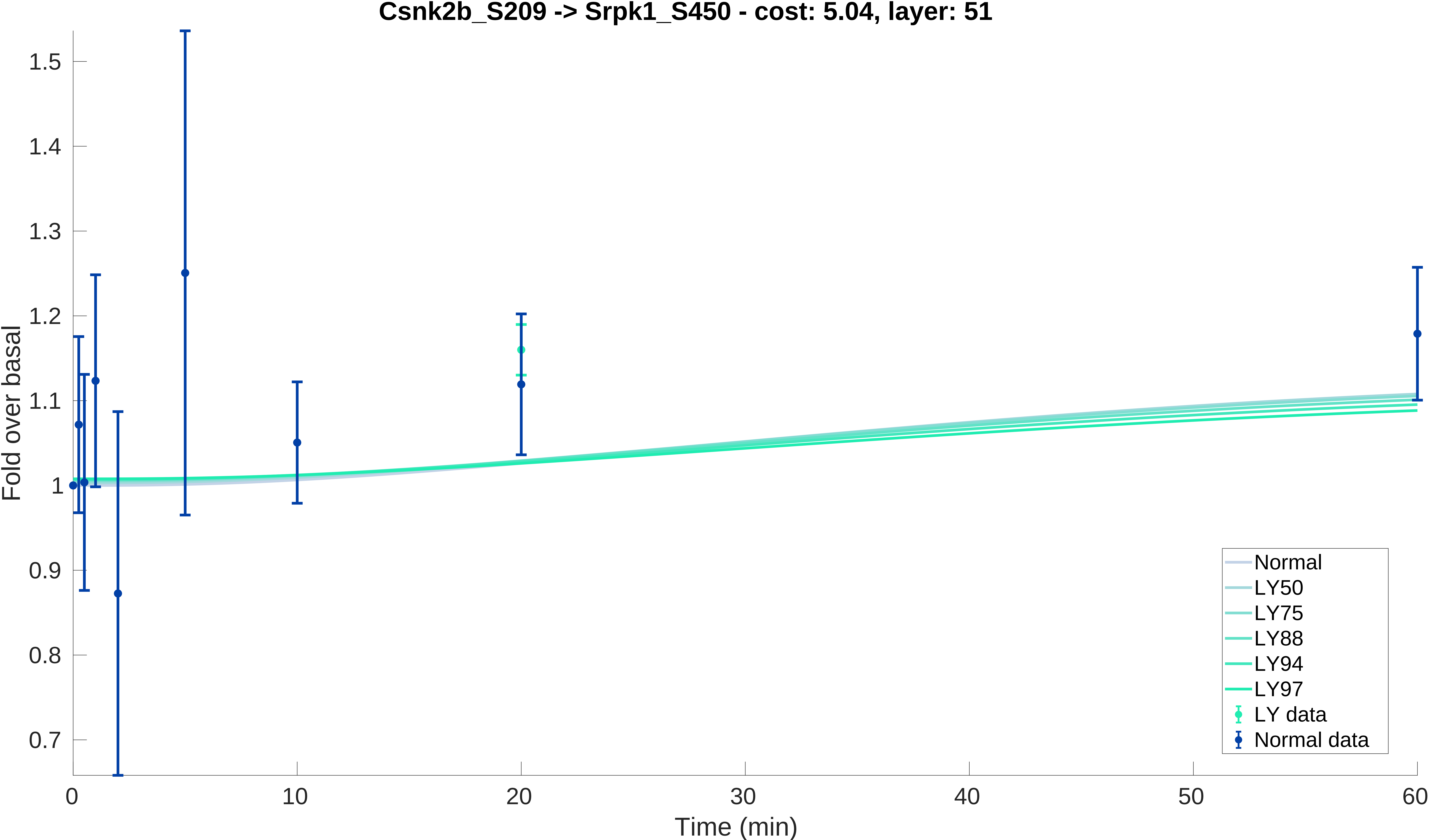
Csnk2b_S209 -> Srpk1_S450 - cost: 5.04, layer: 51
Fold over basal
Time (min)
1.5
1.4
1.3
1.2
1.1
1
0.9
0.8
0.7
0
10
20
30
40
50
60
Normal
LY50
LY75
LY88
LY94
LY97
LY data
Normal data

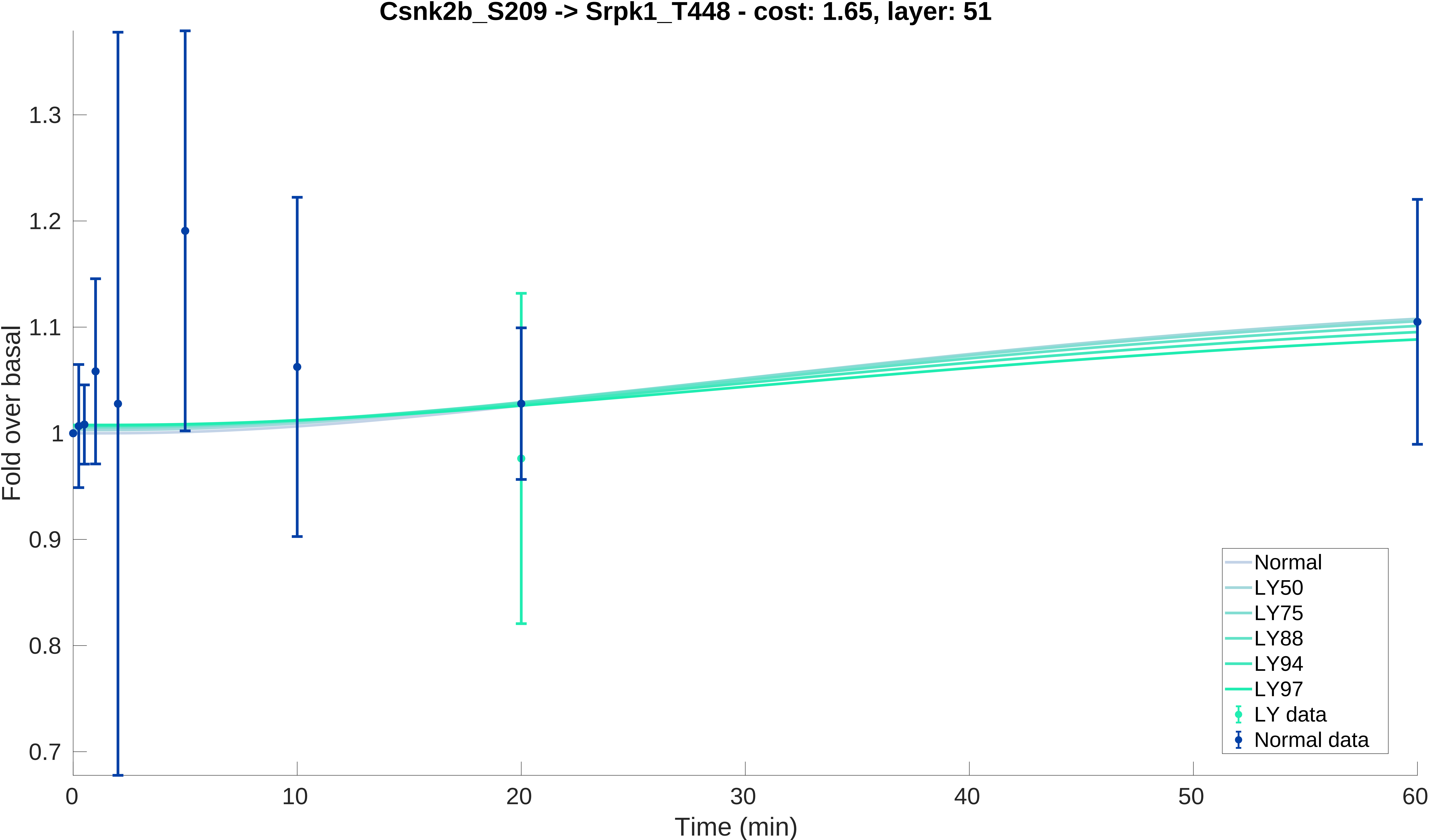
Csnk2b_S209 -> Srpk1_T448 - cost: 1.65, layer: 51
Fold over basal
1.3
1.2
1.1
1
0.9
0.8
0.7
0
10
20
30
40
50
60
Time (min)
Normal
LY50
LY75
LY88
LY94
LY97
LY data
Normal data

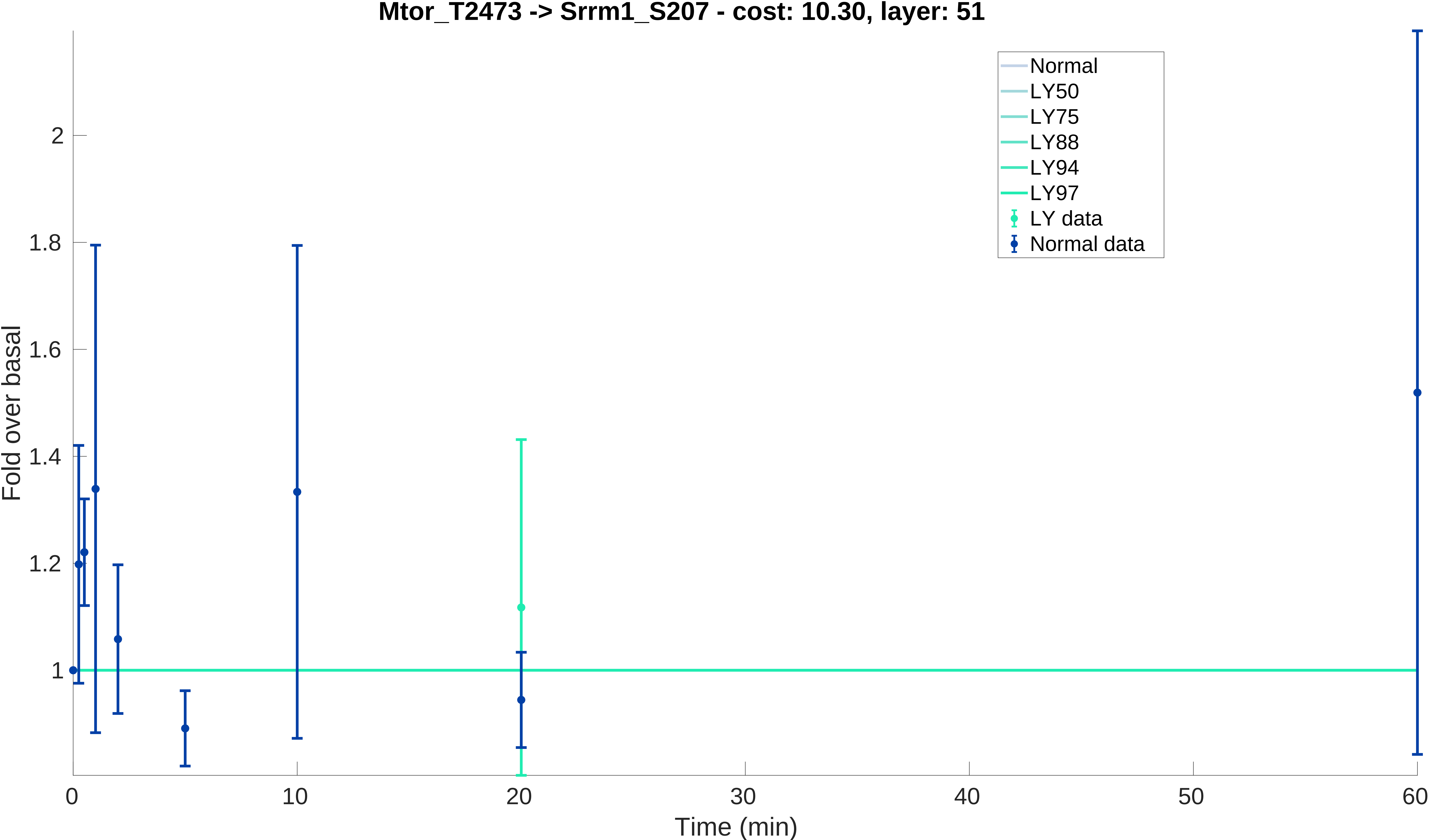

Mtor_T2473 -> Srrm1_S207 - cost: 10.30, layer: 51
Fold over basal
Time (min)
Normal
LY50
LY75
LY88
LY94
LY97
LY data
Normal data
2
1.8
1.6
1.4
1.2
1
0
10
20
30
40
50
60

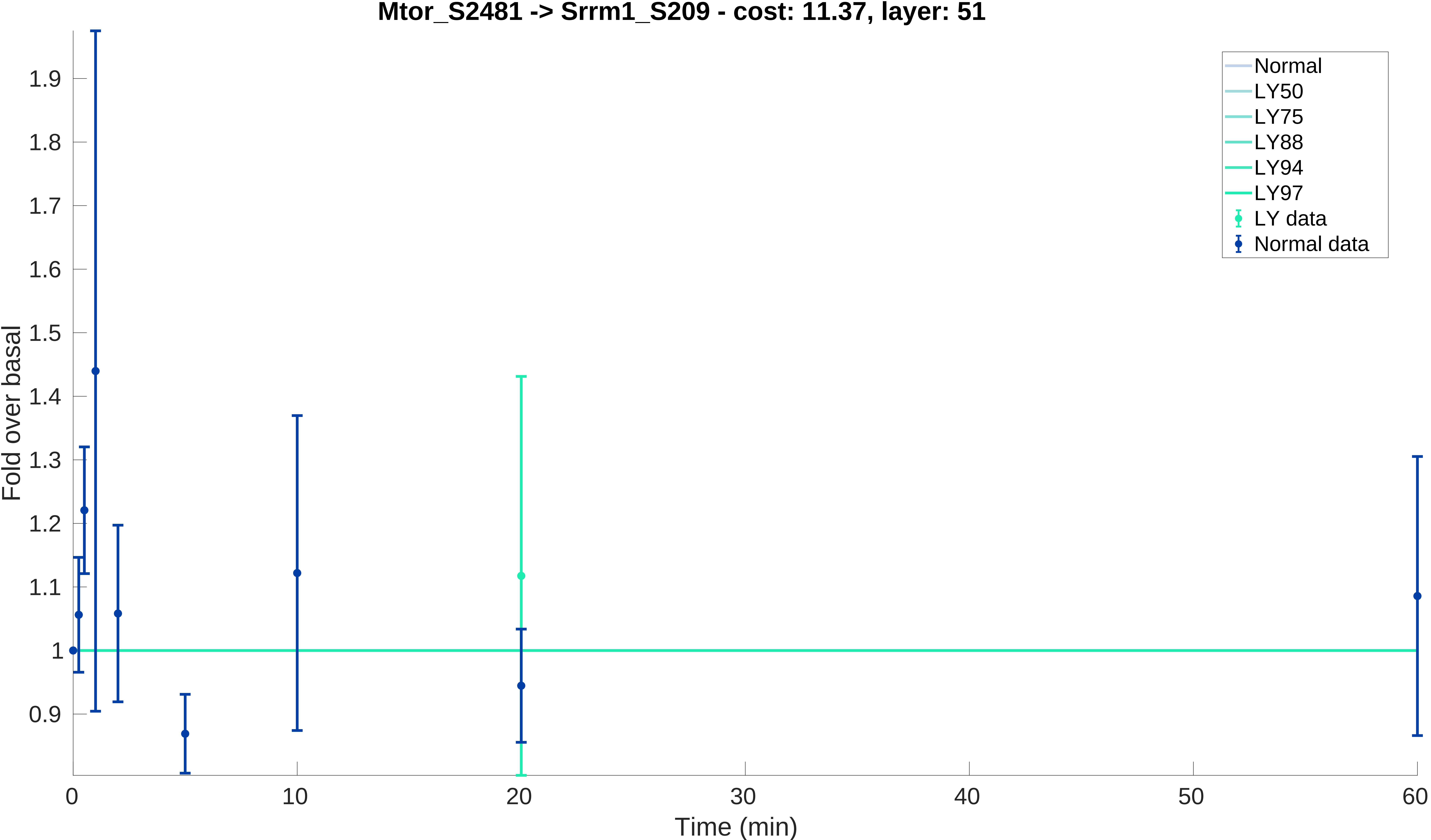

Mtor_S2481 -> Srrm1_S209 - cost: 11.37, layer: 51
Normal
LY50
LY75
LY88
LY94
LY97
LY data
Normal data
Fold over basal
1.9
1.8
1.7
1.6
1.5
1.4
1.3
1.2
1.1
1
0.9
0
10
20
30
40
50
60
Time (min)

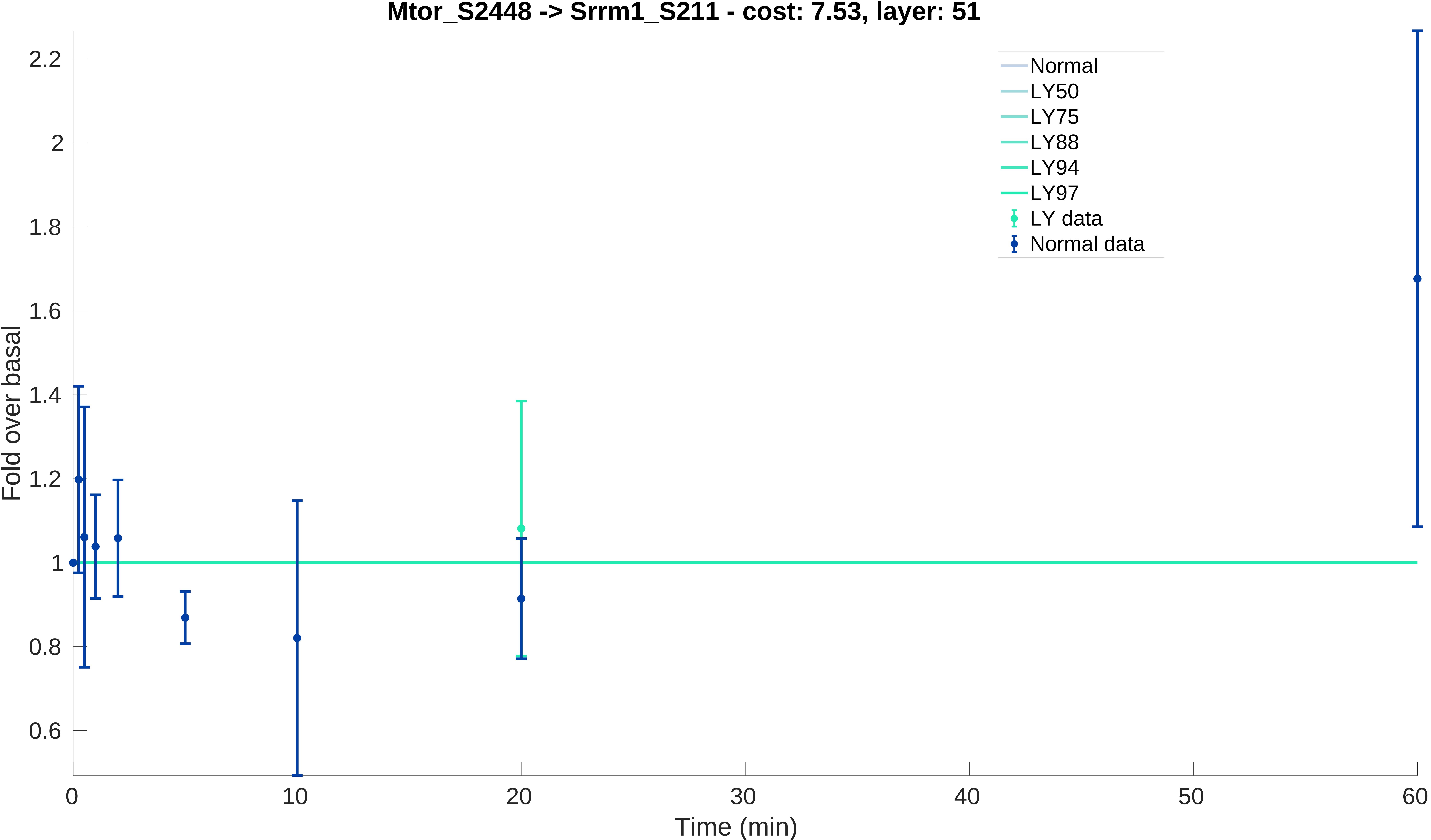

Mtor_S2448 -> Srrm1_S211 - cost: 7.53, layer: 51
Normal
LY50
LY75
LY88
LY94
LY97
LY data
Normal data
Fold over basal
Time (min)
2.2
2
1.8
1.6
1.4
1.2
1
0.8
0.6
0
10
20
30
40
50
60

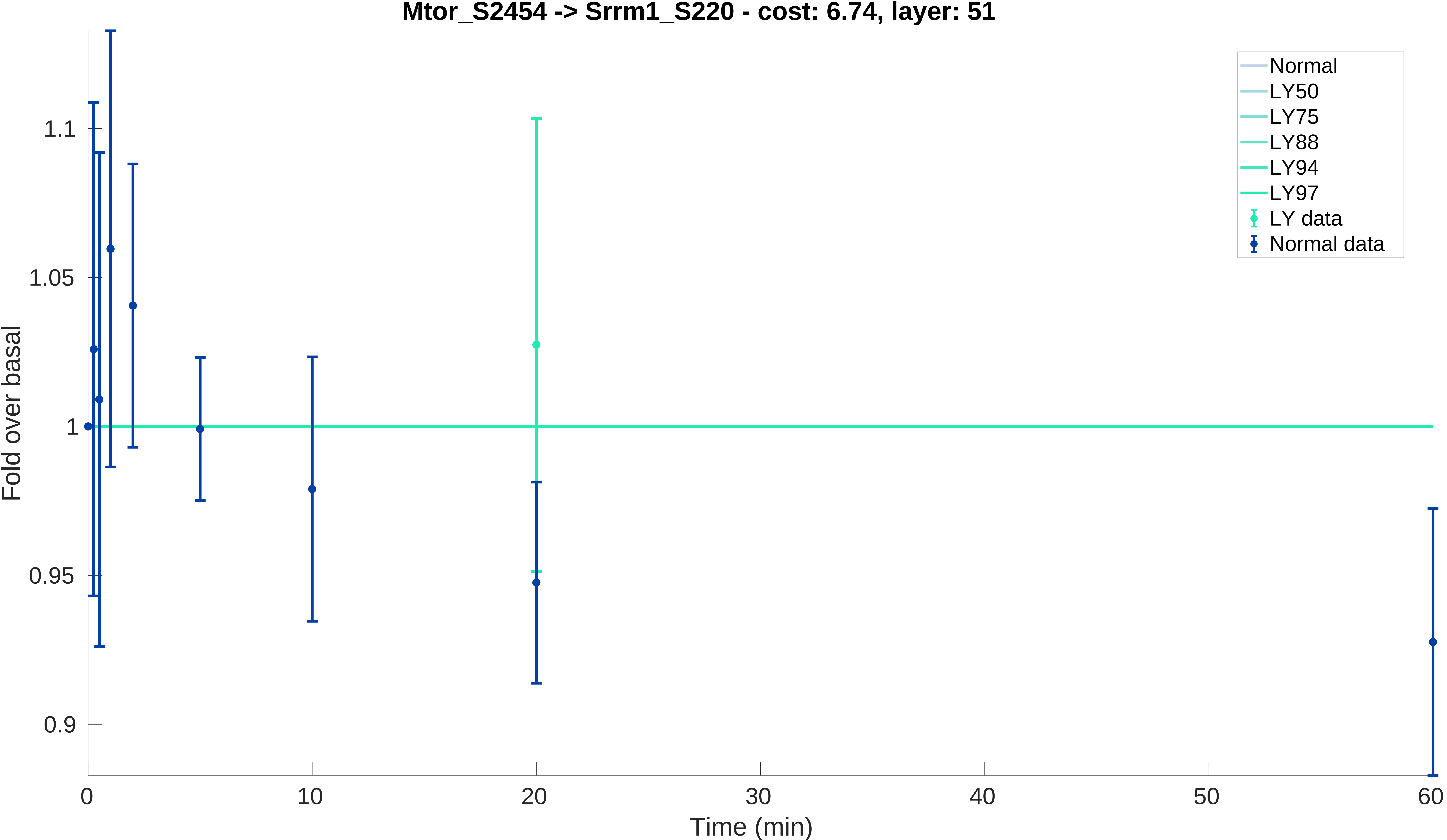

Mtor_S2454 -> Srrm1_S220 - cost: 6.74, layer: 51
Fold over basal
Time (min)
1.1
1.05
1
0.95
0.9
0
10
20
30
40
50
60
Normal
LY50
LY75
LY88
LY94
LY97
LY data
Normal data

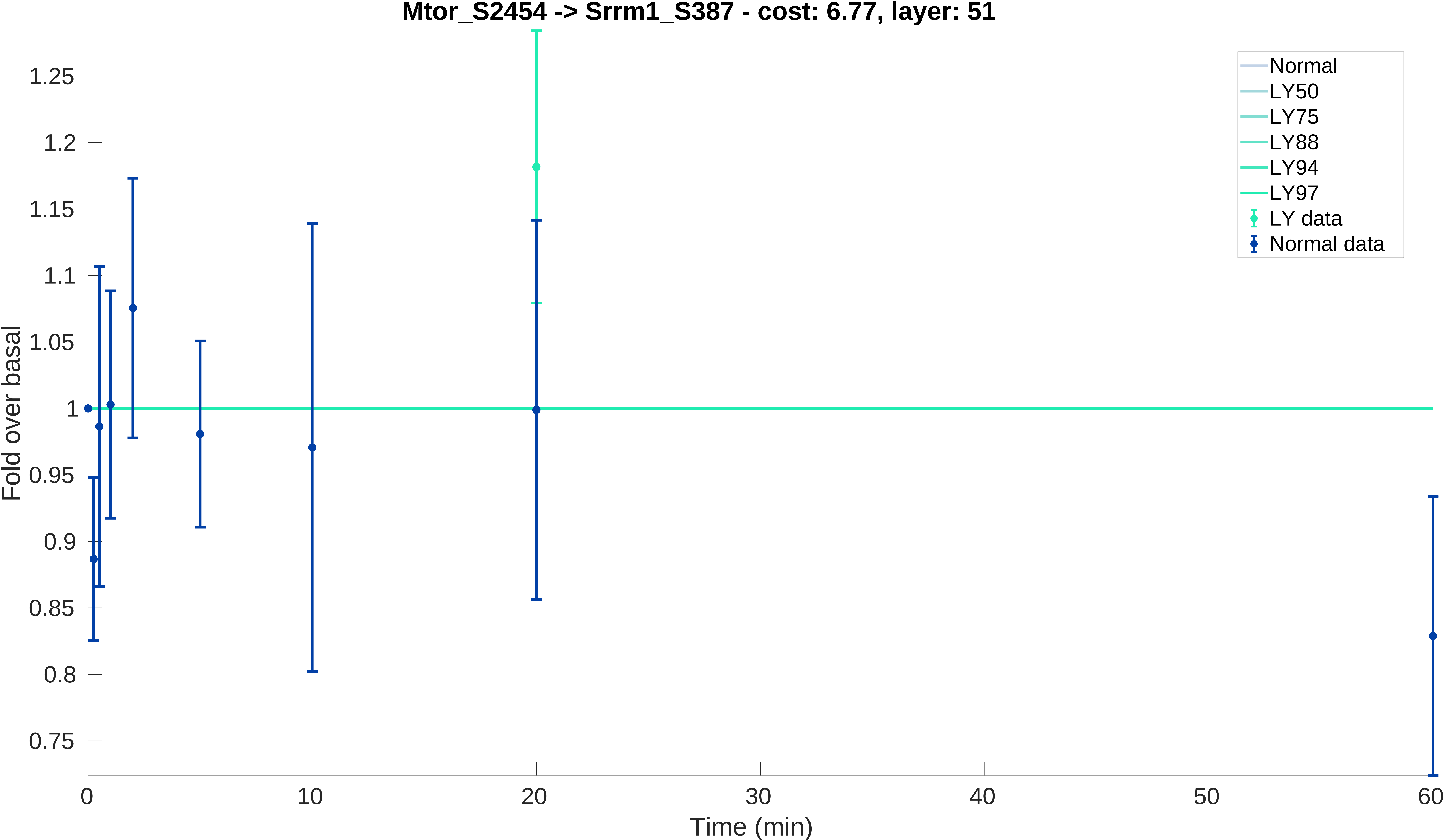
Mtor_S2454 -> Srrm1_S387 - cost: 6.77, layer: 51
Fold over basal
1.25
1.2
1.15
1.1
1.05
1
0.95
0.9
0.85
0.8
0.75
0
10
20
30
40
50
60
Time (min)
Normal
LY50
LY75
LY88
LY94
LY97
LY data
Normal data

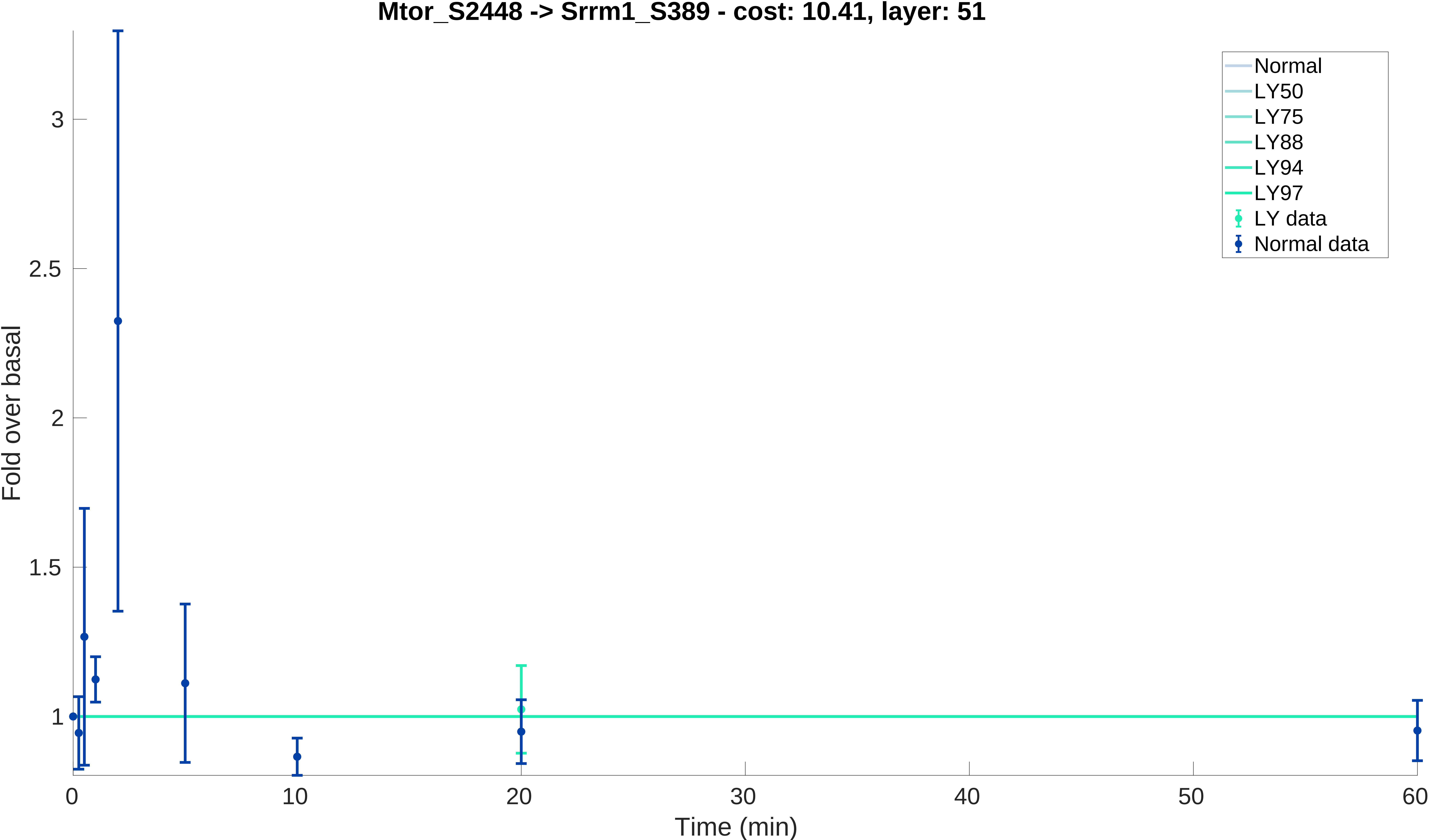
Mtor_S2448 -> Srrm1_S389 - cost: 10.41, layer: 51
Fold over basal
3
2.5
2
1.5
1
Time (min)
0
10
20
30
40
50
60
Normal
LY50
LY75
LY88
LY94
LY97
LY data
Normal data

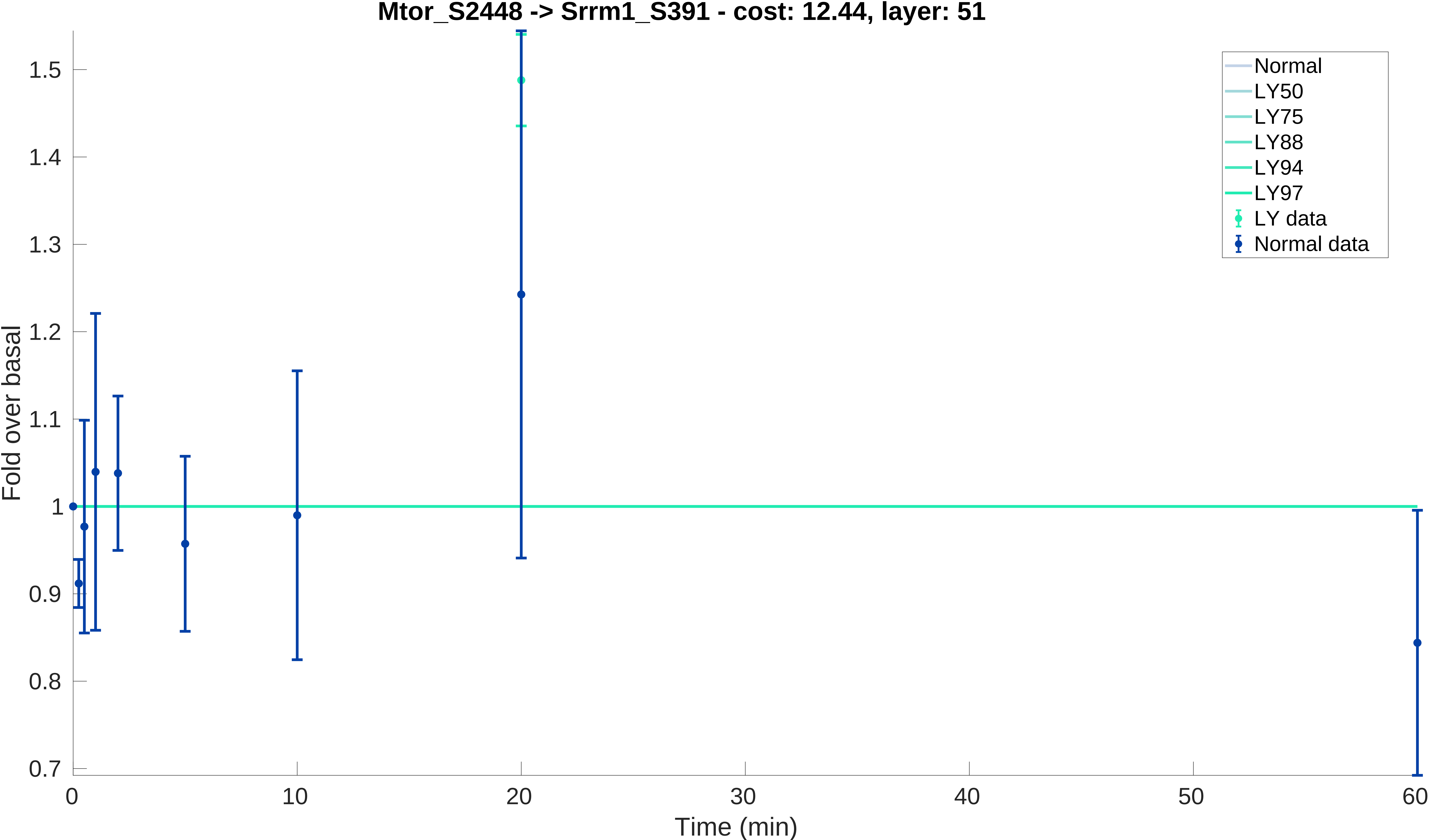
Mtor_S2448 -> Srrm1_S391 - cost: 12.44, layer: 51
Fold over basal
Time (min)
0.7
0.8
0.9
1
1.1
1.2
1.3
1.4
1.5
0
10
20
30
40
50
60
Normal
LY50
LY75
LY88
LY94
LY97
LY data
Normal data

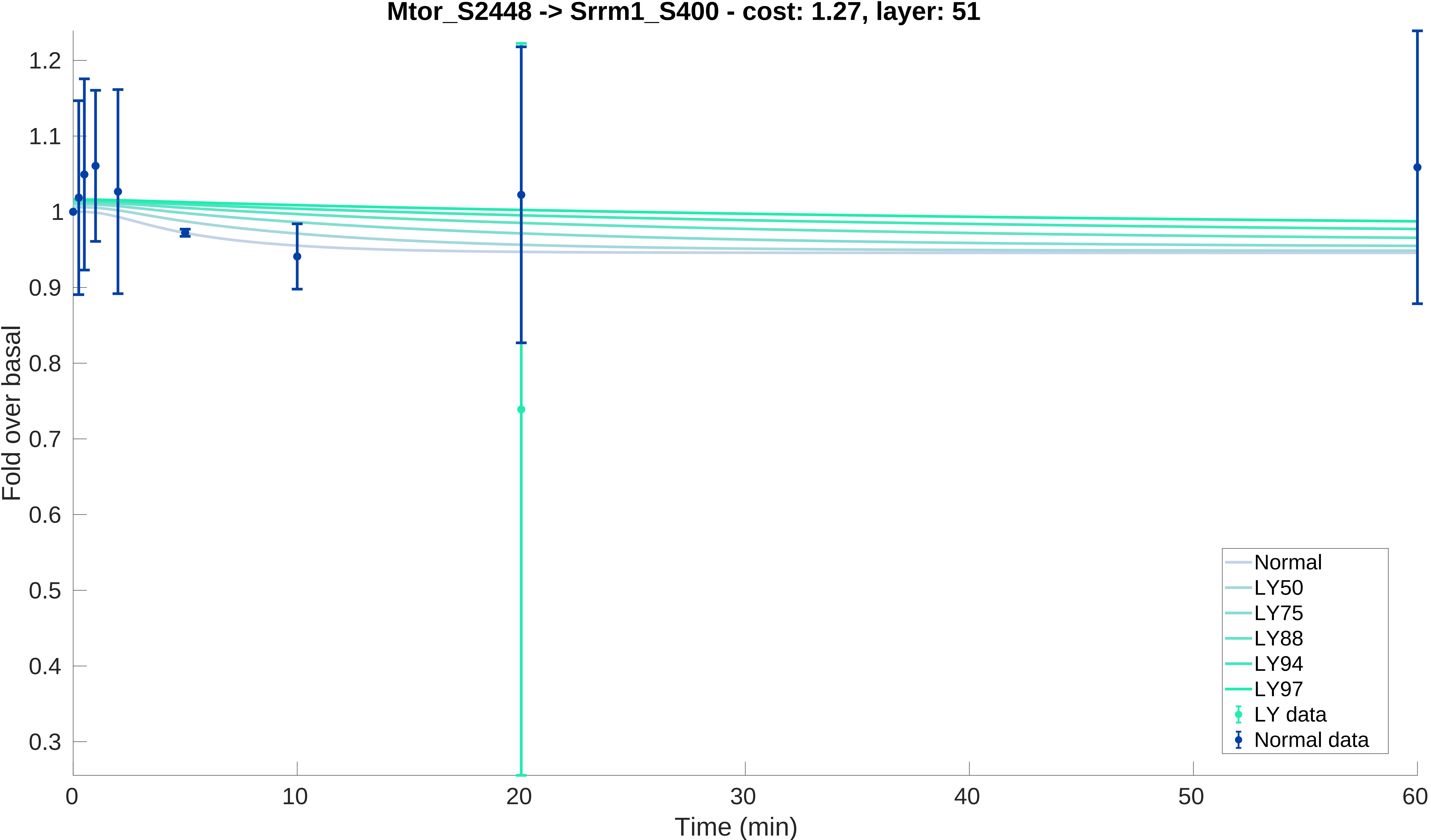
Mtor_S2448 -> Srrm1_S400 - cost: 1.27, layer: 51
Fold over basal
Time (min)
0
10
20
30
40
50
60
1.2
1.1
1
0.9
0.8
0.7
0.6
0.5
0.4
0.3
Normal
LY50
LY75
LY88
LY94
LY97
LY data
Normal data

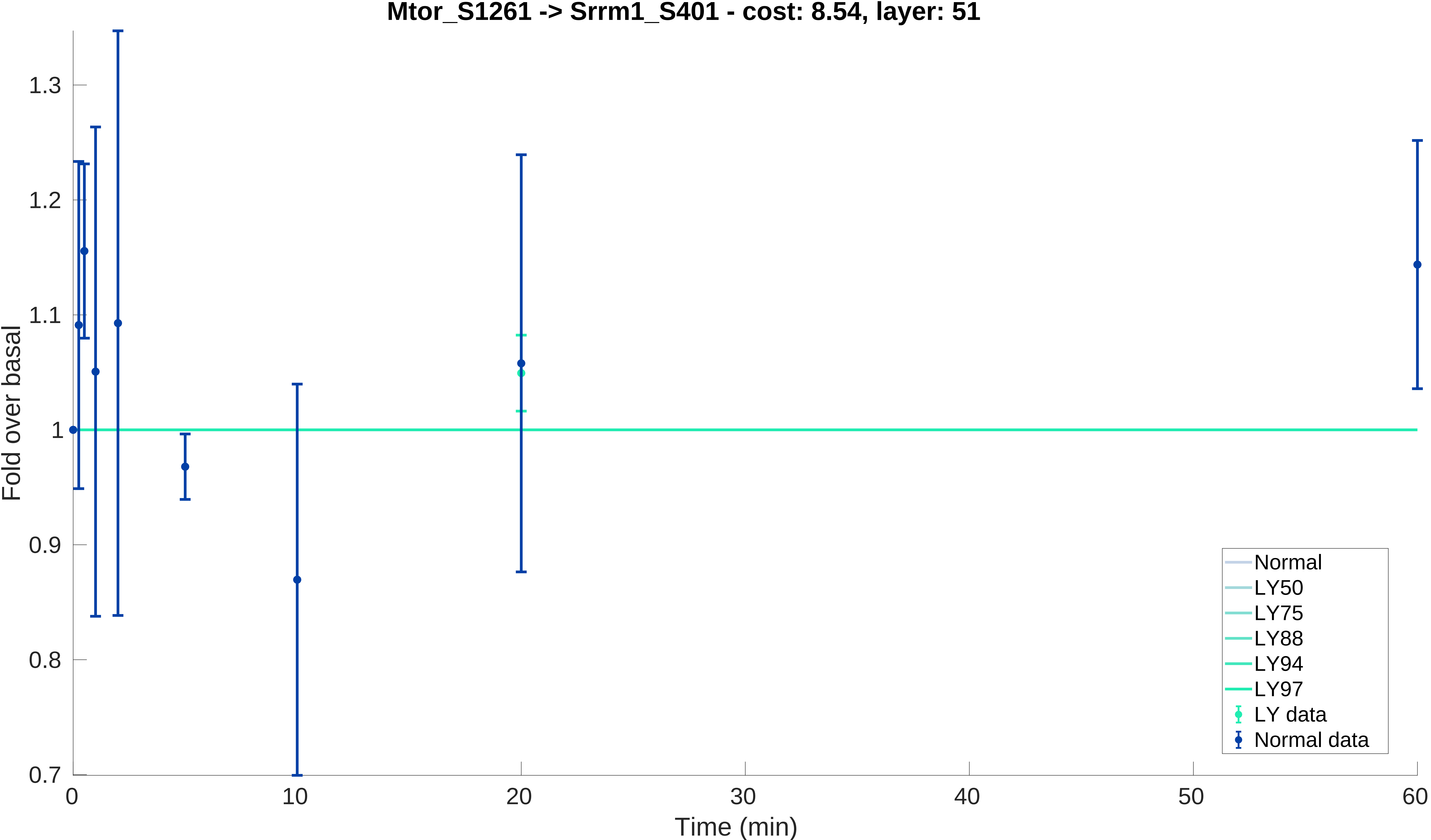

Mtor_S1261 -> Srrm1_S401 - cost: 8.54, layer: 51
Fold over basal
Time (min)
0.7
0.8
0.9
1
1.1
1.2
1.3
0
10
20
30
40
50
60
Normal
LY50
LY75
LY88
LY94
LY97
LY data
Normal data

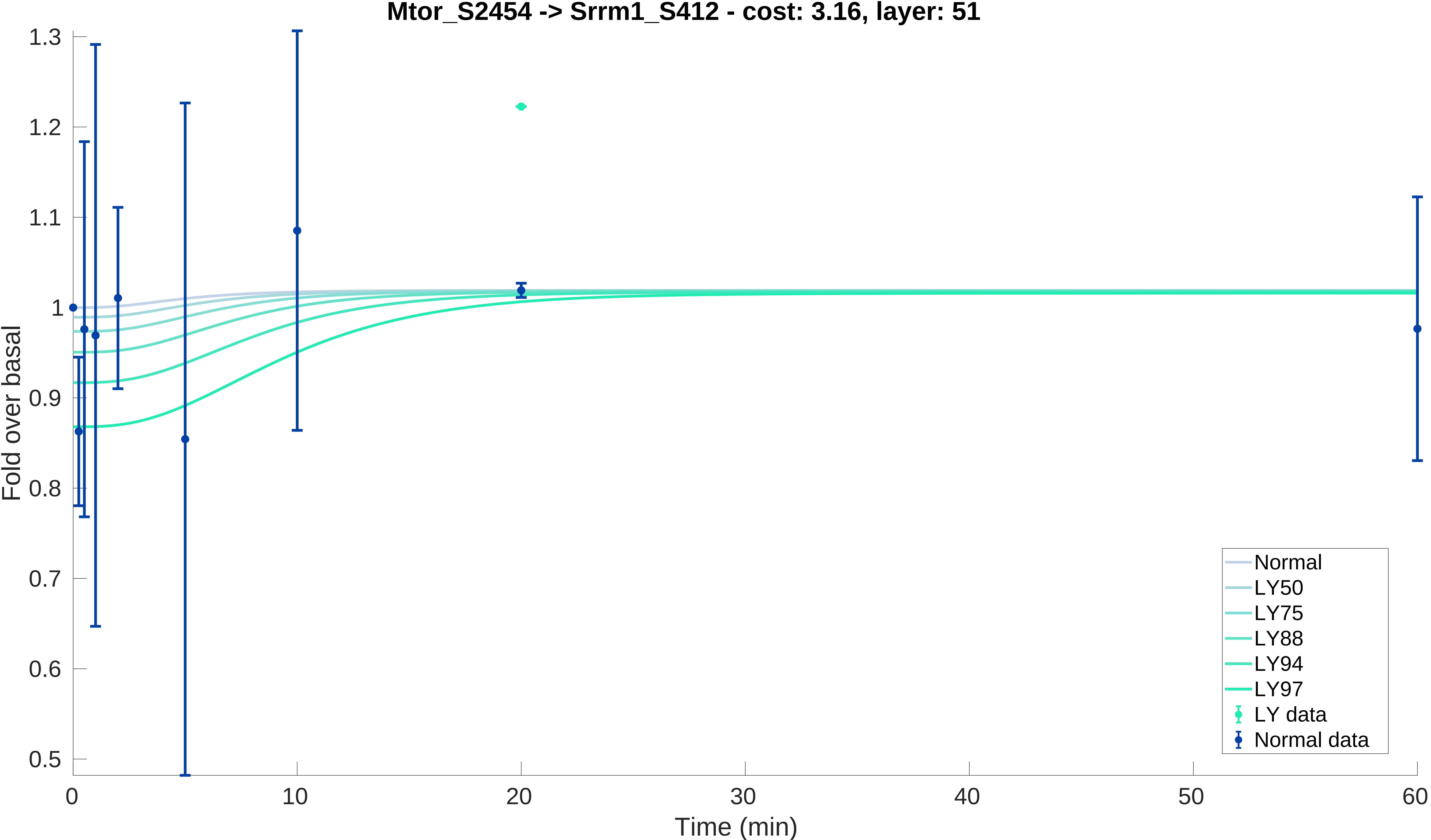

Mtor_S2454 -> Srrm1_S412 - cost: 3.16, layer: 51
1.3
1.2
1.1
1
0.9
0.8
0.7
0.6
0.5
Fold over basal
0
10
20
30
40
50
60
Time (min)
Normal
LY50
LY75
LY88
LY94
LY97
LY data
Normal data

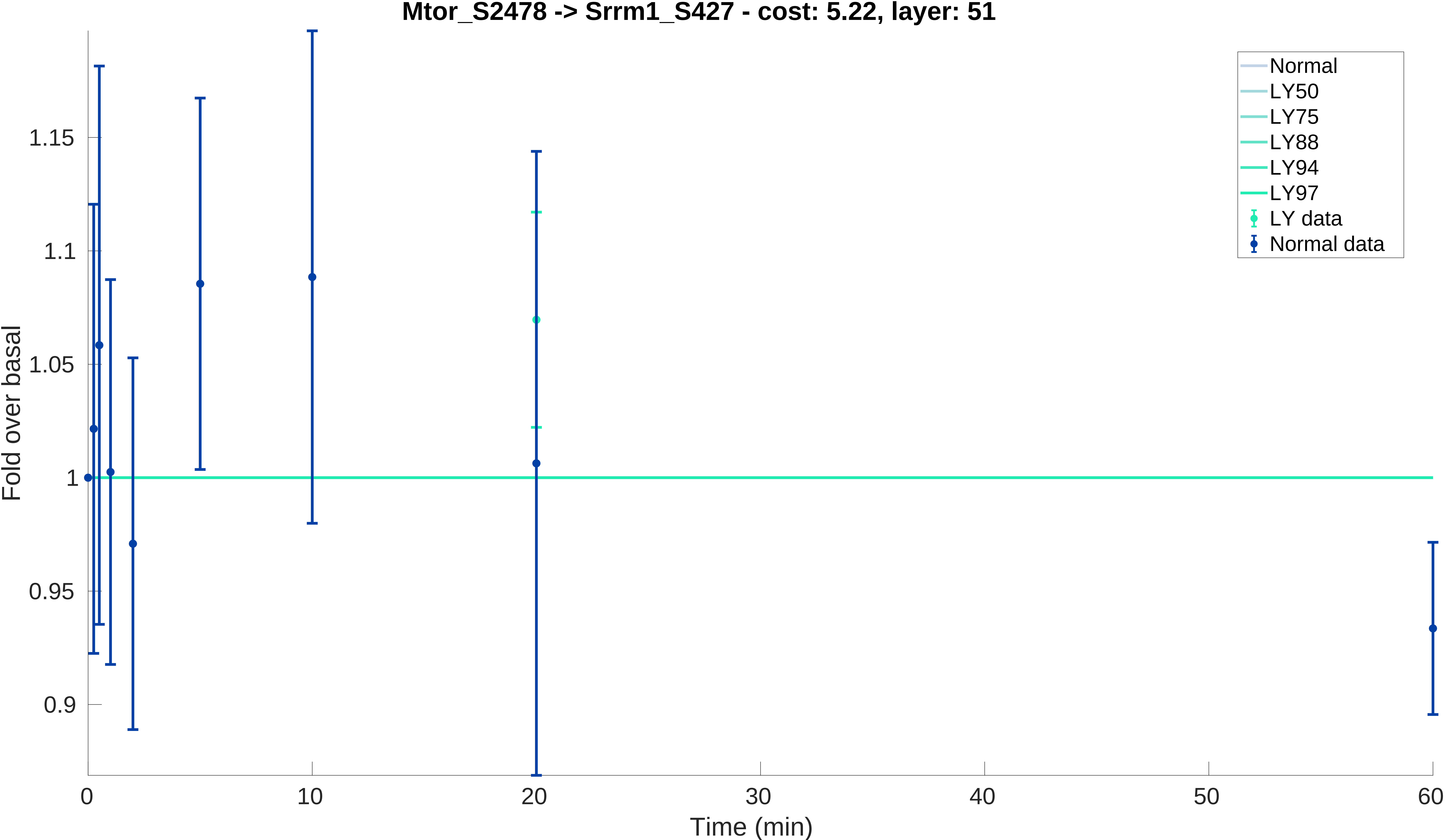

Mtor_S2478 -> Srrm1_S427 - cost: 5.22, layer: 51
Fold over basal
1.15
1.1
1.05
1
0.95
0.9
0
10
20
30
40
50
60
Time (min)
Normal
LY50
LY75
LY88
LY94
LY97
LY data
Normal data

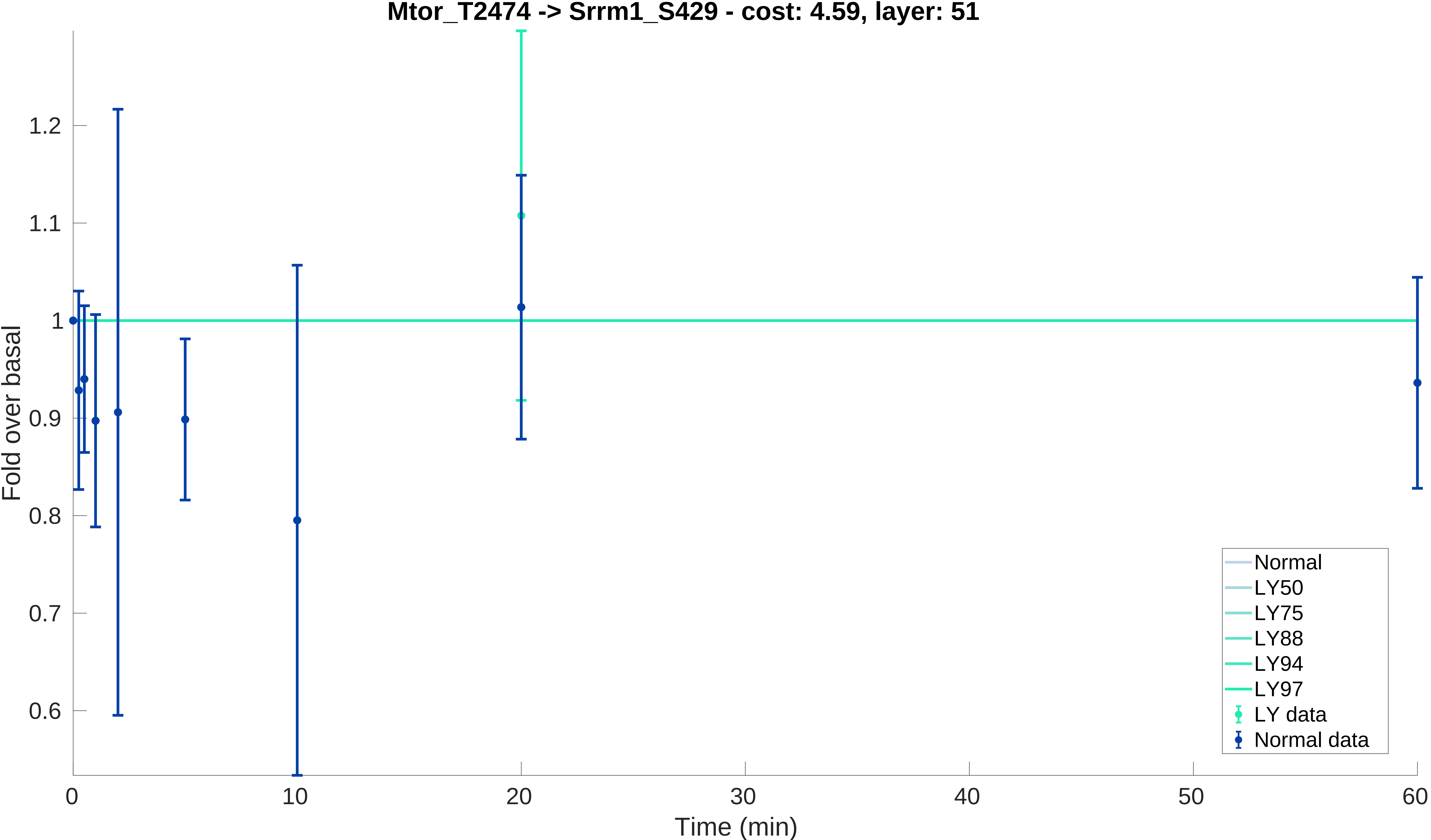

Mtor_T2474 -> Srrm1_S429 - cost: 4.59, layer: 51
Fold over basal
Time (min)
0.6
0.7
0.8
0.9
1
1.1
1.2
0
10
20
30
40
50
60
Normal
LY50
LY75
LY88
LY94
LY97
LY data
Normal data

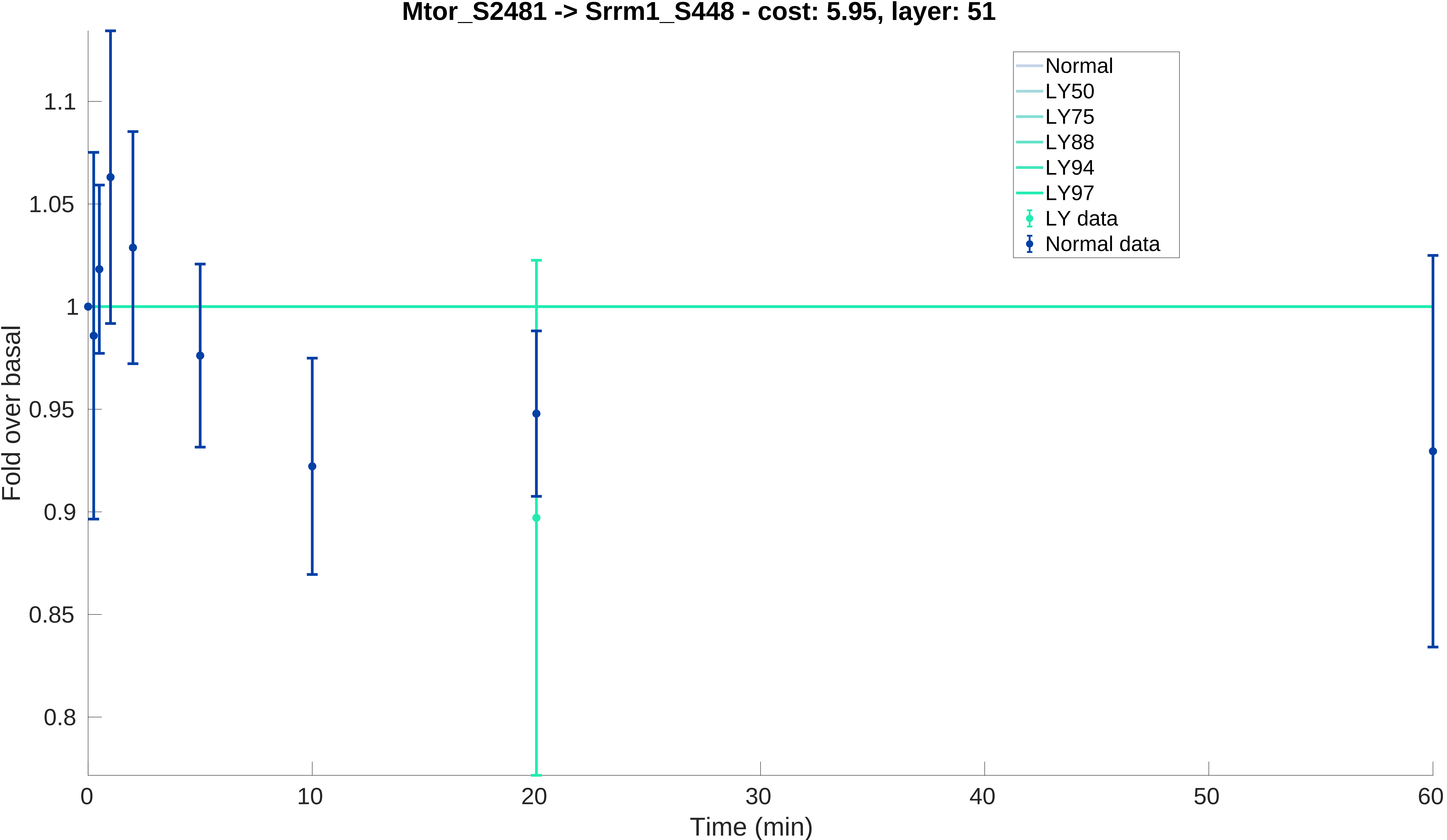

Mtor_S2481 -> Srrm1_S448 - cost: 5.95, layer: 51
Normal
LY50
LY75
LY88
LY94
LY97
LY data
Normal data
1.1
1.05
1
0.95
0.9
0.85
0.8
Fold over basal
0
10
20
30
40
50
60
Time (min)

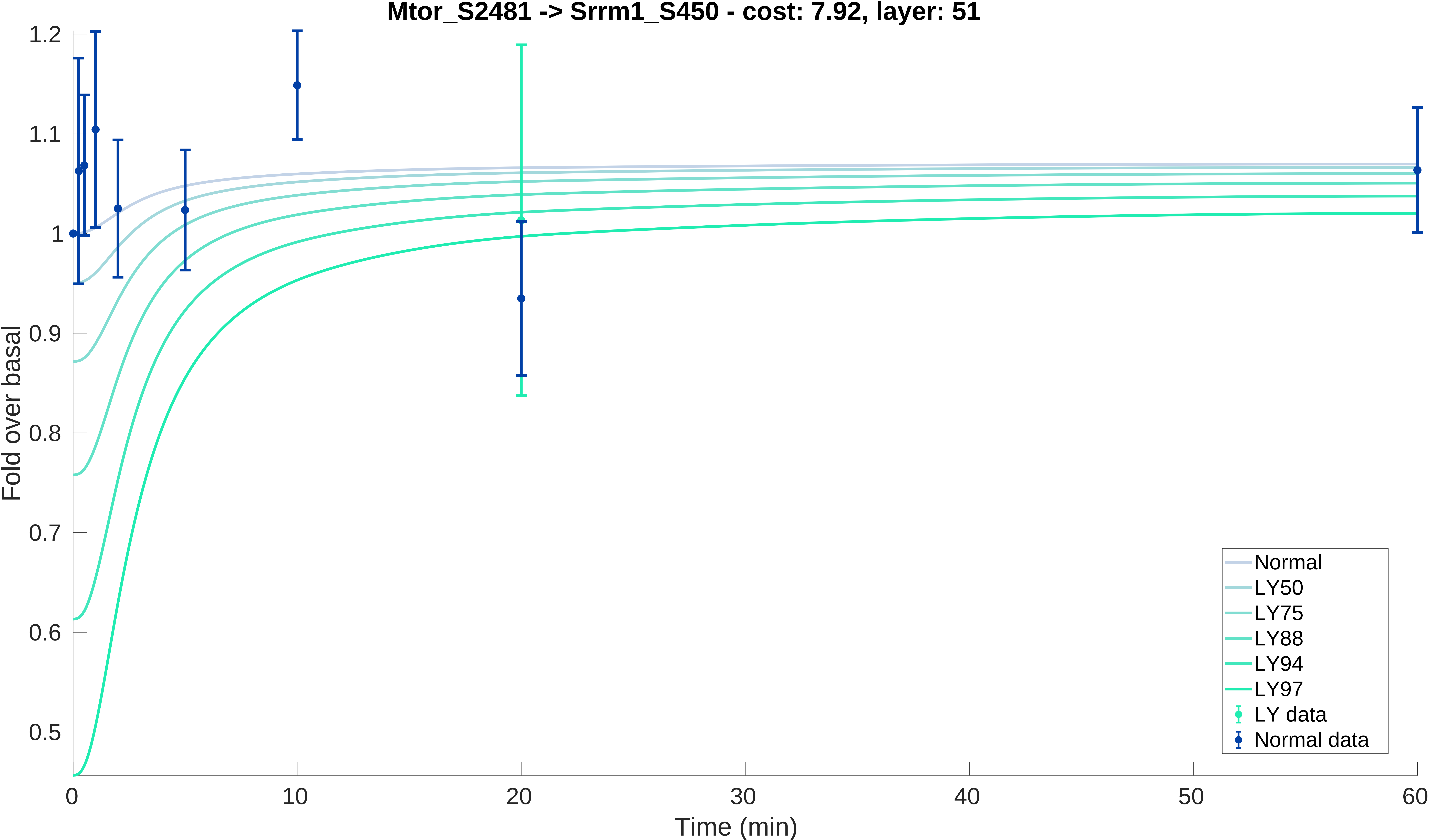

Mtor_S2481 -> Srrm1_S450 - cost: 7.92, layer: 51
Fold over basal
Time (min)
0.5
0.6
0.7
0.8
0.9
1
1.1
1.2
0
10
20
30
40
50
60
Normal
LY50
LY75
LY88
LY94
LY97
LY data
Normal data

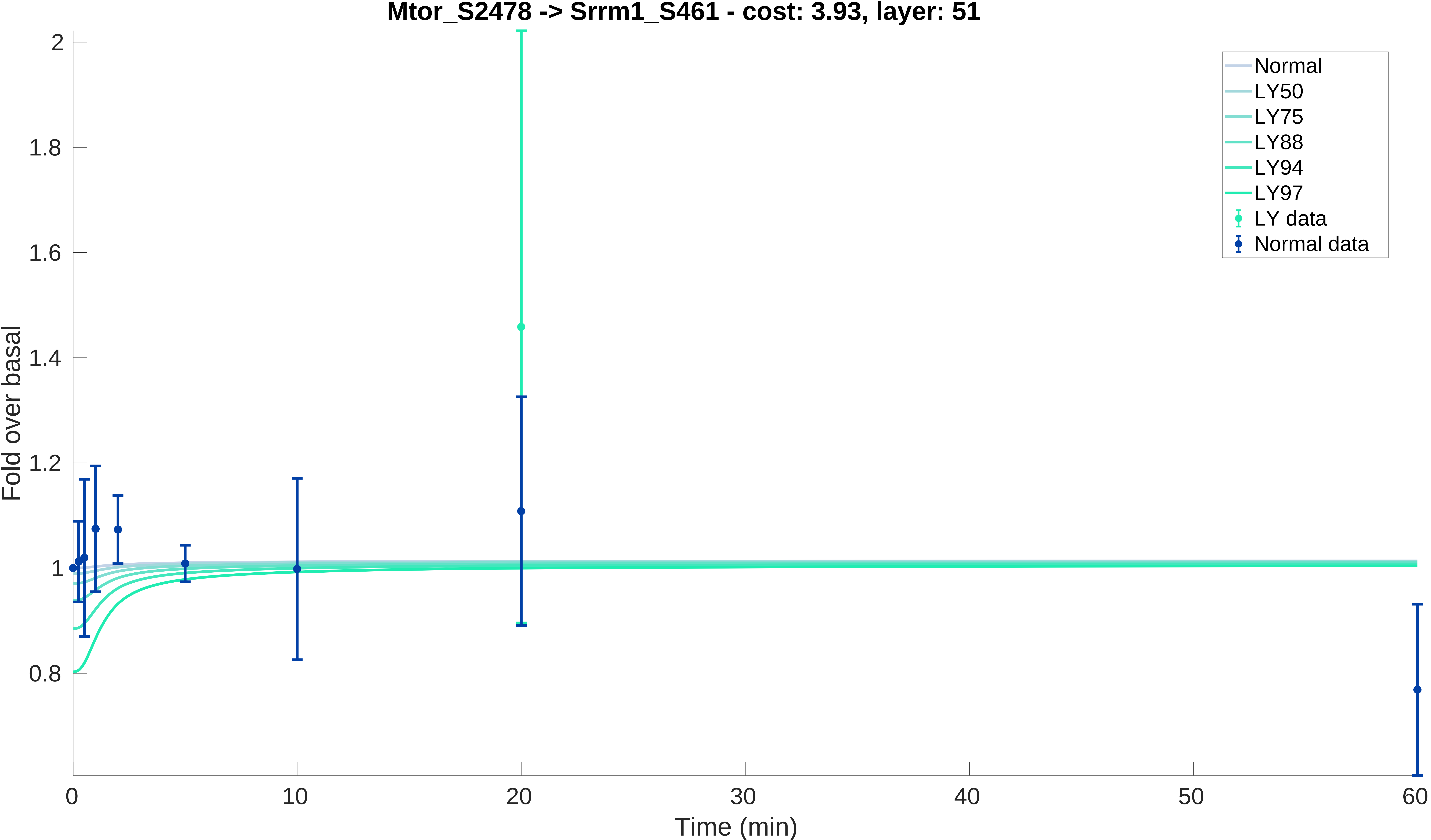

Mtor_S2478 -> Srrm1_S461 - cost: 3.93, layer: 51
Fold over basal
Time (min)
0.8
1
1.2
1.4
1.6
1.8
2
0
10
20
30
40
50
60
Normal
LY50
LY75
LY88
LY94
LY97
LY data
Normal data

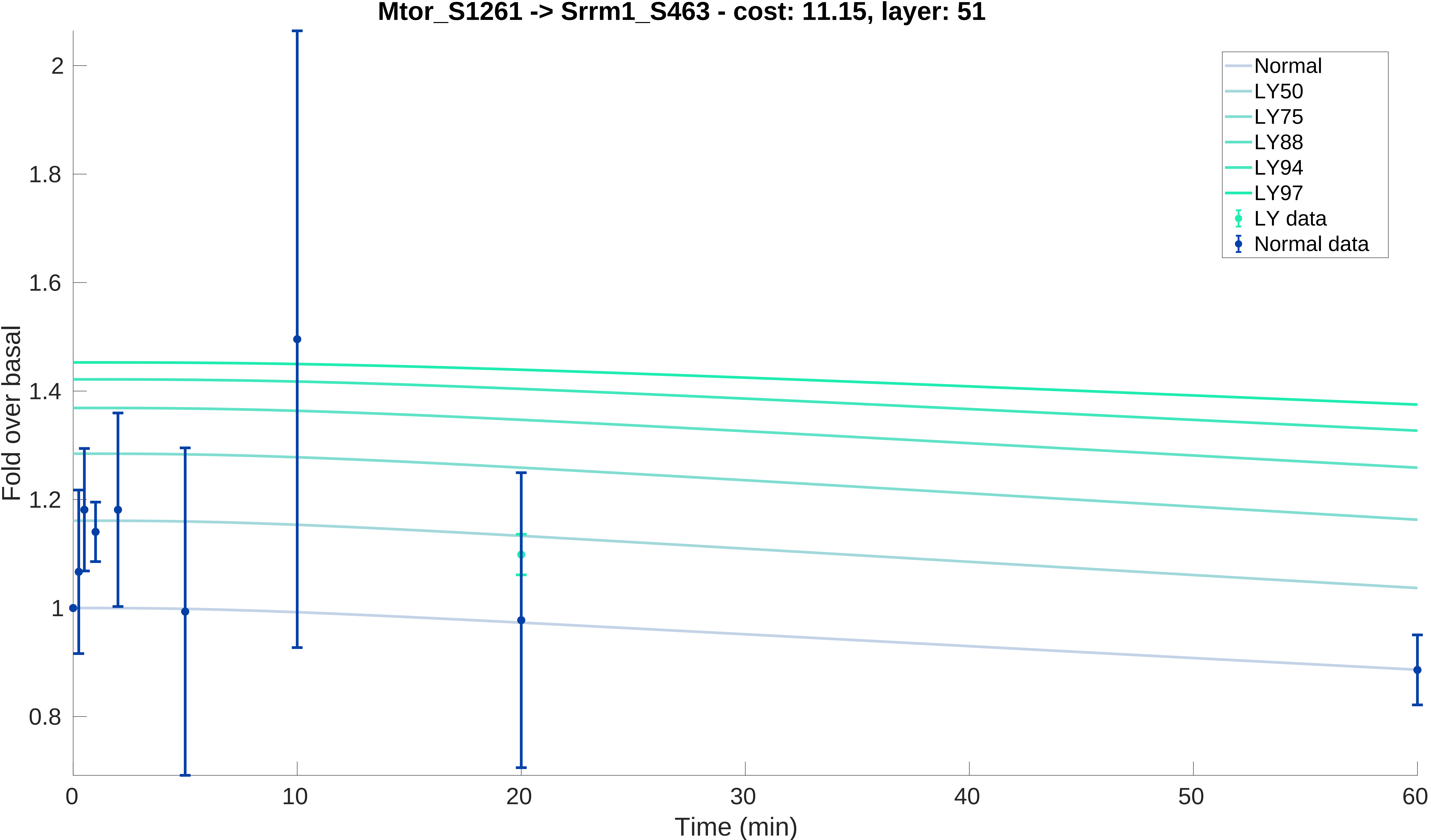
Mtor_S1261 -> Srrm1_S463 - cost: 11.15, layer: 51
Fold over basal
Time (min)
0.8
1
1.2
1.4
1.6
1.8
2
0
10
20
30
40
50
60
Normal
LY50
LY75
LY88
LY94
LY97
LY data
Normal data

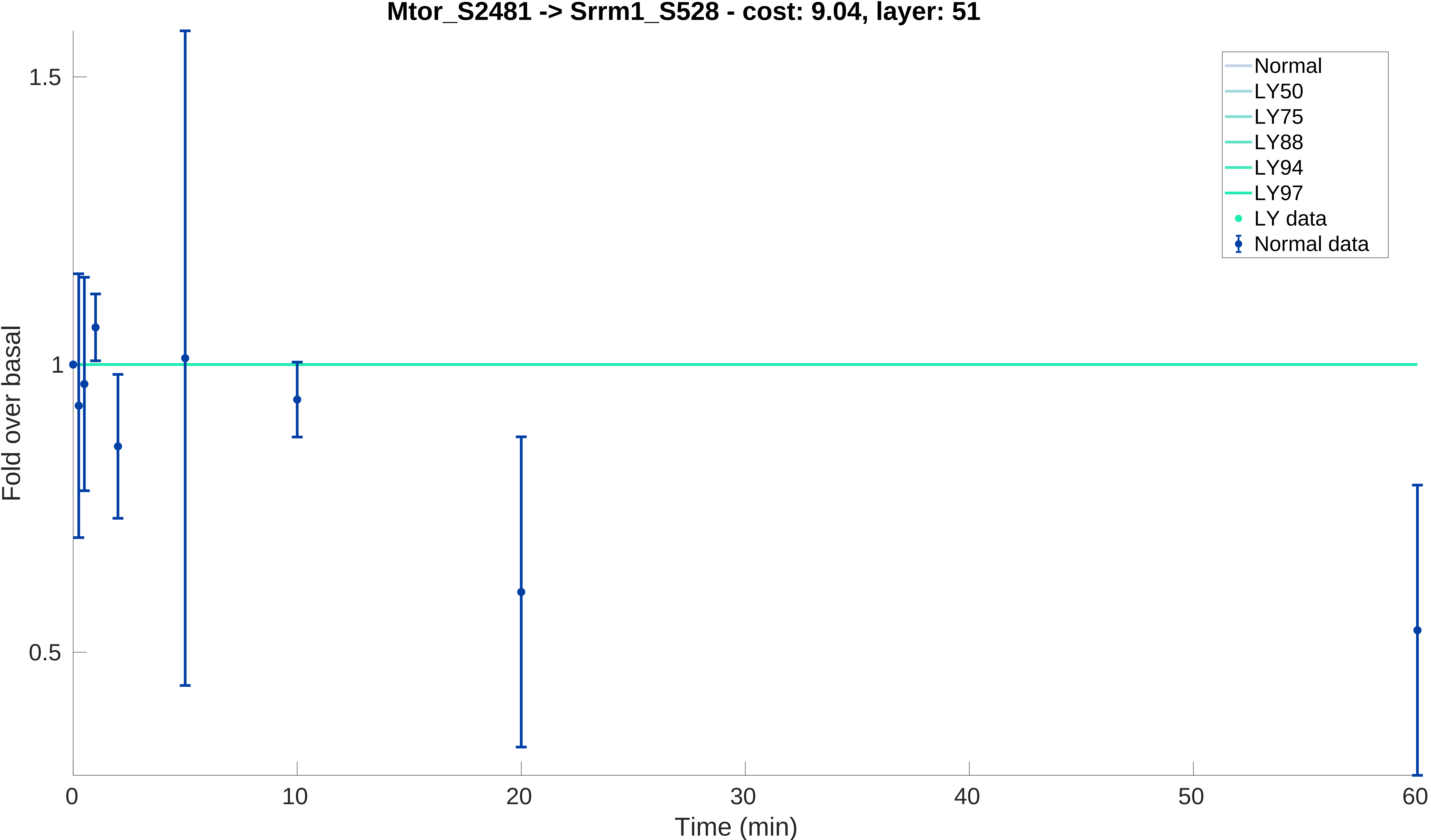
Mtor_S2481 -> Srrm1_S528 - cost: 9.04, layer: 51
Fold over basal
1.5
1
0.5
0
10
20
30
40
50
60
Time (min)
Normal
LY50
LY75
LY88
LY94
LY97
LY data
Normal data

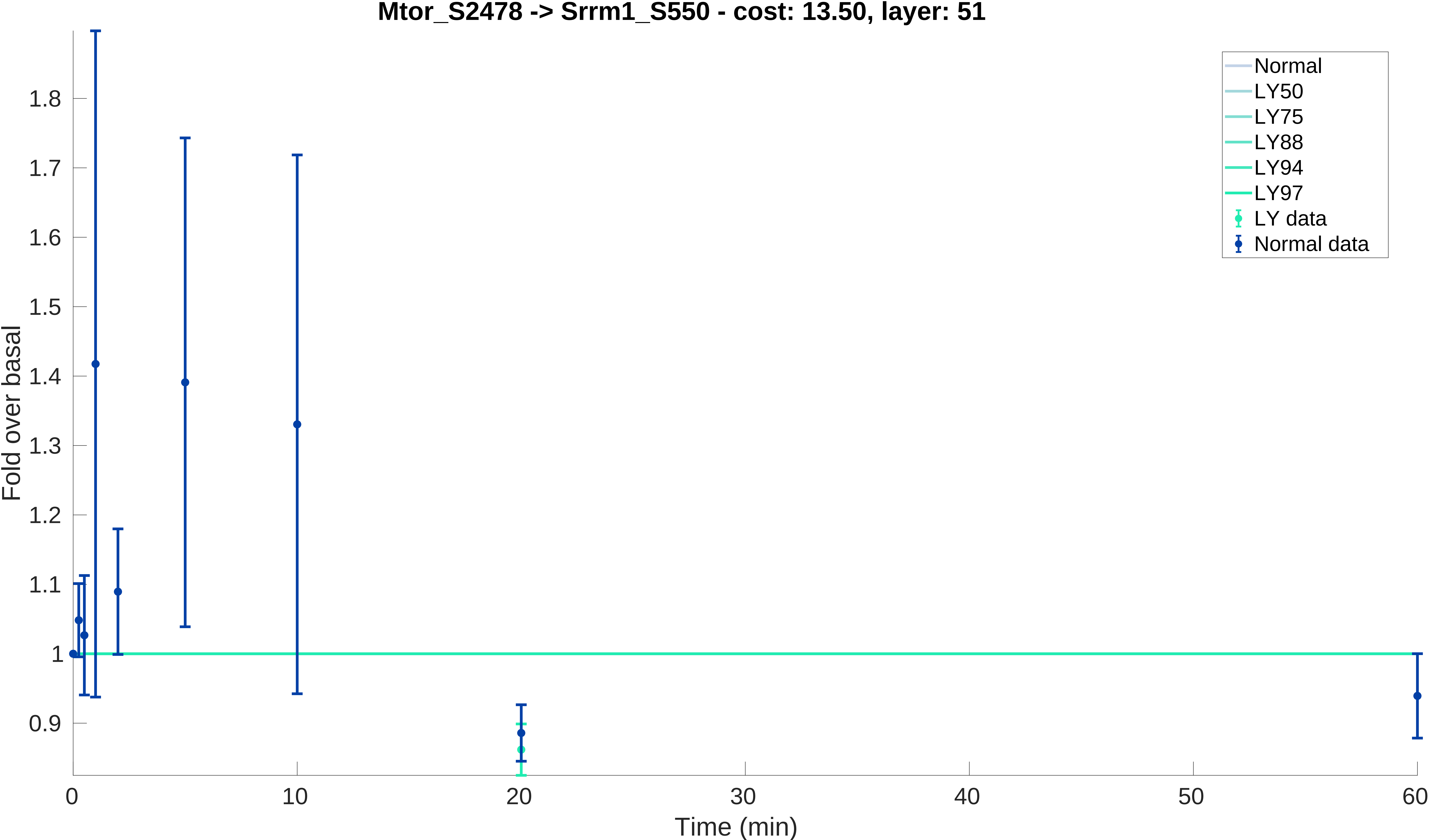

Mtor_S2478 -> Srrm1_S550 - cost: 13.50, layer: 51
Fold over basal
Time (min)
0.9
1
1.1
1.2
1.3
1.4
1.5
1.6
1.7
1.8
0
10
20
30
40
50
60
Normal
LY50
LY75
LY88
LY94
LY97
LY data
Normal data

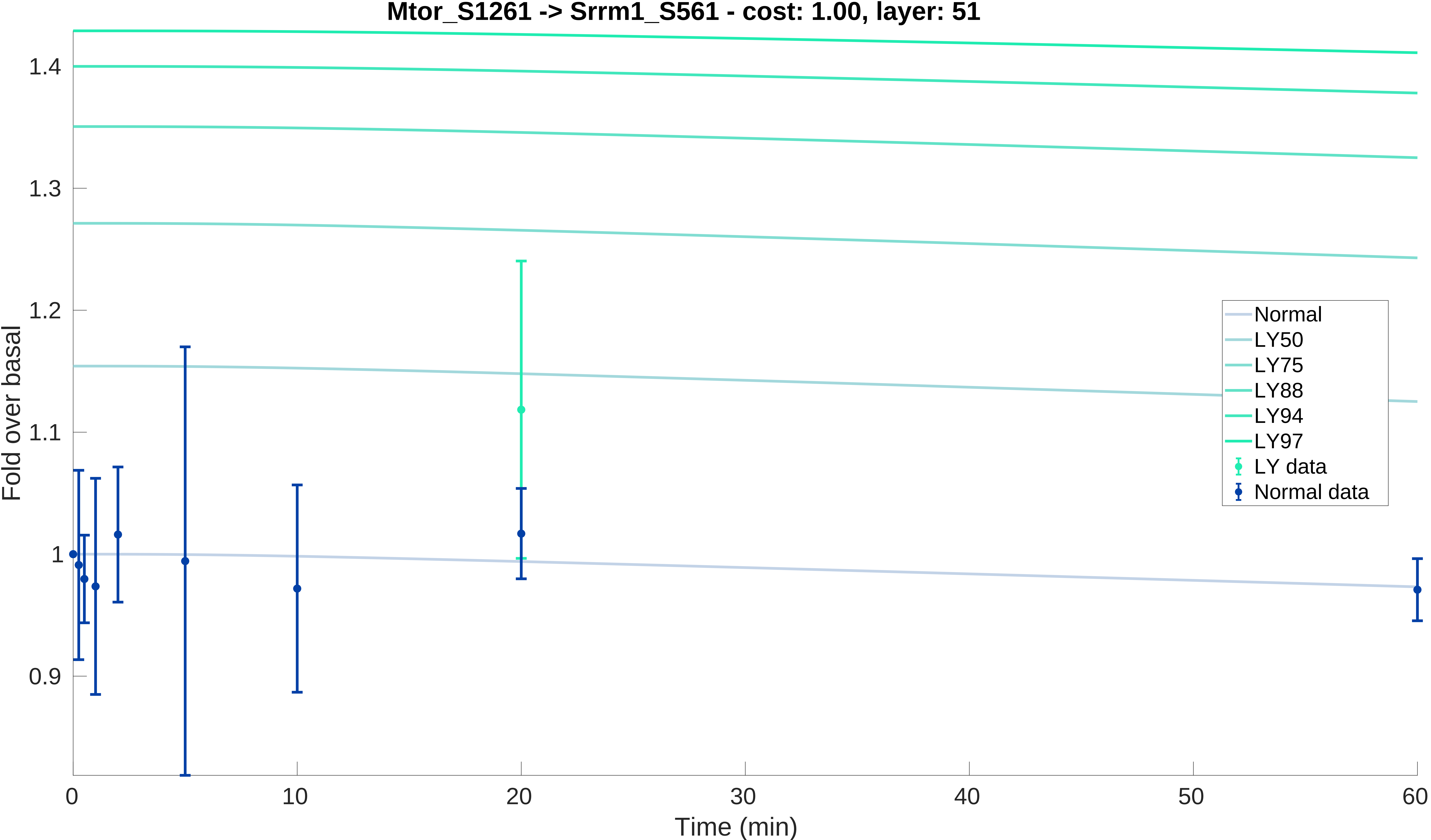

Mtor_S1261 -> Srrm1_S561 - cost: 1.00, layer: 51
Fold over basal
Time (min)
1.4
1.3
1.2
1.1
1
0.9
0
10
20
30
40
50
60
Normal
LY50
LY75
LY88
LY94
LY97
LY data
Normal data

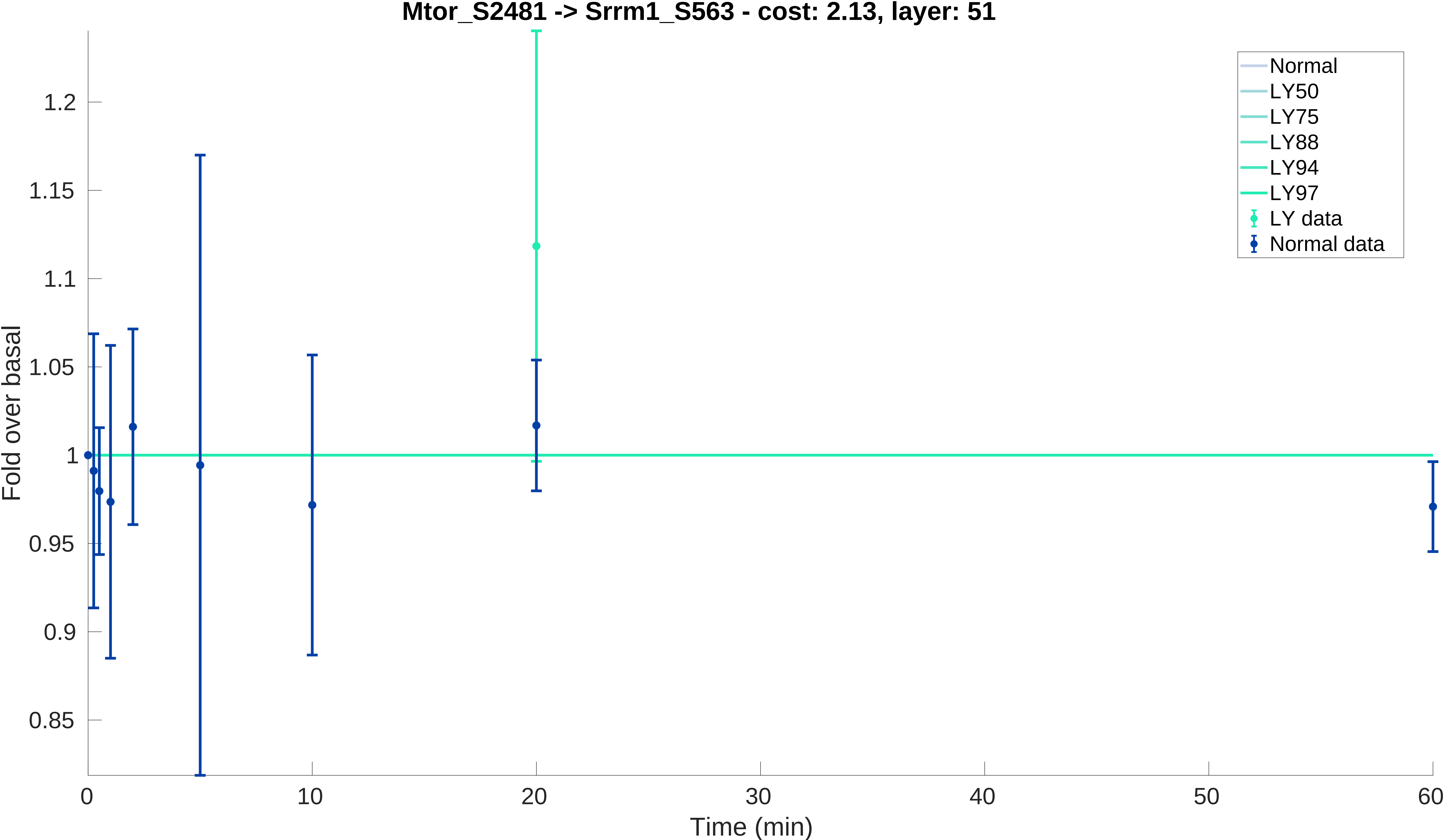
Mtor_S2481 -> Srrm1_S563 - cost: 2.13, layer: 51
Fold over basal
Time (min)
1.2
1.15
1.1
1.05
1
0.95
0.9
0.85
0
10
20
30
40
50
60
Normal
LY50
LY75
LY88
LY94
LY97
LY data
Normal data

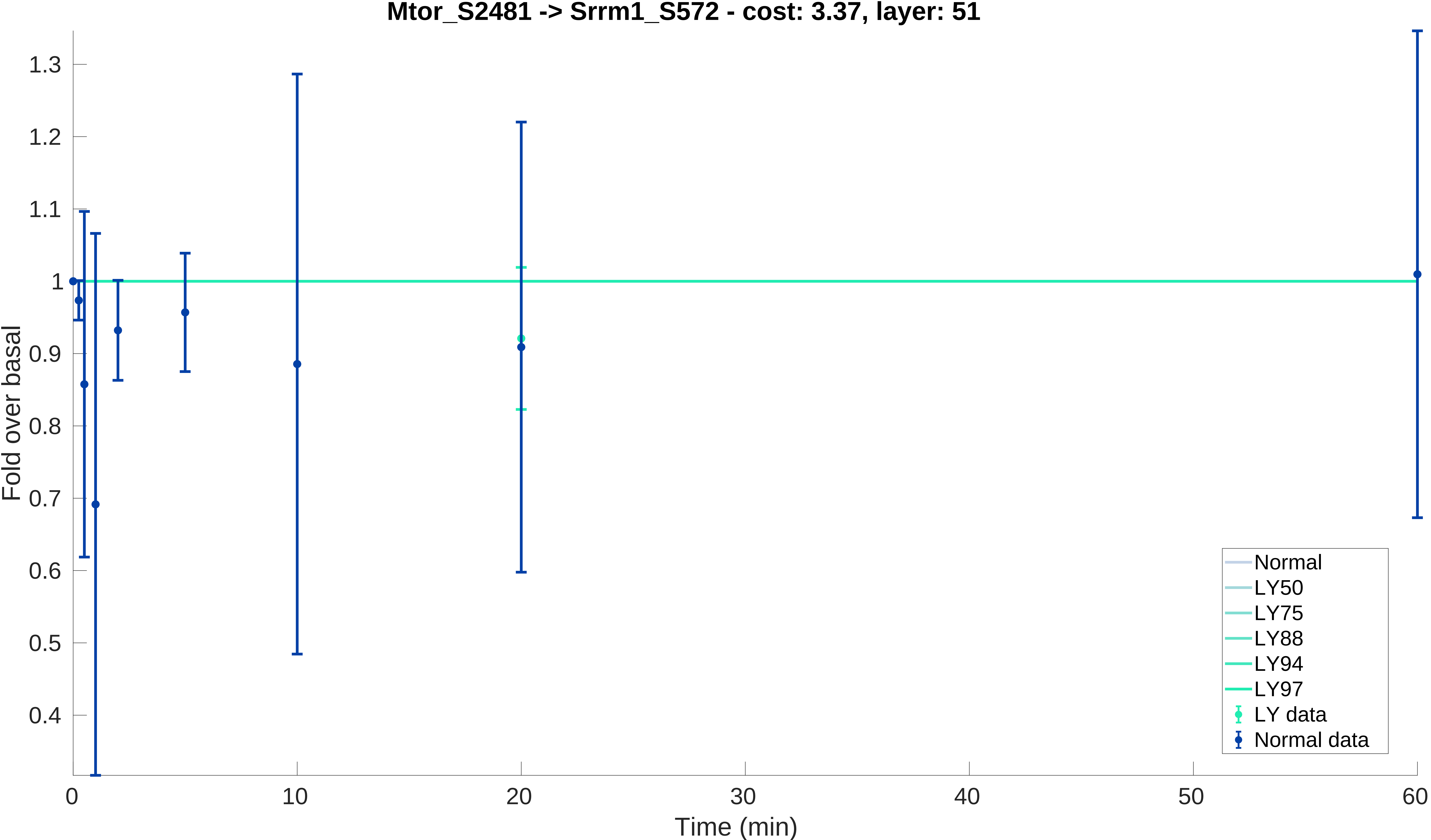

Mtor_S2481 -> Srrm1_S572 - cost: 3.37, layer: 51
1.3
1.2
1.1
1
0.9
0.8
0.7
0.6
0.5
0.4
Fold over basal
0
10
20
30
40
50
60
Time (min)
Normal
LY50
LY75
LY88
LY94
LY97
LY data
Normal data

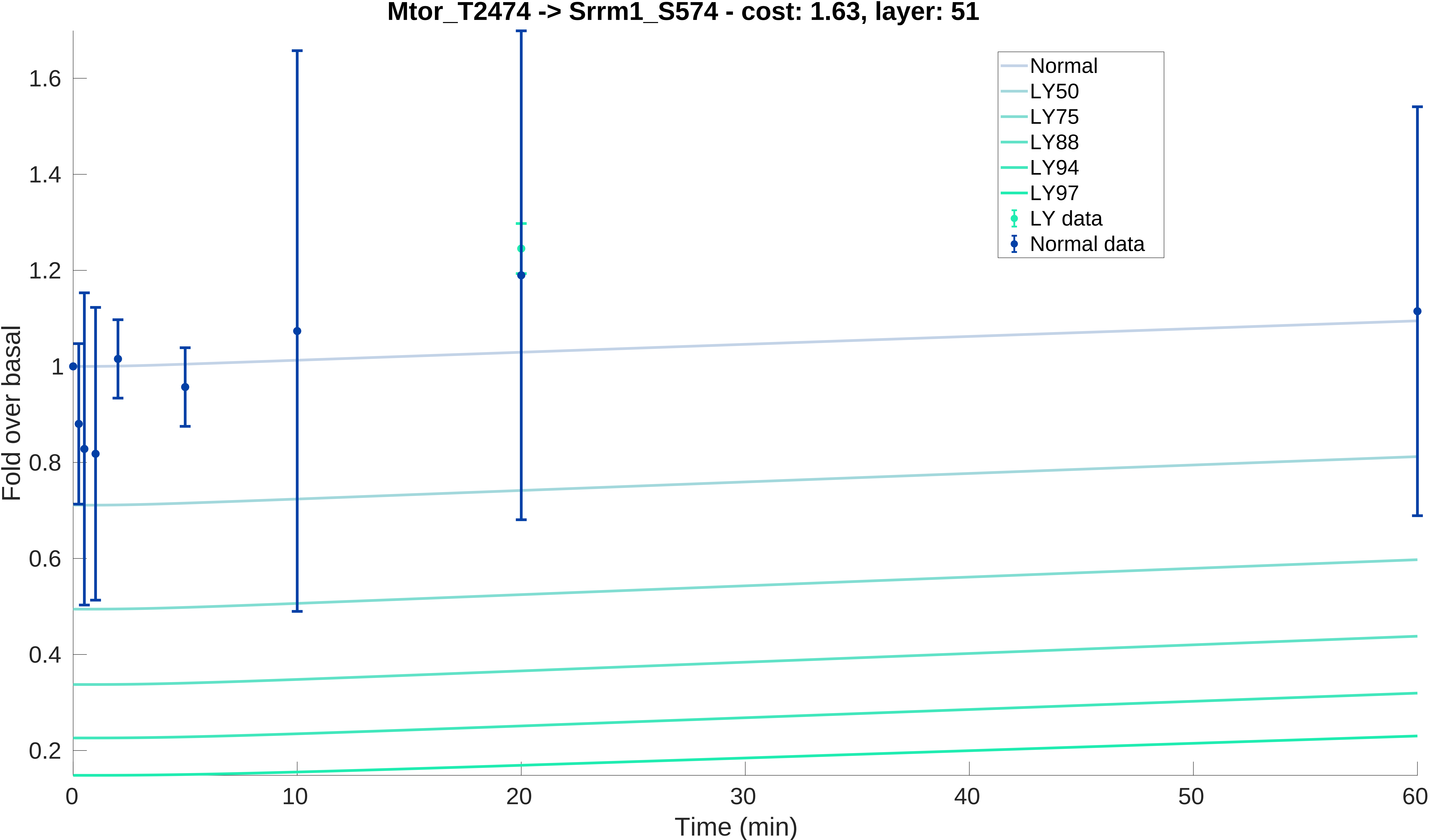
Mtor_T2474 -> Srrm1_S574 - cost: 1.63, layer: 51
Fold over basal
Time (min)
Normal
LY50
LY75
LY88
LY94
LY97
LY data
Normal data
1.6
1.4
1.2
1
0.8
0.6
0.4
0.2
0
10
20
30
40
50
60

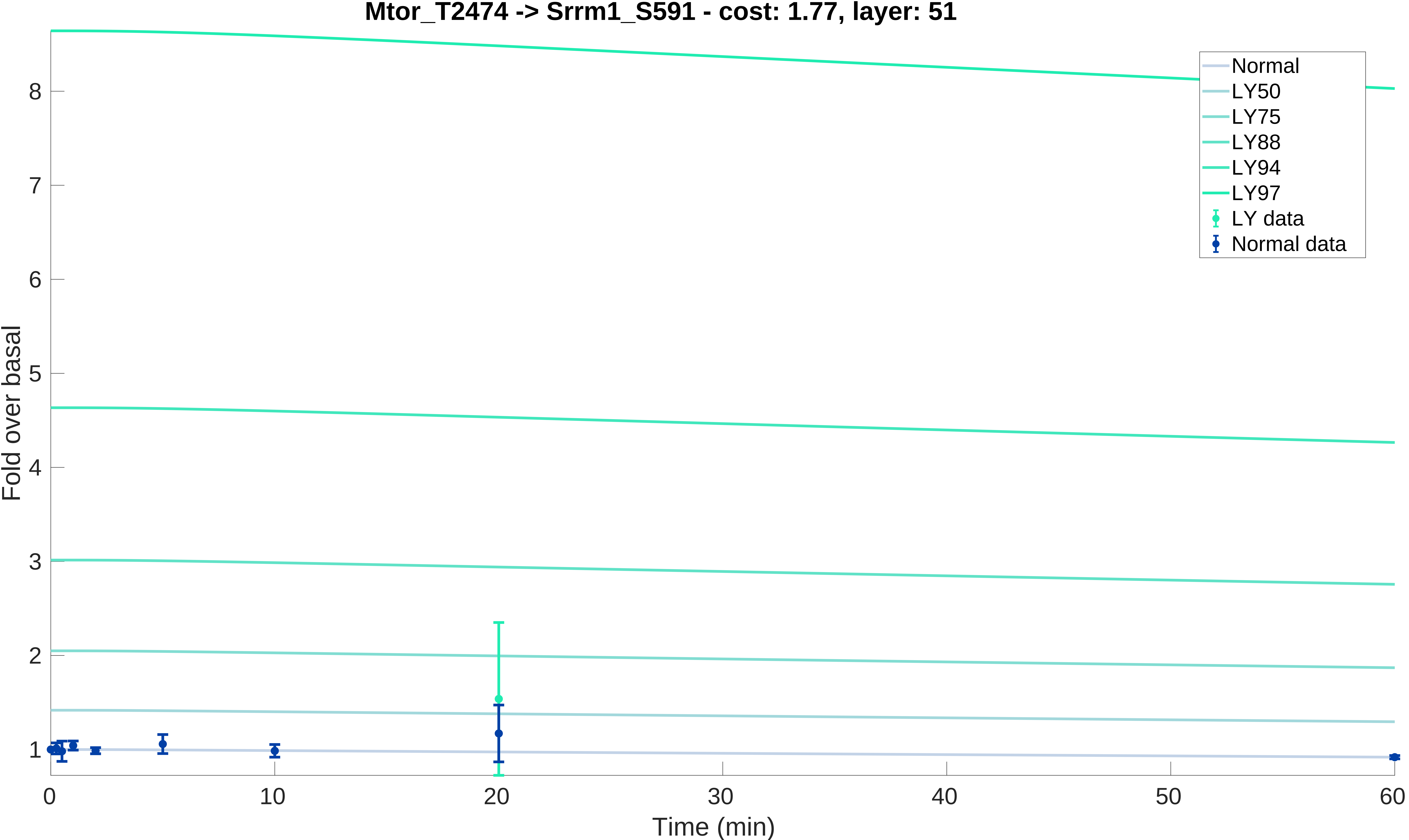

Mtor_T2474 -> Srrm1_S591 - cost: 1.77, layer: 51
Normal
LY50
LY75
LY88
LY94
LY97
LY data
Normal data
Fold over basal
Time (min)
1
2
3
4
5
6
7
8
0
10
20
30
40
50
60

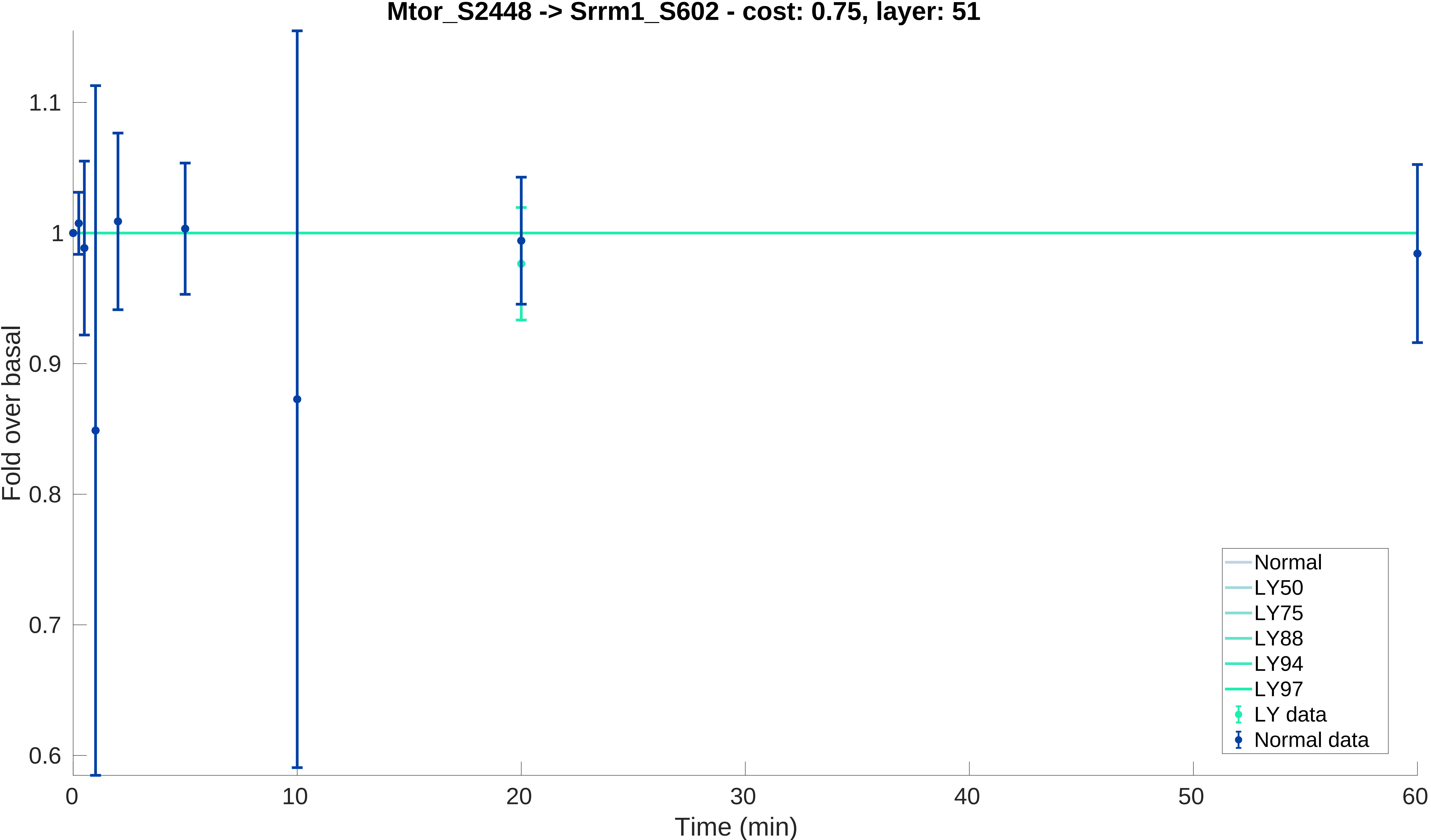

Mtor_S2448 -> Srrm1_S602 - cost: 0.75, layer: 51
Fold over basal
1.1
1
0.9
0.8
0.7
0.6
0
10
20
30
40
50
60
Time (min)
Normal
LY50
LY75
LY88
LY94
LY97
LY data
Normal data

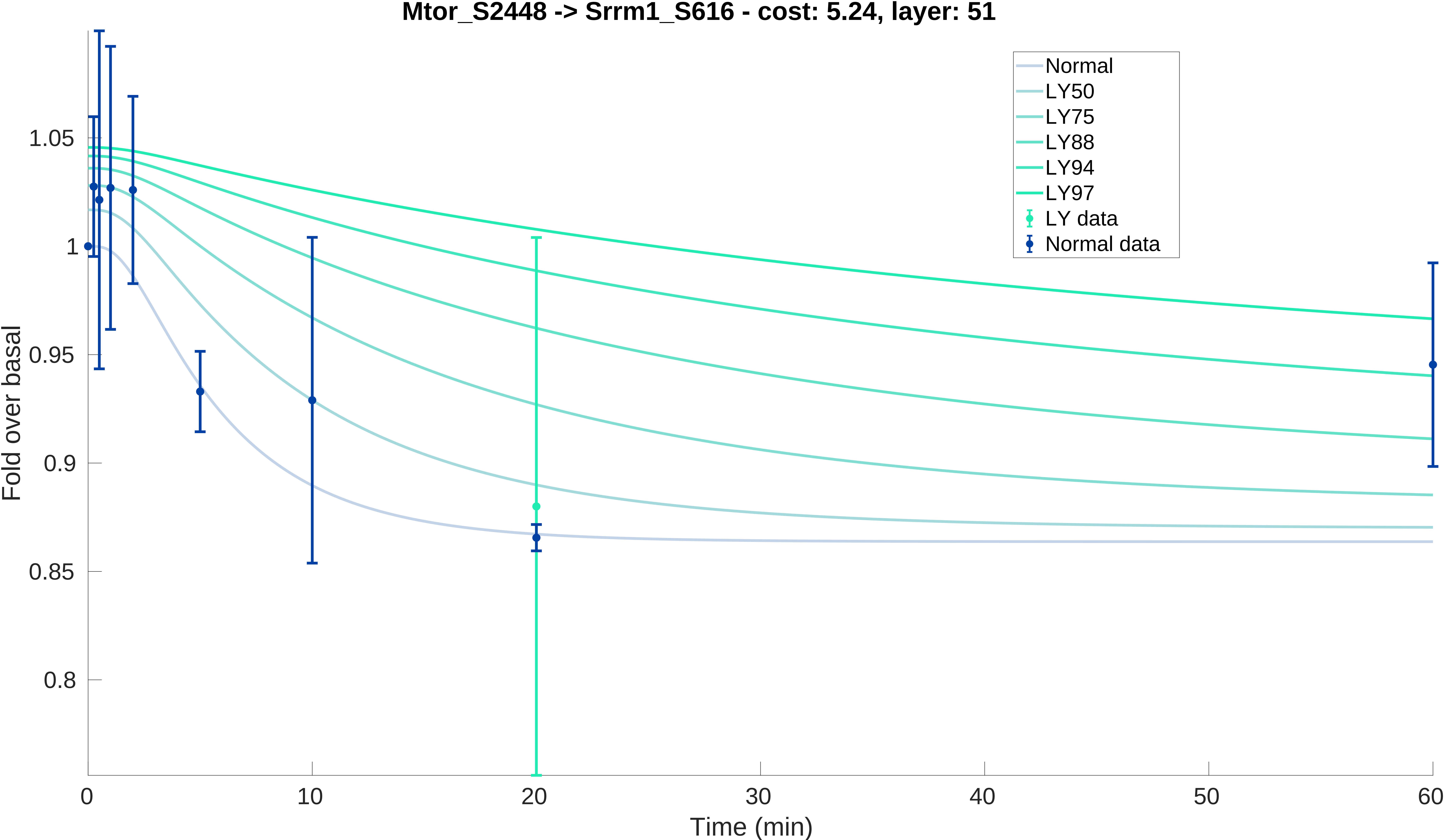

Mtor_S2448 -> Srrm1_S616 - cost: 5.24, layer: 51
Normal
LY50
LY75
LY88
LY94
LY97
LY data
Normal data
Fold over basal
1.05
1
0.95
0.9
0.85
0.8
0
10
20
30
40
50
60
Time (min)

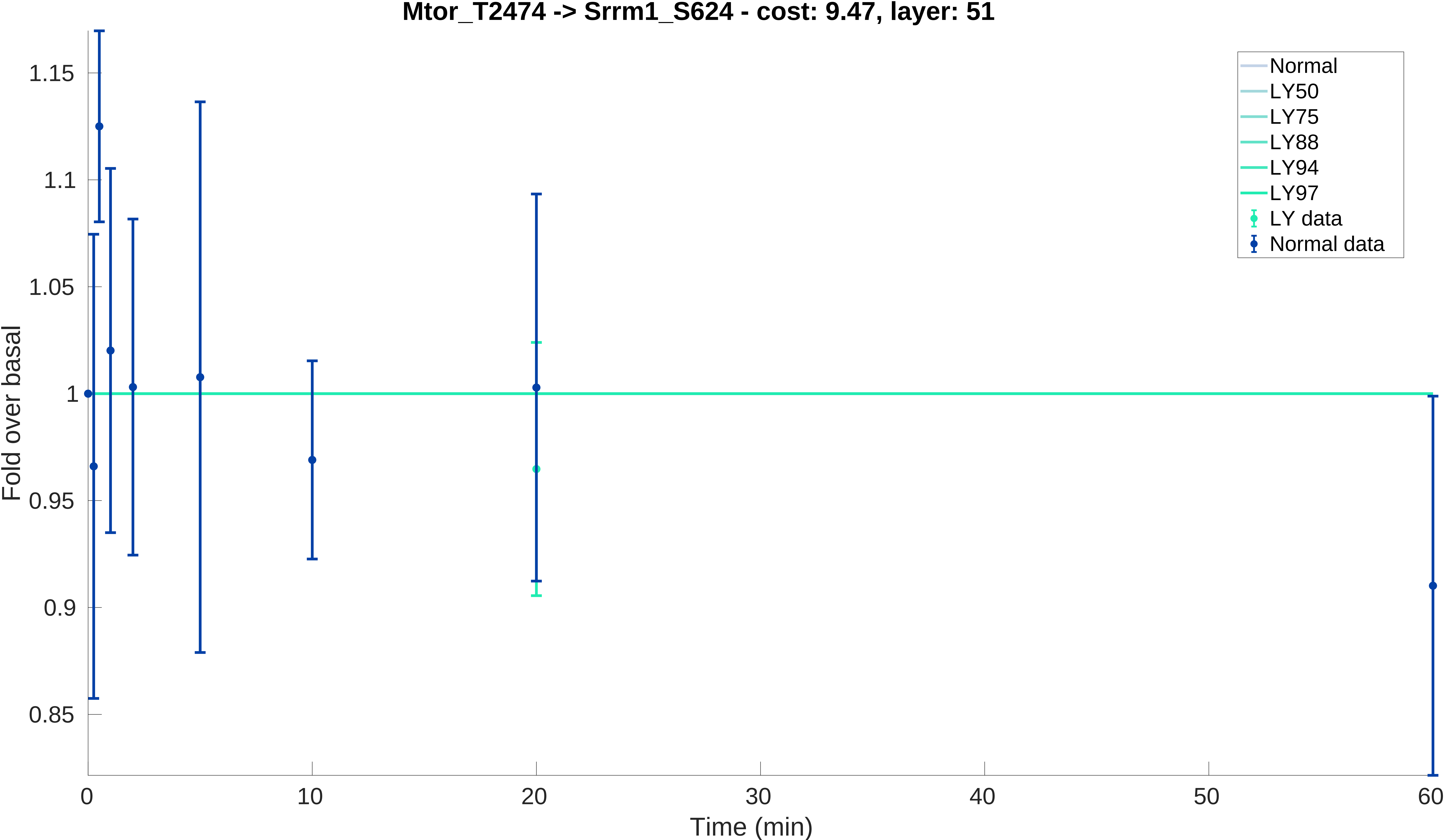

Mtor_T2474 -> Srrm1_S624 - cost: 9.47, layer: 51
Fold over basal
Time (min)
1.15
1.1
1.05
1
0.95
0.9
0.85
0
10
20
30
40
50
60
Normal
LY50
LY75
LY88
LY94
LY97
LY data
Normal data

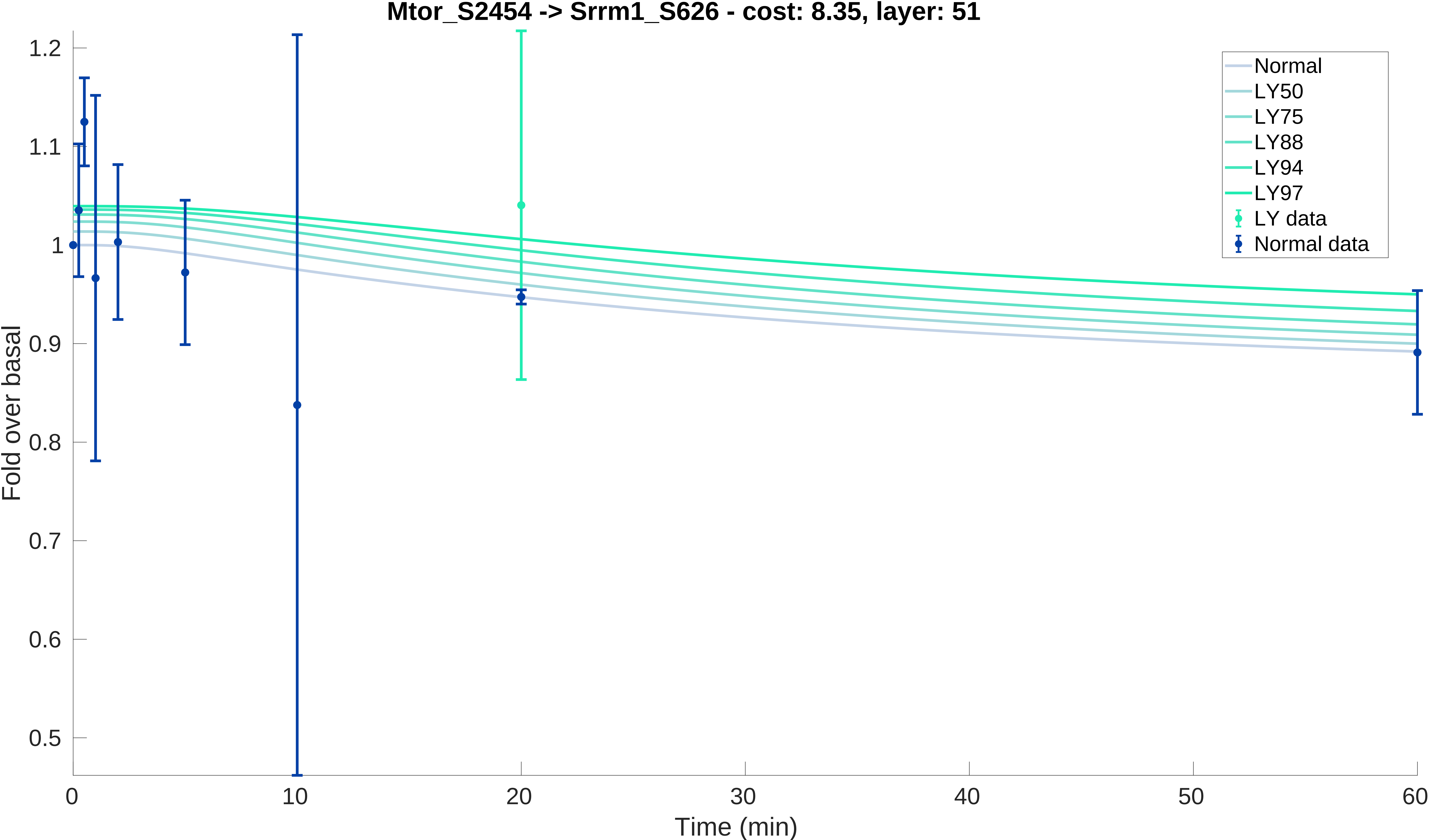

Mtor_S2454 -> Srrm1_S626 - cost: 8.35, layer: 51
Fold over basal
1.2
1.1
1
0.9
0.8
0.7
0.6
0.5
0
10
20
30
40
50
60
Time (min)
Normal
LY50
LY75
LY88
LY94
LY97
LY data
Normal data

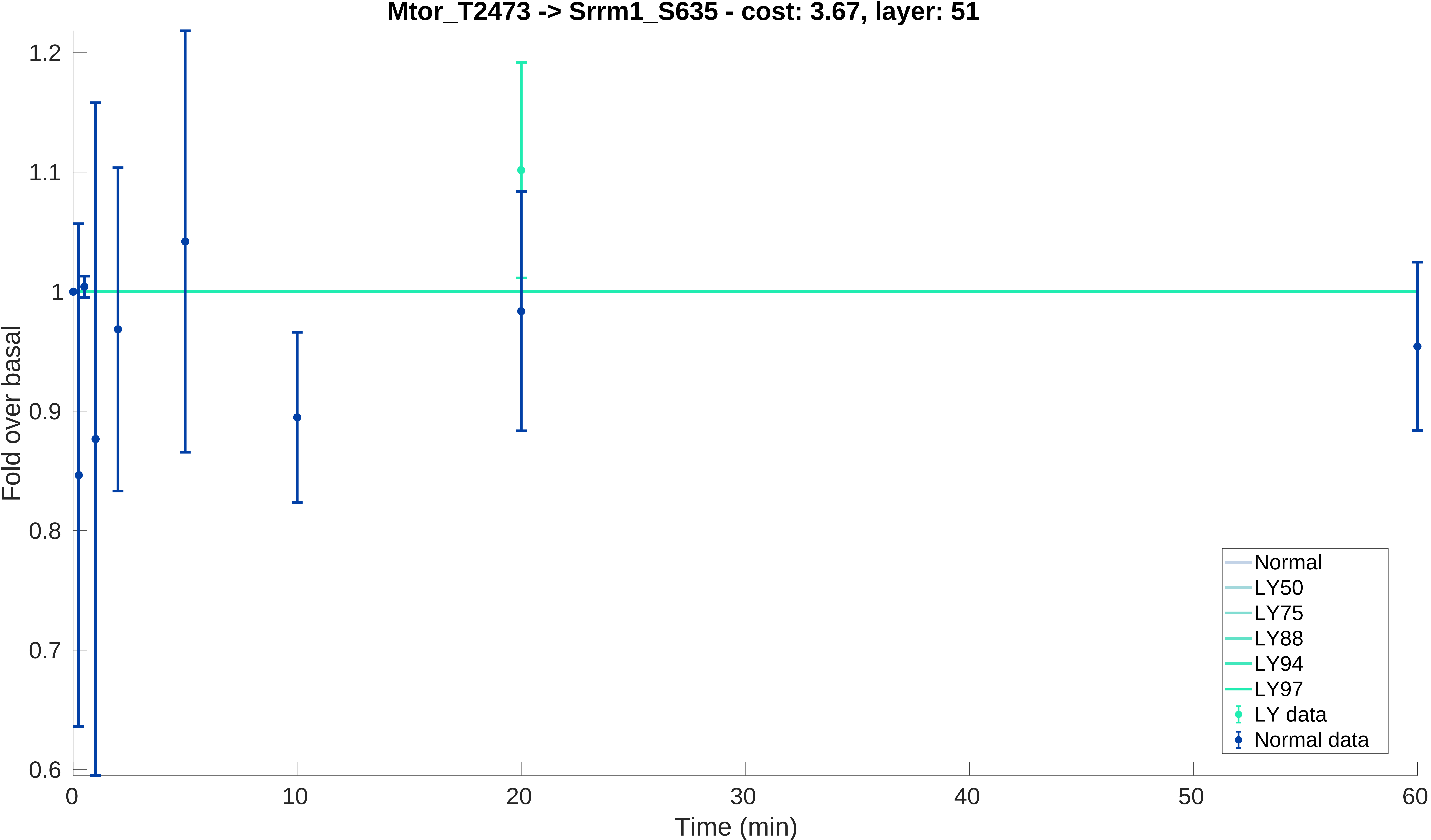

Mtor_T2473 -> Srrm1_S635 - cost: 3.67, layer: 51
Fold over basal
Time (min)
1.2
1.1
1
0.9
0.8
0.7
0.6
0
10
20
30
40
50
60
Normal
LY50
LY75
LY88
LY94
LY97
LY data
Normal data

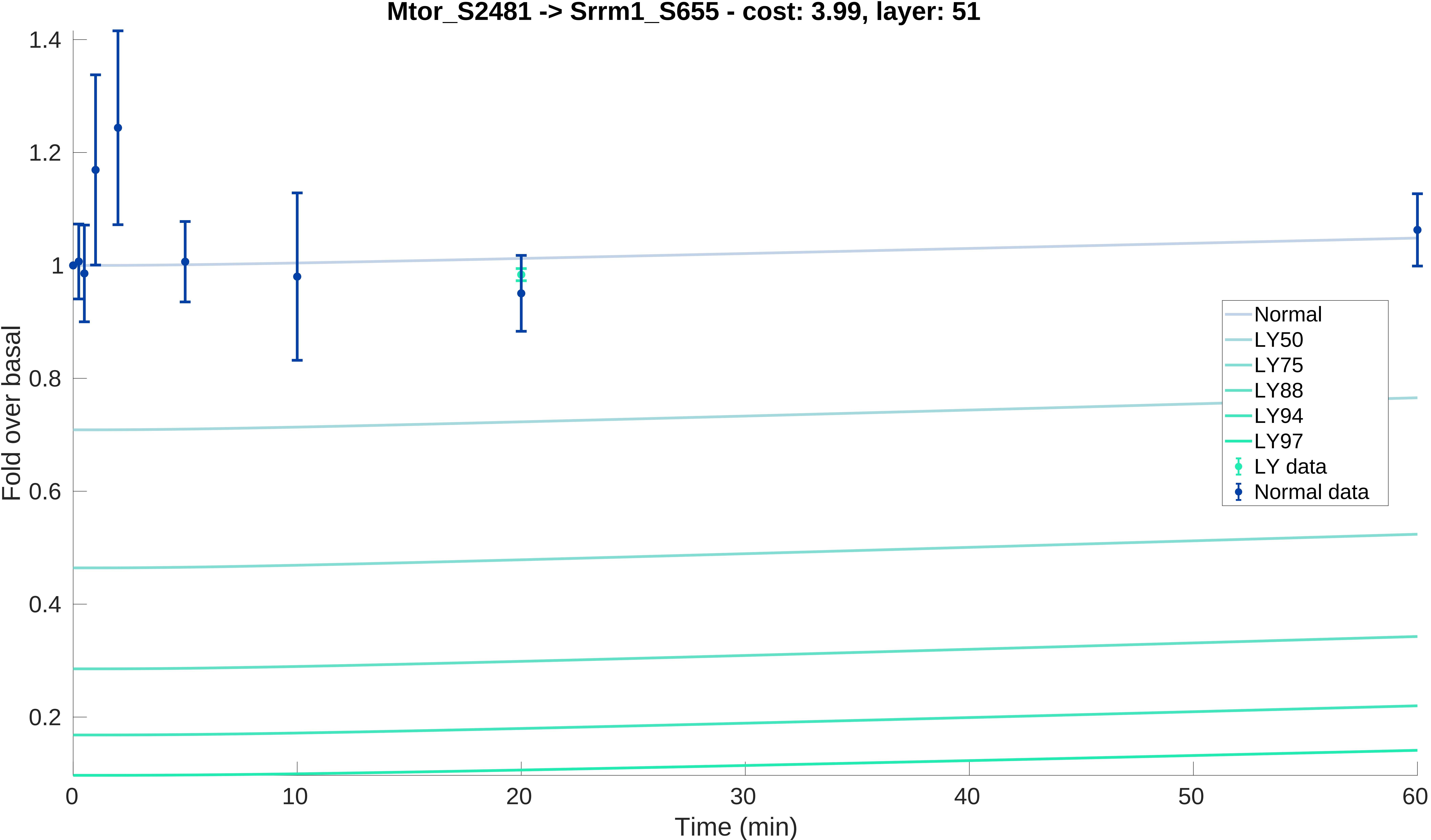
Mtor_S2481 -> Srrm1_S655 - cost: 3.99, layer: 51
Fold over basal
Time (min)
1.4
1.2
1
0.8
0.6
0.4
0.2
0
10
20
30
40
50
60
Normal
LY50
LY75
LY88
LY94
LY97
LY data
Normal data

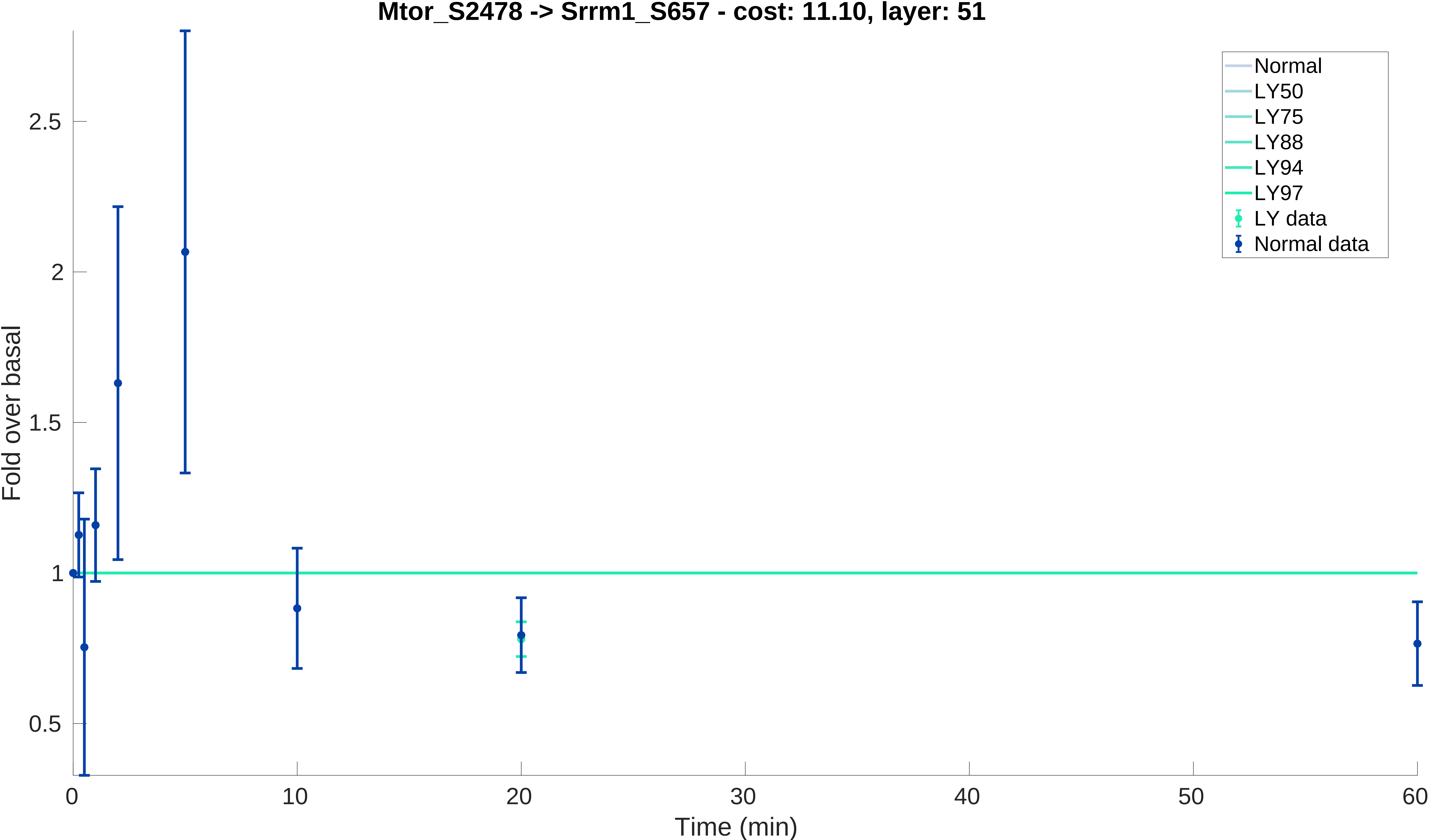

Mtor_S2478 -> Srrm1_S657 - cost: 11.10, layer: 51
Normal
LY50
LY75
LY88
LY94
LY97
LY data
Normal data
Fold over basal
0.5
1
1.5
2
2.5
0
10
20
30
40
50
60
Time (min)

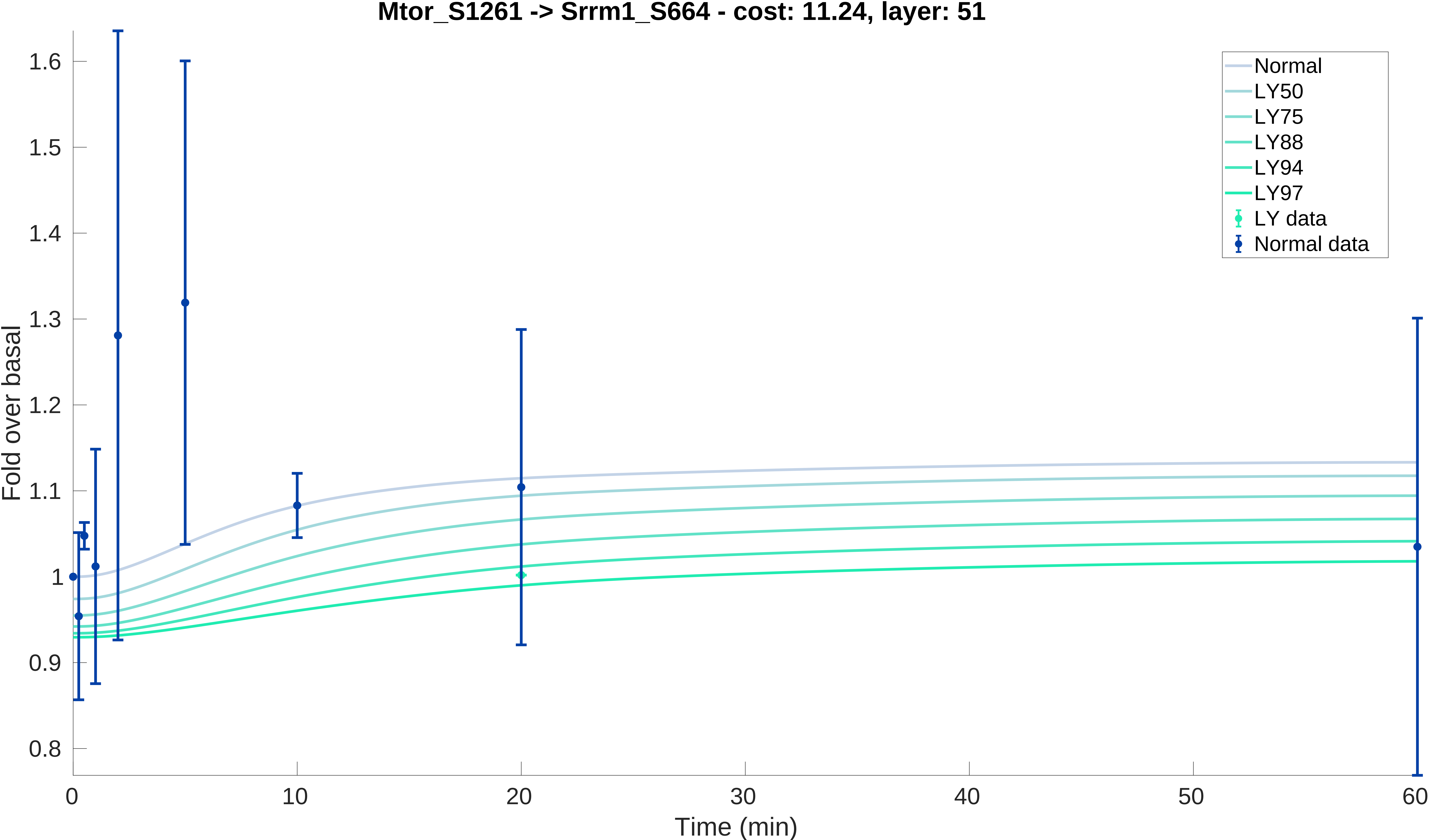

Mtor_S1261 -> Srrm1_S664 - cost: 11.24, layer: 51
Fold over basal
Time (min)
Normal
LY50
LY75
LY88
LY94
LY97
LY data
Normal data
1.6
1.5
1.4
1.3
1.2
1.1
1
0.9
0.8
0
10
20
30
40
50
60

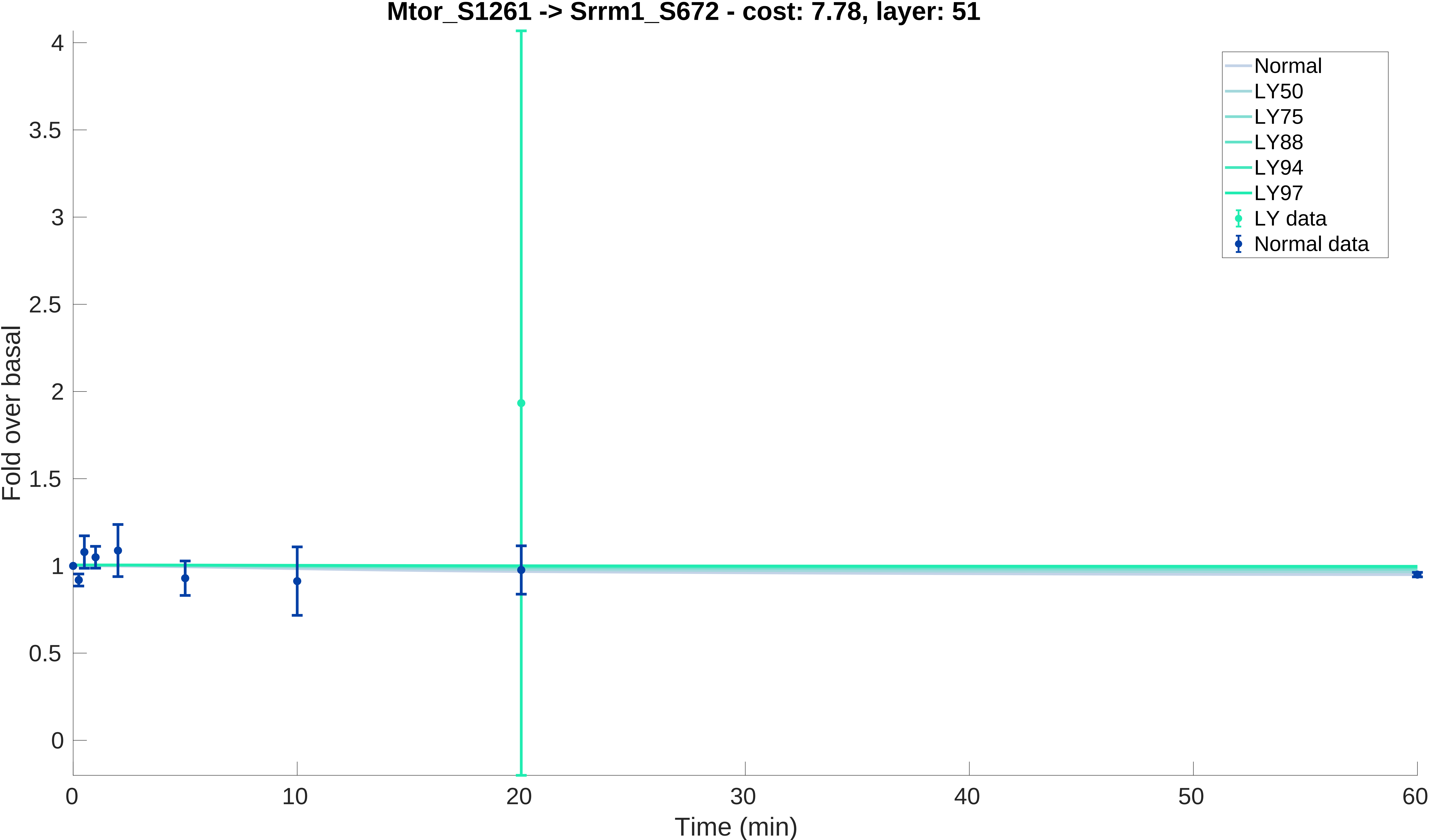

Mtor_S1261 -> Srrm1_S672 - cost: 7.78, layer: 51
Fold over basal
Time (min)
0
0.5
1
1.5
2
2.5
3
3.5
4
0
10
20
30
40
50
60
Normal
LY50
LY75
LY88
LY94
LY97
LY data
Normal data

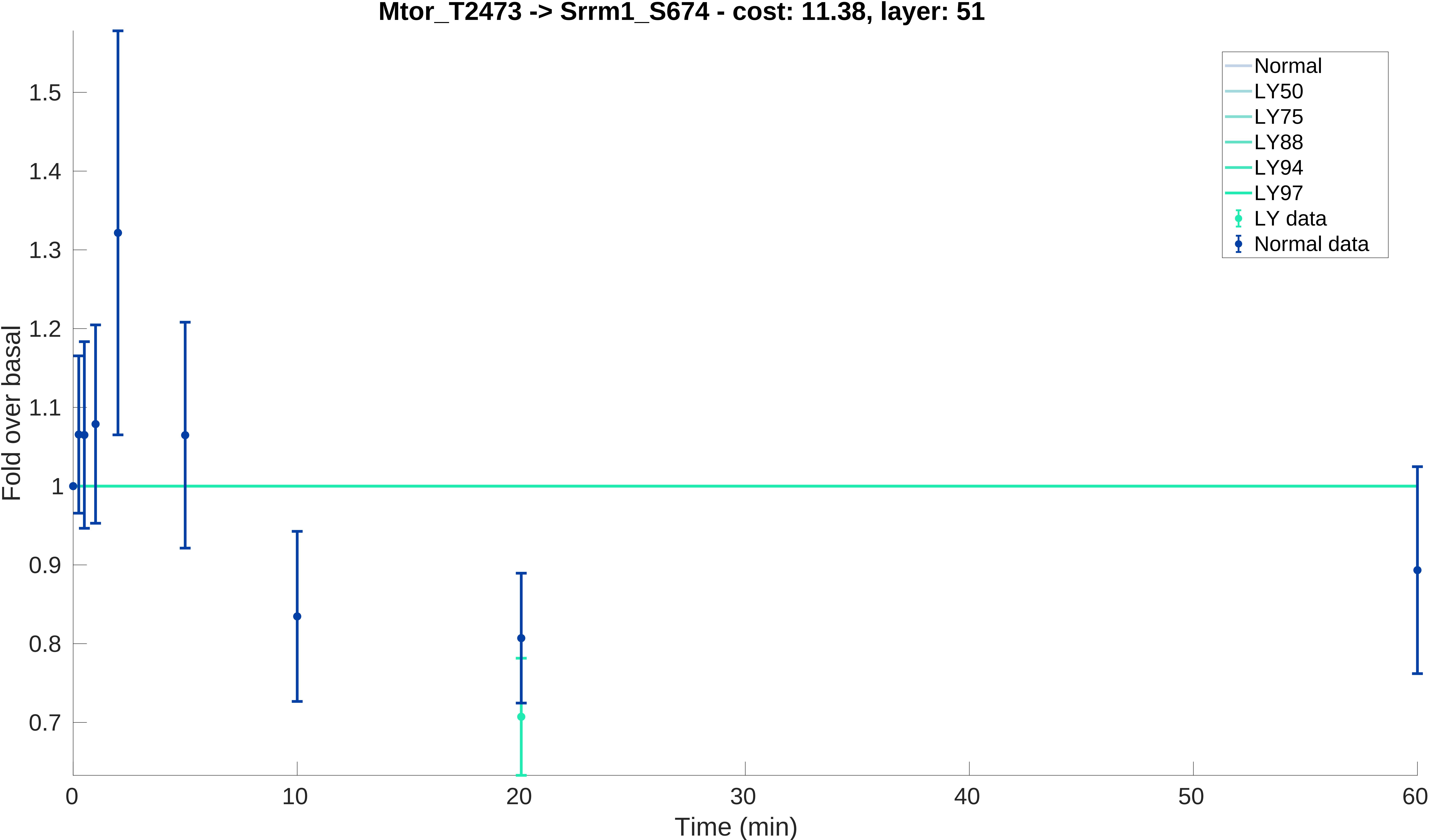
Mtor_T2473 -> Srrm1_S674 - cost: 11.38, layer: 51
Normal
LY50
LY75
LY88
LY94
LY97
LY data
Normal data
Fold over basal
1.5
1.4
1.3
1.2
1.1
1
0.9
0.8
0.7
0
10
20
30
40
50
60
Time (min)

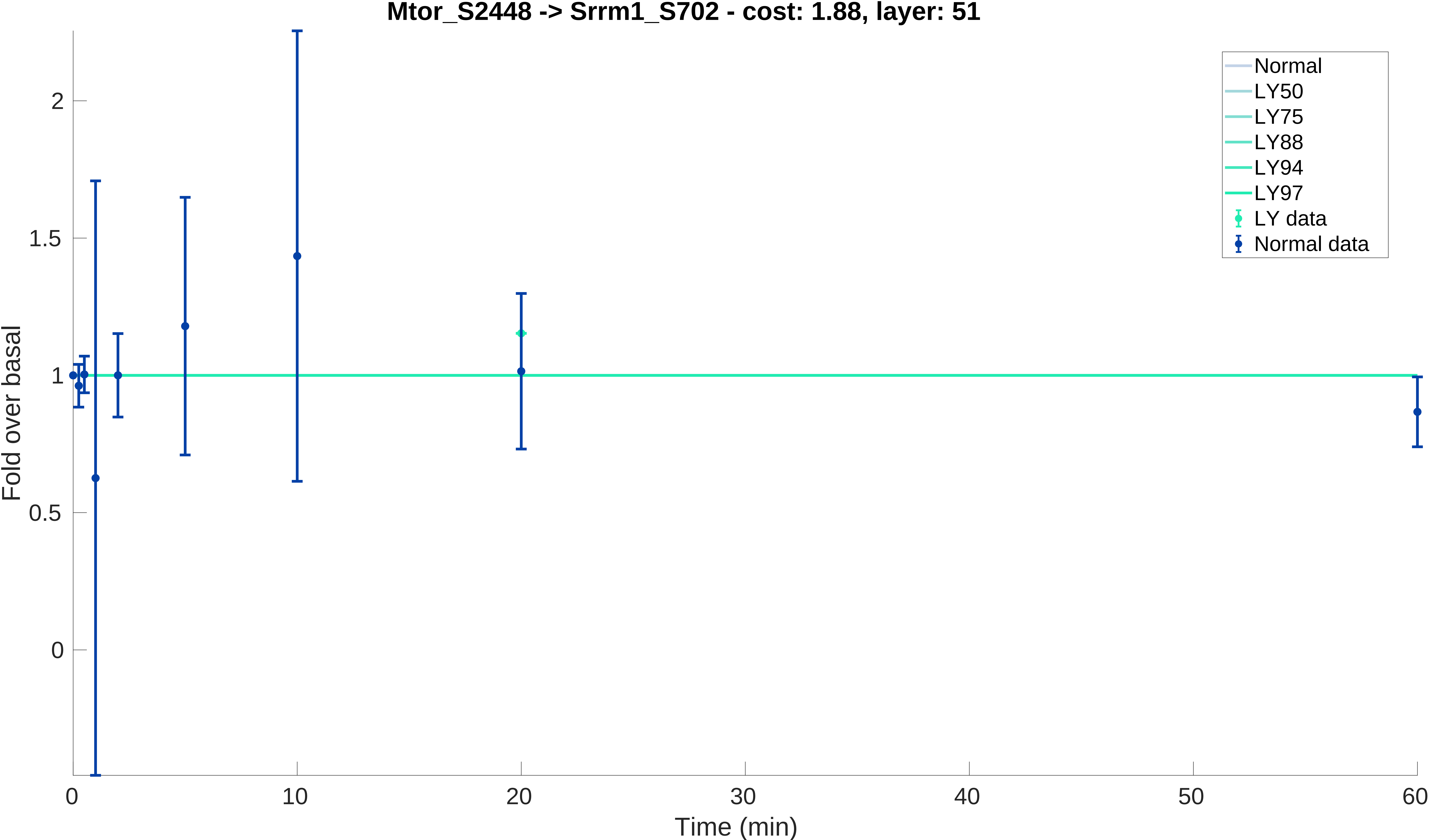

Mtor_S2448 -> Srrm1_S702 - cost: 1.88, layer: 51
Normal
LY50
LY75
LY88
LY94
LY97
LY data
Normal data
Fold over basal
2
1.5
1
0.5
0
Time (min)
0
10
20
30
40
50
60

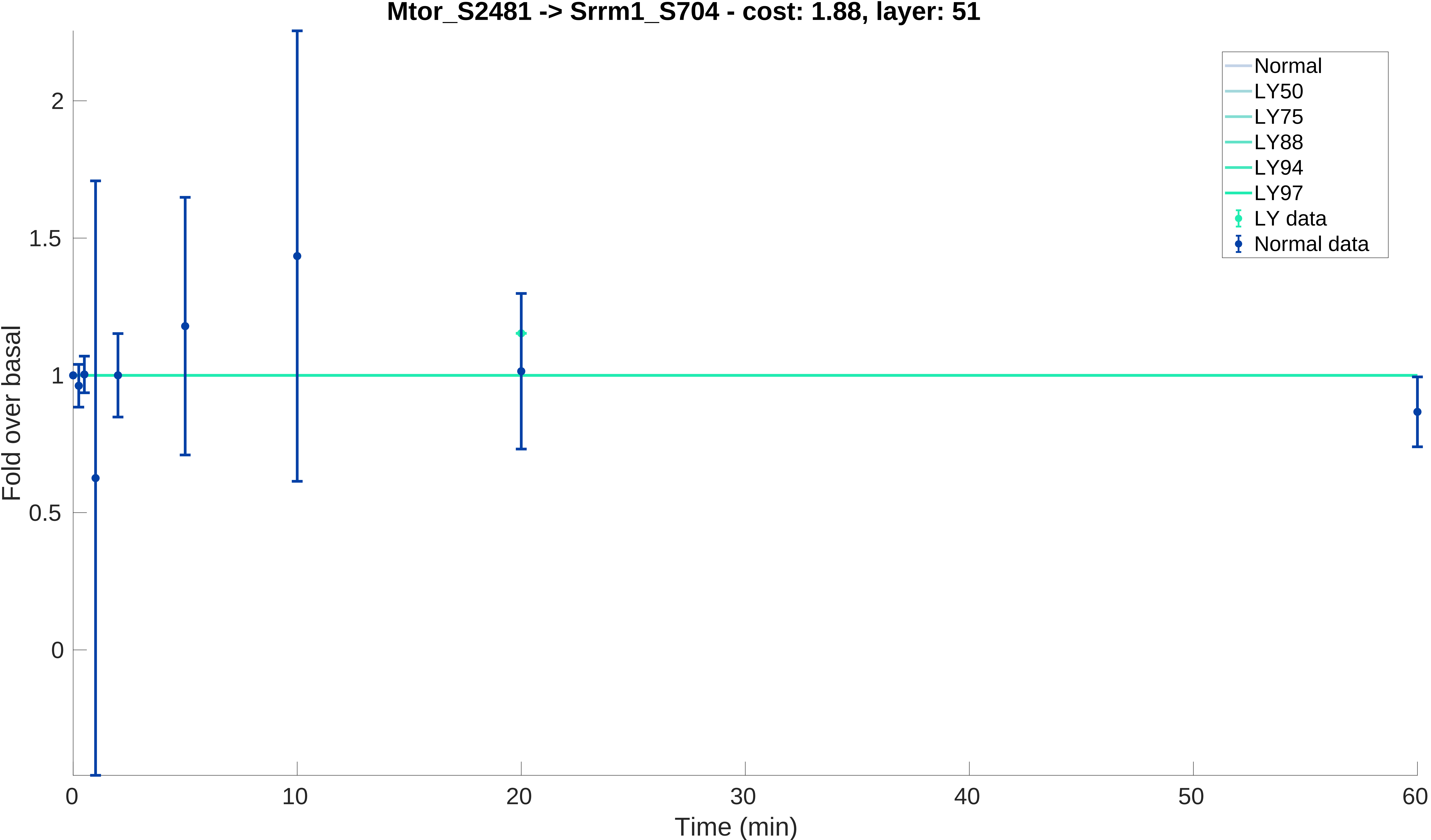

Mtor_S2481 -> Srrm1_S704 - cost: 1.88, layer: 51
Normal
LY50
LY75
LY88
LY94
LY97
LY data
Normal data
Fold over basal
2
1.5
1
0.5
0
0
10
20
30
40
50
60
Time (min)

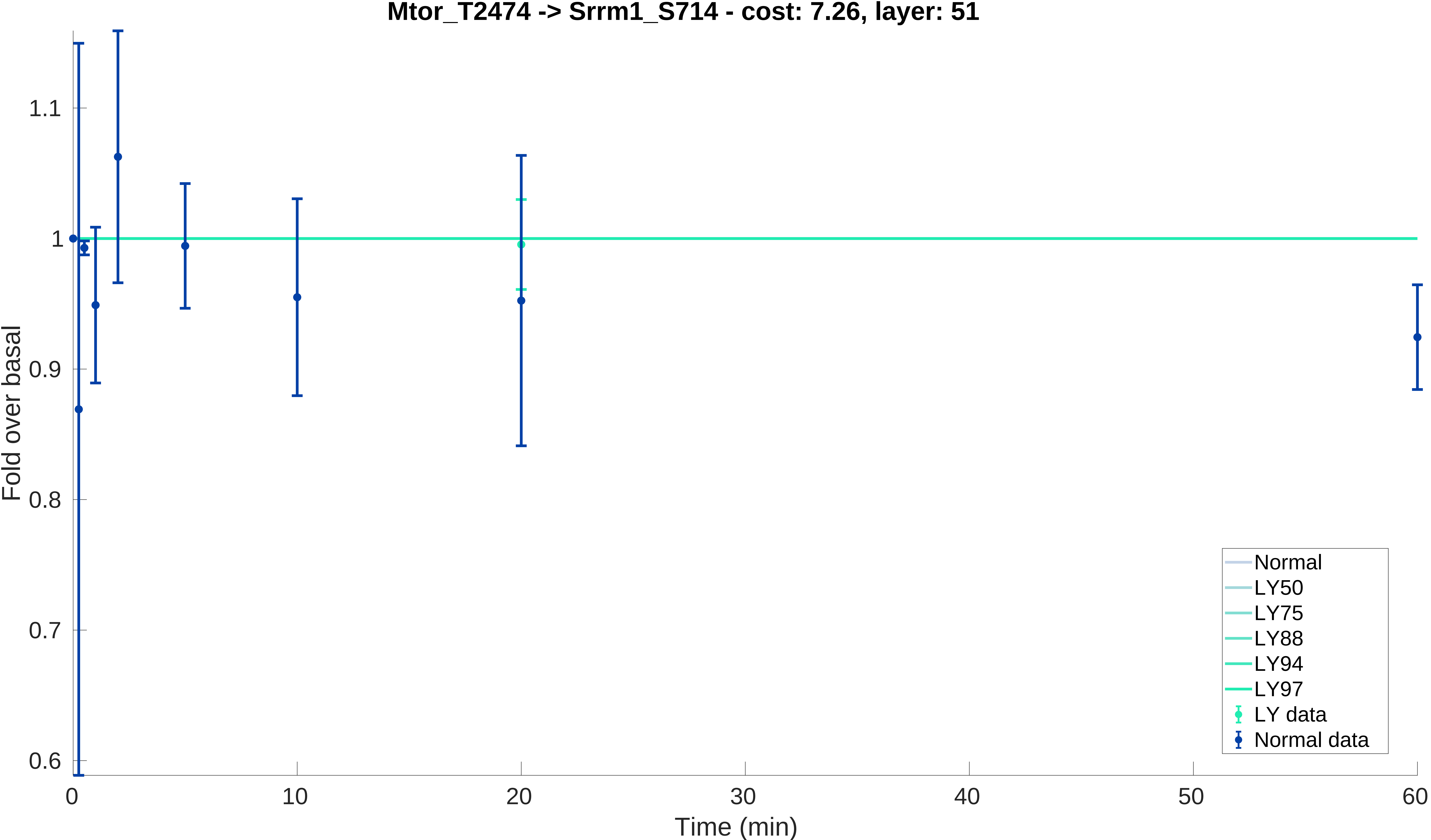

Mtor_T2474 -> Srrm1_S714 - cost: 7.26, layer: 51
Fold over basal
1.1
1
0.9
0.8
0.7
0.6
0
10
20
30
40
50
60
Time (min)
Normal
LY50
LY75
LY88
LY94
LY97
LY data
Normal data

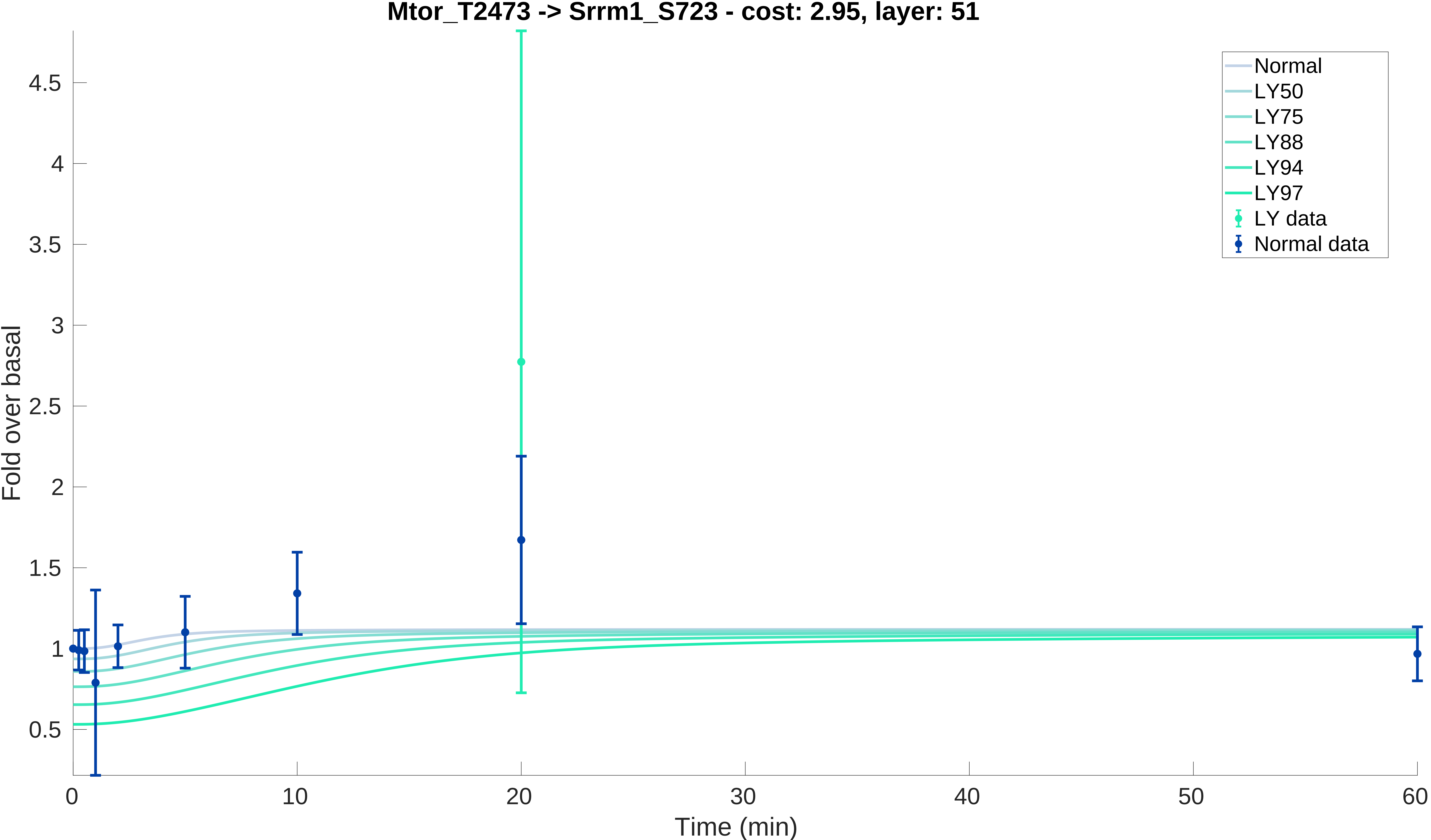

Mtor_T2473 -> Srrm1_S723 - cost: 2.95, layer: 51
Normal
LY50
LY75
LY88
LY94
LY97
LY data
Normal data
Fold over basal
Time (min)
0
10
20
30
40
50
60
0.5
1
1.5
2
2.5
3
3.5
4
4.5

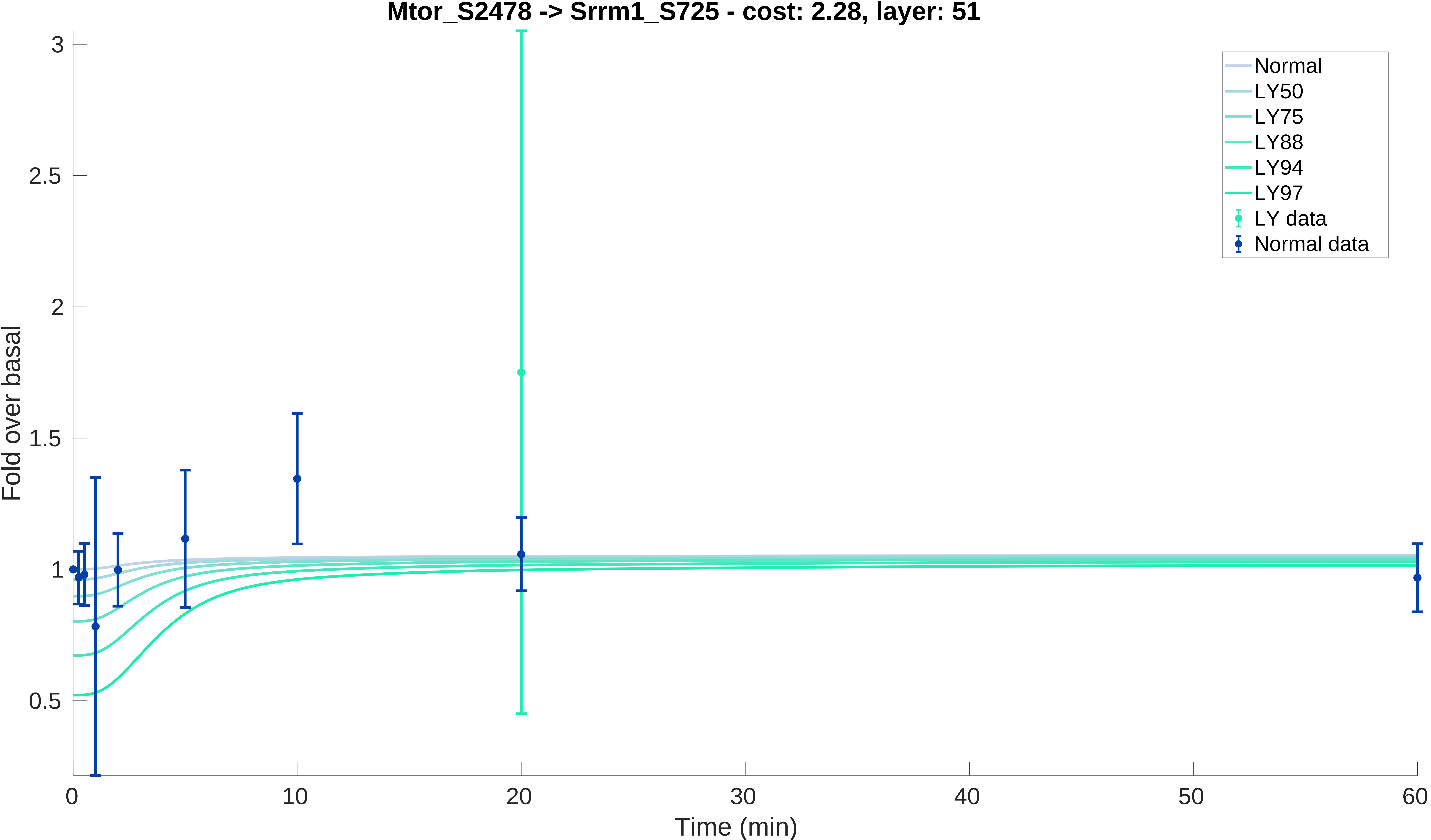

Mtor_S2478 -> Srrm1_S725 - cost: 2.28, layer: 51
Fold over basal
Time (min)
0.5
1
1.5
2
2.5
3
0
10
20
30
40
50
60
Normal
LY50
LY75
LY88
LY94
LY97
LY data
Normal data

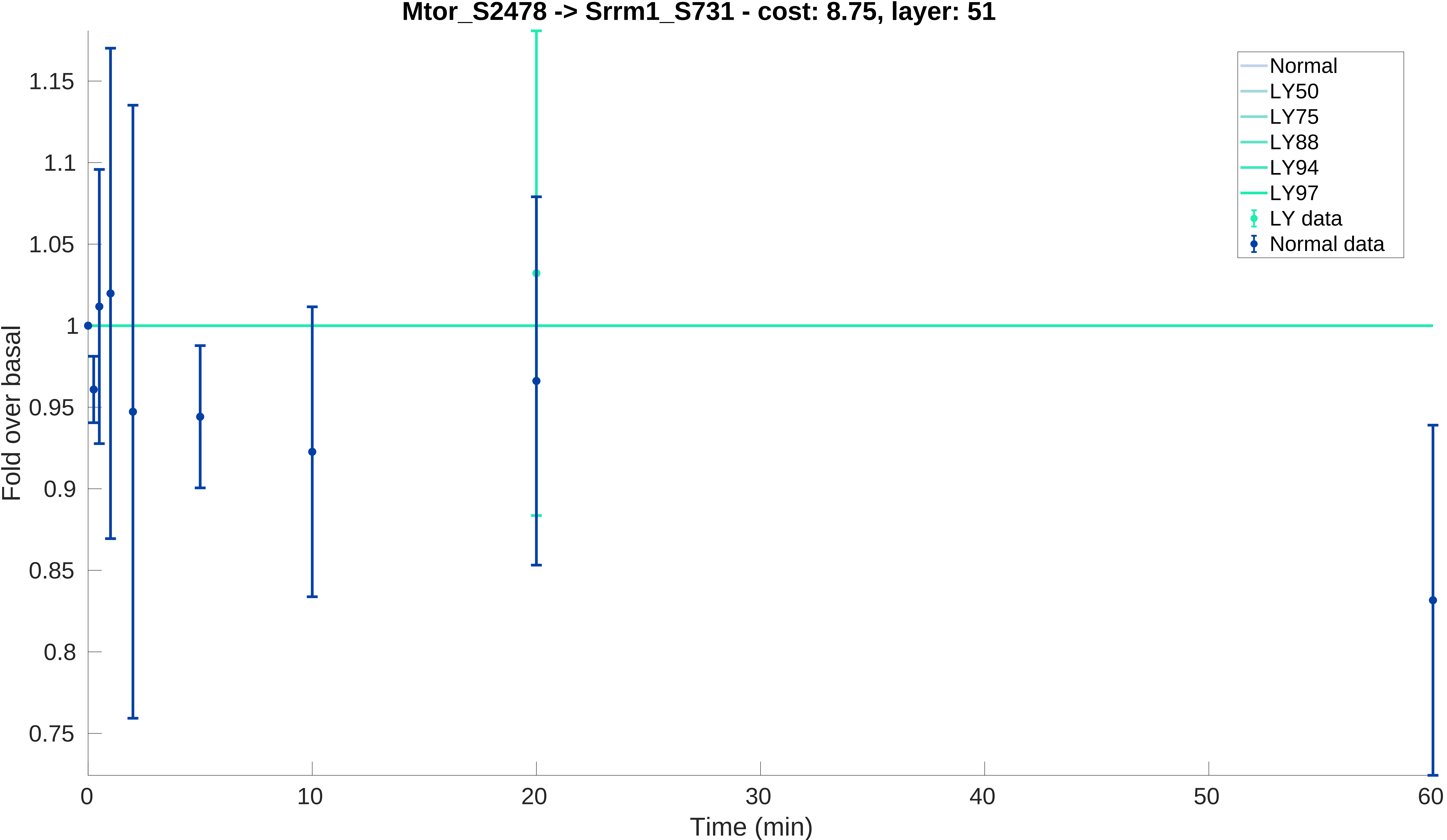

Mtor_S2478 -> Srrm1_S731 - cost: 8.75, layer: 51
Fold over basal
Time (min)
0
10
20
30
40
50
60
0.75
0.8
0.85
0.9
0.95
1
1.05
1.1
1.15
Normal
LY50
LY75
LY88
LY94
LY97
LY data
Normal data

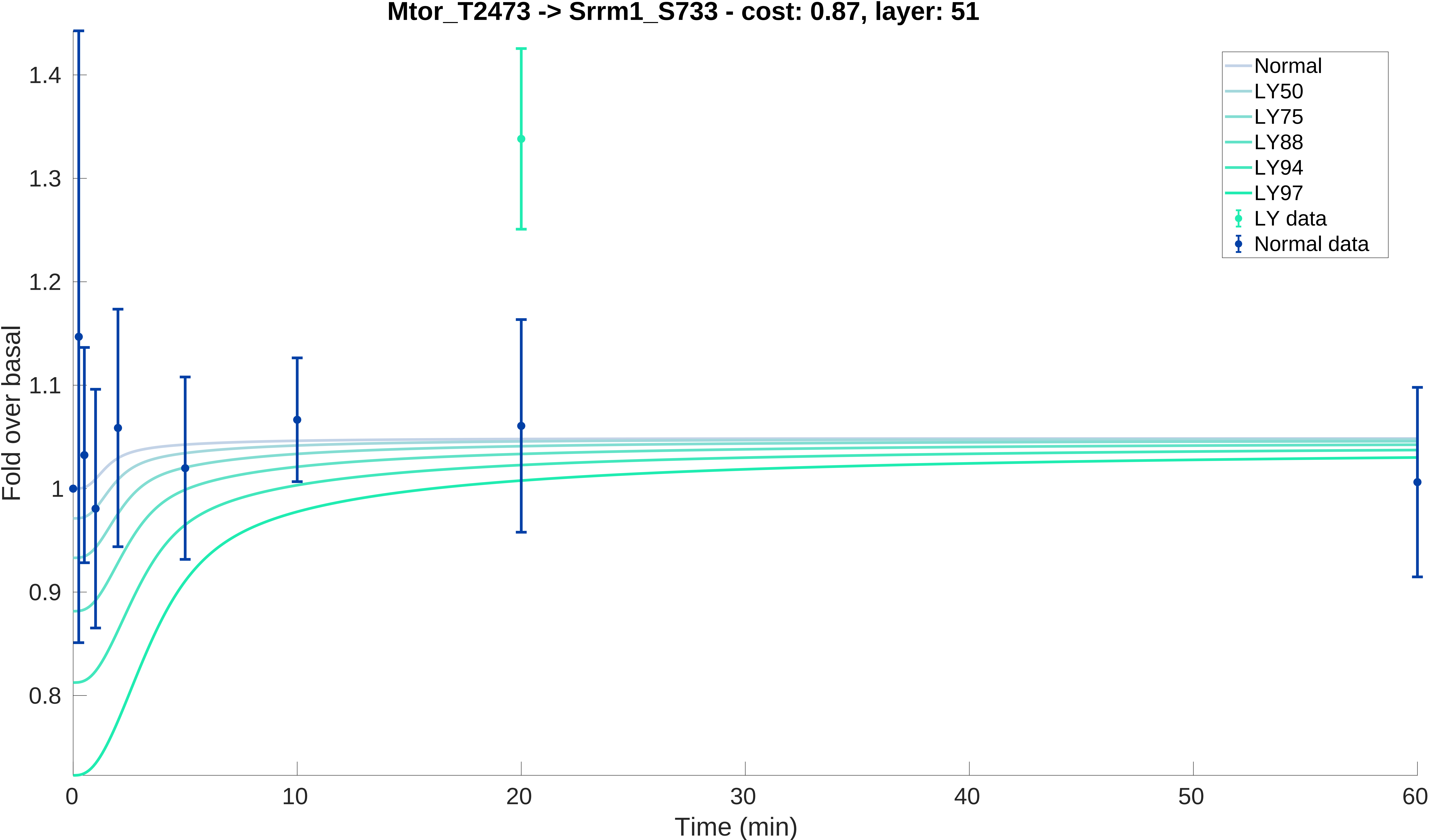

Mtor_T2473 -> Srrm1_S733 - cost: 0.87, layer: 51
Fold over basal
Time (min)
0
10
20
30
40
50
60
0.8
0.9
1
1.1
1.2
1.3
1.4
Normal
LY50
LY75
LY88
LY94
LY97
LY data
Normal data

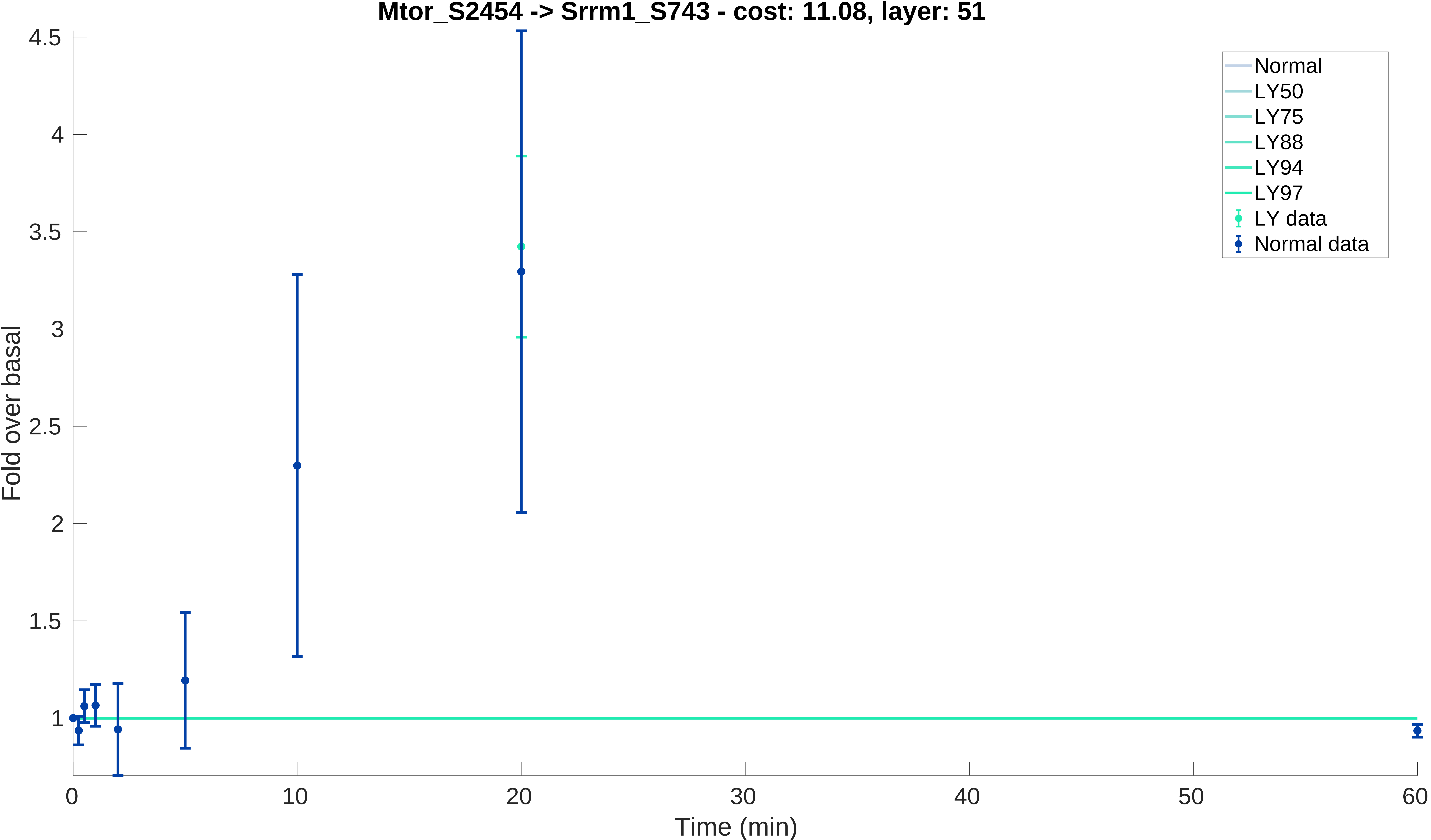

Mtor_S2454 -> Srrm1_S743 - cost: 11.08, layer: 51
Fold over basal
Time (min)
4.5
4
3.5
3
2.5
2
1.5
1
0
10
20
30
40
50
60
Normal
LY50
LY75
LY88
LY94
LY97
LY data
Normal data

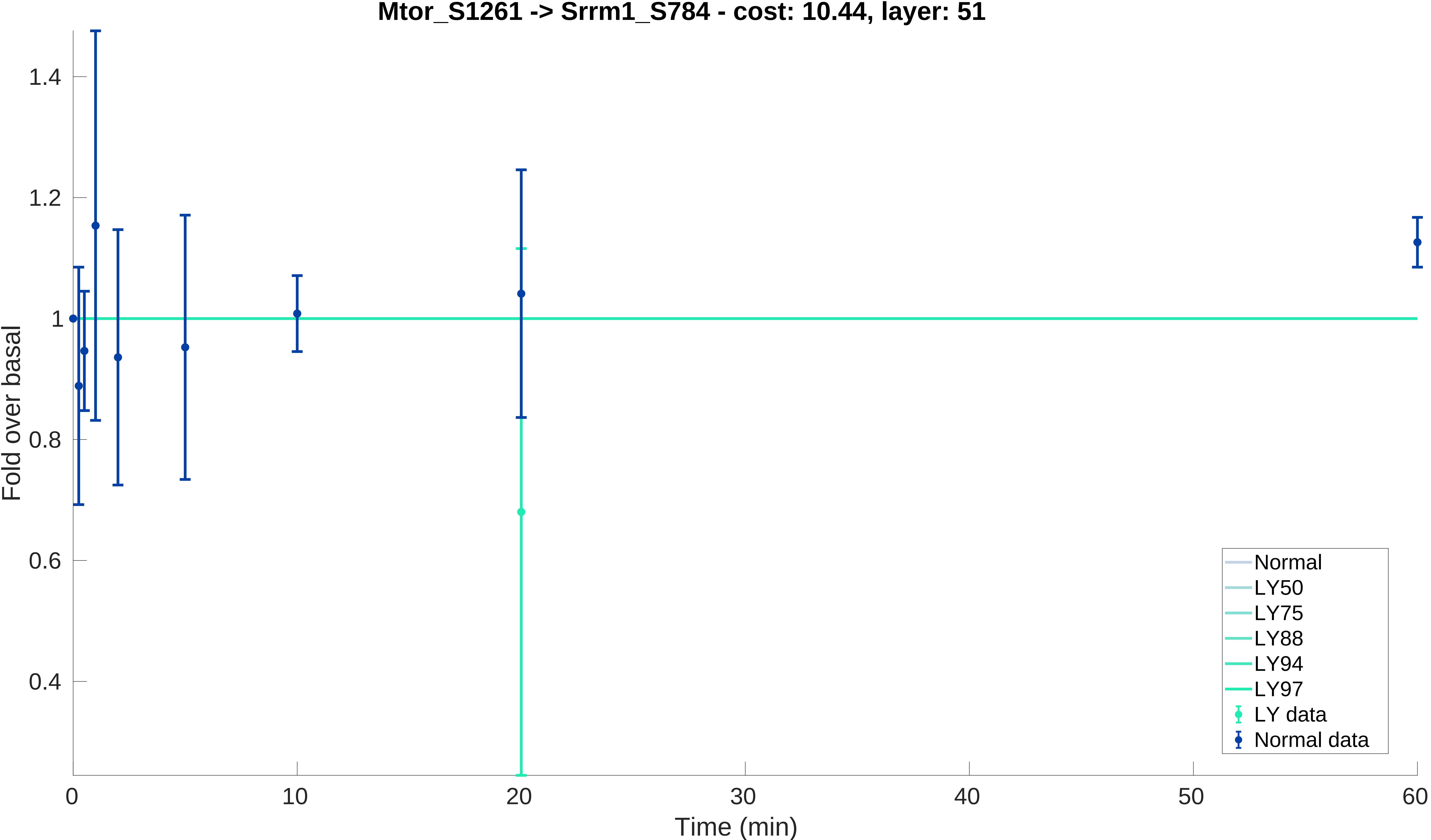
Mtor_S1261 -> Srrm1_S784 - cost: 10.44, layer: 51
Fold over basal
1.4
1.2
1
0.8
0.6
0.4
0
10
20
30
40
50
60
Time (min)
Normal
LY50
LY75
LY88
LY94
LY97
LY data
Normal data

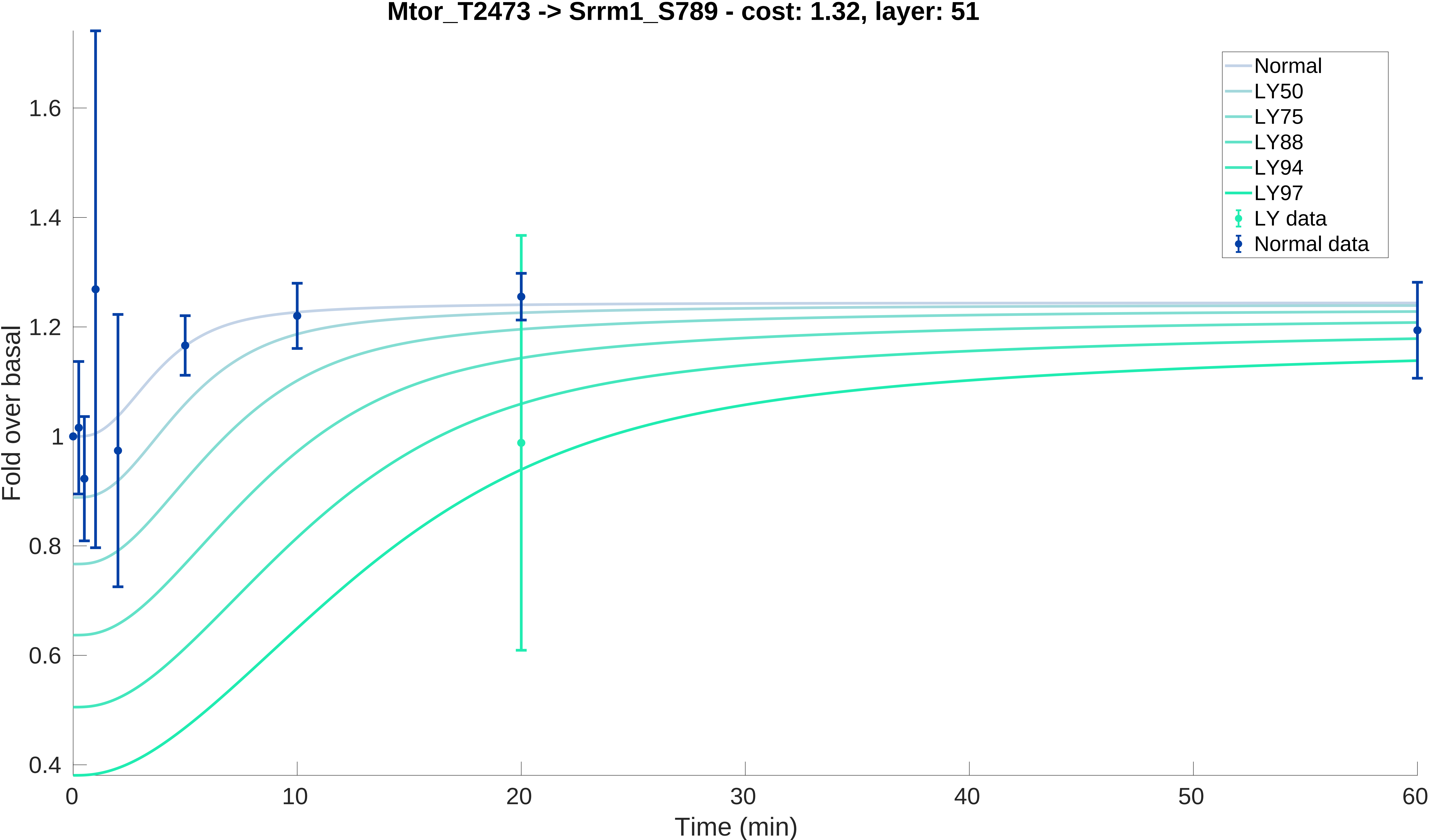
Mtor_T2473 -> Srrm1_S789 - cost: 1.32, layer: 51
Normal
LY50
LY75
LY88
LY94
LY97
LY data
Normal data
Fold over basal
1.6
1.4
1.2
1
0.8
0.6
0.4
0
10
20
30
40
50
60
Time (min)

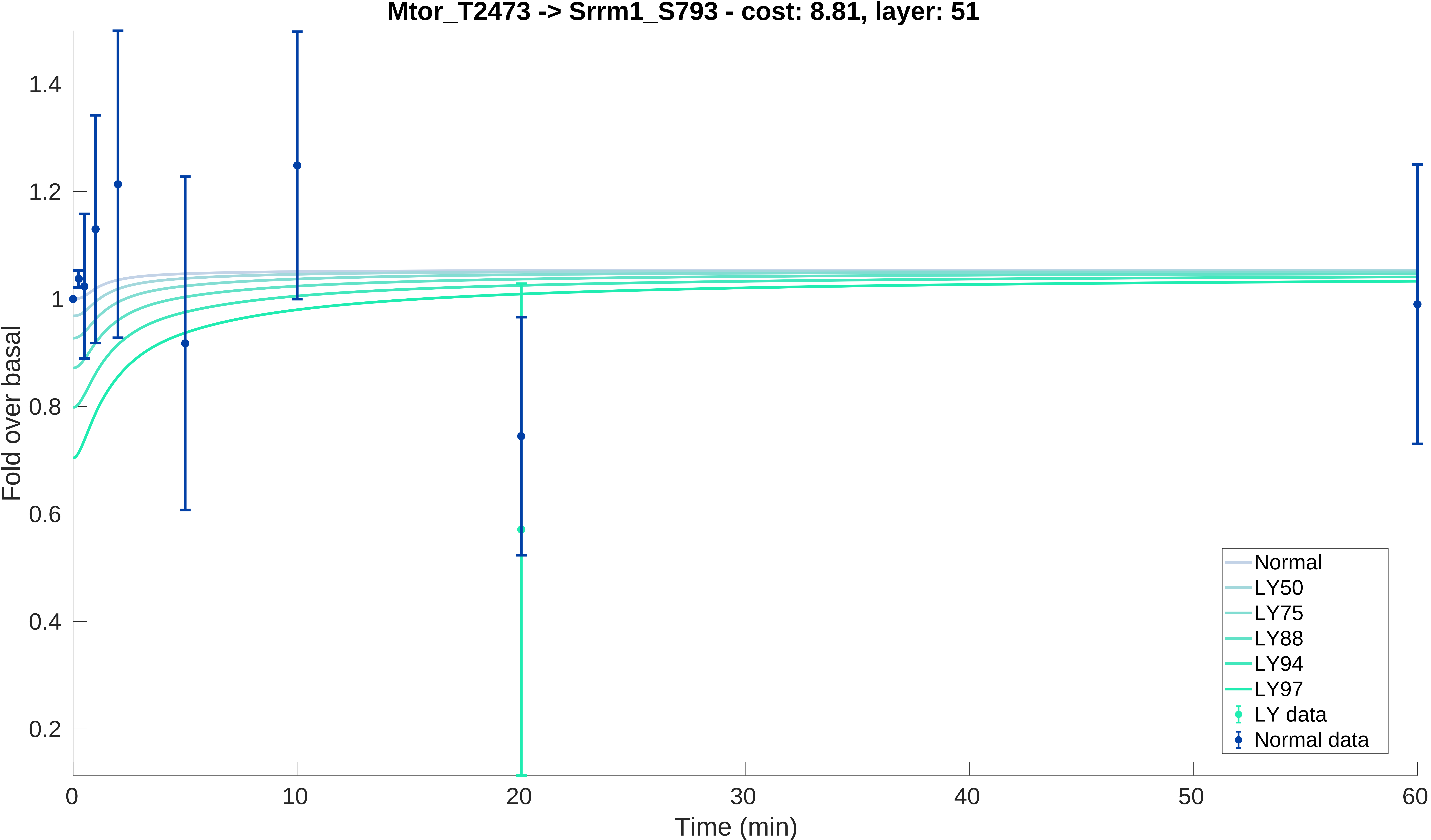

Mtor_T2473 -> Srrm1_S793 - cost: 8.81, layer: 51
Fold over basal
1.4
1.2
1
0.8
0.6
0.4
0.2
0
10
20
30
40
50
60
Time (min)
Normal
LY50
LY75
LY88
LY94
LY97
LY data
Normal data

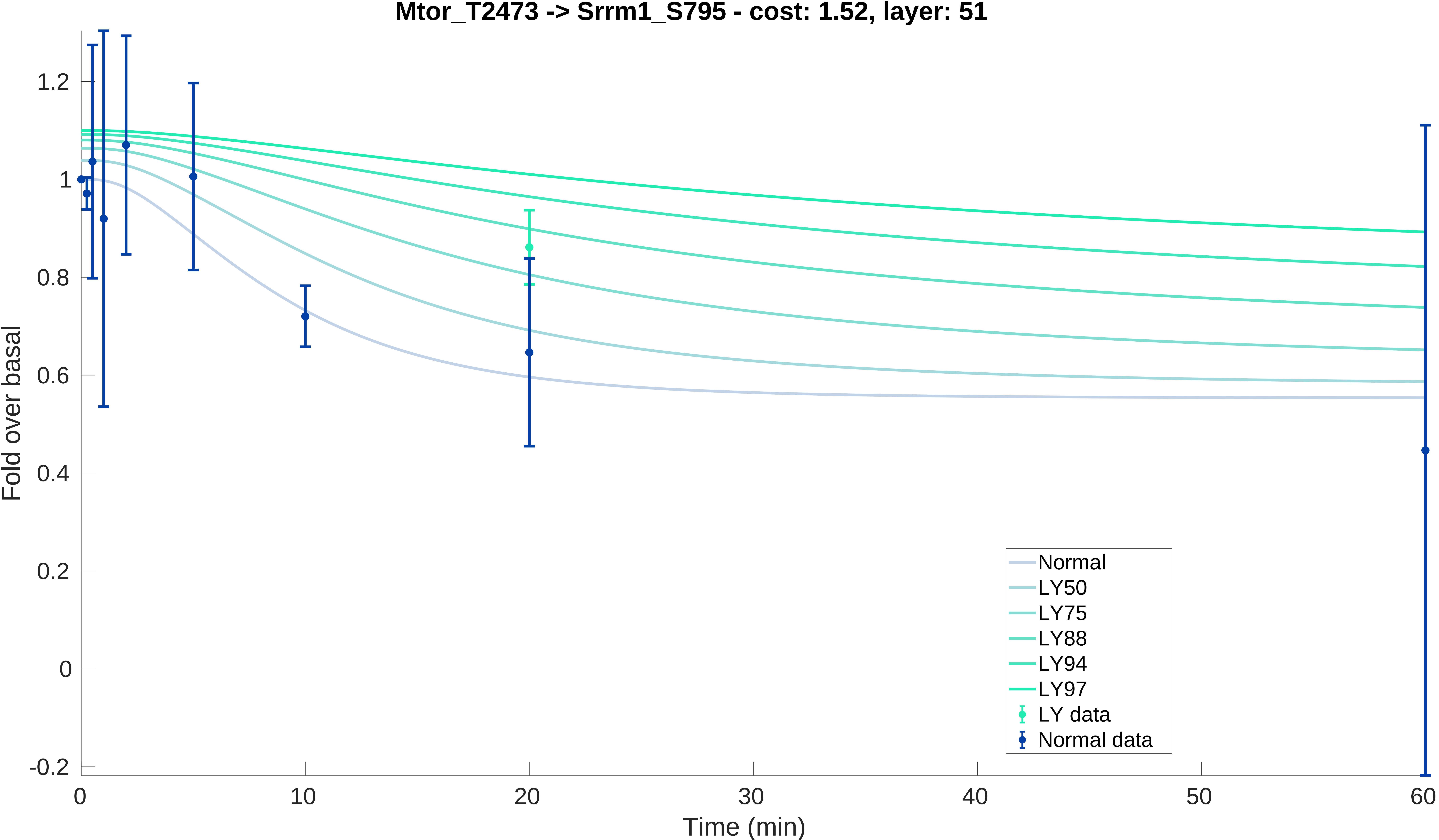

Mtor_T2473 -> Srrm1_S795 - cost: 1.52, layer: 51
Fold over basal
Time (min)
1.2
1
0.8
0.6
0.4
0.2
0
-0.2
0
10
20
30
40
50
60
Normal
LY50
LY75
LY88
LY94
LY97
LY data
Normal data

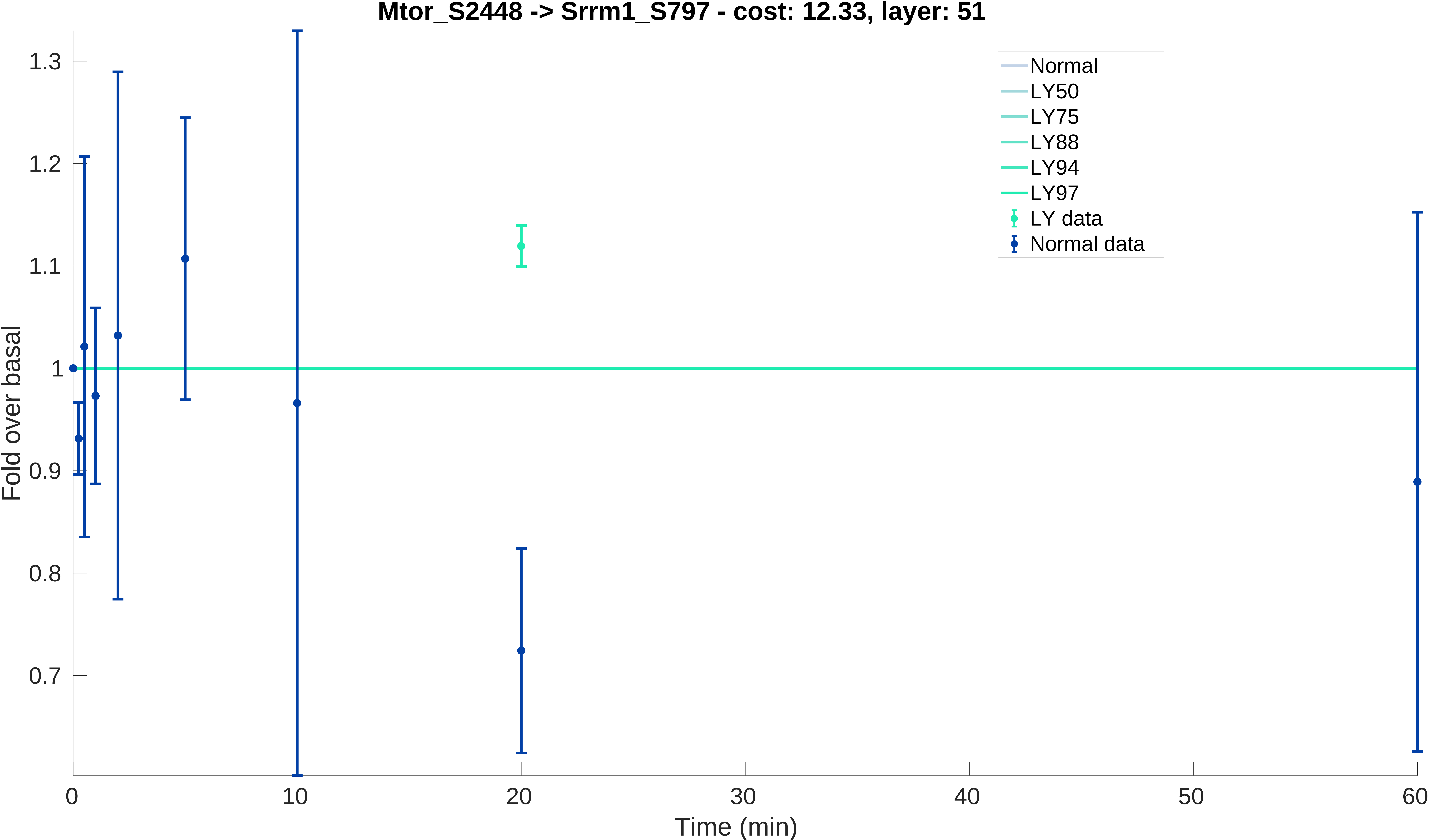

Mtor_S2448 -> Srrm1_S797 - cost: 12.33, layer: 51
Fold over basal
Time (min)
Normal
LY50
LY75
LY88
LY94
LY97
LY data
Normal data
1.3
1.2
1.1
1
0.9
0.8
0.7
0
10
20
30
40
50
60

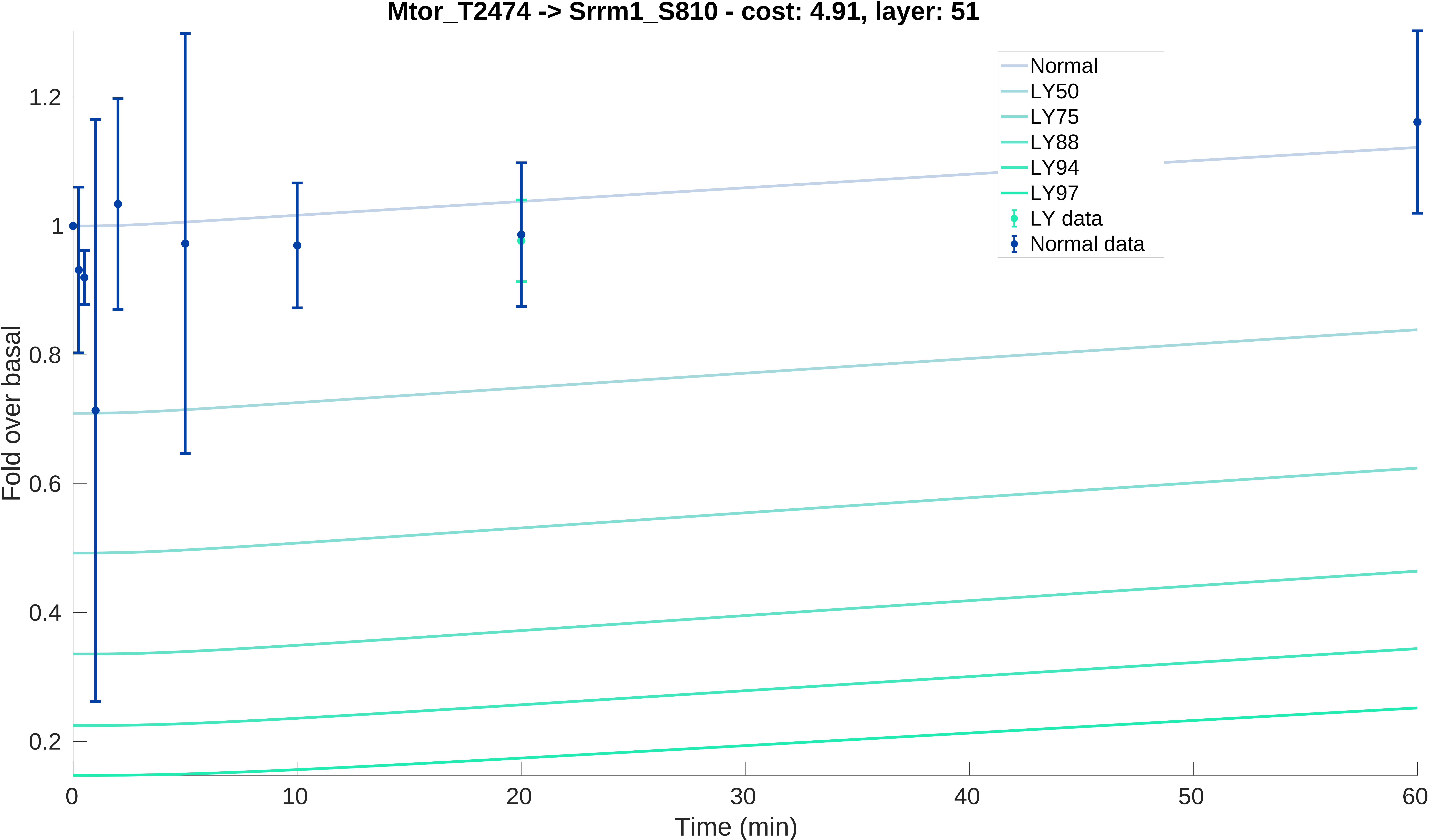

Mtor_T2474 -> Srrm1_S810 - cost: 4.91, layer: 51
Normal
LY50
LY75
LY88
LY94
LY97
LY data
Normal data
Fold over basal
Time (min)
0
10
20
30
40
50
60
0.2
0.4
0.6
0.8
1
1.2

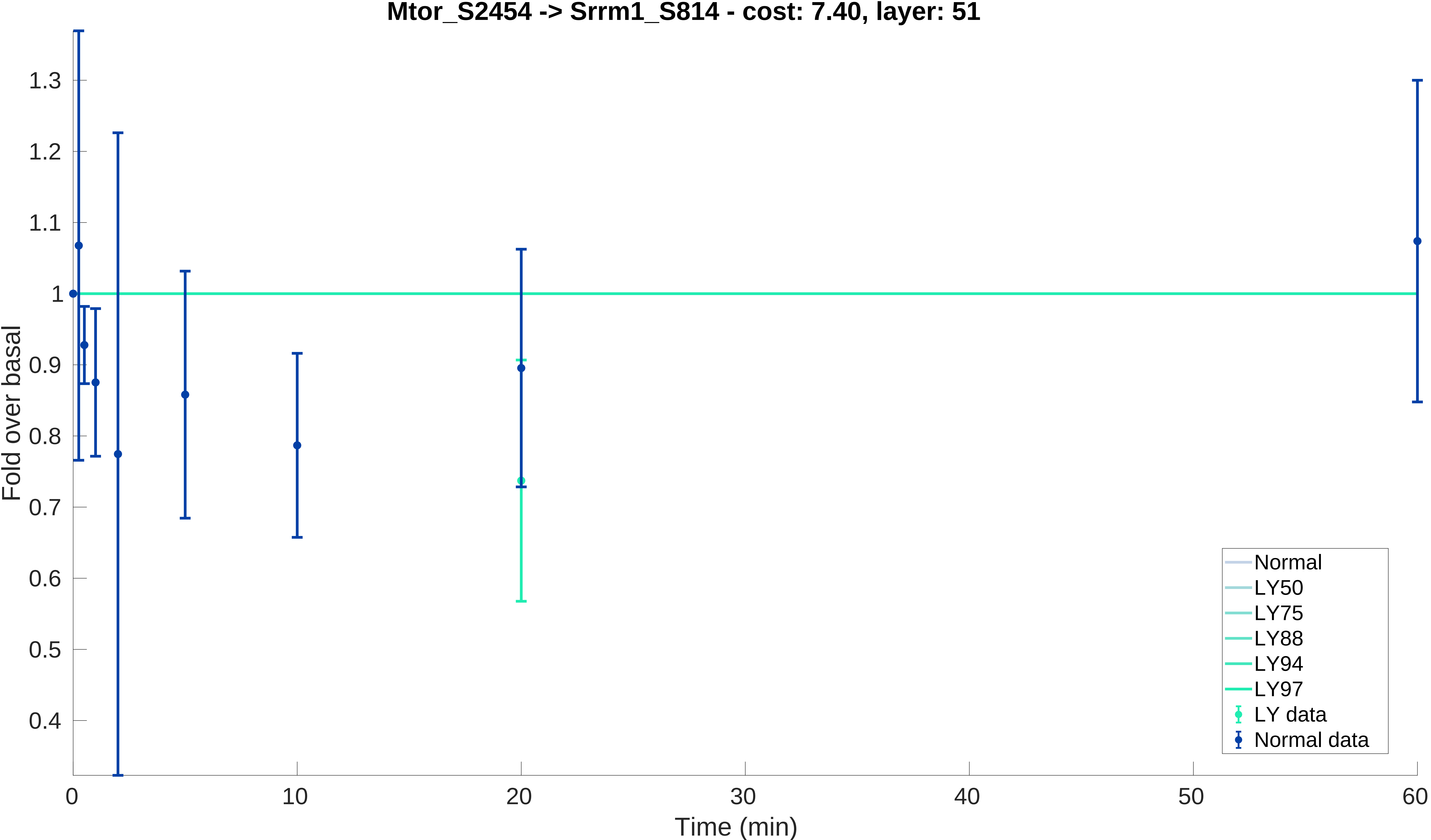
Mtor_S2454 -> Srrm1_S814 - cost: 7.40, layer: 51
Fold over basal
1.3
1.2
1.1
1
0.9
0.8
0.7
0.6
0.5
0.4
0
10
20
30
40
50
60
Time (min)
Normal
LY50
LY75
LY88
LY94
LY97
LY data
Normal data

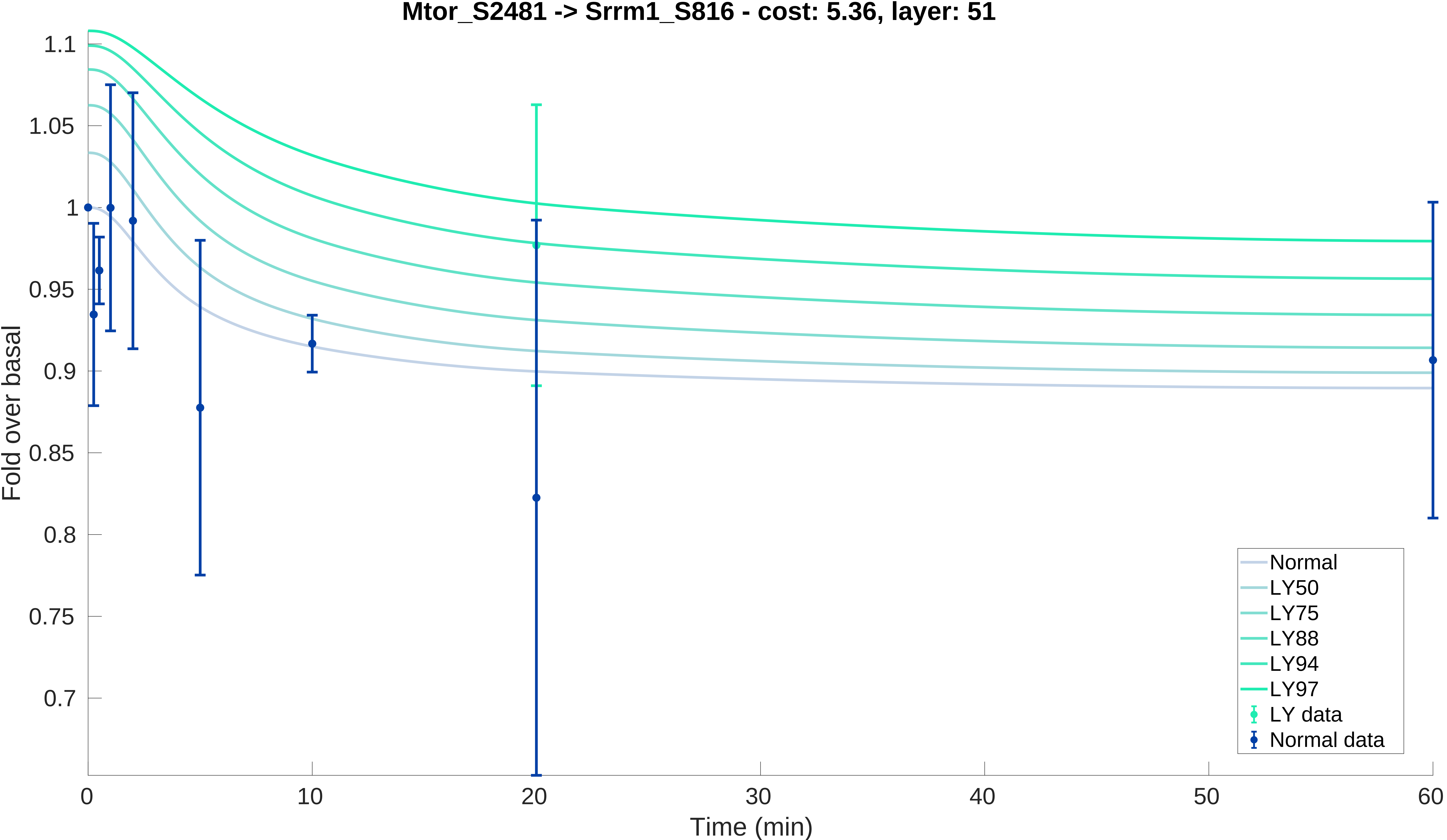

Mtor_S2481 -> Srrm1_S816 - cost: 5.36, layer: 51
Fold over basal
1.1
1.05
1
0.95
0.9
0.85
0.8
0.75
0.7
0
10
20
30
40
50
60
Time (min)
Normal
LY50
LY75
LY88
LY94
LY97
LY data
Normal data

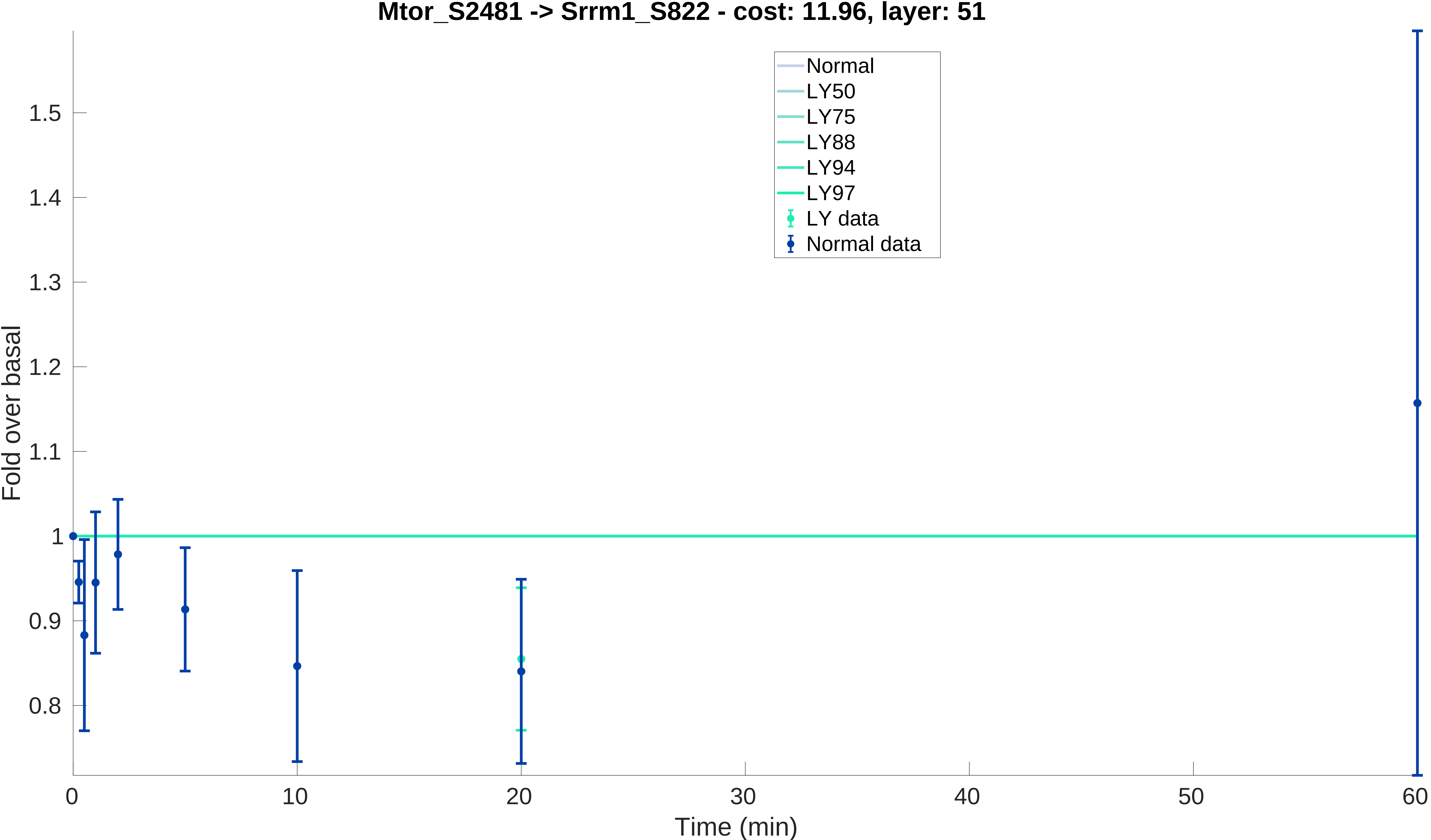

Mtor_S2481 -> Srrm1_S822 - cost: 11.96, layer: 51
Normal
LY50
LY75
LY88
LY94
LY97
LY data
Normal data
Fold over basal
1.5
1.4
1.3
1.2
1.1
1
0.9
0.8
0
10
20
30
40
50
60
Time (min)

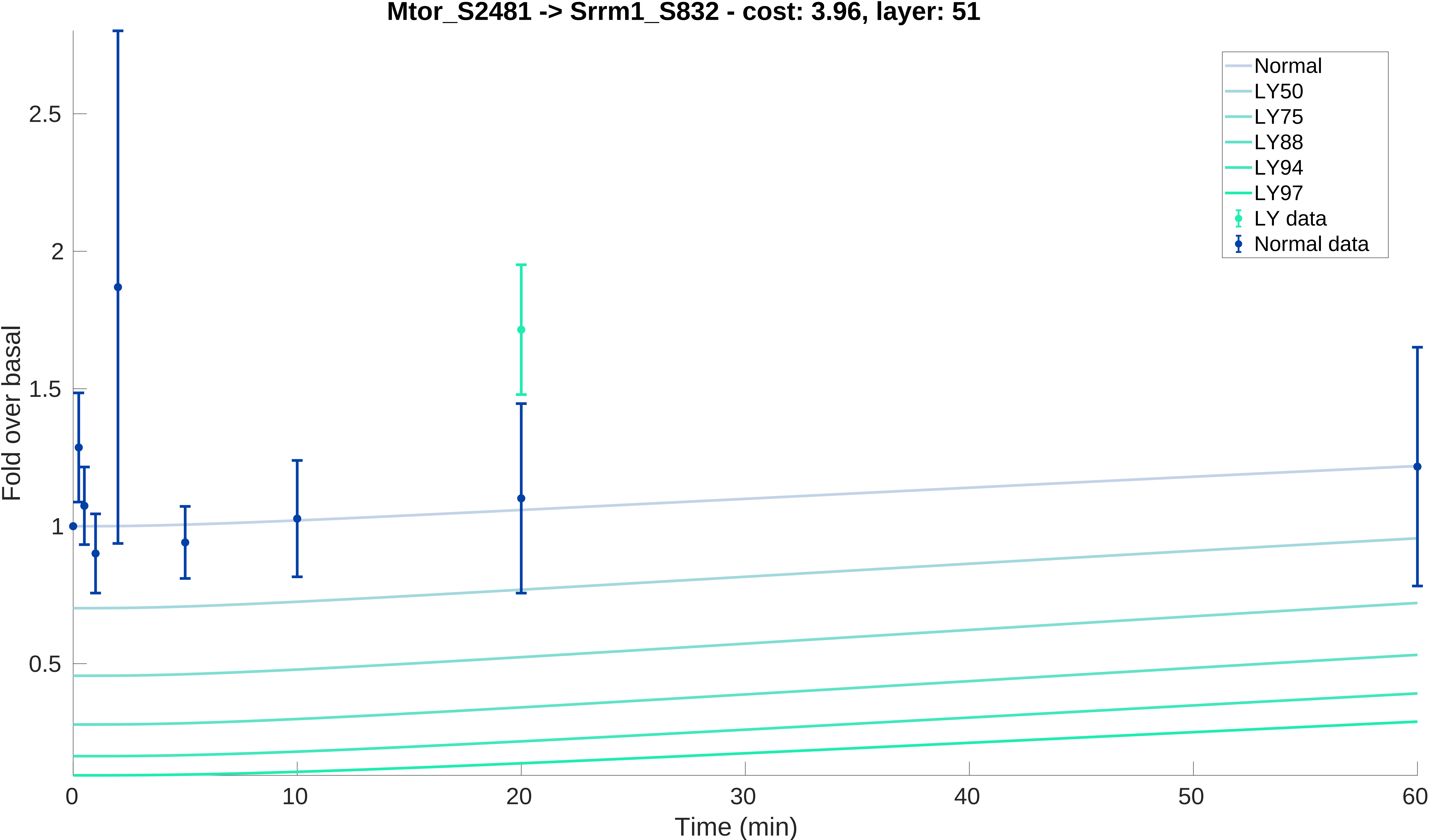

Mtor_S2481 -> Srrm1_S832 - cost: 3.96, layer: 51
Normal
LY50
LY75
LY88
LY94
LY97
LY data
Normal data
Fold over basal
2.5
2
1.5
1
0.5
0
10
20
30
40
50
60
Time (min)

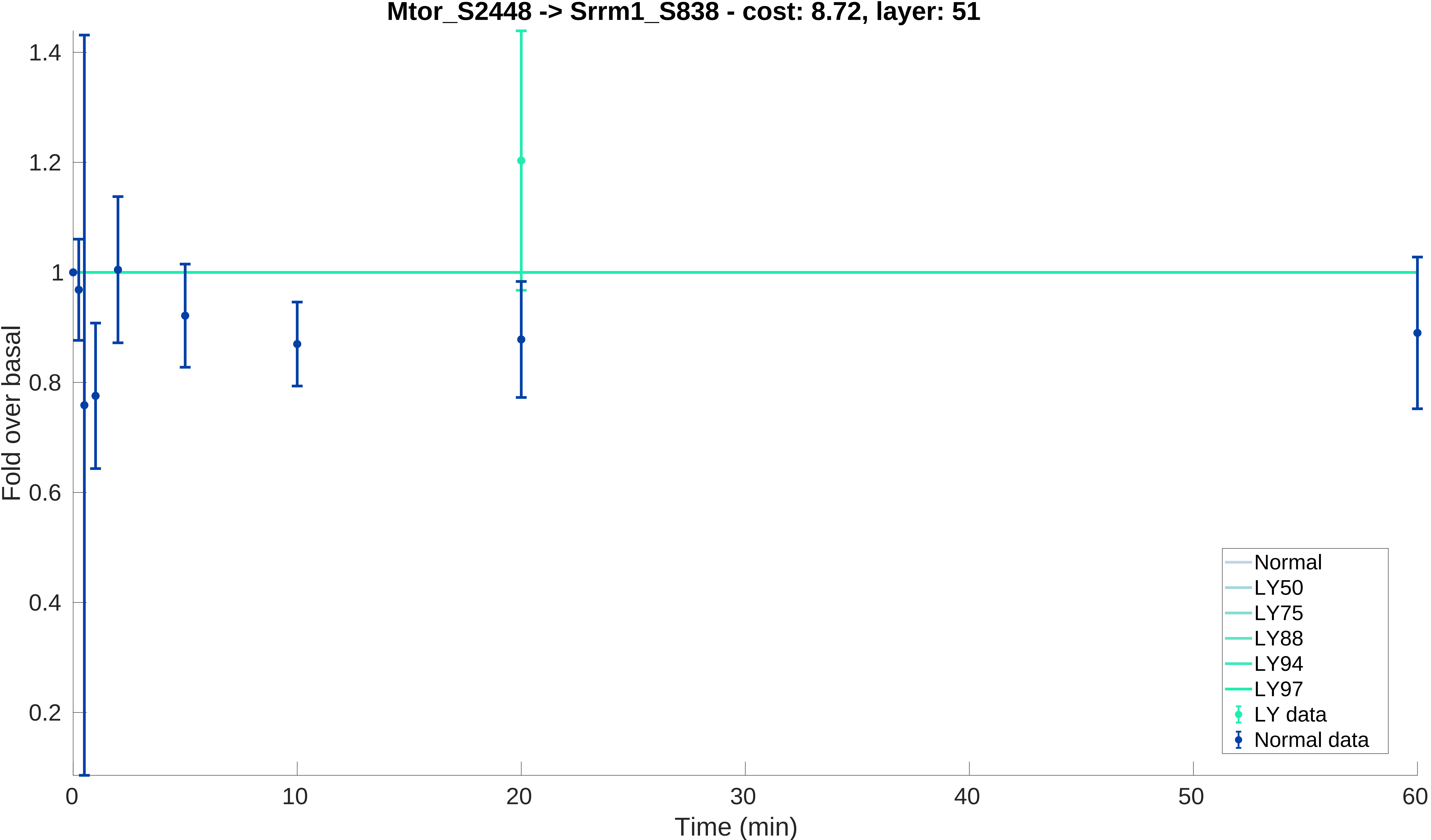

Mtor_S2448 -> Srrm1_S838 - cost: 8.72, layer: 51
Fold over basal
1.4
1.2
1
0.8
0.6
0.4
0.2
0
10
20
30
40
50
60
Time (min)
Normal
LY50
LY75
LY88
LY94
LY97
LY data
Normal data

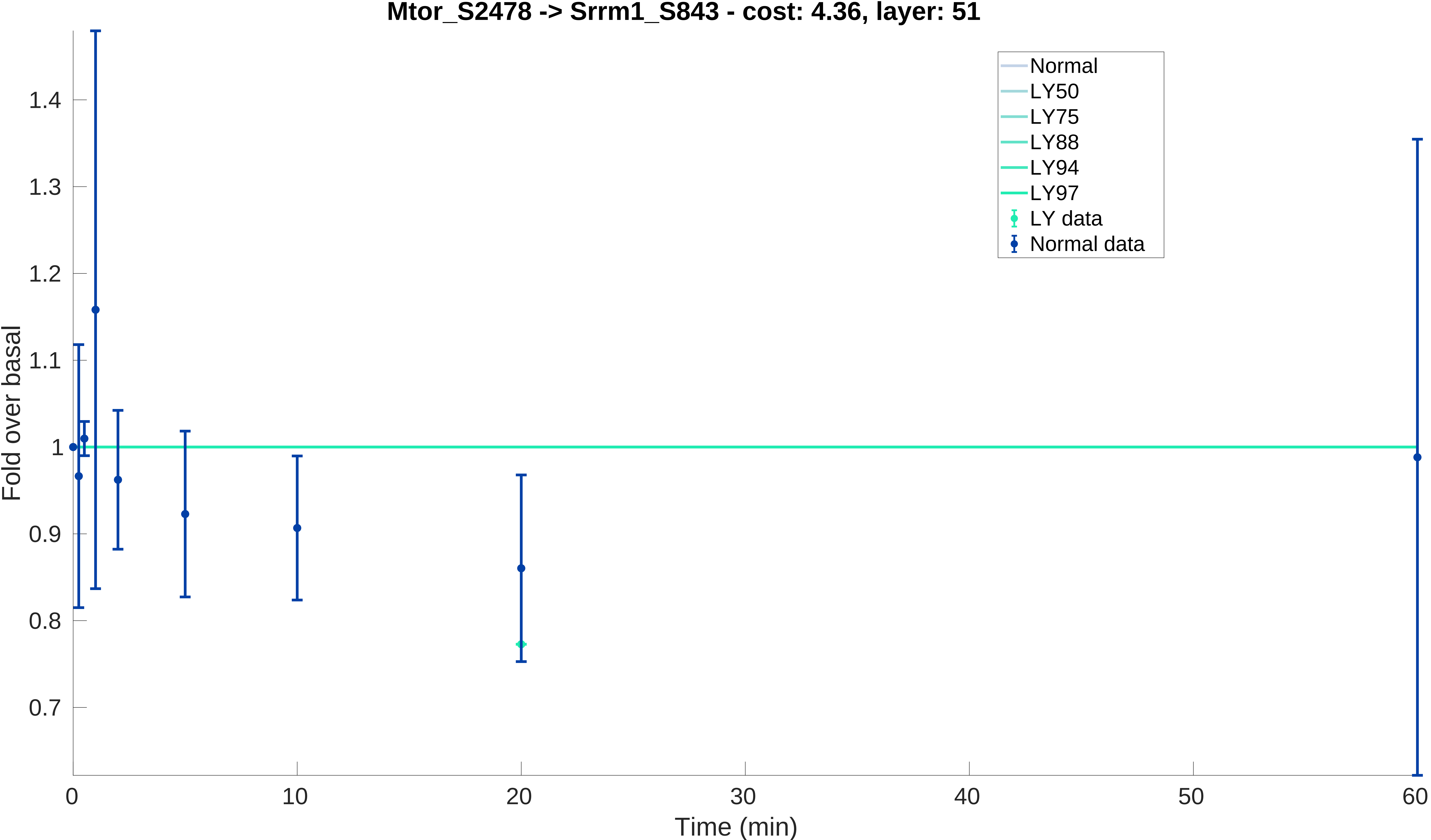

Mtor_S2478 -> Srrm1_S843 - cost: 4.36, layer: 51
Fold over basal
Time (min)
0
10
20
30
40
50
60
0.7
0.8
0.9
1
1.1
1.2
1.3
1.4
Normal
LY50
LY75
LY88
LY94
LY97
LY data
Normal data

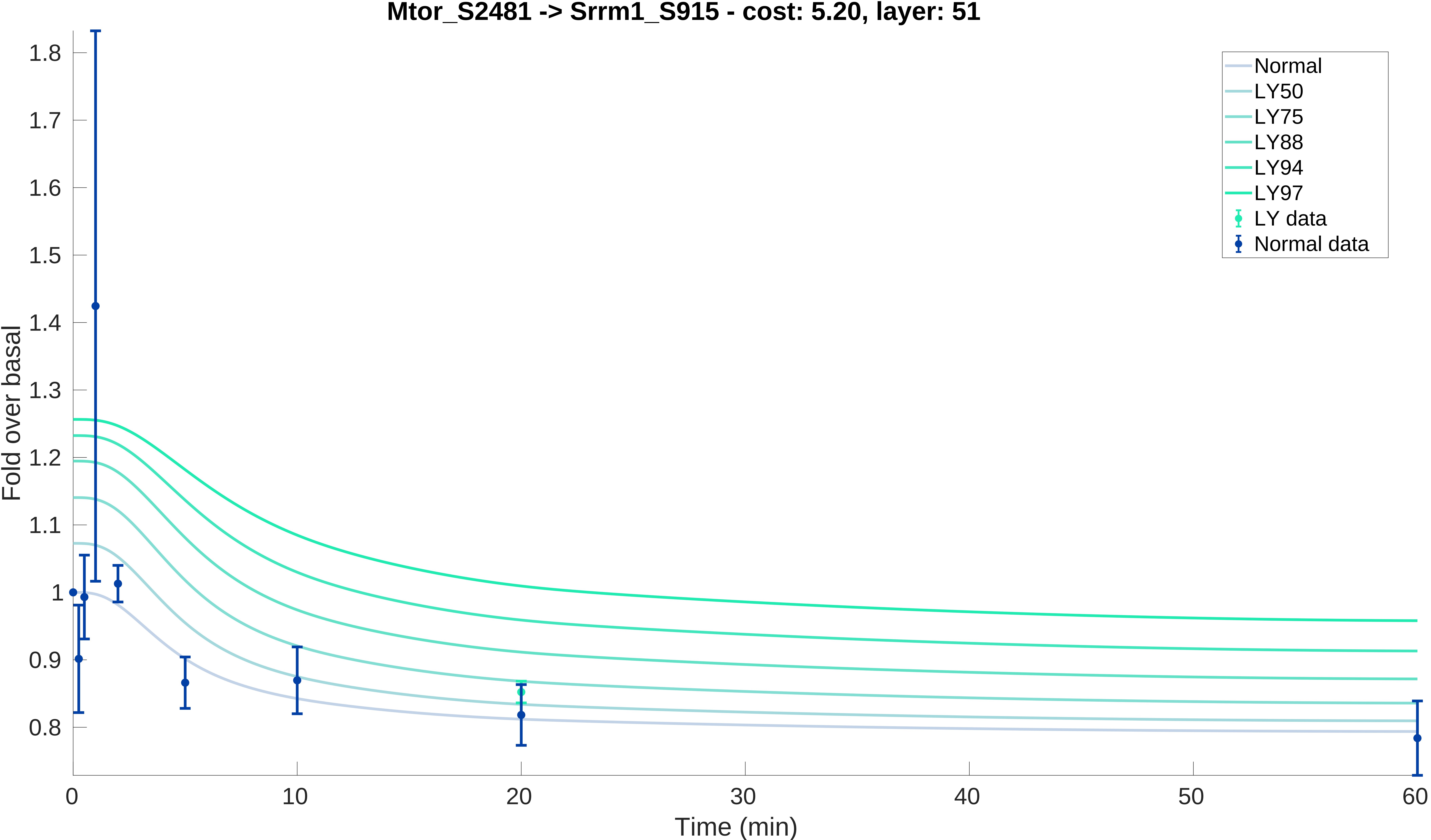

Mtor_S2481 -> Srrm1_S915 - cost: 5.20, layer: 51
Fold over basal
1.8
1.7
1.6
1.5
1.4
1.3
1.2
1.1
1
0.9
0.8
0
10
20
30
40
50
60
Time (min)
Normal
LY50
LY75
LY88
LY94
LY97
LY data
Normal data

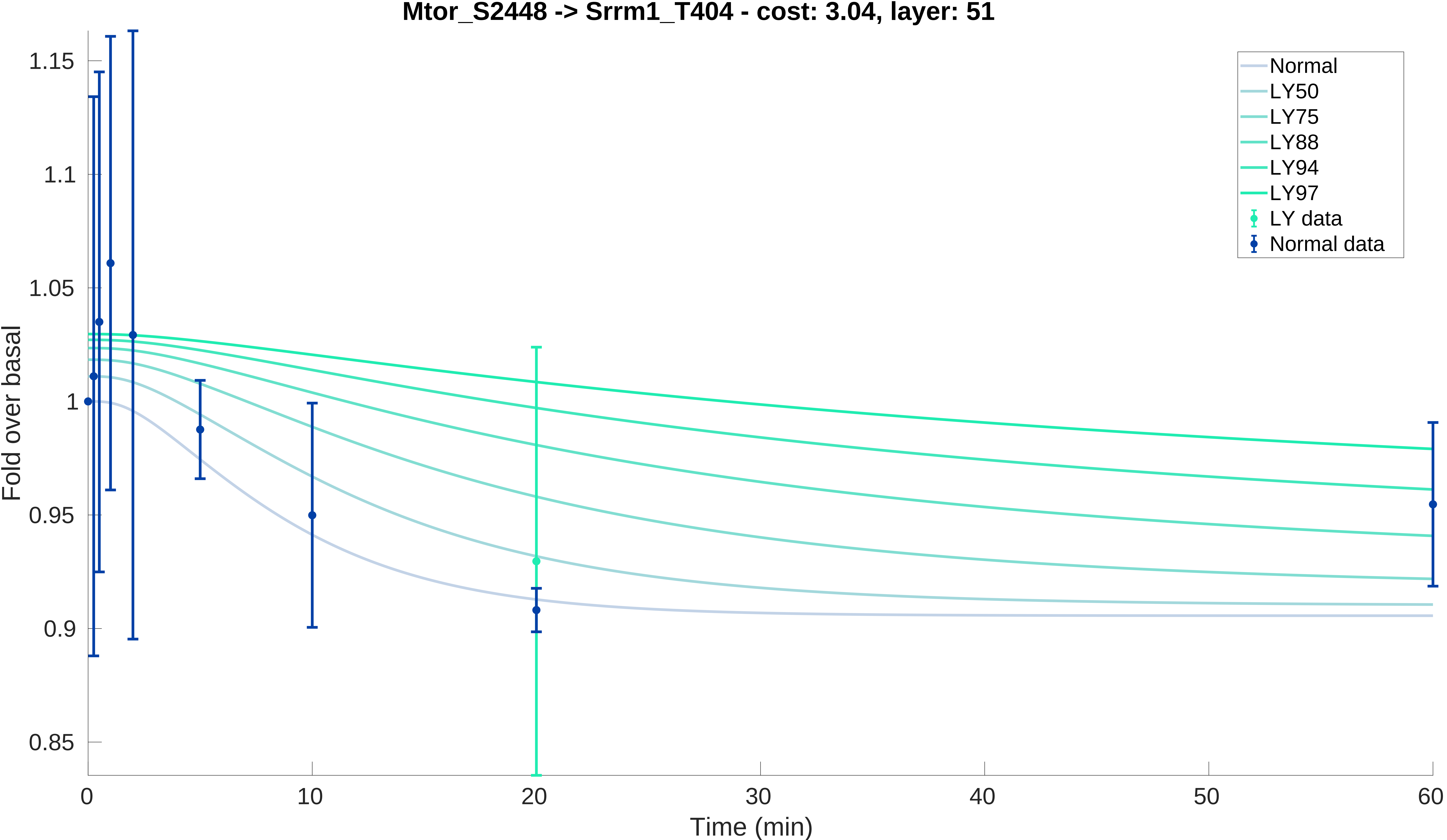

Mtor_S2448 -> Srrm1_T404 - cost: 3.04, layer: 51
Fold over basal
1.15
1.1
1.05
1
0.95
0.9
0.85
0
10
20
30
40
50
60
Time (min)
Normal
LY50
LY75
LY88
LY94
LY97
LY data
Normal data

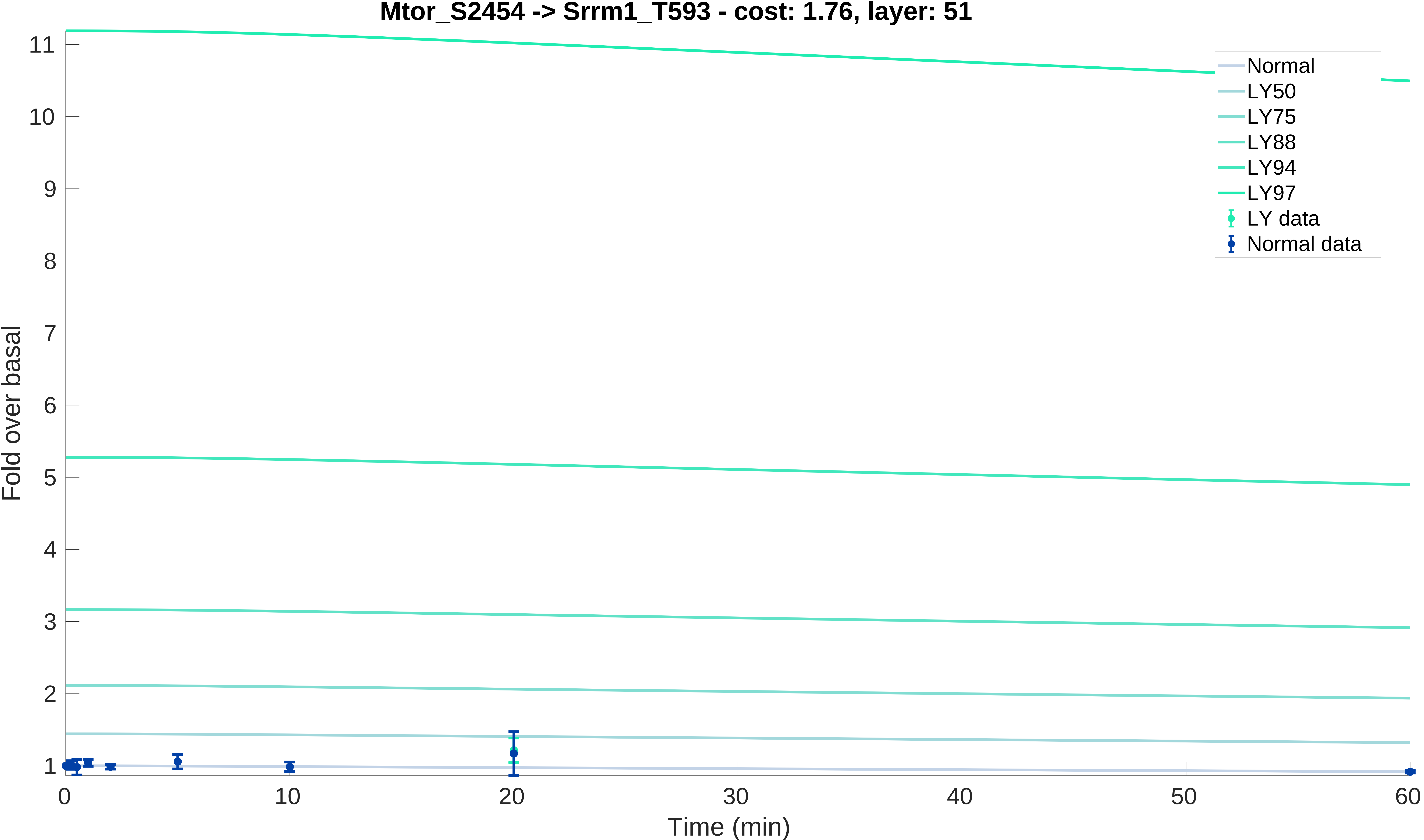

Mtor_S2454 -> Srrm1_T593 - cost: 1.76, layer: 51
Fold over basal
Time (min)
0
10
20
30
40
50
60
1
2
3
4
5
6
7
8
9
10
11
Normal
LY50
LY75
LY88
LY94
LY97
LY data
Normal data

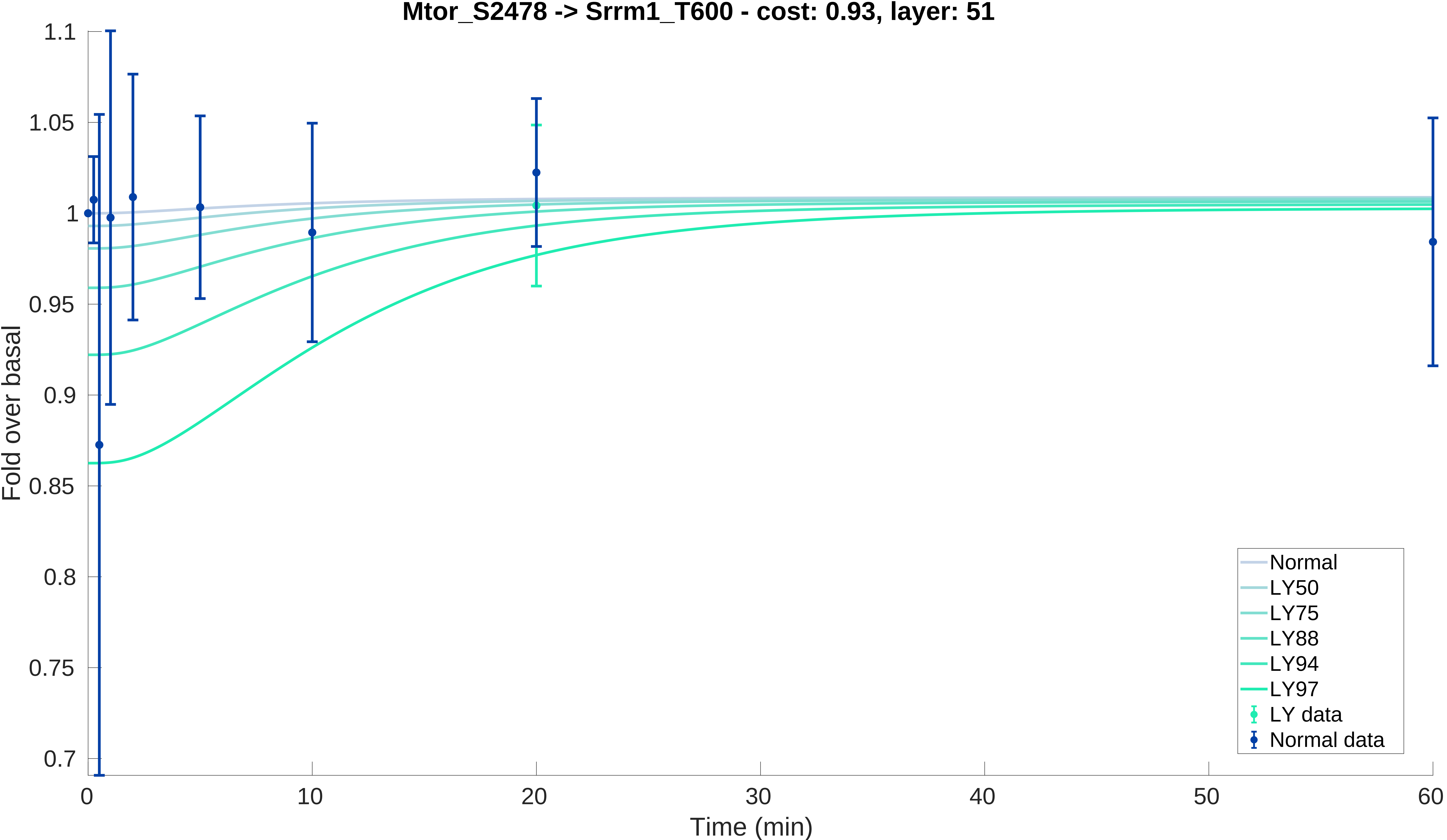

Mtor_S2478 -> Srrm1_T600 - cost: 0.93, layer: 51
Fold over basal
Time (min)
1.1
1.05
1
0.95
0.9
0.85
0.8
0.75
0.7
0
10
20
30
40
50
60
Normal
LY50
LY75
LY88
LY94
LY97
LY data
Normal data

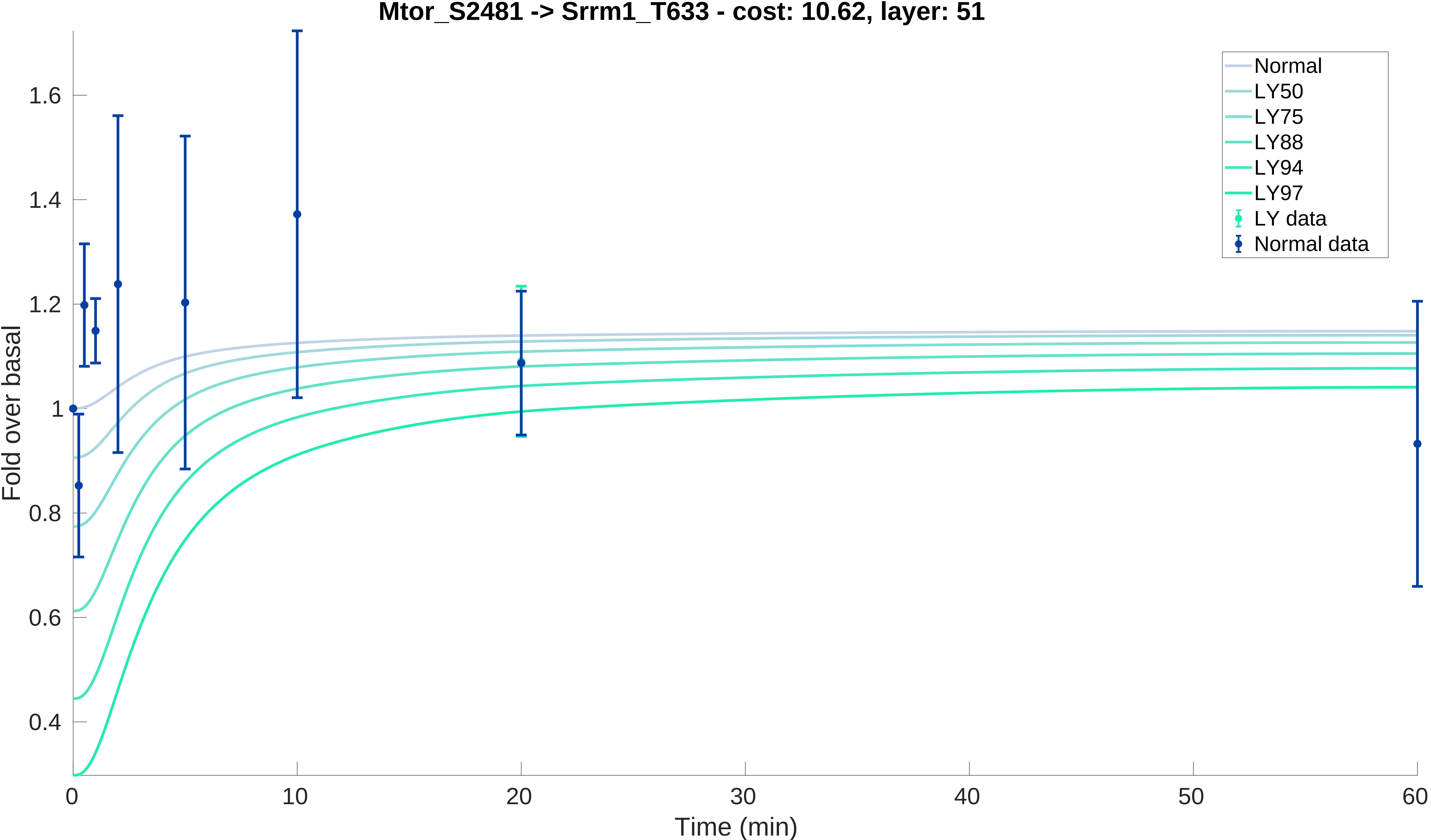

Mtor_S2481 -> Srrm1_T633 - cost: 10.62, layer: 51
Normal
LY50
LY75
LY88
LY94
LY97
LY data
Normal data
Fold over basal
1.6
1.4
1.2
1
0.8
0.6
0.4
0
10
20
30
40
50
60
Time (min)

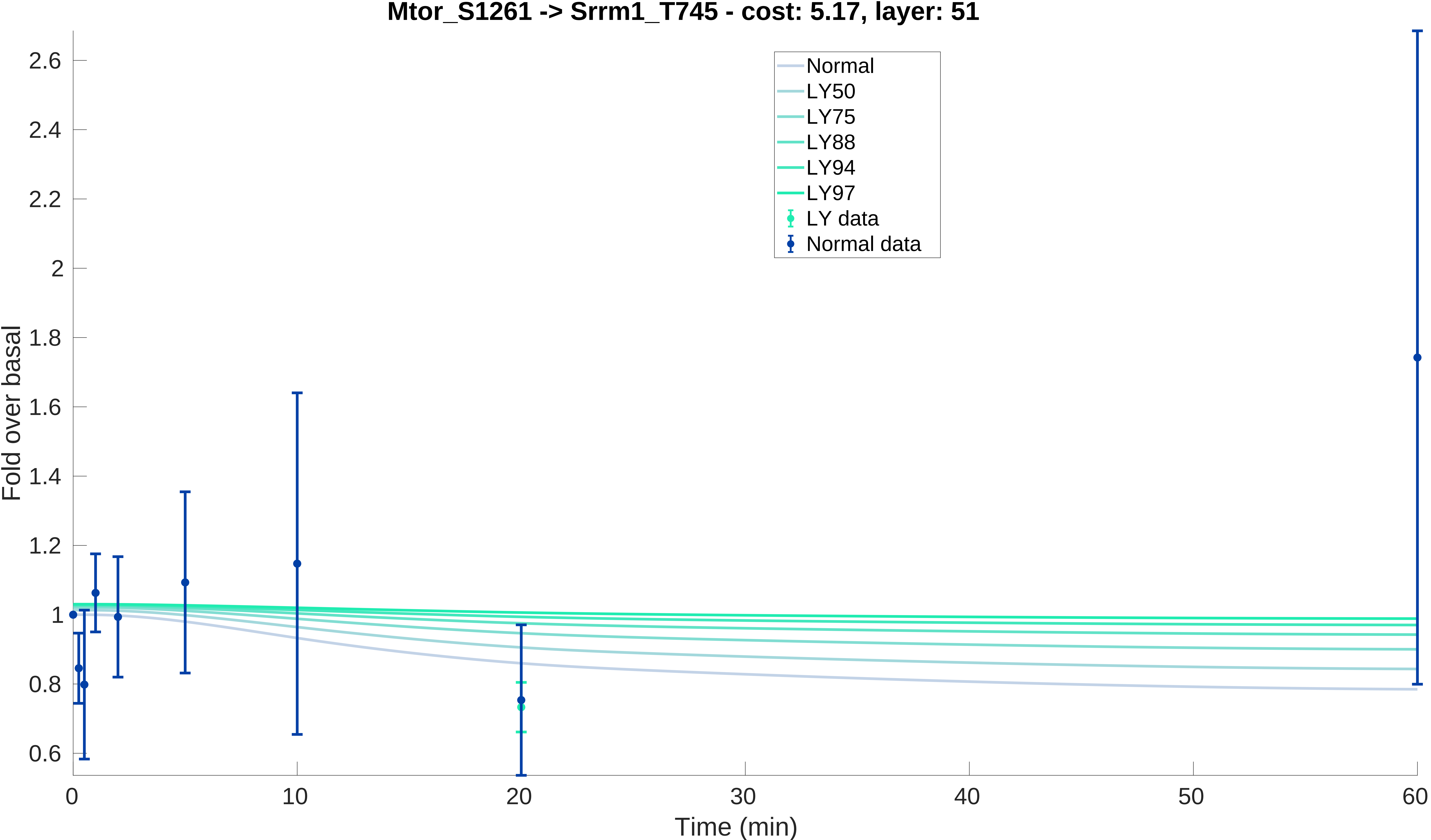
Mtor_S1261 -> Srrm1_T745 - cost: 5.17, layer: 51
Fold over basal
Time (min)
Normal
LY50
LY75
LY88
LY94
LY97
LY data
Normal data
0.6
0.8
1
1.2
1.4
1.6
1.8
2
2.2
2.4
2.6
0
10
20
30
40
50
60

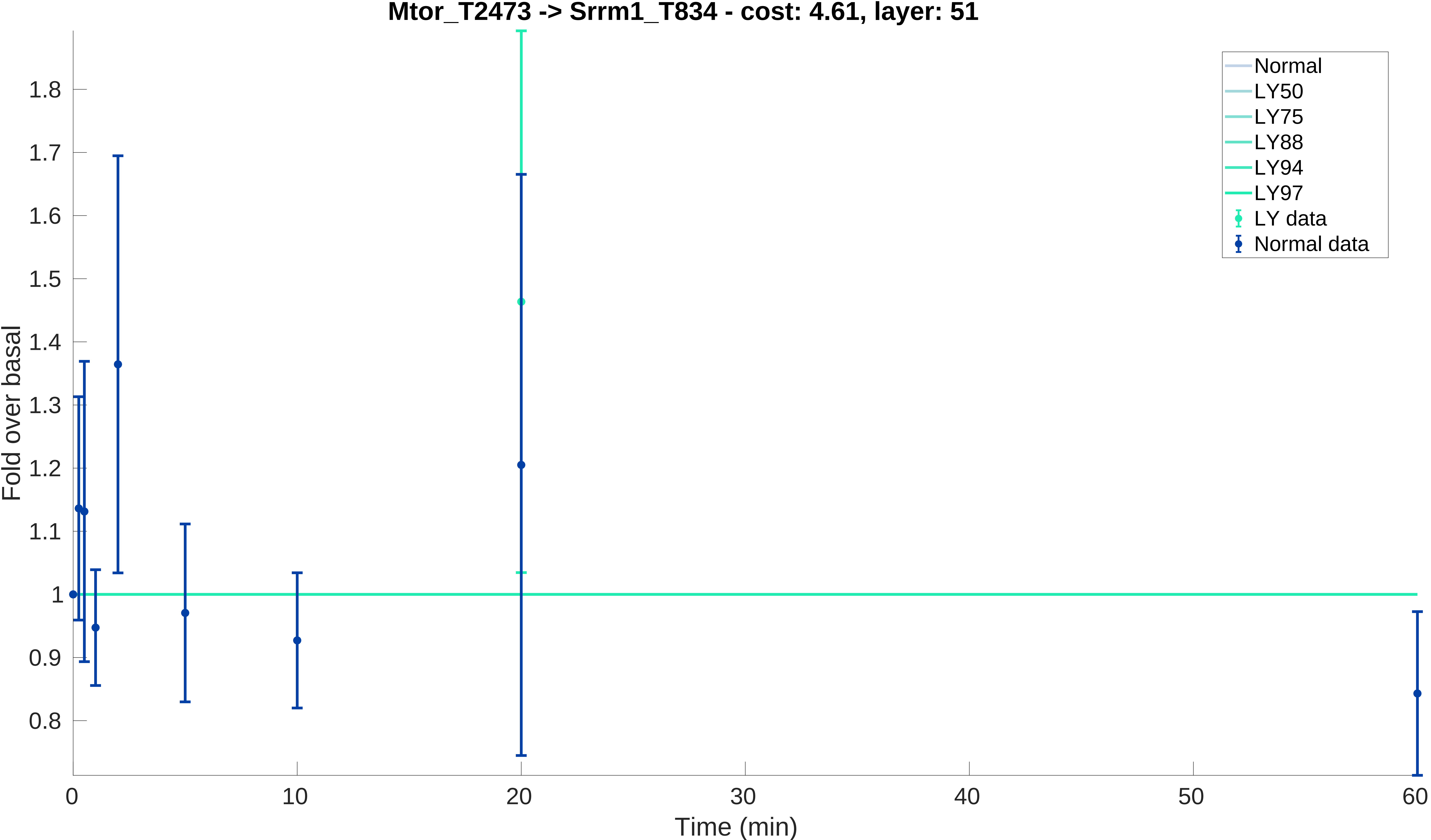

Mtor_T2473 -> Srrm1_T834 - cost: 4.61, layer: 51
Fold over basal
Time (min)
Normal
LY50
LY75
LY88
LY94
LY97
LY data
Normal data
1.8
1.7
1.6
1.5
1.4
1.3
1.2
1.1
1
0.9
0.8
0
10
20
30
40
50
60

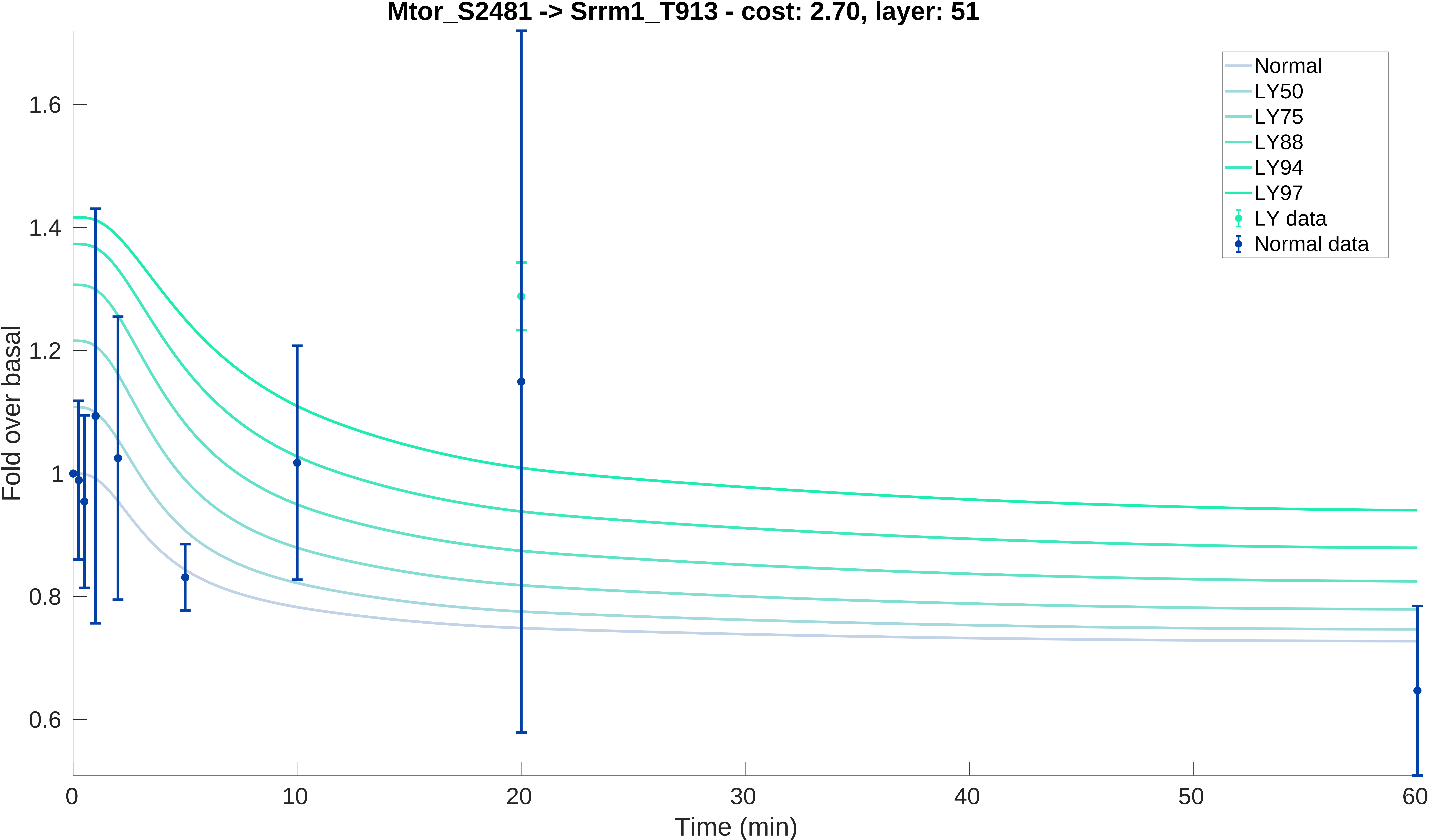

Mtor_S2481 -> Srrm1_T913 - cost: 2.70, layer: 51
Normal
LY50
LY75
LY88
LY94
LY97
LY data
Normal data
Fold over basal
1.6
1.4
1.2
1
0.8
0.6
0
10
20
30
40
50
60
Time (min)

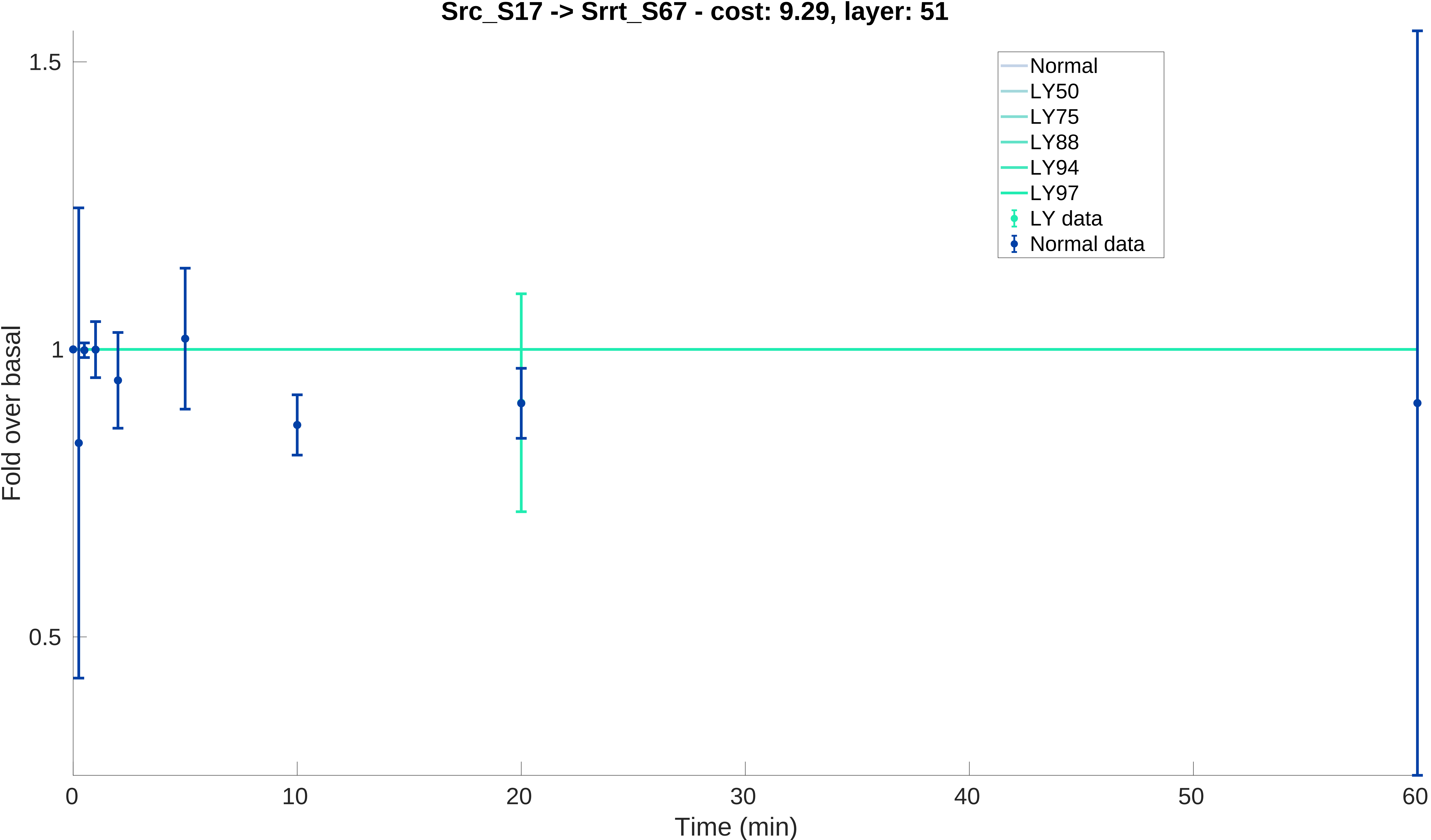
Src_S17 -> Srrt_S67 - cost: 9.29, layer: 51
Normal
LY50
LY75
LY88
LY94
LY97
LY data
Normal data
Fold over basal
1.5
1
0.5
0
10
20
30
40
50
60
Time (min)

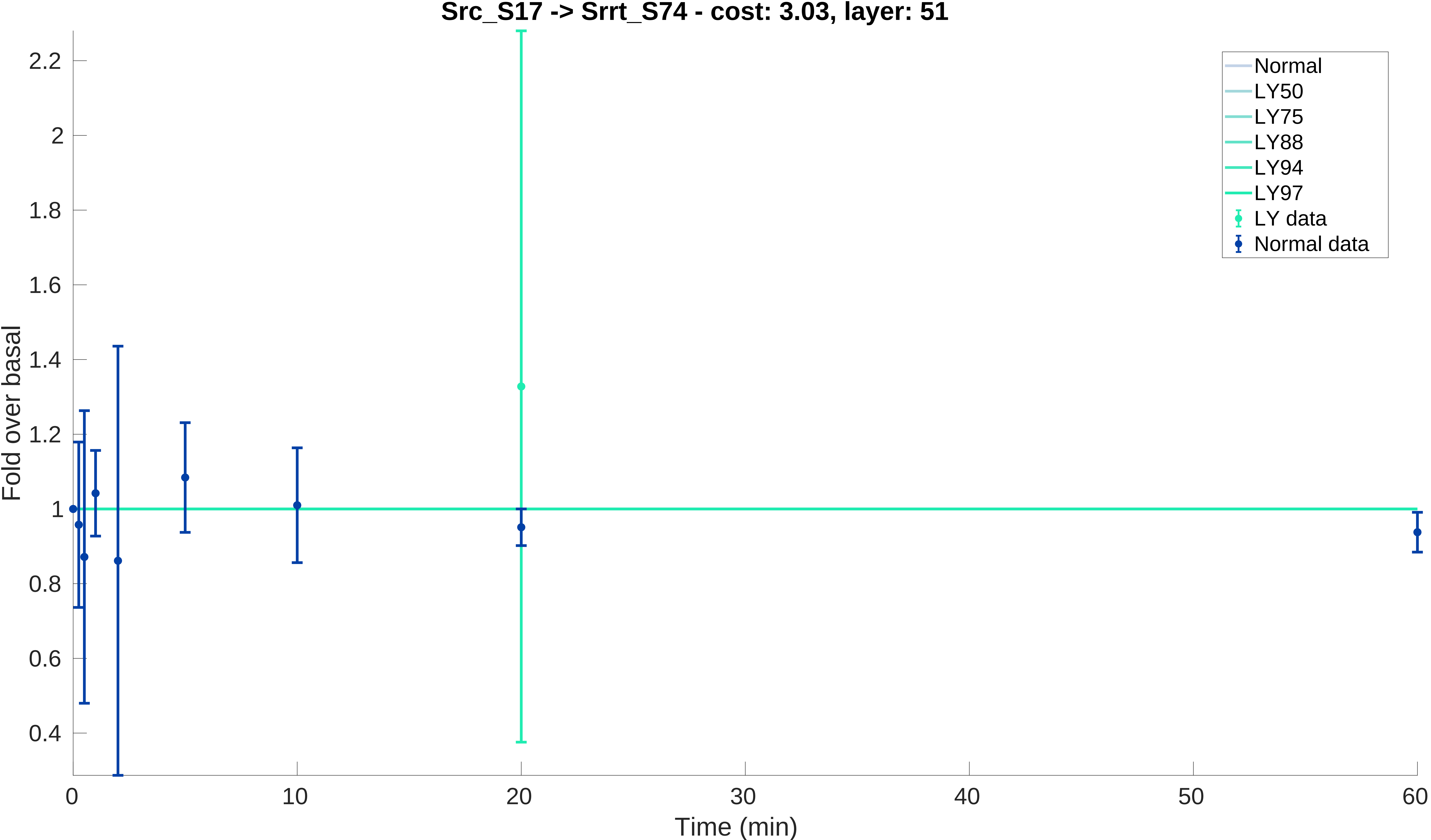

Src_S17 -> Srrt_S74 - cost: 3.03, layer: 51
Normal
LY50
LY75
LY88
LY94
LY97
LY data
Normal data
Fold over basal
Time (min)
0
10
20
30
40
50
60
0.4
0.6
0.8
1
1.2
1.4
1.6
1.8
2
2.2

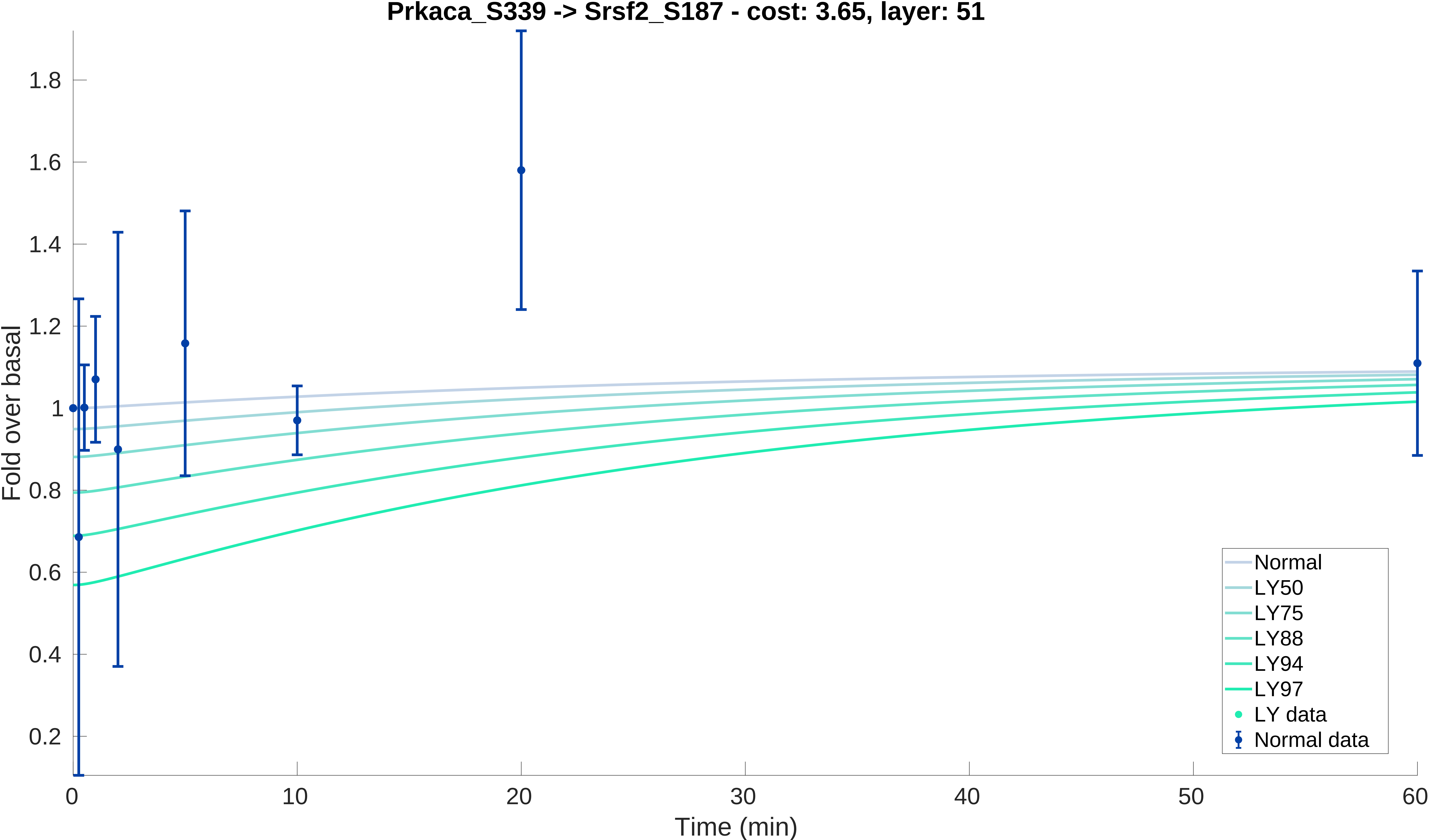

Prkaca_S339 -> Srsf2_S187 - cost: 3.65, layer: 51
Fold over basal
1.8
1.6
1.4
1.2
1
0.8
0.6
0.4
0.2
0
10
20
30
40
50
60
Time (min)
Normal
LY50
LY75
LY88
LY94
LY97
LY data
Normal data

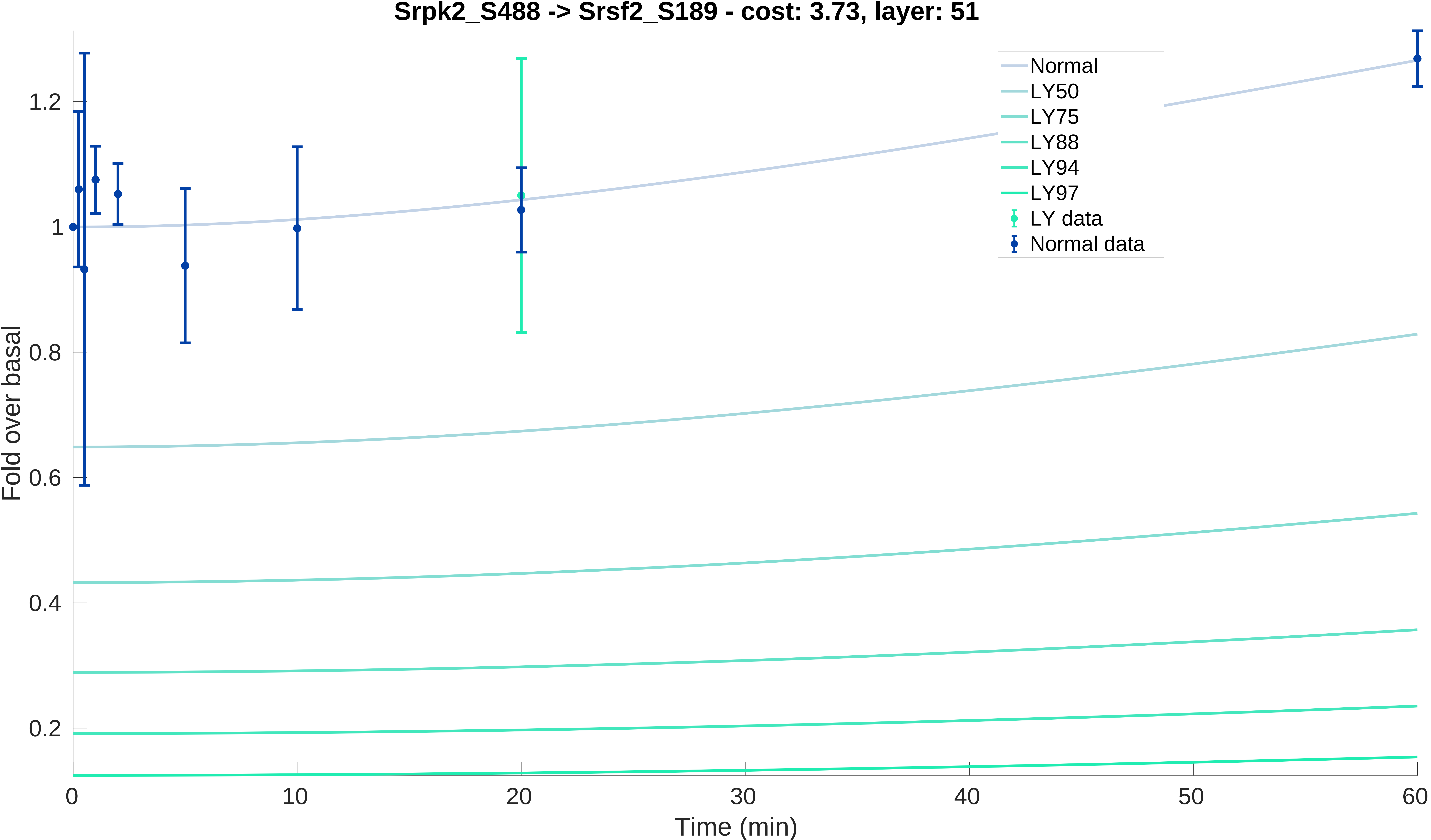

Srpk2_S488 -> Srsf2_S189 - cost: 3.73, layer: 51
Fold over basal
Time (min)
0
10
20
30
40
50
60
0.2
0.4
0.6
0.8
1
1.2
Normal
LY50
LY75
LY88
LY94
LY97
LY data
Normal data

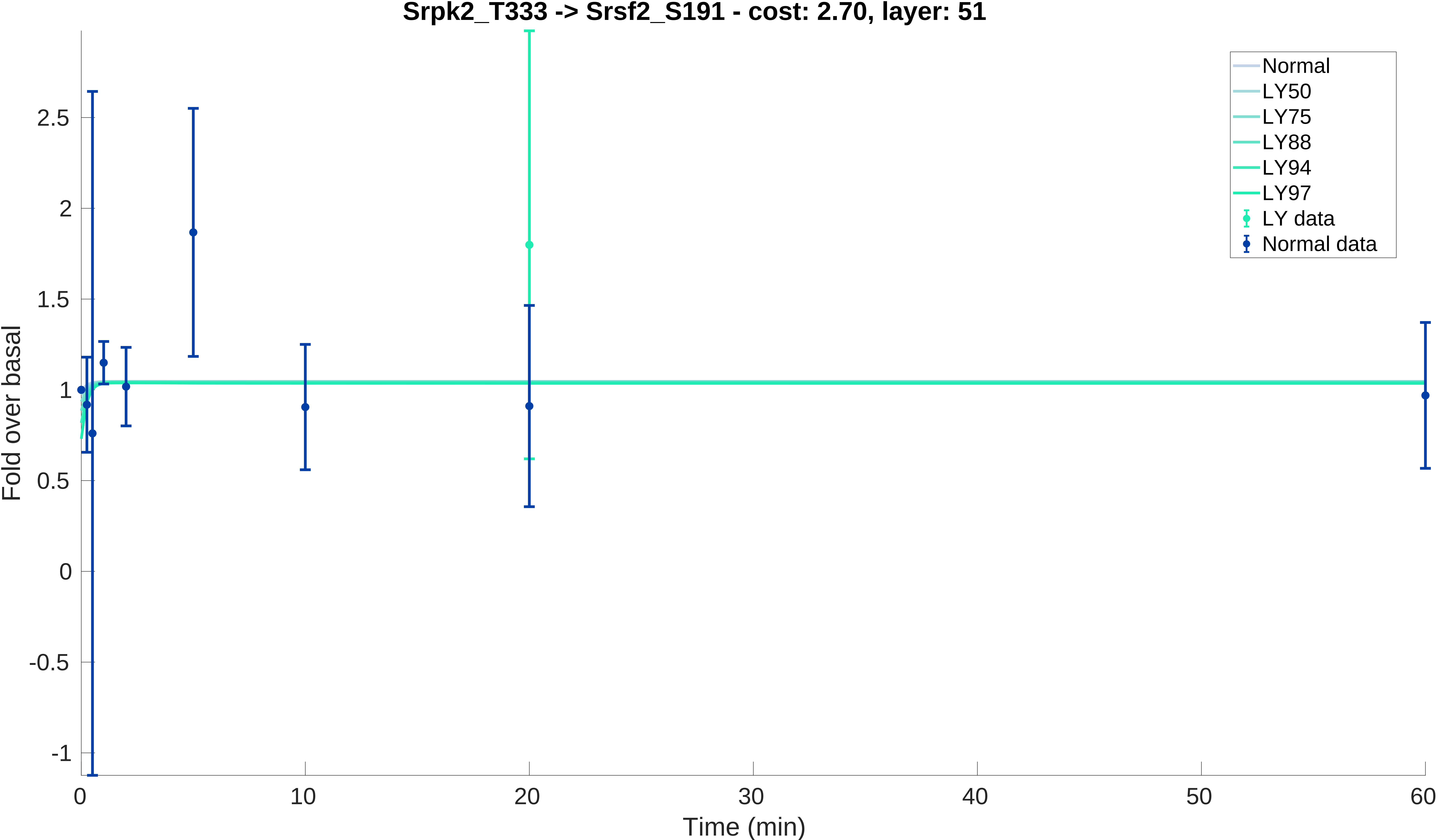

Srpk2_T333 -> Srsf2_S191 - cost: 2.70, layer: 51
Fold over basal
Time (min)
2.5
2
1.5
1
0.5
0
-0.5
-1
0
10
20
30
40
50
60
Normal
LY50
LY75
LY88
LY94
LY97
LY data
Normal data

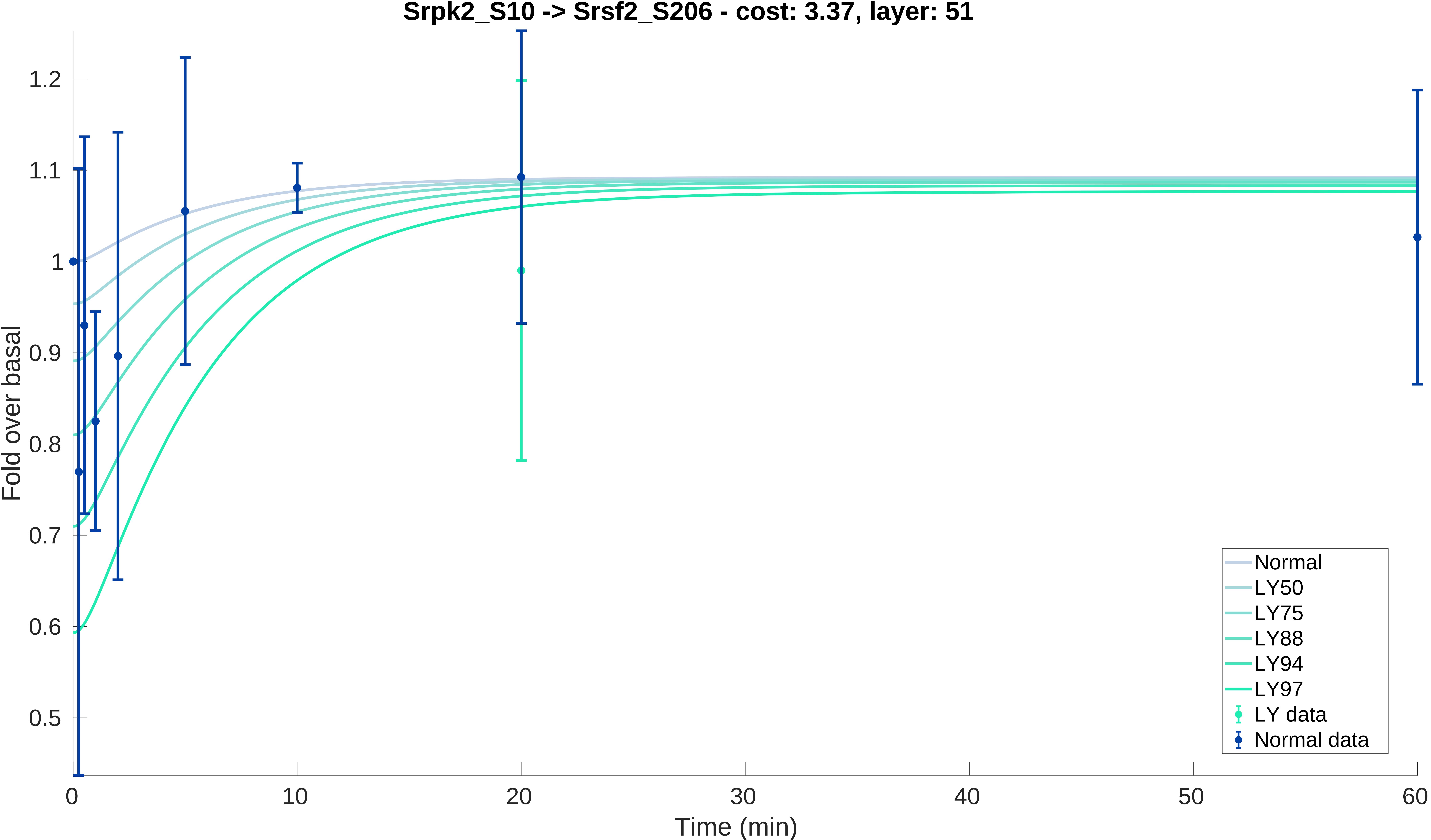

Srpk2_S10 -> Srsf2_S206 - cost: 3.37, layer: 51
Fold over basal
Time (min)
0.5
0.6
0.7
0.8
0.9
1
1.1
1.2
0
10
20
30
40
50
60
Normal
LY50
LY75
LY88
LY94
LY97
LY data
Normal data

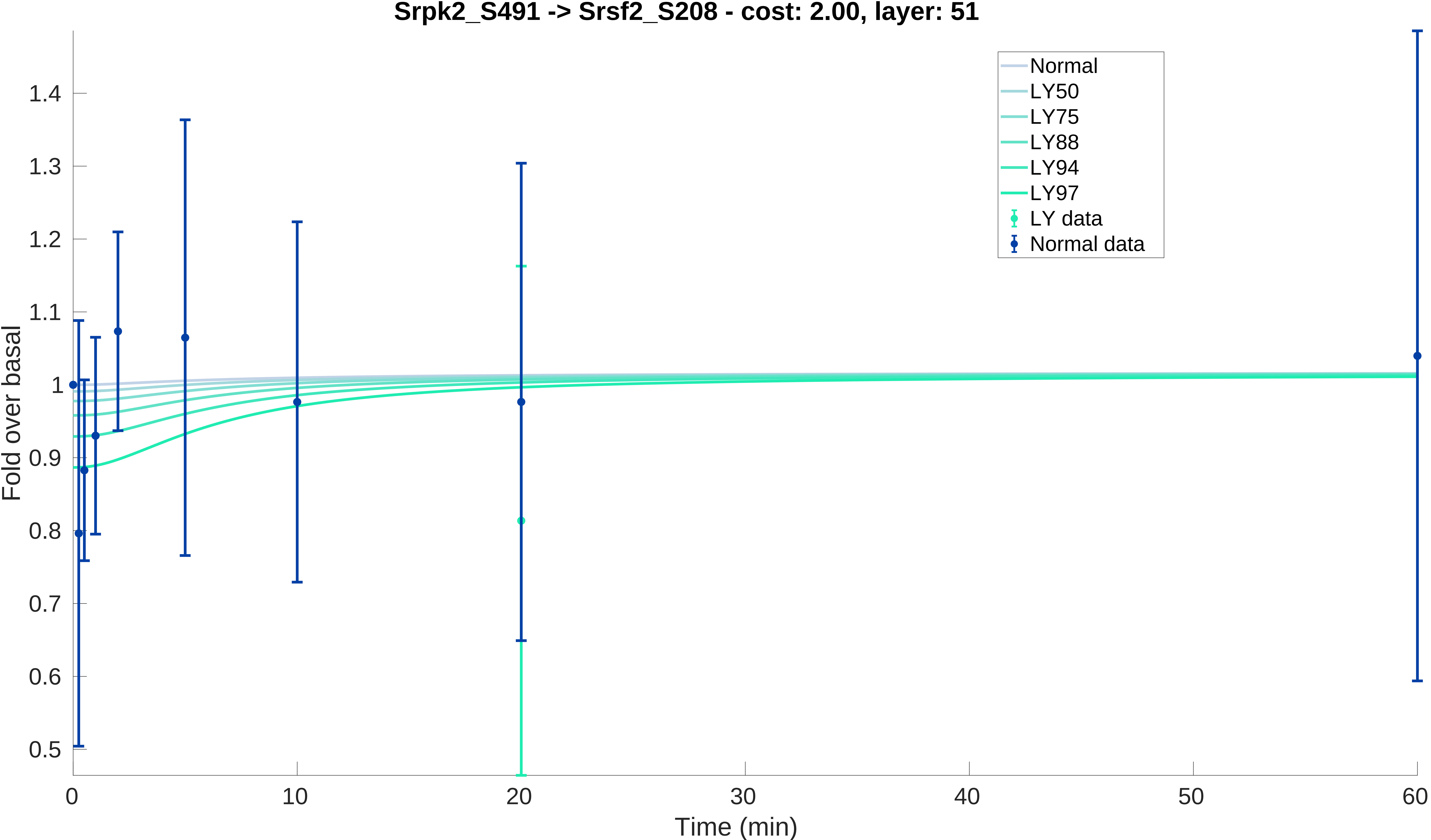
Srpk2_S491 -> Srsf2_S208 - cost: 2.00, layer: 51
Normal
LY50
LY75
LY88
LY94
LY97
LY data
Normal data
Fold over basal
Time (min)
1.4
1.3
1.2
1.1
1
0.9
0.8
0.7
0.6
0.5
0
10
20
30
40
50
60

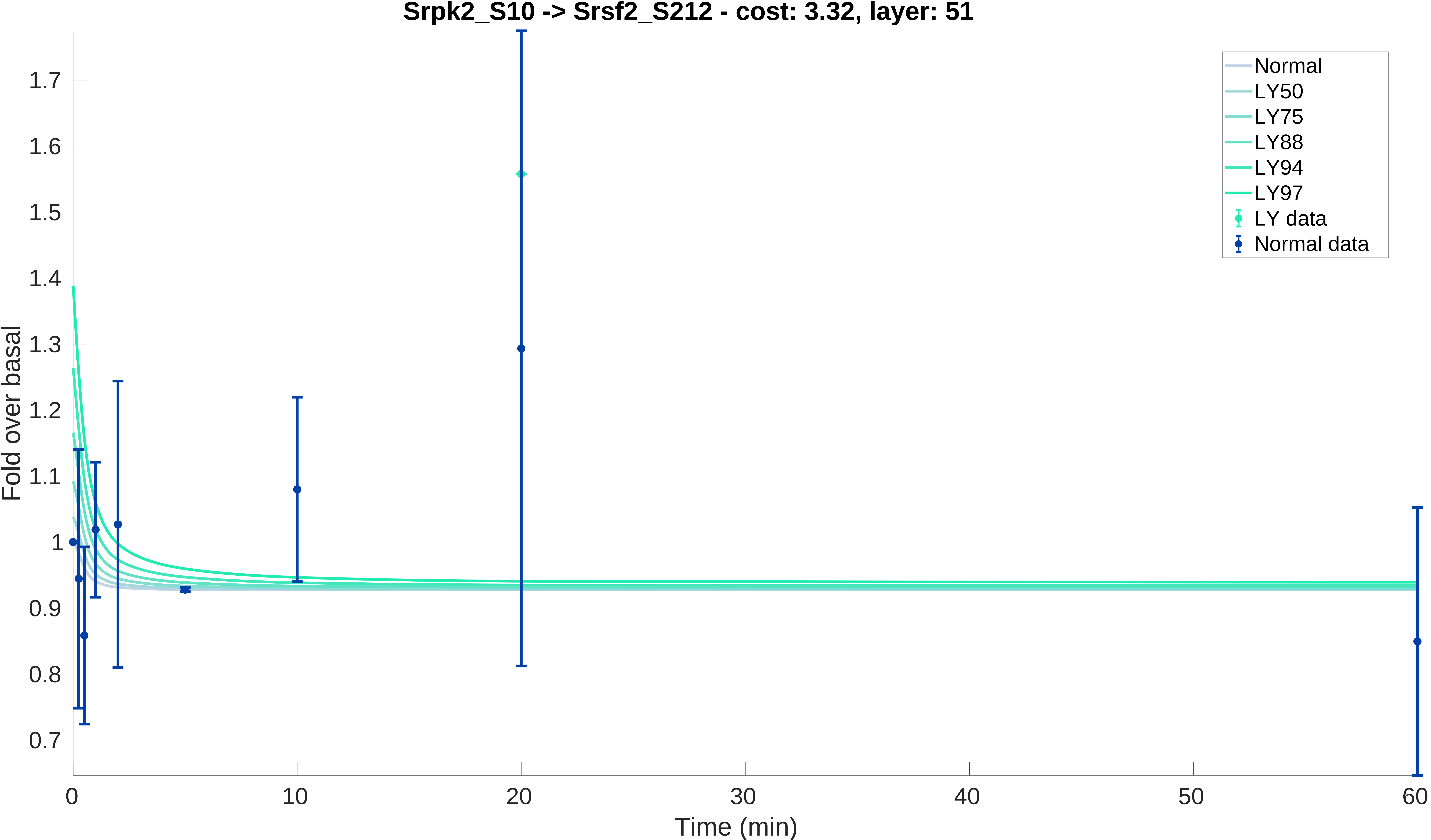

Srpk2_S10 -> Srsf2_S212 - cost: 3.32, layer: 51
Fold over basal
1.7
1.6
1.5
1.4
1.3
1.2
1.1
1
0.9
0.8
0.7
0
10
20
30
40
50
60
Time (min)
Normal
LY50
LY75
LY88
LY94
LY97
LY data
Normal data

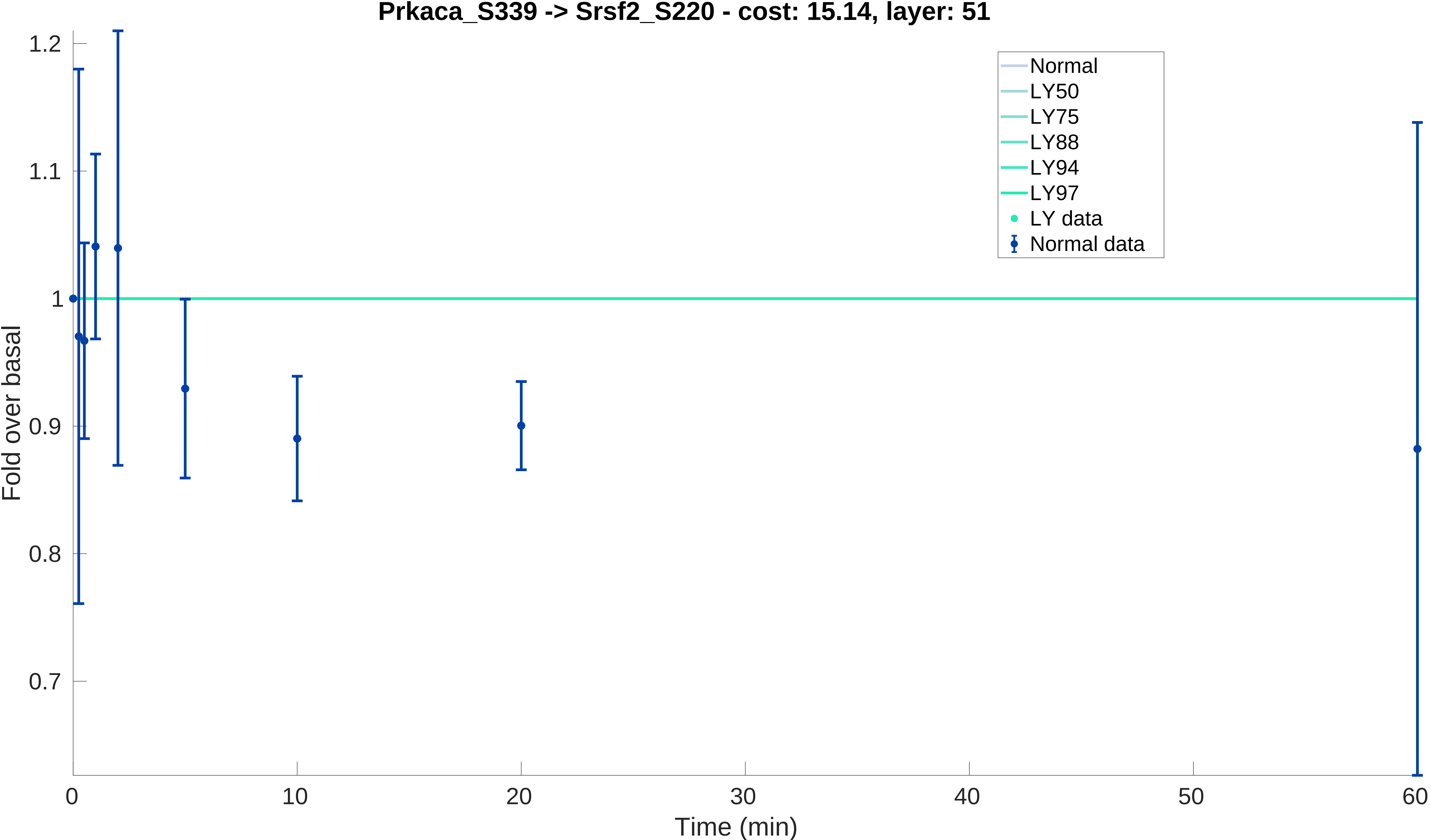

Prkaca_S339 -> Srsf2_S220 - cost: 15.14, layer: 51
Fold over basal
Time (min)
1.2
1.1
1
0.9
0.8
0.7
0
10
20
30
40
50
60
Normal
LY50
LY75
LY88
LY94
LY97
LY data
Normal data

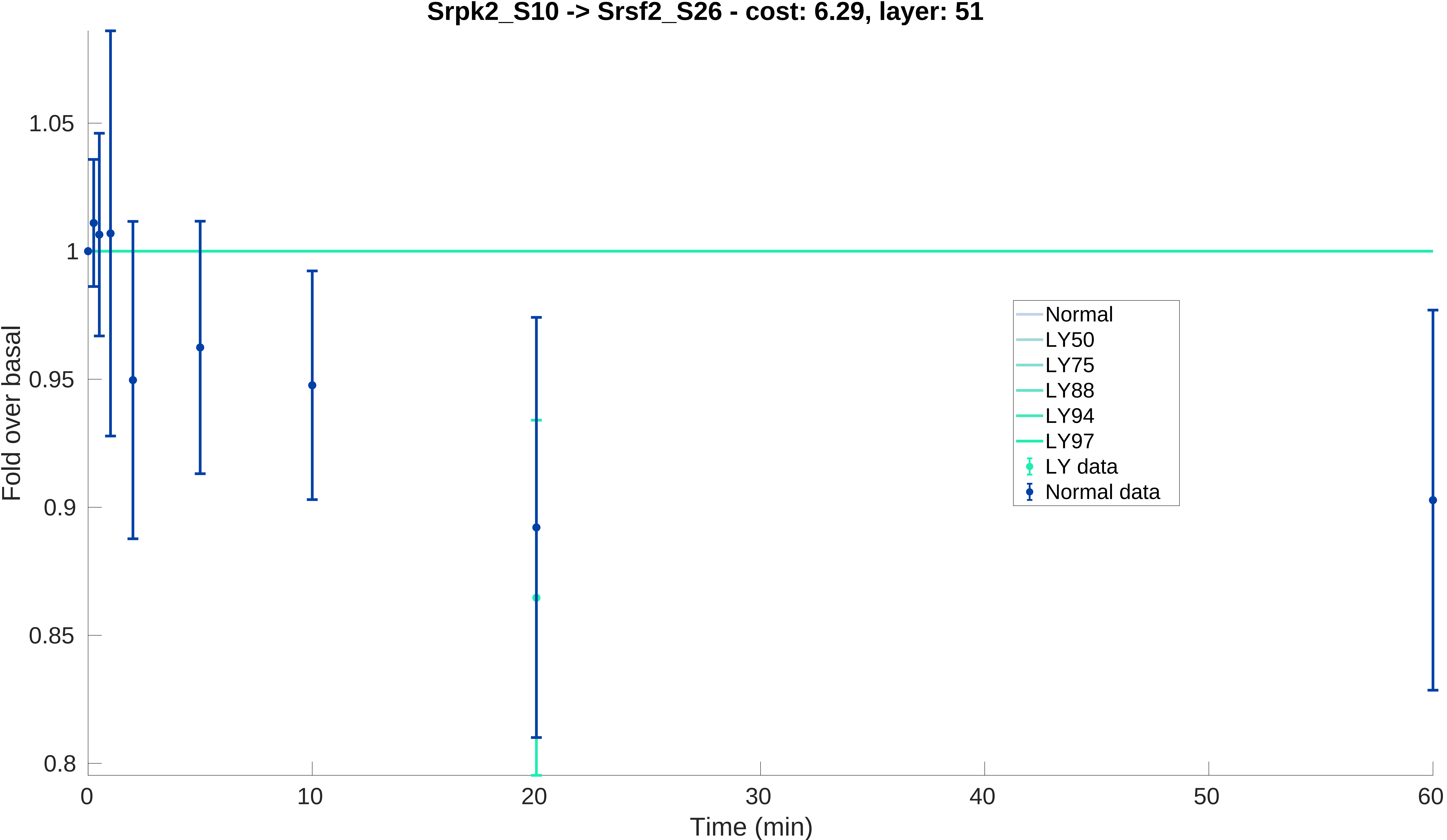

Srpk2_S10 -> Srsf2_S26 - cost: 6.29, layer: 51
Fold over basal
Time (min)
1.05
1
0.95
0.9
0.85
0.8
0
10
20
30
40
50
60
Normal
LY50
LY75
LY88
LY94
LY97
LY data
Normal data

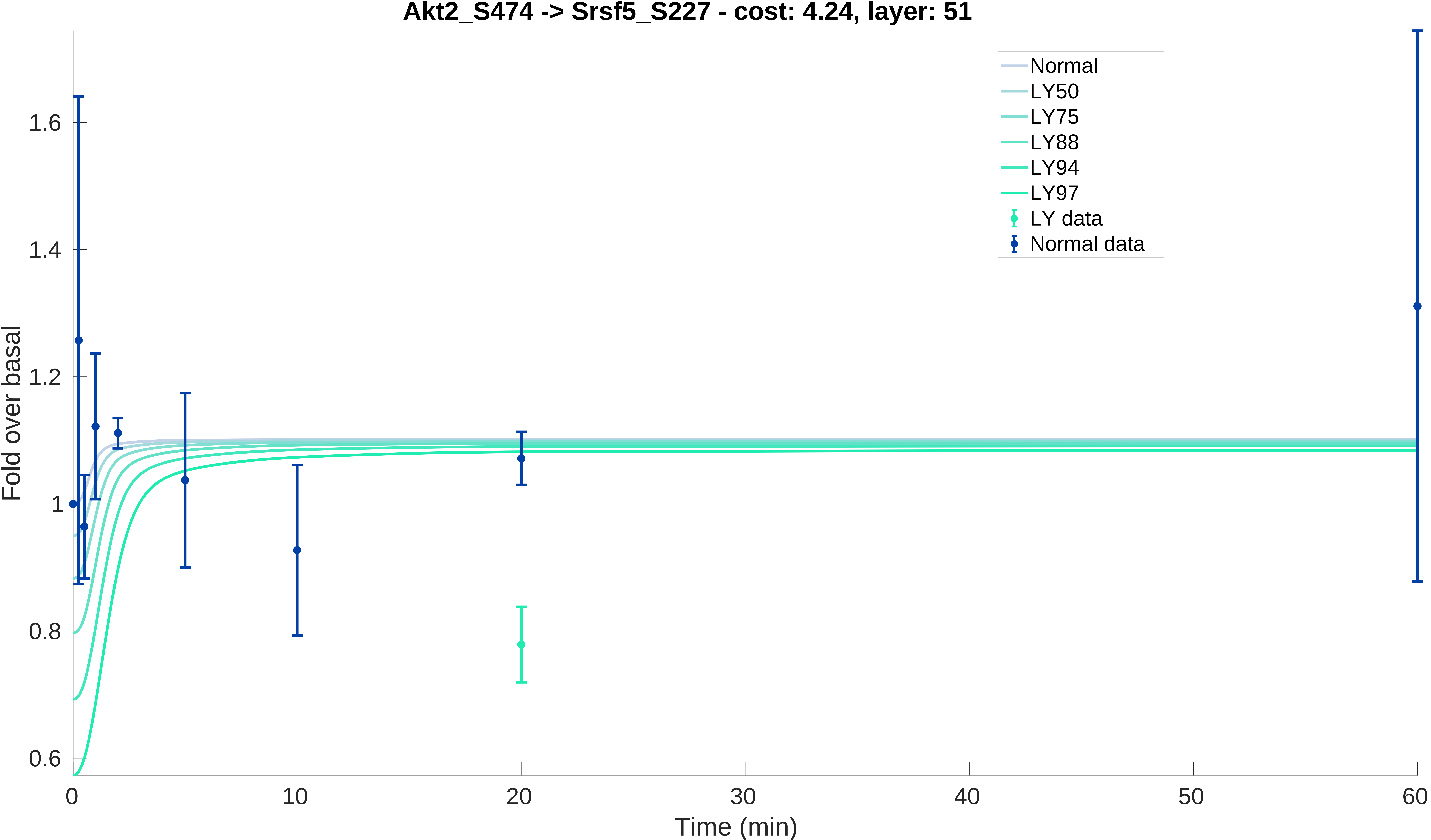

Akt2_S474 -> Srsf5_S227 - cost: 4.24, layer: 51
Normal
LY50
LY75
LY88
LY94
LY97
LY data
Normal data
Fold over basal
0.6
0.8
1
1.2
1.4
1.6
0
10
20
30
40
50
60
Time (min)

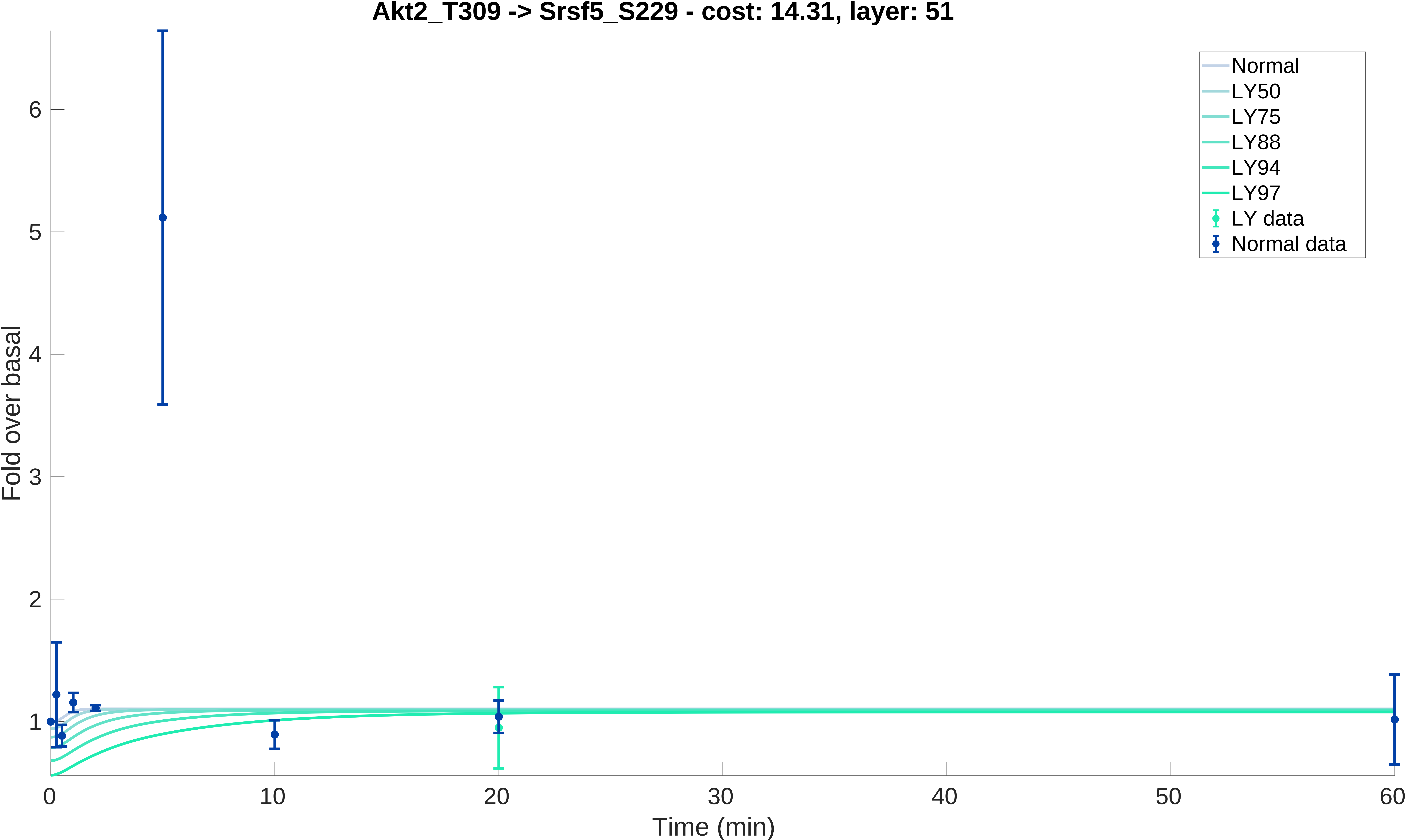

Akt2_T309 -> Srsf5_S229 - cost: 14.31, layer: 51
Fold over basal
Time (min)
Normal
LY50
LY75
LY88
LY94
LY97
LY data
Normal data
1
2
3
4
5
6
0
10
20
30
40
50
60

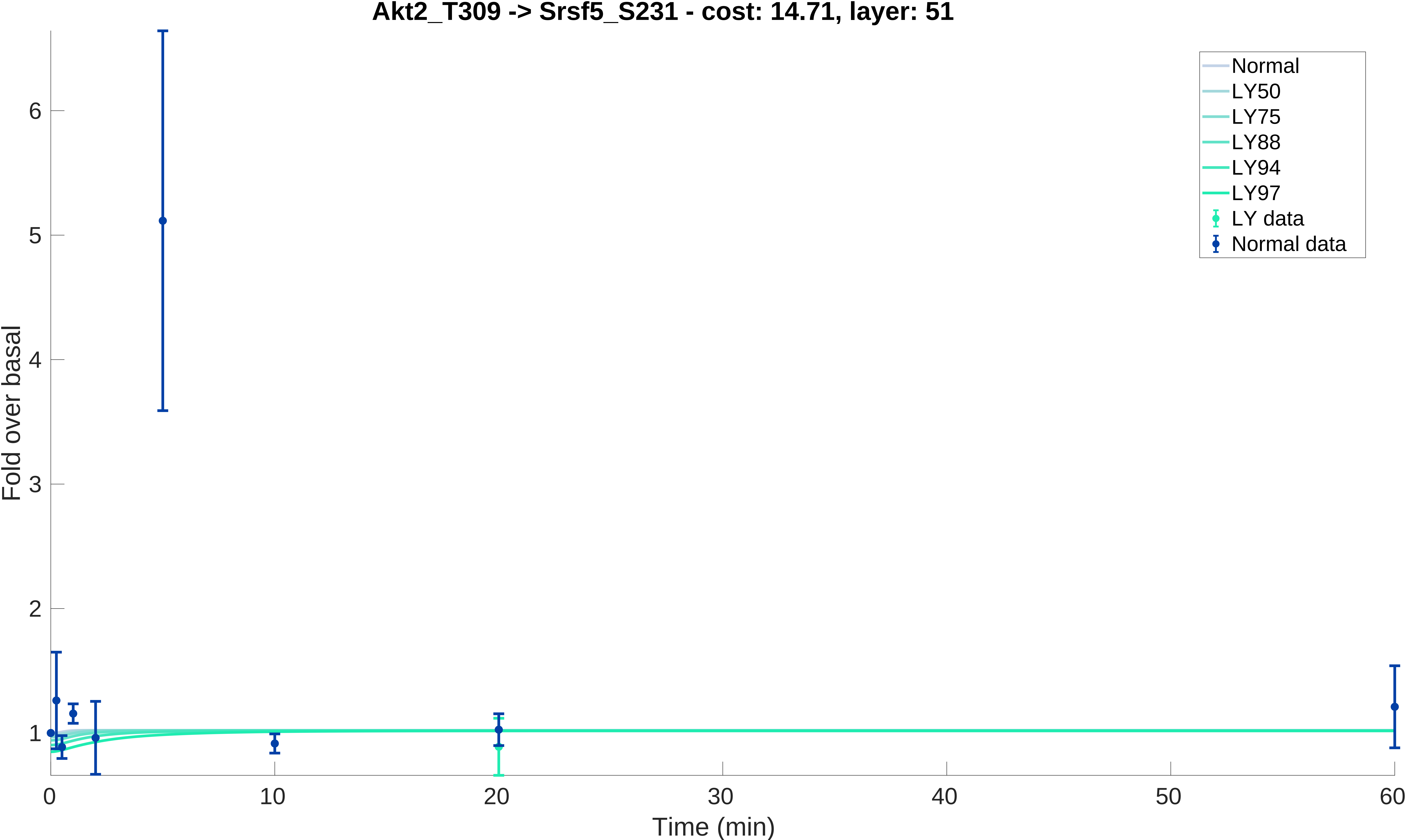

Akt2_T309 -> Srsf5_S231 - cost: 14.71, layer: 51
Fold over basal
Time (min)
0
10
20
30
40
50
60
1
2
3
4
5
6
Normal
LY50
LY75
LY88
LY94
LY97
LY data
Normal data

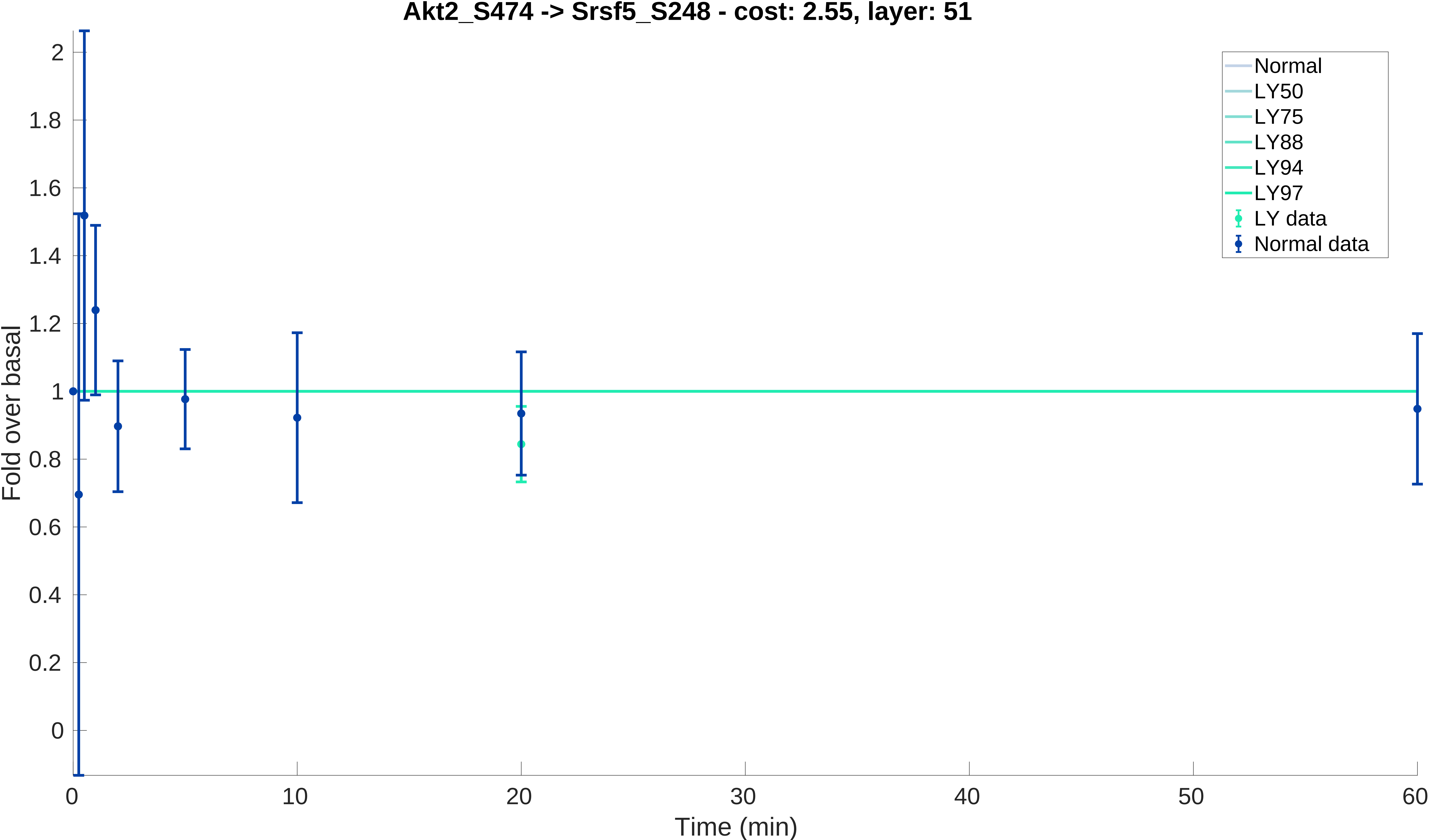

Akt2_S474 -> Srsf5_S248 - cost: 2.55, layer: 51
Normal
LY50
LY75
LY88
LY94
LY97
LY data
Normal data
Fold over basal
Time (min)
0
10
20
30
40
50
60
2
1.8
1.6
1.4
1.2
1
0.8
0.6
0.4
0.2
0

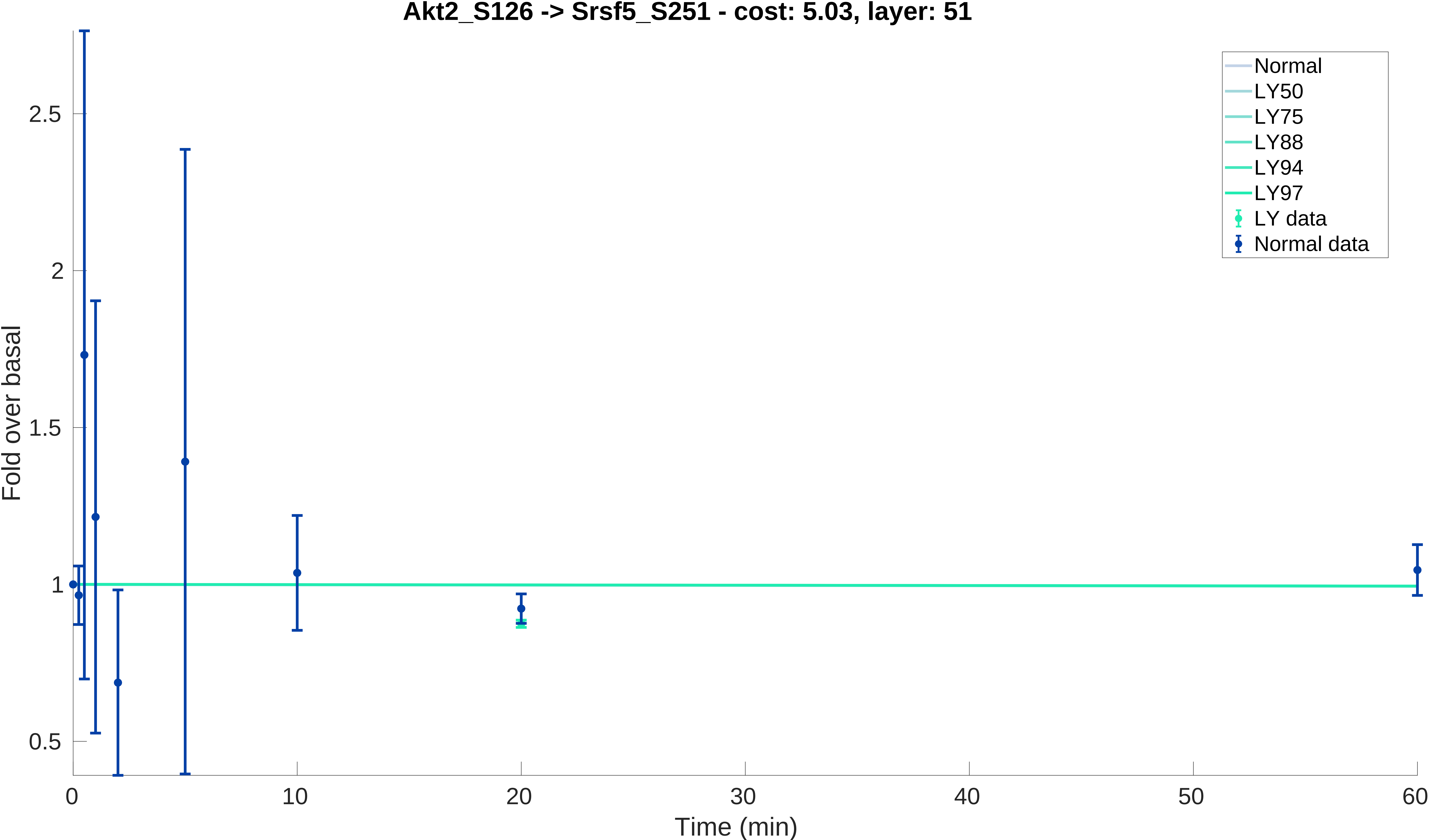

Akt2_S126 -> Srsf5_S251 - cost: 5.03, layer: 51
Fold over basal
Time (min)
0.5
1
1.5
2
2.5
0
10
20
30
40
50
60
Normal
LY50
LY75
LY88
LY94
LY97
LY data
Normal data

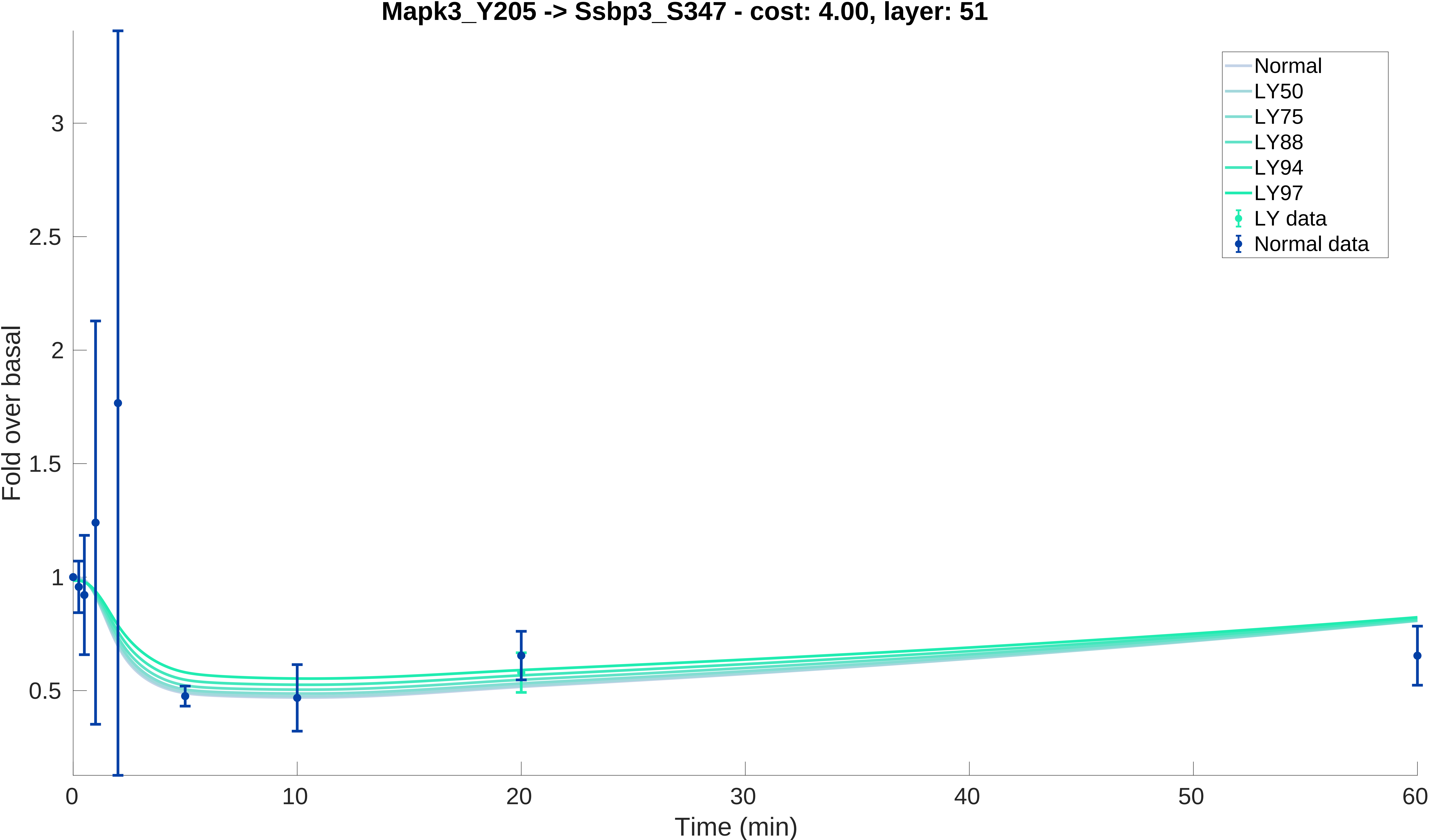

Mapk3_Y205 -> Ssbp3_S347 - cost: 4.00, layer: 51
Normal
LY50
LY75
LY88
LY94
LY97
LY data
Normal data
Fold over basal
3
2.5
2
1.5
1
0.5
0
10
20
30
40
50
60
Time (min)

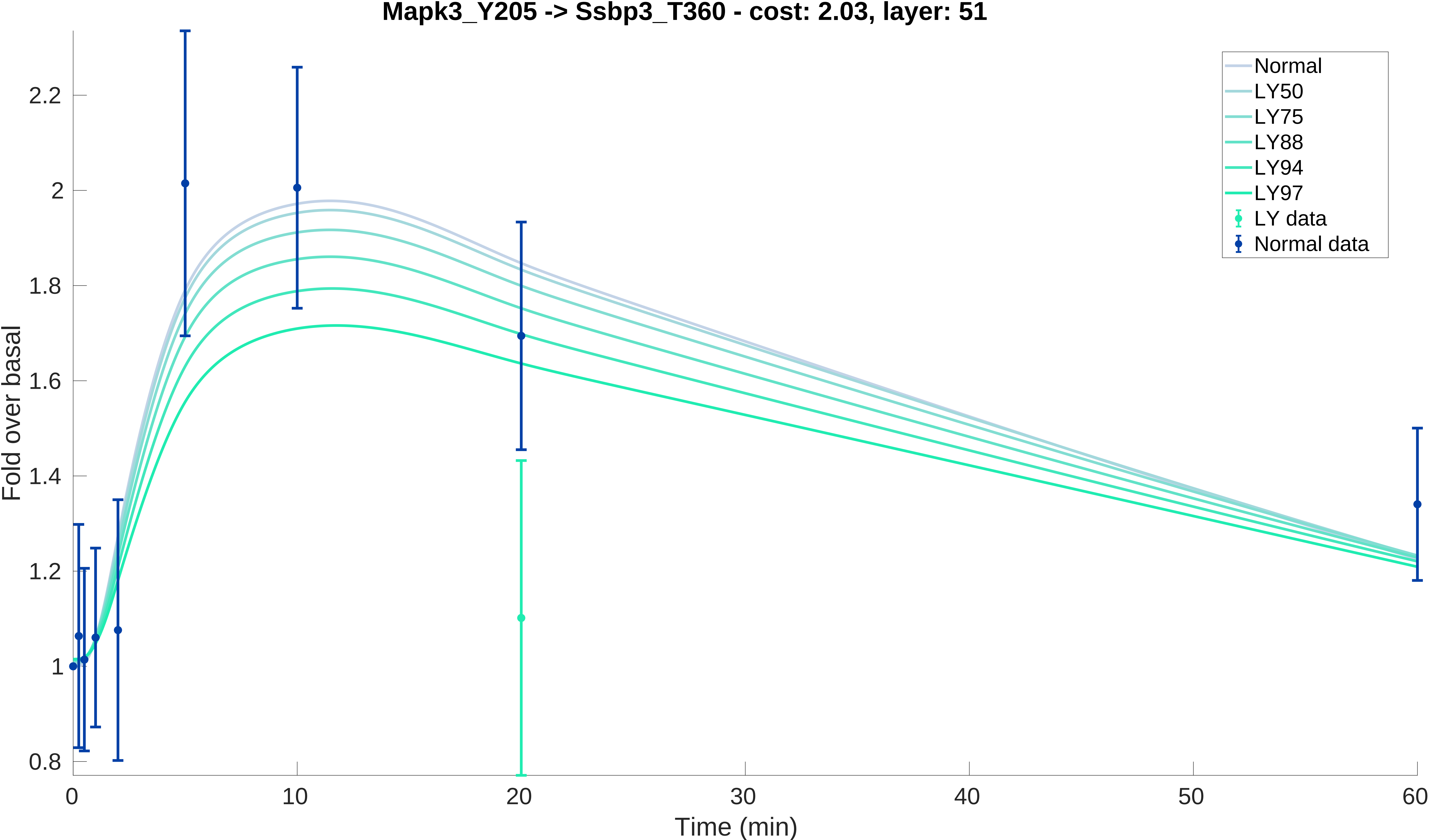

Mapk3_Y205 -> Ssbp3_T360 - cost: 2.03, layer: 51
Fold over basal
Time (min)
0.8
1
1.2
1.4
1.6
1.8
2
2.2
0
10
20
30
40
50
60
Normal
LY50
LY75
LY88
LY94
LY97
LY data
Normal data

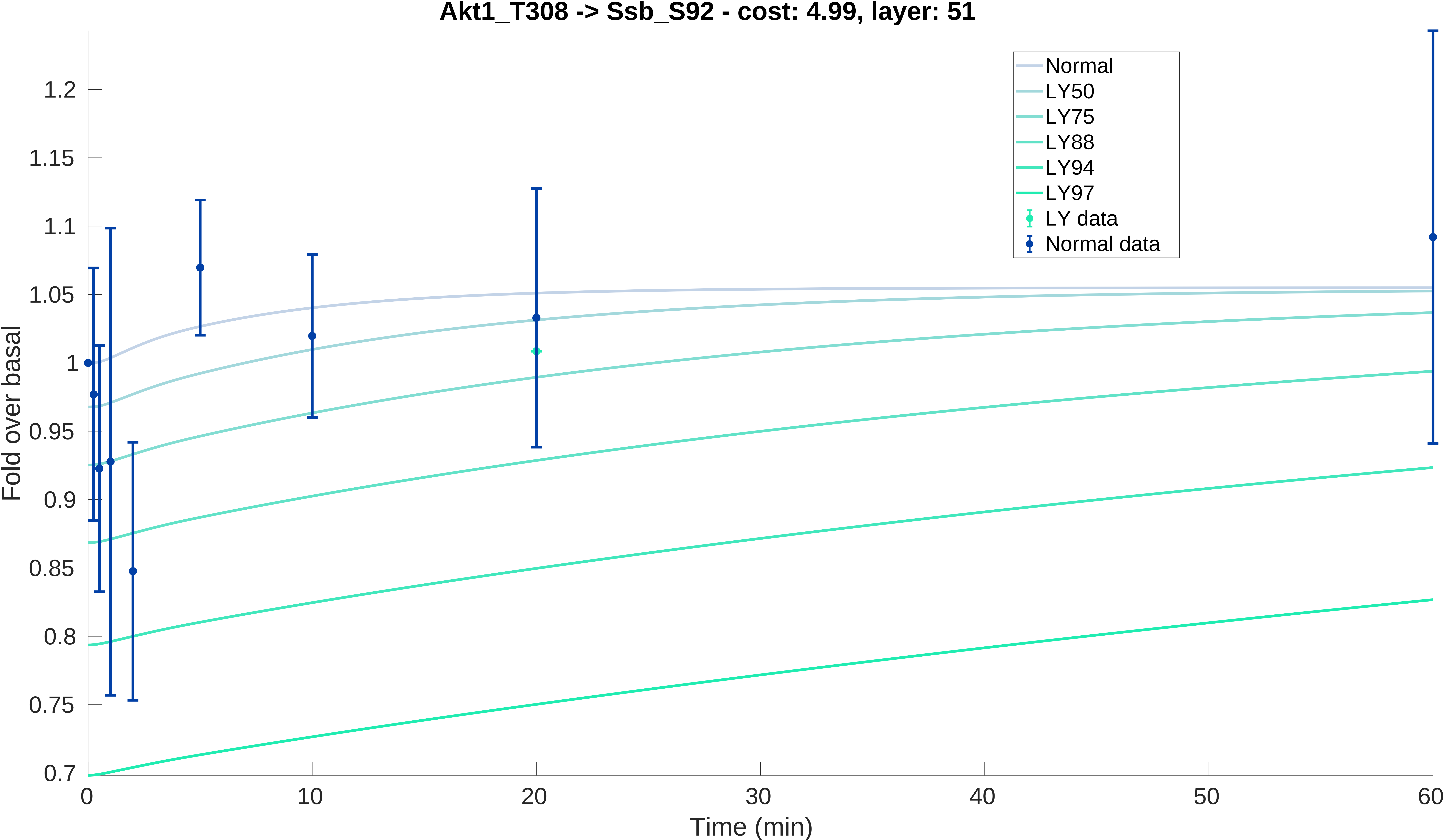

Akt1_T308 -> Ssb_S92 - cost: 4.99, layer: 51
Normal
LY50
LY75
LY88
LY94
LY97
LY data
Normal data
Fold over basal
Time (min)
1.2
1.15
1.1
1.05
1
0.95
0.9
0.85
0.8
0.75
0.7
0
10
20
30
40
50
60

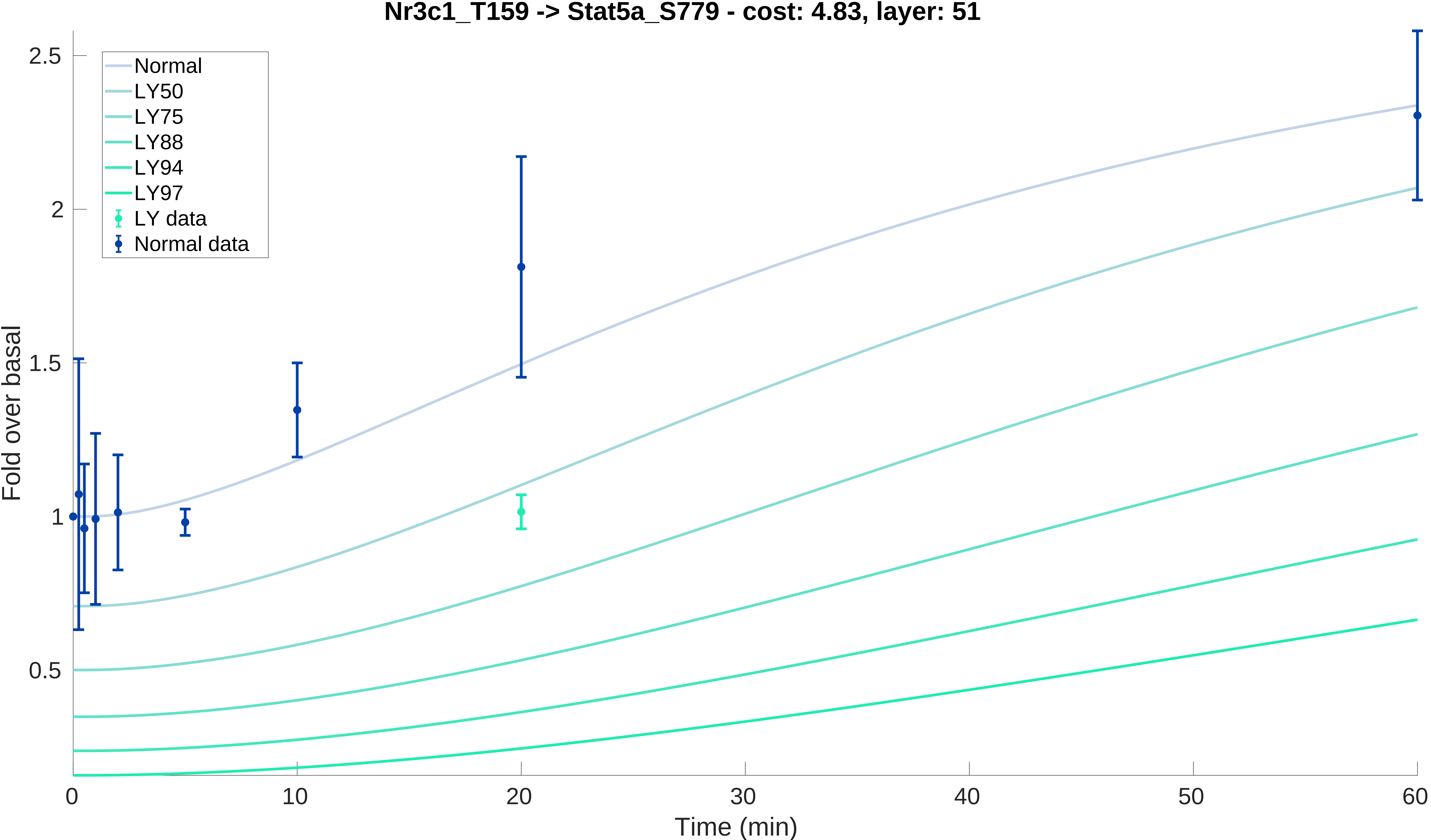

Nr3c1_T159 -> Stat5a_S779 - cost: 4.83, layer: 51
Normal
LY50
LY75
LY88
LY94
LY97
LY data
Normal data
Fold over basal
0.5
1
1.5
2
2.5
0
10
20
30
40
50
60
Time (min)

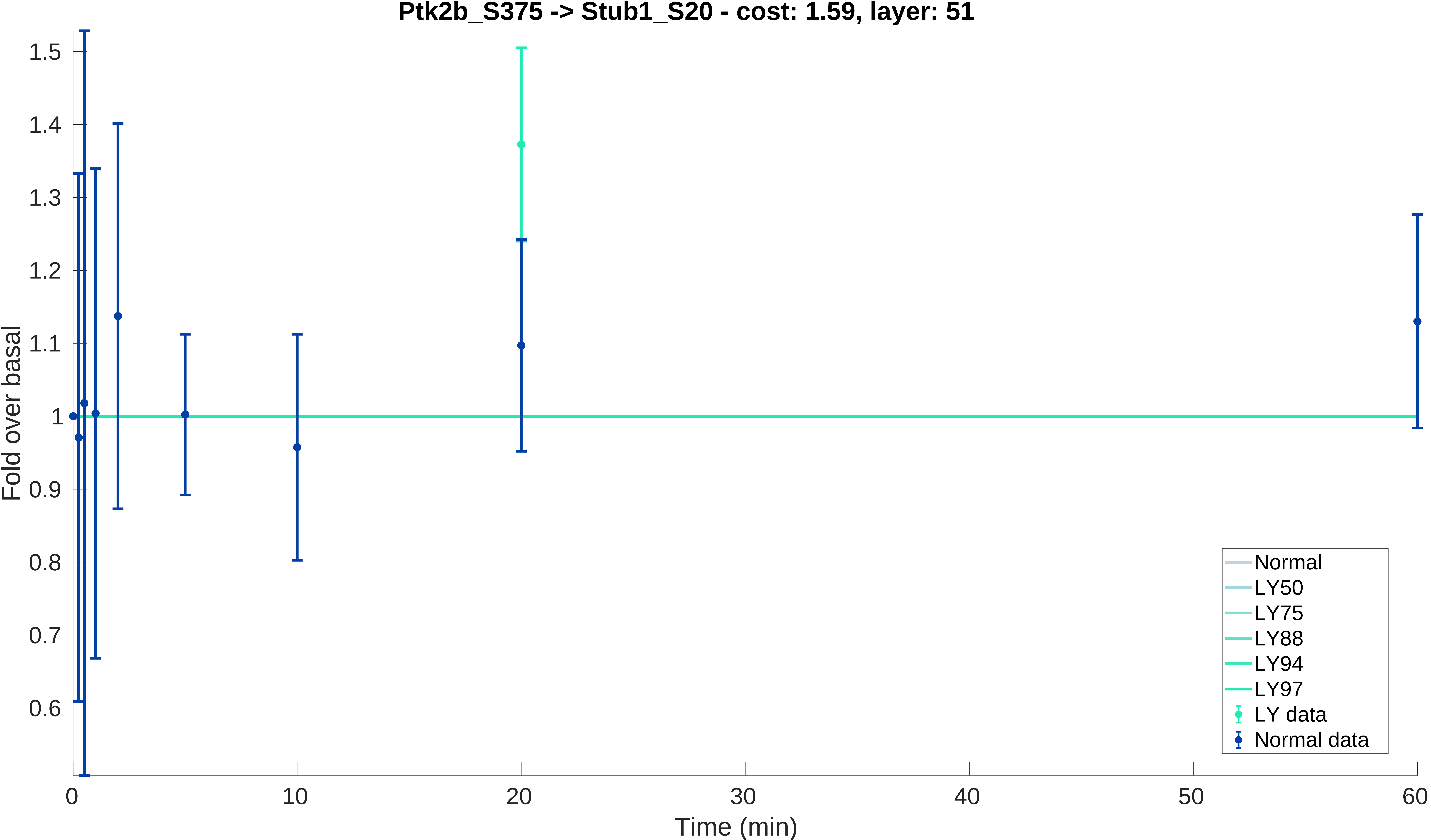

Ptk2b_S375 -> Stub1_S20 - cost: 1.59, layer: 51
Fold over basal
1.5
1.4
1.3
1.2
1.1
1
0.9
0.8
0.7
0.6
0
10
20
30
40
50
60
Time (min)
Normal
LY50
LY75
LY88
LY94
LY97
LY data
Normal data

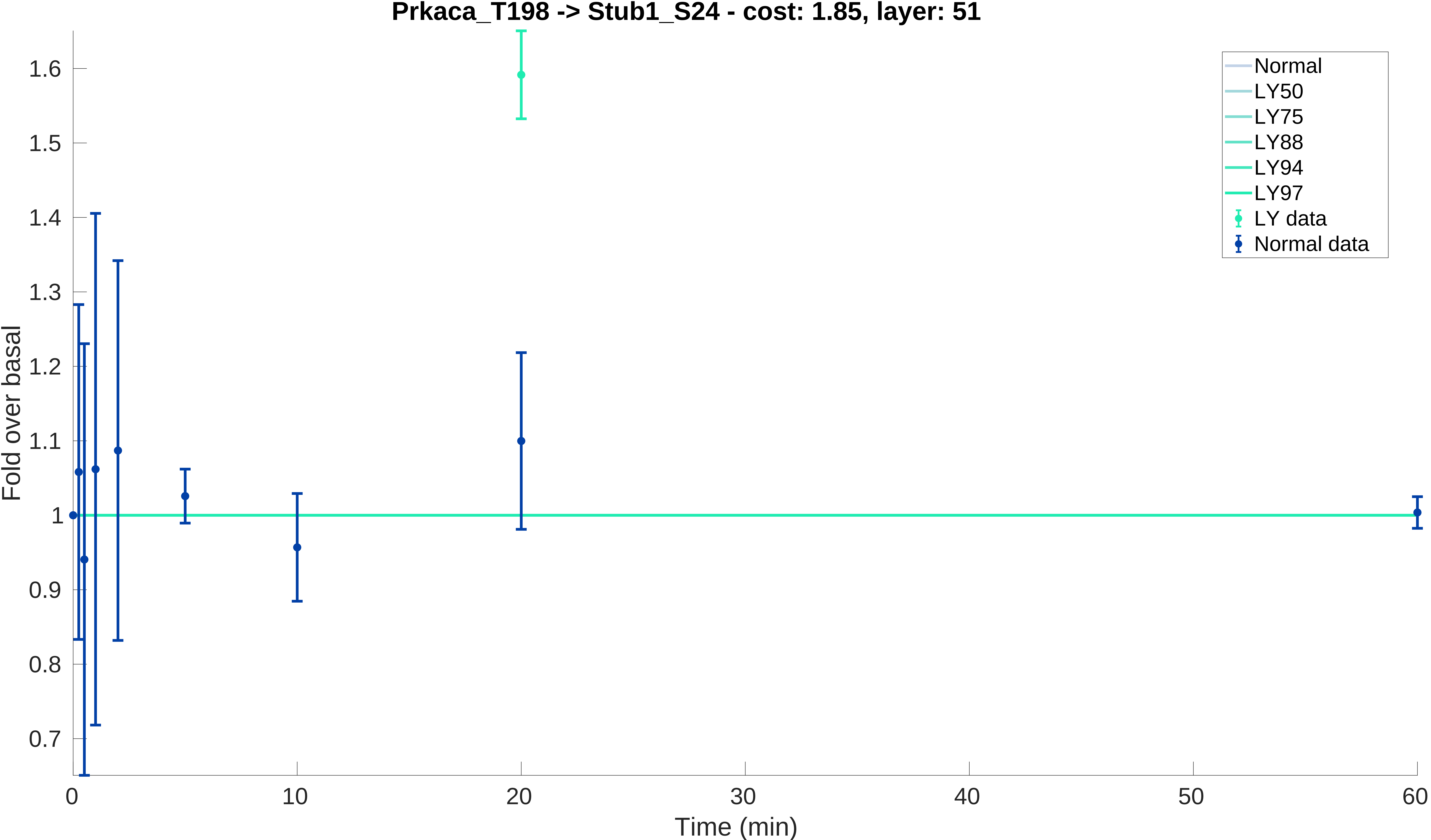
Prkaca_T198 -> Stub1_S24 - cost: 1.85, layer: 51
Fold over basal
Time (min)
1.6
1.5
1.4
1.3
1.2
1.1
1
0.9
0.8
0.7
0
10
20
30
40
50
60
Normal
LY50
LY75
LY88
LY94
LY97
LY data
Normal data

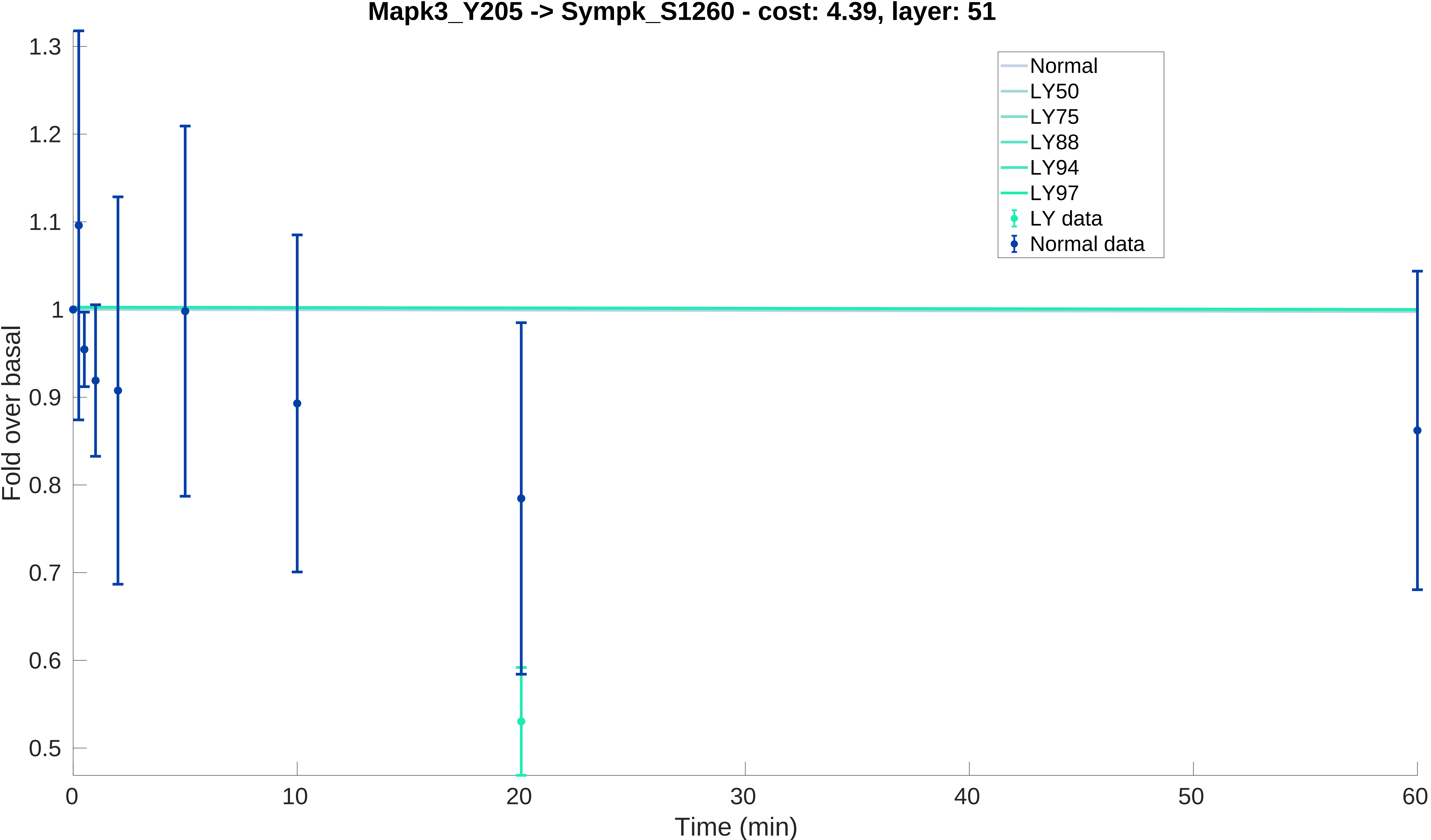

Mapk3_Y205 -> Sympk_S1260 - cost: 4.39, layer: 51
Normal
LY50
LY75
LY88
LY94
LY97
LY data
Normal data
Fold over basal
Time (min)
1.3
1.2
1.1
1
0.9
0.8
0.7
0.6
0.5
0
10
20
30
40
50
60

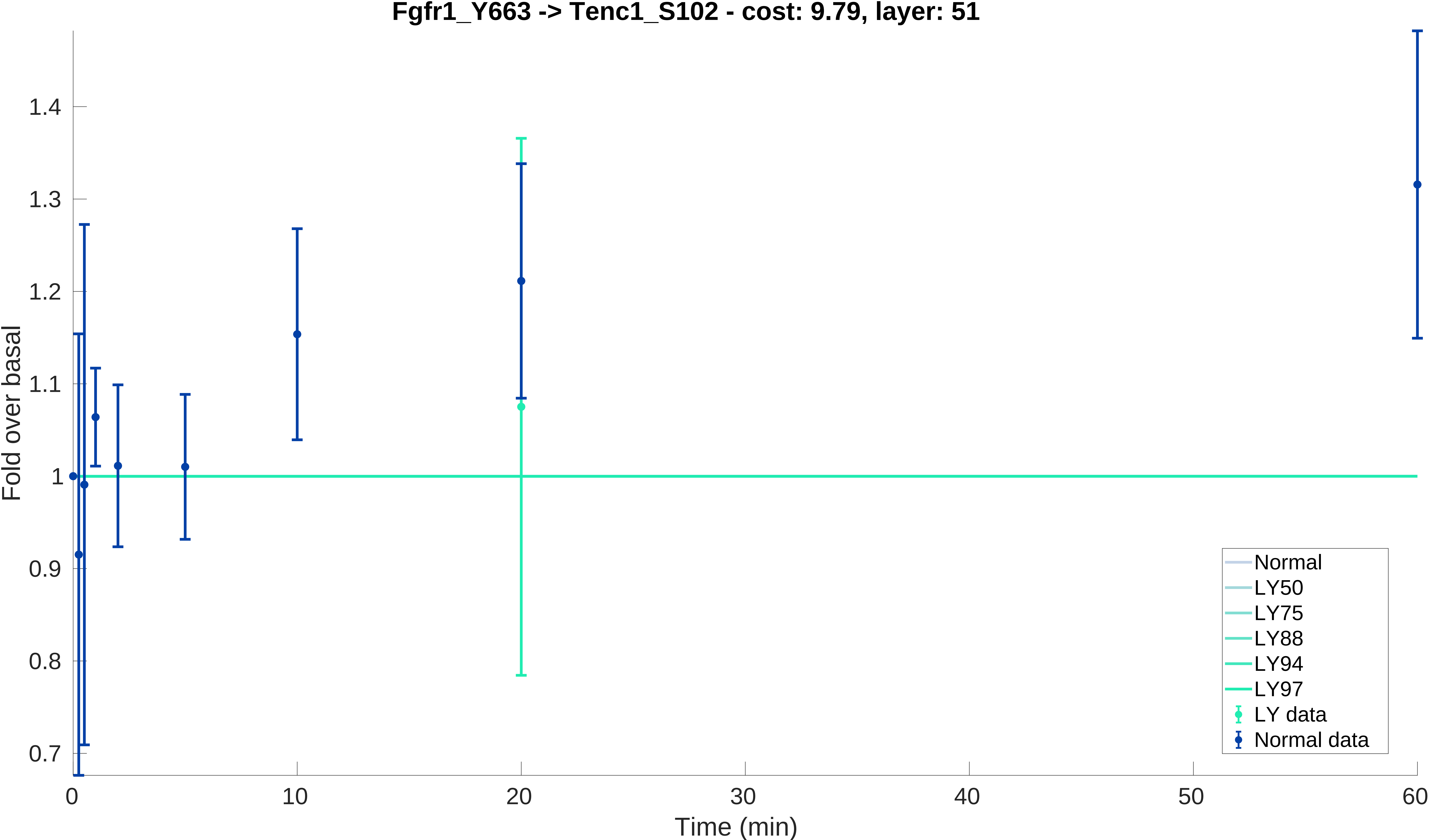

Fgfr1_Y663 -> Tenc1_S102 - cost: 9.79, layer: 51
Fold over basal
1.4
1.3
1.2
1.1
1
0.9
0.8
0.7
0
10
20
30
40
50
60
Time (min)
Normal
LY50
LY75
LY88
LY94
LY97
LY data
Normal data

Fgfr1_Y663 -> Tenc1_S1087 - cost: 12.26, layer: 51

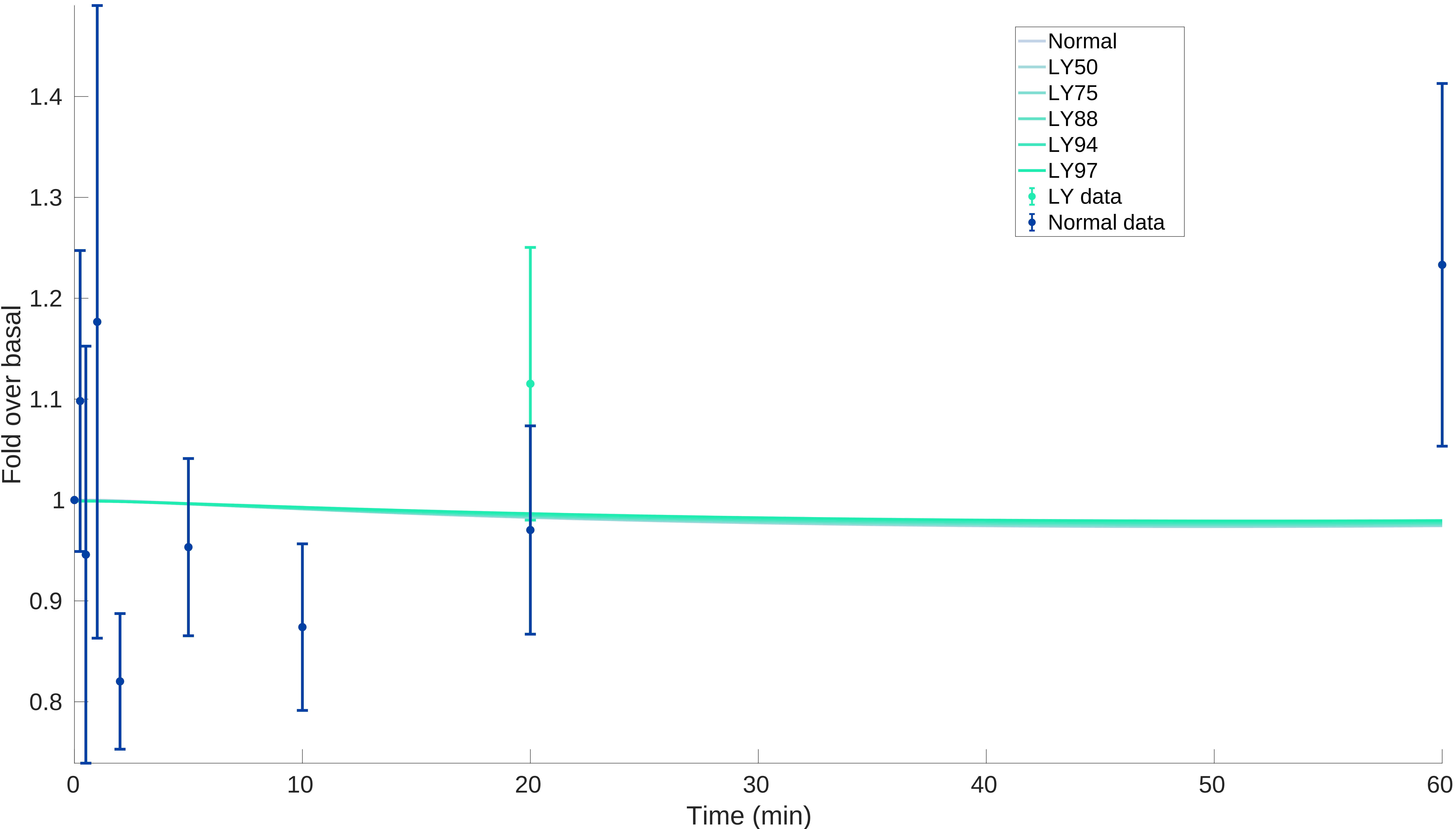

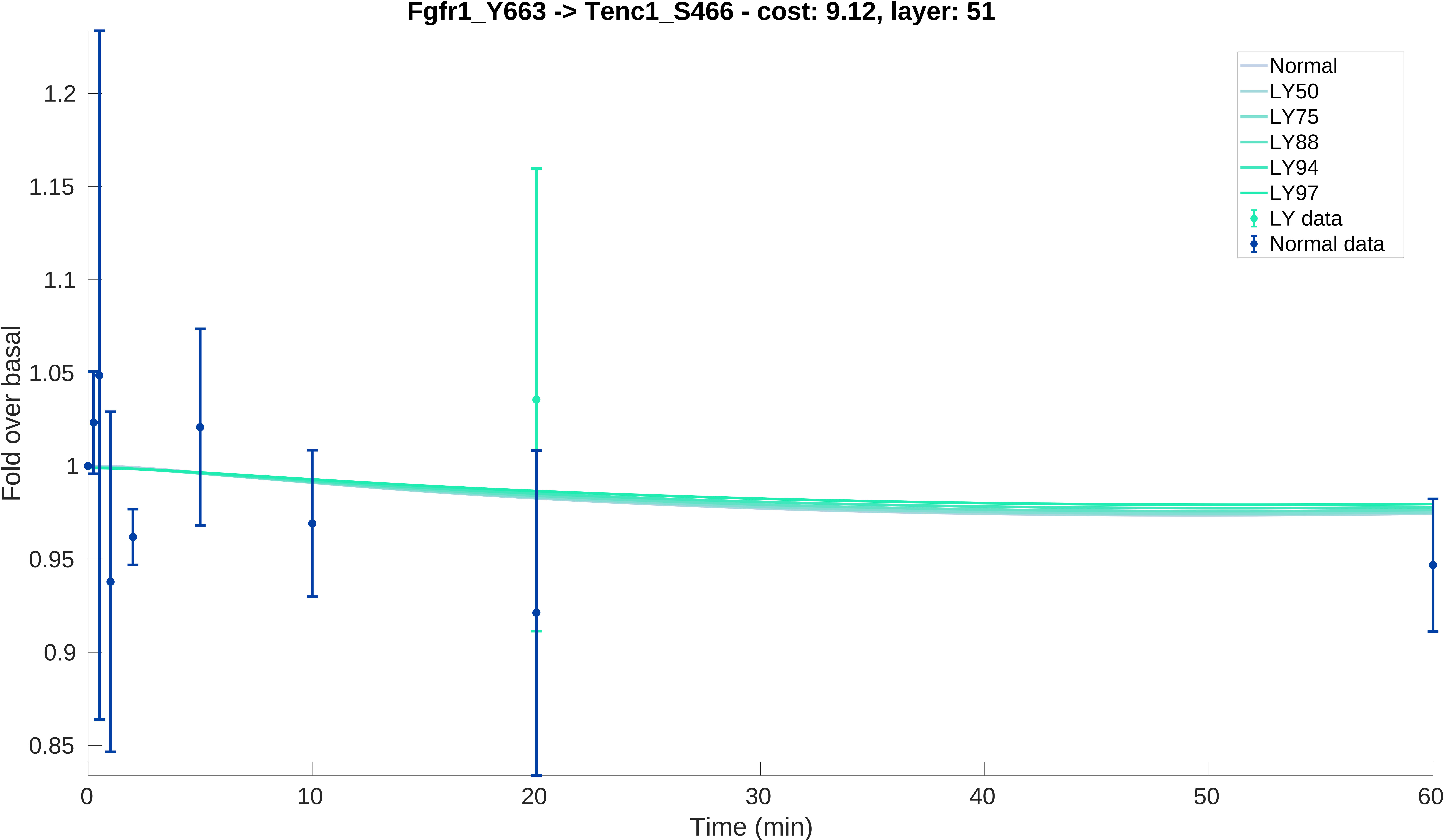

Fgfr1_Y663 -> Tenc1_S466 - cost: 9.12, layer: 51
Normal
LY50
LY75
LY88
LY94
LY97
LY data
Normal data
Fold over basal
1.2
1.15
1.1
1.05
1
0.95
0.9
0.85
0
10
20
30
40
50
60
Time (min)

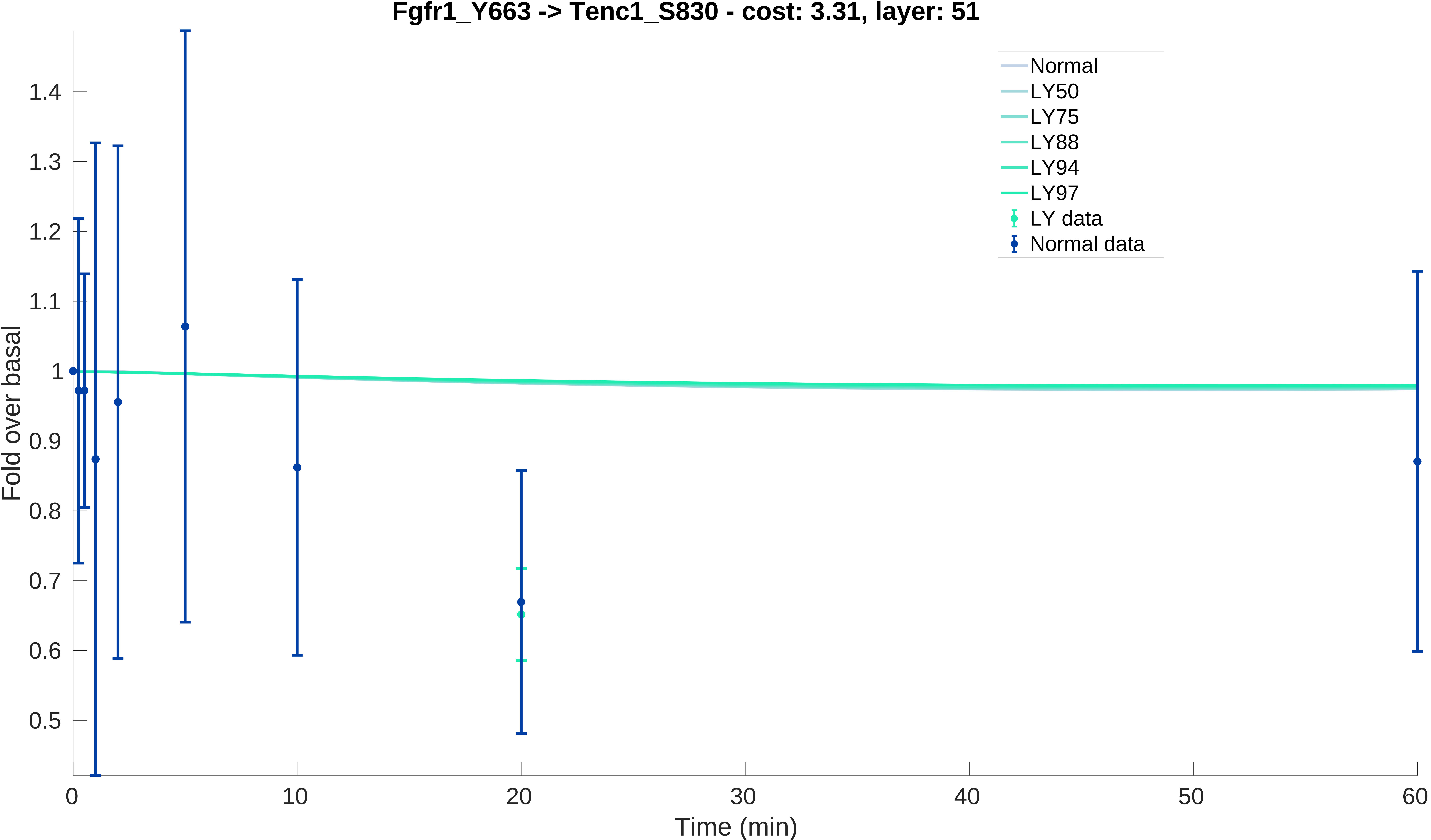

Fgfr1_Y663 -> Tenc1_S830 - cost: 3.31, layer: 51
Fold over basal
1.4
1.3
1.2
1.1
1
0.9
0.8
0.7
0.6
0.5
0
10
20
30
40
50
60
Time (min)
Normal
LY50
LY75
LY88
LY94
LY97
LY data
Normal data

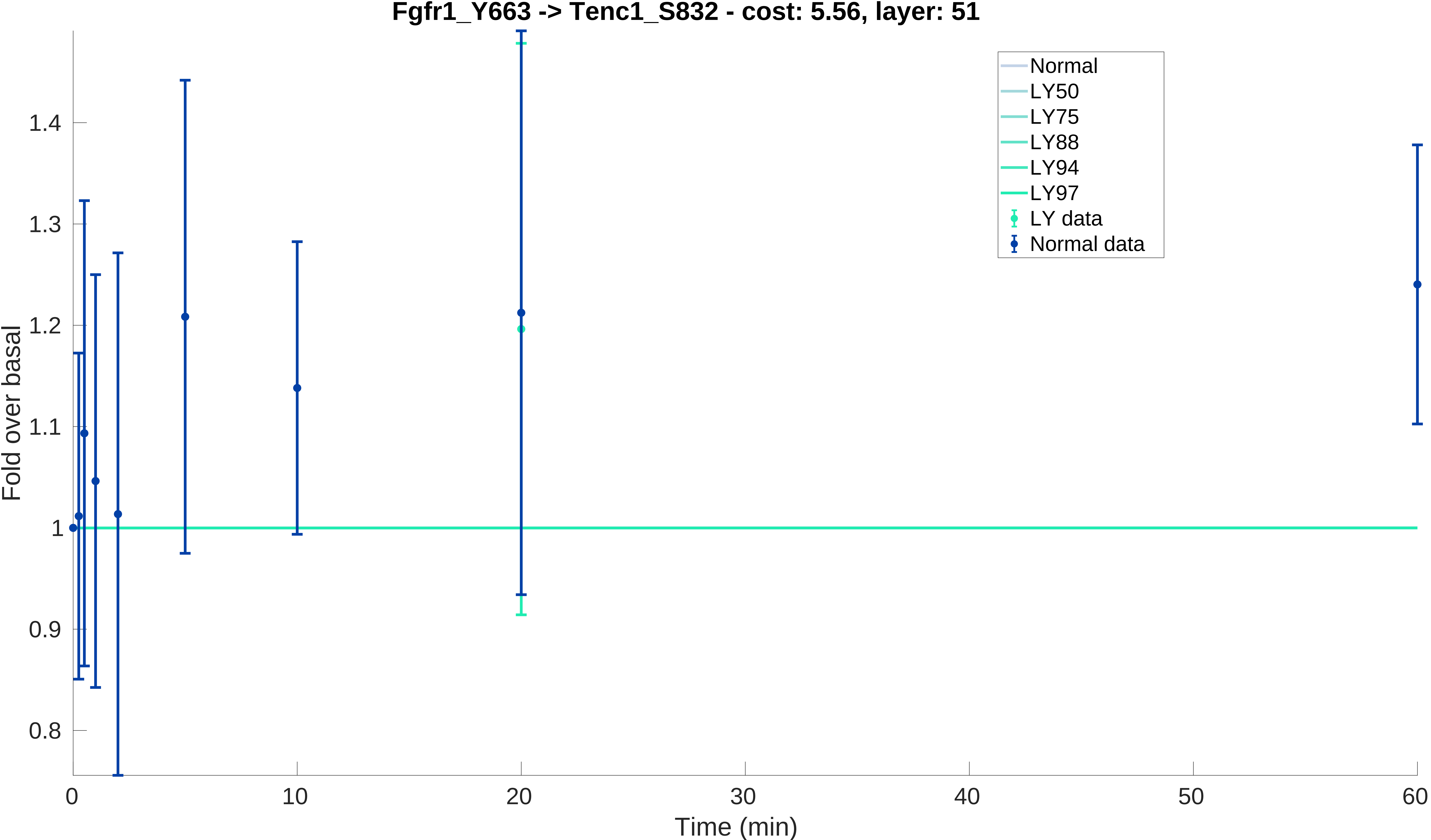

Fgfr1_Y663 -> Tenc1_S832 - cost: 5.56, layer: 51
Normal
LY50
LY75
LY88
LY94
LY97
LY data
Normal data
1.4
1.3
1.2
1.1
1
0.9
0.8
Fold over basal
0
10
20
30
40
50
60
Time (min)

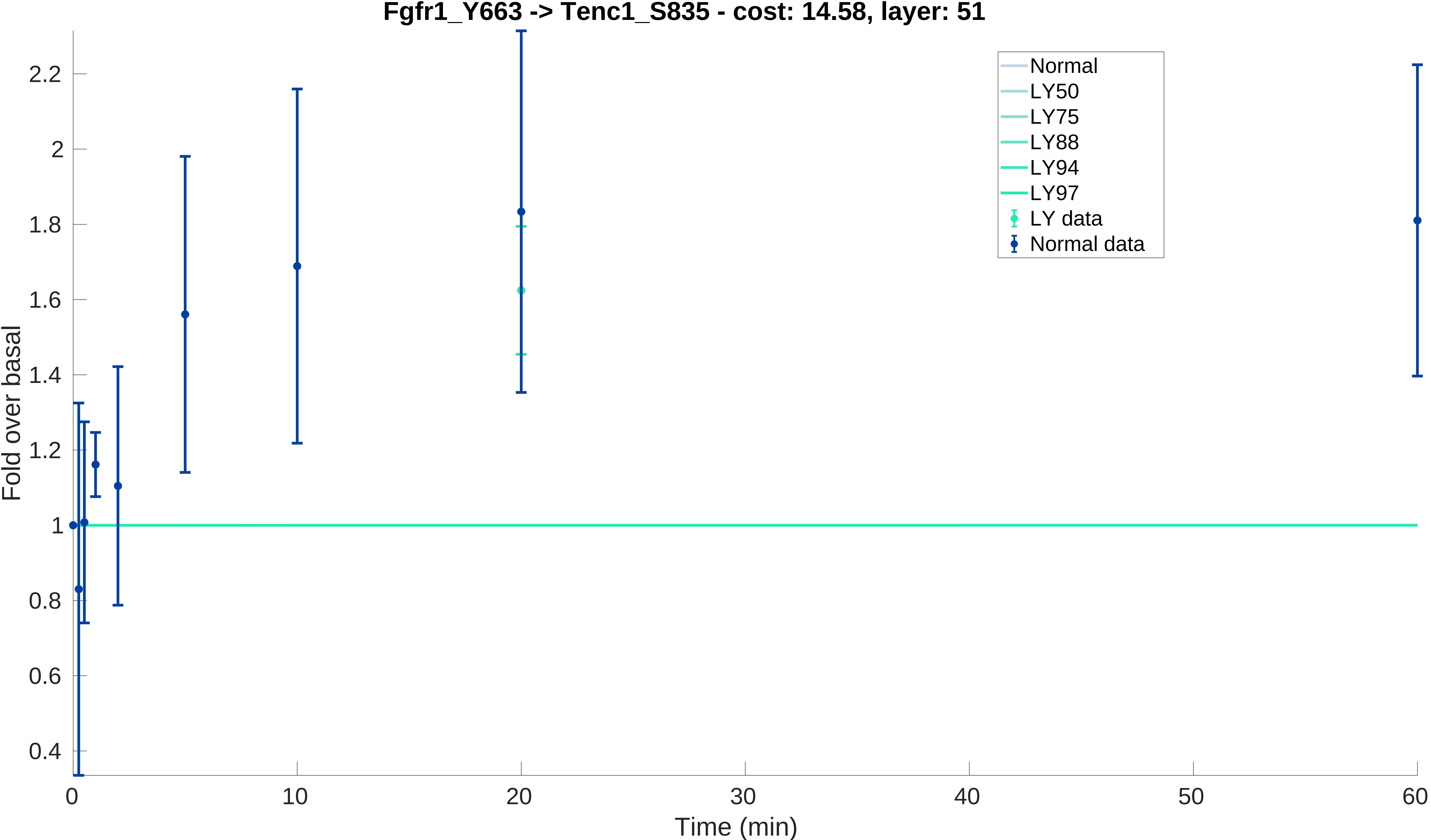

Fgfr1_Y663 -> Tenc1_S835 - cost: 14.58, layer: 51
Normal
LY50
LY75
LY88
LY94
LY97
LY data
Normal data
Fold over basal
Time (min)
0.4
0.6
0.8
1
1.2
1.4
1.6
1.8
2
2.2
0
10
20
30
40
50
60

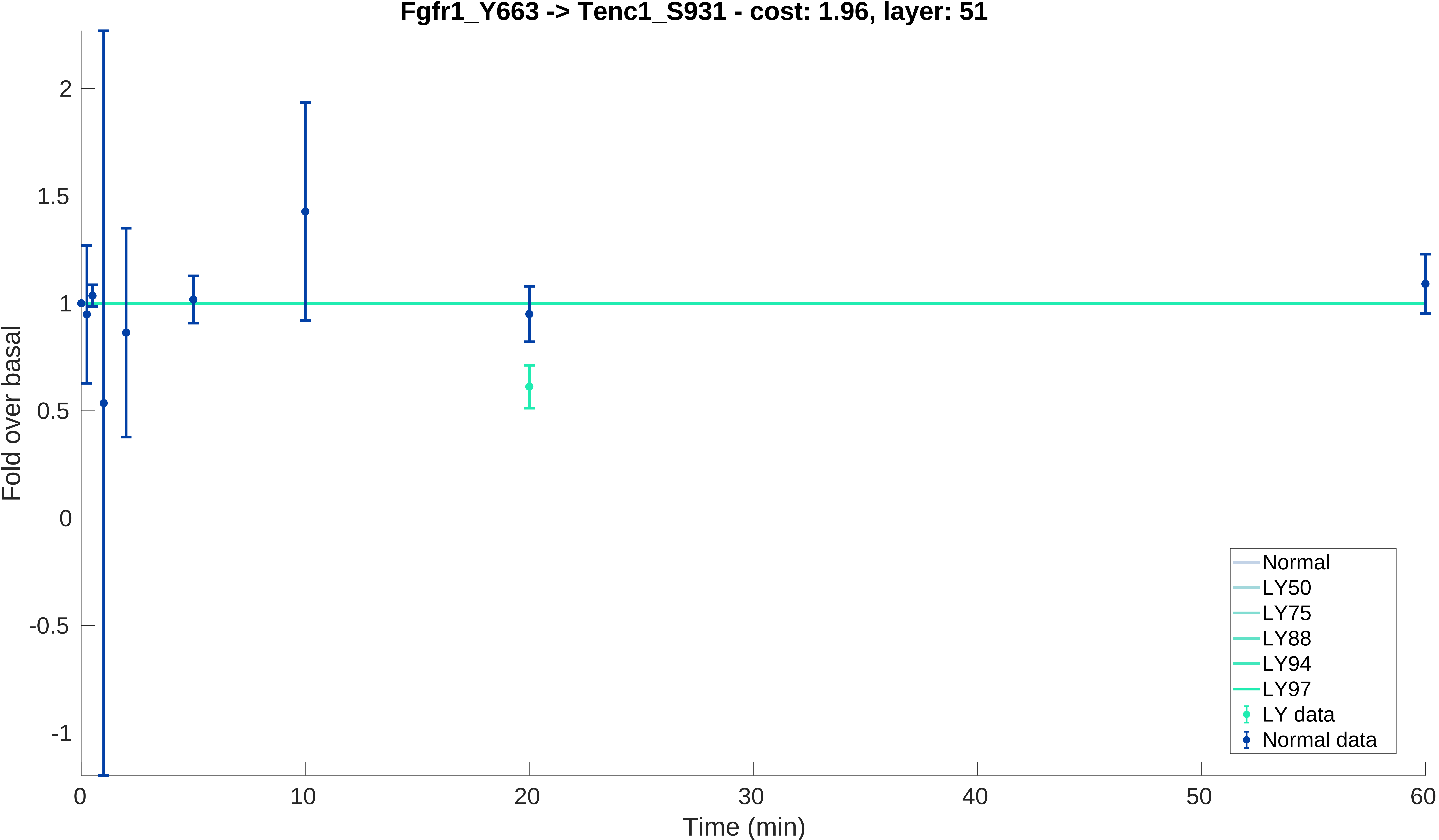

Fgfr1_Y663 -> Tenc1_S931 - cost: 1.96, layer: 51
Fold over basal
Time (min)
2
1.5
1
0.5
0
-0.5
-1
0
10
20
30
40
50
60
Normal
LY50
LY75
LY88
LY94
LY97
LY data
Normal data

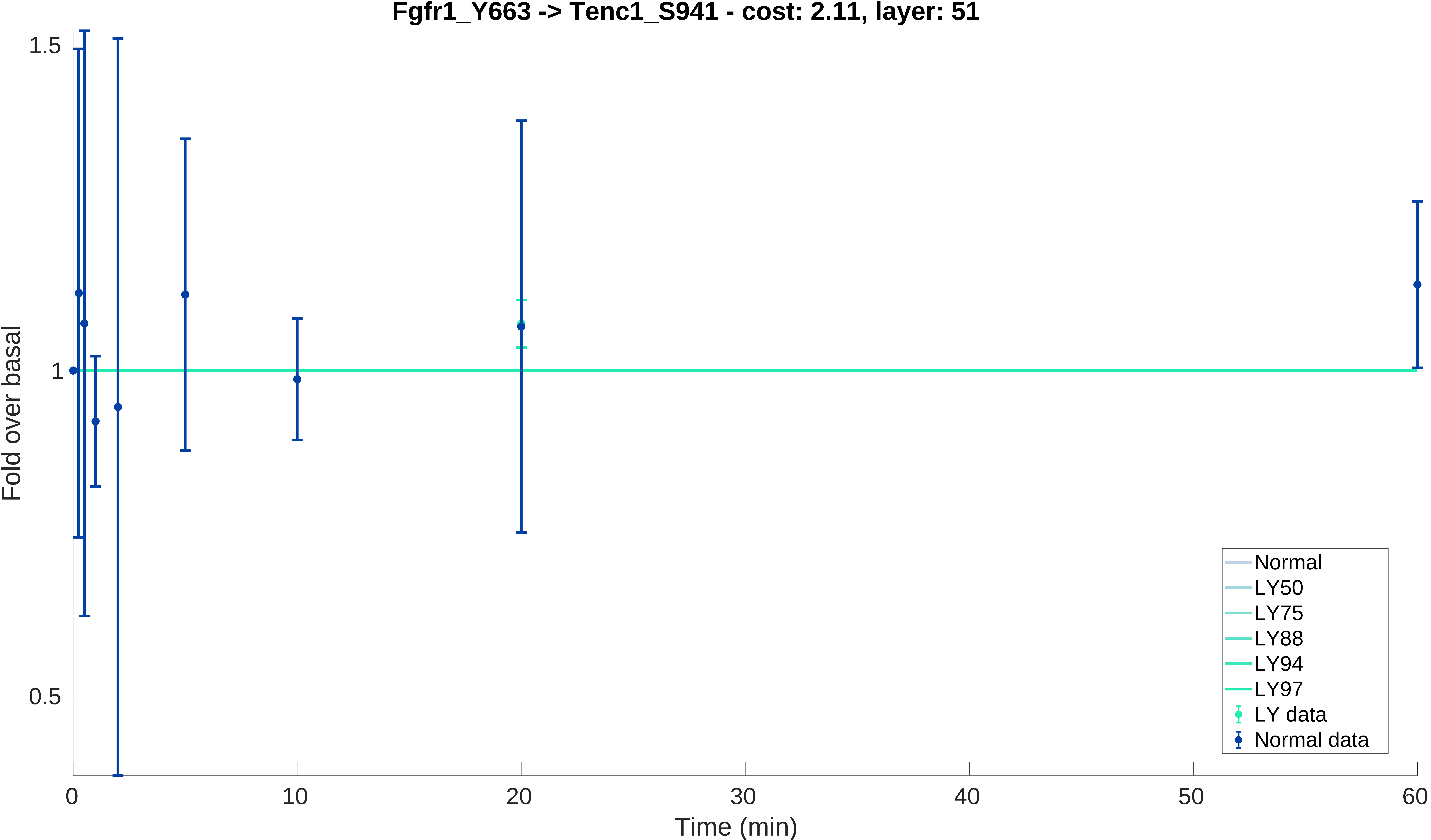

Fgfr1_Y663 -> Tenc1_S941 - cost: 2.11, layer: 51
Fold over basal
1.5
1
0.5
0
10
20
30
40
50
60
Time (min)
Normal
LY50
LY75
LY88
LY94
LY97
LY data
Normal data

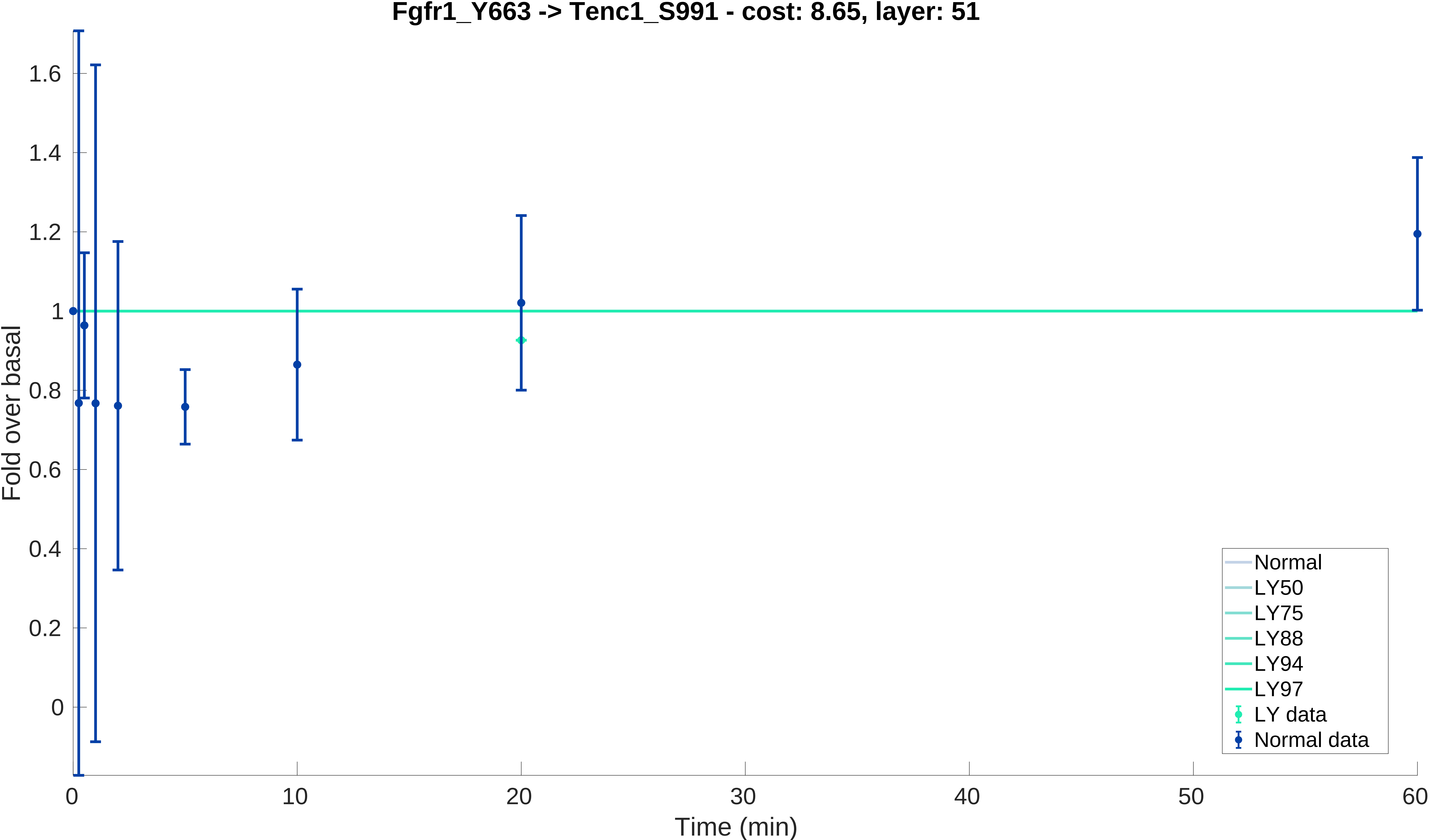

Fgfr1_Y663 -> Tenc1_S991 - cost: 8.65, layer: 51
Fold over basal
1.6
1.4
1.2
1
0.8
0.6
0.4
0.2
0
0
10
20
30
40
50
60
Time (min)
Normal
LY50
LY75
LY88
LY94
LY97
LY data
Normal data

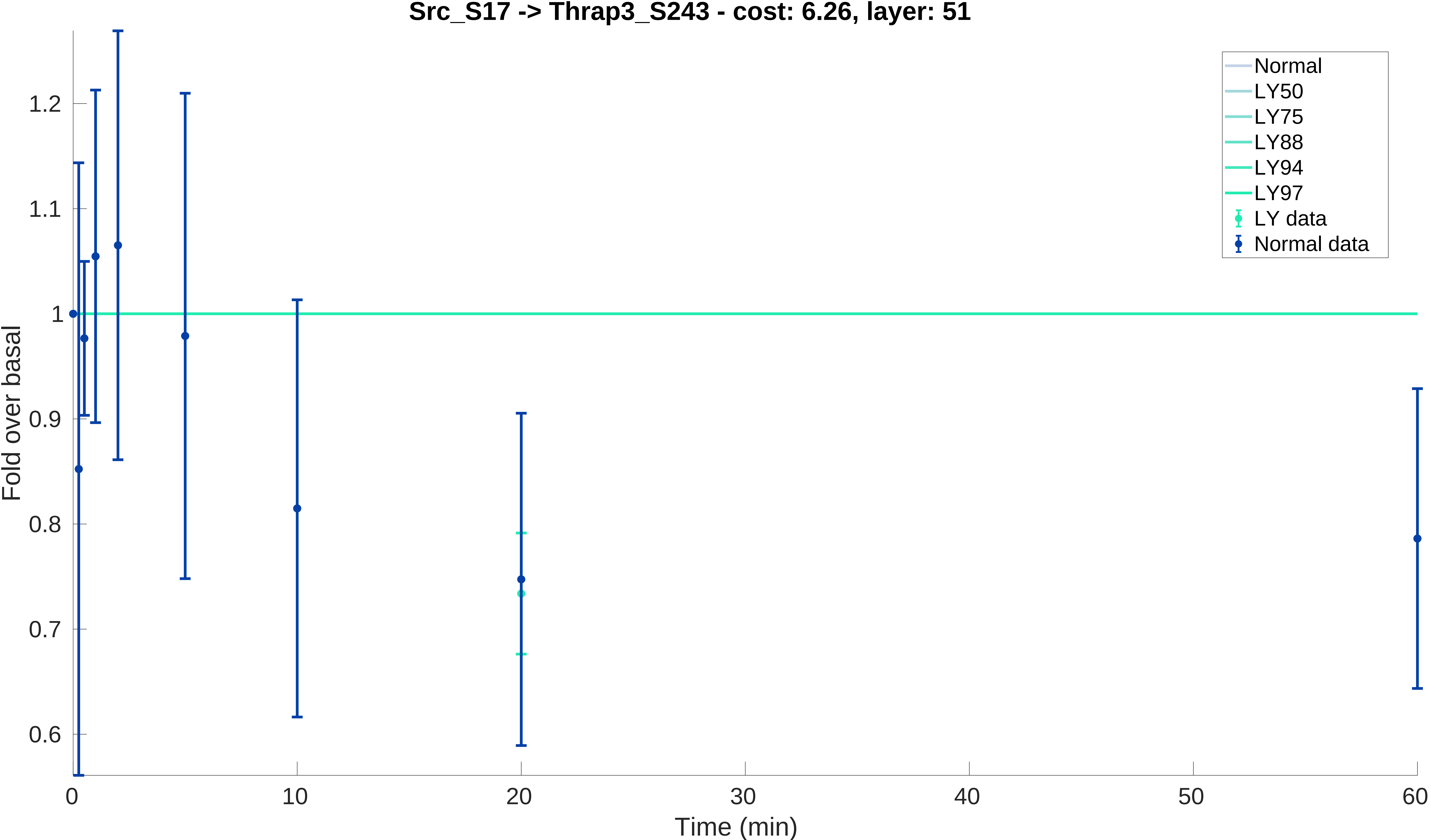

Src_S17 -> Thrap3_S243 - cost: 6.26, layer: 51
Normal
LY50
LY75
LY88
LY94
LY97
LY data
Normal data
Fold over basal
1.2
1.1
1
0.9
0.8
0.7
0.6
0
10
20
30
40
50
60
Time (min)

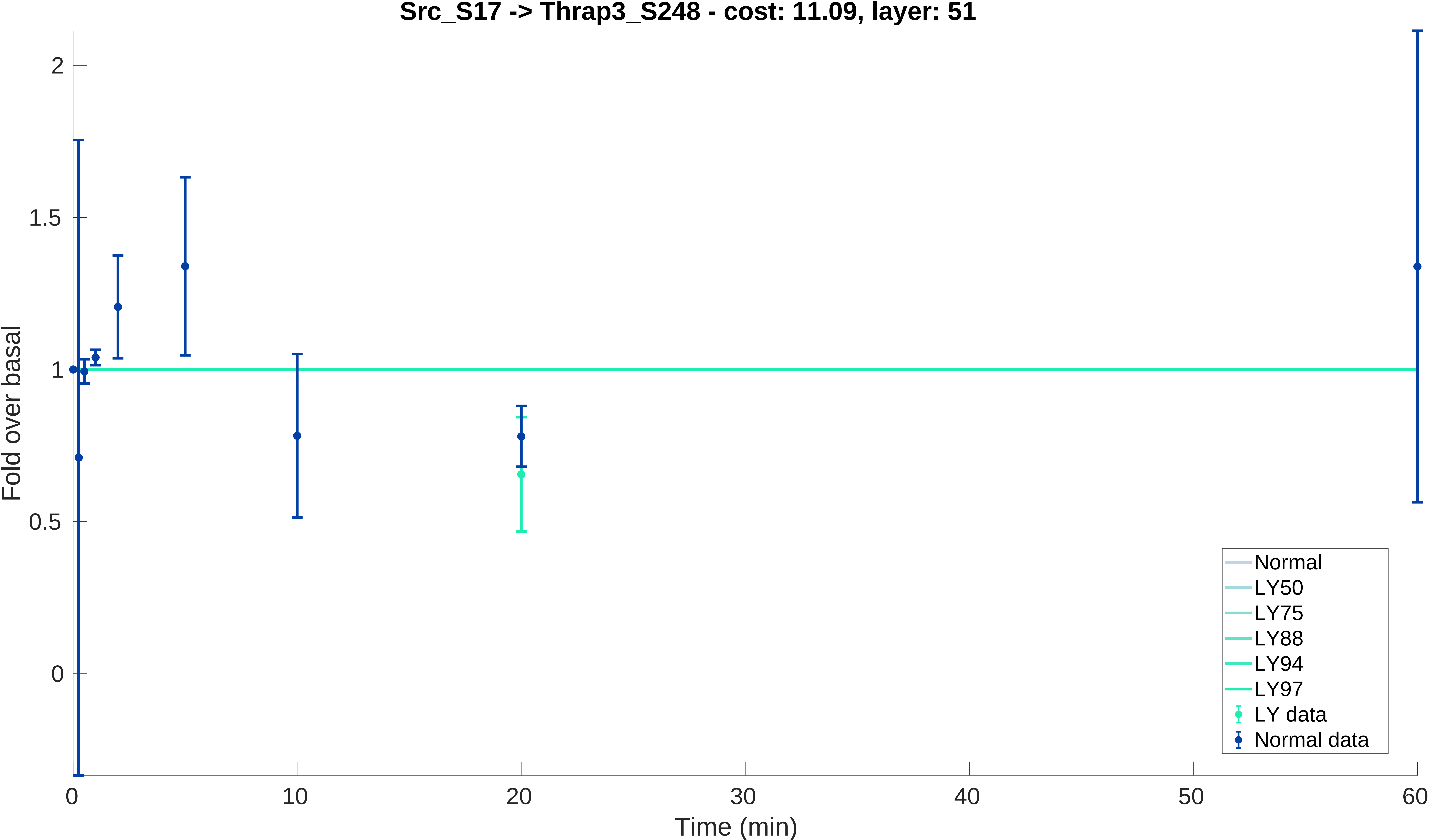
Src_S17 -> Thrap3_S248 - cost: 11.09, layer: 51
Fold over basal
Time (min)
0
10
20
30
40
50
60
0
0.5
1
1.5
2
Normal
LY50
LY75
LY88
LY94
LY97
LY data
Normal data

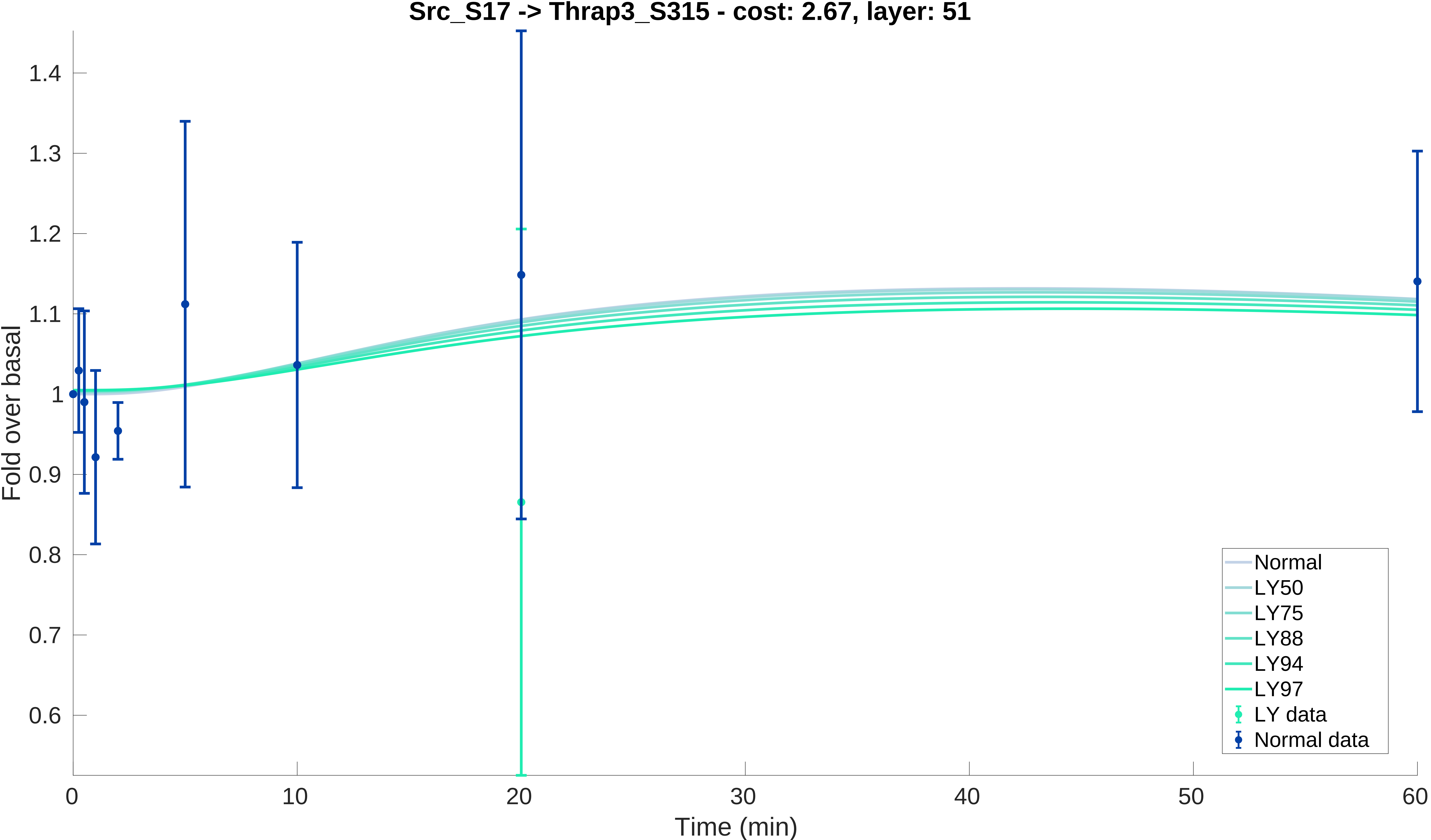

Src_S17 -> Thrap3_S315 - cost: 2.67, layer: 51
Fold over basal
1.4
1.3
1.2
1.1
1
0.9
0.8
0.7
0.6
0
10
20
30
40
50
60
Time (min)
Normal
LY50
LY75
LY88
LY94
LY97
LY data
Normal data

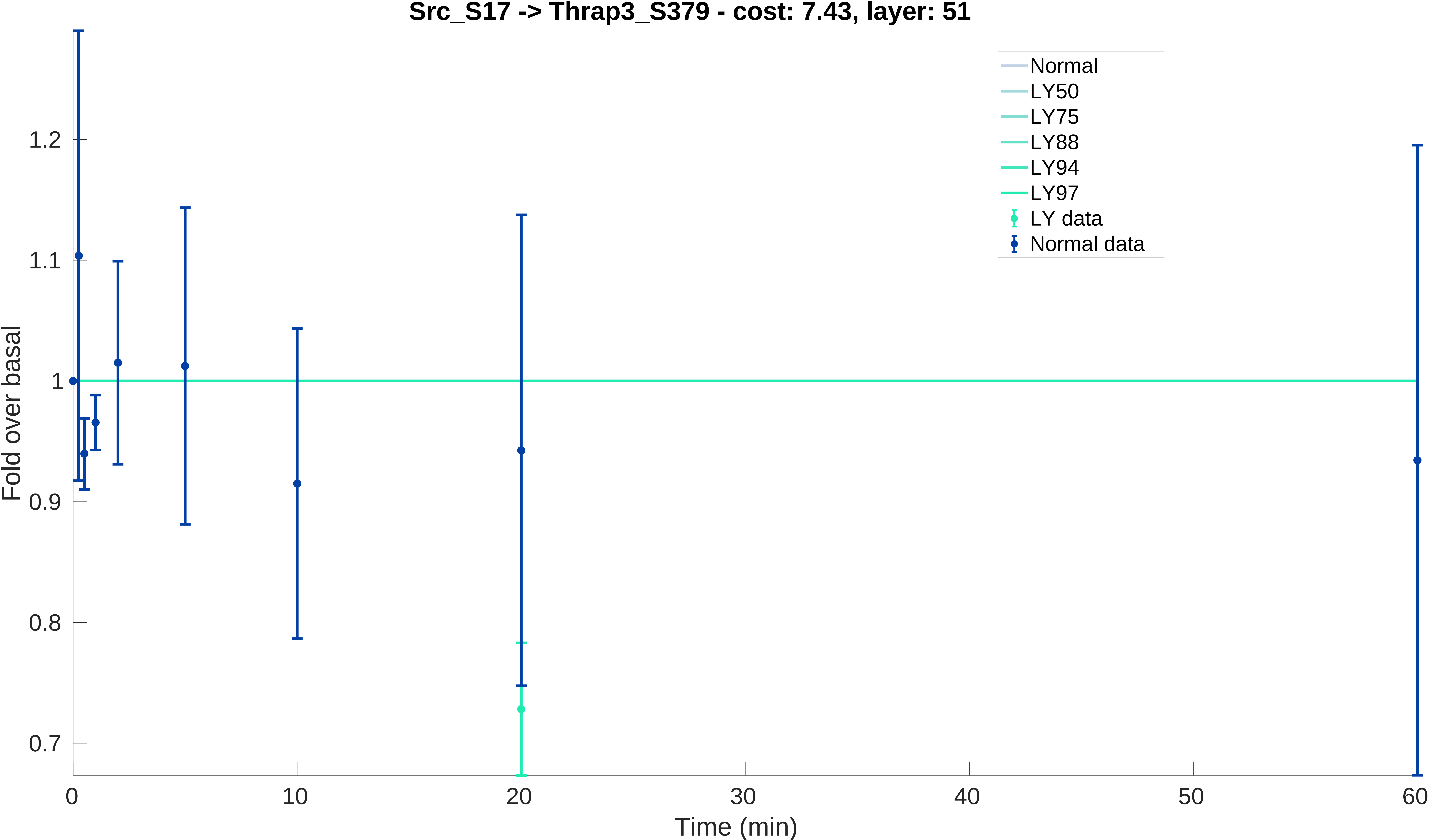
Src_S17 -> Thrap3_S379 - cost: 7.43, layer: 51
Fold over basal
Time (min)
0
10
20
30
40
50
60
0.7
0.8
0.9
1
1.1
1.2
Normal
LY50
LY75
LY88
LY94
LY97
LY data
Normal data

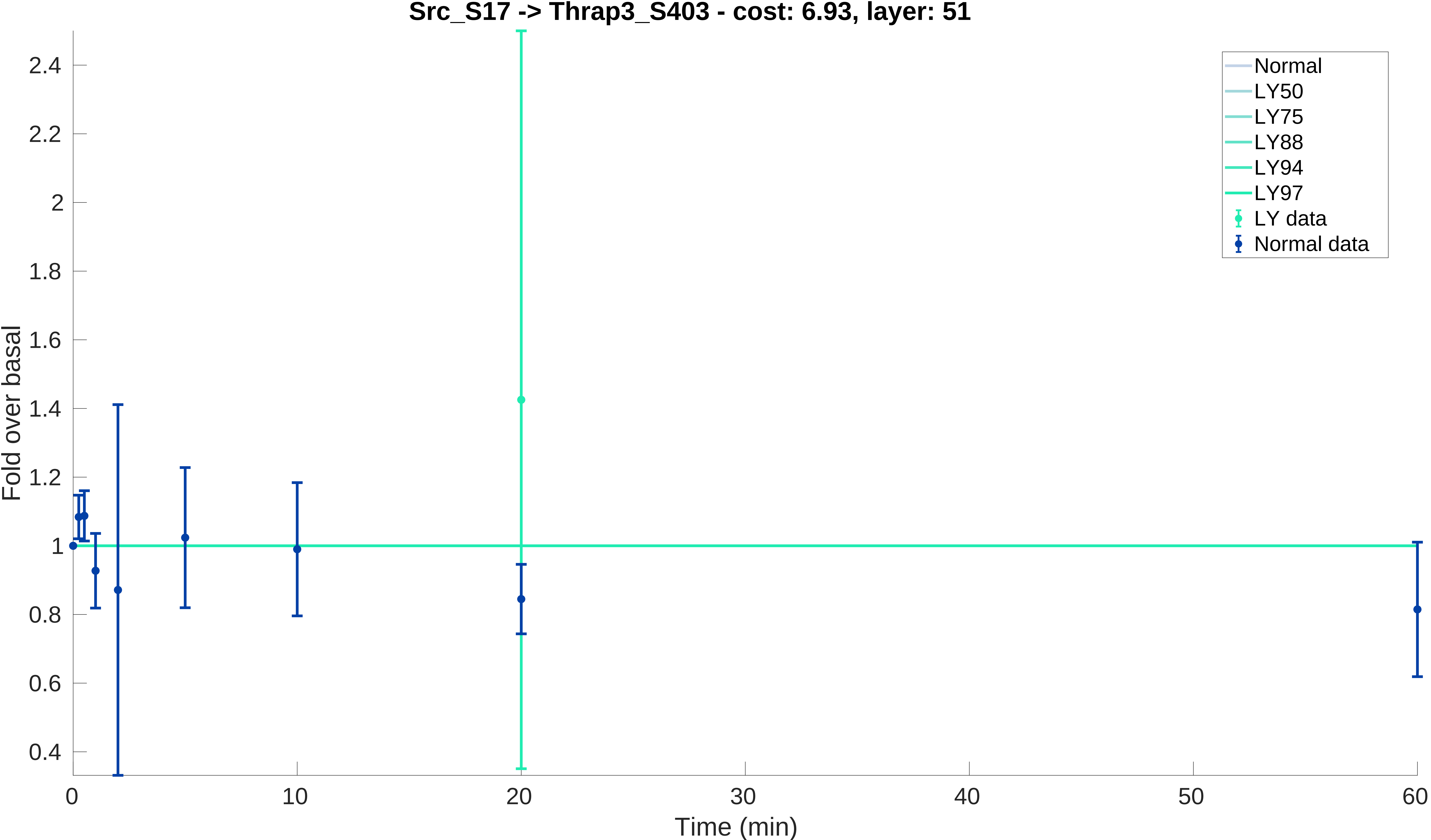
Src_S17 -> Thrap3_S403 - cost: 6.93, layer: 51
Fold over basal
Time (min)
0.4
0.6
0.8
1
1.2
1.4
1.6
1.8
2
2.2
2.4
0
10
20
30
40
50
60
Normal
LY50
LY75
LY88
LY94
LY97
LY data
Normal data

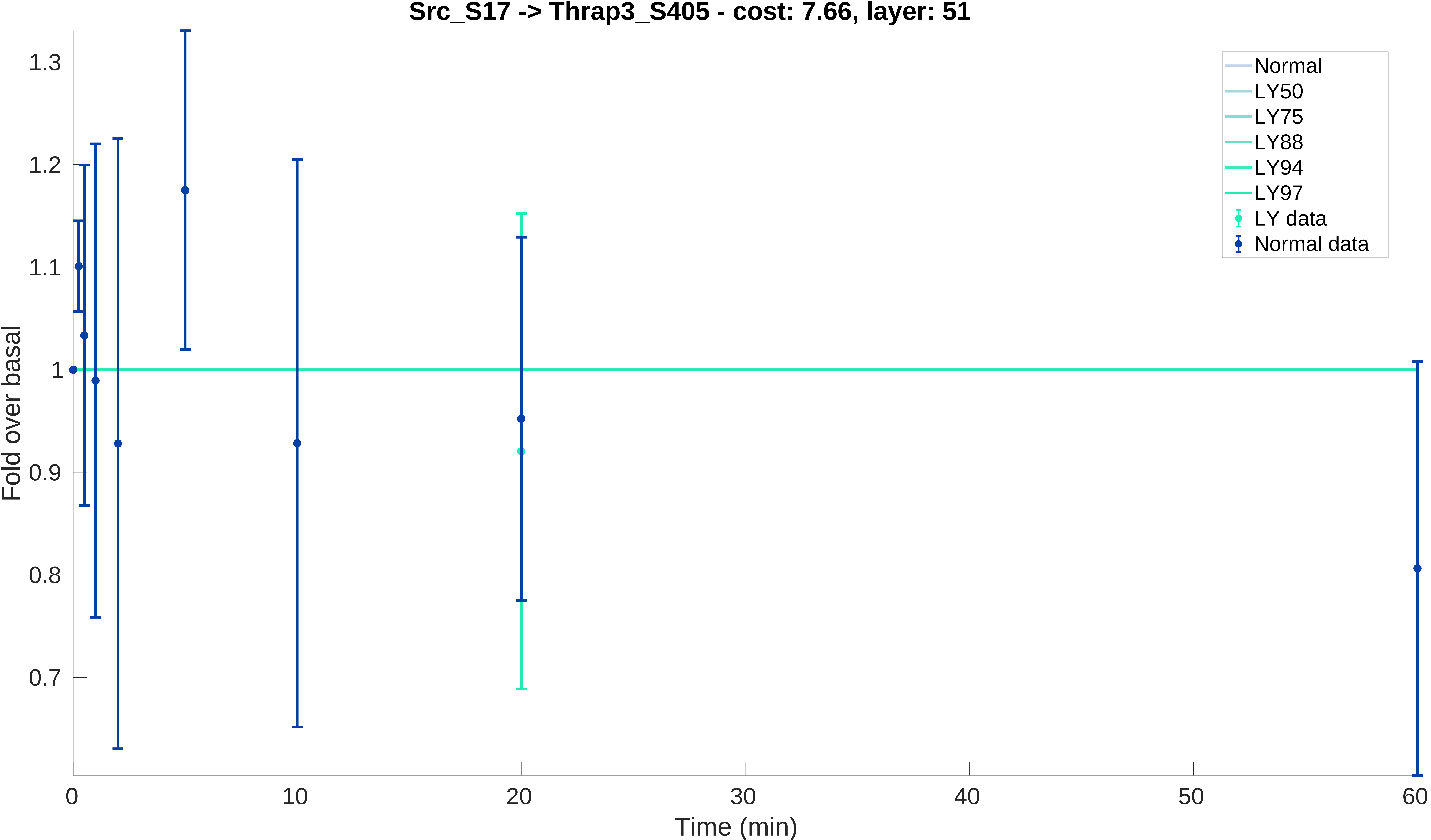
Src_S17 -> Thrap3_S405 - cost: 7.66, layer: 51
Fold over basal
1.3
1.2
1.1
1
0.9
0.8
0.7
0
10
20
30
40
50
60
Time (min)
Normal
LY50
LY75
LY88
LY94
LY97
LY data
Normal data

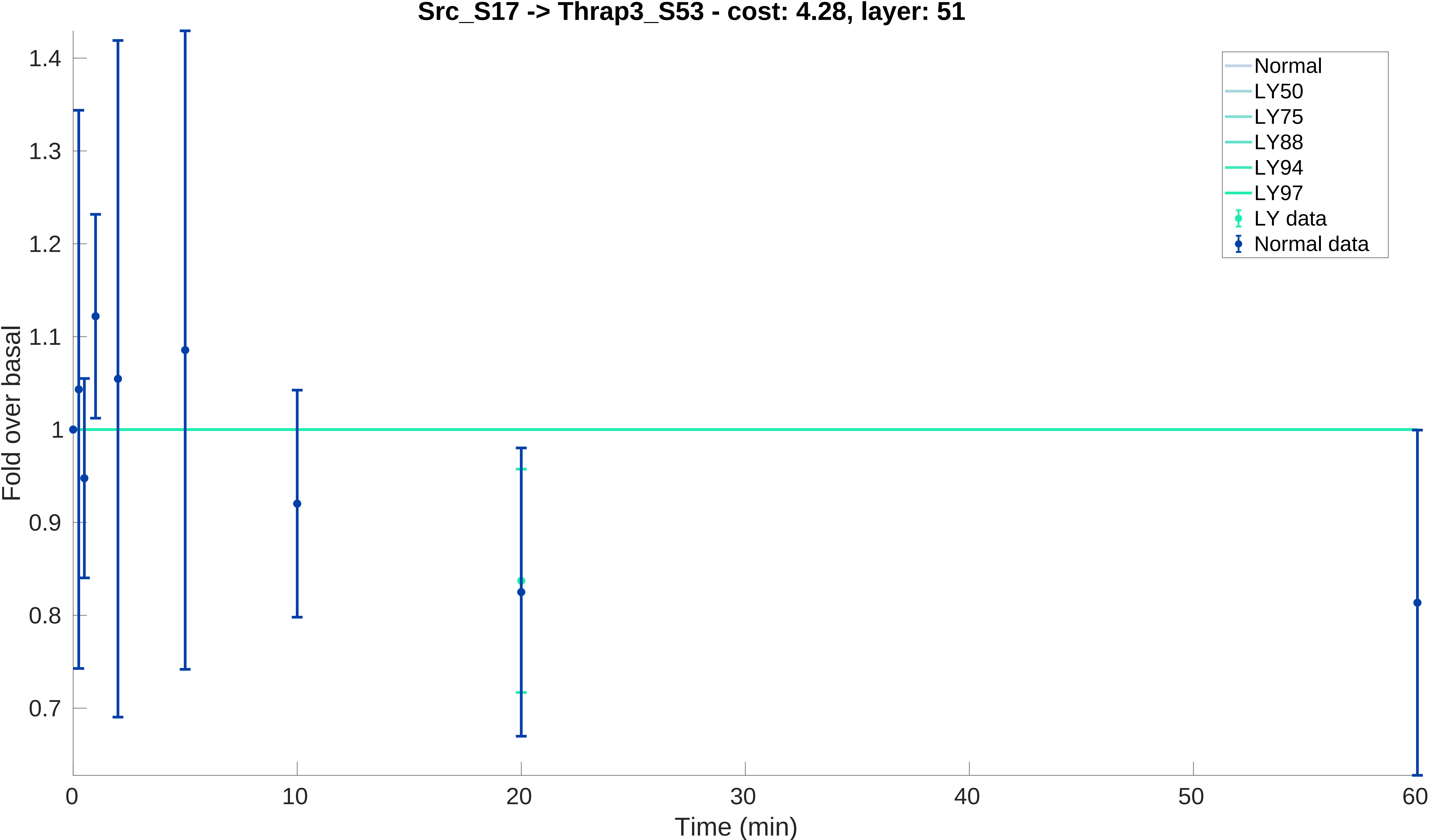

Src_S17 -> Thrap3_S53 - cost: 4.28, layer: 51
Fold over basal
1.4
1.3
1.2
1.1
1
0.9
0.8
0.7
0
10
20
30
40
50
60
Time (min)
Normal
LY50
LY75
LY88
LY94
LY97
LY data
Normal data

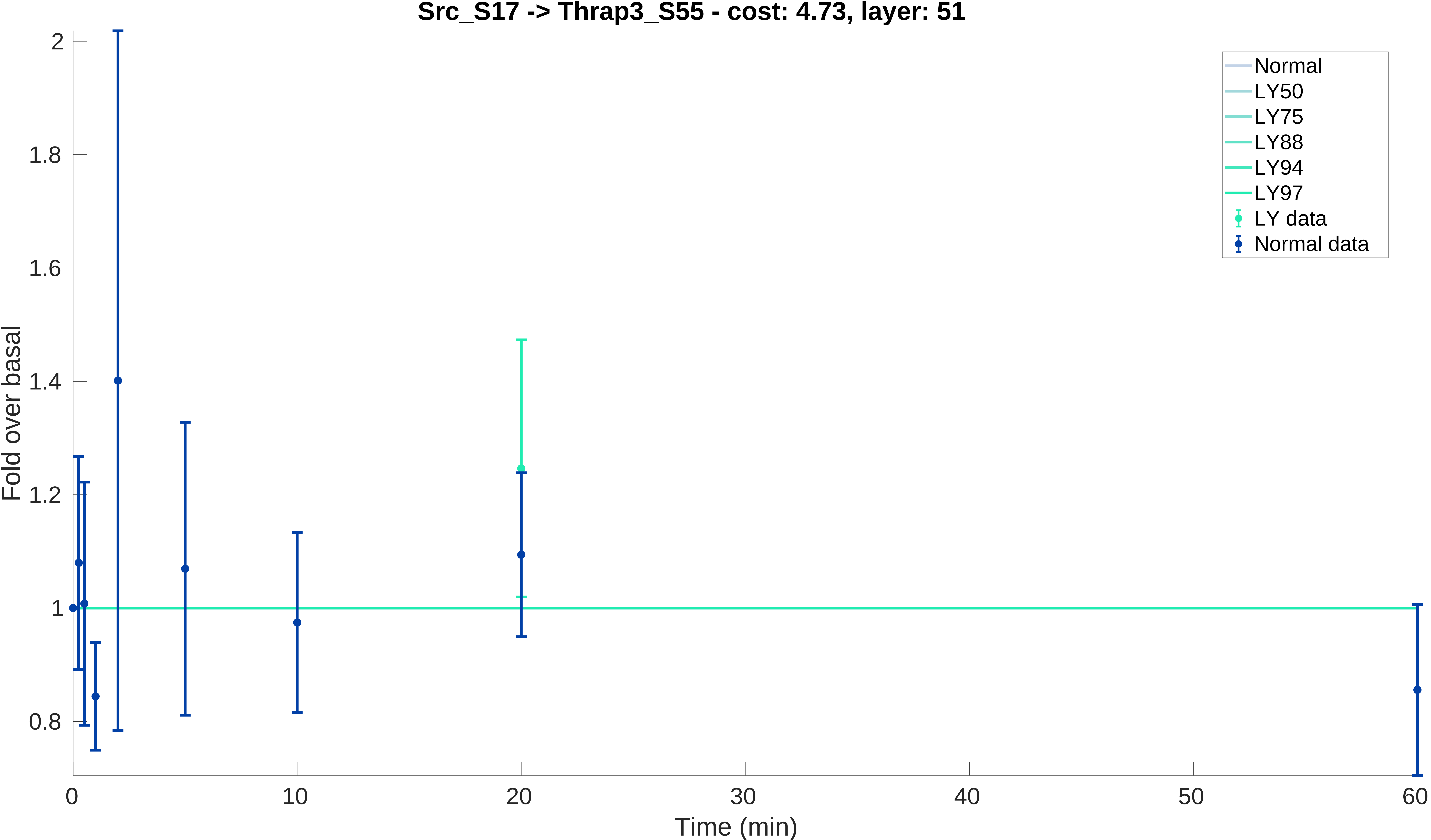
Src_S17 -> Thrap3_S55 - cost: 4.73, layer: 51
Fold over basal
Time (min)
0
10
20
30
40
50
60
0.8
1
1.2
1.4
1.6
1.8
2
Normal
LY50
LY75
LY88
LY94
LY97
LY data
Normal data

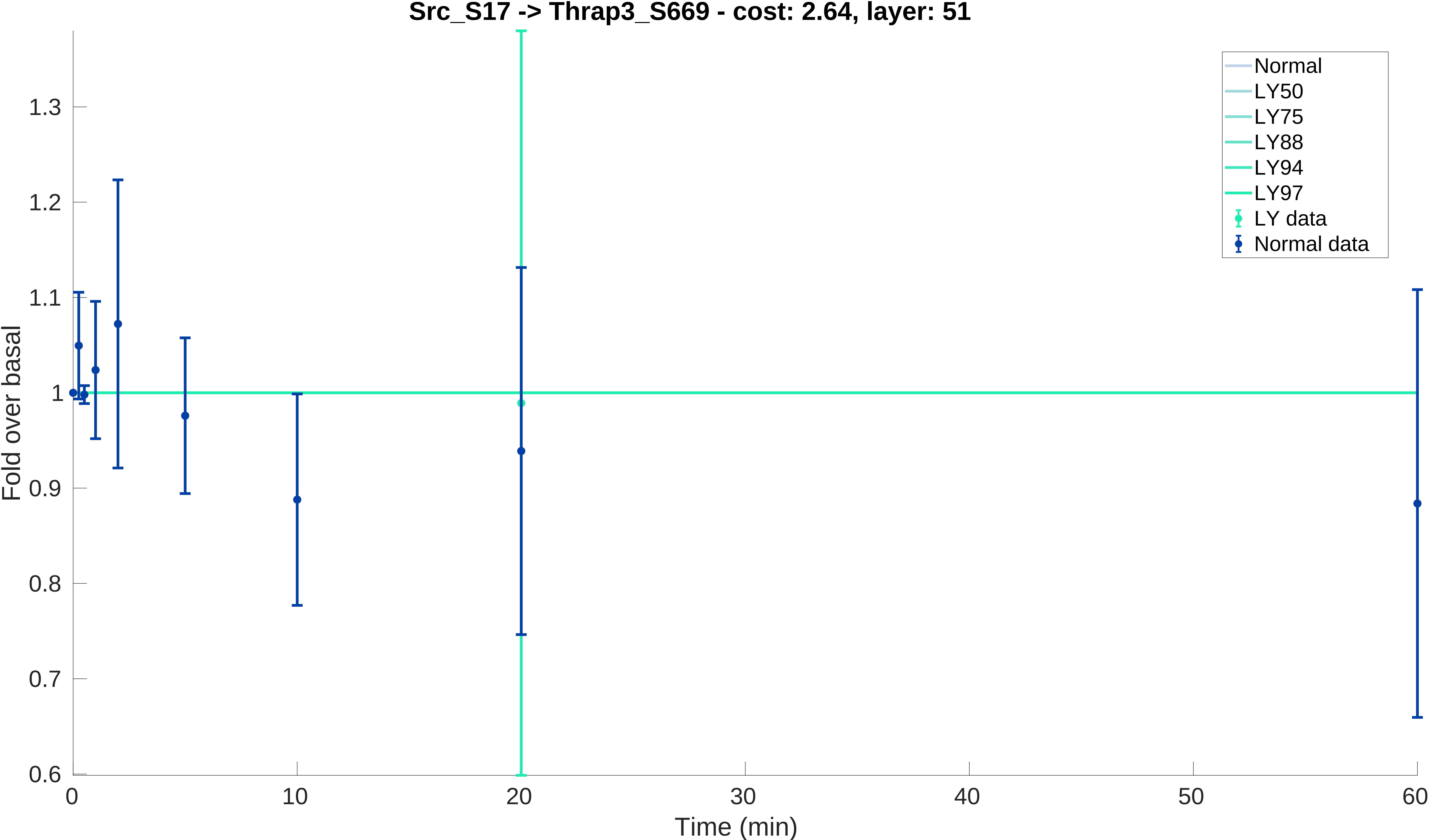

Src_S17 -> Thrap3_S669 - cost: 2.64, layer: 51
Fold over basal
Time (min)
1.3
1.2
1.1
1
0.9
0.8
0.7
0.6
0
10
20
30
40
50
60
Normal
LY50
LY75
LY88
LY94
LY97
LY data
Normal data

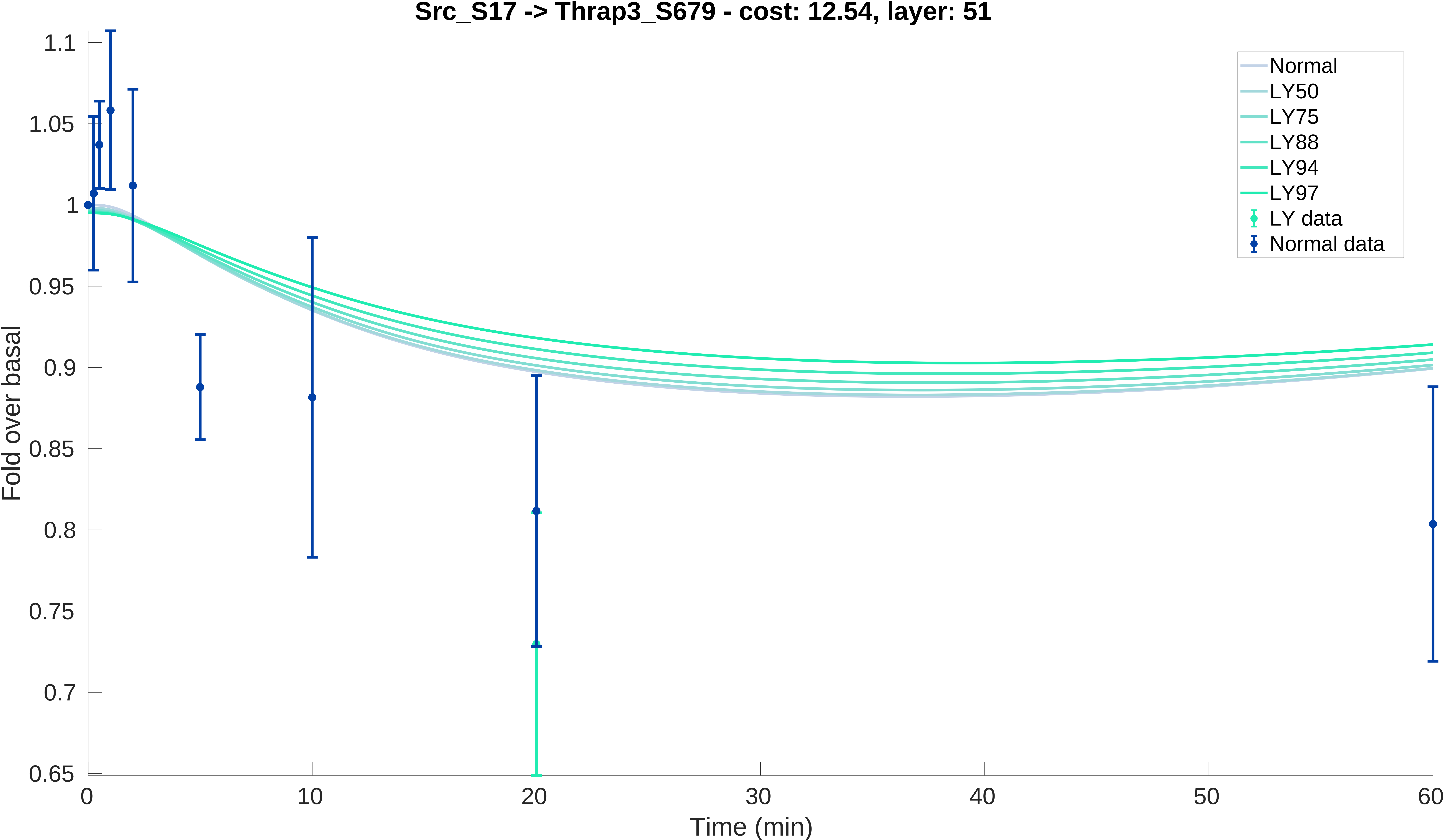

Src_S17 -> Thrap3_S679 - cost: 12.54, layer: 51
Fold over basal
Time (min)
1.1
1.05
1
0.95
0.9
0.85
0.8
0.75
0.7
0.65
0
10
20
30
40
50
60
Normal
LY50
LY75
LY88
LY94
LY97
LY data
Normal data

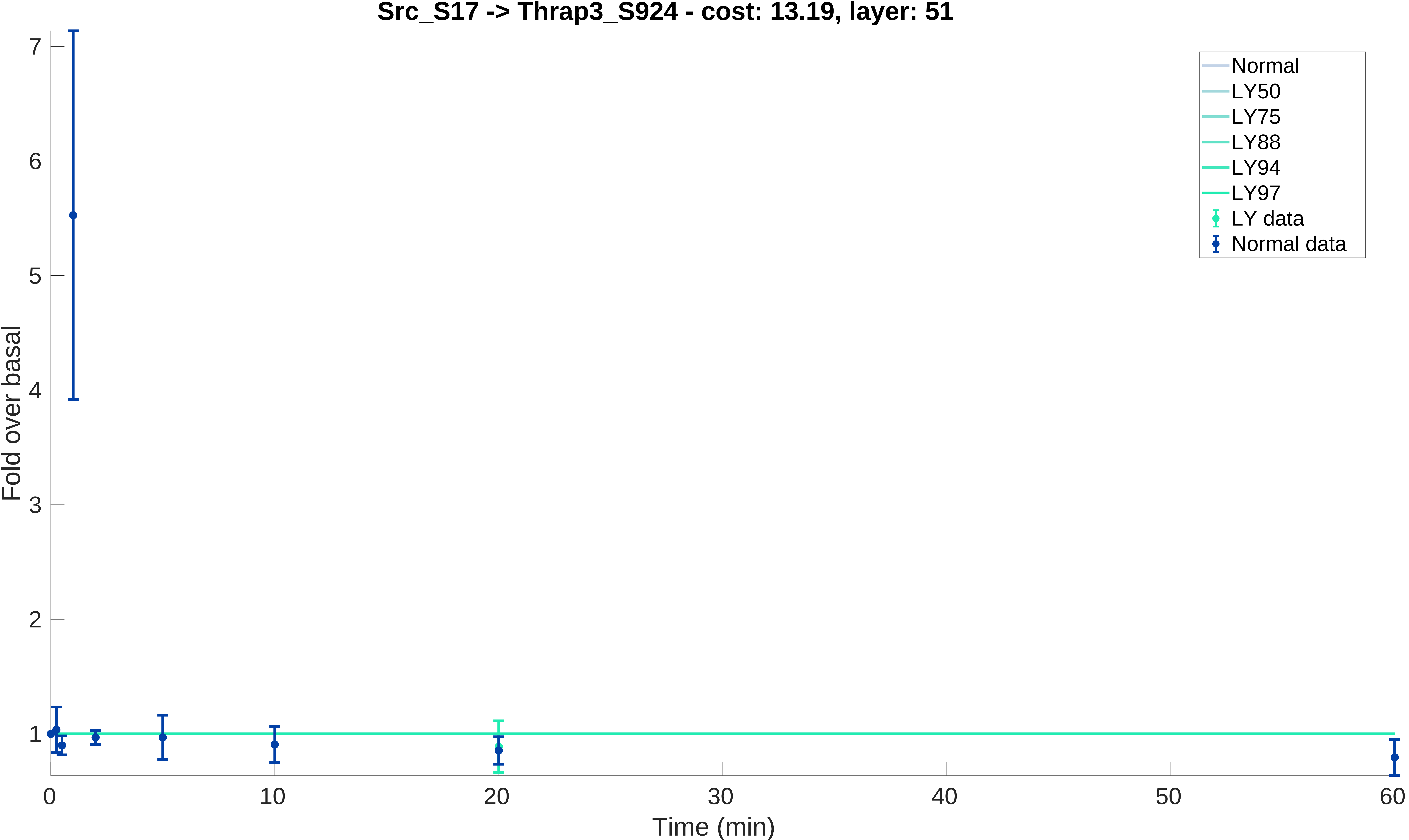

Src_S17 -> Thrap3_S924 - cost: 13.19, layer: 51
Fold over basal
Time (min)
Normal
LY50
LY75
LY88
LY94
LY97
LY data
Normal data
0
10
20
30
40
50
60
1
2
3
4
5
6
7

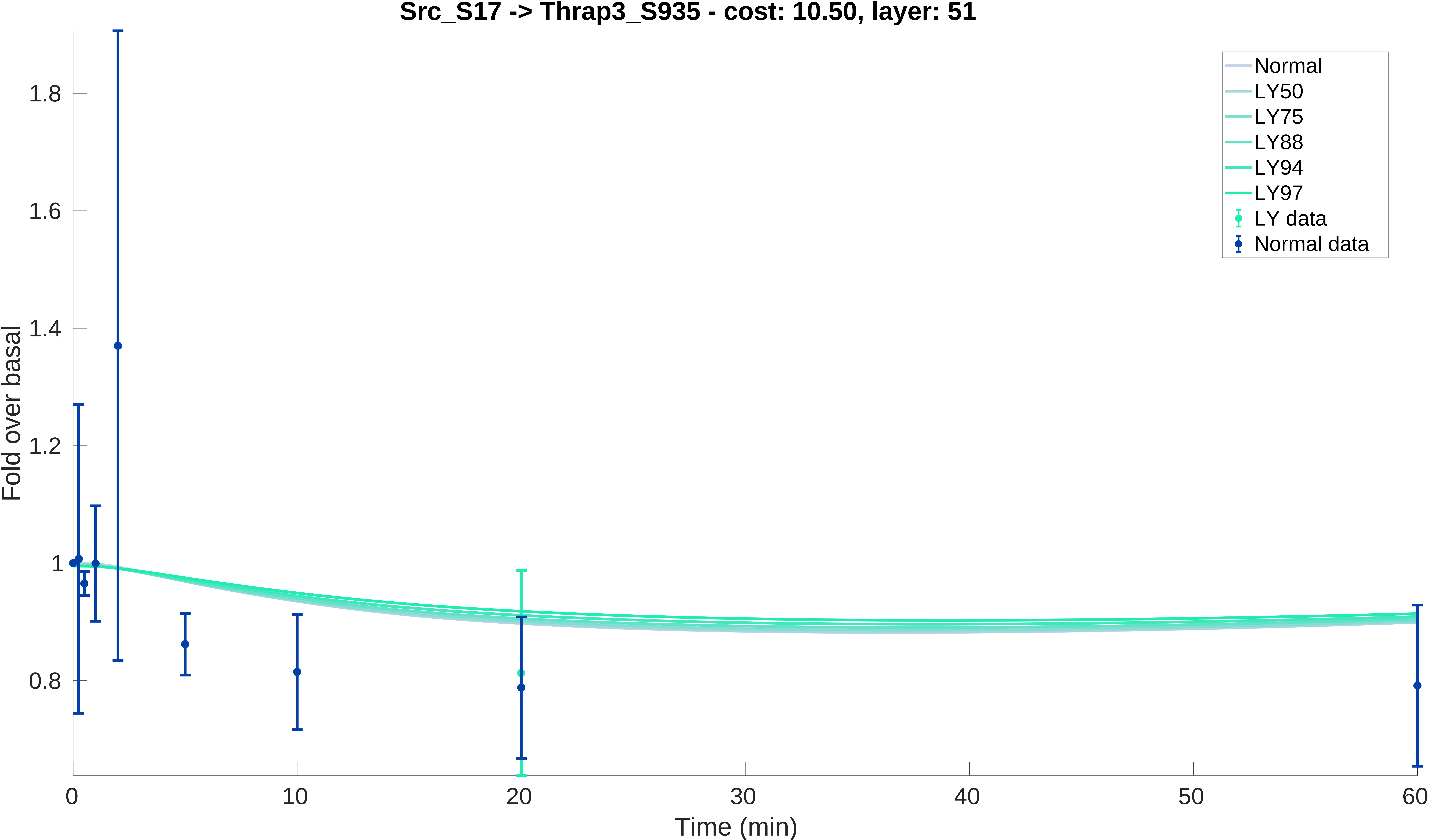

Src_S17 -> Thrap3_S935 - cost: 10.50, layer: 51
Normal
LY50
LY75
LY88
LY94
LY97
LY data
Normal data
Fold over basal
1.8
1.6
1.4
1.2
1
0.8
0
10
20
30
40
50
60
Time (min)

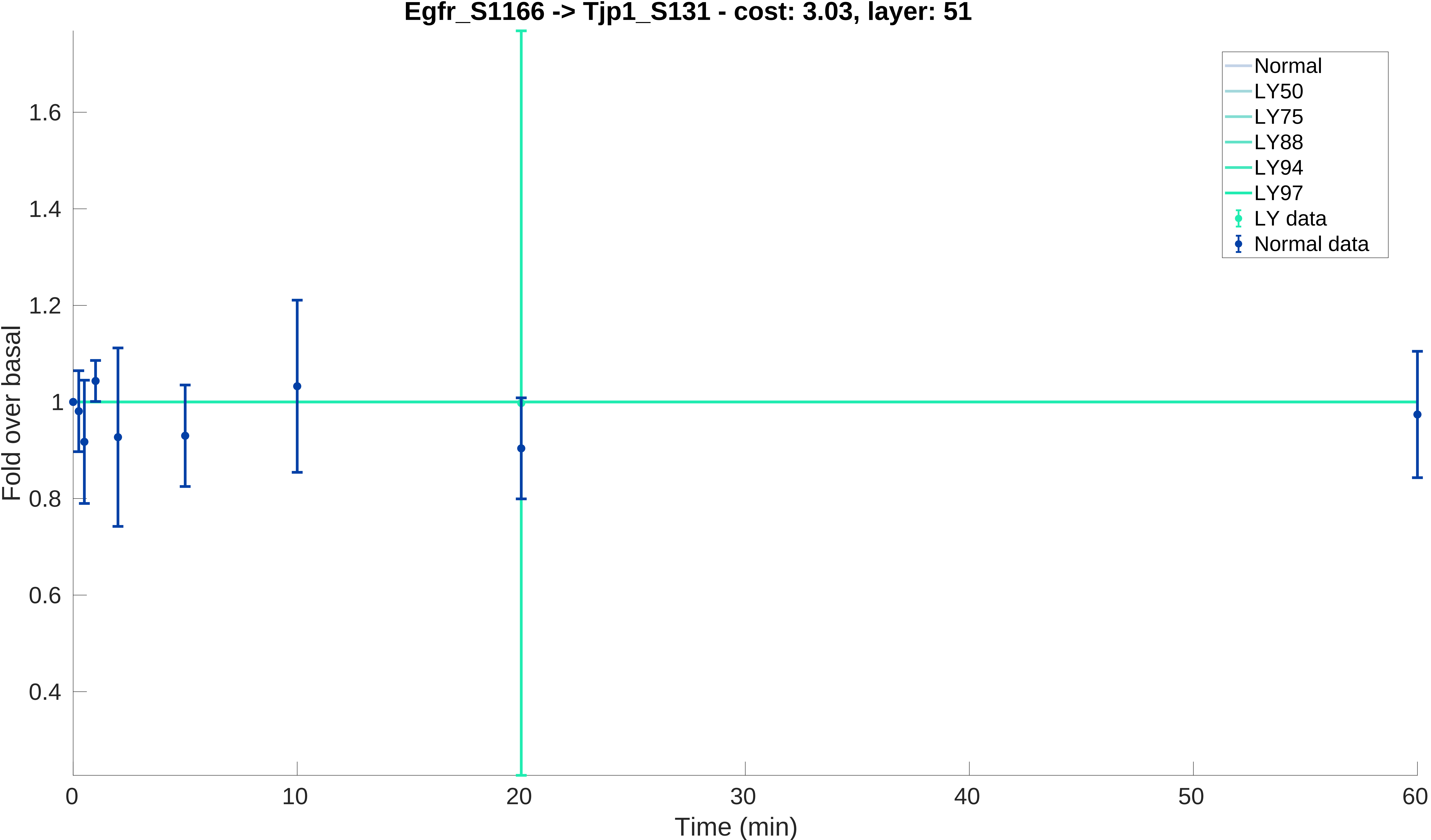

Egfr_S1166 -> Tjp1_S131 - cost: 3.03, layer: 51
Fold over basal
Time (min)
0.4
0.6
0.8
1
1.2
1.4
1.6
0
10
20
30
40
50
60
Normal
LY50
LY75
LY88
LY94
LY97
LY data
Normal data

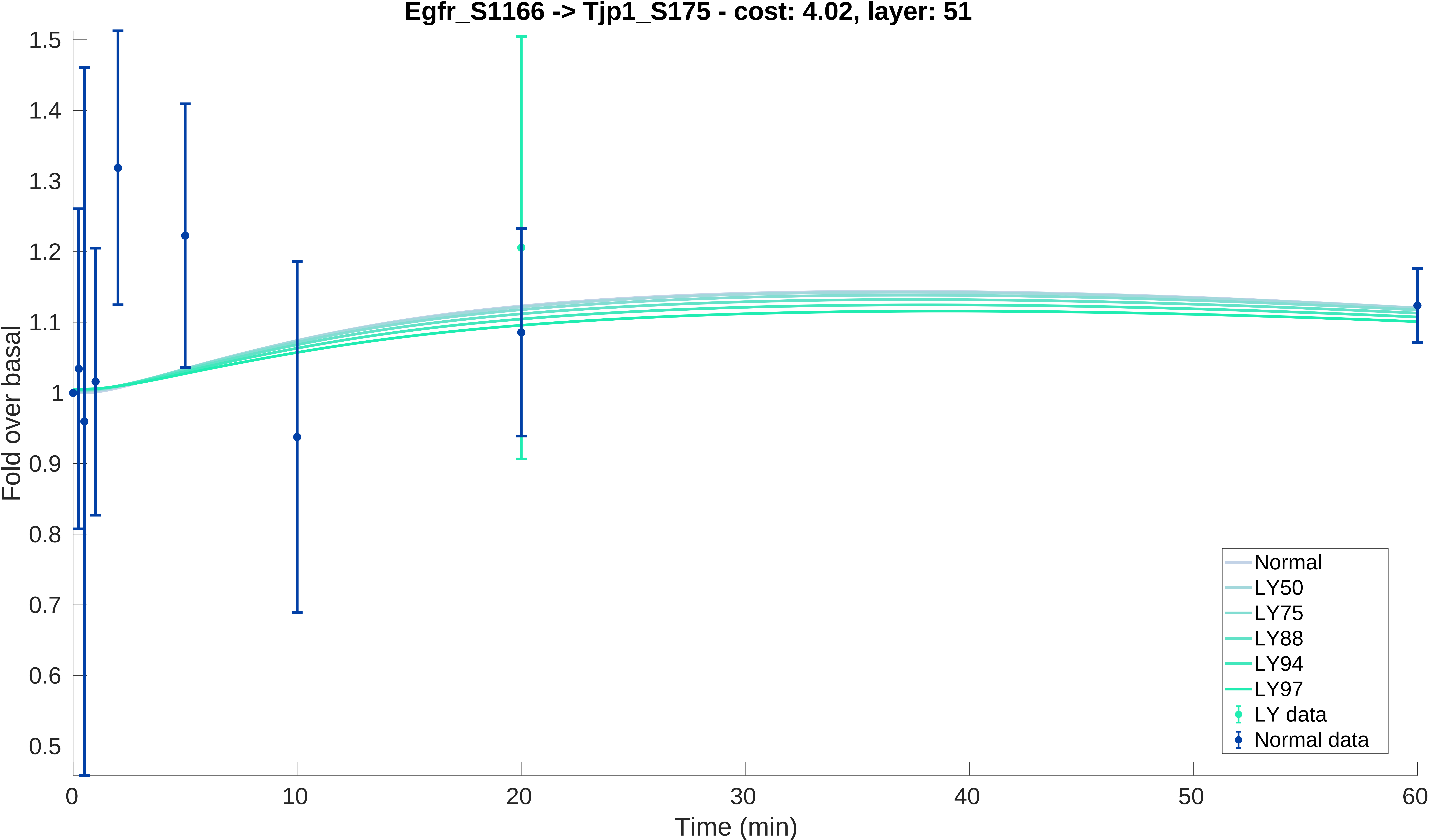

Egfr_S1166 -> Tjp1_S175 - cost: 4.02, layer: 51
Fold over basal
Time (min)
0
10
20
30
40
50
60
0.5
0.6
0.7
0.8
0.9
1
1.1
1.2
1.3
1.4
1.5
Normal
LY50
LY75
LY88
LY94
LY97
LY data
Normal data

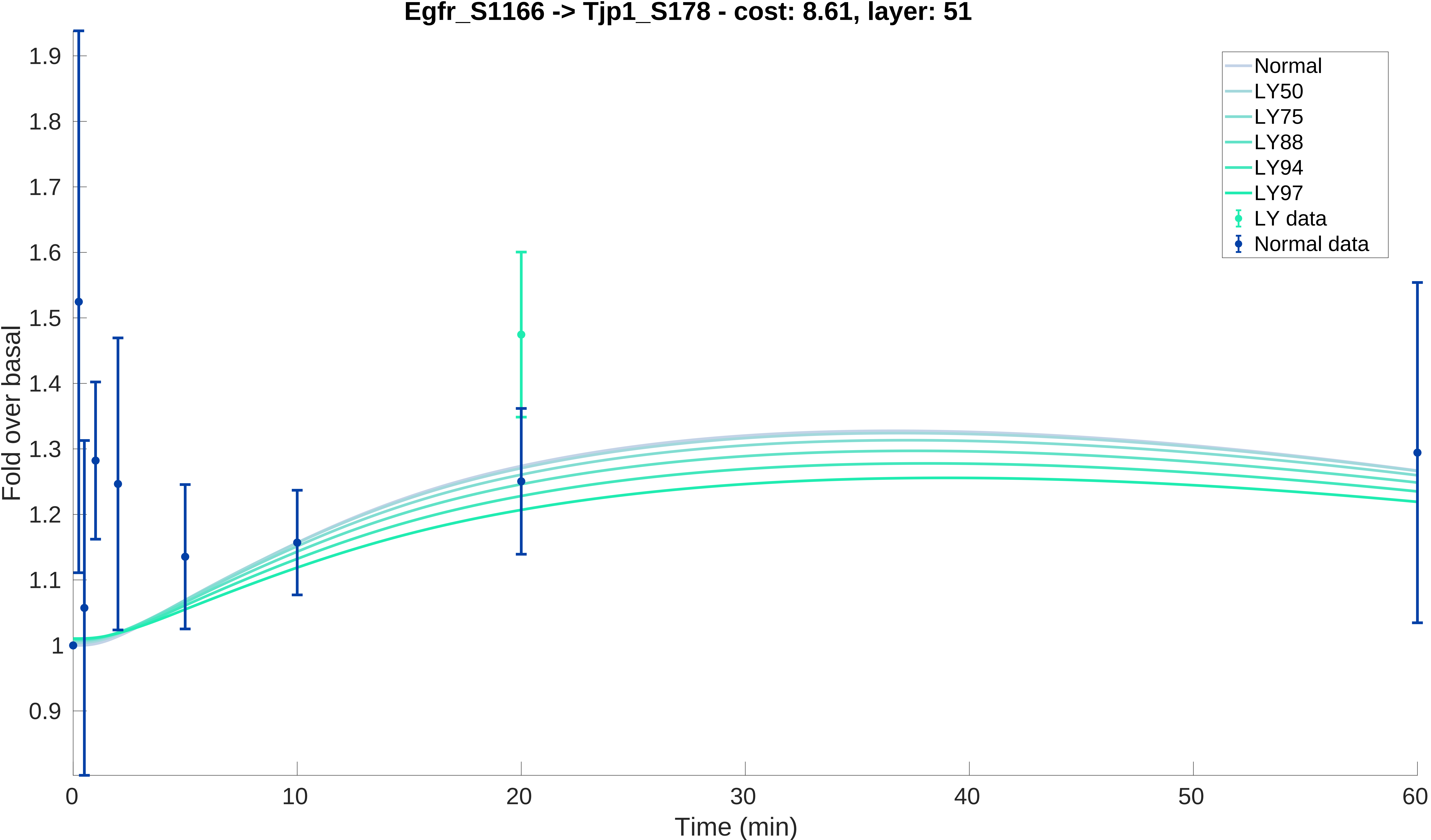

Egfr_S1166 -> Tjp1_S178 - cost: 8.61, layer: 51
Fold over basal
Time (min)
1.9
1.8
1.7
1.6
1.5
1.4
1.3
1.2
1.1
1
0.9
0
10
20
30
40
50
60
Normal
LY50
LY75
LY88
LY94
LY97
LY data
Normal data

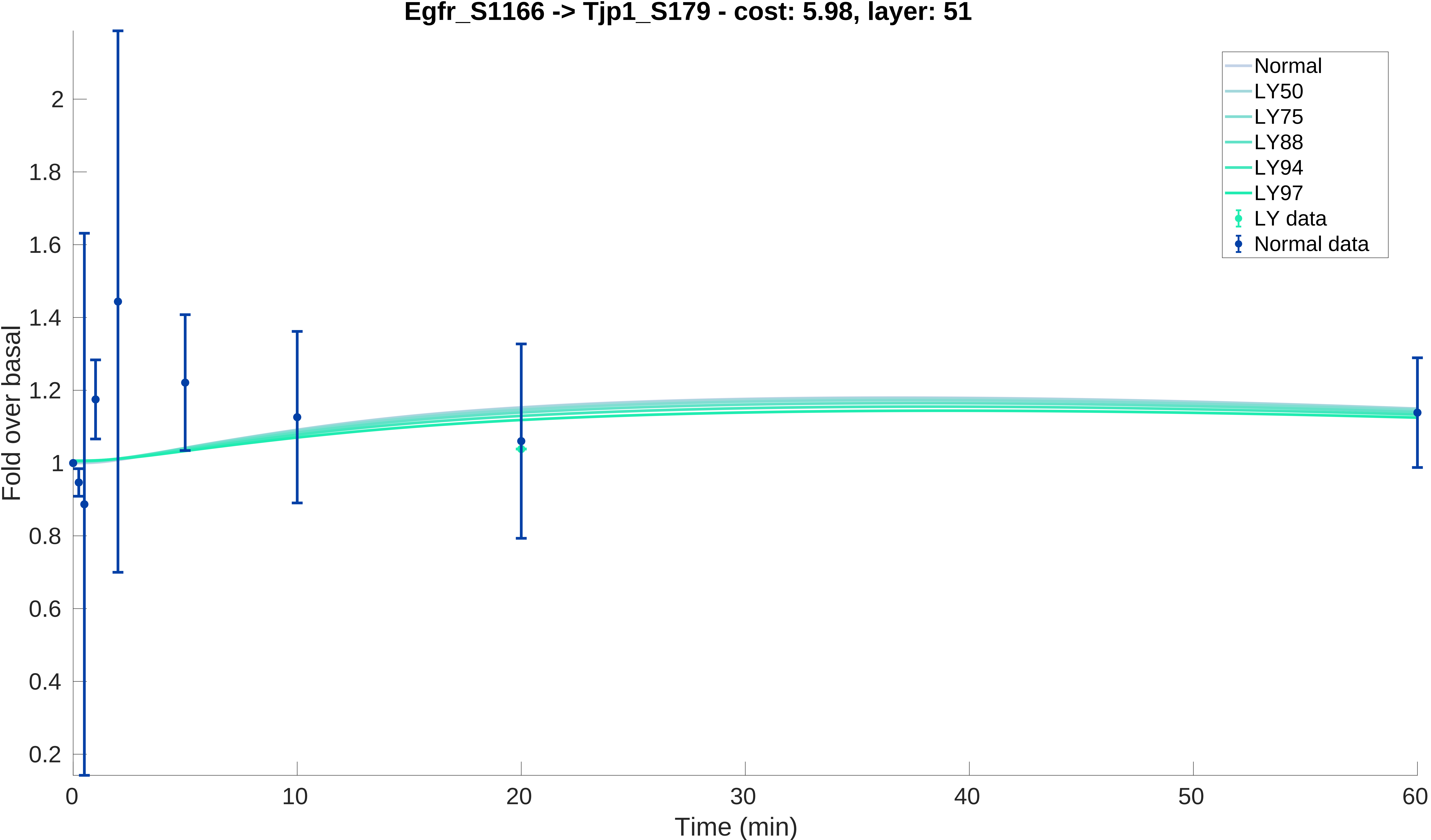

Egfr_S1166 -> Tjp1_S179 - cost: 5.98, layer: 51
Fold over basal
Time (min)
Normal
LY50
LY75
LY88
LY94
LY97
LY data
Normal data
0.2
0.4
0.6
0.8
1
1.2
1.4
1.6
1.8
2
0
10
20
30
40
50
60

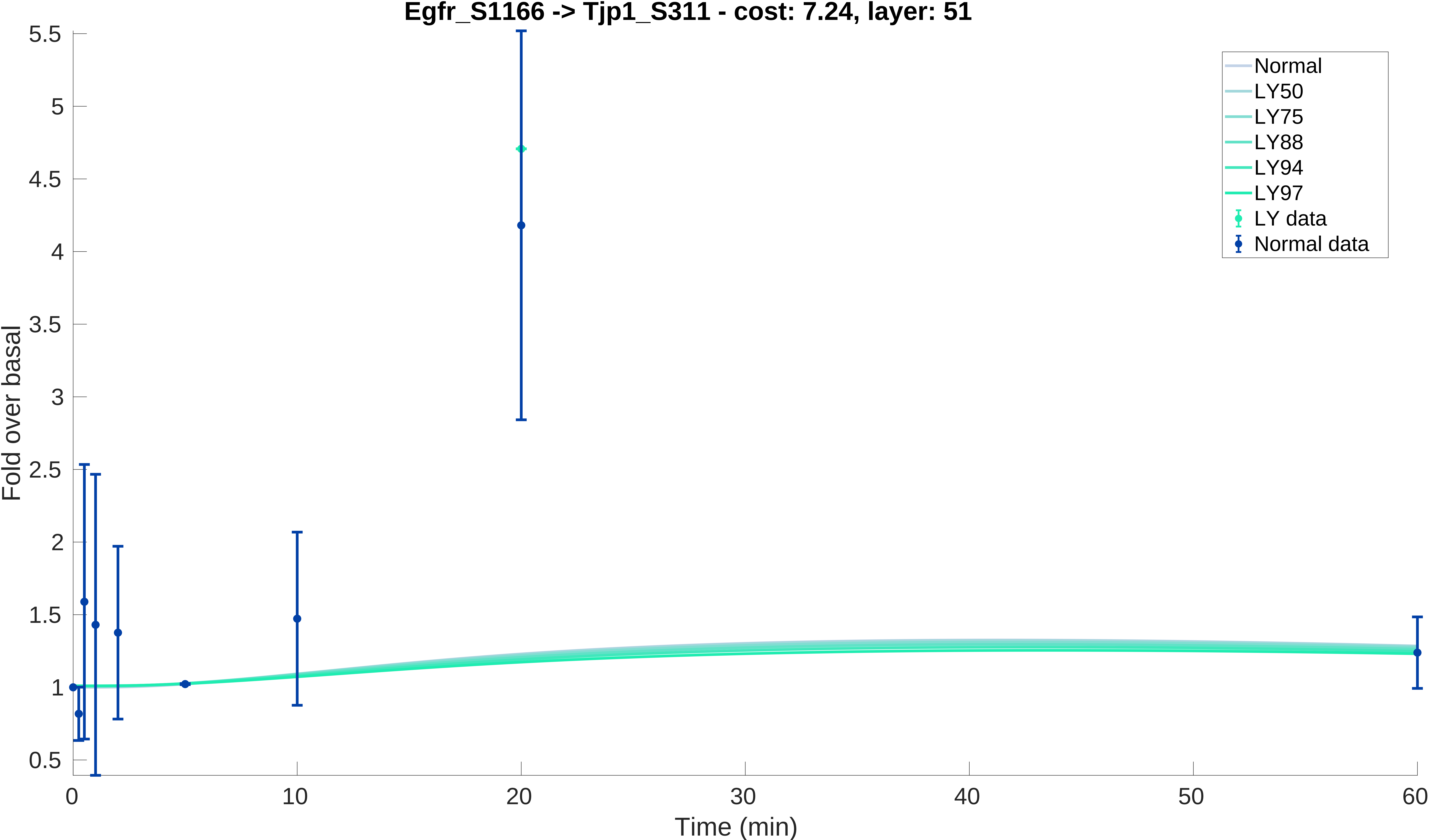
Egfr_S1166 -> Tjp1_S311 - cost: 7.24, layer: 51
Fold over basal
Time (min)
0.5
1
1.5
2
2.5
3
3.5
4
4.5
5
5.5
0
10
20
30
40
50
60
Normal
LY50
LY75
LY88
LY94
LY97
LY data
Normal data

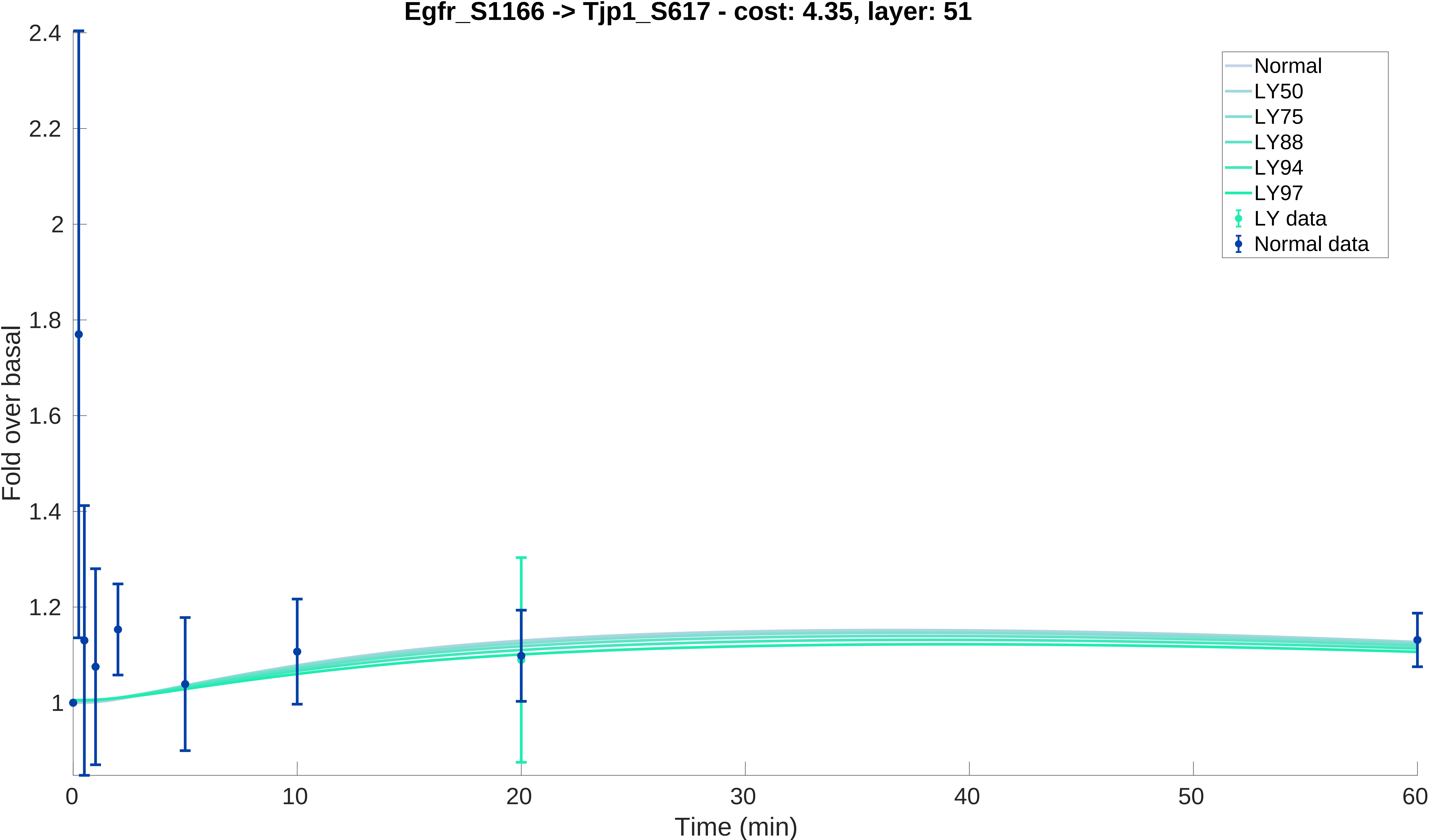

Egfr_S1166 -> Tjp1_S617 - cost: 4.35, layer: 51
Fold over basal
Time (min)
Normal
LY50
LY75
LY88
LY94
LY97
LY data
Normal data
2.4
2.2
2
1.8
1.6
1.4
1.2
1
0
10
20
30
40
50
60

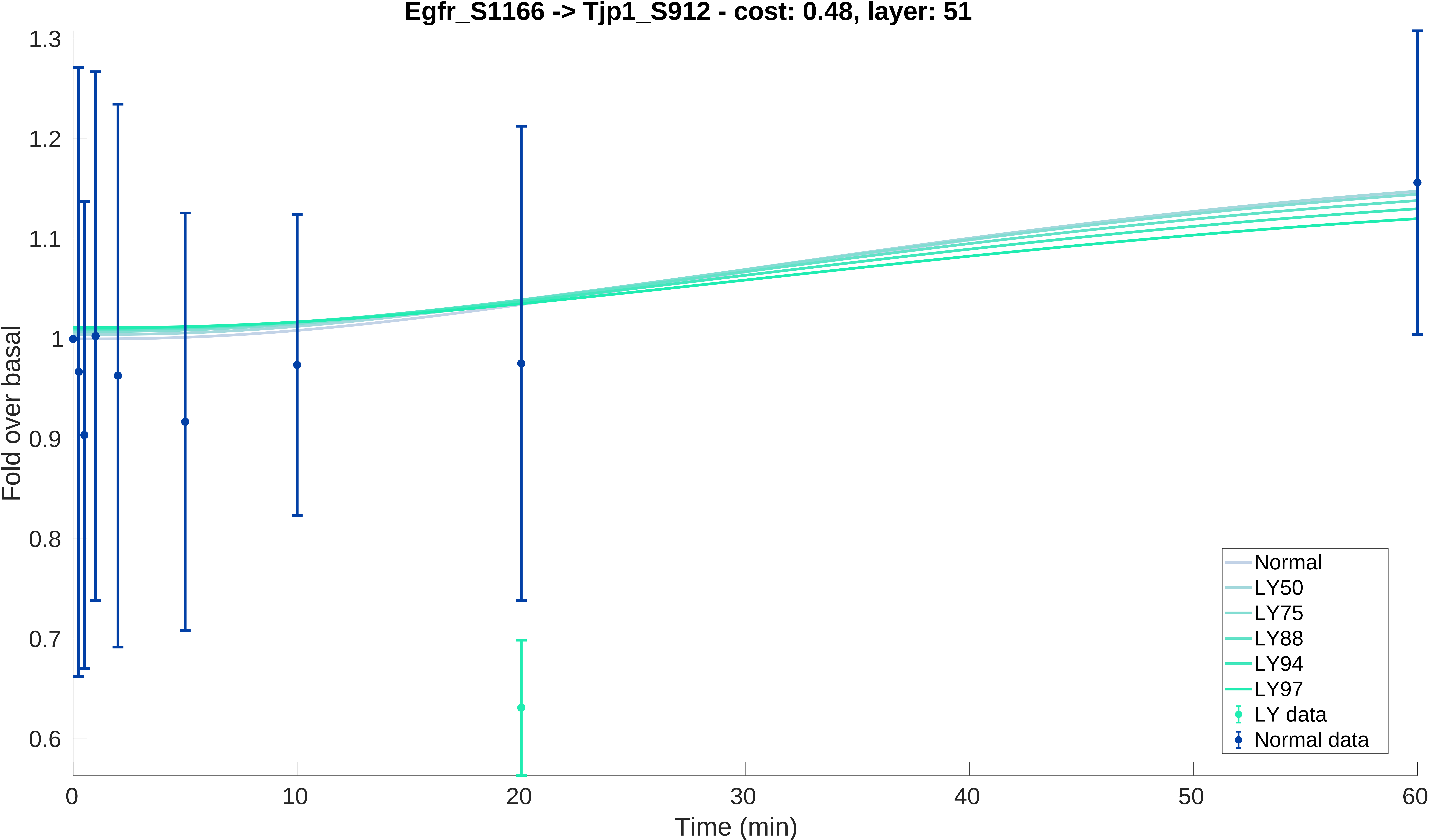

Egfr_S1166 -> Tjp1_S912 - cost: 0.48, layer: 51
1.3
1.2
1.1
1
0.9
0.8
0.7
0.6
Fold over basal
0
10
20
30
40
50
60
Time (min)
Normal
LY50
LY75
LY88
LY94
LY97
LY data
Normal data

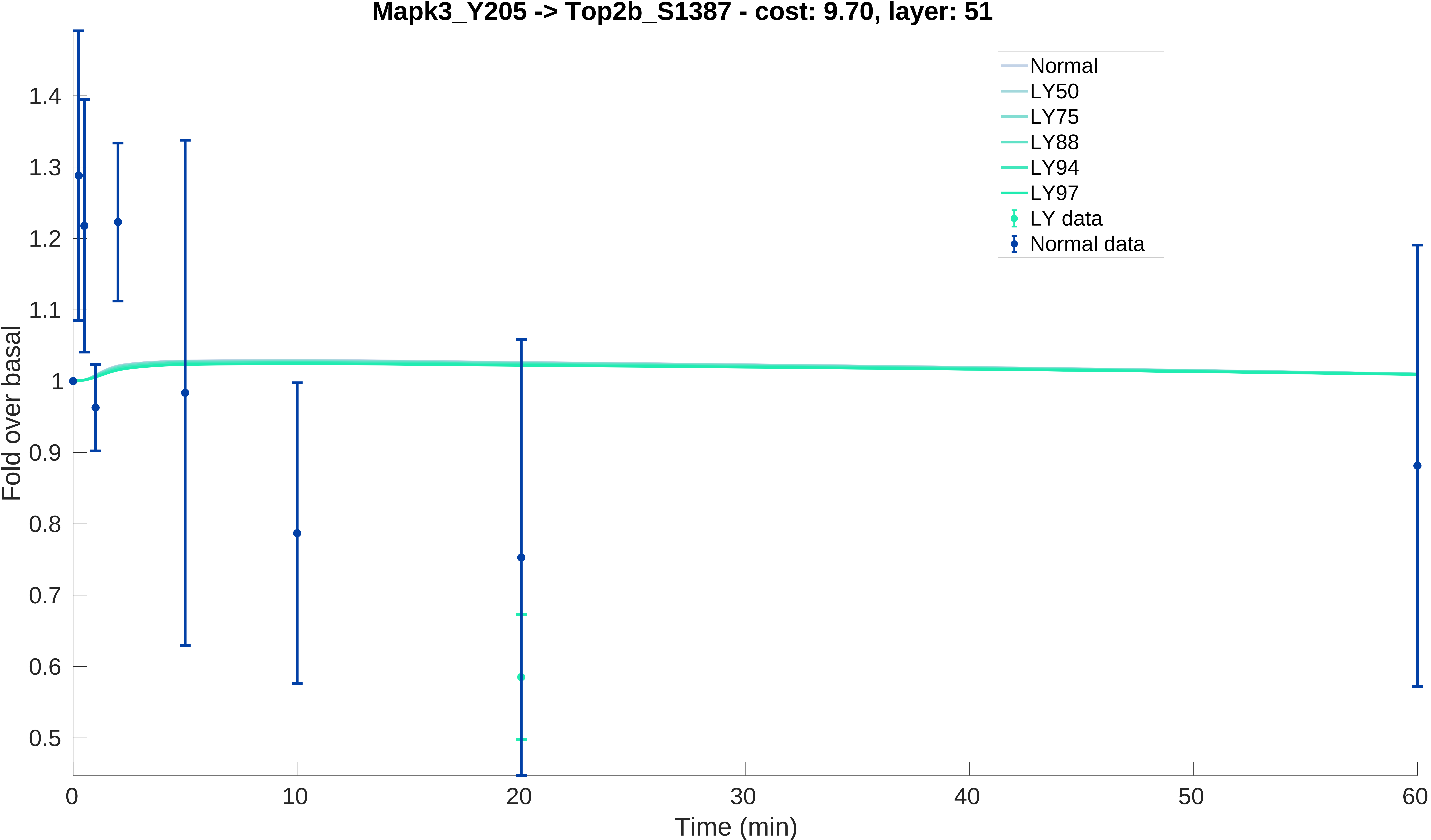

Mapk3_Y205 -> Top2b_S1387 - cost: 9.70, layer: 51
Normal
LY50
LY75
LY88
LY94
LY97
LY data
Normal data
Fold over basal
Time (min)
0
10
20
30
40
50
60
0.5
0.6
0.7
0.8
0.9
1
1.1
1.2
1.3
1.4

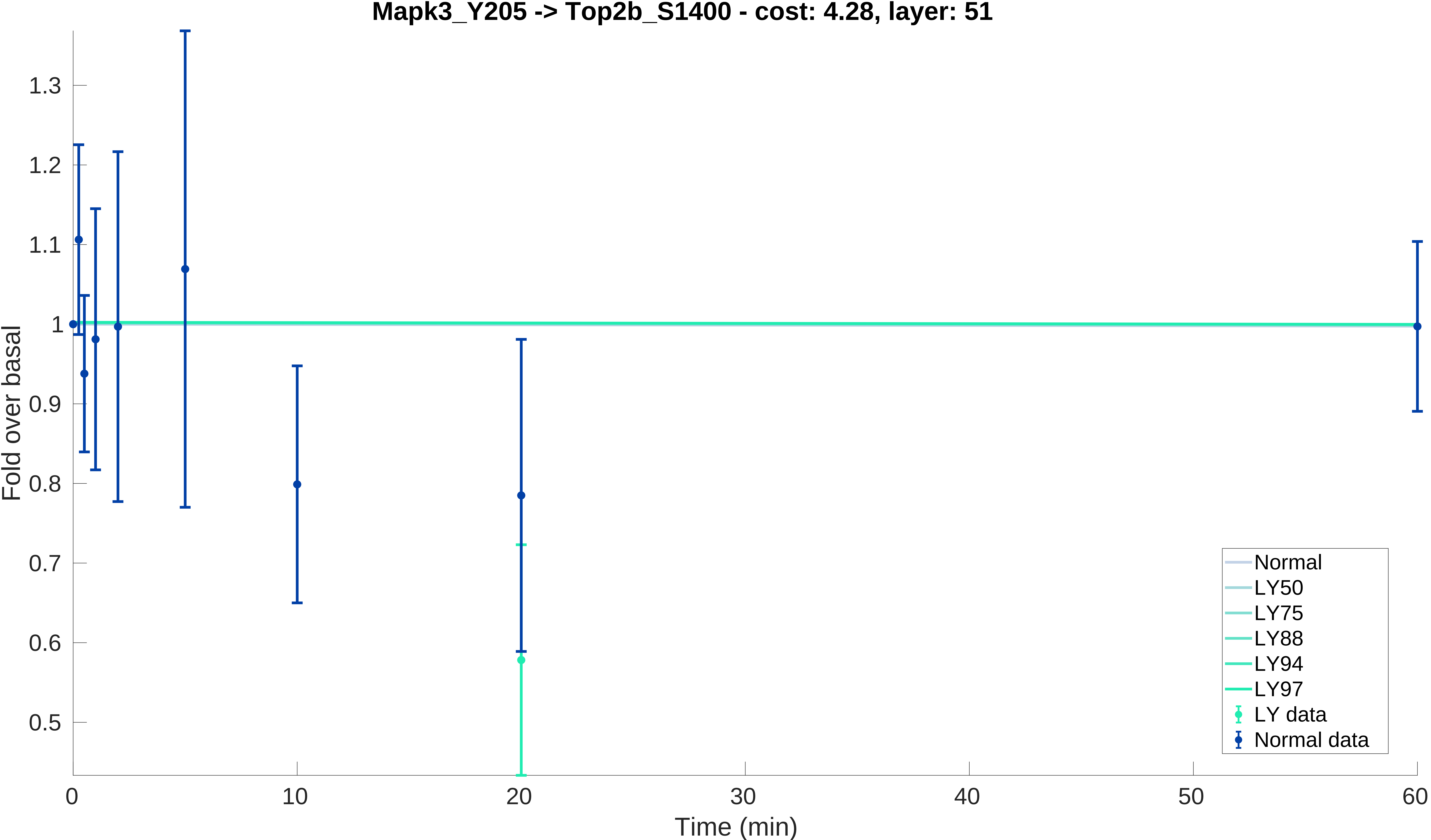

Mapk3_Y205 -> Top2b_S1400 - cost: 4.28, layer: 51
Fold over basal
1.3
1.2
1.1
1
0.9
0.8
0.7
0.6
0.5
0
10
20
30
40
50
60
Time (min)
Normal
LY50
LY75
LY88
LY94
LY97
LY data
Normal data

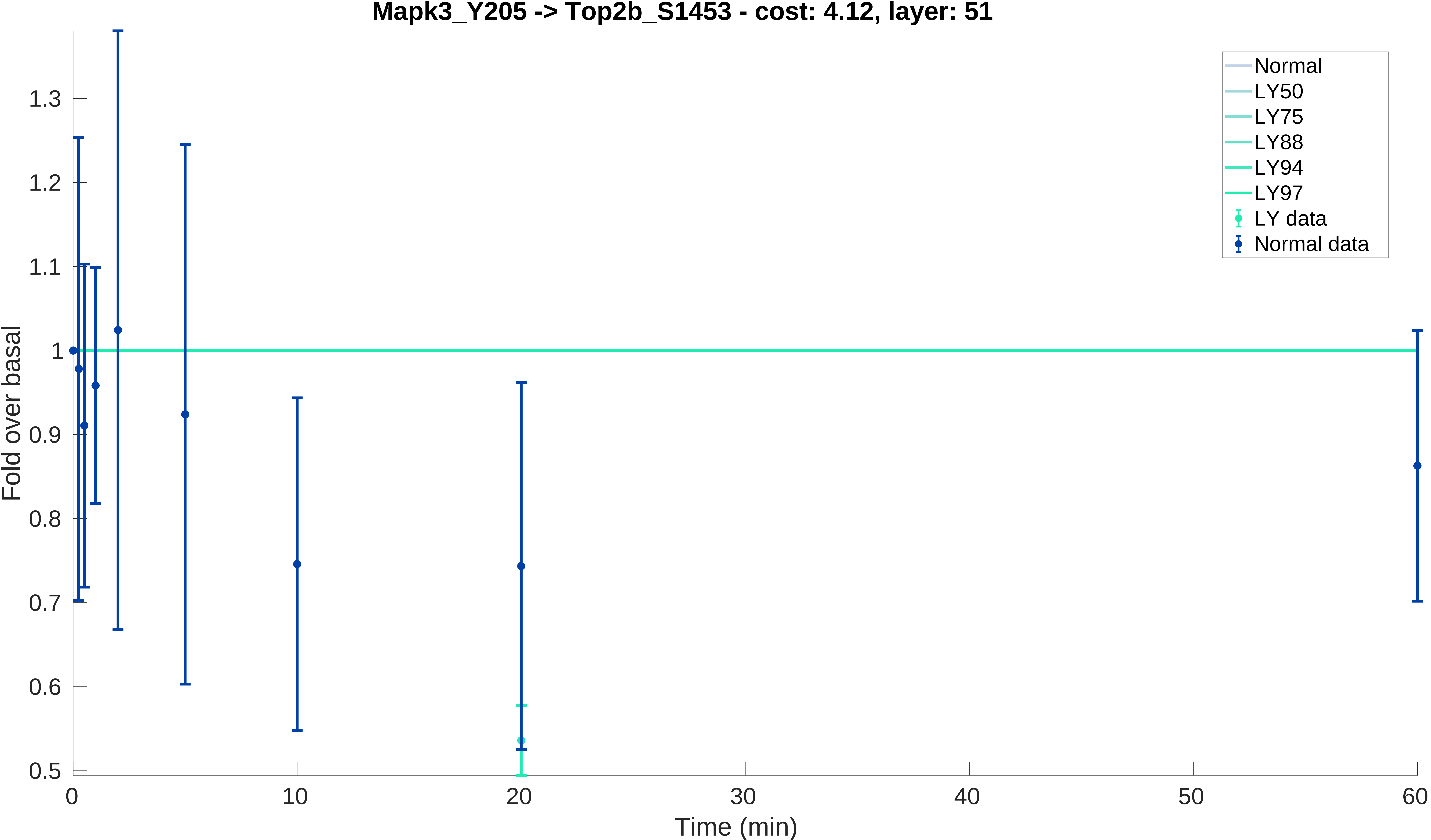

Mapk3_Y205 -> Top2b_S1453 - cost: 4.12, layer: 51
Normal
LY50
LY75
LY88
LY94
LY97
LY data
Normal data
Fold over basal
1.3
1.2
1.1
1
0.9
0.8
0.7
0.6
0.5
0
10
20
30
40
50
60
Time (min)

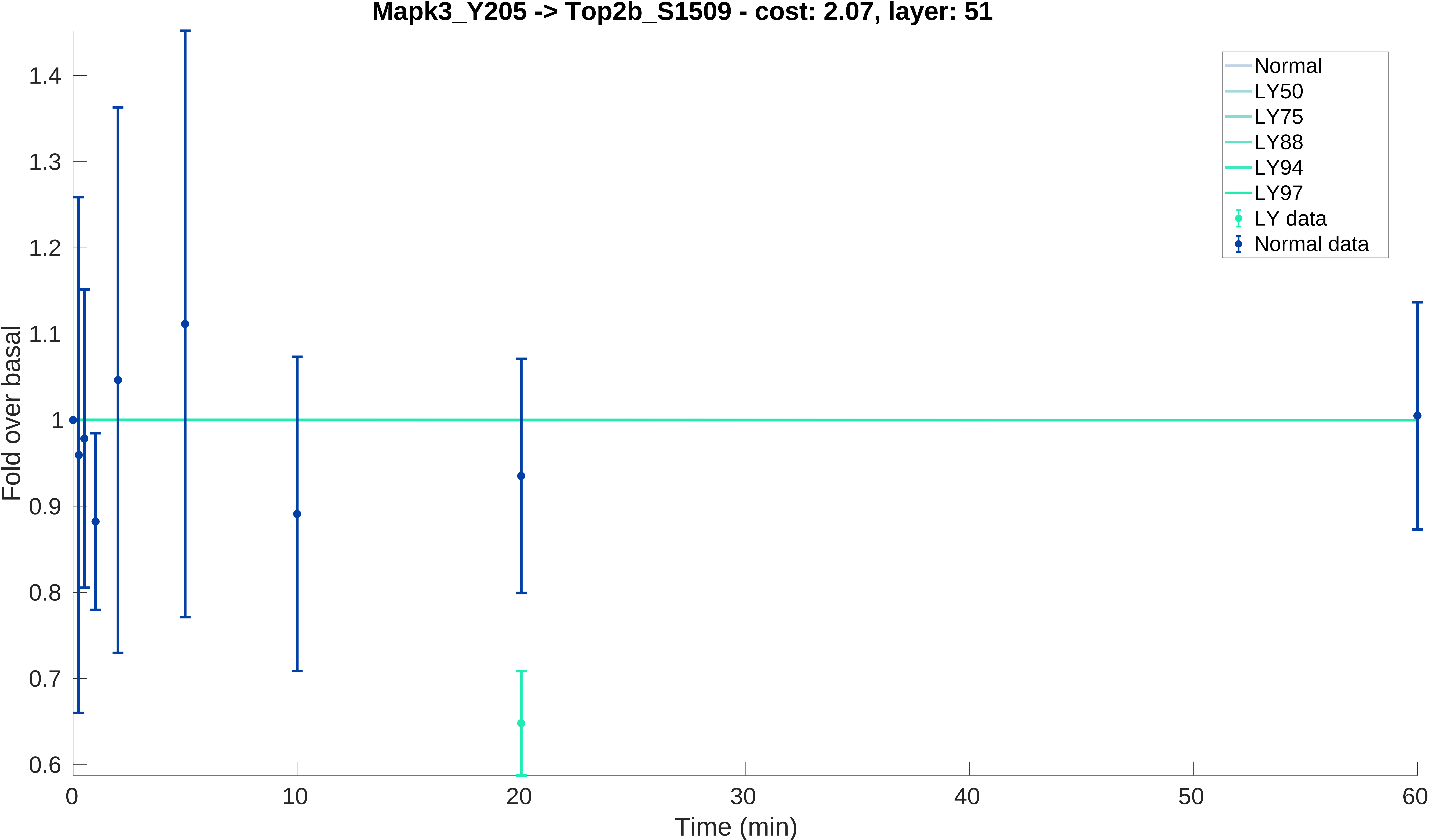

Mapk3_Y205 -> Top2b_S1509 - cost: 2.07, layer: 51
Normal
LY50
LY75
LY88
LY94
LY97
LY data
Normal data
Fold over basal
Time (min)
0.6
0.7
0.8
0.9
1
1.1
1.2
1.3
1.4
0
10
20
30
40
50
60

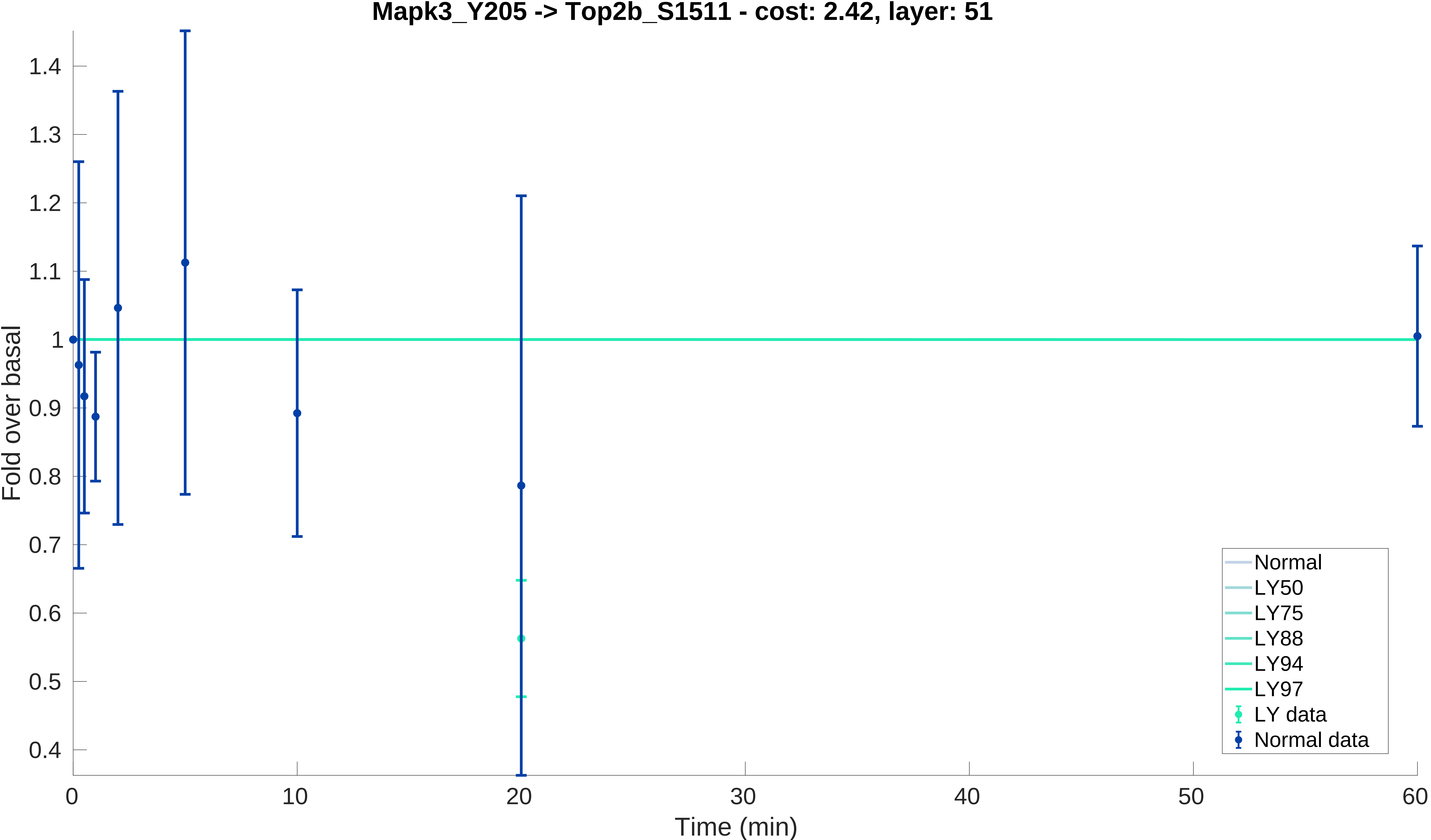
Mapk3_Y205 -> Top2b_S1511 - cost: 2.42, layer: 51
Fold over basal
Time (min)
1.4
1.3
1.2
1.1
1
0.9
0.8
0.7
0.6
0.5
0.4
0
10
20
30
40
50
60
Normal
LY50
LY75
LY88
LY94
LY97
LY data
Normal data

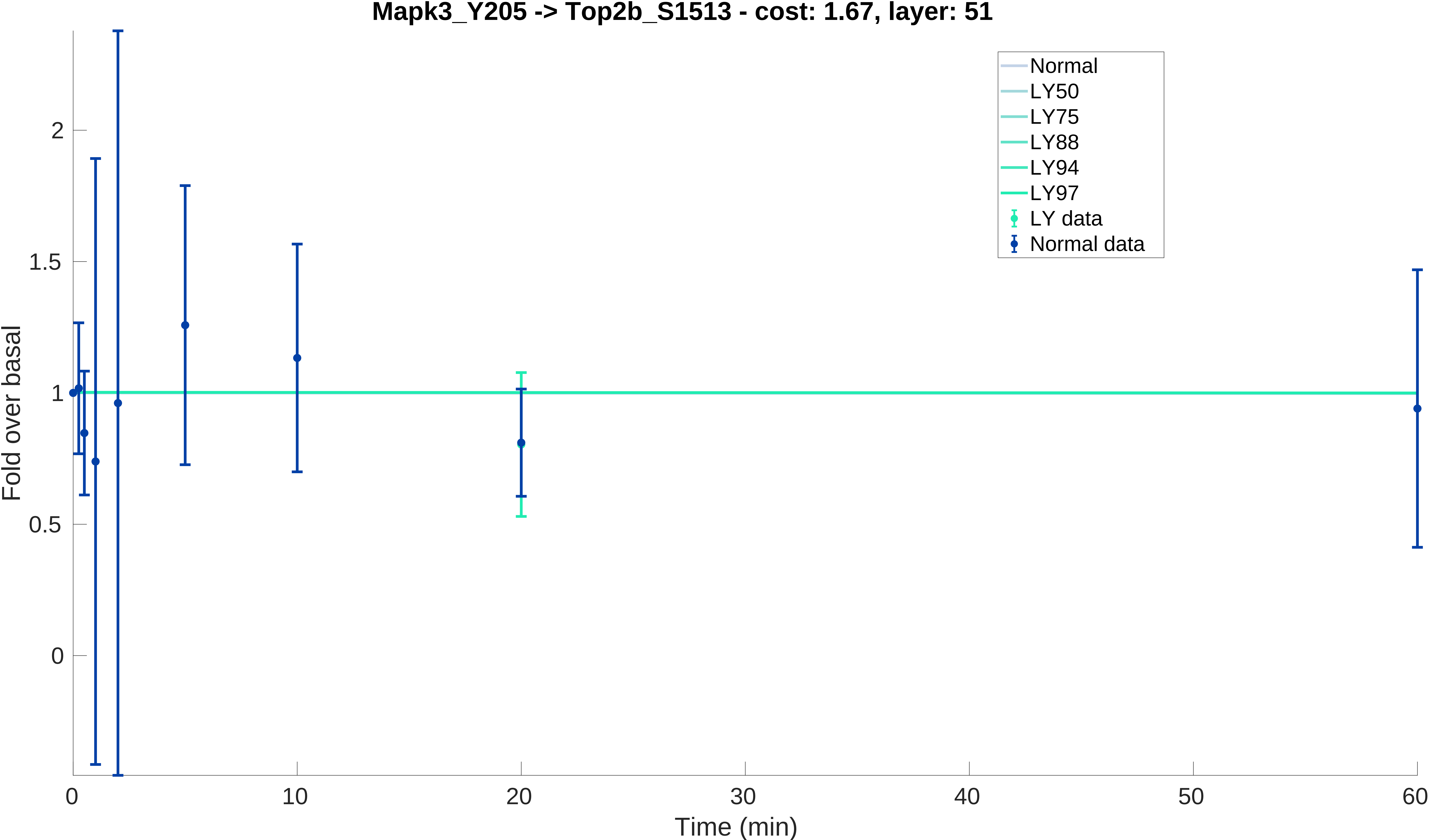

Mapk3_Y205 -> Top2b_S1513 - cost: 1.67, layer: 51
Normal
LY50
LY75
LY88
LY94
LY97
LY data
Normal data
Fold over basal
2
1.5
1
0.5
0
0
10
20
30
40
50
60
Time (min)

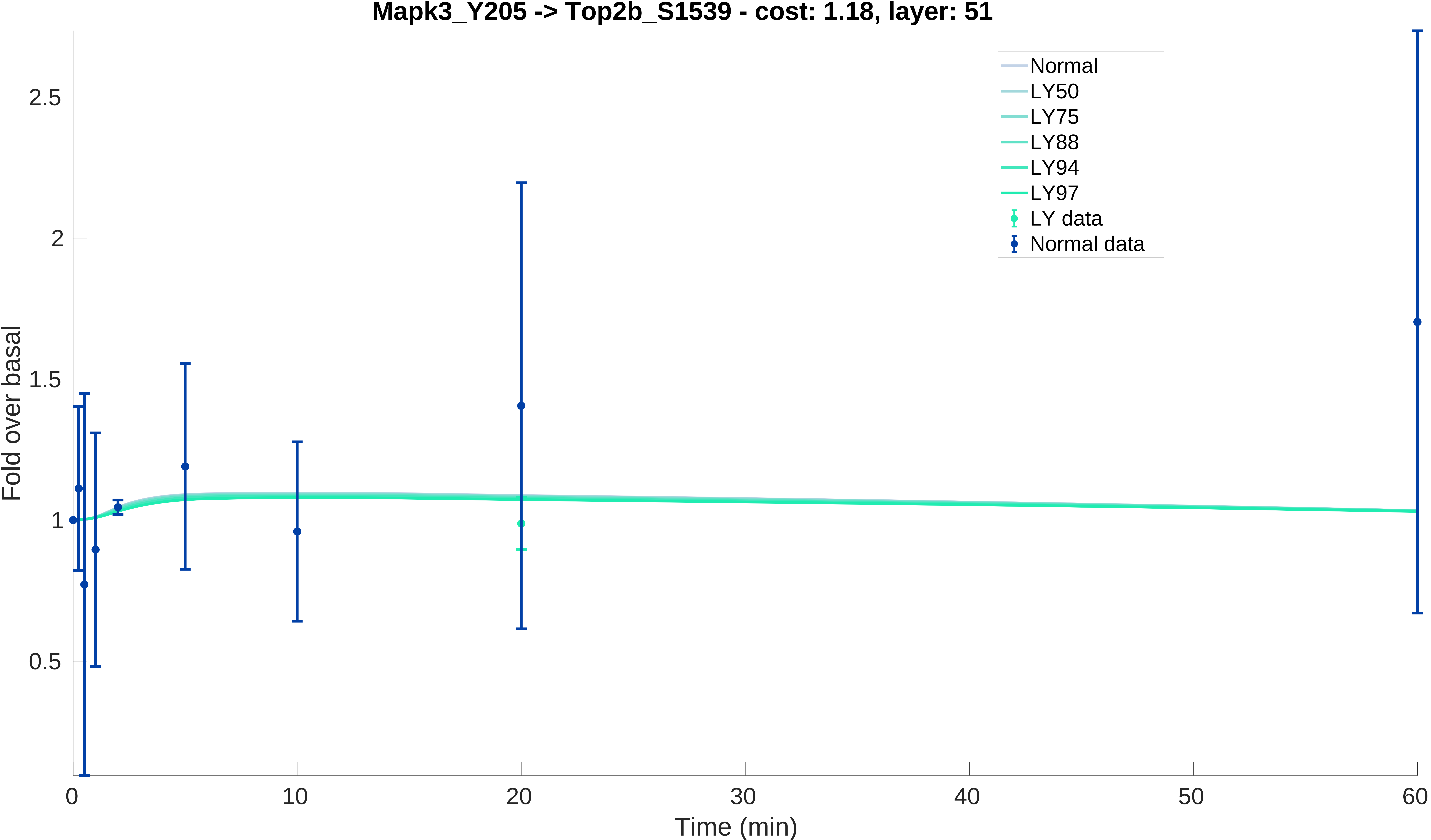

Mapk3_Y205 -> Top2b_S1539 - cost: 1.18, layer: 51
Normal
LY50
LY75
LY88
LY94
LY97
LY data
Normal data
Fold over basal
2.5
2
1.5
1
0.5
0
10
20
30
40
50
60
Time (min)

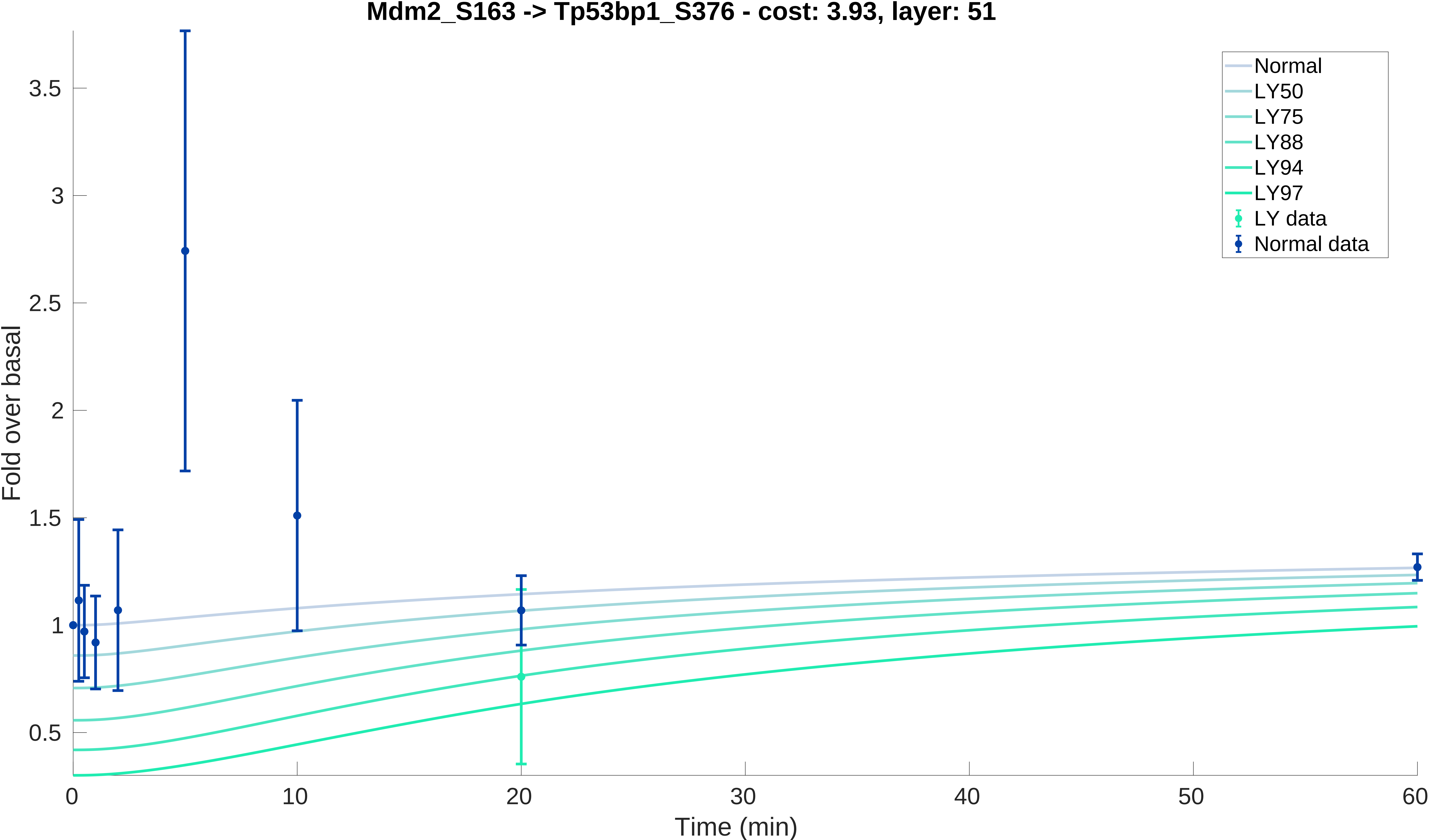

Mdm2_S163 -> Tp53bp1_S376 - cost: 3.93, layer: 51
Fold over basal
Time (min)
0.5
1
1.5
2
2.5
3
3.5
0
10
20
30
40
50
60
Normal
LY50
LY75
LY88
LY94
LY97
LY data
Normal data

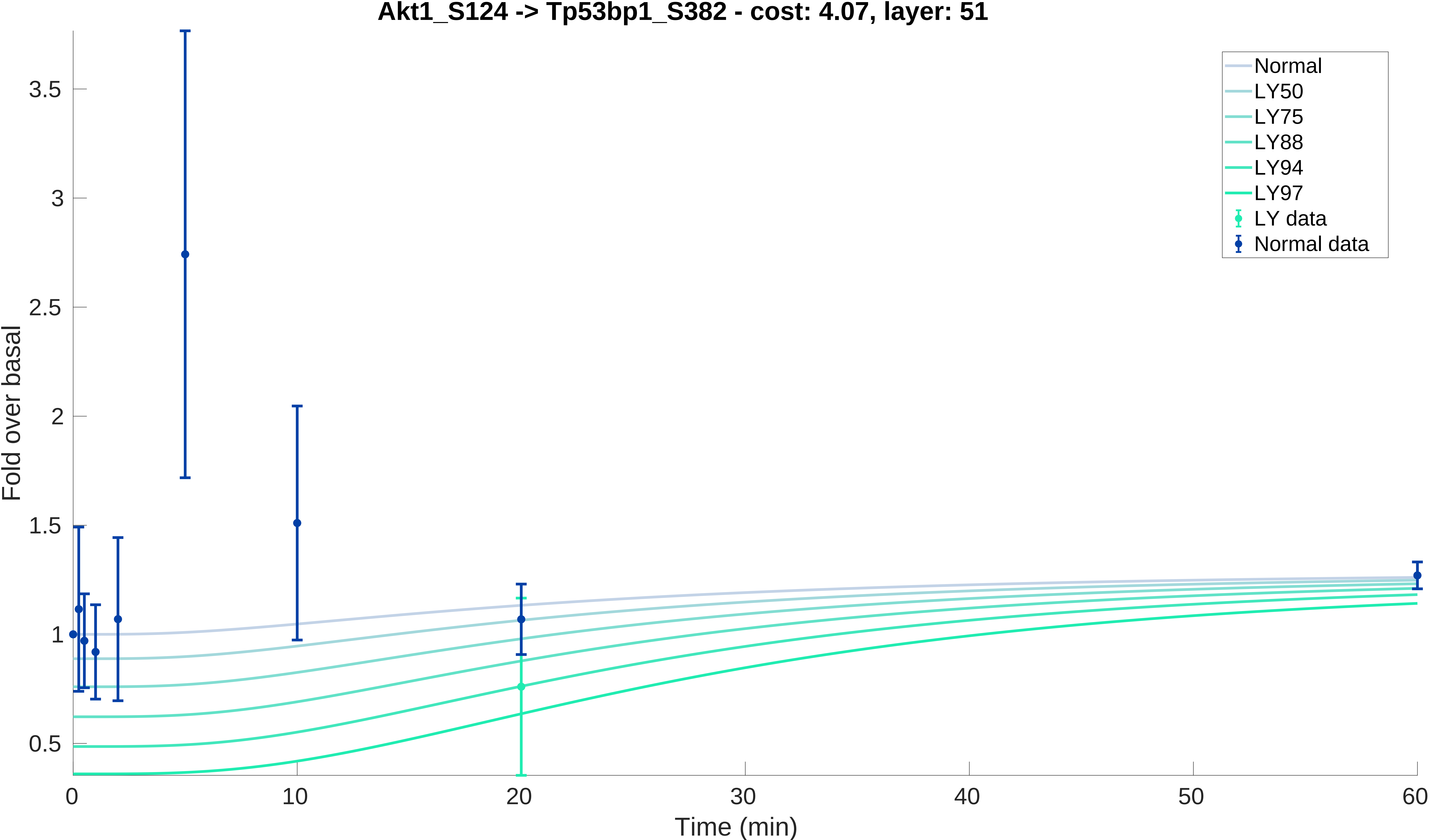

Akt1_S124 -> Tp53bp1_S382 - cost: 4.07, layer: 51
Fold over basal
Time (min)
0.5
1
1.5
2
2.5
3
3.5
0
10
20
30
40
50
60
Normal
LY50
LY75
LY88
LY94
LY97
LY data
Normal data

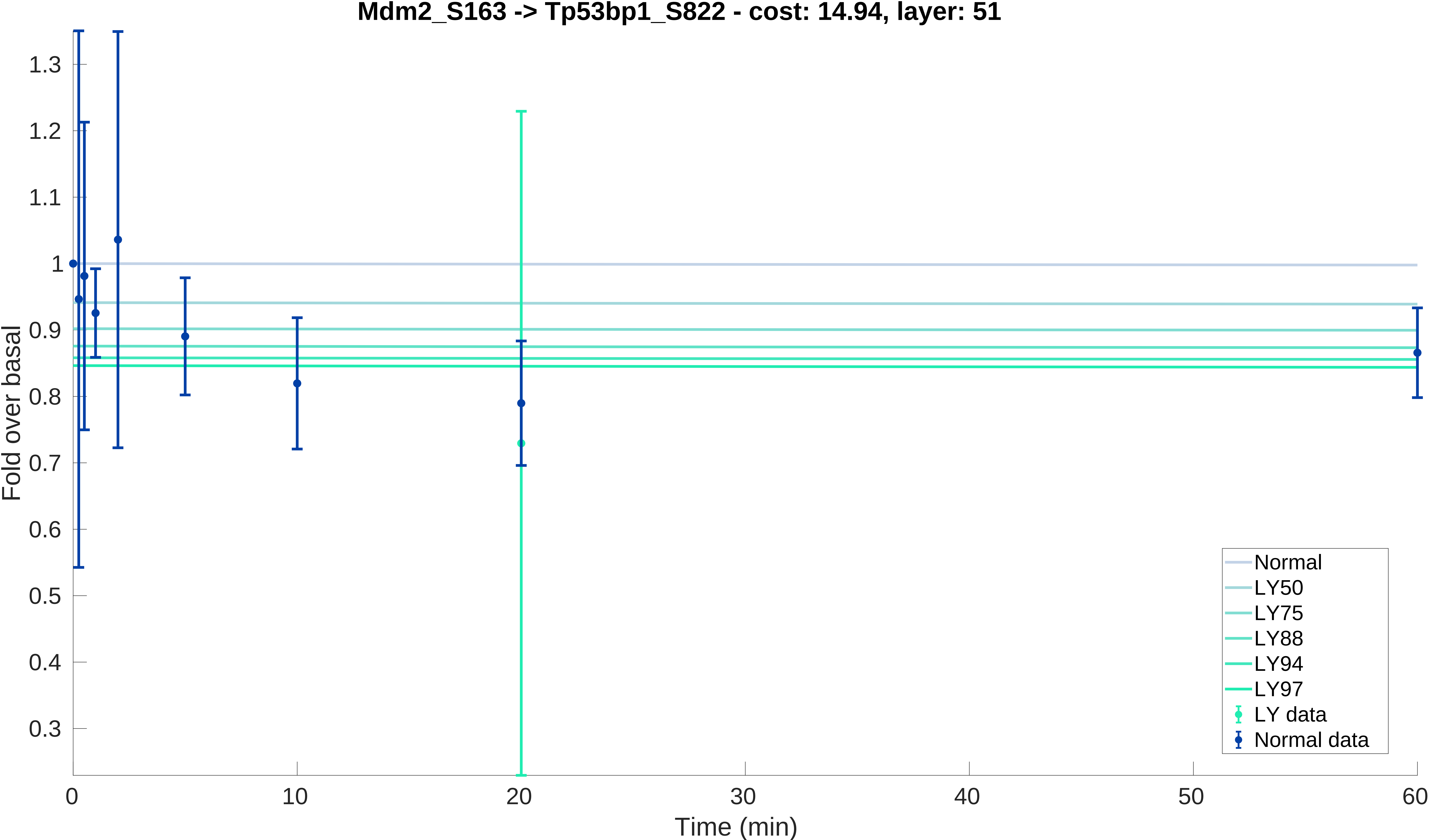

Mdm2_S163 -> Tp53bp1_S822 - cost: 14.94, layer: 51
Fold over basal
1.3
1.2
1.1
1
0.9
0.8
0.7
0.6
0.5
0.4
0.3
0
10
20
30
40
50
60
Time (min)
Normal
LY50
LY75
LY88
LY94
LY97
LY data
Normal data

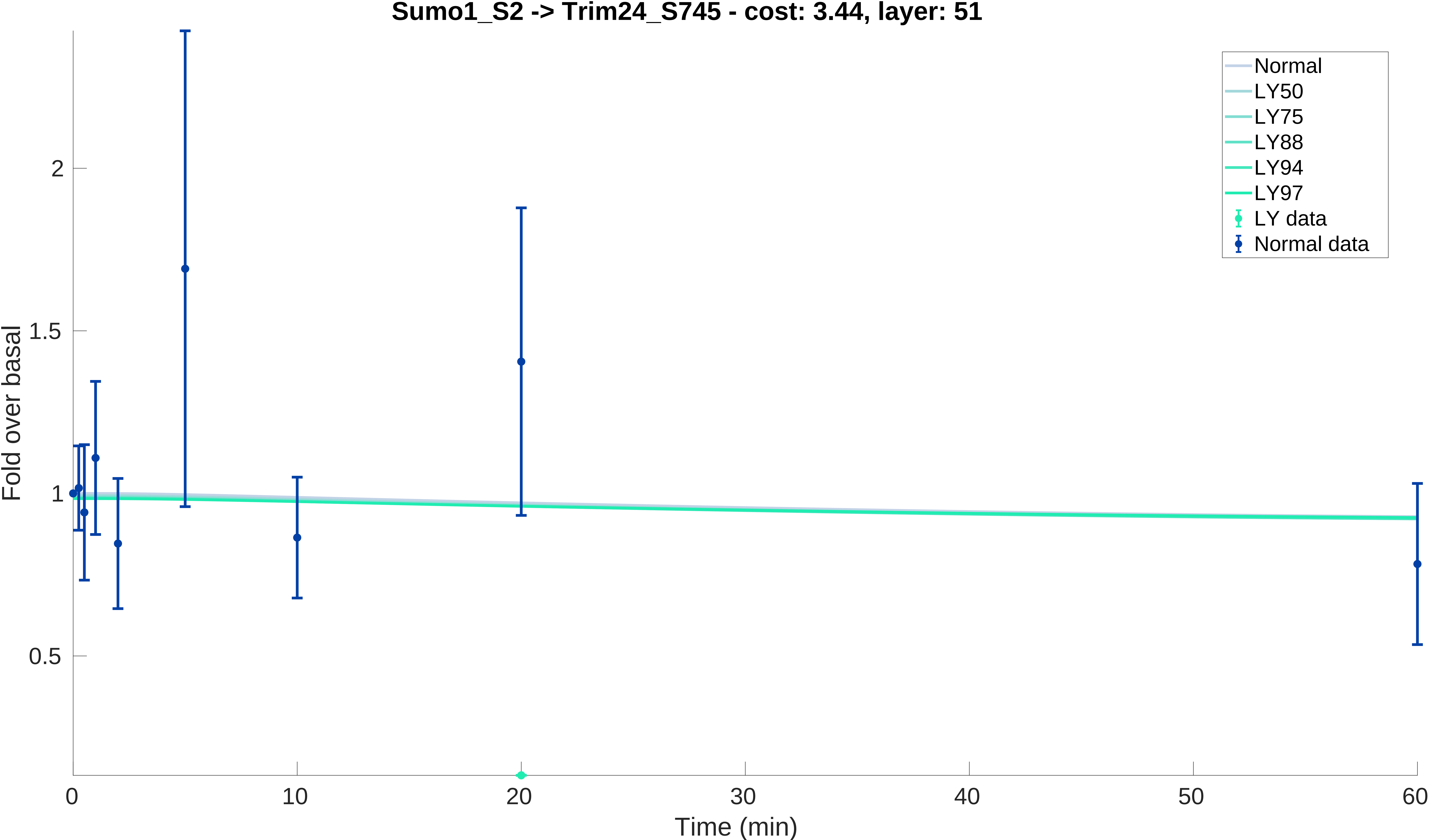
Sumo1_S2 -> Trim24_S745 - cost: 3.44, layer: 51
Fold over basal
Time (min)
0
10
20
30
40
50
60
0.5
1
1.5
2
Normal
LY50
LY75
LY88
LY94
LY97
LY data
Normal data

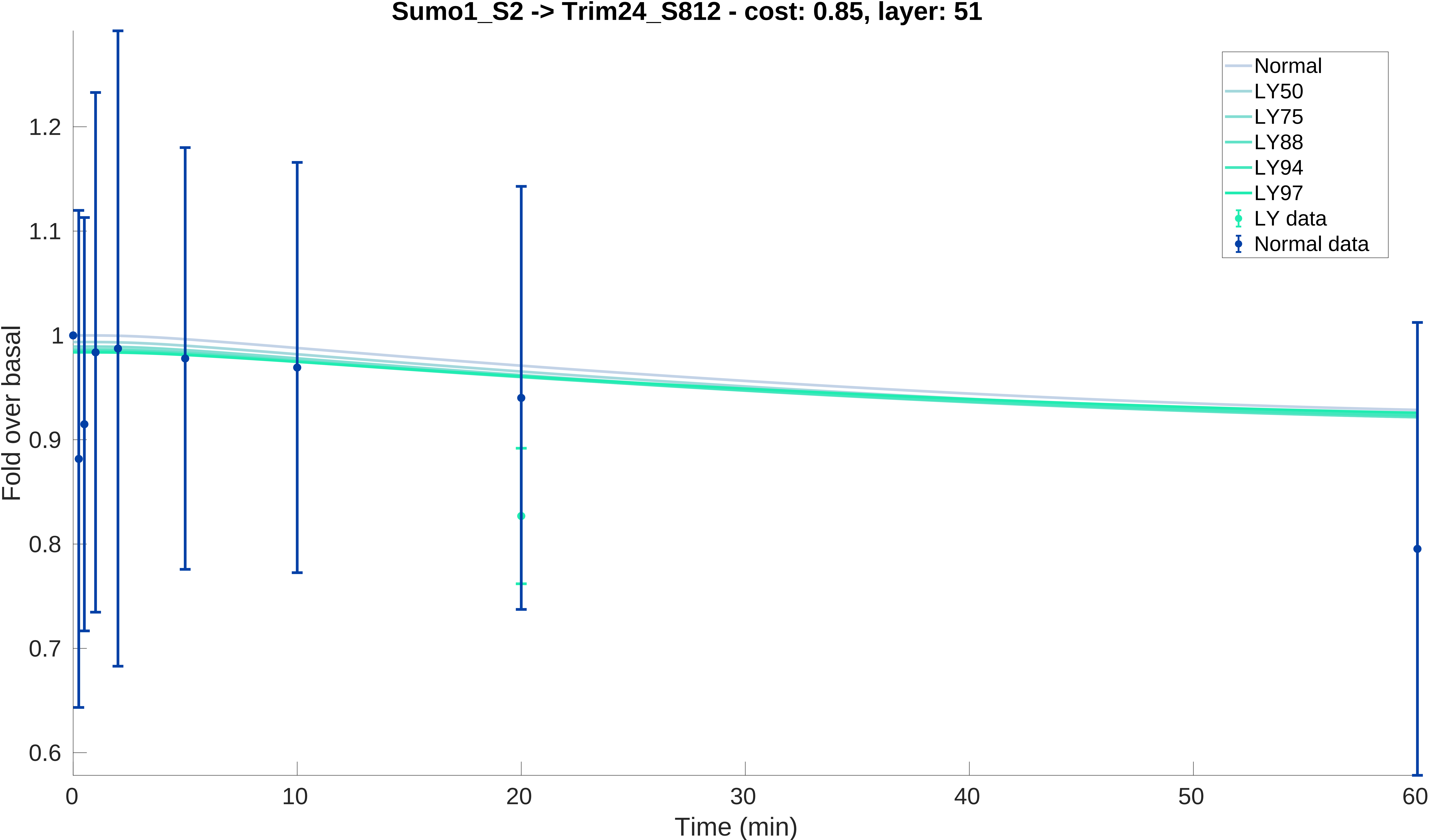

Sumo1_S2 -> Trim24_S812 - cost: 0.85, layer: 51
Normal
LY50
LY75
LY88
LY94
LY97
LY data
Normal data
Fold over basal
1.2
1.1
1
0.9
0.8
0.7
0.6
0
10
20
30
40
50
60
Time (min)

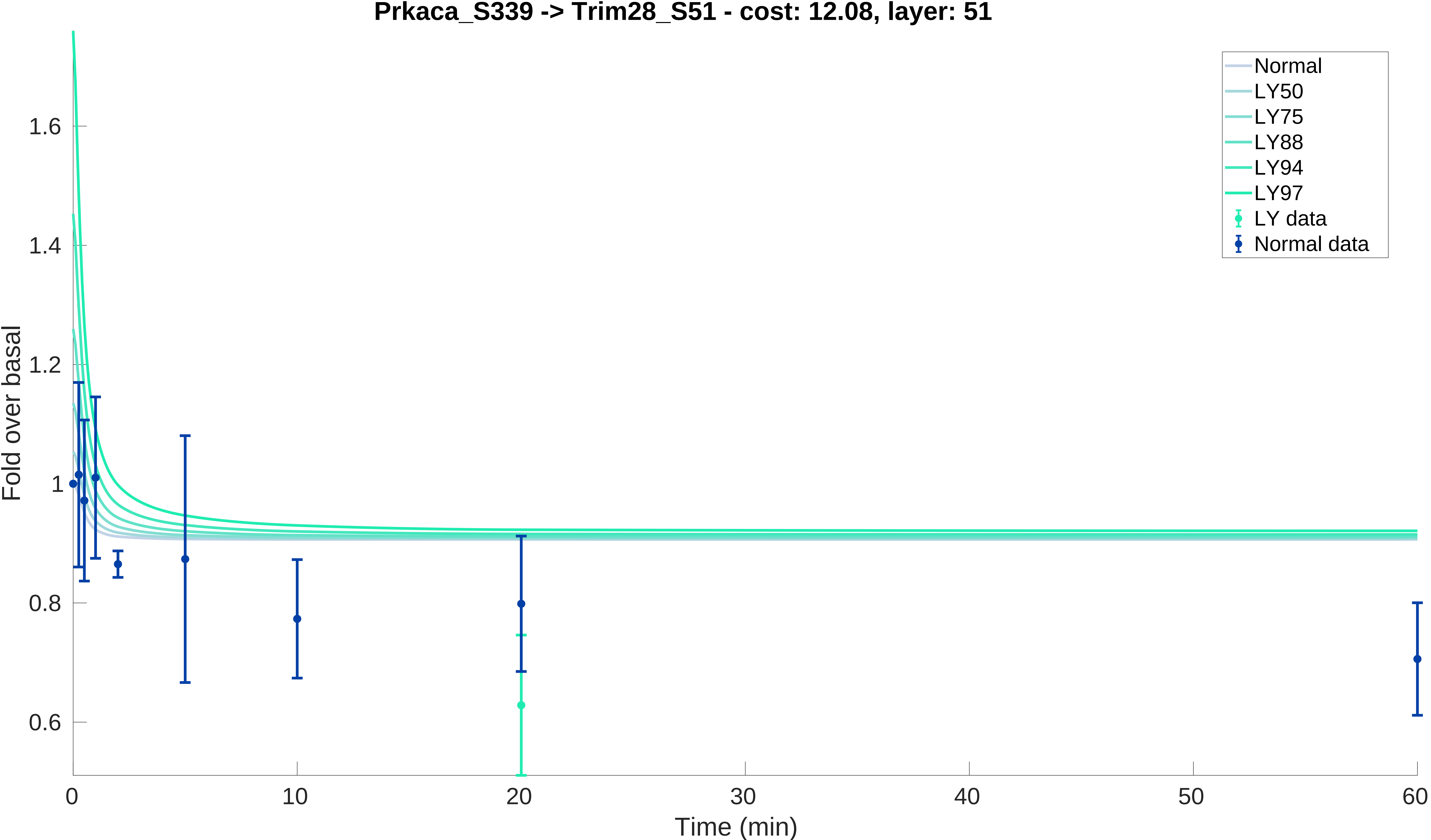

Prrkaca_S339 -> Trim28_S51 - cost: 12.08, layer: 51
Normal
LY50
LY75
LY88
LY94
LY97
LY data
Normal data
Fold over basal
Time (min)
1.6
1.4
1.2
1
0.8
0.6
0
10
20
30
40
50
60

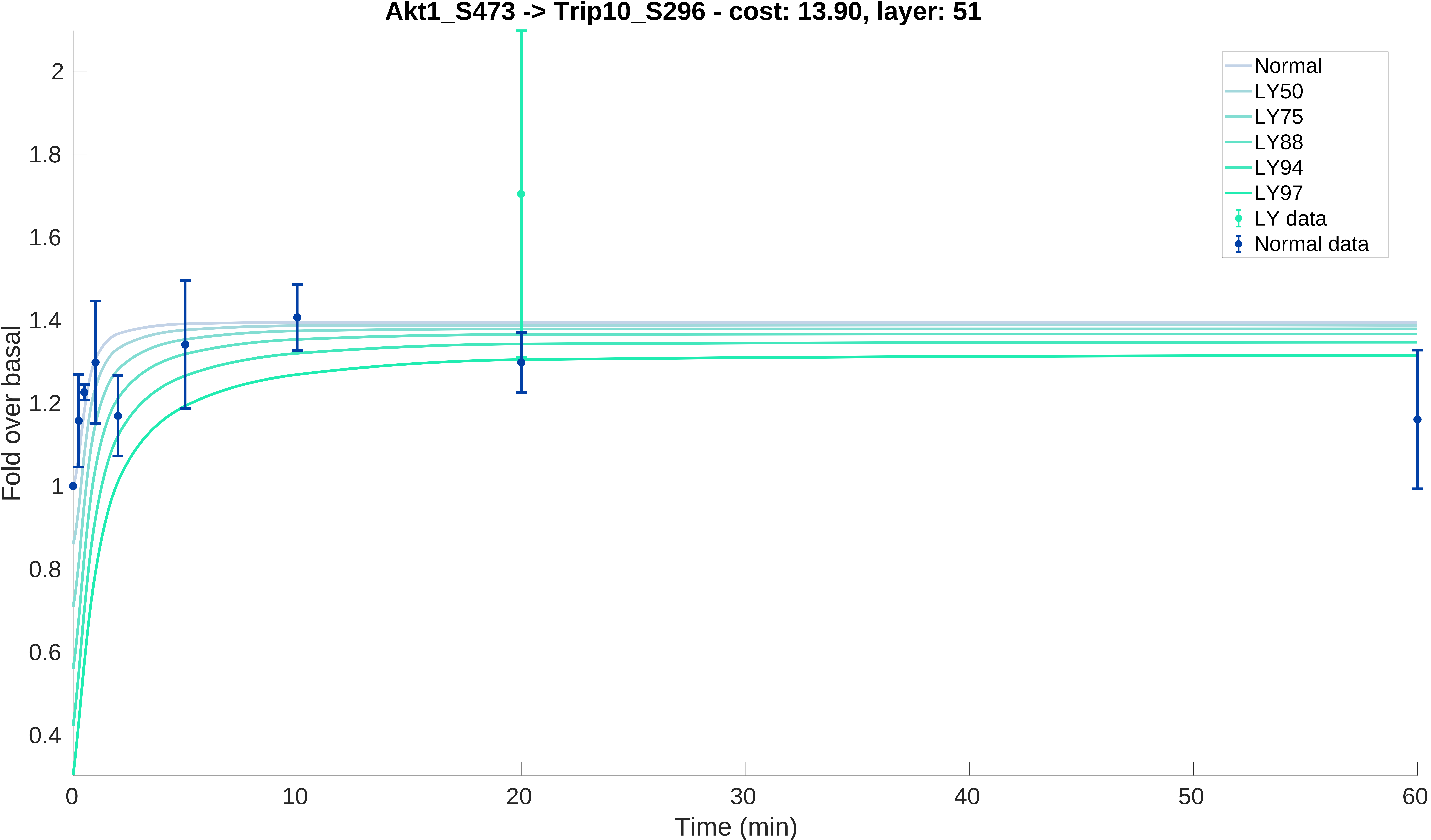

Akt1_S473 -> Trip10_S296 - cost: 13.90, layer: 51
Fold over basal
Time (min)
0.4
0.6
0.8
1
1.2
1.4
1.6
1.8
2
0
10
20
30
40
50
60
Normal
LY50
LY75
LY88
LY94
LY97
LY data
Normal data

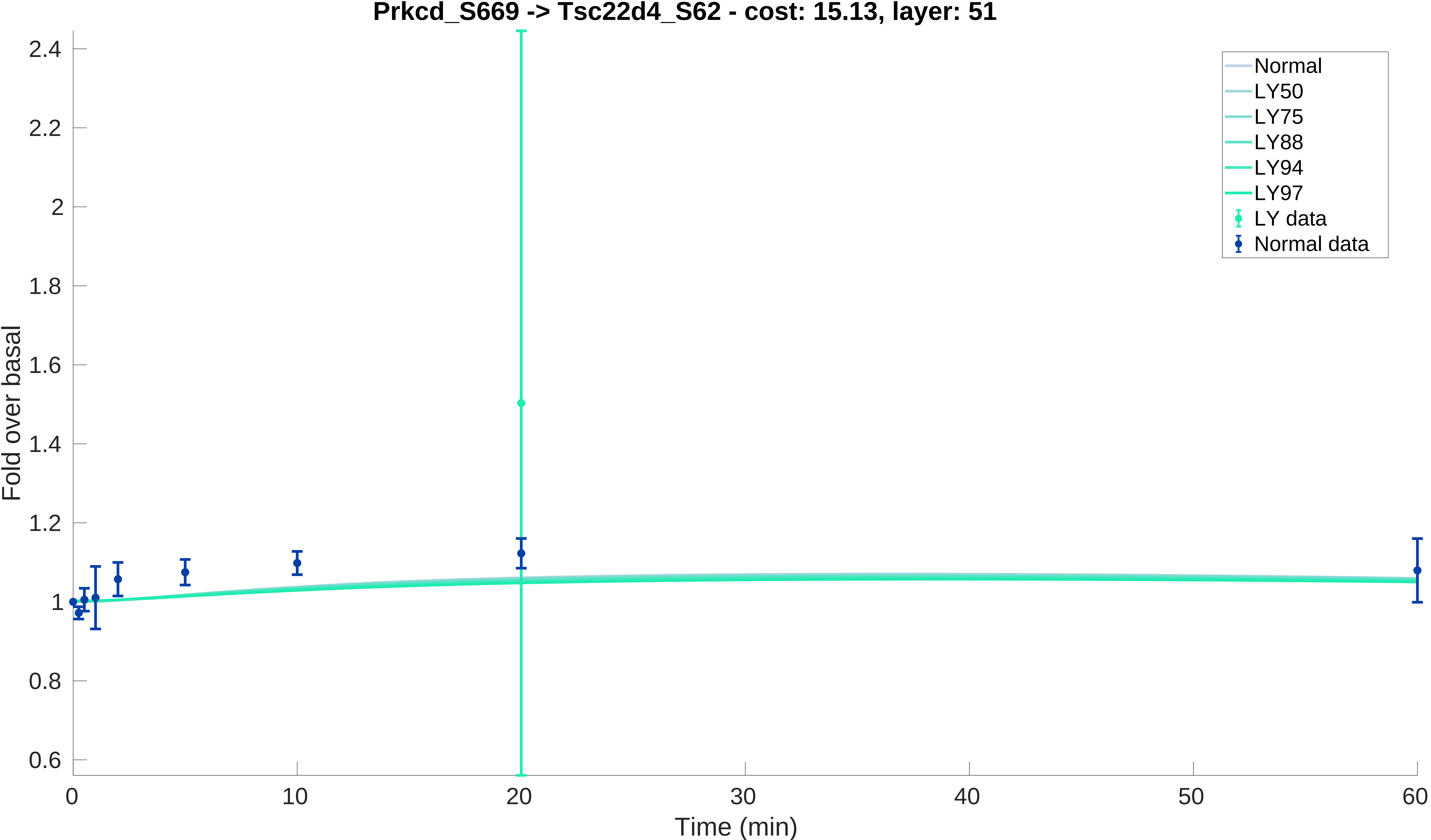

Prkcd_S669 -> Tsc22d4_S62 - cost: 15.13, layer: 51
Fold over basal
Time (min)
0.6
0.8
1
1.2
1.4
1.6
1.8
2
2.2
2.4
0
10
20
30
40
50
60
Normal
LY50
LY75
LY88
LY94
LY97
LY data
Normal data

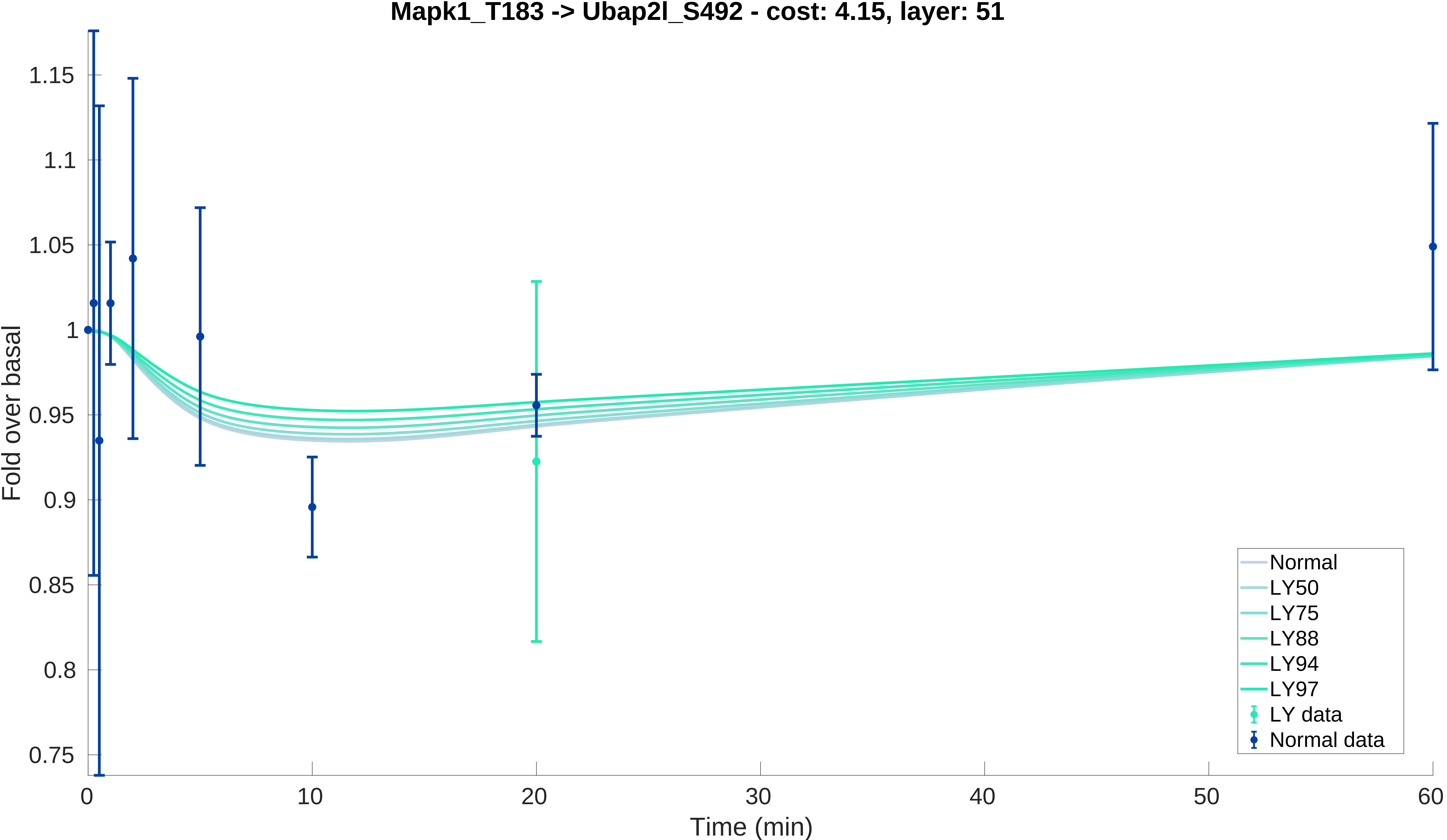
Mapk1_T183 -> Ubap2l_S492 - cost: 4.15, layer: 51
Fold over basal
Time (min)
1.15
1.1
1.05
1
0.95
0.9
0.85
0.8
0.75
0
10
20
30
40
50
60
Normal
LY50
LY75
LY88
LY94
LY97
LY data
Normal data

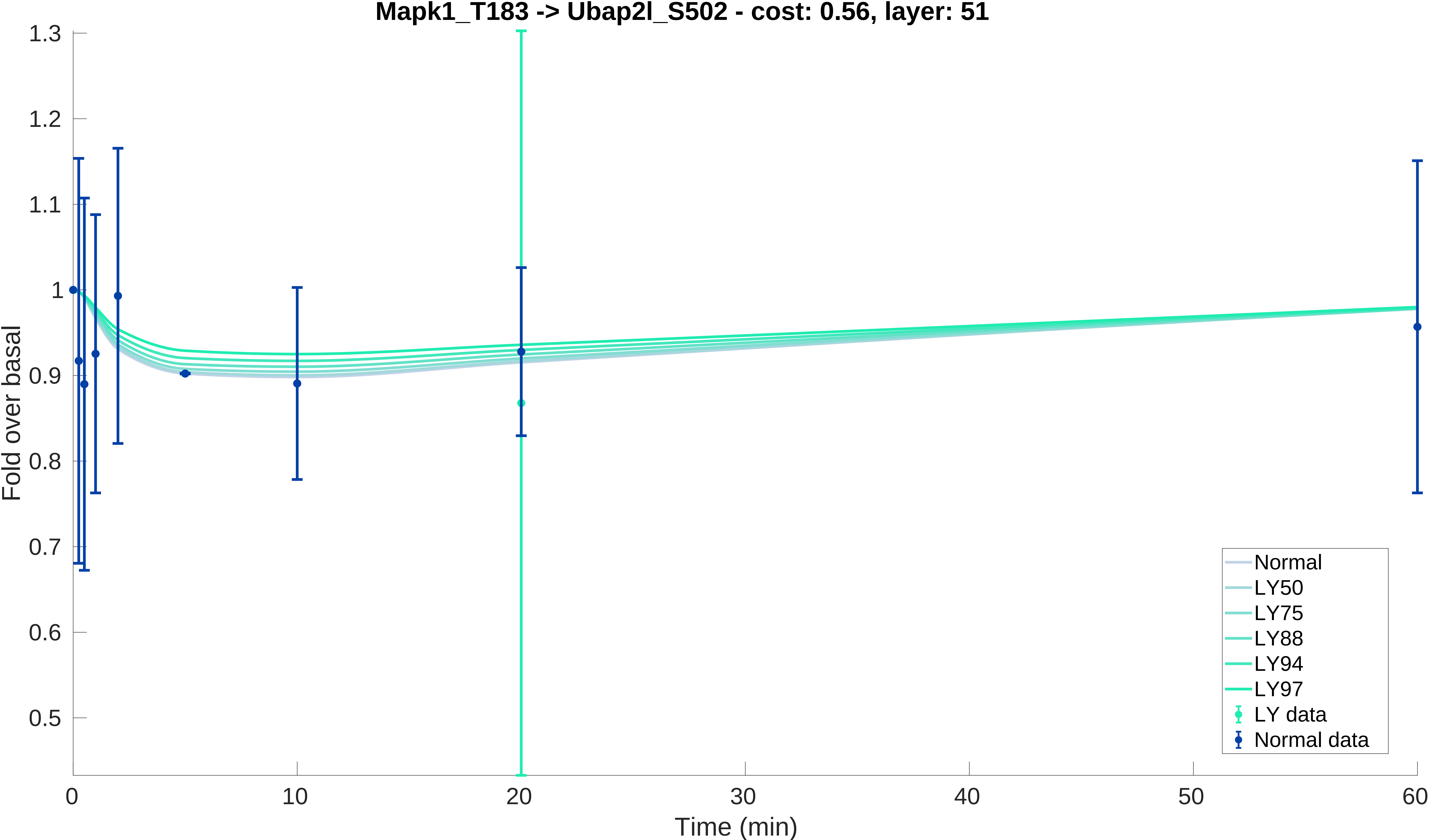

Mapk1_T183 -> Ubap2l_S502 - cost: 0.56, layer: 51
Fold over basal
Time (min)
1.3
1.2
1.1
1
0.9
0.8
0.7
0.6
0.5
0
10
20
30
40
50
60
Normal
LY50
LY75
LY88
LY94
LY97
LY data
Normal data

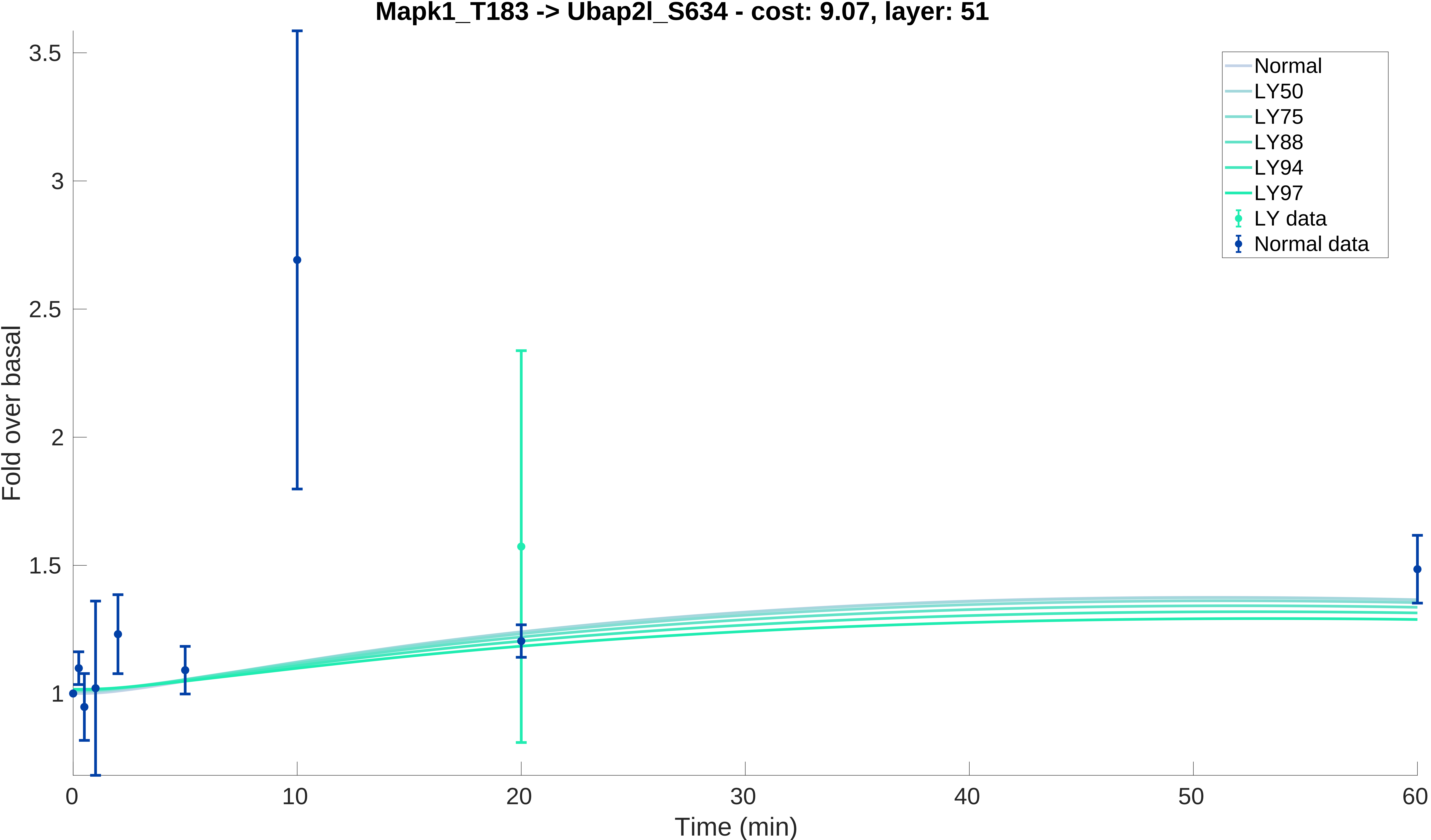
Mapk1_T183 -> Ubap2l_S634 - cost: 9.07, layer: 51
Fold over basal
Time (min)
3.5
3
2.5
2
1.5
1
0
10
20
30
40
50
60
Normal
LY50
LY75
LY88
LY94
LY97
LY data
Normal data

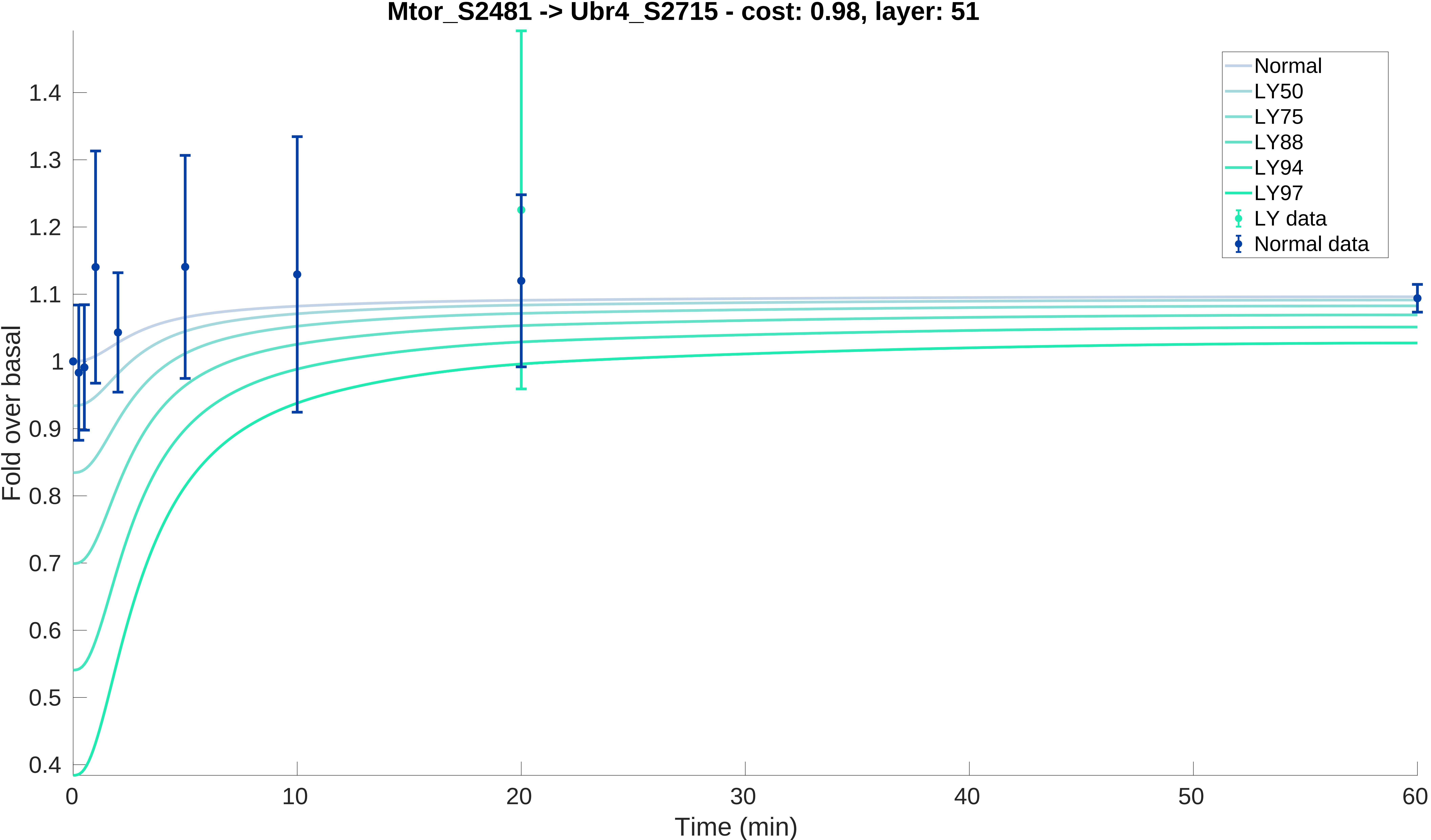

Mtor_S2481 -> Ubr4_S2715 - cost: 0.98, layer: 51
Normal
LY50
LY75
LY88
LY94
LY97
LY data
Normal data
Fold over basal
1.4
1.3
1.2
1.1
1
0.9
0.8
0.7
0.6
0.5
0.4
0
10
20
30
40
50
60
Time (min)

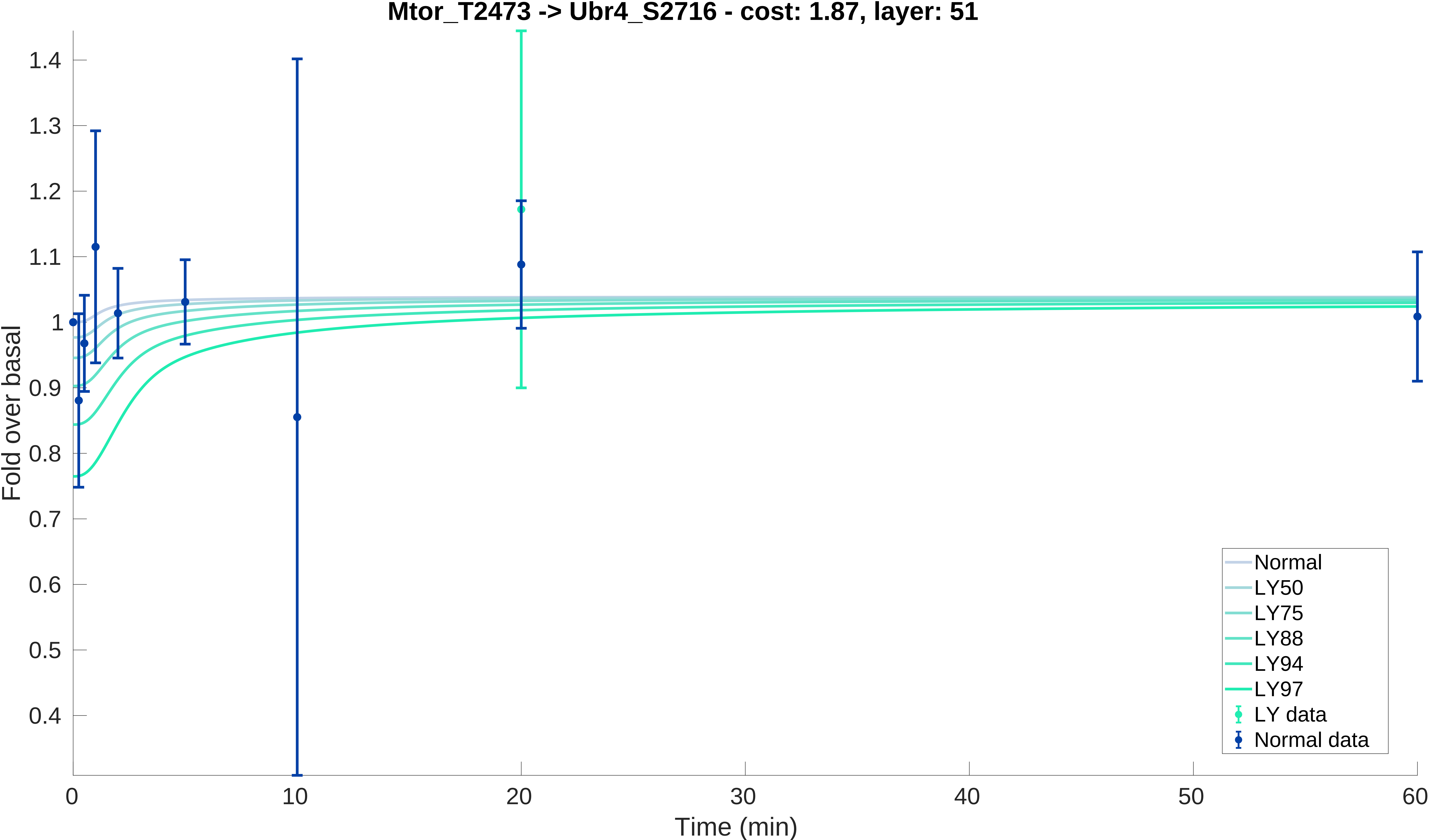

Mtor_T2473 -> Ubr4_S2716 - cost: 1.87, layer: 51
Fold over basal
Time (min)
0
10
20
30
40
50
60
0.4
0.5
0.6
0.7
0.8
0.9
1
1.1
1.2
1.3
1.4
Normal
LY50
LY75
LY88
LY94
LY97
LY data
Normal data

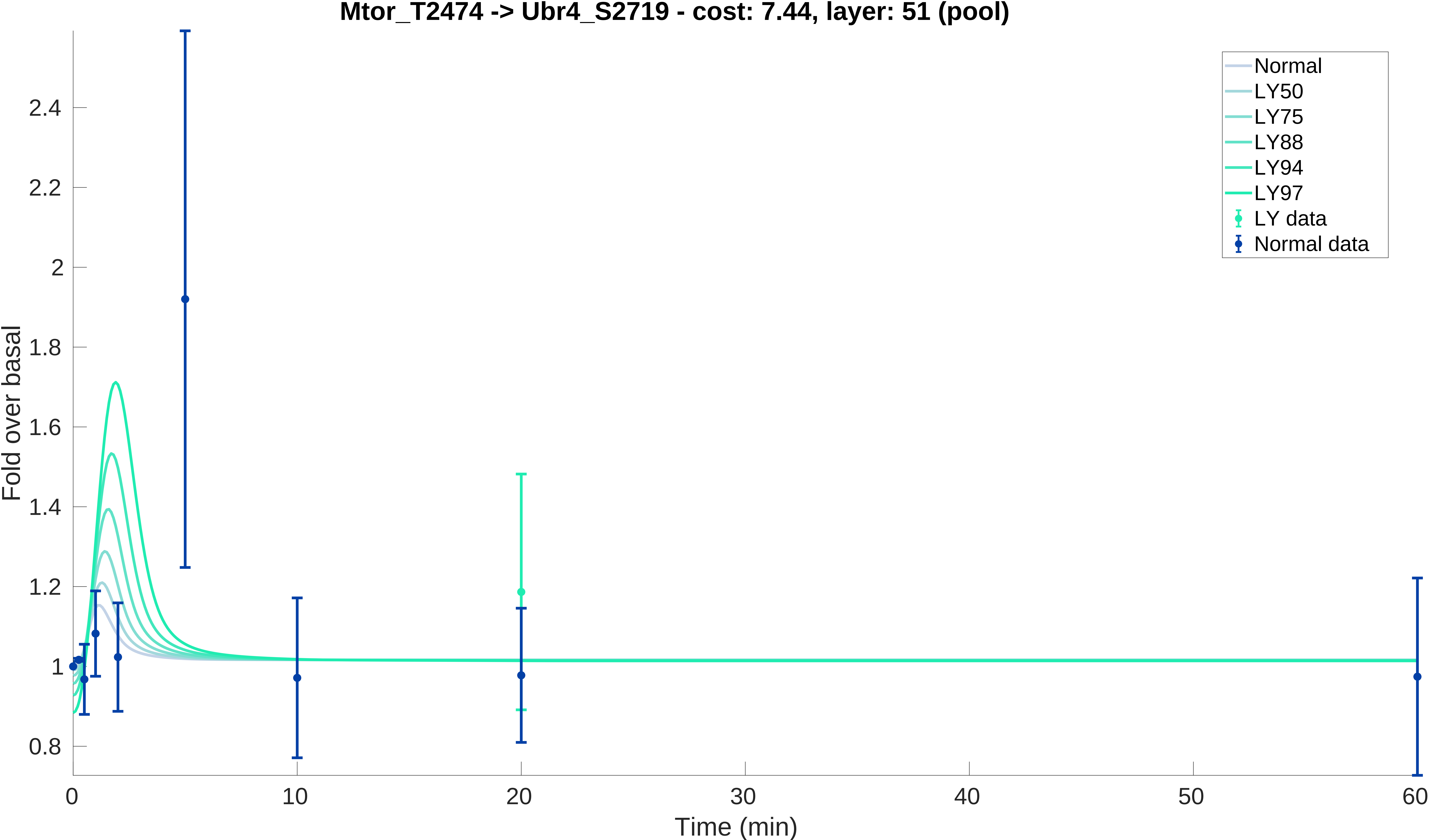

Mtor_T2474 -> Ubr4_S2719 - cost: 7.44, layer: 51 (pool)
Fold over basal
Time (min)
0.8
1
1.2
1.4
1.6
1.8
2
2.2
2.4
0
10
20
30
40
50
60
Normal
LY50
LY75
LY88
LY94
LY97
LY data
Normal data

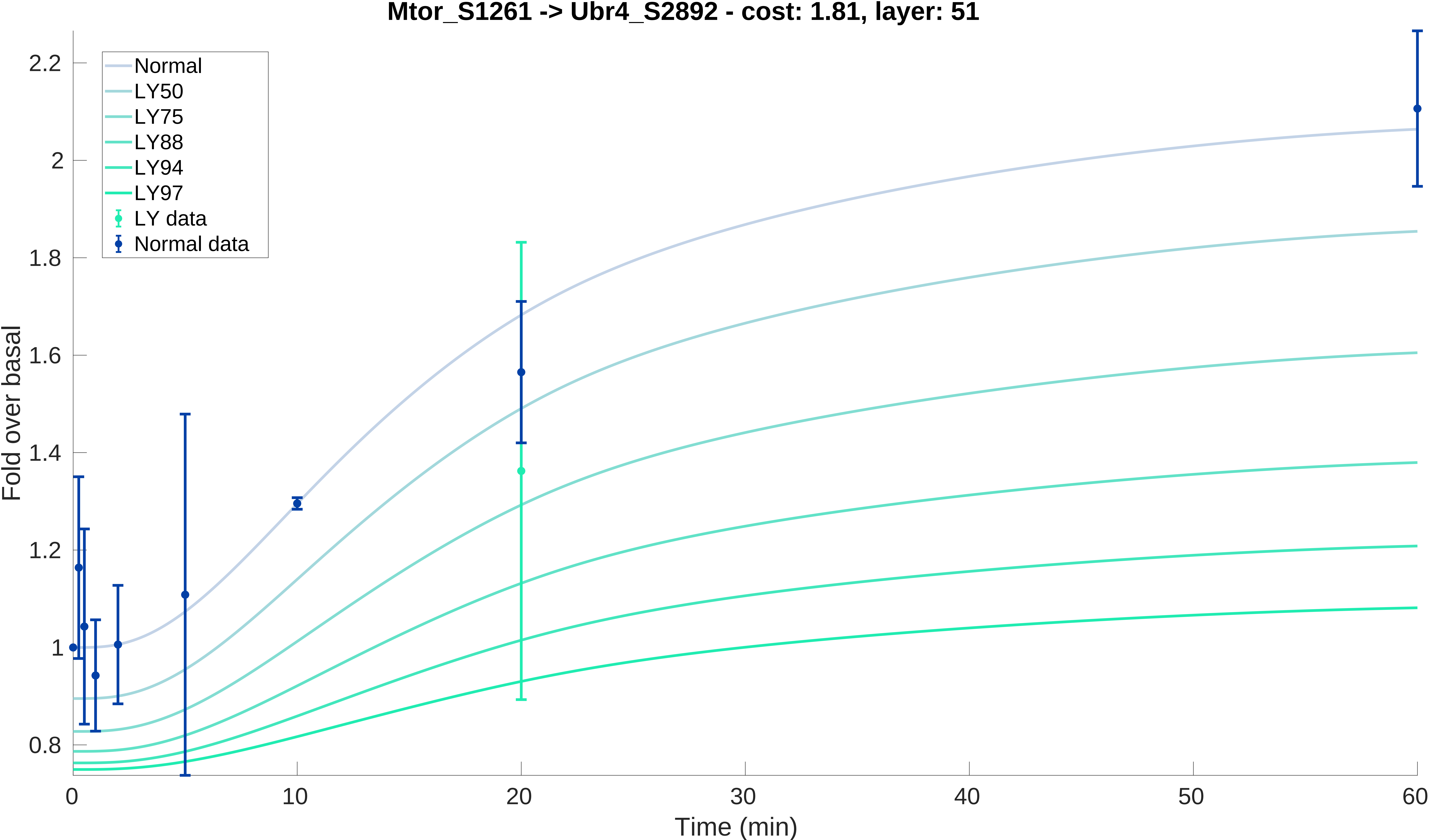

Mtor_S1261 -> Ubr4_S2892 - cost: 1.81, layer: 51
Normal
LY50
LY75
LY88
LY94
LY97
LY data
Normal data
Fold over basal
Time (min)
0.8
1
1.2
1.4
1.6
1.8
2
2.2
0
10
20
30
40
50
60

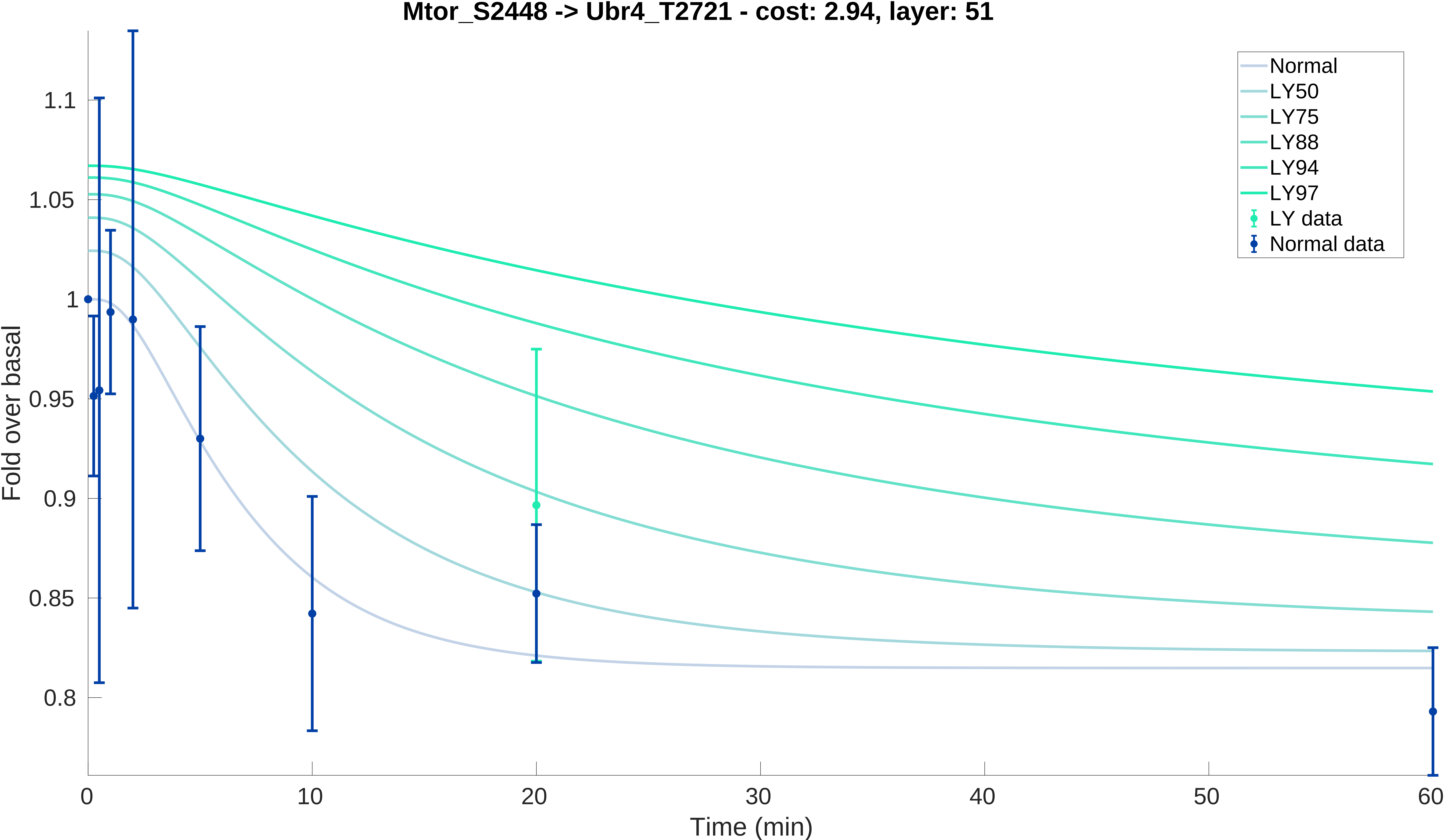
Mtor_S2448 -> Ubr4_T2721 - cost: 2.94, layer: 51
Fold over basal
Time (min)
1.1
1.05
1
0.95
0.9
0.85
0.8
0
10
20
30
40
50
60
Normal
LY50
LY75
LY88
LY94
LY97
LY data
Normal data

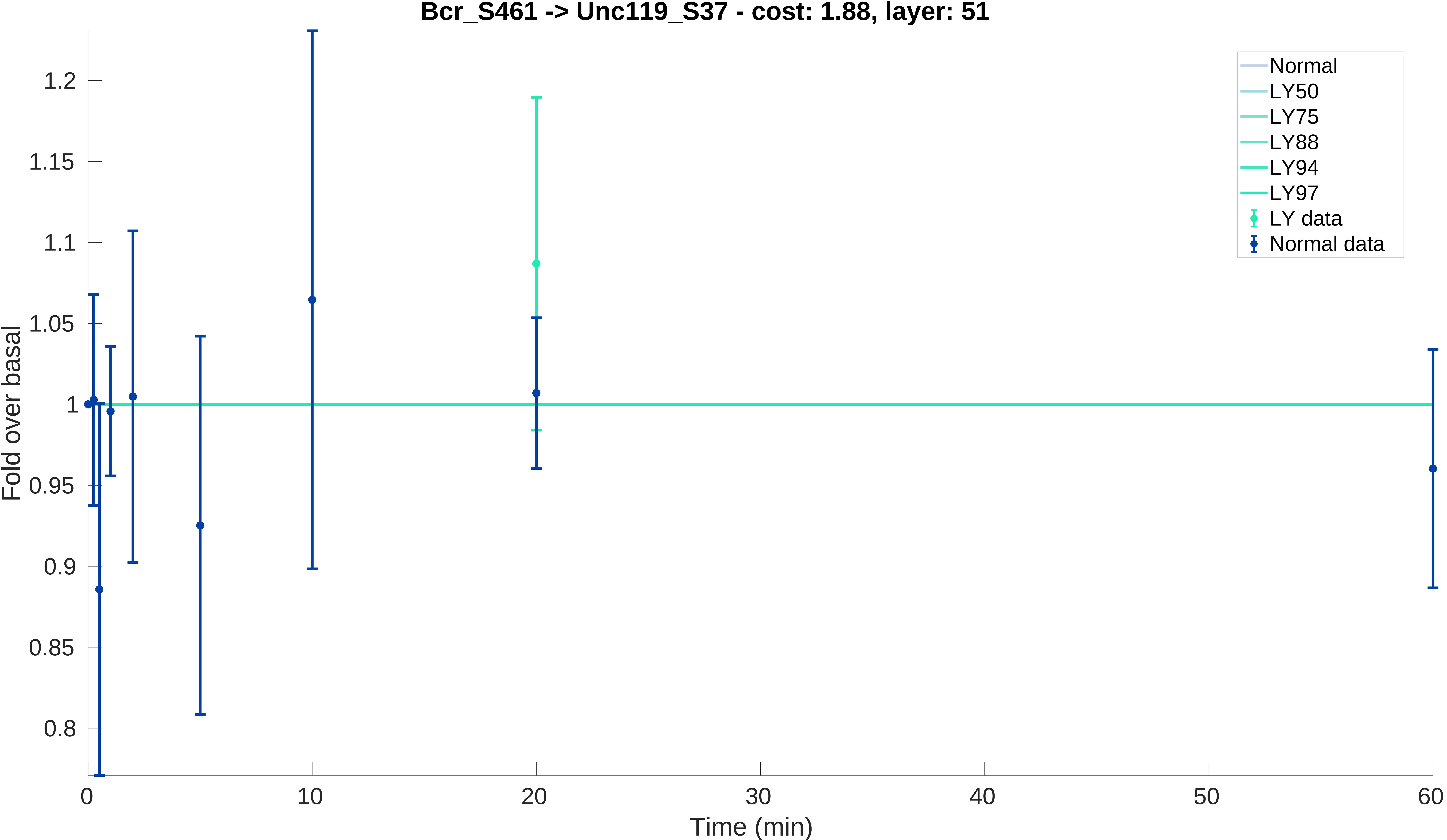
Bcr_S461 -> Unc119_S37 - cost: 1.88, layer: 51
Fold over basal
Time (min)
1.2
1.15
1.1
1.05
1
0.95
0.9
0.85
0.8
0
10
20
30
40
50
60
Normal
LY50
LY75
LY88
LY94
LY97
LY data
Normal data

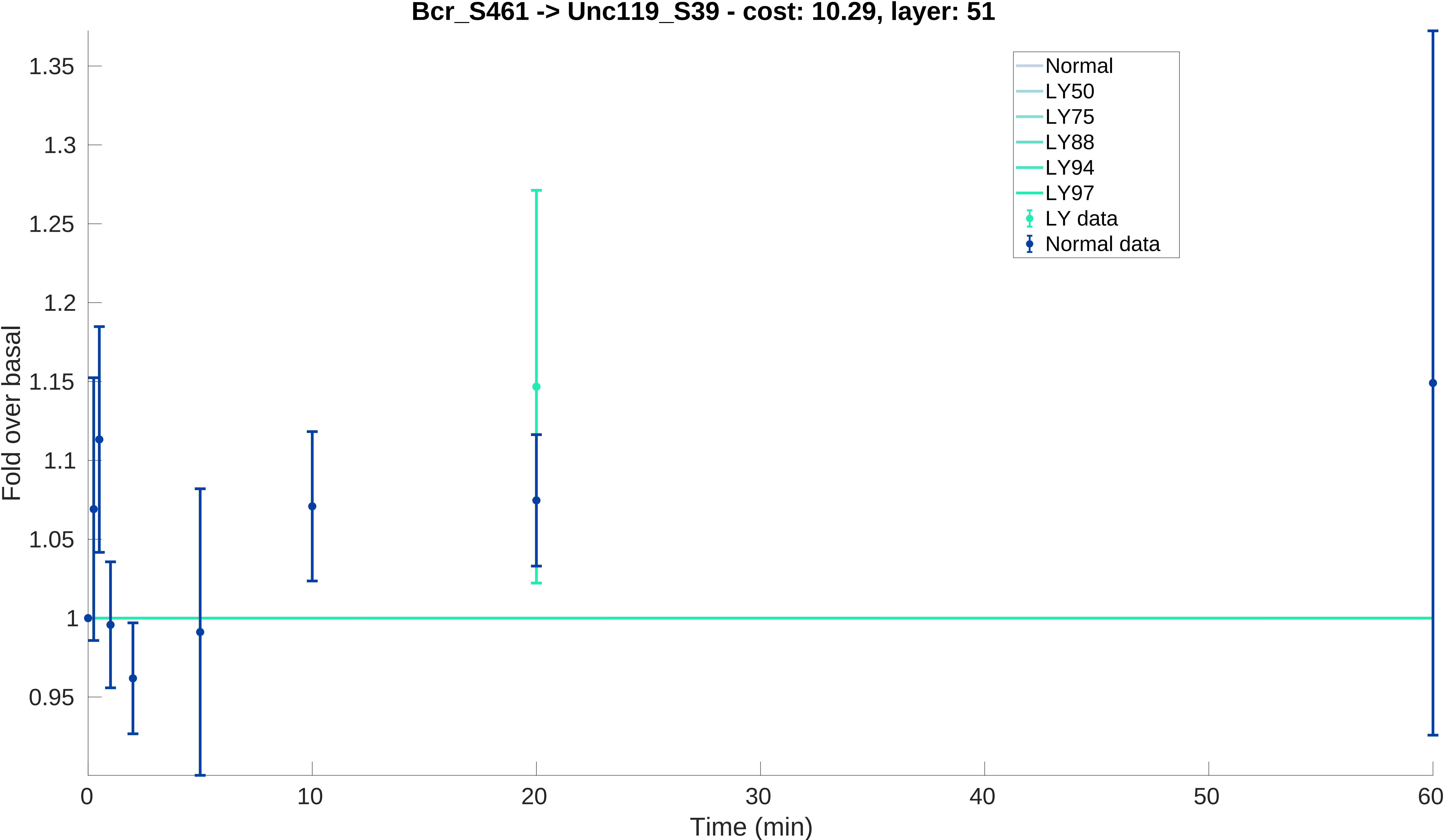
Bcr_S461 -> Unc119_S39 - cost: 10.29, layer: 51
Normal
LY50
LY75
LY88
LY94
LY97
LY data
Normal data
Fold over basal
Time (min)
1.35
1.3
1.25
1.2
1.15
1.1
1.05
1
0.95
0
10
20
30
40
50
60

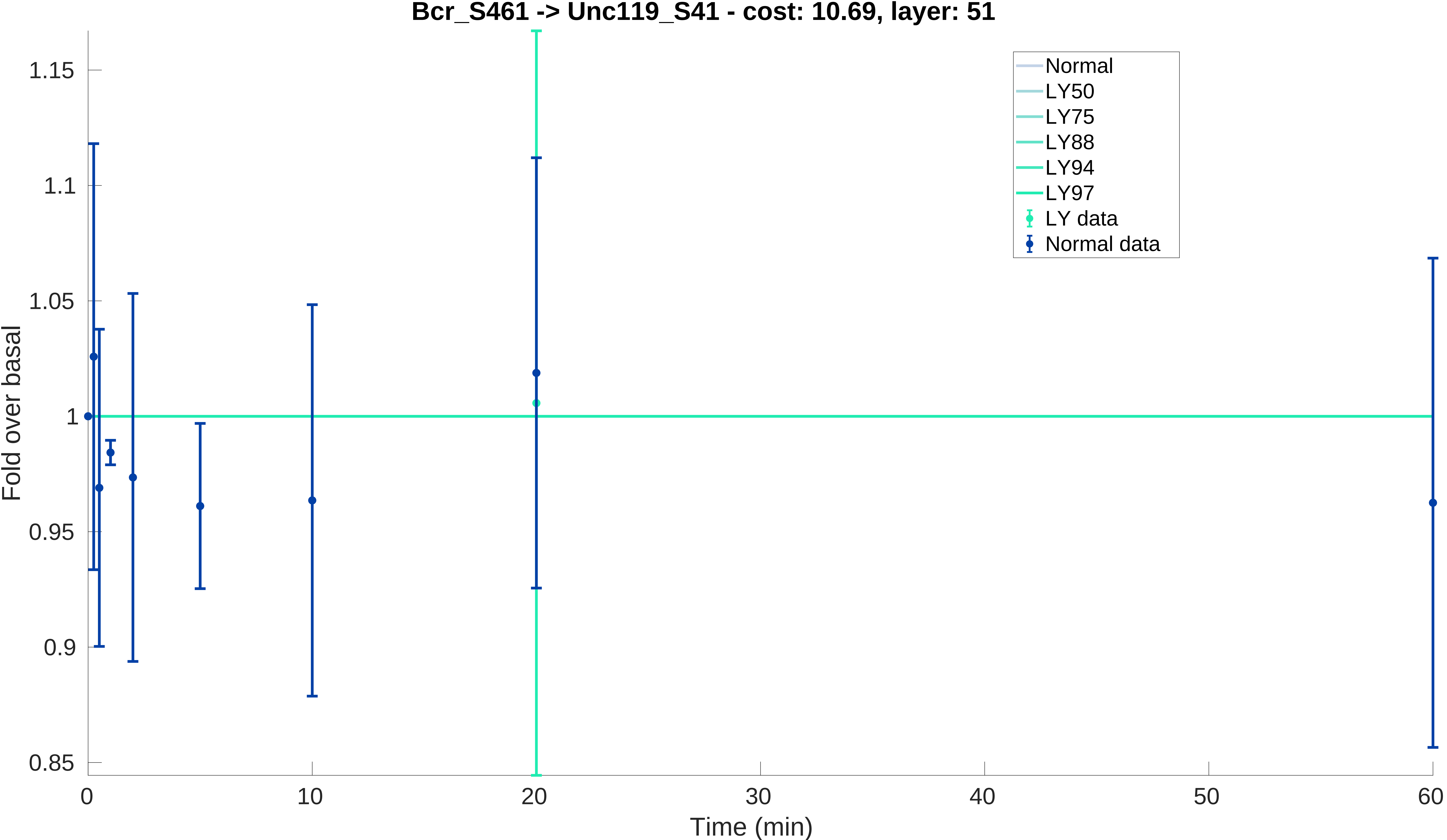

Bcr_S461 -> Unc119_S41 - cost: 10.69, layer: 51
Normal
LY50
LY75
LY88
LY94
LY97
LY data
Normal data
Fold over basal
1.15
1.1
1.05
1
0.95
0.9
0.85
0
10
20
30
40
50
60
Time (min)

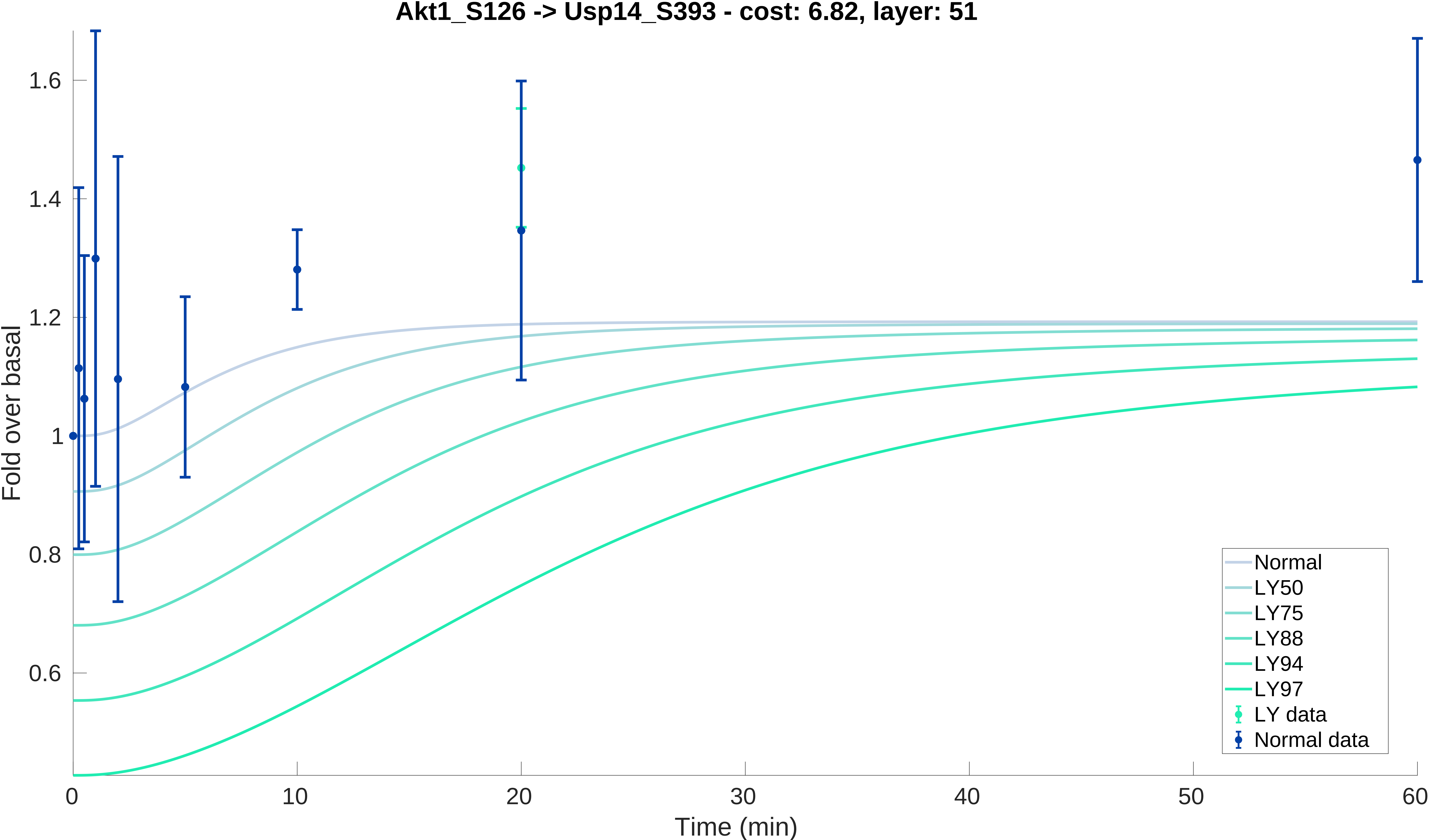

Akt1_S126 -> Usp14_S393 - cost: 6.82, layer: 51
1.6
1.4
1.2
1
0.8
0.6
Fold over basal
0
10
20
30
40
50
60
Time (min)
Normal
LY50
LY75
LY88
LY94
LY97
LY data
Normal data

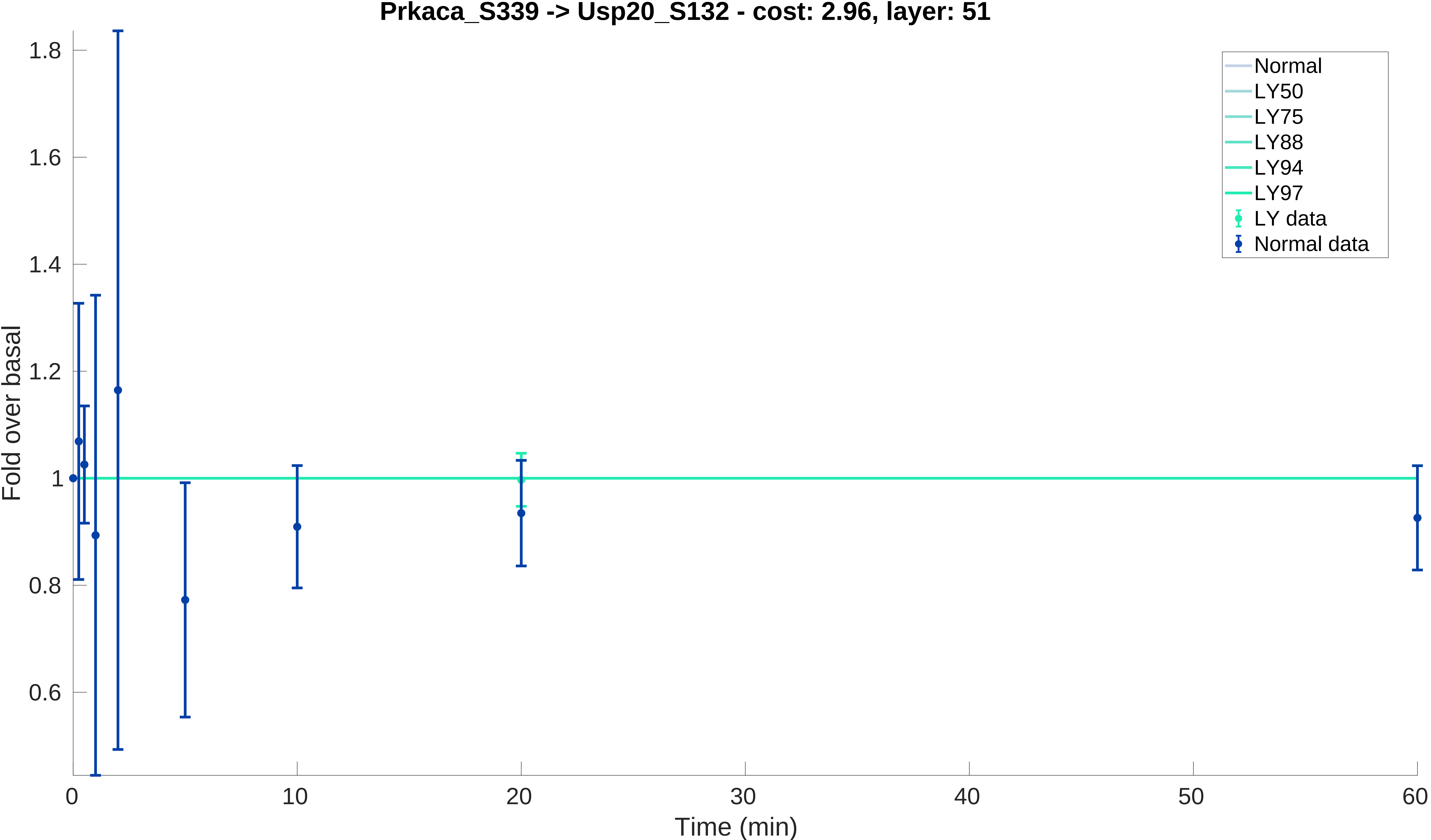
Prkaca_S339 -> Usp20_S132 - cost: 2.96, layer: 51
Normal
LY50
LY75
LY88
LY94
LY97
LY data
Normal data
Fold over basal
Time (min)
0
10
20
30
40
50
60
0.6
0.8
1
1.2
1.4
1.6
1.8

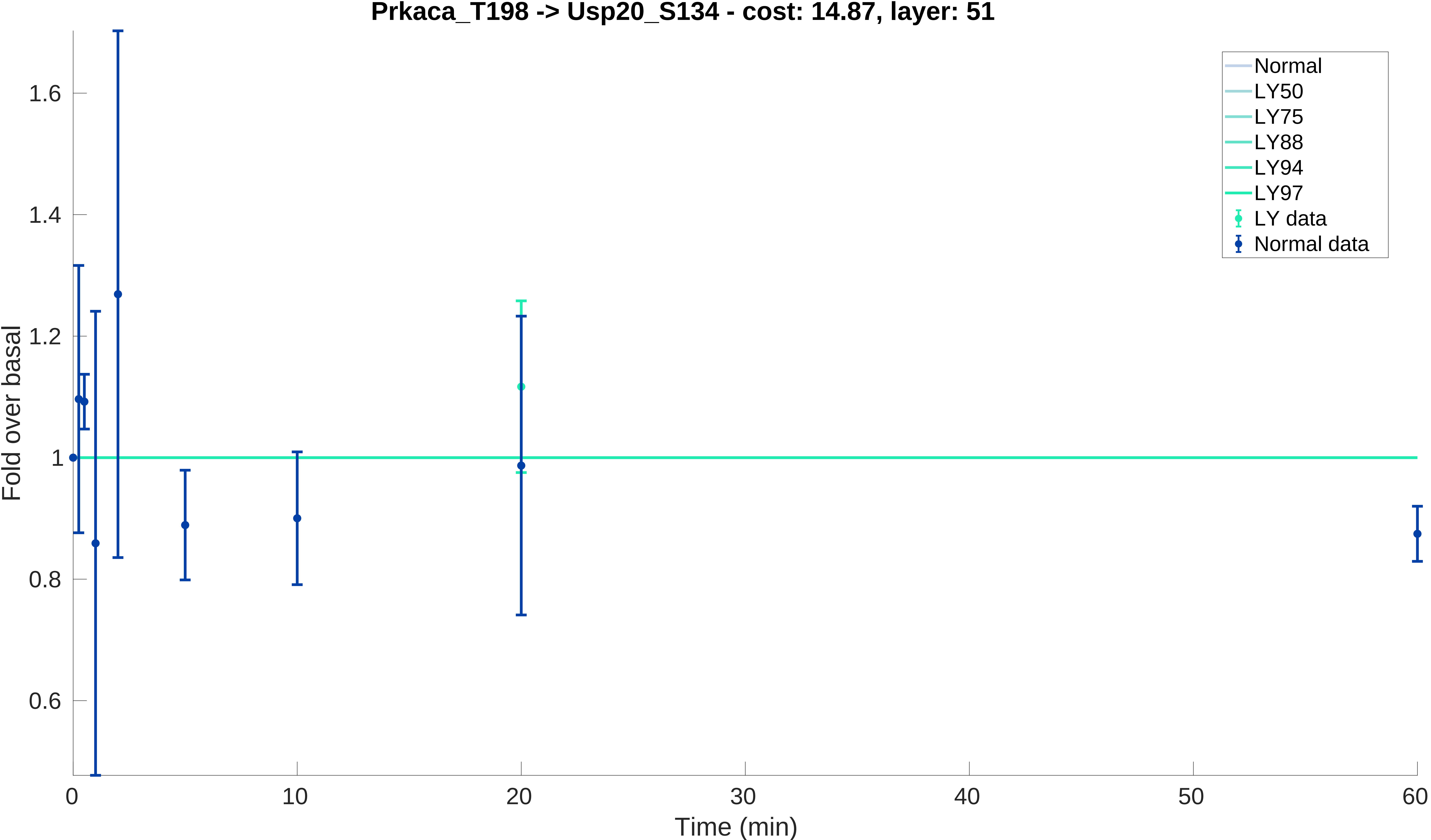

Prkaca_T198 -> Usp20_S134 - cost: 14.87, layer: 51
Fold over basal
Time (min)
0.6
0.8
1
1.2
1.4
1.6
0
10
20
30
40
50
60
Normal
LY50
LY75
LY88
LY94
LY97
LY data
Normal data

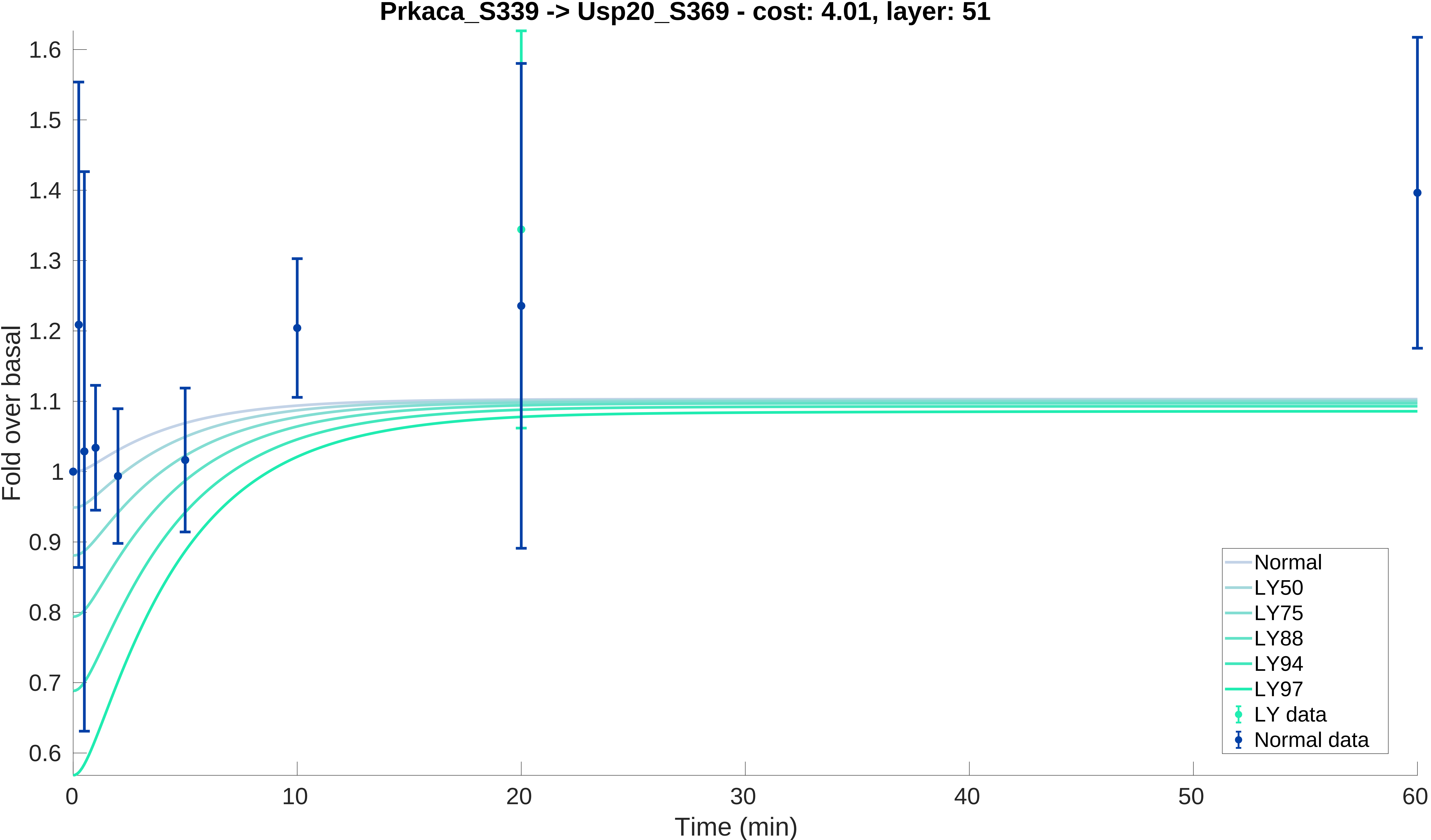

Prkaca_S339 -> Usp20_S369 - cost: 4.01, layer: 51
Fold over basal
1.6
1.5
1.4
1.3
1.2
1.1
1
0.9
0.8
0.7
0.6
0
10
20
30
40
50
60
Time (min)
Normal
LY50
LY75
LY88
LY94
LY97
LY data
Normal data

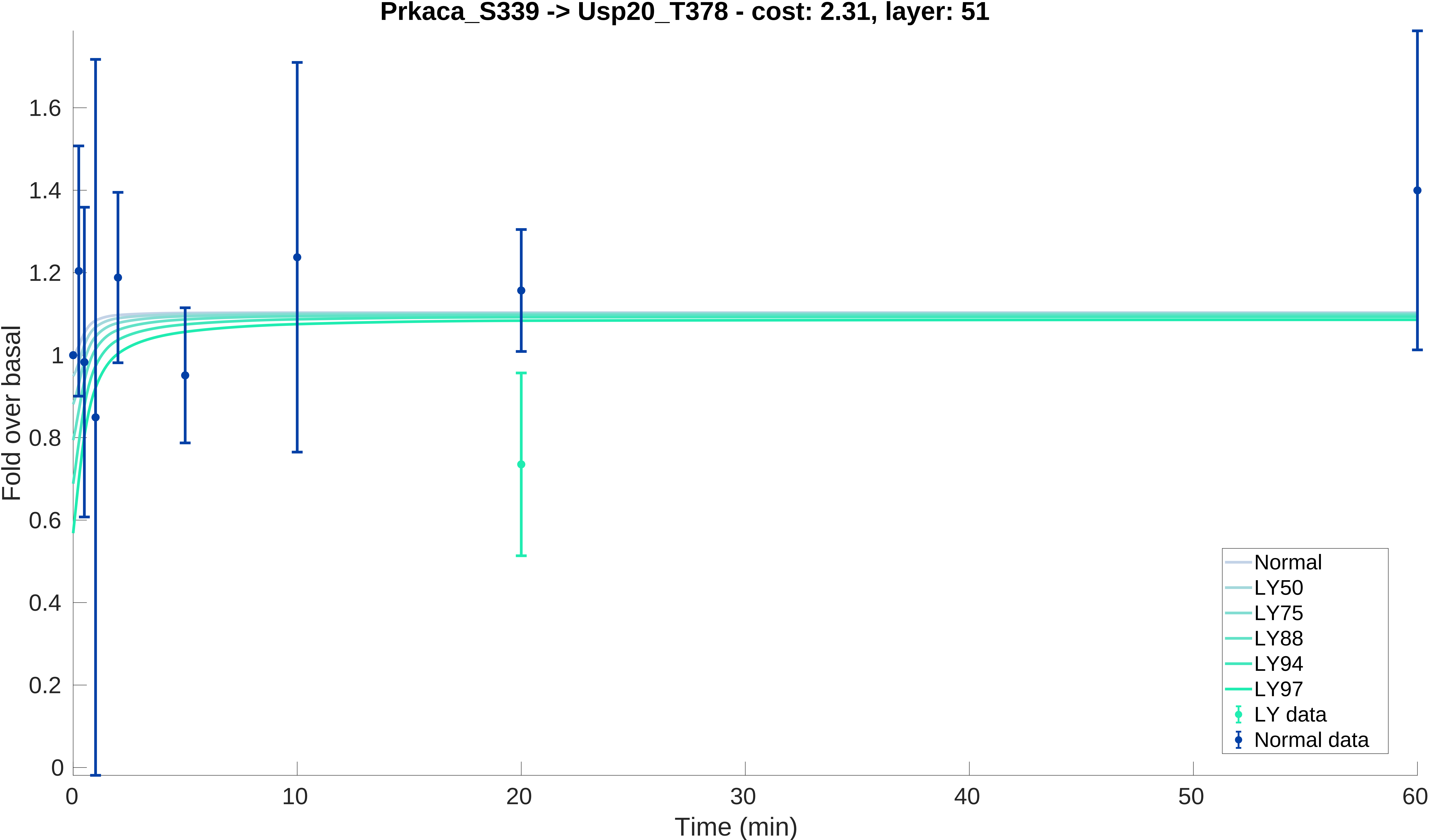

Prkaca_S339 -> Usp20_T378 - cost: 2.31, layer: 51
Fold over basal
Time (min)
0
0.2
0.4
0.6
0.8
1
1.2
1.4
1.6
0
10
20
30
40
50
60
Normal
LY50
LY75
LY88
LY94
LY97
LY data
Normal data

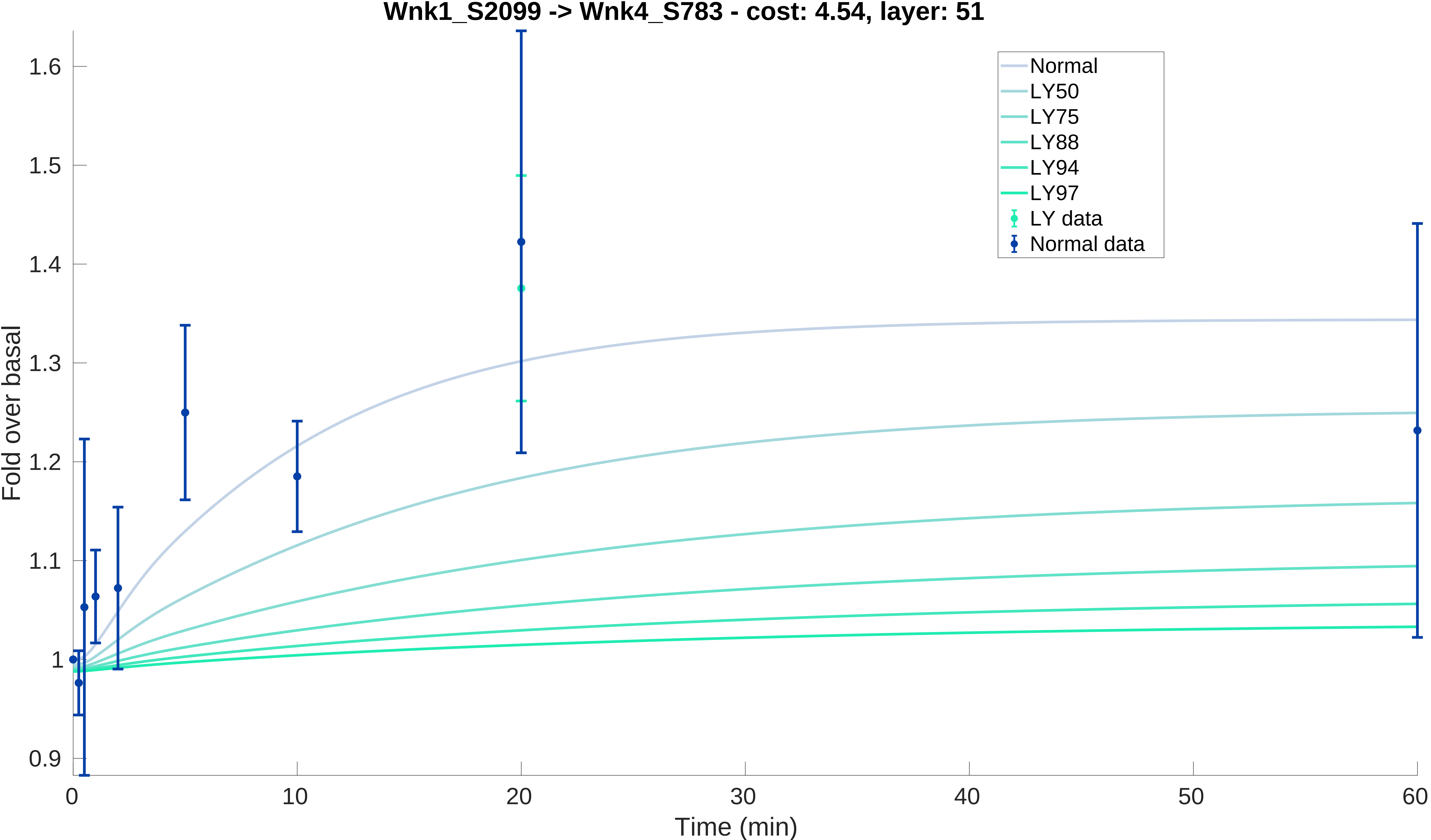
Wnk1_S2099 -> Wnk4_S783 - cost: 4.54, layer: 51
Fold over basal
Time (min)
0.9
1
1.1
1.2
1.3
1.4
1.5
1.6
0
10
20
30
40
50
60
Normal
LY50
LY75
LY88
LY94
LY97
LY data
Normal data

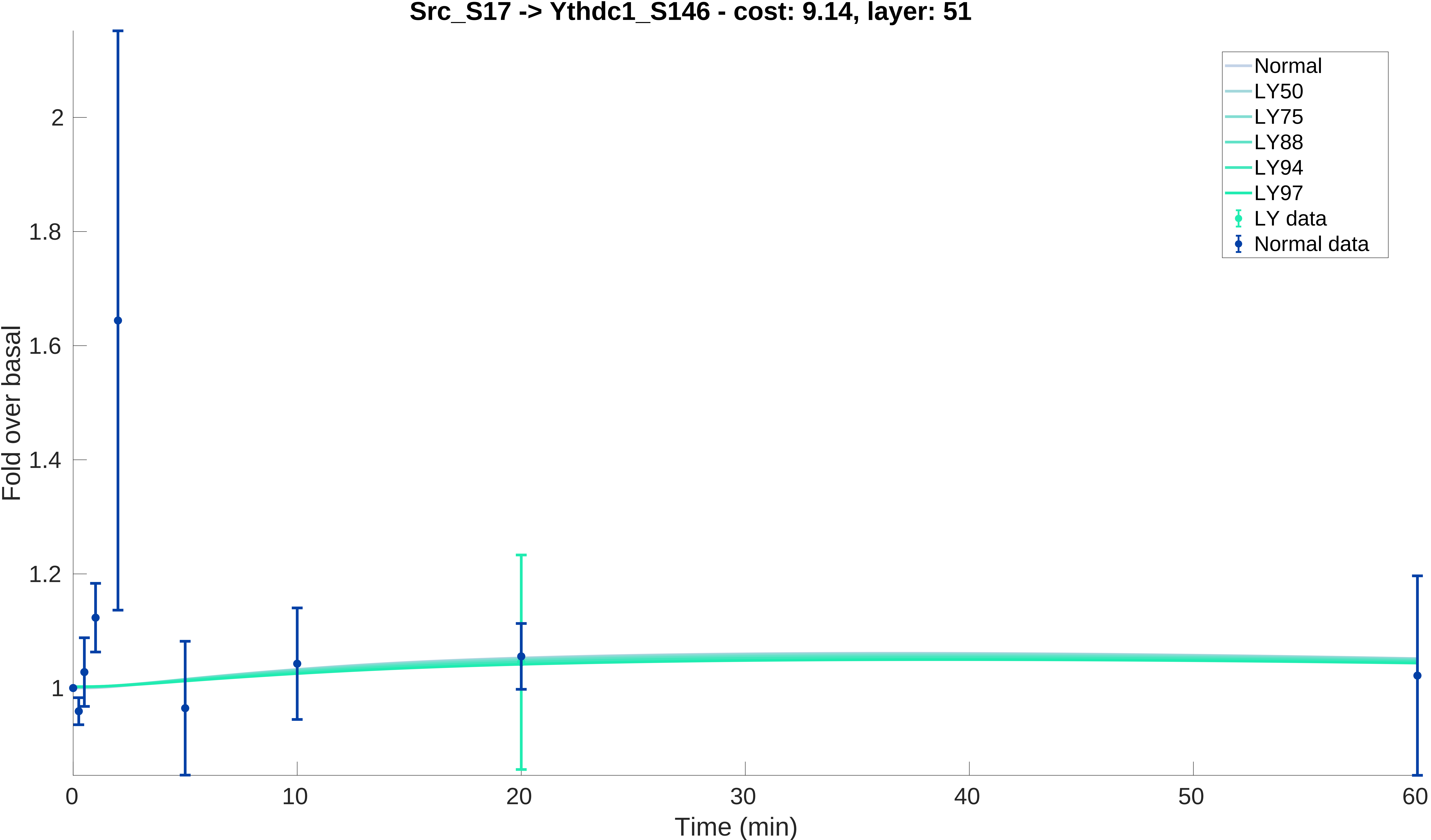

Src_S17 -> Ythdc1_S146 - cost: 9.14, layer: 51
Fold over basal
2
1.8
1.6
1.4
1.2
1
0
10
20
30
40
50
60
Time (min)
Normal
LY50
LY75
LY88
LY94
LY97
LY data
Normal data

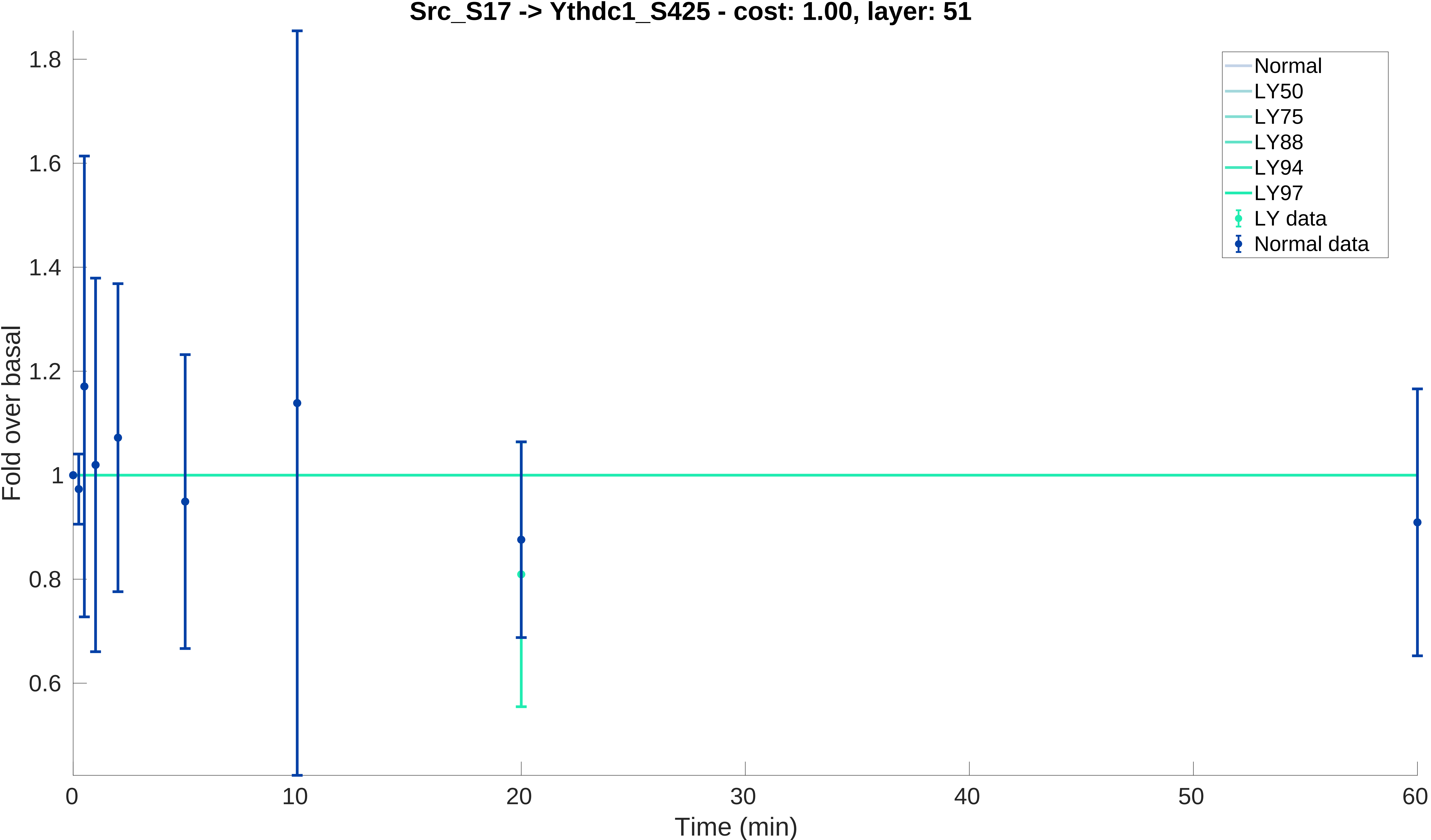

Src_S17 -> Ythdc1_S425 - cost: 1.00, layer: 51
Fold over basal
Time (min)
1.8
1.6
1.4
1.2
1
0.8
0.6
0
10
20
30
40
50
60
Normal
LY50
LY75
LY88
LY94
LY97
LY data
Normal data

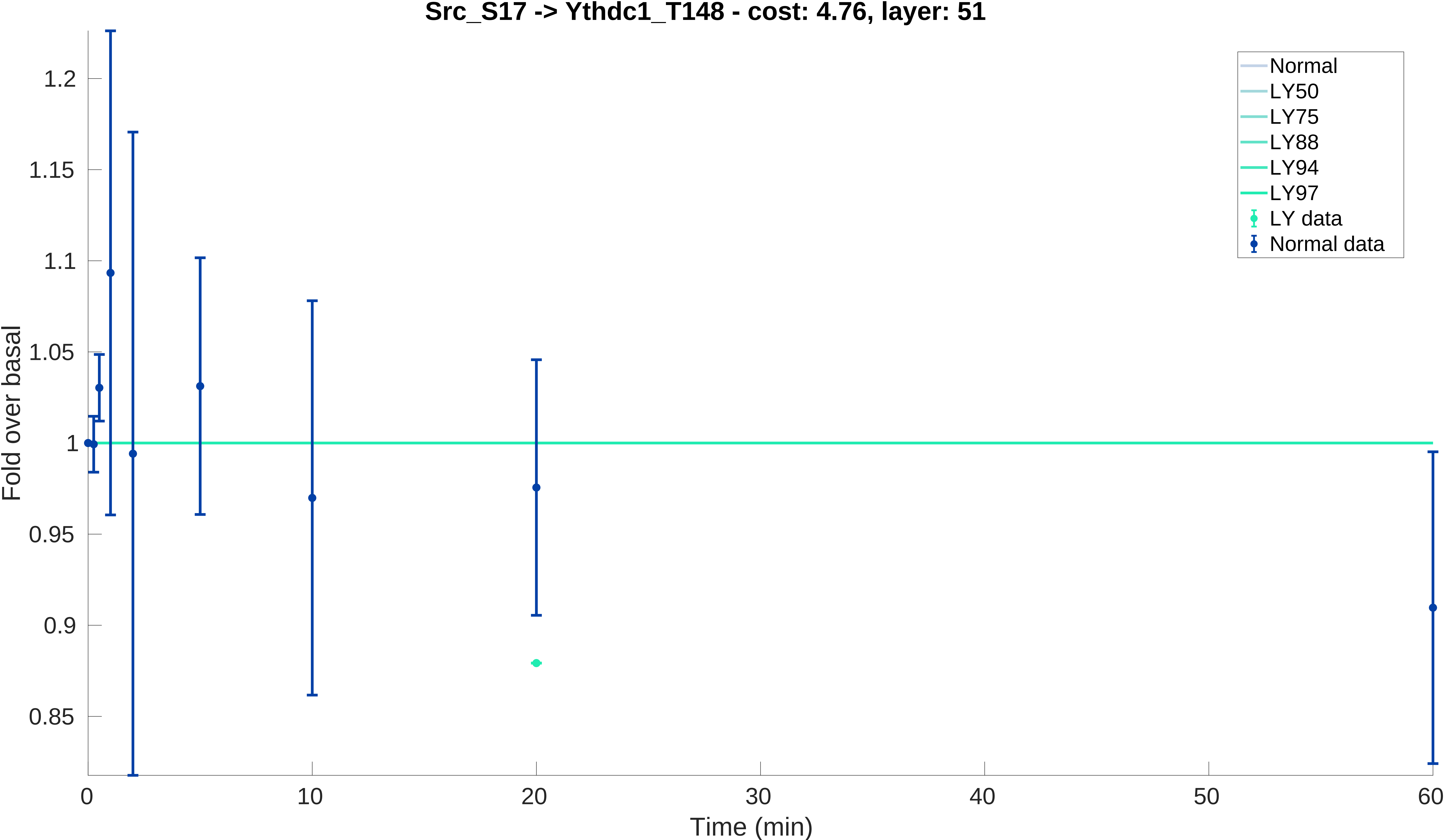

Src_S17 -> Ythdc1_T148 - cost: 4.76, layer: 51
Normal
LY50
LY75
LY88
LY94
LY97
LY data
Normal data
Fold over basal
1.2
1.15
1.1
1.05
1
0.95
0.9
0.85
0
10
20
30
40
50
60
Time (min)

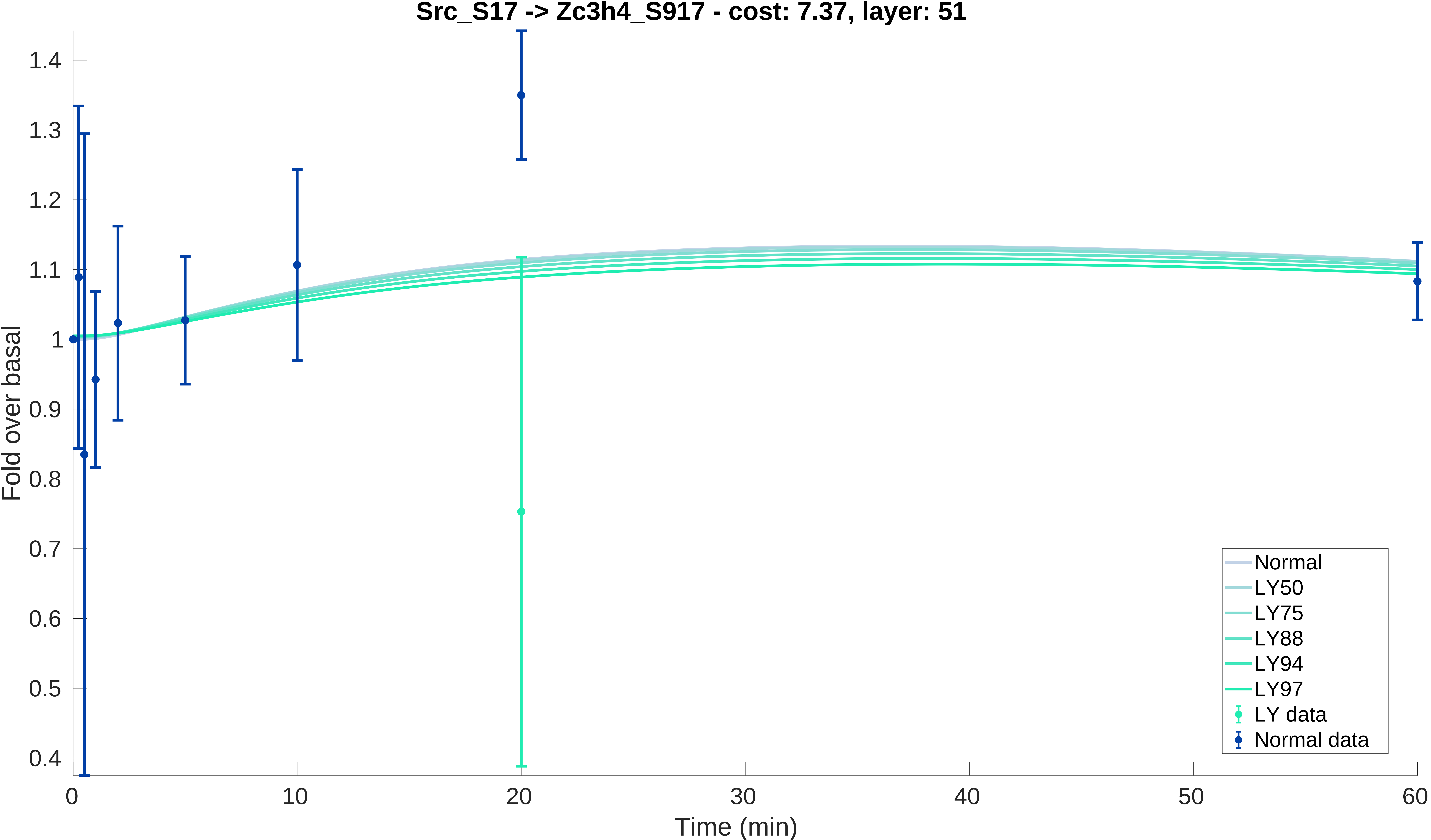

Src_S17 -> Zc3h4_S917 - cost: 7.37, layer: 51
Fold over basal
1.4
1.3
1.2
1.1
1
0.9
0.8
0.7
0.6
0.5
0.4
0
10
20
30
40
50
60
Time (min)
Normal
LY50
LY75
LY88
LY94
LY97
LY data
Normal data

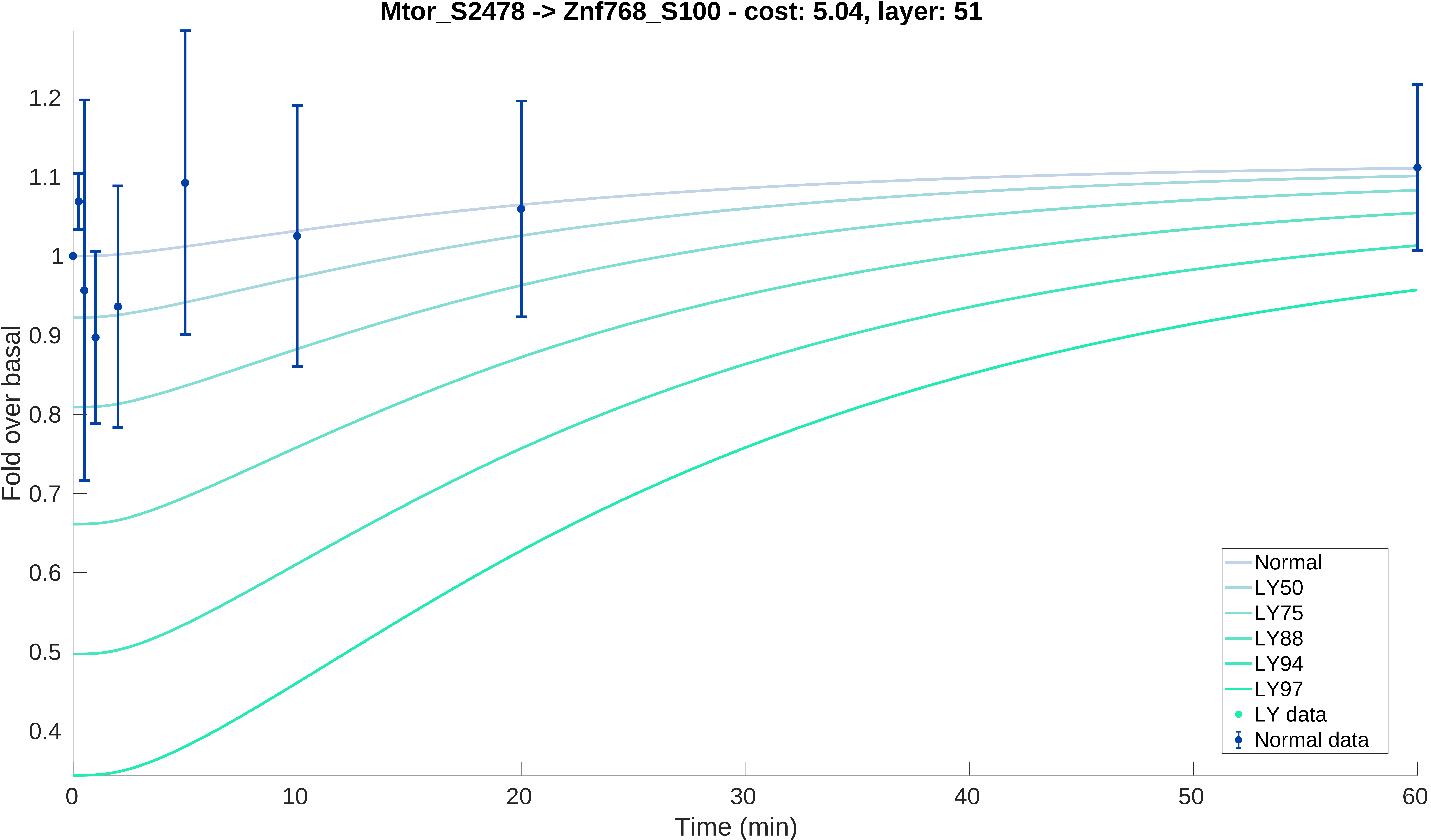

Mtor_S2478 -> Znf768_S100 - cost: 5.04, layer: 51
Fold over basal
1.2
1.1
1
0.9
0.8
0.7
0.6
0.5
0.4
0
10
20
30
40
50
60
Time (min)
Normal
LY50
LY75
LY88
LY94
LY97
LY data
Normal data

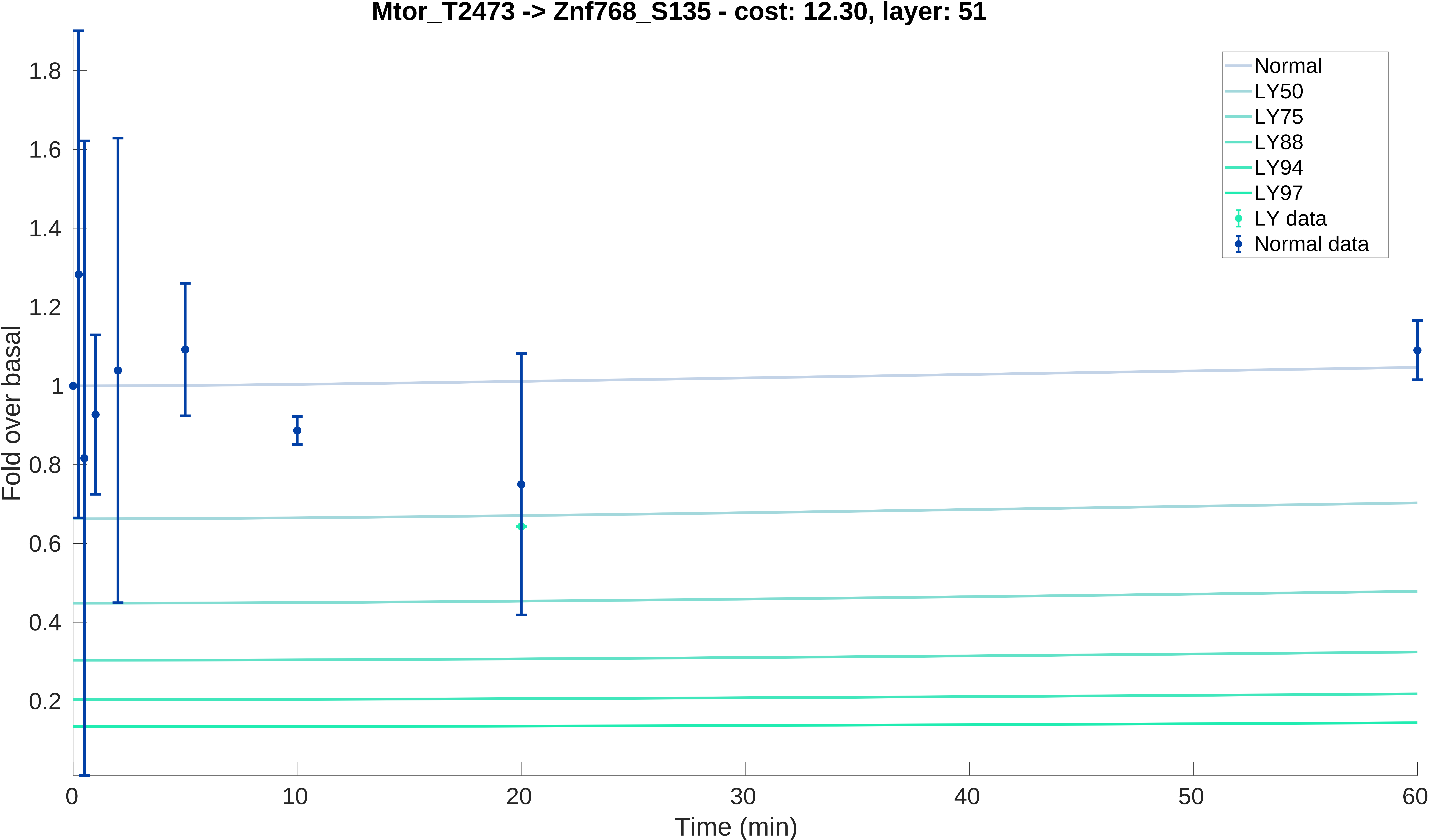

Mtor_T2473 -> Znf768_S135 - cost: 12.30, layer: 51
Fold over basal
Time (min)
Normal
LY50
LY75
LY88
LY94
LY97
LY data
Normal data
0.2
0.4
0.6
0.8
1
1.2
1.4
1.6
1.8
0
10
20
30
40
50
60

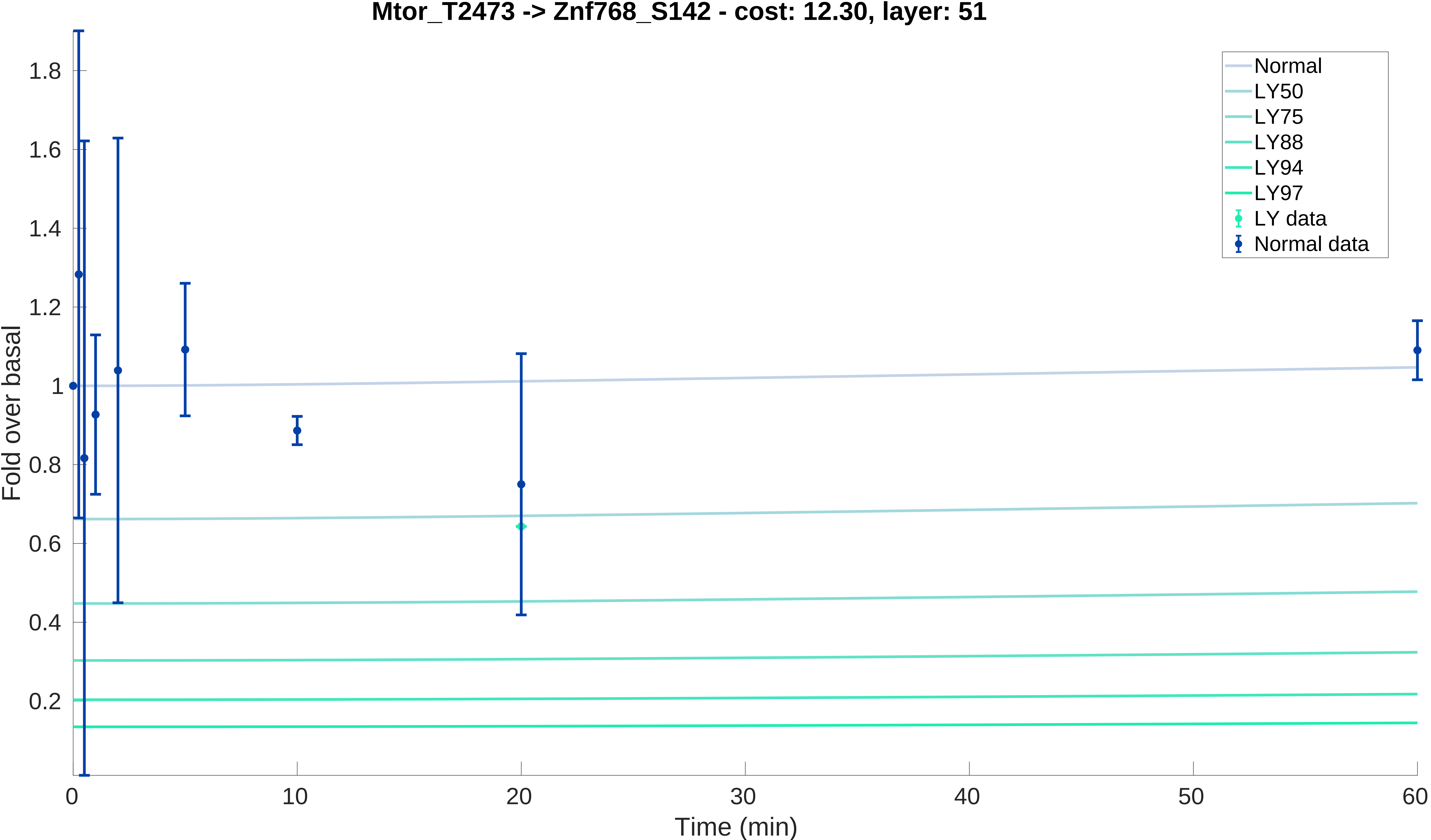

Mtor_T2473 -> Znf768_S142 - cost: 12.30, layer: 51
Fold over basal
Time (min)
Normal
LY50
LY75
LY88
LY94
LY97
LY data
Normal data
1.8
1.6
1.4
1.2
1
0.8
0.6
0.4
0.2
0
10
20
30
40
50
60

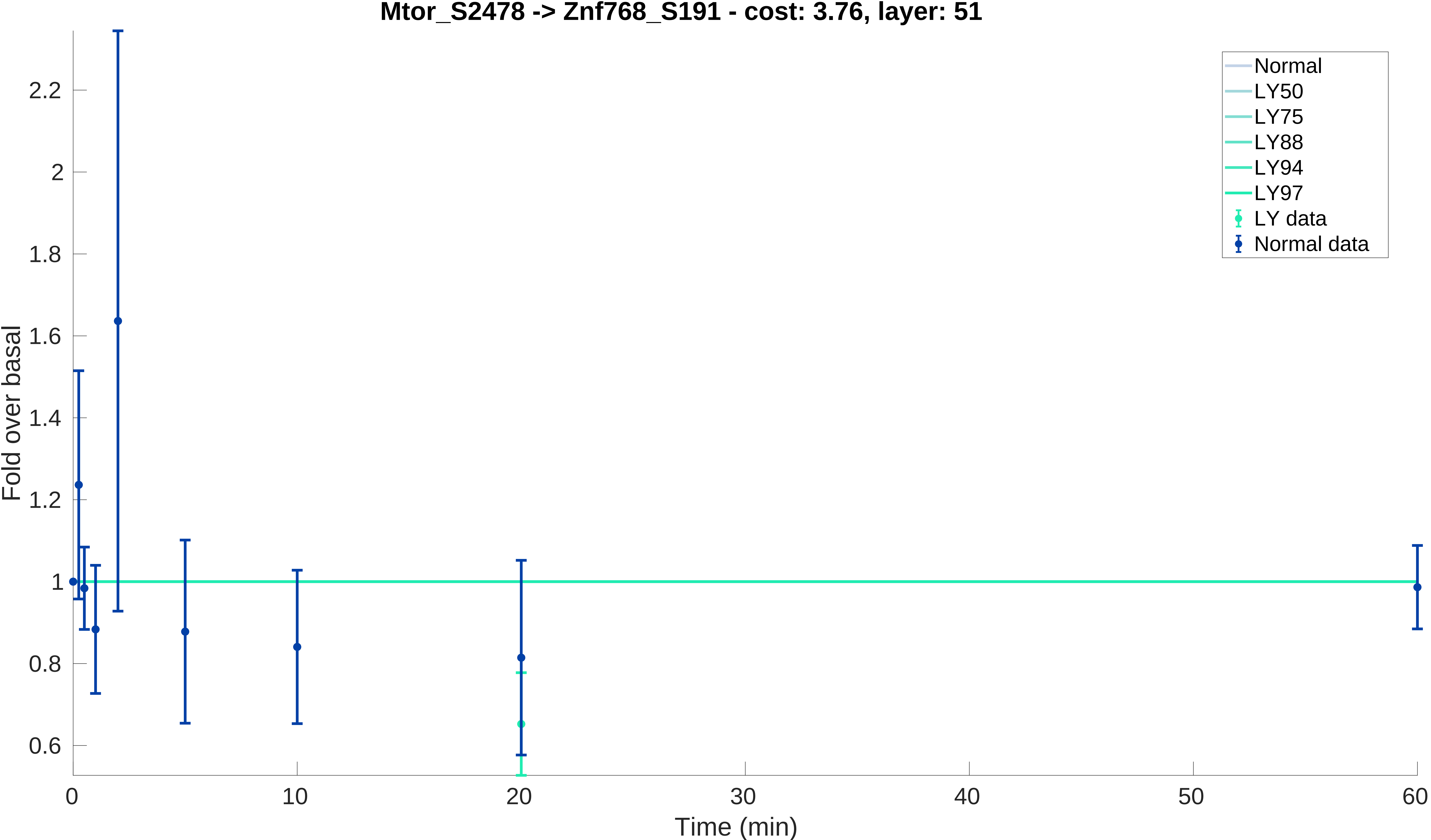

Mtor_S2478 -> Znf768_S191 - cost: 3.76, layer: 51
Fold over basal
Time (min)
0.6
0.8
1
1.2
1.4
1.6
1.8
2
2.2
0
10
20
30
40
50
60
Normal
LY50
LY75
LY88
LY94
LY97
LY data
Normal data

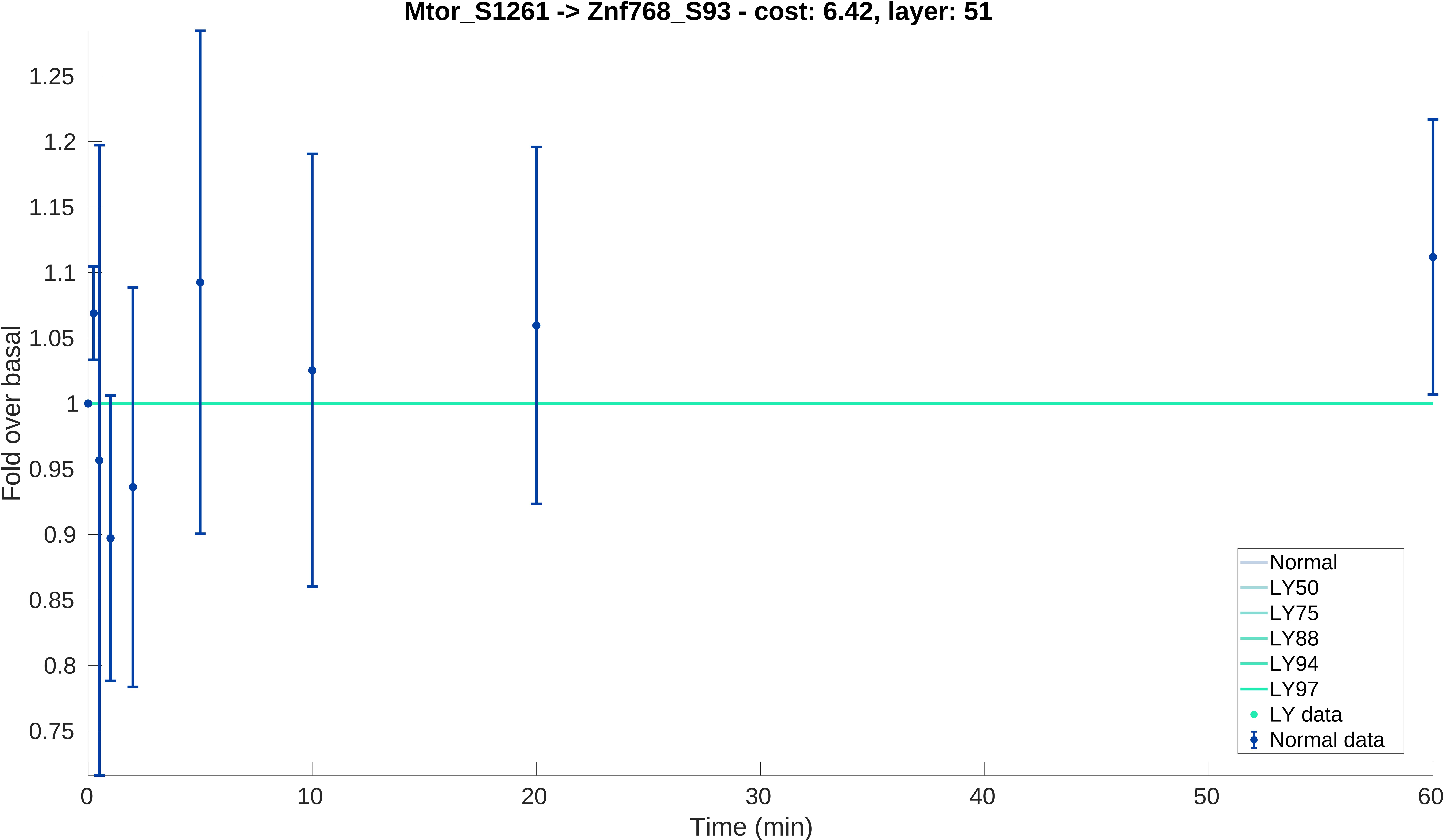

Mtor_S1261 -> Znf768_S93 - cost: 6.42, layer: 51
Fold over basal
Time (min)
1.25
1.2
1.15
1.1
1.05
1
0.95
0.9
0.85
0.8
0.75
0
10
20
30
40
50
60
Normal
LY50
LY75
LY88
LY94
LY97
LY data
Normal data

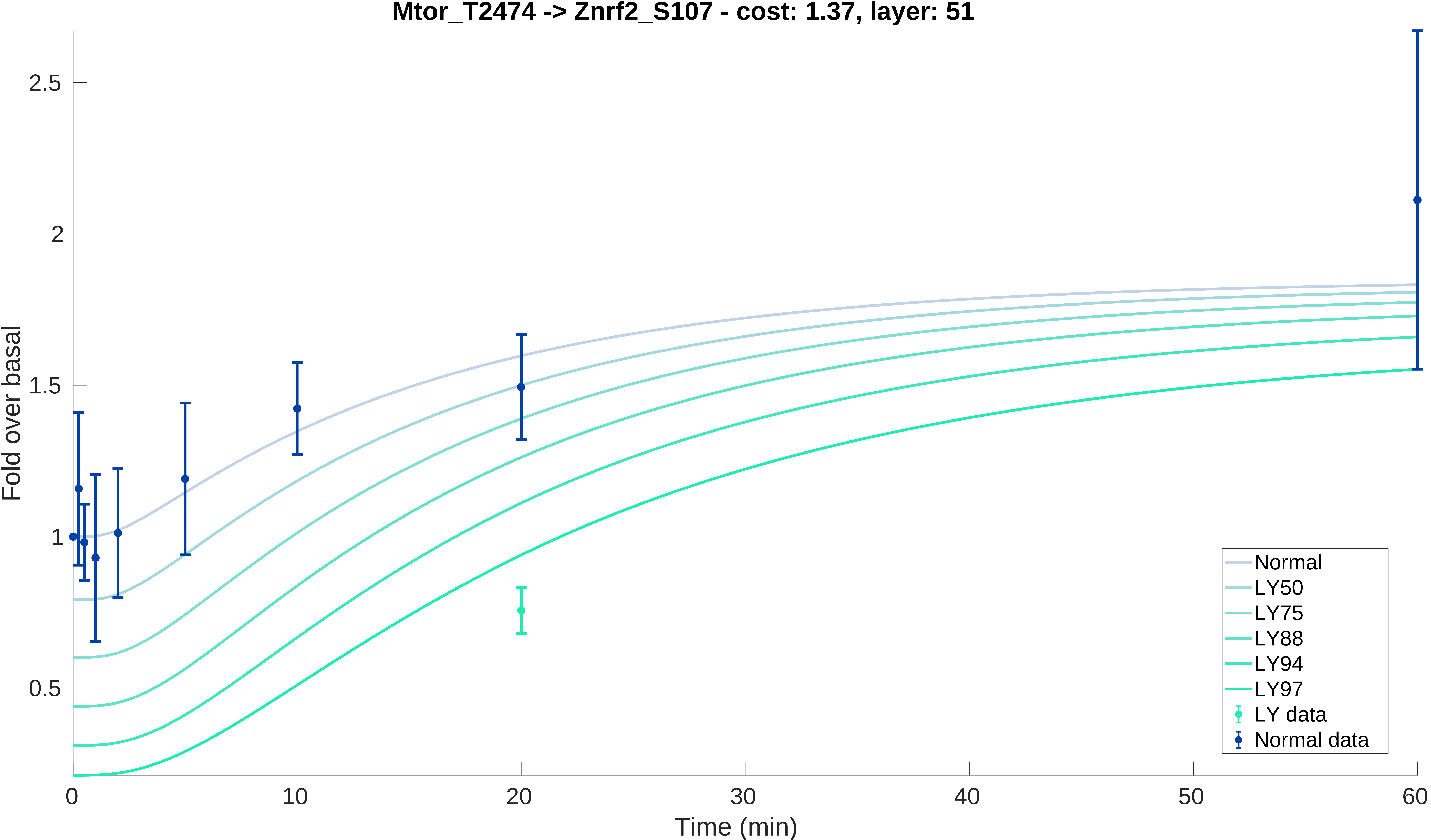
Mtor_T2474 -> Znrf2_S107 - cost: 1.37, layer: 51
Fold over basal
Time (min)
0
10
20
30
40
50
60
0.5
1
1.5
2
2.5
Normal
LY50
LY75
LY88
LY94
LY97
LY data
Normal data

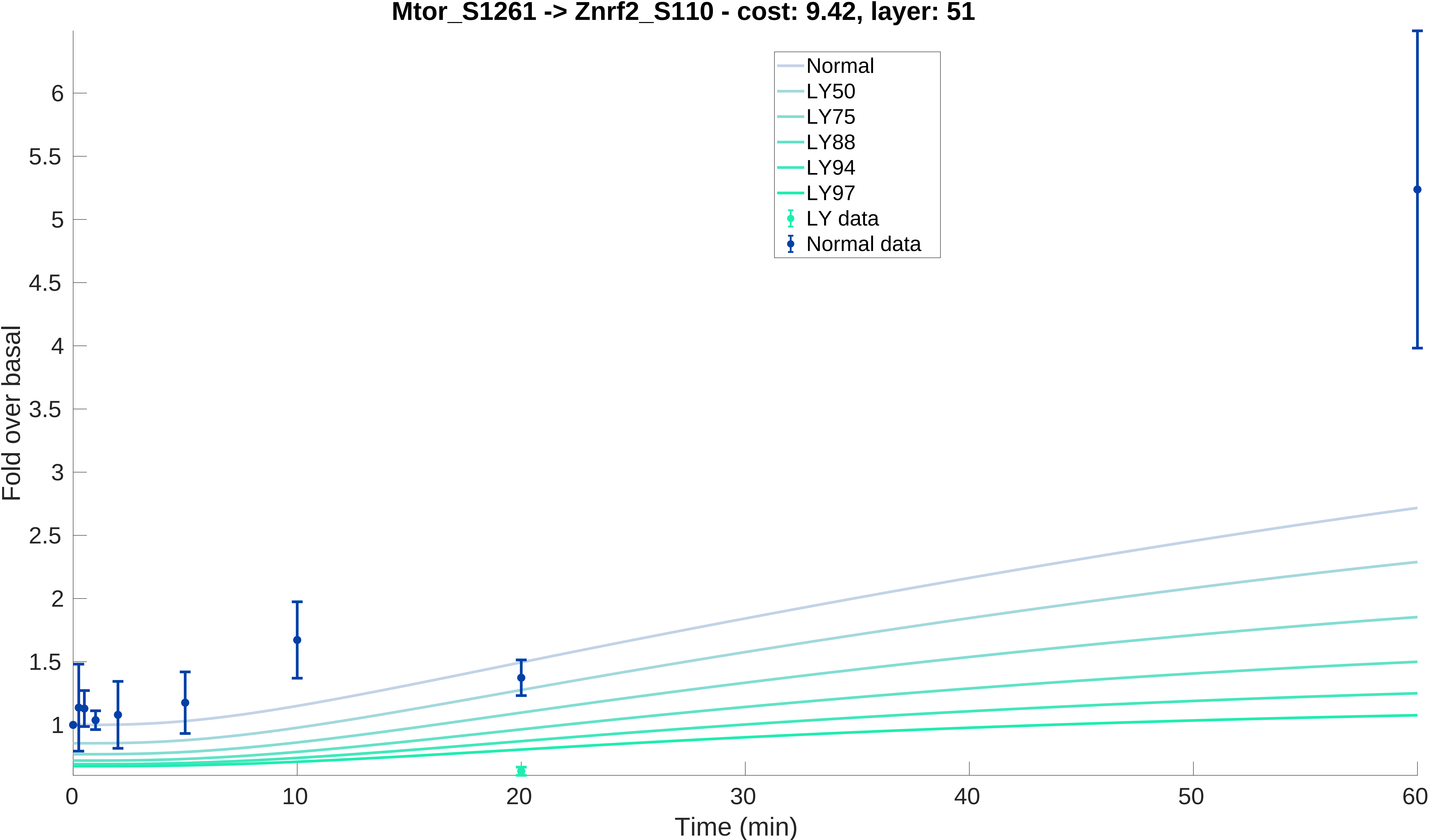
Mtor_S1261 -> Znrf2_S110 - cost: 9.42, layer: 51
Fold over basal
Time (min)
Normal
LY50
LY75
LY88
LY94
LY97
LY data
Normal data
6
5.5
5
4.5
4
3.5
3
2.5
2
1.5
1
0
10
20
30
40
50
60

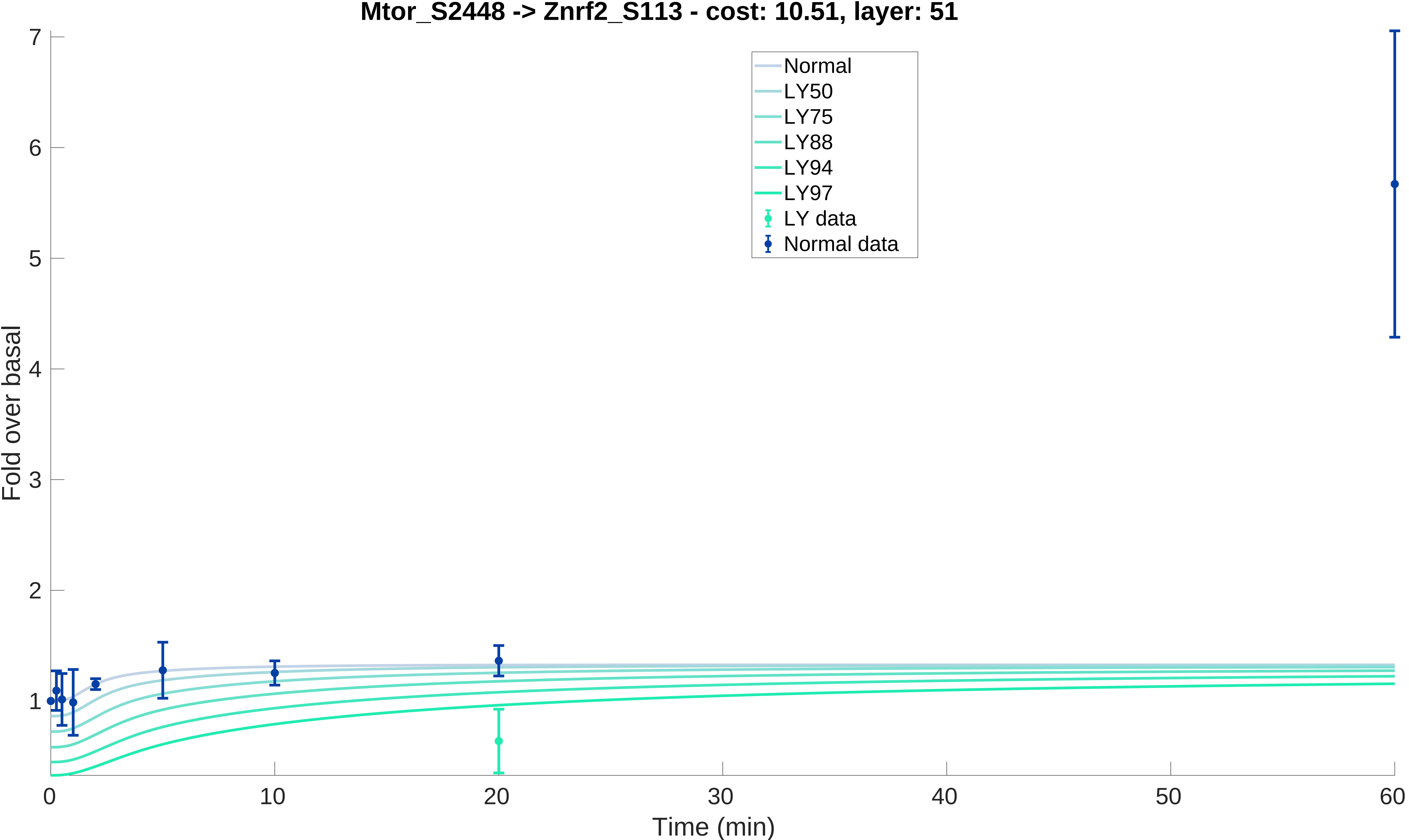

Mtor_S2448 -> Znrf2_S113 - cost: 10.51, layer: 51
Normal
LY50
LY75
LY88
LY94
LY97
LY data
Normal data
Fold over basal
Time (min)
0
10
20
30
40
50
60
1
2
3
4
5
6
7

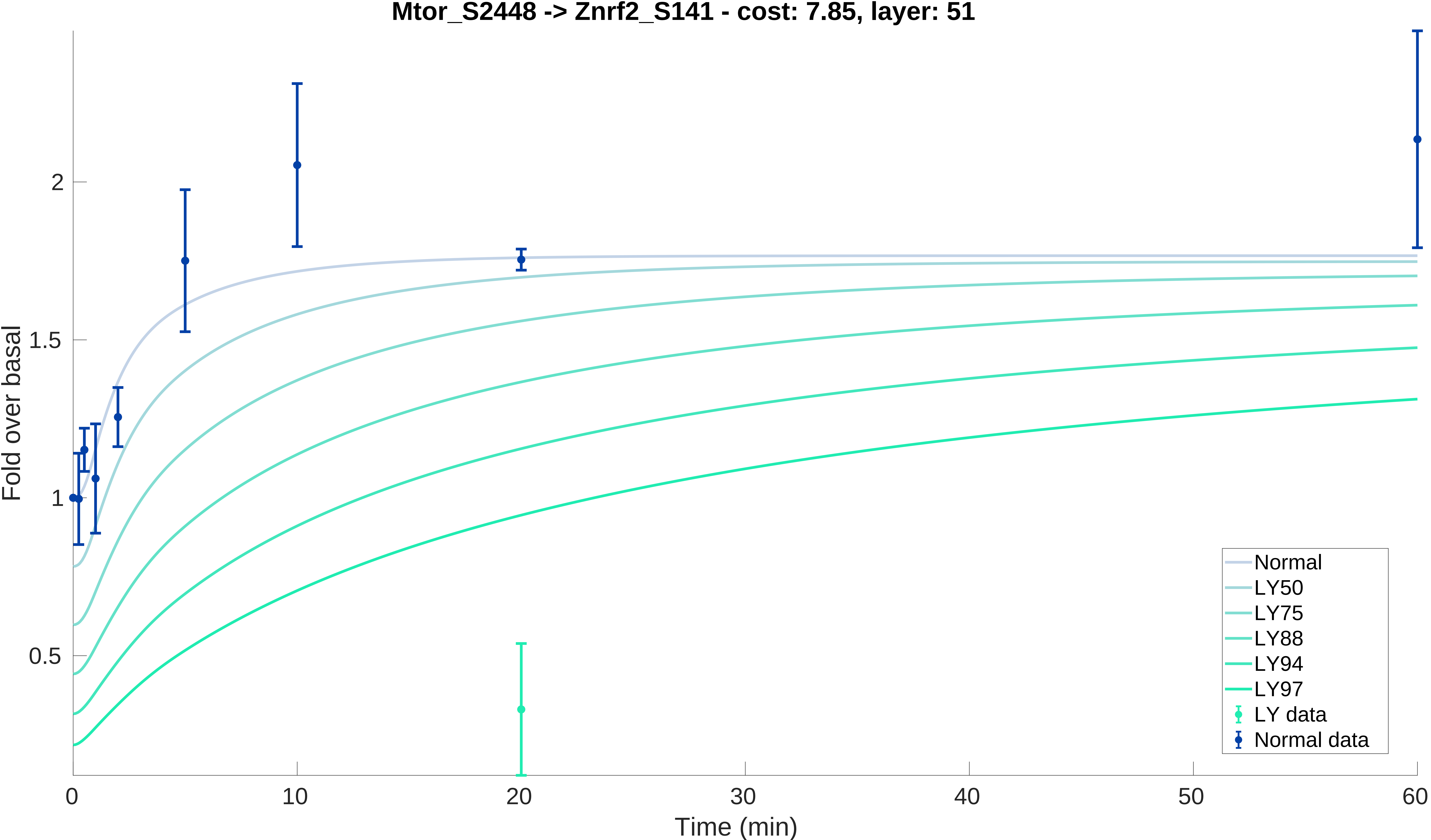

Mtor_S2448 -> Znrf2_S141 - cost: 7.85, layer: 51
Fold over basal
Time (min)
0.5
1
1.5
2
0
10
20
30
40
50
60
Normal
LY50
LY75
LY88
LY94
LY97
LY data
Normal data

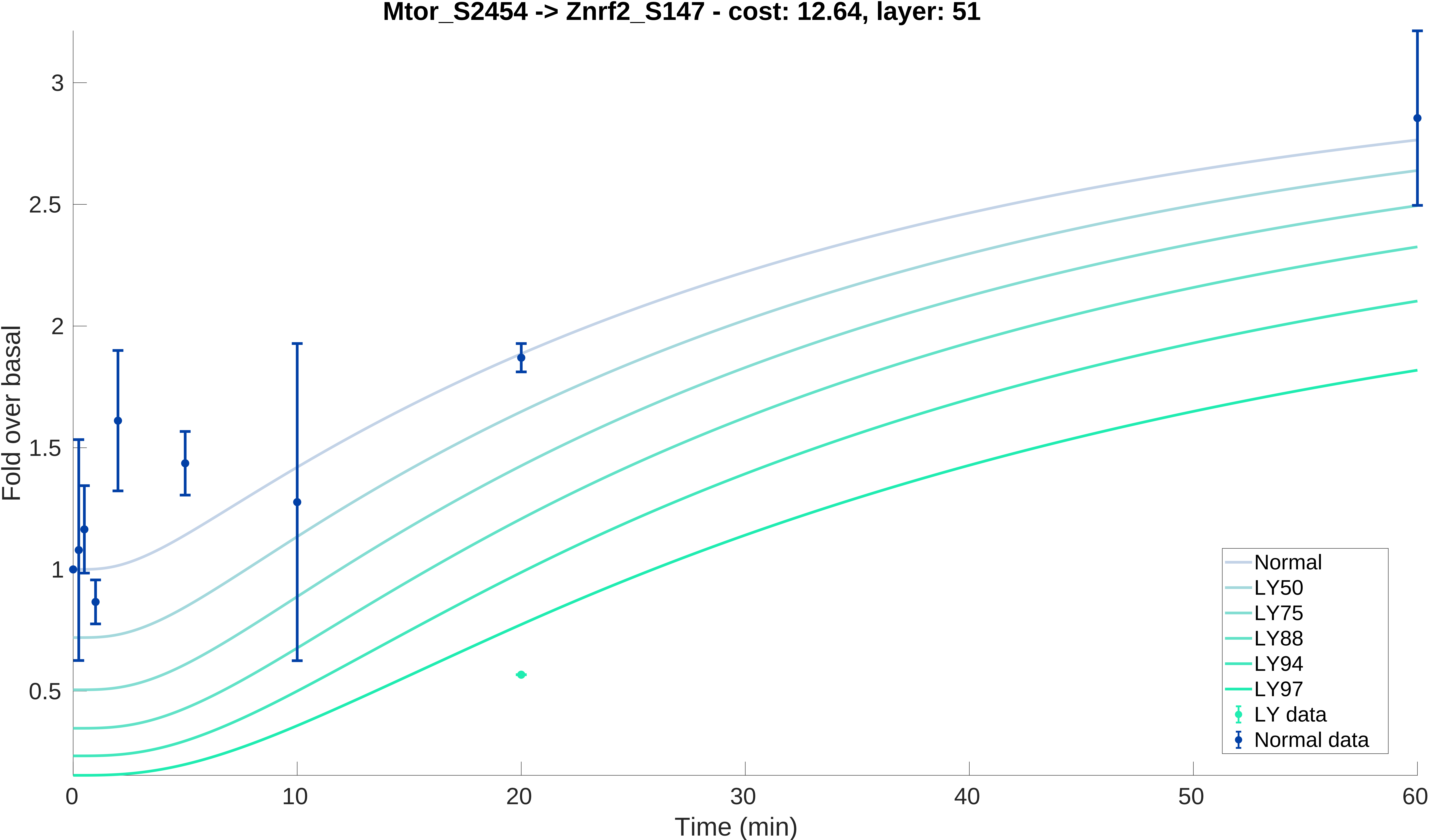

Mtor_S2454 -> Znrf2_S147 - cost: 12.64, layer: 51
Fold over basal
Time (min)
0.5
1
1.5
2
2.5
3
0
10
20
30
40
50
60
Normal
LY50
LY75
LY88
LY94
LY97
LY data
Normal data

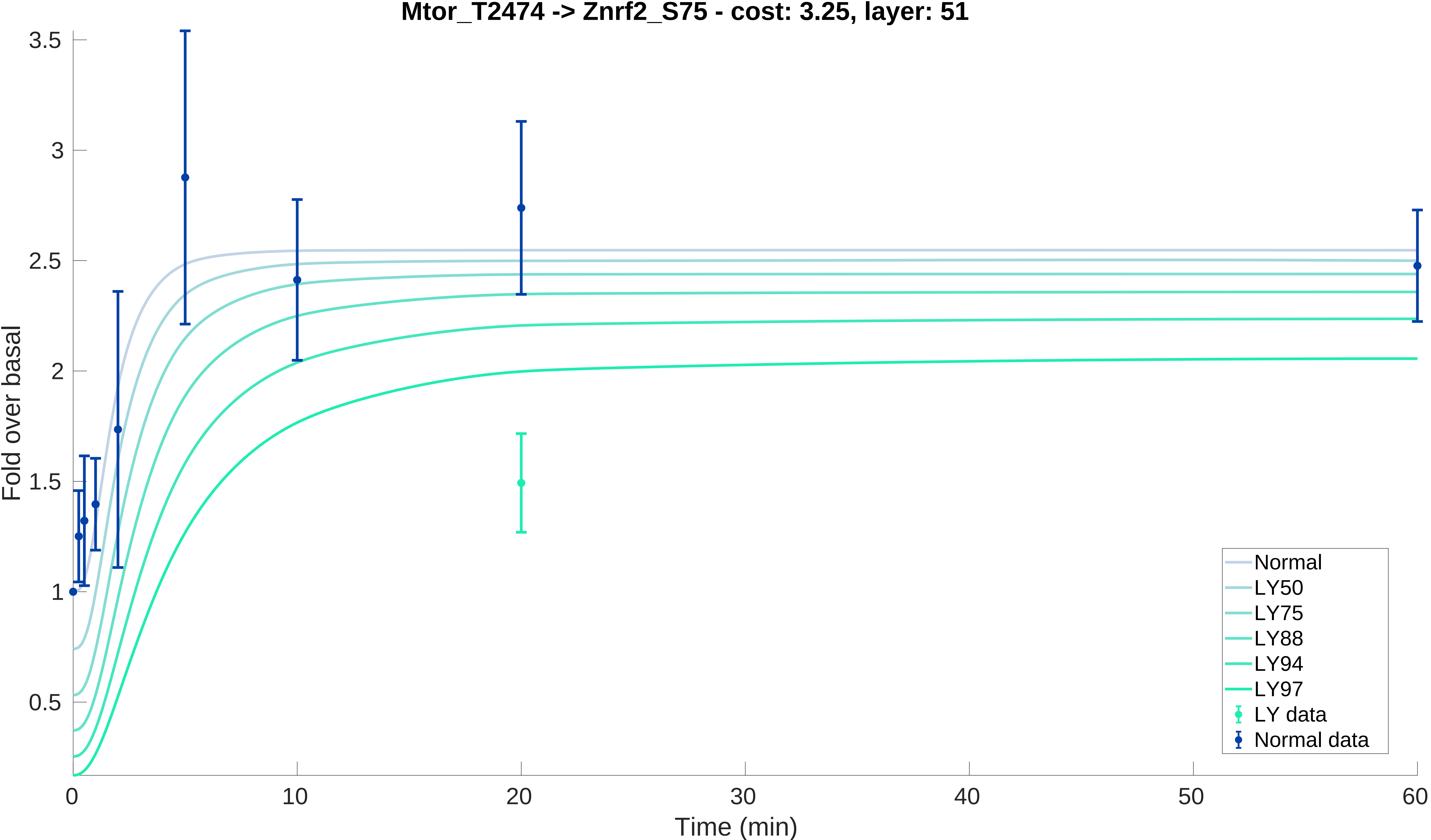

Mtor_T2474 -> Znrf2_S75 - cost: 3.25, layer: 51
Fold over basal
3.5
3
2.5
2
1.5
1
0.5
0
10
20
30
40
50
60
Time (min)
Normal
LY50
LY75
LY88
LY94
LY97
LY data
Normal data

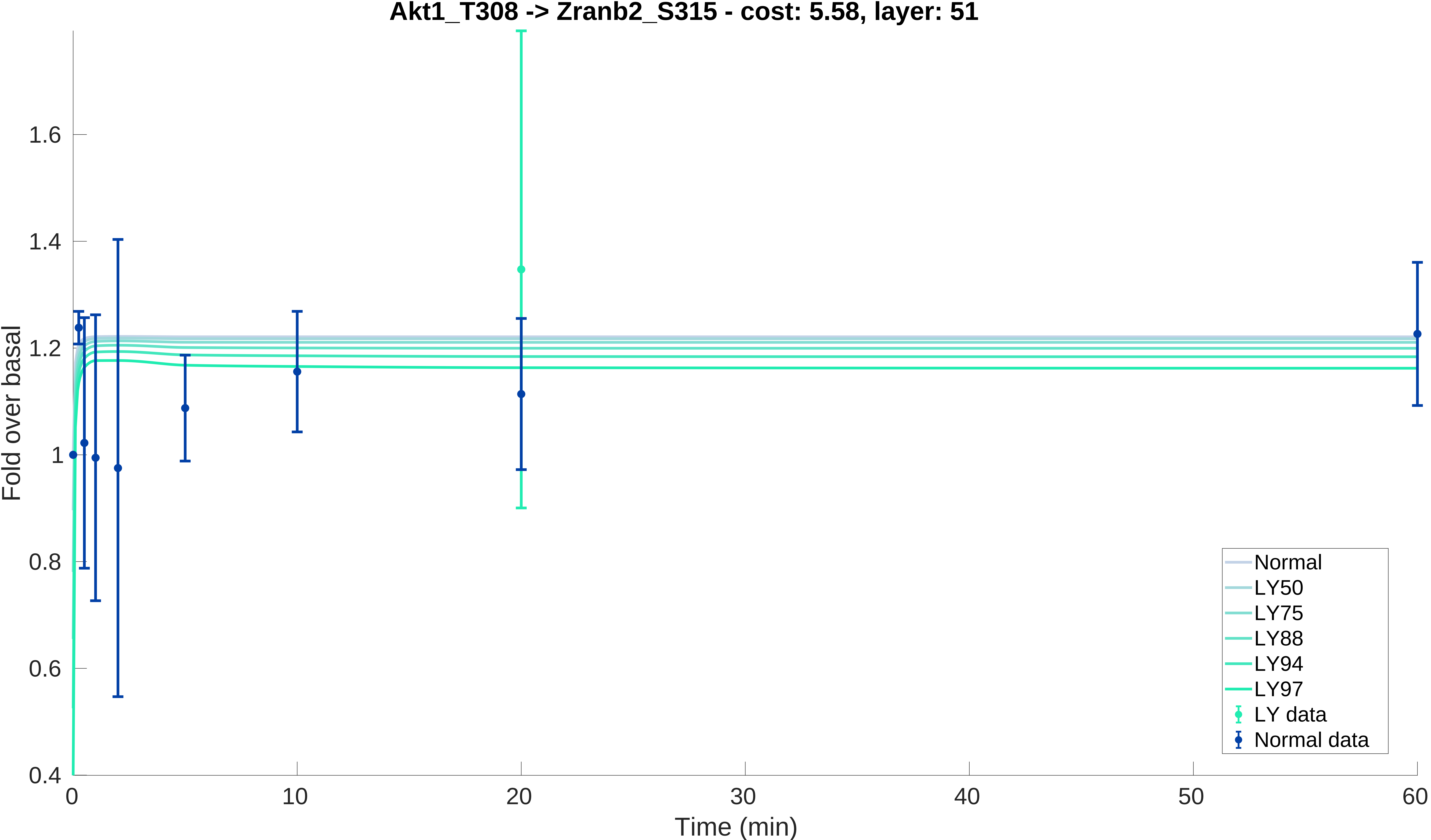

Akt1_T308 -> Zranb2_S315 - cost: 5.58, layer: 51
Fold over basal
1.6
1.4
1.2
1
0.8
0.6
0.4
0
10
20
30
40
50
60
Time (min)
Normal
LY50
LY75
LY88
LY94
LY97
LY data
Normal data

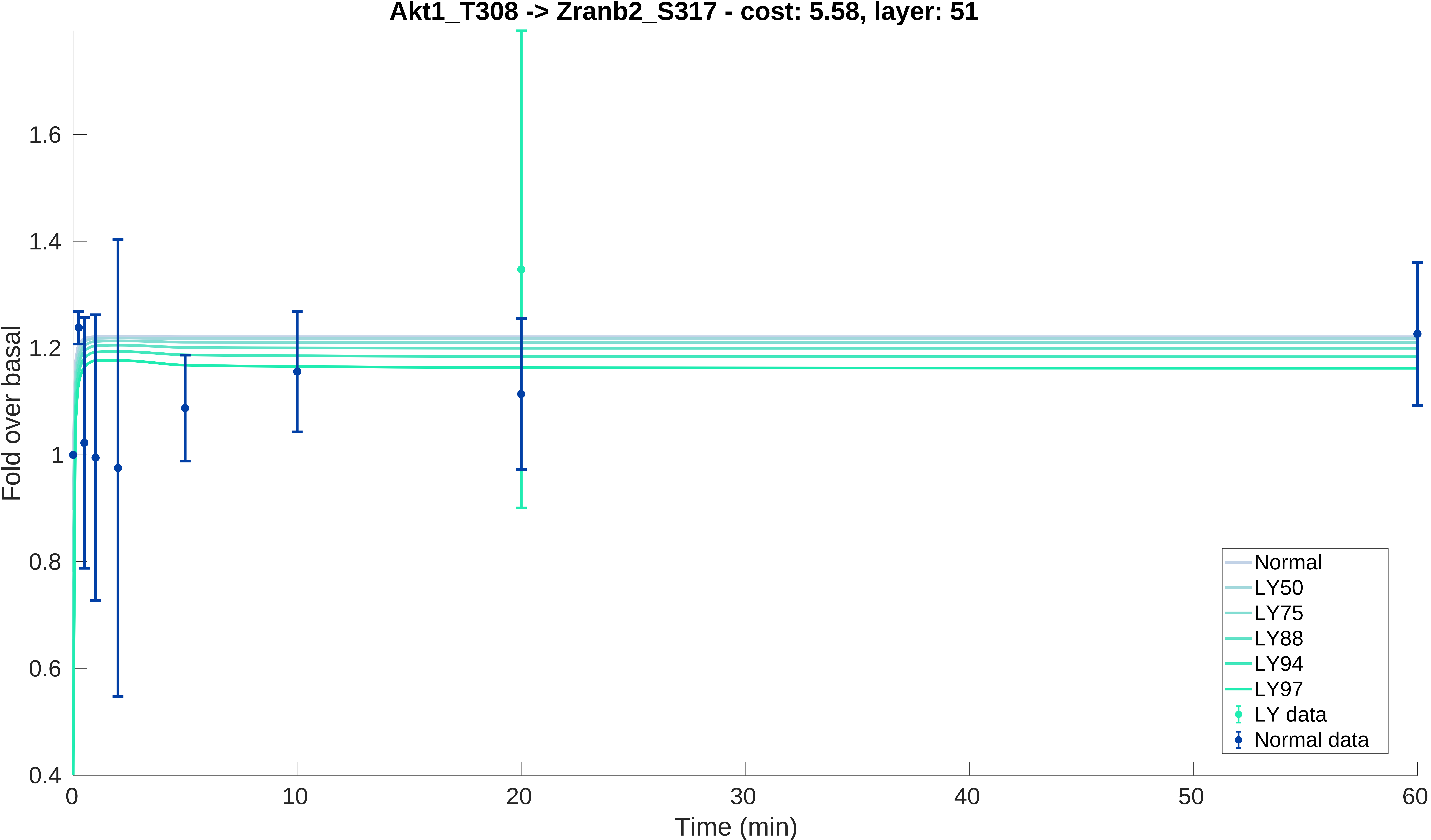

Akt1_T308 -> Zranb2_S317 - cost: 5.58, layer: 51
Fold over basal
Time (min)
0.4
0.6
0.8
1
1.2
1.4
1.6
0
10
20
30
40
50
60
Normal
LY50
LY75
LY88
LY94
LY97
LY data
Normal data

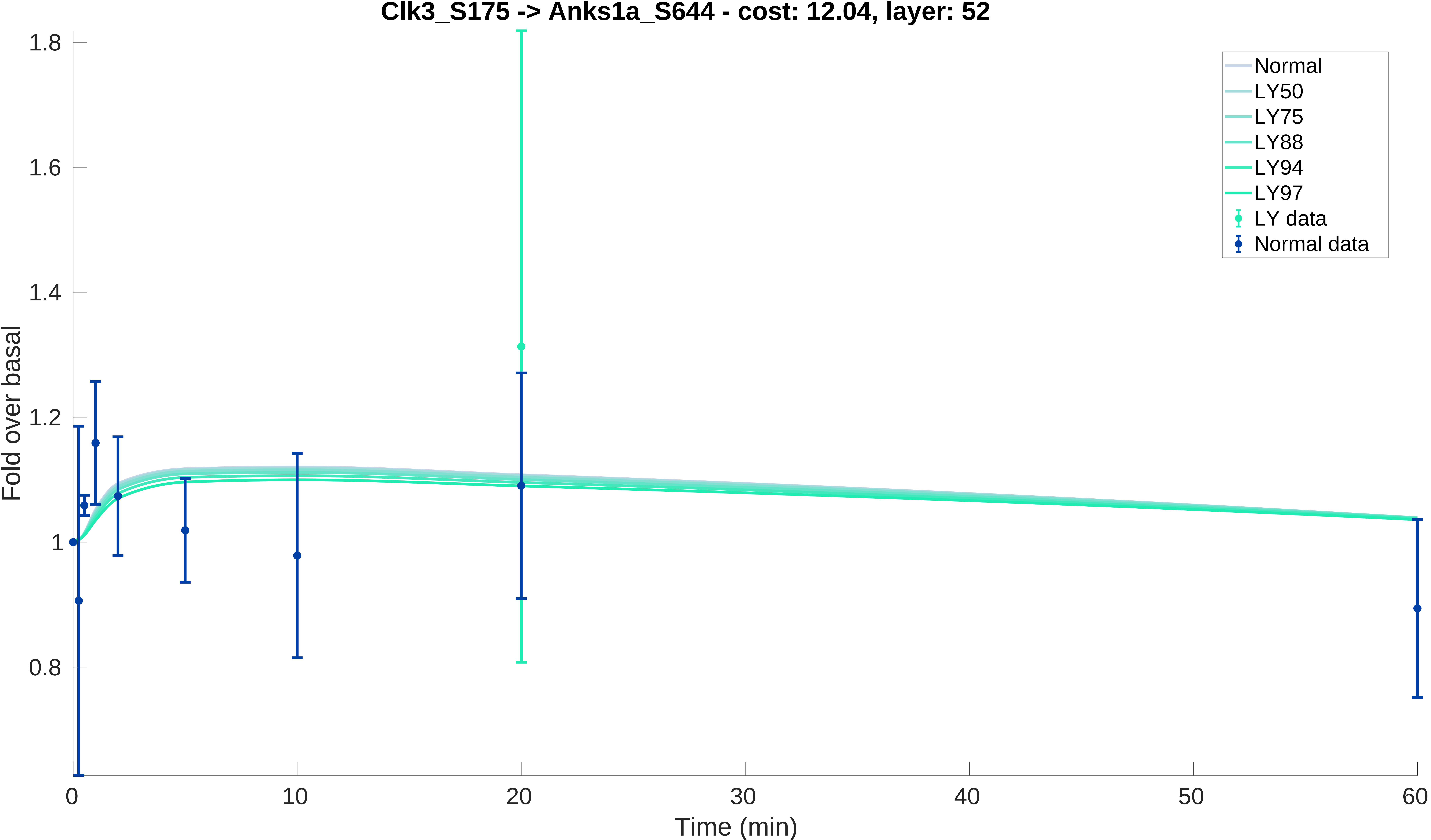

Clk3_S175 -> Anks1a_S644 - cost: 12.04, layer: 52
Fold over basal
Time (min)
1.8
1.6
1.4
1.2
1
0.8
0
10
20
30
40
50
60
Normal
LY50
LY75
LY88
LY94
LY97
LY data
Normal data

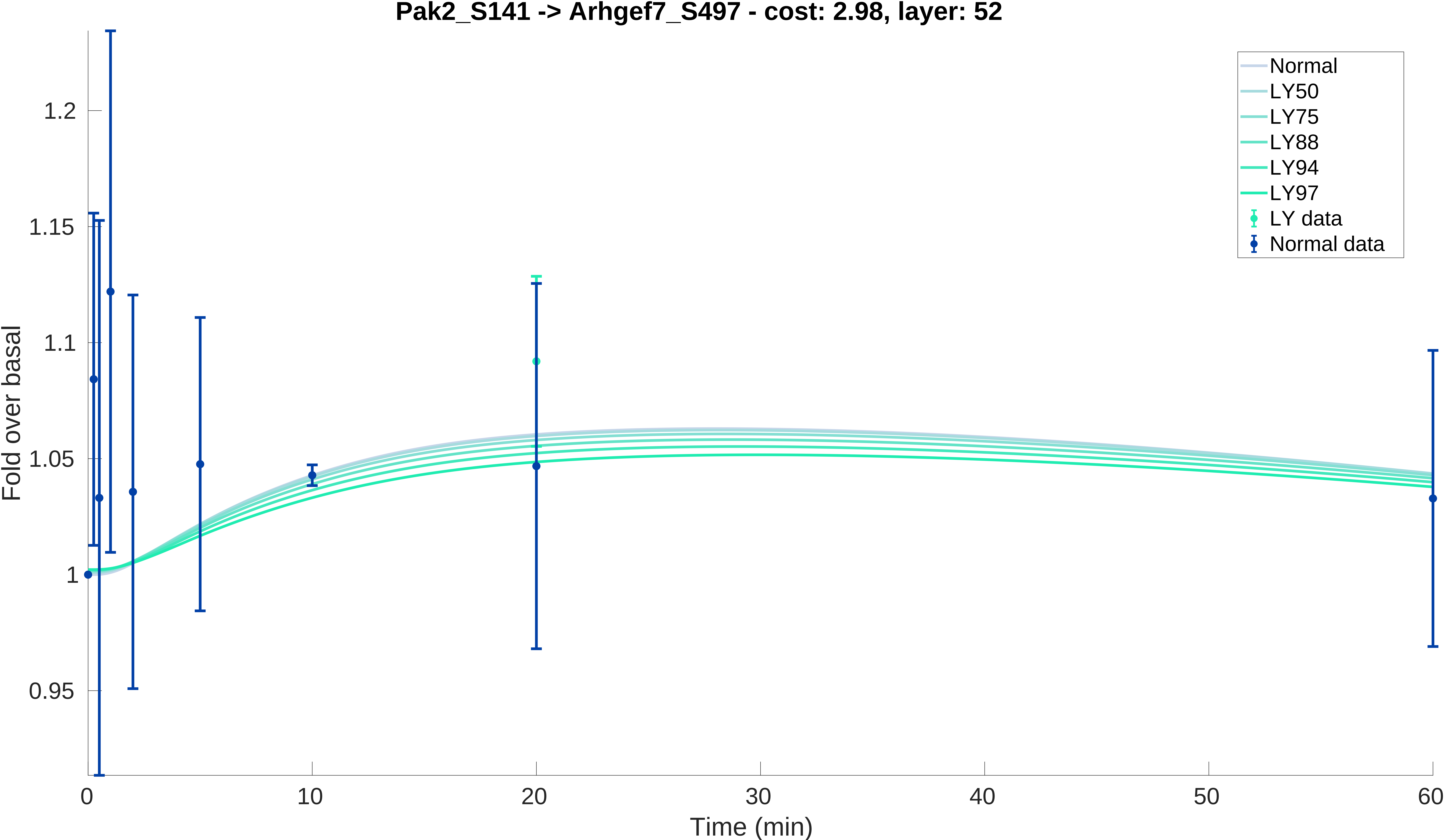

Pak2_S141 -> Arhgef7_S497 - cost: 2.98, layer: 52
Normal
LY50
LY75
LY88
LY94
LY97
LY data
Normal data
Fold over basal
Time (min)
0.95
1
1.05
1.1
1.15
1.2
0
10
20
30
40
50
60

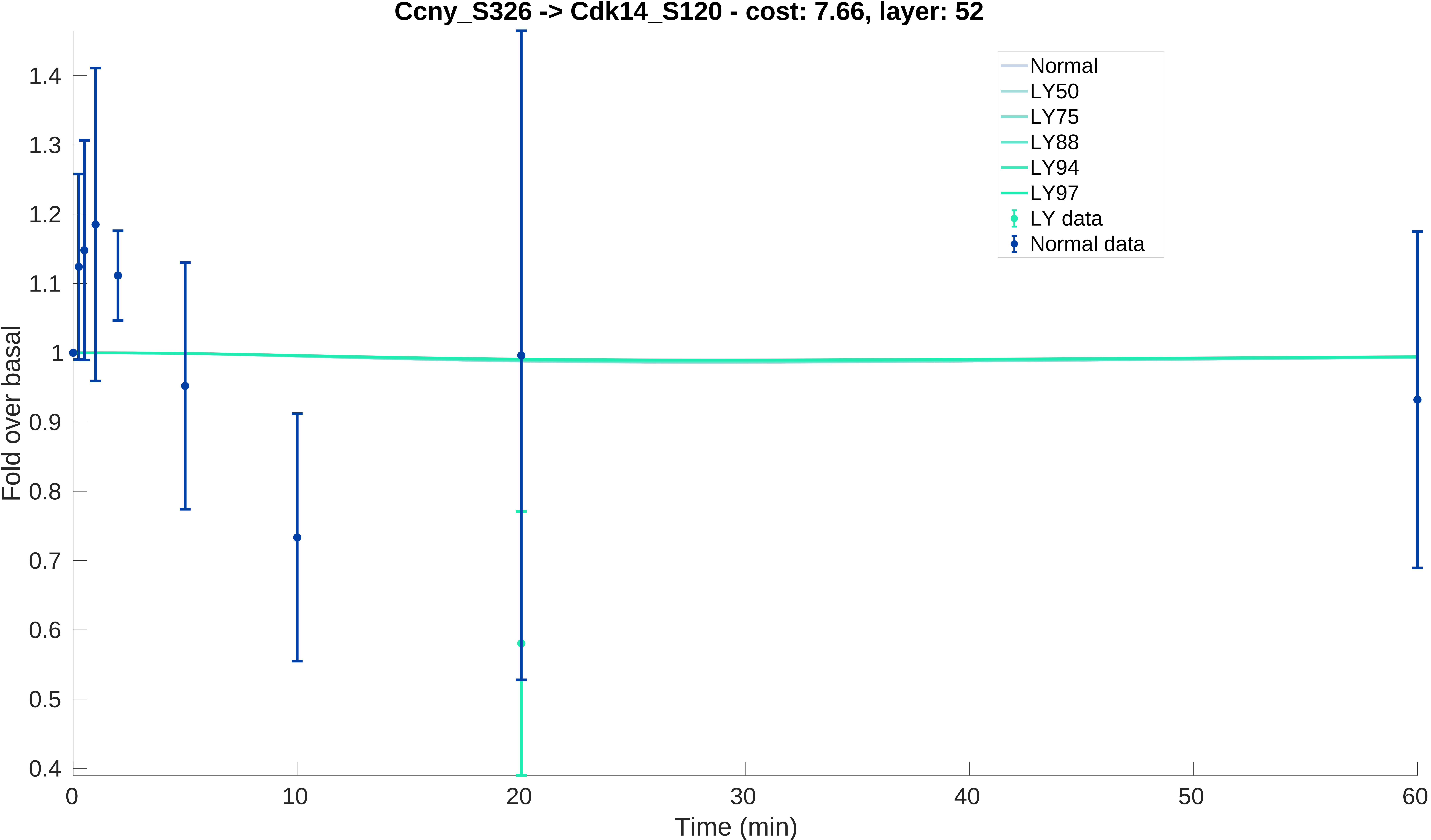
Ccny_S326 -> Cdk14_S120 - cost: 7.66, layer: 52
Normal
LY50
LY75
LY88
LY94
LY97
LY data
Normal data
Fold over basal
Time (min)
1.4
1.3
1.2
1.1
1
0.9
0.8
0.7
0.6
0.5
0.4
0
10
20
30
40
50
60

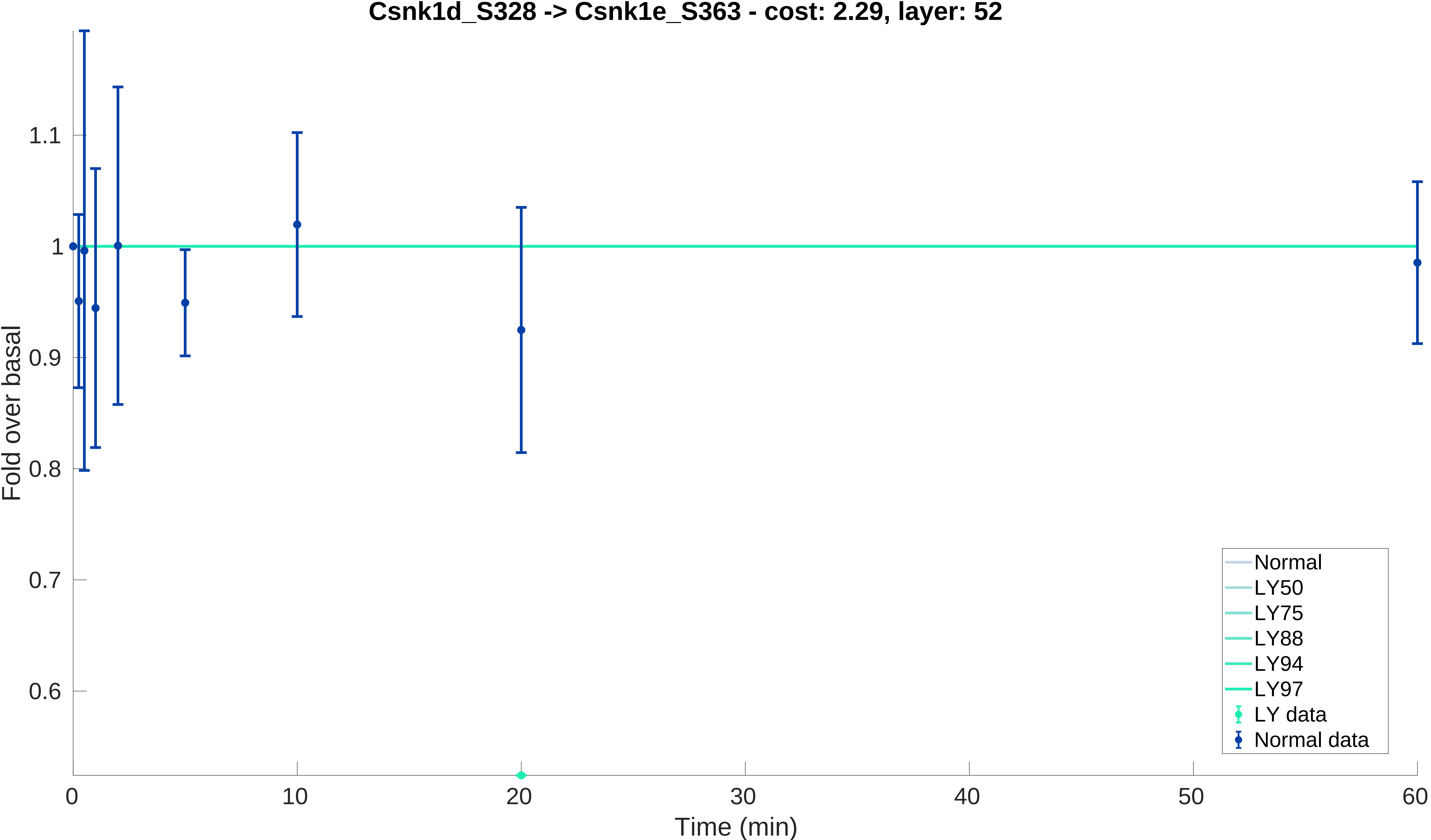

Csnk1d_S328 -> Csnk1e_S363 - cost: 2.29, layer: 52
Fold over basal
1.1
1
0.9
0.8
0.7
0.6
0
10
20
30
40
50
60
Time (min)
Normal
LY50
LY75
LY88
LY94
LY97
LY data
Normal data

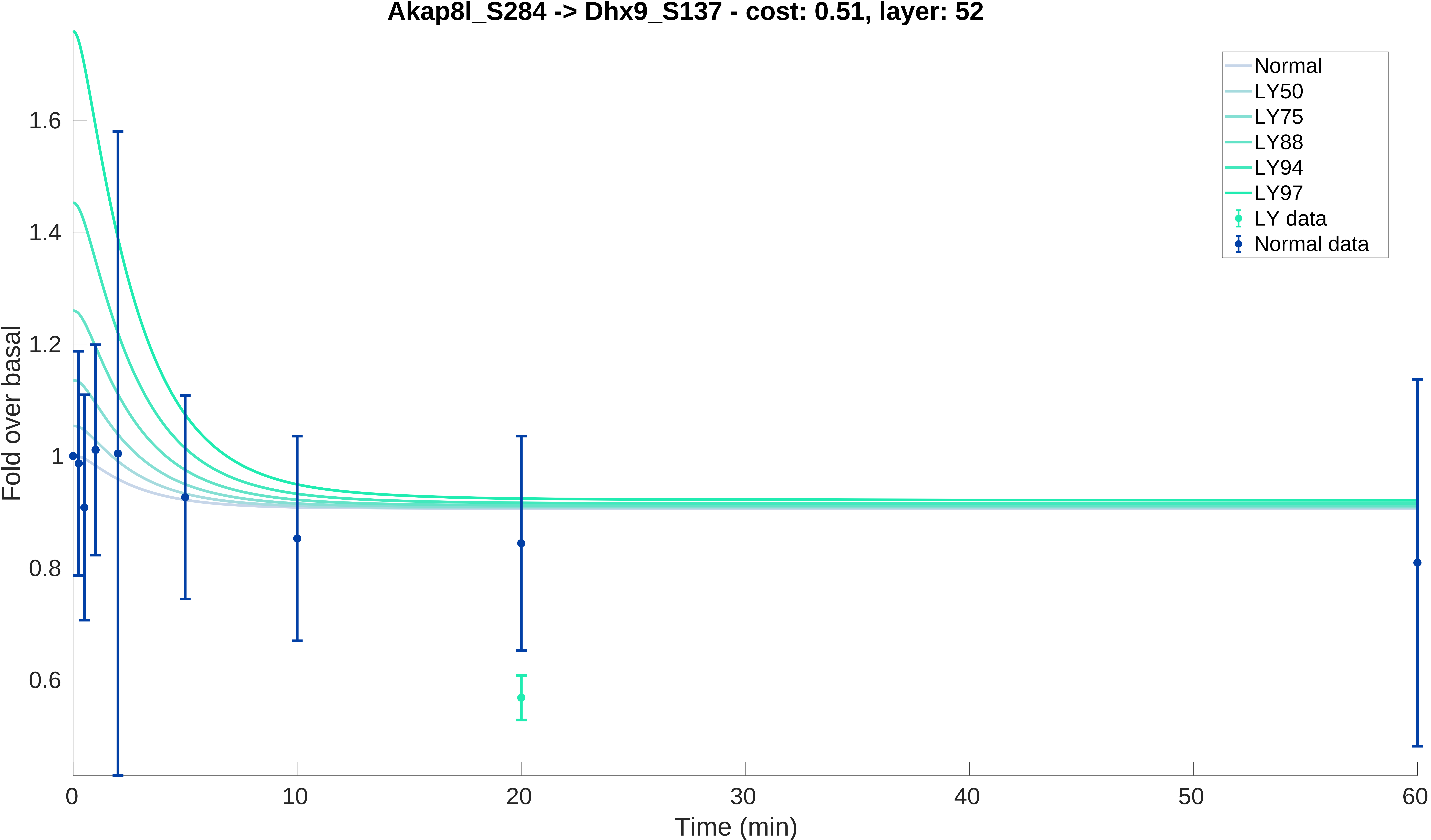

Akap8l_S284 -> Dhx9_S137 - cost: 0.51, layer: 52
Normal
LY50
LY75
LY88
LY94
LY97
LY data
Normal data
Fold over basal
1.6
1.4
1.2
1
0.8
0.6
0
10
20
30
40
50
60
Time (min)

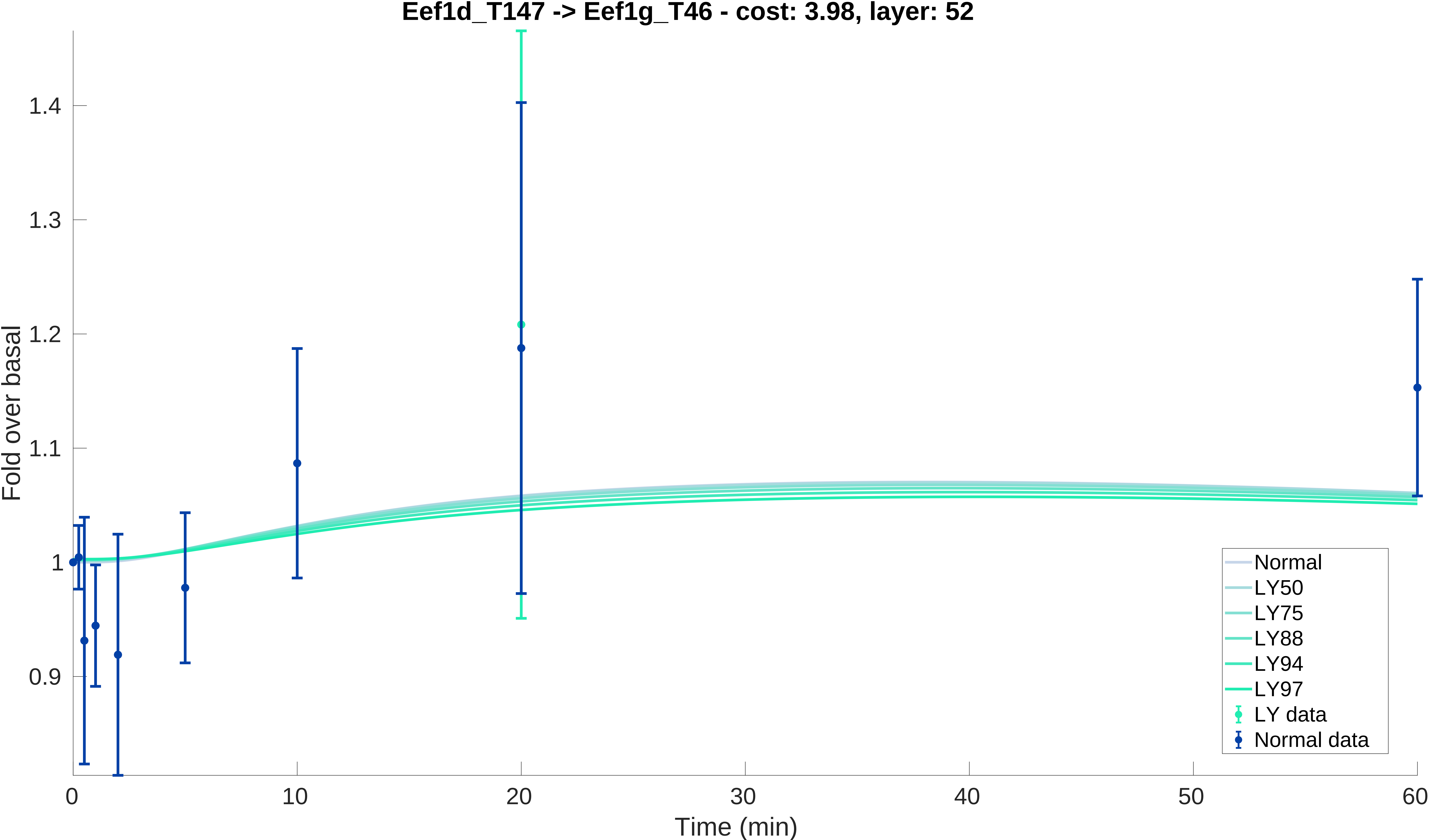
Eef1d_T147 -> Eef1g_T46 - cost: 3.98, layer: 52
Fold over basal
1.4
1.3
1.2
1.1
1
0.9
0
10
20
30
40
50
60
Time (min)
Normal
LY50
LY75
LY88
LY94
LY97
LY data
Normal data

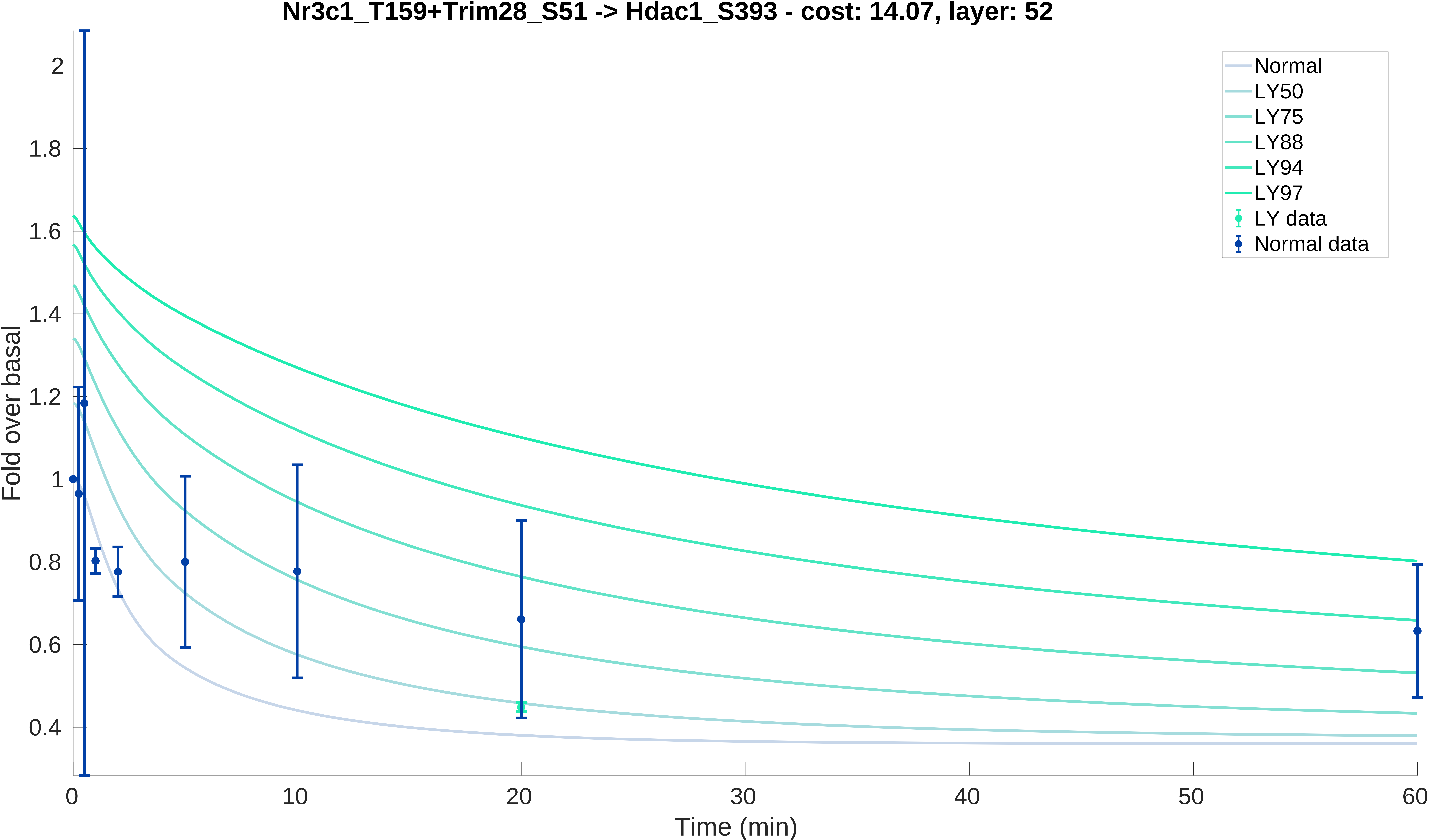
Nr3c1_T159+Trim28_S51 -> Hdac1_S393 - cost: 14.07, layer: 52
Fold over basal
Time (min)
Normal
LY50
LY75
LY88
LY94
LY97
LY data
Normal data
0.4
0.6
0.8
1
1.2
1.4
1.6
1.8
2
0
10
20
30
40
50
60

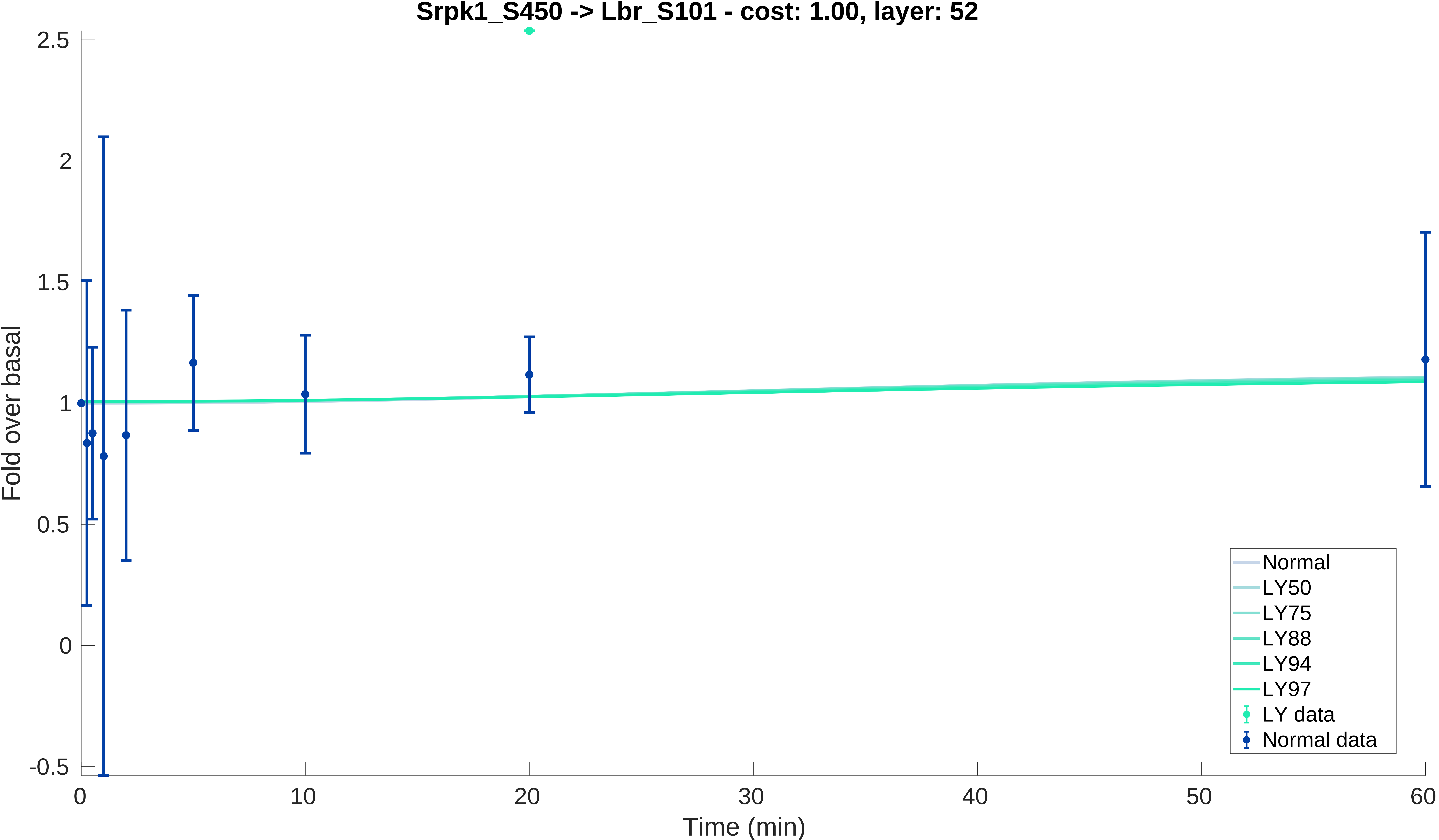

Srpk1_S450 -> Lbr_S101 - cost: 1.00, layer: 52
2.5
2
1.5
1
0.5
0
-0.5
Fold over basal
0
10
20
30
40
50
60
Time (min)
Normal
LY50
LY75
LY88
LY94
LY97
LY data
Normal data

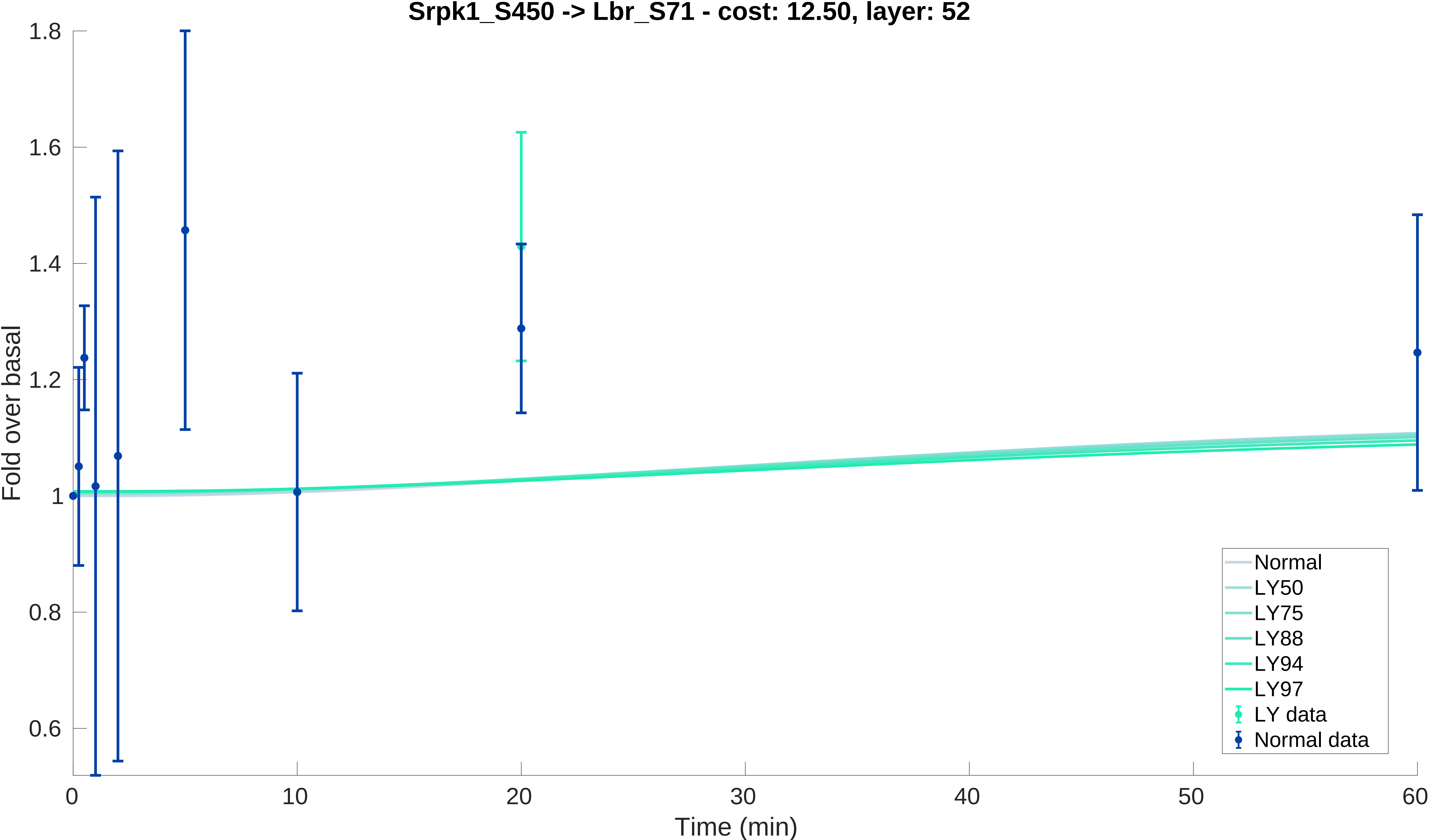

Srpk1_S450 -> Lbr_S71 - cost: 12.50, layer: 52
Fold over basal
1.8
1.6
1.4
1.2
1
0.8
0.6
0
10
20
30
40
50
60
Time (min)
Normal
LY50
LY75
LY88
LY94
LY97
LY data
Normal data

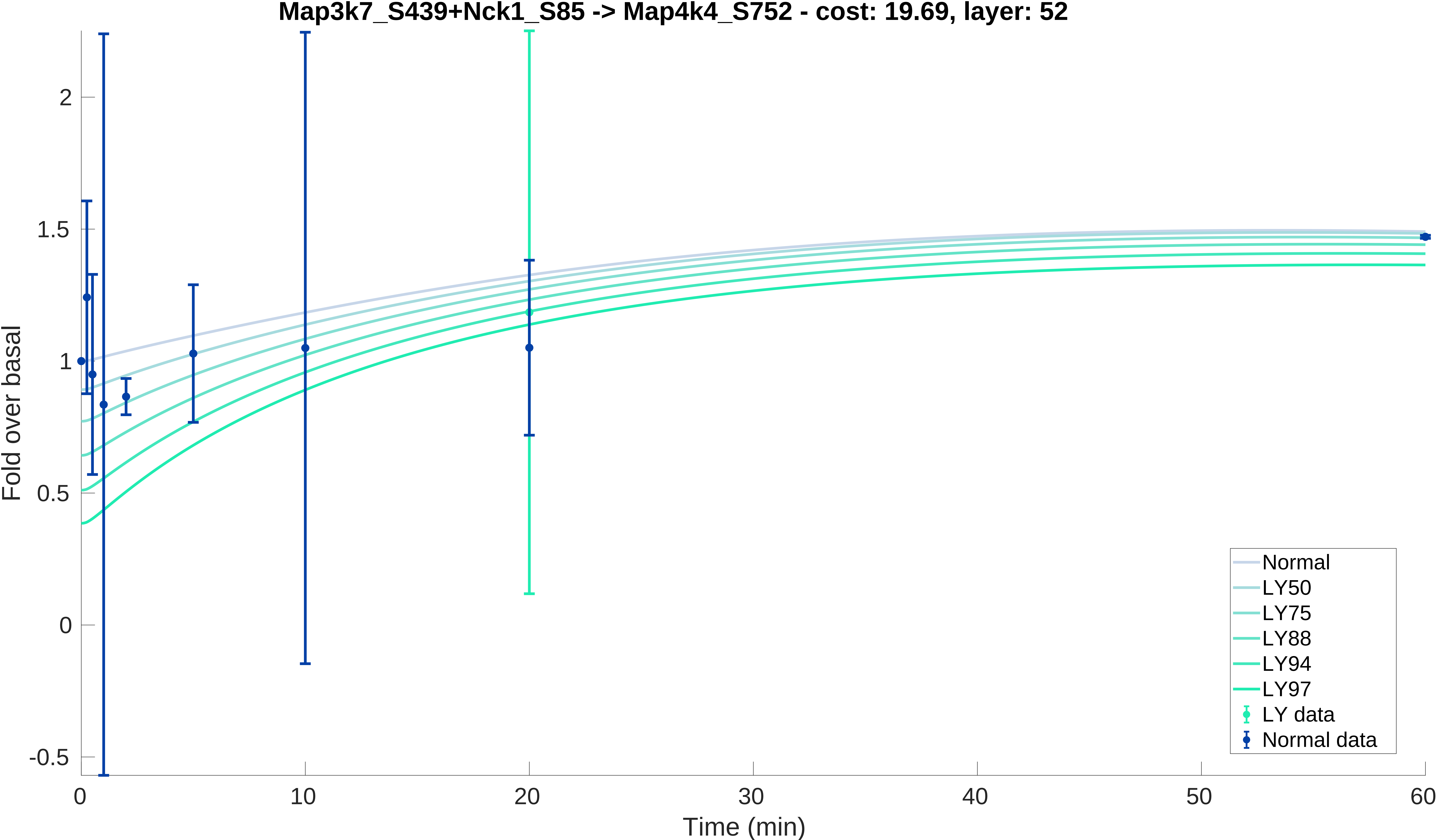

Map3k7_S439+Nck1_S85 -> Map4k4_S752 - cost: 19.69, layer: 52
Fold over basal
2
1.5
1
0.5
0
-0.5
0
10
20
30
40
50
60
Time (min)
Normal
LY50
LY75
LY88
LY94
LY97
LY data
Normal data

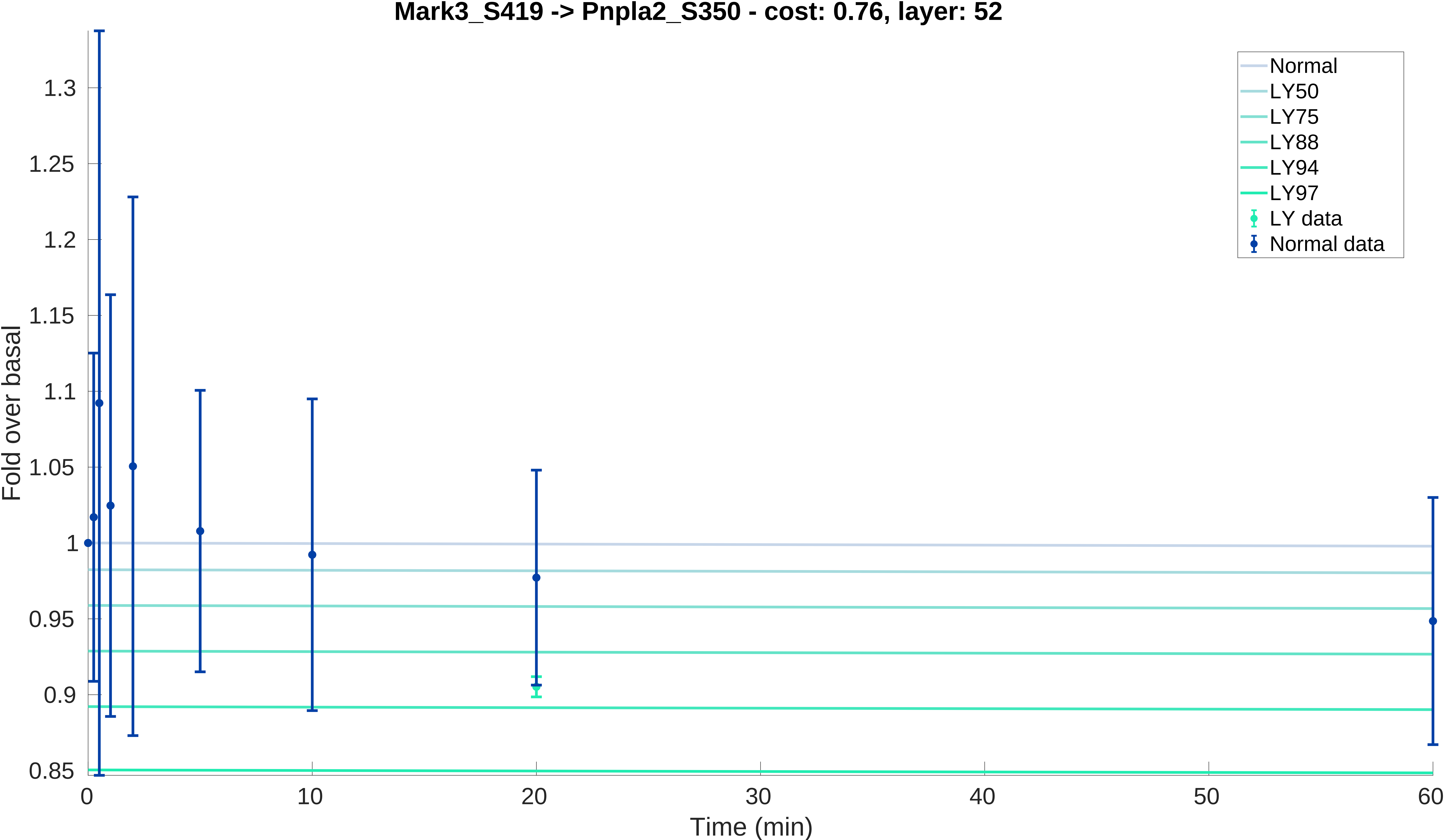

Mark3_S419 -> Pnpla2_S350 - cost: 0.76, layer: 52
Fold over basal
Time (min)
1.3
1.25
1.2
1.15
1.1
1.05
1
0.95
0.9
0.85
0
10
20
30
40
50
60
Normal
LY50
LY75
LY88
LY94
LY97
LY data
Normal data

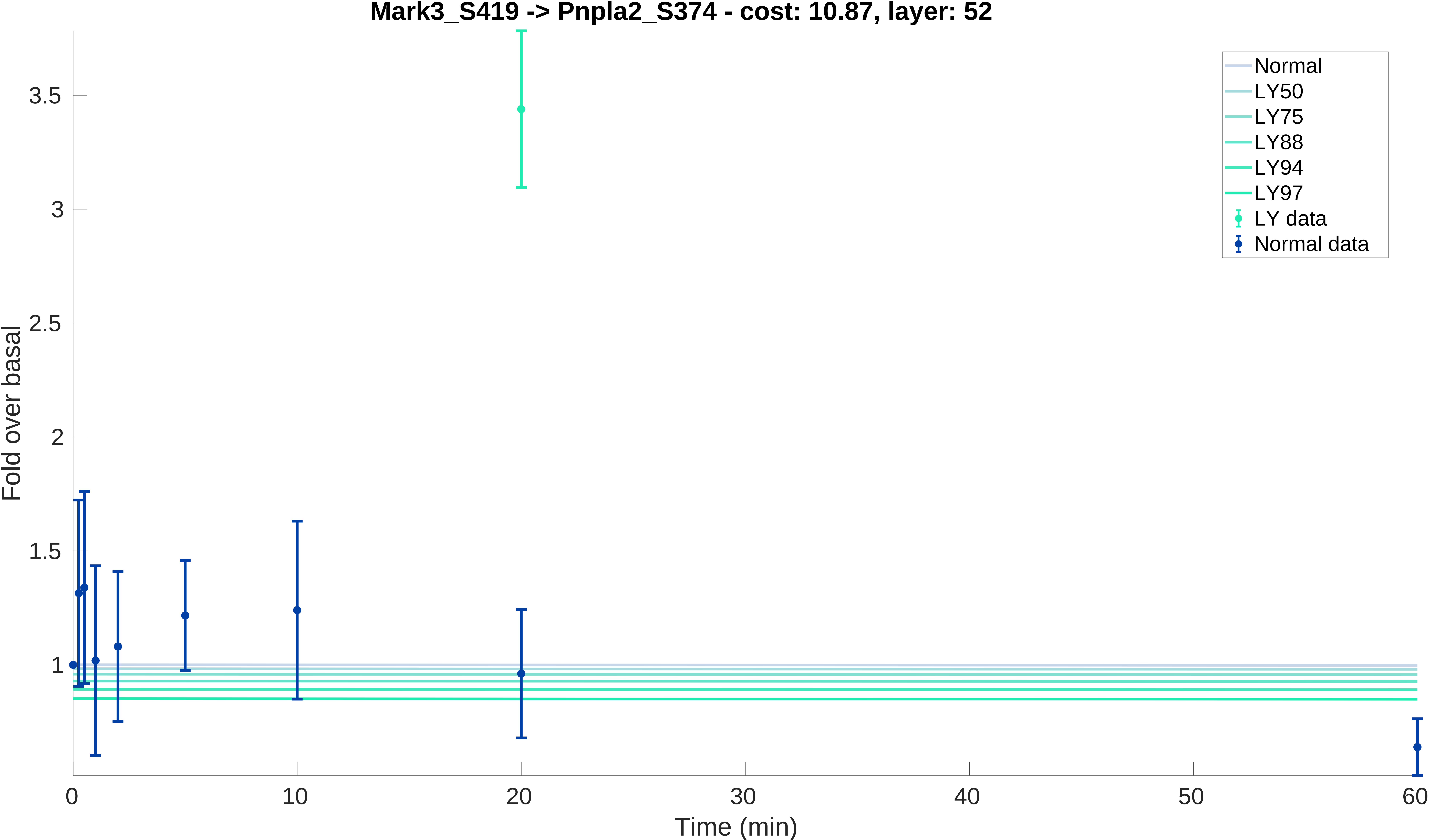
Mark3_S419 -> Pnpla2_S374 - cost: 10.87, layer: 52
Fold over basal
Time (min)
Normal
LY50
LY75
LY88
LY94
LY97
LY data
Normal data
3.5
3
2.5
2
1.5
1
0
10
20
30
40
50
60

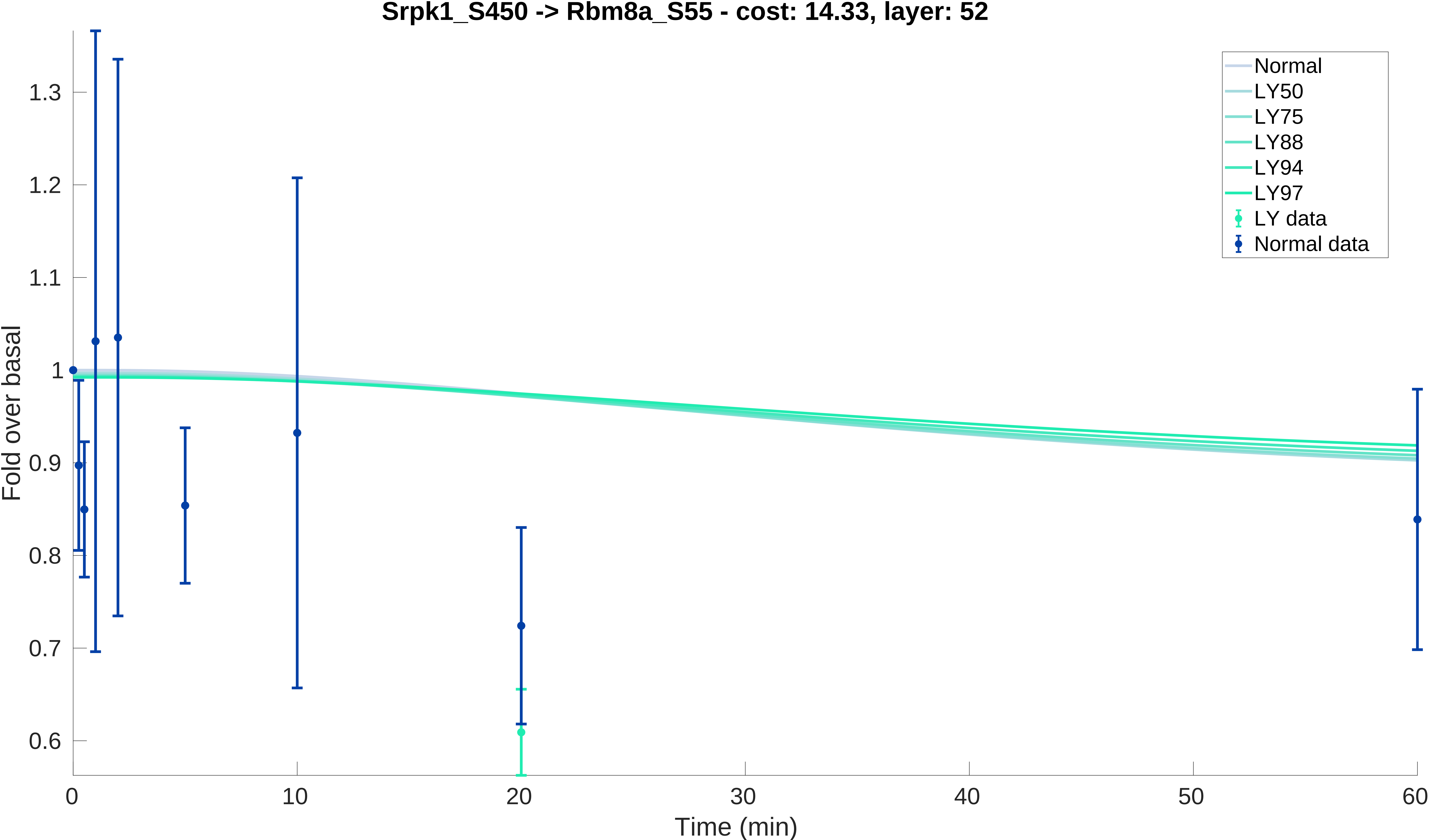

Srpk1_S450 -> Rbm8a_S55 - cost: 14.33, layer: 52
Fold over basal
Time (min)
0
10
20
30
40
50
60
0.6
0.7
0.8
0.9
1
1.1
1.2
1.3
Normal
LY50
LY75
LY88
LY94
LY97
LY data
Normal data

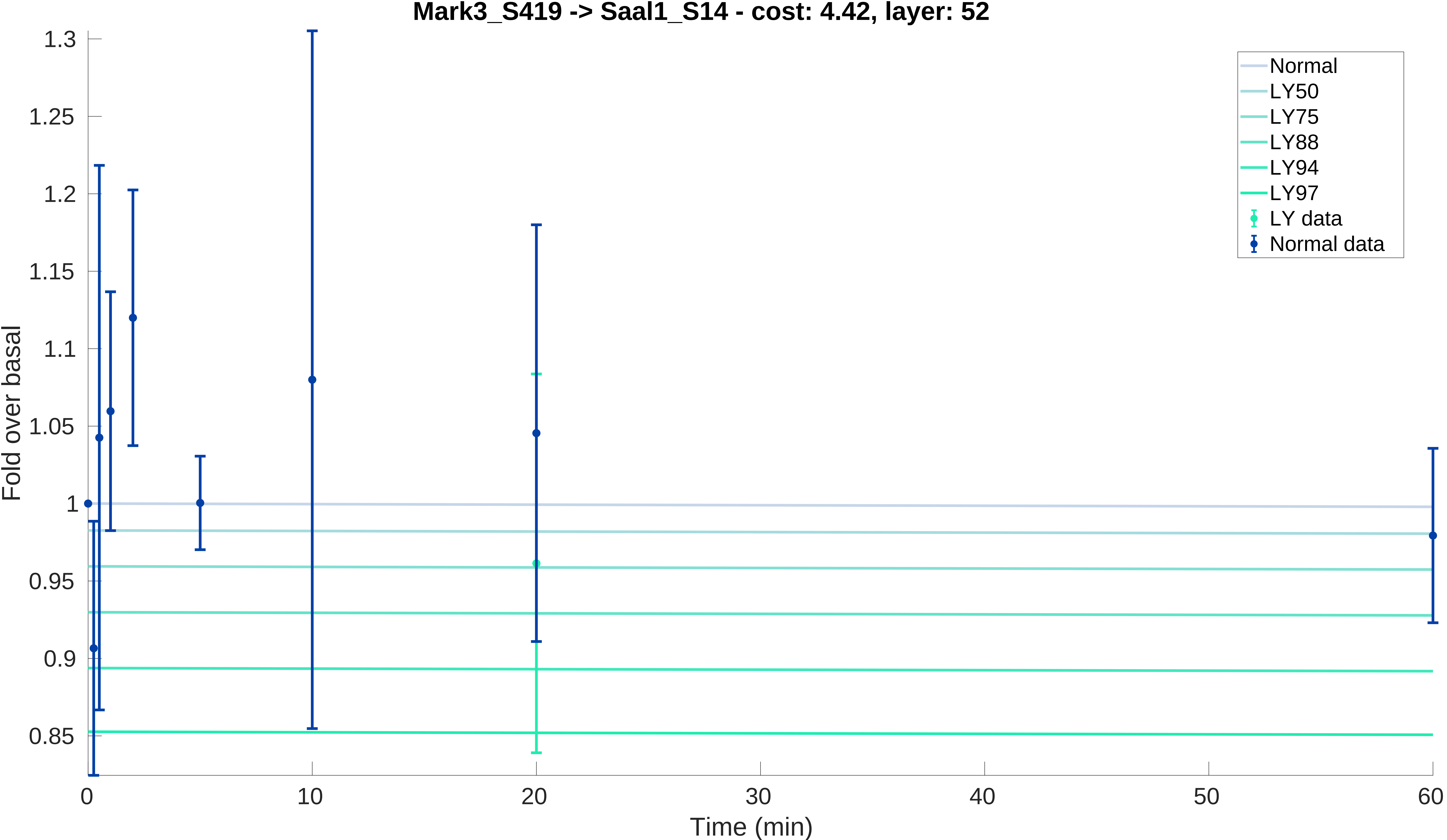
Mark3_S419 -> Saal1_S14 - cost: 4.42, layer: 52
Fold over basal
1.3
1.25
1.2
1.15
1.1
1.05
1
0.95
0.9
0.85
0
10
20
30
40
50
60
Time (min)
Normal
LY50
LY75
LY88
LY94
LY97
LY data
Normal data

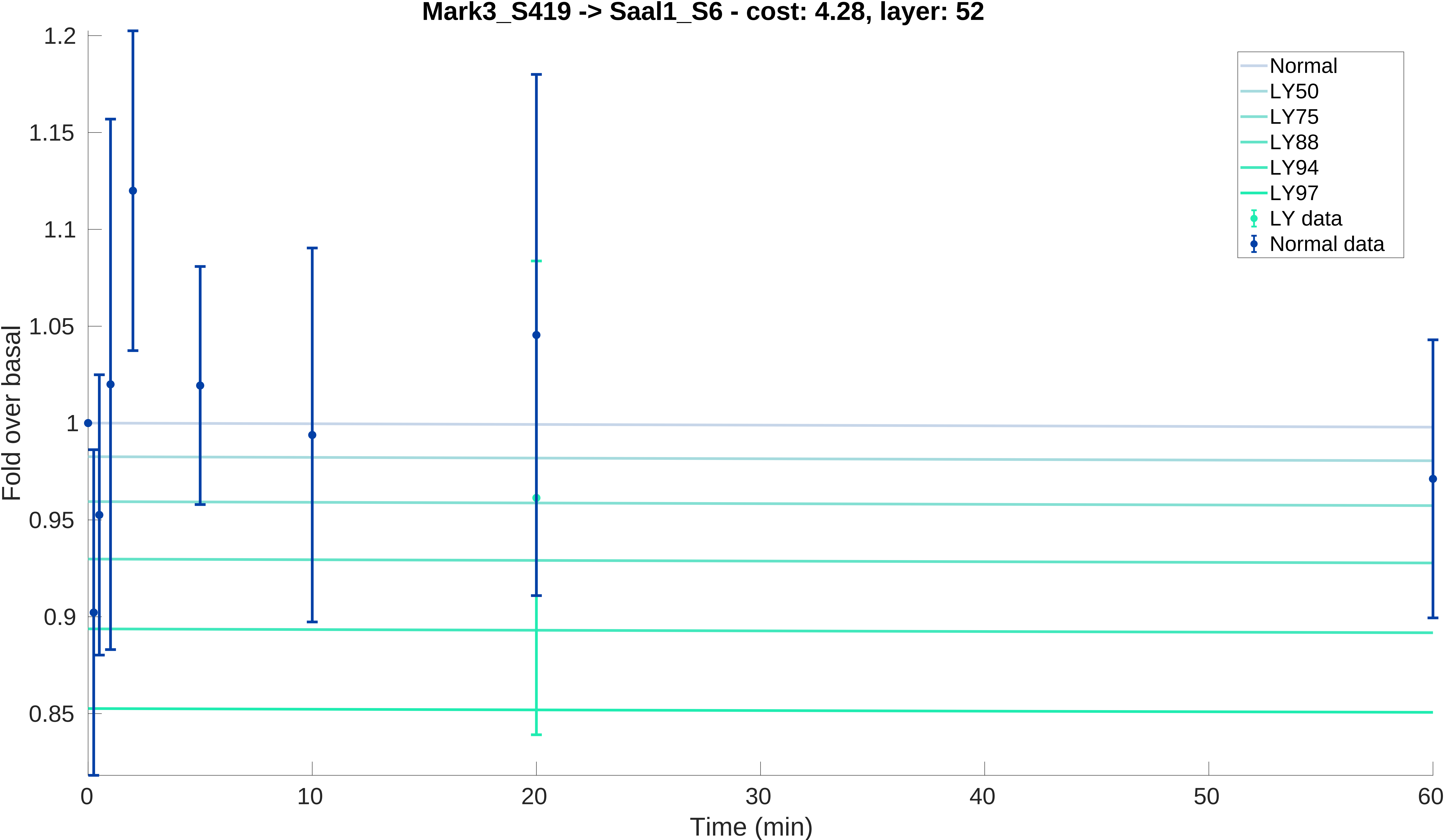

Mark3_S419 -> Saal1_S6 - cost: 4.28, layer: 52
Fold over basal
1.2
1.15
1.1
1.05
1
0.95
0.9
0.85
0
10
20
30
40
50
60
Time (min)
Normal
LY50
LY75
LY88
LY94
LY97
LY data
Normal data

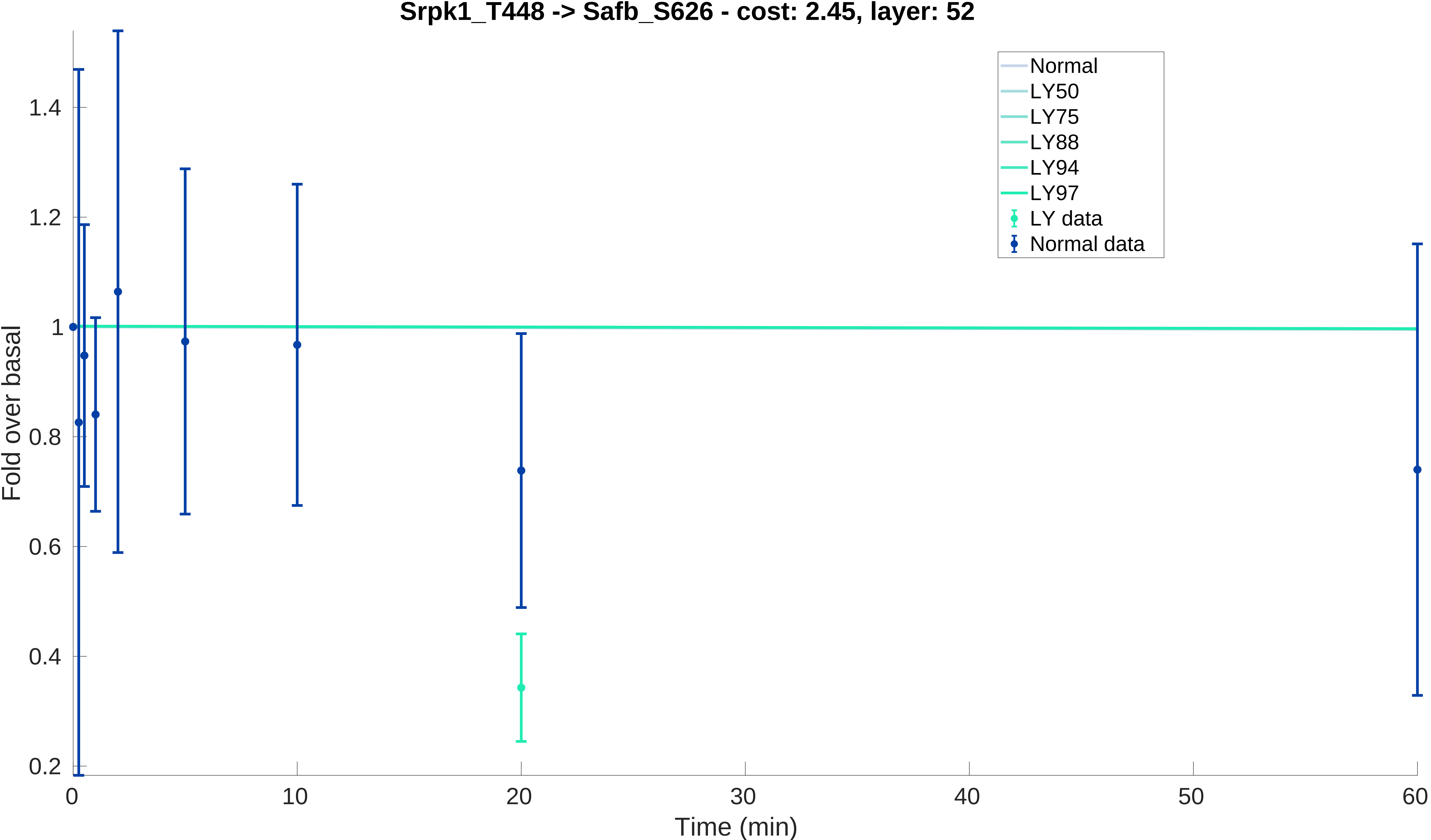

Srpk1_T448 -> Safb_S626 - cost: 2.45, layer: 52
Normal
LY50
LY75
LY88
LY94
LY97
LY data
Normal data
Fold over basal
Time (min)
0.2
0.4
0.6
0.8
1
1.2
1.4
0
10
20
30
40
50
60

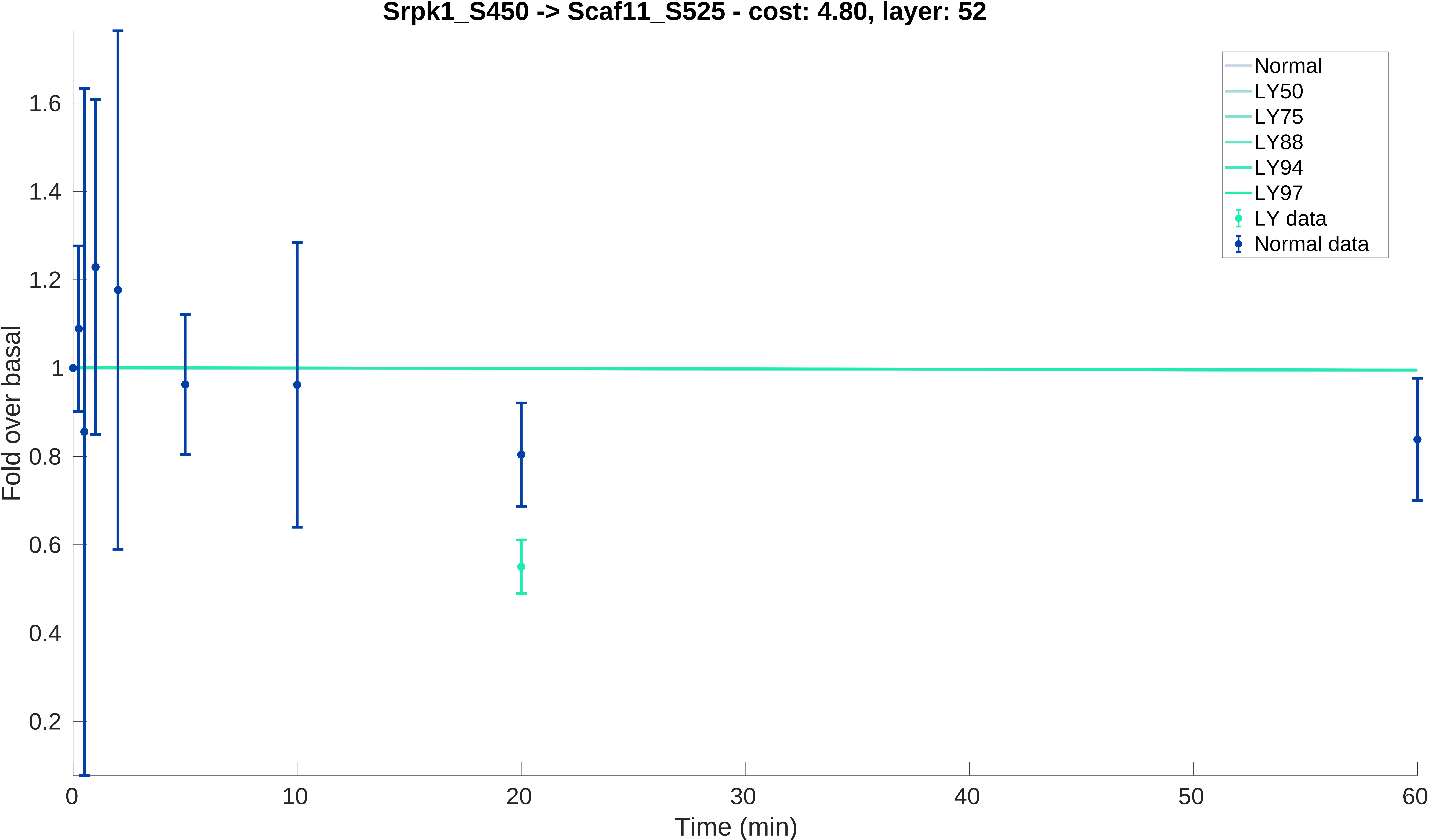

Srpk1_S450 -> Scaf11_S525 - cost: 4.80, layer: 52
Fold over basal
Time (min)
1.6
1.4
1.2
1
0.8
0.6
0.4
0.2
0
10
20
30
40
50
60
Normal
LY50
LY75
LY88
LY94
LY97
LY data
Normal data

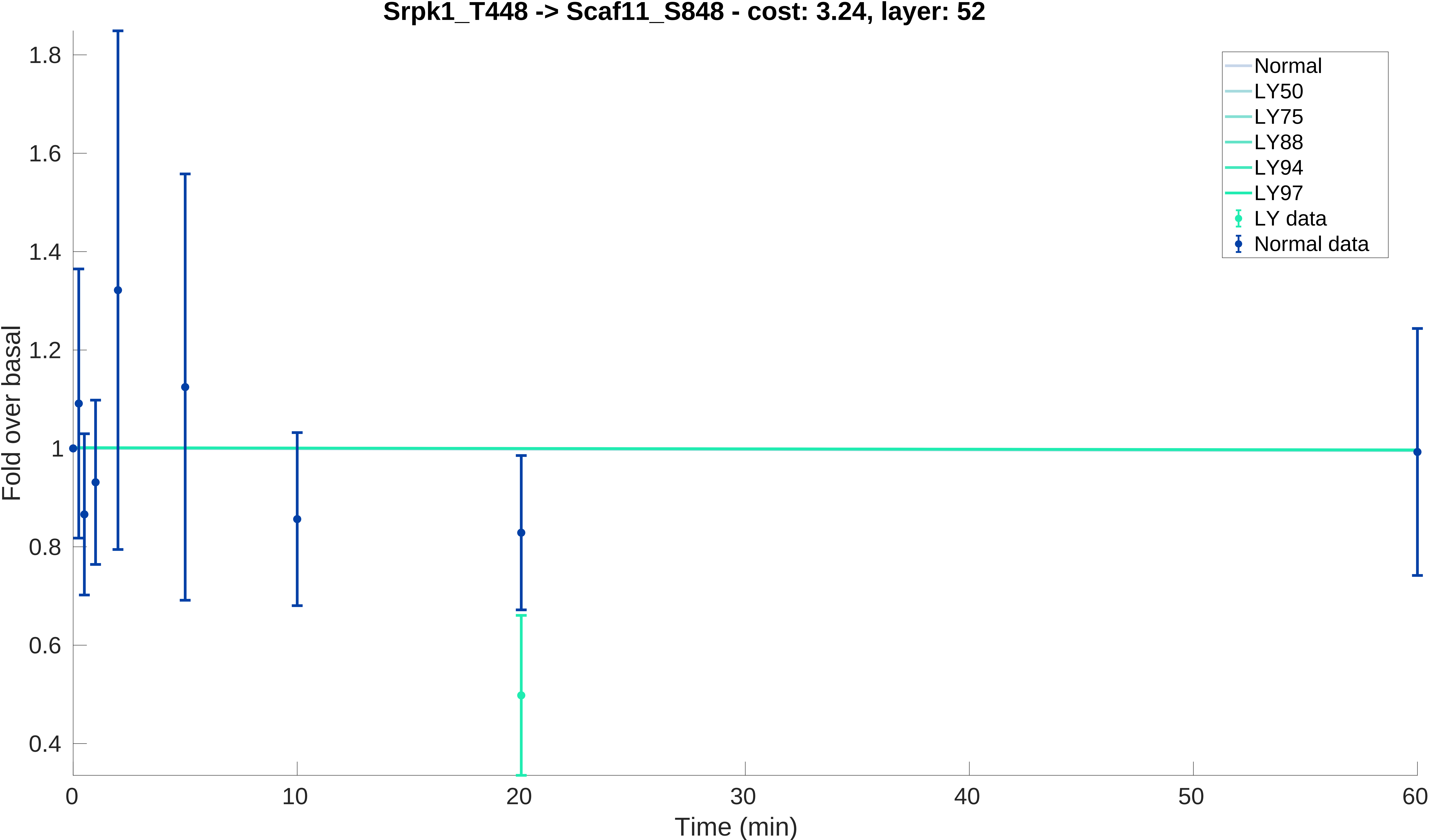

Srpk1_T448 -> Scaf11_S848 - cost: 3.24, layer: 52
Fold over basal
Time (min)
1.8
1.6
1.4
1.2
1
0.8
0.6
0.4
0
10
20
30
40
50
60
Normal
LY50
LY75
LY88
LY94
LY97
LY data
Normal data

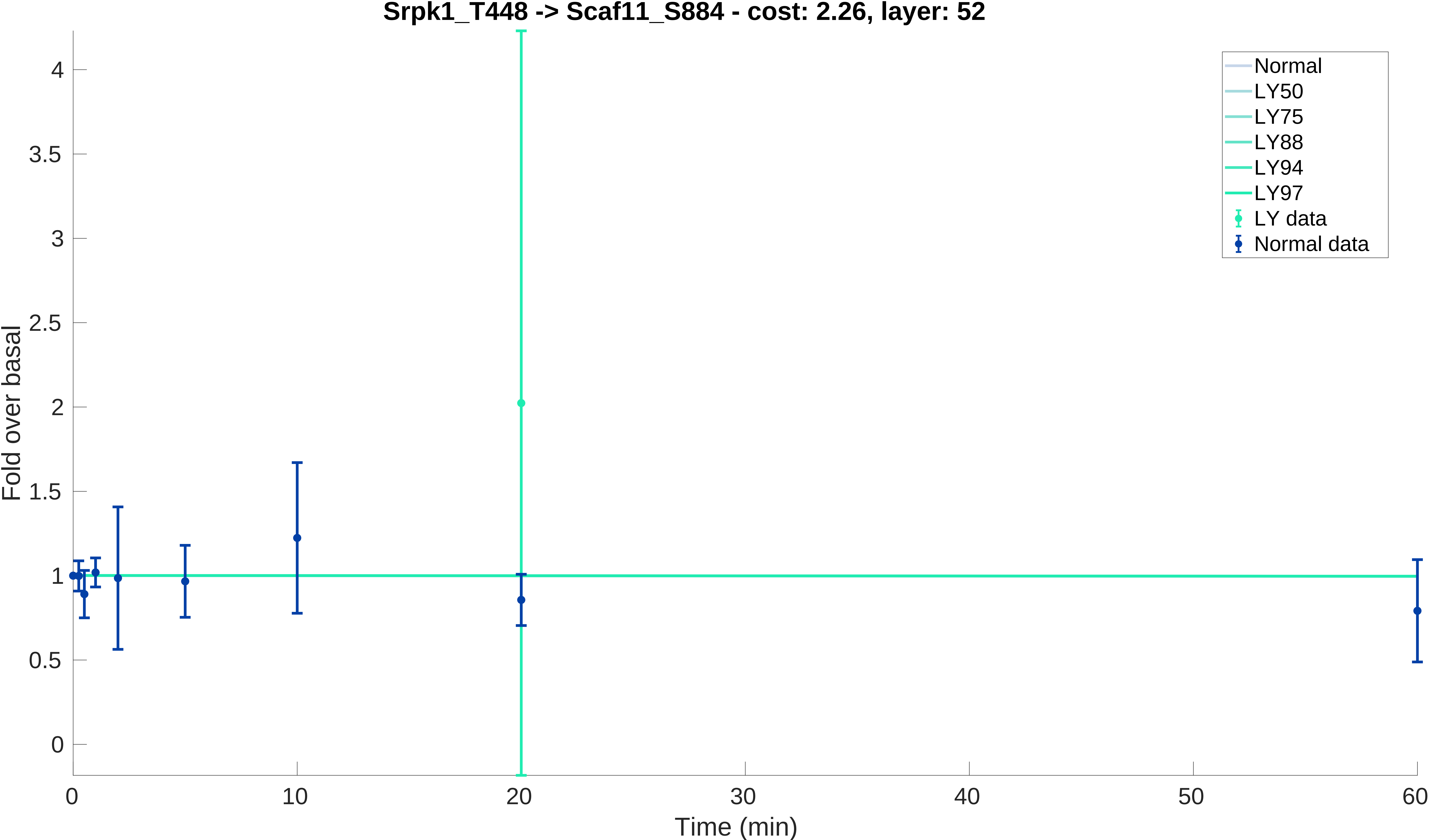

Srpk1_T448 -> Scaf11_S884 - cost: 2.26, layer: 52
Fold over basal
Time (min)
0
0.5
1
1.5
2
2.5
3
3.5
4
0
10
20
30
40
50
60
Normal
LY50
LY75
LY88
LY94
LY97
LY data
Normal data

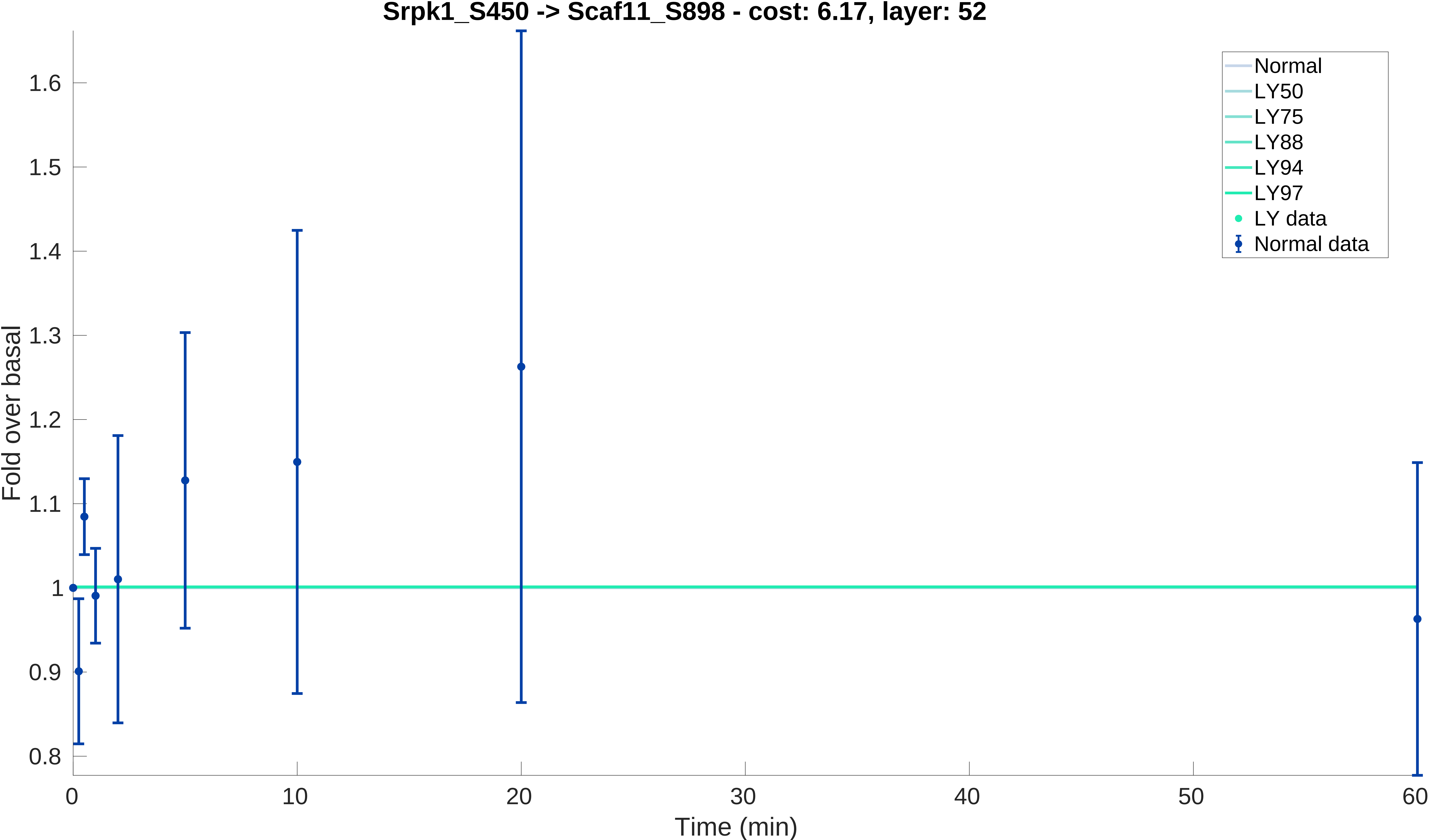
Srpk1_S450 -> Scaf11_S898 - cost: 6.17, layer: 52
Fold over basal
Time (min)
Normal
LY50
LY75
LY88
LY94
LY97
LY data
Normal data
1.6
1.5
1.4
1.3
1.2
1.1
1
0.9
0.8
0
10
20
30
40
50
60

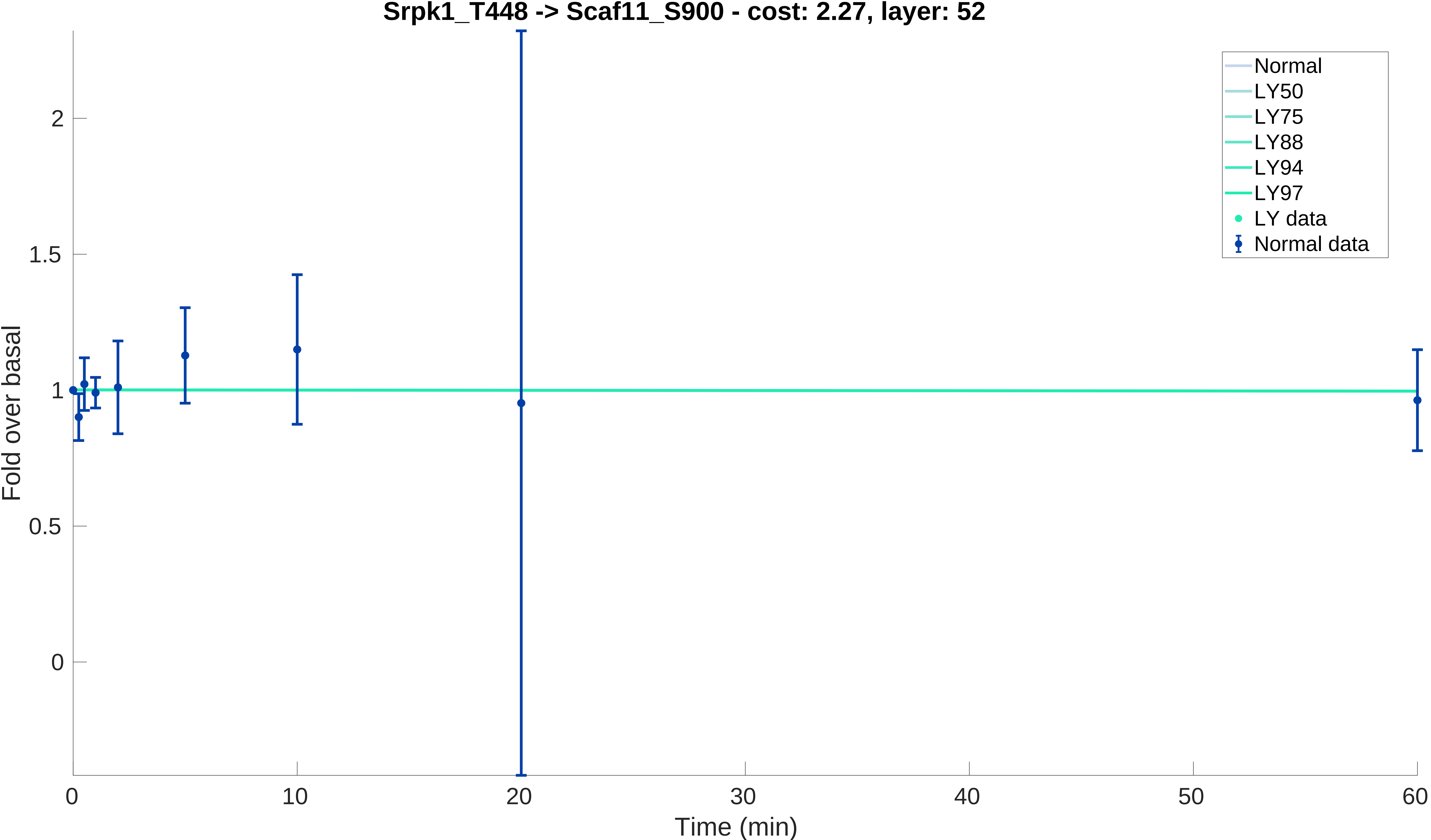

Srpk1_T448 -> Scaf11_S900 - cost: 2.27, layer: 52
Fold over basal
Time (min)
0
0.5
1
1.5
2
0
10
20
30
40
50
60
Normal
LY50
LY75
LY88
LY94
LY97
LY data
Normal data

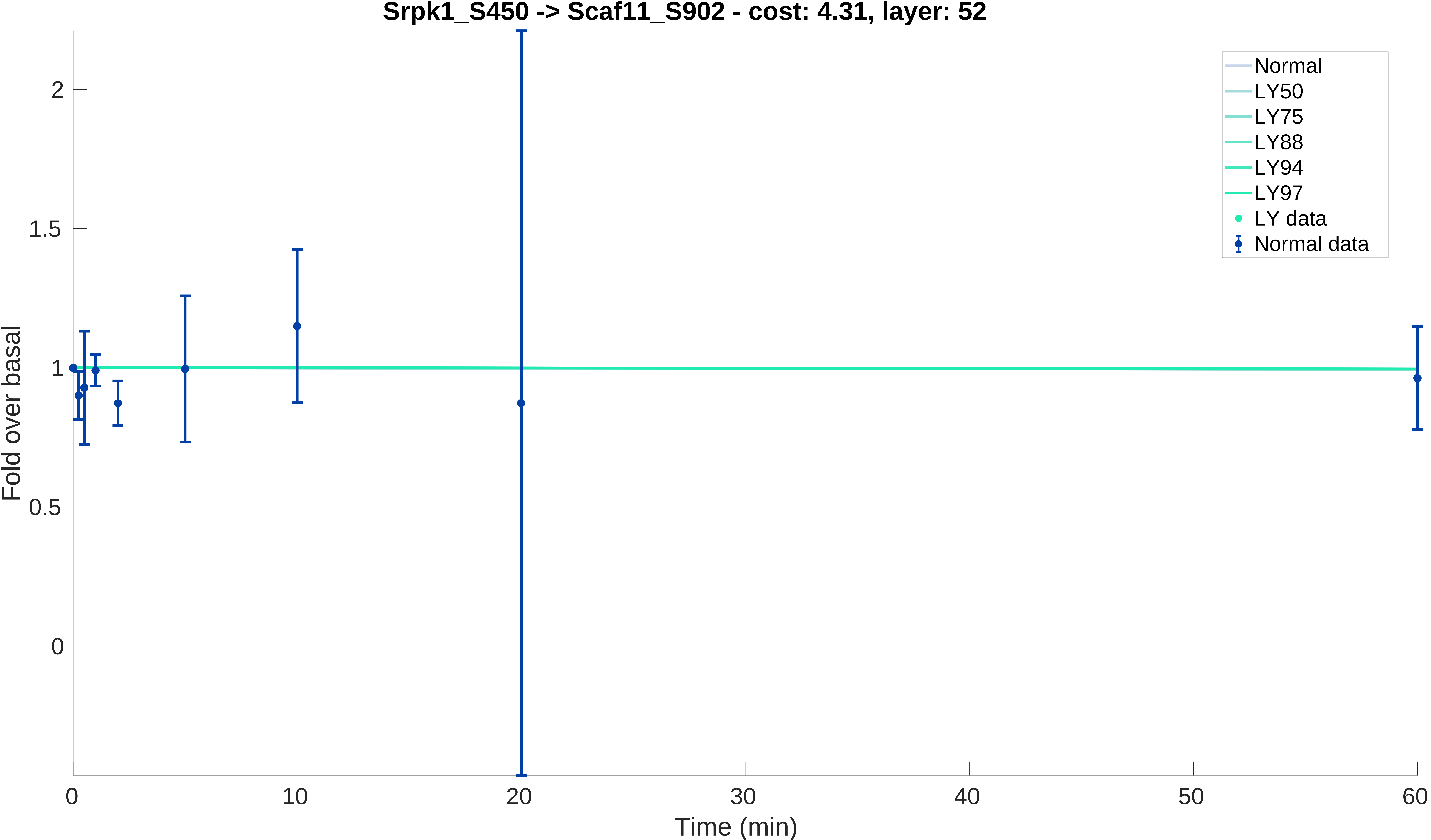

Srpk1_S450 -> Scaf11_S902 - cost: 4.31, layer: 52
Fold over basal
Time (min)
0
10
20
30
40
50
60
-0
0.5
1
1.5
2
Normal
LY50
LY75
LY88
LY94
LY97
LY data
Normal data

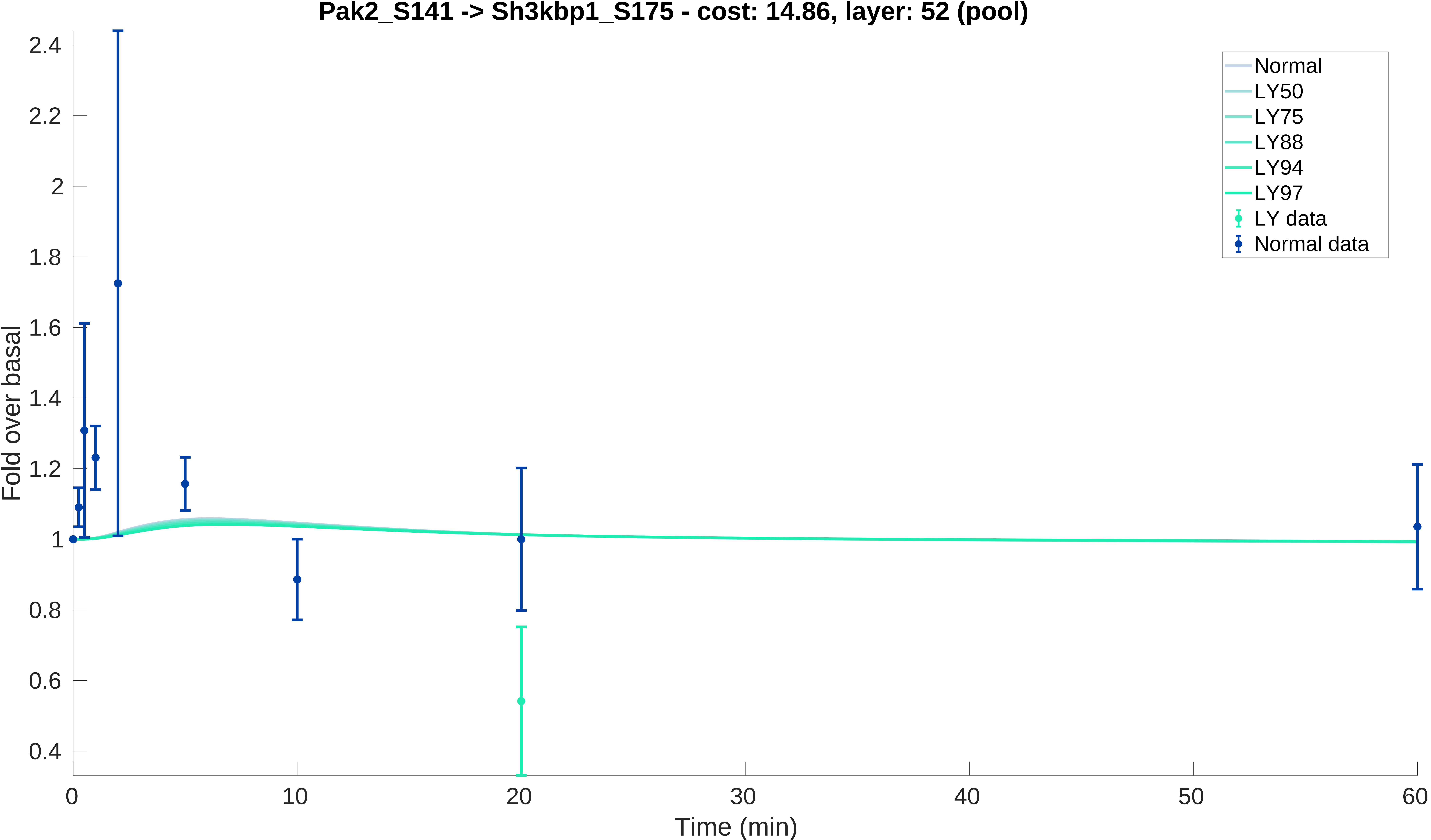

Pak2_S141 -> Sh3kbp1_S175 - cost: 14.86, layer: 52 (pool)
Fold over basal
Time (min)
2.4
2.2
2
1.8
1.6
1.4
1.2
1
0.8
0.6
0.4
0
10
20
30
40
50
60
Normal
LY50
LY75
LY88
LY94
LY97
LY data
Normal data

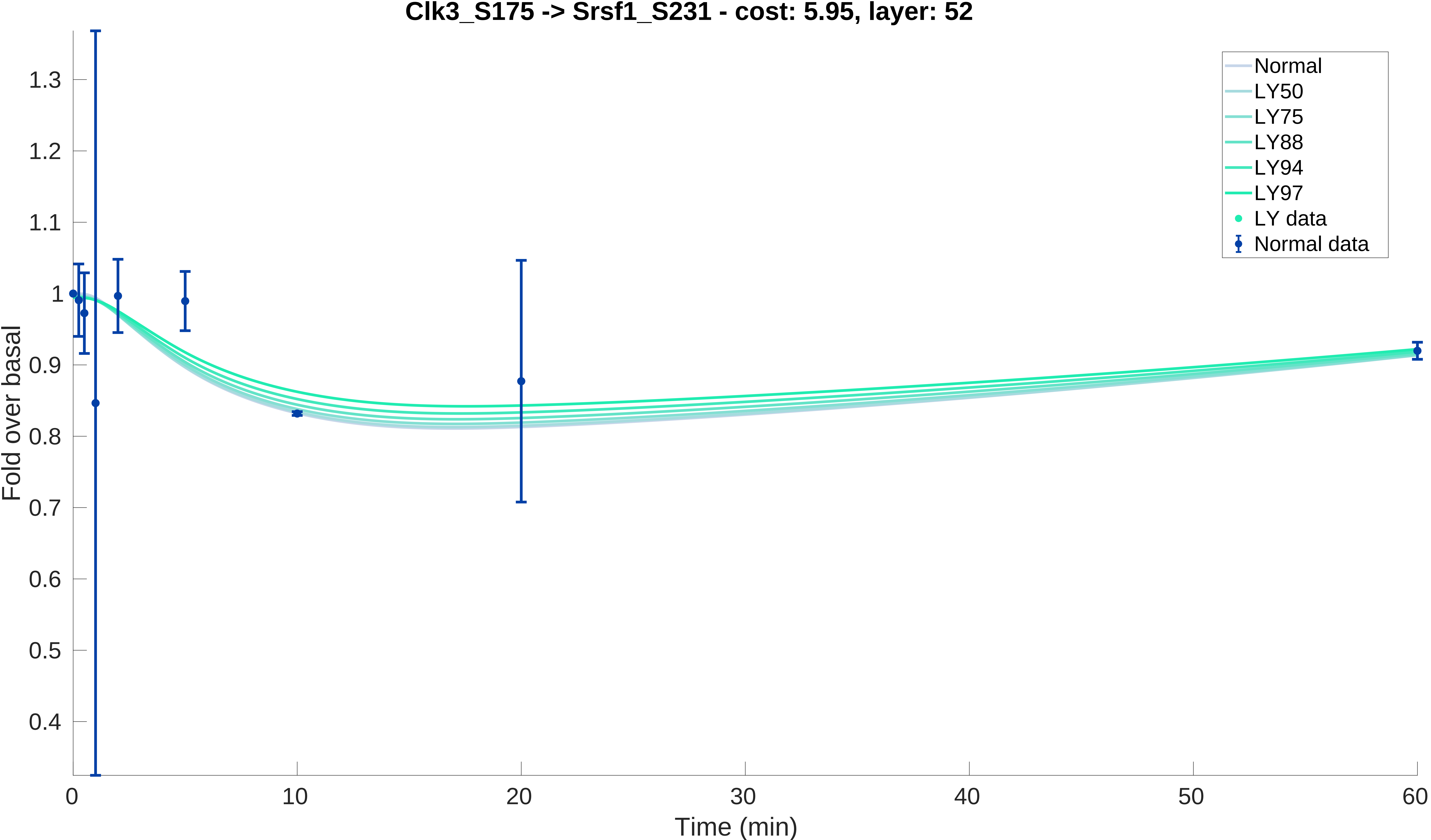

Clk3_S175 -> Srsf1_S231 - cost: 5.95, layer: 52
Fold over basal
Time (min)
1.3
1.2
1.1
1
0.9
0.8
0.7
0.6
0.5
0.4
0
10
20
30
40
50
60
Normal
LY50
LY75
LY88
LY94
LY97
LY data
Normal data

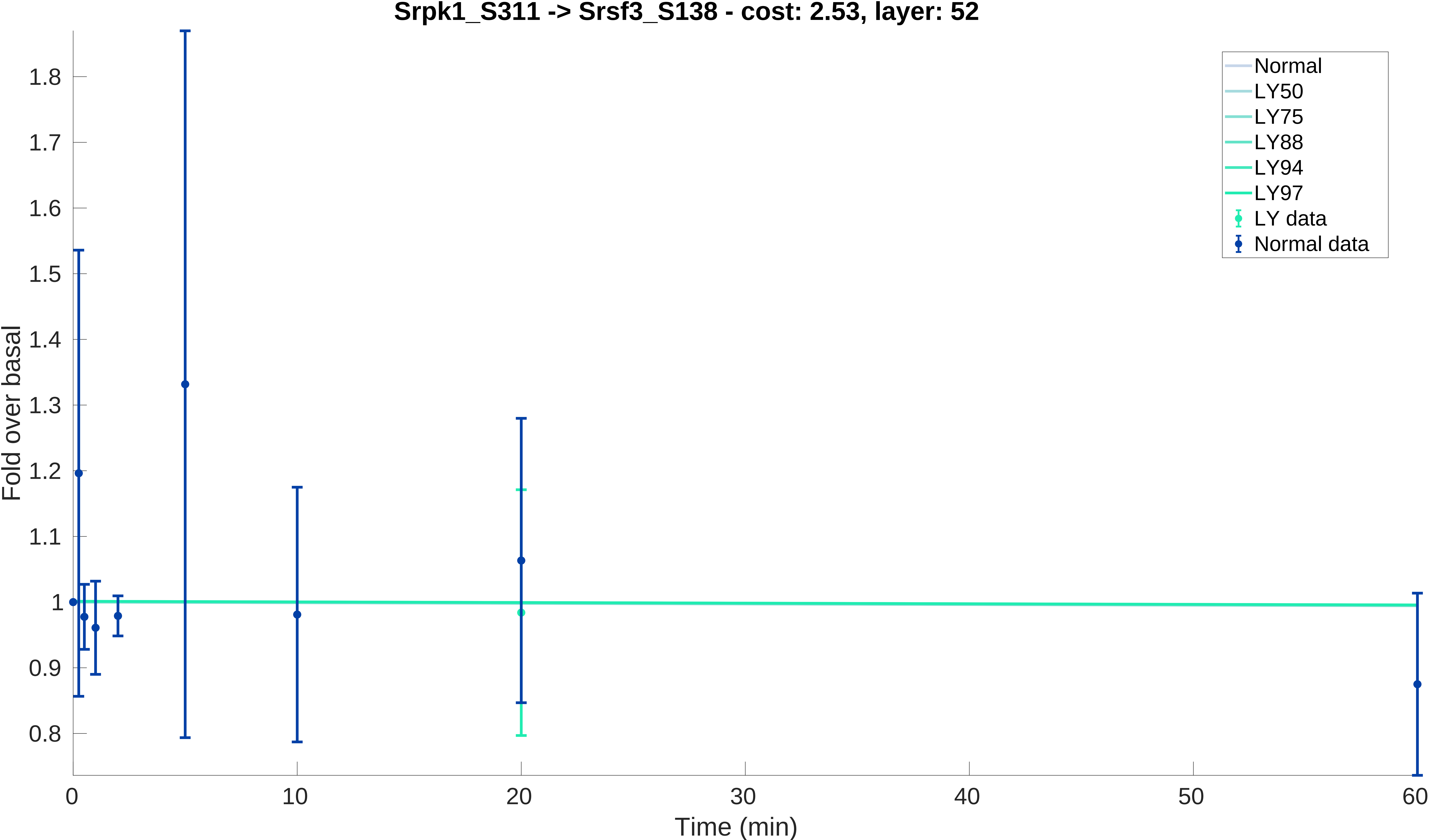
Srpk1_S311 -> Srsf3_S138 - cost: 2.53, layer: 52
Fold over basal
1.8
1.7
1.6
1.5
1.4
1.3
1.2
1.1
1
0.9
0.8
0
10
20
30
40
50
60
Time (min)
Normal
LY50
LY75
LY88
LY94
LY97
LY data
Normal data

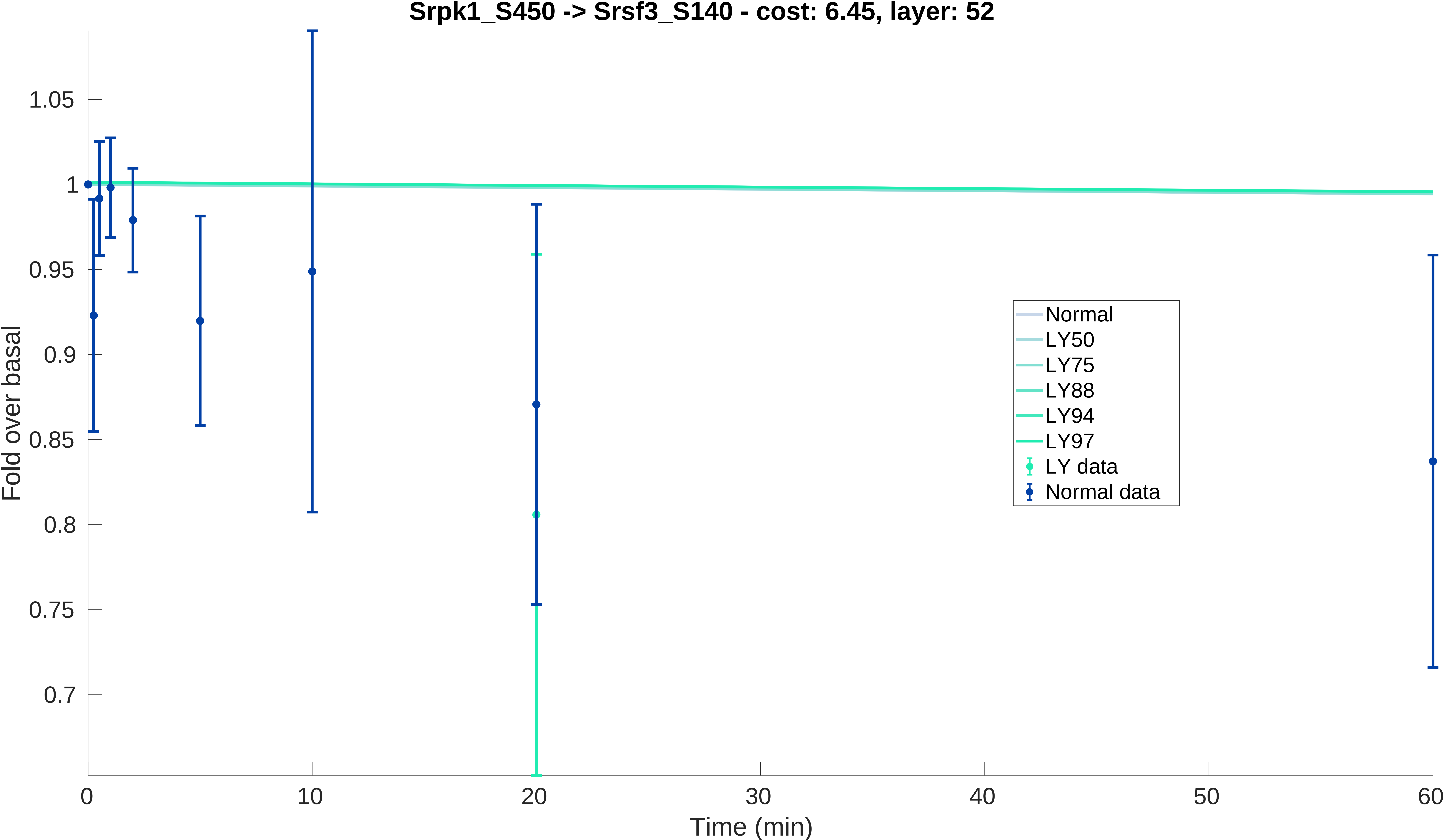

Srpk1_S450 -> Srsf3_S140 - cost: 6.45, layer: 52
Fold over basal
1.05
1
0.95
0.9
0.85
0.8
0.75
0.7
0
10
20
30
40
50
60
Time (min)
Normal
LY50
LY75
LY88
LY94
LY97
LY data
Normal data

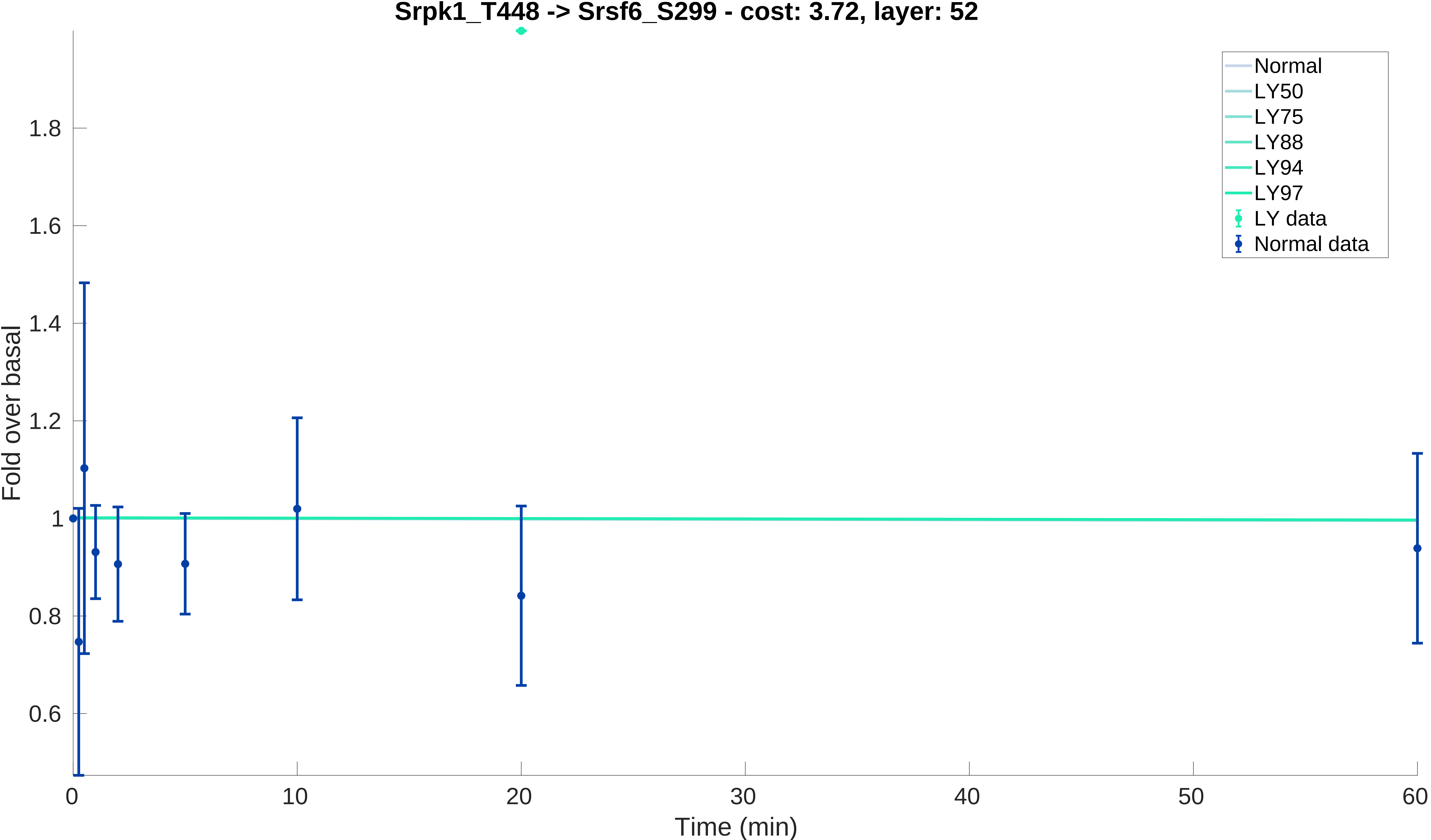

Srpk1_T448 -> Srsf6_S299 - cost: 3.72, layer: 52
Fold over basal
Time (min)
Normal
LY50
LY75
LY88
LY94
LY97
LY data
Normal data
1.8
1.6
1.4
1.2
1
0.8
0.6
0
10
20
30
40
50
60

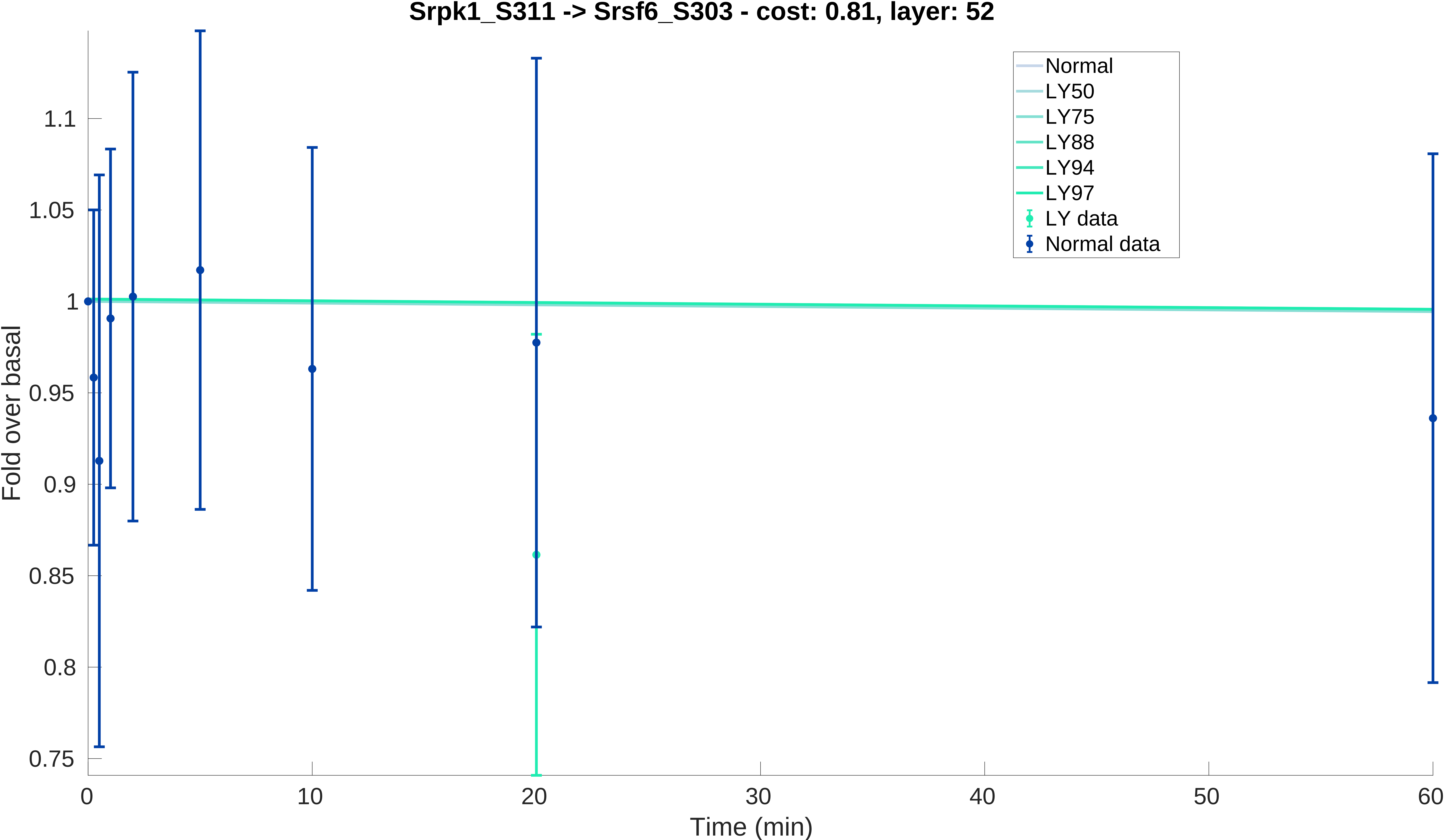

Srpk1_S311 -> Srsf6_S303 - cost: 0.81, layer: 52
Normal
LY50
LY75
LY88
LY94
LY97
LY data
Normal data
Fold over basal
1.1
1.05
1
0.95
0.9
0.85
0.8
0.75
0
10
20
30
40
50
60
Time (min)

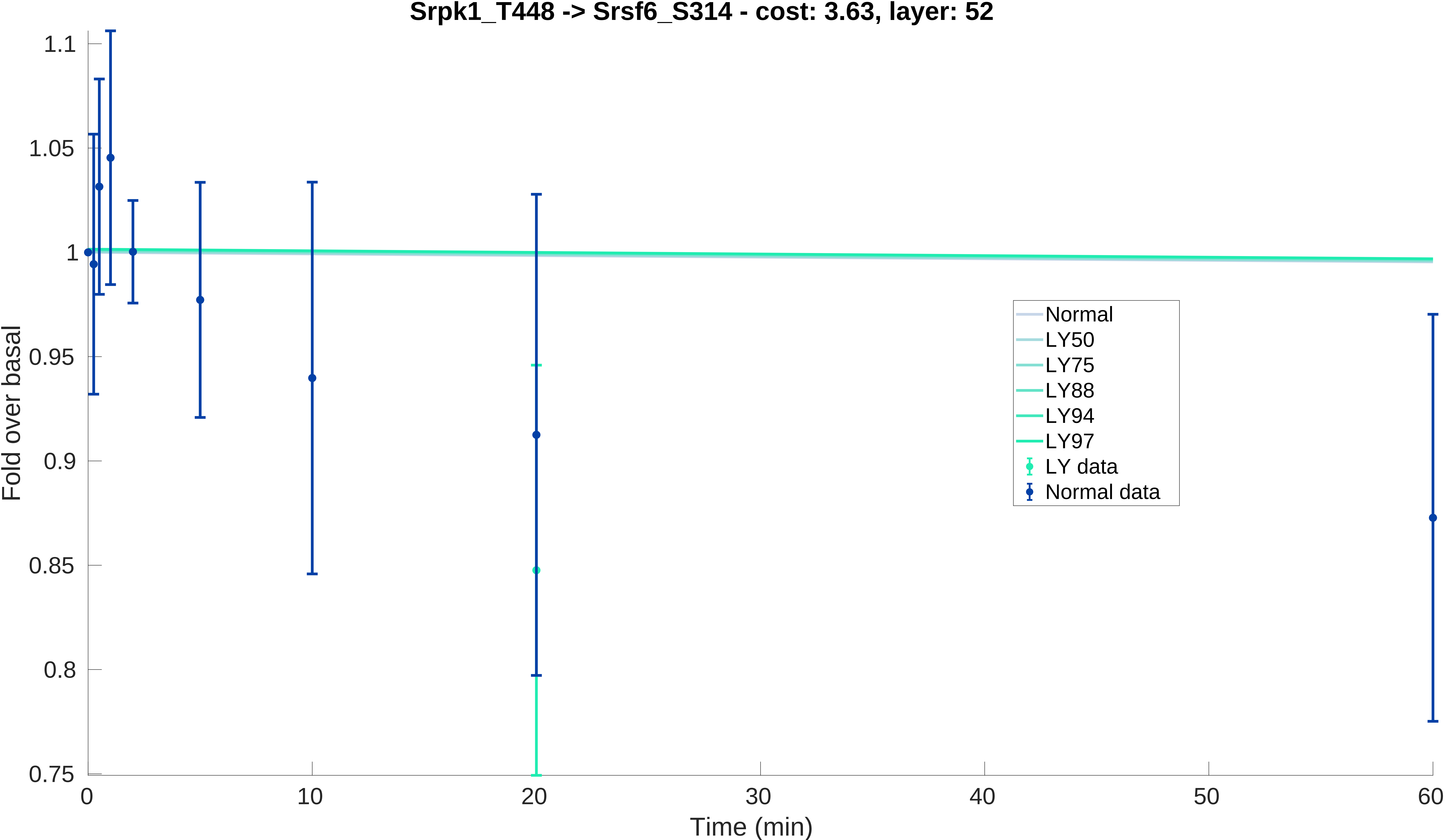

Srpk1_T448 -> Srsf6_S314 - cost: 3.63, layer: 52
Fold over basal
1.1
1.05
1
0.95
0.9
0.85
0.8
0.75
0
10
20
30
40
50
60
Time (min)
Normal
LY50
LY75
LY88
LY94
LY97
LY data
Normal data

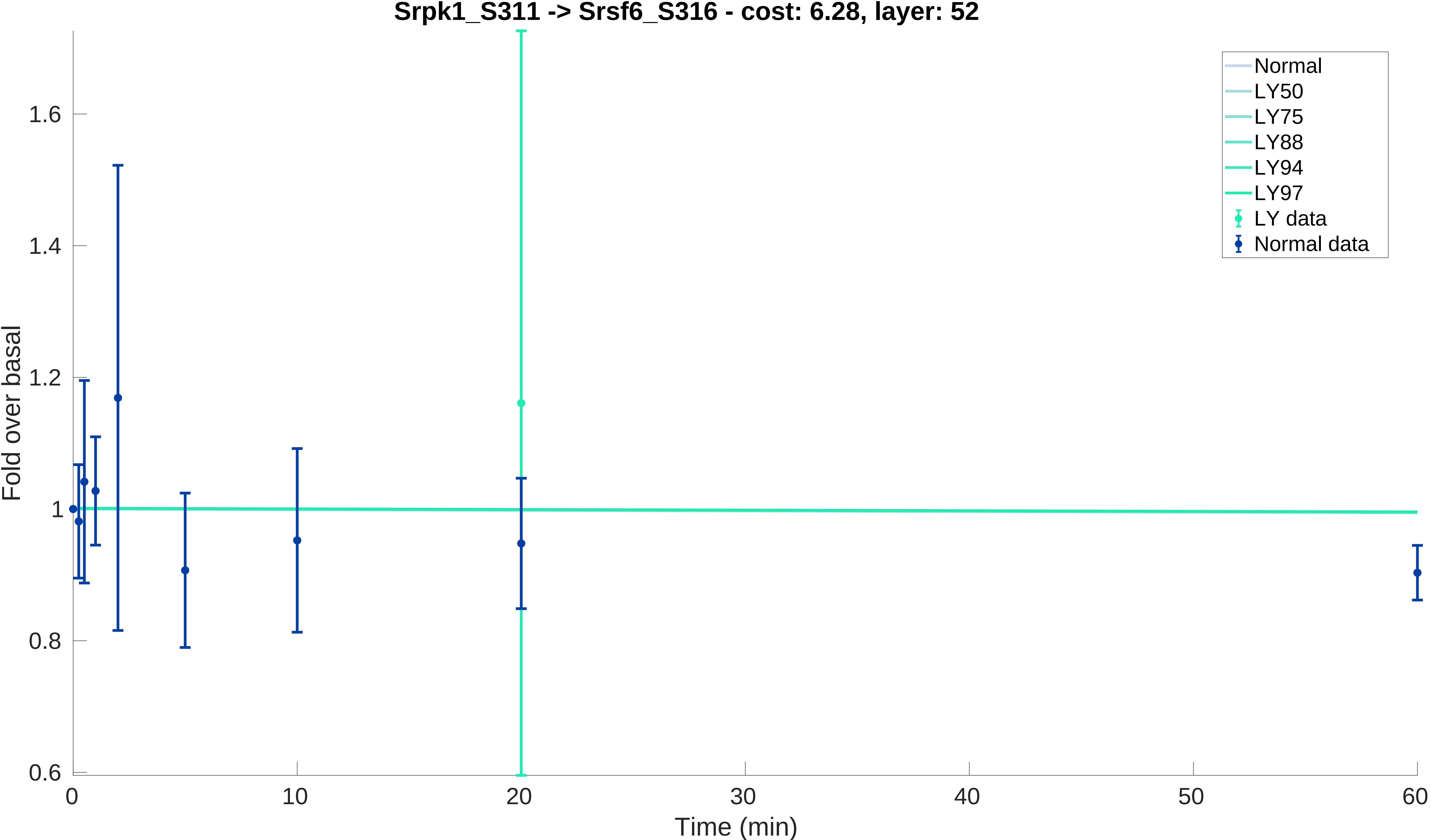

Srpk1_S311 -> Srsf6_S316 - cost: 6.28, layer: 52
Normal
LY50
LY75
LY88
LY94
LY97
LY data
Normal data
Fold over basal
1.6
1.4
1.2
1
0.8
0.6
0
10
20
30
40
50
60
Time (min)

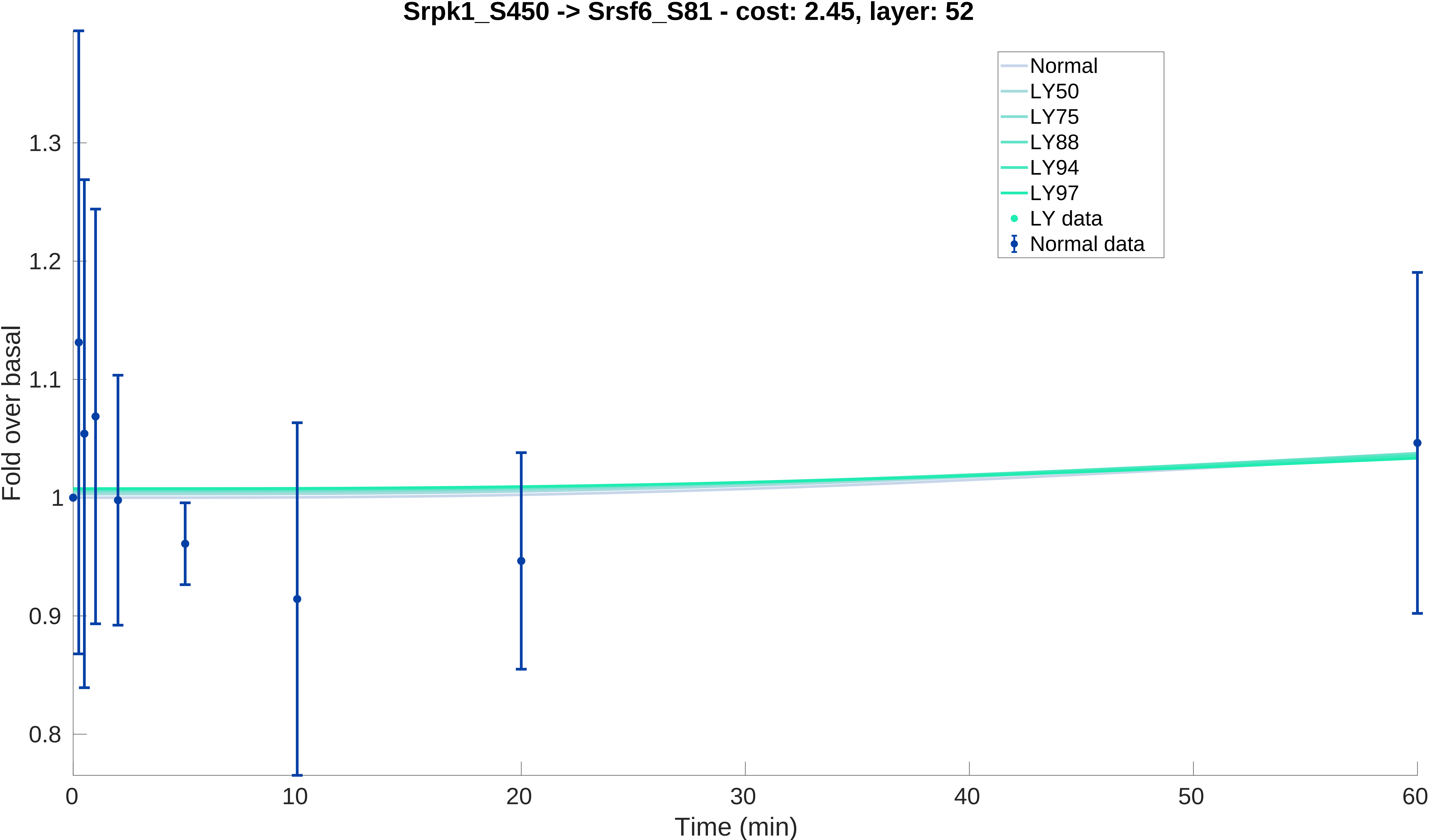

Srpk1_S450 -> Srsf6_S81 - cost: 2.45, layer: 52
Normal
LY50
LY75
LY88
LY94
LY97
LY data
Normal data
Fold over basal
1.3
1.2
1.1
1
0.9
0.8
0
10
20
30
40
50
60
Time (min)

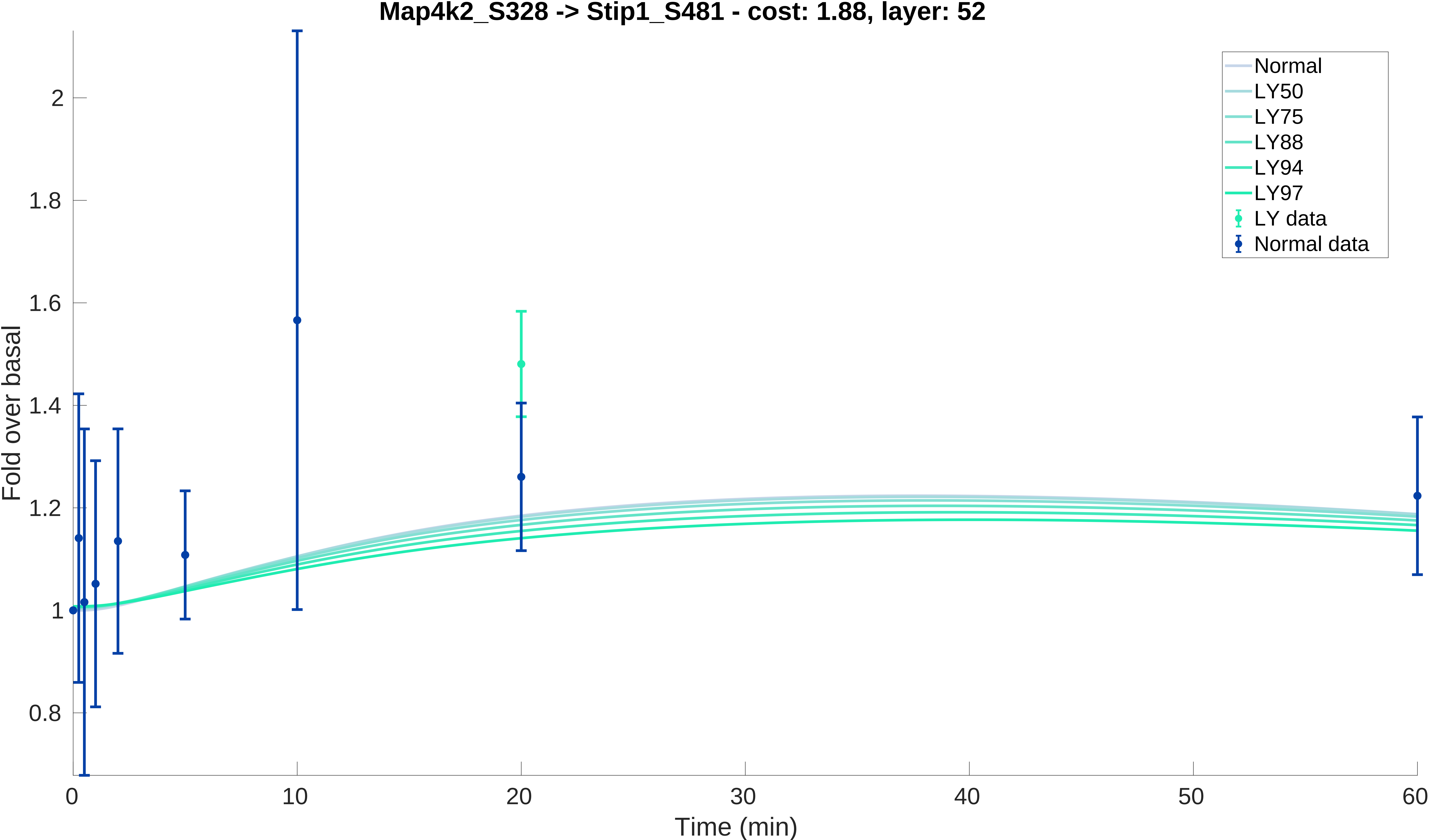

Map4k2_S328 -> Stip1_S481 - cost: 1.88, layer: 52
Normal
LY50
LY75
LY88
LY94
LY97
LY data
Normal data
Fold over basal
0.8
1
1.2
1.4
1.6
1.8
2
0
10
20
30
40
50
60
Time (min)

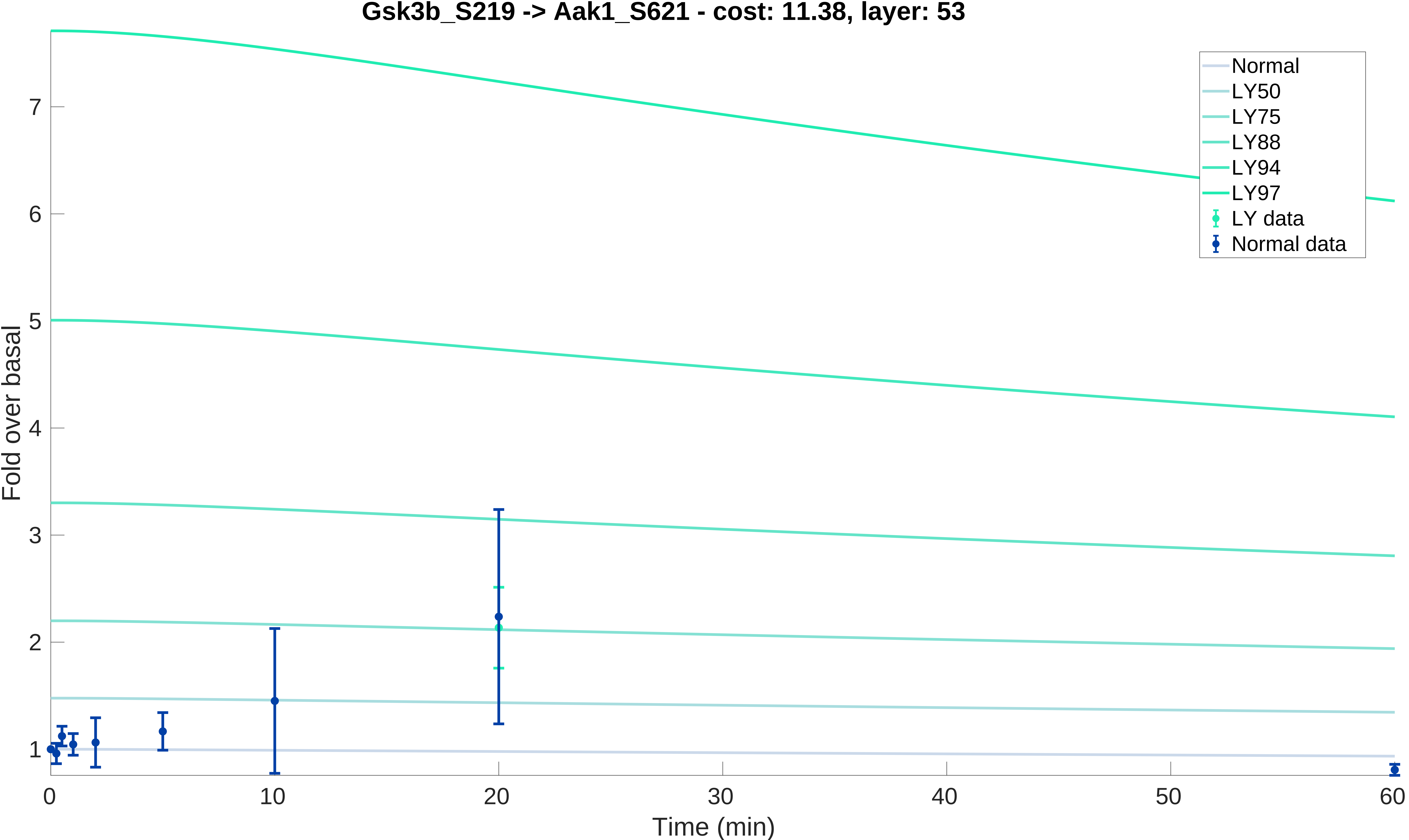
Gsk3b_S219 -> Aak1_S621 - cost: 11.38, layer: 53
Fold over basal
Time (min)
0
10
20
30
40
50
60
1
2
3
4
5
6
7
Normal
LY50
LY75
LY88
LY94
LY97
LY data
Normal data

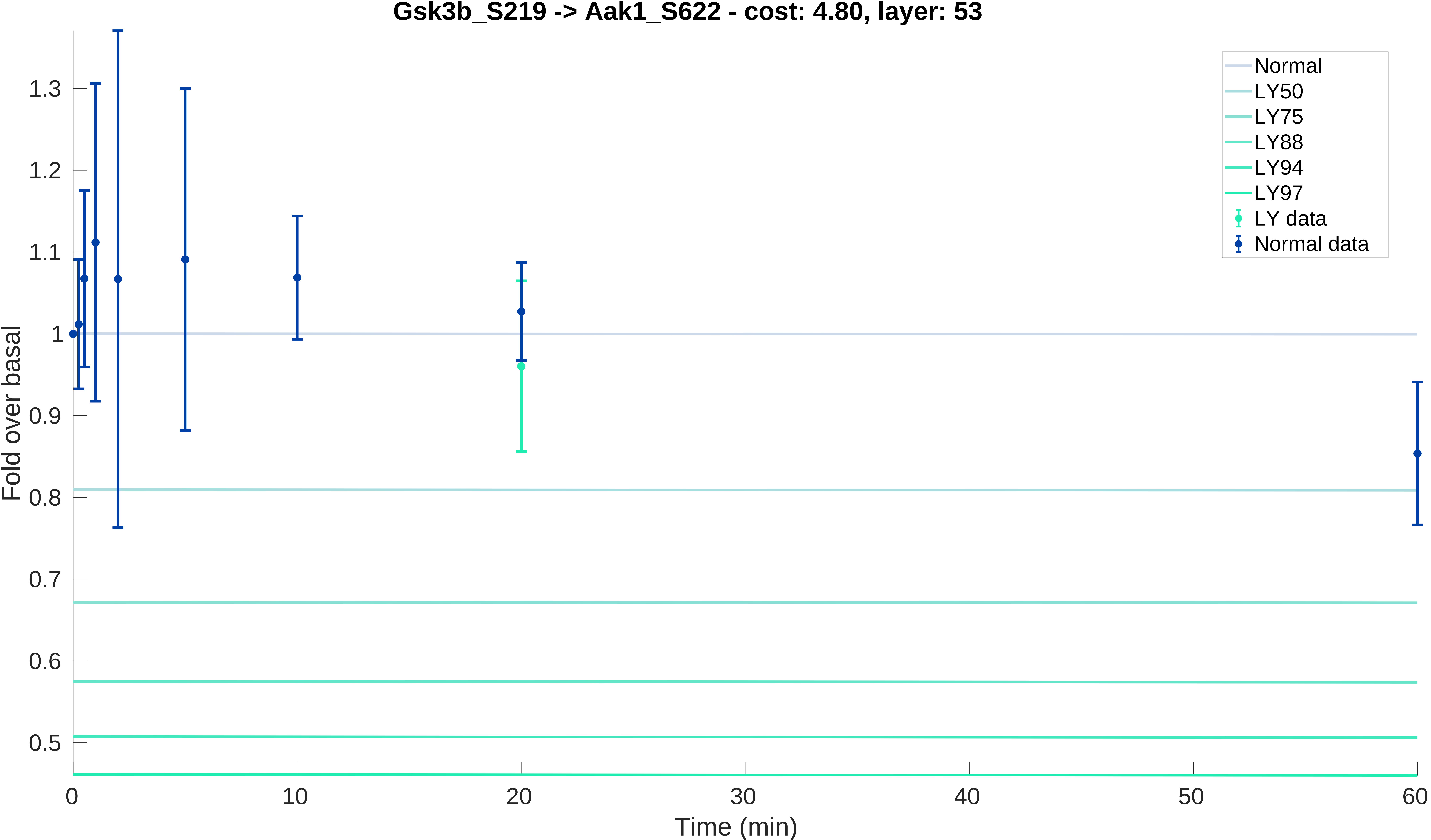

Gsk3b_S219 -> Aak1_S622 - cost: 4.80, layer: 53
Fold over basal
Time (min)
1.3
1.2
1.1
1
0.9
0.8
0.7
0.6
0.5
0
10
20
30
40
50
60
Normal
LY50
LY75
LY88
LY94
LY97
LY data
Normal data

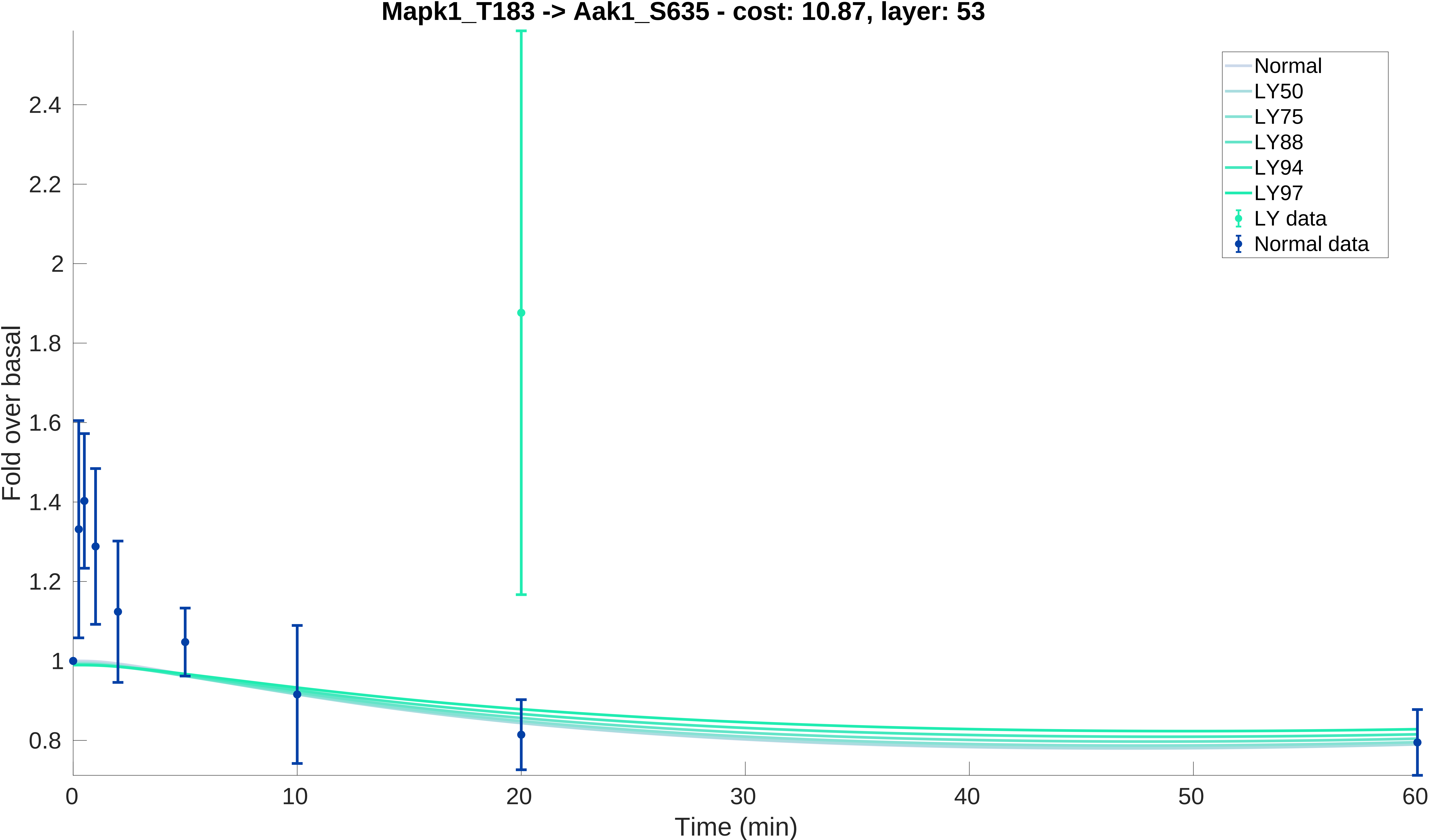

Mapk1_T183 -> Aak1_S635 - cost: 10.87, layer: 53
Normal
LY50
LY75
LY88
LY94
LY97
LY data
Normal data
Fold over basal
2.4
2.2
2
1.8
1.6
1.4
1.2
1
0.8
0
10
20
30
40
50
60
Time (min)

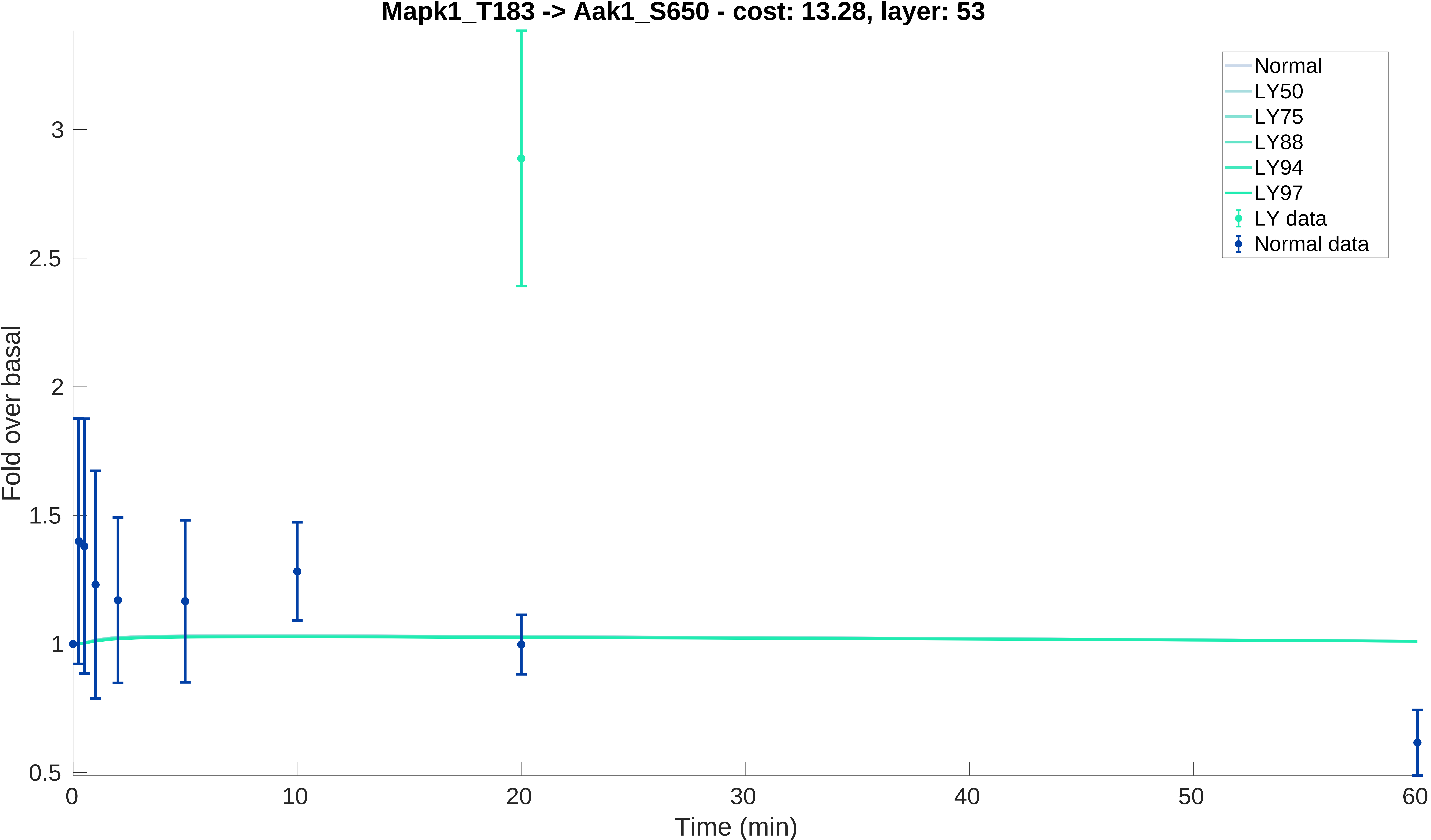

Mapk1_T183 -> Aak1_S650 - cost: 13.28, layer: 53
Fold over basal
Time (min)
0.5
1
1.5
2
2.5
3
0
10
20
30
40
50
60
Normal
LY50
LY75
LY88
LY94
LY97
LY data
Normal data

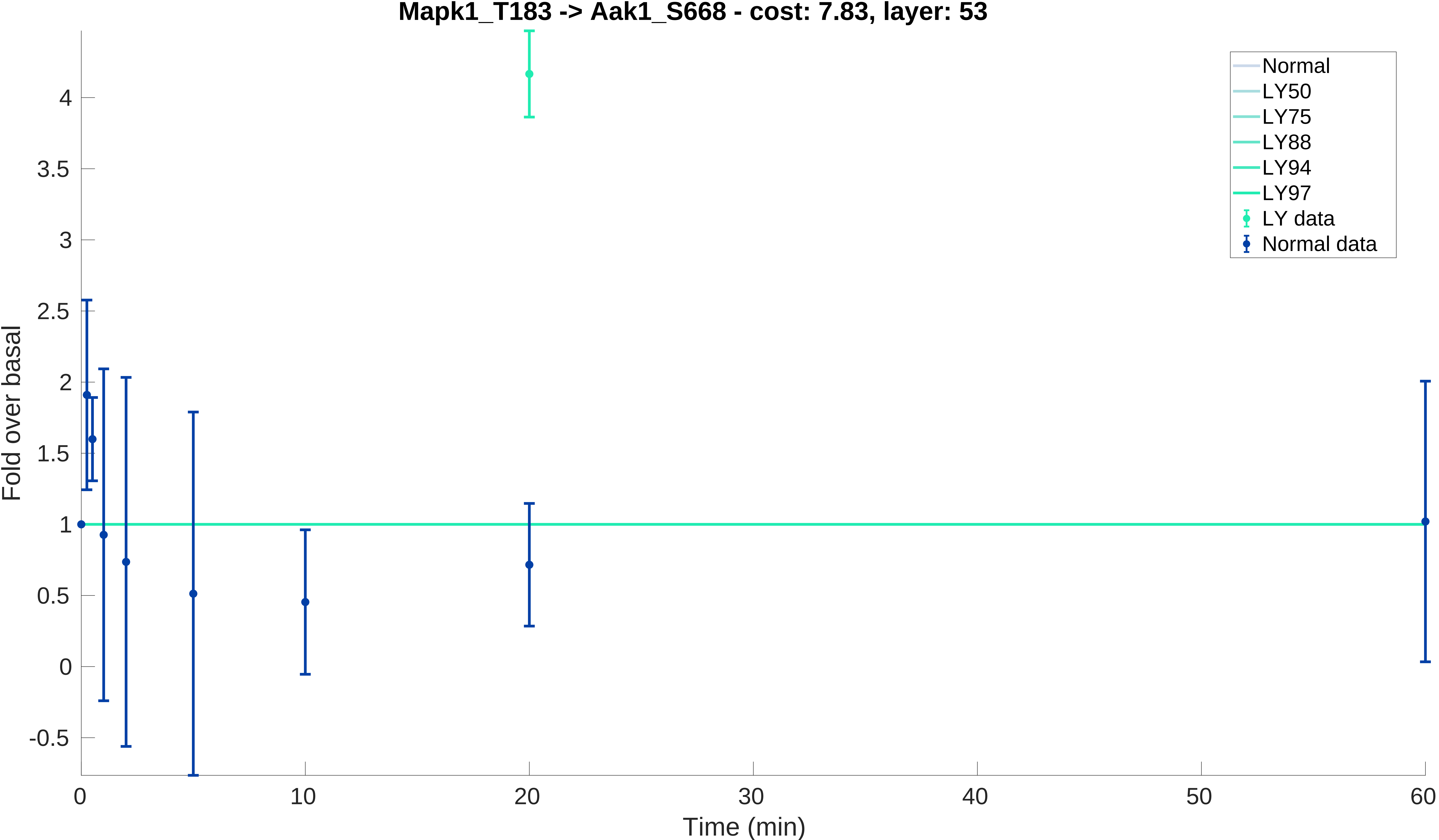

Mapk1_T183 -> Aak1_S668 - cost: 7.83, layer: 53
Fold over basal
Time (min)
Normal
LY50
LY75
LY88
LY94
LY97
LY data
Normal data
4
3.5
3
2.5
2
1.5
1
0.5
0
-0.5
0
10
20
30
40
50
60

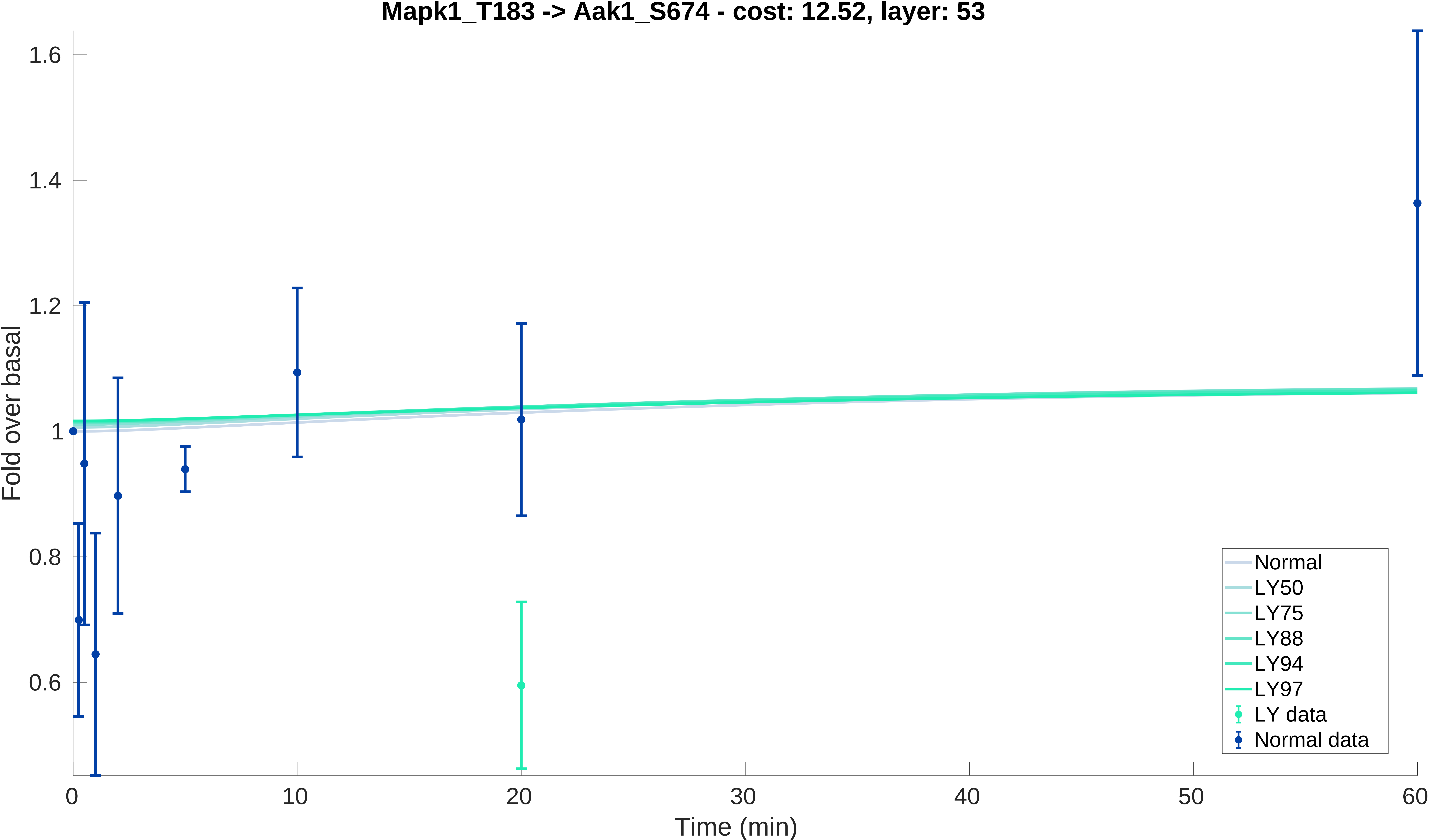

Mapk1_T183 -> Aak1_S674 - cost: 12.52, layer: 53
Fold over basal
1.6
1.4
1.2
1
0.8
0.6
0
10
20
30
40
50
60
Time (min)
Normal
LY50
LY75
LY88
LY94
LY97
LY data
Normal data

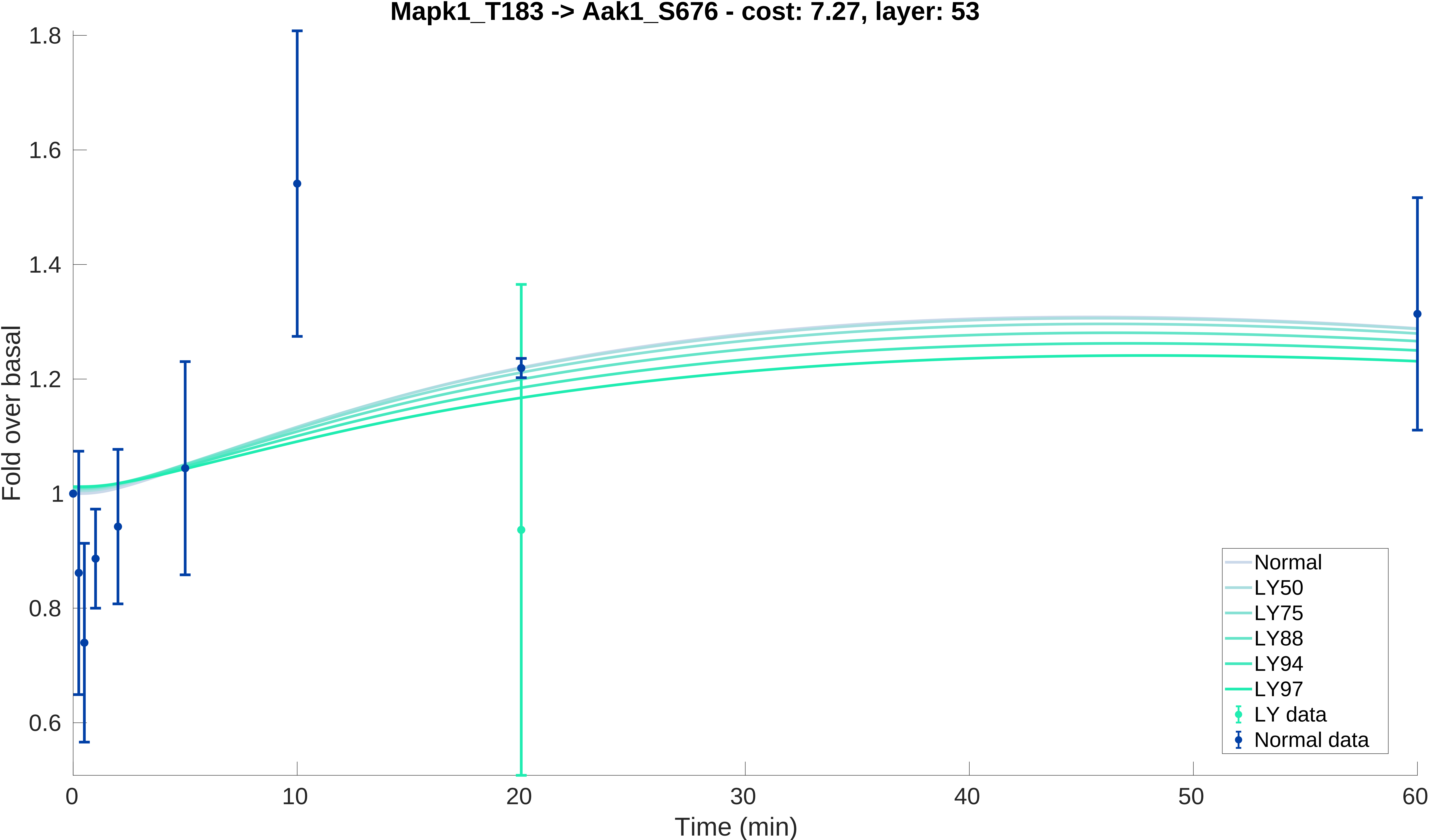

Mapk1_T183 -> Aak1_S676 - cost: 7.27, layer: 53
Fold over basal
Time (min)
1.8
1.6
1.4
1.2
1
0.8
0.6
0
10
20
30
40
50
60
Normal
LY50
LY75
LY88
LY94
LY97
LY data
Normal data

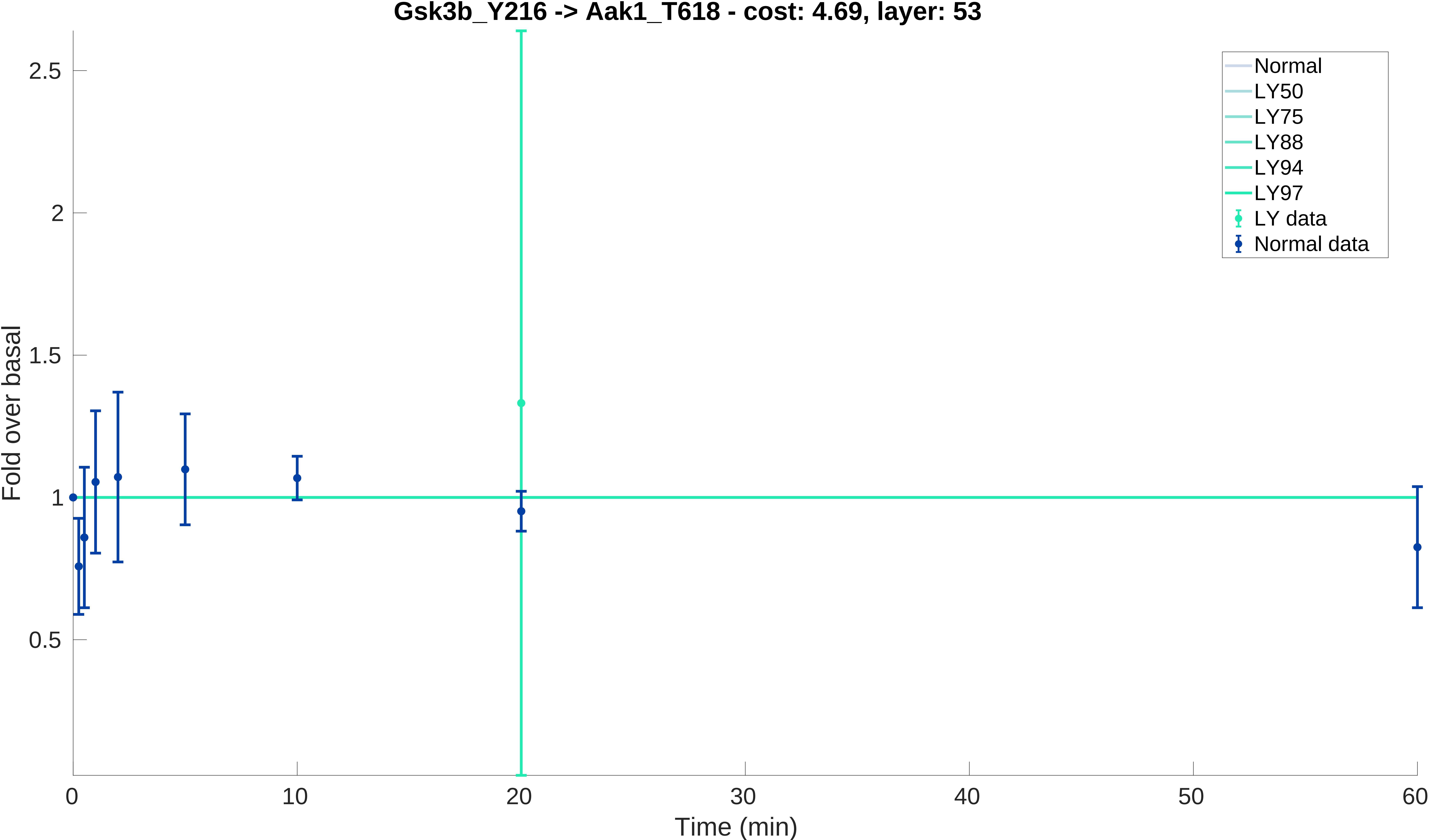
Gsk3b_Y216 -> Aak1_T618 - cost: 4.69, layer: 53
Fold over basal
Time (min)
0
10
20
30
40
50
60
0.5
1
1.5
2
2.5
Normal
LY50
LY75
LY88
LY94
LY97
LY data
Normal data

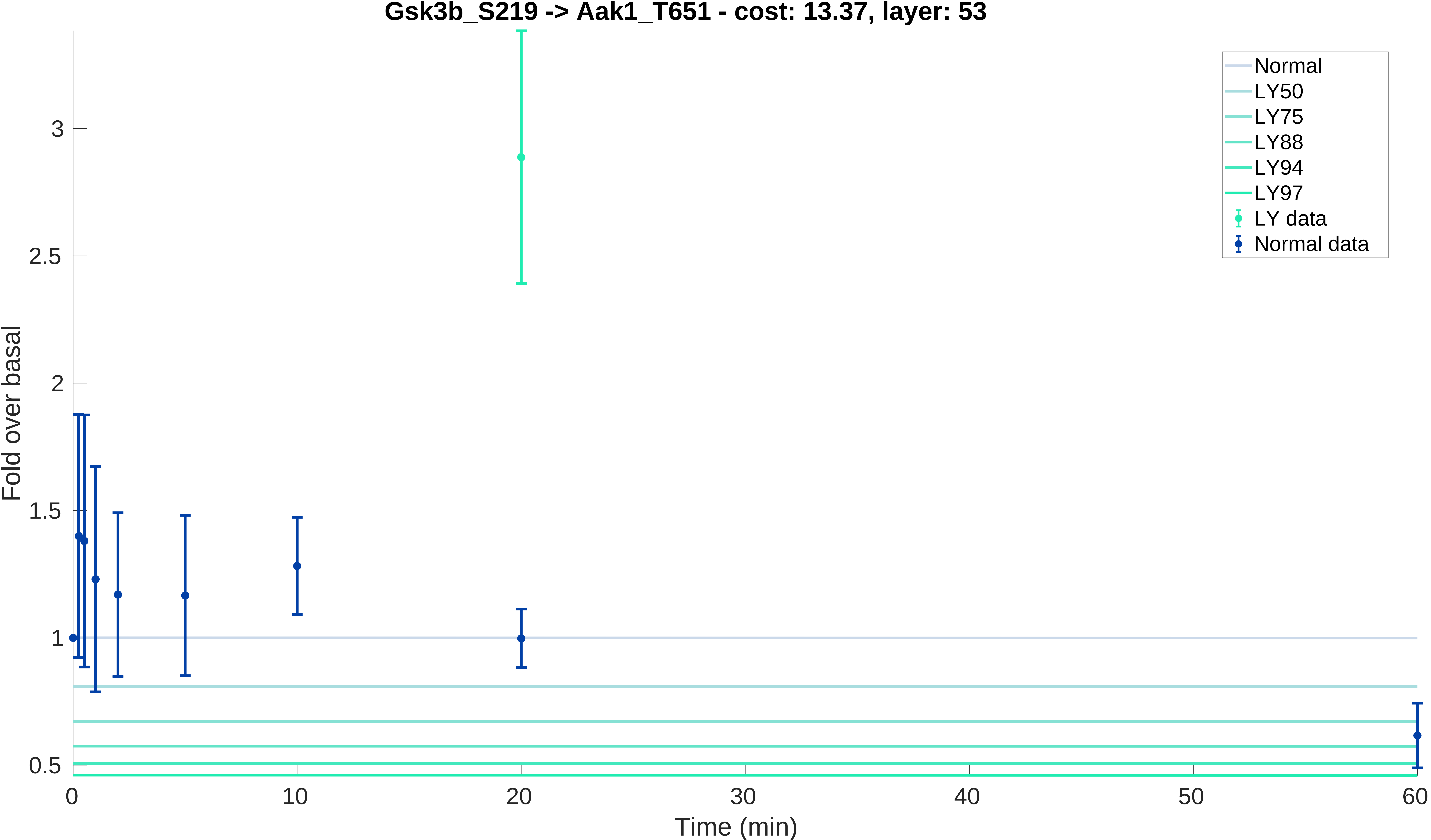

Gsk3b_S219 -> Aak1_T651 - cost: 13.37, layer: 53
Fold over basal
Time (min)
Normal
LY50
LY75
LY88
LY94
LY97
LY data
Normal data
0.5
1
1.5
2
2.5
3
0
10
20
30
40
50
60

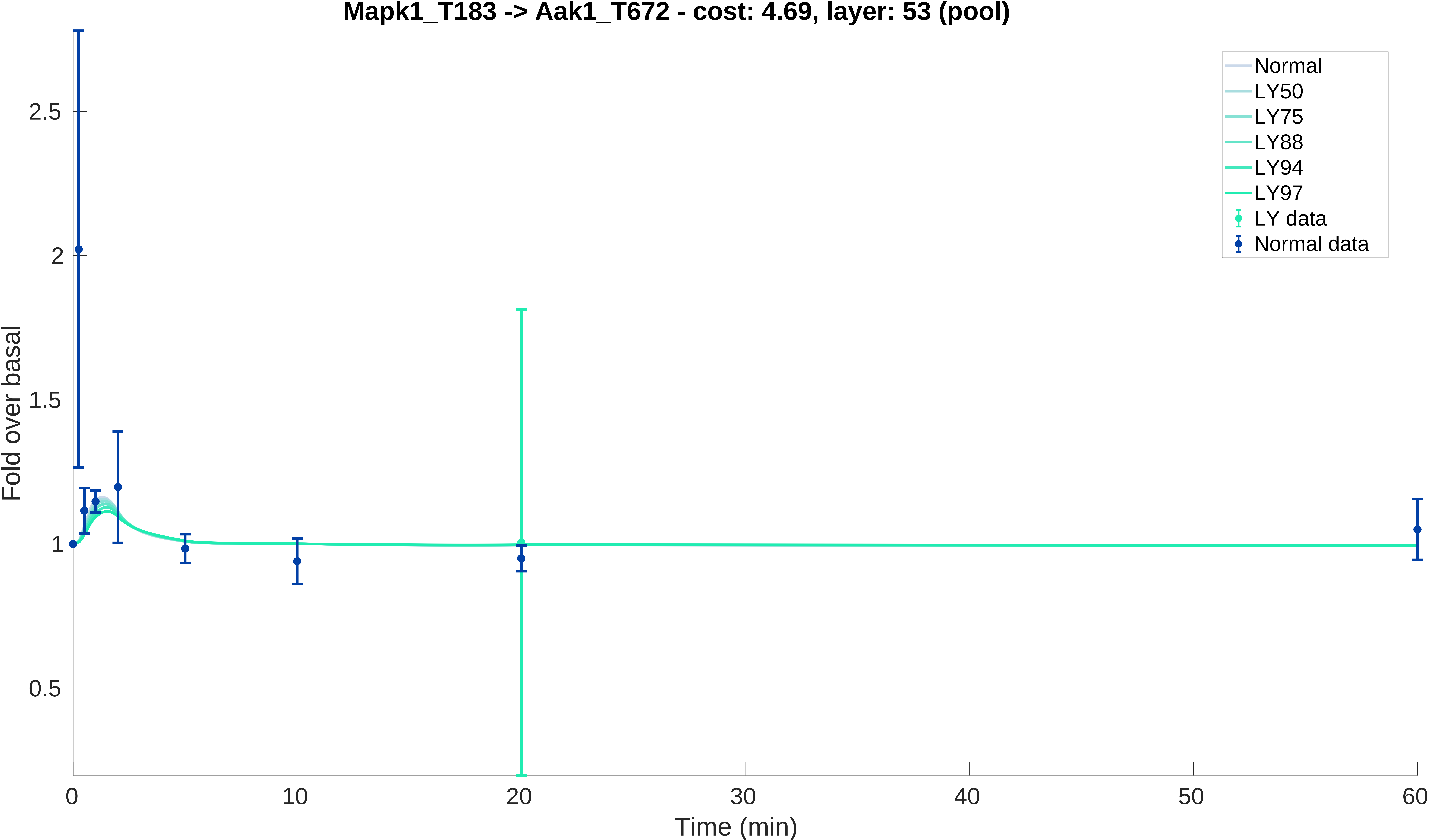

Mapk1_T183 -> Aak1_T672 - cost: 4.69, layer: 53 (pool)
Normal
LY50
LY75
LY88
LY94
LY97
LY data
Normal data
Fold over basal
2.5
2
1.5
1
0.5
0
10
20
30
40
50
60
Time (min)

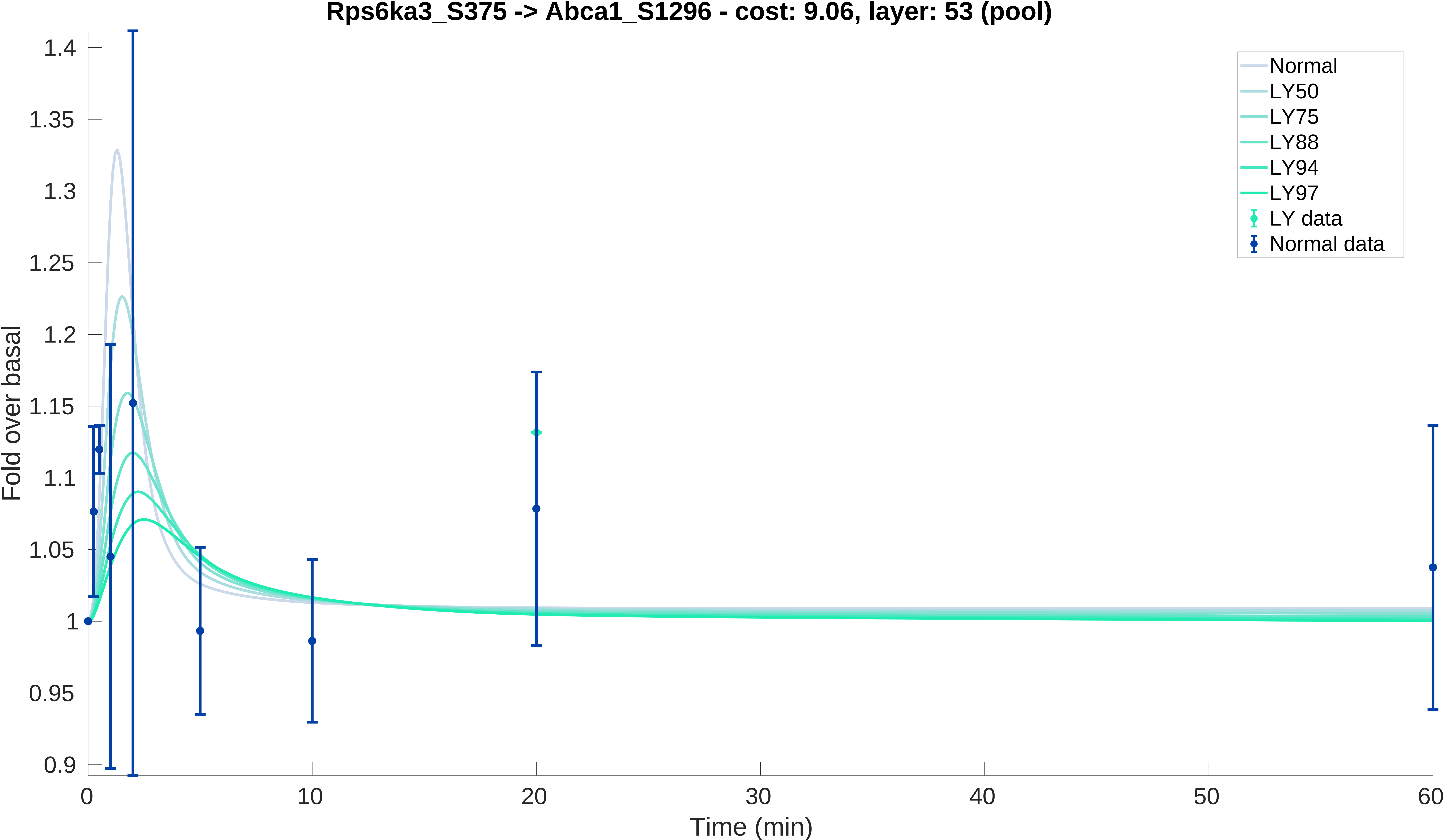

Rps6ka3_S375 -> Abca1_S1296 - cost: 9.06, layer: 53 (pool)
Fold over basal
Time (min)
Normal
LY50
LY75
LY88
LY94
LY97
LY data
Normal data
1.4
1.35
1.3
1.25
1.2
1.15
1.1
1.05
1
0.95
0.9
0
10
20
30
40
50
60

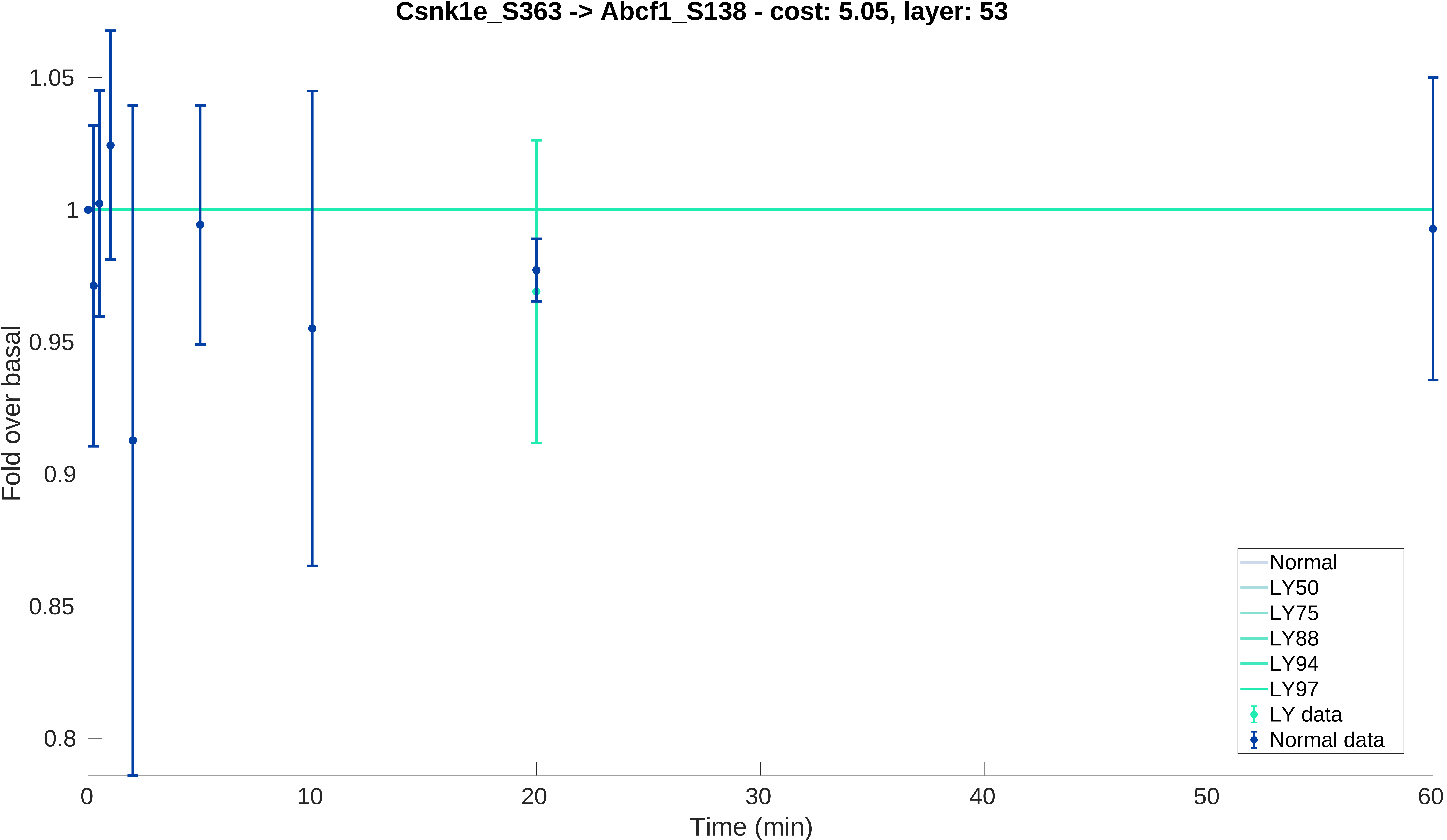

Csnk1e_S363 -> Abcf1_S138 - cost: 5.05, layer: 53
Fold over basal
1.05
1
0.95
0.9
0.85
0.8
0
10
20
30
40
50
60
Time (min)
Normal
LY50
LY75
LY88
LY94
LY97
LY data
Normal data

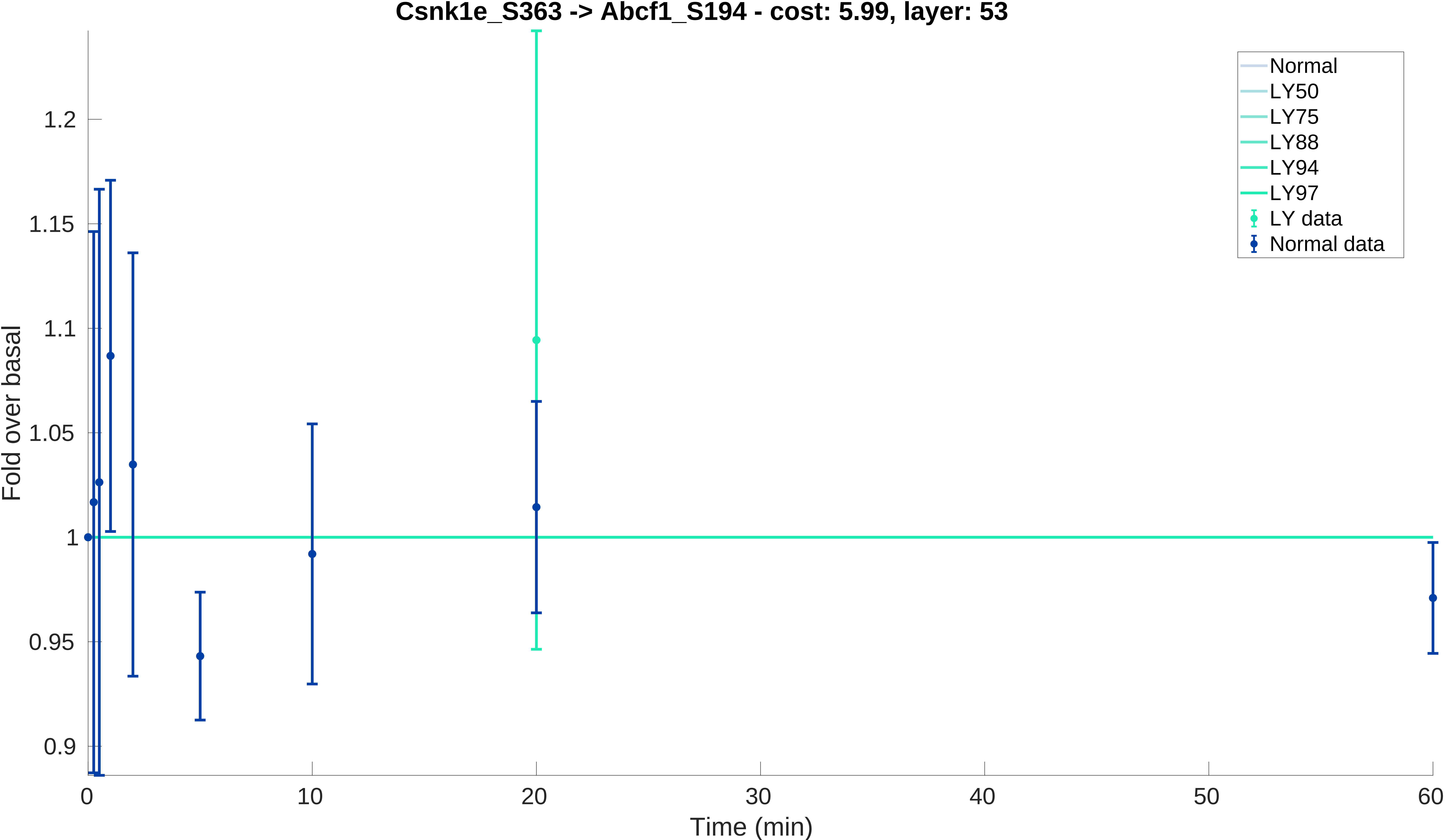

Csnk1e_S363 -> Abcf1_S194 - cost: 5.99, layer: 53
Normal
LY50
LY75
LY88
LY94
LY97
LY data
Normal data
Fold over basal
1.2
1.15
1.1
1.05
1
0.95
0.9
0
10
20
30
40
50
60
Time (min)

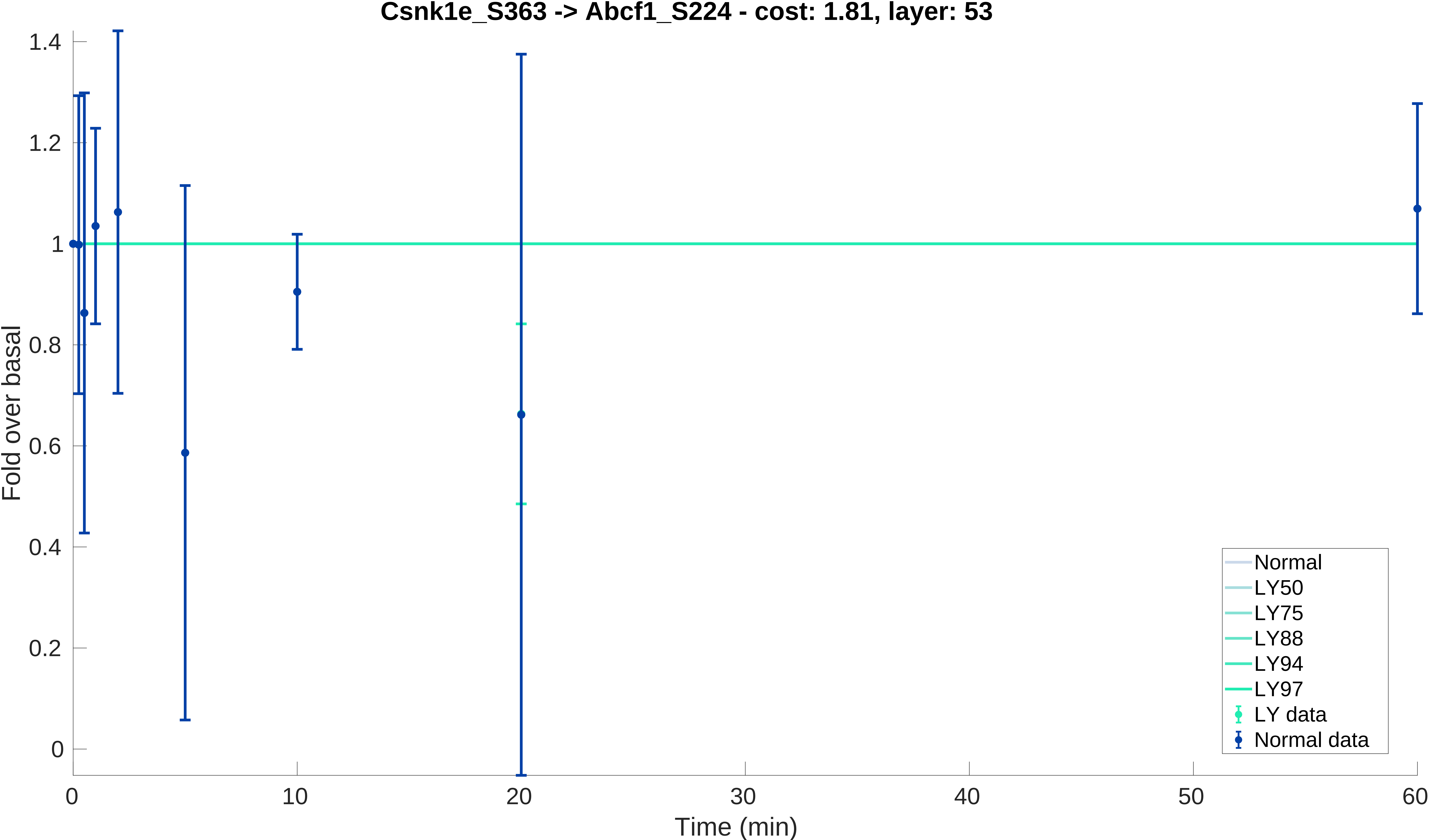

Csnk1e_S363 -> Abcf1_S224 - cost: 1.81, layer: 53
Fold over basal
Time (min)
0
0.2
0.4
0.6
0.8
1
1.2
1.4
0
10
20
30
40
50
60
Normal
LY50
LY75
LY88
LY94
LY97
LY data
Normal data

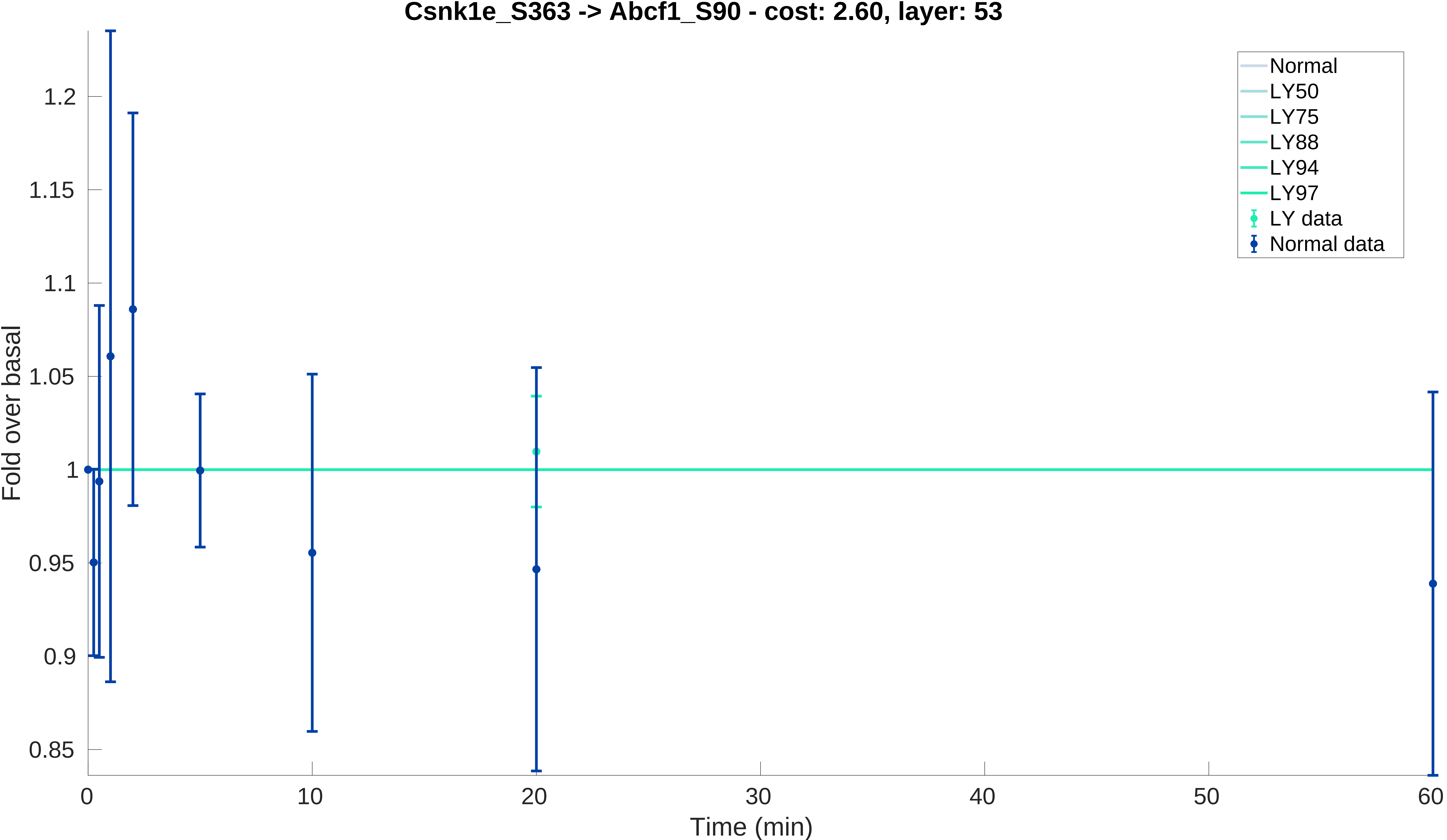

Csnk1e_S363 -> Abcf1_S90 - cost: 2.60, layer: 53
Fold over basal
Time (min)
1.2
1.15
1.1
1.05
1
0.95
0.9
0.85
0
10
20
30
40
50
60
Normal
LY50
LY75
LY88
LY94
LY97
LY data
Normal data

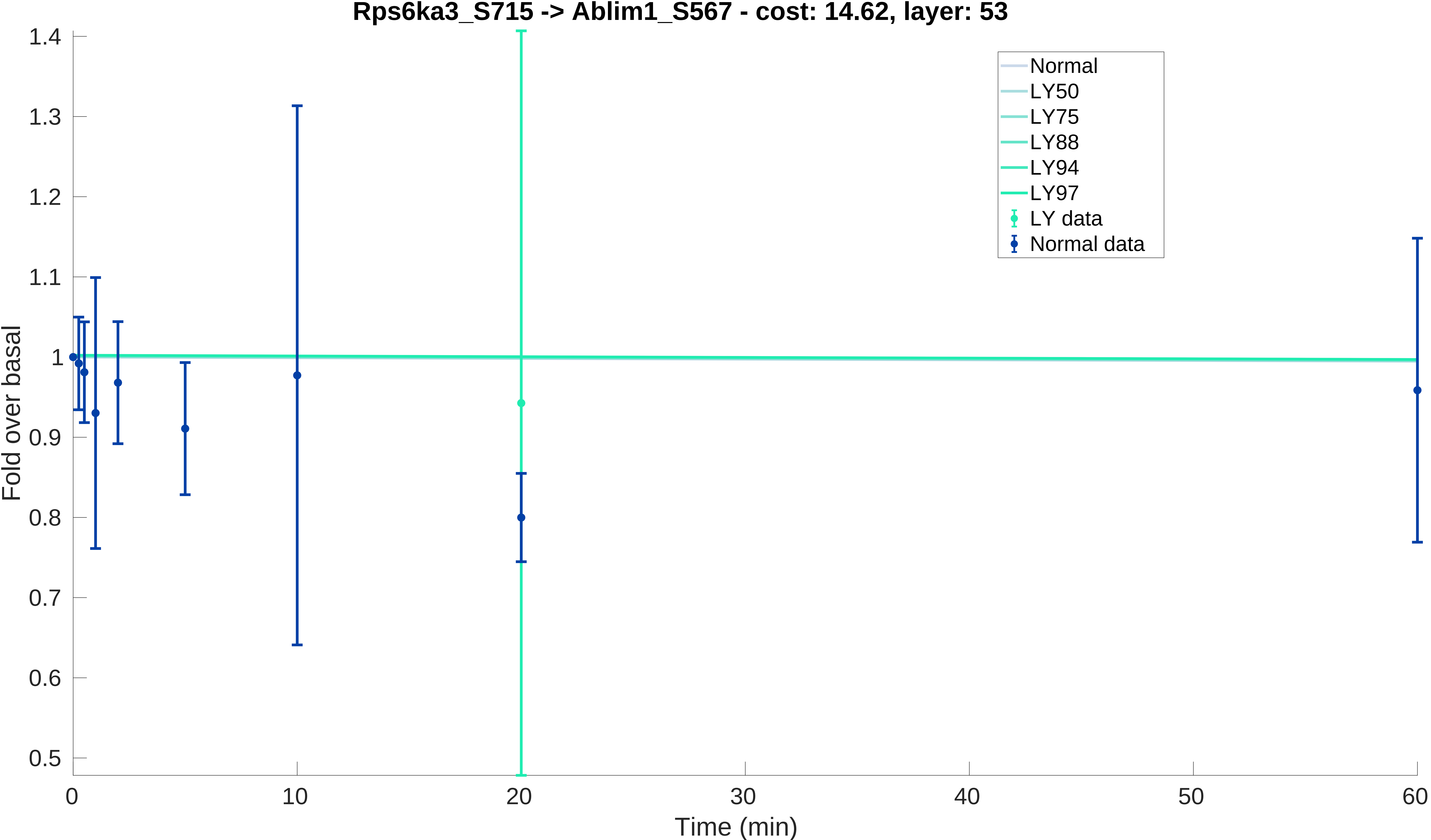
Rps6ka3_S715 -> Ablim1_S567 - cost: 14.62, layer: 53
Fold over basal
Time (min)
1.4
1.3
1.2
1.1
1
0.9
0.8
0.7
0.6
0.5
0
10
20
30
40
50
60
Normal
LY50
LY75
LY88
LY94
LY97
LY data
Normal data

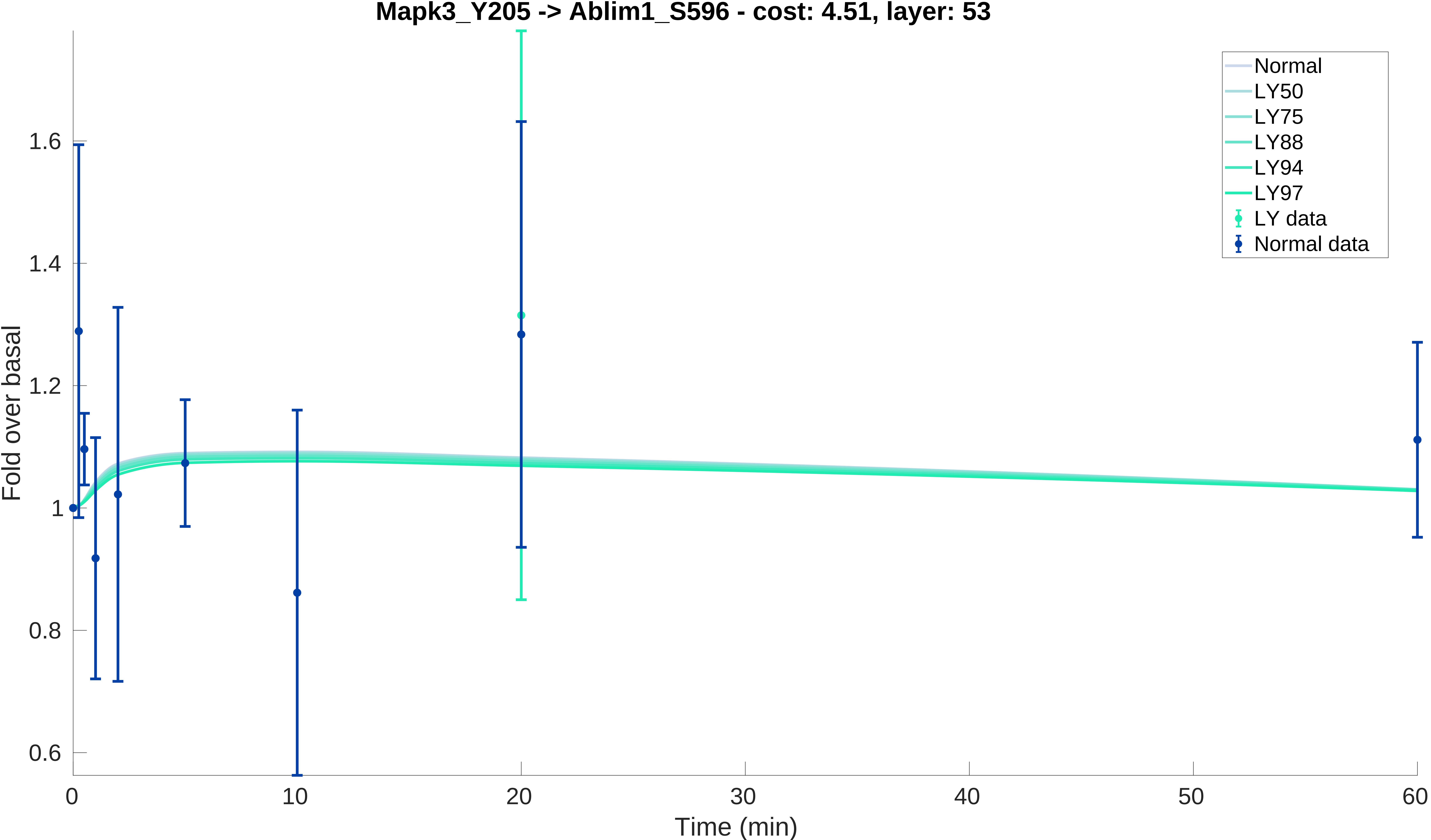

Mapk3_Y205 -> Ablim1_S596 - cost: 4.51, layer: 53
Fold over basal
Time (min)
0.6
0.8
1
1.2
1.4
1.6
0
10
20
30
40
50
60
Normal
LY50
LY75
LY88
LY94
LY97
LY data
Normal data

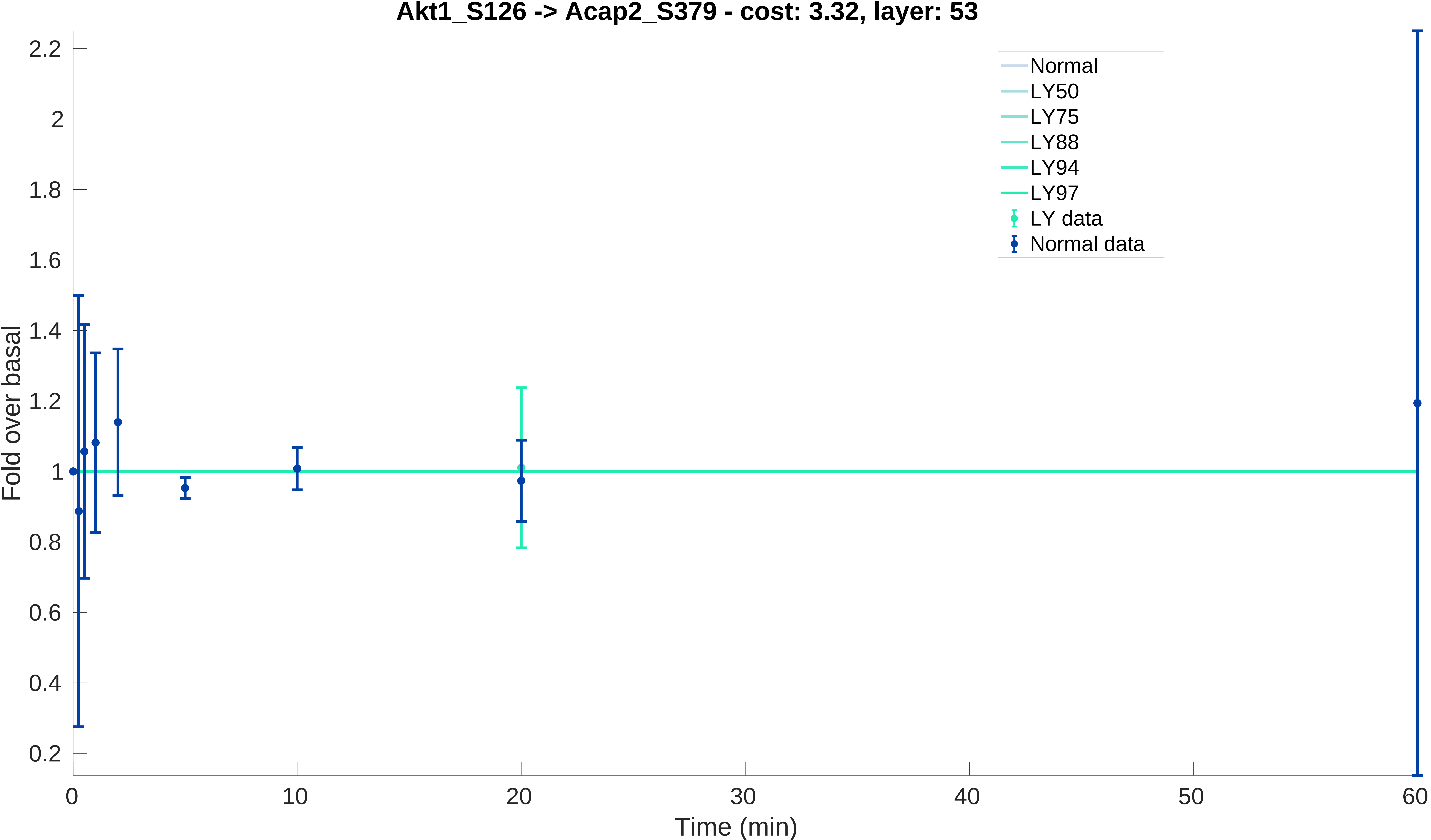

Akt1_S126 -> Acap2_S379 - cost: 3.32, layer: 53
Normal
LY50
LY75
LY88
LY94
LY97
LY data
Normal data
Fold over basal
Time (min)
0.2
0.4
0.6
0.8
1
1.2
1.4
1.6
1.8
2
2.2
0
10
20
30
40
50
60

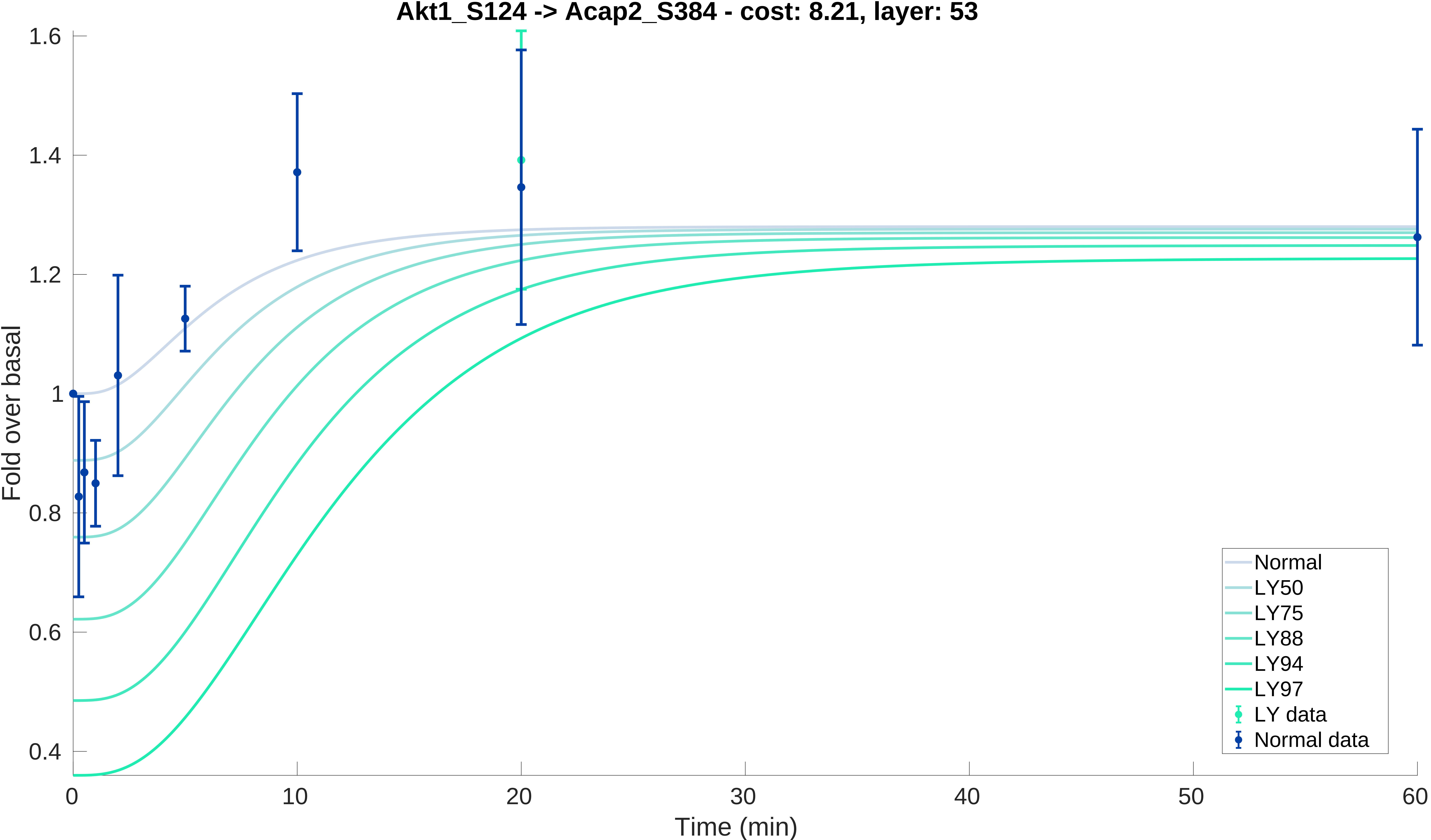

Akt1_S124 -> Acap2_S384 - cost: 8.21, layer: 53
Fold over basal
1.6
1.4
1.2
1
0.8
0.6
0.4
0
10
20
30
40
50
60
Time (min)
Normal
LY50
LY75
LY88
LY94
LY97
LY data
Normal data

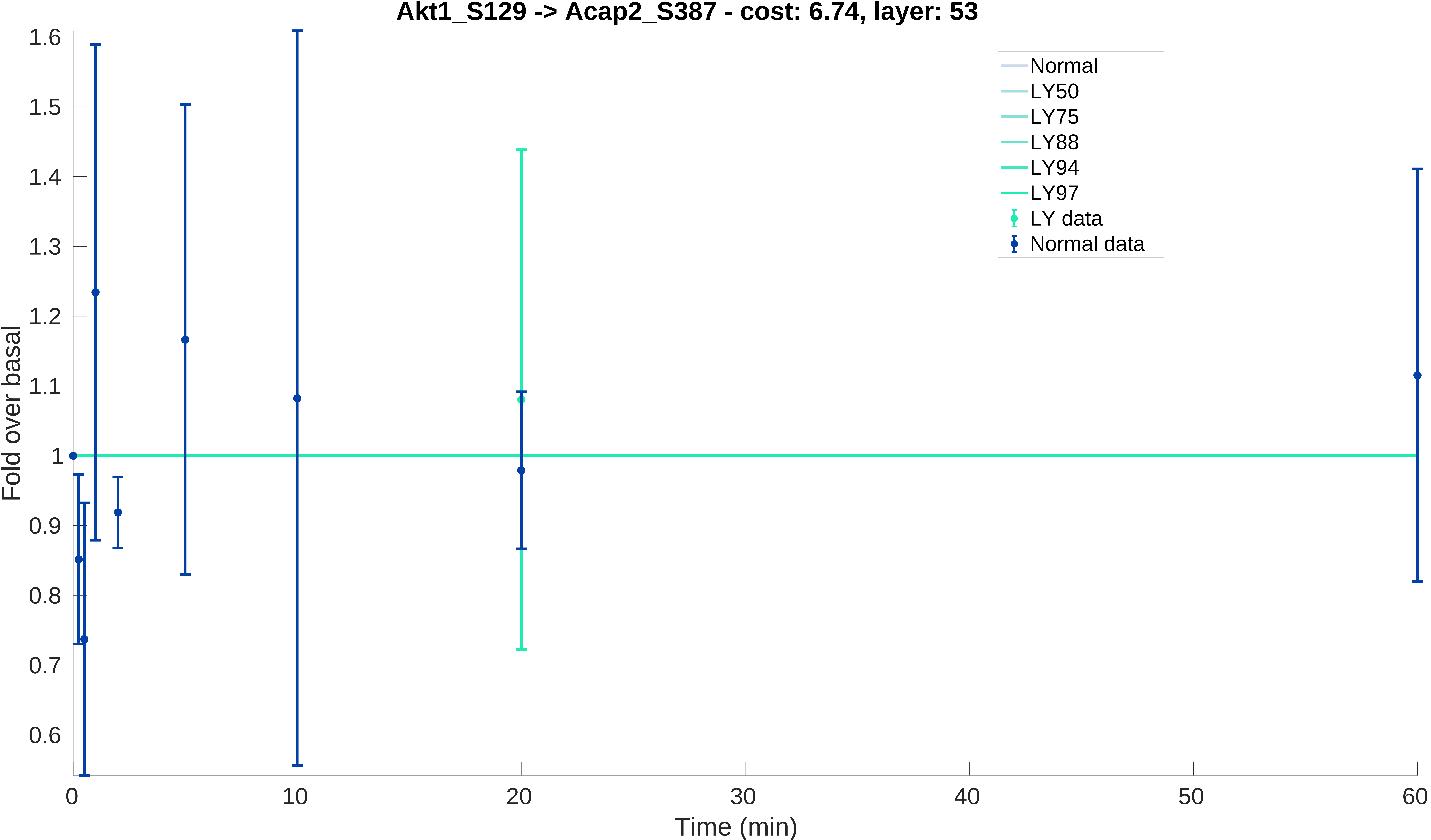

Akt1_S129 -> Acap2_S387 - cost: 6.74, layer: 53
Fold over basal
Time (min)
Normal
LY50
LY75
LY88
LY94
LY97
LY data
Normal data
0.6
0.7
0.8
0.9
1
1.1
1.2
1.3
1.4
1.5
1.6
0
10
20
30
40
50
60

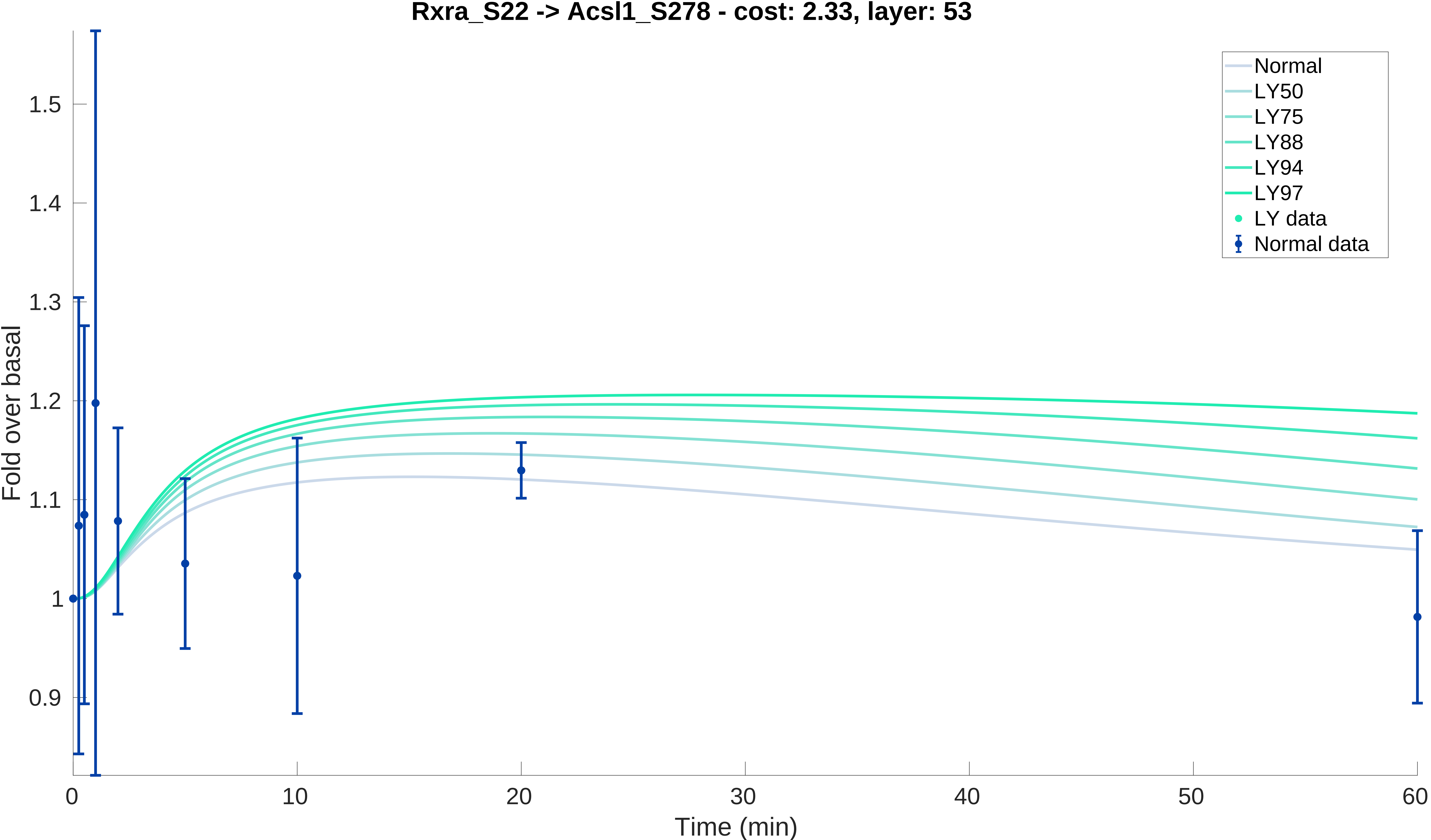

Rxra_S22 -> Acsl1_S278 - cost: 2.33, layer: 53
Normal
LY50
LY75
LY88
LY94
LY97
LY data
Normal data
Fold over basal
1.5
1.4
1.3
1.2
1.1
1
0.9
0
10
20
30
40
50
60
Time (min)

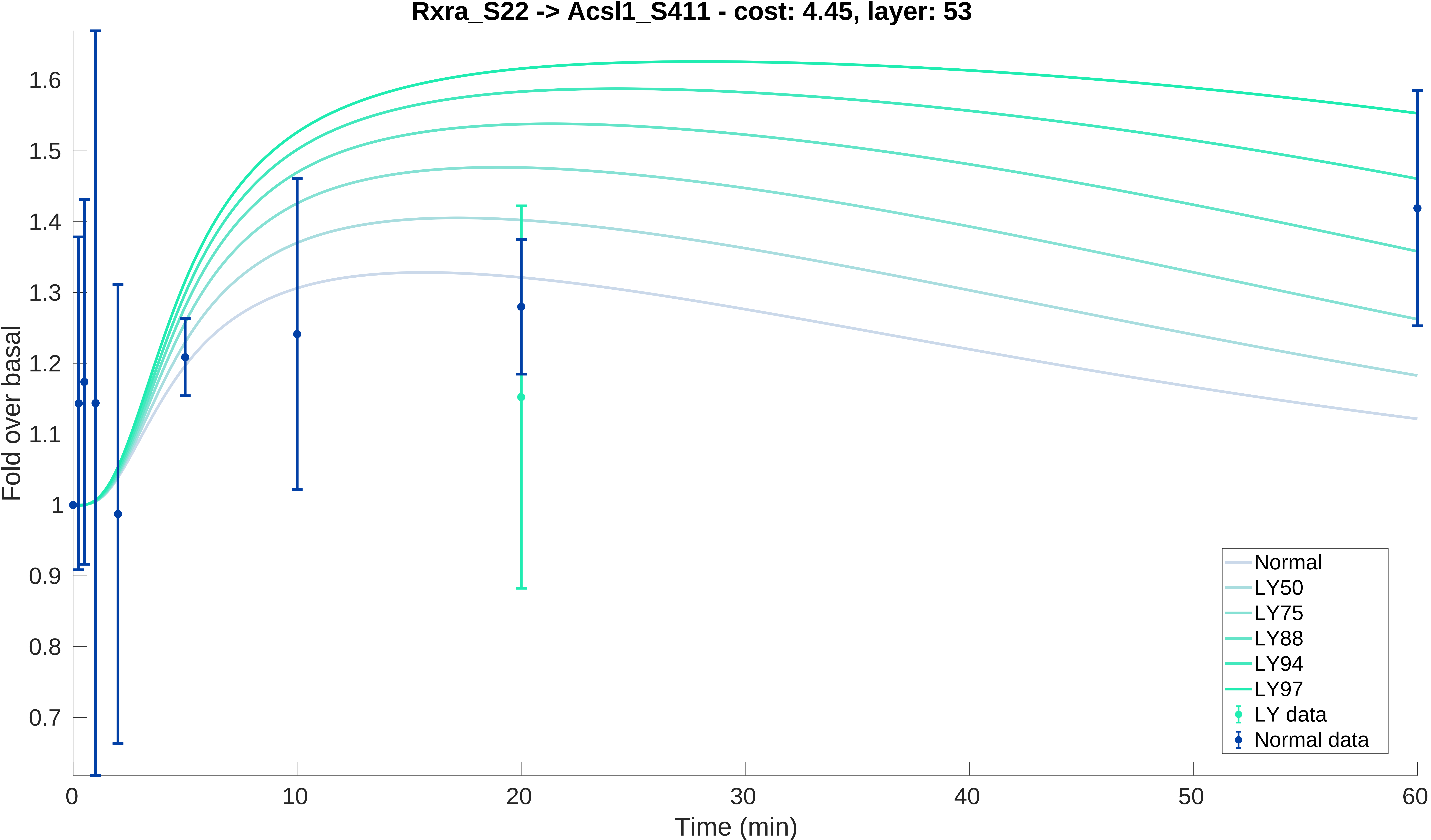
Rxra_S22 -> Acsl1_S411 - cost: 4.45, layer: 53
Fold over basal
Time (min)
0.7
0.8
0.9
1
1.1
1.2
1.3
1.4
1.5
1.6
0
10
20
30
40
50
60
Normal
LY50
LY75
LY88
LY94
LY97
LY data
Normal data

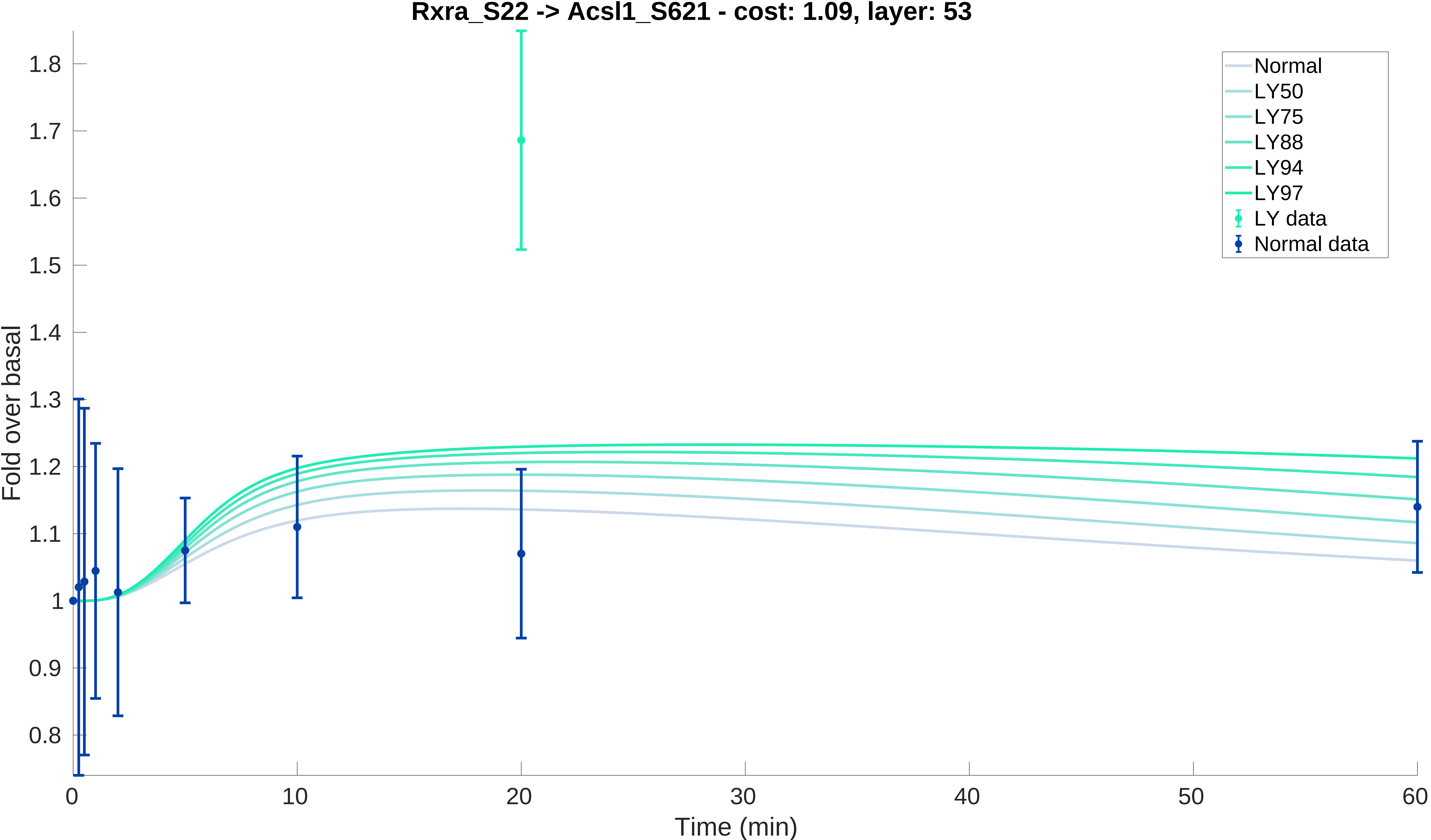

Rxra_S22 -> Acsl1_S621 - cost: 1.09, layer: 53
Fold over basal
Time (min)
1.8
1.7
1.6
1.5
1.4
1.3
1.2
1.1
1
0.9
0.8
0
10
20
30
40
50
60
Normal
LY50
LY75
LY88
LY94
LY97
LY data
Normal data

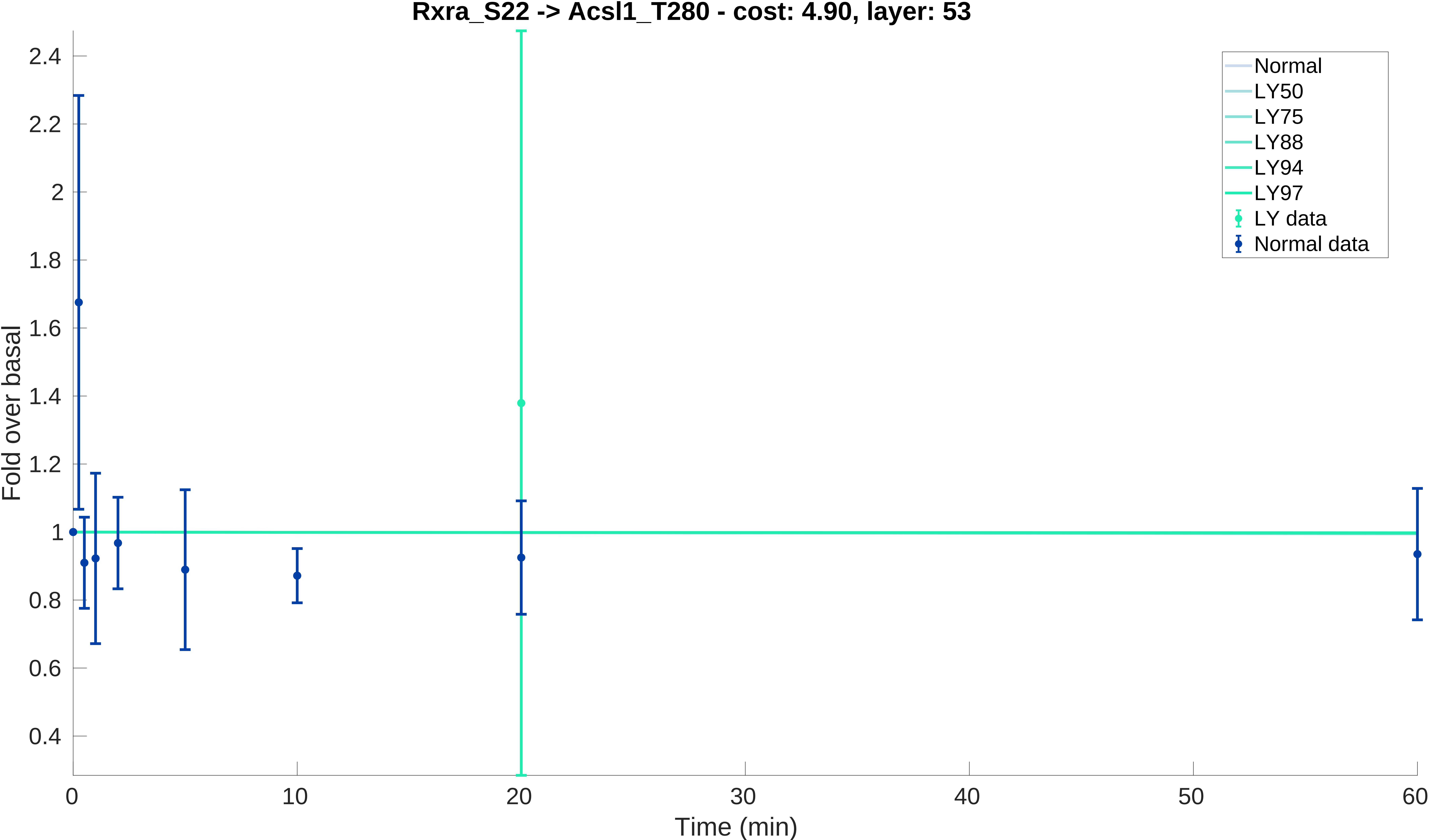

Rxra_S22 -> Acsl1_T280 - cost: 4.90, layer: 53
Fold over basal
Time (min)
2.4
2.2
2
1.8
1.6
1.4
1.2
1
0.8
0.6
0.4
0
10
20
30
40
50
60
Normal
LY50
LY75
LY88
LY94
LY97
LY data
Normal data

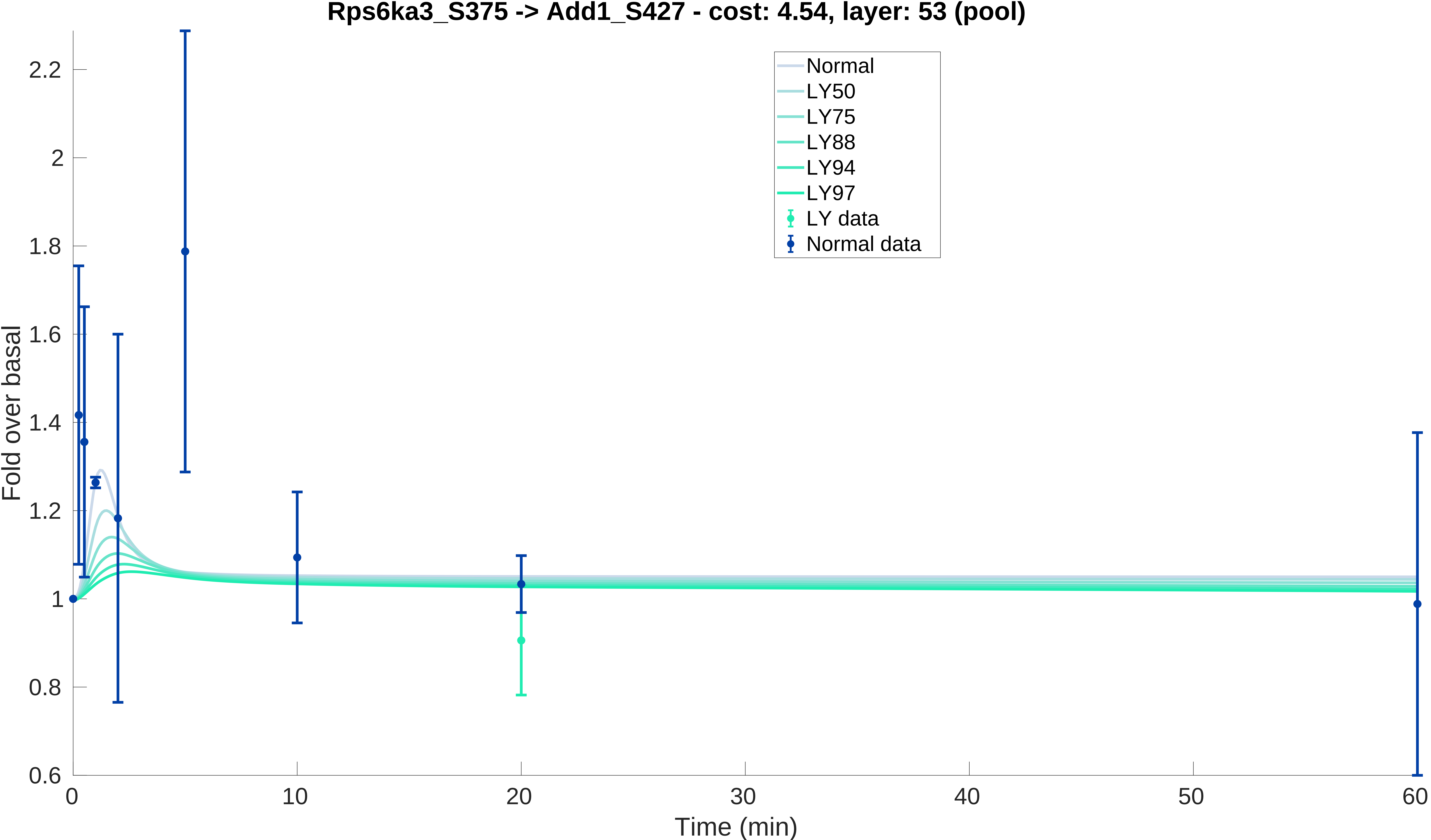

Rps6ka3_S375 -> Add1_S427 - cost: 4.54, layer: 53 (pool)
Normal
LY50
LY75
LY88
LY94
LY97
LY data
Normal data
Fold over basal
Time (min)
2.2
2
1.8
1.6
1.4
1.2
1
0.8
0.6
0
10
20
30
40
50
60

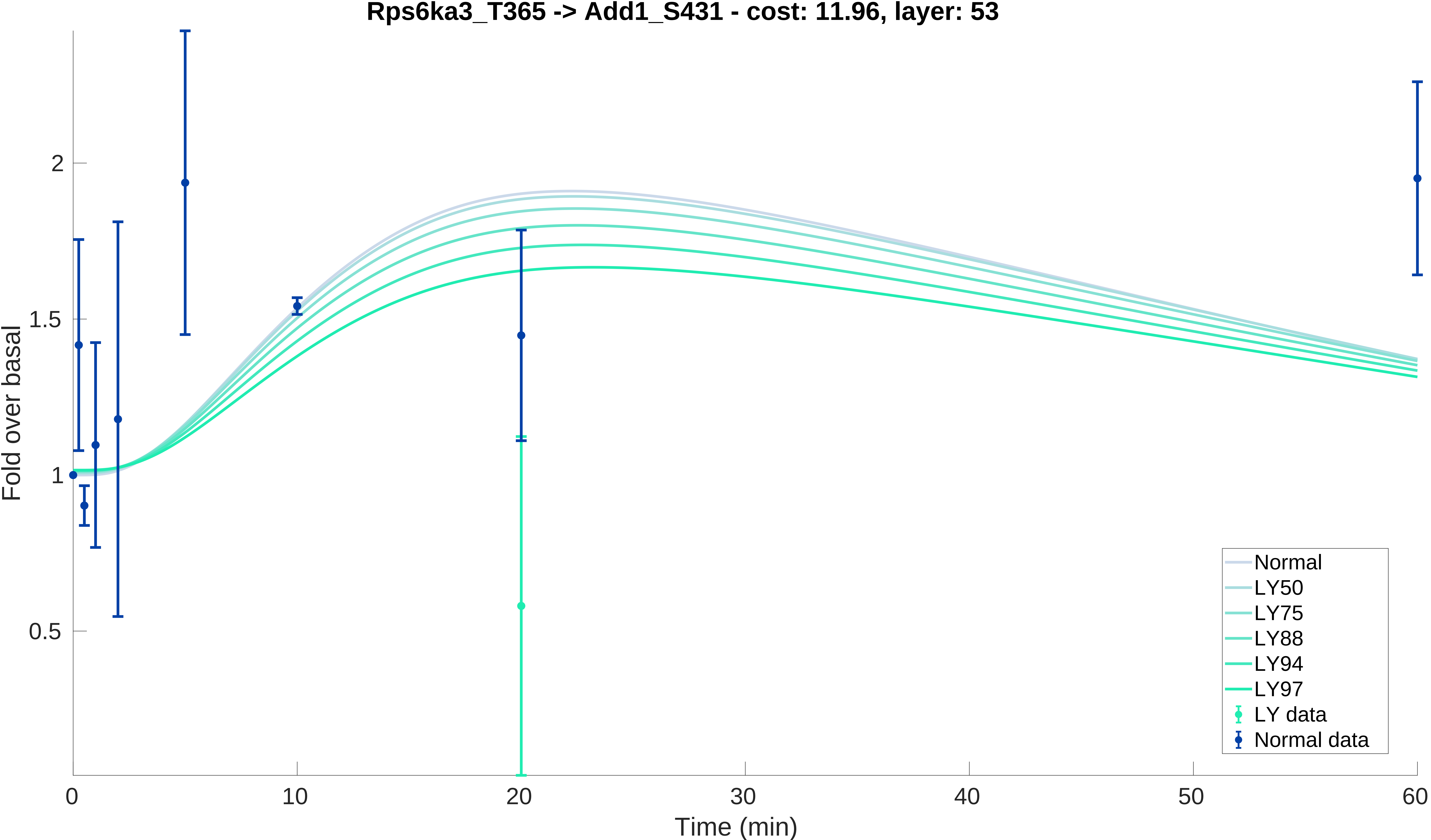

Rps6ka3_T365 -> Add1_S431 - cost: 11.96, layer: 53
Fold over basal
Time (min)
0
10
20
30
40
50
60
0.5
1
1.5
2
Normal
LY50
LY75
LY88
LY94
LY97
LY data
Normal data

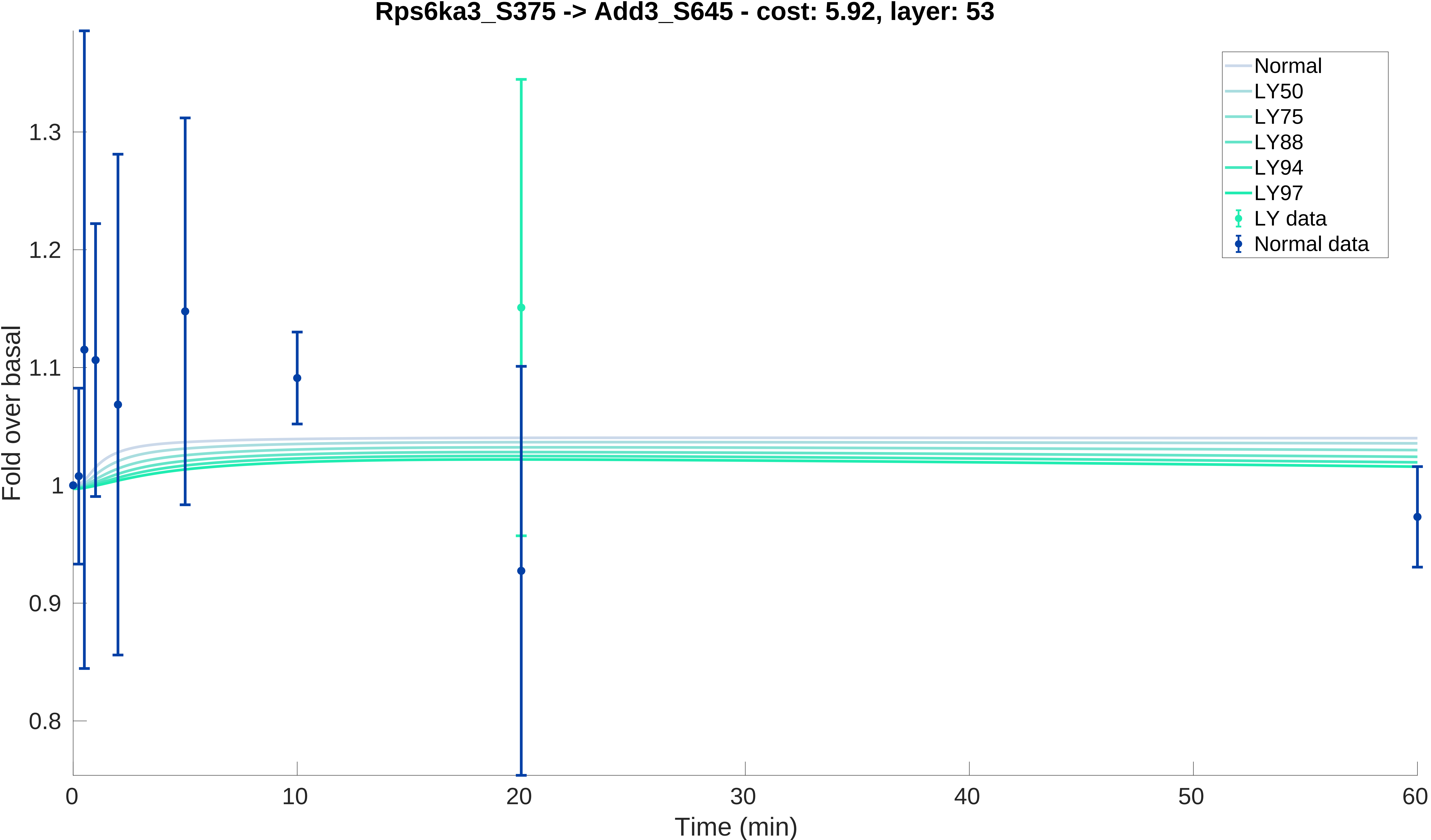
Rps6ka3_S375 -> Add3_S645 - cost: 5.92, layer: 53
Normal
LY50
LY75
LY88
LY94
LY97
LY data
Normal data
1.3
1.2
1.1
1
0.9
0.8
Fold over basal
0
10
20
30
40
50
60
Time (min)

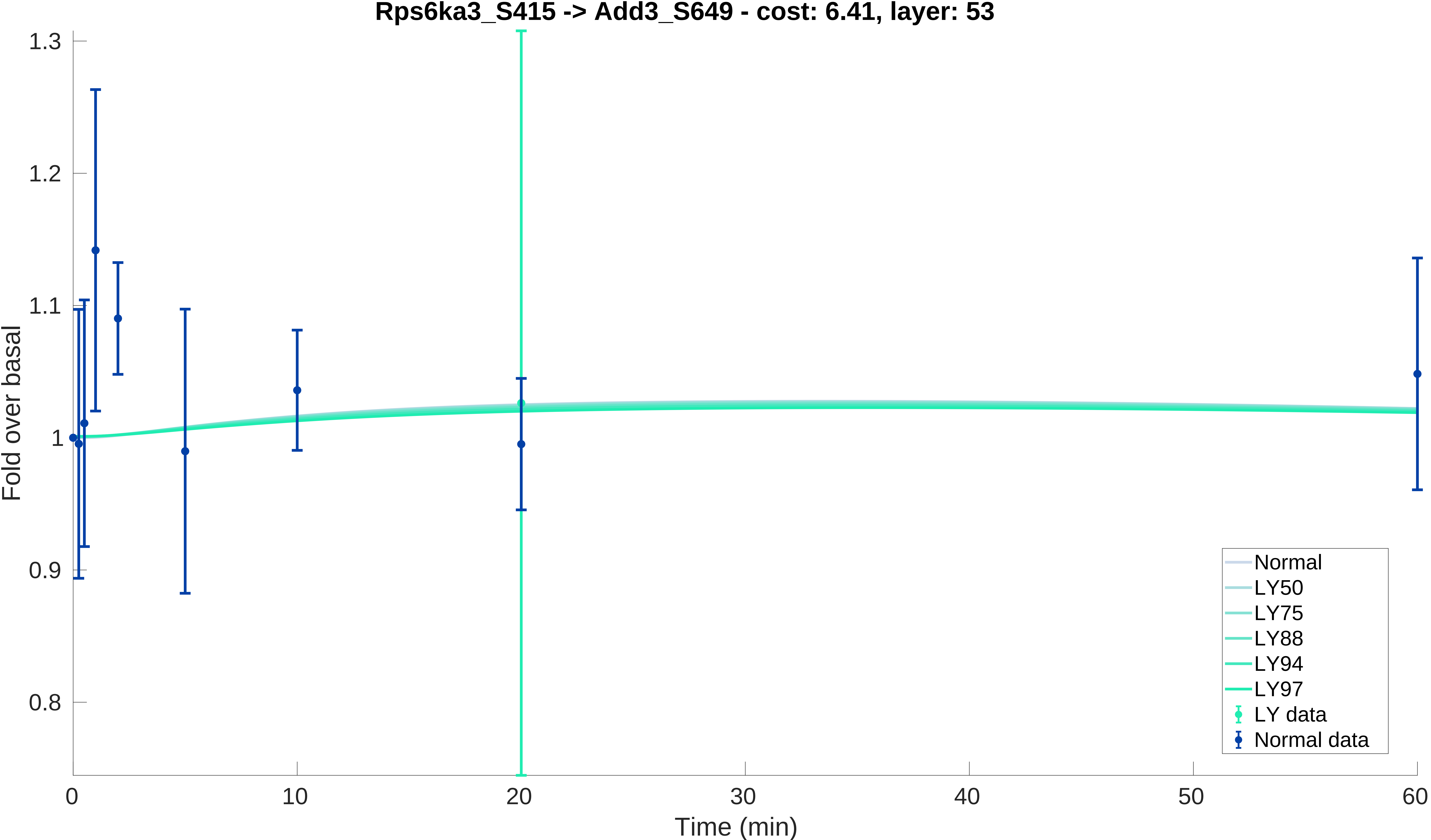

Rps6ka3_S415 -> Add3_S649 - cost: 6.41, layer: 53
Fold over basal
1.3
1.2
1.1
1
0.9
0.8
0
10
20
30
40
50
60
Time (min)
Normal
LY50
LY75
LY88
LY94
LY97
LY data
Normal data

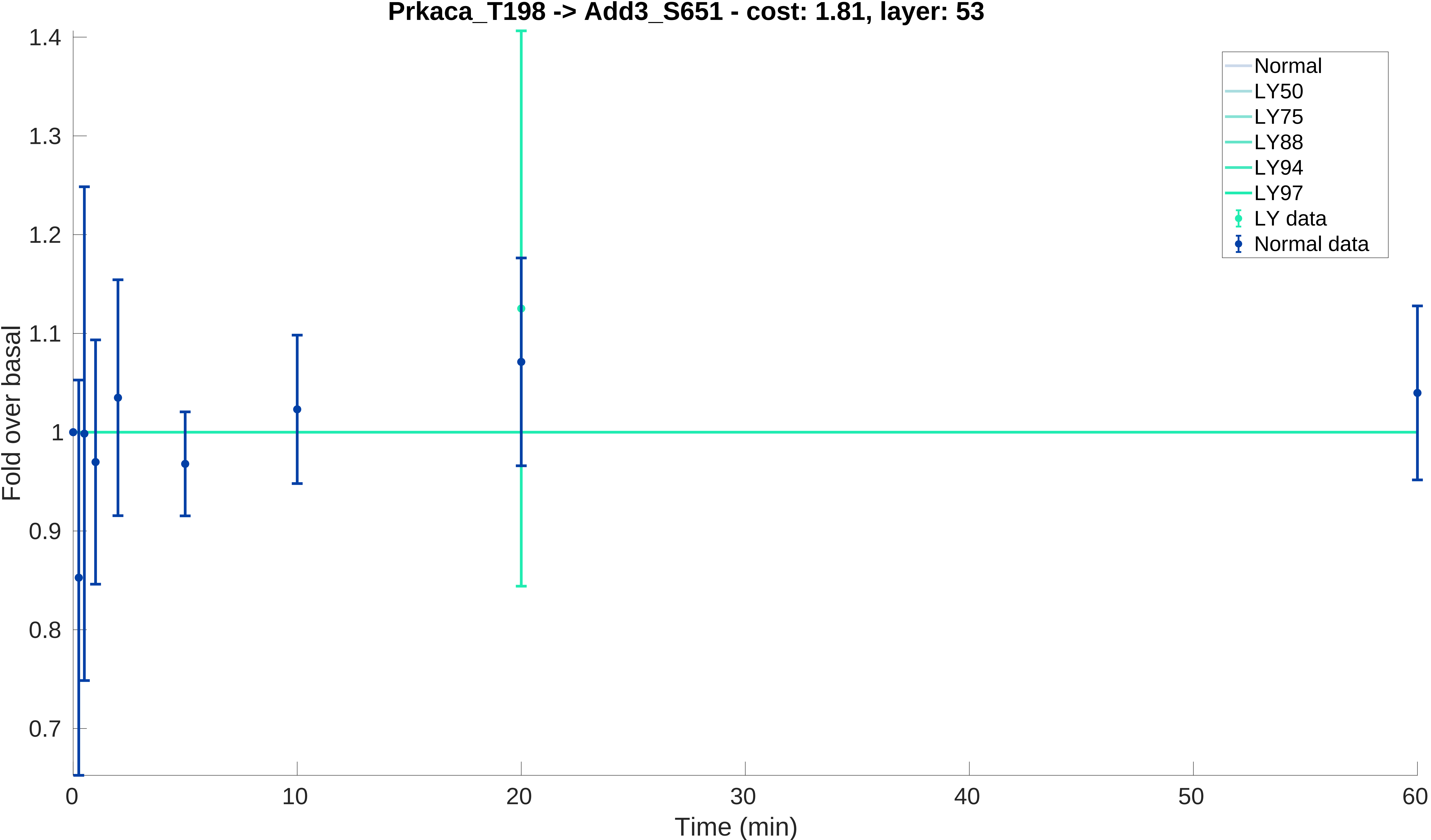

Prkaca_T198 -> Add3_S651 - cost: 1.81, layer: 53
Fold over basal
Time (min)
1.4
1.3
1.2
1.1
1
0.9
0.8
0.7
0
10
20
30
40
50
60
Normal
LY50
LY75
LY88
LY94
LY97
LY data
Normal data

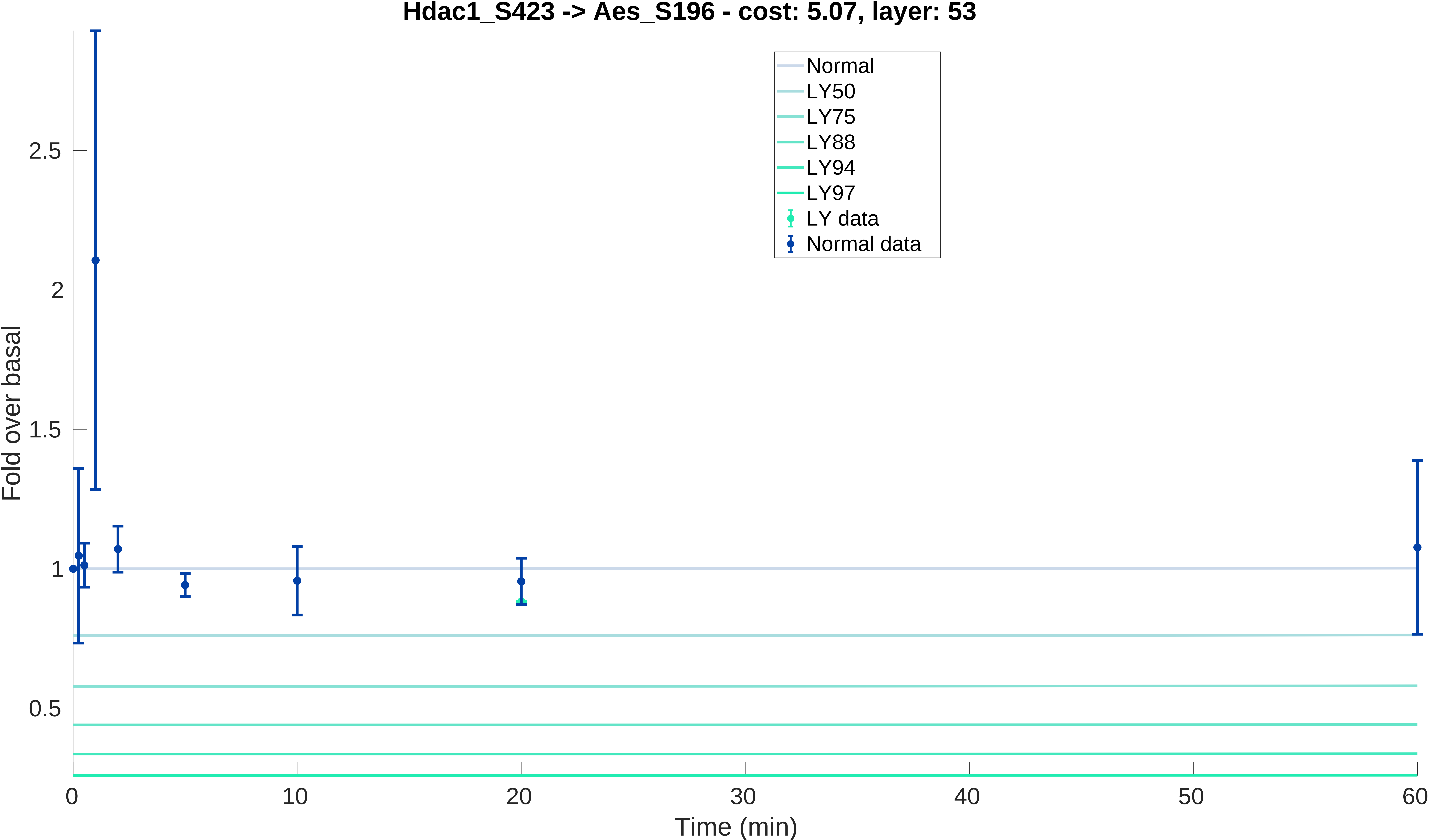

Hdac1_S423 -> Aes_S196 - cost: 5.07, layer: 53
Normal
LY50
LY75
LY88
LY94
LY97
LY data
Normal data
Fold over basal
2.5
2
1.5
1
0.5
0
10
20
30
40
50
60
Time (min)

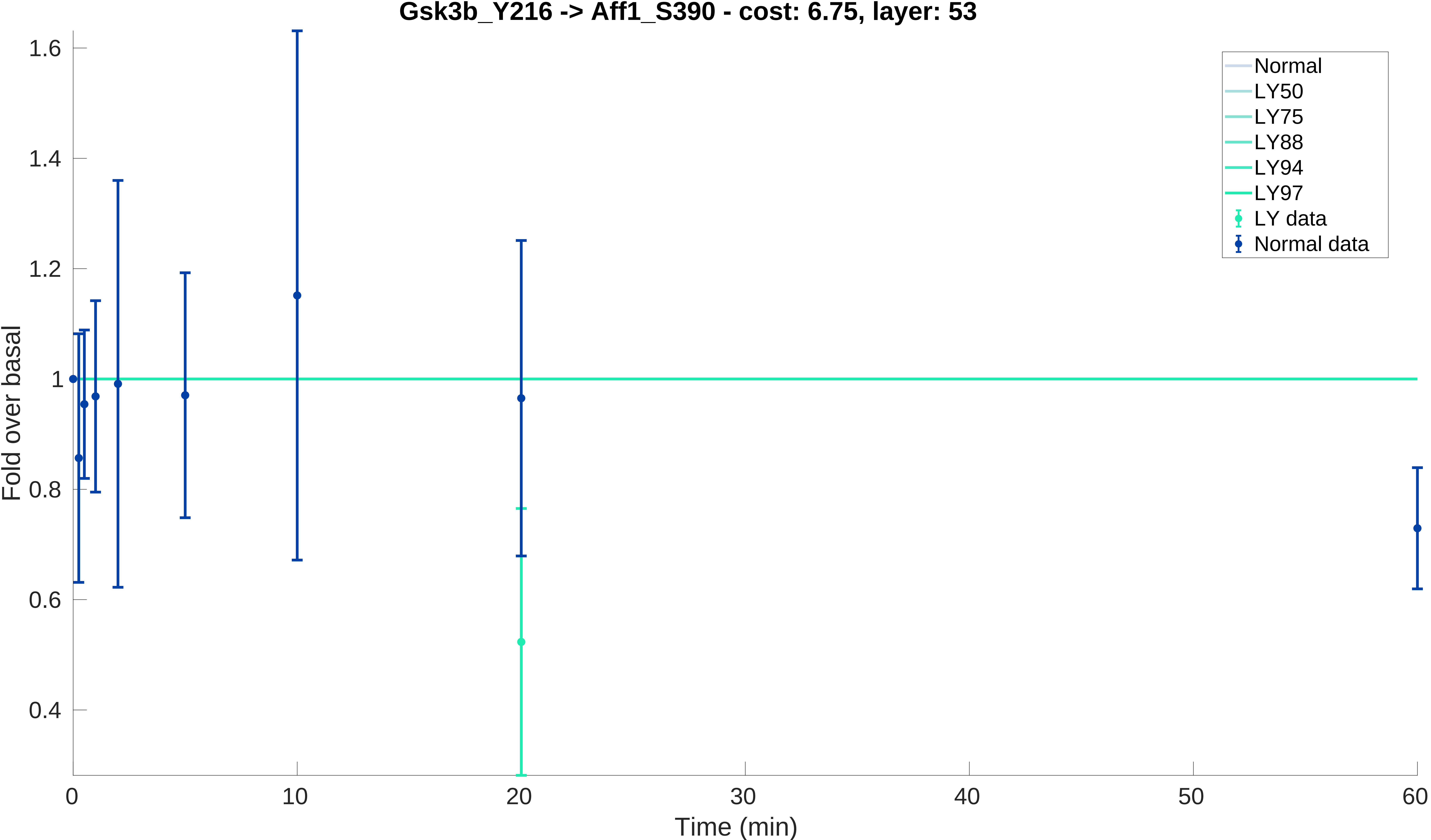

Gsk3b_Y216 -> Aff1_S390 - cost: 6.75, layer: 53
Normal
LY50
LY75
LY88
LY94
LY97
LY data
Normal data
Fold over basal
1.6
1.4
1.2
1
0.8
0.6
0.4
0
10
20
30
40
50
60
Time (min)

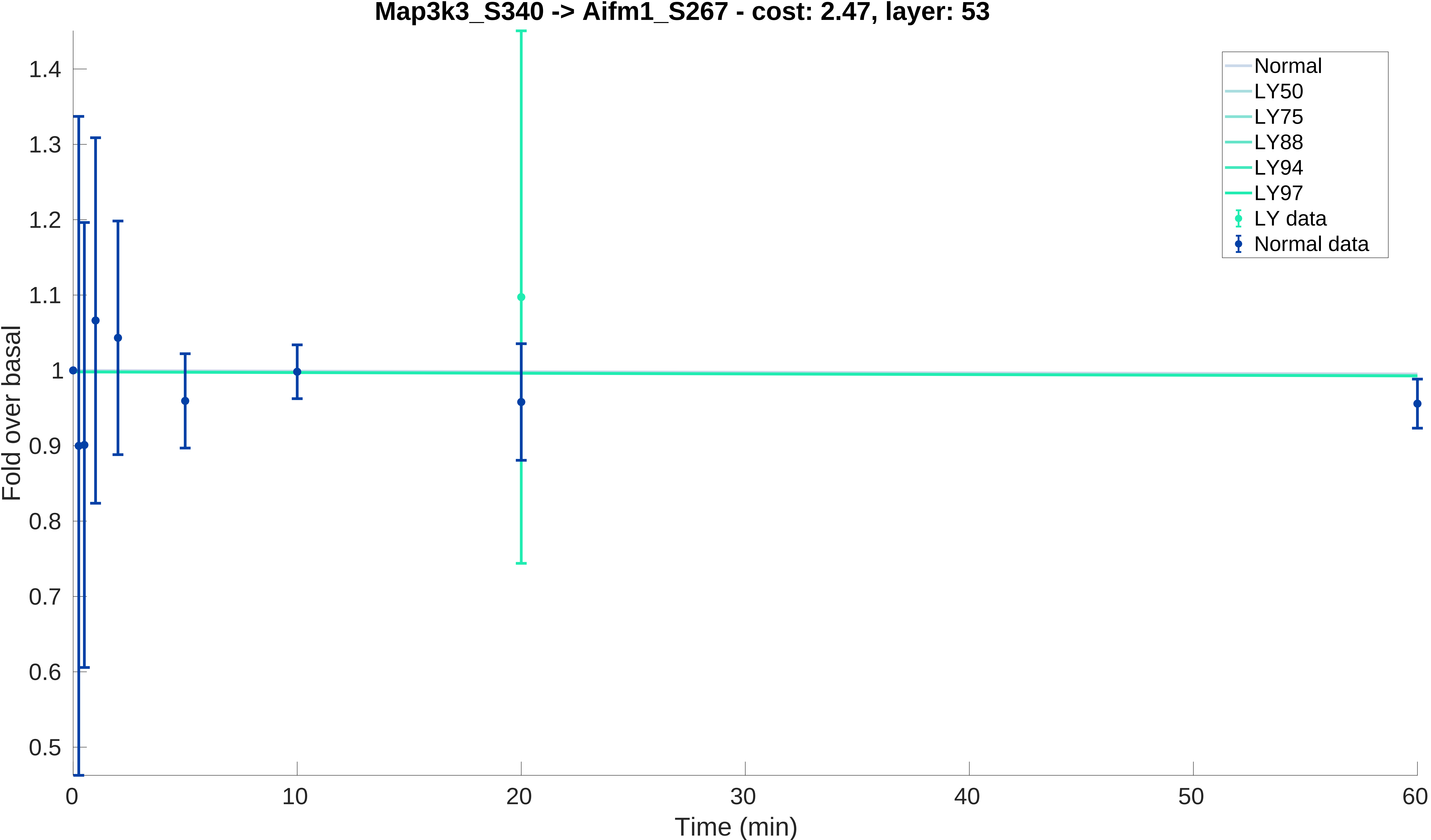
Map3k3_S340 -> Aifm1_S267 - cost: 2.47, layer: 53
Fold over basal
Time (min)
Normal
LY50
LY75
LY88
LY94
LY97
LY data
Normal data
1.4
1.3
1.2
1.1
1
0.9
0.8
0.7
0.6
0.5
0
10
20
30
40
50
60

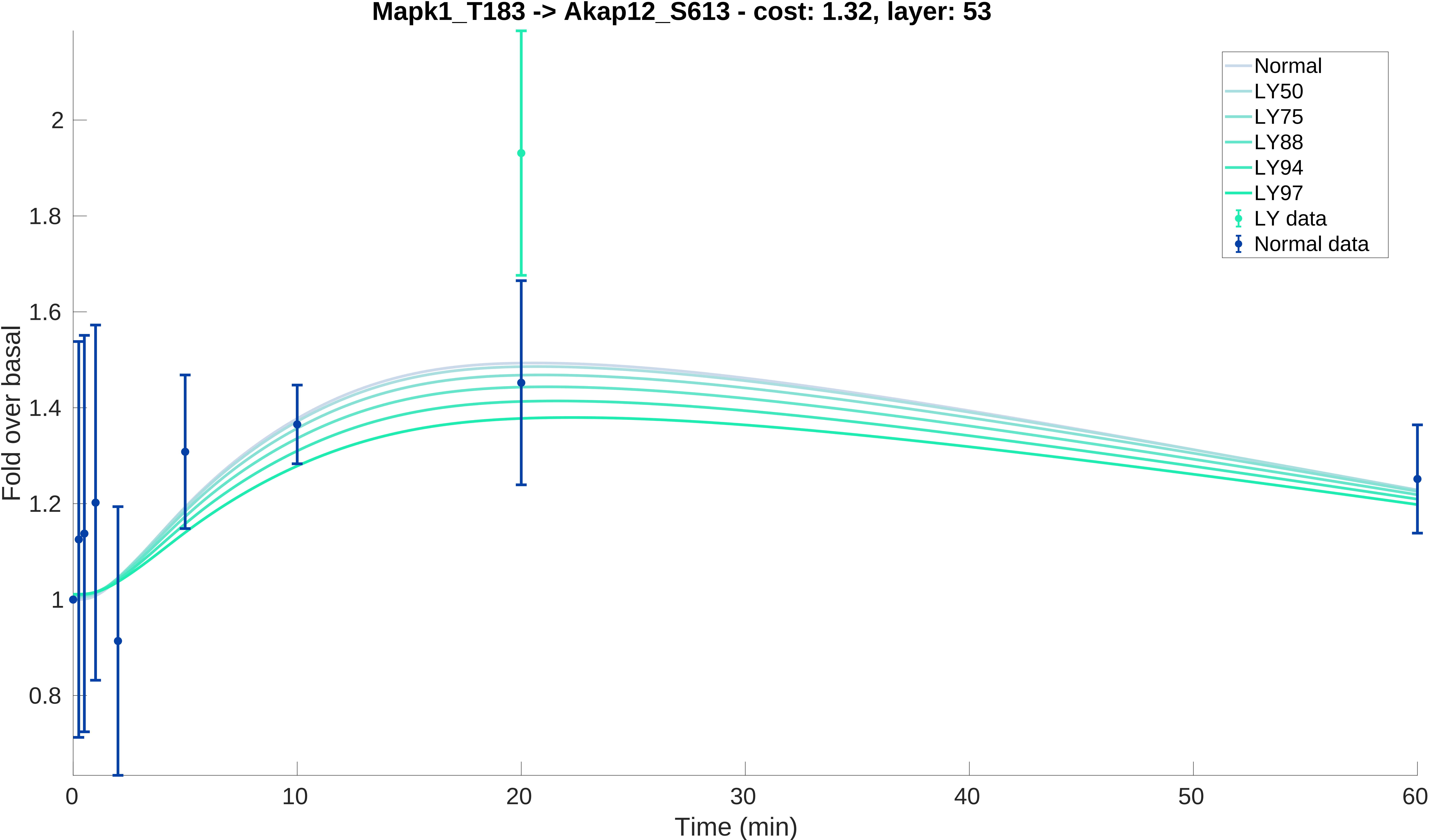

Mapk1_T183 -> Akap12_S613 - cost: 1.32, layer: 53
Fold over basal
Time (min)
0.8
1
1.2
1.4
1.6
1.8
2
0
10
20
30
40
50
60
Normal
LY50
LY75
LY88
LY94
LY97
LY data
Normal data

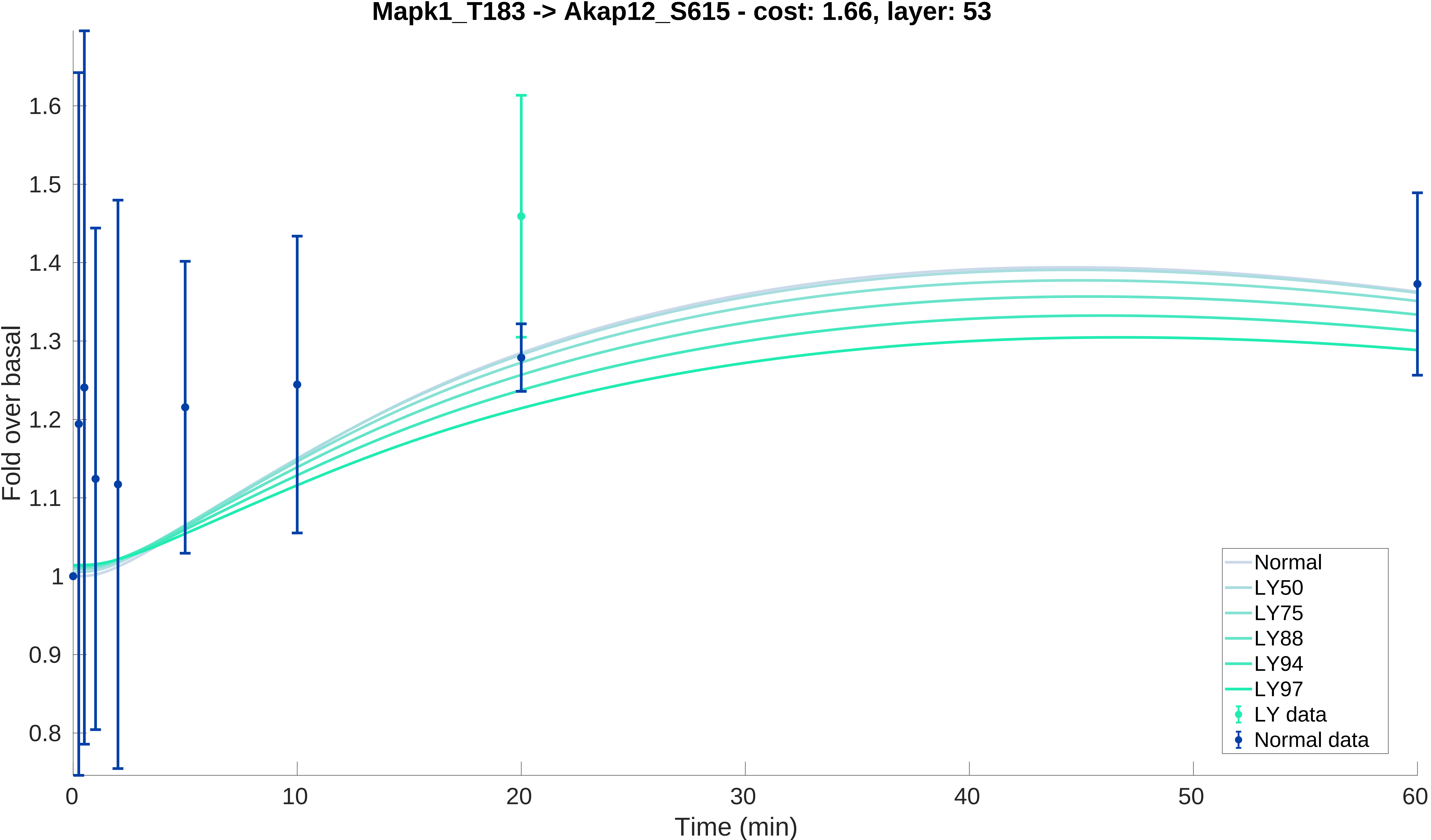

Mapk1_T183 -> Akap12_S615 - cost: 1.66, layer: 53
Fold over basal
1.6
1.5
1.4
1.3
1.2
1.1
1
0.9
0.8
0
10
20
30
40
50
60
Time (min)
Normal
LY50
LY75
LY88
LY94
LY97
LY data
Normal data

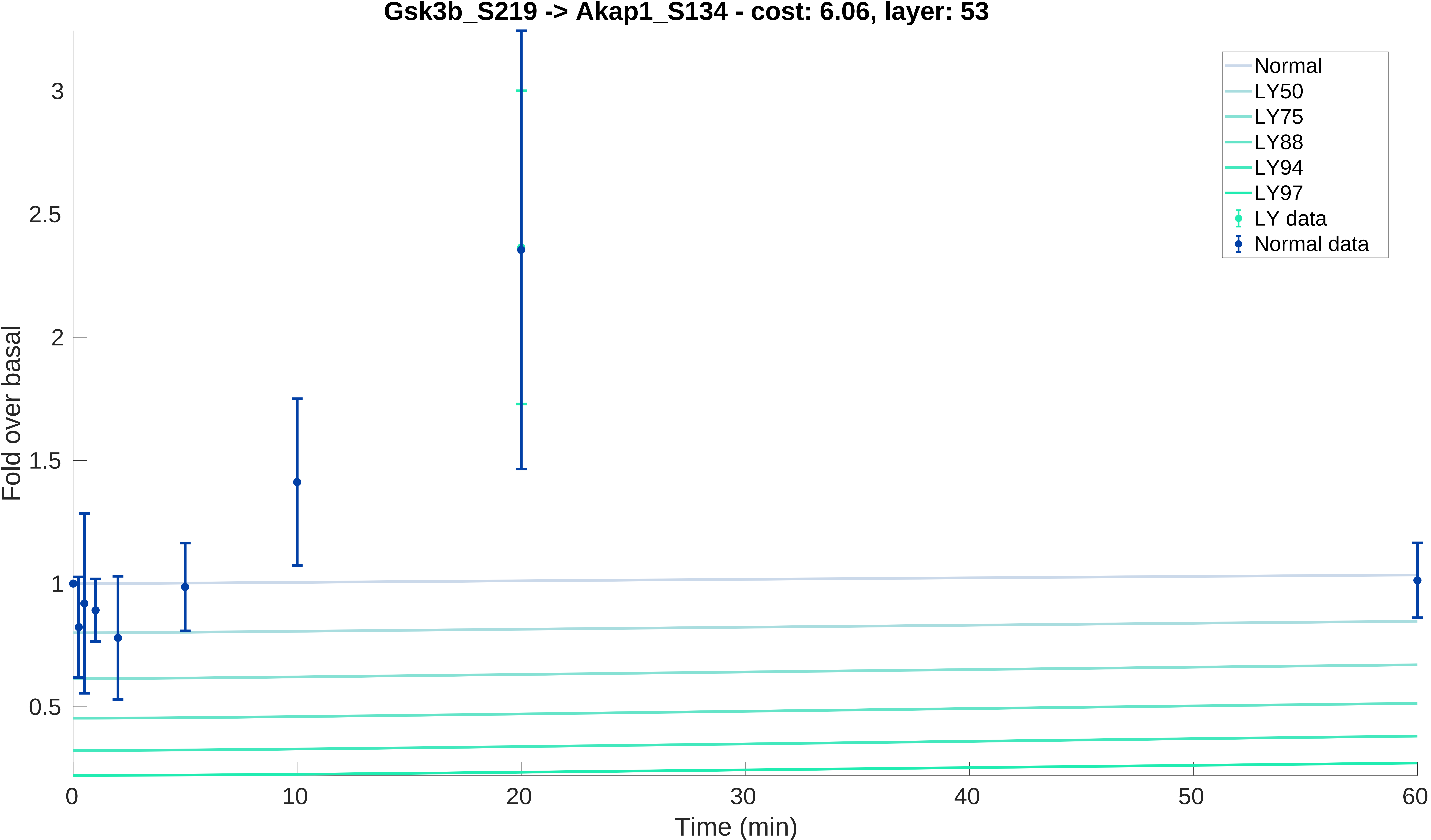
Gsk3b_S219 -> Akap1_S134 - cost: 6.06, layer: 53
Fold over basal
Time (min)
0
10
20
30
40
50
60
0.5
1
1.5
2
2.5
3
Normal
LY50
LY75
LY88
LY94
LY97
LY data
Normal data

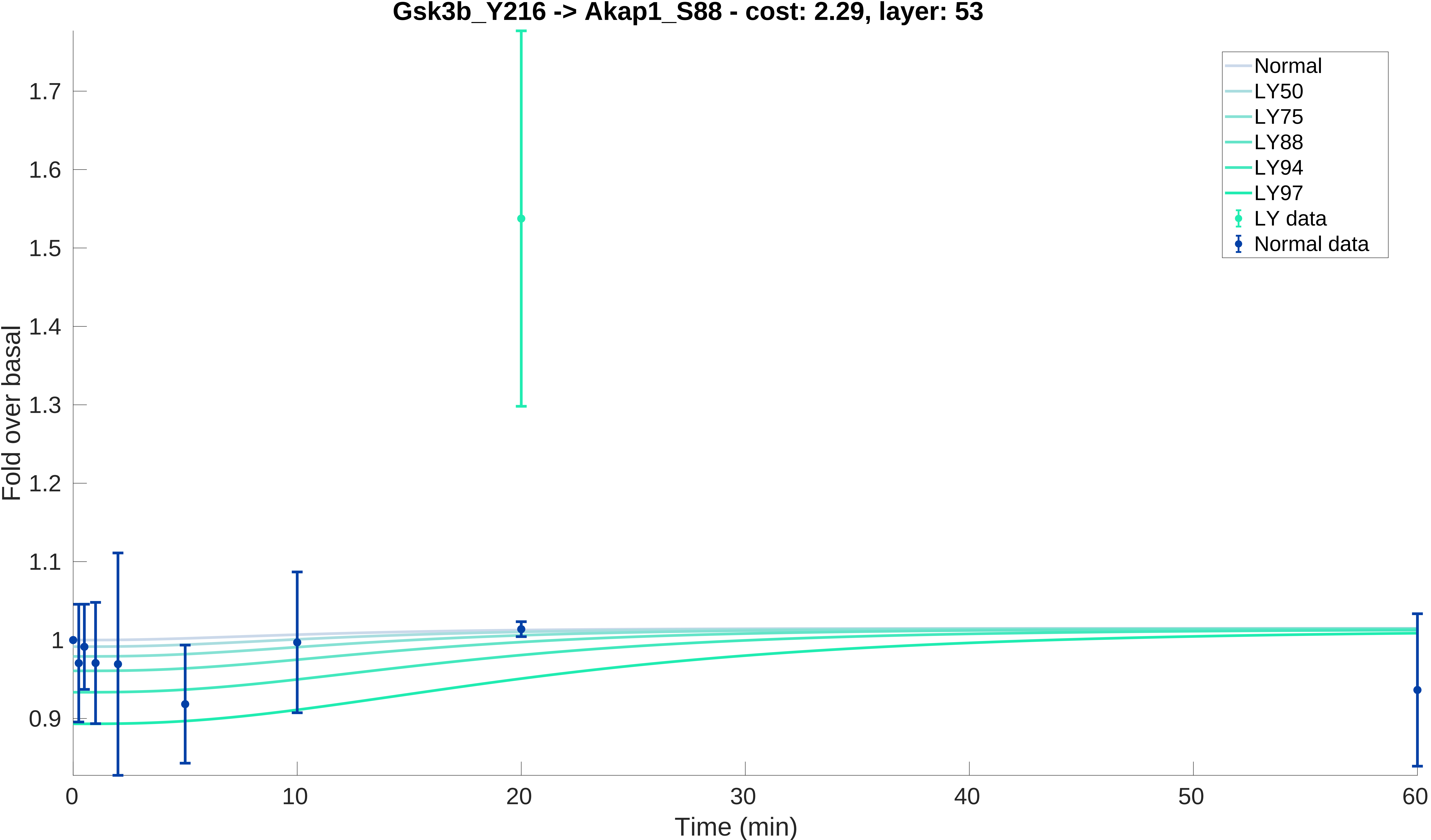

Gsk3b_Y216 -> Akap1_S88 - cost: 2.29, layer: 53
Fold over basal
Time (min)
0
10
20
30
40
50
60
0.9
1
1.1
1.2
1.3
1.4
1.5
1.6
1.7
Normal
LY50
LY75
LY88
LY94
LY97
LY data
Normal data

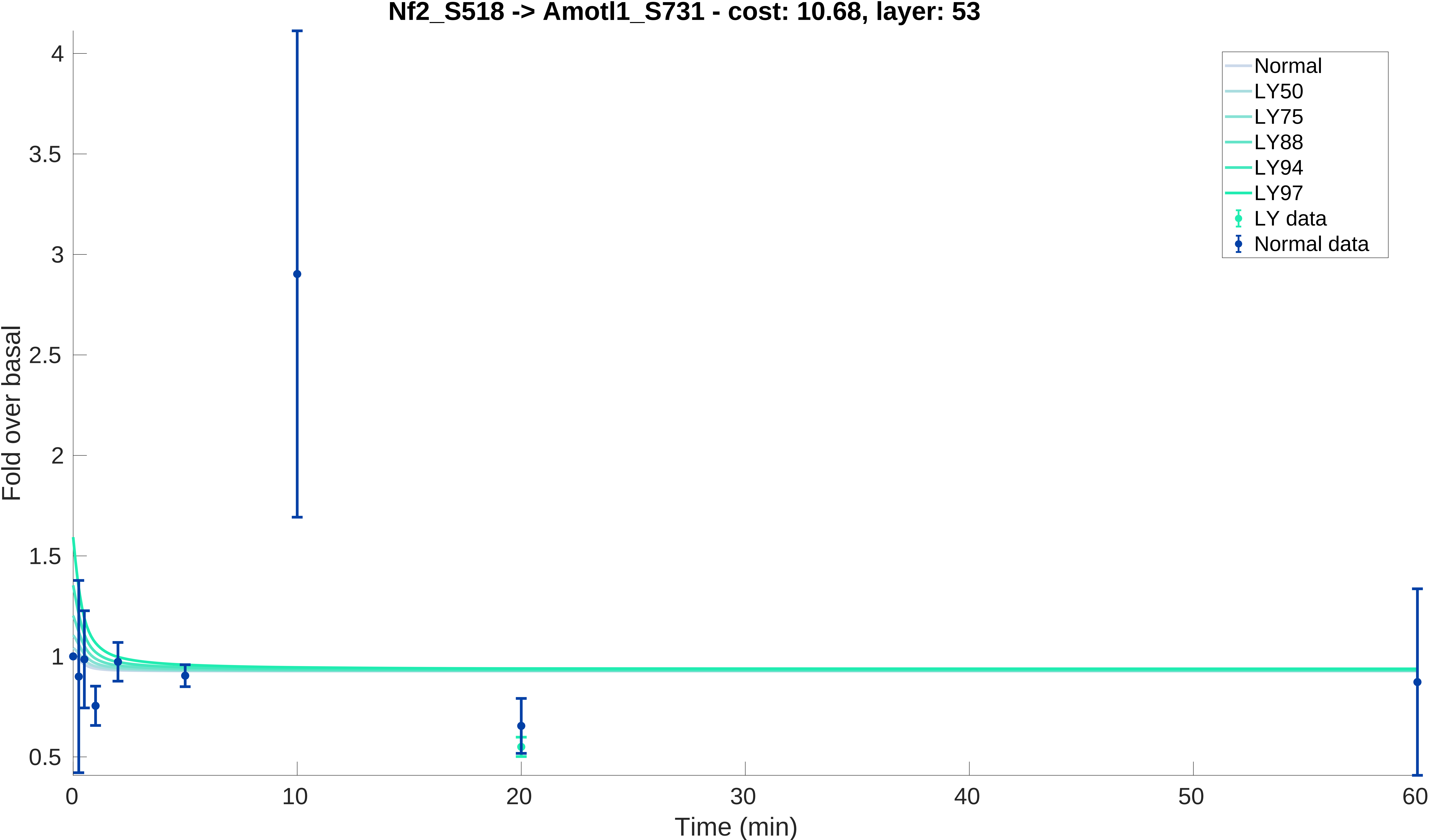

Nf2_S518 -> Amotl1_S731 - cost: 10.68, layer: 53
Fold over basal
Time (min)
0.5
1
1.5
2
2.5
3
3.5
4
0
10
20
30
40
50
60
Normal
LY50
LY75
LY88
LY94
LY97
LY data
Normal data

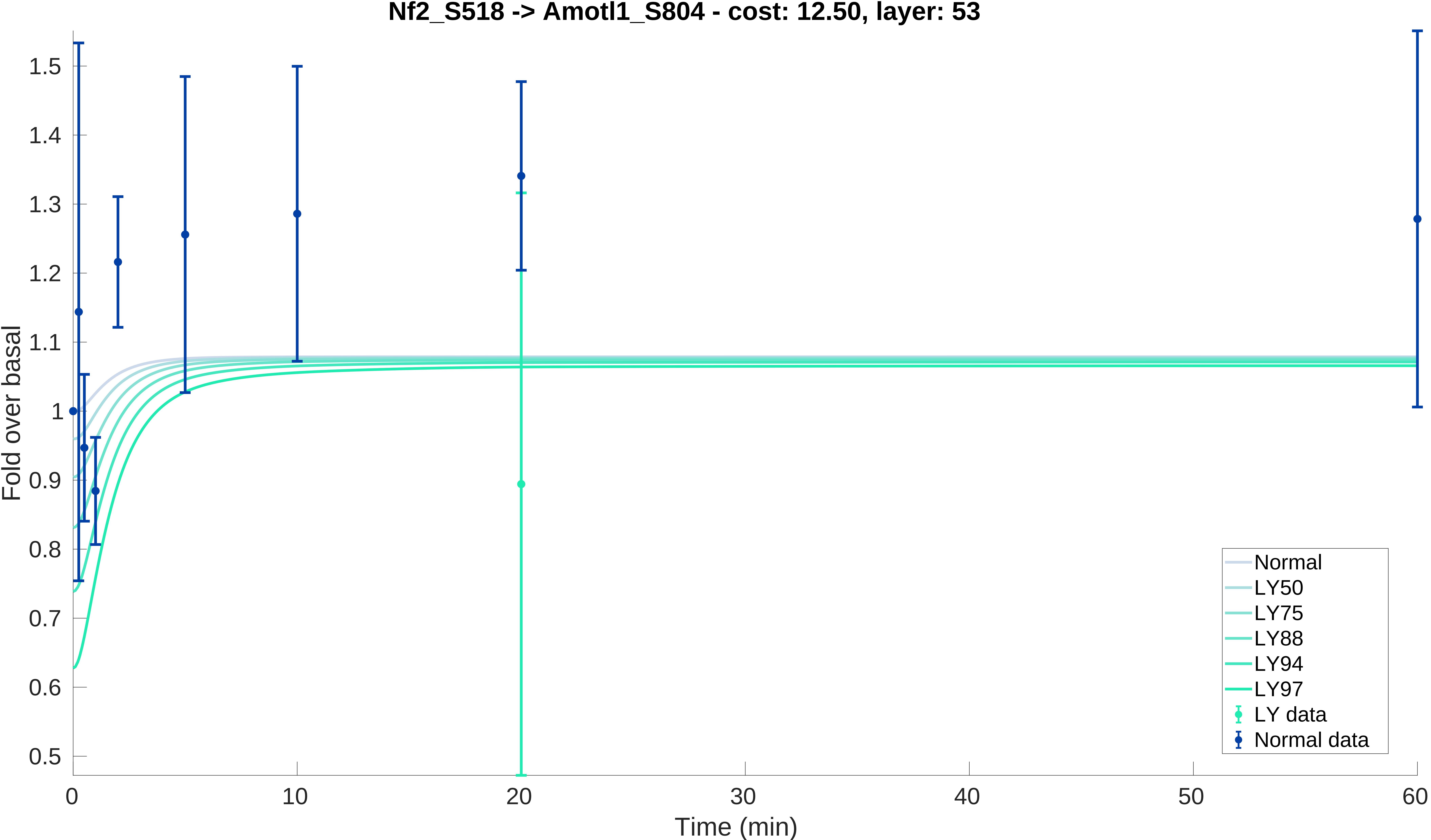

Nf2_S518 -> Amotl1_S804 - cost: 12.50, layer: 53
Fold over basal
Time (min)
1.5
1.4
1.3
1.2
1.1
1
0.9
0.8
0.7
0.6
0.5
0
10
20
30
40
50
60
Normal
LY50
LY75
LY88
LY94
LY97
LY data
Normal data

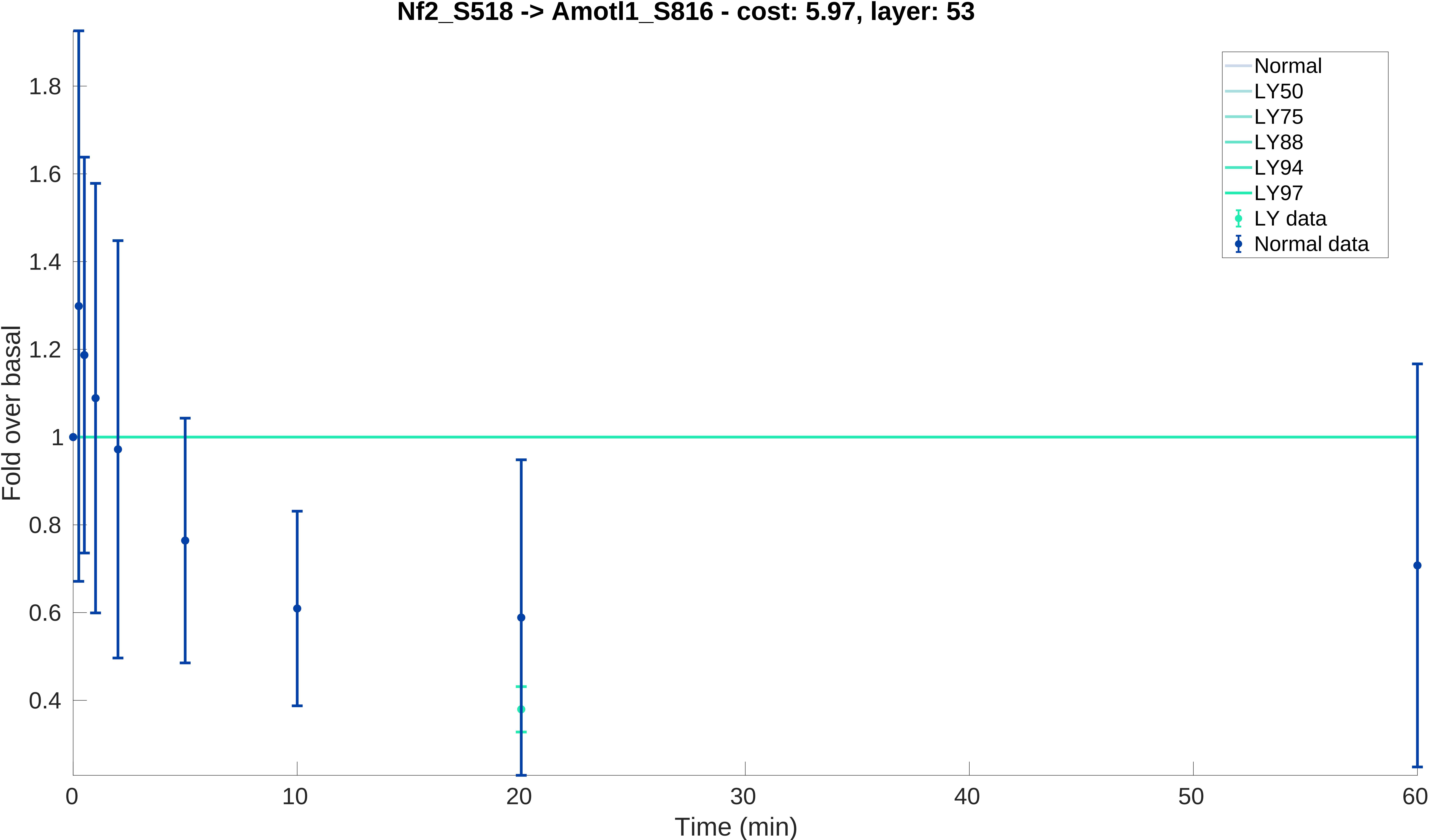

Nf2_S518 -> Amotl1_S816 - cost: 5.97, layer: 53
Normal
LY50
LY75
LY88
LY94
LY97
LY data
Normal data
Fold over basal
1.8
1.6
1.4
1.2
1
0.8
0.6
0.4
0
10
20
30
40
50
60
Time (min)

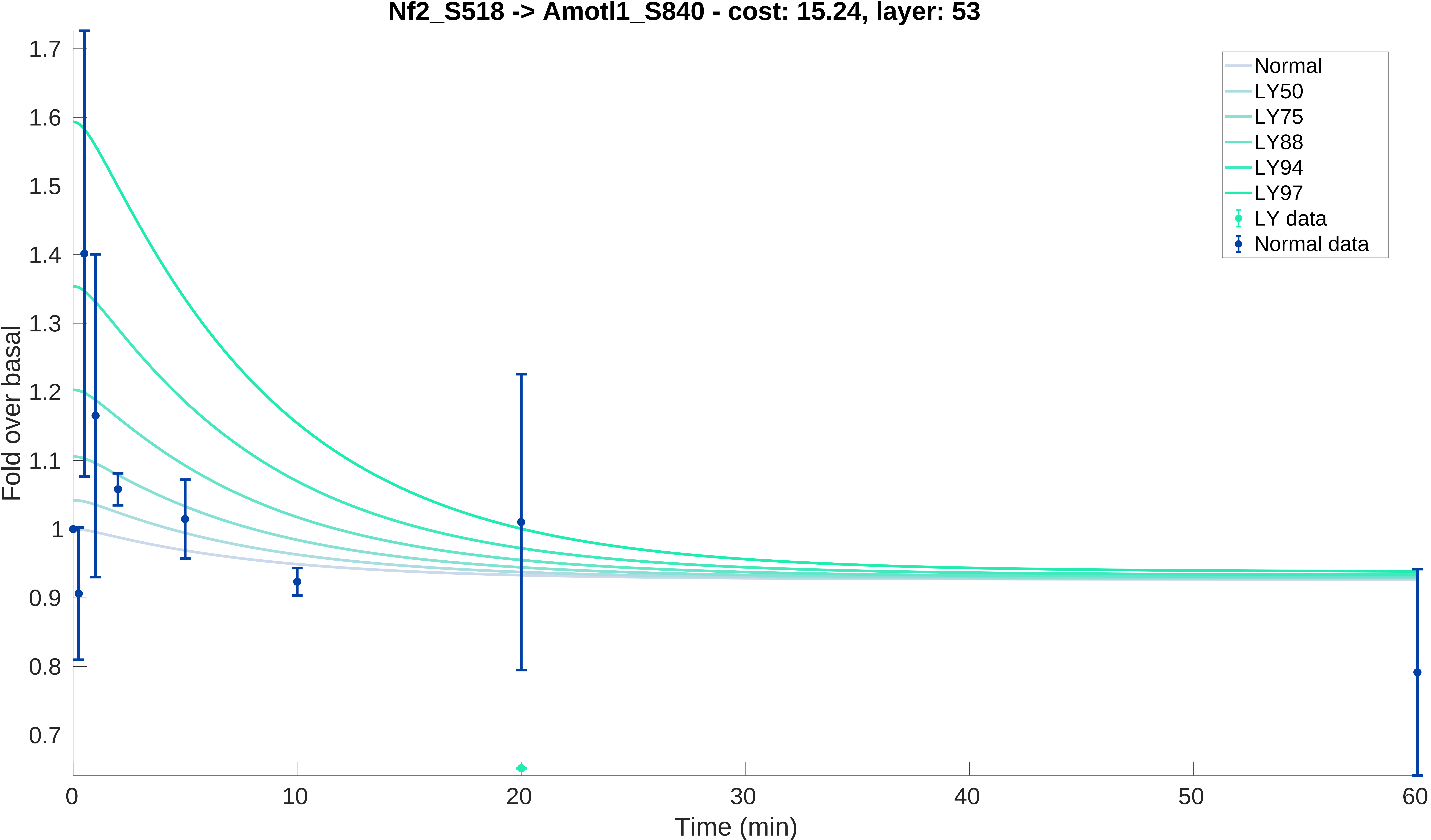
Nf2_S518 -> Amotl1_S840 - cost: 15.24, layer: 53
Fold over basal
Time (min)
0
10
20
30
40
50
60
0.7
0.8
0.9
1
1.1
1.2
1.3
1.4
1.5
1.6
1.7
Normal
LY50
LY75
LY88
LY94
LY97
LY data
Normal data

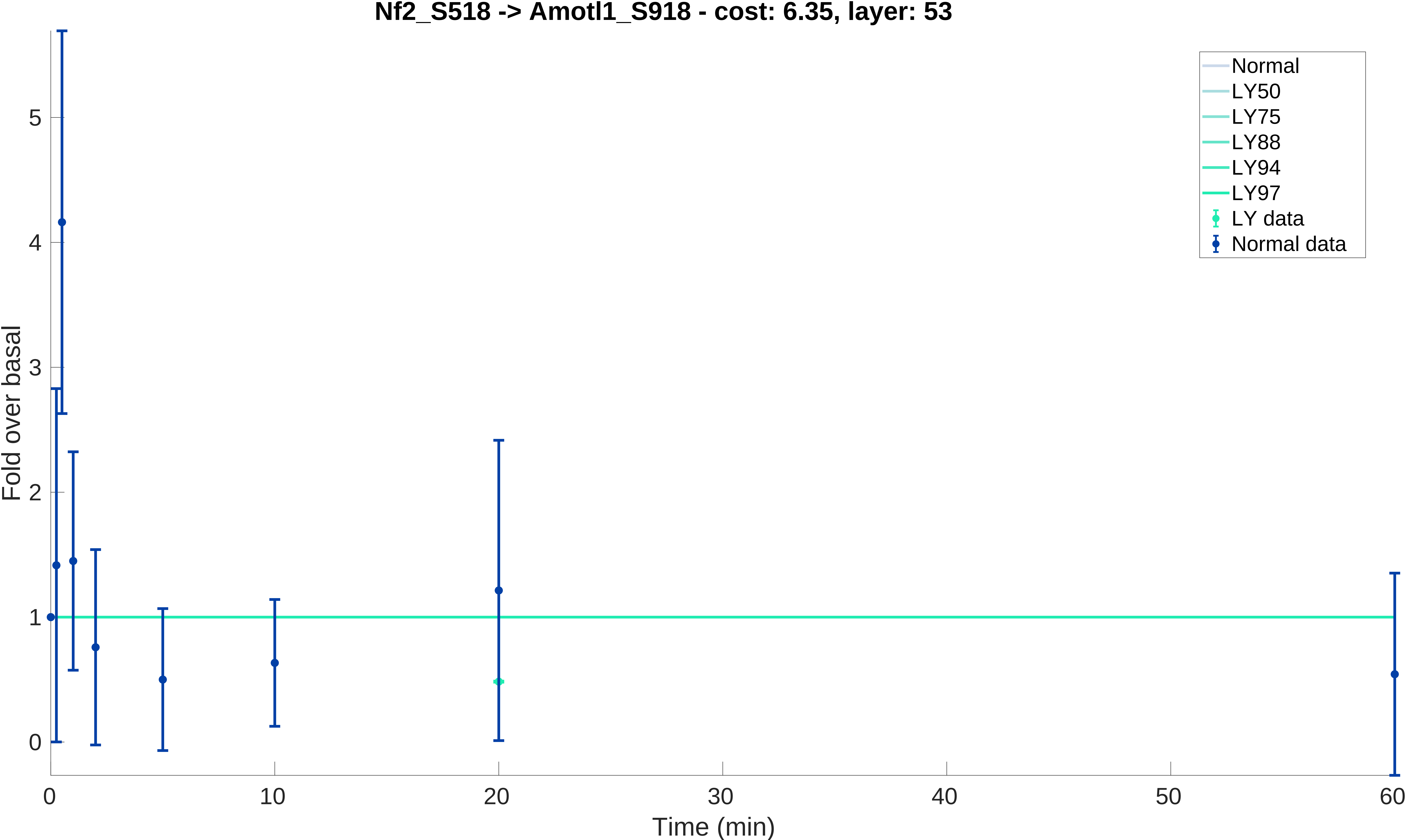

Nf2_S518 -> Amotl1_S918 - cost: 6.35, layer: 53
Fold over basal
Time (min)
0
1
2
3
4
5
0
10
20
30
40
50
60
Normal
LY50
LY75
LY88
LY94
LY97
LY data
Normal data

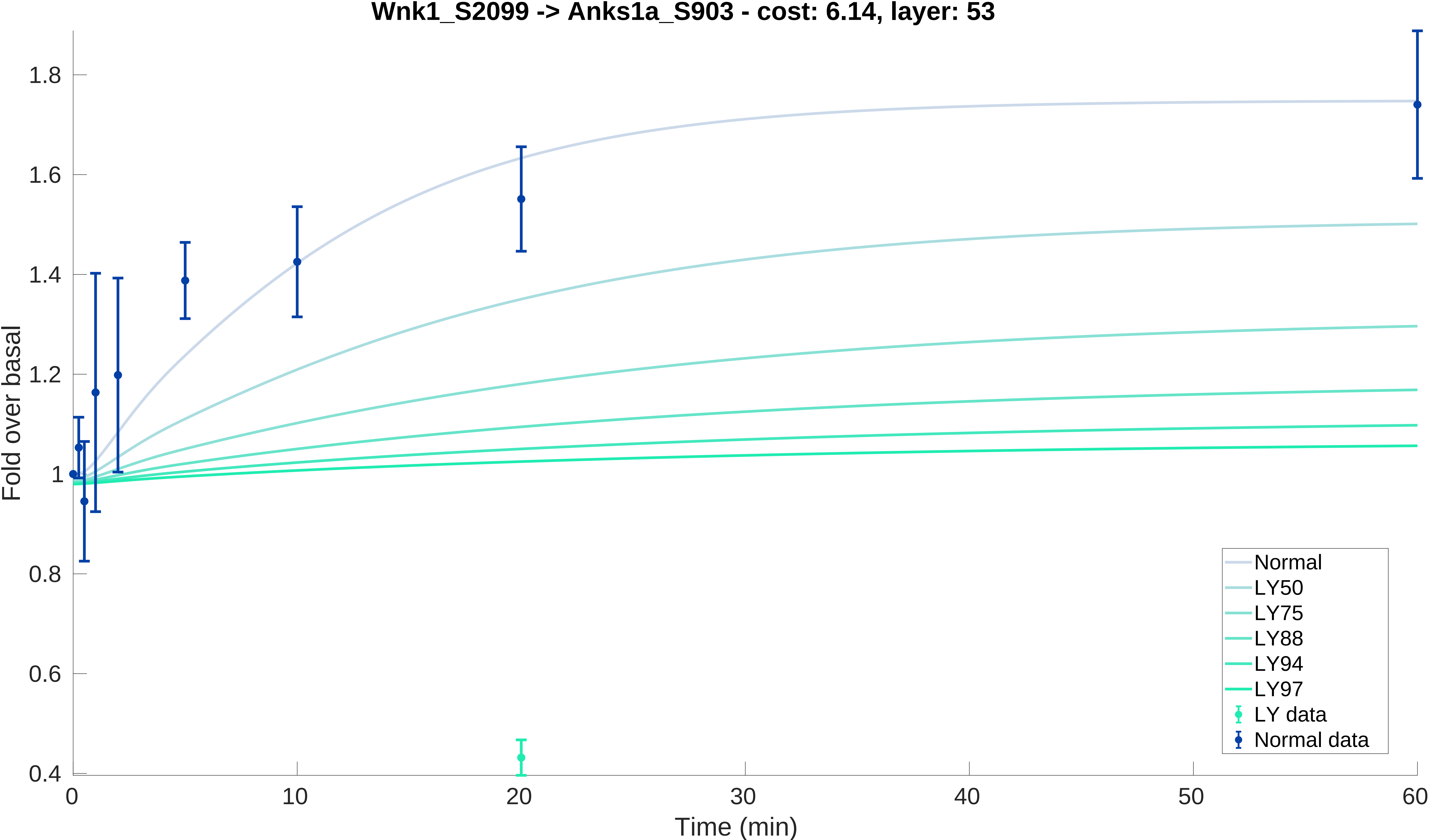

Wnk1_S2099 -> Anks1a_S903 - cost: 6.14, layer: 53
Fold over basal
1.8
1.6
1.4
1.2
1
0.8
0.6
0.4
0
10
20
30
40
50
60
Time (min)
Normal
LY50
LY75
LY88
LY94
LY97
LY data
Normal data

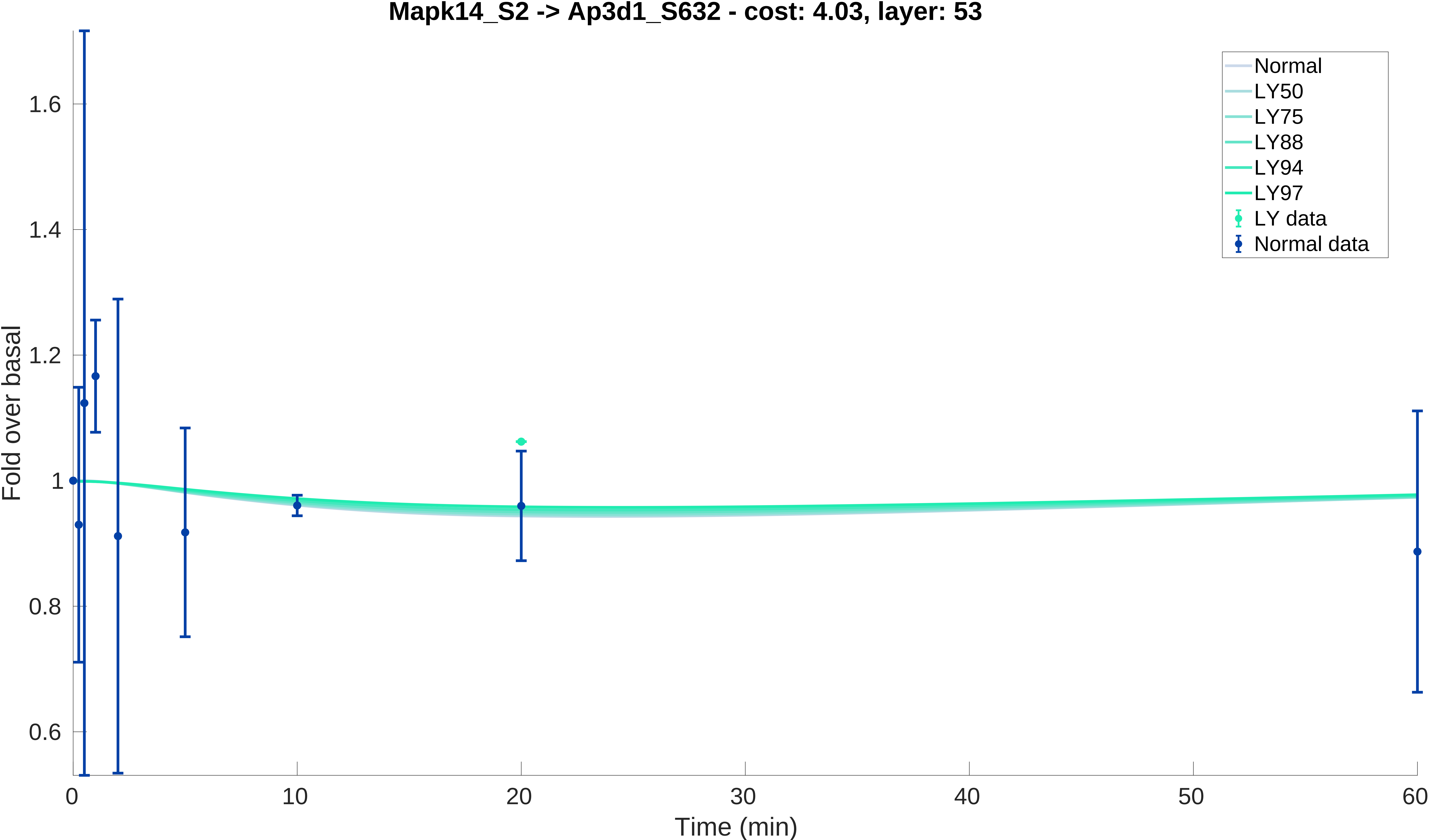

Mapk14_S2 -> Ap3d1_S632 - cost: 4.03, layer: 53
Fold over basal
Time (min)
1.6
1.4
1.2
1
0.8
0.6
0
10
20
30
40
50
60
Normal
LY50
LY75
LY88
LY94
LY97
LY data
Normal data

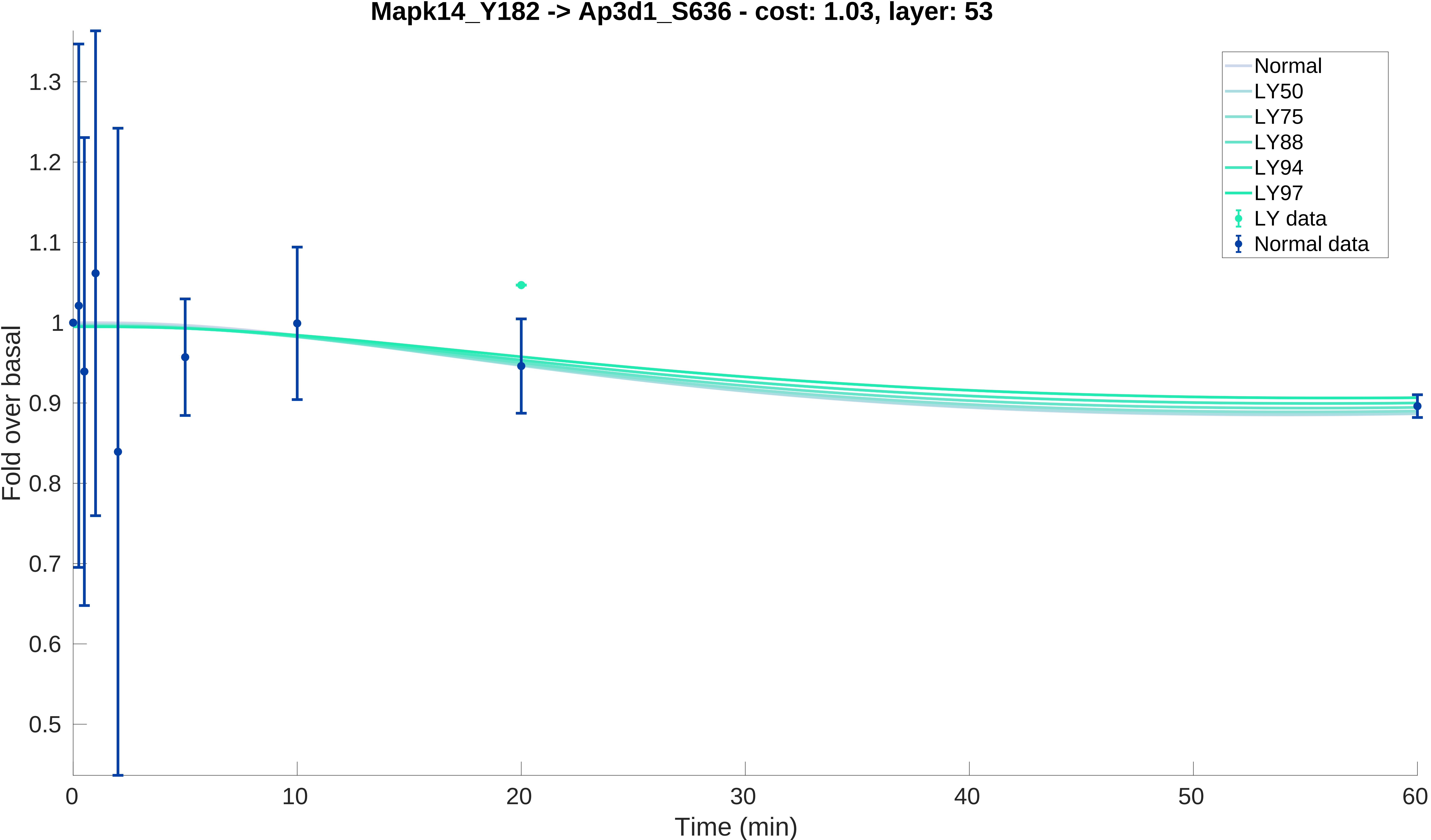

Mapk14_Y182 -> Ap3d1_S636 - cost: 1.03, layer: 53
Normal
LY50
LY75
LY88
LY94
LY97
LY data
Normal data
Fold over basal
Time (min)
0
10
20
30
40
50
60
0.5
0.6
0.7
0.8
0.9
1
1.1
1.2
1.3

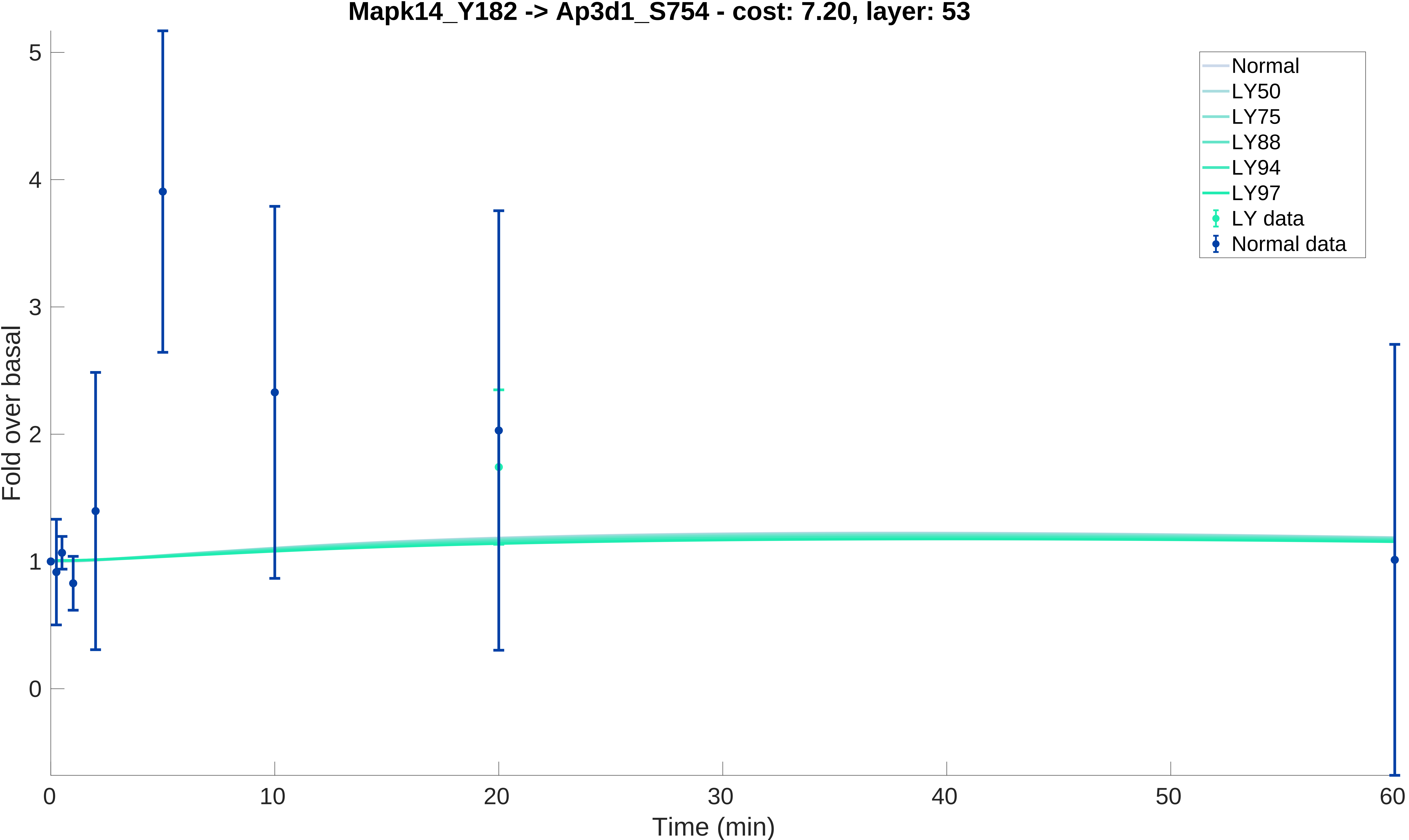

Mapk14_Y182 -> Ap3d1_S754 - cost: 7.20, layer: 53
Fold over basal
Time (min)
Normal
LY50
LY75
LY88
LY94
LY97
LY data
Normal data
0
1
2
3
4
5
10
20
30
40
50
60

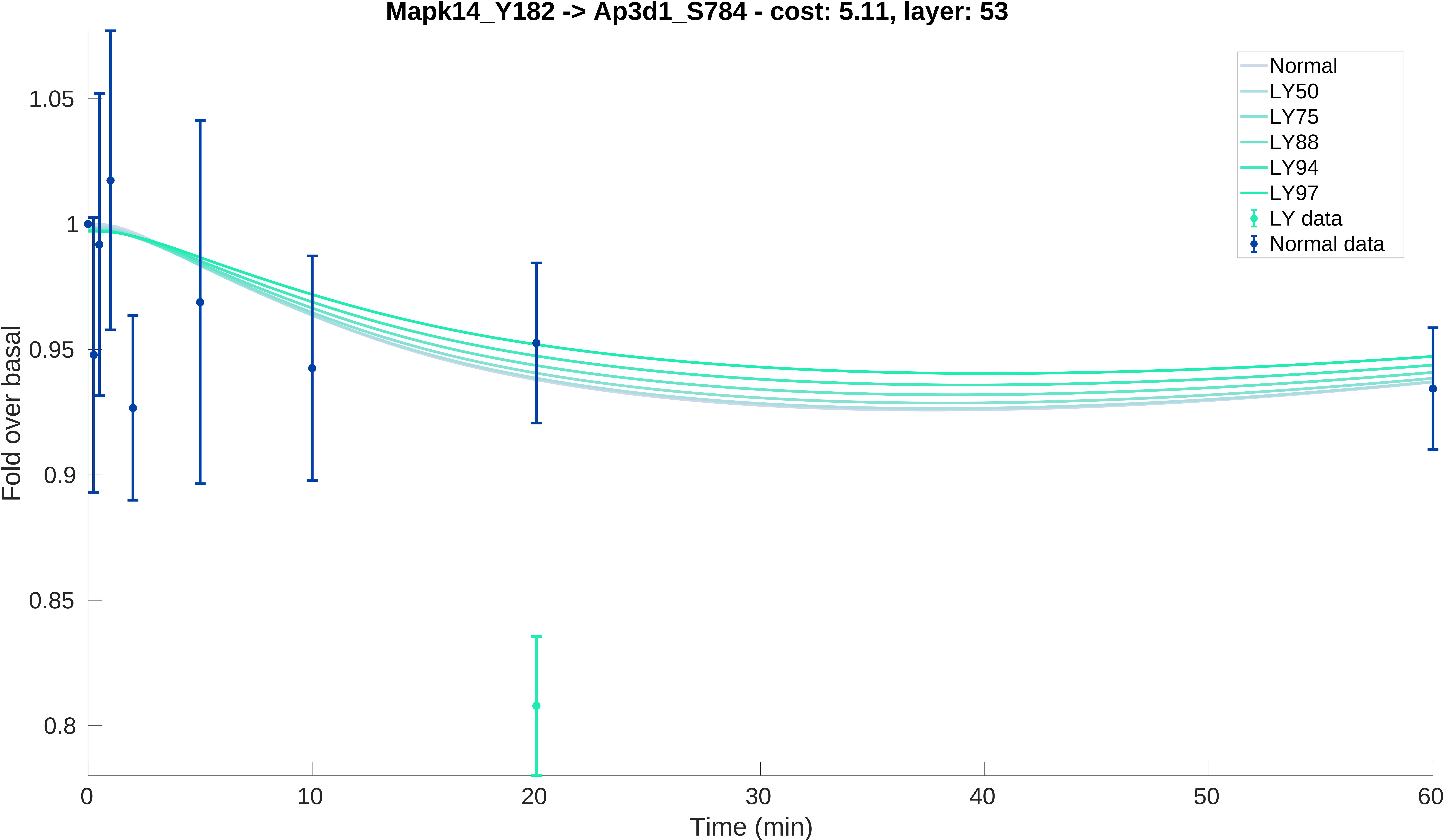
Mapk14_Y182 -> Ap3d1_S784 - cost: 5.11, layer: 53
Fold over basal
Time (min)
0.8
0.85
0.9
0.95
1
1.05
0
10
20
30
40
50
60
Normal
LY50
LY75
LY88
LY94
LY97
LY data
Normal data

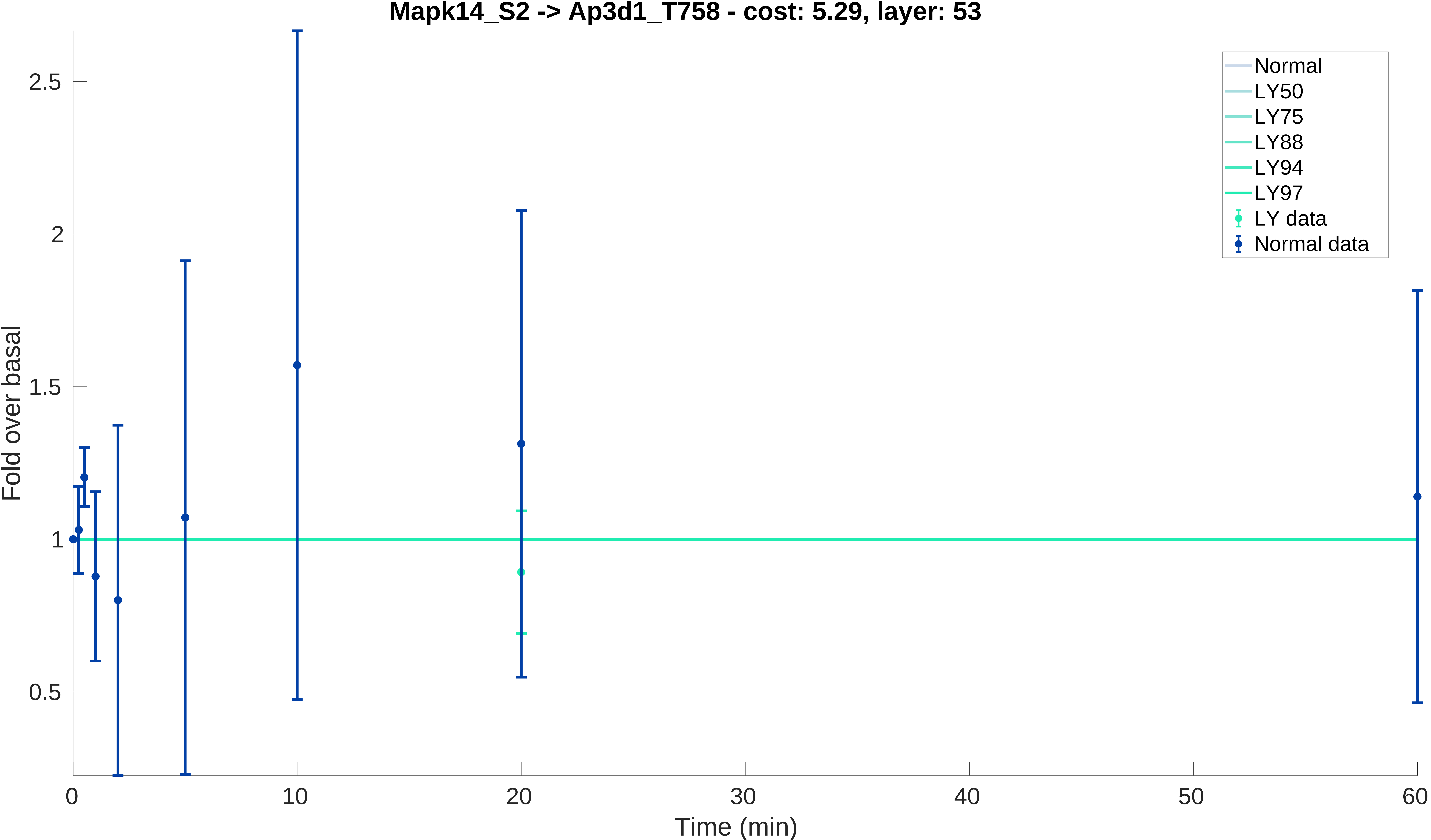

Mapk14_S2 -> Ap3d1_T758 - cost: 5.29, layer: 53
Fold over basal
Time (min)
0.5
1
1.5
2
2.5
0
10
20
30
40
50
60
Normal
LY50
LY75
LY88
LY94
LY97
LY data
Normal data

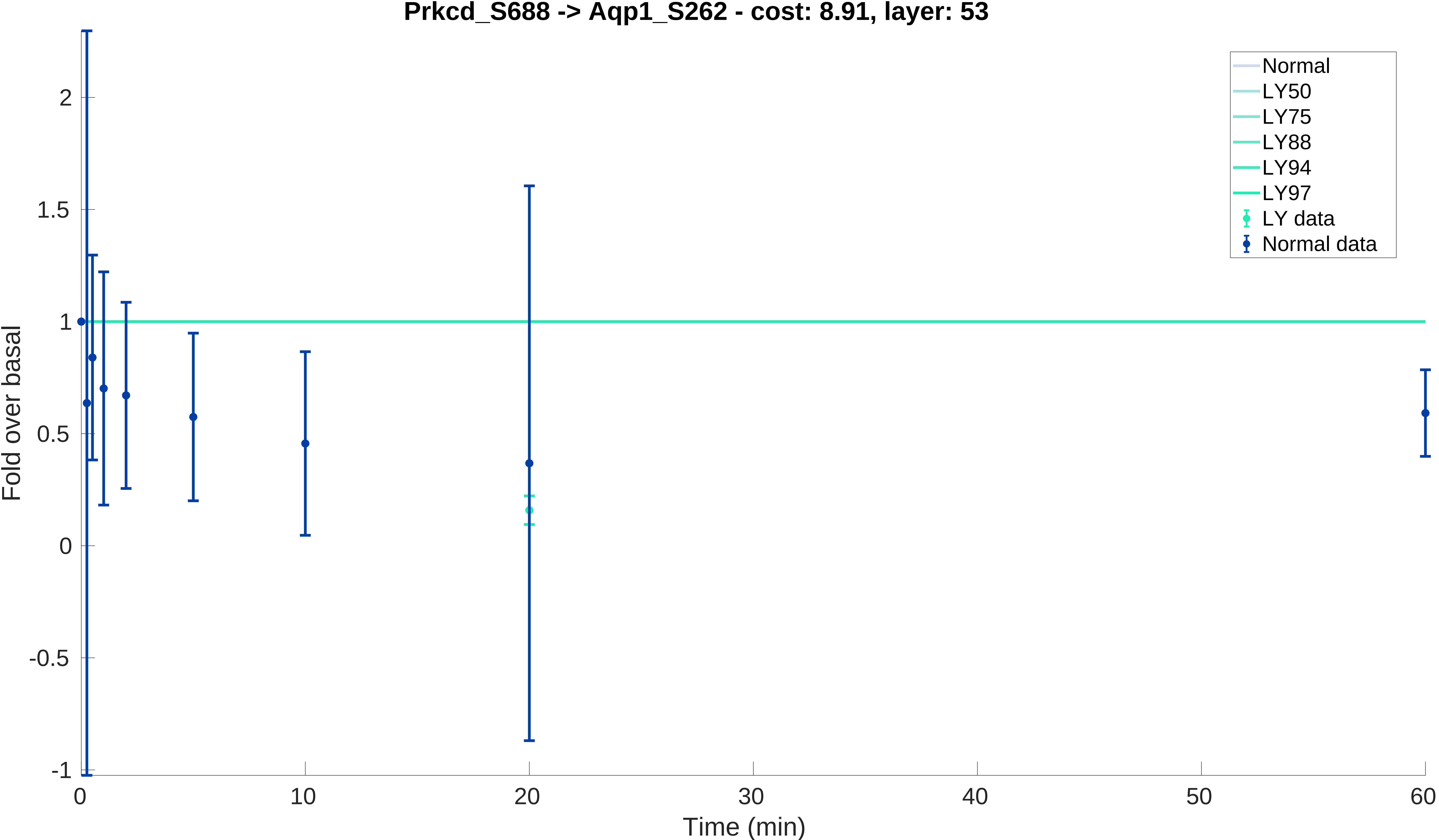

Prkcd_S688 -> Aqp1_S262 - cost: 8.91, layer: 53
Fold over basal
Time (min)
-1
-0.5
0
0.5
1
1.5
2
0
10
20
30
40
50
60
Normal
LY50
LY75
LY88
LY94
LY97
LY data
Normal data

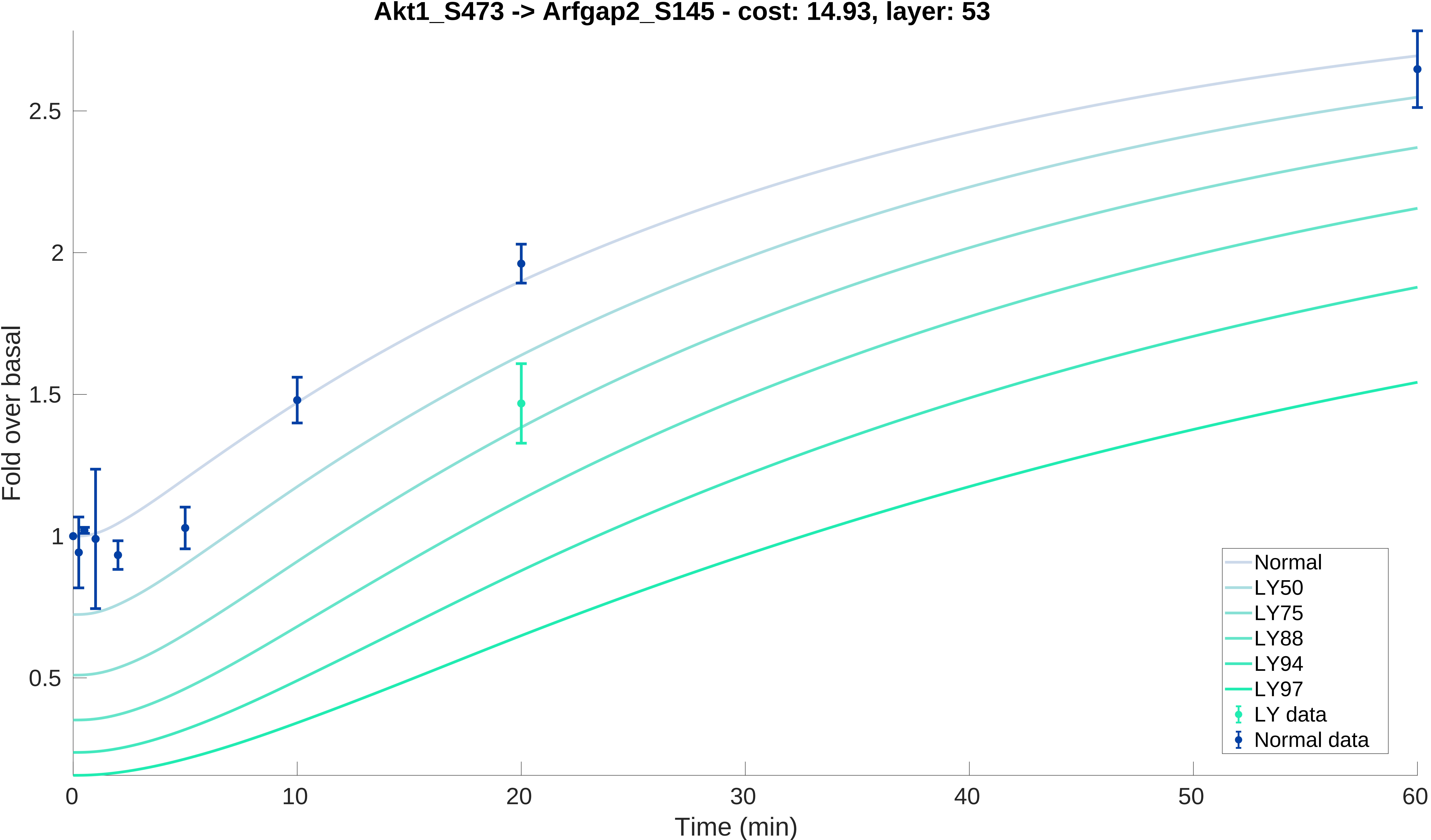
Akt1_S473 -> Arfgap2_S145 - cost: 14.93, layer: 53
Fold over basal
2.5
2
1.5
1
0.5
0
10
20
30
40
50
60
Time (min)
Normal
LY50
LY75
LY88
LY94
LY97
LY data
Normal data

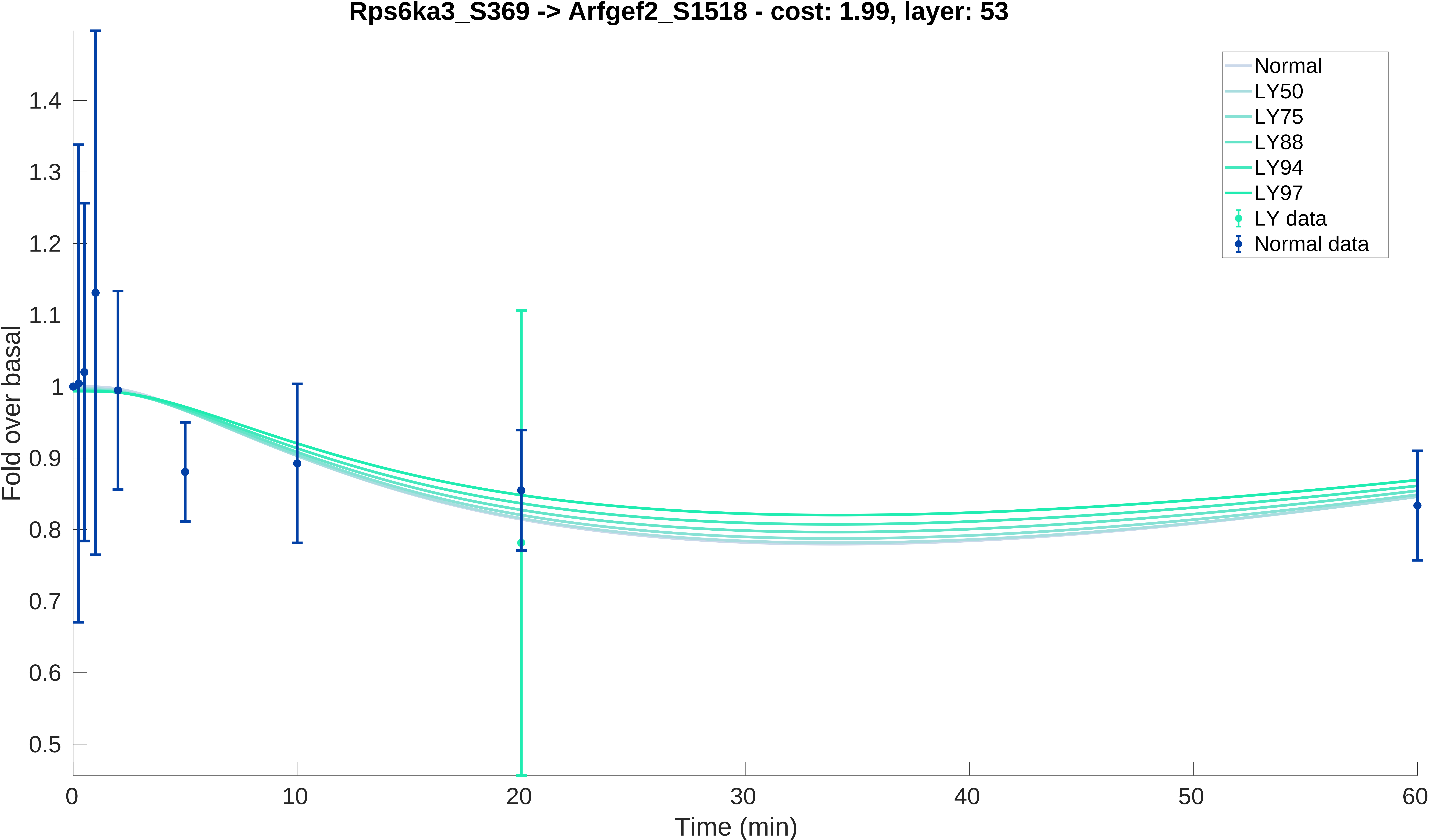

Rps6ka3_S369 -> Arfgef2_S1518 - cost: 1.99, layer: 53
Fold over basal
Time (min)
0
10
20
30
40
50
60
0.5
0.6
0.7
0.8
0.9
1
1.1
1.2
1.3
1.4
Normal
LY50
LY75
LY88
LY94
LY97
LY data
Normal data

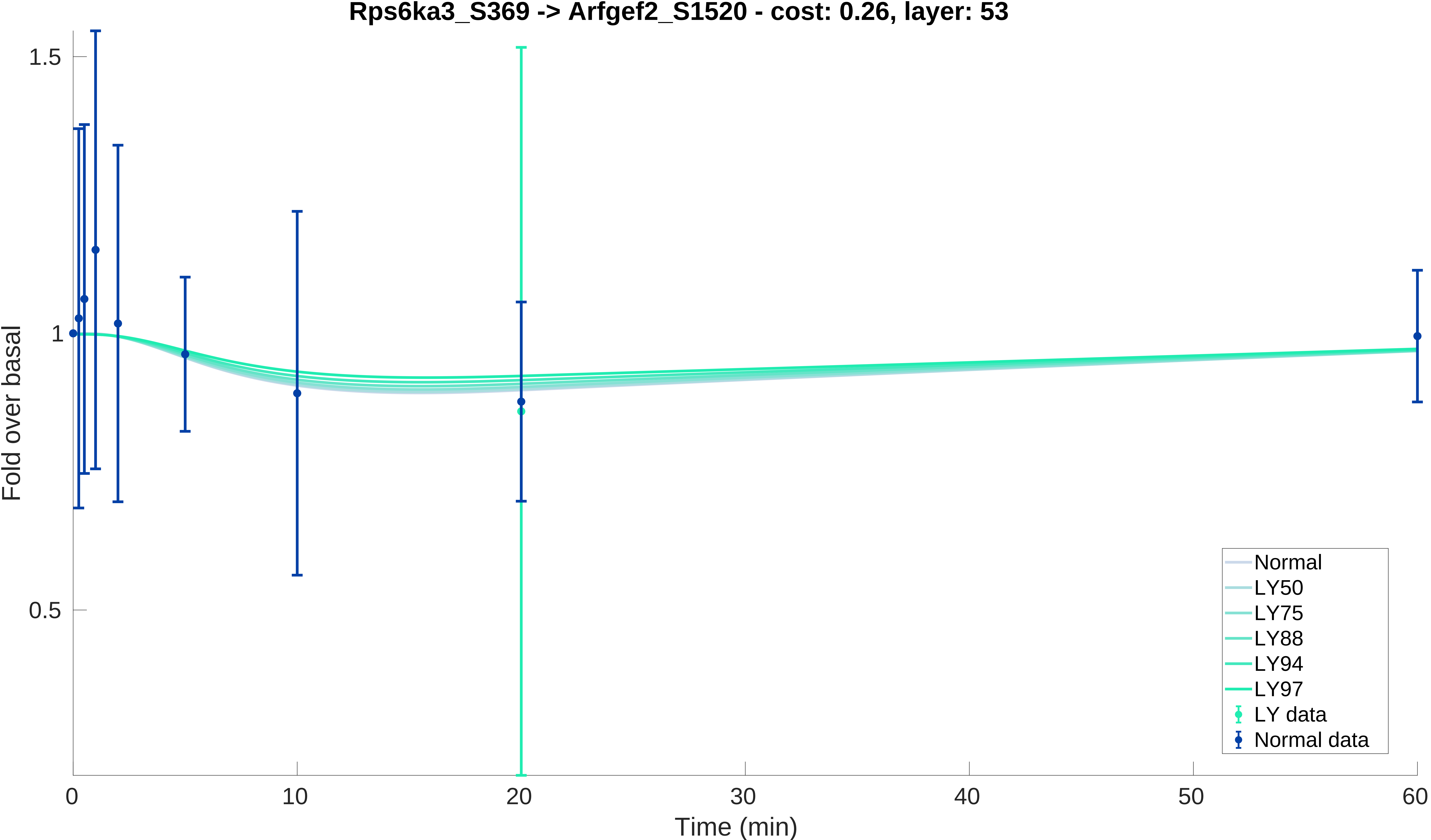

Rps6ka3_S369 -> Arfgef2_S1520 - cost: 0.26, layer: 53
Fold over basal
1.5
1
0.5
0
10
20
30
40
50
60
Time (min)
Normal
LY50
LY75
LY88
LY94
LY97
LY data
Normal data

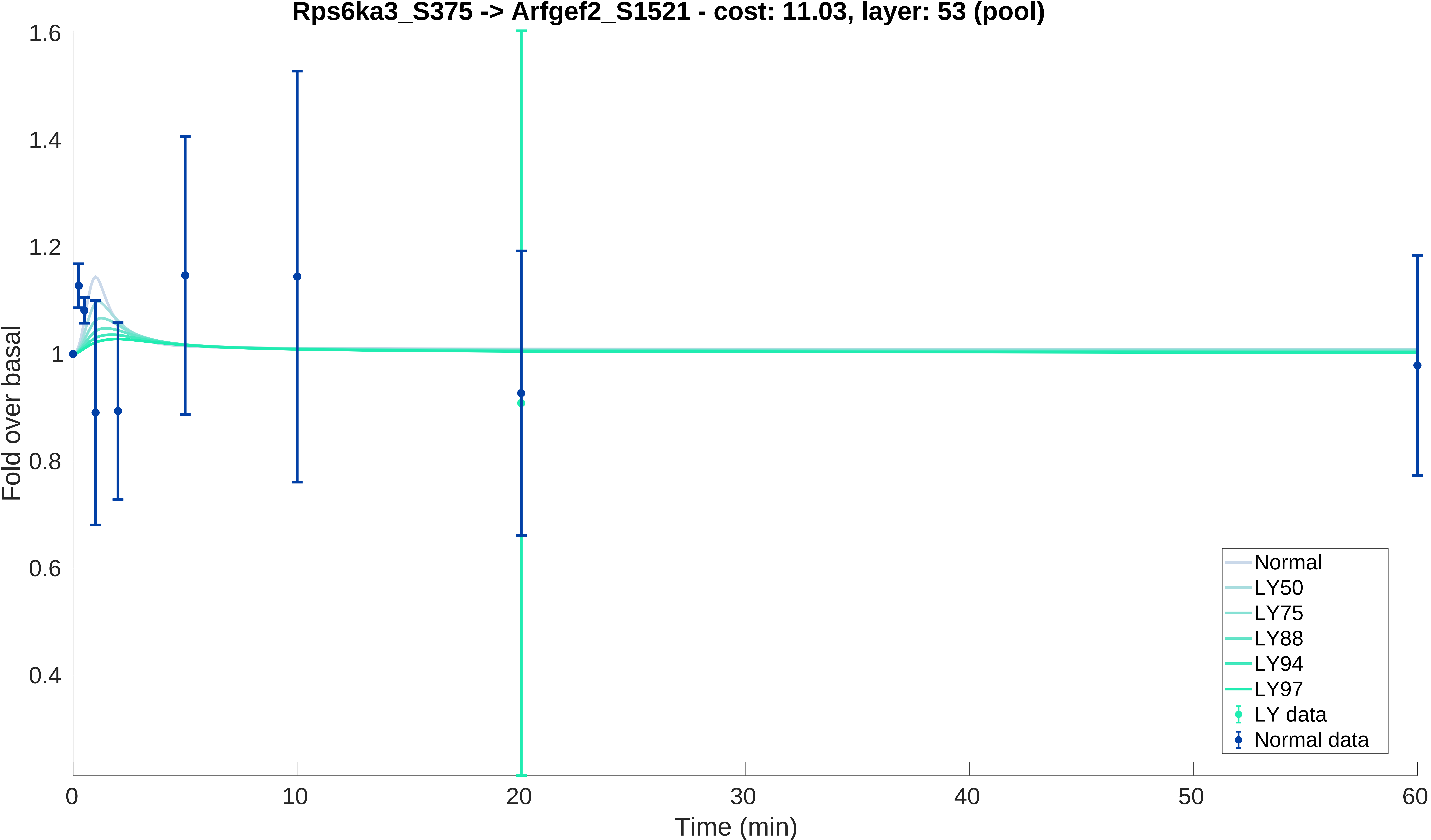

Rps6ka3_S375 -> Arfgef2_S1521 - cost: 11.03, layer: 53 (pool)
Fold over basal
1.6
1.4
1.2
1
0.8
0.6
0.4
0
10
20
30
40
50
60
Time (min)
Normal
LY50
LY75
LY88
LY94
LY97
LY data
Normal data

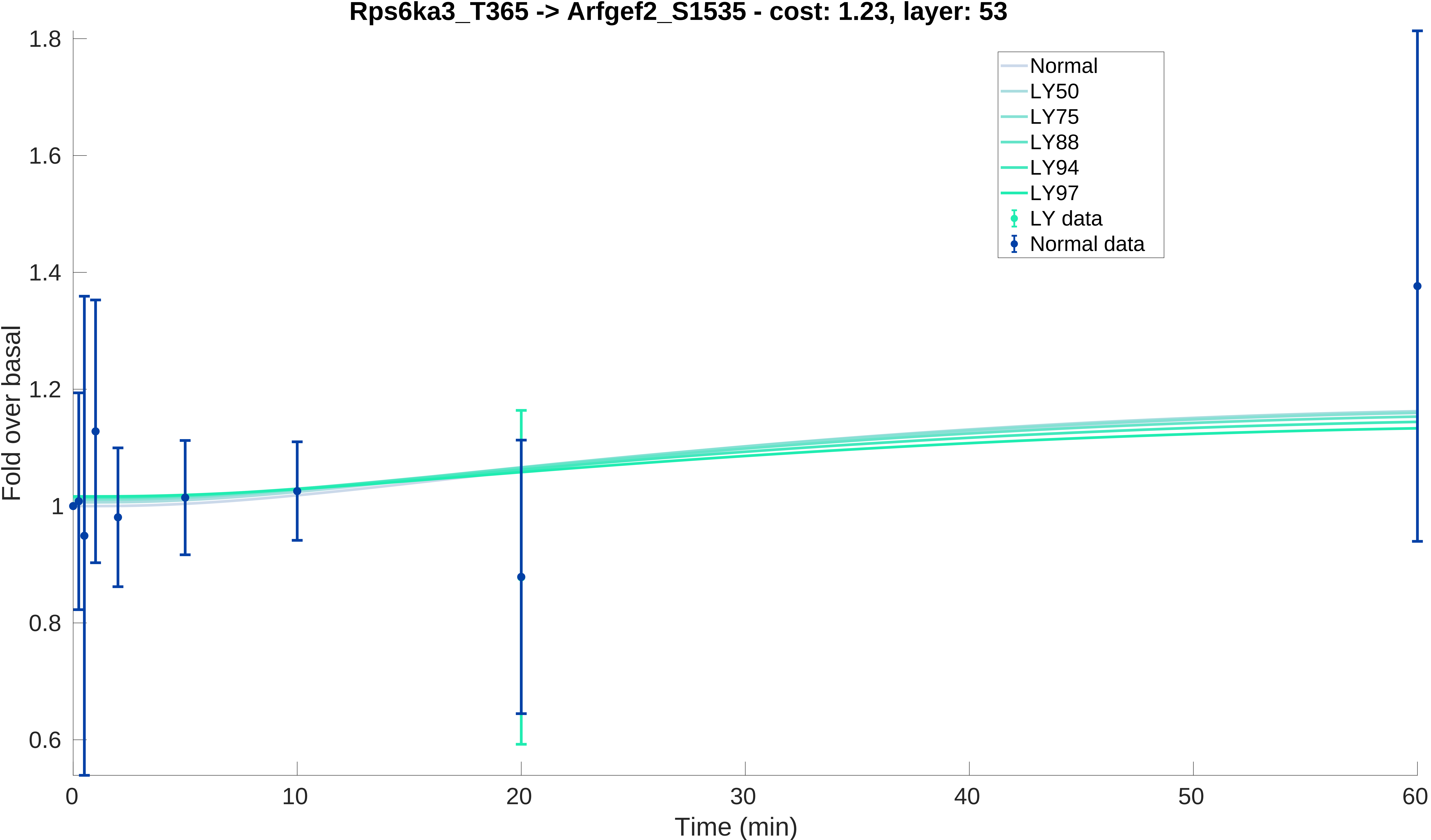
Rps6ka3_T365 -> Arfgef2_S1535 - cost: 1.23, layer: 53
Fold over basal
1.8
1.6
1.4
1.2
1
0.8
0.6
Normal
LY50
LY75
LY88
LY94
LY97
LY data
Normal data
0
10
20
30
40
50
60
Time (min)

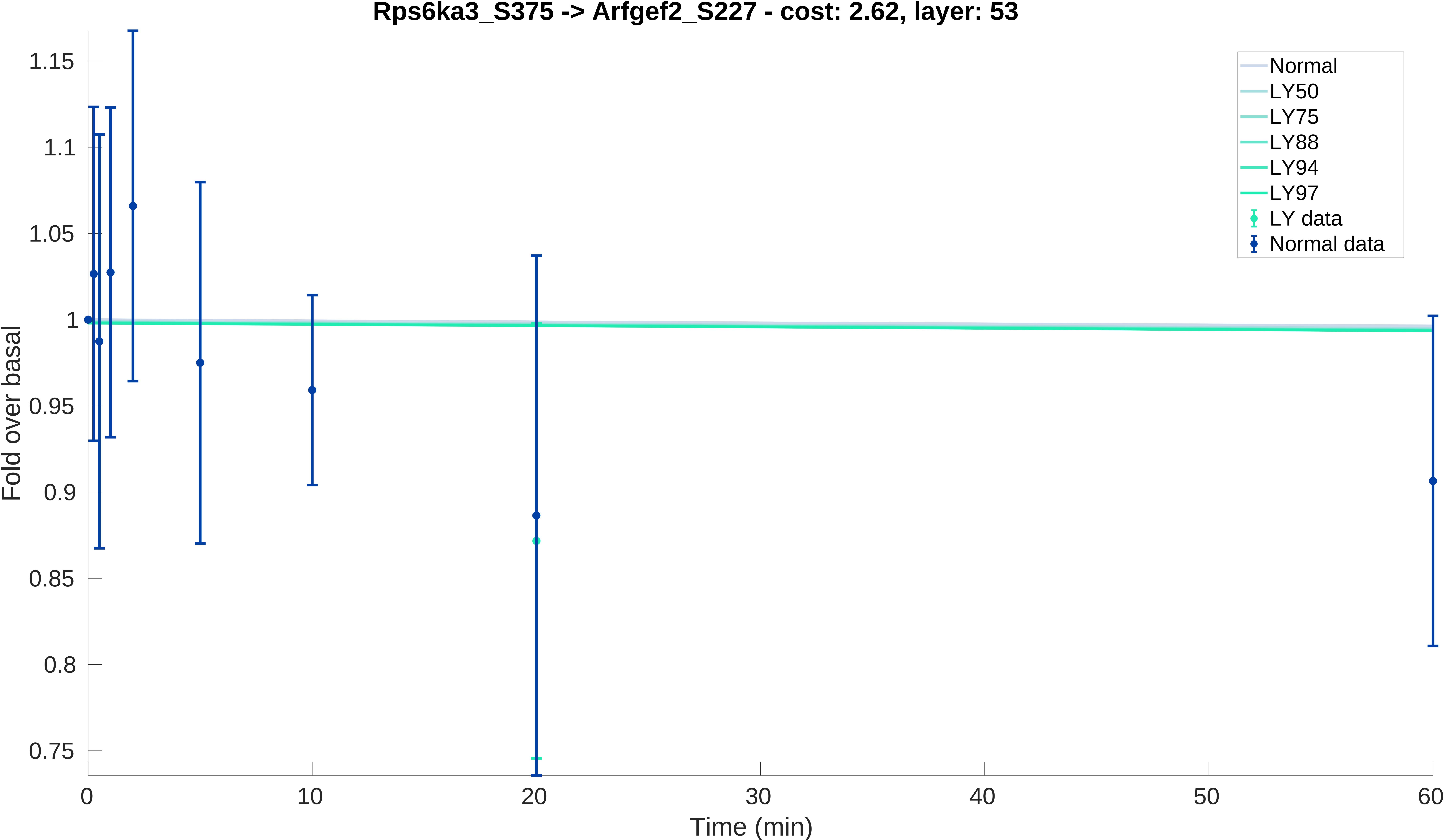

Rps6ka3_S375 -> Arfgef2_S227 - cost: 2.62, layer: 53
Fold over basal
1.15
1.1
1.05
1
0.95
0.9
0.85
0.8
0.75
0
10
20
30
40
50
60
Time (min)
Normal
LY50
LY75
LY88
LY94
LY97
LY data
Normal data

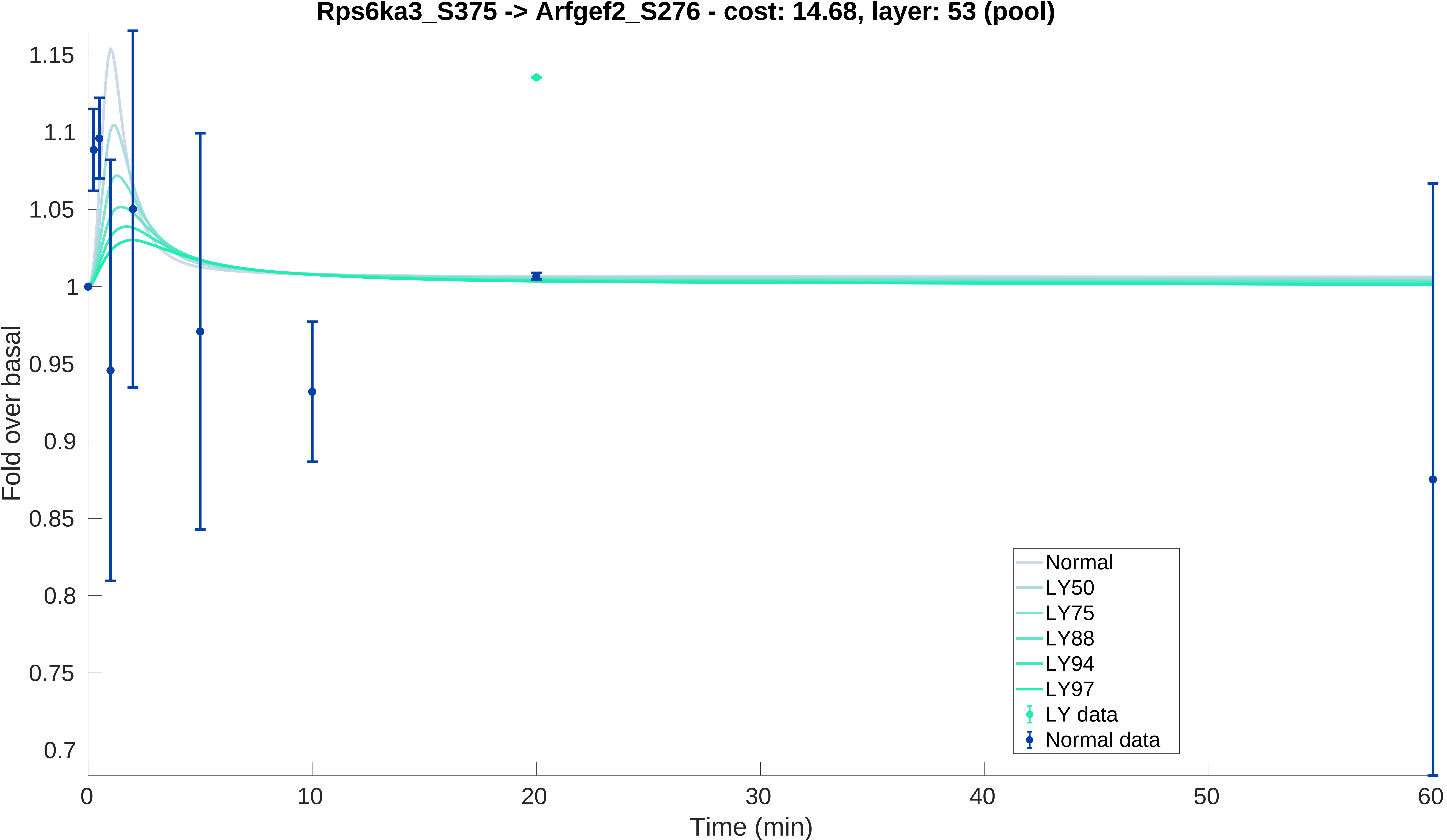

Rps6ka3_S375 -> Arfgef2_S276 - cost: 14.68, layer: 53 (pool)
Fold over basal
1.15
1.1
1.05
1
0.95
0.9
0.85
0.8
0.75
0.7
0
10
20
30
40
50
60
Time (min)
Normal
LY50
LY75
LY88
LY94
LY97
LY data
Normal data

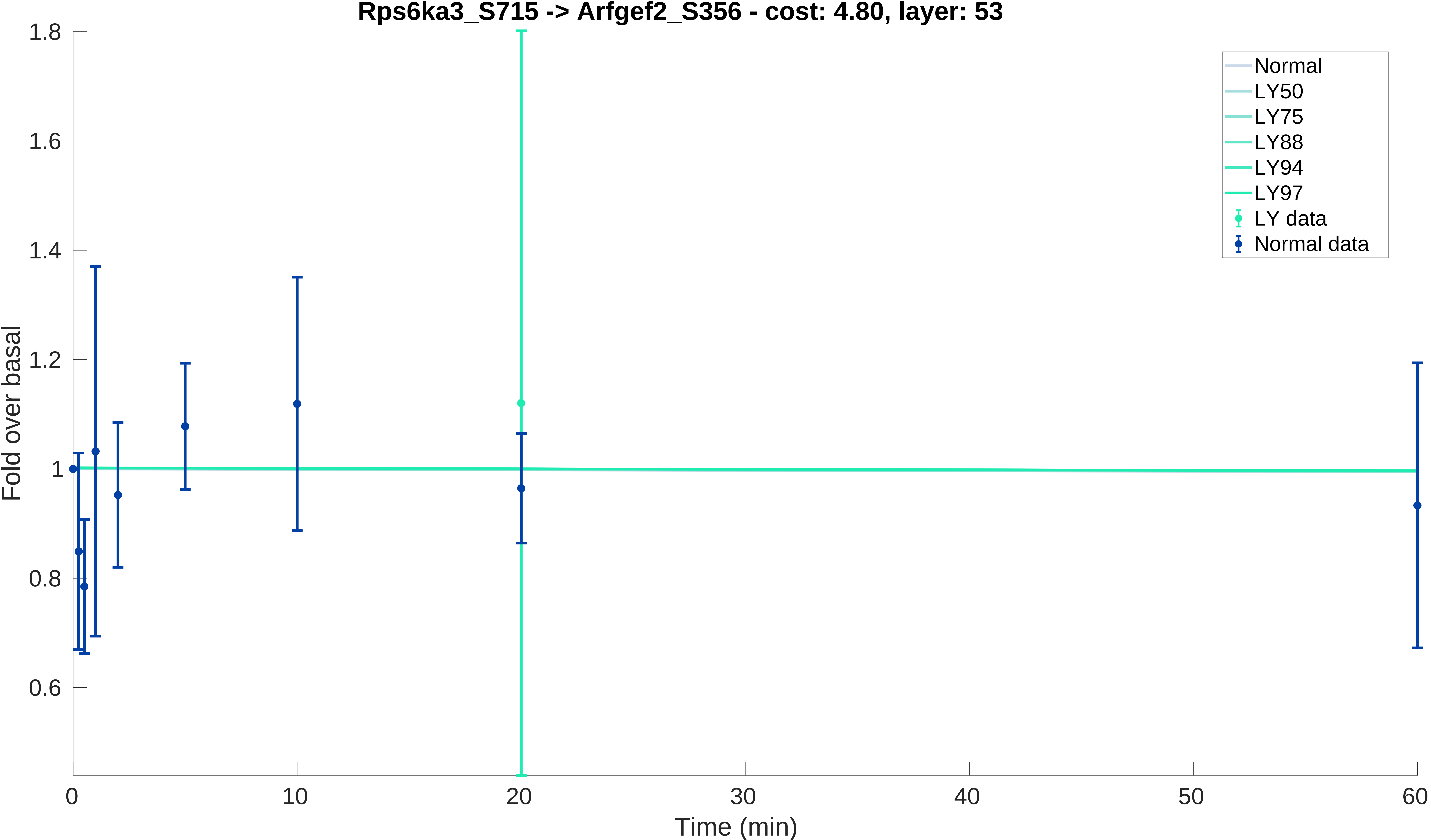

Rps6ka3_S715 -> Arfgef2_S356 - cost: 4.80, layer: 53
Fold over basal
Time (min)
1.8
1.6
1.4
1.2
1
0.8
0.6
0
10
20
30
40
50
60
Normal
LY50
LY75
LY88
LY94
LY97
LY data
Normal data

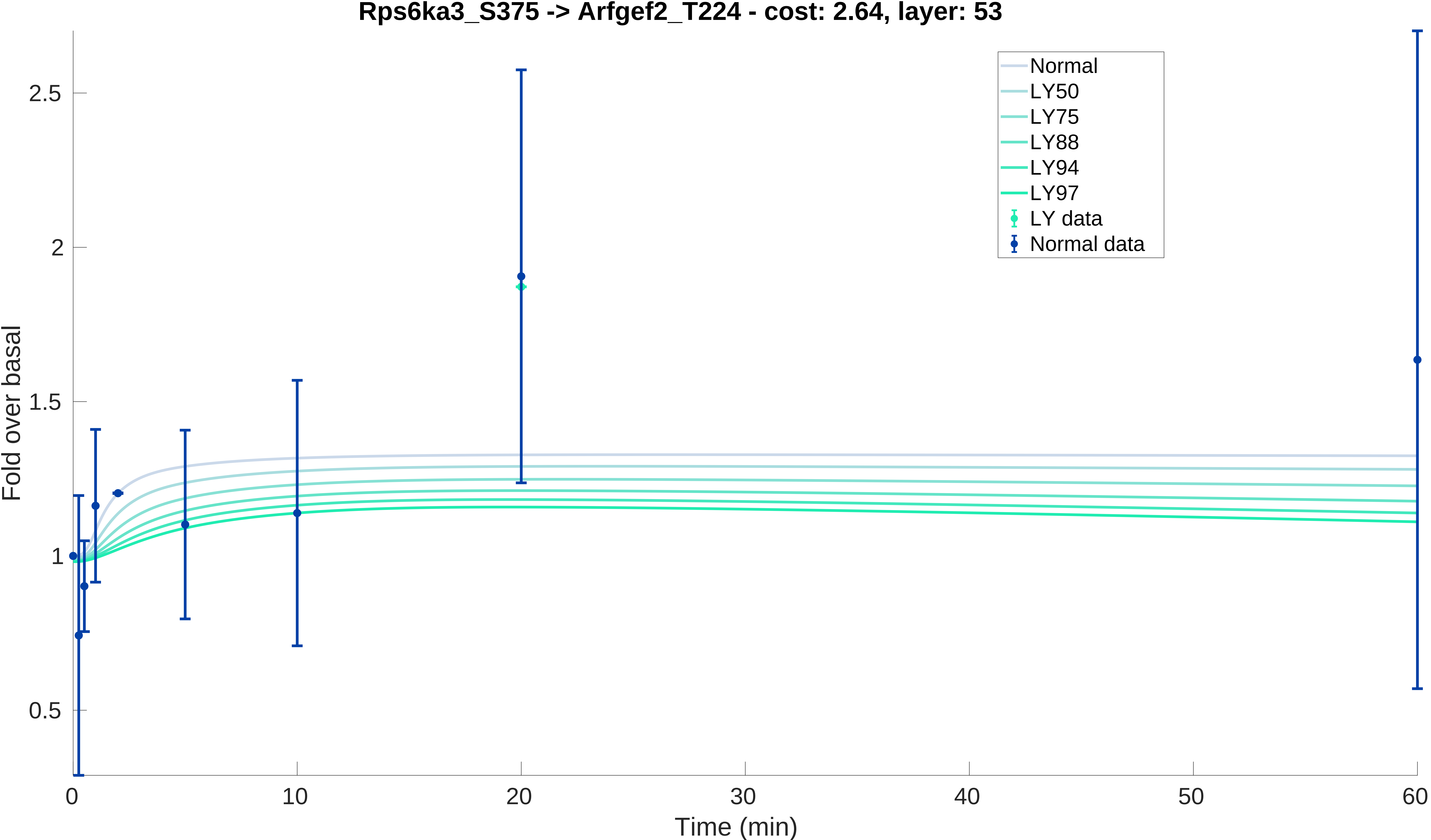

Rps6ka3_S375 -> Arfgef2_T224 - cost: 2.64, layer: 53
Fold over basal
Time (min)
0.5
1
1.5
2
2.5
0
10
20
30
40
50
60
Normal
LY50
LY75
LY88
LY94
LY97
LY data
Normal data

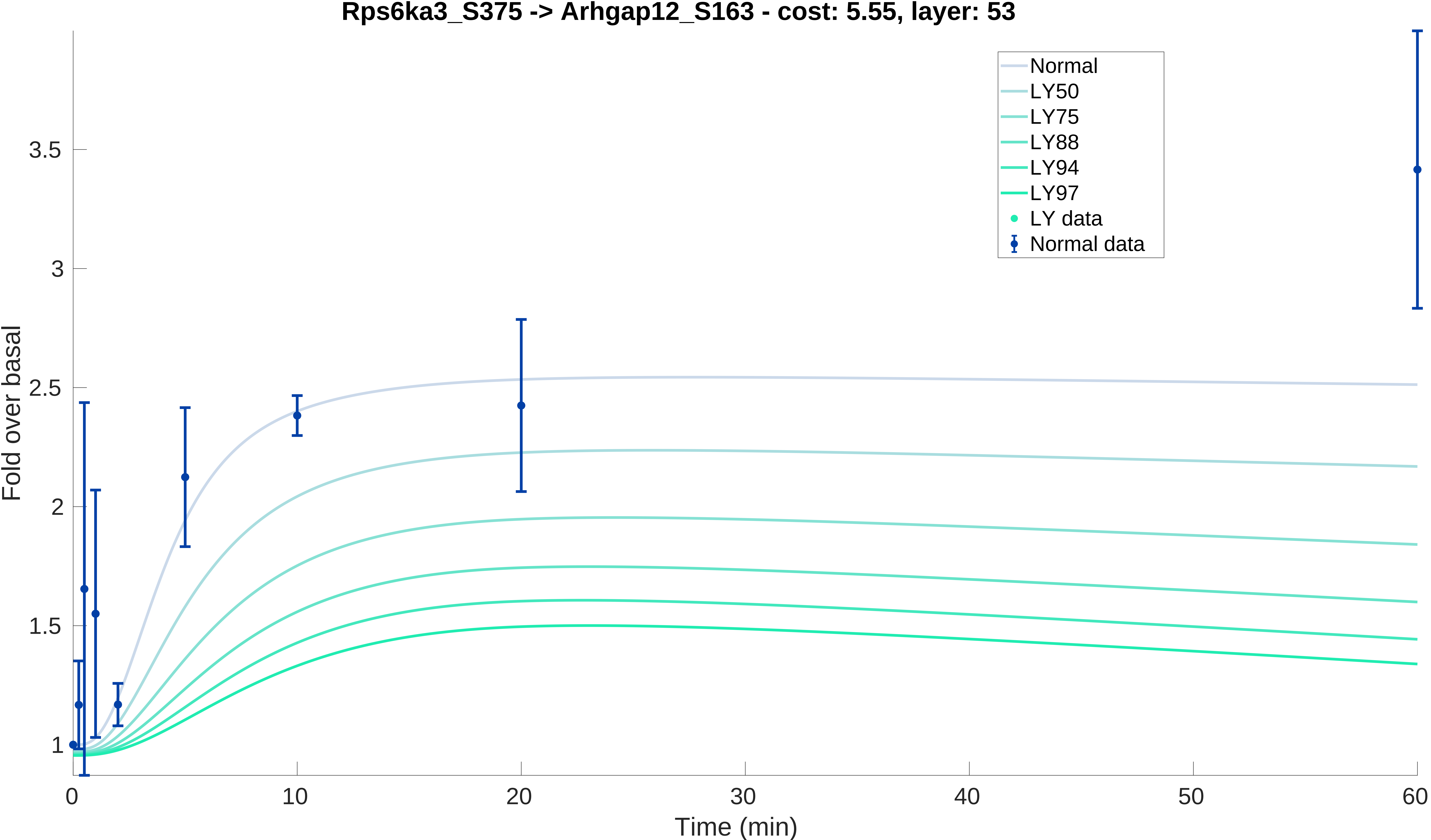

Rps6ka3_S375 -> Arhgap12_S163 - cost: 5.55, layer: 53
Normal
LY50
LY75
LY88
LY94
LY97
LY data
Normal data
Fold over basal
1
1.5
2
2.5
3
3.5
0
10
20
30
40
50
60
Time (min)

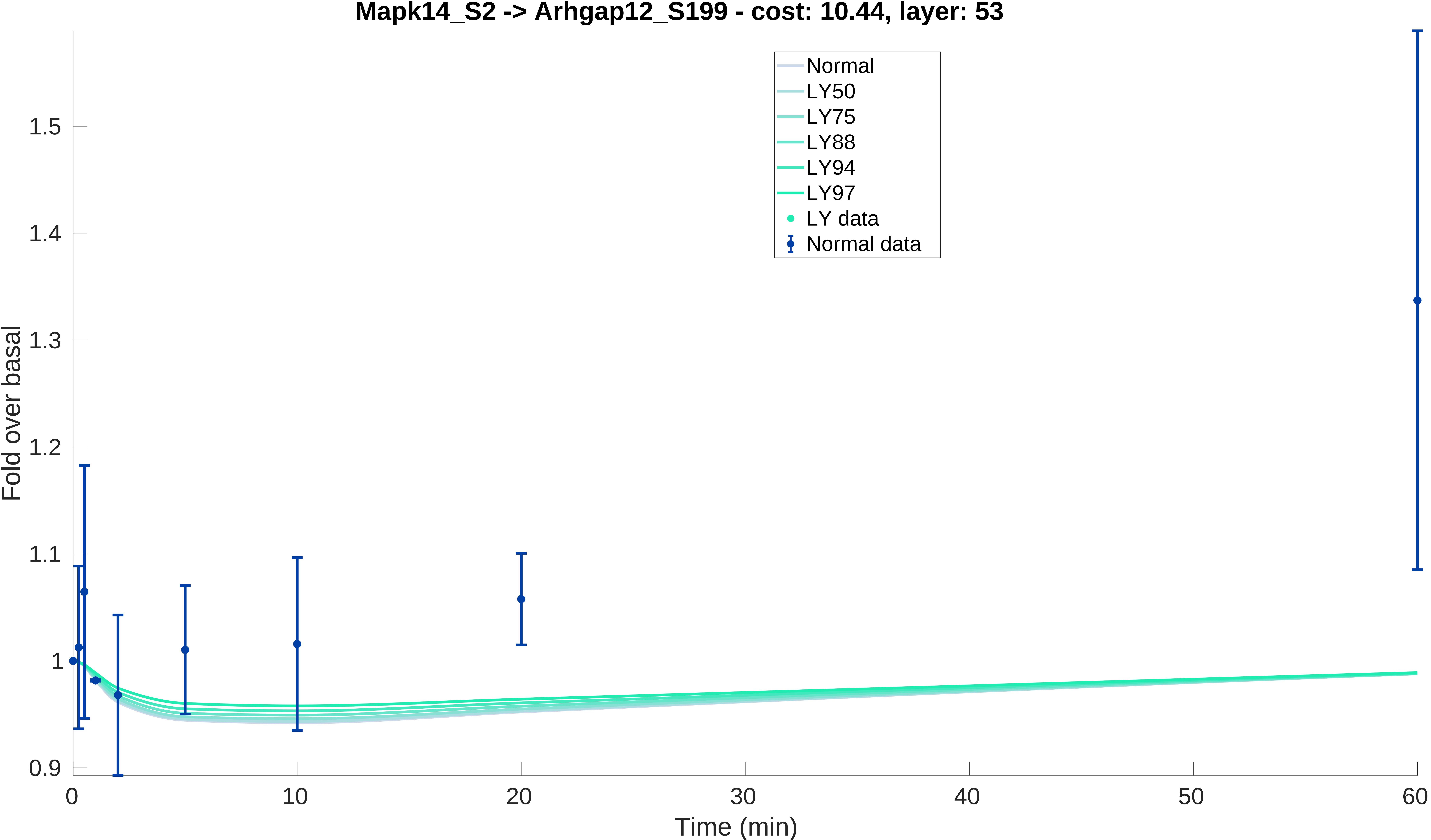

Mapk14_S2 -> Arhgap12_S199 - cost: 10.44, layer: 53
Normal
LY50
LY75
LY88
LY94
LY97
LY data
Normal data
Fold over basal
1.5
1.4
1.3
1.2
1.1
1
0.9
0
10
20
30
40
50
60
Time (min)

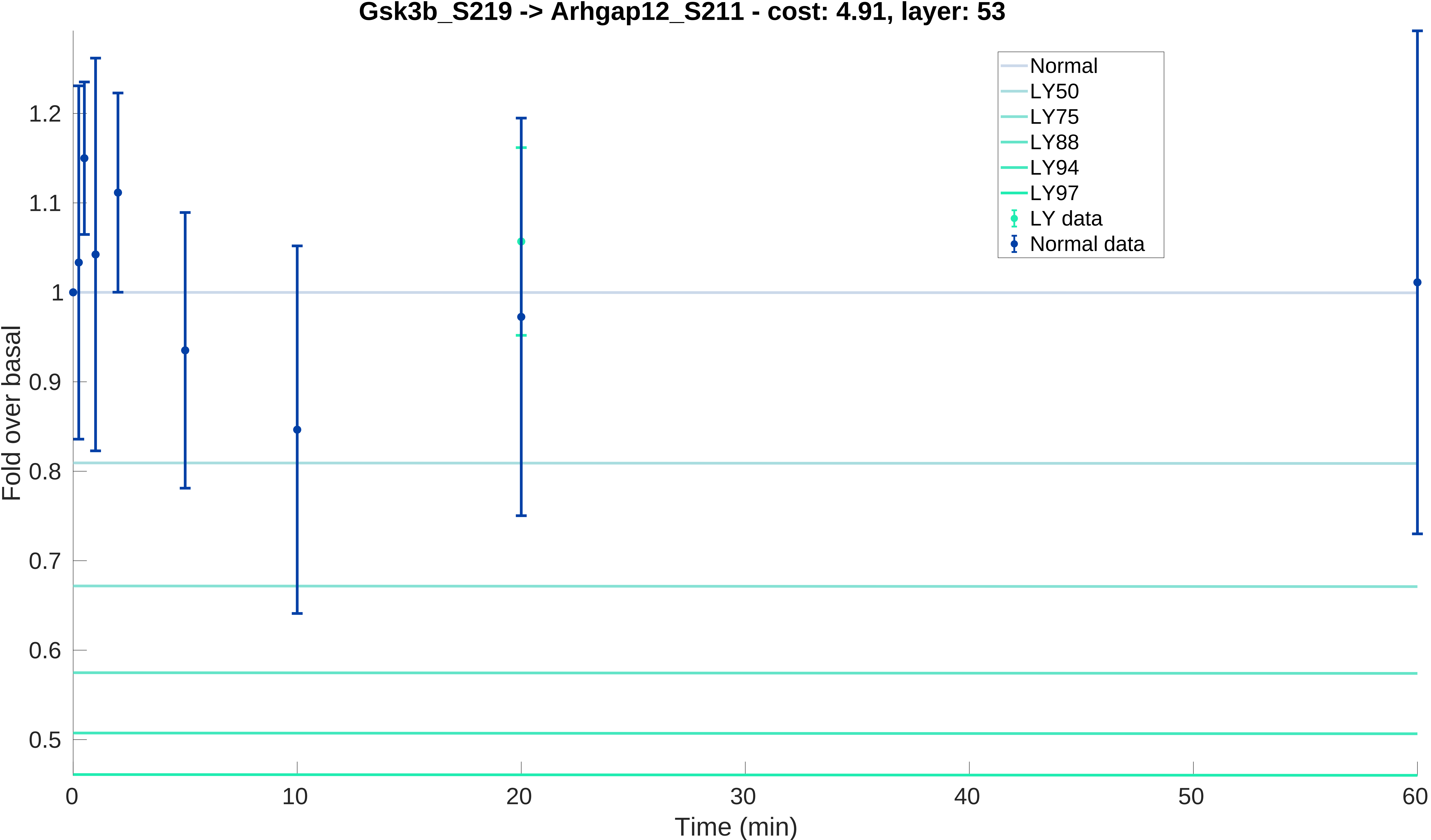

Gsk3b_S219 -> Arhgap12_S211 - cost: 4.91, layer: 53
Normal
LY50
LY75
LY88
LY94
LY97
LY data
Normal data
Fold over basal
Time (min)
0
10
20
30
40
50
60
0.5
0.6
0.7
0.8
0.9
1
1.1
1.2

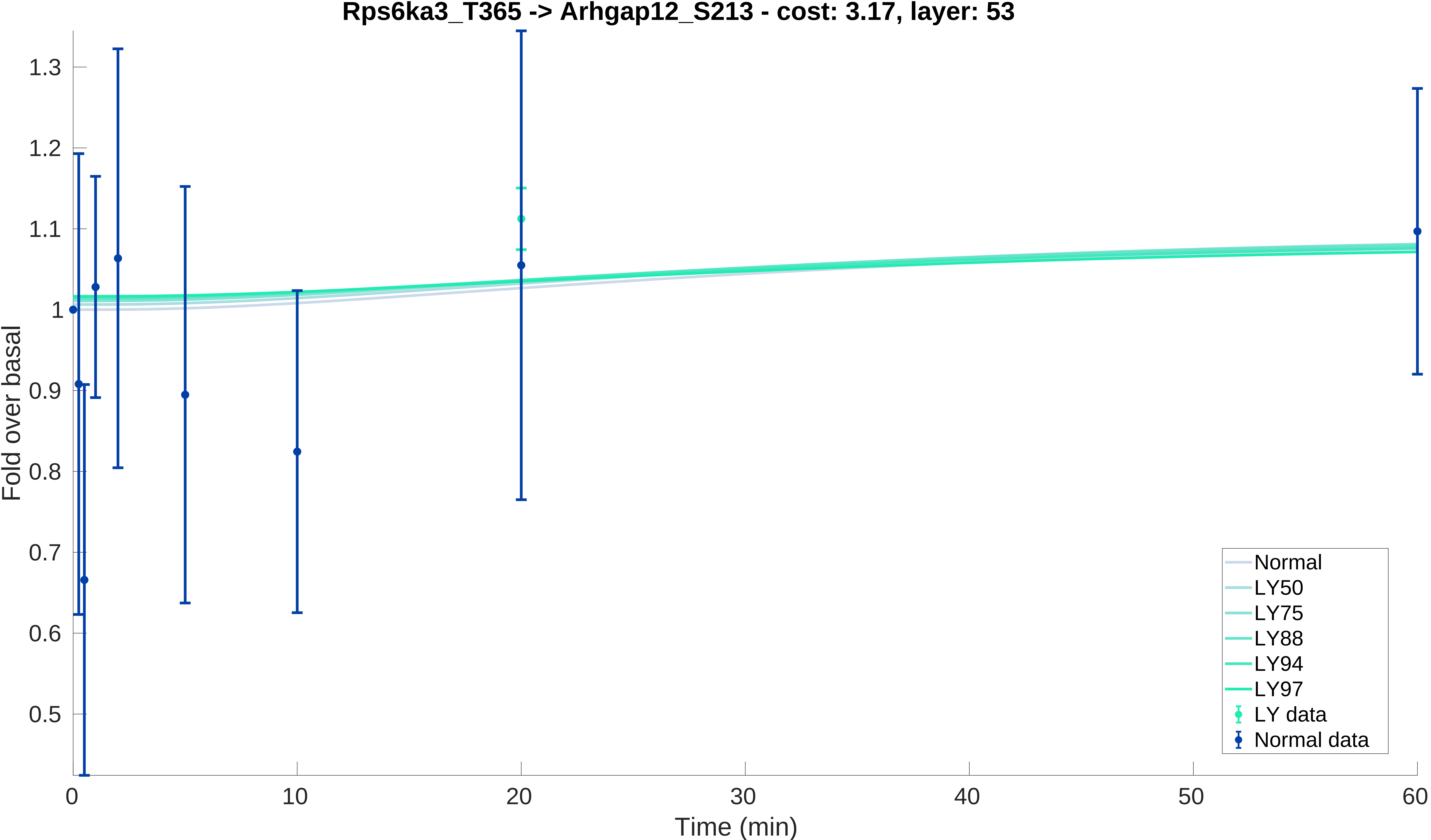

Rps6ka3_T365 -> Arhgap12_S213 - cost: 3.17, layer: 53
Fold over basal
1.3
1.2
1.1
1
0.9
0.8
0.7
0.6
0.5
0
10
20
30
40
50
60
Time (min)
Normal
LY50
LY75
LY88
LY94
LY97
LY data
Normal data

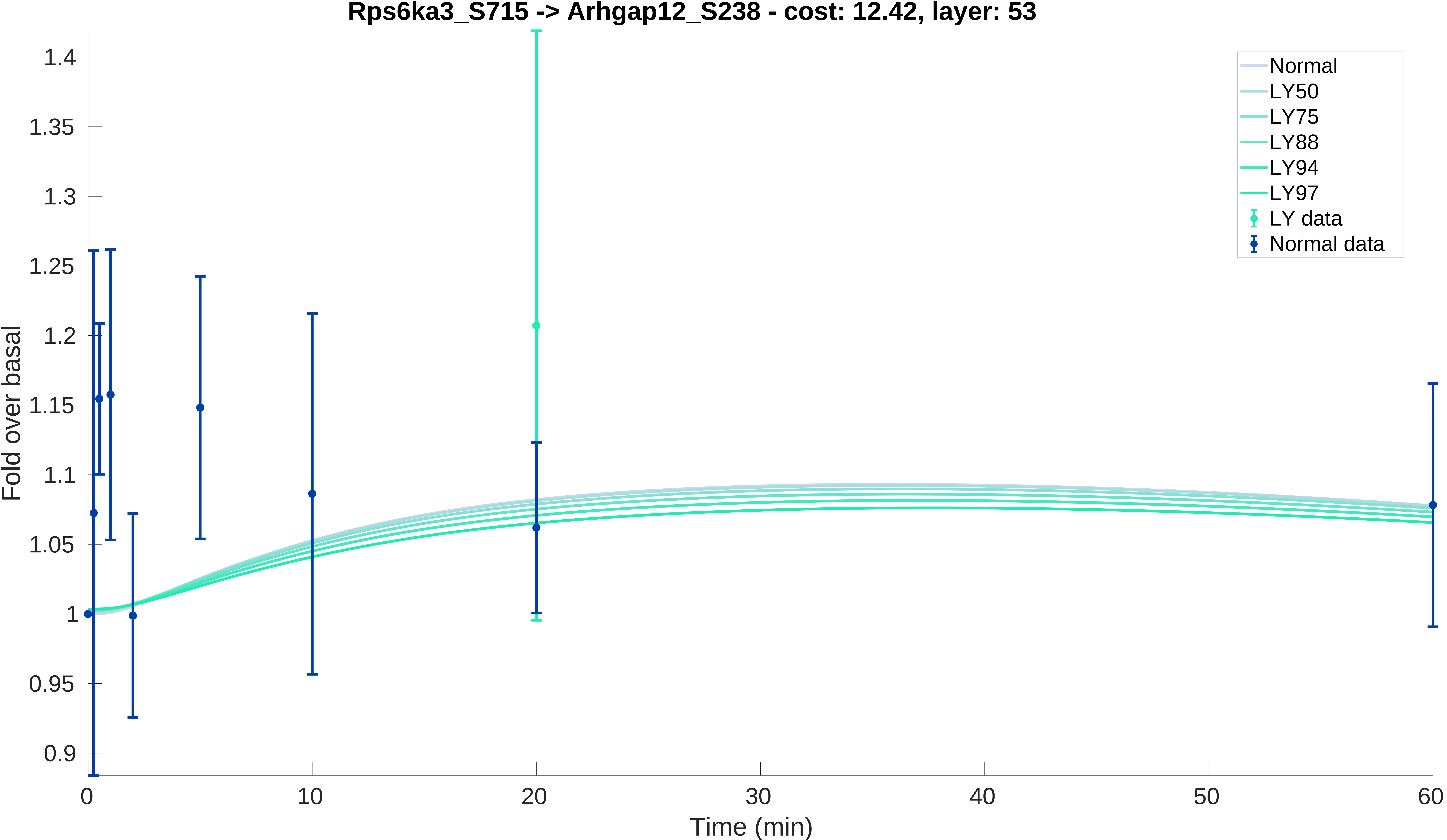

Rps6ka3_S715 -> Arhgap12_S238 - cost: 12.42, layer: 53
Fold over basal
1.4
1.35
1.3
1.25
1.2
1.15
1.1
1.05
1
0.95
0.9
0
10
20
30
40
50
60
Time (min)
Normal
LY50
LY75
LY88
LY94
LY97
LY data
Normal data

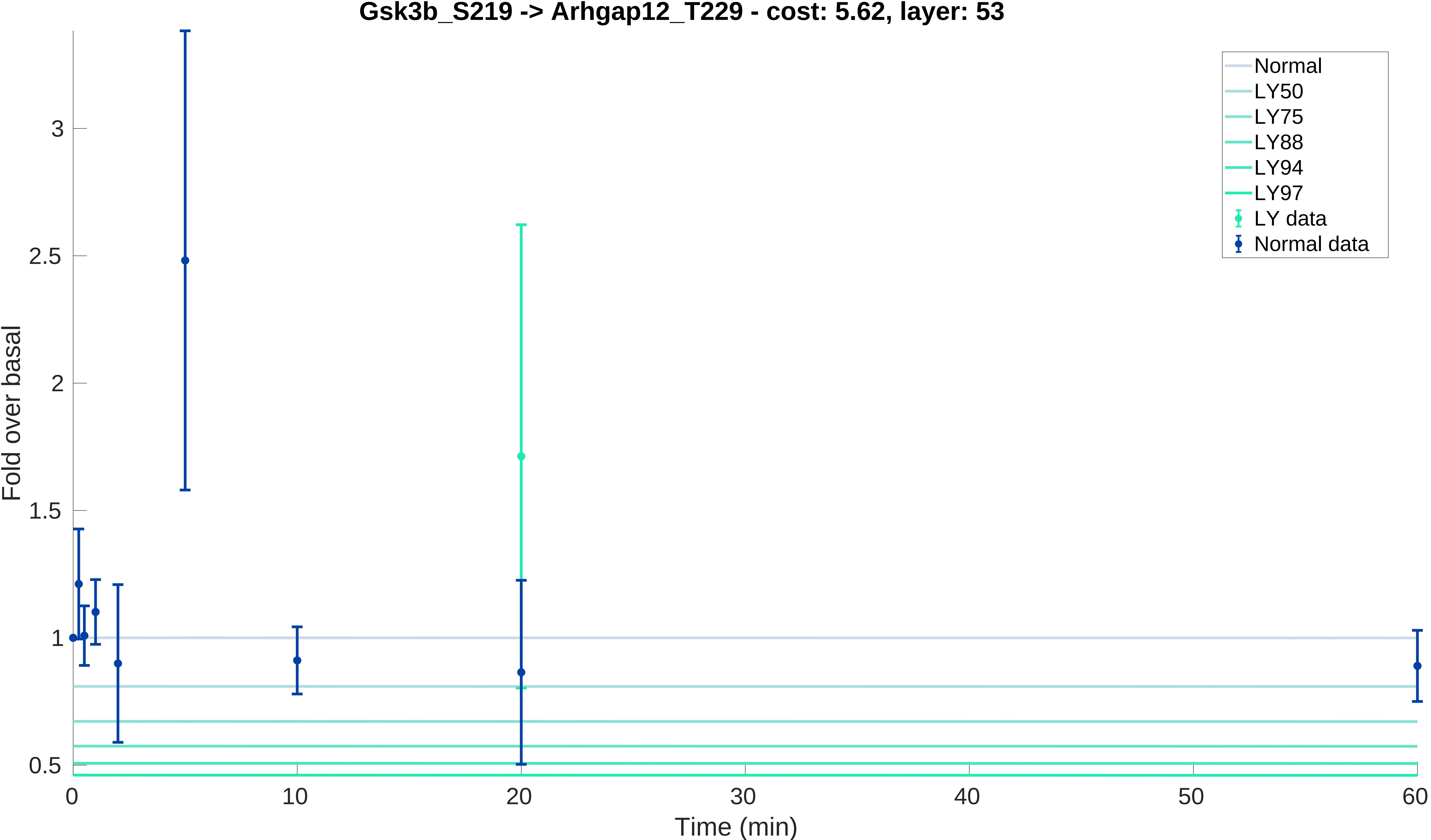

Gsk3b_S219 -> Arhgap12_T229 - cost: 5.62, layer: 53
Fold over basal
Time (min)
0.5
1
1.5
2
2.5
3
0
10
20
30
40
50
60
Normal
LY50
LY75
LY88
LY94
LY97
LY data
Normal data

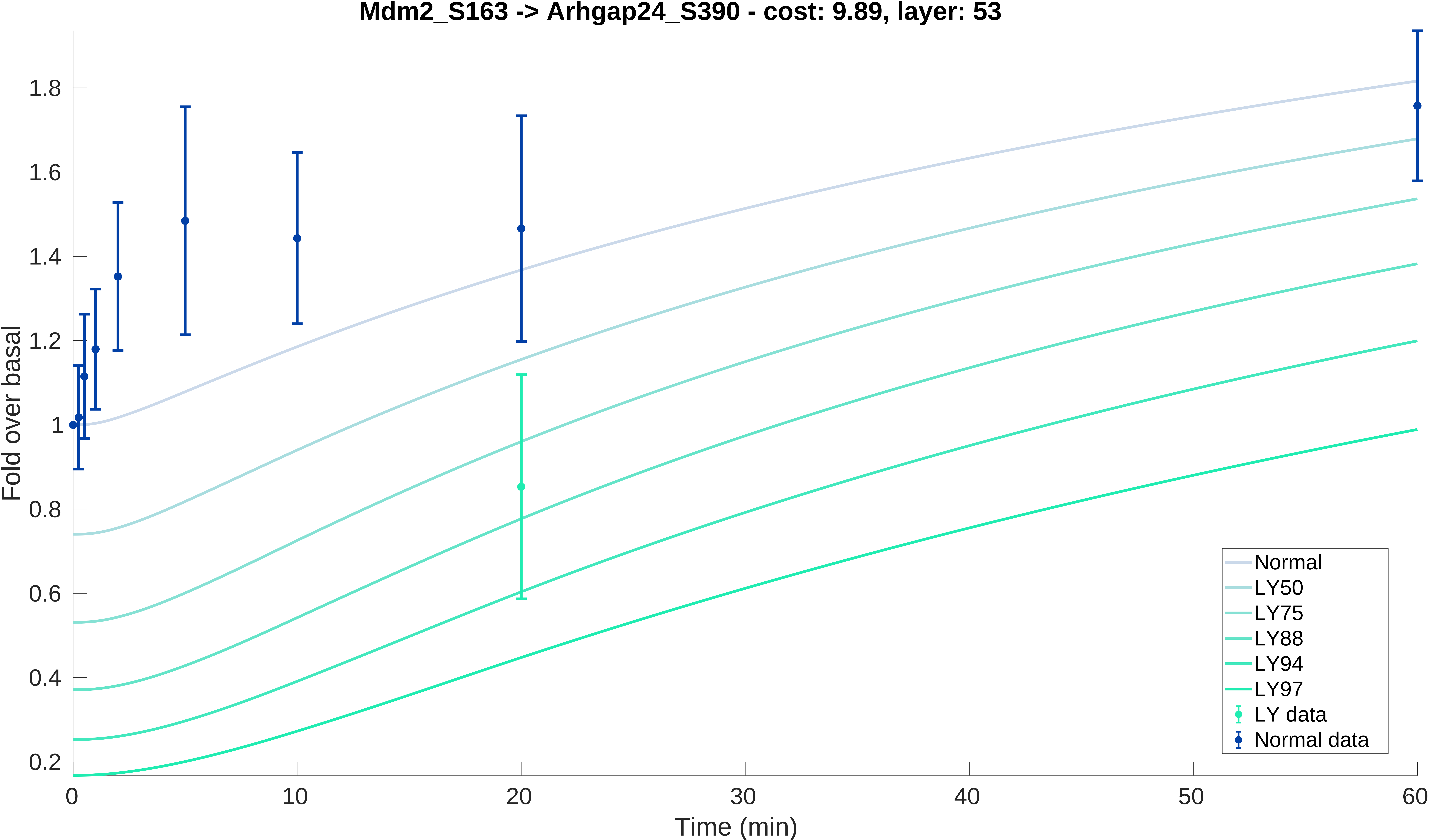

Mdm2_S163 -> Arhgap24_S390 - cost: 9.89, layer: 53
Fold over basal
Time (min)
0.2
0.4
0.6
0.8
1
1.2
1.4
1.6
1.8
0
10
20
30
40
50
60
Normal
LY50
LY75
LY88
LY94
LY97
LY data
Normal data

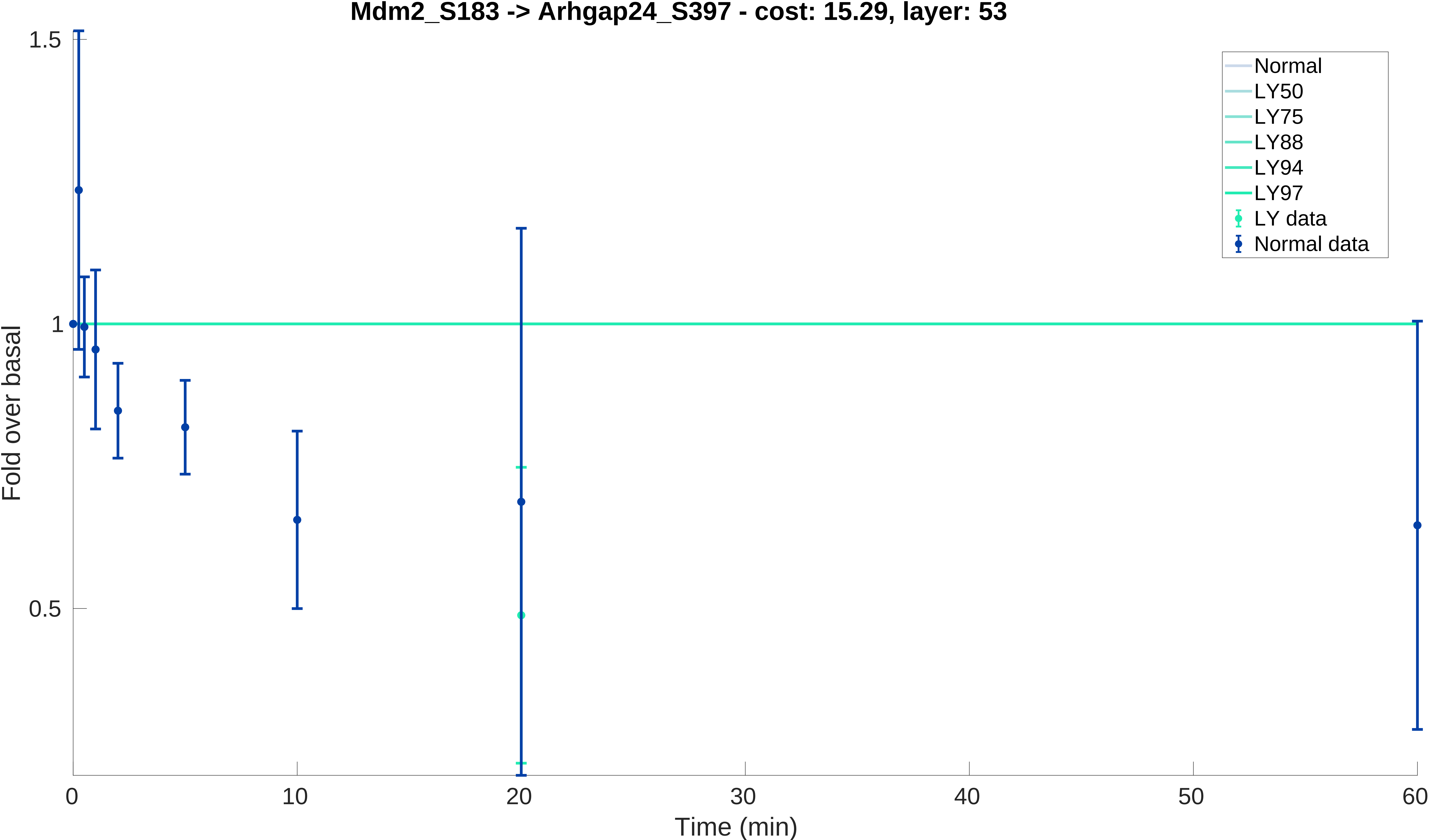
Mdm2_S183 -> Arhgap24_S397 - cost: 15.29, layer: 53
Fold over basal
1.5
1
0.5
0
10
20
30
40
50
60
Time (min)
Normal
LY50
LY75
LY88
LY94
LY97
LY data
Normal data

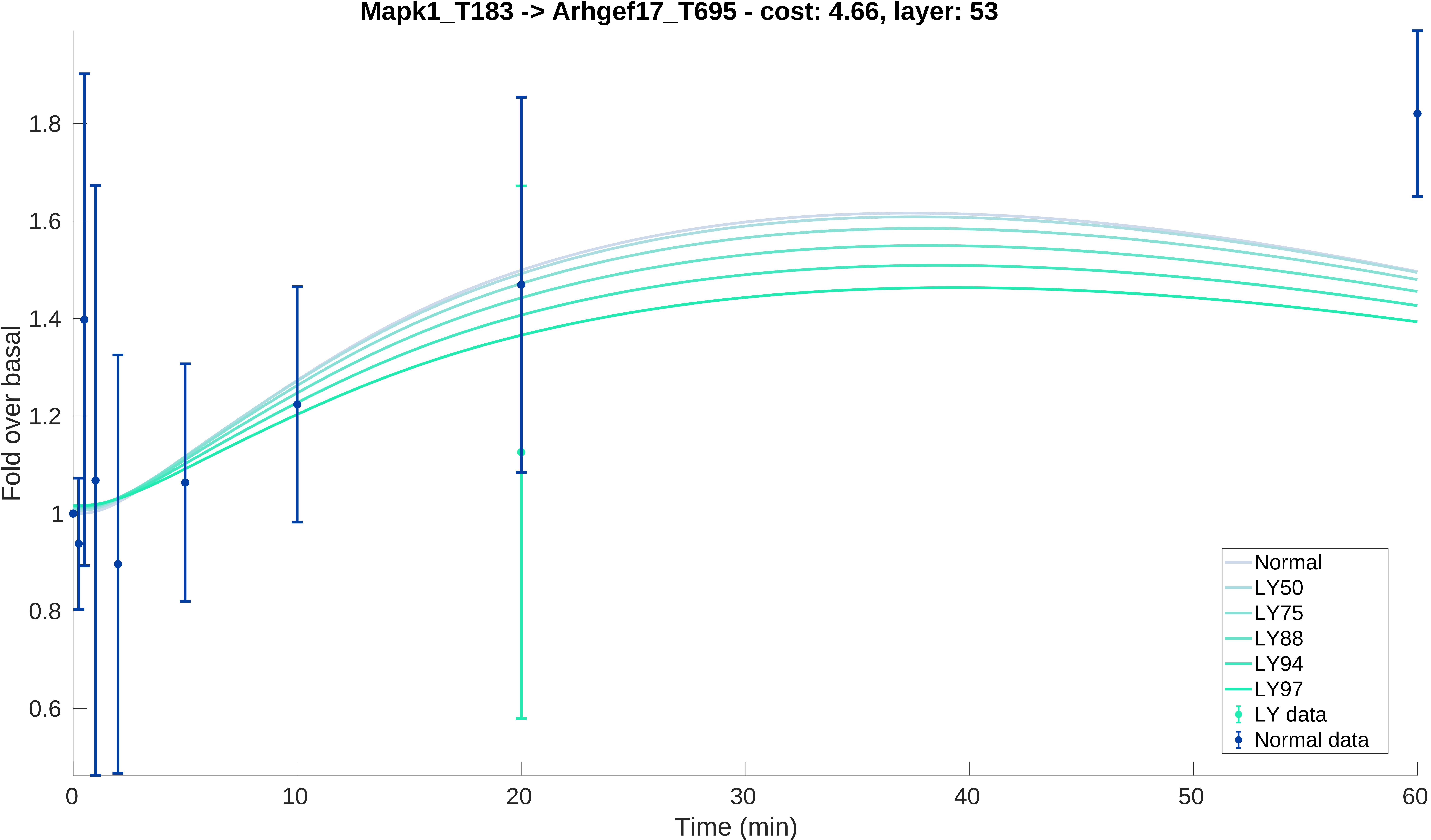

Mapk1_T183 -> Arhgef17_T695 - cost: 4.66, layer: 53
Fold over basal
Time (min)
0.6
0.8
1
1.2
1.4
1.6
1.8
0
10
20
30
40
50
60
Normal
LY50
LY75
LY88
LY94
LY97
LY data
Normal data

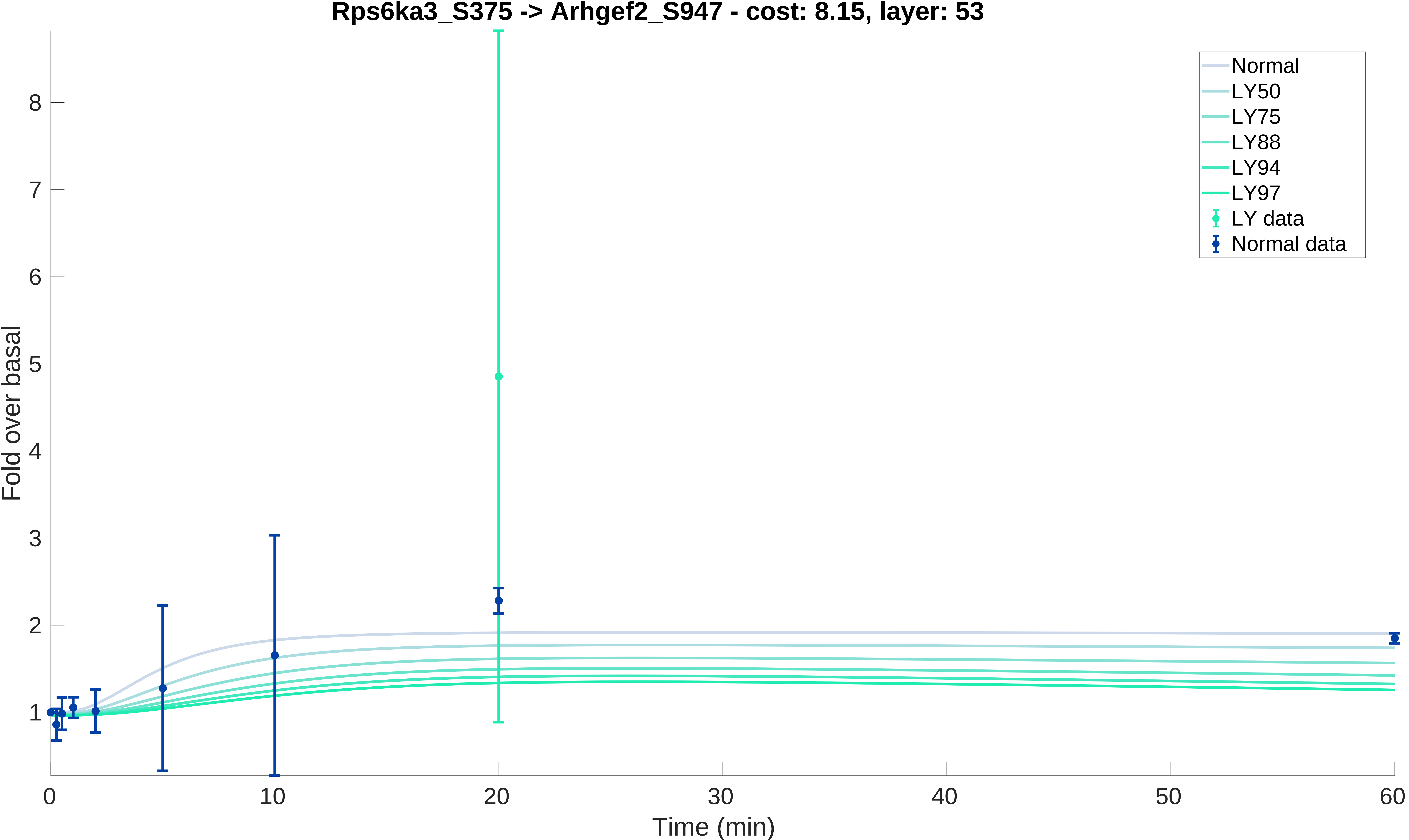

Rps6ka3_S375 -> Arhgef2_S947 - cost: 8.15, layer: 53
Fold over basal
Time (min)
0
10
20
30
40
50
60
1
2
3
4
5
6
7
8
Normal
LY50
LY75
LY88
LY94
LY97
LY data
Normal data

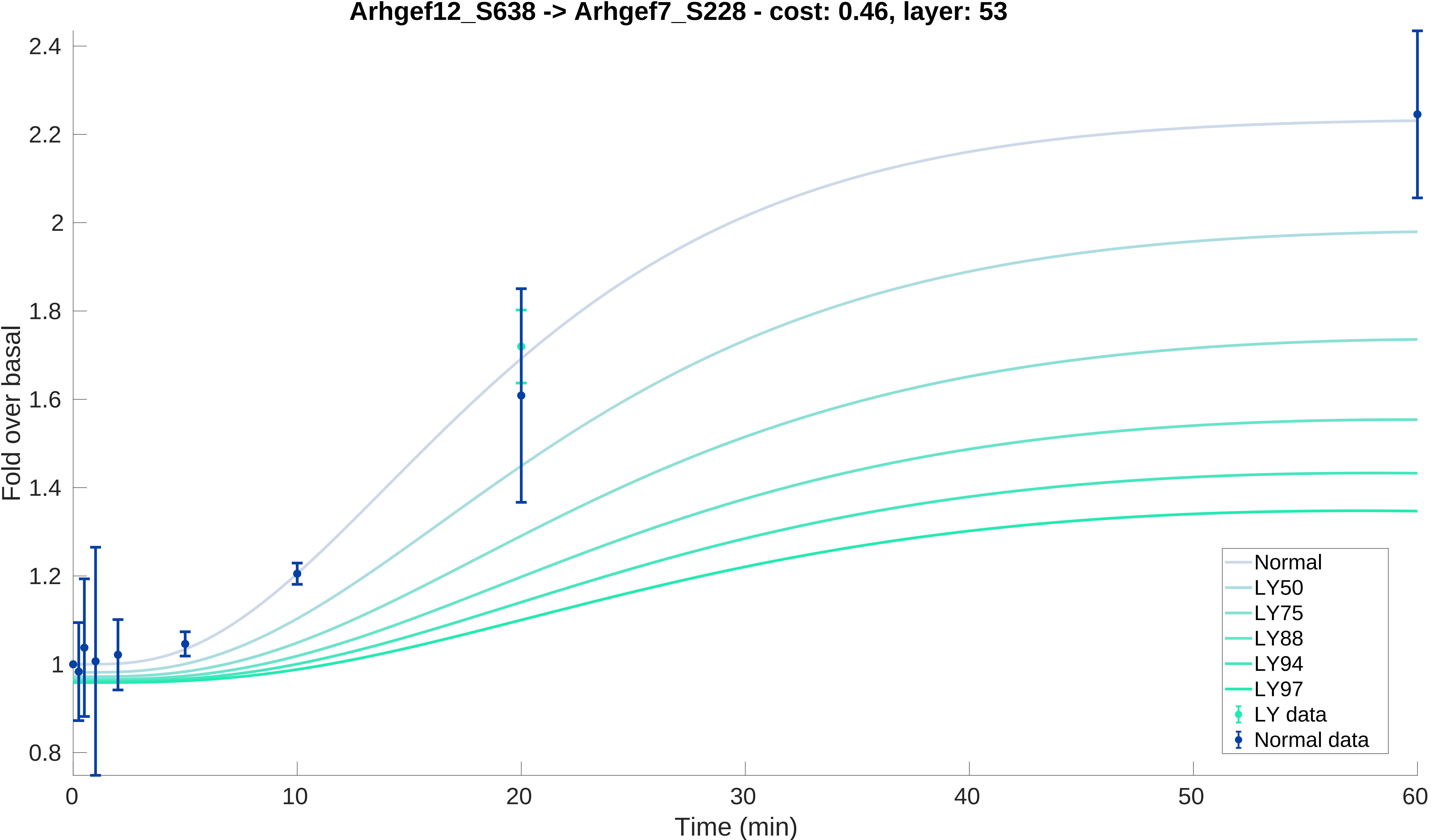

Arhgef12_S638 -> Arhgef7_S228 - cost: 0.46, layer: 53
Fold over basal
2.4
2.2
2
1.8
1.6
1.4
1.2
1
0.8
0
10
20
30
40
50
60
Time (min)
Normal
LY50
LY75
LY88
LY94
LY97
LY data
Normal data

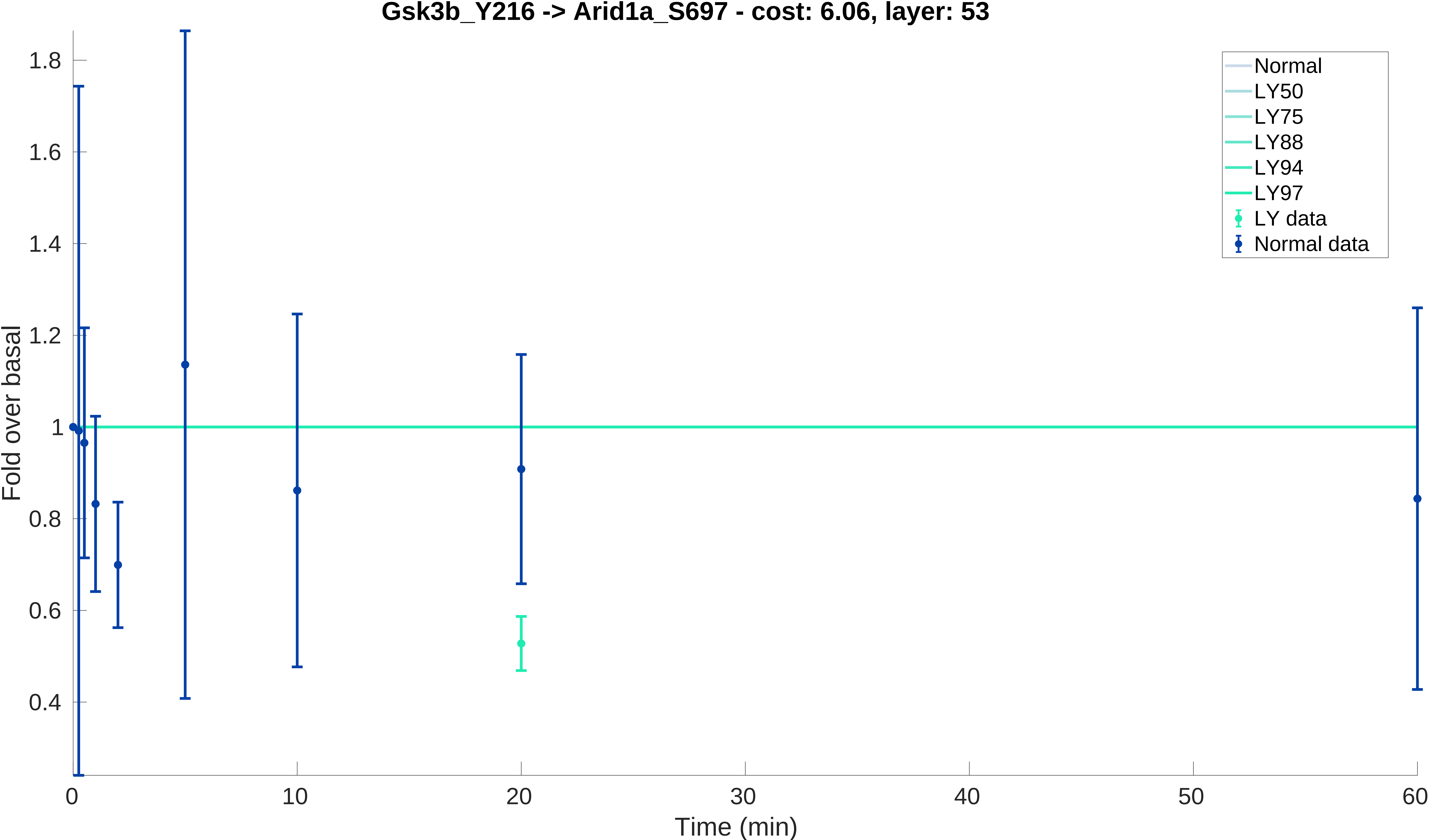
Gsk3b_Y216 -> Arid1a_S697 - cost: 6.06, layer: 53
Fold over basal
1.8
1.6
1.4
1.2
1
0.8
0.6
0.4
0
10
20
30
40
50
60
Time (min)
Normal
LY50
LY75
LY88
LY94
LY97
LY data
Normal data

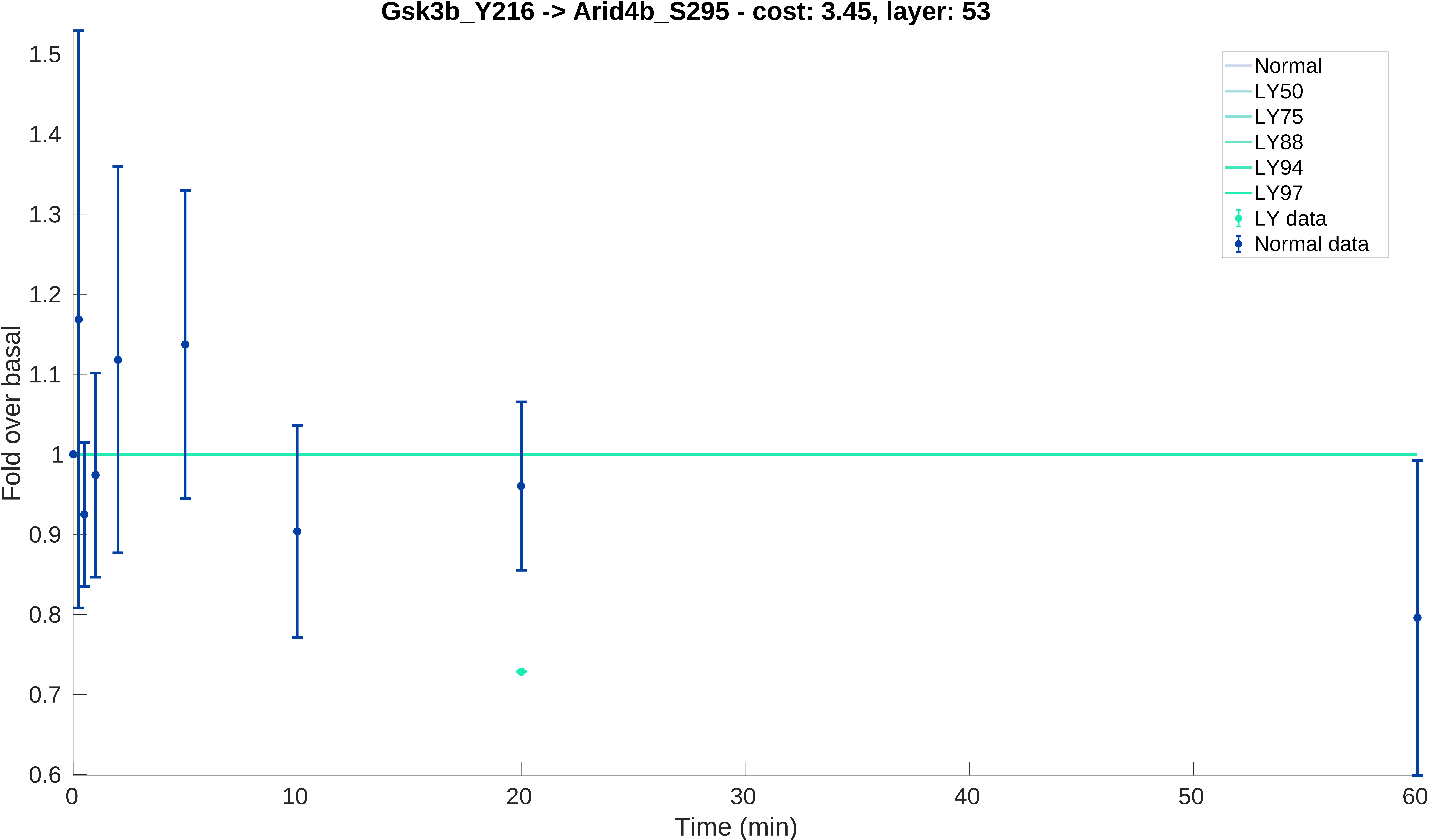

Gsk3b_Y216 -> Arid4b_S295 - cost: 3.45, layer: 53
Normal
LY50
LY75
LY88
LY94
LY97
LY data
Normal data
Fold over basal
1.5
1.4
1.3
1.2
1.1
1
0.9
0.8
0.7
0.6
0
10
20
30
40
50
60
Time (min)

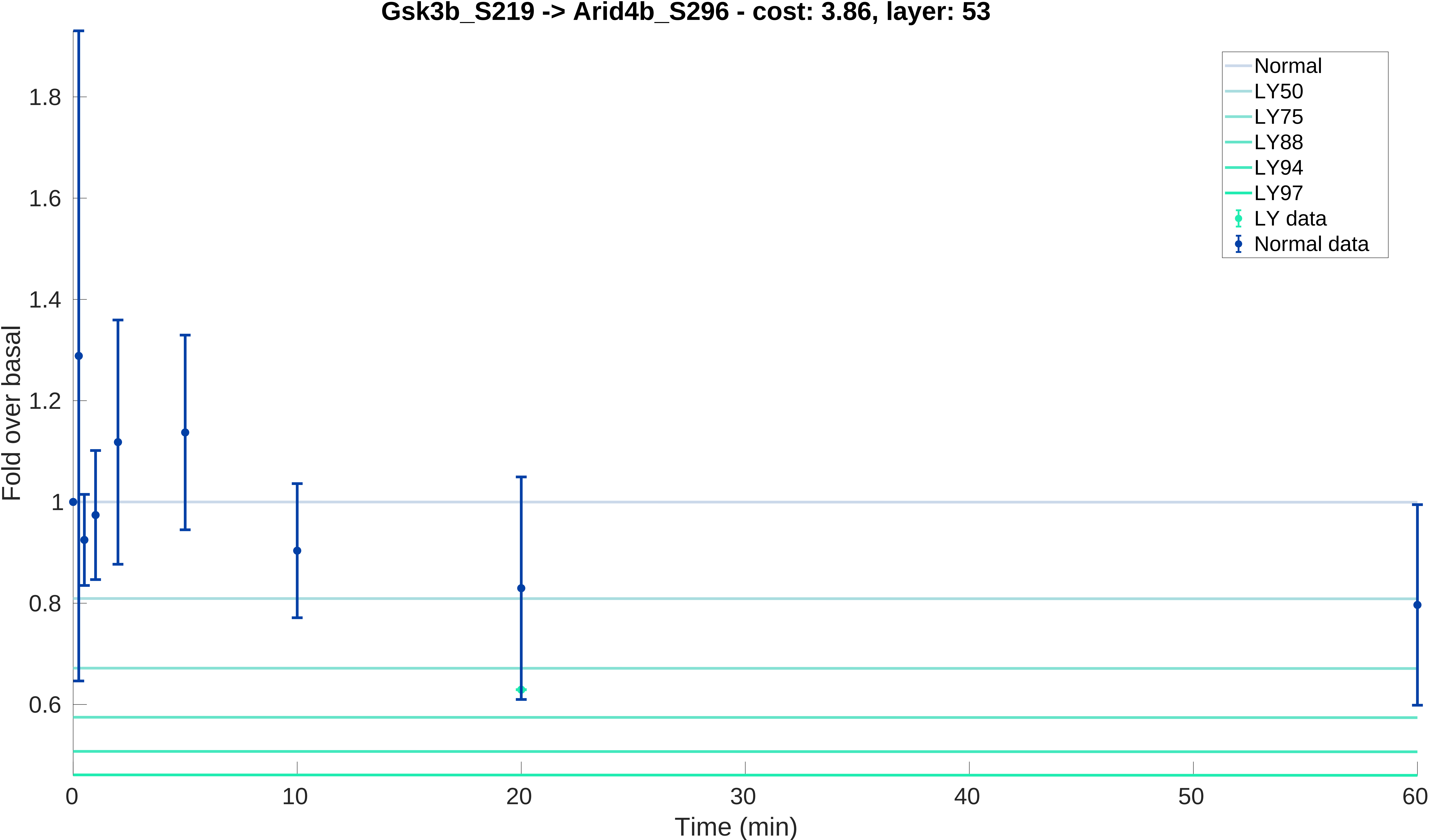

Gsk3b_S219 -> Arid4b_S296 - cost: 3.86, layer: 53
Fold over basal
1.8
1.6
1.4
1.2
1
0.8
0.6
Time (min)
0
10
20
30
40
50
60
Normal
LY50
LY75
LY88
LY94
LY97
LY data
Normal data

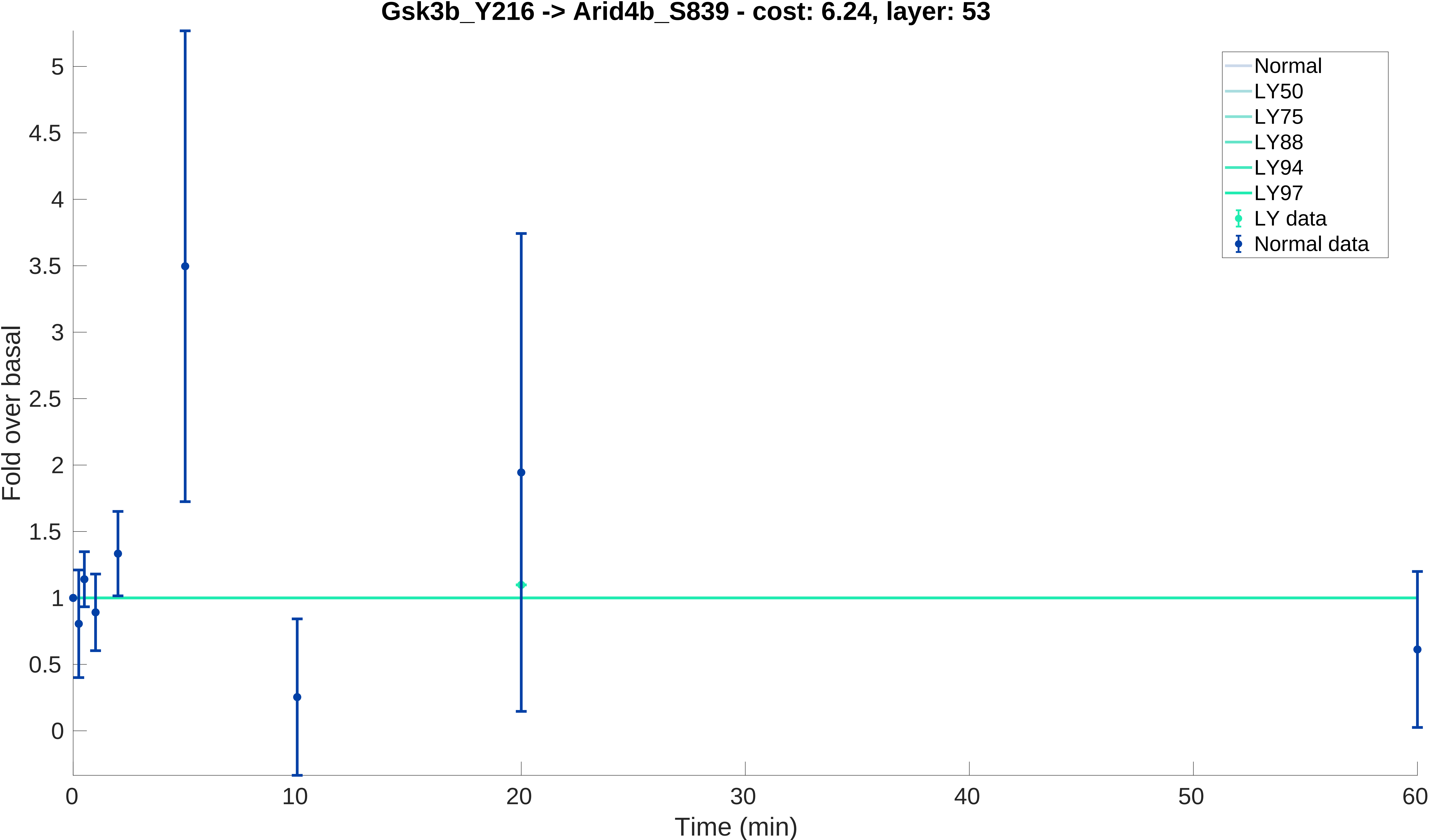
Gsk3b_Y216 -> Arid4b_S839 - cost: 6.24, layer: 53
Fold over basal
Time (min)
0
0.5
1
1.5
2
2.5
3
3.5
4
4.5
5
0
10
20
30
40
50
60
Normal
LY50
LY75
LY88
LY94
LY97
LY data
Normal data

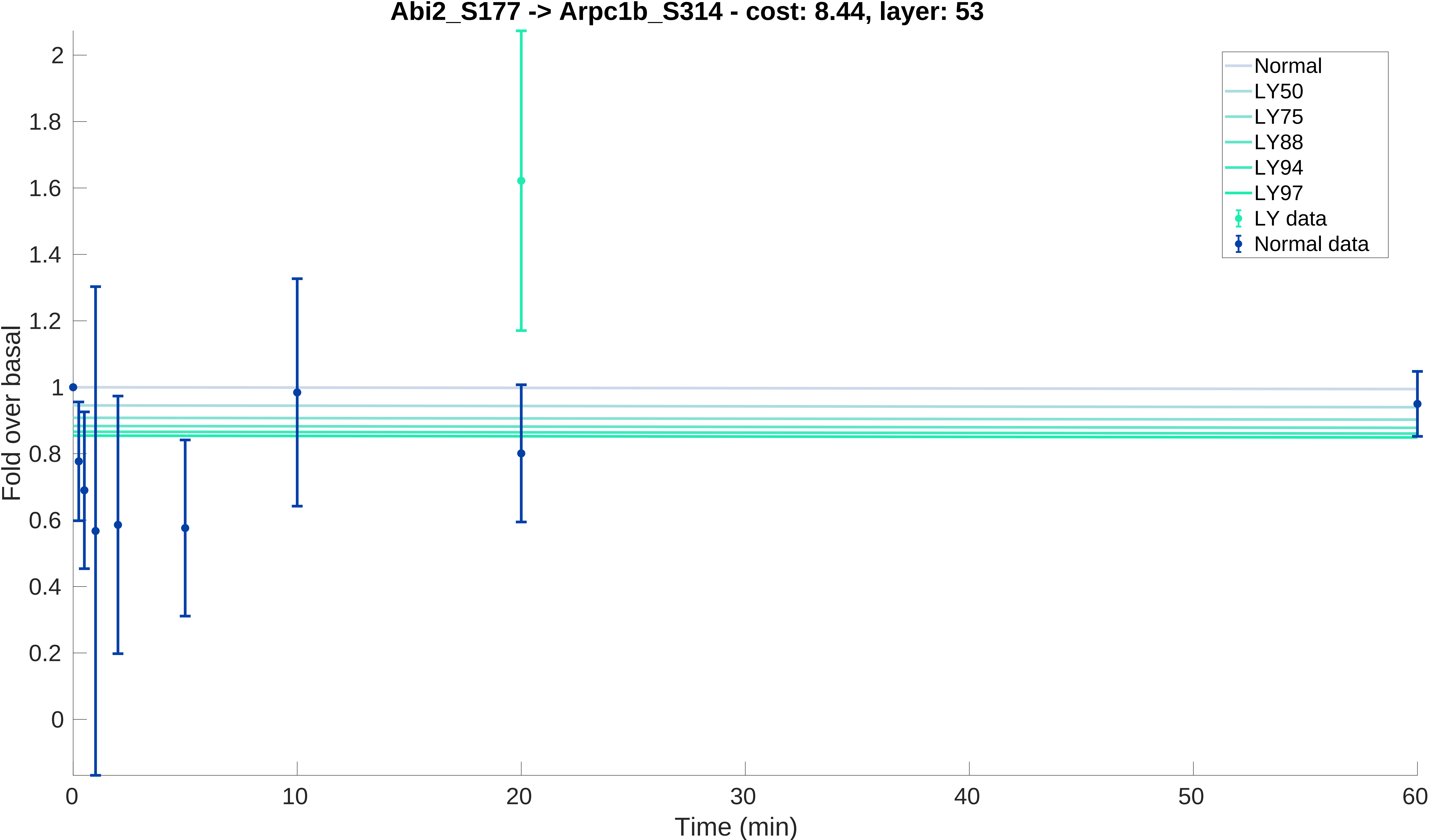
Abi2_S177 -> Arpc1b_S314 - cost: 8.44, layer: 53
Normal
LY50
LY75
LY88
LY94
LY97
LY data
Normal data
Fold over basal
Time (min)
0
0.2
0.4
0.6
0.8
1
1.2
1.4
1.6
1.8
2
0
10
20
30
40
50
60

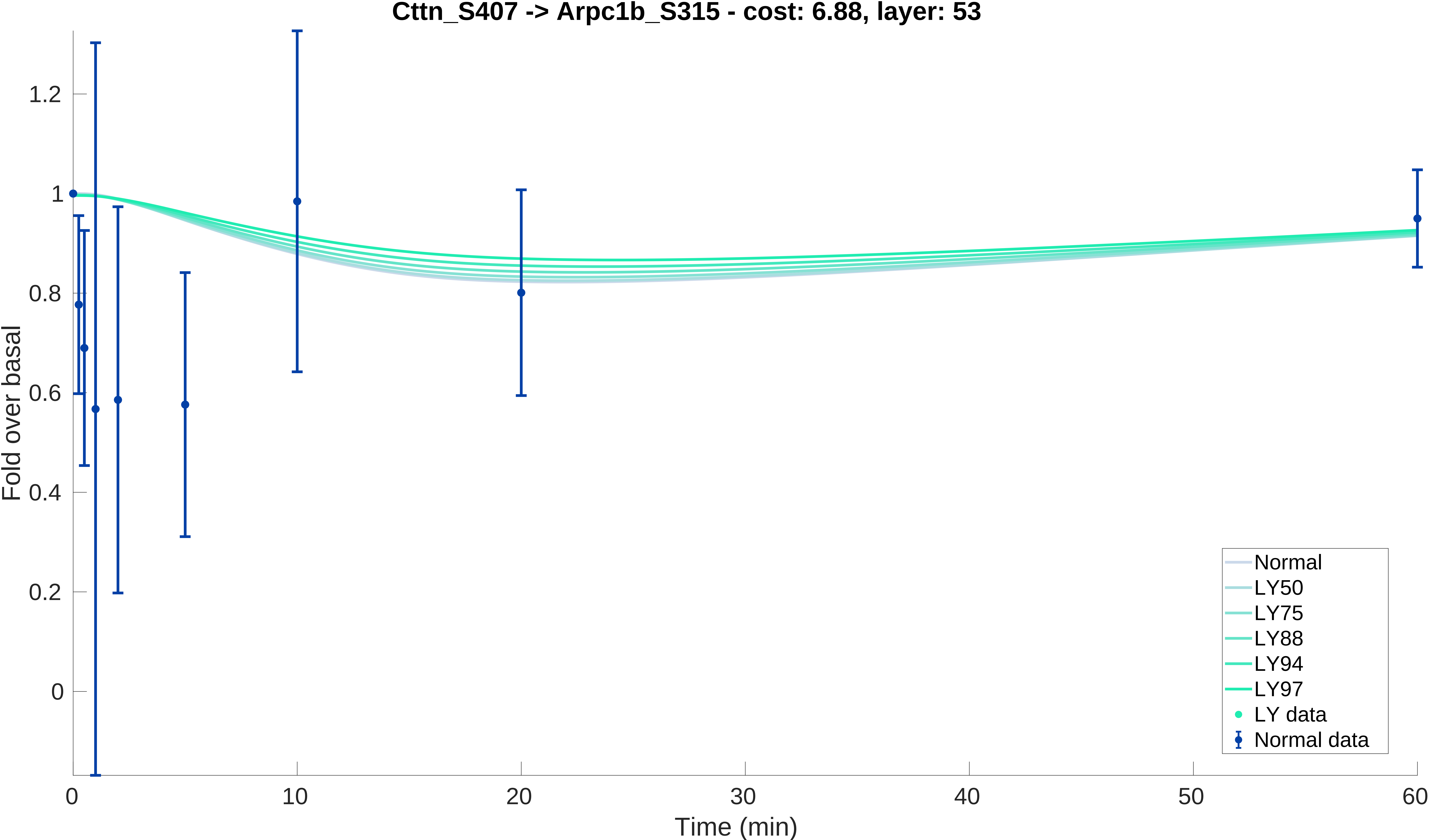

Cttn_S407 -> Arpc1b_S315 - cost: 6.88, layer: 53
Fold over basal
Time (min)
0
0.2
0.4
0.6
0.8
1
1.2
0
10
20
30
40
50
60
Normal
LY50
LY75
LY88
LY94
LY97
LY data
Normal data

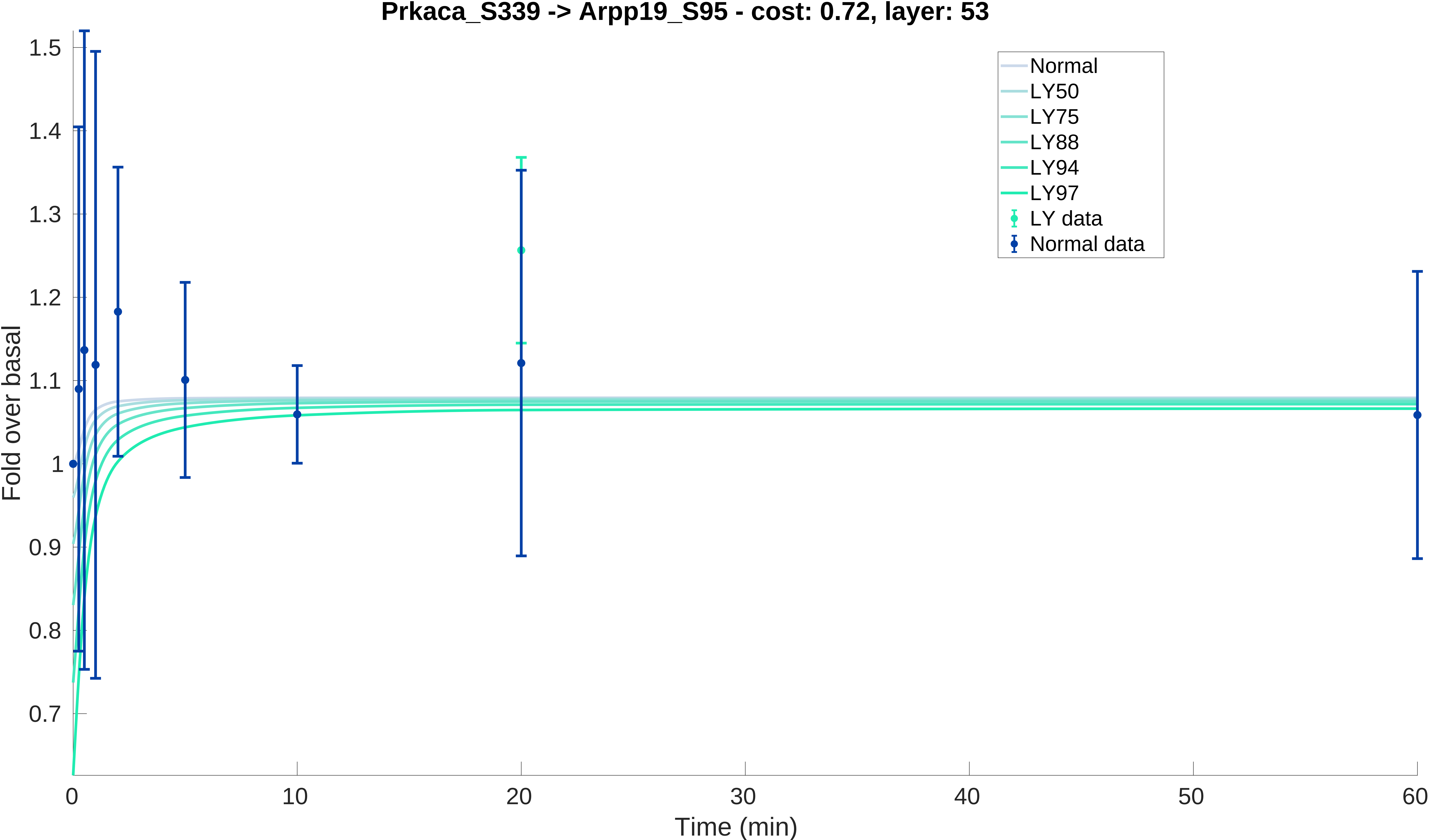

Prkaca_S339 -> Arpp19_S95 - cost: 0.72, layer: 53
Fold over basal
Time (min)
Normal
LY50
LY75
LY88
LY94
LY97
LY data
Normal data
1.5
1.4
1.3
1.2
1.1
1
0.9
0.8
0.7
0
10
20
30
40
50
60

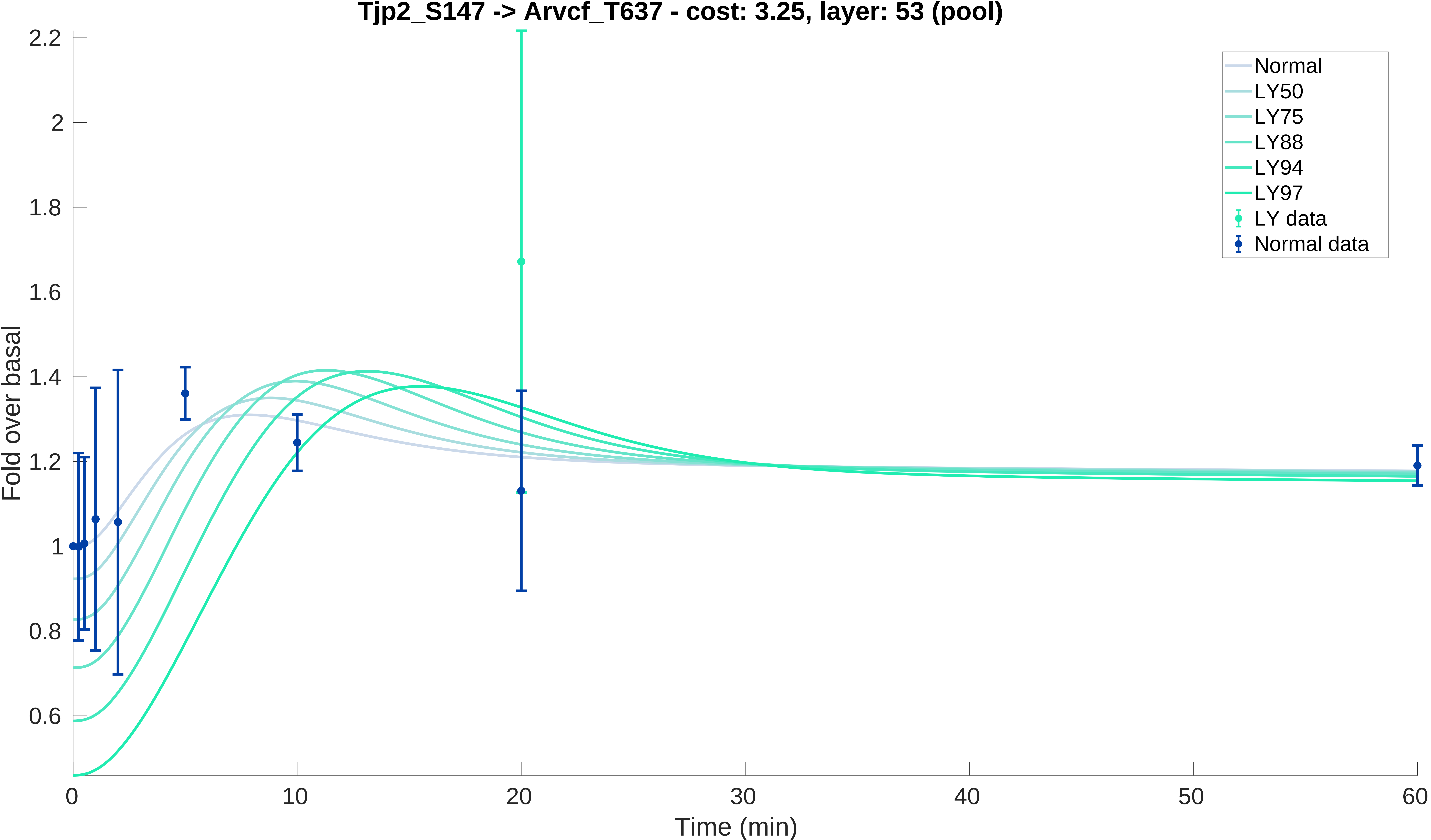

Tjp2_S147 -> Arvcf_T637 - cost: 3.25, layer: 53 (pool)
Fold over basal
Time (min)
2.2
2
1.8
1.6
1.4
1.2
1
0.8
0.6
0
10
20
30
40
50
60
Normal
LY50
LY75
LY88
LY94
LY97
LY data
Normal data

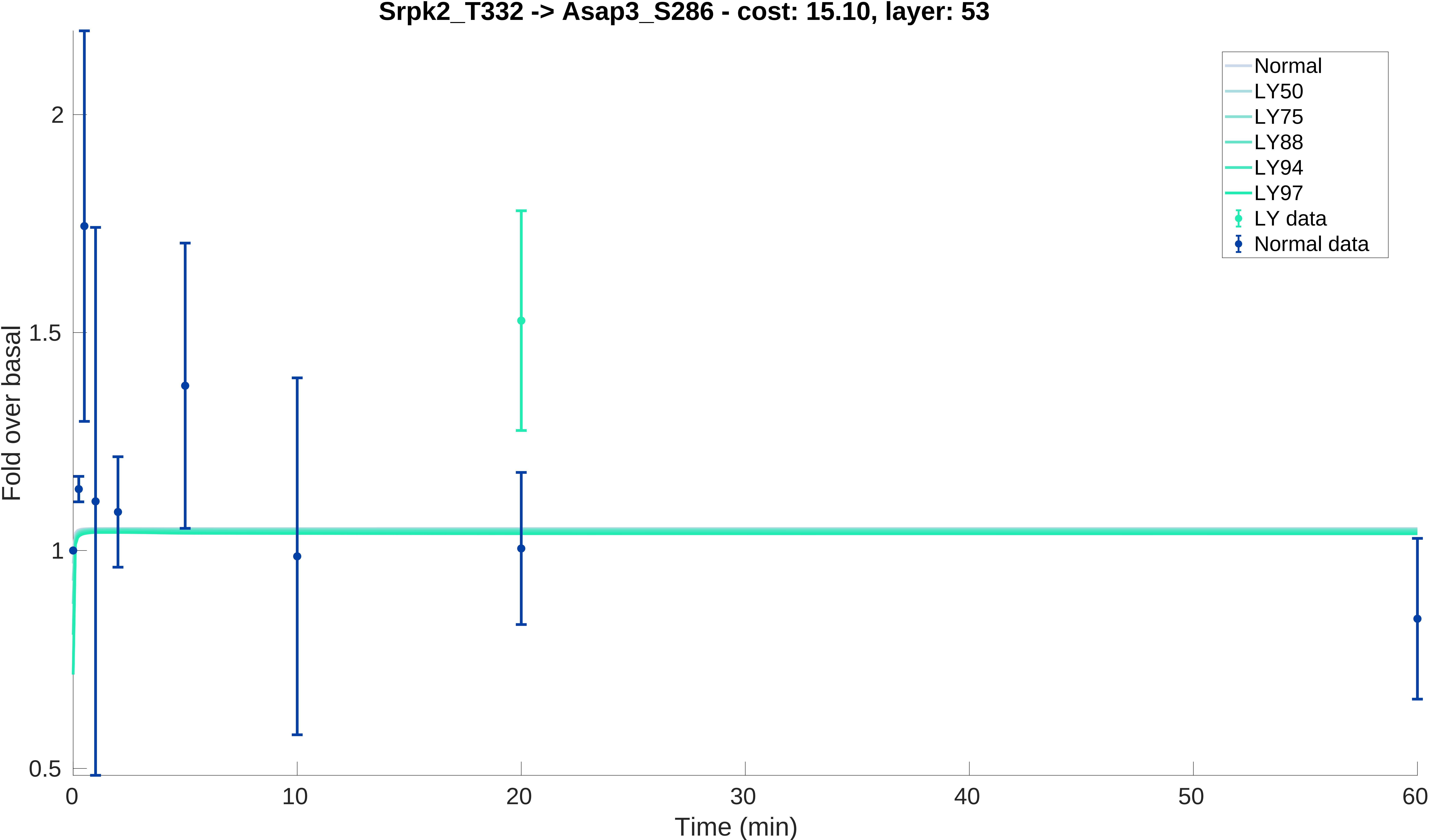

Srpk2_T332 -> Asap3_S286 - cost: 15.10, layer: 53
Fold over basal
Time (min)
0.5
1
1.5
2
0
10
20
30
40
50
60
Normal
LY50
LY75
LY88
LY94
LY97
LY data
Normal data

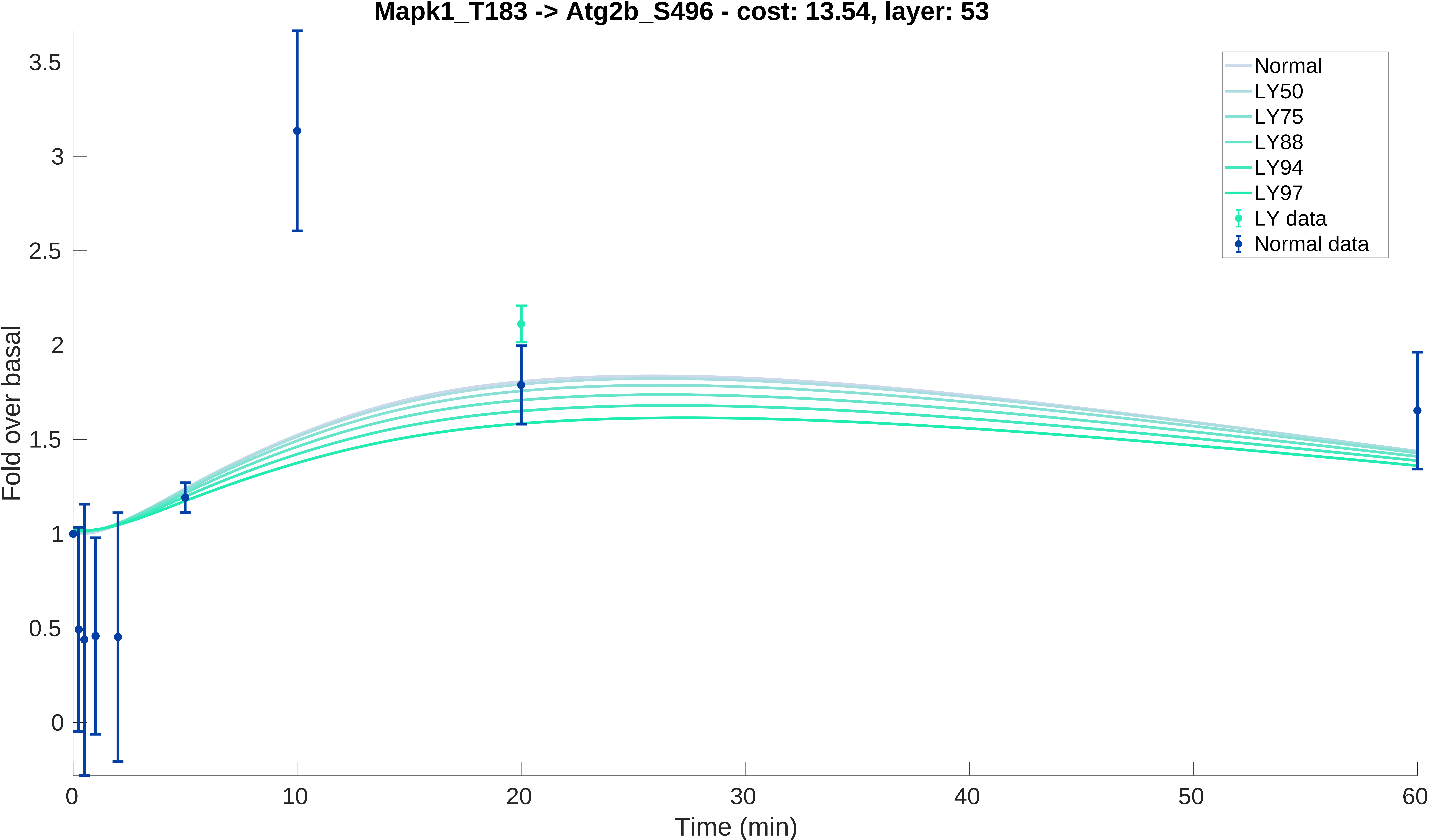

Mapk1_T183 -> Atg2b_S496 - cost: 13.54, layer: 53
Normal
LY50
LY75
LY88
LY94
LY97
LY data
Normal data
Fold over basal
3.5
3
2.5
2
1.5
1
0.5
0
0
10
20
30
40
50
60
Time (min)

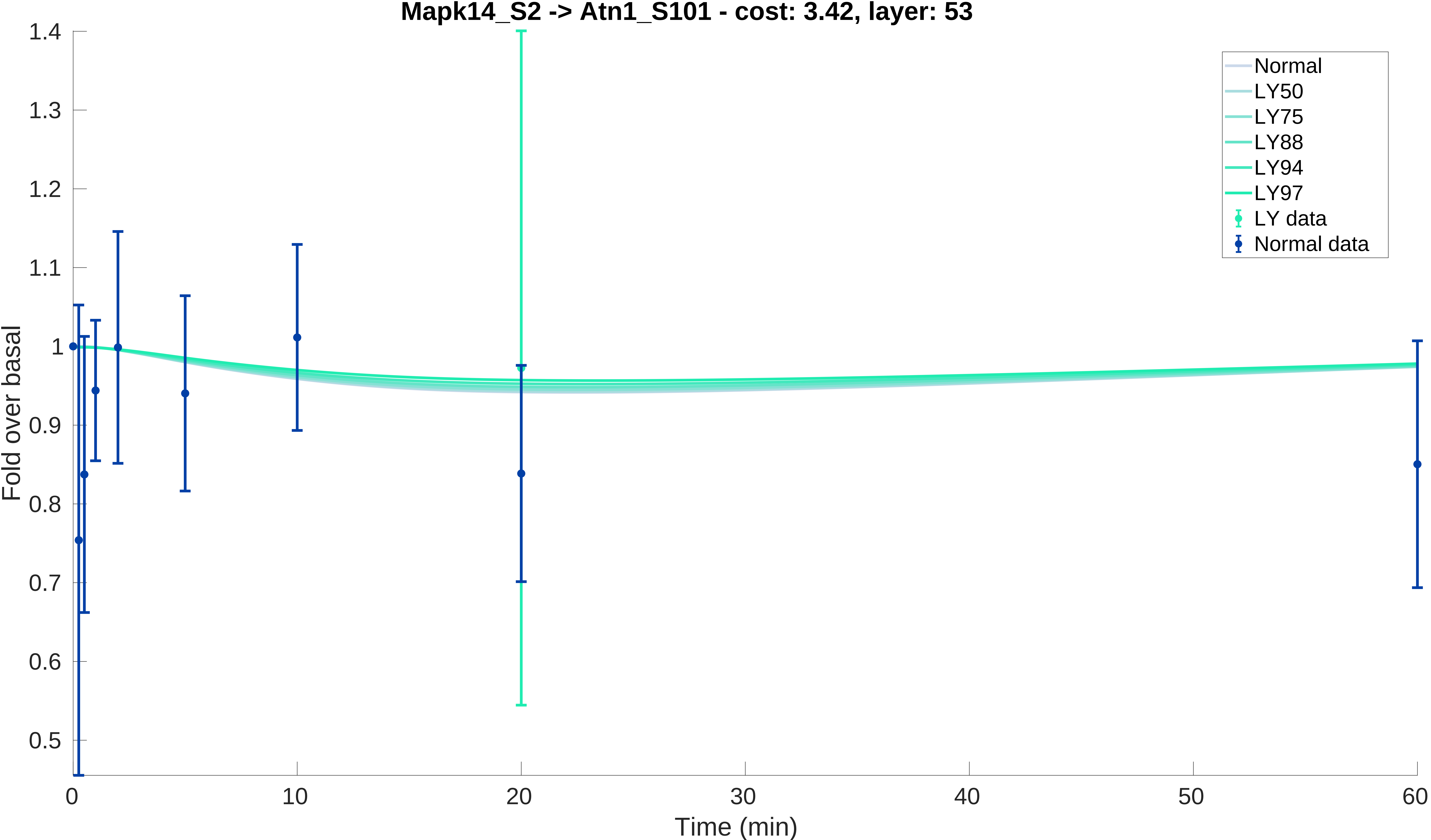

Mapk14_S2 -> Atn1_S101 - cost: 3.42, layer: 53
Fold over basal
Time (min)
0.5
0.6
0.7
0.8
0.9
1
1.1
1.2
1.3
1.4
0
10
20
30
40
50
60
Normal
LY50
LY75
LY88
LY94
LY97
LY data
Normal data

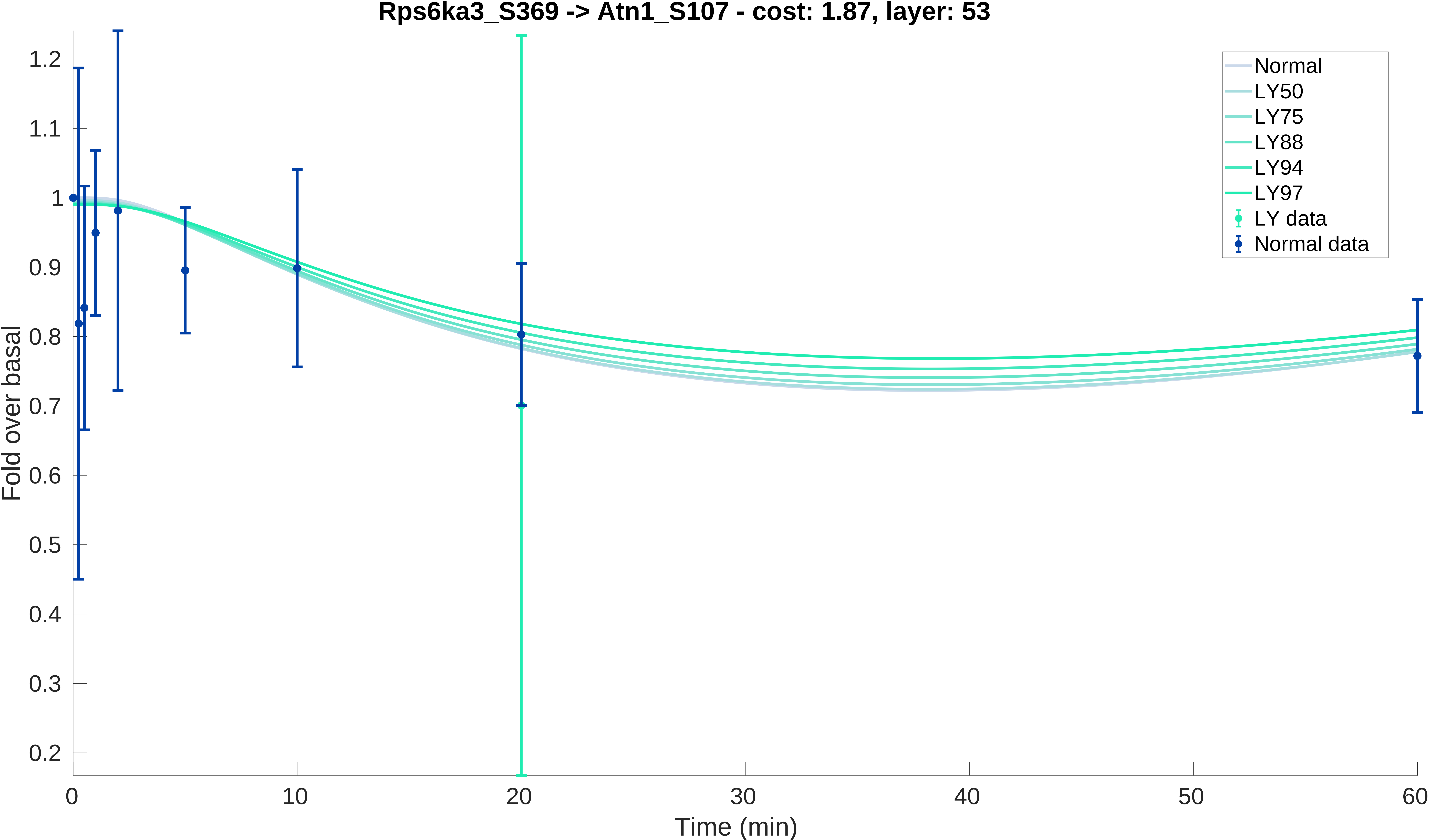

Rps6ka3_S369 -> Atn1_S107 - cost: 1.87, layer: 53
Fold over basal
Time (min)
0
10
20
30
40
50
60
0.2
0.3
0.4
0.5
0.6
0.7
0.8
0.9
1
1.1
1.2
Normal
LY50
LY75
LY88
LY94
LY97
LY data
Normal data

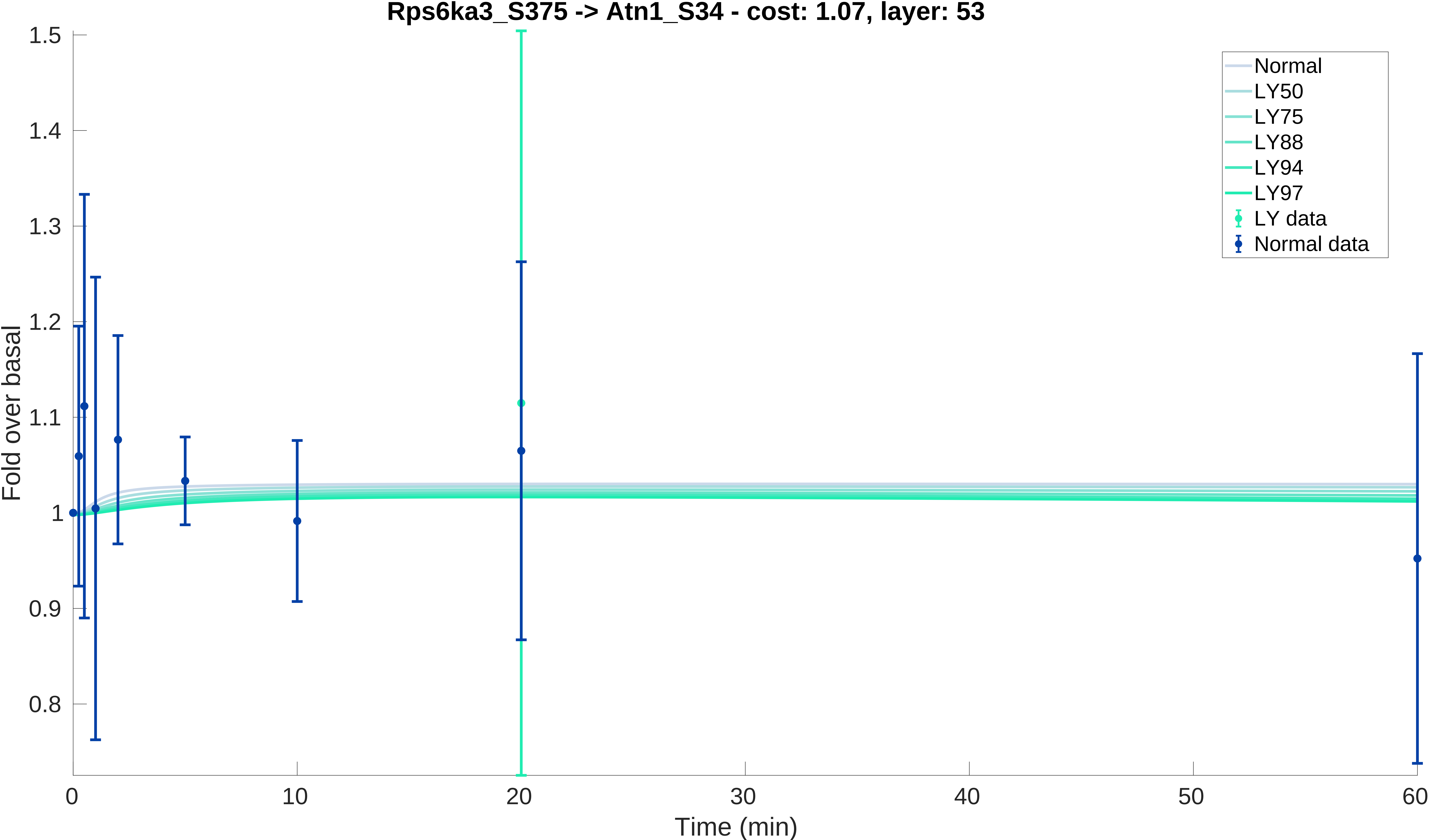

Rps6ka3_S375 -> Atn1_S34 - cost: 1.07, layer: 53
Fold over basal
Time (min)
Normal
LY50
LY75
LY88
LY94
LY97
LY data
Normal data
1.5
1.4
1.3
1.2
1.1
1
0.9
0.8
0
10
20
30
40
50
60

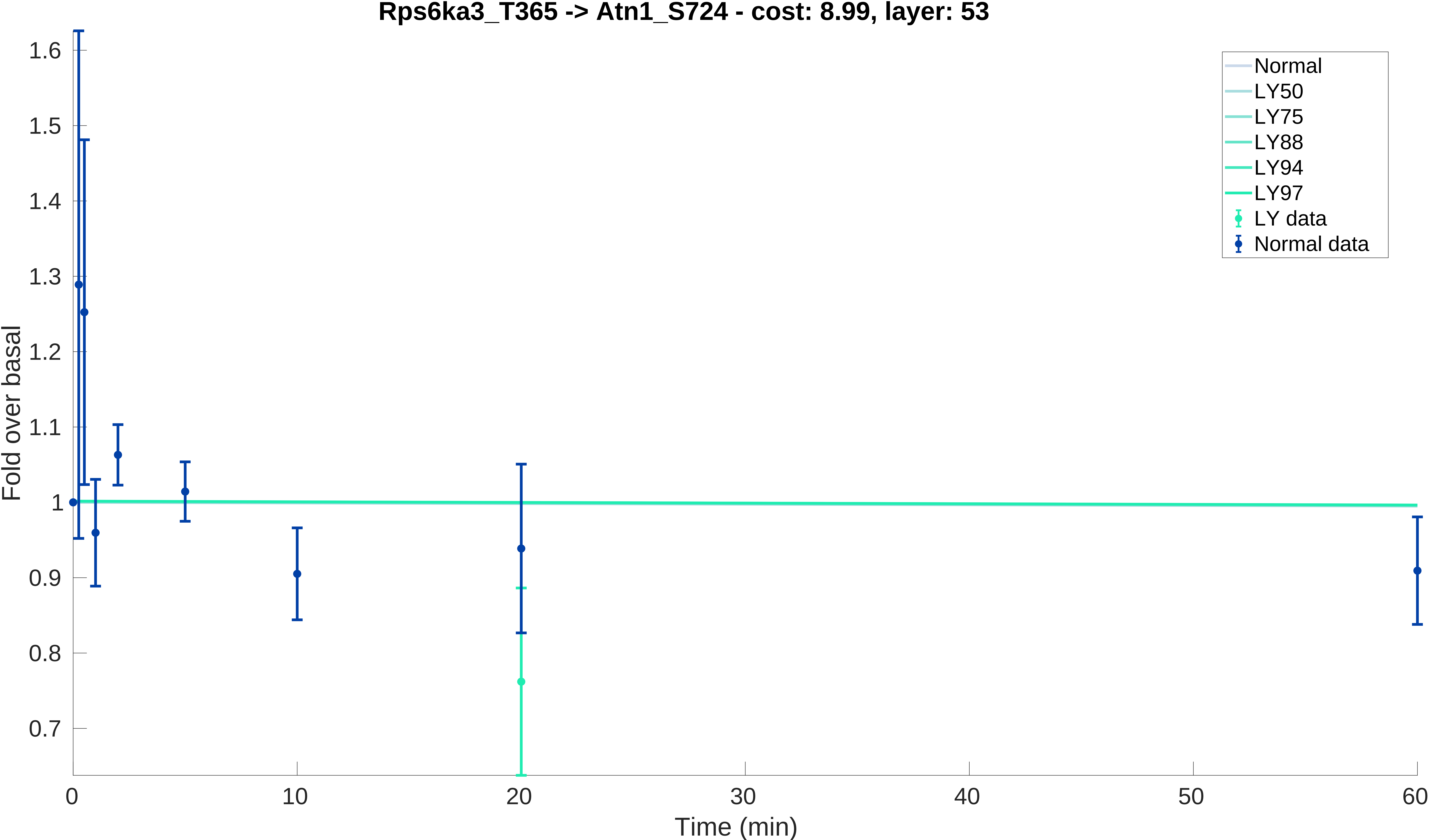

Rps6ka3_T365 -> Atn1_S724 - cost: 8.99, layer: 53
Fold over basal
1.6
1.5
1.4
1.3
1.2
1.1
1
0.9
0.8
0.7
0
10
20
30
40
50
60
Time (min)
Normal
LY50
LY75
LY88
LY94
LY97
LY data
Normal data

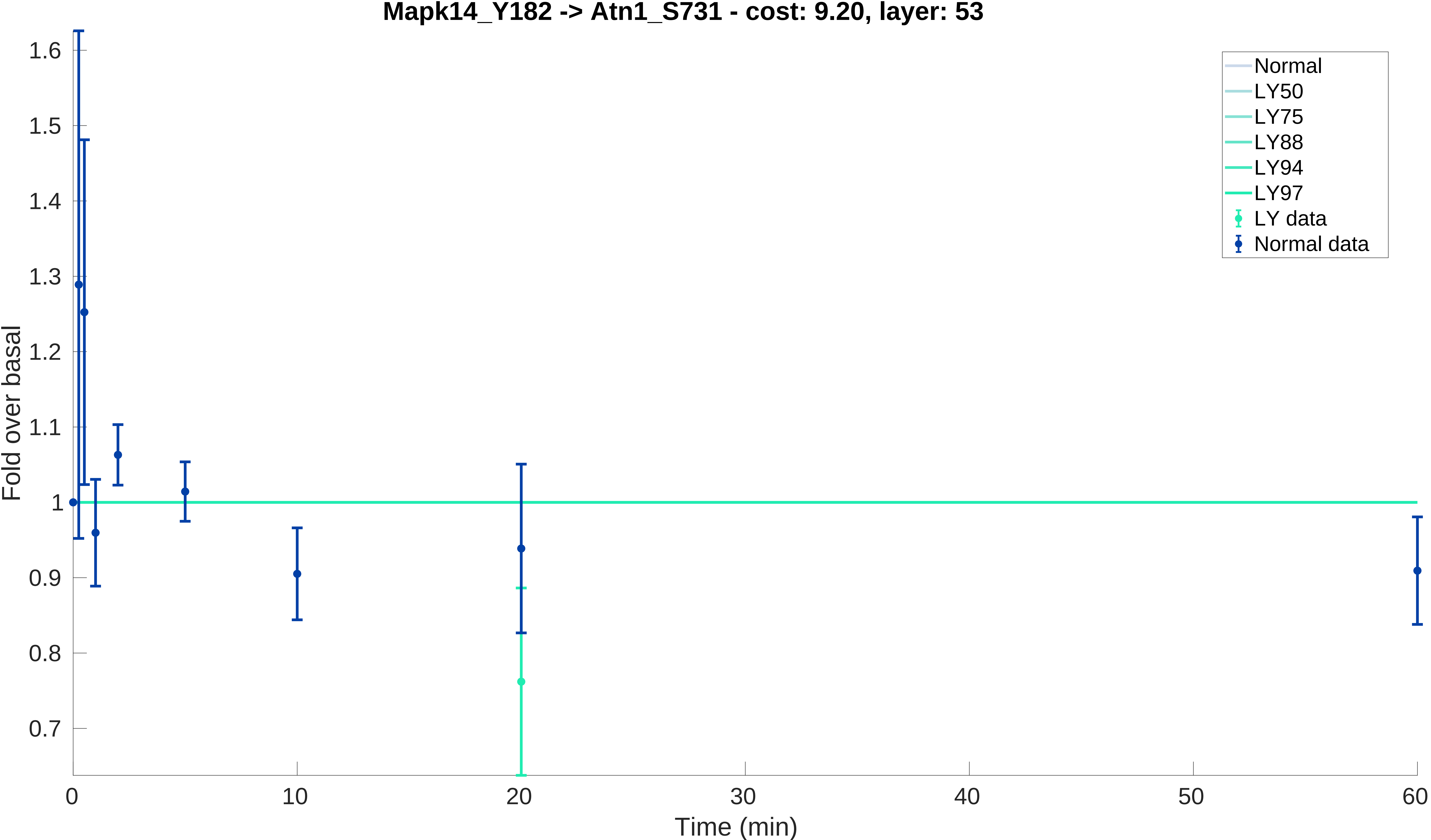

Mapk14_Y182 -> Atn1_S731 - cost: 9.20, layer: 53
Fold over basal
1.6
1.5
1.4
1.3
1.2
1.1
1
0.9
0.8
0.7
0
10
20
30
40
50
60
Time (min)
Normal
LY50
LY75
LY88
LY94
LY97
LY data
Normal data

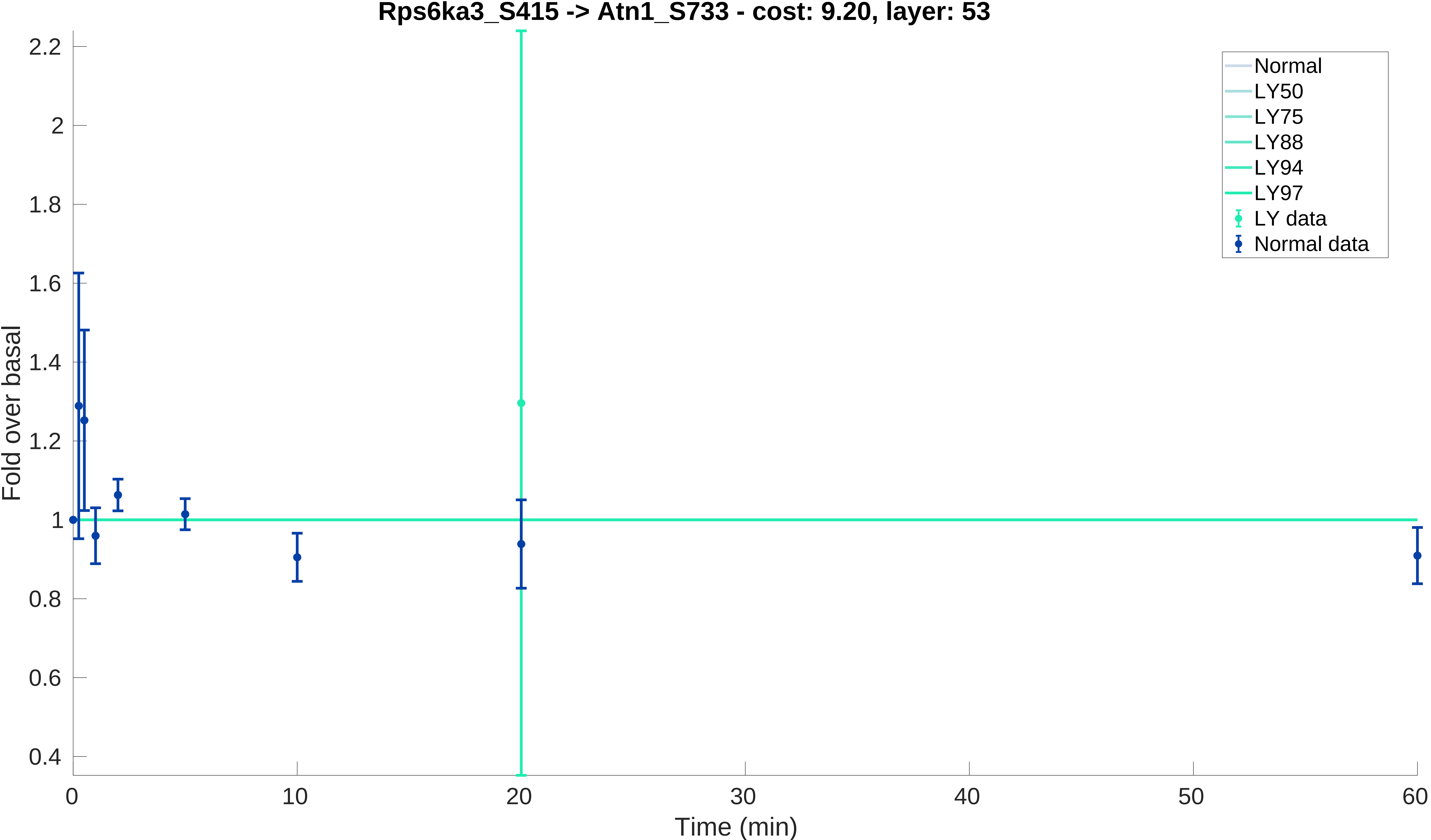

Rps6ka3_S415 -> Atn1_S733 - cost: 9.20, layer: 53
Fold over basal
Time (min)
2.2
2
1.8
1.6
1.4
1.2
1
0.8
0.6
0.4
0
10
20
30
40
50
60
Normal
LY50
LY75
LY88
LY94
LY97
LY data
Normal data

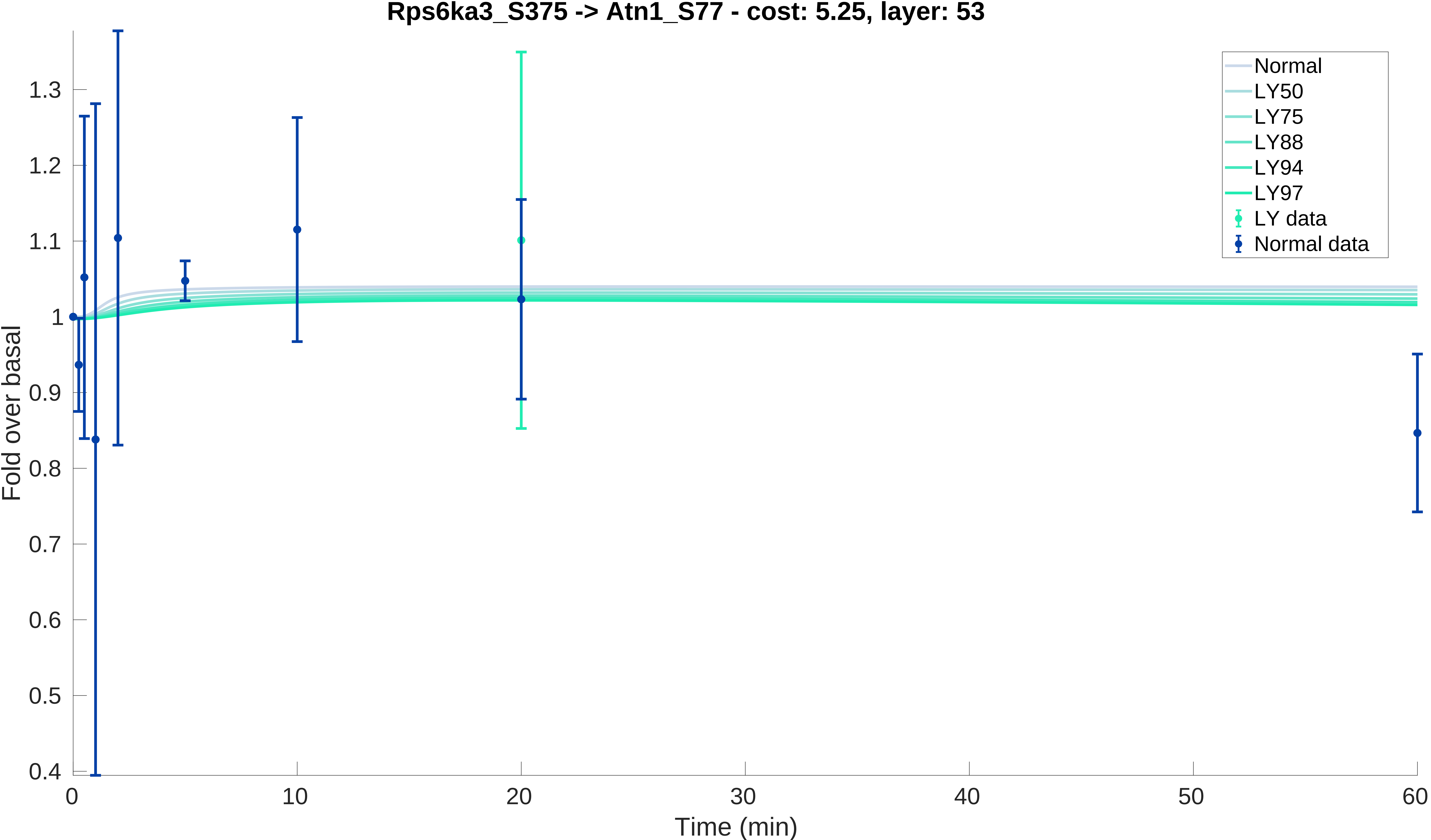

Rps6ka3_S375 -> Atn1_S77 - cost: 5.25, layer: 53
Fold over basal
1.3
1.2
1.1
1
0.9
0.8
0.7
0.6
0.5
0.4
0
10
20
30
40
50
60
Time (min)
Normal
LY50
LY75
LY88
LY94
LY97
LY data
Normal data

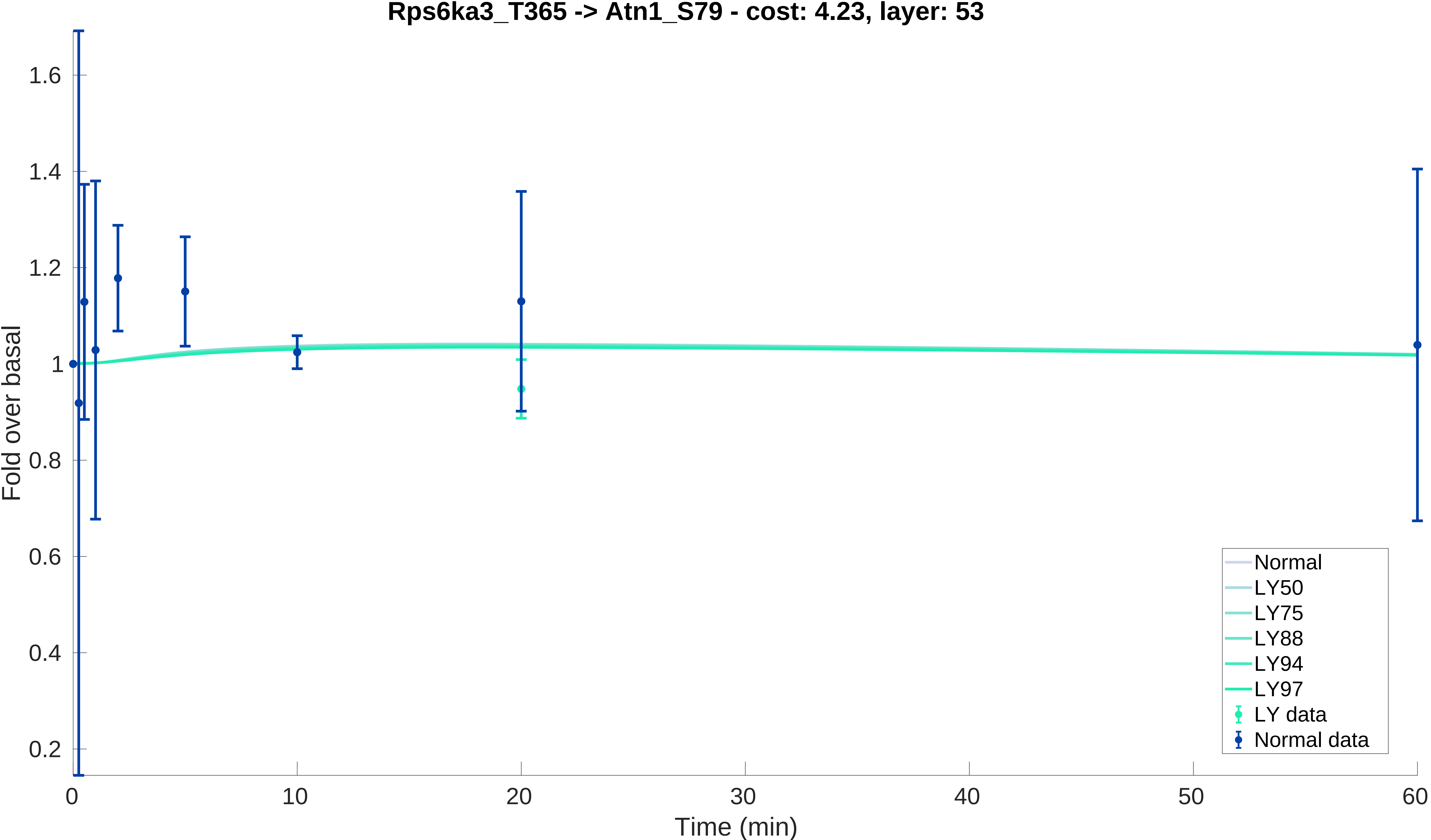

Rps6ka3_T365 -> Atn1_S79 - cost: 4.23, layer: 53
Fold over basal
Time (min)
0.2
0.4
0.6
0.8
1
1.2
1.4
1.6
0
10
20
30
40
50
60
Normal
LY50
LY75
LY88
LY94
LY97
LY data
Normal data

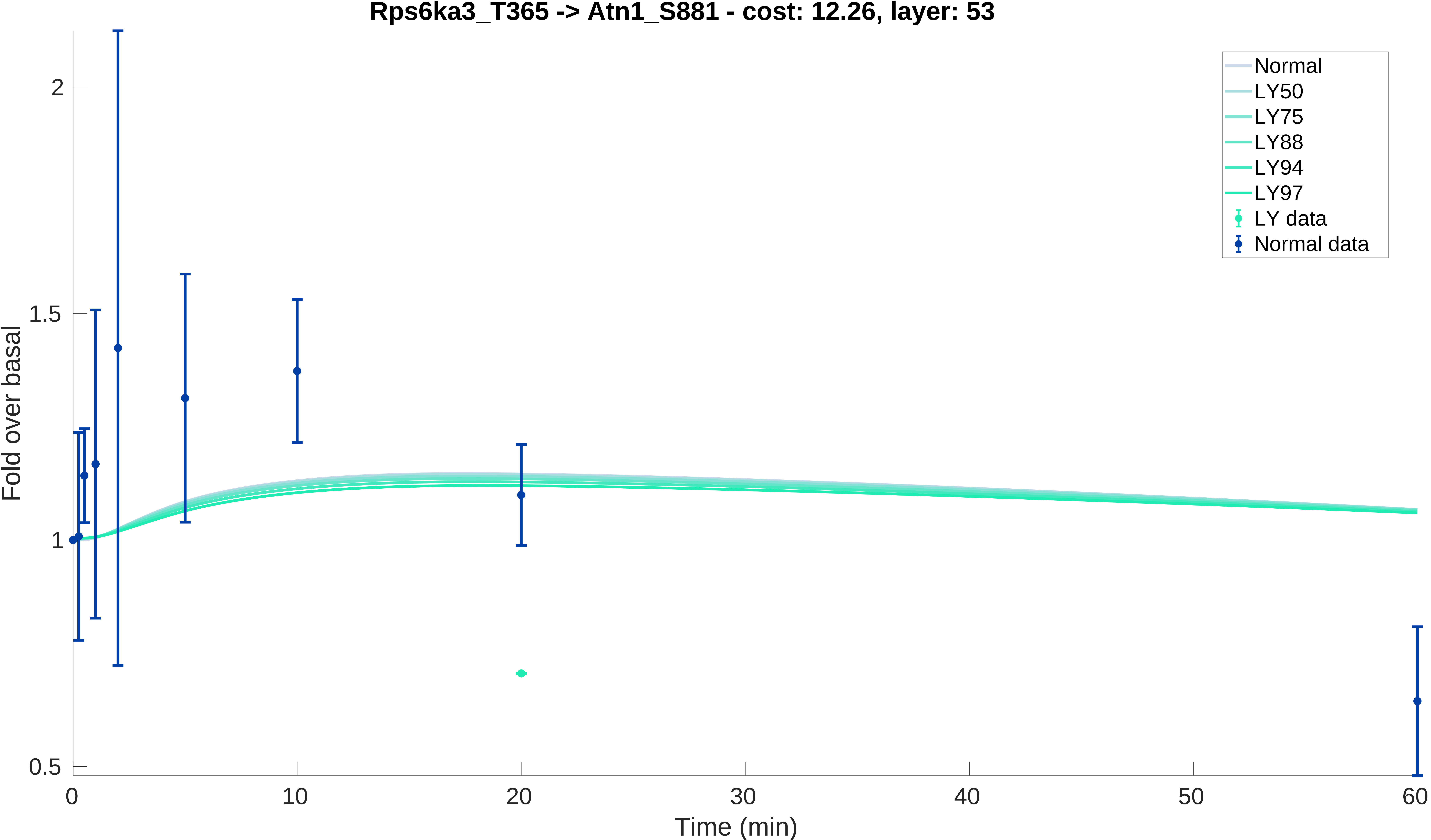
Rps6ka3_T365 -> Atn1_S881 - cost: 12.26, layer: 53
Fold over basal
Time (min)
2
1.5
1
0.5
0
10
20
30
40
50
60
Normal
LY50
LY75
LY88
LY94
LY97
LY data
Normal data

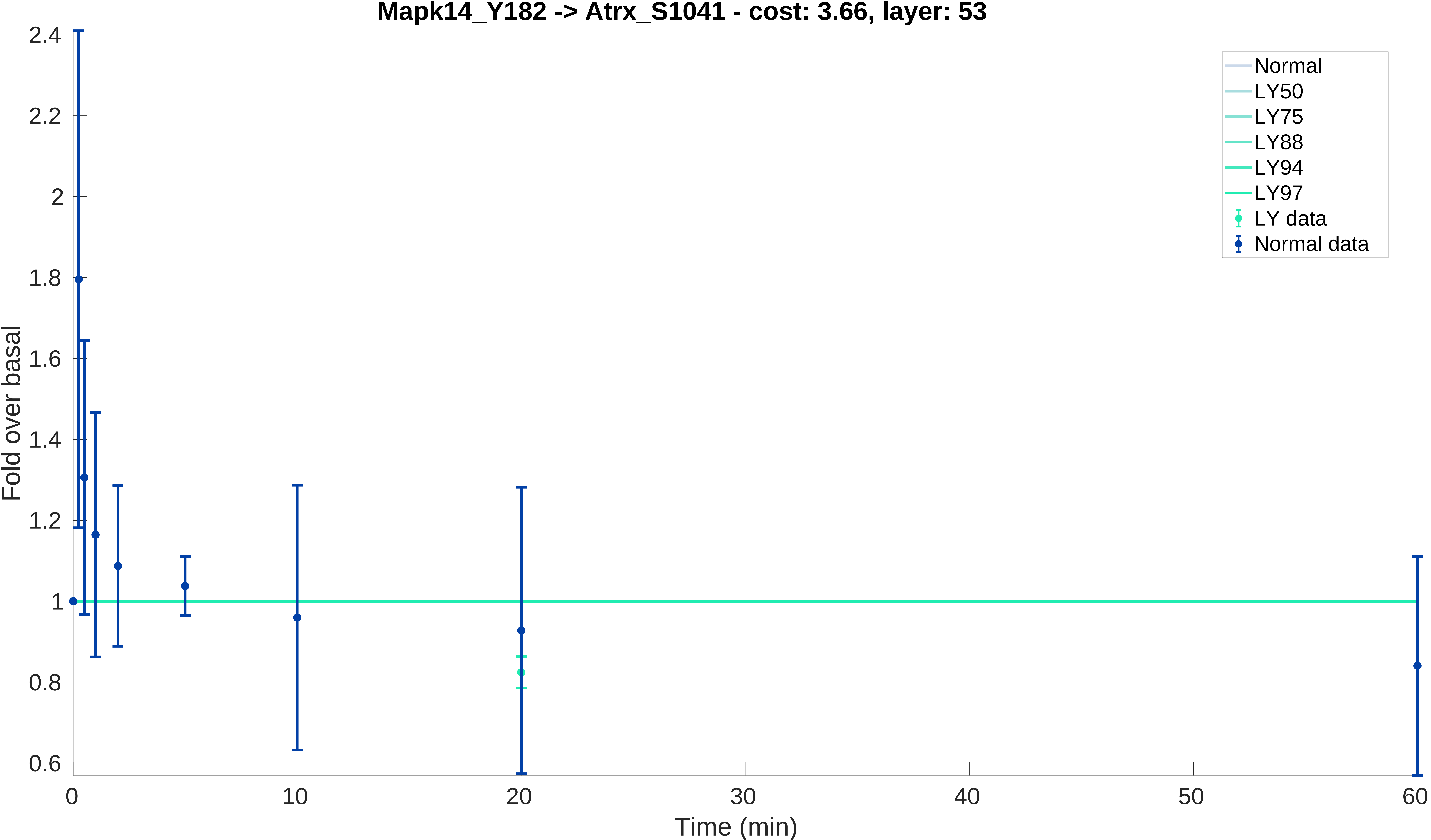

Mapk14_Y182 -> Atrx_S1041 - cost: 3.66, layer: 53
Fold over basal
Time (min)
2.4
2.2
2
1.8
1.6
1.4
1.2
1
0.8
0.6
0
10
20
30
40
50
60
Normal
LY50
LY75
LY88
LY94
LY97
LY data
Normal data

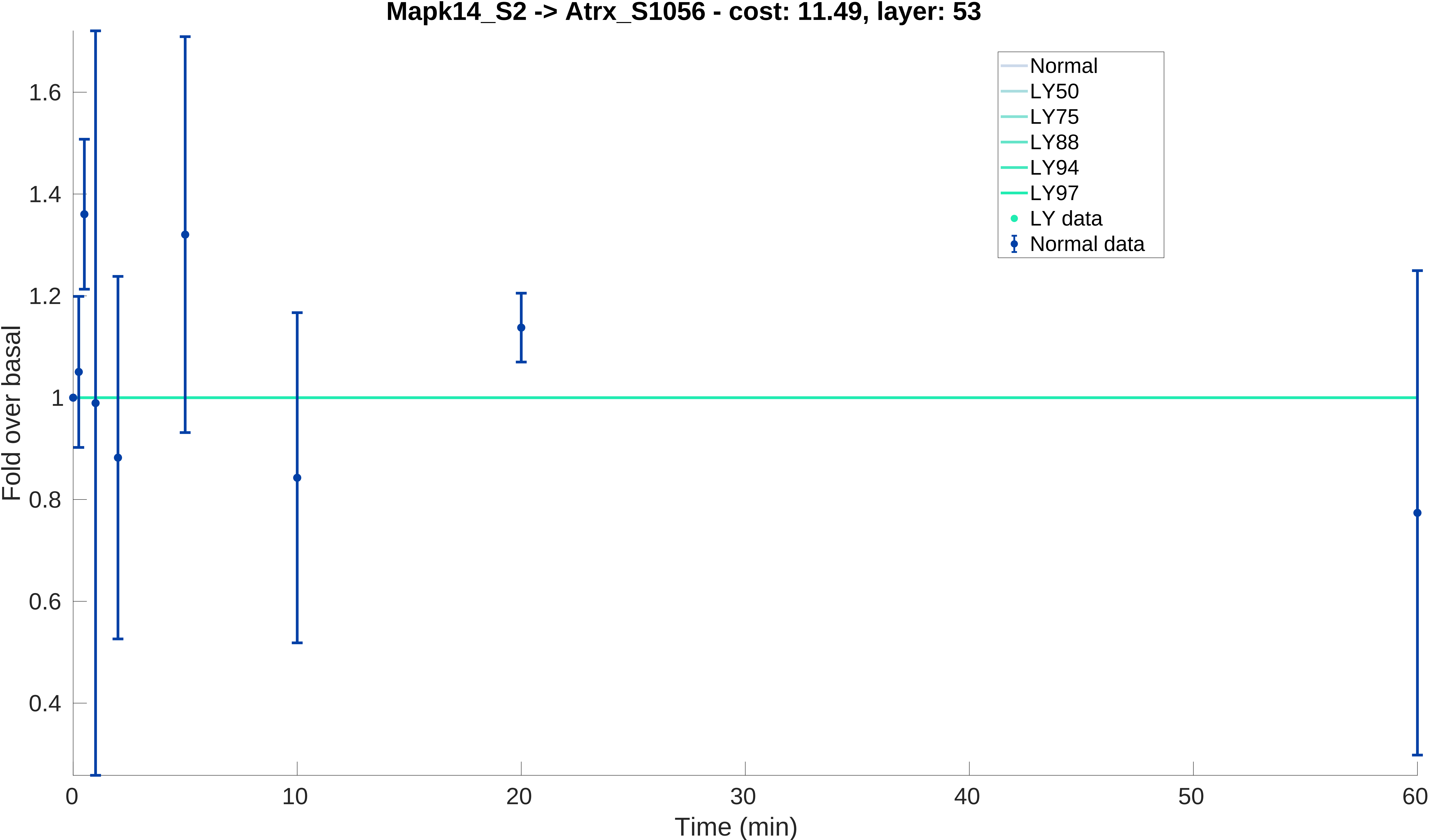

Mapk14_S2 -> Atrx_S1056 - cost: 11.49, layer: 53
Normal
LY50
LY75
LY88
LY94
LY97
LY data
Normal data
Fold over basal
1.6
1.4
1.2
1
0.8
0.6
0.4
0
10
20
30
40
50
60
Time (min)

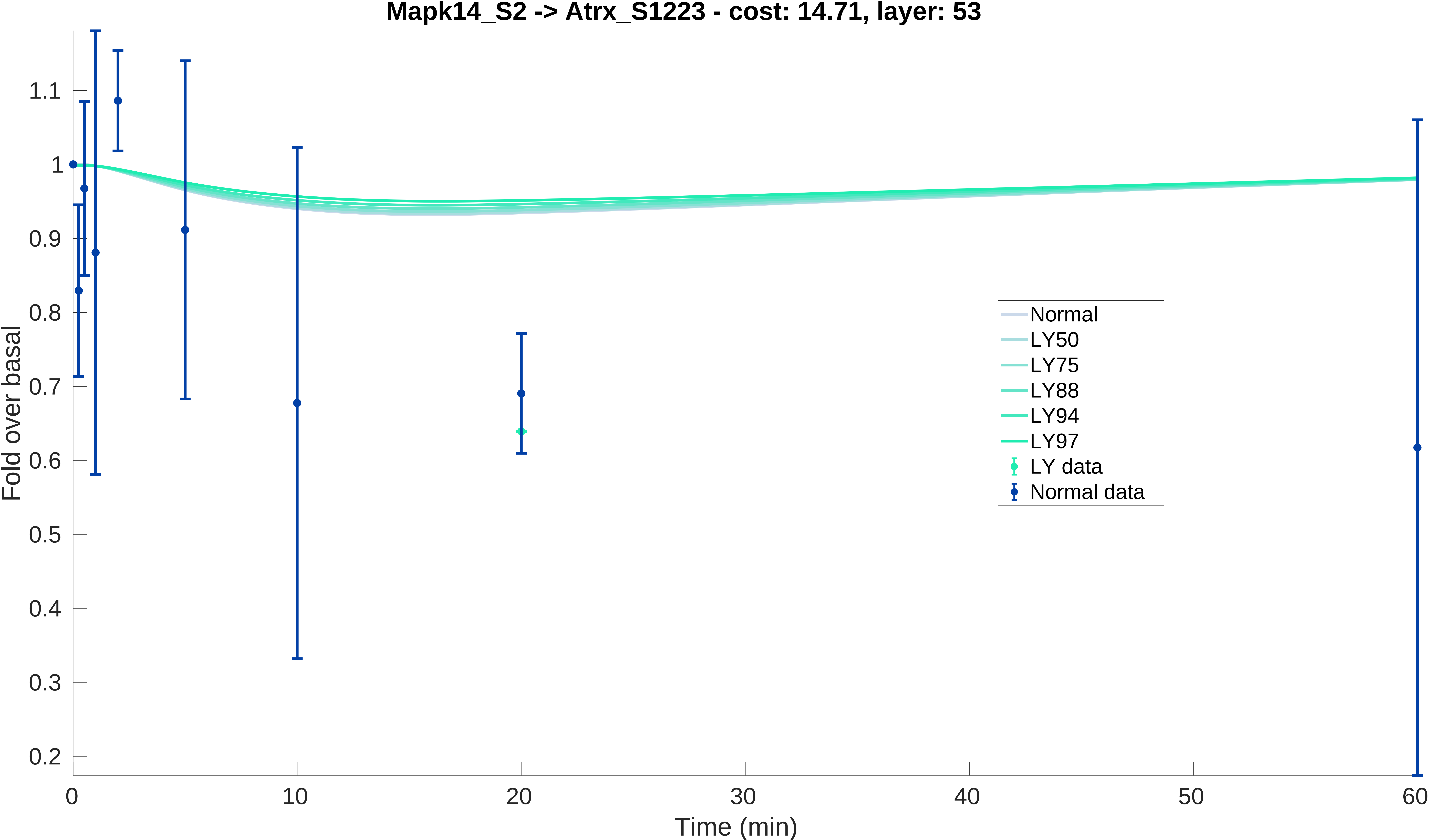

Mapk14_S2 -> Atrx_S1223 - cost: 14.71, layer: 53
Fold over basal
1.1
1
0.9
0.8
0.7
0.6
0.5
0.4
0.3
0.2
0
10
20
30
40
50
60
Time (min)
Normal
LY50
LY75
LY88
LY94
LY97
LY data
Normal data

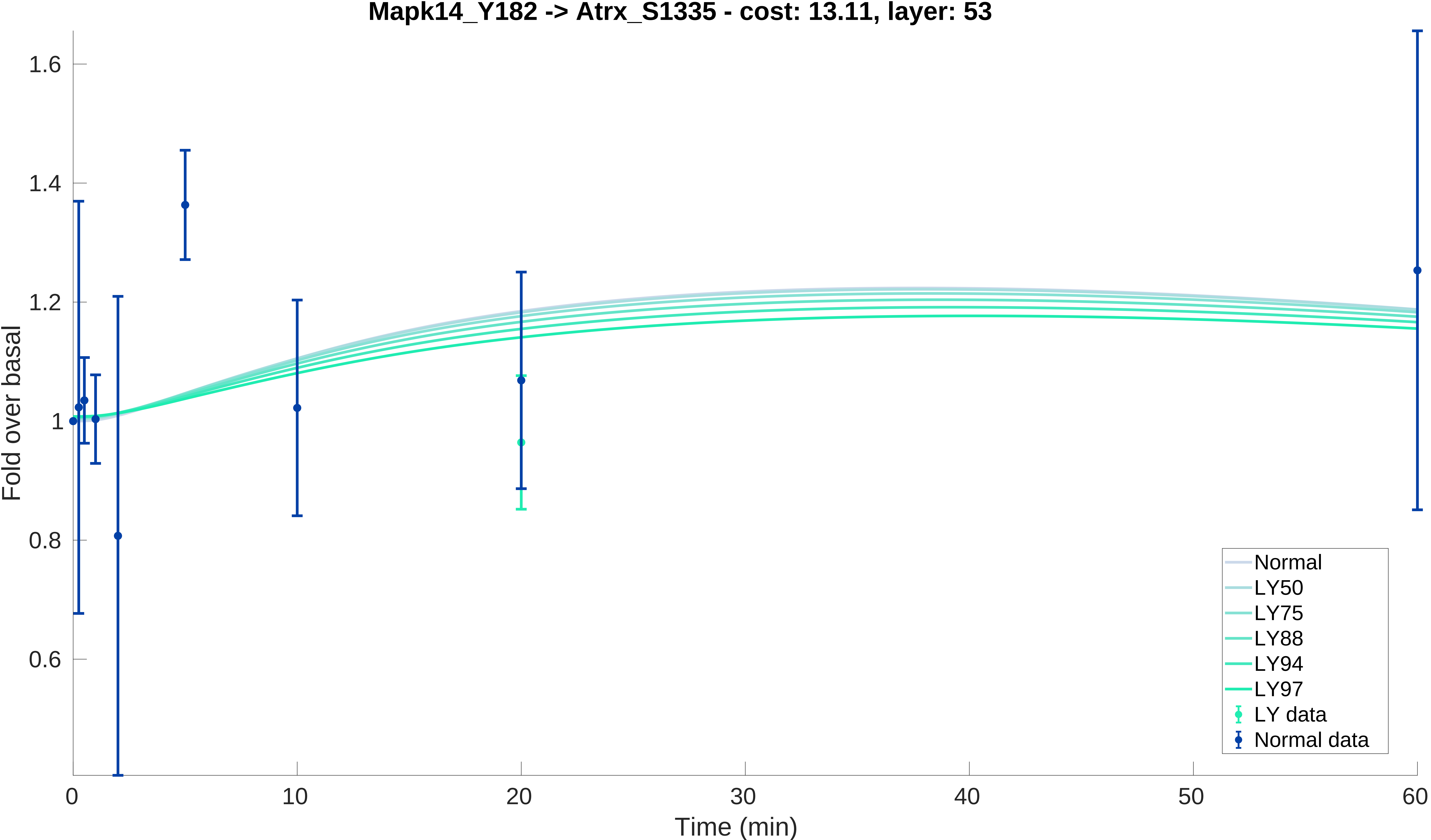

Mapk14_Y182 -> Atrx_S1335 - cost: 13.11, layer: 53
Fold over basal
1.6
1.4
1.2
1
0.8
0.6
0
10
20
30
40
50
60
Time (min)
Normal
LY50
LY75
LY88
LY94
LY97
LY data
Normal data

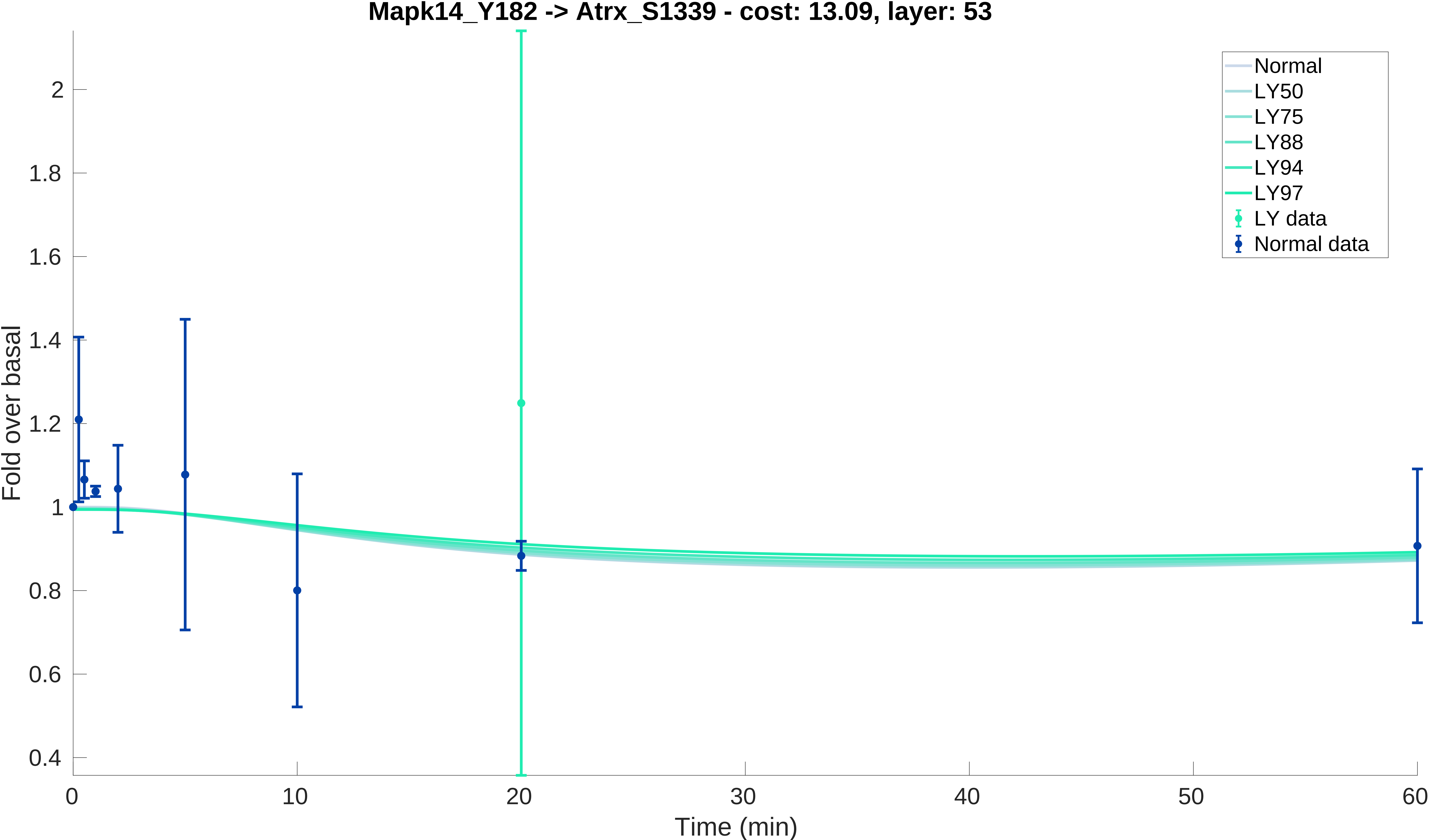

Mapk14_Y182 -> Atrx_S1339 - cost: 13.09, layer: 53
Fold over basal
Time (min)
0.4
0.6
0.8
1
1.2
1.4
1.6
1.8
2
0
10
20
30
40
50
60
Normal
LY50
LY75
LY88
LY94
LY97
LY data
Normal data

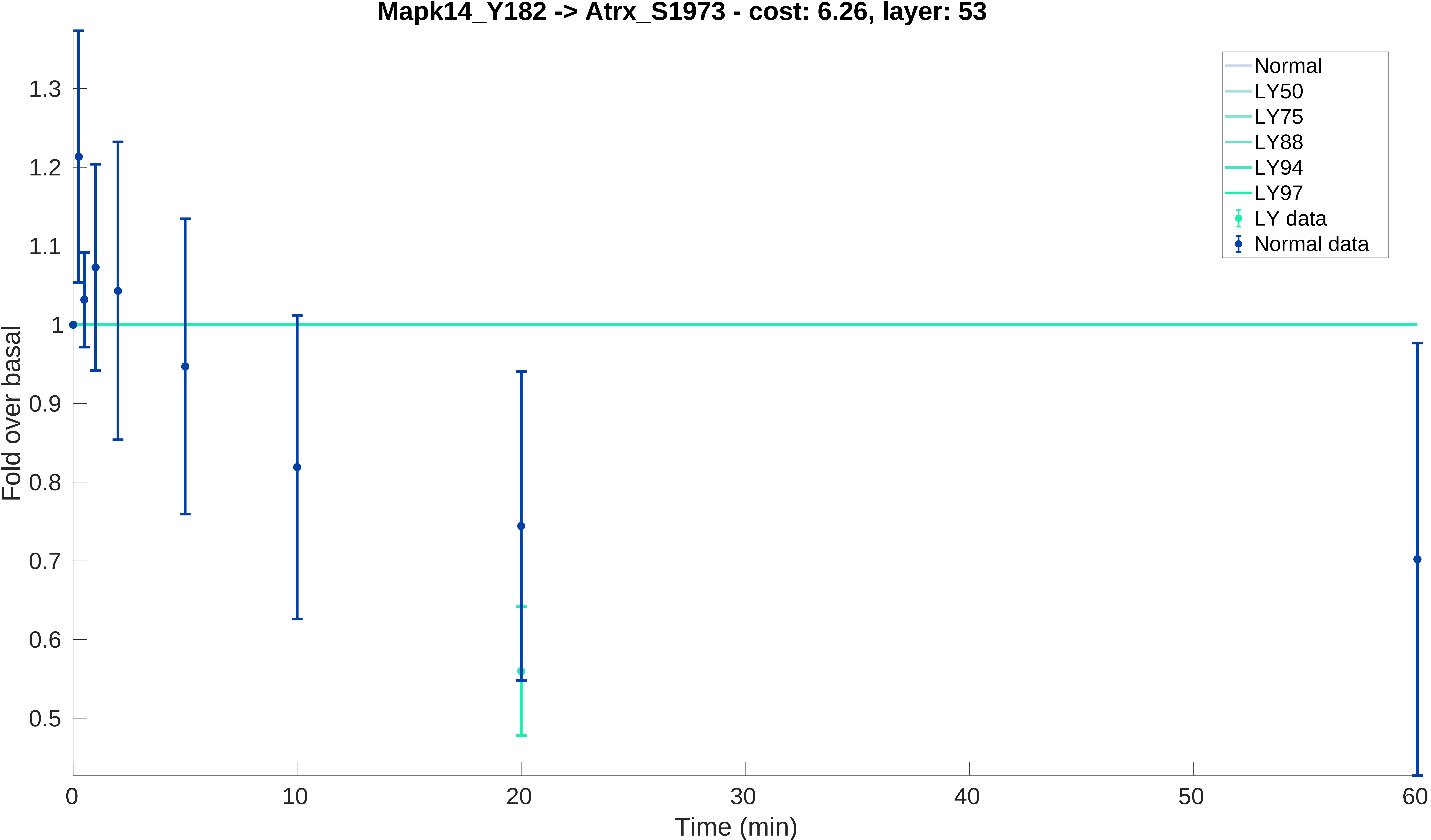
Mapk14_Y182 -> Atrx_S1973 - cost: 6.26, layer: 53
Fold over basal
Time (min)
1.3
1.2
1.1
1
0.9
0.8
0.7
0.6
0.5
0
10
20
30
40
50
60
Normal
LY50
LY75
LY88
LY94
LY97
LY data
Normal data

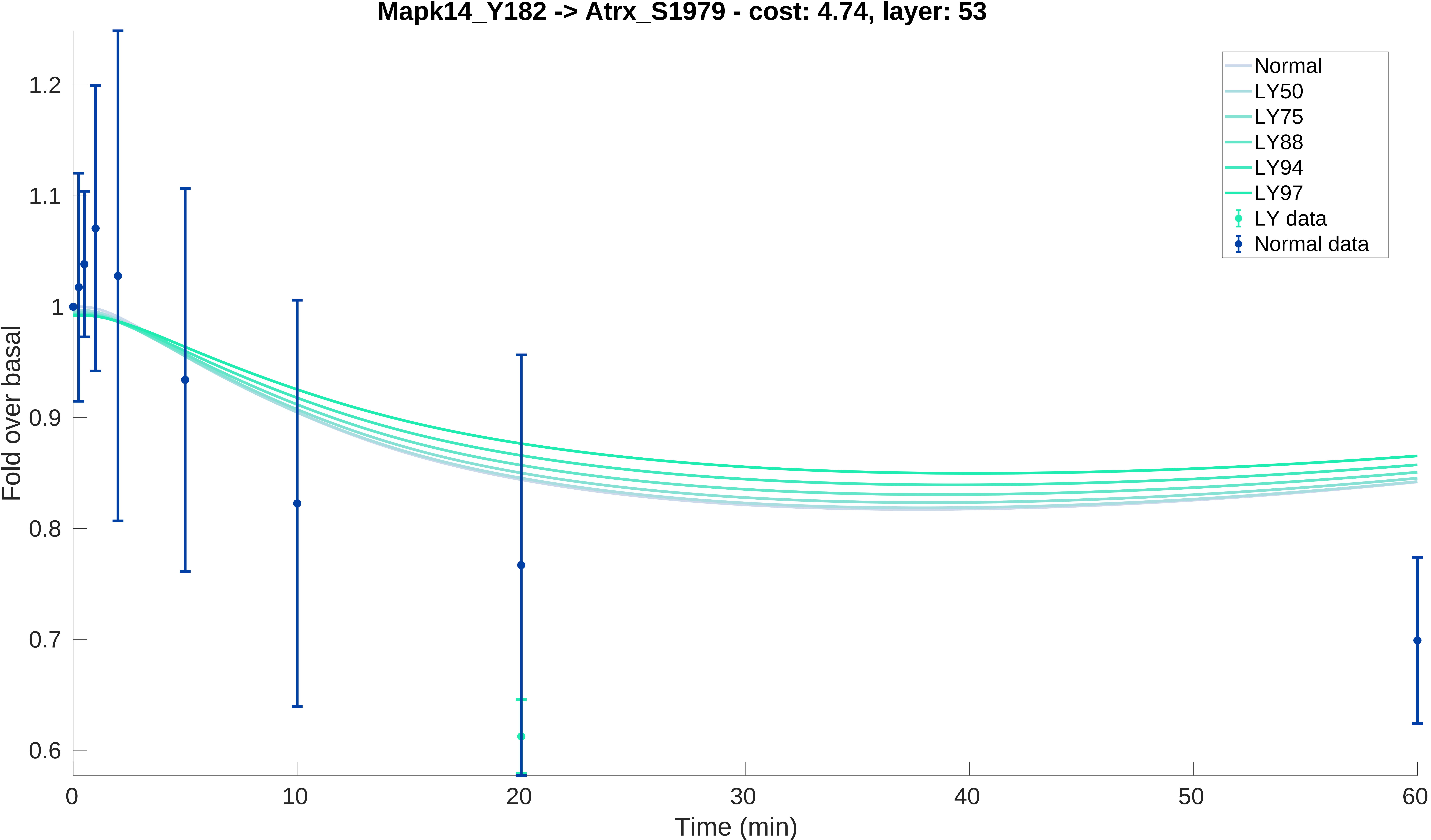

Mapk14_Y182 -> Atrx_S1979 - cost: 4.74, layer: 53
Fold over basal
Time (min)
0.6
0.7
0.8
0.9
1
1.1
1.2
0
10
20
30
40
50
60
Normal
LY50
LY75
LY88
LY94
LY97
LY data
Normal data

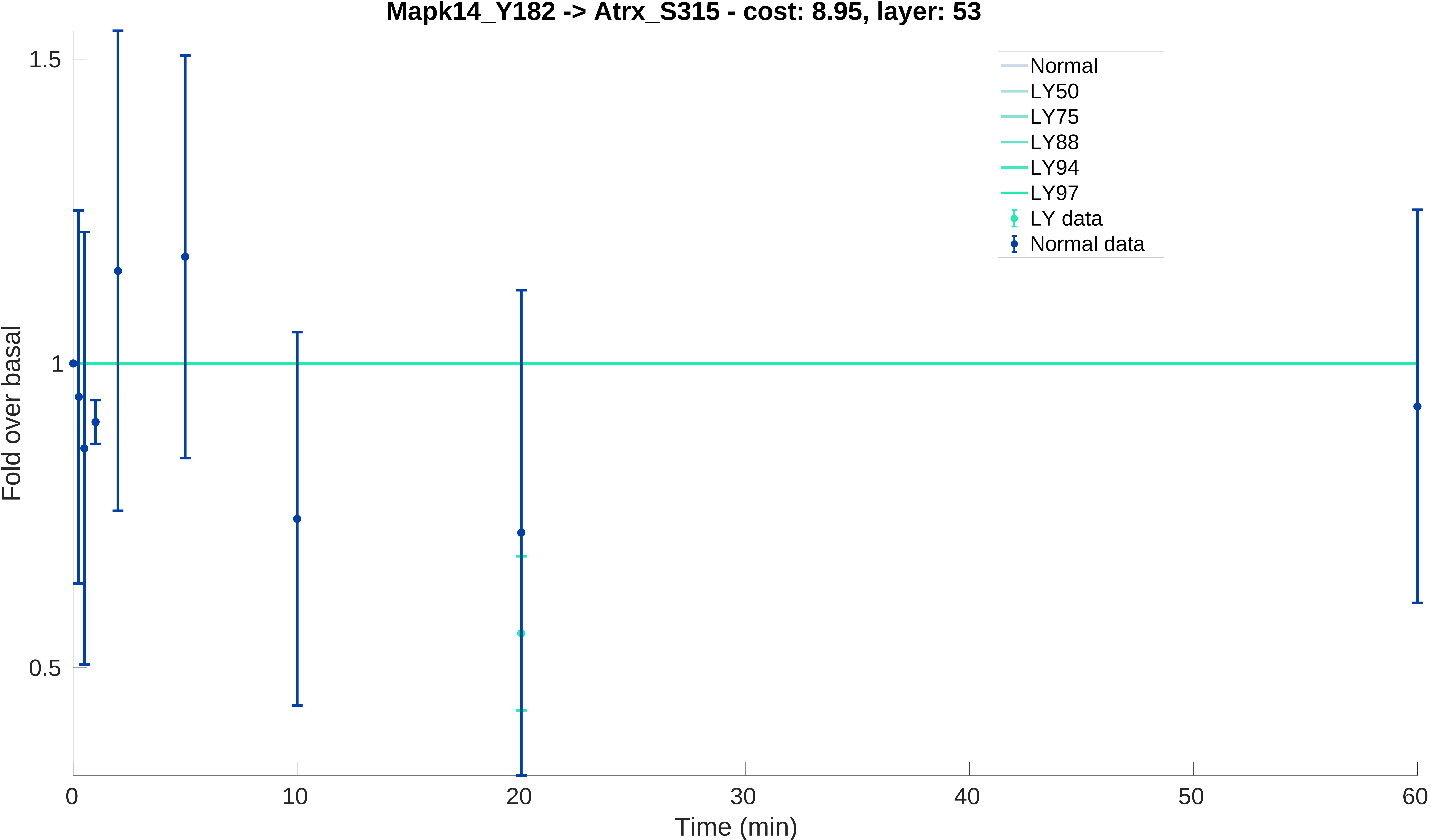

Mapk14_Y182 -> Atrx_S315 - cost: 8.95, layer: 53
Normal
LY50
LY75
LY88
LY94
LY97
LY data
Normal data
Fold over basal
1.5
1
0.5
0
10
20
30
40
50
60
Time (min)

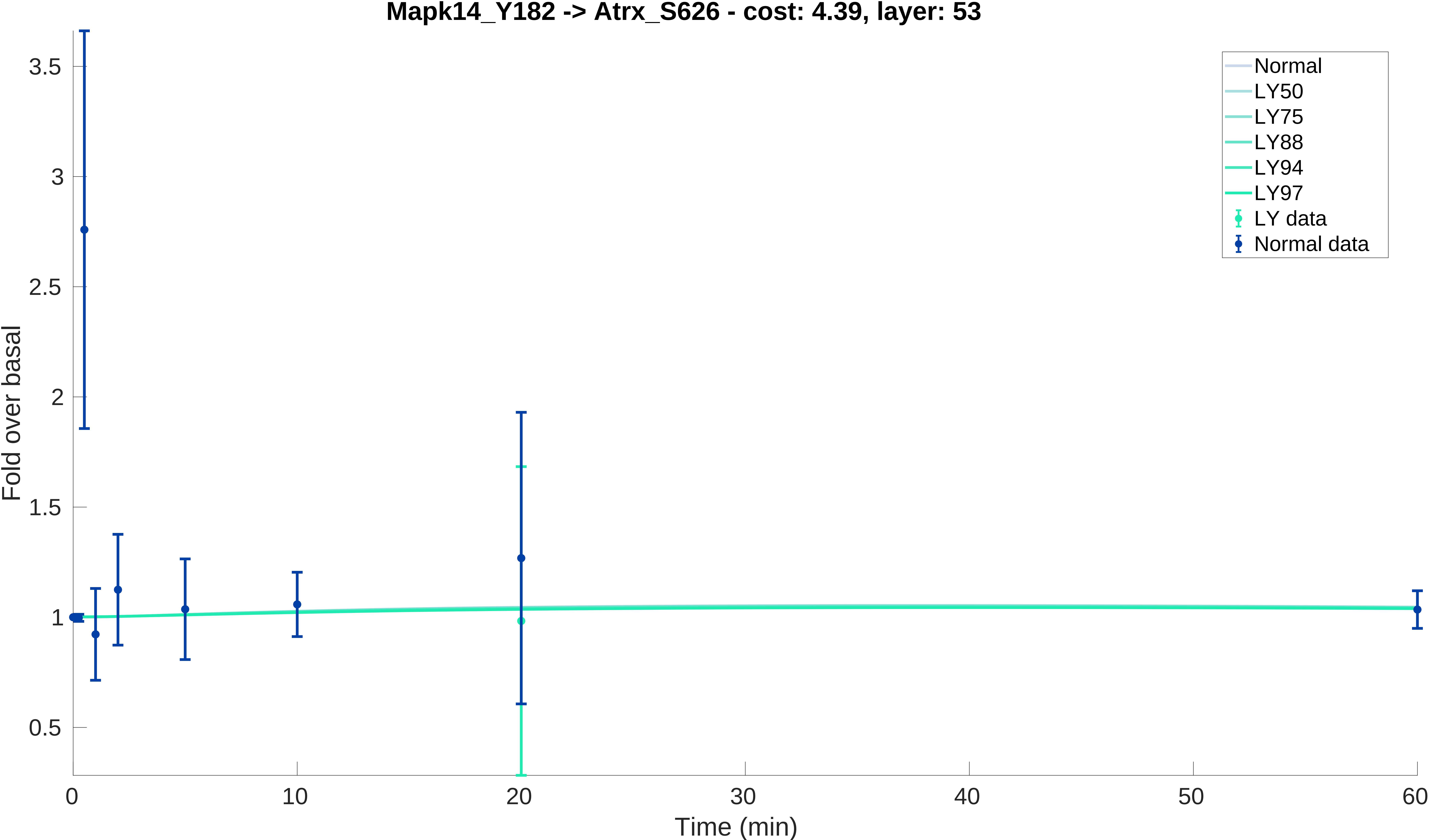

Mapk14_Y182 -> Atrx_S626 - cost: 4.39, layer: 53
Normal
LY50
LY75
LY88
LY94
LY97
LY data
Normal data
Fold over basal
Time (min)
0.5
1
1.5
2
2.5
3
3.5
0
10
20
30
40
50
60

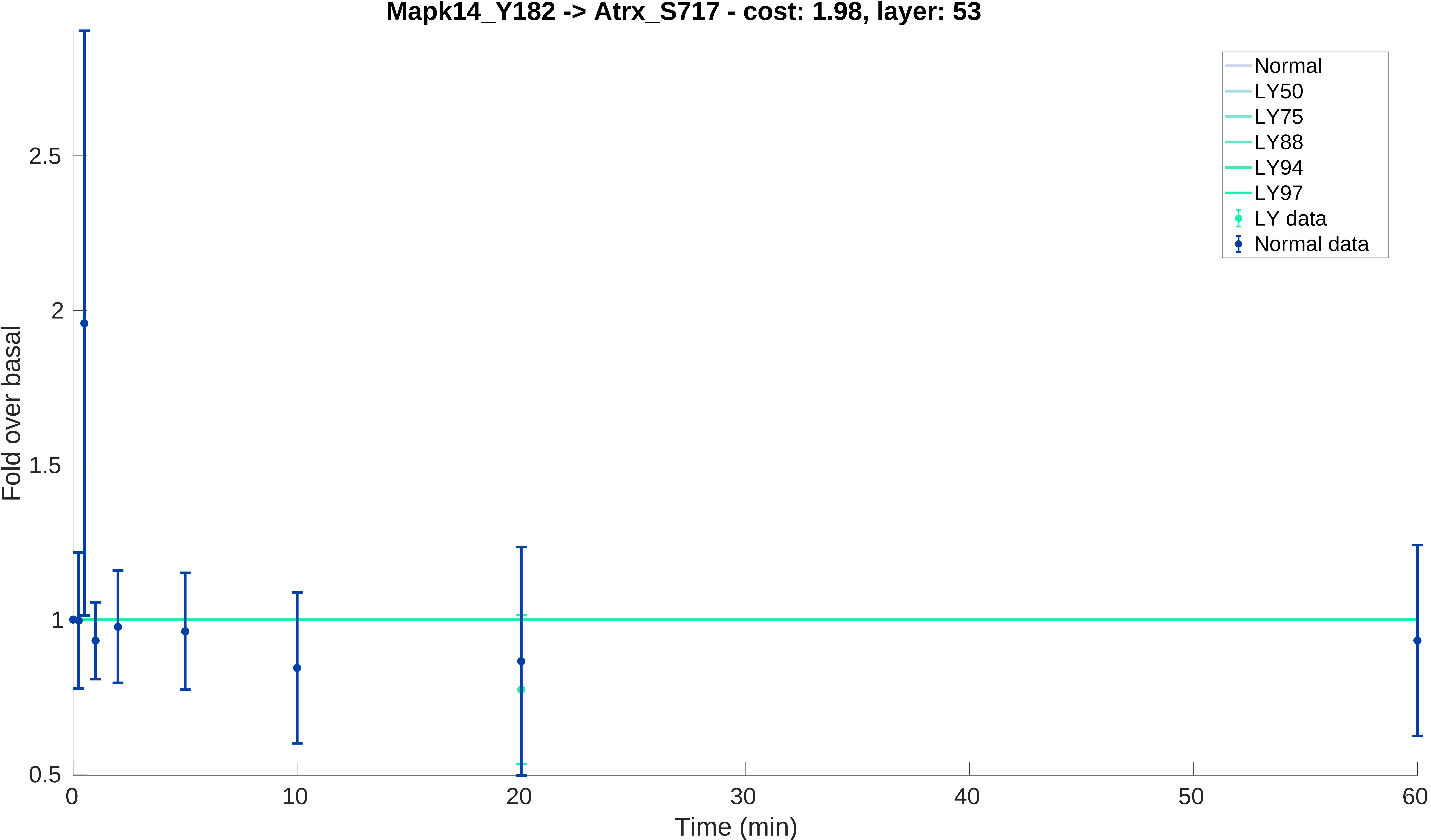

Mapk14_Y182 -> Atrx_S717 - cost: 1.98, layer: 53
Fold over basal
2.5
2
1.5
1
0.5
0
10
20
30
40
50
60
Time (min)
Normal
LY50
LY75
LY88
LY94
LY97
LY data
Normal data

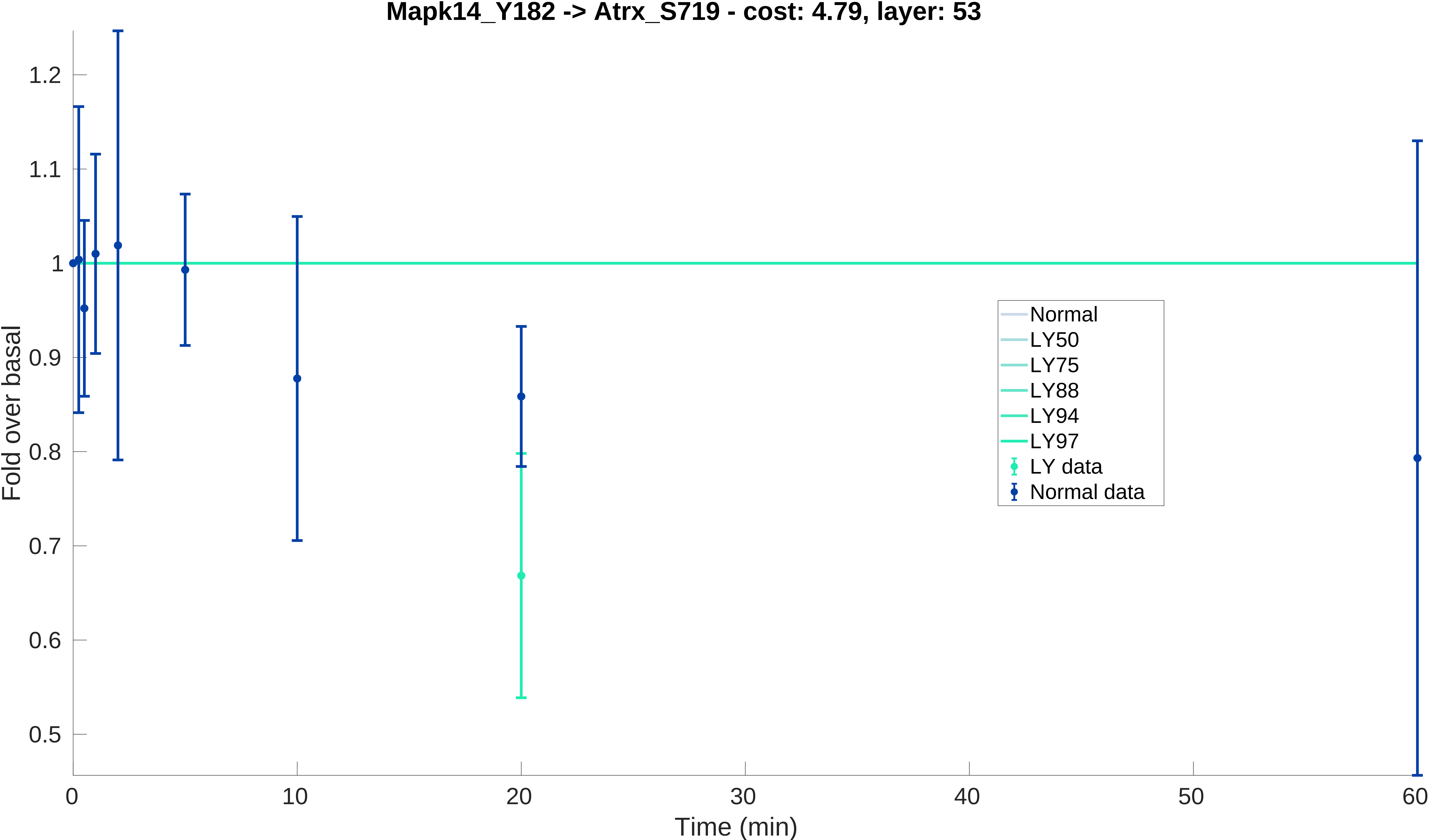
Mapk14_Y182 -> Atrx_S719 - cost: 4.79, layer: 53
Fold over basal
1.2
1.1
1
0.9
0.8
0.7
0.6
0.5
0
10
20
30
40
50
60
Time (min)
Normal
LY50
LY75
LY88
LY94
LY97
LY data
Normal data

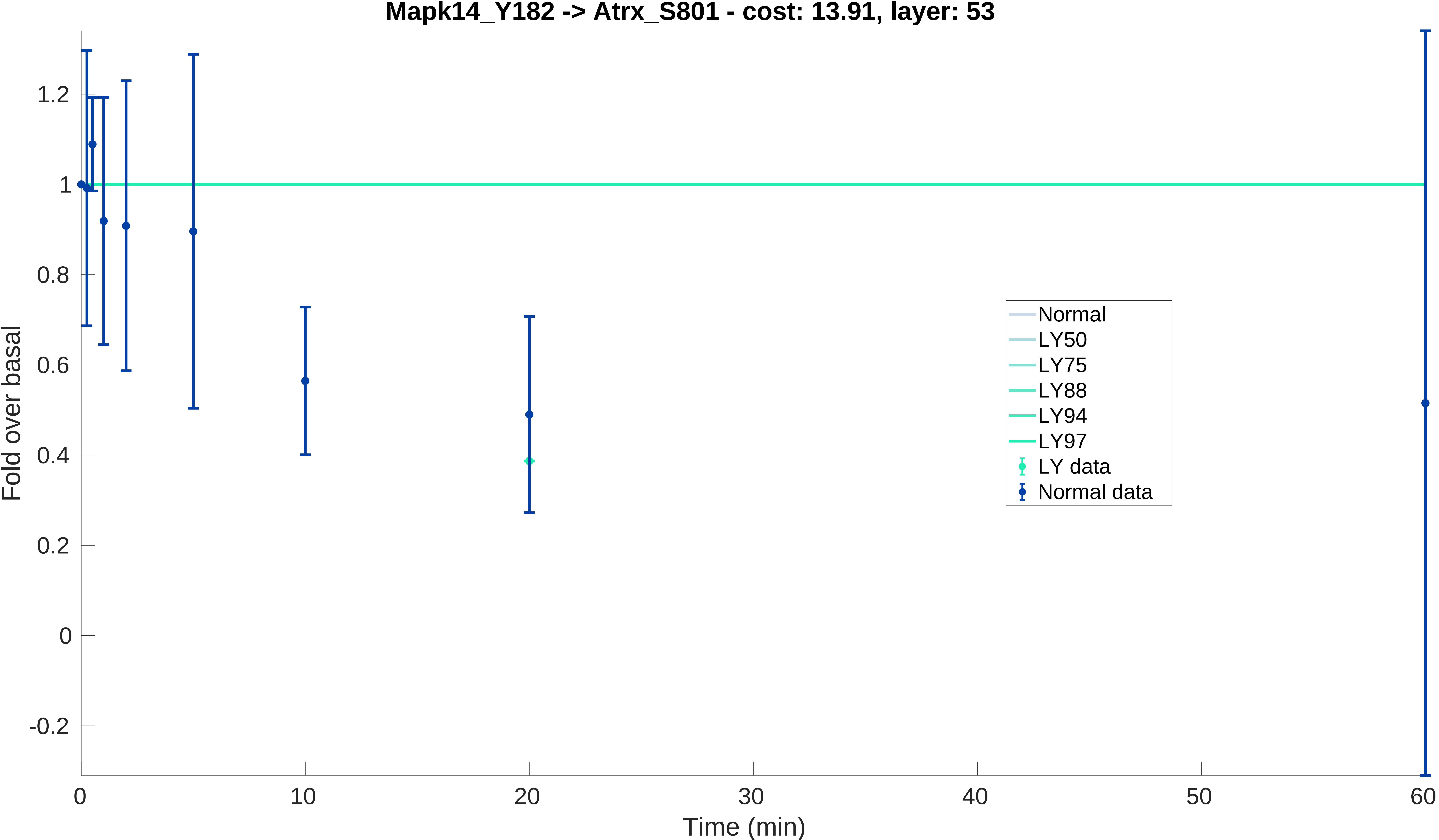

Mapk14_Y182 -> Atrx_S801 - cost: 13.91, layer: 53
1.2
1
0.8
0.6
0.4
0.2
0
-0.2
Fold over basal
0
10
20
30
40
50
60
Time (min)
Normal
LY50
LY75
LY88
LY94
LY97
LY data
Normal data

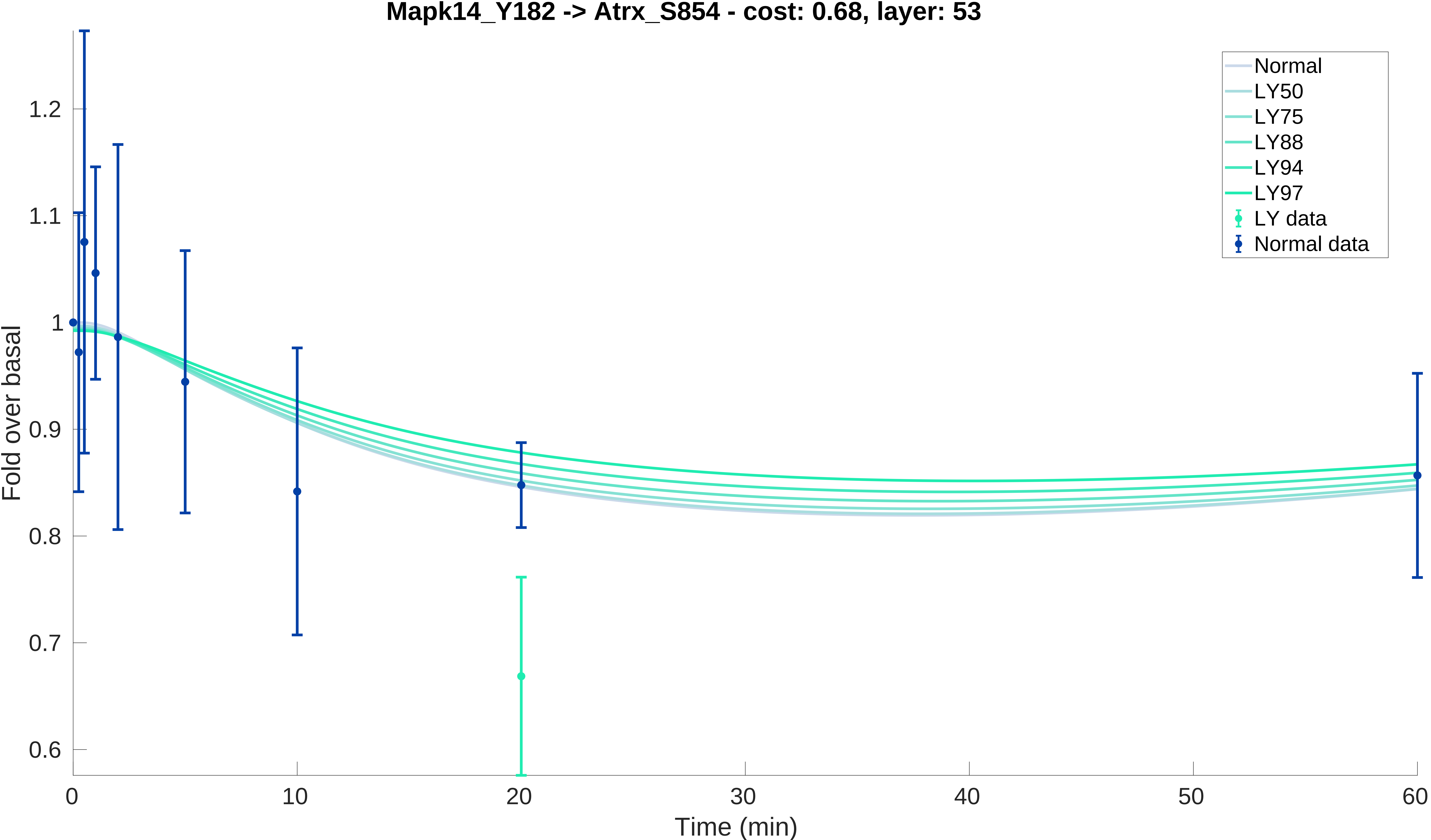
Mapk14_Y182 -> Atrx_S854 - cost: 0.68, layer: 53
Normal
LY50
LY75
LY88
LY94
LY97
LY data
Normal data
Fold over basal
1.2
1.1
1
0.9
0.8
0.7
0.6
0
10
20
30
40
50
60
Time (min)

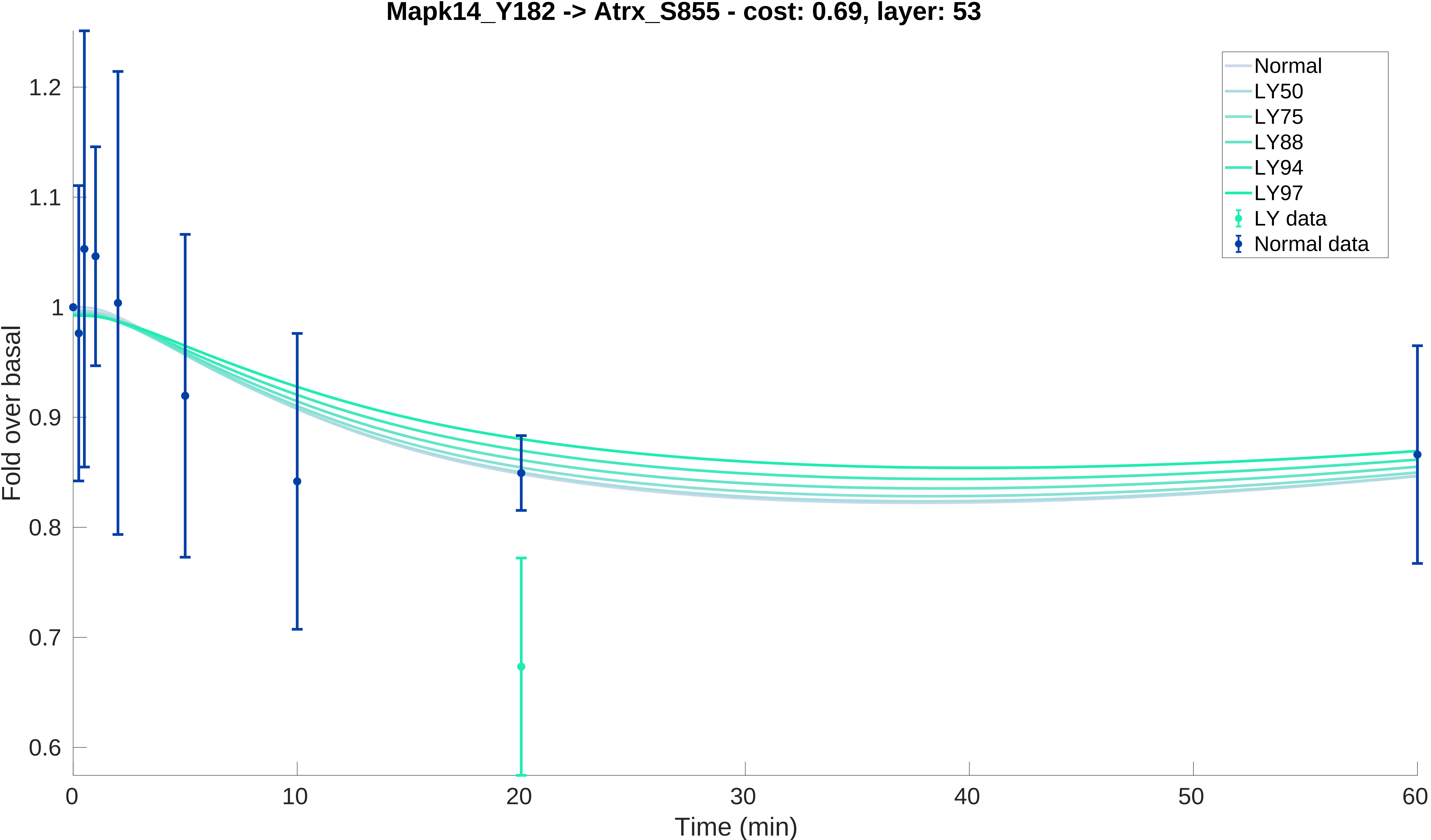

Mapk14_Y182 -> Atrx_S855 - cost: 0.69, layer: 53
Fold over basal
Time (min)
0.6
0.7
0.8
0.9
1
1.1
1.2
0
10
20
30
40
50
60
Normal
LY50
LY75
LY88
LY94
LY97
LY data
Normal data

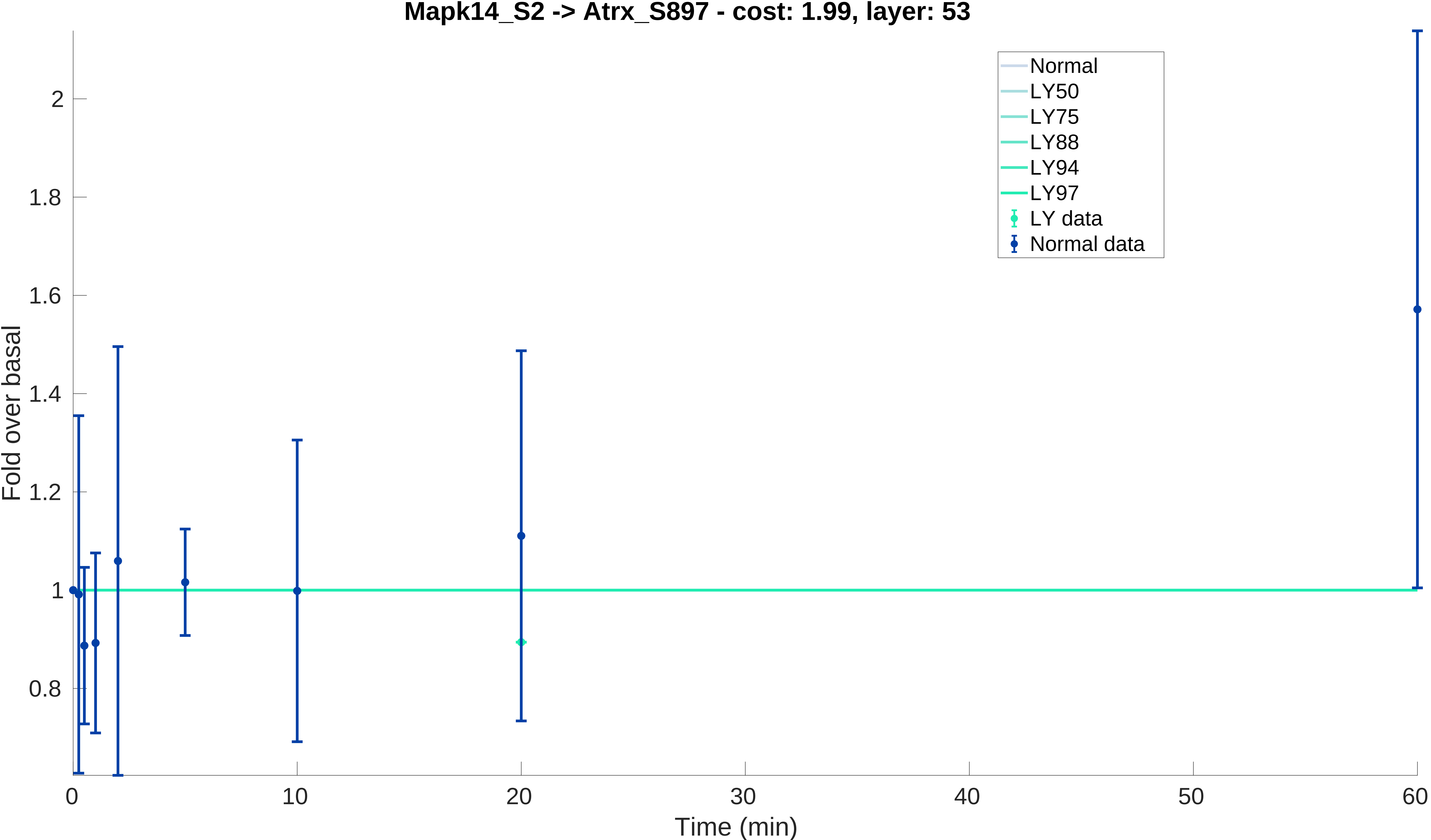

Mapk14_S2 -> Atrx_S897 - cost: 1.99, layer: 53
Normal
LY50
LY75
LY88
LY94
LY97
LY data
Normal data
Fold over basal
0.8
1
1.2
1.4
1.6
1.8
2
0
10
20
30
40
50
60
Time (min)

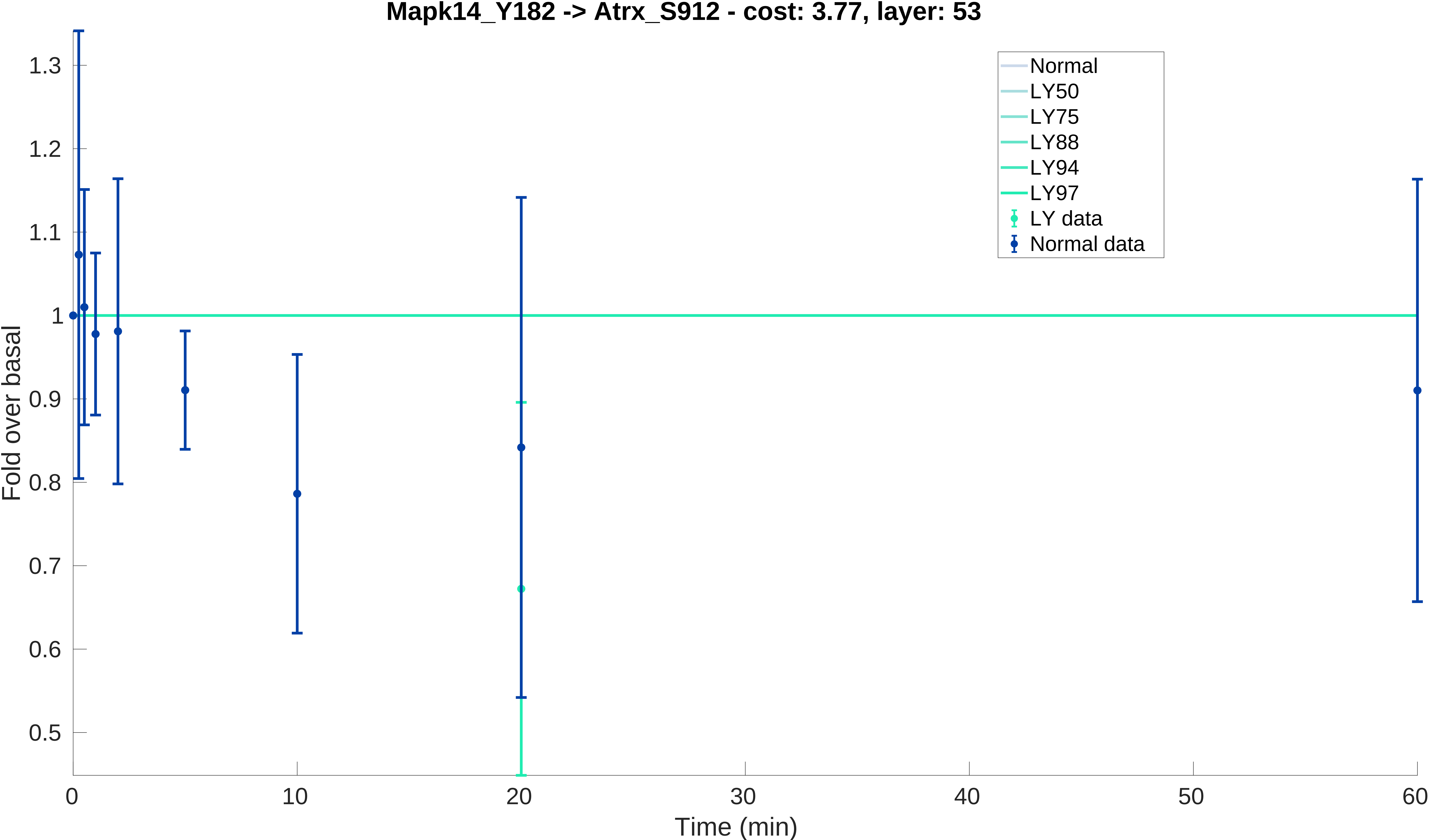

Mapk14_Y182 -> Atrx_S912 - cost: 3.77, layer: 53
Normal
LY50
LY75
LY88
LY94
LY97
LY data
Normal data
Fold over basal
1.3
1.2
1.1
1
0.9
0.8
0.7
0.6
0.5
0
10
20
30
40
50
60
Time (min)

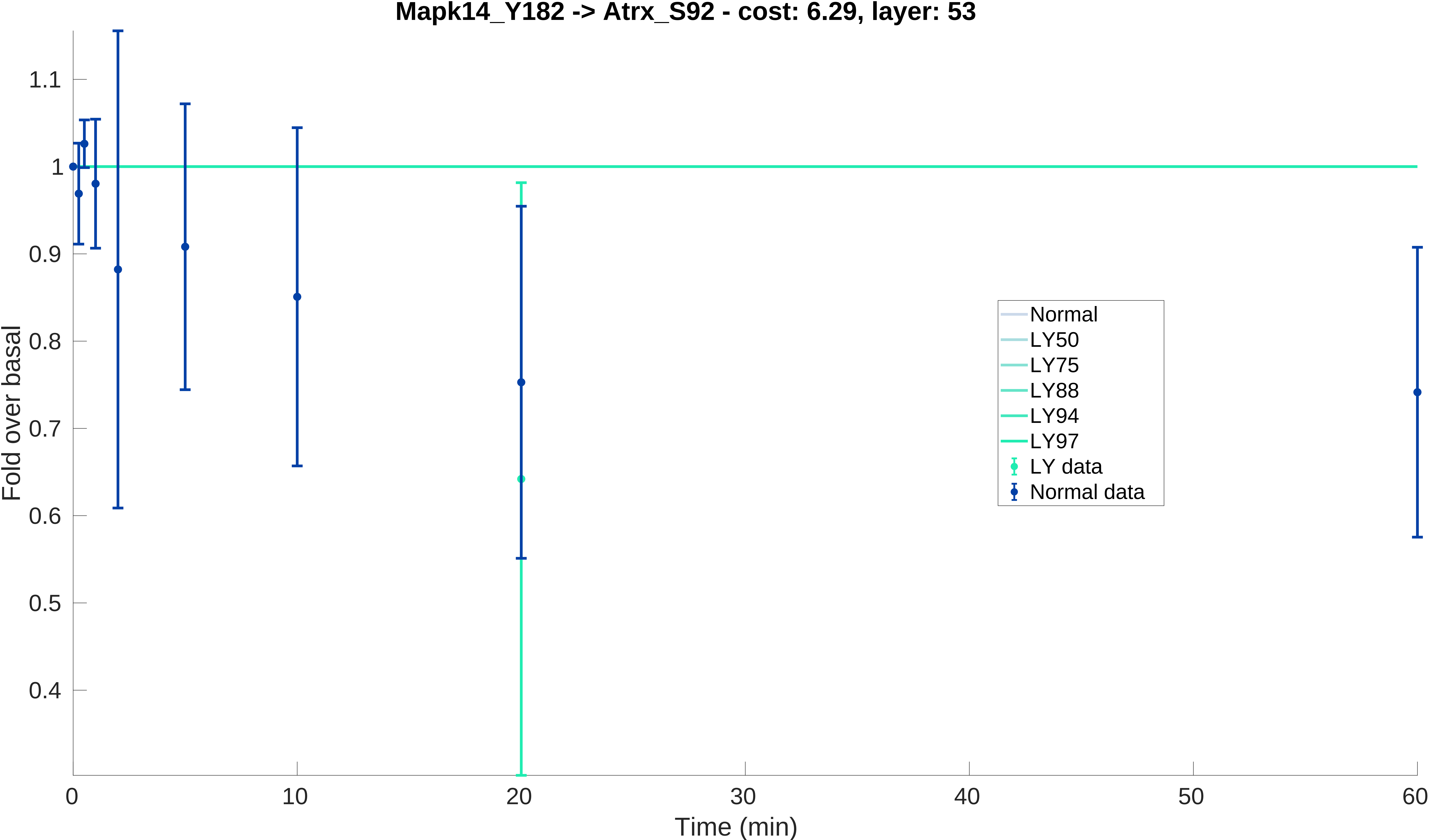

Mapk14_Y182 -> Atrx_S92 - cost: 6.29, layer: 53
Fold over basal
1.1
1
0.9
0.8
0.7
0.6
0.5
0.4
0
10
20
30
40
50
60
Time (min)
Normal
LY50
LY75
LY88
LY94
LY97
LY data
Normal data

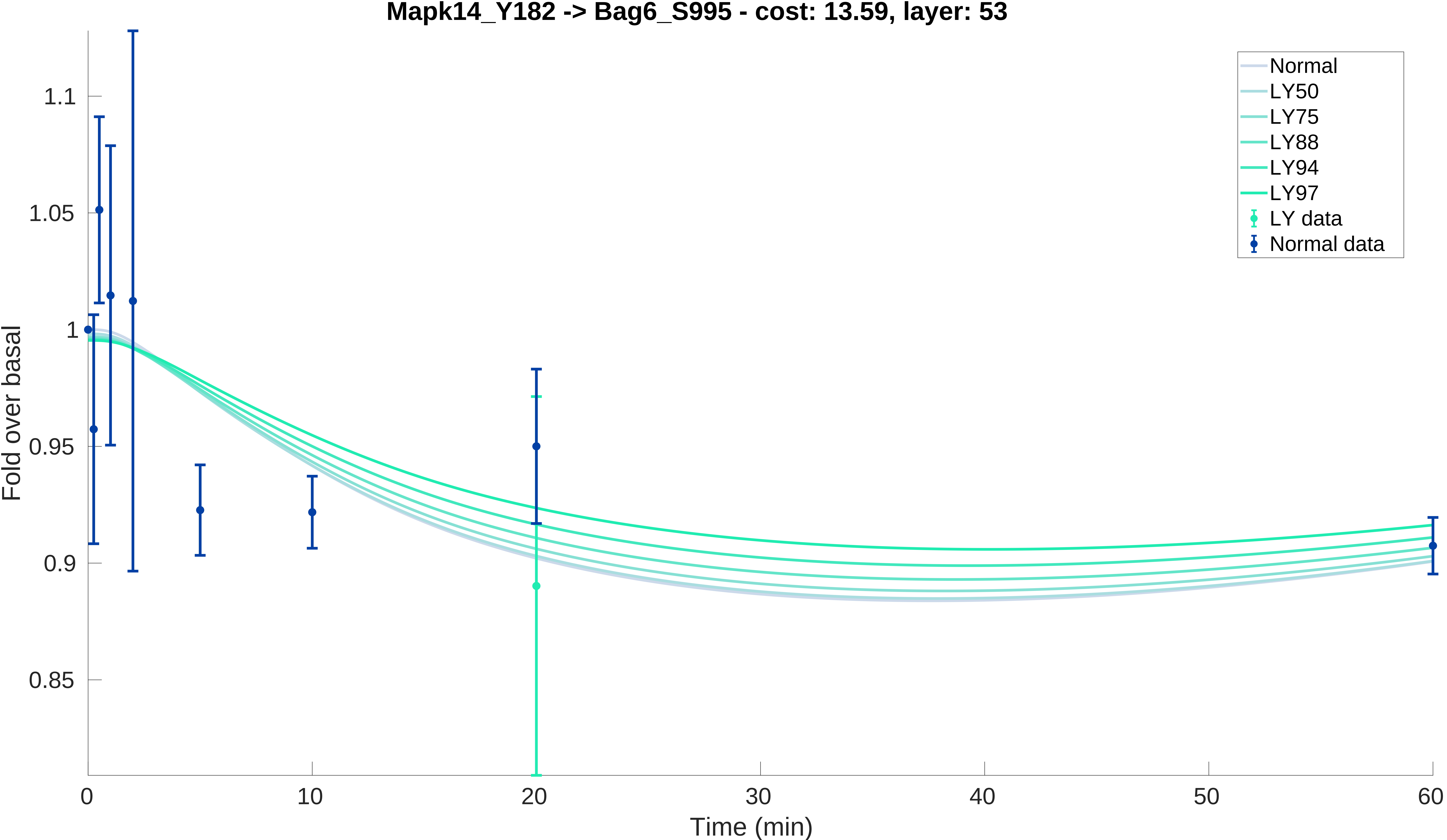

Mapk14_Y182 -> Bag6_S995 - cost: 13.59, layer: 53
Fold over basal
Time (min)
1.1
1.05
1
0.95
0.9
0.85
0
10
20
30
40
50
60
Normal
LY50
LY75
LY88
LY94
LY97
LY data
Normal data

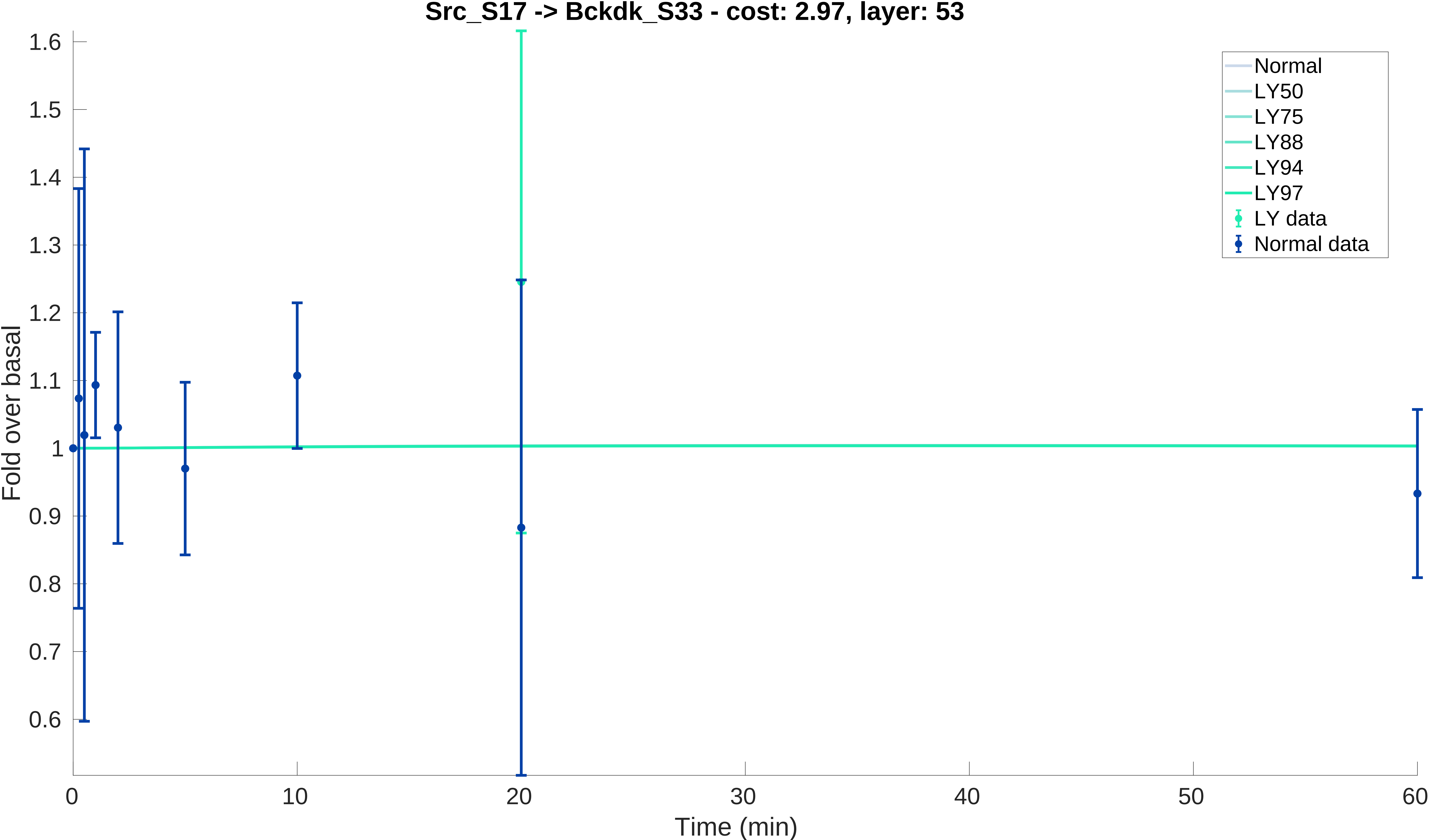

Src_S17 -> Bckdk_S33 - cost: 2.97, layer: 53
Normal
LY50
LY75
LY88
LY94
LY97
LY data
Normal data
Fold over basal
Time (min)
1.6
1.5
1.4
1.3
1.2
1.1
1
0.9
0.8
0.7
0.6
0
10
20
30
40
50
60

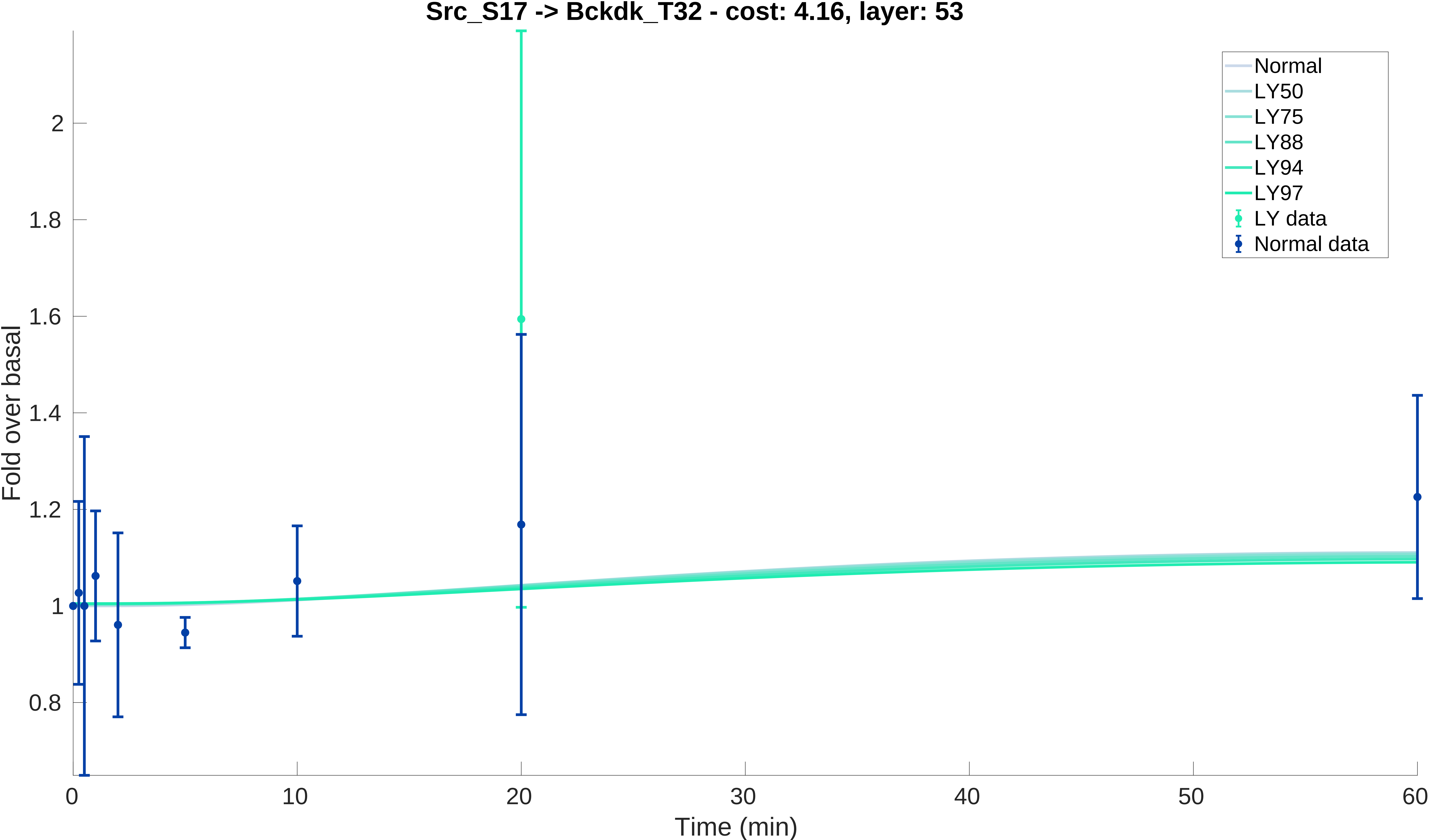

Src_S17 -> Bckdk_T32 - cost: 4.16, layer: 53
Normal
LY50
LY75
LY88
LY94
LY97
LY data
Normal data
Fold over basal
Time (min)
0.8
1
1.2
1.4
1.6
1.8
2
0
10
20
30
40
50
60

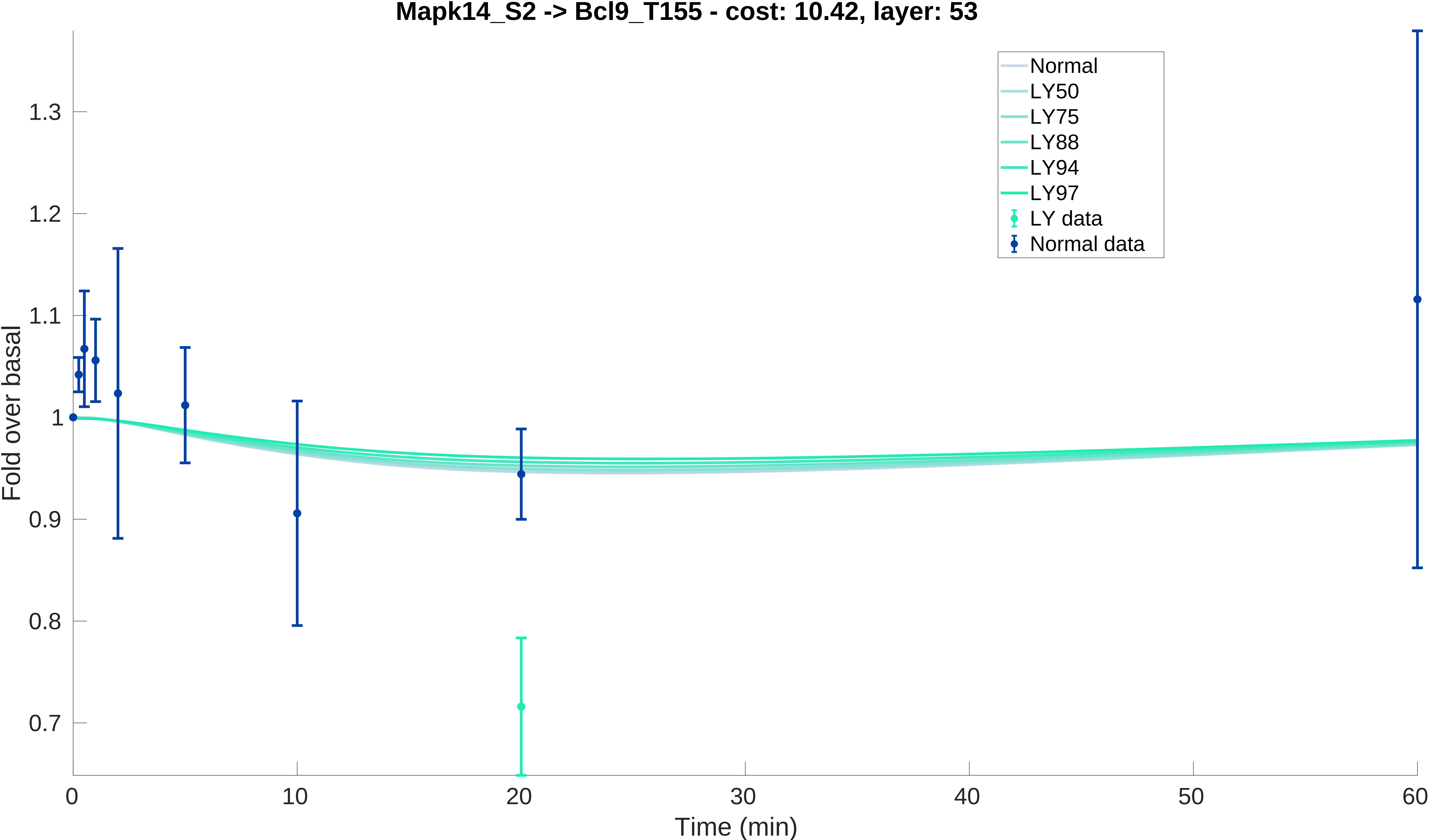

Mapk14_S2 -> Bcl9_T155 - cost: 10.42, layer: 53
Normal
LY50
LY75
LY88
LY94
LY97
LY data
Normal data
Fold over basal
Time (min)
1.3
1.2
1.1
1
0.9
0.8
0.7
0
10
20
30
40
50
60

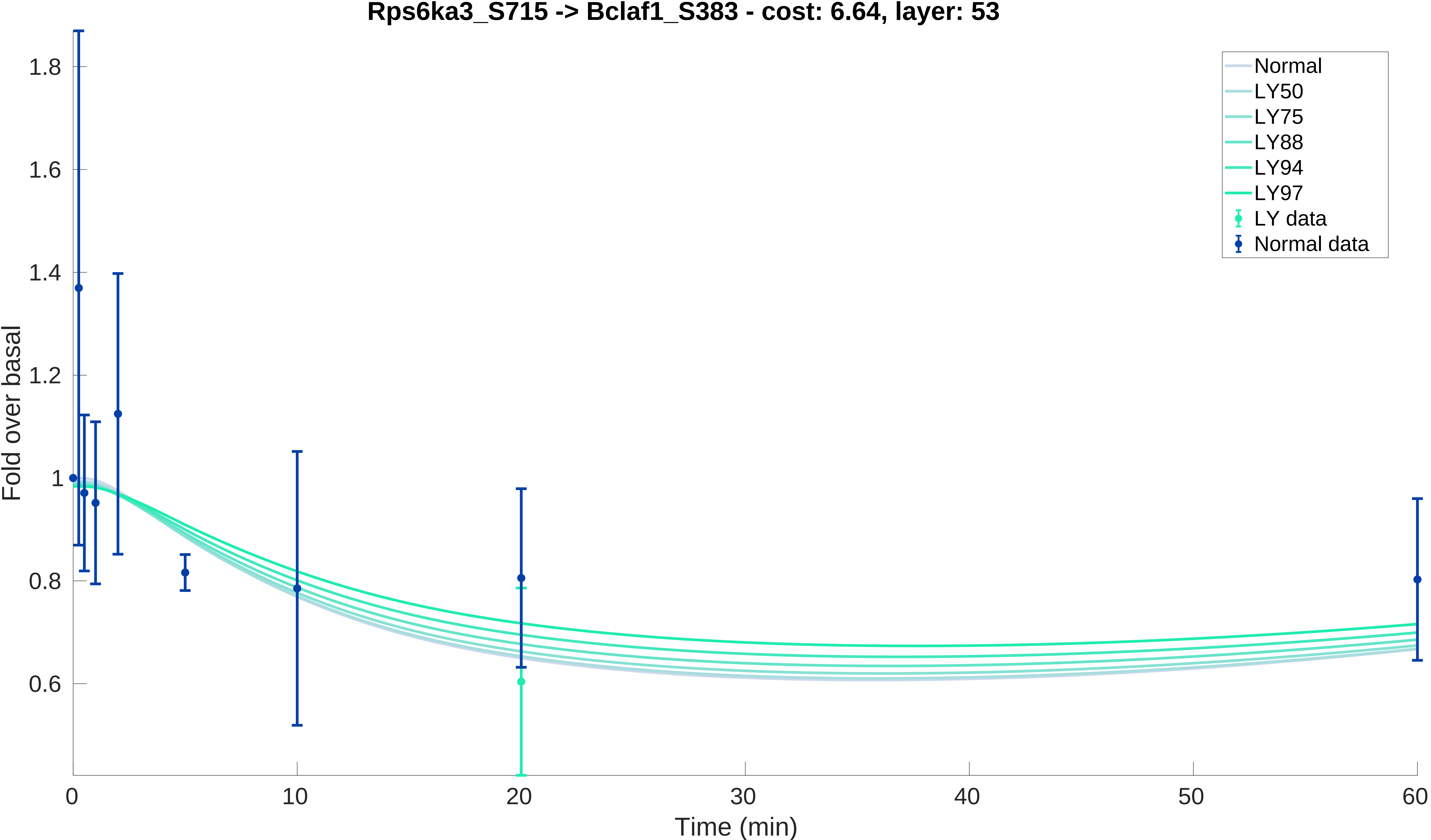

Rps6ka3_S715 -> Bclaf1_S383 - cost: 6.64, layer: 53
Fold over basal
Time (min)
1.8
1.6
1.4
1.2
1
0.8
0.6
0
10
20
30
40
50
60
Normal
LY50
LY75
LY88
LY94
LY97
LY data
Normal data

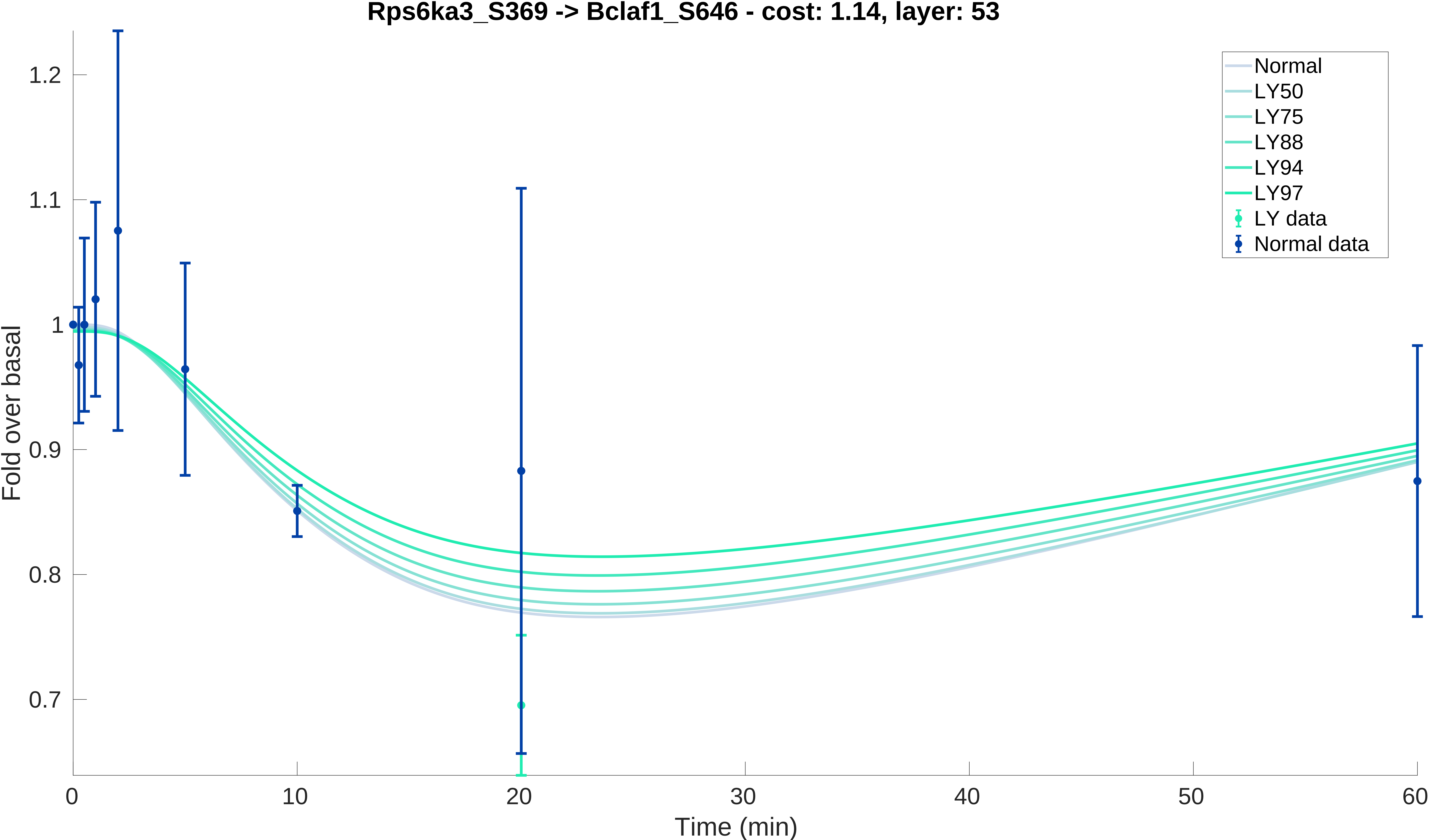

Rps6ka3_S369 -> Bclaf1_S646 - cost: 1.14, layer: 53
Fold over basal
Time (min)
1.2
1.1
1
0.9
0.8
0.7
0
10
20
30
40
50
60
Normal
LY50
LY75
LY88
LY94
LY97
LY data
Normal data

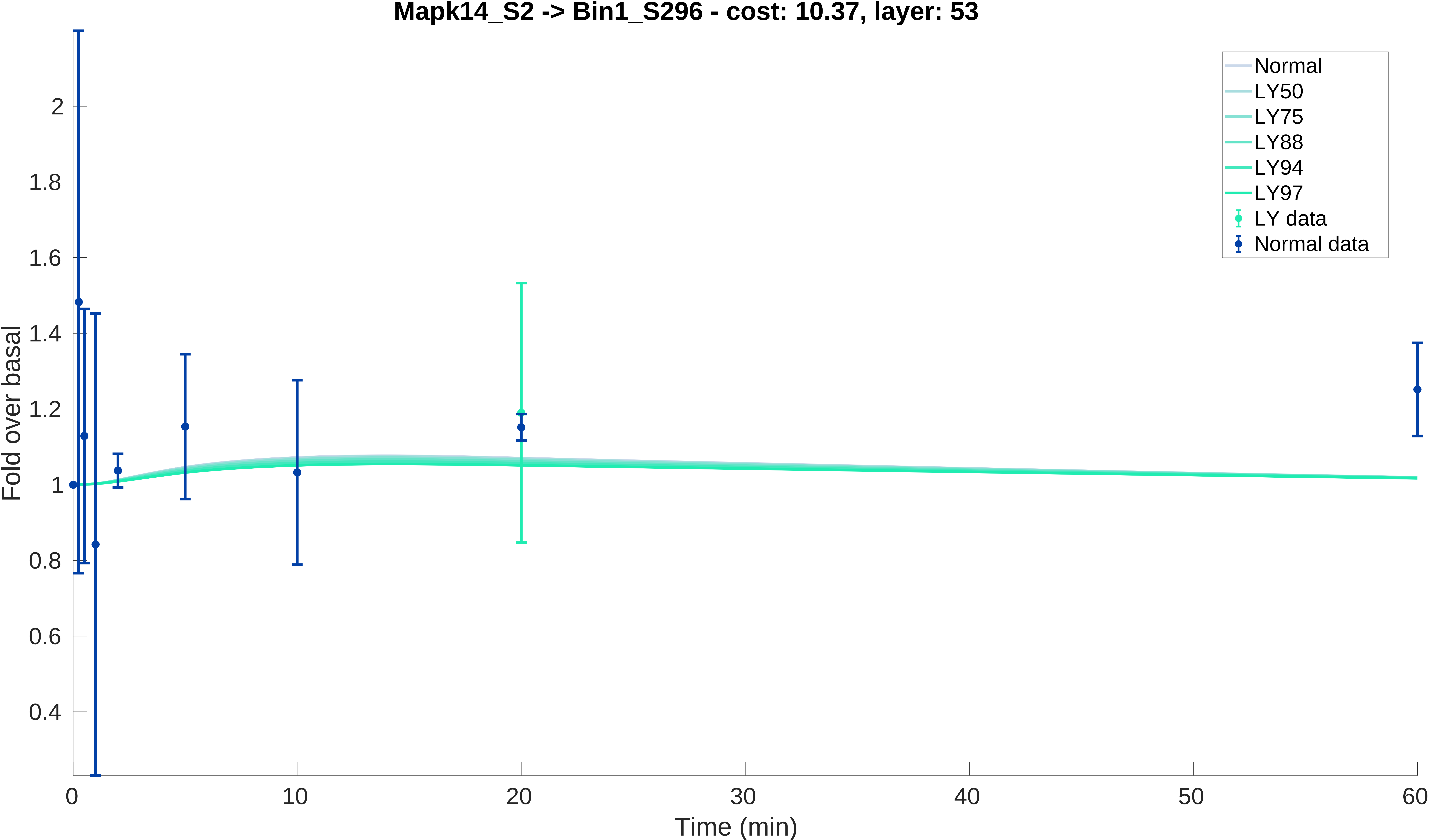

Mapk14_S2 -> Bin1_S296 - cost: 10.37, layer: 53
Normal
LY50
LY75
LY88
LY94
LY97
LY data
Normal data
Fold over basal
Time (min)
0.4
0.6
0.8
1
1.2
1.4
1.6
1.8
2
0
10
20
30
40
50
60

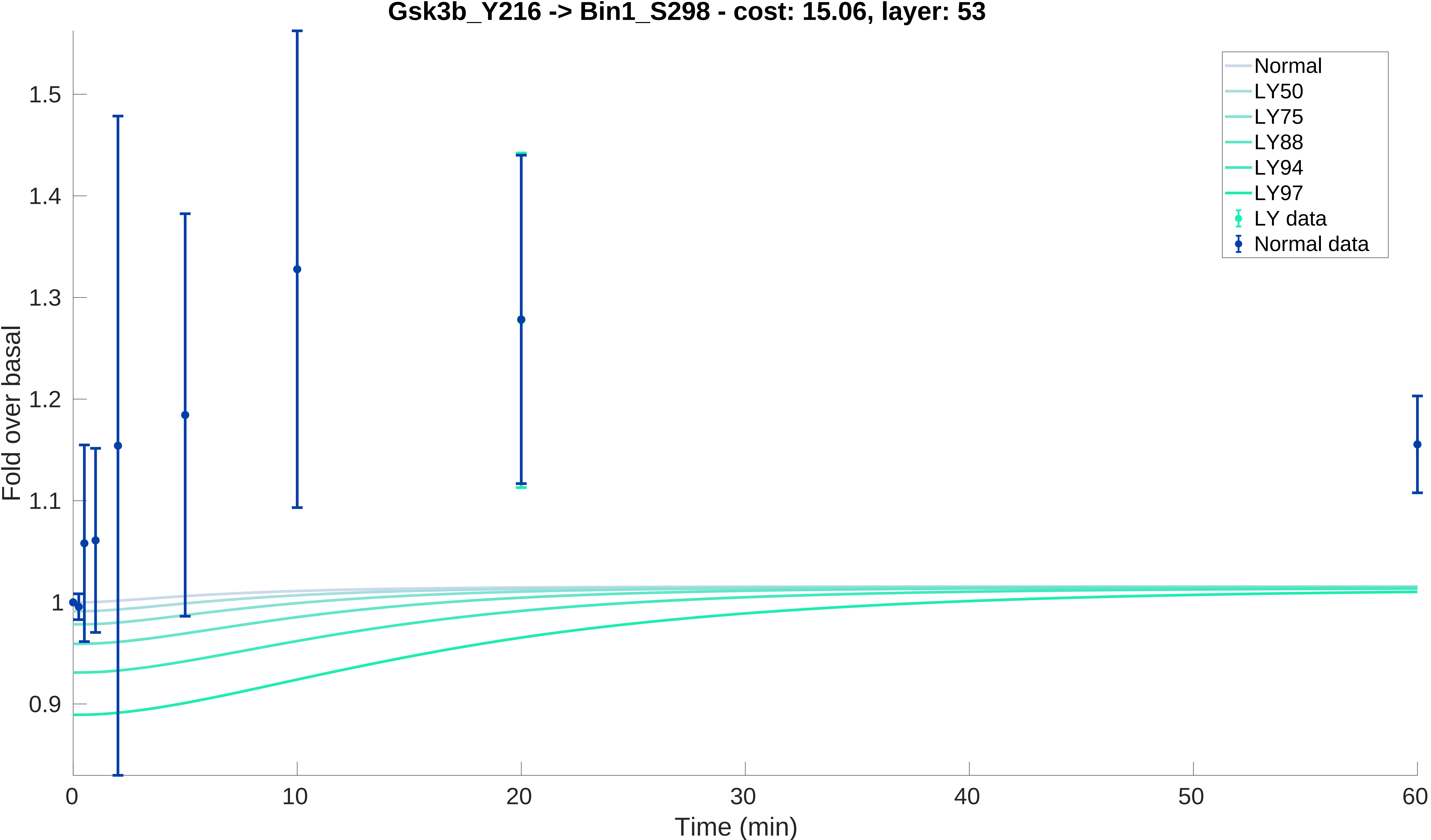

Gsk3b_Y216 -> Bin1_S298 - cost: 15.06, layer: 53
Fold over basal
Time (min)
1.5
1.4
1.3
1.2
1.1
1
0.9
0
10
20
30
40
50
60
Normal
LY50
LY75
LY88
LY94
LY97
LY data
Normal data

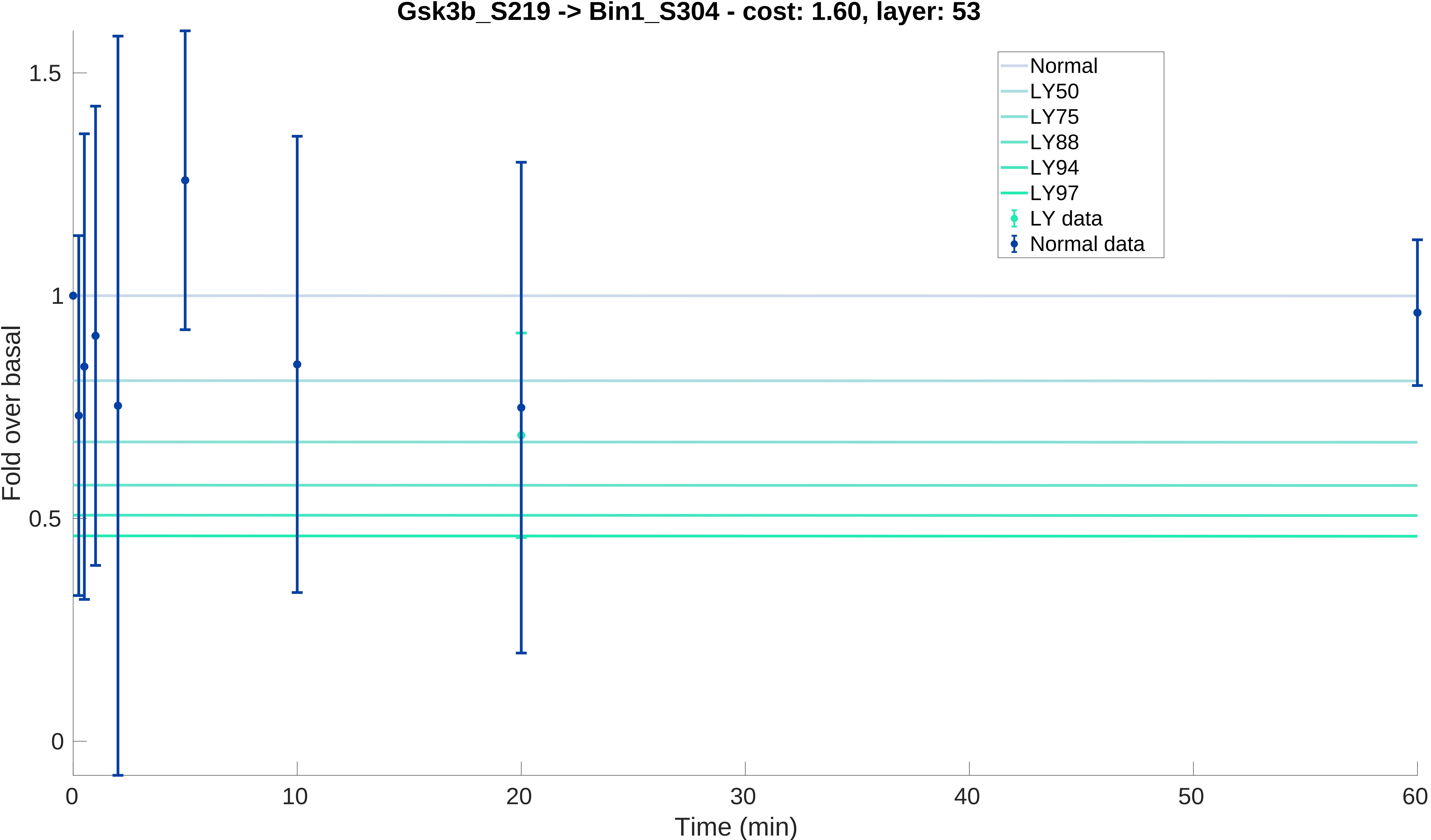

Gsk3b_S219 -> Bin1_S304 - cost: 1.60, layer: 53
Normal
LY50
LY75
LY88
LY94
LY97
LY data
Normal data
Fold over basal
1.5
1
0.5
0
Time (min)
0
10
20
30
40
50
60

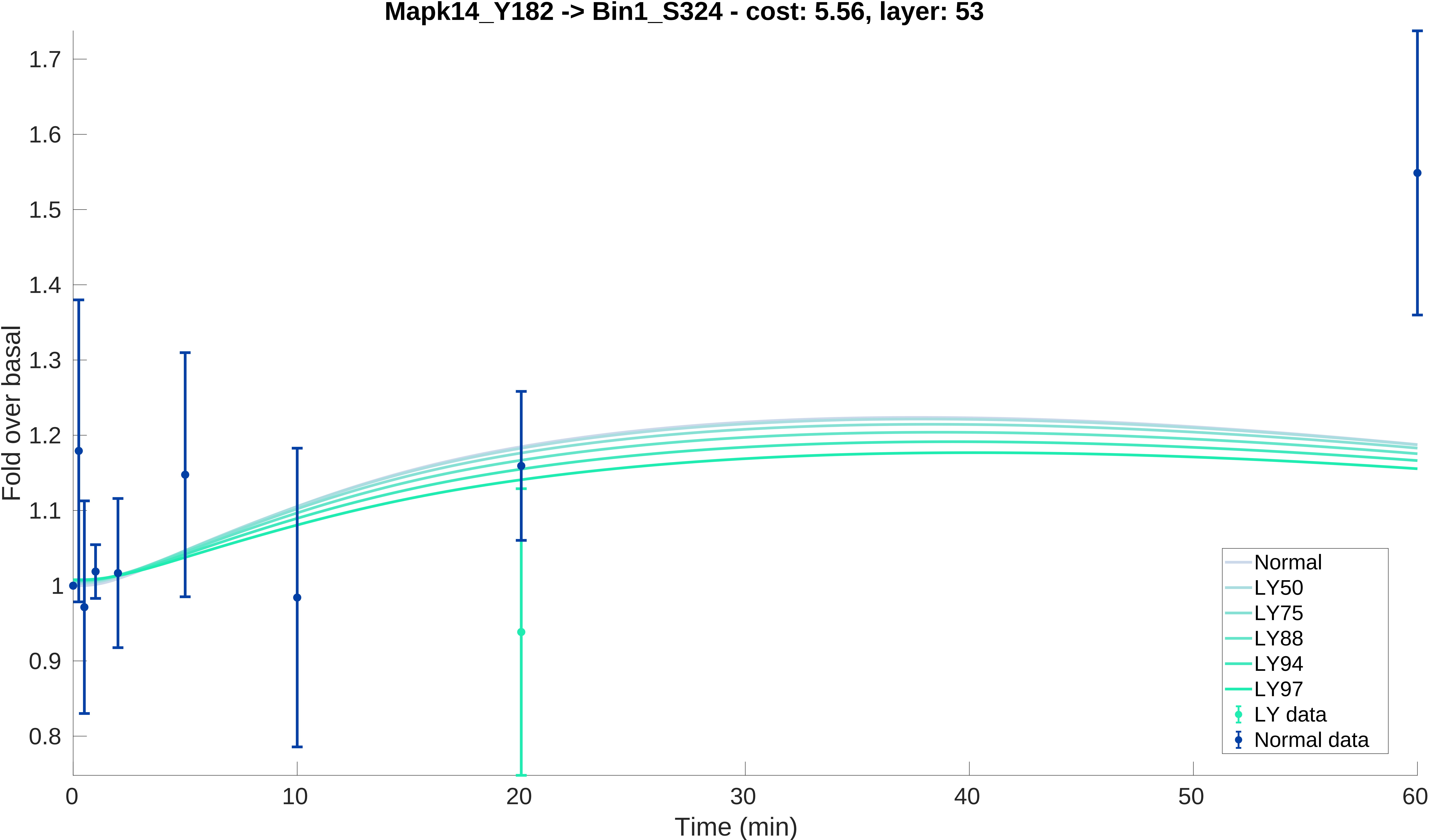

Mapk14_Y182 -> Bin1_S324 - cost: 5.56, layer: 53
Fold over basal
1.7
1.6
1.5
1.4
1.3
1.2
1.1
1
0.9
0.8
0
10
20
30
40
50
60
Time (min)
Normal
LY50
LY75
LY88
LY94
LY97
LY data
Normal data

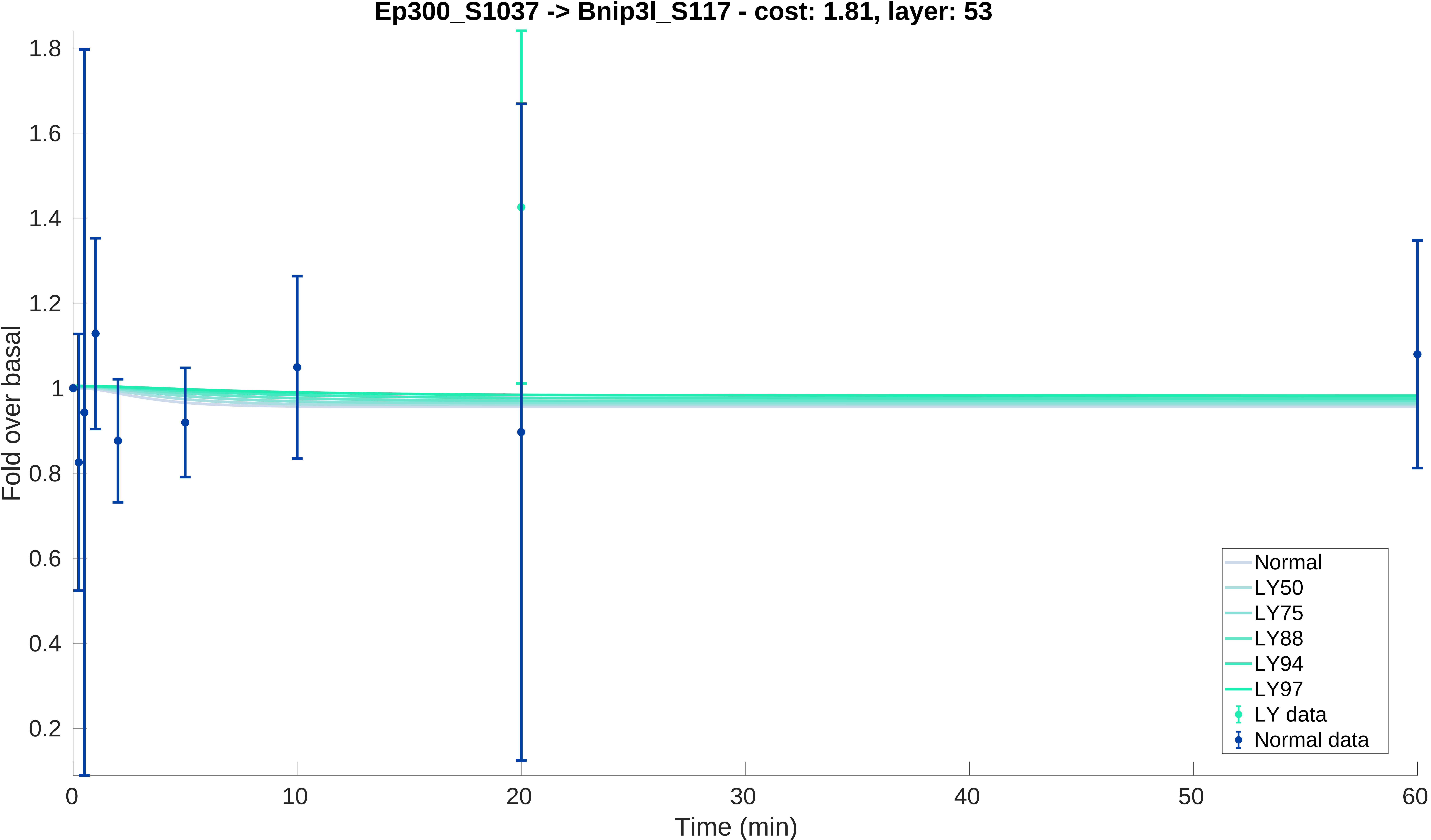
Ep300_S1037 -> Bnip3l_S117 - cost: 1.81, layer: 53
Fold over basal
Time (min)
0
10
20
30
40
50
60
0.2
0.4
0.6
0.8
1
1.2
1.4
1.6
1.8
Normal
LY50
LY75
LY88
LY94
LY97
LY data
Normal data

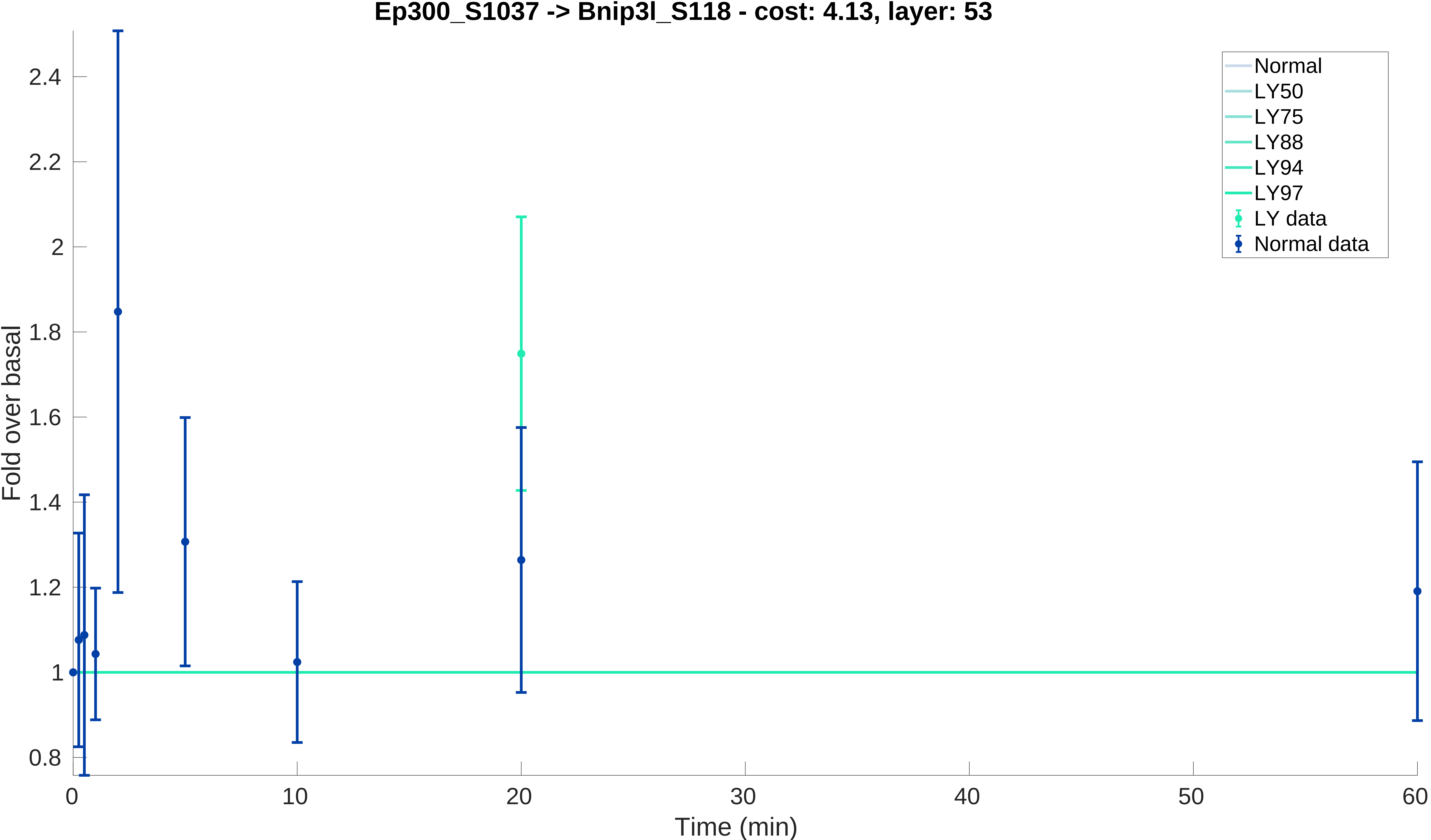

Ep300_S1037 -> Bnip3l_S118 - cost: 4.13, layer: 53
Normal
LY50
LY75
LY88
LY94
LY97
LY data
Normal data
Fold over basal
Time (min)
0.8
1
1.2
1.4
1.6
1.8
2
2.2
2.4
0
10
20
30
40
50
60

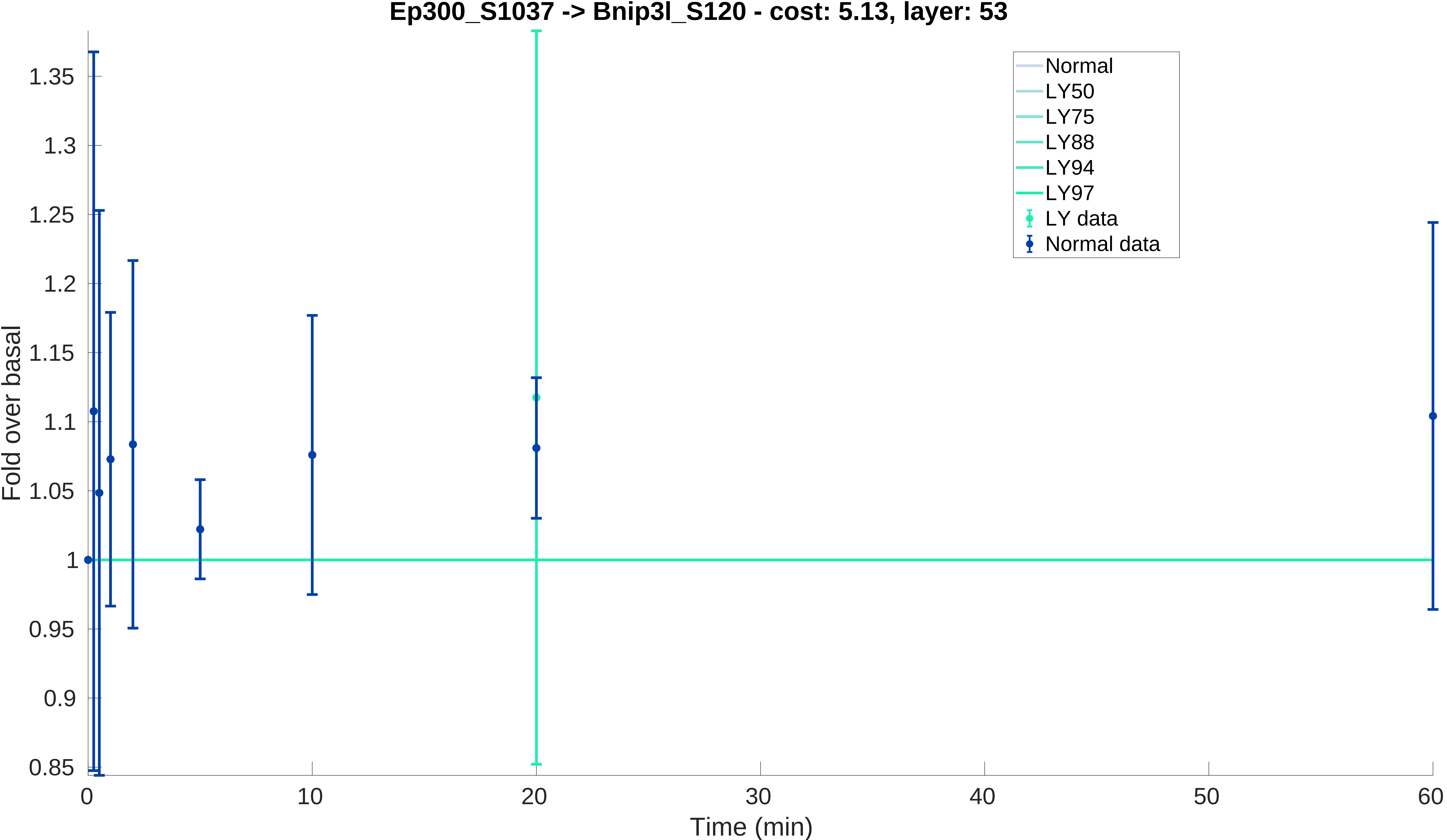

Ep300_S1037 -> Bnip3l_S120 - cost: 5.13, layer: 53
Normal
LY50
LY75
LY88
LY94
LY97
LY data
Normal data
Fold over basal
1.35
1.3
1.25
1.2
1.15
1.1
1.05
1
0.95
0.9
0.85
0
10
20
30
40
50
60
Time (min)

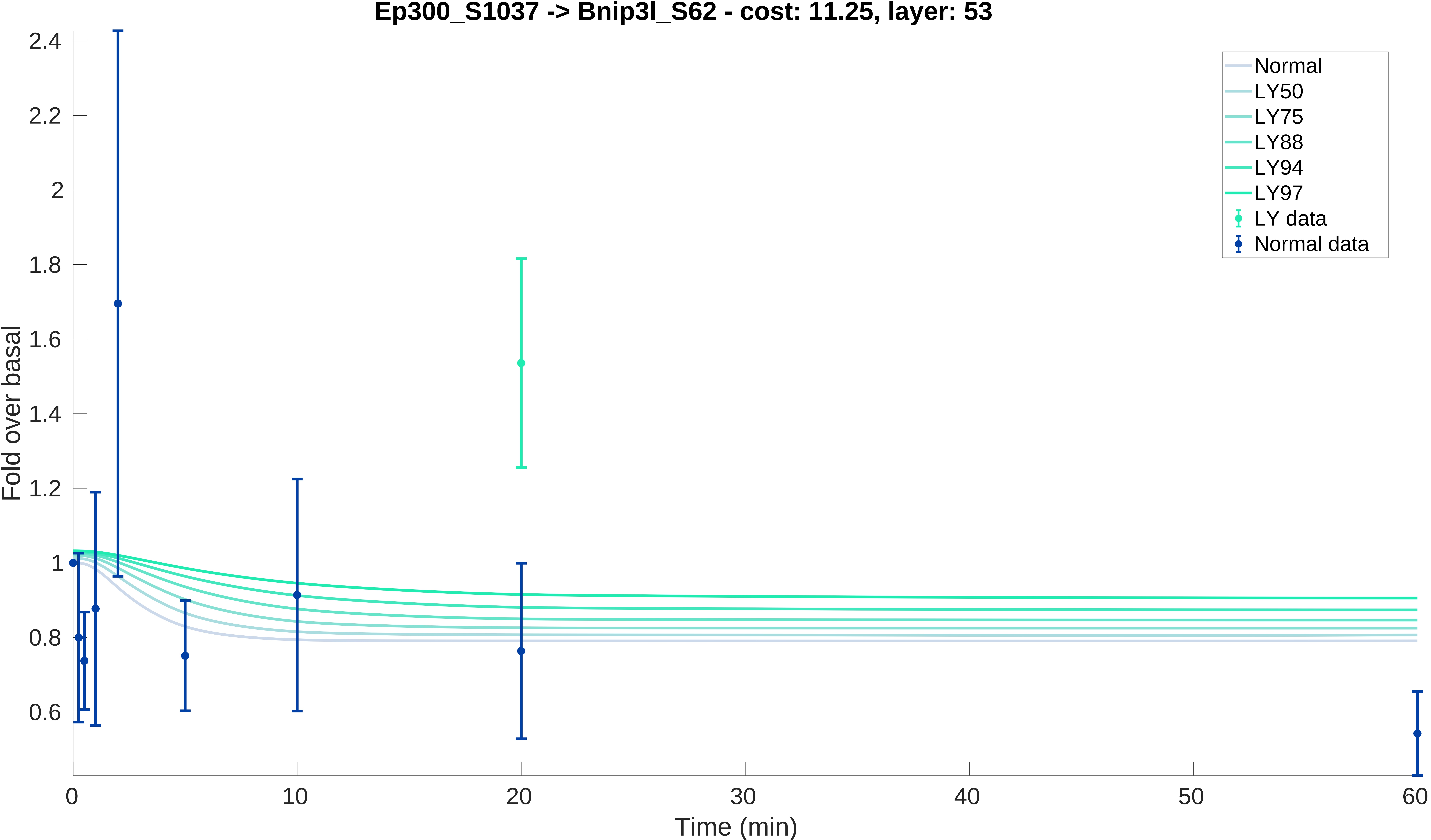

Ep300_S1037 -> Bnip3l_S62 - cost: 11.25, layer: 53
Fold over basal
Time (min)
Normal
LY50
LY75
LY88
LY94
LY97
LY data
Normal data
2.4
2.2
2
1.8
1.6
1.4
1.2
1
0.8
0.6
0
10
20
30
40
50
60

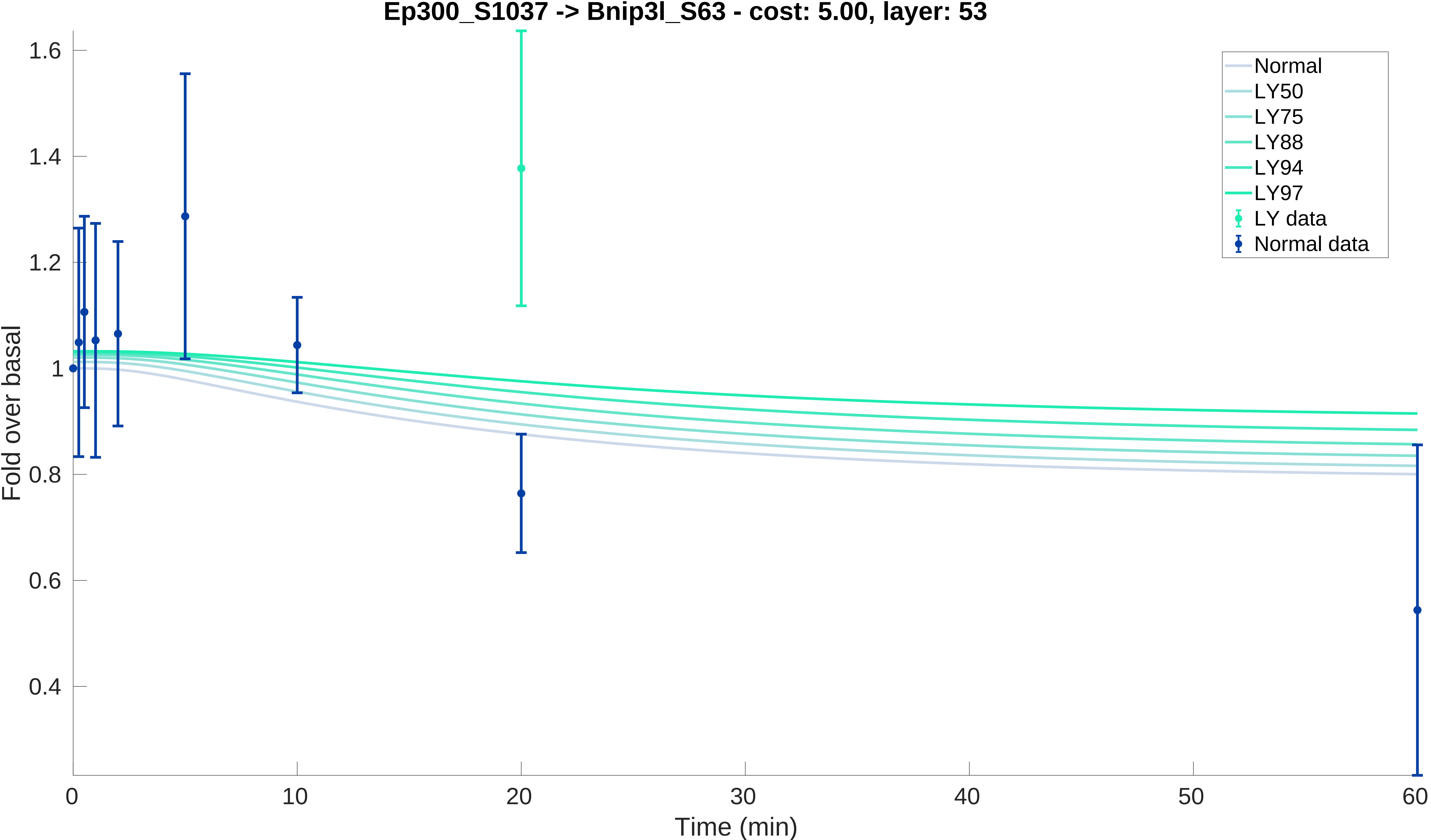

Ep300_S1037 -> Bnip3l_S63 - cost: 5.00, layer: 53
Fold over basal
Time (min)
0.4
0.6
0.8
1
1.2
1.4
1.6
0
10
20
30
40
50
60
Normal
LY50
LY75
LY88
LY94
LY97
LY data
Normal data

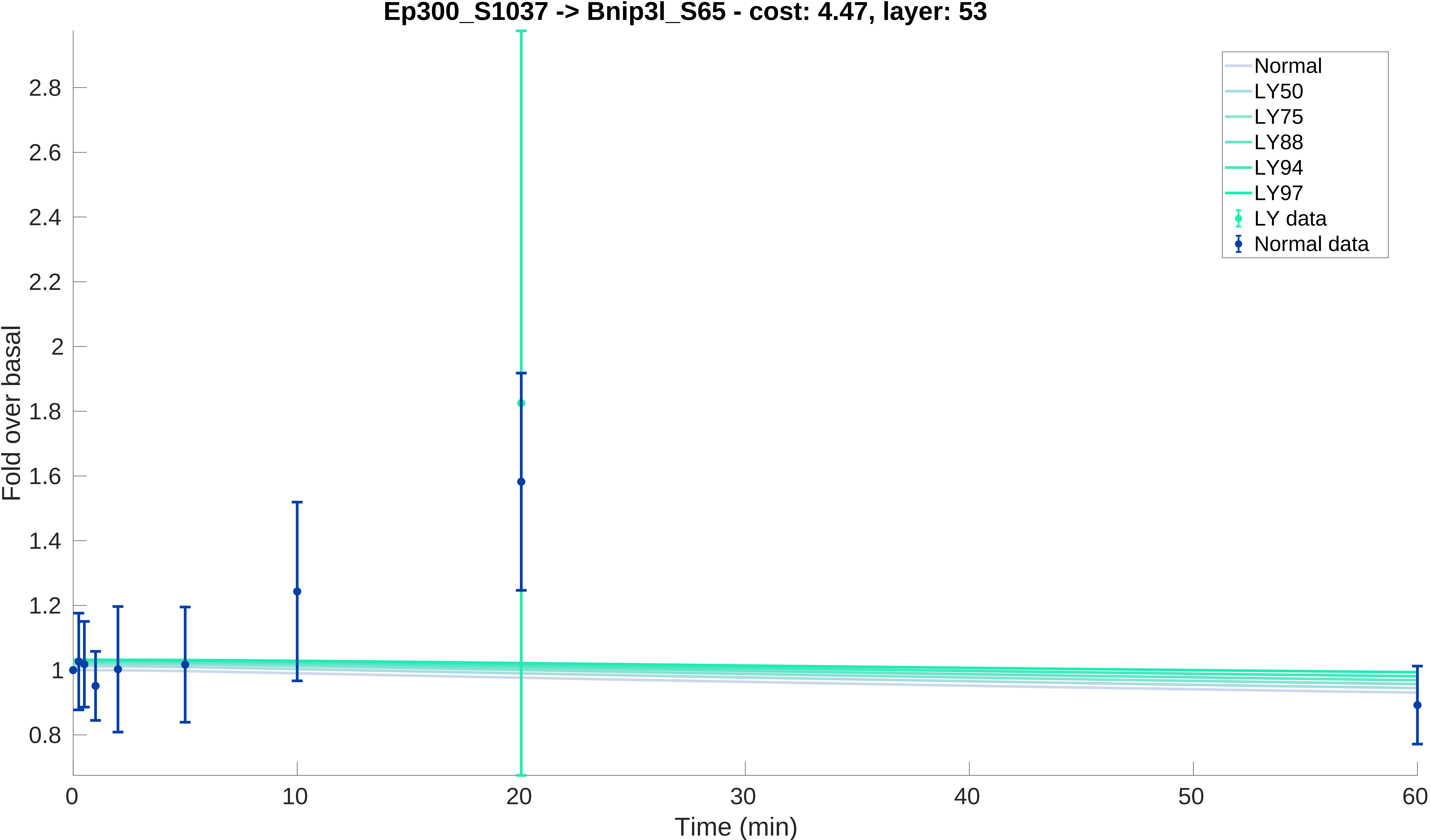

Ep300_S1037 -> Bnip3l_S65 - cost: 4.47, layer: 53
Fold over basal
Time (min)
Normal
LY50
LY75
LY88
LY94
LY97
LY data
Normal data
2.8
2.6
2.4
2.2
2
1.8
1.6
1.4
1.2
1
0.8
0
10
20
30
40
50
60

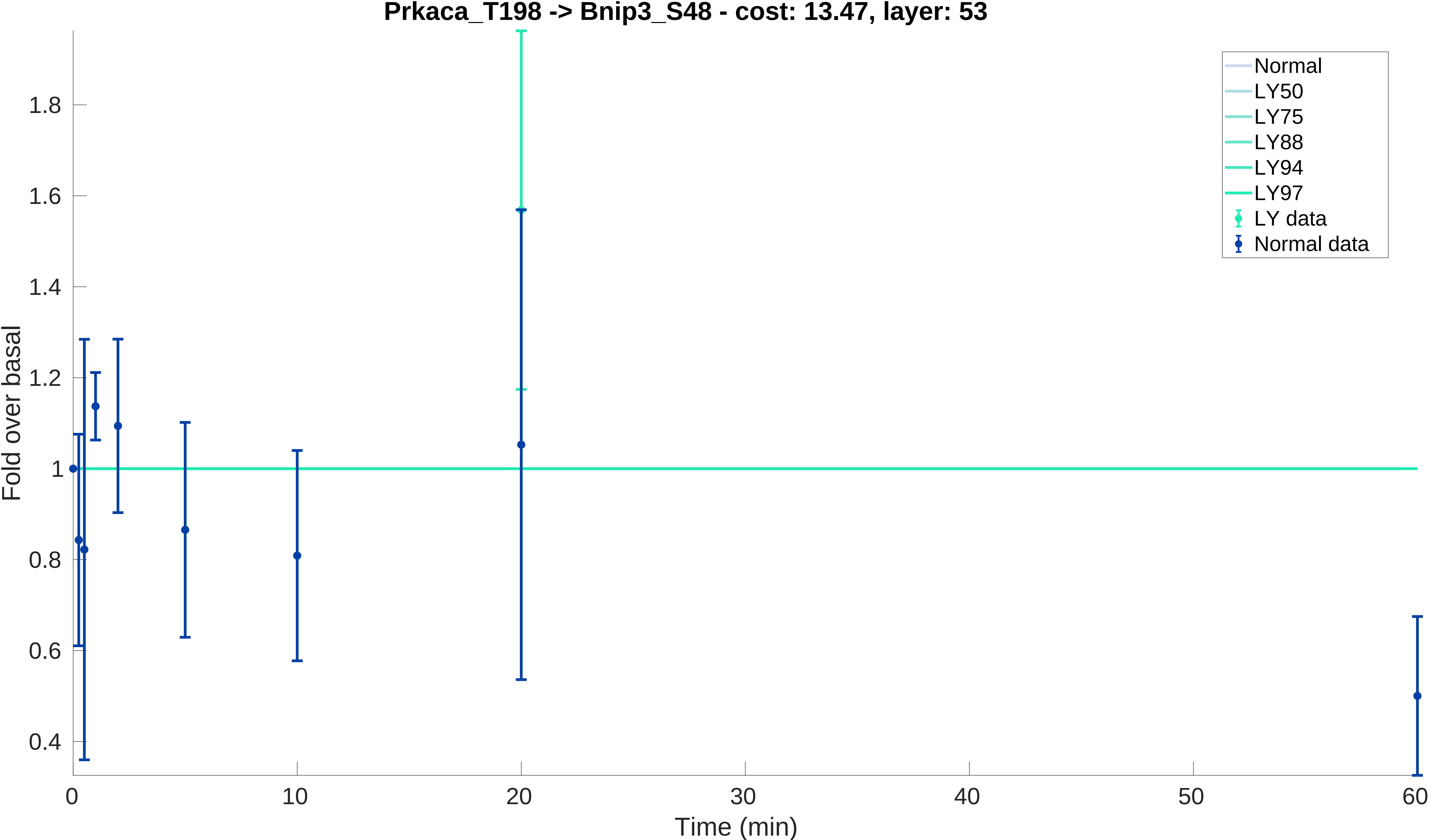
Prrkaca_T198 -> Bnip3_S48 - cost: 13.47, layer: 53
Fold over basal
Time (min)
1.8
1.6
1.4
1.2
1
0.8
0.6
0.4
0
10
20
30
40
50
60
Normal
LY50
LY75
LY88
LY94
LY97
LY data
Normal data

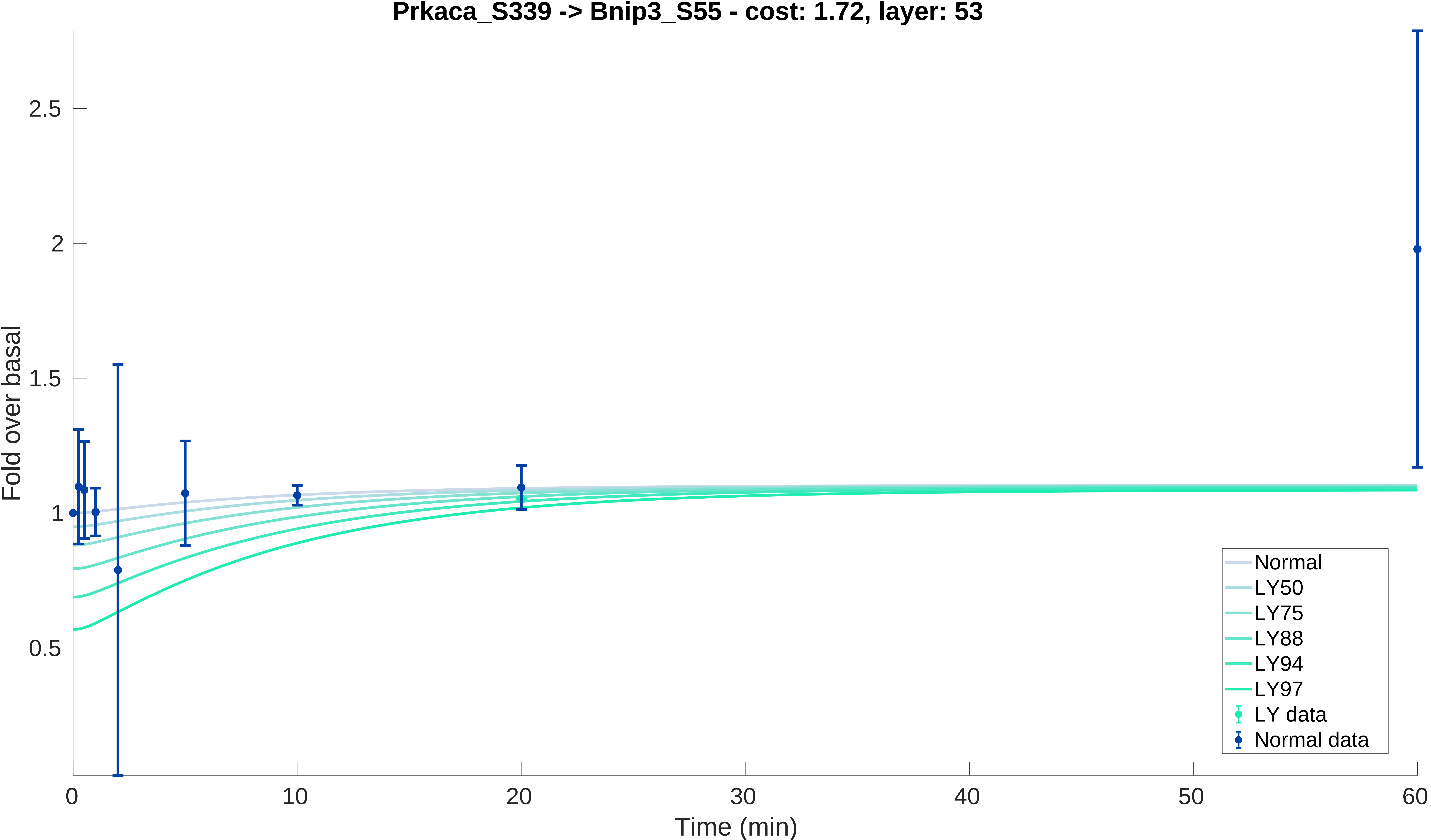

Prrkaca_S339 -> Bnip3_S55 - cost: 1.72, layer: 53
Fold over basal
Time (min)
0.5
1
1.5
2
2.5
0
10
20
30
40
50
60
Normal
LY50
LY75
LY88
LY94
LY97
LY data
Normal data

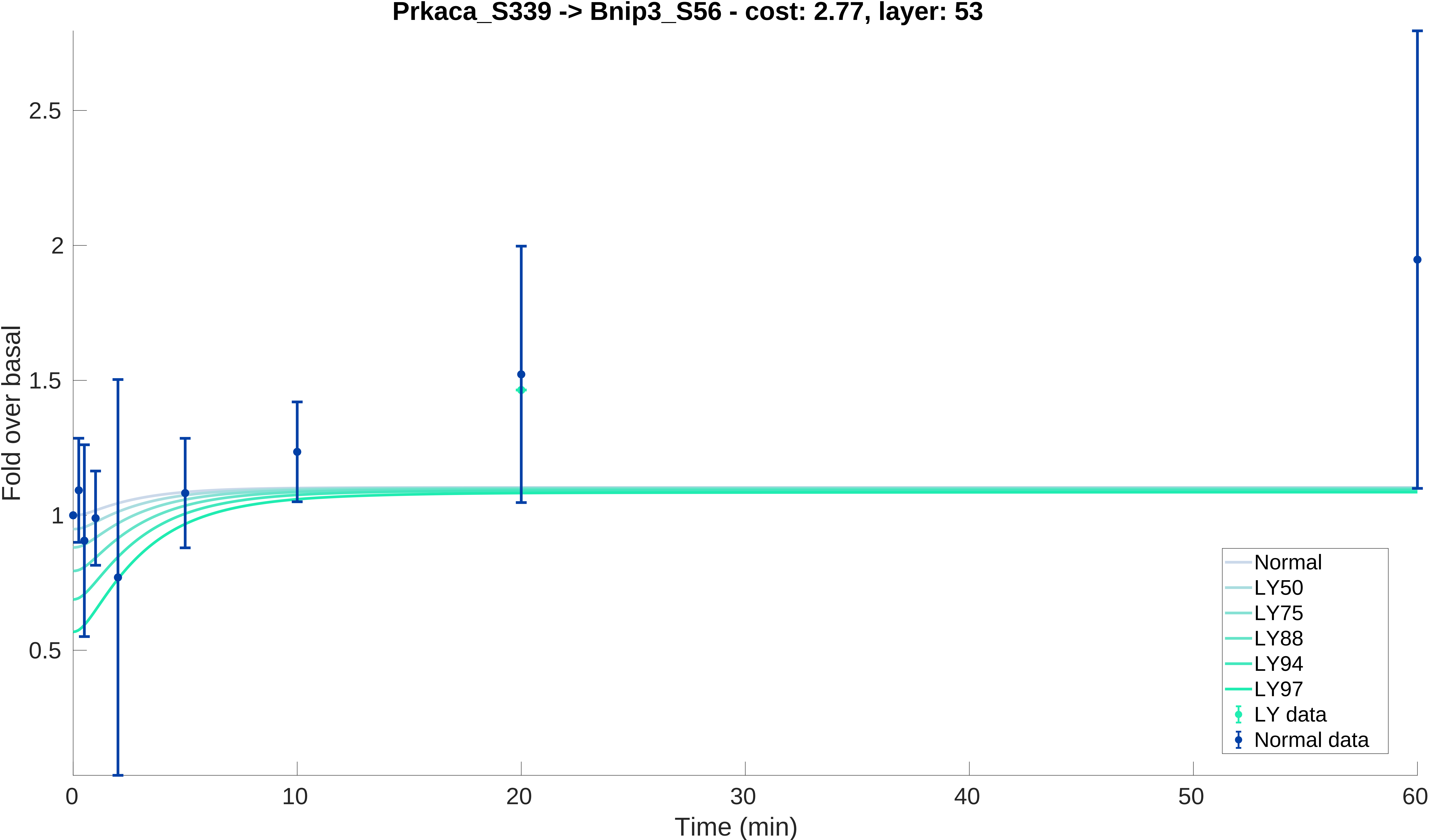
Prkaca_S339 -> Bnip3_S56 - cost: 2.77, layer: 53
Fold over basal
0.5
1
1.5
2
2.5
0
10
20
30
40
50
60
Time (min)
Normal
LY50
LY75
LY88
LY94
LY97
LY data
Normal data

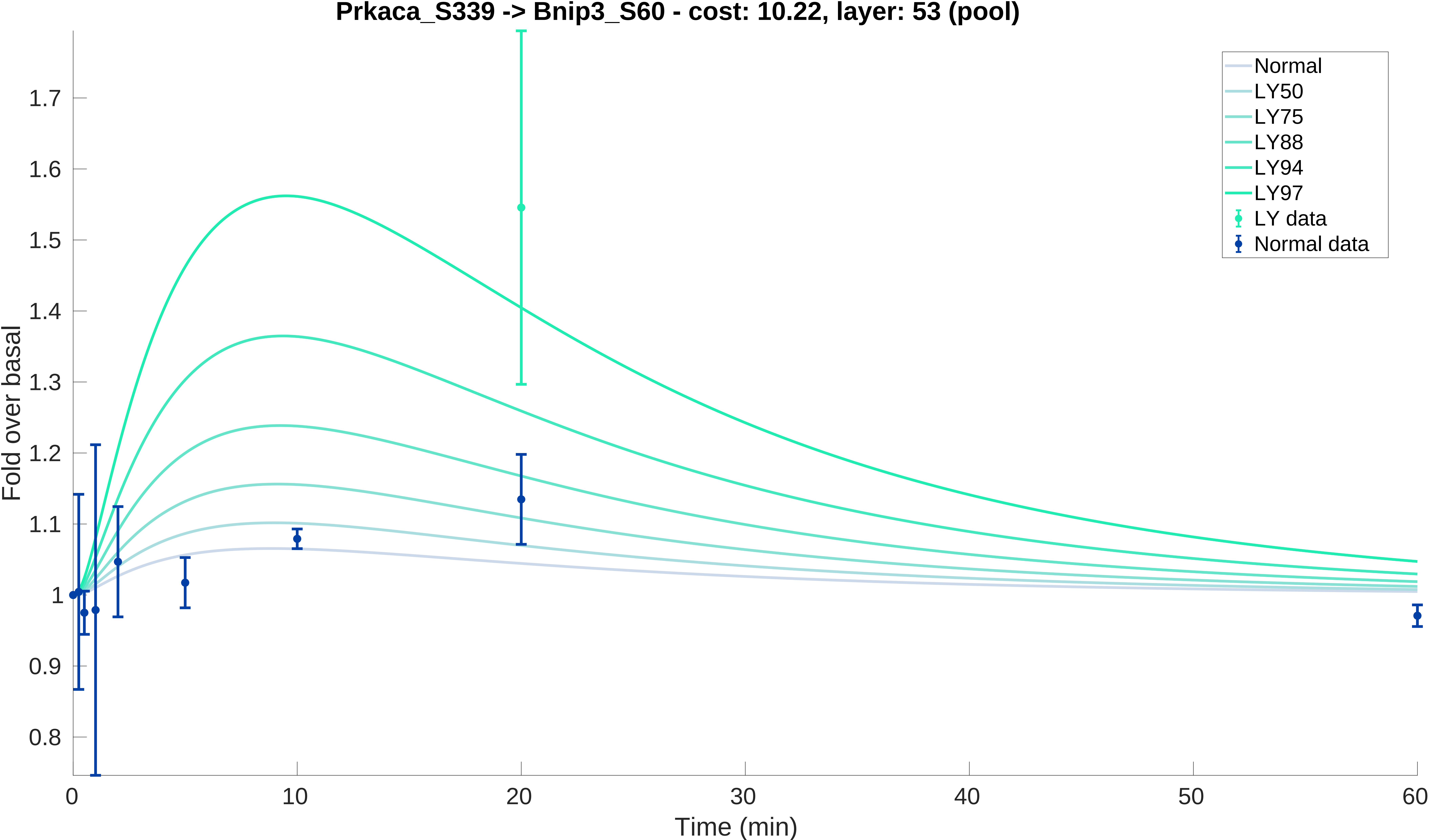

Prkaca_S339 -> Bnip3_S60 - cost: 10.22, layer: 53 (pool)
Normal
LY50
LY75
LY88
LY94
LY97
LY data
Normal data
Fold over basal
Time (min)
0
10
20
30
40
50
60
0.8
0.9
1
1.1
1.2
1.3
1.4
1.5
1.6
1.7

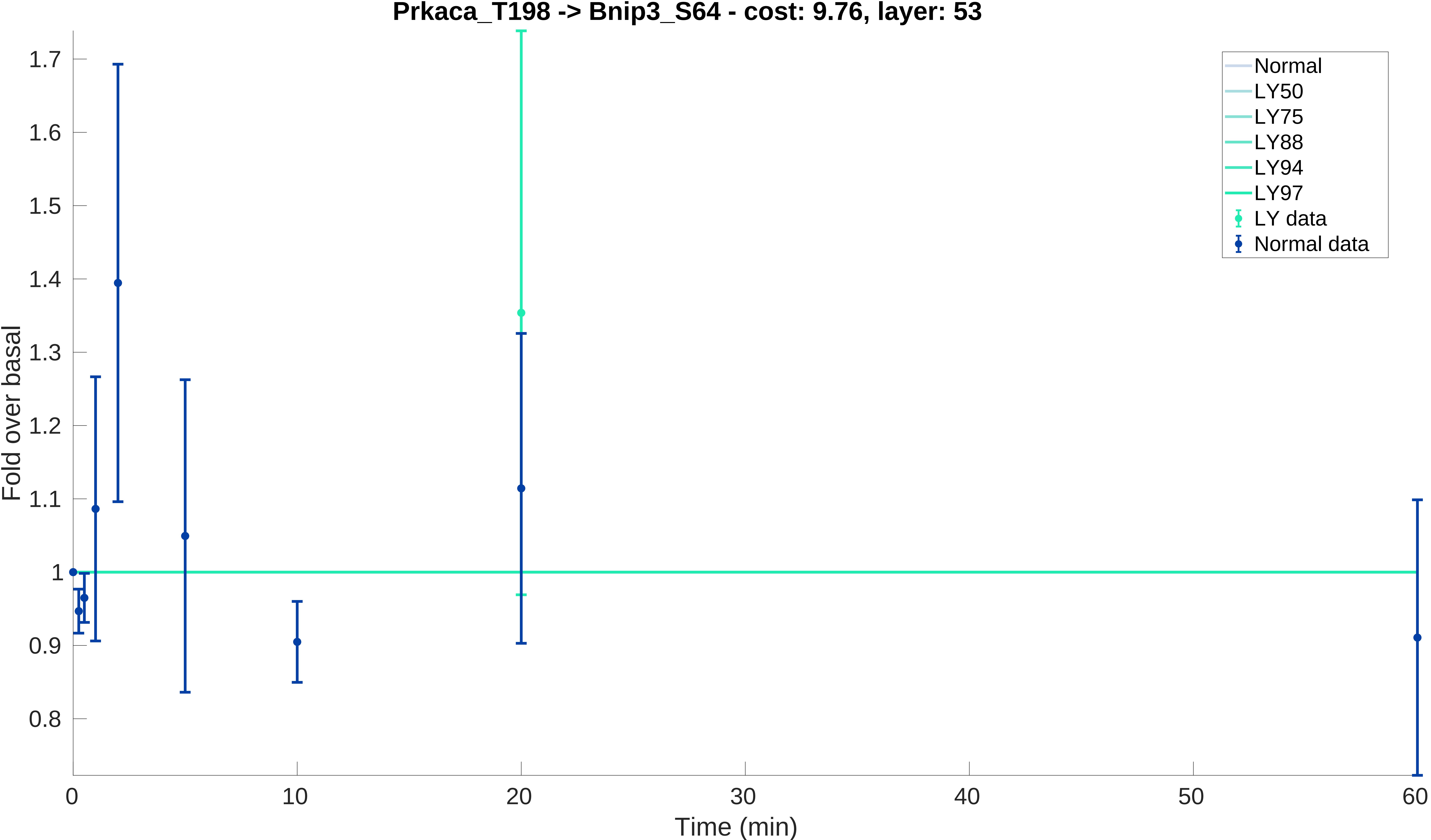
Prrkaca_T198 -> Bnip3_S64 - cost: 9.76, layer: 53
Fold over basal
Time (min)
0.8
0.9
1
1.1
1.2
1.3
1.4
1.5
1.6
1.7
0
10
20
30
40
50
60
Normal
LY50
LY75
LY88
LY94
LY97
LY data
Normal data

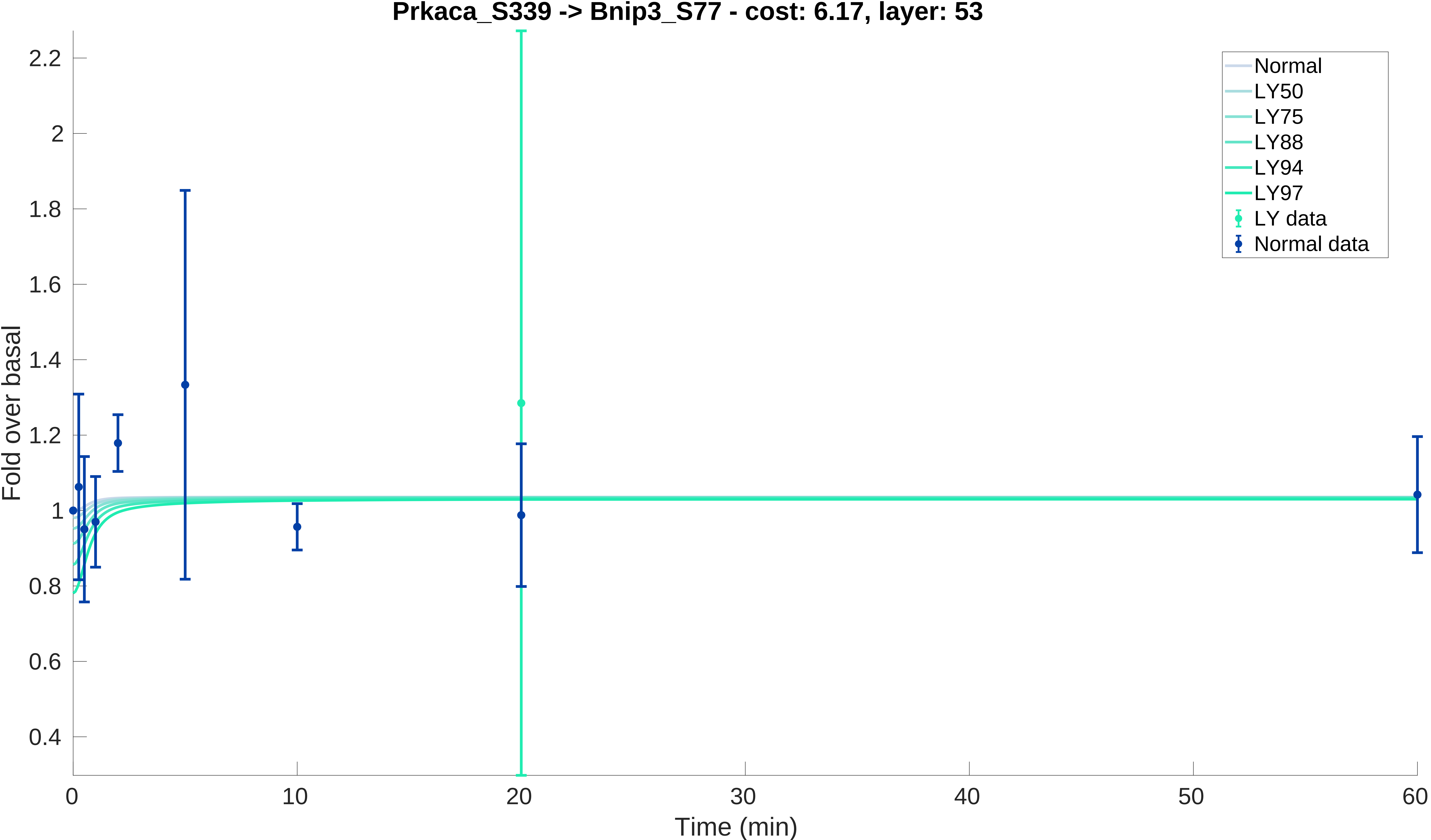

Prkaca_S339 -> Bnip3_S77 - cost: 6.17, layer: 53
Fold over basal
Time (min)
Normal
LY50
LY75
LY88
LY94
LY97
LY data
Normal data
0.4
0.6
0.8
1
1.2
1.4
1.6
1.8
2
2.2
0
10
20
30
40
50
60

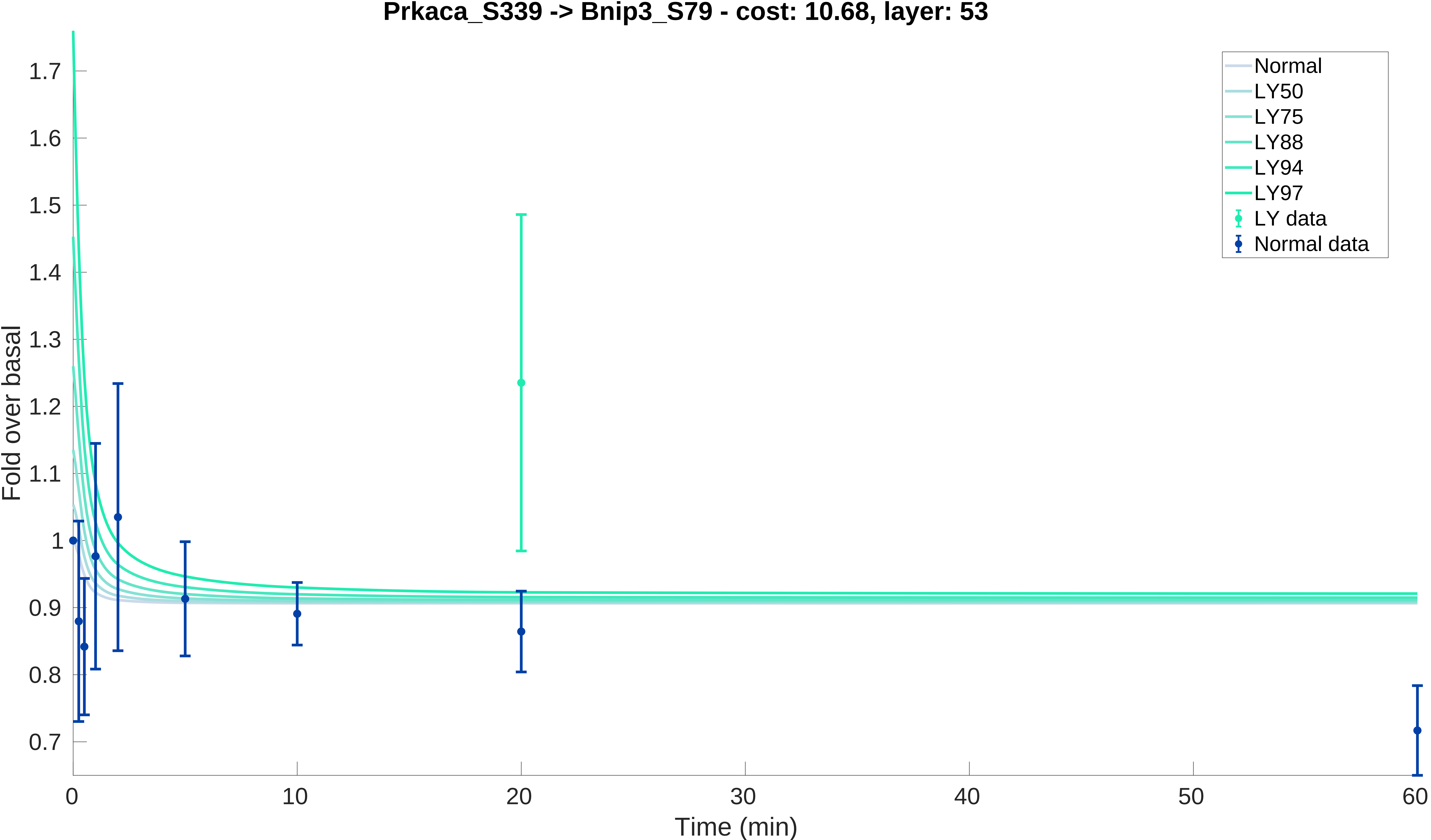
Prkaca_S339 -> Bnip3_S79 - cost: 10.68, layer: 53
Fold over basal
Time (min)
0.7
0.8
0.9
1
1.1
1.2
1.3
1.4
1.5
1.6
1.7
0
10
20
30
40
50
60
Normal
LY50
LY75
LY88
LY94
LY97
LY data
Normal data

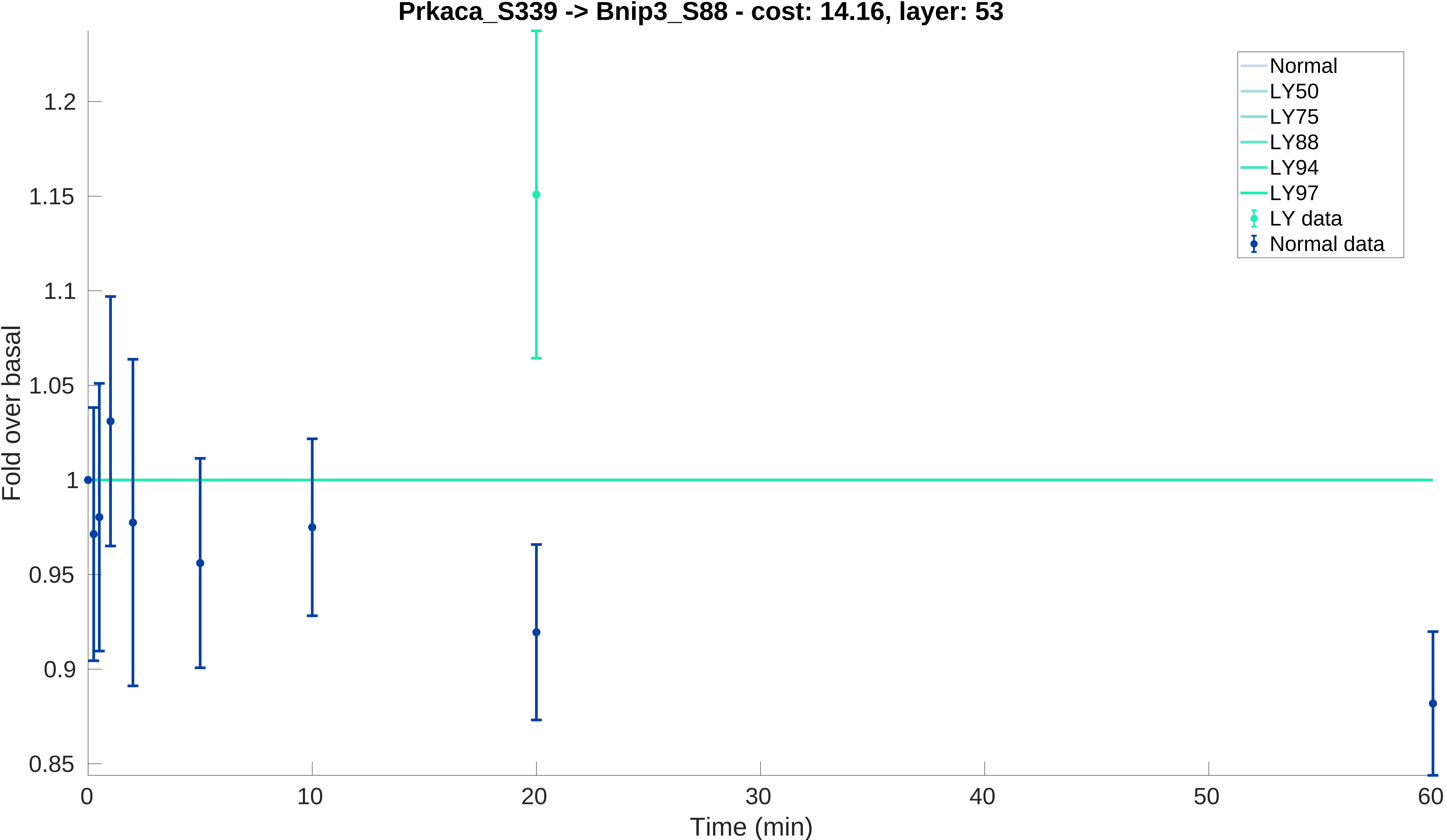
Prkaca_S339 -> Bnip3_S88 - cost: 14.16, layer: 53
Fold over basal
Time (min)
1.2
1.15
1.1
1.05
1
0.95
0.9
0.85
0
10
20
30
40
50
60
Normal
LY50
LY75
LY88
LY94
LY97
LY data
Normal data

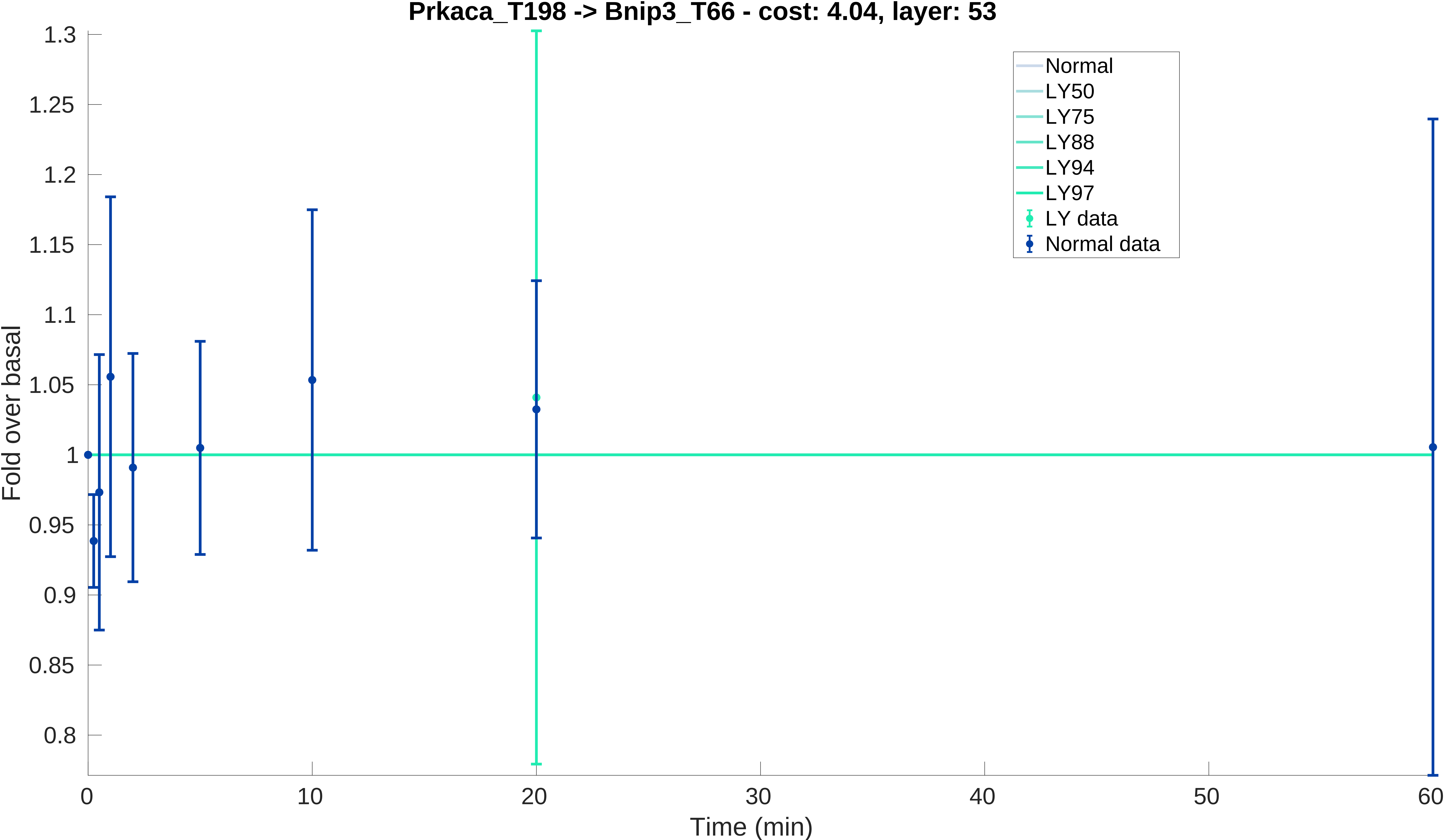

Prkaca_T198 -> Bnip3_T66 - cost: 4.04, layer: 53
Fold over basal
Time (min)
Normal
LY50
LY75
LY88
LY94
LY97
LY data
Normal data
1.3
1.25
1.2
1.15
1.1
1.05
1
0.95
0.9
0.85
0.8
0
10
20
30
40
50
60

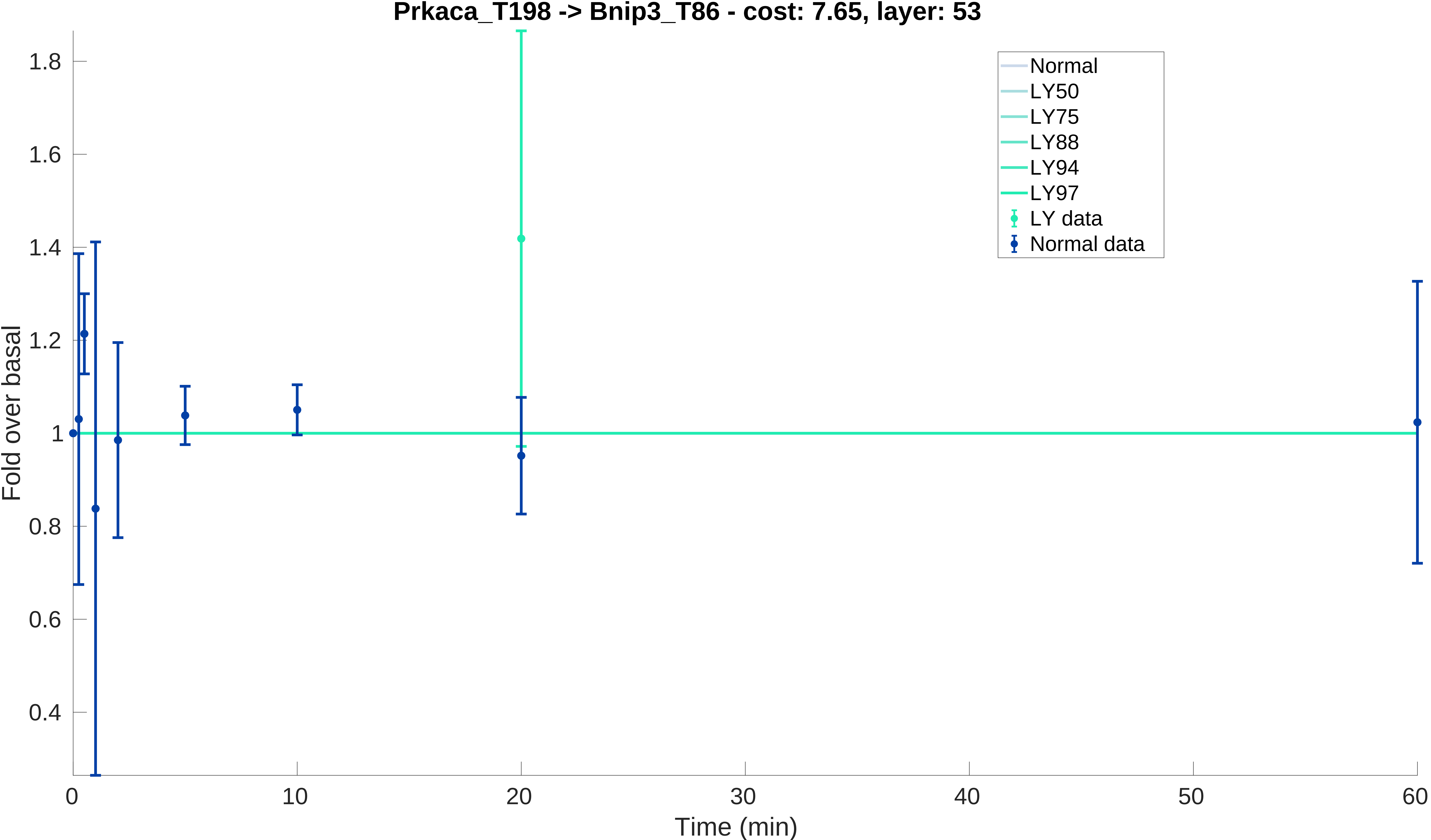

Prrkaca_T198 -> Bnip3_T86 - cost: 7.65, layer: 53
Normal
LY50
LY75
LY88
LY94
LY97
LY data
Normal data
Fold over basal
Time (min)
1.8
1.6
1.4
1.2
1
0.8
0.6
0.4
0
10
20
30
40
50
60

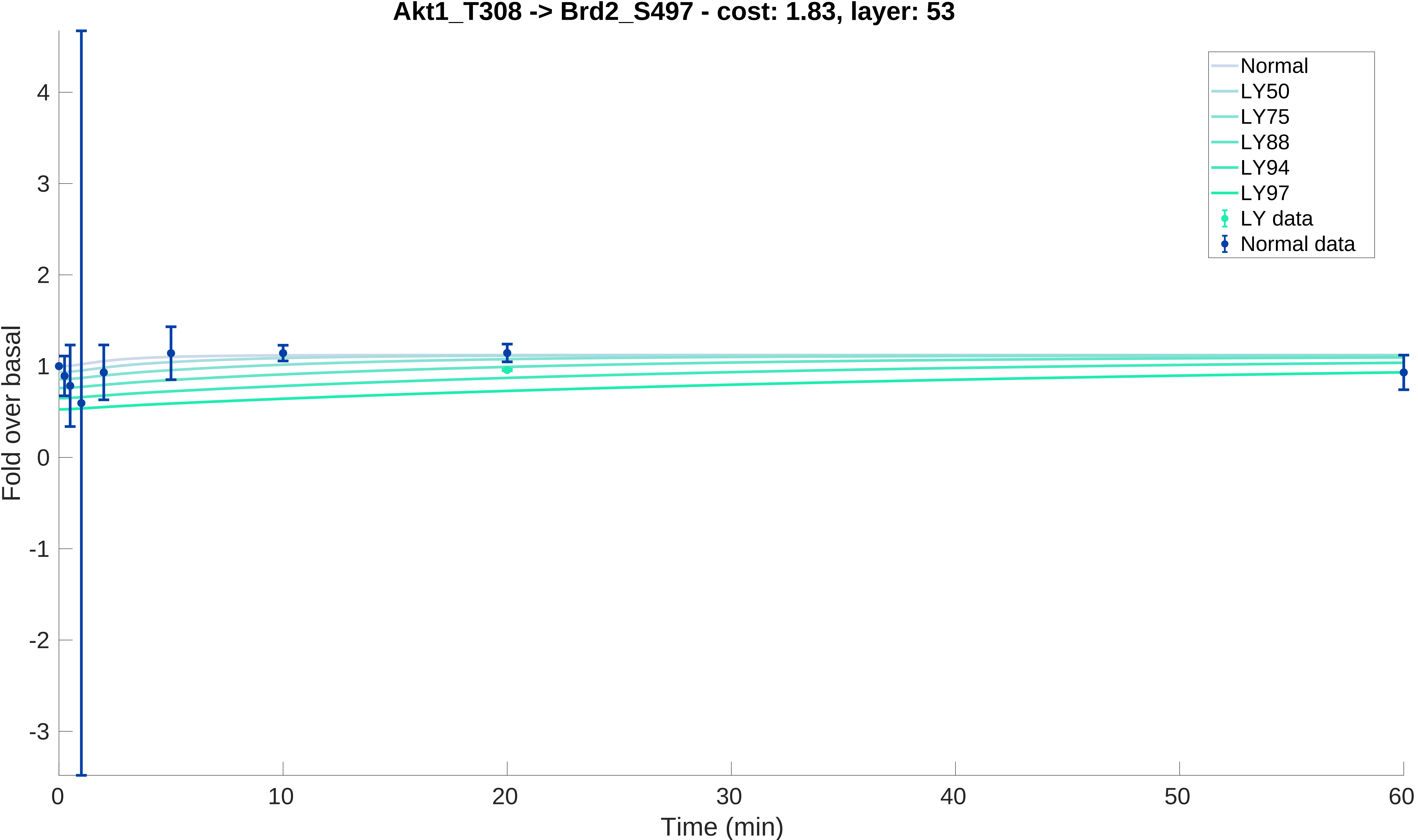

Akt1_T308 -> Brd2_S497 - cost: 1.83, layer: 53
Normal
LY50
LY75
LY88
LY94
LY97
LY data
Normal data
Fold over basal
4
3
2
1
0
-1
-2
-3
0
10
20
30
40
50
60
Time (min)

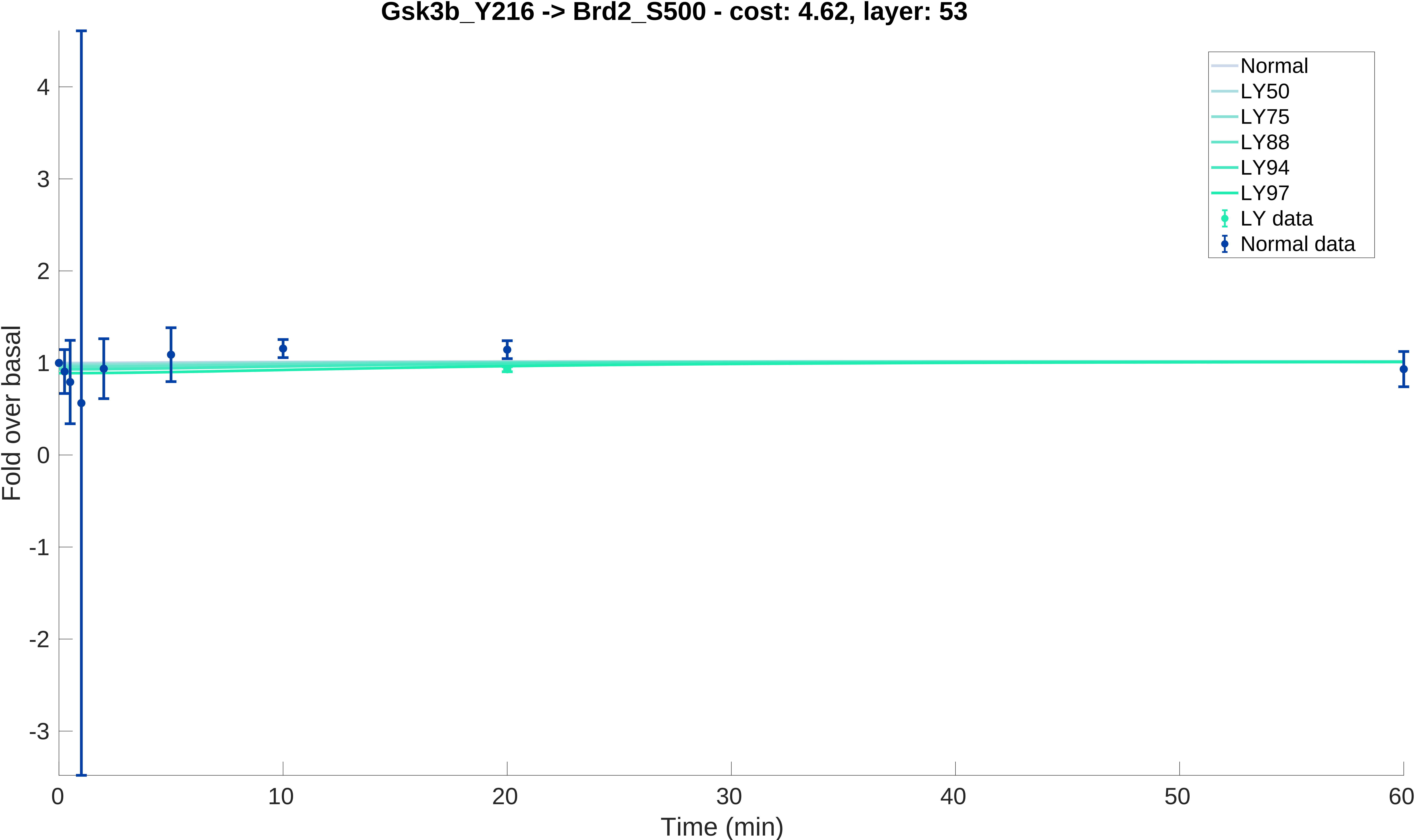

Gsk3b_Y216 -> Brd2_S500 - cost: 4.62, layer: 53
Normal
LY50
LY75
LY88
LY94
LY97
LY data
Normal data
Fold over basal
Time (min)
4
3
2
1
0
-1
-2
-3
0
10
20
30
40
50
60

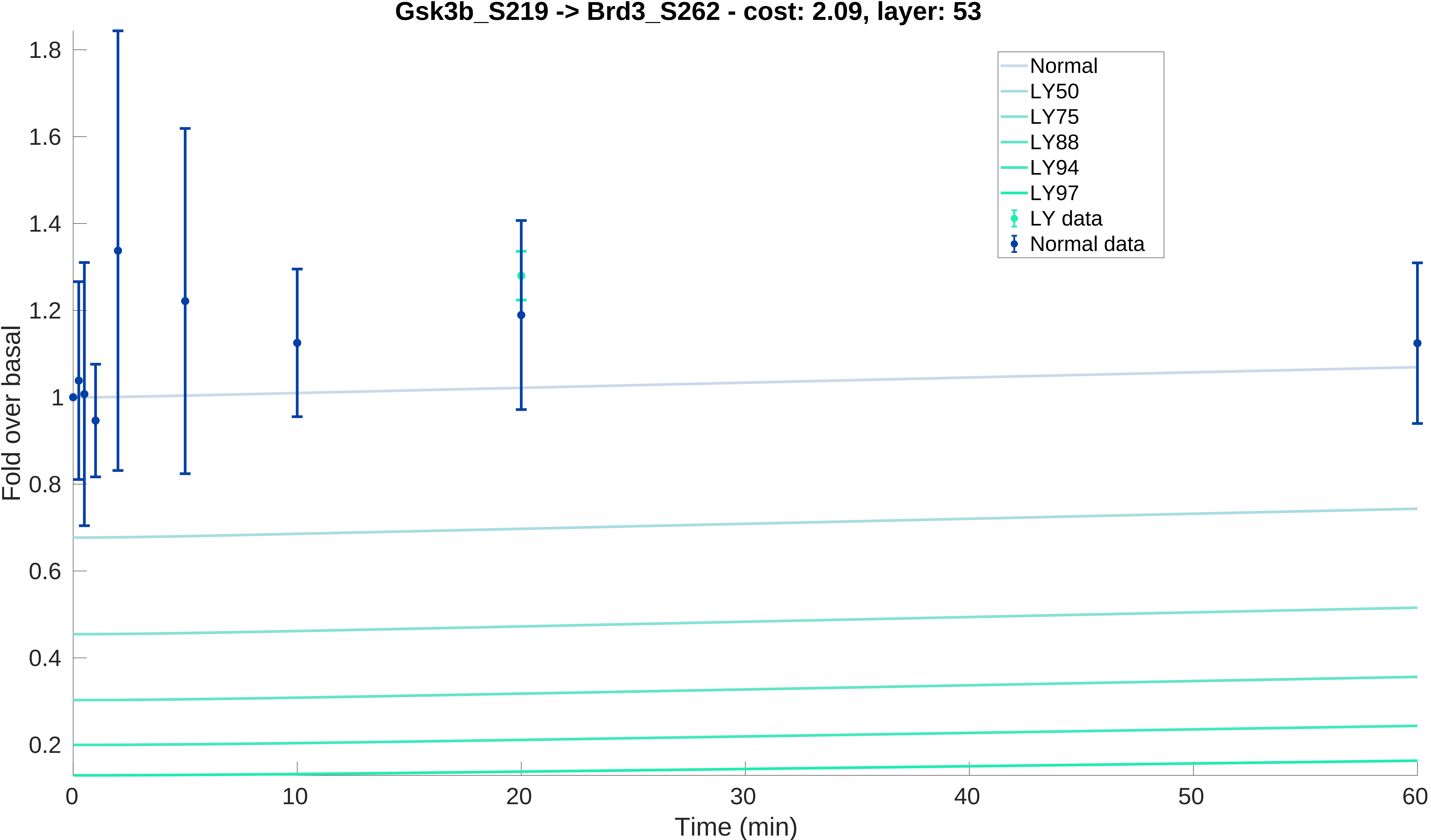

Gsk3b_S219 -> Brd3_S262 - cost: 2.09, layer: 53
Fold over basal
1.8
1.6
1.4
1.2
1
0.8
0.6
0.4
0.2
0
10
20
30
40
50
60
Time (min)
Normal
LY50
LY75
LY88
LY94
LY97
LY data
Normal data

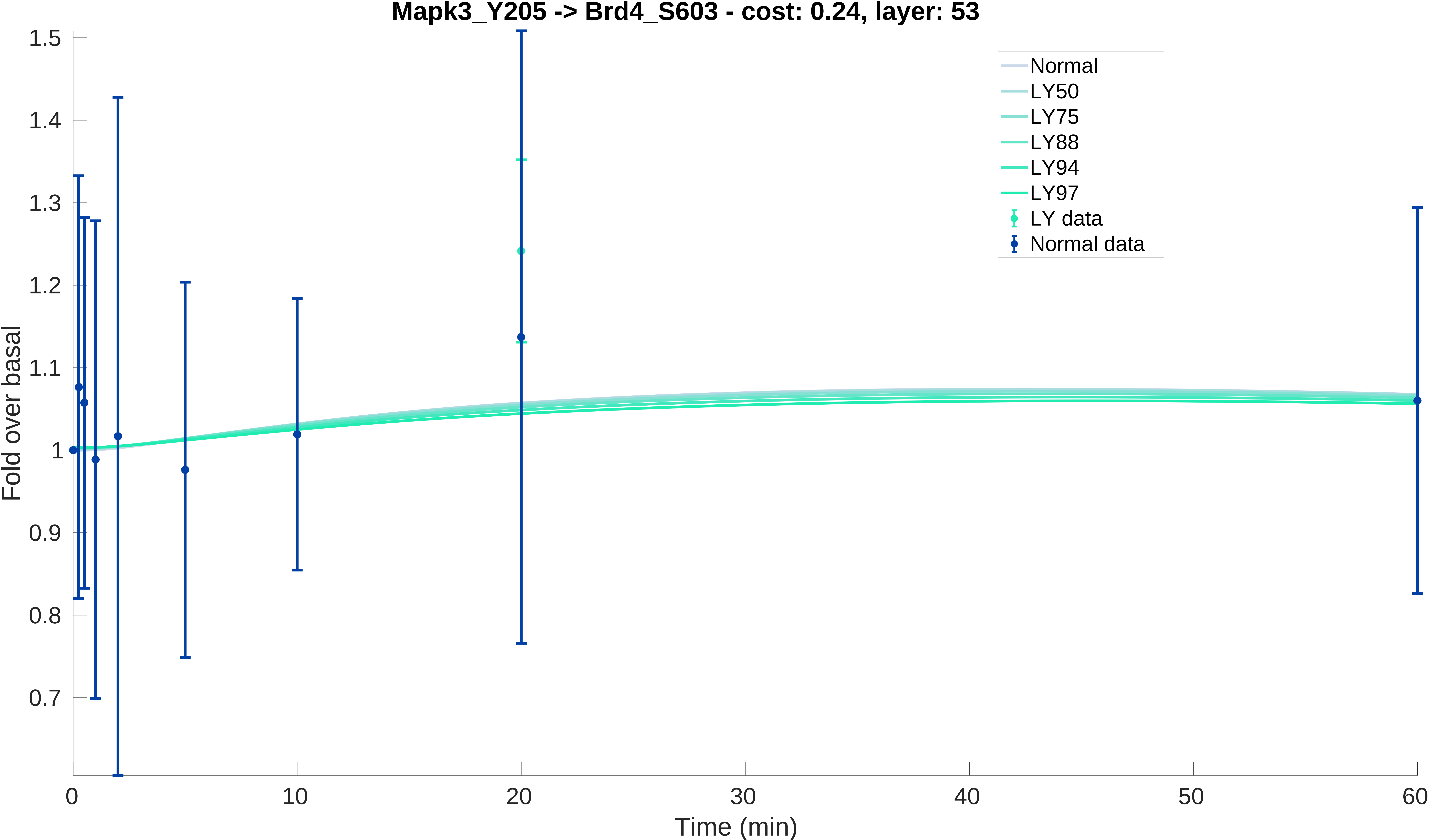

Mapk3_Y205 -> Brd4_S603 - cost: 0.24, layer: 53
Fold over basal
1.5
1.4
1.3
1.2
1.1
1
0.9
0.8
0.7
0
10
20
30
40
50
60
Time (min)
Normal
LY50
LY75
LY88
LY94
LY97
LY data
Normal data

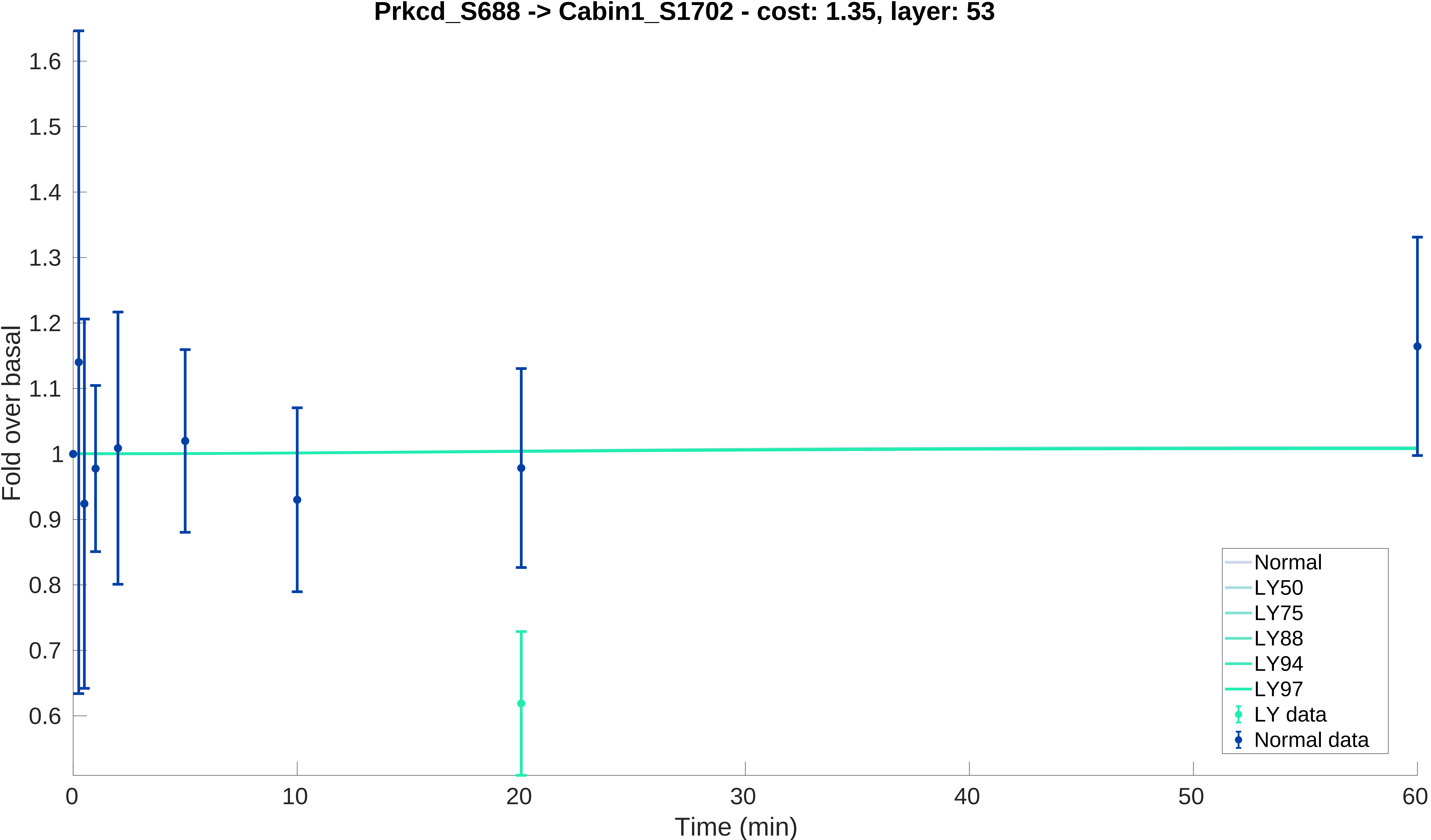

Prkcd_S688 -> Cabin1_S1702 - cost: 1.35, layer: 53
Fold over basal
1.6
1.5
1.4
1.3
1.2
1.1
1
0.9
0.8
0.7
0.6
0
10
20
30
40
50
60
Time (min)
Normal
LY50
LY75
LY88
LY94
LY97
LY data
Normal data

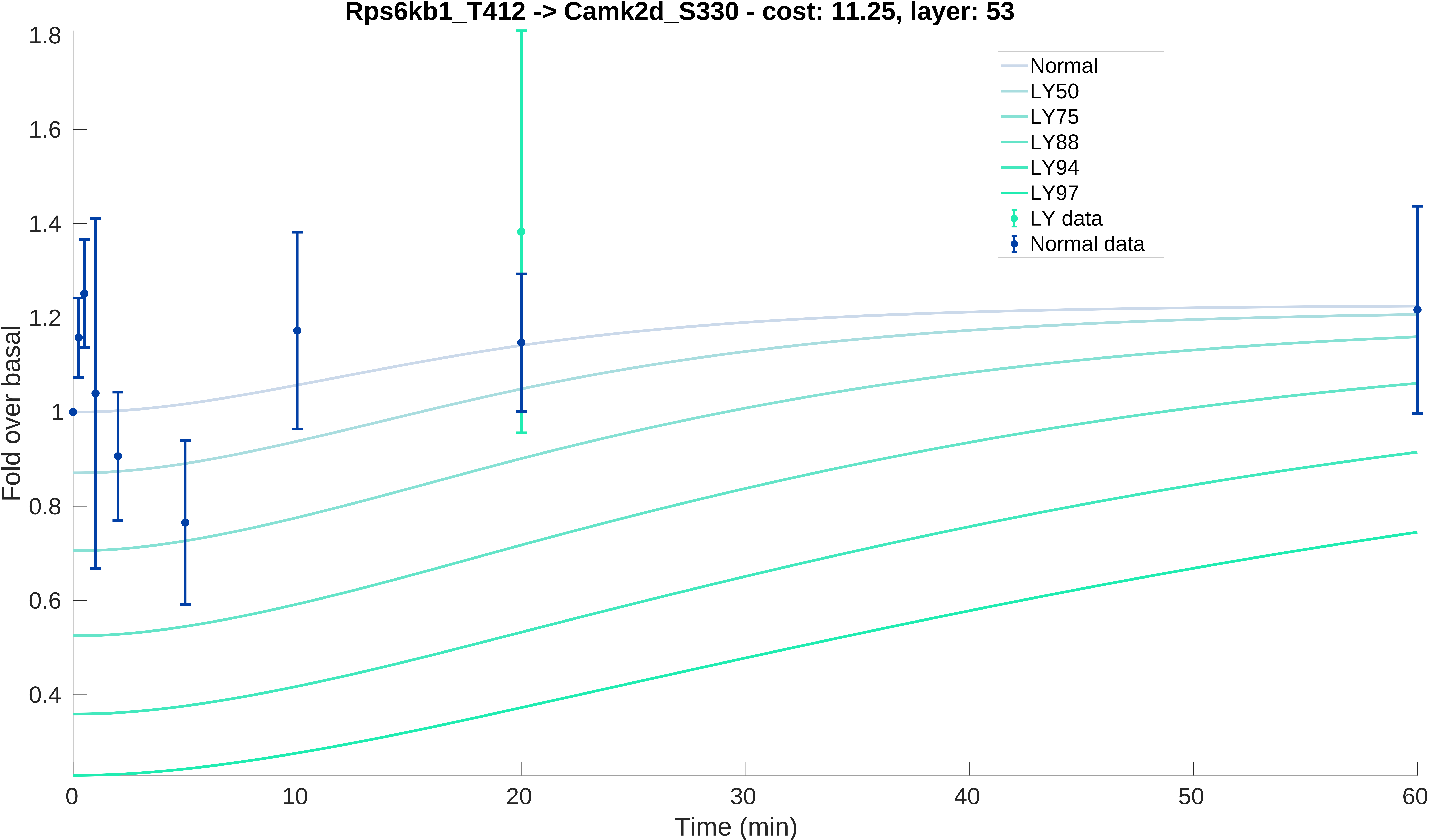

Rps6kb1_T412 -> Camk2d_S330 - cost: 11.25, layer: 53
Fold over basal
Time (min)
1.8
1.6
1.4
1.2
1
0.8
0.6
0.4
0
10
20
30
40
50
60
Normal
LY50
LY75
LY88
LY94
LY97
LY data
Normal data

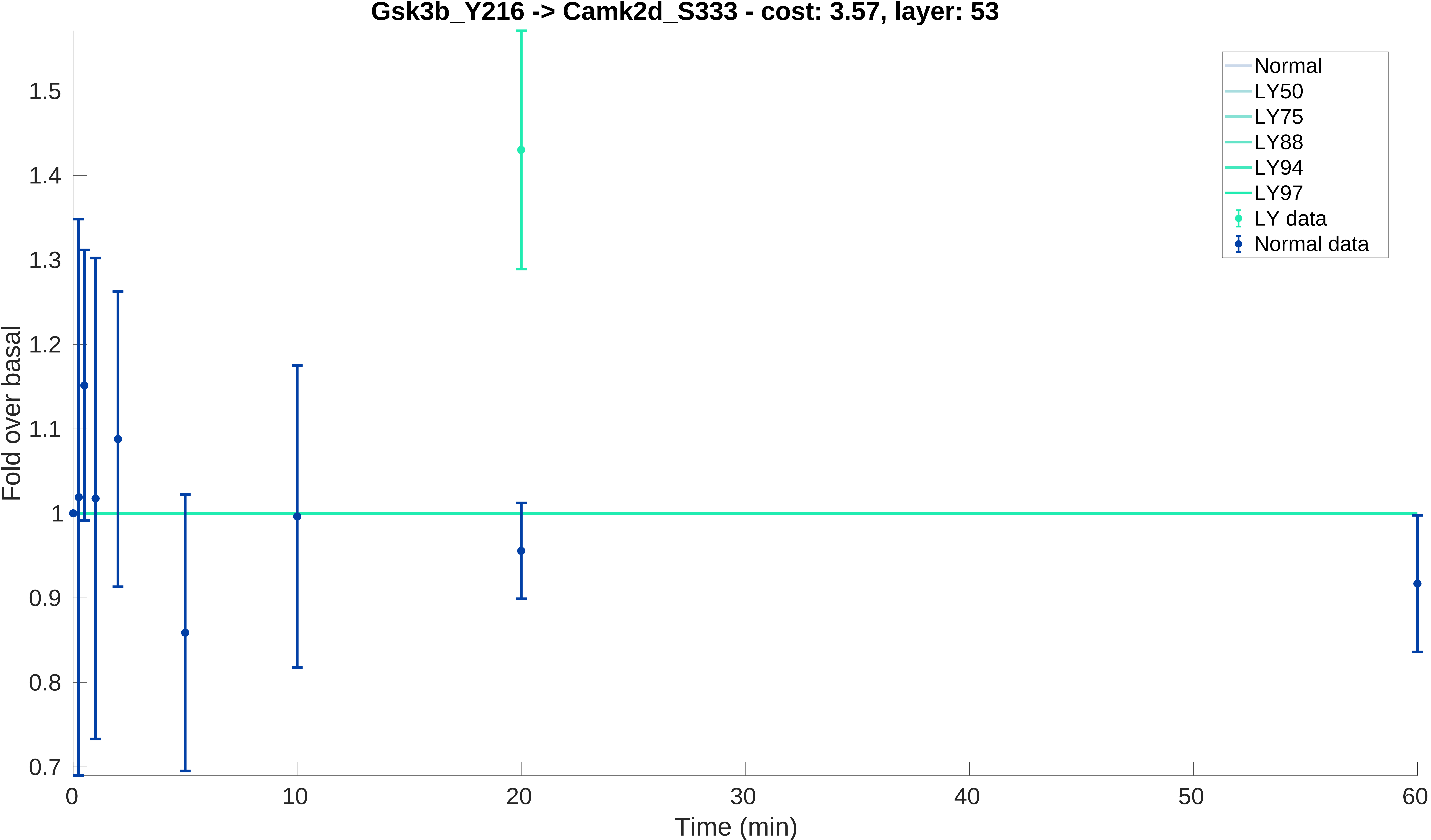

Gsk3b_Y216 -> Camk2d_S333 - cost: 3.57, layer: 53
Fold over basal
Time (min)
0.7
0.8
0.9
1
1.1
1.2
1.3
1.4
1.5
0
10
20
30
40
50
60
Normal
LY50
LY75
LY88
LY94
LY97
LY data
Normal data

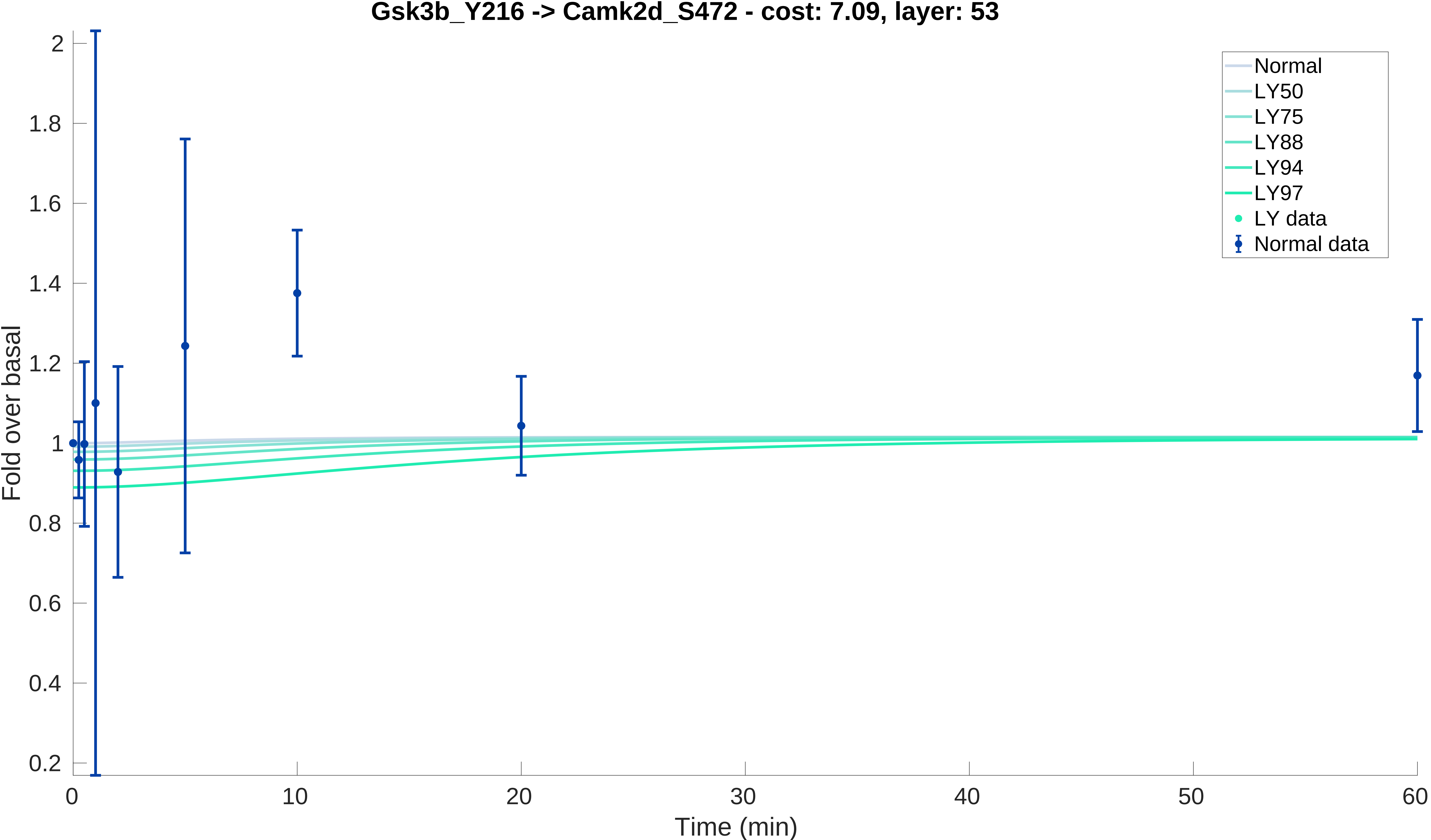
Gsk3b_Y216 -> Camk2d_S472 - cost: 7.09, layer: 53
Fold over basal
Time (min)
Normal
LY50
LY75
LY88
LY94
LY97
LY data
Normal data
0.2
0.4
0.6
0.8
1
1.2
1.4
1.6
1.8
2
0
10
20
30
40
50
60

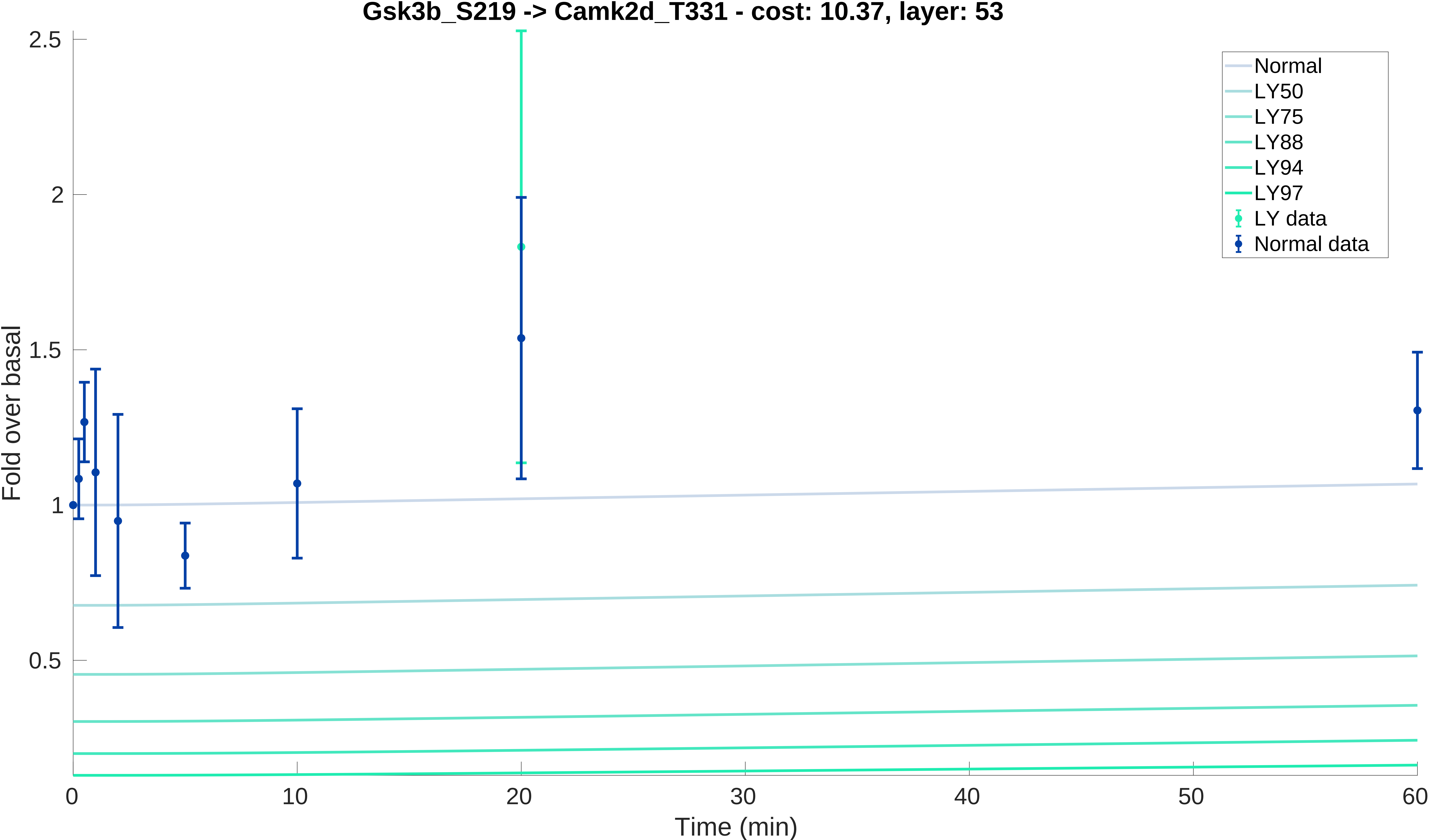

Gsk3b_S219 -> Camk2d_T331 - cost: 10.37, layer: 53
Fold over basal
Time (min)
0
10
20
30
40
50
60
0.5
1
1.5
2
2.5
Normal
LY50
LY75
LY88
LY94
LY97
LY data
Normal data

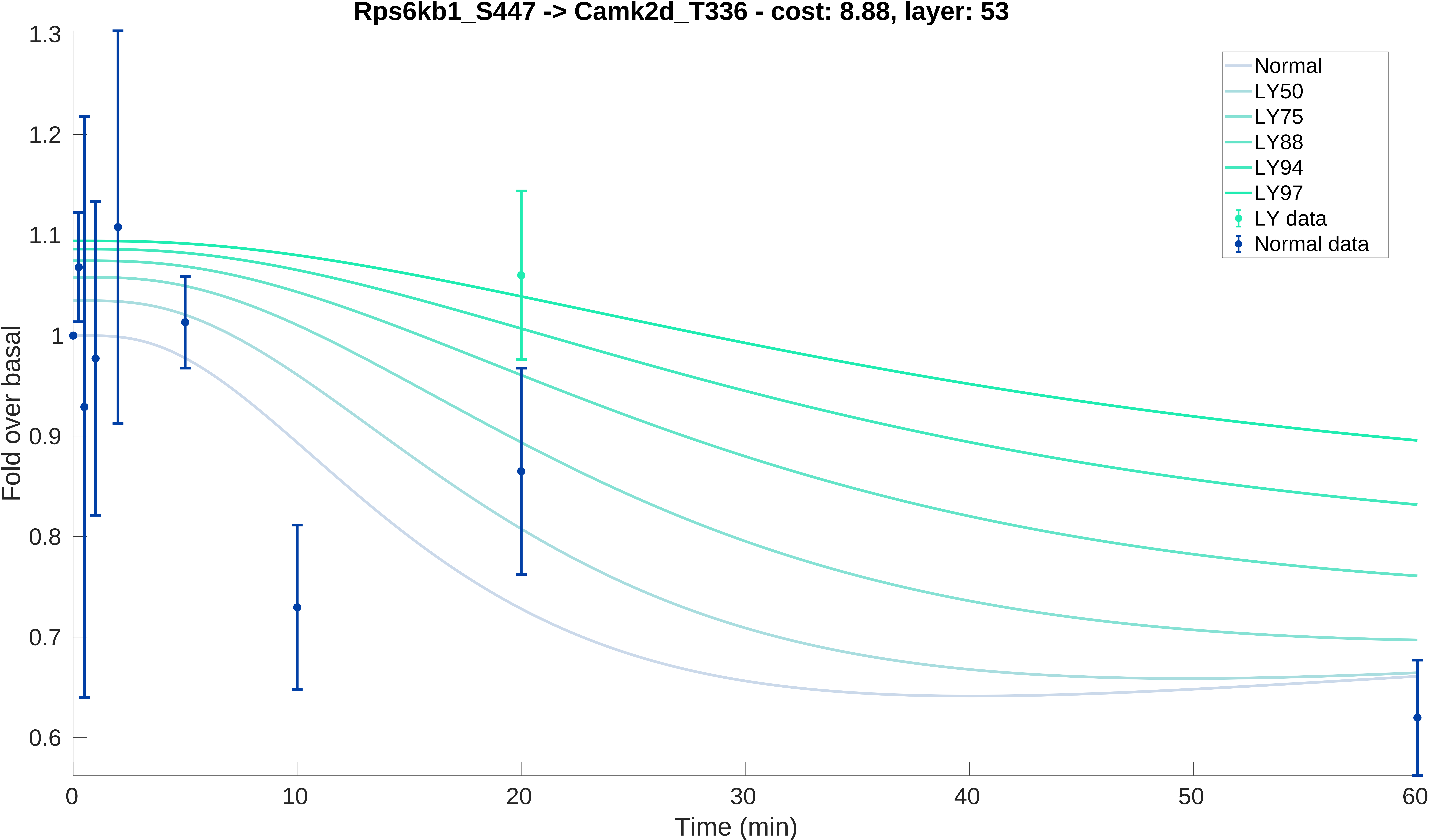

Rps6kb1_S447 -> Camk2d_T336 - cost: 8.88, layer: 53
Fold over basal
Time (min)
1.3
1.2
1.1
1
0.9
0.8
0.7
0.6
0
10
20
30
40
50
60
Normal
LY50
LY75
LY88
LY94
LY97
LY data
Normal data

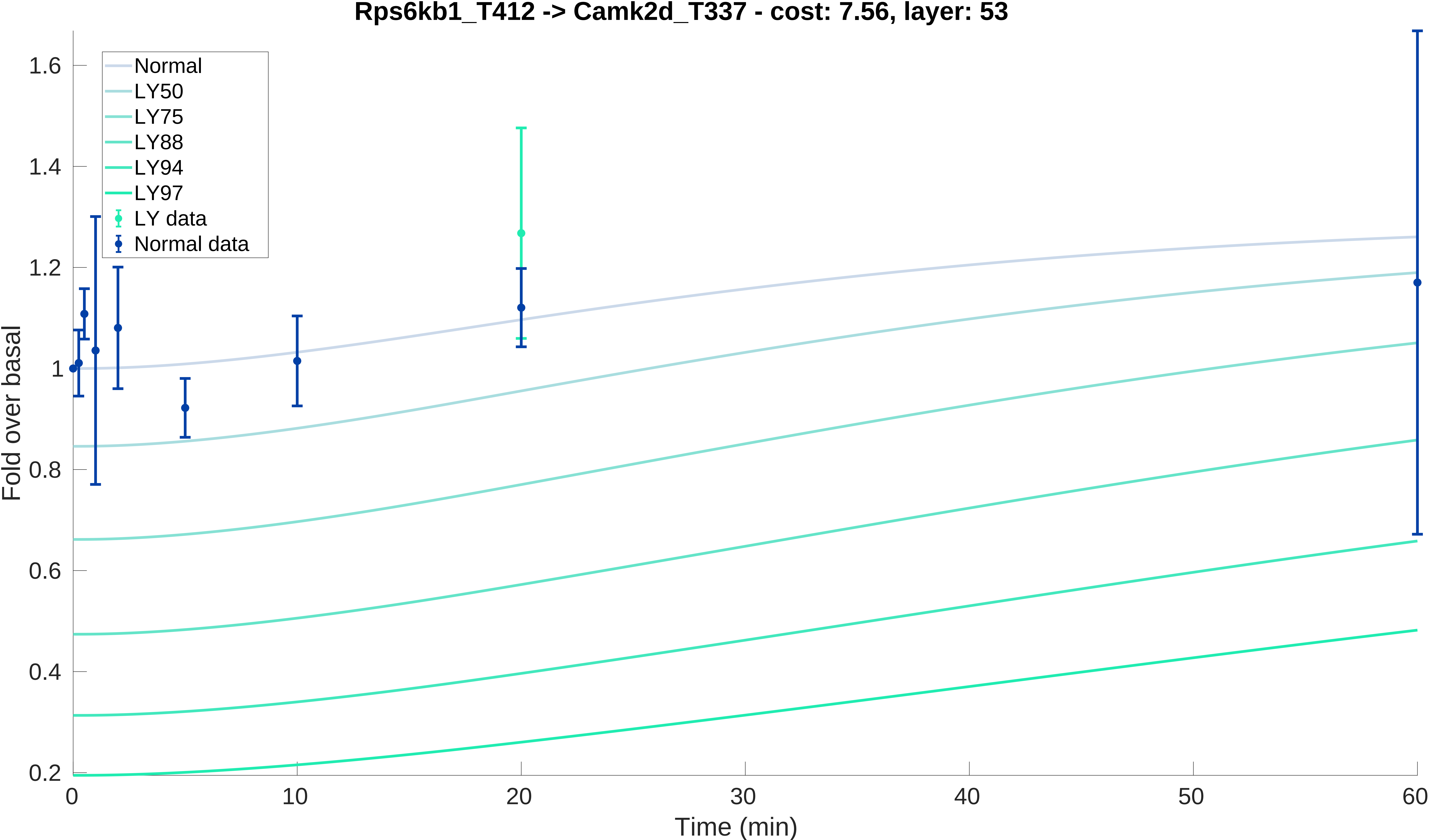

Rps6kb1_T412 -> Camk2d_T337 - cost: 7.56, layer: 53
Normal
LY50
LY75
LY88
LY94
LY97
LY data
Normal data
Fold over basal
Time (min)
1.6
1.4
1.2
1
0.8
0.6
0.4
0.2
0
10
20
30
40
50
60

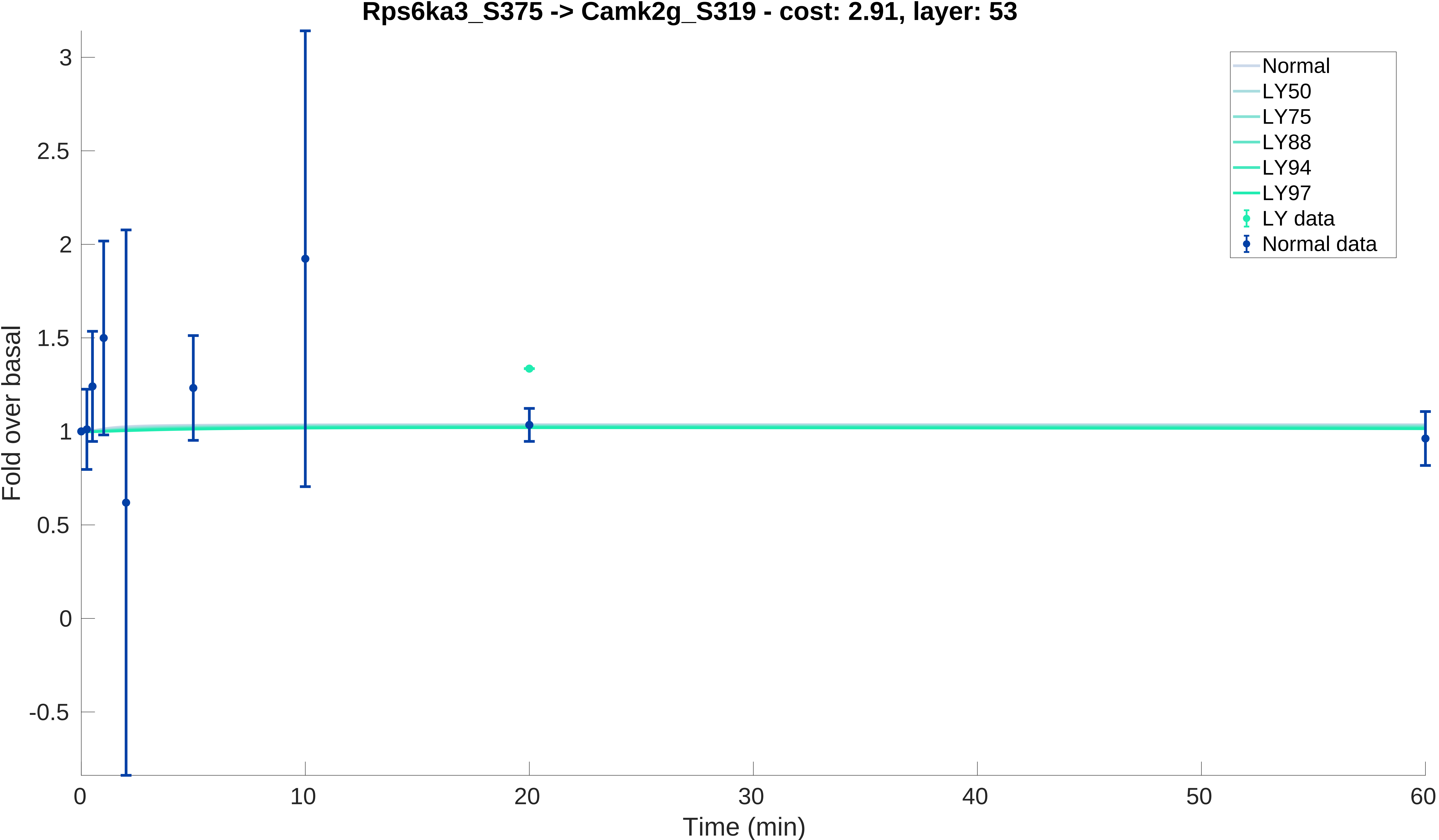

Rps6ka3_S375 -> Camk2g_S319 - cost: 2.91, layer: 53
Fold over basal
Time (min)
3
2.5
2
1.5
1
0.5
0
-0.5
0
10
20
30
40
50
60
Normal
LY50
LY75
LY88
LY94
LY97
LY data
Normal data

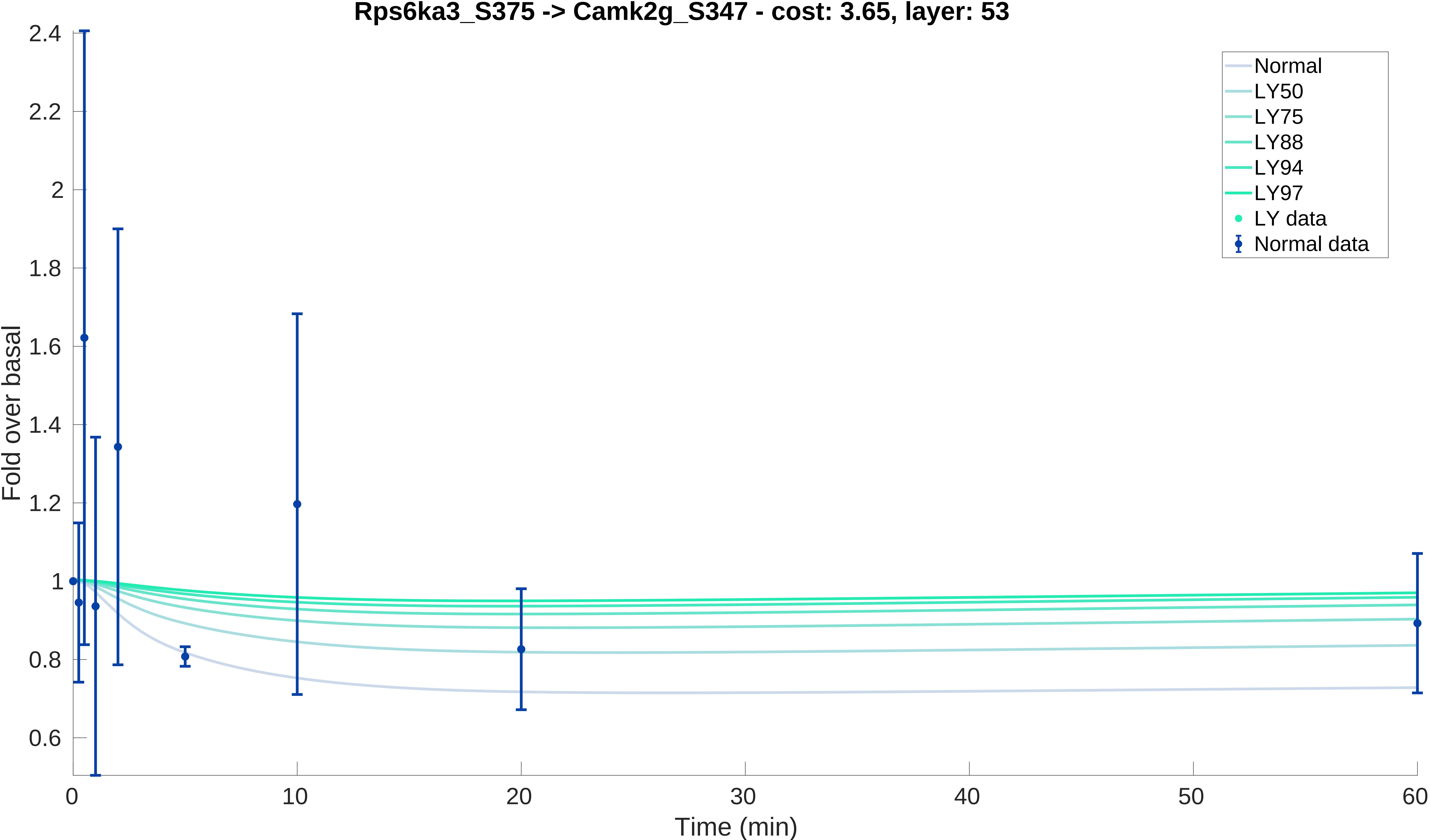

Rps6ka3_S375 -> Camk2g_S347 - cost: 3.65, layer: 53
Fold over basal
Time (min)
2.4
2.2
2
1.8
1.6
1.4
1.2
1
0.8
0.6
0
10
20
30
40
50
60
Normal
LY50
LY75
LY88
LY94
LY97
LY data
Normal data

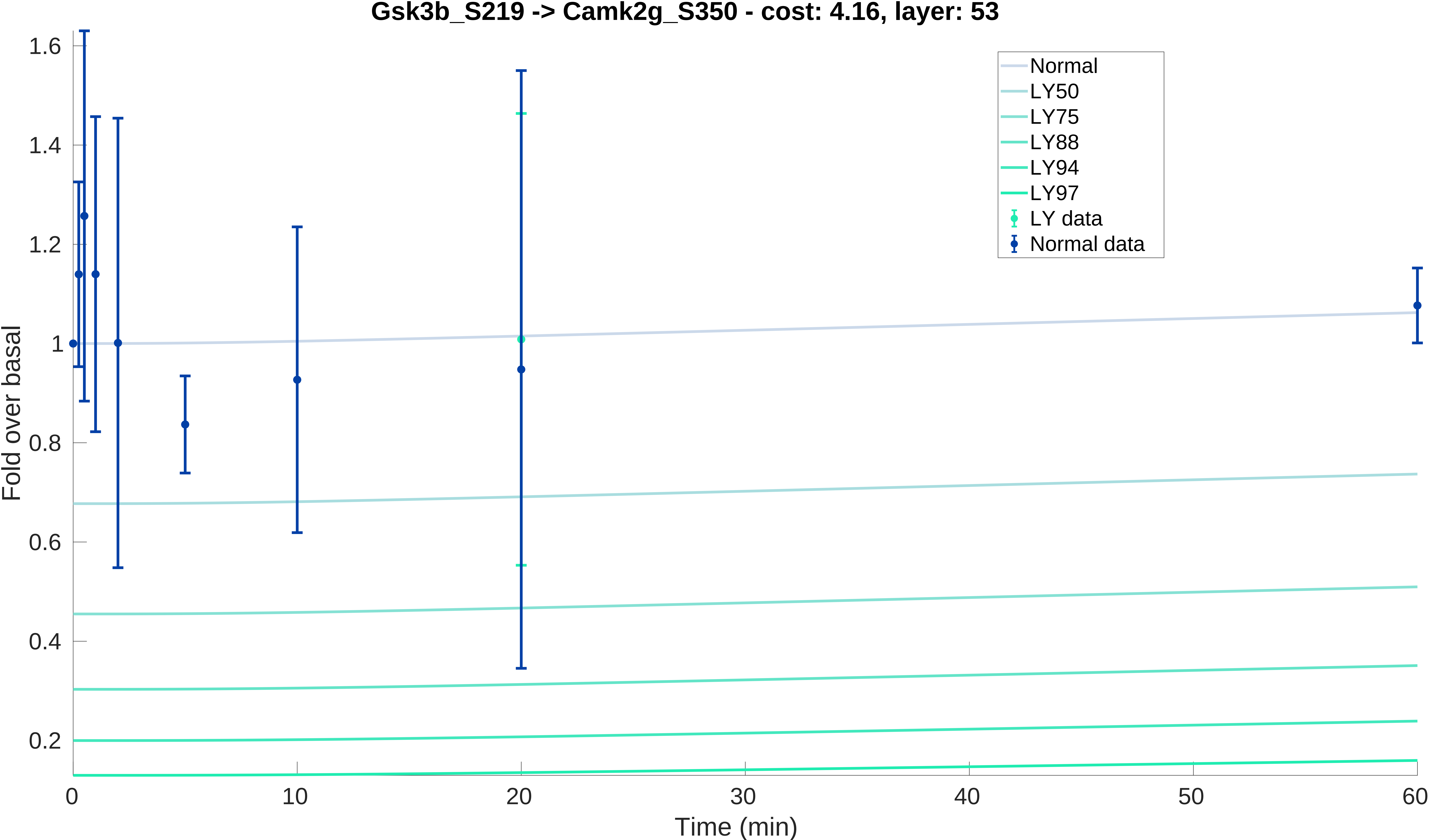

Gsk3b_S219 -> Camk2g_S350 - cost: 4.16, layer: 53
Fold over basal
Time (min)
0
10
20
30
40
50
60
0.2
0.4
0.6
0.8
1
1.2
1.4
1.6
Normal
LY50
LY75
LY88
LY94
LY97
LY data
Normal data

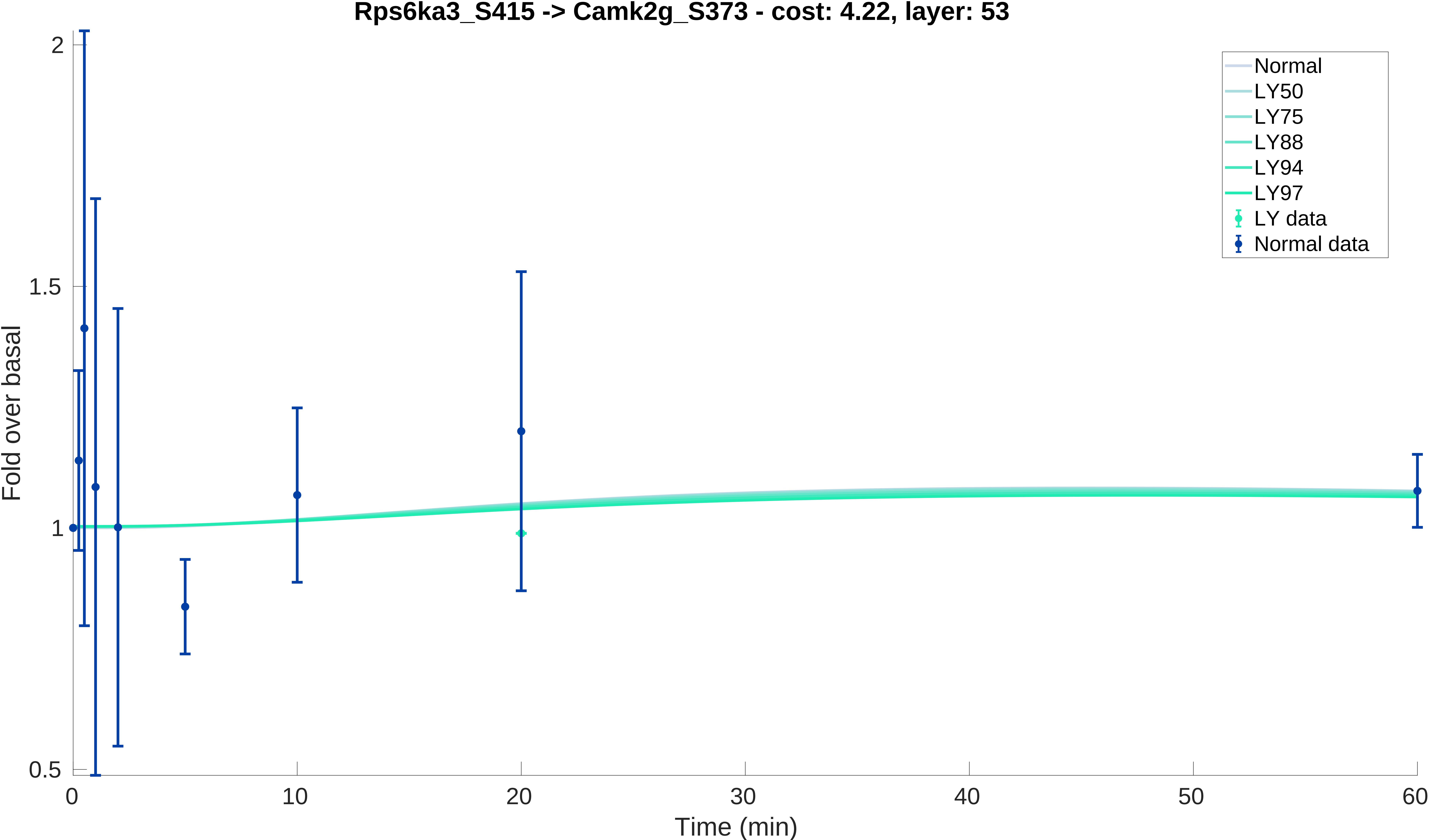

Rps6ka3_S415 -> Camk2g_S373 - cost: 4.22, layer: 53
Fold over basal
Time (min)
2
1.5
1
0.5
0
10
20
30
40
50
60
Normal
LY50
LY75
LY88
LY94
LY97
LY data
Normal data

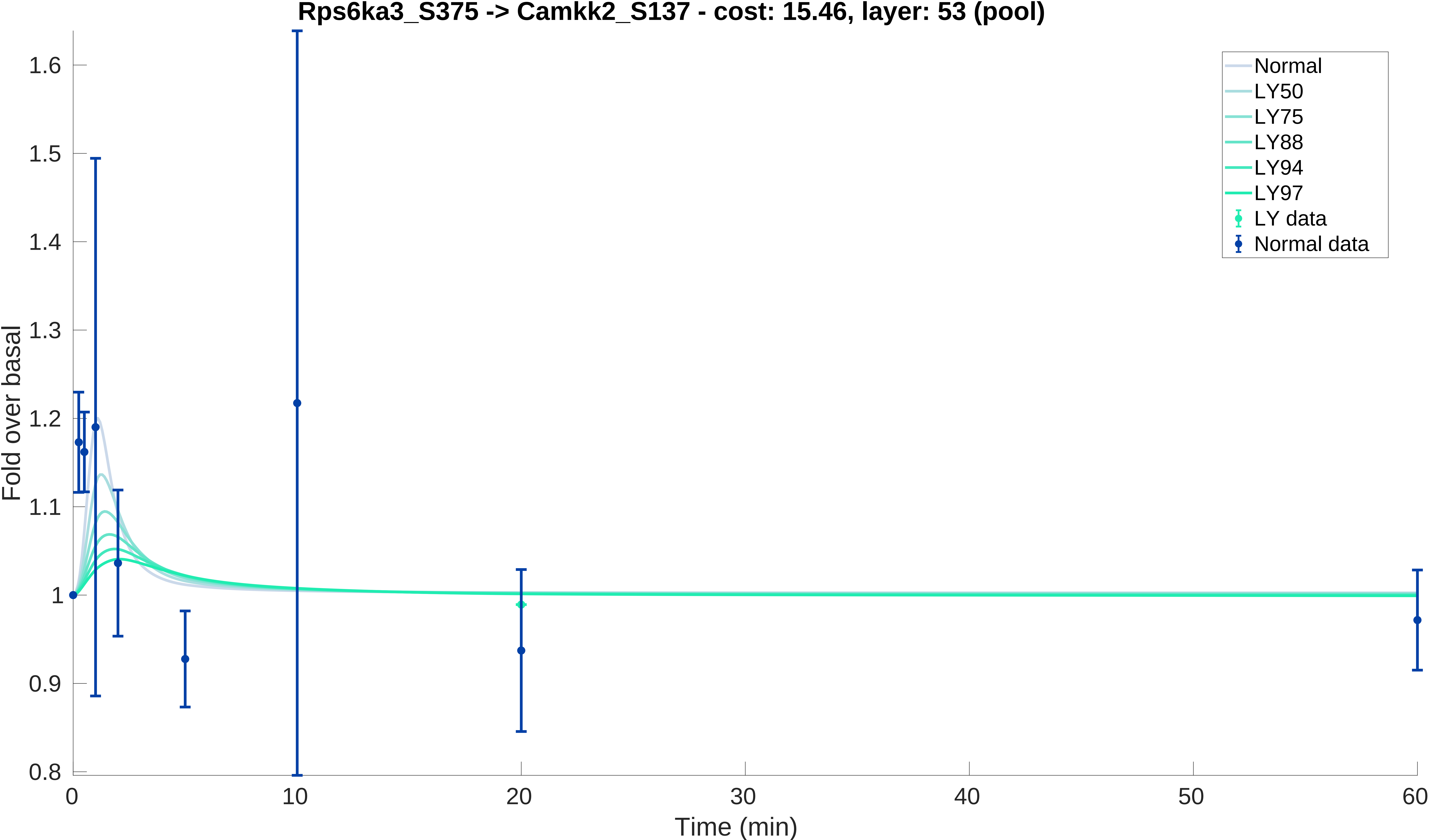

Rps6ka3_S375 -> Camkk2_S137 - cost: 15.46, layer: 53 (pool)
Fold over basal
1.6
1.5
1.4
1.3
1.2
1.1
1
0.9
0.8
0
10
20
30
40
50
60
Time (min)
Normal
LY50
LY75
LY88
LY94
LY97
LY data
Normal data

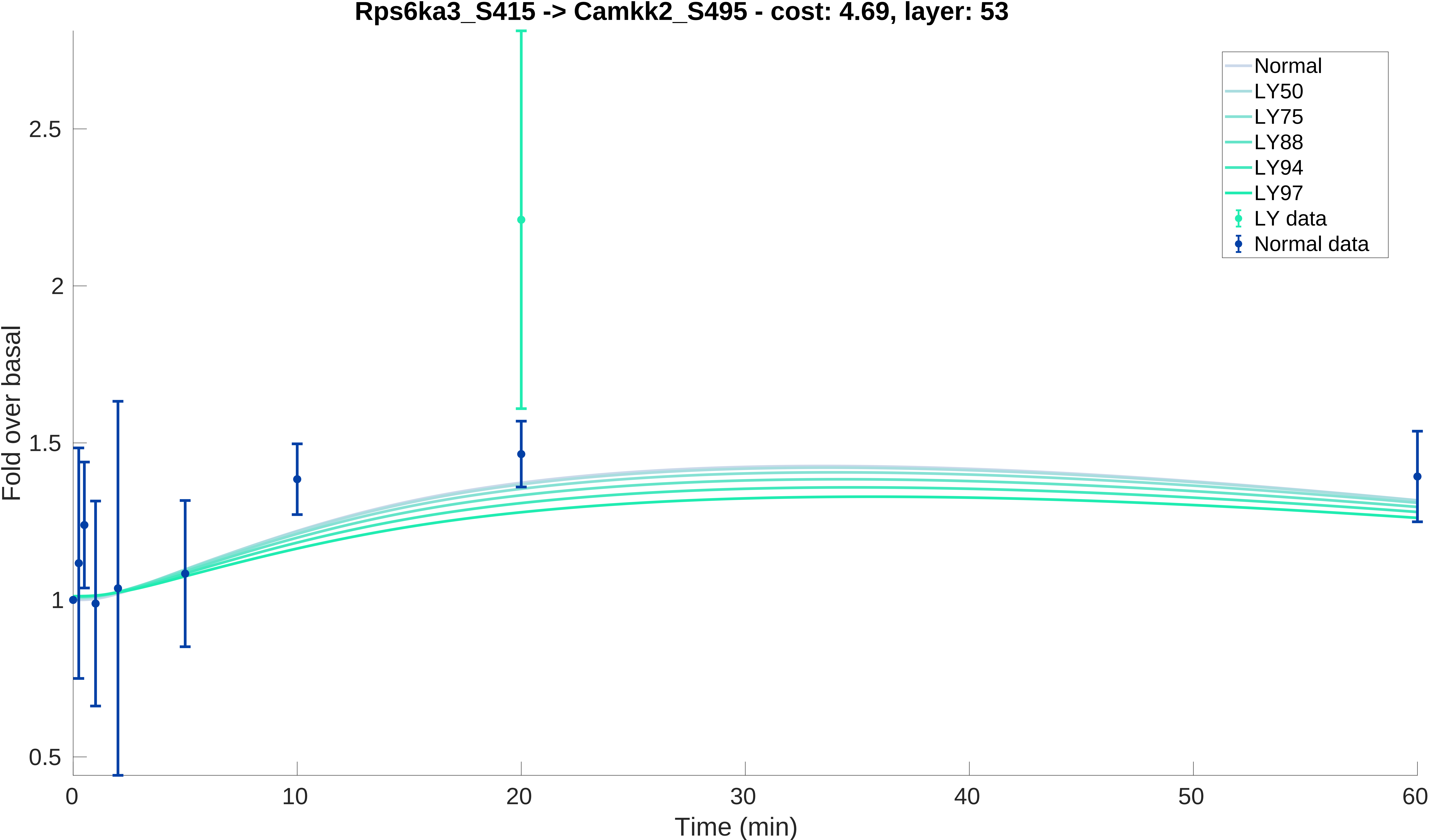

Rps6ka3_S415 -> Camkk2_S495 - cost: 4.69, layer: 53
Normal
LY50
LY75
LY88
LY94
LY97
LY data
Normal data
Fold over basal
2.5
2
1.5
1
0.5
0
10
20
30
40
50
60
Time (min)

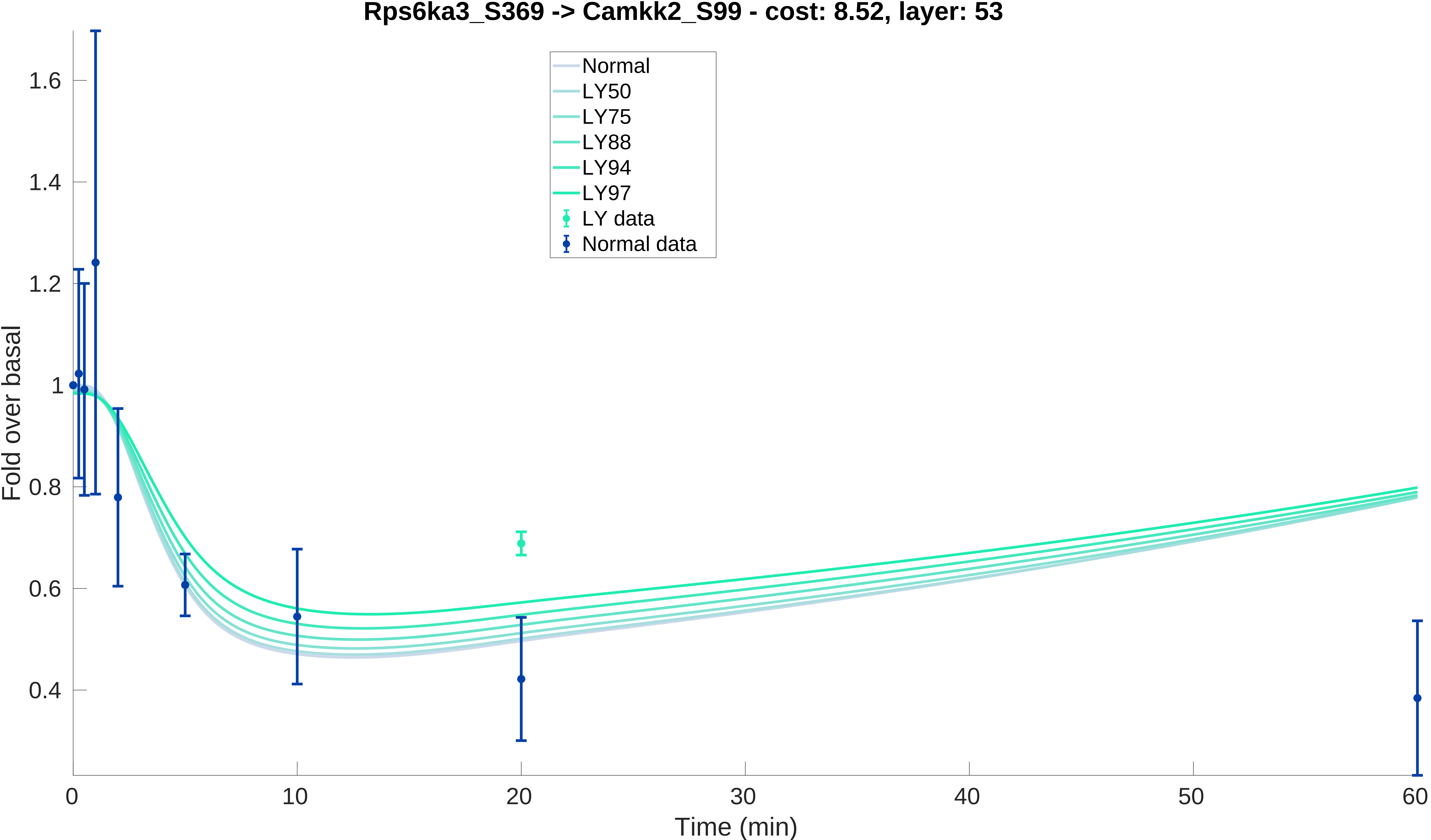

Rps6ka3_S369 -> Camkk2_S99 - cost: 8.52, layer: 53
Normal
LY50
LY75
LY88
LY94
LY97
LY data
Normal data
Fold over basal
Time (min)
0.4
0.6
0.8
1
1.2
1.4
1.6
0
10
20
30
40
50
60

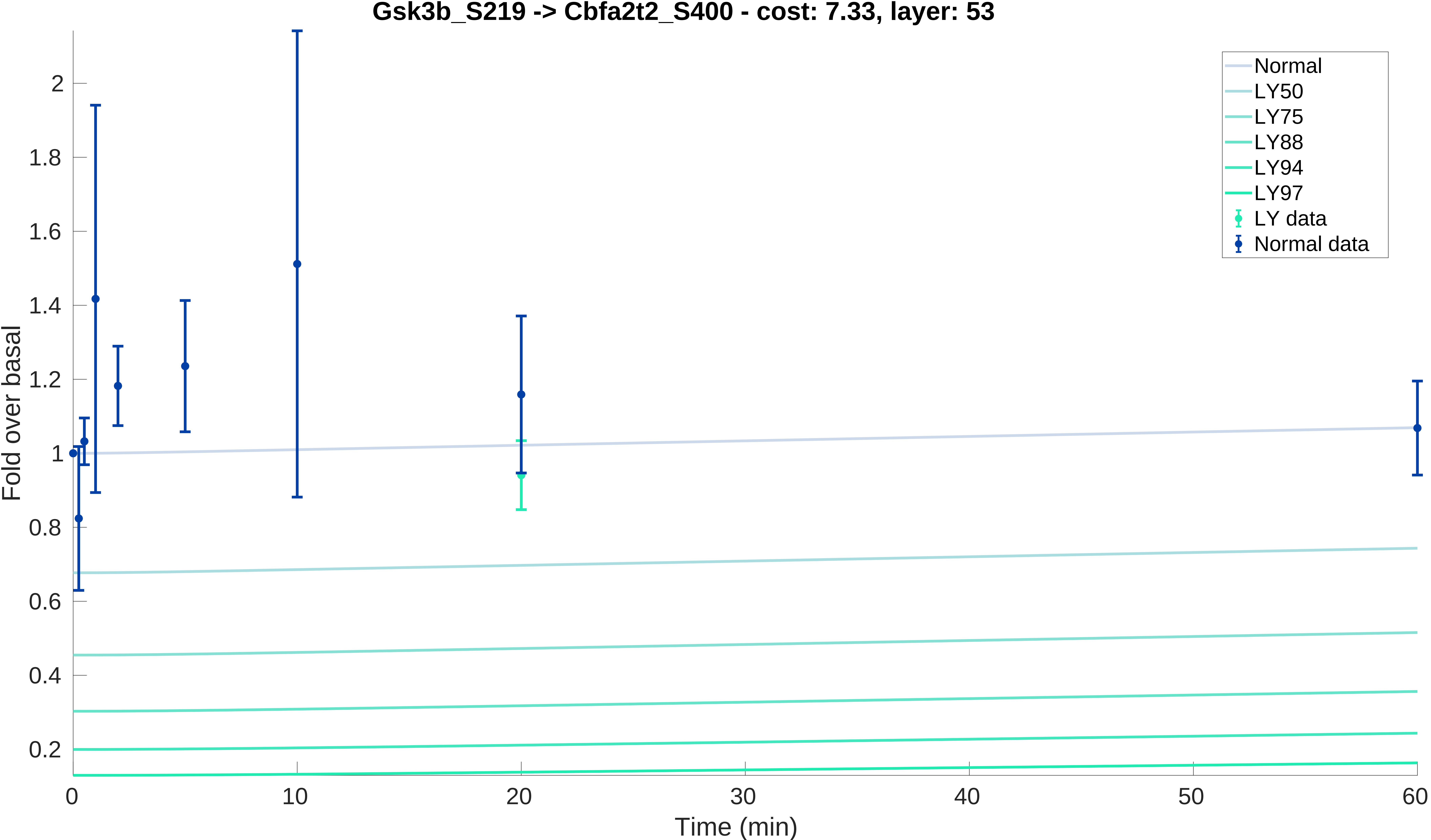

Gsk3b_S219 -> Cbfa2t2_S400 - cost: 7.33, layer: 53
Normal
LY50
LY75
LY88
LY94
LY97
LY data
Normal data
Fold over basal
2
1.8
1.6
1.4
1.2
1
0.8
0.6
0.4
0.2
0
10
20
30
40
50
60
Time (min)

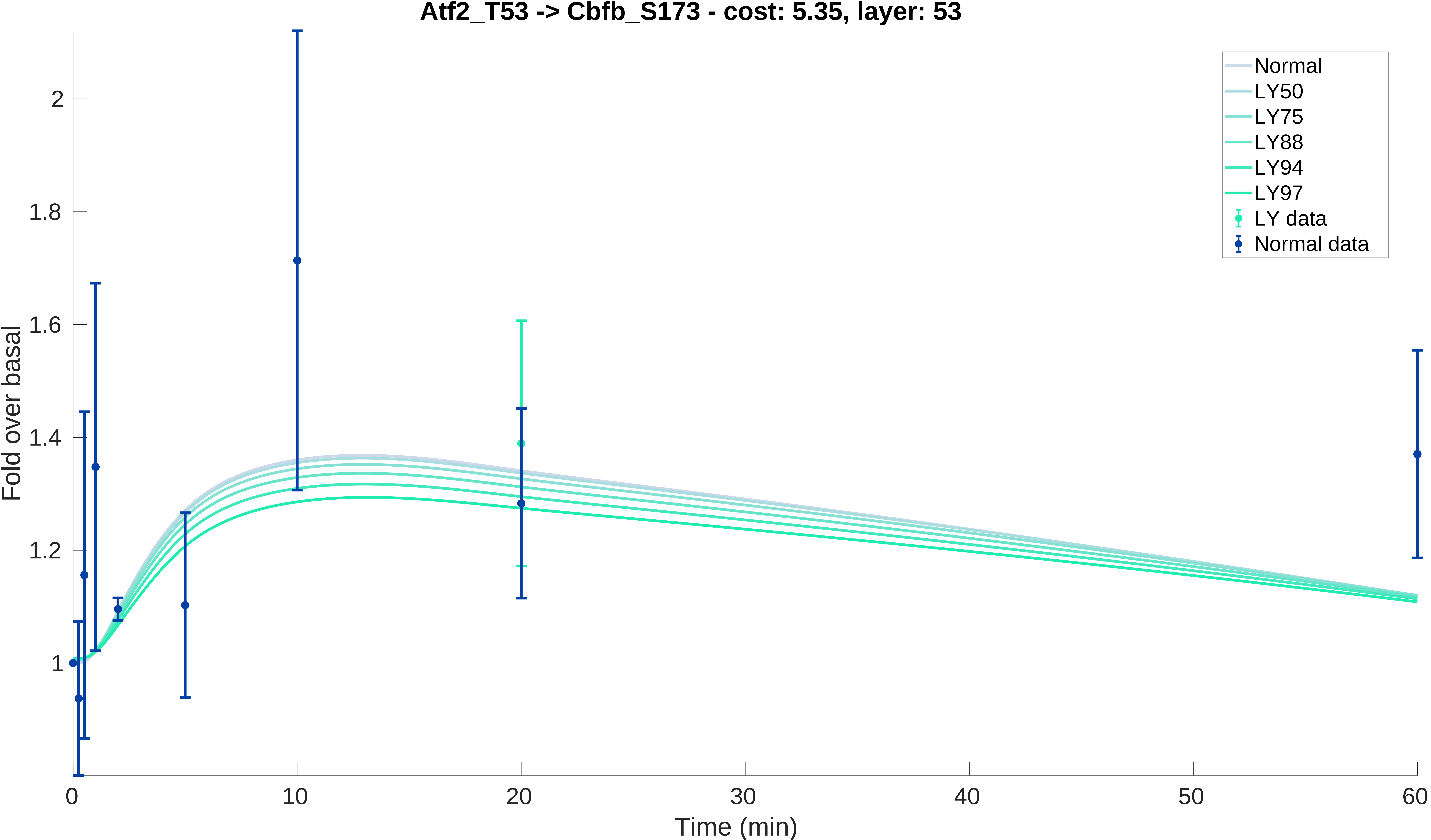
Atf2_T53 -> Cbfb_S173 - cost: 5.35, layer: 53
Fold over basal
Time (min)
2
1.8
1.6
1.4
1.2
1
0
10
20
30
40
50
60
Normal
LY50
LY75
LY88
LY94
LY97
LY data
Normal data

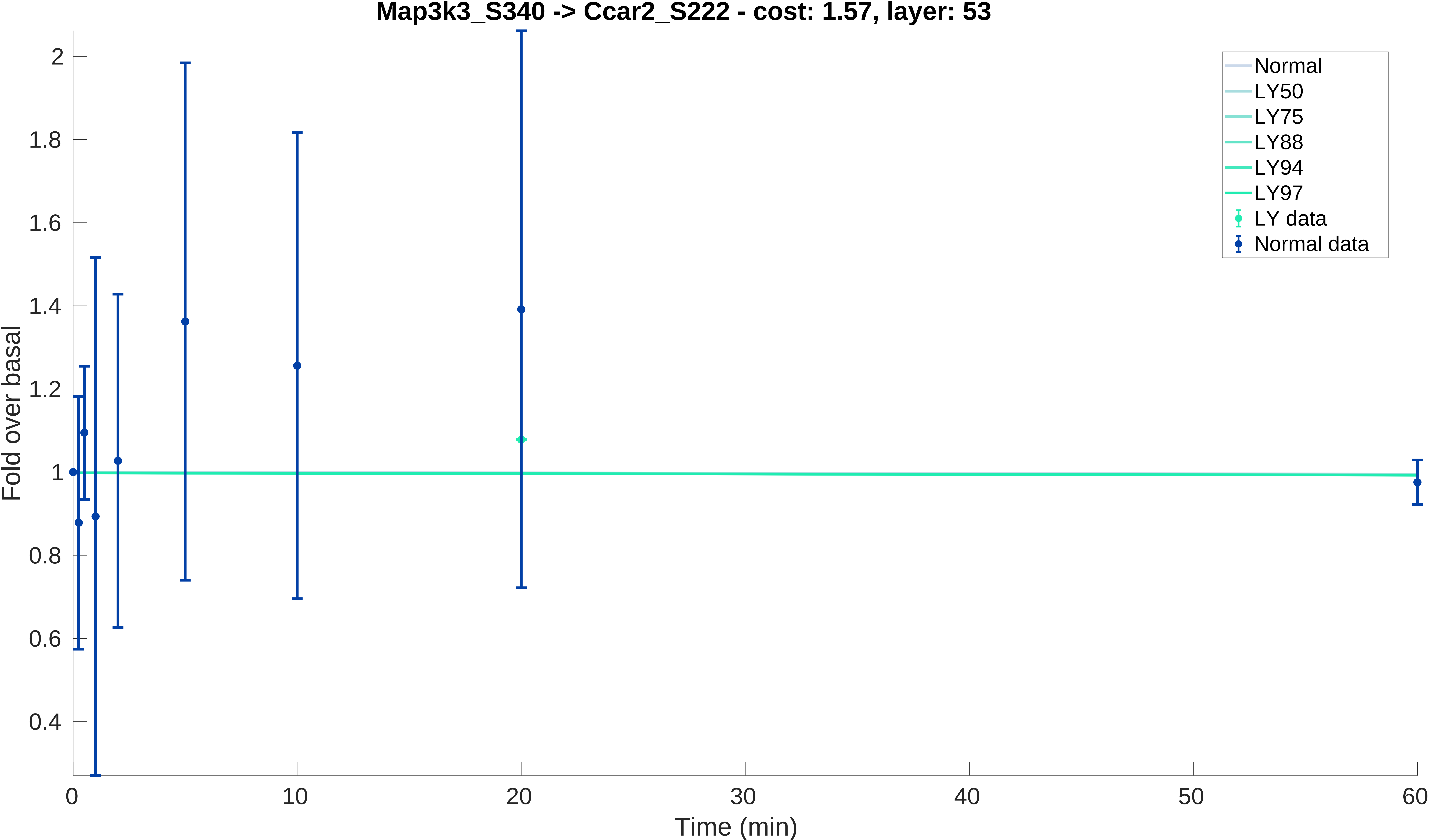

Map3k3_S340 -> Ccar2_S222 - cost: 1.57, layer: 53
Fold over basal
Time (min)
2
1.8
1.6
1.4
1.2
1
0.8
0.6
0.4
0
10
20
30
40
50
60
Normal
LY50
LY75
LY88
LY94
LY97
LY data
Normal data

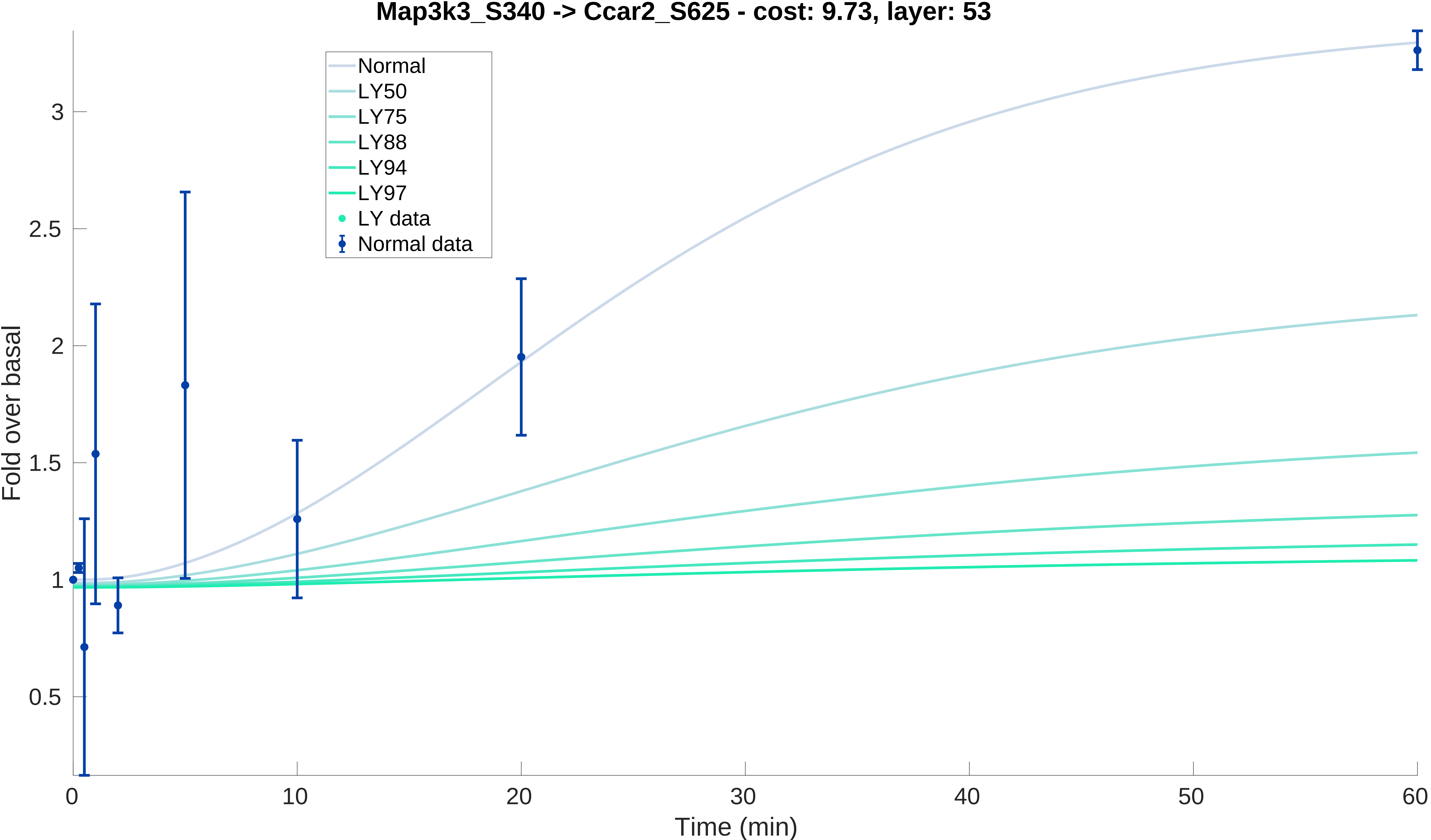

Map3k3_S340 -> Ccar2_S625 - cost: 9.73, layer: 53
Normal
LY50
LY75
LY88
LY94
LY97
LY data
Normal data
Fold over basal
Time (min)
0.5
1
1.5
2
2.5
3
0
10
20
30
40
50
60

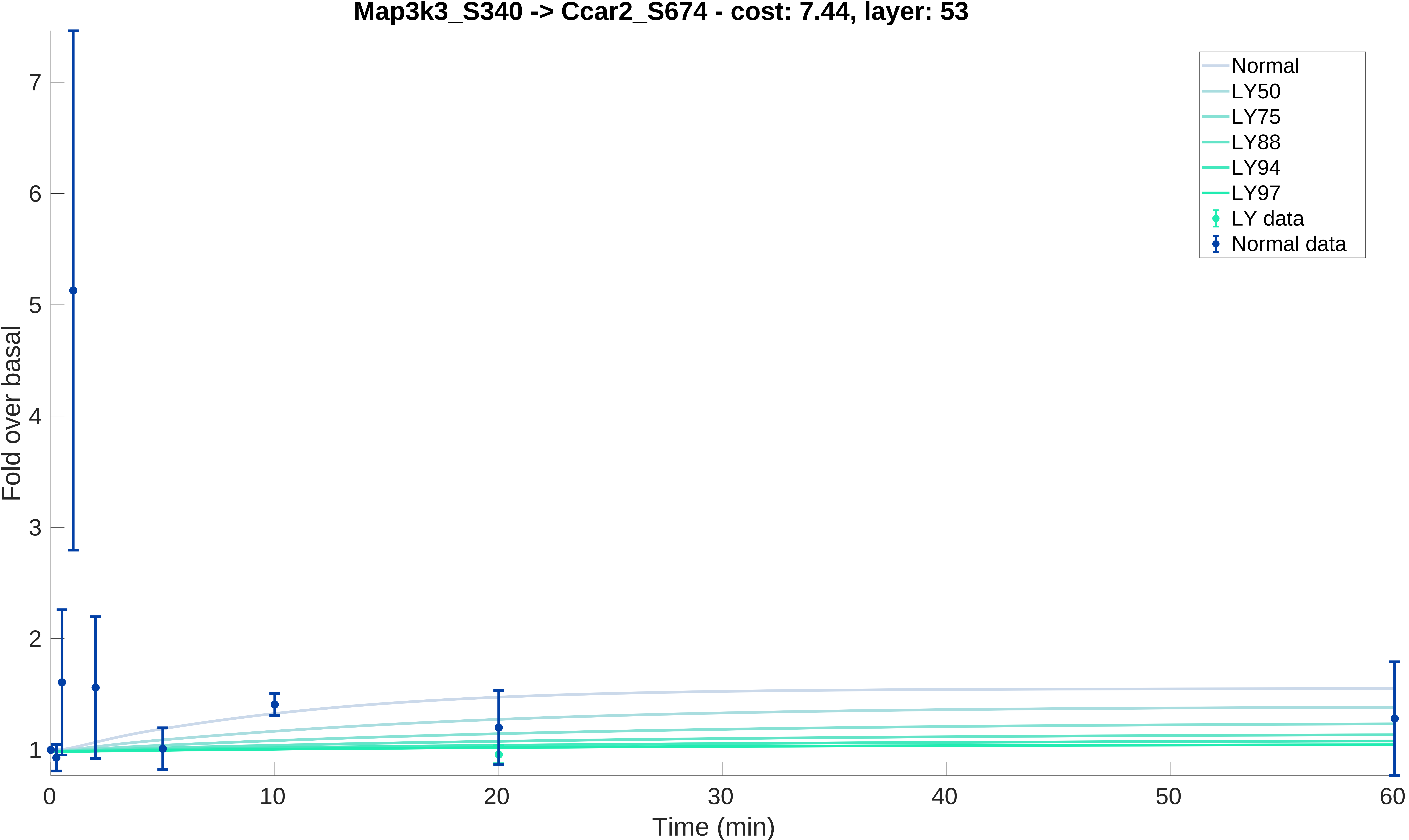

Map3k3_S340 -> Ccar2_S674 - cost: 7.44, layer: 53
Fold over basal
Time (min)
0
10
20
30
40
50
60
1
2
3
4
5
6
7
Normal
LY50
LY75
LY88
LY94
LY97
LY data
Normal data

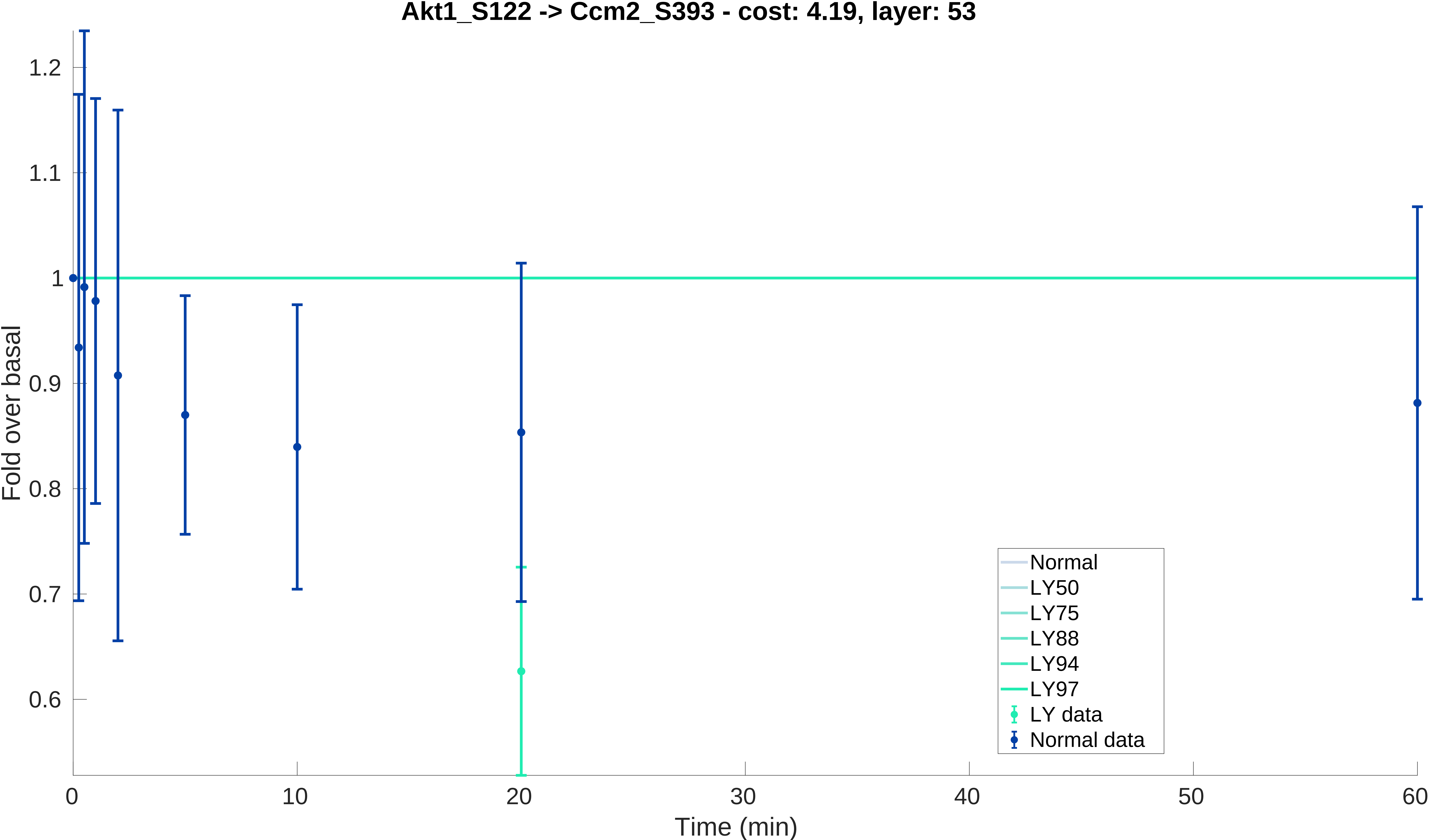

Akt1_S122 -> Ccm2_S393 - cost: 4.19, layer: 53
Fold over basal
1.2
1.1
1
0.9
0.8
0.7
0.6
0
10
20
30
40
50
60
Time (min)
Normal
LY50
LY75
LY88
LY94
LY97
LY data
Normal data

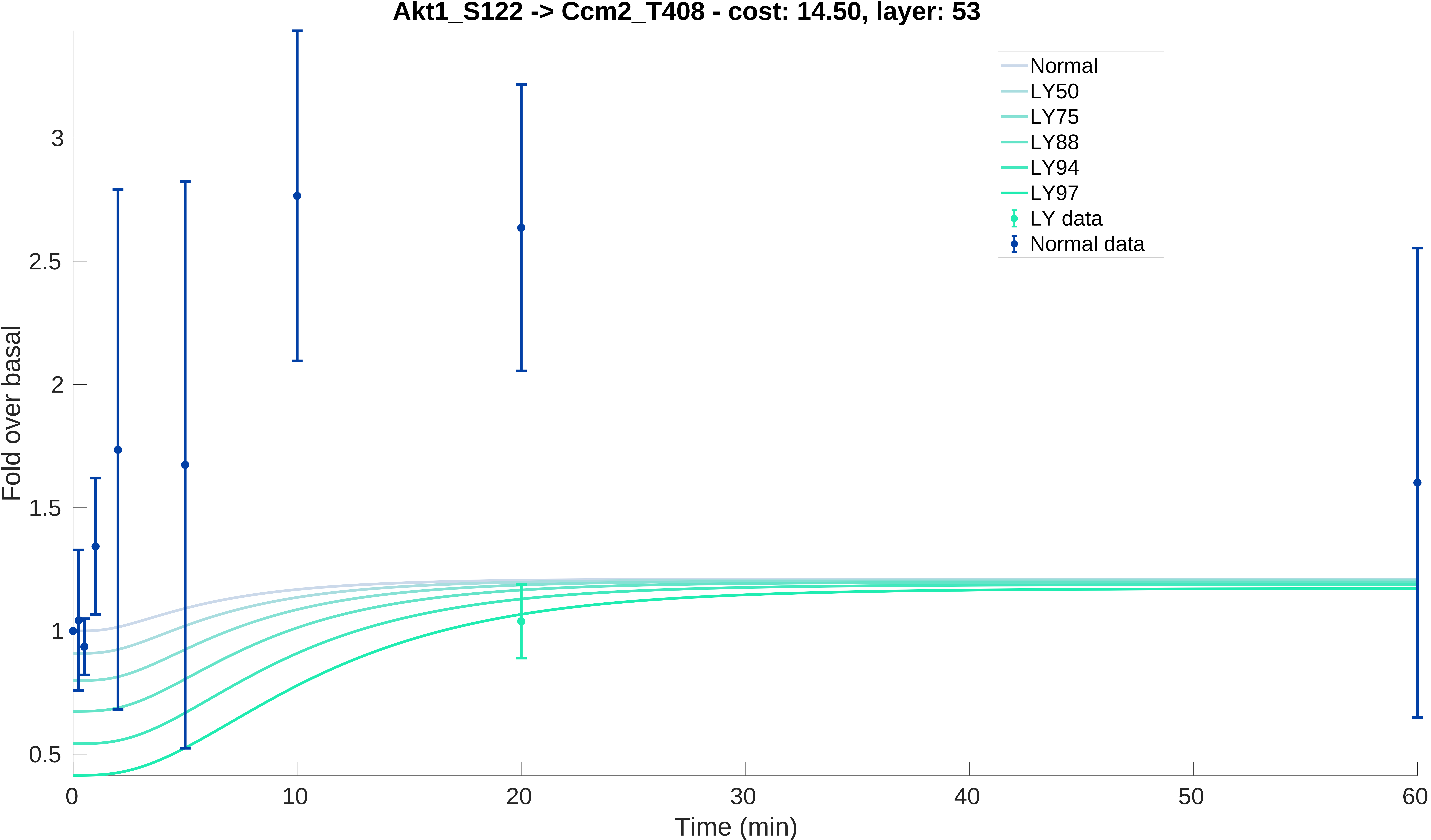

Akt1_S122 -> Ccm2_T408 - cost: 14.50, layer: 53
Normal
LY50
LY75
LY88
LY94
LY97
LY data
Normal data
Fold over basal
Time (min)
0.5
1
1.5
2
2.5
3
0
10
20
30
40
50
60

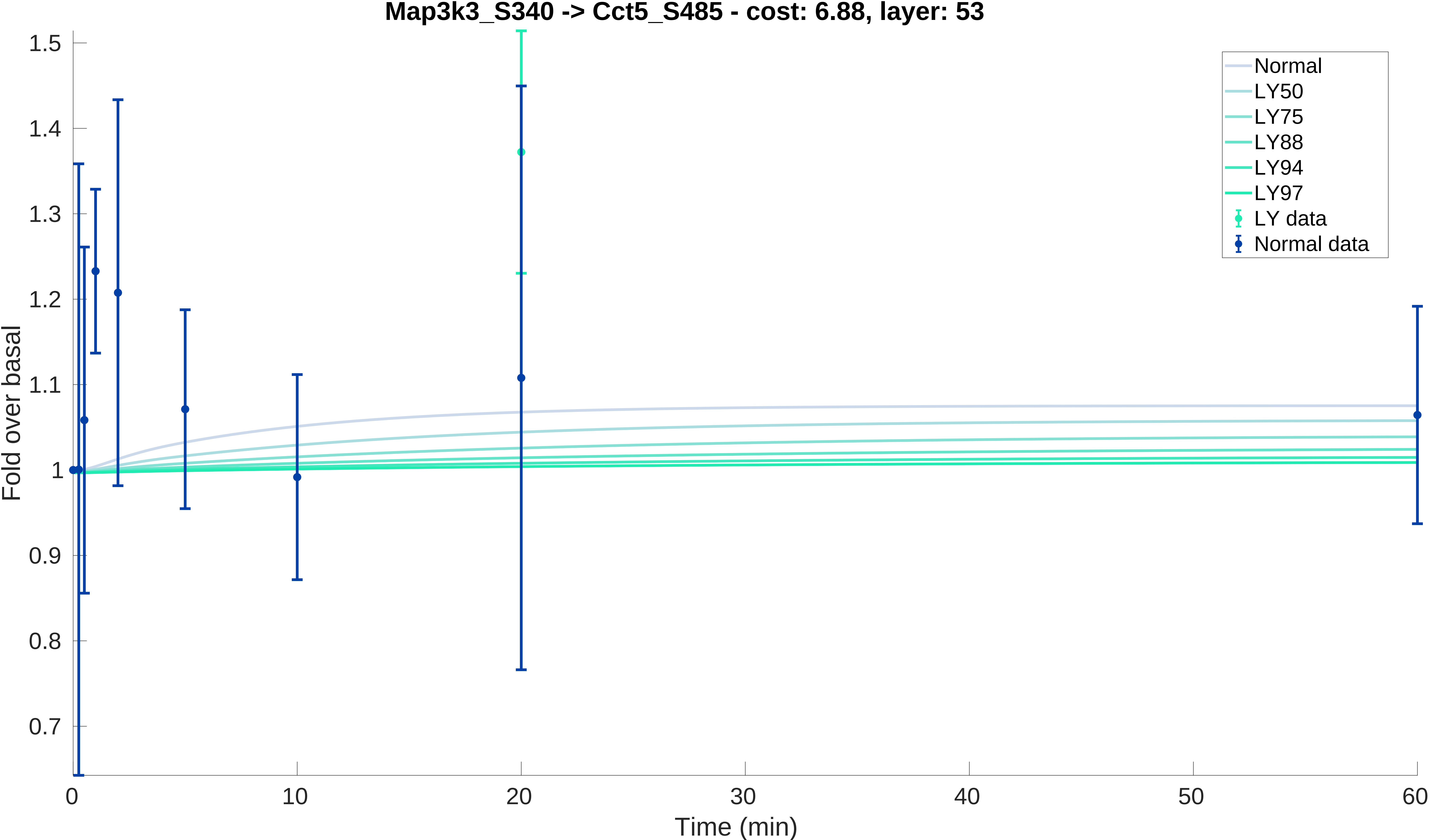

Map3k3_S340 -> Cct5_S485 - cost: 6.88, layer: 53
Fold over basal
Time (min)
1.5
1.4
1.3
1.2
1.1
1
0.9
0.8
0.7
0
10
20
30
40
50
60
Normal
LY50
LY75
LY88
LY94
LY97
LY data
Normal data

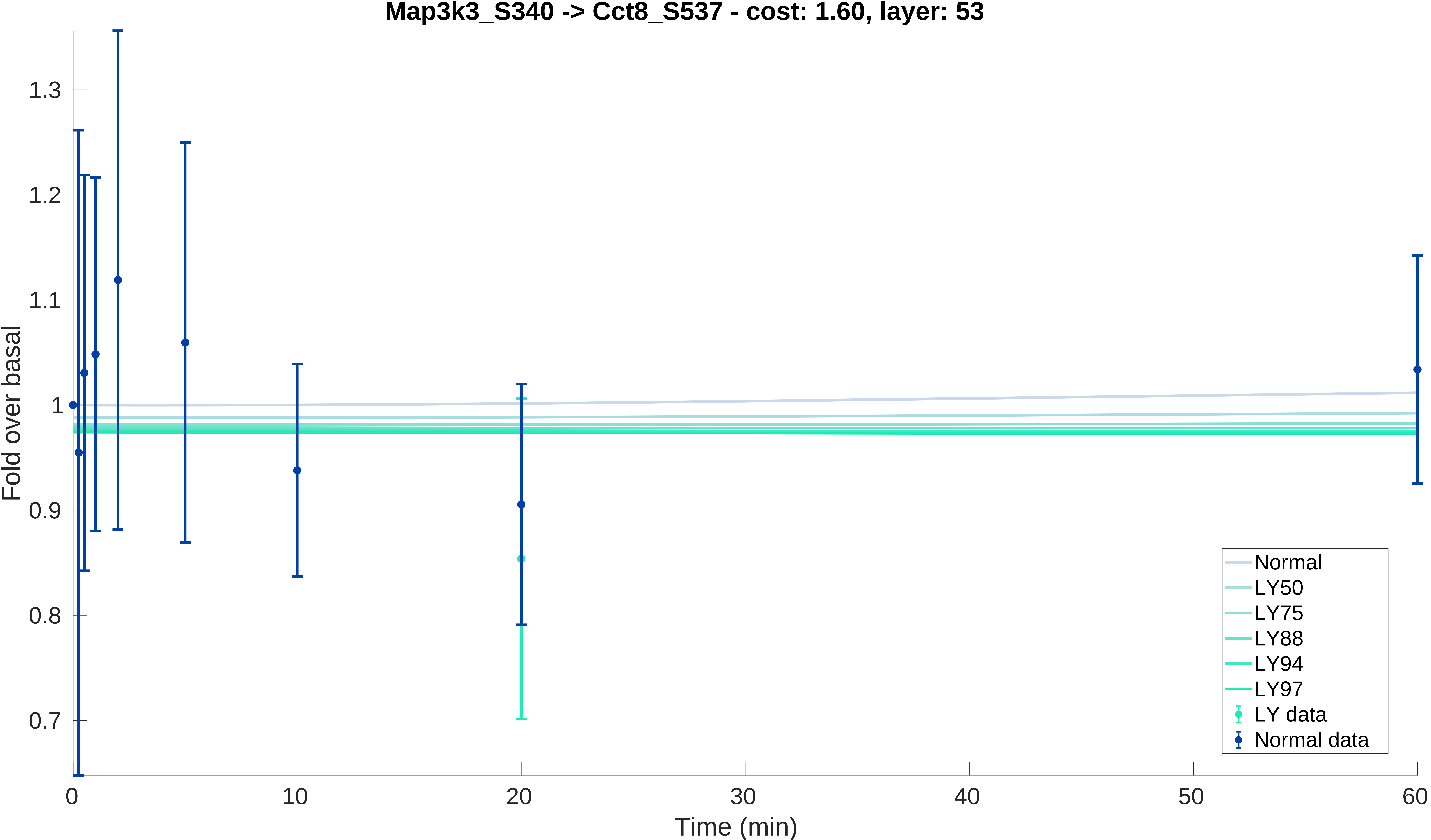

Map3k3_S340 -> Cct8_S537 - cost: 1.60, layer: 53
Fold over basal
Time (min)
1.3
1.2
1.1
1
0.9
0.8
0.7
0
10
20
30
40
50
60
Normal
LY50
LY75
LY88
LY94
LY97
LY data
Normal data

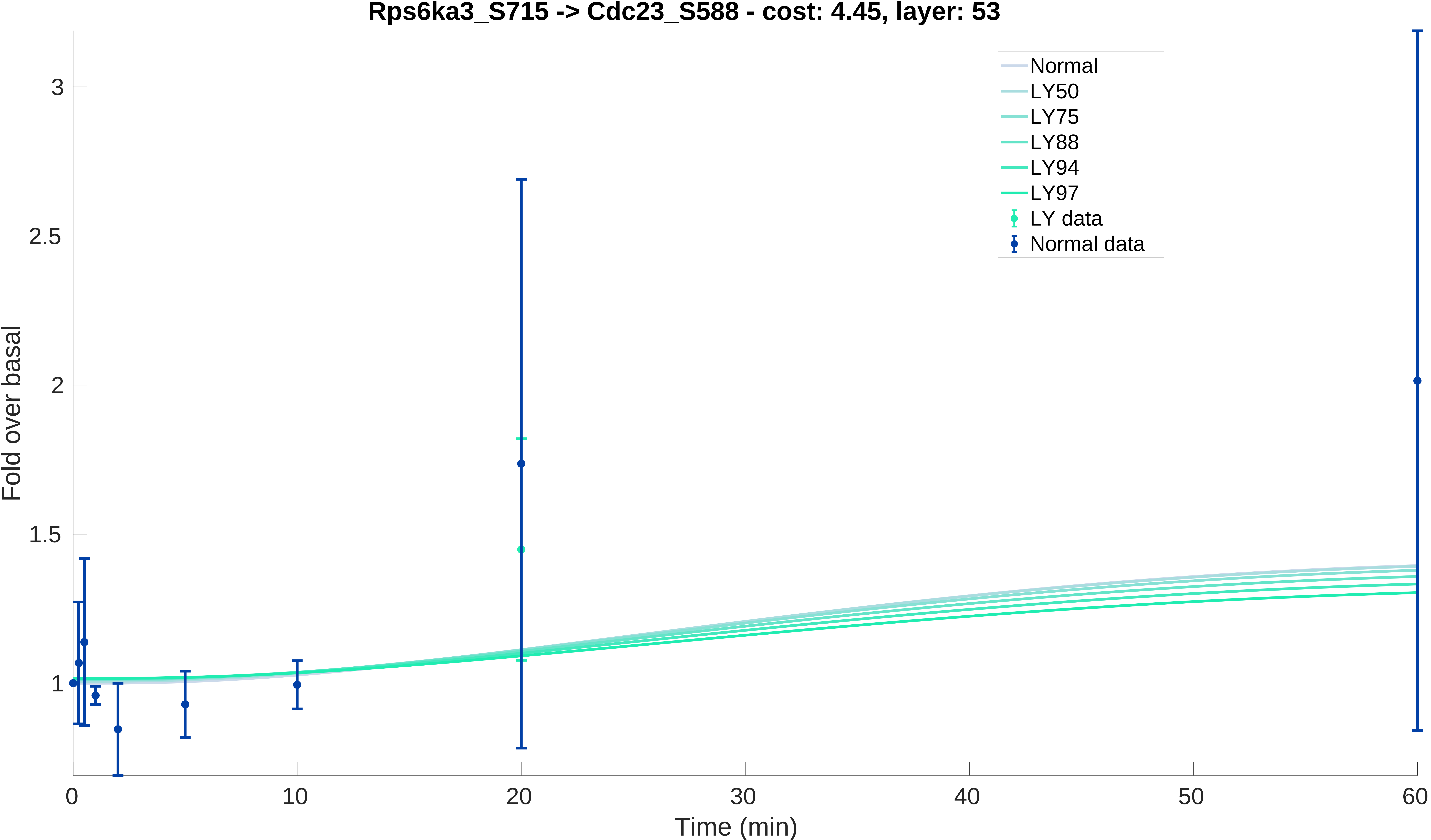

Rps6ka3_S715 -> Cdc23_S588 - cost: 4.45, layer: 53
Normal
LY50
LY75
LY88
LY94
LY97
LY data
Normal data
Fold over basal
Time (min)
3
2.5
2
1.5
1
0
10
20
30
40
50
60

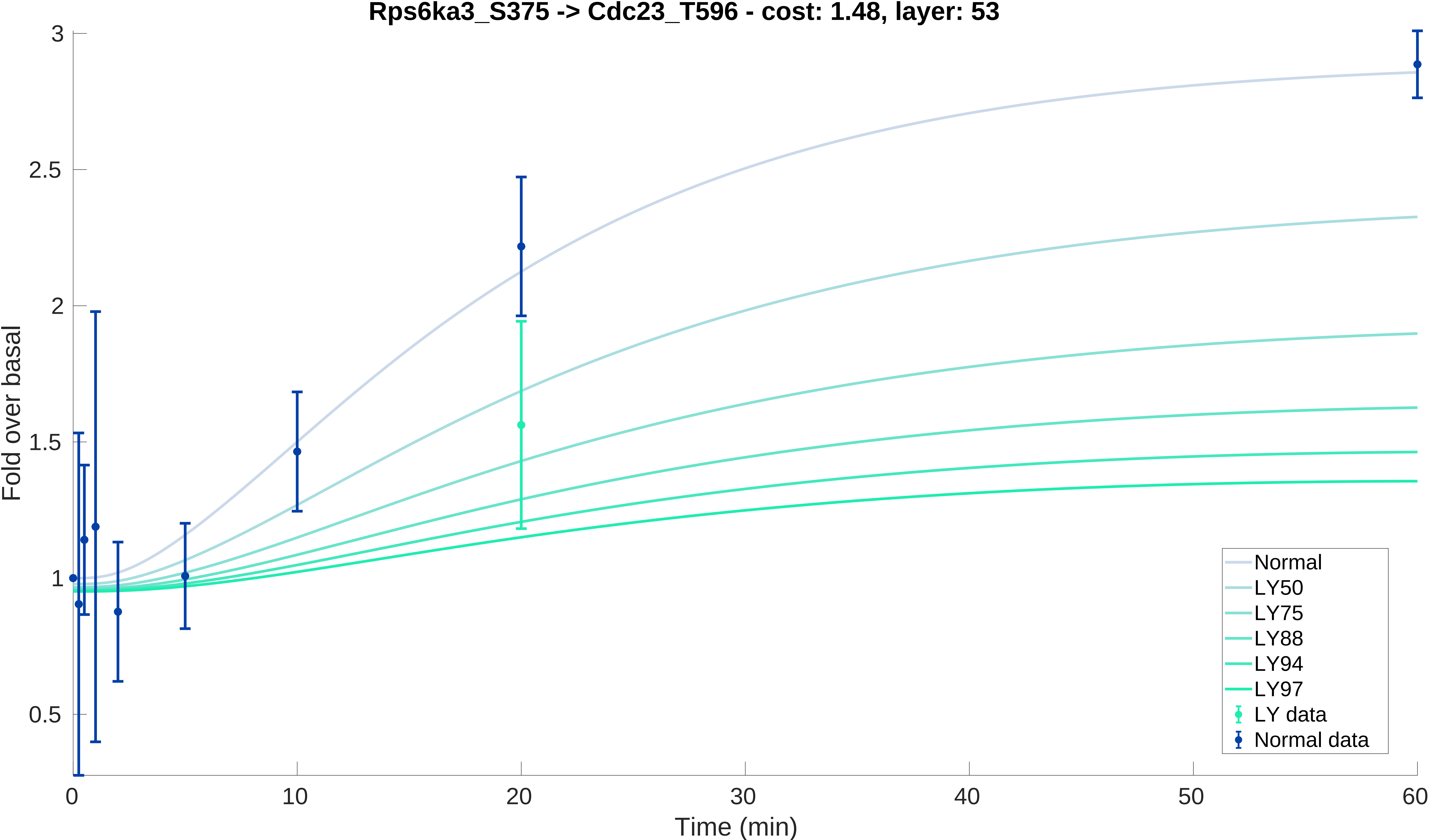

Rps6ka3_S375 -> Cdc23_T596 - cost: 1.48, layer: 53
Fold over basal
Time (min)
0.5
1
1.5
2
2.5
3
0
10
20
30
40
50
60
Normal
LY50
LY75
LY88
LY94
LY97
LY data
Normal data

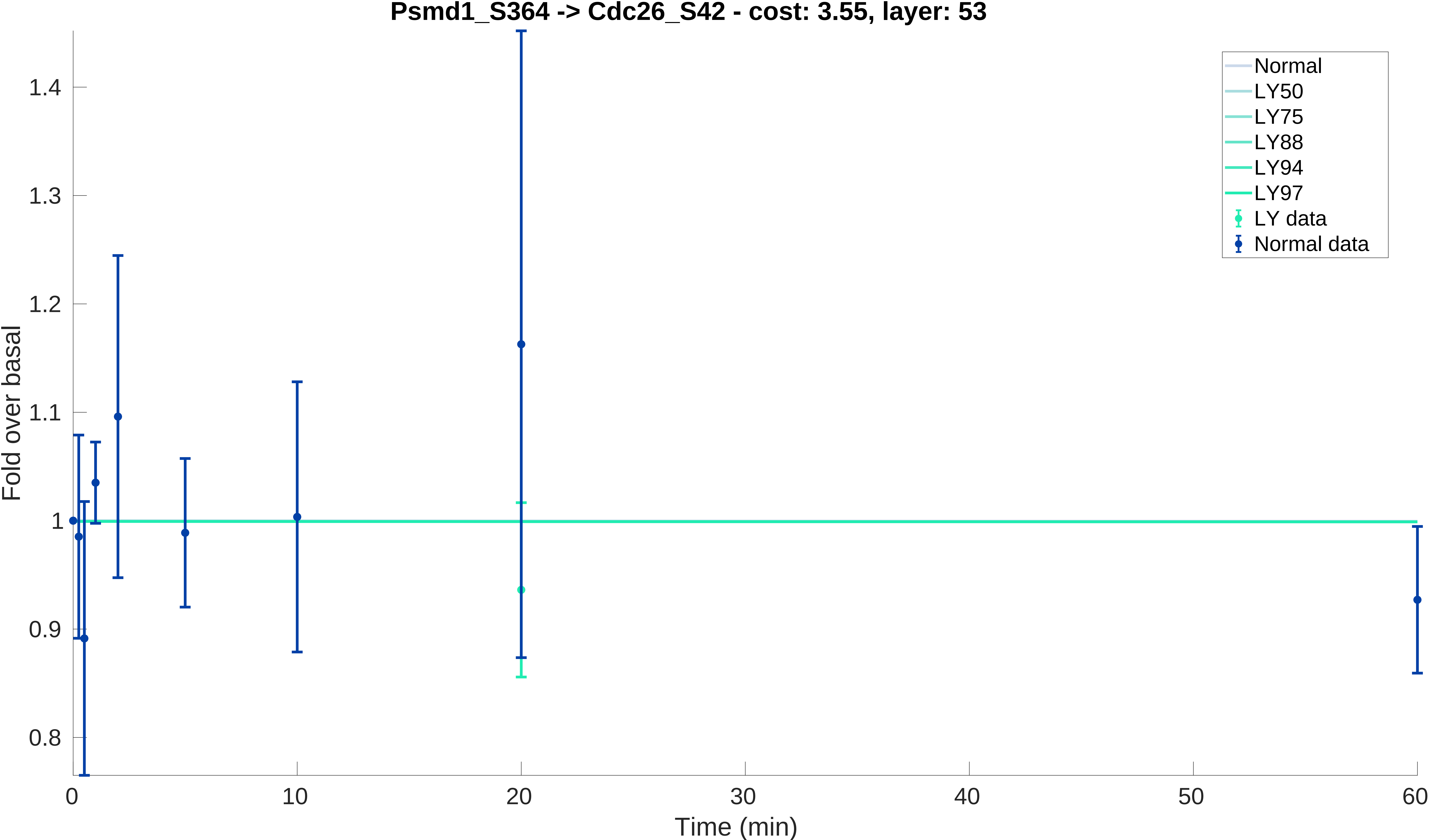

Psmd1_S364 -> Cdc26_S42 - cost: 3.55, layer: 53
Fold over basal
Time (min)
0.8
0.9
1
1.1
1.2
1.3
1.4
0
10
20
30
40
50
60
Normal
LY50
LY75
LY88
LY94
LY97
LY data
Normal data

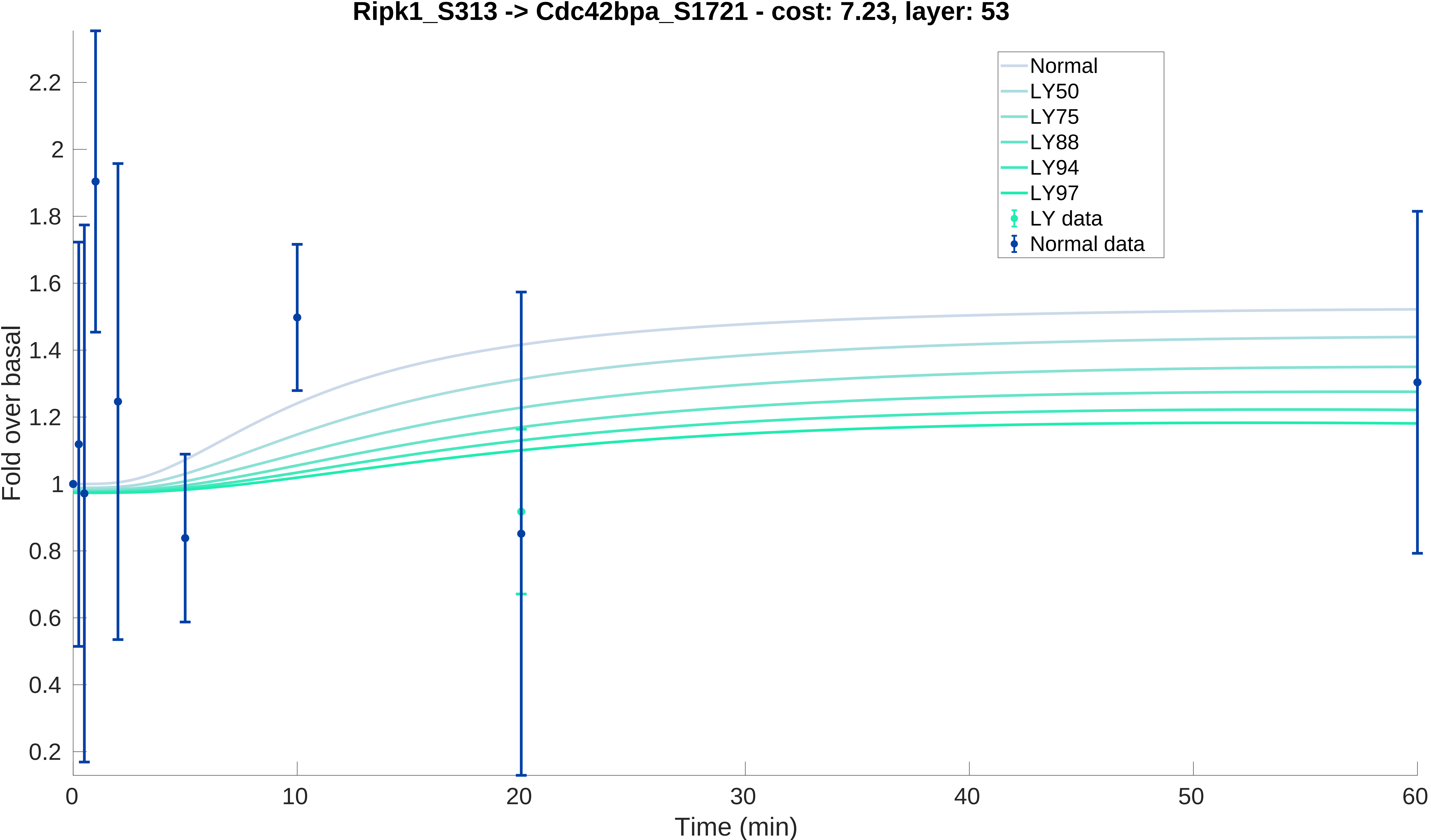

Ripk1_S313 -> Cdc42bpa_S1721 - cost: 7.23, layer: 53
Normal
LY50
LY75
LY88
LY94
LY97
LY data
Normal data
Fold over basal
2.2
2
1.8
1.6
1.4
1.2
1
0.8
0.6
0.4
0.2
0
10
20
30
40
50
60
Time (min)

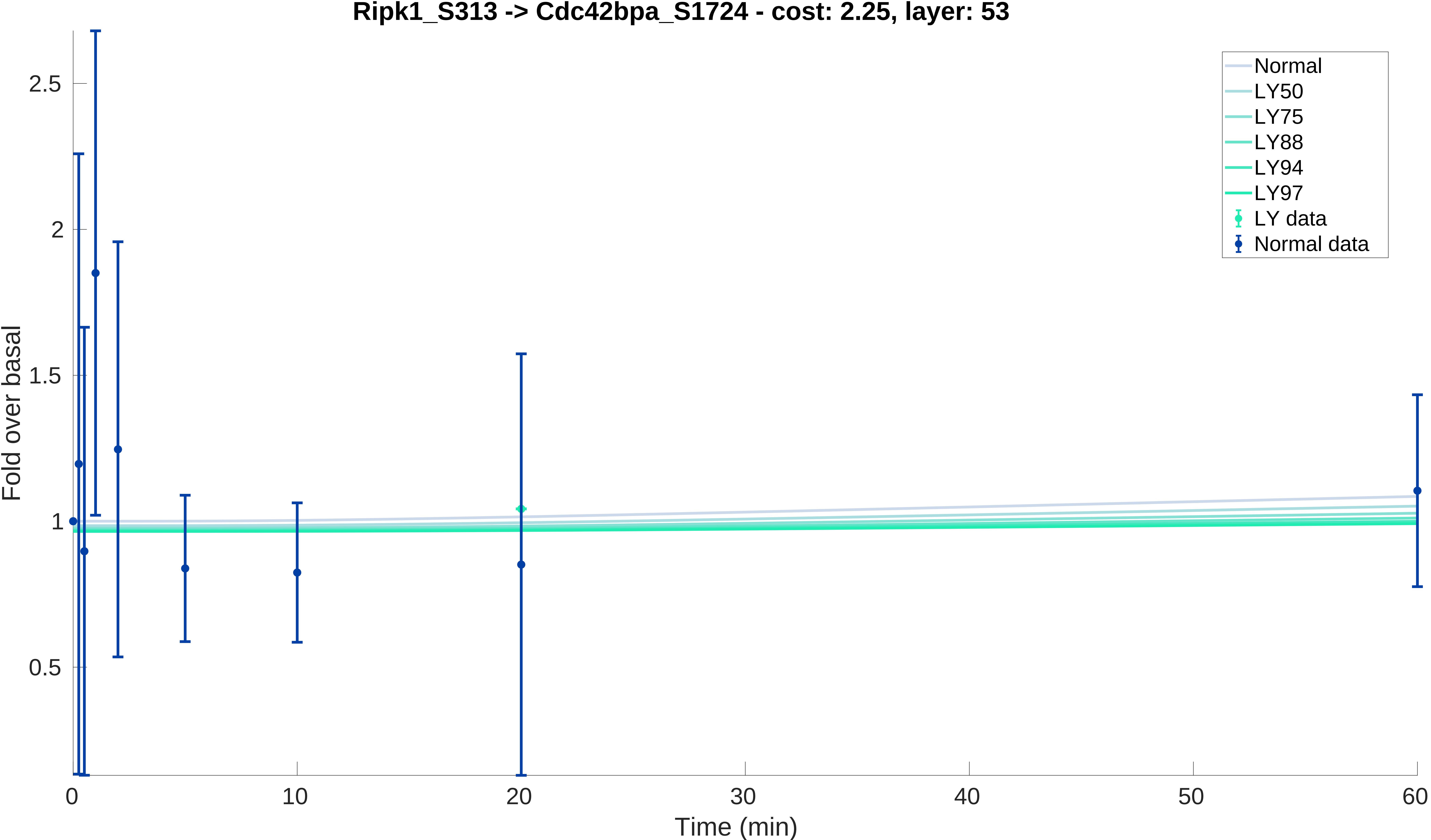

Ripk1_S313 -> Cdc42bpa_S1724 - cost: 2.25, layer: 53
Normal
LY50
LY75
LY88
LY94
LY97
LY data
Normal data
Fold over basal
Time (min)
0.5
1
1.5
2
2.5
0
10
20
30
40
50
60

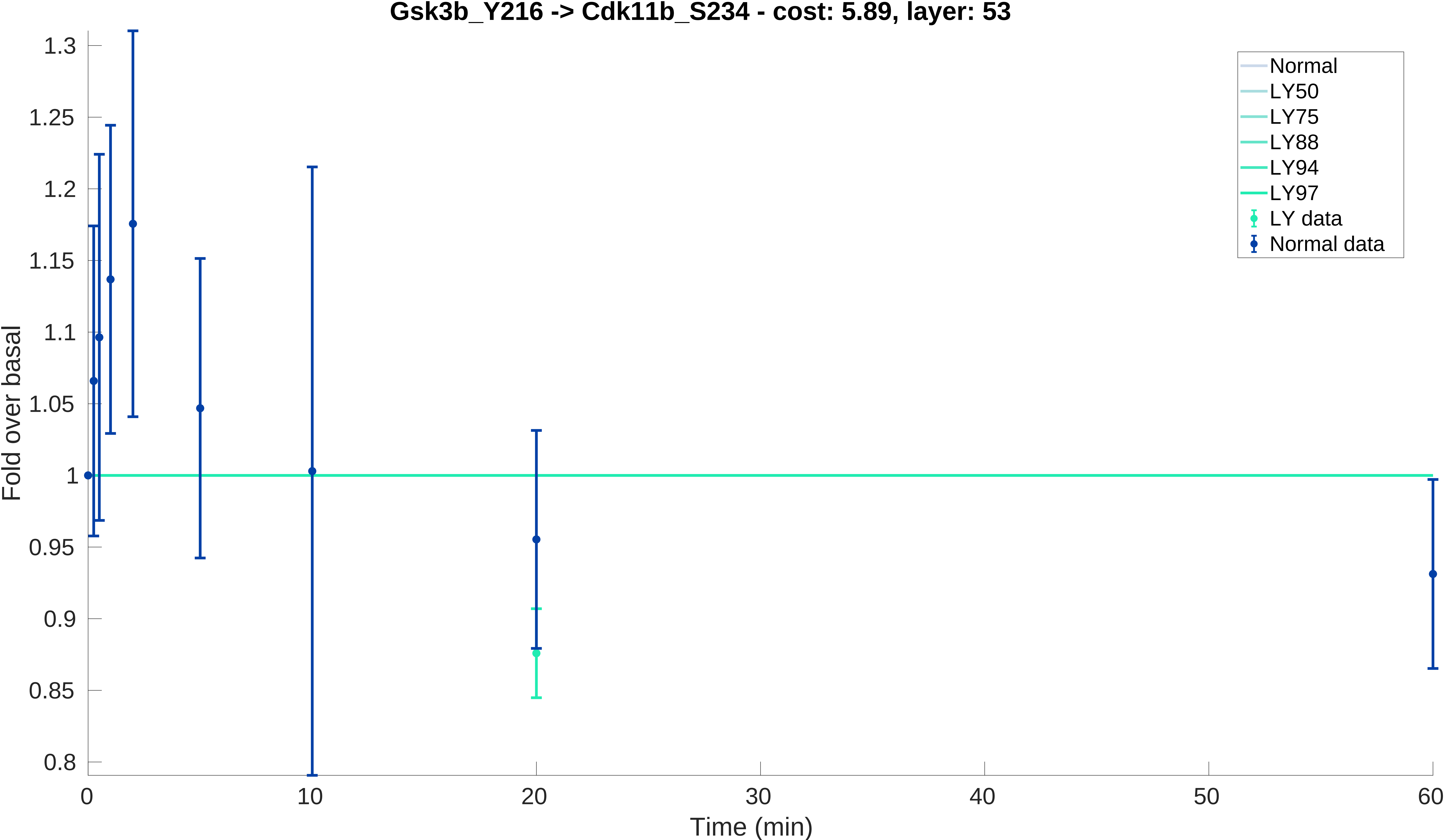

Gsk3b_Y216 -> Cdk11b_S234 - cost: 5.89, layer: 53
Fold over basal
Time (min)
1.3
1.25
1.2
1.15
1.1
1.05
1
0.95
0.9
0.85
0.8
0
10
20
30
40
50
60
Normal
LY50
LY75
LY88
LY94
LY97
LY data
Normal data

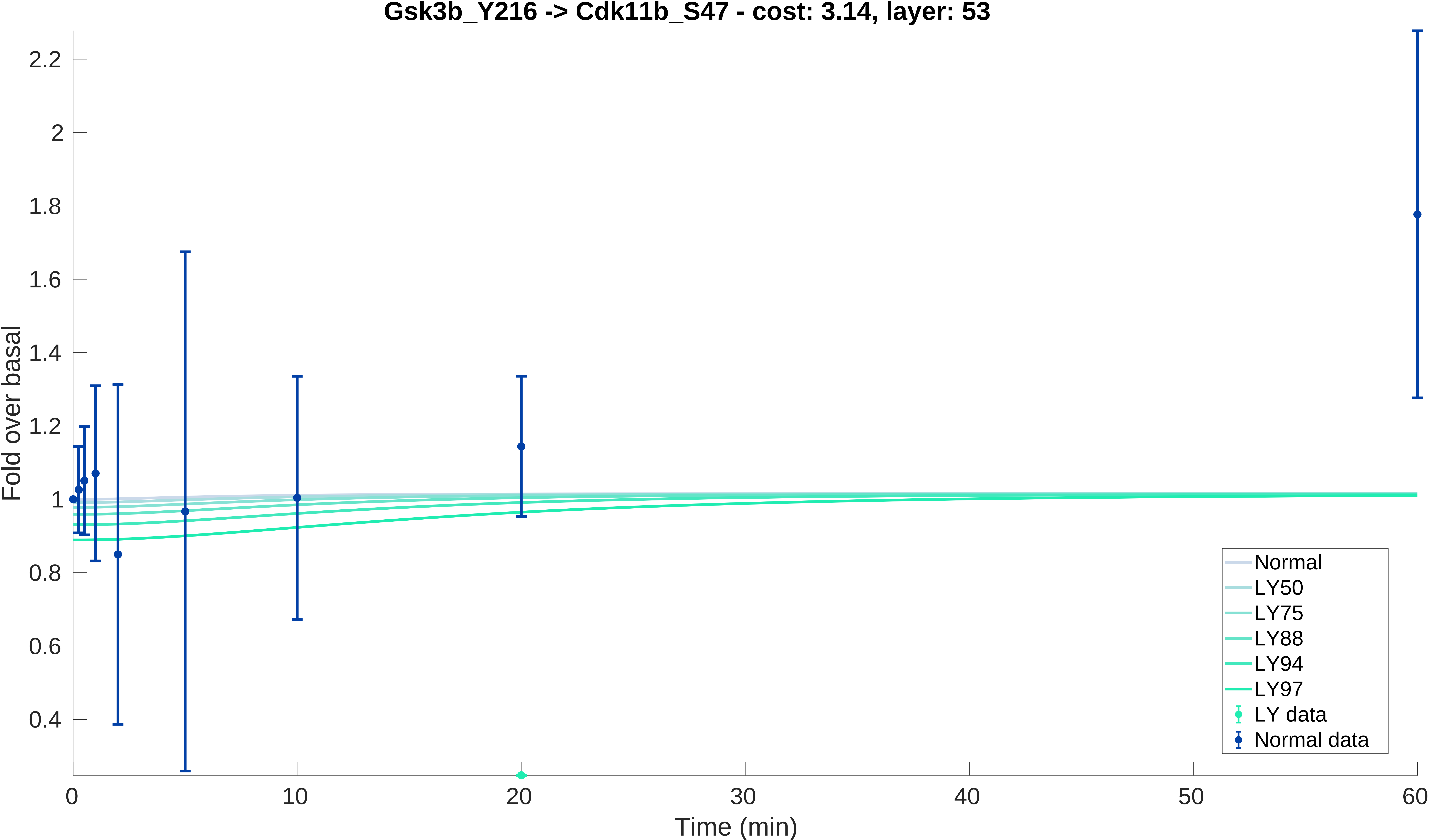

Gsk3b_Y216 -> Cdk11b_S47 - cost: 3.14, layer: 53
Fold over basal
2.2
2
1.8
1.6
1.4
1.2
1
0.8
0.6
0.4
0
10
20
30
40
50
60
Time (min)
Normal
LY50
LY75
LY88
LY94
LY97
LY data
Normal data

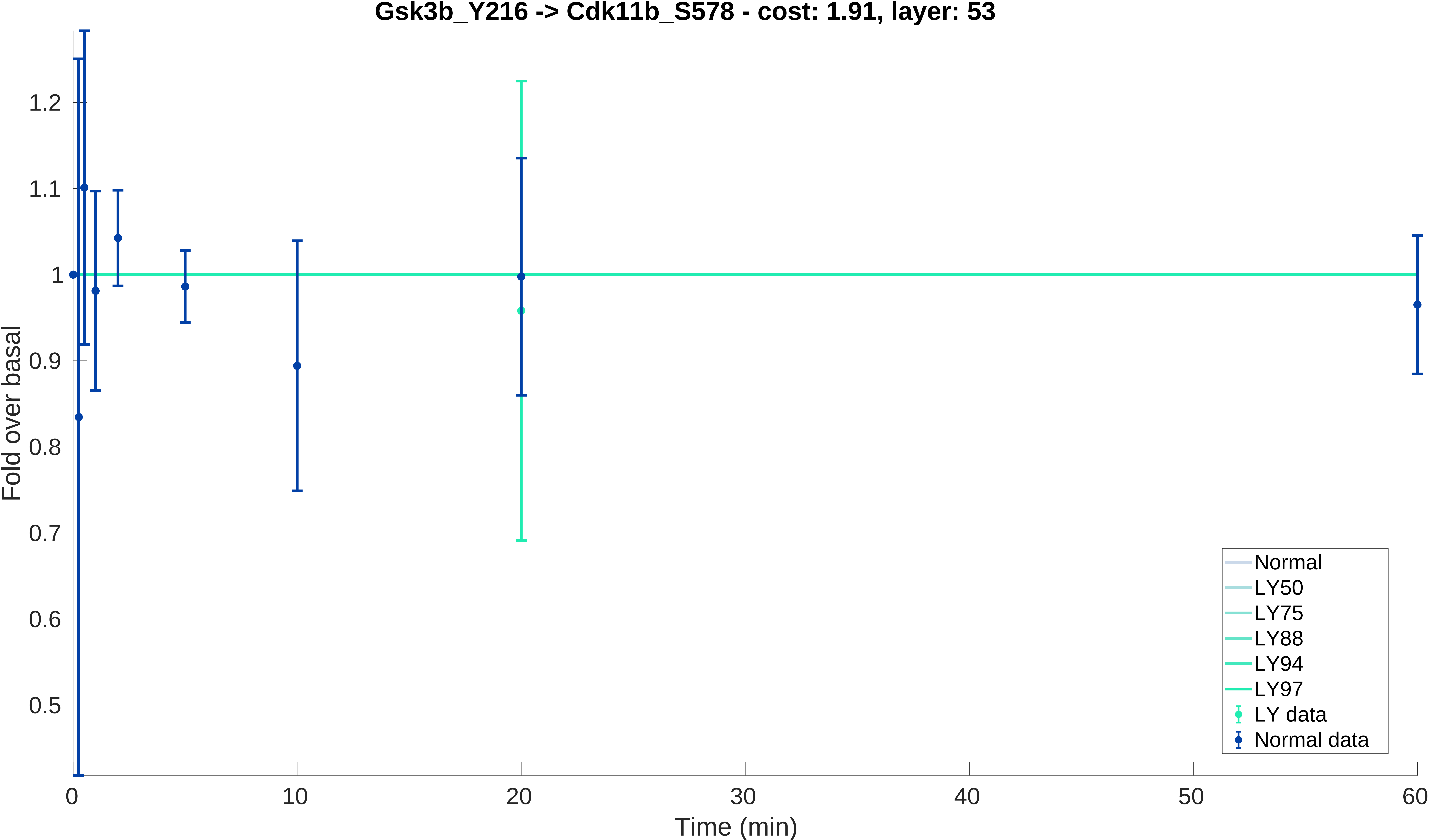

Gsk3b_Y216 -> Cdk11b_S578 - cost: 1.91, layer: 53
Fold over basal
1.2
1.1
1
0.9
0.8
0.7
0.6
0.5
0
10
20
30
40
50
60
Time (min)
Normal
LY50
LY75
LY88
LY94
LY97
LY data
Normal data

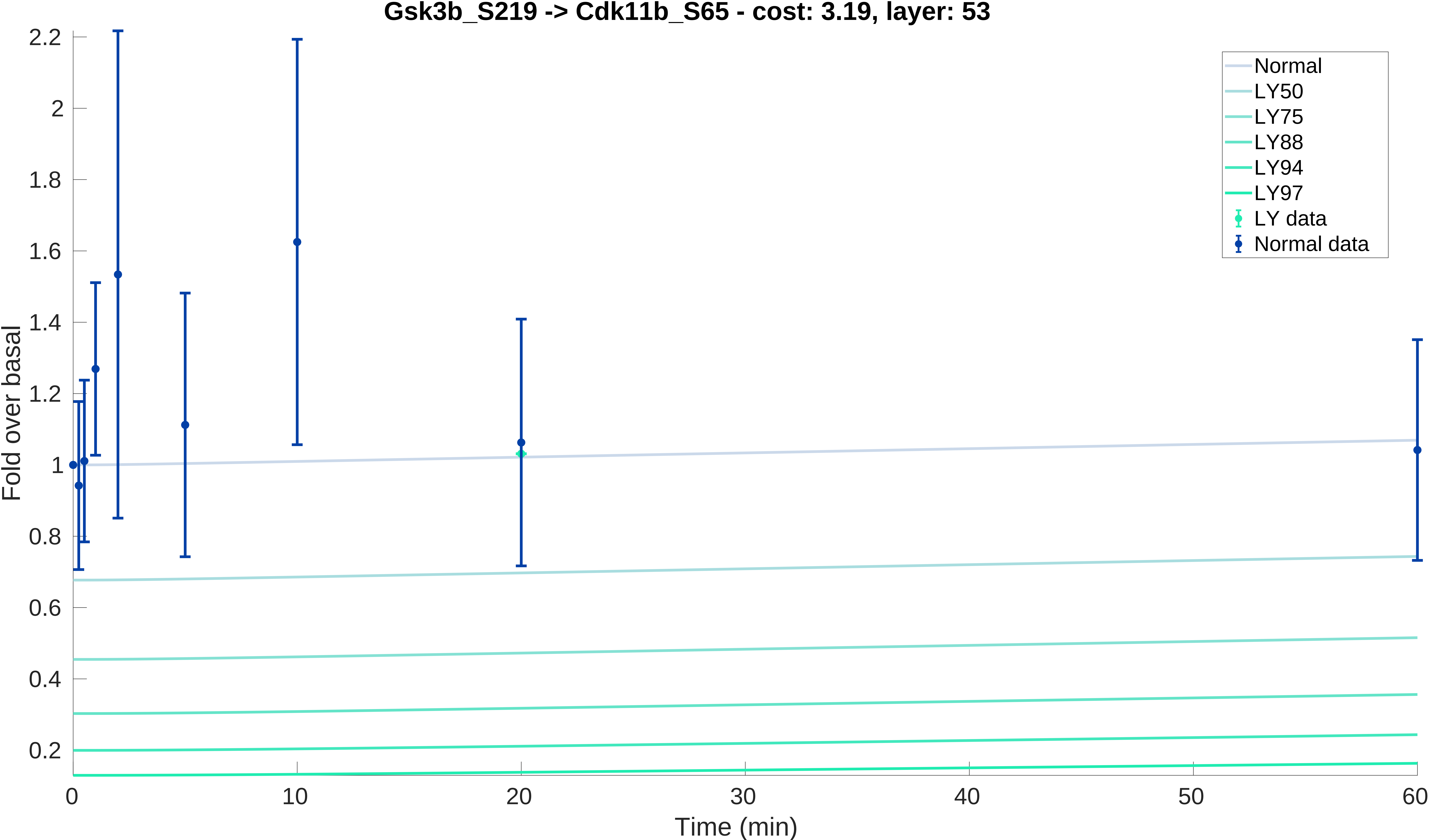
Gsk3b_S219 -> Cdk11b_S65 - cost: 3.19, layer: 53
Fold over basal
Time (min)
0
10
20
30
40
50
60
0.2
0.4
0.6
0.8
1
1.2
1.4
1.6
1.8
2
2.2
Normal
LY50
LY75
LY88
LY94
LY97
LY data
Normal data

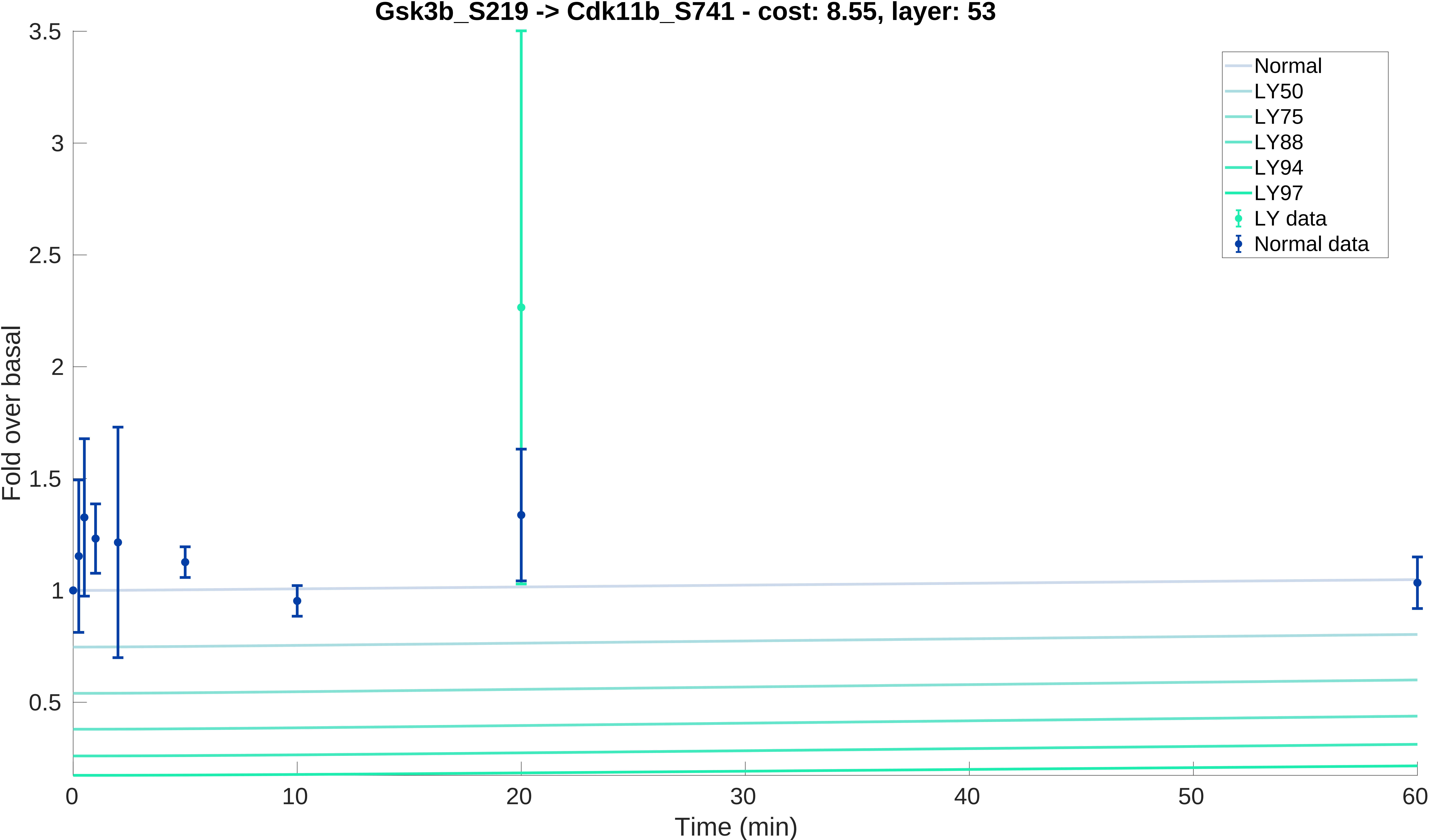

Gsk3b_S219 -> Cdk11b_S741 - cost: 8.55, layer: 53
Fold over basal
Time (min)
0.5
1
1.5
2
2.5
3
3.5
0
10
20
30
40
50
60
Normal
LY50
LY75
LY88
LY94
LY97
LY data
Normal data

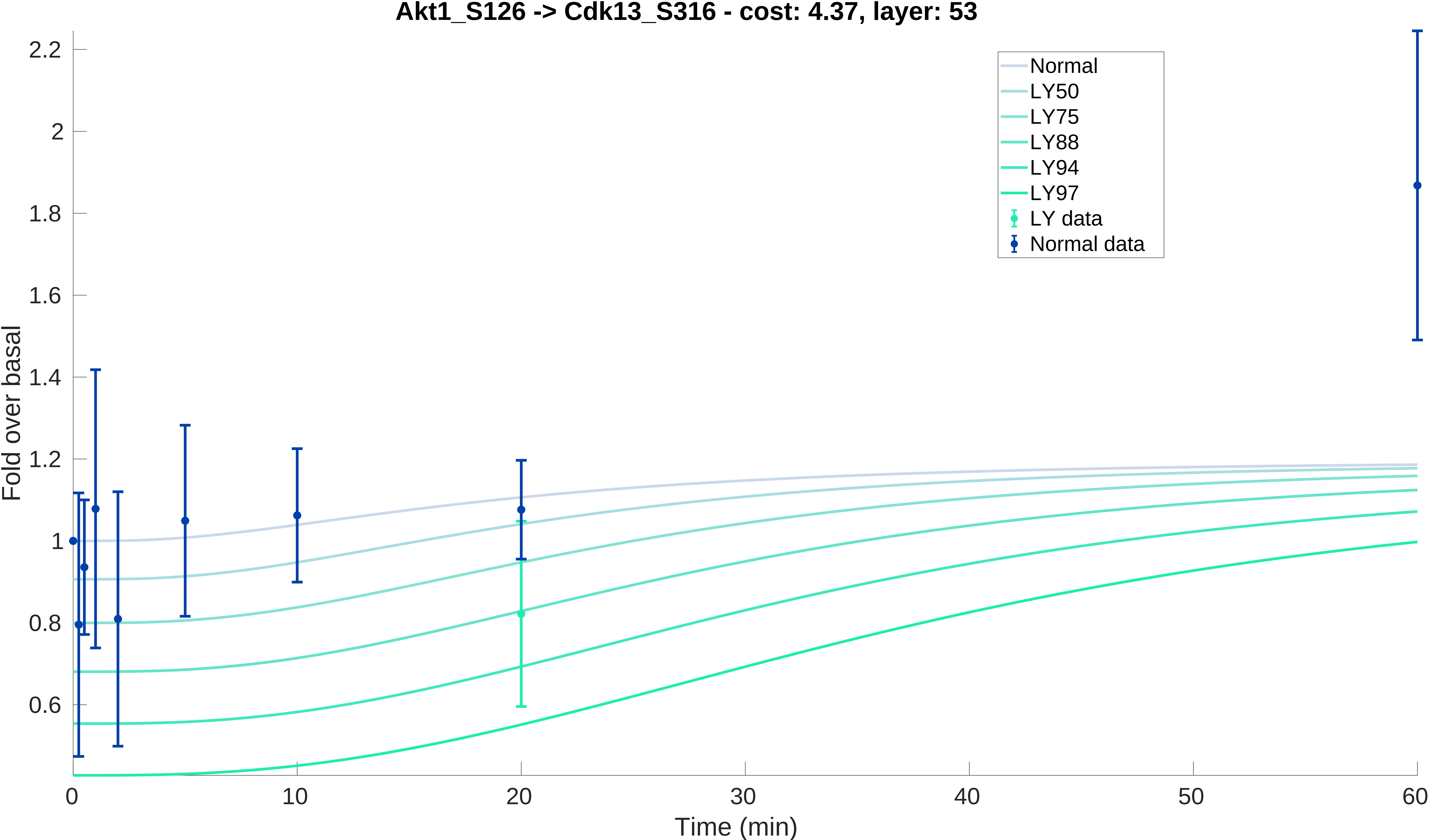

Akt1_S126 -> Cdk13_S316 - cost: 4.37, layer: 53
Fold over basal
2.2
2
1.8
1.6
1.4
1.2
1
0.8
0.6
0
10
20
30
40
50
60
Time (min)
Normal
LY50
LY75
LY88
LY94
LY97
LY data
Normal data

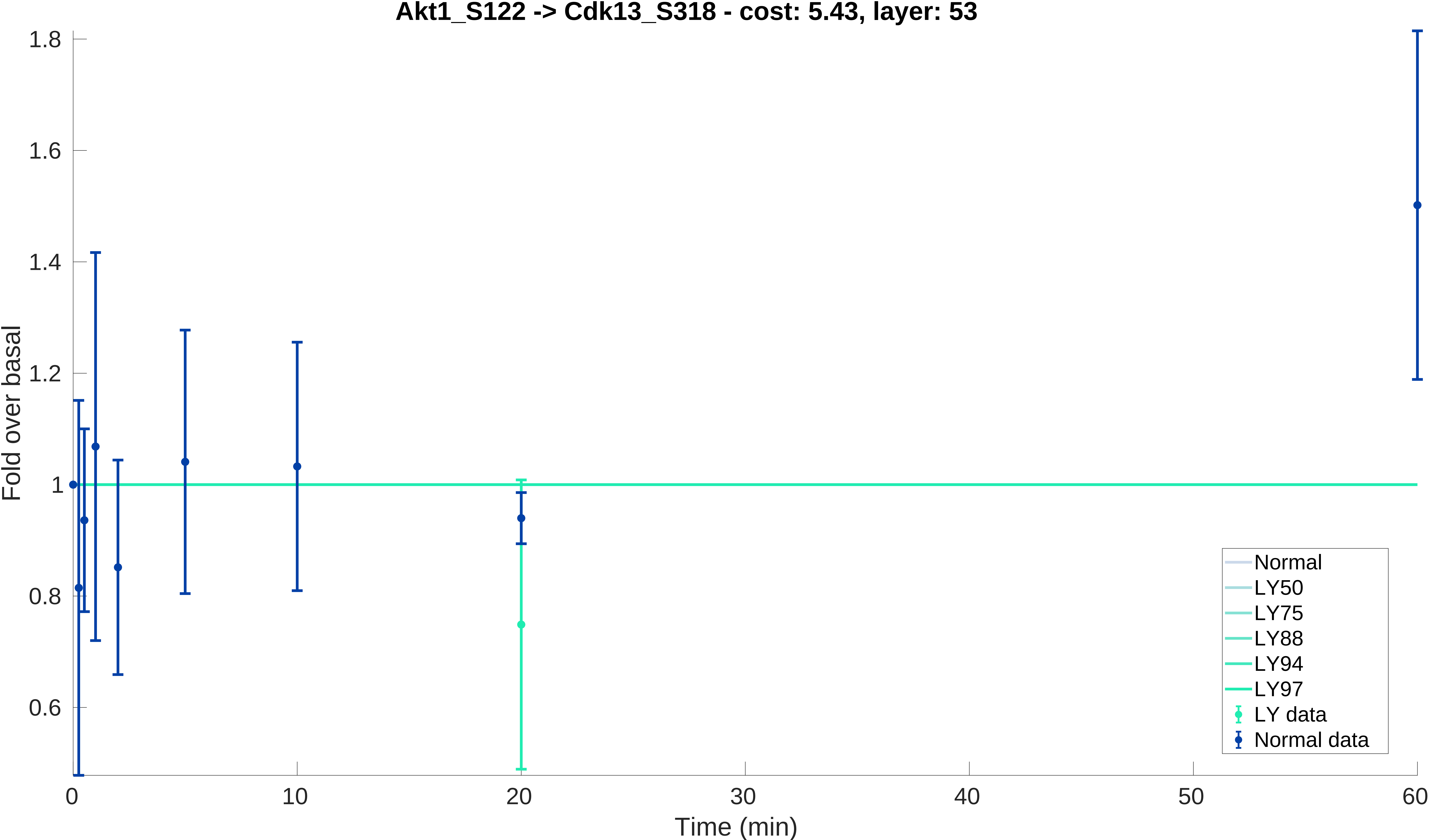

Akt1_S122 -> Cdk13_S318 - cost: 5.43, layer: 53
Fold over basal
Time (min)
1.8
1.6
1.4
1.2
1
0.8
0.6
0
10
20
30
40
50
60
Normal
LY50
LY75
LY88
LY94
LY97
LY data
Normal data

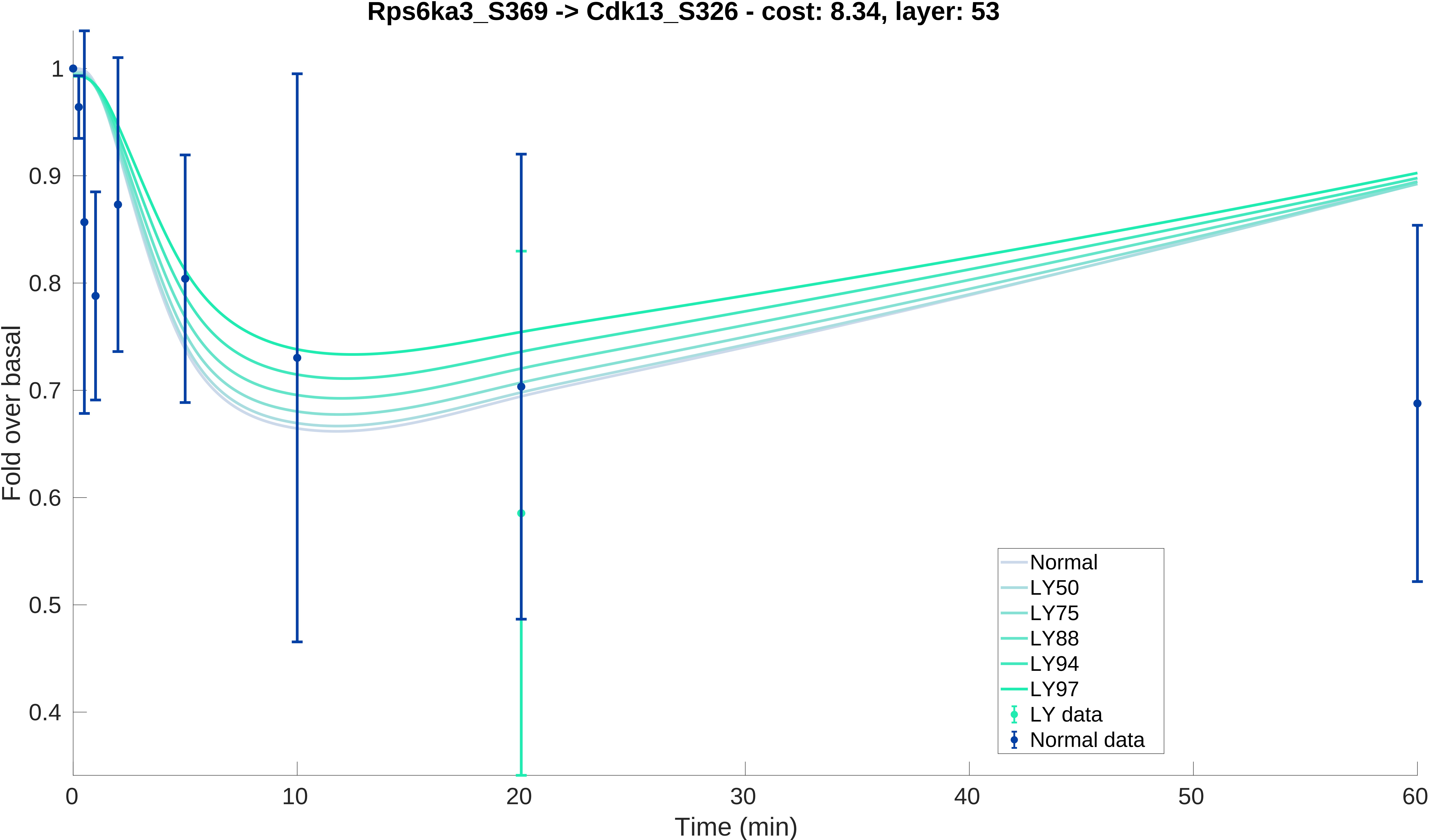
Rps6ka3_S369 -> Cdk13_S326 - cost: 8.34, layer: 53
Fold over basal
1
0.9
0.8
0.7
0.6
0.5
0.4
0
10
20
30
40
50
60
Time (min)
Normal
LY50
LY75
LY88
LY94
LY97
LY data
Normal data

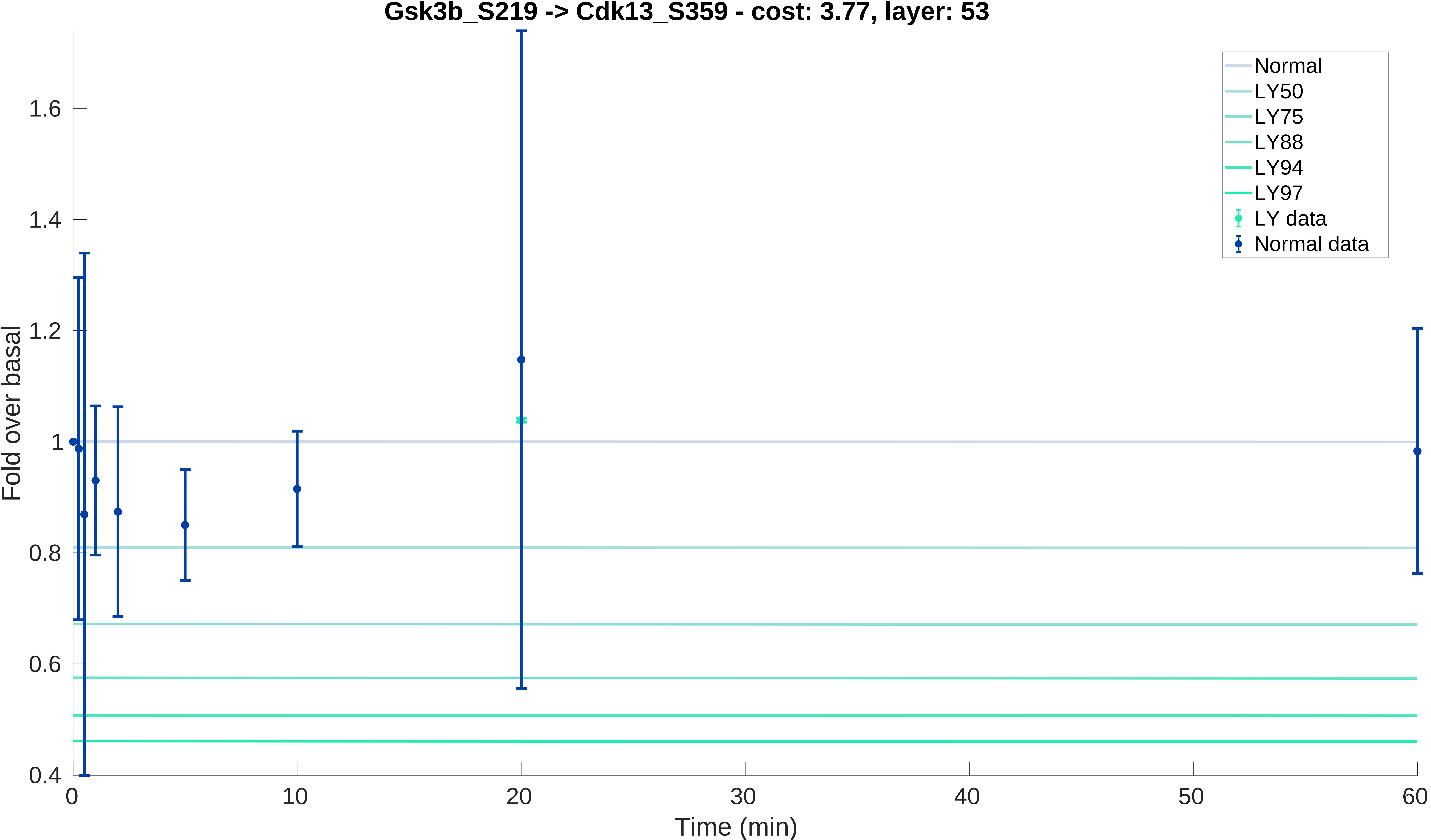
Gsk3b_S219 -> Cdk13_S359 - cost: 3.77, layer: 53
Fold over basal
Time (min)
0.4
0.6
0.8
1
1.2
1.4
1.6
0
10
20
30
40
50
60
Normal
LY50
LY75
LY88
LY94
LY97
LY data
Normal data

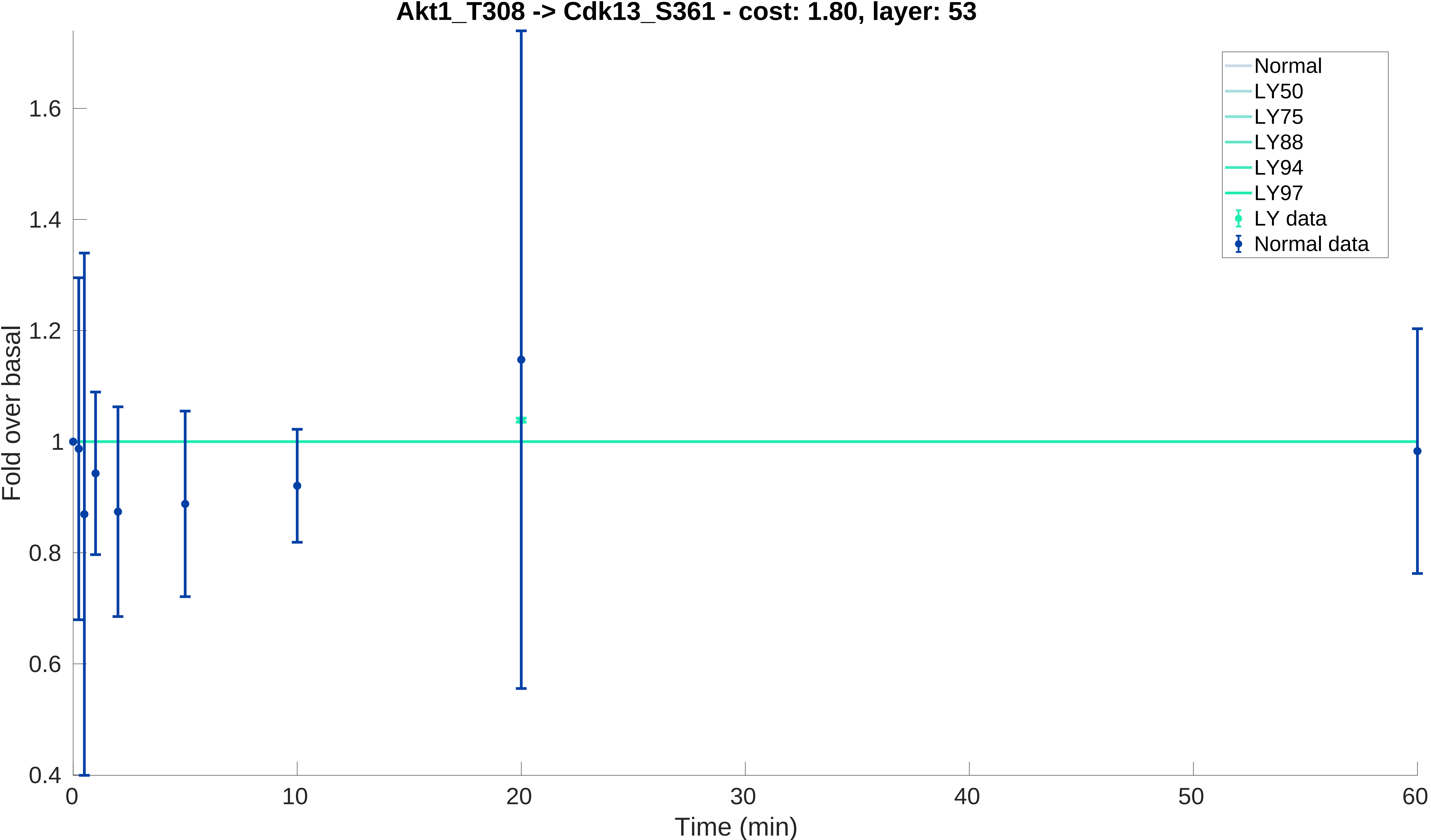
Akt1_T308 -> Cdk13_S361 - cost: 1.80, layer: 53
Fold over basal
Time (min)
1.6
1.4
1.2
1
0.8
0.6
0.4
0
10
20
30
40
50
60
Normal
LY50
LY75
LY88
LY94
LY97
LY data
Normal data

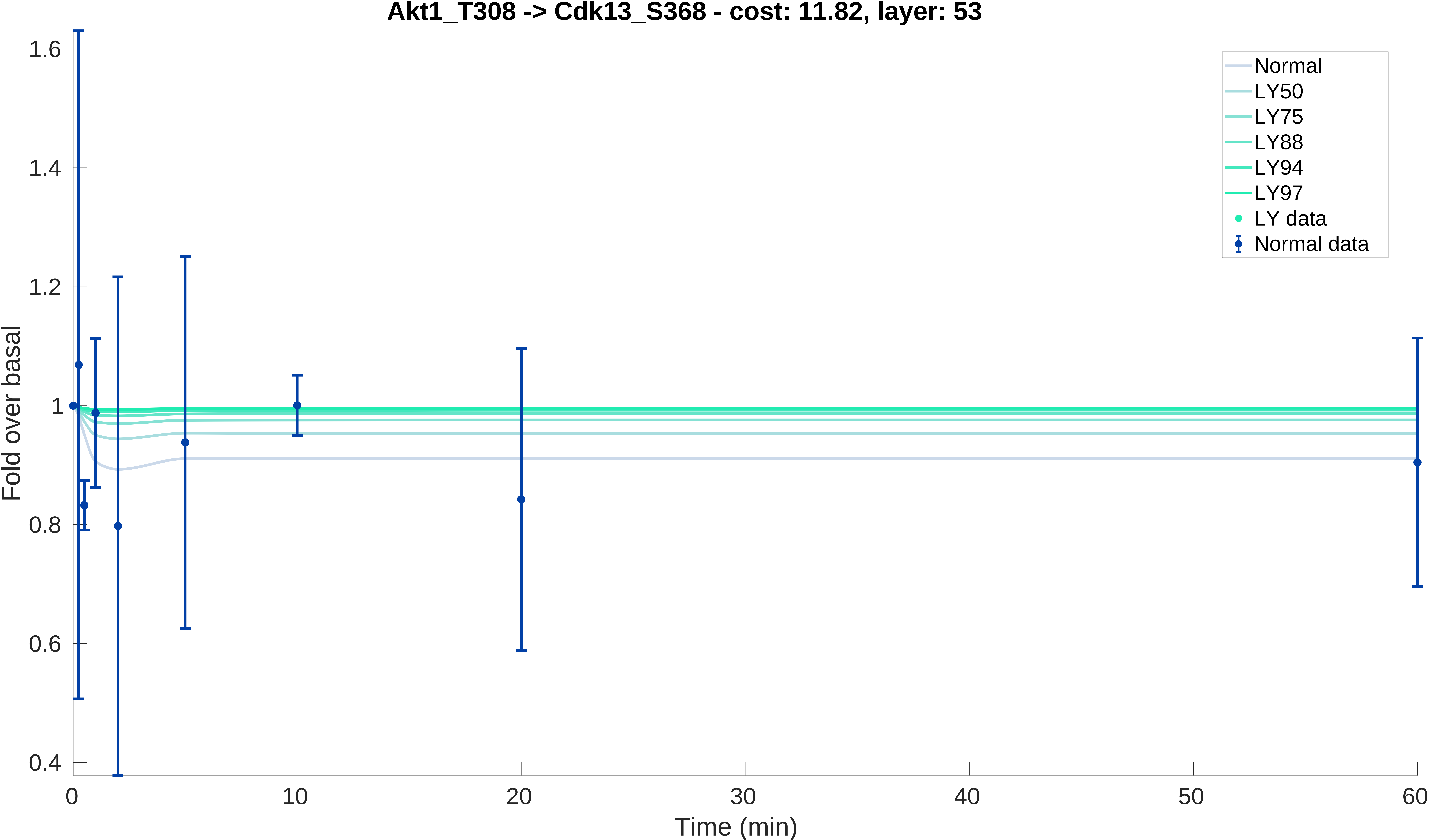

Akt1_T308 -> Cdk13_S368 - cost: 11.82, layer: 53
Fold over basal
Time (min)
1.6
1.4
1.2
1
0.8
0.6
0.4
0
10
20
30
40
50
60
Normal
LY50
LY75
LY88
LY94
LY97
LY data
Normal data

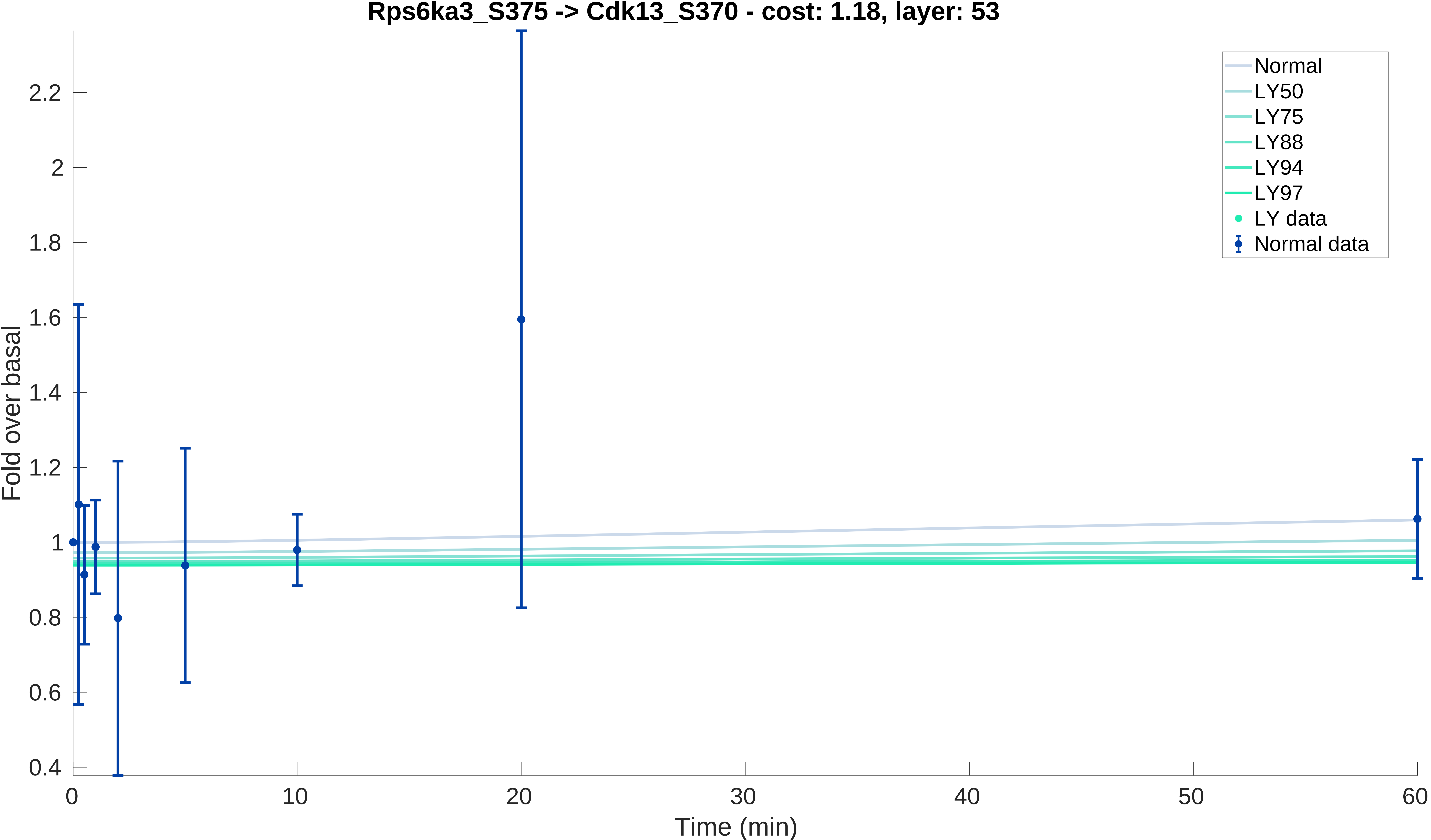

Rps6ka3_S375 -> Cdk13_S370 - cost: 1.18, layer: 53
Normal
LY50
LY75
LY88
LY94
LY97
LY data
Normal data
Fold over basal
Time (min)
0
10
20
30
40
50
60
0.4
0.6
0.8
1
1.2
1.4
1.6
1.8
2
2.2

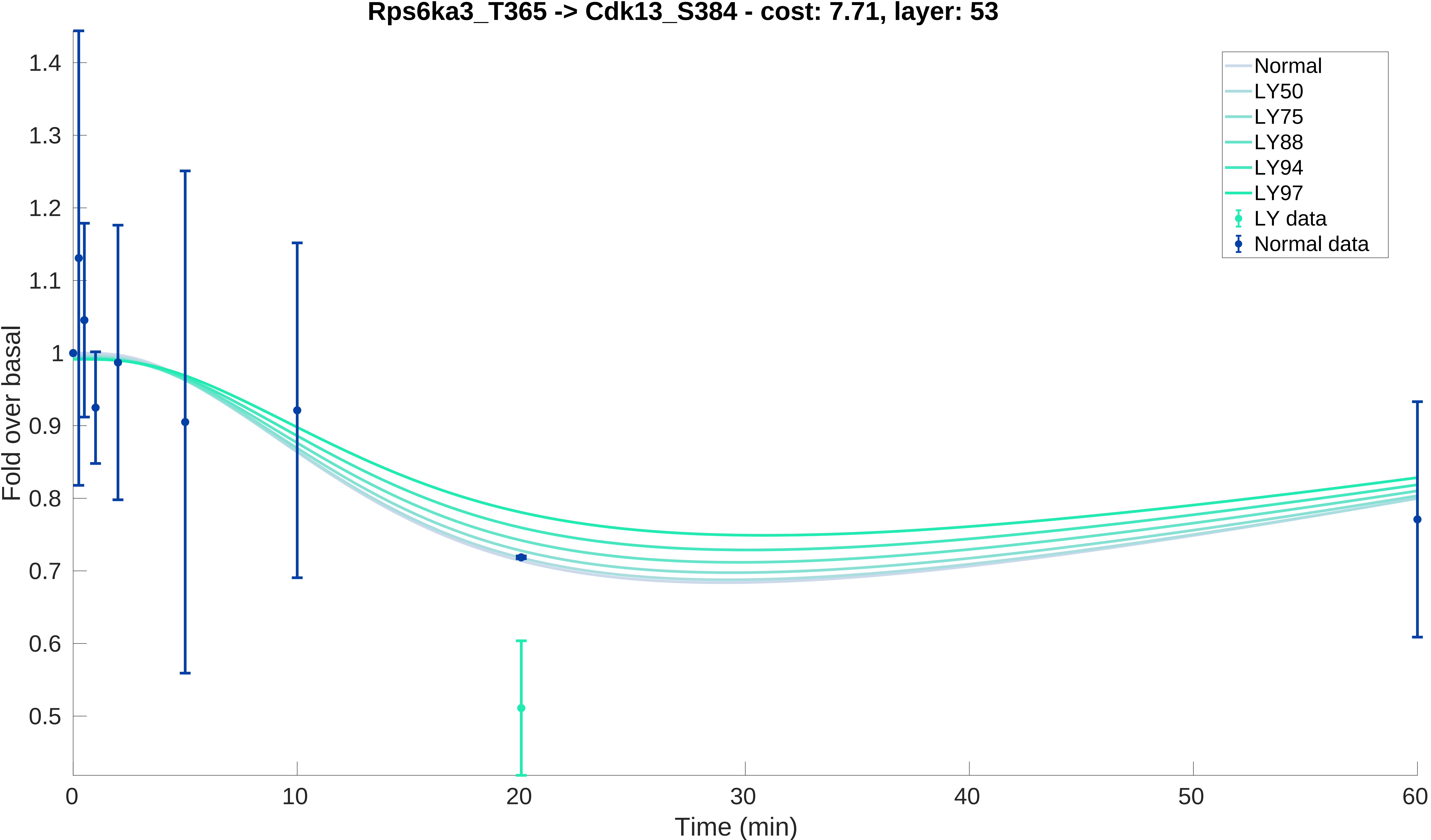

Rps6ka3_T365 -> Cdk13_S384 - cost: 7.71, layer: 53
Normal
LY50
LY75
LY88
LY94
LY97
LY data
Normal data
Fold over basal
1.4
1.3
1.2
1.1
1
0.9
0.8
0.7
0.6
0.5
0
10
20
30
40
50
60
Time (min)

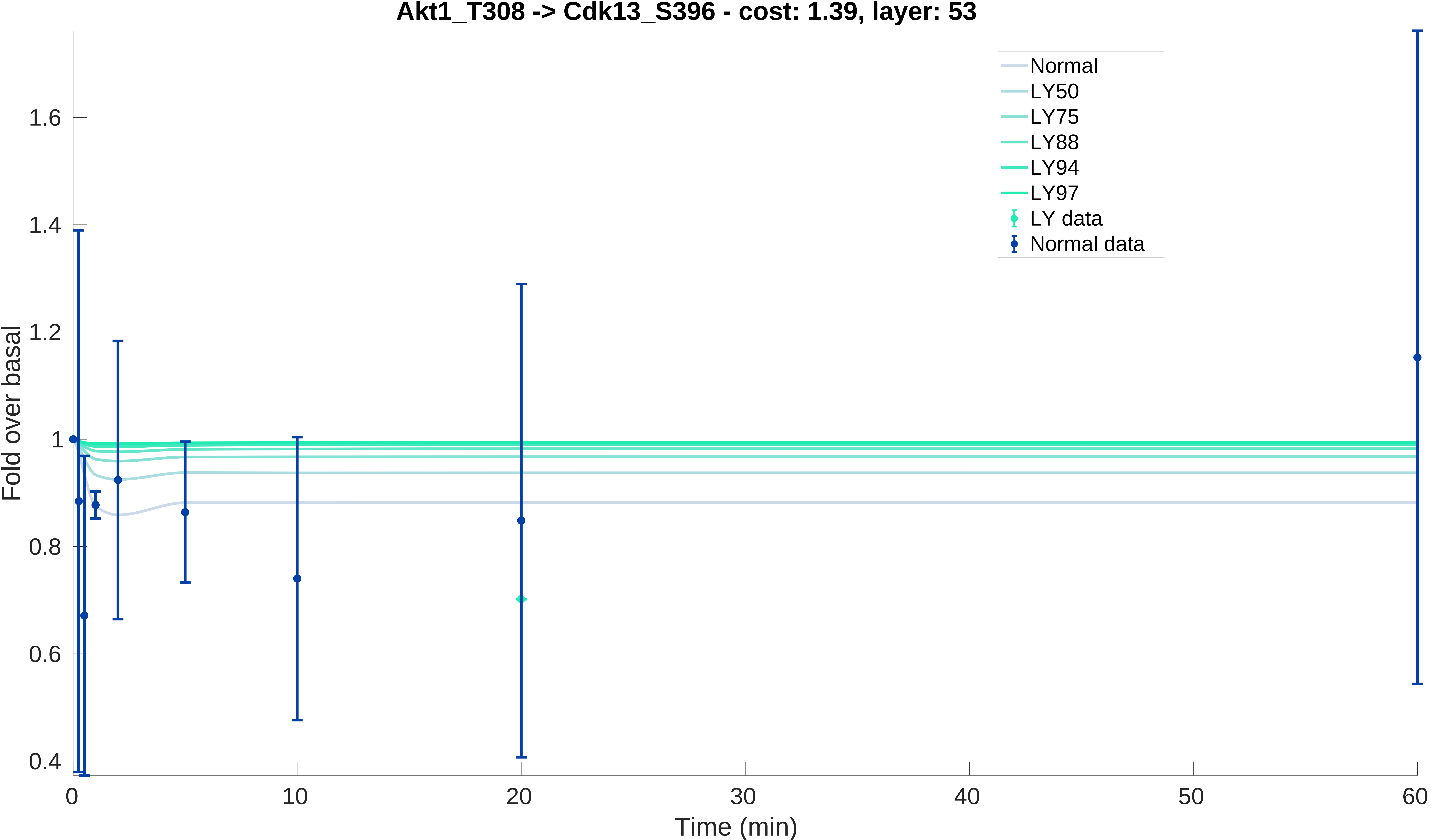

Akt1_T308 -> Cdk13_S396 - cost: 1.39, layer: 53
Normal
LY50
LY75
LY88
LY94
LY97
LY data
Normal data
1.6
1.4
1.2
1
0.8
0.6
0.4
Fold over basal
0
10
20
30
40
50
60
Time (min)

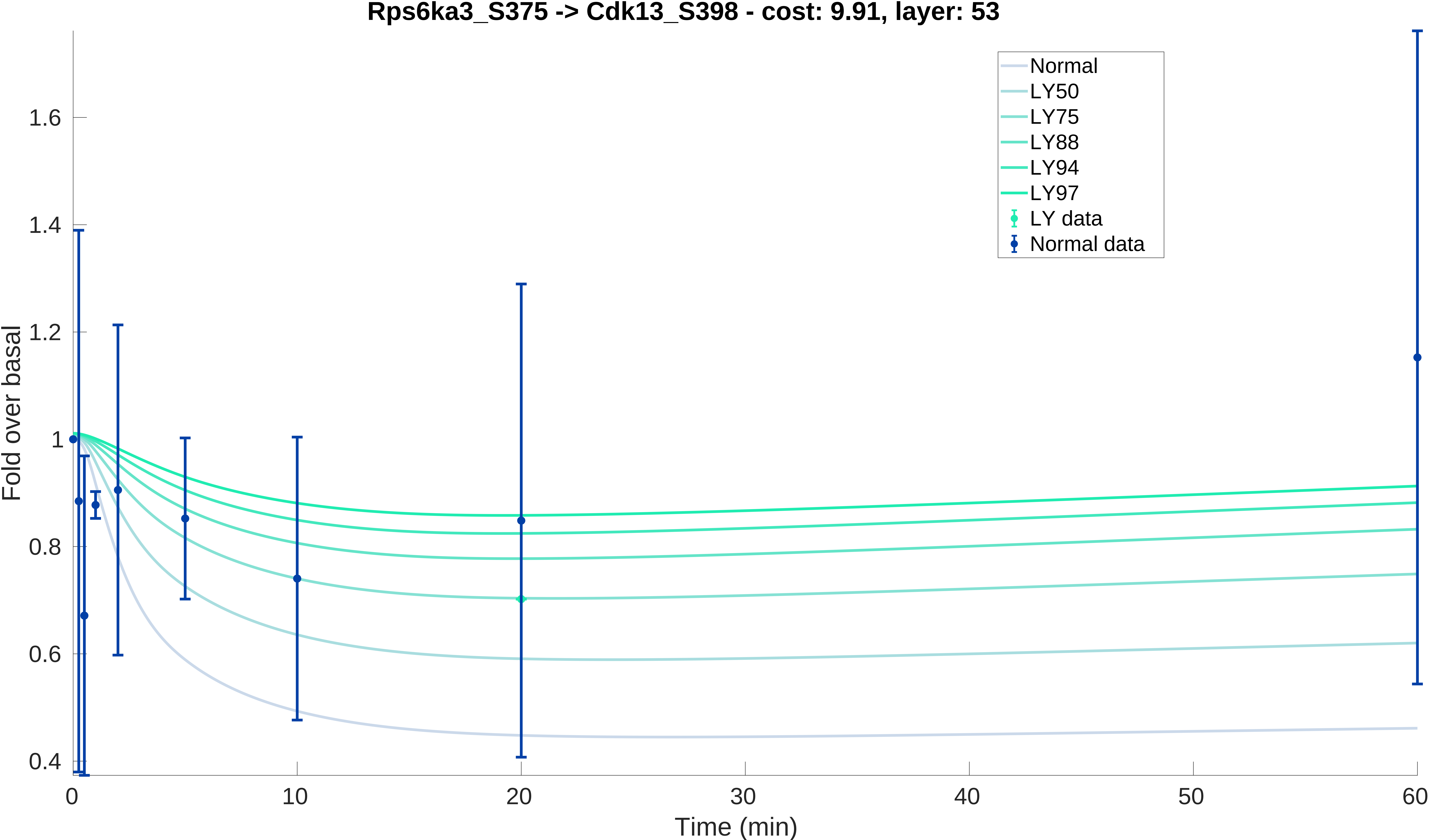
Rps6ka3_S375 -> Cdk13_S398 - cost: 9.91, layer: 53
Fold over basal
Time (min)
0.4
0.6
0.8
1
1.2
1.4
1.6
0
10
20
30
40
50
60
Normal
LY50
LY75
LY88
LY94
LY97
LY data
Normal data

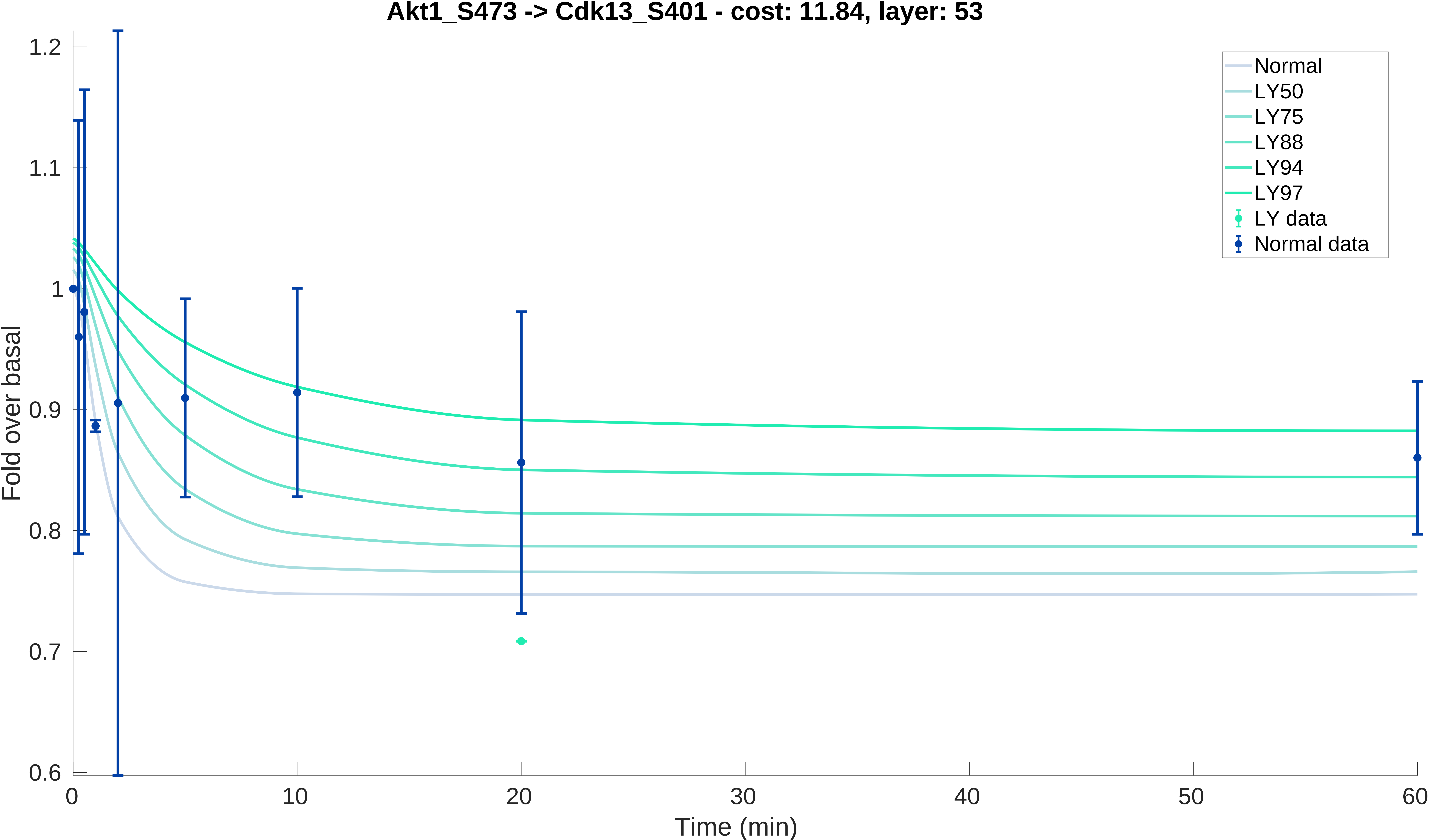
Akt1_S473 -> Cdk13_S401 - cost: 11.84, layer: 53
Normal
LY50
LY75
LY88
LY94
LY97
LY data
Normal data
Fold over basal
Time (min)
0.6
0.7
0.8
0.9
1
1.1
1.2
0
10
20
30
40
50
60

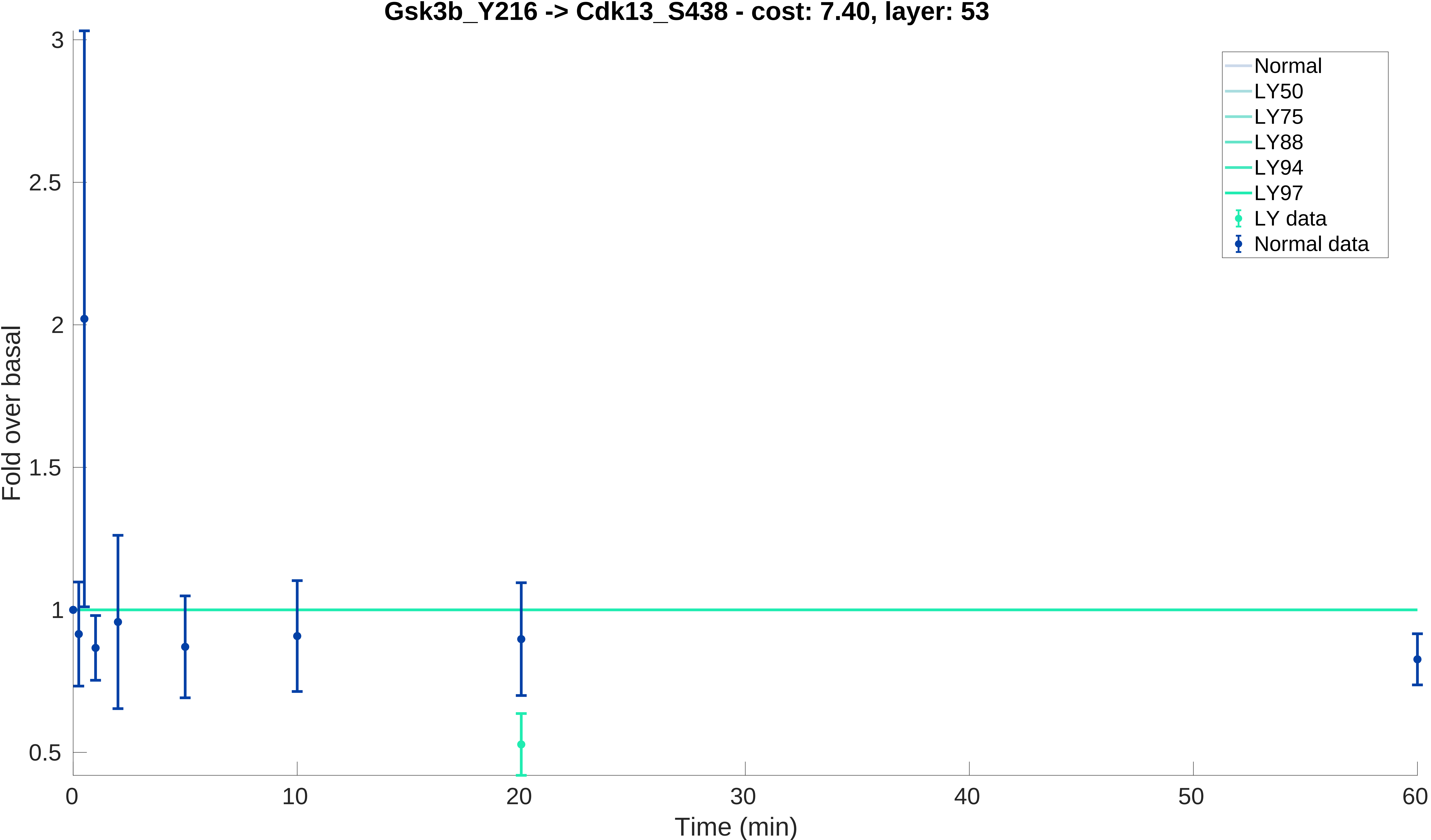

Gsk3b_Y216 -> Cdk13_S438 - cost: 7.40, layer: 53
Fold over basal
3
2.5
2
1.5
1
0.5
0
10
20
30
40
50
60
Time (min)
Normal
LY50
LY75
LY88
LY94
LY97
LY data
Normal data

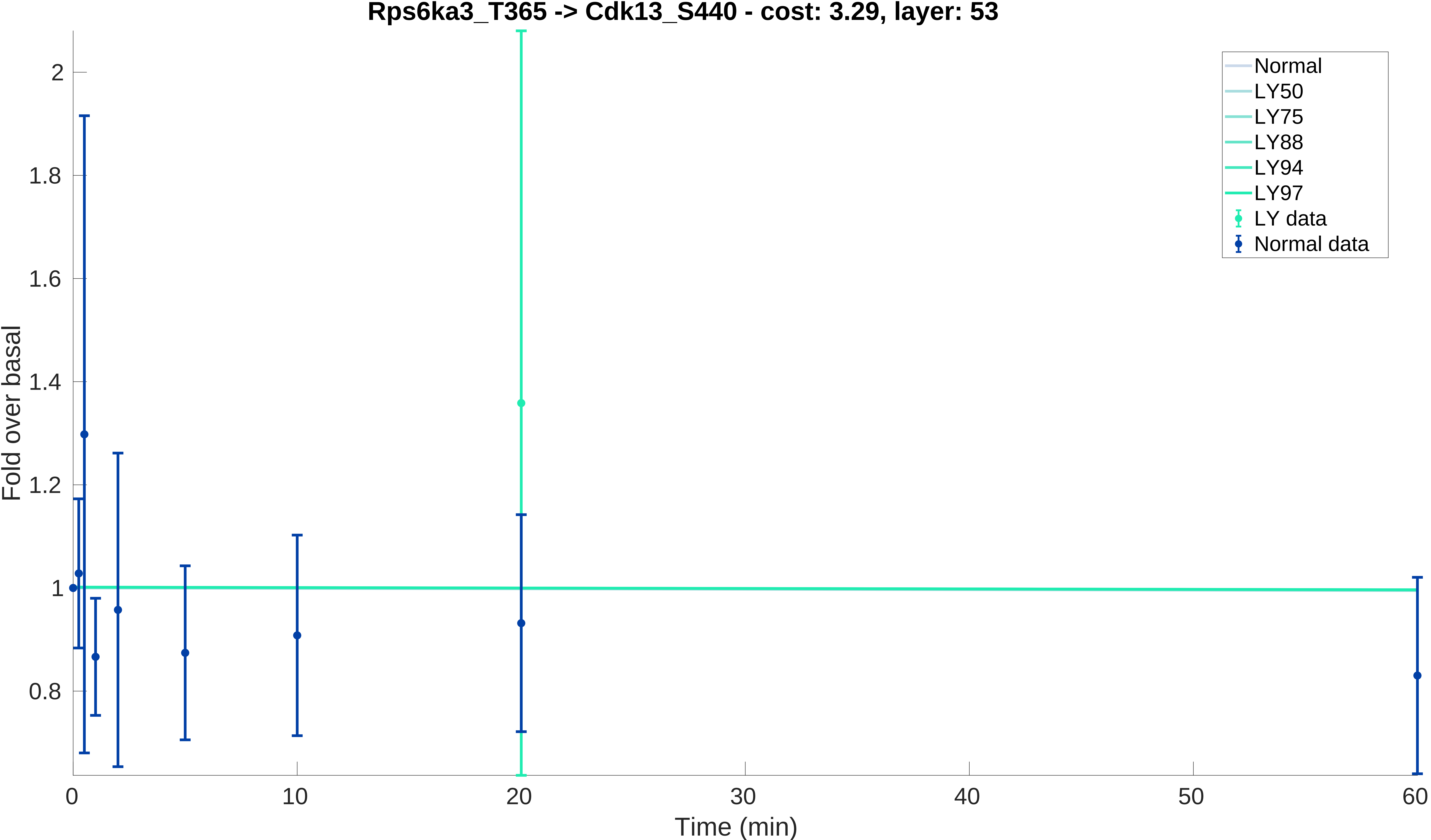

Rps6ka3_T365 -> Cdk13_S440 - cost: 3.29, layer: 53
Normal
LY50
LY75
LY88
LY94
LY97
LY data
Normal data
Fold over basal
2
1.8
1.6
1.4
1.2
1
0.8
0
10
20
30
40
50
60
Time (min)

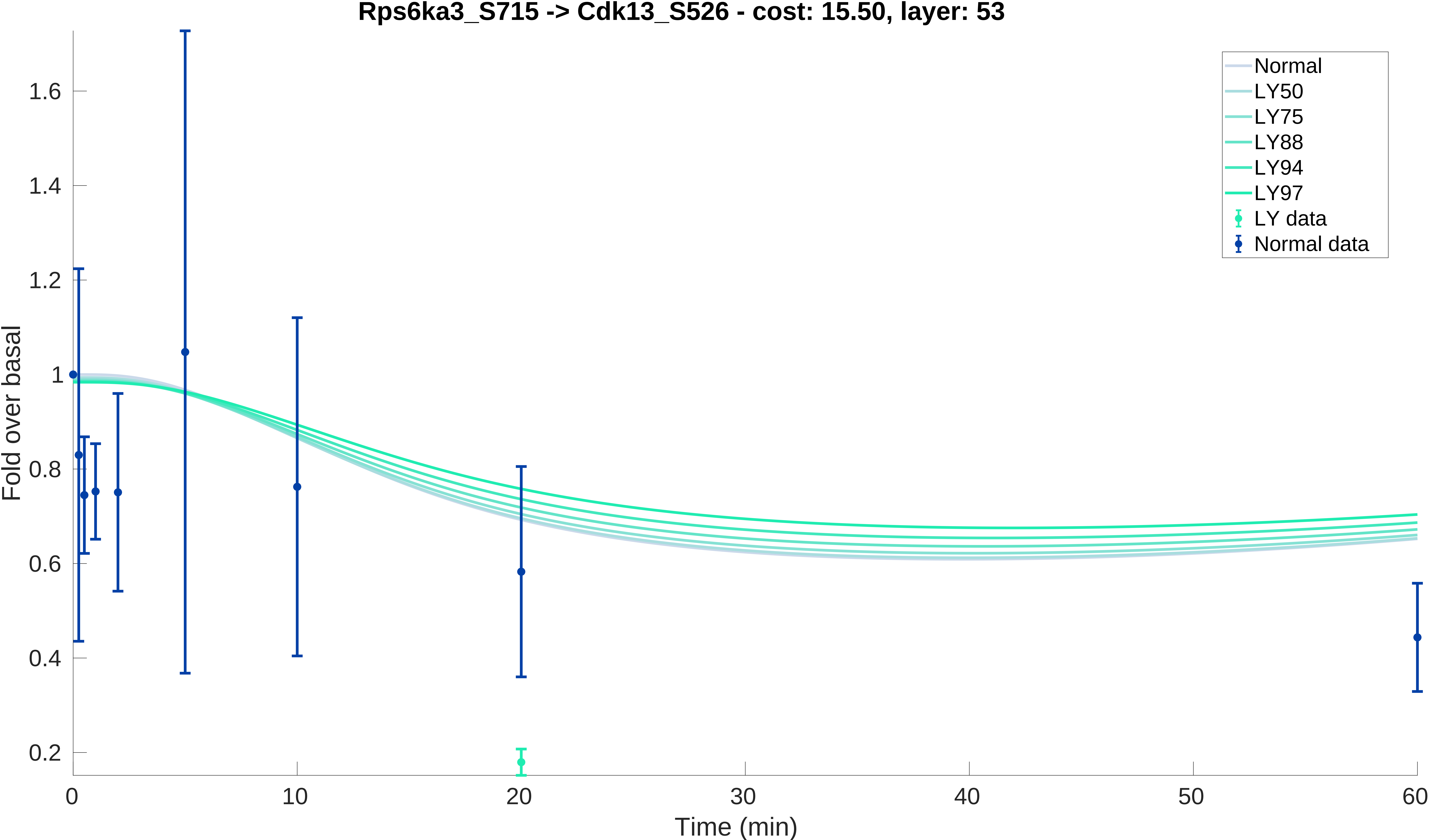

Rps6ka3_S715 -> Cdk13_S526 - cost: 15.50, layer: 53
Normal
LY50
LY75
LY88
LY94
LY97
LY data
Normal data
Fold over basal
1.6
1.4
1.2
1
0.8
0.6
0.4
0.2
0
10
20
30
40
50
60
Time (min)

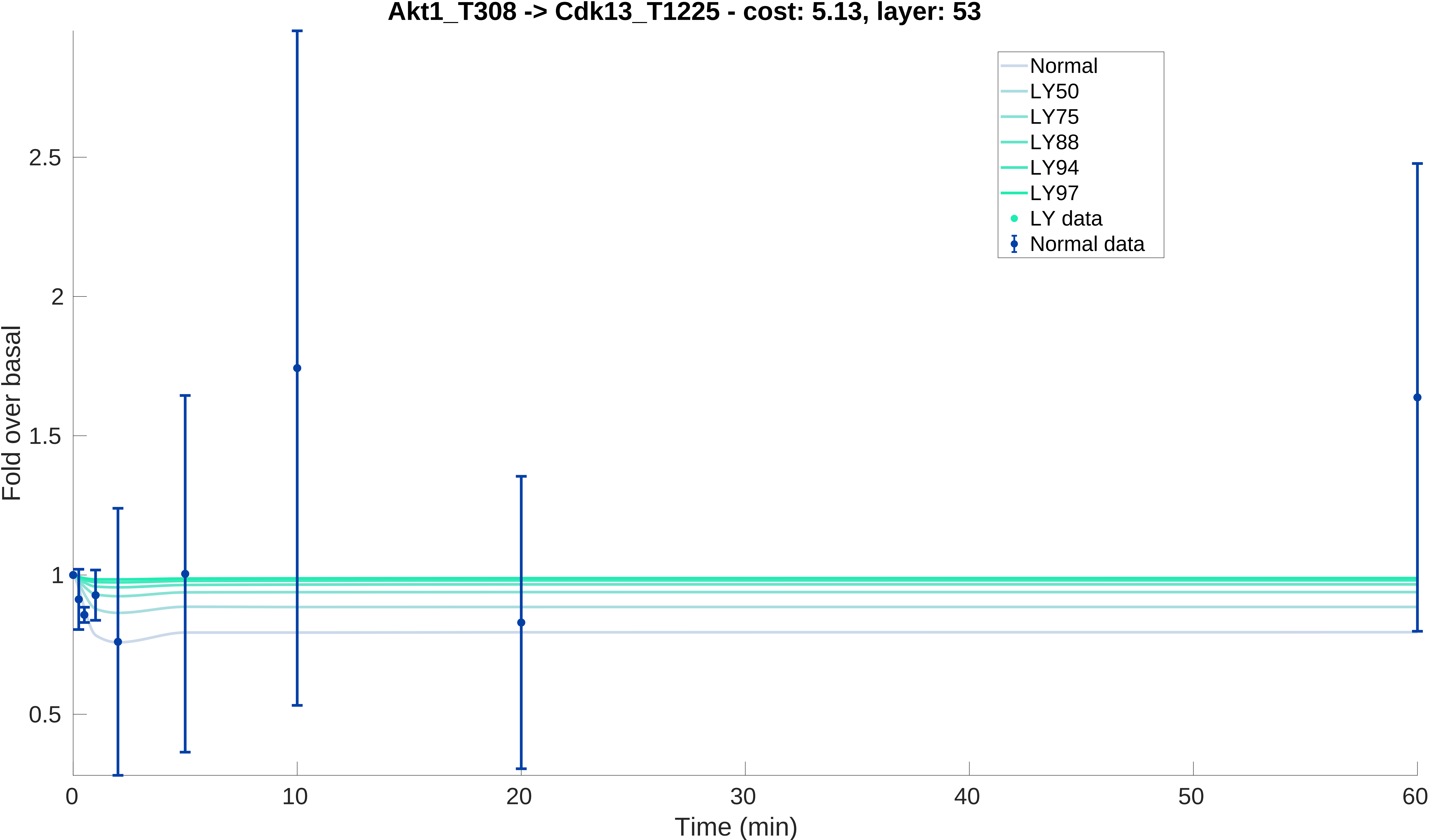

Akt1_T308 -> Cdk13_T1225 - cost: 5.13, layer: 53
Normal
LY50
LY75
LY88
LY94
LY97
LY data
Normal data
Fold over basal
Time (min)
0.5
1
1.5
2
2.5
0
10
20
30
40
50
60

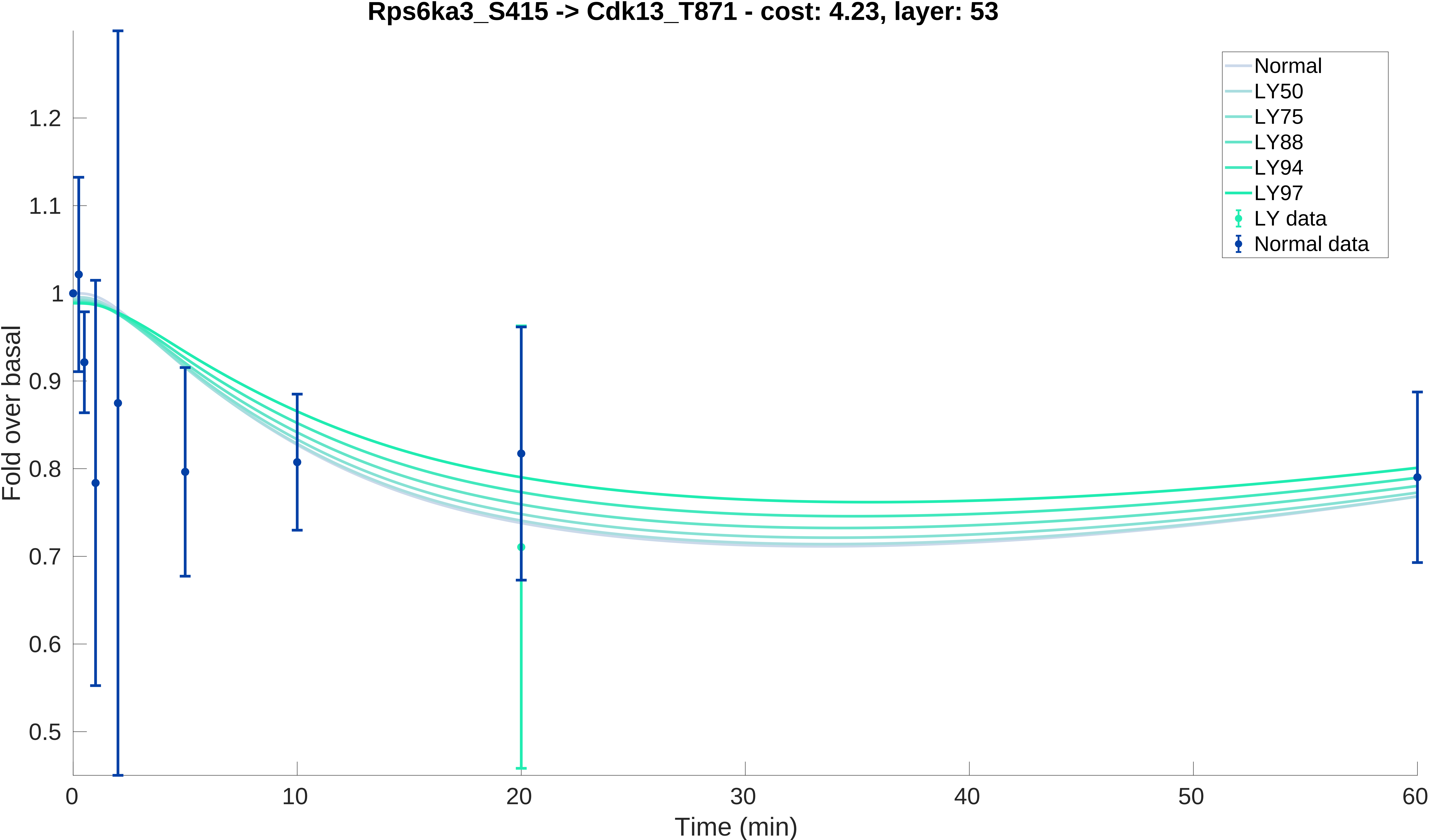
Rps6ka3_S415 -> Cdk13_T871 - cost: 4.23, layer: 53
Fold over basal
Time (min)
Normal
LY50
LY75
LY88
LY94
LY97
LY data
Normal data
1.2
1.1
1
0.9
0.8
0.7
0.6
0.5
0
10
20
30
40
50
60

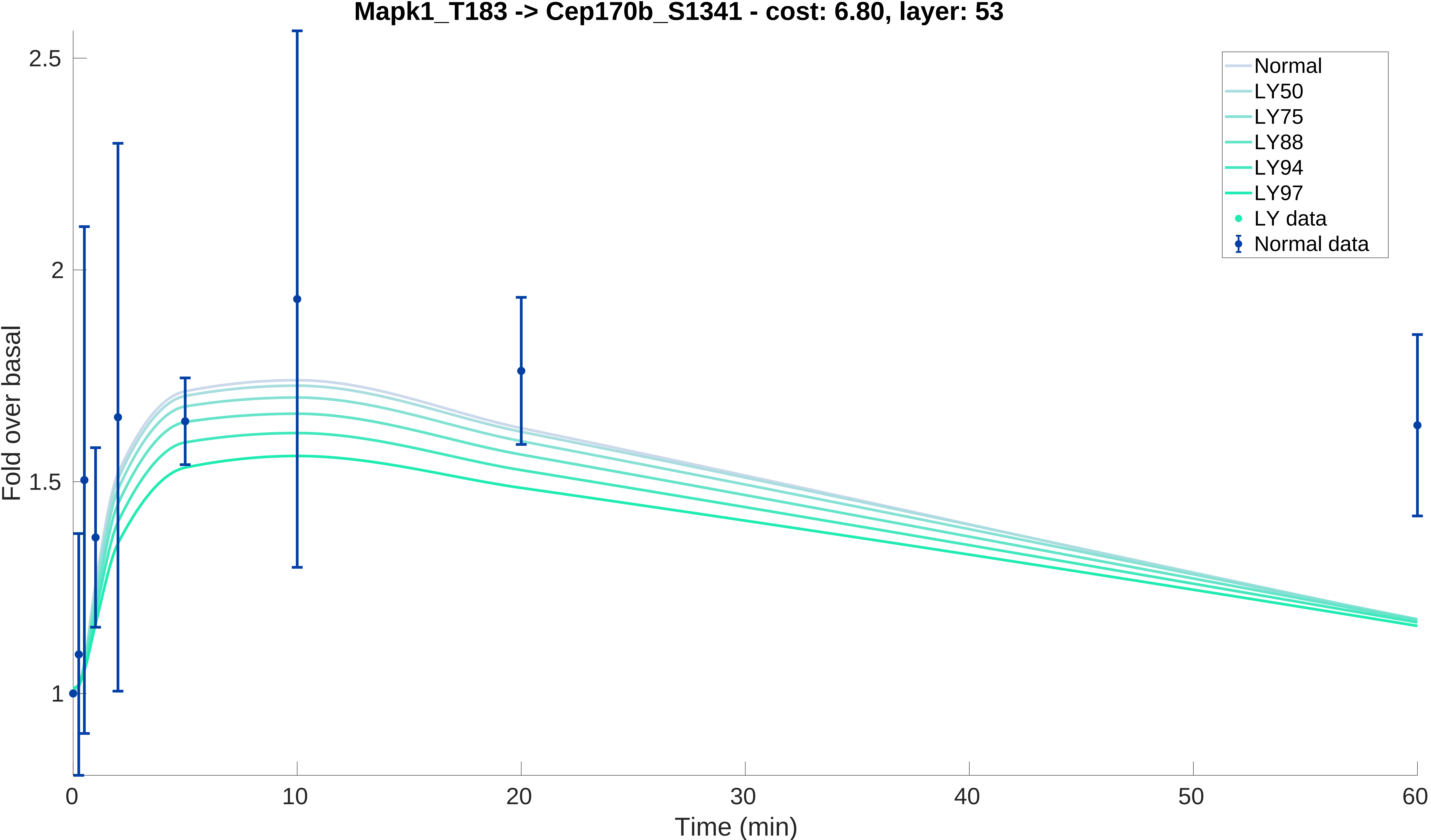

Mapk1_T183 -> Cep170b_S1341 - cost: 6.80, layer: 53
Normal
LY50
LY75
LY88
LY94
LY97
LY data
Normal data
Fold over basal
2.5
2
1.5
1
0
10
20
30
40
50
60
Time (min)

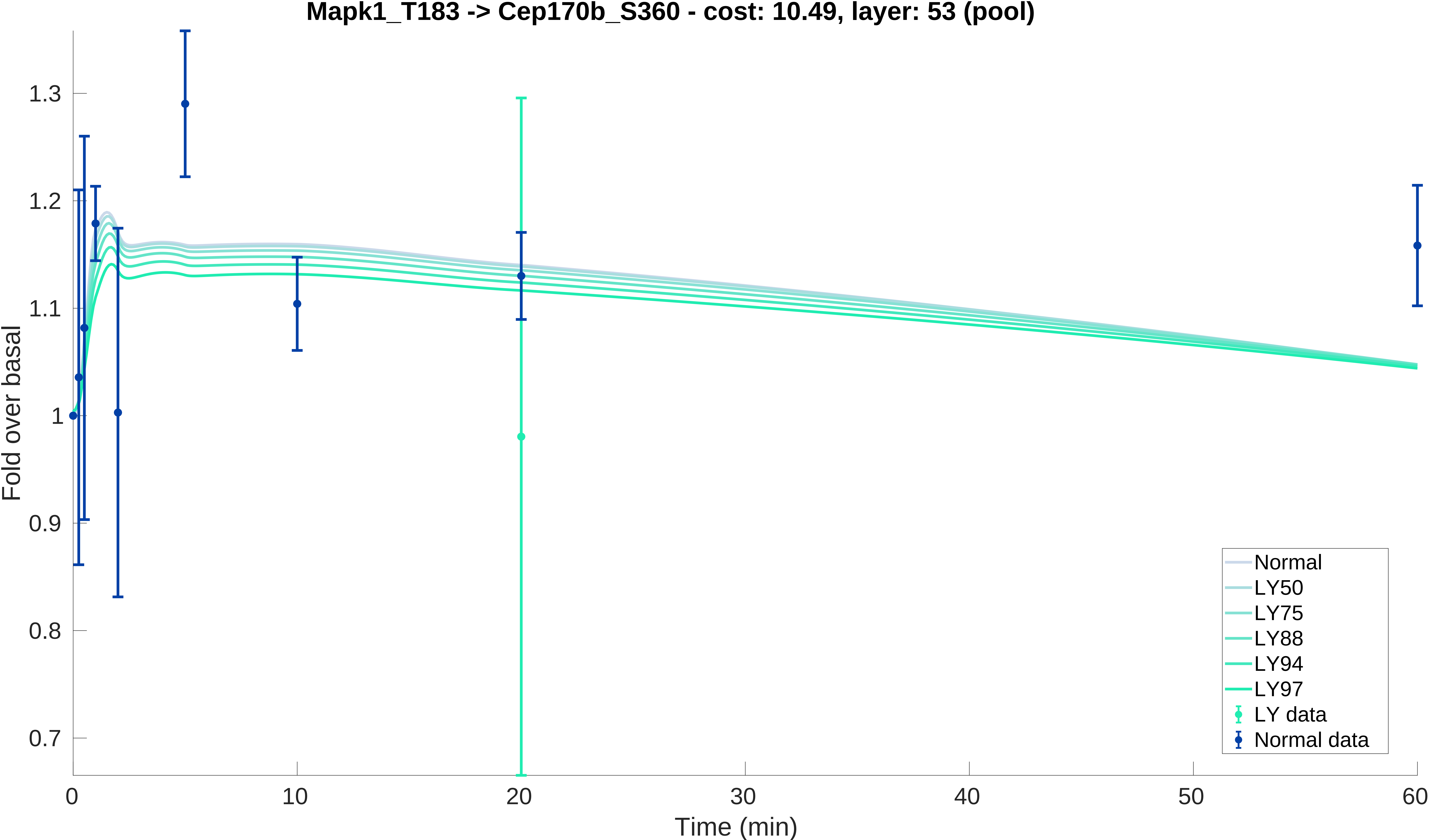

Mapk1_T183 -> Cep170b_S360 - cost: 10.49, layer: 53 (pool)
1.3
1.2
1.1
1
0.9
0.8
0.7
Fold over basal
0
10
20
30
40
50
60
Time (min)
Normal
LY50
LY75
LY88
LY94
LY97
LY data
Normal data

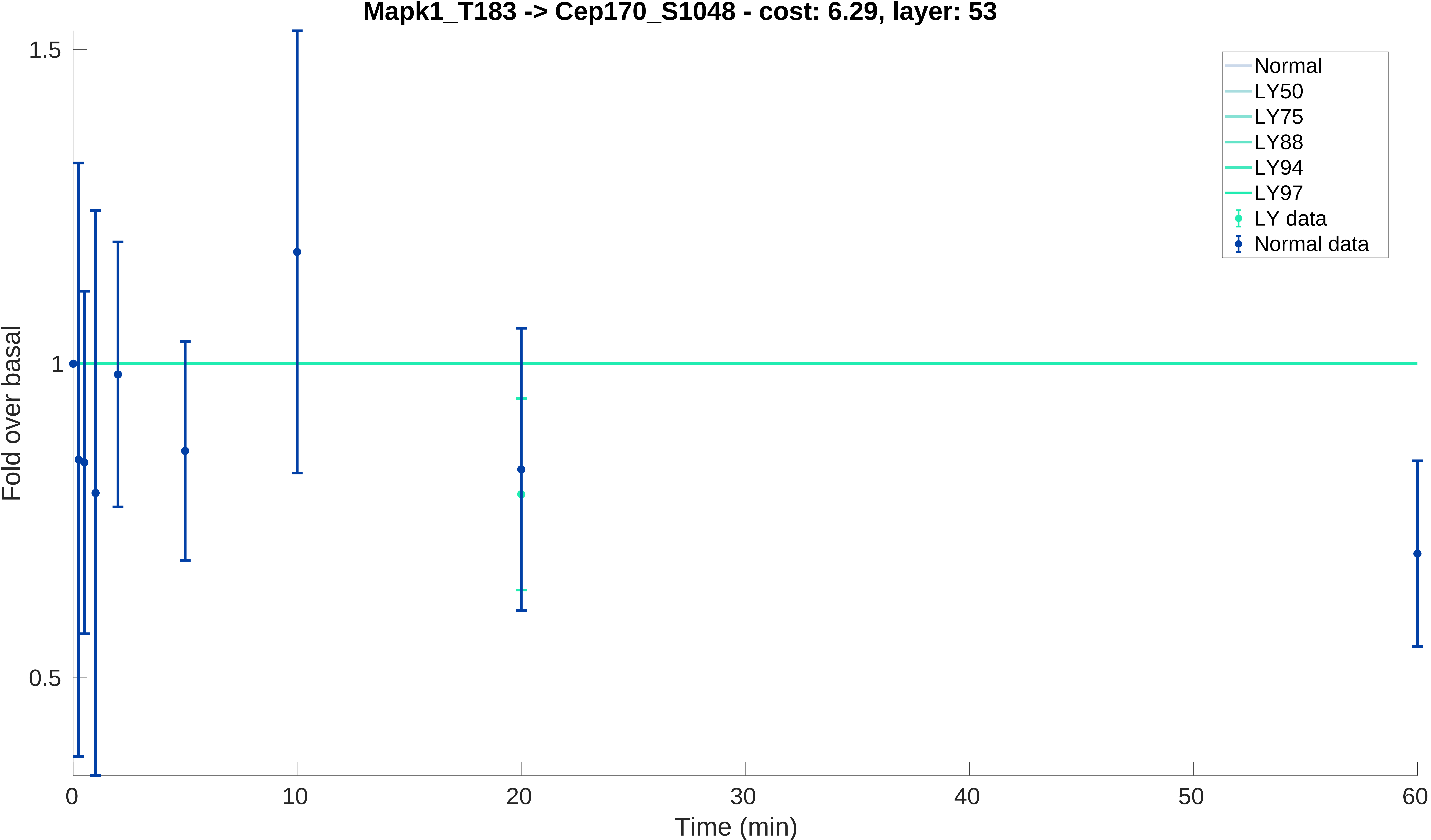

Mapk1_T183 -> Cep170_S1048 - cost: 6.29, layer: 53
1.5
1
0.5
Fold over basal
0
10
20
30
40
50
60
Time (min)
Normal
LY50
LY75
LY88
LY94
LY97
LY data
Normal data

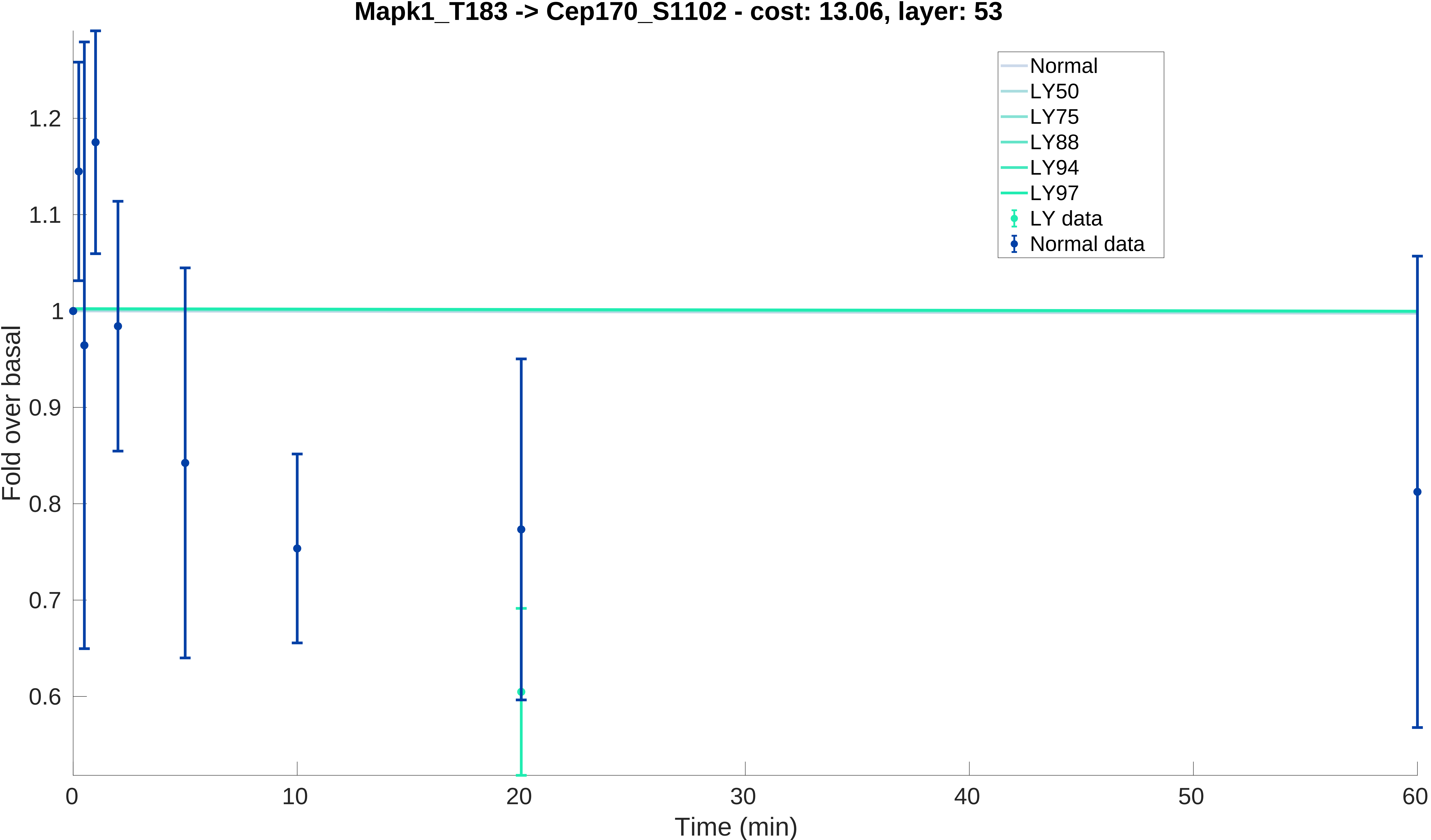

Mapk1_T183 -> Cep170_S1102 - cost: 13.06, layer: 53
Fold over basal
1.2
1.1
1
0.9
0.8
0.7
0.6
0
10
20
30
40
50
60
Time (min)
Normal
LY50
LY75
LY88
LY94
LY97
LY data
Normal data

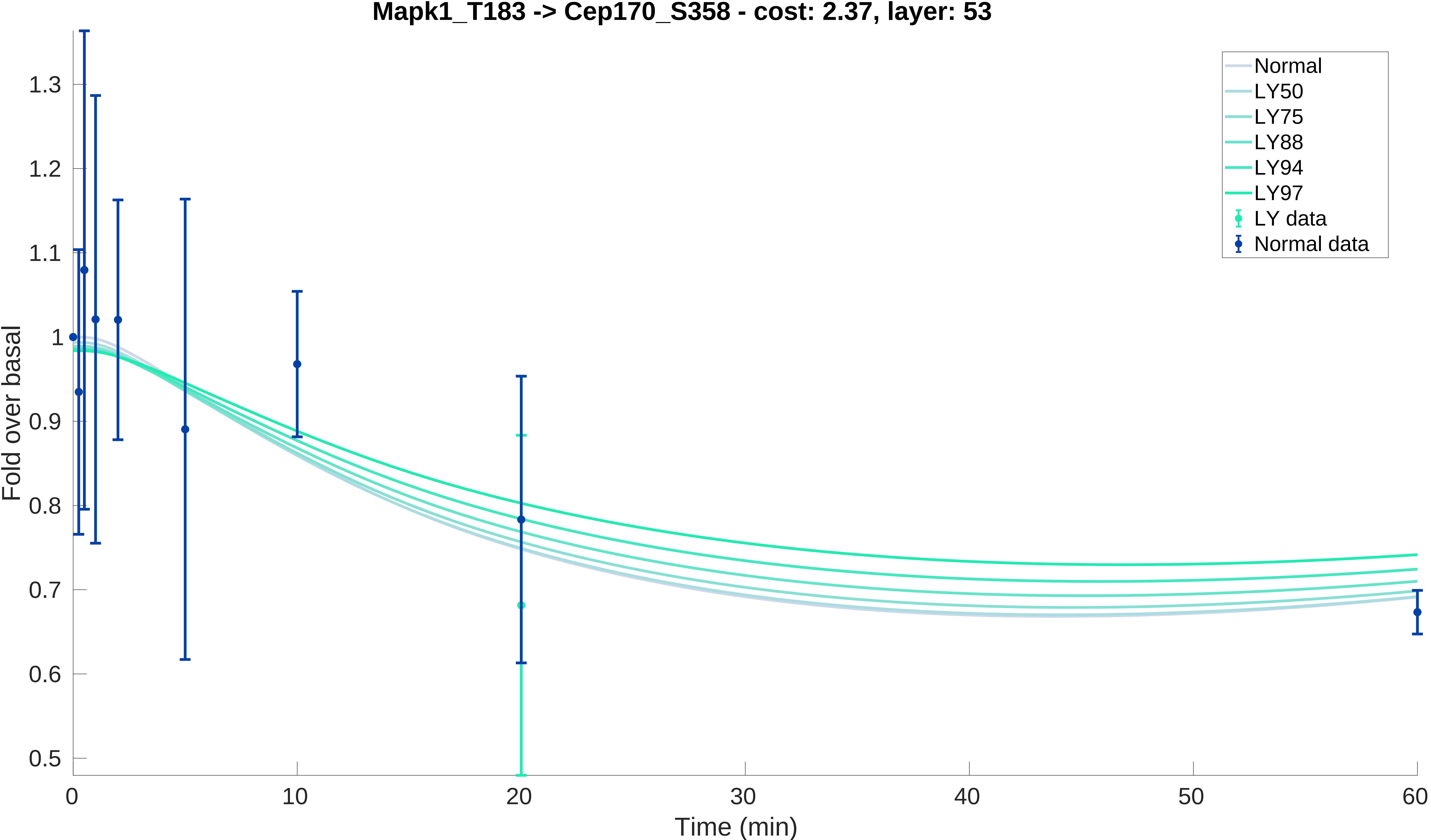
Mapk1_T183 -> Cep170_S358 - cost: 2.37, layer: 53
Fold over basal
Time (min)
0.5
0.6
0.7
0.8
0.9
1
1.1
1.2
1.3
0
10
20
30
40
50
60
Normal
LY50
LY75
LY88
LY94
LY97
LY data
Normal data

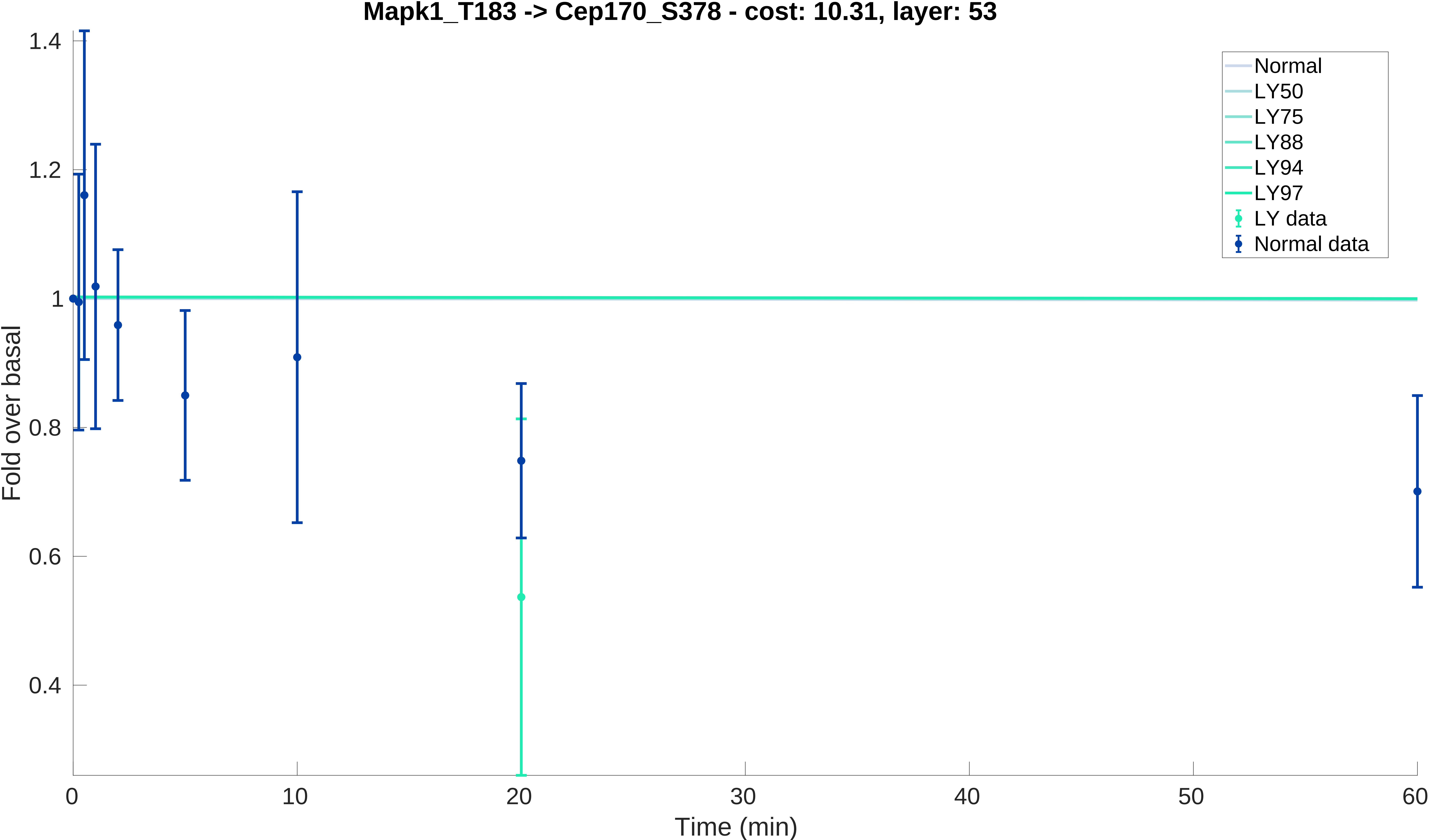

Mapk1_T183 -> Cep170_S378 - cost: 10.31, layer: 53
1.4
1.2
1
0.8
0.6
0.4
Fold over basal
0
10
20
30
40
50
60
Time (min)
Normal
LY50
LY75
LY88
LY94
LY97
LY data
Normal data

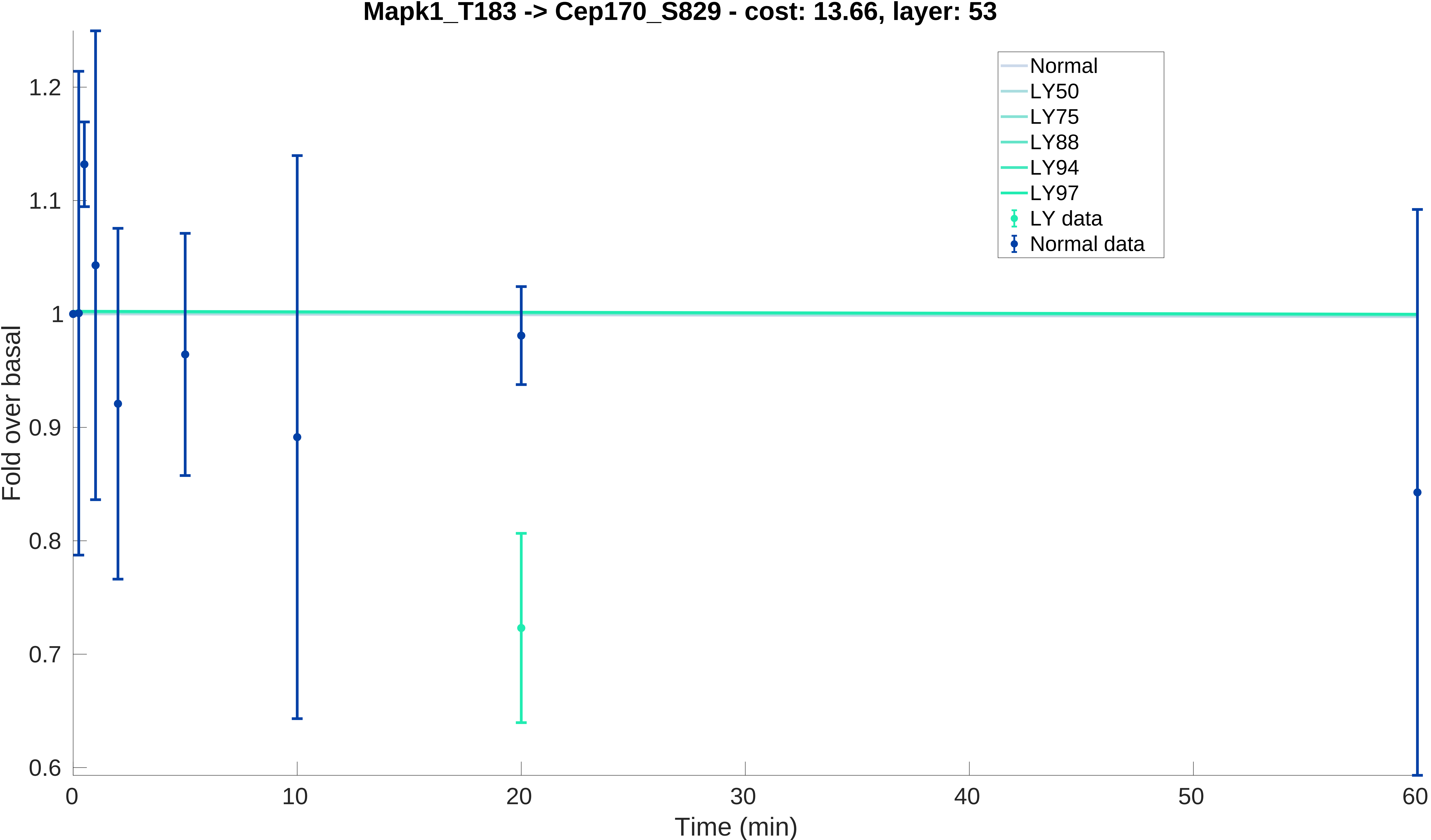

Mapk1_T183 -> Cep170_S829 - cost: 13.66, layer: 53
Normal
LY50
LY75
LY88
LY94
LY97
LY data
Normal data
Fold over basal
1.2
1.1
1
0.9
0.8
0.7
0.6
0
10
20
30
40
50
60
Time (min)

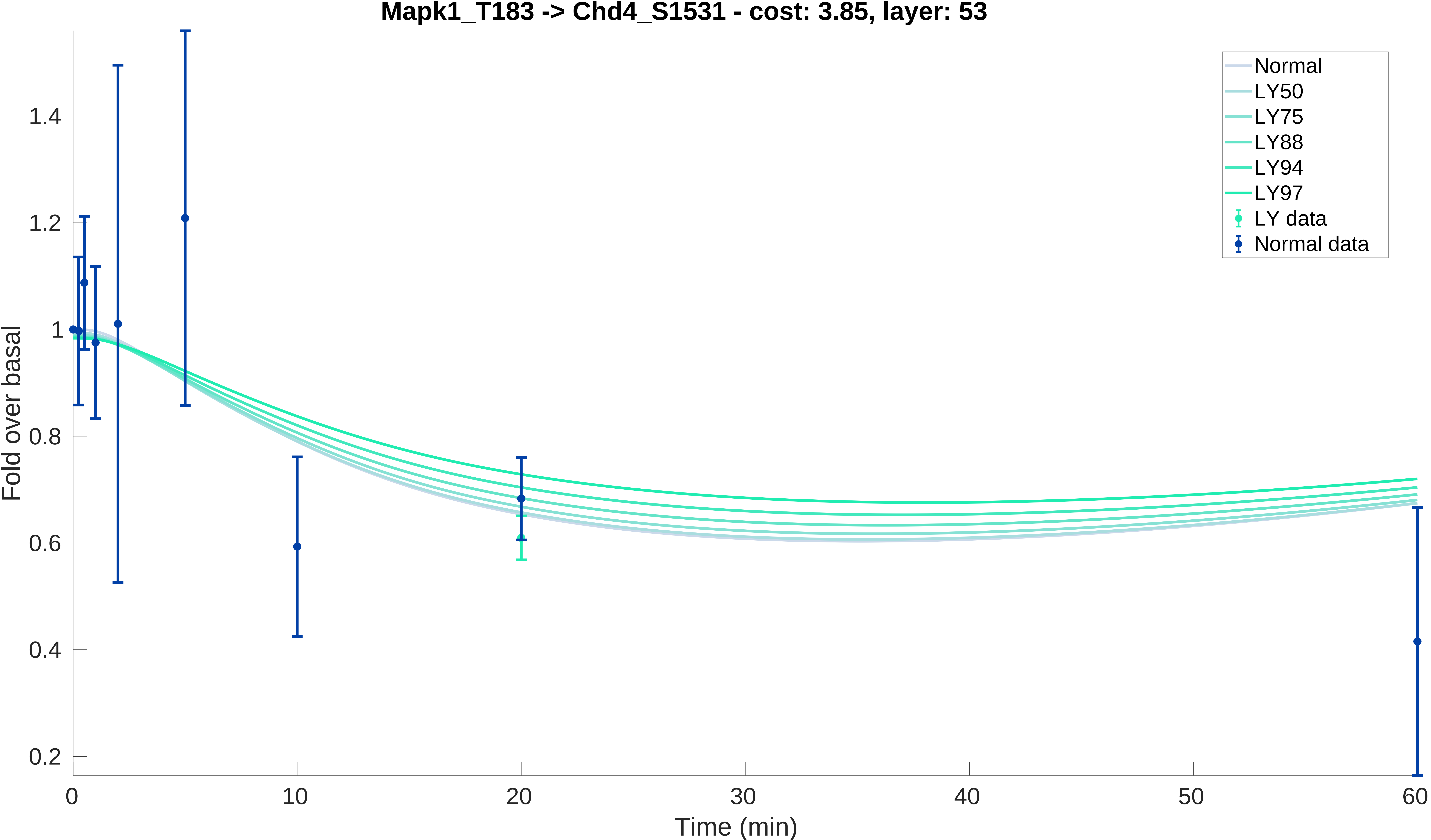

Mapk1_T183 -> Chd4_S1531 - cost: 3.85, layer: 53
Fold over basal
Time (min)
0.2
0.4
0.6
0.8
1
1.2
1.4
0
10
20
30
40
50
60
Normal
LY50
LY75
LY88
LY94
LY97
LY data
Normal data

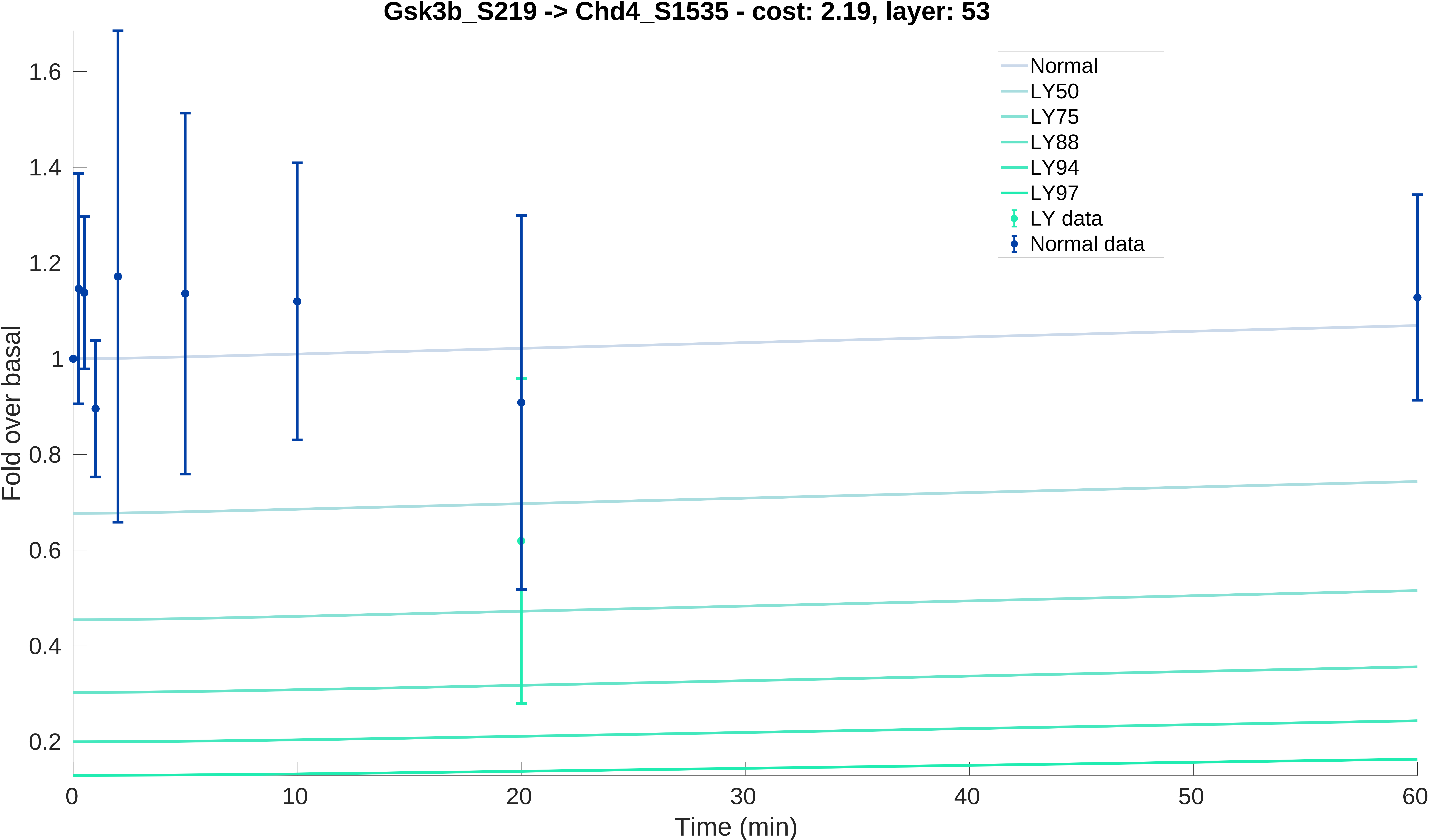

Gsk3b_S219 -> Chd4_S1535 - cost: 2.19, layer: 53
Fold over basal
Time (min)
1.6
1.4
1.2
1
0.8
0.6
0.4
0.2
0
10
20
30
40
50
60
Normal
LY50
LY75
LY88
LY94
LY97
LY data
Normal data

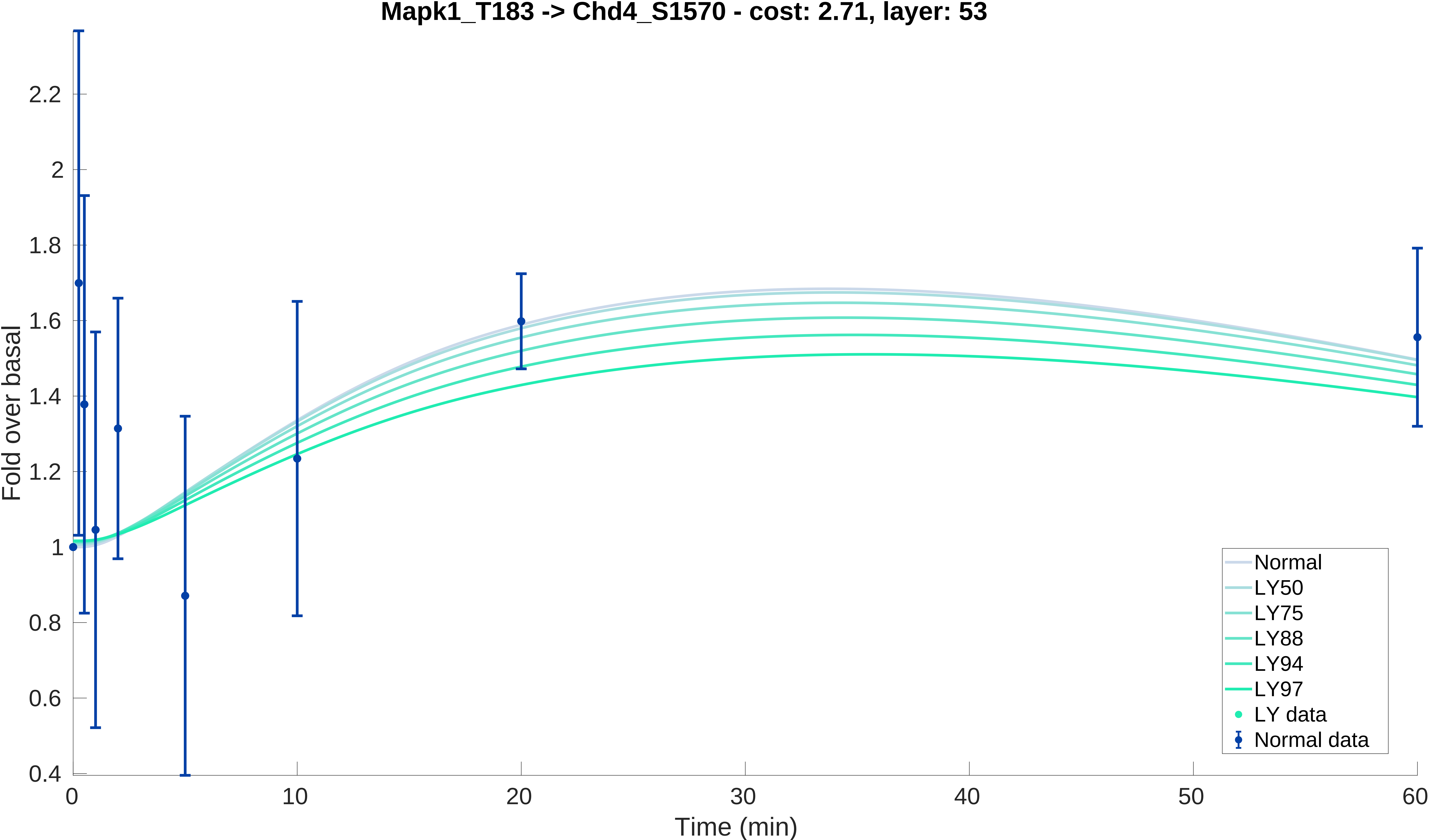

Mapk1_T183 -> Chd4_S1570 - cost: 2.71, layer: 53
Fold over basal
Time (min)
0.4
0.6
0.8
1
1.2
1.4
1.6
1.8
2
2.2
0
10
20
30
40
50
60
Normal
LY50
LY75
LY88
LY94
LY97
LY data
Normal data

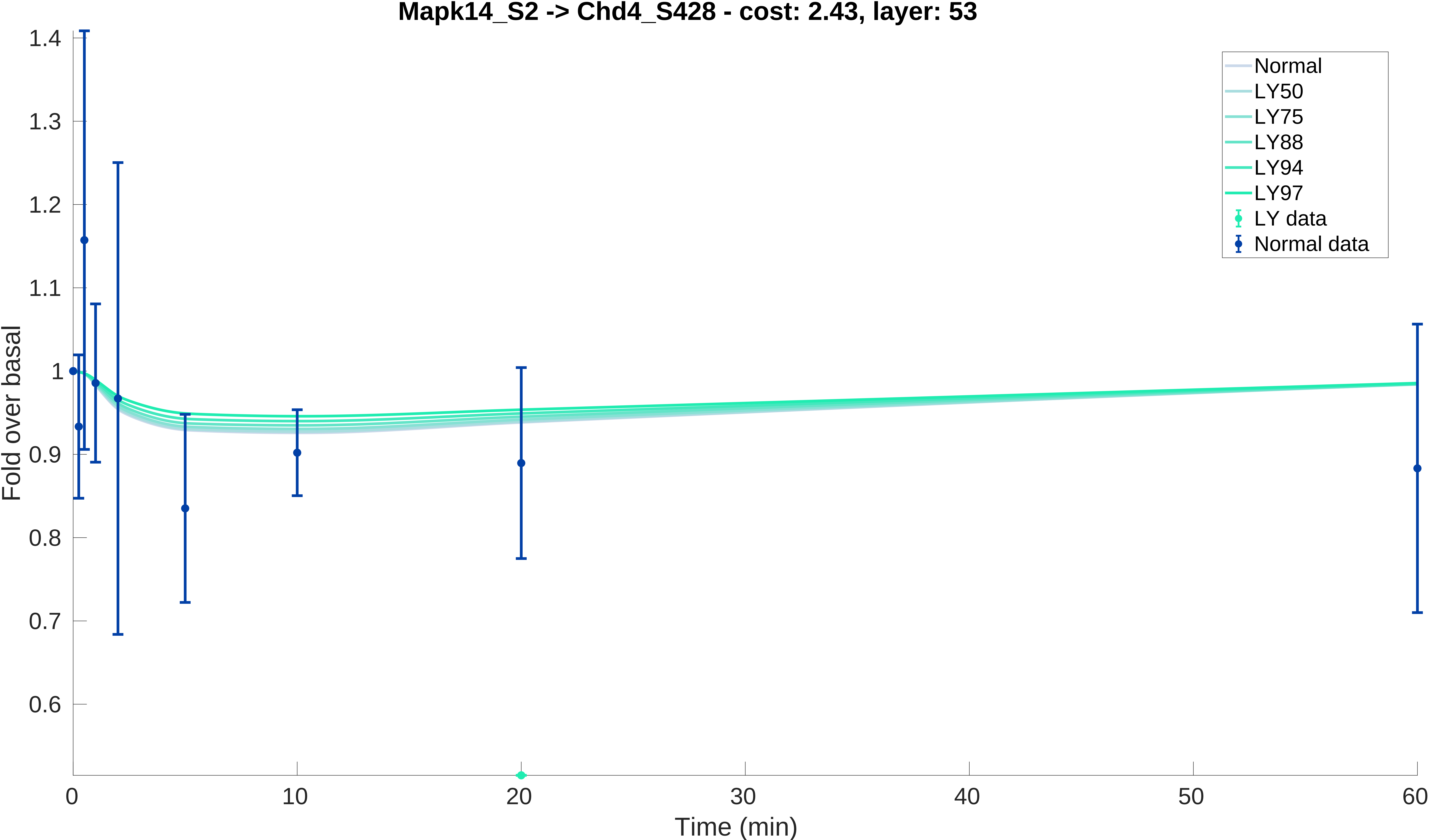

Mapk14_S2 -> Chd4_S428 - cost: 2.43, layer: 53
Fold over basal
Time (min)
0.6
0.7
0.8
0.9
1
1.1
1.2
1.3
1.4
0
10
20
30
40
50
60
Normal
LY50
LY75
LY88
LY94
LY97
LY data
Normal data

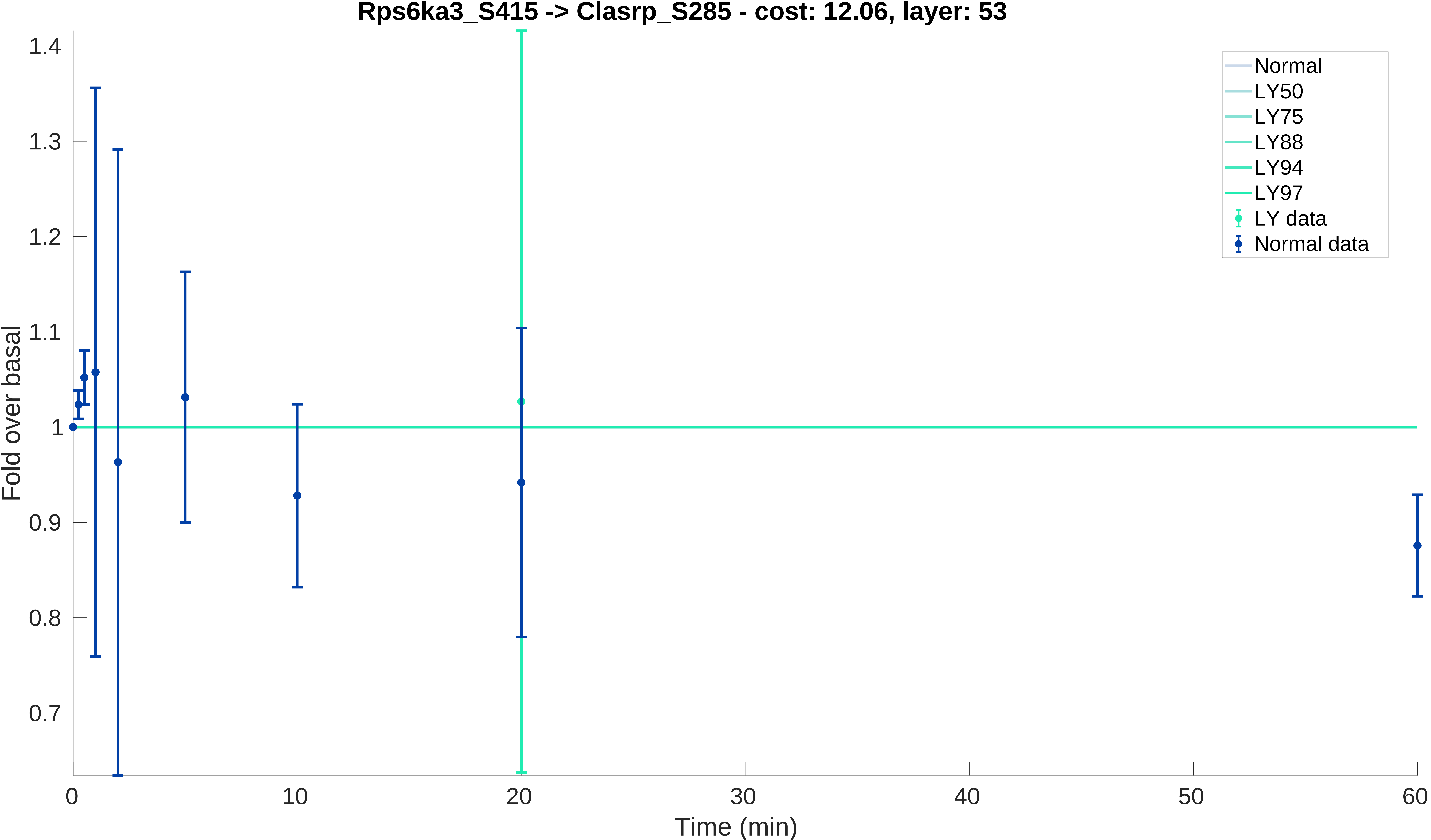

Rps6ka3_S415 -> Clasrp_S285 - cost: 12.06, layer: 53
Fold over basal
Time (min)
1.4
1.3
1.2
1.1
1
0.9
0.8
0.7
0
10
20
30
40
50
60
Normal
LY50
LY75
LY88
LY94
LY97
LY data
Normal data

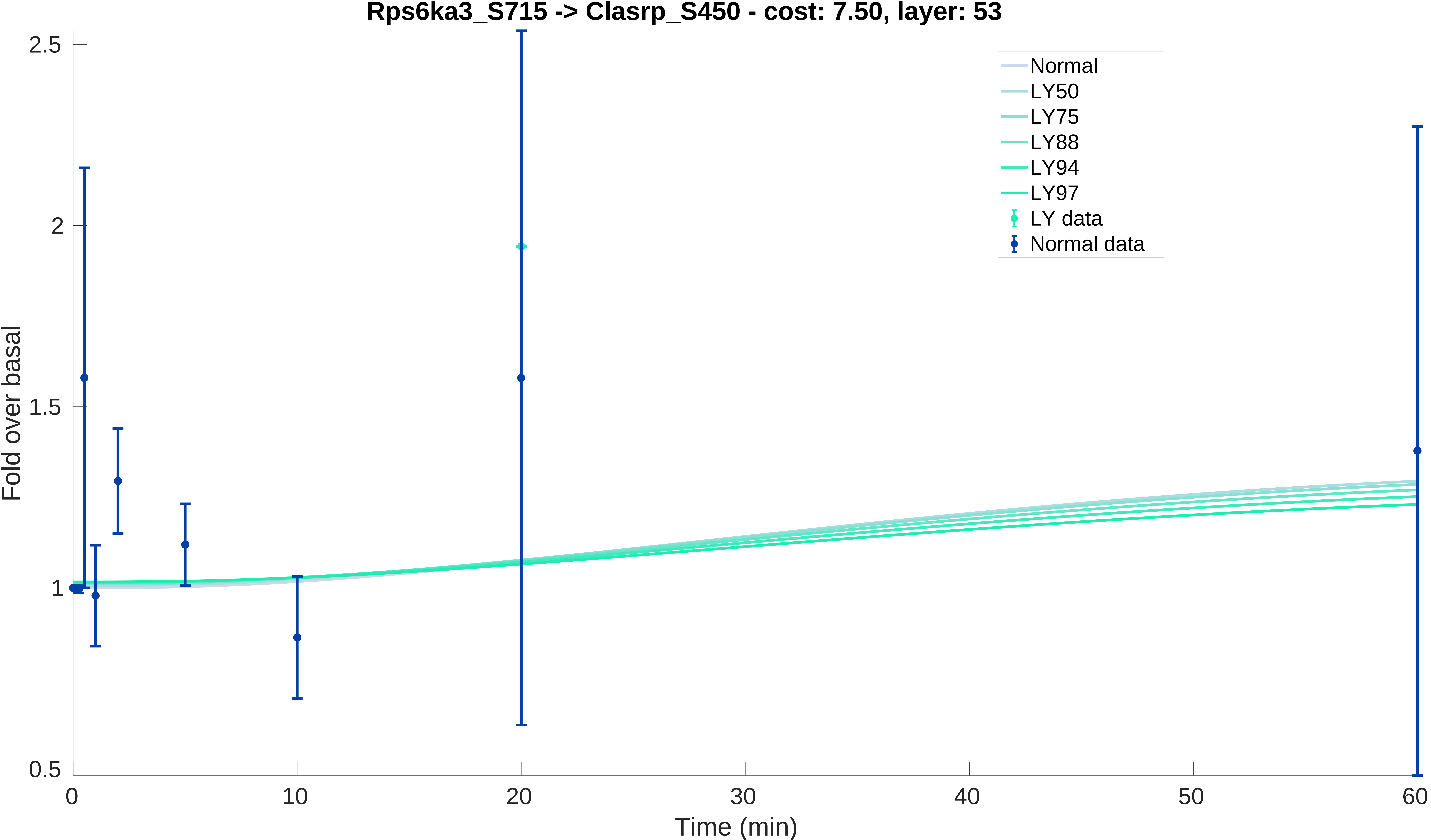

Rps6ka3_S715 -> Clasrp_S450 - cost: 7.50, layer: 53
Fold over basal
Time (min)
0.5
1
1.5
2
2.5
0
10
20
30
40
50
60
Normal
LY50
LY75
LY88
LY94
LY97
LY data
Normal data

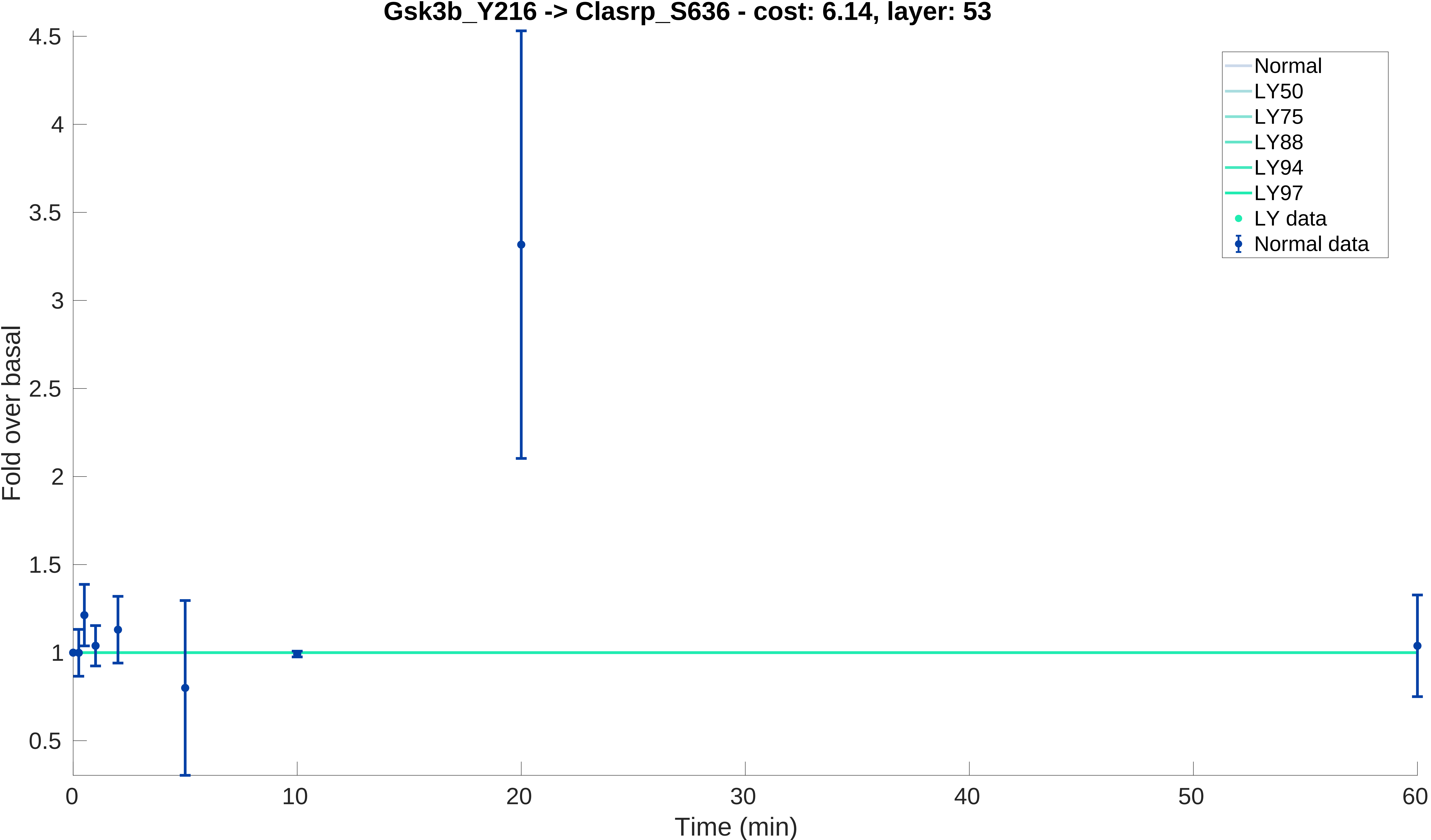

Gsk3b_Y216 -> Clasrp_S636 - cost: 6.14, layer: 53
Fold over basal
Time (min)
4.5
4
3.5
3
2.5
2
1.5
1
0.5
0
10
20
30
40
50
60
Normal
LY50
LY75
LY88
LY94
LY97
LY data
Normal data

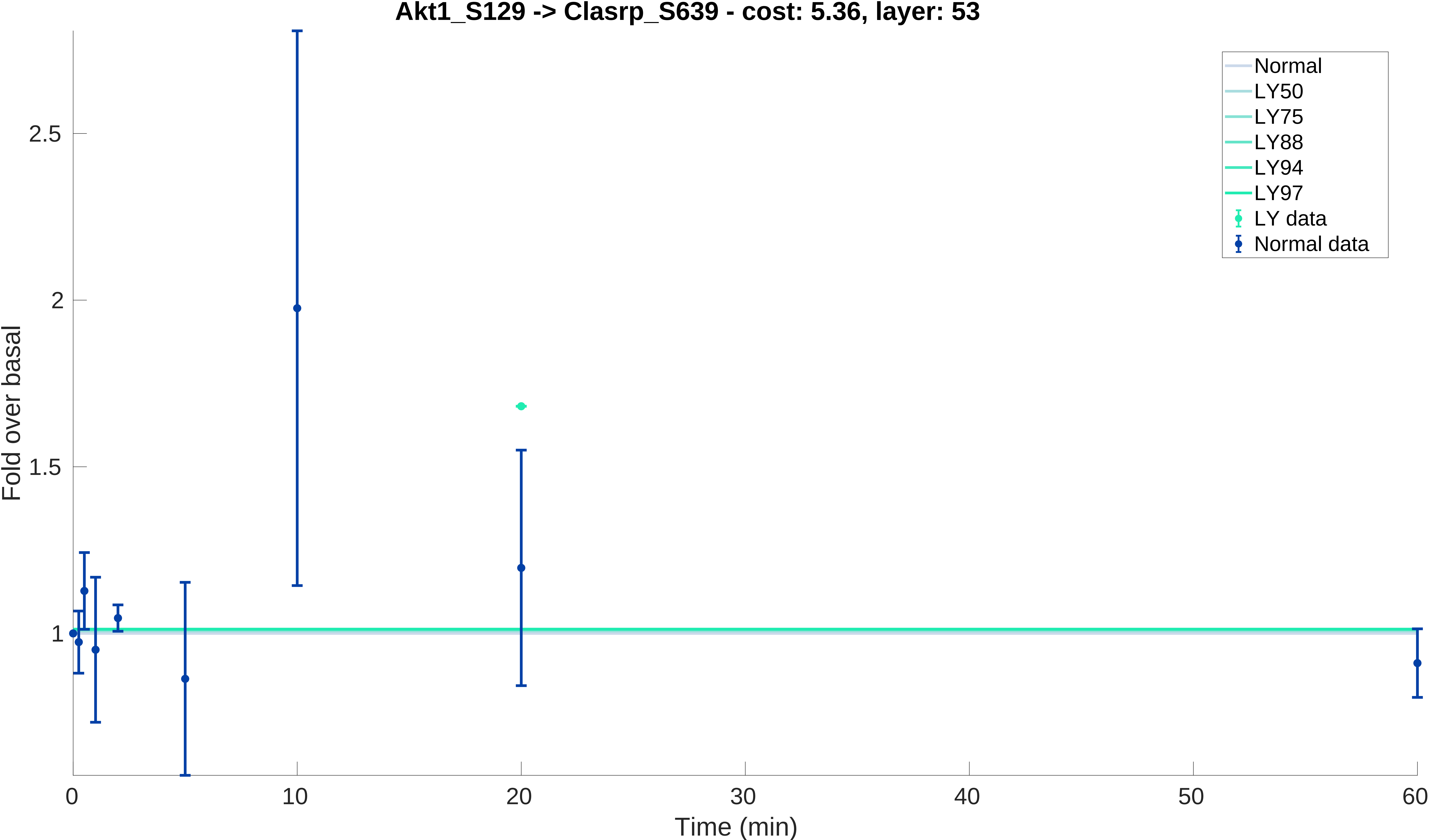

Akt1_S129 -> Clasrp_S639 - cost: 5.36, layer: 53
Normal
LY50
LY75
LY88
LY94
LY97
LY data
Normal data
Fold over basal
2.5
2
1.5
1
0
10
20
30
40
50
60
Time (min)

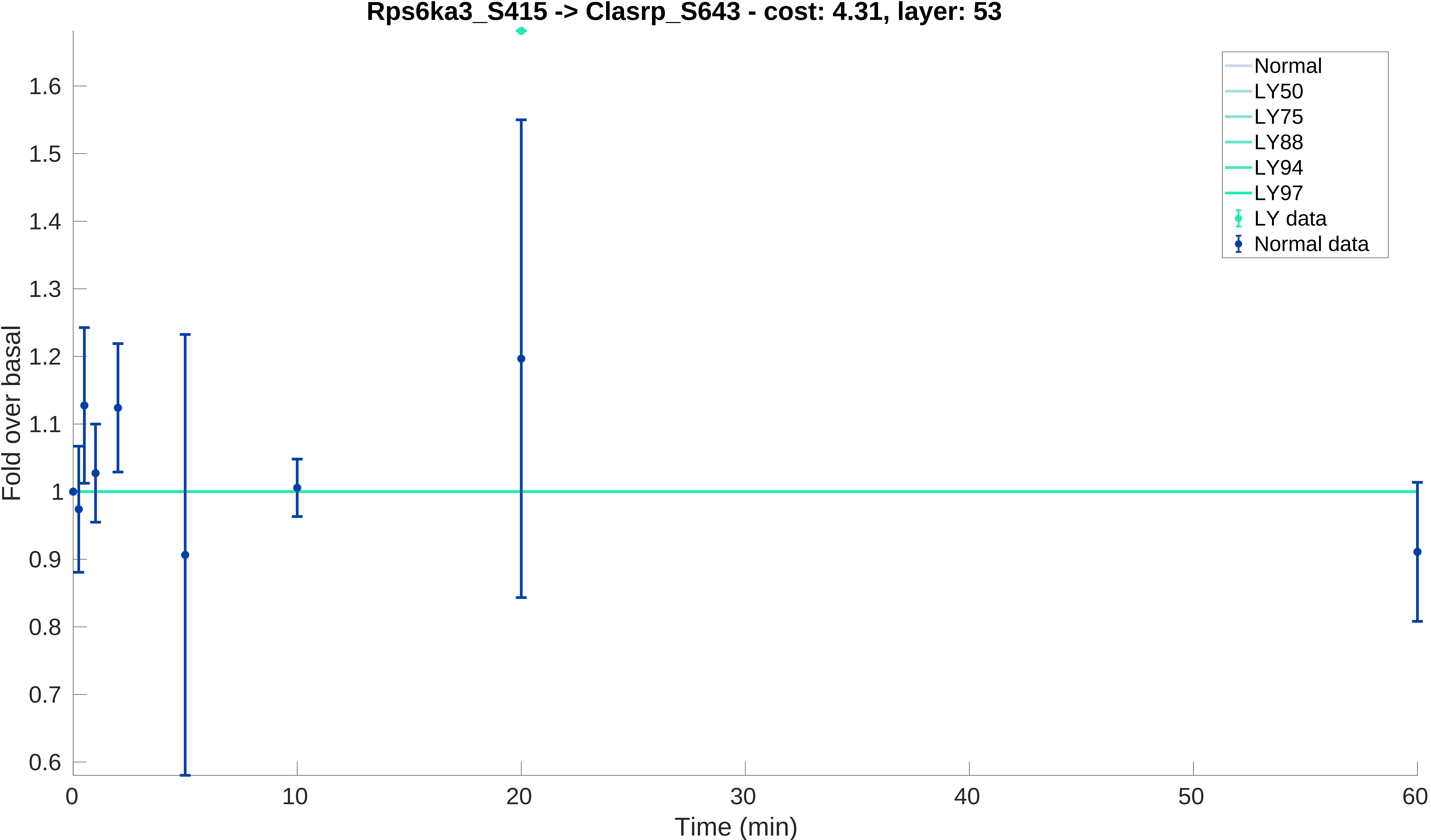

Rps6ka3_S415 -> Clasrp_S643 - cost: 4.31, layer: 53
Fold over basal
Time (min)
0.6
0.7
0.8
0.9
1
1.1
1.2
1.3
1.4
1.5
1.6
0
10
20
30
40
50
60
Normal
LY50
LY75
LY88
LY94
LY97
LY data
Normal data

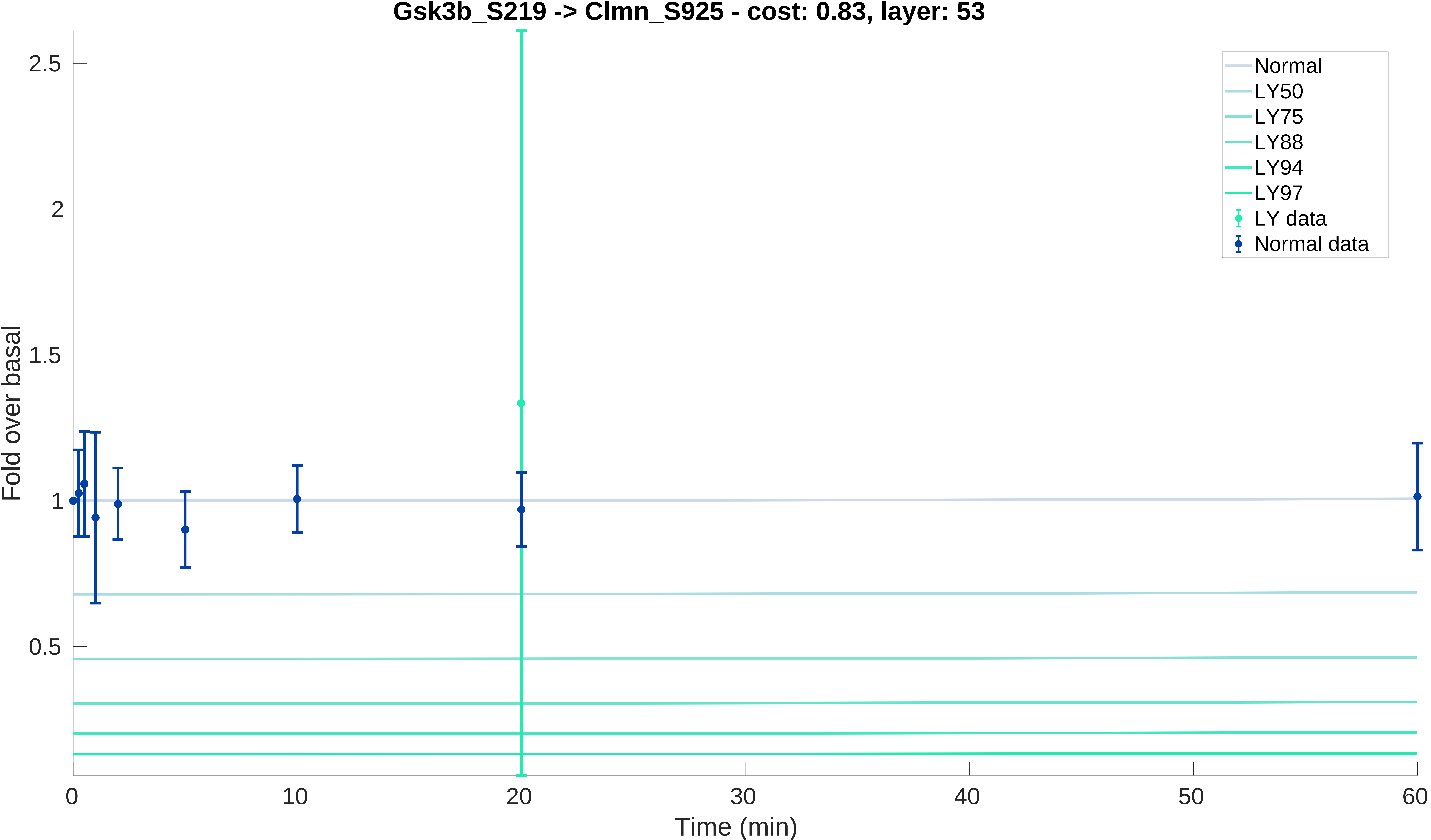

Gsk3b_S219 -> Clmn_S925 - cost: 0.83, layer: 53
Fold over basal
Time (min)
0
10
20
30
40
50
60
0.5
1
1.5
2
2.5
Normal
LY50
LY75
LY88
LY94
LY97
LY data
Normal data

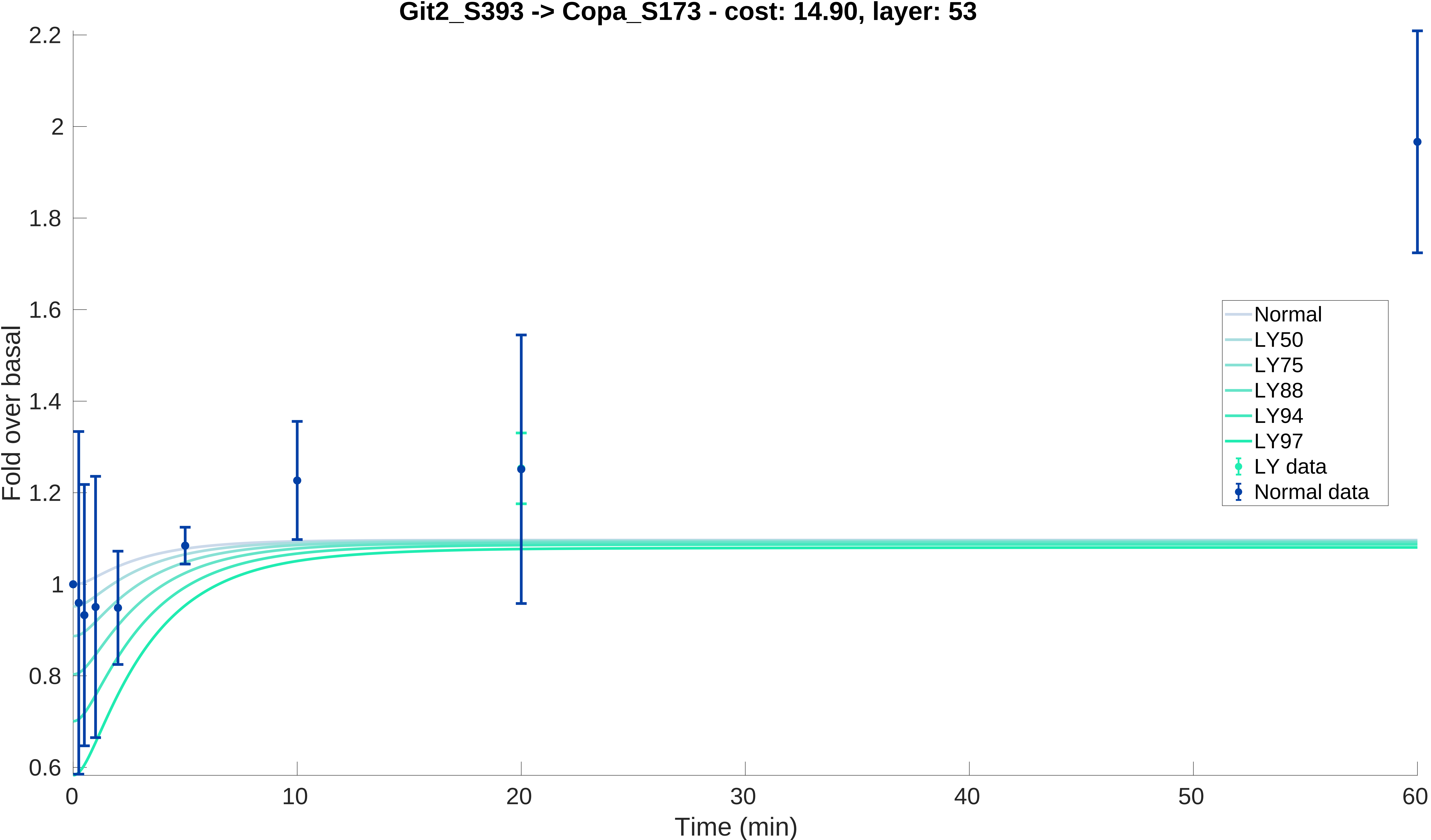

Git2_S393 -> Copa_S173 - cost: 14.90, layer: 53
2.2
2
1.8
1.6
1.4
1.2
1
0.8
0.6
Fold over basal
0
10
20
30
40
50
60
Time (min)
Normal
LY50
LY75
LY88
LY94
LY97
LY data
Normal data

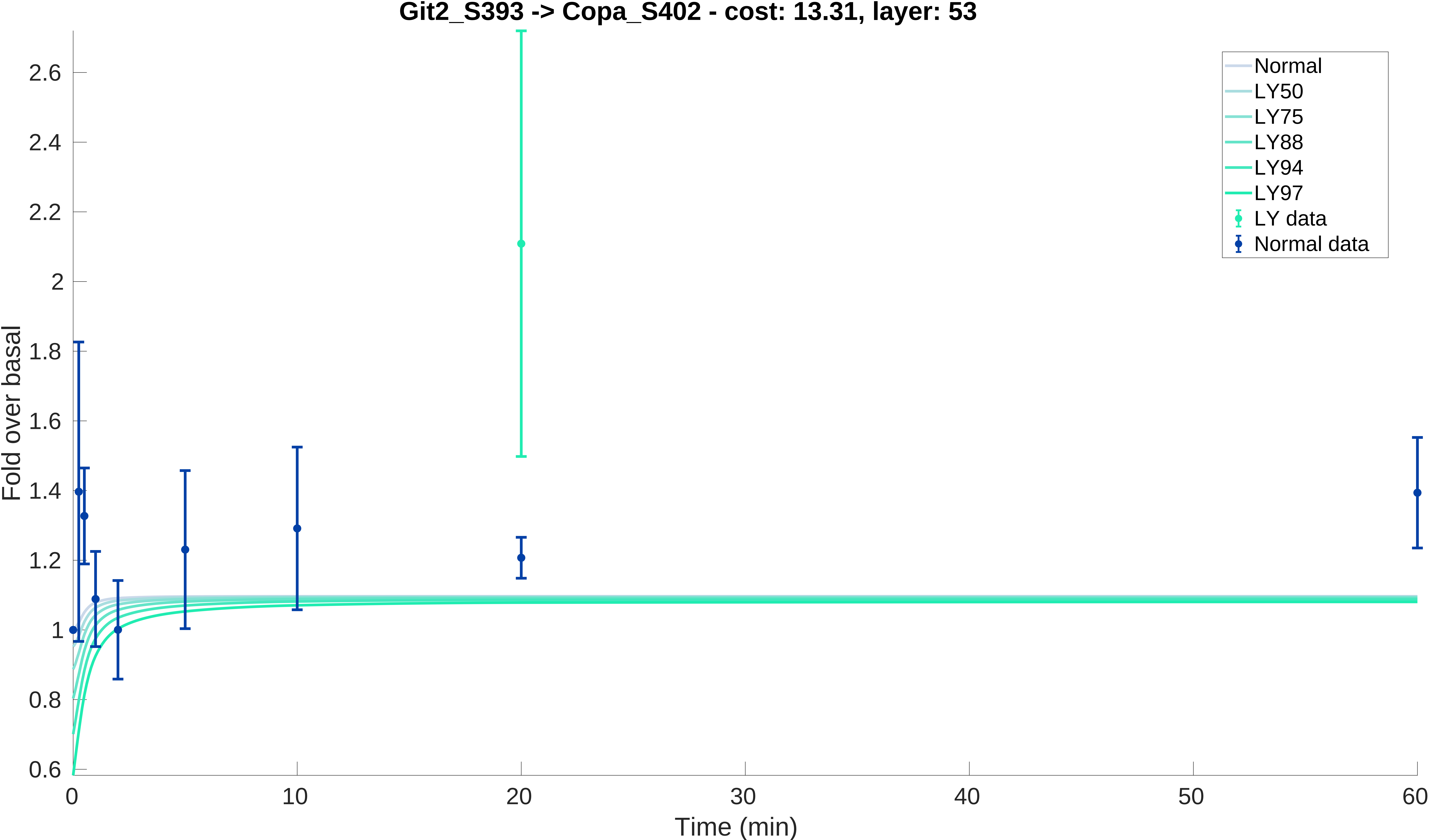
Git2_S393 -> Copa_S402 - cost: 13.31, layer: 53
Fold over basal
Time (min)
0.6
0.8
1
1.2
1.4
1.6
1.8
2
2.2
2.4
2.6
0
10
20
30
40
50
60
Normal
LY50
LY75
LY88
LY94
LY97
LY data
Normal data

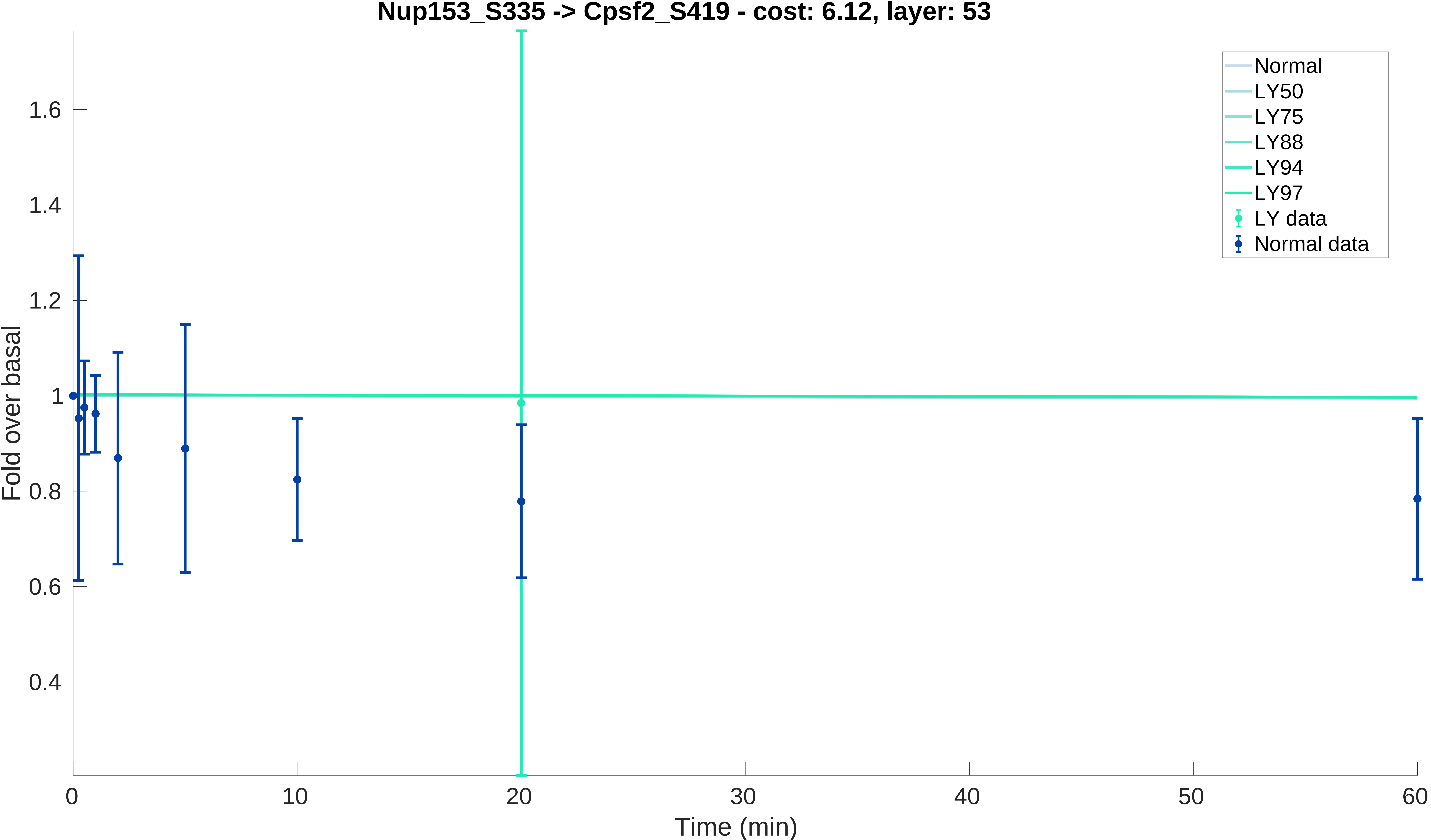

Nup153_S335 -> Cpsf2_S419 - cost: 6.12, layer: 53
Normal
LY50
LY75
LY88
LY94
LY97
LY data
Normal data
Fold over basal
1.6
1.4
1.2
1
0.8
0.6
0.4
0
10
20
30
40
50
60
Time (min)

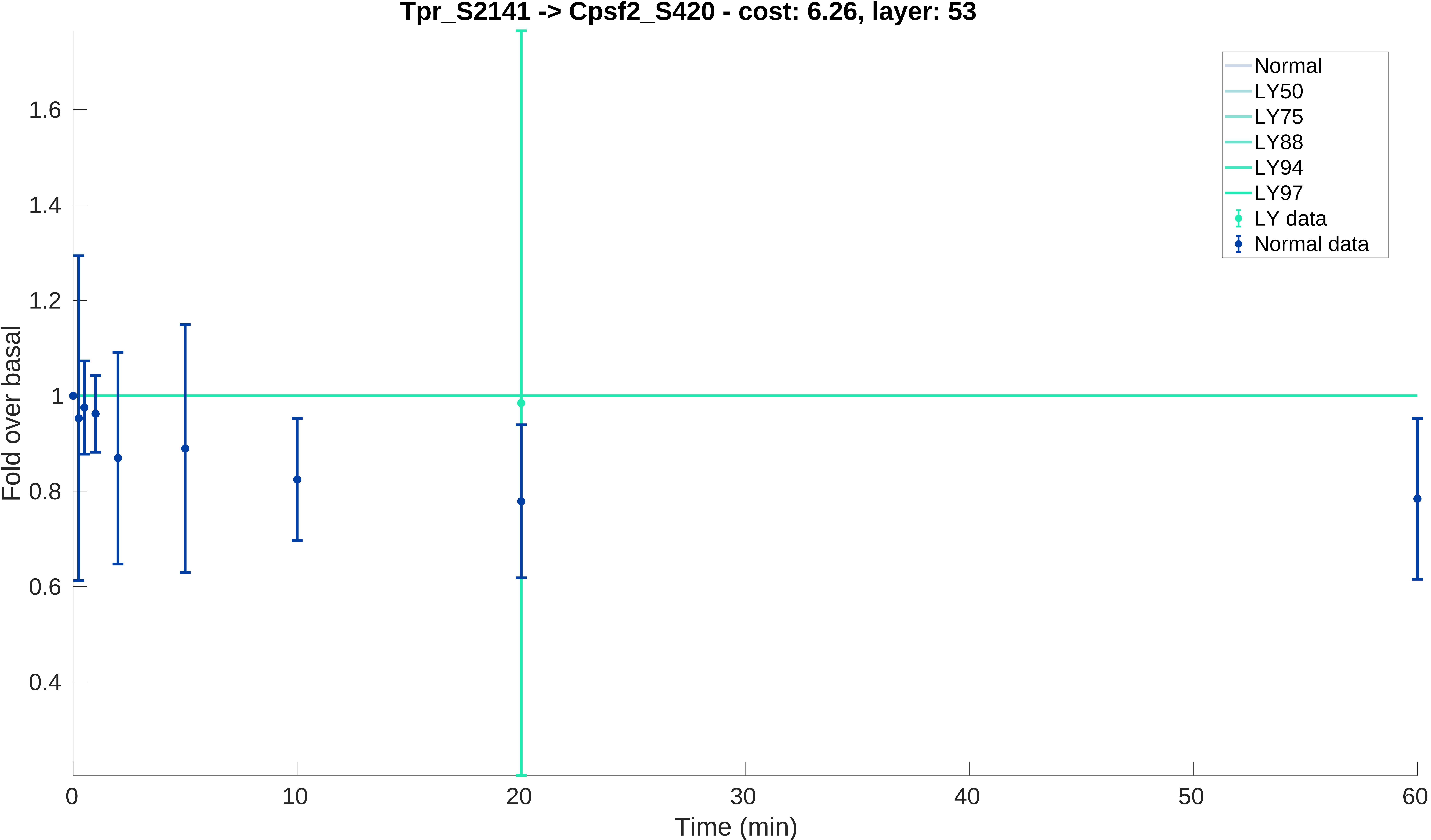

Tpr_S2141 -> Cpsf2_S420 - cost: 6.26, layer: 53
Fold over basal
Time (min)
1.6
1.4
1.2
1
0.8
0.6
0.4
0
10
20
30
40
50
60
Normal
LY50
LY75
LY88
LY94
LY97
LY data
Normal data

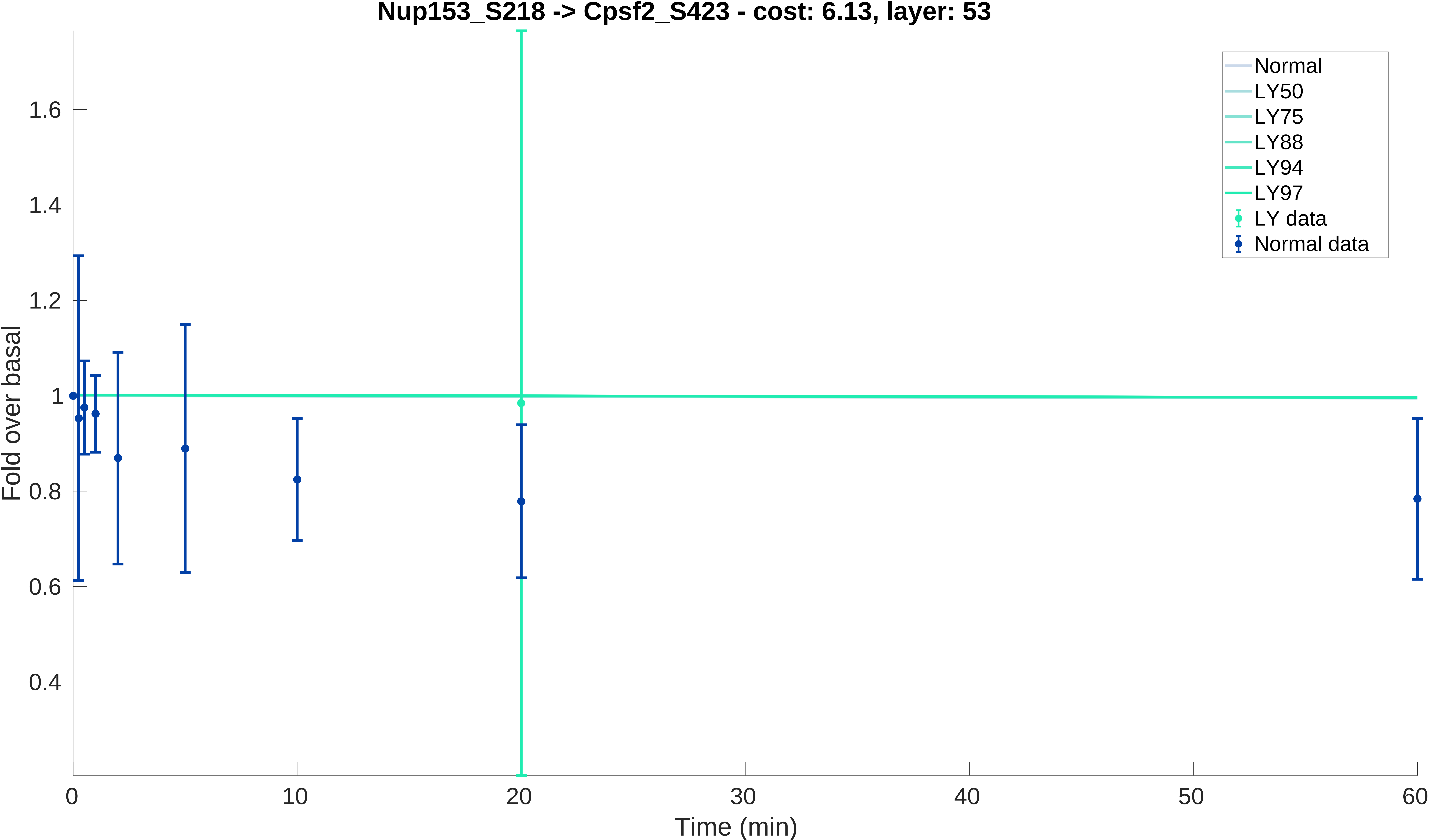
Nup153_S218 -> Cpsf2_S423 - cost: 6.13, layer: 53
Normal
LY50
LY75
LY88
LY94
LY97
LY data
Normal data
Fold over basal
1.6
1.4
1.2
1
0.8
0.6
0.4
0
10
20
30
40
50
60
Time (min)

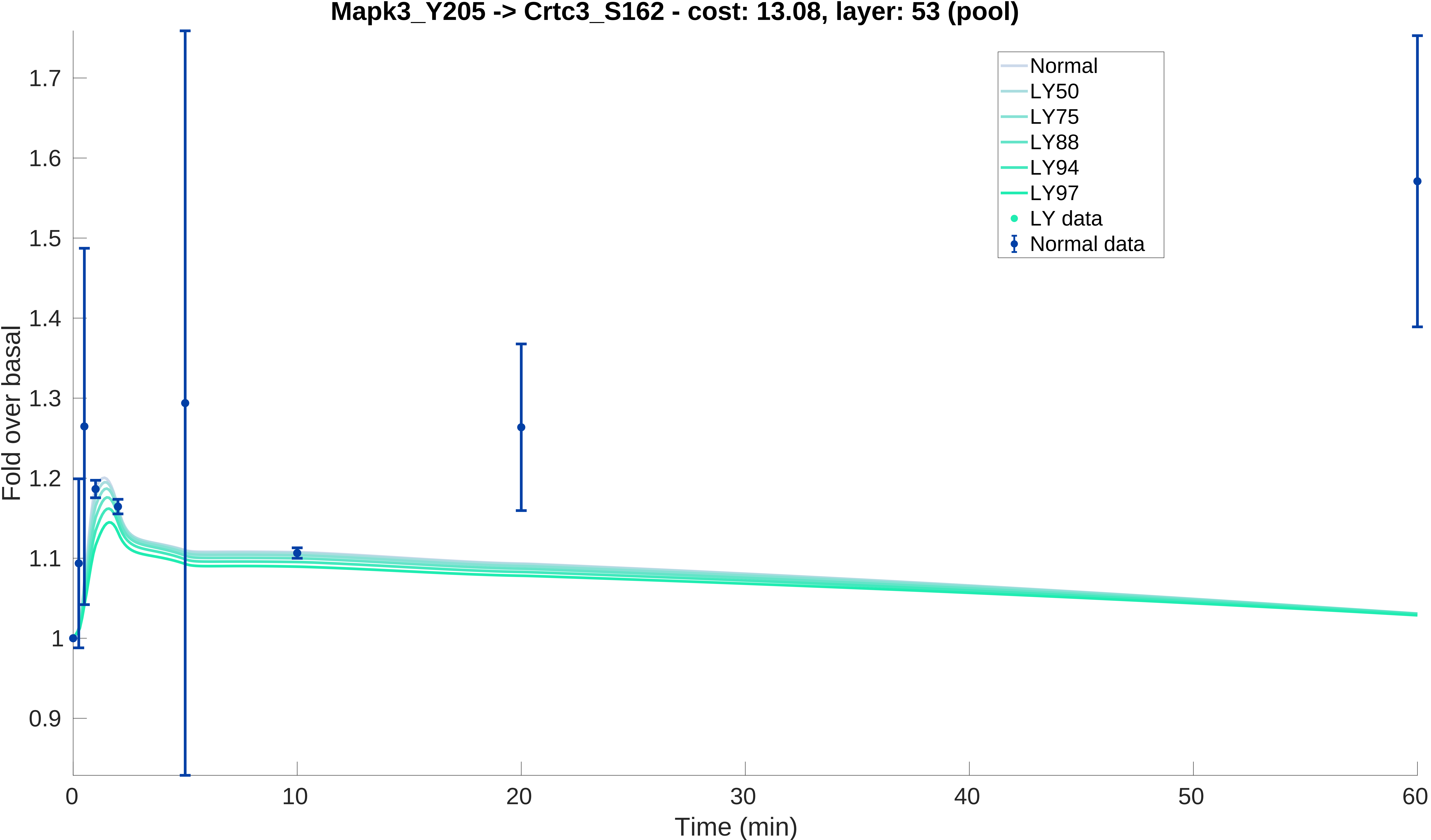
Mapk3_Y205 -> Crtc3_S162 - cost: 13.08, layer: 53 (pool)
Fold over basal
Time (min)
Normal
LY50
LY75
LY88
LY94
LY97
LY data
Normal data
0.9
1
1.1
1.2
1.3
1.4
1.5
1.6
1.7
0
10
20
30
40
50
60

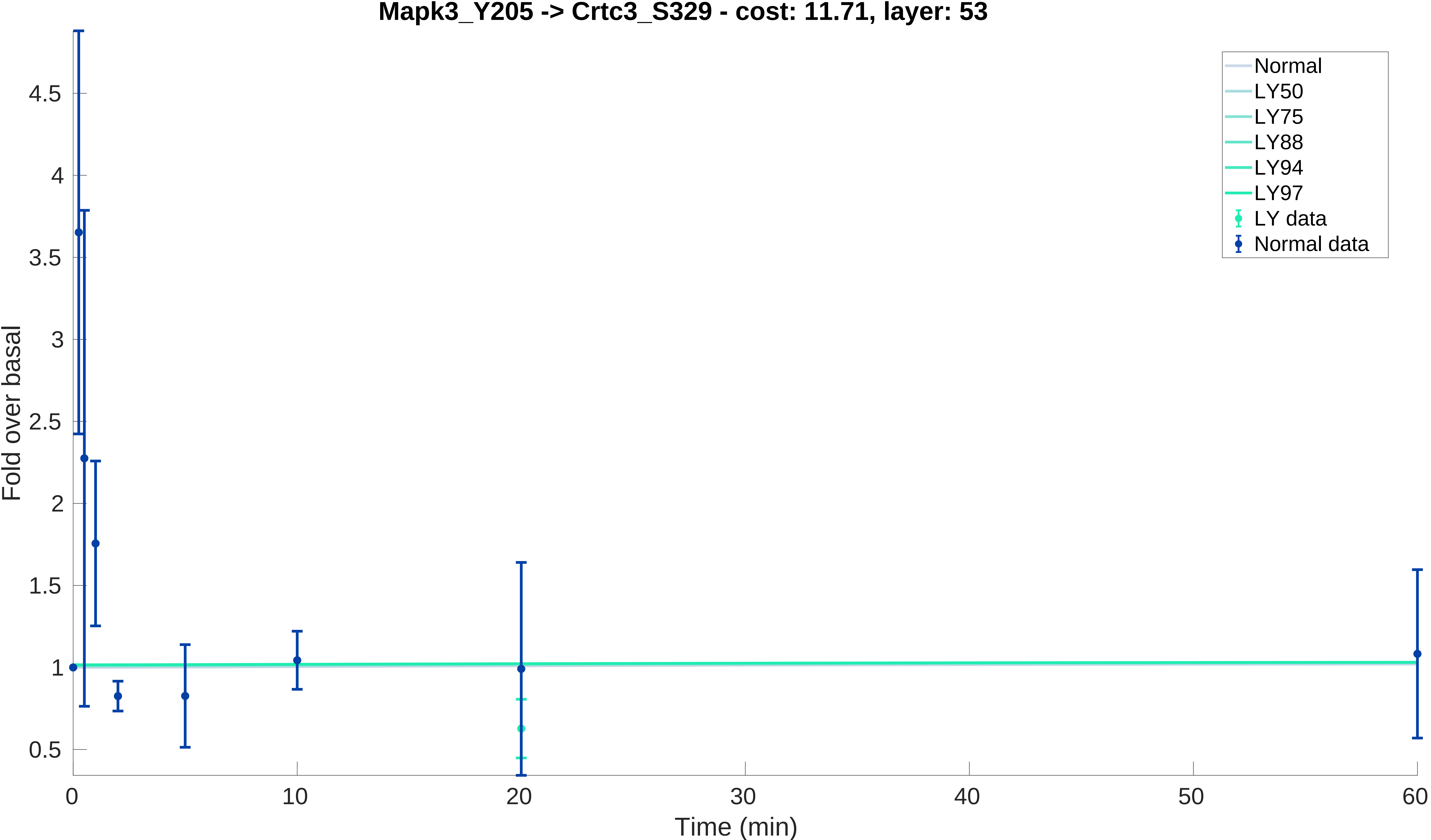
Mapk3_Y205 -> Crtc3_S329 - cost: 11.71, layer: 53
Normal
LY50
LY75
LY88
LY94
LY97
LY data
Normal data
Fold over basal
4.5
4
3.5
3
2.5
2
1.5
1
0.5
0
10
20
30
40
50
60
Time (min)

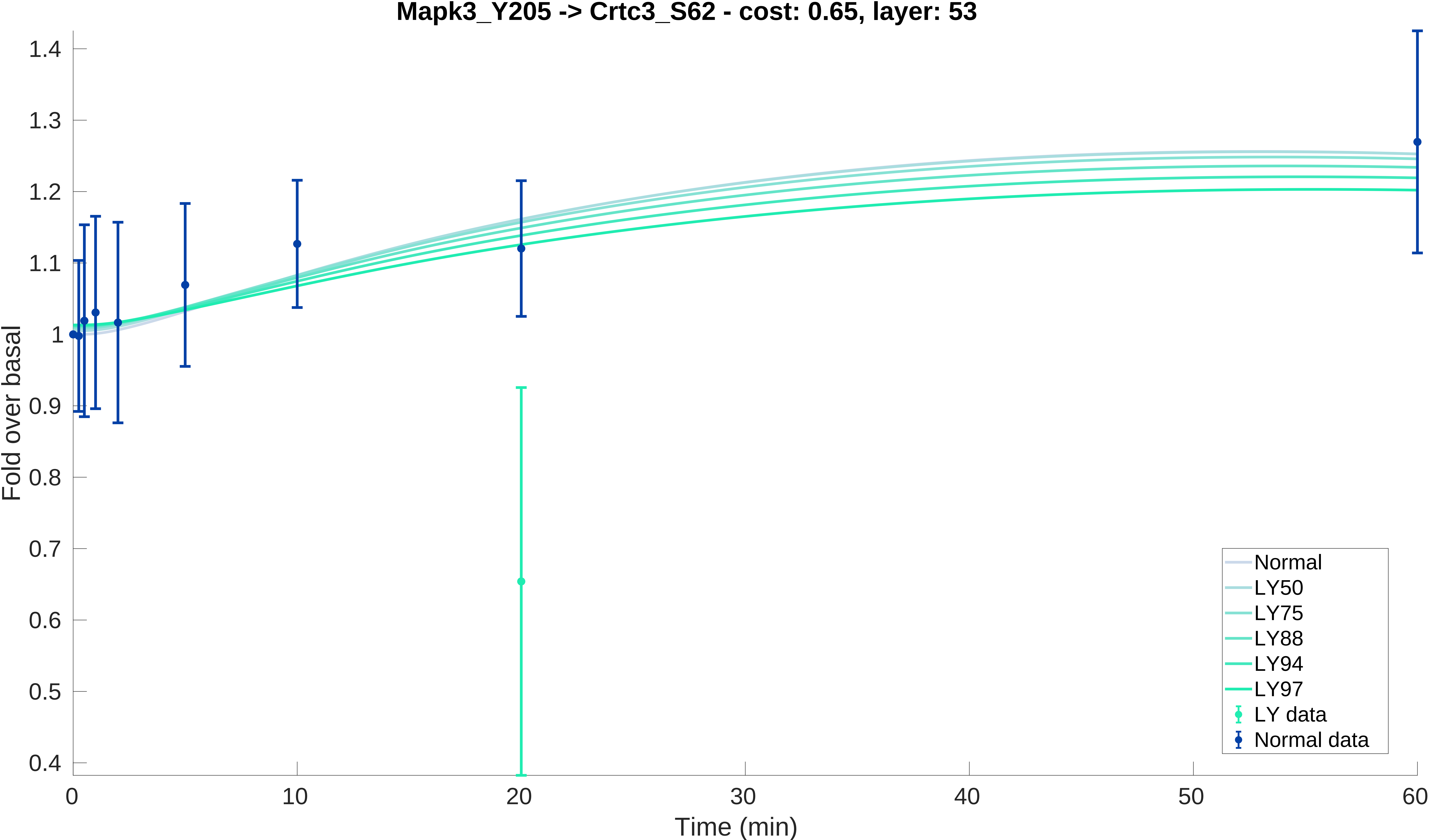

Mapk3_Y205 -> Crtc3_S62 - cost: 0.65, layer: 53
Fold over basal
Time (min)
1.4
1.3
1.2
1.1
1
0.9
0.8
0.7
0.6
0.5
0.4
0
10
20
30
40
50
60
Normal
LY50
LY75
LY88
LY94
LY97
LY data
Normal data

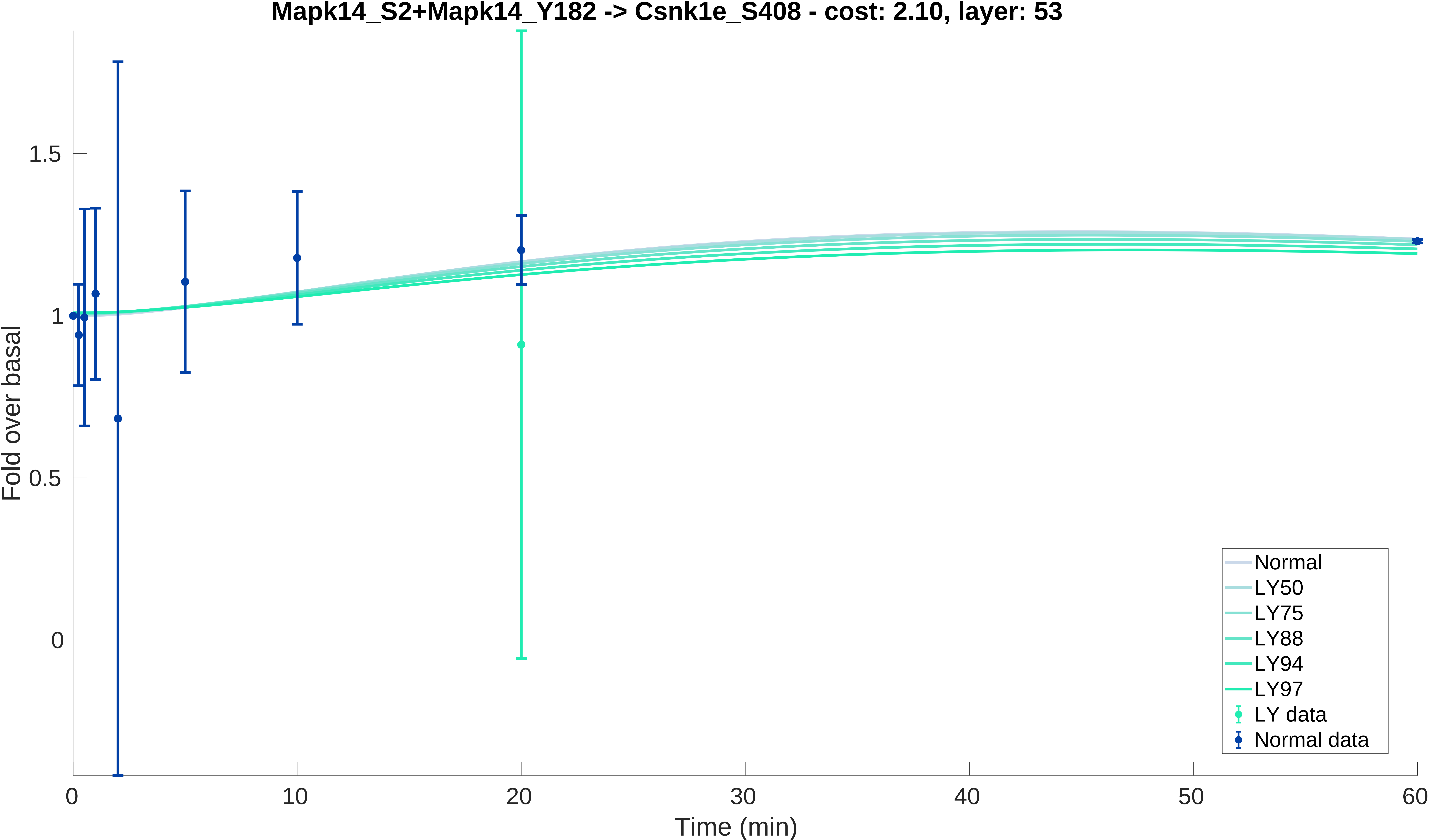
Mapk14_S2+Mapk14_Y182 -> Csnk1e_S408 - cost: 2.10, layer: 53
Fold over basal
1.5
1
0.5
0
0
10
20
30
40
50
60
Time (min)
Normal
LY50
LY75
LY88
LY94
LY97
LY data
Normal data

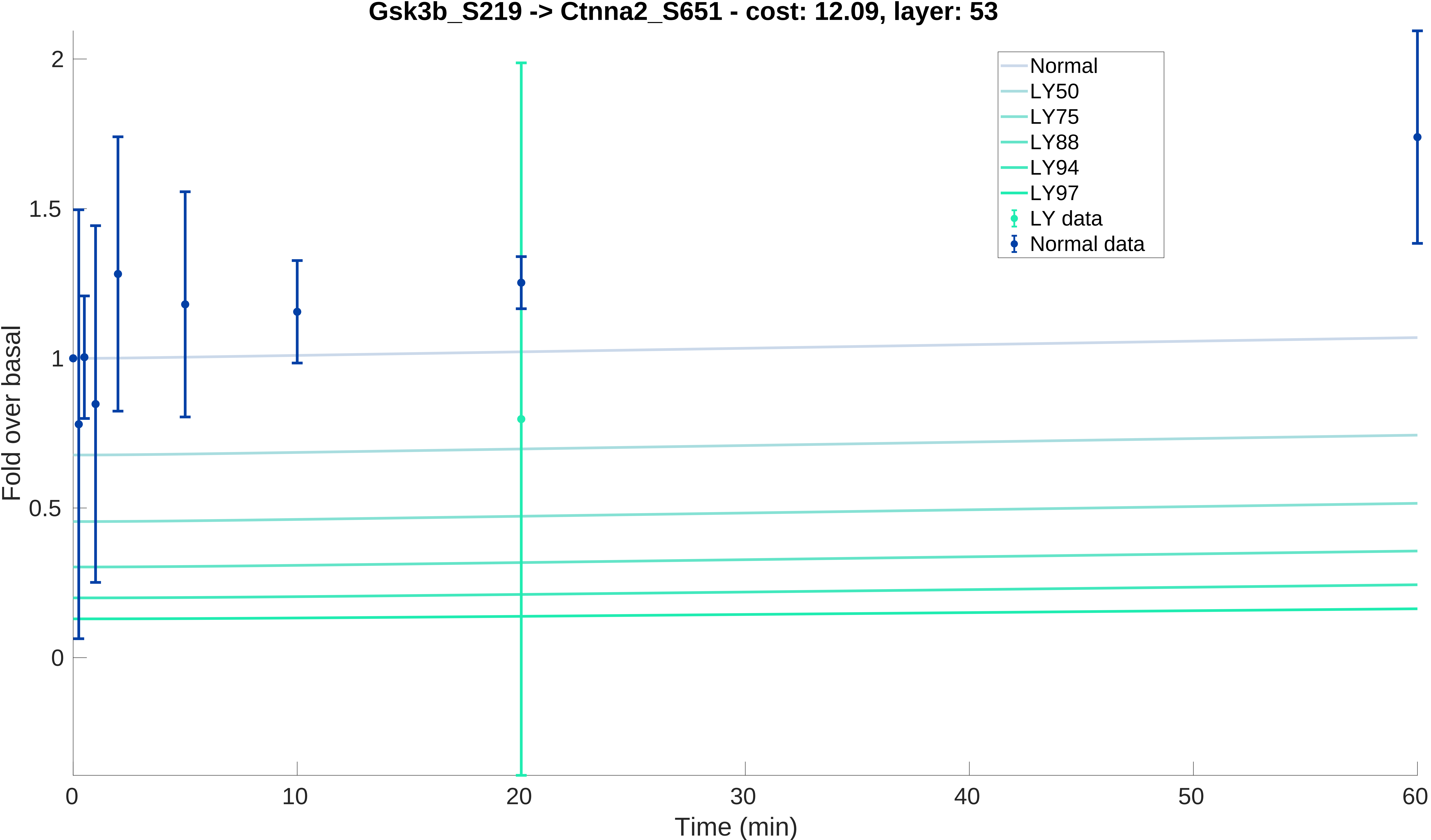

Gsk3b_S219 -> Ctnna2_S651 - cost: 12.09, layer: 53
Normal
LY50
LY75
LY88
LY94
LY97
LY data
Normal data
Fold over basal
Time (min)
0
0.5
1
1.5
2
0
10
20
30
40
50
60

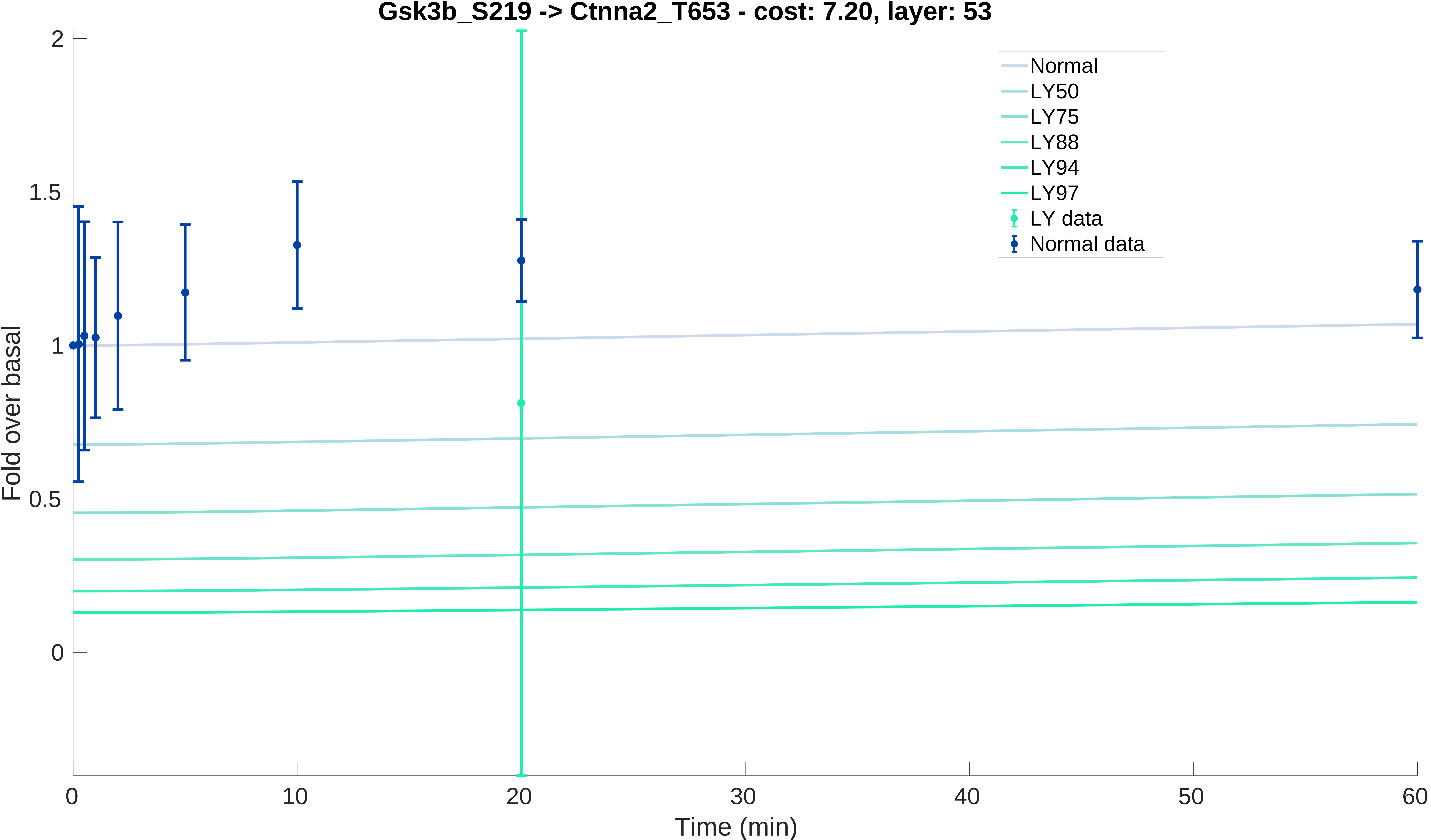
Gsk3b_S219 -> Ctnna2_T653 - cost: 7.20, layer: 53
Normal
LY50
LY75
LY88
LY94
LY97
LY data
Normal data
Fold over basal
Time (min)
0
0.5
1
1.5
2
0
10
20
30
40
50
60

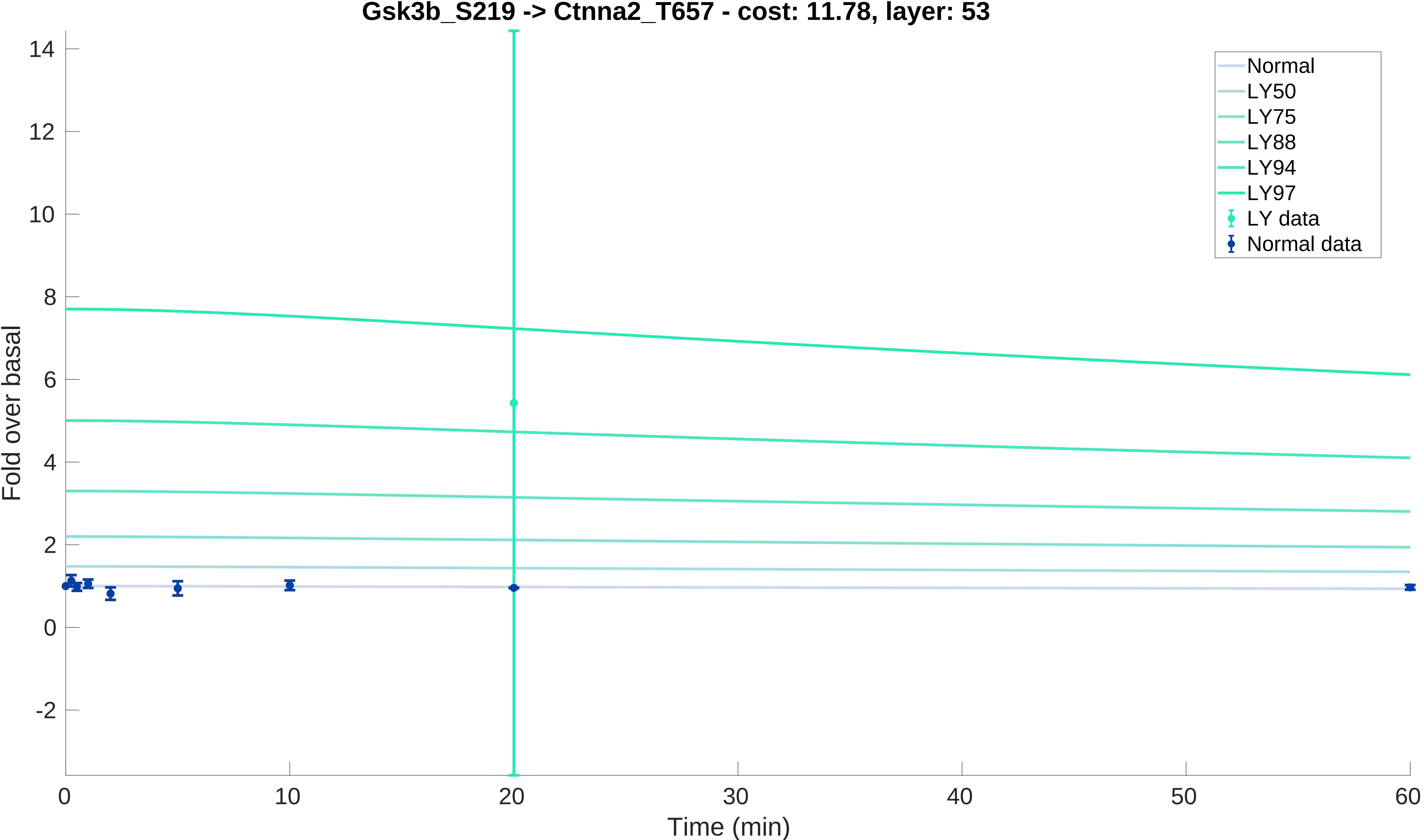

Gsk3b_S219 -> Ctnna2_T657 - cost: 11.78, layer: 53
Fold over basal
Time (min)
14
12
10
8
6
4
2
0
-2
0
10
20
30
40
50
60
Normal
LY50
LY75
LY88
LY94
LY97
LY data
Normal data

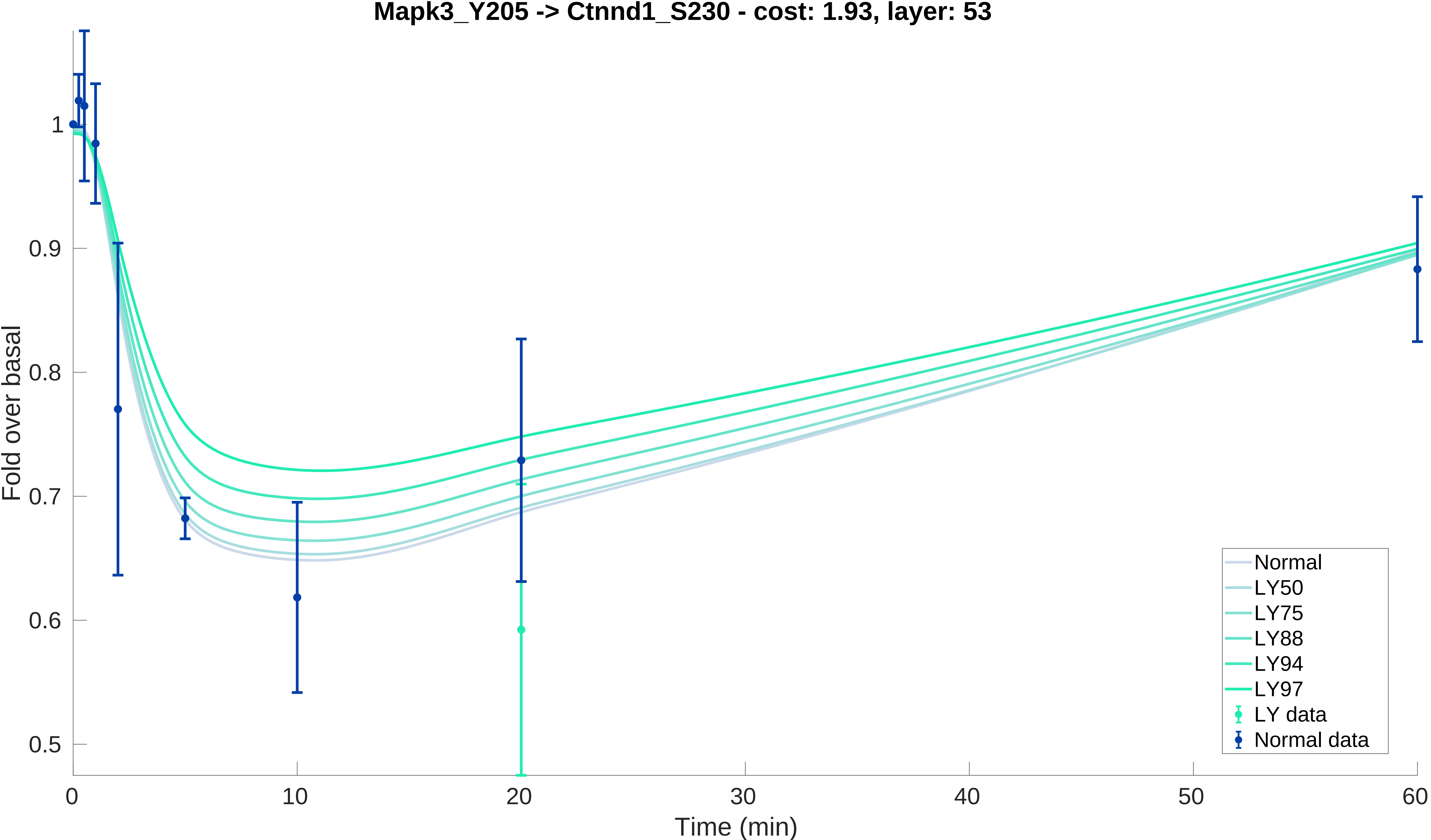

Mapk3_Y205 -> Ctnnd1_S230 - cost: 1.93, layer: 53
Fold over basal
1
0.9
0.8
0.7
0.6
0.5
0
10
20
30
40
50
60
Time (min)
Normal
LY50
LY75
LY88
LY94
LY97
LY data
Normal data

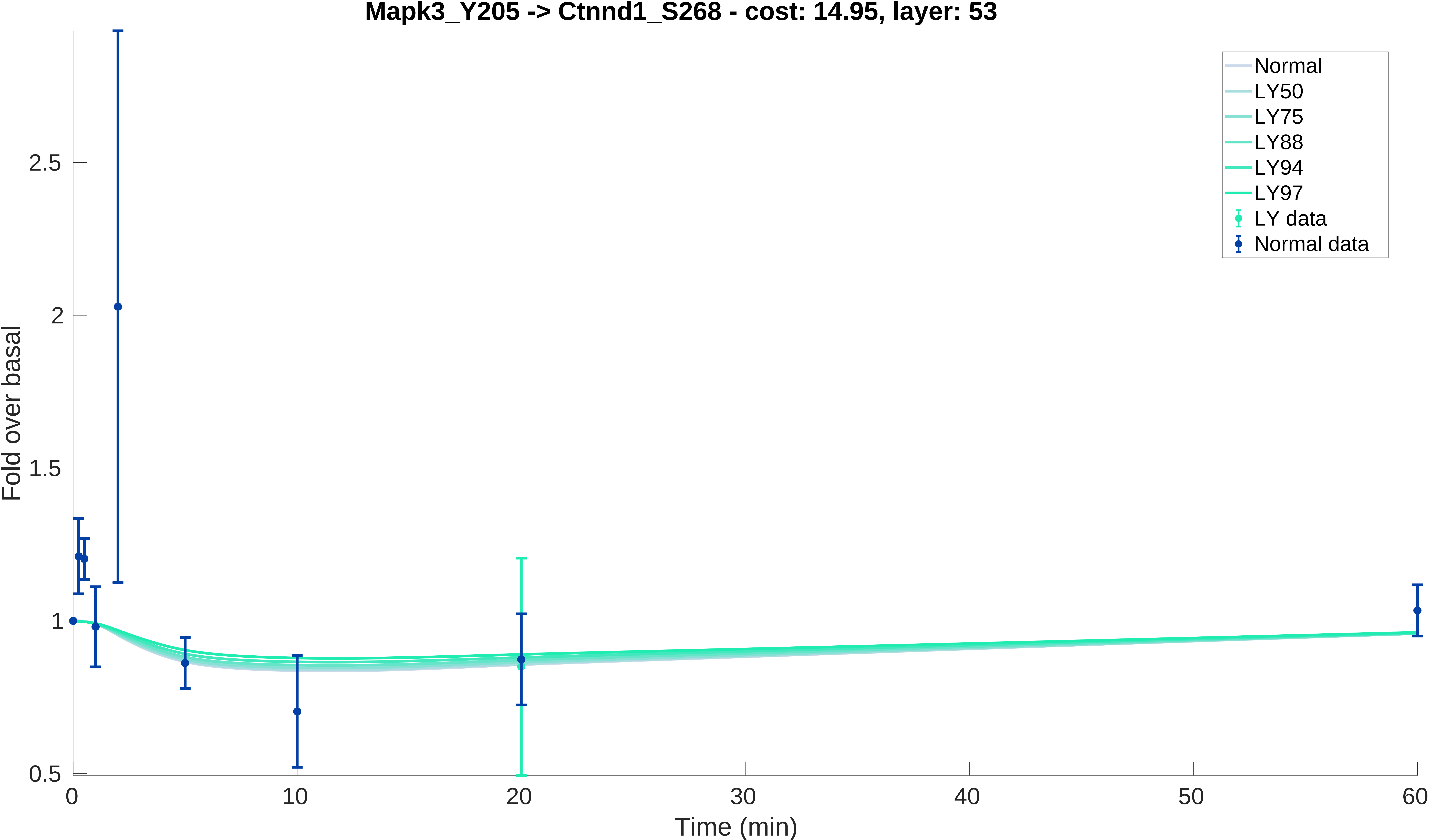

Mapk3_Y205 -> Ctnnd1_S268 - cost: 14.95, layer: 53
Normal
LY50
LY75
LY88
LY94
LY97
LY data
Normal data
Fold over basal
Time (min)
0.5
1
1.5
2
2.5
0
10
20
30
40
50
60

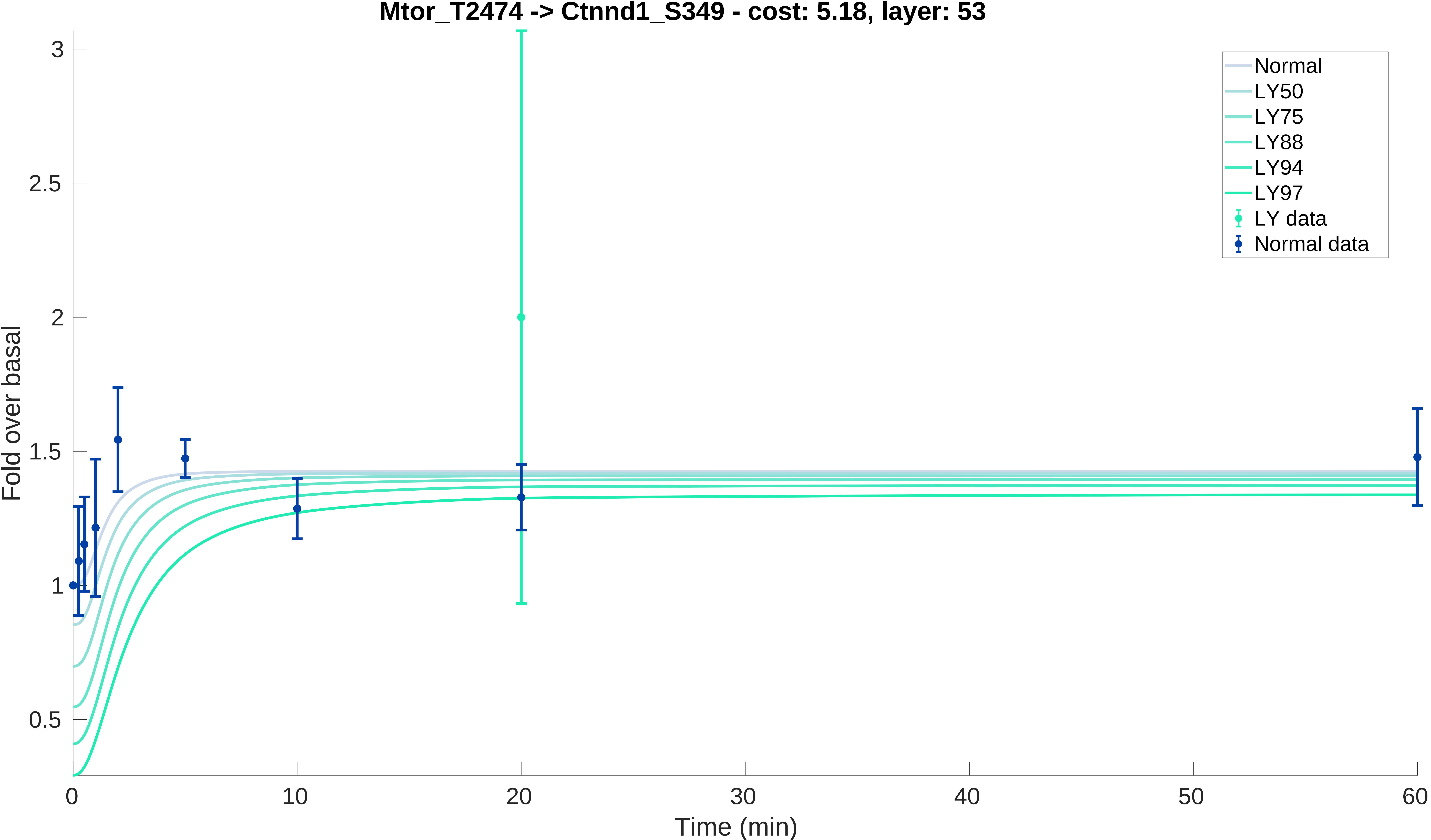

Mtor_T2474 -> Ctnnd1_S349 - cost: 5.18, layer: 53
Fold over basal
Time (min)
0
10
20
30
40
50
60
0.5
1
1.5
2
2.5
3
Normal
LY50
LY75
LY88
LY94
LY97
LY data
Normal data

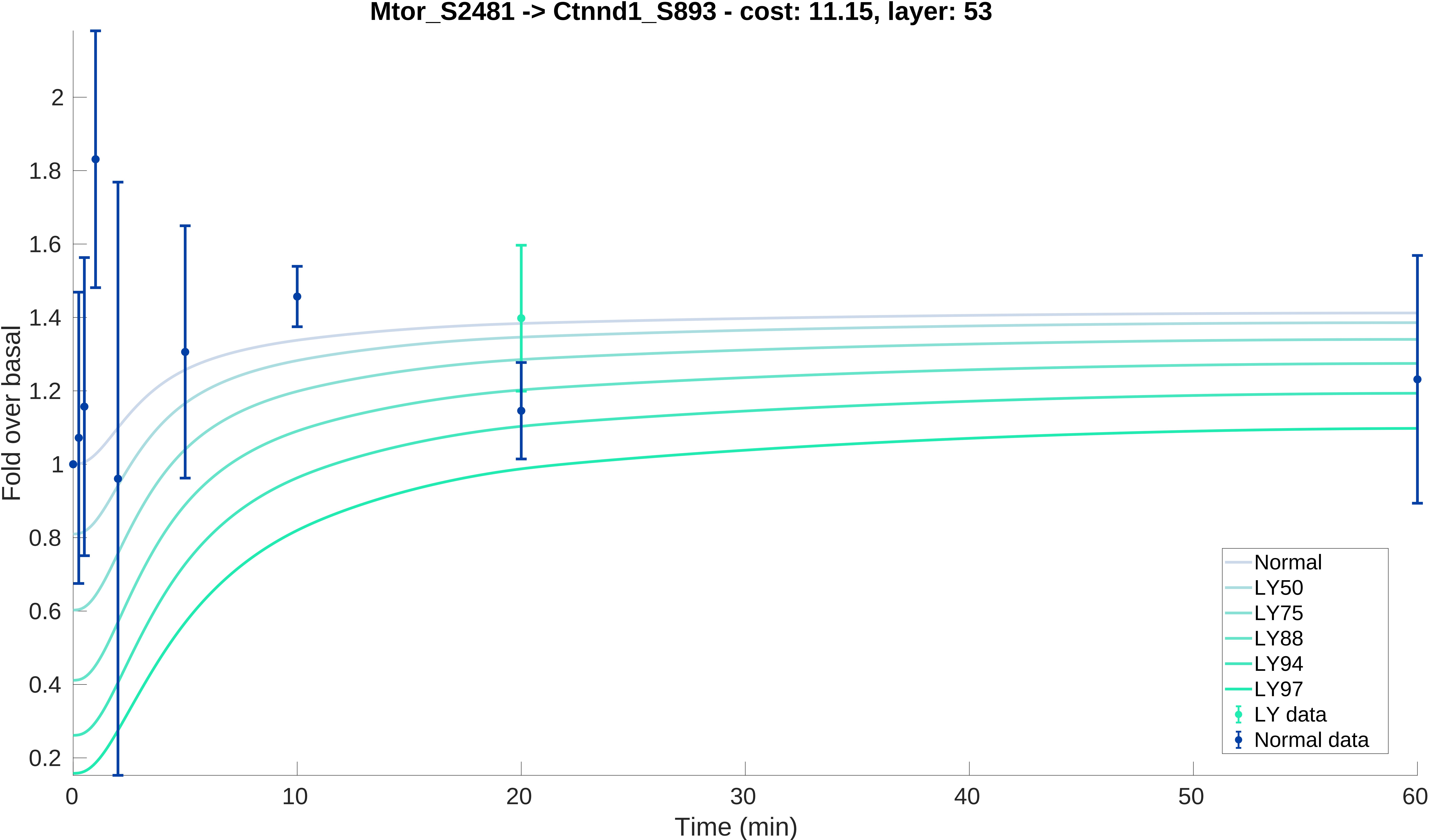

Mtor_S2481 -> Ctnnd1_S893 - cost: 11.15, layer: 53
Fold over basal
Time (min)
0.2
0.4
0.6
0.8
1
1.2
1.4
1.6
1.8
2
0
10
20
30
40
50
60
Normal
LY50
LY75
LY88
LY94
LY97
LY data
Normal data

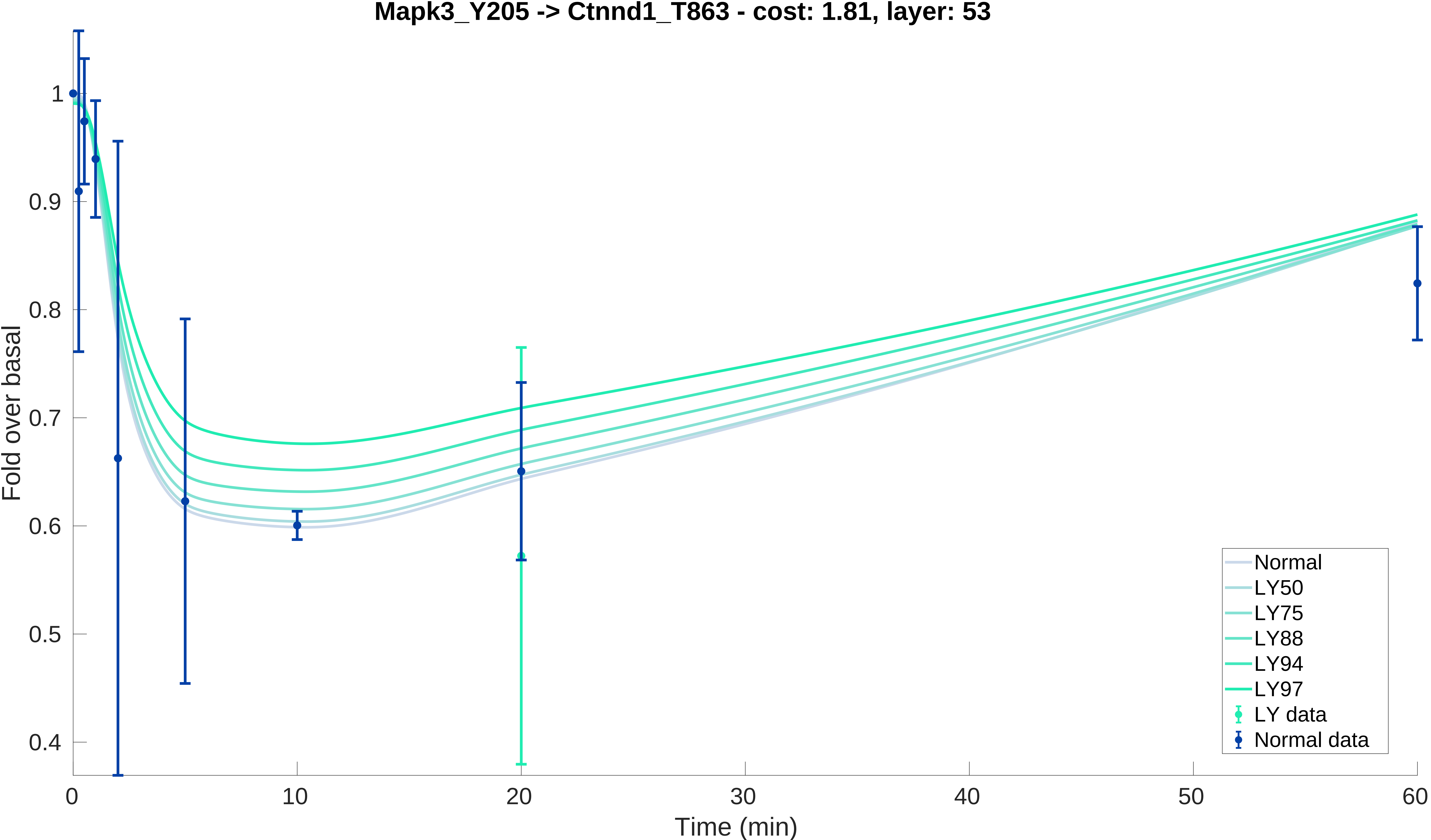
Mapk3_Y205 -> Ctnnd1_T863 - cost: 1.81, layer: 53
Fold over basal
1
0.9
0.8
0.7
0.6
0.5
0.4
0
10
20
30
40
50
60
Time (min)
Normal
LY50
LY75
LY88
LY94
LY97
LY data
Normal data

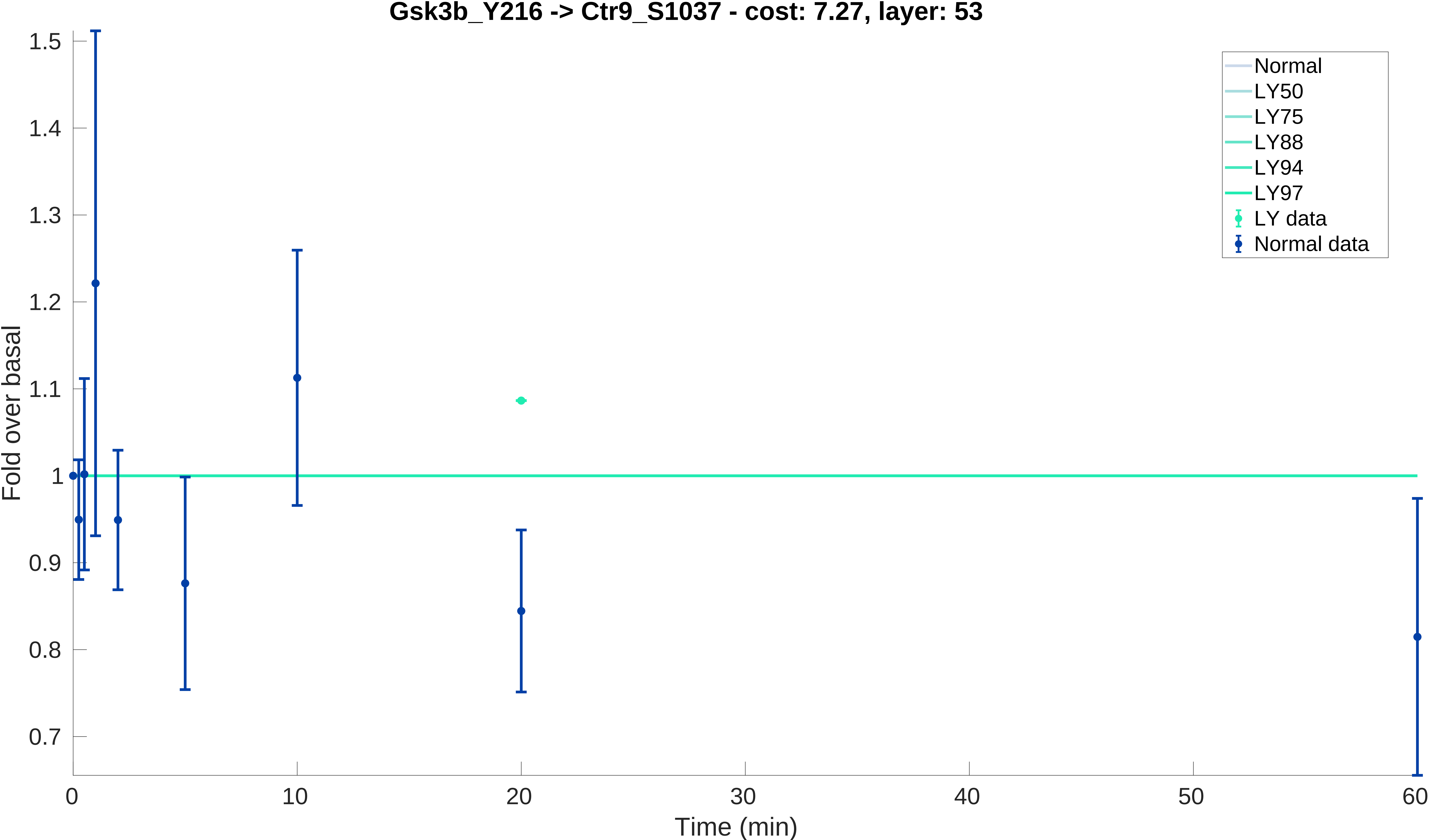

Gsk3b_Y216 -> Ctr9_S1037 - cost: 7.27, layer: 53
Fold over basal
1.5
1.4
1.3
1.2
1.1
1
0.9
0.8
0.7
0
10
20
30
40
50
60
Time (min)
Normal
LY50
LY75
LY88
LY94
LY97
LY data
Normal data

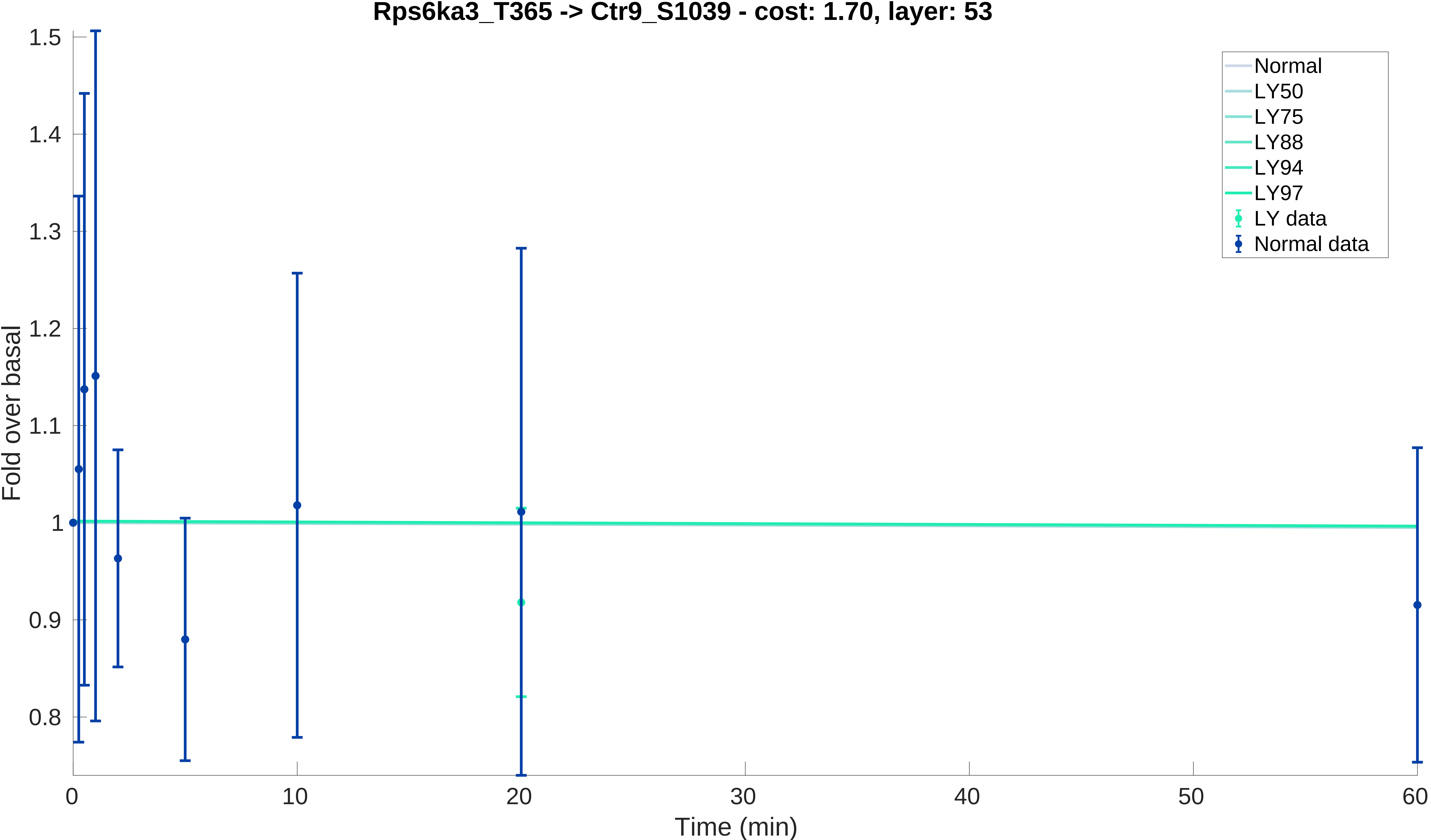
Rps6ka3_T365 -> Ctr9_S1039 - cost: 1.70, layer: 53
Fold over basal
Time (min)
0.8
0.9
1
1.1
1.2
1.3
1.4
1.5
0
10
20
30
40
50
60
Normal
LY50
LY75
LY88
LY94
LY97
LY data
Normal data

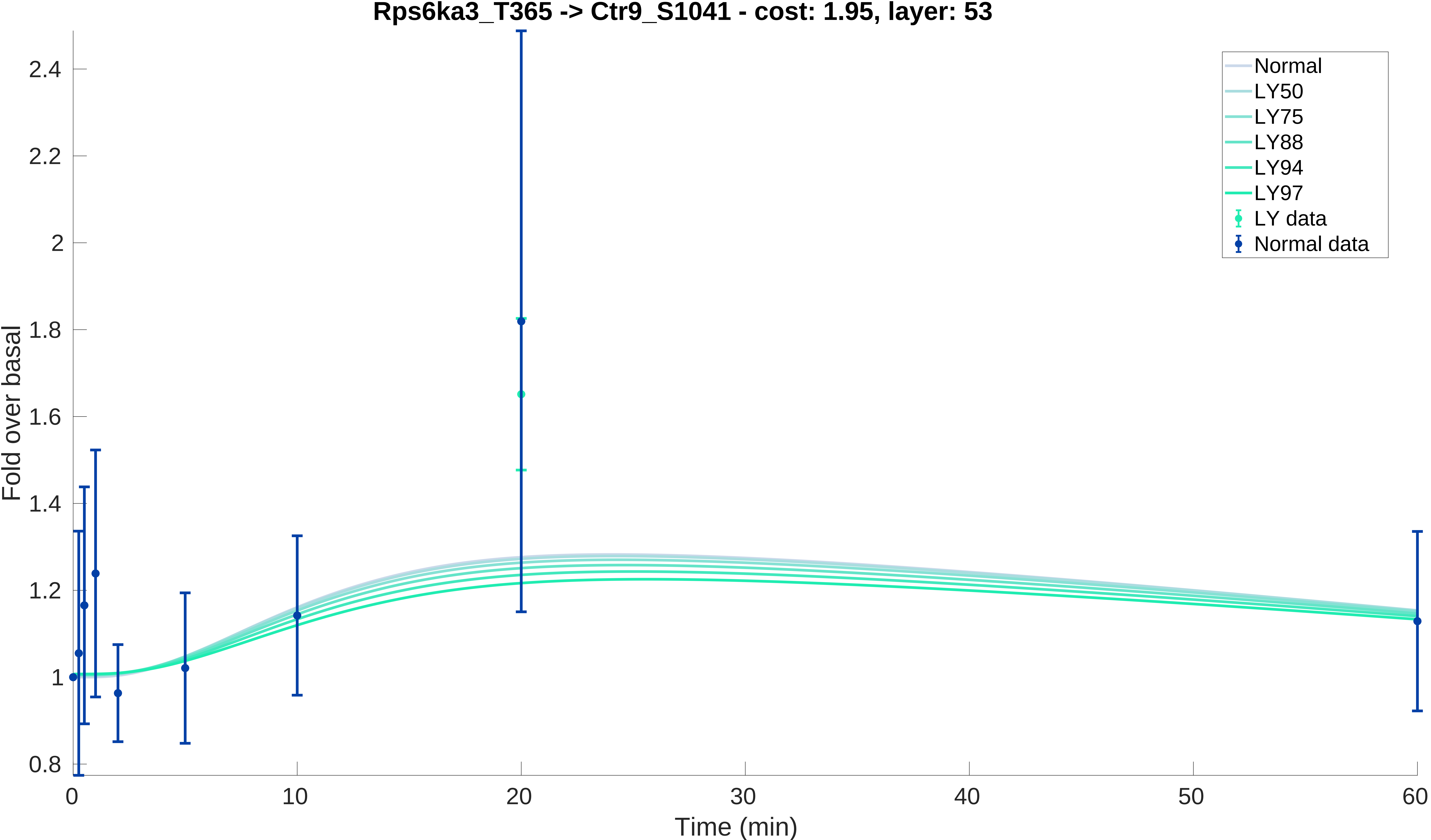

Rps6ka3_T365 -> Ctr9_S1041 - cost: 1.95, layer: 53
Fold over basal
Time (min)
0.8
1
1.2
1.4
1.6
1.8
2
2.2
2.4
0
10
20
30
40
50
60
Normal
LY50
LY75
LY88
LY94
LY97
LY data
Normal data

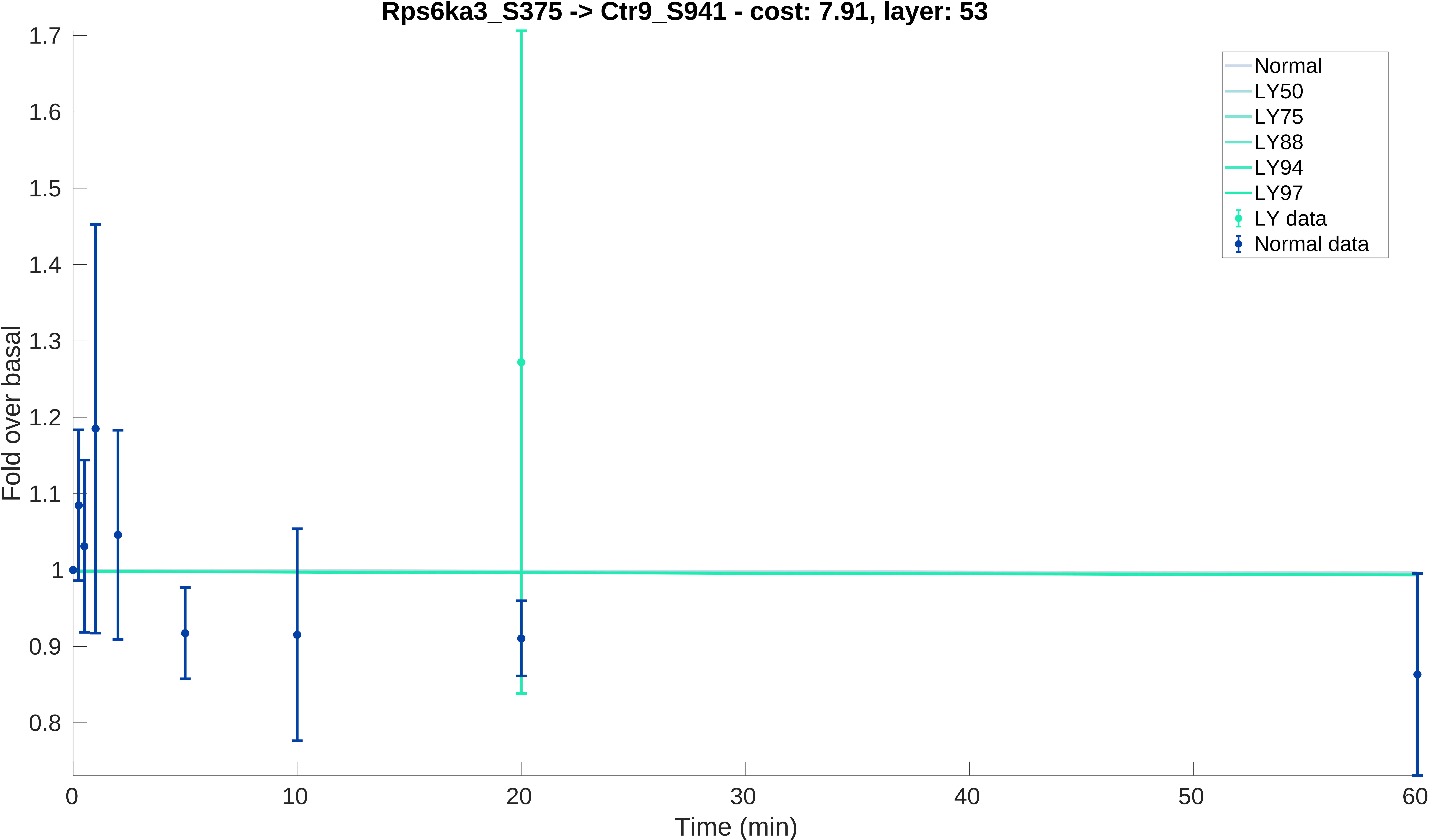

Rps6ka3_S375 -> Ctr9_S941 - cost: 7.91, layer: 53
Fold over basal
Time (min)
Normal
LY50
LY75
LY88
LY94
LY97
LY data
Normal data
0.8
0.9
1
1.1
1.2
1.3
1.4
1.5
1.6
1.7
0
10
20
30
40
50
60

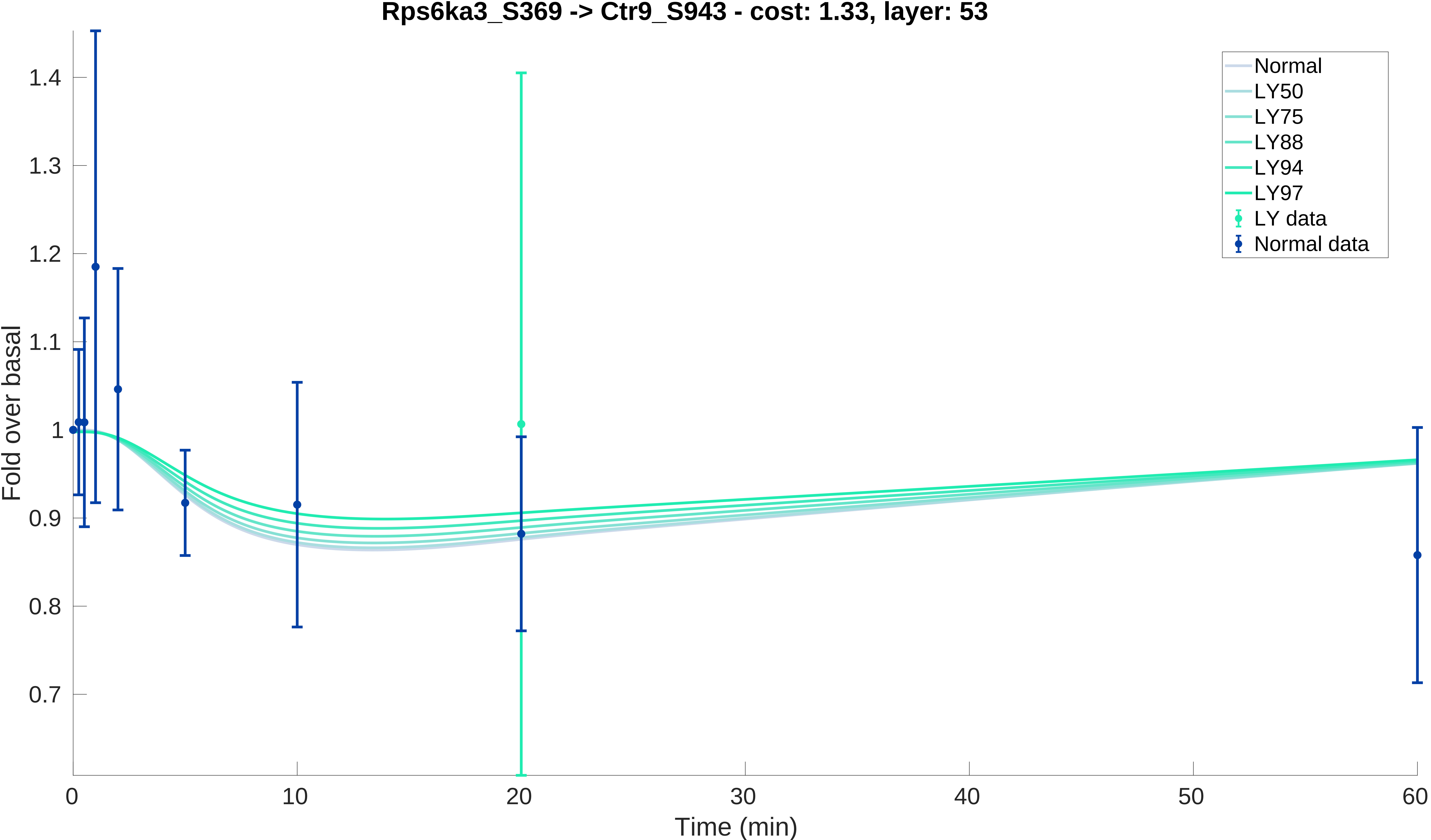

Rps6ka3_S369 -> Ctr9_S943 - cost: 1.33, layer: 53
Fold over basal
Time (min)
0.7
0.8
0.9
1
1.1
1.2
1.3
1.4
0
10
20
30
40
50
60
Normal
LY50
LY75
LY88
LY94
LY97
LY data
Normal data

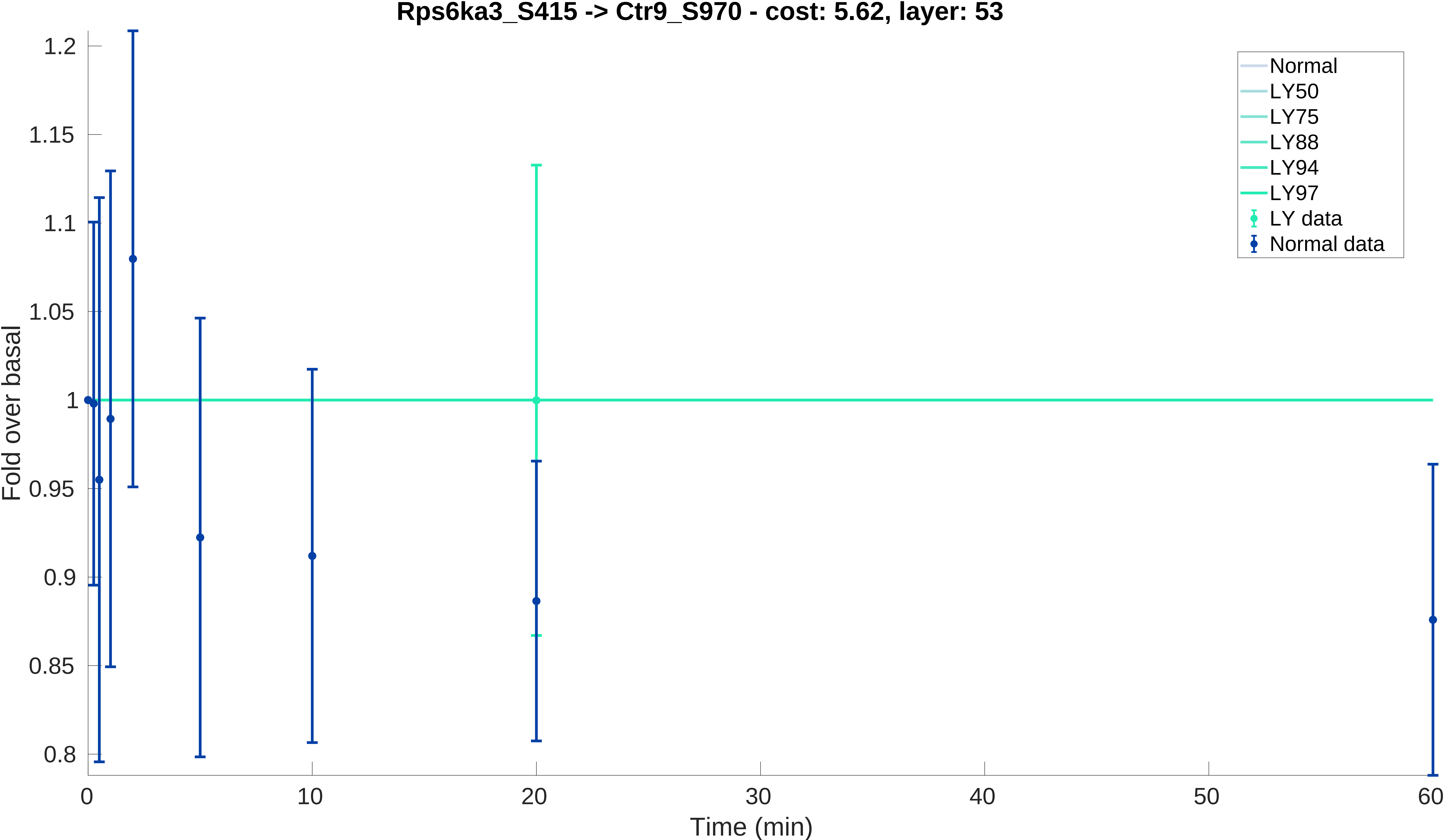

Rps6ka3_S415 -> Ctr9_S970 - cost: 5.62, layer: 53
Fold over basal
1.2
1.15
1.1
1.05
1
0.95
0.9
0.85
0.8
0
10
20
30
40
50
60
Time (min)
Normal
LY50
LY75
LY88
LY94
LY97
LY data
Normal data

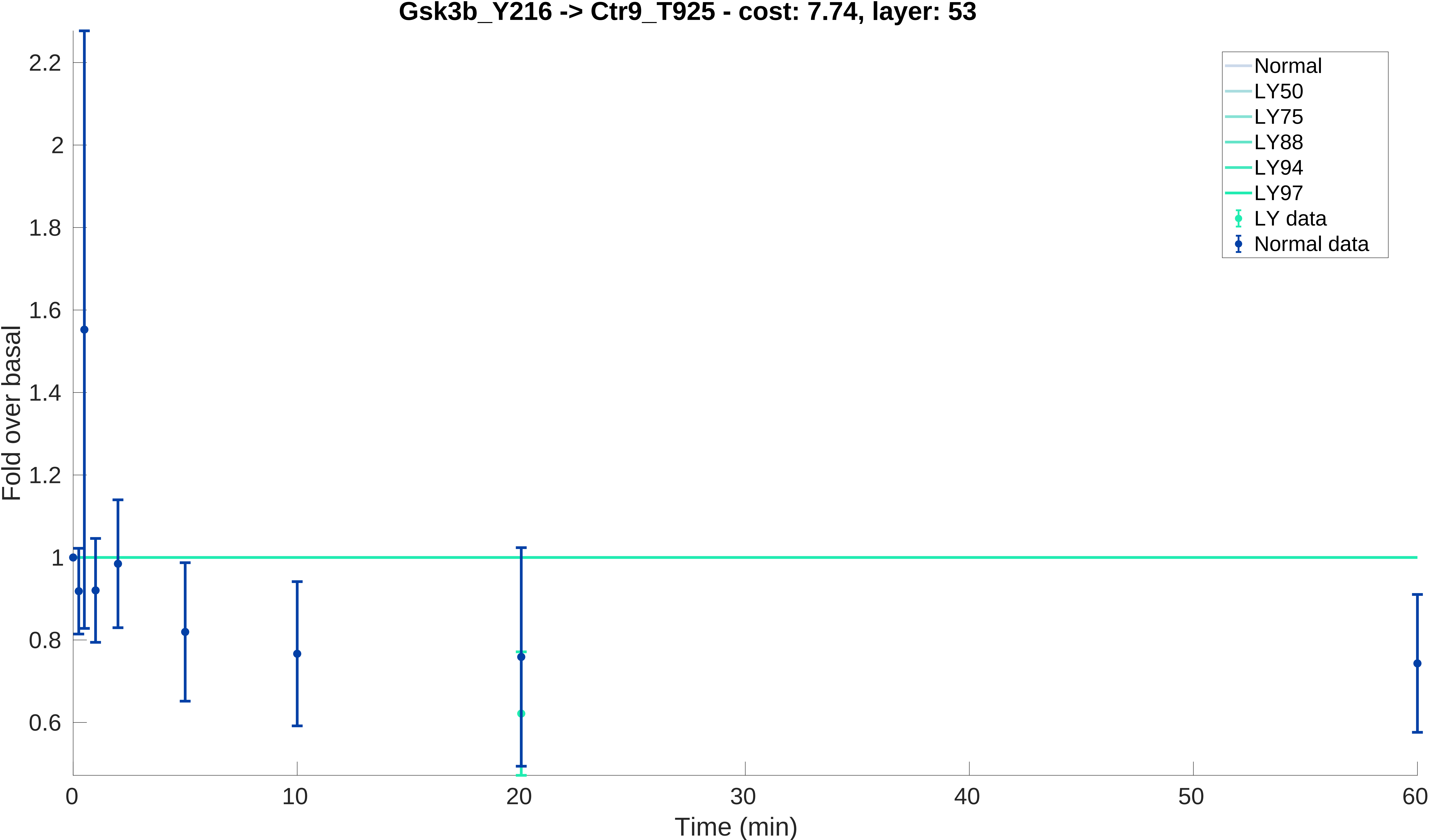

Gsk3b_Y216 -> Ctr9_T925 - cost: 7.74, layer: 53
Normal
LY50
LY75
LY88
LY94
LY97
LY data
Normal data
Fold over basal
Time (min)
0.6
0.8
1
1.2
1.4
1.6
1.8
2
2.2
0
10
20
30
40
50
60

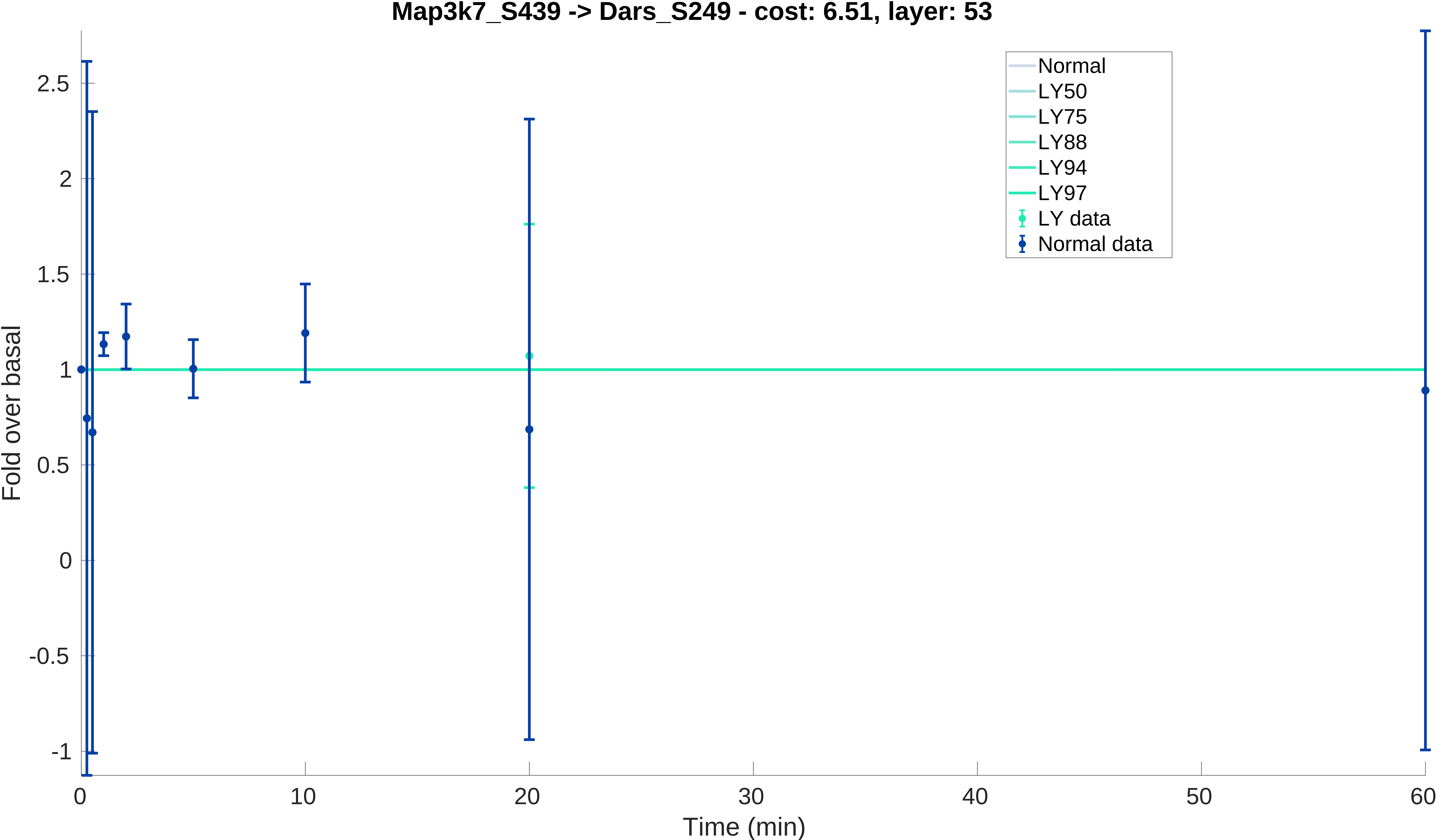

Map3k7_S439 -> Dars_S249 - cost: 6.51, layer: 53
Fold over basal
Time (min)
2.5
2
1.5
1
0.5
0
-0.5
-1
0
10
20
30
40
50
60
Normal
LY50
LY75
LY88
LY94
LY97
LY data
Normal data

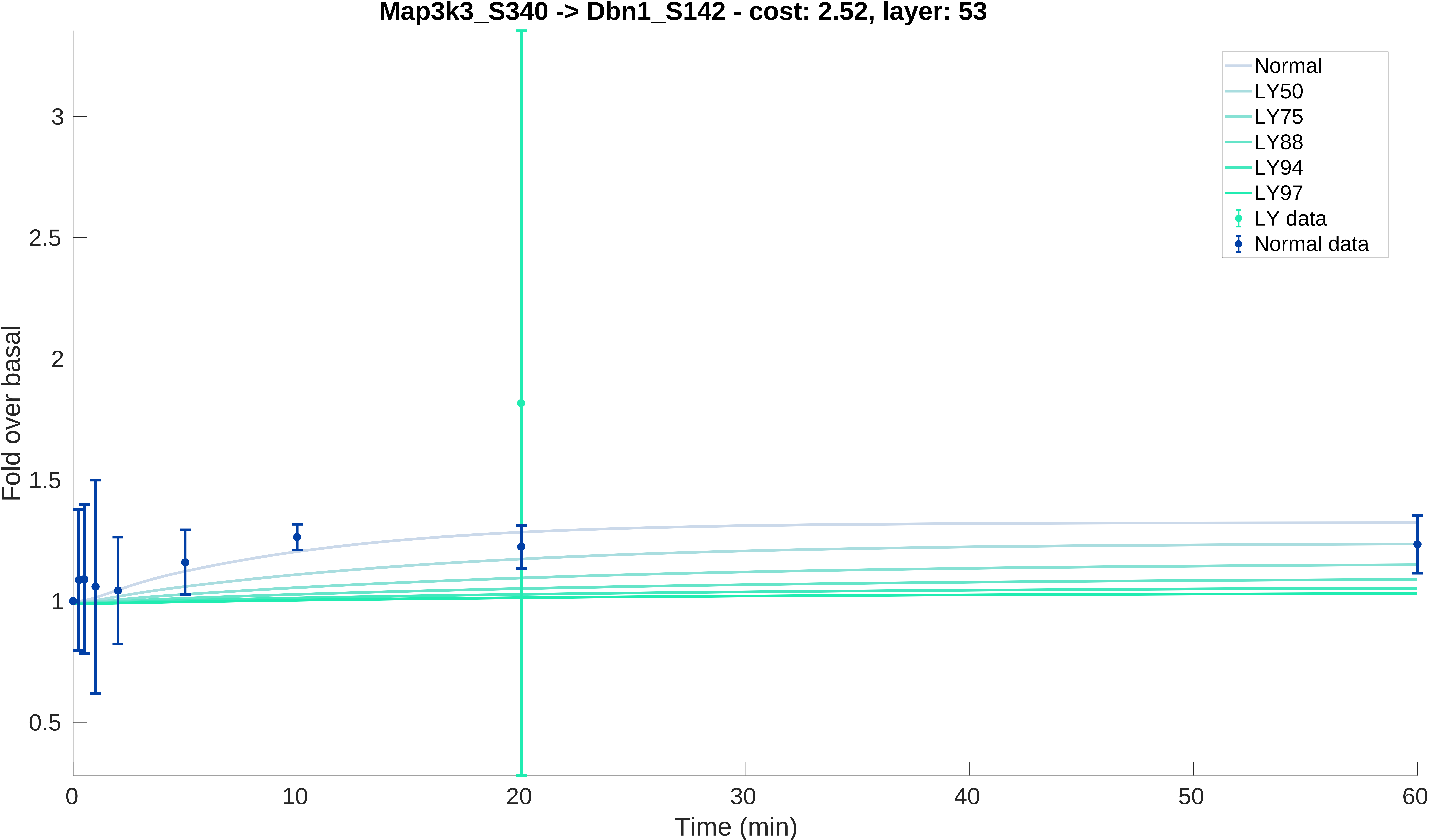
Map3k3_S340 -> Dbn1_S142 - cost: 2.52, layer: 53
Fold over basal
Time (min)
0.5
1
1.5
2
2.5
3
0
10
20
30
40
50
60
Normal
LY50
LY75
LY88
LY94
LY97
LY data
Normal data

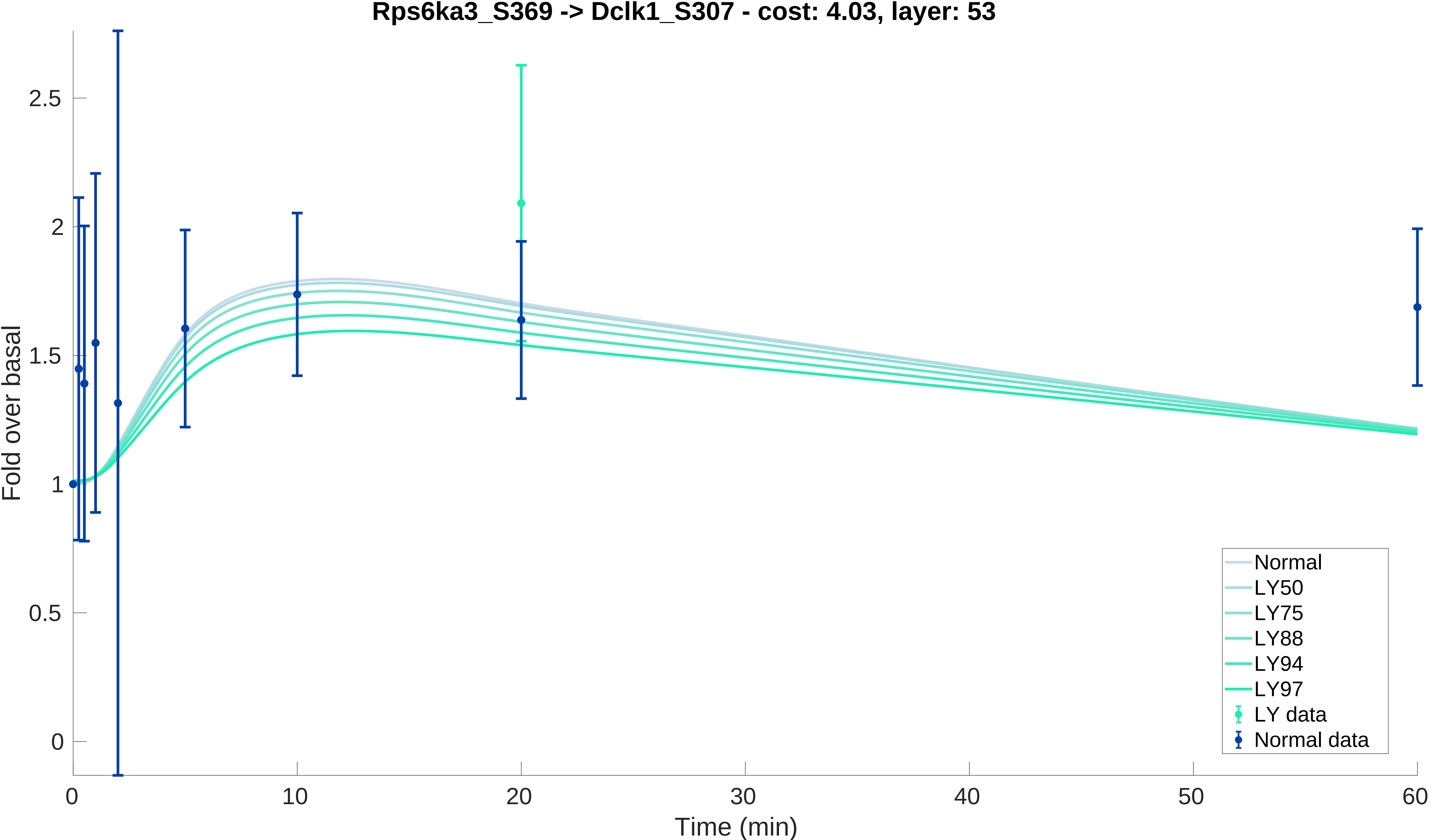

Rps6ka3_S369 -> Dclk1_S307 - cost: 4.03, layer: 53
Fold over basal
Time (min)
0
0.5
1
1.5
2
2.5
0
10
20
30
40
50
60
Normal
LY50
LY75
LY88
LY94
LY97
LY data
Normal data

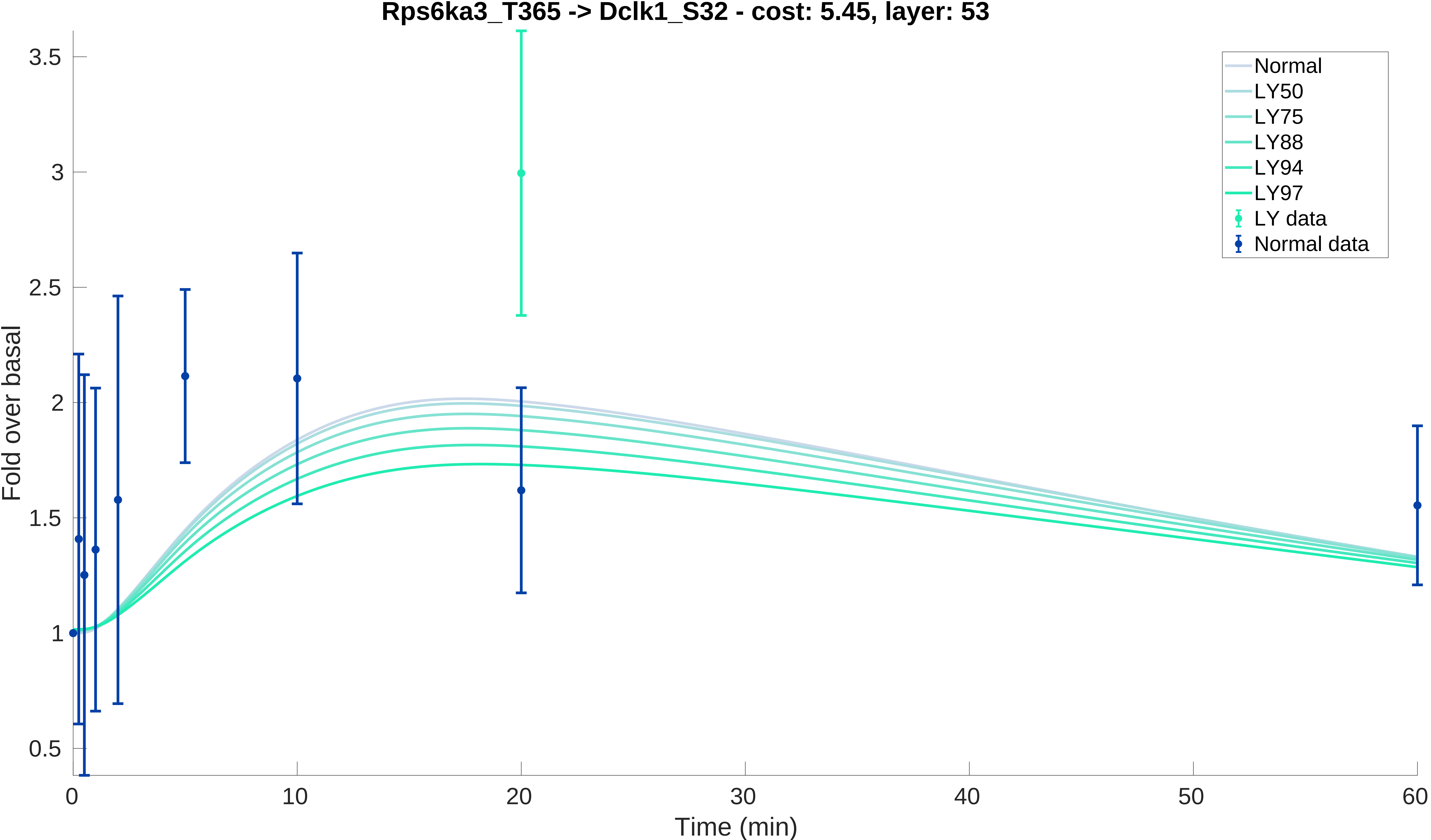
Rps6ka3_T365 -> Dclk1_S32 - cost: 5.45, layer: 53
Fold over basal
Time (min)
Normal
LY50
LY75
LY88
LY94
LY97
LY data
Normal data
0.5
1
1.5
2
2.5
3
3.5
0
10
20
30
40
50
60

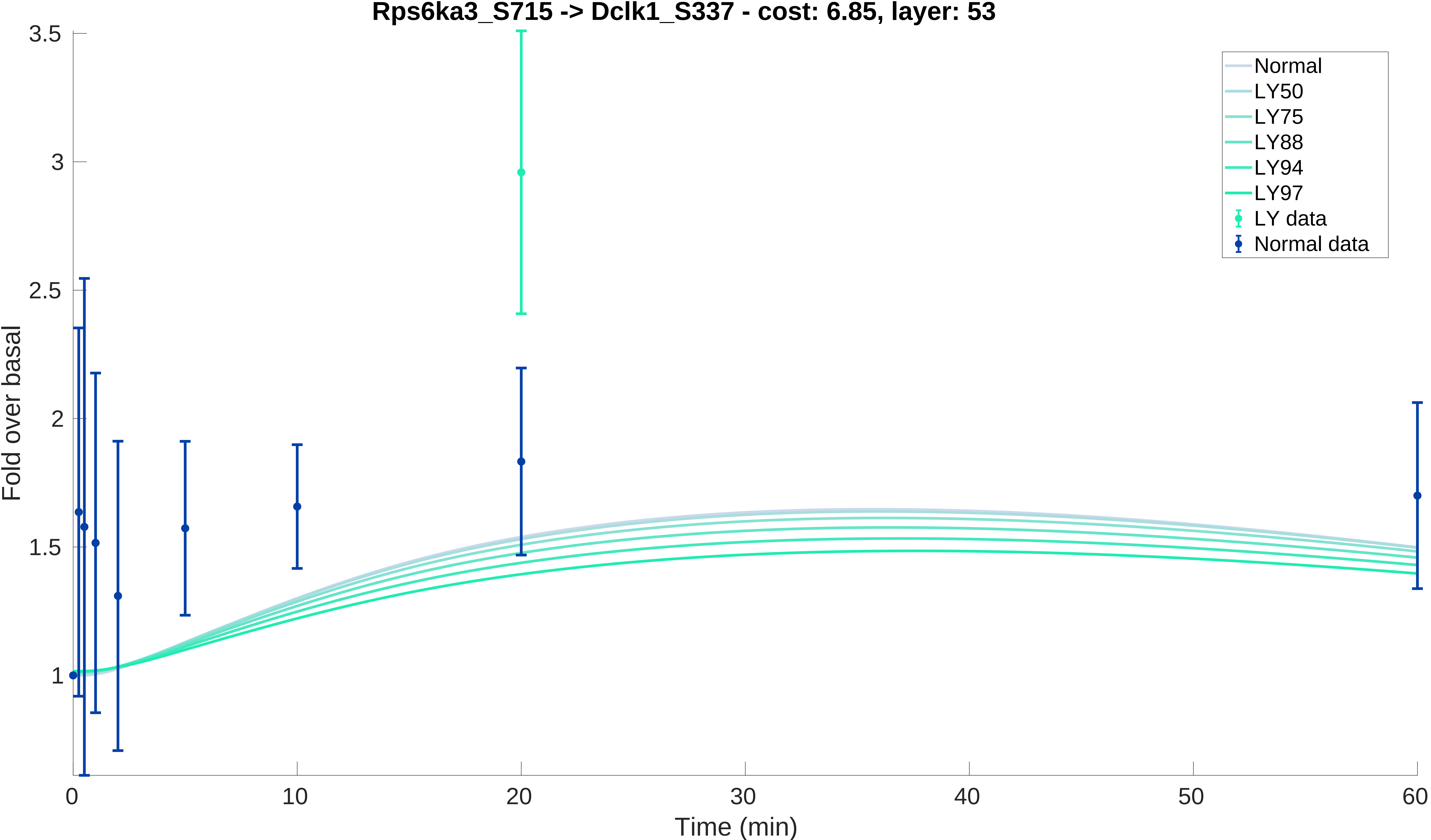

Rps6ka3_S715 -> Dclk1_S337 - cost: 6.85, layer: 53
3.5
3
2.5
2
1.5
1
0
10
20
30
40
50
60
Fold over basal
Time (min)
Normal
LY50
LY75
LY88
LY94
LY97
LY data
Normal data

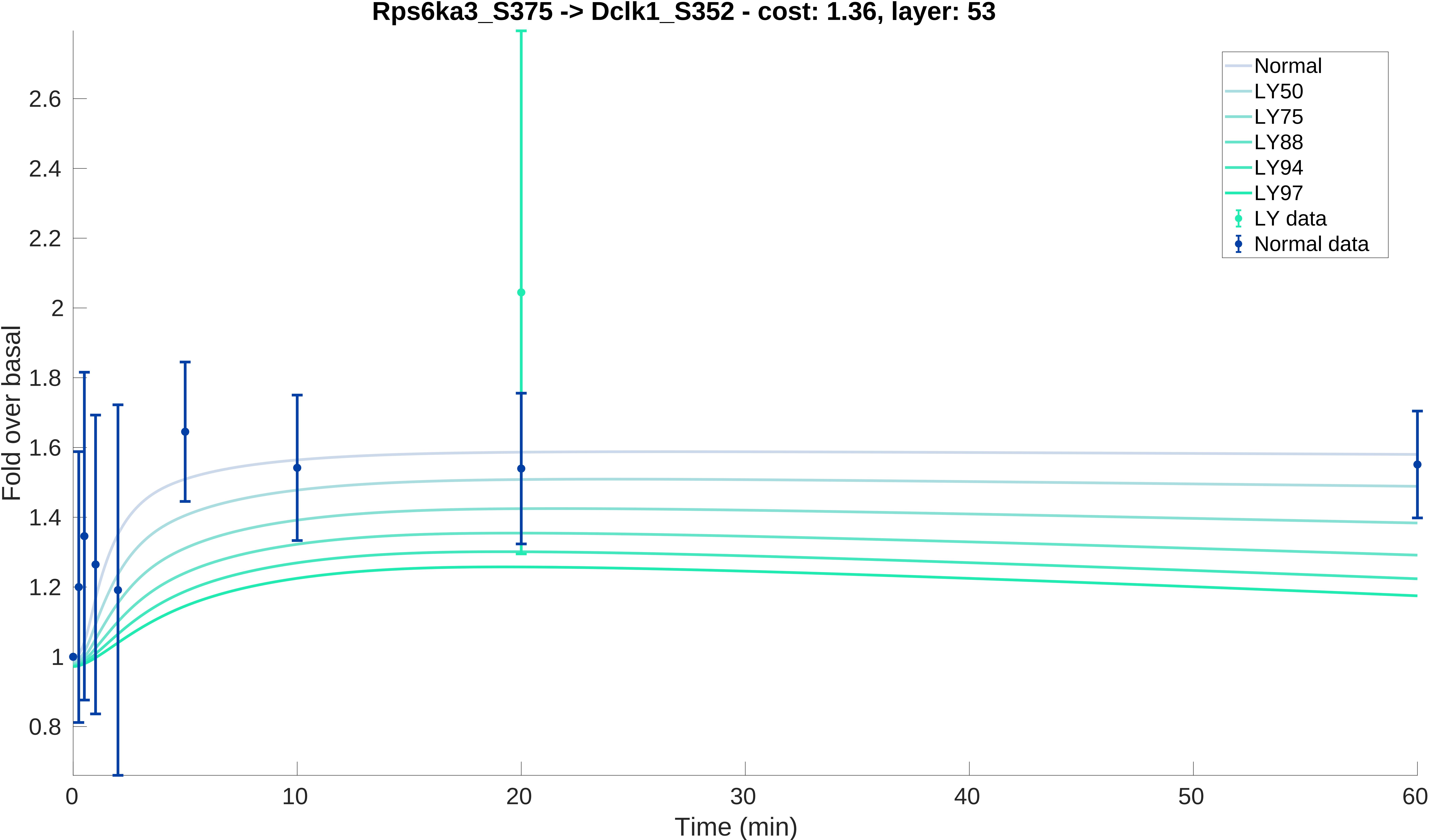

Rps6ka3_S375 -> Dclk1_S352 - cost: 1.36, layer: 53
Normal
LY50
LY75
LY88
LY94
LY97
LY data
Normal data
Fold over basal
Time (min)
0.8
1
1.2
1.4
1.6
1.8
2
2.2
2.4
2.6
0
10
20
30
40
50
60

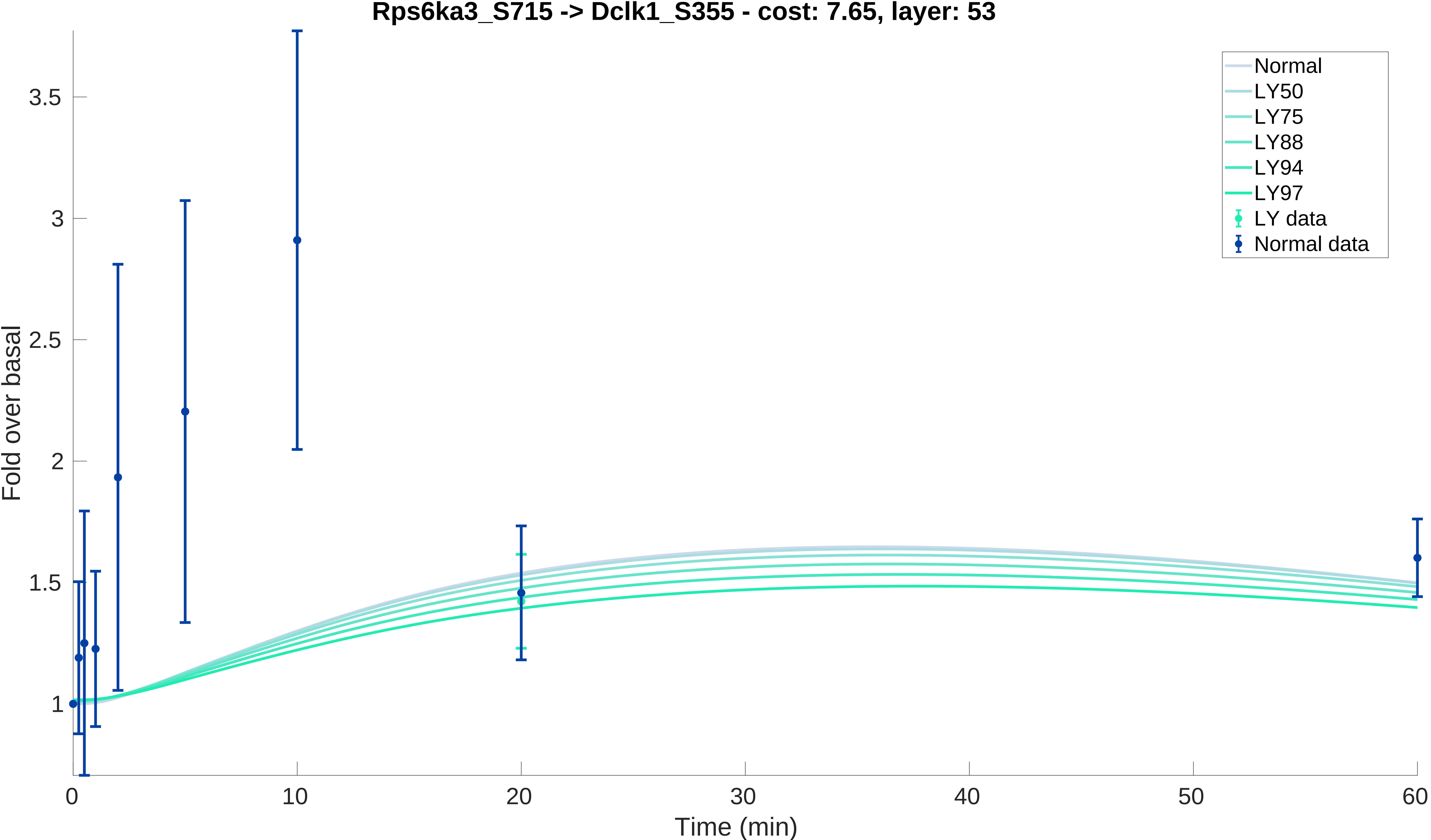
Rps6ka3_S715 -> Dclk1_S355 - cost: 7.65, layer: 53
Fold over basal
Time (min)
3.5
3
2.5
2
1.5
1
0
10
20
30
40
50
60
Normal
LY50
LY75
LY88
LY94
LY97
LY data
Normal data

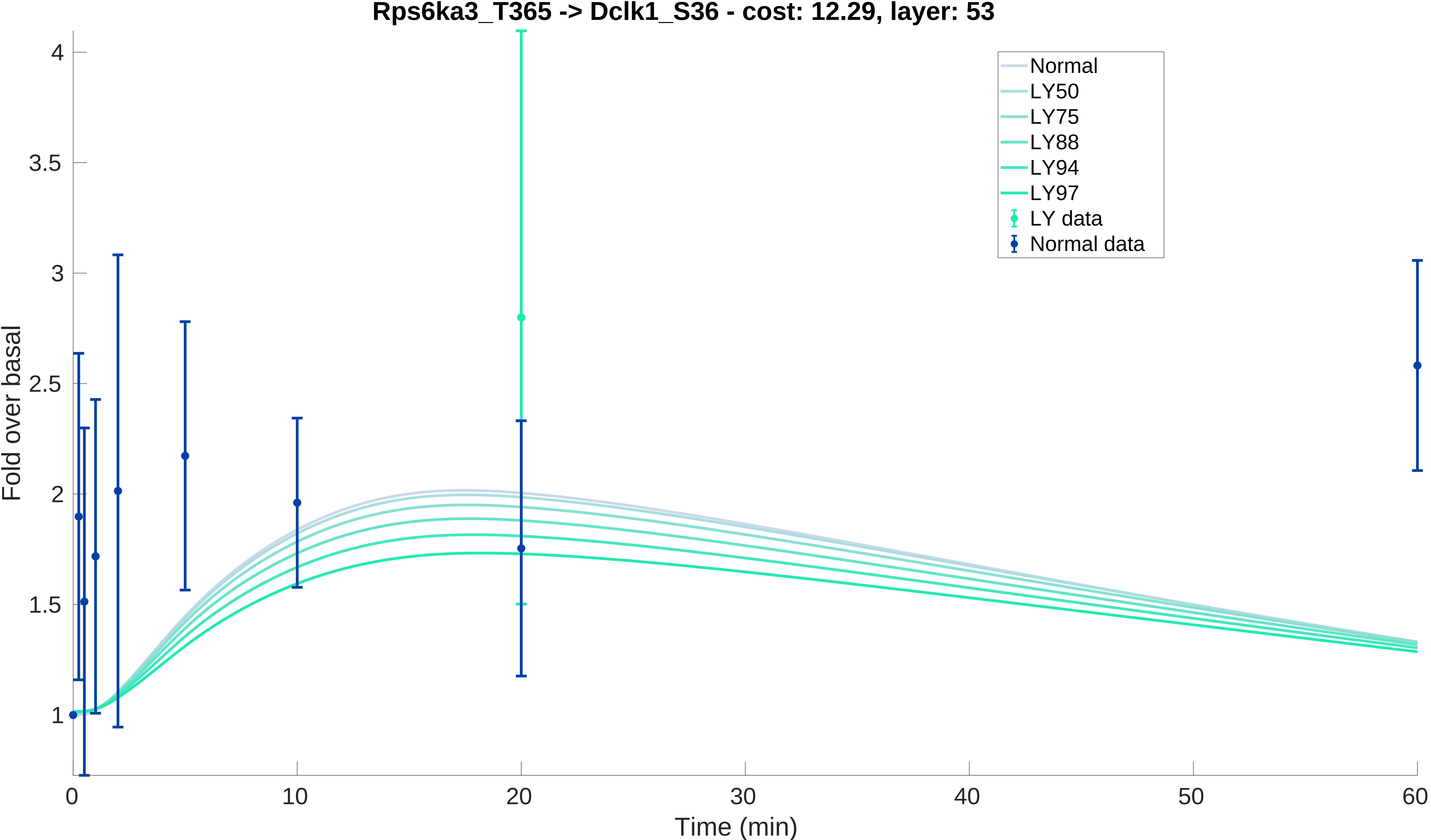

Rps6ka3_T365 -> Dclk1_S36 - cost: 12.29, layer: 53
Fold over basal
Time (min)
4
3.5
3
2.5
2
1.5
1
0
10
20
30
40
50
60
Normal
LY50
LY75
LY88
LY94
LY97
LY data
Normal data

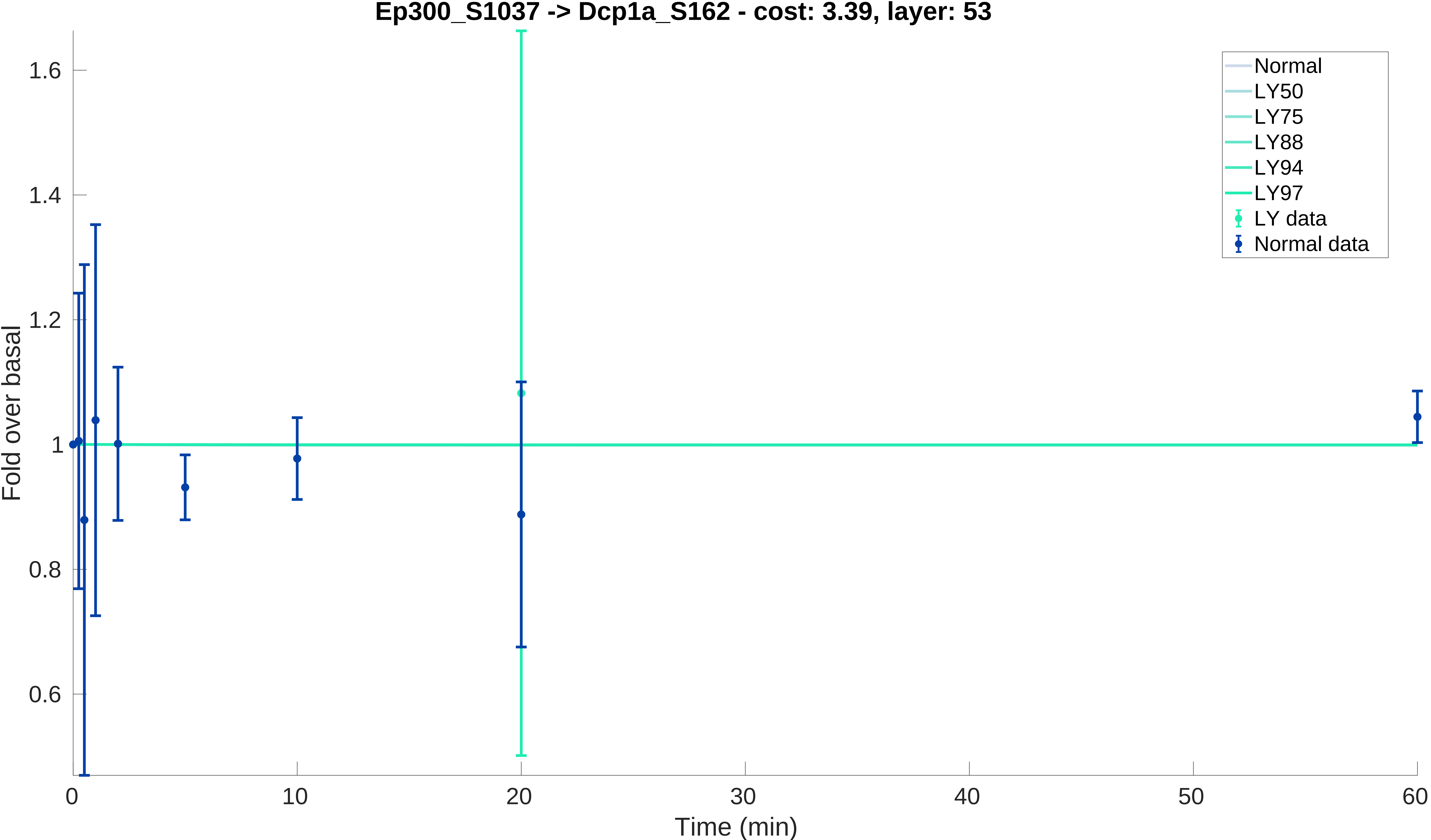

Ep300_S1037 -> Dcp1a_S162 - cost: 3.39, layer: 53
Fold over basal
1.6
1.4
1.2
1
0.8
0.6
0
10
20
30
40
50
60
Time (min)
Normal
LY50
LY75
LY88
LY94
LY97
LY data
Normal data

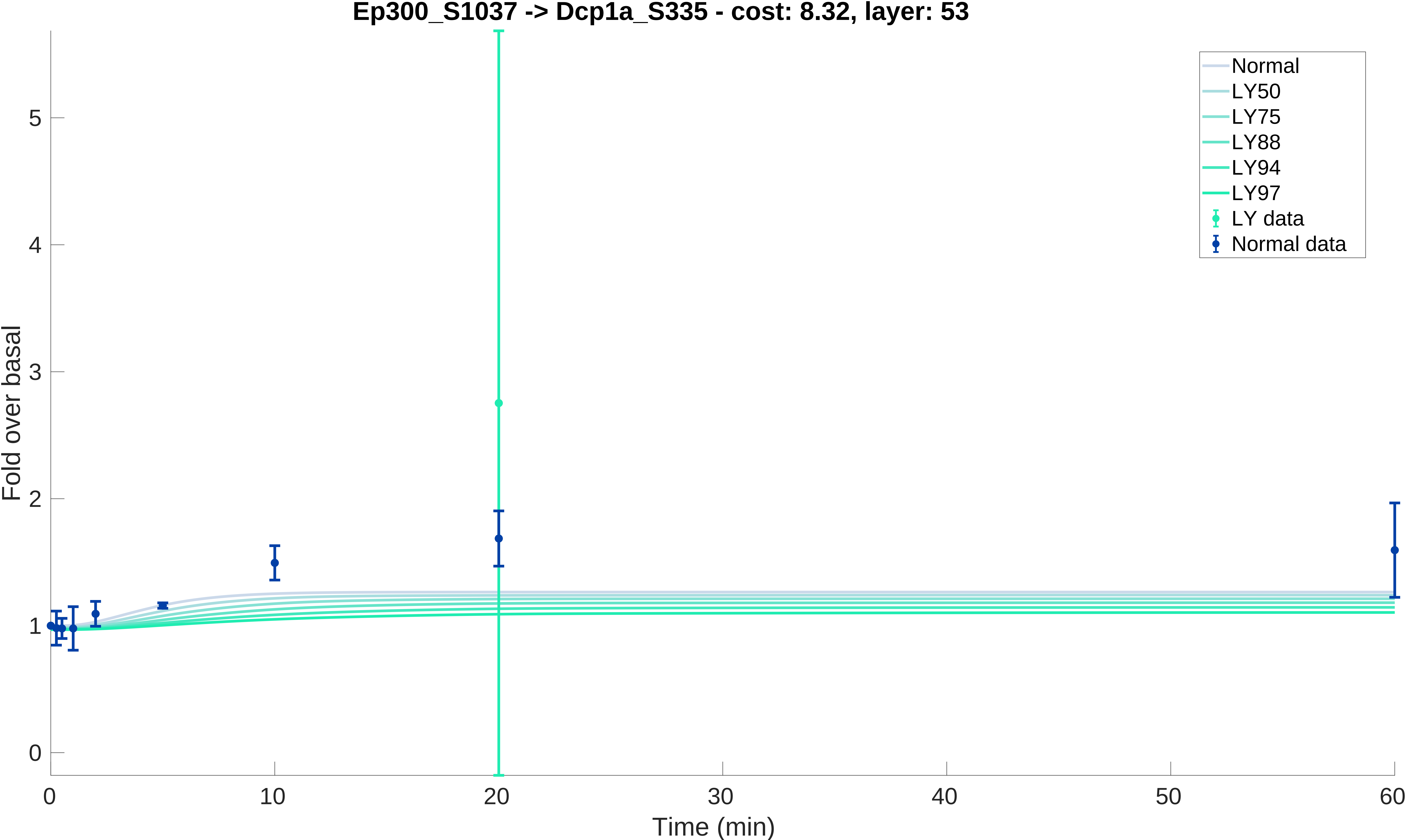
Ep300_S1037 -> Dcp1a_S335 - cost: 8.32, layer: 53
Fold over basal
Time (min)
0
1
2
3
4
5
0
10
20
30
40
50
60
Normal
LY50
LY75
LY88
LY94
LY97
LY data
Normal data

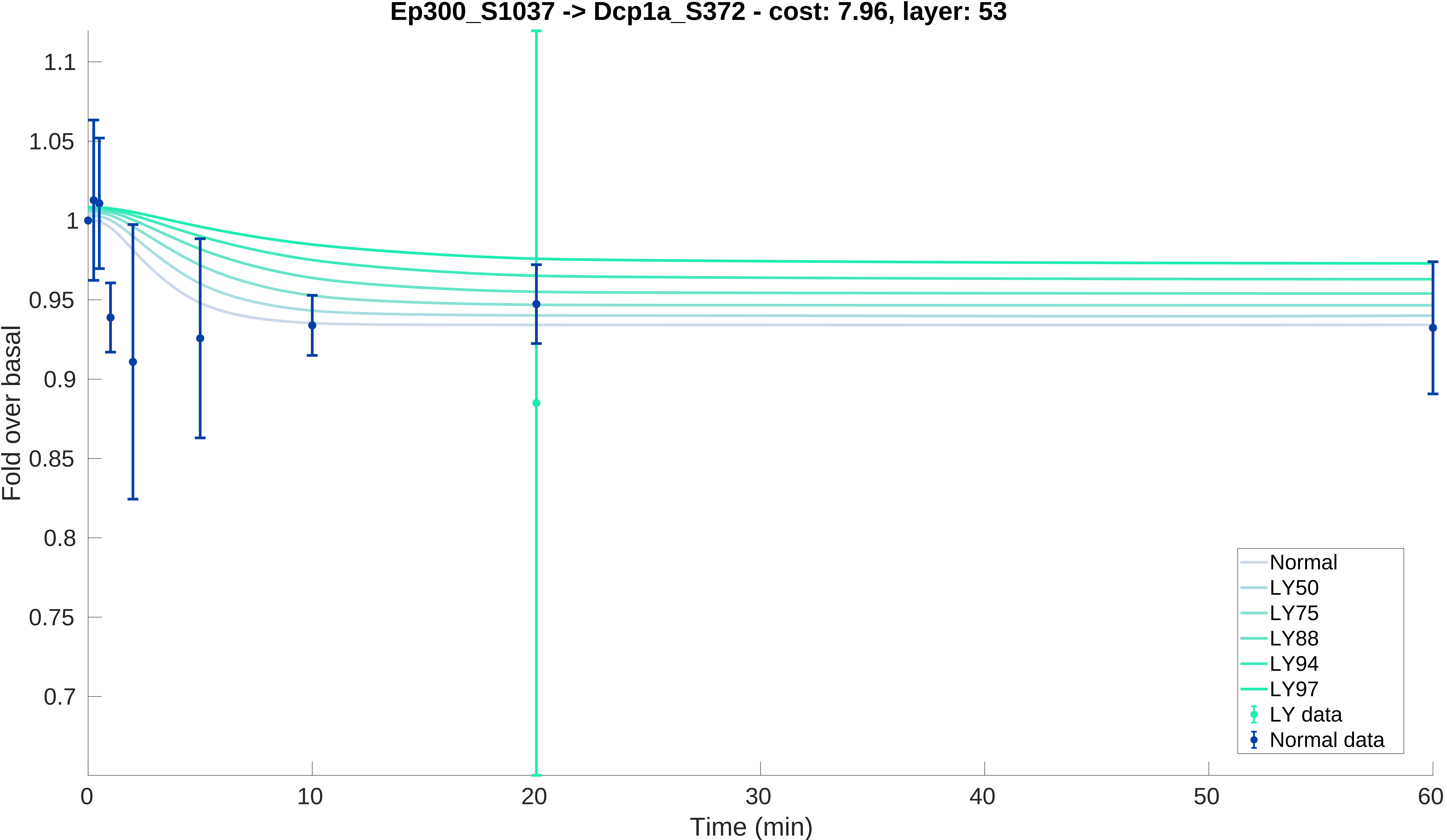

Ep300_S1037 -> Dcp1a_S372 - cost: 7.96, layer: 53
Fold over basal
1.1
1.05
1
0.95
0.9
0.85
0.8
0.75
0.7
0
10
20
30
40
50
60
Time (min)
Normal
LY50
LY75
LY88
LY94
LY97
LY data
Normal data

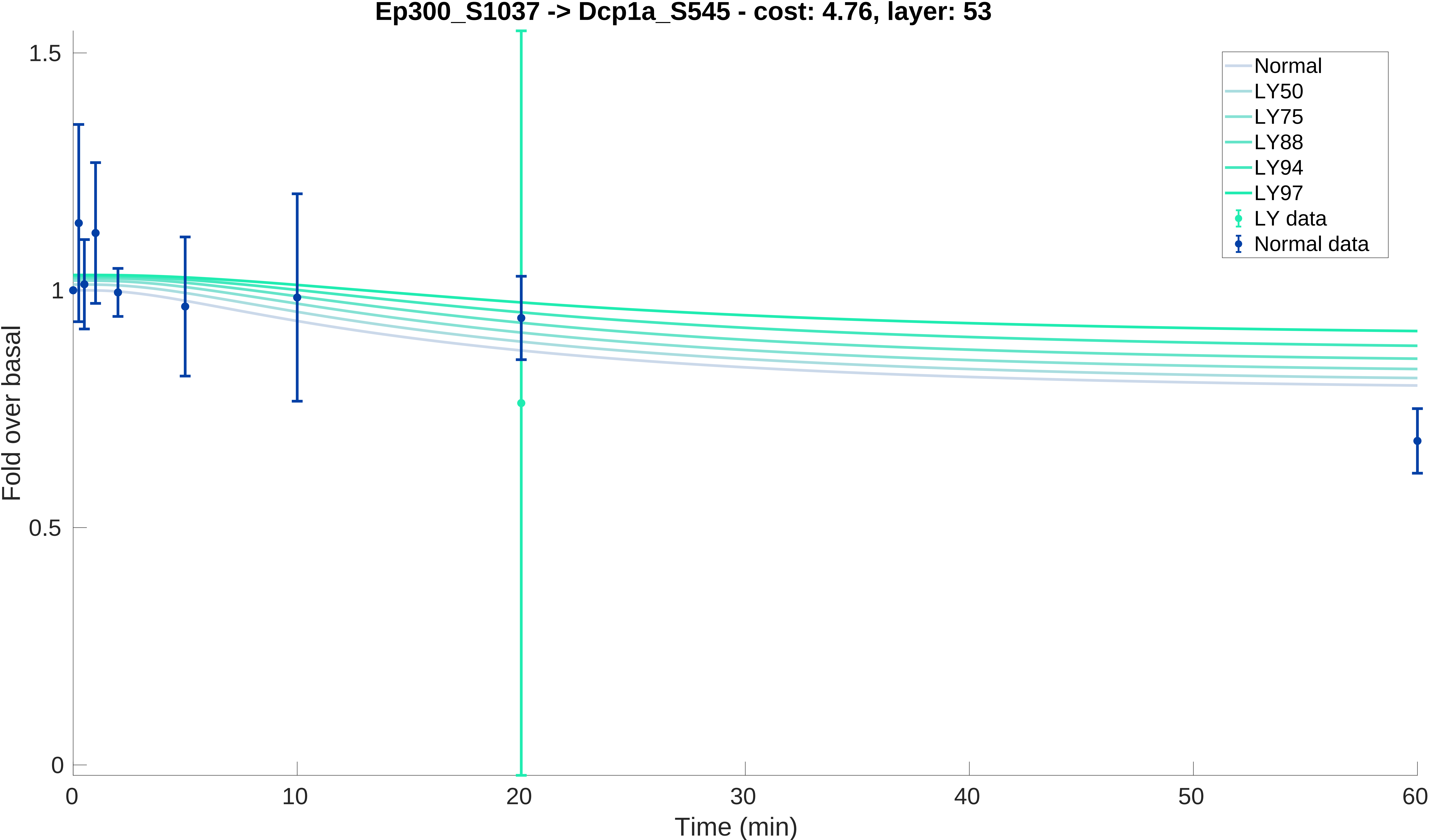

Ep300_S1037 -> Dcp1a_S545 - cost: 4.76, layer: 53
Fold over basal
Time (min)
1.5
1
0.5
0
0
10
20
30
40
50
60
Normal
LY50
LY75
LY88
LY94
LY97
LY data
Normal data

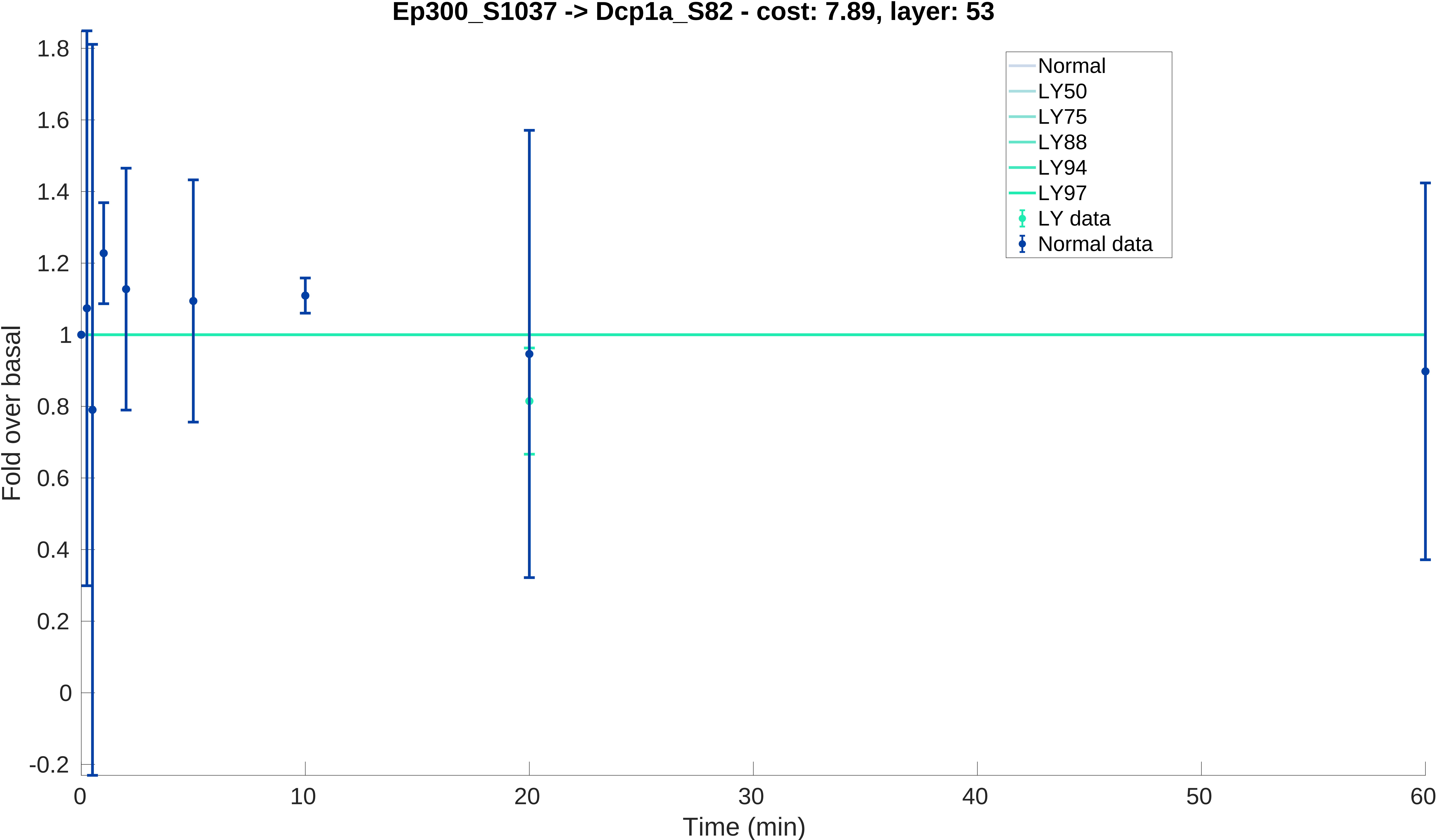

Ep300_S1037 -> Dcp1a_S82 - cost: 7.89, layer: 53
Normal
LY50
LY75
LY88
LY94
LY97
LY data
Normal data
Fold over basal
Time (min)
1.8
1.6
1.4
1.2
1
0.8
0.6
0.4
0.2
0
-0.2
0
10
20
30
40
50
60

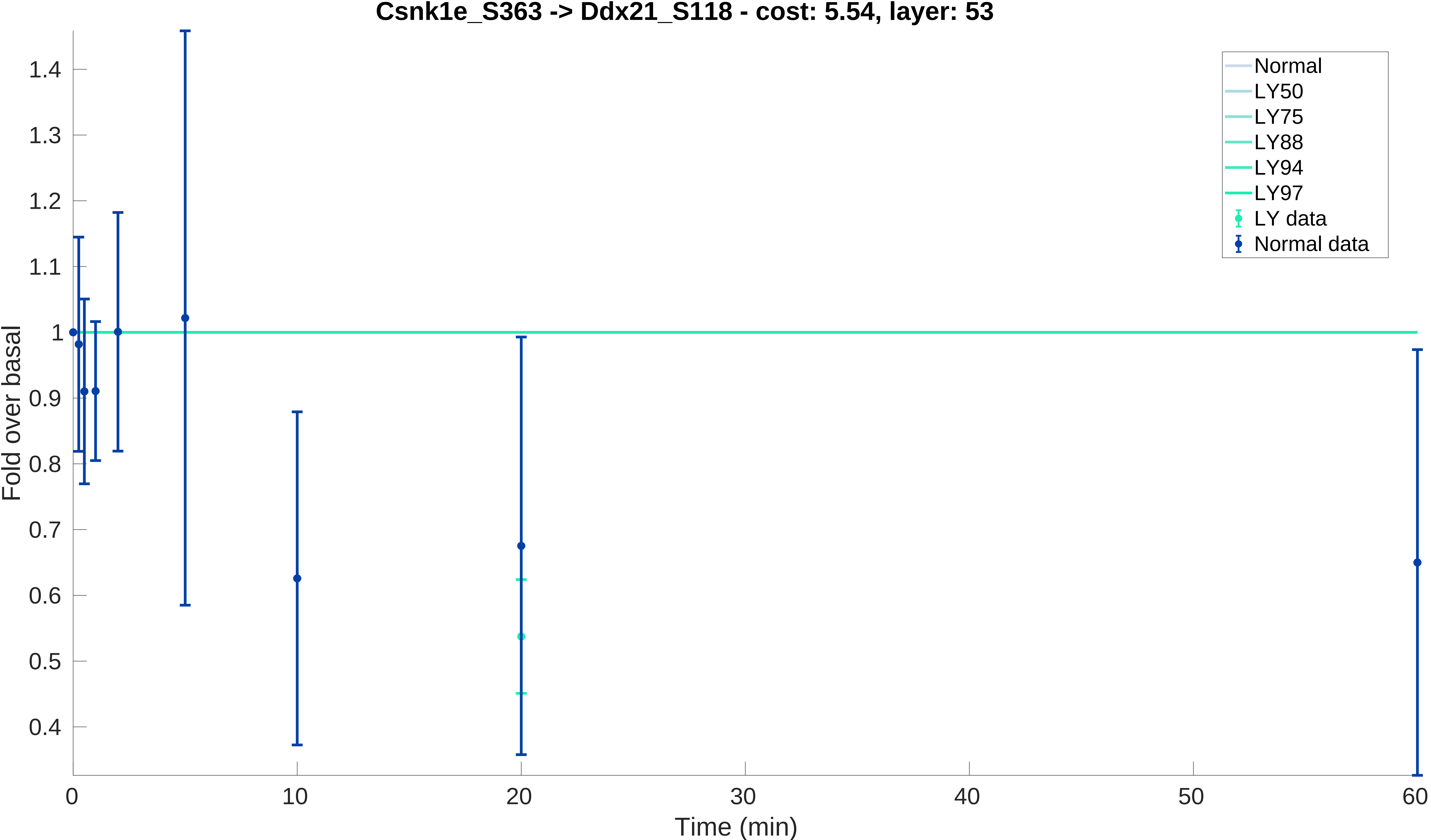

Csnk1e_S363 -> Ddx21_S118 - cost: 5.54, layer: 53
Fold over basal
Time (min)
1.4
1.3
1.2
1.1
1
0.9
0.8
0.7
0.6
0.5
0.4
0
10
20
30
40
50
60
Normal
LY50
LY75
LY88
LY94
LY97
LY data
Normal data

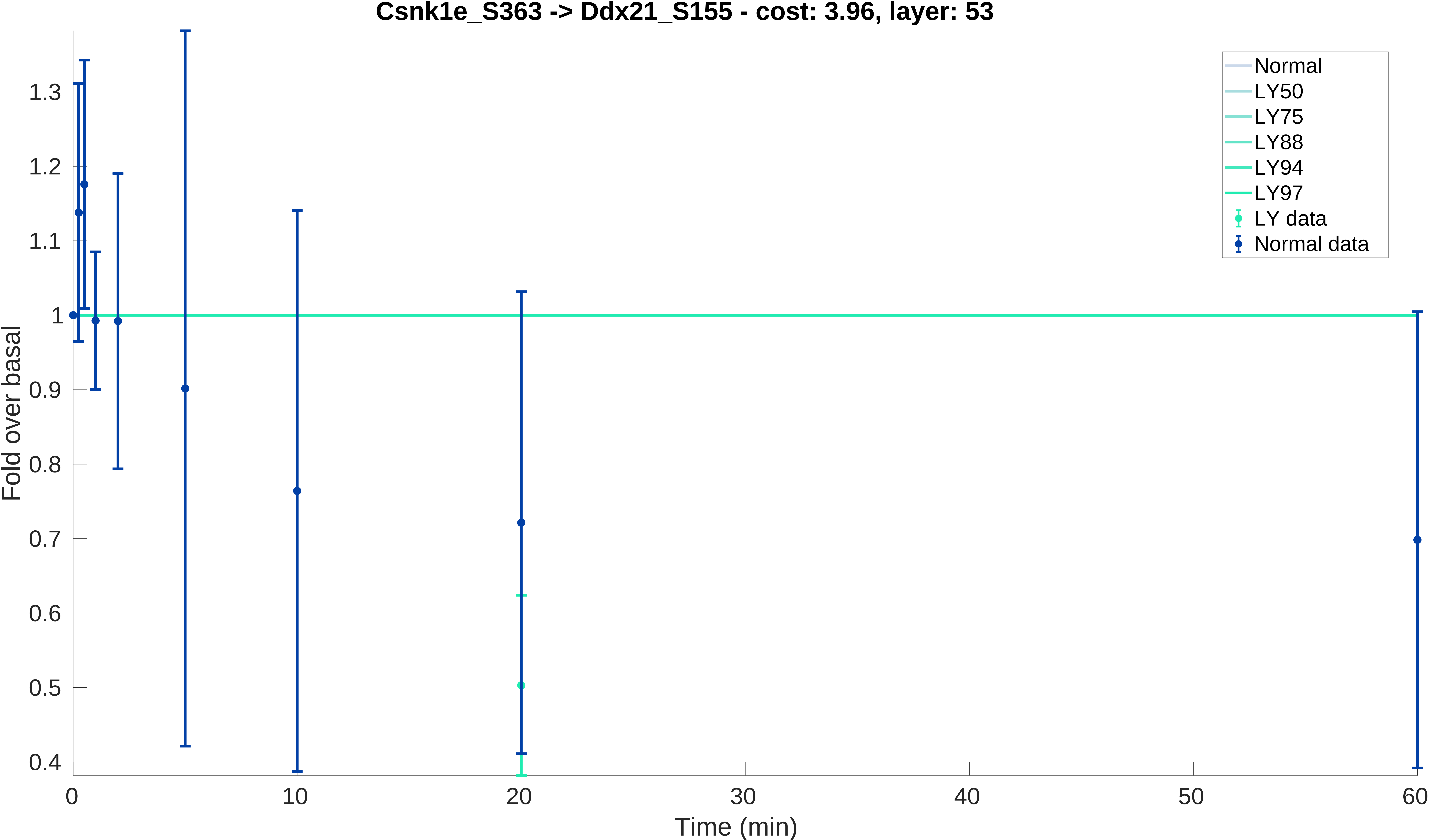

Csnk1e_S363 -> Ddx21_S155 - cost: 3.96, layer: 53
Fold over basal
Time (min)
1.3
1.2
1.1
1
0.9
0.8
0.7
0.6
0.5
0.4
0
10
20
30
40
50
60
Normal
LY50
LY75
LY88
LY94
LY97
LY data
Normal data

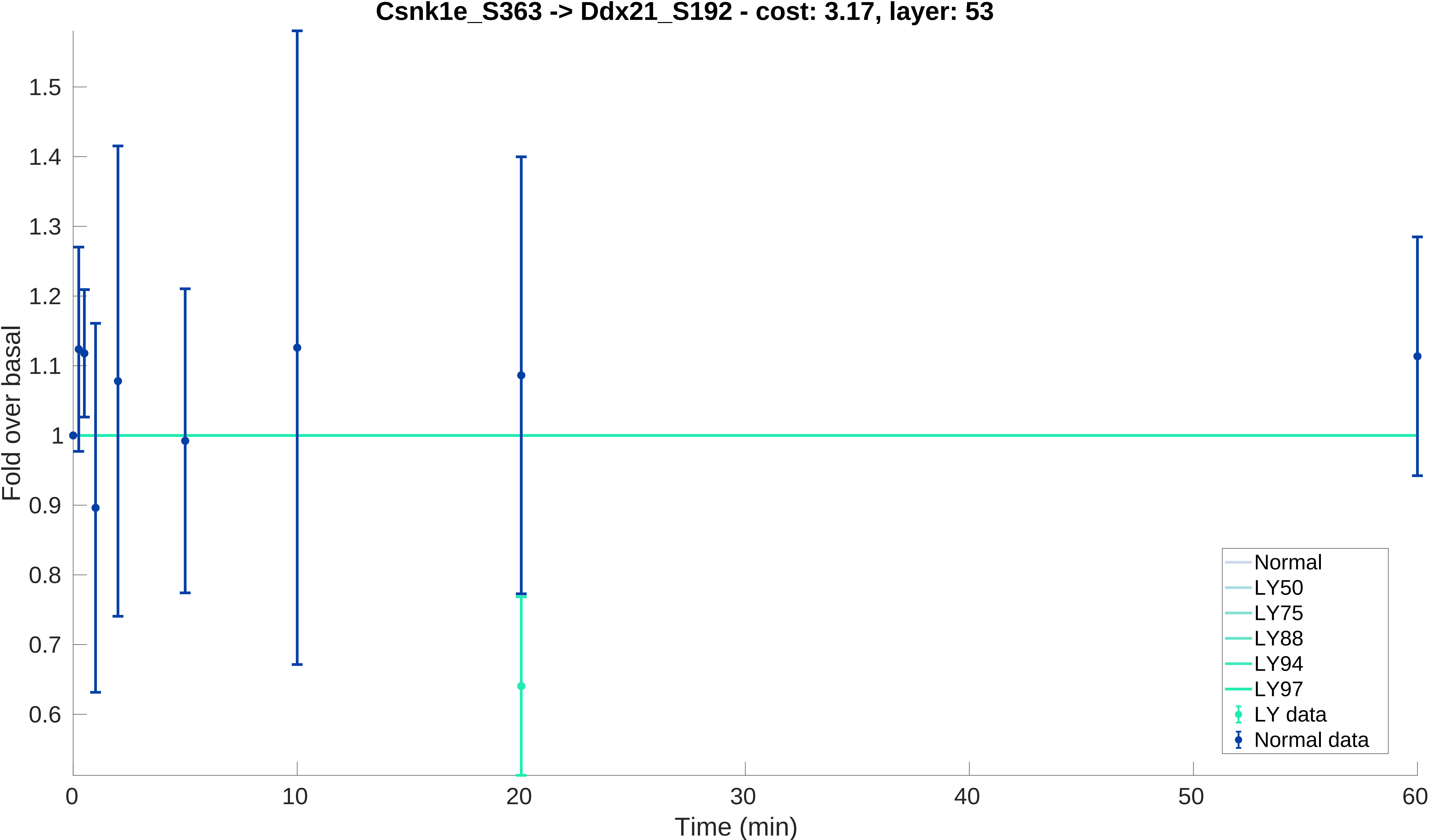
Csnk1e_S363 -> Ddx21_S192 - cost: 3.17, layer: 53
Fold over basal
Time (min)
0.6
0.7
0.8
0.9
1
1.1
1.2
1.3
1.4
1.5
0
10
20
30
40
50
60
Normal
LY50
LY75
LY88
LY94
LY97
LY data
Normal data

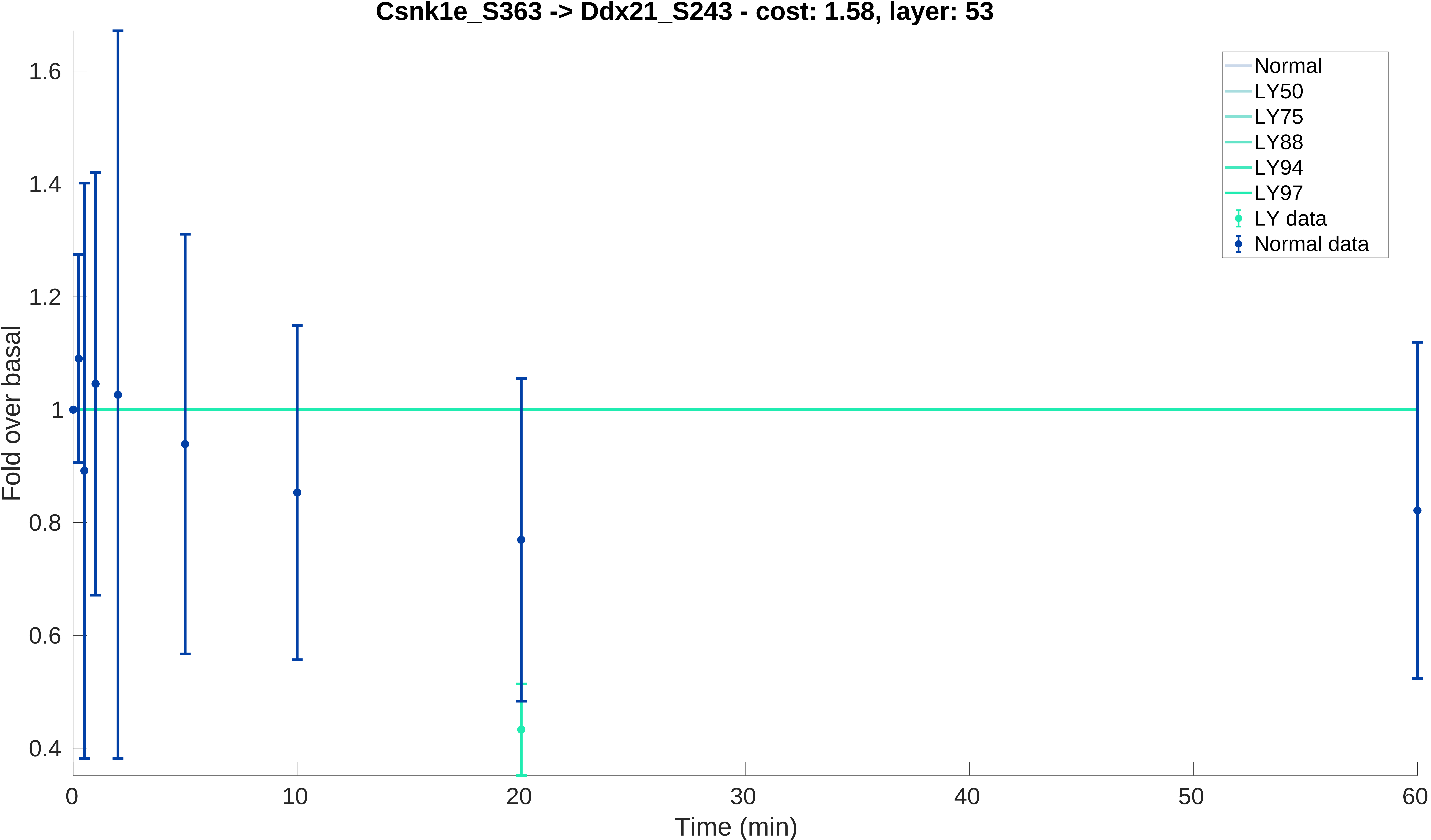

Csnk1e_S363 -> Ddx21_S243 - cost: 1.58, layer: 53
Fold over basal
Time (min)
0.4
0.6
0.8
1
1.2
1.4
1.6
0
10
20
30
40
50
60
Normal
LY50
LY75
LY88
LY94
LY97
LY data
Normal data

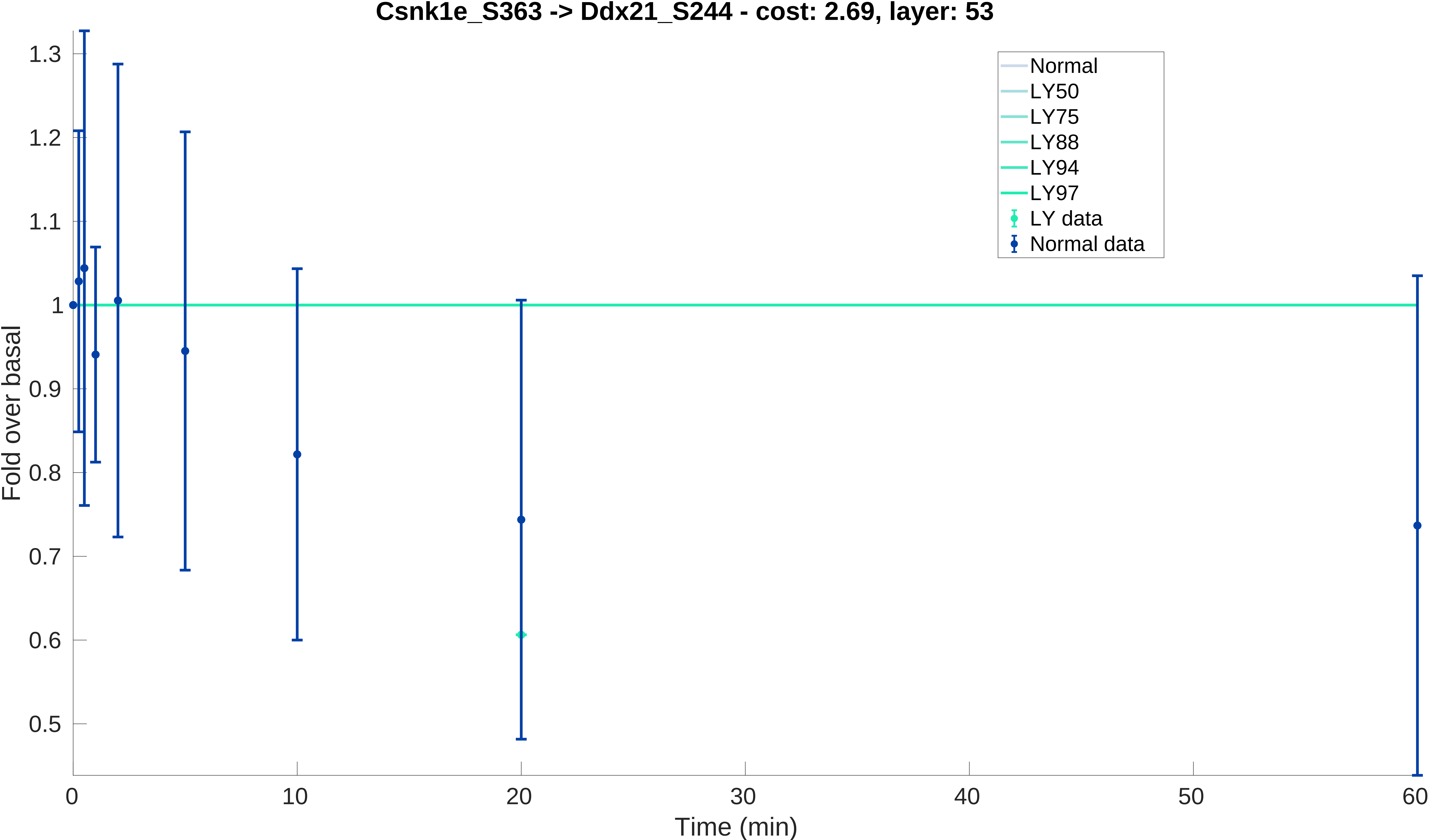
Csnk1e_S363 -> Ddx21_S244 - cost: 2.69, layer: 53
Fold over basal
Time (min)
Normal
LY50
LY75
LY88
LY94
LY97
LY data
Normal data
1.3
1.2
1.1
1
0.9
0.8
0.7
0.6
0.5
0
10
20
30
40
50
60

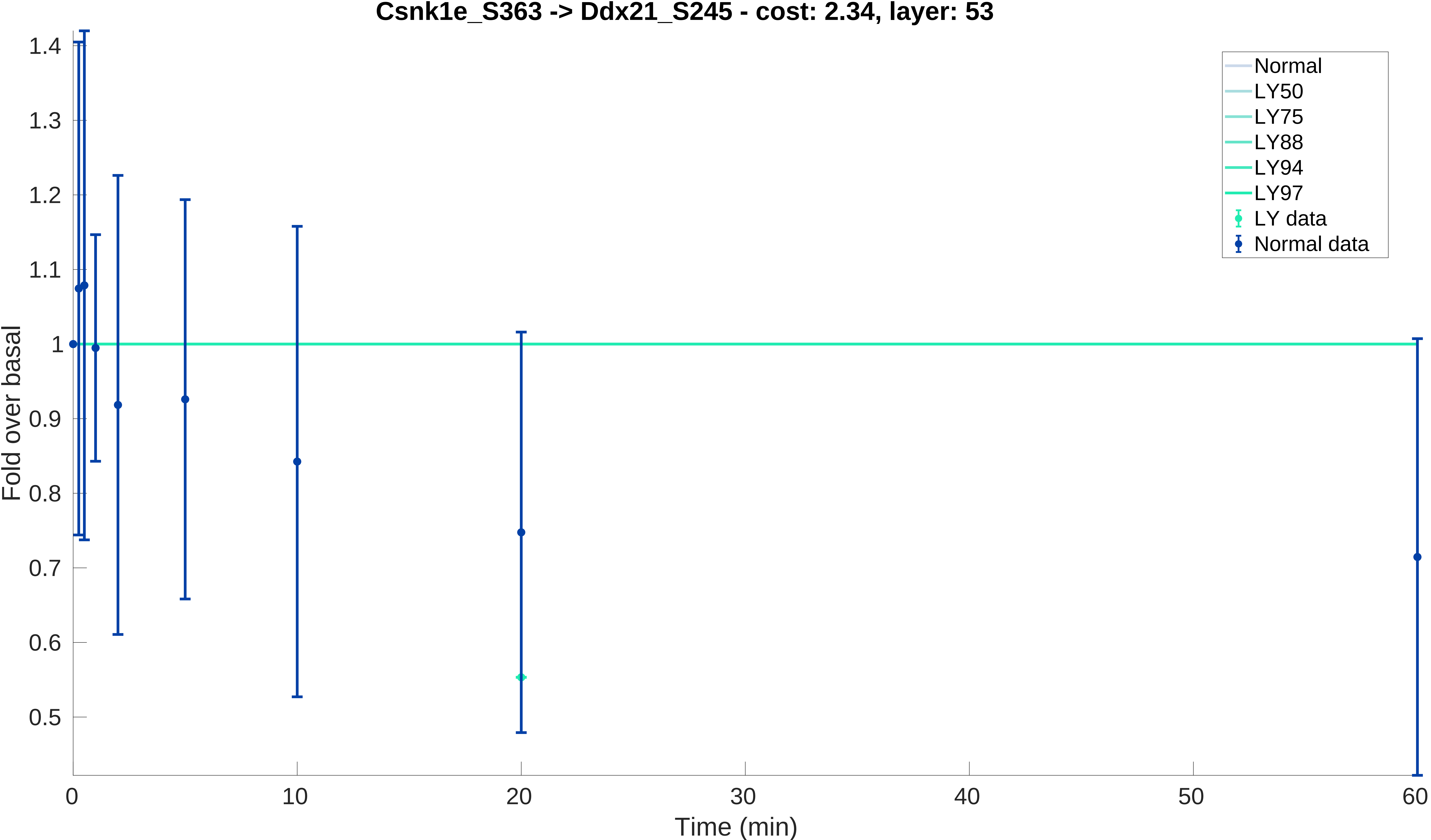

Csnk1e_S363 -> Ddx21_S245 - cost: 2.34, layer: 53
Fold over basal
Time (min)
1.4
1.3
1.2
1.1
1
0.9
0.8
0.7
0.6
0.5
0
10
20
30
40
50
60
Normal
LY50
LY75
LY88
LY94
LY97
LY data
Normal data

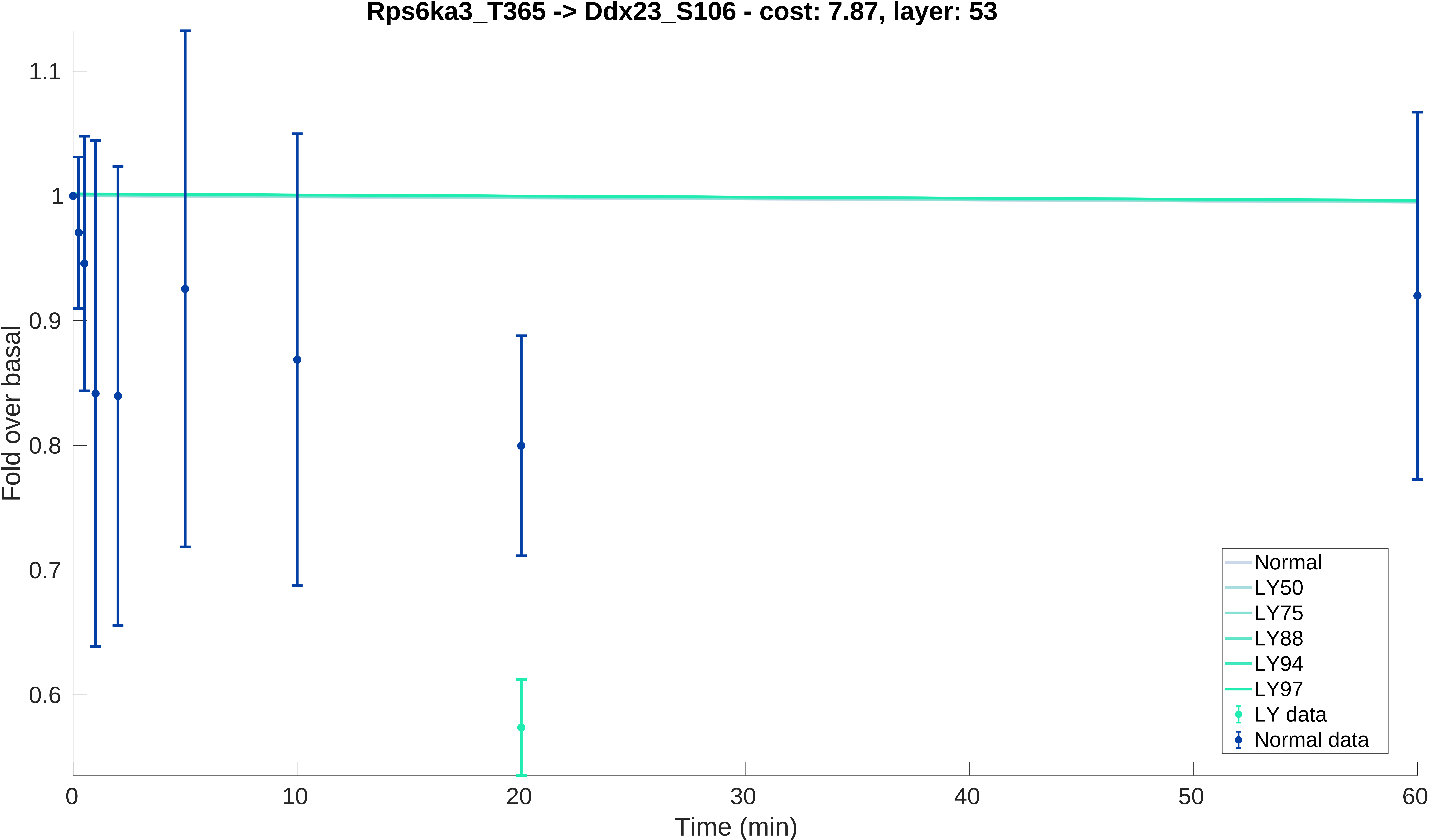

Rps6ka3_T365 -> Ddx23_S106 - cost: 7.87, layer: 53
Fold over basal
1.1
1
0.9
0.8
0.7
0.6
0
10
20
30
40
50
60
Time (min)
Normal
LY50
LY75
LY88
LY94
LY97
LY data
Normal data

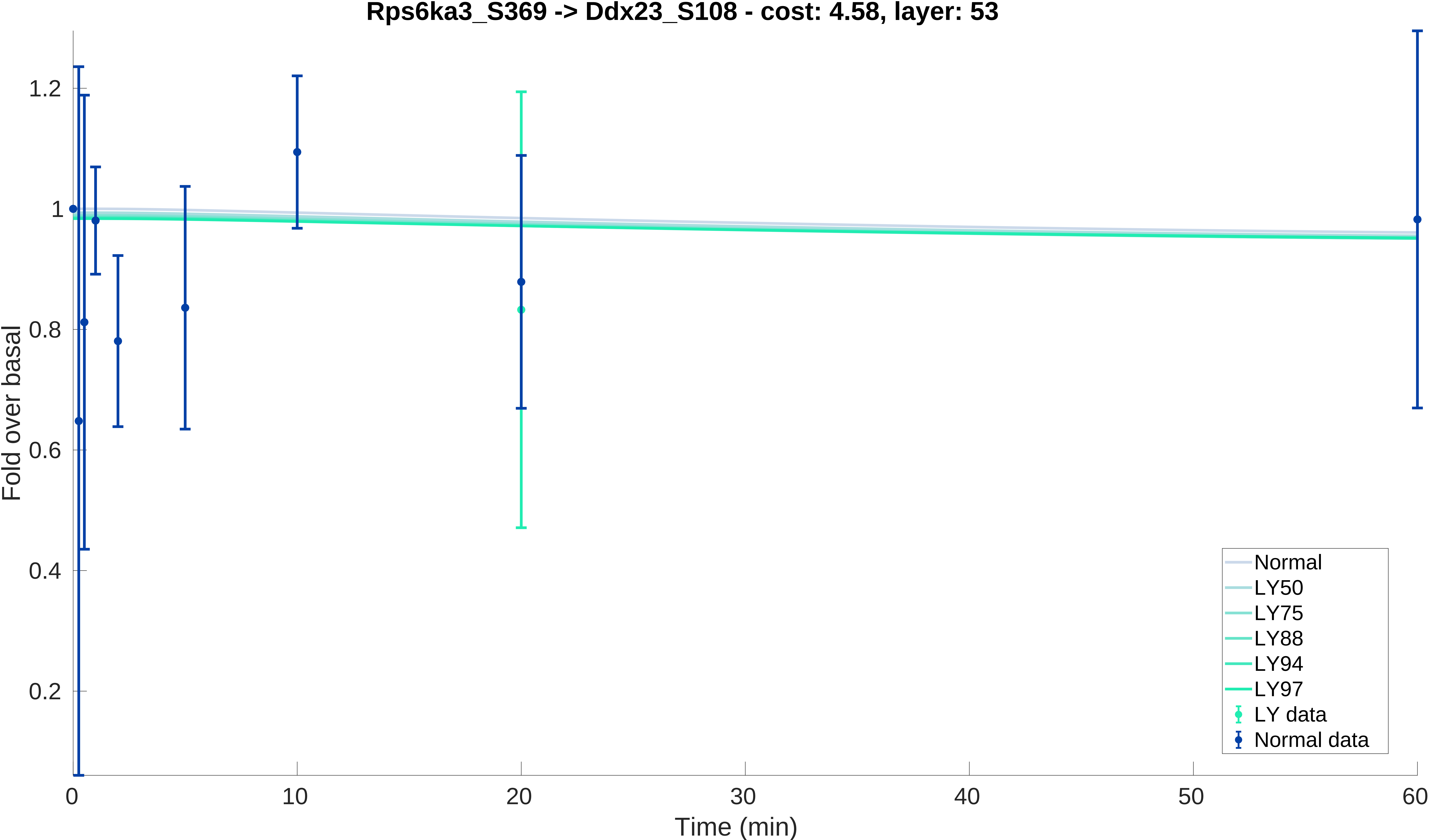

Rps6ka3_S369 -> Ddx23_S108 - cost: 4.58, layer: 53
Fold over basal
Time (min)
1.2
1
0.8
0.6
0.4
0.2
0
10
20
30
40
50
60
Normal
LY50
LY75
LY88
LY94
LY97
LY data
Normal data

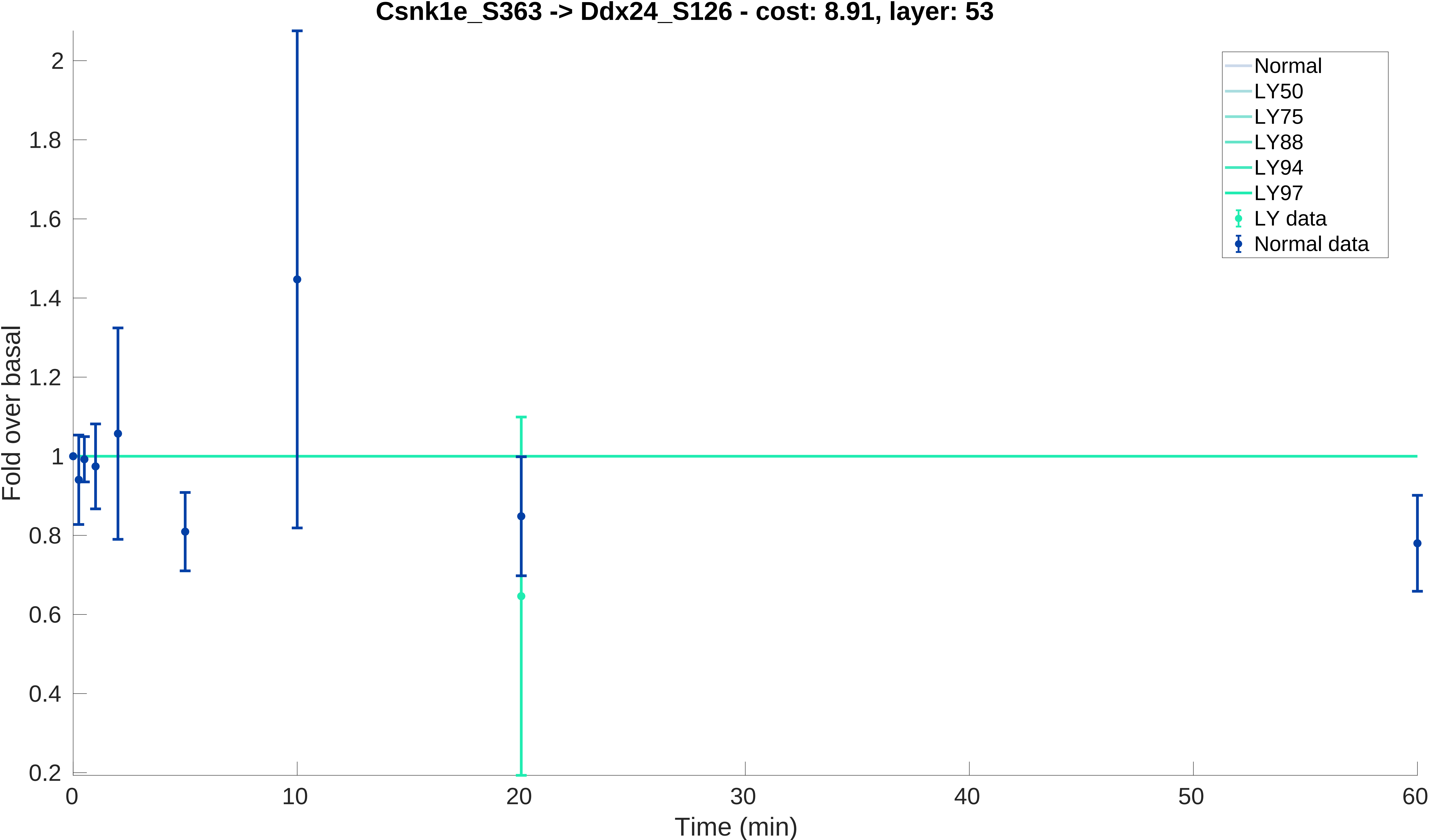

Csnk1e_S363 -> Ddx24_S126 - cost: 8.91, layer: 53
Normal
LY50
LY75
LY88
LY94
LY97
LY data
Normal data
Fold over basal
Time (min)
0
10
20
30
40
50
60
0.2
0.4
0.6
0.8
1
1.2
1.4
1.6
1.8
2

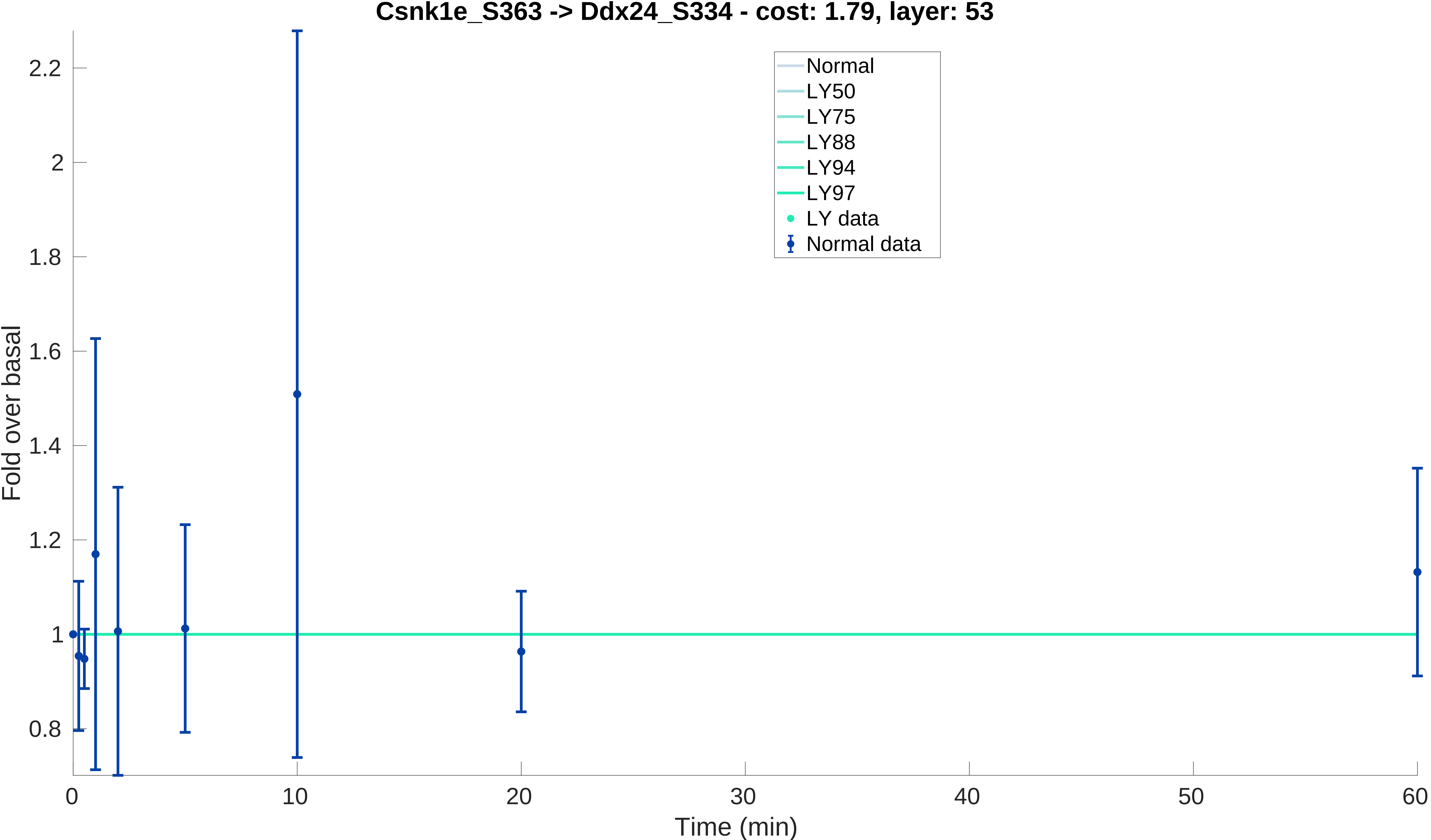

Csnk1e_S363 -> Ddx24_S334 - cost: 1.79, layer: 53
Normal
LY50
LY75
LY88
LY94
LY97
LY data
Normal data
Fold over basal
Time (min)
0.8
1
1.2
1.4
1.6
1.8
2
2.2
0
10
20
30
40
50
60

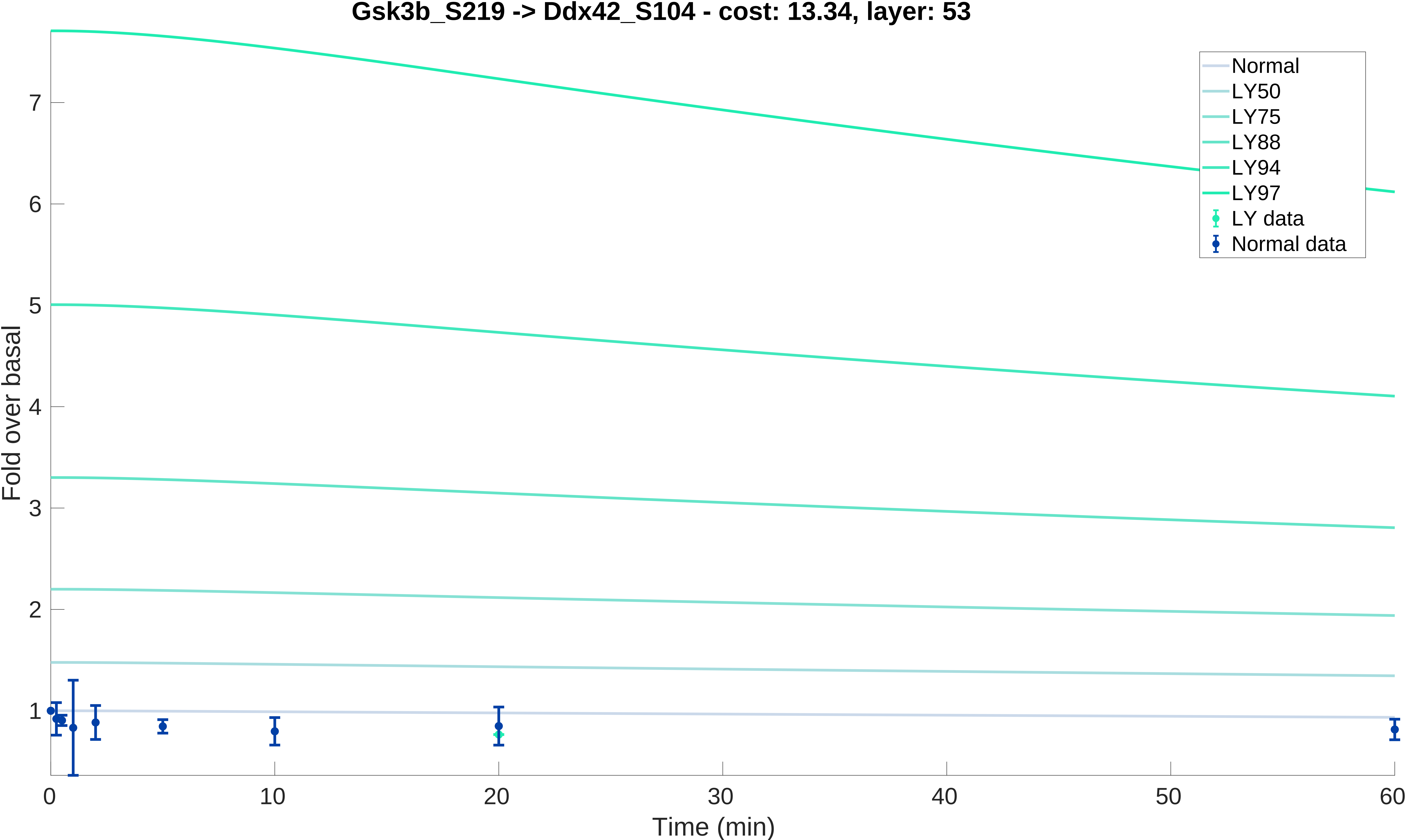
Gsk3b_S219 -> Ddx42_S104 - cost: 13.34, layer: 53
Fold over basal
Time (min)
0
10
20
30
40
50
60
1
2
3
4
5
6
7
Normal
LY50
LY75
LY88
LY94
LY97
LY data
Normal data

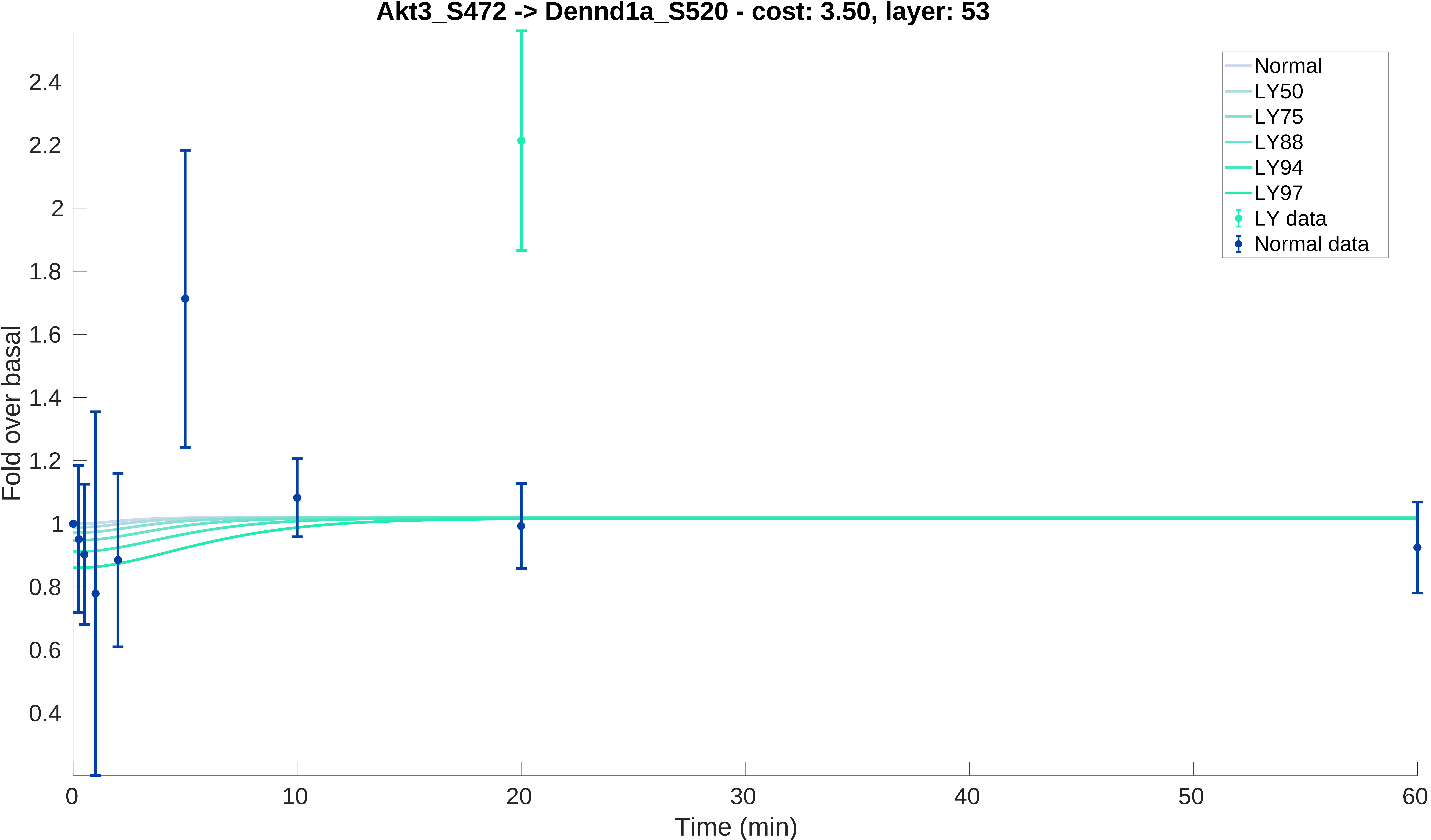

Akt3_S472 -> Dennd1a_S520 - cost: 3.50, layer: 53
Normal
LY50
LY75
LY88
LY94
LY97
LY data
Normal data
Fold over basal
Time (min)
2.4
2.2
2
1.8
1.6
1.4
1.2
1
0.8
0.6
0.4
0
10
20
30
40
50
60

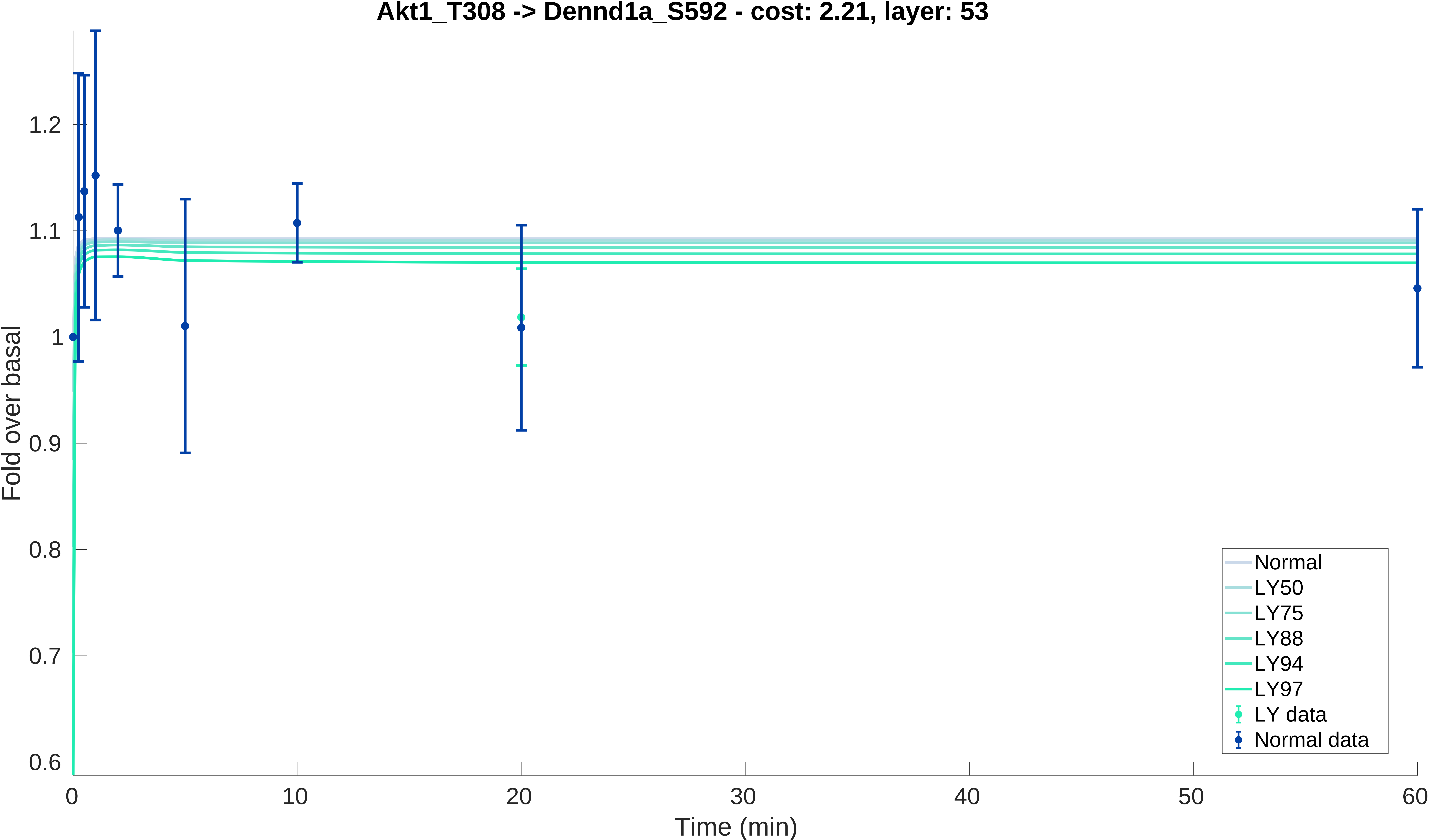
Akt1_T308 -> Dennd1a_S592 - cost: 2.21, layer: 53
Fold over basal
Time (min)
0.6
0.7
0.8
0.9
1
1.1
1.2
0
10
20
30
40
50
60
Normal
LY50
LY75
LY88
LY94
LY97
LY data
Normal data

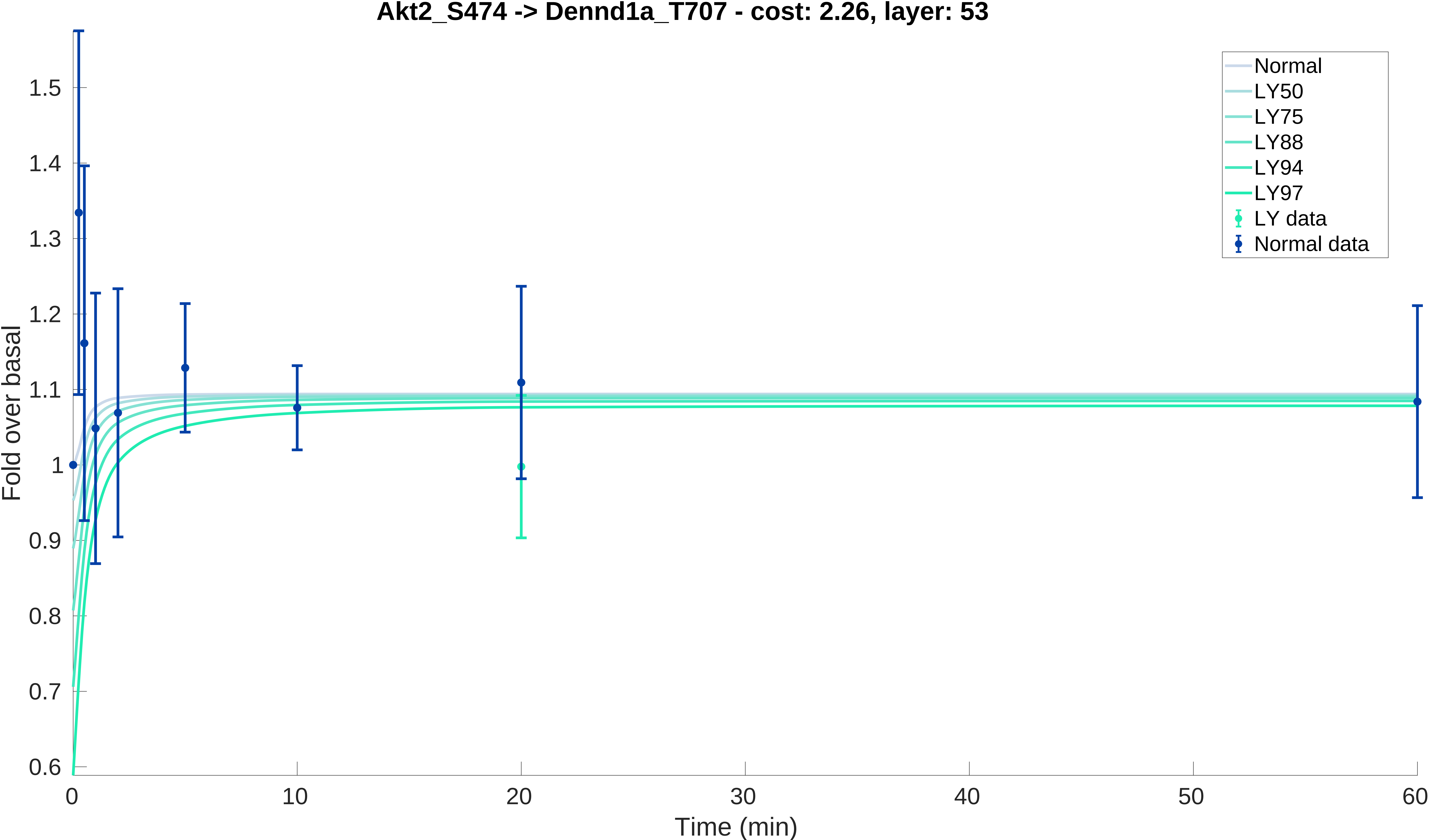

Akt2_S474 -> Dennd1a_T707 - cost: 2.26, layer: 53
Fold over basal
Time (min)
0.6
0.7
0.8
0.9
1
1.1
1.2
1.3
1.4
1.5
0
10
20
30
40
50
60
Normal
LY50
LY75
LY88
LY94
LY97
LY data
Normal data

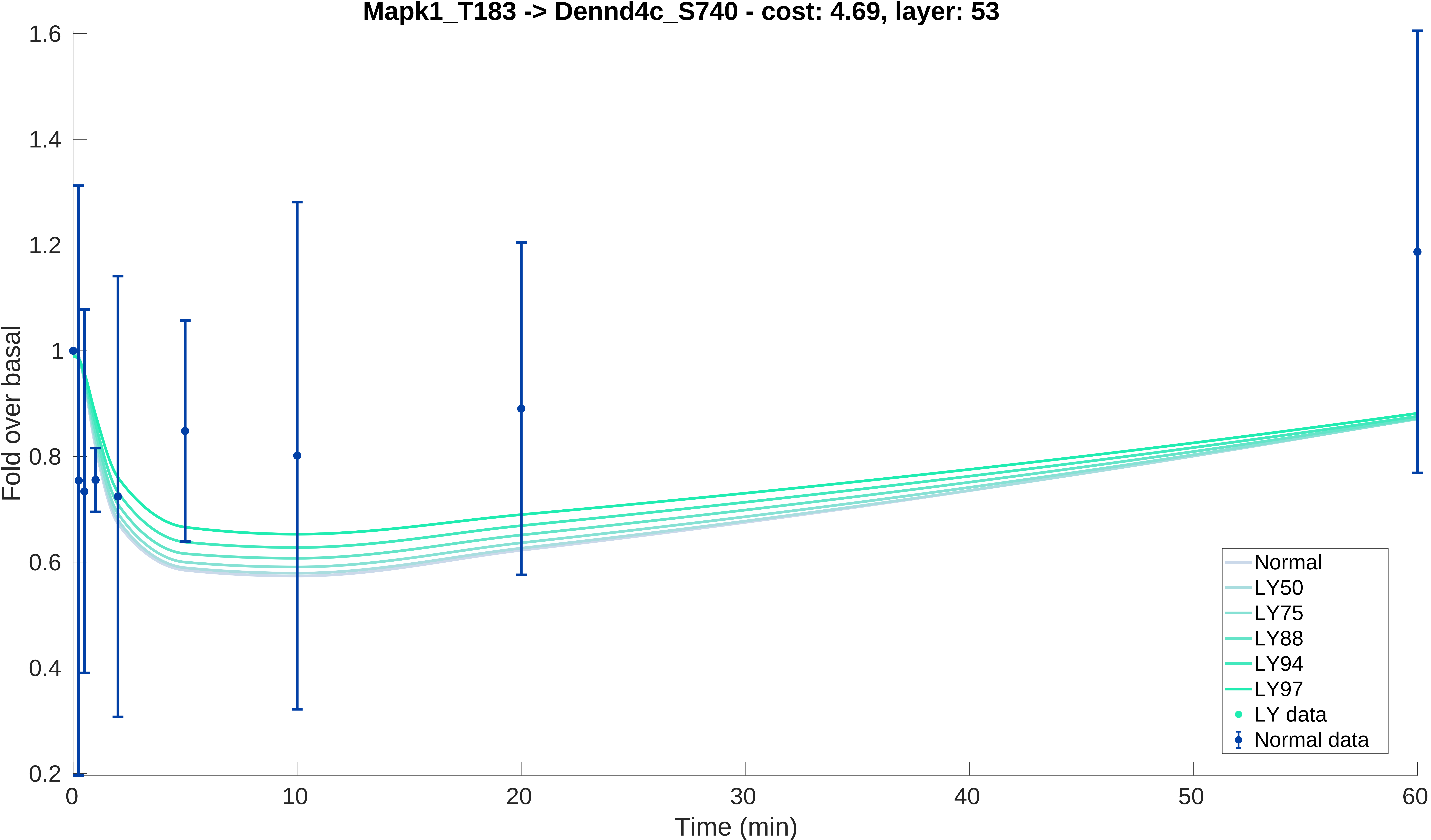

Mapk1_T183 -> Dennd4c_S740 - cost: 4.69, layer: 53
Fold over basal
Time (min)
1.6
1.4
1.2
1
0.8
0.6
0.4
0.2
0
10
20
30
40
50
60
Normal
LY50
LY75
LY88
LY94
LY97
LY data
Normal data

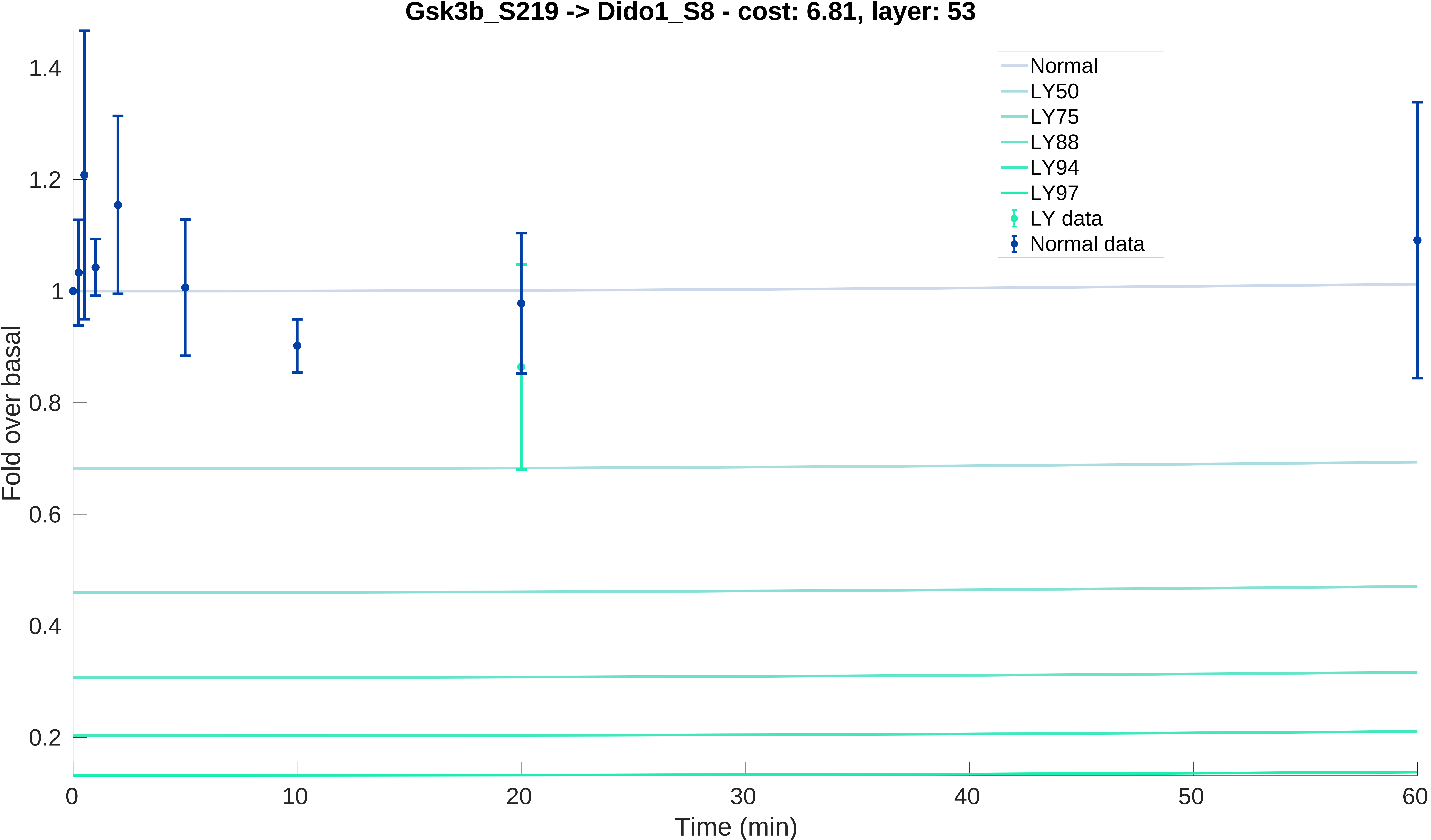
Gsk3b_S219 -> Dido1_S8 - cost: 6.81, layer: 53
Fold over basal
Time (min)
0
10
20
30
40
50
60
0.2
0.4
0.6
0.8
1
1.2
1.4
Normal
LY50
LY75
LY88
LY94
LY97
LY data
Normal data

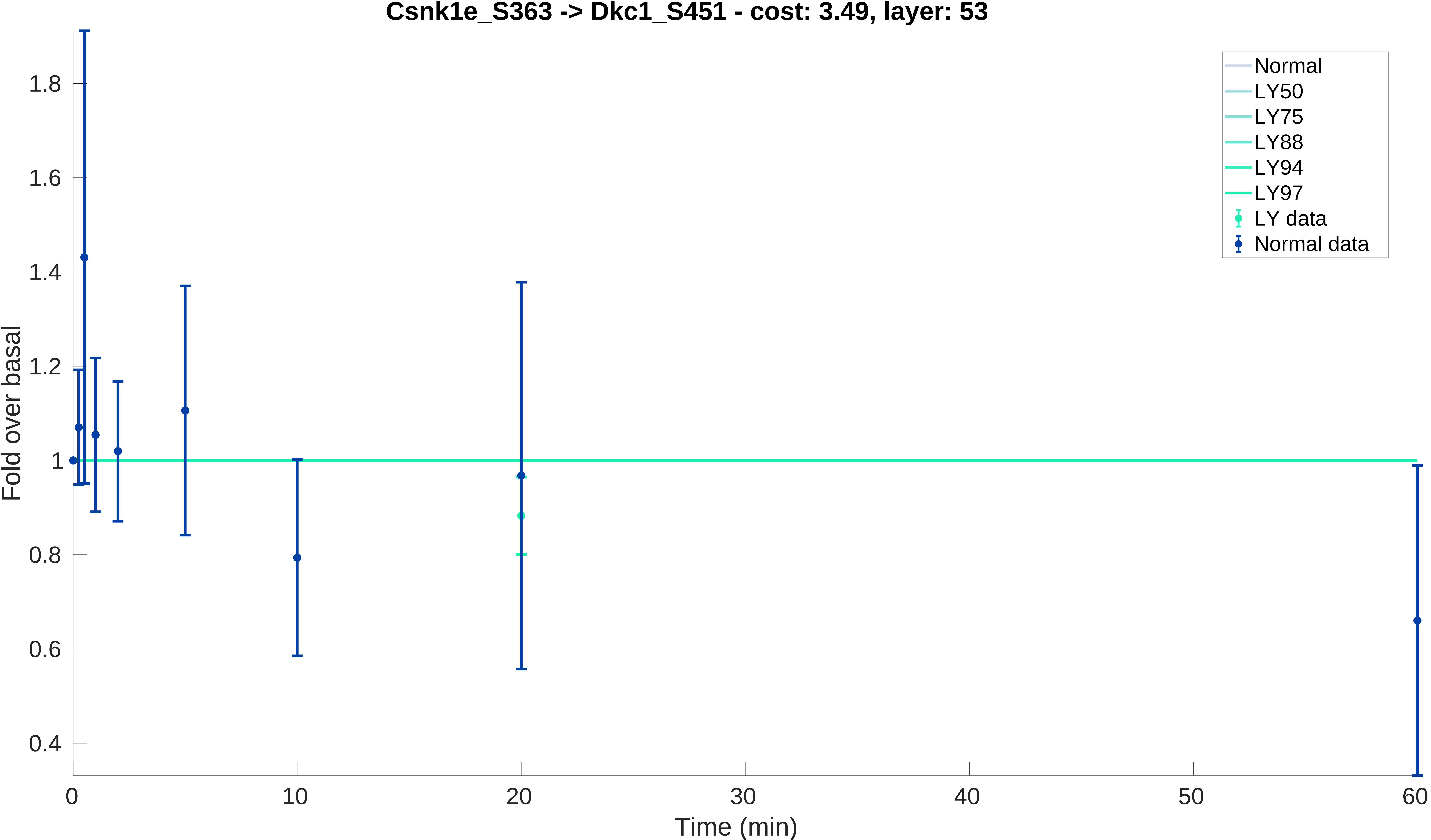

Csnk1e_S363 -> Dkc1_S451 - cost: 3.49, layer: 53
Normal
LY50
LY75
LY88
LY94
LY97
LY data
Normal data
Fold over basal
1.8
1.6
1.4
1.2
1
0.8
0.6
0.4
0
10
20
30
40
50
60
Time (min)

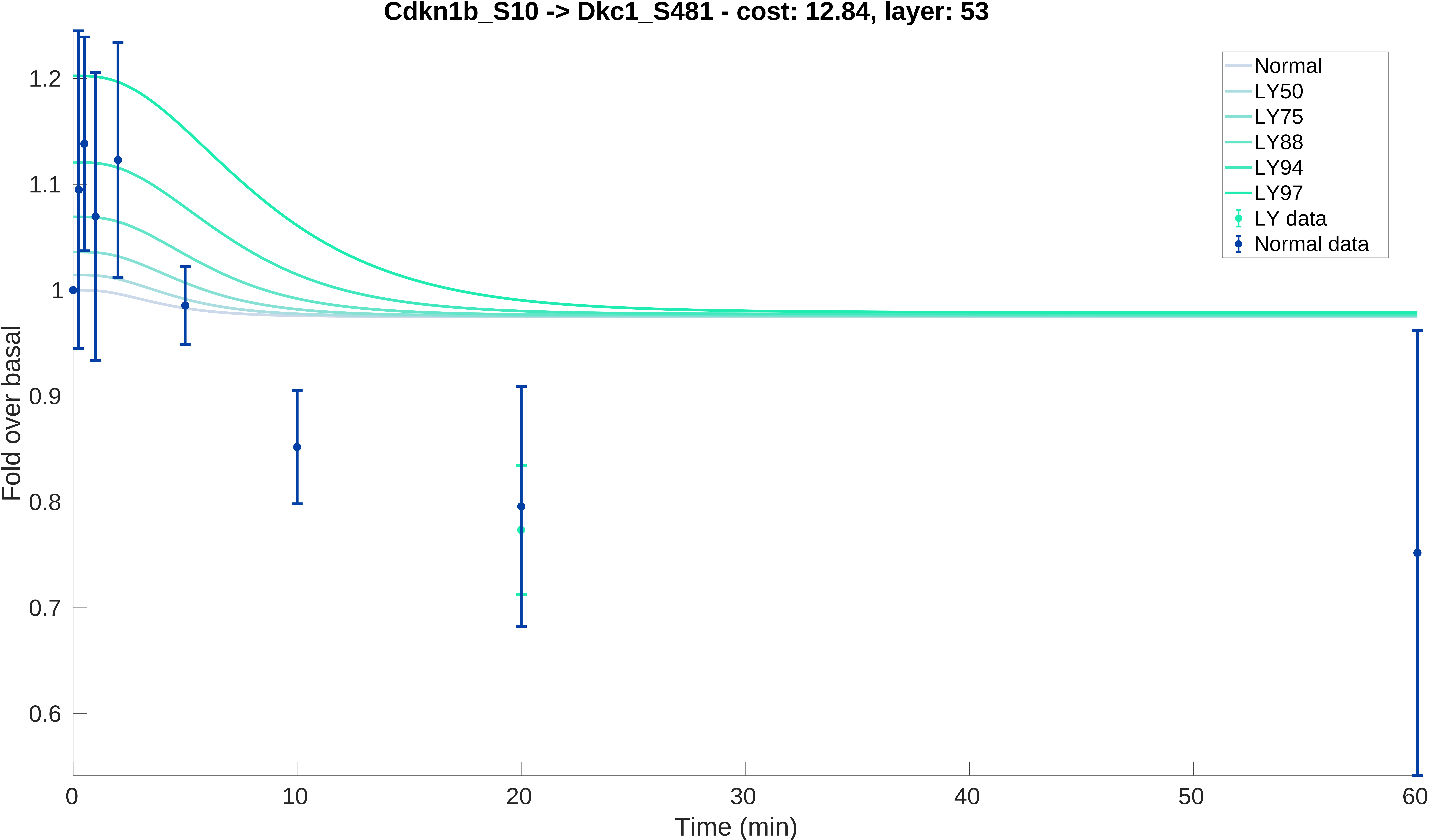

Cdkn1b_S10 -> Dkc1_S481 - cost: 12.84, layer: 53
Normal
LY50
LY75
LY88
LY94
LY97
LY data
Normal data
Fold over basal
Time (min)
1.2
1.1
1
0.9
0.8
0.7
0.6
0
10
20
30
40
50
60

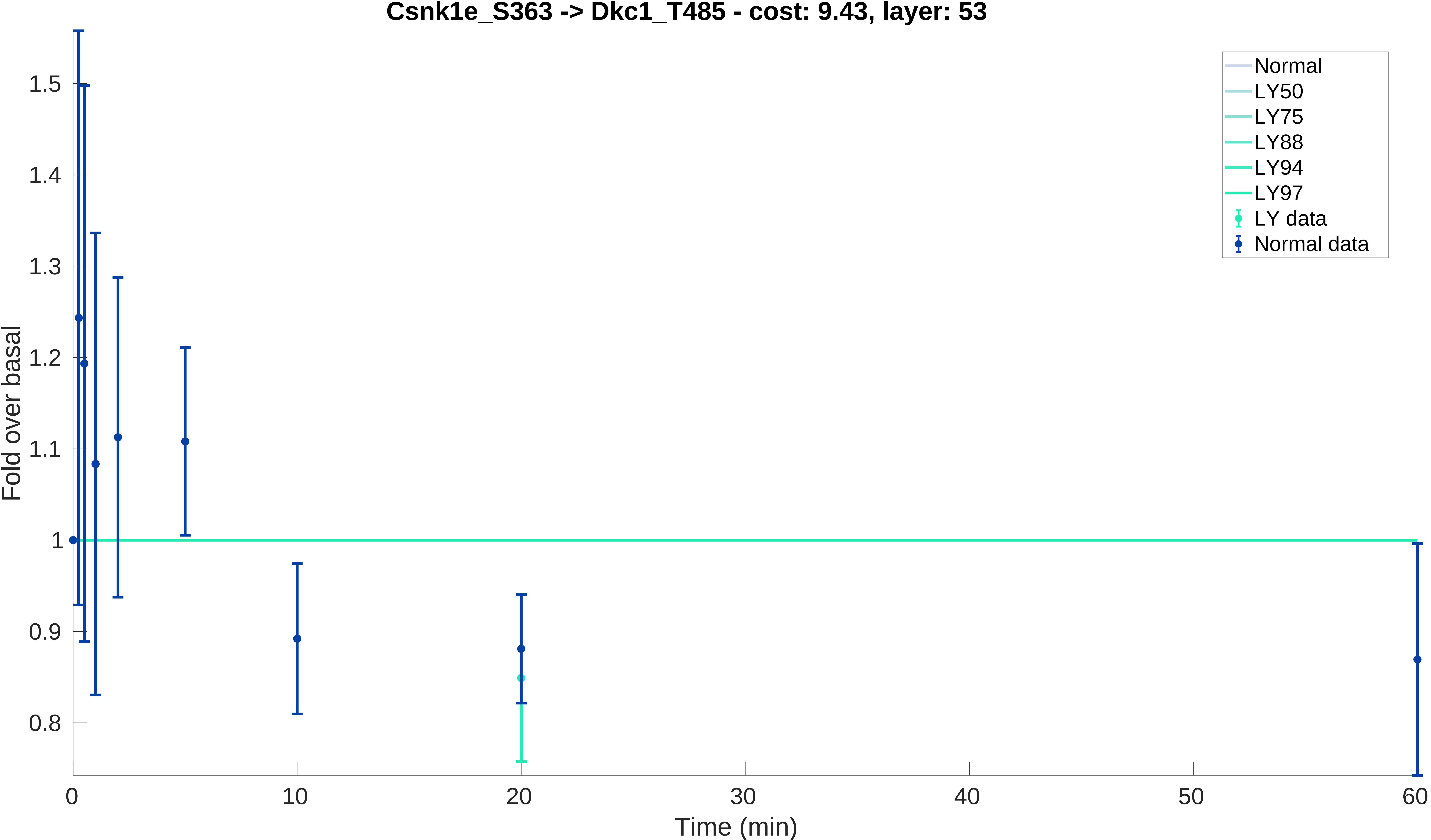

Csnk1e_S363 -> Dkc1_T485 - cost: 9.43, layer: 53
Fold over basal
Time (min)
0
10
20
30
40
50
60
0.8
0.9
1
1.1
1.2
1.3
1.4
1.5
Normal
LY50
LY75
LY88
LY94
LY97
LY data
Normal data

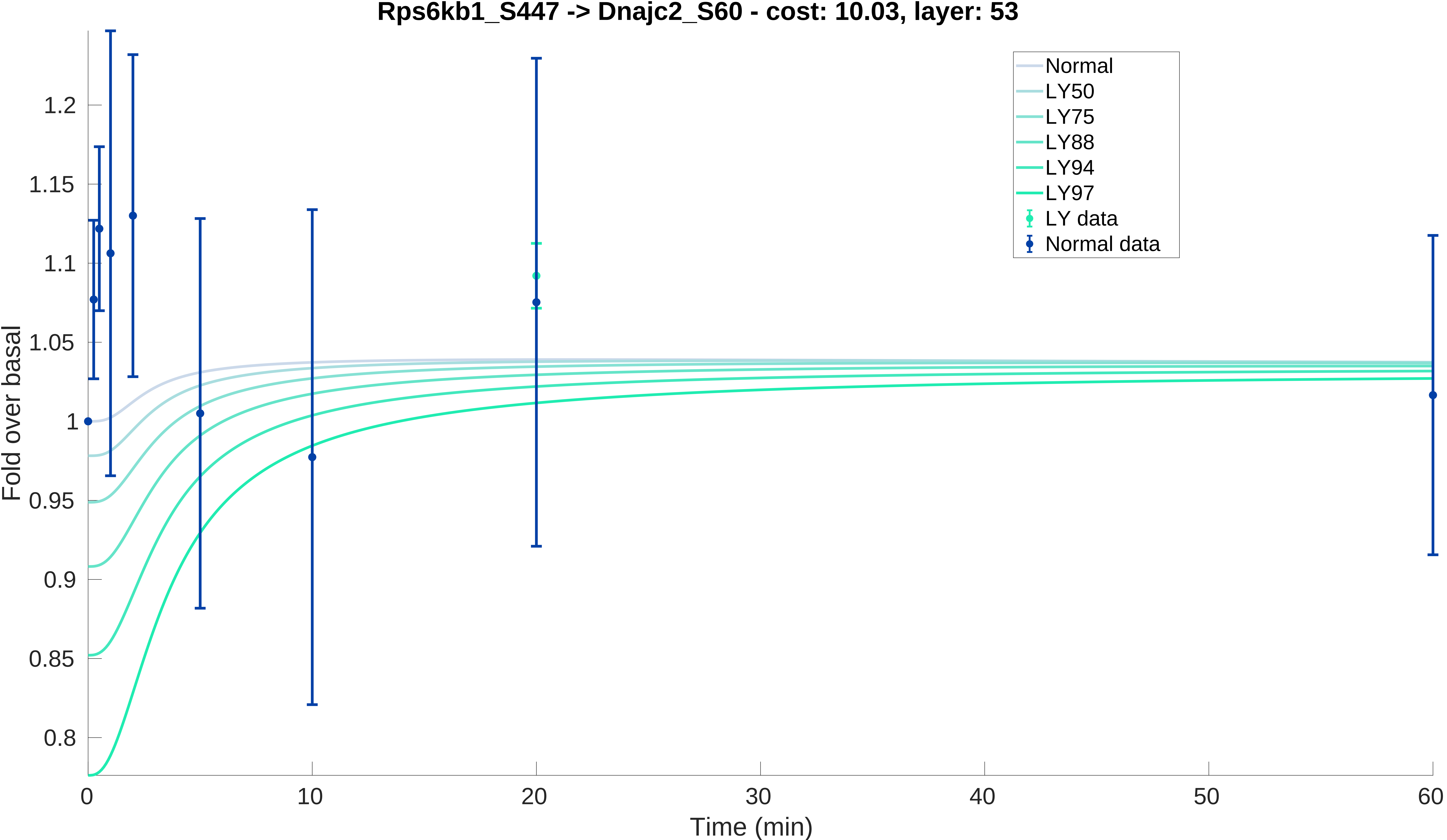

Rps6kb1_S447 -> Dnajc2_S60 - cost: 10.03, layer: 53
Fold over basal
Time (min)
Normal
LY50
LY75
LY88
LY94
LY97
LY data
Normal data
1.2
1.15
1.1
1.05
1
0.95
0.9
0.85
0.8
0
10
20
30
40
50
60

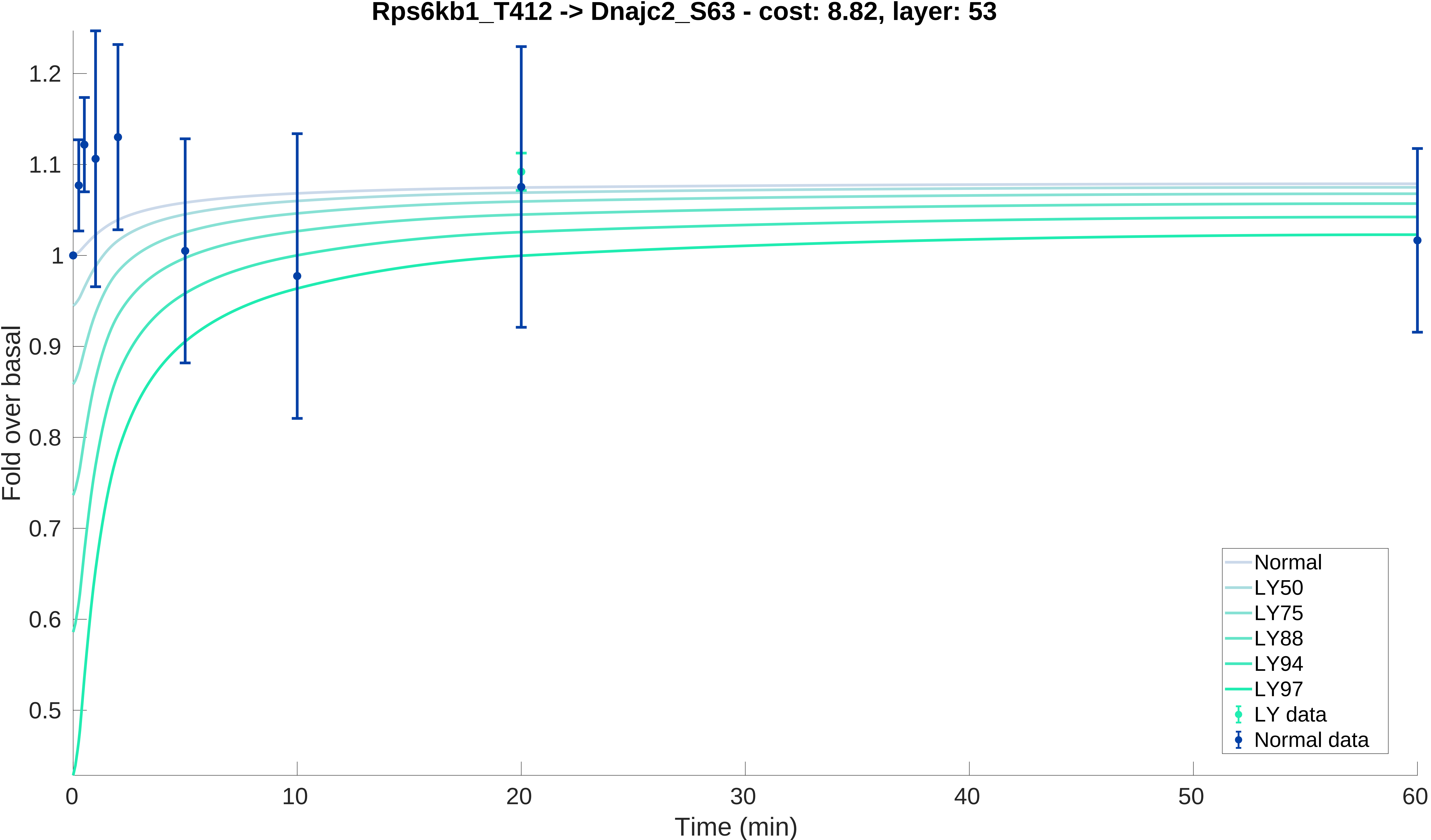

Rps6kb1_T412 -> Dnajc2_S63 - cost: 8.82, layer: 53
Fold over basal
Time (min)
1.2
1.1
1
0.9
0.8
0.7
0.6
0.5
0
10
20
30
40
50
60
Normal
LY50
LY75
LY88
LY94
LY97
LY data
Normal data

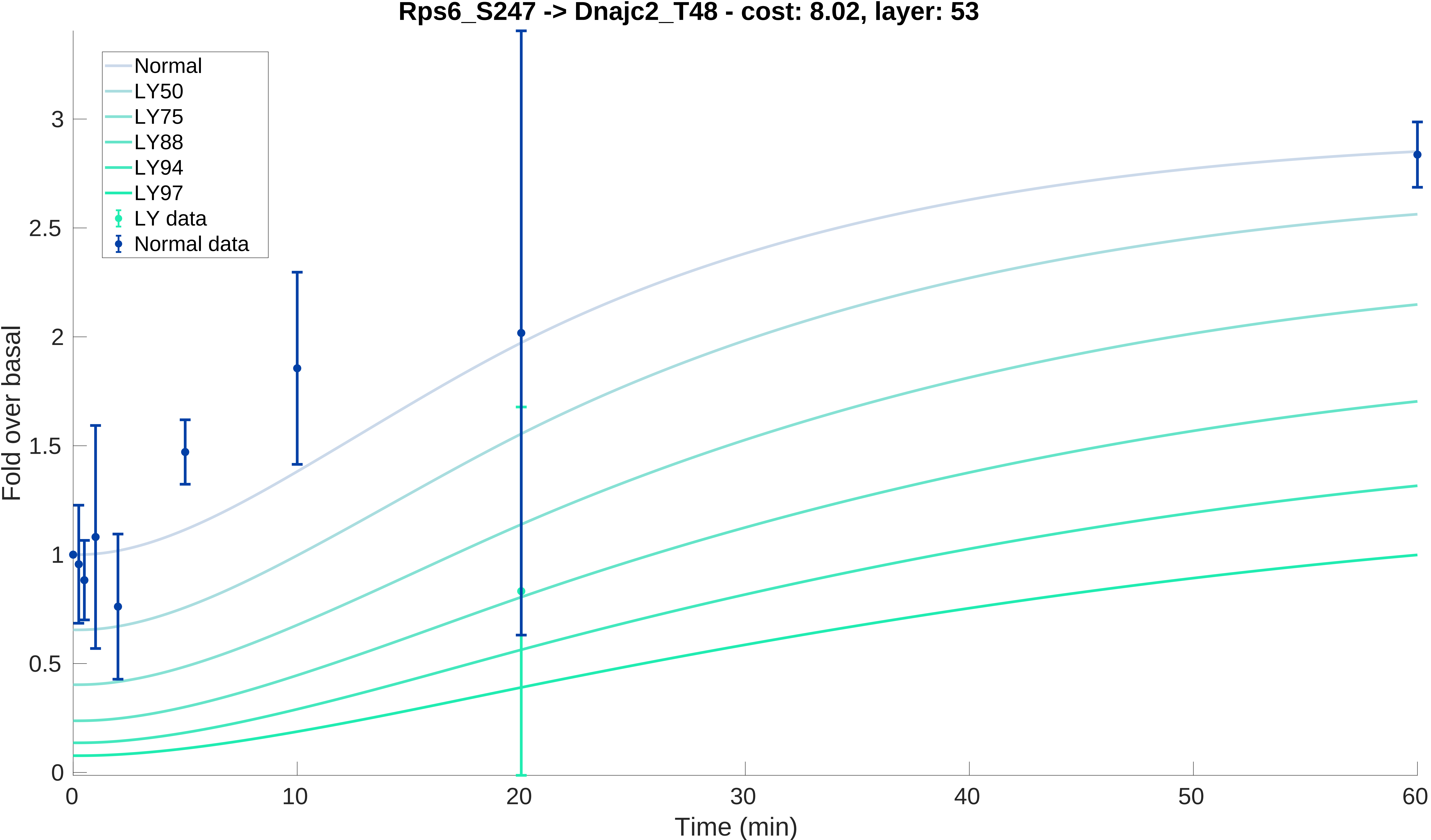

Rps6_S247 -> Dnajc2_T48 - cost: 8.02, layer: 53
Normal
LY50
LY75
LY88
LY94
LY97
LY data
Normal data
Fold over basal
Time (min)
0
0.5
1
1.5
2
2.5
3
0
10
20
30
40
50
60

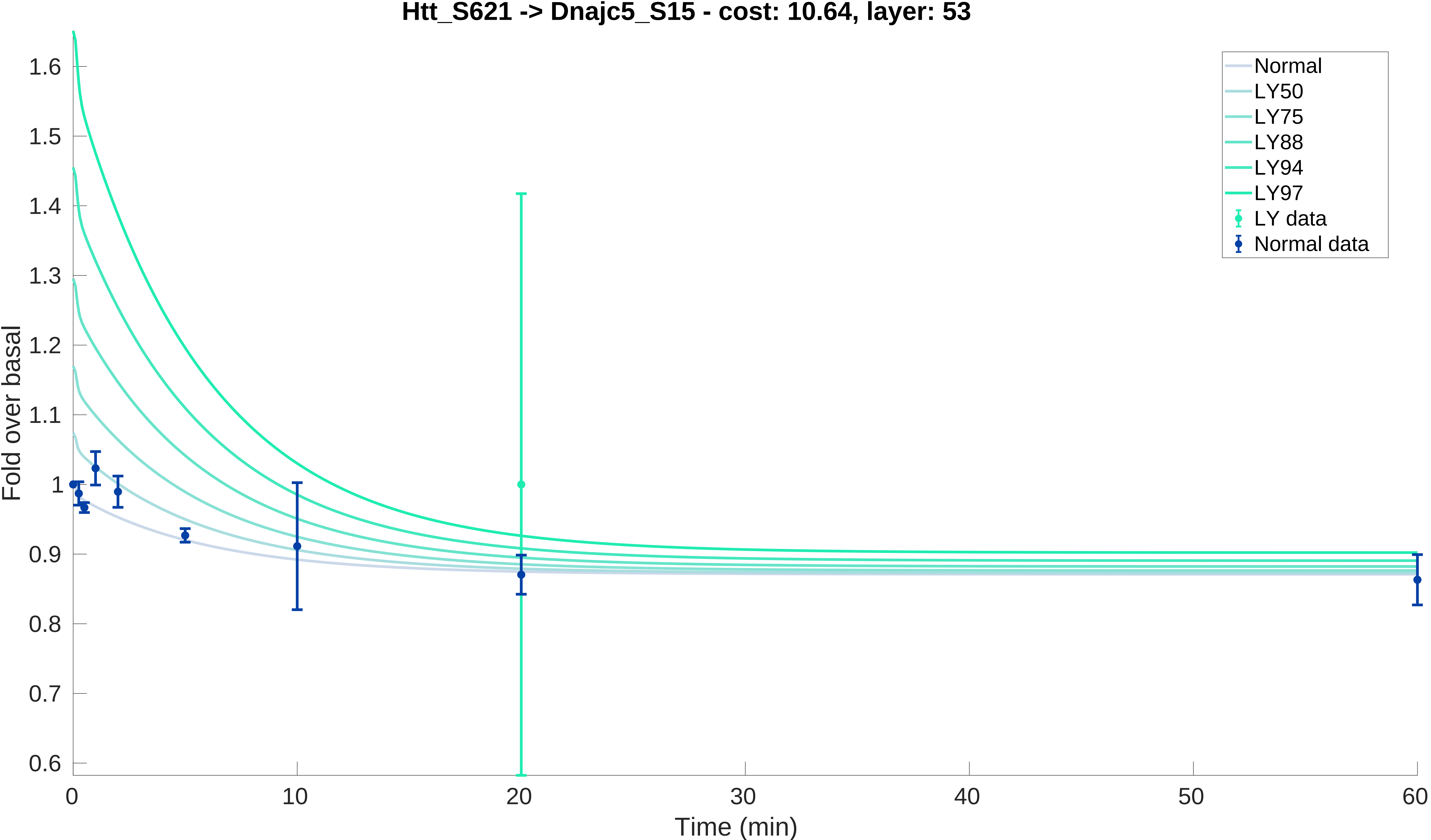

Htt_S621 -> Dnajc5_S15 - cost: 10.64, layer: 53
Fold over basal
Time (min)
0.6
0.7
0.8
0.9
1
1.1
1.2
1.3
1.4
1.5
1.6
0
10
20
30
40
50
60
Normal
LY50
LY75
LY88
LY94
LY97
LY data
Normal data

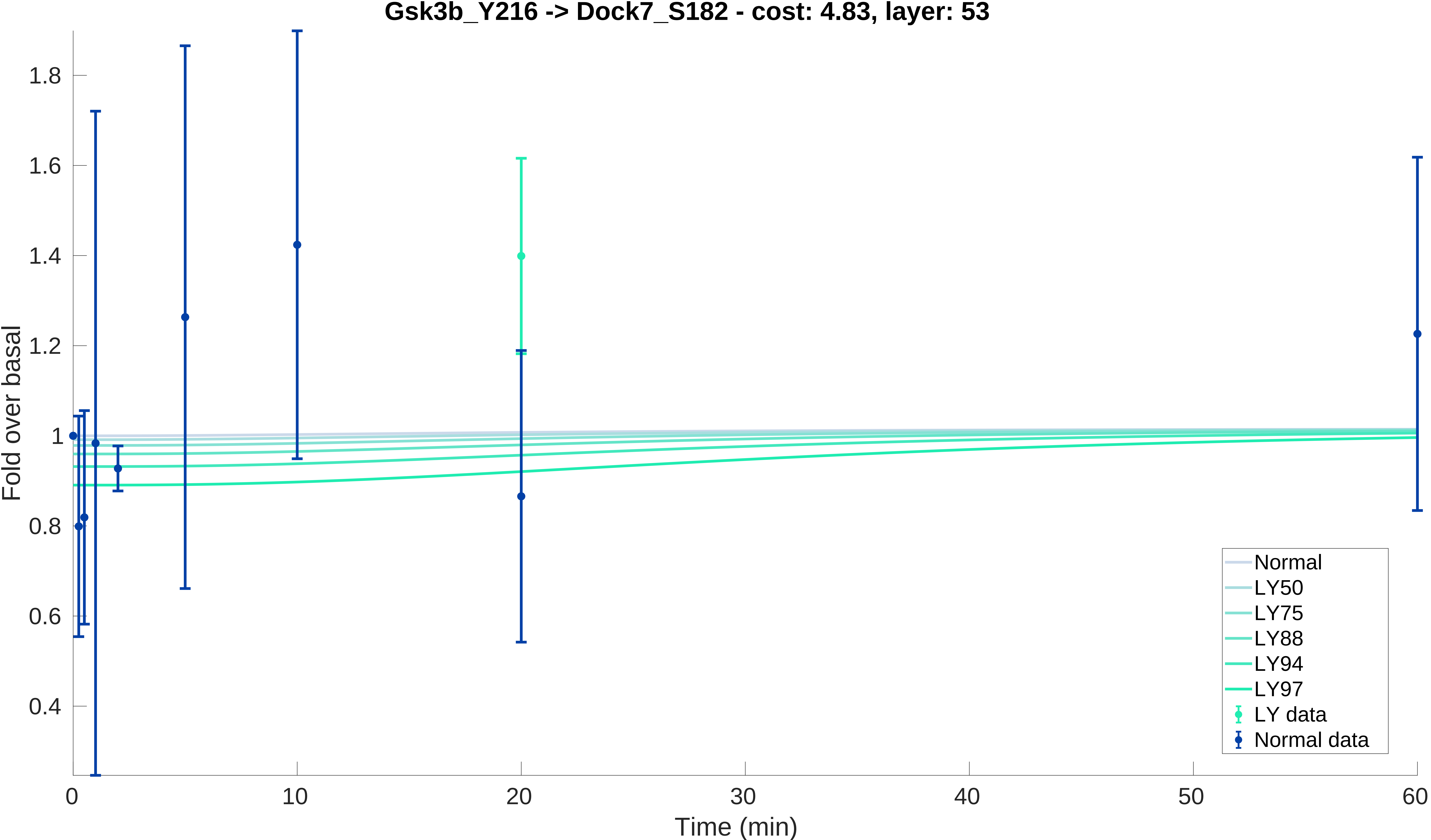

Gsk3b_Y216 -> Dock7_S182 - cost: 4.83, layer: 53
Fold over basal
Time (min)
1.8
1.6
1.4
1.2
1
0.8
0.6
0.4
0
10
20
30
40
50
60
Normal
LY50
LY75
LY88
LY94
LY97
LY data
Normal data

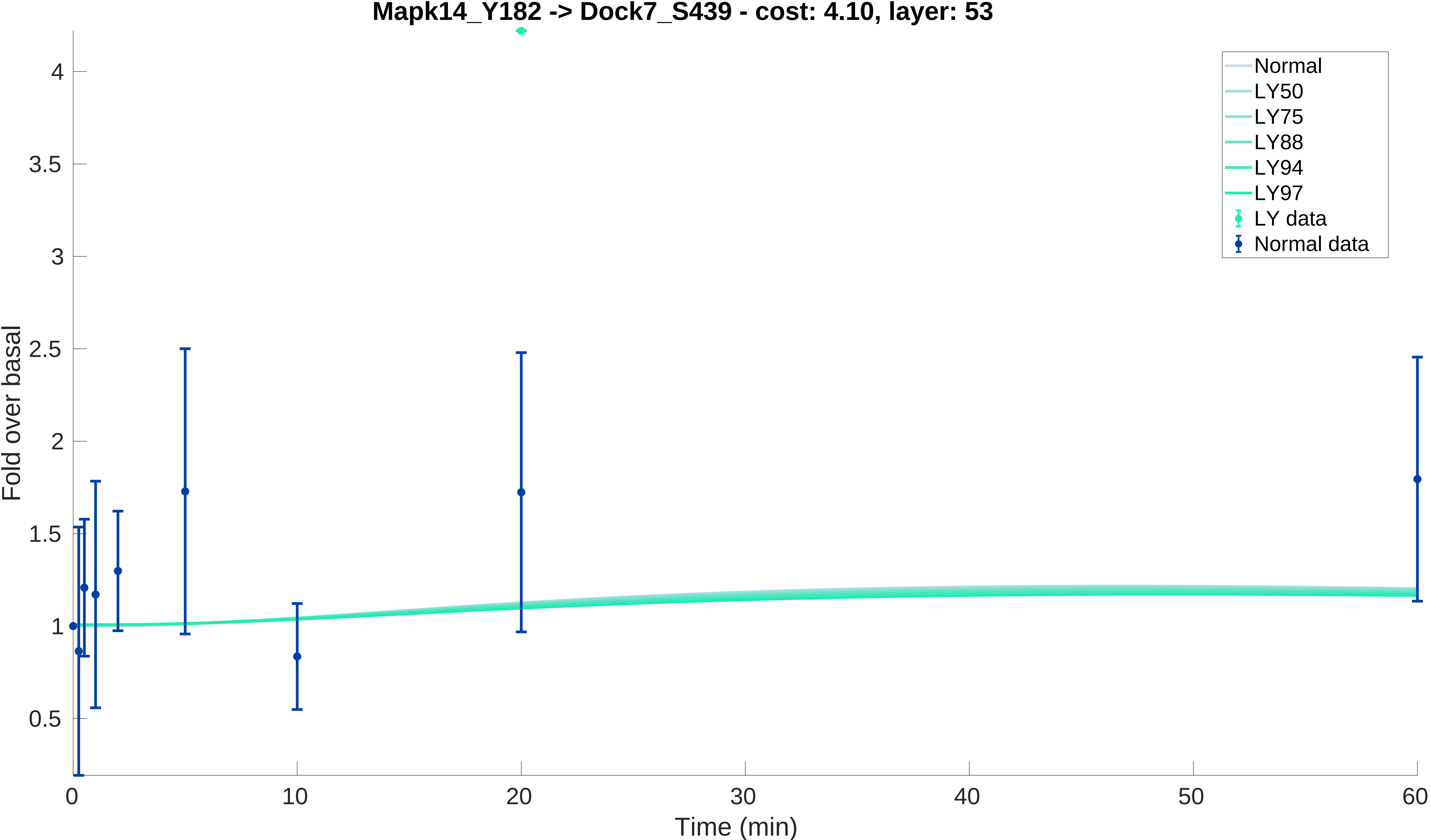
Mapk14_Y182 -> Dock7_S439 - cost: 4.10, layer: 53
Fold over basal
Time (min)
0.5
1
1.5
2
2.5
3
3.5
4
0
10
20
30
40
50
60
Normal
LY50
LY75
LY88
LY94
LY97
LY data
Normal data

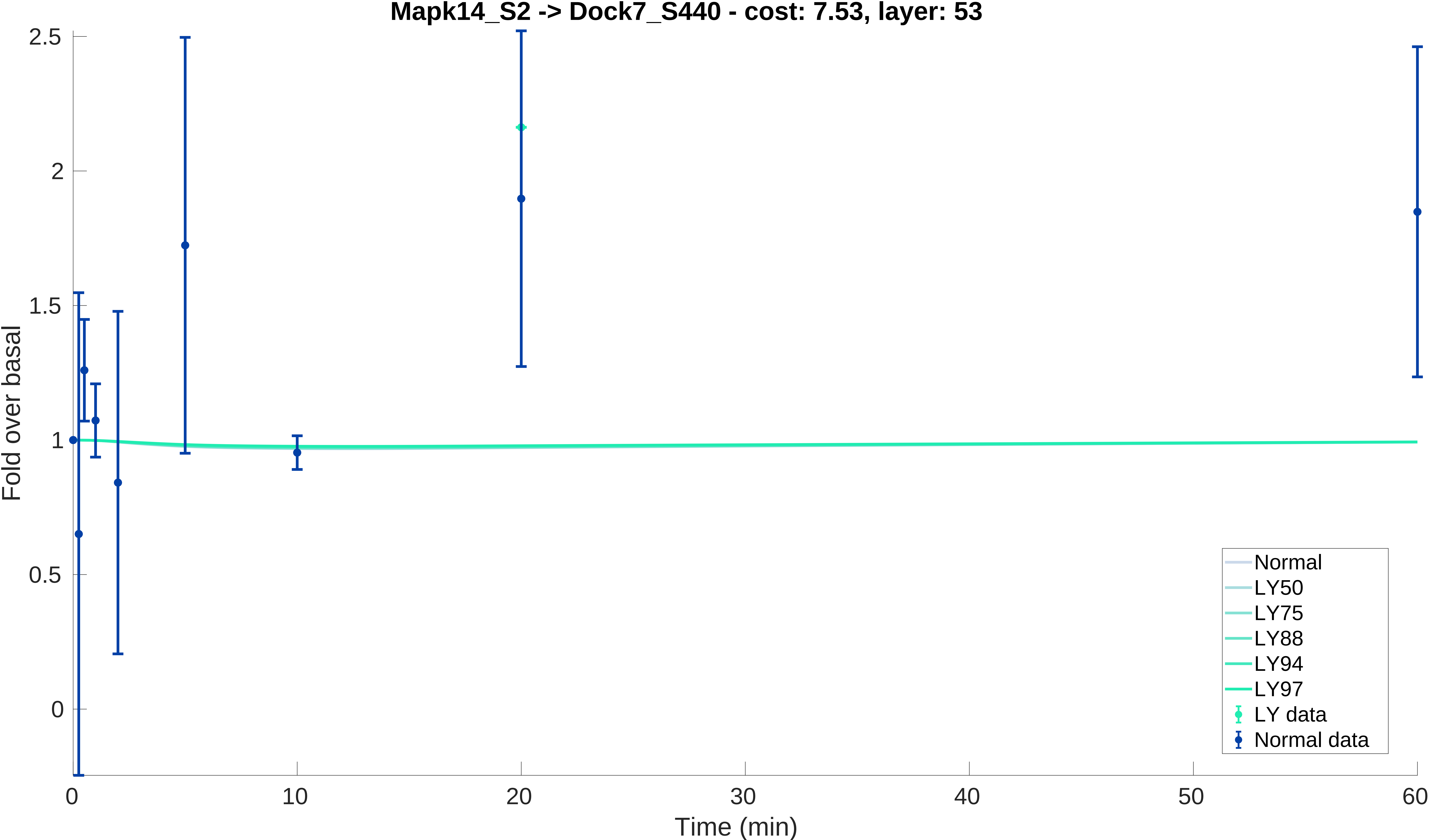
Mapk14_S2 -> Dock7_S440 - cost: 7.53, layer: 53
2.5
2
1.5
1
0.5
0
Fold over basal
0
10
20
30
40
50
60
Time (min)
Normal
LY50
LY75
LY88
LY94
LY97
LY data
Normal data

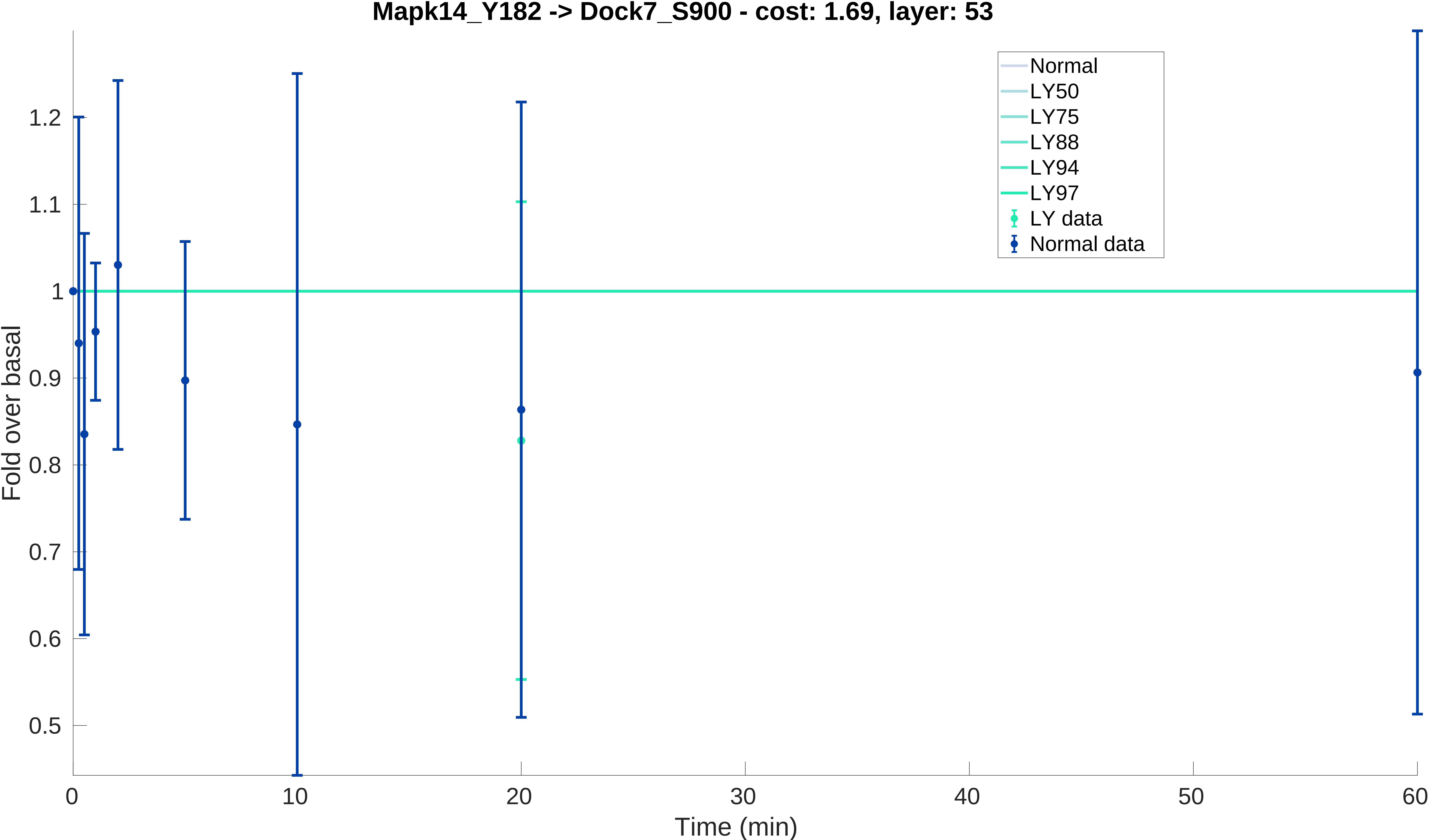

Mapk14_Y182 -> Dock7_S900 - cost: 1.69, layer: 53
Normal
LY50
LY75
LY88
LY94
LY97
LY data
Normal data
Fold over basal
1.2
1.1
1
0.9
0.8
0.7
0.6
0.5
0
10
20
30
40
50
60
Time (min)

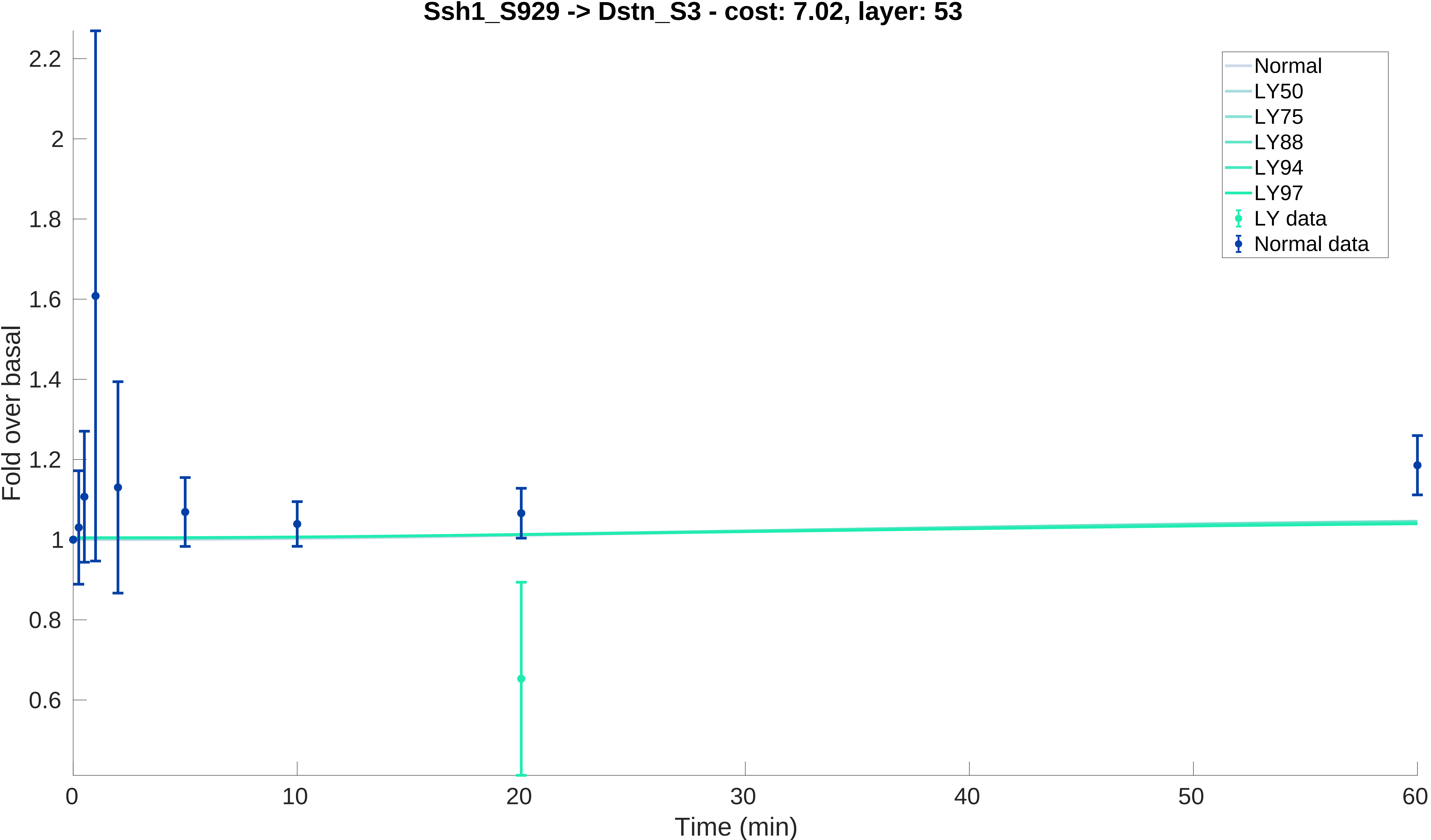

Ssh1_S929 -> Dstn_S3 - cost: 7.02, layer: 53
Normal
LY50
LY75
LY88
LY94
LY97
LY data
Normal data
Fold over basal
2.2
2
1.8
1.6
1.4
1.2
1
0.8
0.6
0
10
20
30
40
50
60
Time (min)

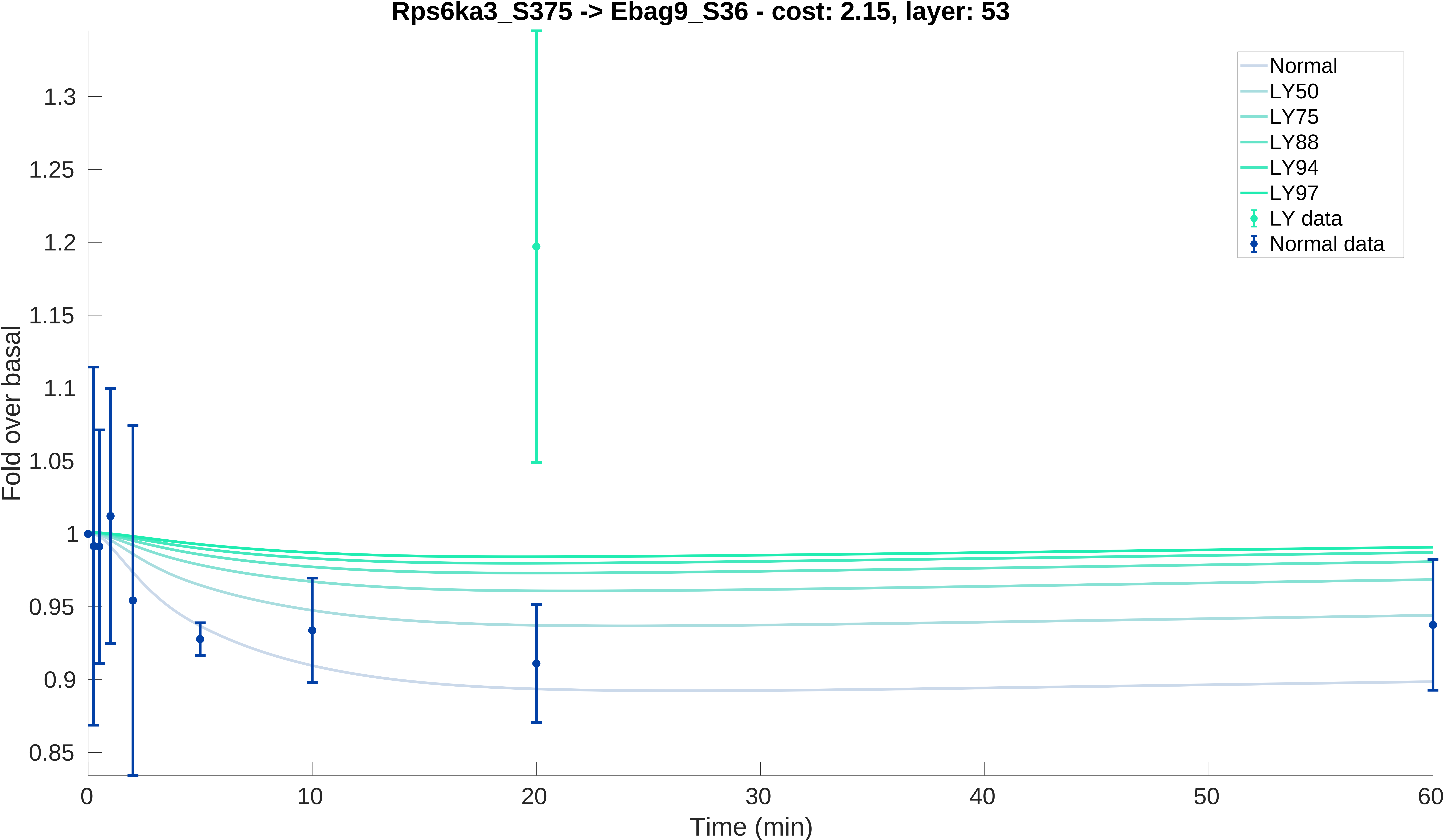

Rps6ka3_S375 -> Ebag9_S36 - cost: 2.15, layer: 53
Fold over basal
Time (min)
0.85
0.9
0.95
1
1.05
1.1
1.15
1.2
1.25
1.3
0
10
20
30
40
50
60
Normal
LY50
LY75
LY88
LY94
LY97
LY data
Normal data

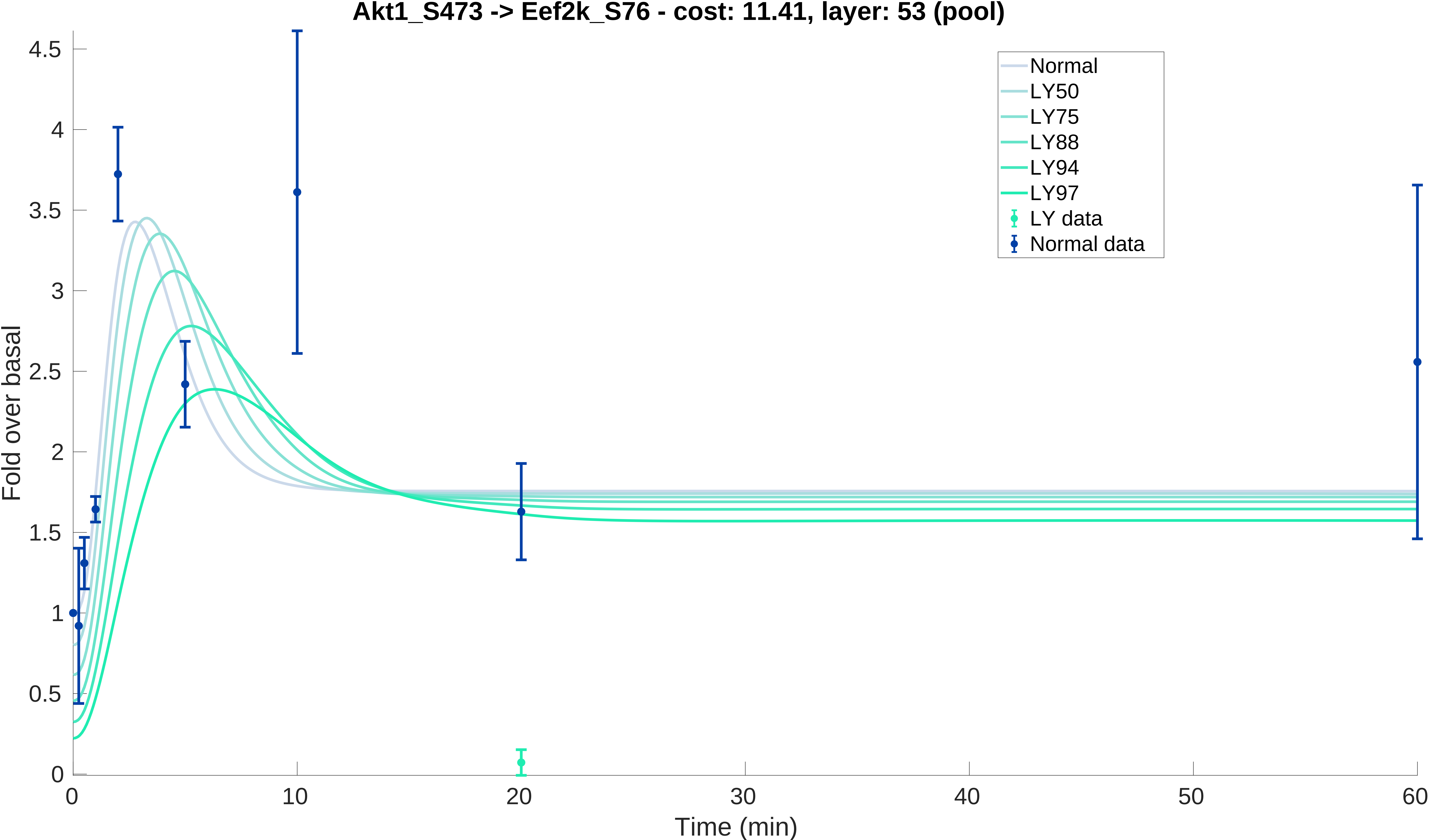

Akt1_S473 -> Eef2k_S76 - cost: 11.41, layer: 53 (pool)
Normal
LY50
LY75
LY88
LY94
LY97
LY data
Normal data
Fold over basal
Time (min)
0
0.5
1
1.5
2
2.5
3
3.5
4
4.5
0
10
20
30
40
50
60

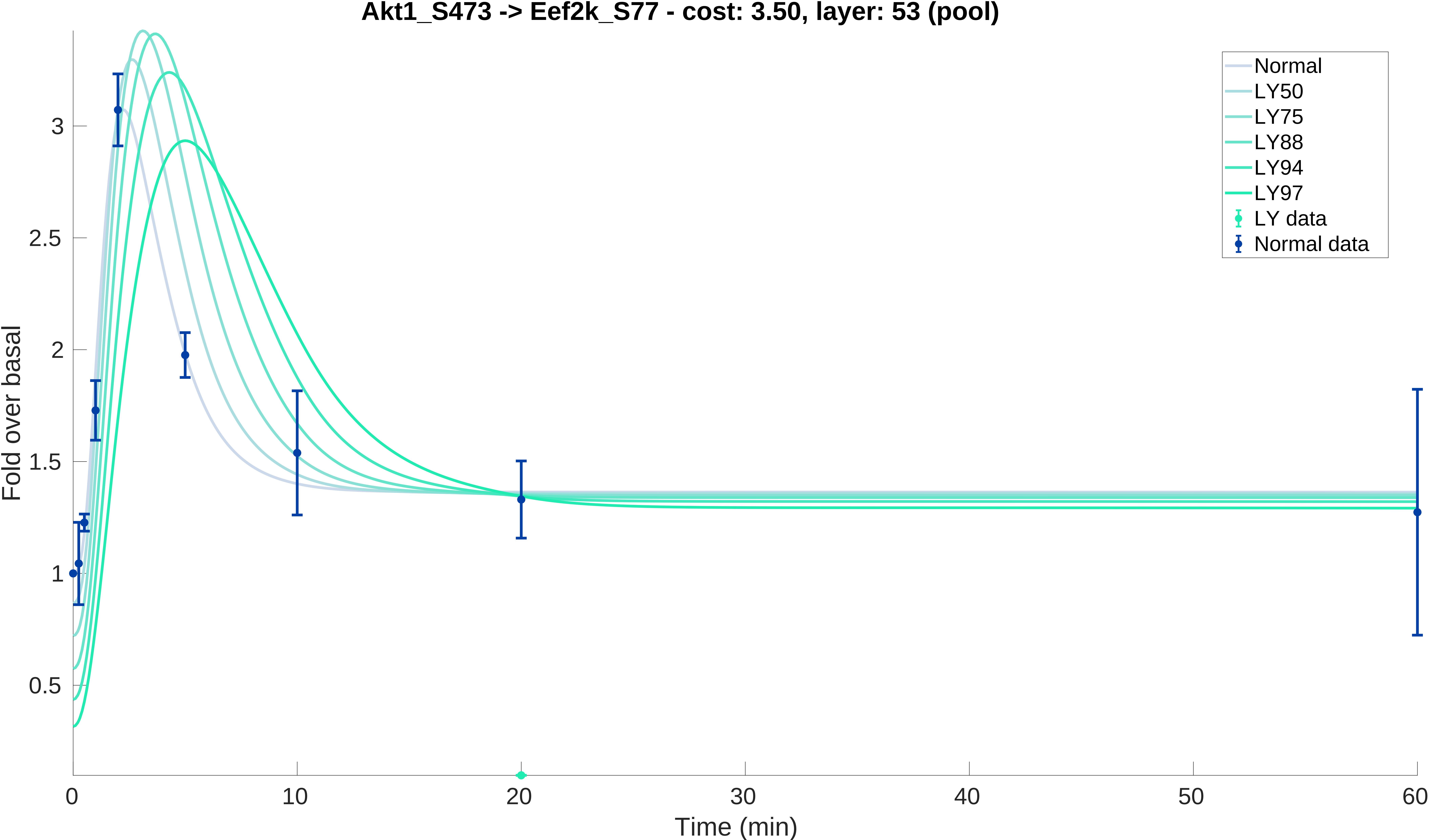
Akt1_S473 -> Eef2k_S77 - cost: 3.50, layer: 53 (pool)
Fold over basal
Time (min)
0
10
20
30
40
50
60
0.5
1
1.5
2
2.5
3
Normal
LY50
LY75
LY88
LY94
LY97
LY data
Normal data

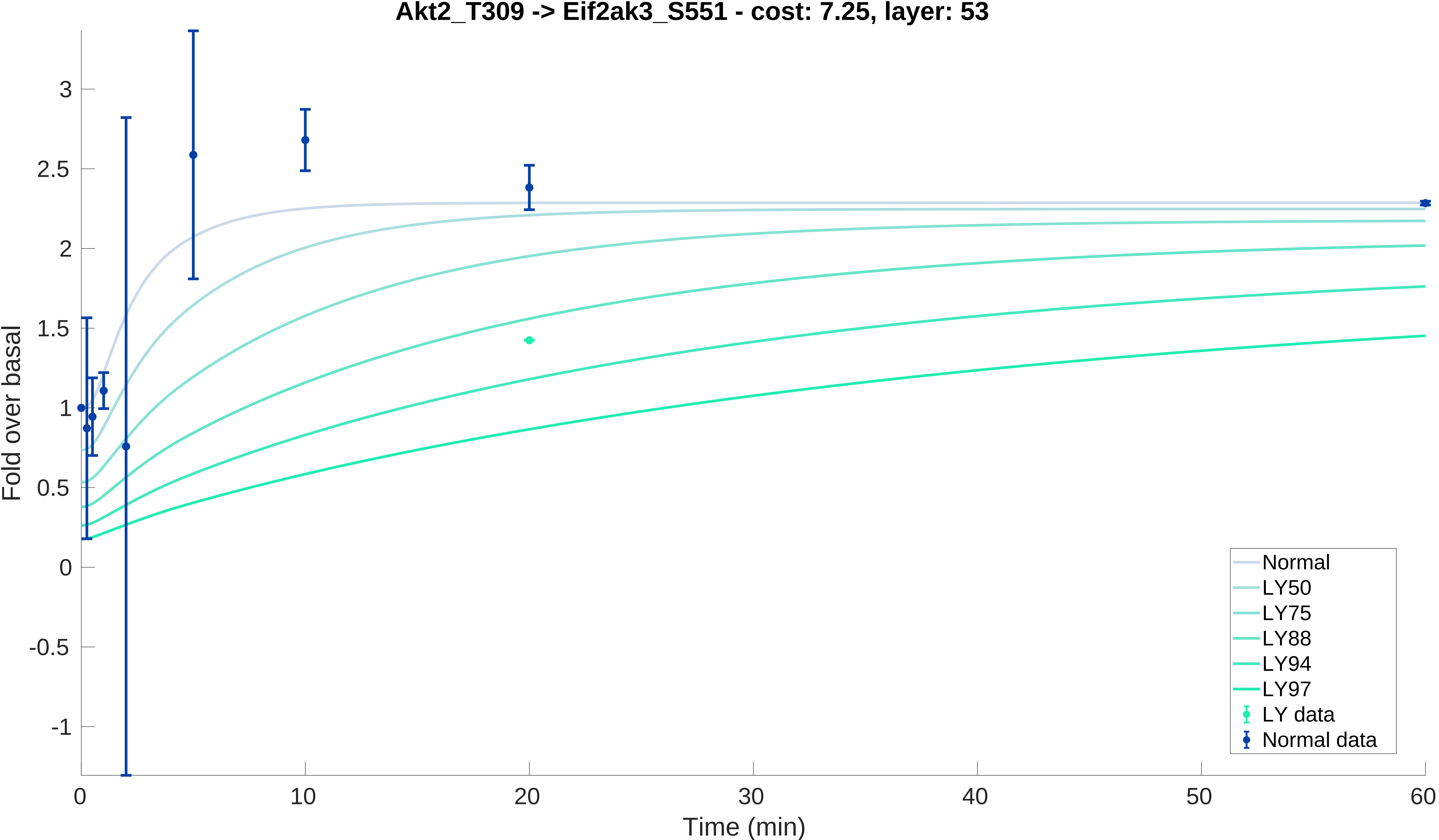

Akt2_T309 -> Eif2ak3_S551 - cost: 7.25, layer: 53
Fold over basal
Time (min)
3
2.5
2
1.5
1
0.5
0
-0.5
-1
0
10
20
30
40
50
60
Normal
LY50
LY75
LY88
LY94
LY97
LY data
Normal data

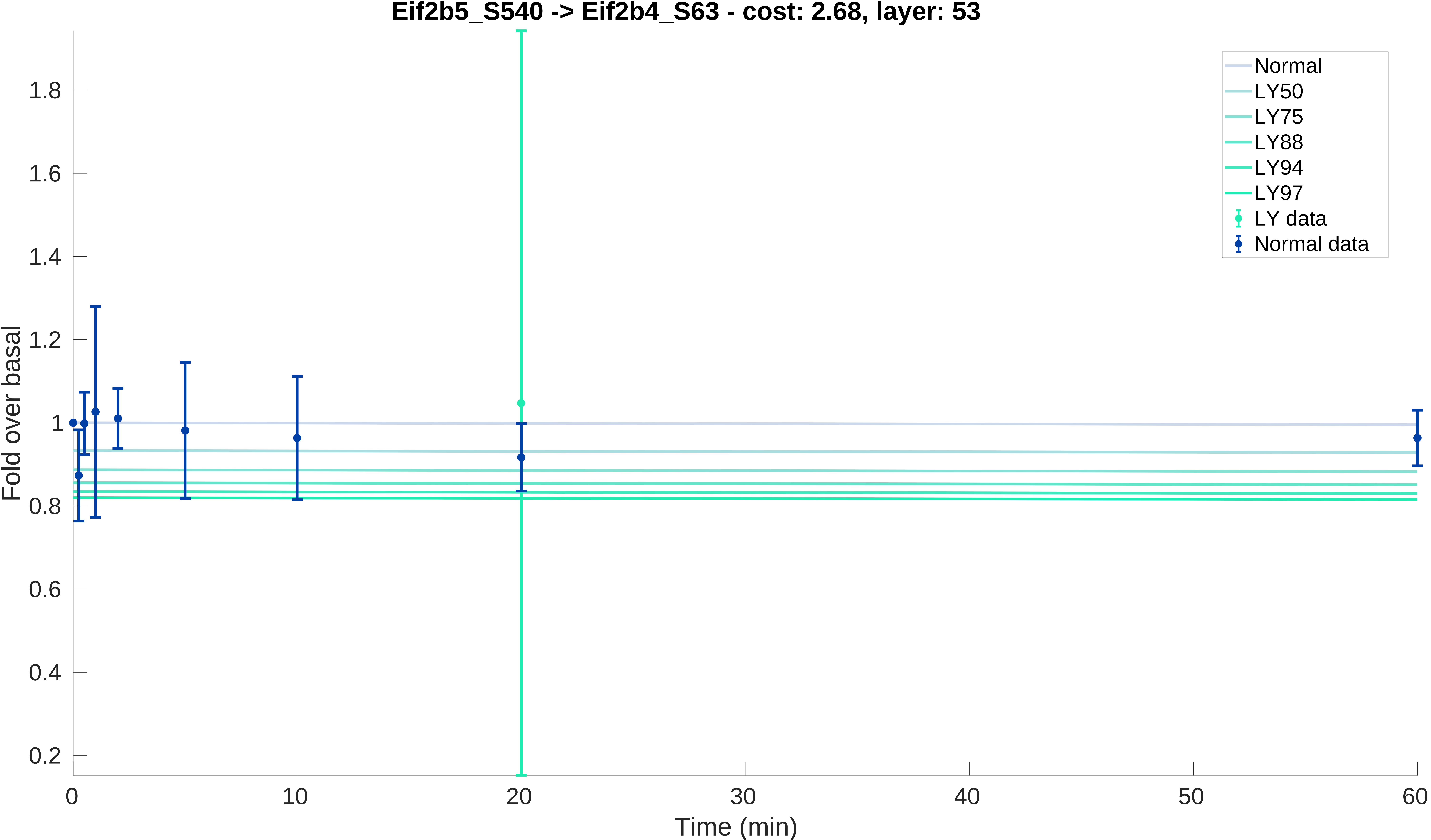

Eif2b5_S540 -> Eif2b4_S63 - cost: 2.68, layer: 53
Normal
LY50
LY75
LY88
LY94
LY97
LY data
Normal data
Fold over basal
1.8
1.6
1.4
1.2
1
0.8
0.6
0.4
0.2
0
10
20
30
40
50
60
Time (min)

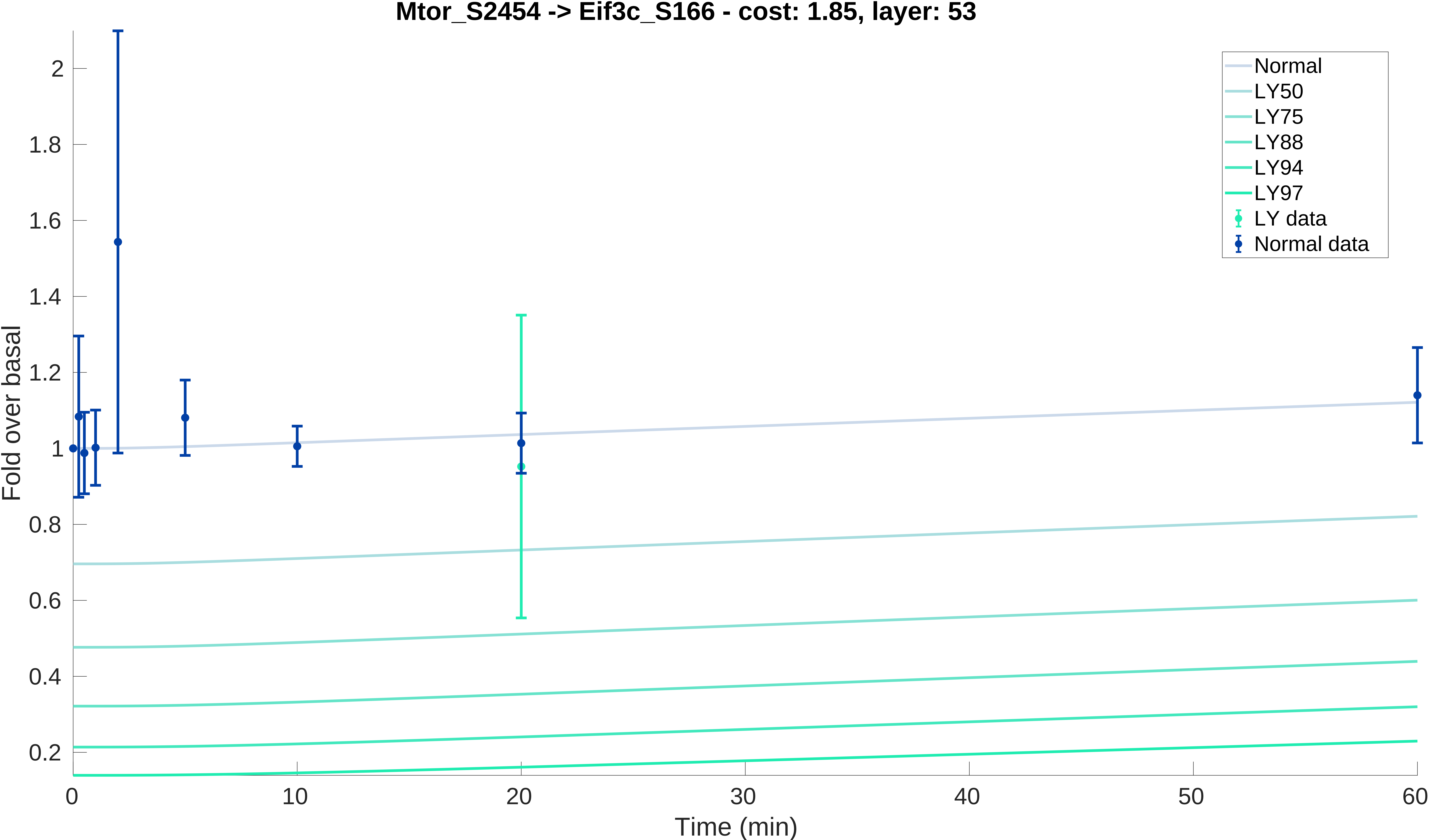

Mtor_S2454 -> Eif3c_S166 - cost: 1.85, layer: 53
Fold over basal
Time (min)
Normal
LY50
LY75
LY88
LY94
LY97
LY data
Normal data
0
10
20
30
40
50
60
0.2
0.4
0.6
0.8
1
1.2
1.4
1.6
1.8
2

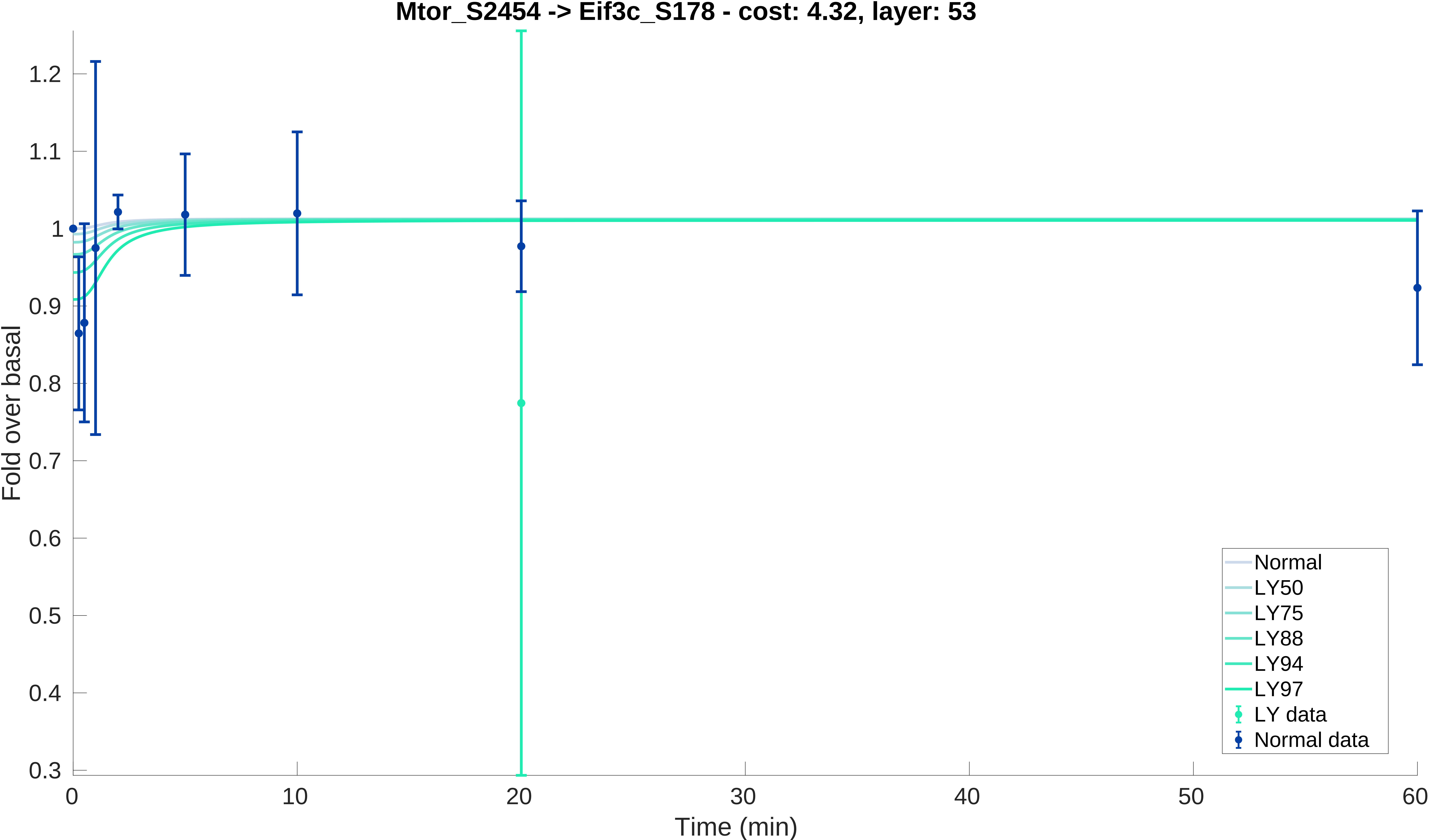

Mtor_S2454 -> Eif3c_S178 - cost: 4.32, layer: 53
Fold over basal
1.2
1.1
1
0.9
0.8
0.7
0.6
0.5
0.4
0.3
0
10
20
30
40
50
60
Time (min)
Normal
LY50
LY75
LY88
LY94
LY97
LY data
Normal data

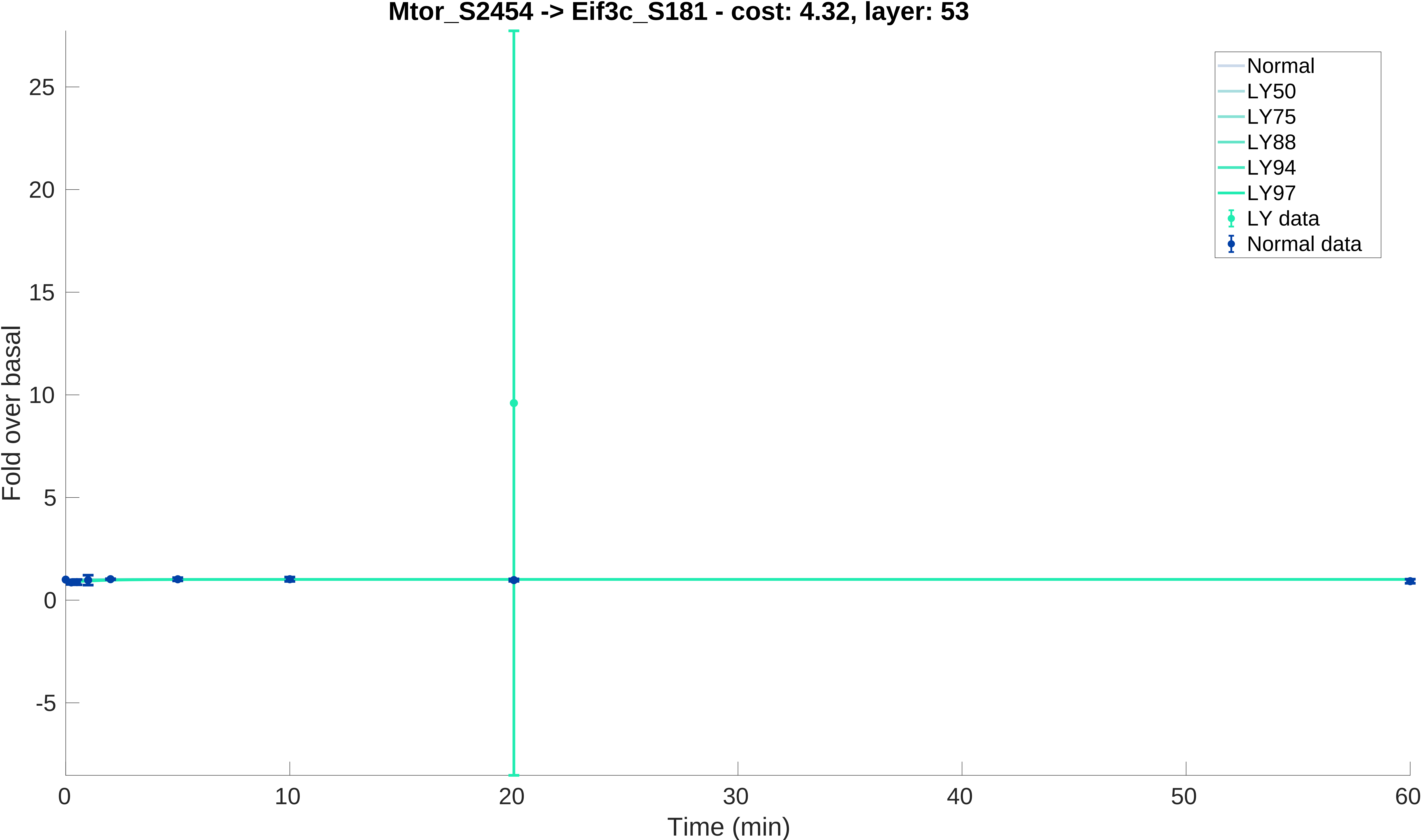

Mtor_S2454 -> Eif3c_S181 - cost: 4.32, layer: 53
Fold over basal
Time (min)
25
20
15
10
5
0
-5
0
10
20
30
40
50
60
Normal
LY50
LY75
LY88
LY94
LY97
LY data
Normal data

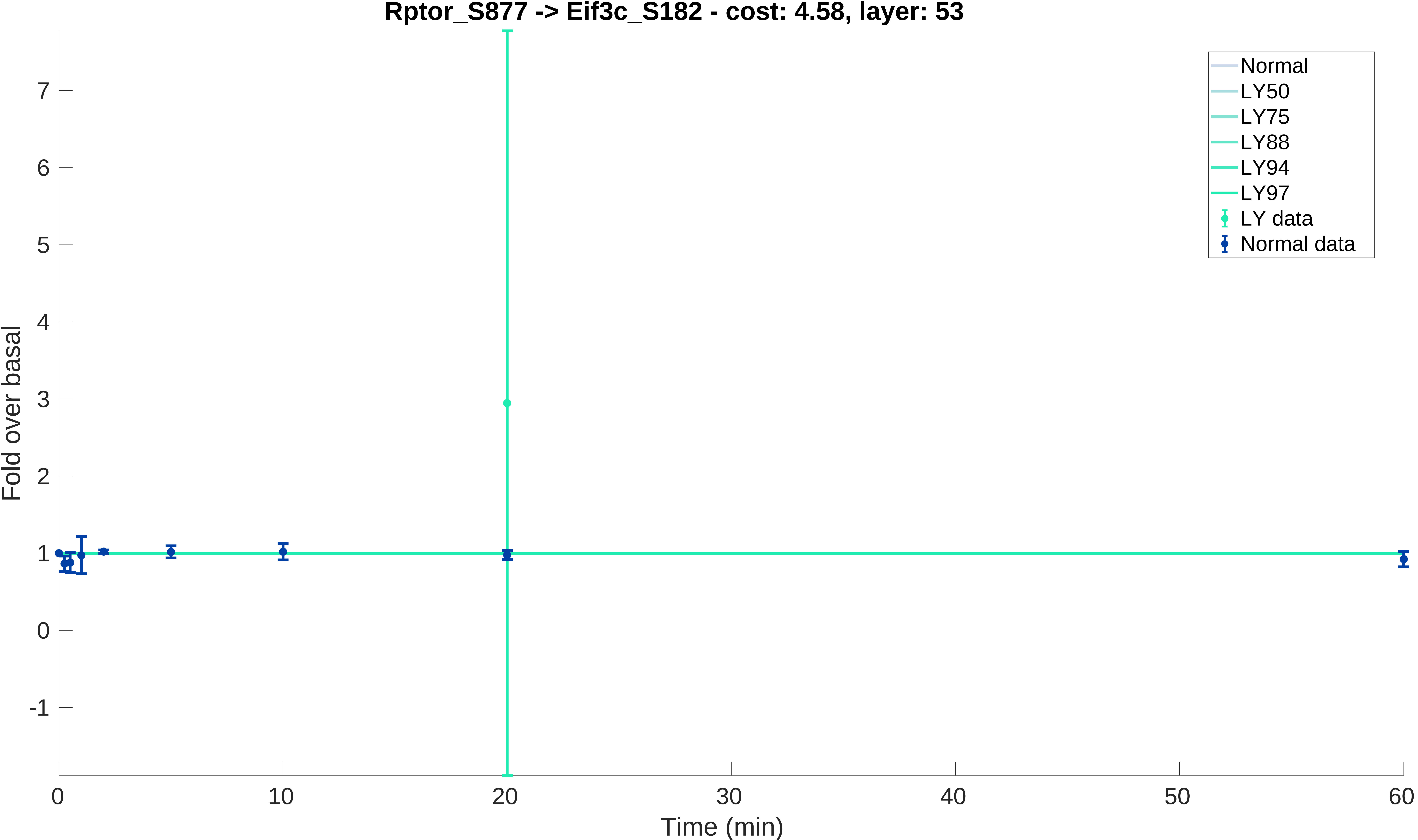
Rptor_S877 -> Eif3c_S182 - cost: 4.58, layer: 53
Normal
LY50
LY75
LY88
LY94
LY97
LY data
Normal data
Fold over basal
Time (min)
7
6
5
4
3
2
1
0
-1
0
10
20
30
40
50
60

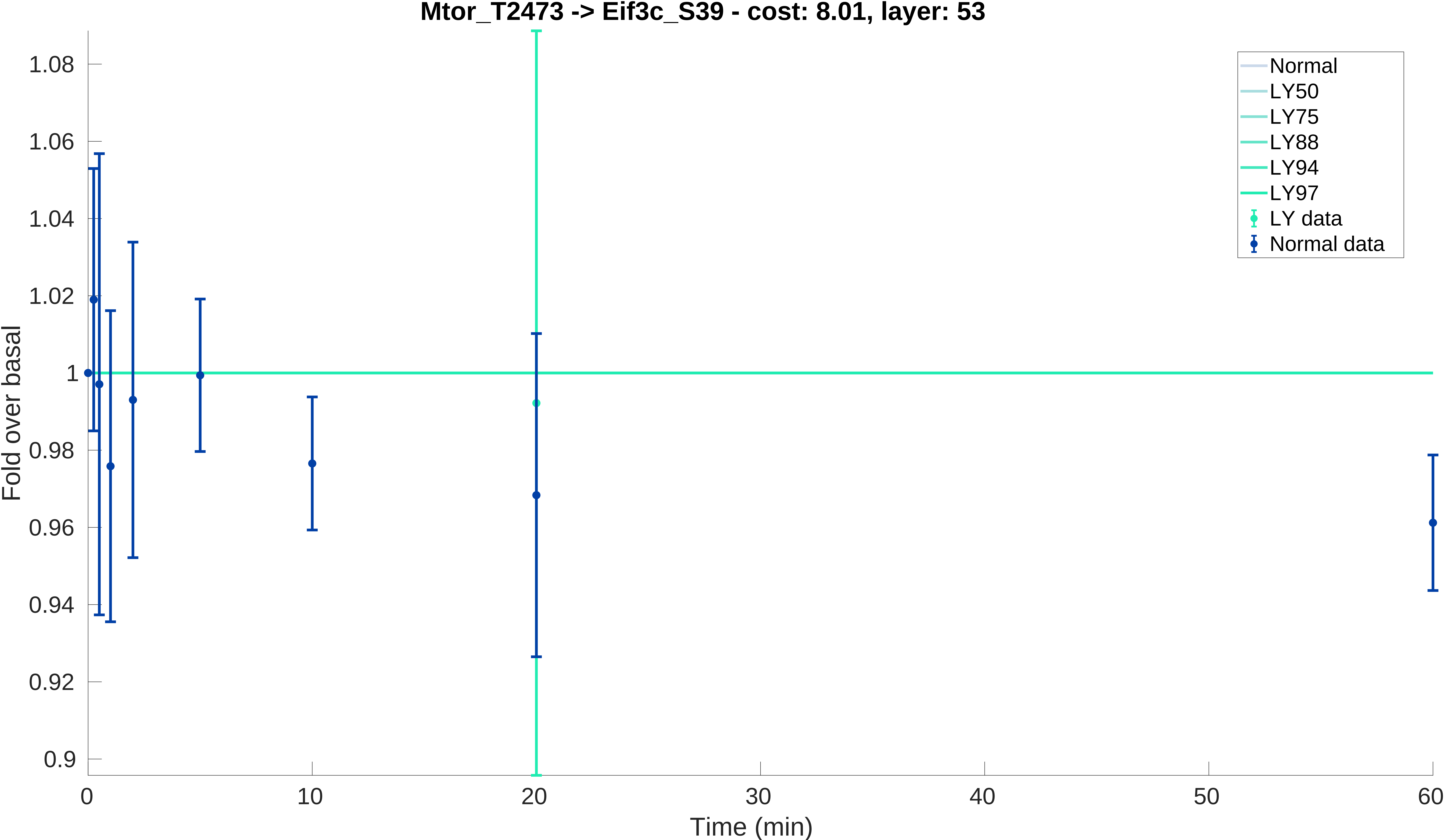
Mtor_T2473 -> Eif3c_S39 - cost: 8.01, layer: 53
Fold over basal
1.08
1.06
1.04
1.02
1
0.98
0.96
0.94
0.92
0.9
0
10
20
30
40
50
60
Time (min)
Normal
LY50
LY75
LY88
LY94
LY97
LY data
Normal data

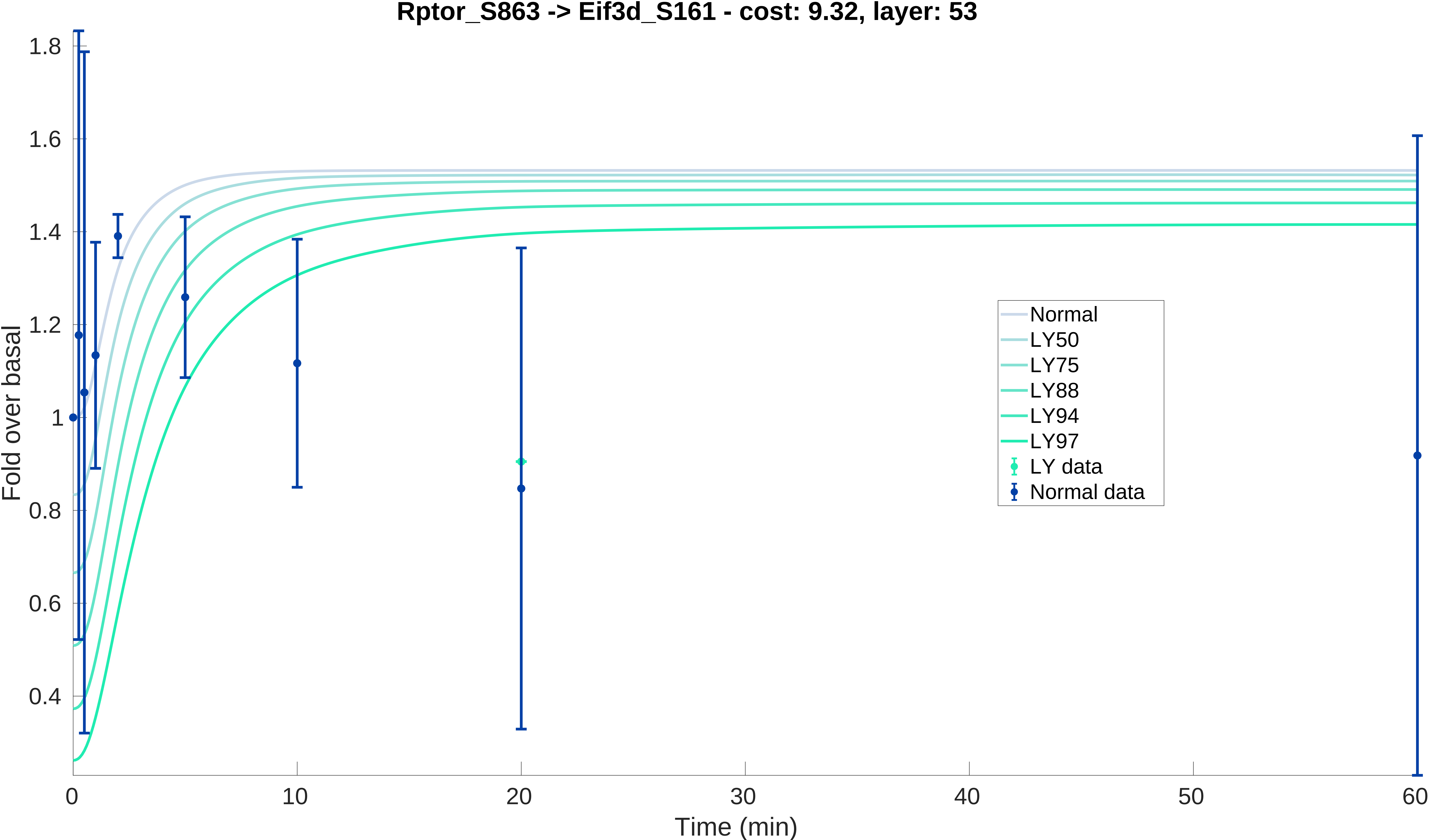

Rptor_S863 -> Eif3d_S161 - cost: 9.32, layer: 53
Fold over basal
1.8
1.6
1.4
1.2
1
0.8
0.6
0.4
0
10
20
30
40
50
60
Time (min)
Normal
LY50
LY75
LY88
LY94
LY97
LY data
Normal data

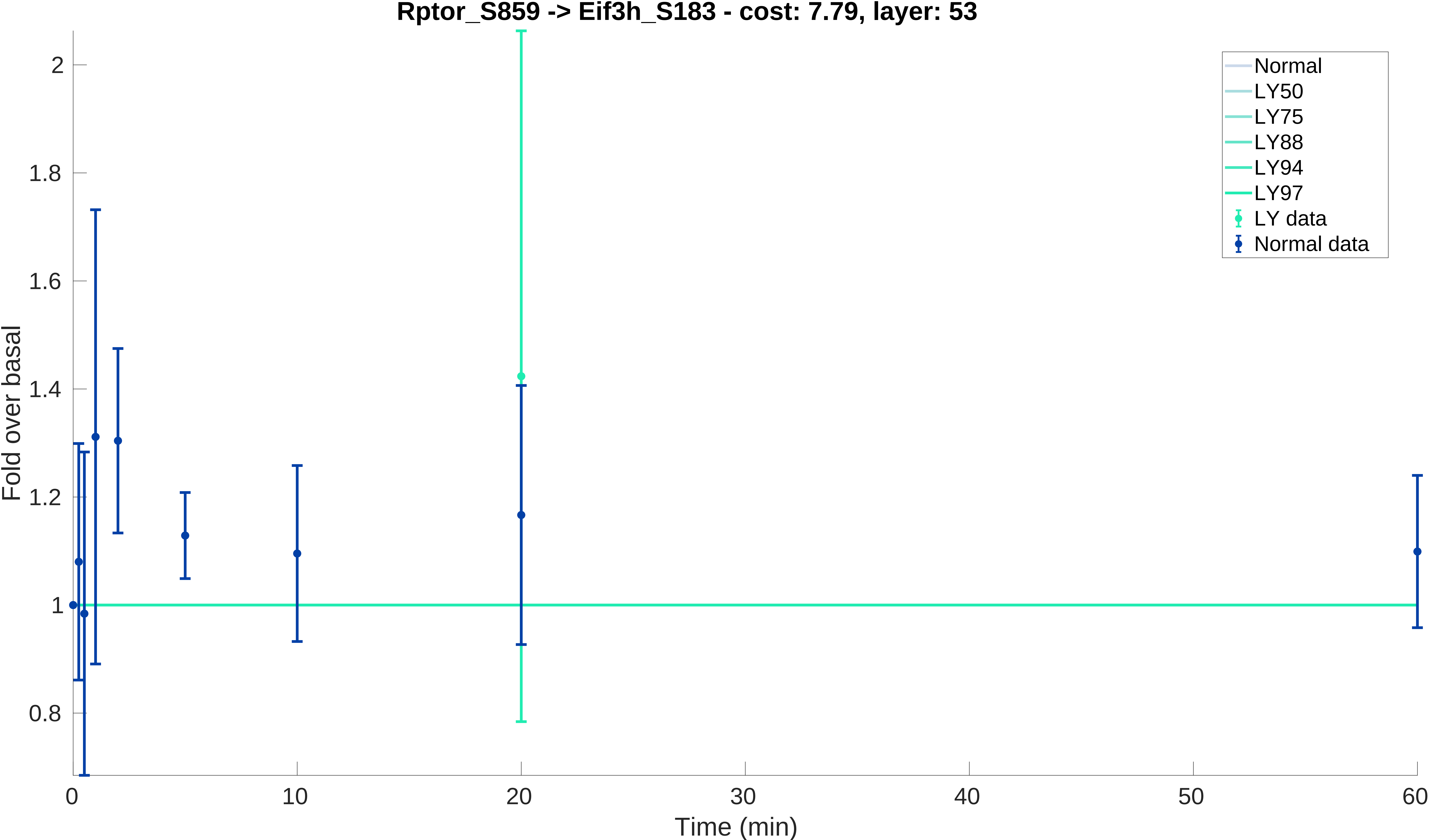

Rptor_S859 -> Eif3h_S183 - cost: 7.79, layer: 53
Fold over basal
Time (min)
0.8
1
1.2
1.4
1.6
1.8
2
0
10
20
30
40
50
60
Normal
LY50
LY75
LY88
LY94
LY97
LY data
Normal data

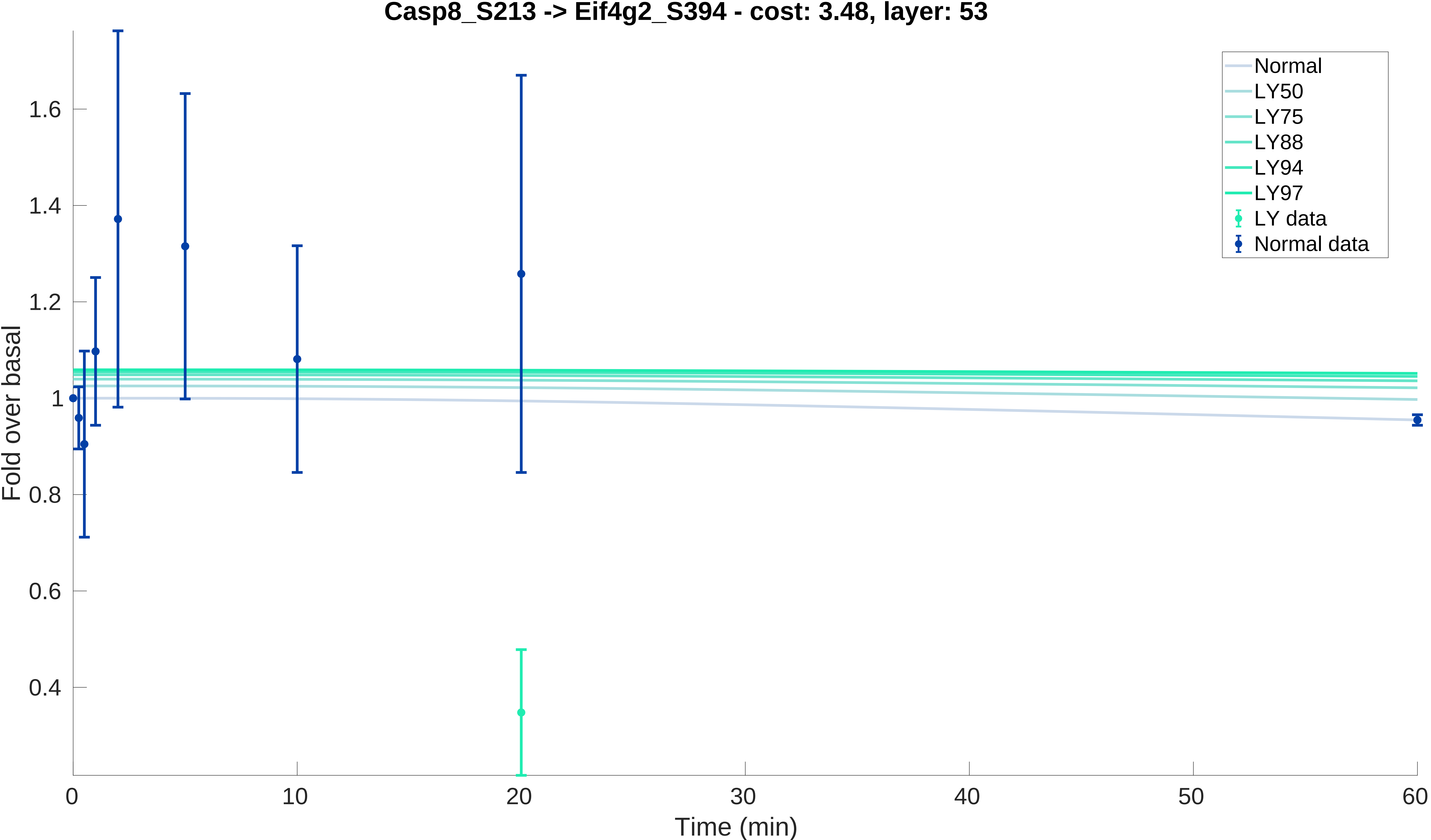

Casp8_S213 -> Eif4g2_S394 - cost: 3.48, layer: 53
Normal
LY50
LY75
LY88
LY94
LY97
LY data
Normal data
Fold over basal
1.6
1.4
1.2
1
0.8
0.6
0.4
0
10
20
30
40
50
60
Time (min)

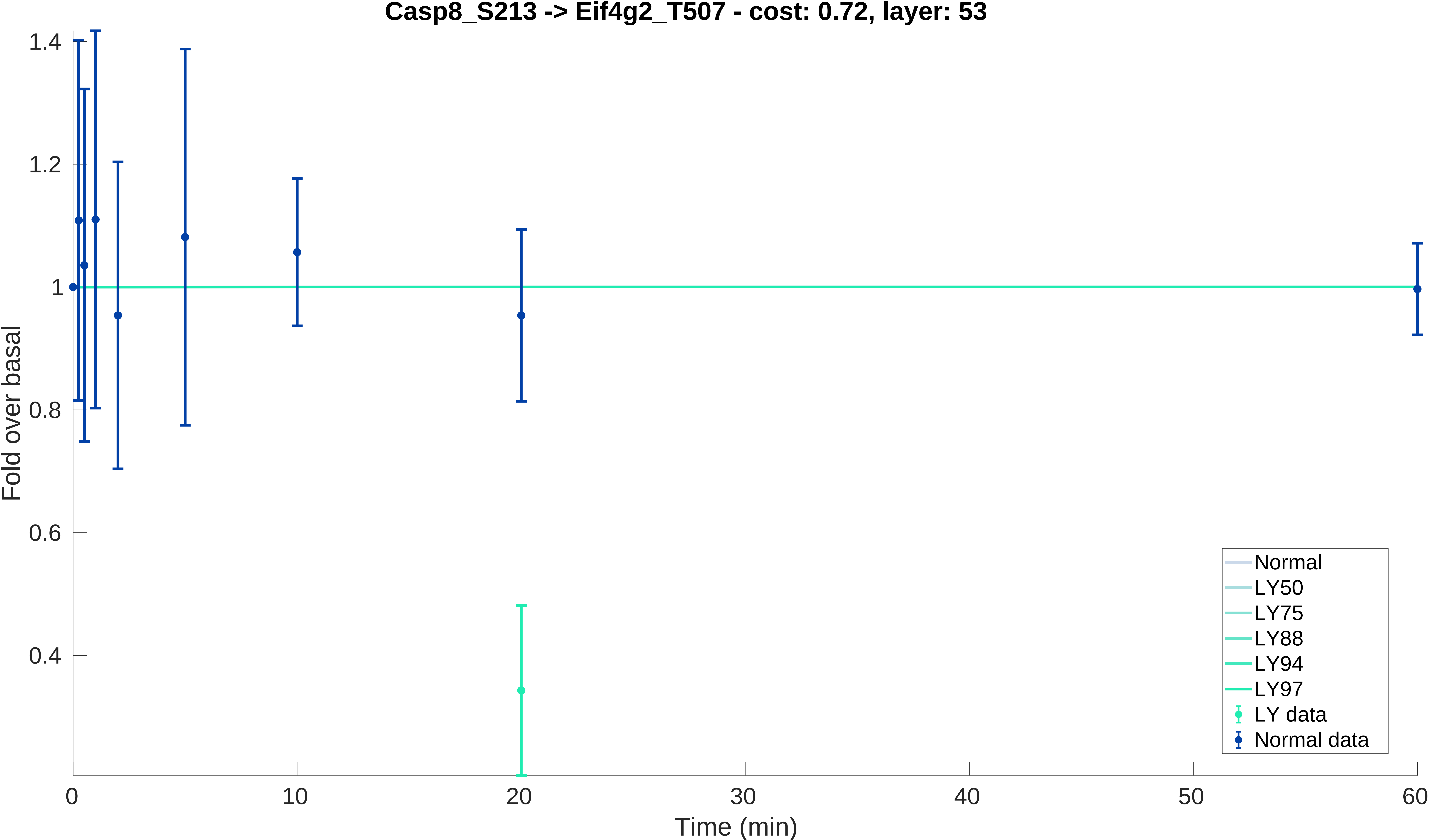

Casp8_S213 -> Eif4g2_T507 - cost: 0.72, layer: 53
Fold over basal
1.4
1.2
1
0.8
0.6
0.4
0
10
20
30
40
50
60
Time (min)
Normal
LY50
LY75
LY88
LY94
LY97
LY data
Normal data

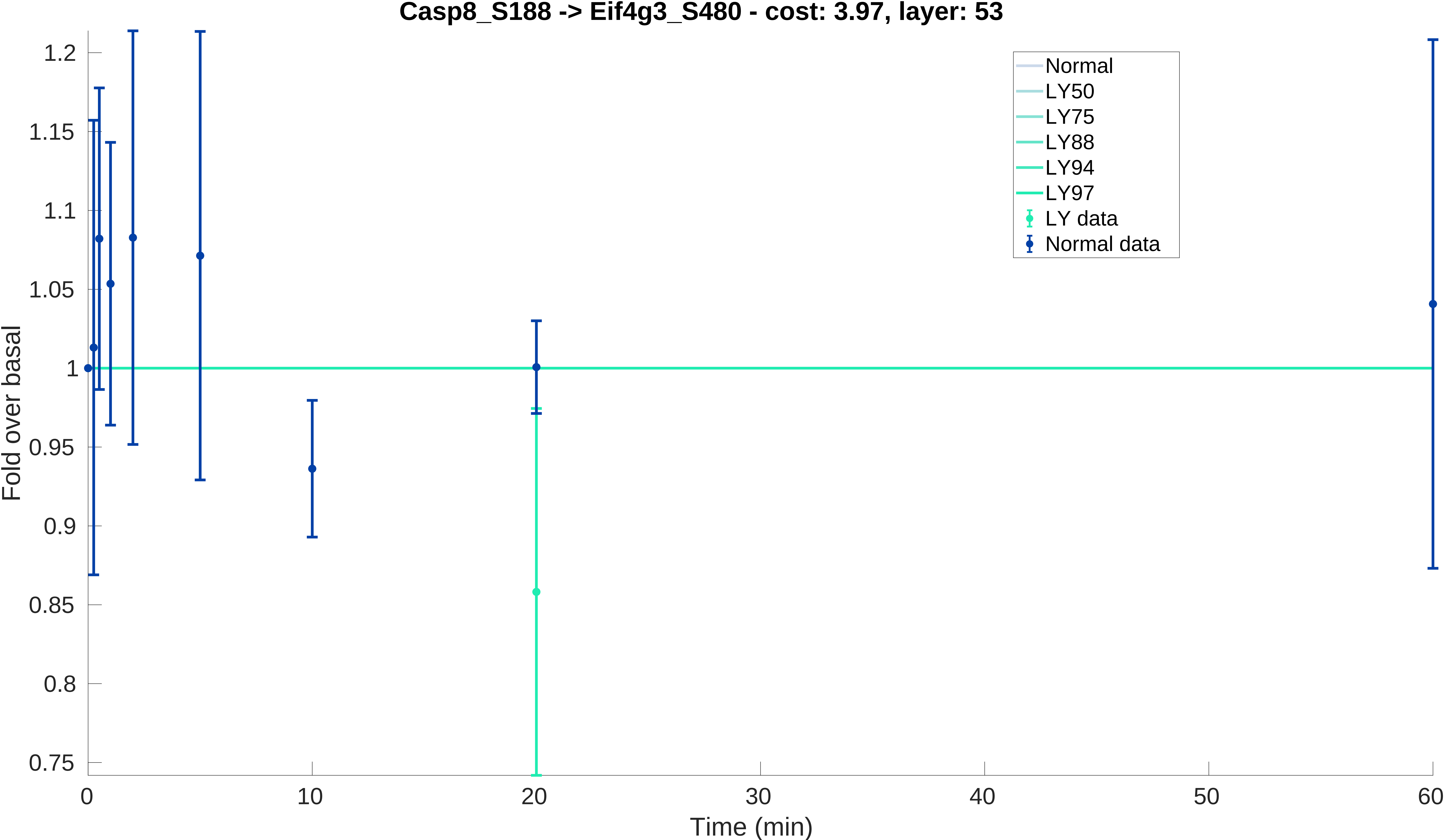

Casp8_S188 -> Eif4g3_S480 - cost: 3.97, layer: 53
Fold over basal
Time (min)
1.2
1.15
1.1
1.05
1
0.95
0.9
0.85
0.8
0.75
0
10
20
30
40
50
60
Normal
LY50
LY75
LY88
LY94
LY97
LY data
Normal data

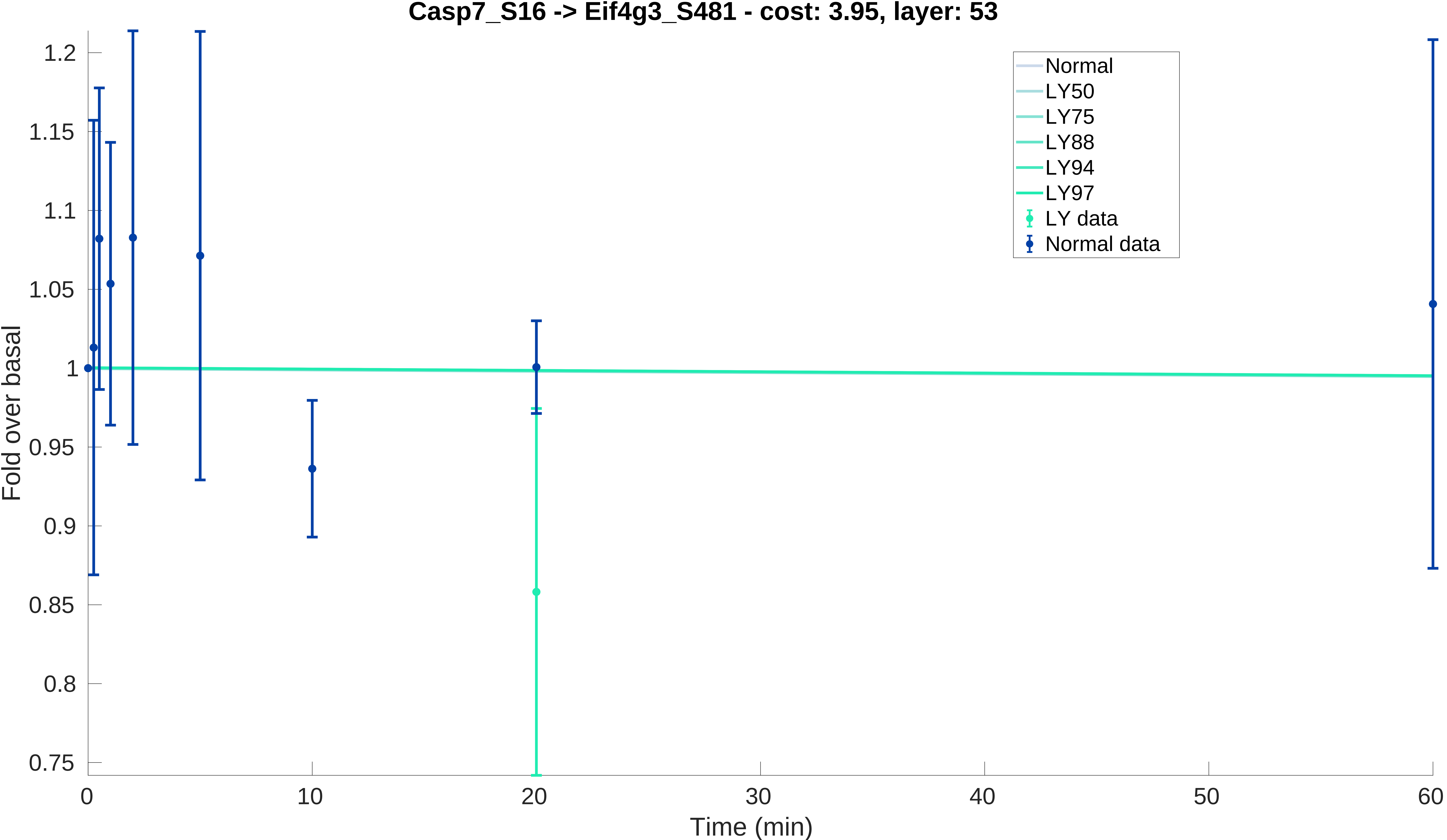

Casp7_S16 -> Eif4g3_S481 - cost: 3.95, layer: 53
Fold over basal
Time (min)
1.2
1.15
1.1
1.05
1
0.95
0.9
0.85
0.8
0.75
0
10
20
30
40
50
60
Normal
LY50
LY75
LY88
LY94
LY97
LY data
Normal data

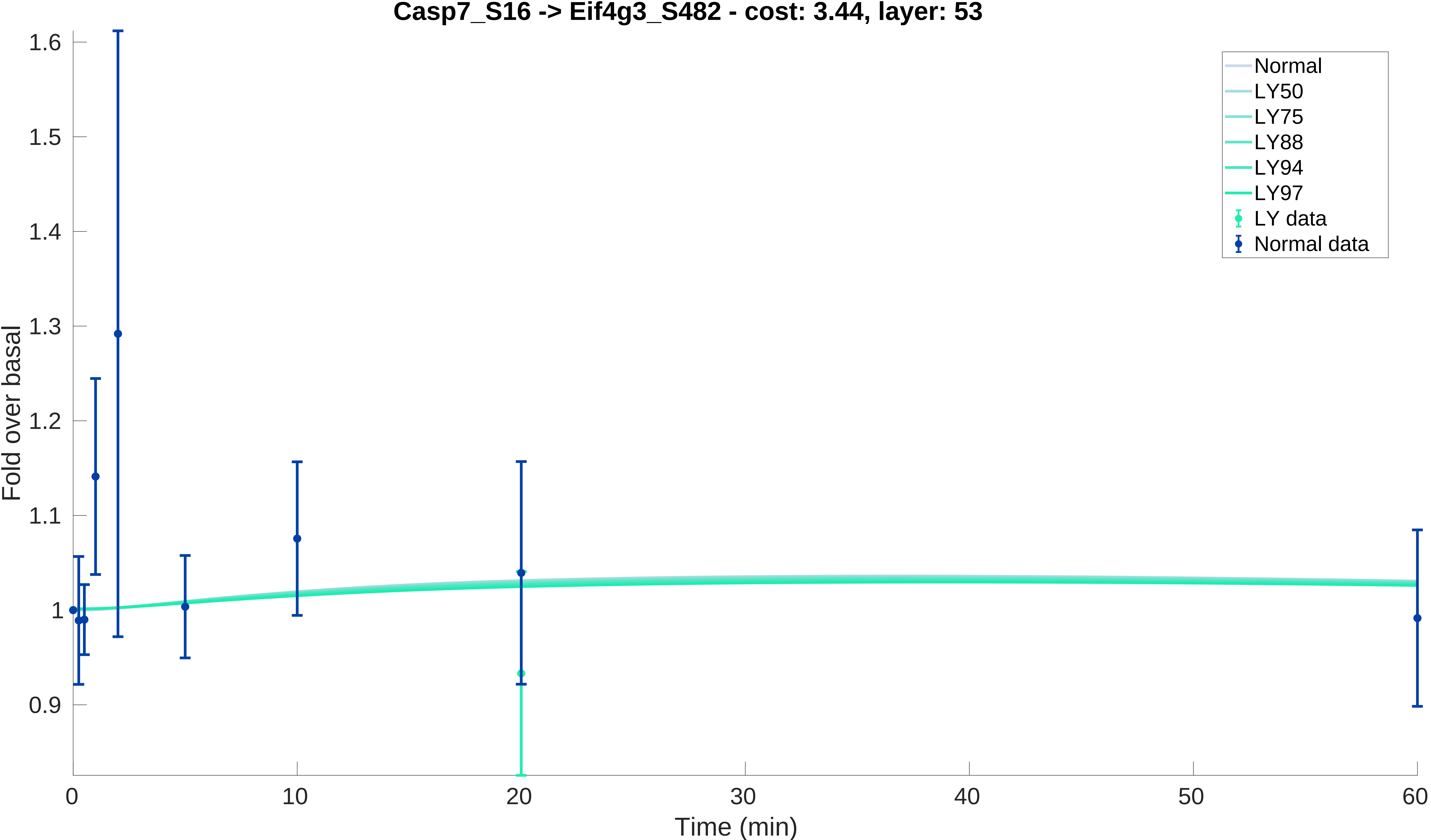

Casp7_S16 -> Eif4g3_S482 - cost: 3.44, layer: 53
Fold over basal
1.6
1.5
1.4
1.3
1.2
1.1
1
0.9
0
10
20
30
40
50
60
Time (min)
Normal
LY50
LY75
LY88
LY94
LY97
LY data
Normal data

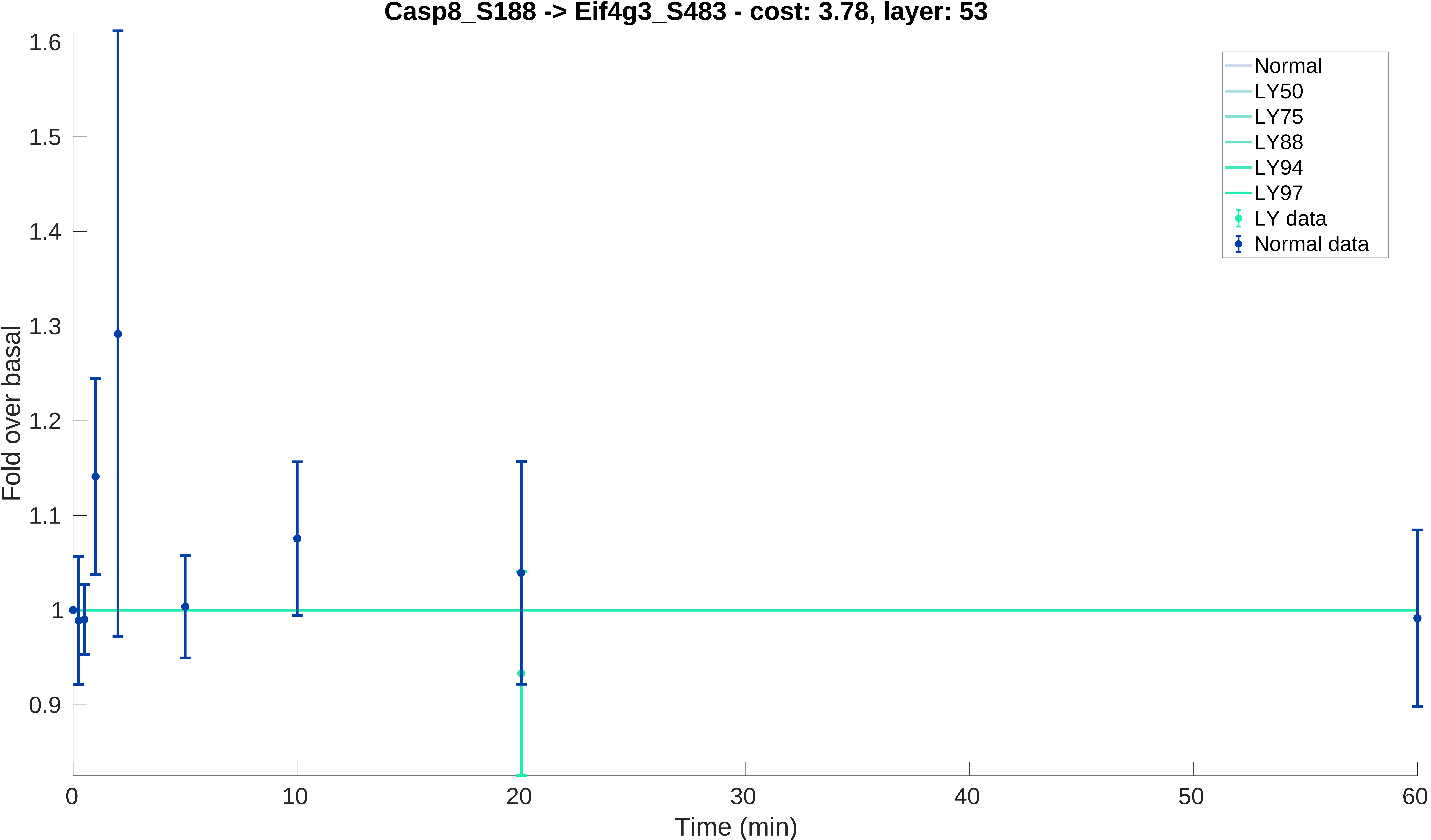

Casp8_S188 -> Eif4g3_S483 - cost: 3.78, layer: 53
Fold over basal
1.6
1.5
1.4
1.3
1.2
1.1
1
0.9
Time (min)
0
10
20
30
40
50
60
Normal
LY50
LY75
LY88
LY94
LY97
LY data
Normal data

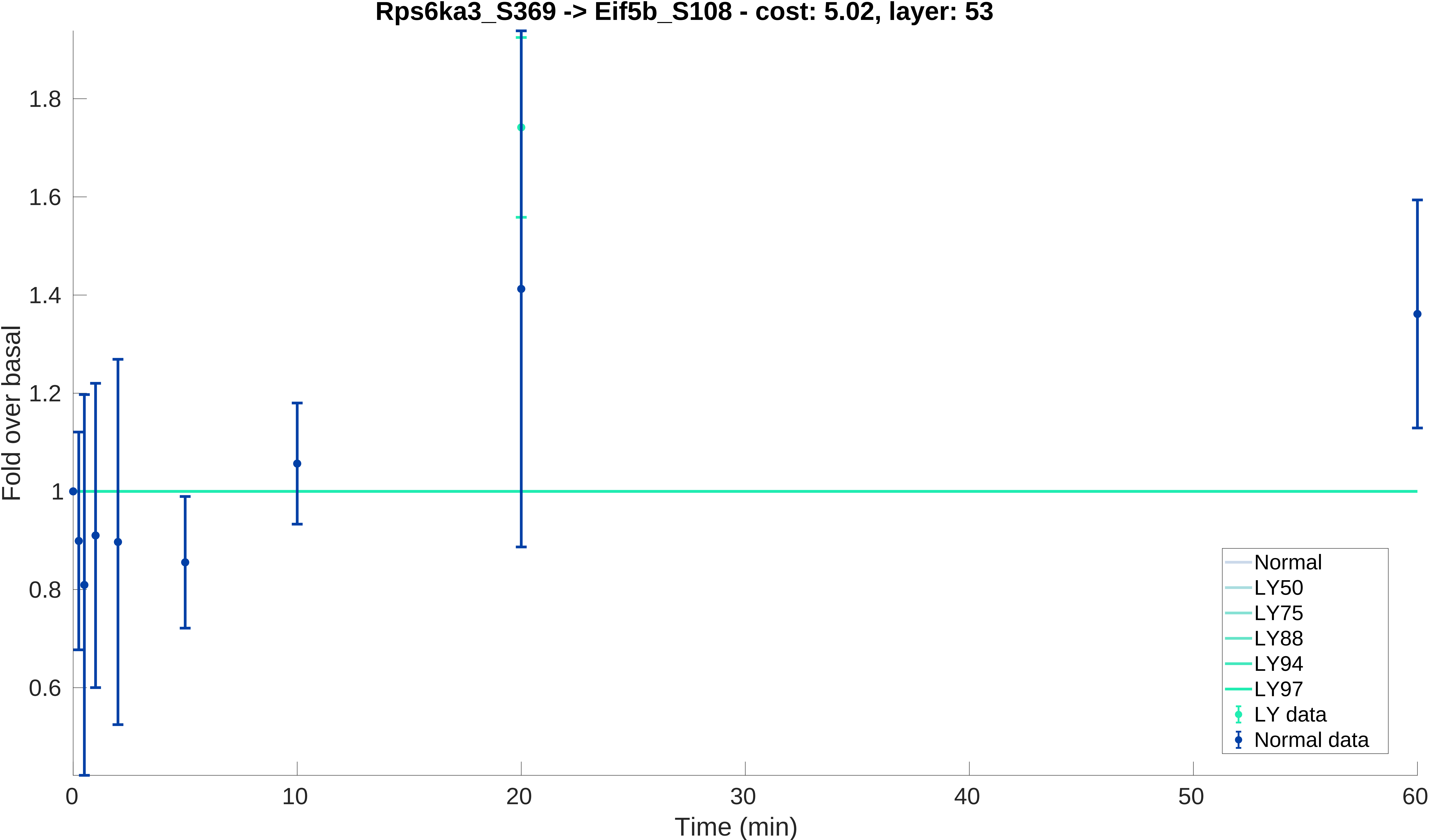

Rps6ka3_S369 -> Eif5b_S108 - cost: 5.02, layer: 53
Fold over basal
1.8
1.6
1.4
1.2
1
0.8
0.6
0
10
20
30
40
50
60
Time (min)
Normal
LY50
LY75
LY88
LY94
LY97
LY data
Normal data

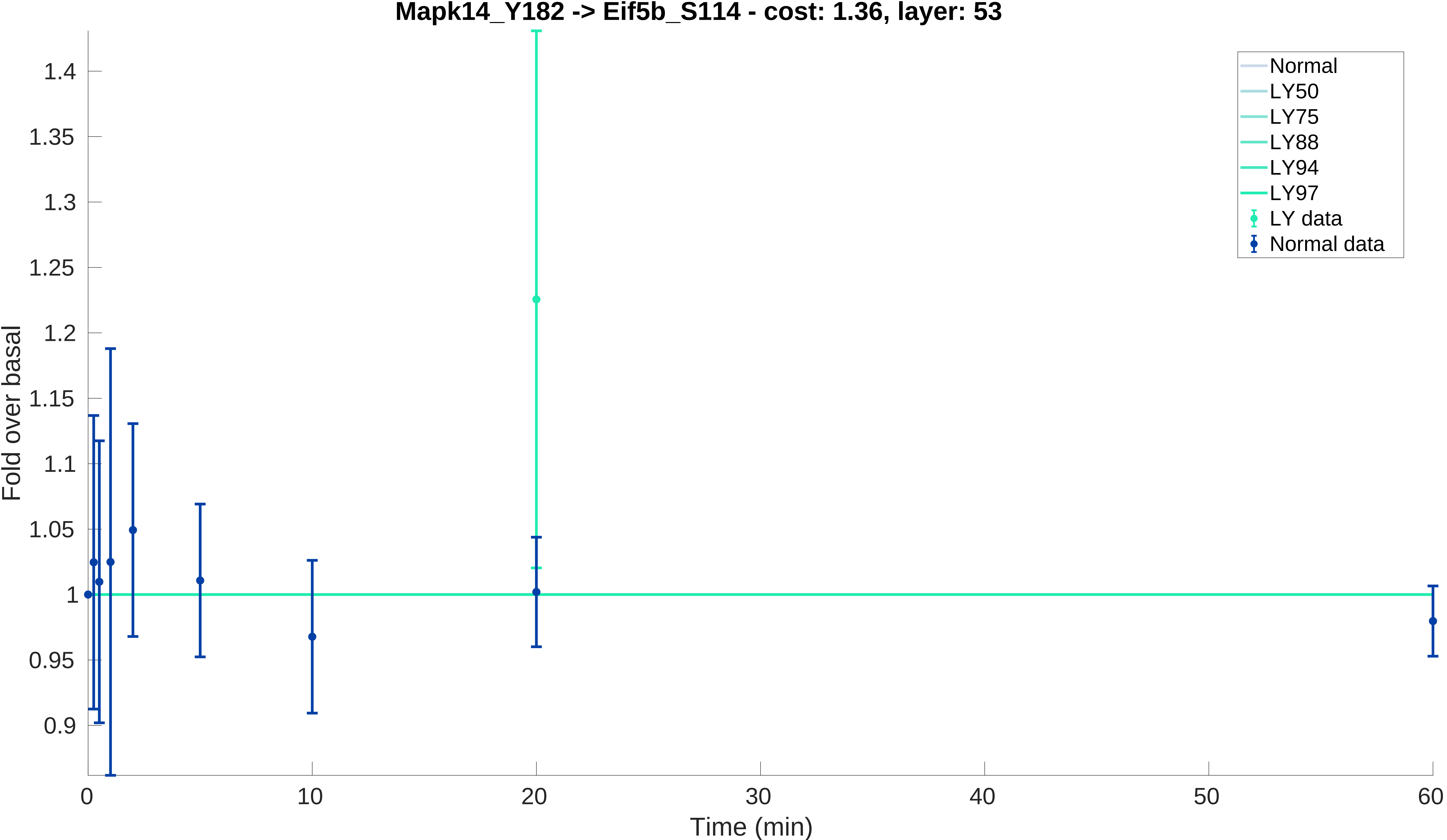

Mapk14_Y182 -> Eif5b_S114 - cost: 1.36, layer: 53
Fold over basal
Time (min)
1.4
1.35
1.3
1.25
1.2
1.15
1.1
1.05
1
0.95
0.9
0
10
20
30
40
50
60
Normal
LY50
LY75
LY88
LY94
LY97
LY data
Normal data

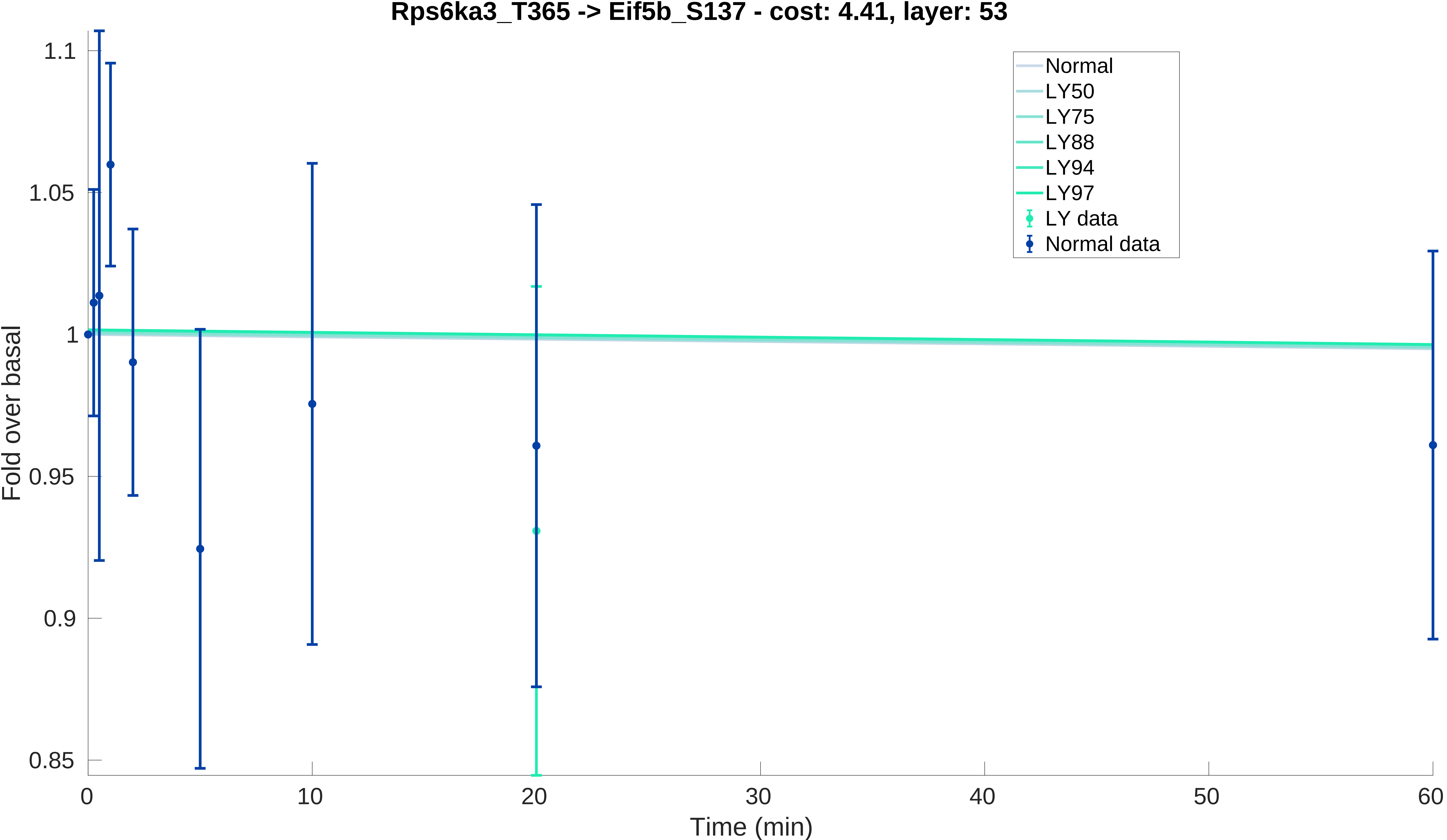

Rps6ka3_T365 -> Eif5b_S137 - cost: 4.41, layer: 53
Normal
LY50
LY75
LY88
LY94
LY97
LY data
Normal data
Fold over basal
1.1
1.05
1
0.95
0.9
0.85
0
10
20
30
40
50
60
Time (min)

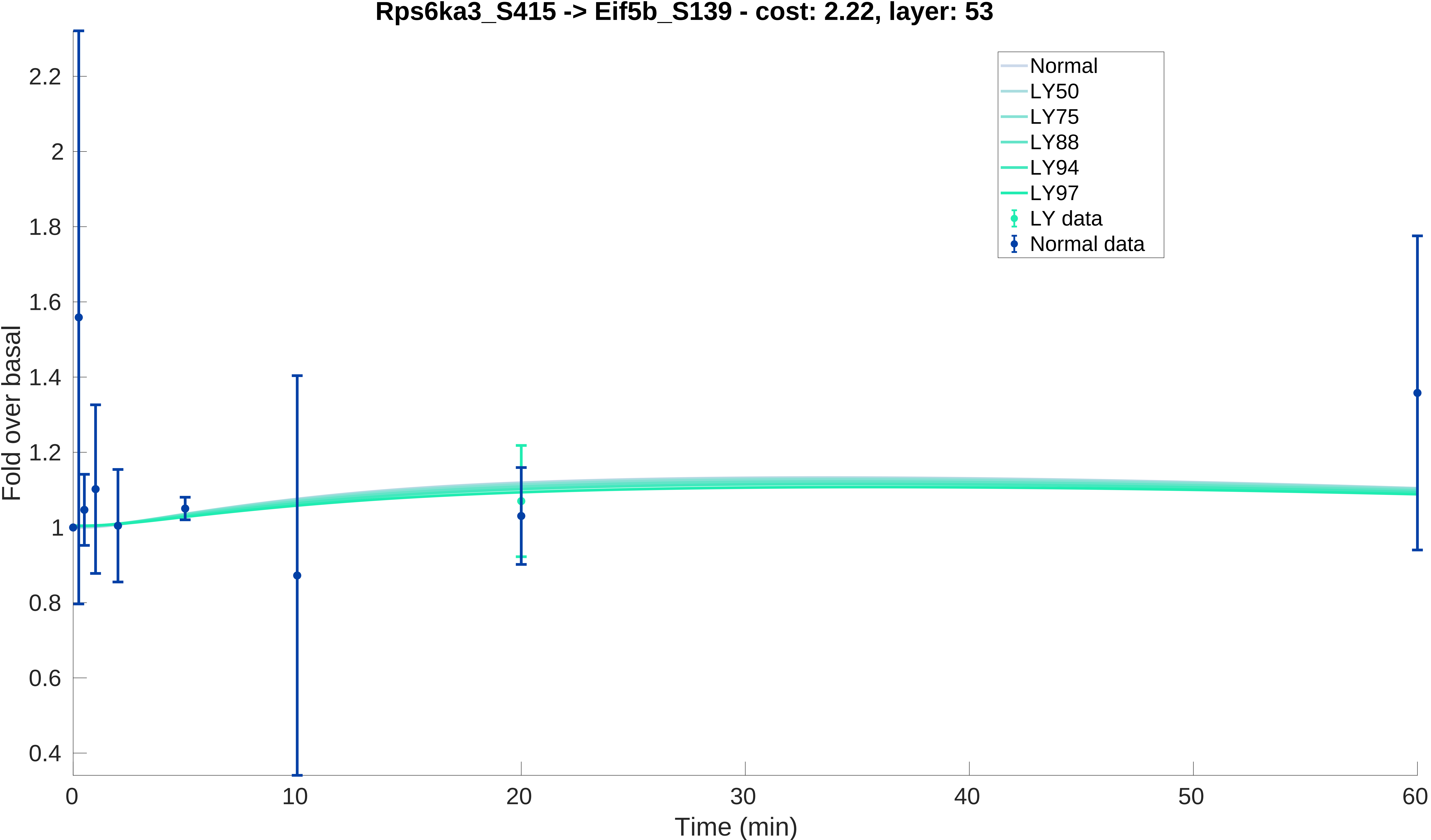

Rps6ka3_S415 -> Eif5b_S139 - cost: 2.22, layer: 53
Fold over basal
Time (min)
Normal
LY50
LY75
LY88
LY94
LY97
LY data
Normal data
0.4
0.6
0.8
1
1.2
1.4
1.6
1.8
2
2.2
0
10
20
30
40
50
60

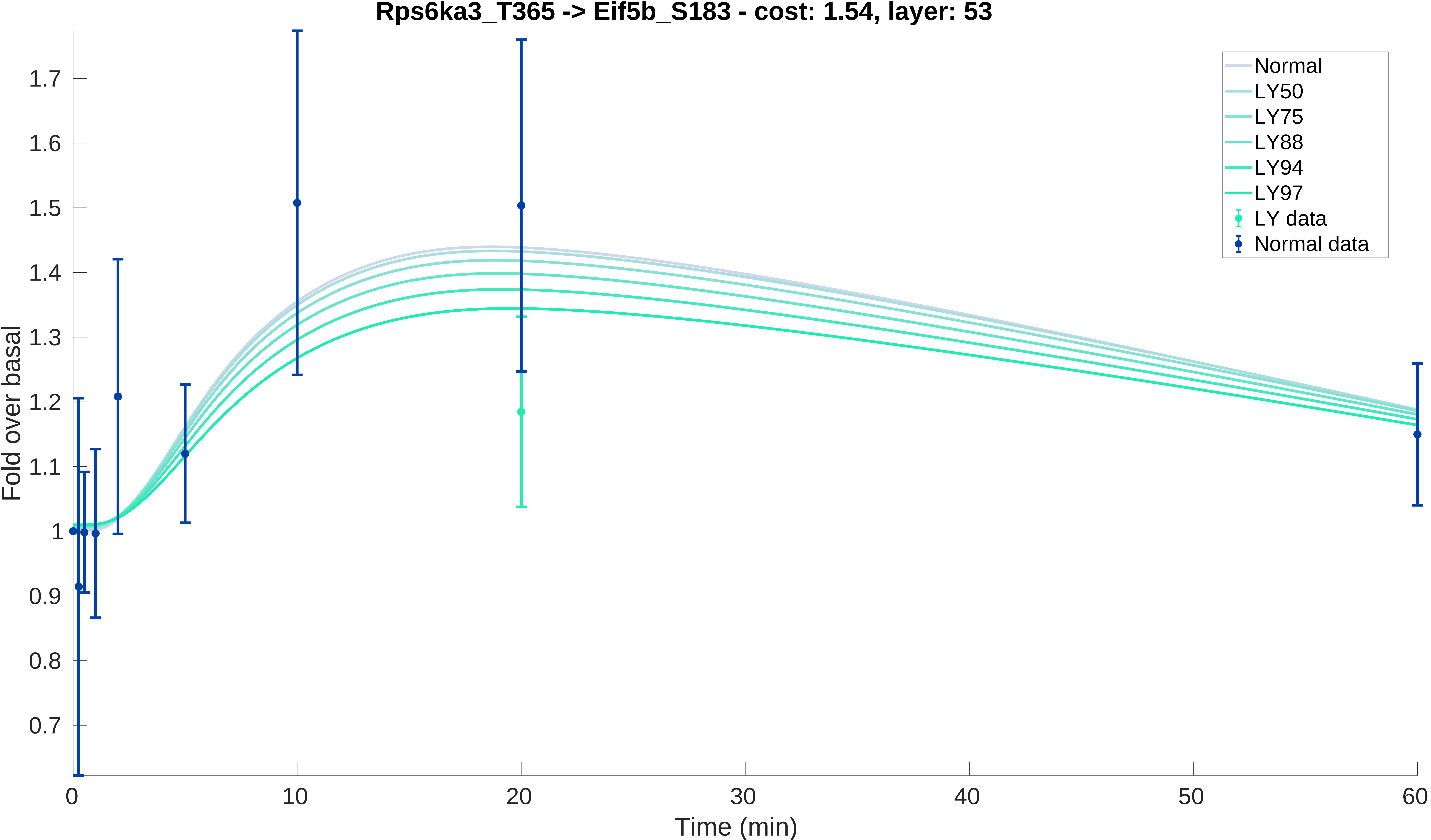

Rps6ka3_T365 -> Eif5b_S183 - cost: 1.54, layer: 53
Fold over basal
Time (min)
0.7
0.8
0.9
1
1.1
1.2
1.3
1.4
1.5
1.6
1.7
0
10
20
30
40
50
60
Normal
LY50
LY75
LY88
LY94
LY97
LY data
Normal data

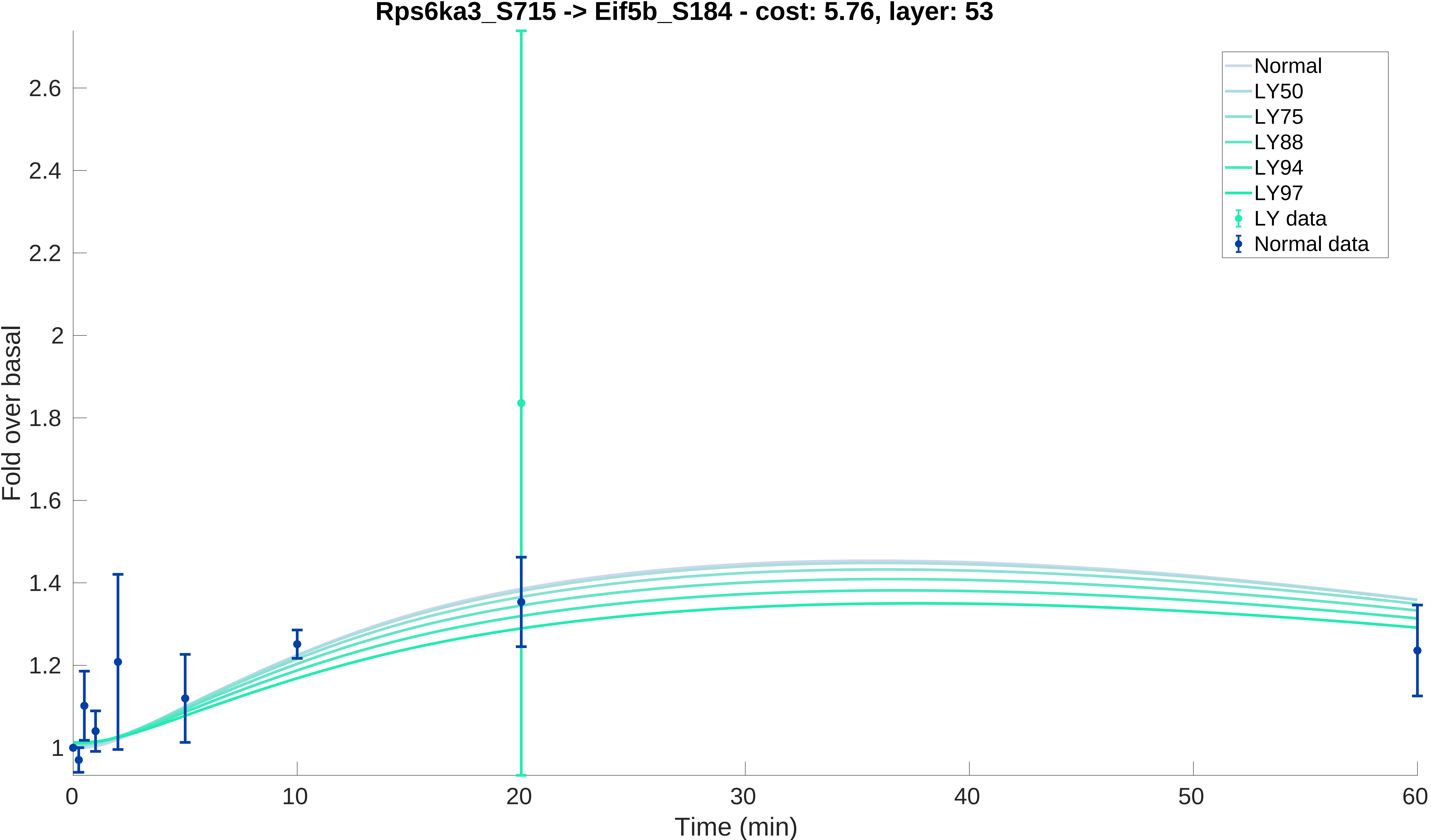
Rps6ka3_S715 -> Eif5b_S184 - cost: 5.76, layer: 53
Fold over basal
2.6
2.4
2.2
2
1.8
1.6
1.4
1.2
1
0
10
20
30
40
50
60
Time (min)
Normal
LY50
LY75
LY88
LY94
LY97
LY data
Normal data

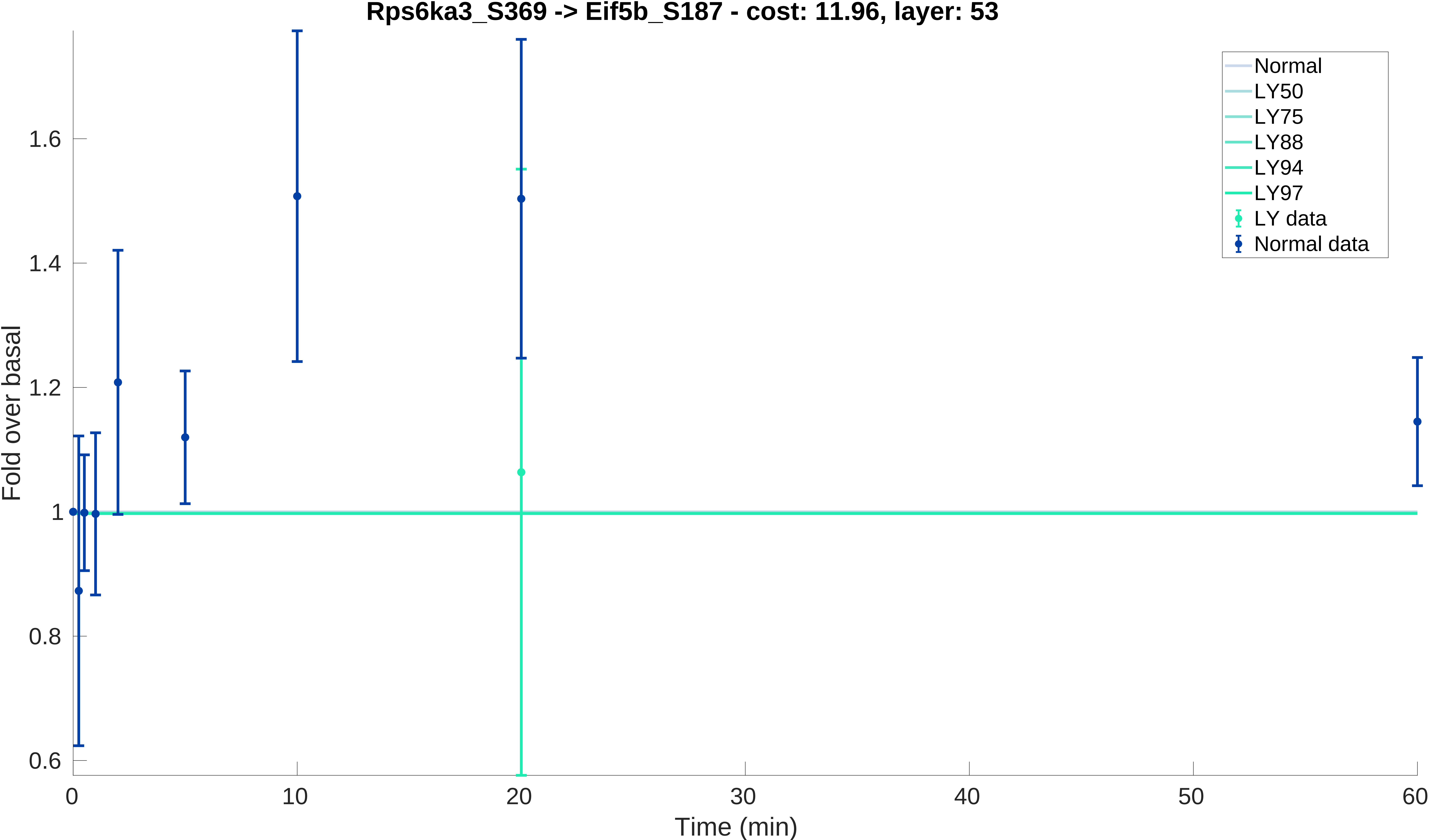
Rps6ka3_S369 -> Eif5b_S187 - cost: 11.96, layer: 53
Fold over basal
Time (min)
0.6
0.8
1
1.2
1.4
1.6
0
10
20
30
40
50
60
Normal
LY50
LY75
LY88
LY94
LY97
LY data
Normal data

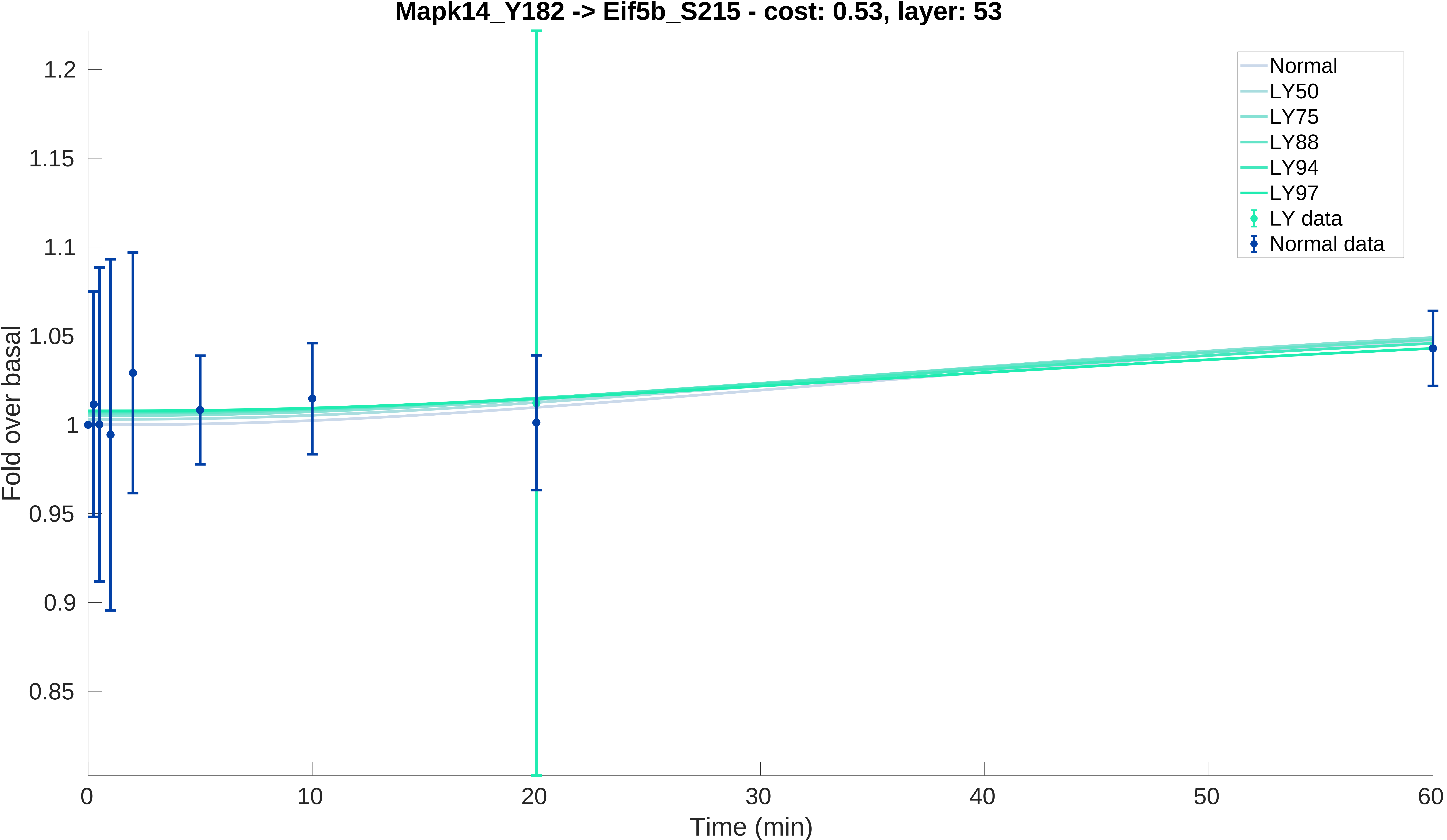

Mapk14_Y182 -> Eif5b_S215 - cost: 0.53, layer: 53
Normal
LY50
LY75
LY88
LY94
LY97
LY data
Normal data
Fold over basal
1.2
1.15
1.1
1.05
1
0.95
0.9
0.85
0
10
20
30
40
50
60
Time (min)

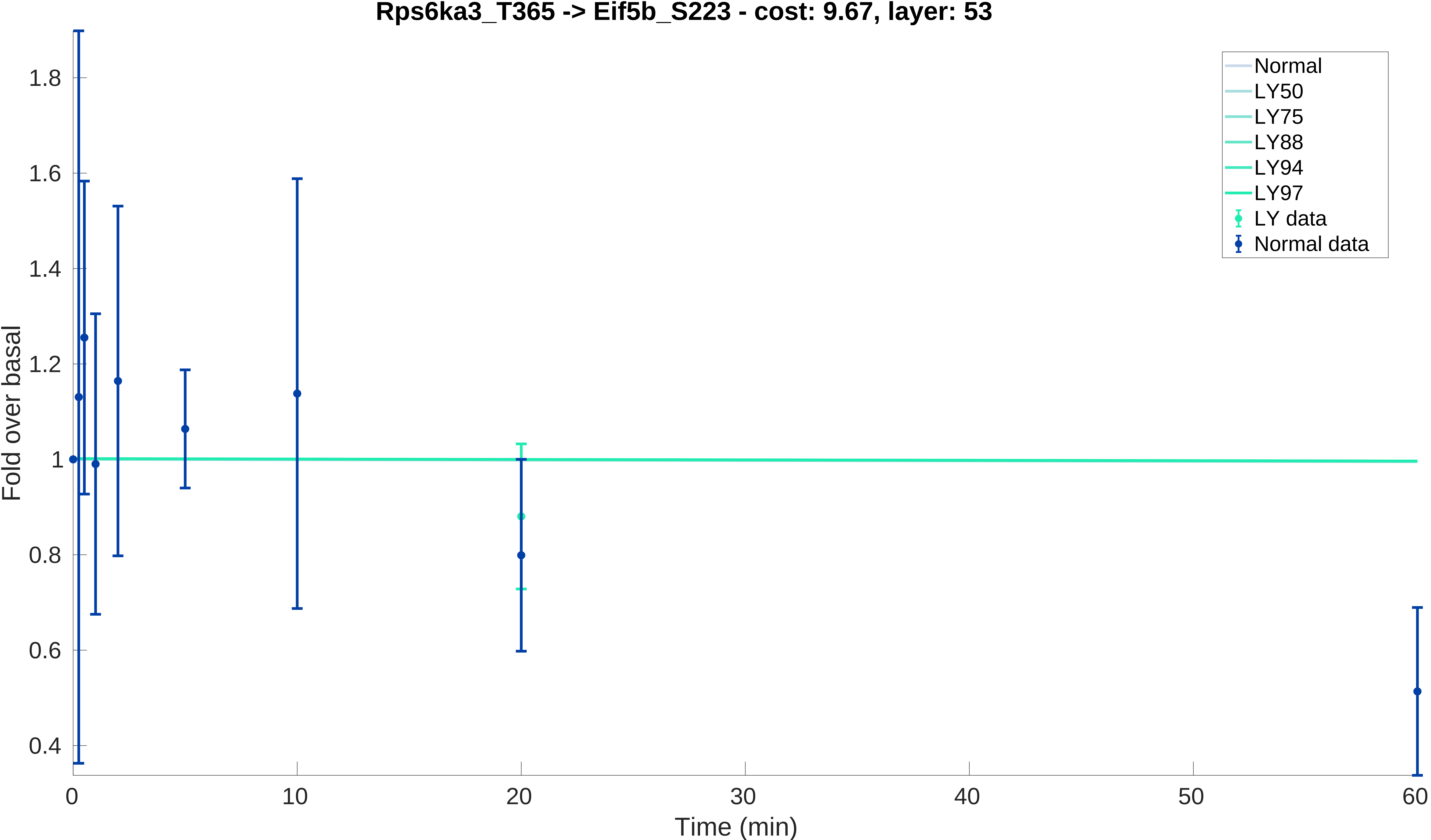

Rps6ka3_T365 -> Eif5b_S223 - cost: 9.67, layer: 53
Normal
LY50
LY75
LY88
LY94
LY97
LY data
Normal data
Fold over basal
1.8
1.6
1.4
1.2
1
0.8
0.6
0.4
0
10
20
30
40
50
60
Time (min)

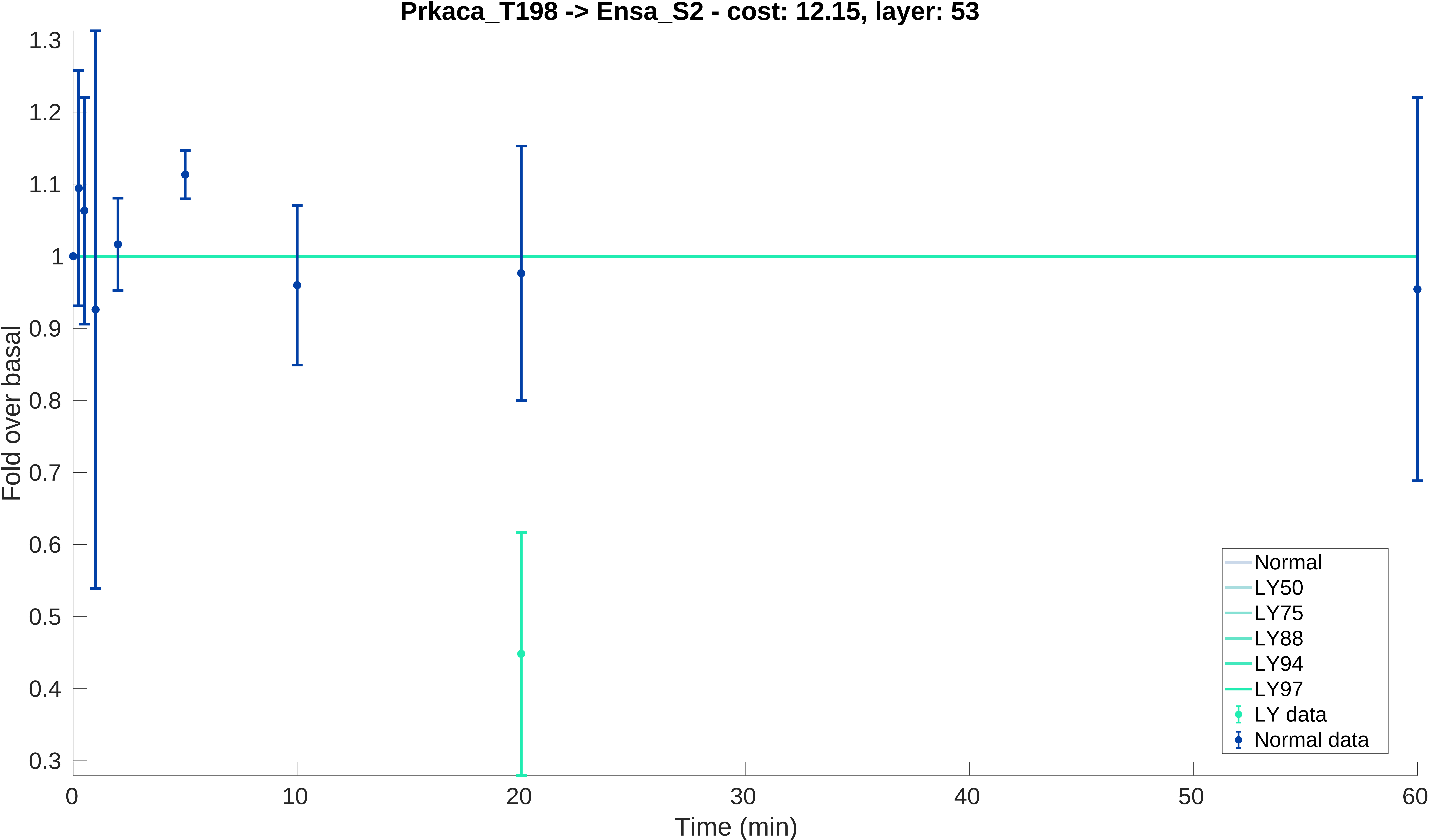

Prkaca_T198 -> Ensa_S2 - cost: 12.15, layer: 53
Fold over basal
Time (min)
1.3
1.2
1.1
1
0.9
0.8
0.7
0.6
0.5
0.4
0.3
0
10
20
30
40
50
60
Normal
LY50
LY75
LY88
LY94
LY97
LY data
Normal data

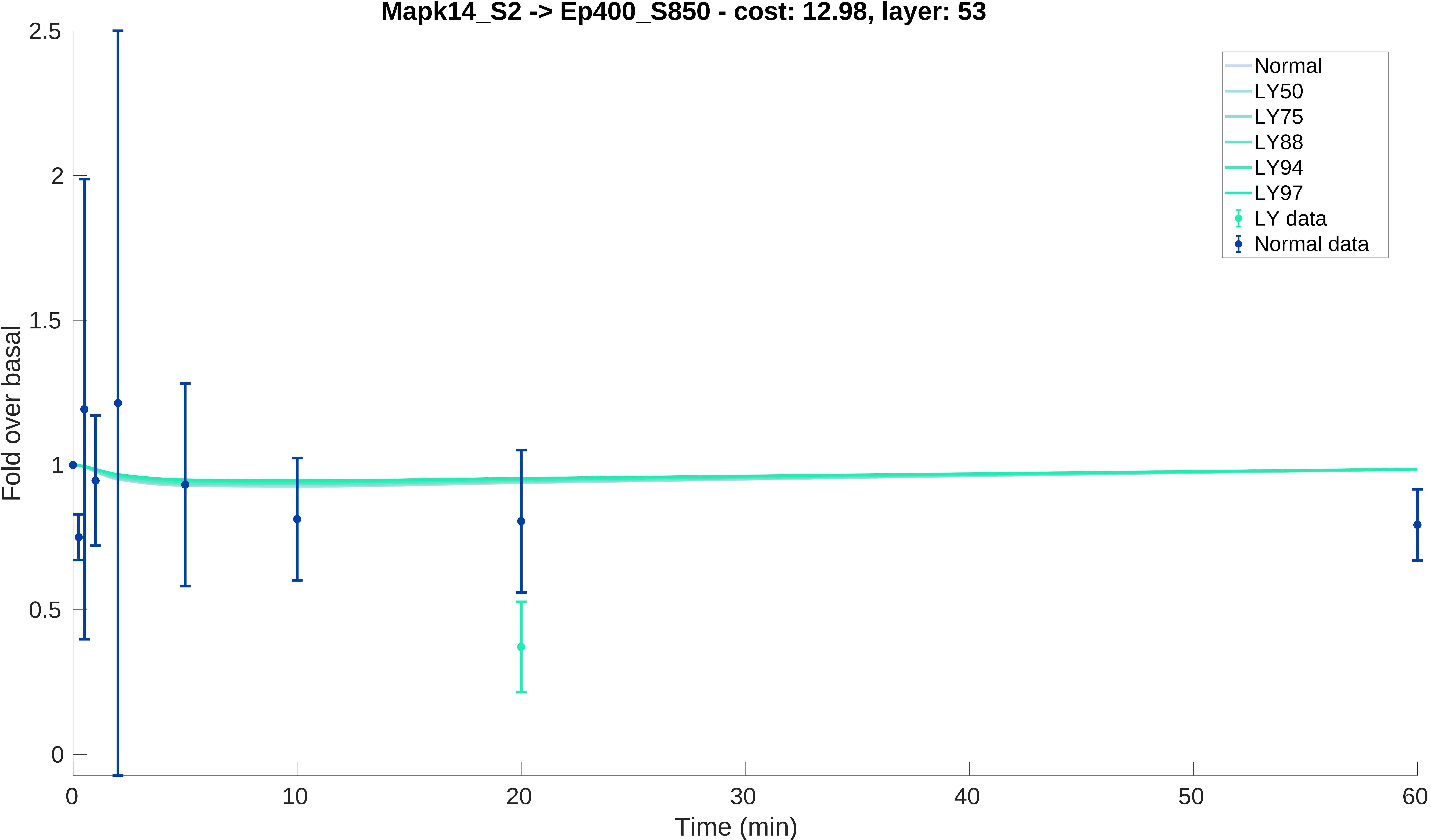

Mapk14_S2 -> Ep400_S850 - cost: 12.98, layer: 53
Fold over basal
Time (min)
0
0.5
1
1.5
2
2.5
0
10
20
30
40
50
60
Normal
LY50
LY75
LY88
LY94
LY97
LY data
Normal data

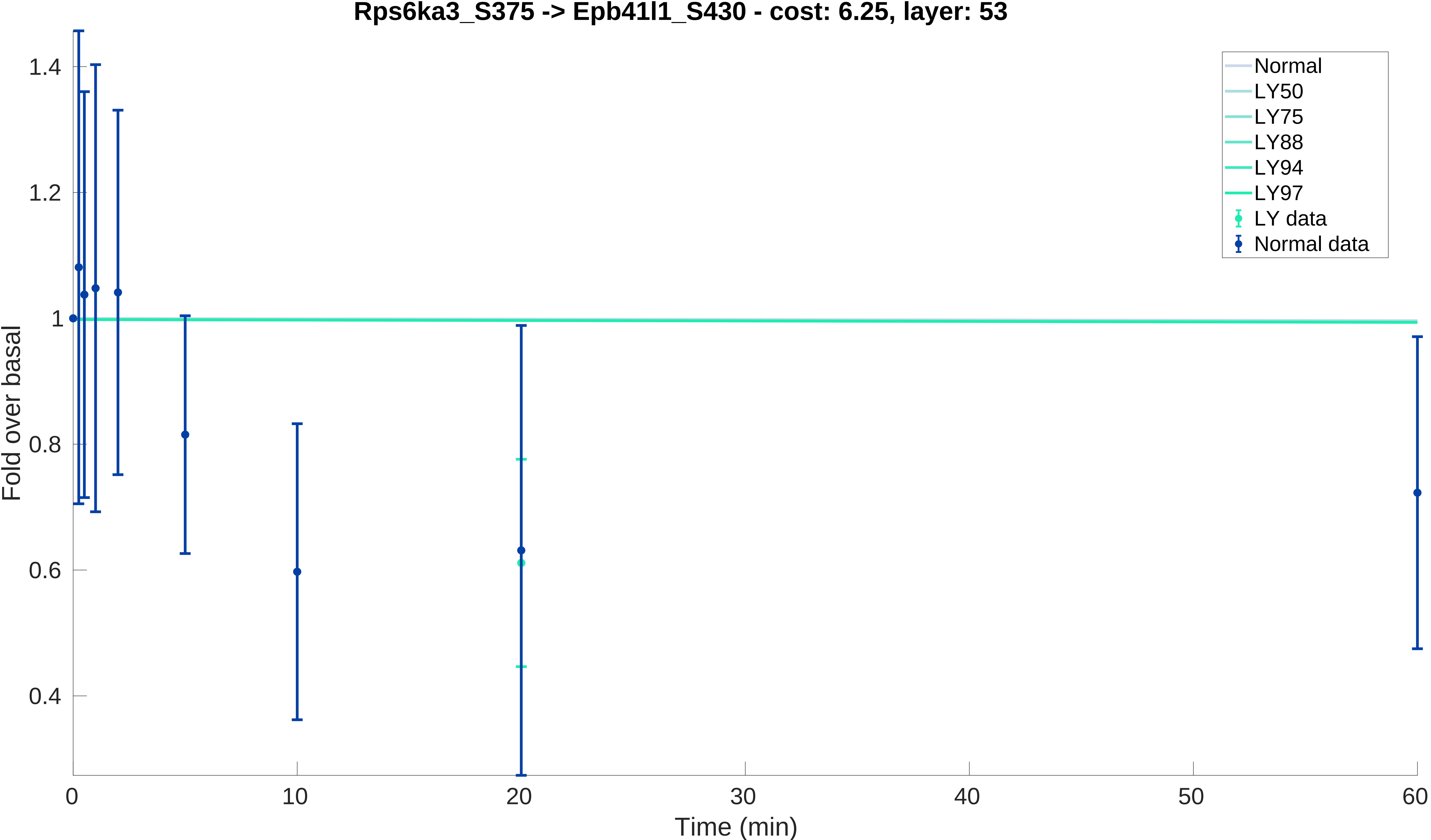

Rps6ka3_S375 -> Epb41l1_S430 - cost: 6.25, layer: 53
Normal
LY50
LY75
LY88
LY94
LY97
LY data
Normal data
Fold over basal
1.4
1.2
1
0.8
0.6
0.4
0
10
20
30
40
50
60
Time (min)

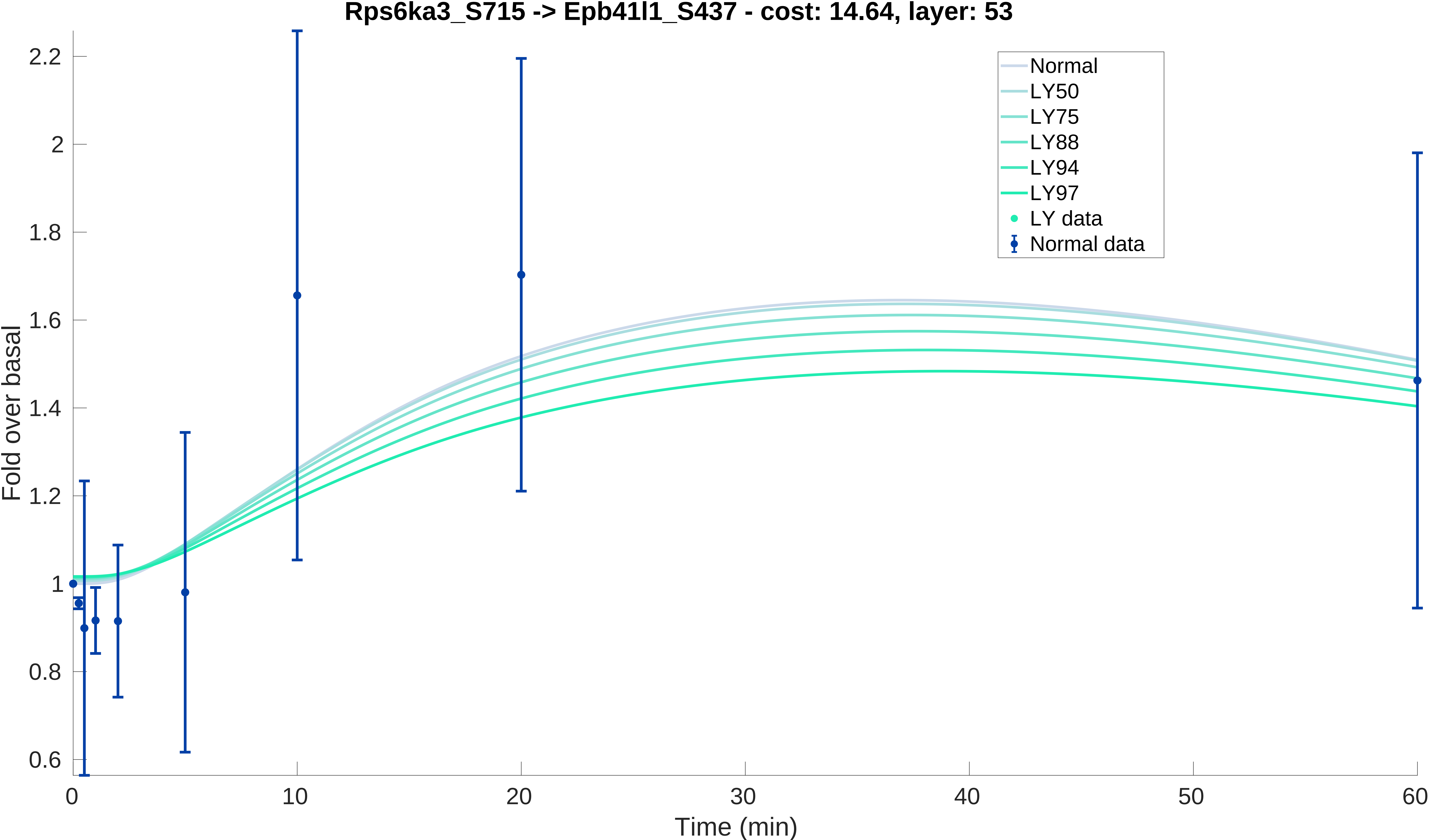
Rps6ka3_S715 -> Epb41l1_S437 - cost: 14.64, layer: 53
Normal
LY50
LY75
LY88
LY94
LY97
LY data
Normal data
Fold over basal
2.2
2
1.8
1.6
1.4
1.2
1
0.8
0.6
0
10
20
30
40
50
60
Time (min)

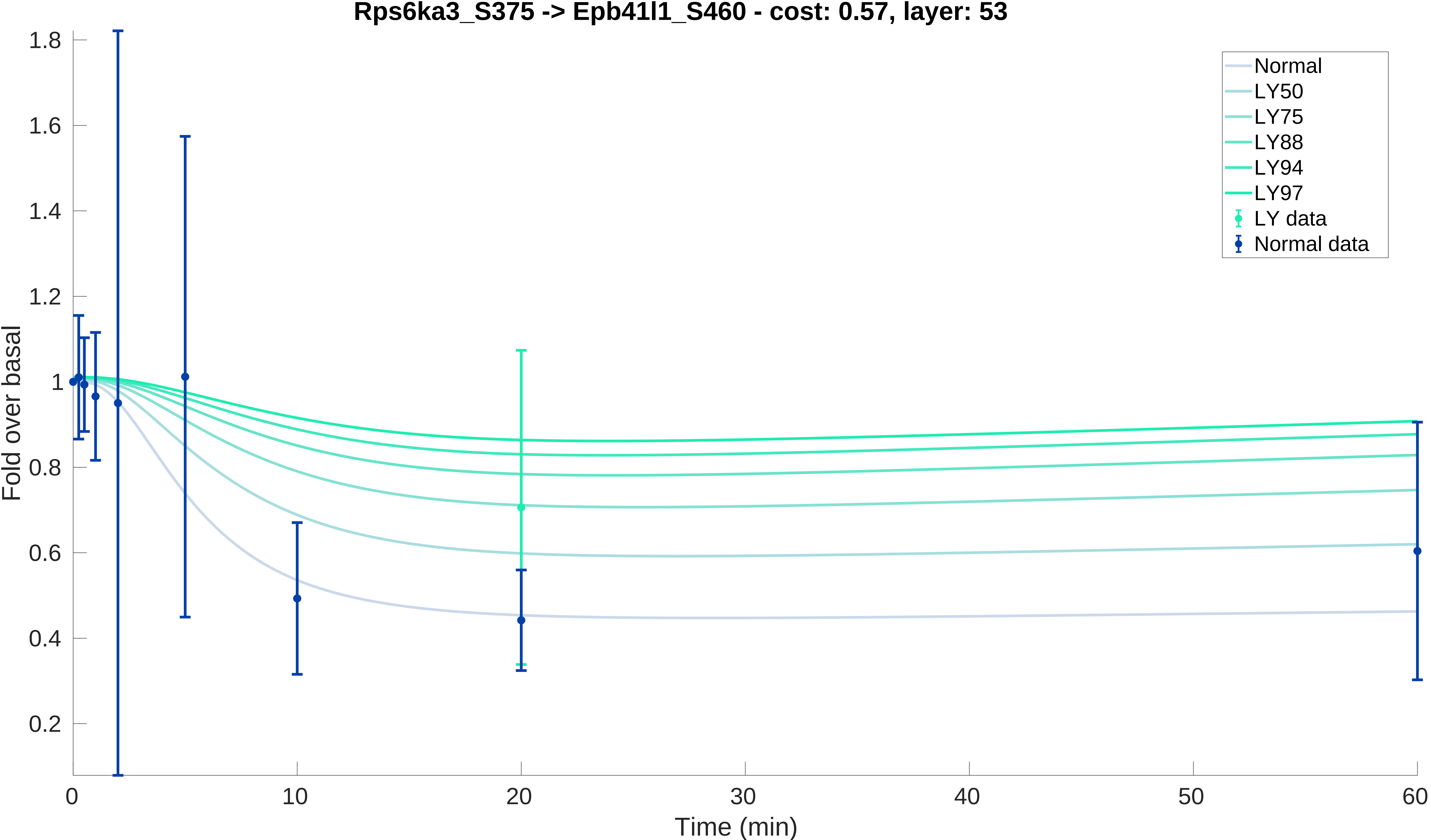

Rps6ka3_S375 -> Epb41l1_S460 - cost: 0.57, layer: 53
Fold over basal
1.8
1.6
1.4
1.2
1
0.8
0.6
0.4
0.2
0
10
20
30
40
50
60
Time (min)
Normal
LY50
LY75
LY88
LY94
LY97
LY data
Normal data

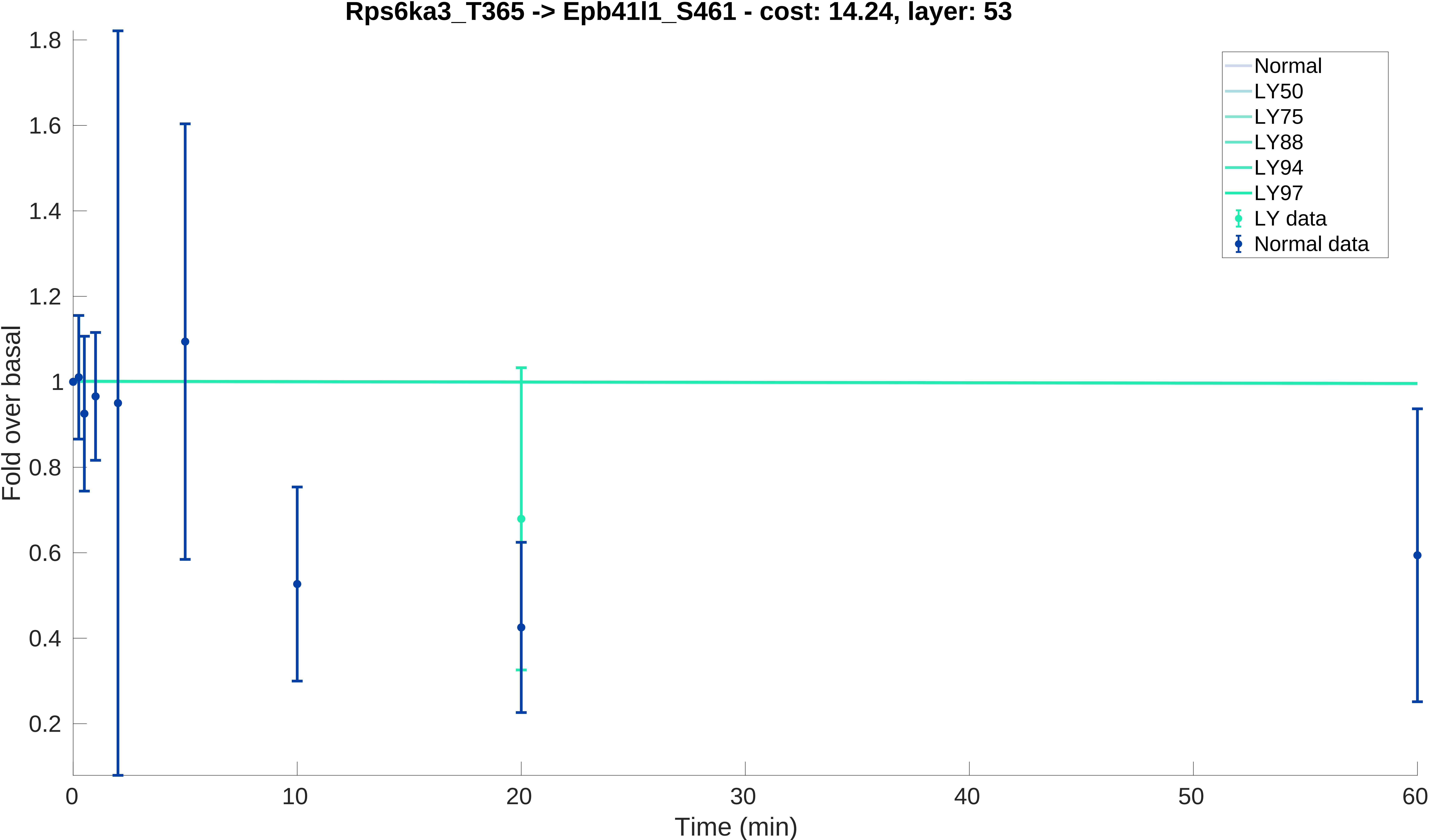

Rps6ka3_T365 -> Epb41l1_S461 - cost: 14.24, layer: 53
Fold over basal
Time (min)
1.8
1.6
1.4
1.2
1
0.8
0.6
0.4
0.2
0
10
20
30
40
50
60
Normal
LY50
LY75
LY88
LY94
LY97
LY data
Normal data

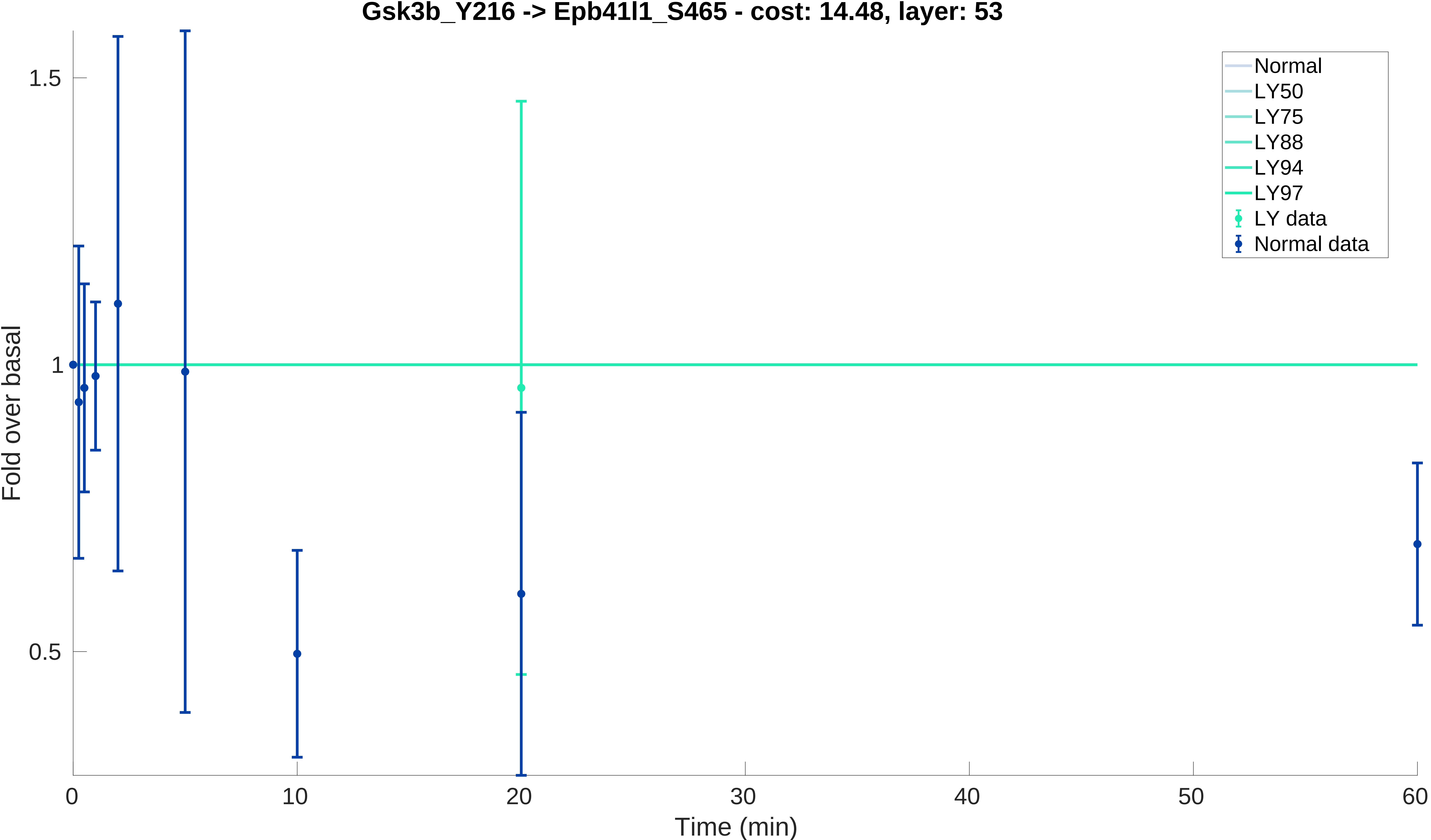

Gsk3b_Y216 -> Epb41l1_S465 - cost: 14.48, layer: 53
Normal
LY50
LY75
LY88
LY94
LY97
LY data
Normal data
Fold over basal
1.5
1
0.5
0
10
20
30
40
50
60
Time (min)

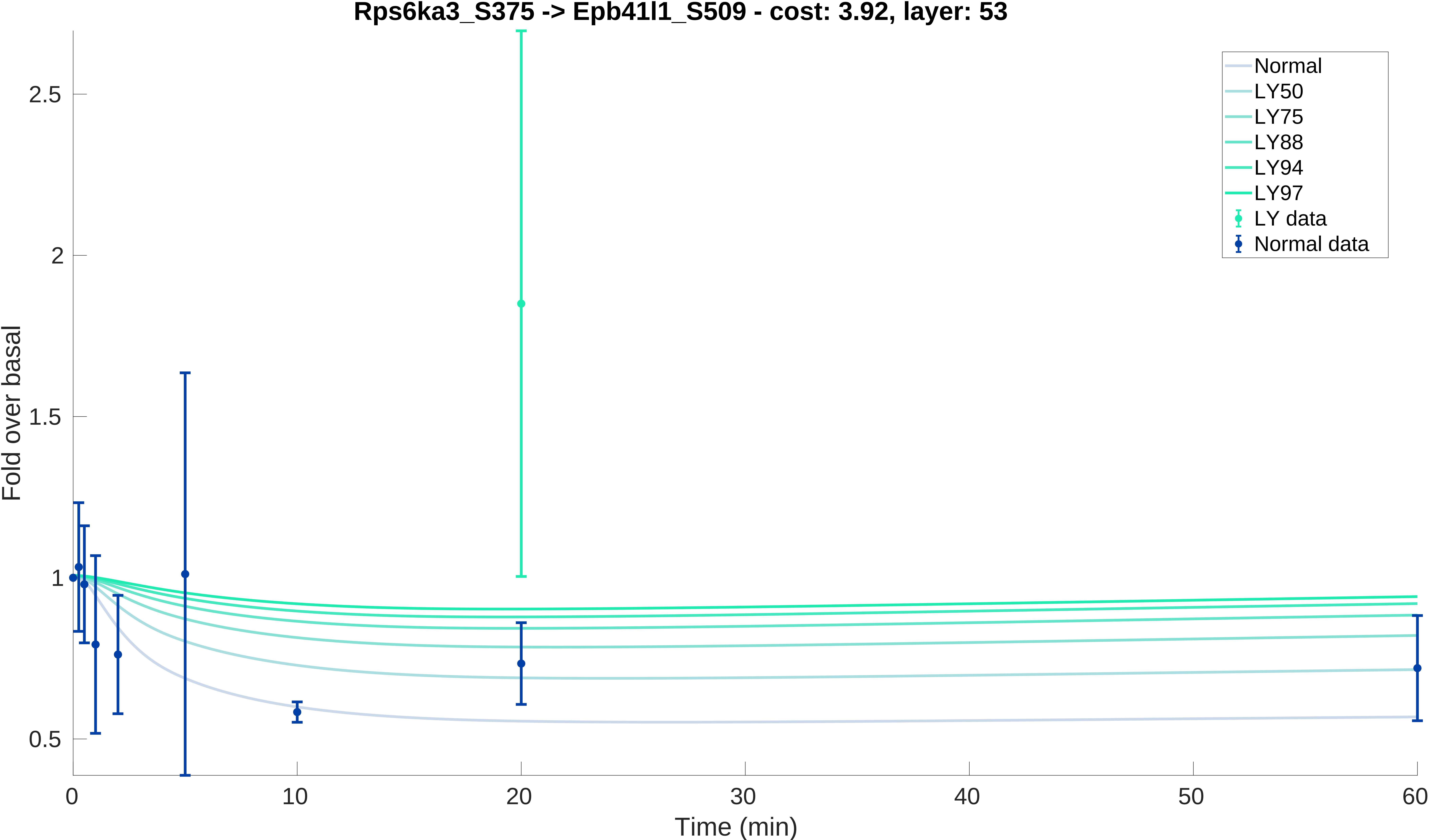
Rps6ka3_S375 -> Epb41l1_S509 - cost: 3.92, layer: 53
Fold over basal
2.5
2
1.5
1
0.5
0
10
20
30
40
50
60
Time (min)
Normal
LY50
LY75
LY88
LY94
LY97
LY data
Normal data

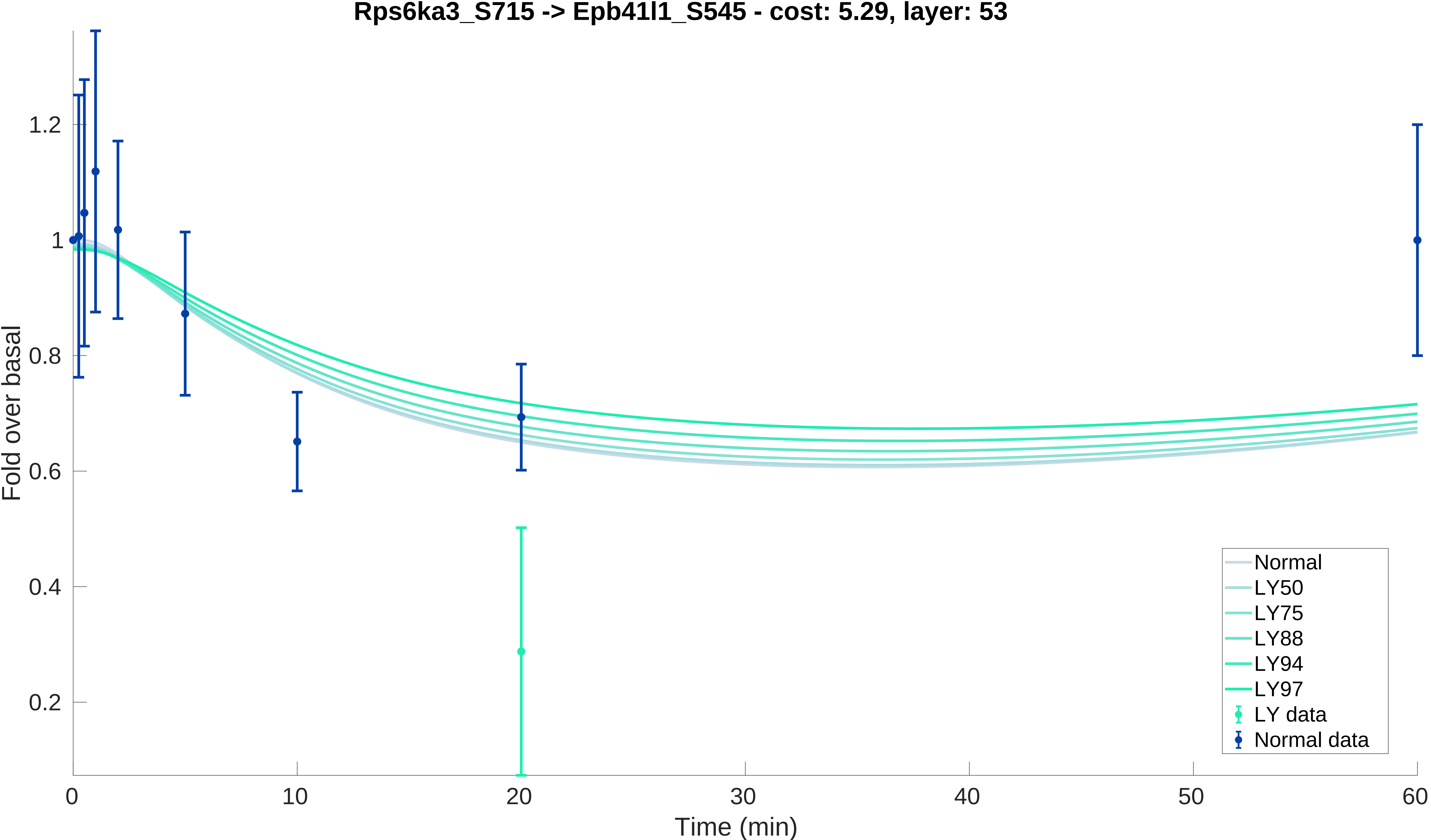
Rps6ka3_S715 -> Epb41l1_S545 - cost: 5.29, layer: 53
Fold over basal
1.2
1
0.8
0.6
0.4
0.2
0
10
20
30
40
50
60
Time (min)
Normal
LY50
LY75
LY88
LY94
LY97
LY data
Normal data

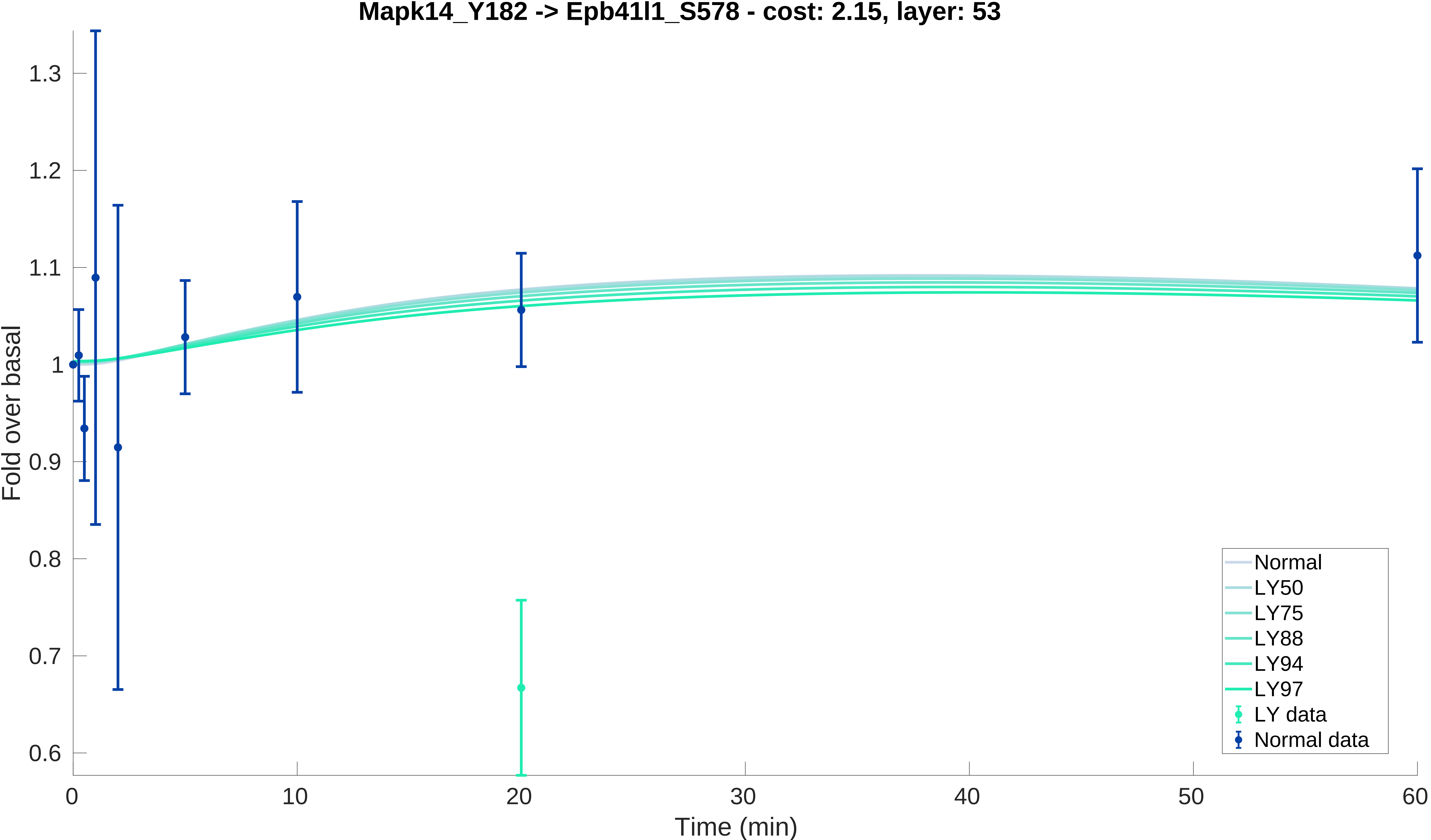

Mapk14_Y182 -> Epb41l1_S578 - cost: 2.15, layer: 53
Fold over basal
1.3
1.2
1.1
1
0.9
0.8
0.7
0.6
0
10
20
30
40
50
60
Time (min)
Normal
LY50
LY75
LY88
LY94
LY97
LY data
Normal data

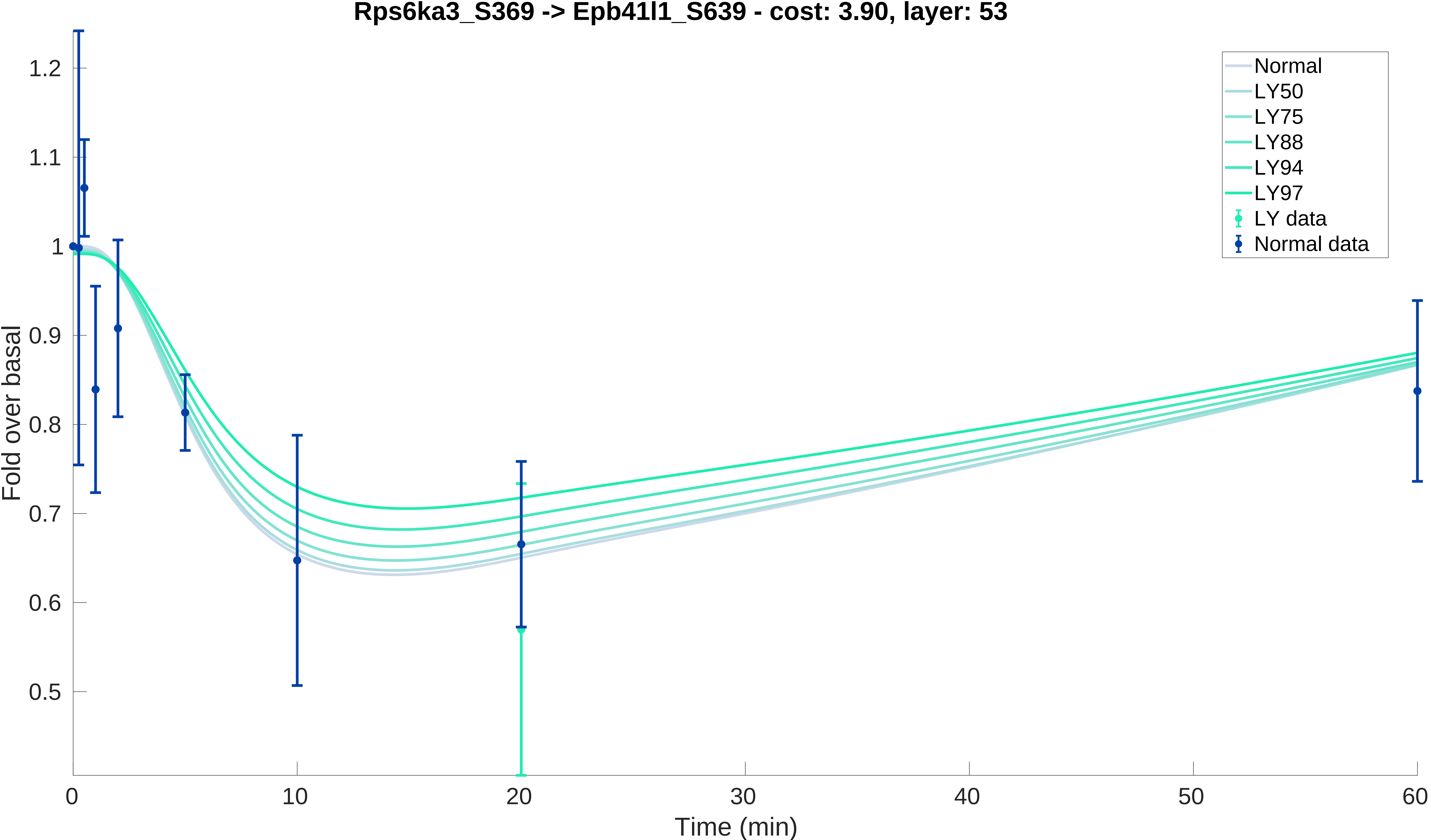

Rps6ka3_S369 -> Epb41l1_S639 - cost: 3.90, layer: 53
Normal
LY50
LY75
LY88
LY94
LY97
LY data
Normal data
Fold over basal
1.2
1.1
1
0.9
0.8
0.7
0.6
0.5
0
10
20
30
40
50
60
Time (min)

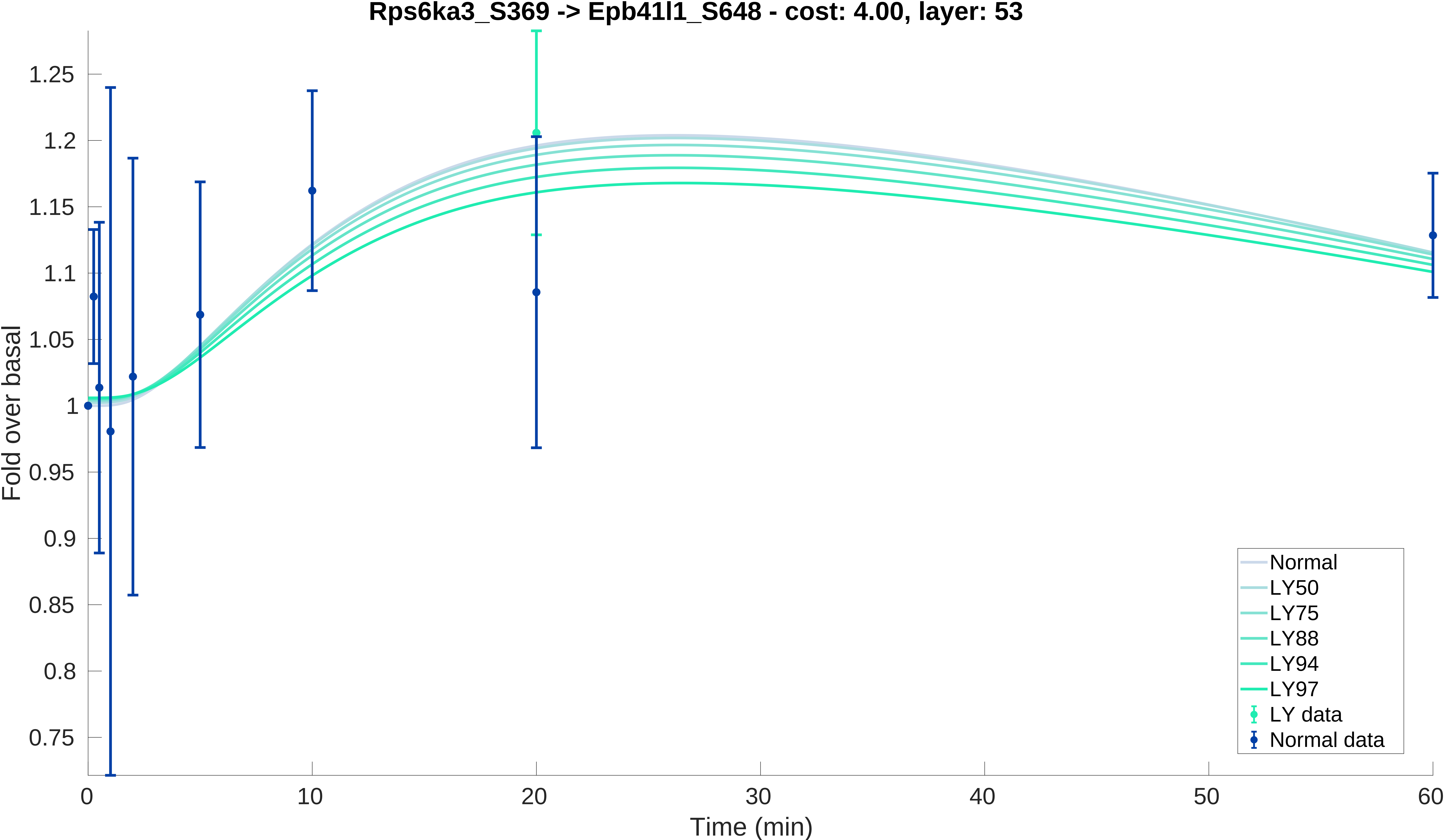

Rps6ka3_S369 -> Epb41l1_S648 - cost: 4.00, layer: 53
Fold over basal
Time (min)
1.25
1.2
1.15
1.1
1.05
1
0.95
0.9
0.85
0.8
0.75
0
10
20
30
40
50
60
Normal
LY50
LY75
LY88
LY94
LY97
LY data
Normal data

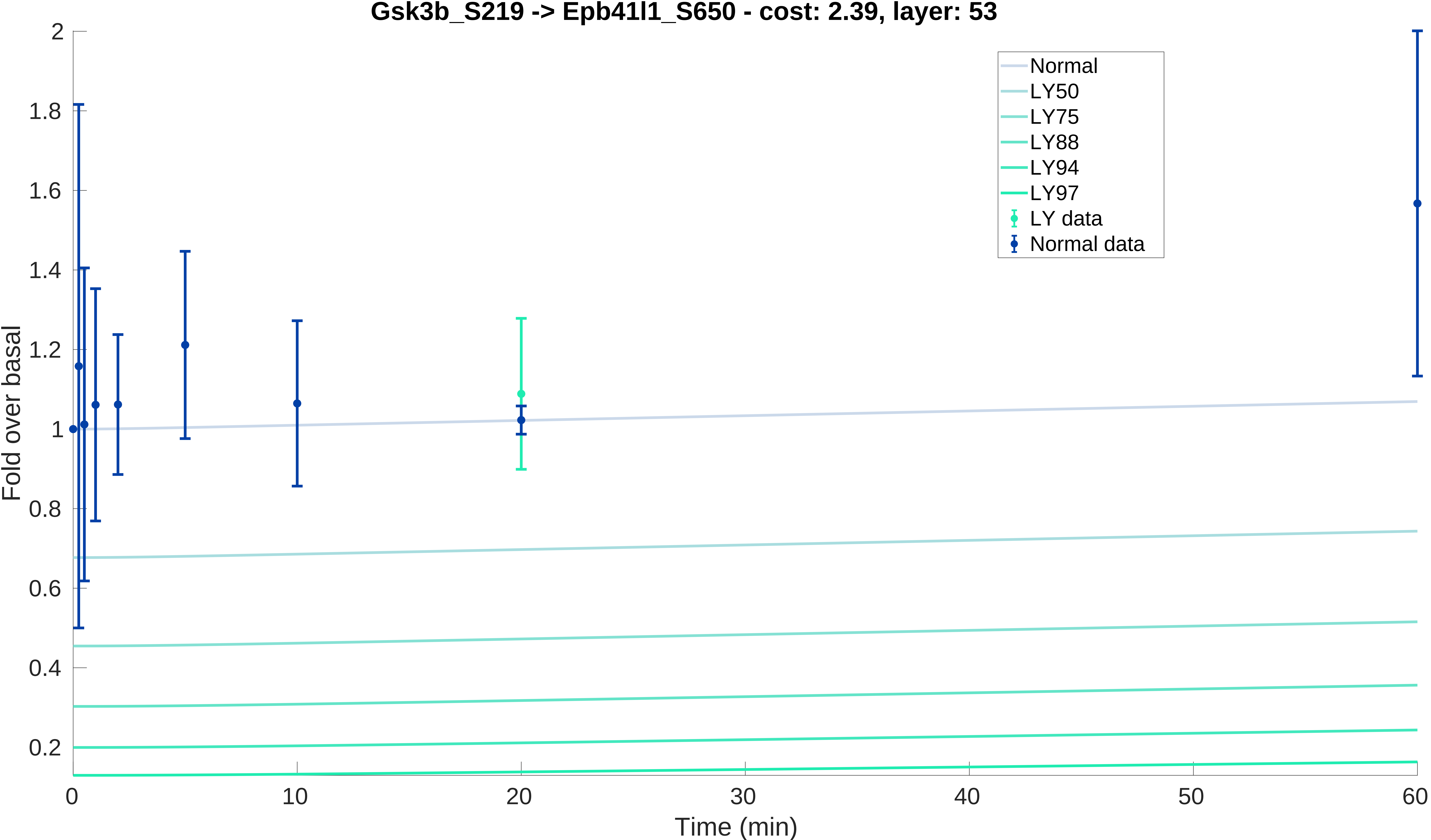

Gsk3b_S219 -> Epb41l1_S650 - cost: 2.39, layer: 53
Fold over basal
Time (min)
2
1.8
1.6
1.4
1.2
1
0.8
0.6
0.4
0.2
0
10
20
30
40
50
60
Normal
LY50
LY75
LY88
LY94
LY97
LY data
Normal data

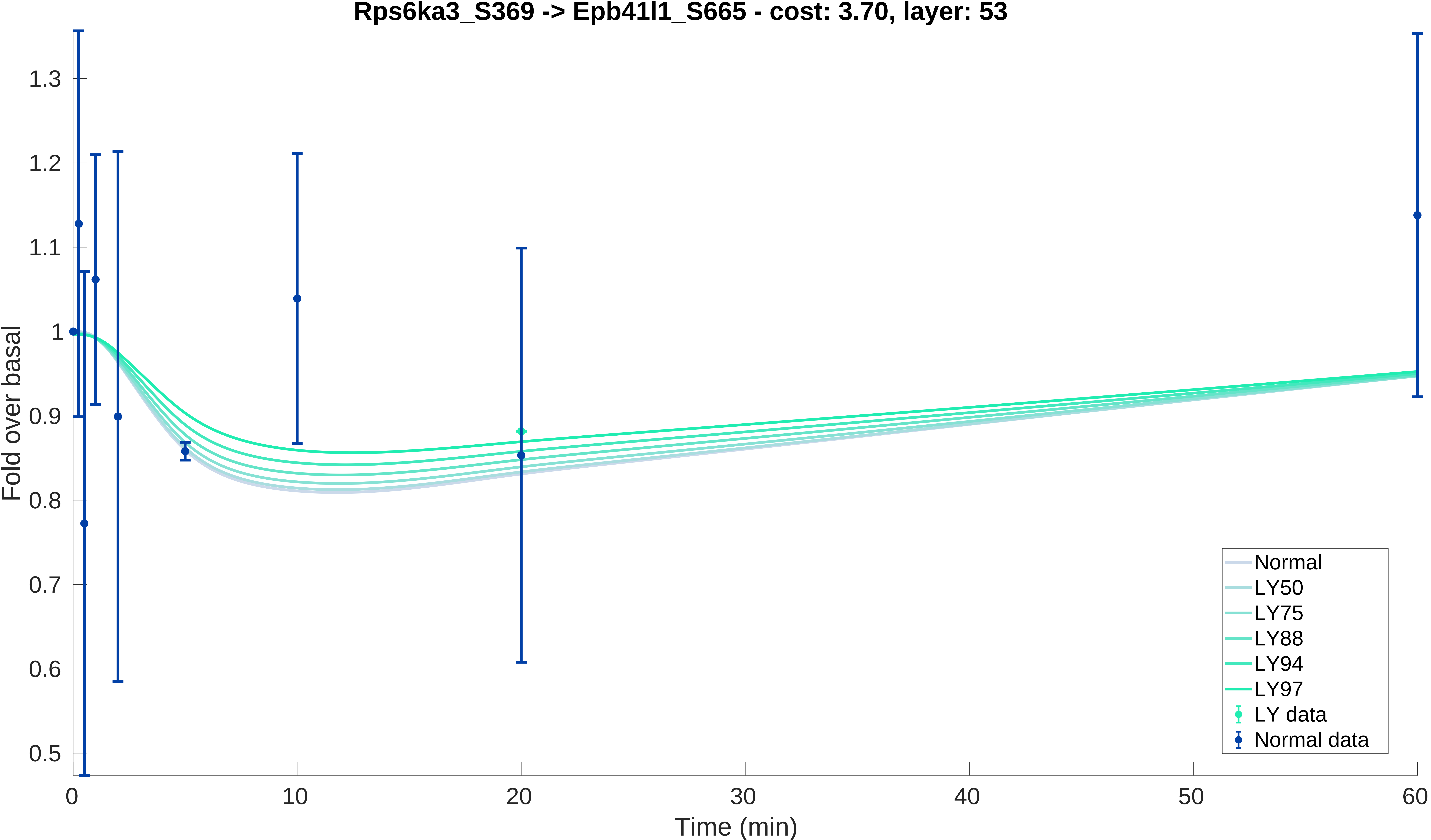

Rps6ka3_S369 -> Epb41l1_S665 - cost: 3.70, layer: 53
Fold over basal
1.3
1.2
1.1
1
0.9
0.8
0.7
0.6
0.5
0
10
20
30
40
50
60
Time (min)
Normal
LY50
LY75
LY88
LY94
LY97
LY data
Normal data

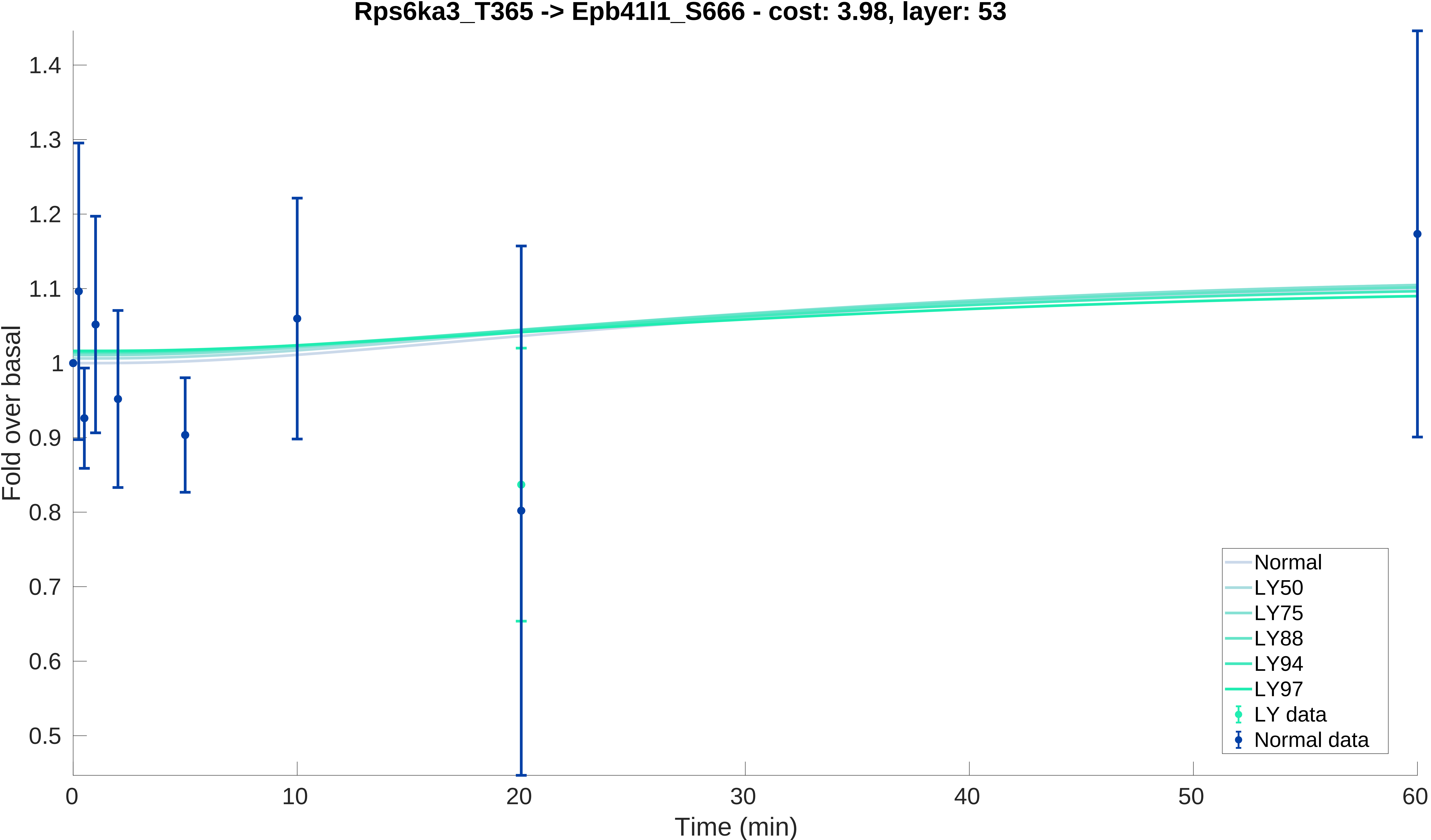

Rps6ka3_T365 -> Epb41l1_S666 - cost: 3.98, layer: 53
Fold over basal
Time (min)
0
10
20
30
40
50
60
0.5
0.6
0.7
0.8
0.9
1
1.1
1.2
1.3
1.4
Normal
LY50
LY75
LY88
LY94
LY97
LY data
Normal data

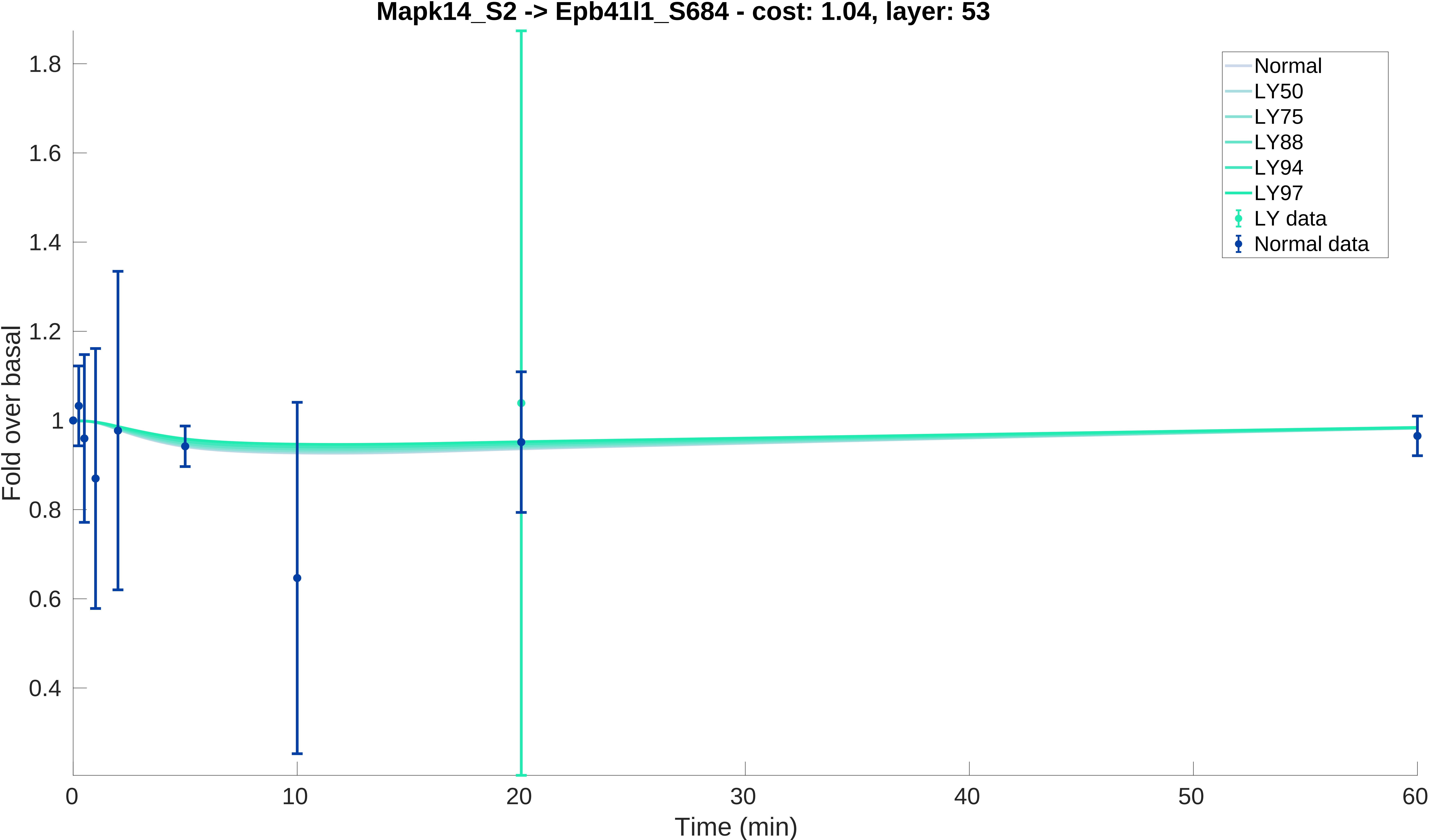
Mapk14_S2 -> Epb41l1_S684 - cost: 1.04, layer: 53
Fold over basal
Time (min)
1.8
1.6
1.4
1.2
1
0.8
0.6
0.4
0
10
20
30
40
50
60
Normal
LY50
LY75
LY88
LY94
LY97
LY data
Normal data

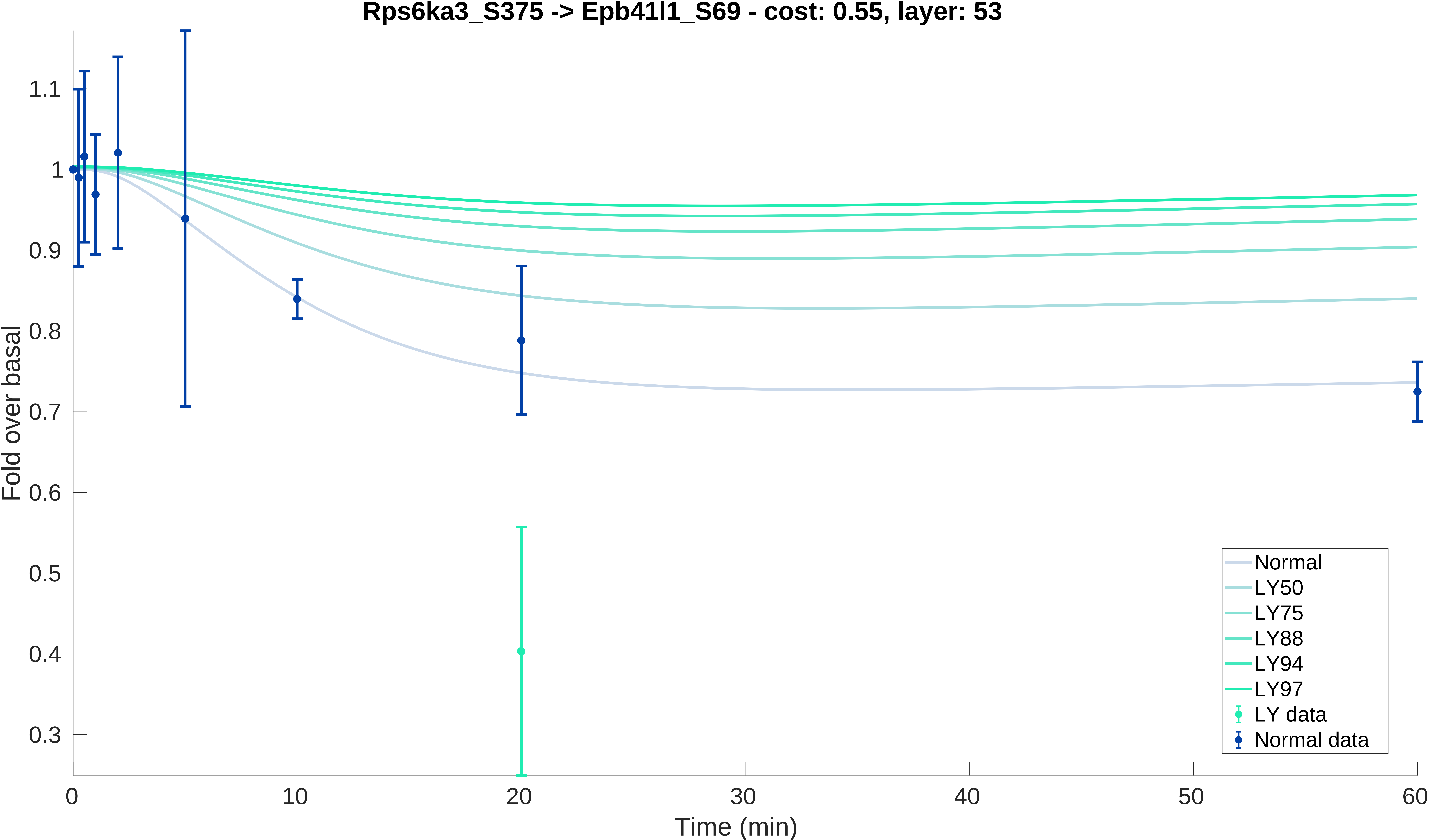

Rps6ka3_S375 -> Epb41l1_S69 - cost: 0.55, layer: 53
Fold over basal
1.1
1
0.9
0.8
0.7
0.6
0.5
0.4
0.3
0
10
20
30
40
50
60
Time (min)
Normal
LY50
LY75
LY88
LY94
LY97
LY data
Normal data

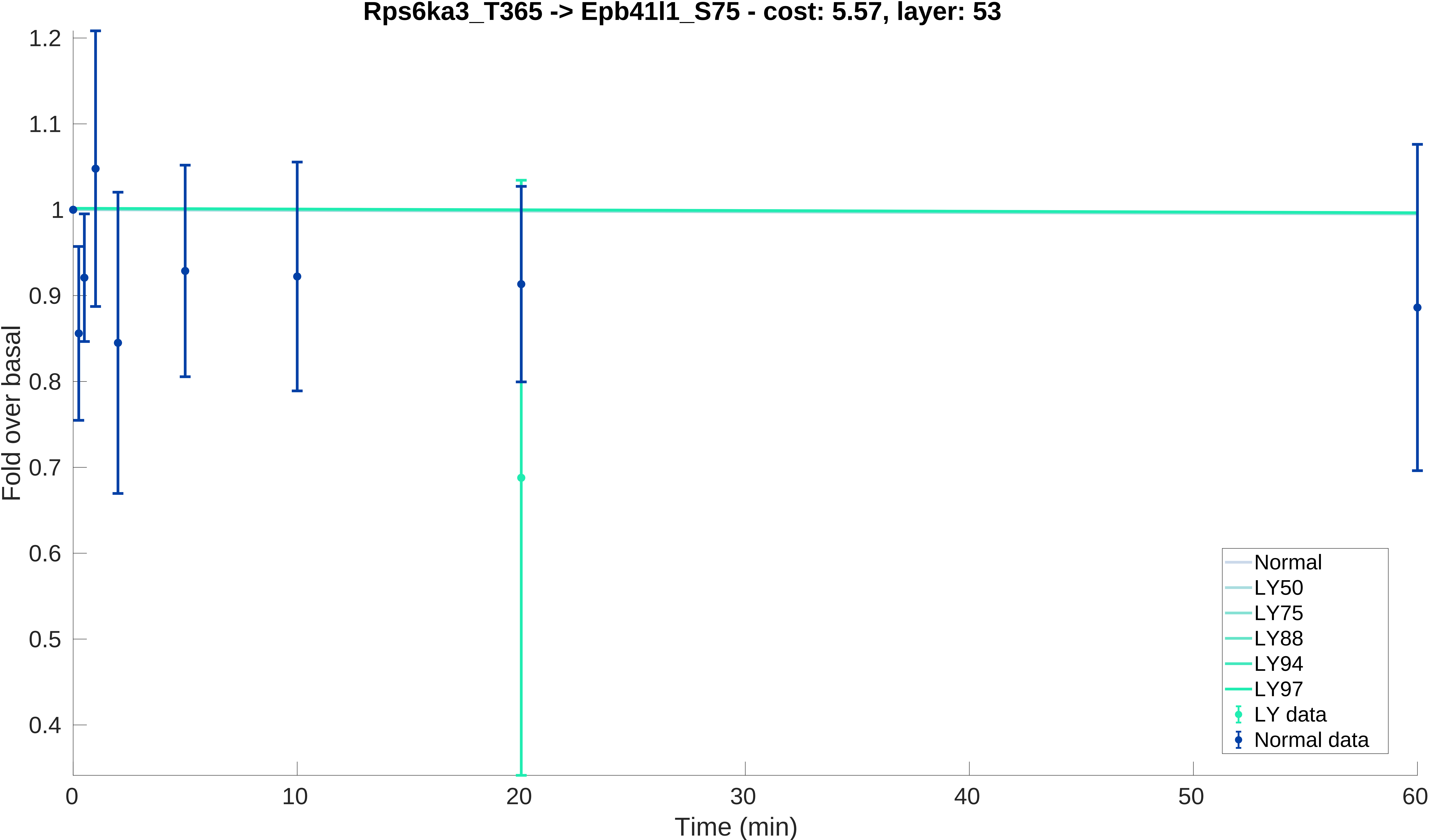
Rps6ka3_T365 -> Epb41l1_S75 - cost: 5.57, layer: 53
Fold over basal
Time (min)
1.2
1.1
1
0.9
0.8
0.7
0.6
0.5
0.4
0
10
20
30
40
50
60
Normal
LY50
LY75
LY88
LY94
LY97
LY data
Normal data

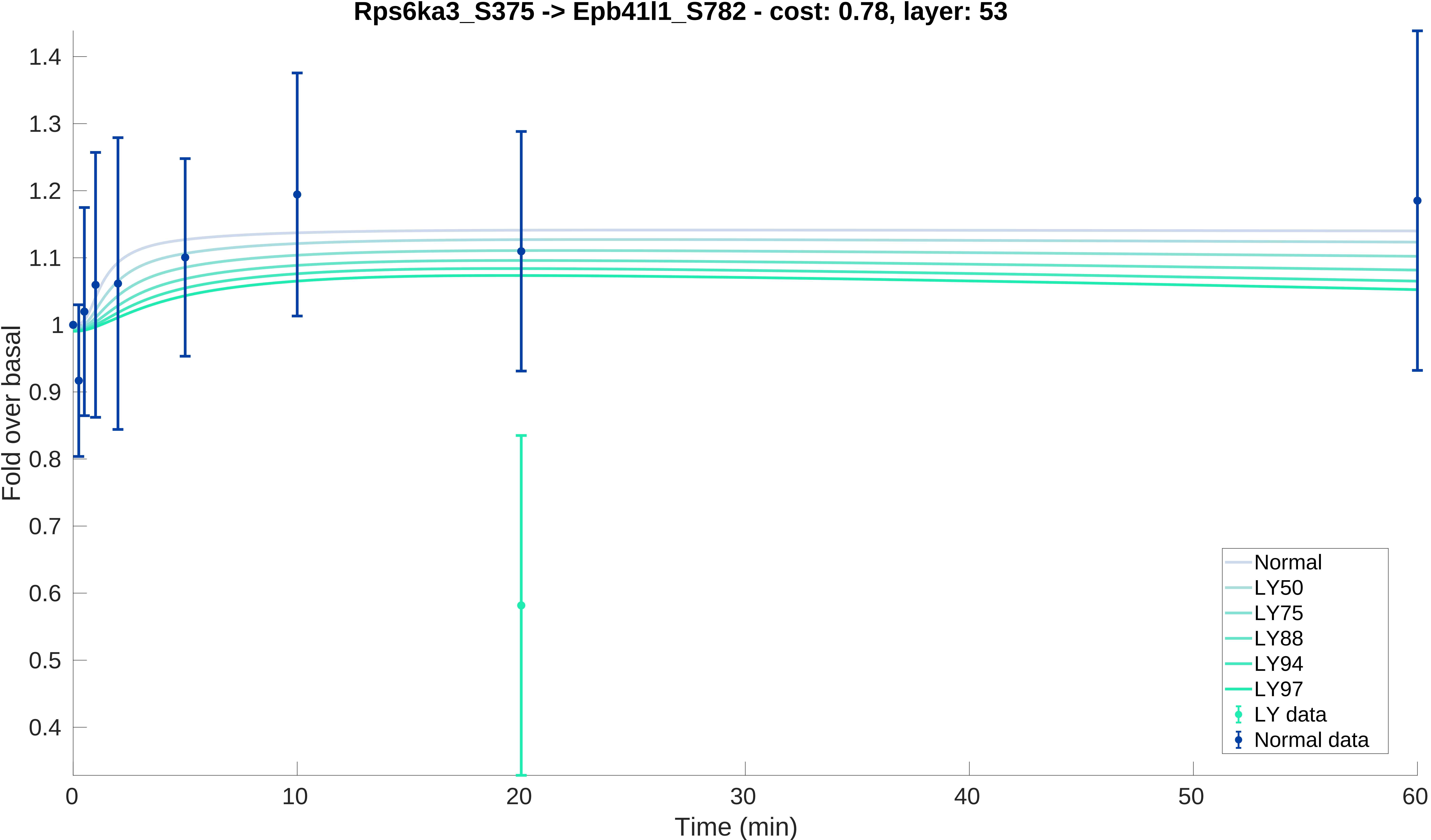

Rps6ka3_S375 -> Epb41l1_S782 - cost: 0.78, layer: 53
Fold over basal
1.4
1.3
1.2
1.1
1
0.9
0.8
0.7
0.6
0.5
0.4
0
10
20
30
40
50
60
Time (min)
Normal
LY50
LY75
LY88
LY94
LY97
LY data
Normal data

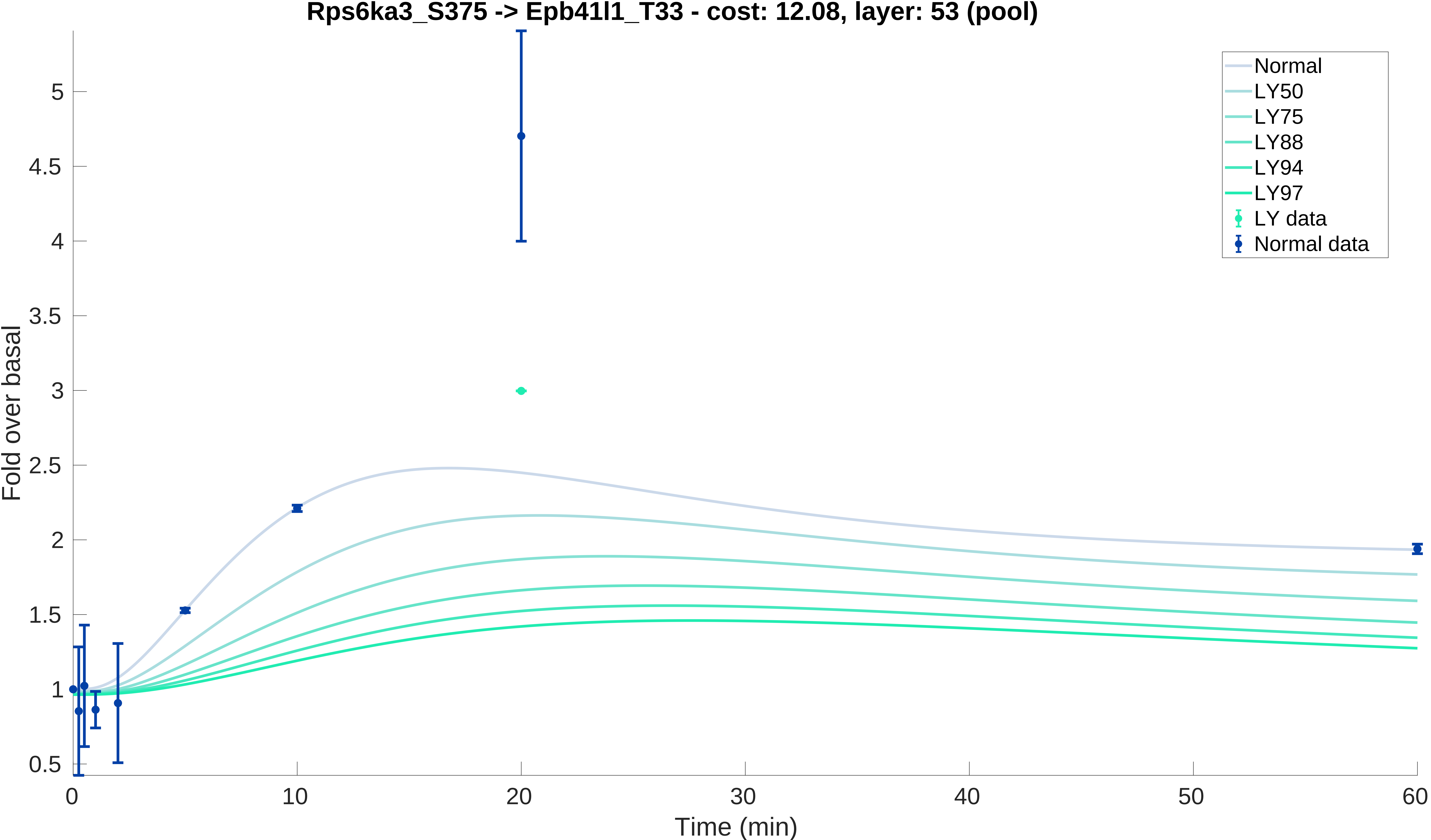

Rps6ka3_S375 -> Epb41l1_T33 - cost: 12.08, layer: 53 (pool)
Fold over basal
Time (min)
0.5
1
1.5
2
2.5
3
3.5
4
4.5
5
0
10
20
30
40
50
60
Normal
LY50
LY75
LY88
LY94
LY97
LY data
Normal data

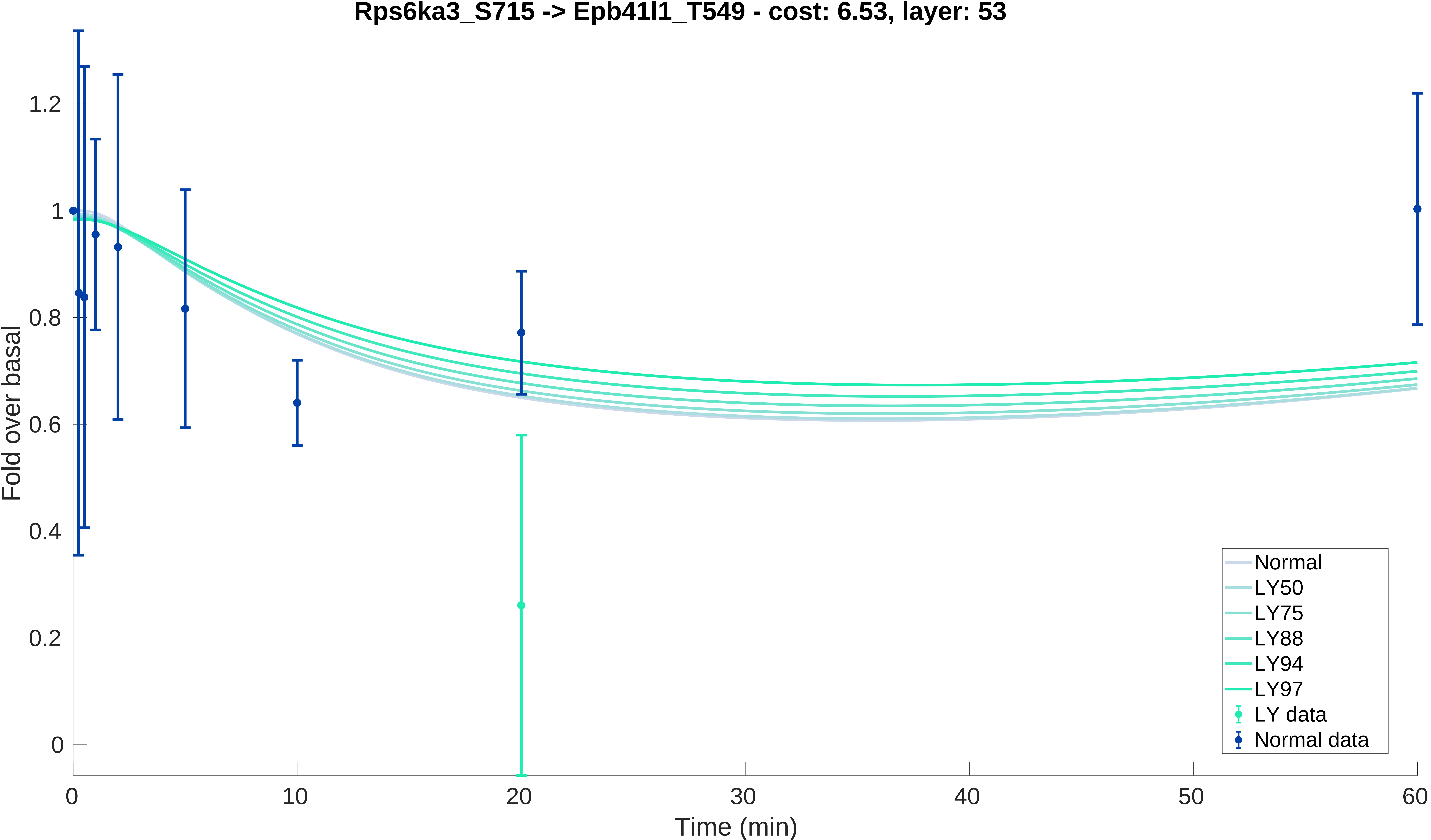

Rps6ka3_S715 -> Epb41l1_T549 - cost: 6.53, layer: 53
Fold over basal
Time (min)
0
0.2
0.4
0.6
0.8
1
1.2
0
10
20
30
40
50
60
Normal
LY50
LY75
LY88
LY94
LY97
LY data
Normal data

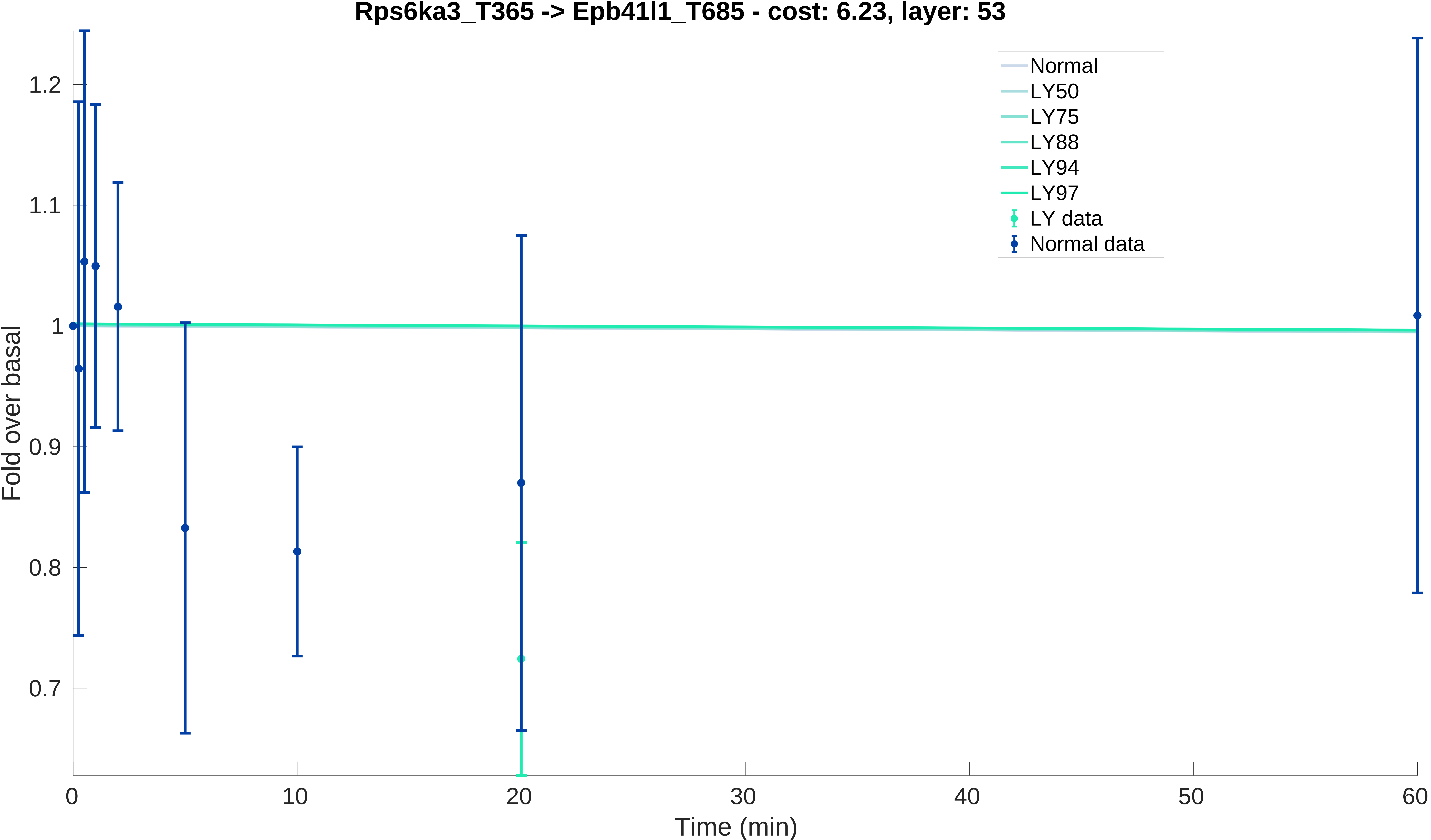
Rps6ka3_T365 -> Epb41l1_T685 - cost: 6.23, layer: 53
Normal
LY50
LY75
LY88
LY94
LY97
LY data
Normal data
Fold over basal
1.2
1.1
1
0.9
0.8
0.7
0
10
20
30
40
50
60
Time (min)

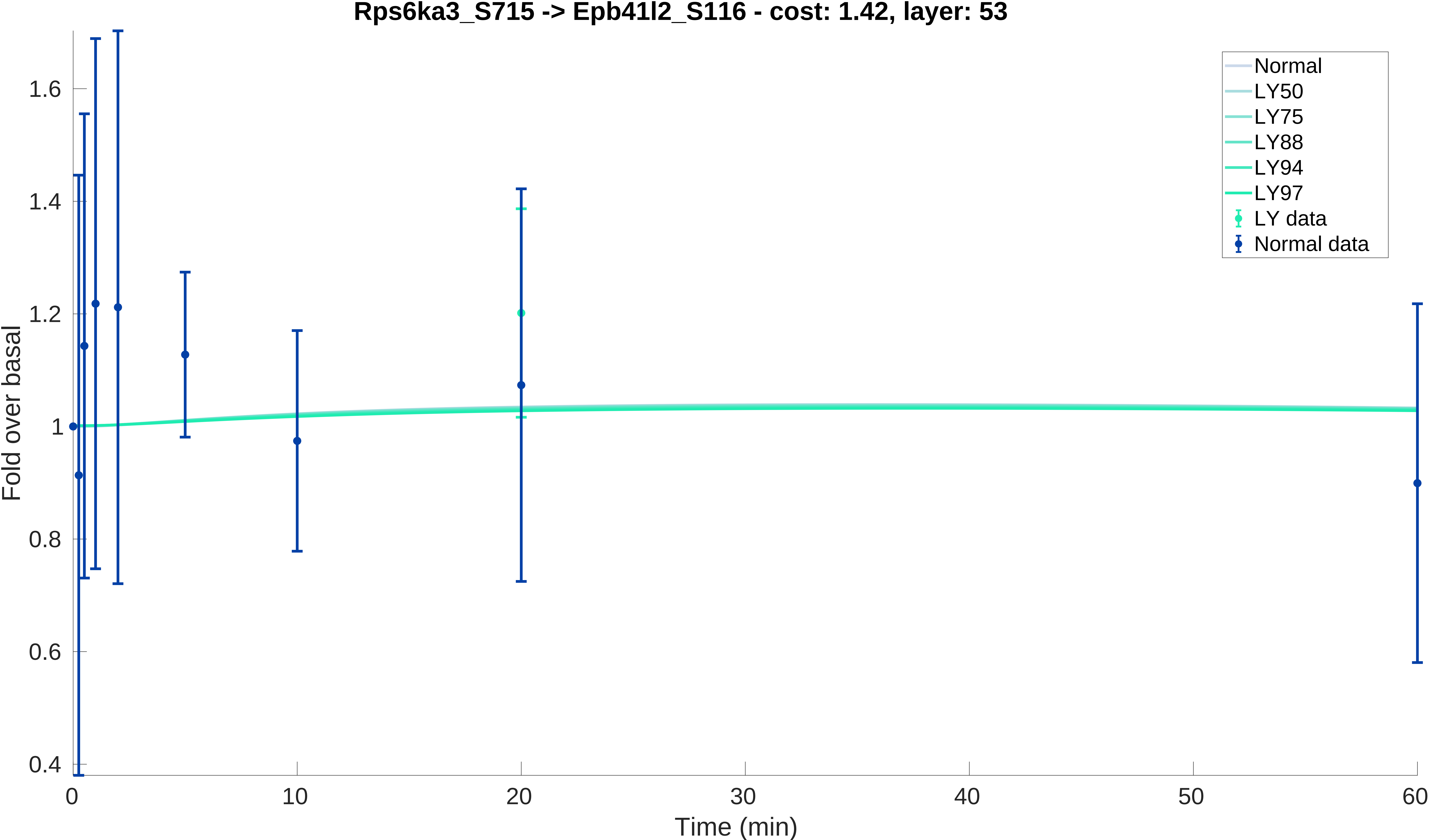
Rps6ka3_S715 -> Epb41l2_S116 - cost: 1.42, layer: 53
Normal
LY50
LY75
LY88
LY94
LY97
LY data
Normal data
Fold over basal
1.6
1.4
1.2
1
0.8
0.6
0.4
0
10
20
30
40
50
60
Time (min)

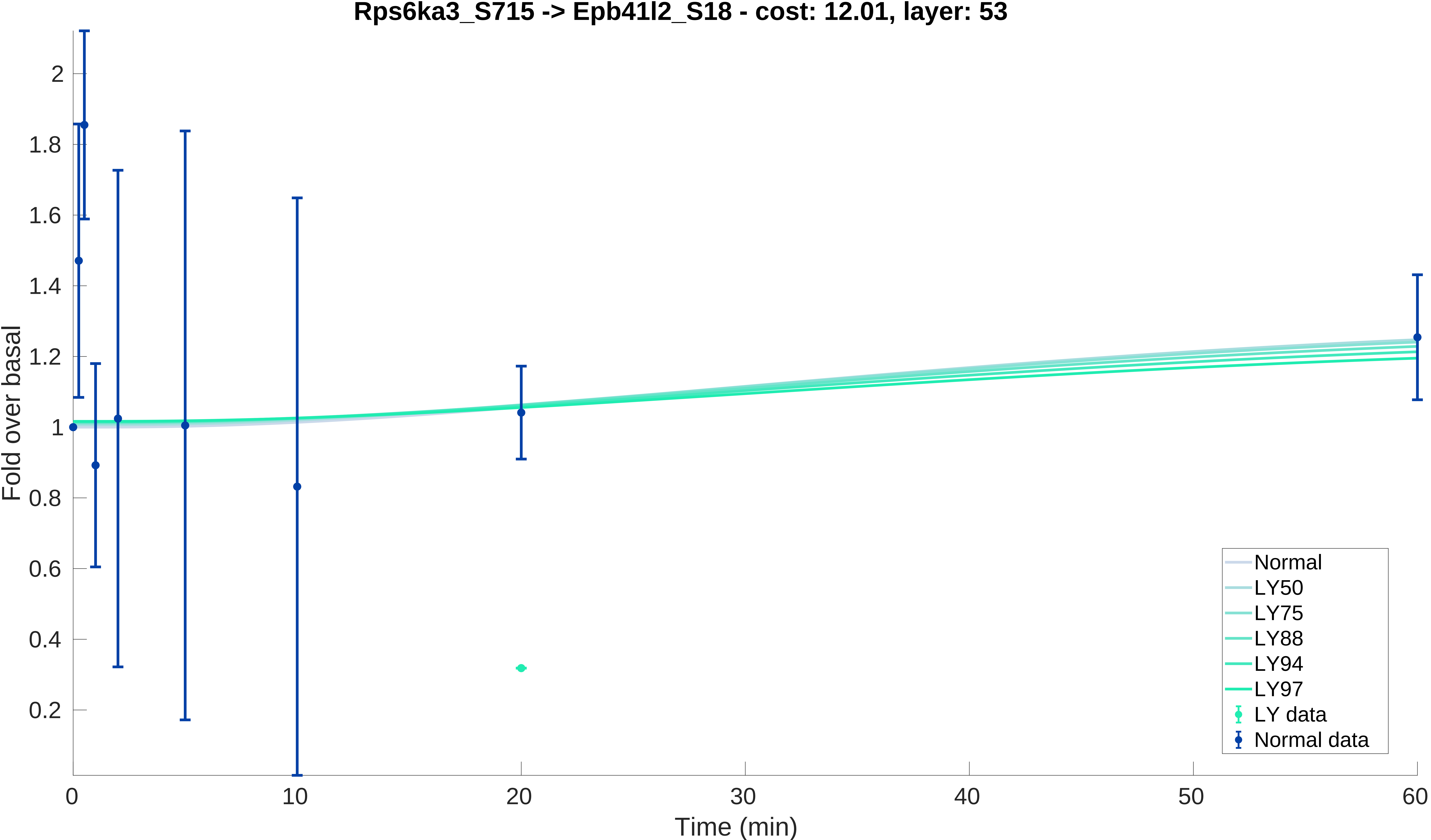

Rps6ka3_S715 -> Epb41l2_S18 - cost: 12.01, layer: 53
Fold over basal
Time (min)
0
10
20
30
40
50
60
0.2
0.4
0.6
0.8
1
1.2
1.4
1.6
1.8
2
Normal
LY50
LY75
LY88
LY94
LY97
LY data
Normal data

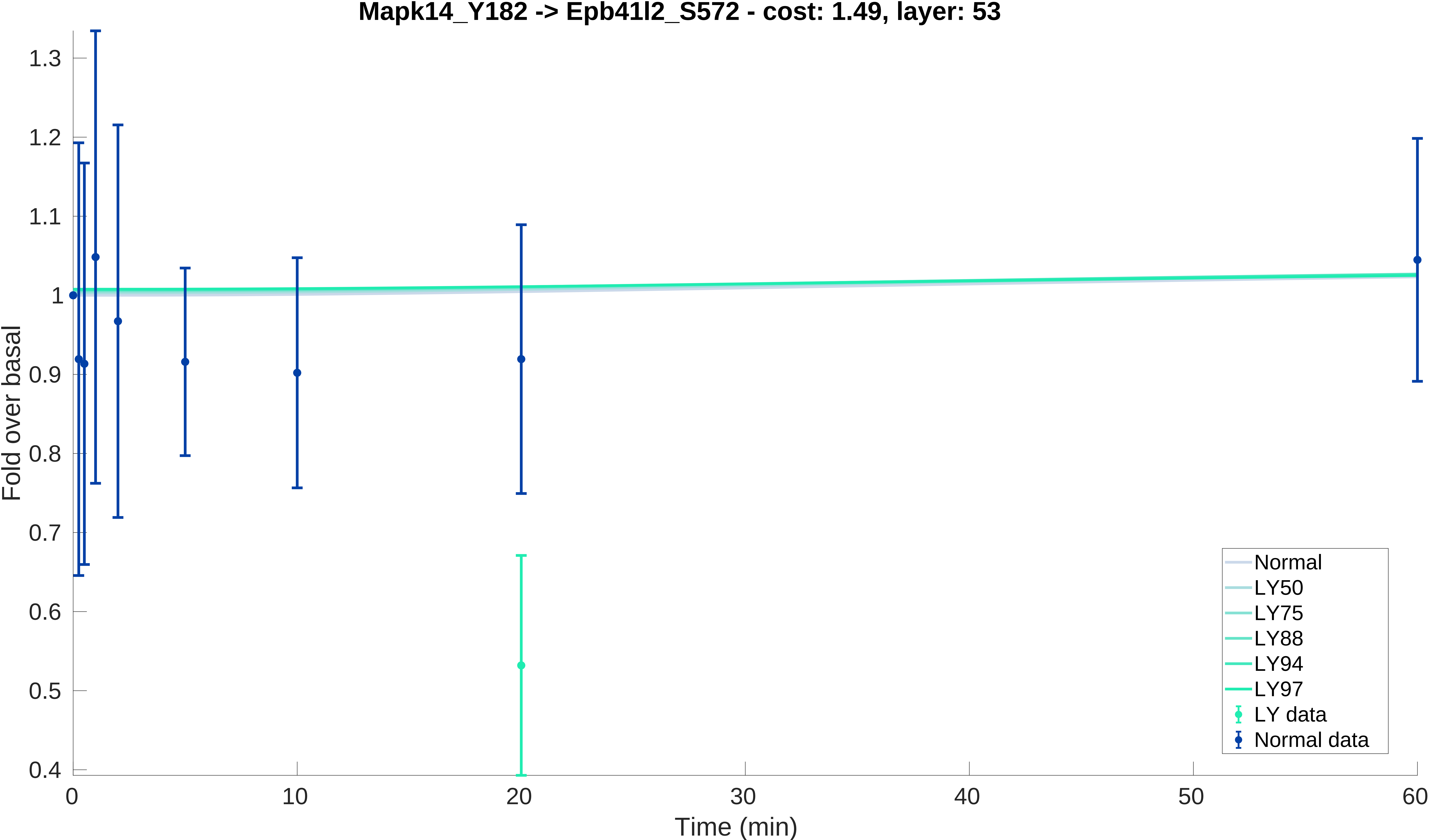

Mapk14_Y182 -> Epb41l2_S572 - cost: 1.49, layer: 53
Fold over basal
Time (min)
0
10
20
30
40
50
60
0.4
0.5
0.6
0.7
0.8
0.9
1
1.1
1.2
1.3
Normal
LY50
LY75
LY88
LY94
LY97
LY data
Normal data

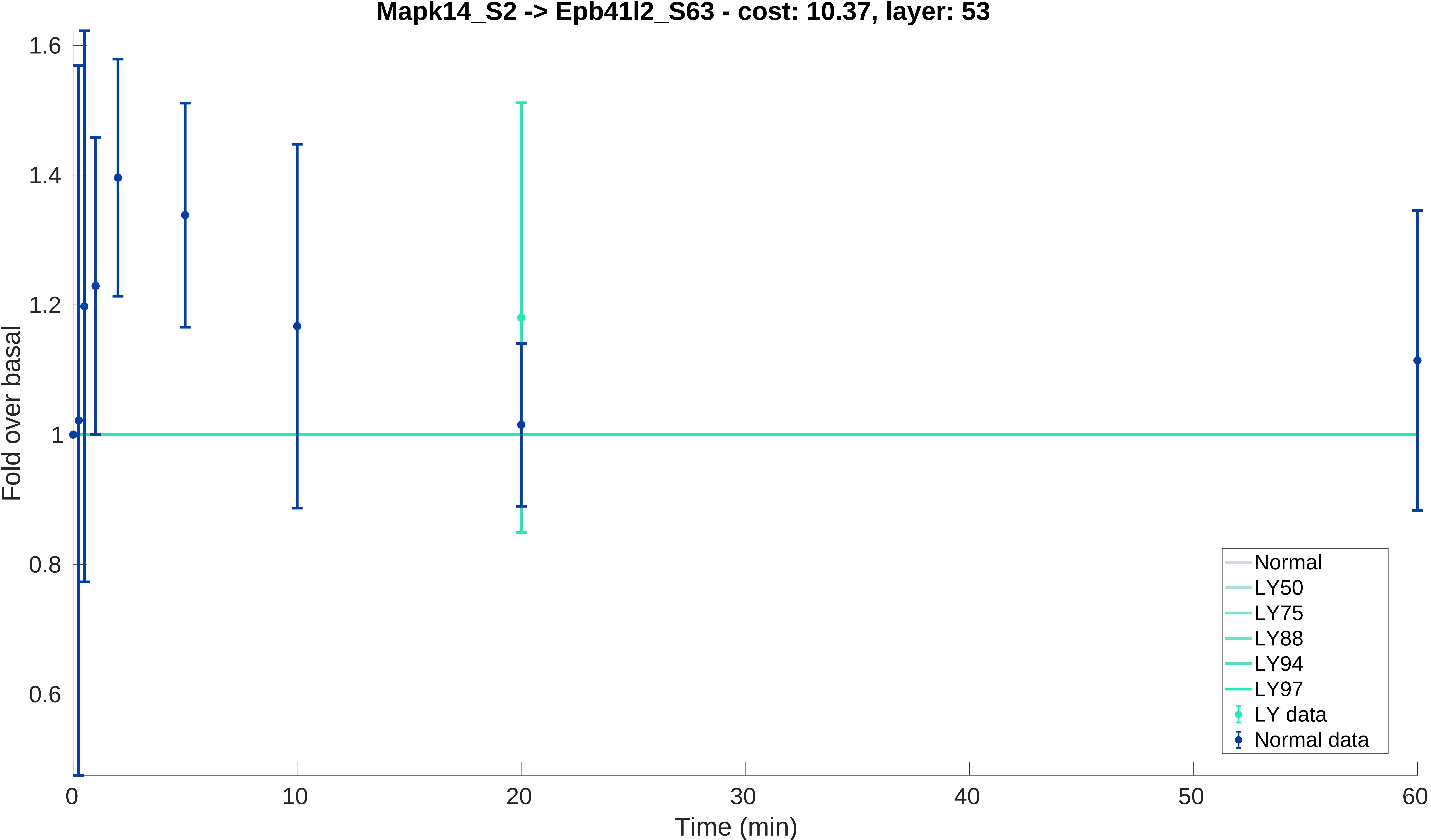

Mapk14_S2 -> Epb41l2_S63 - cost: 10.37, layer: 53
Fold over basal
1.6
1.4
1.2
1
0.8
0.6
0
10
20
30
40
50
60
Time (min)
Normal
LY50
LY75
LY88
LY94
LY97
LY data
Normal data

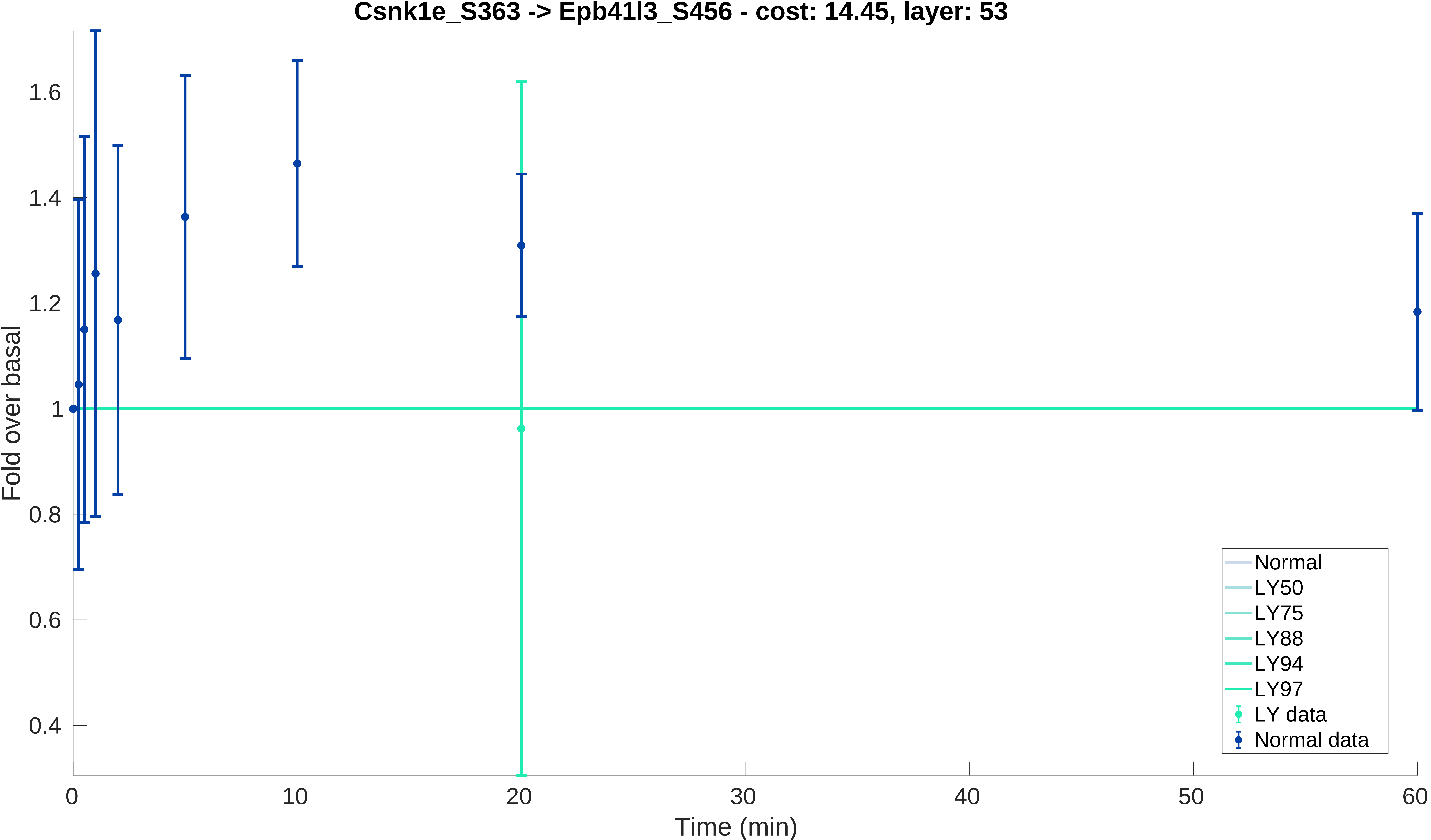

Csnk1e_S363 -> Epb41l3_S456 - cost: 14.45, layer: 53
Fold over basal
1.6
1.4
1.2
1
0.8
0.6
0.4
0
10
20
30
40
50
60
Time (min)
Normal
LY50
LY75
LY88
LY94
LY97
LY data
Normal data

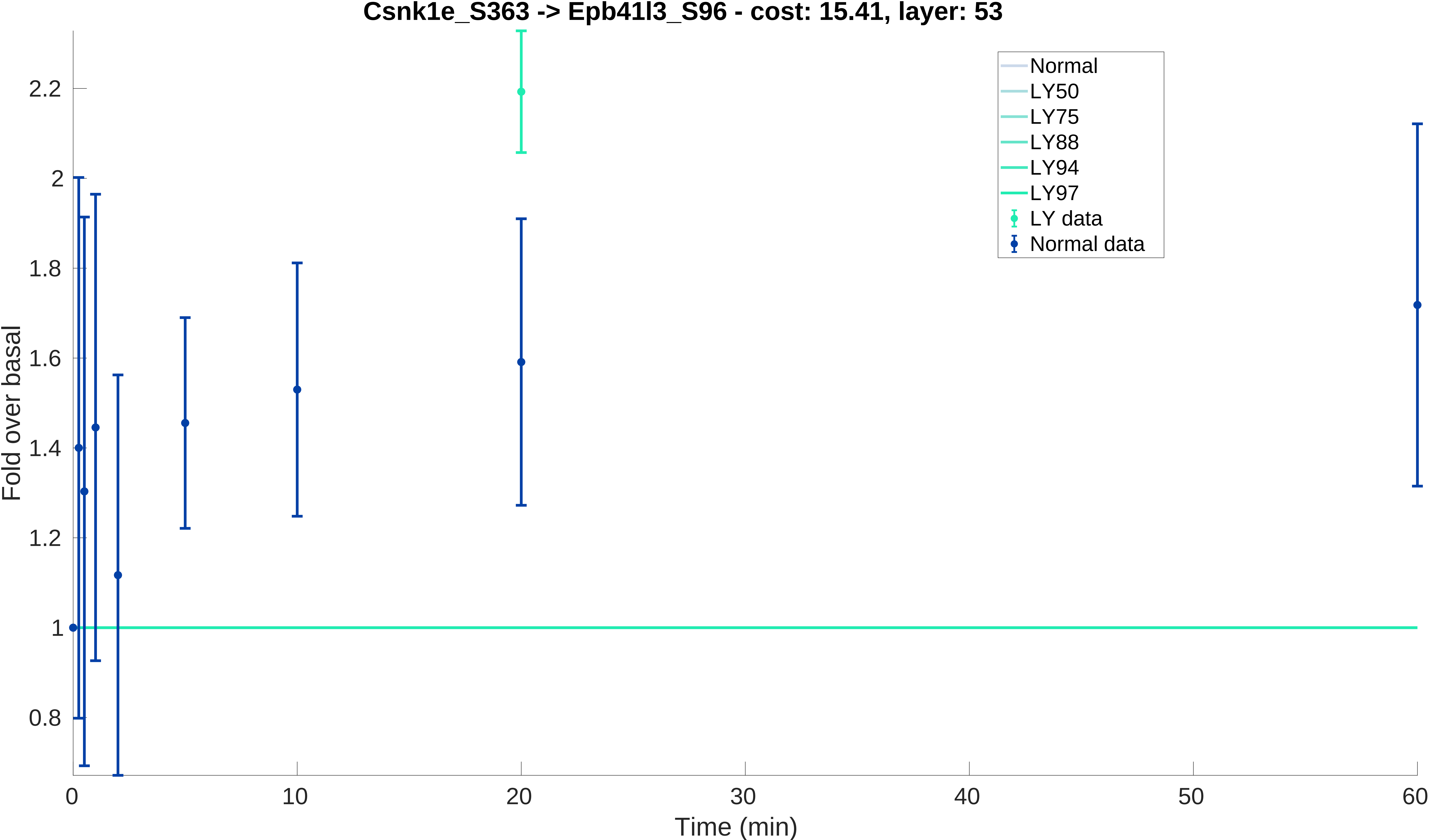

Csnk1e_S363 -> Epb41l3_S96 - cost: 15.41, layer: 53
Normal
LY50
LY75
LY88
LY94
LY97
LY data
Normal data
Fold over basal
Time (min)
2.2
2
1.8
1.6
1.4
1.2
1
0.8
0
10
20
30
40
50
60

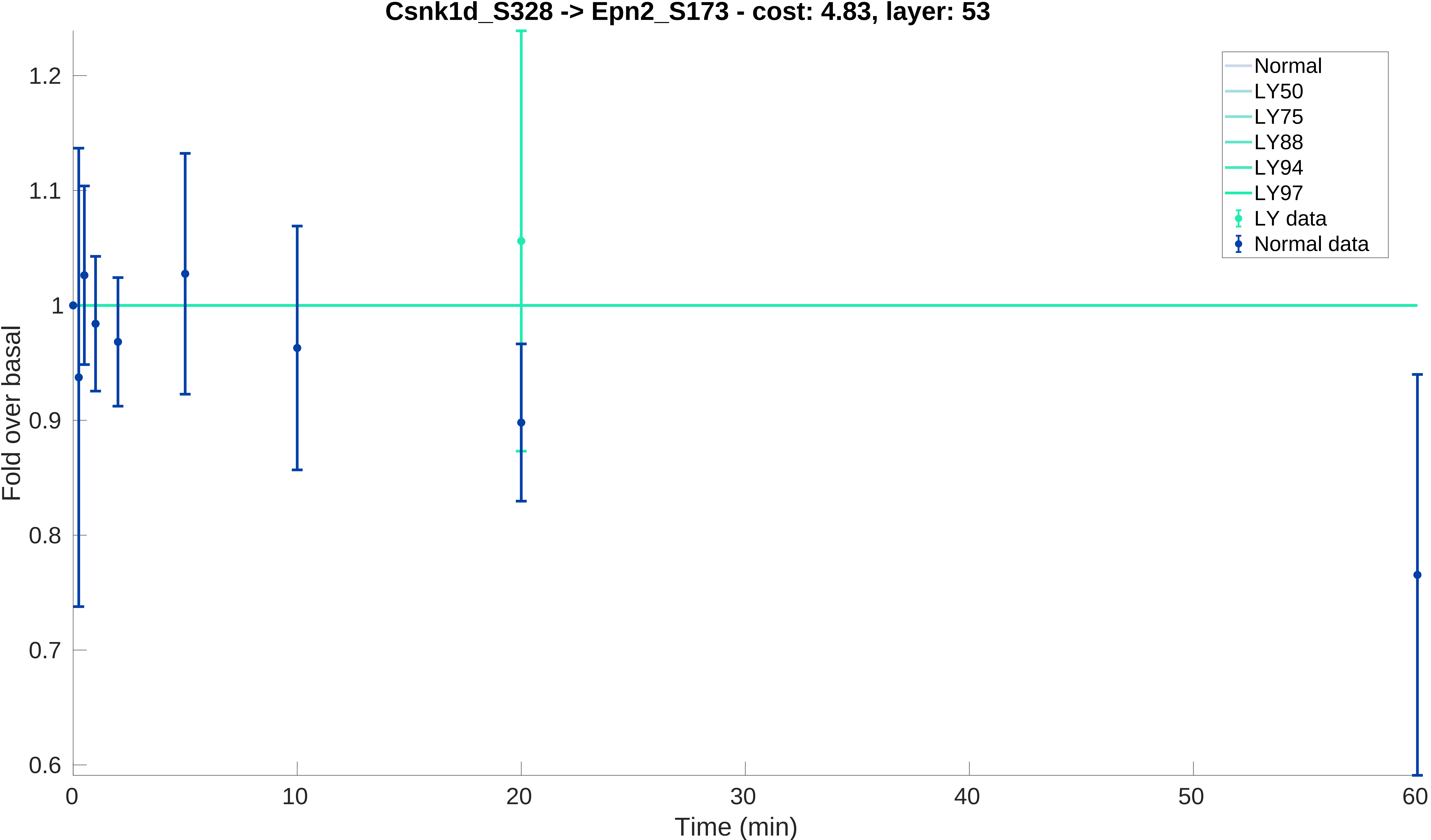

Csnk1d_S328 -> Epn2_S173 - cost: 4.83, layer: 53
Fold over basal
Time (min)
0.6
0.7
0.8
0.9
1
1.1
1.2
0
10
20
30
40
50
60
Normal
LY50
LY75
LY88
LY94
LY97
LY data
Normal data

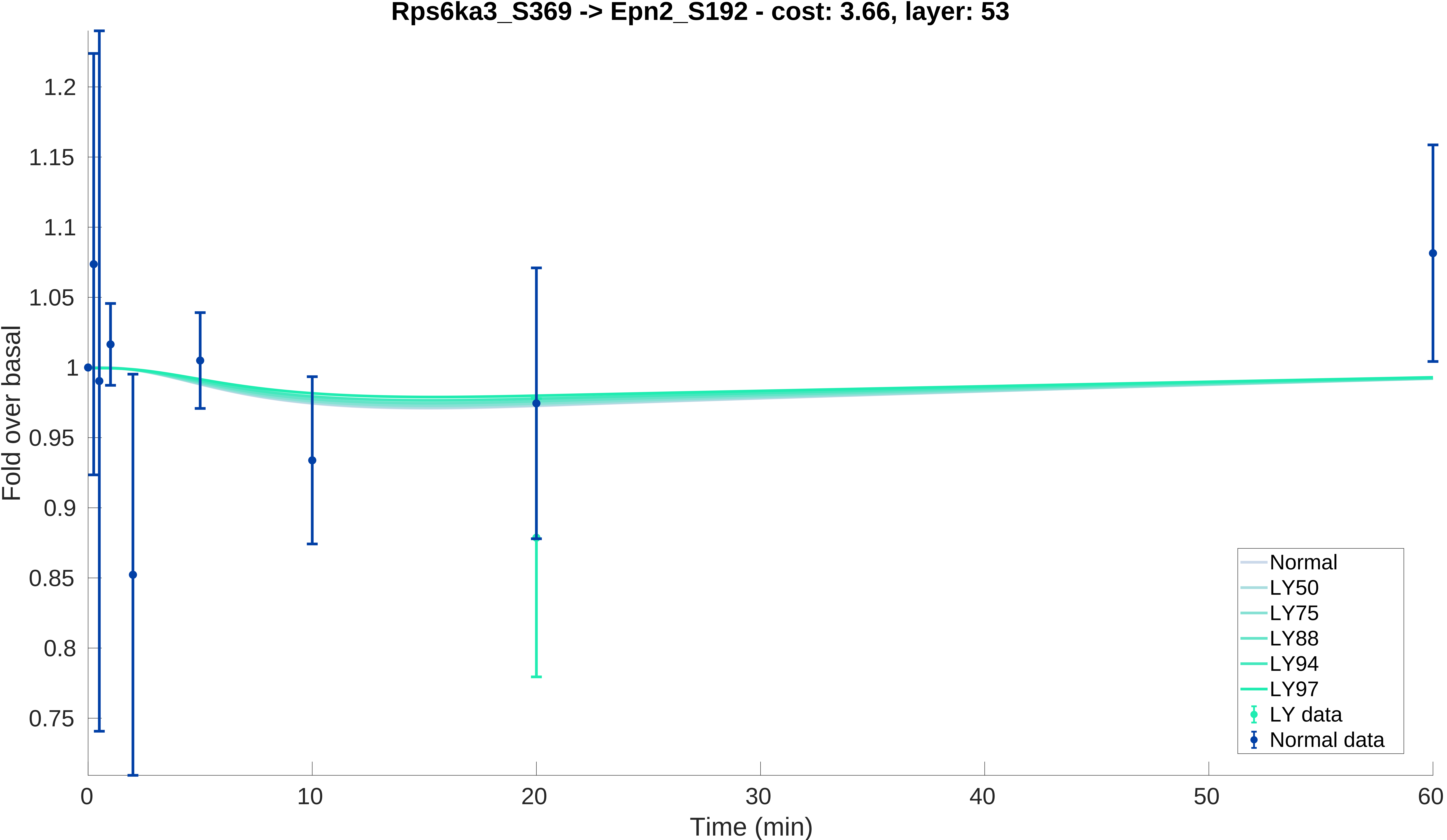

Rps6ka3_S369 -> Epn2_S192 - cost: 3.66, layer: 53
Fold over basal
1.2
1.15
1.1
1.05
1
0.95
0.9
0.85
0.8
0.75
0
10
20
30
40
50
60
Time (min)
Normal
LY50
LY75
LY88
LY94
LY97
LY data
Normal data

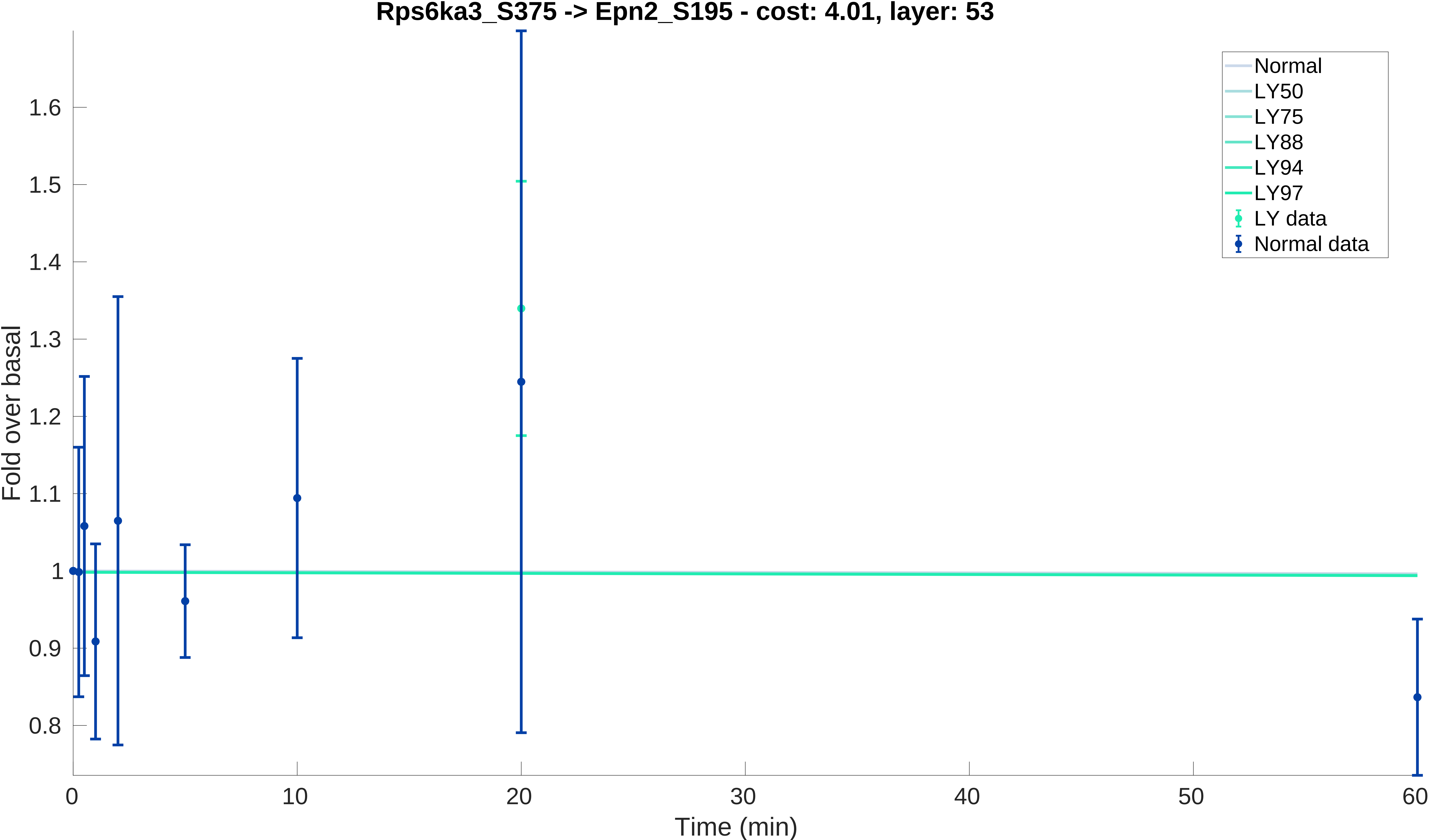

Rps6ka3_S375 -> Epn2_S195 - cost: 4.01, layer: 53
Fold over basal
Time (min)
0.8
0.9
1
1.1
1.2
1.3
1.4
1.5
1.6
0
10
20
30
40
50
60
Normal
LY50
LY75
LY88
LY94
LY97
LY data
Normal data

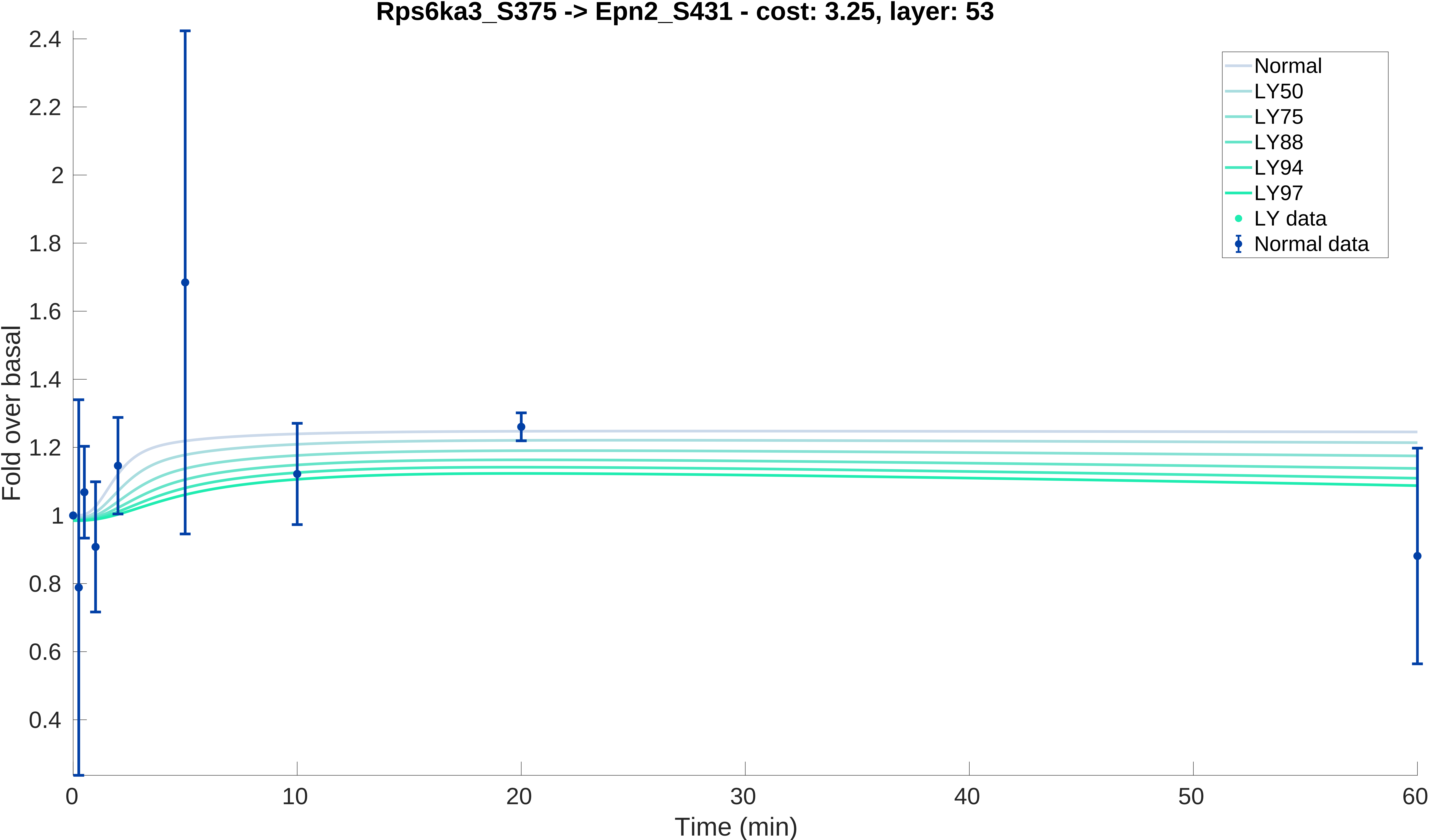
Rps6ka3_S375 -> Epn2_S431 - cost: 3.25, layer: 53
Fold over basal
2.4
2.2
2
1.8
1.6
1.4
1.2
1
0.8
0.6
0.4
0
10
20
30
40
50
60
Time (min)
Normal
LY50
LY75
LY88
LY94
LY97
LY data
Normal data

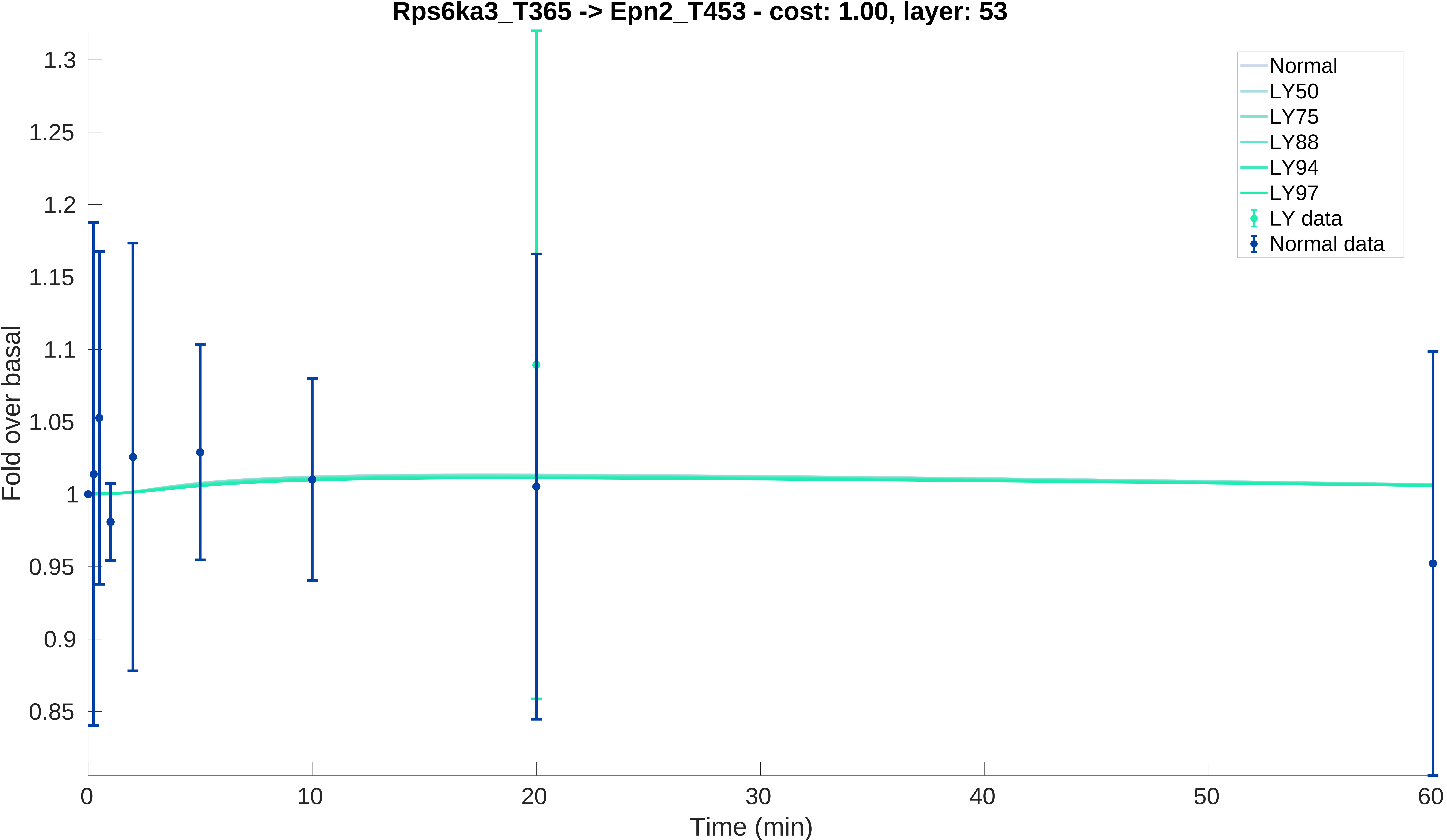
Rps6ka3_T365 -> Epn2_T453 - cost: 1.00, layer: 53
Fold over basal
Time (min)
1.3
1.25
1.2
1.15
1.1
1.05
1
0.95
0.9
0.85
0
10
20
30
40
50
60
Normal
LY50
LY75
LY88
LY94
LY97
LY data
Normal data

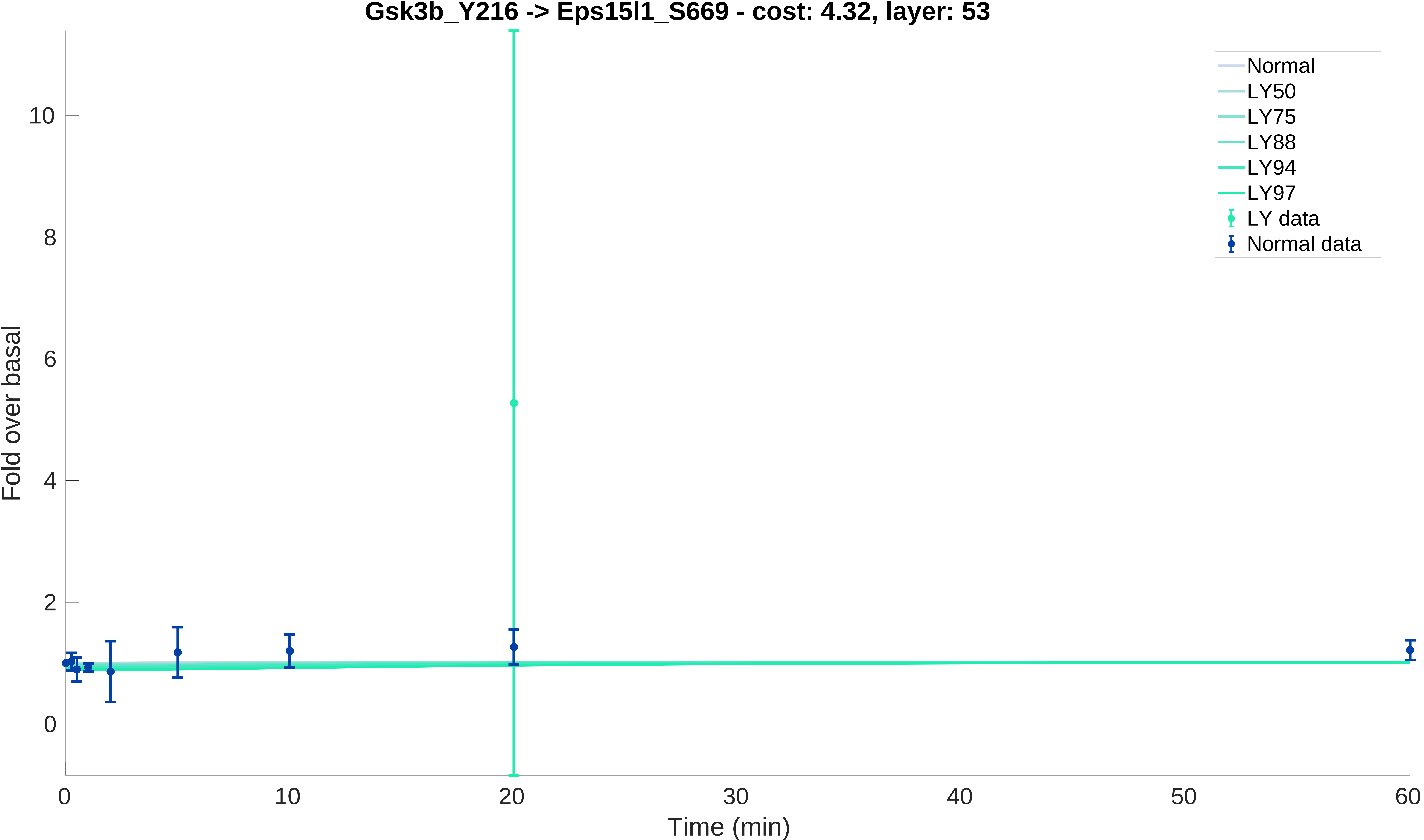

Gsk3b_Y216 -> Eps15l1_S669 - cost: 4.32, layer: 53
Normal
LY50
LY75
LY88
LY94
LY97
LY data
Normal data
Fold over basal
Time (min)
0
2
4
6
8
10
0
10
20
30
40
50
60

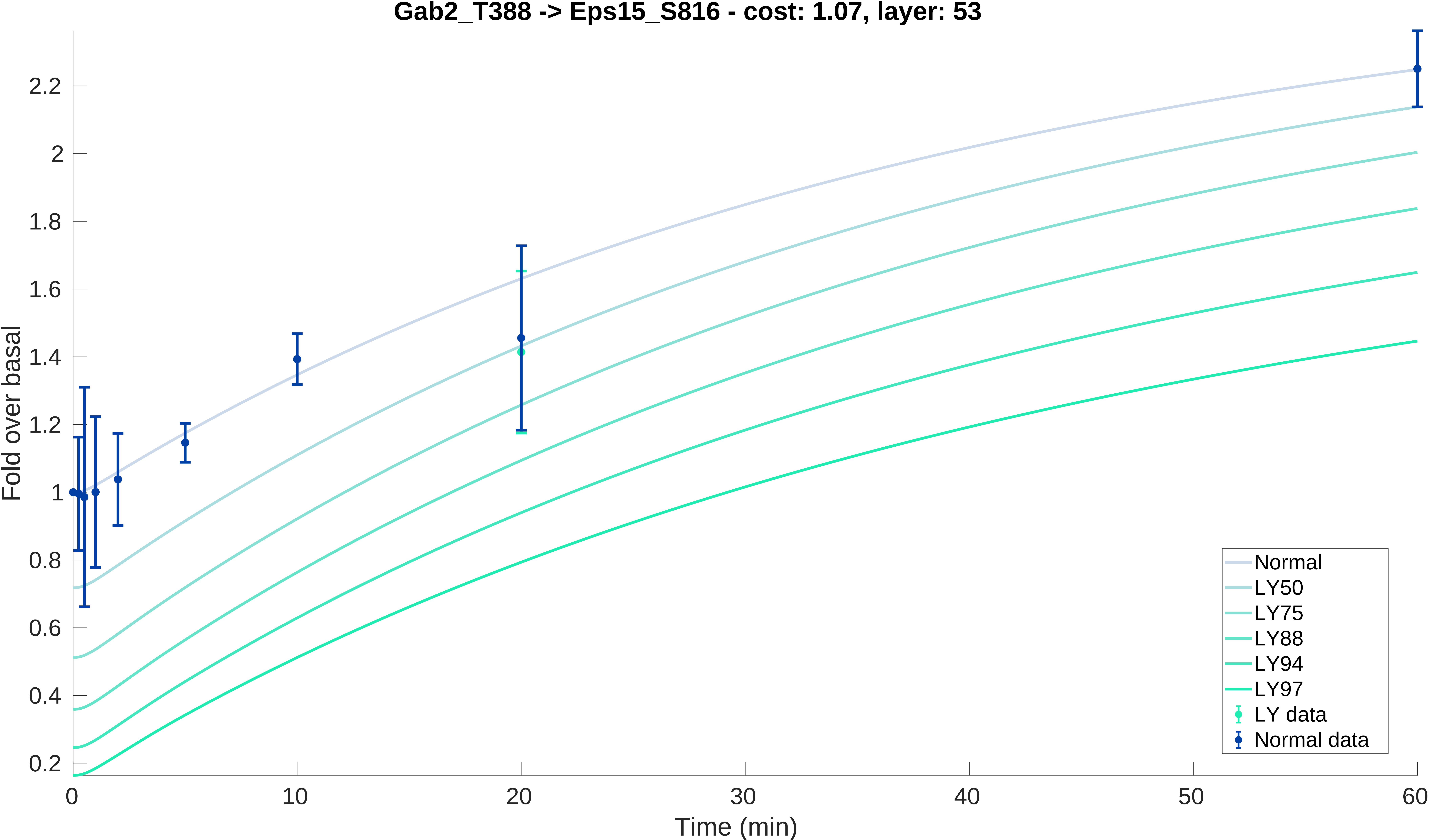

Gab2_T388 -> Eps15_S816 - cost: 1.07, layer: 53
Fold over basal
Time (min)
0
10
20
30
40
50
60
0.2
0.4
0.6
0.8
1
1.2
1.4
1.6
1.8
2
2.2
Normal
LY50
LY75
LY88
LY94
LY97
LY data
Normal data

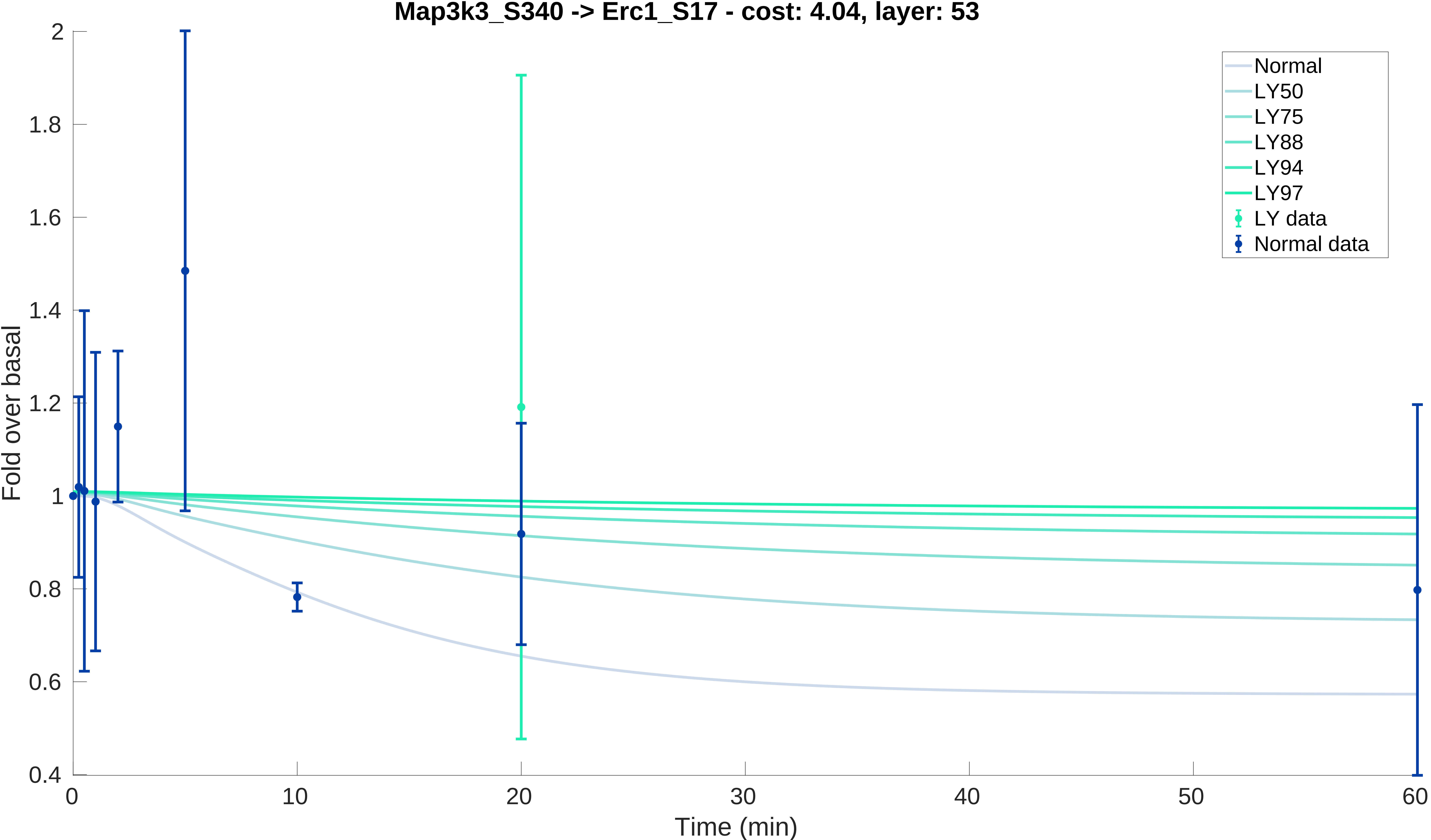

Map3k3_S340 -> Erc1_S17 - cost: 4.04, layer: 53
Fold over basal
Time (min)
0.4
0.6
0.8
1
1.2
1.4
1.6
1.8
2
0
10
20
30
40
50
60
Normal
LY50
LY75
LY88
LY94
LY97
LY data
Normal data

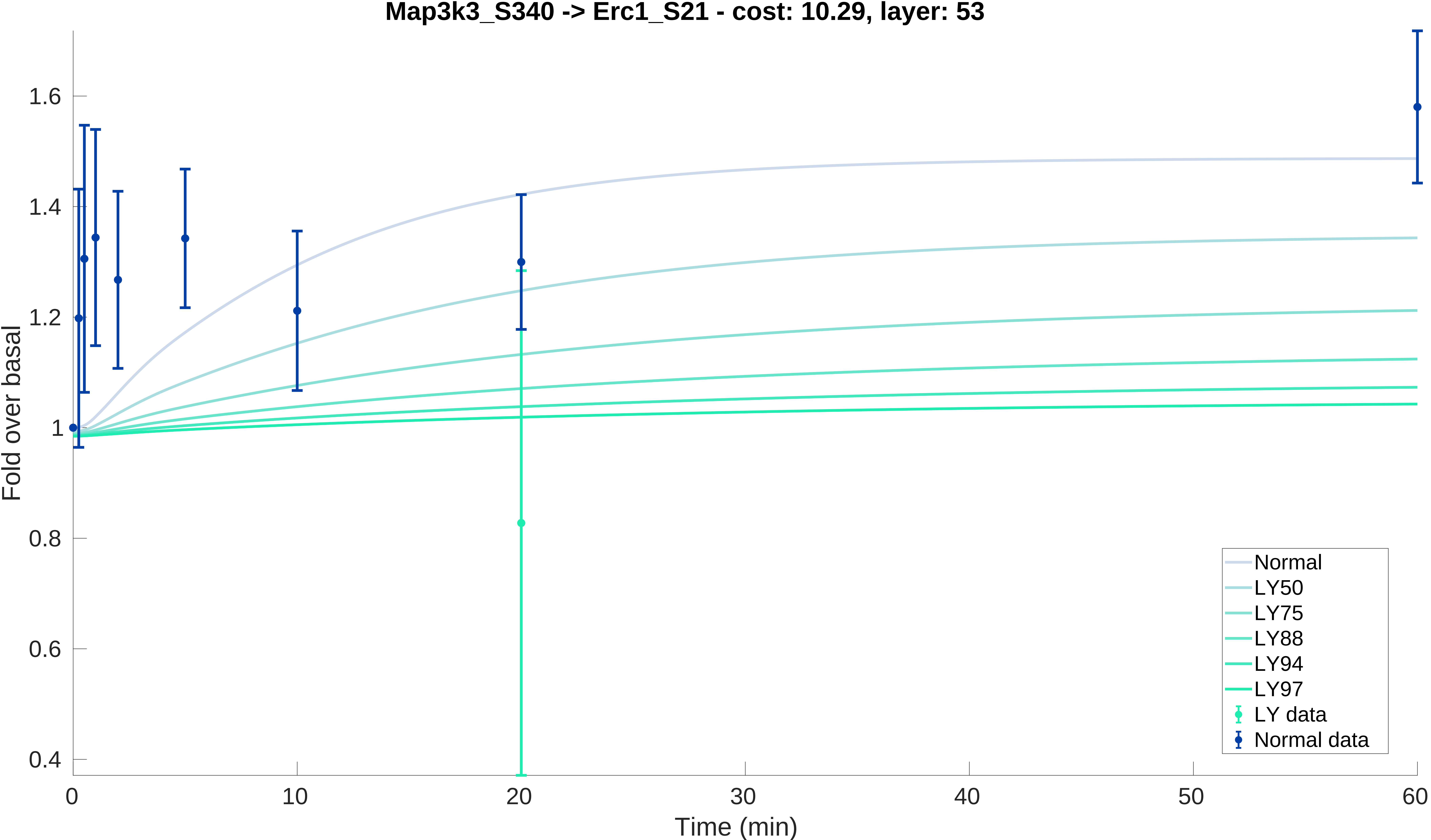

Map3k3_S340 -> Erc1_S21 - cost: 10.29, layer: 53
Fold over basal
Time (min)
0.4
0.6
0.8
1
1.2
1.4
1.6
0
10
20
30
40
50
60
Normal
LY50
LY75
LY88
LY94
LY97
LY data
Normal data

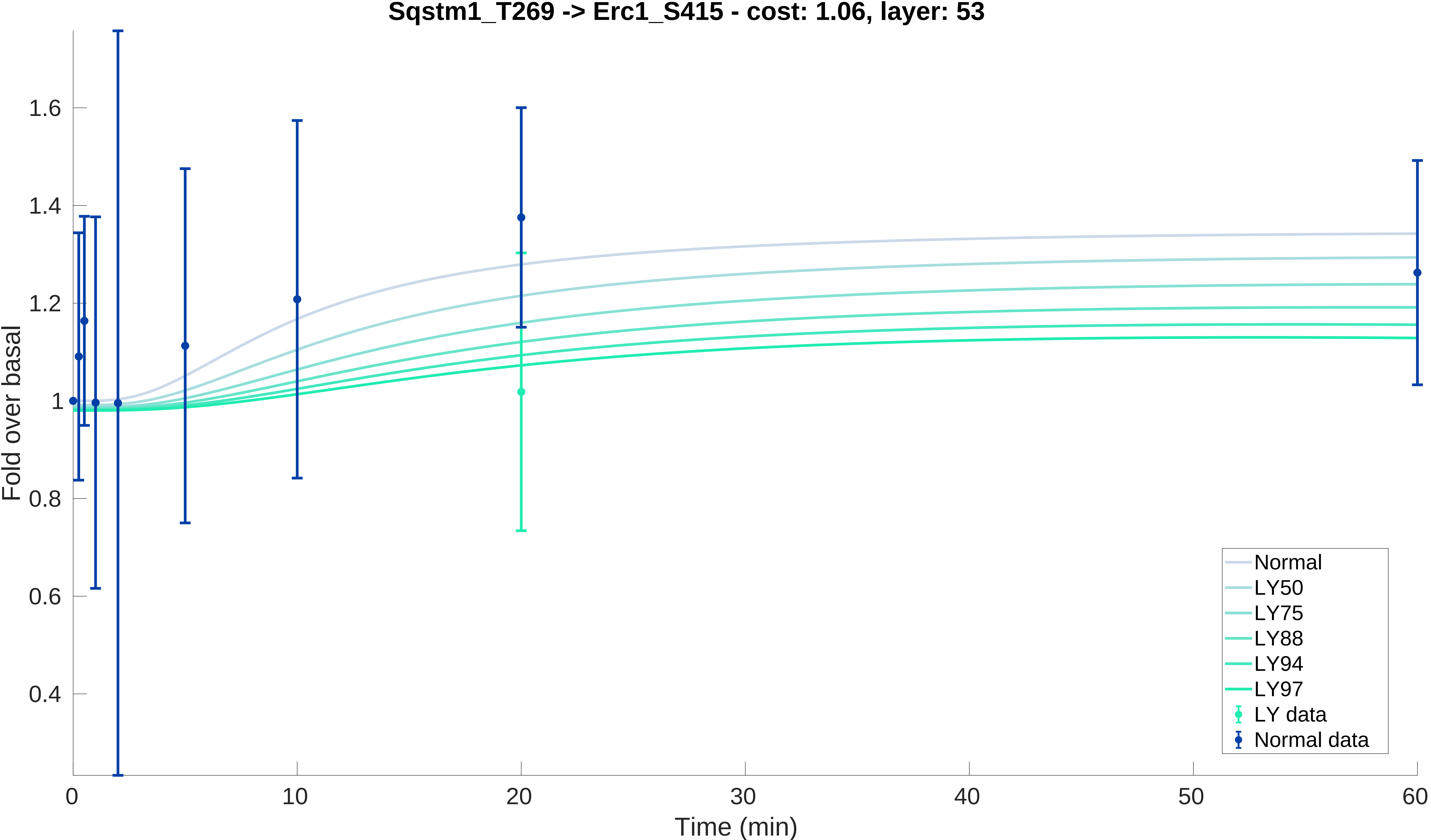

Sqstm1_T269 -> Erc1_S415 - cost: 1.06, layer: 53
Fold over basal
Time (min)
0.4
0.6
0.8
1
1.2
1.4
1.6
0
10
20
30
40
50
60
Normal
LY50
LY75
LY88
LY94
LY97
LY data
Normal data

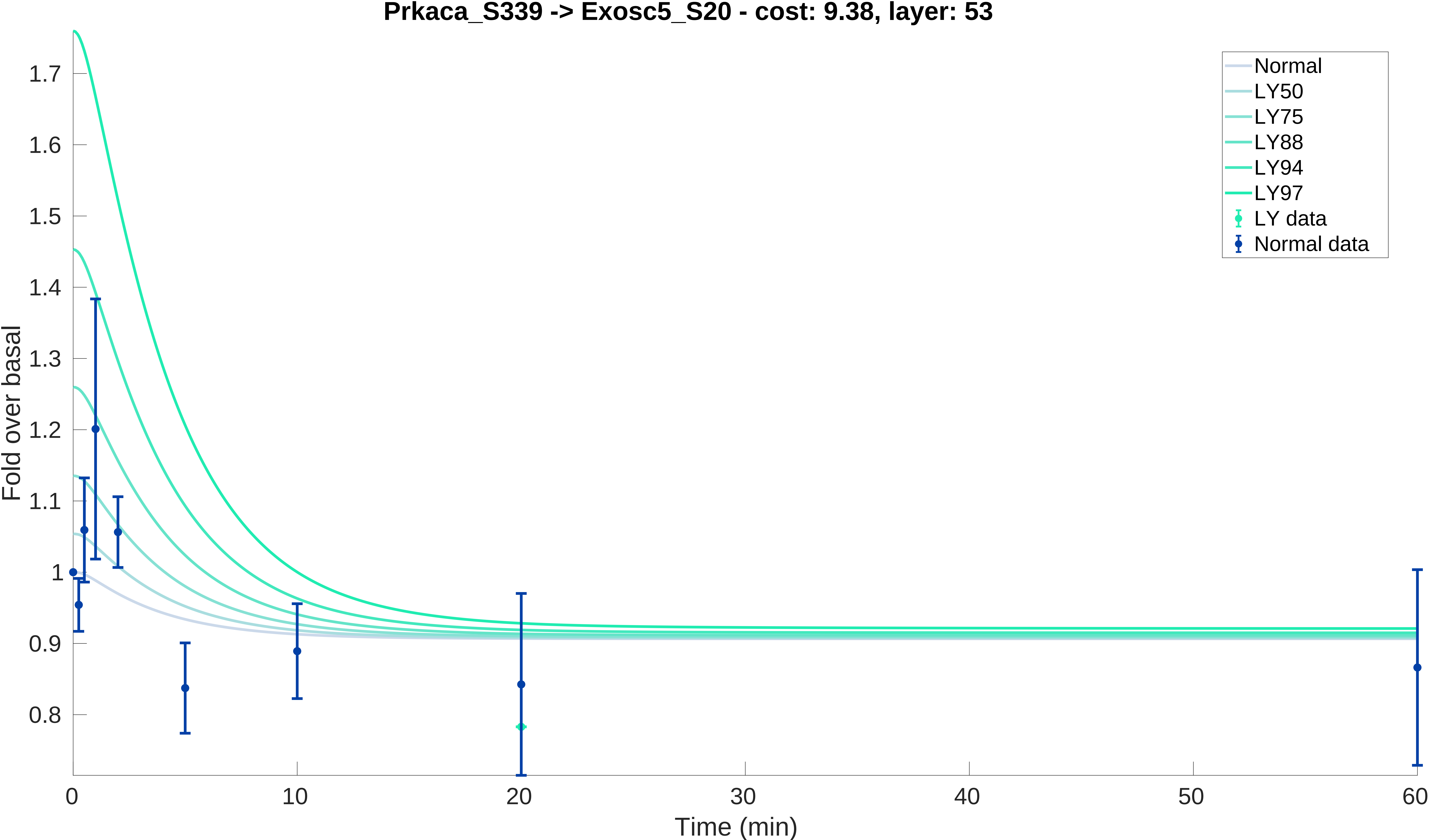
Prkaca_S339 -> Exosc5_S20 - cost: 9.38, layer: 53
Normal
LY50
LY75
LY88
LY94
LY97
LY data
Normal data
Fold over basal
Time (min)
1.7
1.6
1.5
1.4
1.3
1.2
1.1
1
0.9
0.8
0
10
20
30
40
50
60

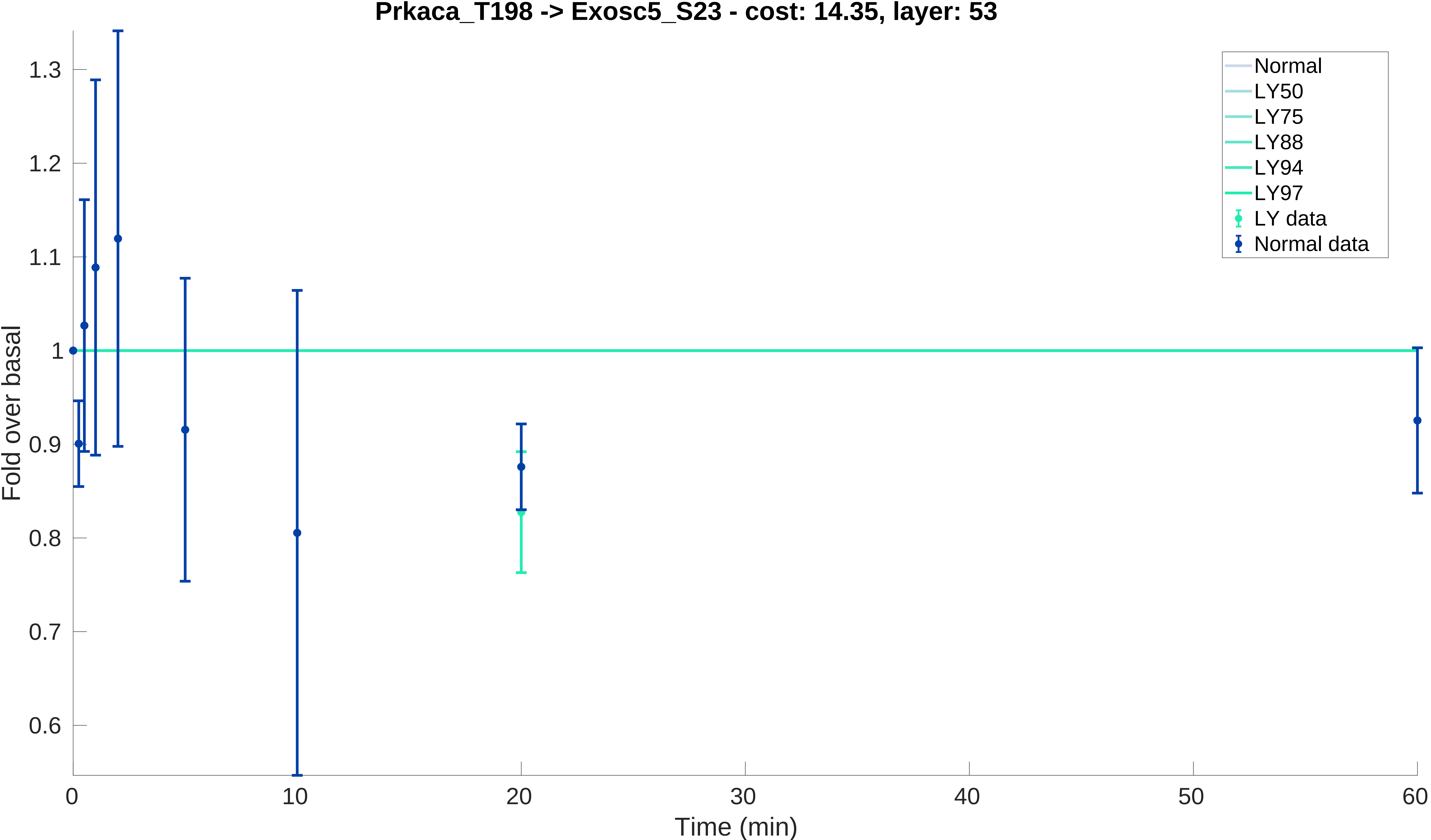

Prkaca_T198 -> Exosc5_S23 - cost: 14.35, layer: 53
Normal
LY50
LY75
LY88
LY94
LY97
LY data
Normal data
Fold over basal
1.3
1.2
1.1
1
0.9
0.8
0.7
0.6
0
10
20
30
40
50
60
Time (min)

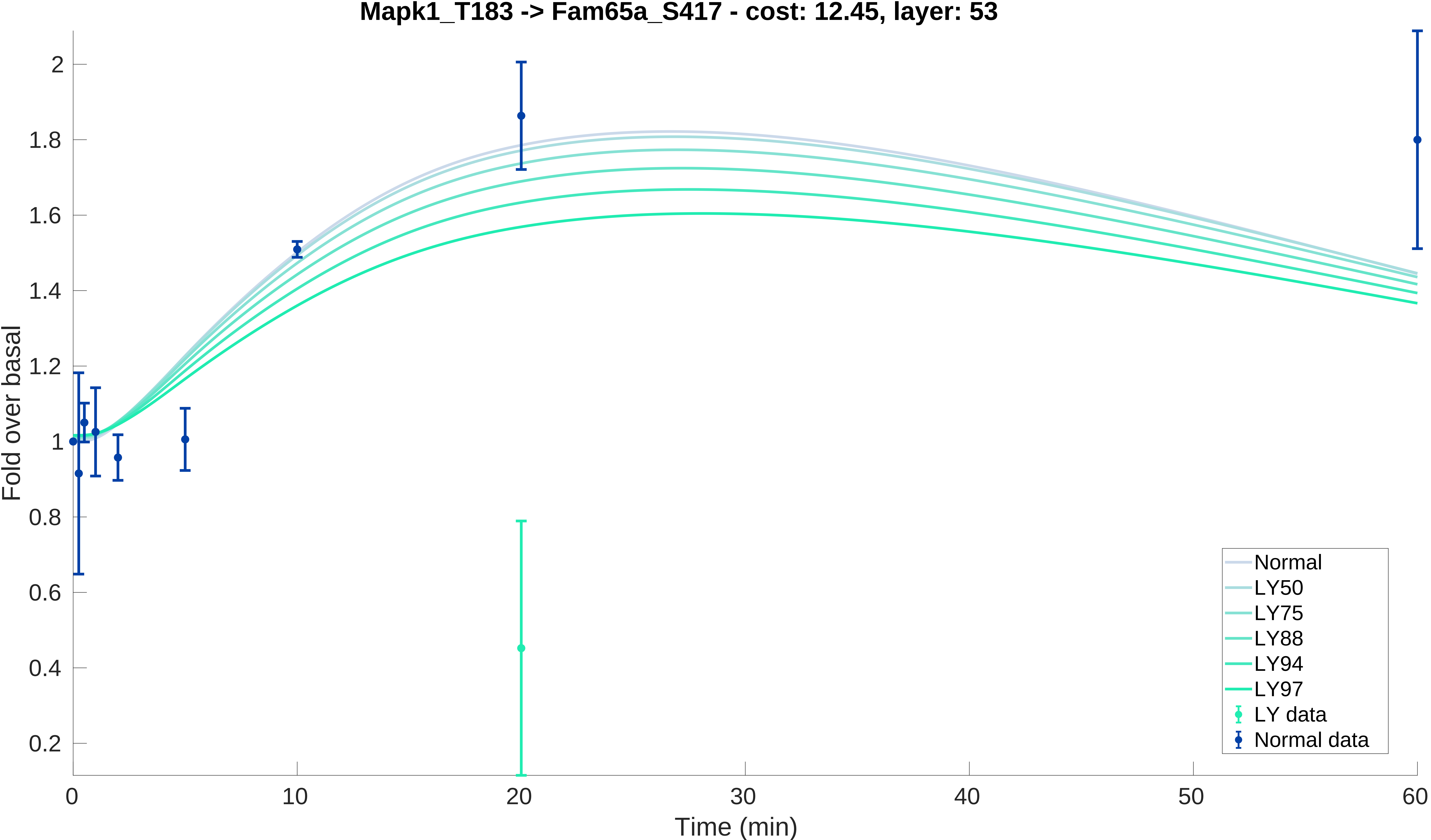
Mapk1_T183 -> Fam65a_S417 - cost: 12.45, layer: 53
Fold over basal
Time (min)
0.2
0.4
0.6
0.8
1
1.2
1.4
1.6
1.8
2
0
10
20
30
40
50
60
Normal
LY50
LY75
LY88
LY94
LY97
LY data
Normal data

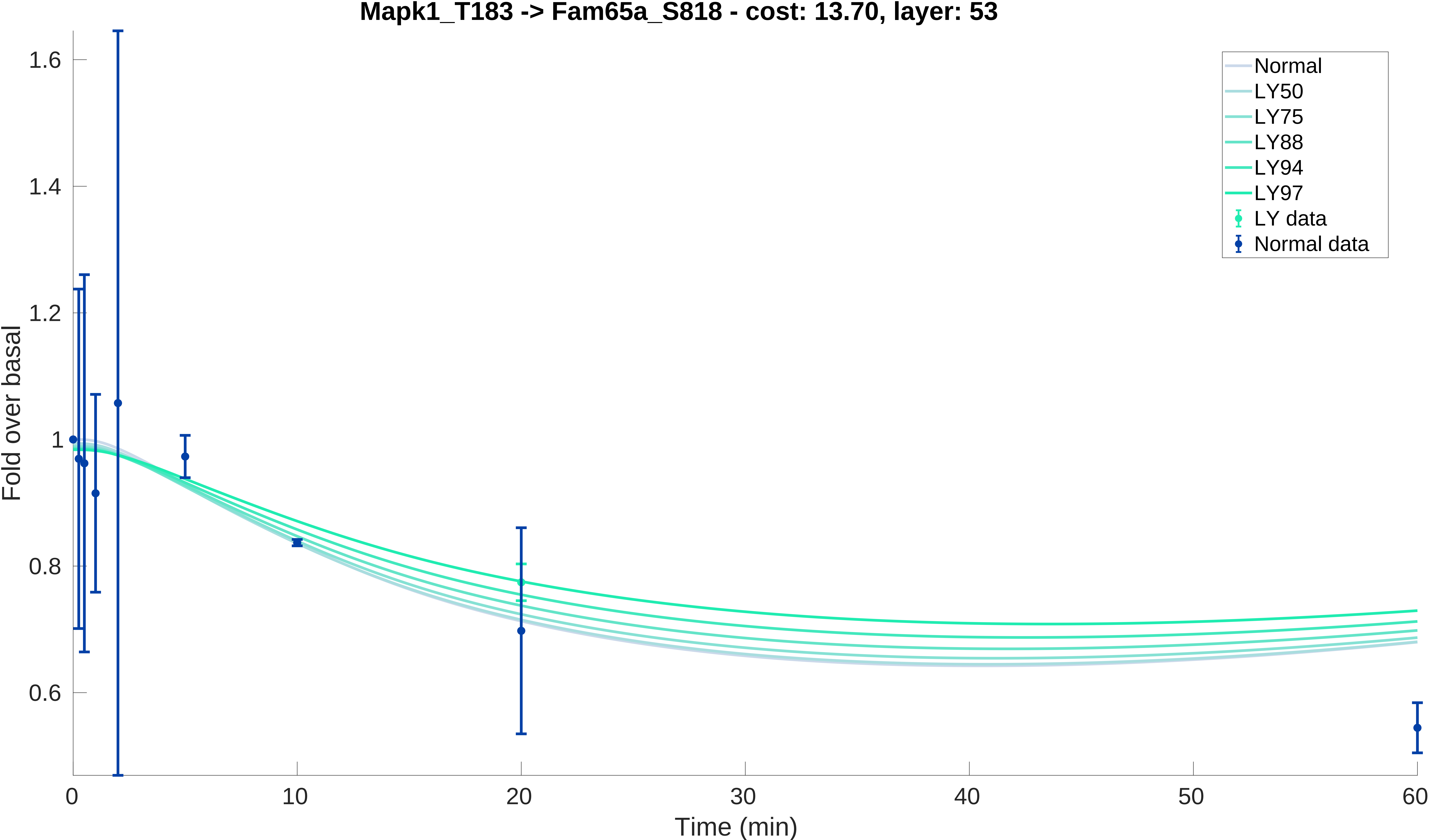
Mapk1_T183 -> Fam65a_S818 - cost: 13.70, layer: 53
Fold over basal
1.6
1.4
1.2
1
0.8
0.6
0
10
20
30
40
50
60
Time (min)
Normal
LY50
LY75
LY88
LY94
LY97
LY data
Normal data

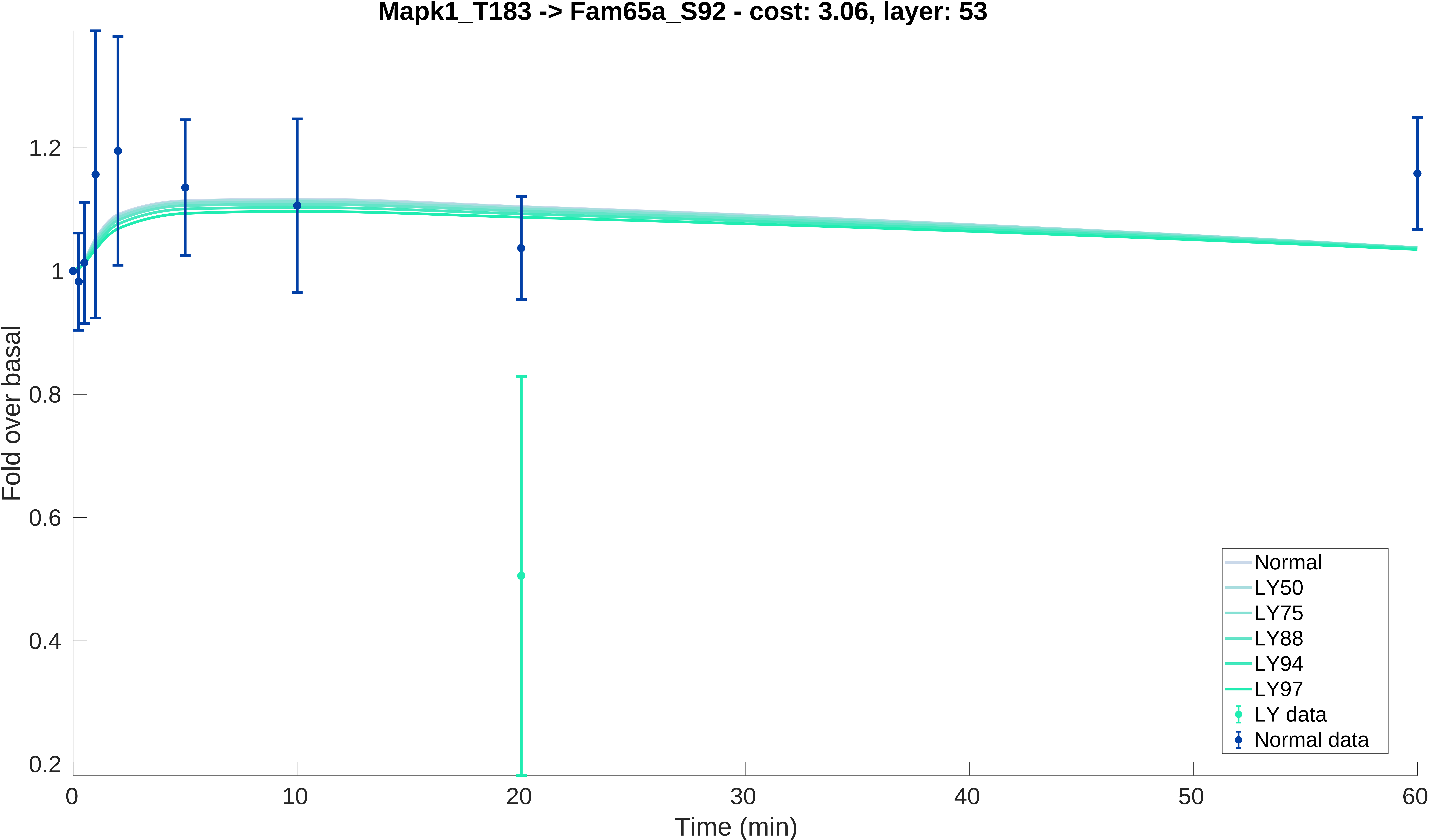
Mapk1_T183 -> Fam65a_S92 - cost: 3.06, layer: 53
Fold over basal
Time (min)
1.2
1
0.8
0.6
0.4
0.2
0
10
20
30
40
50
60
Normal
LY50
LY75
LY88
LY94
LY97
LY data
Normal data

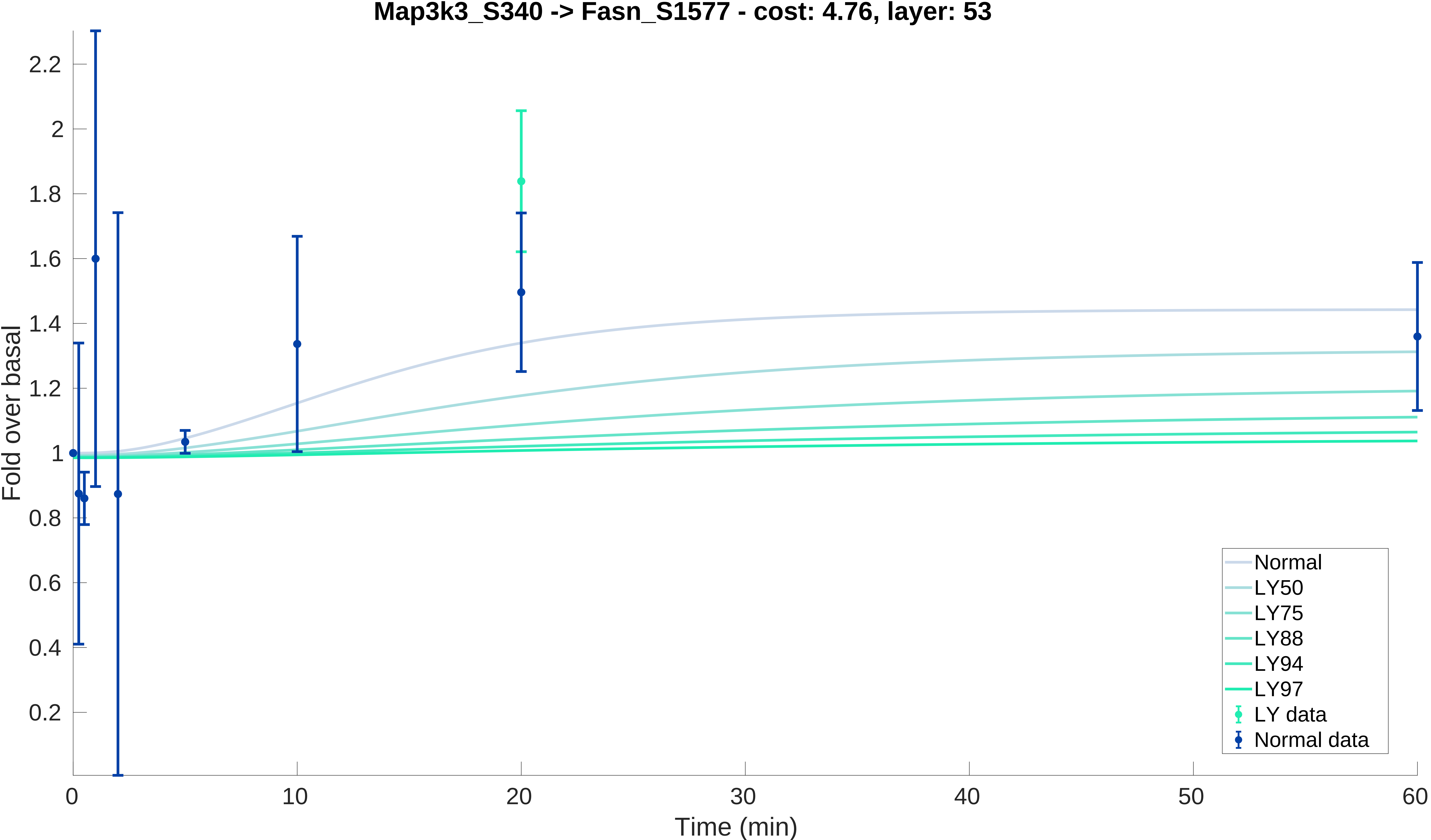

Map3k3_S340 -> Fasn_S1577 - cost: 4.76, layer: 53
Fold over basal
Time (min)
2.2
2
1.8
1.6
1.4
1.2
1
0.8
0.6
0.4
0.2
0
10
20
30
40
50
60
Normal
LY50
LY75
LY88
LY94
LY97
LY data
Normal data

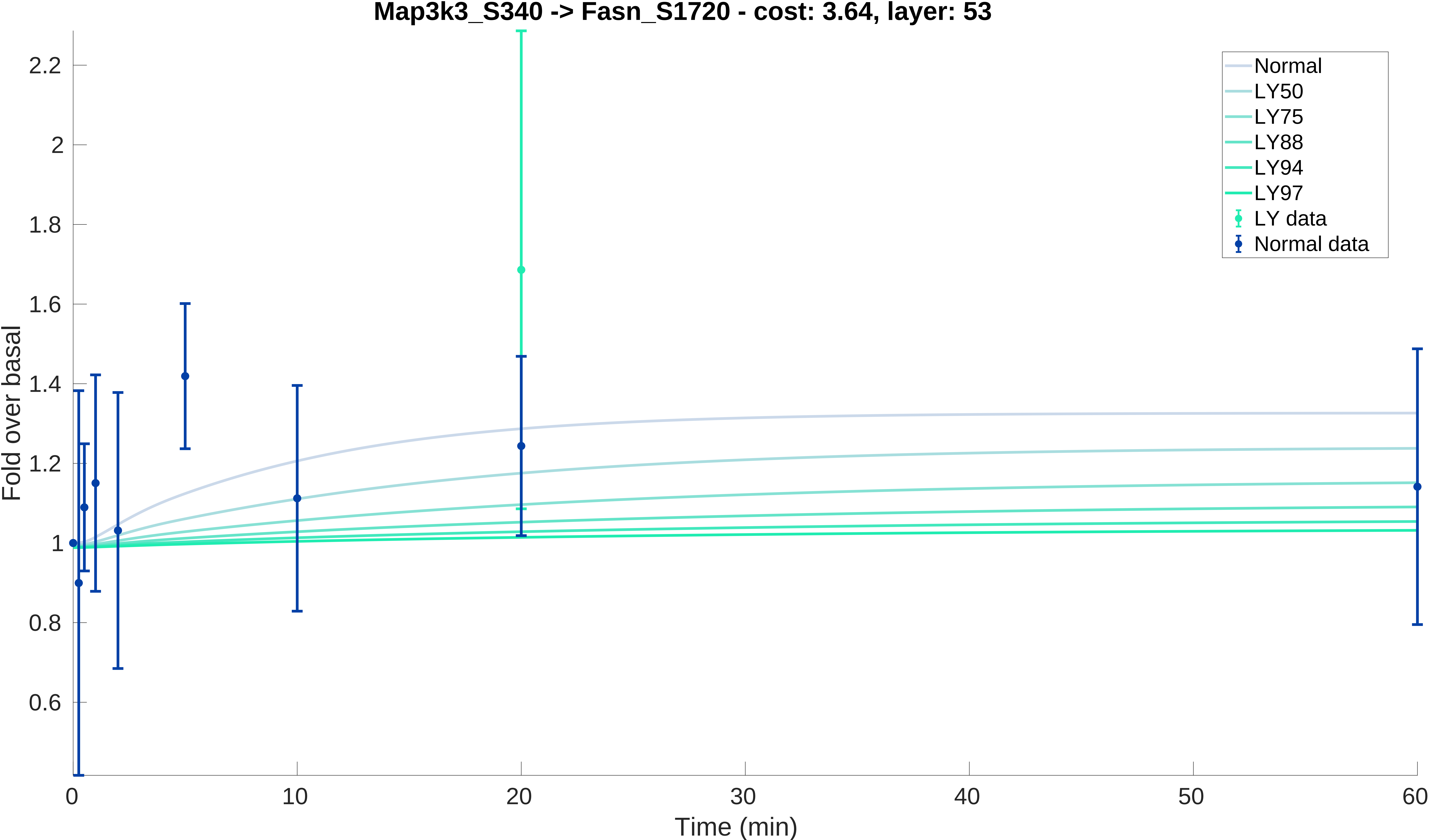

Map3k3_S340 -> Fasn_S1720 - cost: 3.64, layer: 53
Fold over basal
Time (min)
0.6
0.8
1
1.2
1.4
1.6
1.8
2
2.2
0
10
20
30
40
50
60
Normal
LY50
LY75
LY88
LY94
LY97
LY data
Normal data

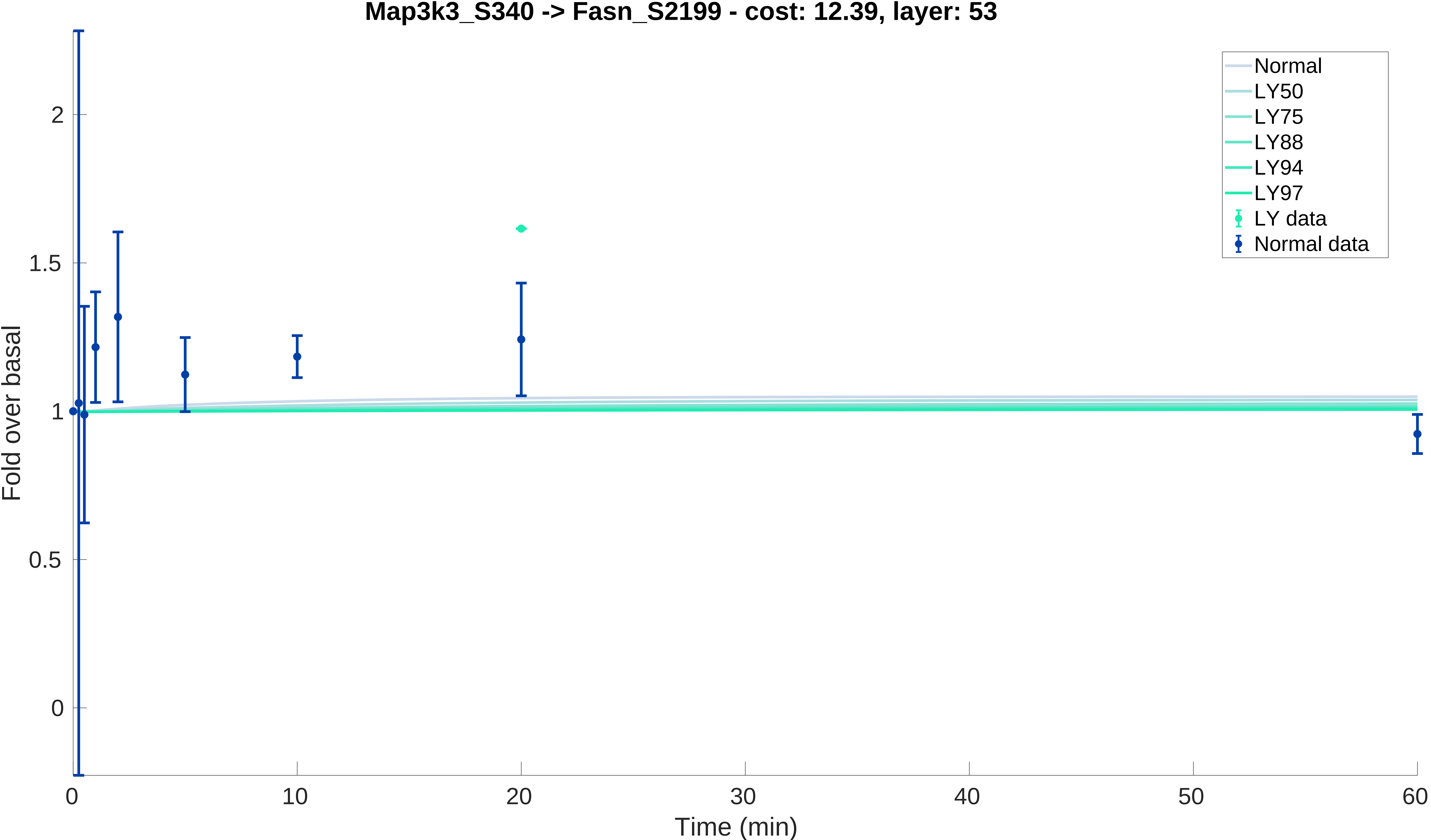

Map3k3_S340 -> Fasn_S2199 - cost: 12.39, layer: 53
Normal
LY50
LY75
LY88
LY94
LY97
LY data
Normal data
Fold over basal
2
1.5
1
0.5
0
0
10
20
30
40
50
60
Time (min)

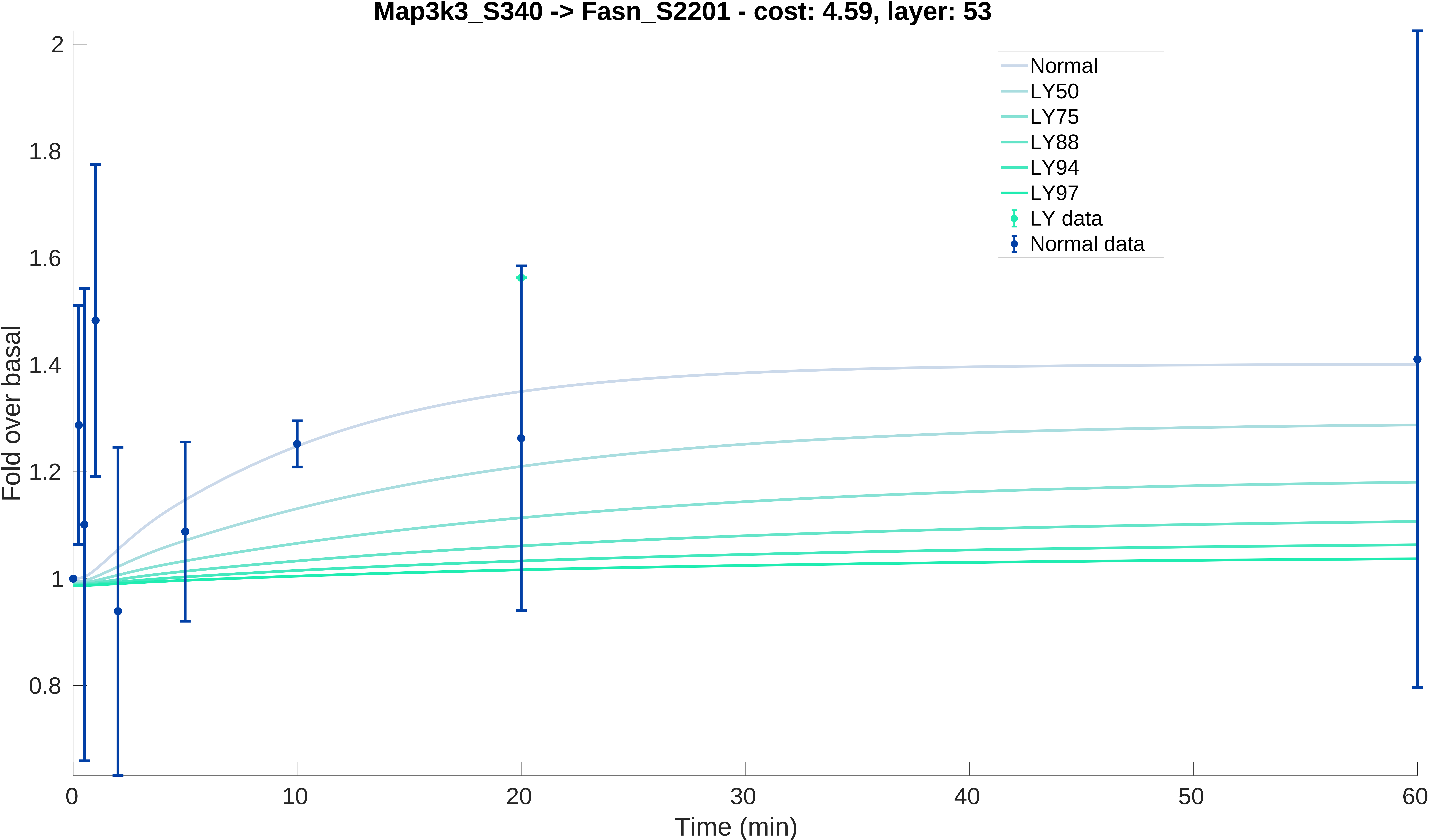

Map3k3_S340 -> Fasn_S2201 - cost: 4.59, layer: 53
Normal
LY50
LY75
LY88
LY94
LY97
LY data
Normal data
Fold over basal
Time (min)
2
1.8
1.6
1.4
1.2
1
0.8
0
10
20
30
40
50
60

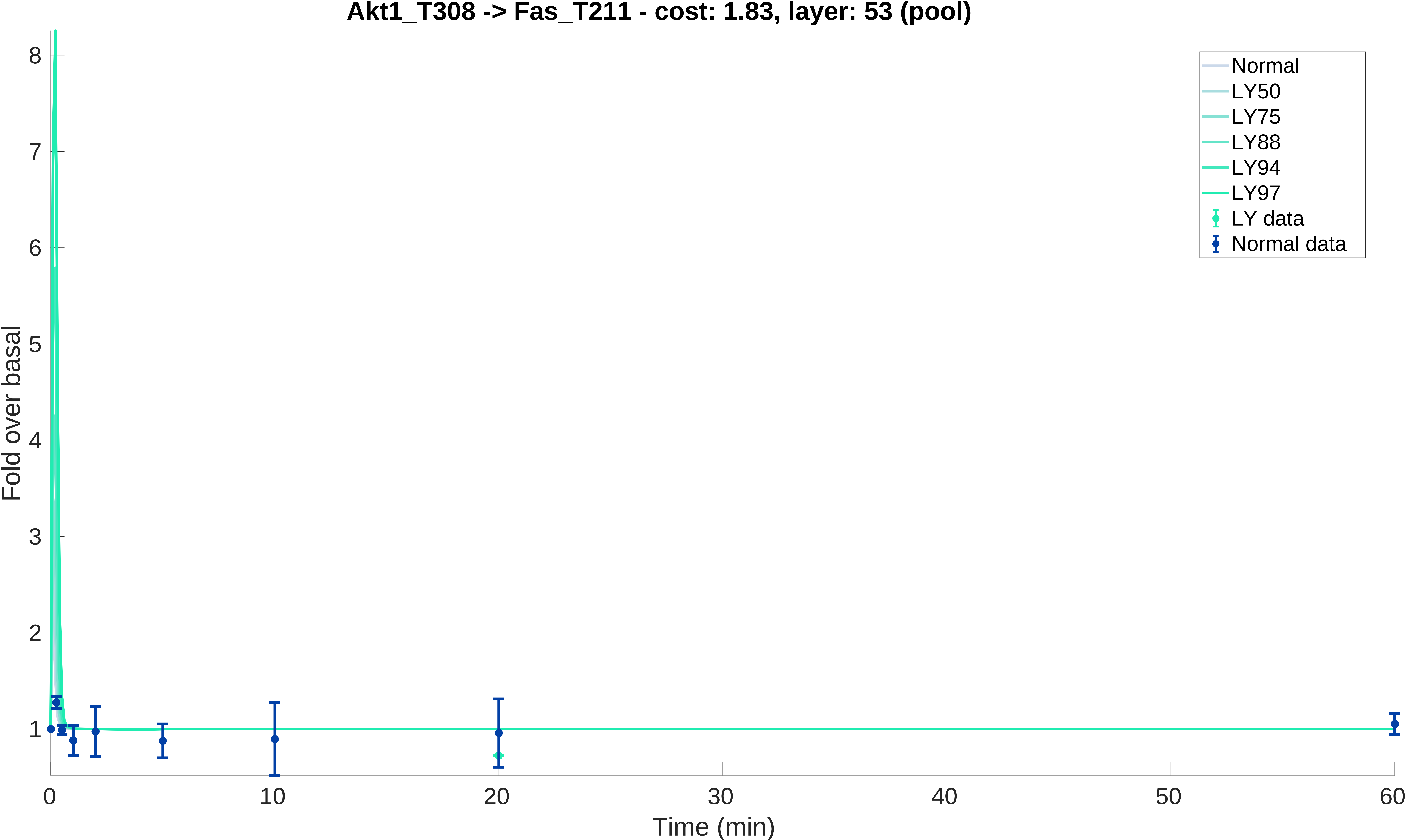

Akt1_T308 -> Fas_T211 - cost: 1.83, layer: 53 (pool)
Fold over basal
Time (min)
Normal
LY50
LY75
LY88
LY94
LY97
LY data
Normal data
1
2
3
4
5
6
7
8
0
10
20
30
40
50
60

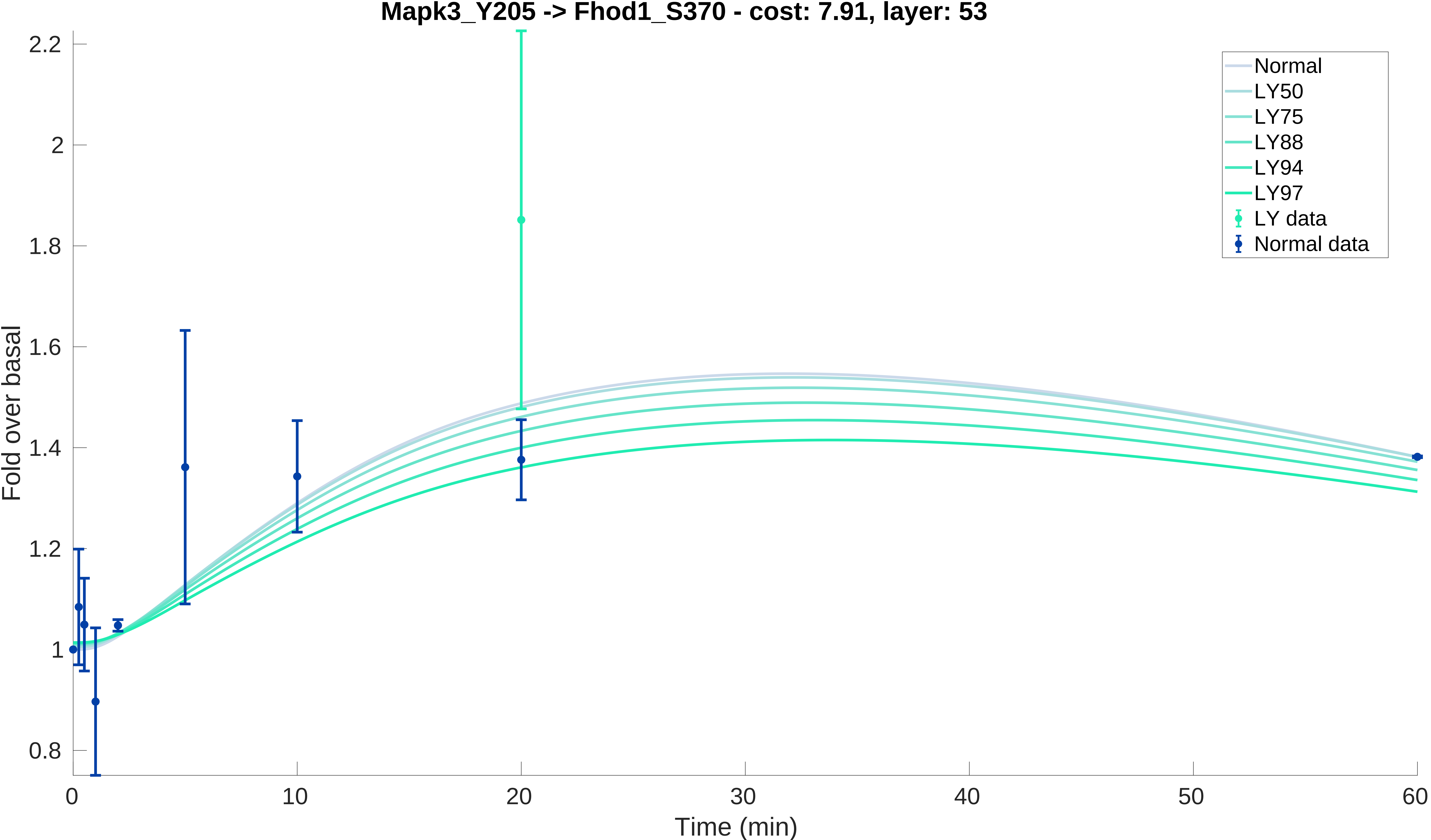

Mapk3_Y205 -> Fhod1_S370 - cost: 7.91, layer: 53
Fold over basal
Time (min)
0.8
1
1.2
1.4
1.6
1.8
2
2.2
0
10
20
30
40
50
60
Normal
LY50
LY75
LY88
LY94
LY97
LY data
Normal data

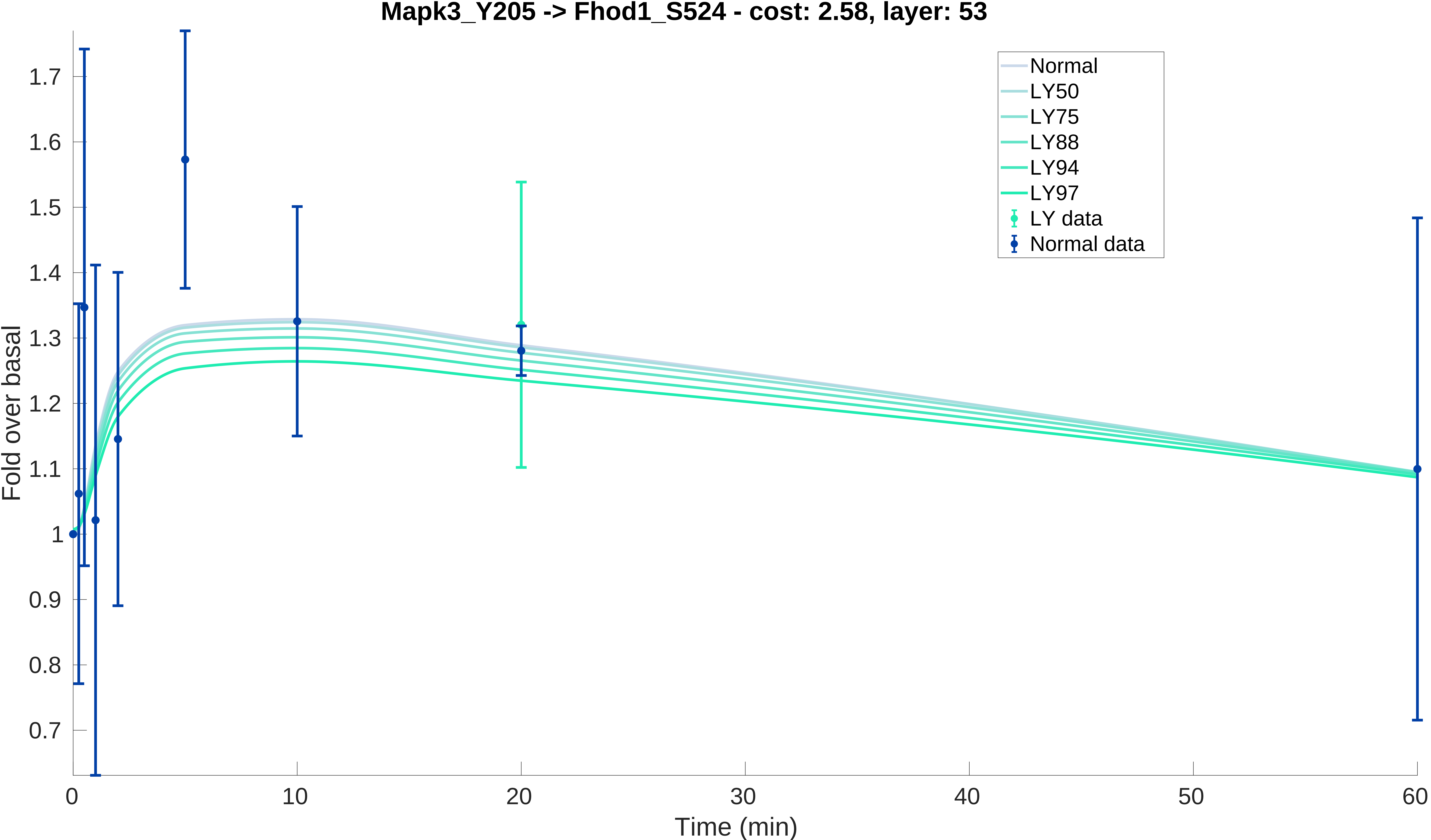

Mapk3_Y205 -> Fhod1_S524 - cost: 2.58, layer: 53
Normal
LY50
LY75
LY88
LY94
LY97
LY data
Normal data
Fold over basal
1.7
1.6
1.5
1.4
1.3
1.2
1.1
1
0.9
0.8
0.7
0
10
20
30
40
50
60
Time (min)

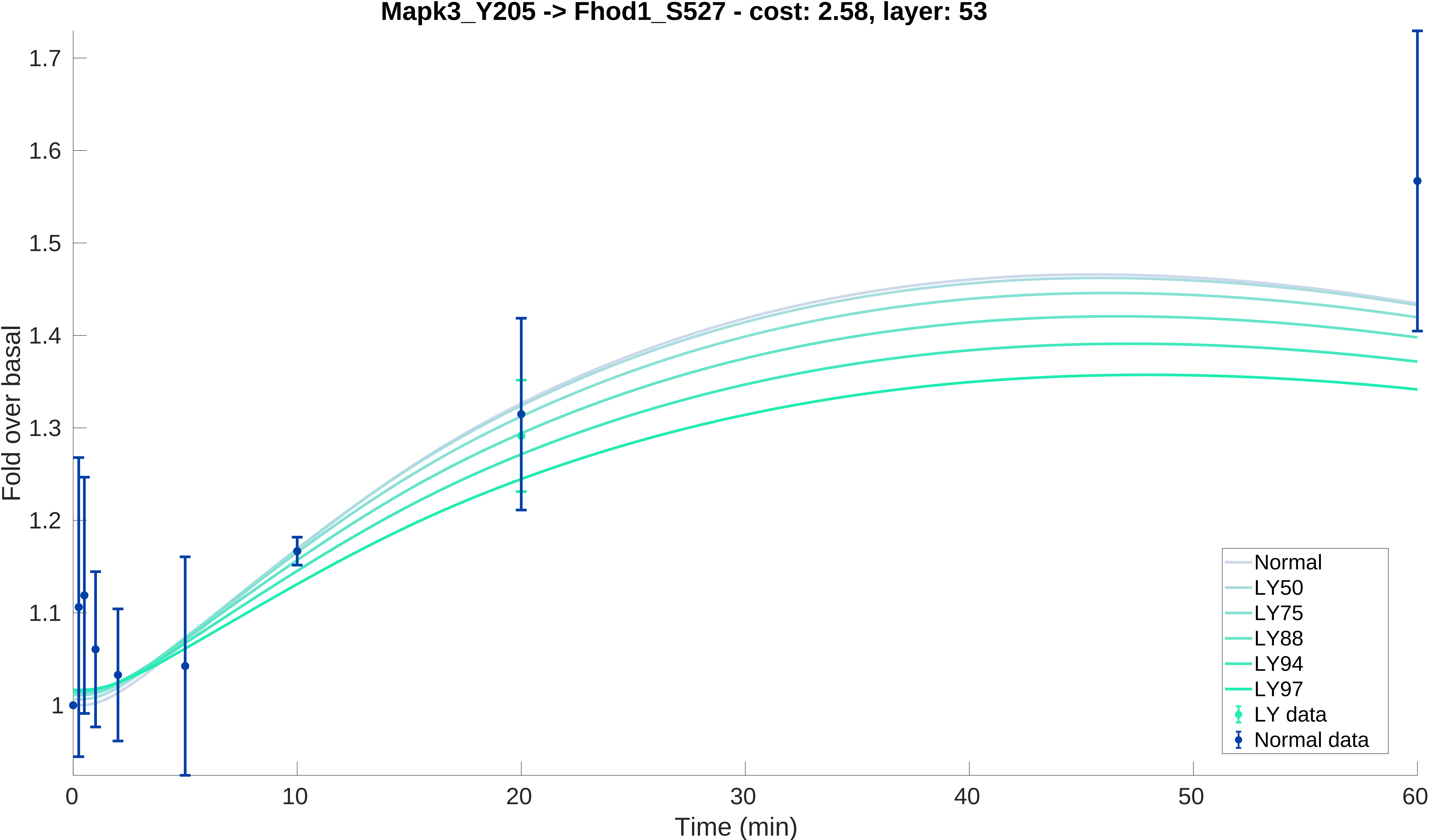
Mapk3_Y205 -> Fhod1_S527 - cost: 2.58, layer: 53
Fold over basal
1.7
1.6
1.5
1.4
1.3
1.2
1.1
1
Time (min)
0
10
20
30
40
50
60
Normal
LY50
LY75
LY88
LY94
LY97
LY data
Normal data

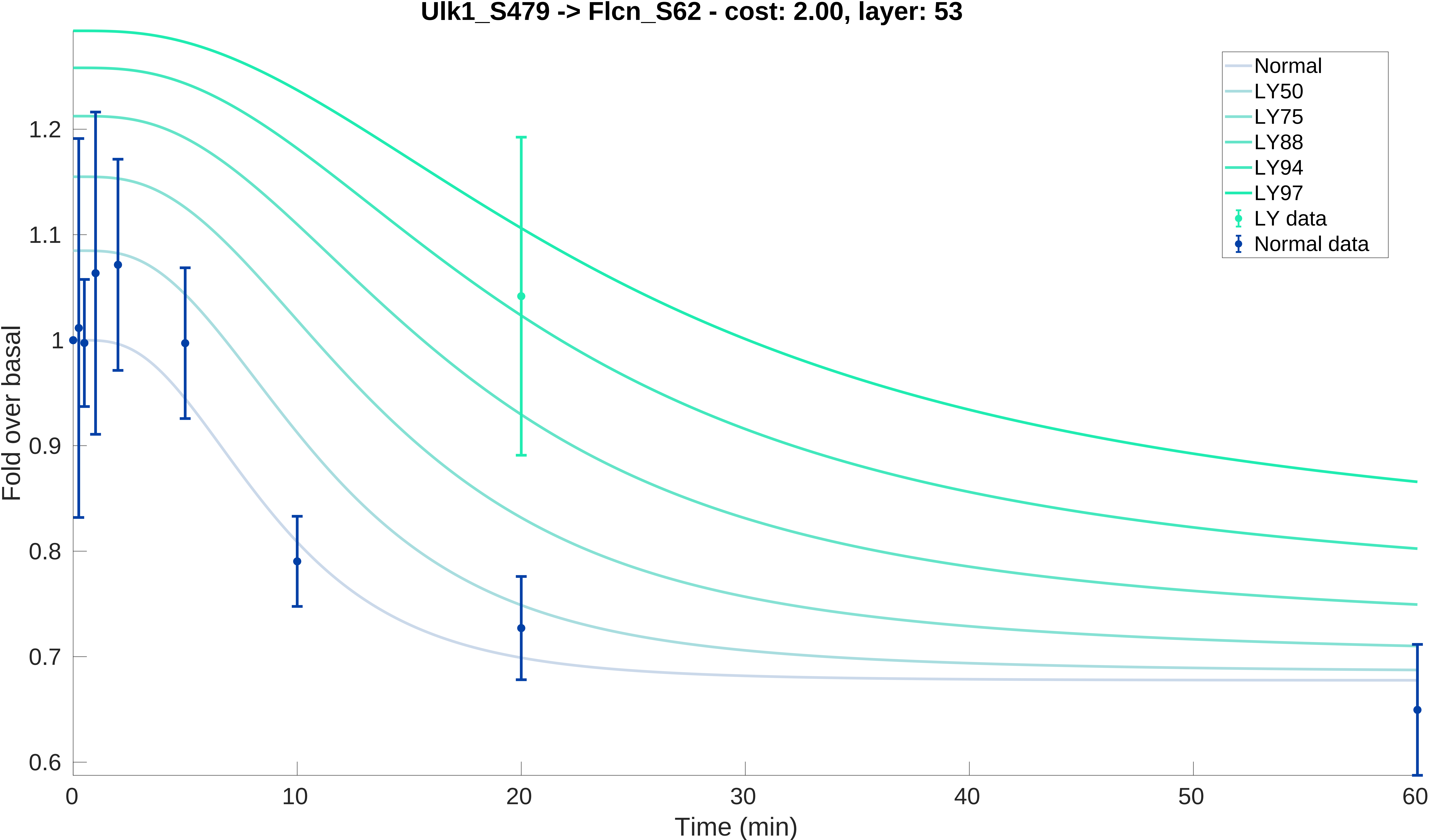

Ulk1_S479 -> Flcn_S62 - cost: 2.00, layer: 53
Normal
LY50
LY75
LY88
LY94
LY97
LY data
Normal data
Fold over basal
1.2
1.1
1
0.9
0.8
0.7
0.6
0
10
20
30
40
50
60
Time (min)

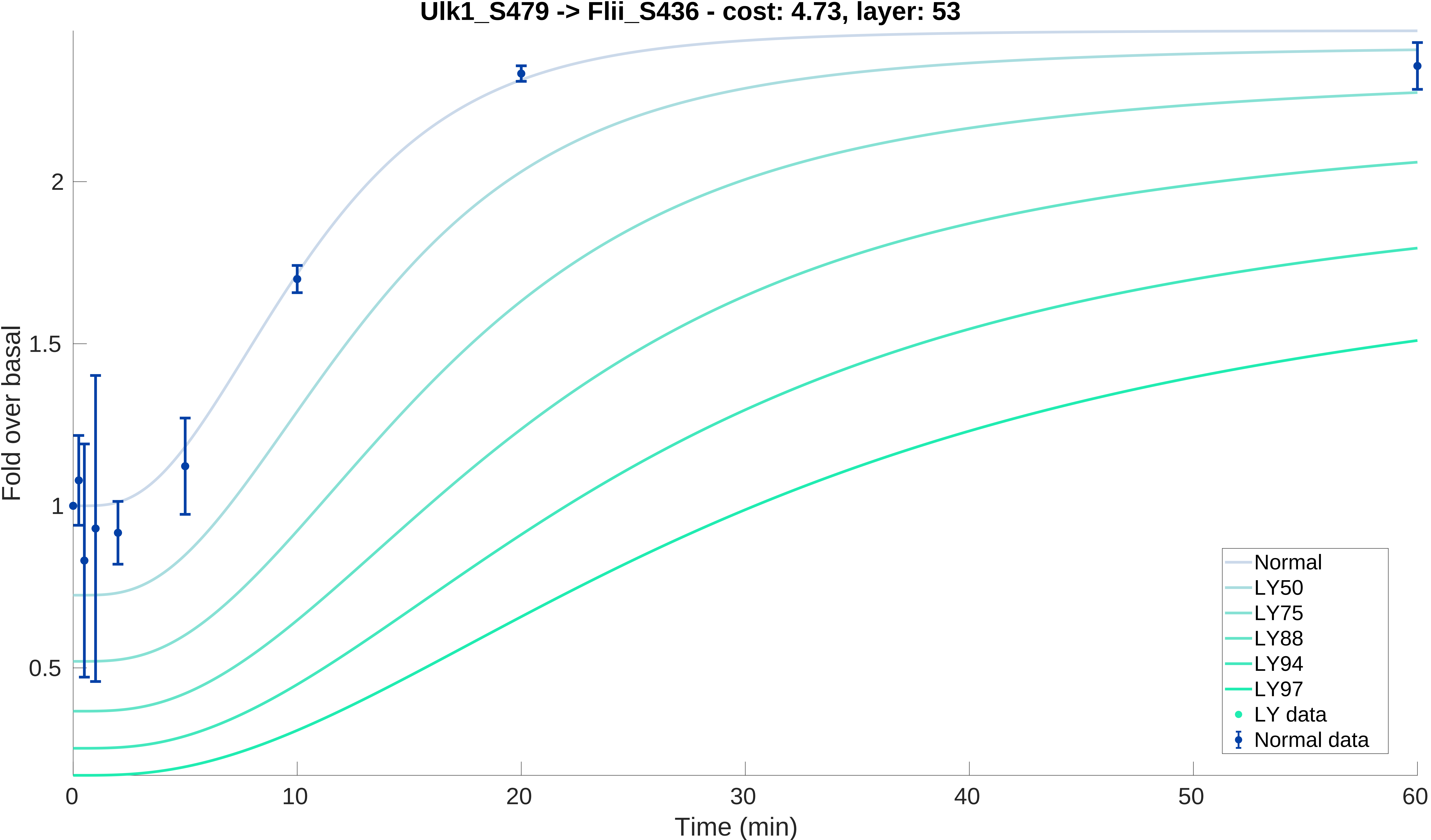

Ulk1_S479 -> Flii_S436 - cost: 4.73, layer: 53
Fold over basal
Time (min)
0
10
20
30
40
50
60
0.5
1
1.5
2
Normal
LY50
LY75
LY88
LY94
LY97
LY data
Normal data

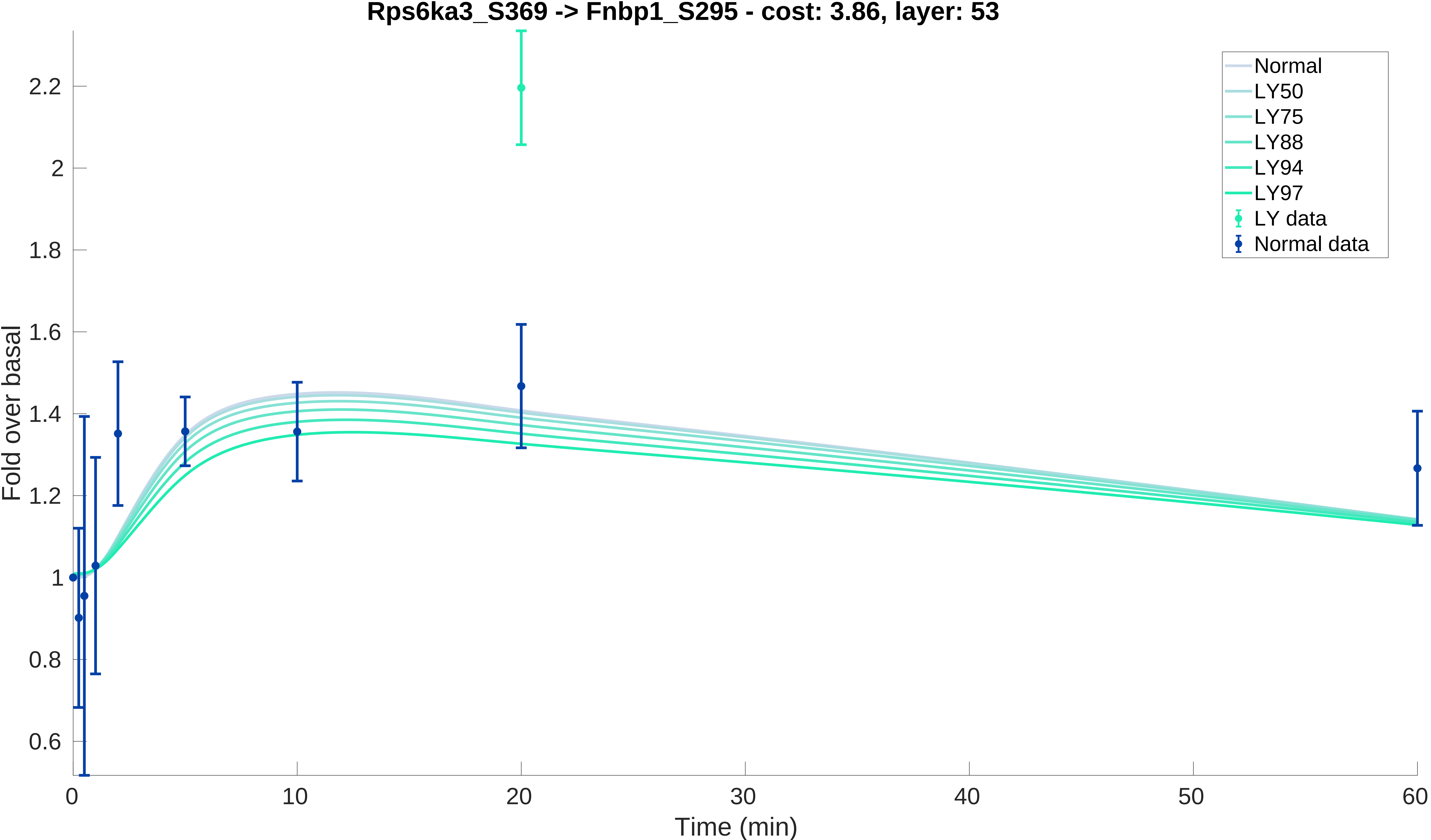
Rps6ka3_S369 -> Fnbp1_S295 - cost: 3.86, layer: 53
Normal
LY50
LY75
LY88
LY94
LY97
LY data
Normal data
Fold over basal
2.2
2
1.8
1.6
1.4
1.2
1
0.8
0.6
0
10
20
30
40
50
60
Time (min)

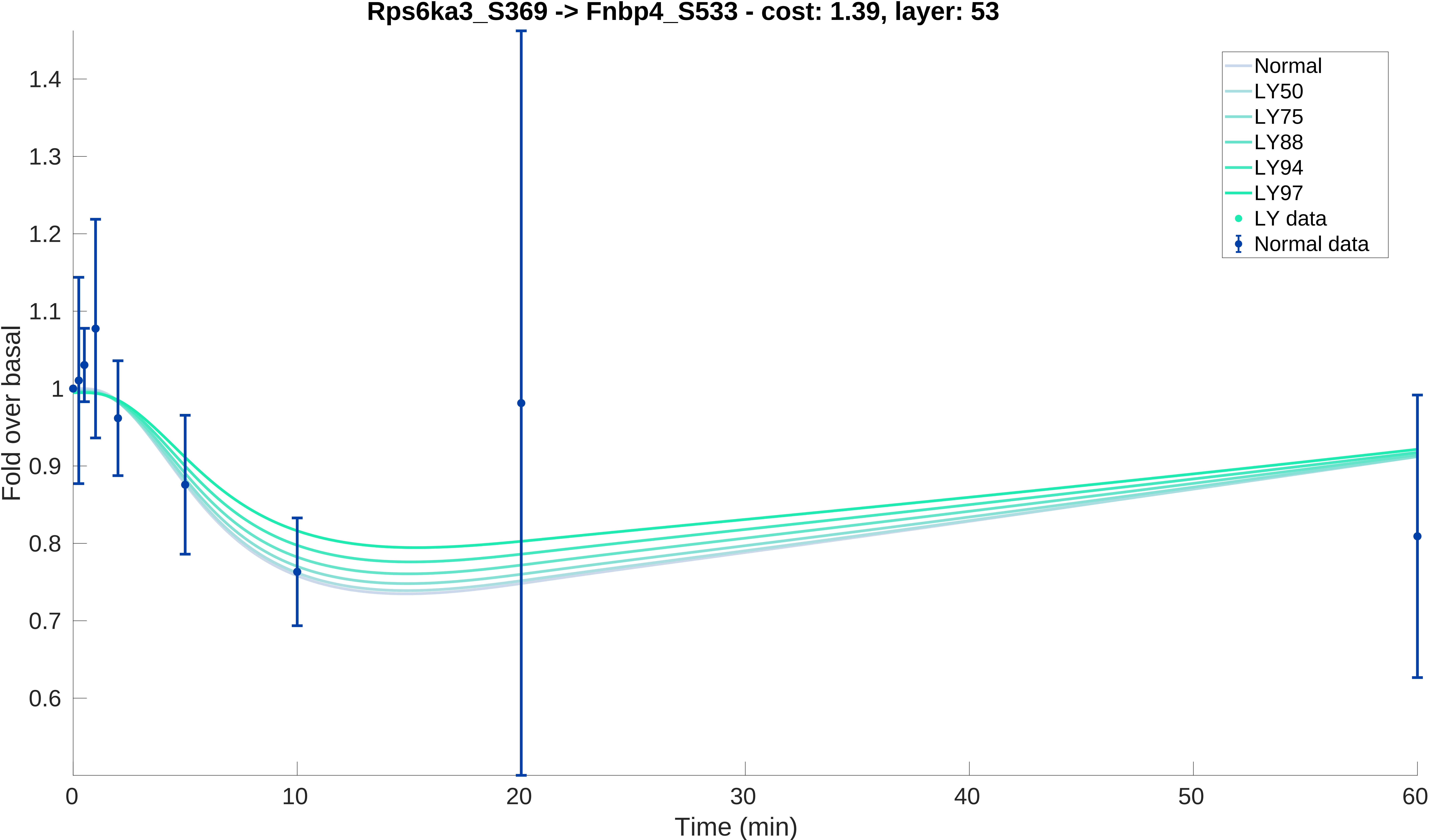

Rps6ka3_S369 -> Fnbp4_S533 - cost: 1.39, layer: 53
Fold over basal
Time (min)
1.4
1.3
1.2
1.1
1
0.9
0.8
0.7
0.6
0
10
20
30
40
50
60
Normal
LY50
LY75
LY88
LY94
LY97
LY data
Normal data

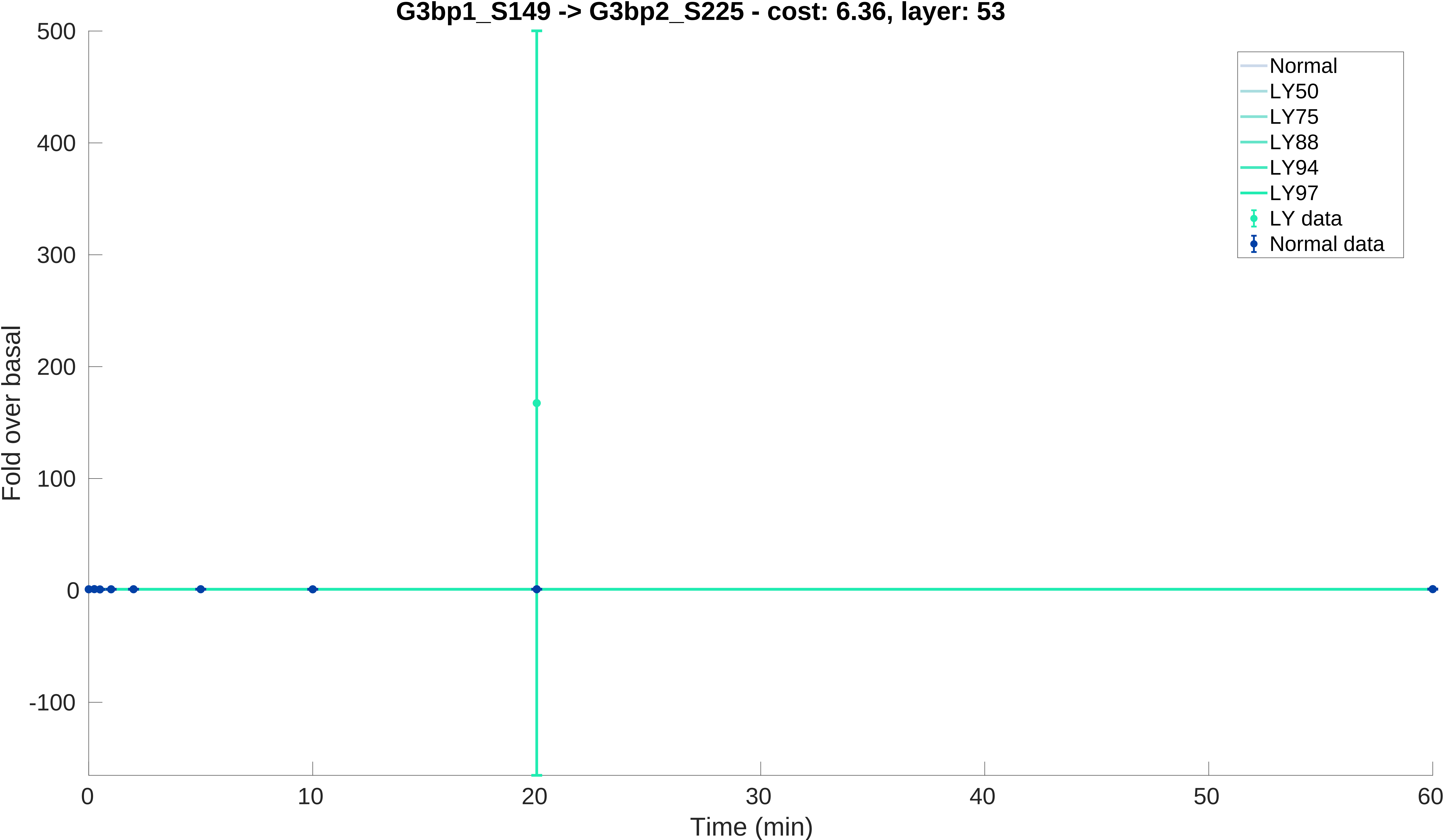

G3bp1_S149 -> G3bp2_S225 - cost: 6.36, layer: 53
Fold over basal
Time (min)
500
400
300
200
100
0
-100
0
10
20
30
40
50
60
Normal
LY50
LY75
LY88
LY94
LY97
LY data
Normal data

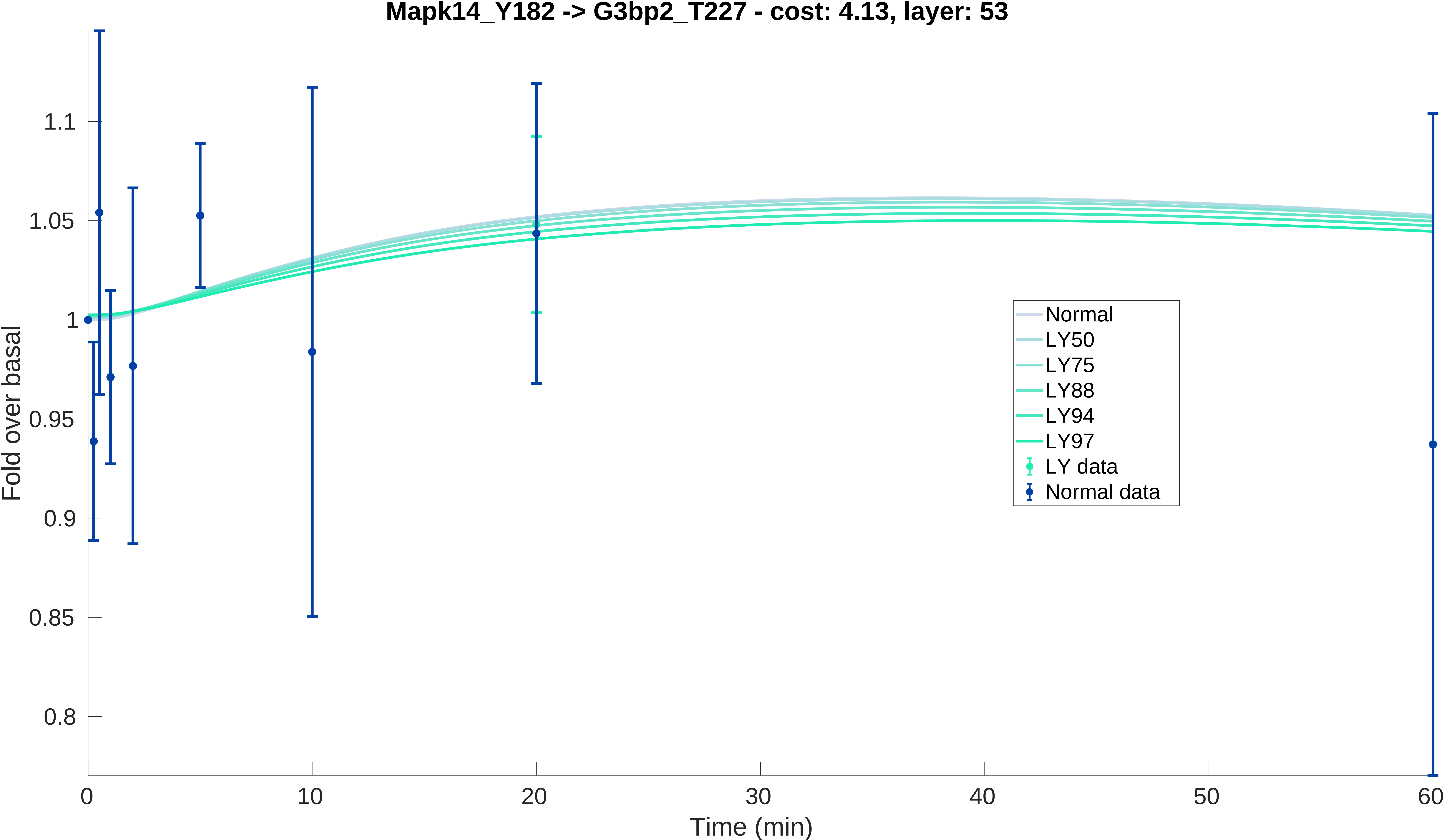

Mapk14_Y182 -> G3bp2_T227 - cost: 4.13, layer: 53
Fold over basal
1.1
1.05
1
0.95
0.9
0.85
0.8
0
10
20
30
40
50
60
Time (min)
Normal
LY50
LY75
LY88
LY94
LY97
LY data
Normal data

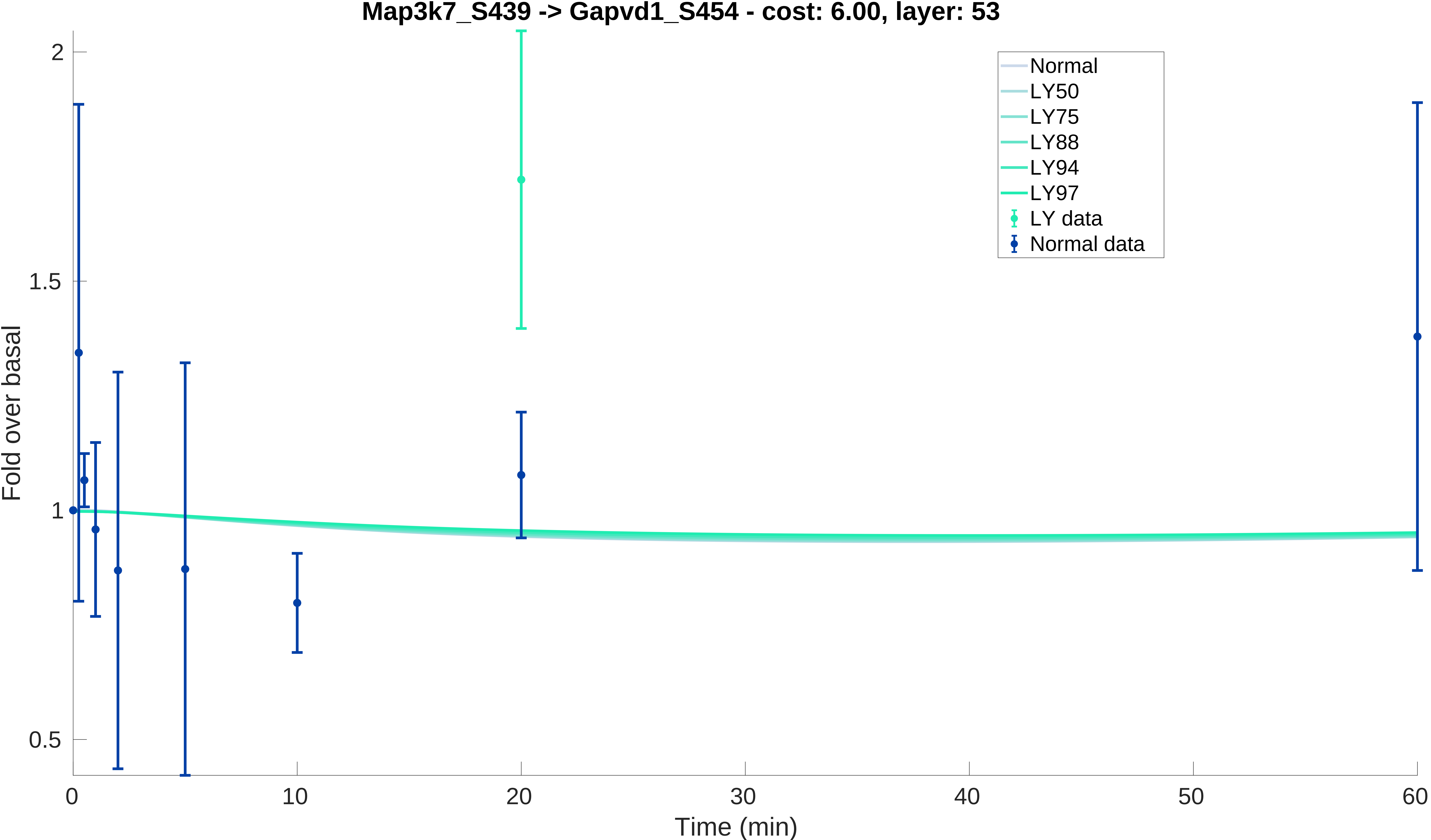

Map3k7_S439 -> Gapvd1_S454 - cost: 6.00, layer: 53
Fold over basal
Time (min)
0.5
1
1.5
2
0
10
20
30
40
50
60
Normal
LY50
LY75
LY88
LY94
LY97
LY data
Normal data

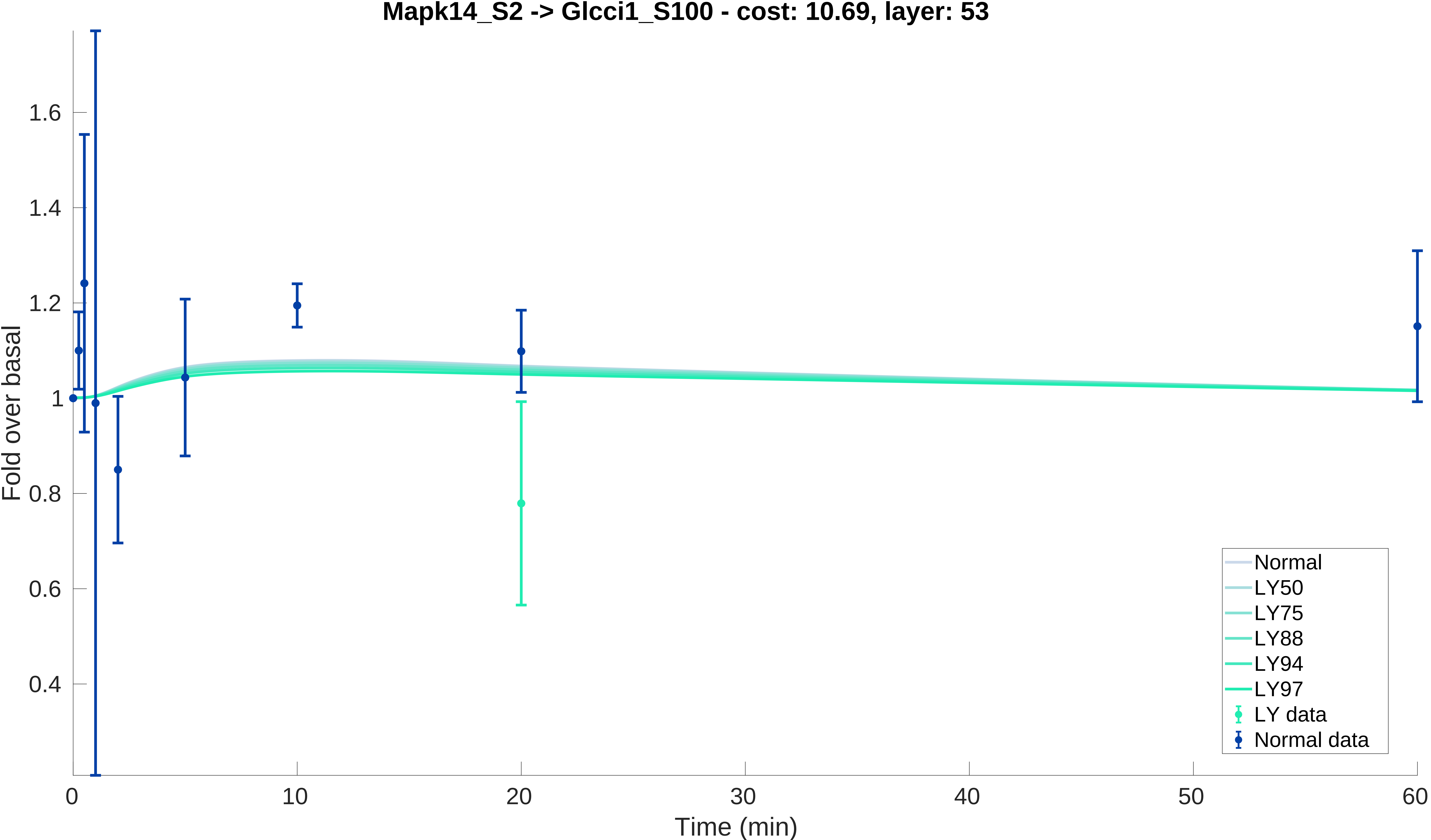

Mapk14_S2 -> Glcci1_S100 - cost: 10.69, layer: 53
Fold over basal
1.6
1.4
1.2
1
0.8
0.6
0.4
0
10
20
30
40
50
60
Time (min)
Normal
LY50
LY75
LY88
LY94
LY97
LY data
Normal data

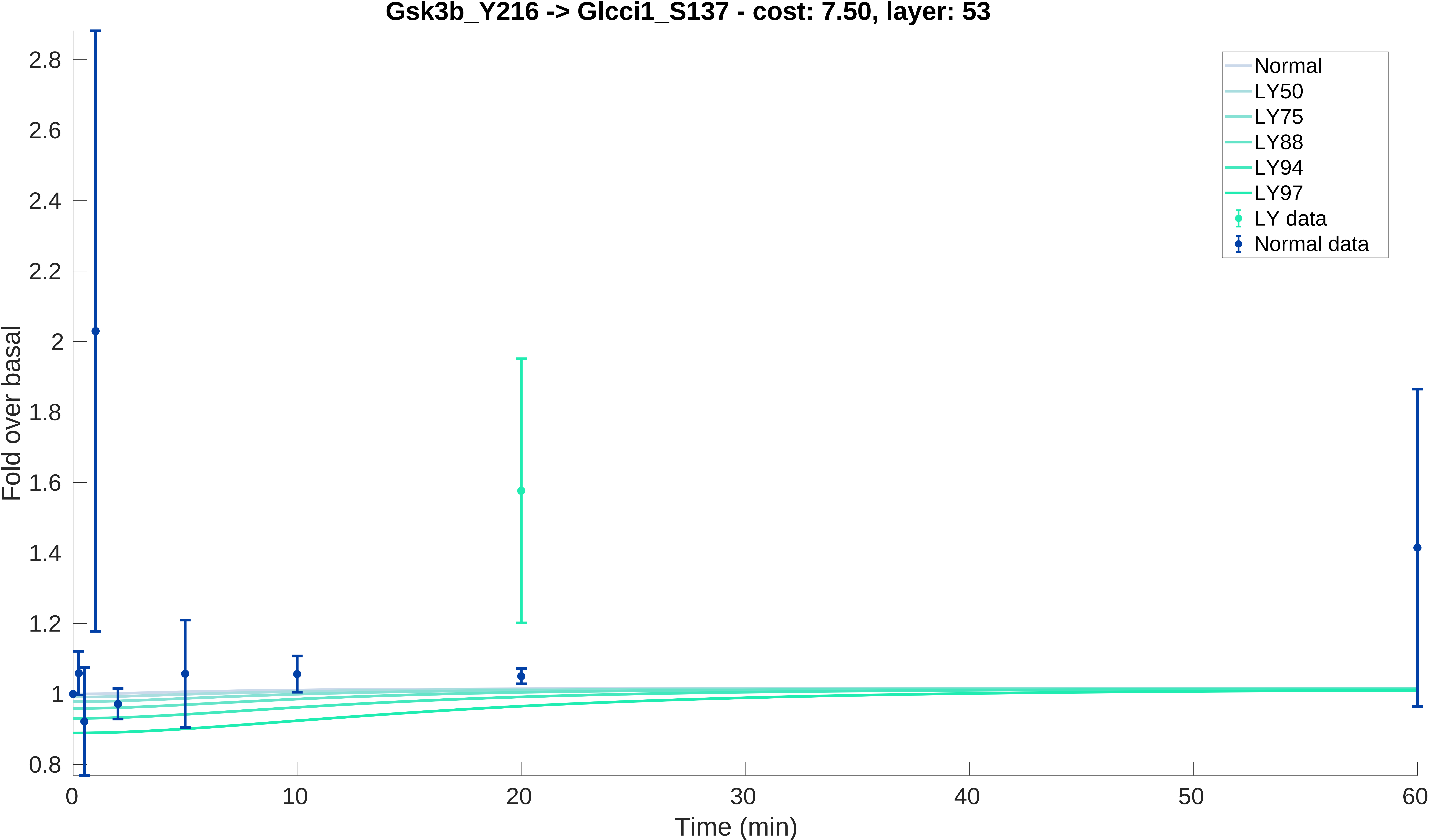
Gsk3b_Y216 -> Glcci1_S137 - cost: 7.50, layer: 53
Fold over basal
2.8
2.6
2.4
2.2
2
1.8
1.6
1.4
1.2
1
0.8
0
10
20
30
40
50
60
Time (min)
Normal
LY50
LY75
LY88
LY94
LY97
LY data
Normal data

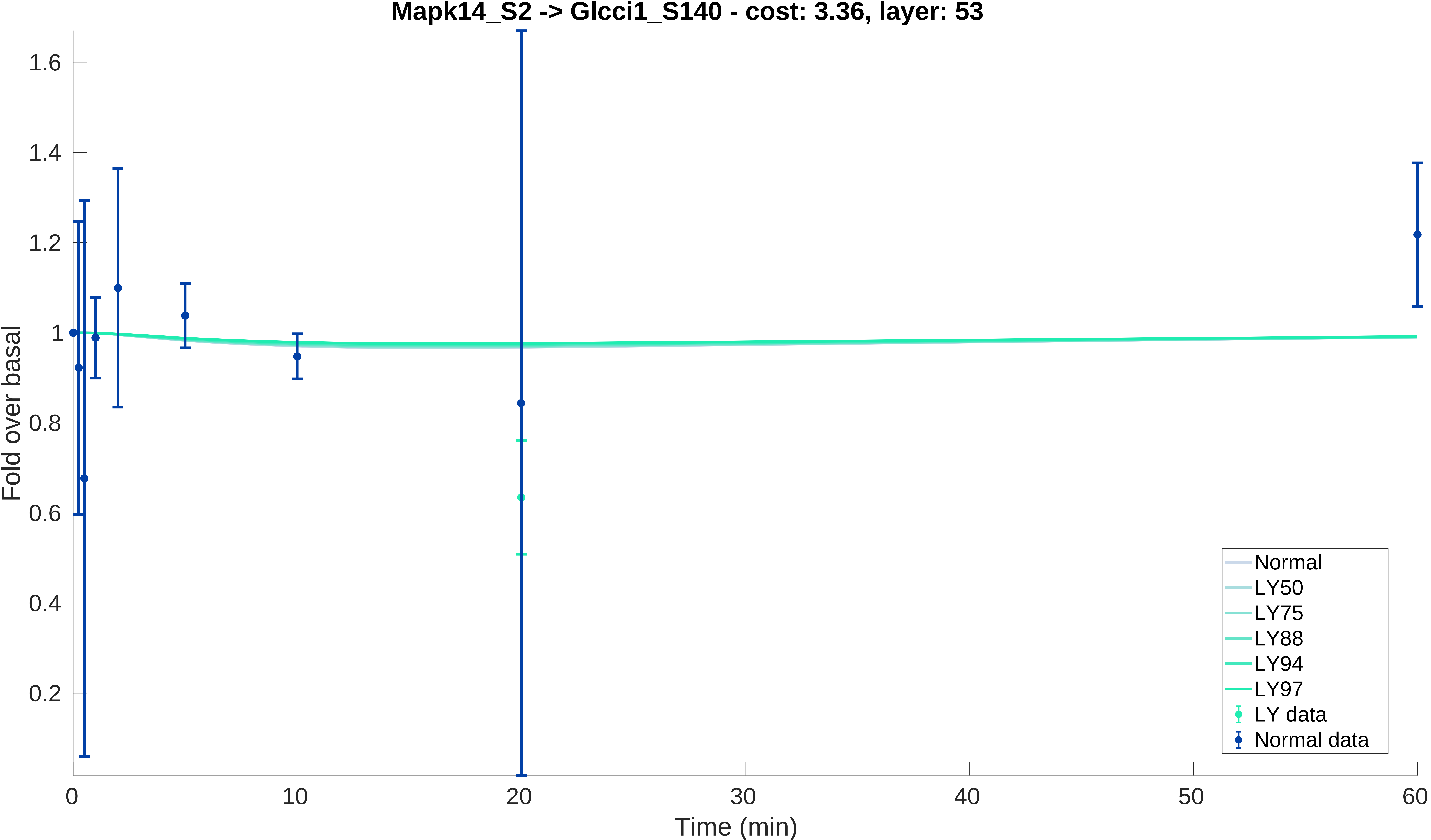

Mapk14_S2 -> Glcci1_S140 - cost: 3.36, layer: 53
Fold over basal
1.6
1.4
1.2
1
0.8
0.6
0.4
0.2
0
10
20
30
40
50
60
Time (min)
Normal
LY50
LY75
LY88
LY94
LY97
LY data
Normal data

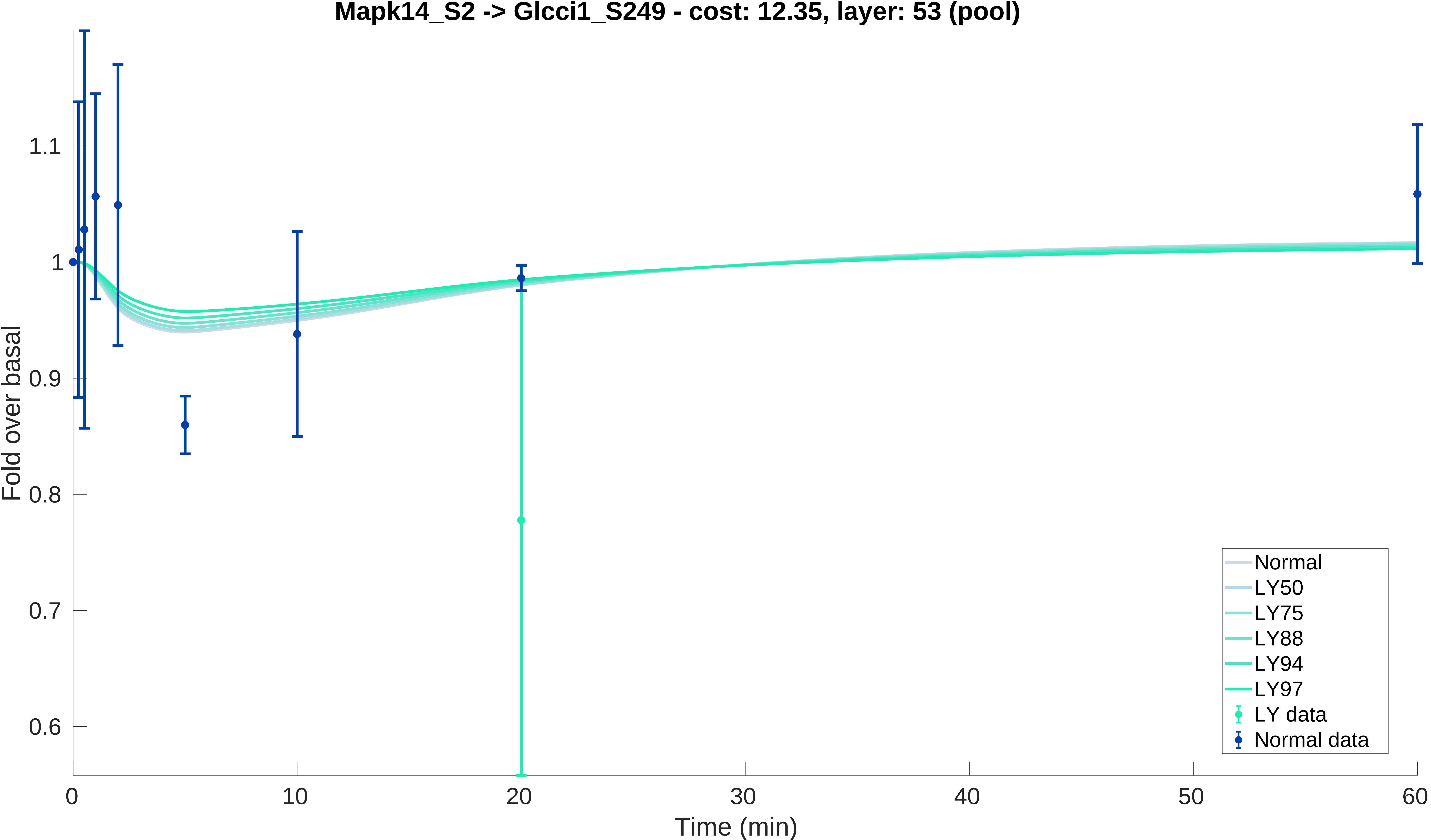
Mapk14_S2 -> Glcci1_S249 - cost: 12.35, layer: 53 (pool)
Fold over basal
Time (min)
1.1
1
0.9
0.8
0.7
0.6
0
10
20
30
40
50
60
Normal
LY50
LY75
LY88
LY94
LY97
LY data
Normal data

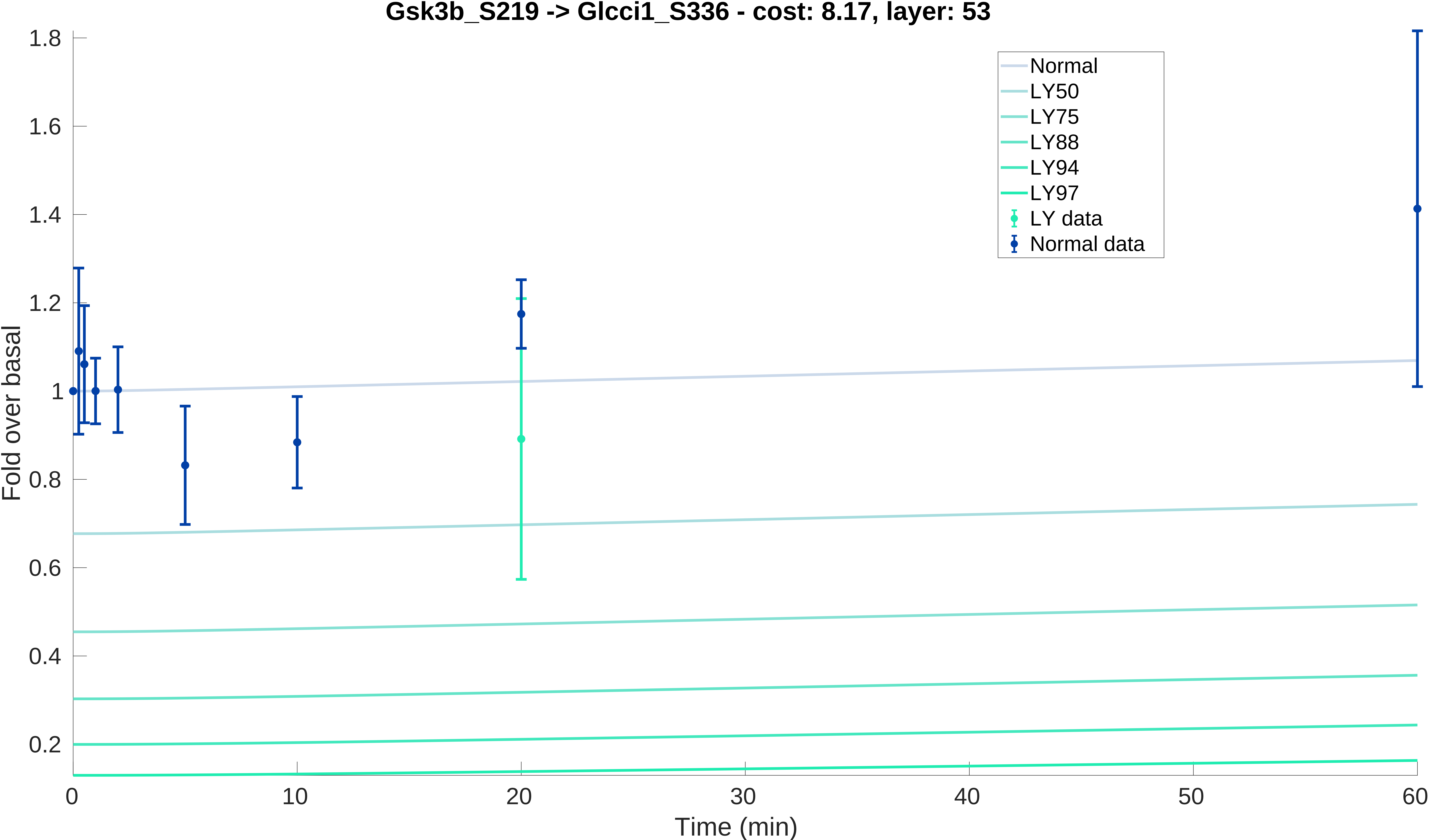

Gsk3b_S219 -> Glcci1_S336 - cost: 8.17, layer: 53
Fold over basal
Time (min)
0.2
0.4
0.6
0.8
1
1.2
1.4
1.6
1.8
0
10
20
30
40
50
60
Normal
LY50
LY75
LY88
LY94
LY97
LY data
Normal data

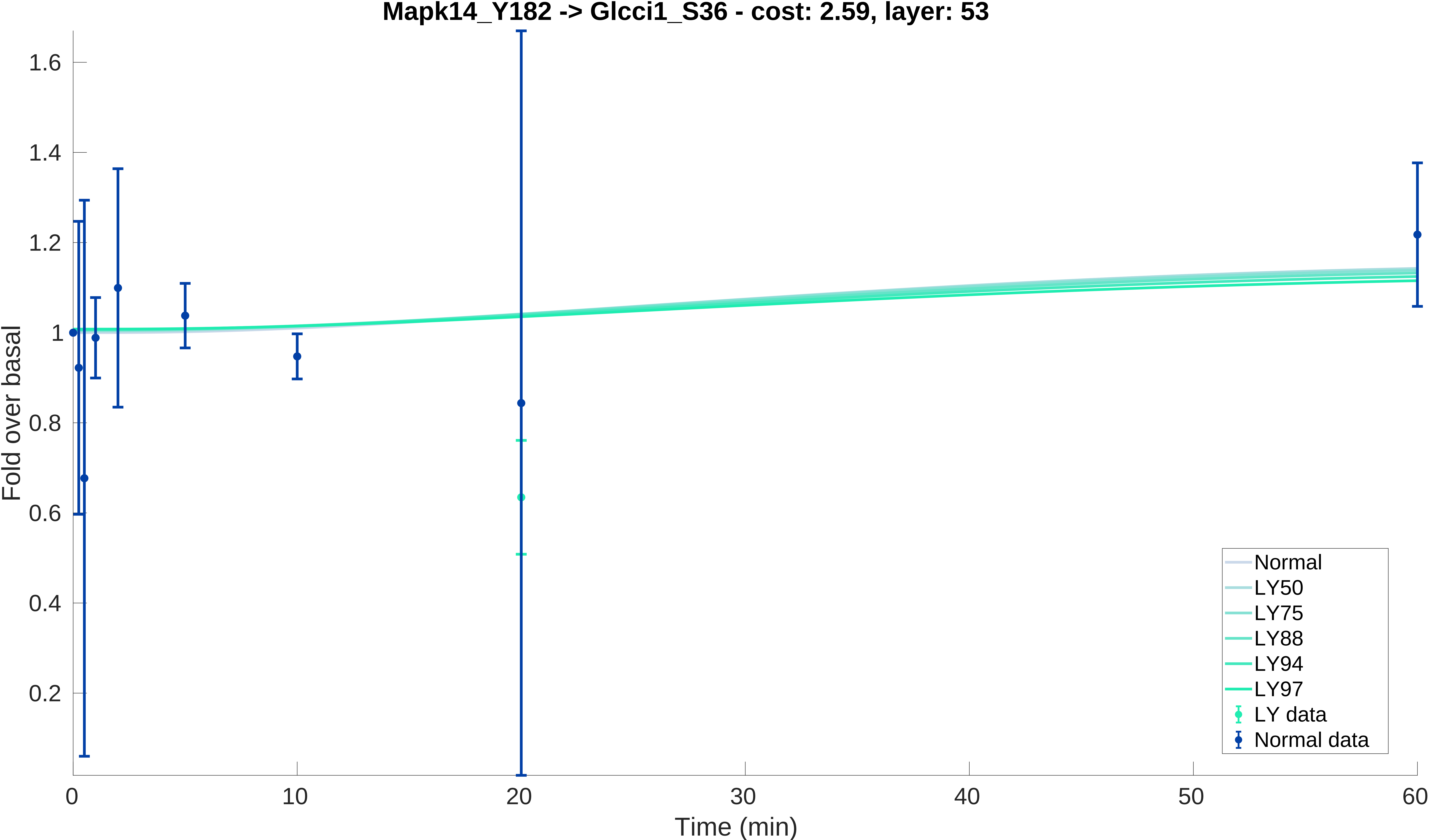

Mapk14_Y182 -> Glcci1_S36 - cost: 2.59, layer: 53
Fold over basal
Time (min)
0
10
20
30
40
50
60
0.2
0.4
0.6
0.8
1
1.2
1.4
1.6
Normal
LY50
LY75
LY88
LY94
LY97
LY data
Normal data

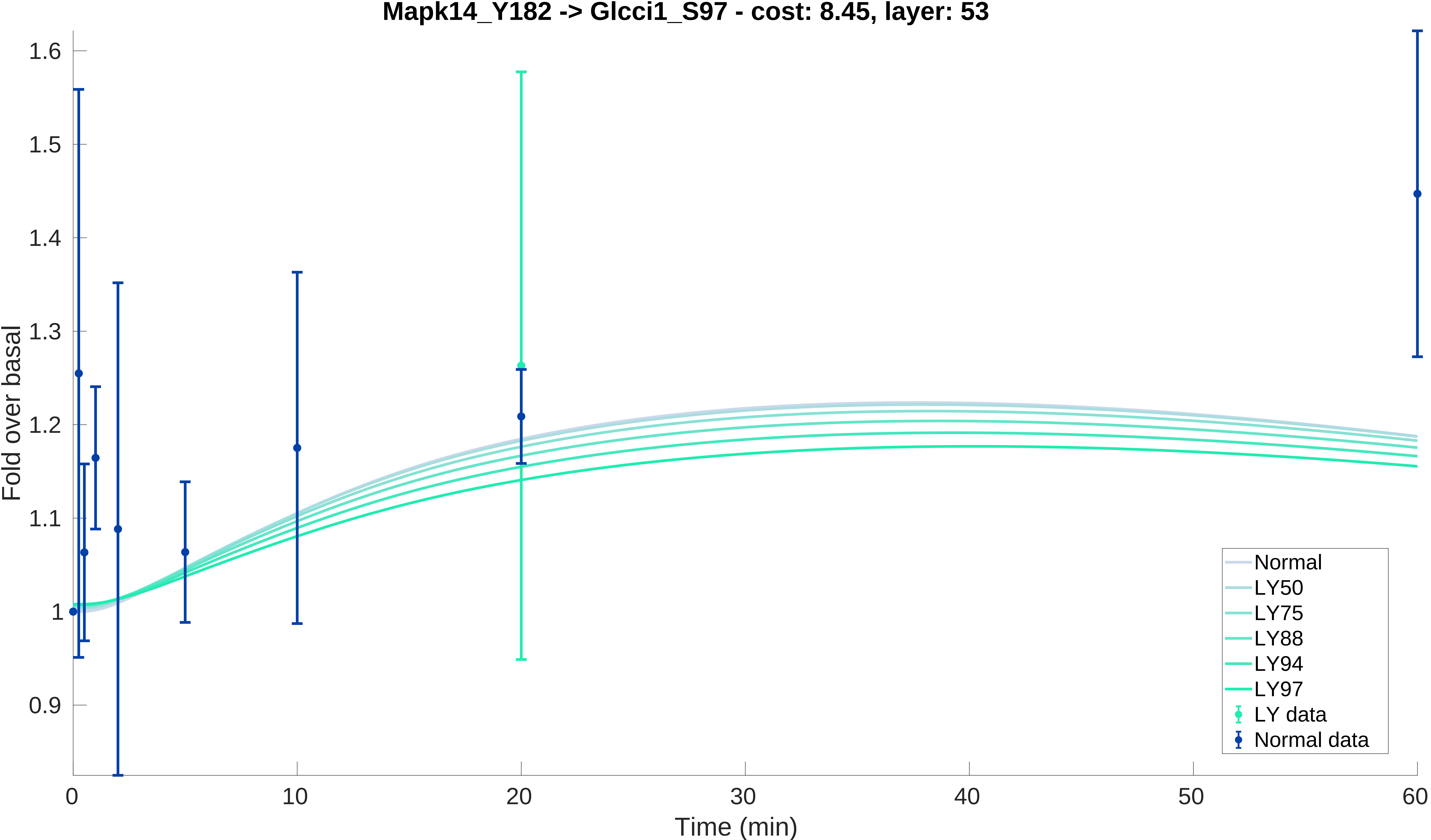

Mapk14_Y182 -> Glcci1_S97 - cost: 8.45, layer: 53
Fold over basal
1.6
1.5
1.4
1.3
1.2
1.1
1
0.9
0
10
20
30
40
50
60
Time (min)
Normal
LY50
LY75
LY88
LY94
LY97
LY data
Normal data

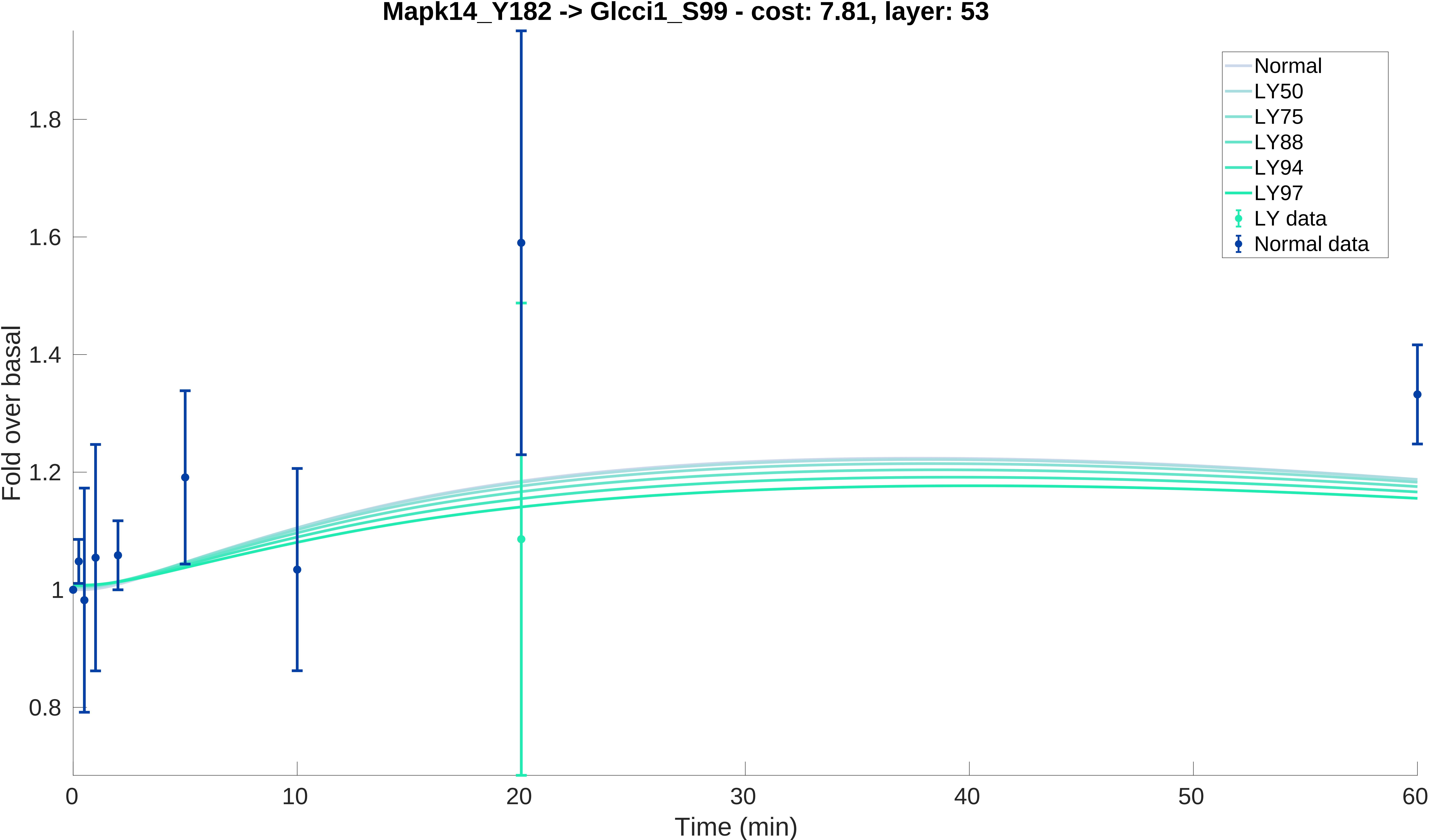

Mapk14_Y182 -> Glcci1_S99 - cost: 7.81, layer: 53
Fold over basal
Time (min)
0.8
1
1.2
1.4
1.6
1.8
0
10
20
30
40
50
60
Normal
LY50
LY75
LY88
LY94
LY97
LY data
Normal data

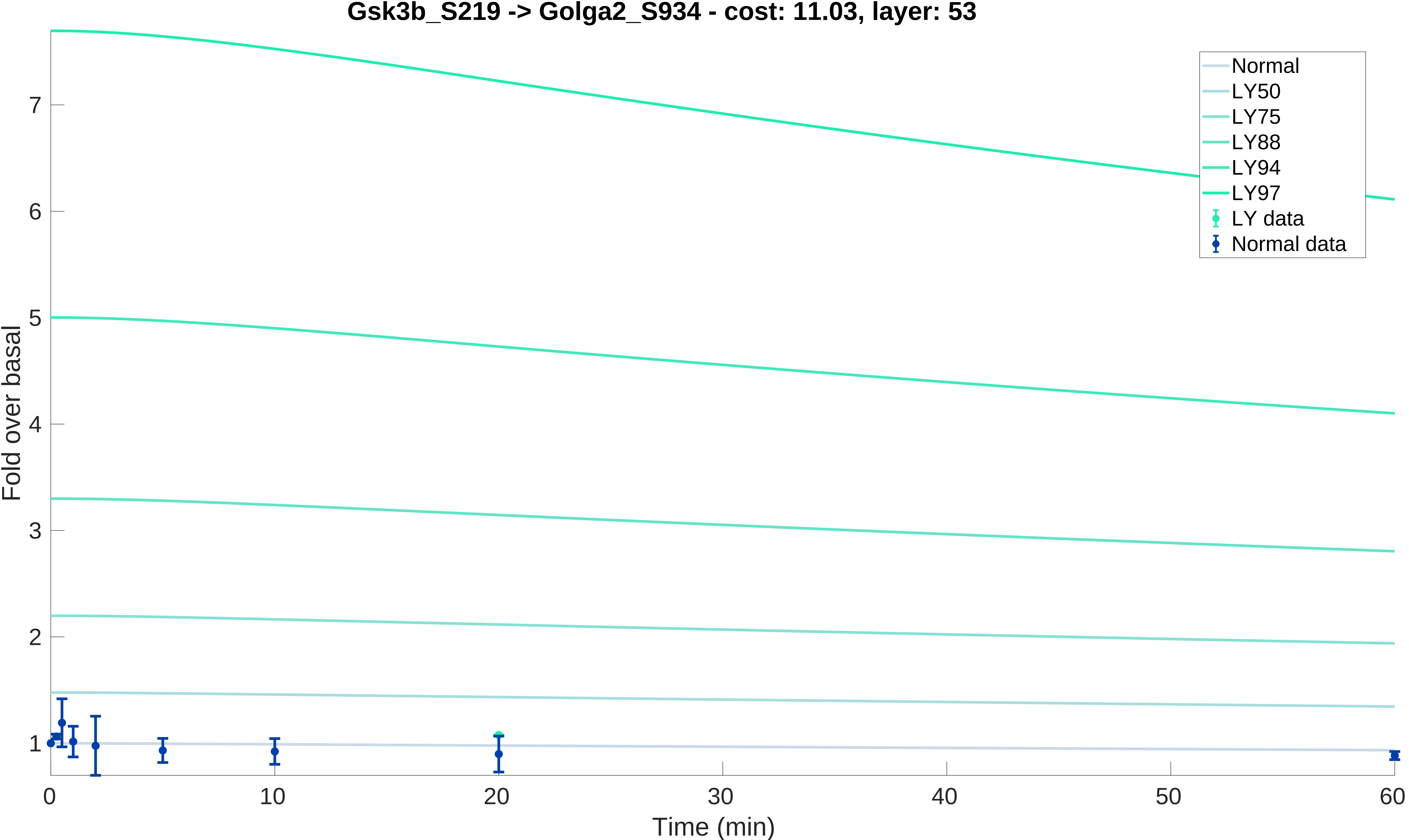
Gsk3b_S219 -> Golga2_S934 - cost: 11.03, layer: 53
Fold over basal
Time (min)
Normal
LY50
LY75
LY88
LY94
LY97
LY data
Normal data
0
10
20
30
40
50
60
1
2
3
4
5
6
7

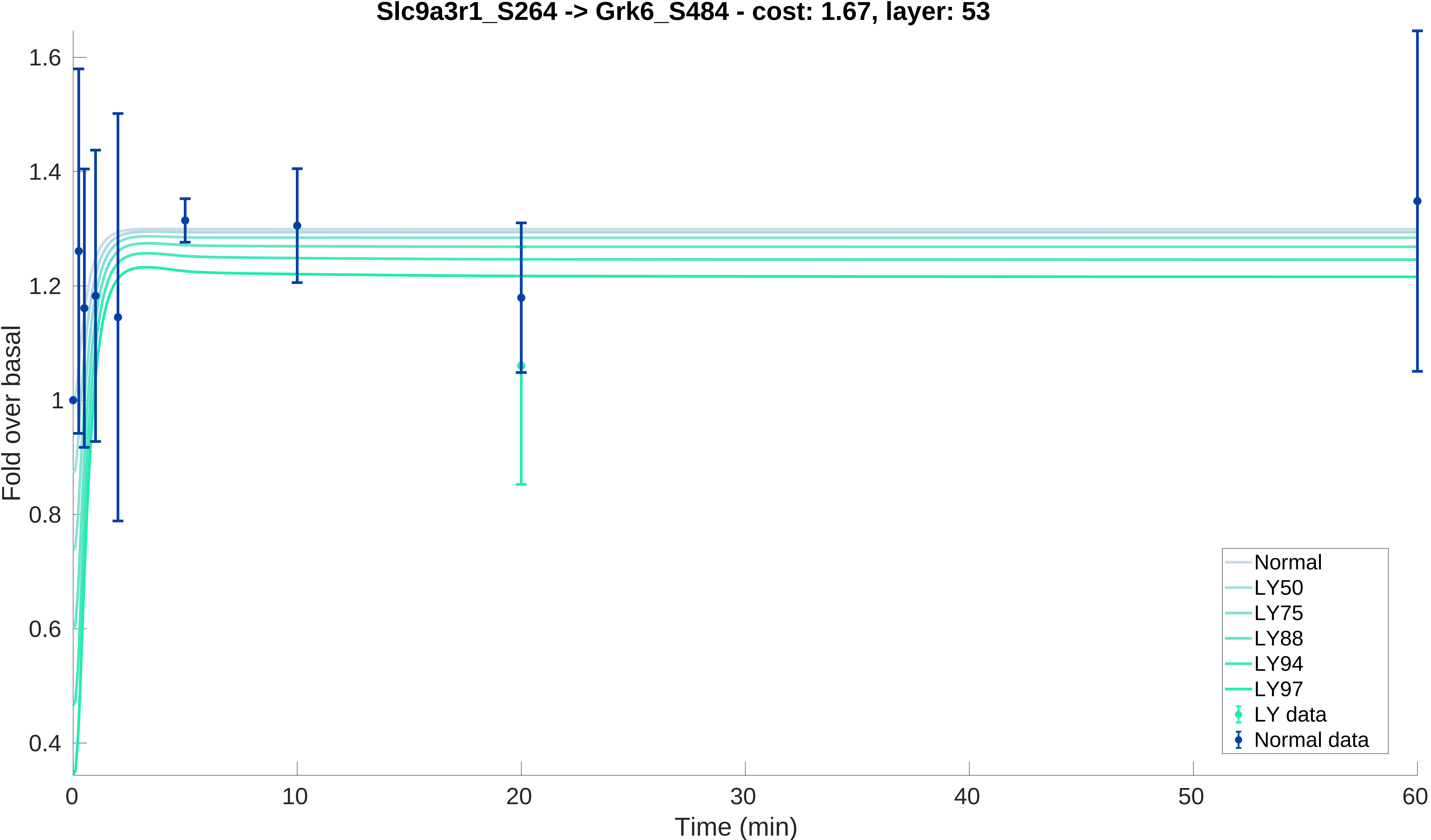

Slc9a3r1_S264 -> Grk6_S484 - cost: 1.67, layer: 53
Fold over basal
Time (min)
1.6
1.4
1.2
1
0.8
0.6
0.4
0
10
20
30
40
50
60
Normal
LY50
LY75
LY88
LY94
LY97
LY data
Normal data

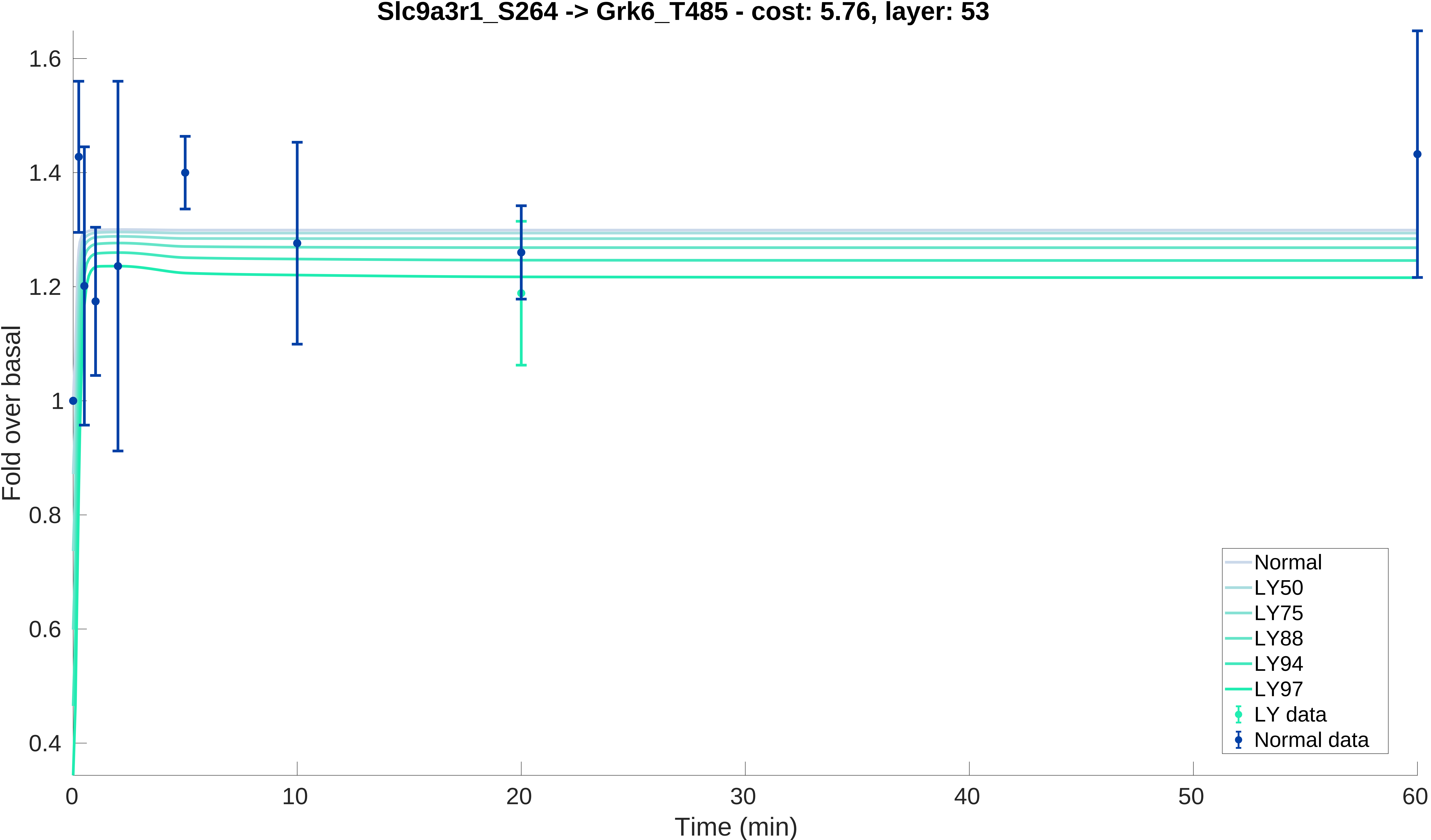

Slc9a3r1_S264 -> Grk6_T485 - cost: 5.76, layer: 53
Fold over basal
Time (min)
1.6
1.4
1.2
1
0.8
0.6
0.4
0
10
20
30
40
50
60
Normal
LY50
LY75
LY88
LY94
LY97
LY data
Normal data

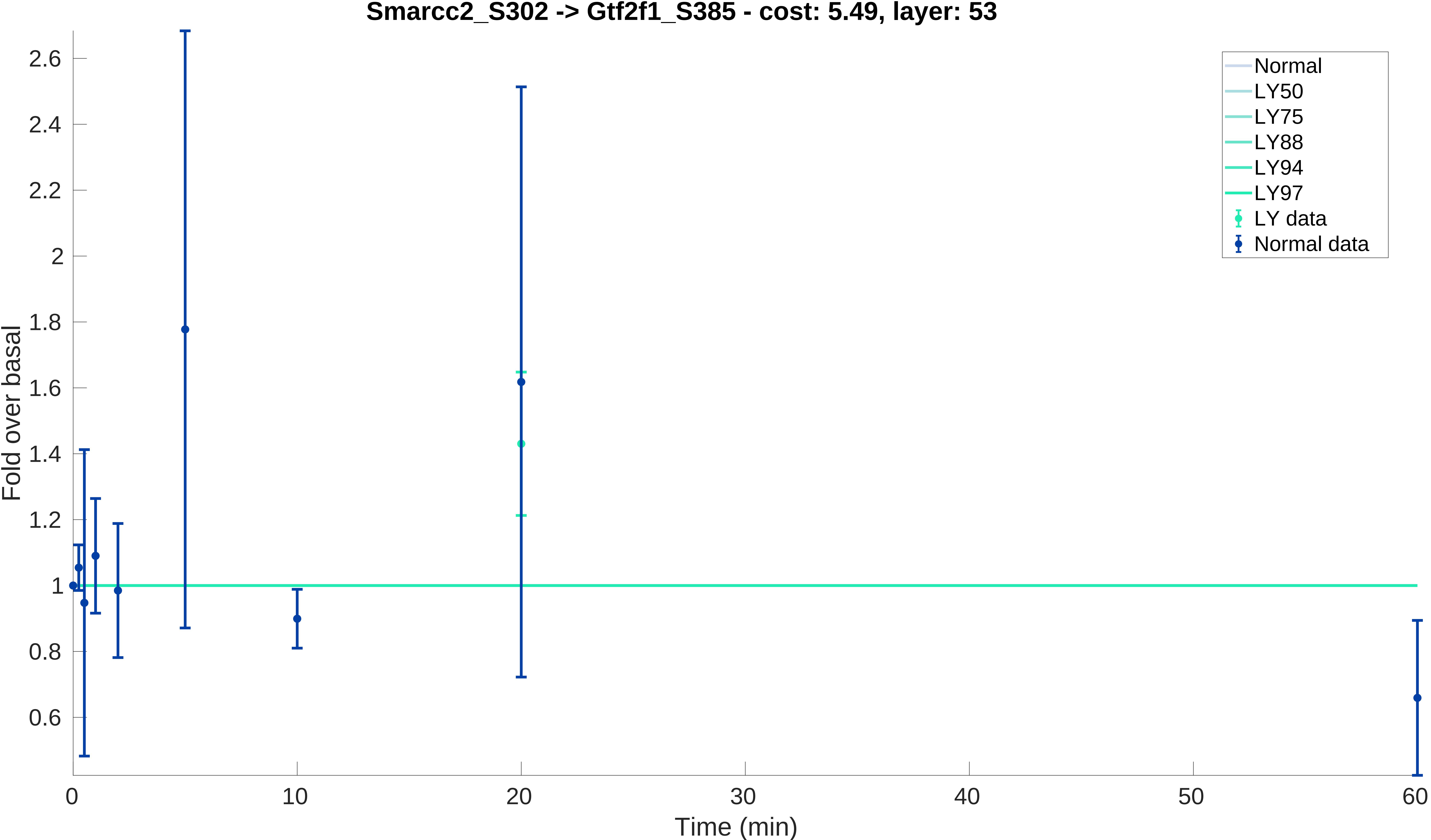

Smarcc2_S302 -> Gtf2f1_S385 - cost: 5.49, layer: 53
Fold over basal
Time (min)
2.6
2.4
2.2
2
1.8
1.6
1.4
1.2
1
0.8
0.6
0
10
20
30
40
50
60
Normal
LY50
LY75
LY88
LY94
LY97
LY data
Normal data

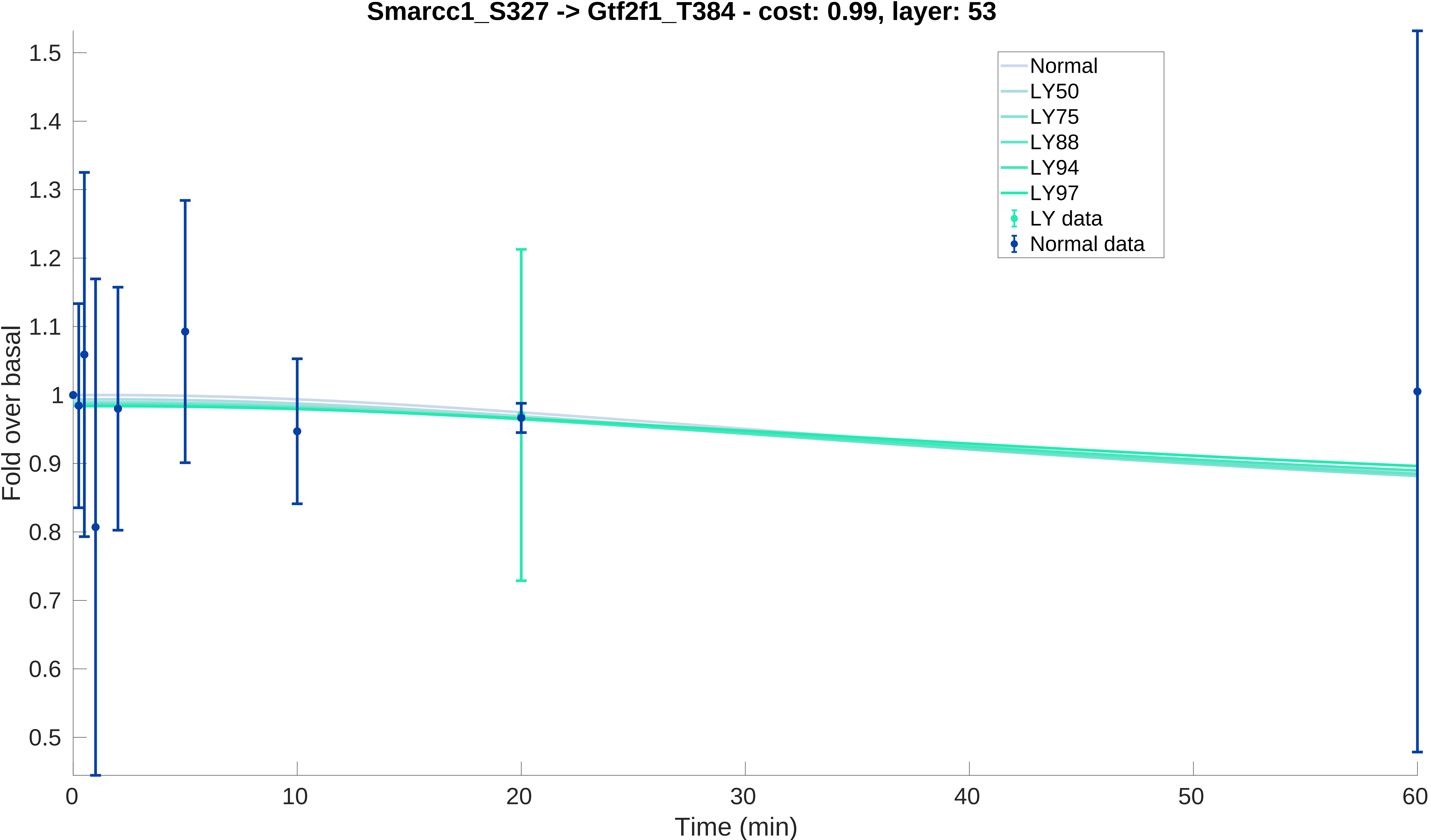
Smarcc1_S327 -> Gtf2f1_T384 - cost: 0.99, layer: 53
Fold over basal
Time (min)
Normal
LY50
LY75
LY88
LY94
LY97
LY data
Normal data
1.5
1.4
1.3
1.2
1.1
1
0.9
0.8
0.7
0.6
0.5
0
10
20
30
40
50
60

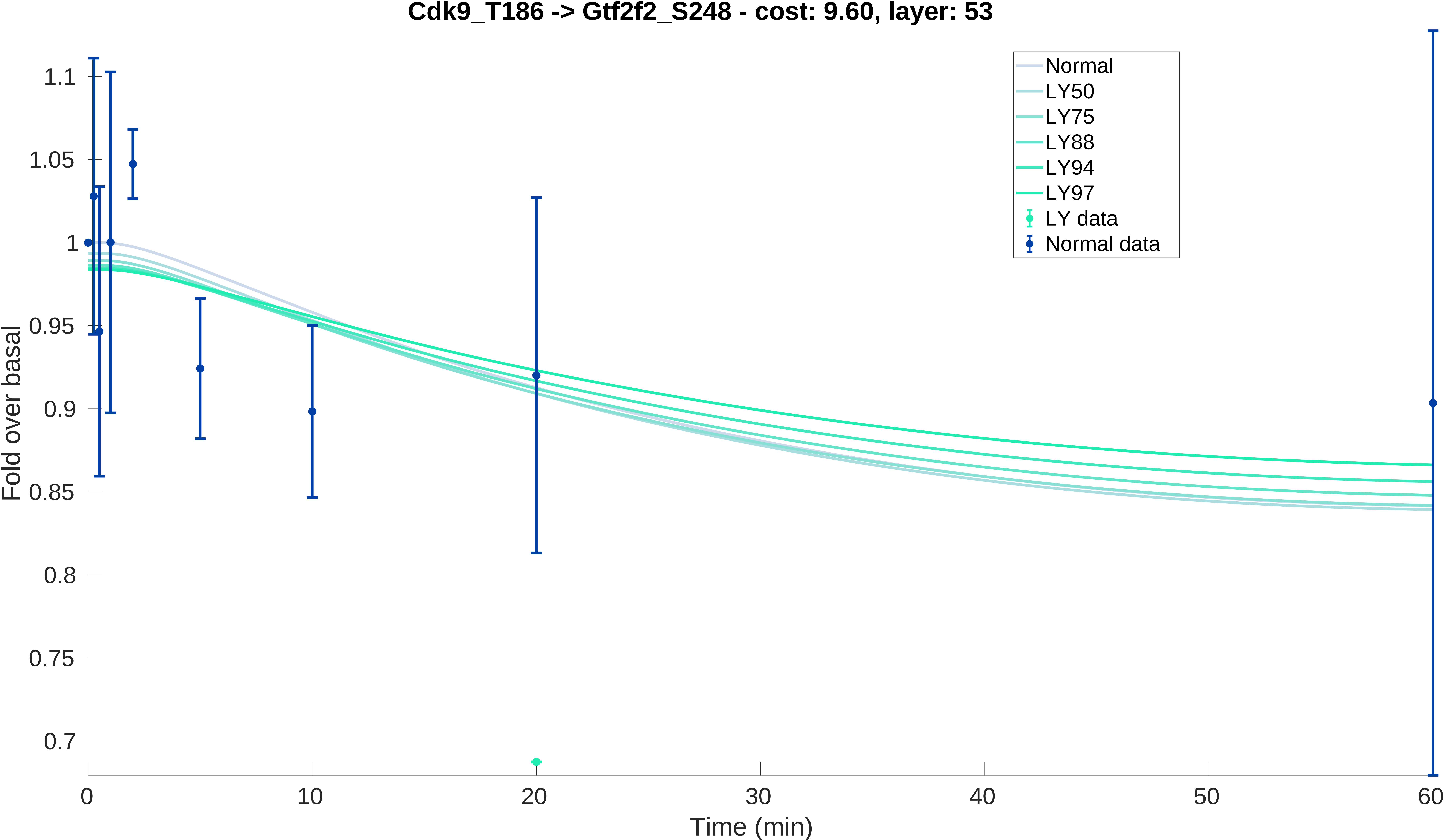

Cdk9_T186 -> Gtf2f2_S248 - cost: 9.60, layer: 53
Normal
LY50
LY75
LY88
LY94
LY97
LY data
Normal data
Fold over basal
Time (min)
1.1
1.05
1
0.95
0.9
0.85
0.8
0.75
0.7
0
10
20
30
40
50
60

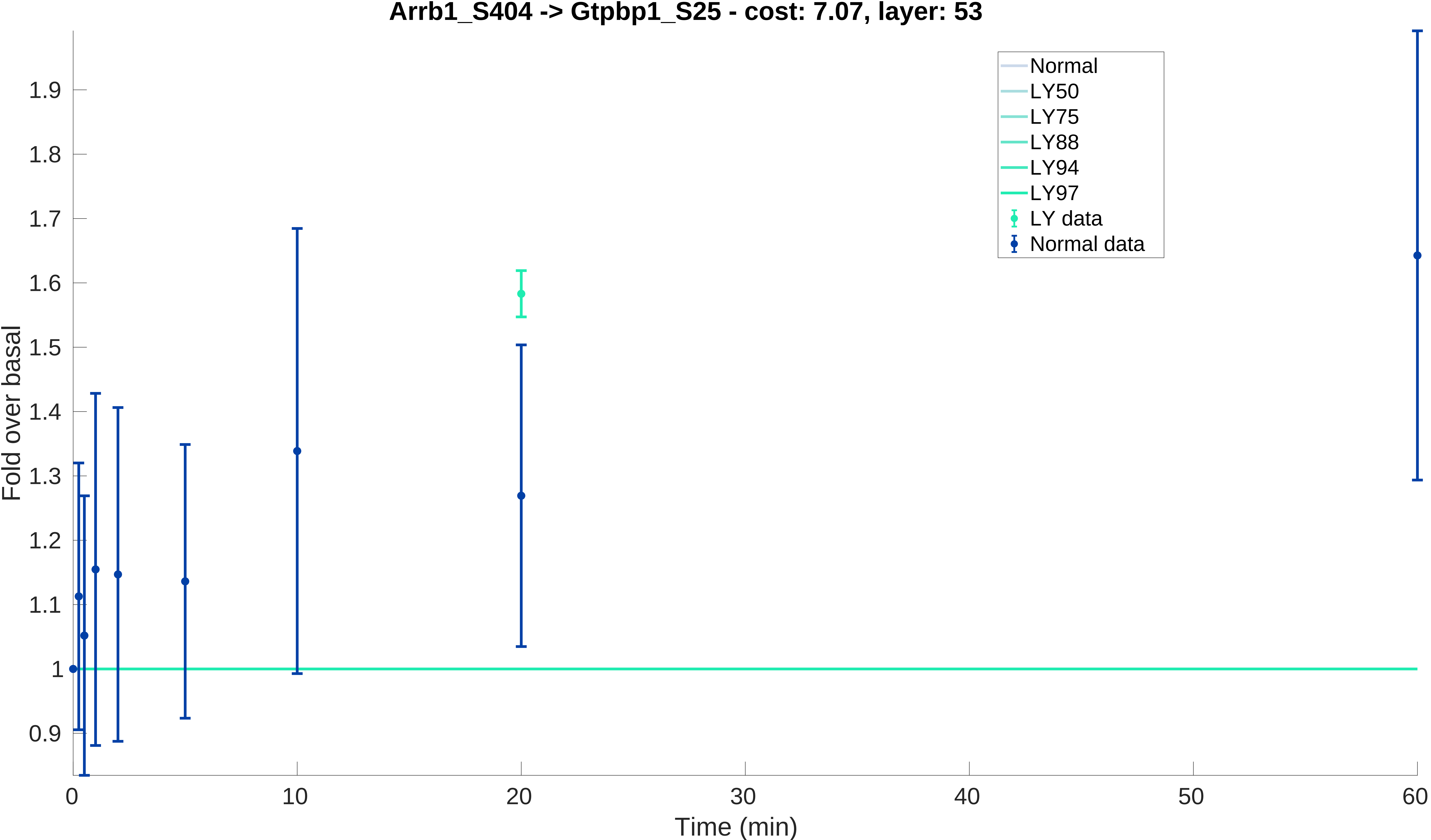
Arrb1_S404 -> Gtpbp1_S25 - cost: 7.07, layer: 53
Fold over basal
Time (min)
Normal
LY50
LY75
LY88
LY94
LY97
LY data
Normal data
1.9
1.8
1.7
1.6
1.5
1.4
1.3
1.2
1.1
1
0.9
0
10
20
30
40
50
60

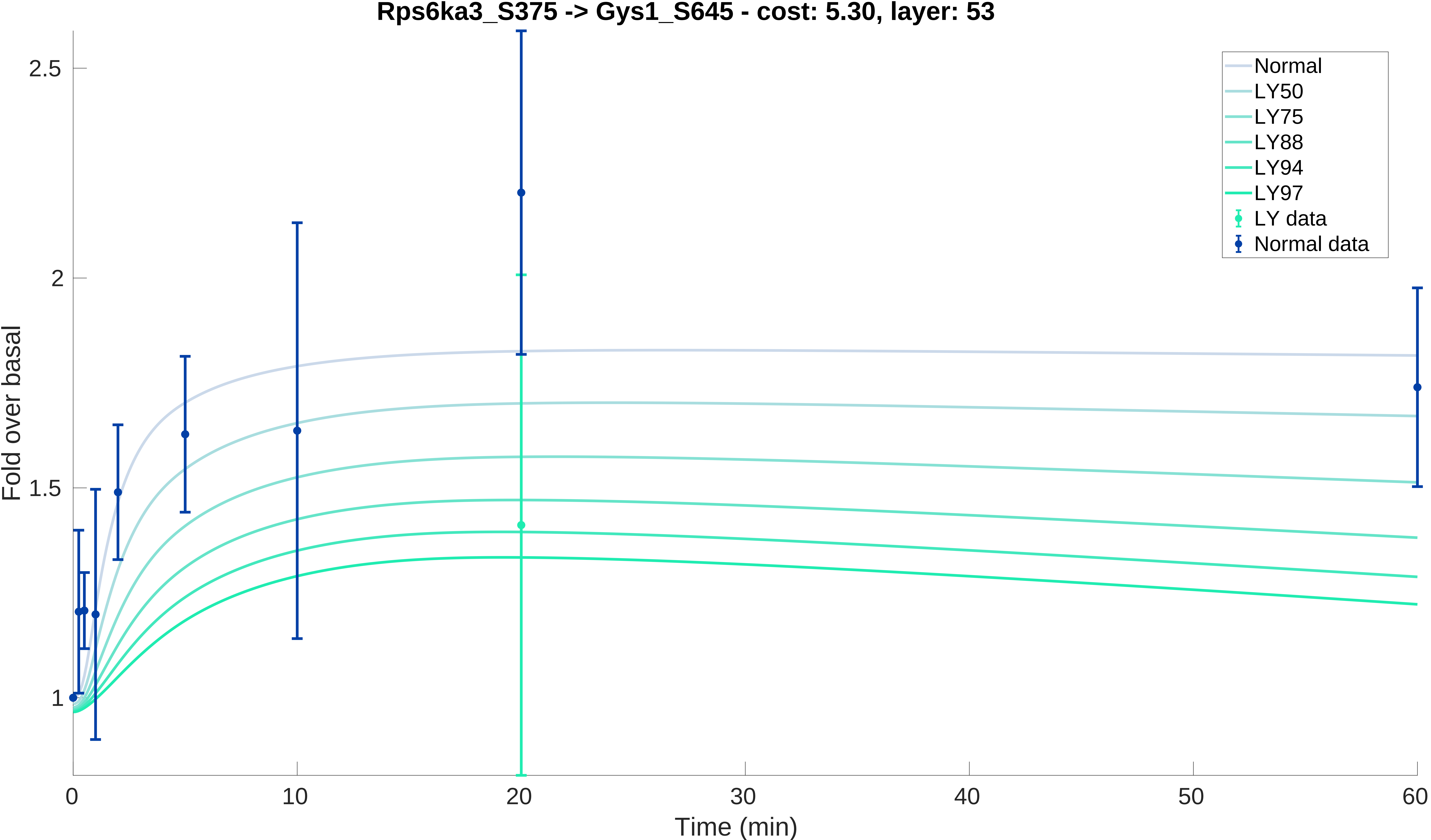

Rps6ka3_S375 -> Gys1_S645 - cost: 5.30, layer: 53
Fold over basal
Time (min)
2.5
2
1.5
1
0
10
20
30
40
50
60
Normal
LY50
LY75
LY88
LY94
LY97
LY data
Normal data

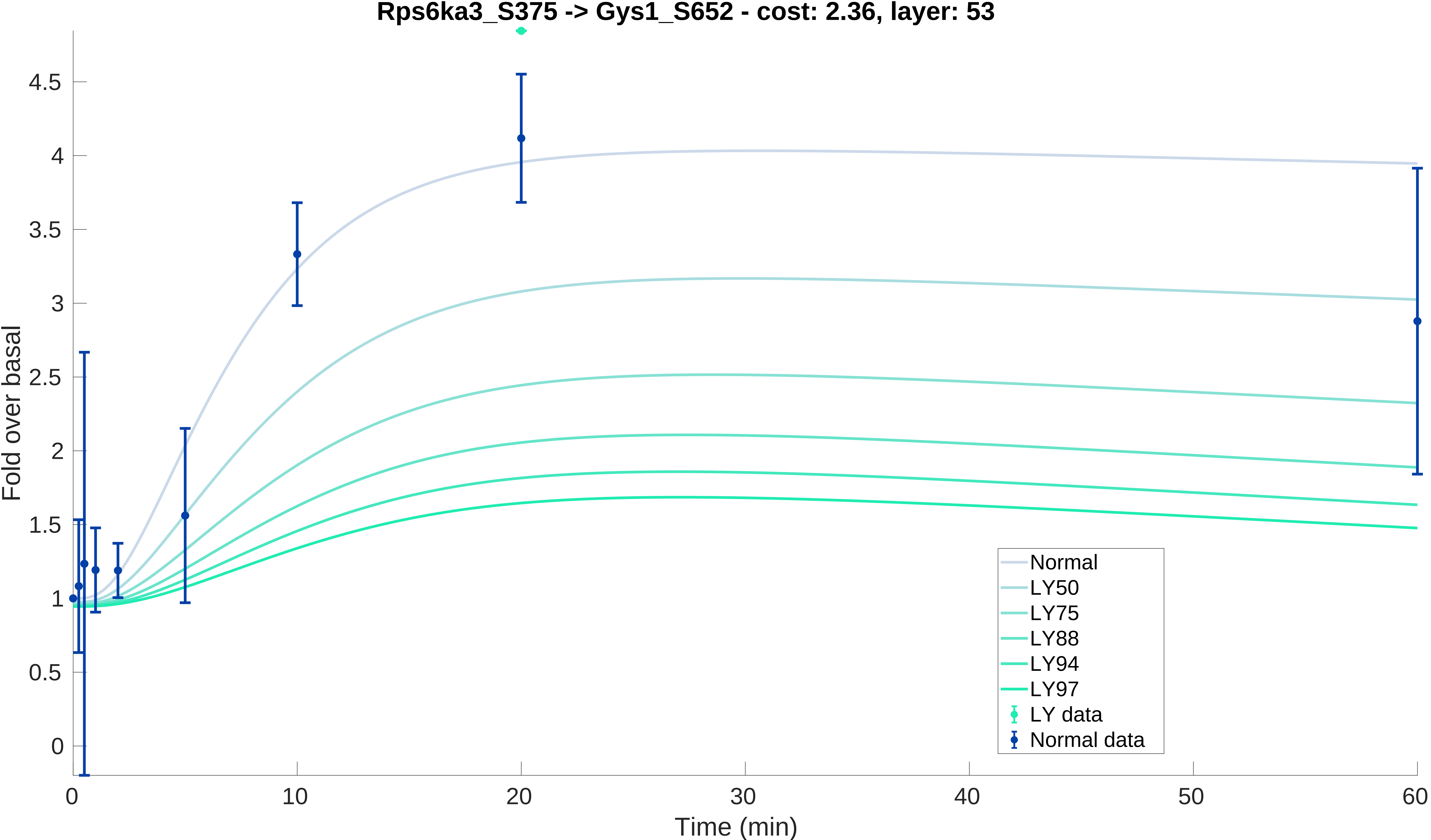

Rps6ka3_S375 -> Gys1_S652 - cost: 2.36, layer: 53
Fold over basal
Time (min)
0
0.5
1
1.5
2
2.5
3
3.5
4
4.5
0
10
20
30
40
50
60
Normal
LY50
LY75
LY88
LY94
LY97
LY data
Normal data

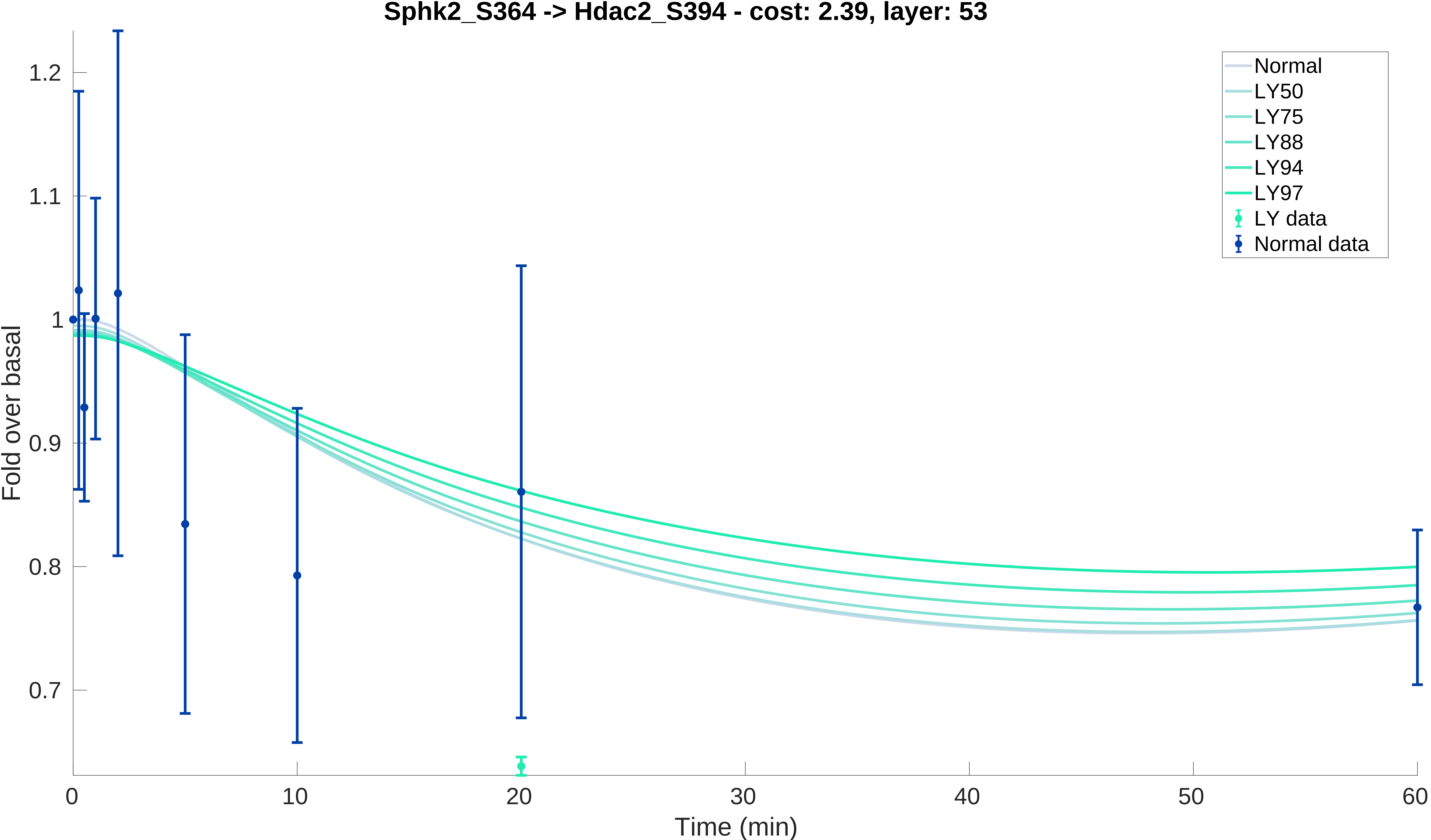
Sphk2_S364 -> Hdac2_S394 - cost: 2.39, layer: 53
Fold over basal
1.2
1.1
1
0.9
0.8
0.7
0
10
20
30
40
50
60
Time (min)
Normal
LY50
LY75
LY88
LY94
LY97
LY data
Normal data

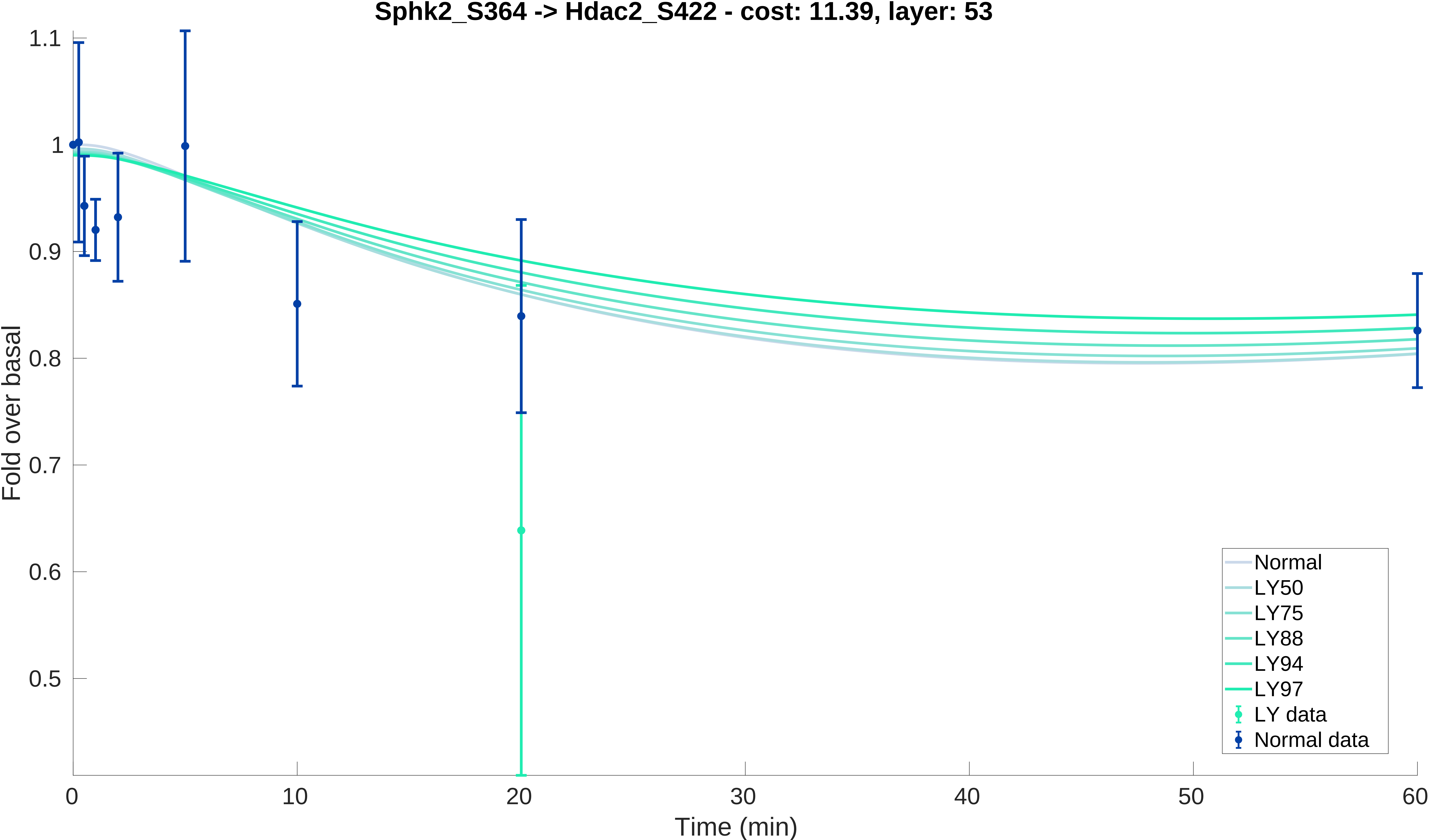

Sphk2_S364 -> Hdac2_S422 - cost: 11.39, layer: 53
Fold over basal
1.1
1
0.9
0.8
0.7
0.6
0.5
0
10
20
30
40
50
60
Time (min)
Normal
LY50
LY75
LY88
LY94
LY97
LY data
Normal data

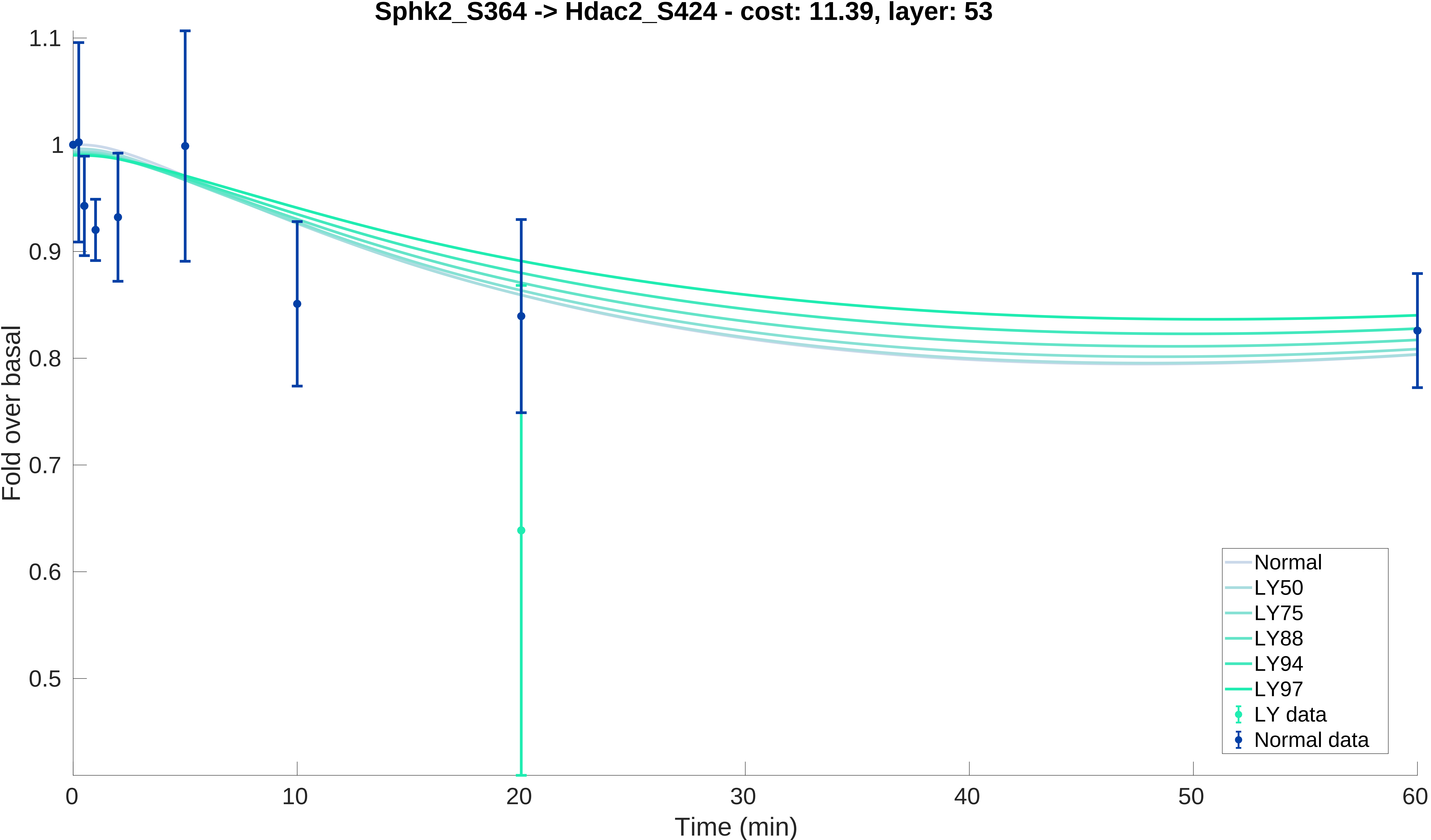
Sphk2_S364 -> Hdac2_S424 - cost: 11.39, layer: 53
Fold over basal
1.1
1
0.9
0.8
0.7
0.6
0.5
0
10
20
30
40
50
60
Time (min)
Normal
LY50
LY75
LY88
LY94
LY97
LY data
Normal data

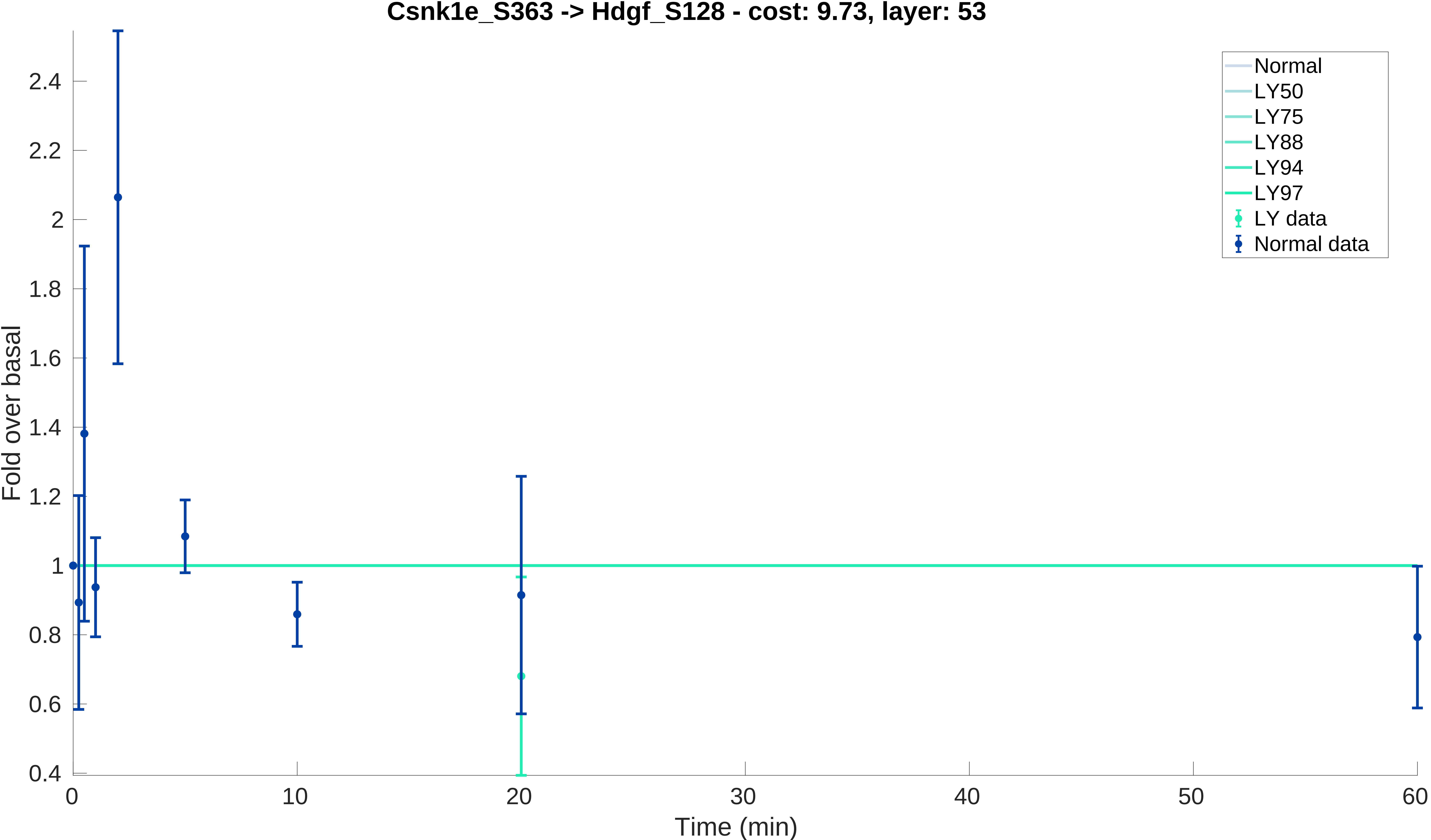
Csnk1e_S363 -> Hdgf_S128 - cost: 9.73, layer: 53
Fold over basal
Time (min)
0.4
0.6
0.8
1
1.2
1.4
1.6
1.8
2
2.2
2.4
0
10
20
30
40
50
60
Normal
LY50
LY75
LY88
LY94
LY97
LY data
Normal data

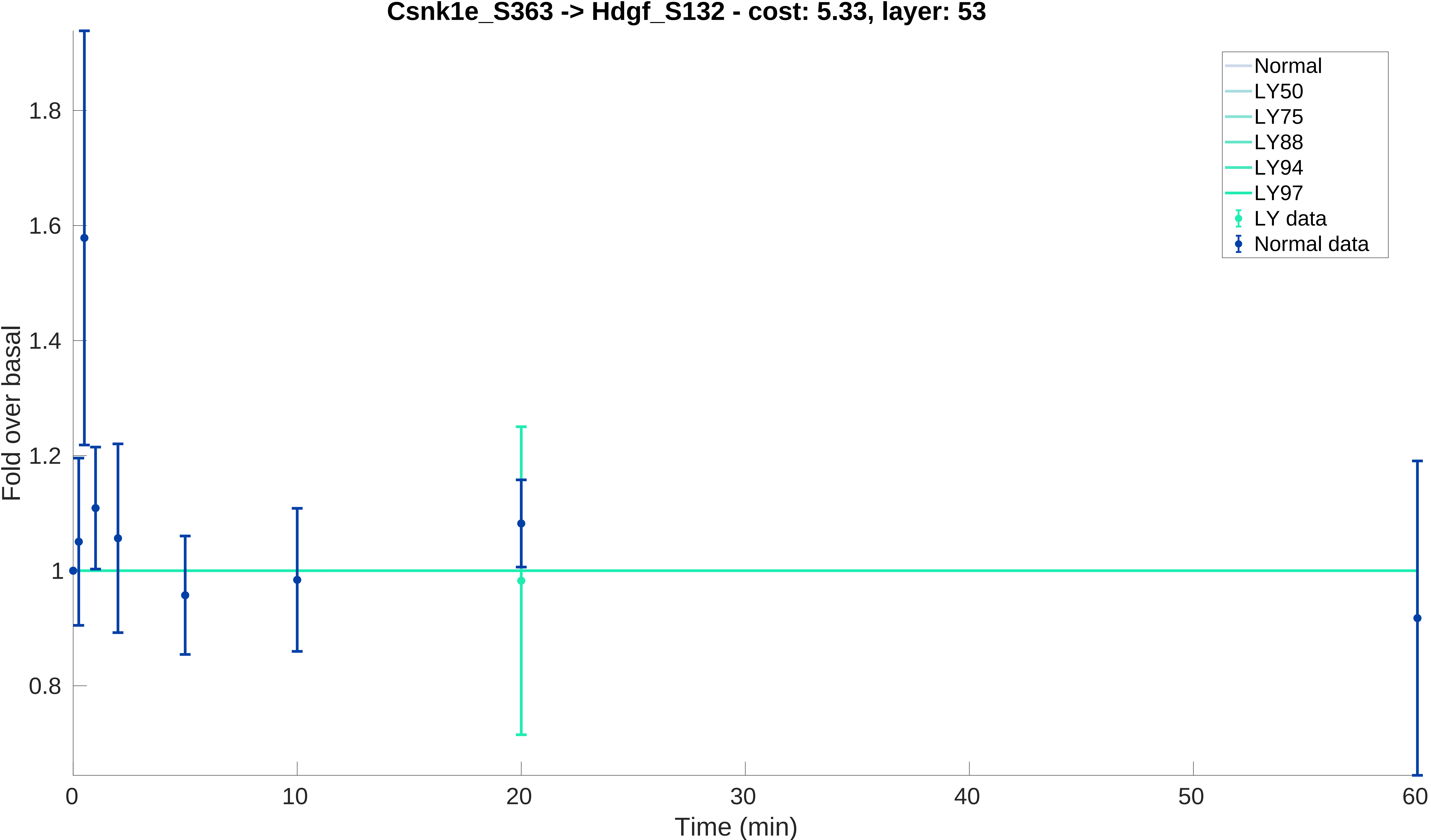

Csnk1e_S363 -> Hdgf_S132 - cost: 5.33, layer: 53
Fold over basal
1.8
1.6
1.4
1.2
1
0.8
0
10
20
30
40
50
60
Time (min)
Normal
LY50
LY75
LY88
LY94
LY97
LY data
Normal data

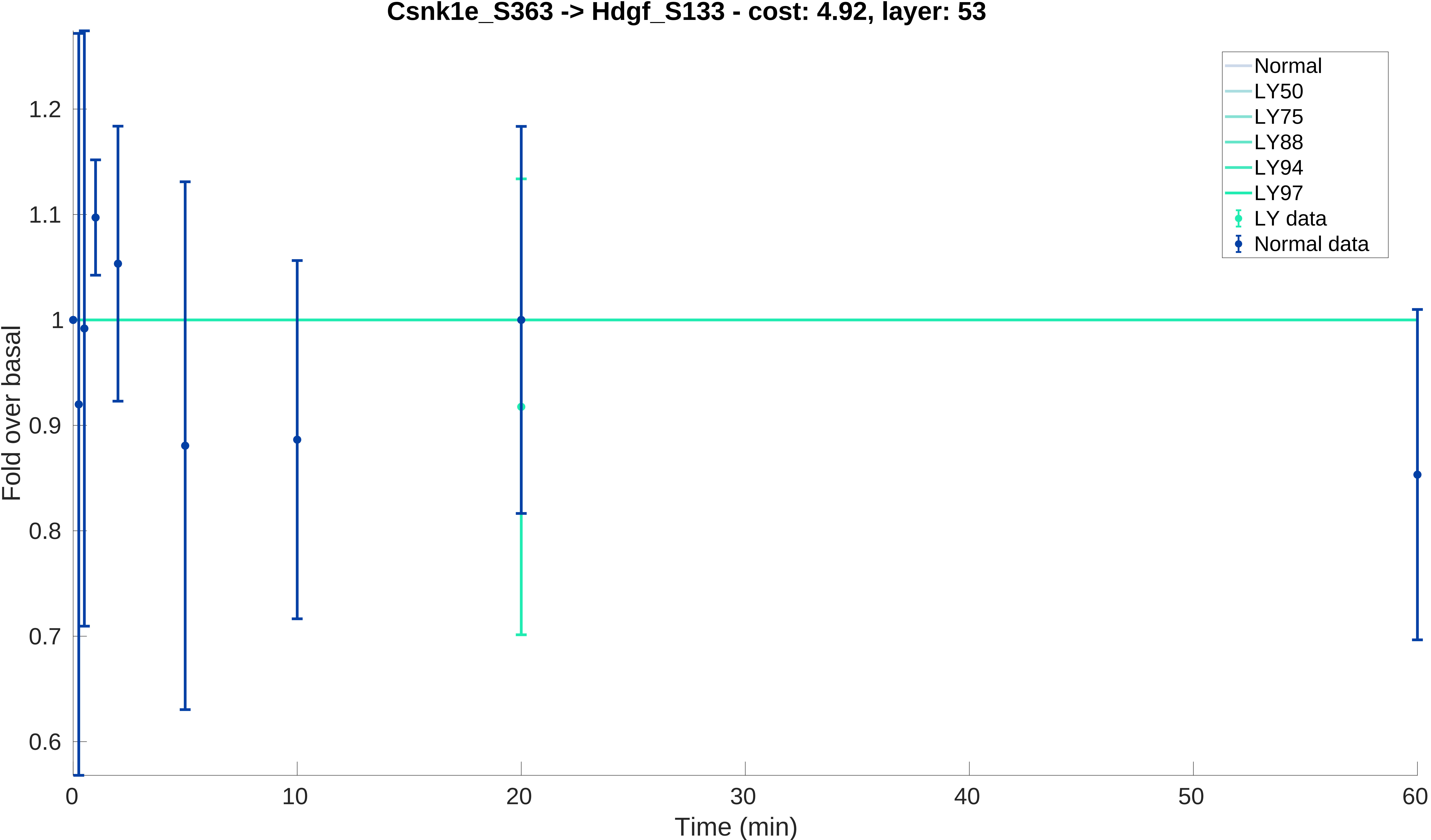

Csnk1e_S363 -> Hdgf_S133 - cost: 4.92, layer: 53
Fold over basal
Time (min)
0.6
0.7
0.8
0.9
1
1.1
1.2
0
10
20
30
40
50
60
Normal
LY50
LY75
LY88
LY94
LY97
LY data
Normal data

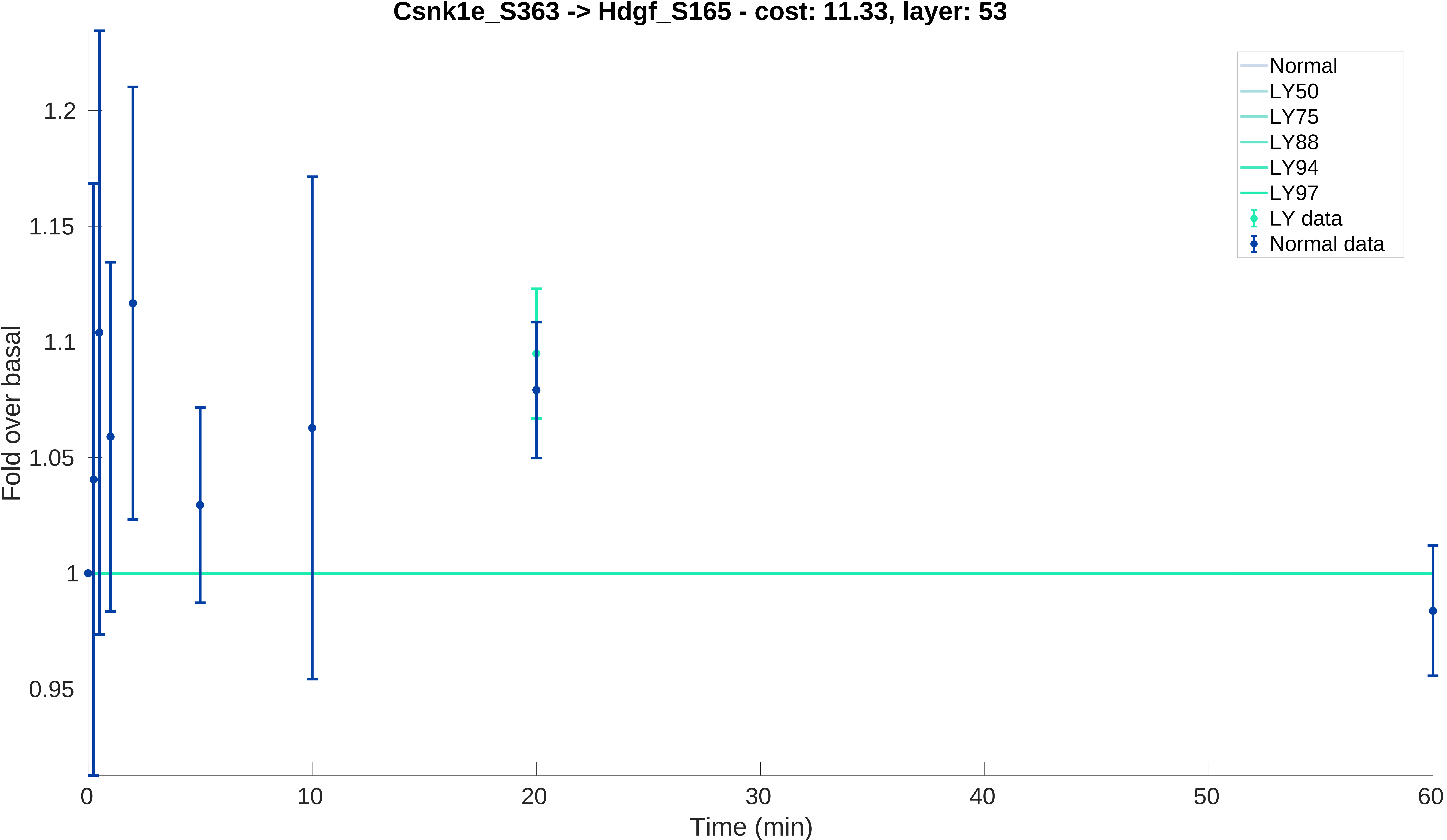

Csnk1e_S363 -> Hdgf_S165 - cost: 11.33, layer: 53
Normal
LY50
LY75
LY88
LY94
LY97
LY data
Normal data
Fold over basal
1.2
1.15
1.1
1.05
1
0.95
0
10
20
30
40
50
60
Time (min)

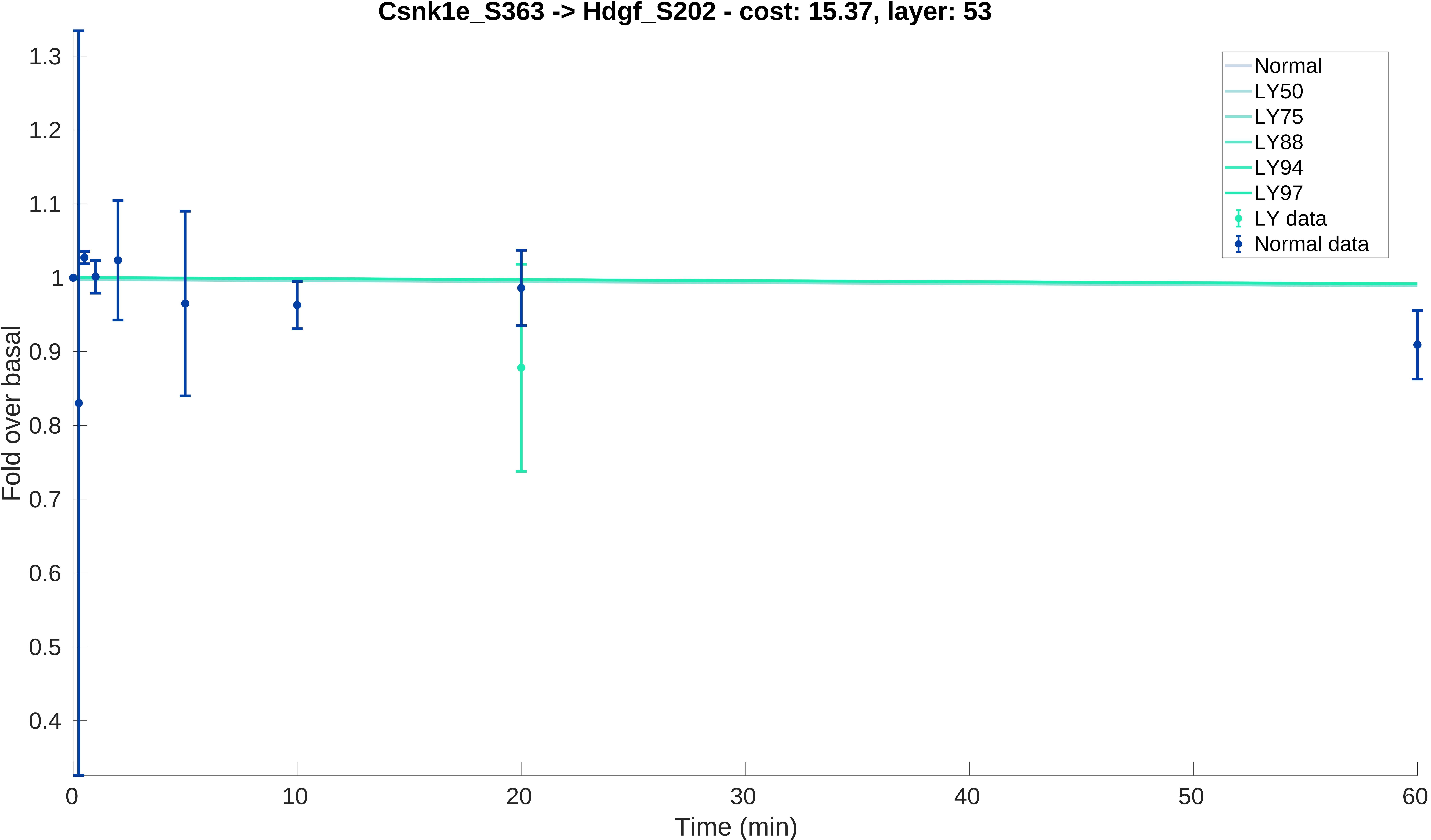
Csnk1e_S363 -> Hdgf_S202 - cost: 15.37, layer: 53
Fold over basal
1.3
1.2
1.1
1
0.9
0.8
0.7
0.6
0.5
0.4
0
10
20
30
40
50
60
Time (min)
Normal
LY50
LY75
LY88
LY94
LY97
LY data
Normal data

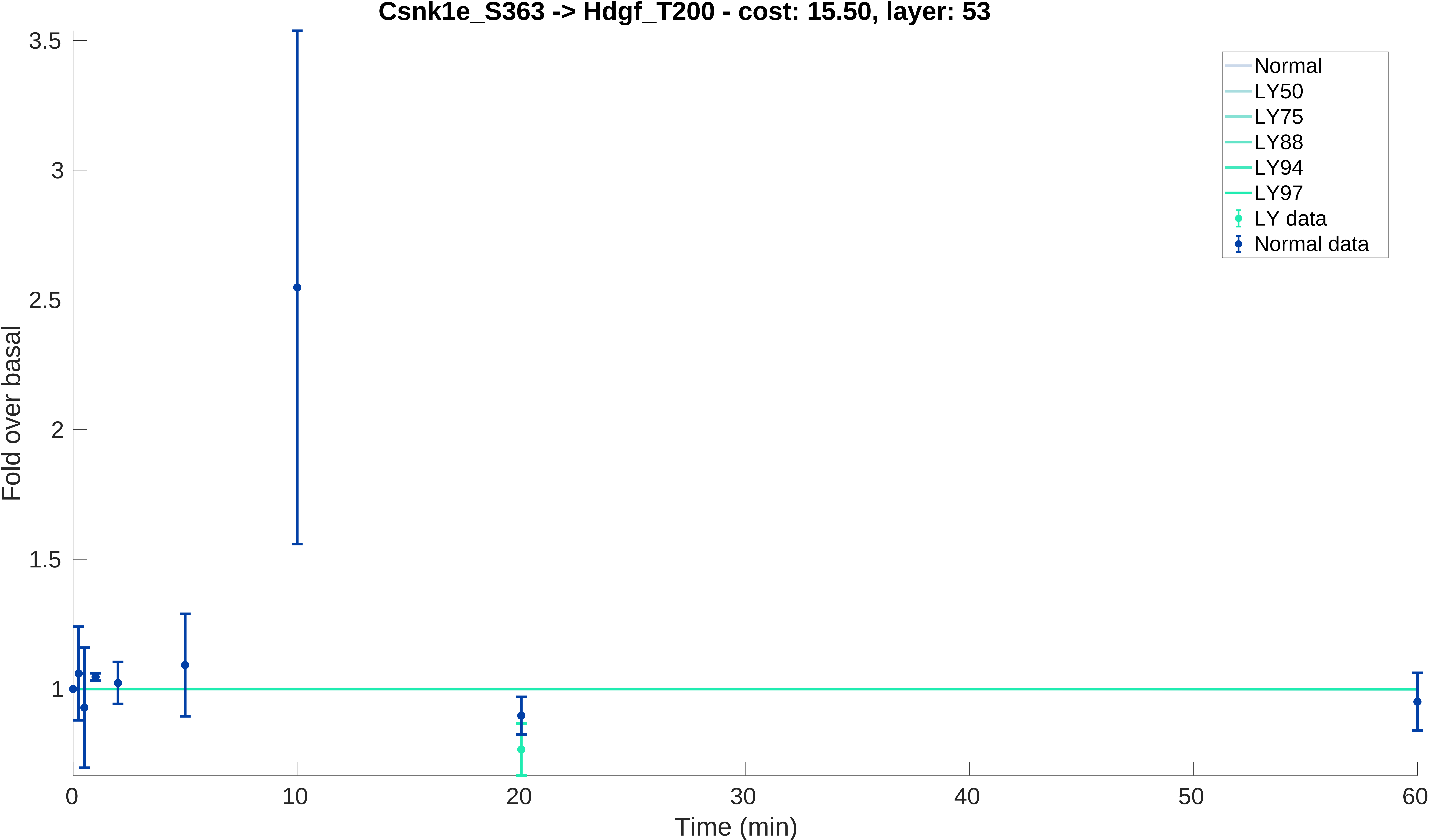

Csnk1e_S363 -> Hdgf_T200 - cost: 15.50, layer: 53
Fold over basal
Time (min)
3.5
3
2.5
2
1.5
1
0
10
20
30
40
50
60
Normal
LY50
LY75
LY88
LY94
LY97
LY data
Normal data

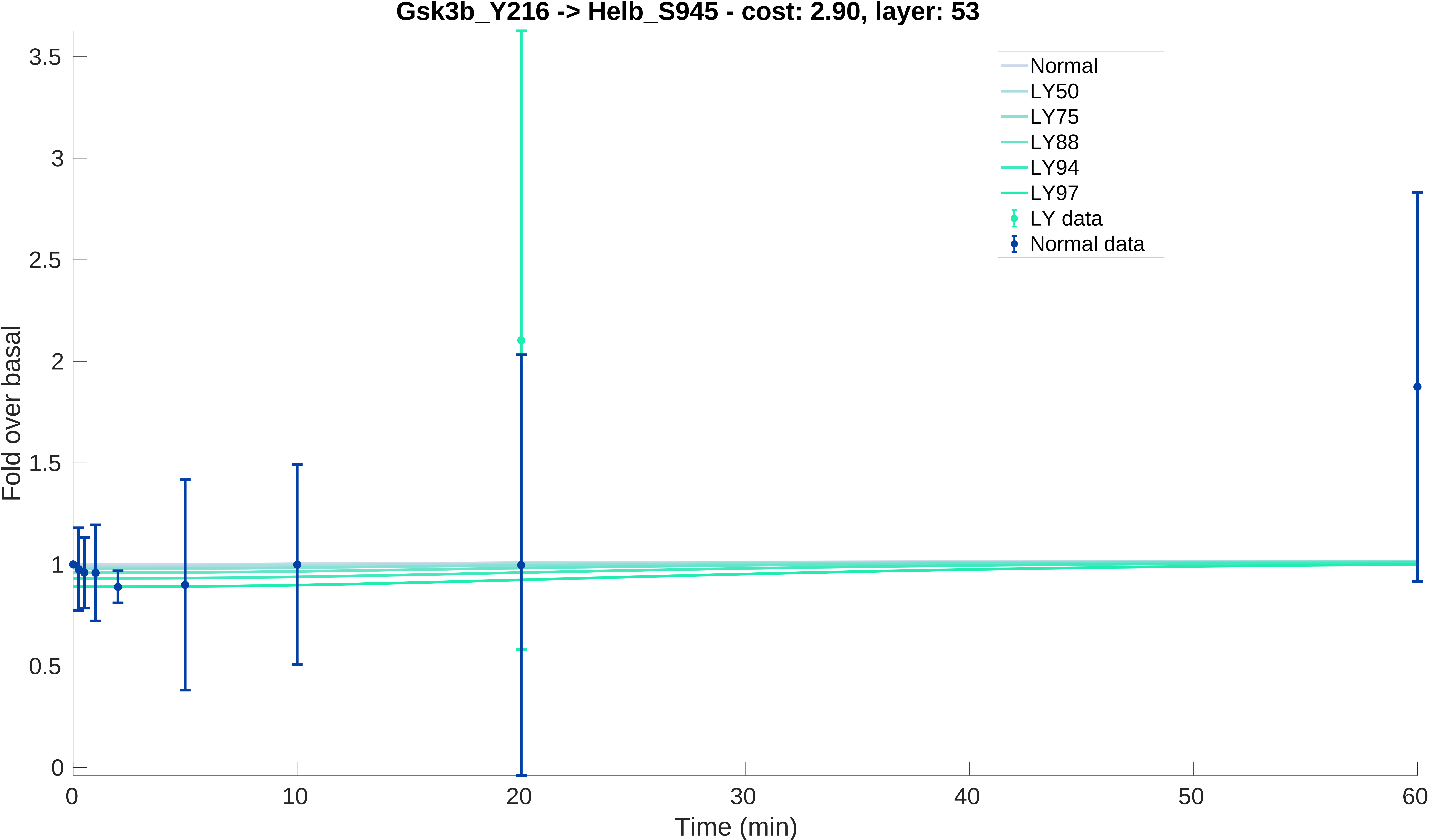

Gsk3b_Y216 -> Helb_S945 - cost: 2.90, layer: 53
Normal
LY50
LY75
LY88
LY94
LY97
LY data
Normal data
Fold over basal
0
0.5
1
1.5
2
2.5
3
3.5
0
10
20
30
40
50
60
Time (min)

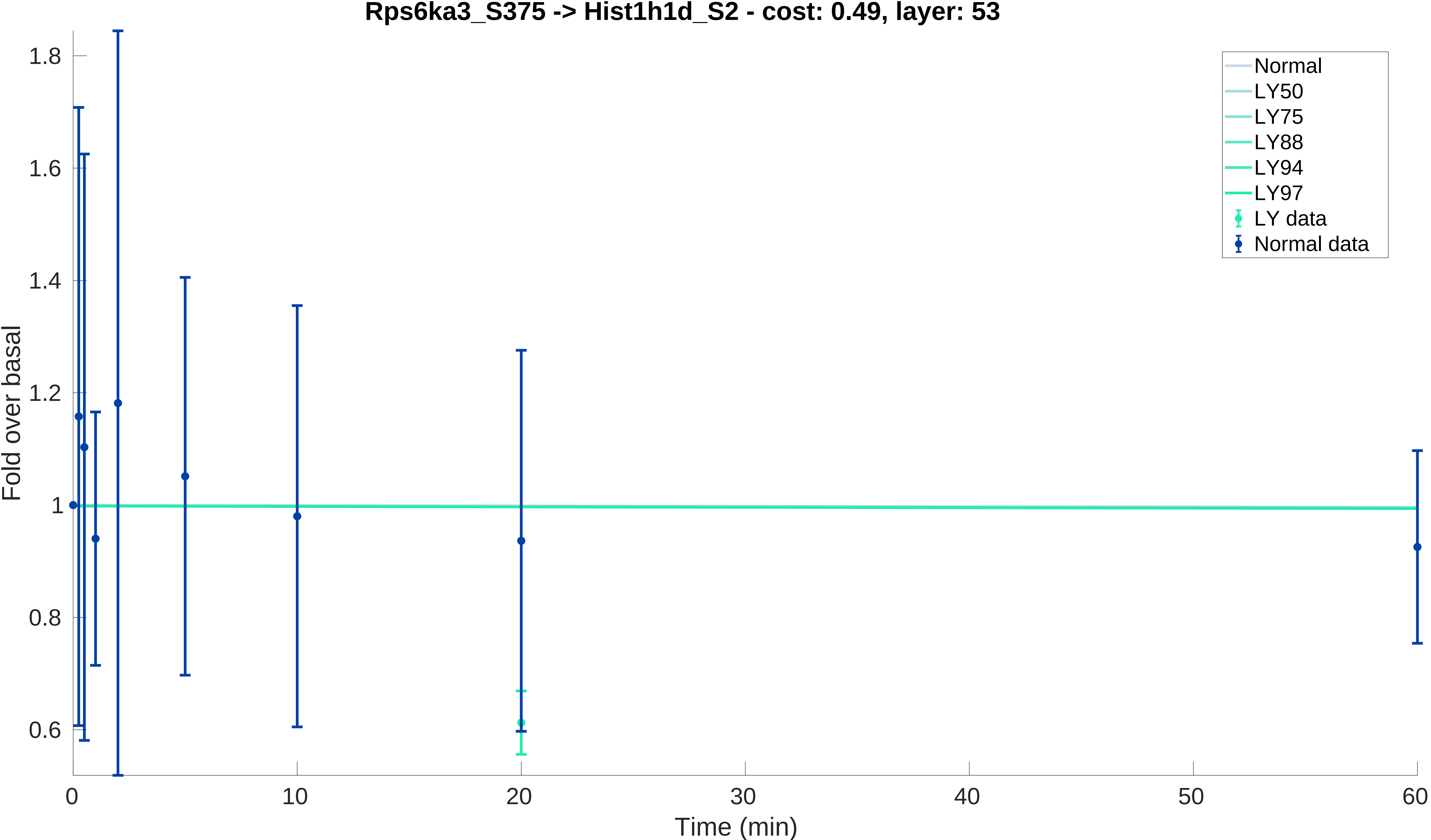
Rps6ka3_S375 -> Hist1h1d_S2 - cost: 0.49, layer: 53
Fold over basal
1.8
1.6
1.4
1.2
1
0.8
0.6
0
10
20
30
40
50
60
Time (min)
Normal
LY50
LY75
LY88
LY94
LY97
LY data
Normal data

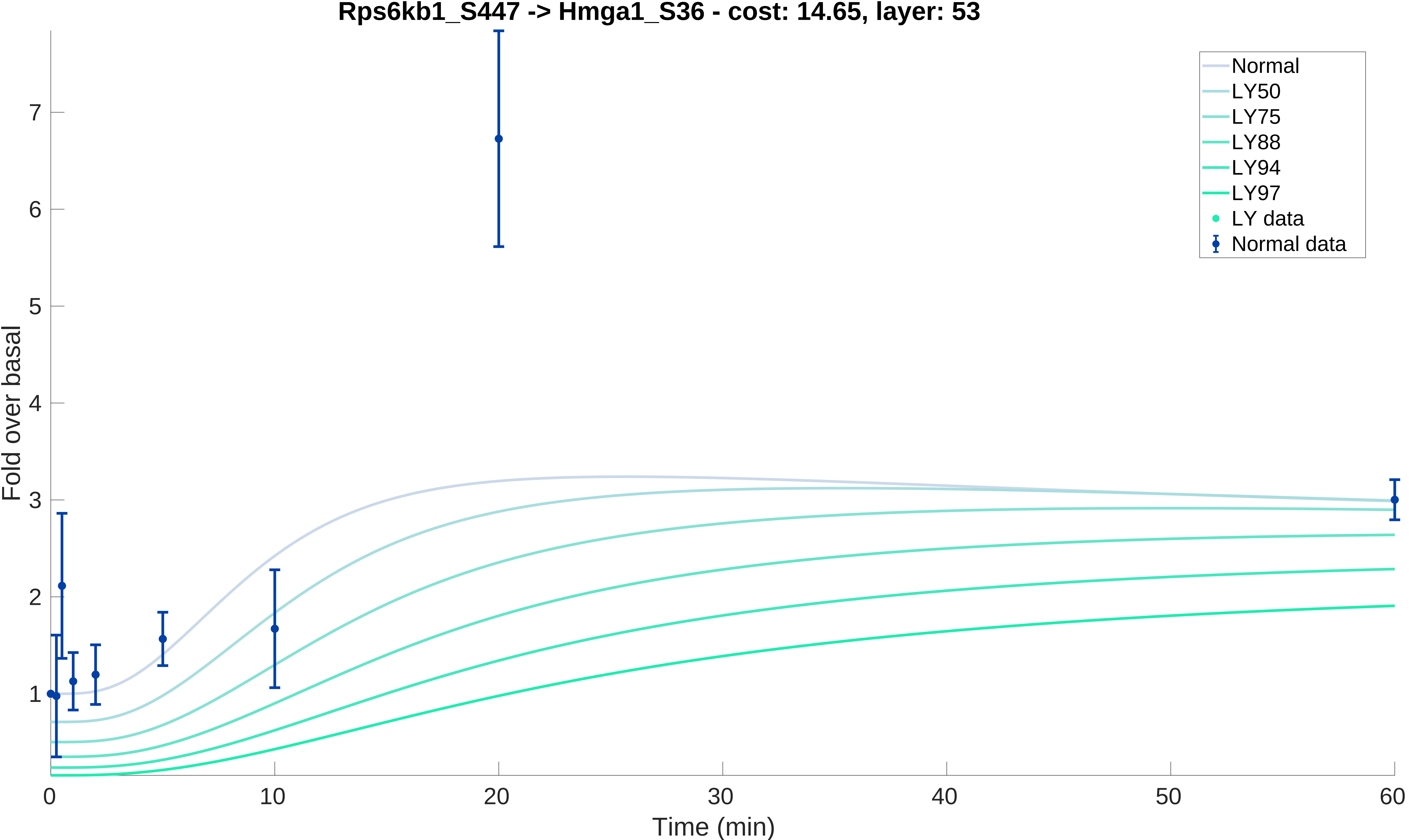
Rps6kb1_S447 -> Hmga1_S36 - cost: 14.65, layer: 53
Fold over basal
Time (min)
Normal
LY50
LY75
LY88
LY94
LY97
LY data
Normal data
0
10
20
30
40
50
60
1
2
3
4
5
6
7

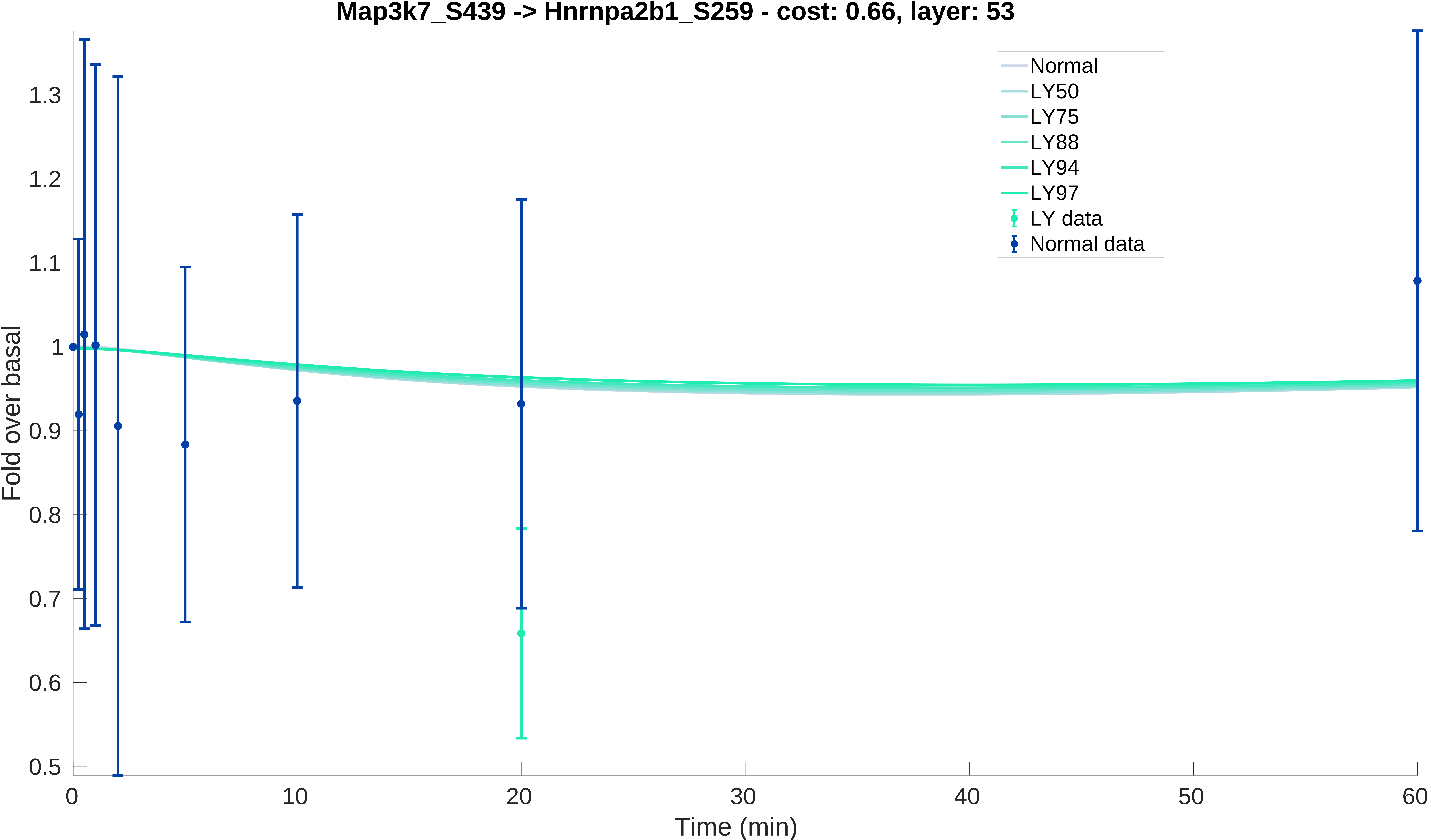

Map3k7_S439 -> Hnrnpa2b1_S259 - cost: 0.66, layer: 53
Normal
LY50
LY75
LY88
LY94
LY97
LY data
Normal data
Fold over basal
Time (min)
0.5
0.6
0.7
0.8
0.9
1
1.1
1.2
1.3
0
10
20
30
40
50
60

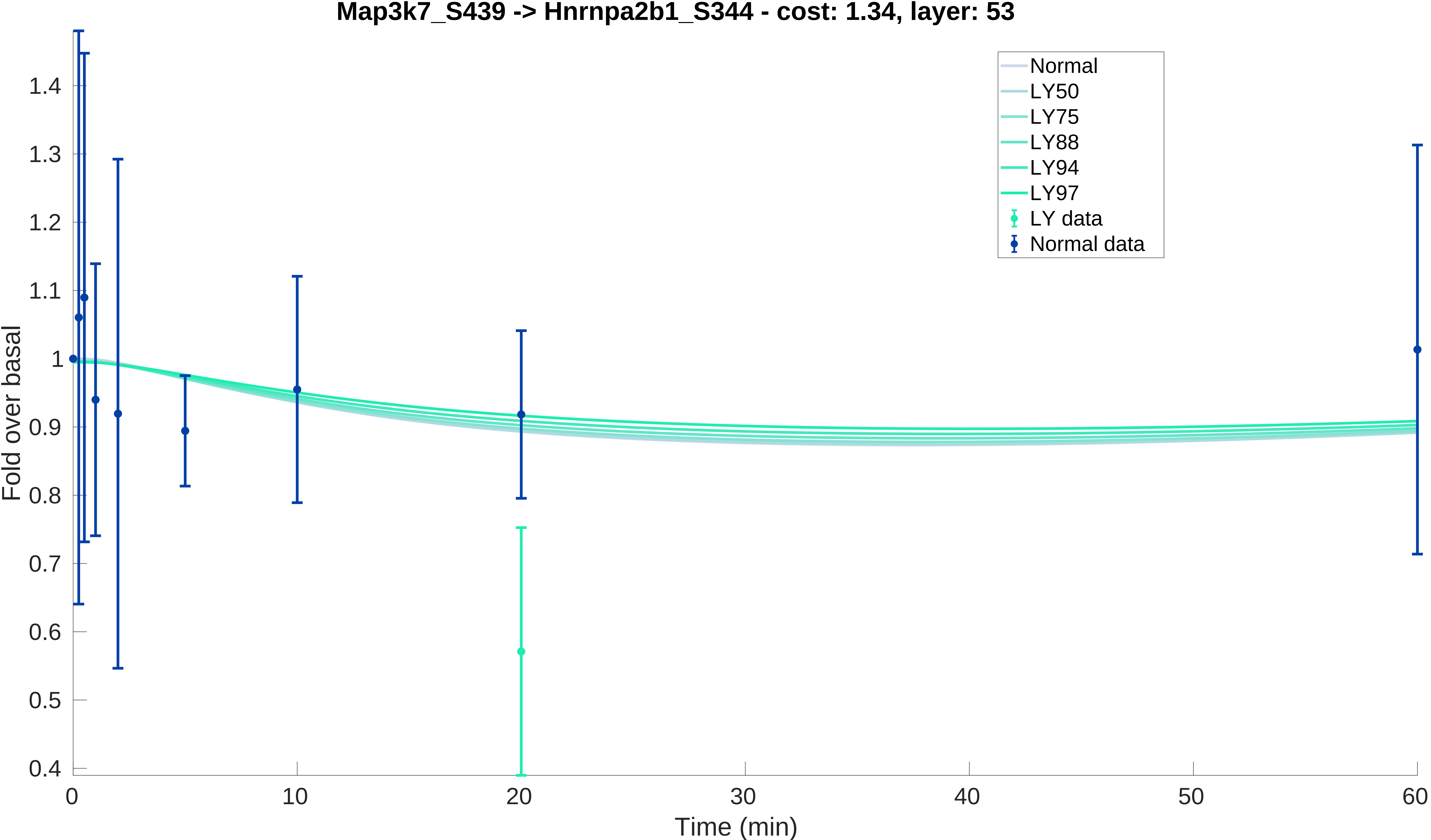

Map3k7_S439 -> Hnrnpa2b1_S344 - cost: 1.34, layer: 53
Normal
LY50
LY75
LY88
LY94
LY97
LY data
Normal data
Fold over basal
1.4
1.3
1.2
1.1
1
0.9
0.8
0.7
0.6
0.5
0.4
0
10
20
30
40
50
60
Time (min)

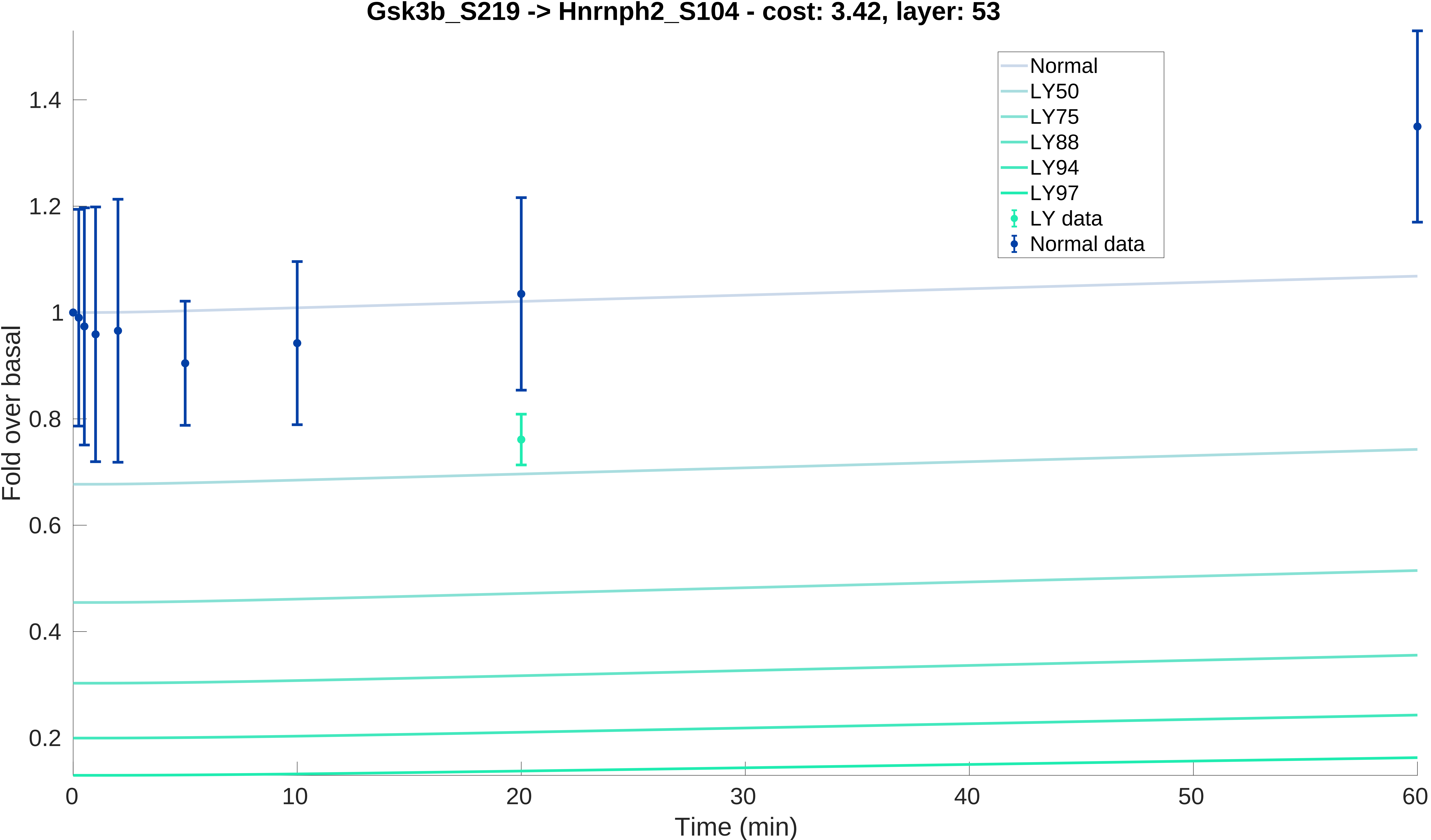
Gsk3b_S219 -> Hnrnph2_S104 - cost: 3.42, layer: 53
Normal
LY50
LY75
LY88
LY94
LY97
LY data
Normal data
Fold over basal
Time (min)
0.2
0.4
0.6
0.8
1
1.2
1.4
0
10
20
30
40
50
60

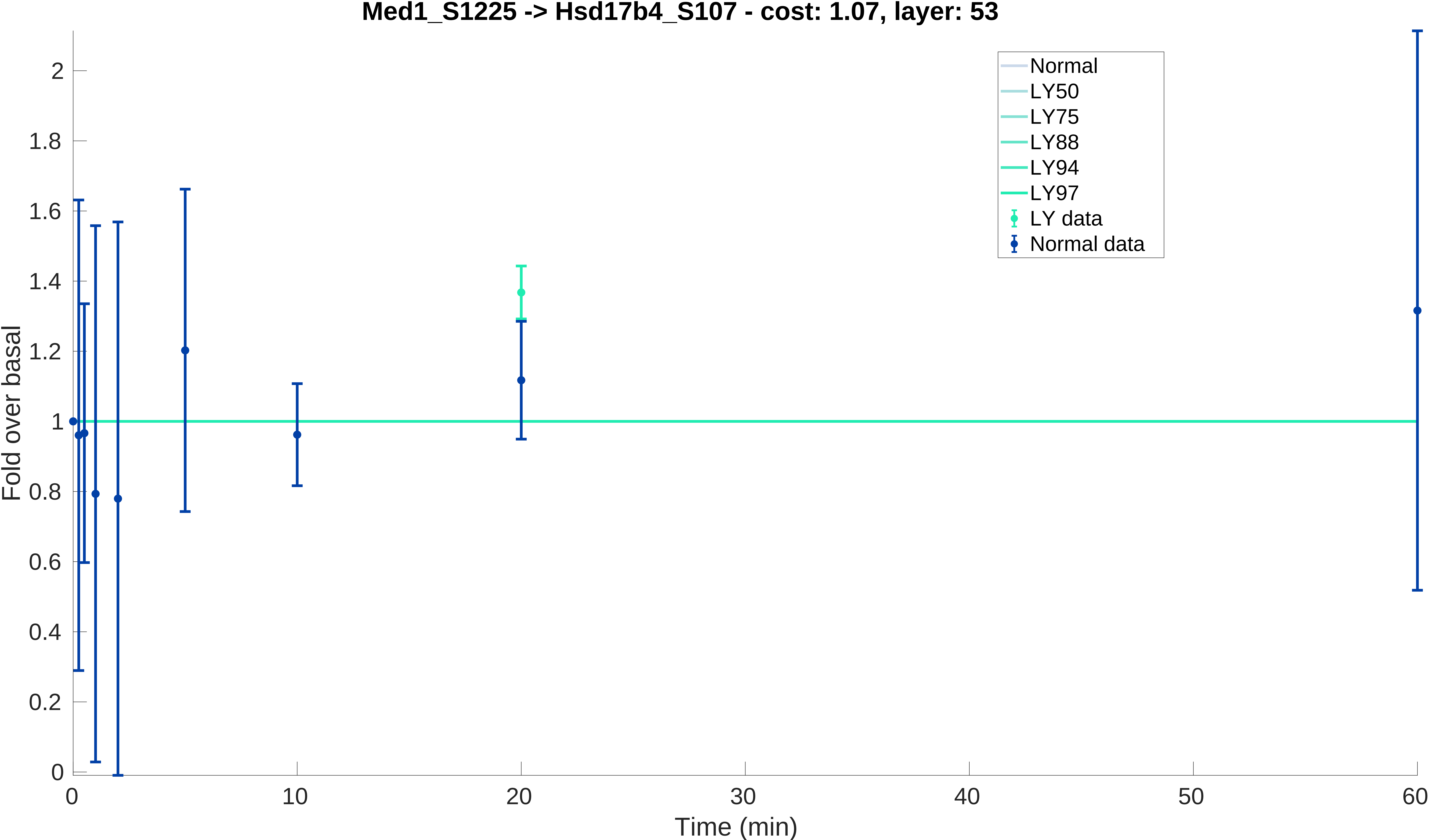

Med1_S1225 -> Hsd17b4_S107 - cost: 1.07, layer: 53
Normal
LY50
LY75
LY88
LY94
LY97
LY data
Normal data
Fold over basal
0
0.2
0.4
0.6
0.8
1
1.2
1.4
1.6
1.8
2
0
10
20
30
40
50
60
Time (min)

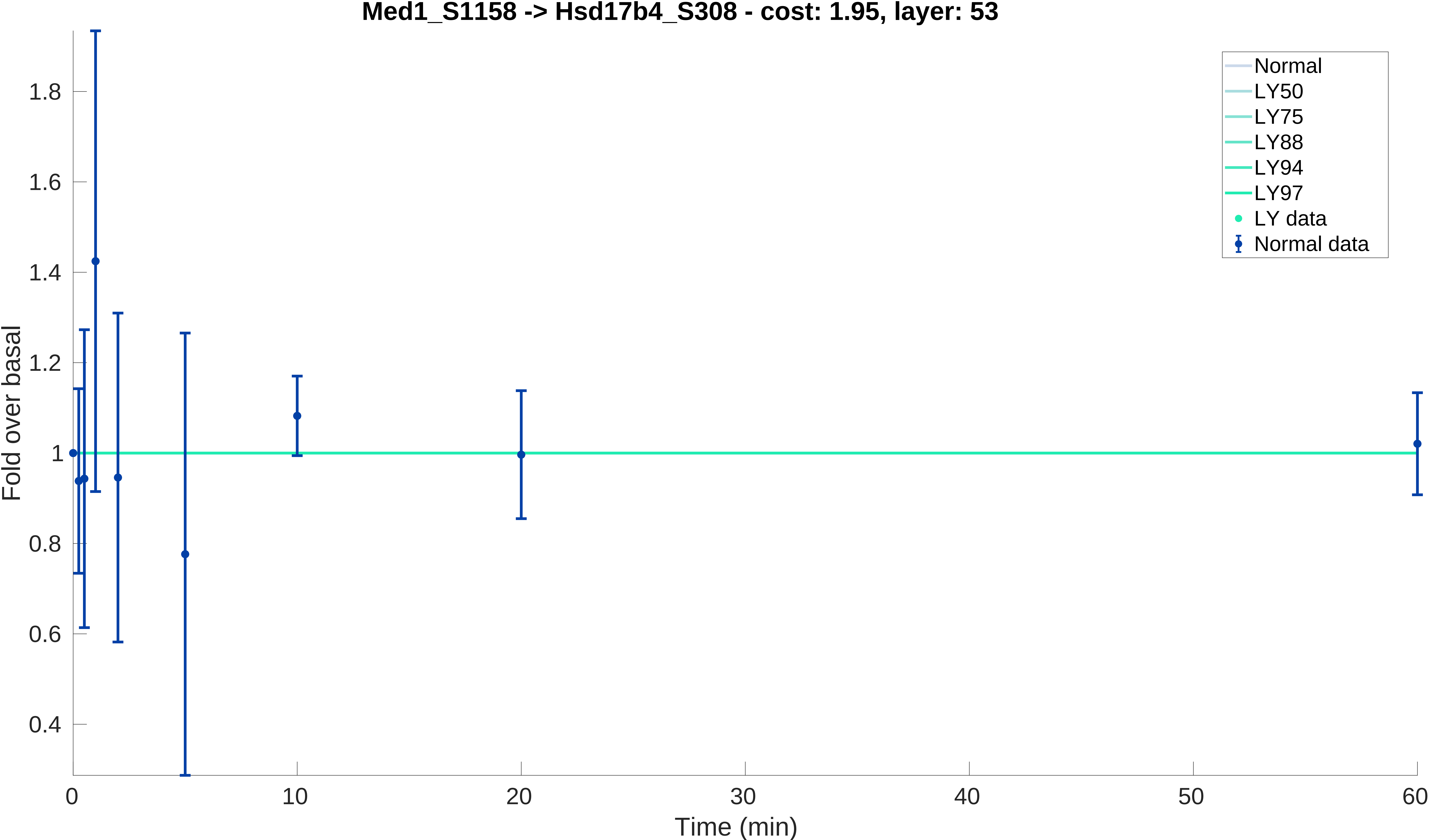
Med1_S1158 -> Hsd17b4_S308 - cost: 1.95, layer: 53
Fold over basal
Time (min)
0.4
0.6
0.8
1
1.2
1.4
1.6
1.8
0
10
20
30
40
50
60
Normal
LY50
LY75
LY88
LY94
LY97
LY data
Normal data

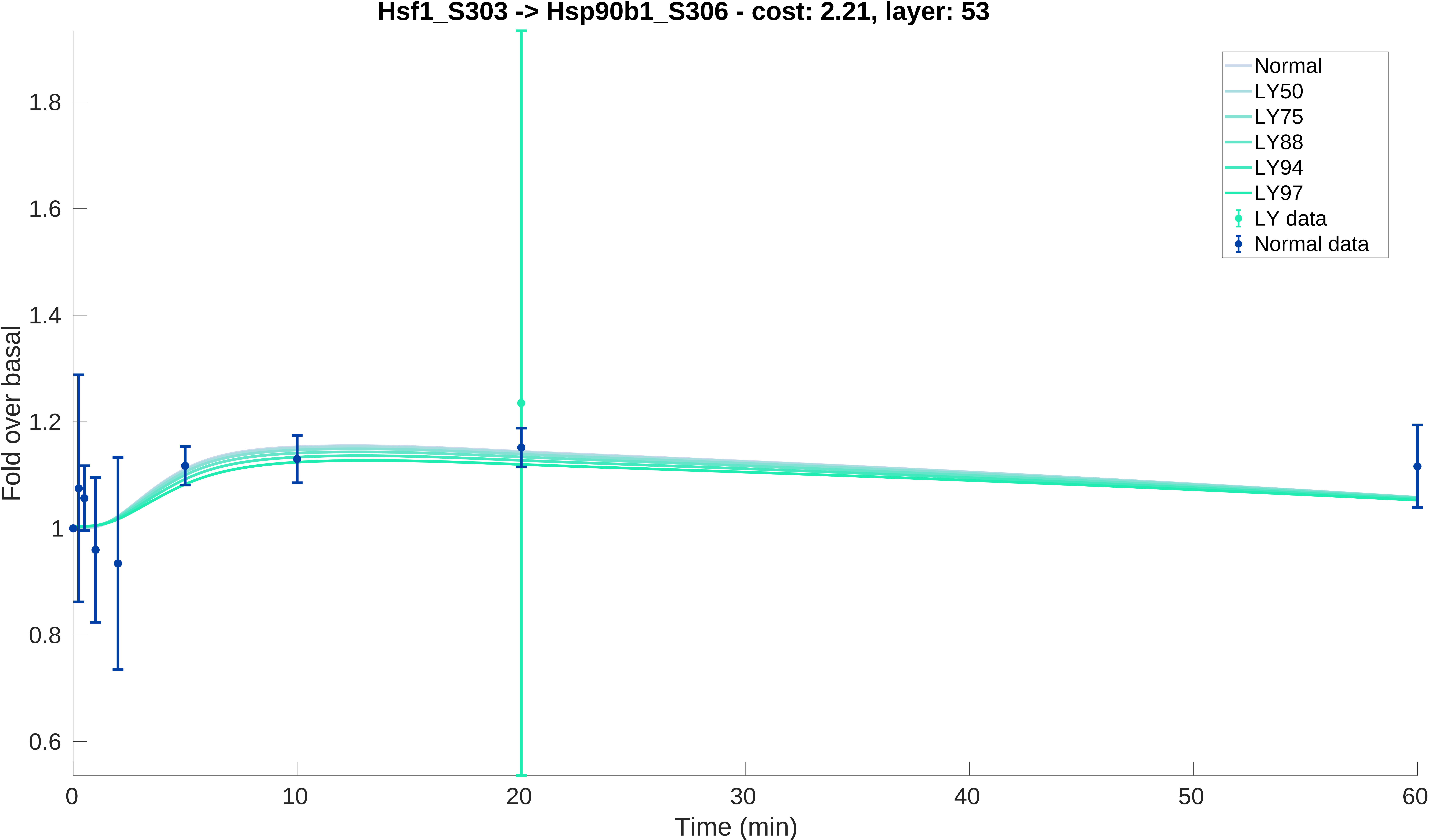
Hsf1_S303 -> Hsp90b1_S306 - cost: 2.21, layer: 53
Fold over basal
1.8
1.6
1.4
1.2
1
0.8
0.6
0
10
20
30
40
50
60
Time (min)
Normal
LY50
LY75
LY88
LY94
LY97
LY data
Normal data

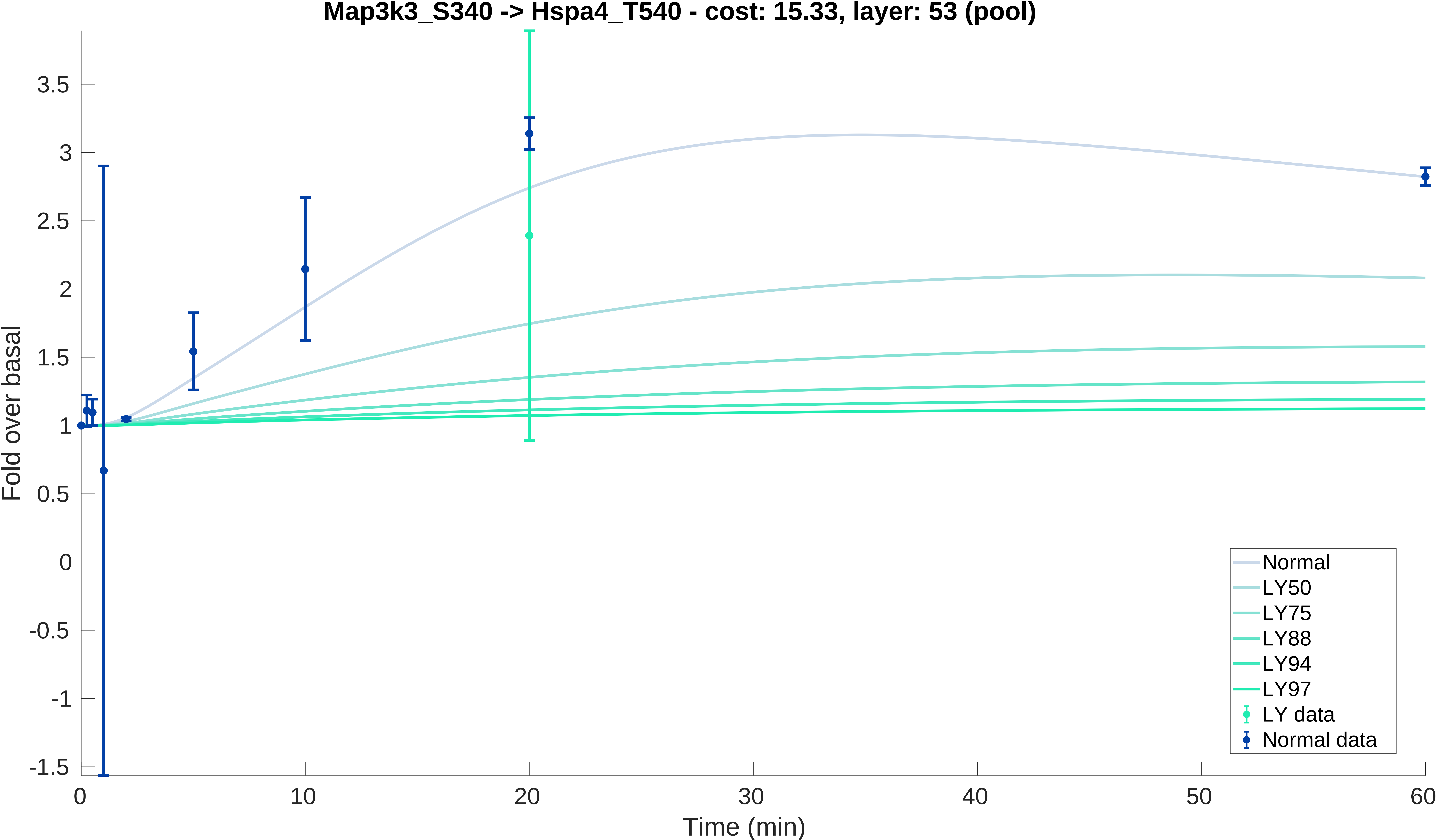
Map3k3_S340 -> Hspa4_T540 - cost: 15.33, layer: 53 (pool)
Fold over basal
Time (min)
3.5
3
2.5
2
1.5
1
0.5
0
-0.5
-1
-1.5
0
10
20
30
40
50
60
Normal
LY50
LY75
LY88
LY94
LY97
LY data
Normal data

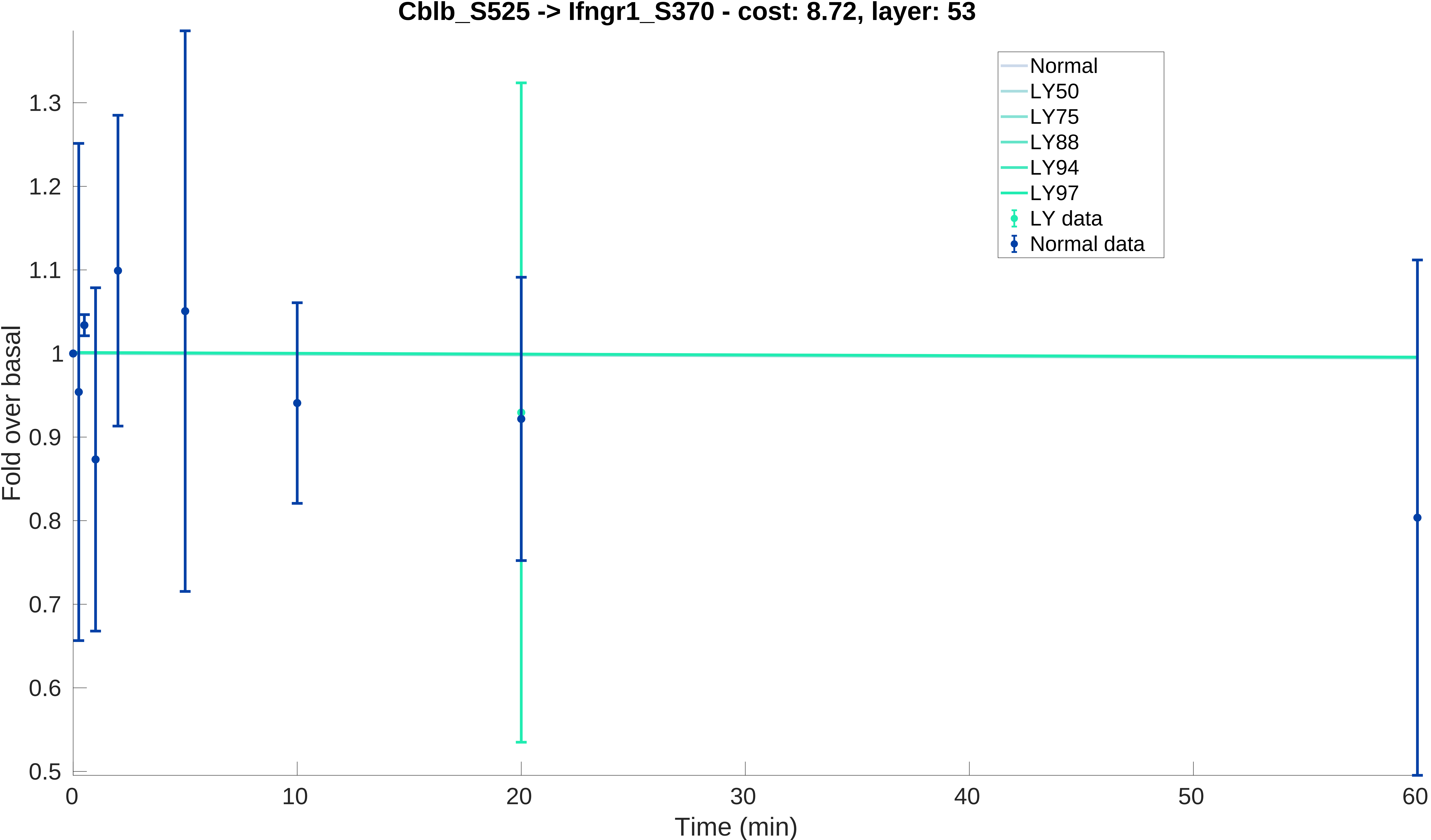

Cblb_S525 -> Ifngr1_S370 - cost: 8.72, layer: 53
Normal
LY50
LY75
LY88
LY94
LY97
LY data
Normal data
Fold over basal
Time (min)
0.5
0.6
0.7
0.8
0.9
1
1.1
1.2
1.3
0
10
20
30
40
50
60

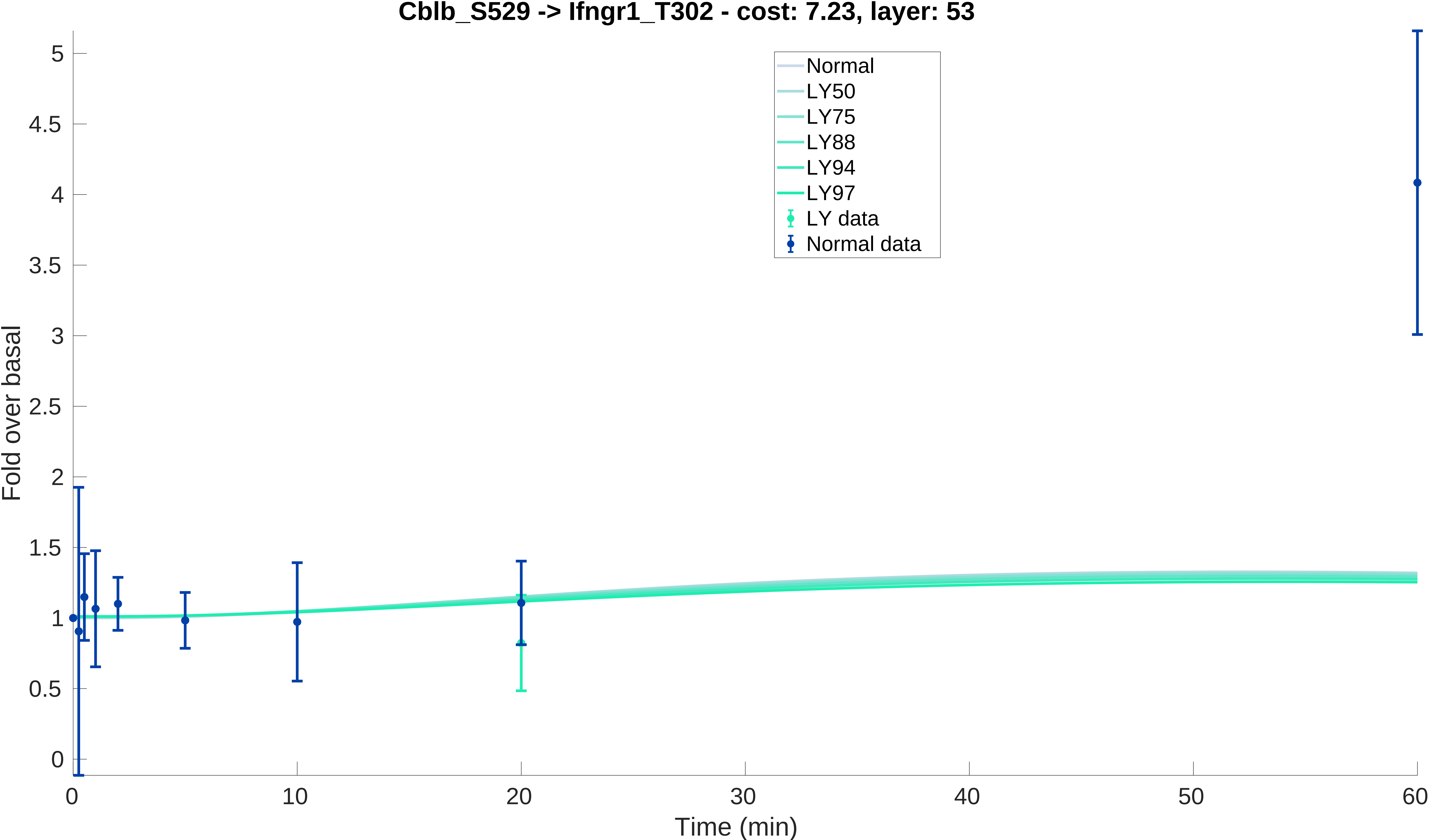
Cblb_S529 -> Ifngr1_T302 - cost: 7.23, layer: 53
Normal
LY50
LY75
LY88
LY94
LY97
LY data
Normal data
Fold over basal
Time (min)
0
0.5
1
1.5
2
2.5
3
3.5
4
4.5
5
0
10
20
30
40
50
60

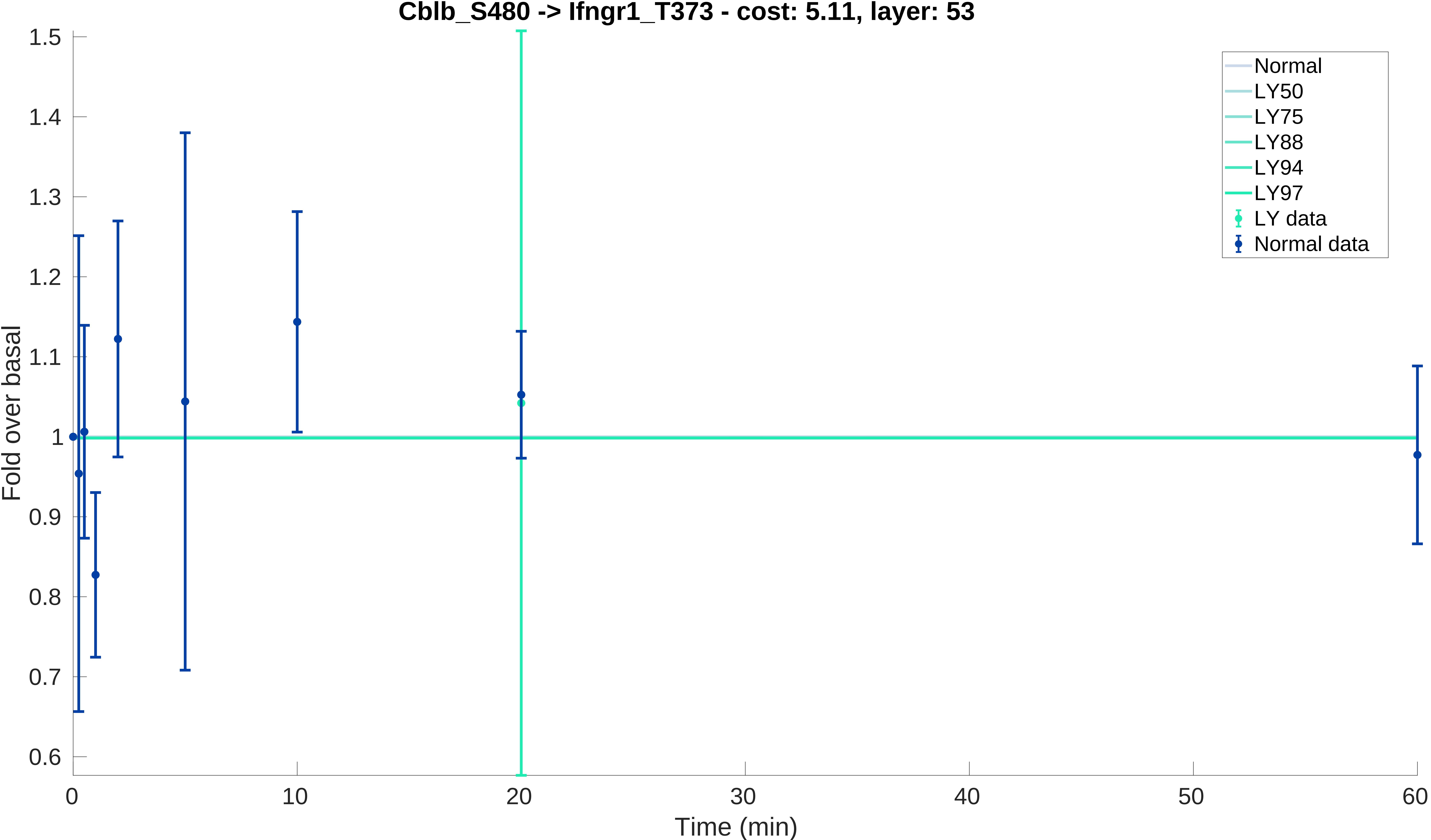

Cblb_S480 -> Ifngr1_T373 - cost: 5.11, layer: 53
Fold over basal
1.5
1.4
1.3
1.2
1.1
1
0.9
0.8
0.7
0.6
0
10
20
30
40
50
60
Time (min)
Normal
LY50
LY75
LY88
LY94
LY97
LY data
Normal data

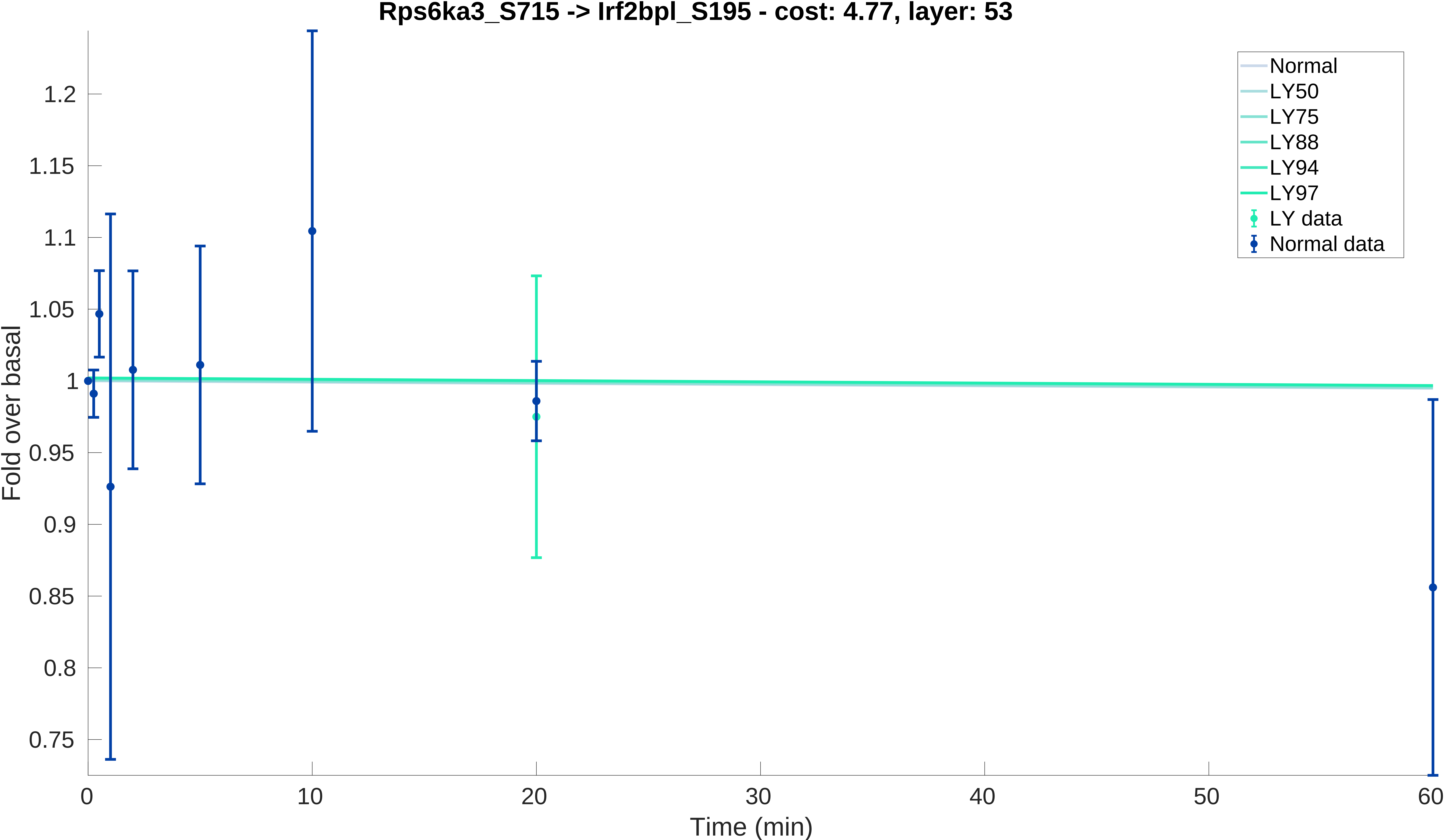
Rps6ka3_S715 -> Irf2bpl_S195 - cost: 4.77, layer: 53
Fold over basal
Time (min)
1.2
1.15
1.1
1.05
1
0.95
0.9
0.85
0.8
0.75
0
10
20
30
40
50
60
Normal
LY50
LY75
LY88
LY94
LY97
LY data
Normal data

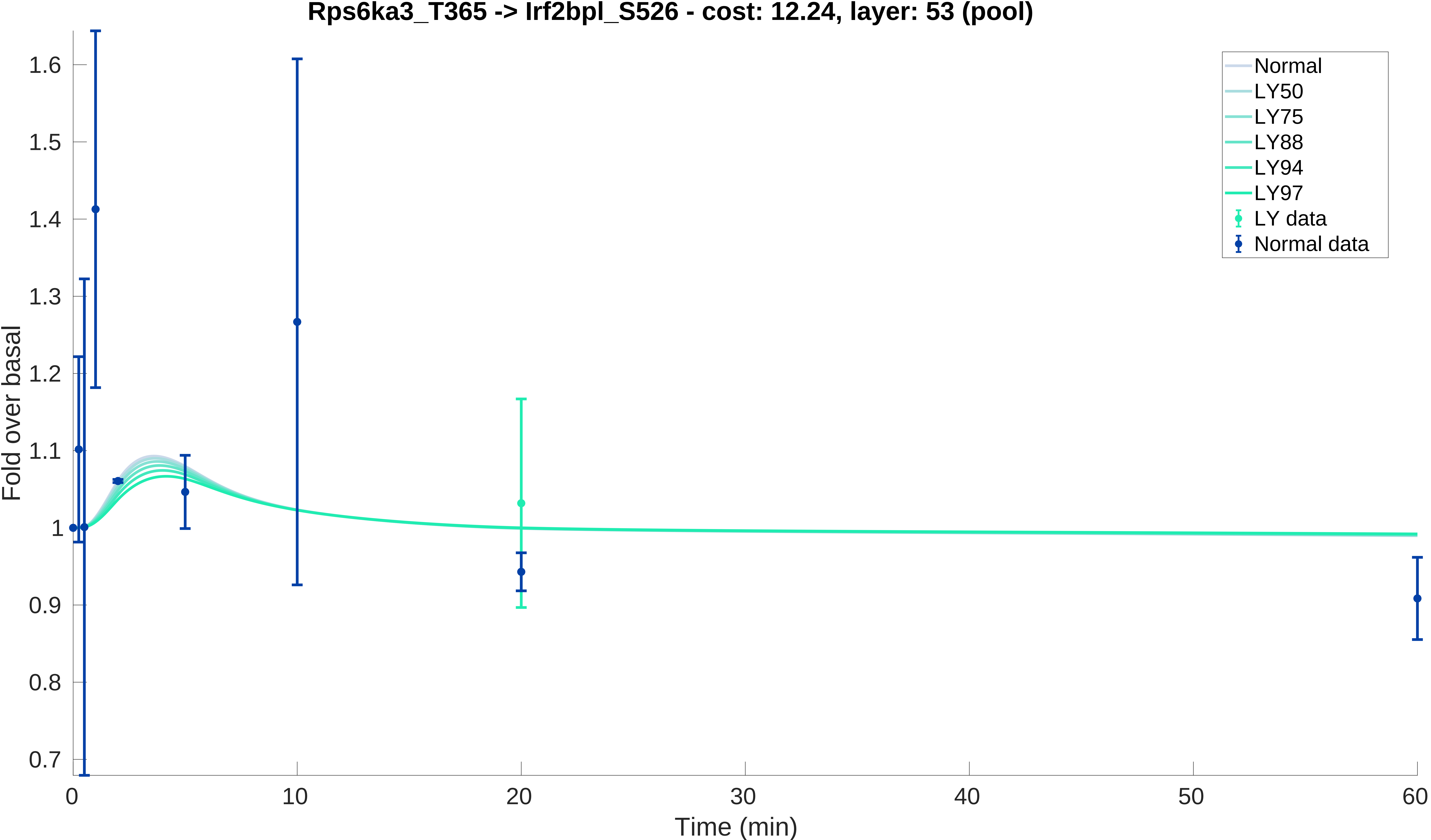

Rps6ka3_T365 -> Irf2bpl_S526 - cost: 12.24, layer: 53 (pool)
Fold over basal
Time (min)
1.6
1.5
1.4
1.3
1.2
1.1
1
0.9
0.8
0.7
0
10
20
30
40
50
60
Normal
LY50
LY75
LY88
LY94
LY97
LY data
Normal data

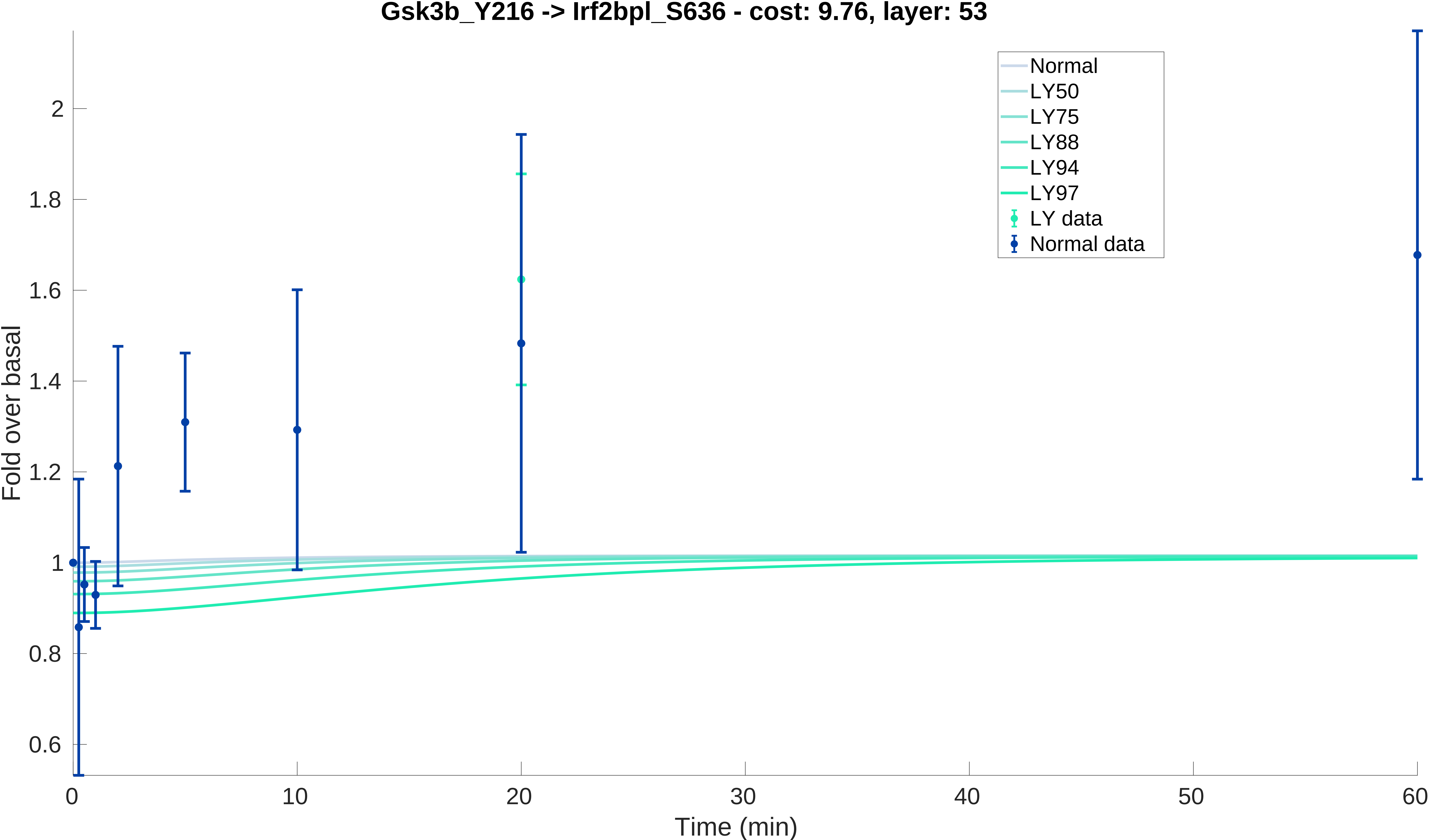

Gsk3b_Y216 -> Irf2bpl_S636 - cost: 9.76, layer: 53
Normal
LY50
LY75
LY88
LY94
LY97
LY data
Normal data
Fold over basal
Time (min)
0.6
0.8
1
1.2
1.4
1.6
1.8
2
0
10
20
30
40
50
60

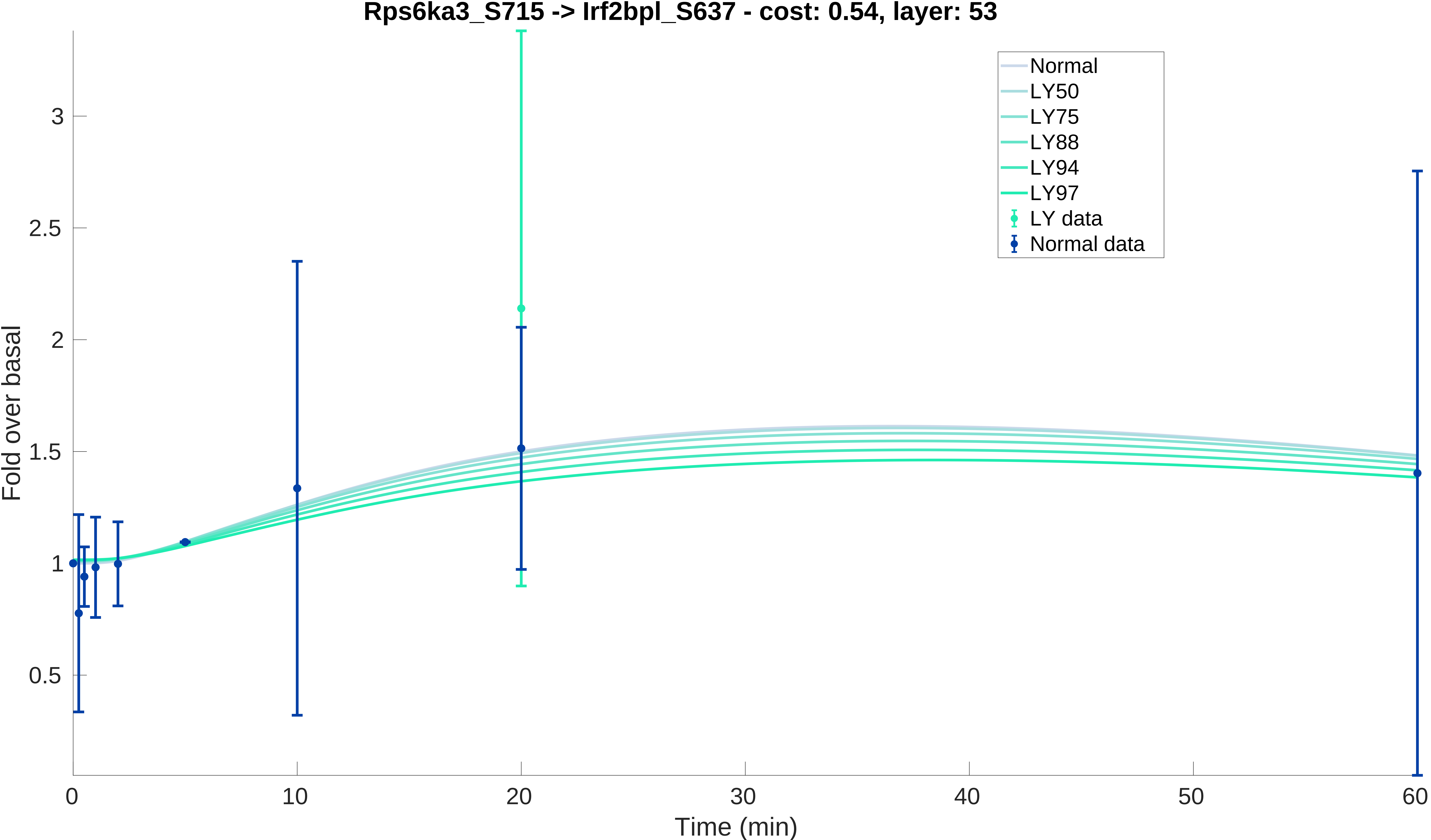
Rps6ka3_S715 -> Irf2bpl_S637 - cost: 0.54, layer: 53
Normal
LY50
LY75
LY88
LY94
LY97
LY data
Normal data
Fold over basal
Time (min)
0.5
1
1.5
2
2.5
3
0
10
20
30
40
50
60

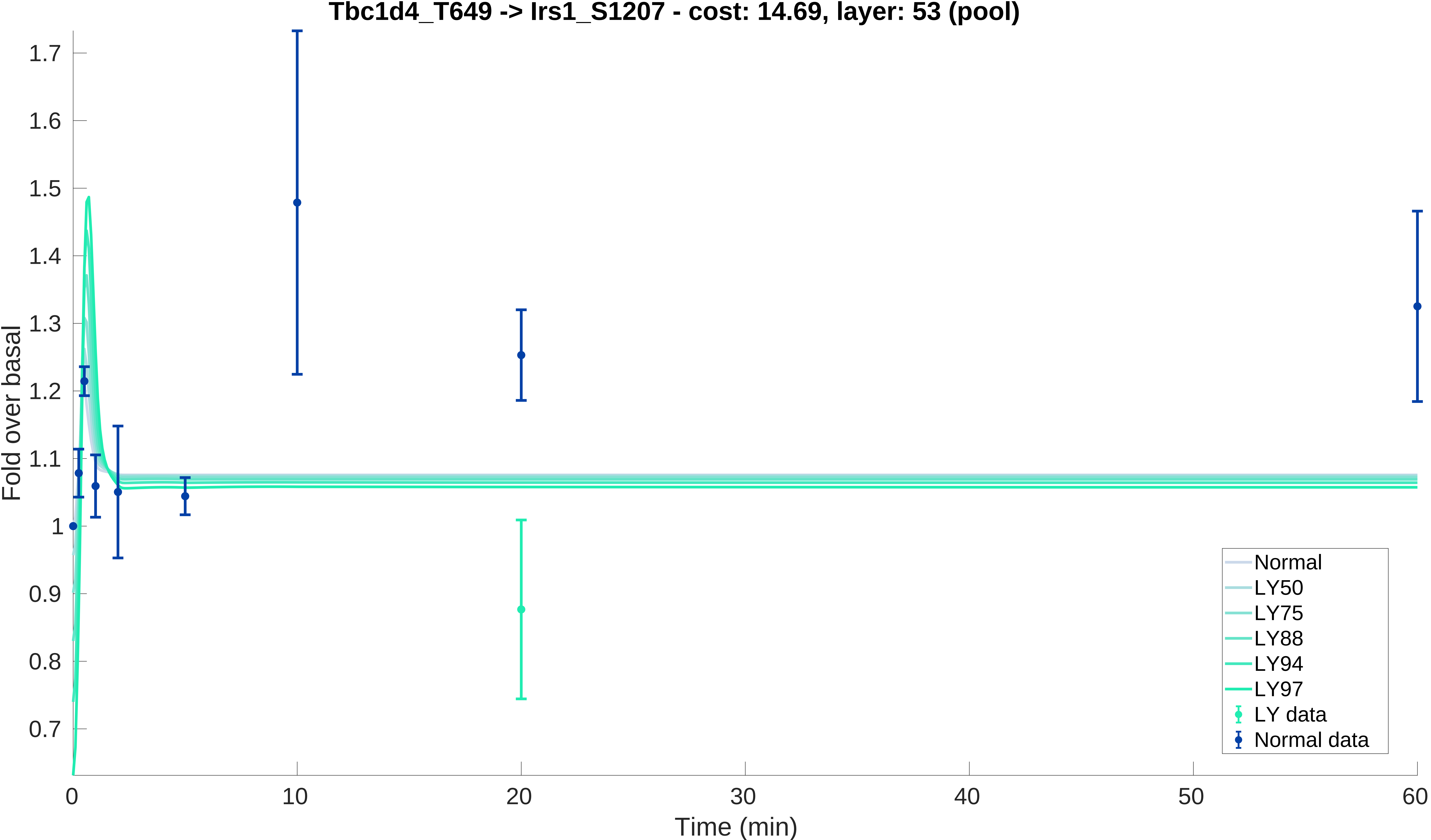
Tbc1d4_T649 -> Irs1_S1207 - cost: 14.69, layer: 53 (pool)
Fold over basal
Time (min)
1.7
1.6
1.5
1.4
1.3
1.2
1.1
1
0.9
0.8
0.7
0
10
20
30
40
50
60
Normal
LY50
LY75
LY88
LY94
LY97
LY data
Normal data

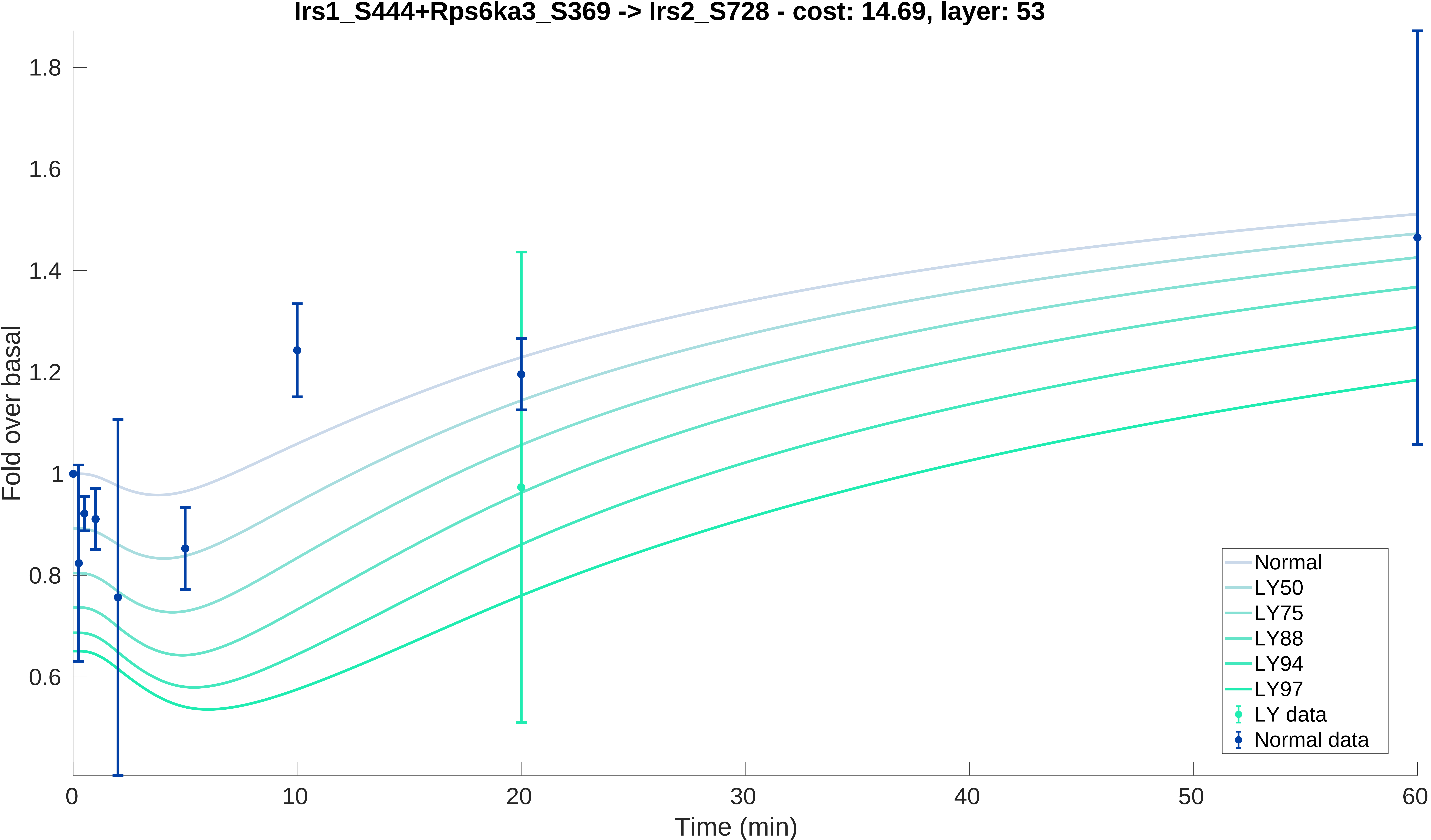

Irs1_S444+Rps6ka3_S369 -> Irs2_S728 - cost: 14.69, layer: 53
Fold over basal
1.8
1.6
1.4
1.2
1
0.8
0.6
0
10
20
30
40
50
60
Time (min)
Normal
LY50
LY75
LY88
LY94
LY97
LY data
Normal data

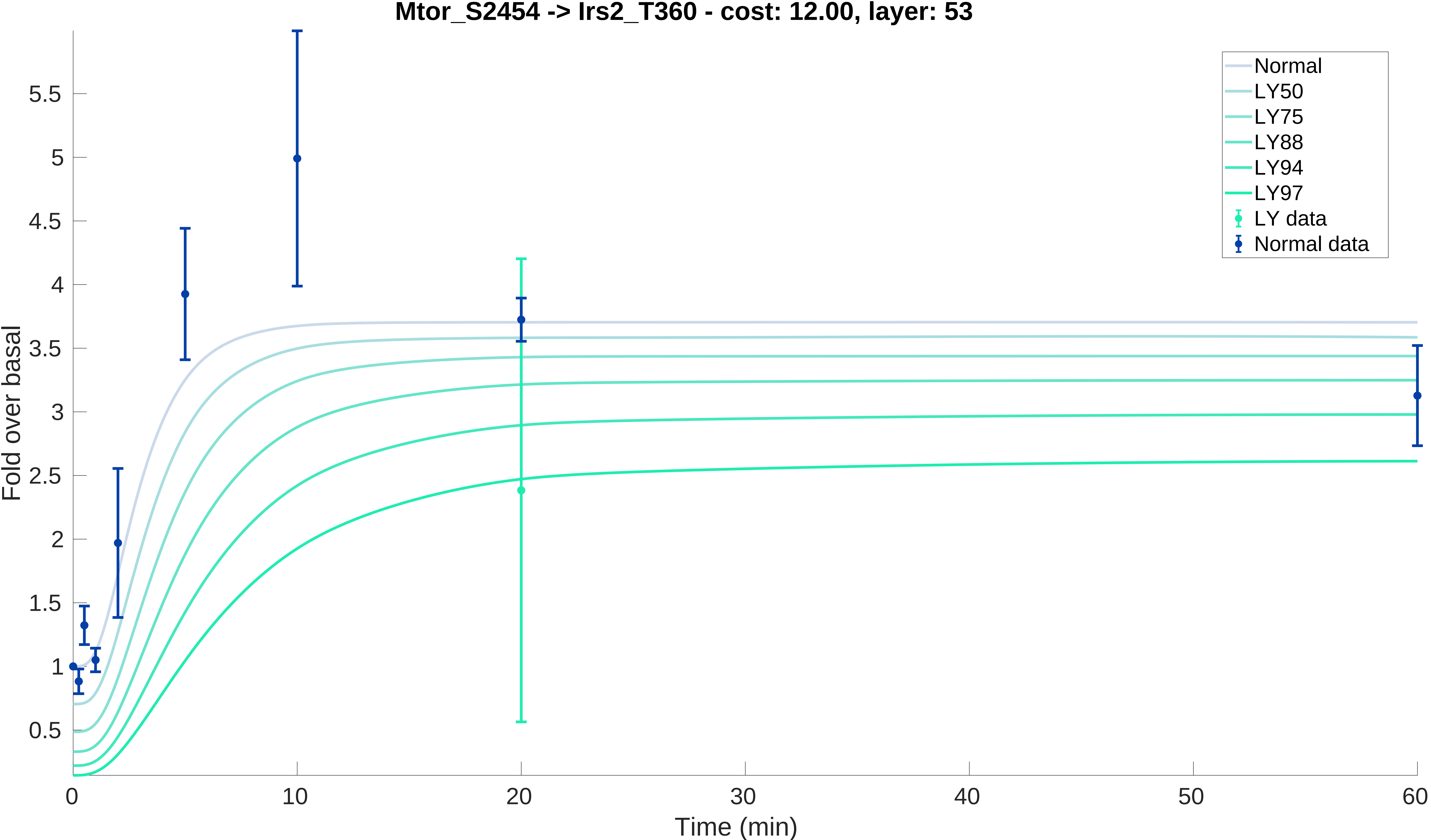

Mtor_S2454 -> Irs2_T360 - cost: 12.00, layer: 53
Normal
LY50
LY75
LY88
LY94
LY97
LY data
Normal data
Fold over basal
Time (min)
0
10
20
30
40
50
60
0.5
1
1.5
2
2.5
3
3.5
4
4.5
5
5.5

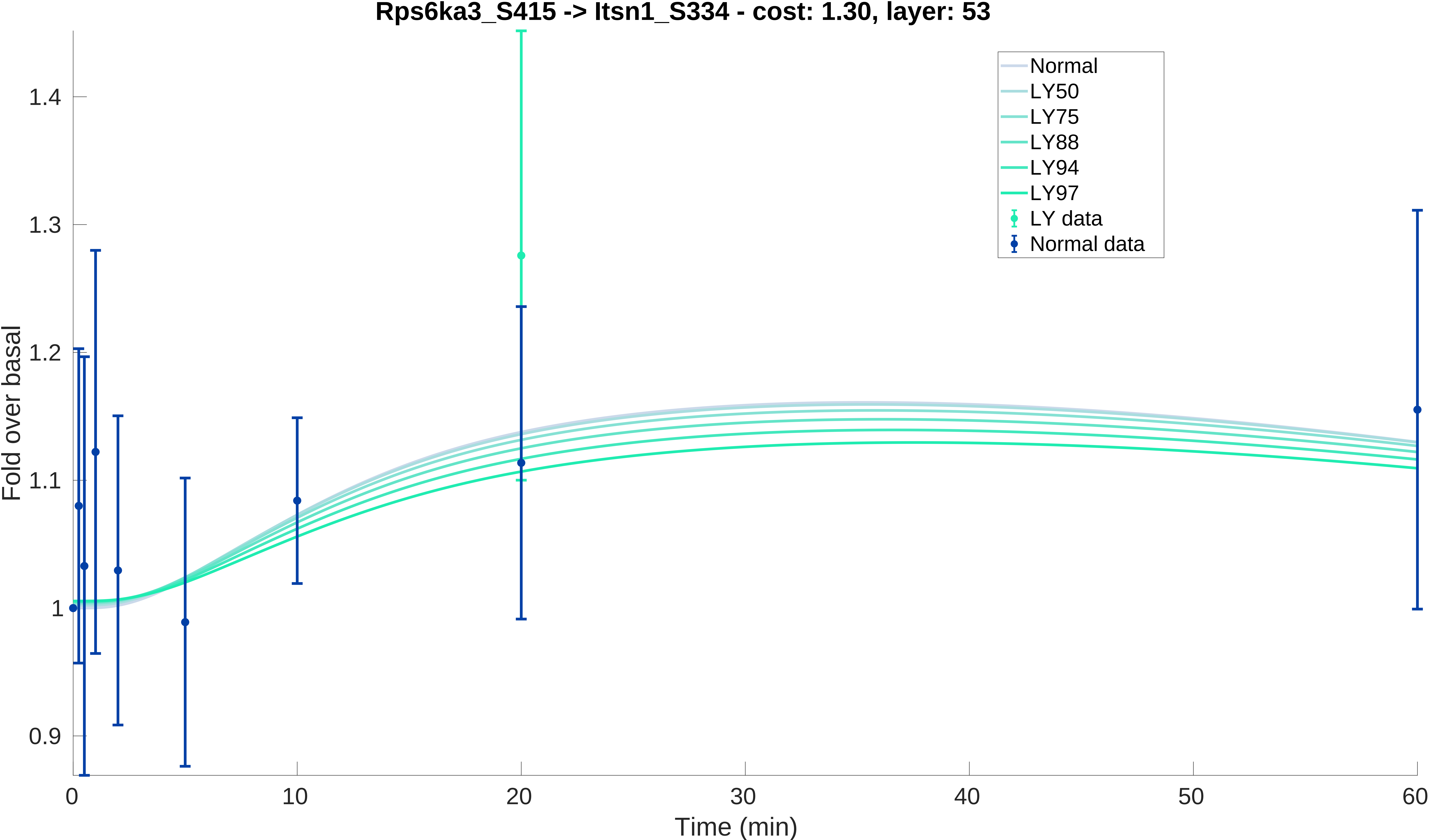

Rps6ka3_S415 -> Itsn1_S334 - cost: 1.30, layer: 53
Fold over basal
Time (min)
0.9
1
1.1
1.2
1.3
1.4
0
10
20
30
40
50
60
Normal
LY50
LY75
LY88
LY94
LY97
LY data
Normal data

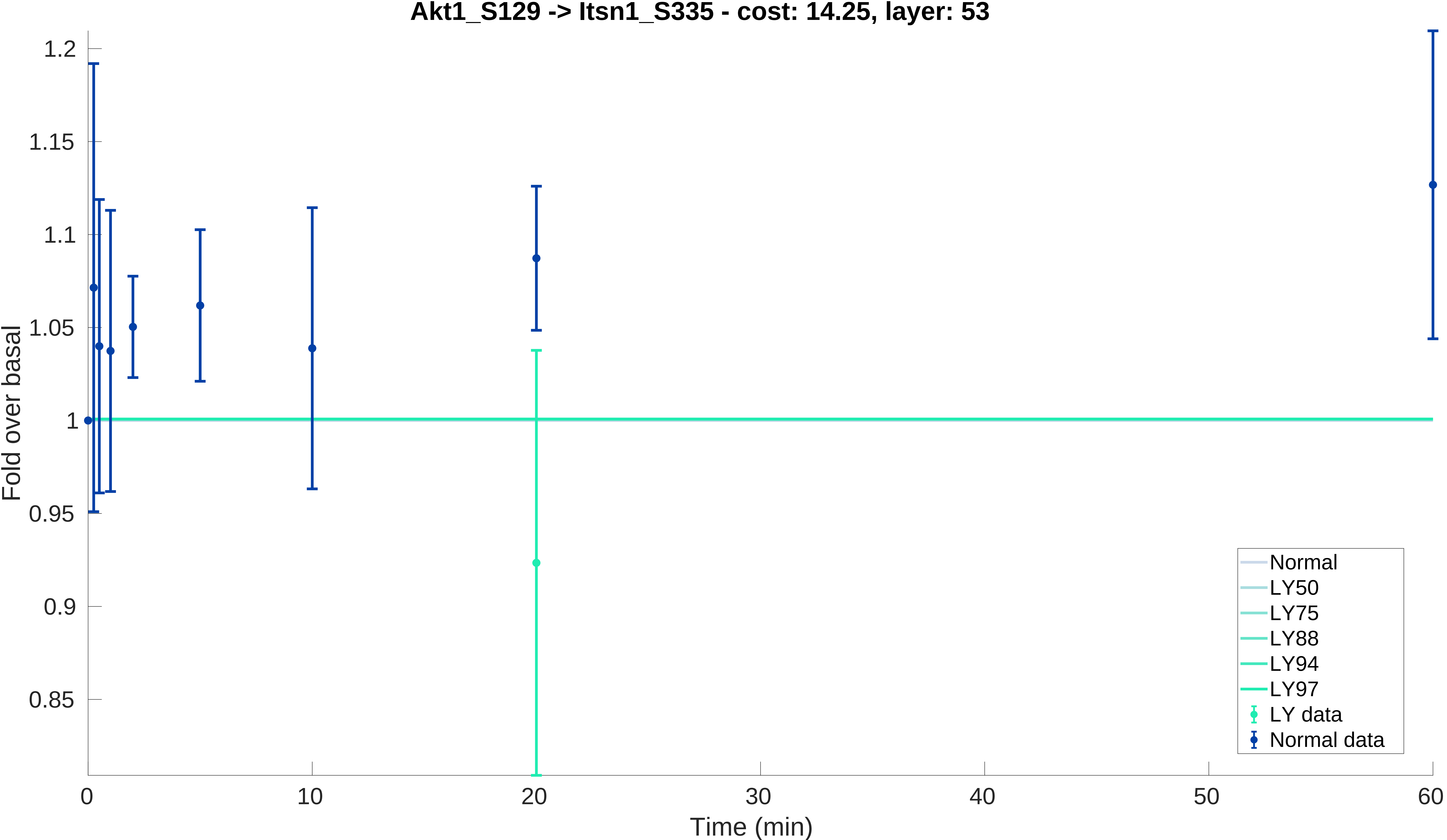

Akt1_S129 -> Itsn1_S335 - cost: 14.25, layer: 53
Fold over basal
1.2
1.15
1.1
1.05
1
0.95
0.9
0.85
0
10
20
30
40
50
60
Time (min)
Normal
LY50
LY75
LY88
LY94
LY97
LY data
Normal data

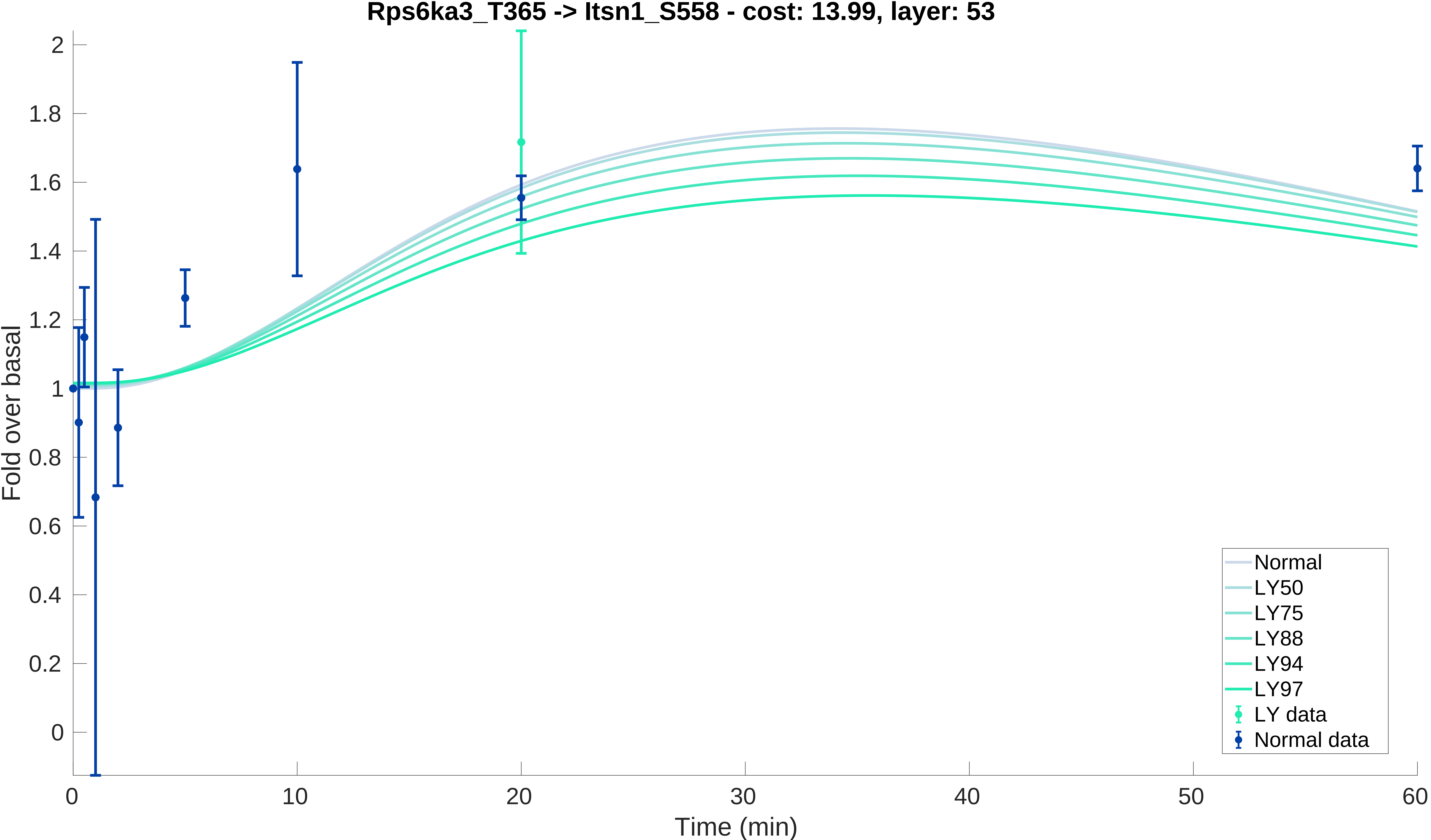
Rps6ka3_T365 -> Itsn1_S558 - cost: 13.99, layer: 53
Fold over basal
Time (min)
0
0.2
0.4
0.6
0.8
1
1.2
1.4
1.6
1.8
2
0
10
20
30
40
50
60
Normal
LY50
LY75
LY88
LY94
LY97
LY data
Normal data

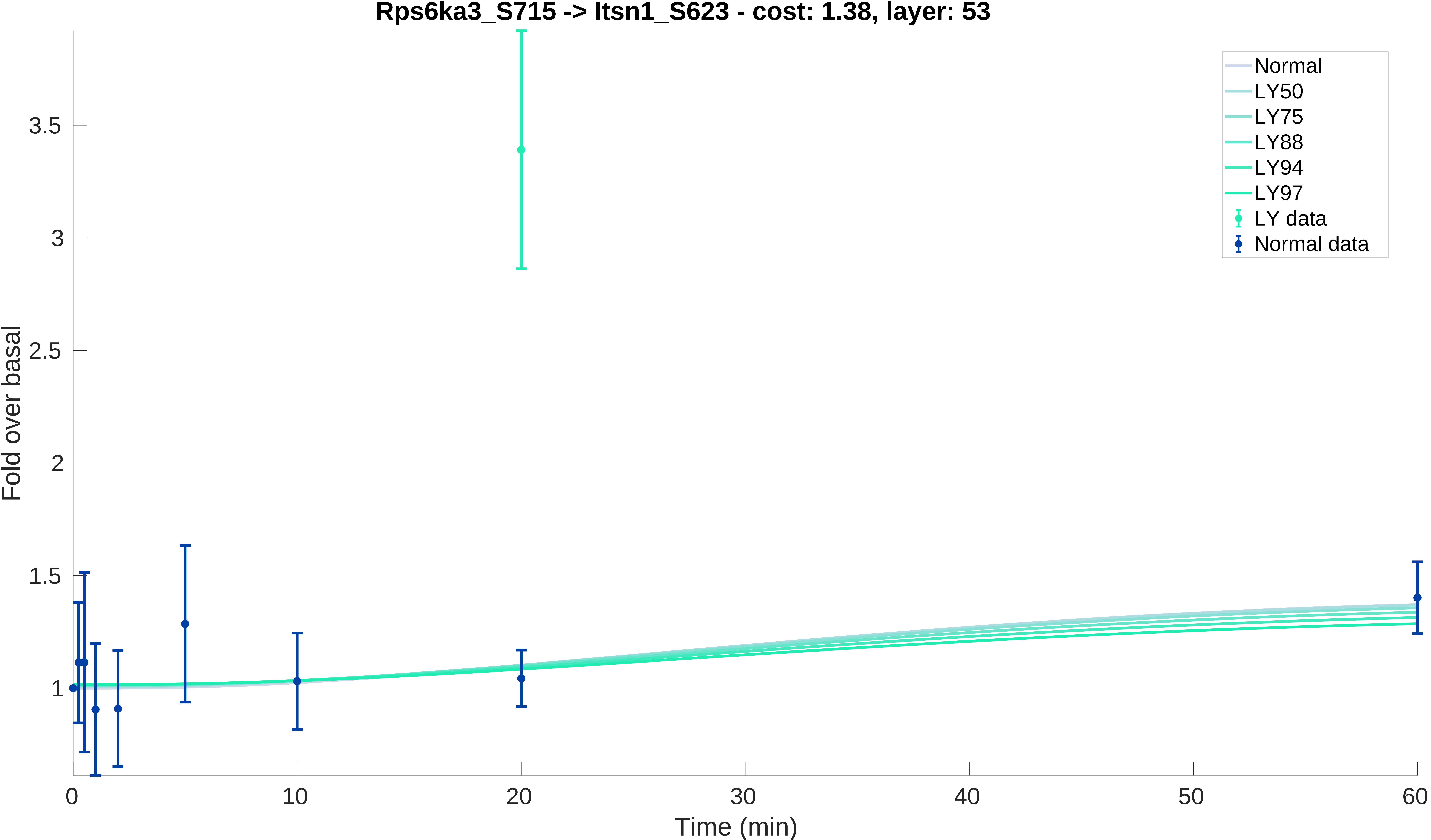

Rps6ka3_S715 -> Itsn1_S623 - cost: 1.38, layer: 53
Fold over basal
Time (min)
3.5
3
2.5
2
1.5
1
0
10
20
30
40
50
60
Normal
LY50
LY75
LY88
LY94
LY97
LY data
Normal data

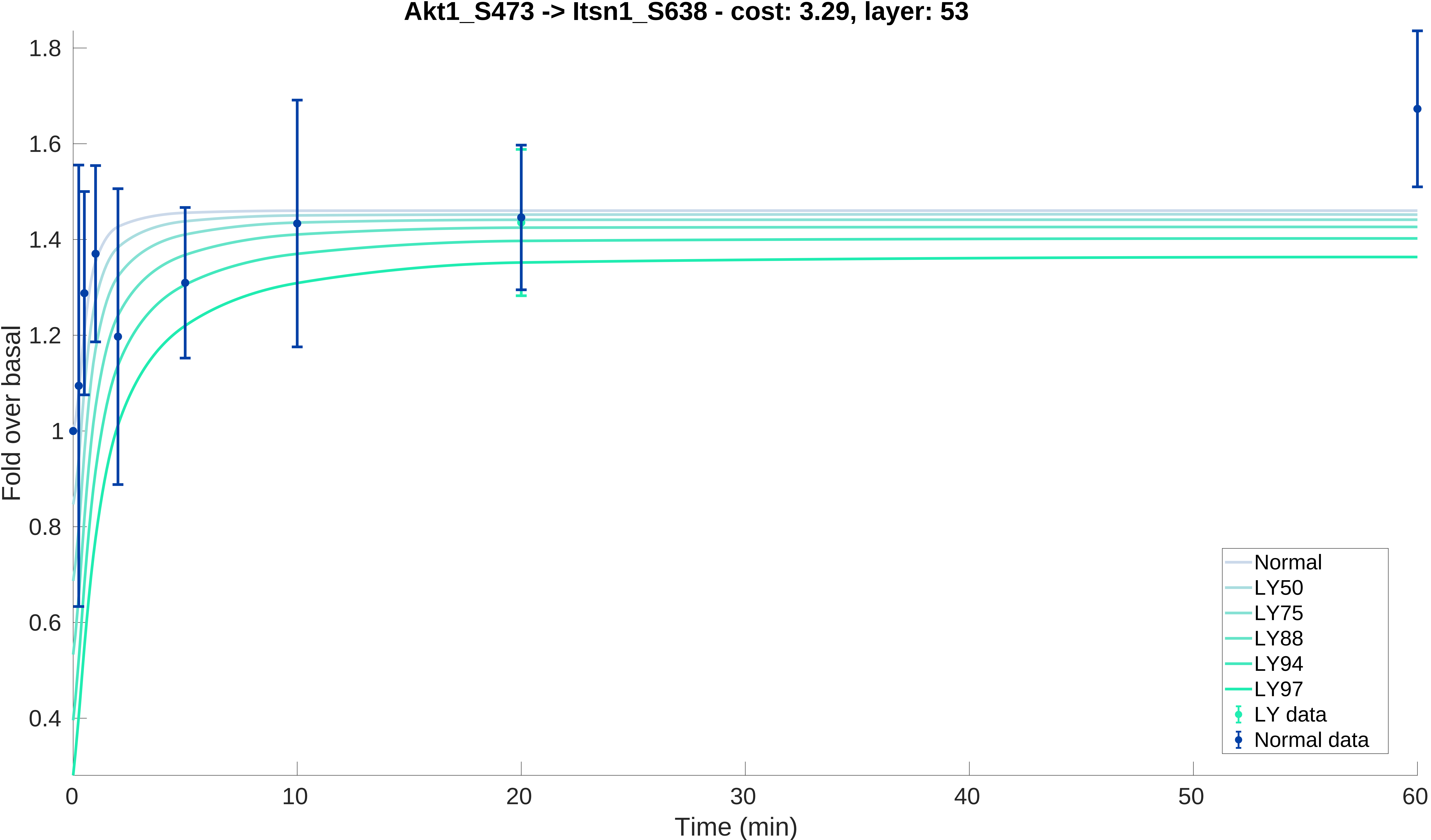
Akt1_S473 -> Itsn1_S638 - cost: 3.29, layer: 53
Fold over basal
Time (min)
1.8
1.6
1.4
1.2
1
0.8
0.6
0.4
0
10
20
30
40
50
60
Normal
LY50
LY75
LY88
LY94
LY97
LY data
Normal data

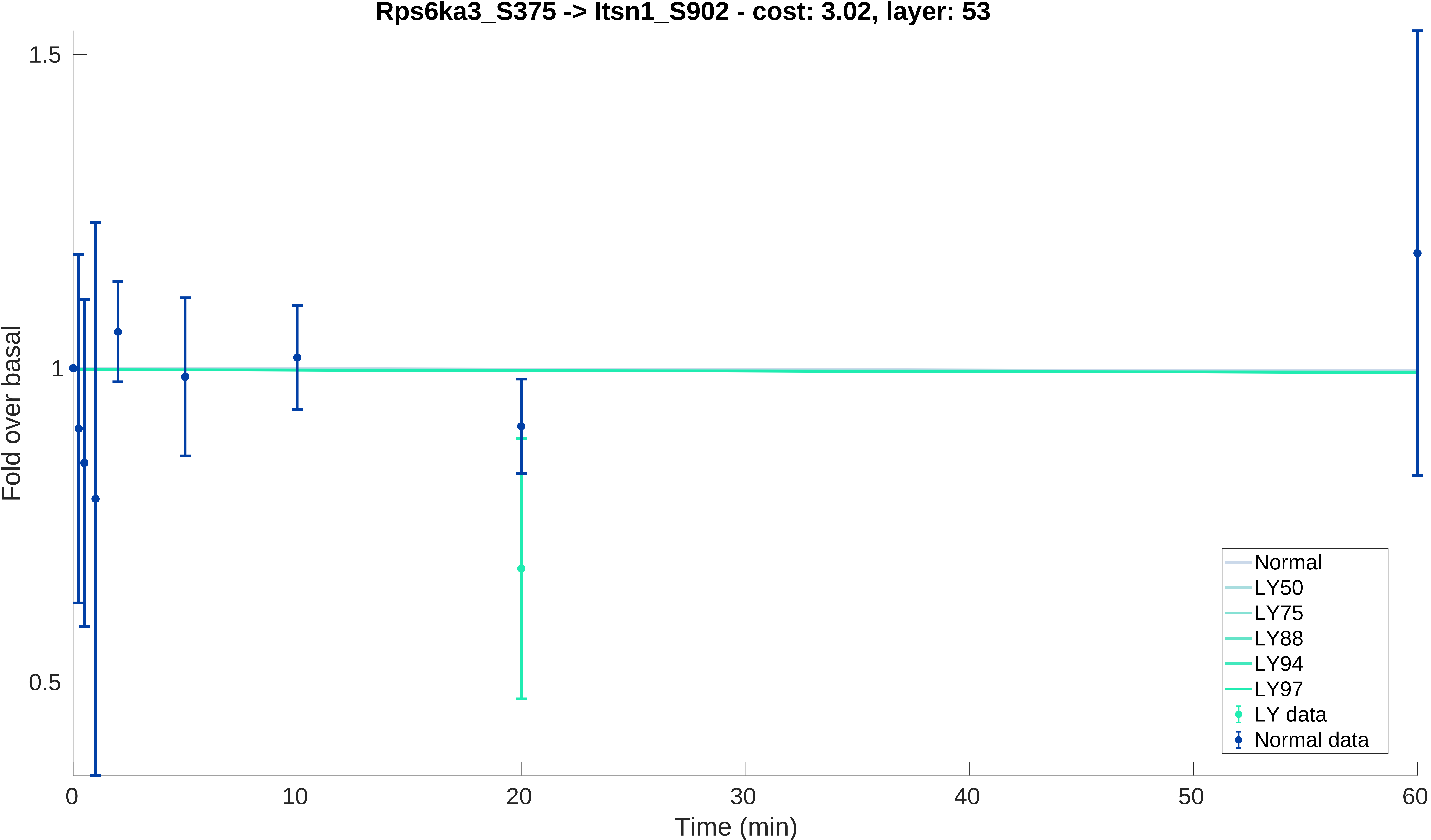

Rps6ka3_S375 -> Itsn1_S902 - cost: 3.02, layer: 53
Fold over basal
1.5
1
0.5
0
10
20
30
40
50
60
Time (min)
Normal
LY50
LY75
LY88
LY94
LY97
LY data
Normal data

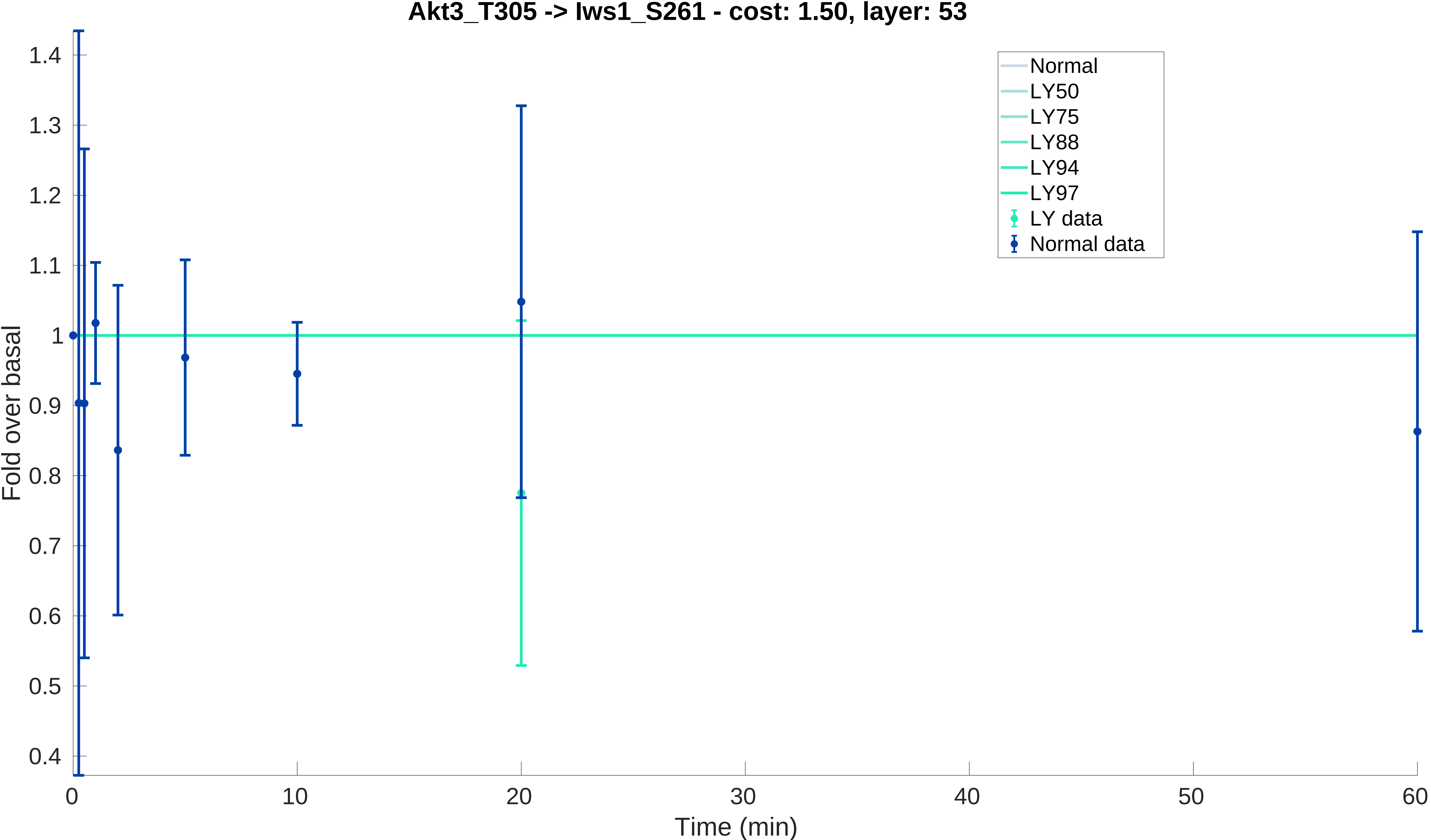

Akt3_T305 -> Iws1_S261 - cost: 1.50, layer: 53
Normal
LY50
LY75
LY88
LY94
LY97
LY data
Normal data
Fold over basal
1.4
1.3
1.2
1.1
1
0.9
0.8
0.7
0.6
0.5
0.4
0
10
20
30
40
50
60
Time (min)

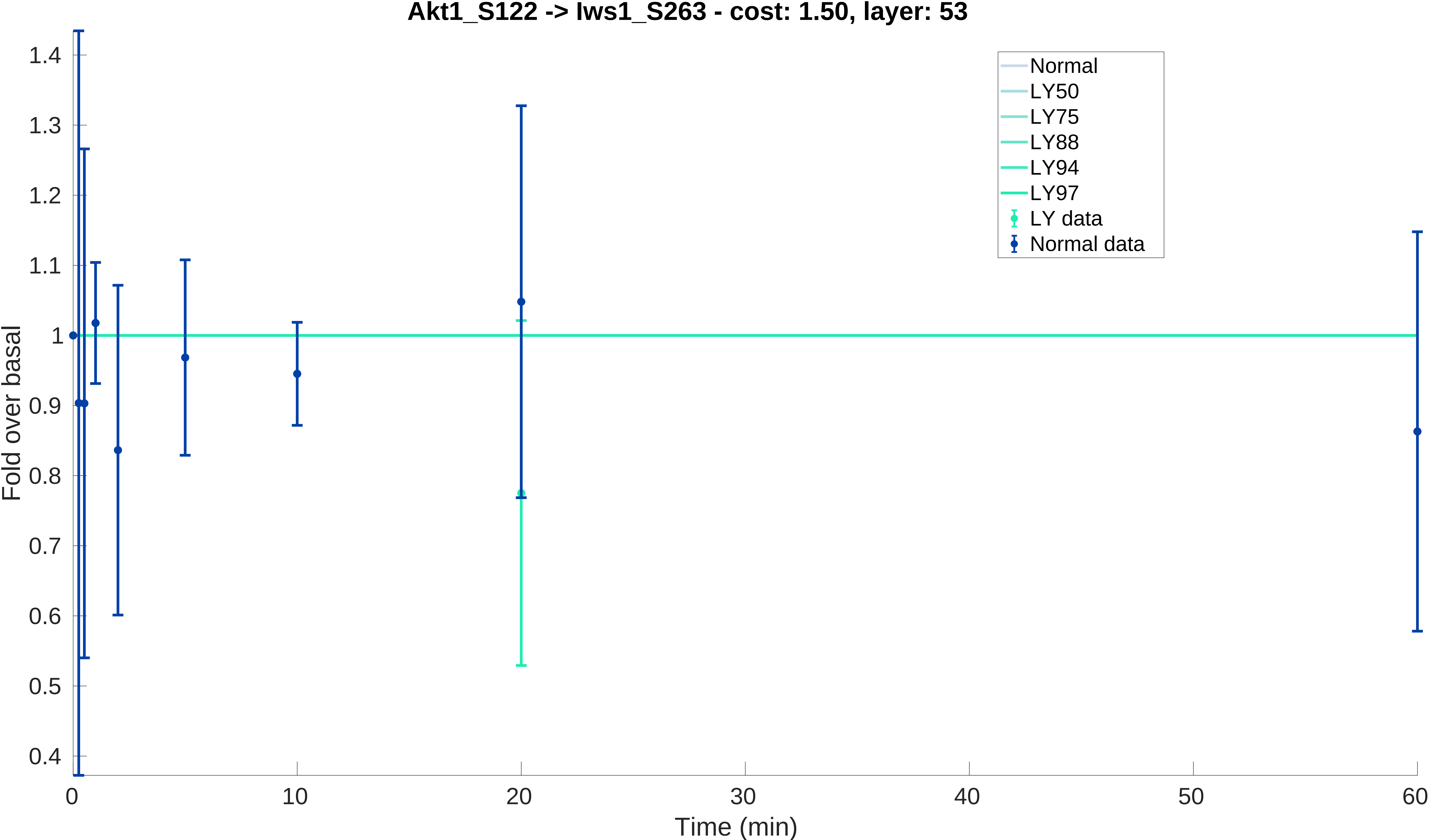

Akt1_S122 -> Iws1_S263 - cost: 1.50, layer: 53
Fold over basal
Time (min)
Normal
LY50
LY75
LY88
LY94
LY97
LY data
Normal data
0.4
0.5
0.6
0.7
0.8
0.9
1
1.1
1.2
1.3
1.4
0
10
20
30
40
50
60

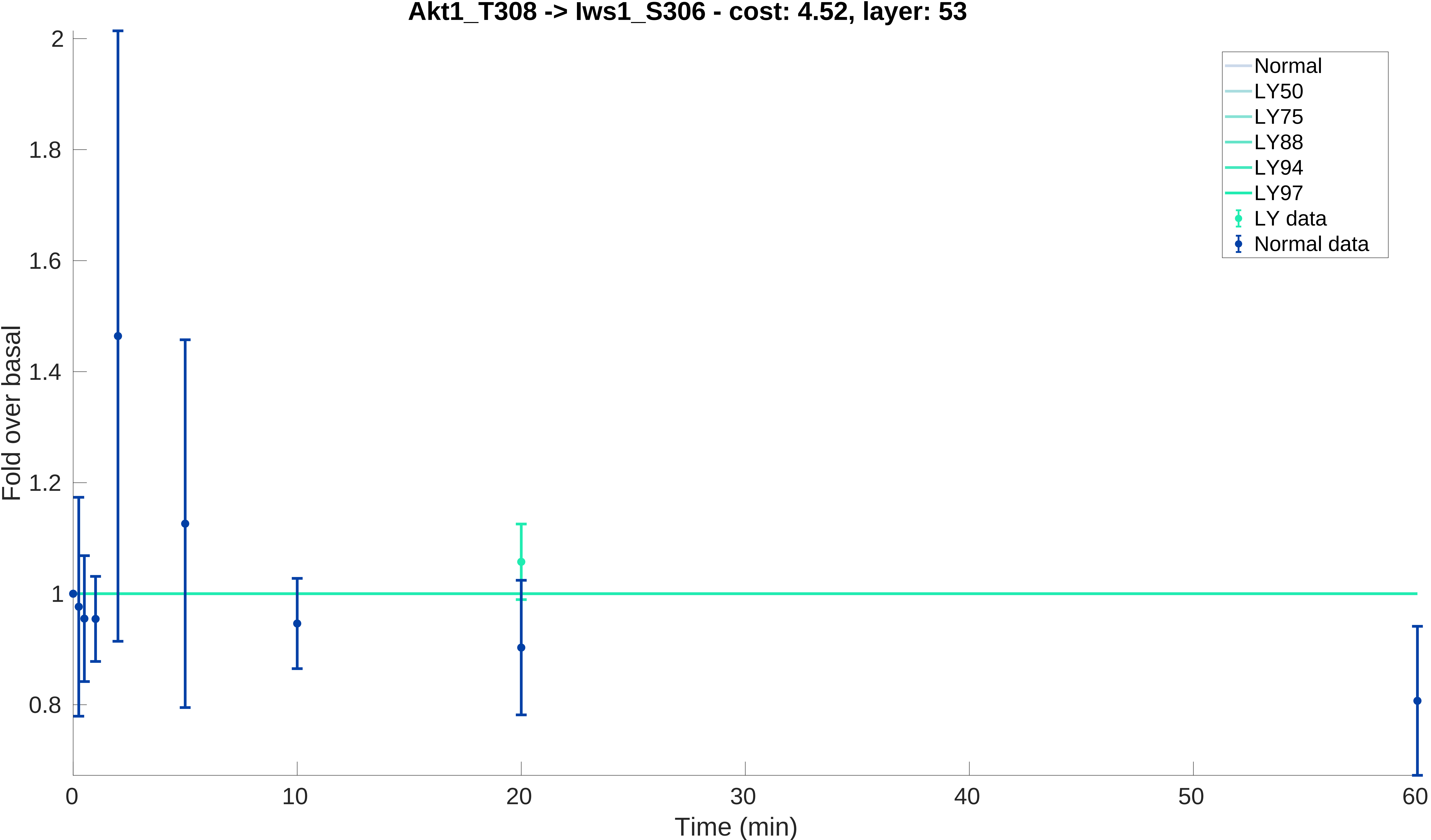

Akt1_T308 -> Iws1_S306 - cost: 4.52, layer: 53
Fold over basal
2
1.8
1.6
1.4
1.2
1
0.8
0
10
20
30
40
50
60
Time (min)
Normal
LY50
LY75
LY88
LY94
LY97
LY data
Normal data

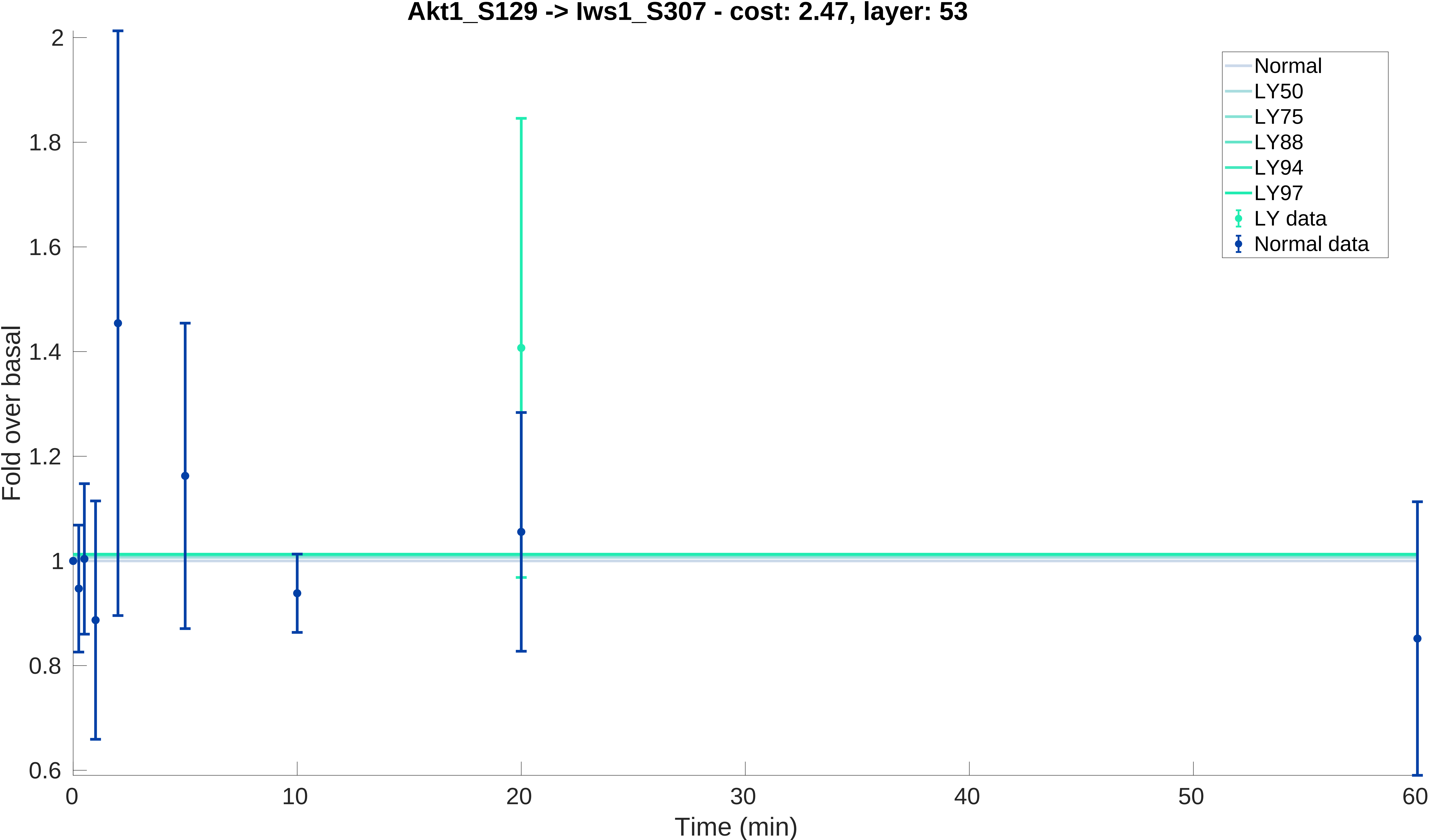

Akt1_S129 -> Iws1_S307 - cost: 2.47, layer: 53
Fold over basal
Time (min)
0.6
0.8
1
1.2
1.4
1.6
1.8
2
0
10
20
30
40
50
60
Normal
LY50
LY75
LY88
LY94
LY97
LY data
Normal data

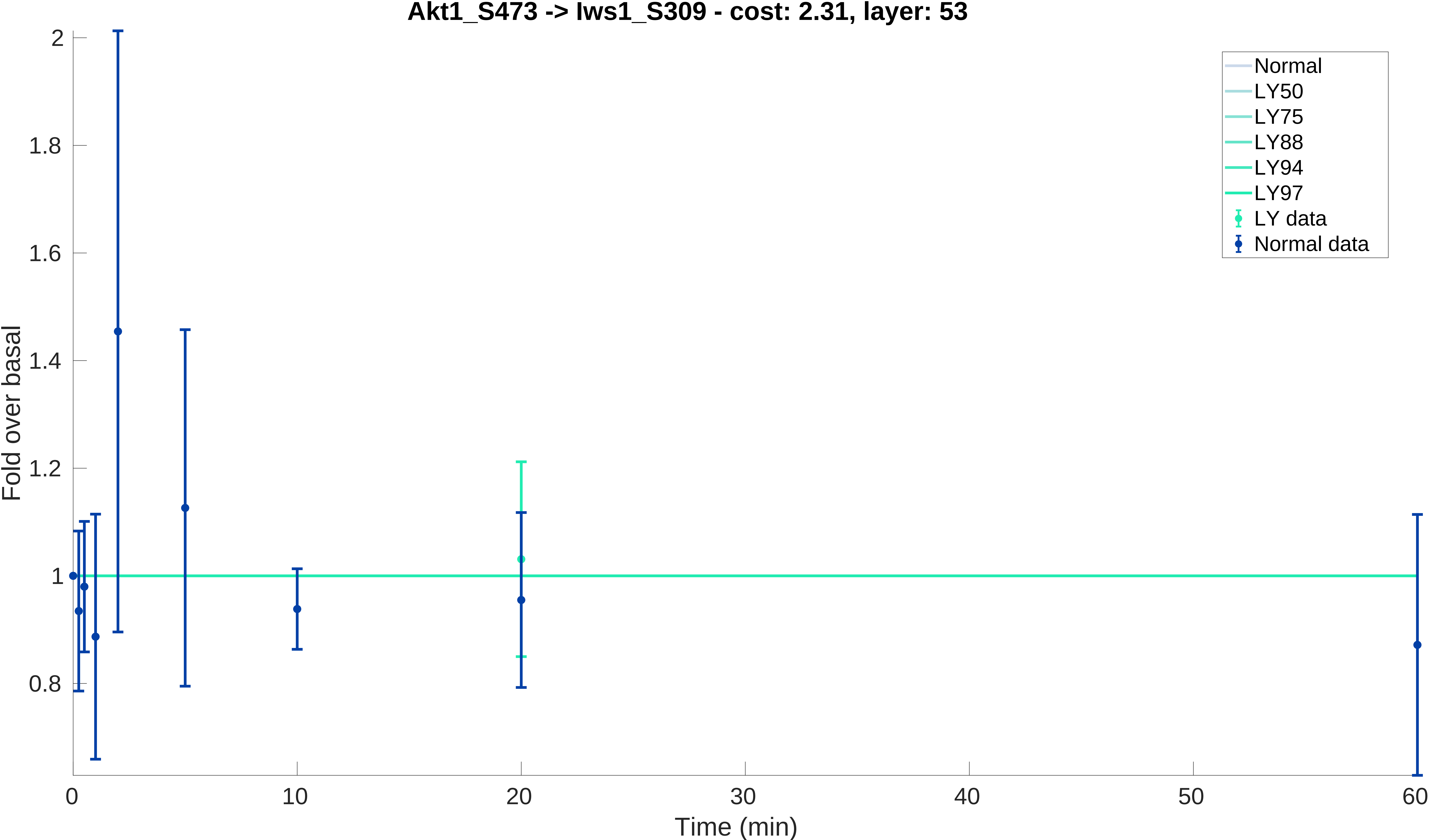

Akt1_S473 -> Iws1_S309 - cost: 2.31, layer: 53
Fold over basal
Time (min)
Normal
LY50
LY75
LY88
LY94
LY97
LY data
Normal data
0.8
1
1.2
1.4
1.6
1.8
2
0
10
20
30
40
50
60

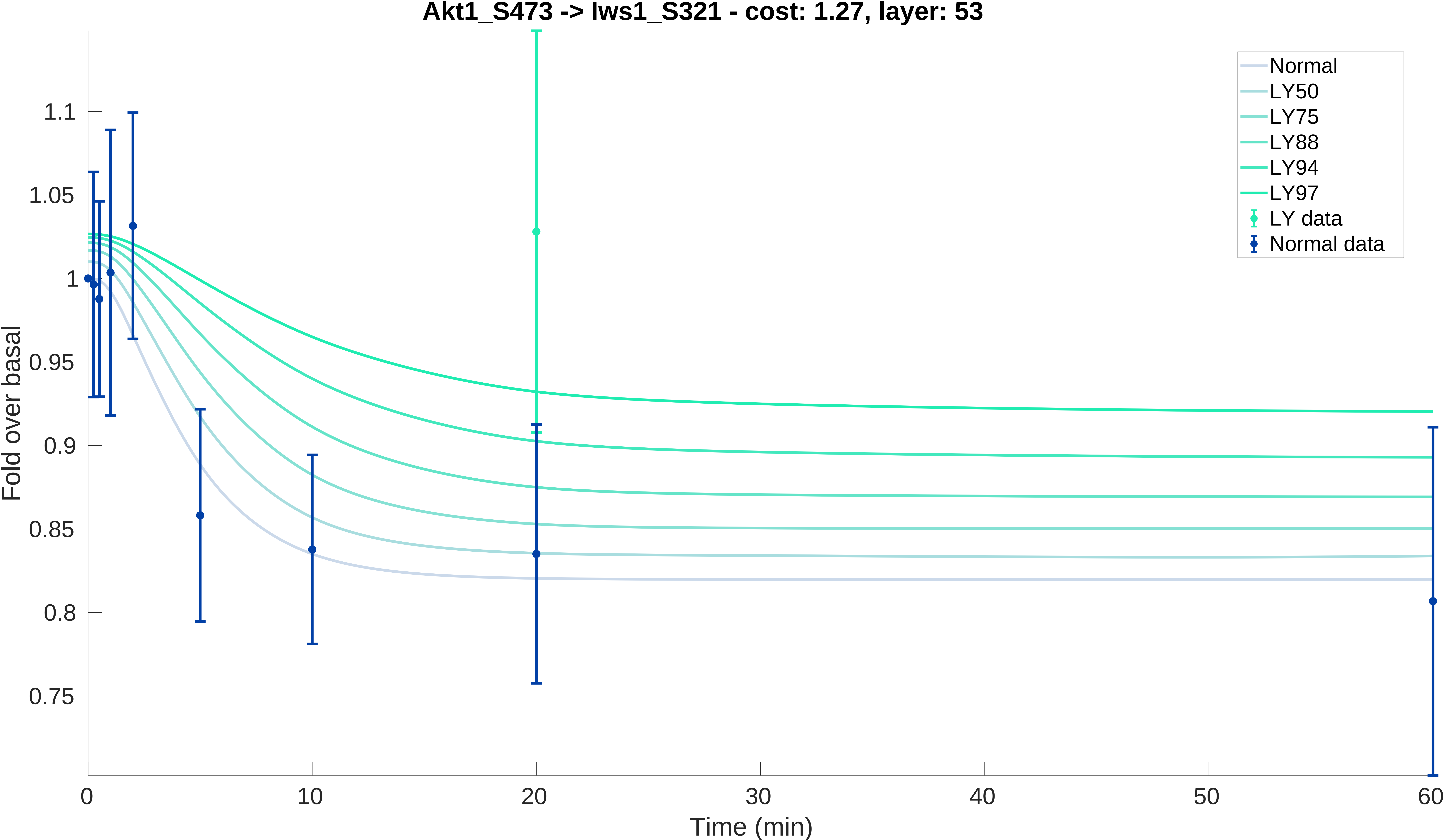

Akt1_S473 -> Iws1_S321 - cost: 1.27, layer: 53
Fold over basal
1.1
1.05
1
0.95
0.9
0.85
0.8
0.75
0
10
20
30
40
50
60
Time (min)
Normal
LY50
LY75
LY88
LY94
LY97
LY data
Normal data

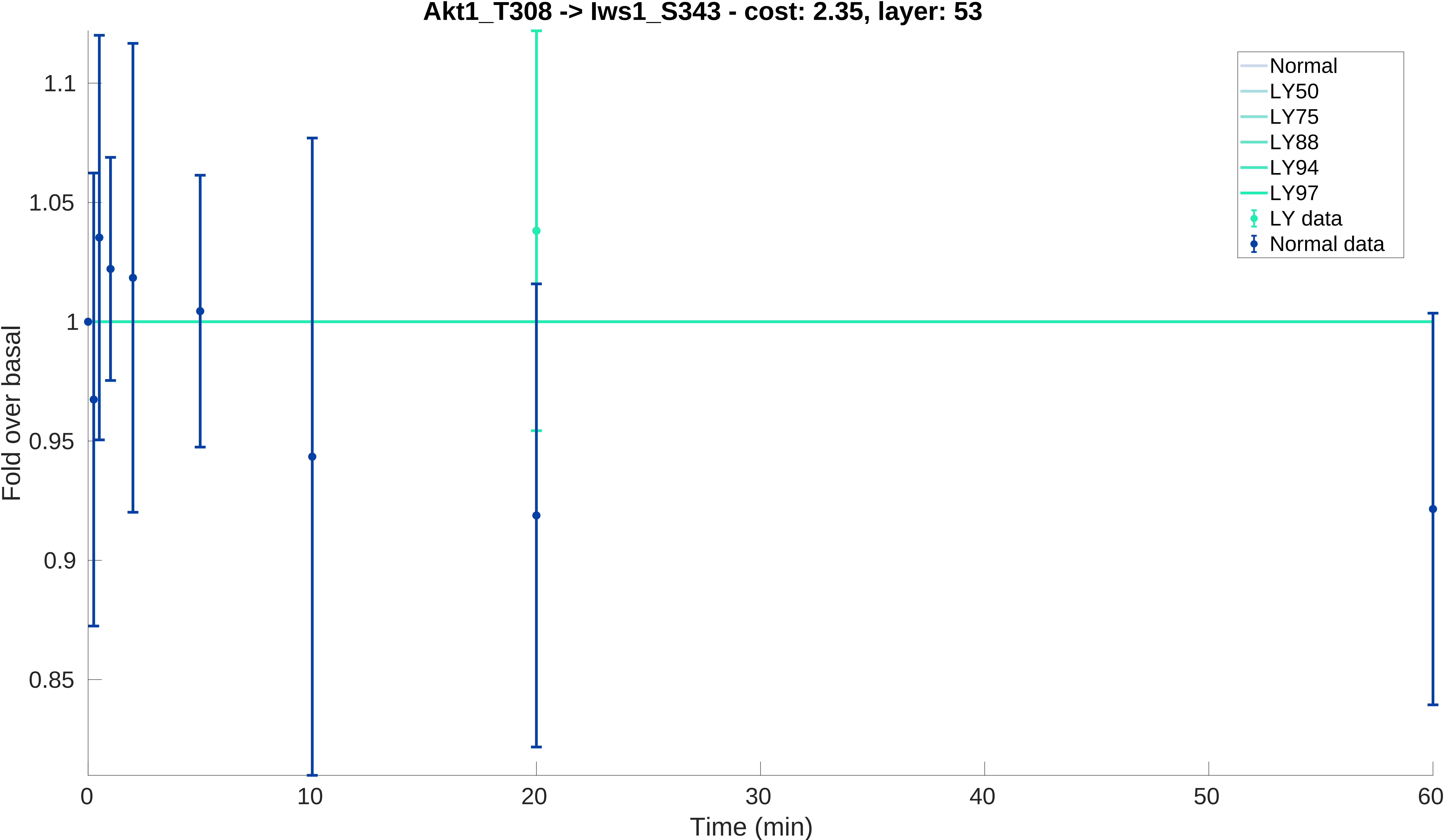
Akt1_T308 -> Iws1_S343 - cost: 2.35, layer: 53
Fold over basal
1.1
1.05
1
0.95
0.9
0.85
0
10
20
30
40
50
60
Time (min)
Normal
LY50
LY75
LY88
LY94
LY97
LY data
Normal data

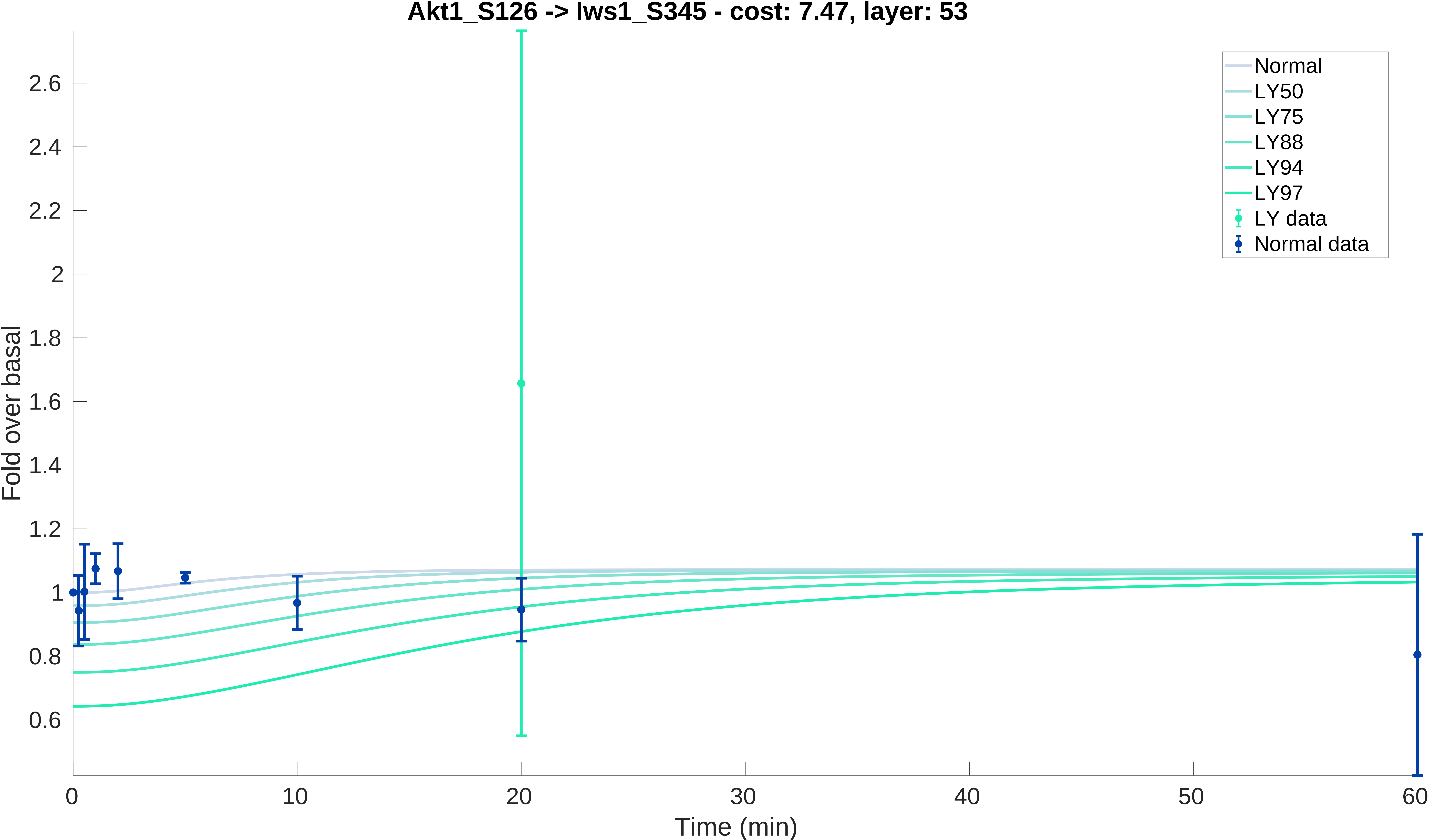

Akt1_S126 -> Iws1_S345 - cost: 7.47, layer: 53
Fold over basal
Time (min)
0.6
0.8
1
1.2
1.4
1.6
1.8
2
2.2
2.4
2.6
0
10
20
30
40
50
60
Normal
LY50
LY75
LY88
LY94
LY97
LY data
Normal data

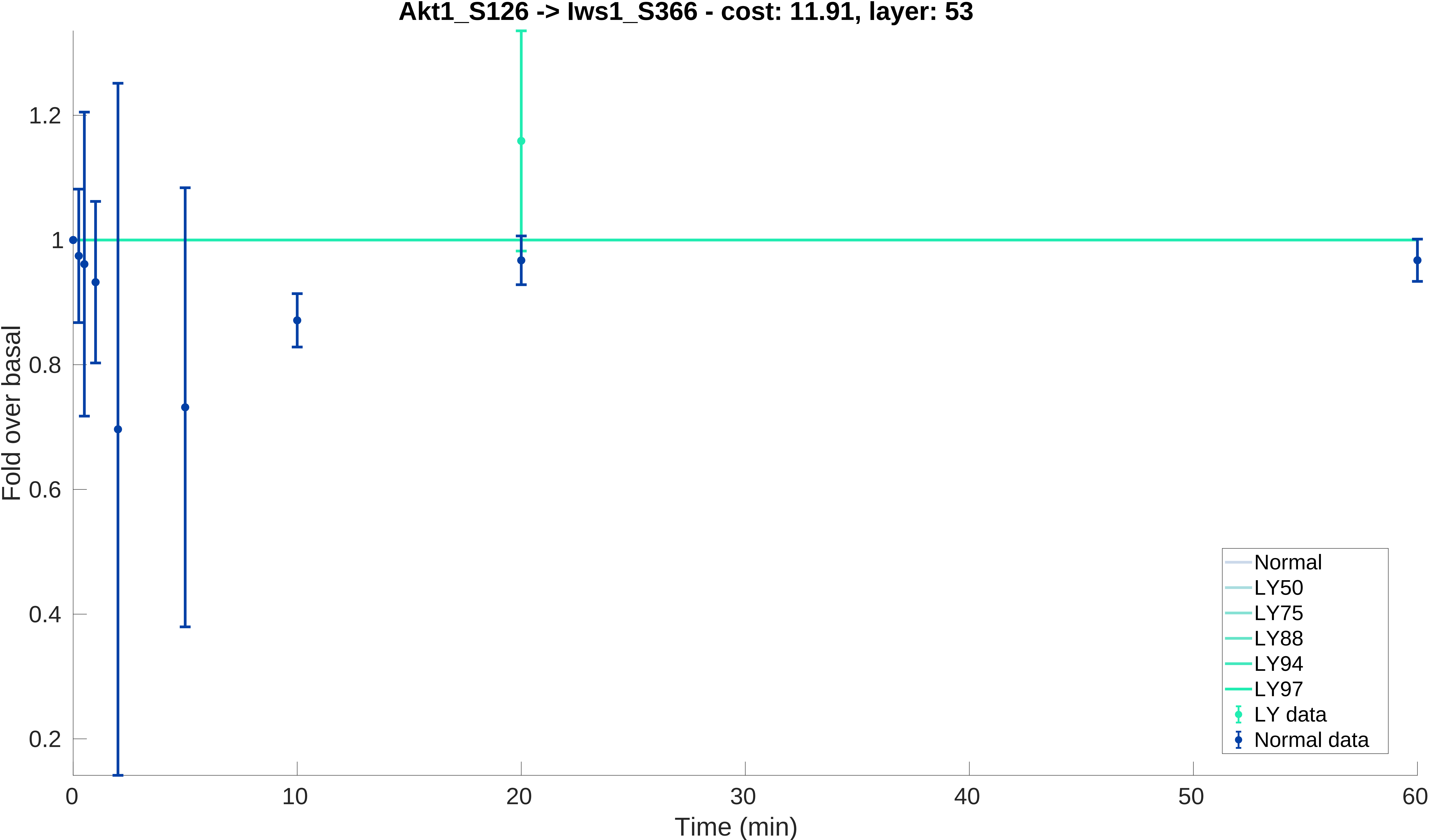
Akt1_S126 -> Iws1_S366 - cost: 11.91, layer: 53
Fold over basal
1.2
1
0.8
0.6
0.4
0.2
0
10
20
30
40
50
60
Time (min)
Normal
LY50
LY75
LY88
LY94
LY97
LY data
Normal data

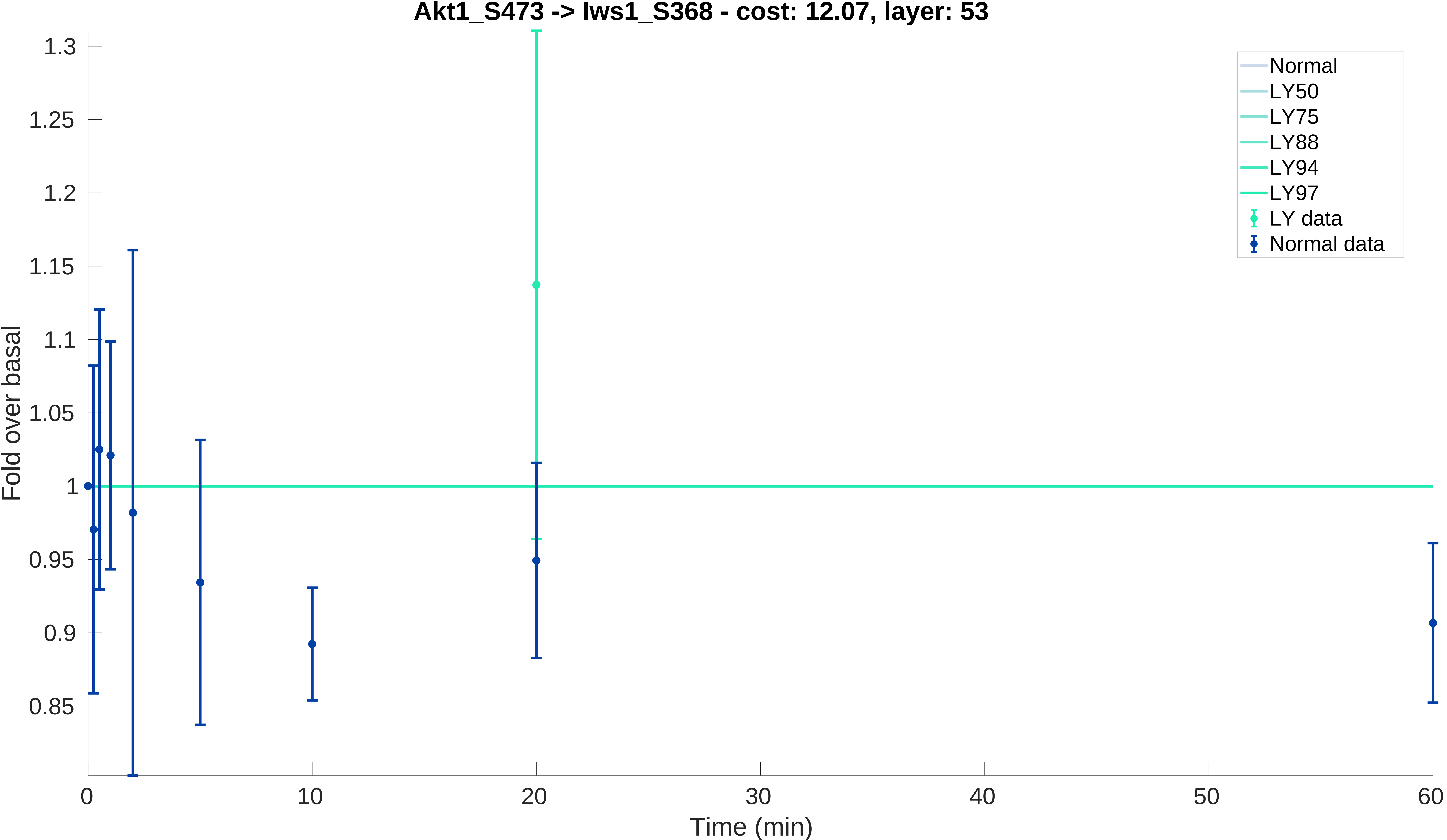

Akt1_S473 -> Iws1_S368 - cost: 12.07, layer: 53
Fold over basal
Time (min)
1.3
1.25
1.2
1.15
1.1
1.05
1
0.95
0.9
0.85
0
10
20
30
40
50
60
Normal
LY50
LY75
LY88
LY94
LY97
LY data
Normal data

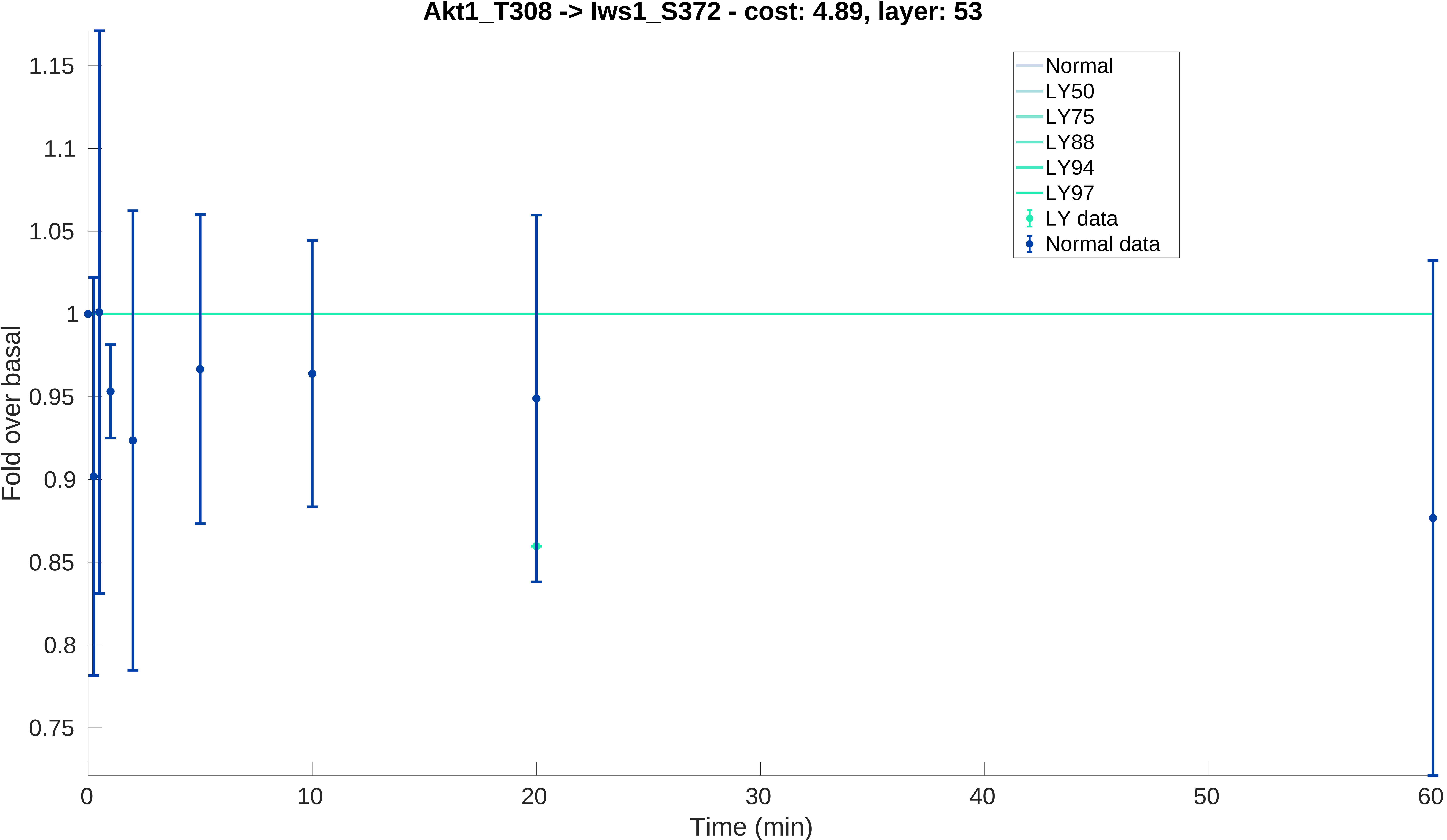

Akt1_T308 -> Iws1_S372 - cost: 4.89, layer: 53
Normal
LY50
LY75
LY88
LY94
LY97
LY data
Normal data
Fold over basal
1.15
1.1
1.05
1
0.95
0.9
0.85
0.8
0.75
0
10
20
30
40
50
60
Time (min)

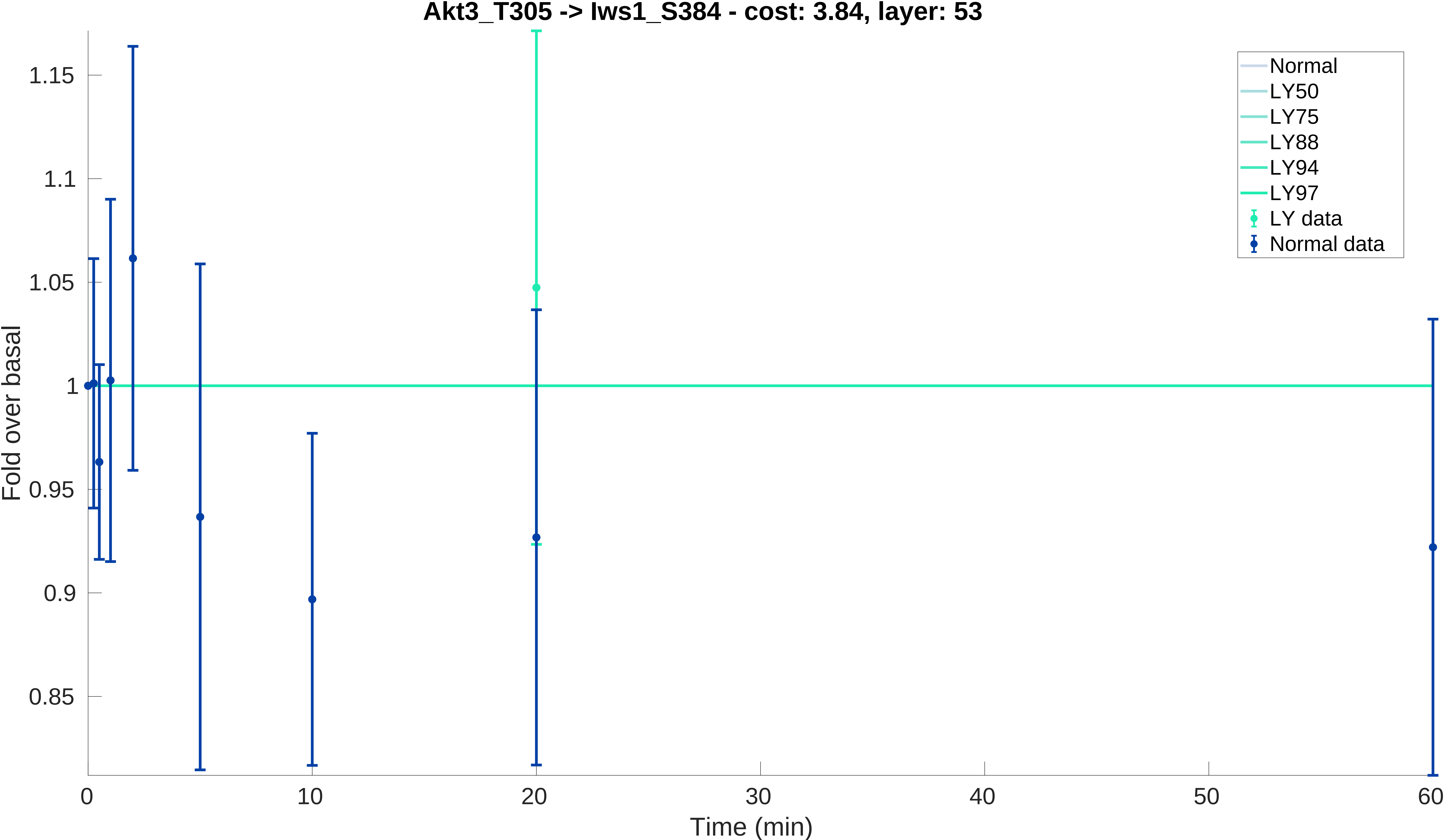
Akt3_T305 -> Iws1_S384 - cost: 3.84, layer: 53
Fold over basal
1.15
1.1
1.05
1
0.95
0.9
0.85
0
10
20
30
40
50
60
Time (min)
Normal
LY50
LY75
LY88
LY94
LY97
LY data
Normal data

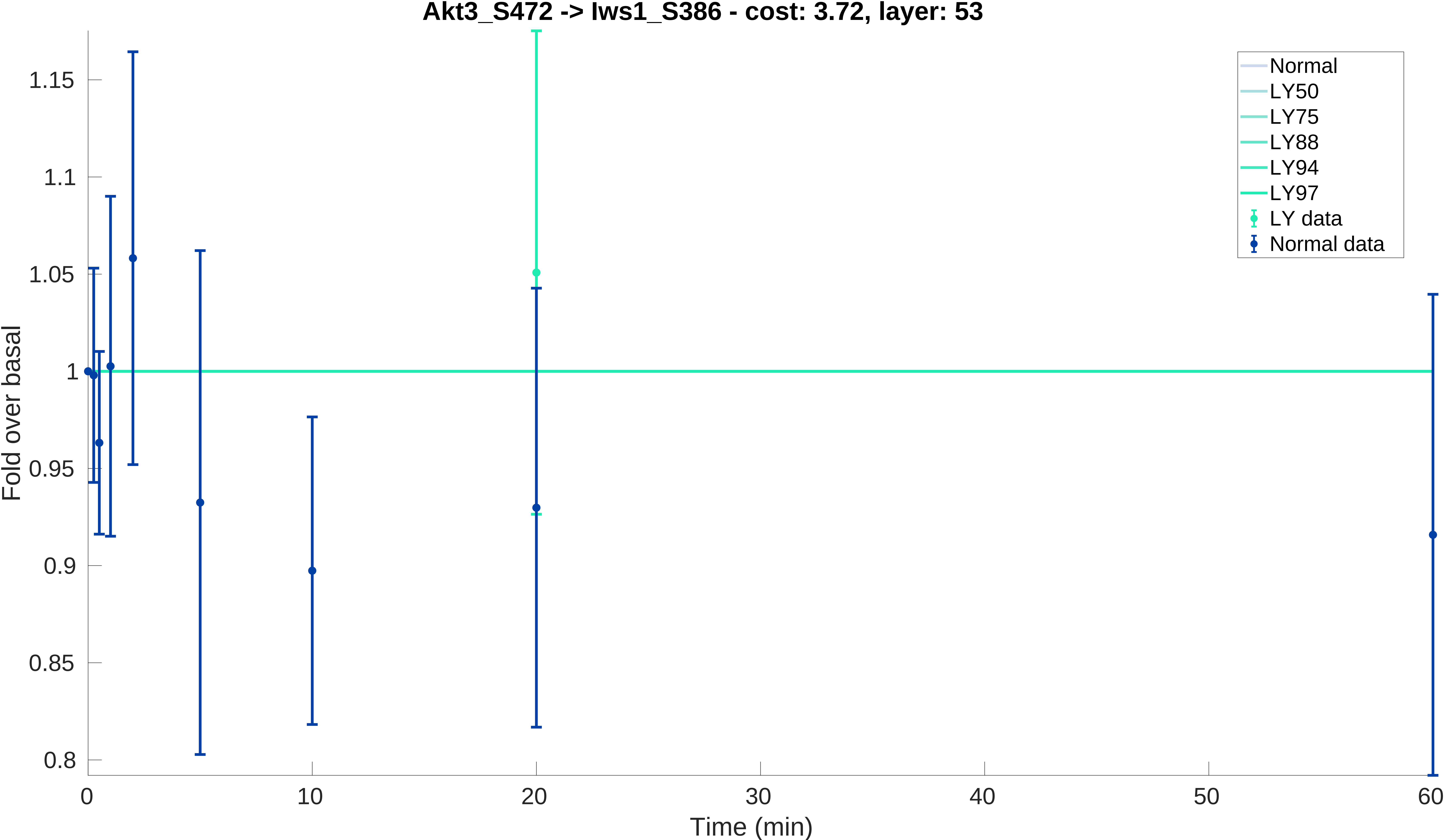

Akt3_S472 -> Iws1_S386 - cost: 3.72, layer: 53
Fold over basal
1.15
1.1
1.05
1
0.95
0.9
0.85
0.8
0
10
20
30
40
50
60
Time (min)
Normal
LY50
LY75
LY88
LY94
LY97
LY data
Normal data

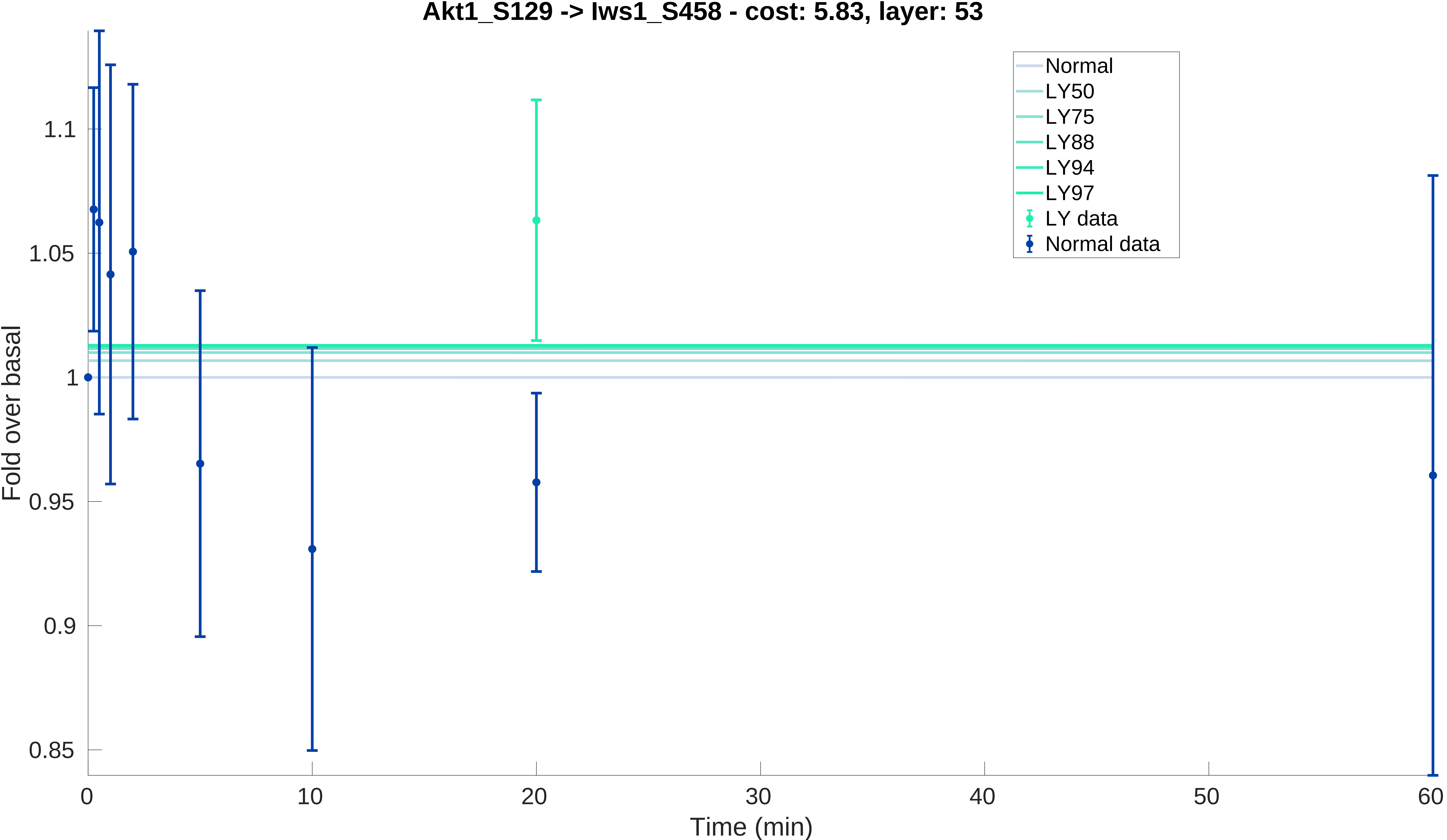

Akt1_S129 -> Iws1_S458 - cost: 5.83, layer: 53
Normal
LY50
LY75
LY88
LY94
LY97
LY data
Normal data
Fold over basal
1.1
1.05
1
0.95
0.9
0.85
0
10
20
30
40
50
60
Time (min)

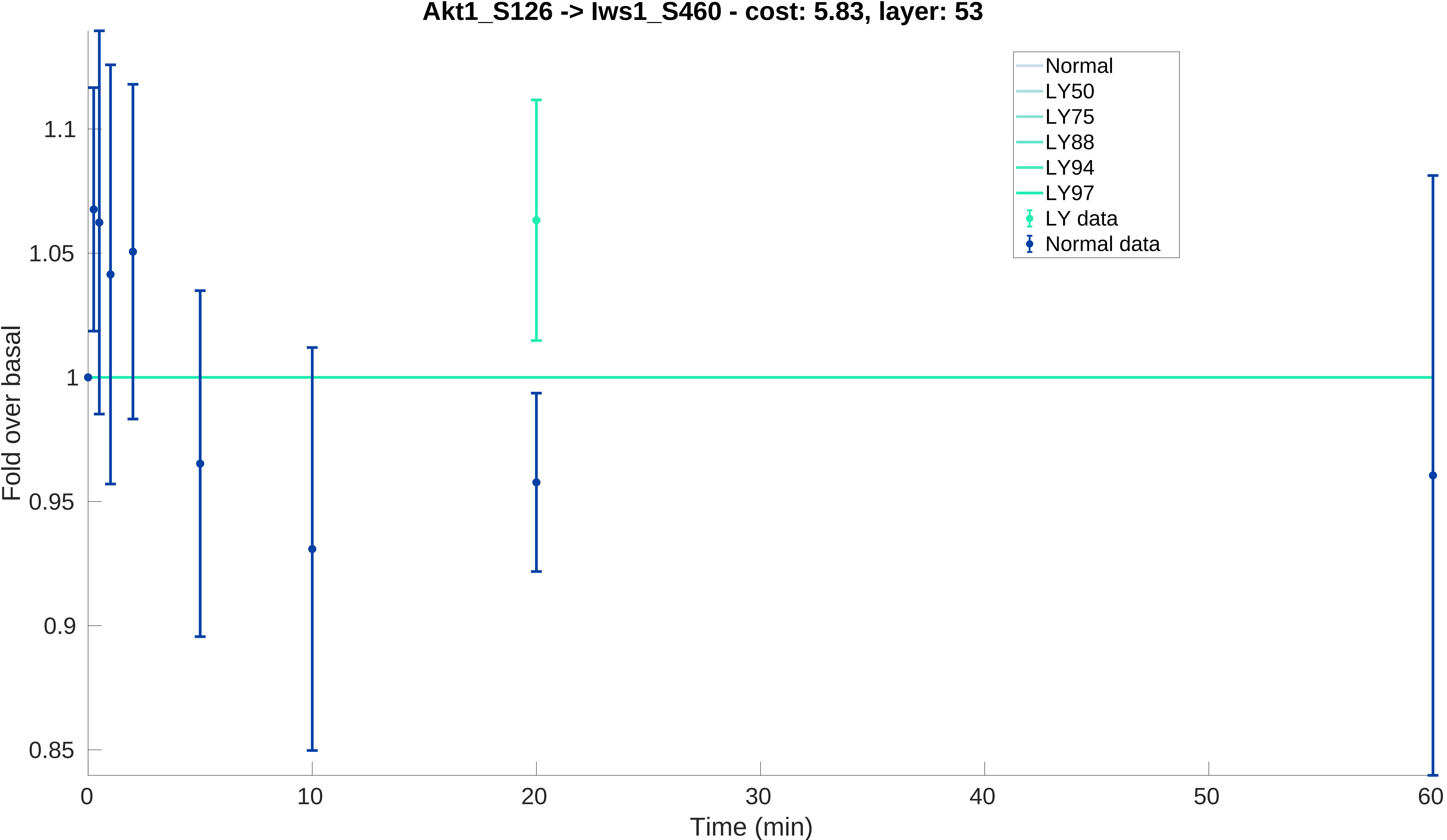

Akt1_S126 -> Iws1_S460 - cost: 5.83, layer: 53
Normal
LY50
LY75
LY88
LY94
LY97
LY data
Normal data
Fold over basal
1.1
1.05
1
0.95
0.9
0.85
0
10
20
30
40
50
60
Time (min)

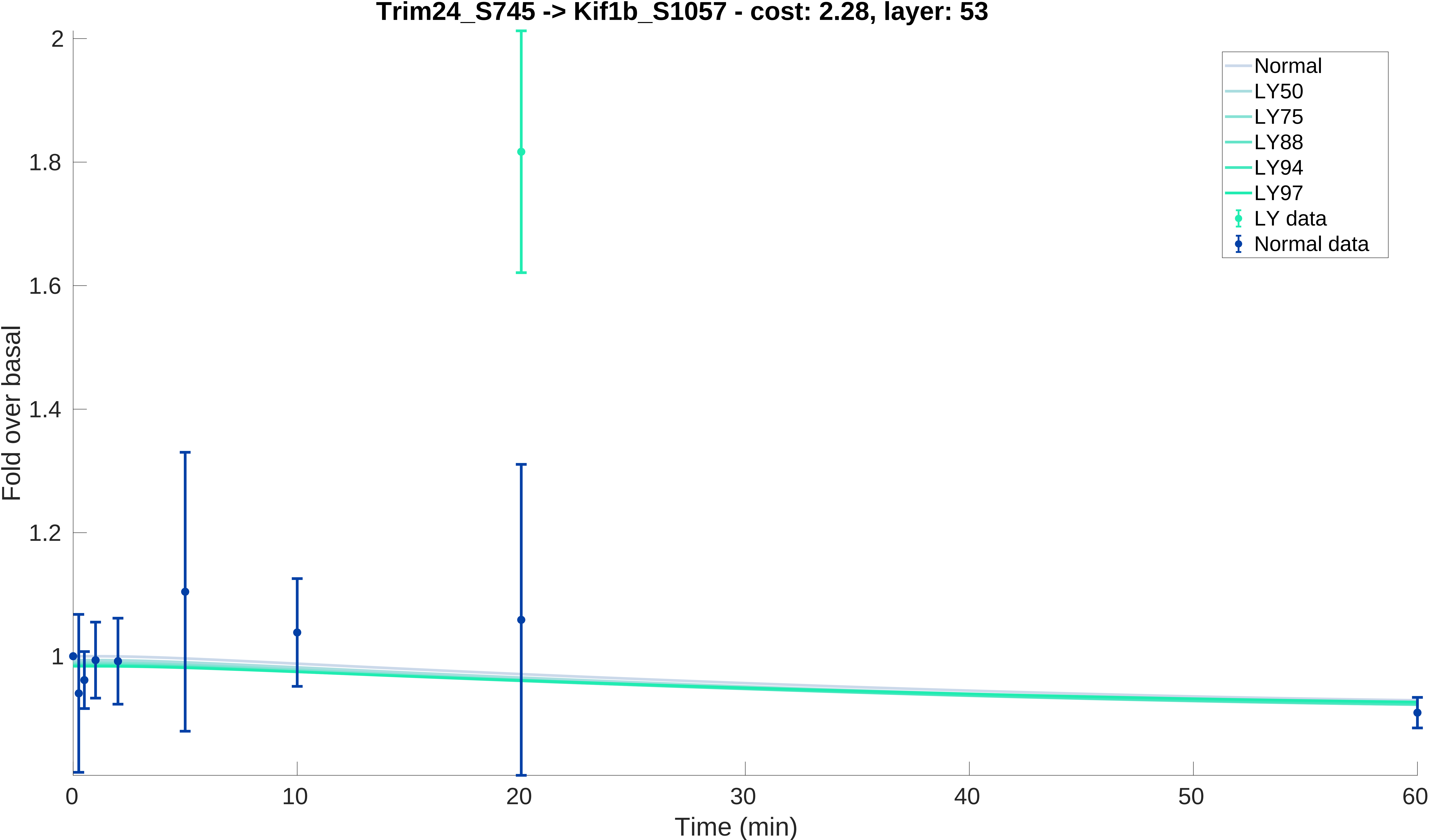
Trim24_S745 -> Kif1b_S1057 - cost: 2.28, layer: 53
Fold over basal
Time (min)
2
1.8
1.6
1.4
1.2
1
0
10
20
30
40
50
60
Normal
LY50
LY75
LY88
LY94
LY97
LY data
Normal data

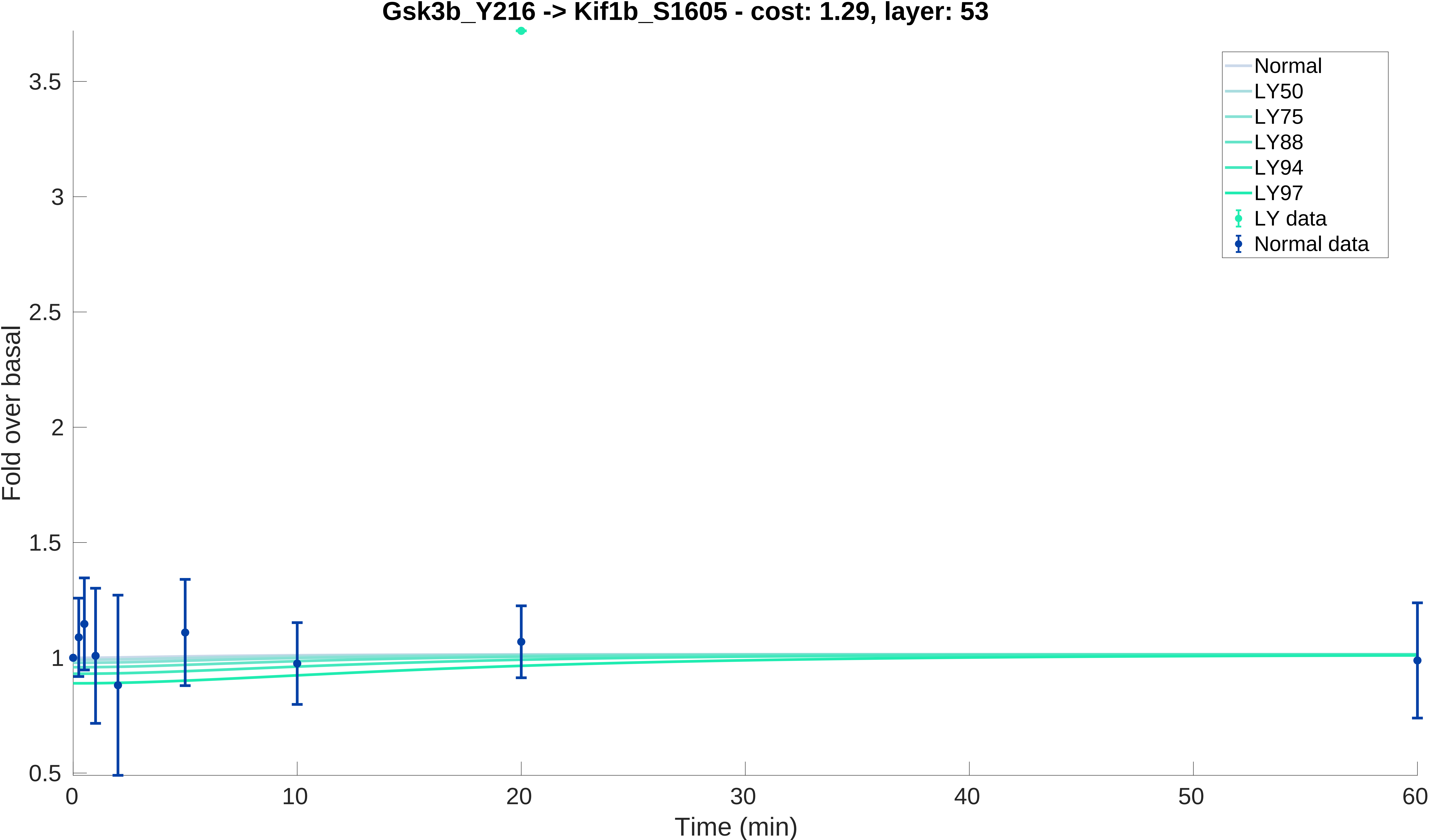

Gsk3b_Y216 -> Kif1b_S1605 - cost: 1.29, layer: 53
Normal
LY50
LY75
LY88
LY94
LY97
LY data
Normal data
Fold over basal
Time (min)
0.5
1
1.5
2
2.5
3
3.5
0
10
20
30
40
50
60

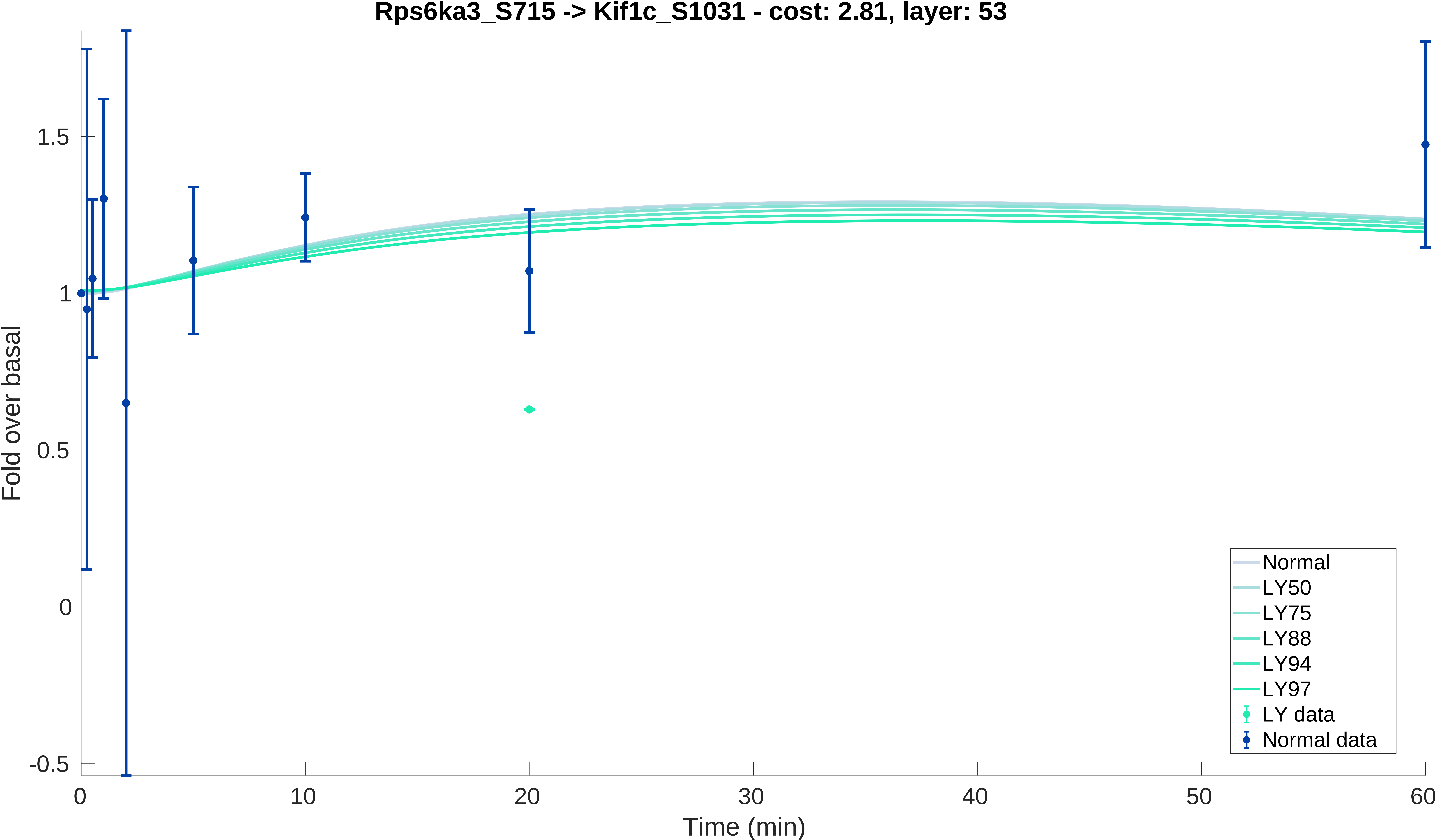

Rps6ka3_S715 -> Kif1c_S1031 - cost: 2.81, layer: 53
Fold over basal
1.5
1
0.5
0
-0.5
0
10
20
30
40
50
60
Time (min)
Normal
LY50
LY75
LY88
LY94
LY97
LY data
Normal data

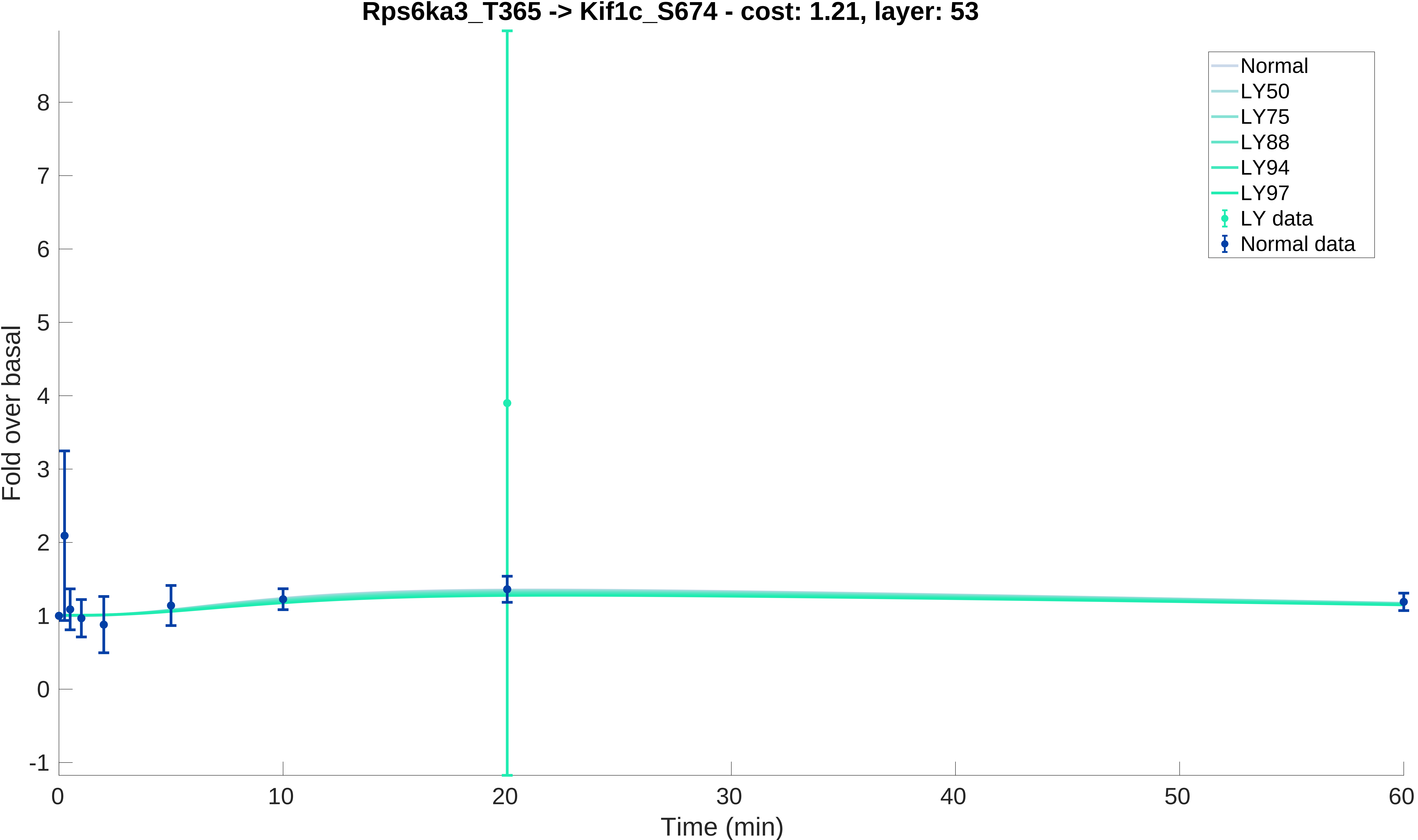

Rps6ka3_T365 -> Kif1c_S674 - cost: 1.21, layer: 53
Normal
LY50
LY75
LY88
LY94
LY97
LY data
Normal data
Fold over basal
Time (min)
-1
0
1
2
3
4
5
6
7
8
0
10
20
30
40
50
60

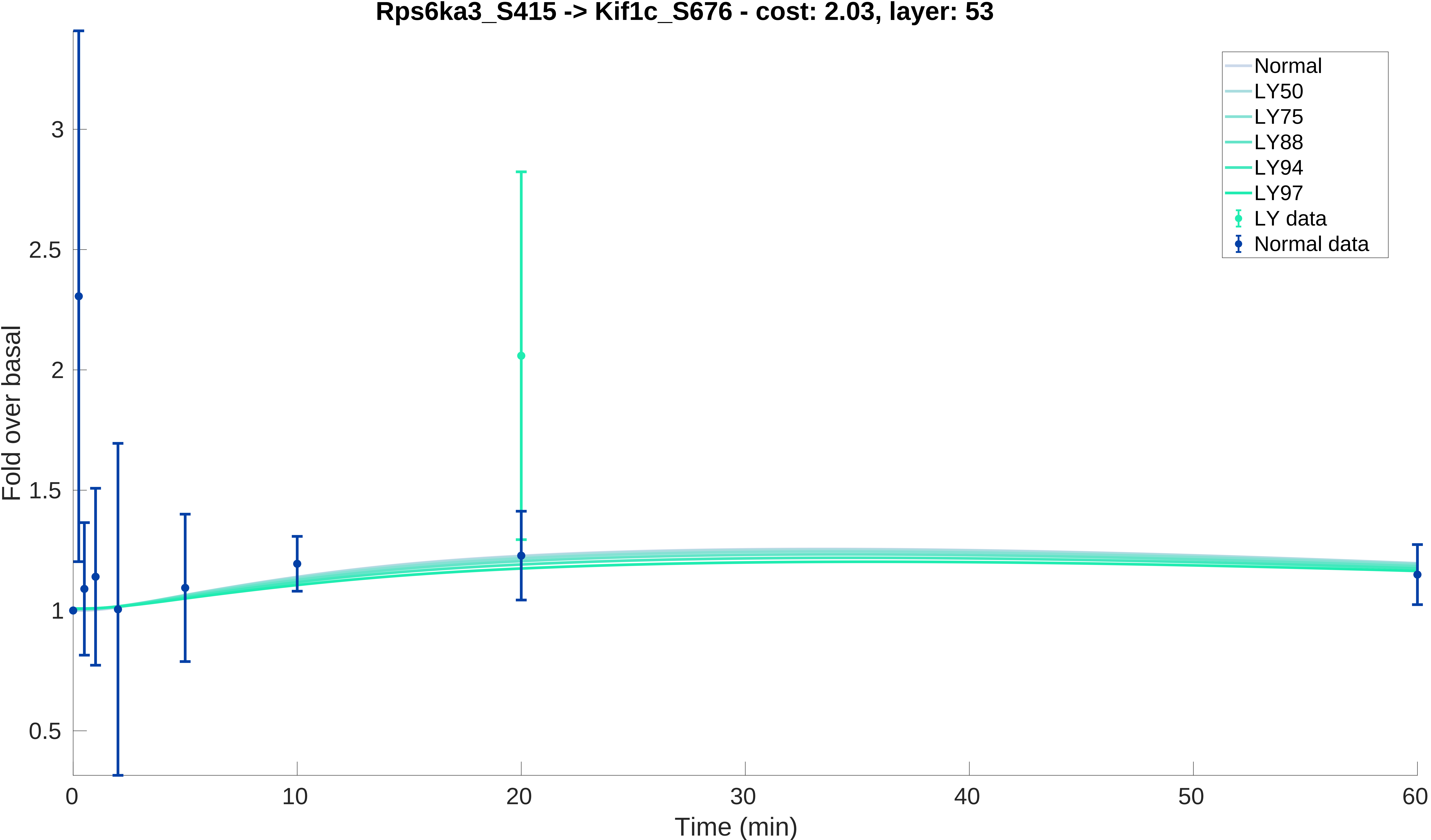
Rps6ka3_S415 -> Kif1c_S676 - cost: 2.03, layer: 53
Fold over basal
Time (min)
0.5
1
1.5
2
2.5
3
0
10
20
30
40
50
60
Normal
LY50
LY75
LY88
LY94
LY97
LY data
Normal data

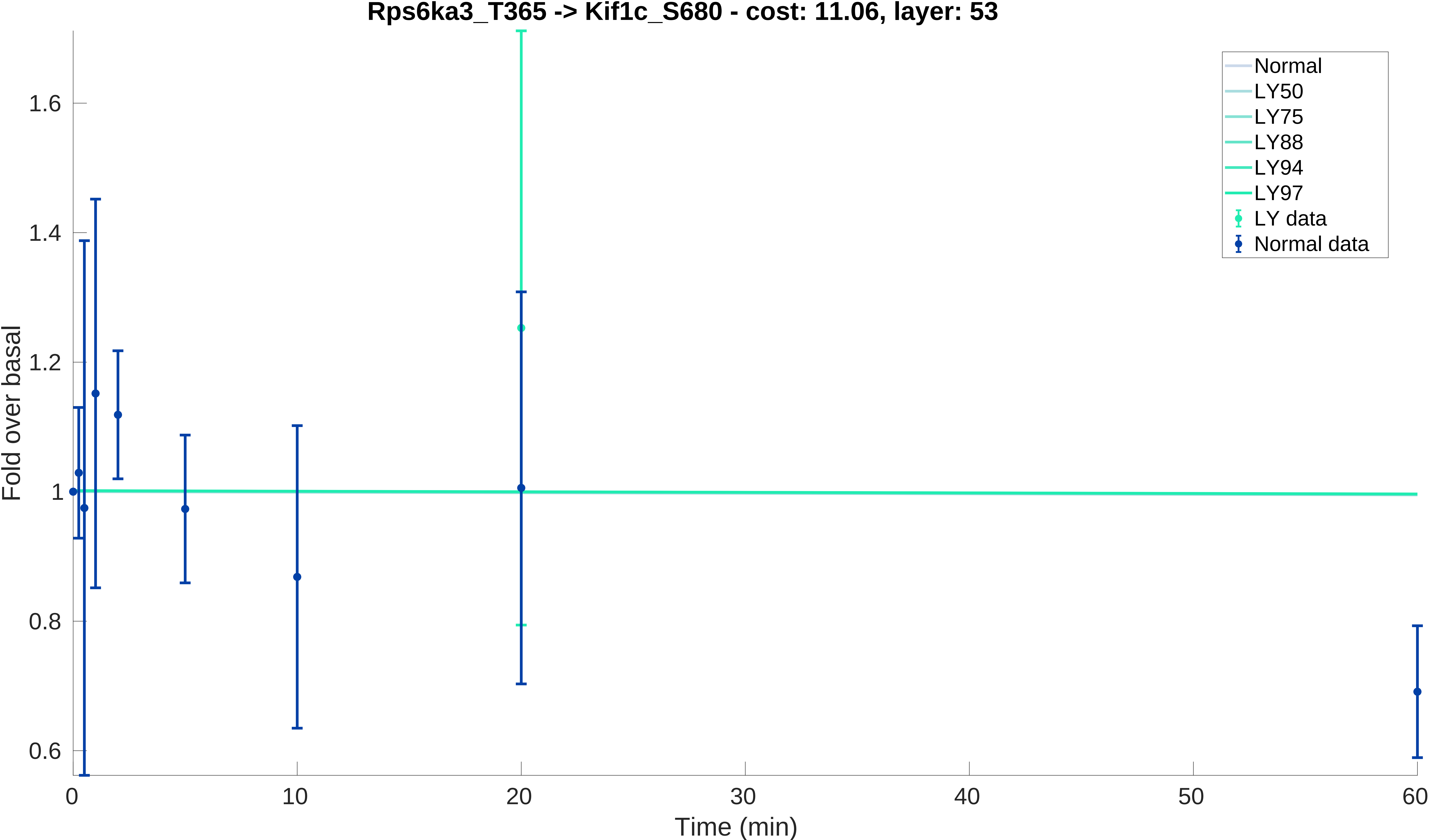

Rps6ka3_T365 -> Kif1c_S680 - cost: 11.06, layer: 53
Fold over basal
1.6
1.4
1.2
1
0.8
0.6
0
10
20
30
40
50
60
Time (min)
Normal
LY50
LY75
LY88
LY94
LY97
LY data
Normal data

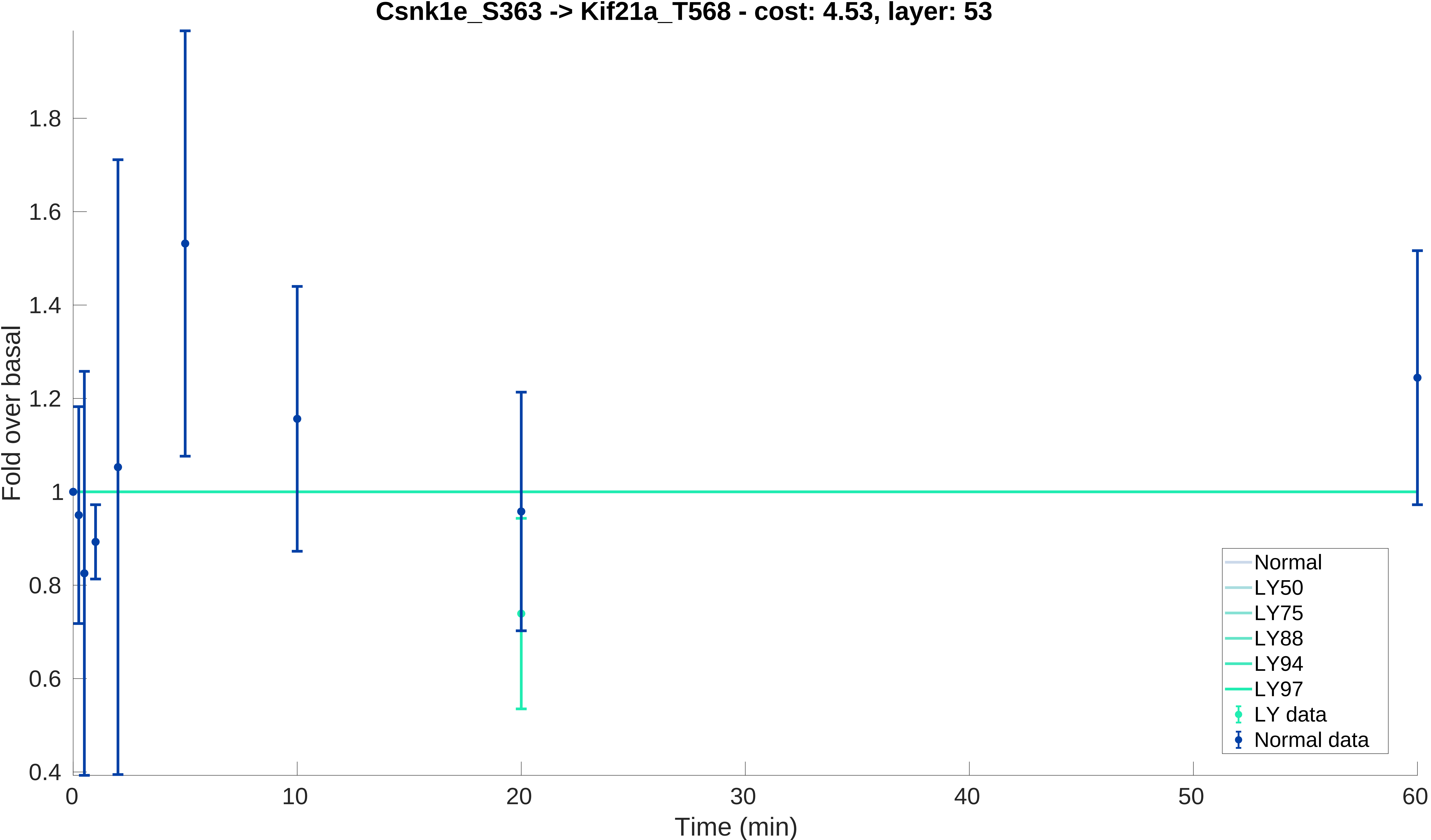

Csnk1e_S363 -> Kif21a_T568 - cost: 4.53, layer: 53
Fold over basal
Time (min)
1.8
1.6
1.4
1.2
1
0.8
0.6
0.4
0
10
20
30
40
50
60
Normal
LY50
LY75
LY88
LY94
LY97
LY data
Normal data

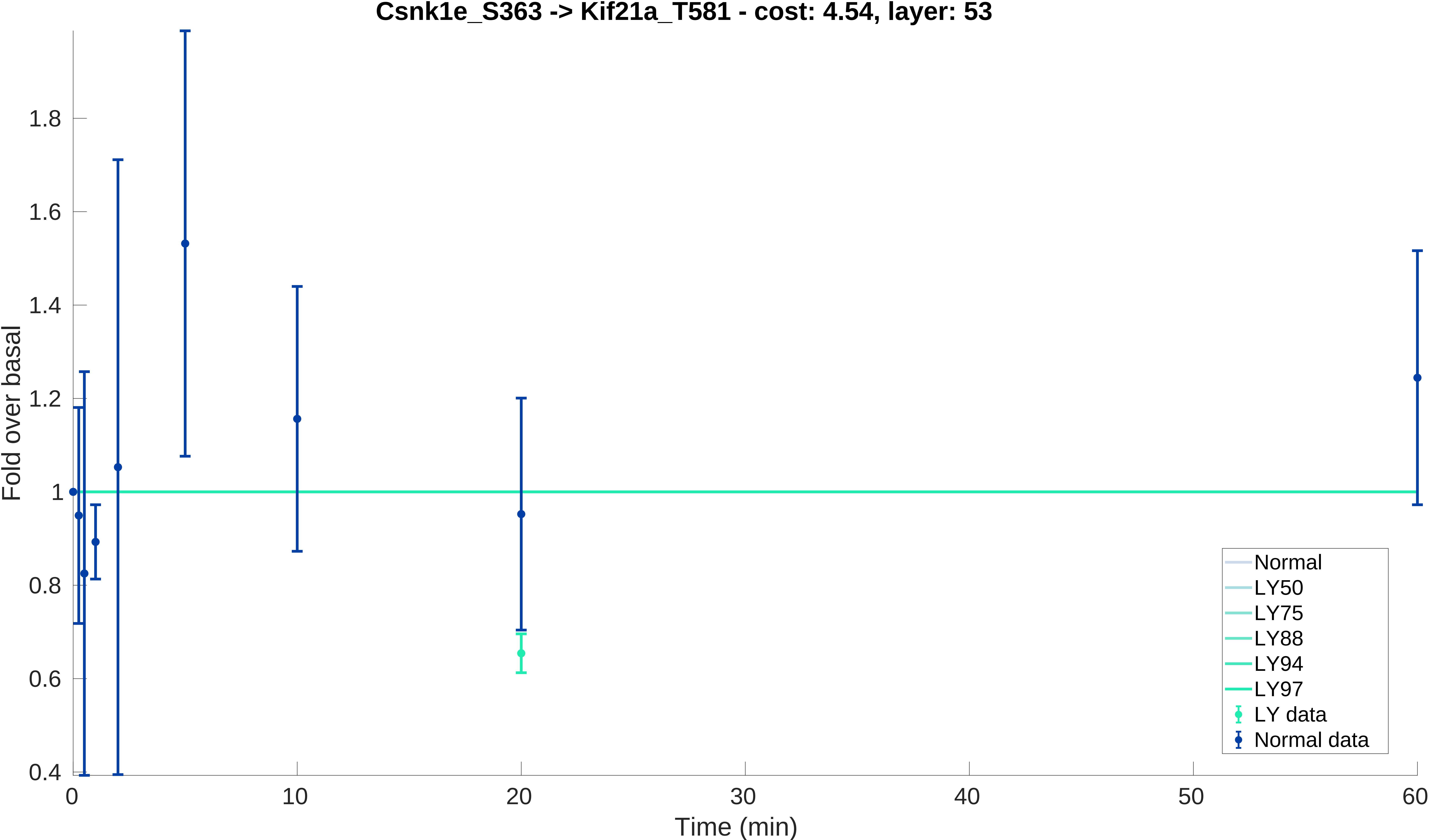
Csnk1e_S363 -> Kif21a_T581 - cost: 4.54, layer: 53
Fold over basal
1.8
1.6
1.4
1.2
1
0.8
0.6
0.4
0
10
20
30
40
50
60
Time (min)
Normal
LY50
LY75
LY88
LY94
LY97
LY data
Normal data

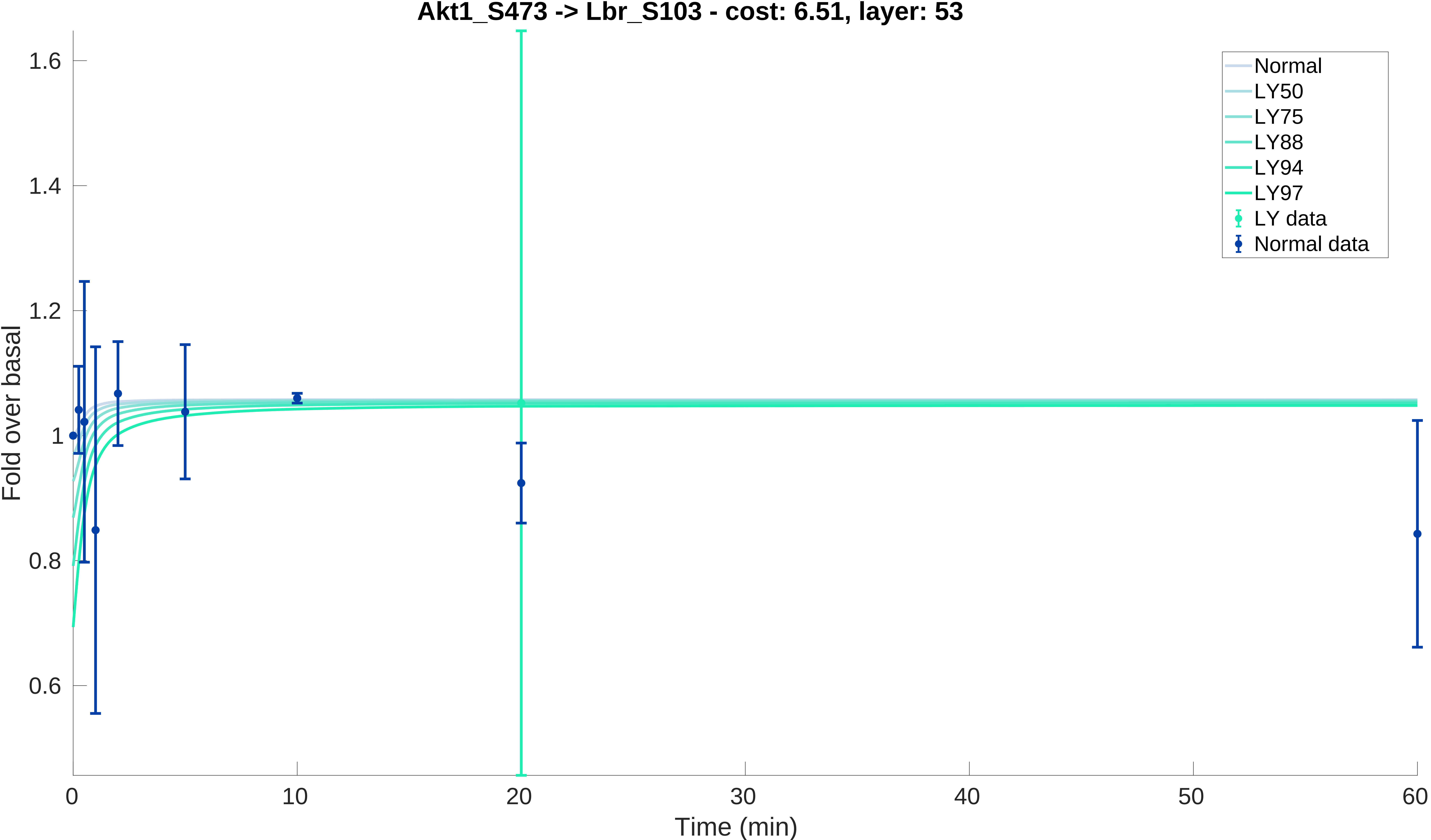
Akt1_S473 -> Lbr_S103 - cost: 6.51, layer: 53
Fold over basal
Time (min)
0.6
0.8
1
1.2
1.4
1.6
0
10
20
30
40
50
60
Normal
LY50
LY75
LY88
LY94
LY97
LY data
Normal data

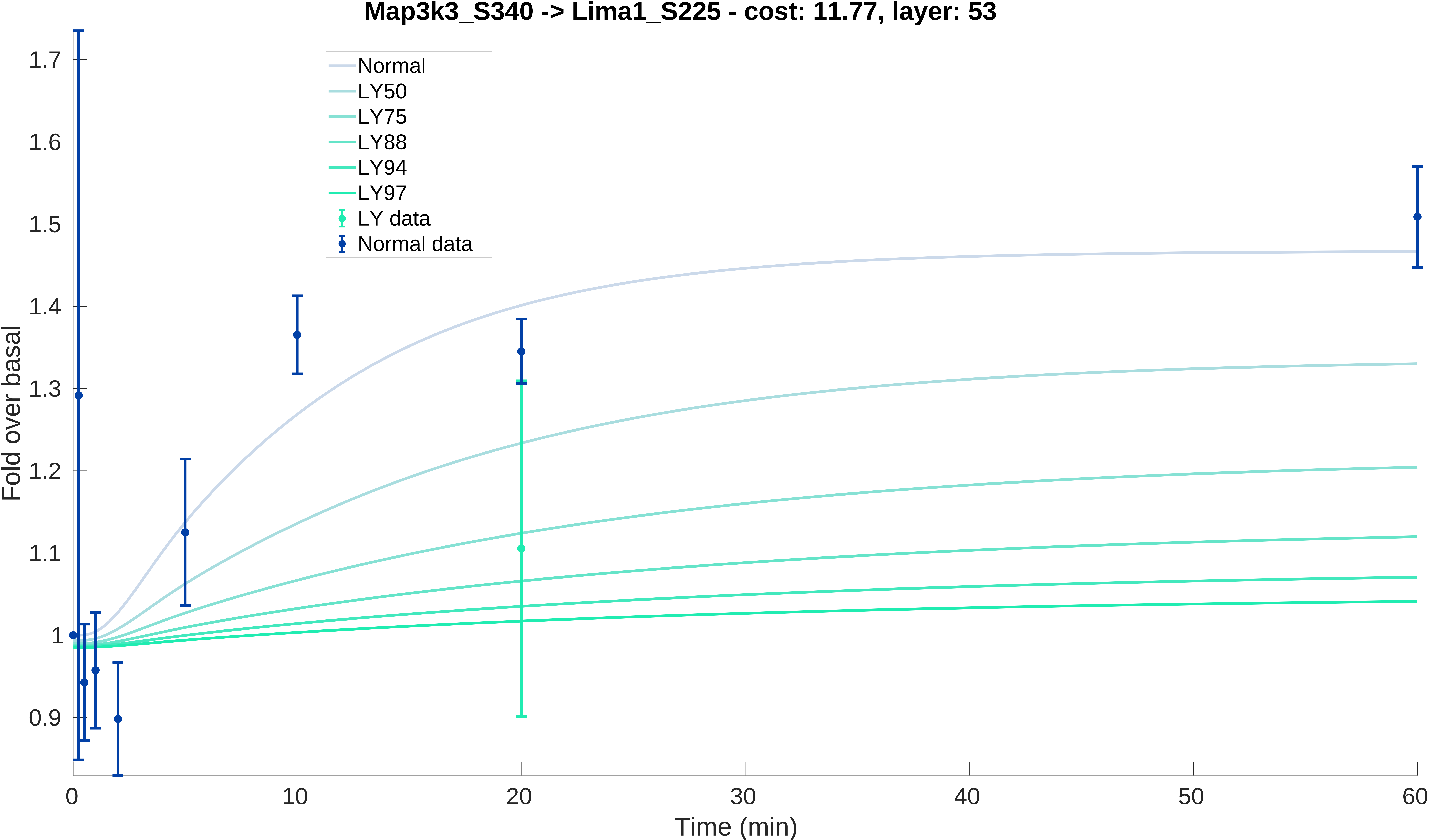

Map3k3_S340 -> Lima1_S225 - cost: 11.77, layer: 53
Normal
LY50
LY75
LY88
LY94
LY97
LY data
Normal data
Fold over basal
1.7
1.6
1.5
1.4
1.3
1.2
1.1
1
0.9
0
10
20
30
40
50
60
Time (min)

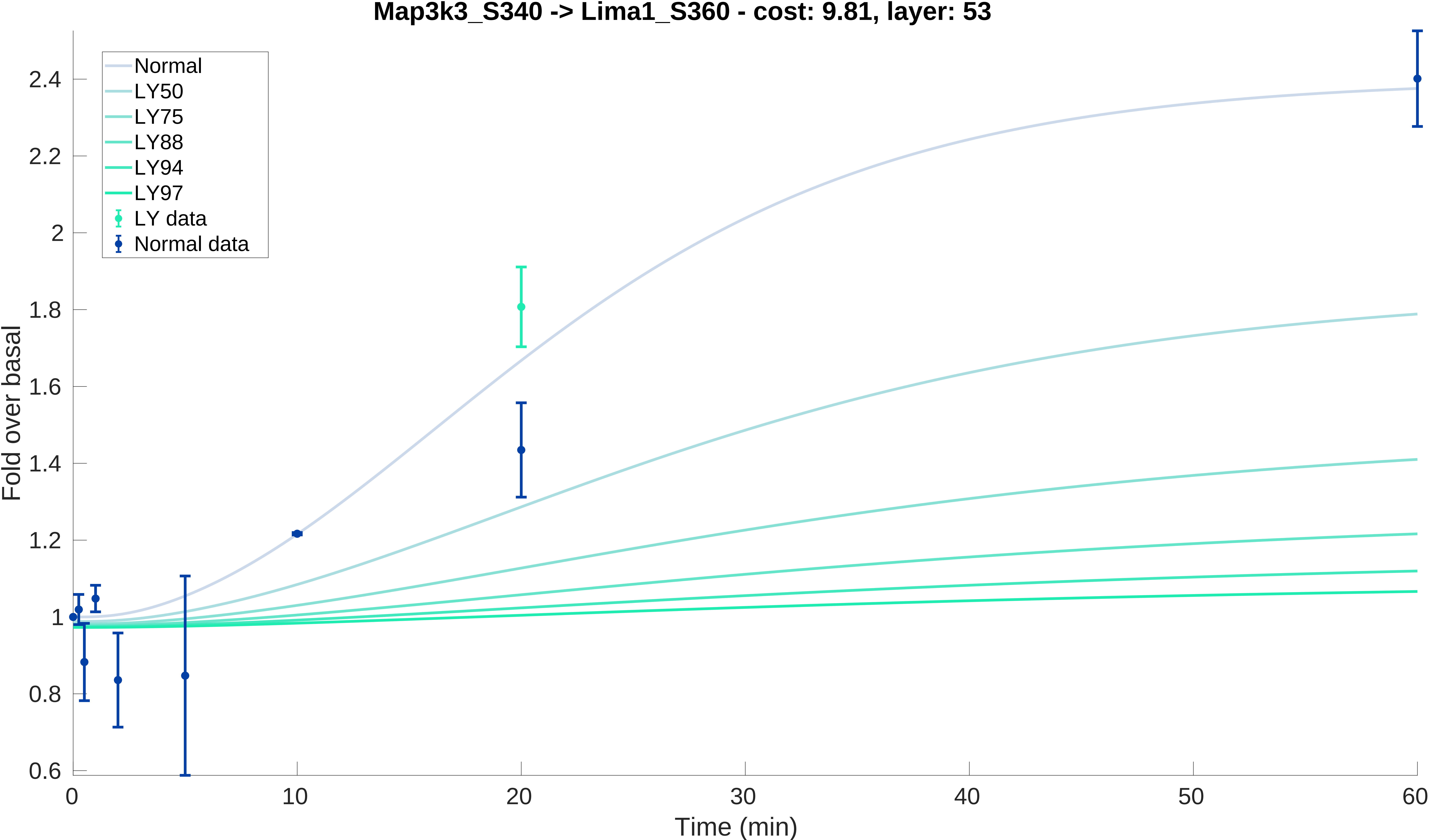
Map3k3_S340 -> Lima1_S360 - cost: 9.81, layer: 53
Normal
LY50
LY75
LY88
LY94
LY97
LY data
Normal data
Fold over basal
Time (min)
0.6
0.8
1
1.2
1.4
1.6
1.8
2
2.2
2.4
0
10
20
30
40
50
60

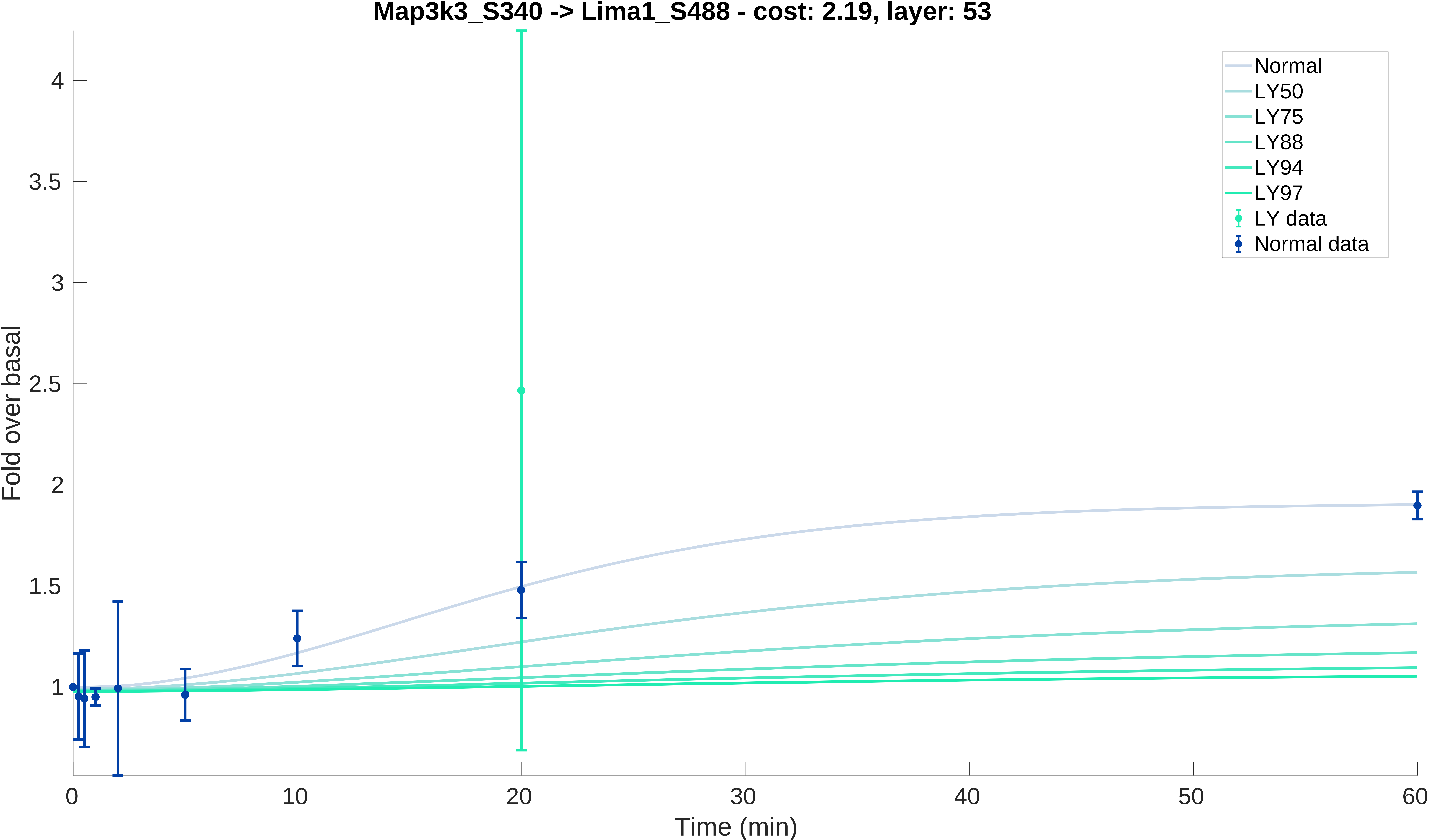

Map3k3_S340 -> Lima1_S488 - cost: 2.19, layer: 53
Normal
LY50
LY75
LY88
LY94
LY97
LY data
Normal data
Fold over basal
Time (min)
4
3.5
3
2.5
2
1.5
1
0
10
20
30
40
50
60

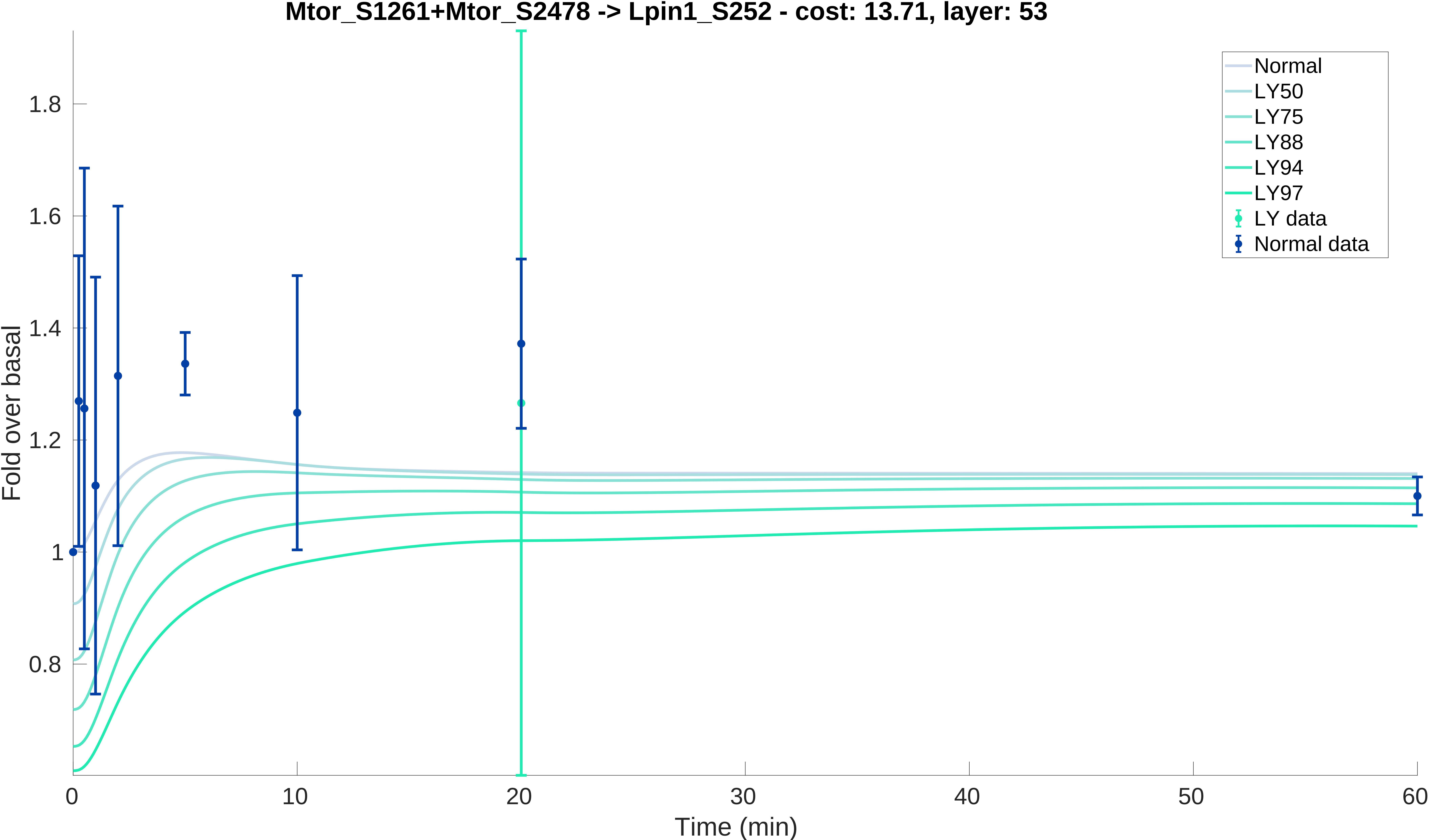

Mtor_S1261+Mtor_S2478 -> Lpin1_S252 - cost: 13.71, layer: 53
Normal
LY50
LY75
LY88
LY94
LY97
LY data
Normal data
Fold over basal
1.8
1.6
1.4
1.2
1
0.8
0
10
20
30
40
50
60
Time (min)

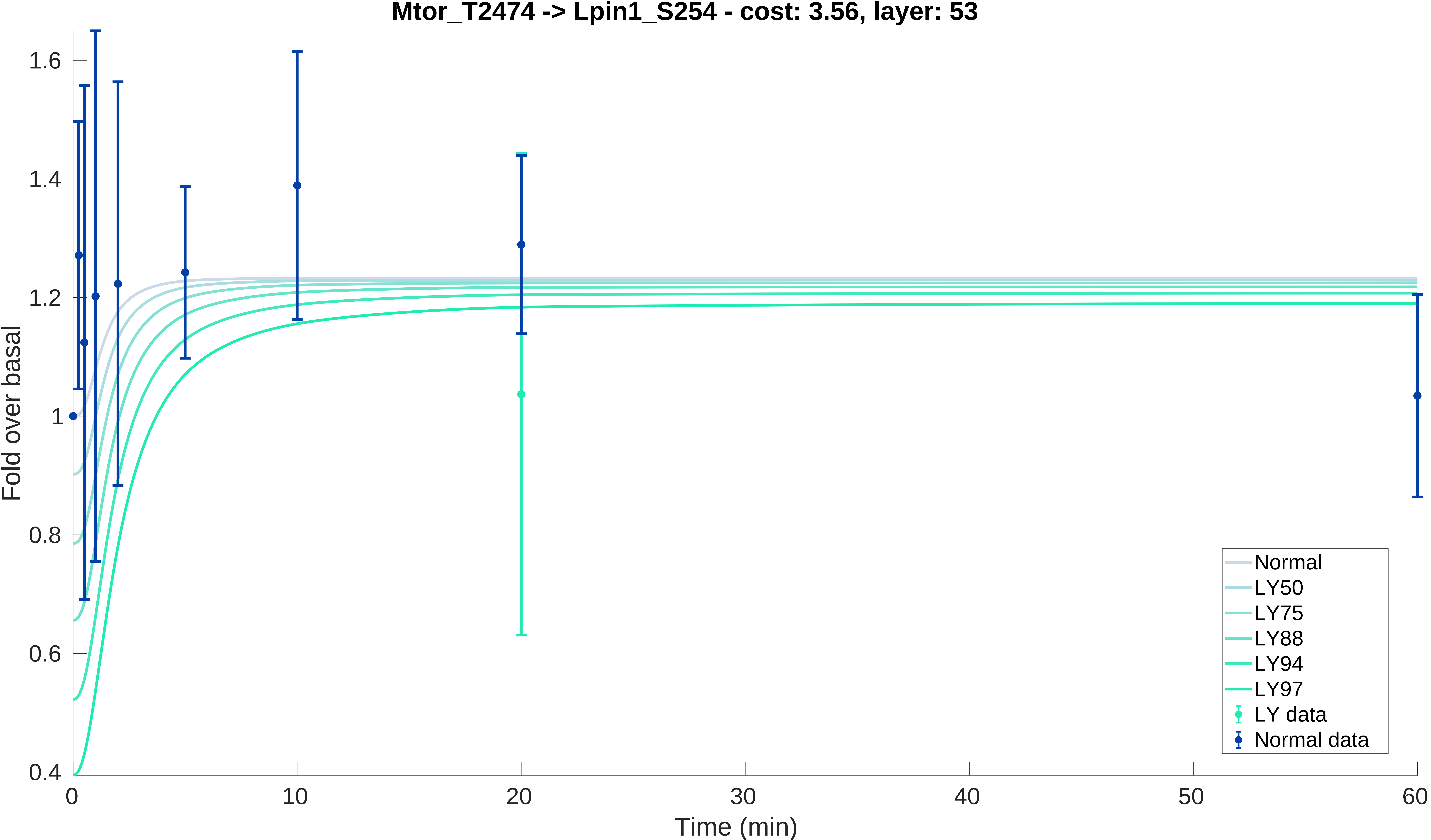

Mtor_T2474 -> Lpin1_S254 - cost: 3.56, layer: 53
Fold over basal
1.6
1.4
1.2
1
0.8
0.6
0.4
0
10
20
30
40
50
60
Time (min)
Normal
LY50
LY75
LY88
LY94
LY97
LY data
Normal data

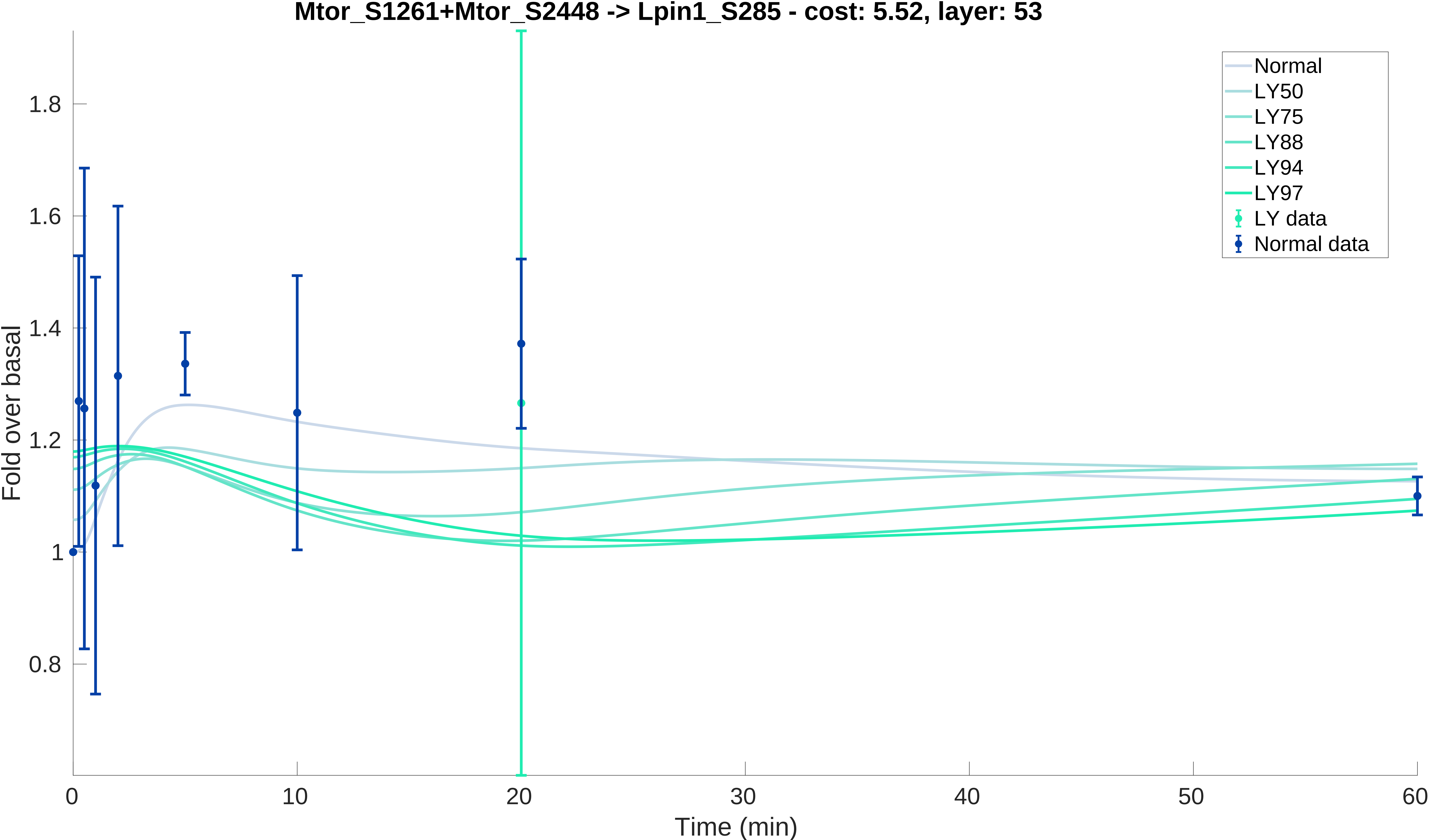

Mtor_S1261+Mtor_S2448 -> Lpin1_S285 - cost: 5.52, layer: 53
Fold over basal
Time (min)
Normal
LY50
LY75
LY88
LY94
LY97
LY data
Normal data
0.8
1
1.2
1.4
1.6
1.8
0
10
20
30
40
50
60

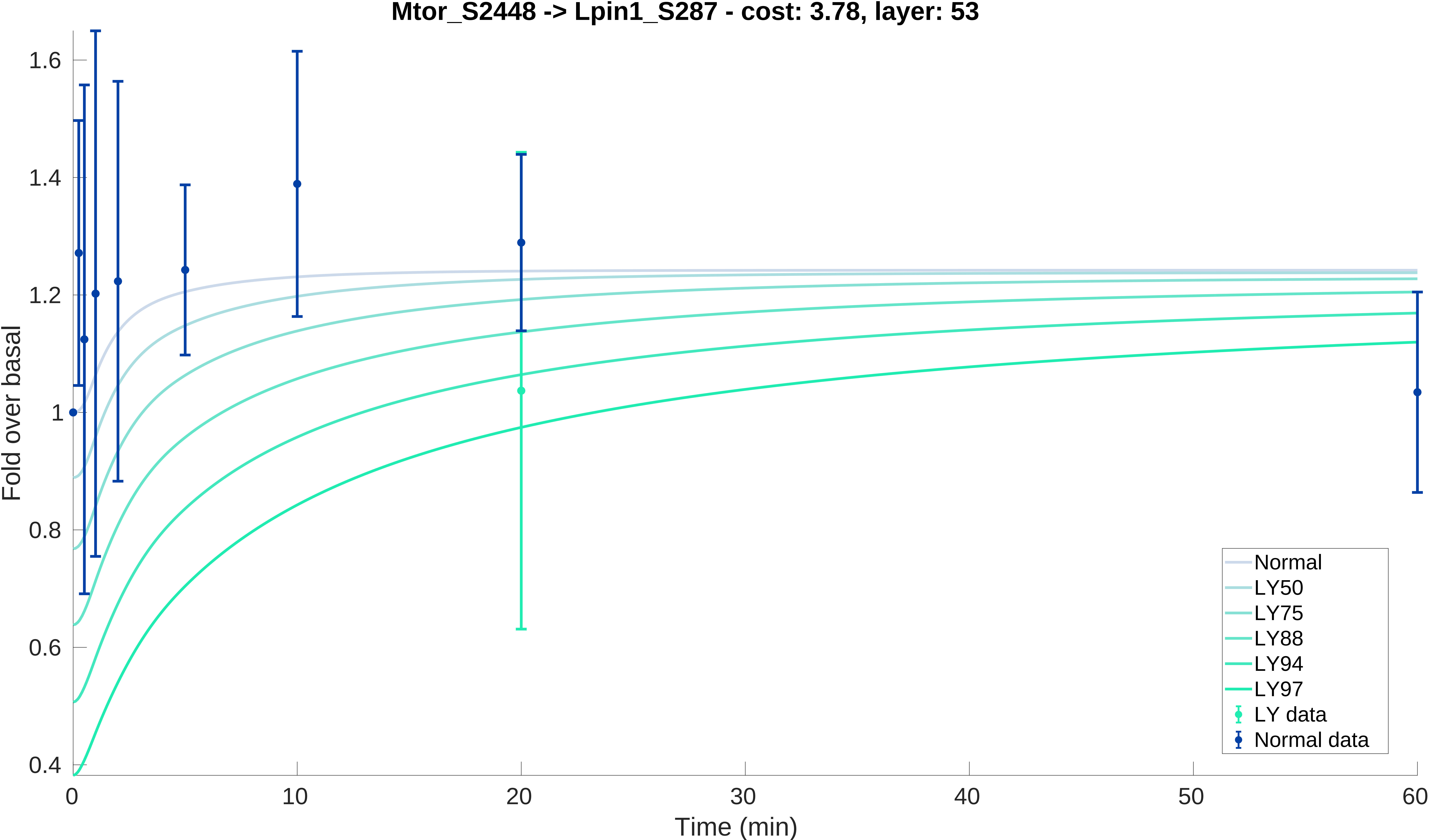

Mtor_S2448 -> Lpin1_S287 - cost: 3.78, layer: 53
Fold over basal
1.6
1.4
1.2
1
0.8
0.6
0.4
0
10
20
30
40
50
60
Time (min)
Normal
LY50
LY75
LY88
LY94
LY97
LY data
Normal data

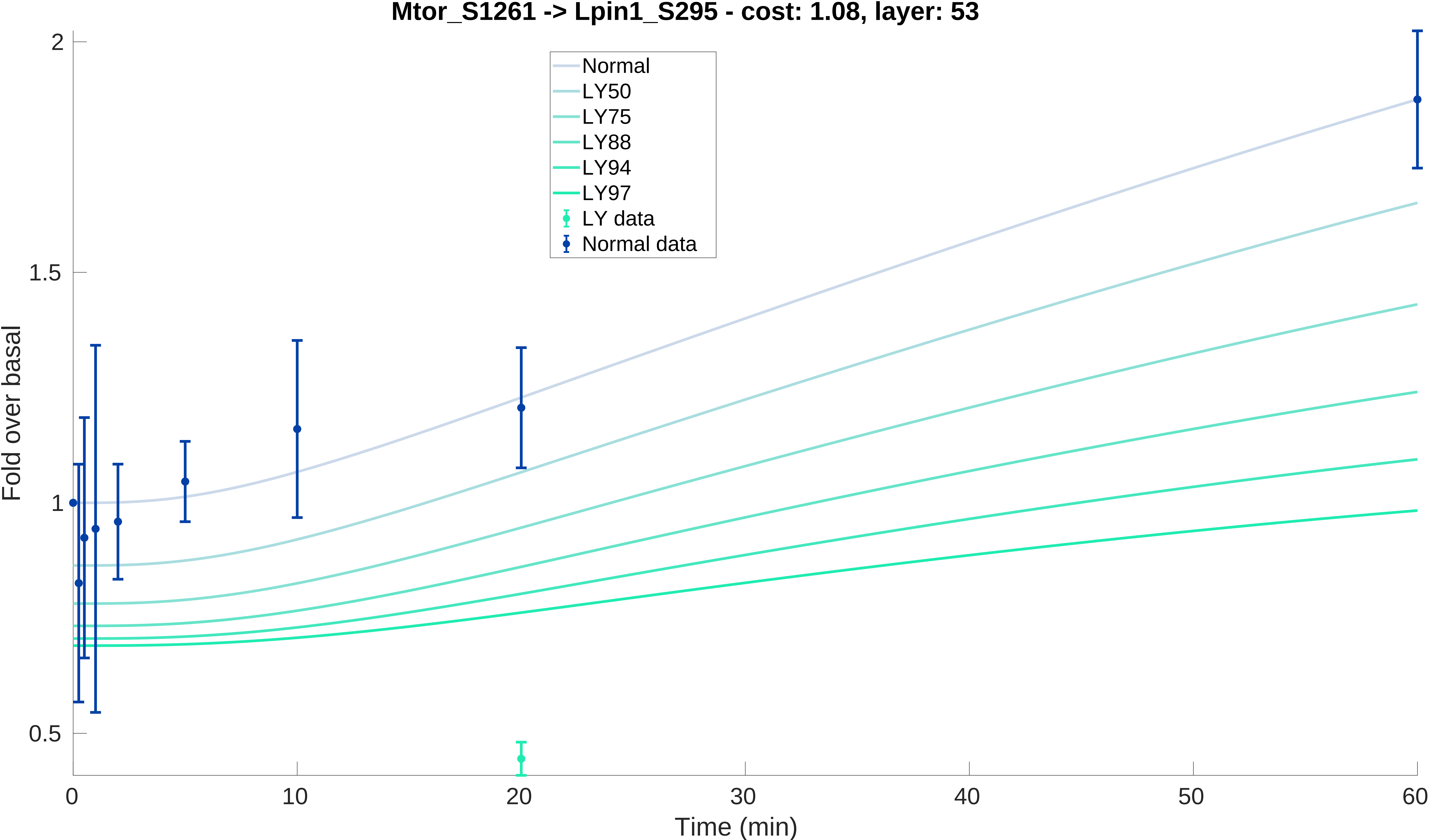
Mtor_S1261 -> Lpin1_S295 - cost: 1.08, layer: 53
Normal
LY50
LY75
LY88
LY94
LY97
LY data
Normal data
Fold over basal
Time (min)
0.5
1
1.5
2
0
10
20
30
40
50
60

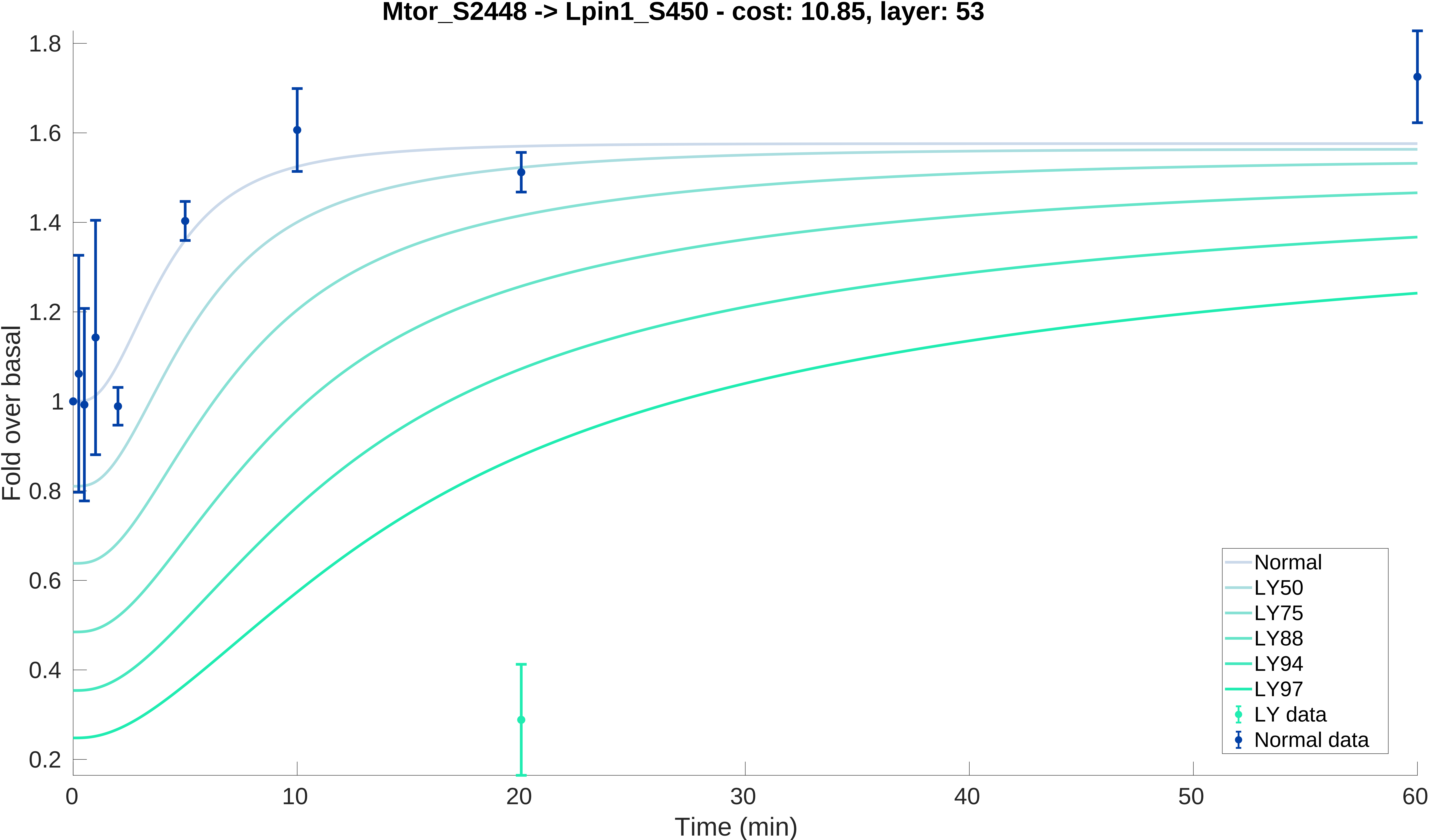
Mtor_S2448 -> Lpin1_S450 - cost: 10.85, layer: 53
Fold over basal
Time (min)
0.2
0.4
0.6
0.8
1
1.2
1.4
1.6
1.8
0
10
20
30
40
50
60
Normal
LY50
LY75
LY88
LY94
LY97
LY data
Normal data

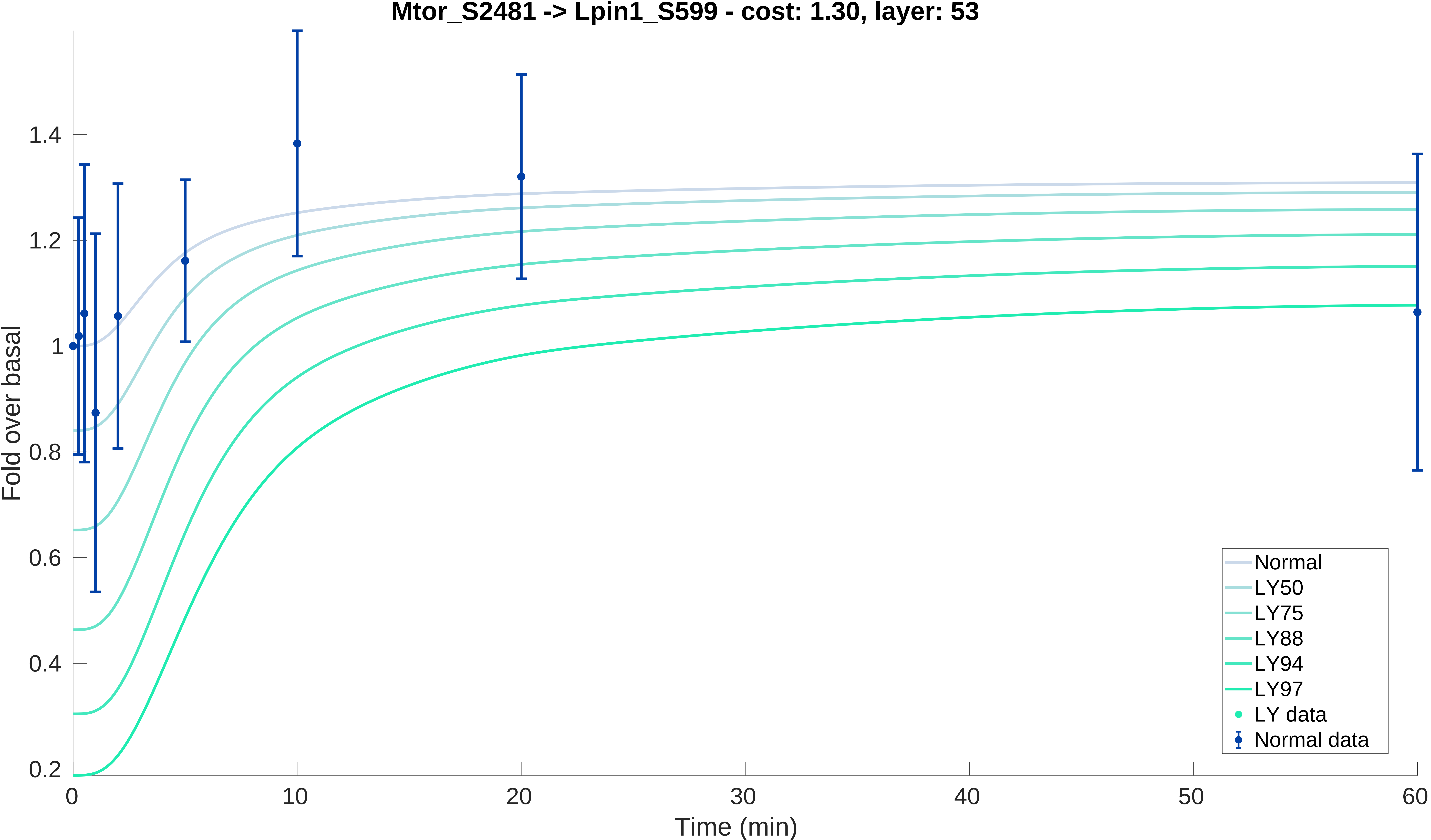

Mtor_S2481 -> Lpin1_S599 - cost: 1.30, layer: 53
Fold over basal
1.4
1.2
1
0.8
0.6
0.4
0.2
0
10
20
30
40
50
60
Time (min)
Normal
LY50
LY75
LY88
LY94
LY97
LY data
Normal data

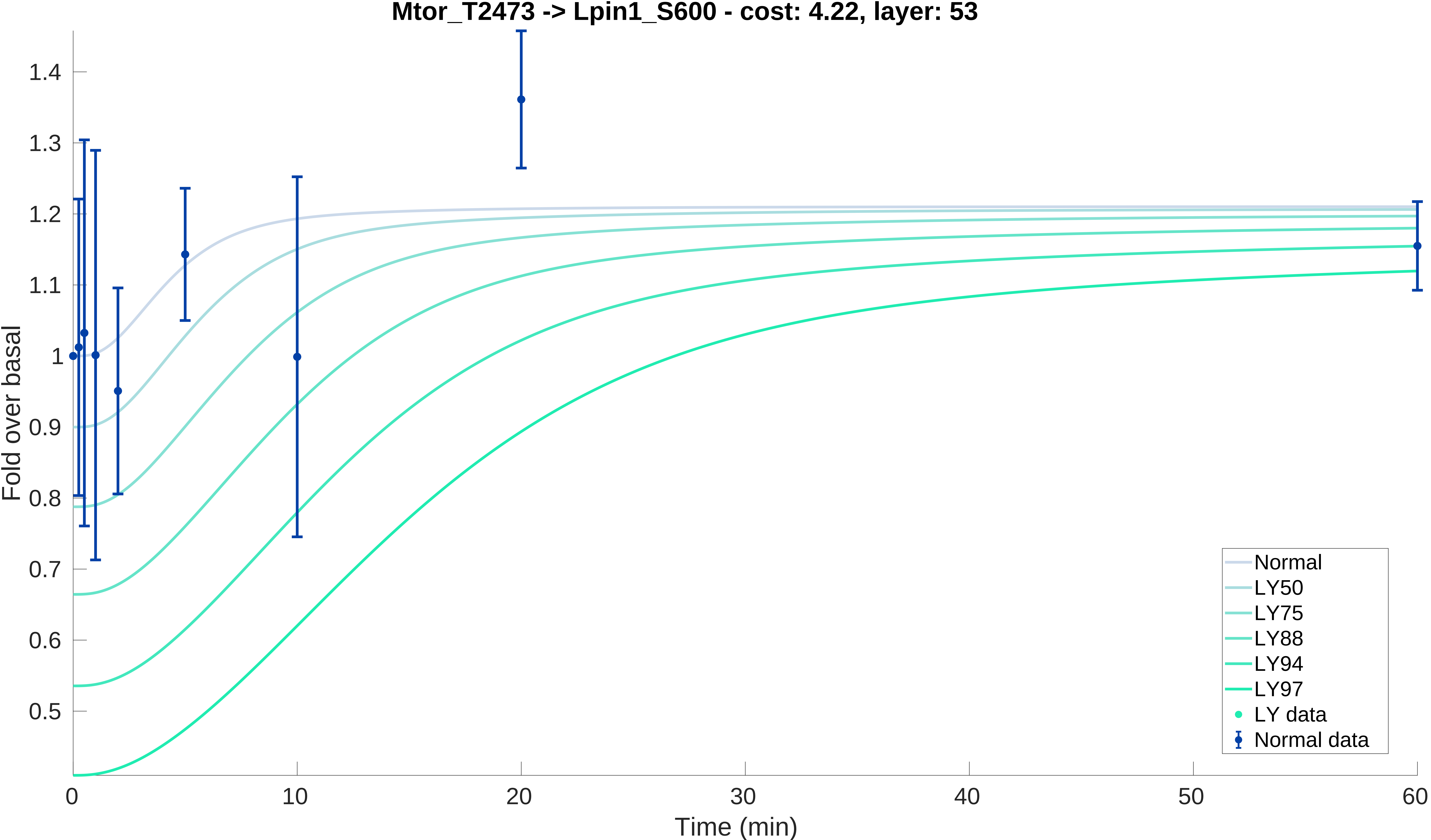
Mtor_T2473 -> Lpin1_S600 - cost: 4.22, layer: 53
Fold over basal
1.4
1.3
1.2
1.1
1
0.9
0.8
0.7
0.6
0.5
0
10
20
30
40
50
60
Time (min)
Normal
LY50
LY75
LY88
LY94
LY97
LY data
Normal data

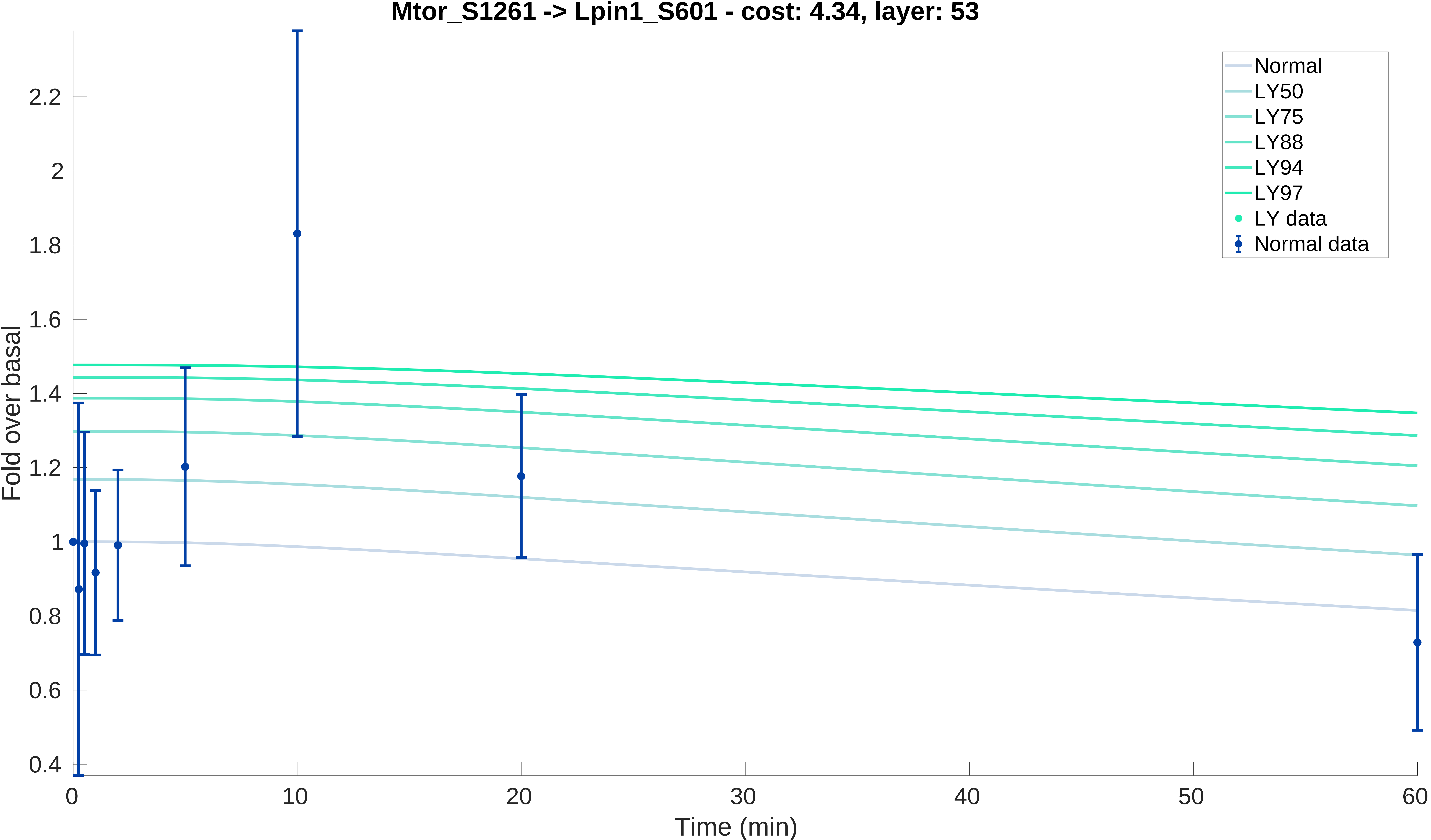

Mtor_S1261 -> Lpin1_S601 - cost: 4.34, layer: 53
Fold over basal
Time (min)
2.2
2
1.8
1.6
1.4
1.2
1
0.8
0.6
0.4
0
10
20
30
40
50
60
Normal
LY50
LY75
LY88
LY94
LY97
LY data
Normal data

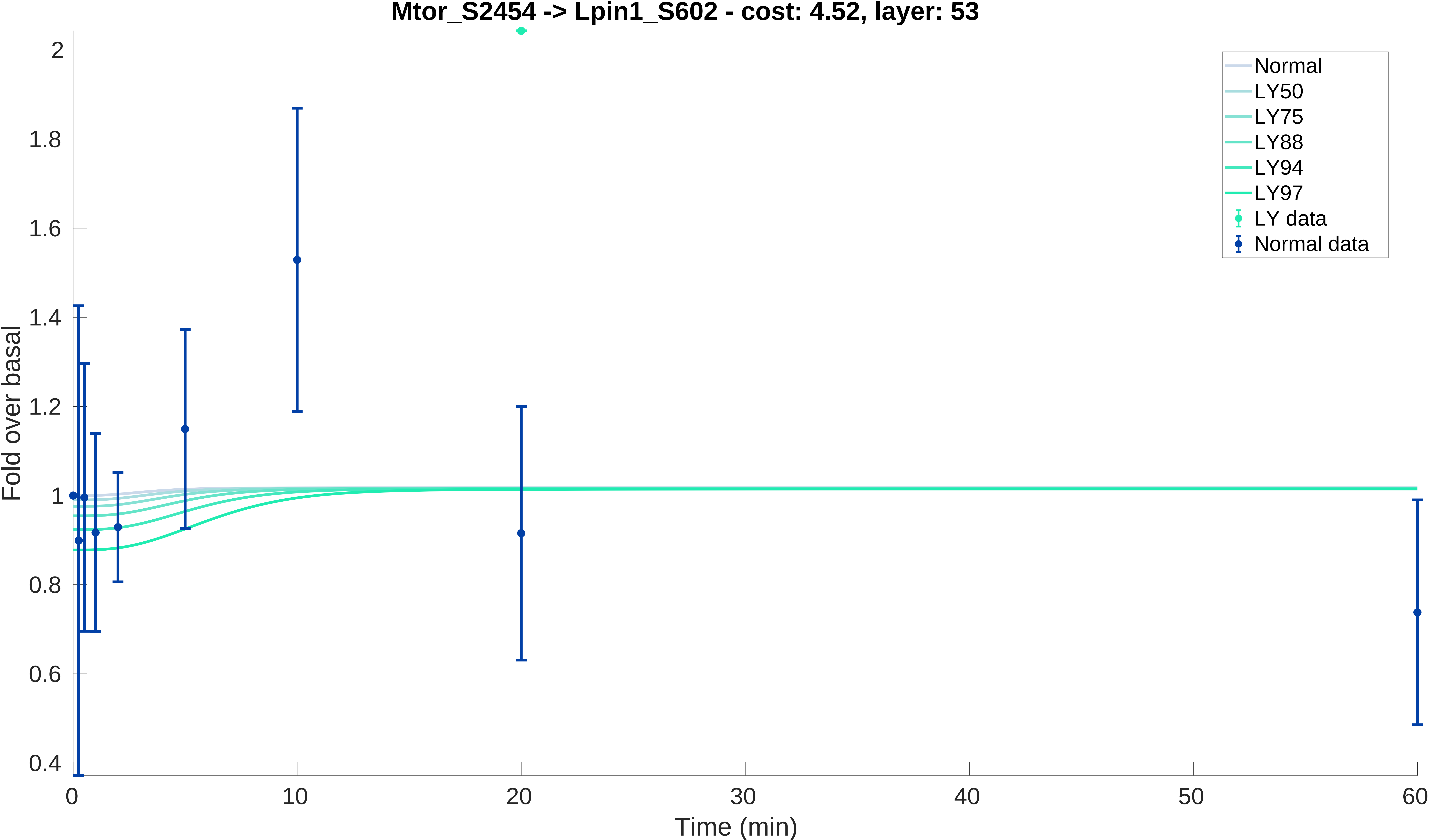

Mtor_S2454 -> Lpin1_S602 - cost: 4.52, layer: 53
Fold over basal
Time (min)
0.4
0.6
0.8
1
1.2
1.4
1.6
1.8
2
0
10
20
30
40
50
60
Normal
LY50
LY75
LY88
LY94
LY97
LY data
Normal data

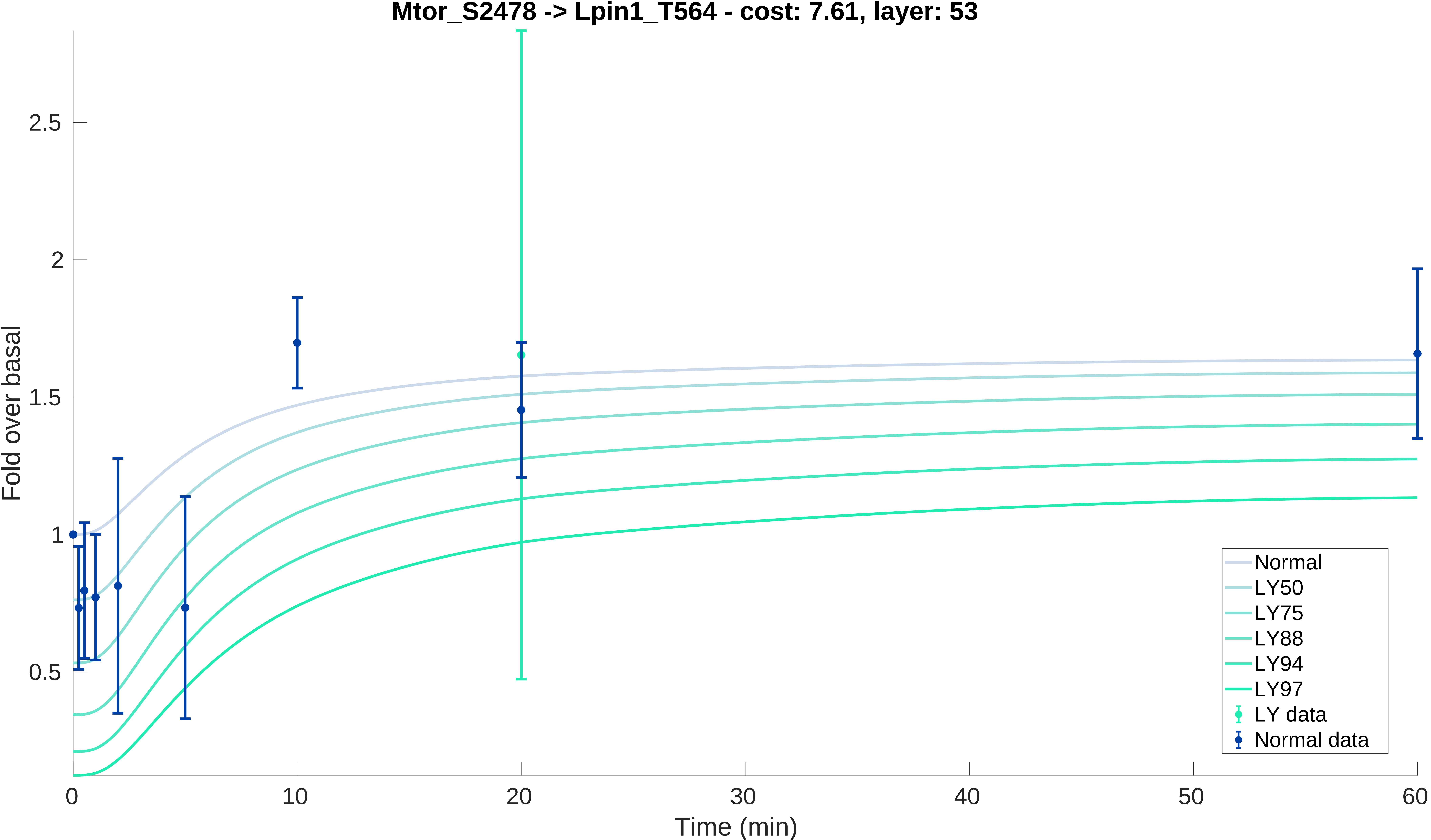

Mtor_S2478 -> Lpin1_T564 - cost: 7.61, layer: 53
Fold over basal
2.5
2
1.5
1
0.5
0
10
20
30
40
50
60
Time (min)
Normal
LY50
LY75
LY88
LY94
LY97
LY data
Normal data

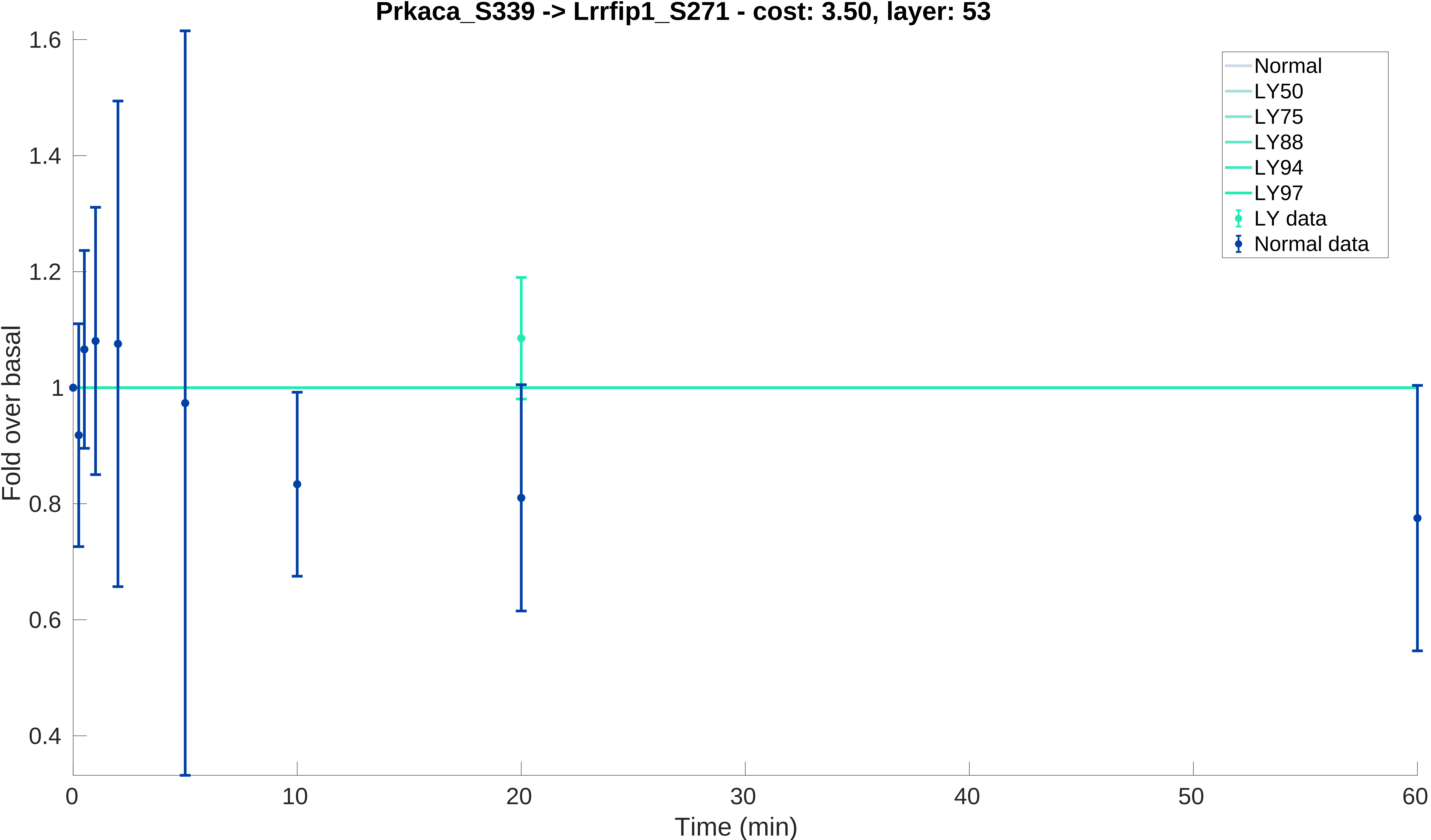
Prkaca_S339 -> Lrrfip1_S271 - cost: 3.50, layer: 53
Fold over basal
Time (min)
1.6
1.4
1.2
1
0.8
0.6
0.4
0
10
20
30
40
50
60
Normal
LY50
LY75
LY88
LY94
LY97
LY data
Normal data

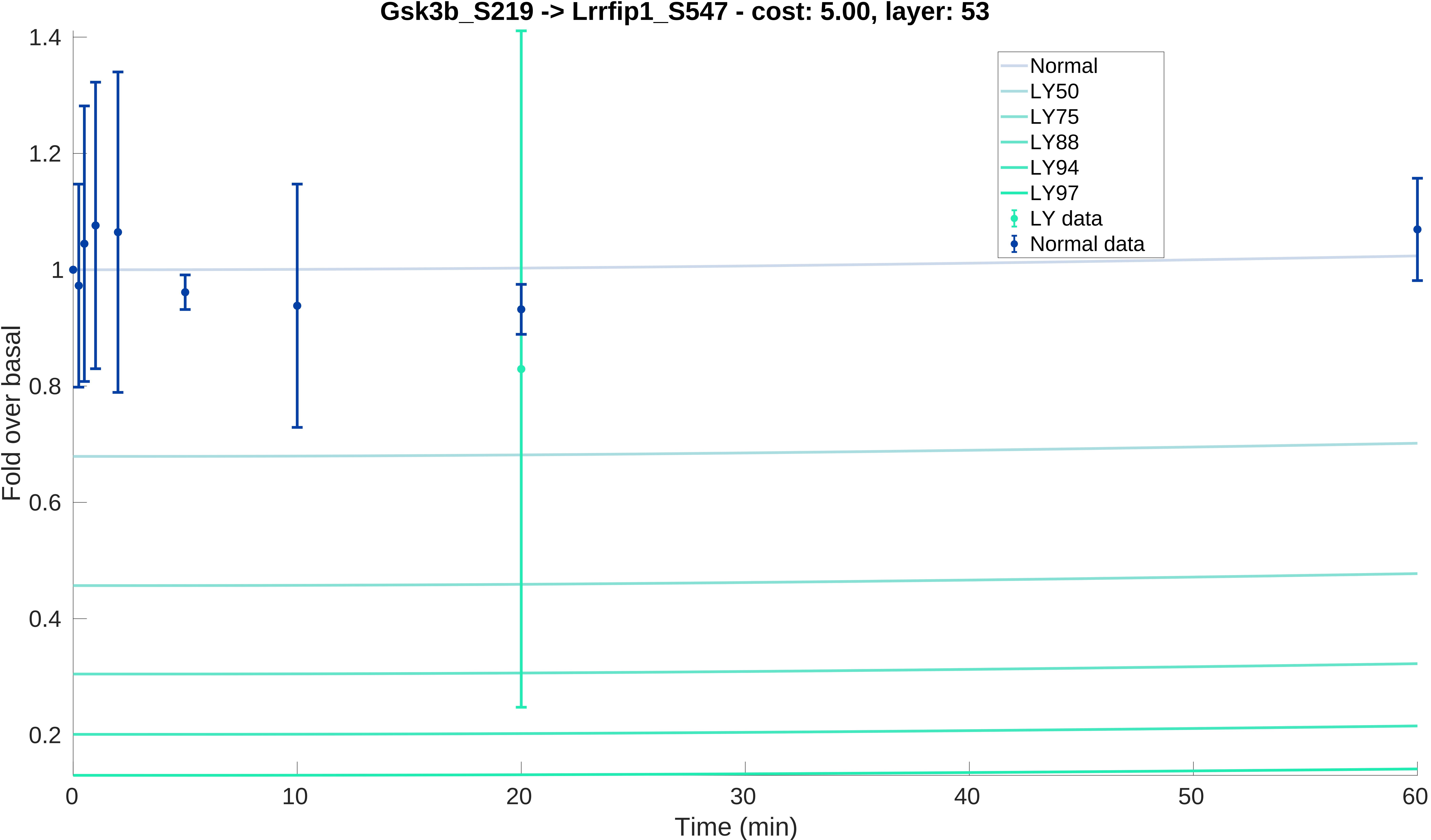

Gsk3b_S219 -> Lrrfip1_S547 - cost: 5.00, layer: 53
Fold over basal
Time (min)
1.4
1.2
1
0.8
0.6
0.4
0.2
0
10
20
30
40
50
60
Normal
LY50
LY75
LY88
LY94
LY97
LY data
Normal data

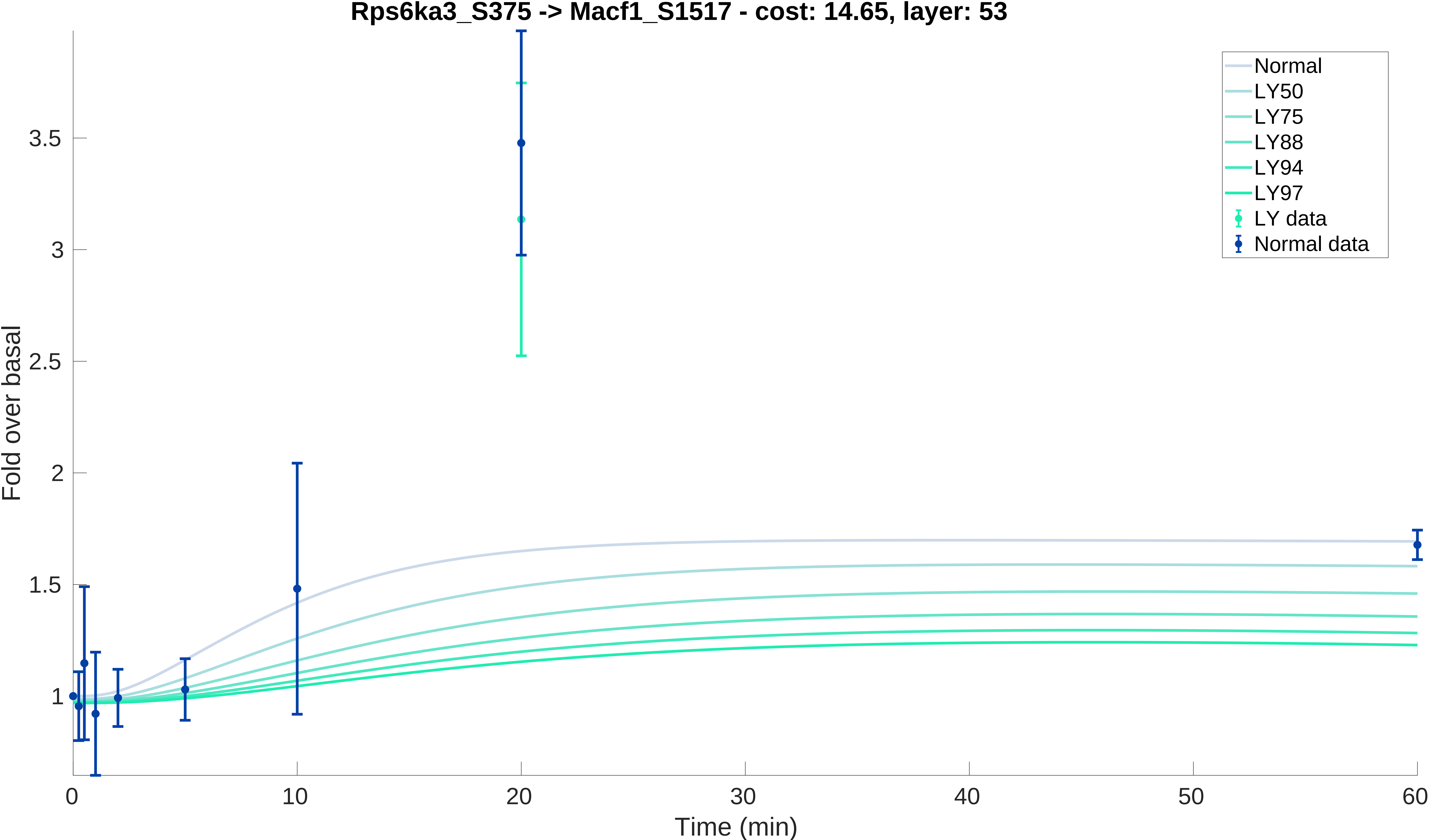
Rps6ka3_S375 -> Macf1_S1517 - cost: 14.65, layer: 53
Normal
LY50
LY75
LY88
LY94
LY97
LY data
Normal data
Fold over basal
1
1.5
2
2.5
3
3.5
0
10
20
30
40
50
60
Time (min)

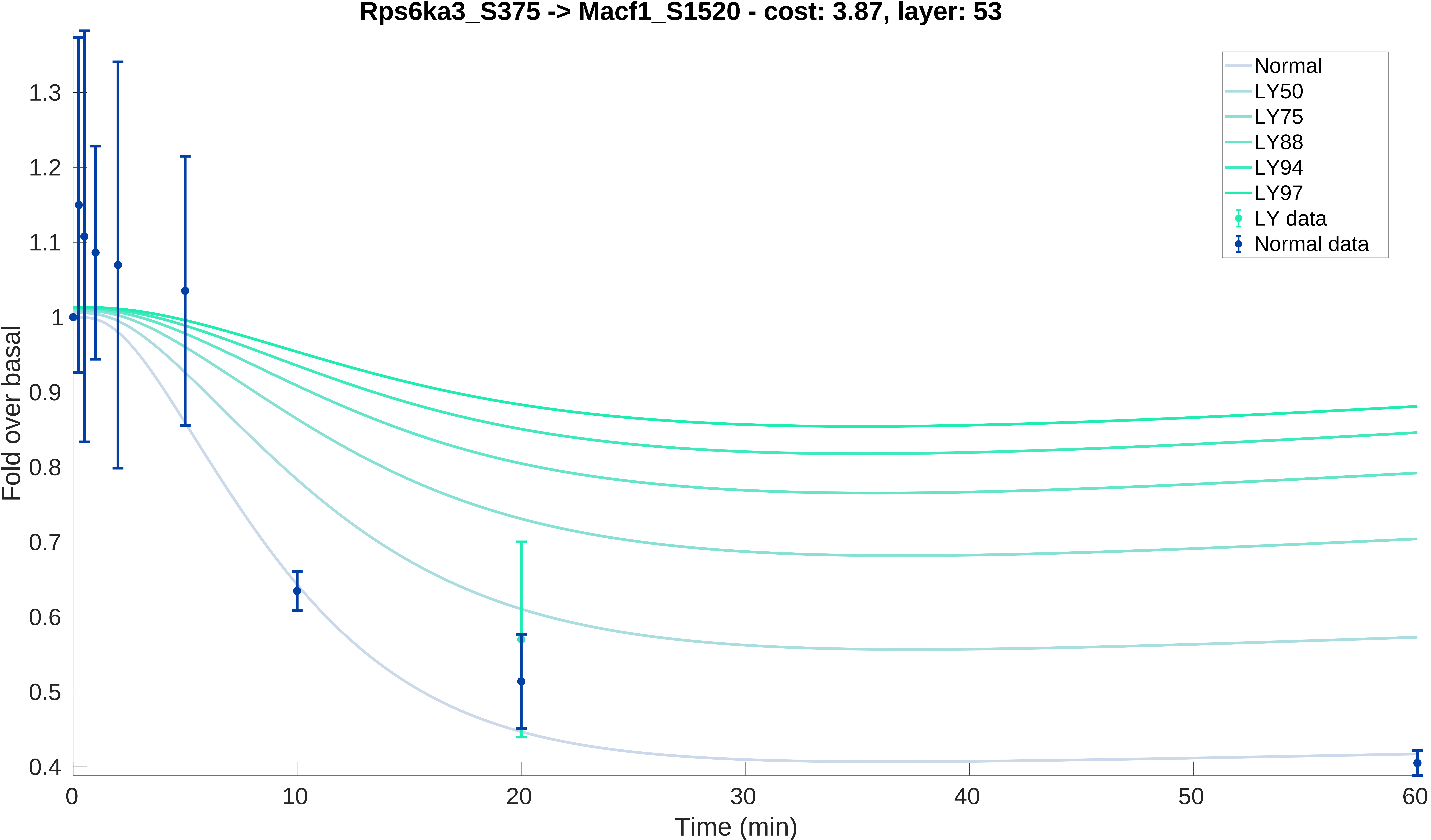

Rps6ka3_S375 -> Macf1_S1520 - cost: 3.87, layer: 53
Fold over basal
Time (min)
0
10
20
30
40
50
60
0.4
0.5
0.6
0.7
0.8
0.9
1
1.1
1.2
1.3
Normal
LY50
LY75
LY88
LY94
LY97
LY data
Normal data

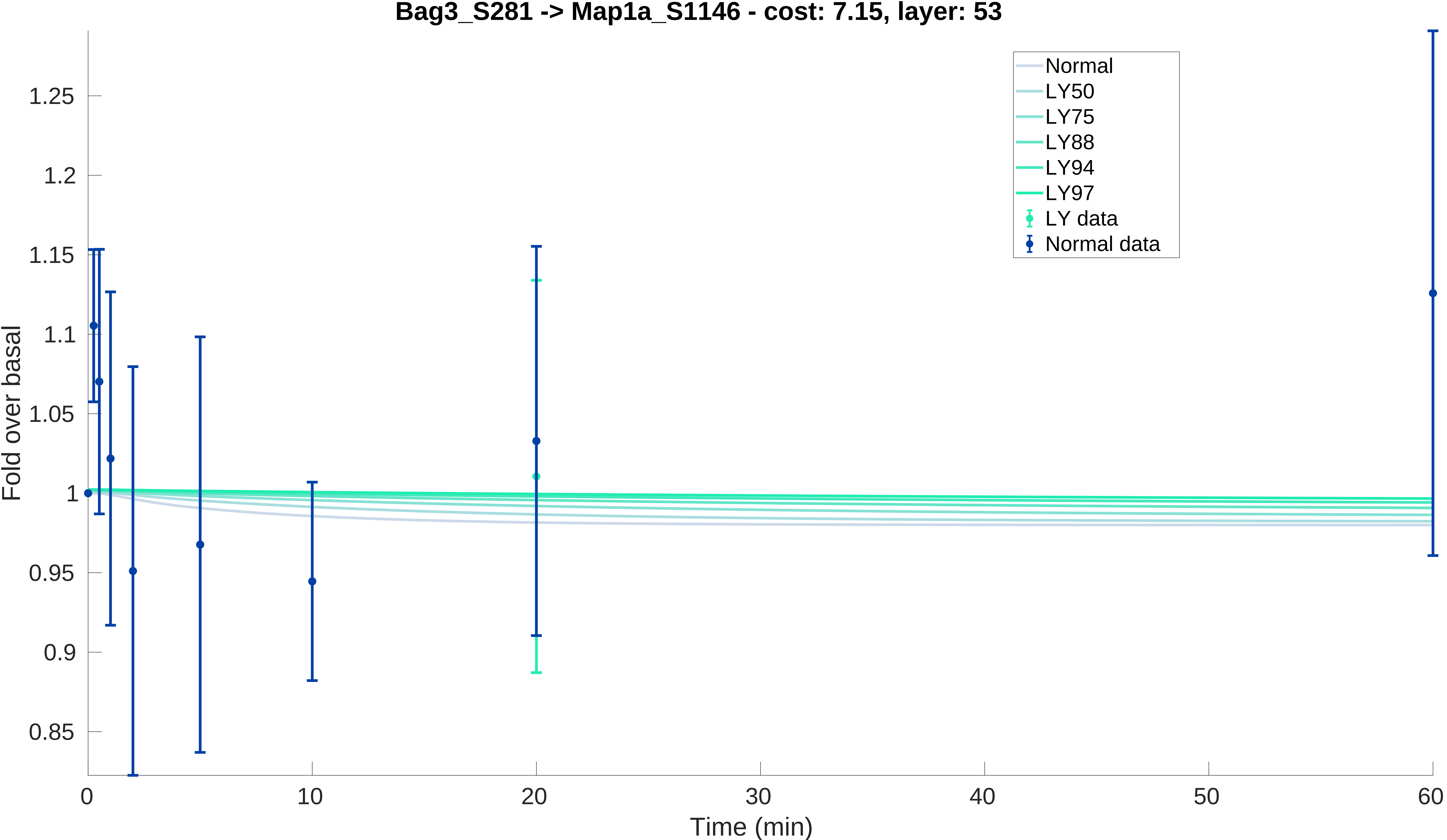

Bag3_S281 -> Map1a_S1146 - cost: 7.15, layer: 53
Fold over basal
Time (min)
1.25
1.2
1.15
1.1
1.05
1
0.95
0.9
0.85
0
10
20
30
40
50
60
Normal
LY50
LY75
LY88
LY94
LY97
LY data
Normal data

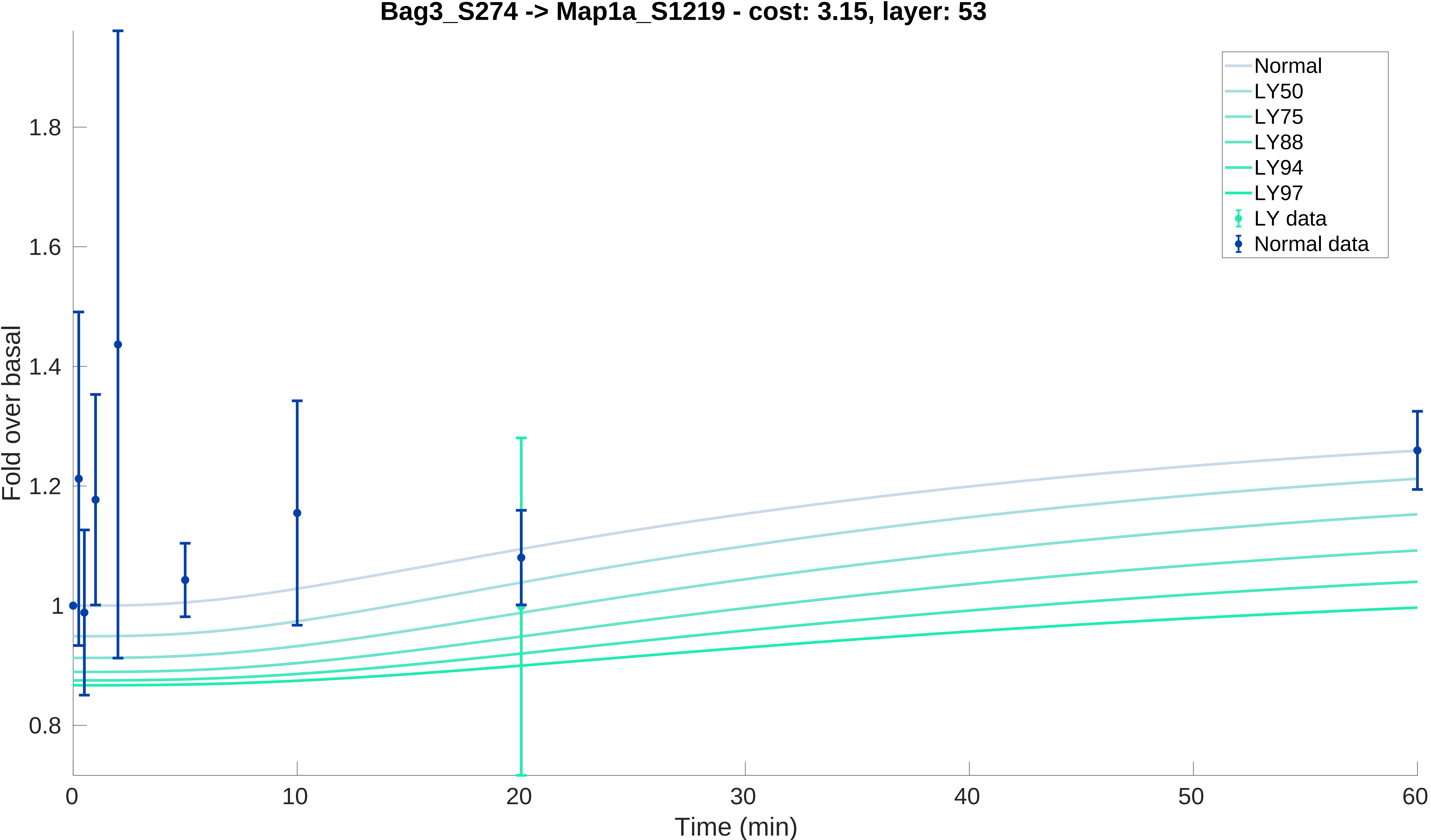
Bag3_S274 -> Map1a_S1219 - cost: 3.15, layer: 53
Normal
LY50
LY75
LY88
LY94
LY97
LY data
Normal data
Fold over basal
1.8
1.6
1.4
1.2
1
0.8
0
10
20
30
40
50
60
Time (min)

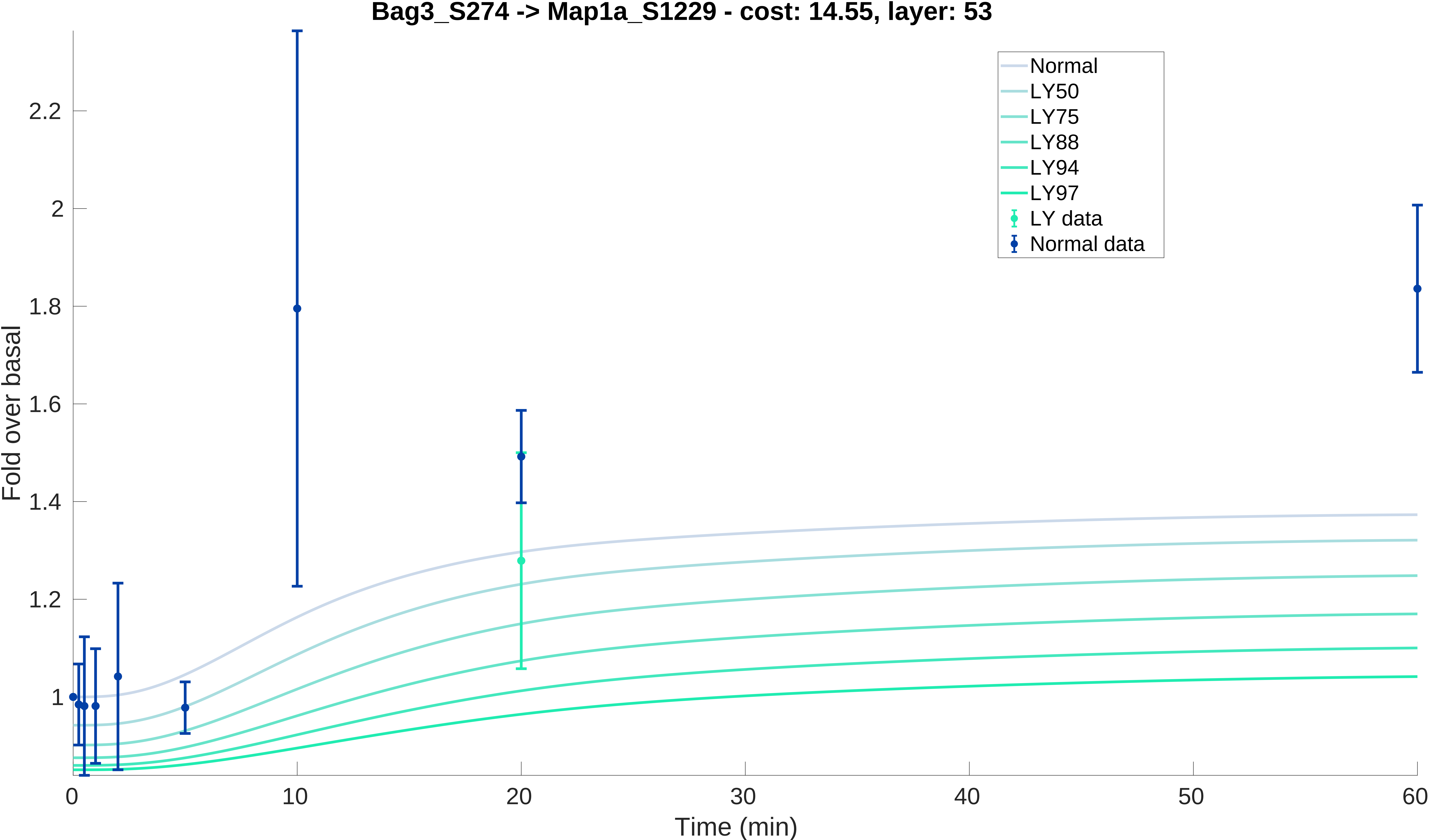
Bag3_S274 -> Map1a_S1229 - cost: 14.55, layer: 53
Normal
LY50
LY75
LY88
LY94
LY97
LY data
Normal data
Fold over basal
2.2
2
1.8
1.6
1.4
1.2
1
0
10
20
30
40
50
60
Time (min)

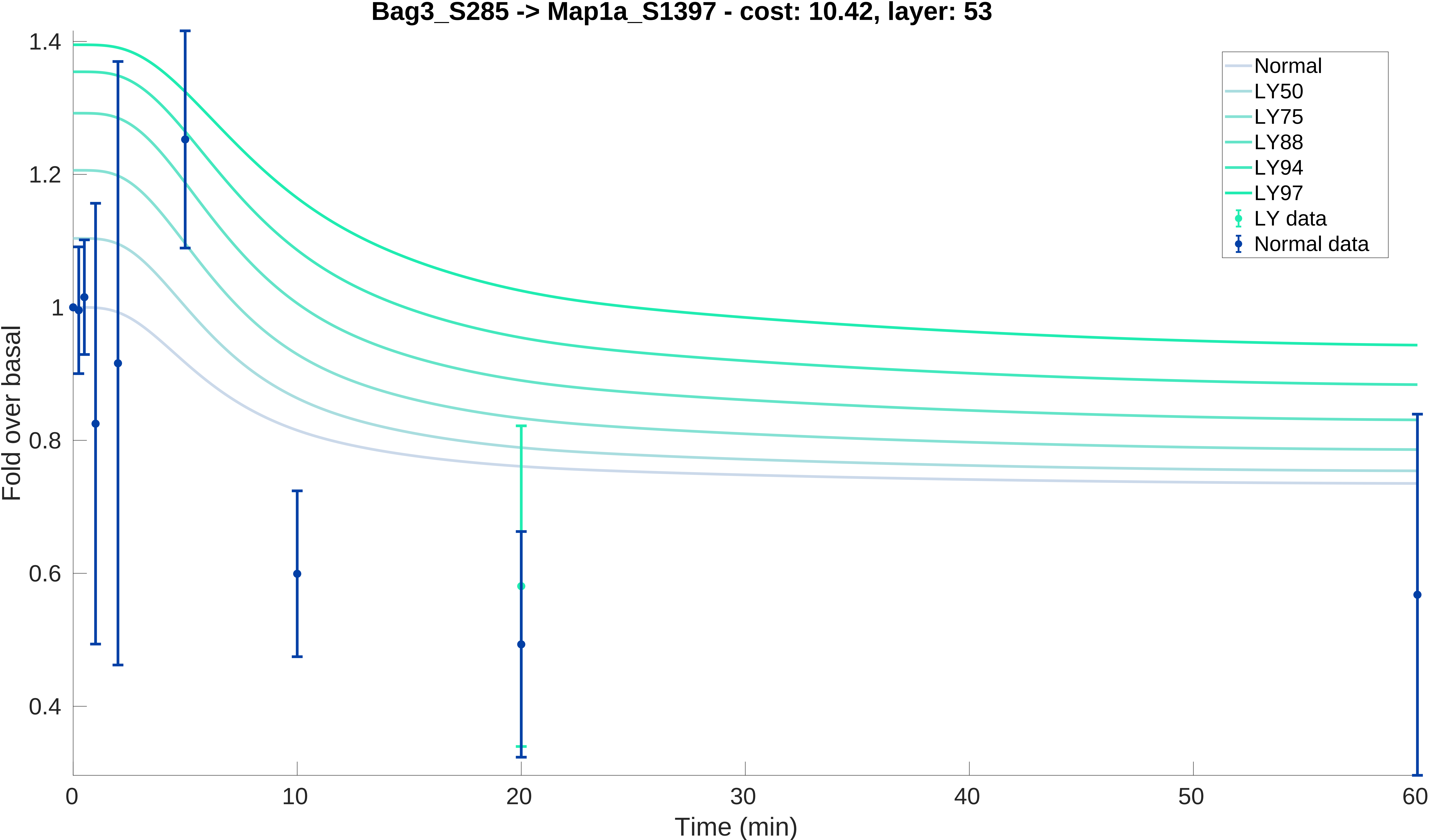

Bag3_S285 -> Map1a_S1397 - cost: 10.42, layer: 53
Fold over basal
Time (min)
0.4
0.6
0.8
1
1.2
1.4
0
10
20
30
40
50
60
Normal
LY50
LY75
LY88
LY94
LY97
LY data
Normal data

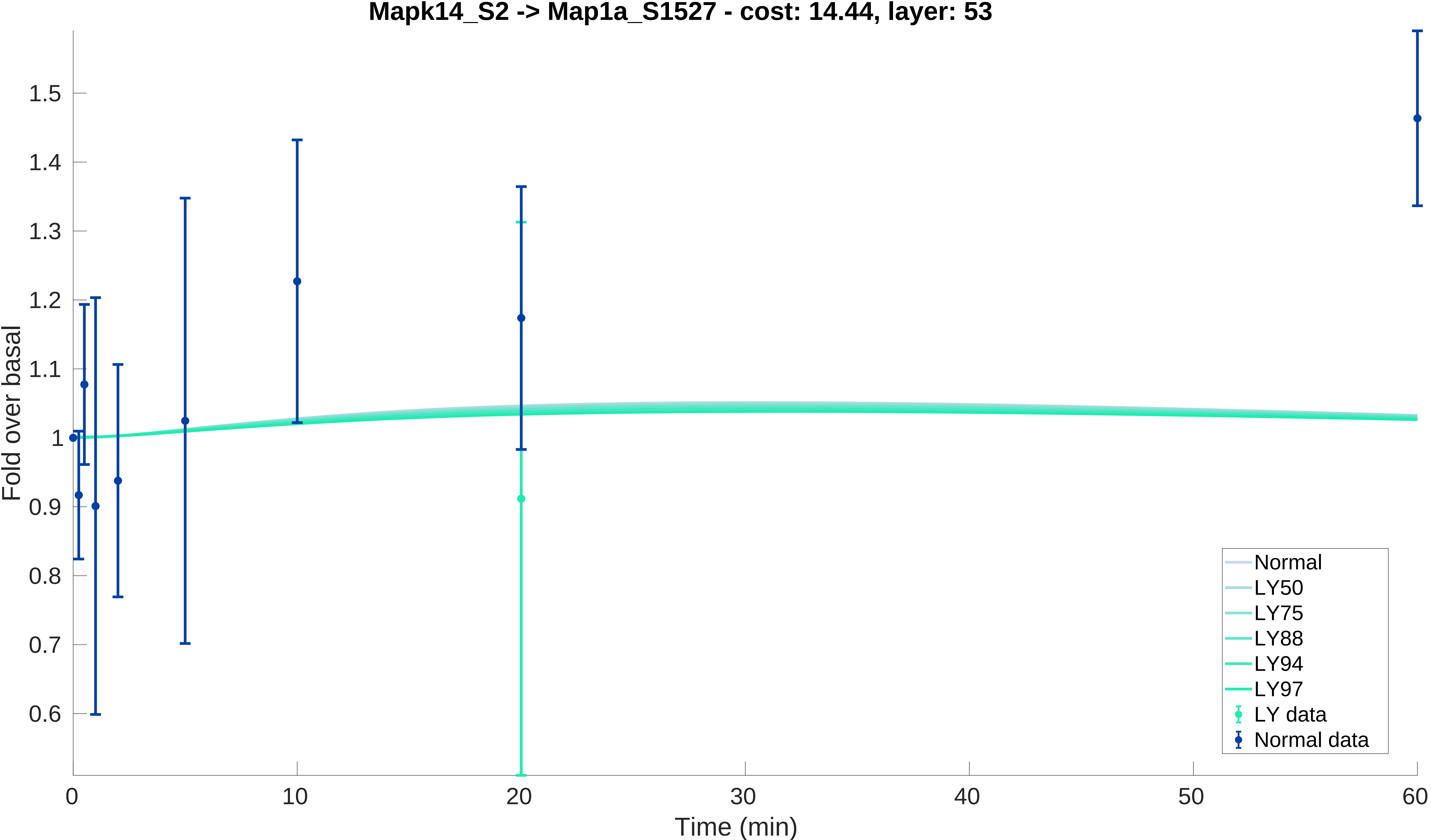

Mapk14_S2 -> Map1a_S1527 - cost: 14.44, layer: 53
Fold over basal
1.5
1.4
1.3
1.2
1.1
1
0.9
0.8
0.7
0.6
0
10
20
30
40
50
60
Time (min)
Normal
LY50
LY75
LY88
LY94
LY97
LY data
Normal data

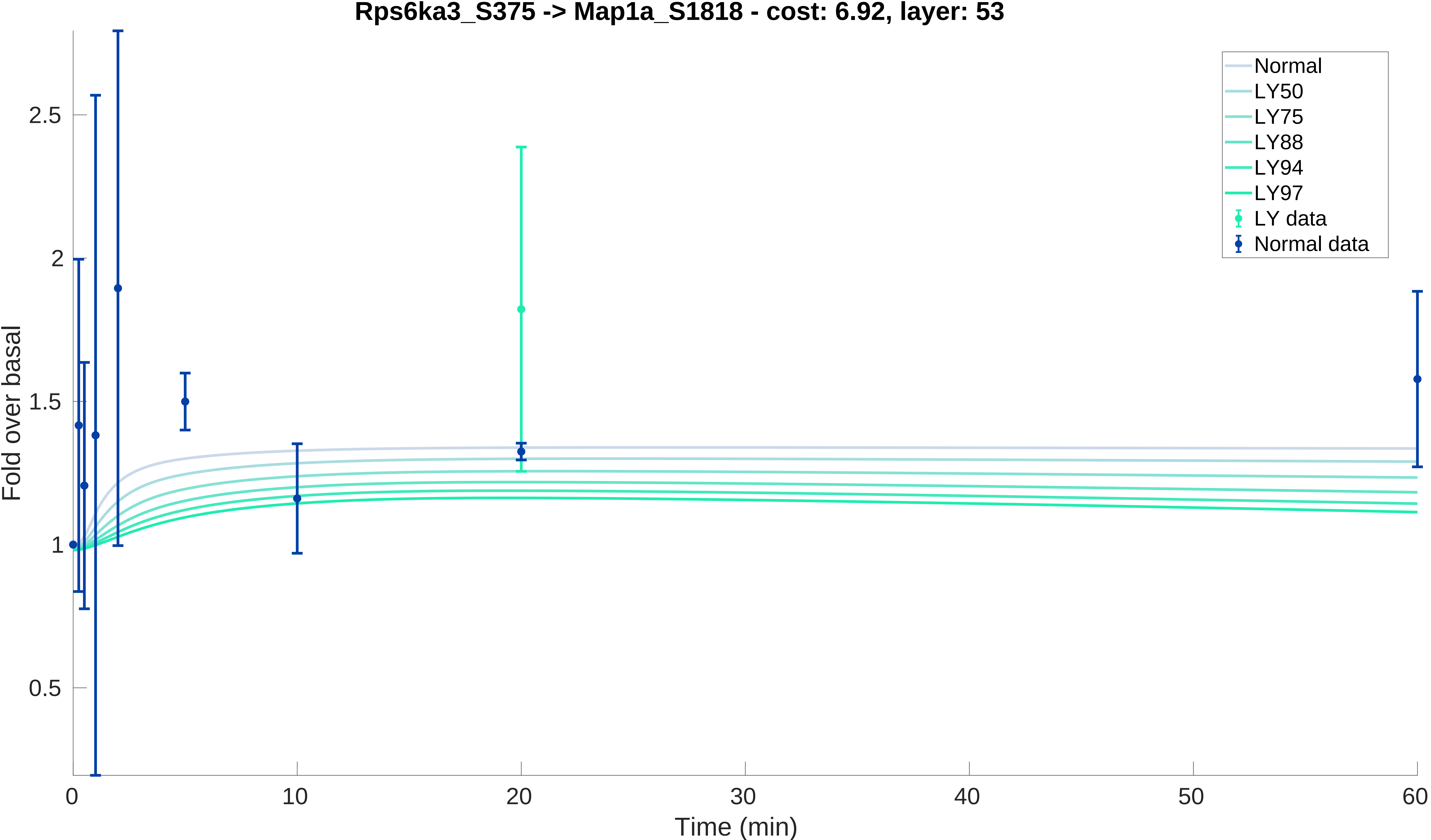

Rps6ka3_S375 -> Map1a_S1818 - cost: 6.92, layer: 53
Fold over basal
0.5
1
1.5
2
2.5
0
10
20
30
40
50
60
Time (min)
Normal
LY50
LY75
LY88
LY94
LY97
LY data
Normal data

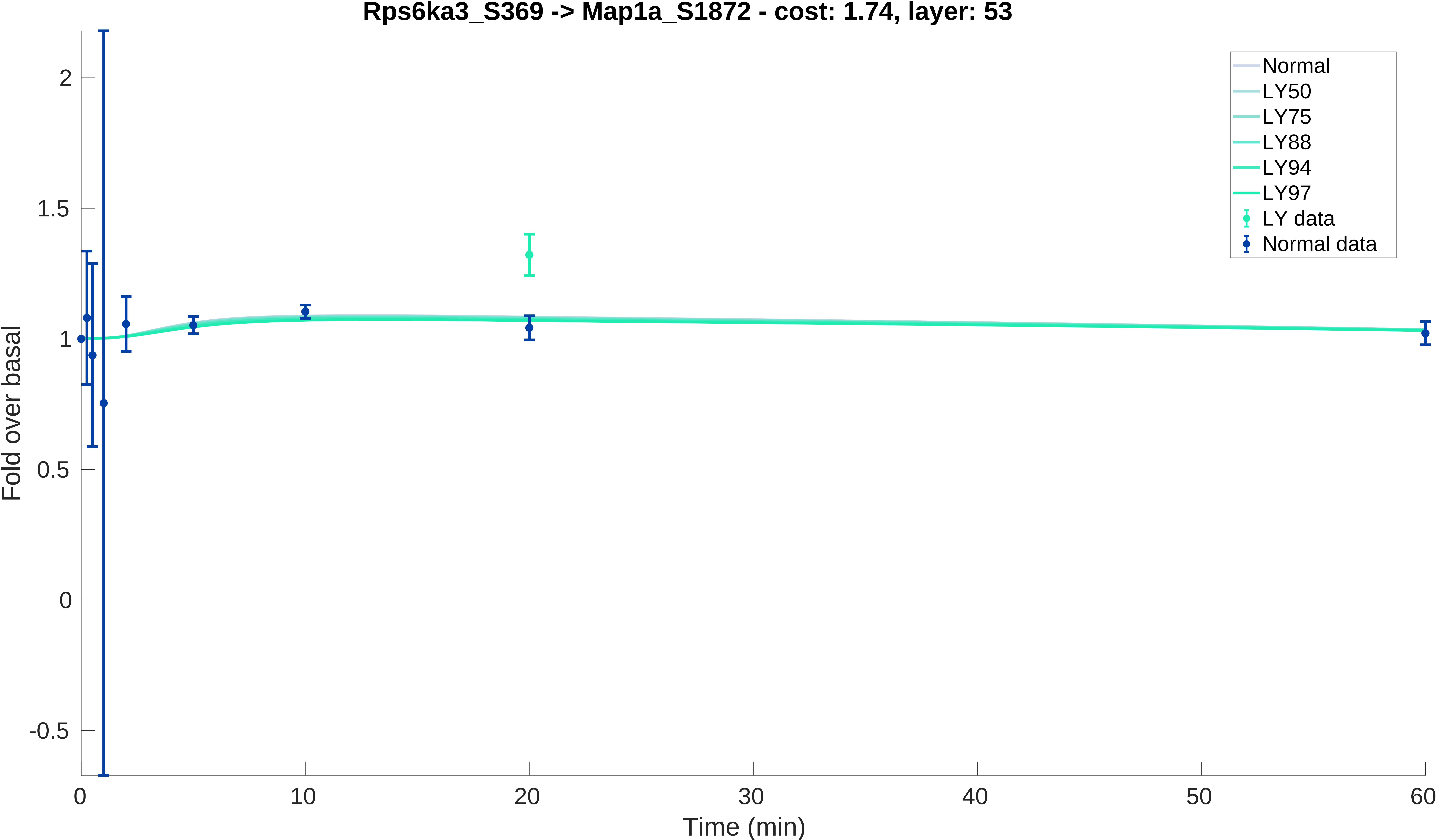

Rps6ka3_S369 -> Map1a_S1872 - cost: 1.74, layer: 53
Fold over basal
Time (min)
2
1.5
1
0.5
0
-0.5
0
10
20
30
40
50
60
Normal
LY50
LY75
LY88
LY94
LY97
LY data
Normal data

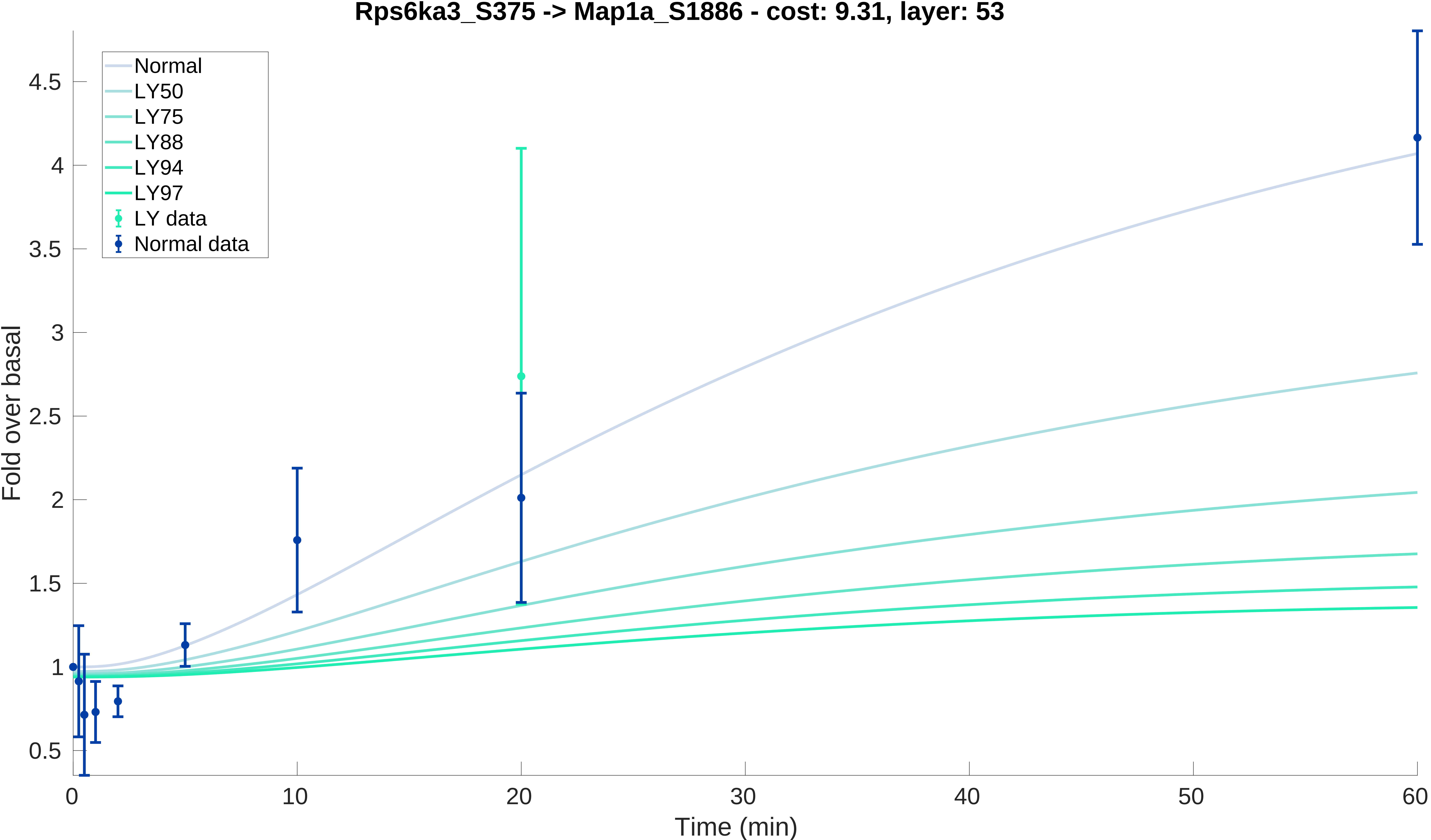

Rps6ka3_S375 -> Map1a_S1886 - cost: 9.31, layer: 53
Normal
LY50
LY75
LY88
LY94
LY97
LY data
Normal data
Fold over basal
Time (min)
0.5
1
1.5
2
2.5
3
3.5
4
4.5
0
10
20
30
40
50
60

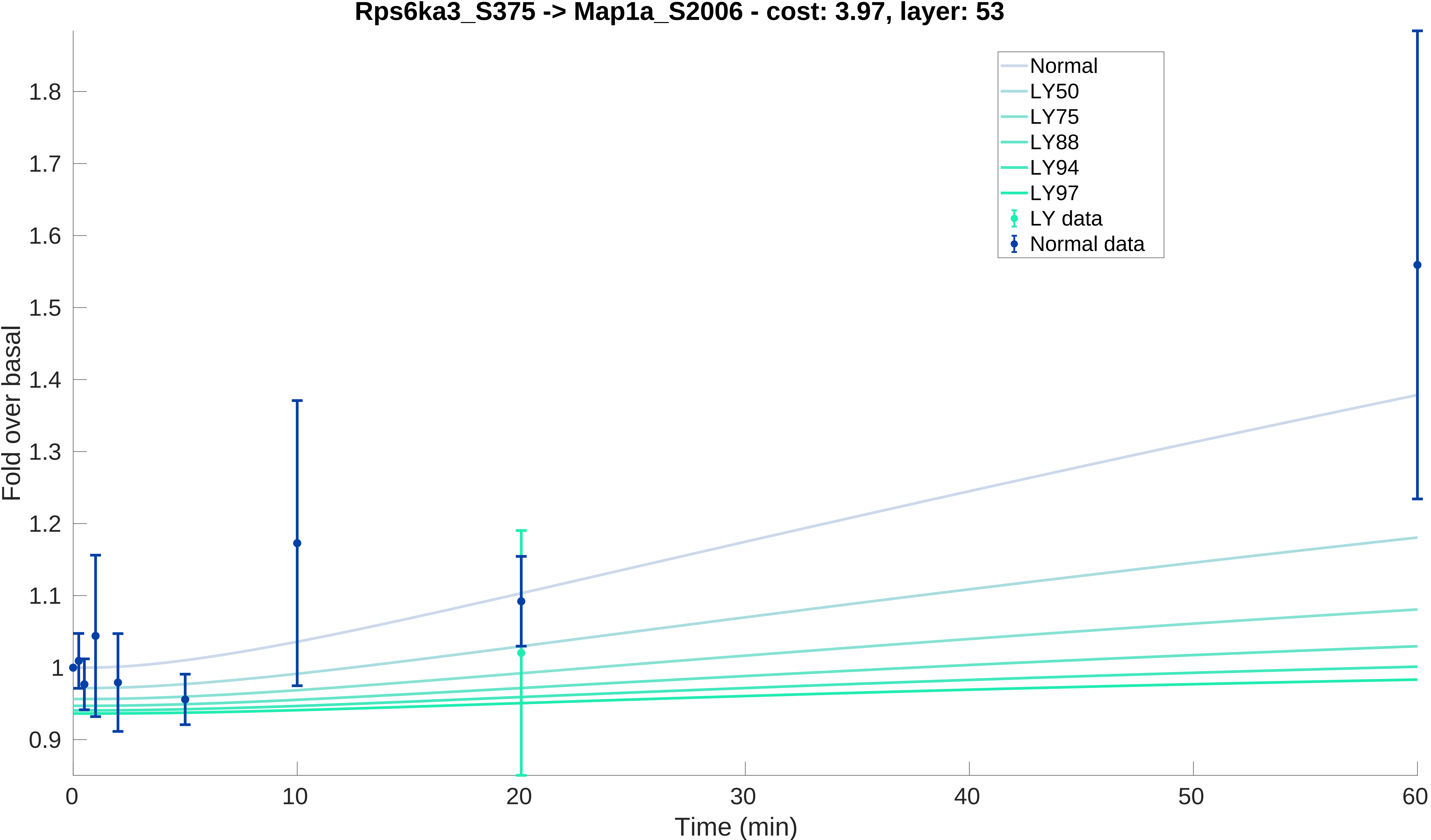
Rps6ka3_S375 -> Map1a_S2006 - cost: 3.97, layer: 53
Fold over basal
Time (min)
0.9
1
1.1
1.2
1.3
1.4
1.5
1.6
1.7
1.8
0
10
20
30
40
50
60
Normal
LY50
LY75
LY88
LY94
LY97
LY data
Normal data

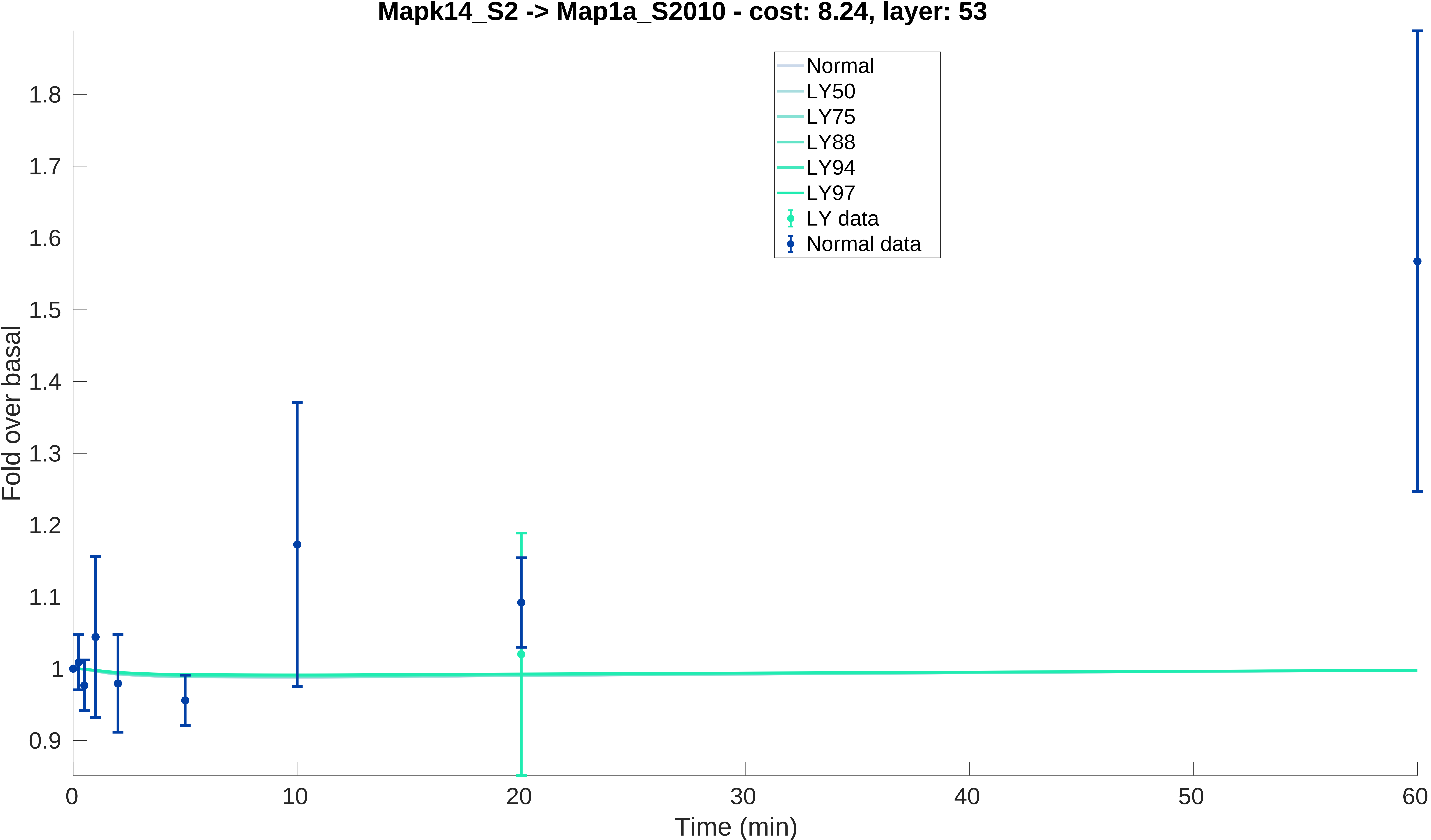

Mapk14_S2 -> Map1a_S2010 - cost: 8.24, layer: 53
Fold over basal
Time (min)
0.9
1
1.1
1.2
1.3
1.4
1.5
1.6
1.7
1.8
0
10
20
30
40
50
60
Normal
LY50
LY75
LY88
LY94
LY97
LY data
Normal data

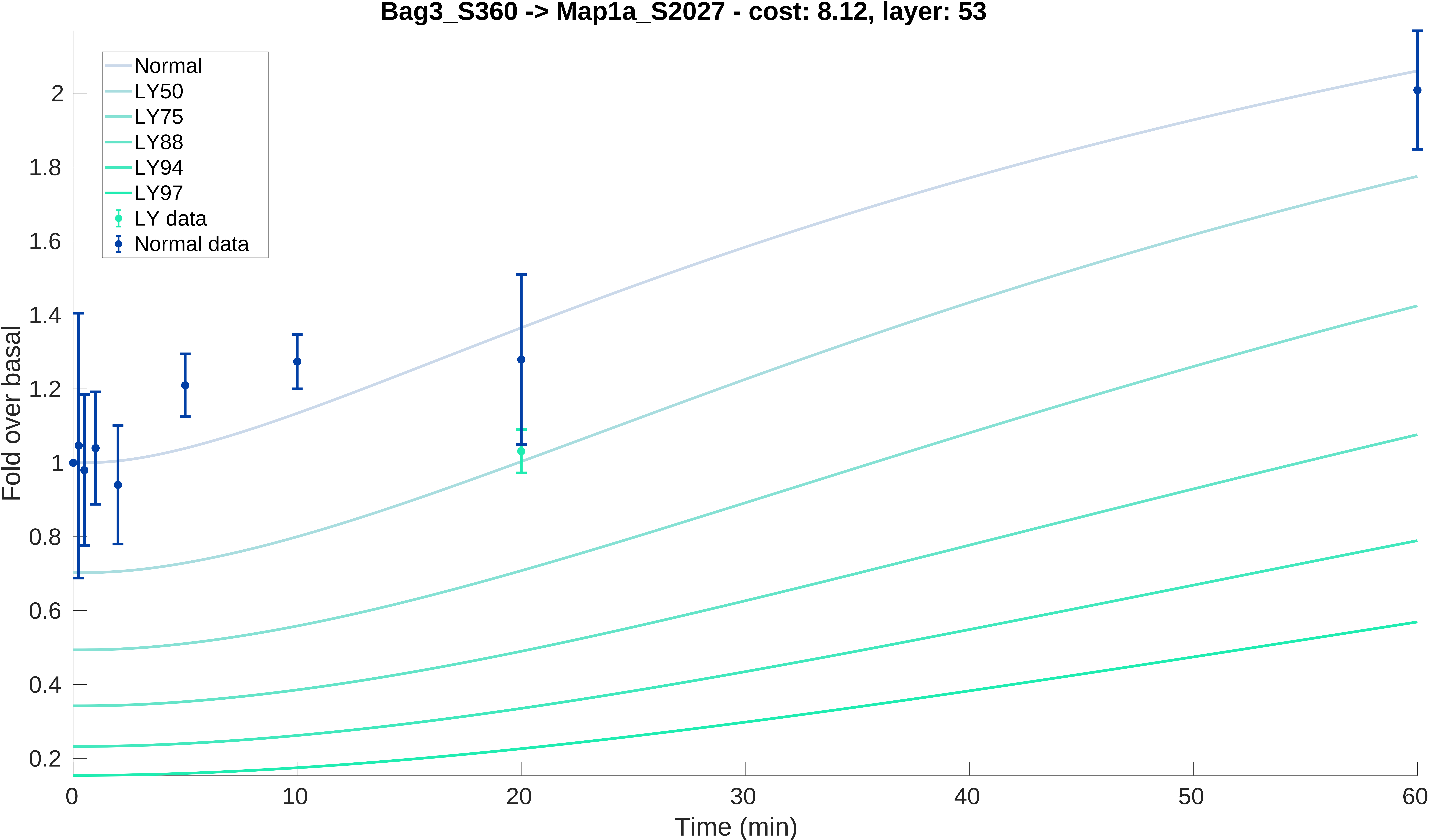

Bag3_S360 -> Map1a_S2027 - cost: 8.12, layer: 53
Normal
LY50
LY75
LY88
LY94
LY97
LY data
Normal data
Fold over basal
Time (min)
0
10
20
30
40
50
60
0.2
0.4
0.6
0.8
1
1.2
1.4
1.6
1.8
2

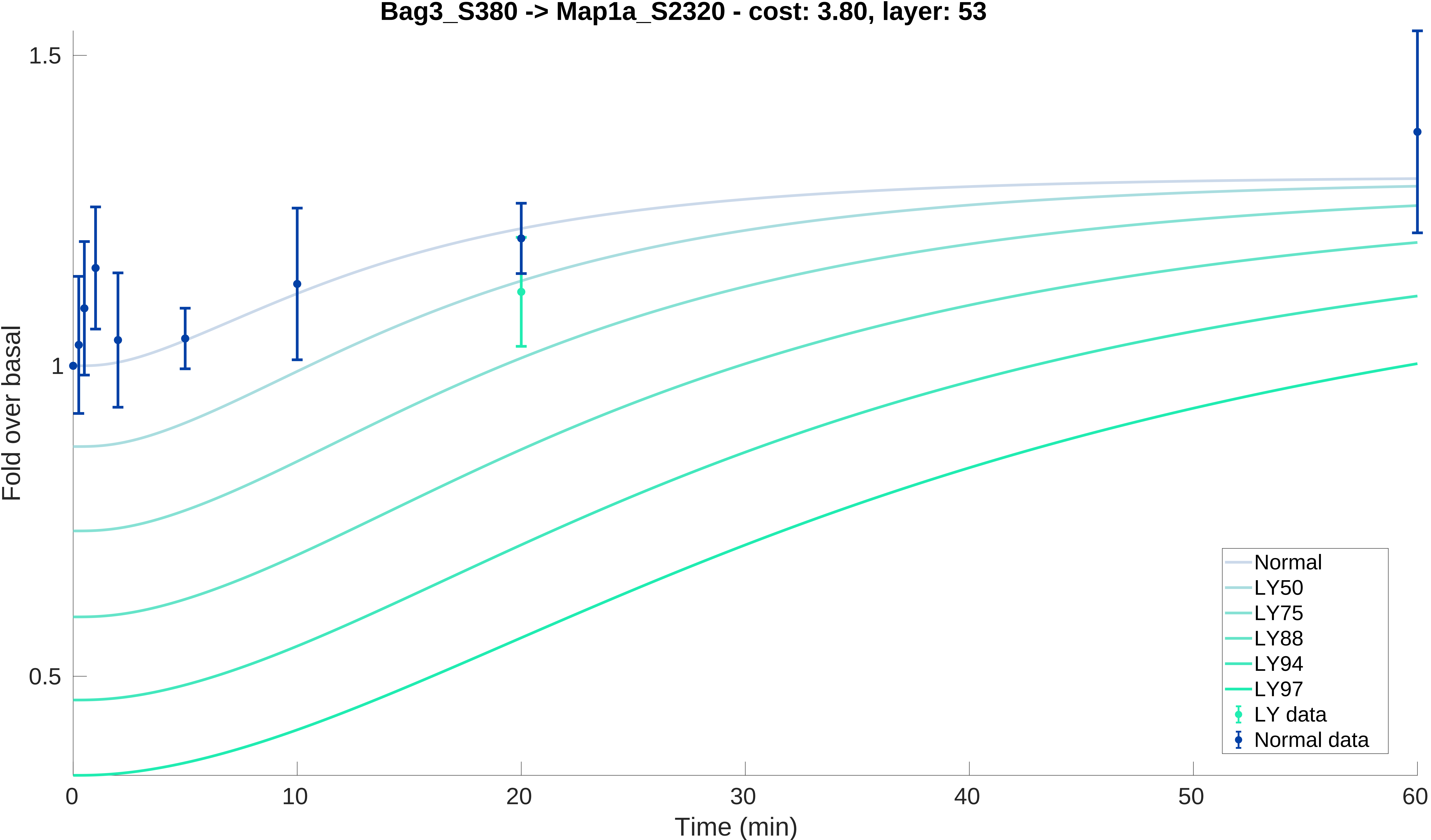

Bag3_S380 -> Map1a_S2320 - cost: 3.80, layer: 53
Fold over basal
1.5
1
0.5
0
10
20
30
40
50
60
Time (min)
Normal
LY50
LY75
LY88
LY94
LY97
LY data
Normal data

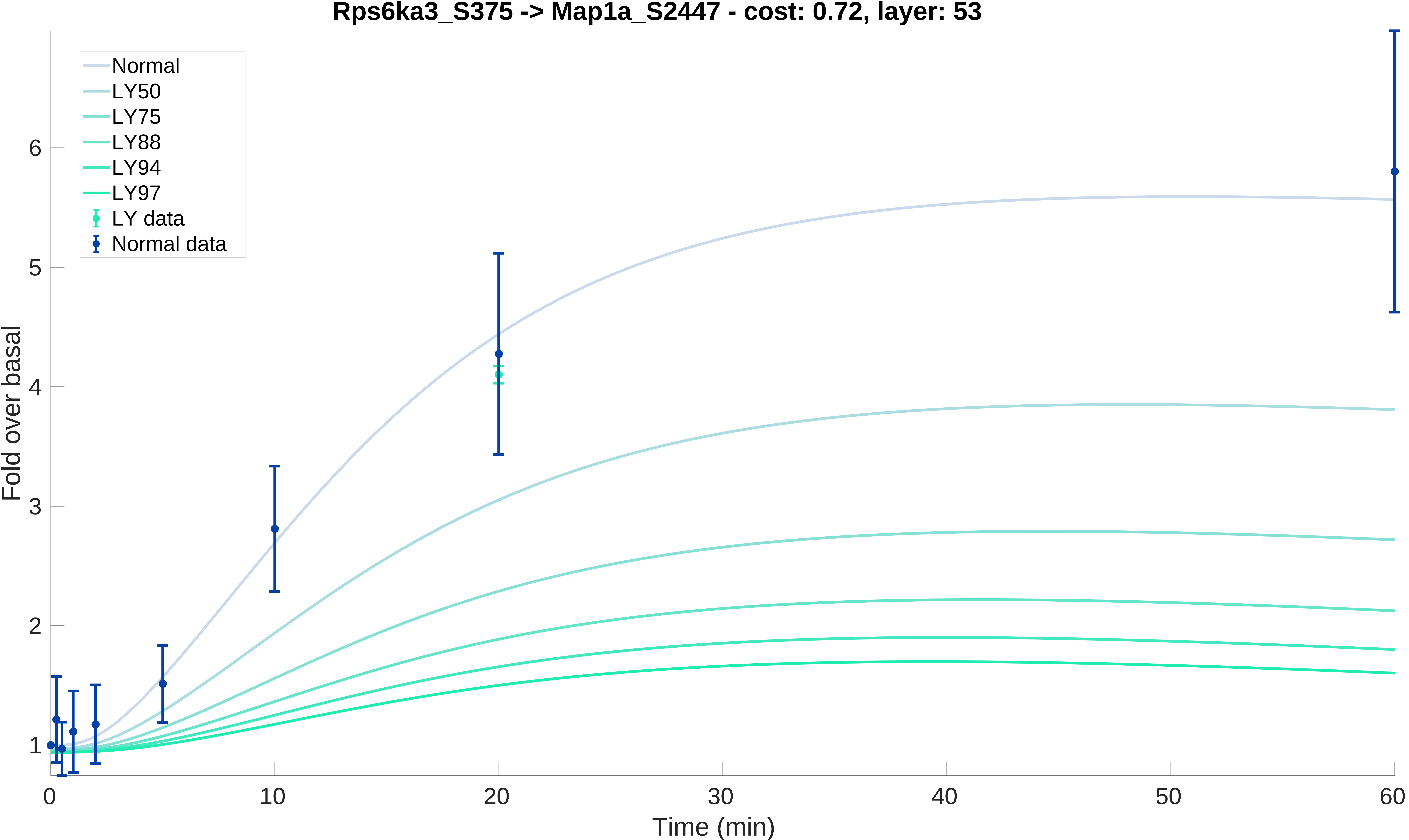
Rps6ka3_S375 -> Map1a_S2447 - cost: 0.72, layer: 53
Normal
LY50
LY75
LY88
LY94
LY97
LY data
Normal data
Fold over basal
1
2
3
4
5
6
0
10
20
30
40
50
60
Time (min)

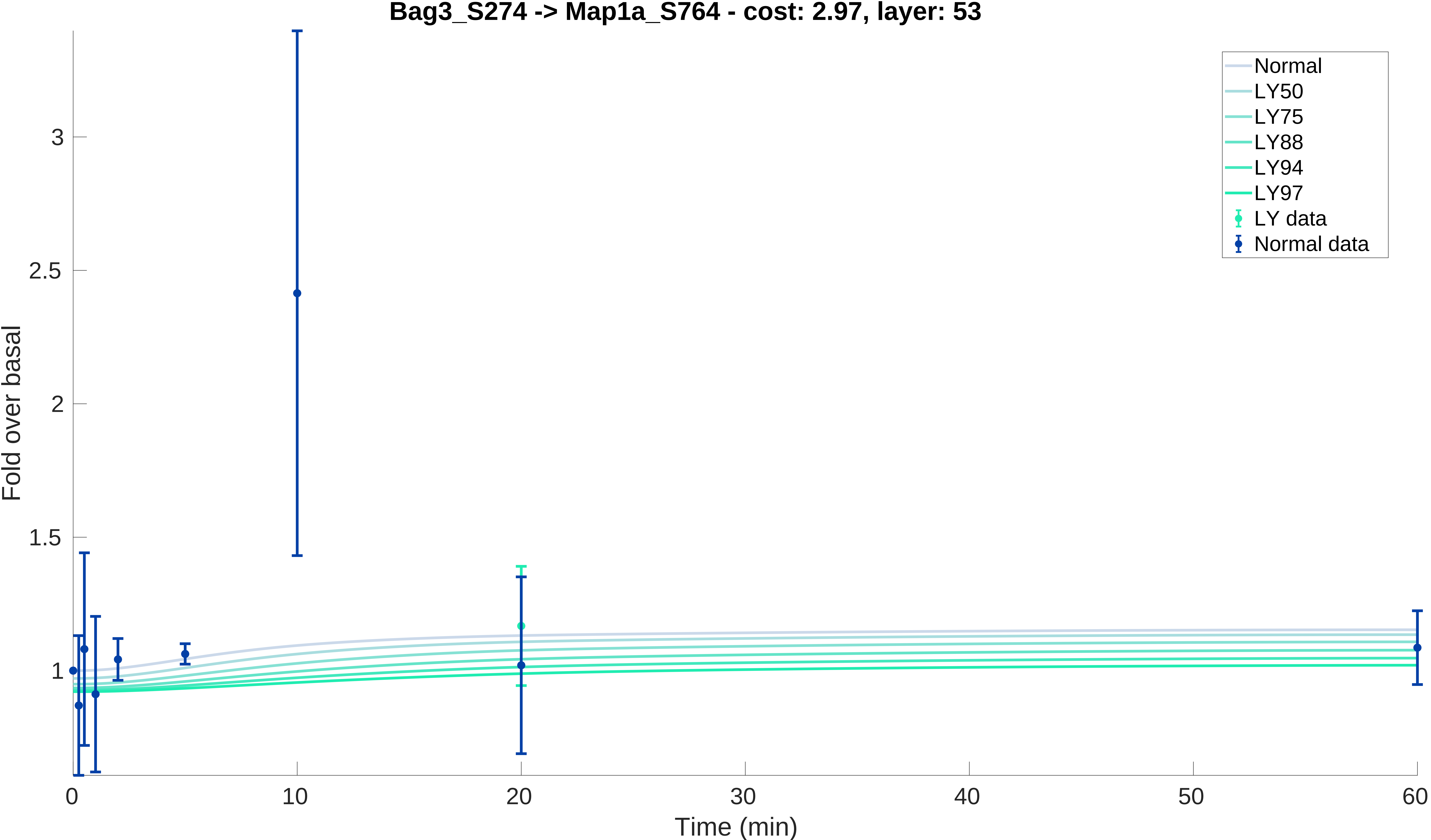

Bag3_S274 -> Map1a_S764 - cost: 2.97, layer: 53
Fold over basal
Time (min)
3
2.5
2
1.5
1
0
10
20
30
40
50
60
Normal
LY50
LY75
LY88
LY94
LY97
LY data
Normal data

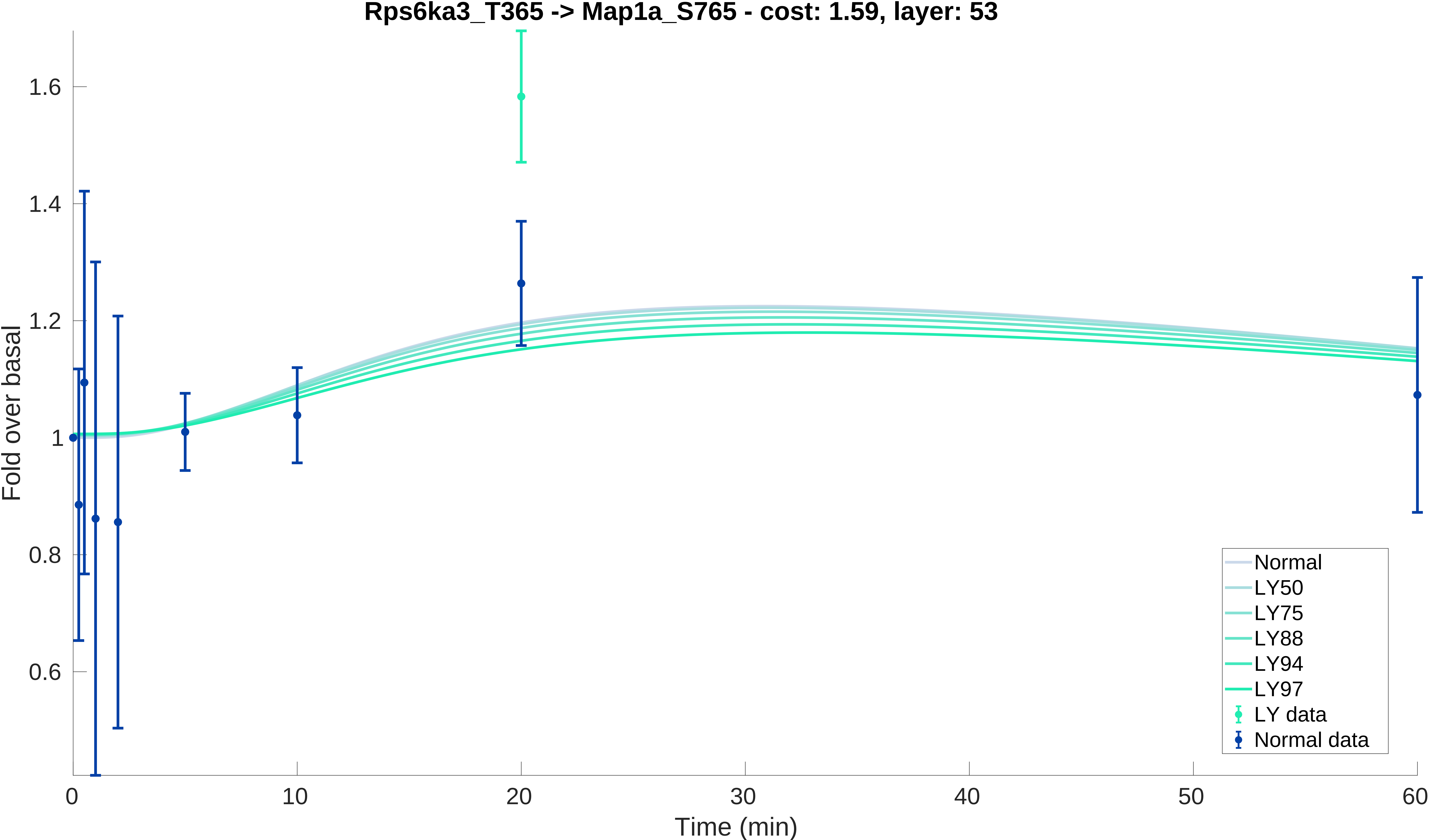
Rps6ka3_T365 -> Map1a_S765 - cost: 1.59, layer: 53
Fold over basal
1.6
1.4
1.2
1
0.8
0.6
0
10
20
30
40
50
60
Time (min)
Normal
LY50
LY75
LY88
LY94
LY97
LY data
Normal data

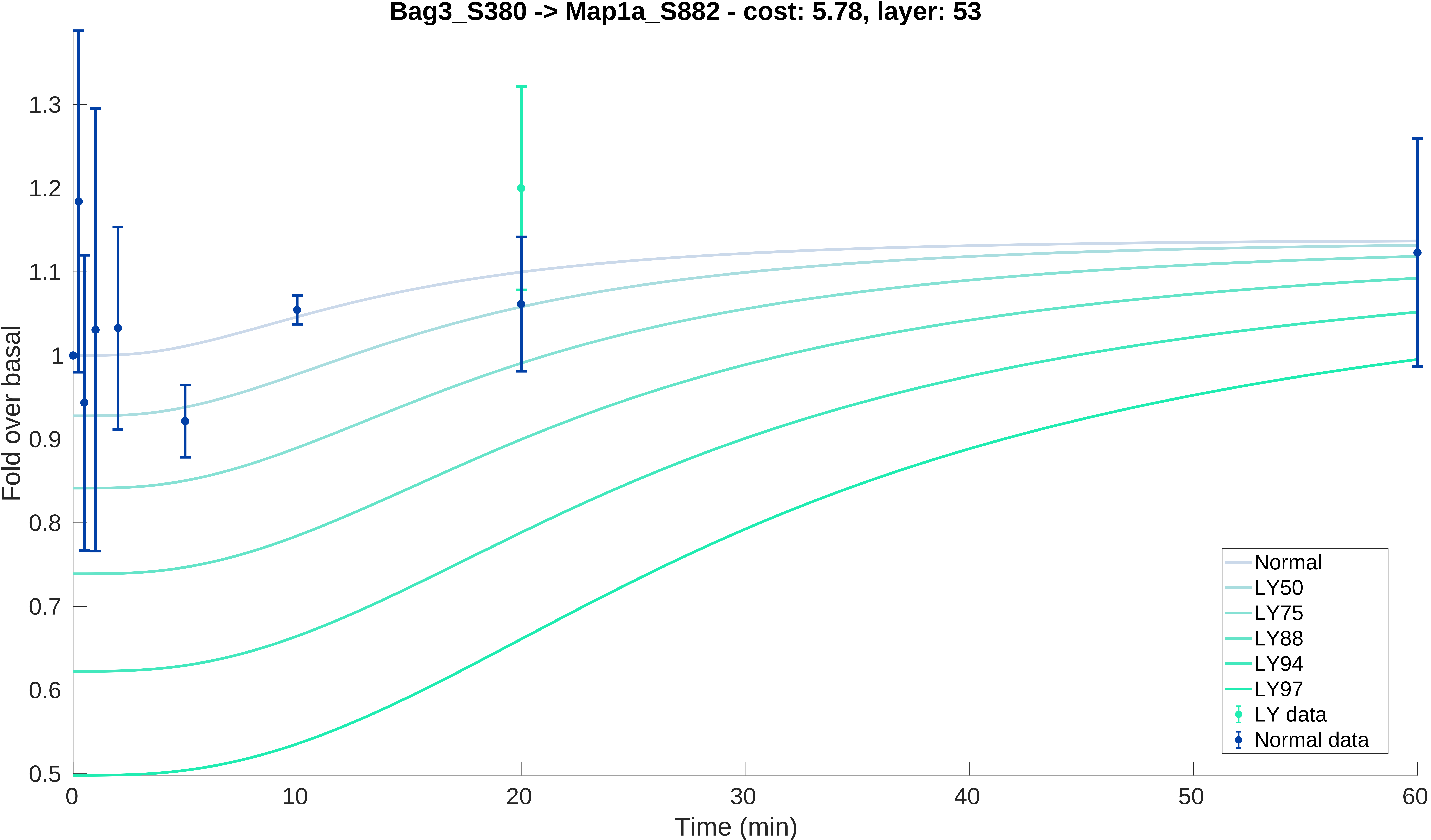

Bag3_S380 -> Map1a_S882 - cost: 5.78, layer: 53
Fold over basal
Time (min)
0.5
0.6
0.7
0.8
0.9
1
1.1
1.2
1.3
0
10
20
30
40
50
60
Normal
LY50
LY75
LY88
LY94
LY97
LY data
Normal data

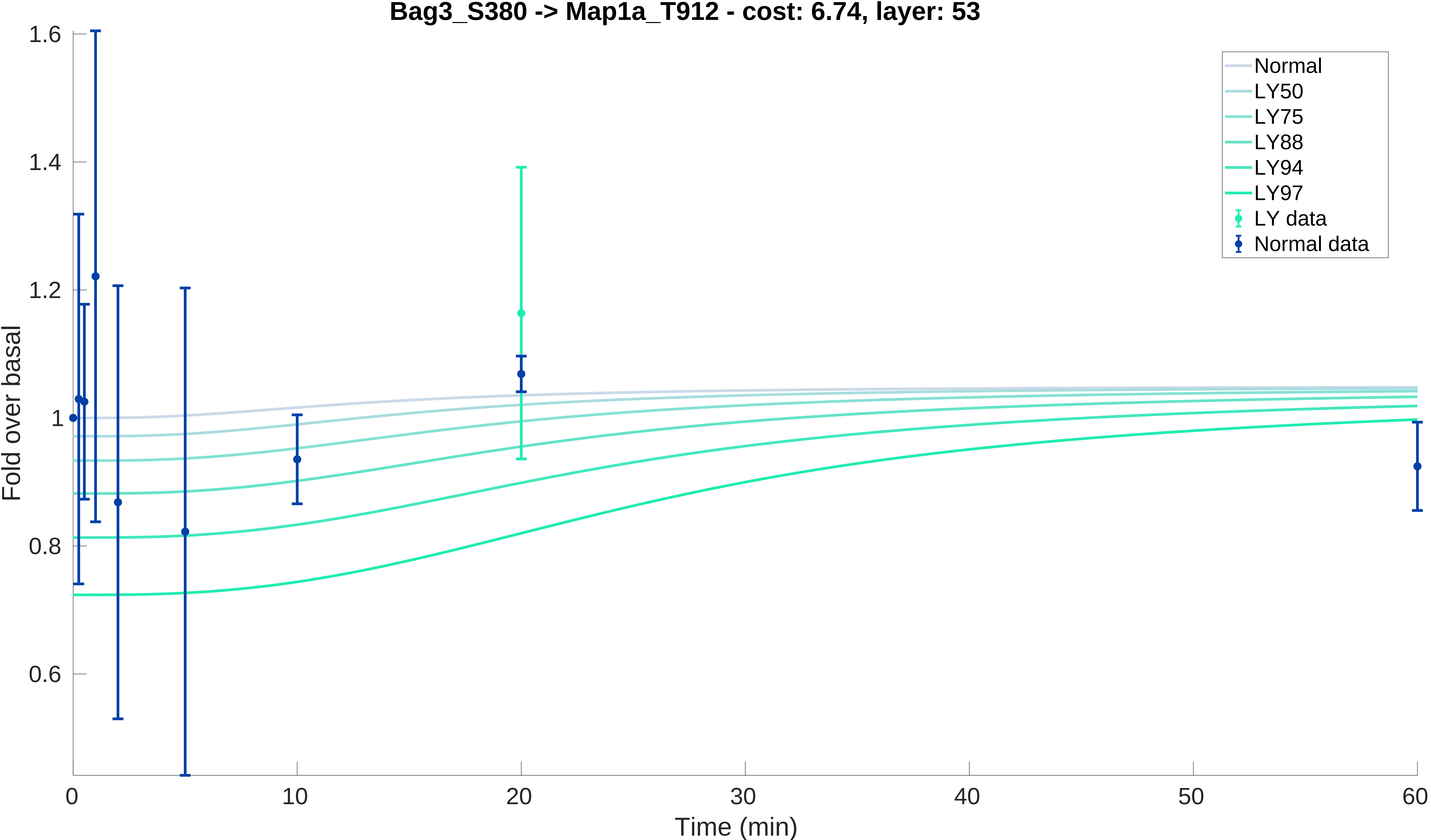
Bag3_S380 -> Map1a_T912 - cost: 6.74, layer: 53
Fold over basal
1.6
1.4
1.2
1
0.8
0.6
0
10
20
30
40
50
60
Time (min)
Normal
LY50
LY75
LY88
LY94
LY97
LY data
Normal data

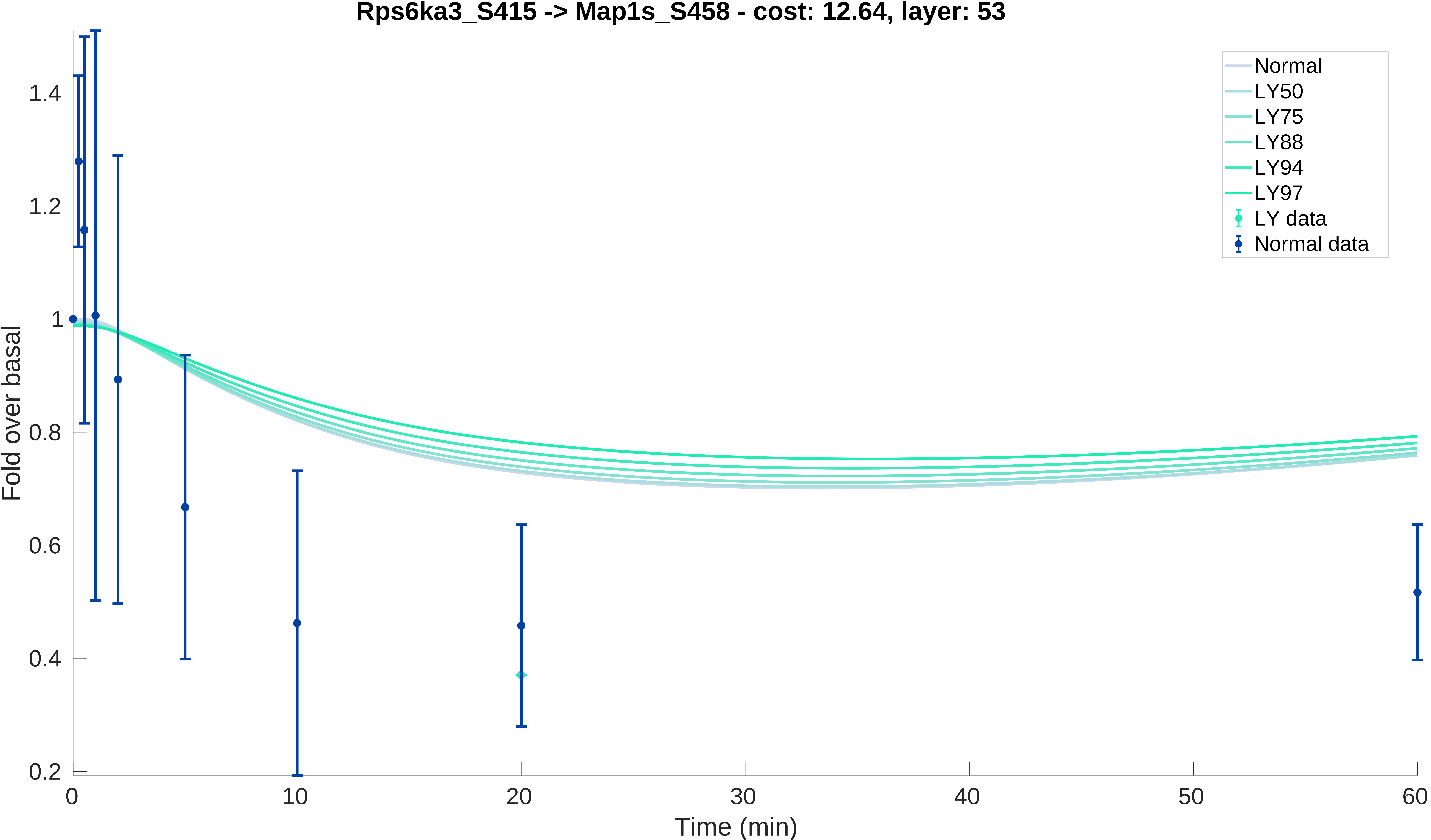

Rps6ka3_S415 -> Map1s_S458 - cost: 12.64, layer: 53
Normal
LY50
LY75
LY88
LY94
LY97
LY data
Normal data
Fold over basal
Time (min)
0.2
0.4
0.6
0.8
1
1.2
1.4
0
10
20
30
40
50
60

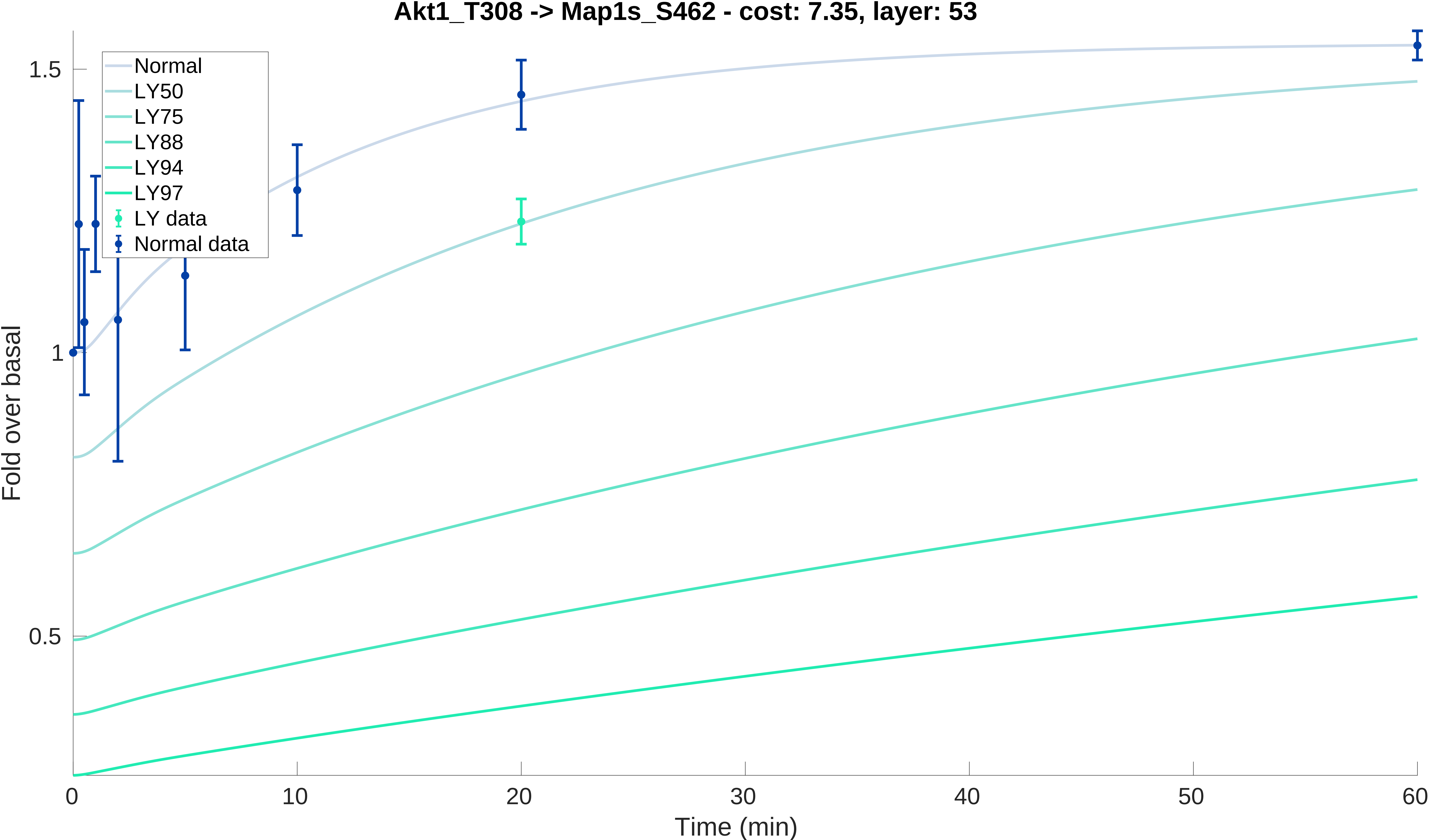

Akt1_T308 -> Map1s_S462 - cost: 7.35, layer: 53
Normal
LY50
LY75
LY88
LY94
LY97
LY data
Normal data
Fold over basal
1.5
1
0.5
0
10
20
30
40
50
60
Time (min)

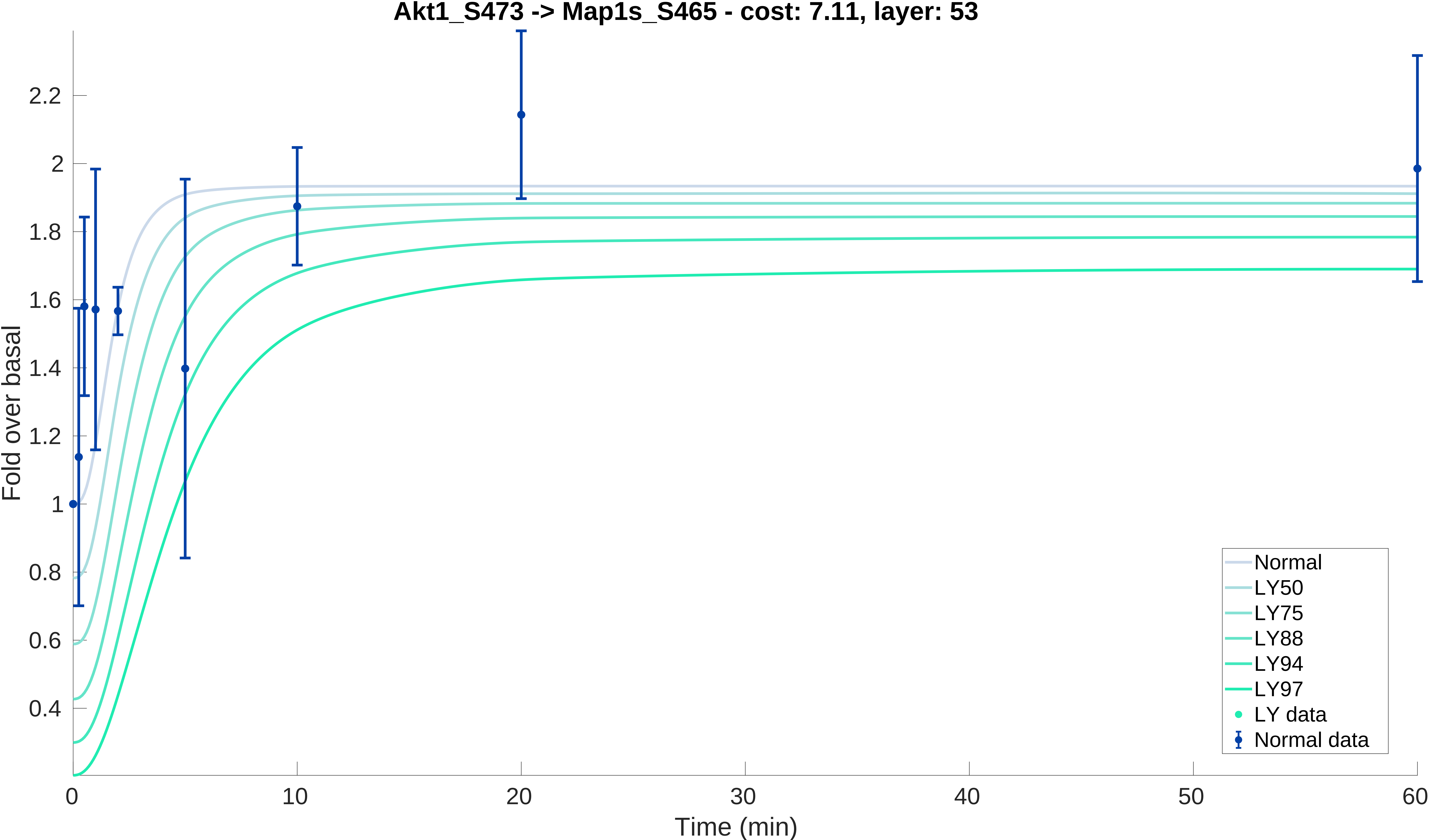

Akt1_S473 -> Map1s_S465 - cost: 7.11, layer: 53
Fold over basal
0.4
0.6
0.8
1
1.2
1.4
1.6
1.8
2
2.2
Time (min)
0
10
20
30
40
50
60
Normal
LY50
LY75
LY88
LY94
LY97
LY data
Normal data

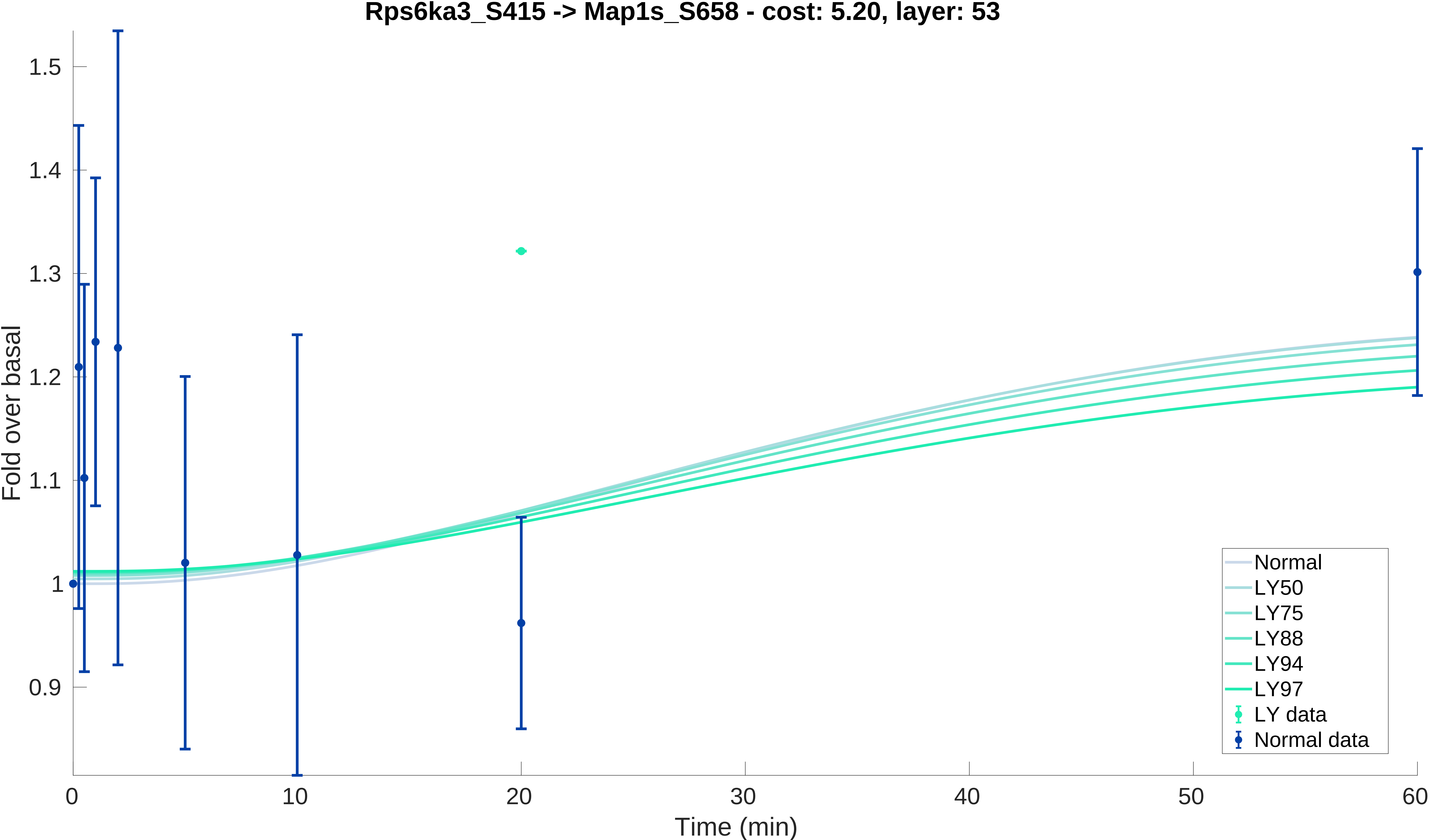
Rps6ka3_S415 -> Map1s_S658 - cost: 5.20, layer: 53
Fold over basal
Time (min)
1.5
1.4
1.3
1.2
1.1
1
0.9
0
10
20
30
40
50
60
Normal
LY50
LY75
LY88
LY94
LY97
LY data
Normal data

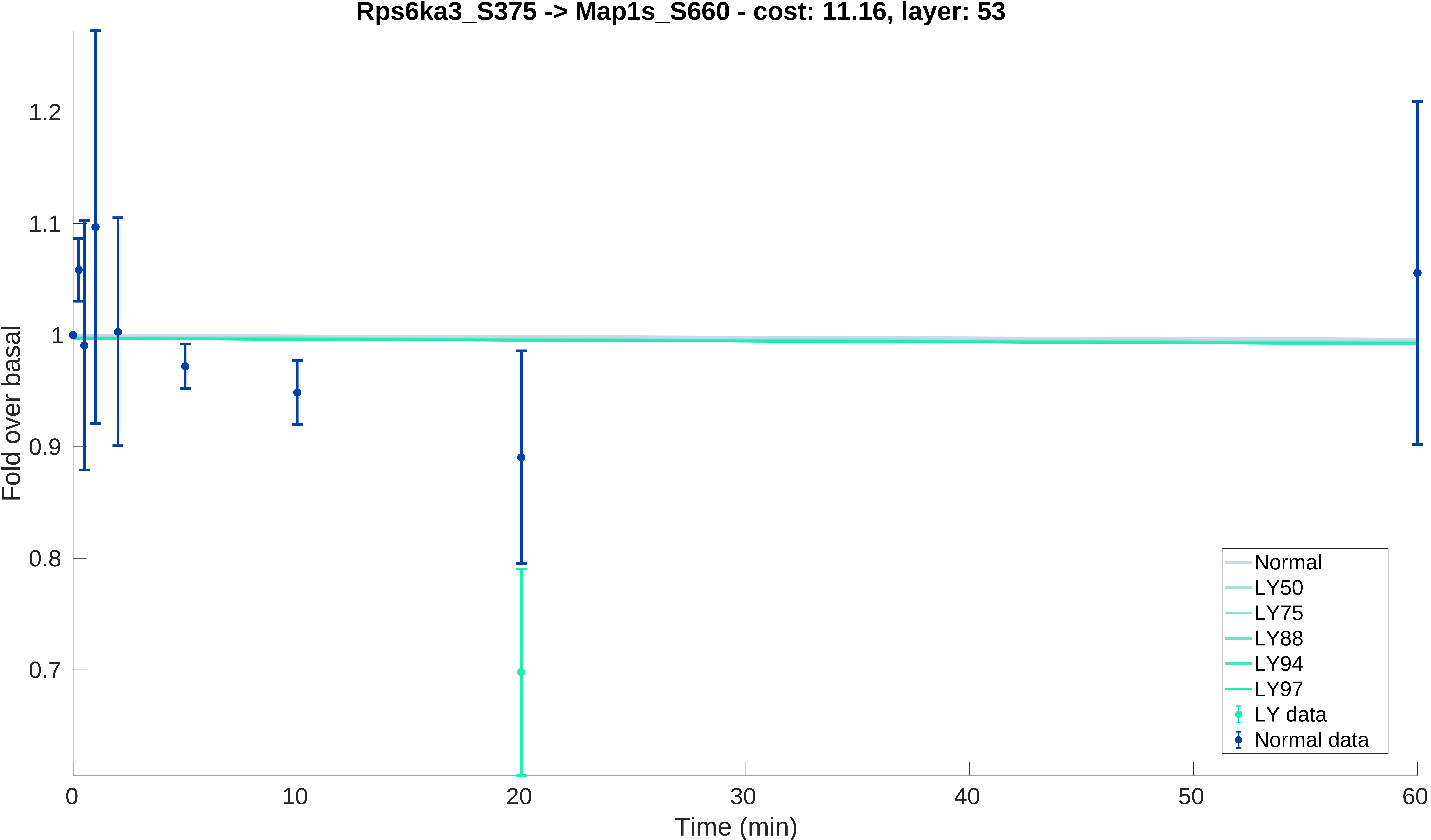

Rps6ka3_S375 -> Map1s_S660 - cost: 11.16, layer: 53
Fold over basal
1.2
1.1
1
0.9
0.8
0.7
0
10
20
30
40
50
60
Time (min)
Normal
LY50
LY75
LY88
LY94
LY97
LY data
Normal data

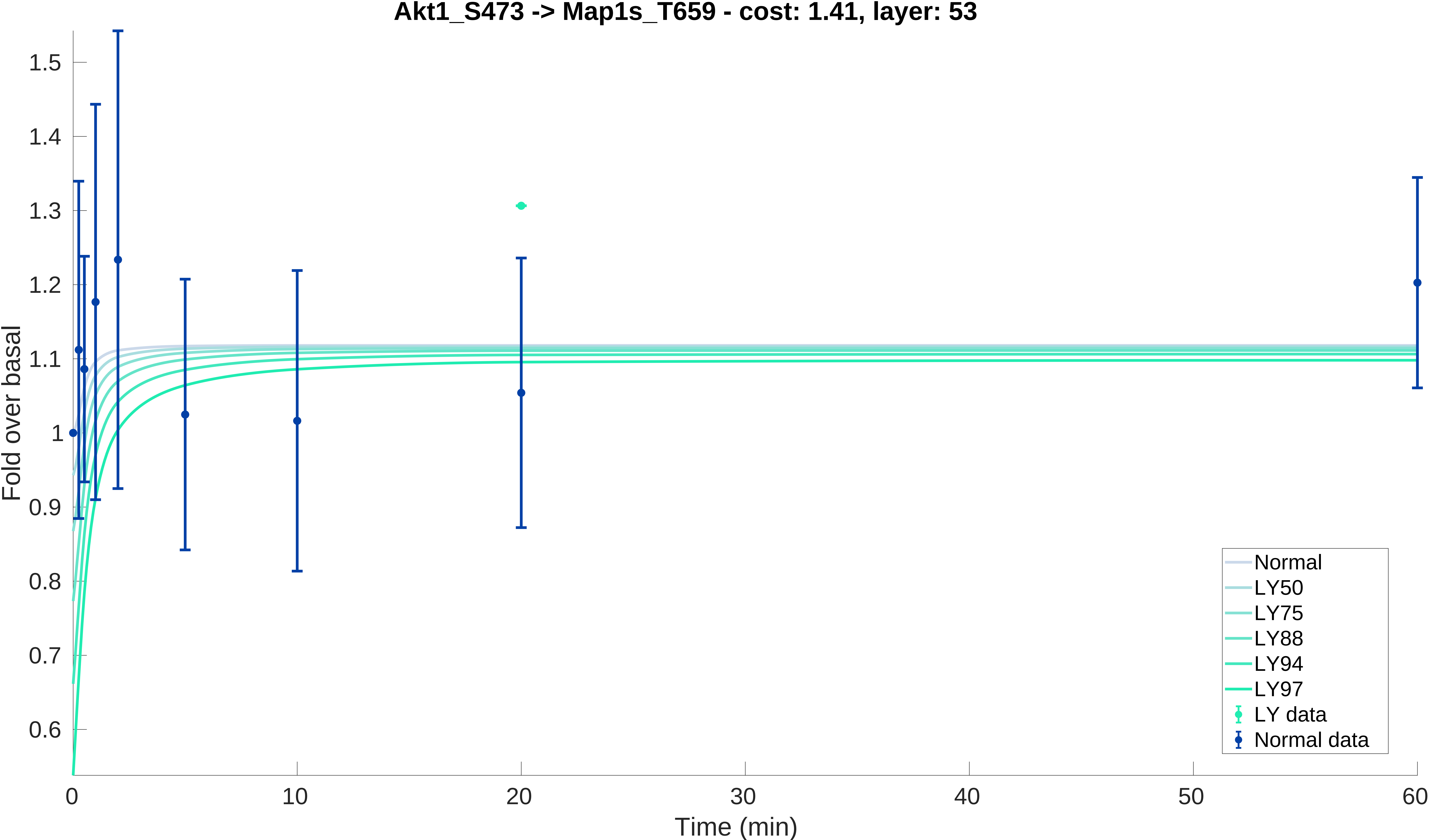

Akt1_S473 -> Map1s_T659 - cost: 1.41, layer: 53
Fold over basal
1.5
1.4
1.3
1.2
1.1
1
0.9
0.8
0.7
0.6
0
10
20
30
40
50
60
Time (min)
Normal
LY50
LY75
LY88
LY94
LY97
LY data
Normal data

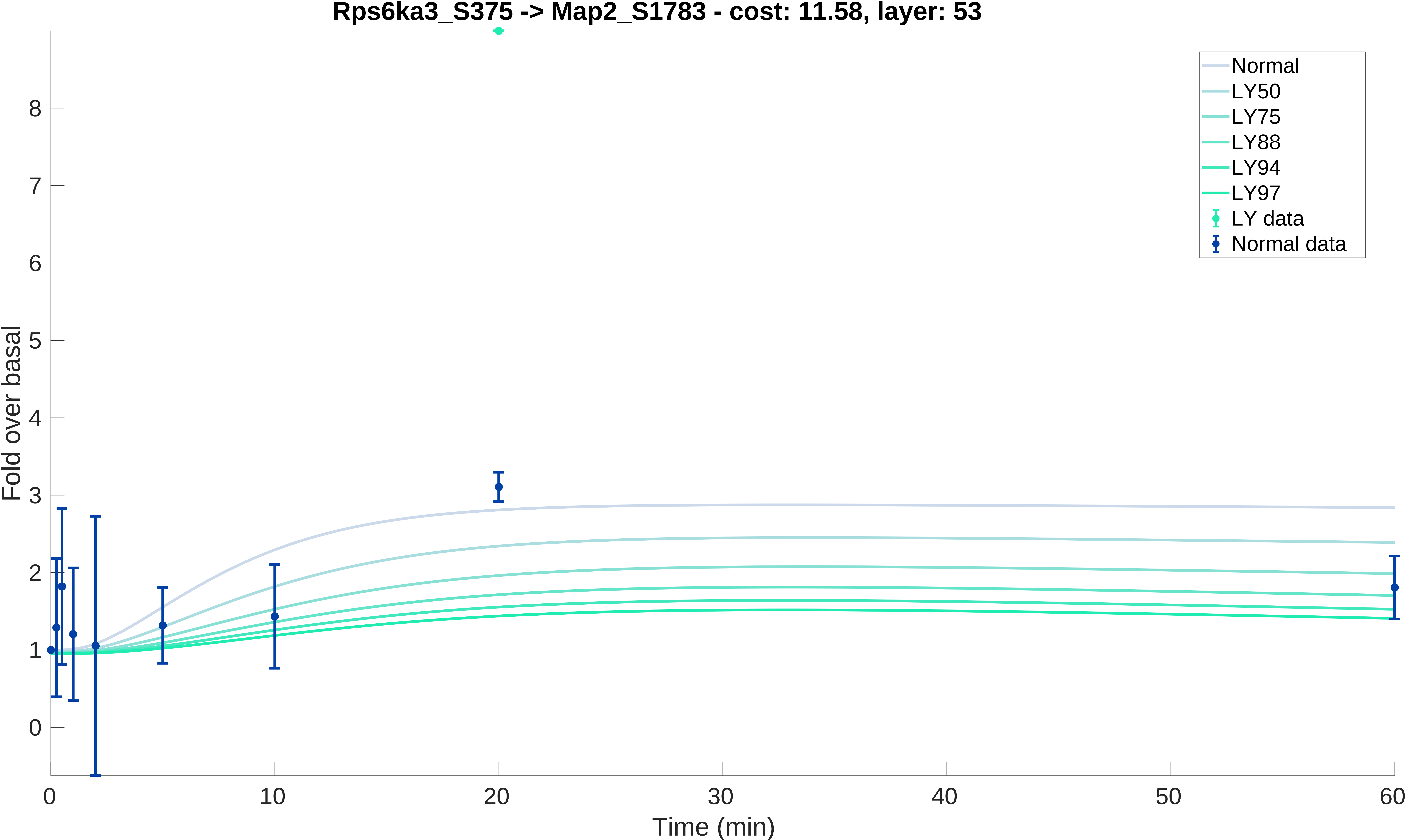

Rps6ka3_S375 -> Map2_S1783 - cost: 11.58, layer: 53
Fold over basal
Time (min)
0
1
2
3
4
5
6
7
8
10
20
30
40
50
60
Normal
LY50
LY75
LY88
LY94
LY97
LY data
Normal data

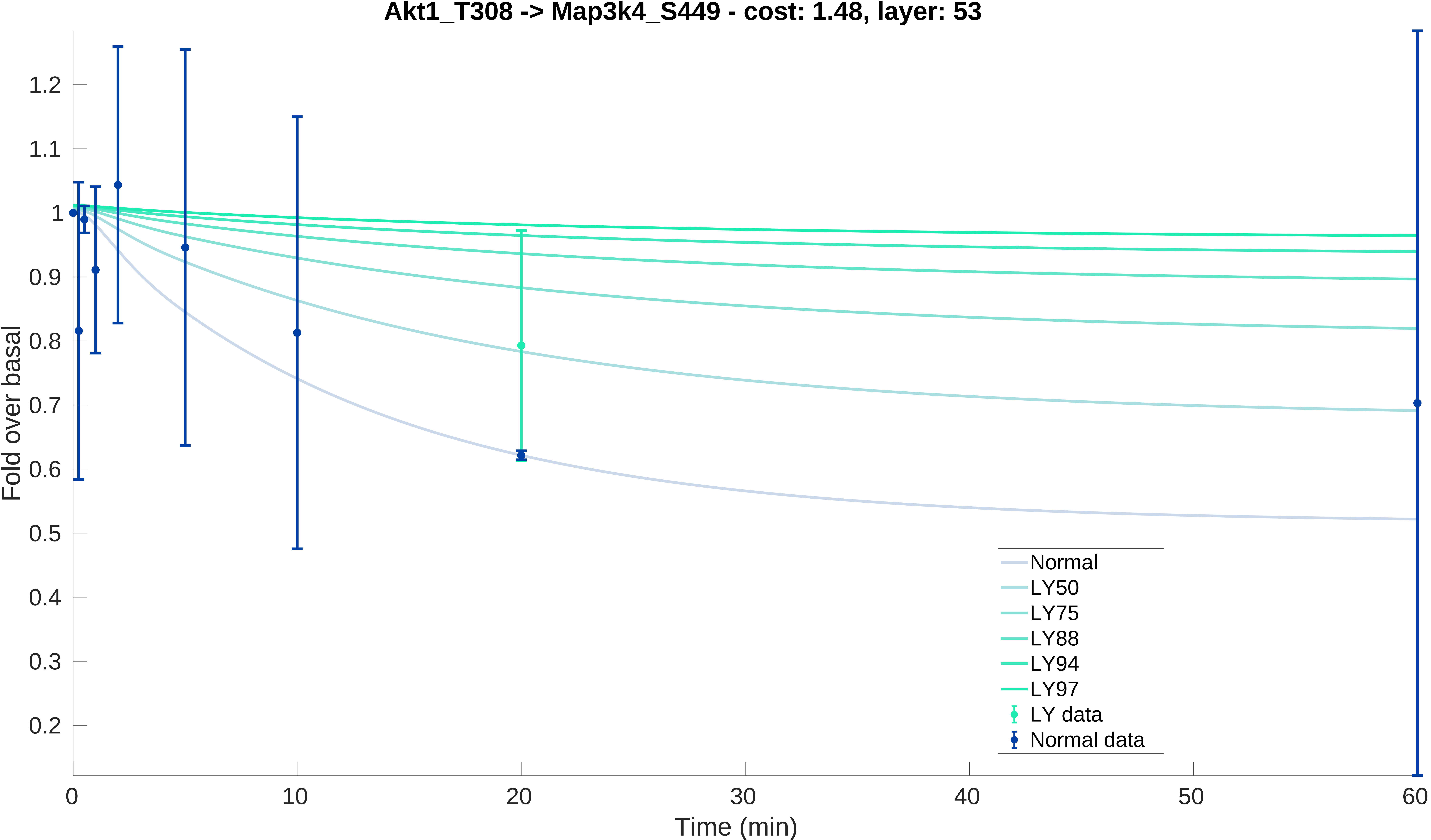
Akt1_T308 -> Map3k4_S449 - cost: 1.48, layer: 53
Fold over basal
Time (min)
0
10
20
30
40
50
60
0.2
0.3
0.4
0.5
0.6
0.7
0.8
0.9
1
1.1
1.2
Normal
LY50
LY75
LY88
LY94
LY97
LY data
Normal data

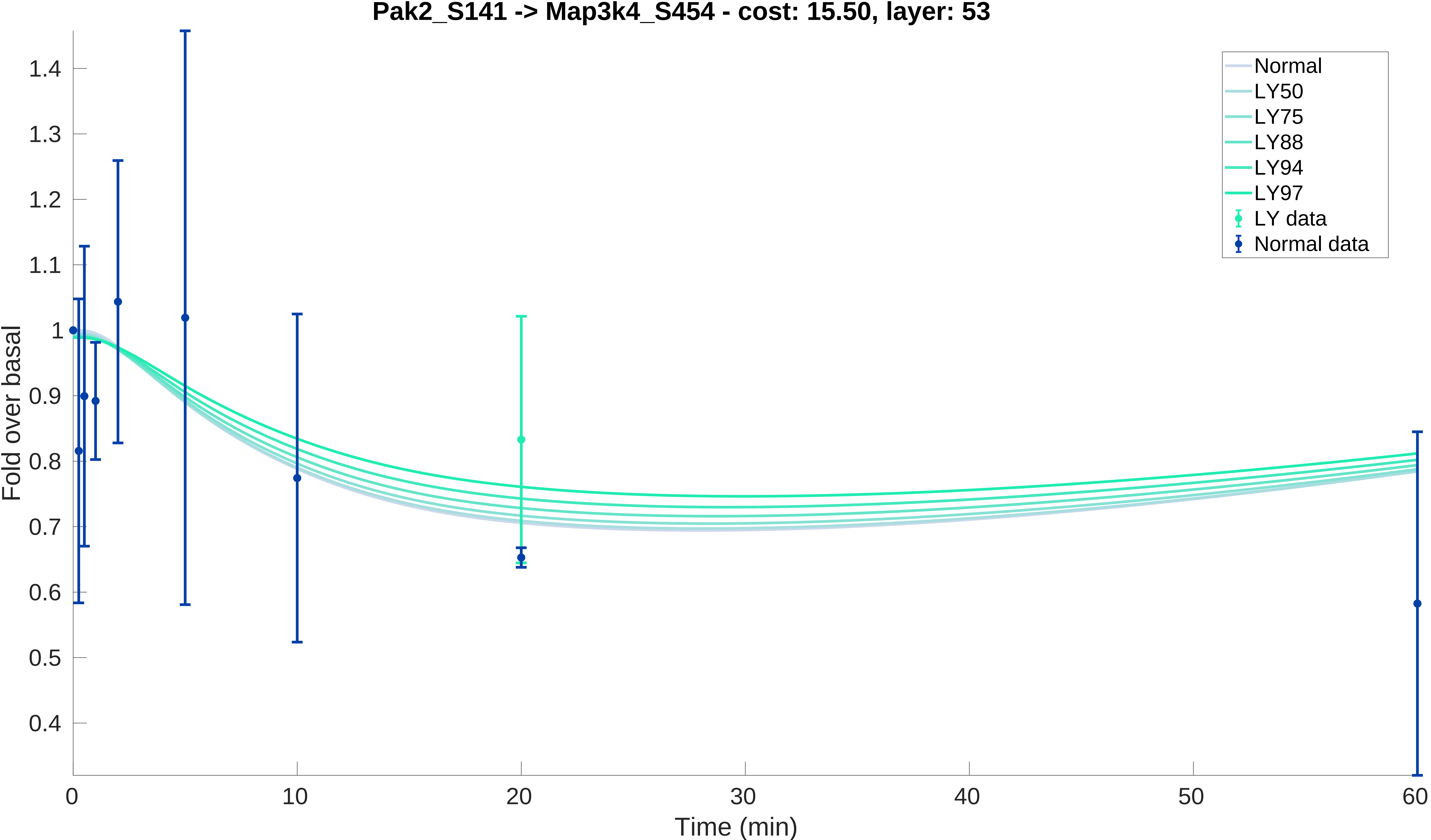

Pak2_S141 -> Map3k4_S454 - cost: 15.50, layer: 53
Fold over basal
Time (min)
1.4
1.3
1.2
1.1
1
0.9
0.8
0.7
0.6
0.5
0.4
0
10
20
30
40
50
60
Normal
LY50
LY75
LY88
LY94
LY97
LY data
Normal data

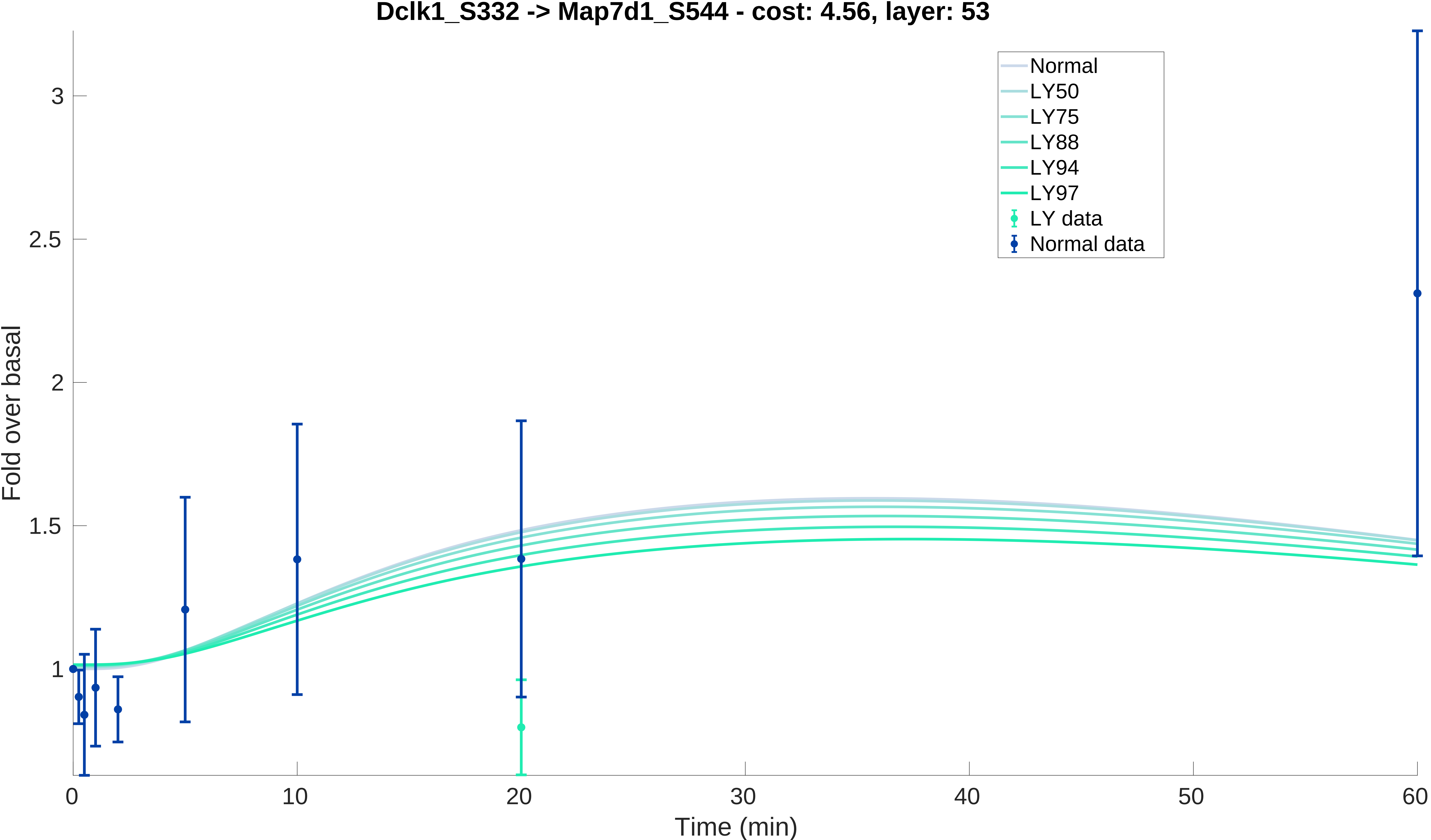

Dclk1_S332 -> Map7d1_S544 - cost: 4.56, layer: 53
Fold over basal
Time (min)
Normal
LY50
LY75
LY88
LY94
LY97
LY data
Normal data
3
2.5
2
1.5
1
0
10
20
30
40
50
60

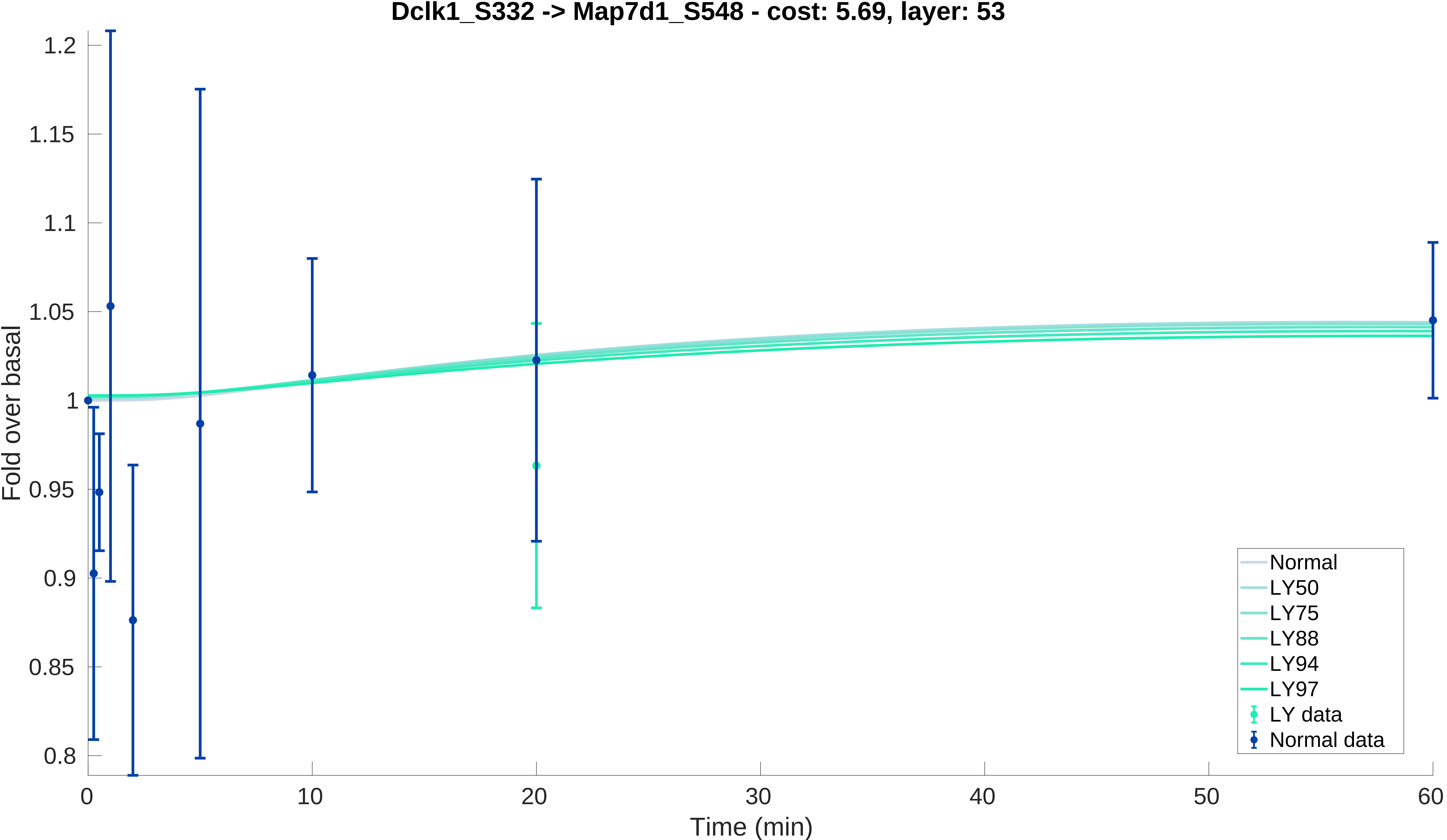

Dclk1_S332 -> Map7d1_S548 - cost: 5.69, layer: 53
Fold over basal
Time (min)
1.2
1.15
1.1
1.05
1
0.95
0.9
0.85
0.8
0
10
20
30
40
50
60
Normal
LY50
LY75
LY88
LY94
LY97
LY data
Normal data

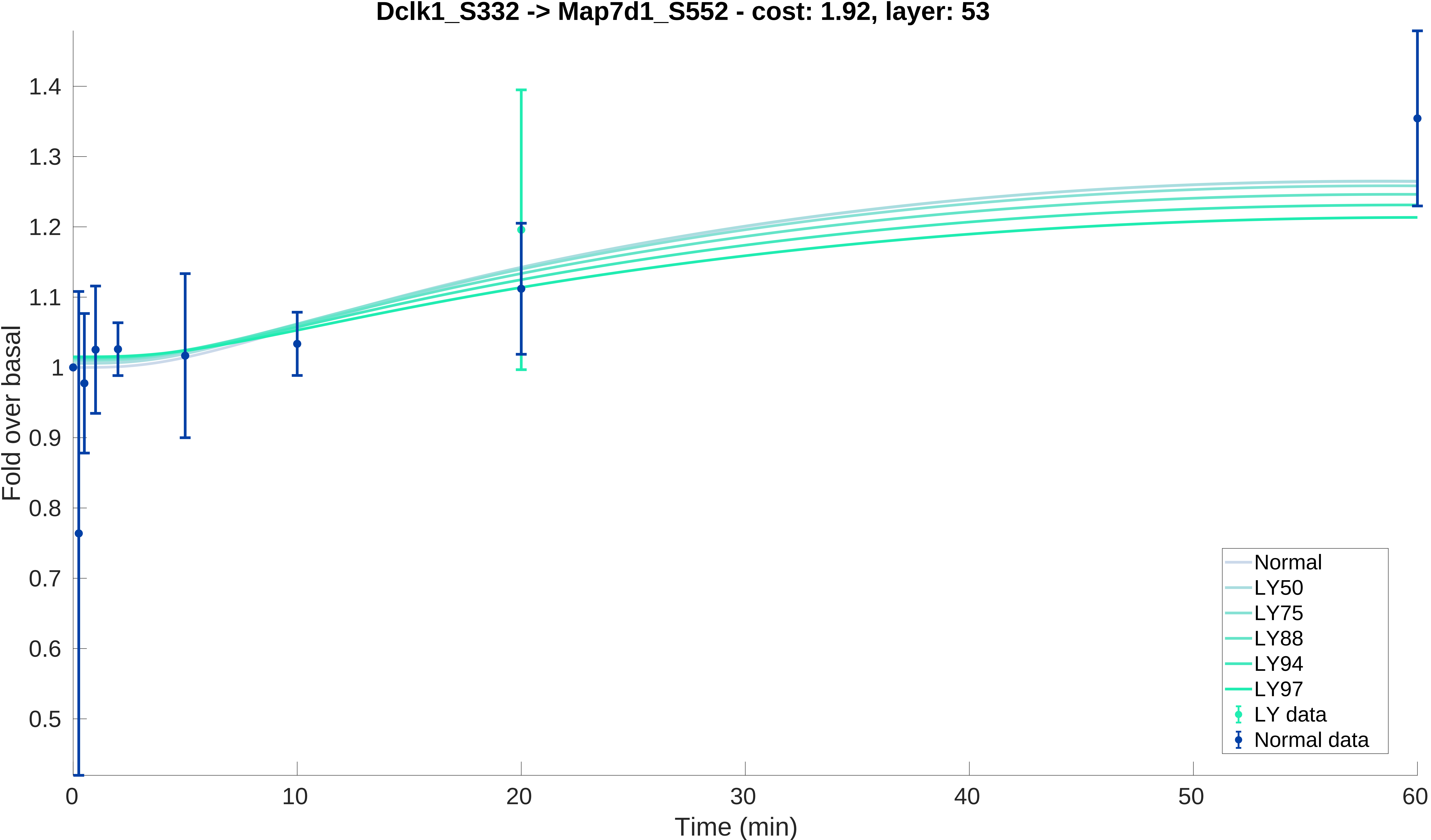

Dclk1_S332 -> Map7d1_S552 - cost: 1.92, layer: 53
Fold over basal
Time (min)
1.4
1.3
1.2
1.1
1
0.9
0.8
0.7
0.6
0.5
0
10
20
30
40
50
60
Normal
LY50
LY75
LY88
LY94
LY97
LY data
Normal data

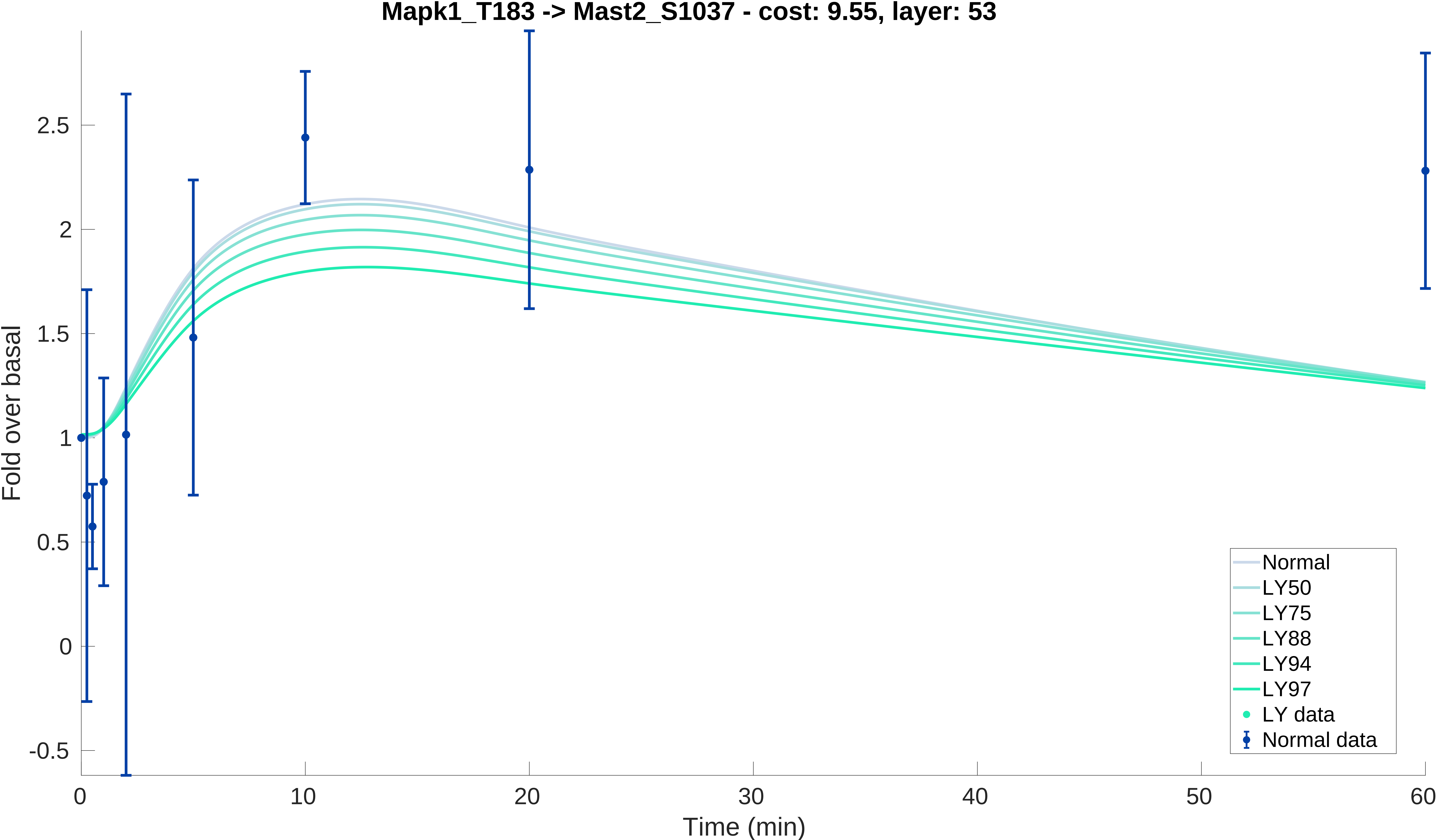

Mapk1_T183 -> Mast2_S1037 - cost: 9.55, layer: 53
Fold over basal
Time (min)
2.5
2
1.5
1
0.5
0
-0.5
0
10
20
30
40
50
60
Normal
LY50
LY75
LY88
LY94
LY97
LY data
Normal data

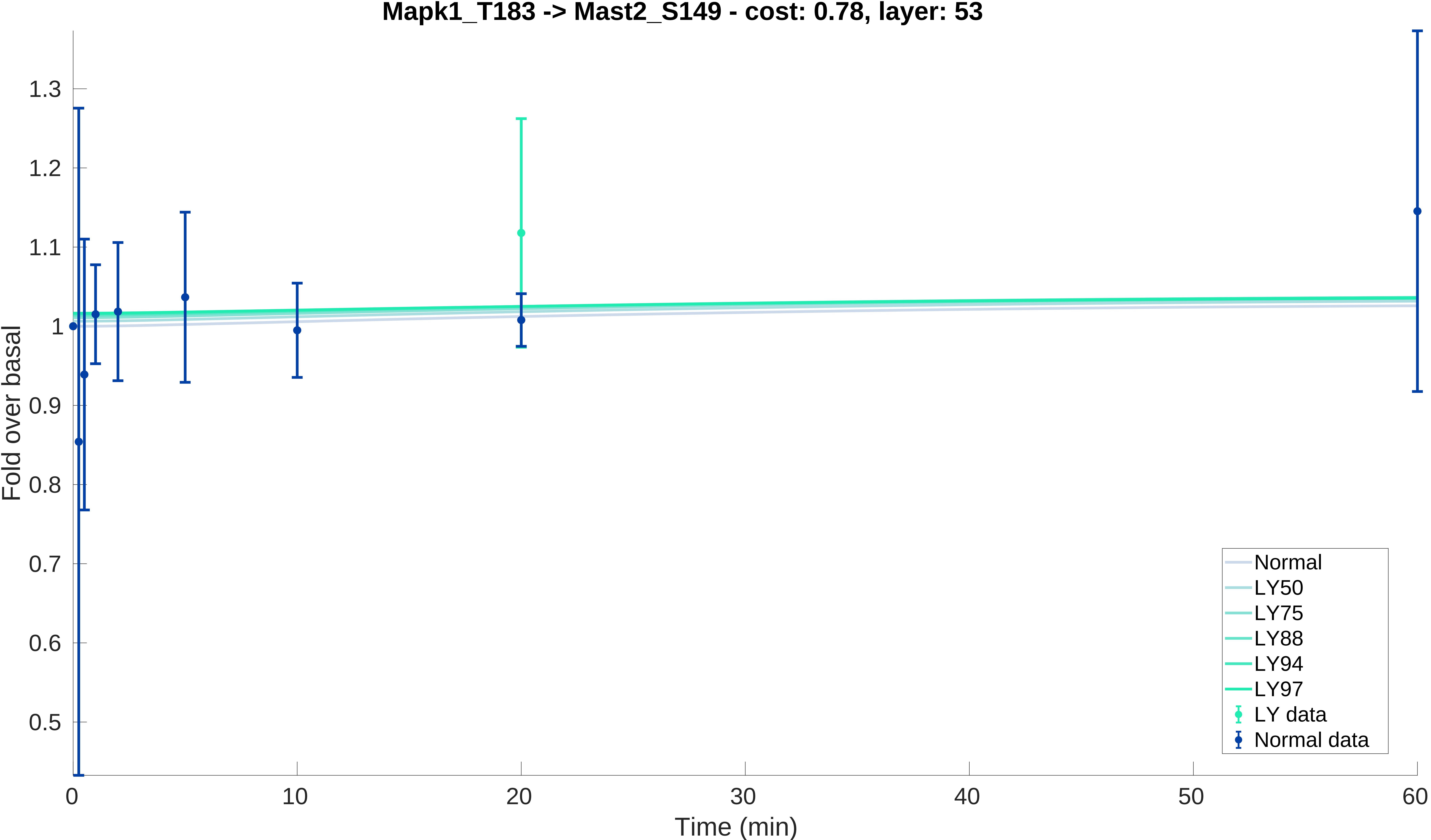

Mapk1_T183 -> Mast2_S149 - cost: 0.78, layer: 53
Fold over basal
Time (min)
1.3
1.2
1.1
1
0.9
0.8
0.7
0.6
0.5
0
10
20
30
40
50
60
Normal
LY50
LY75
LY88
LY94
LY97
LY data
Normal data

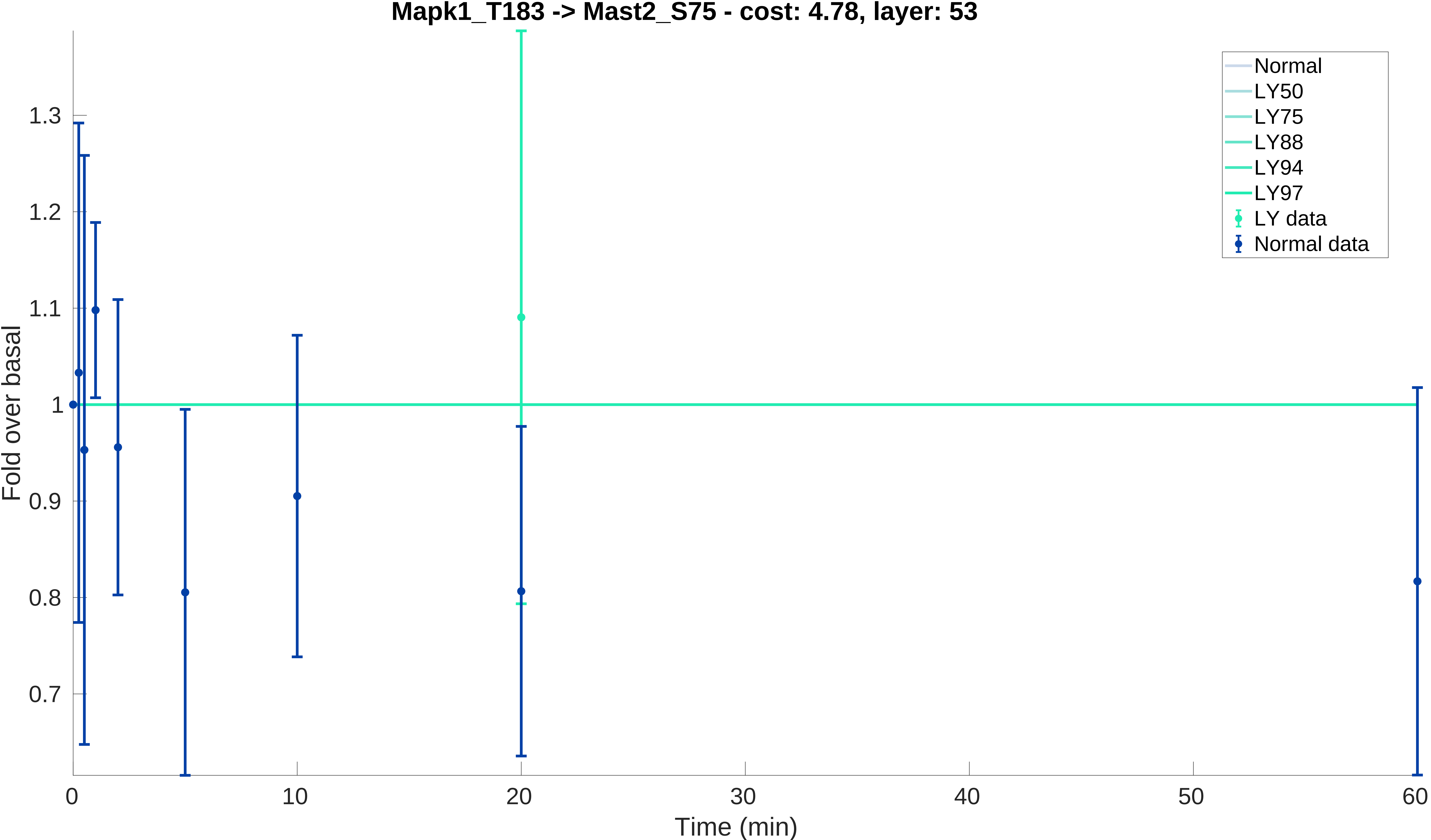

Mapk1_T183 -> Mast2_S75 - cost: 4.78, layer: 53
Fold over basal
1.3
1.2
1.1
1
0.9
0.8
0.7
0
10
20
30
40
50
60
Time (min)
Normal
LY50
LY75
LY88
LY94
LY97
LY data
Normal data

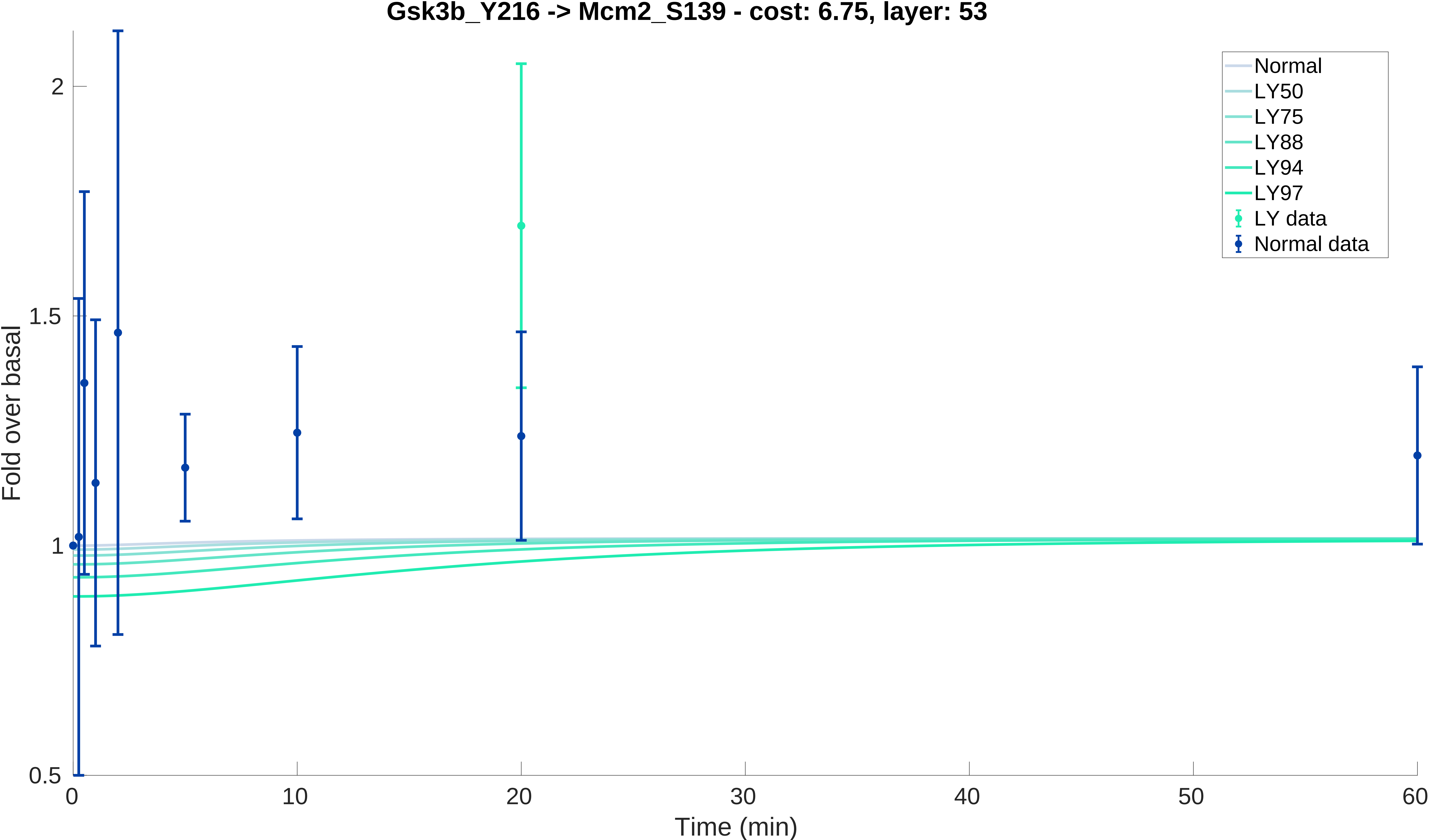
Gsk3b_Y216 -> Mcm2_S139 - cost: 6.75, layer: 53
Fold over basal
Time (min)
0.5
1
1.5
2
0
10
20
30
40
50
60
Normal
LY50
LY75
LY88
LY94
LY97
LY data
Normal data

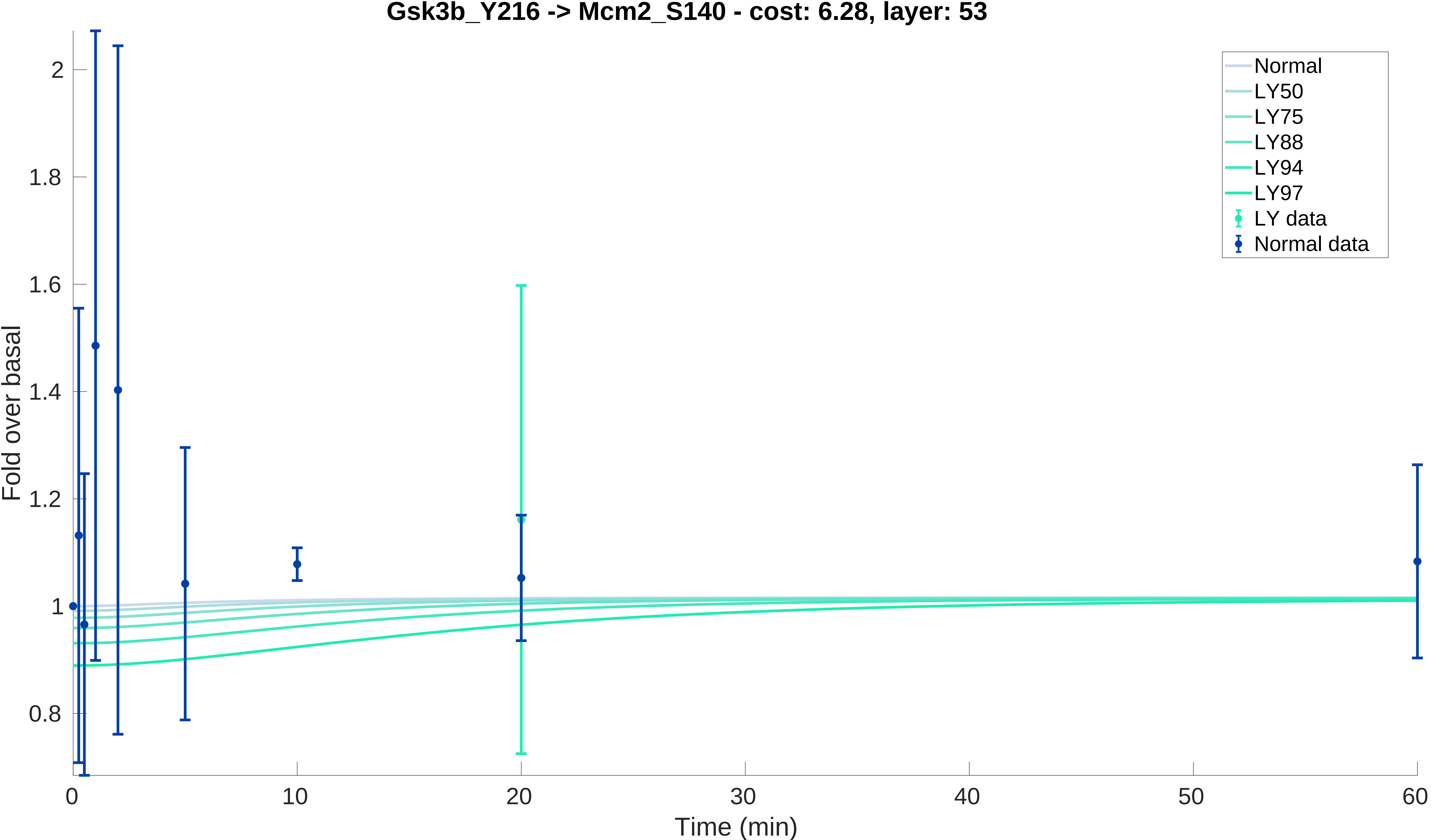

Gsk3b_Y216 -> Mcm2_S140 - cost: 6.28, layer: 53
Fold over basal
Time (min)
0.8
1
1.2
1.4
1.6
1.8
2
0
10
20
30
40
50
60
Normal
LY50
LY75
LY88
LY94
LY97
LY data
Normal data

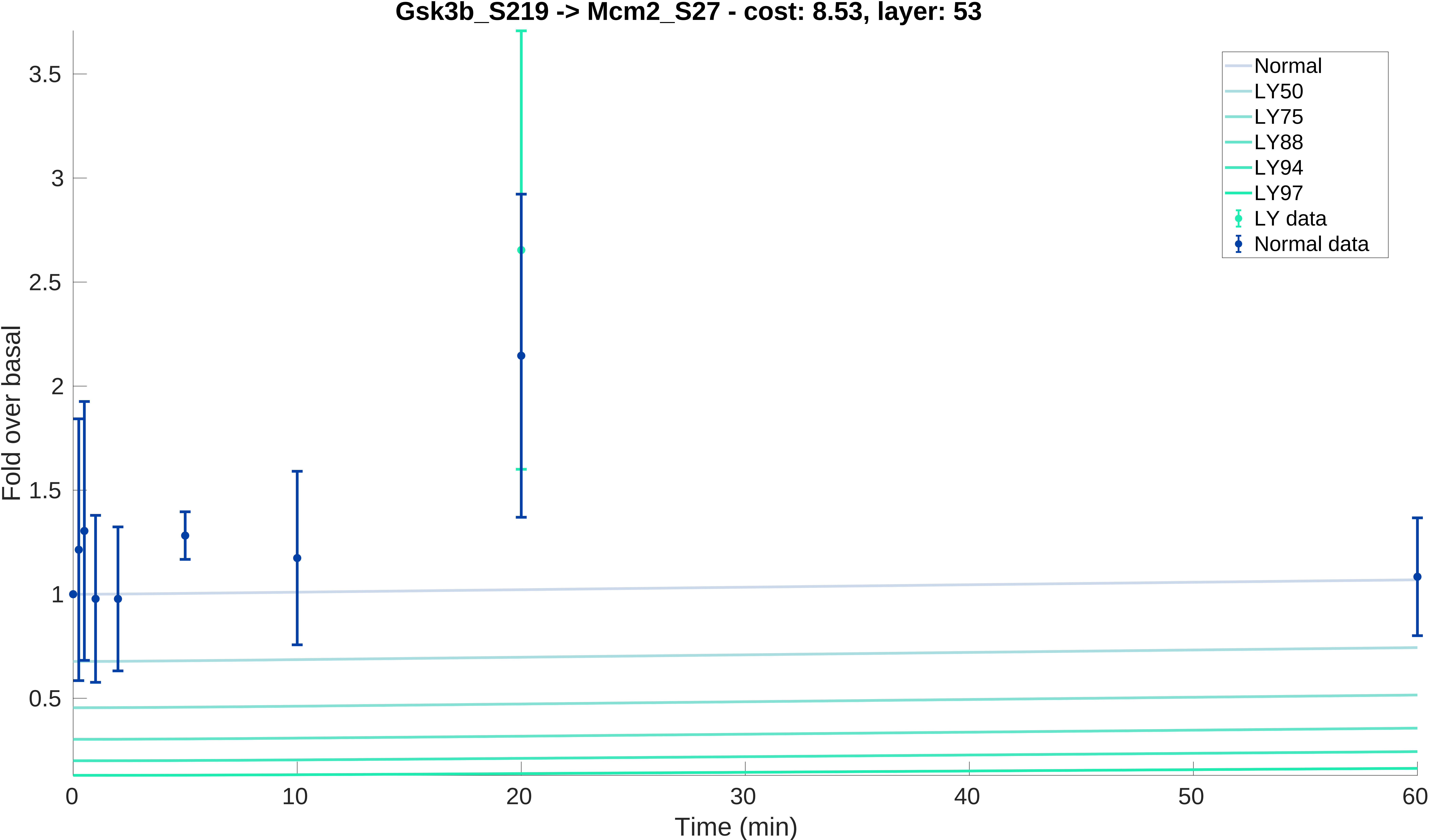

Gsk3b_S219 -> Mcm2_S27 - cost: 8.53, layer: 53
Fold over basal
Time (min)
Normal
LY50
LY75
LY88
LY94
LY97
LY data
Normal data
0.5
1
1.5
2
2.5
3
3.5
0
10
20
30
40
50
60

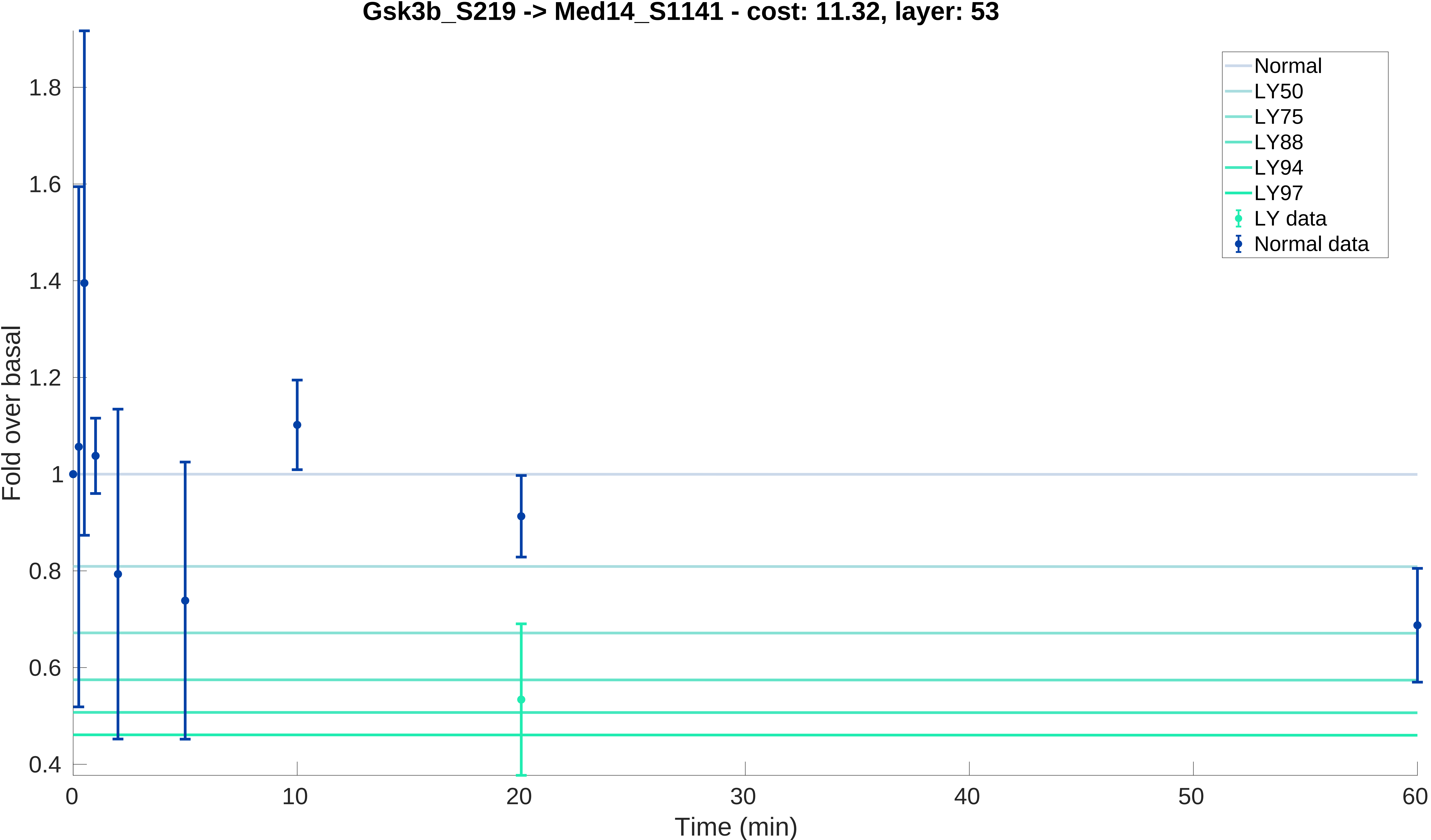
Gsk3b_S219 -> Med14_S1141 - cost: 11.32, layer: 53
Fold over basal
Time (min)
0.4
0.6
0.8
1
1.2
1.4
1.6
1.8
0
10
20
30
40
50
60
Normal
LY50
LY75
LY88
LY94
LY97
LY data
Normal data

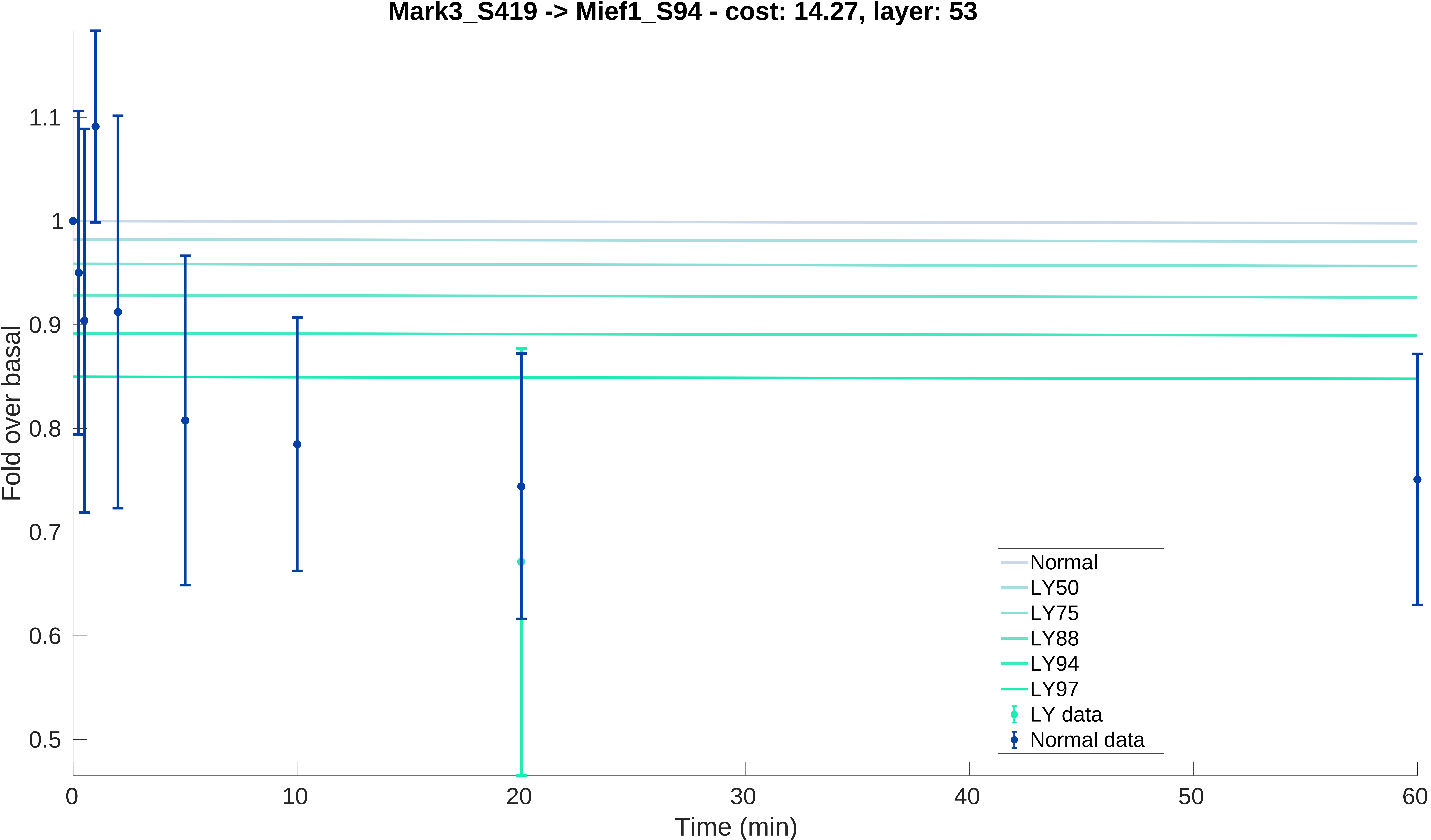
Mark3_S419 -> Mief1_S94 - cost: 14.27, layer: 53
Fold over basal
Time (min)
0.5
0.6
0.7
0.8
0.9
1
1.1
0
10
20
30
40
50
60
Normal
LY50
LY75
LY88
LY94
LY97
LY data
Normal data

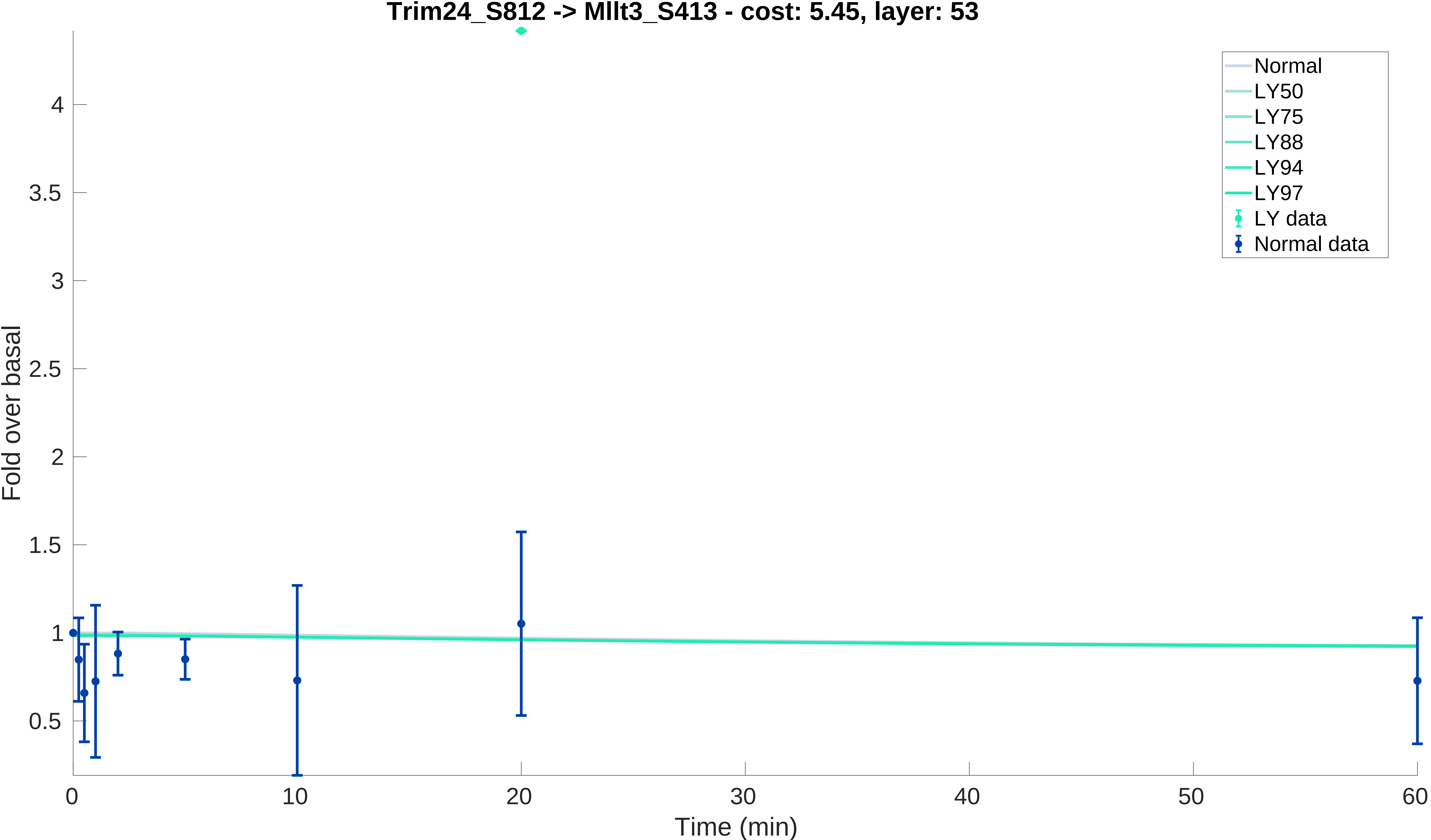

Trim24_S812 -> Mllt3_S413 - cost: 5.45, layer: 53
Fold over basal
Time (min)
Normal
LY50
LY75
LY88
LY94
LY97
LY data
Normal data
4
3.5
3
2.5
2
1.5
1
0.5
0
10
20
30
40
50
60

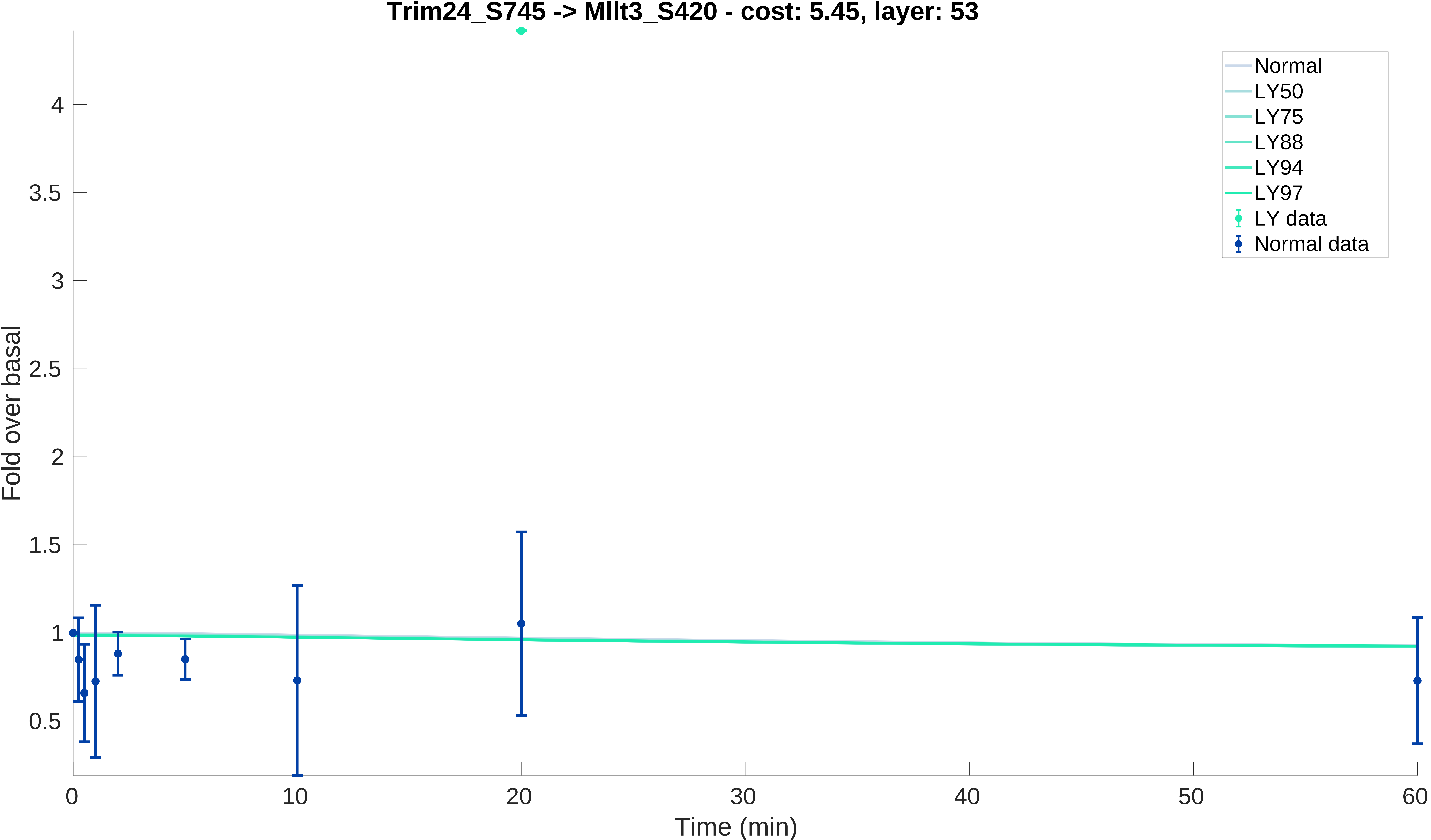

Trim24_S745 -> Mllt3_S420 - cost: 5.45, layer: 53
Fold over basal
Time (min)
Normal
LY50
LY75
LY88
LY94
LY97
LY data
Normal data
0.5
1
1.5
2
2.5
3
3.5
4
0
10
20
30
40
50
60

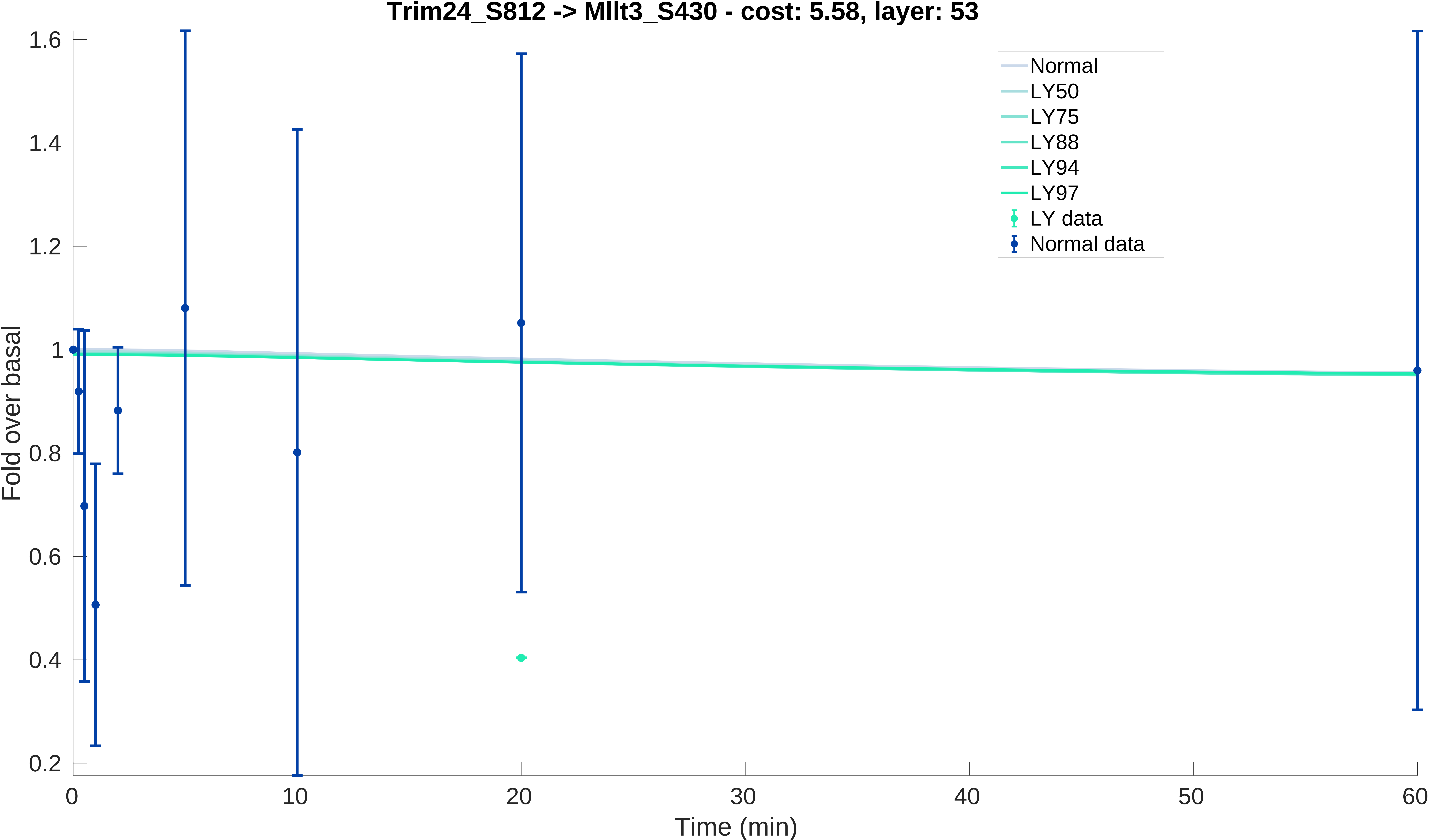

Trim24_S812 -> Mllt3_S430 - cost: 5.58, layer: 53
Fold over basal
Time (min)
Normal
LY50
LY75
LY88
LY94
LY97
LY data
Normal data
1.6
1.4
1.2
1
0.8
0.6
0.4
0.2
0
10
20
30
40
50
60

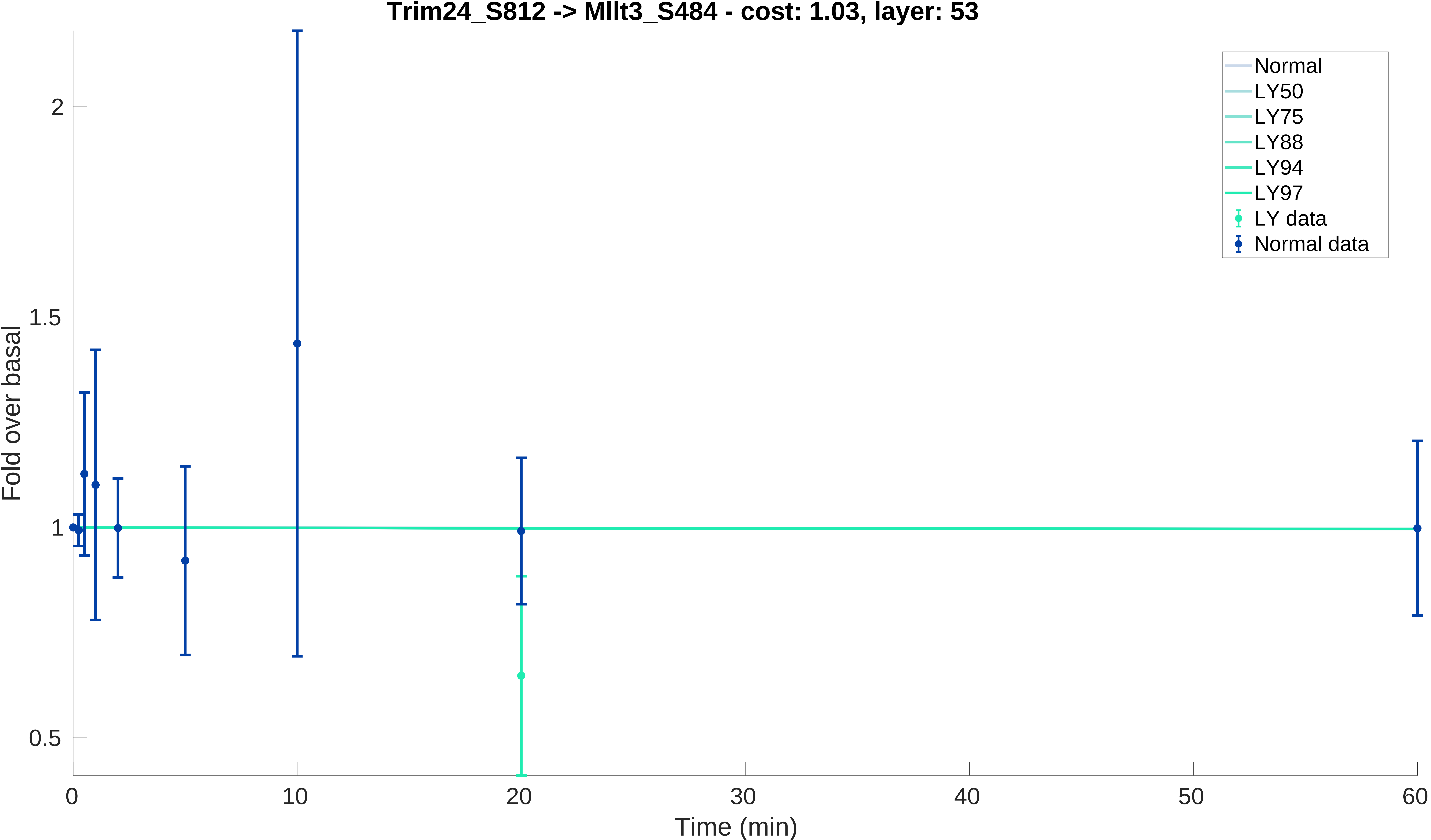

Trim24_S812 -> Mllt3_S484 - cost: 1.03, layer: 53
Fold over basal
0.5
1
1.5
2
0
10
20
30
40
50
60
Time (min)
Normal
LY50
LY75
LY88
LY94
LY97
LY data
Normal data

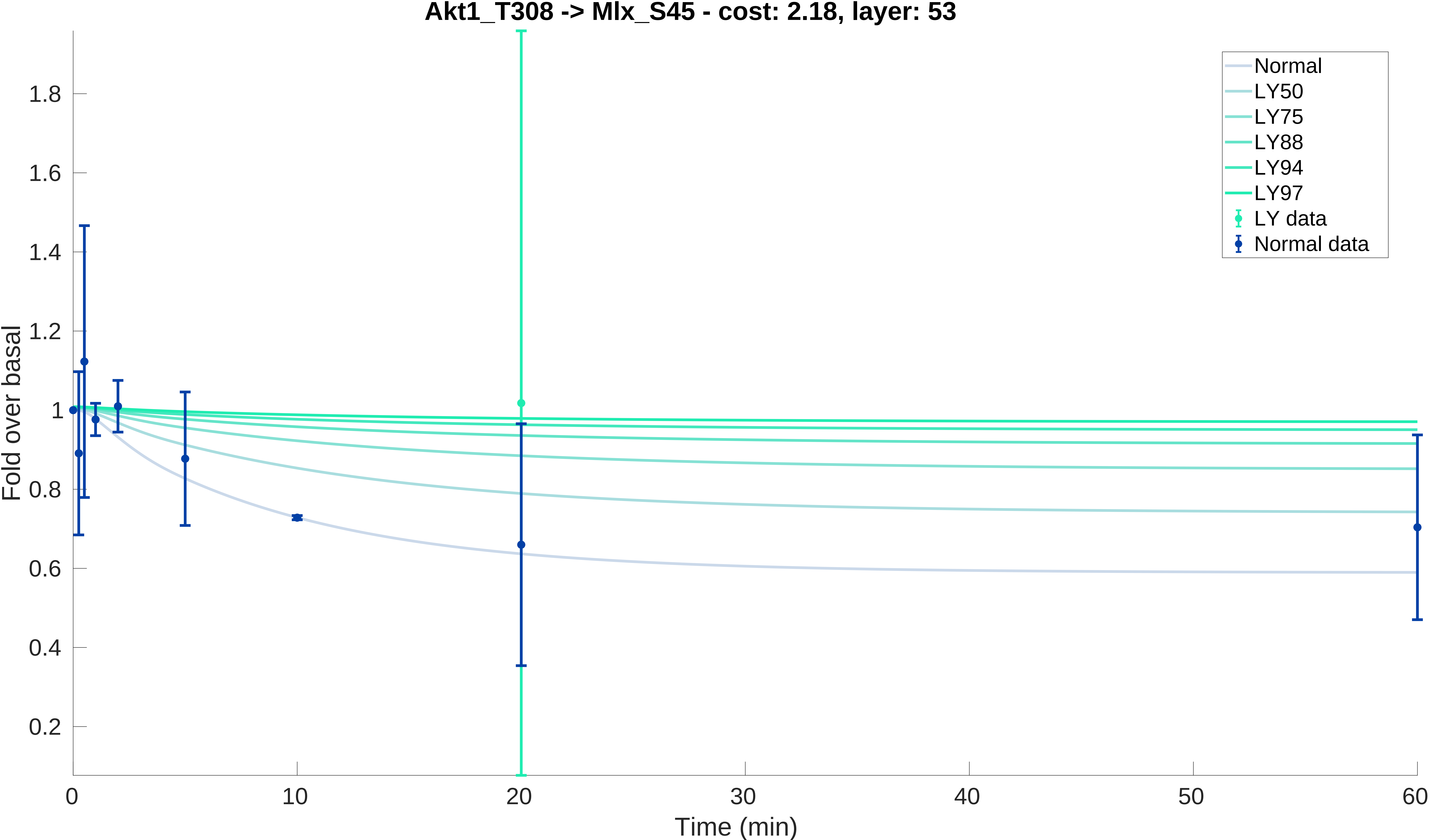
Akt1_T308 -> Mlx_S45 - cost: 2.18, layer: 53
Fold over basal
Time (min)
0.2
0.4
0.6
0.8
1
1.2
1.4
1.6
1.8
0
10
20
30
40
50
60
Normal
LY50
LY75
LY88
LY94
LY97
LY data
Normal data

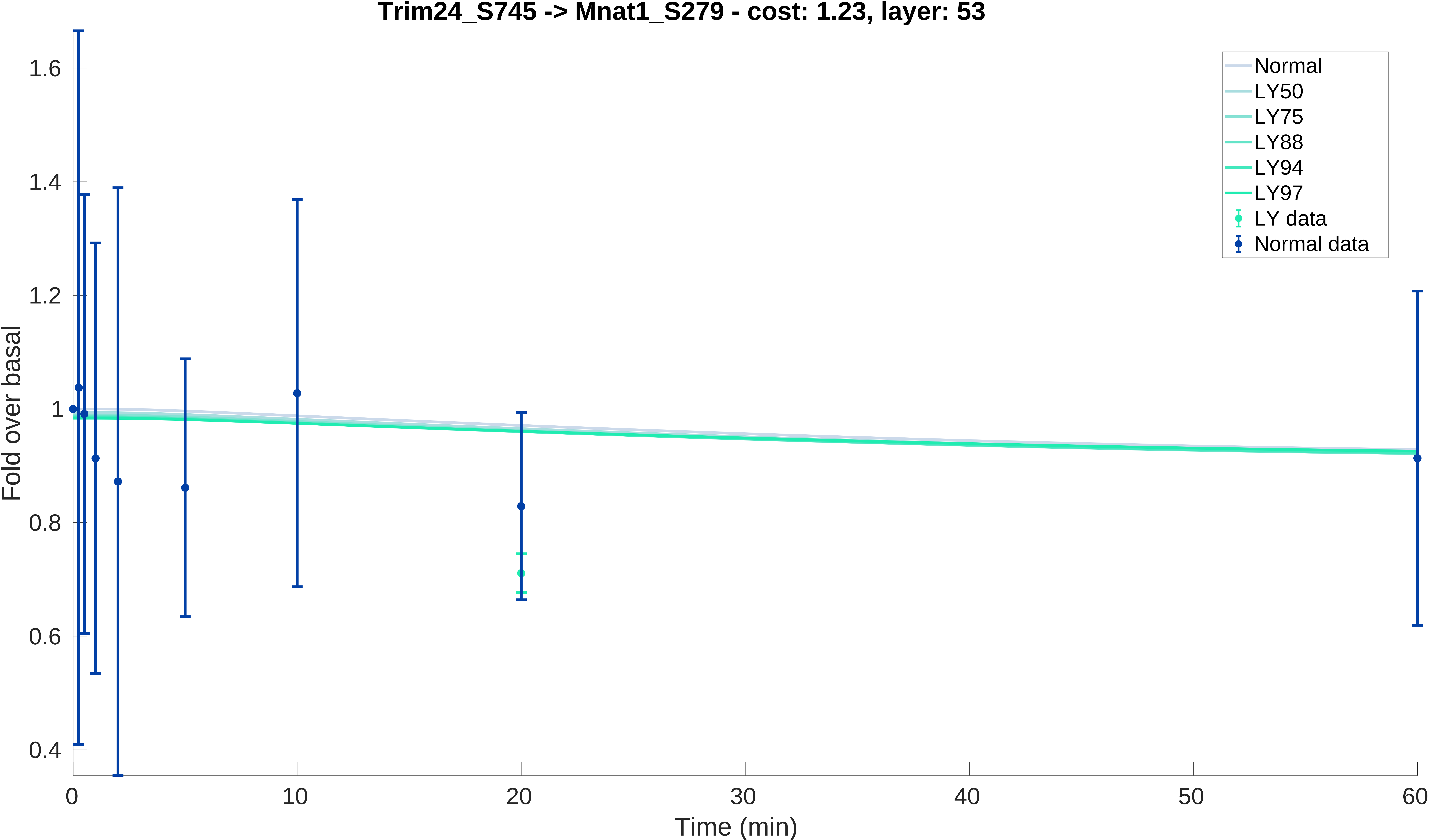

Trim24_S745 -> Mnat1_S279 - cost: 1.23, layer: 53
Normal
LY50
LY75
LY88
LY94
LY97
LY data
Normal data
Fold over basal
1.6
1.4
1.2
1
0.8
0.6
0.4
0
10
20
30
40
50
60
Time (min)

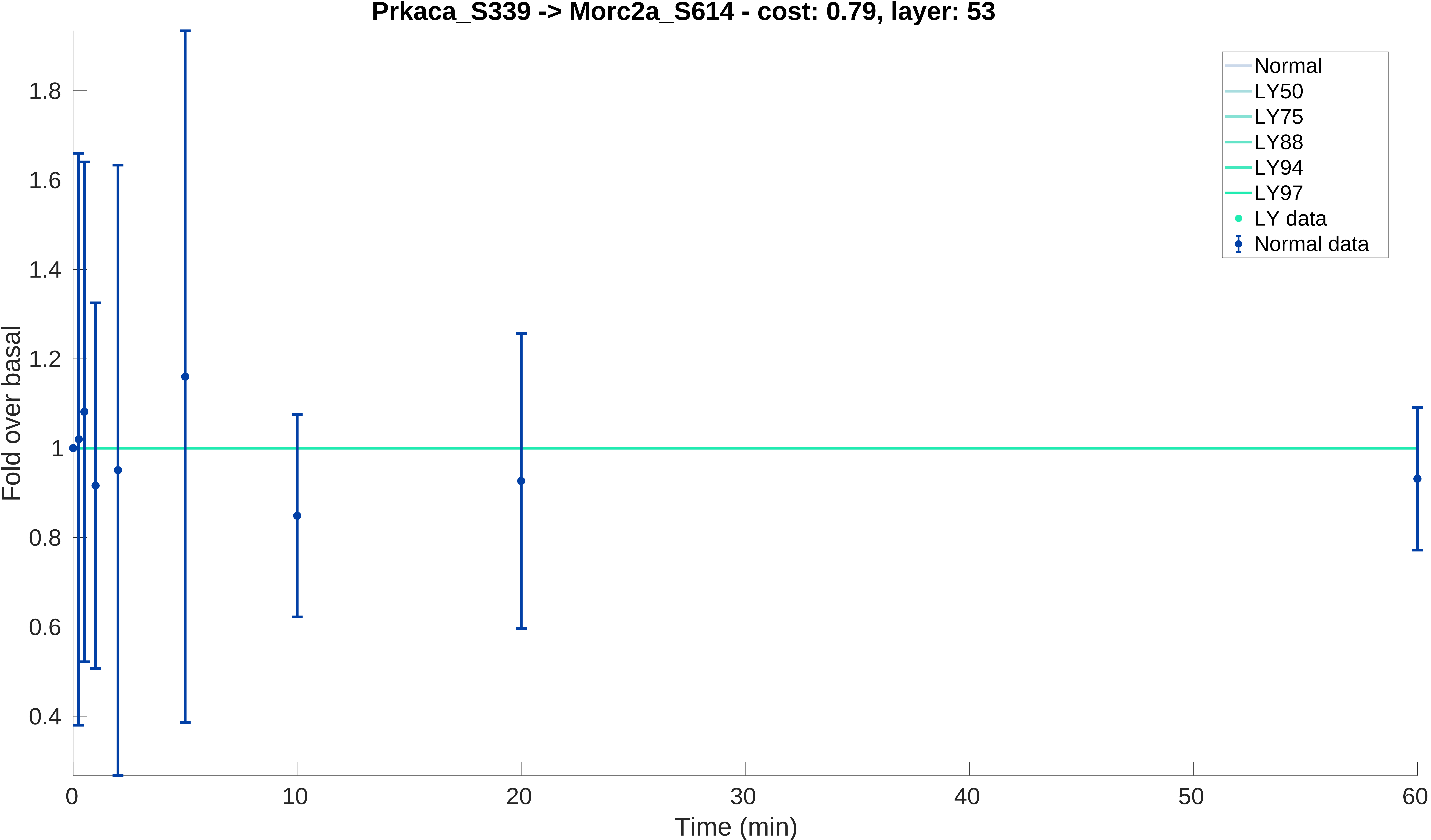

Prkaca_S339 -> Morc2a_S614 - cost: 0.79, layer: 53
Fold over basal
1.8
1.6
1.4
1.2
1
0.8
0.6
0.4
0
10
20
30
40
50
60
Time (min)
Normal
LY50
LY75
LY88
LY94
LY97
LY data
Normal data

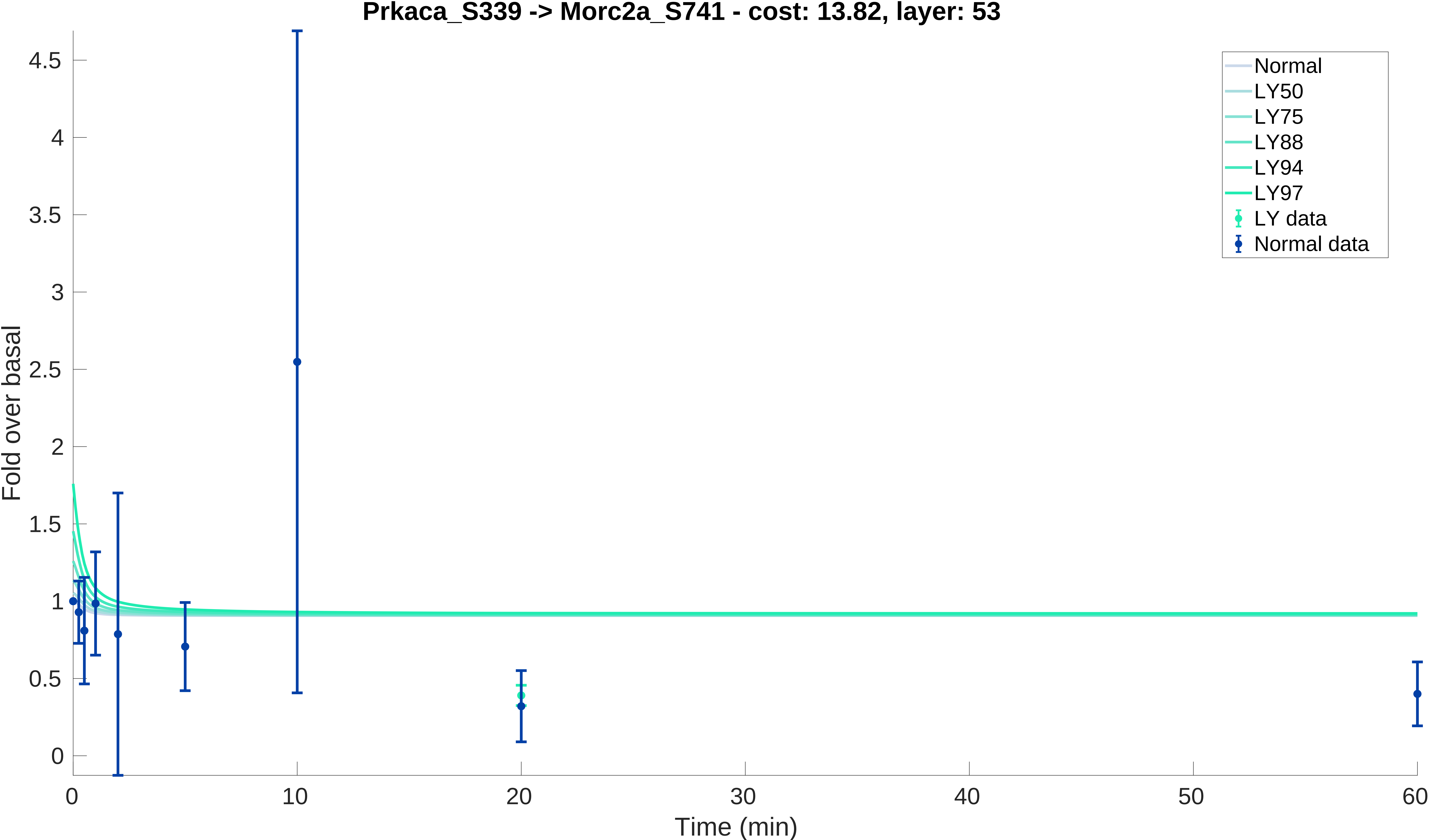

Prrkaca_S339 -> Morc2a_S741 - cost: 13.82, layer: 53
Fold over basal
Time (min)
0
0.5
1
1.5
2
2.5
3
3.5
4
4.5
0
10
20
30
40
50
60
Normal
LY50
LY75
LY88
LY94
LY97
LY data
Normal data

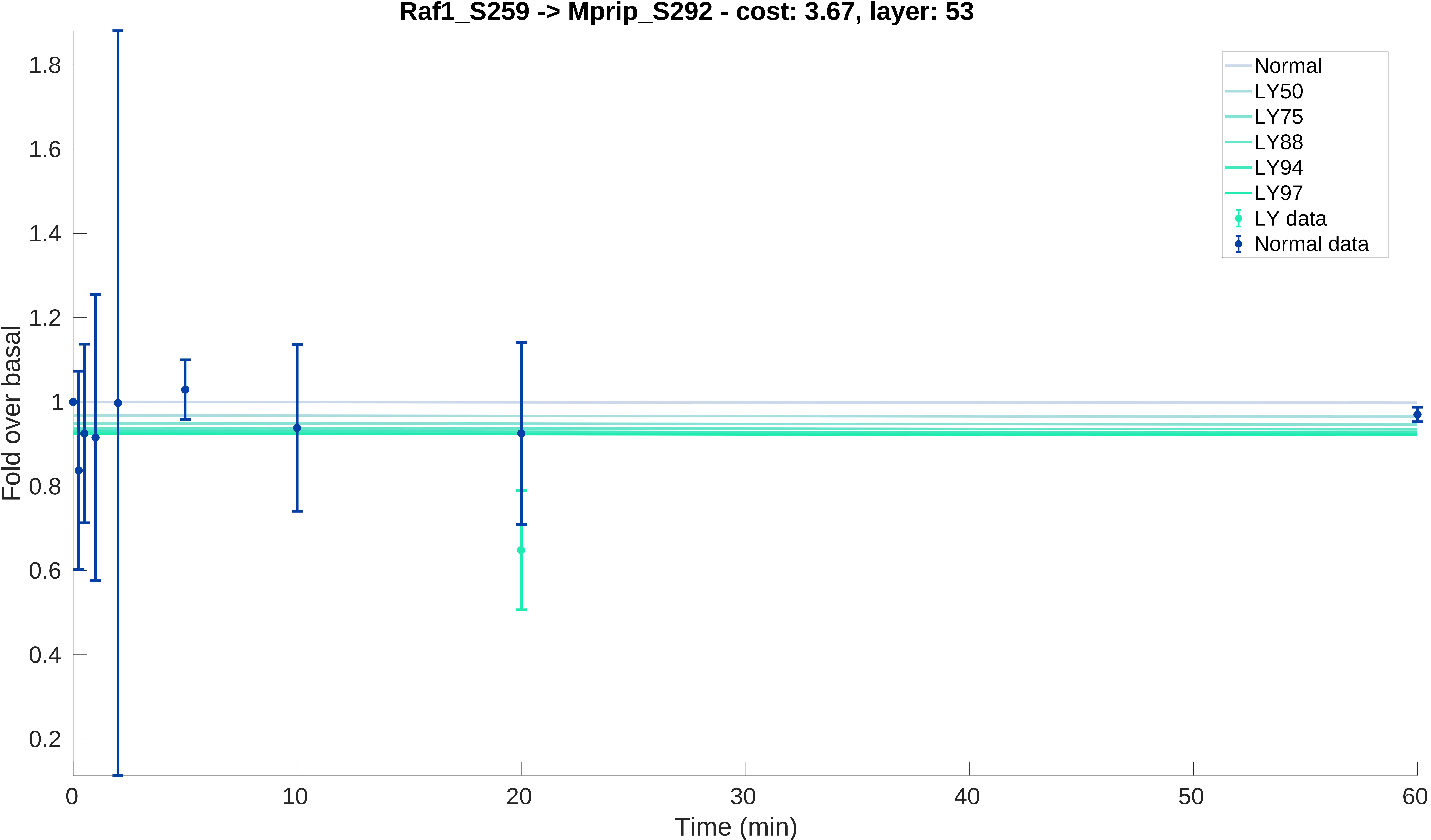

Raf1_S259 -> Mprip_S292 - cost: 3.67, layer: 53
Normal
LY50
LY75
LY88
LY94
LY97
LY data
Normal data
Fold over basal
Time (min)
1.8
1.6
1.4
1.2
1
0.8
0.6
0.4
0.2
0
10
20
30
40
50
60

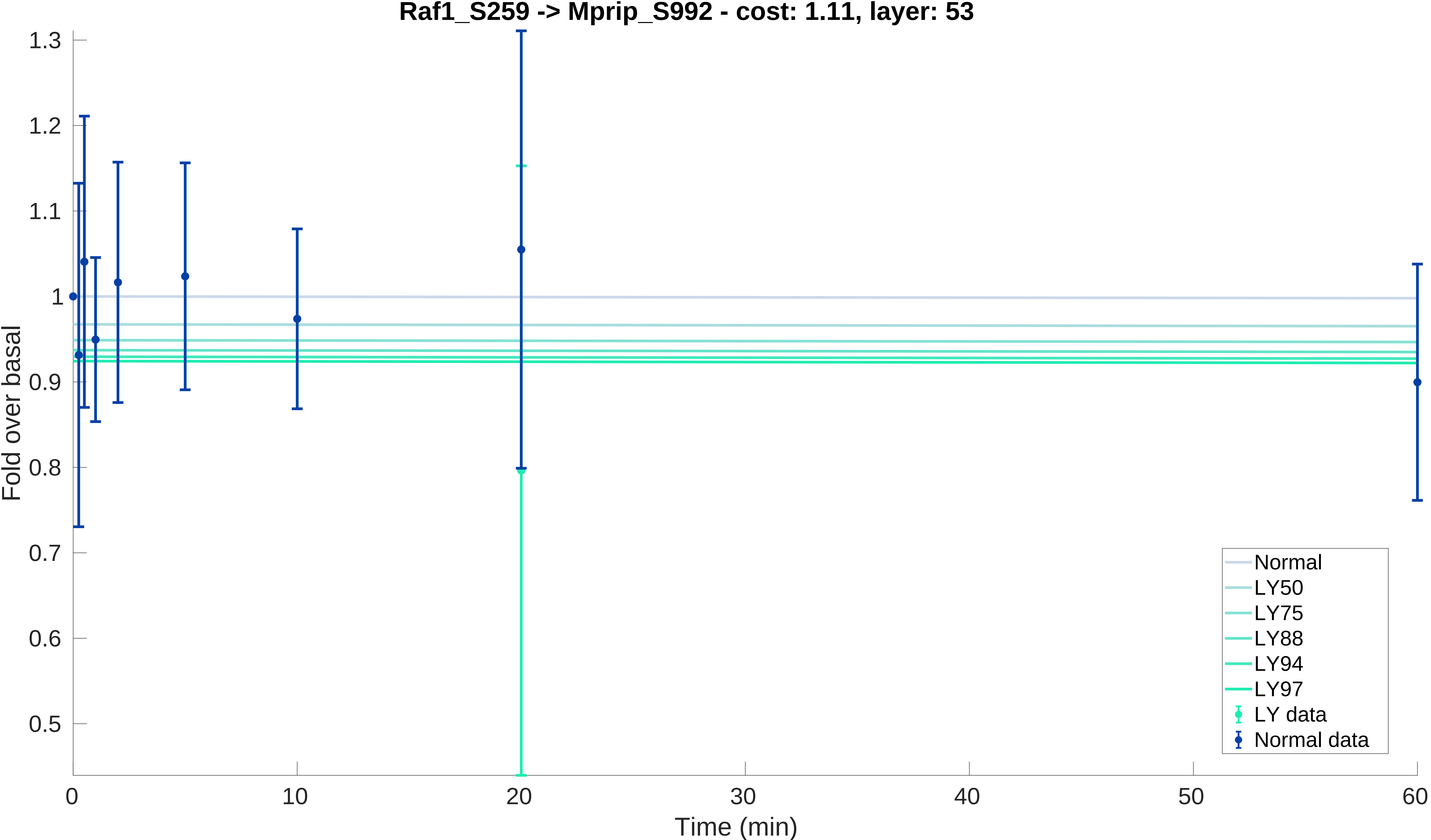

Raf1_S259 -> Mprip_S992 - cost: 1.11, layer: 53
Fold over basal
Time (min)
1.3
1.2
1.1
1
0.9
0.8
0.7
0.6
0.5
0
10
20
30
40
50
60
Normal
LY50
LY75
LY88
LY94
LY97
LY data
Normal data

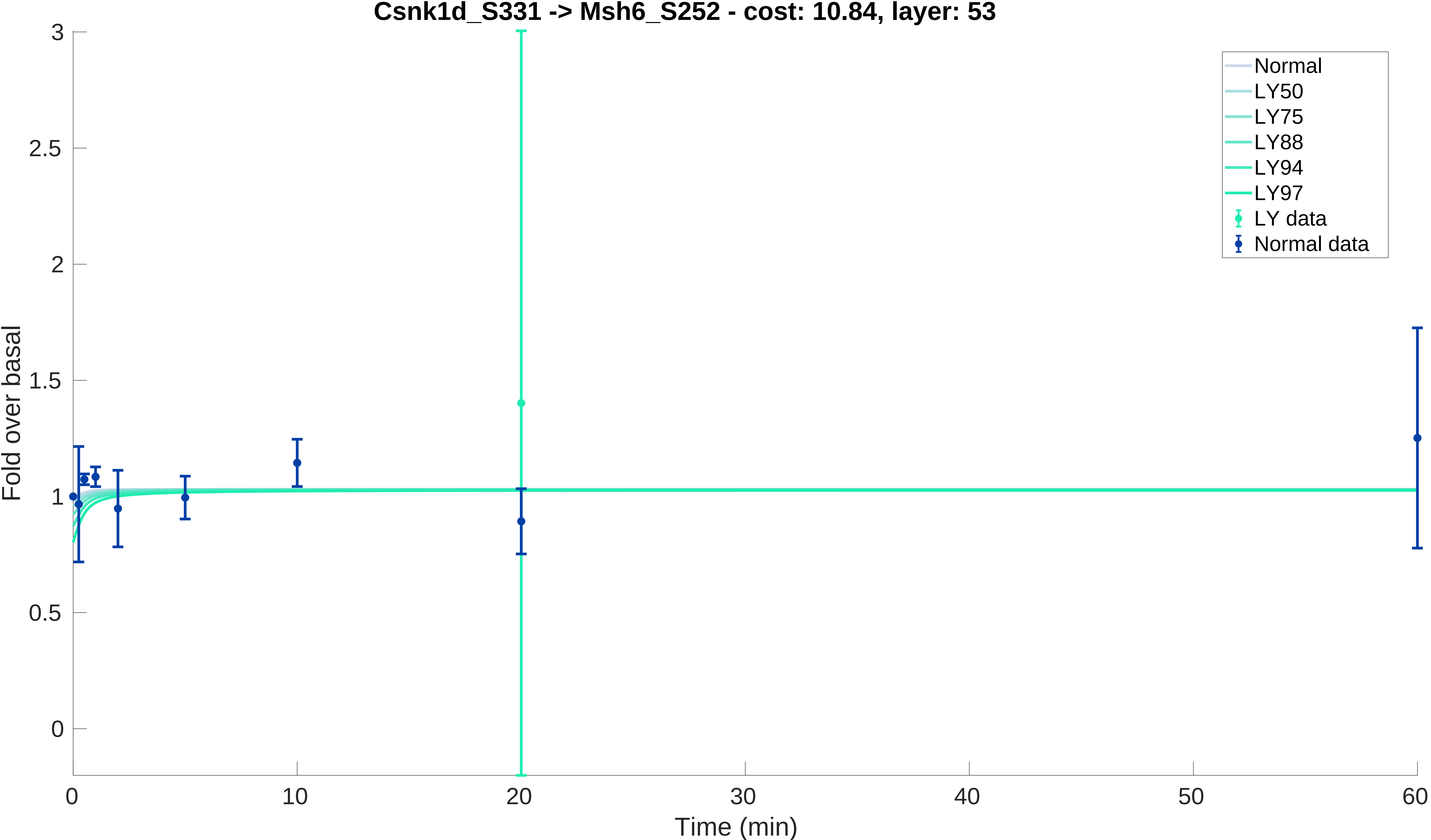

Csnk1d_S331 -> Msh6_S252 - cost: 10.84, layer: 53
Fold over basal
Time (min)
0
0.5
1
1.5
2
2.5
3
0
10
20
30
40
50
60
Normal
LY50
LY75
LY88
LY94
LY97
LY data
Normal data

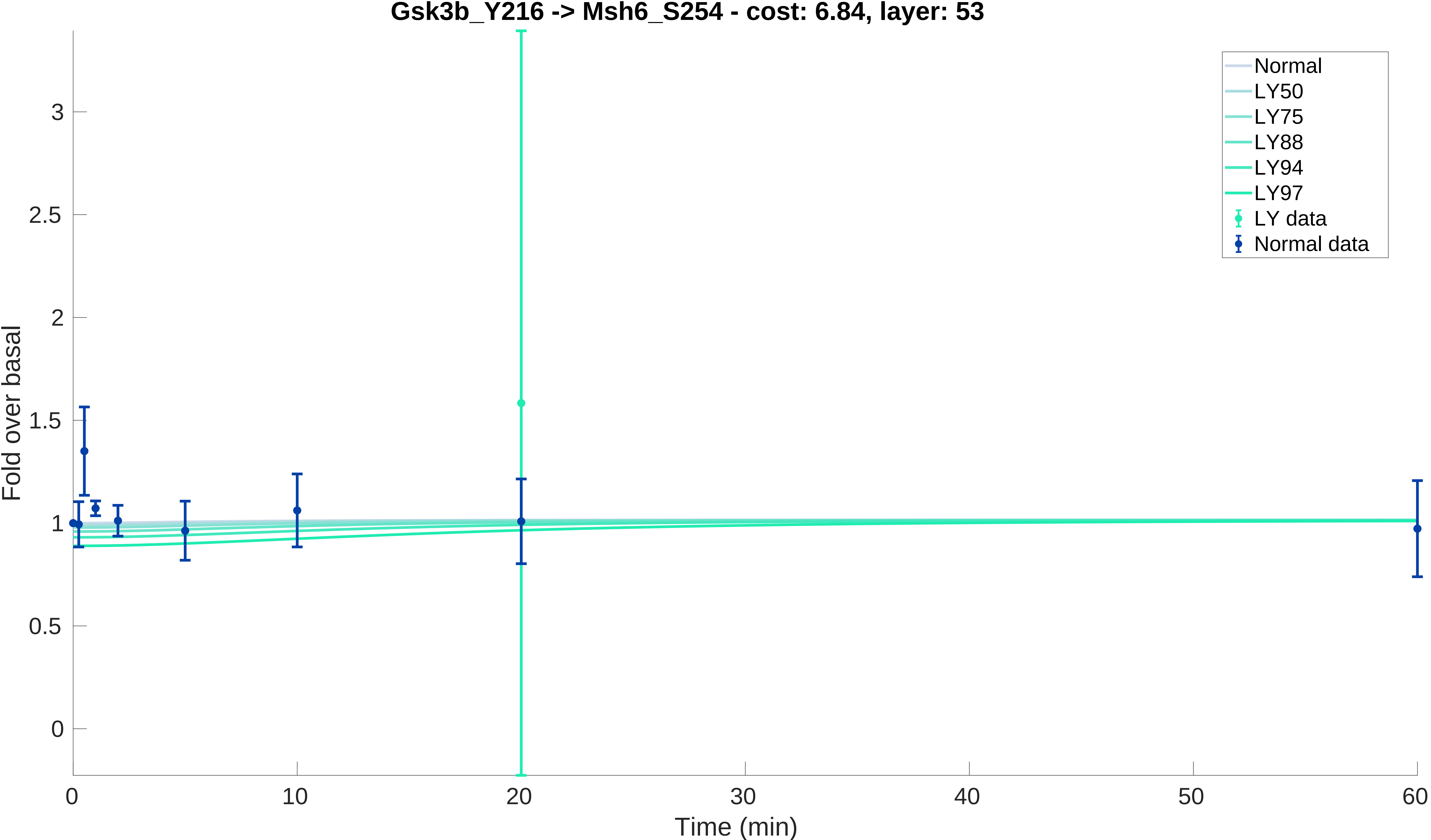

Gsk3b_Y216 -> Msh6_S254 - cost: 6.84, layer: 53
Fold over basal
Time (min)
0
0.5
1
1.5
2
2.5
3
0
10
20
30
40
50
60
Normal
LY50
LY75
LY88
LY94
LY97
LY data
Normal data

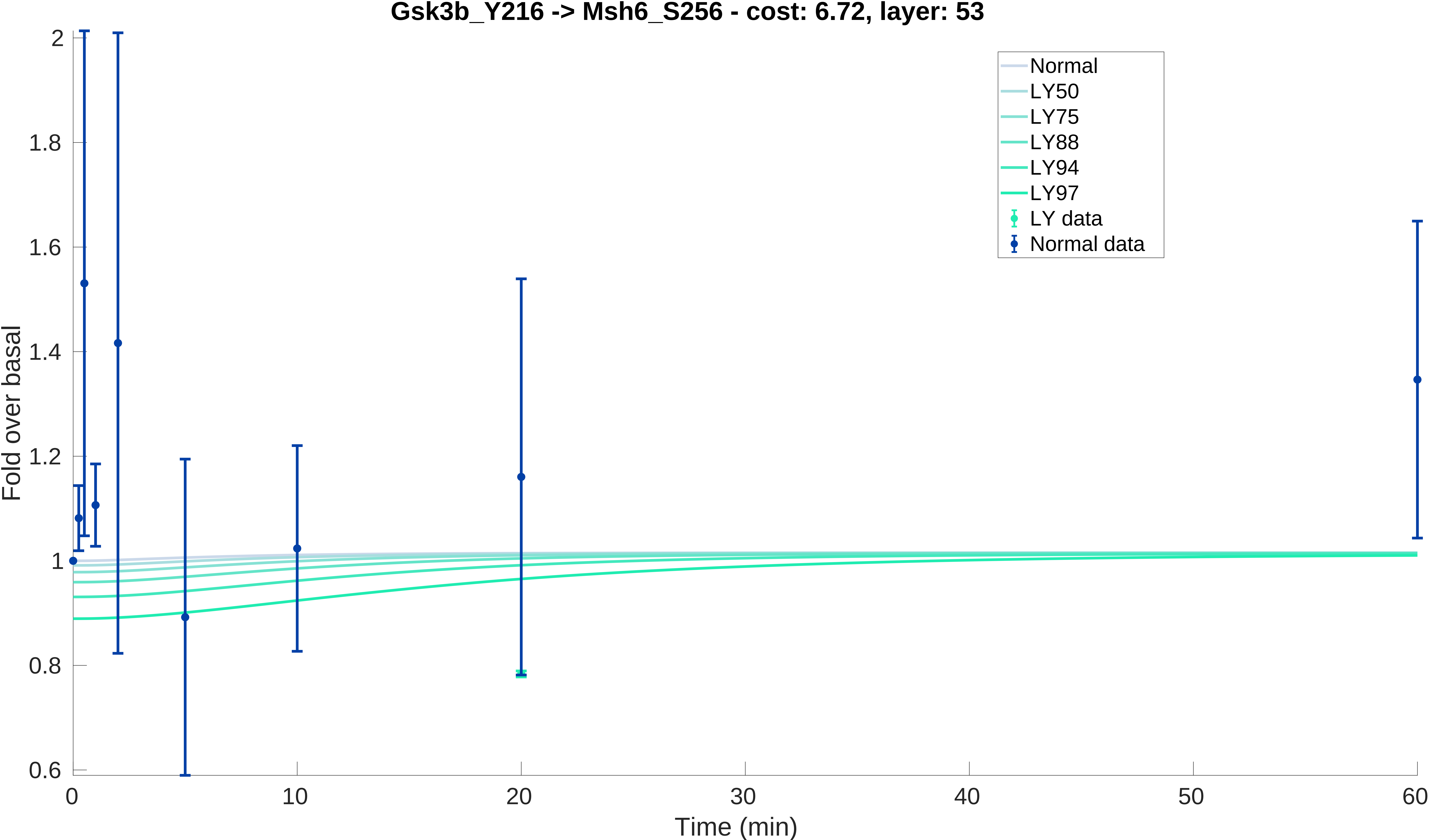

Gsk3b_Y216 -> Msh6_S256 - cost: 6.72, layer: 53
Fold over basal
Time (min)
0.6
0.8
1
1.2
1.4
1.6
1.8
2
0
10
20
30
40
50
60
Normal
LY50
LY75
LY88
LY94
LY97
LY data
Normal data

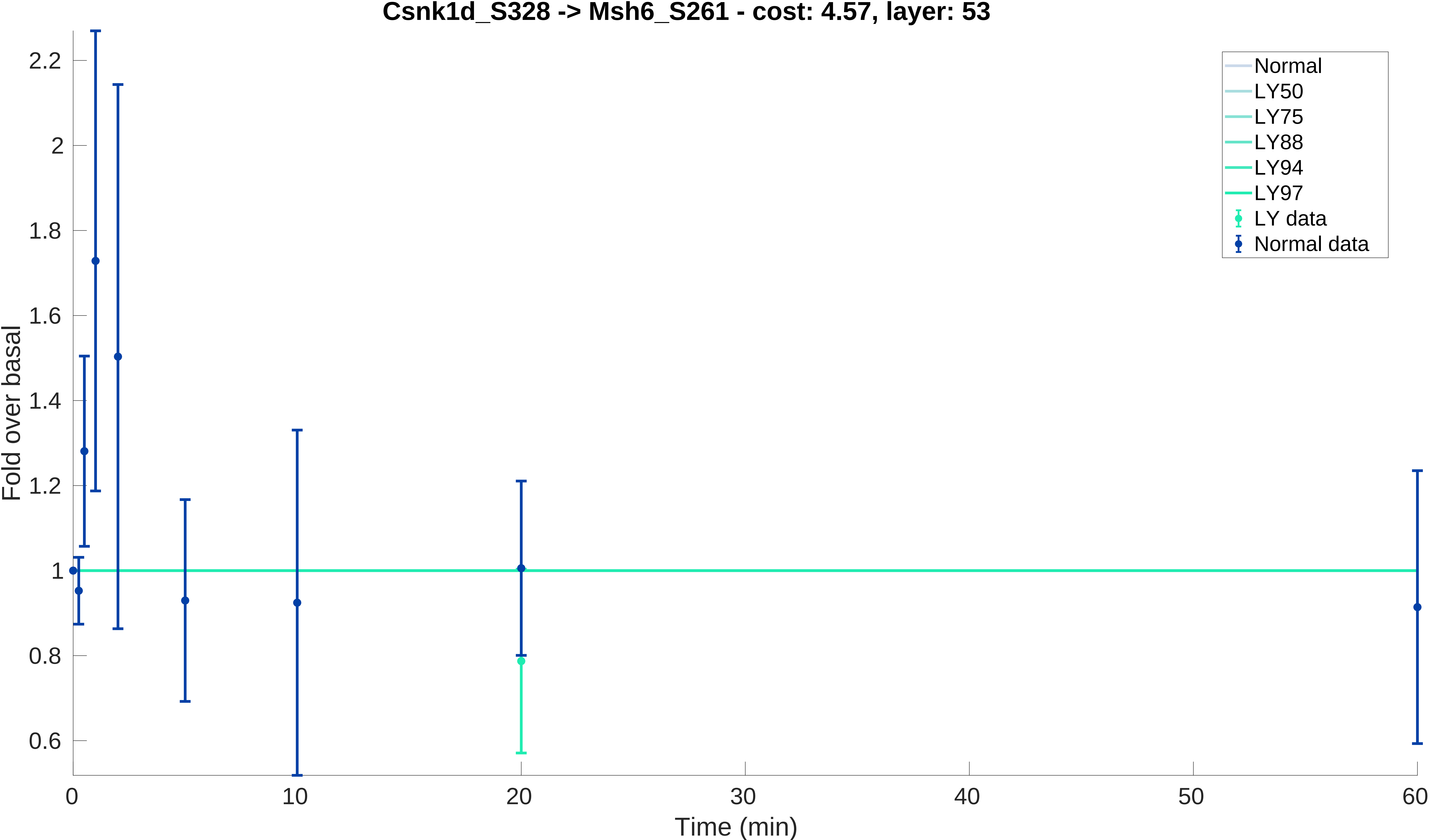

Csnk1d_S328 -> Msh6_S261 - cost: 4.57, layer: 53
Fold over basal
Time (min)
Normal
LY50
LY75
LY88
LY94
LY97
LY data
Normal data
0.6
0.8
1
1.2
1.4
1.6
1.8
2
2.2
0
10
20
30
40
50
60

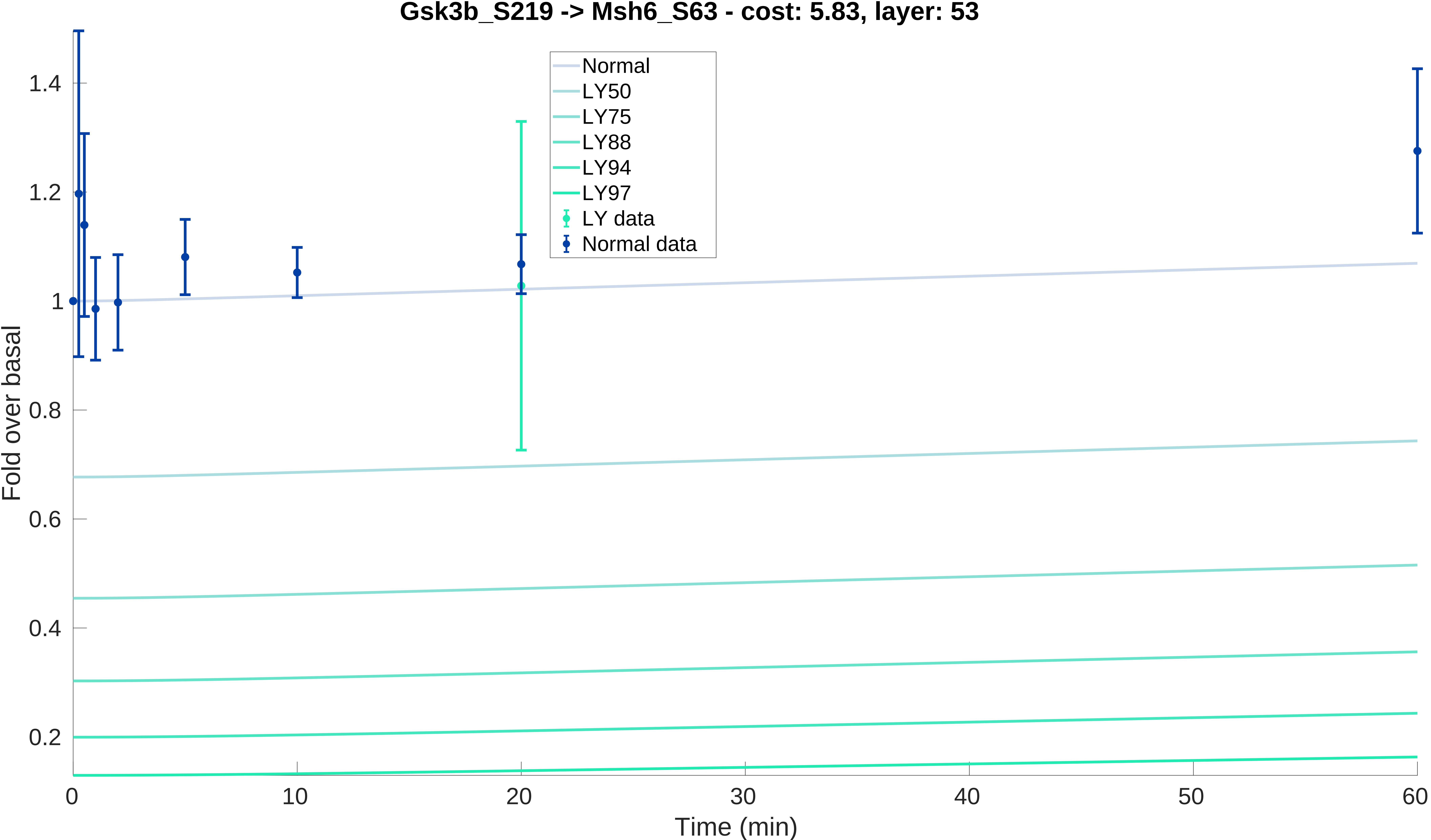

Gsk3b_S219 -> Msh6_S63 - cost: 5.83, layer: 53
Normal
LY50
LY75
LY88
LY94
LY97
LY data
Normal data
Fold over basal
1.4
1.2
1
0.8
0.6
0.4
0.2
0
10
20
30
40
50
60
Time (min)

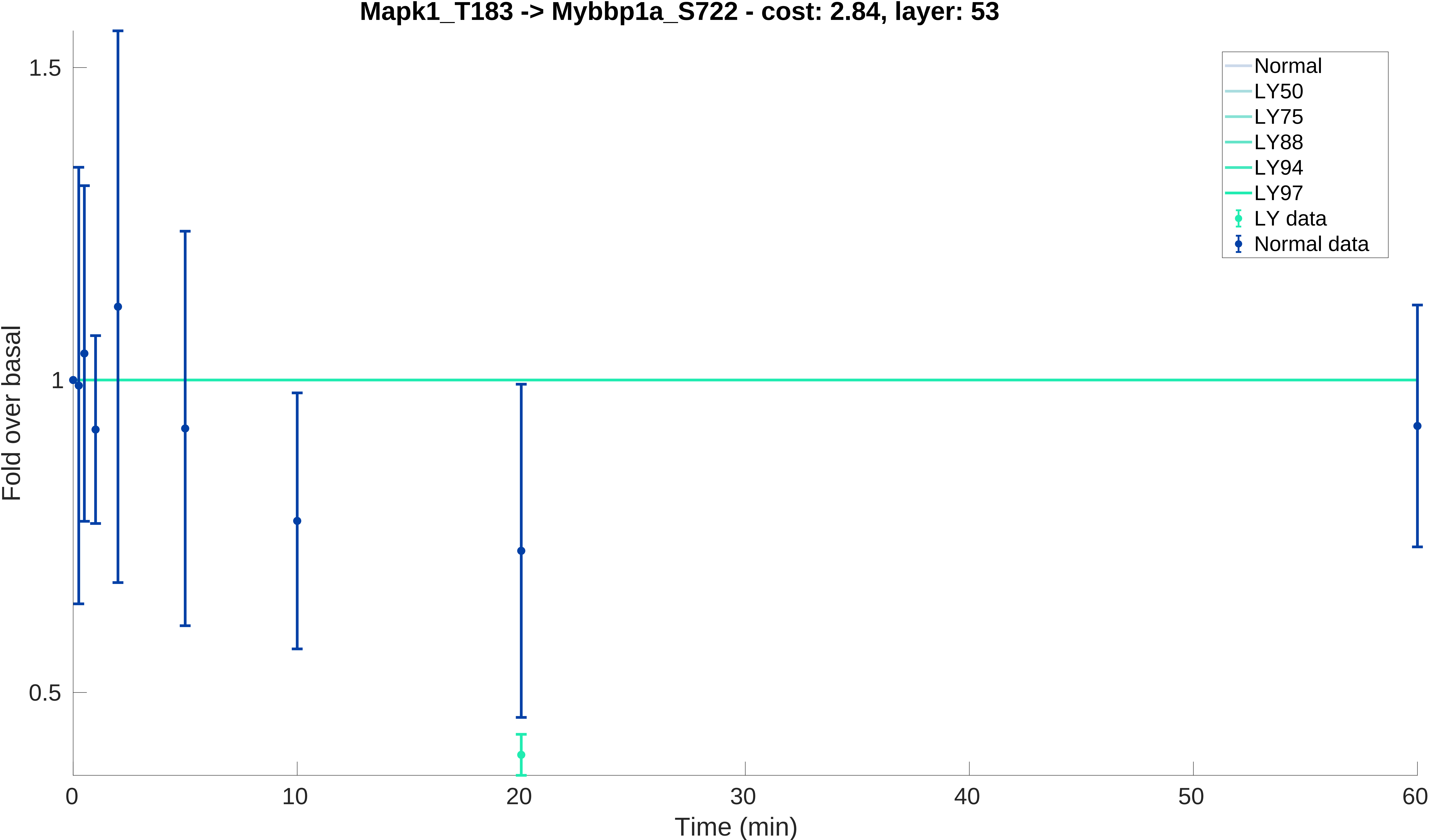

Mapk1_T183 -> Mybbp1a_S722 - cost: 2.84, layer: 53
Normal
LY50
LY75
LY88
LY94
LY97
LY data
Normal data
1.5
1
0.5
Fold over basal
0
10
20
30
40
50
60
Time (min)

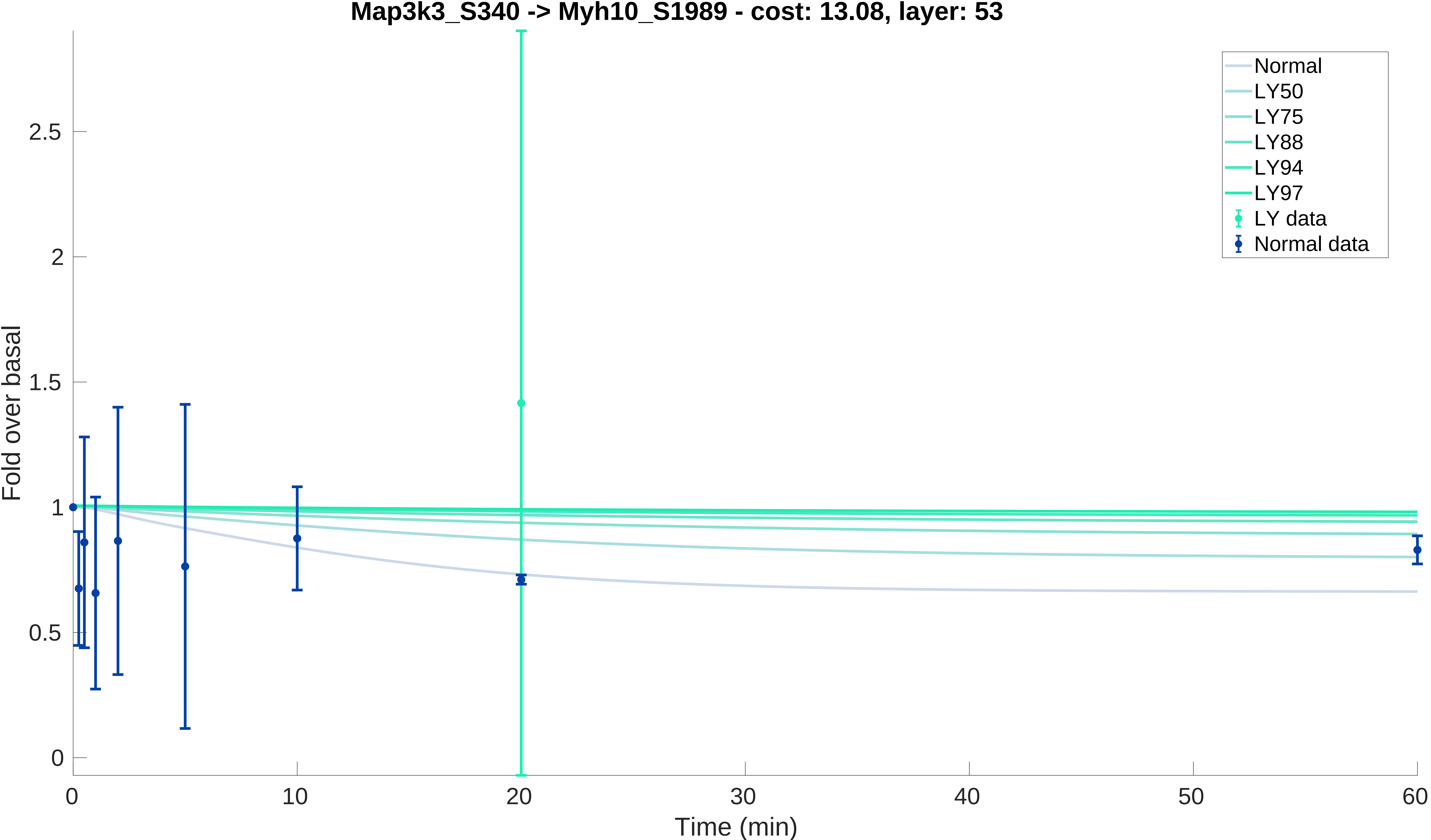

Map3k3_S340 -> Myh10_S1989 - cost: 13.08, layer: 53
Fold over basal
Time (min)
0
0.5
1
1.5
2
2.5
10
20
30
40
50
60
Normal
LY50
LY75
LY88
LY94
LY97
LY data
Normal data

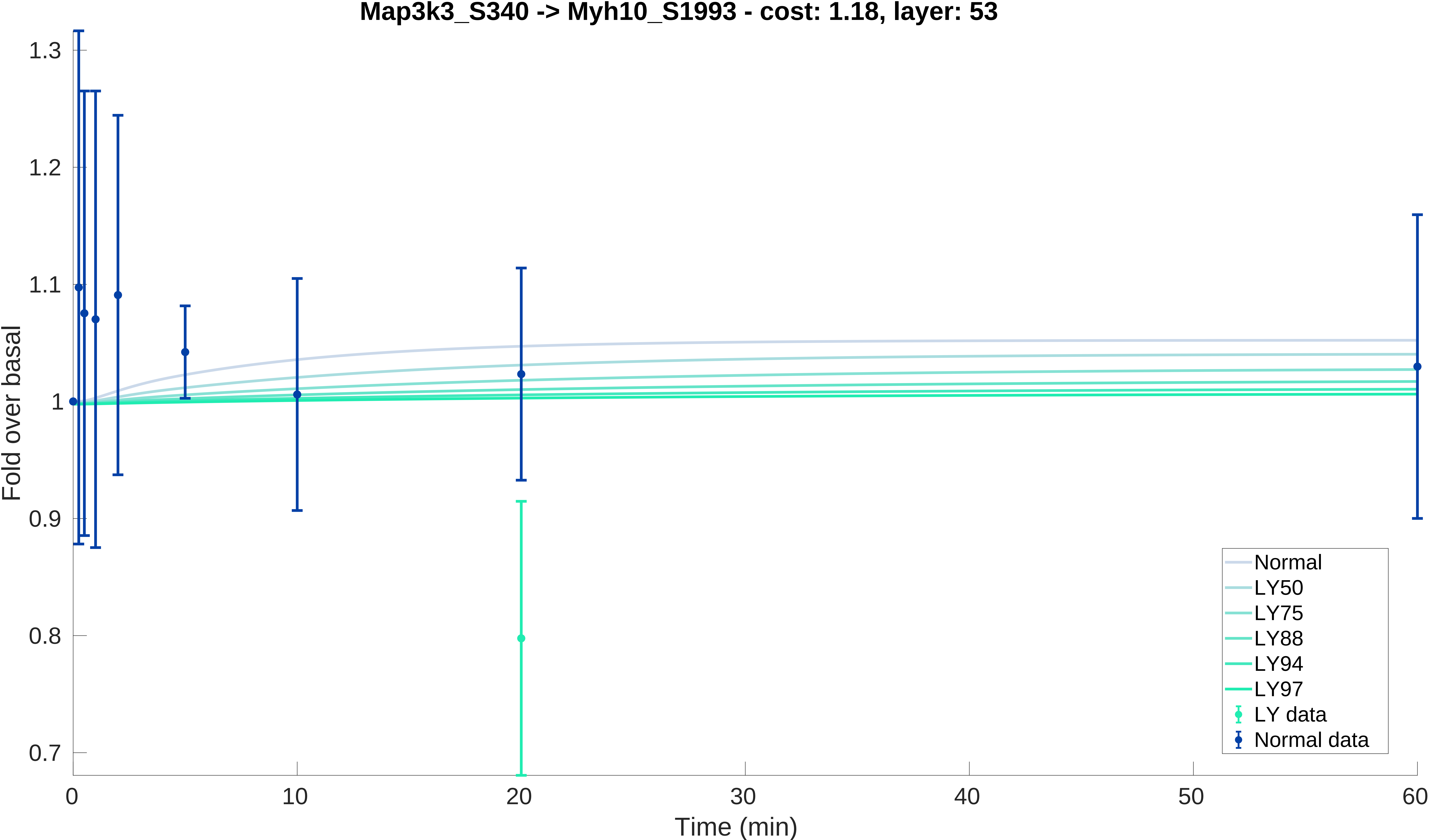
Map3k3_S340 -> Myh10_S1993 - cost: 1.18, layer: 53
Fold over basal
1.3
1.2
1.1
1
0.9
0.8
0.7
0
10
20
30
40
50
60
Time (min)
Normal
LY50
LY75
LY88
LY94
LY97
LY data
Normal data

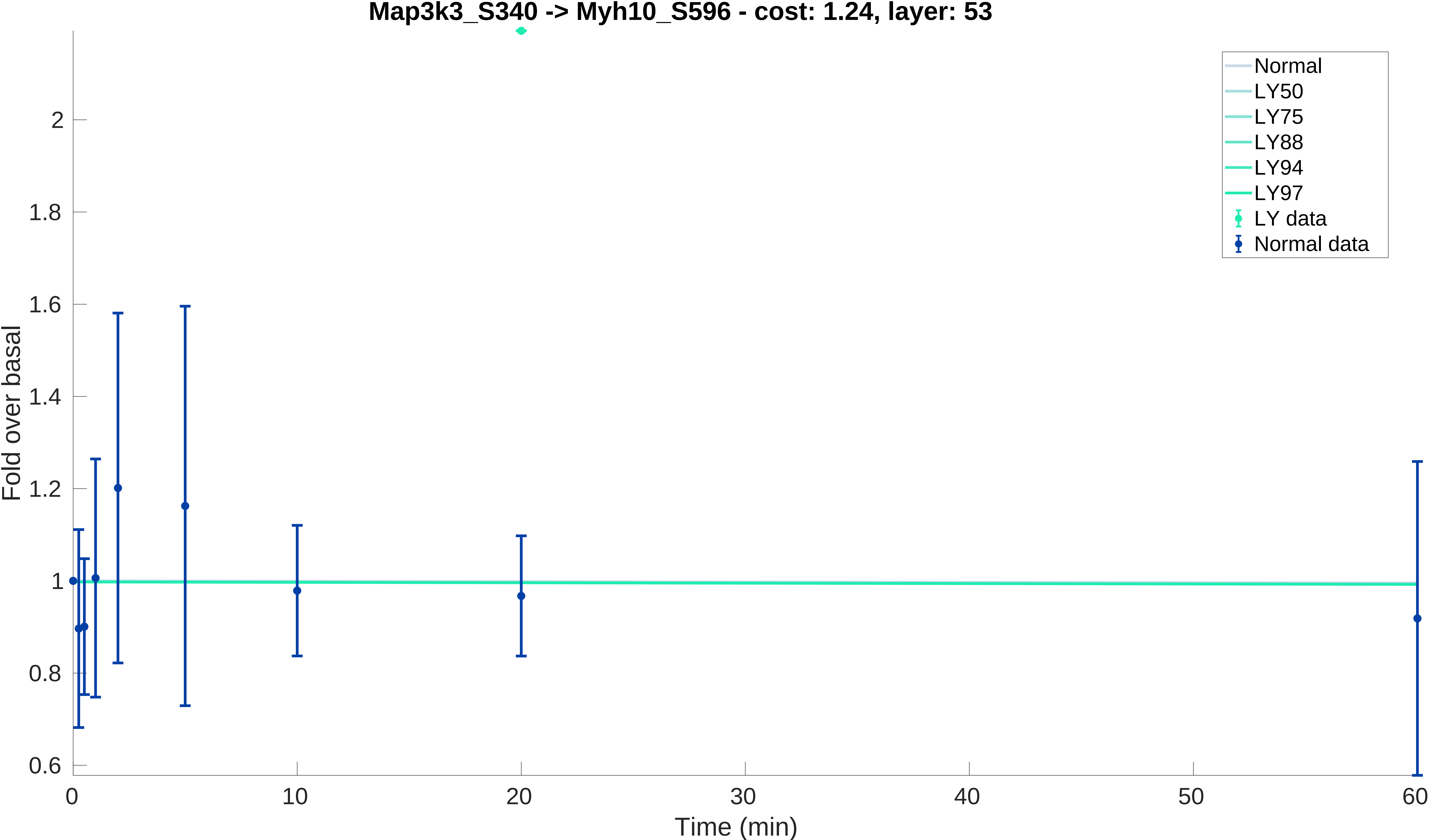
Map3k3_S340 -> Myh10_S596 - cost: 1.24, layer: 53
Fold over basal
Time (min)
0.6
0.8
1
1.2
1.4
1.6
1.8
2
0
10
20
30
40
50
60
Normal
LY50
LY75
LY88
LY94
LY97
LY data
Normal data

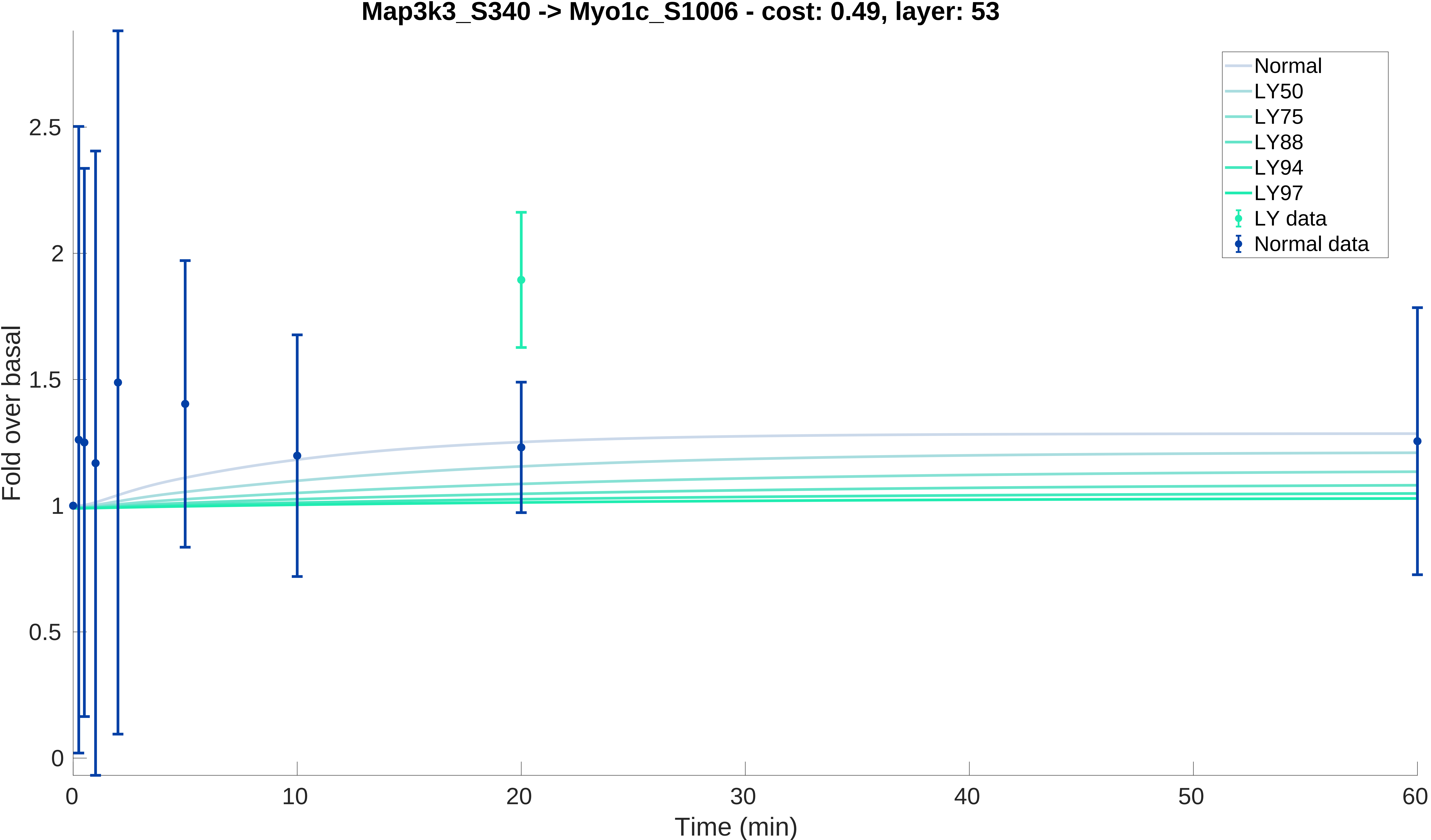

Map3k3_S340 -> Myo1c_S1006 - cost: 0.49, layer: 53
Fold over basal
Time (min)
0
0.5
1
1.5
2
2.5
0
10
20
30
40
50
60
Normal
LY50
LY75
LY88
LY94
LY97
LY data
Normal data

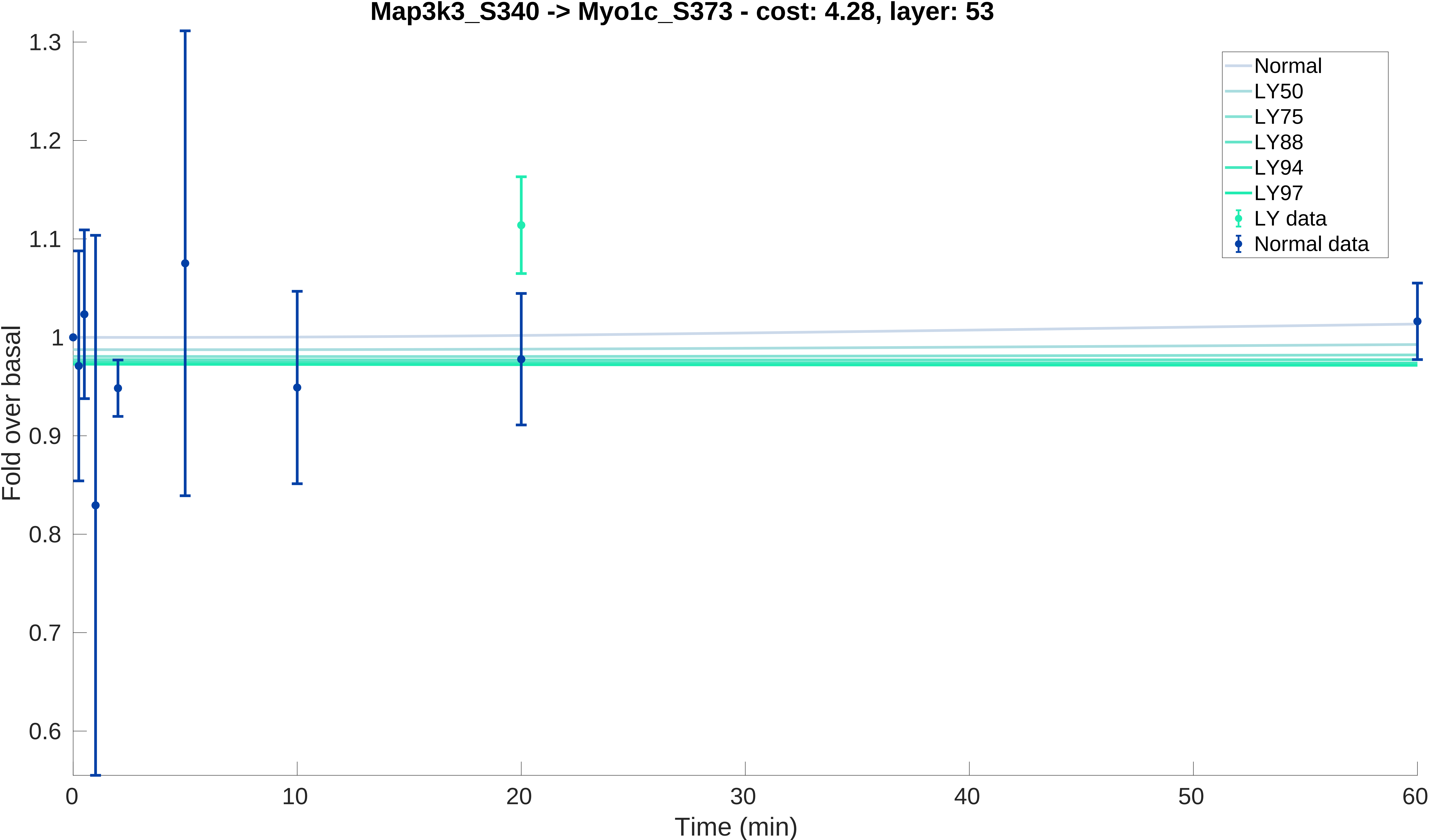
Map3k3_S340 -> Myo1c_S373 - cost: 4.28, layer: 53
Fold over basal
Time (min)
0.6
0.7
0.8
0.9
1
1.1
1.2
1.3
0
10
20
30
40
50
60
Normal
LY50
LY75
LY88
LY94
LY97
LY data
Normal data

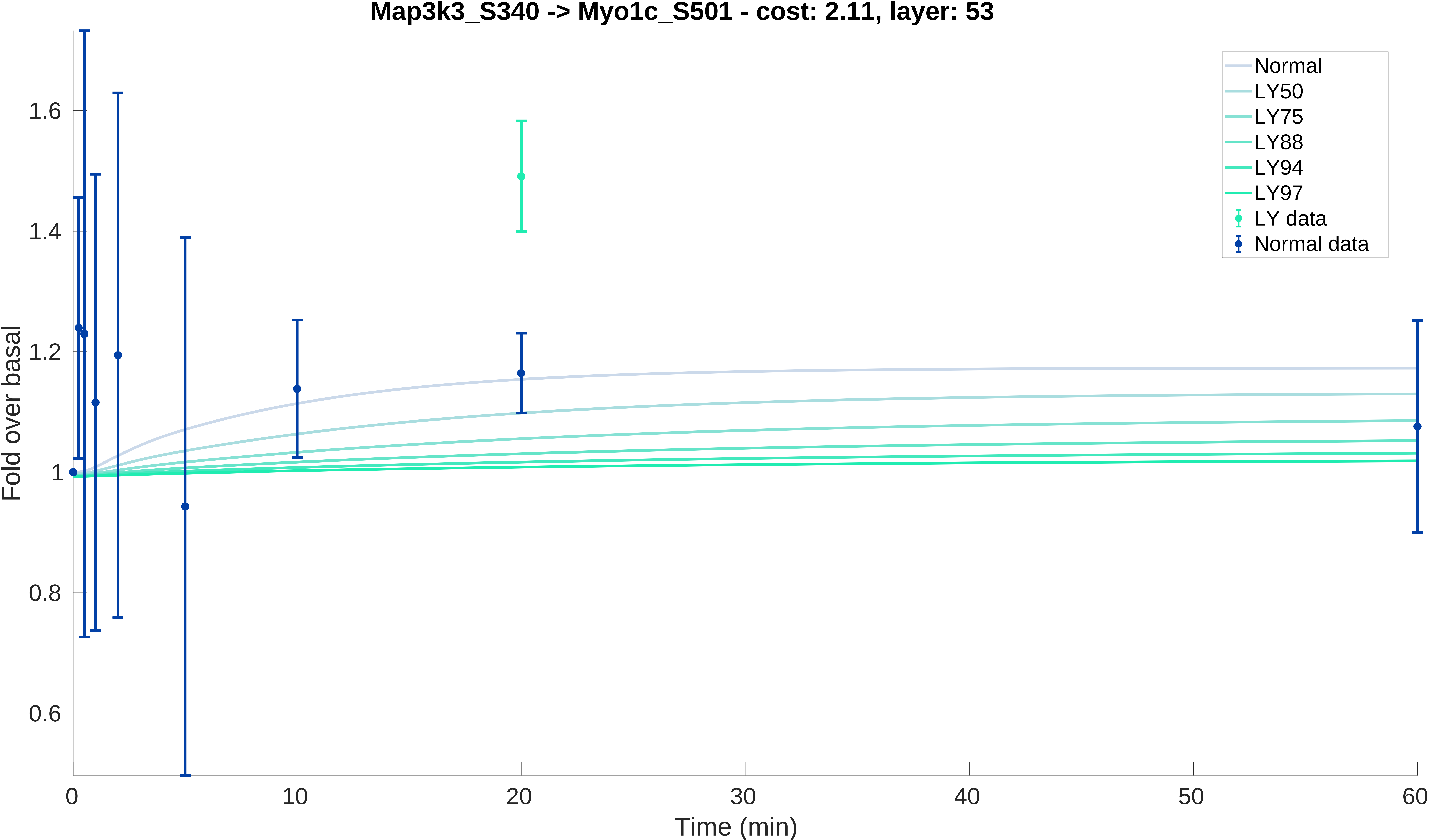

Map3k3_S340 -> Myo1c_S501 - cost: 2.11, layer: 53
Fold over basal
Time (min)
1.6
1.4
1.2
1
0.8
0.6
0
10
20
30
40
50
60
Normal
LY50
LY75
LY88
LY94
LY97
LY data
Normal data

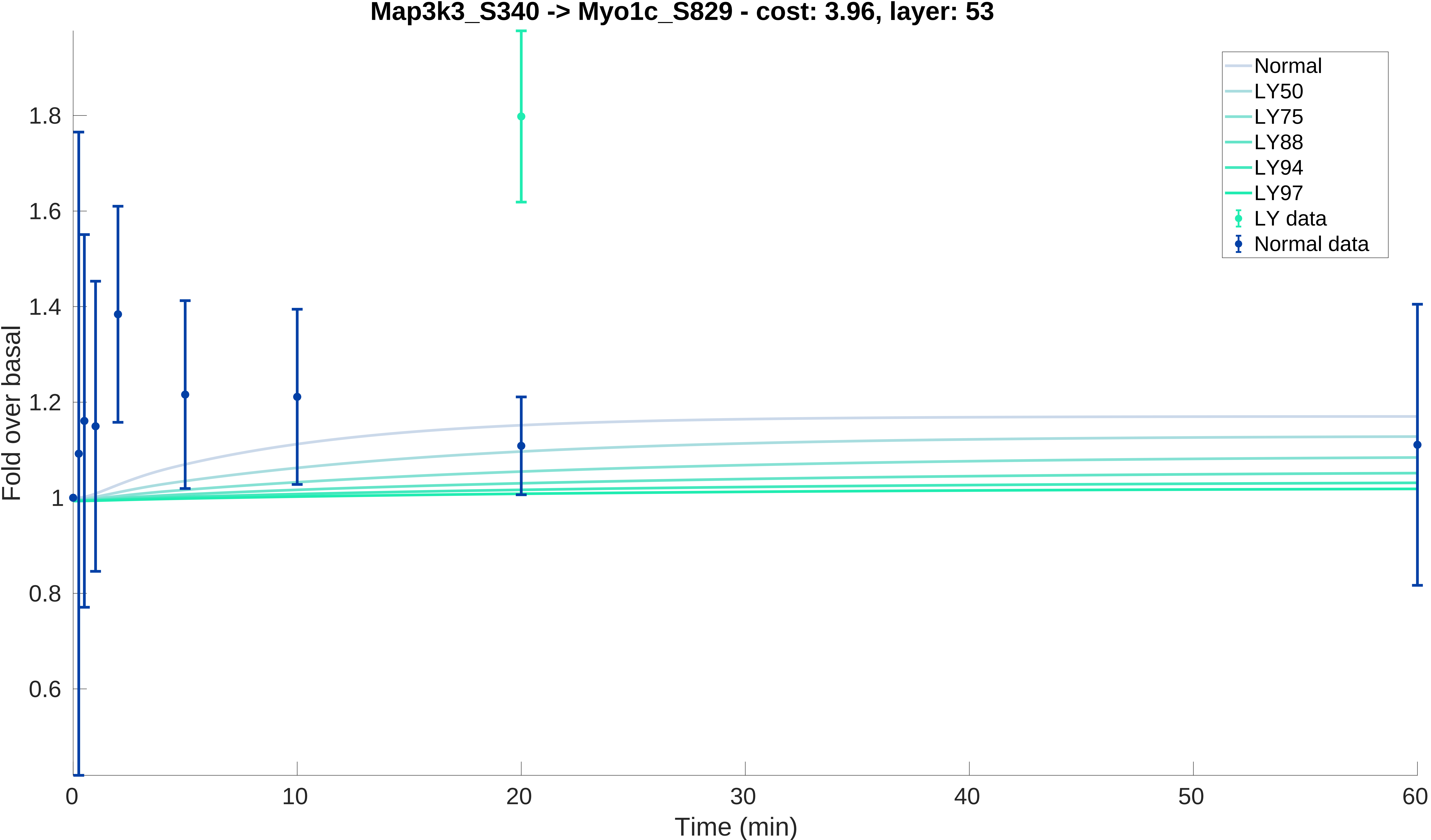

Map3k3_S340 -> Myo1c_S829 - cost: 3.96, layer: 53
Normal
LY50
LY75
LY88
LY94
LY97
LY data
Normal data
Fold over basal
1.8
1.6
1.4
1.2
1
0.8
0.6
0
10
20
30
40
50
60
Time (min)

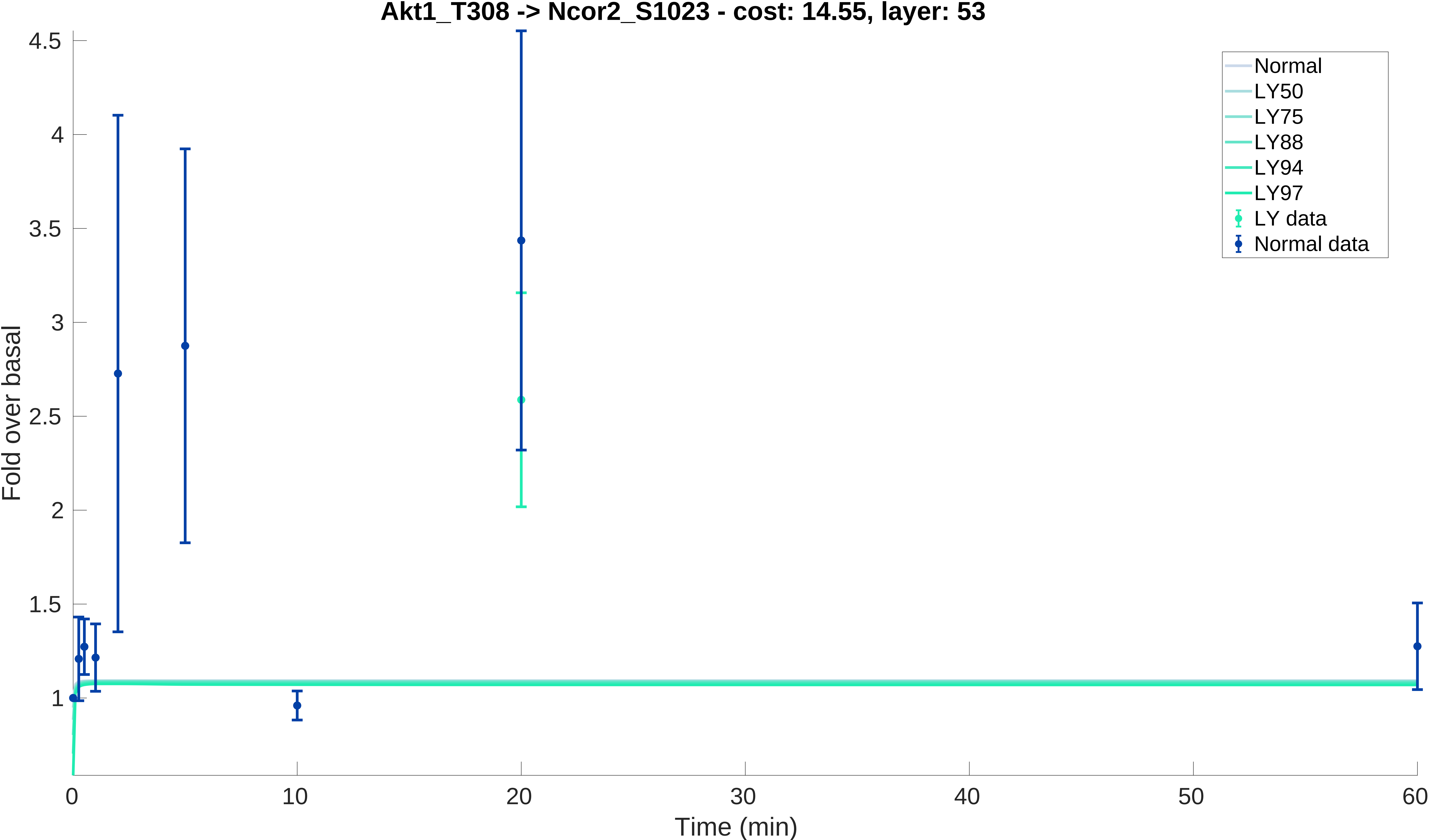

Akt1_T308 -> Ncor2_S1023 - cost: 14.55, layer: 53
Fold over basal
Time (min)
4.5
4
3.5
3
2.5
2
1.5
1
0
10
20
30
40
50
60
Normal
LY50
LY75
LY88
LY94
LY97
LY data
Normal data

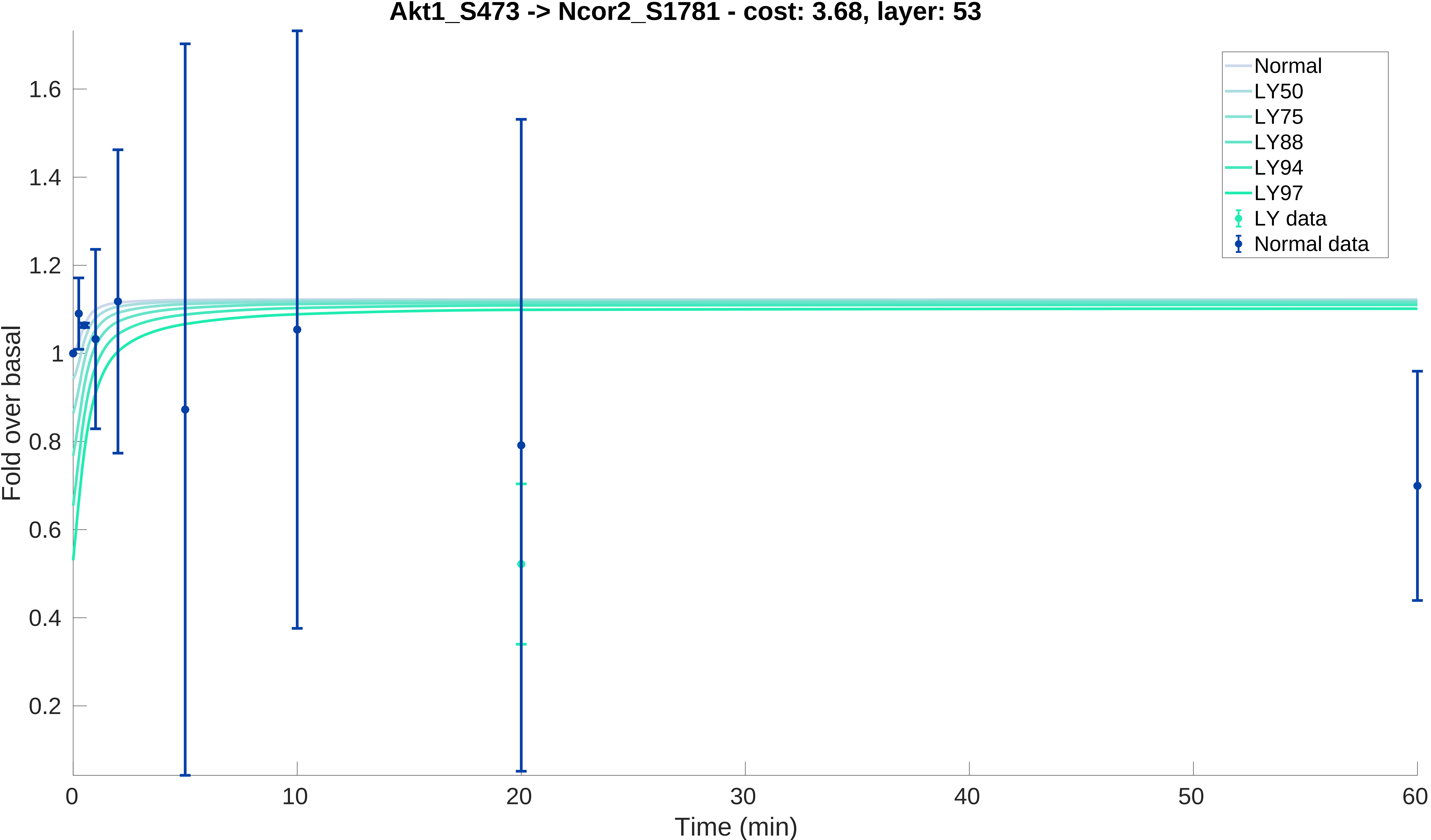

Akt1_S473 -> Ncor2_S1781 - cost: 3.68, layer: 53
Fold over basal
Time (min)
0.2
0.4
0.6
0.8
1
1.2
1.4
1.6
0
10
20
30
40
50
60
Normal
LY50
LY75
LY88
LY94
LY97
LY data
Normal data

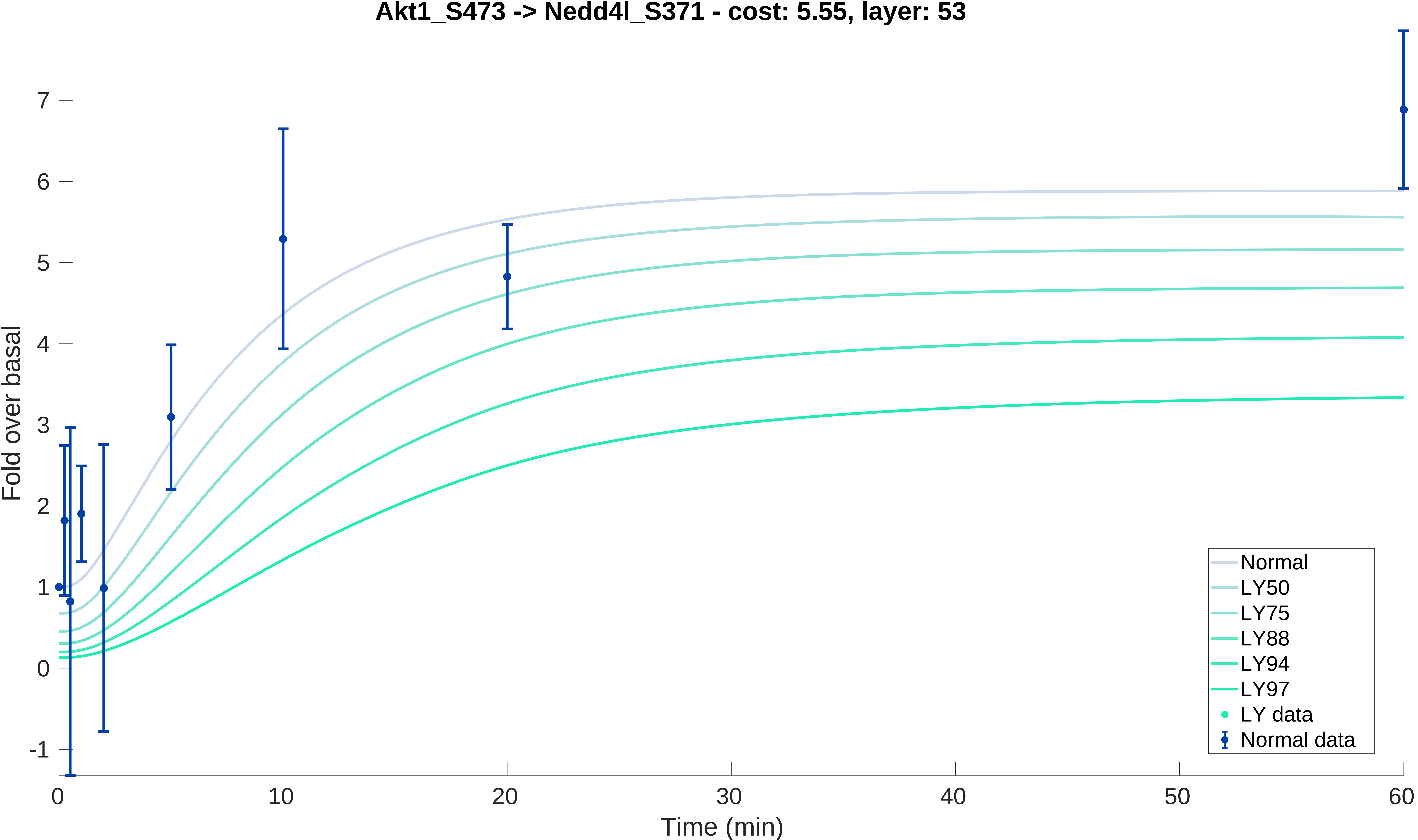

Akt1_S473 -> Nedd4l_S371 - cost: 5.55, layer: 53
Fold over basal
Time (min)
7
6
5
4
3
2
1
0
-1
0
10
20
30
40
50
60
Normal
LY50
LY75
LY88
LY94
LY97
LY data
Normal data

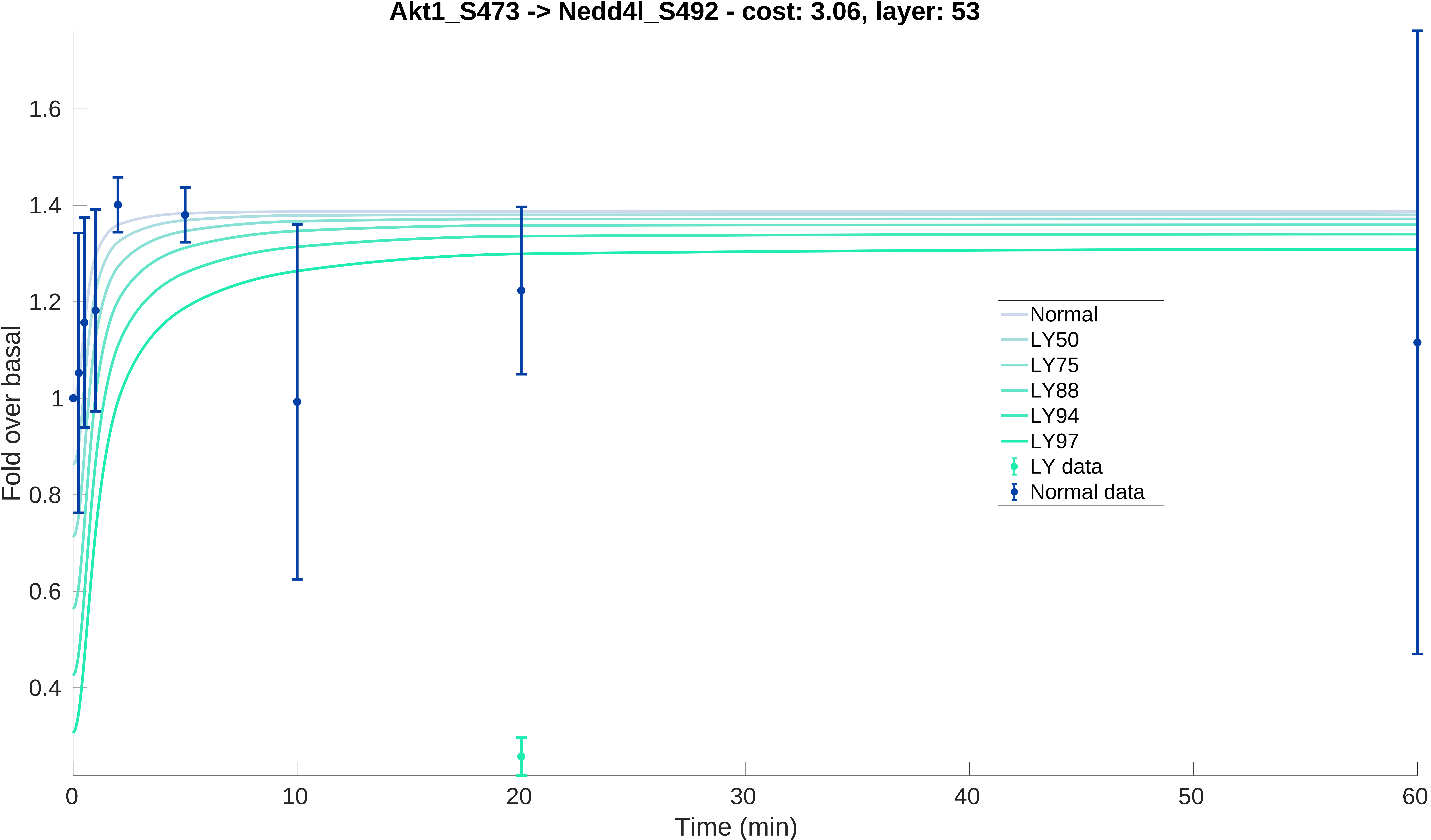

Akt1_S473 -> Nedd4l_S492 - cost: 3.06, layer: 53
Fold over basal
1.6
1.4
1.2
1
0.8
0.6
0.4
0
10
20
30
40
50
60
Time (min)
Normal
LY50
LY75
LY88
LY94
LY97
LY data
Normal data

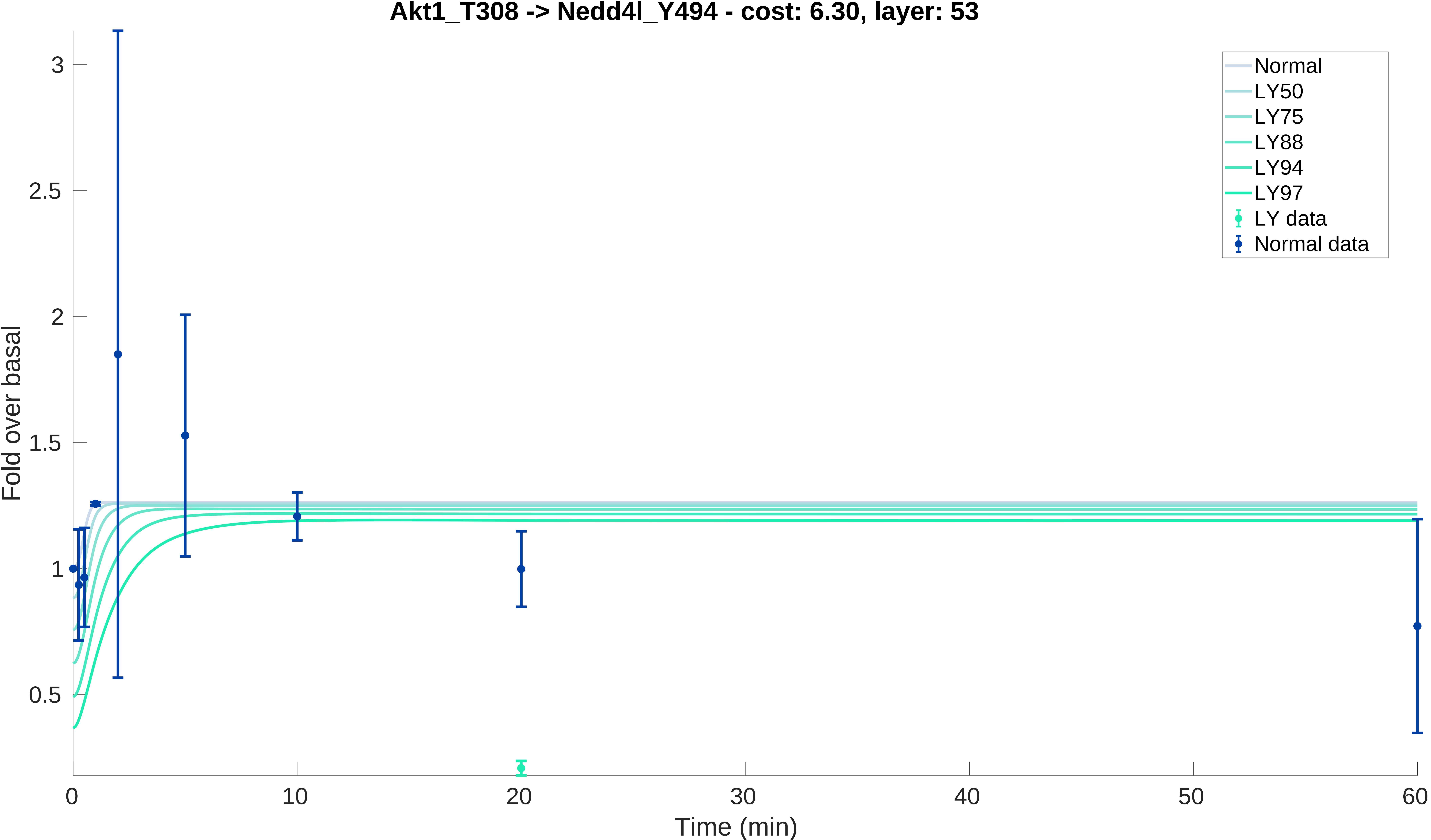

Akt1_T308 -> Nedd4l_Y494 - cost: 6.30, layer: 53
Fold over basal
Time (min)
0
10
20
30
40
50
60
0.5
1
1.5
2
2.5
3
Normal
LY50
LY75
LY88
LY94
LY97
LY data
Normal data

Gsk3b_S219 -> Nfia_S287 - cost: 15.49, layer: 53

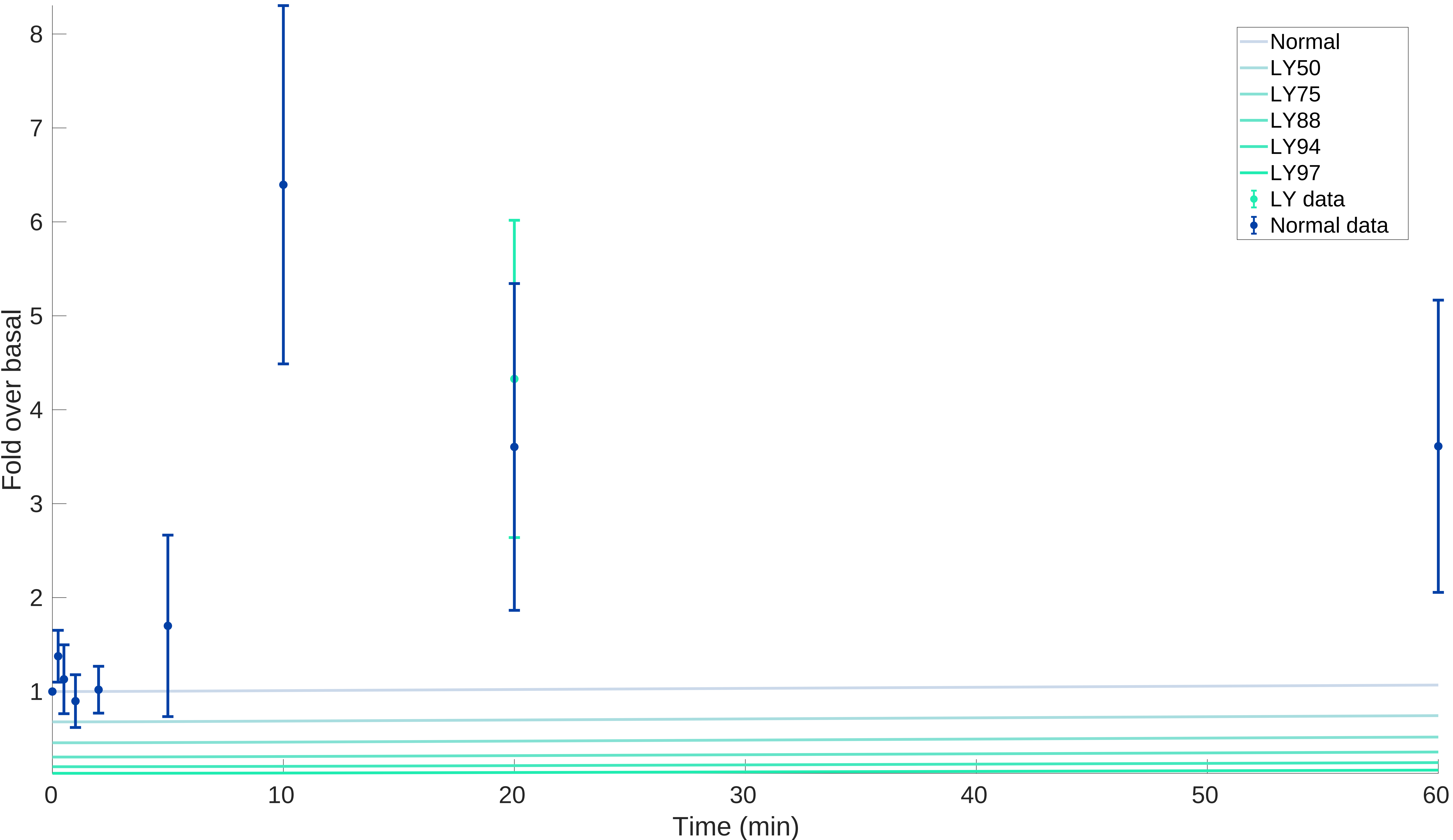

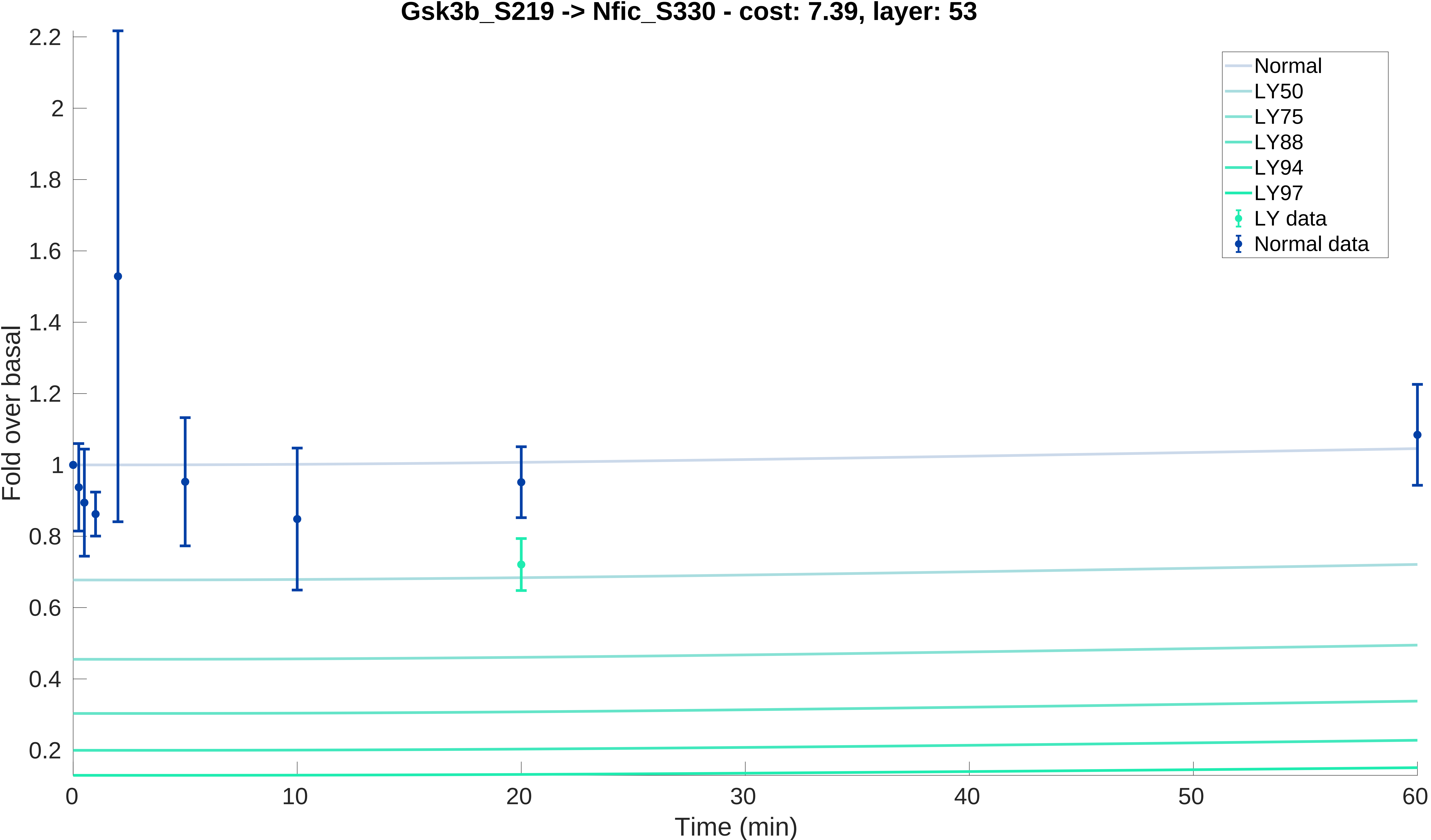

Gsk3b_S219 -> Nfic_S330 - cost: 7.39, layer: 53
Fold over basal
2.2
2
1.8
1.6
1.4
1.2
1
0.8
0.6
0.4
0.2
0
10
20
30
40
50
60
Time (min)
Normal
LY50
LY75
LY88
LY94
LY97
LY data
Normal data

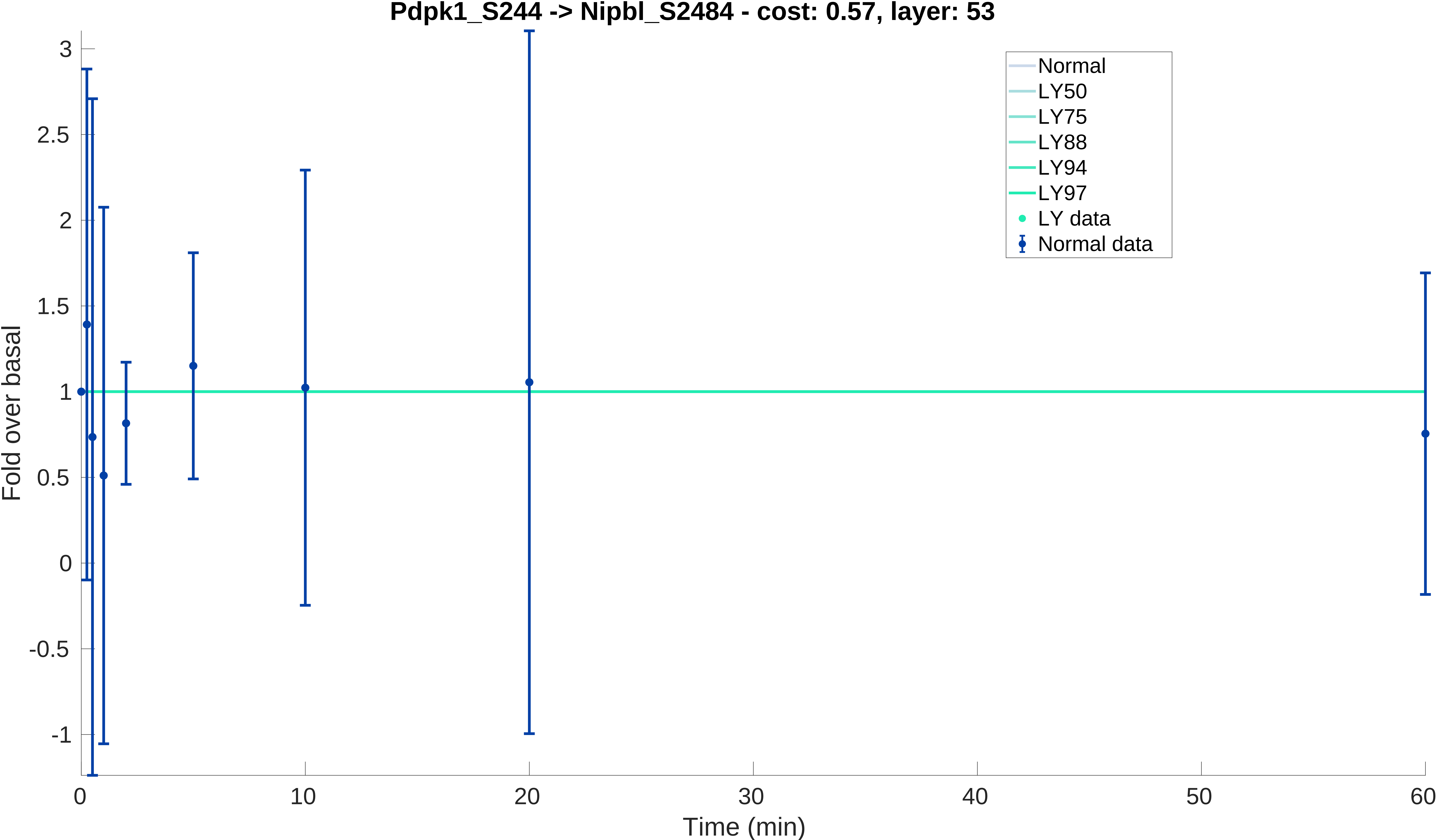

Pdpk1_S244 -> Nipbl_S2484 - cost: 0.57, layer: 53
Normal
LY50
LY75
LY88
LY94
LY97
LY data
Normal data
Fold over basal
Time (min)
3
2.5
2
1.5
1
0.5
0
-0.5
-1
0
10
20
30
40
50
60

Pdpk1_S244 -> Nipbl_S2485 - cost: 0.57, layer: 53

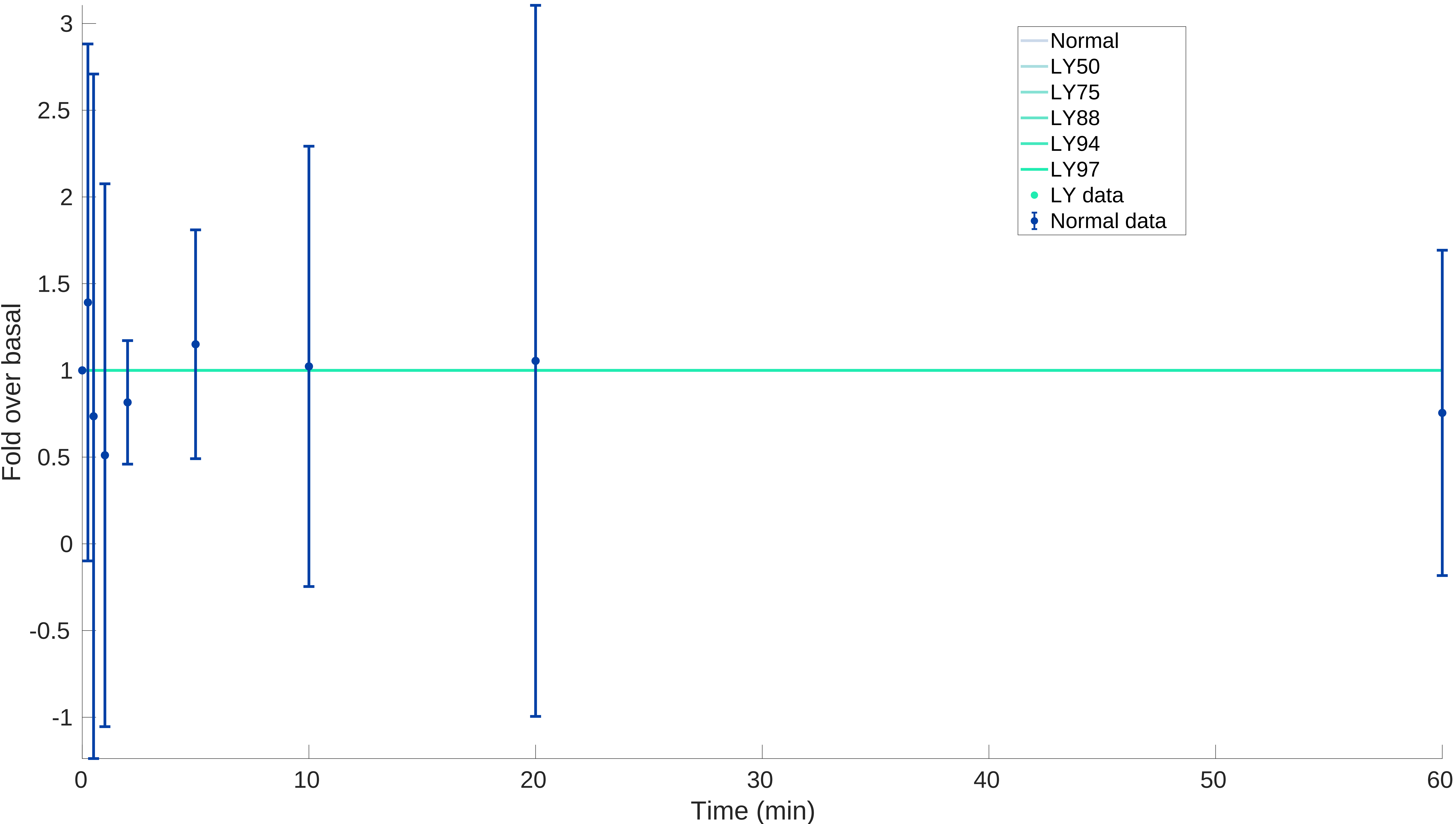

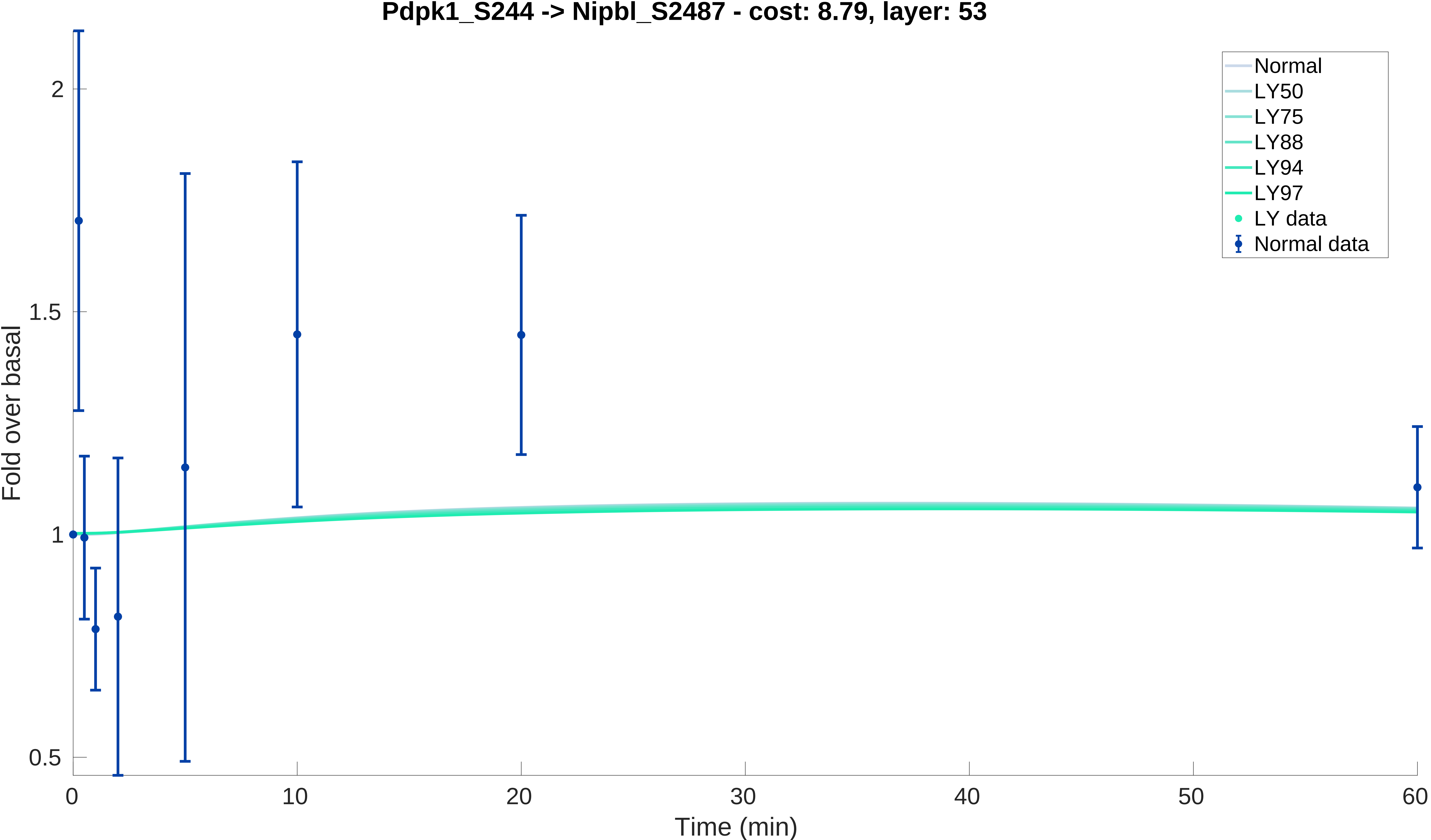
Pdpk1_S244 -> Nipbl_S2487 - cost: 8.79, layer: 53
Fold over basal
Time (min)
0.5
1
1.5
2
0
10
20
30
40
50
60
Normal
LY50
LY75
LY88
LY94
LY97
LY data
Normal data

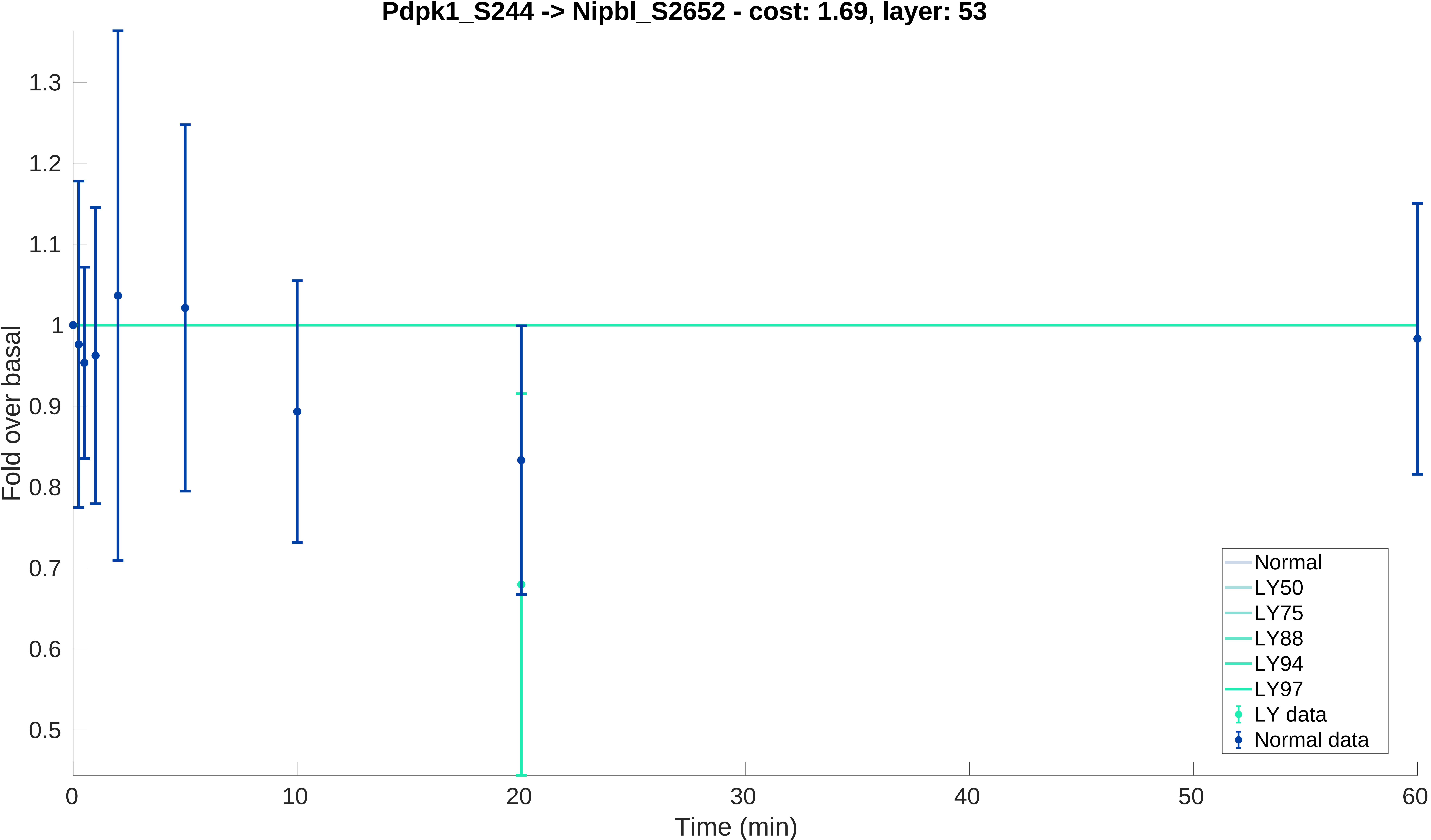
Pdpk1_S244 -> Nipbl_S2652 - cost: 1.69, layer: 53
Fold over basal
Time (min)
1.3
1.2
1.1
1
0.9
0.8
0.7
0.6
0.5
0
10
20
30
40
50
60
Normal
LY50
LY75
LY88
LY94
LY97
LY data
Normal data

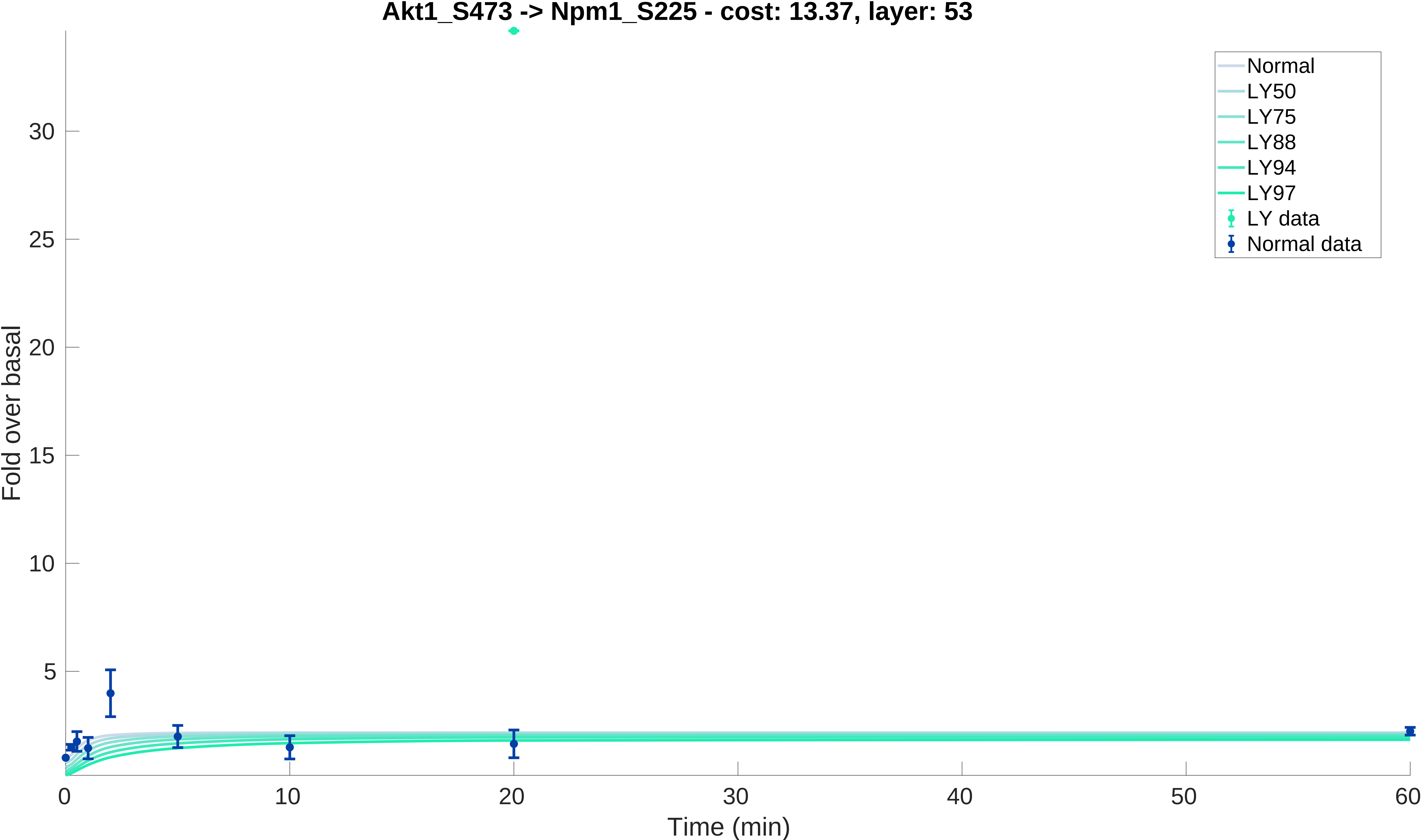

Akt1_S473 -> Npm1_S225 - cost: 13.37, layer: 53
Normal
LY50
LY75
LY88
LY94
LY97
LY data
Normal data
Fold over basal
Time (min)
0
10
20
30
40
50
60
5
10
15
20
25
30

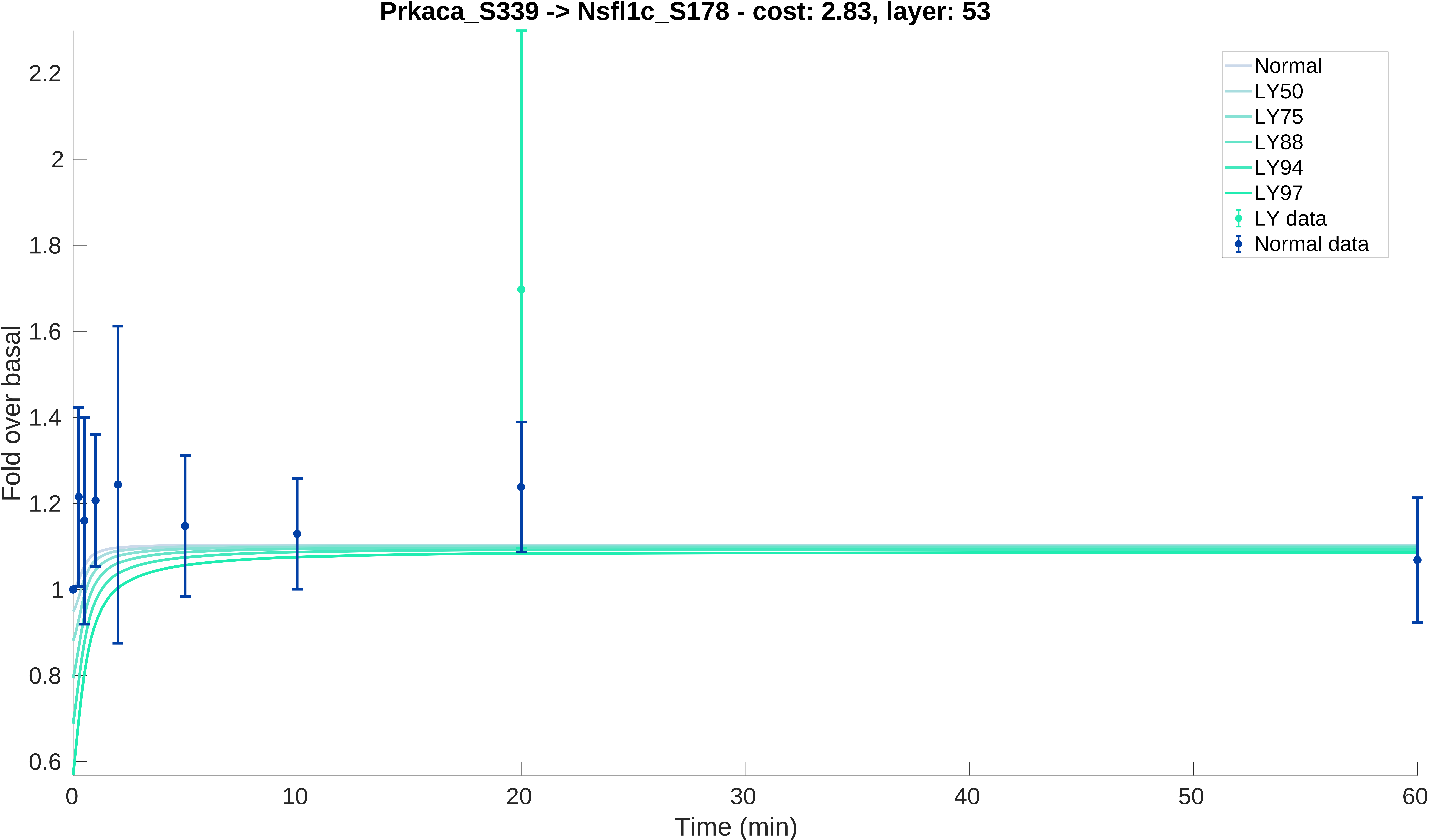
Prkaca_S339 -> Nsfl1c_S178 - cost: 2.83, layer: 53
Fold over basal
Time (min)
0.6
0.8
1
1.2
1.4
1.6
1.8
2
2.2
0
10
20
30
40
50
60
Normal
LY50
LY75
LY88
LY94
LY97
LY data
Normal data

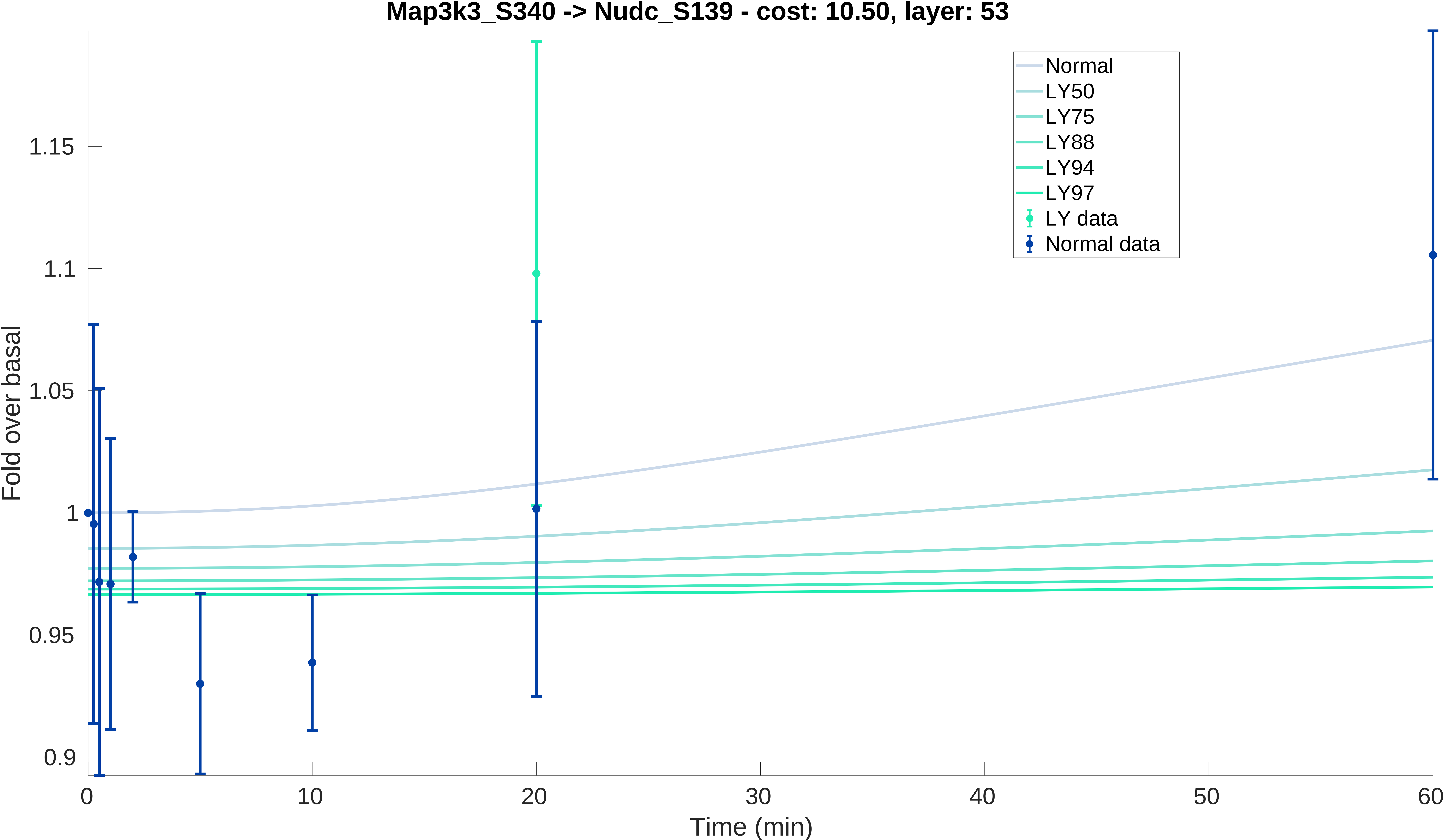

Map3k3_S340 -> Nudc_S139 - cost: 10.50, layer: 53
Normal
LY50
LY75
LY88
LY94
LY97
LY data
Normal data
Fold over basal
1.15
1.1
1.05
1
0.95
0.9
0
10
20
30
40
50
60
Time (min)

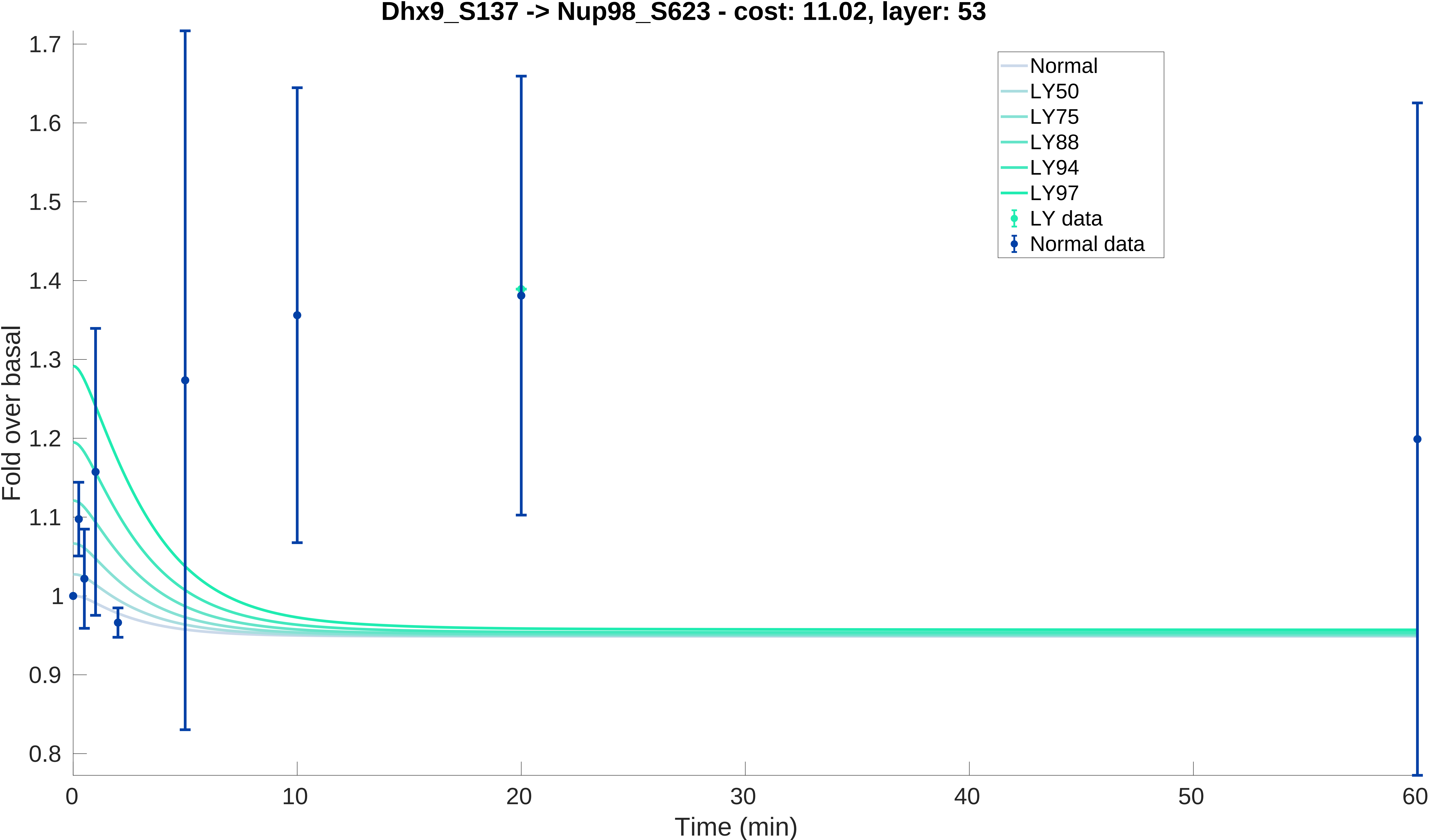

Dhx9_S137 -> Nup98_S623 - cost: 11.02, layer: 53
Fold over basal
Time (min)
1.7
1.6
1.5
1.4
1.3
1.2
1.1
1
0.9
0.8
0
10
20
30
40
50
60
Normal
LY50
LY75
LY88
LY94
LY97
LY data
Normal data

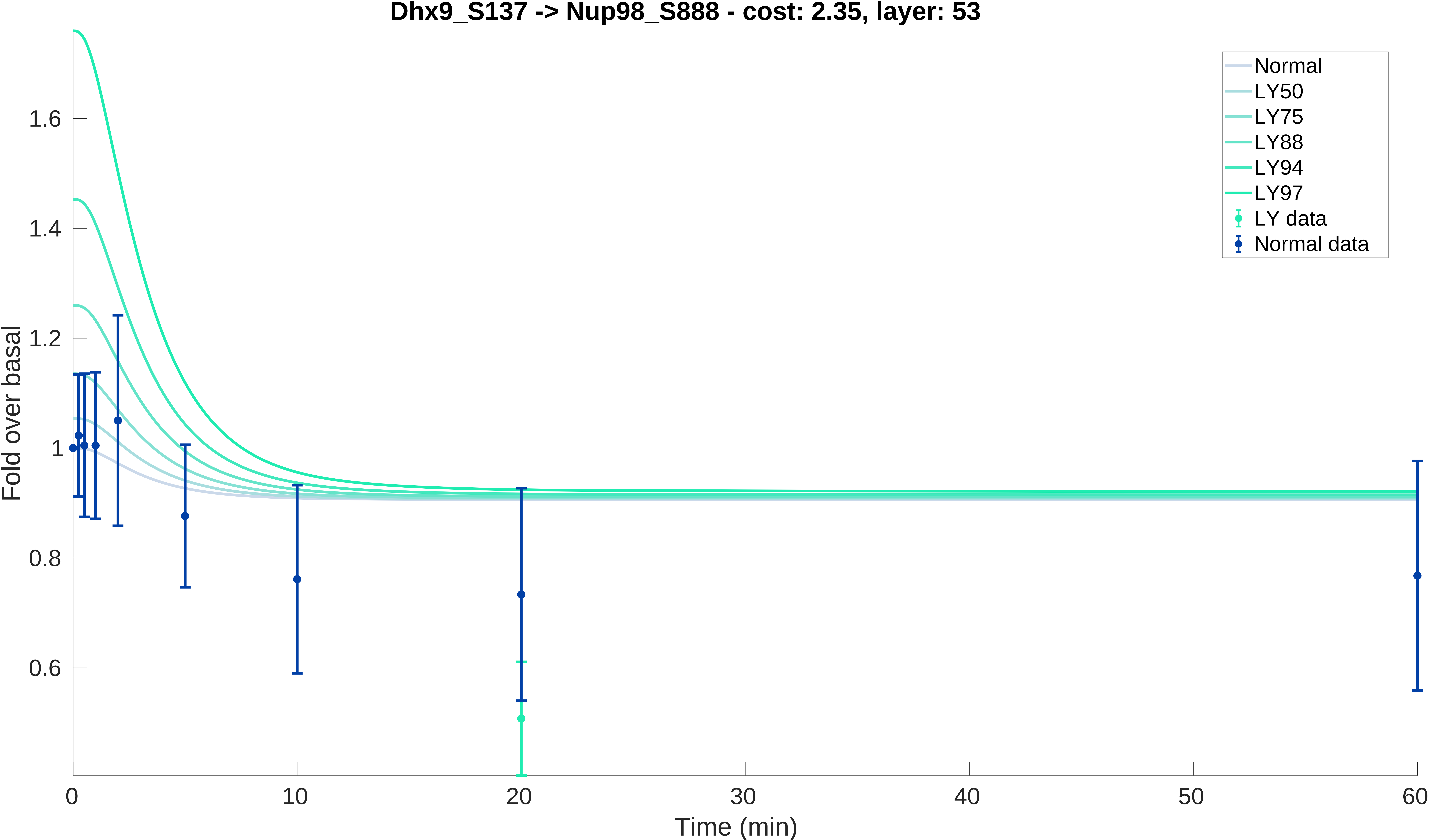

Dhx9_S137 -> Nup98_S888 - cost: 2.35, layer: 53
Normal
LY50
LY75
LY88
LY94
LY97
LY data
Normal data
Fold over basal
1.6
1.4
1.2
1
0.8
0.6
0
10
20
30
40
50
60
Time (min)

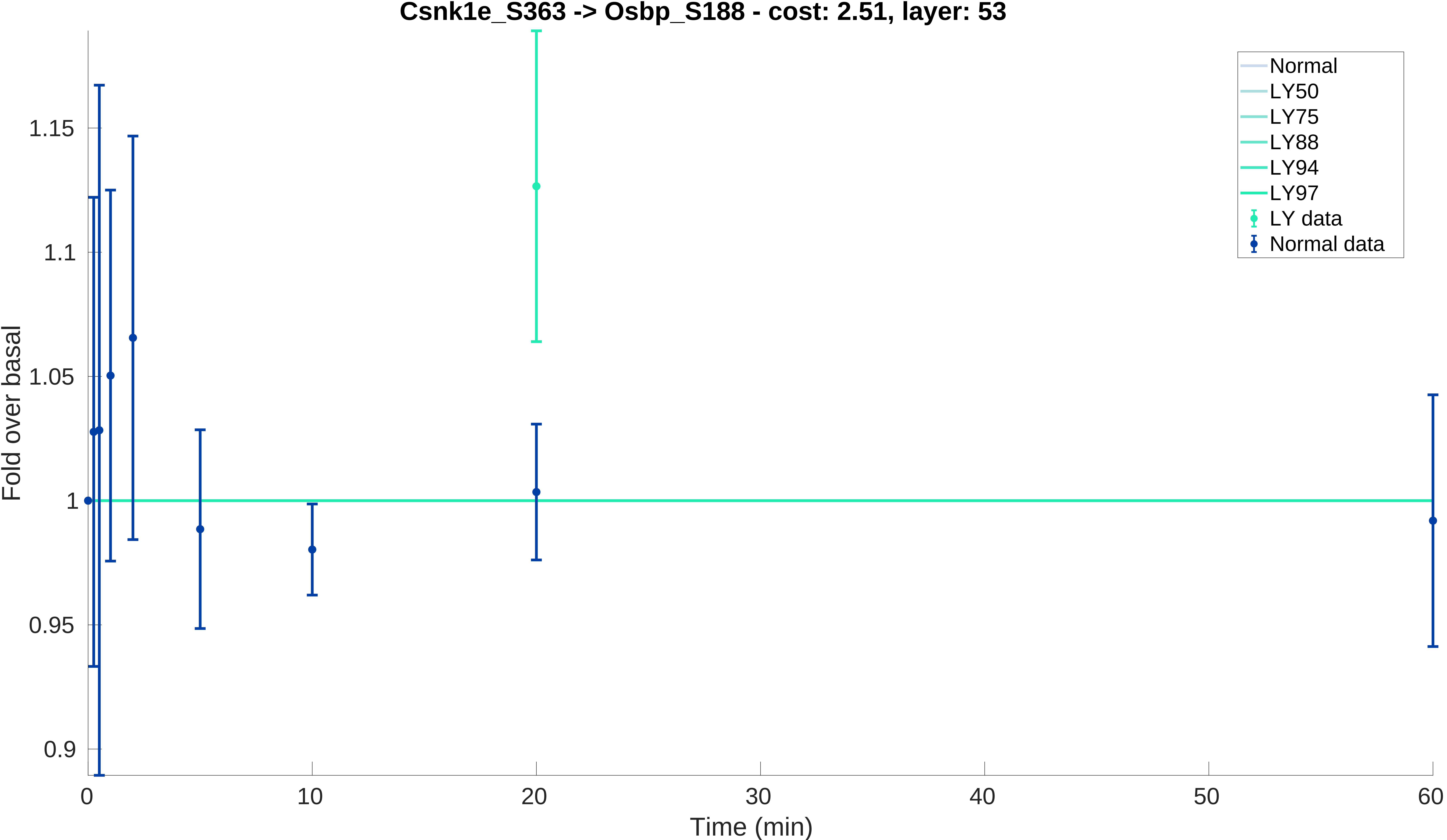

Csnk1e_S363 -> Osbp_S188 - cost: 2.51, layer: 53
Fold over basal
Time (min)
0.9
0.95
1
1.05
1.1
1.15
0
10
20
30
40
50
60
Normal
LY50
LY75
LY88
LY94
LY97
LY data
Normal data

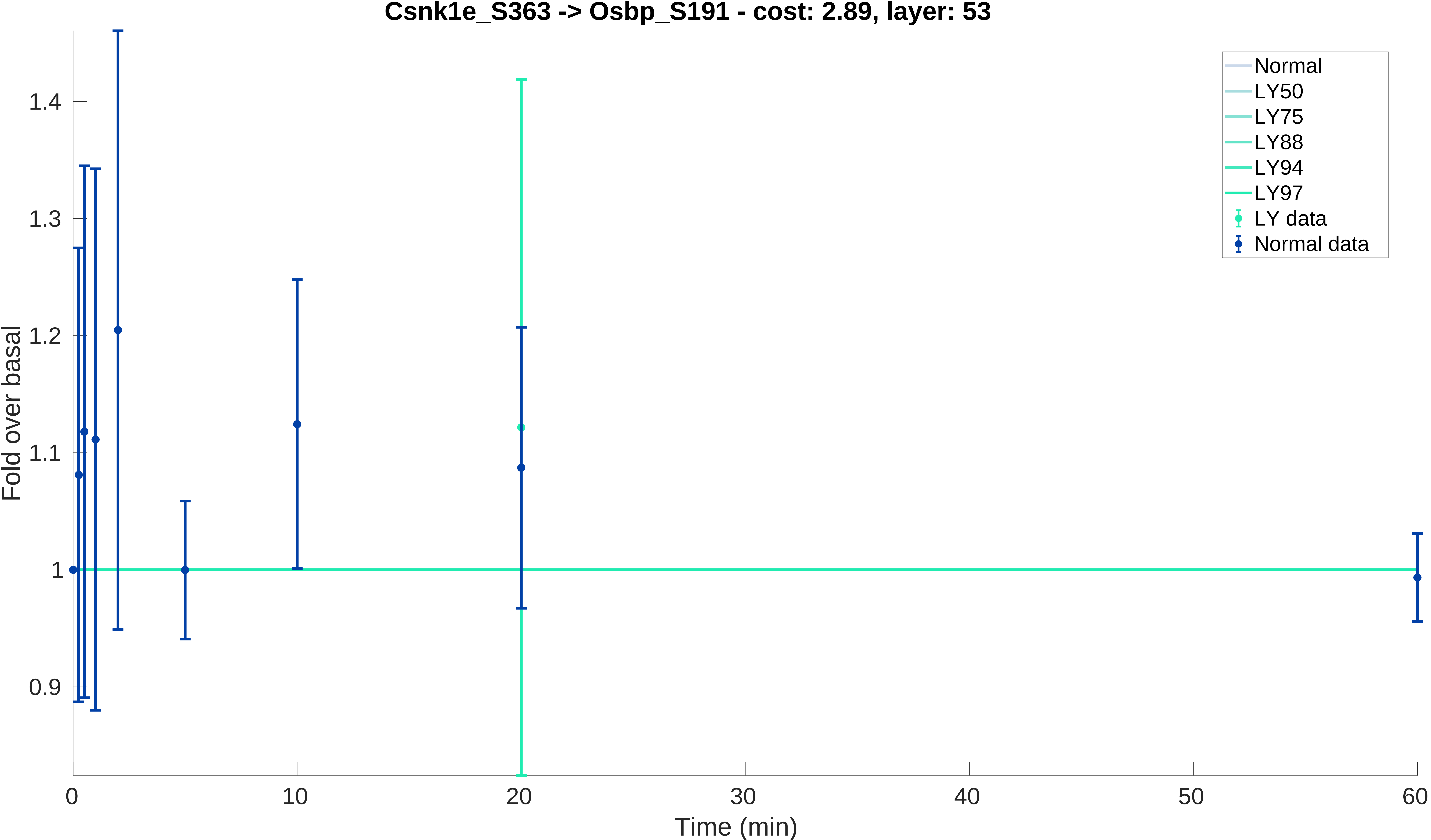
Csnk1e_S363 -> Osbp_S191 - cost: 2.89, layer: 53
Fold over basal
Time (min)
0.9
1
1.1
1.2
1.3
1.4
0
10
20
30
40
50
60
Normal
LY50
LY75
LY88
LY94
LY97
LY data
Normal data

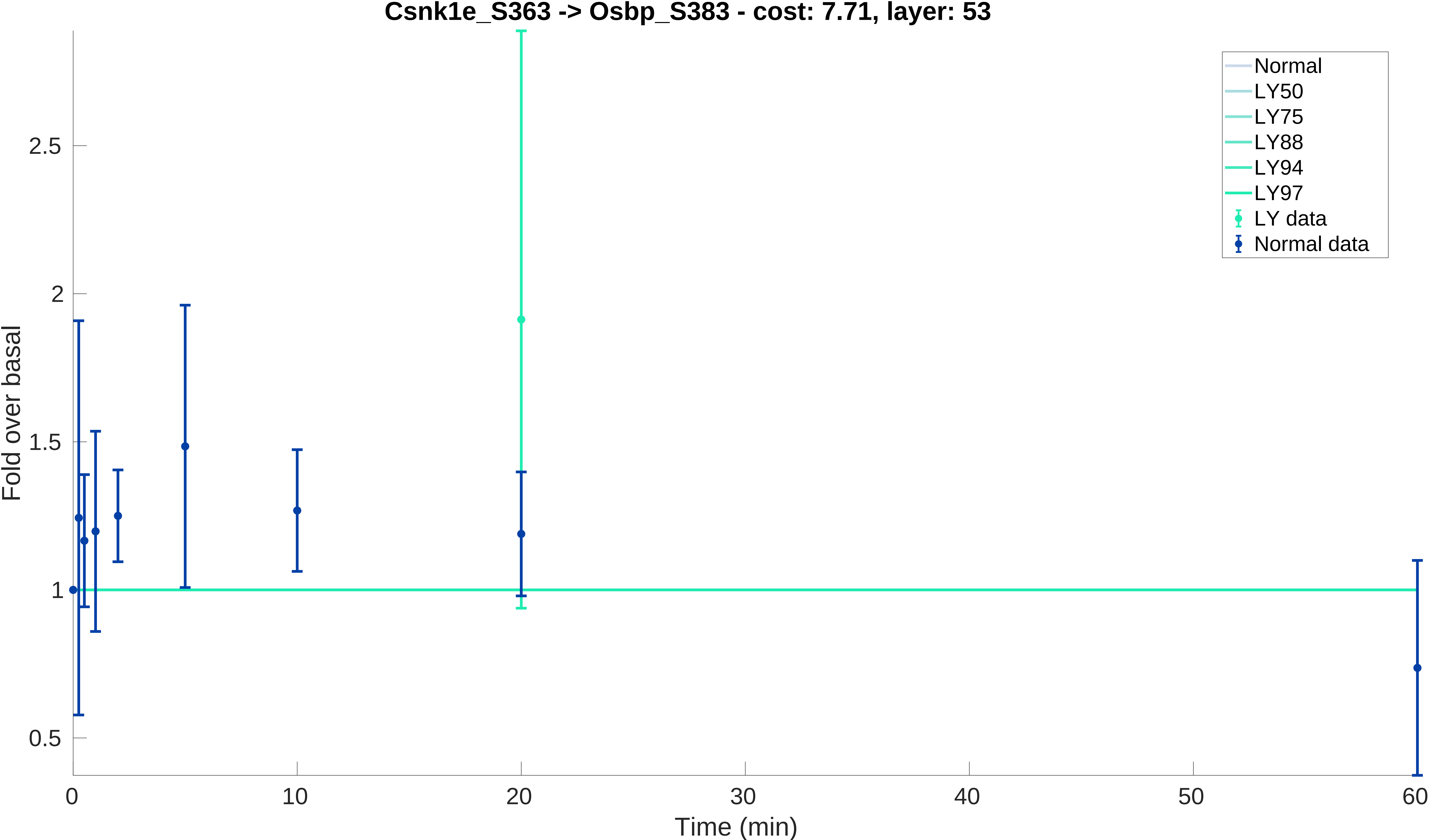

Csnk1e_S363 -> Osbp_S383 - cost: 7.71, layer: 53
Normal
LY50
LY75
LY88
LY94
LY97
LY data
Normal data
Fold over basal
2.5
2
1.5
1
0.5
0
10
20
30
40
50
60
Time (min)

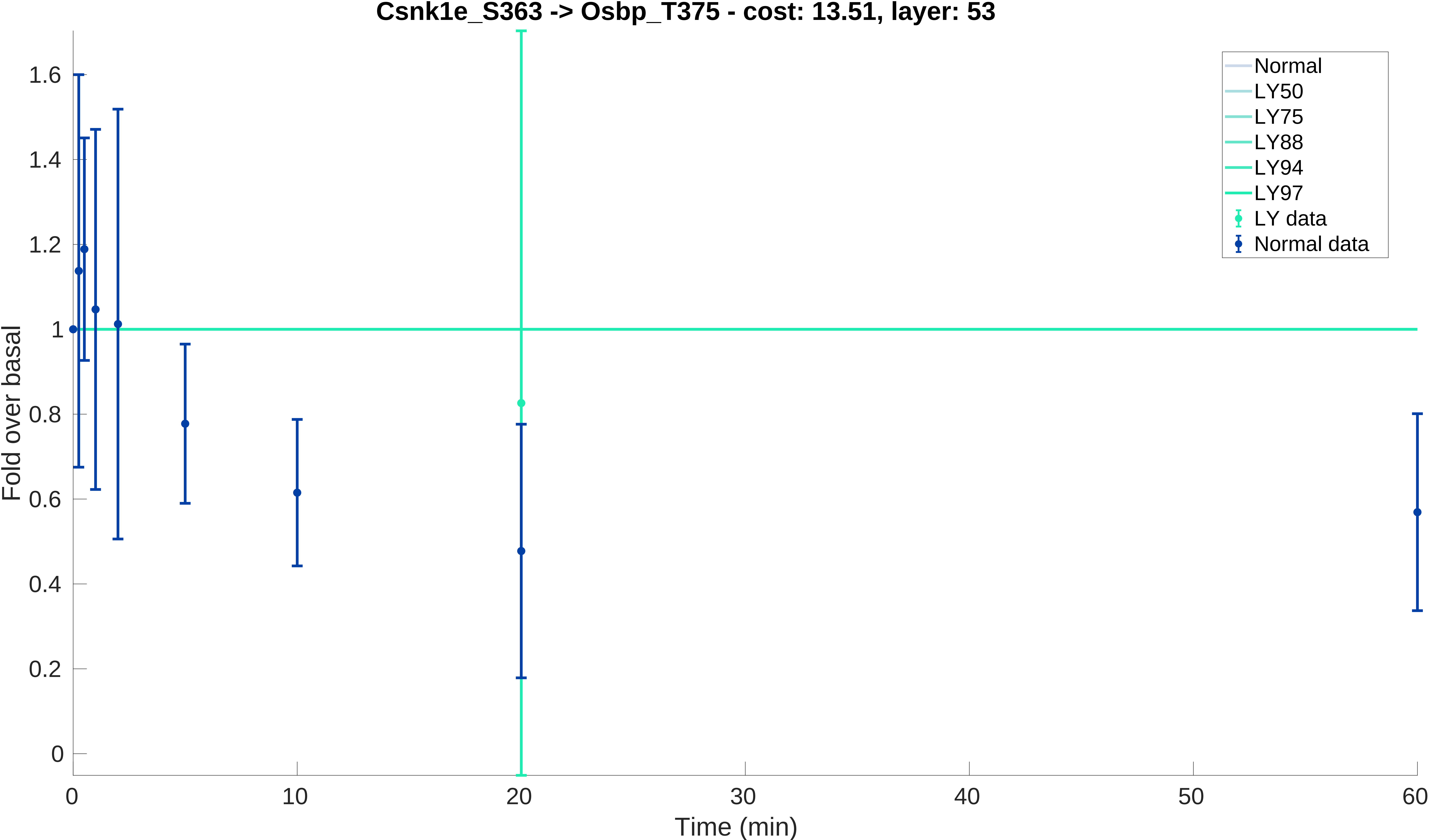

Csnk1e_S363 -> Osbp_T375 - cost: 13.51, layer: 53
Fold over basal
Time (min)
0
0.2
0.4
0.6
0.8
1
1.2
1.4
1.6
0
10
20
30
40
50
60
Normal
LY50
LY75
LY88
LY94
LY97
LY data
Normal data

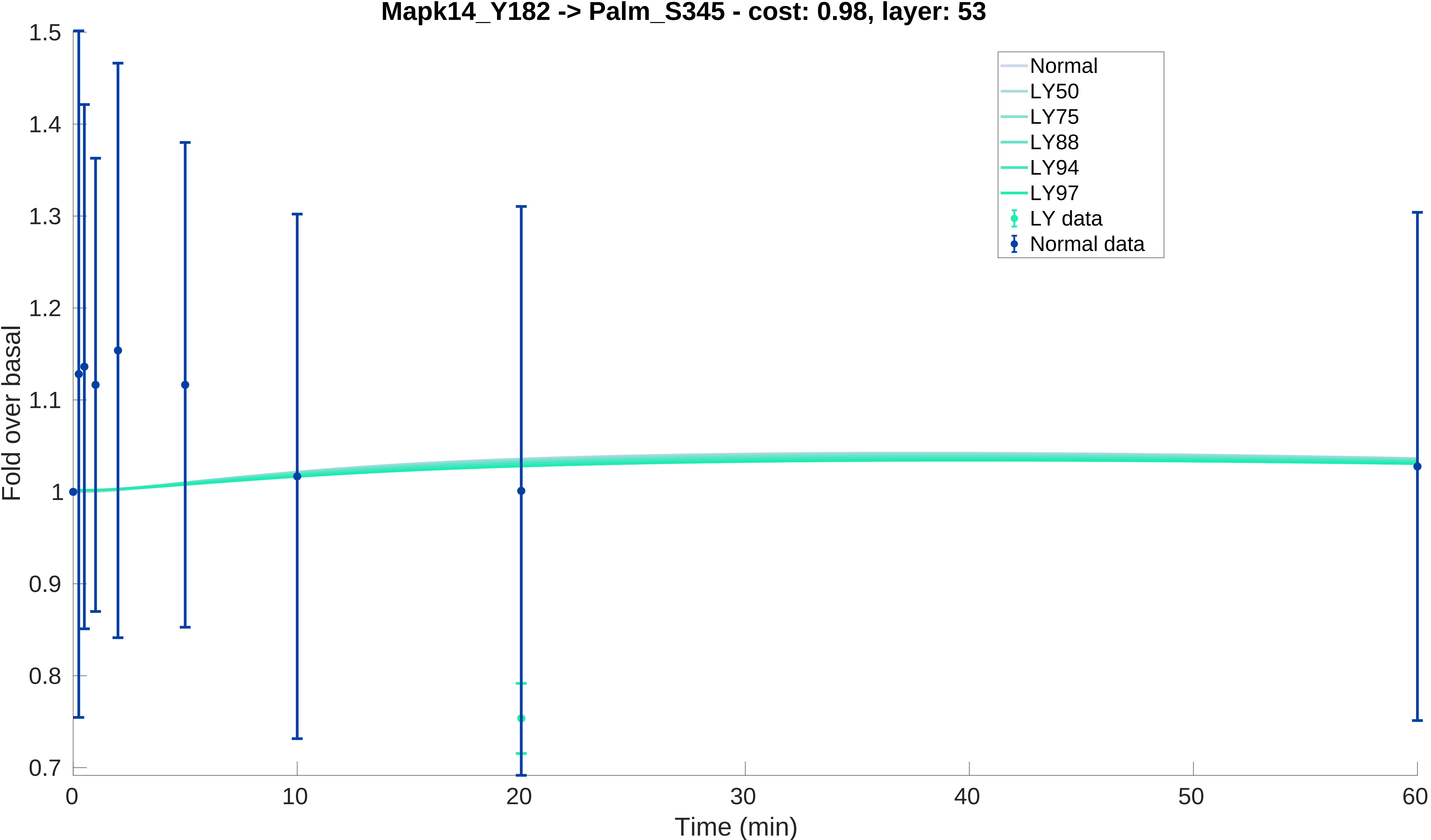

Mapk14_Y182 -> Palm_S345 - cost: 0.98, layer: 53
Fold over basal
Time (min)
Normal
LY50
LY75
LY88
LY94
LY97
LY data
Normal data
1.5
1.4
1.3
1.2
1.1
1
0.9
0.8
0.7
0
10
20
30
40
50
60

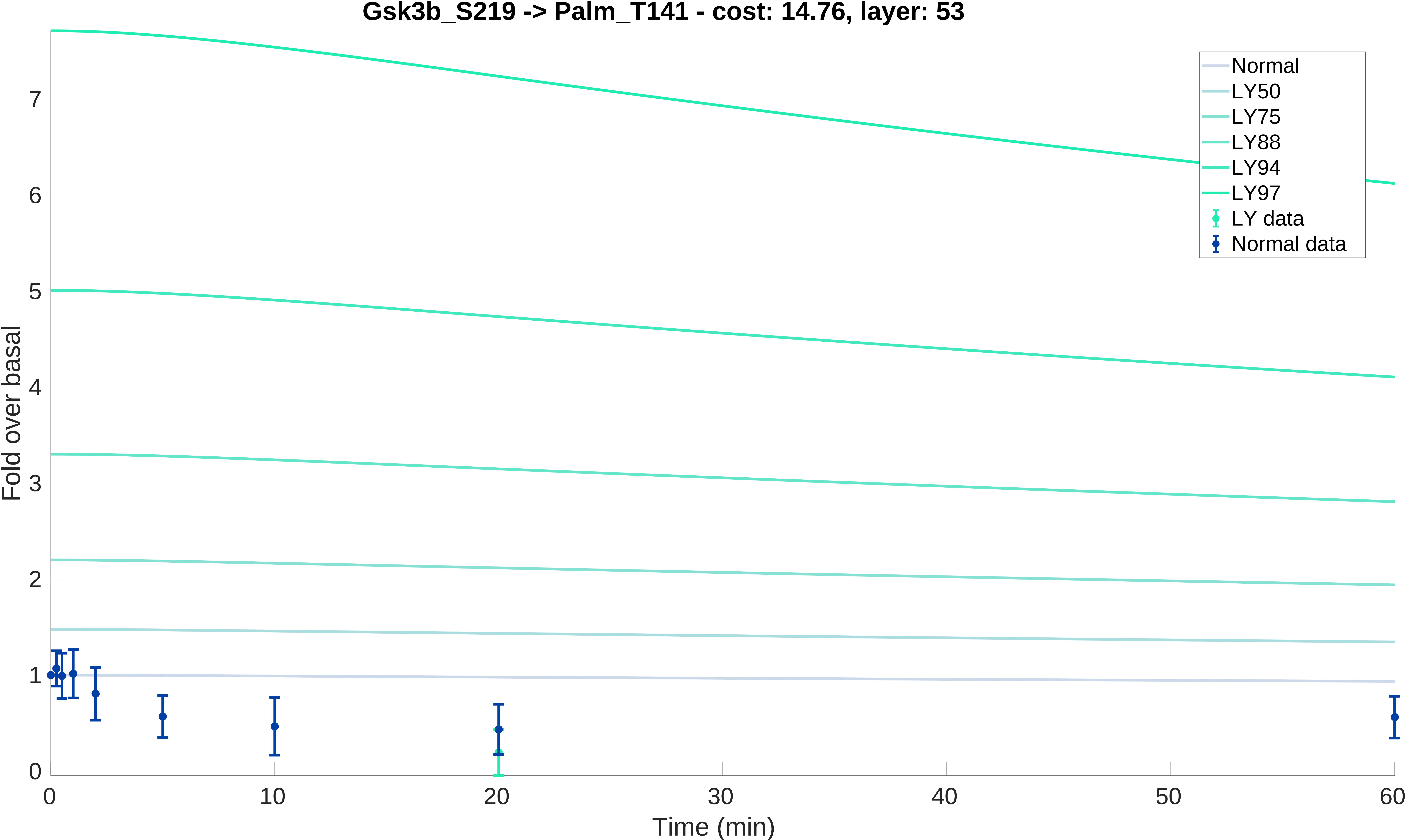

Gsk3b_S219 -> Palm_T141 - cost: 14.76, layer: 53
Normal
LY50
LY75
LY88
LY94
LY97
LY data
Normal data
Fold over basal
Time (min)
0
1
2
3
4
5
6
7
0
10
20
30
40
50
60

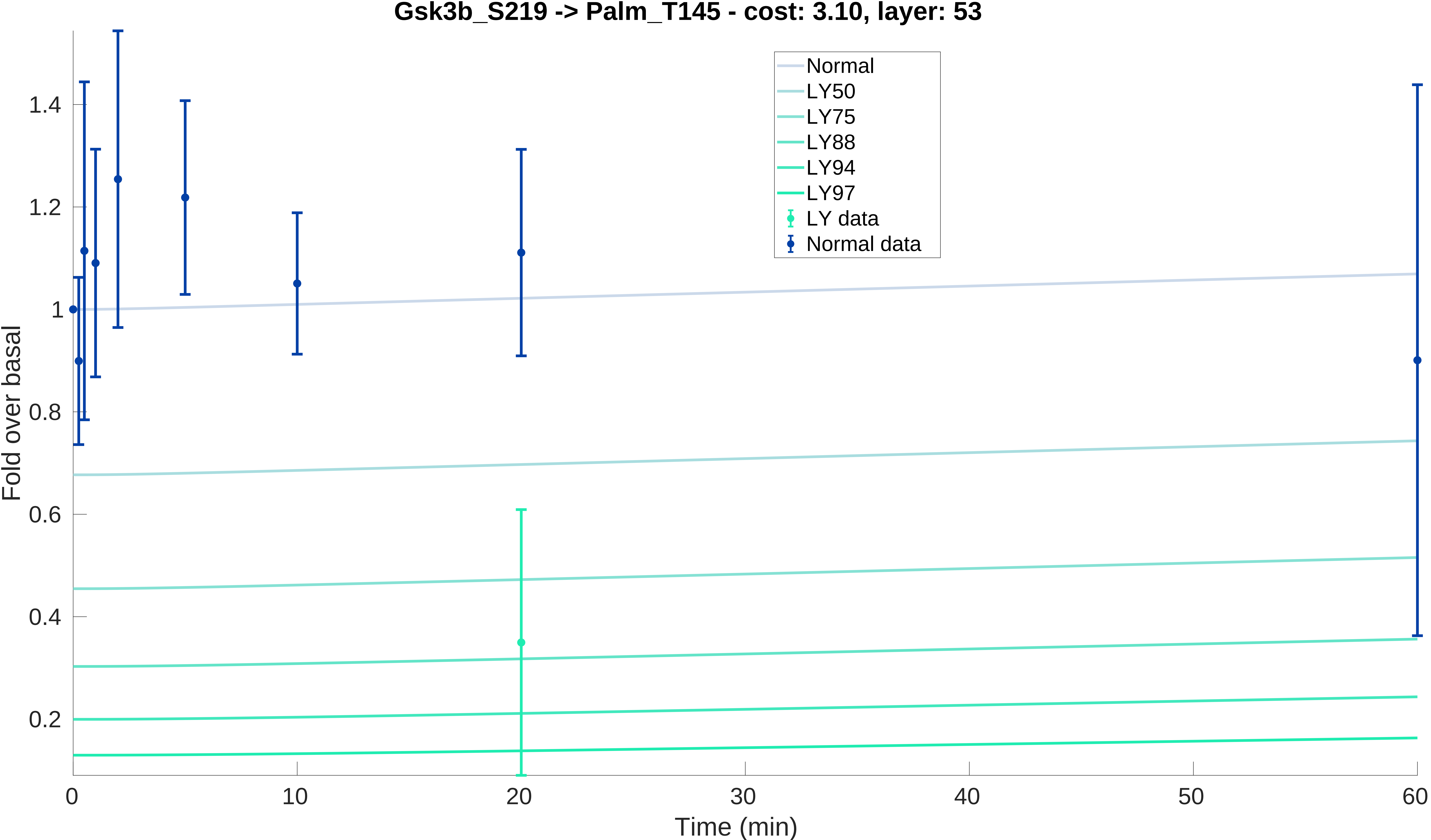

Gsk3b_S219 -> Palm_T145 - cost: 3.10, layer: 53
Normal
LY50
LY75
LY88
LY94
LY97
LY data
Normal data
Fold over basal
1.4
1.2
1
0.8
0.6
0.4
0.2
0
10
20
30
40
50
60
Time (min)

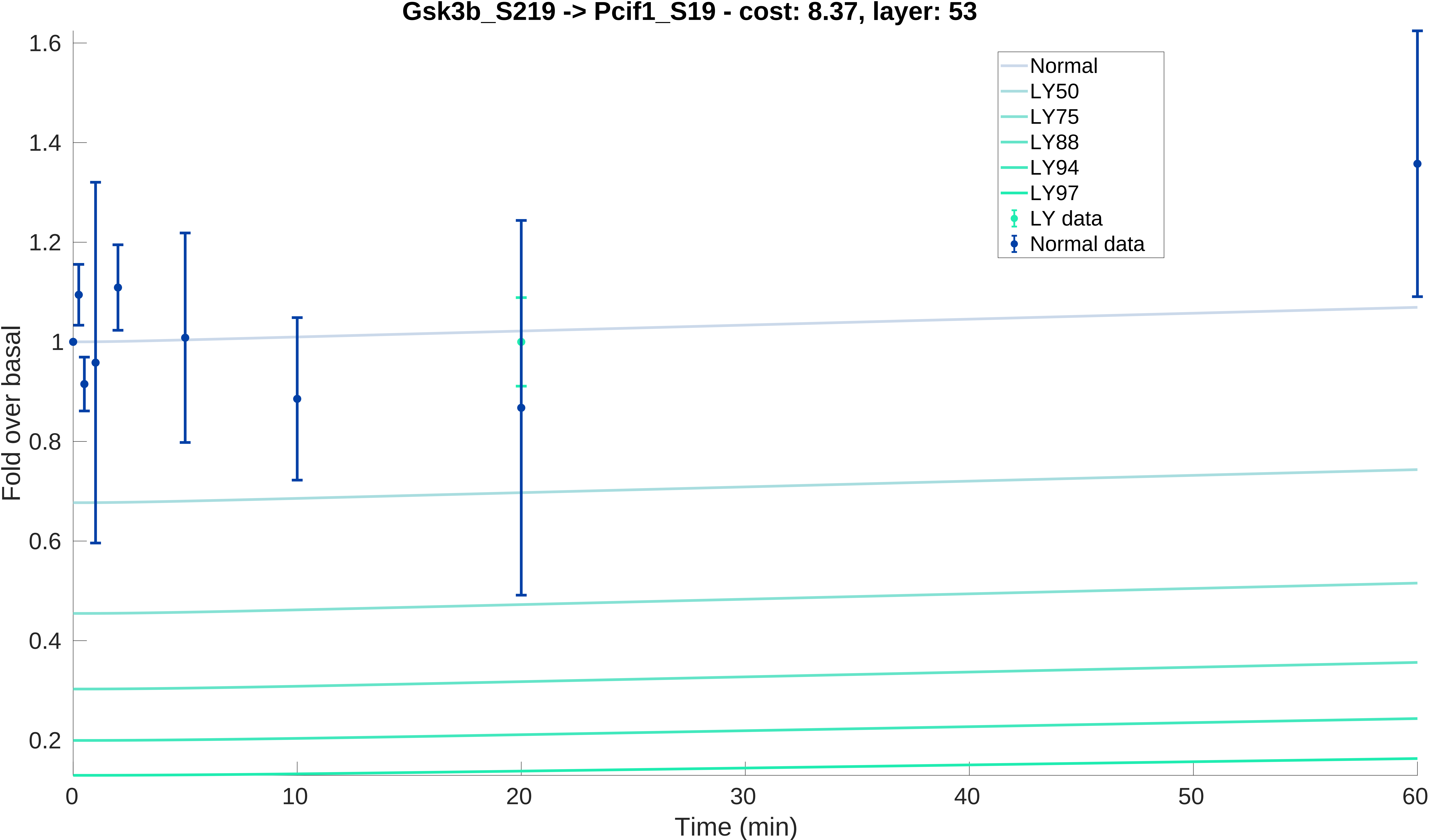

Gsk3b_S219 -> Pcif1_S19 - cost: 8.37, layer: 53
Fold over basal
Time (min)
Normal
LY50
LY75
LY88
LY94
LY97
LY data
Normal data
1.6
1.4
1.2
1
0.8
0.6
0.4
0.2
0
10
20
30
40
50
60

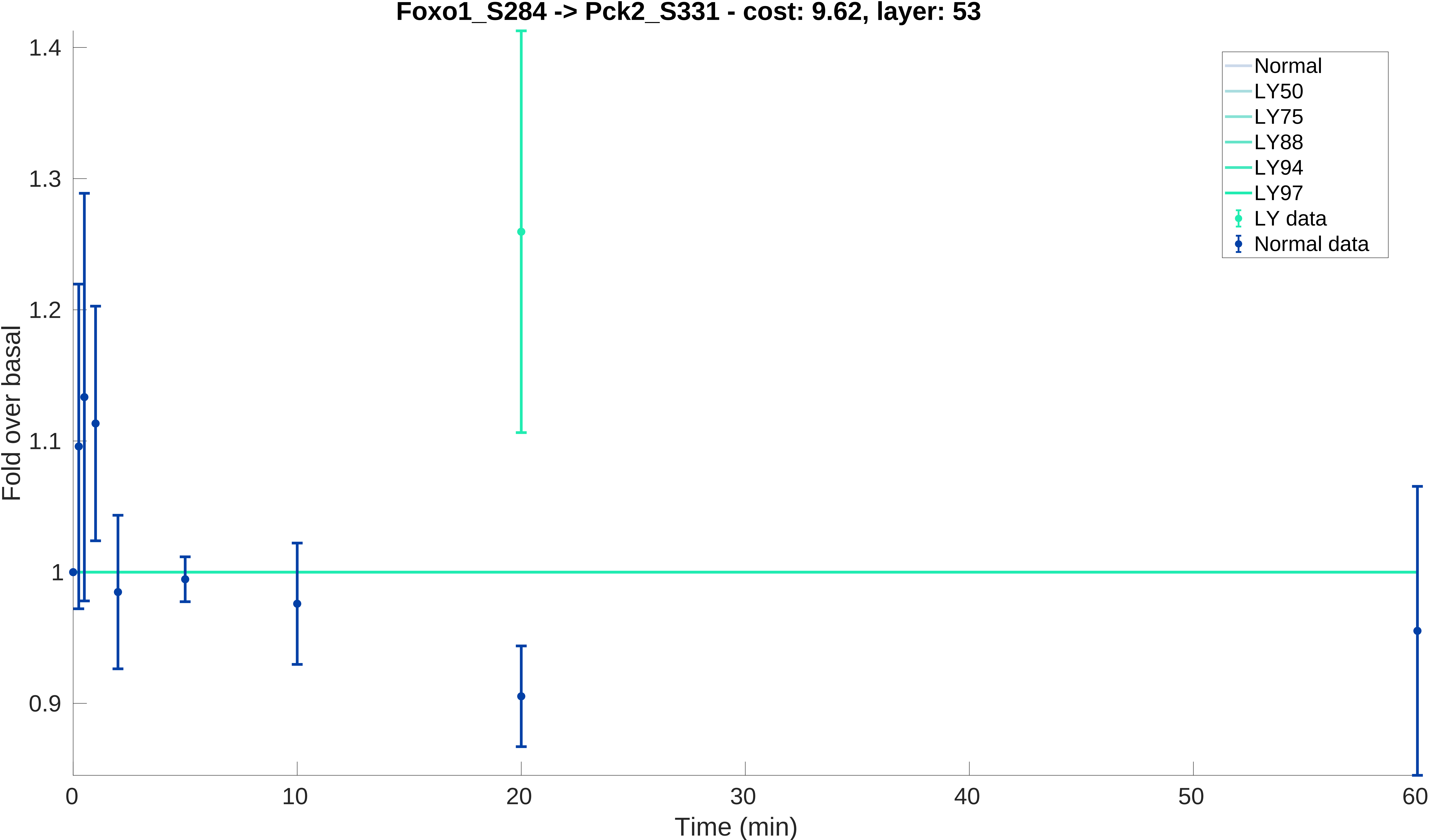

Foxo1_S284 -> Pck2_S331 - cost: 9.62, layer: 53
Fold over basal
Time (min)
1.4
1.3
1.2
1.1
1
0.9
0
10
20
30
40
50
60
Normal
LY50
LY75
LY88
LY94
LY97
LY data
Normal data

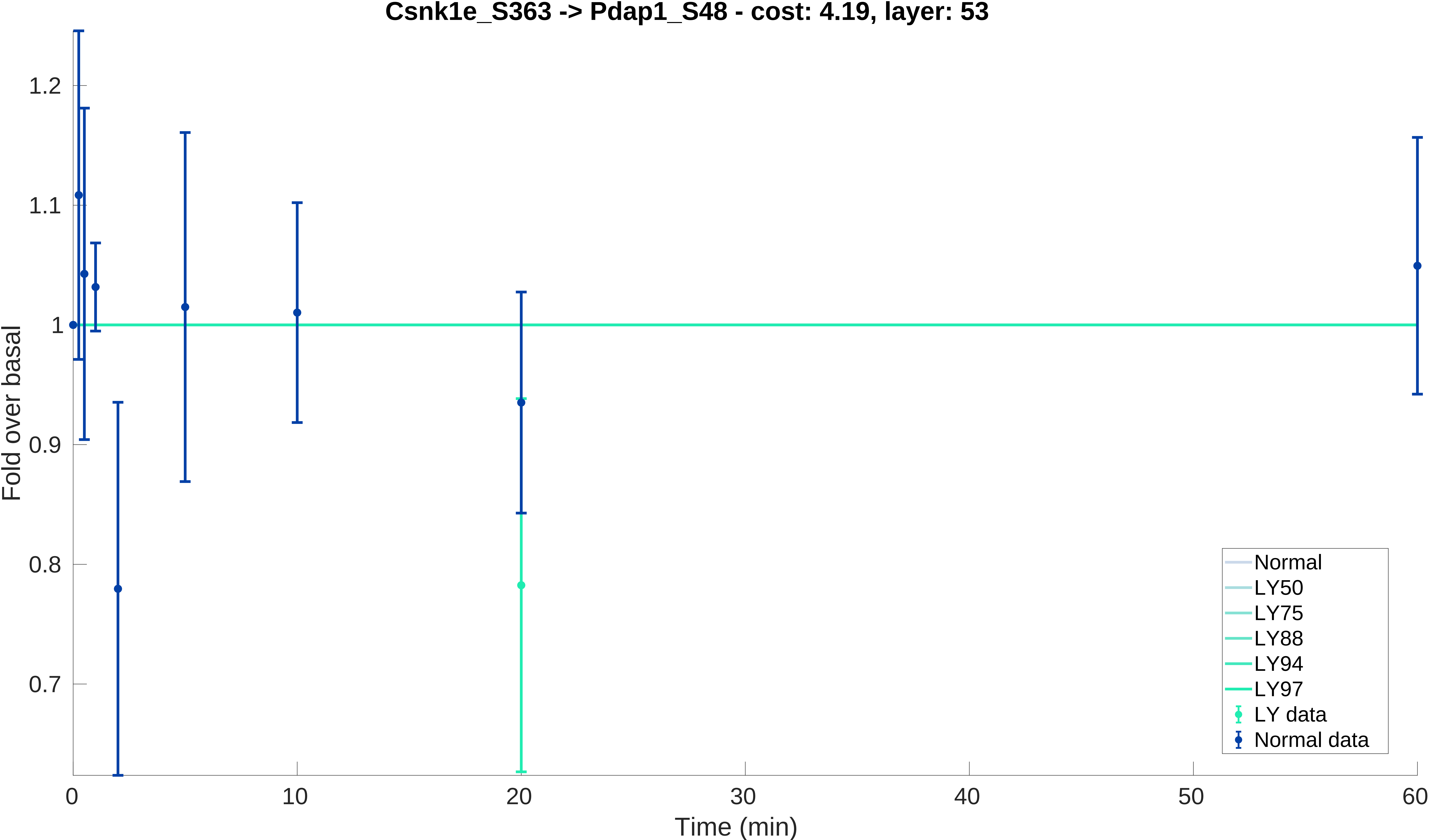
Csnk1e_S363 -> Pdap1_S48 - cost: 4.19, layer: 53
Fold over basal
1.2
1.1
1
0.9
0.8
0.7
0
10
20
30
40
50
60
Time (min)
Normal
LY50
LY75
LY88
LY94
LY97
LY data
Normal data

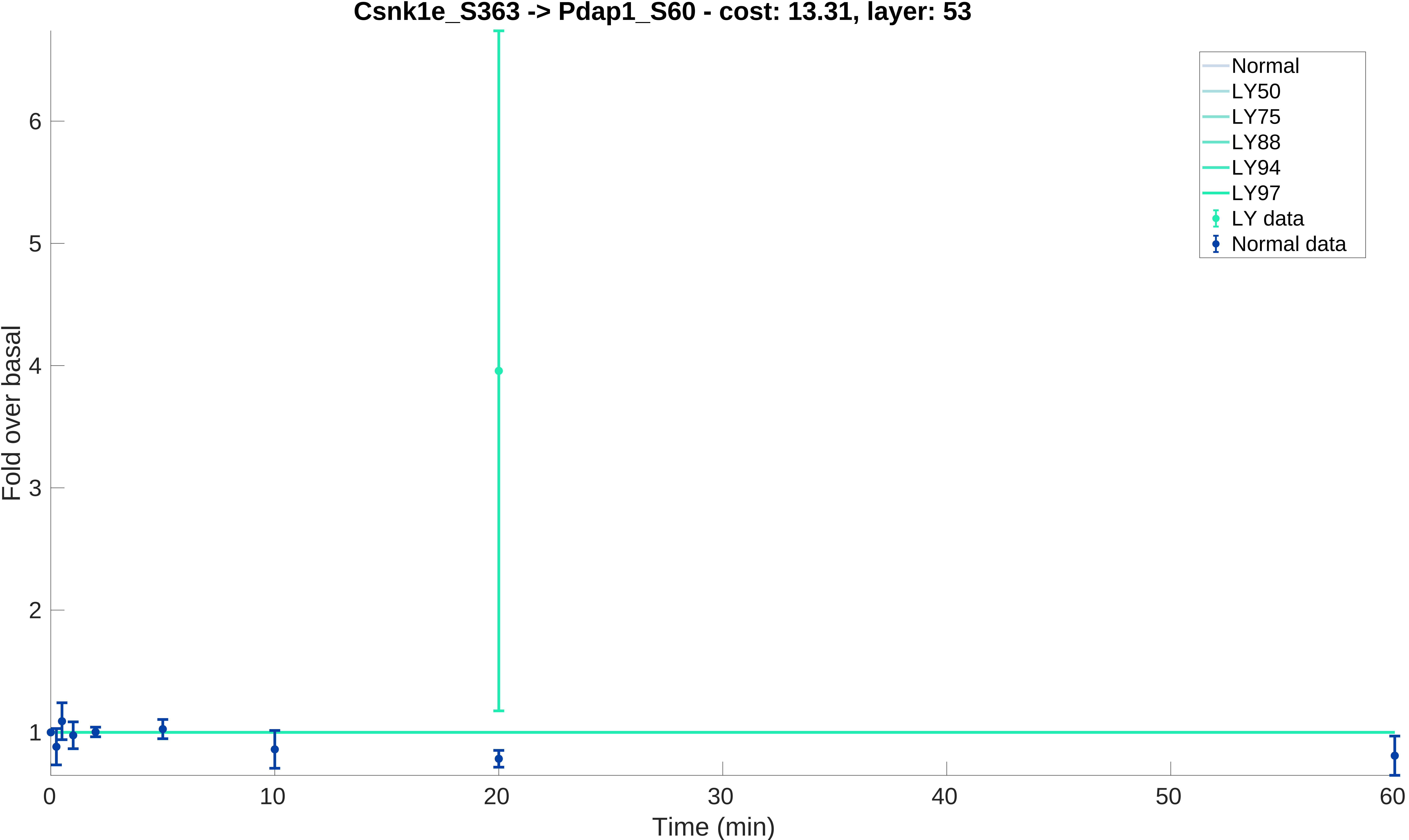

Csnk1e_S363 -> Pdap1_S60 - cost: 13.31, layer: 53
Fold over basal
Time (min)
0
10
20
30
40
50
60
1
2
3
4
5
6
Normal
LY50
LY75
LY88
LY94
LY97
LY data
Normal data

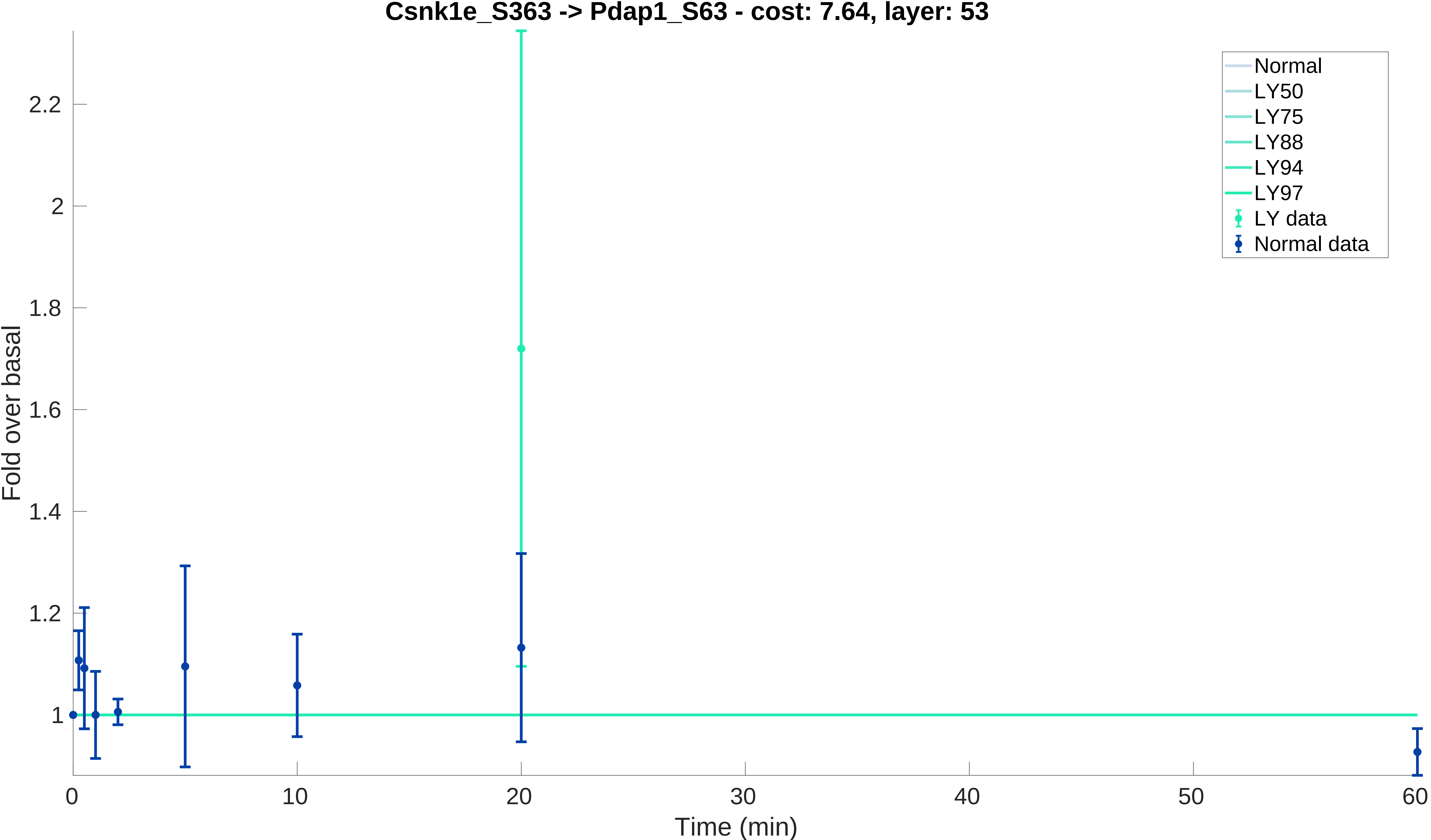

Csnk1e_S363 -> Pdap1_S63 - cost: 7.64, layer: 53
Normal
LY50
LY75
LY88
LY94
LY97
LY data
Normal data
Fold over basal
2.2
2
1.8
1.6
1.4
1.2
1
0
10
20
30
40
50
60
Time (min)

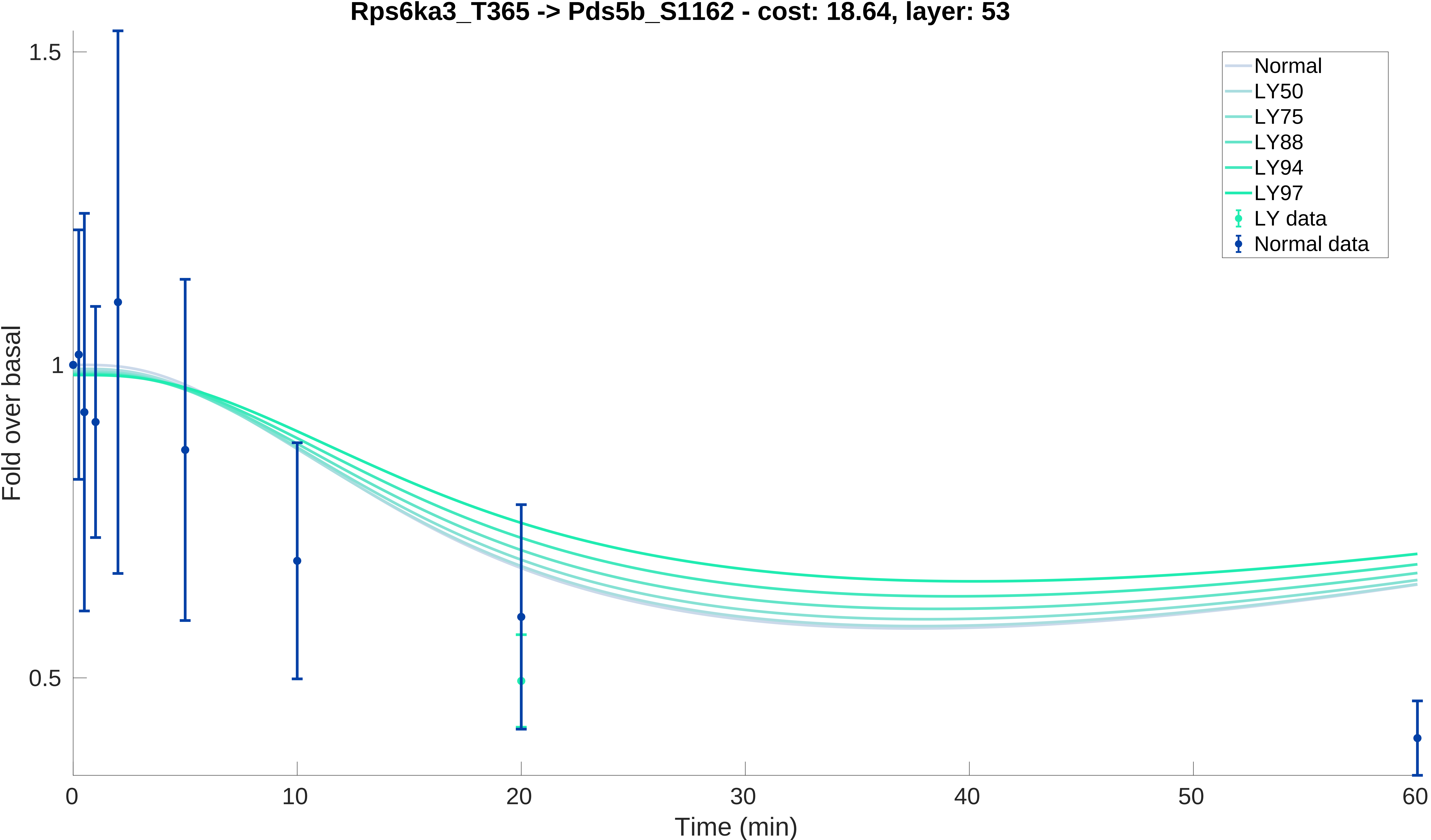
Rps6ka3_T365 -> Pds5b_S1162 - cost: 18.64, layer: 53
Fold over basal
Time (min)
1.5
1
0.5
0
10
20
30
40
50
60
Normal
LY50
LY75
LY88
LY94
LY97
LY data
Normal data

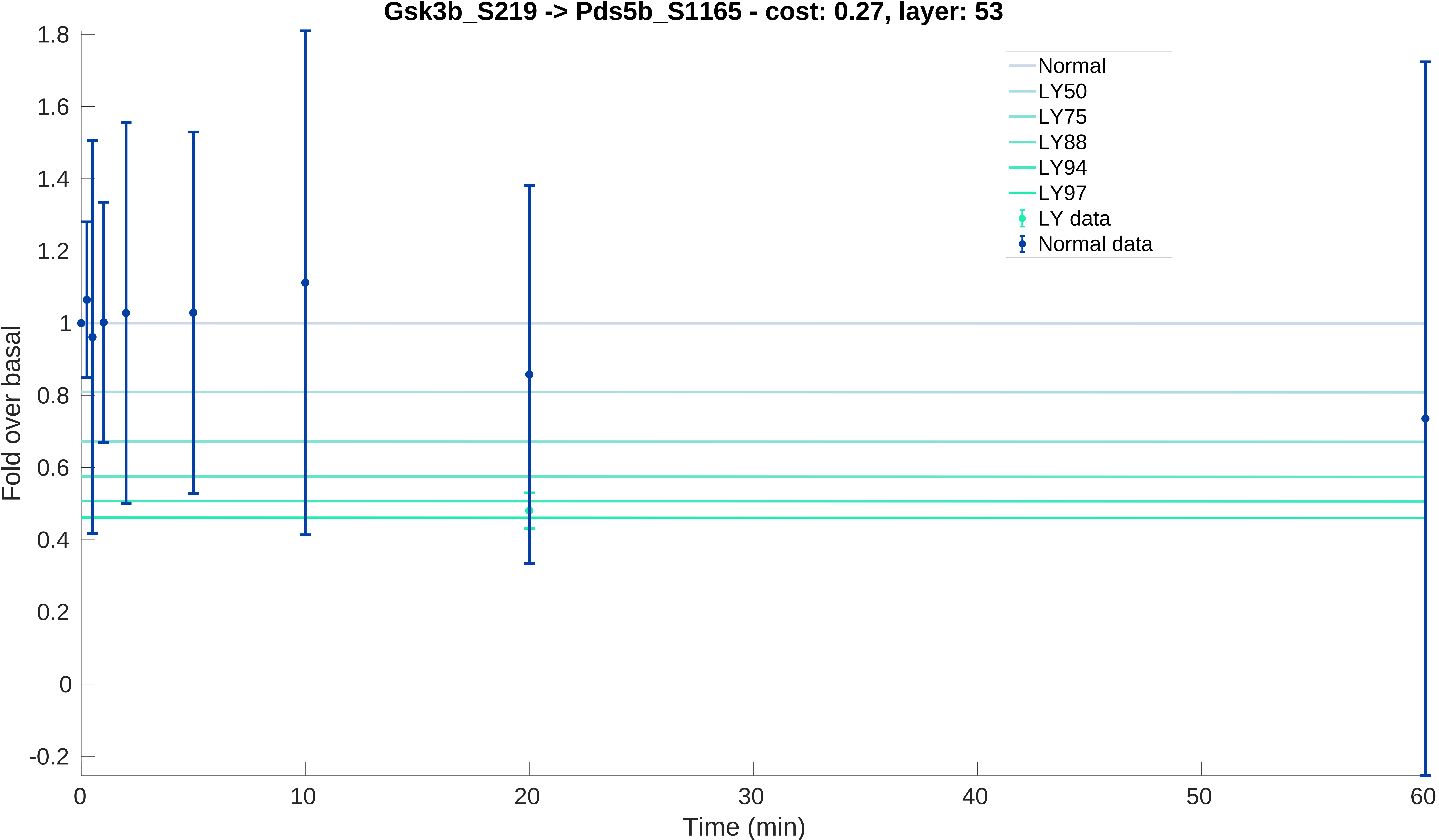

Gsk3b_S219 -> Pds5b_S1165 - cost: 0.27, layer: 53
Normal
LY50
LY75
LY88
LY94
LY97
LY data
Normal data
Fold over basal
Time (min)
1.8
1.6
1.4
1.2
1
0.8
0.6
0.4
0.2
0
-0.2
0
10
20
30
40
50
60

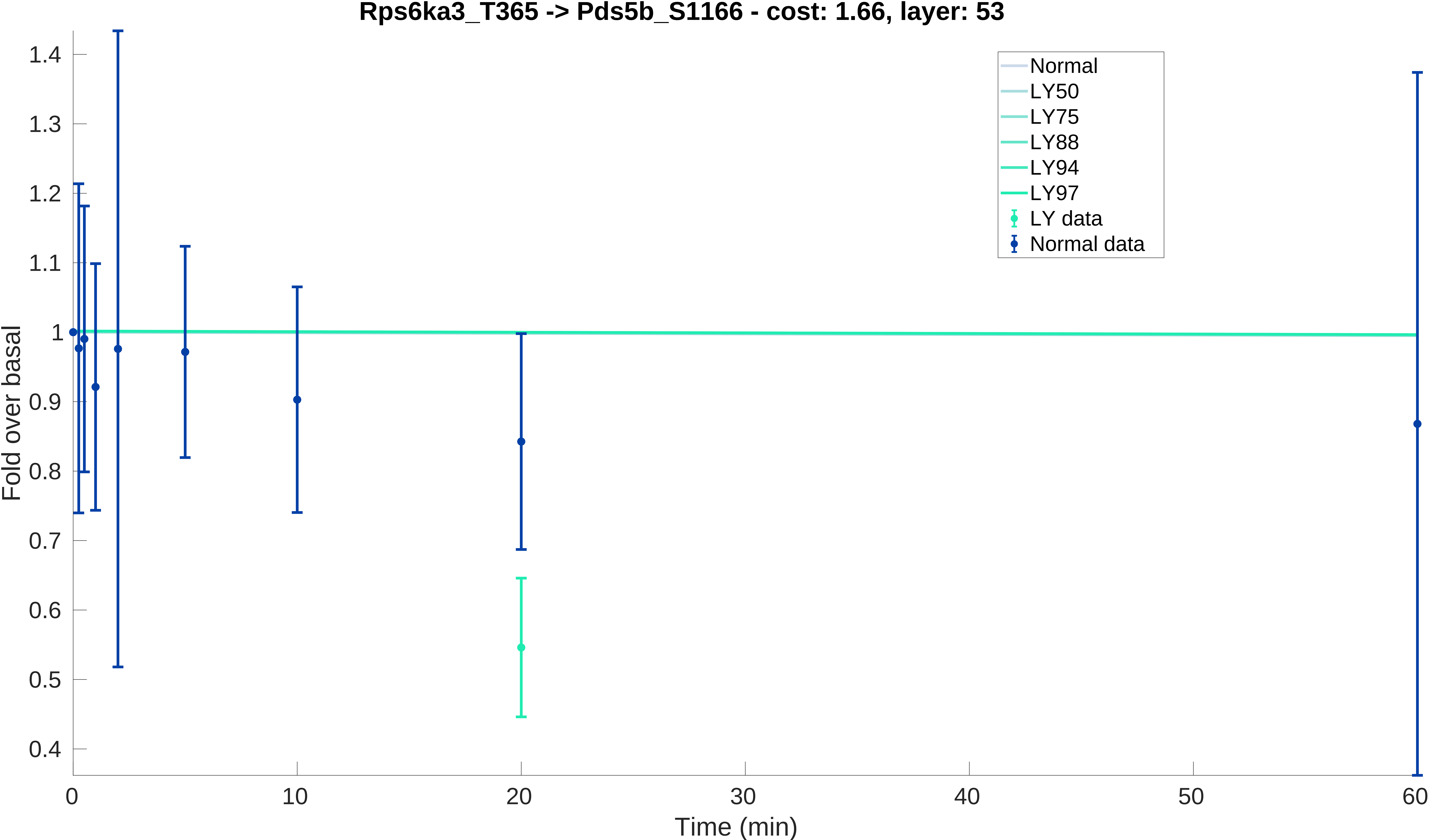

Rps6ka3_T365 -> Pds5b_S1166 - cost: 1.66, layer: 53
Normal
LY50
LY75
LY88
LY94
LY97
LY data
Normal data
Fold over basal
1.4
1.3
1.2
1.1
1
0.9
0.8
0.7
0.6
0.5
0.4
0
10
20
30
40
50
60
Time (min)

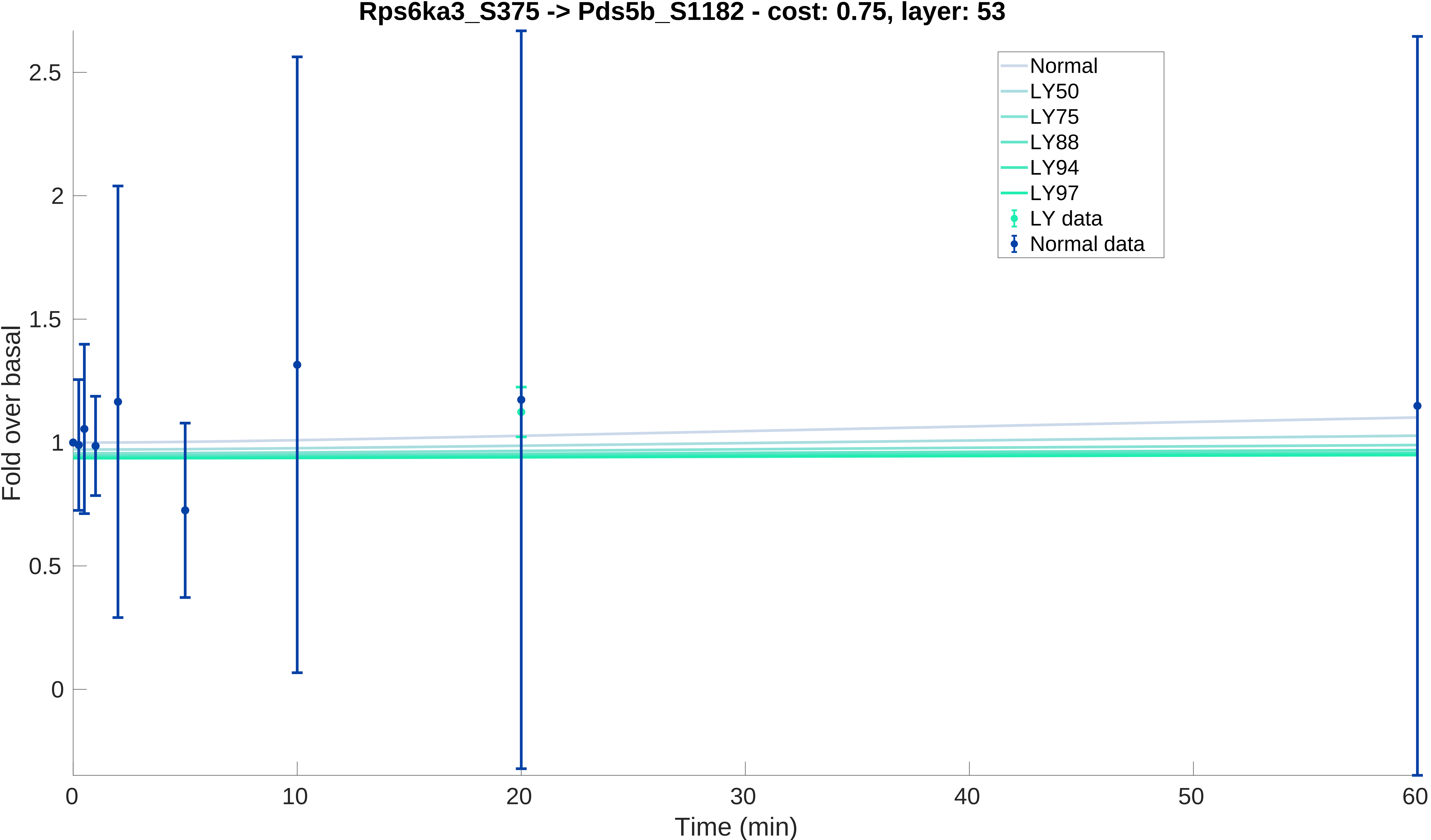

Rps6ka3_S375 -> Pds5b_S1182 - cost: 0.75, layer: 53
Fold over basal
Time (min)
Normal
LY50
LY75
LY88
LY94
LY97
LY data
Normal data
2.5
2
1.5
1
0.5
0
0
10
20
30
40
50
60

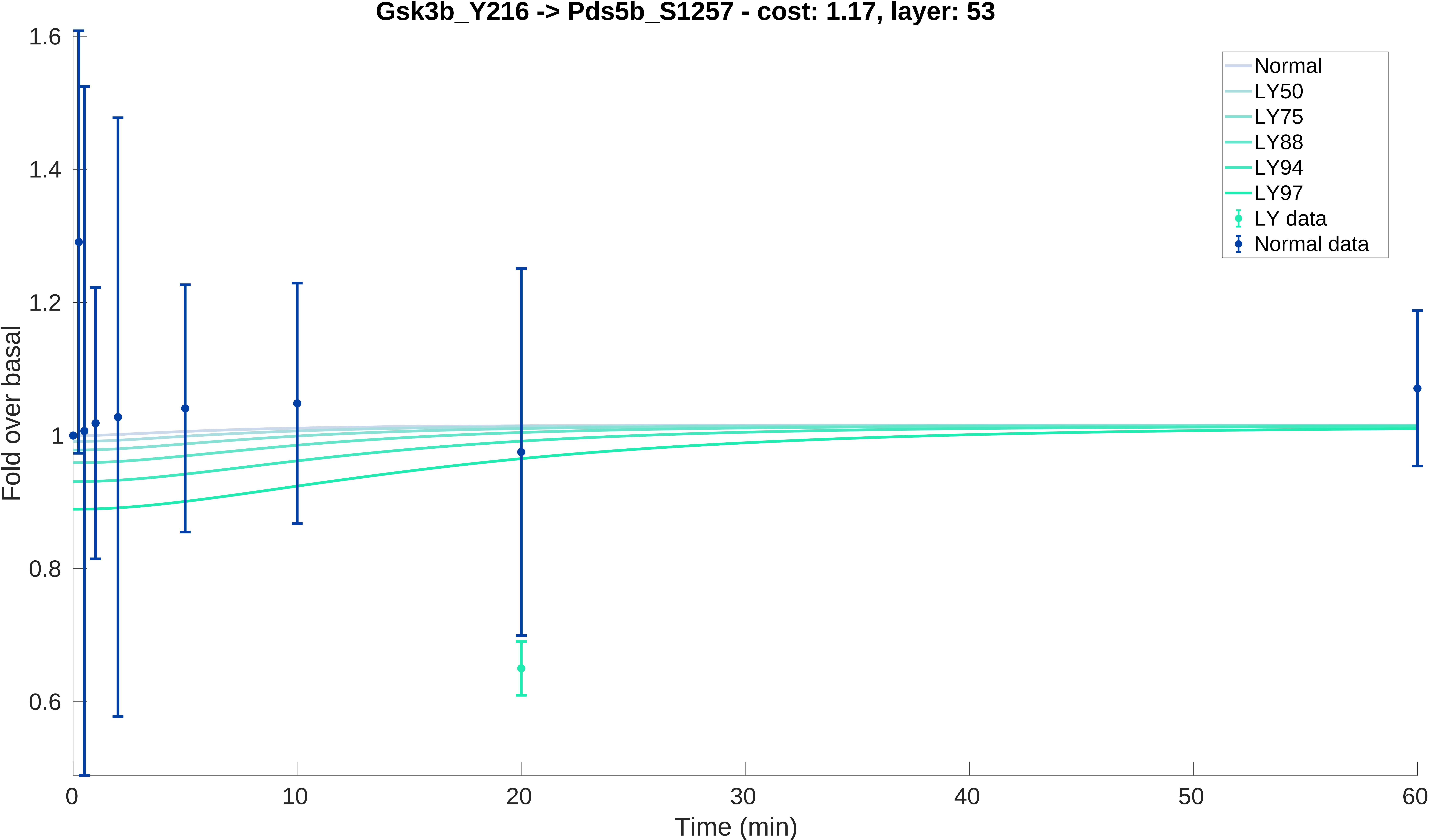

Gsk3b_Y216 -> Pds5b_S1257 - cost: 1.17, layer: 53
Fold over basal
1.6
1.4
1.2
1
0.8
0.6
0
10
20
30
40
50
60
Time (min)
Normal
LY50
LY75
LY88
LY94
LY97
LY data
Normal data

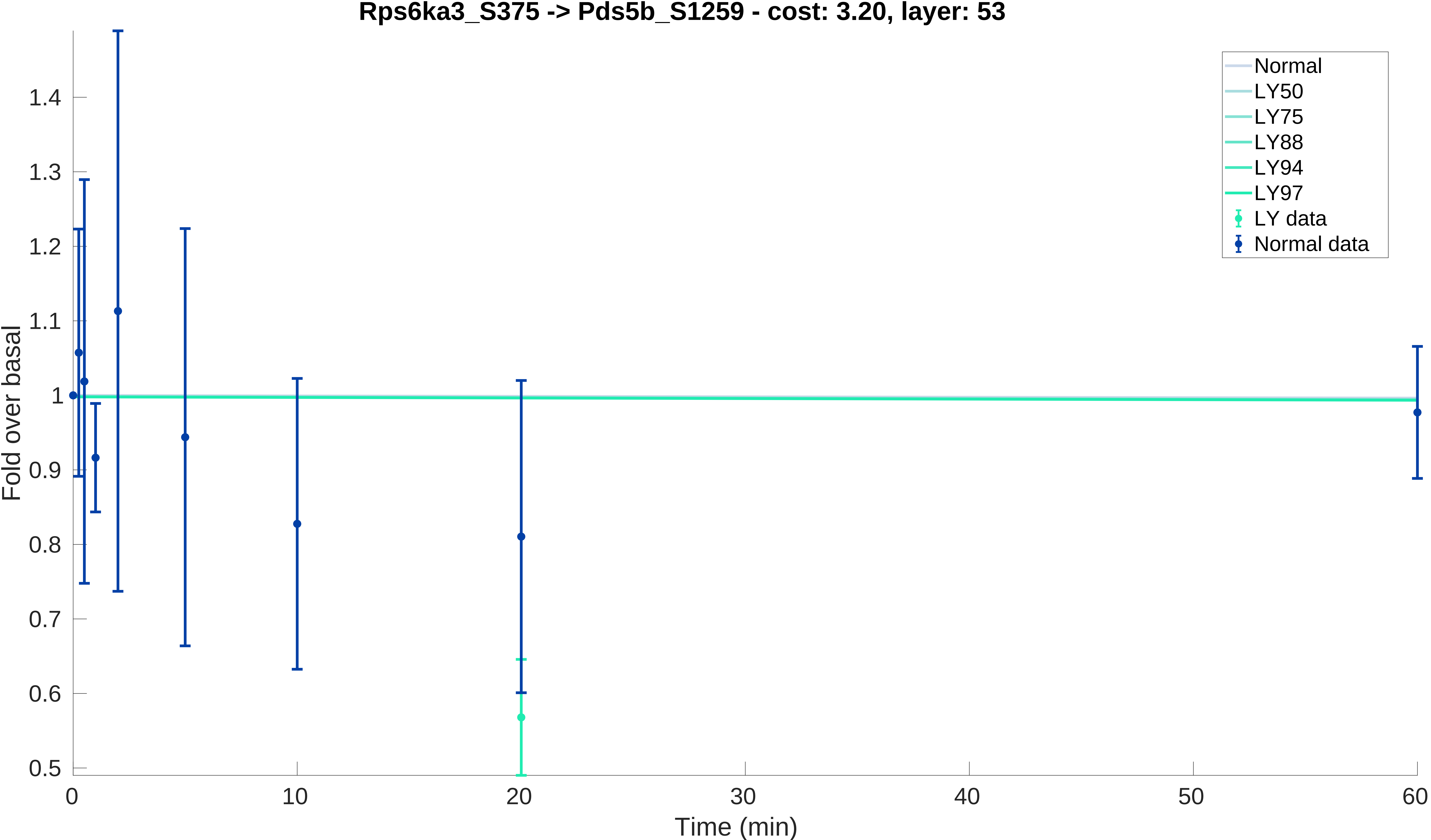
Rps6ka3_S375 -> Pds5b_S1259 - cost: 3.20, layer: 53
Fold over basal
Time (min)
0
10
20
30
40
50
60
0.5
0.6
0.7
0.8
0.9
1
1.1
1.2
1.3
1.4
Normal
LY50
LY75
LY88
LY94
LY97
LY data
Normal data

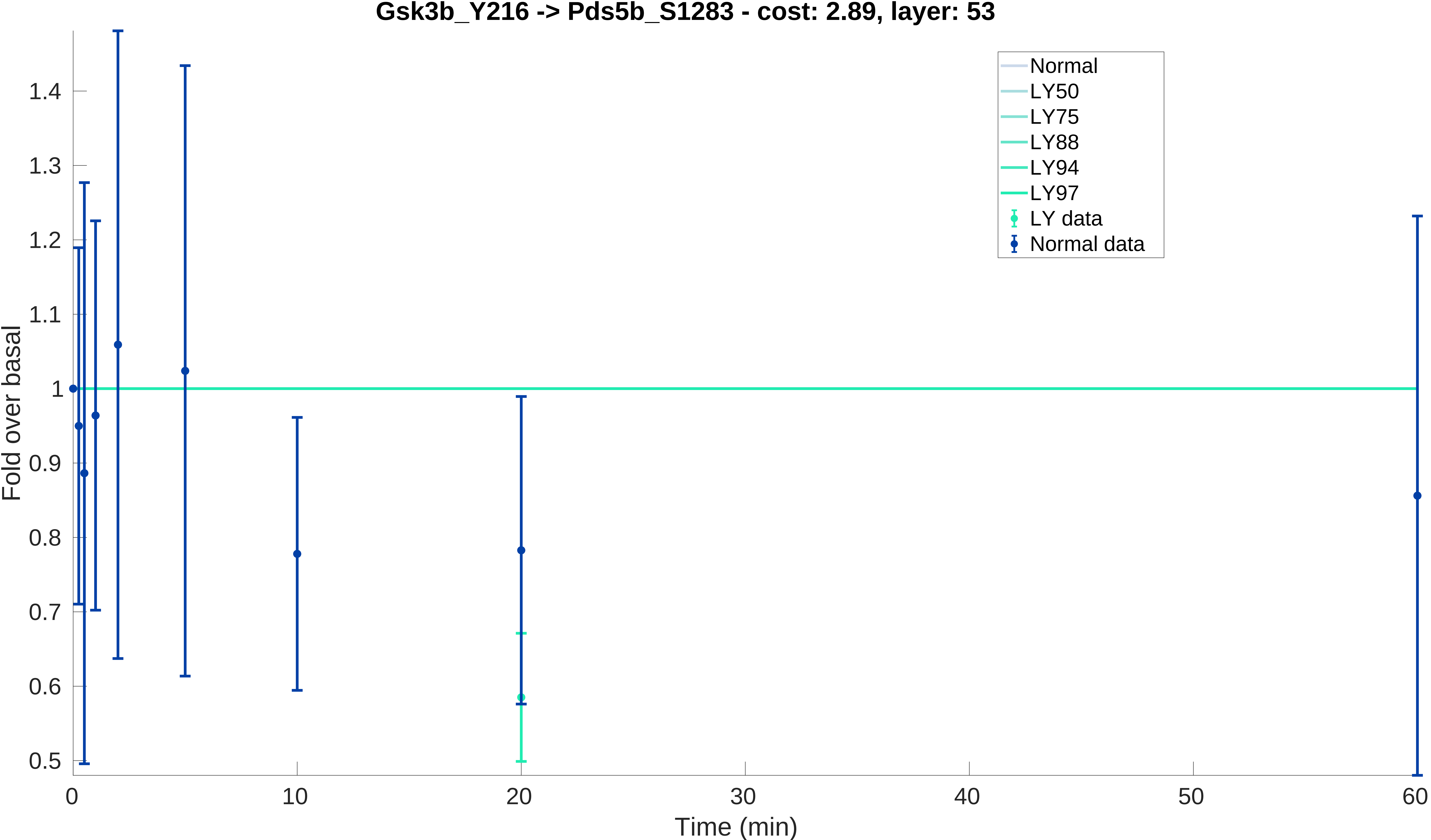

Gsk3b_Y216 -> Pds5b_S1283 - cost: 2.89, layer: 53
Normal
LY50
LY75
LY88
LY94
LY97
LY data
Normal data
Fold over basal
1.4
1.3
1.2
1.1
1
0.9
0.8
0.7
0.6
0.5
0
10
20
30
40
50
60
Time (min)

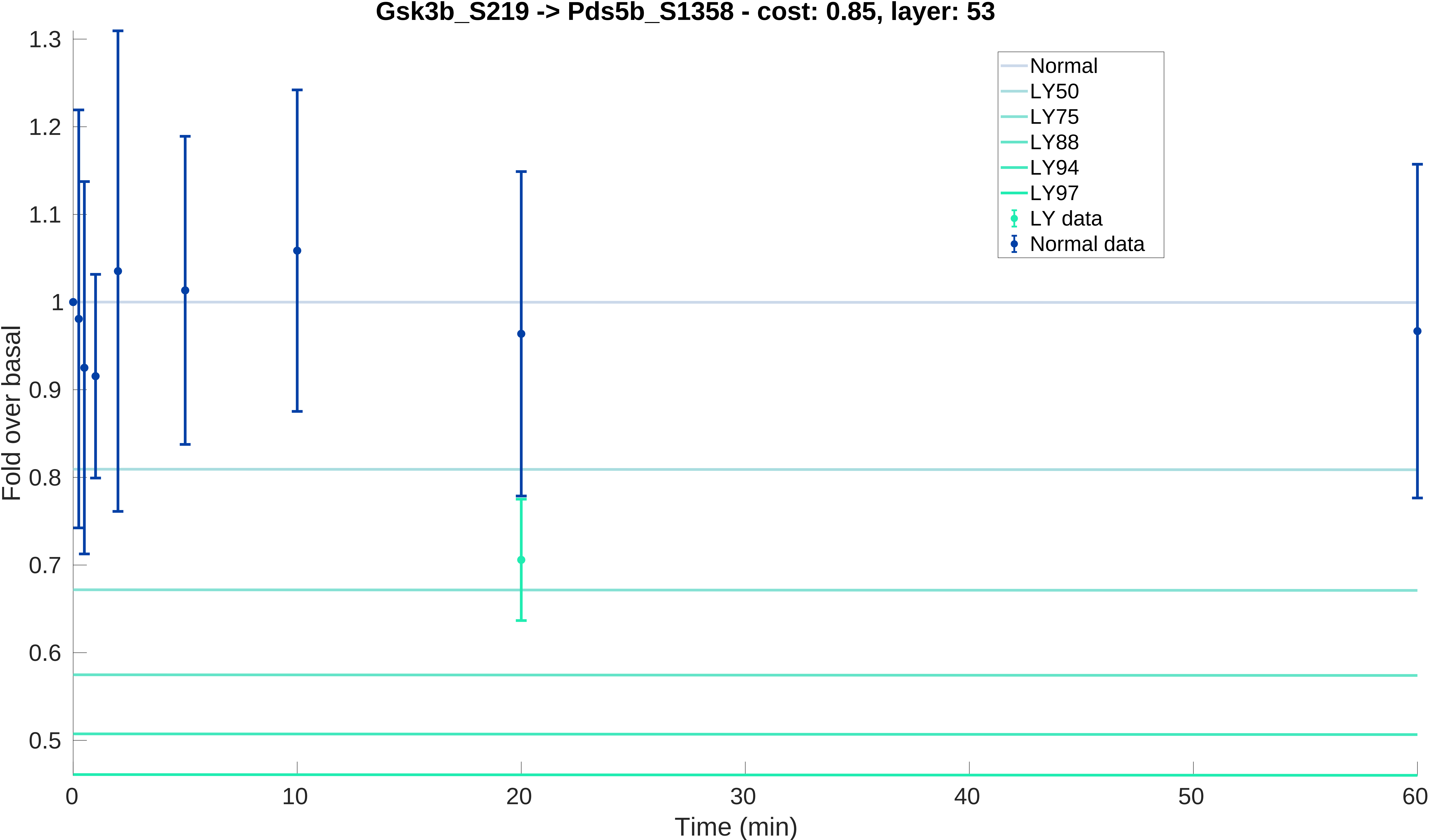

Gsk3b_S219 -> Pds5b_S1358 - cost: 0.85, layer: 53
Fold over basal
Time (min)
Normal
LY50
LY75
LY88
LY94
LY97
LY data
Normal data
1.3
1.2
1.1
1
0.9
0.8
0.7
0.6
0.5
0
10
20
30
40
50
60

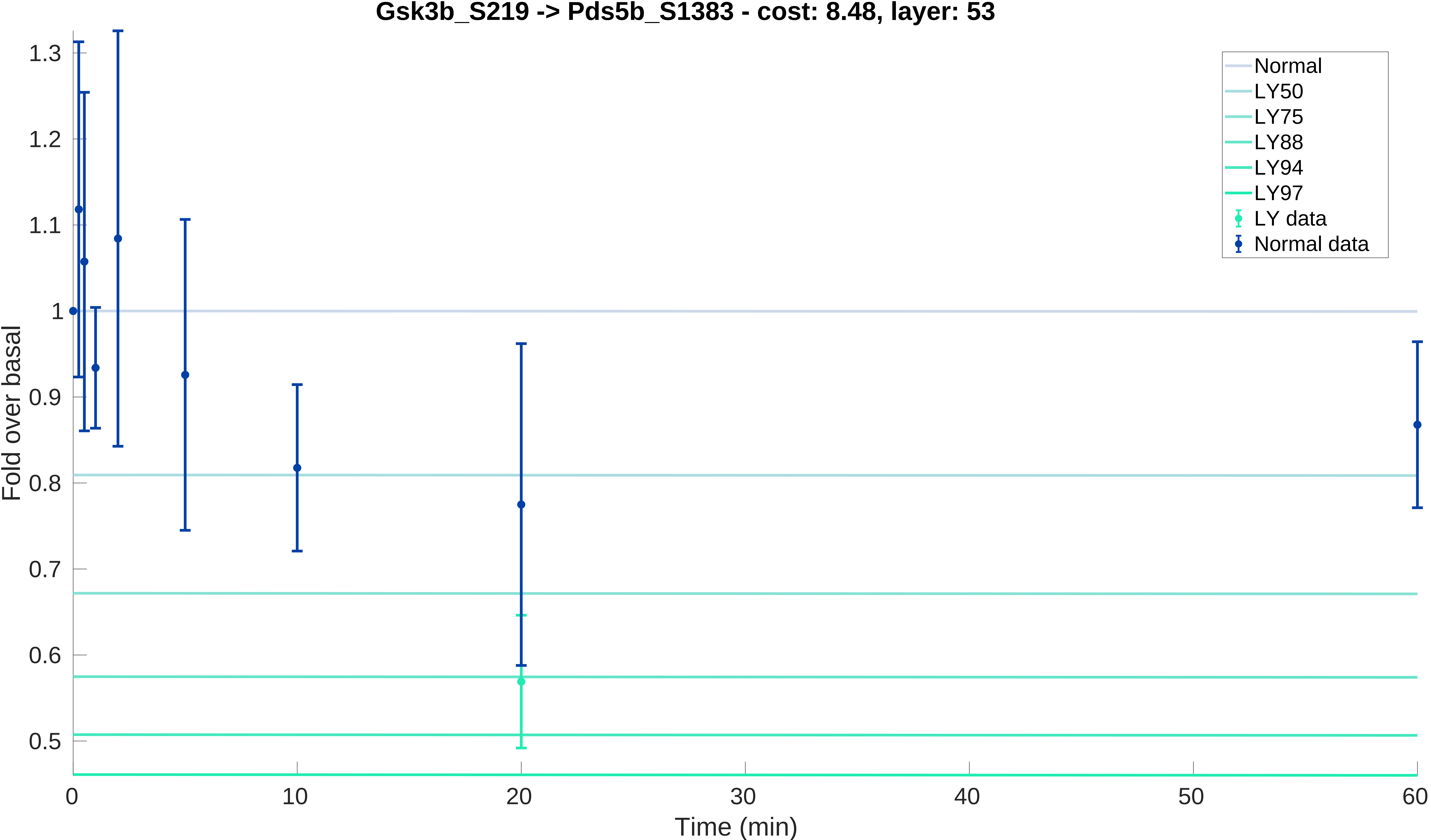

Gsk3b_S219 -> Pds5b_S1383 - cost: 8.48, layer: 53
Fold over basal
Time (min)
1.3
1.2
1.1
1
0.9
0.8
0.7
0.6
0.5
0
10
20
30
40
50
60
Normal
LY50
LY75
LY88
LY94
LY97
LY data
Normal data

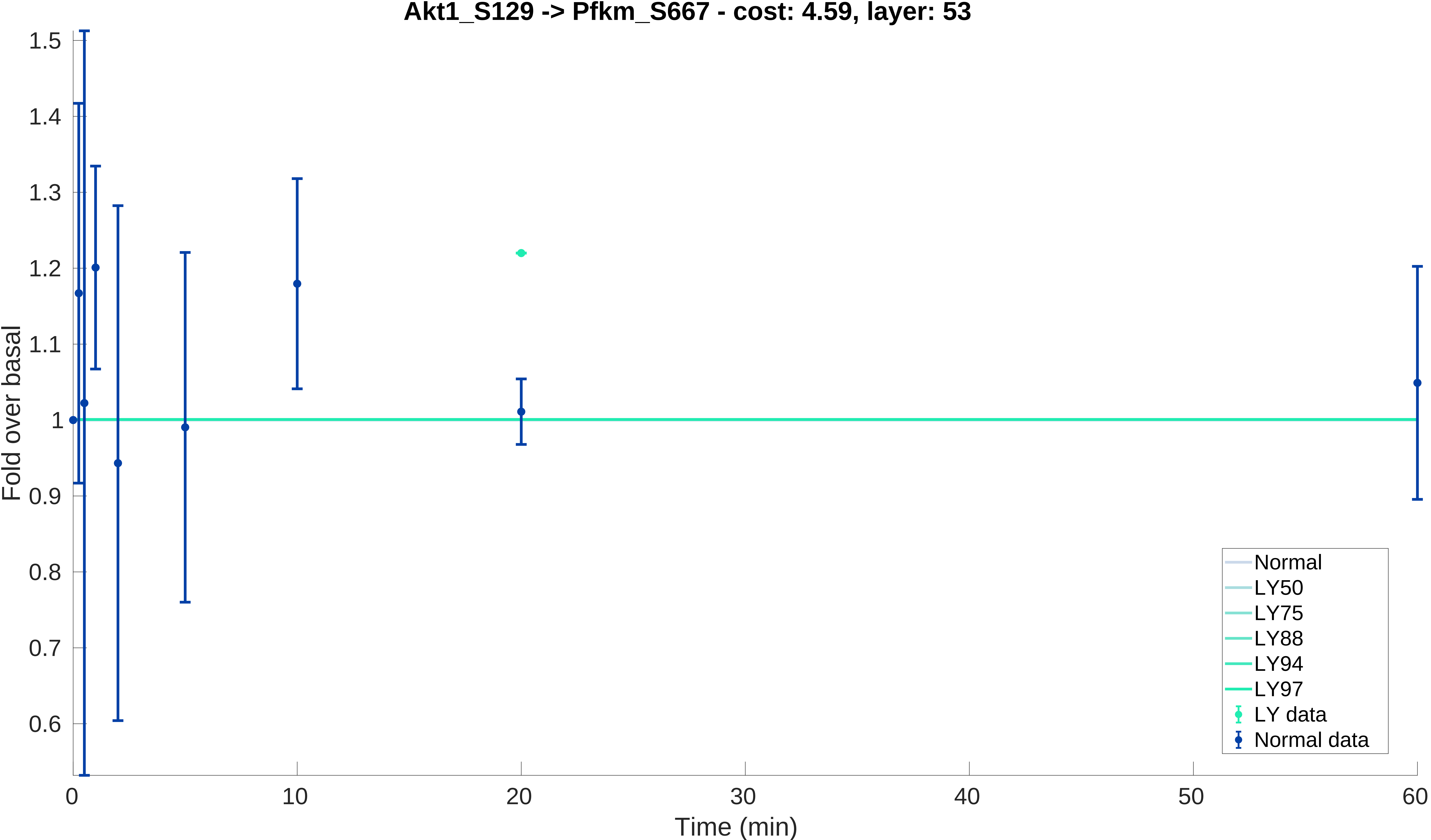

Akt1_S129 -> Pfkm_S667 - cost: 4.59, layer: 53
Fold over basal
1.5
1.4
1.3
1.2
1.1
1
0.9
0.8
0.7
0.6
0
10
20
30
40
50
60
Time (min)
Normal
LY50
LY75
LY88
LY94
LY97
LY data
Normal data

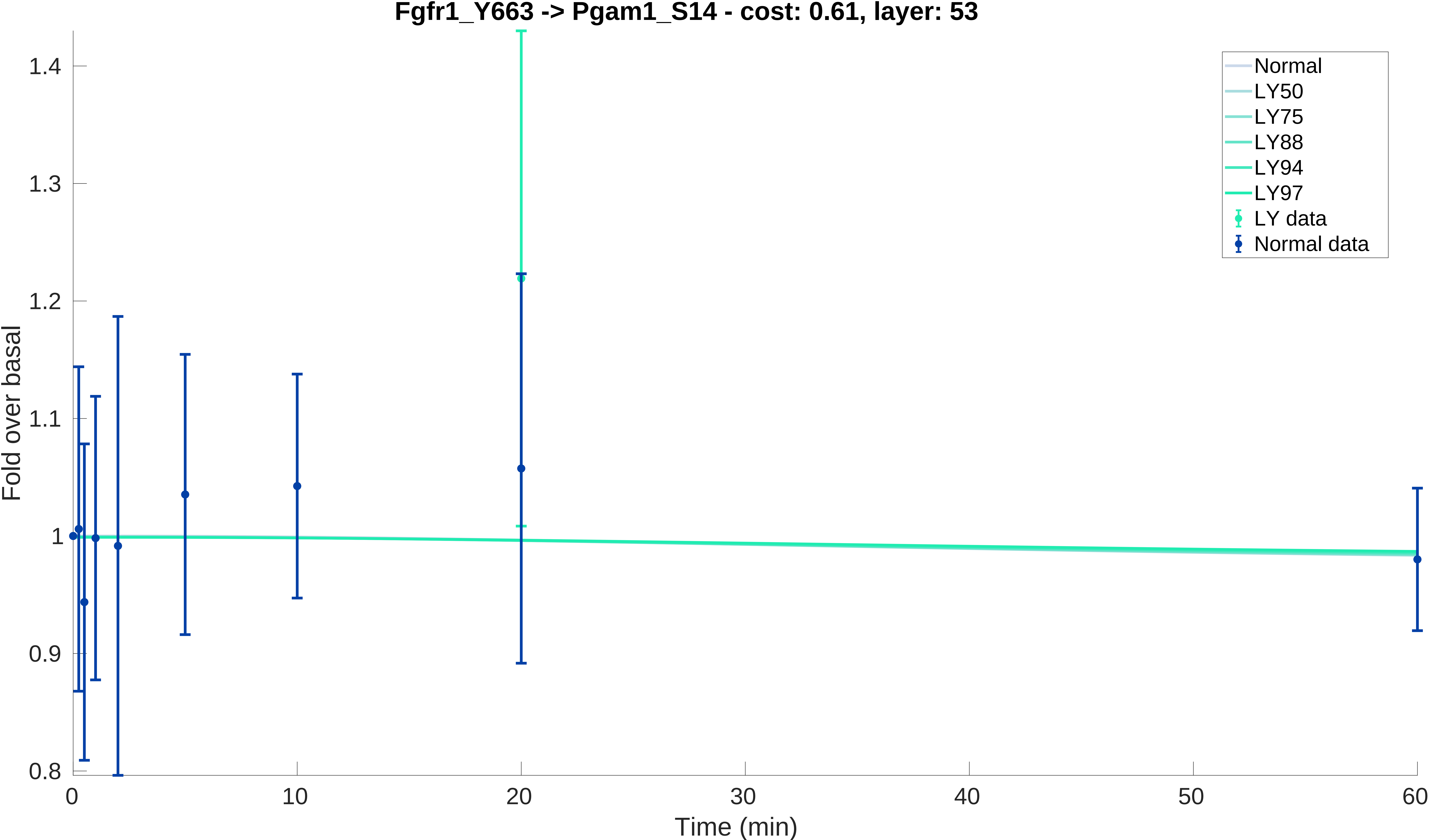

Fgfr1_Y663 -> Pgam1_S14 - cost: 0.61, layer: 53
Fold over basal
Time (min)
1.4
1.3
1.2
1.1
1
0.9
0.8
0
10
20
30
40
50
60
Normal
LY50
LY75
LY88
LY94
LY97
LY data
Normal data

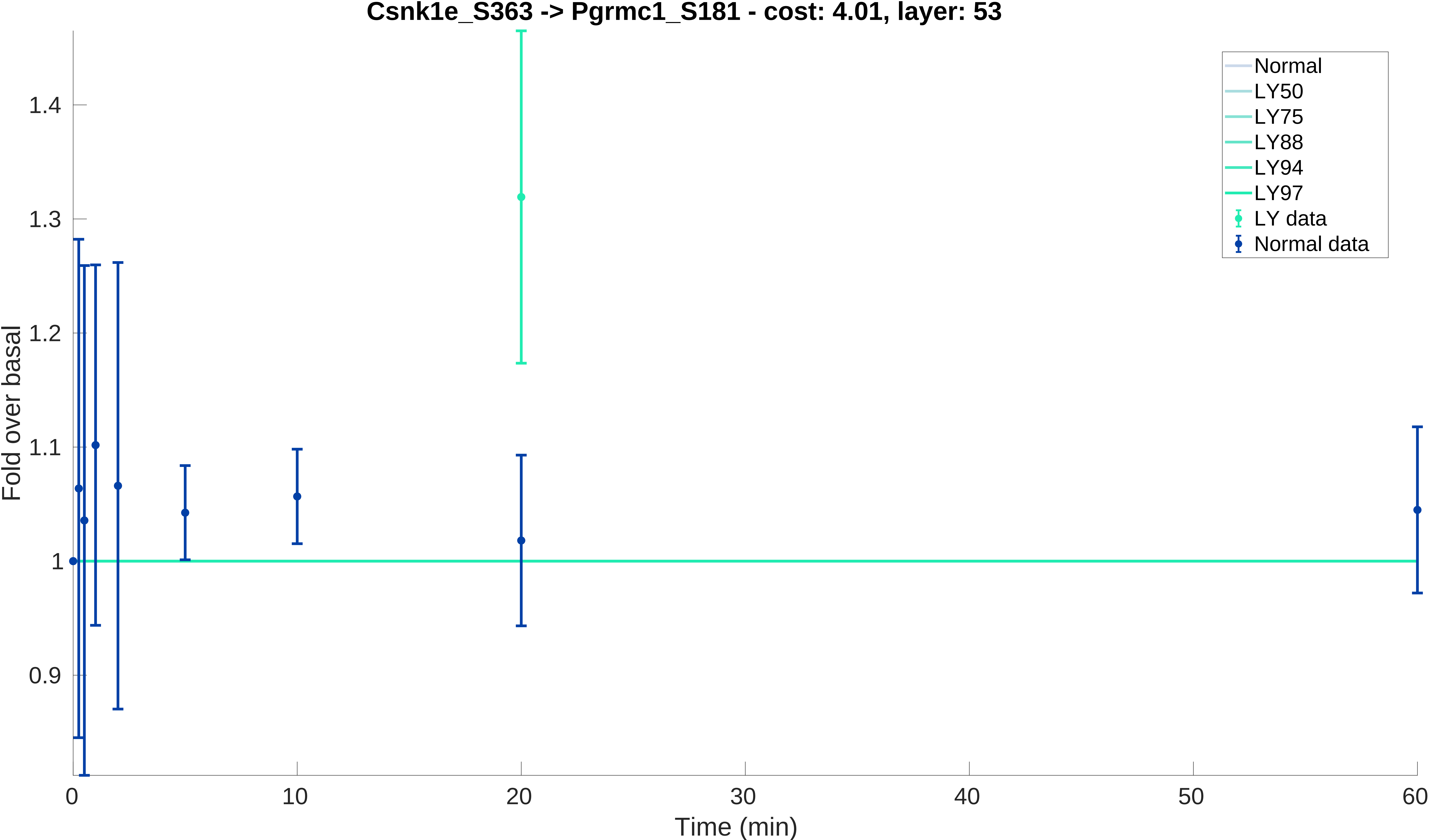

Csnk1e_S363 -> Pgrmc1_S181 - cost: 4.01, layer: 53
Fold over basal
Time (min)
0.9
1
1.1
1.2
1.3
1.4
0
10
20
30
40
50
60
Normal
LY50
LY75
LY88
LY94
LY97
LY data
Normal data

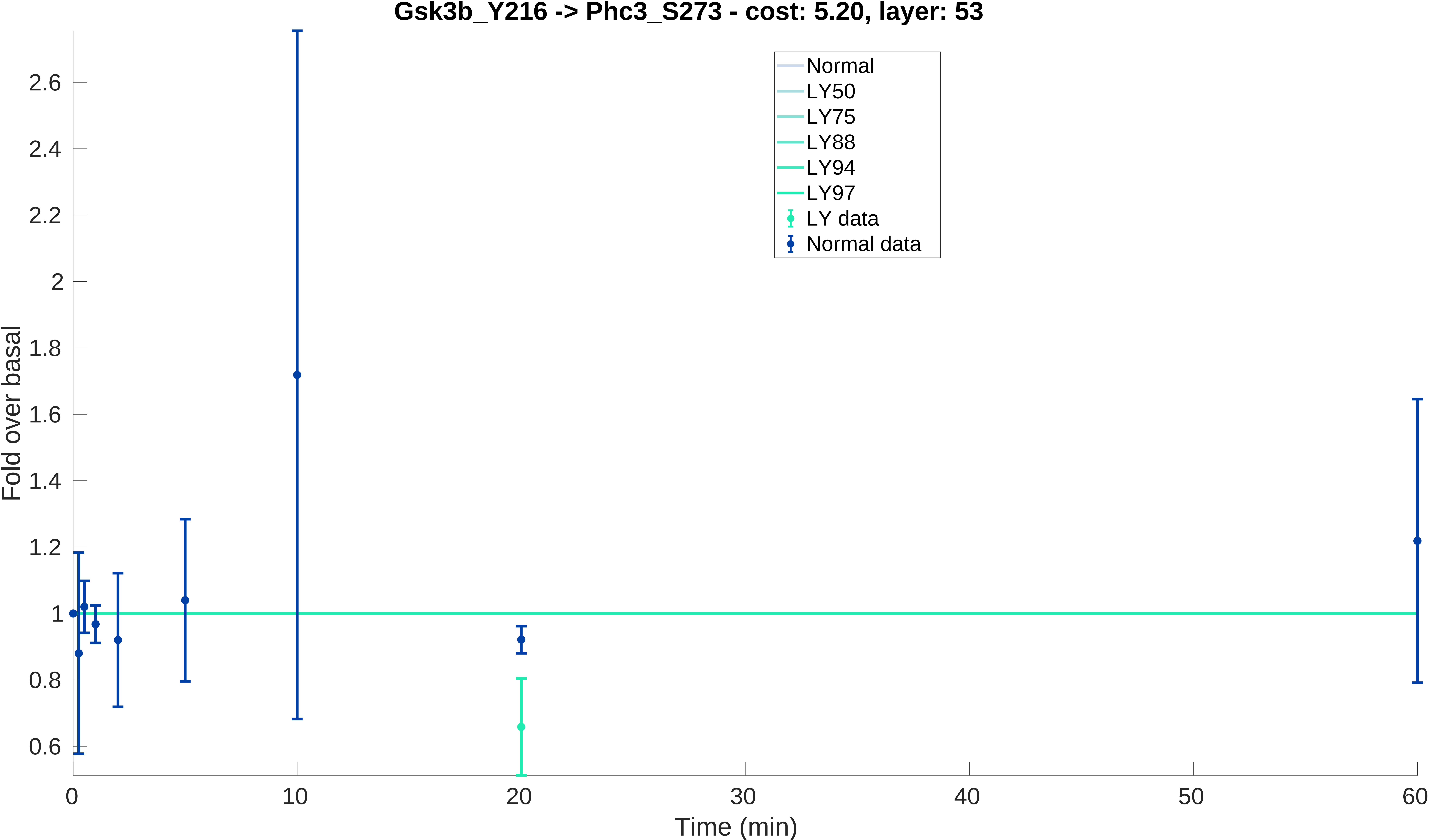

Gsk3b_Y216 -> Phc3_S273 - cost: 5.20, layer: 53
Normal
LY50
LY75
LY88
LY94
LY97
LY data
Normal data
Fold over basal
2.6
2.4
2.2
2
1.8
1.6
1.4
1.2
1
0.8
0.6
0
10
20
30
40
50
60
Time (min)

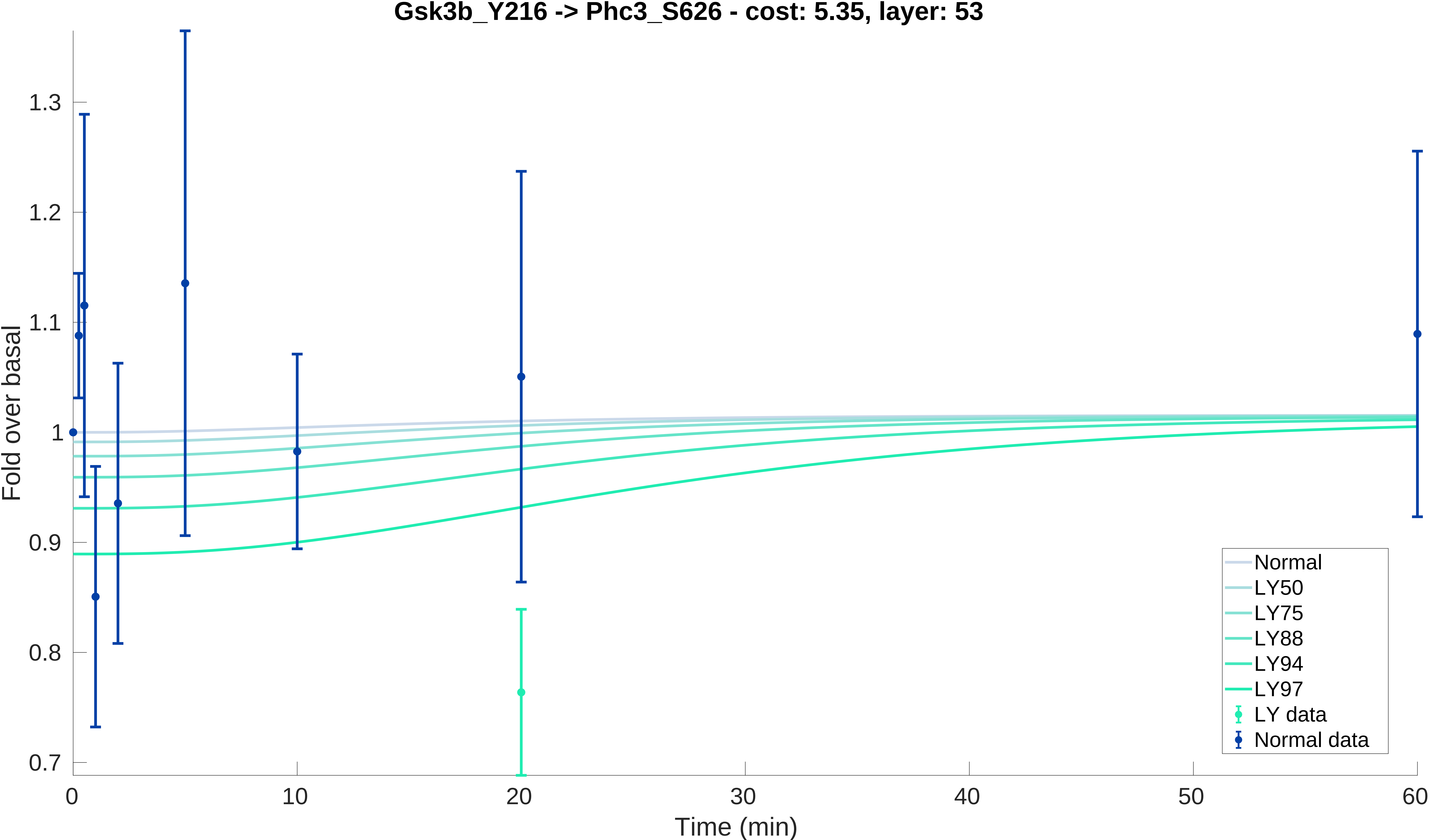

Gsk3b_Y216 -> Phc3_S626 - cost: 5.35, layer: 53
Fold over basal
Time (min)
0.7
0.8
0.9
1
1.1
1.2
1.3
0
10
20
30
40
50
60
Normal
LY50
LY75
LY88
LY94
LY97
LY data
Normal data

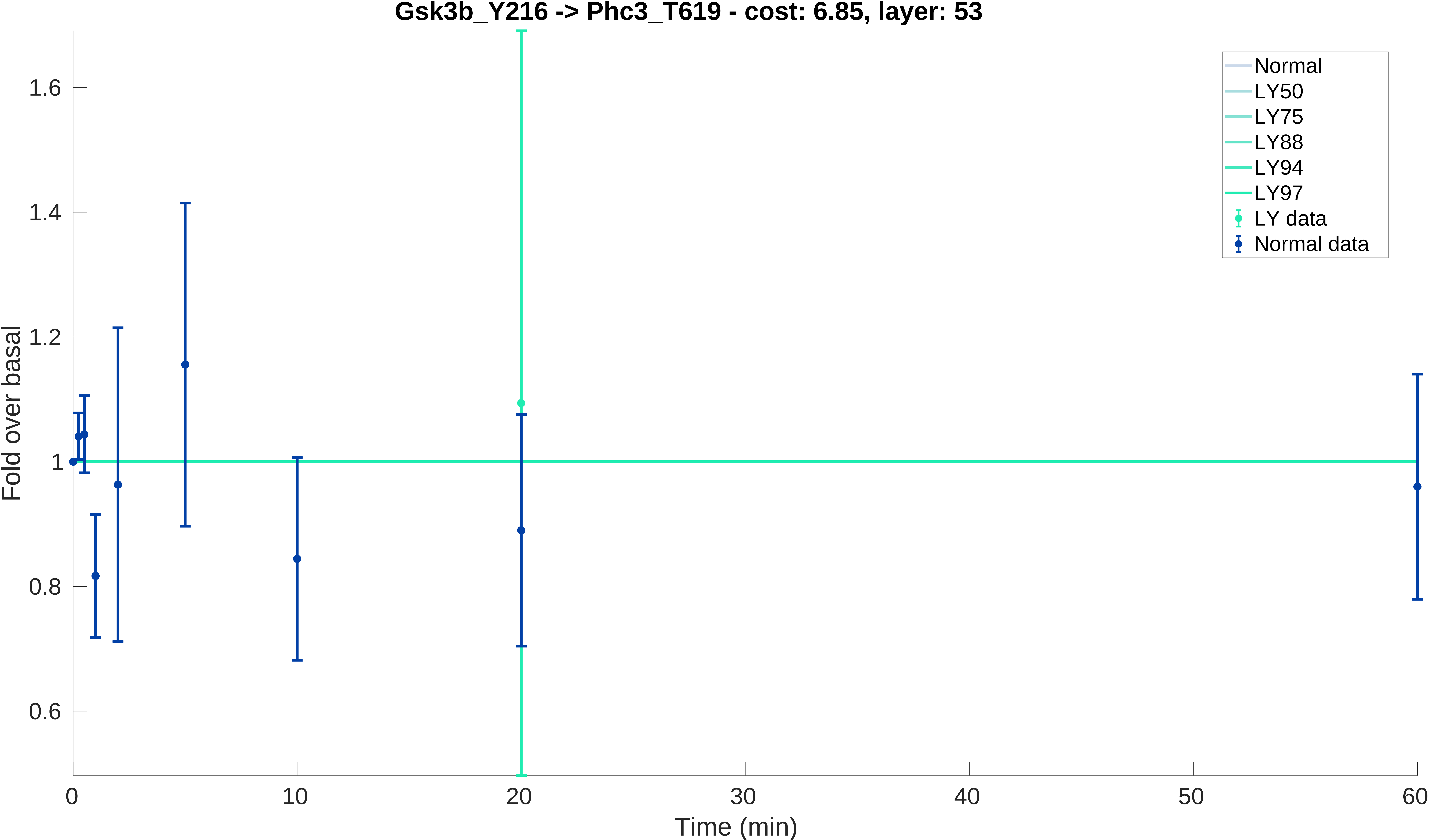

Gsk3b_Y216 -> Phc3_T619 - cost: 6.85, layer: 53
Fold over basal
Time (min)
0.6
0.8
1
1.2
1.4
1.6
0
10
20
30
40
50
60
Normal
LY50
LY75
LY88
LY94
LY97
LY data
Normal data

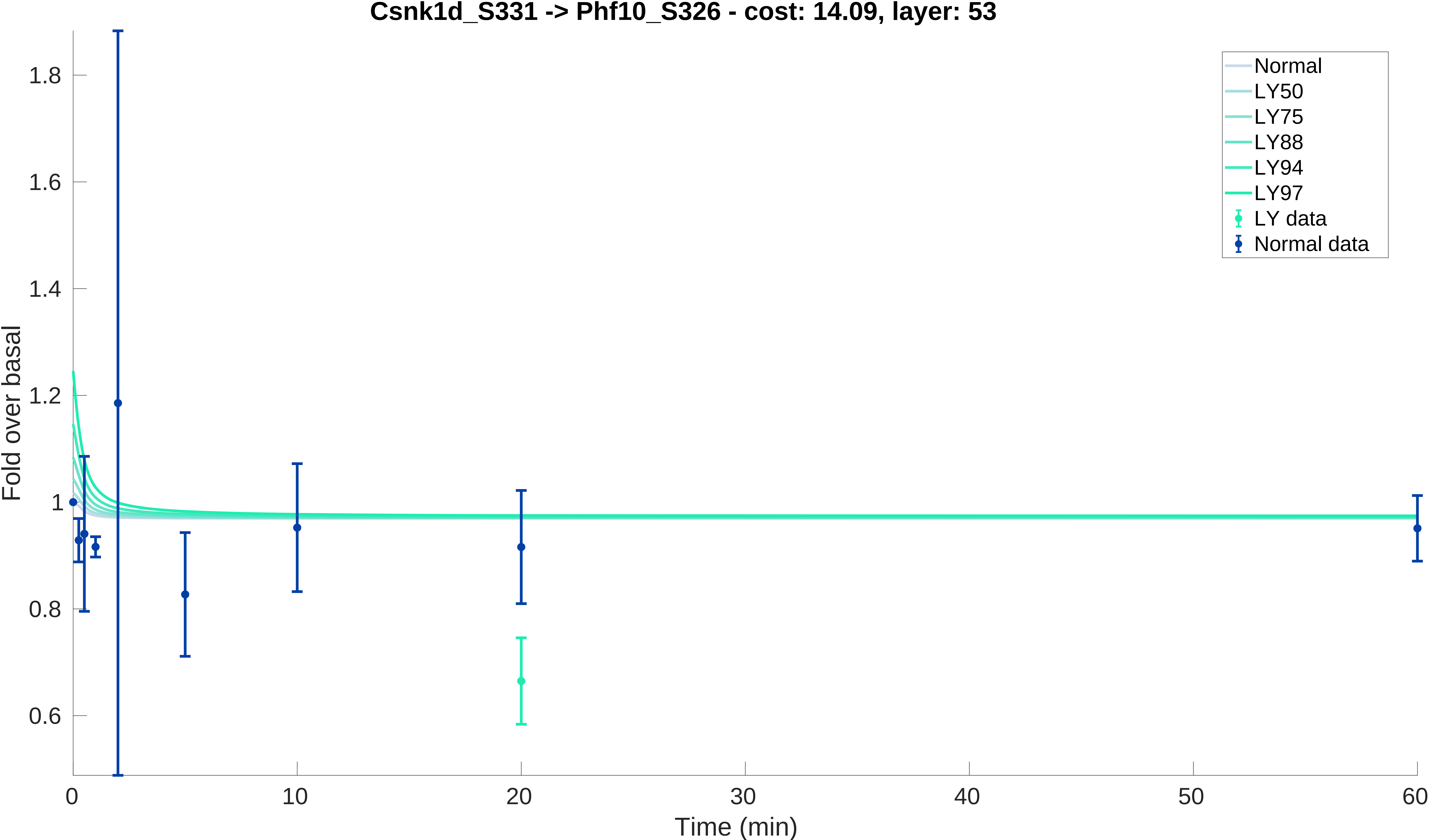

Csnk1d_S331 -> Phf10_S326 - cost: 14.09, layer: 53
Normal
LY50
LY75
LY88
LY94
LY97
LY data
Normal data
Fold over basal
1.8
1.6
1.4
1.2
1
0.8
0.6
0
10
20
30
40
50
60
Time (min)

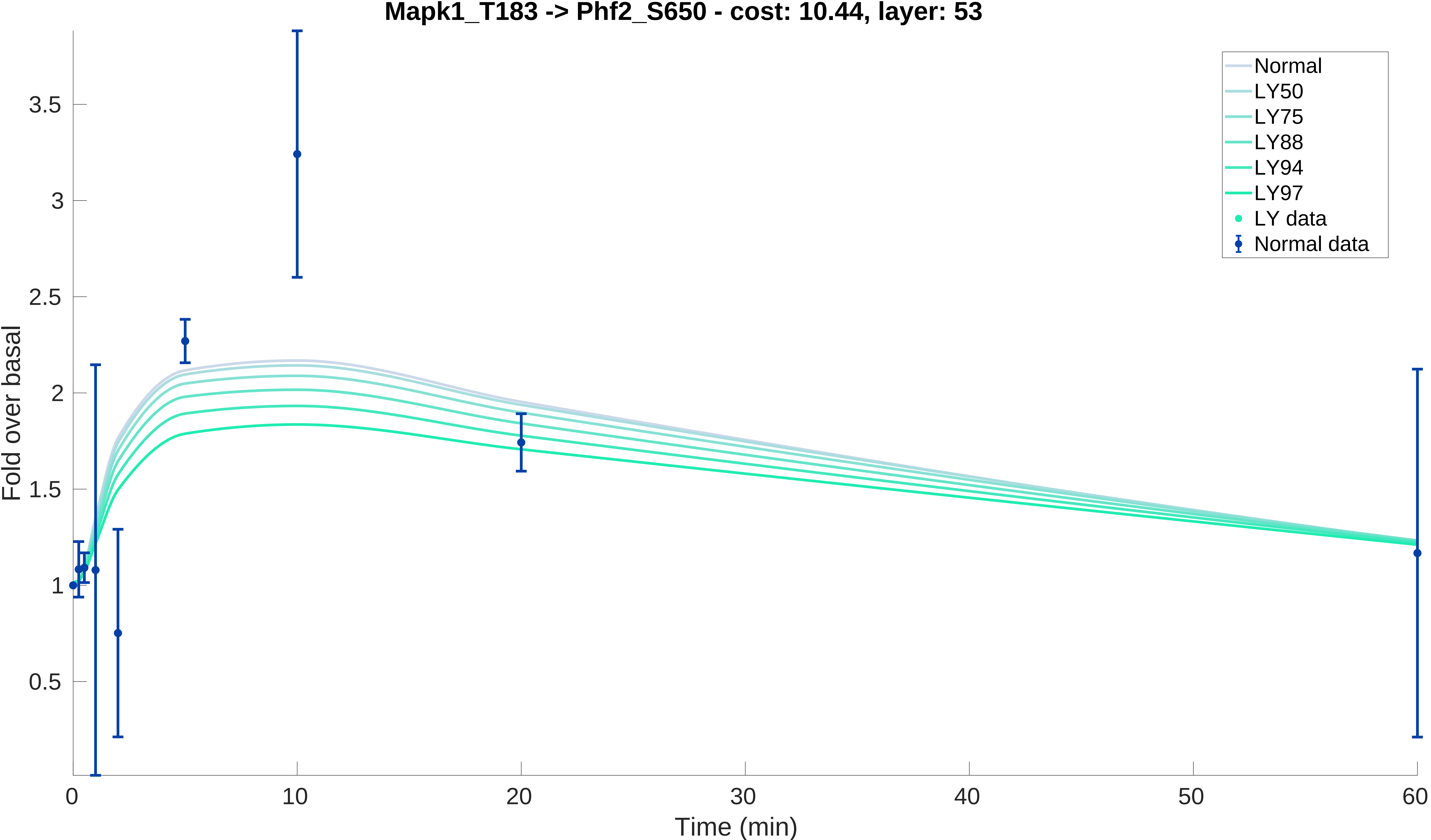
Mapk1_T183 -> Phf2_S650 - cost: 10.44, layer: 53
Fold over basal
Time (min)
0.5
1
1.5
2
2.5
3
3.5
0
10
20
30
40
50
60
Normal
LY50
LY75
LY88
LY94
LY97
LY data
Normal data

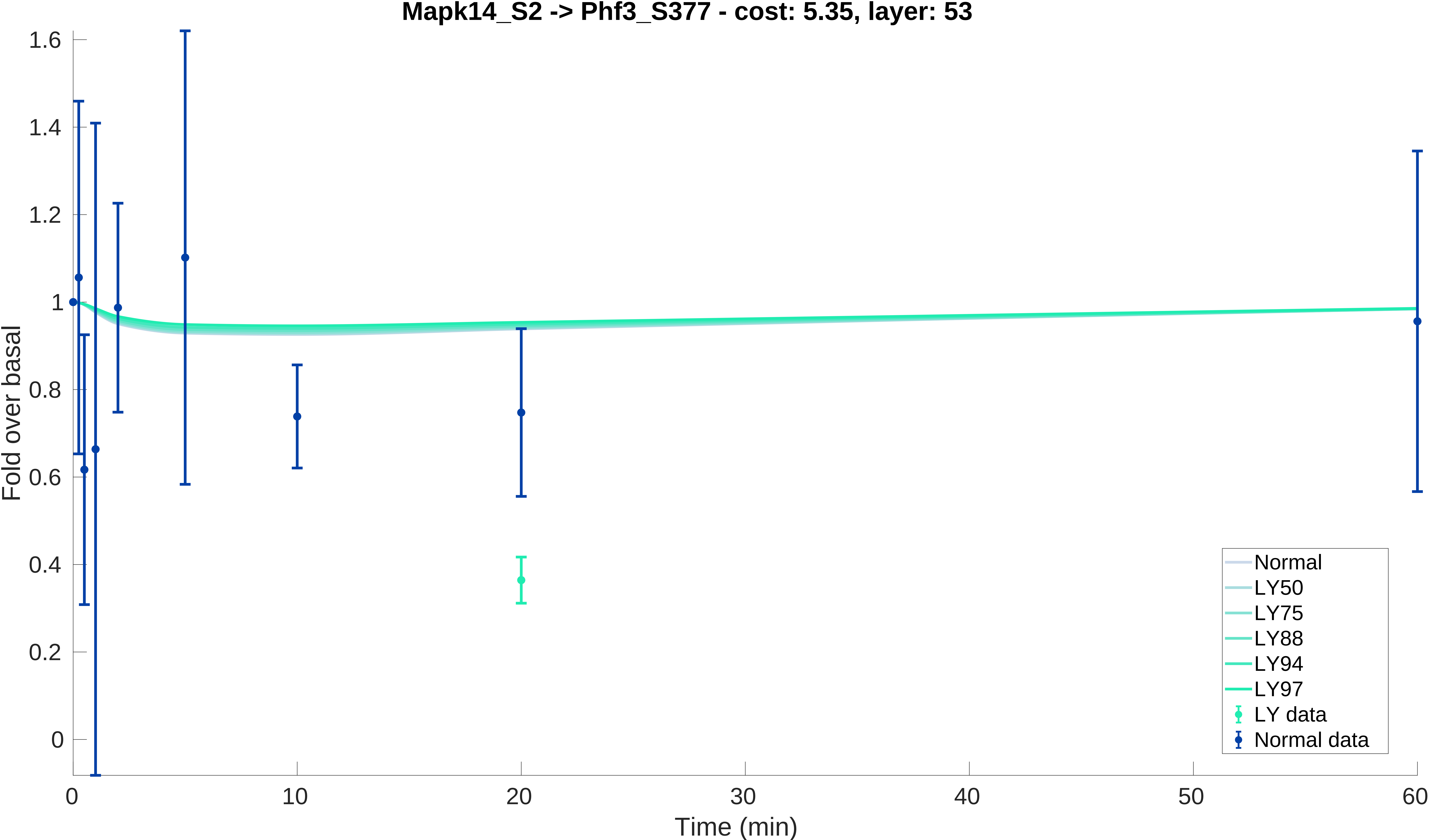

Mapk14_S2 -> Phf3_S377 - cost: 5.35, layer: 53
Fold over basal
1.6
1.4
1.2
1
0.8
0.6
0.4
0.2
0
0
10
20
30
40
50
60
Time (min)
Normal
LY50
LY75
LY88
LY94
LY97
LY data
Normal data

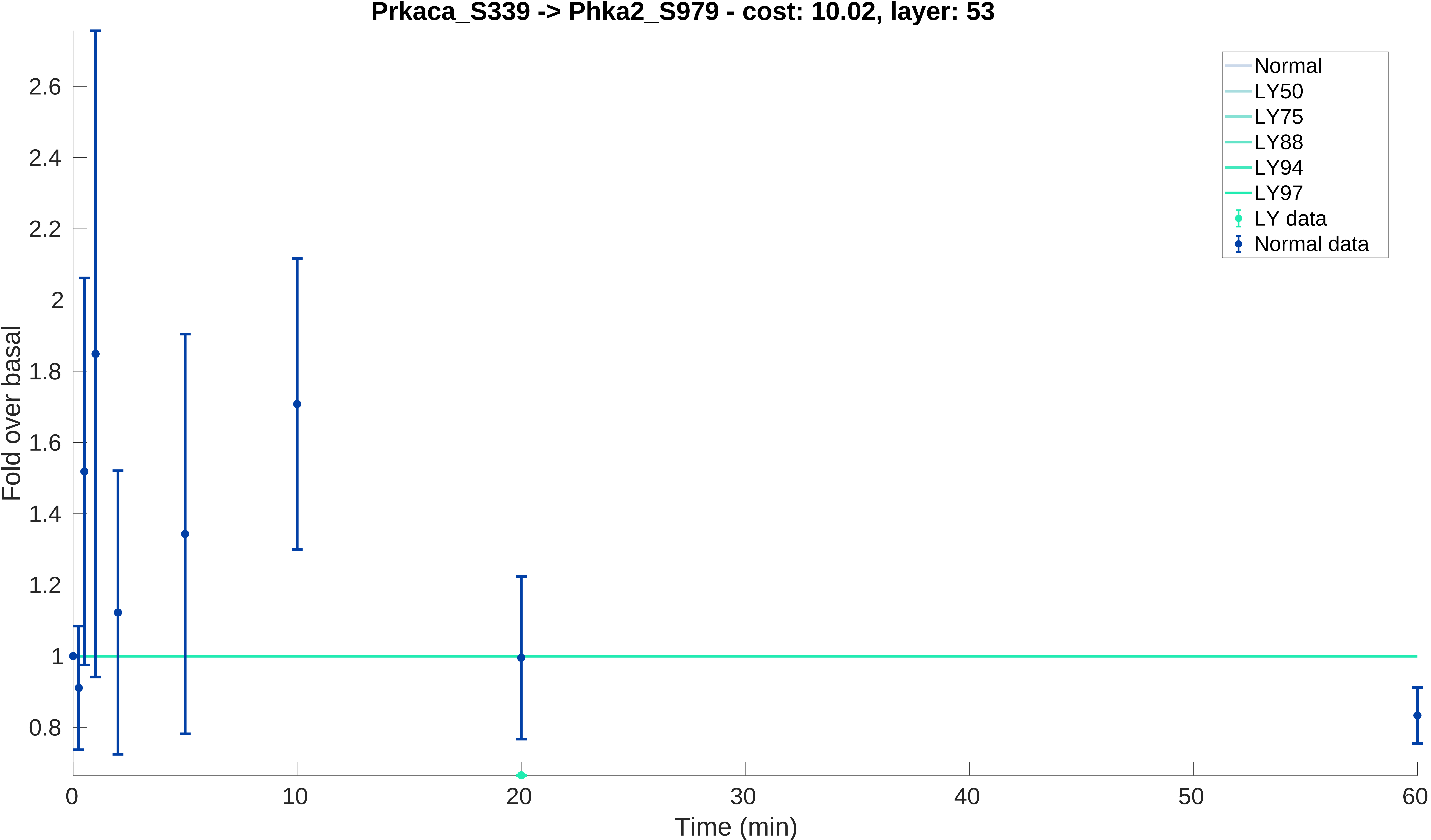
Prkaca_S339 -> Phka2_S979 - cost: 10.02, layer: 53
Fold over basal
Time (min)
Normal
LY50
LY75
LY88
LY94
LY97
LY data
Normal data
0.8
1
1.2
1.4
1.6
1.8
2
2.2
2.4
2.6
0
10
20
30
40
50
60

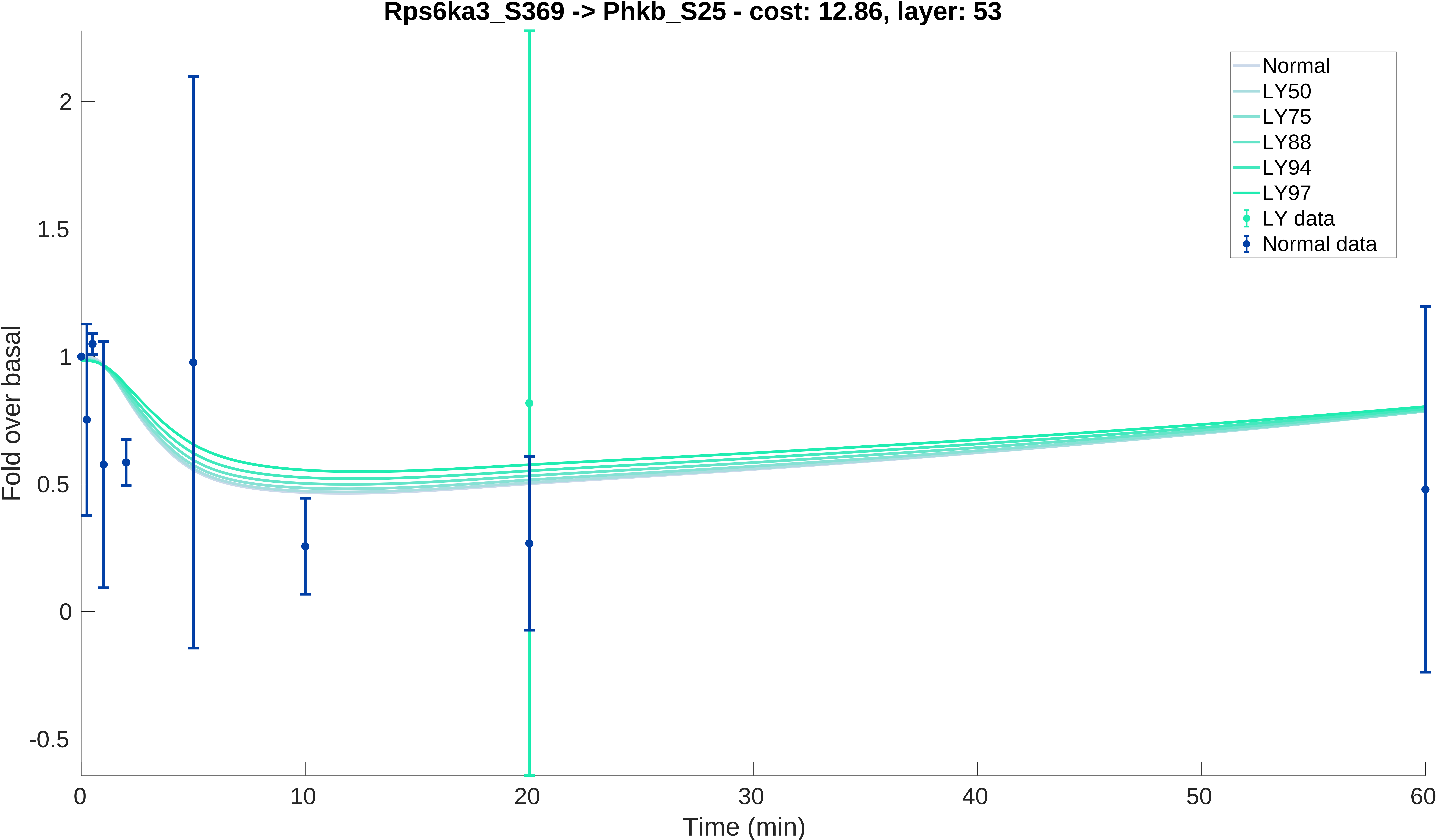

Rps6ka3_S369 -> Phkb_S25 - cost: 12.86, layer: 53
Fold over basal
Time (min)
2
1.5
1
0.5
0
-0.5
0
10
20
30
40
50
60
Normal
LY50
LY75
LY88
LY94
LY97
LY data
Normal data

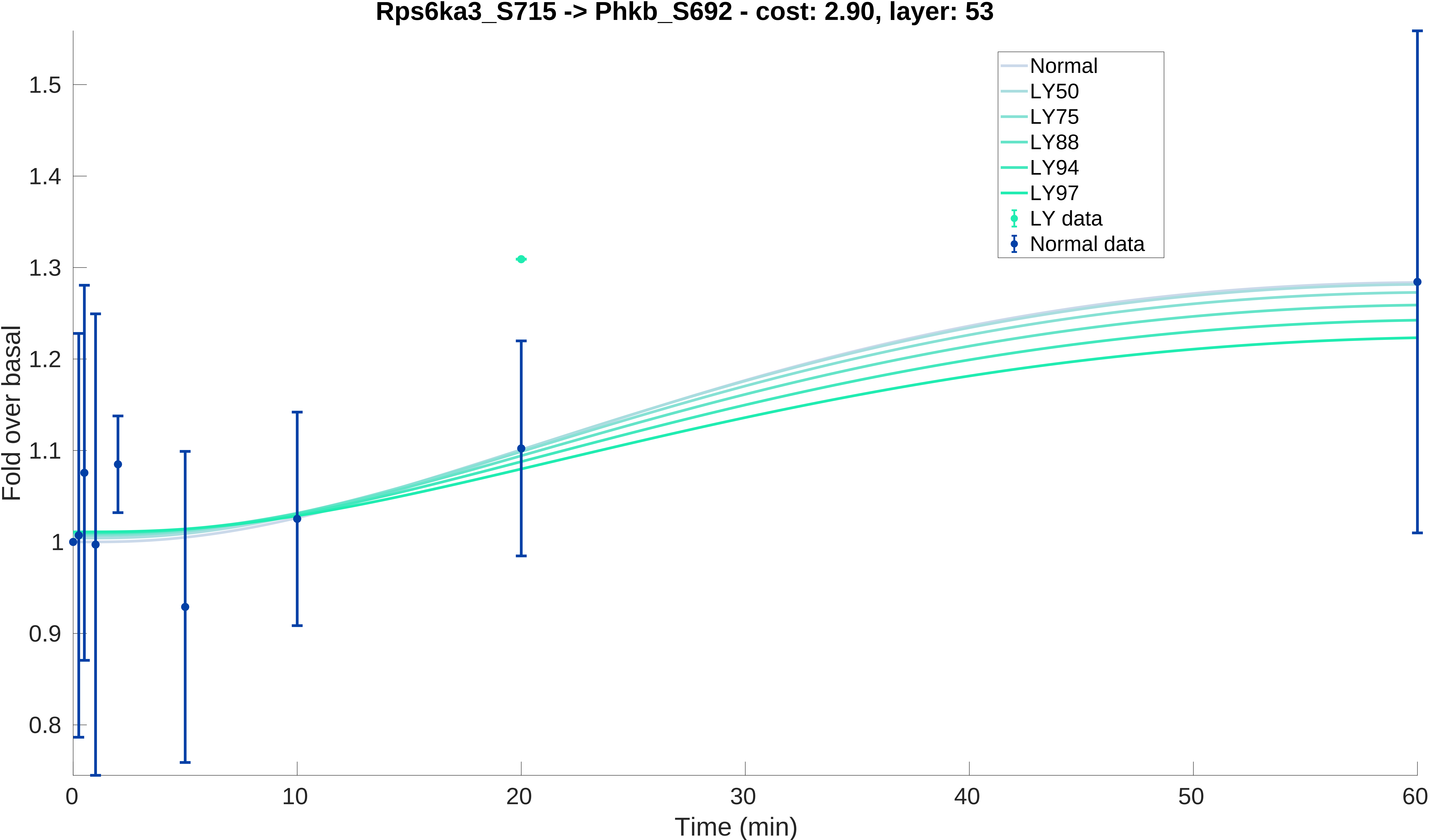

Rps6ka3_S715 -> Phkb_S692 - cost: 2.90, layer: 53
Normal
LY50
LY75
LY88
LY94
LY97
LY data
Normal data
Fold over basal
1.5
1.4
1.3
1.2
1.1
1
0.9
0.8
0
10
20
30
40
50
60
Time (min)

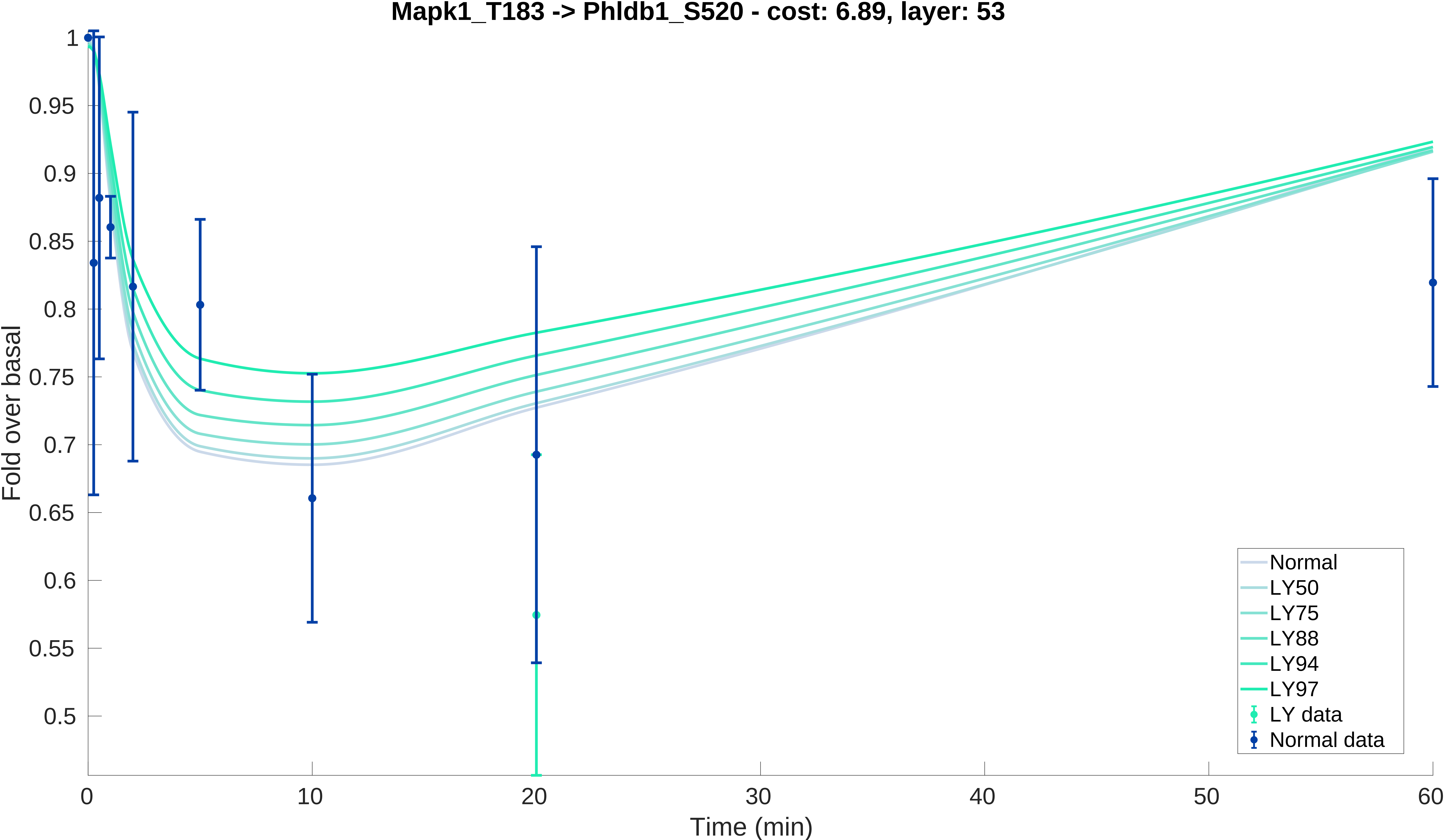

Mapk1_T183 -> Phldb1_S520 - cost: 6.89, layer: 53
Fold over basal
Time (min)
1
0.95
0.9
0.85
0.8
0.75
0.7
0.65
0.6
0.55
0.5
0
10
20
30
40
50
60
Normal
LY50
LY75
LY88
LY94
LY97
LY data
Normal data

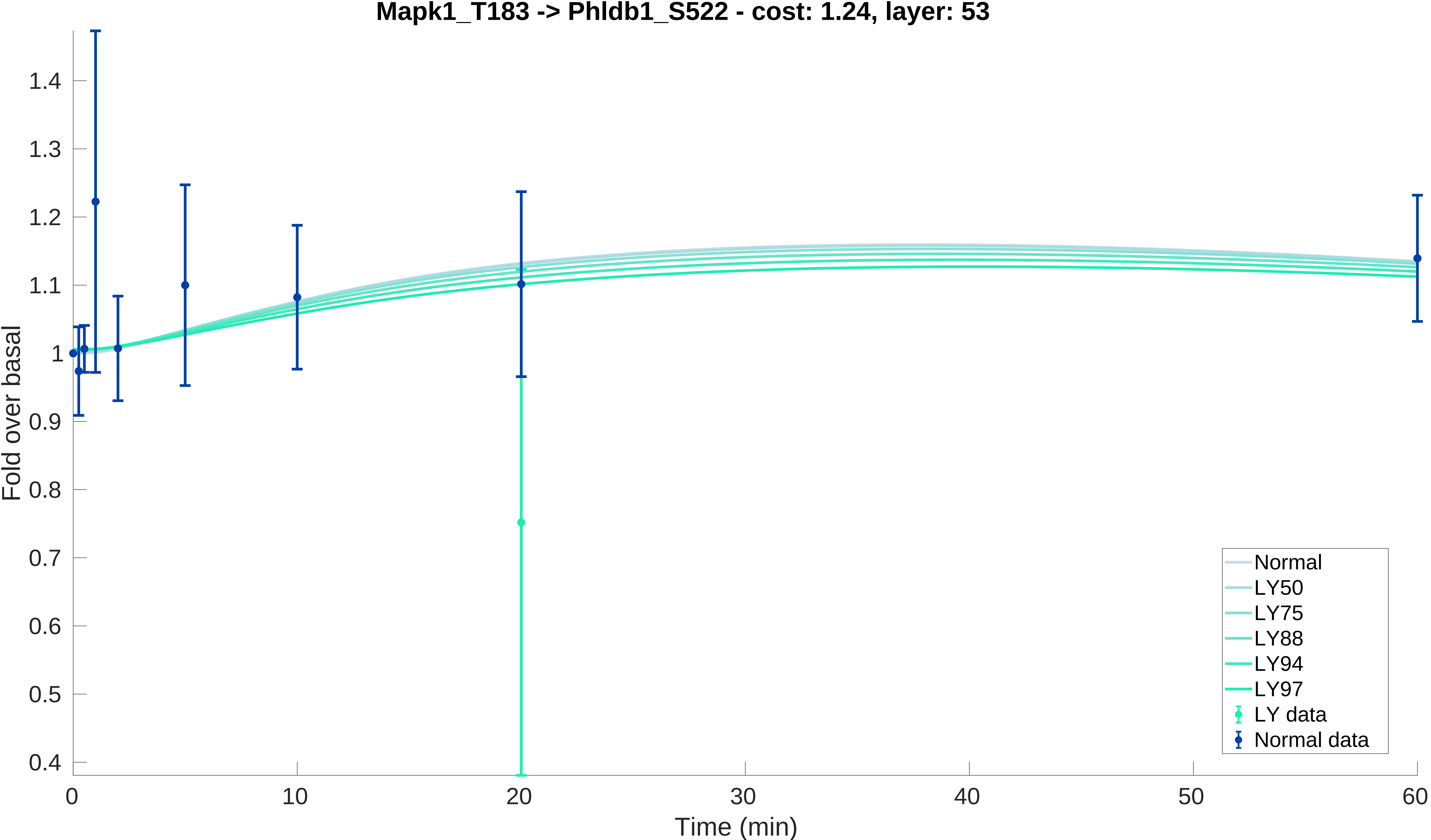
Mapk1_T183 -> Phldb1_S522 - cost: 1.24, layer: 53
Fold over basal
1.4
1.3
1.2
1.1
1
0.9
0.8
0.7
0.6
0.5
0.4
0
10
20
30
40
50
60
Time (min)
Normal
LY50
LY75
LY88
LY94
LY97
LY data
Normal data

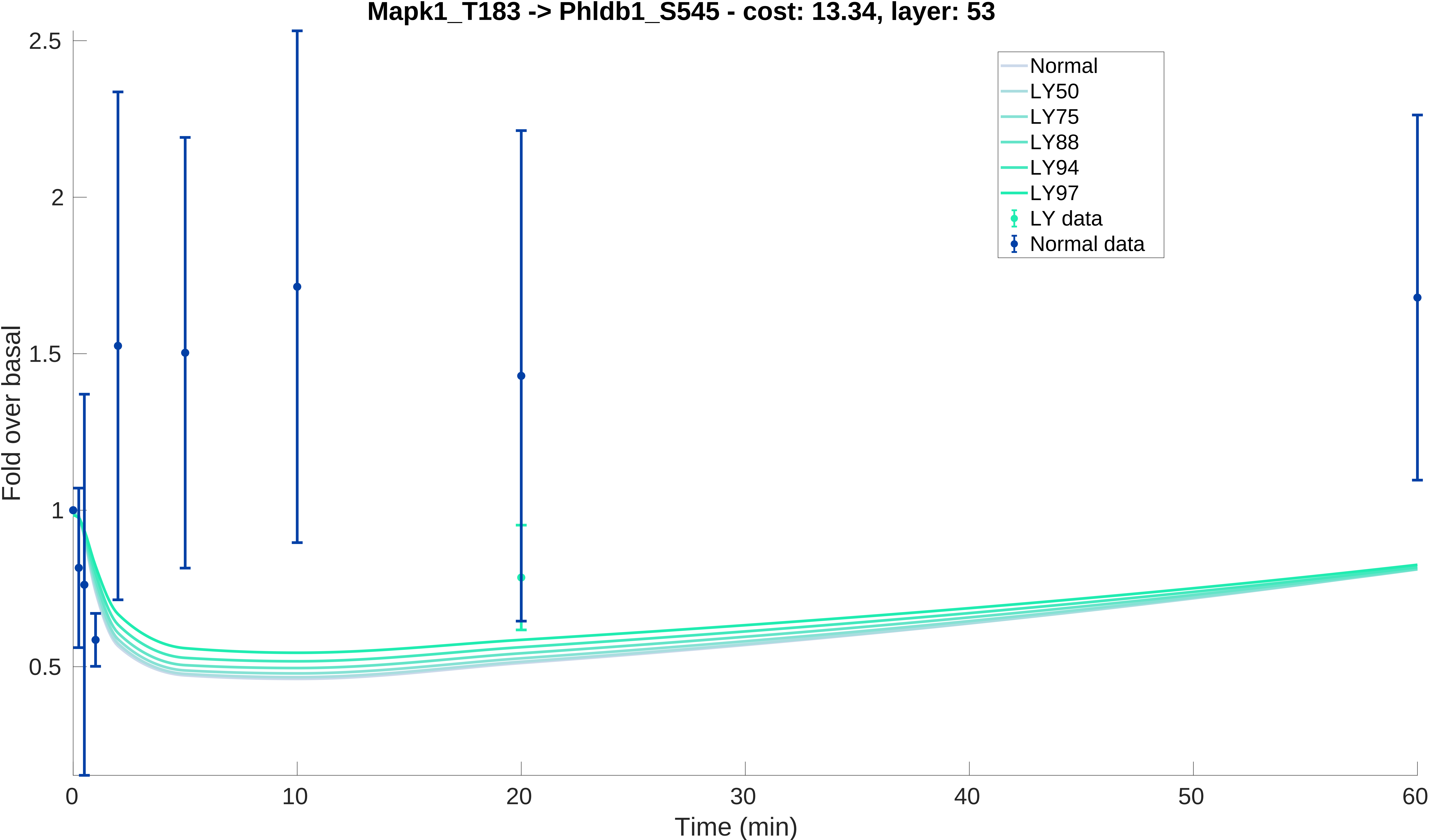

Mapk1_T183 -> Phldb1_S545 - cost: 13.34, layer: 53
Fold over basal
Time (min)
2.5
2
1.5
1
0.5
0
10
20
30
40
50
60
Normal
LY50
LY75
LY88
LY94
LY97
LY data
Normal data

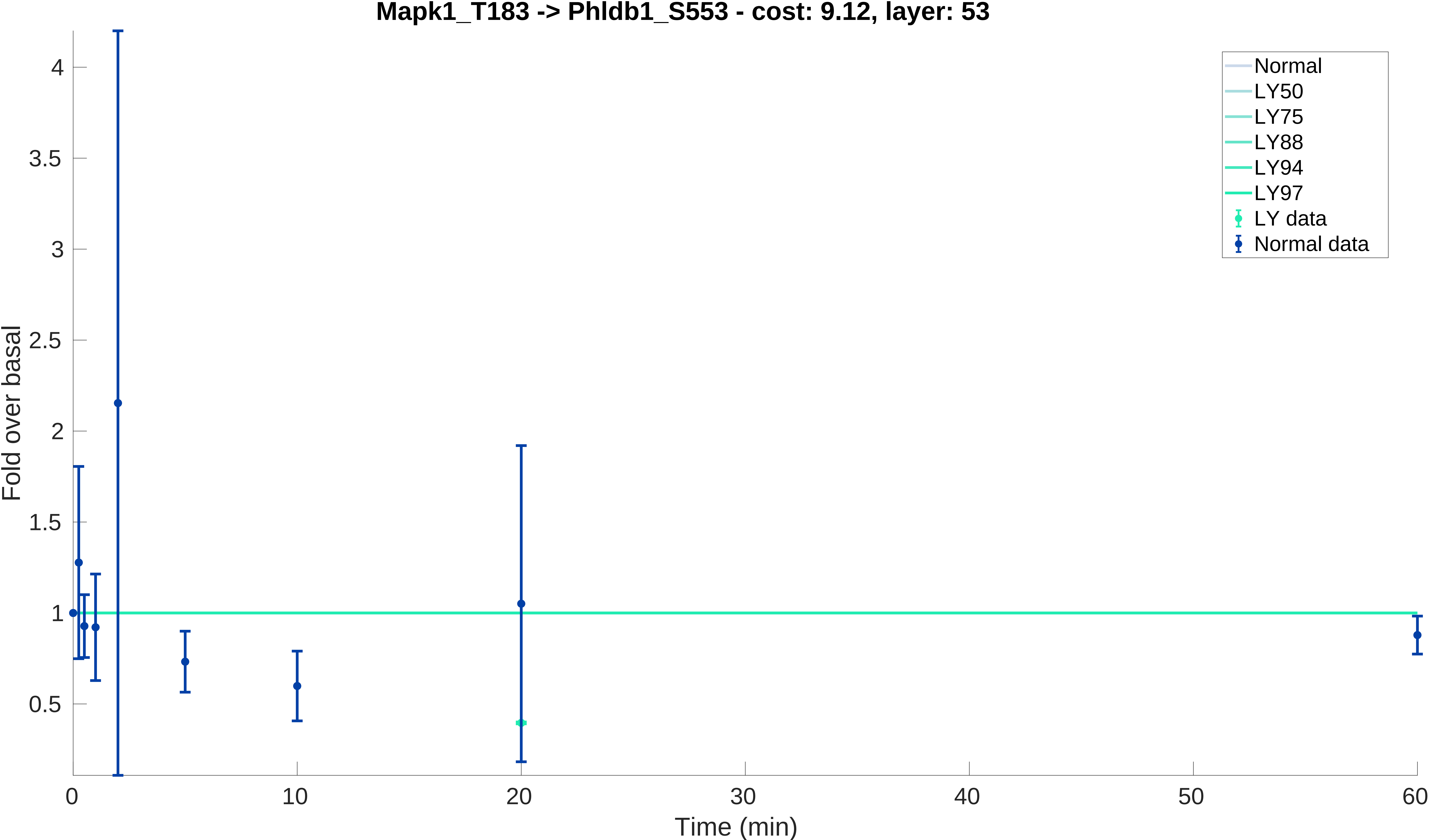

Mapk1_T183 -> Phldb1_S553 - cost: 9.12, layer: 53
Fold over basal
Time (min)
0
10
20
30
40
50
60
0.5
1
1.5
2
2.5
3
3.5
4
Normal
LY50
LY75
LY88
LY94
LY97
LY data
Normal data

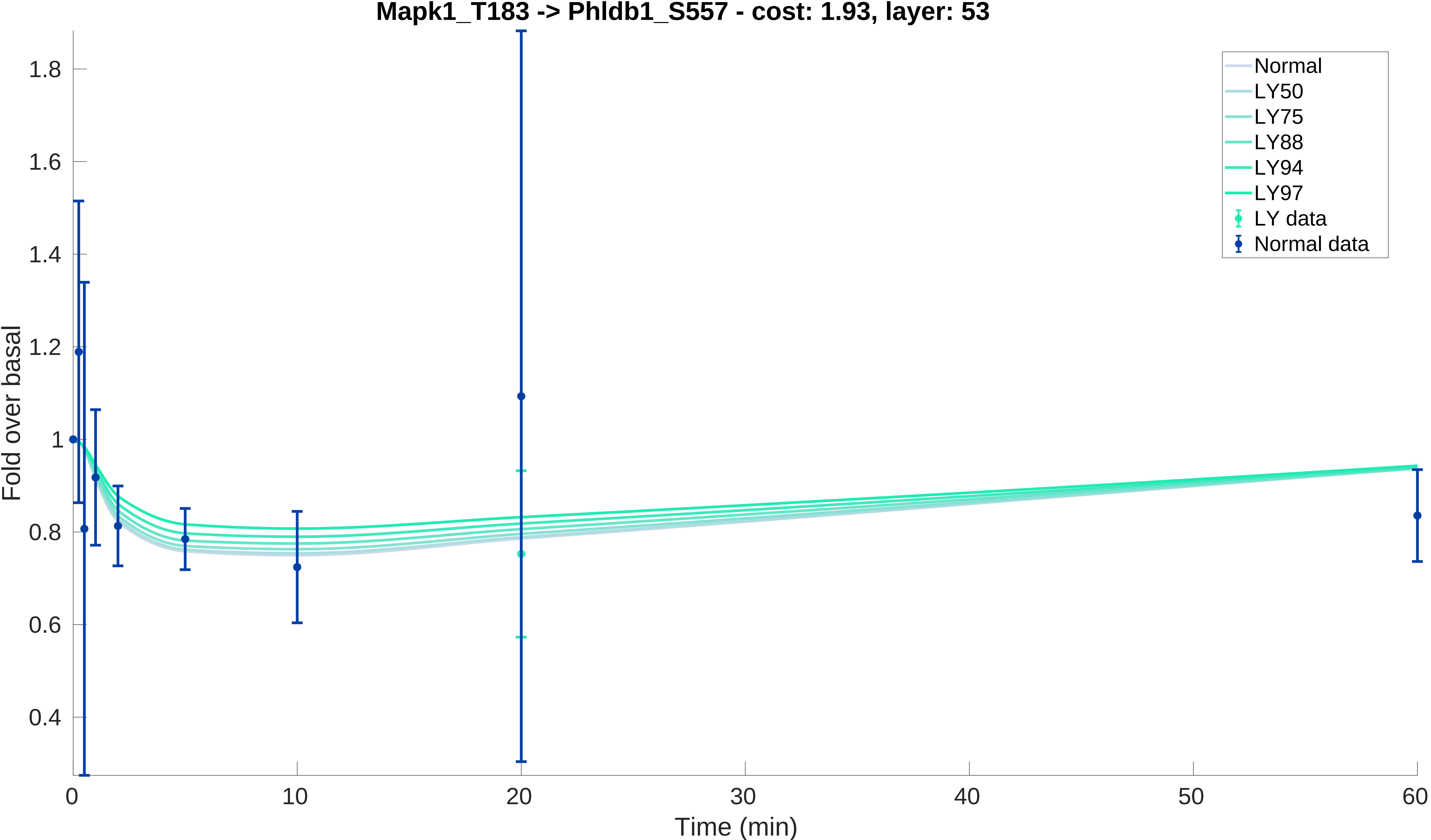

Mapk1_T183 -> Phldb1_S557 - cost: 1.93, layer: 53
Normal
LY50
LY75
LY88
LY94
LY97
LY data
Normal data
Fold over basal
1.8
1.6
1.4
1.2
1
0.8
0.6
0.4
0
10
20
30
40
50
60
Time (min)

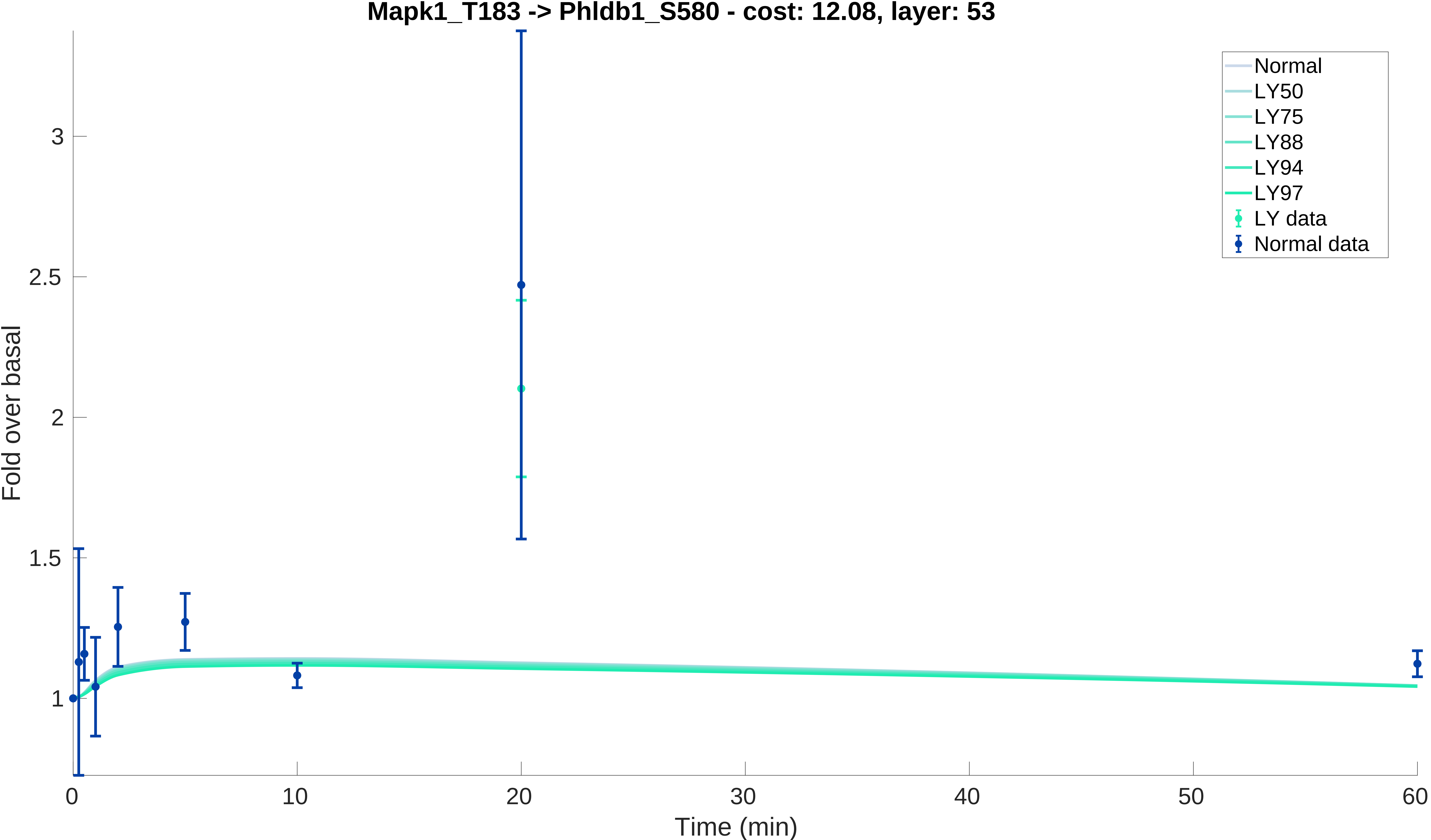

Mapk1_T183 -> Phldb1_S580 - cost: 12.08, layer: 53
Fold over basal
Time (min)
0
10
20
30
40
50
60
1
1.5
2
2.5
3
Normal
LY50
LY75
LY88
LY94
LY97
LY data
Normal data

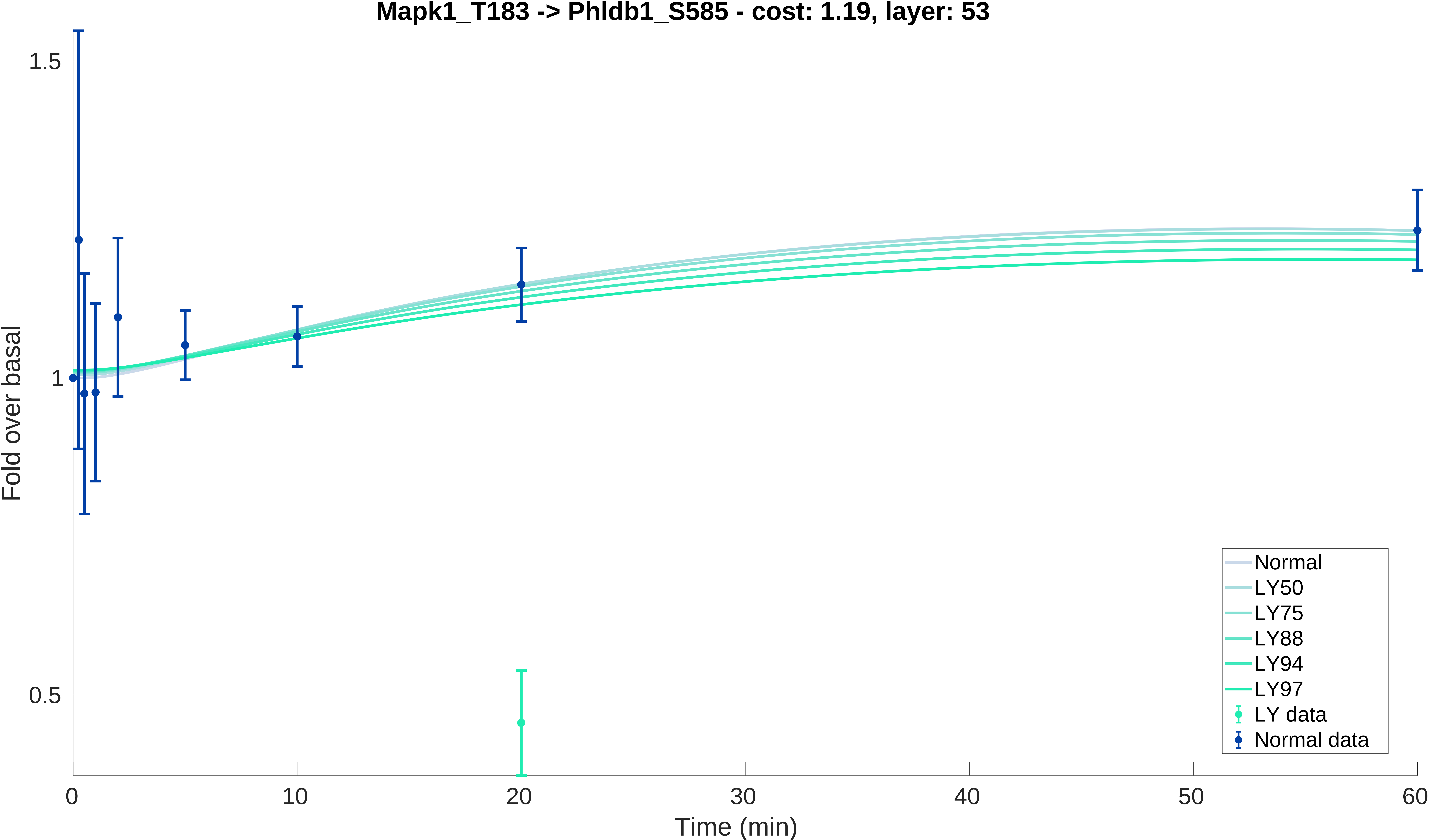

Mapk1_T183 -> Phldb1_S585 - cost: 1.19, layer: 53
1.5
1
0.5
Fold over basal
0
10
20
30
40
50
60
Time (min)
Normal
LY50
LY75
LY88
LY94
LY97
LY data
Normal data

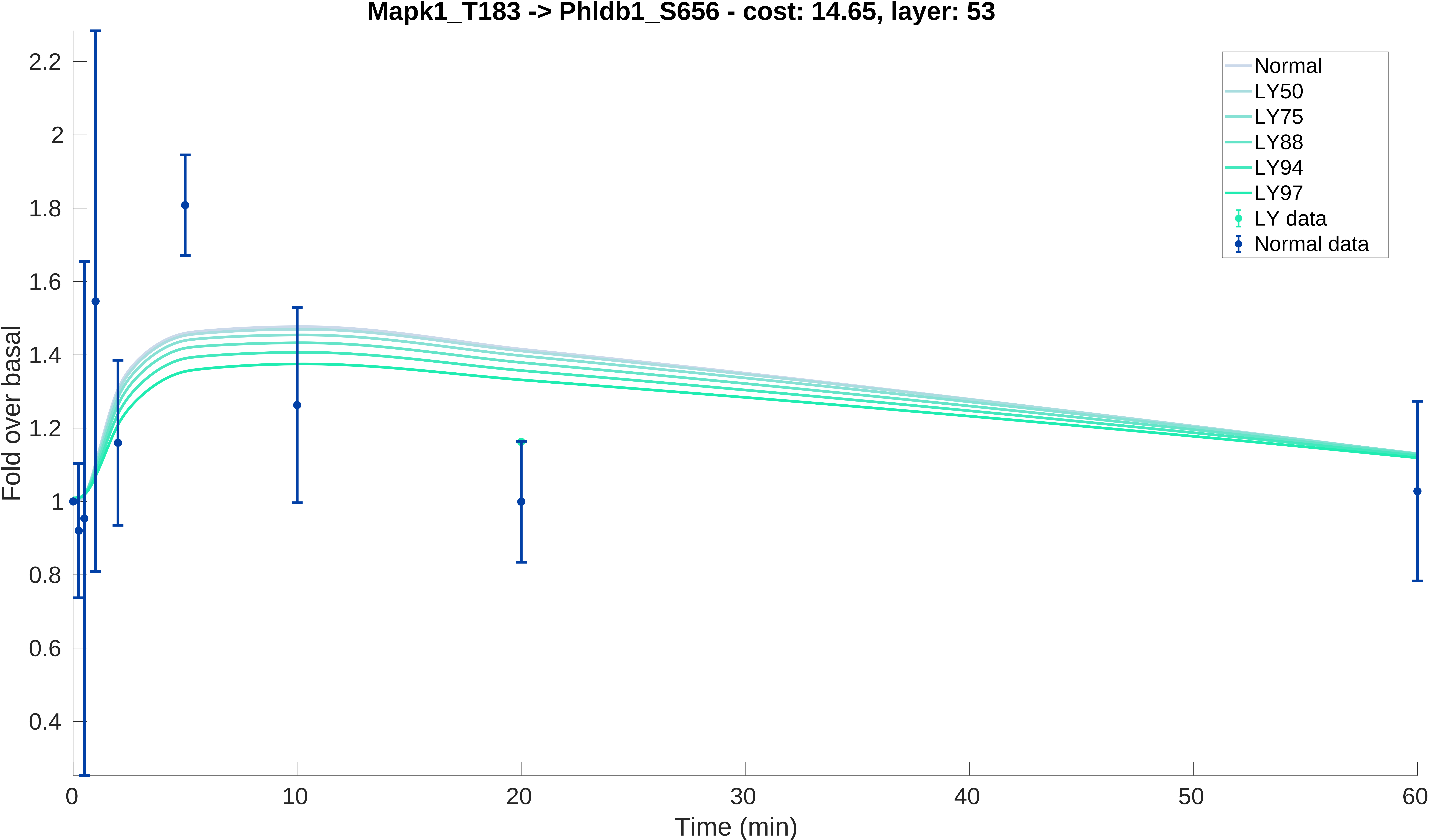

Mapk1_T183 -> Phldb1_S656 - cost: 14.65, layer: 53
Fold over basal
Time (min)
0.4
0.6
0.8
1
1.2
1.4
1.6
1.8
2
2.2
0
10
20
30
40
50
60
Normal
LY50
LY75
LY88
LY94
LY97
LY data
Normal data

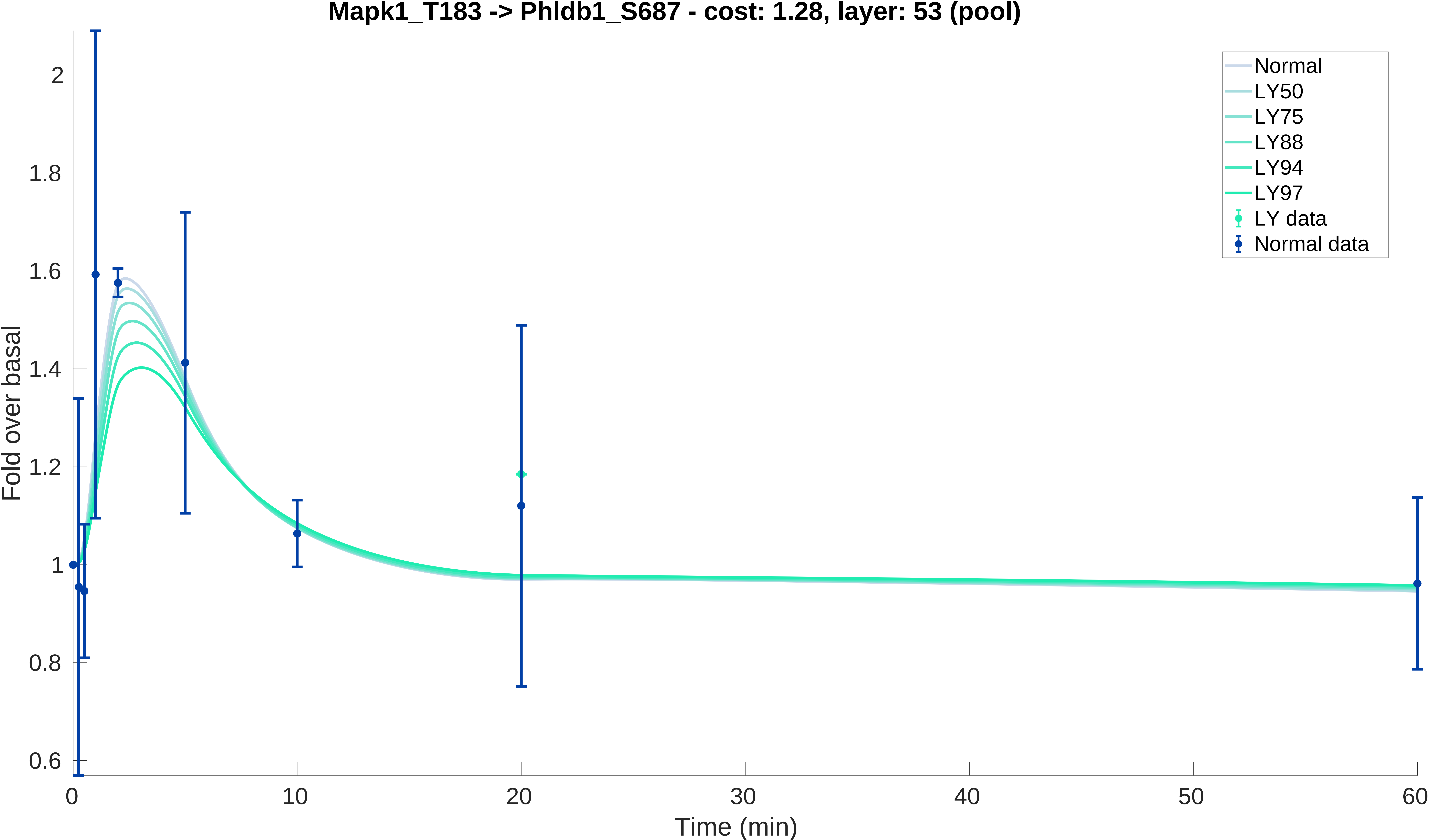

Mapk1_T183 -> Phldb1_S687 - cost: 1.28, layer: 53 (pool)
Fold over basal
Time (min)
0.6
0.8
1
1.2
1.4
1.6
1.8
2
0
10
20
30
40
50
60
Normal
LY50
LY75
LY88
LY94
LY97
LY data
Normal data

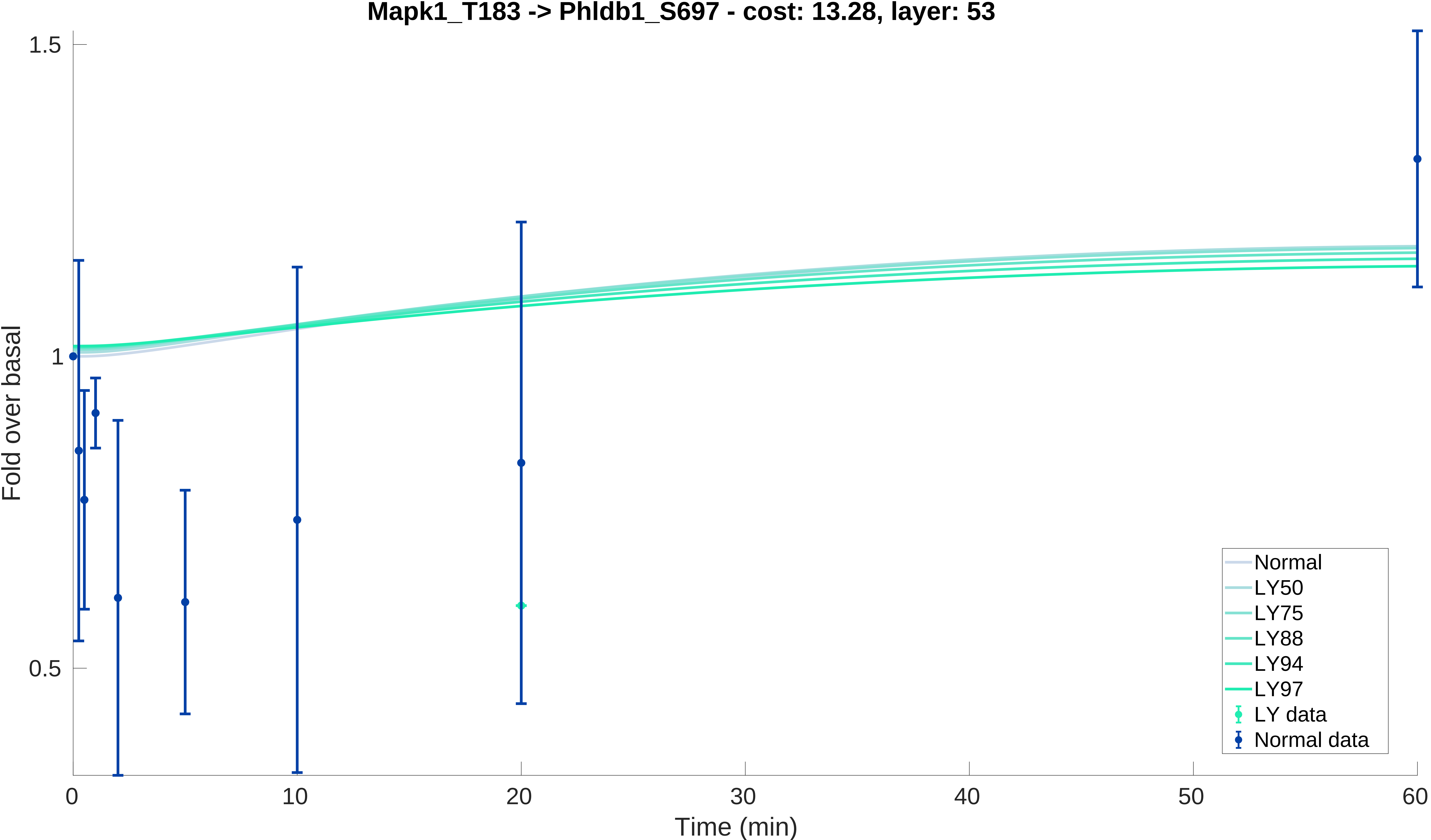
Mapk1_T183 -> Phldb1_S697 - cost: 13.28, layer: 53
1.5
1
0.5
Fold over basal
0
10
20
30
40
50
60
Time (min)
Normal
LY50
LY75
LY88
LY94
LY97
LY data
Normal data

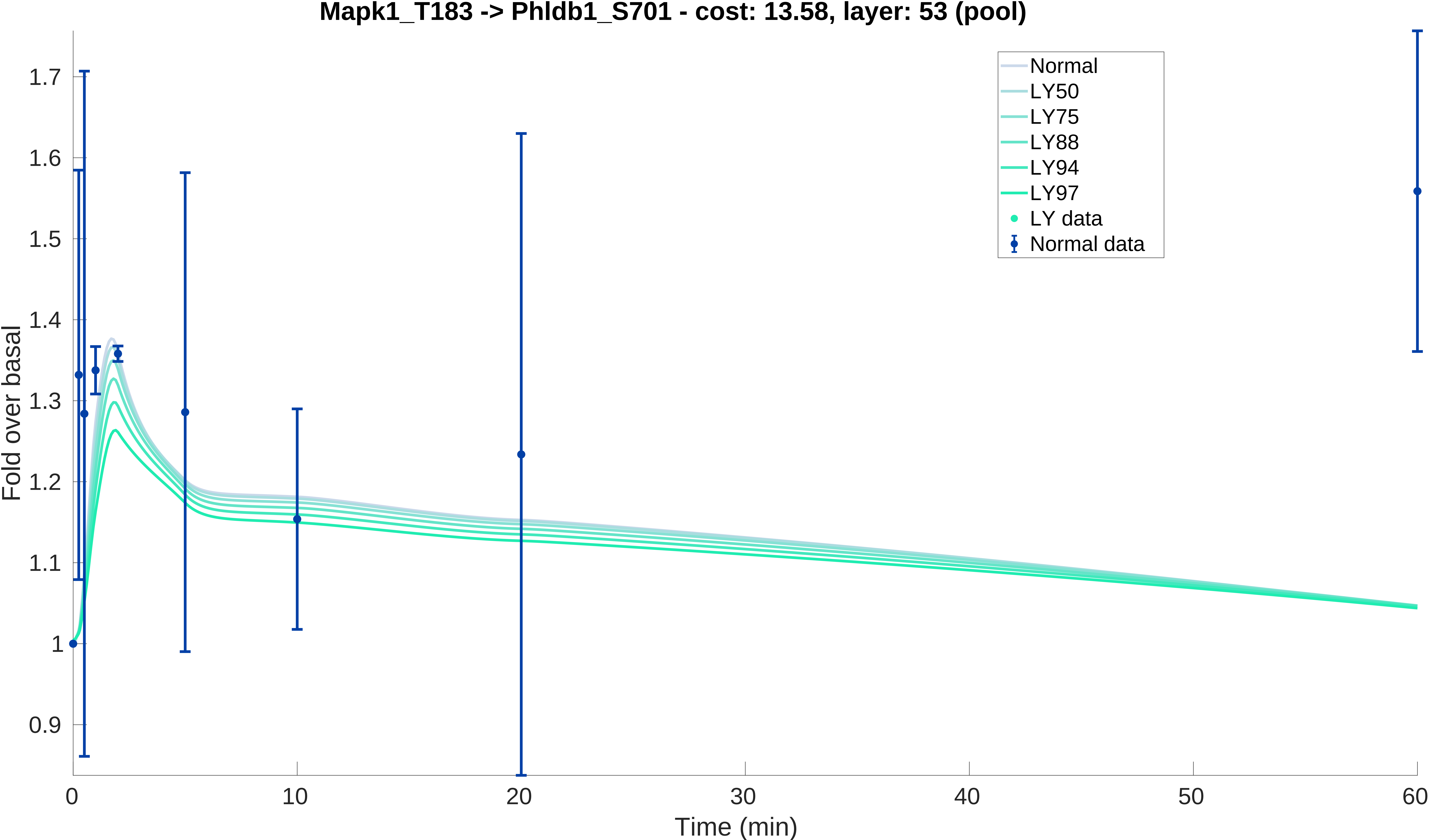
Mapk1_T183 -> Phldb1_S701 - cost: 13.58, layer: 53 (pool)
Normal
LY50
LY75
LY88
LY94
LY97
LY data
Normal data
Fold over basal
Time (min)
0.9
1
1.1
1.2
1.3
1.4
1.5
1.6
1.7
0
10
20
30
40
50
60

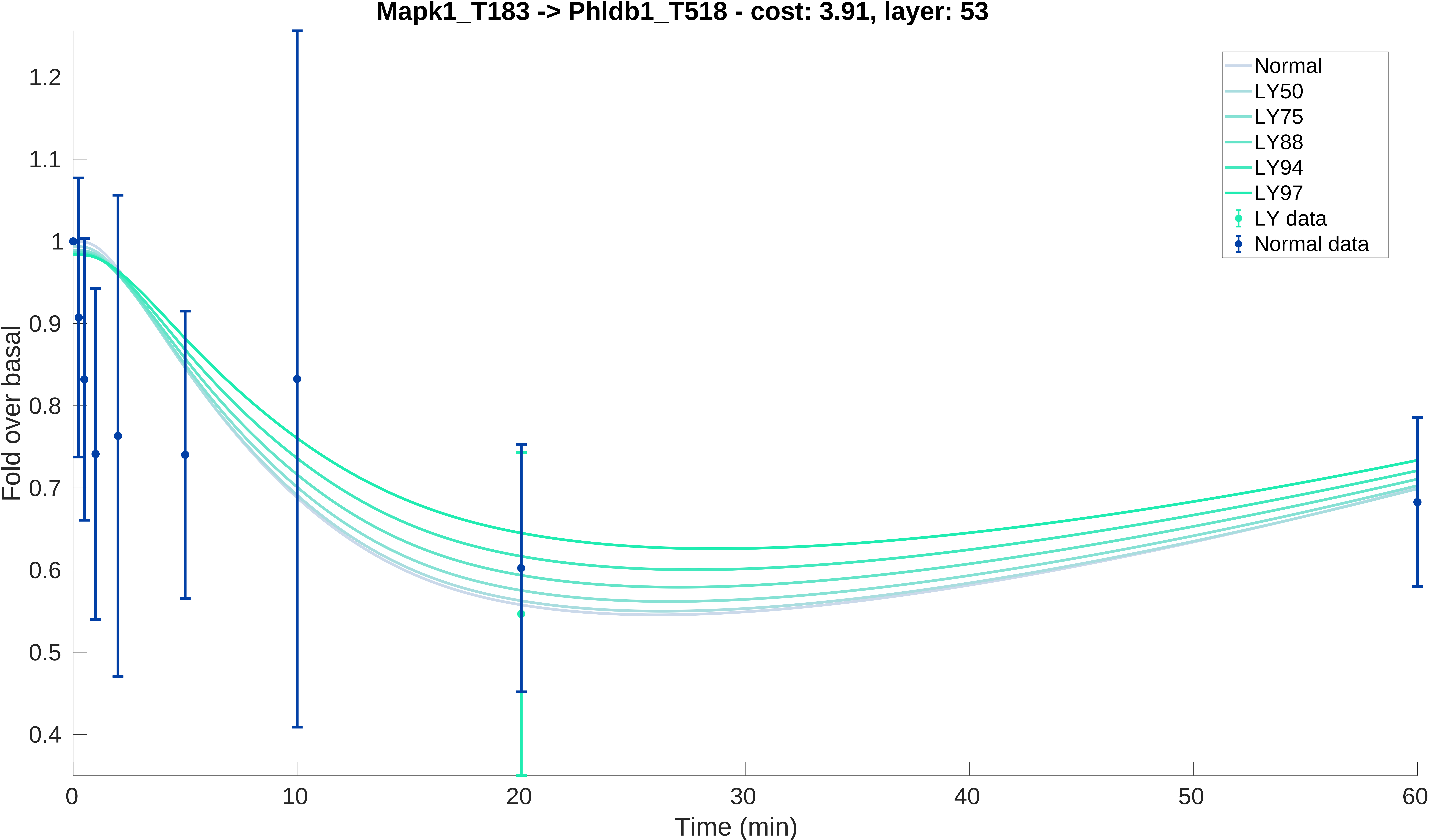

Mapk1_T183 -> Phldb1_T518 - cost: 3.91, layer: 53
Fold over basal
Time (min)
0.4
0.5
0.6
0.7
0.8
0.9
1
1.1
1.2
0
10
20
30
40
50
60
Normal
LY50
LY75
LY88
LY94
LY97
LY data
Normal data

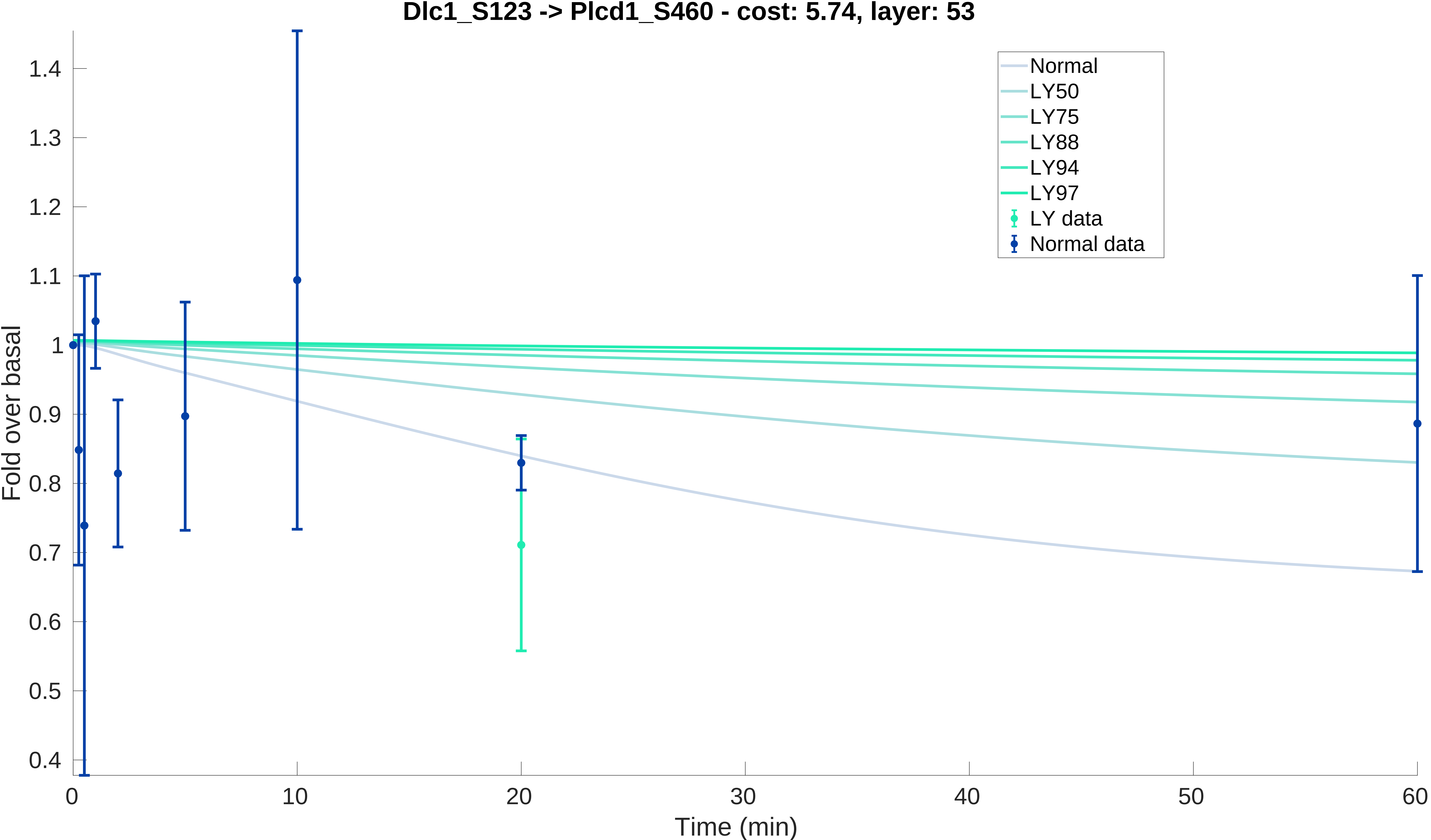

Dlc1_S123 -> Plcd1_S460 - cost: 5.74, layer: 53
Fold over basal
1.4
1.3
1.2
1.1
1
0.9
0.8
0.7
0.6
0.5
0.4
0
10
20
30
40
50
60
Time (min)
Normal
LY50
LY75
LY88
LY94
LY97
LY data
Normal data

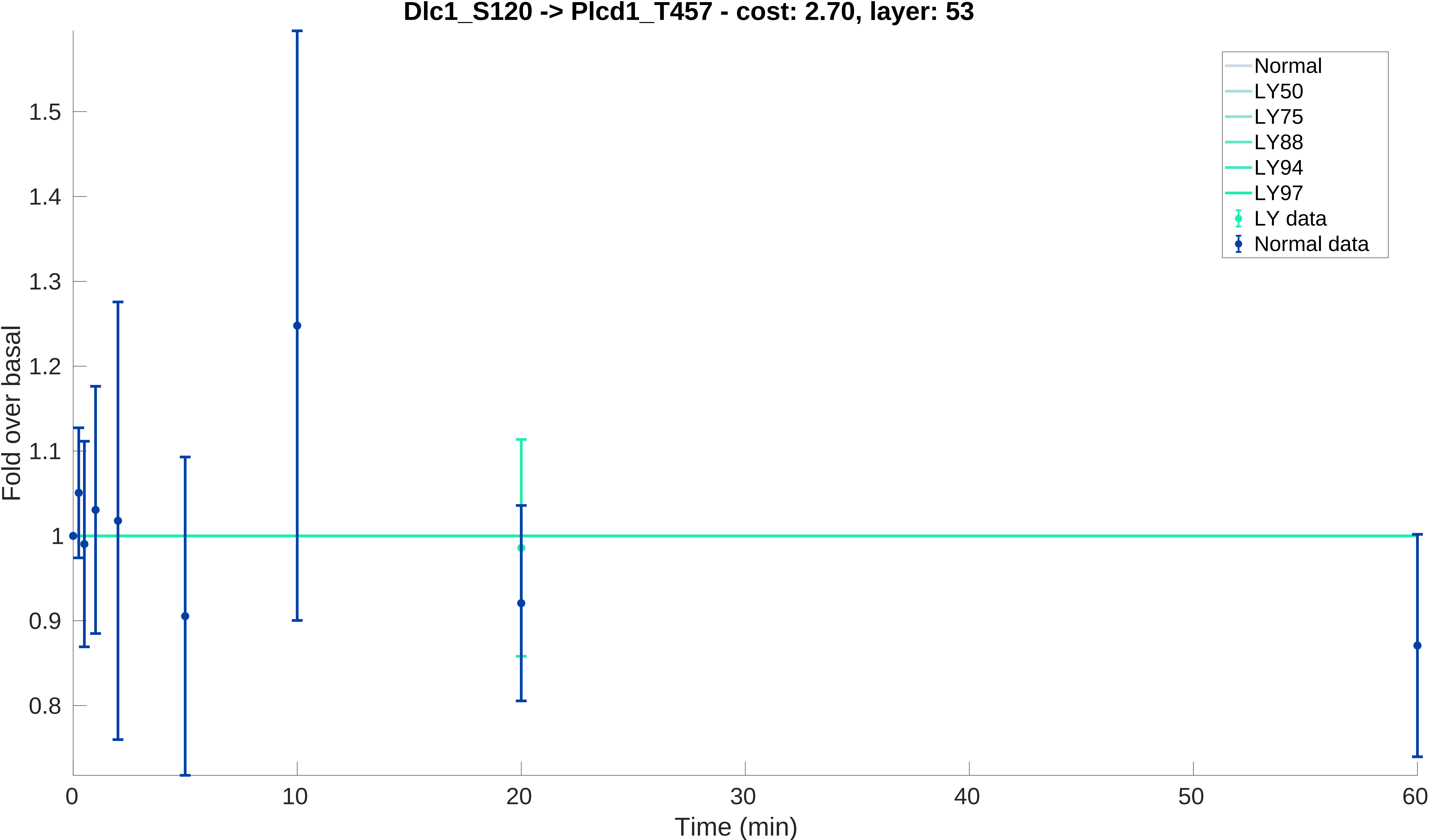

Dlc1_S120 -> Plcd1_T457 - cost: 2.70, layer: 53
Fold over basal
Time (min)
1.5
1.4
1.3
1.2
1.1
1
0.9
0.8
0
10
20
30
40
50
60
Normal
LY50
LY75
LY88
LY94
LY97
LY data
Normal data

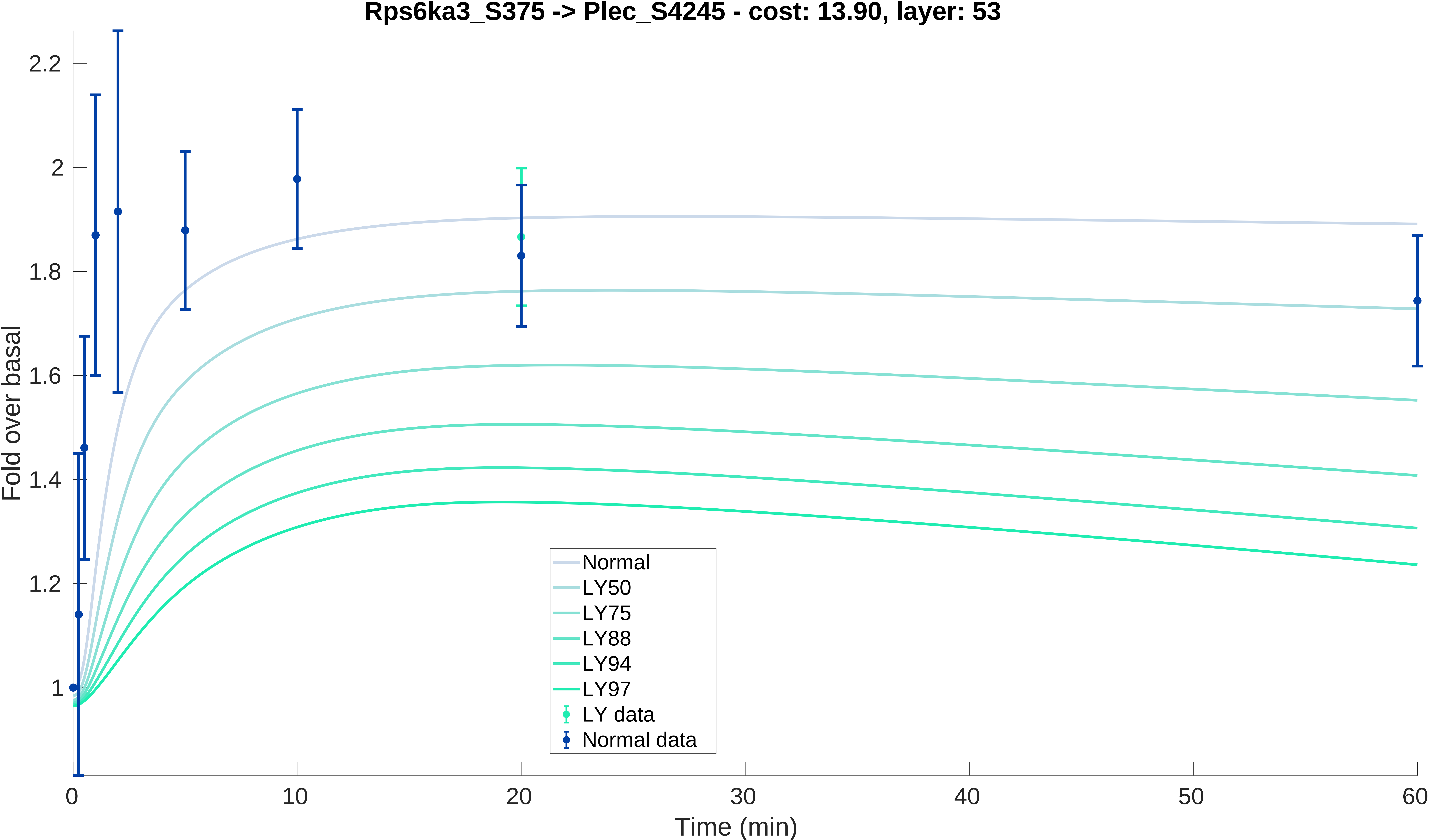
Rps6ka3_S375 -> Plec_S4245 - cost: 13.90, layer: 53
Fold over basal
Time (min)
2.2
2
1.8
1.6
1.4
1.2
1
0
10
20
30
40
50
60
Normal
LY50
LY75
LY88
LY94
LY97
LY data
Normal data

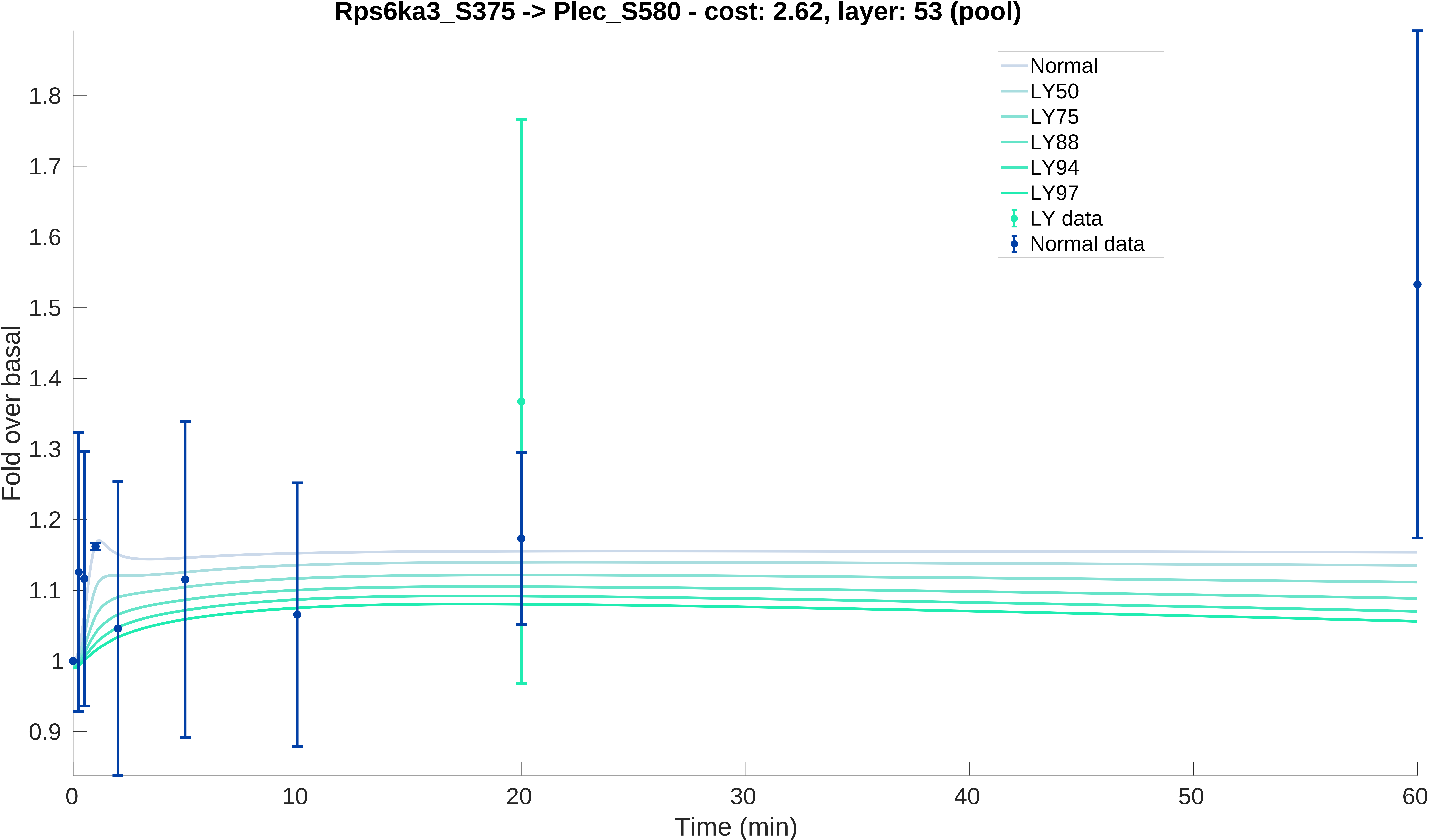

Rps6ka3_S375 -> Plec_S580 - cost: 2.62, layer: 53 (pool)
Normal
LY50
LY75
LY88
LY94
LY97
LY data
Normal data
Fold over basal
1.8
1.7
1.6
1.5
1.4
1.3
1.2
1.1
1
0.9
0
10
20
30
40
50
60
Time (min)

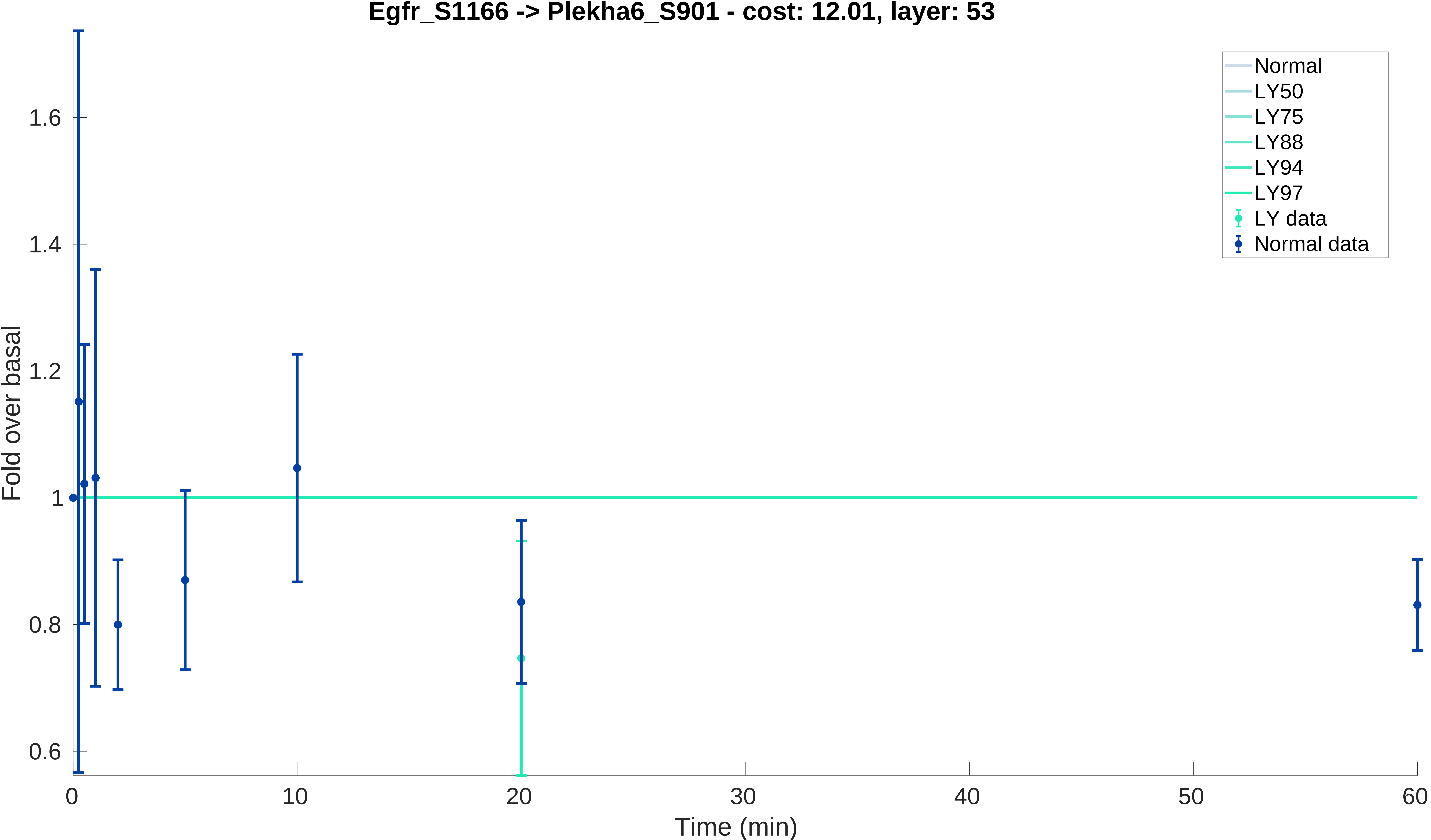

Egfr_S1166 -> Plekha6_S901 - cost: 12.01, layer: 53
Normal
LY50
LY75
LY88
LY94
LY97
LY data
Normal data
Fold over basal
1.6
1.4
1.2
1
0.8
0.6
0
10
20
30
40
50
60
Time (min)

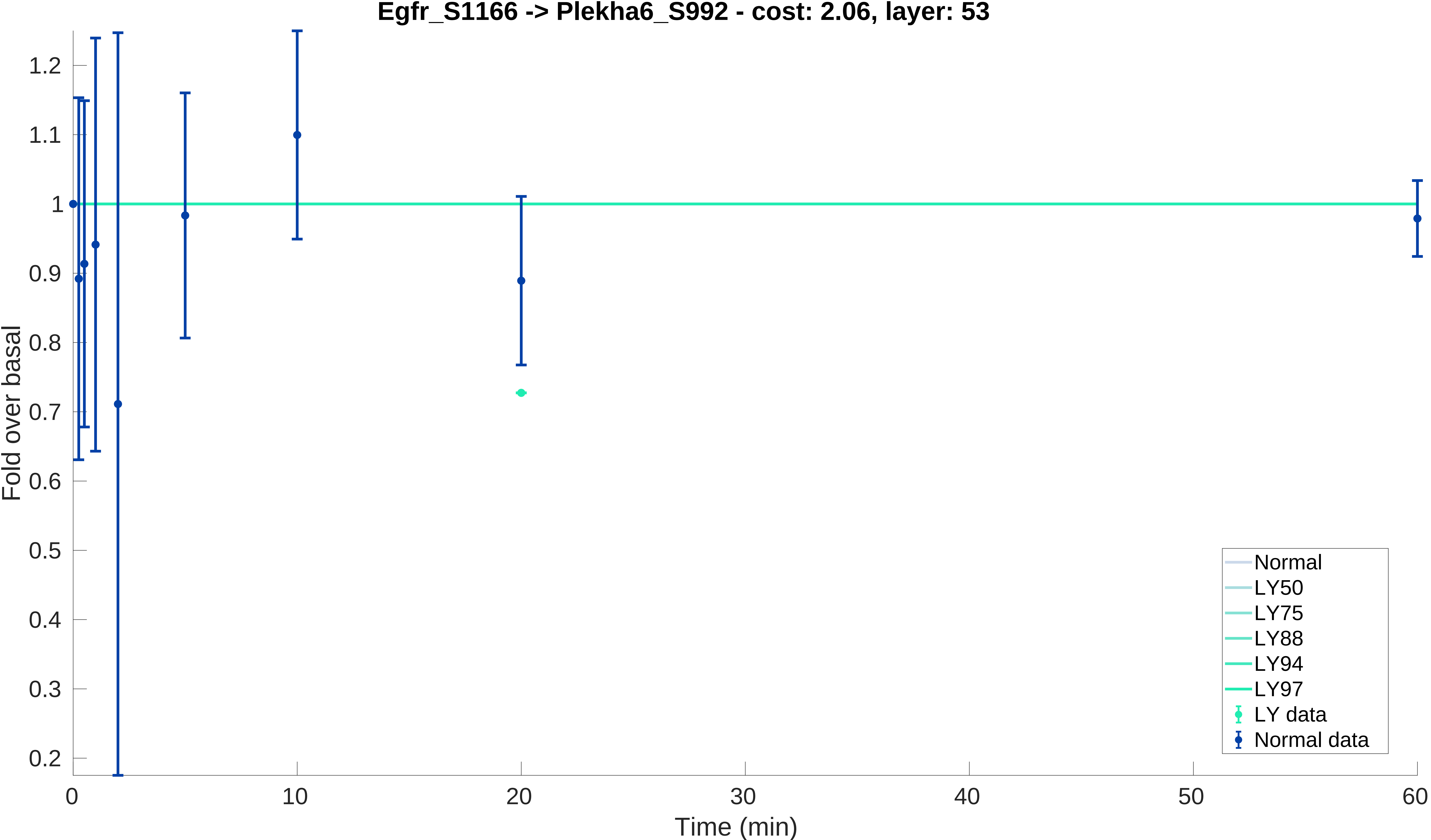
Egfr_S1166 -> Plekha6_S992 - cost: 2.06, layer: 53
Fold over basal
1.2
1.1
1
0.9
0.8
0.7
0.6
0.5
0.4
0.3
0.2
0
10
20
30
40
50
60
Time (min)
Normal
LY50
LY75
LY88
LY94
LY97
LY data
Normal data

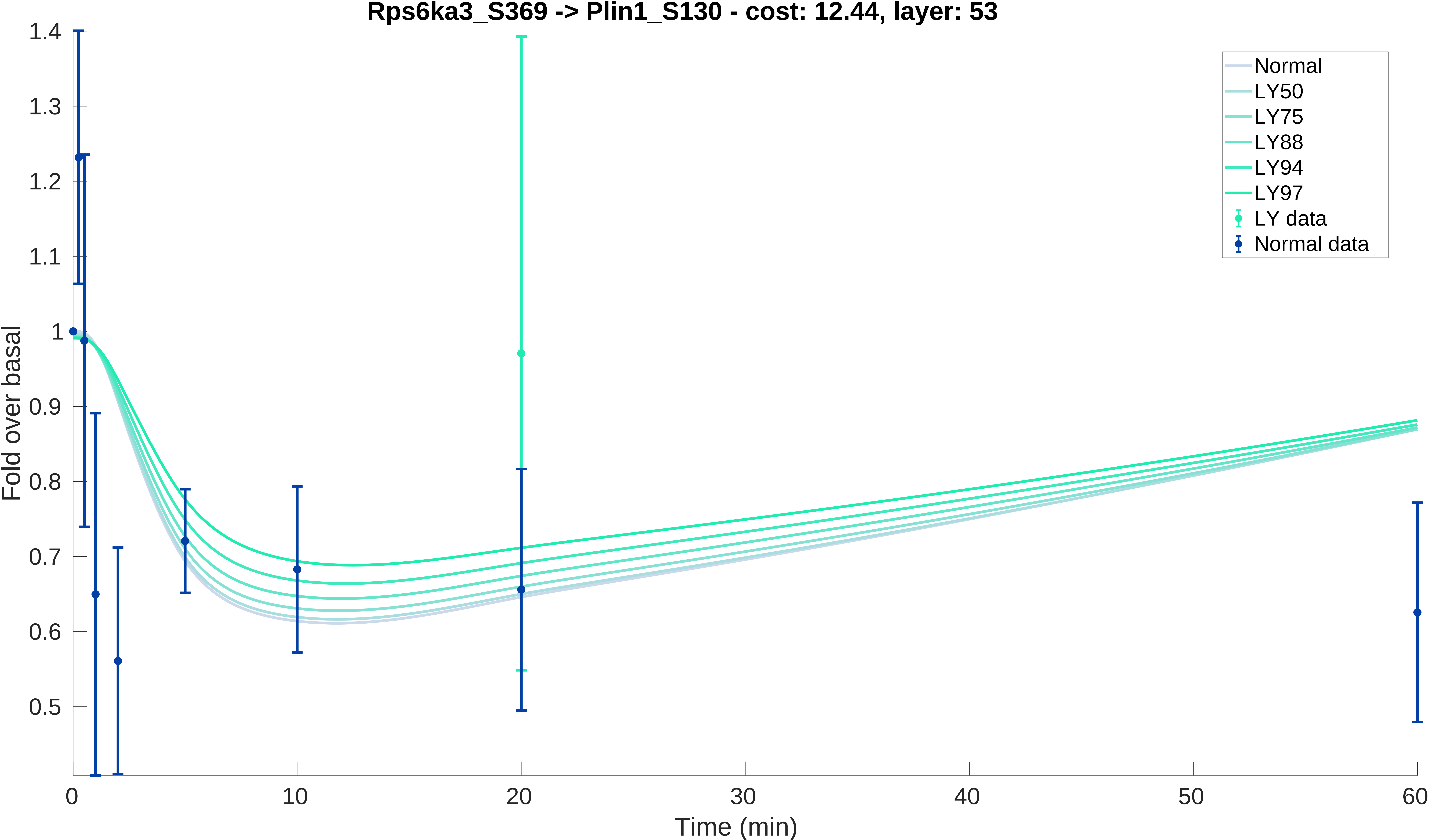
Rps6ka3_S369 -> Plin1_S130 - cost: 12.44, layer: 53
1.4
1.3
1.2
1.1
1
0.9
0.8
0.7
0.6
0.5
Fold over basal
0
10
20
30
40
50
60
Time (min)
Normal
LY50
LY75
LY88
LY94
LY97
LY data
Normal data

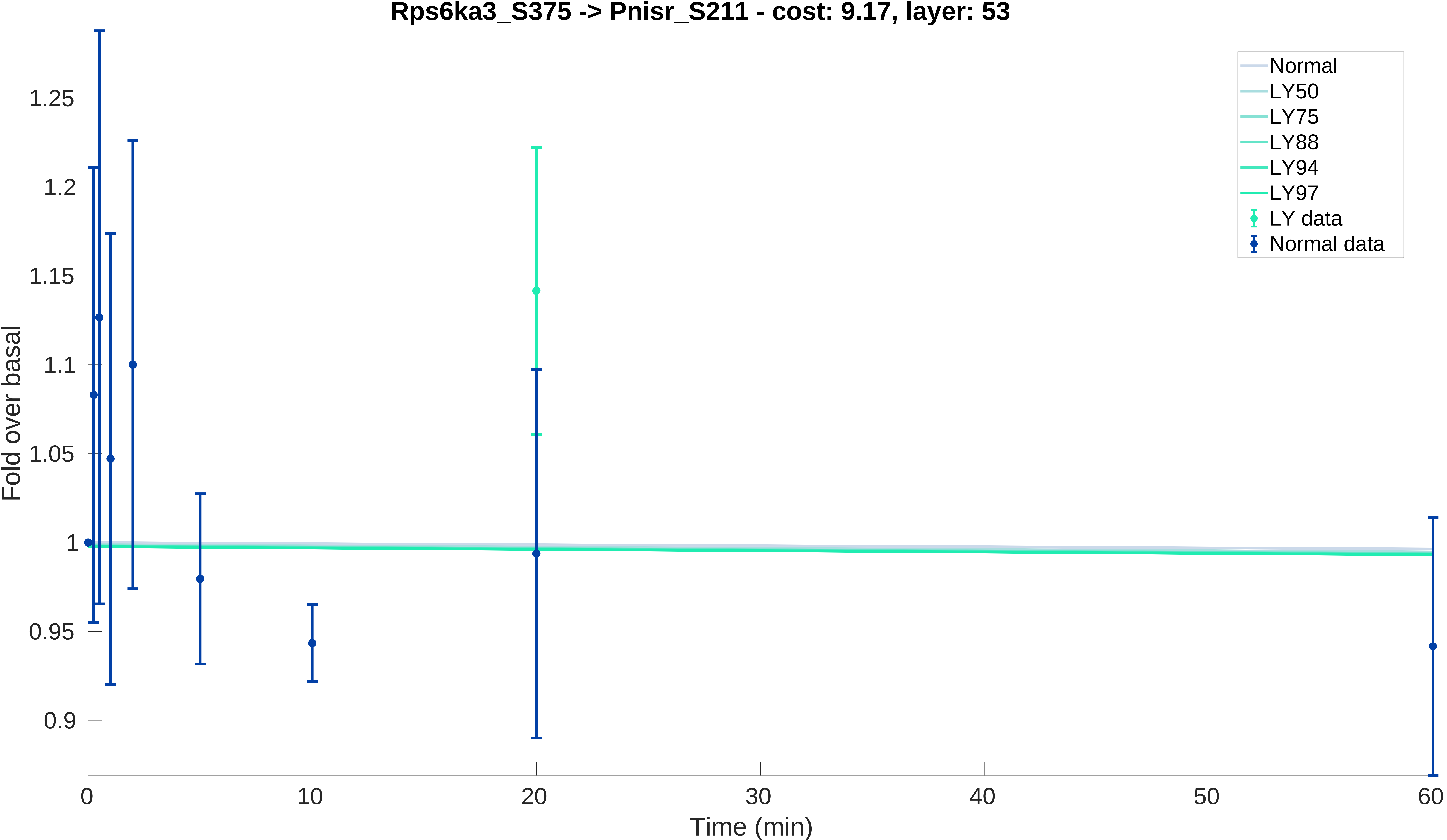

Rps6ka3_S375 -> Pnisr_S211 - cost: 9.17, layer: 53
Fold over basal
Time (min)
Normal
LY50
LY75
LY88
LY94
LY97
LY data
Normal data
1.25
1.2
1.15
1.1
1.05
1
0.95
0.9
0
10
20
30
40
50
60

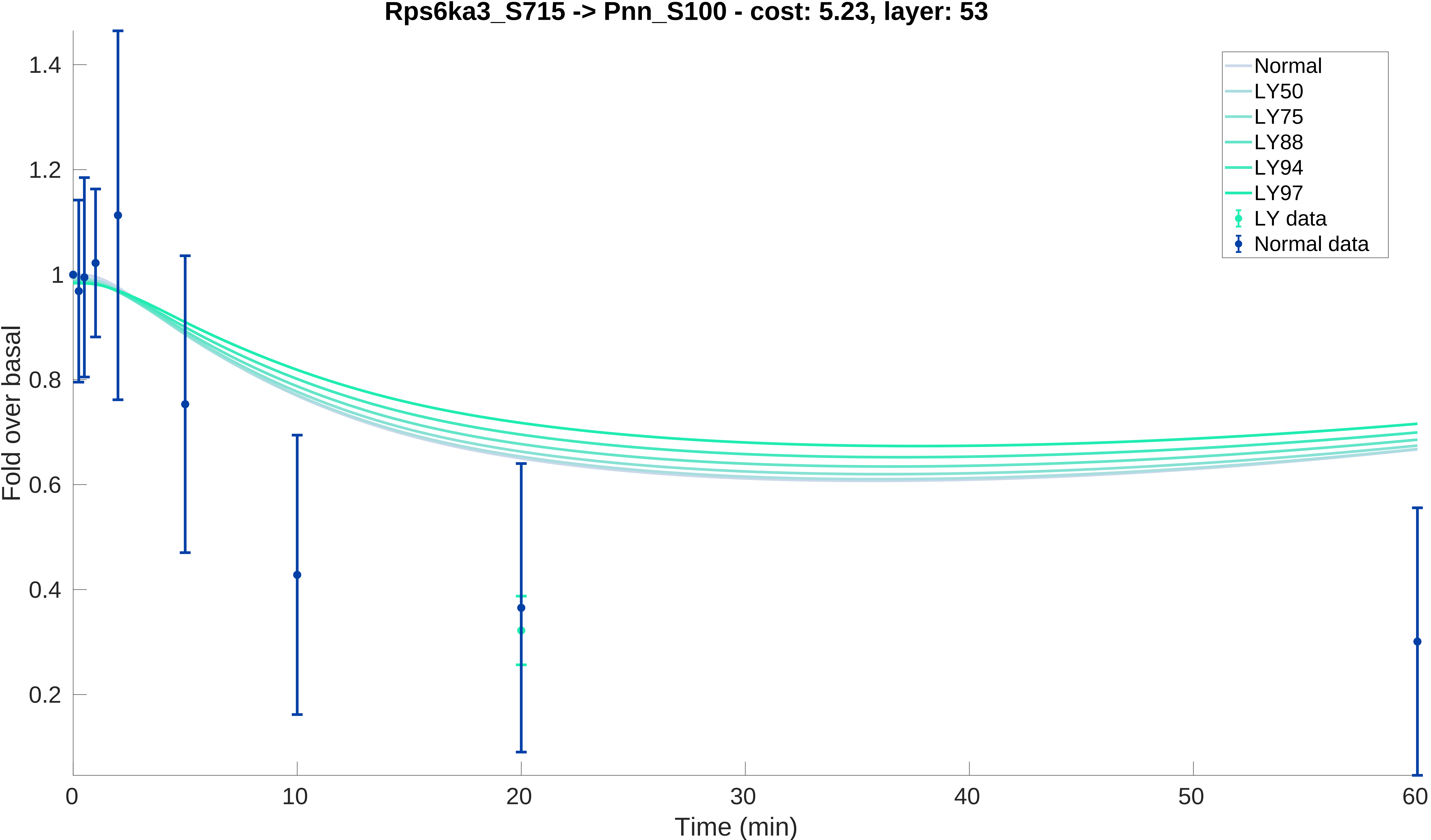

Rps6ka3_S715 -> Pnn_S100 - cost: 5.23, layer: 53
Fold over basal
Time (min)
1.4
1.2
1
0.8
0.6
0.4
0.2
0
10
20
30
40
50
60
Normal
LY50
LY75
LY88
LY94
LY97
LY data
Normal data

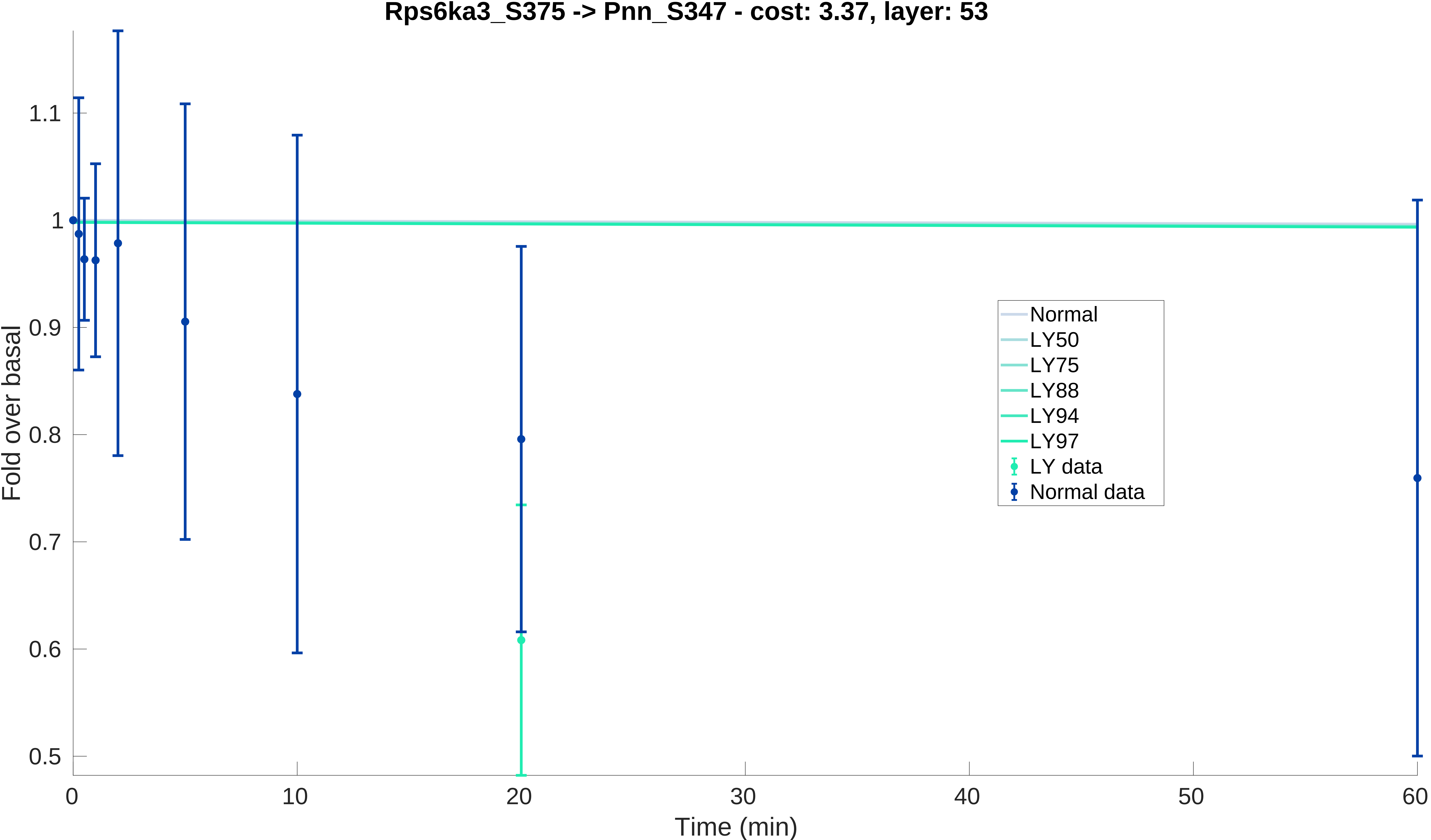

Rps6ka3_S375 -> Pnn_S347 - cost: 3.37, layer: 53
Fold over basal
1.1
1
0.9
0.8
0.7
0.6
0.5
0
10
20
30
40
50
60
Time (min)
Normal
LY50
LY75
LY88
LY94
LY97
LY data
Normal data

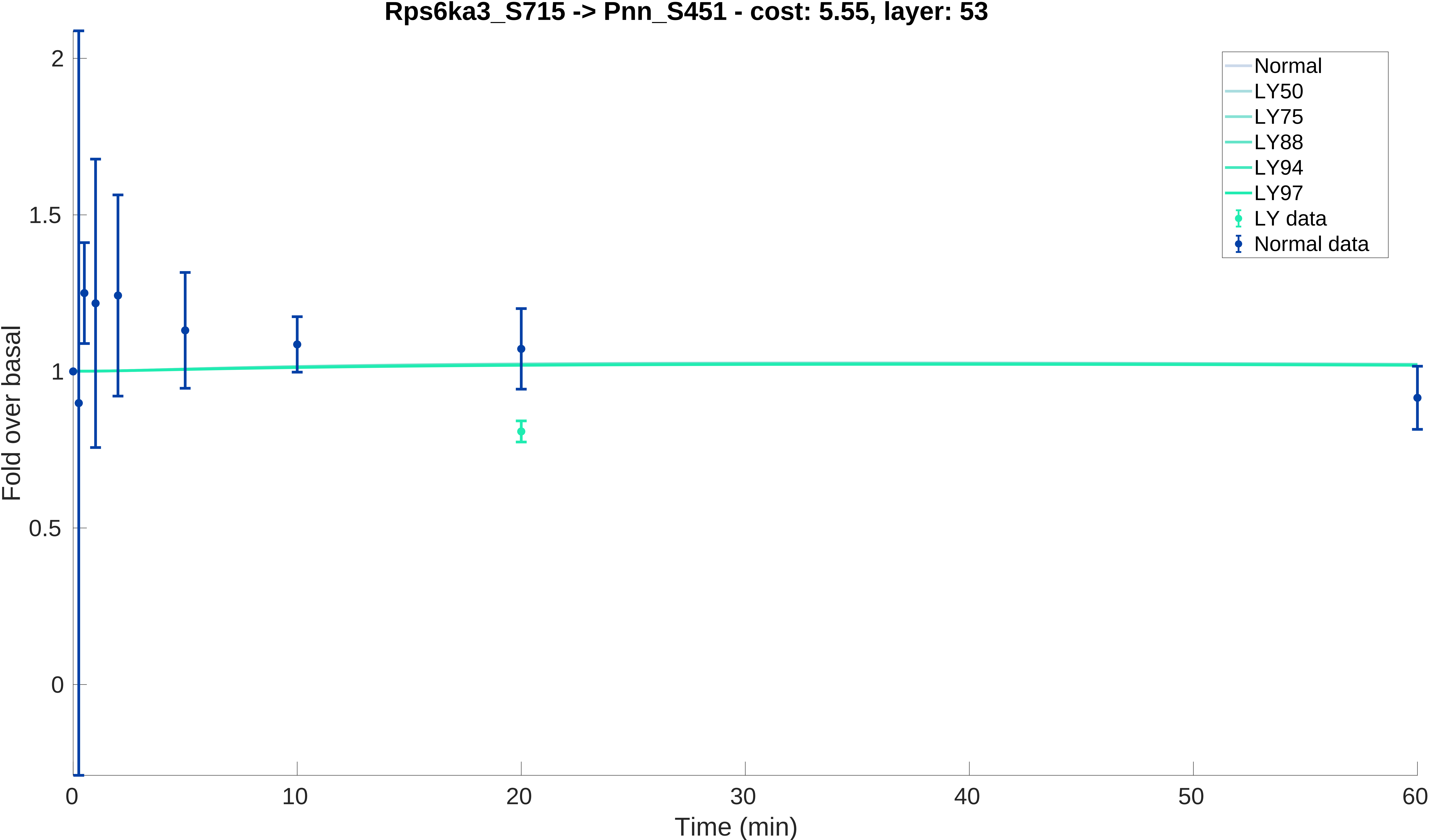
Rps6ka3_S715 -> Pnn_S451 - cost: 5.55, layer: 53
Fold over basal
Time (min)
Normal
LY50
LY75
LY88
LY94
LY97
LY data
Normal data
2
1.5
1
0.5
0
0
10
20
30
40
50
60

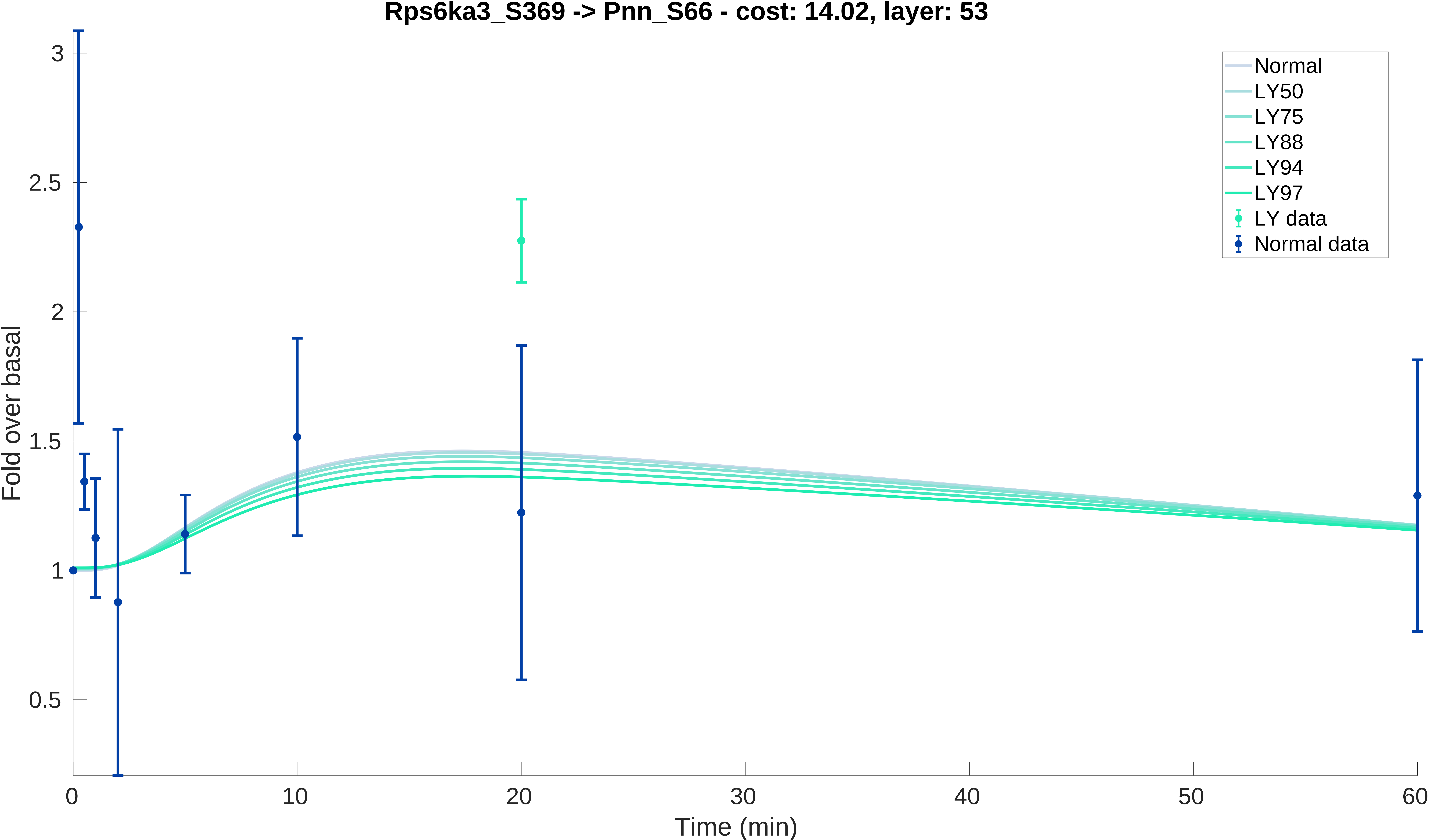

Rps6ka3_S369 -> Pnn_S66 - cost: 14.02, layer: 53
Fold over basal
Time (min)
Normal
LY50
LY75
LY88
LY94
LY97
LY data
Normal data
3
2.5
2
1.5
1
0.5
0
10
20
30
40
50
60

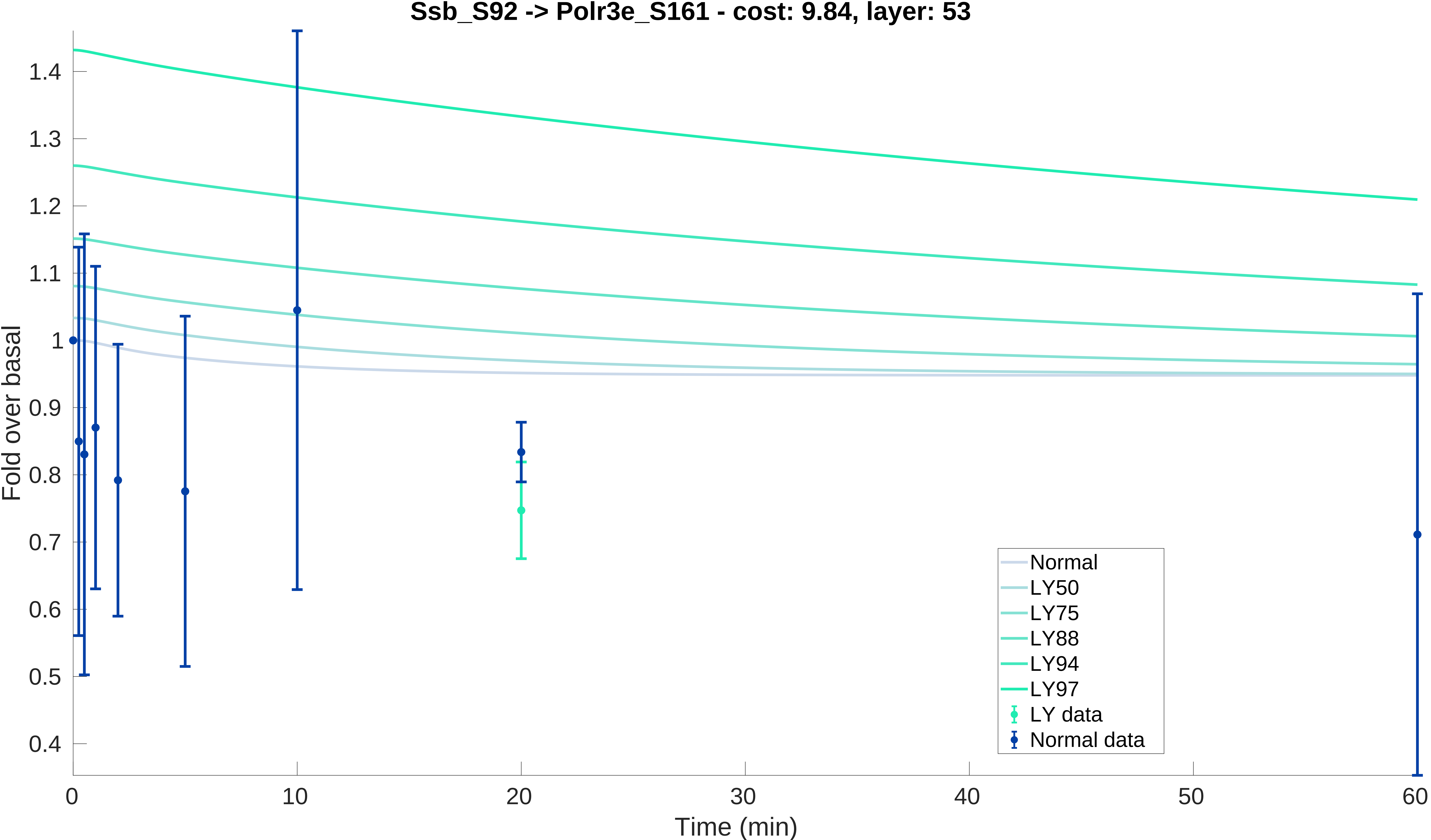

Ssb_S92 -> Polr3e_S161 - cost: 9.84, layer: 53
Fold over basal
1.4
1.3
1.2
1.1
1
0.9
0.8
0.7
0.6
0.5
0.4
0
10
20
30
40
50
60
Time (min)
Normal
LY50
LY75
LY88
LY94
LY97
LY data
Normal data

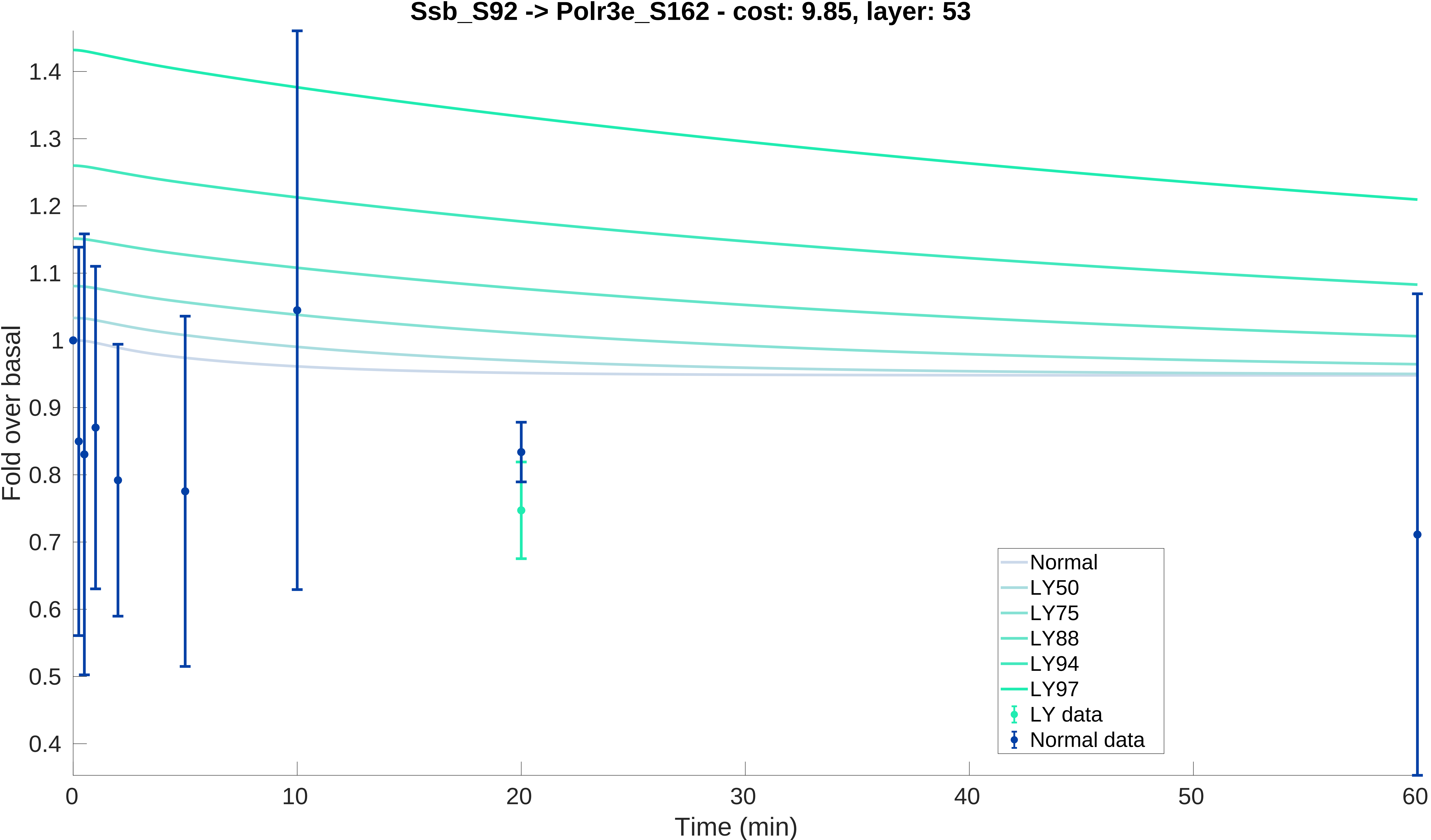

Ssb_S92 -> Polr3e_S162 - cost: 9.85, layer: 53
Fold over basal
Time (min)
0
10
20
30
40
50
60
0.4
0.5
0.6
0.7
0.8
0.9
1
1.1
1.2
1.3
1.4
Normal
LY50
LY75
LY88
LY94
LY97
LY data
Normal data

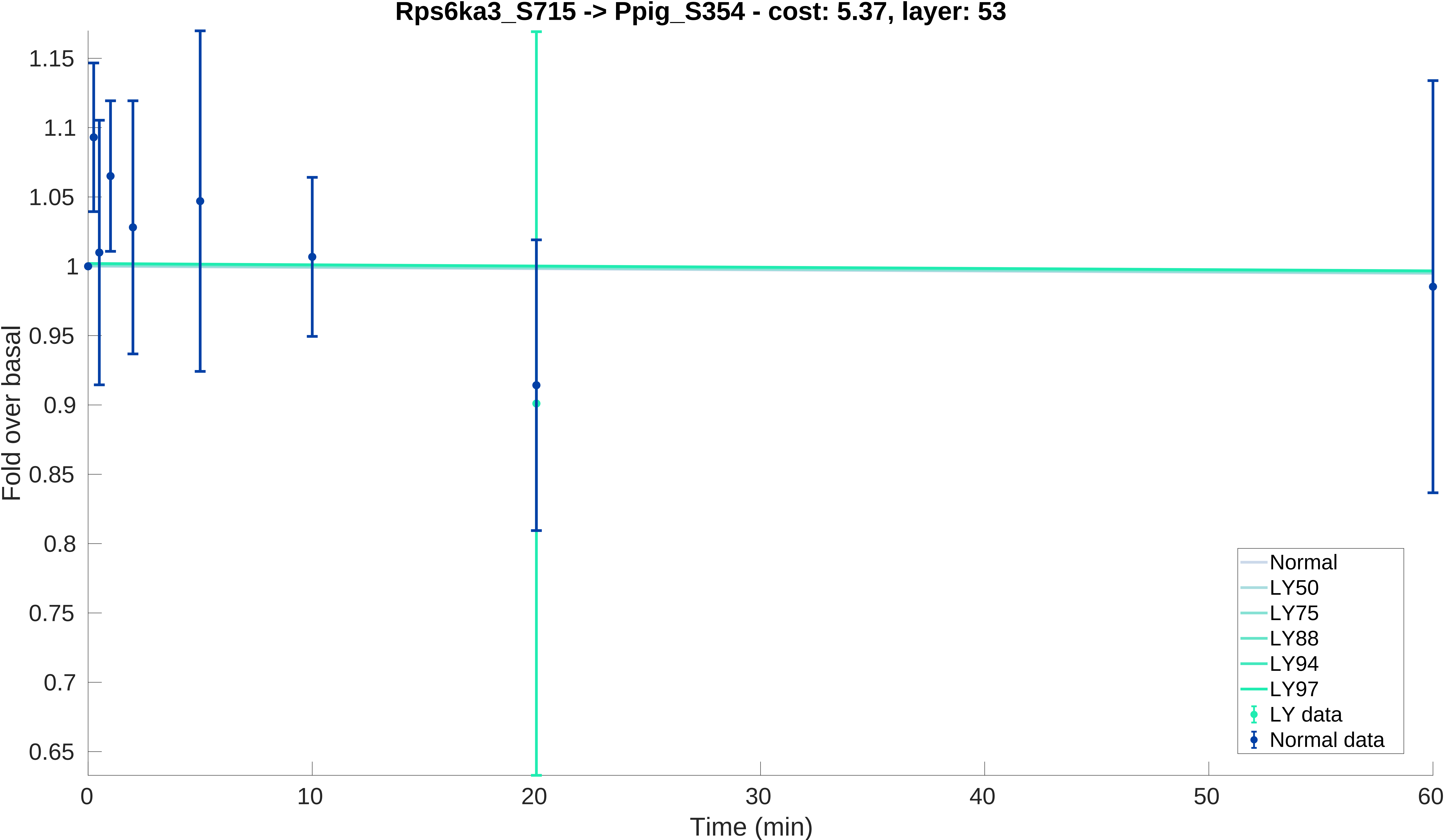

Rps6ka3_S715 -> Ppig_S354 - cost: 5.37, layer: 53
Fold over basal
1.15
1.1
1.05
1
0.95
0.9
0.85
0.8
0.75
0.7
0.65
0
10
20
30
40
50
60
Time (min)
Normal
LY50
LY75
LY88
LY94
LY97
LY data
Normal data

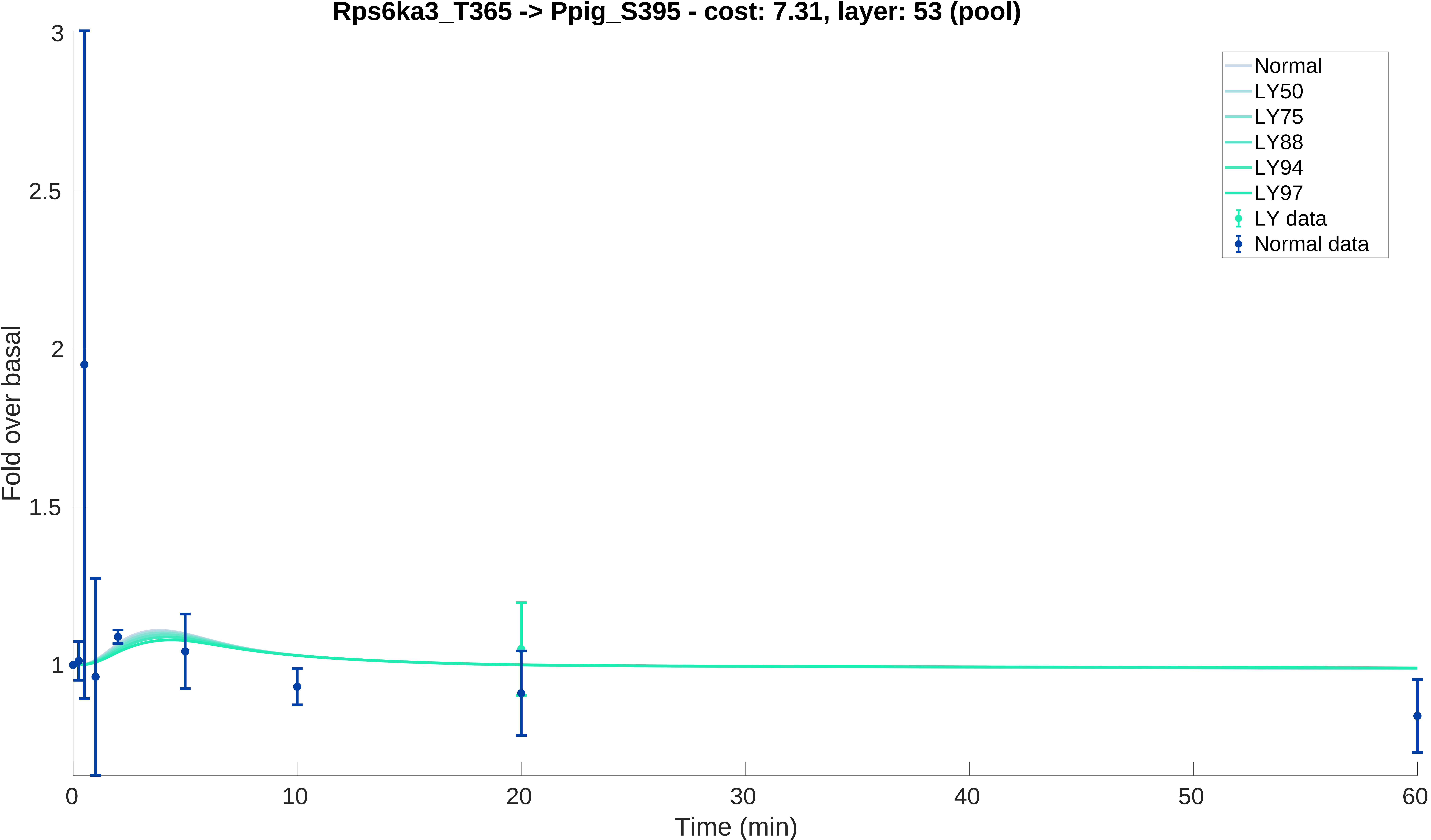

Rps6ka3_T365 -> Ppig_S395 - cost: 7.31, layer: 53 (pool)
Fold over basal
Time (min)
3
2.5
2
1.5
1
0
10
20
30
40
50
60
Normal
LY50
LY75
LY88
LY94
LY97
LY data
Normal data

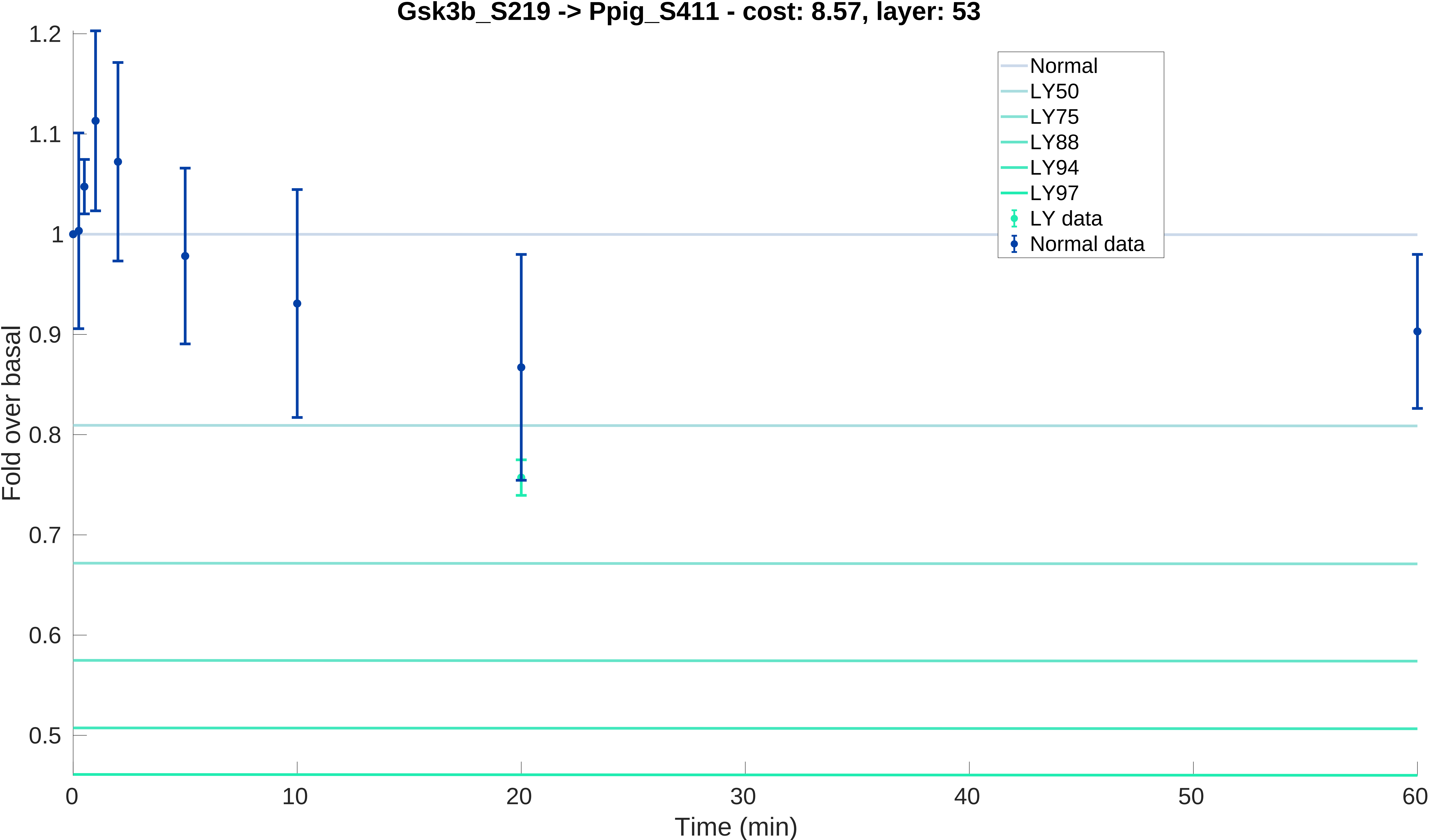

Gsk3b_S219 -> Ppig_S411 - cost: 8.57, layer: 53
Fold over basal
Time (min)
1.2
1.1
1
0.9
0.8
0.7
0.6
0.5
0
10
20
30
40
50
60
Normal
LY50
LY75
LY88
LY94
LY97
LY data
Normal data

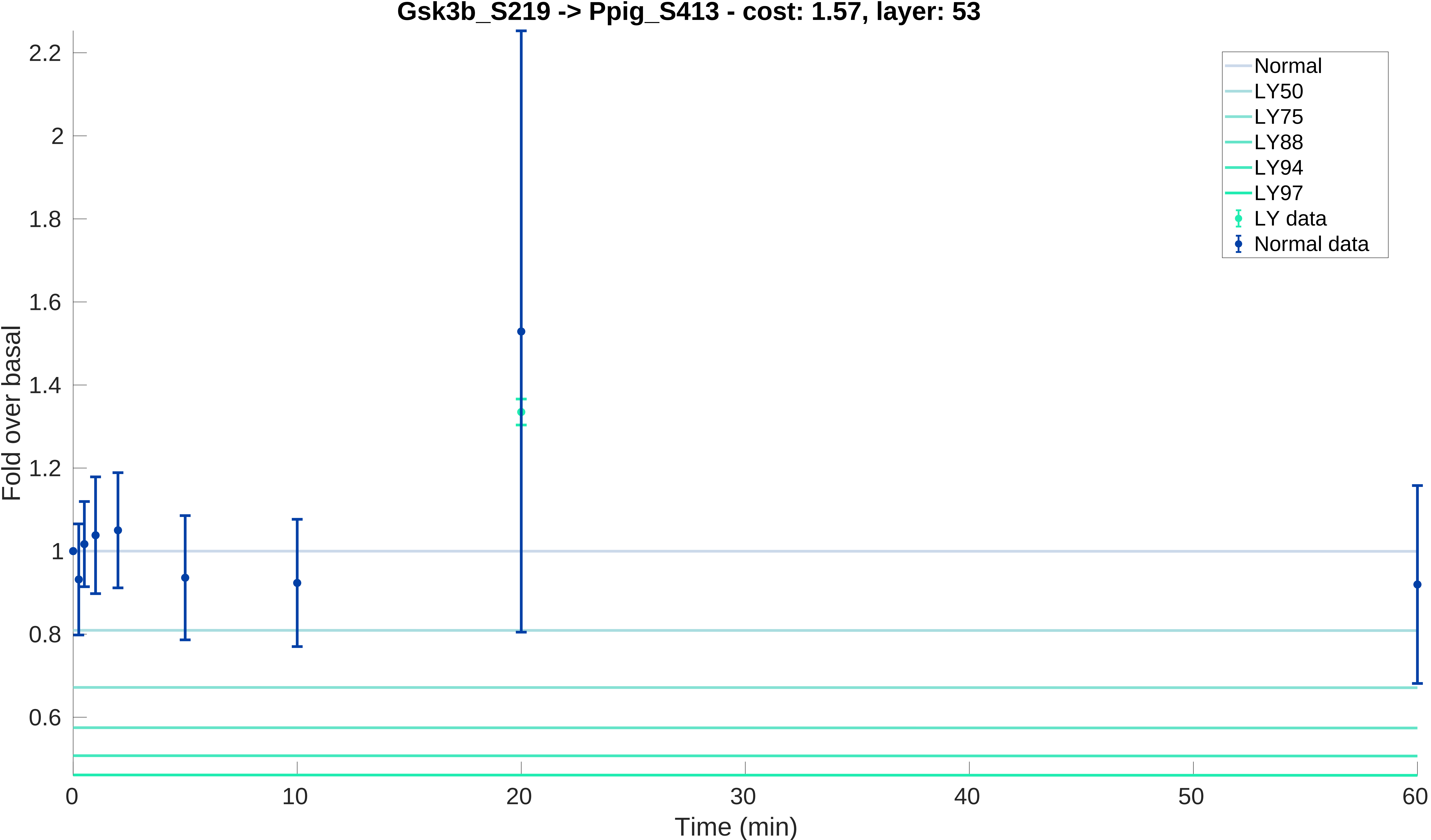
Gsk3b_S219 -> Ppig_S413 - cost: 1.57, layer: 53
Fold over basal
Time (min)
0
10
20
30
40
50
60
0.6
0.8
1
1.2
1.4
1.6
1.8
2
2.2
Normal
LY50
LY75
LY88
LY94
LY97
LY data
Normal data

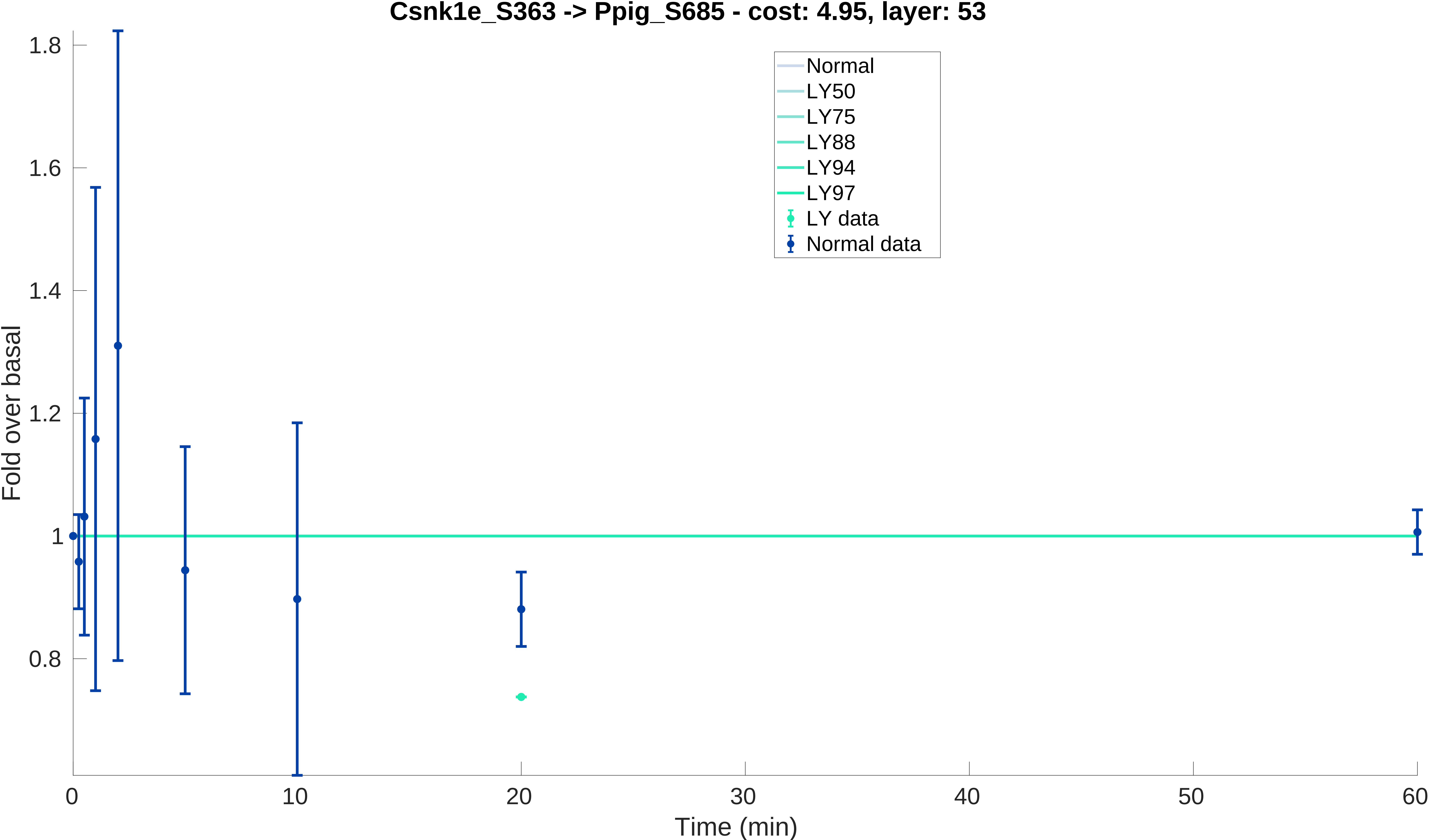

Csnk1e_S363 -> Ppig_S685 - cost: 4.95, layer: 53
Normal
LY50
LY75
LY88
LY94
LY97
LY data
Normal data
Fold over basal
1.8
1.6
1.4
1.2
1
0.8
0
10
20
30
40
50
60
Time (min)

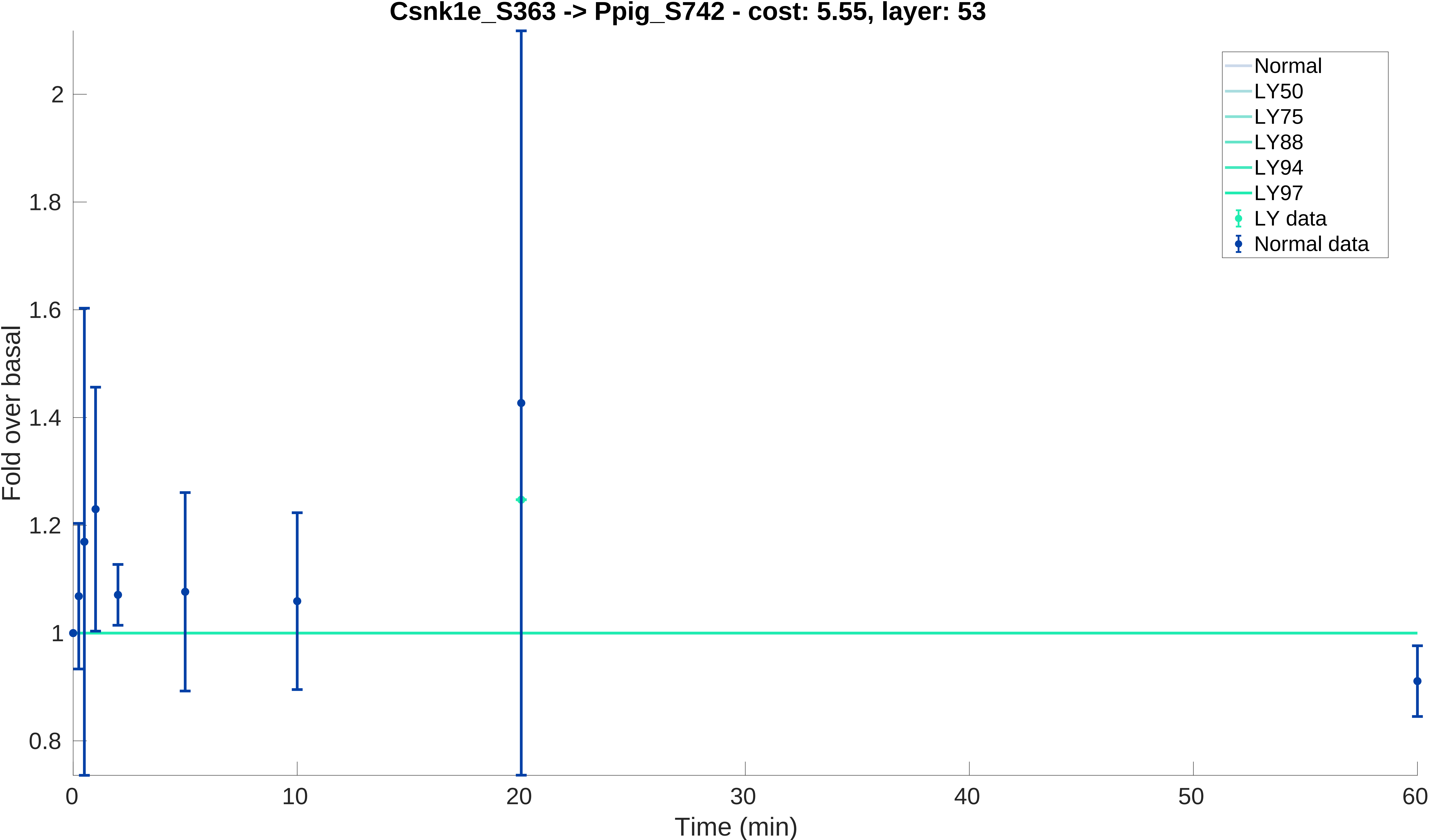

Csnk1e_S363 -> Ppig_S742 - cost: 5.55, layer: 53
Normal
LY50
LY75
LY88
LY94
LY97
LY data
Normal data
Fold over basal
2
1.8
1.6
1.4
1.2
1
0.8
0
10
20
30
40
50
60
Time (min)

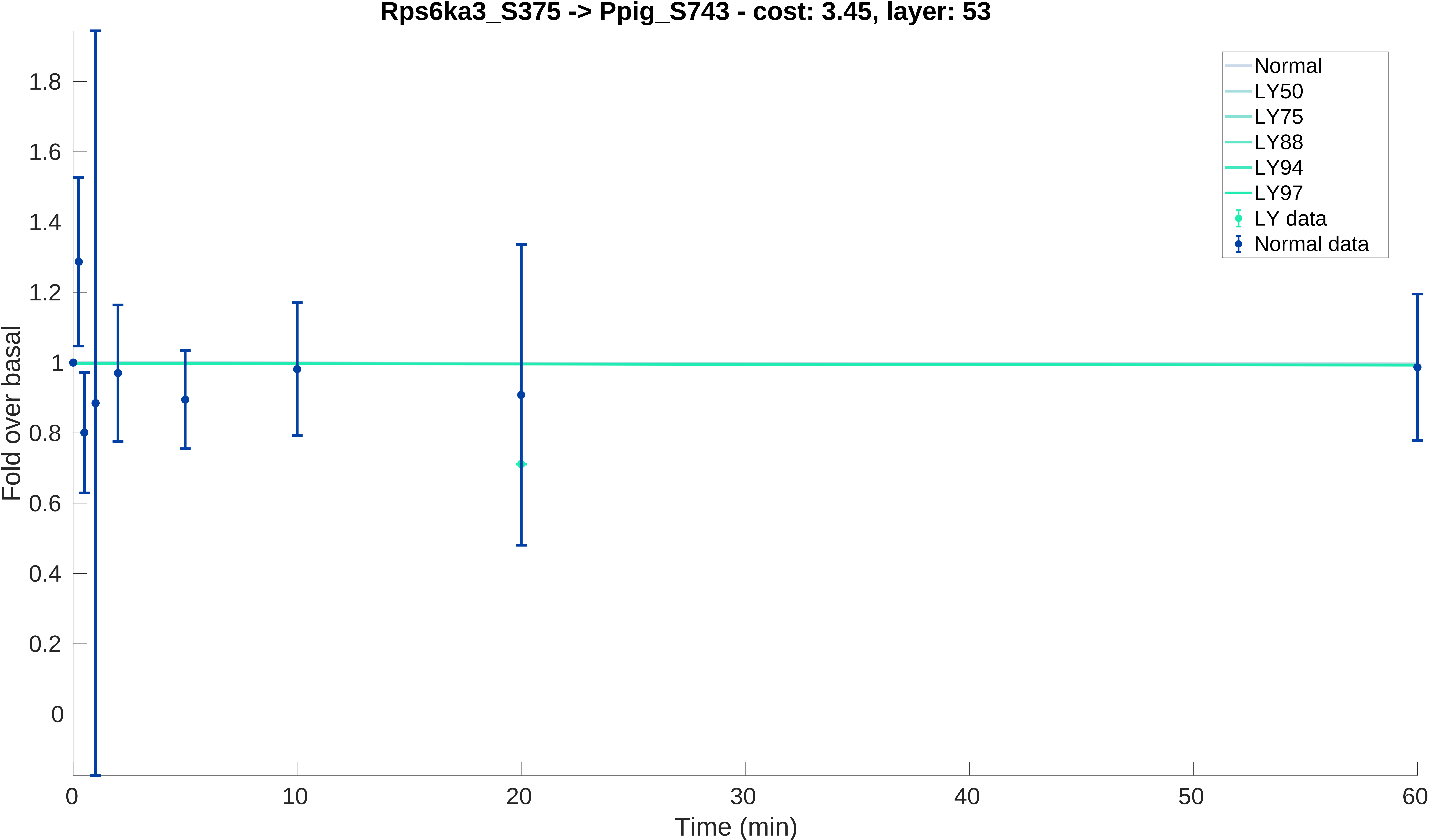

Rps6ka3_S375 -> Ppig_S743 - cost: 3.45, layer: 53
Fold over basal
Time (min)
Normal
LY50
LY75
LY88
LY94
LY97
LY data
Normal data
0
0.2
0.4
0.6
0.8
1
1.2
1.4
1.6
1.8
0
10
20
30
40
50
60

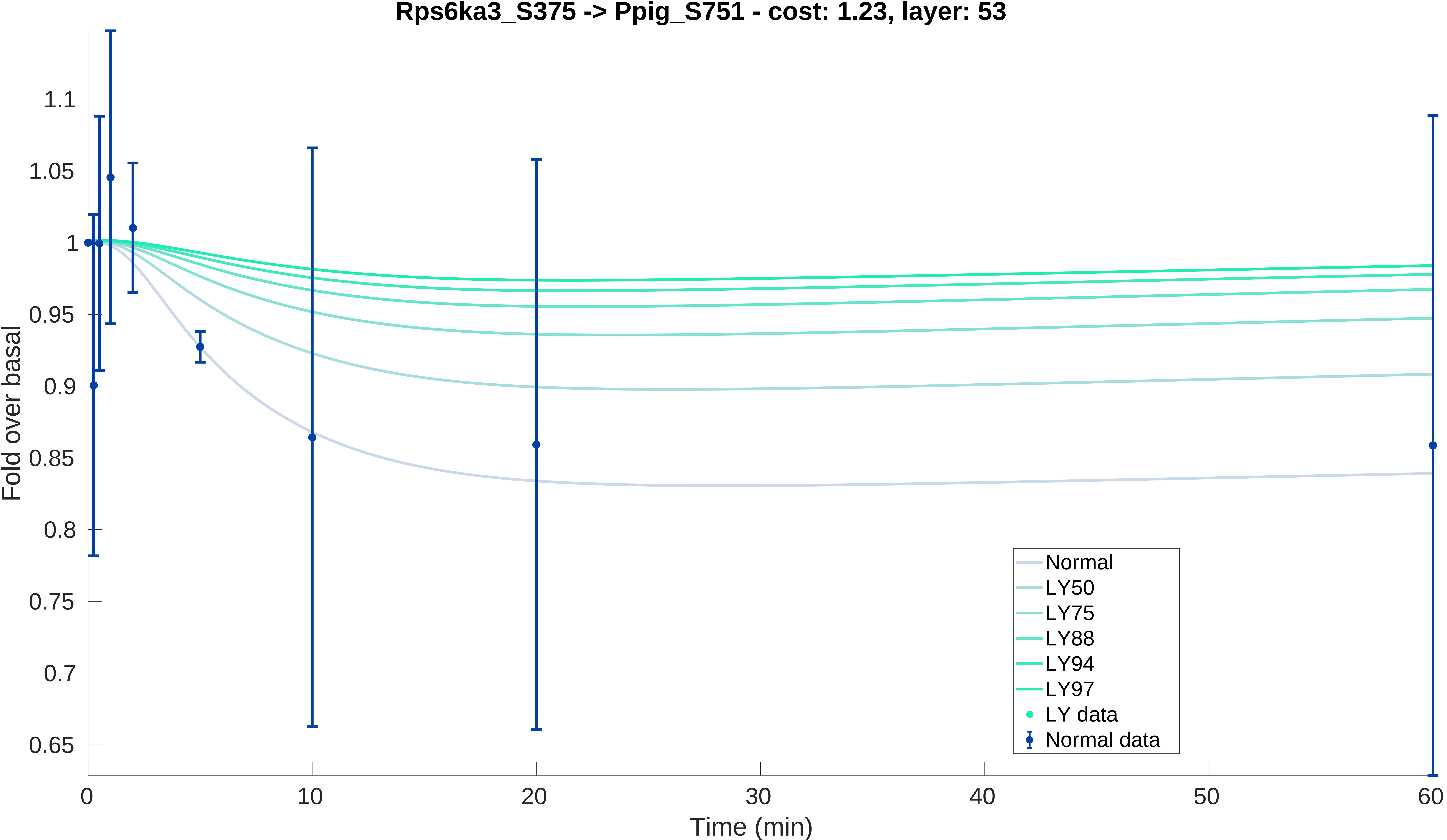

Rps6ka3_S375 -> Ppig_S751 - cost: 1.23, layer: 53
Fold over basal
1.1
1.05
1
0.95
0.9
0.85
0.8
0.75
0.7
0.65
0
10
20
30
40
50
60
Time (min)
Normal
LY50
LY75
LY88
LY94
LY97
LY data
Normal data

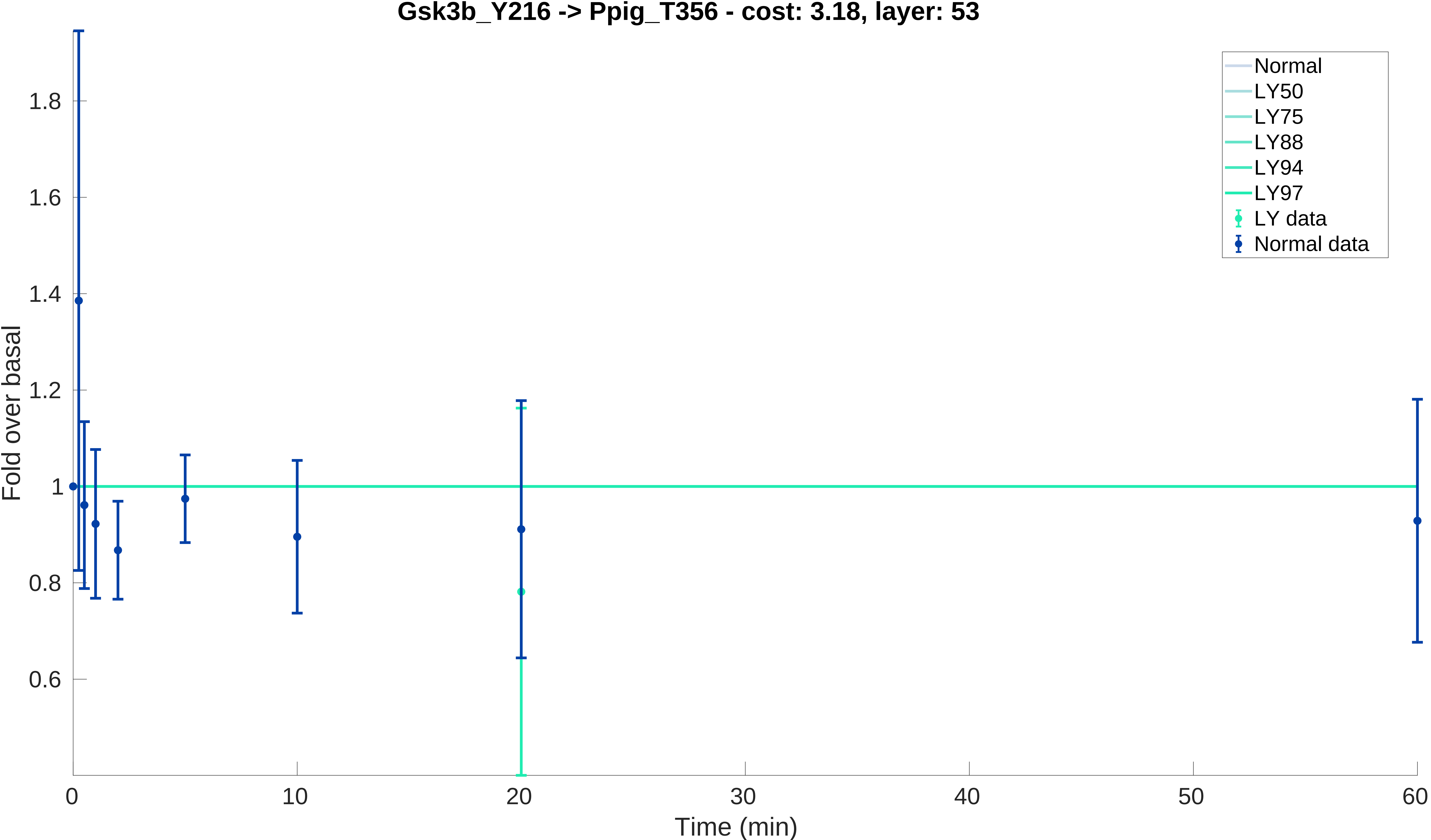

Gsk3b_Y216 -> Ppig_T356 - cost: 3.18, layer: 53
Normal
LY50
LY75
LY88
LY94
LY97
LY data
Normal data
Fold over basal
1.8
1.6
1.4
1.2
1
0.8
0.6
0
10
20
30
40
50
60
Time (min)

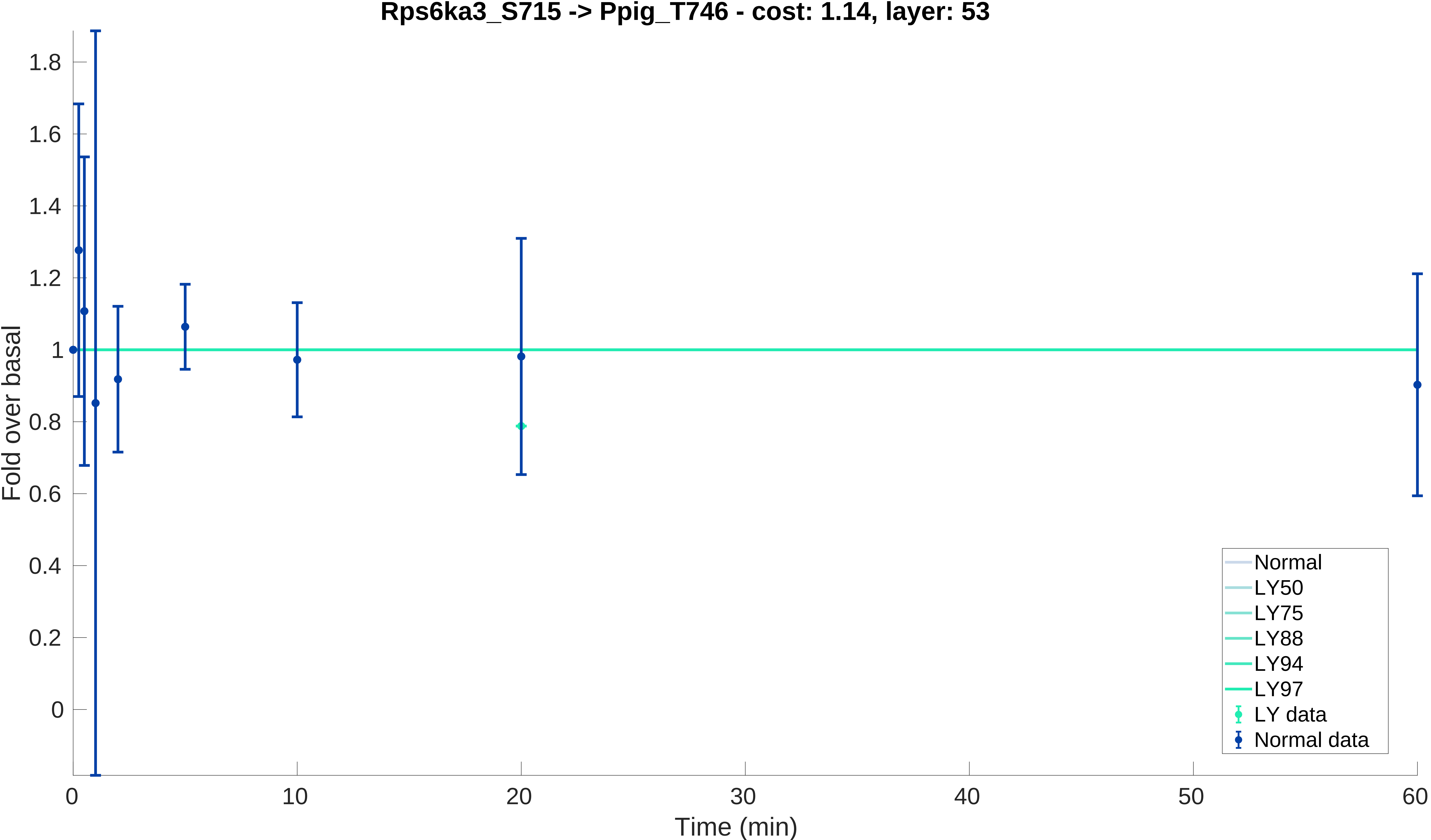
Rps6ka3_S715 -> Ppig_T746 - cost: 1.14, layer: 53
Fold over basal
Time (min)
1.8
1.6
1.4
1.2
1
0.8
0.6
0.4
0.2
0
0
10
20
30
40
50
60
Normal
LY50
LY75
LY88
LY94
LY97
LY data
Normal data

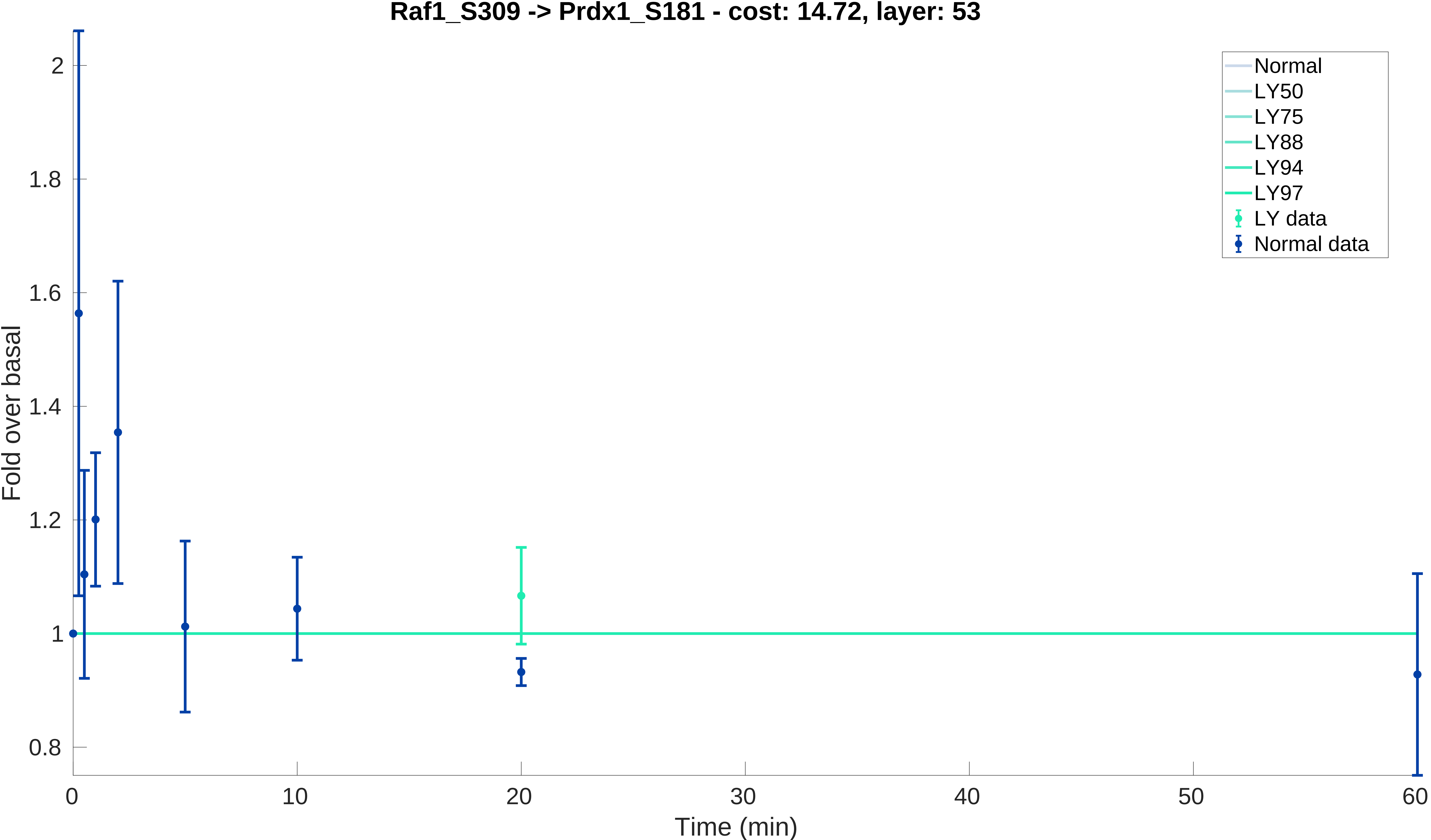

Raf1_S309 -> Prdx1_S181 - cost: 14.72, layer: 53
Fold over basal
Time (min)
2
1.8
1.6
1.4
1.2
1
0.8
0
10
20
30
40
50
60
Normal
LY50
LY75
LY88
LY94
LY97
LY data
Normal data

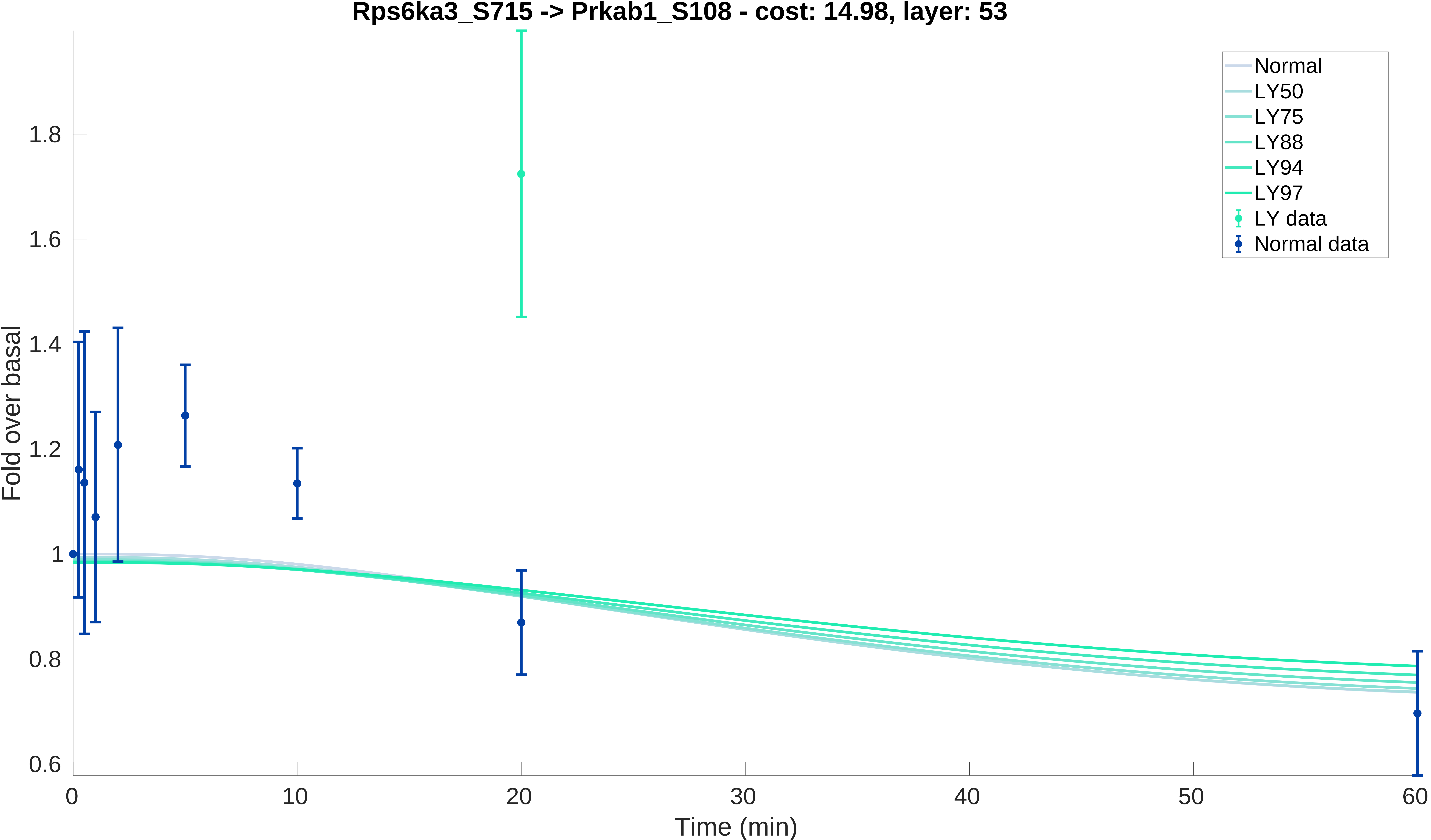

Rps6ka3_S715 -> Prkab1_S108 - cost: 14.98, layer: 53
Normal
LY50
LY75
LY88
LY94
LY97
LY data
Normal data
Fold over basal
1.8
1.6
1.4
1.2
1
0.8
0.6
0
10
20
30
40
50
60
Time (min)

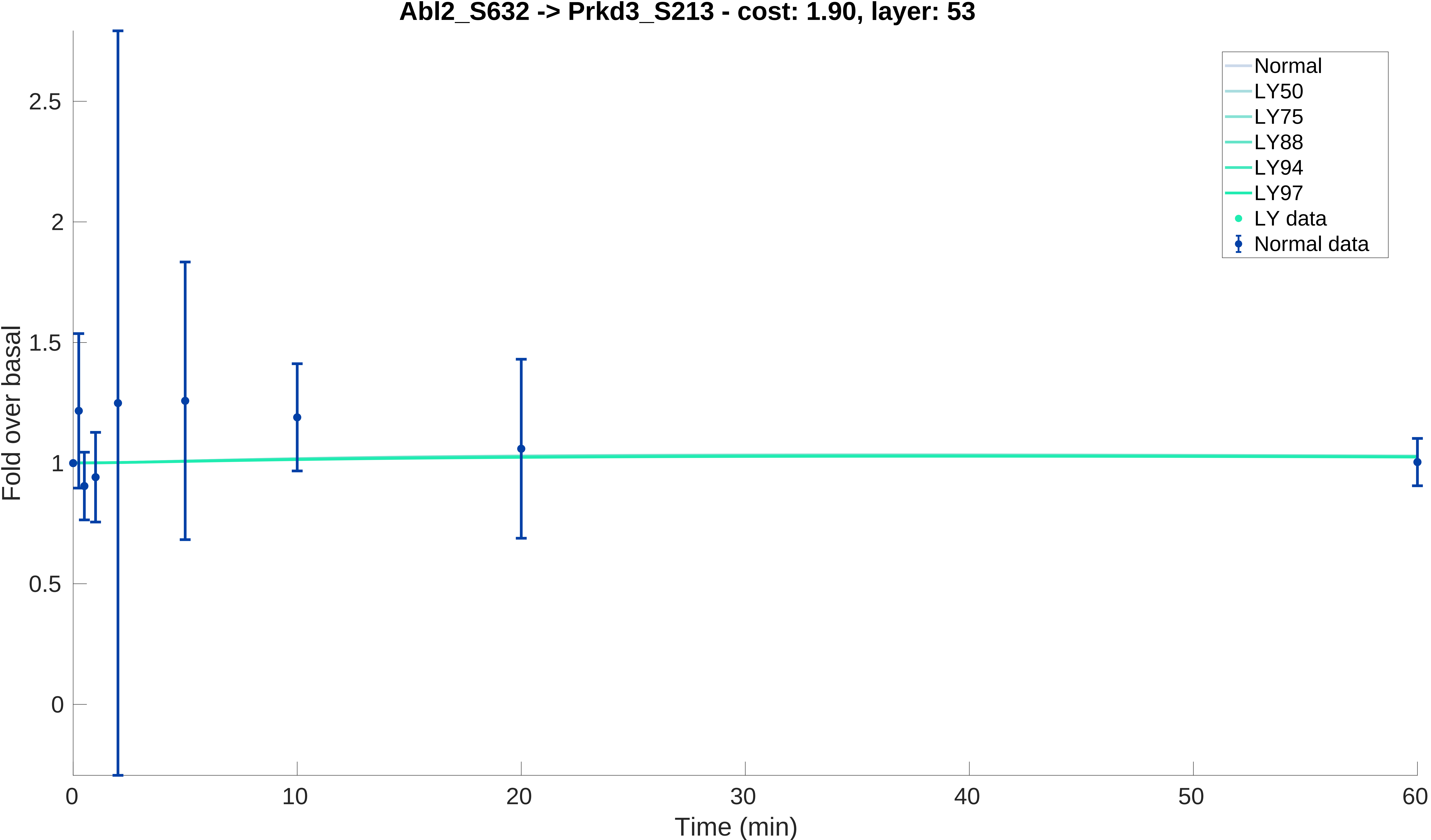

Abl2_S632 -> Prkd3_S213 - cost: 1.90, layer: 53
Fold over basal
Time (min)
0
0.5
1
1.5
2
2.5
0
10
20
30
40
50
60
Normal
LY50
LY75
LY88
LY94
LY97
LY data
Normal data

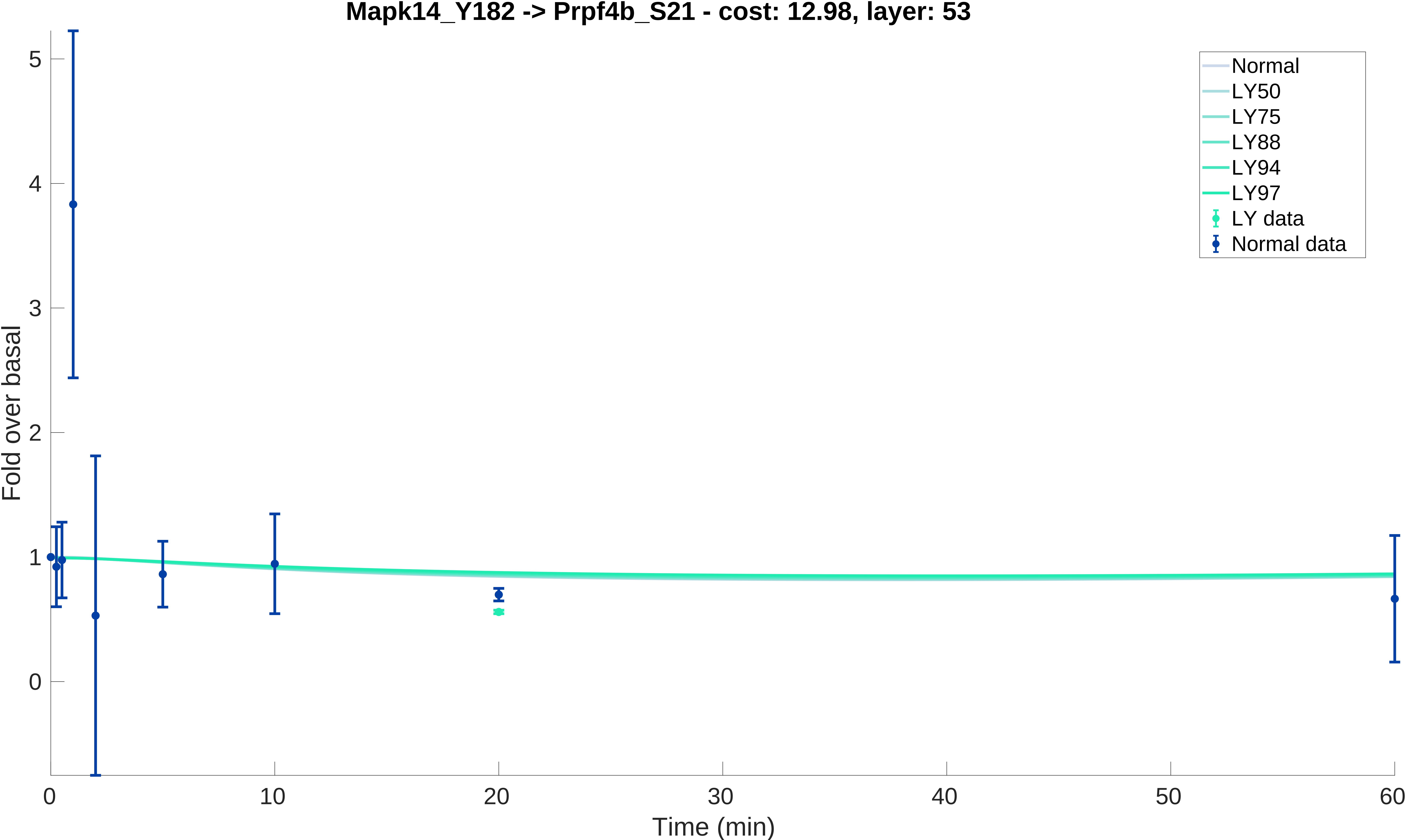

Mapk14_Y182 -> Prpf4b_S21 - cost: 12.98, layer: 53
Fold over basal
Time (min)
0
10
20
30
40
50
60
5
4
3
2
1
0
Normal
LY50
LY75
LY88
LY94
LY97
LY data
Normal data

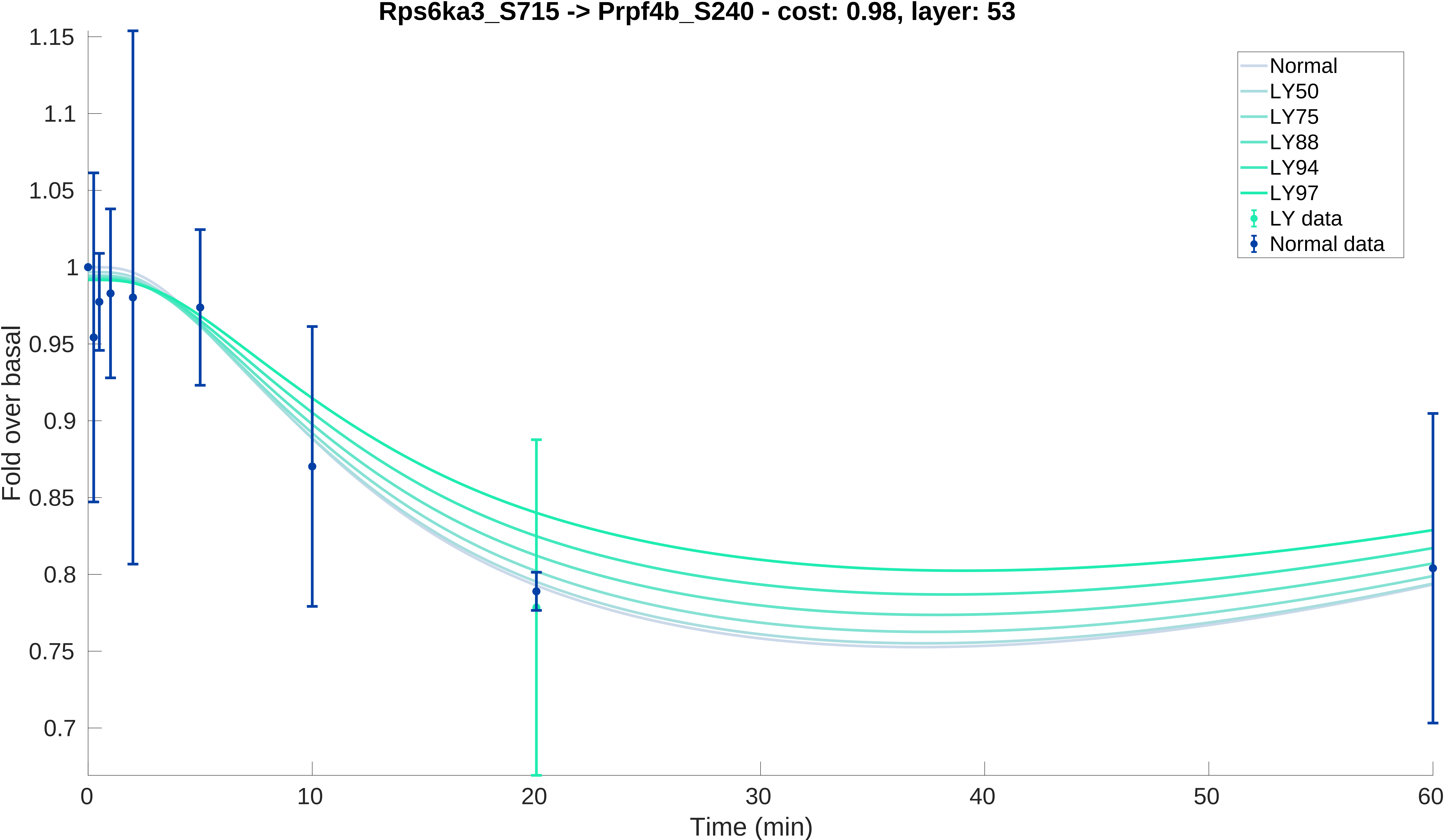
Rps6ka3_S715 -> Prpf4b_S240 - cost: 0.98, layer: 53
Fold over basal
1.15
1.1
1.05
1
0.95
0.9
0.85
0.8
0.75
0.7
0
10
20
30
40
50
60
Time (min)
Normal
LY50
LY75
LY88
LY94
LY97
LY data
Normal data

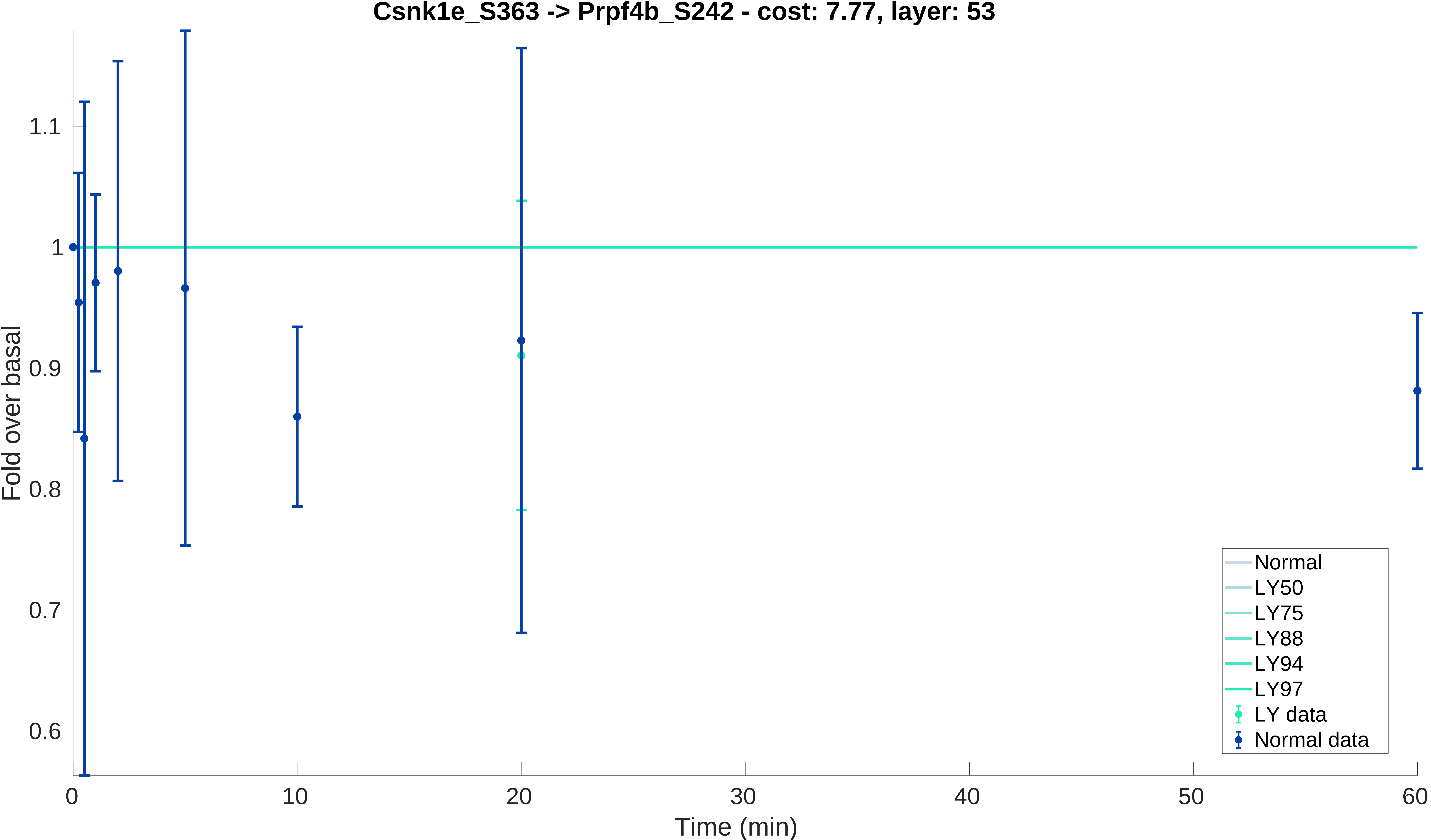

Csnk1e_S363 -> Prpf4b_S242 - cost: 7.77, layer: 53
Fold over basal
1.1
1
0.9
0.8
0.7
0.6
0
10
20
30
40
50
60
Time (min)
Normal
LY50
LY75
LY88
LY94
LY97
LY data
Normal data

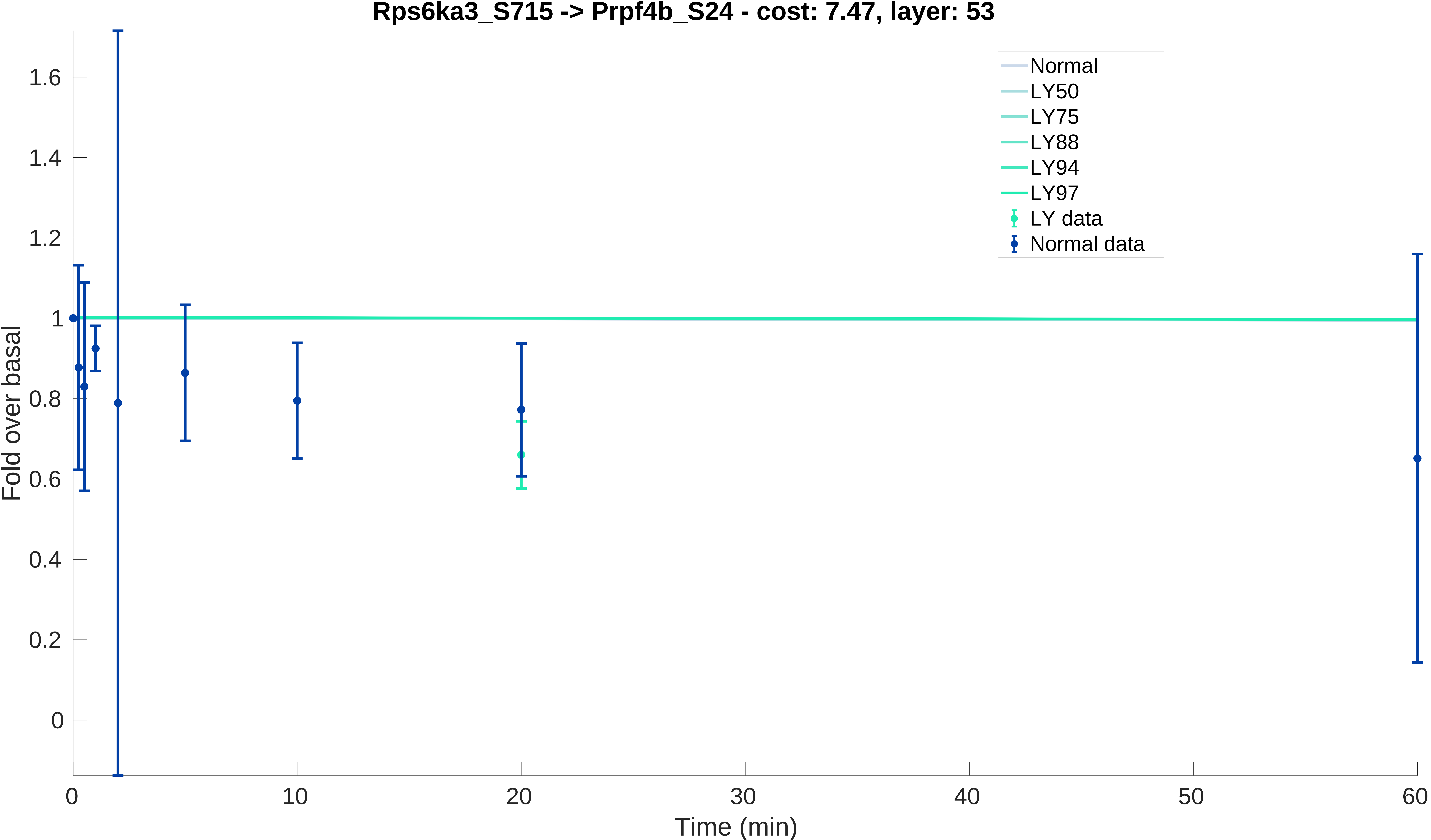
Rps6ka3_S715 -> Prpf4b_S24 - cost: 7.47, layer: 53
Fold over basal
Time (min)
Normal
LY50
LY75
LY88
LY94
LY97
LY data
Normal data
0
0.2
0.4
0.6
0.8
1
1.2
1.4
1.6
0
10
20
30
40
50
60

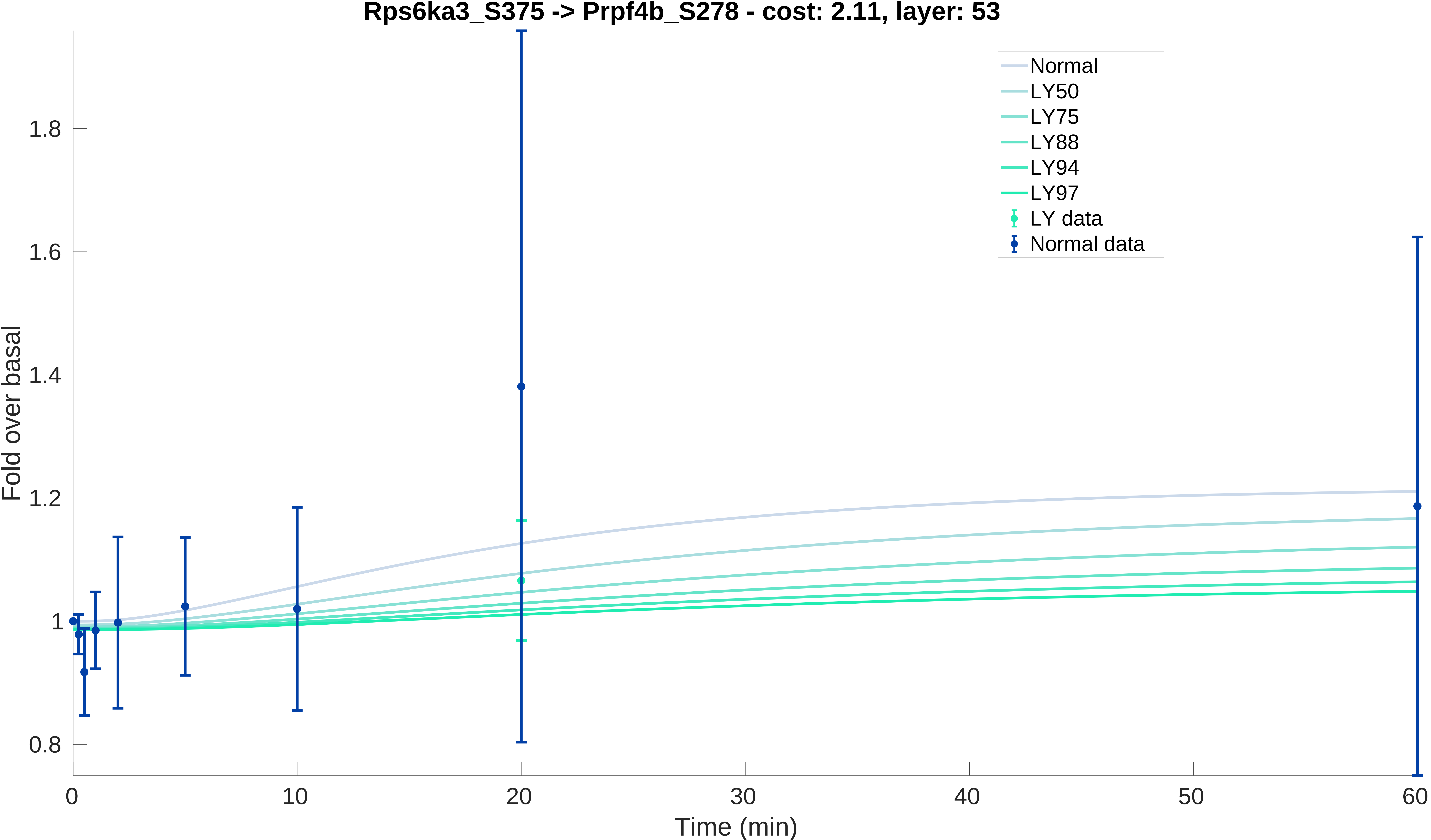

Rps6ka3_S375 -> Prpf4b_S278 - cost: 2.11, layer: 53
Fold over basal
Time (min)
0.8
1
1.2
1.4
1.6
1.8
0
10
20
30
40
50
60
Normal
LY50
LY75
LY88
LY94
LY97
LY data
Normal data

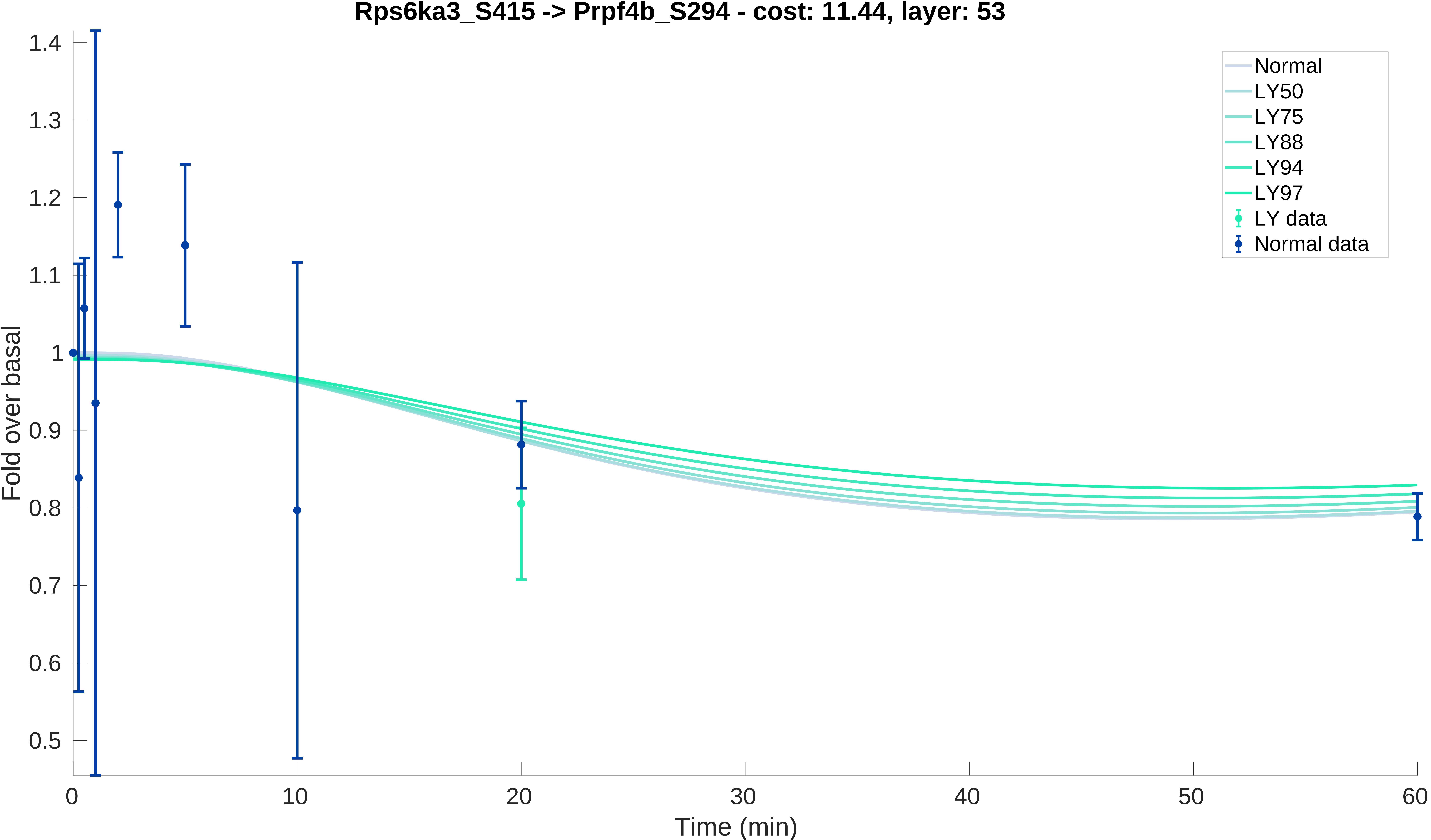

Rps6ka3_S415 -> Prpf4b_S294 - cost: 11.44, layer: 53
Fold over basal
Time (min)
1.4
1.3
1.2
1.1
1
0.9
0.8
0.7
0.6
0.5
0
10
20
30
40
50
60
Normal
LY50
LY75
LY88
LY94
LY97
LY data
Normal data

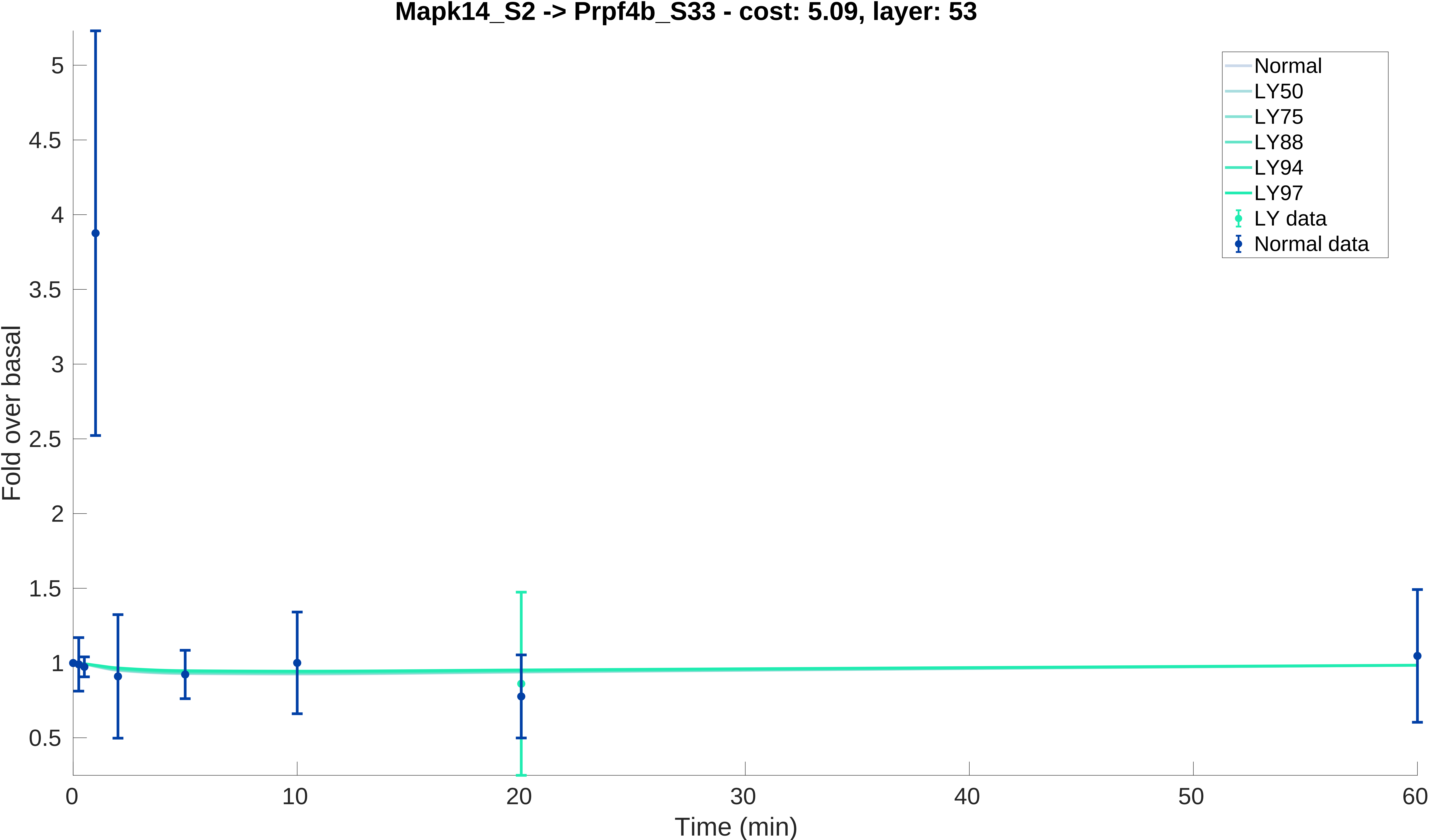

Mapk14_S2 -> Prpf4b_S33 - cost: 5.09, layer: 53
Fold over basal
Time (min)
Normal
LY50
LY75
LY88
LY94
LY97
LY data
Normal data
0.5
1
1.5
2
2.5
3
3.5
4
4.5
5
0
10
20
30
40
50
60

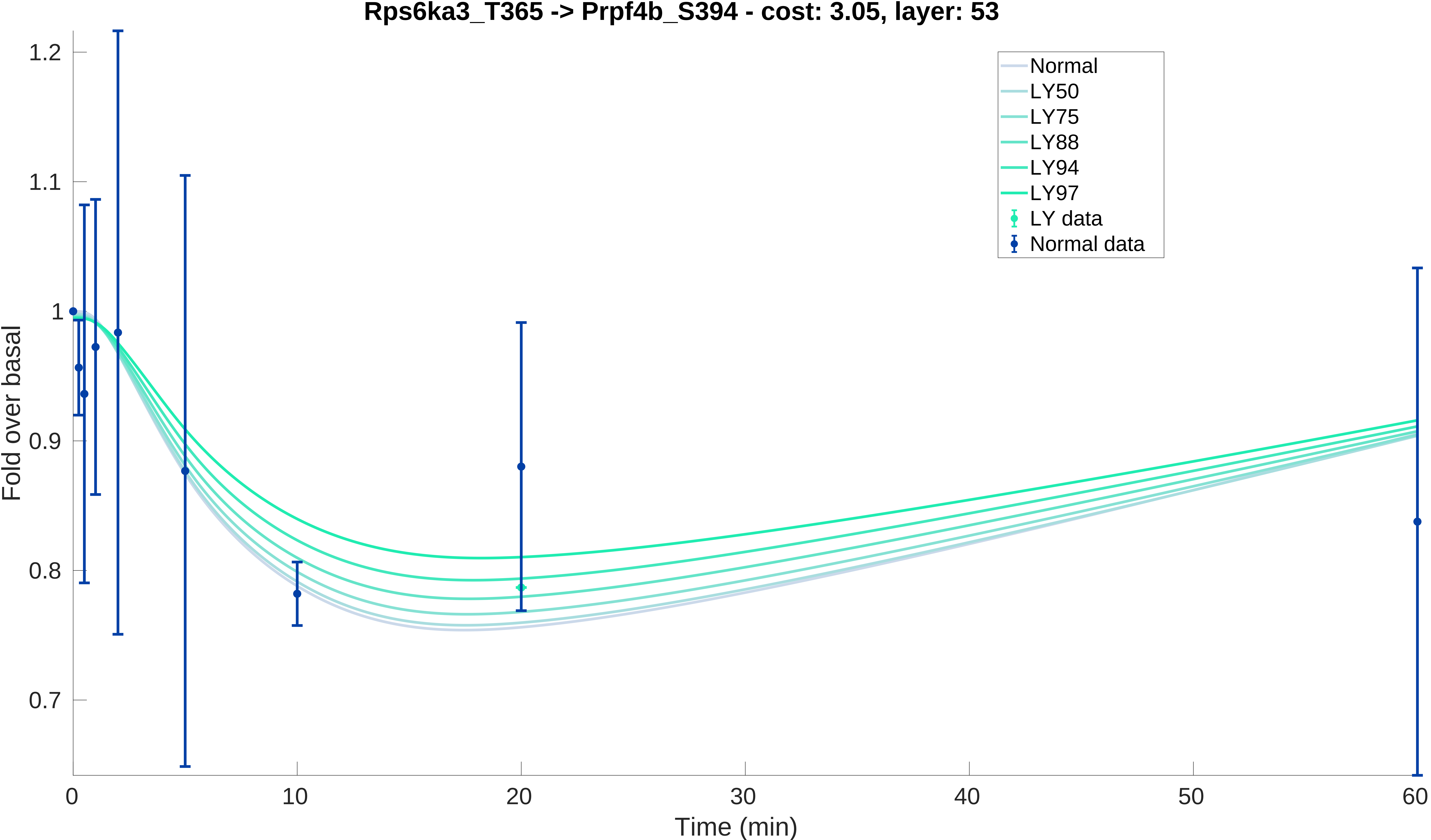

Rps6ka3_T365 -> Prpf4b_S394 - cost: 3.05, layer: 53
Normal
LY50
LY75
LY88
LY94
LY97
LY data
Normal data
1.2
1.1
1
0.9
0.8
0.7
Fold over basal
0
10
20
30
40
50
60
Time (min)

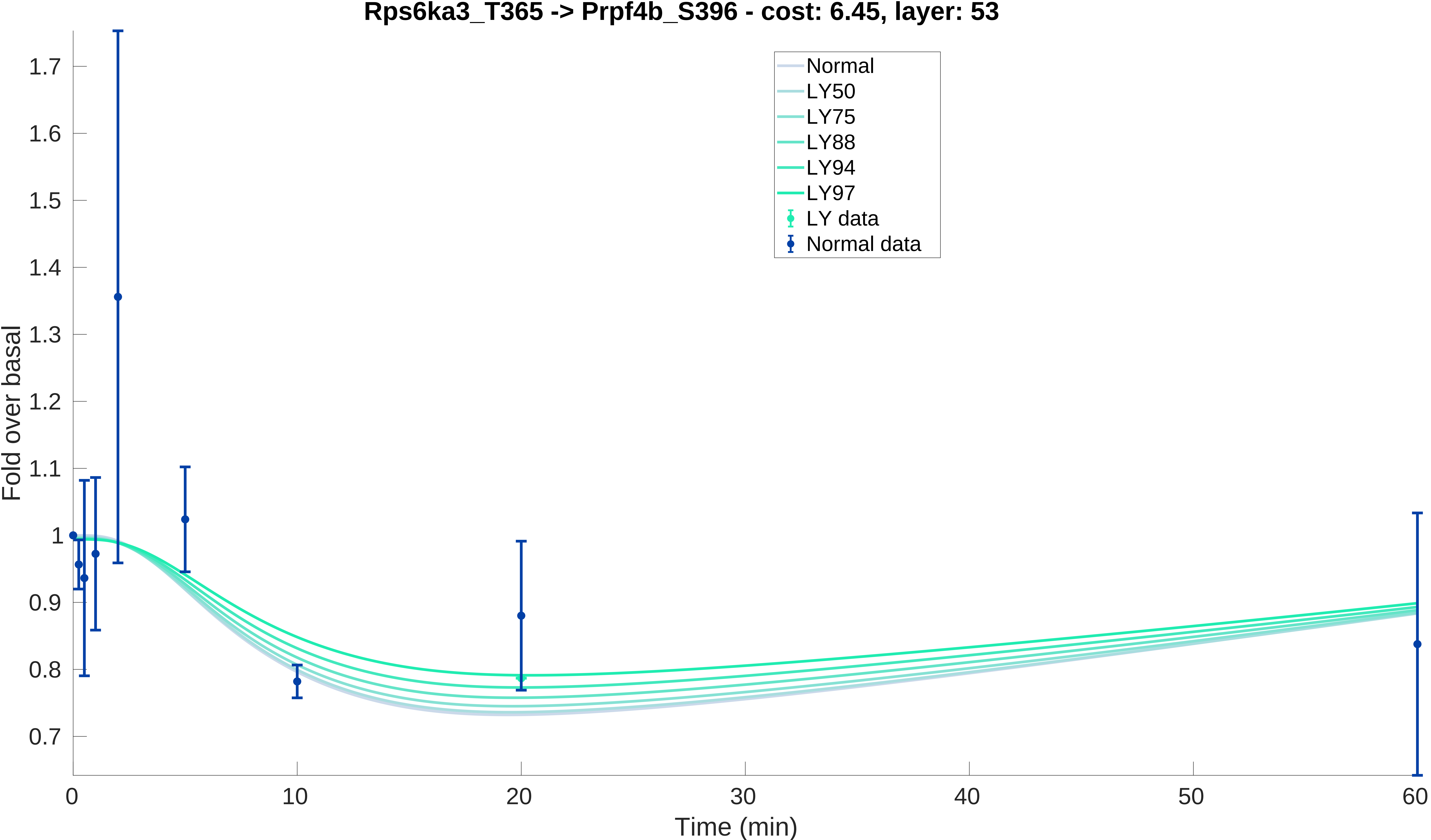
Rps6ka3_T365 -> Prpf4b_S396 - cost: 6.45, layer: 53
Normal
LY50
LY75
LY88
LY94
LY97
LY data
Normal data
Fold over basal
1.7
1.6
1.5
1.4
1.3
1.2
1.1
1
0.9
0.8
0.7
0
10
20
30
40
50
60
Time (min)

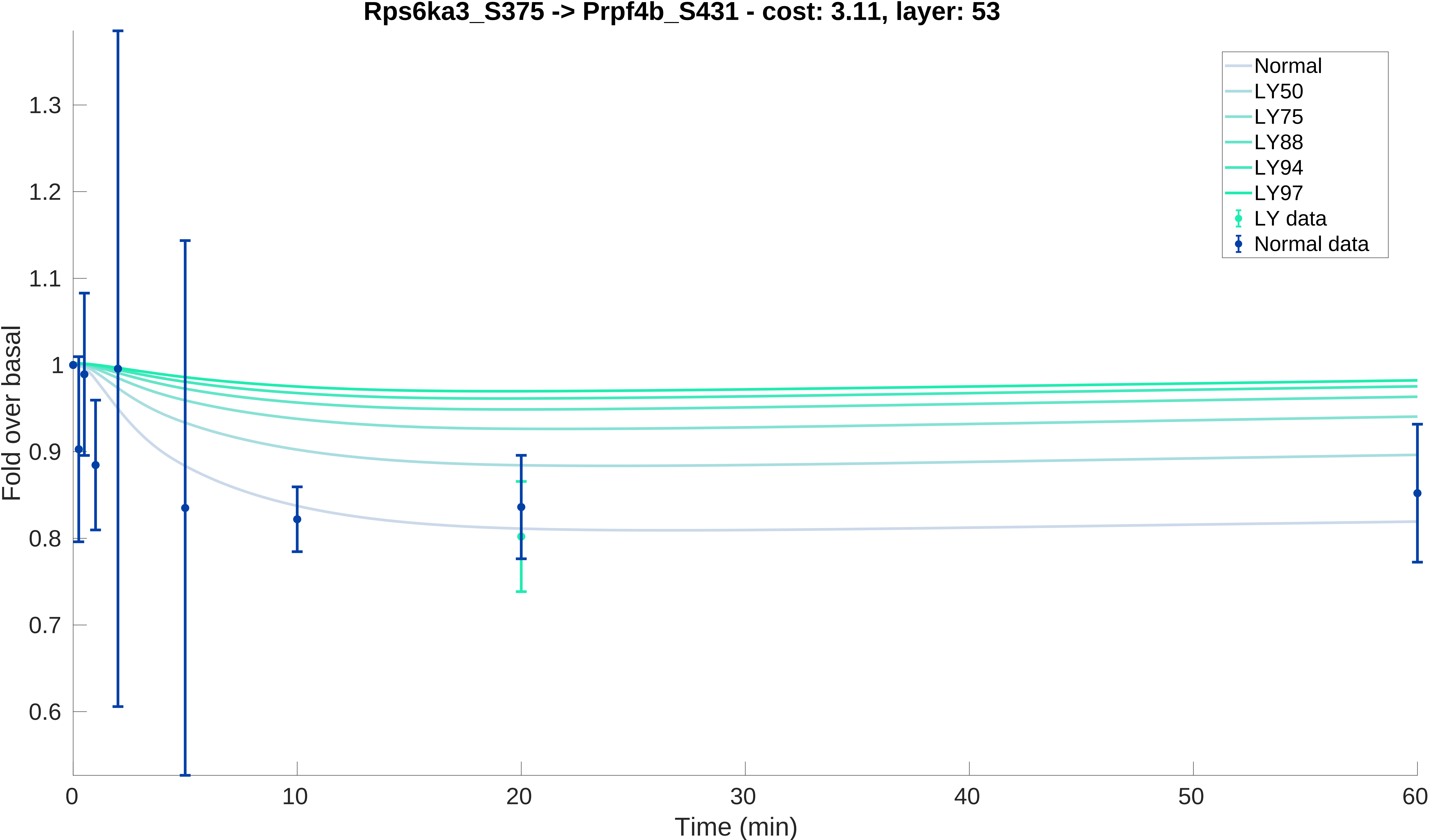
Rps6ka3_S375 -> Prpf4b_S431 - cost: 3.11, layer: 53
Fold over basal
Time (min)
1.3
1.2
1.1
1
0.9
0.8
0.7
0.6
0
10
20
30
40
50
60
Normal
LY50
LY75
LY88
LY94
LY97
LY data
Normal data

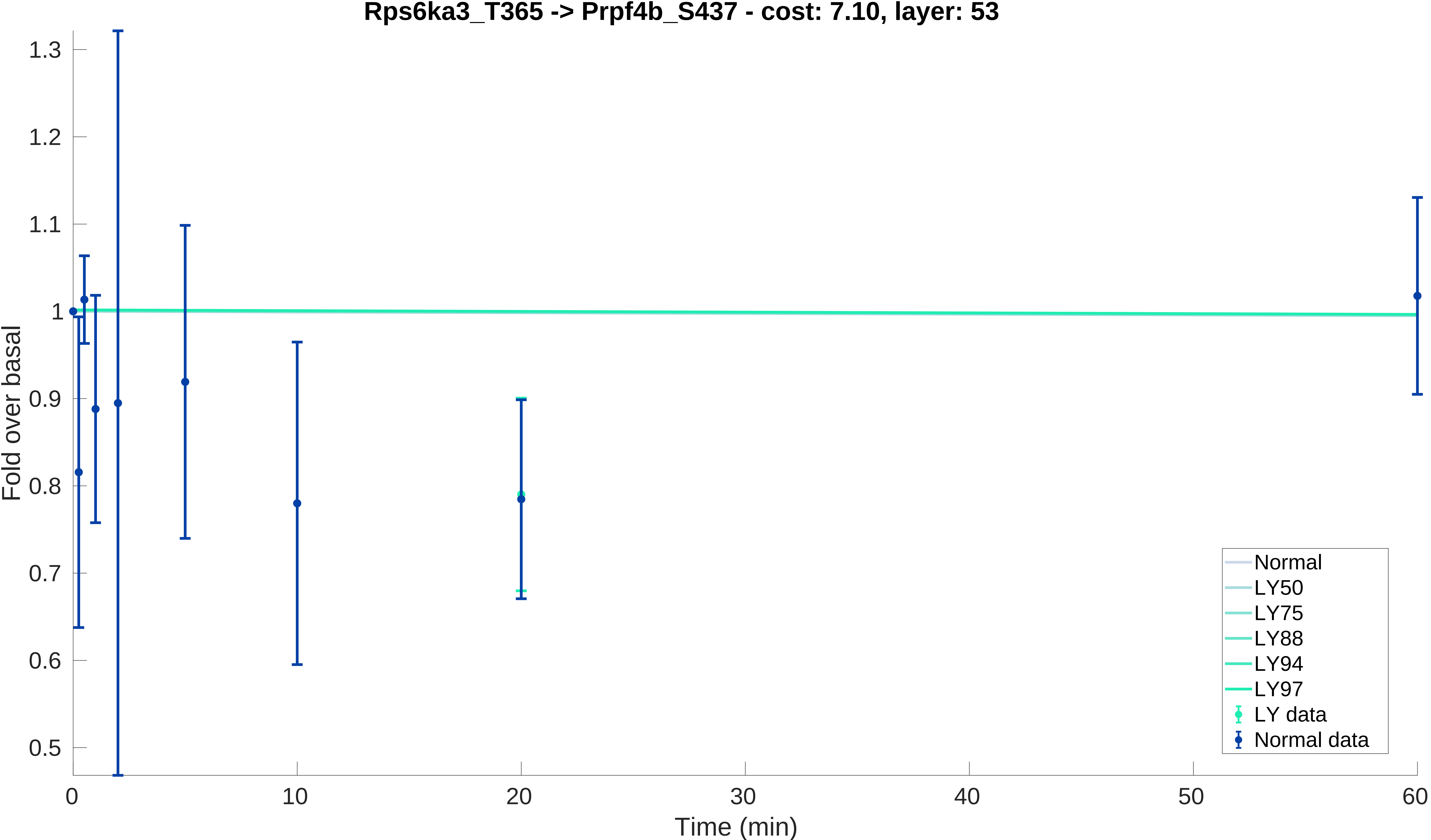

Rps6ka3_T365 -> Prpf4b_S437 - cost: 7.10, layer: 53
Fold over basal
1.3
1.2
1.1
1
0.9
0.8
0.7
0.6
0.5
Time (min)
0
10
20
30
40
50
60
Normal
LY50
LY75
LY88
LY94
LY97
LY data
Normal data

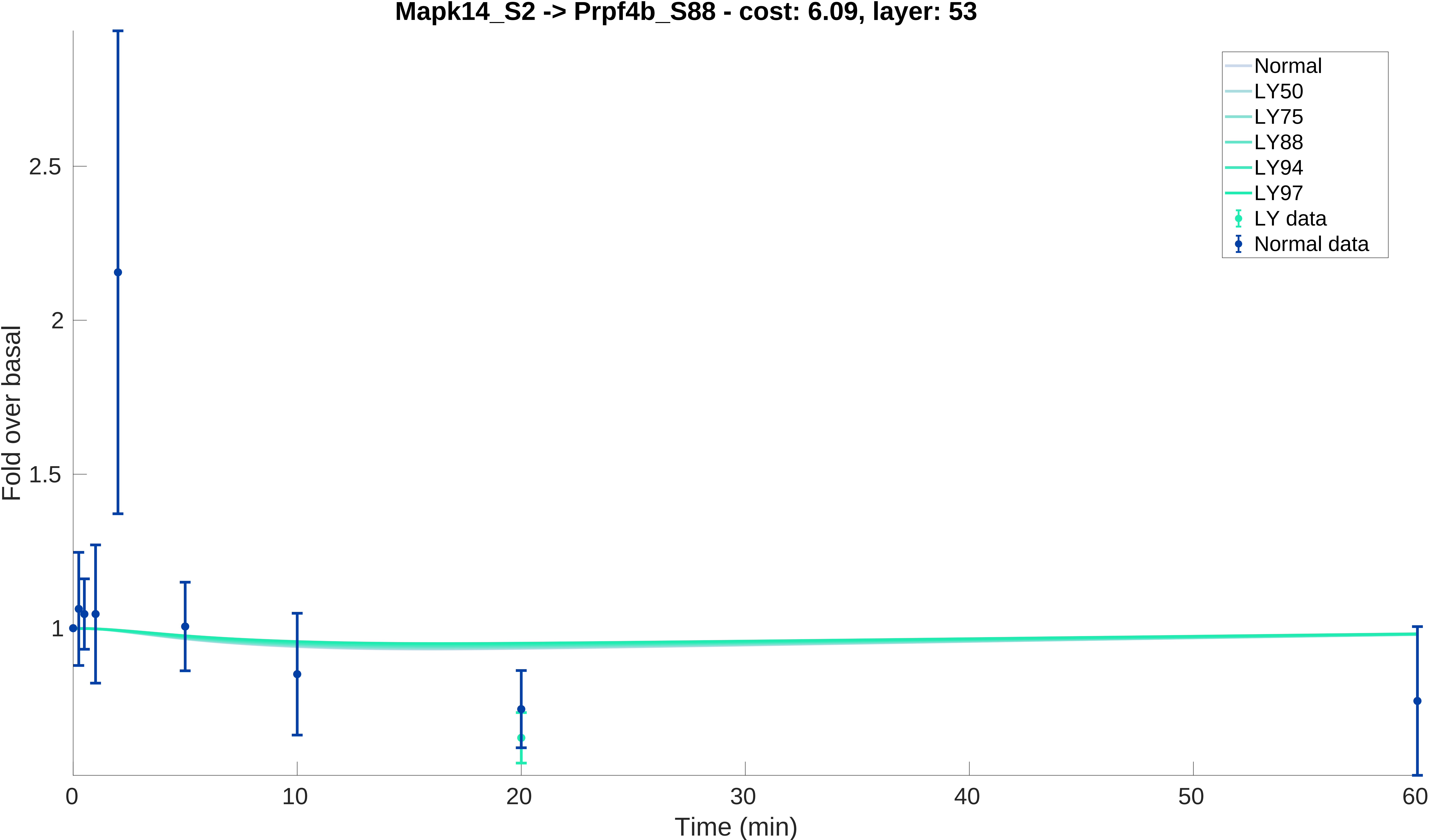

Mapk14_S2 -> Prpf4b_S88 - cost: 6.09, layer: 53
Normal
LY50
LY75
LY88
LY94
LY97
LY data
Normal data
Fold over basal
2.5
2
1.5
1
0
10
20
30
40
50
60
Time (min)

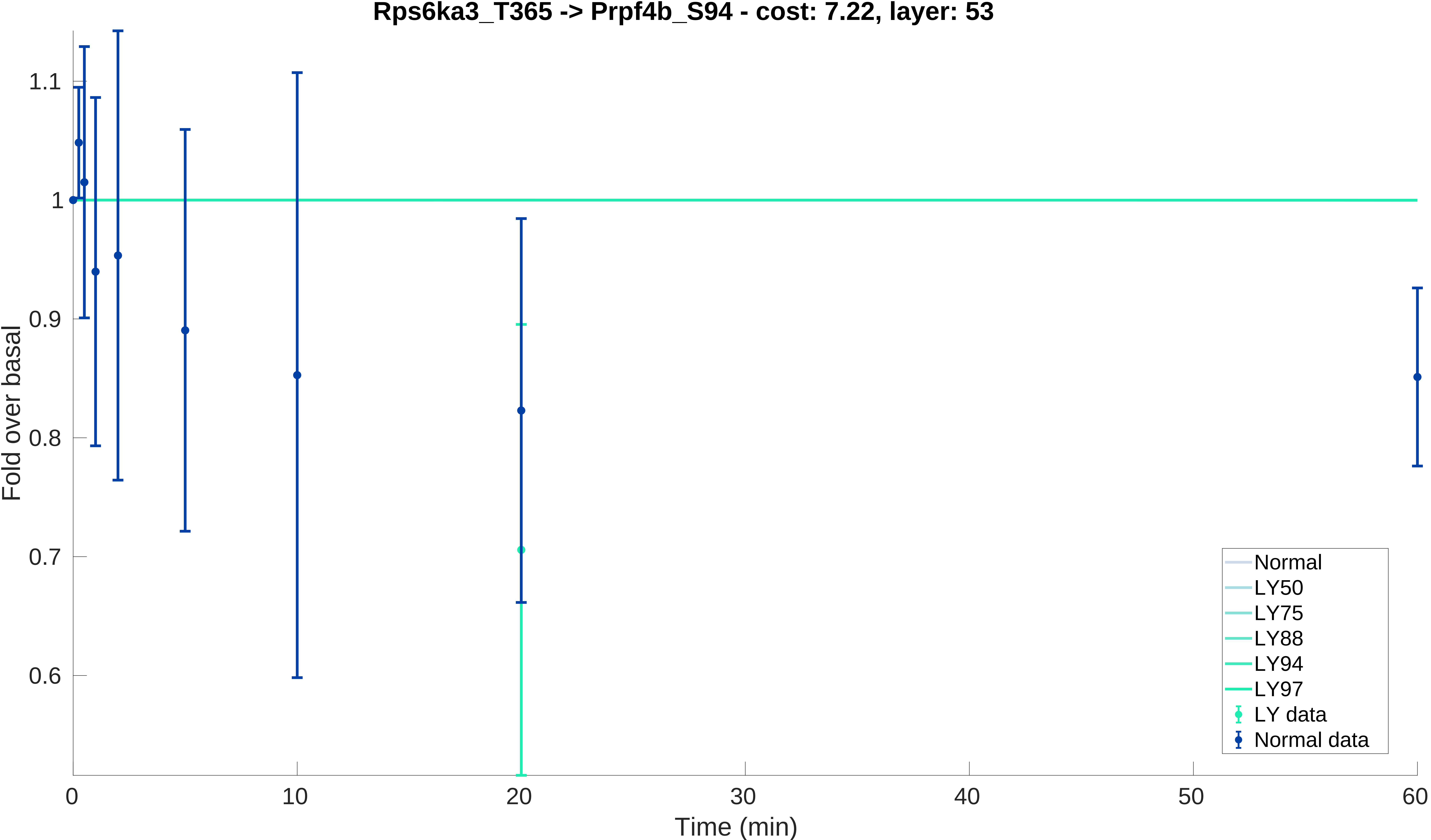

Rps6ka3_T365 -> Prpf4b_S94 - cost: 7.22, layer: 53
Fold over basal
1.1
1
0.9
0.8
0.7
0.6
0
10
20
30
40
50
60
Time (min)
Normal
LY50
LY75
LY88
LY94
LY97
LY data
Normal data

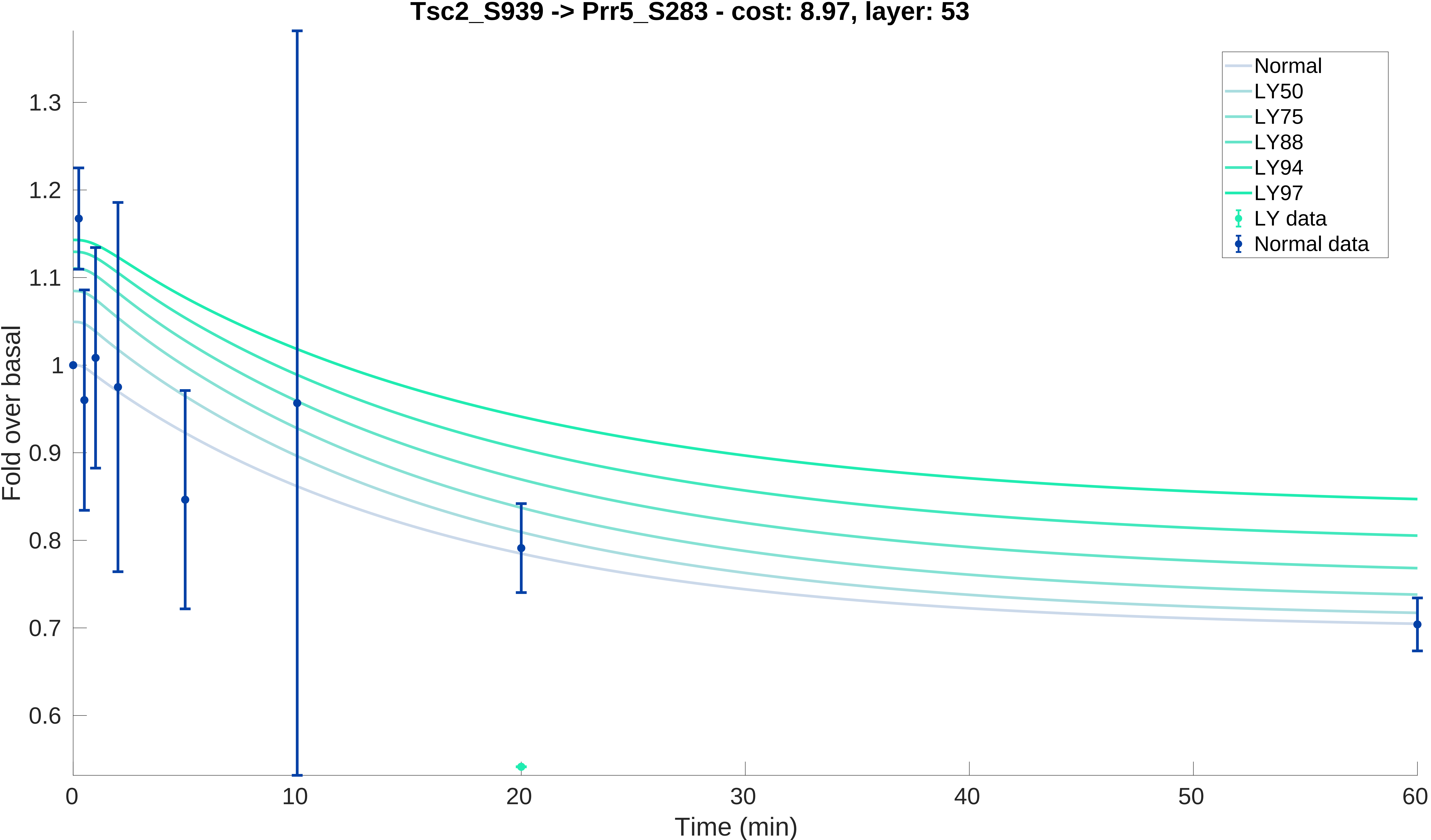

Tsc2_S939 -> Prr5_S283 - cost: 8.97, layer: 53
Normal
LY50
LY75
LY88
LY94
LY97
LY data
Normal data
Fold over basal
1.3
1.2
1.1
1
0.9
0.8
0.7
0.6
0
10
20
30
40
50
60
Time (min)

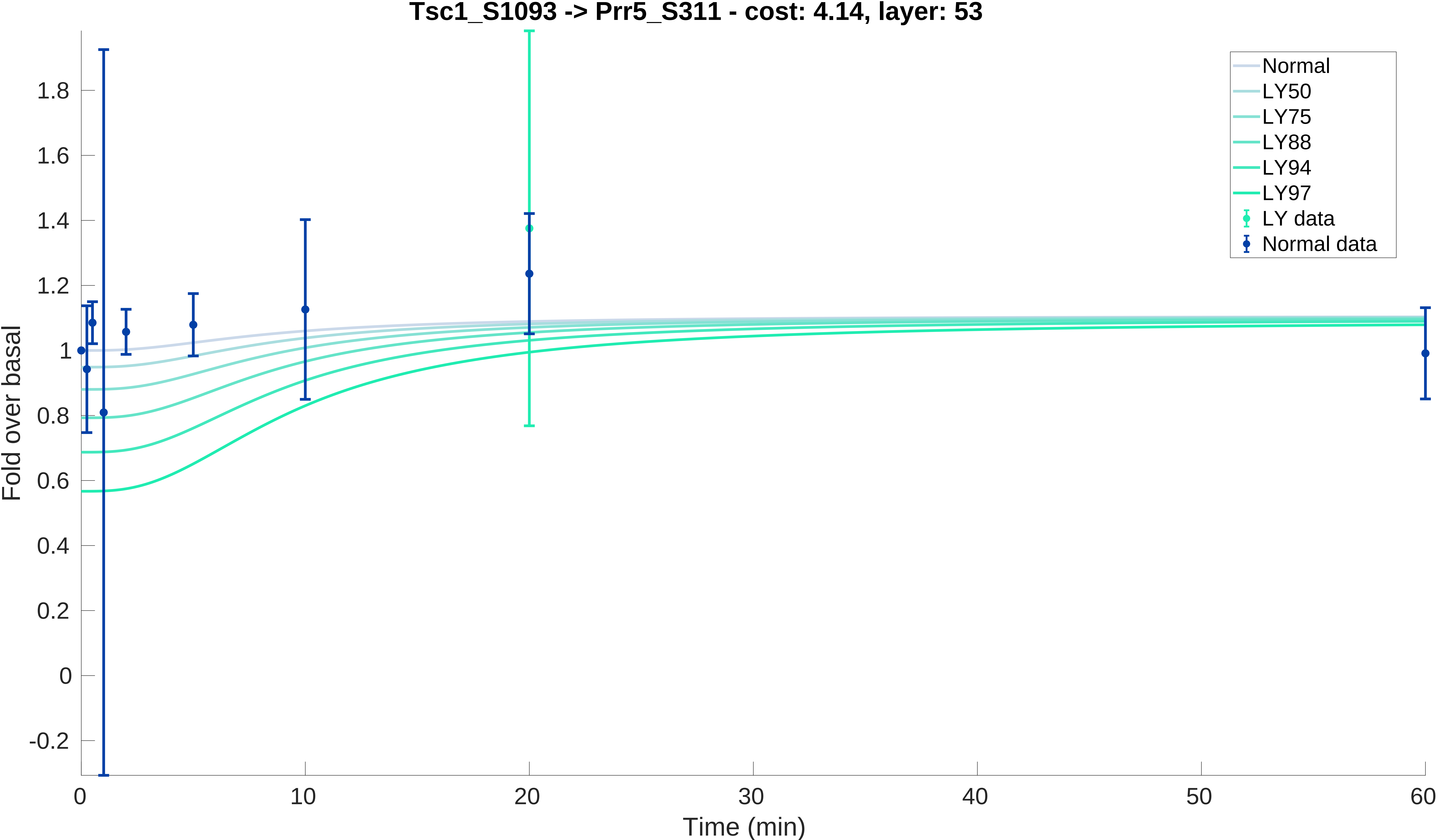
Tsc1_S1093 -> Prr5_S311 - cost: 4.14, layer: 53
Fold over basal
Time (min)
1.8
1.6
1.4
1.2
1
0.8
0.6
0.4
0.2
0
-0.2
0
10
20
30
40
50
60
Normal
LY50
LY75
LY88
LY94
LY97
LY data
Normal data

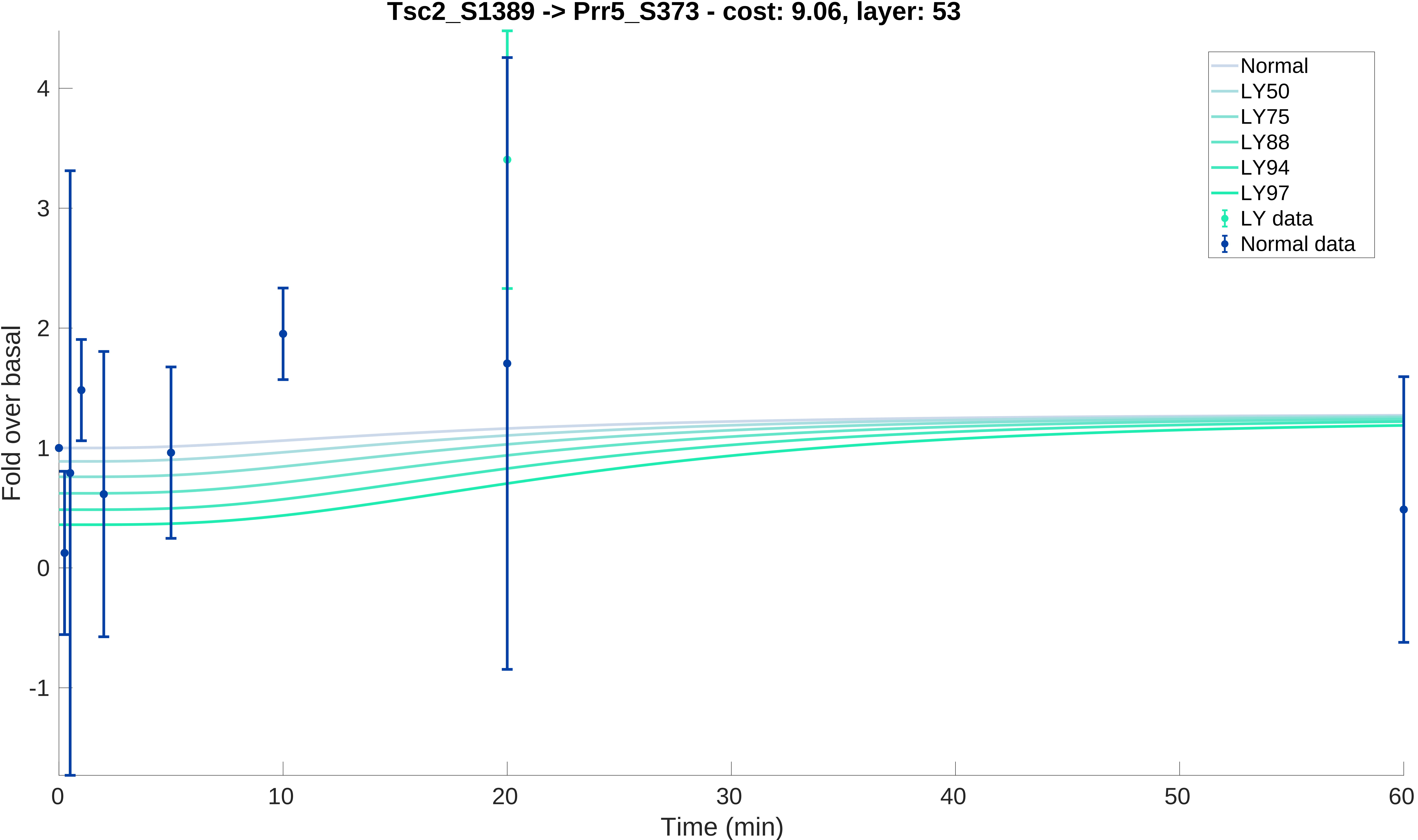
Tsc2_S1389 -> Prr5_S373 - cost: 9.06, layer: 53
Fold over basal
Time (min)
4
3
2
1
0
-1
0
10
20
30
40
50
60
Normal
LY50
LY75
LY88
LY94
LY97
LY data
Normal data

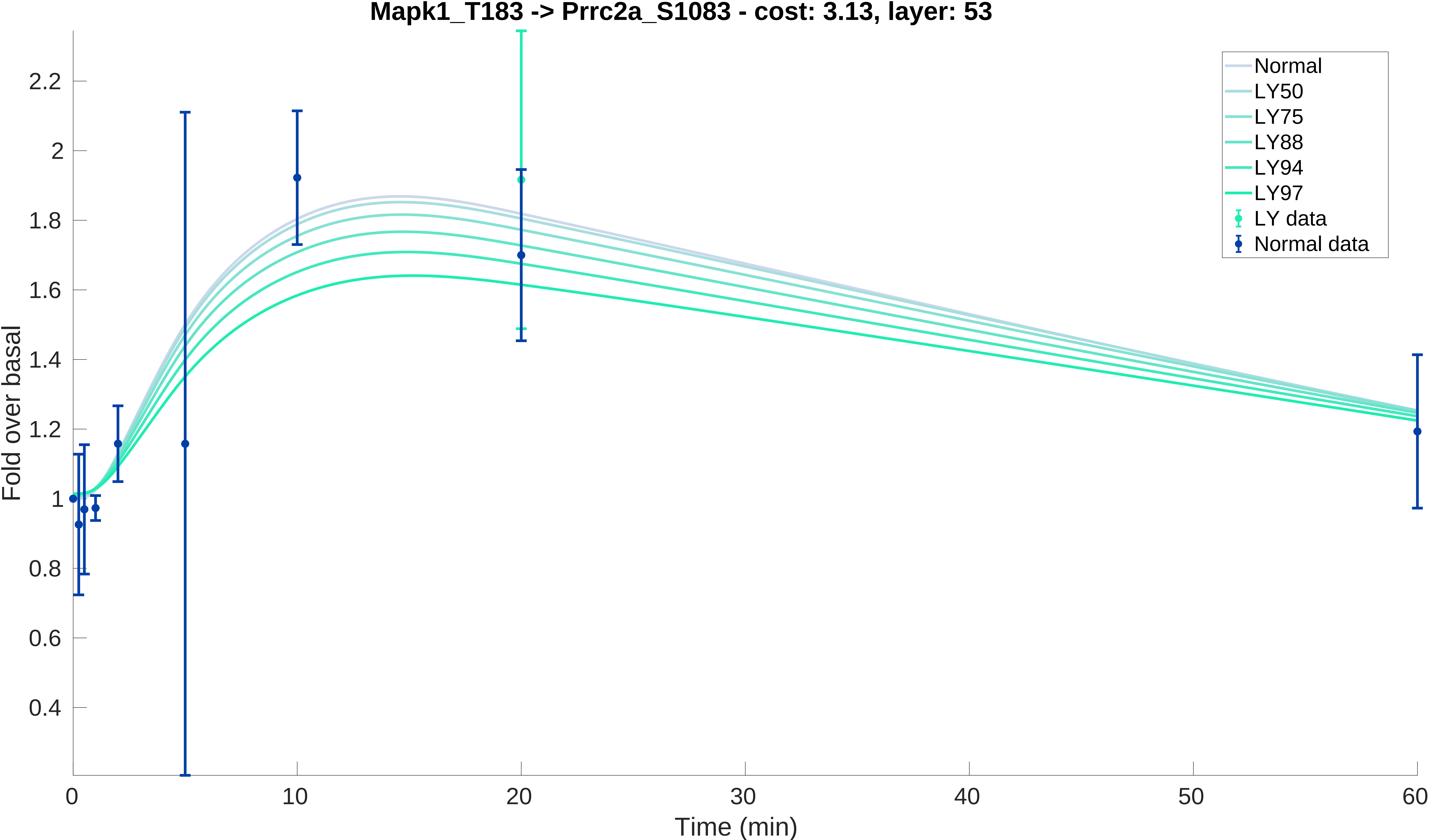

Mapk1_T183 -> Prrc2a_S1083 - cost: 3.13, layer: 53
Fold over basal
Time (min)
Normal
LY50
LY75
LY88
LY94
LY97
LY data
Normal data
2.2
2
1.8
1.6
1.4
1.2
1
0.8
0.6
0.4
0
10
20
30
40
50
60

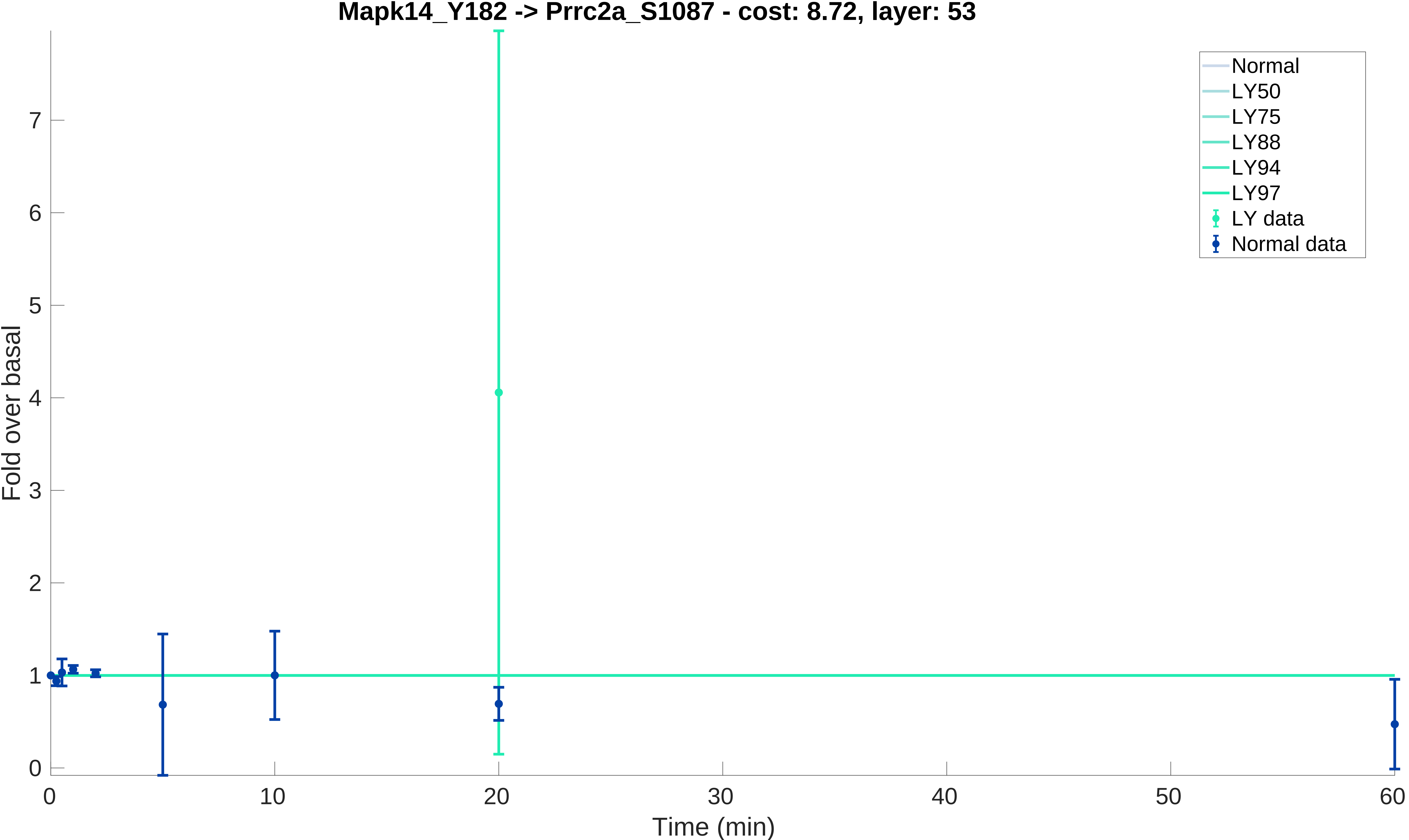

Mapk14_Y182 -> Prrc2a_S1087 - cost: 8.72, layer: 53
Fold over basal
Time (min)
0
1
2
3
4
5
6
7
0
10
20
30
40
50
60
Normal
LY50
LY75
LY88
LY94
LY97
LY data
Normal data

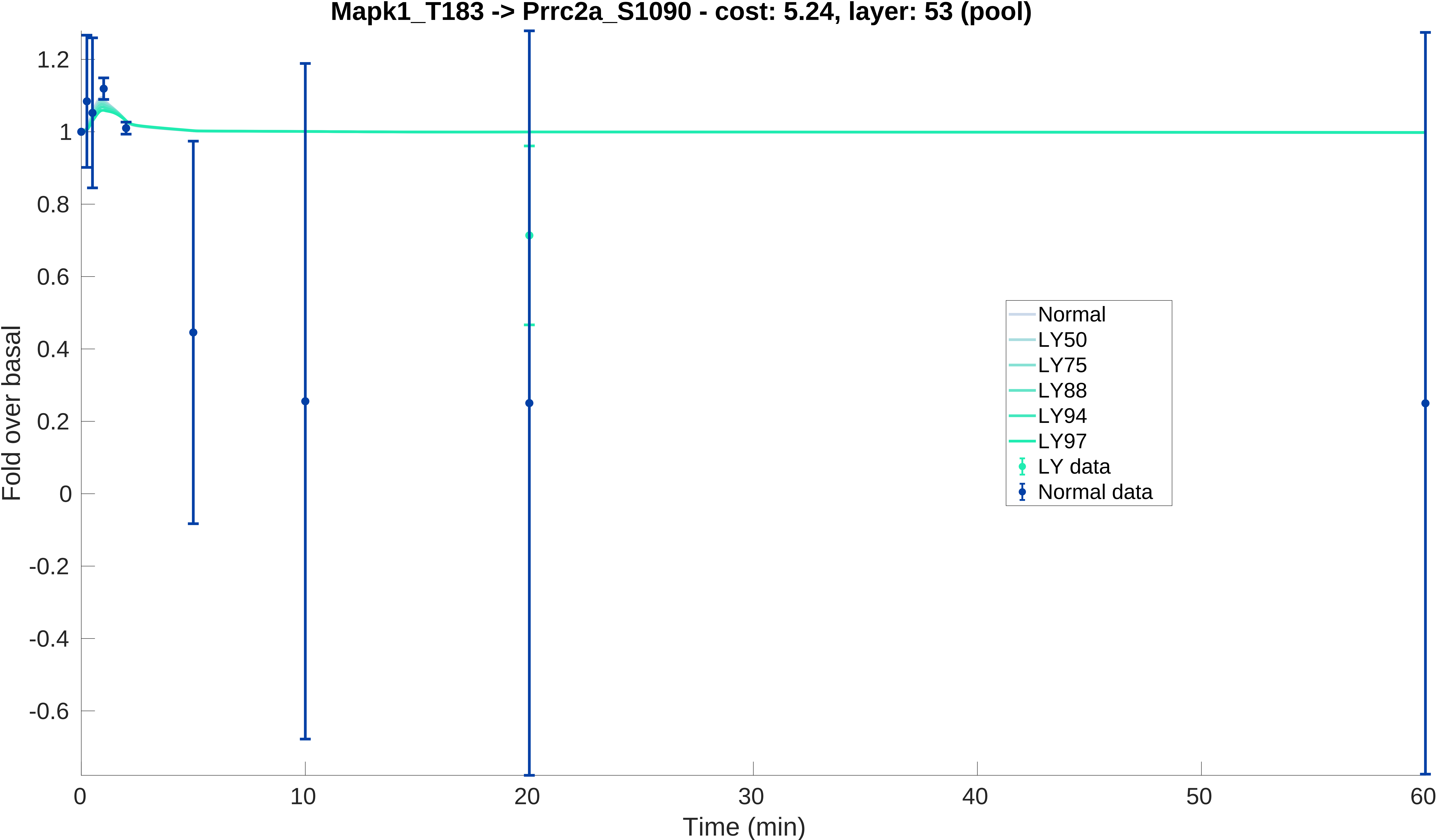

Mapk1_T183 -> Prrc2a_S1090 - cost: 5.24, layer: 53 (pool)
Fold over basal
Time (min)
1.2
1
0.8
0.6
0.4
0.2
0
-0.2
-0.4
-0.6
0
10
20
30
40
50
60
Normal
LY50
LY75
LY88
LY94
LY97
LY data
Normal data

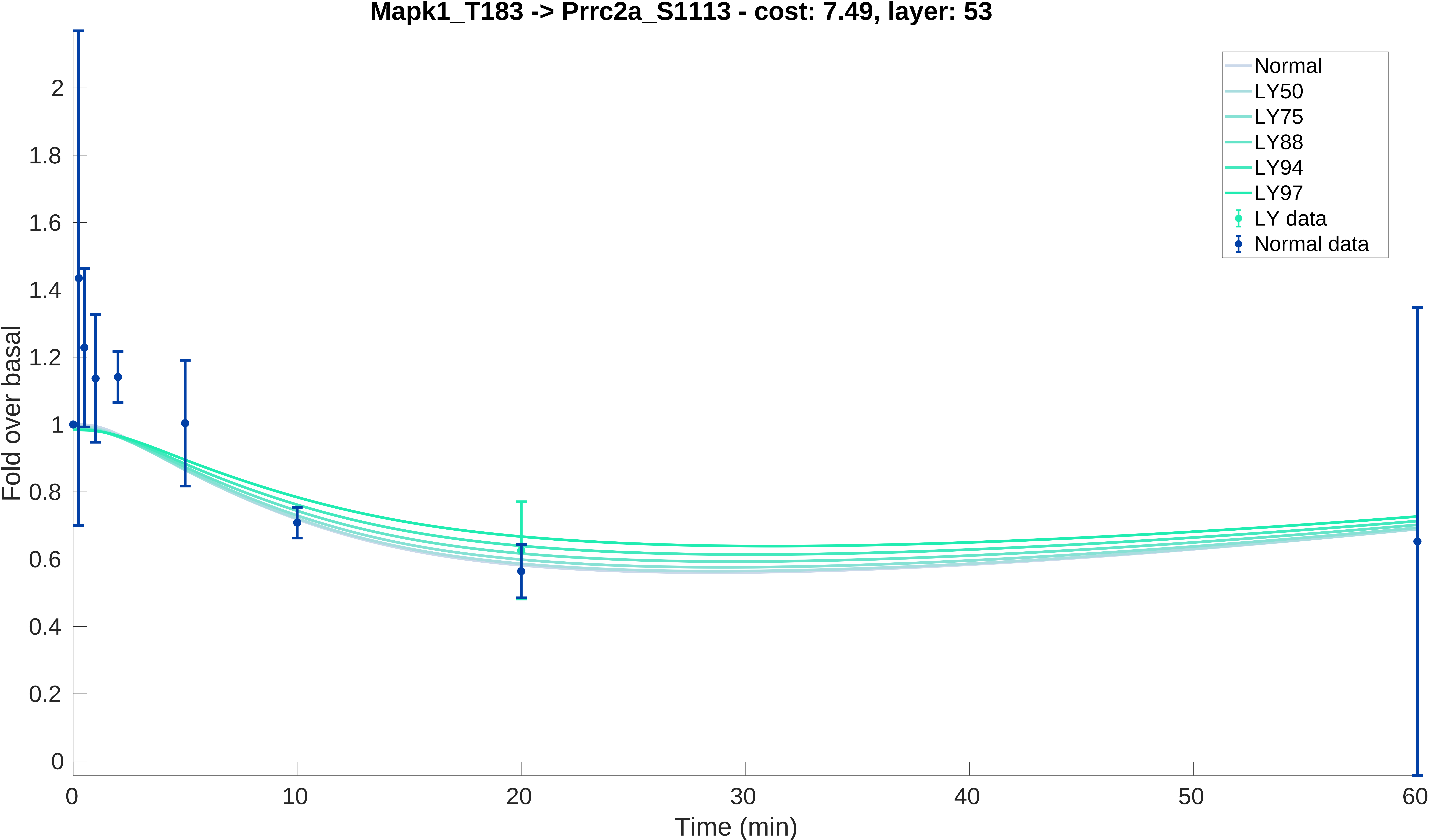
Mapk1_T183 -> Prrc2a_S1113 - cost: 7.49, layer: 53
Fold over basal
Time (min)
0
0.2
0.4
0.6
0.8
1
1.2
1.4
1.6
1.8
2
0
10
20
30
40
50
60
Normal
LY50
LY75
LY88
LY94
LY97
LY data
Normal data

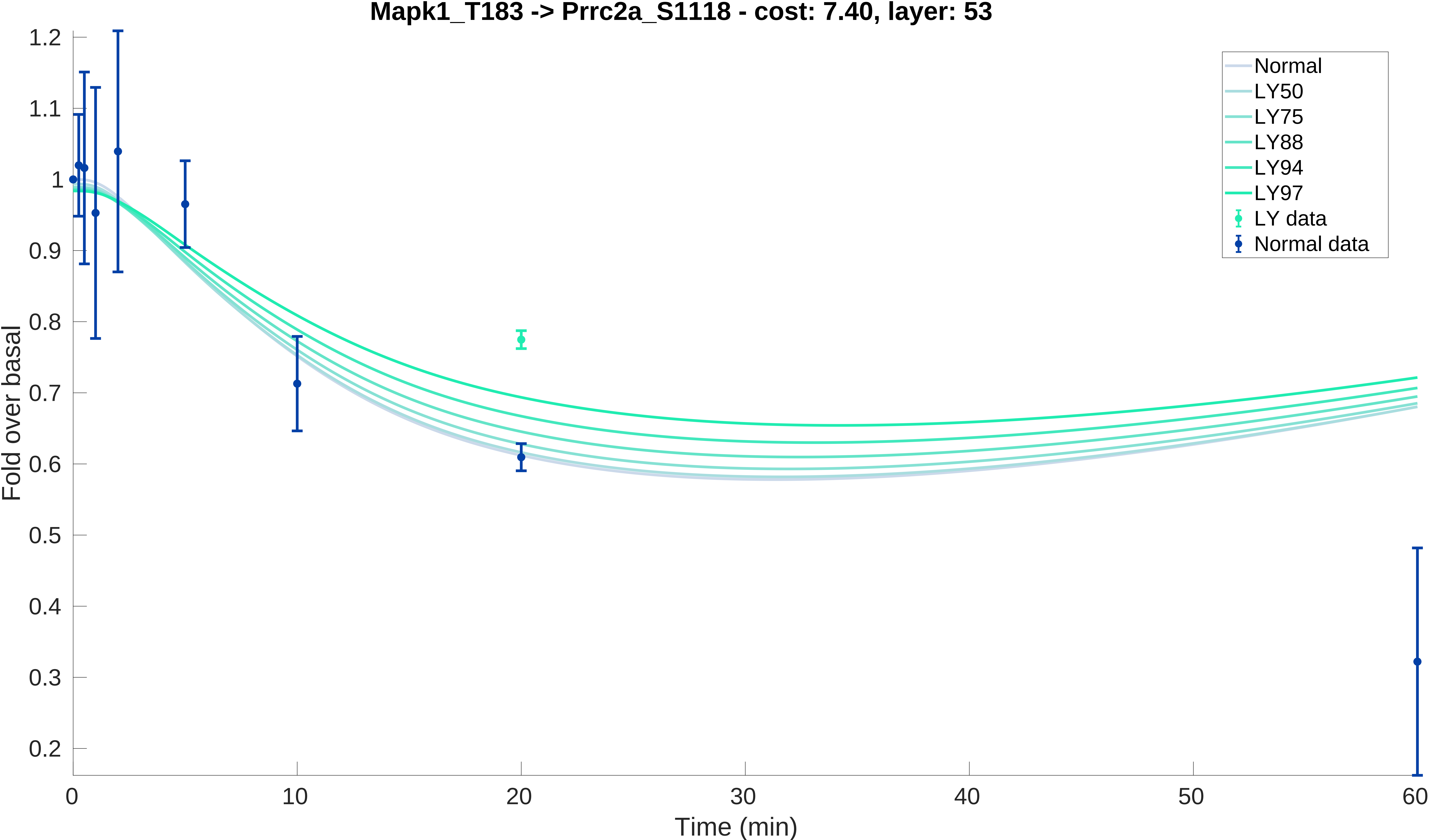
Mapk1_T183 -> Prrc2a_S1118 - cost: 7.40, layer: 53
Fold over basal
Time (min)
1.2
1.1
1
0.9
0.8
0.7
0.6
0.5
0.4
0.3
0.2
0
10
20
30
40
50
60
Normal
LY50
LY75
LY88
LY94
LY97
LY data
Normal data

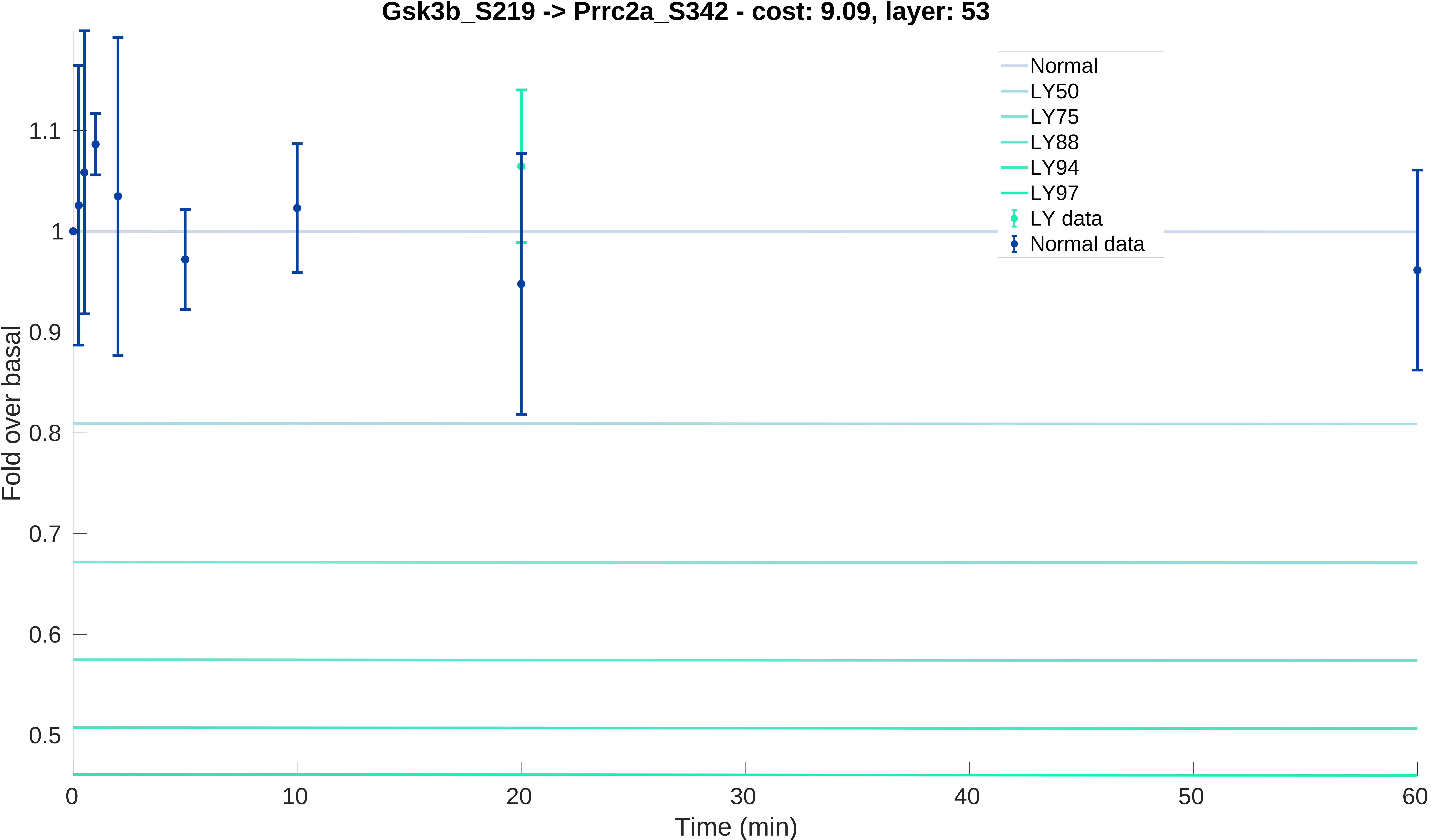

Gsk3b_S219 -> Prrc2a_S342 - cost: 9.09, layer: 53
Normal
LY50
LY75
LY88
LY94
LY97
LY data
Normal data
Fold over basal
1.1
1
0.9
0.8
0.7
0.6
0.5
0
10
20
30
40
50
60
Time (min)

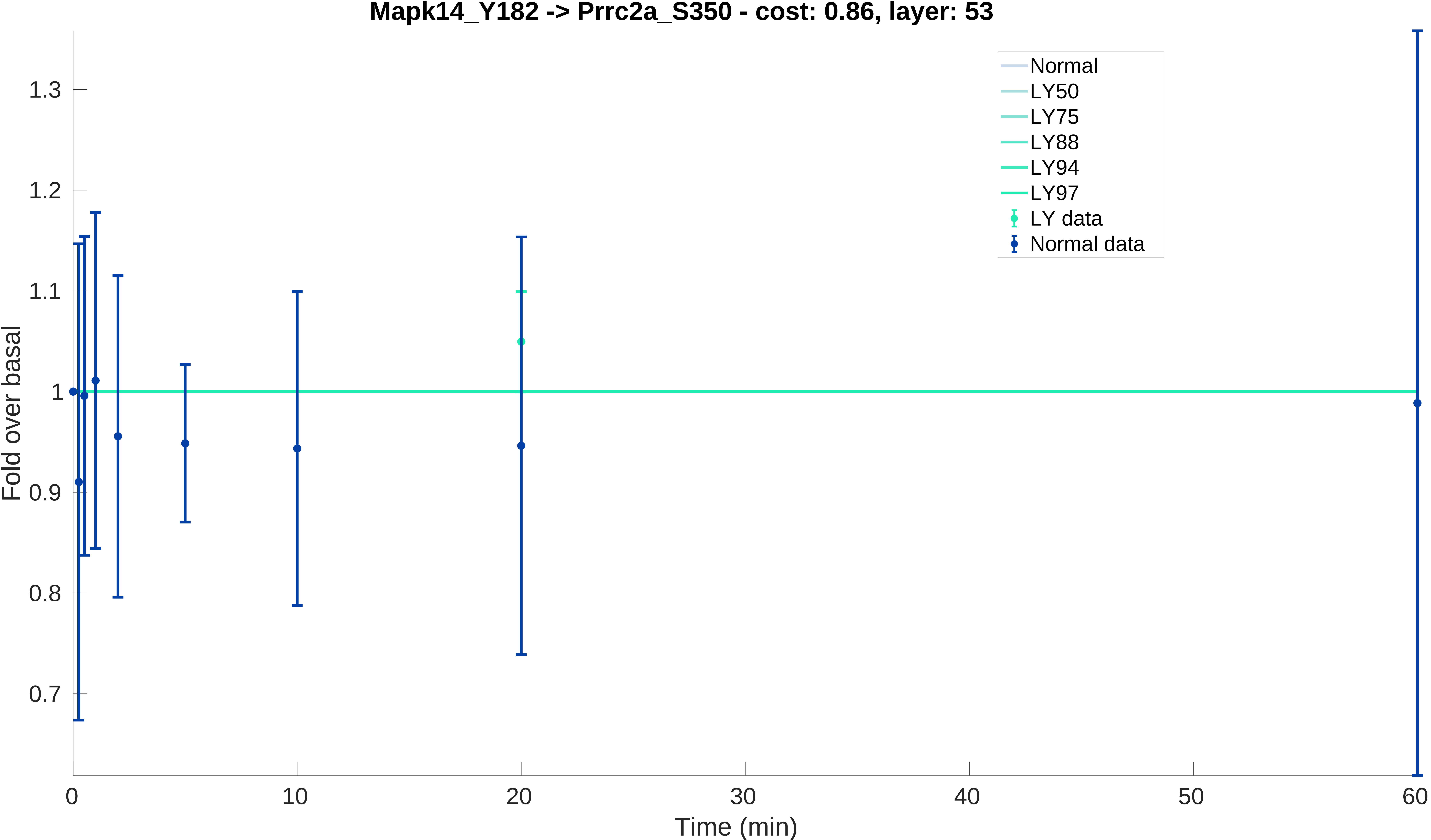

Mapk14_Y182 -> Prrc2a_S350 - cost: 0.86, layer: 53
Normal
LY50
LY75
LY88
LY94
LY97
LY data
Normal data
Fold over basal
1.3
1.2
1.1
1
0.9
0.8
0.7
0
10
20
30
40
50
60
Time (min)

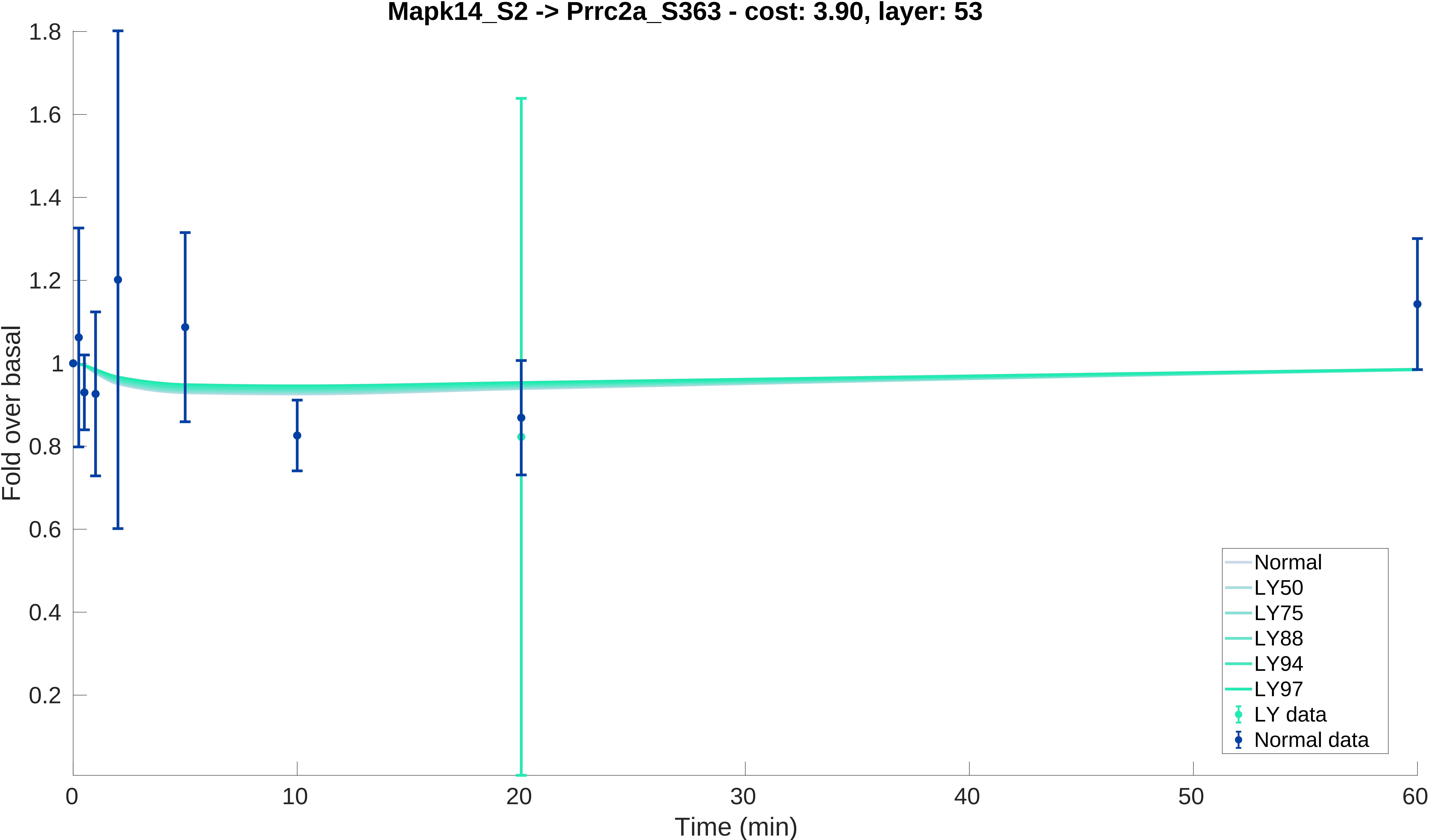

Mapk14_S2 -> Prrc2a_S363 - cost: 3.90, layer: 53
Fold over basal
Time (min)
1.8
1.6
1.4
1.2
1
0.8
0.6
0.4
0.2
0
10
20
30
40
50
60
Normal
LY50
LY75
LY88
LY94
LY97
LY data
Normal data

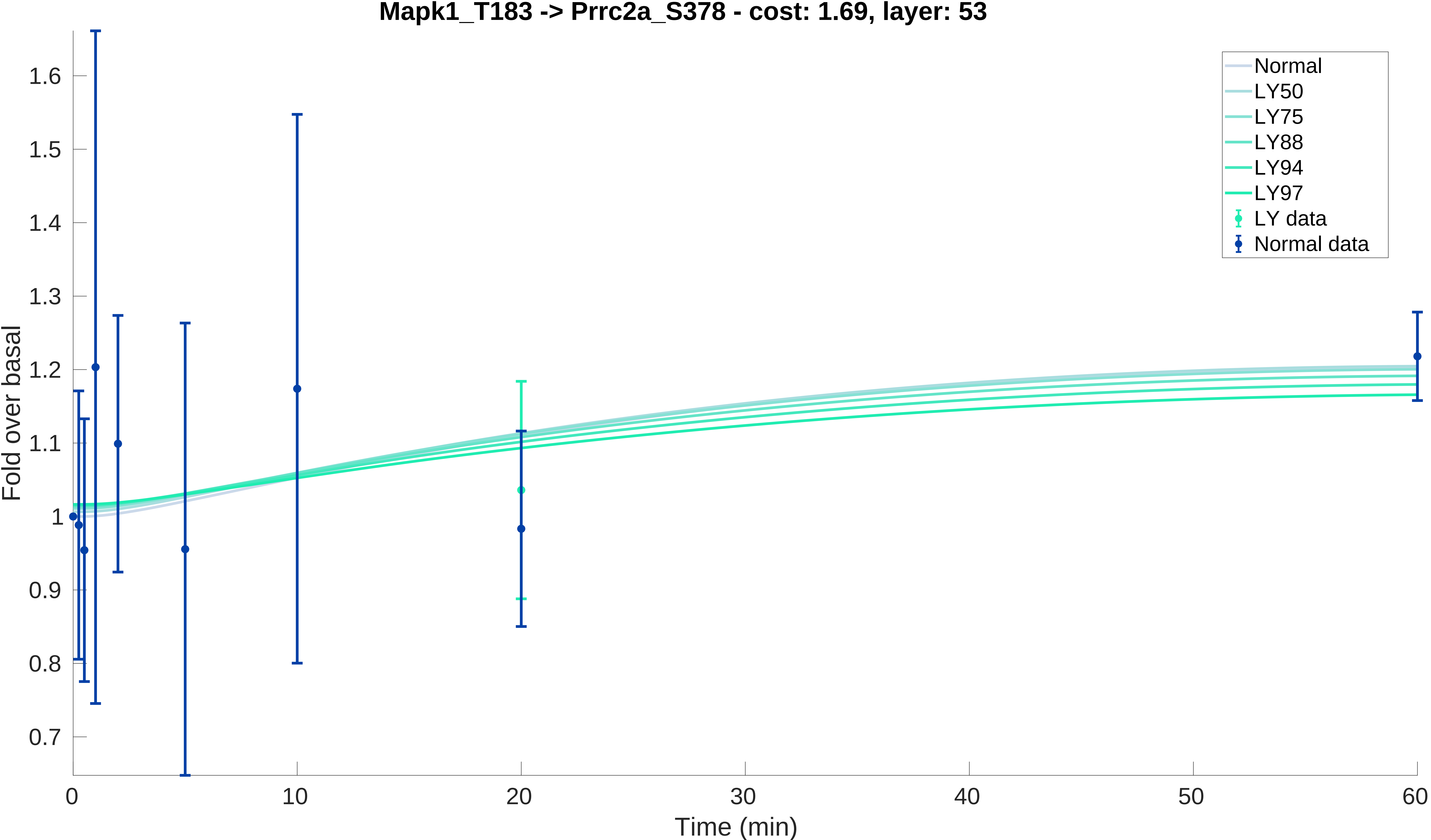
Mapk1_T183 -> Prrc2a_S378 - cost: 1.69, layer: 53
Fold over basal
Time (min)
0.7
0.8
0.9
1
1.1
1.2
1.3
1.4
1.5
1.6
0
10
20
30
40
50
60
Normal
LY50
LY75
LY88
LY94
LY97
LY data
Normal data

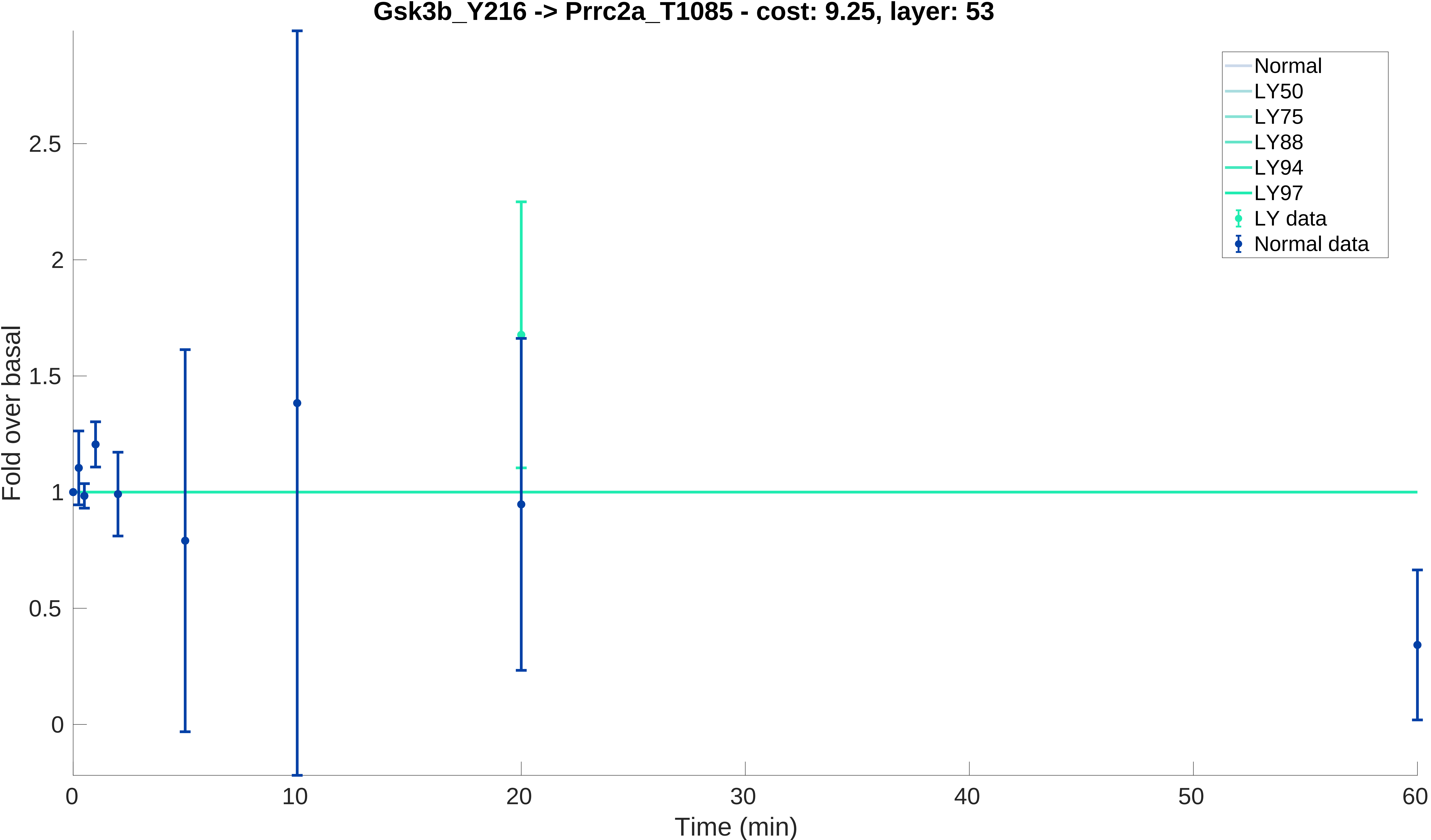

Gsk3b_Y216 -> Prrc2a_T1085 - cost: 9.25, layer: 53
Fold over basal
Time (min)
0
0.5
1
1.5
2
2.5
0
10
20
30
40
50
60
Normal
LY50
LY75
LY88
LY94
LY97
LY data
Normal data

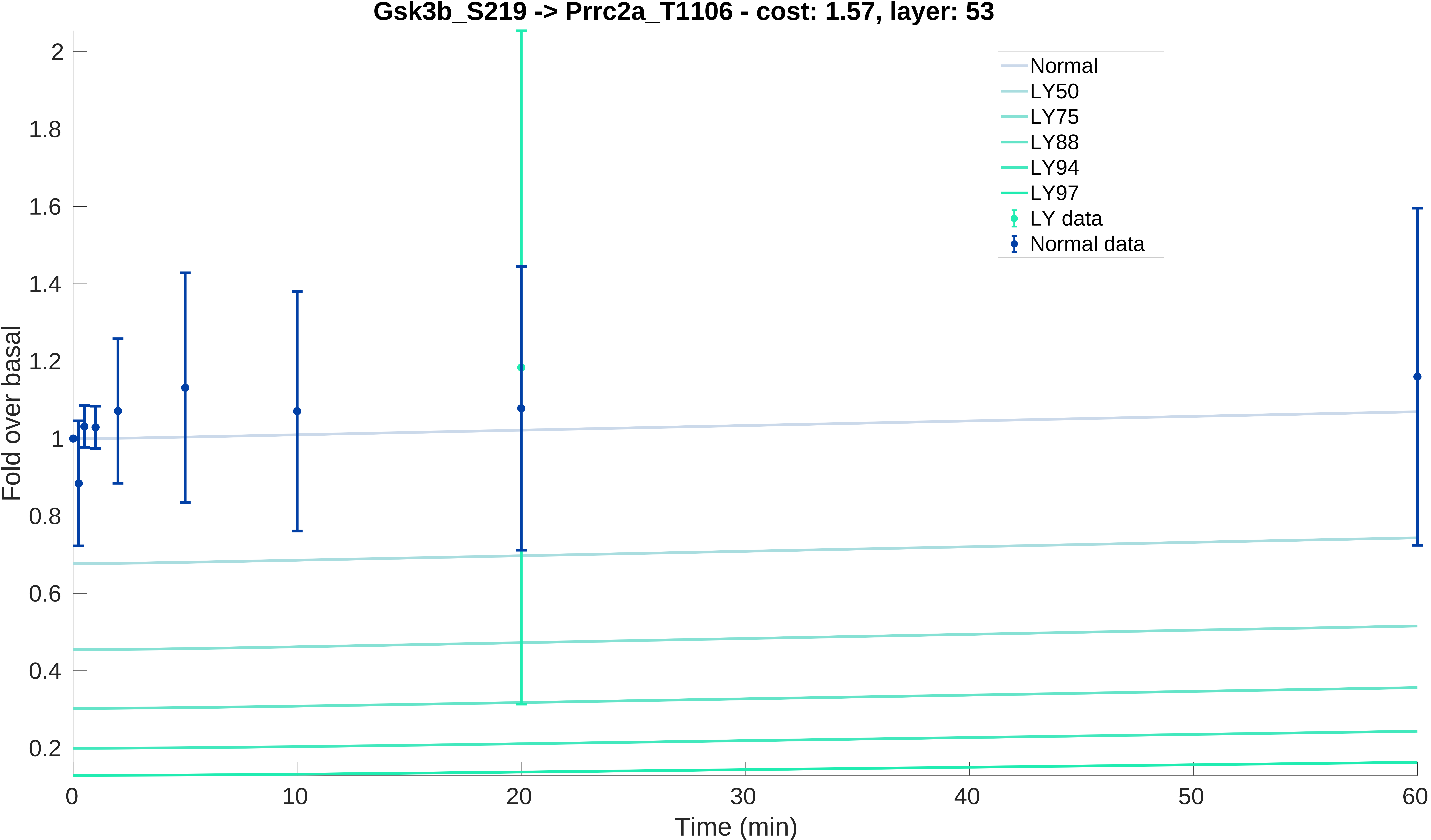
Gsk3b_S219 -> Prrc2a_T1106 - cost: 1.57, layer: 53
Normal
LY50
LY75
LY88
LY94
LY97
LY data
Normal data
Fold over basal
Time (min)
0.2
0.4
0.6
0.8
1
1.2
1.4
1.6
1.8
2
0
10
20
30
40
50
60

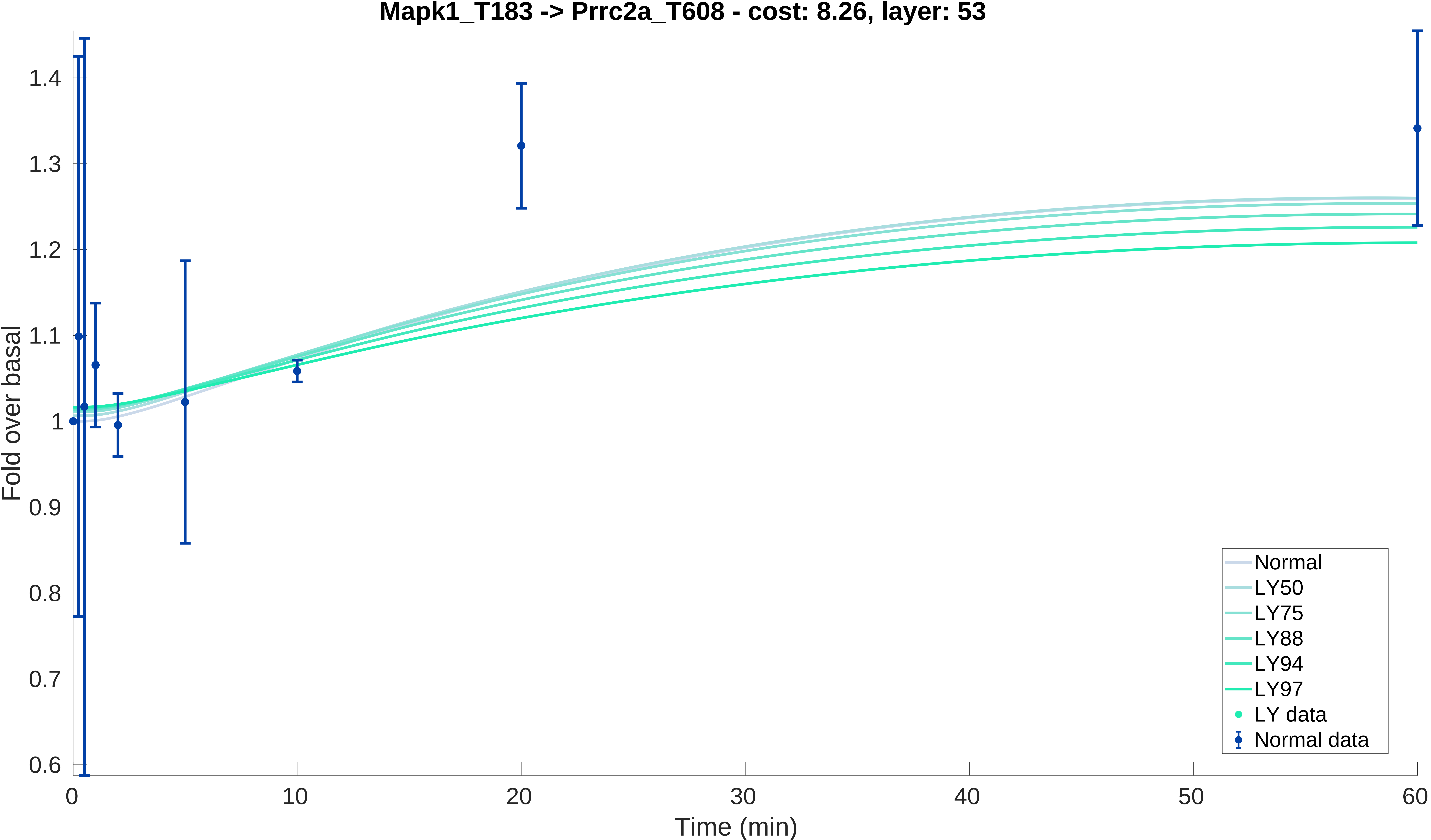

Mapk1_T183 -> Prrc2a_T608 - cost: 8.26, layer: 53
Fold over basal
1.4
1.3
1.2
1.1
1
0.9
0.8
0.7
0.6
0
10
20
30
40
50
60
Time (min)
Normal
LY50
LY75
LY88
LY94
LY97
LY data
Normal data

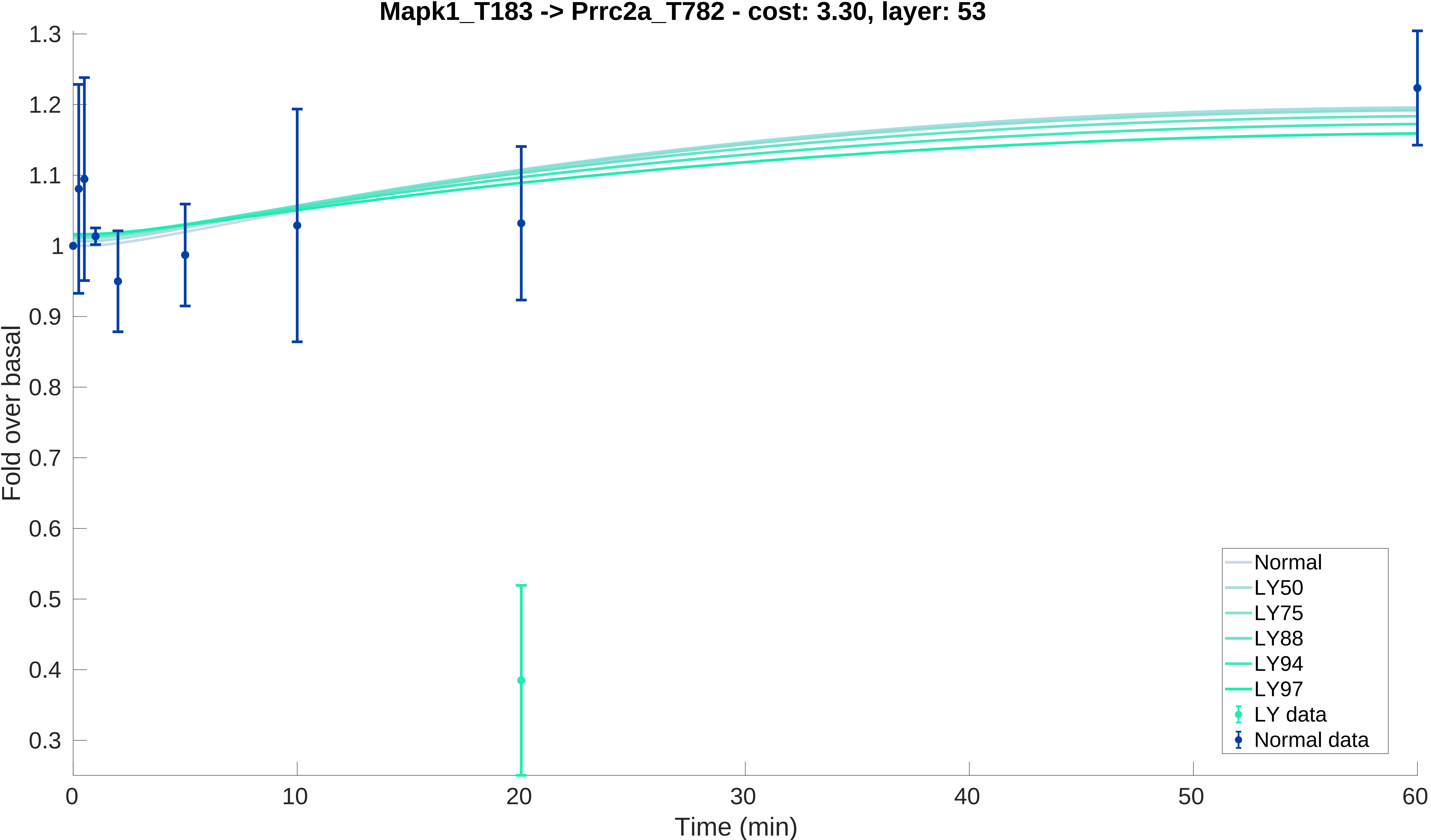

Mapk1_T183 -> Prrc2a_T782 - cost: 3.30, layer: 53
Fold over basal
Time (min)
1.3
1.2
1.1
1
0.9
0.8
0.7
0.6
0.5
0.4
0.3
0
10
20
30
40
50
60
Normal
LY50
LY75
LY88
LY94
LY97
LY data
Normal data

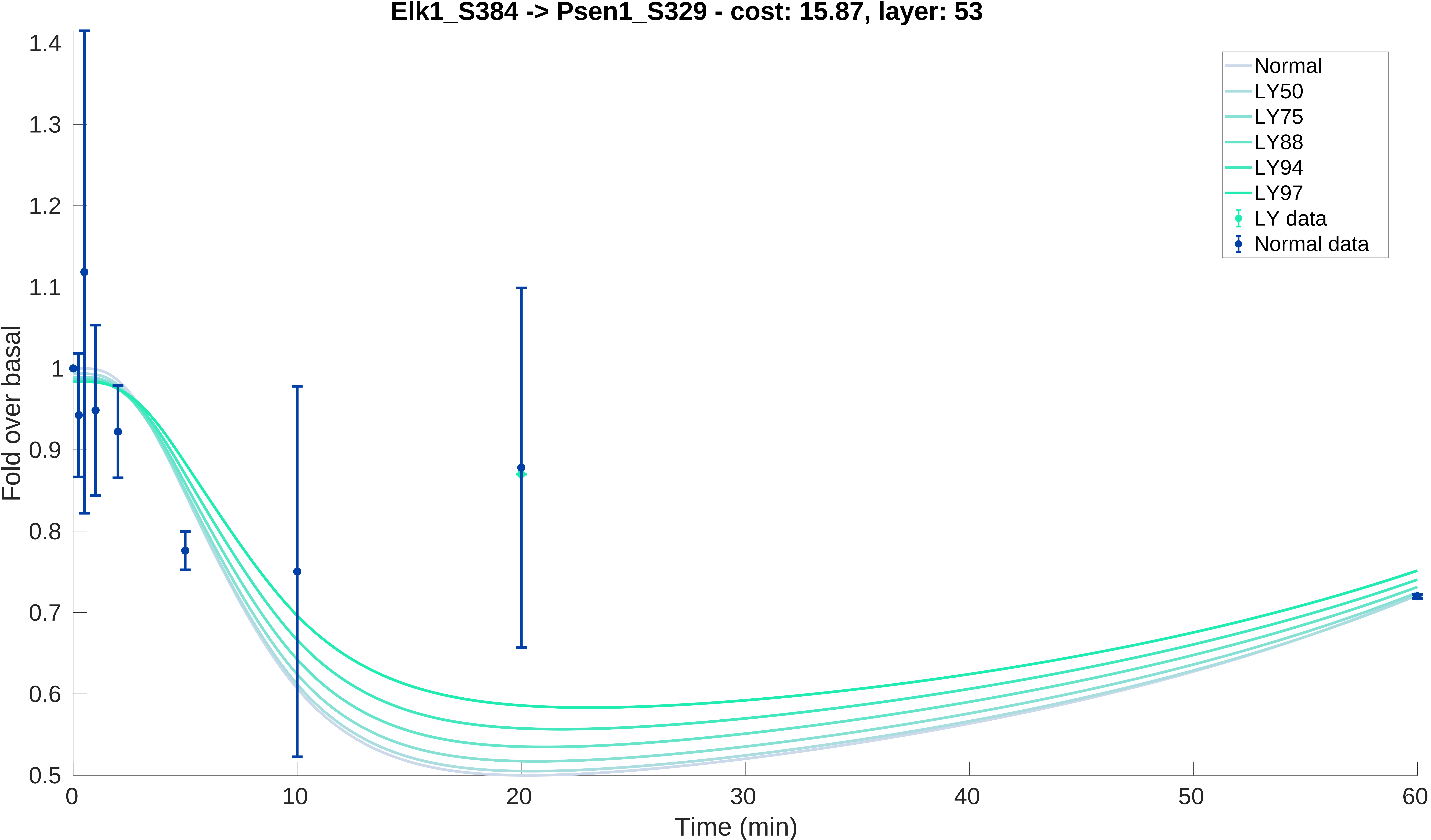
Elk1_S384 -> Psen1_S329 - cost: 15.87, layer: 53
Fold over basal
Time (min)
1.4
1.3
1.2
1.1
1
0.9
0.8
0.7
0.6
0.5
0
10
20
30
40
50
60
Normal
LY50
LY75
LY88
LY94
LY97
LY data
Normal data

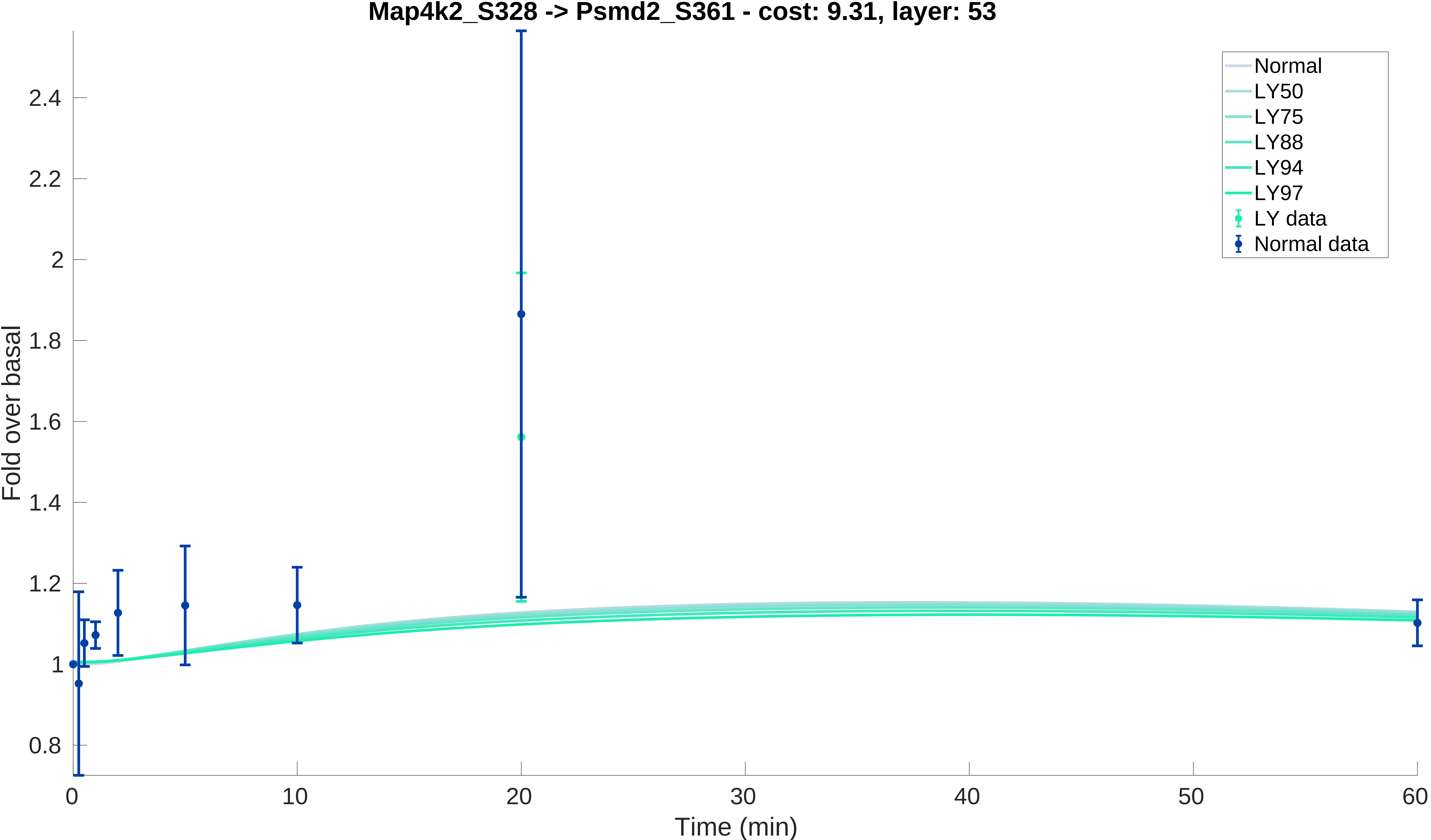

Map4k2_S328 -> Psmd2_S361 - cost: 9.31, layer: 53
Normal
LY50
LY75
LY88
LY94
LY97
LY data
Normal data
Fold over basal
Time (min)
2.4
2.2
2
1.8
1.6
1.4
1.2
1
0.8
0
10
20
30
40
50
60

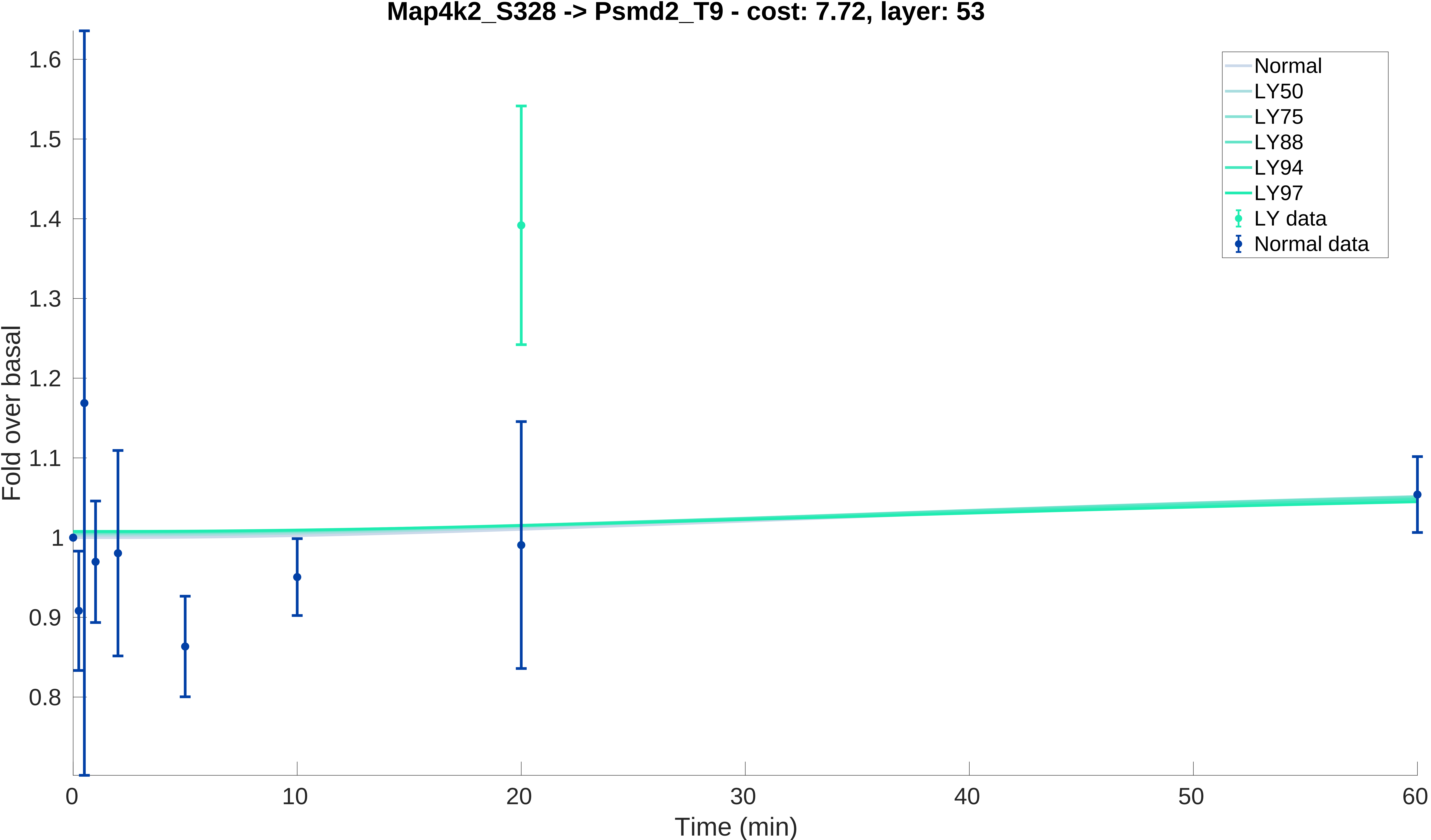

Map4k2_S328 -> Psmd2_T9 - cost: 7.72, layer: 53
Fold over basal
Time (min)
1.6
1.5
1.4
1.3
1.2
1.1
1
0.9
0.8
0
10
20
30
40
50
60
Normal
LY50
LY75
LY88
LY94
LY97
LY data
Normal data

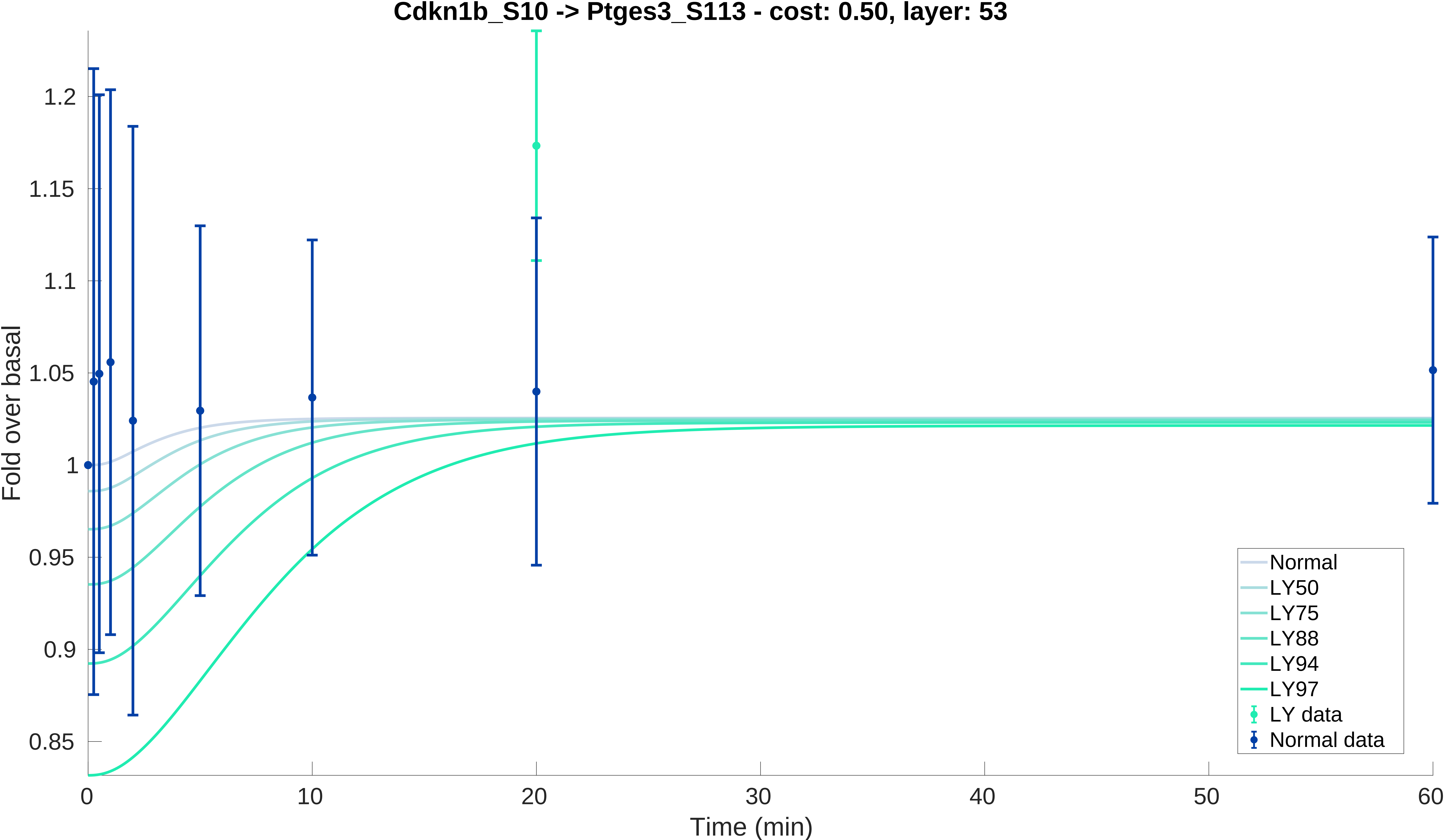

Cdkn1b_S10 -> Ptges3_S113 - cost: 0.50, layer: 53
Fold over basal
Time (min)
0.85
0.9
0.95
1
1.05
1.1
1.15
1.2
0
10
20
30
40
50
60
Normal
LY50
LY75
LY88
LY94
LY97
LY data
Normal data

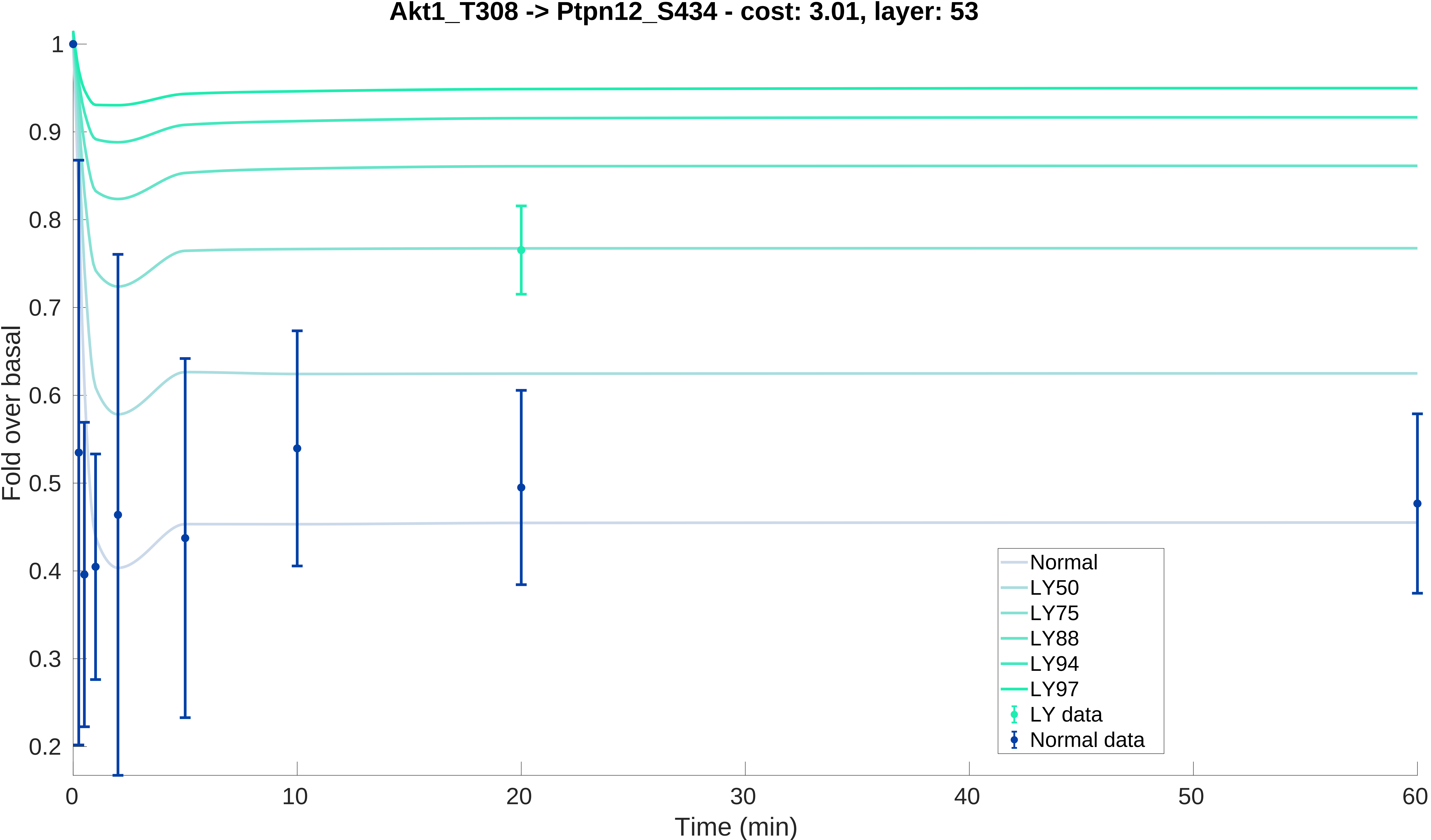

Akt1_T308 -> Ptpn12_S434 - cost: 3.01, layer: 53
Fold over basal
1
0.9
0.8
0.7
0.6
0.5
0.4
0.3
0.2
0
10
20
30
40
50
60
Time (min)
Normal
LY50
LY75
LY88
LY94
LY97
LY data
Normal data

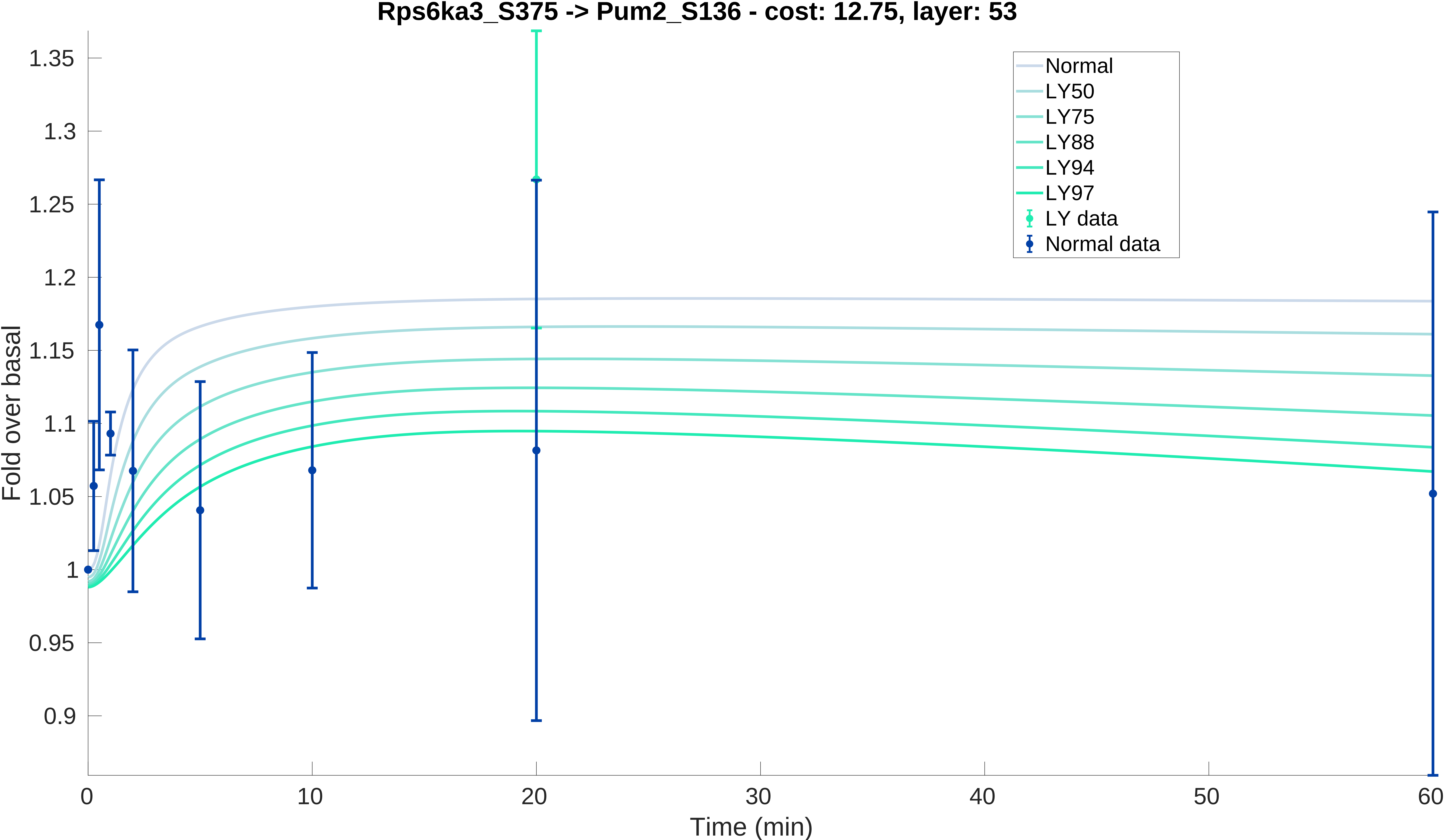

Rps6ka3_S375 -> Pum2_S136 - cost: 12.75, layer: 53
Fold over basal
Time (min)
1.35
1.3
1.25
1.2
1.15
1.1
1.05
1
0.95
0.9
0
10
20
30
40
50
60
Normal
LY50
LY75
LY88
LY94
LY97
LY data
Normal data

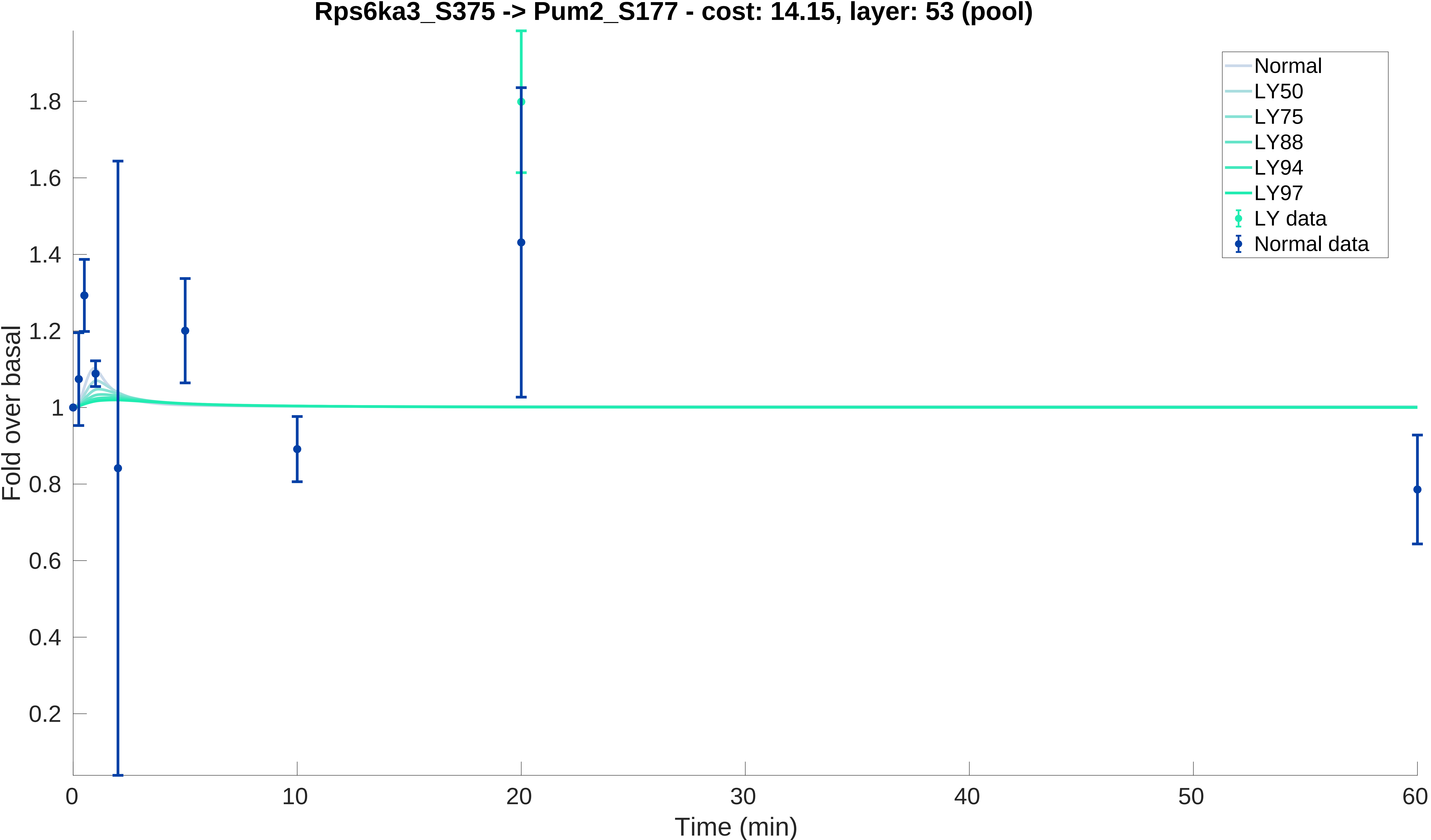

Rps6ka3_S375 -> Pum2_S177 - cost: 14.15, layer: 53 (pool)
Normal
LY50
LY75
LY88
LY94
LY97
LY data
Normal data
Fold over basal
Time (min)
0
10
20
30
40
50
60
0.2
0.4
0.6
0.8
1
1.2
1.4
1.6
1.8

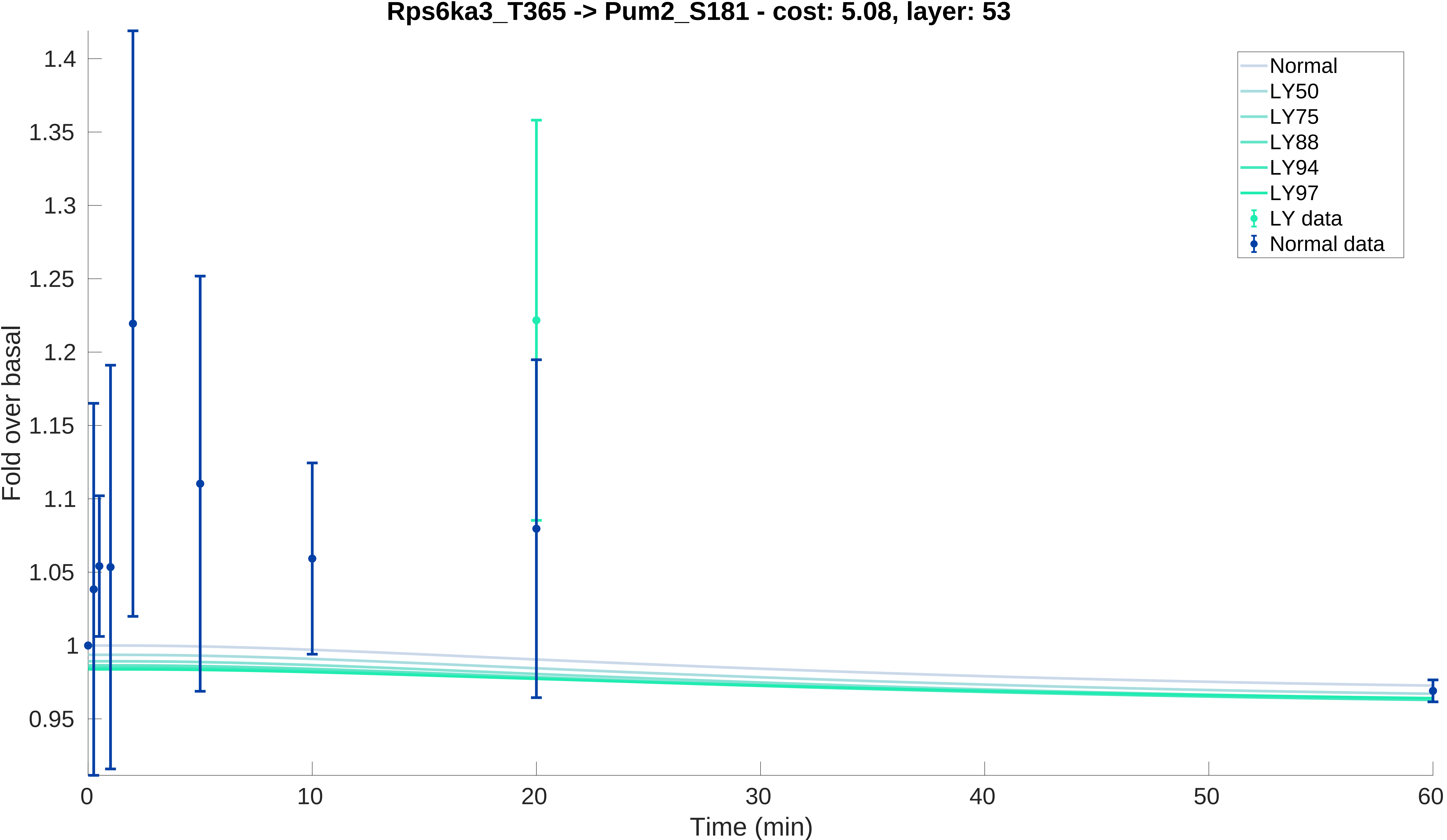
Rps6ka3_T365 -> Pum2_S181 - cost: 5.08, layer: 53
Fold over basal
Time (min)
0
10
20
30
40
50
60
0.95
1
1.05
1.1
1.15
1.2
1.25
1.3
1.35
1.4
Normal
LY50
LY75
LY88
LY94
LY97
LY data
Normal data

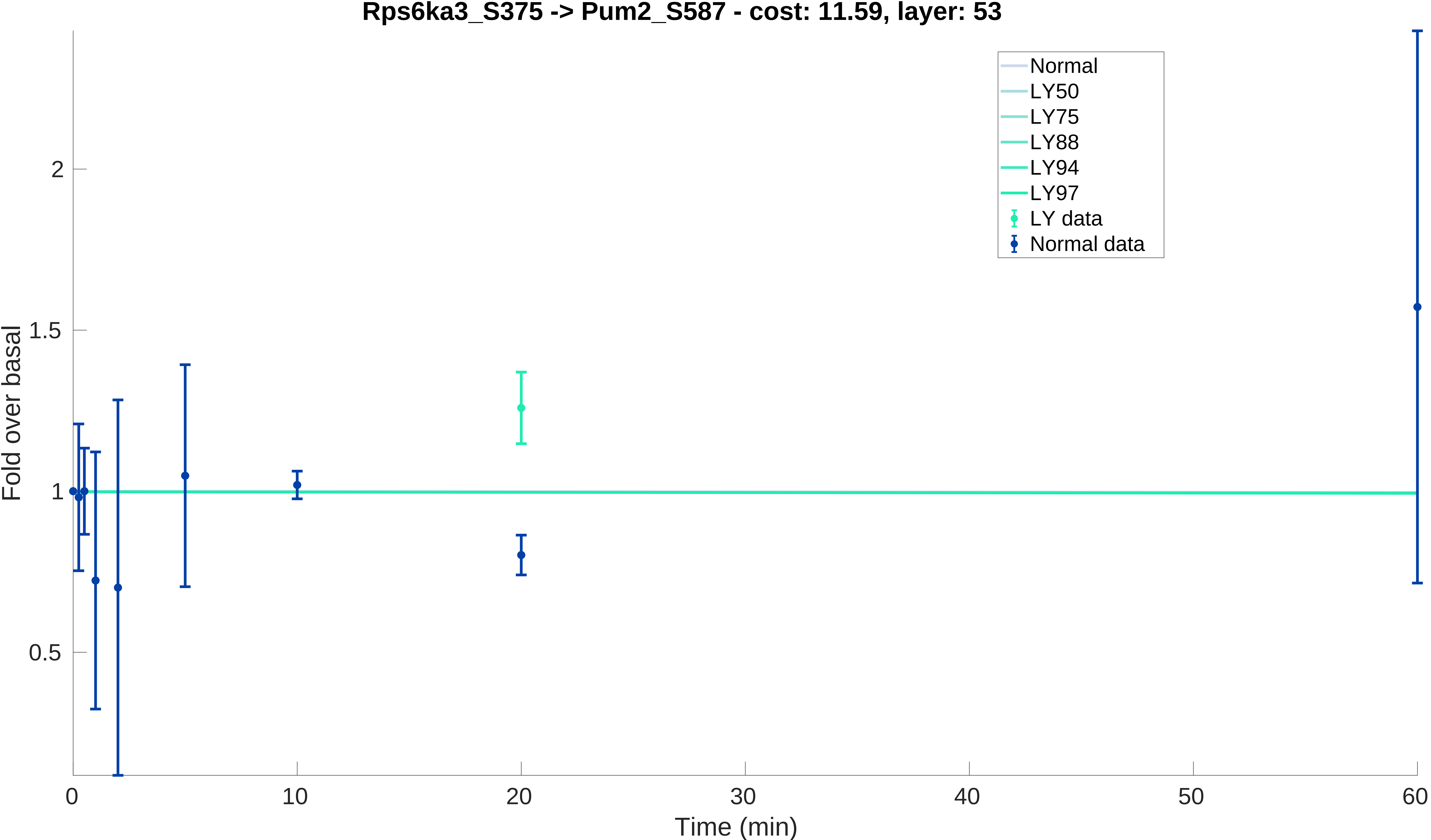

Rps6ka3_S375 -> Pum2_S587 - cost: 11.59, layer: 53
Normal
LY50
LY75
LY88
LY94
LY97
LY data
Normal data
Fold over basal
2
1.5
1
0.5
0
10
20
30
40
50
60
Time (min)

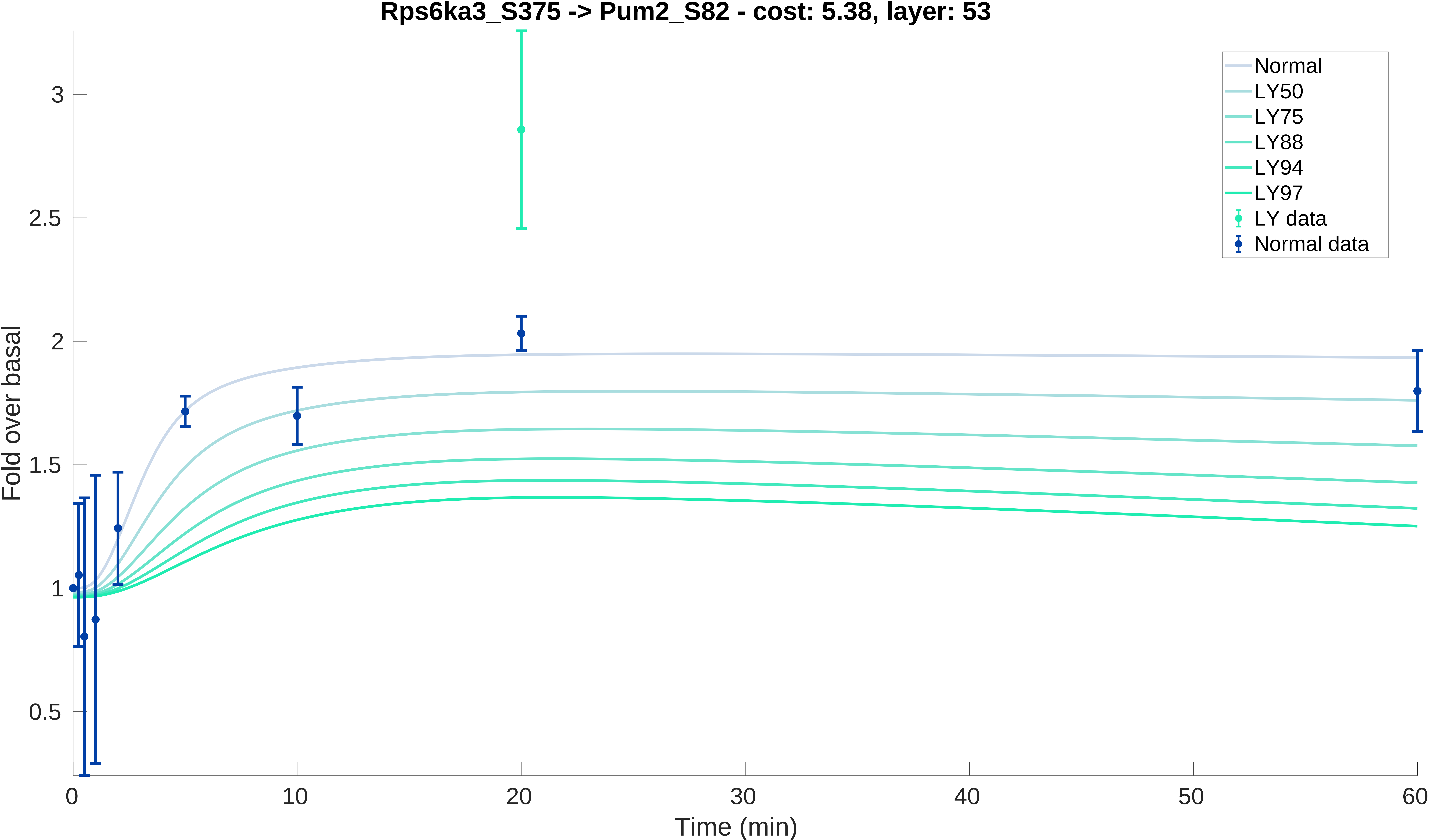

Rps6ka3_S375 -> Pum2_S82 - cost: 5.38, layer: 53
Fold over basal
Time (min)
0.5
1
1.5
2
2.5
3
0
10
20
30
40
50
60
Normal
LY50
LY75
LY88
LY94
LY97
LY data
Normal data

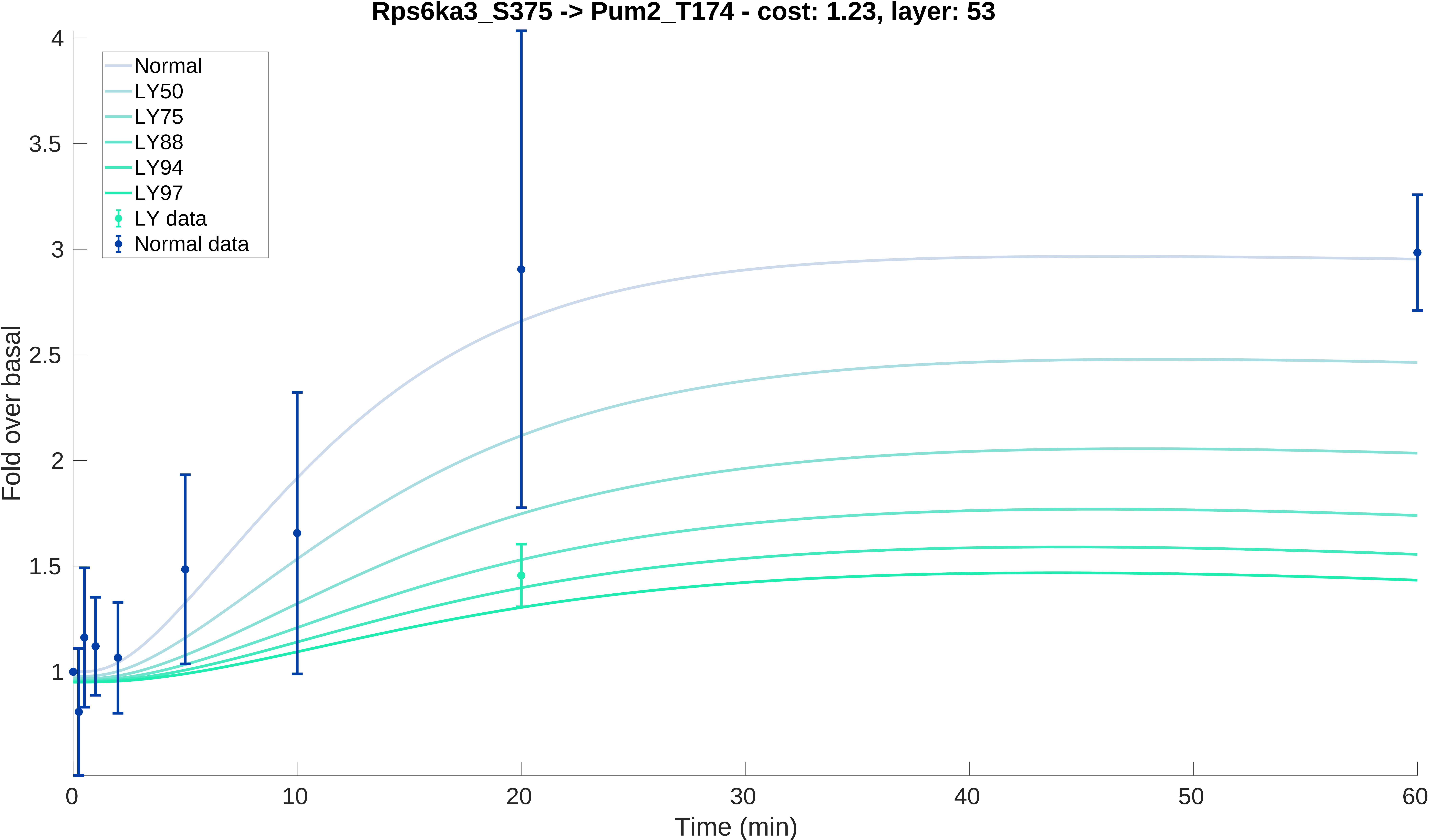

Rps6ka3_S375 -> Pum2_T174 - cost: 1.23, layer: 53
Normal
LY50
LY75
LY88
LY94
LY97
LY data
Normal data
Fold over basal
4
3.5
3
2.5
2
1.5
1
0
10
20
30
40
50
60
Time (min)

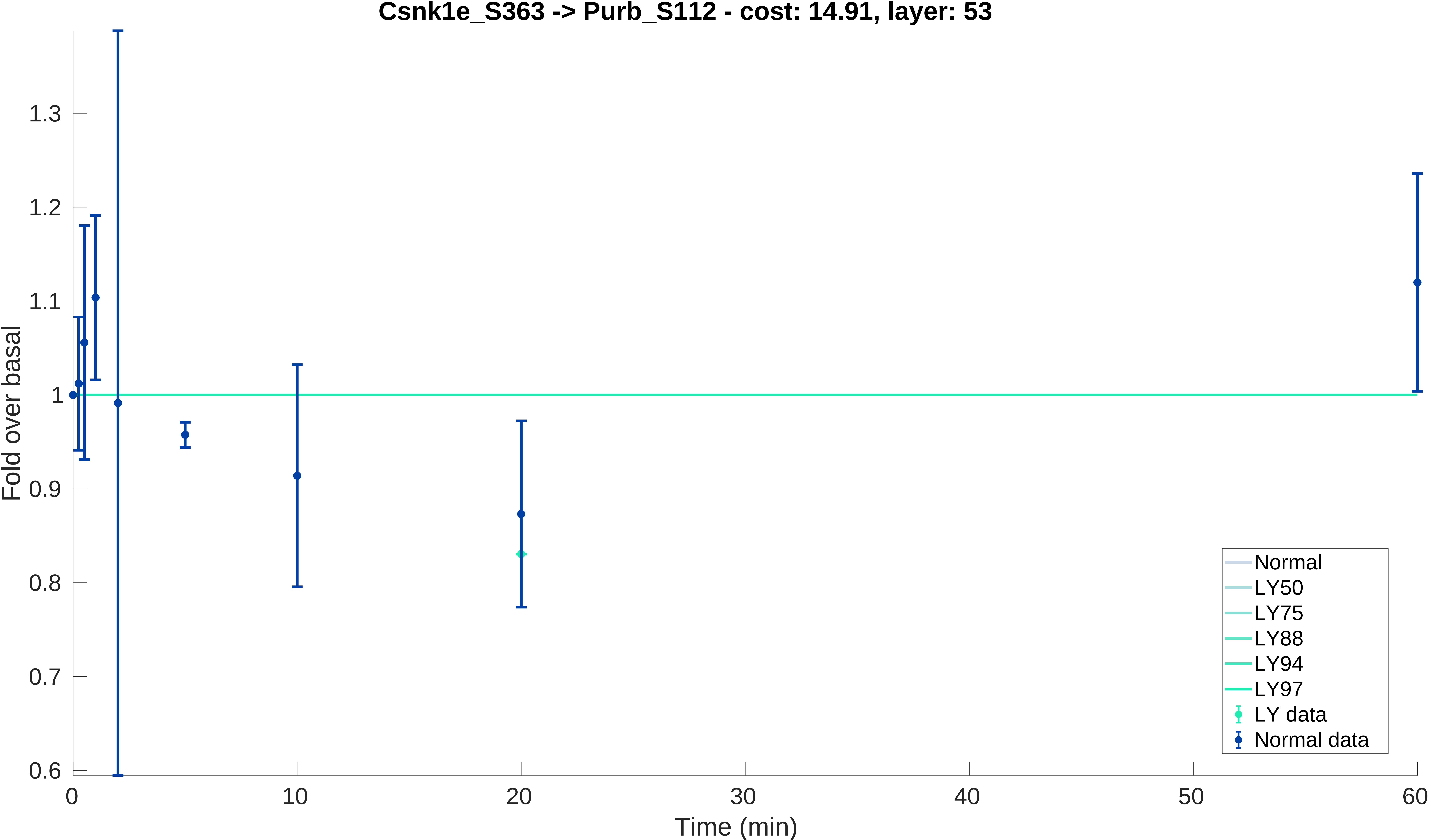
Csnk1e_S363 -> Purb_S112 - cost: 14.91, layer: 53
Fold over basal
Time (min)
1.3
1.2
1.1
1
0.9
0.8
0.7
0.6
0
10
20
30
40
50
60
Normal
LY50
LY75
LY88
LY94
LY97
LY data
Normal data

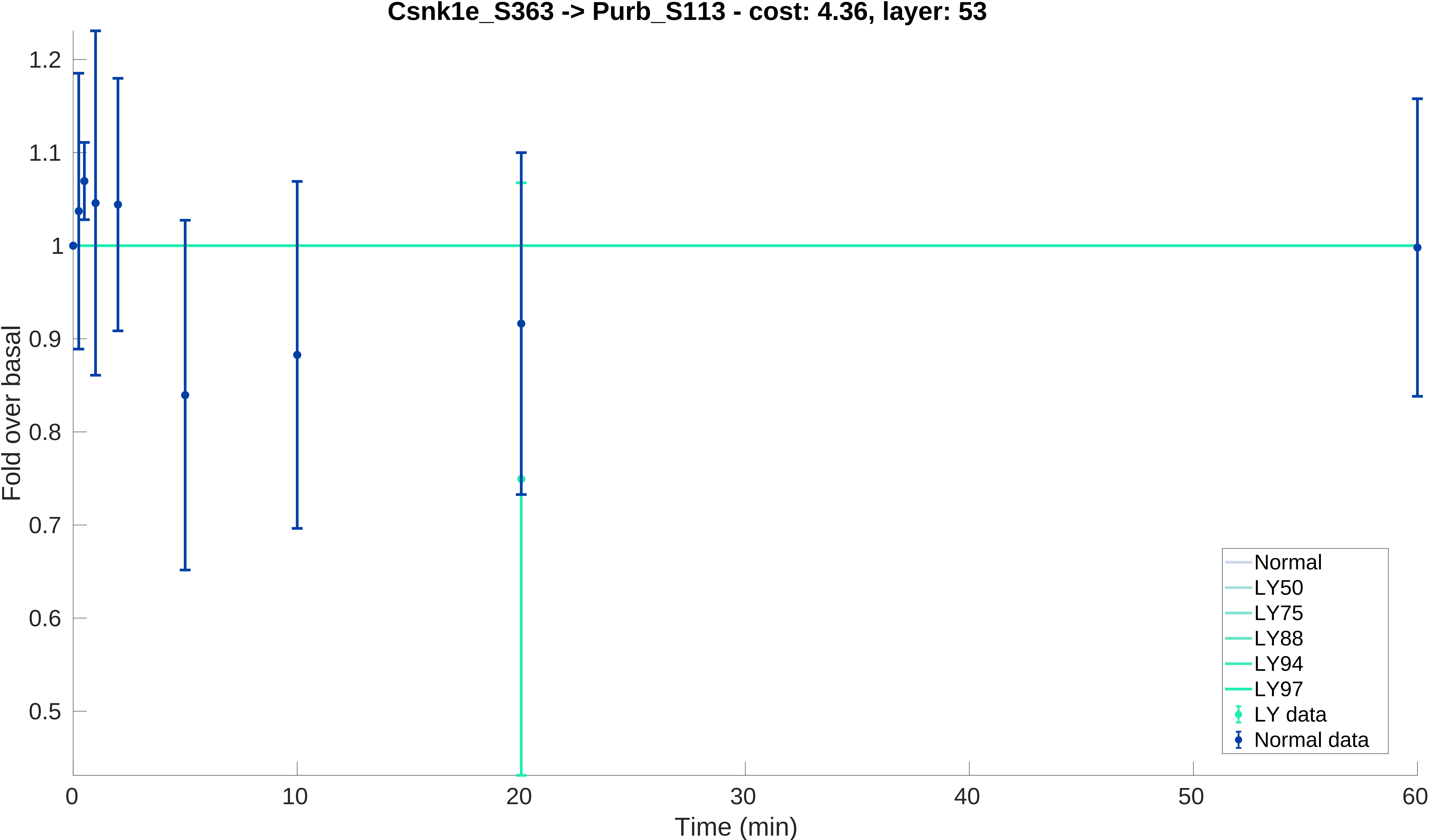

Csnk1e_S363 -> Purb_S113 - cost: 4.36, layer: 53
Fold over basal
1.2
1.1
1
0.9
0.8
0.7
0.6
0.5
0
10
20
30
40
50
60
Time (min)
Normal
LY50
LY75
LY88
LY94
LY97
LY data
Normal data

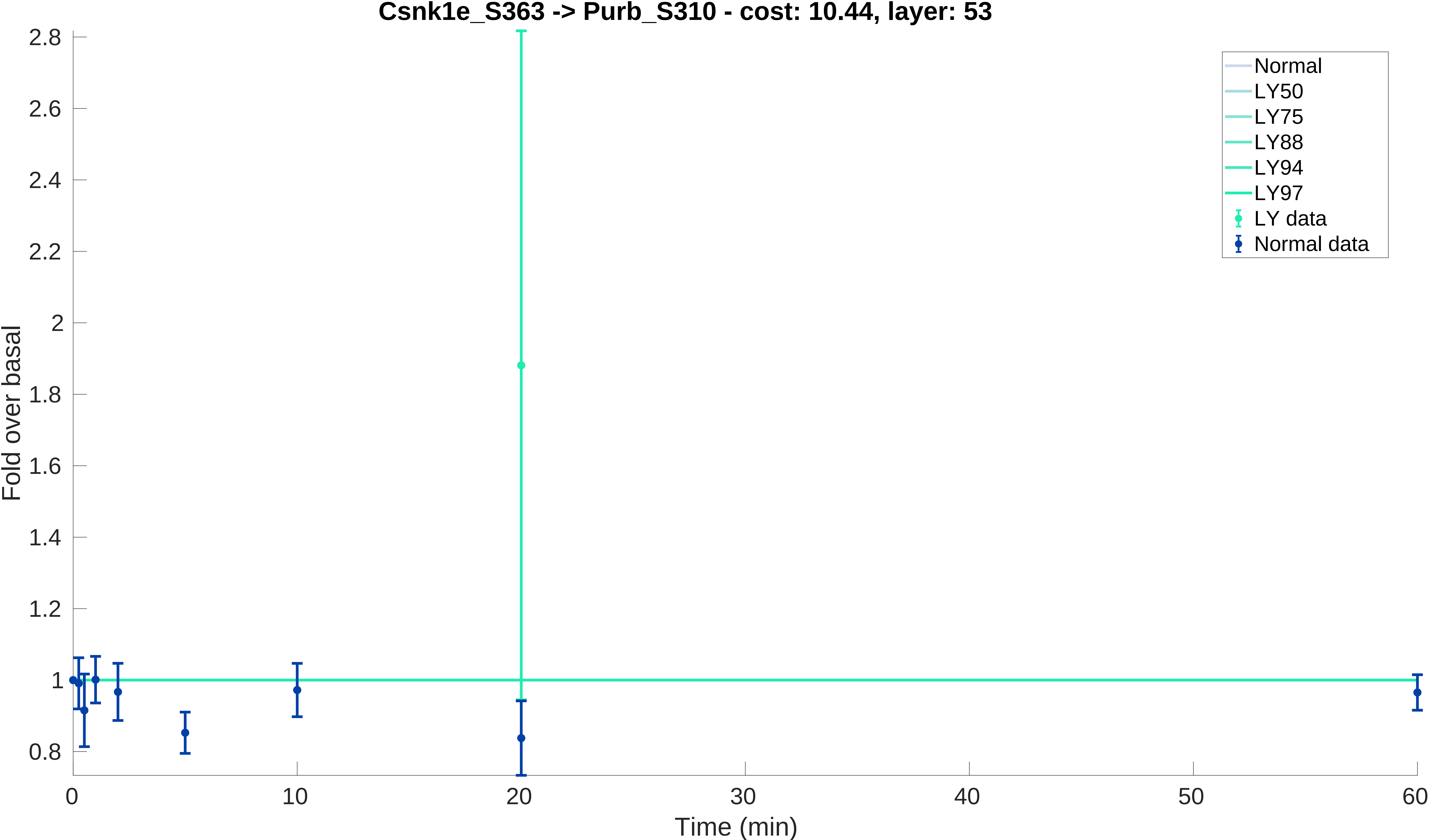

Csnk1e_S363 -> Purb_S310 - cost: 10.44, layer: 53
Fold over basal
2.8
2.6
2.4
2.2
2
1.8
1.6
1.4
1.2
1
0.8
0
10
20
30
40
50
60
Time (min)
Normal
LY50
LY75
LY88
LY94
LY97
LY data
Normal data

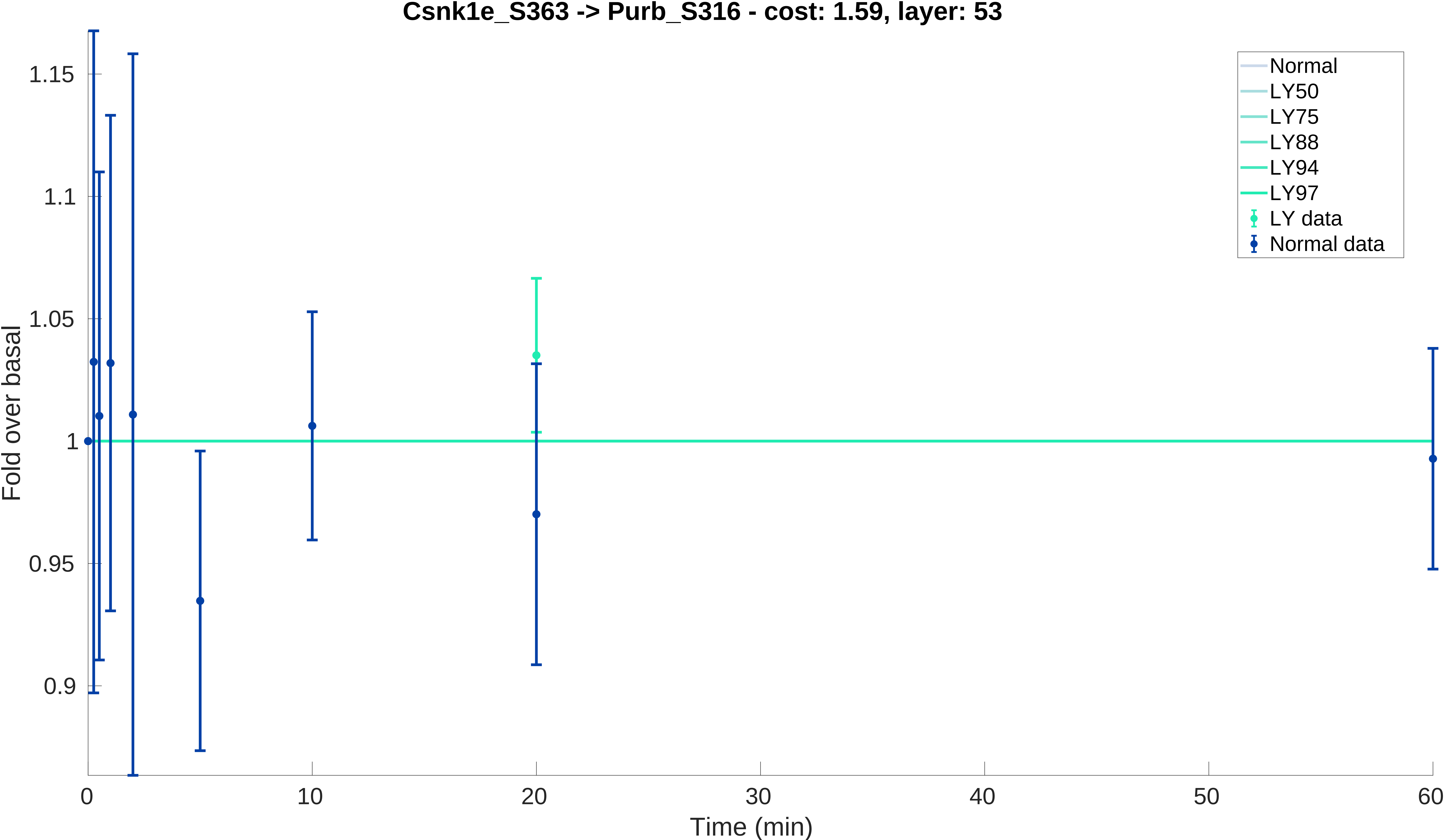

Csnk1e_S363 -> Purb_S316 - cost: 1.59, layer: 53
Normal
LY50
LY75
LY88
LY94
LY97
LY data
Normal data
Fold over basal
1.15
1.1
1.05
1
0.95
0.9
Time (min)
0
10
20
30
40
50
60

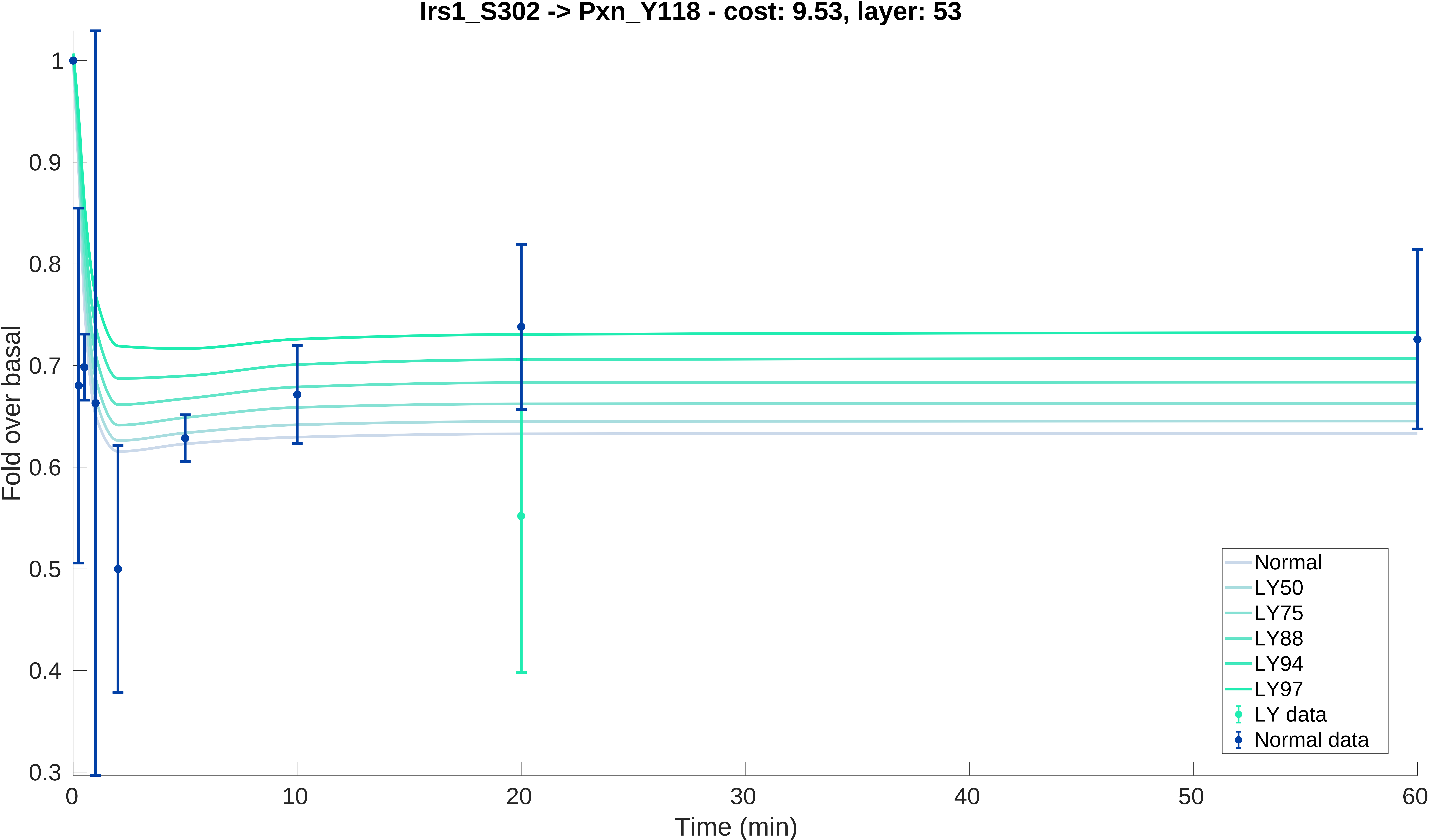

Irs1_S302 -> Pxn_Y118 - cost: 9.53, layer: 53
Fold over basal
1
0.9
0.8
0.7
0.6
0.5
0.4
0.3
0
10
20
30
40
50
60
Time (min)
Normal
LY50
LY75
LY88
LY94
LY97
LY data
Normal data

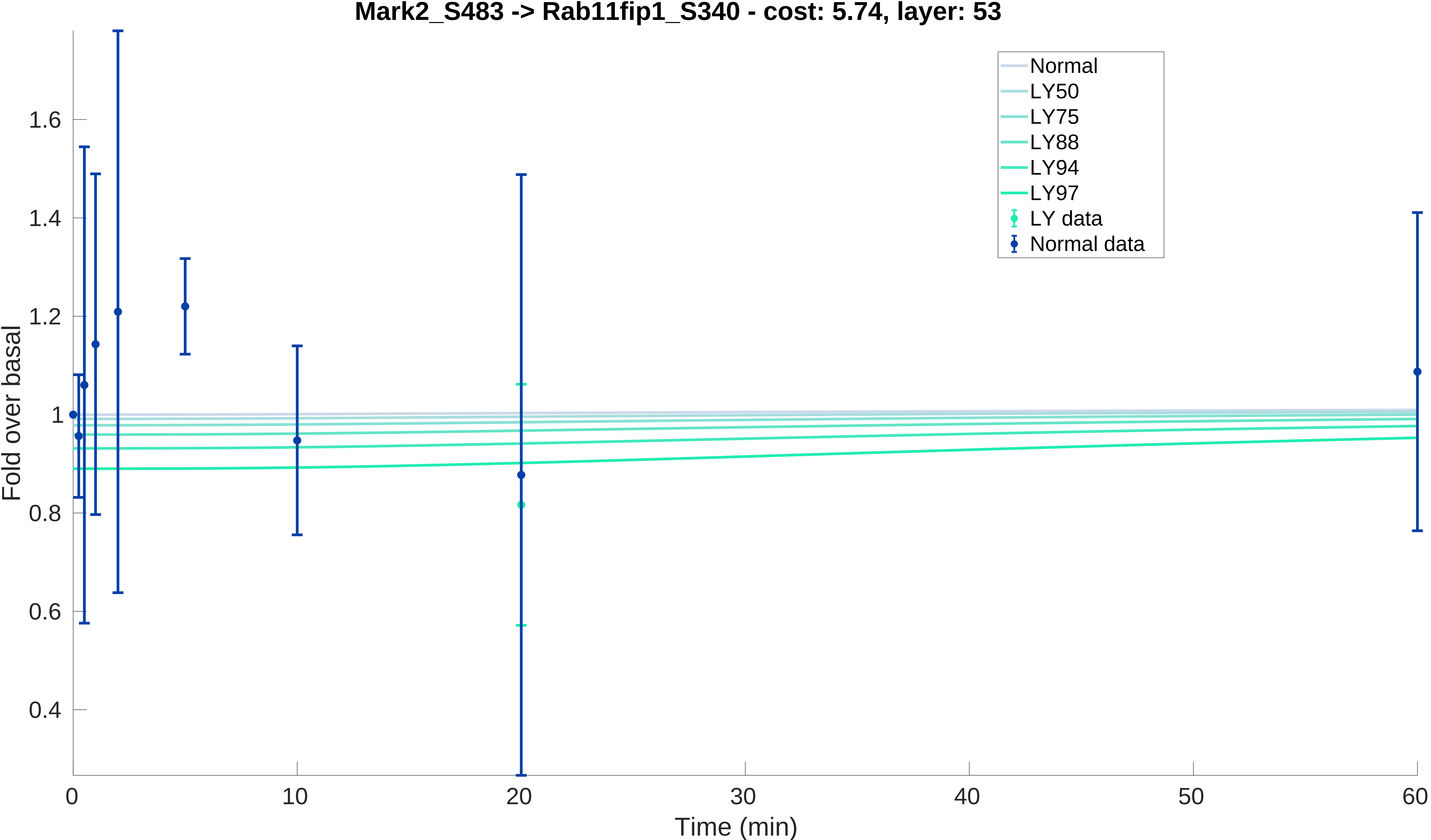

Mark2_S483 -> Rab11fip1_S340 - cost: 5.74, layer: 53
Normal
LY50
LY75
LY88
LY94
LY97
LY data
Normal data
1.6
1.4
1.2
1
0.8
0.6
0.4
Fold over basal
0
10
20
30
40
50
60
Time (min)

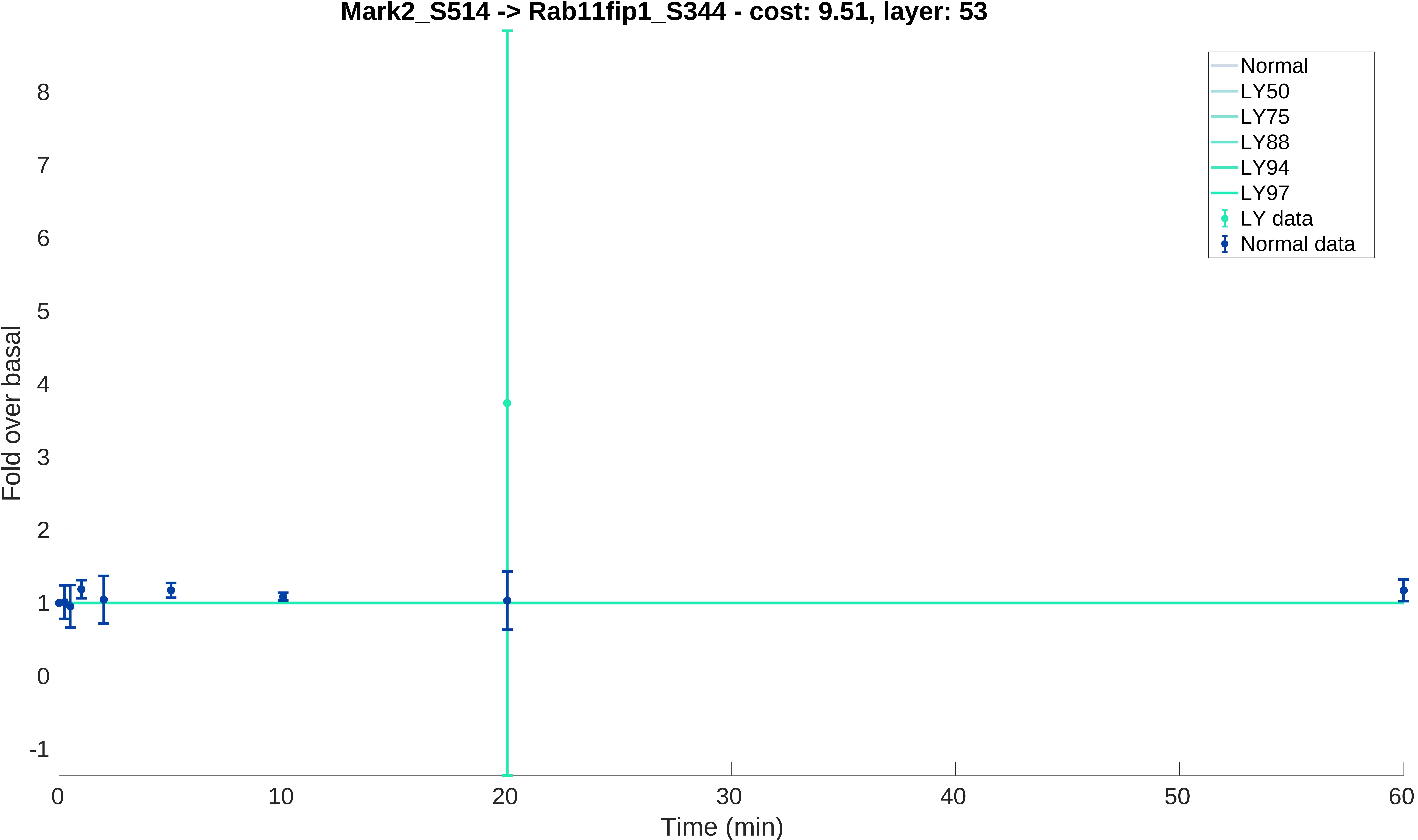

Mark2_S514 -> Rab11fip1_S344 - cost: 9.51, layer: 53
Fold over basal
Time (min)
8
7
6
5
4
3
2
1
0
-1
0
10
20
30
40
50
60
Normal
LY50
LY75
LY88
LY94
LY97
LY data
Normal data

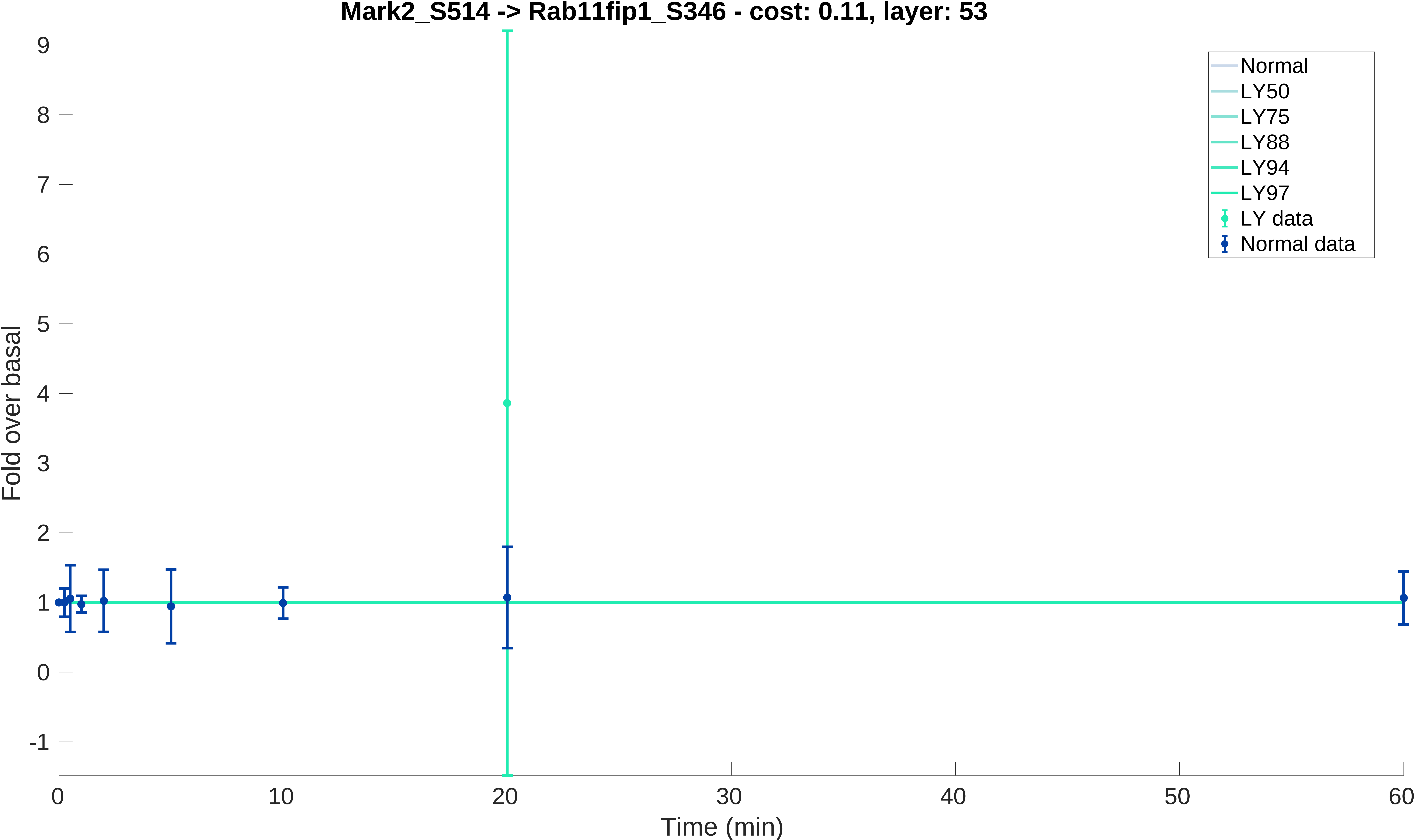
Mark2_S514 -> Rab11fip1_S346 - cost: 0.11, layer: 53
Normal
LY50
LY75
LY88
LY94
LY97
LY data
Normal data
Fold over basal
Time (min)
9
8
7
6
5
4
3
2
1
0
-1
0
10
20
30
40
50
60

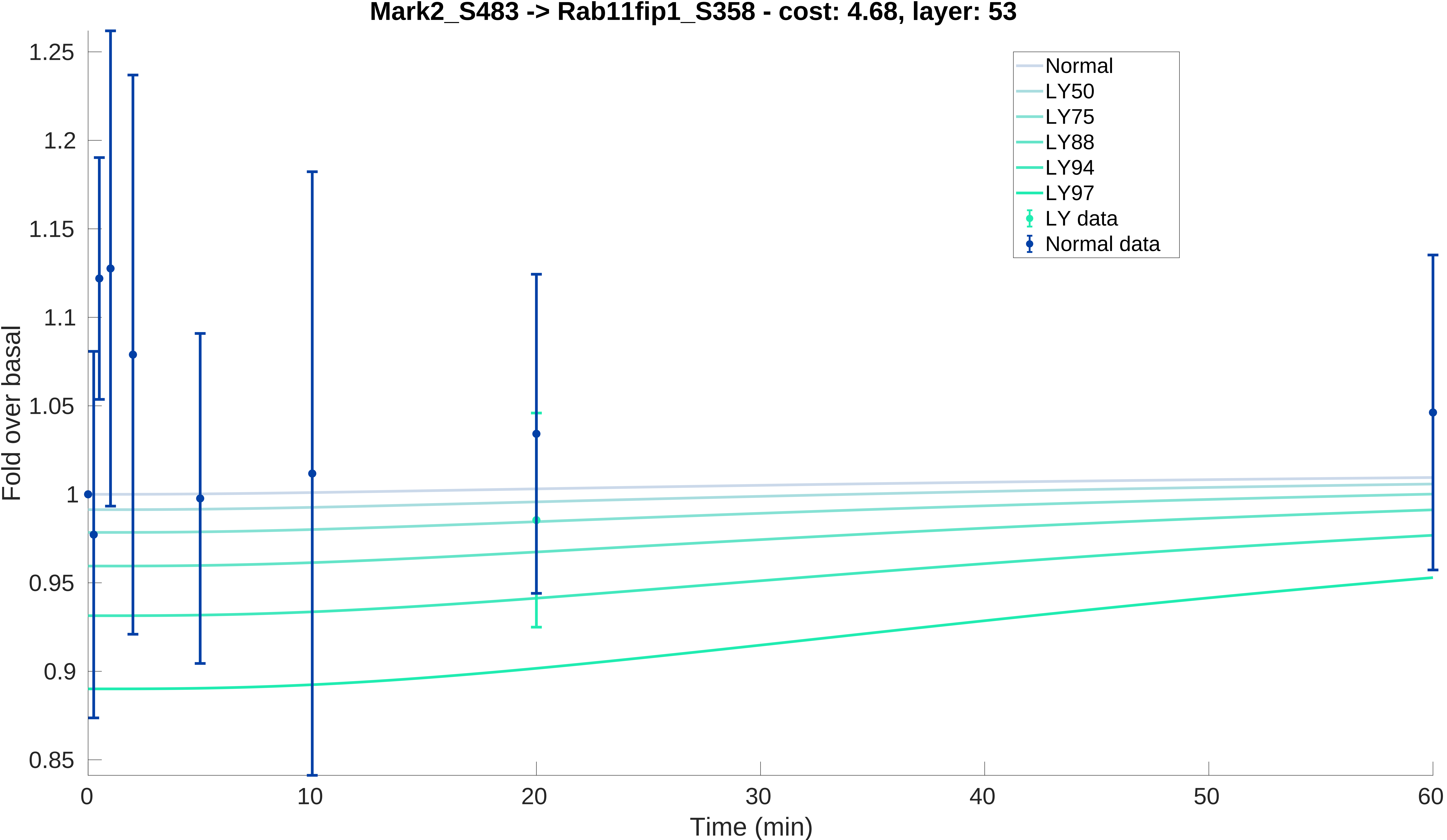

Mark2_S483 -> Rab11fip1_S358 - cost: 4.68, layer: 53
Fold over basal
Time (min)
1.25
1.2
1.15
1.1
1.05
1
0.95
0.9
0.85
0
10
20
30
40
50
60
Normal
LY50
LY75
LY88
LY94
LY97
LY data
Normal data

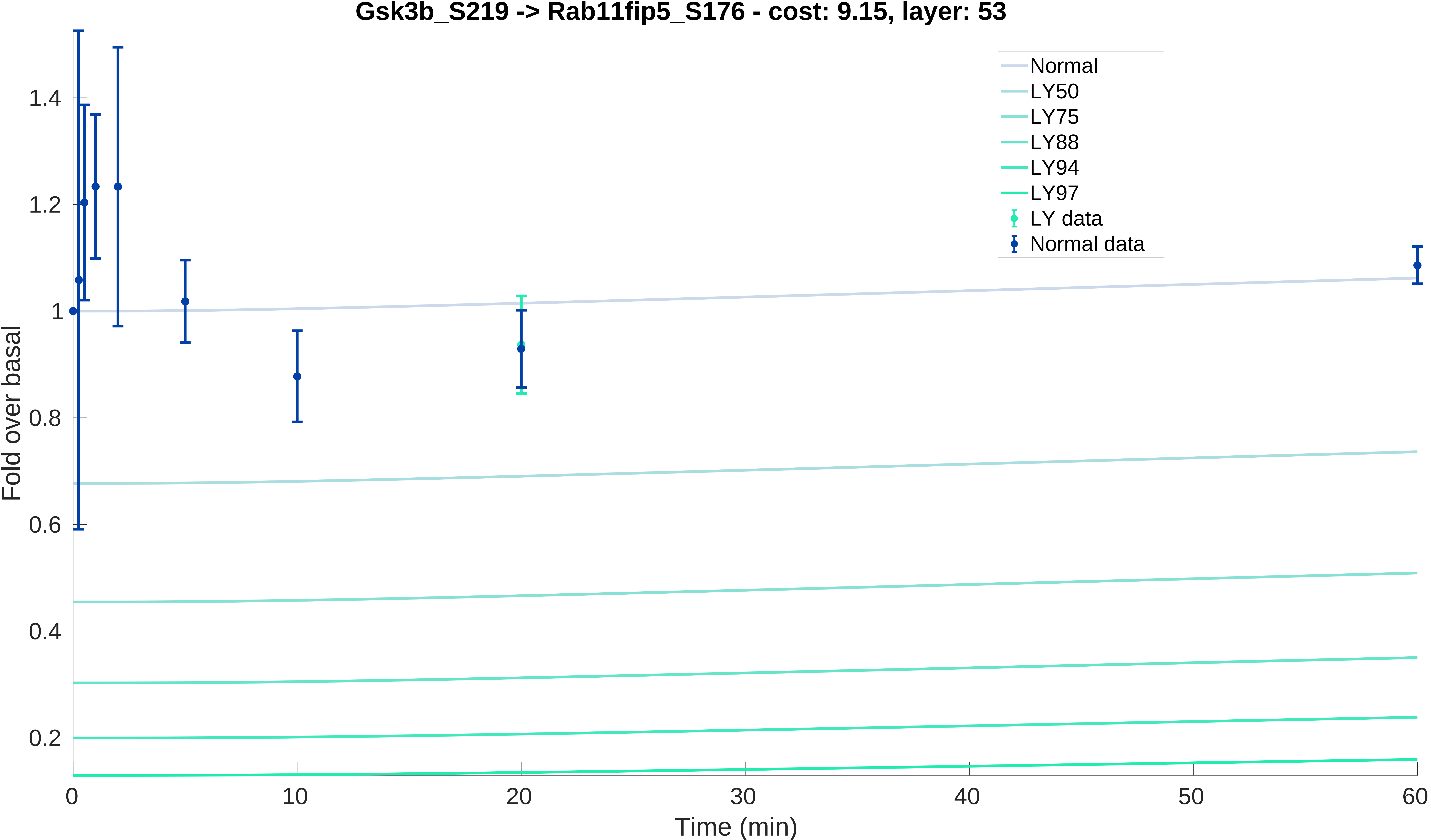

Gsk3b_S219 -> Rab11fip5_S176 - cost: 9.15, layer: 53
Normal
LY50
LY75
LY88
LY94
LY97
LY data
Normal data
Fold over basal
1.4
1.2
1
0.8
0.6
0.4
0.2
0
10
20
30
40
50
60
Time (min)

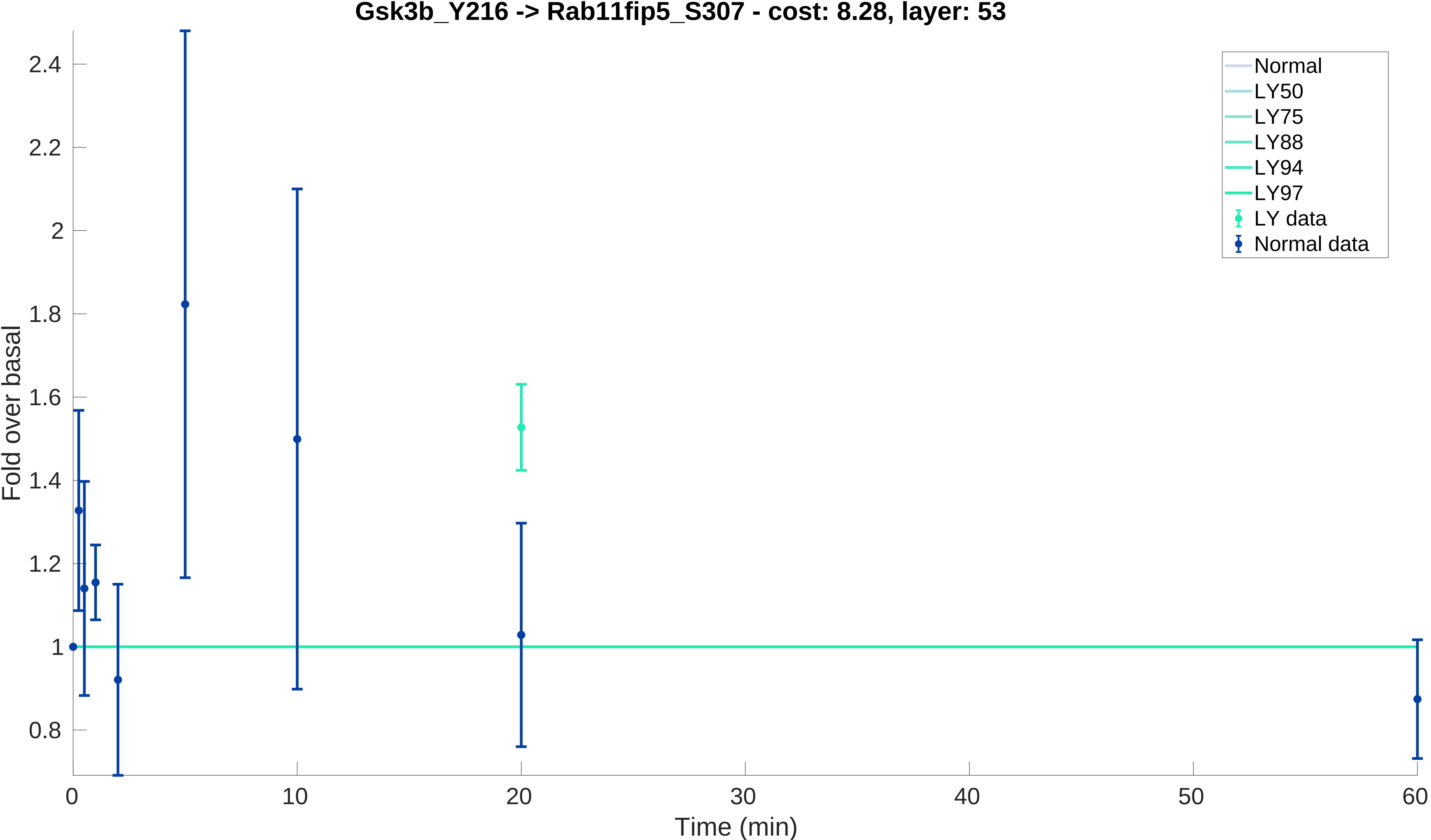
Gsk3b_Y216 -> Rab11fip5_S307 - cost: 8.28, layer: 53
Fold over basal
Time (min)
2.4
2.2
2
1.8
1.6
1.4
1.2
1
0.8
0
10
20
30
40
50
60
Normal
LY50
LY75
LY88
LY94
LY97
LY data
Normal data

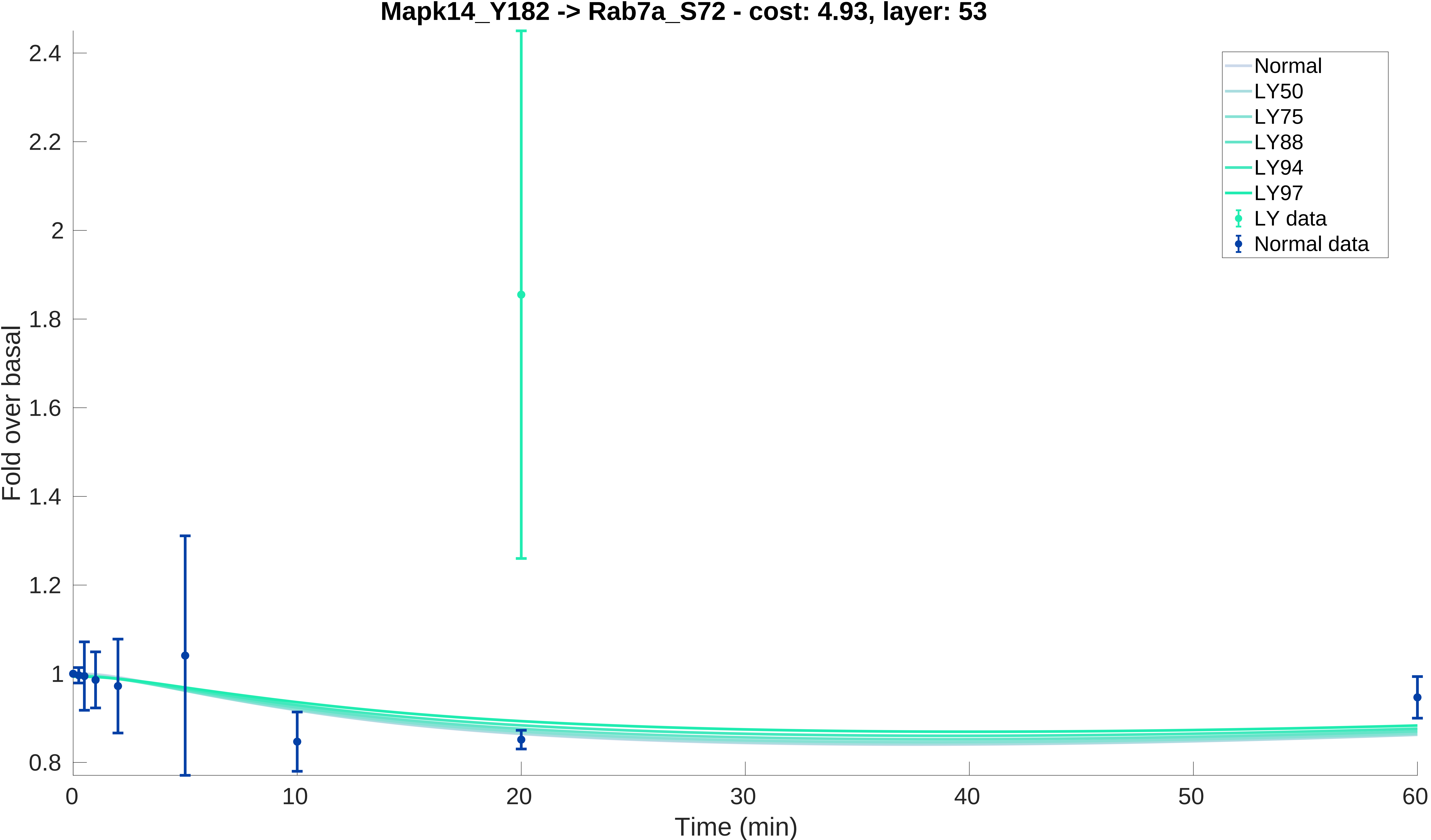
Mapk14_Y182 -> Rab7a_S72 - cost: 4.93, layer: 53
Fold over basal
Time (min)
2.4
2.2
2
1.8
1.6
1.4
1.2
1
0.8
0
10
20
30
40
50
60
Normal
LY50
LY75
LY88
LY94
LY97
LY data
Normal data

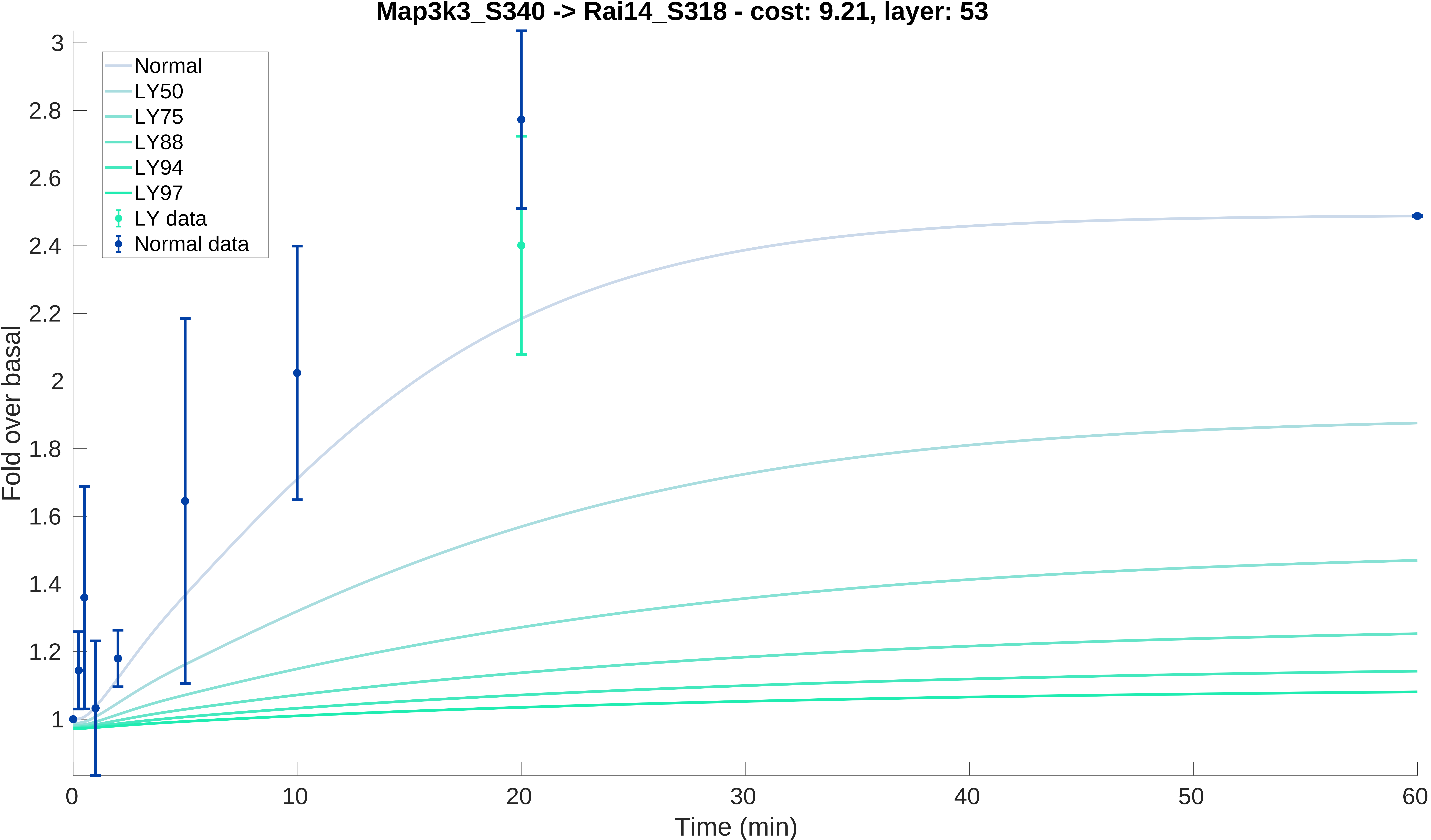

Map3k3_S340 -> Rai14_S318 - cost: 9.21, layer: 53
Normal
LY50
LY75
LY88
LY94
LY97
LY data
Normal data
Fold over basal
Time (min)
3
2.8
2.6
2.4
2.2
2
1.8
1.6
1.4
1.2
1
0
10
20
30
40
50
60

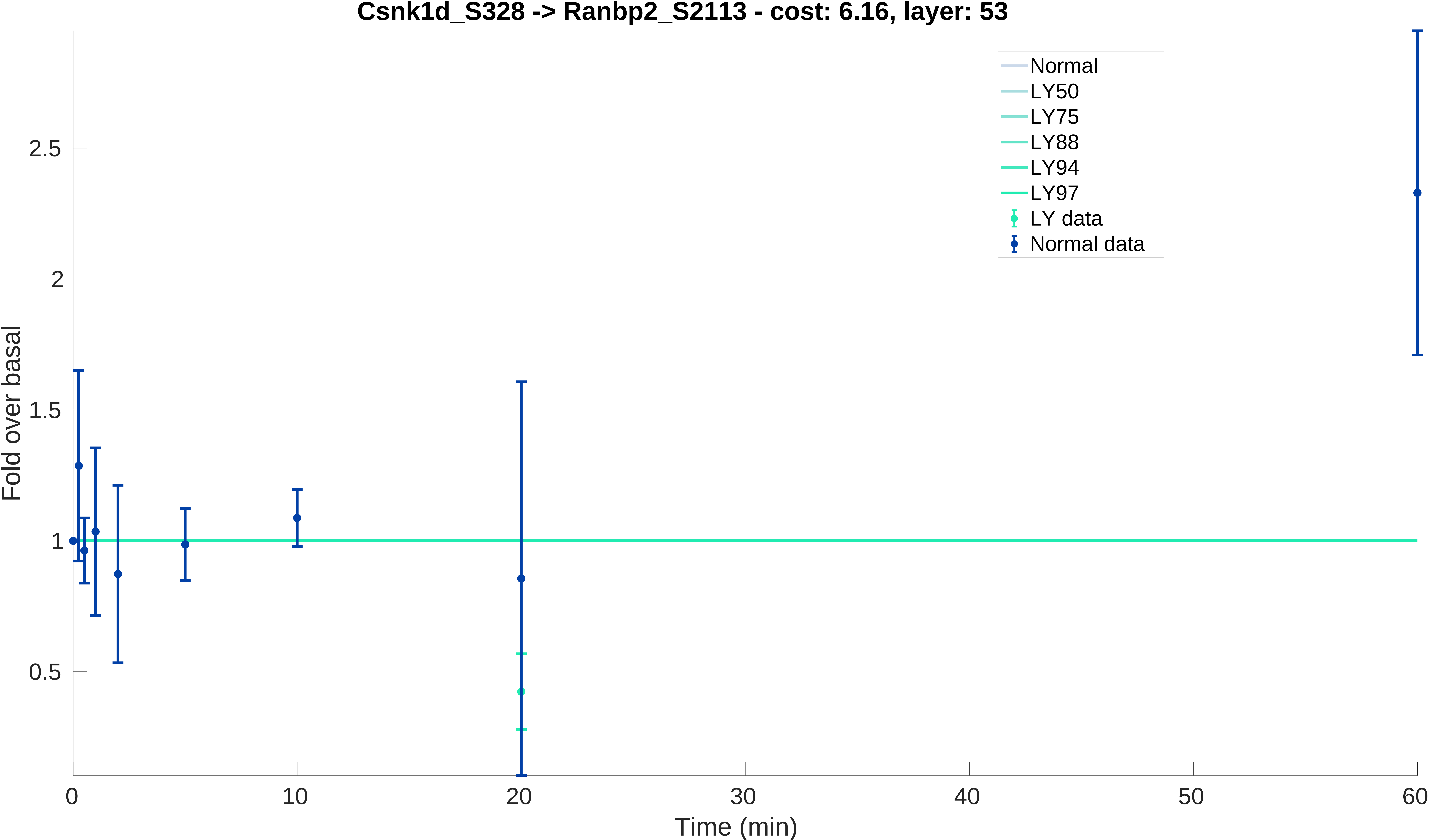

Csnk1d_S328 -> Ranbp2_S2113 - cost: 6.16, layer: 53
Normal
LY50
LY75
LY88
LY94
LY97
LY data
Normal data
2.5
2
1.5
1
0.5
Fold over basal
0
10
20
30
40
50
60
Time (min)

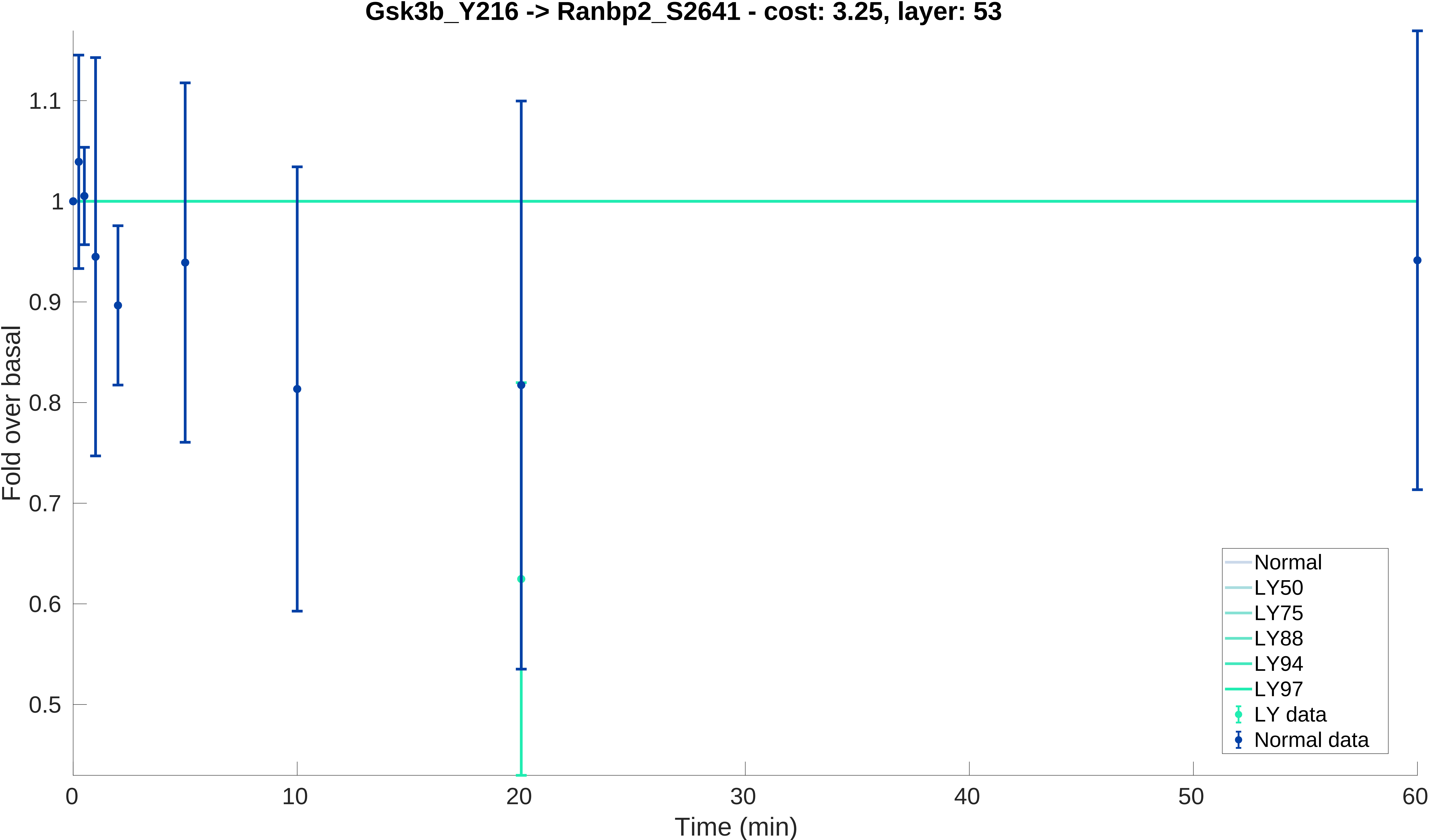
Gsk3b_Y216 -> Ranbp2_S2641 - cost: 3.25, layer: 53
Fold over basal
1.1
1
0.9
0.8
0.7
0.6
0.5
0
10
20
30
40
50
60
Time (min)
Normal
LY50
LY75
LY88
LY94
LY97
LY data
Normal data

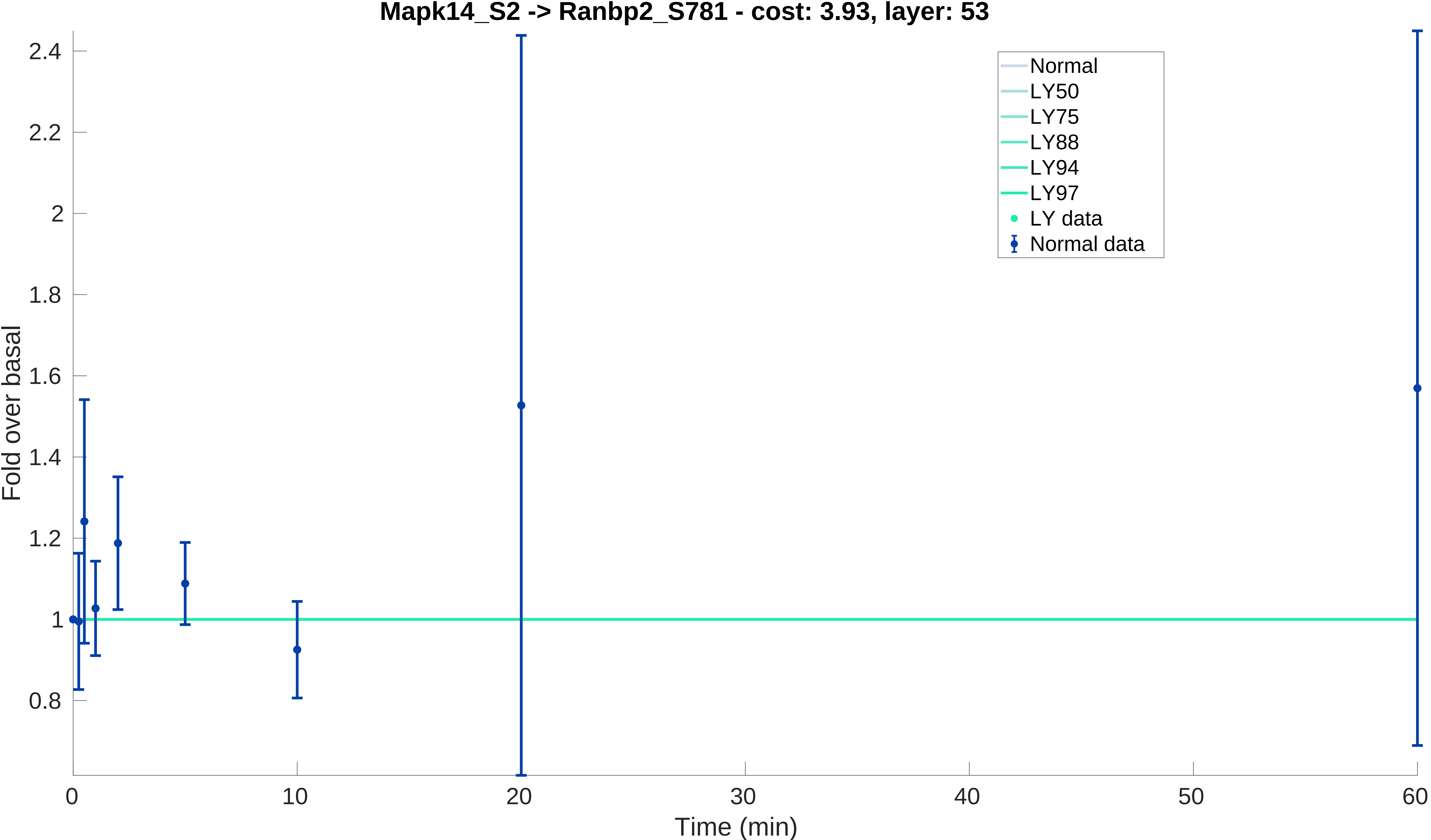

Mapk14_S2 -> Ranbp2_S781 - cost: 3.93, layer: 53
Fold over basal
Time (min)
0.8
1
1.2
1.4
1.6
1.8
2
2.2
2.4
0
10
20
30
40
50
60
Normal
LY50
LY75
LY88
LY94
LY97
LY data
Normal data

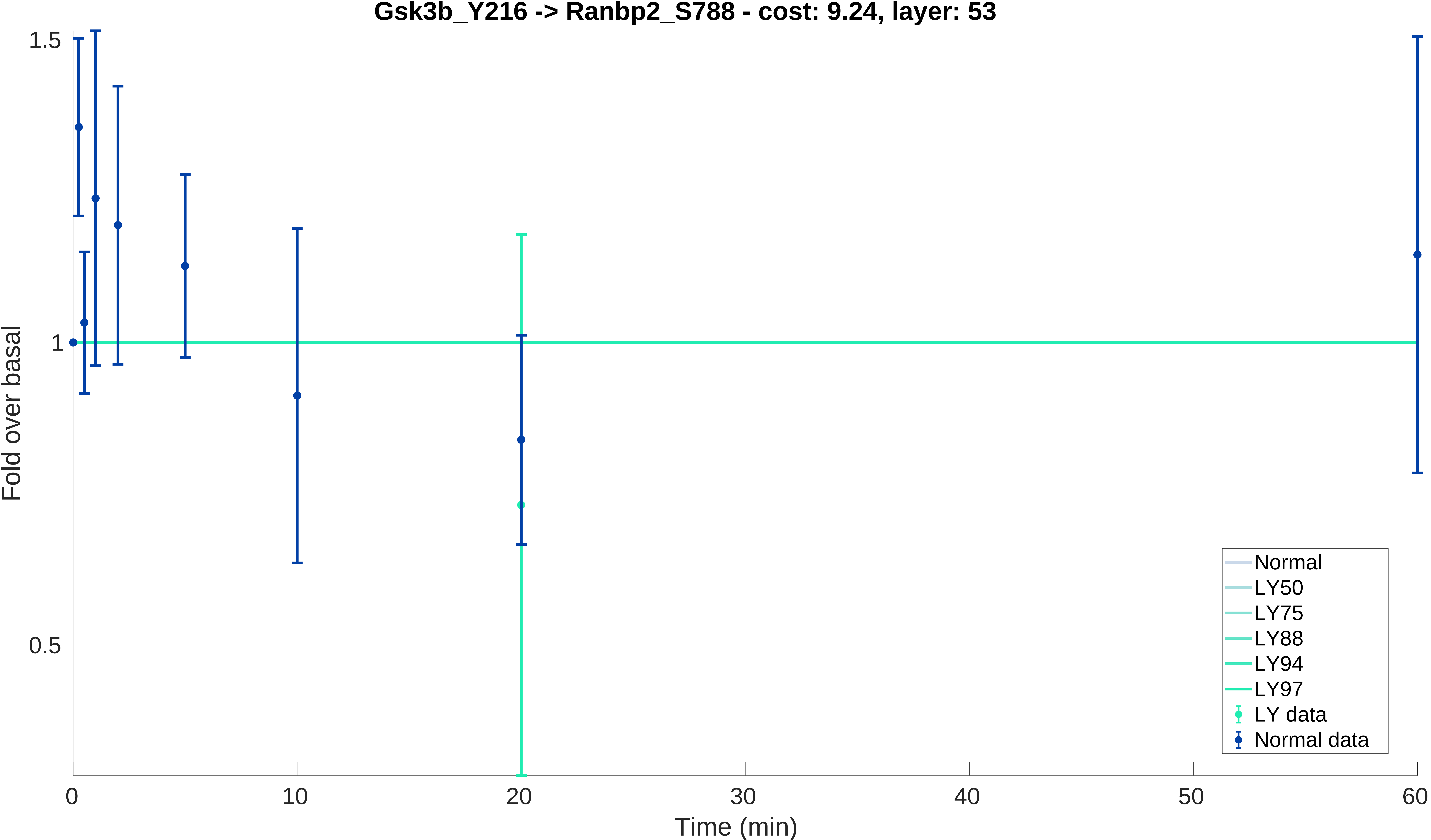

Gsk3b_Y216 -> Ranbp2_S788 - cost: 9.24, layer: 53
1.5
1
0.5
Fold over basal
0
10
20
30
40
50
60
Time (min)
Normal
LY50
LY75
LY88
LY94
LY97
LY data
Normal data

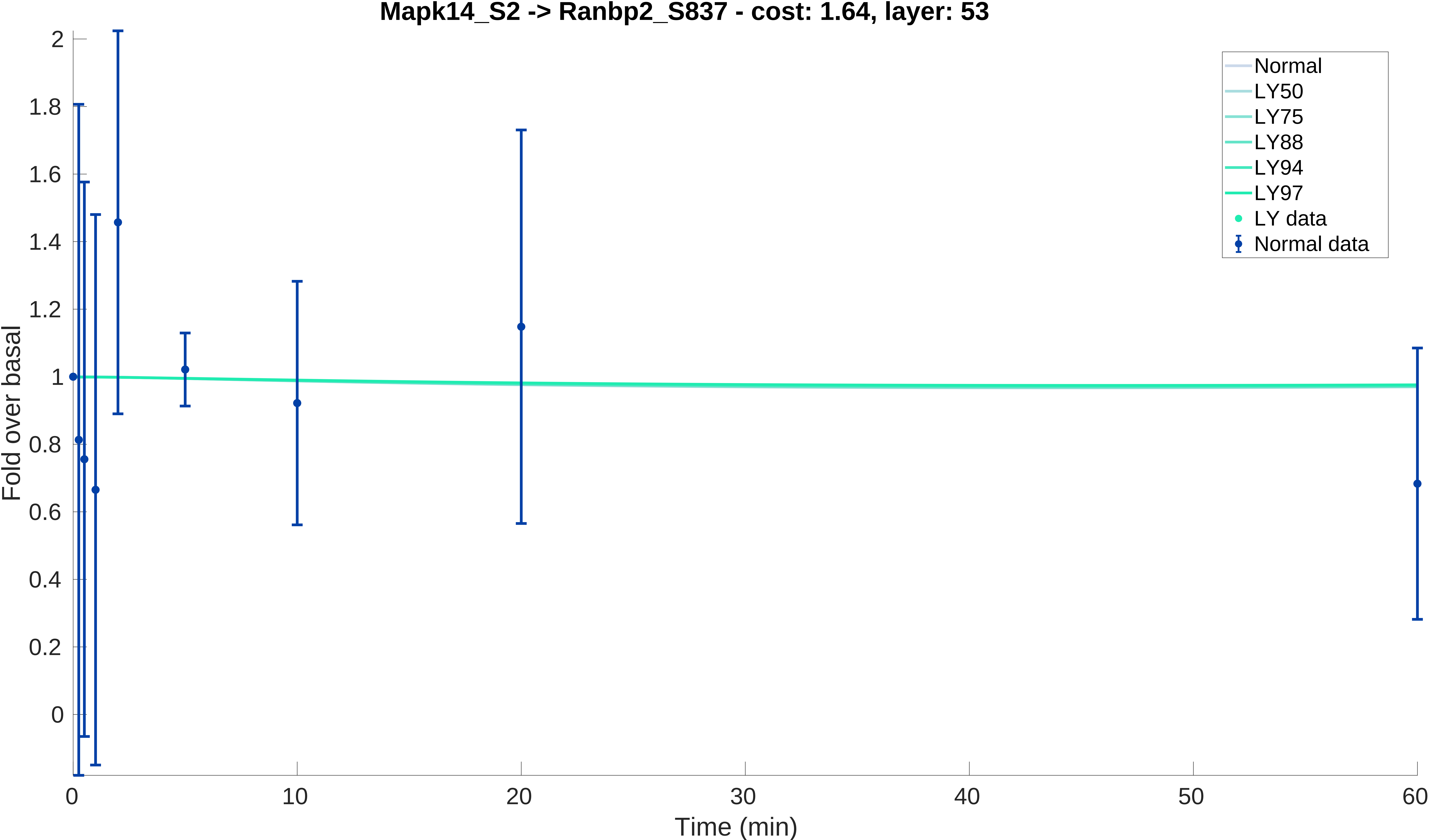

Mapk14_S2 -> Ranbp2_S837 - cost: 1.64, layer: 53
Fold over basal
Time (min)
2
1.8
1.6
1.4
1.2
1
0.8
0.6
0.4
0.2
0
0
10
20
30
40
50
60
Normal
LY50
LY75
LY88
LY94
LY97
LY data
Normal data

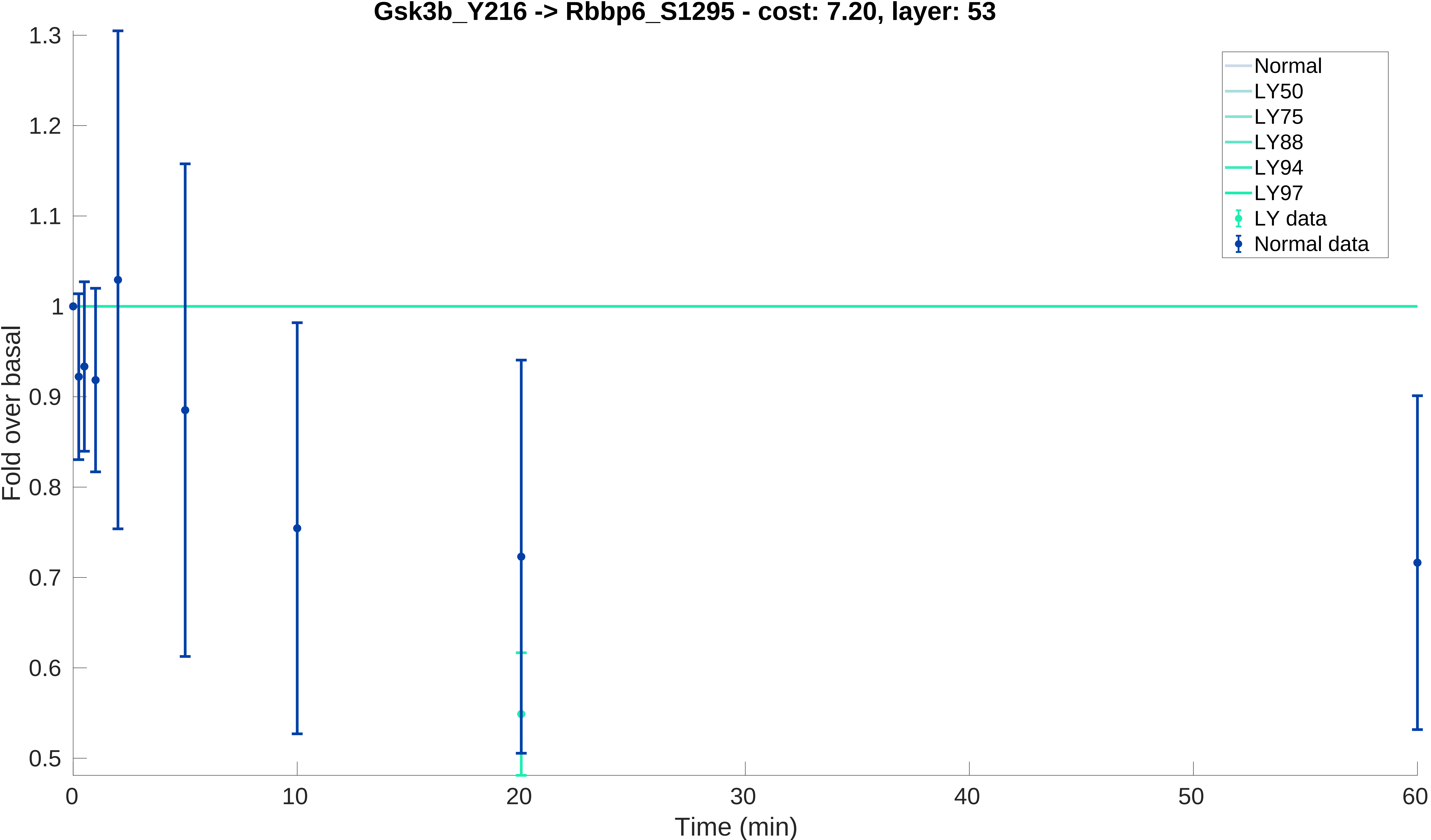

Gsk3b_Y216 -> Rbbp6_S1295 - cost: 7.20, layer: 53
Fold over basal
Time (min)
1.3
1.2
1.1
1
0.9
0.8
0.7
0.6
0.5
0
10
20
30
40
50
60
Normal
LY50
LY75
LY88
LY94
LY97
LY data
Normal data

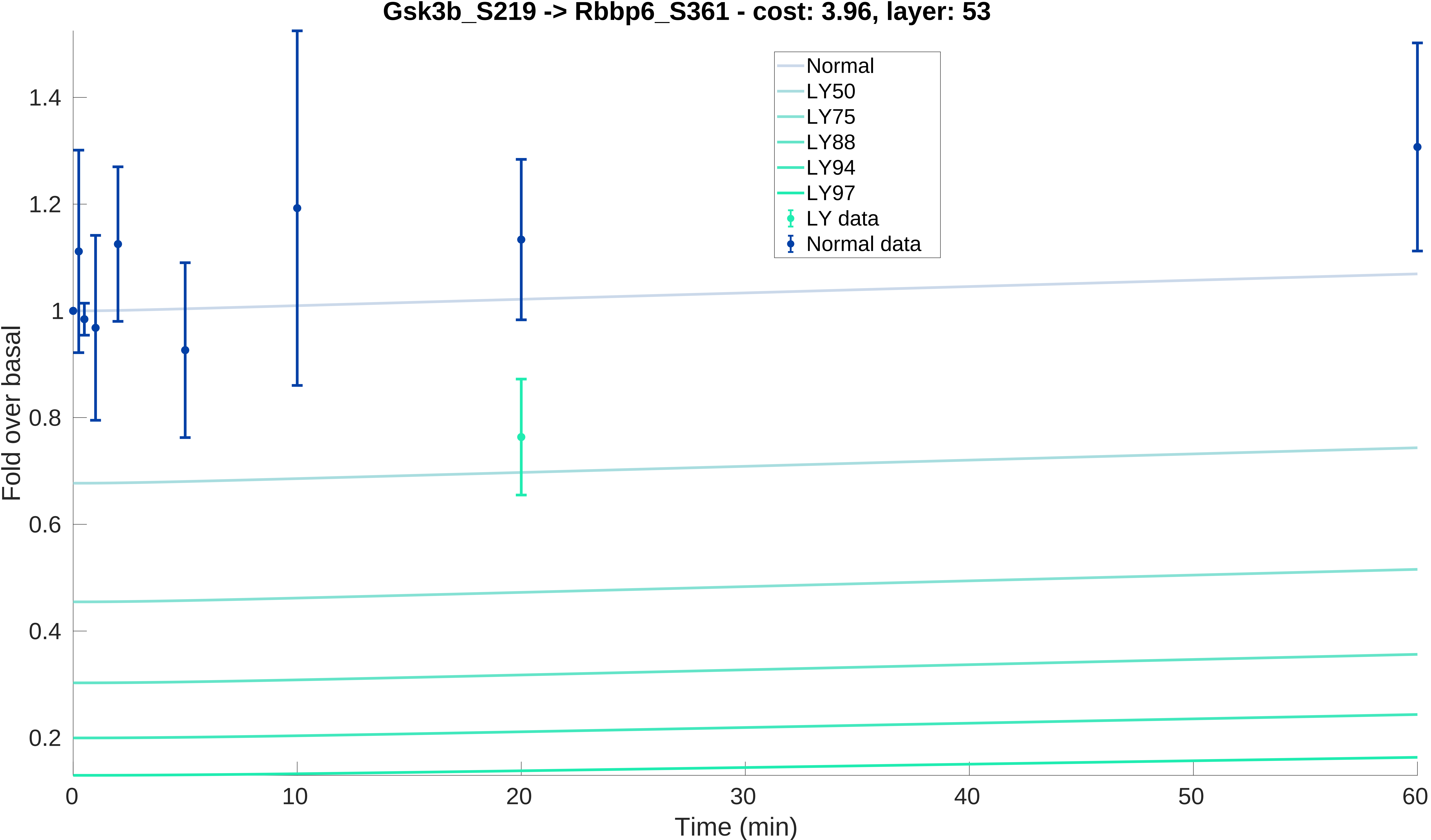
Gsk3b_S219 -> Rbbp6_S361 - cost: 3.96, layer: 53
Normal
LY50
LY75
LY88
LY94
LY97
LY data
Normal data
Fold over basal
Time (min)
1.4
1.2
1
0.8
0.6
0.4
0.2
0
10
20
30
40
50
60

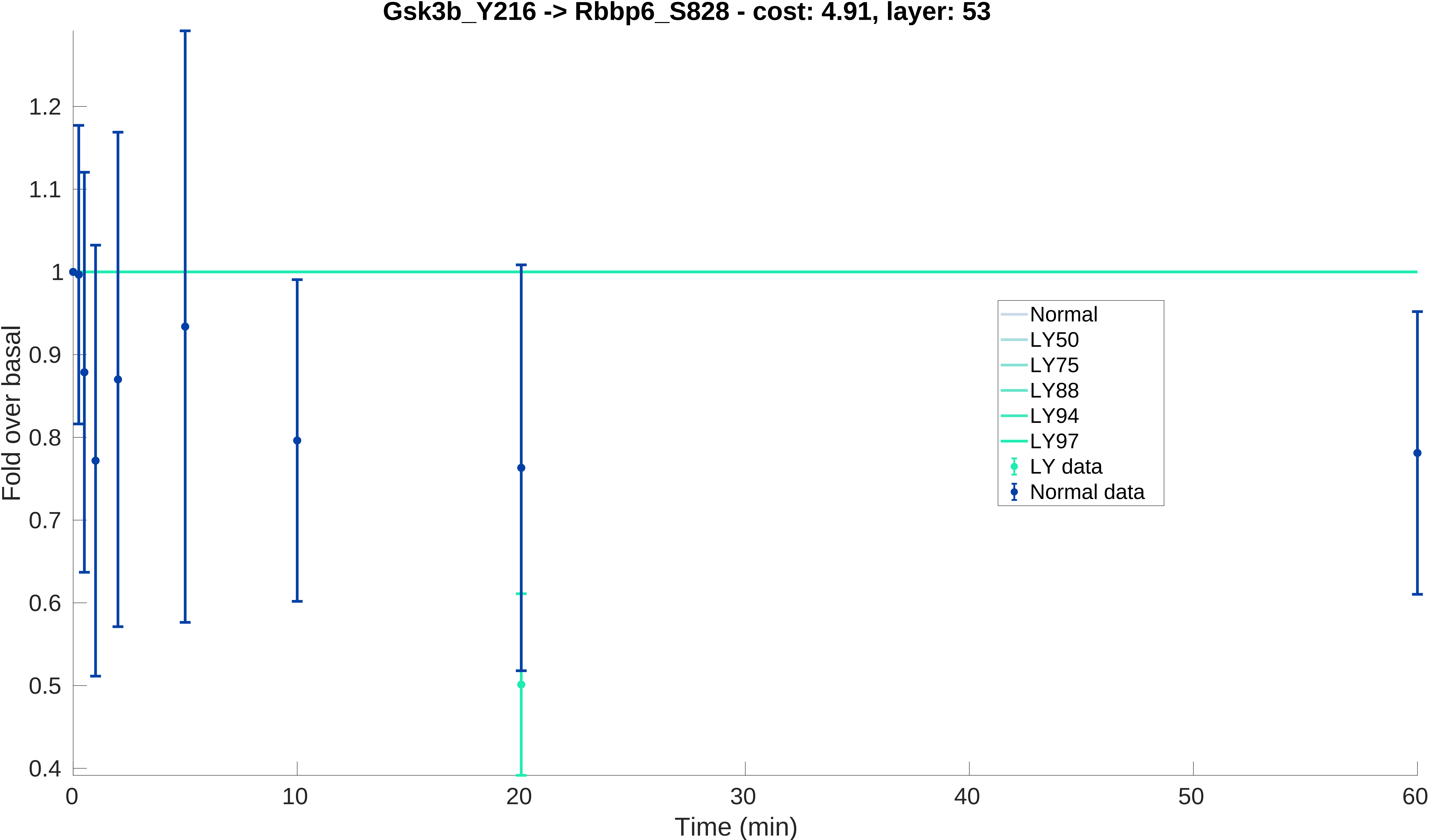

Gsk3b_Y216 -> Rbbp6_S828 - cost: 4.91, layer: 53
Fold over basal
Time (min)
0
10
20
30
40
50
60
0.4
0.5
0.6
0.7
0.8
0.9
1
1.1
1.2
Normal
LY50
LY75
LY88
LY94
LY97
LY data
Normal data

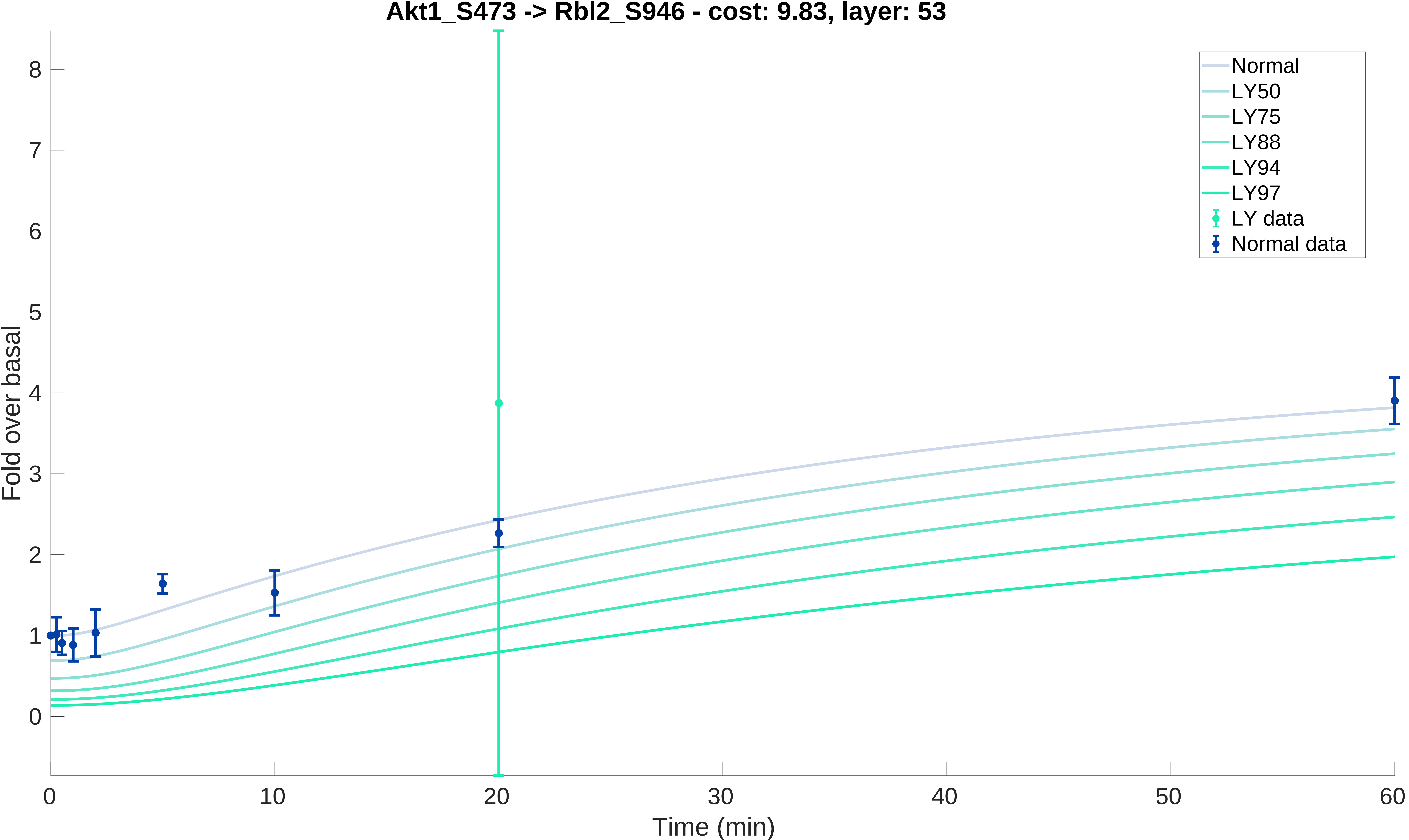
Akt1_S473 -> Rbl2_S946 - cost: 9.83, layer: 53
Fold over basal
Time (min)
0
1
2
3
4
5
6
7
8
0
10
20
30
40
50
60
Normal
LY50
LY75
LY88
LY94
LY97
LY data
Normal data

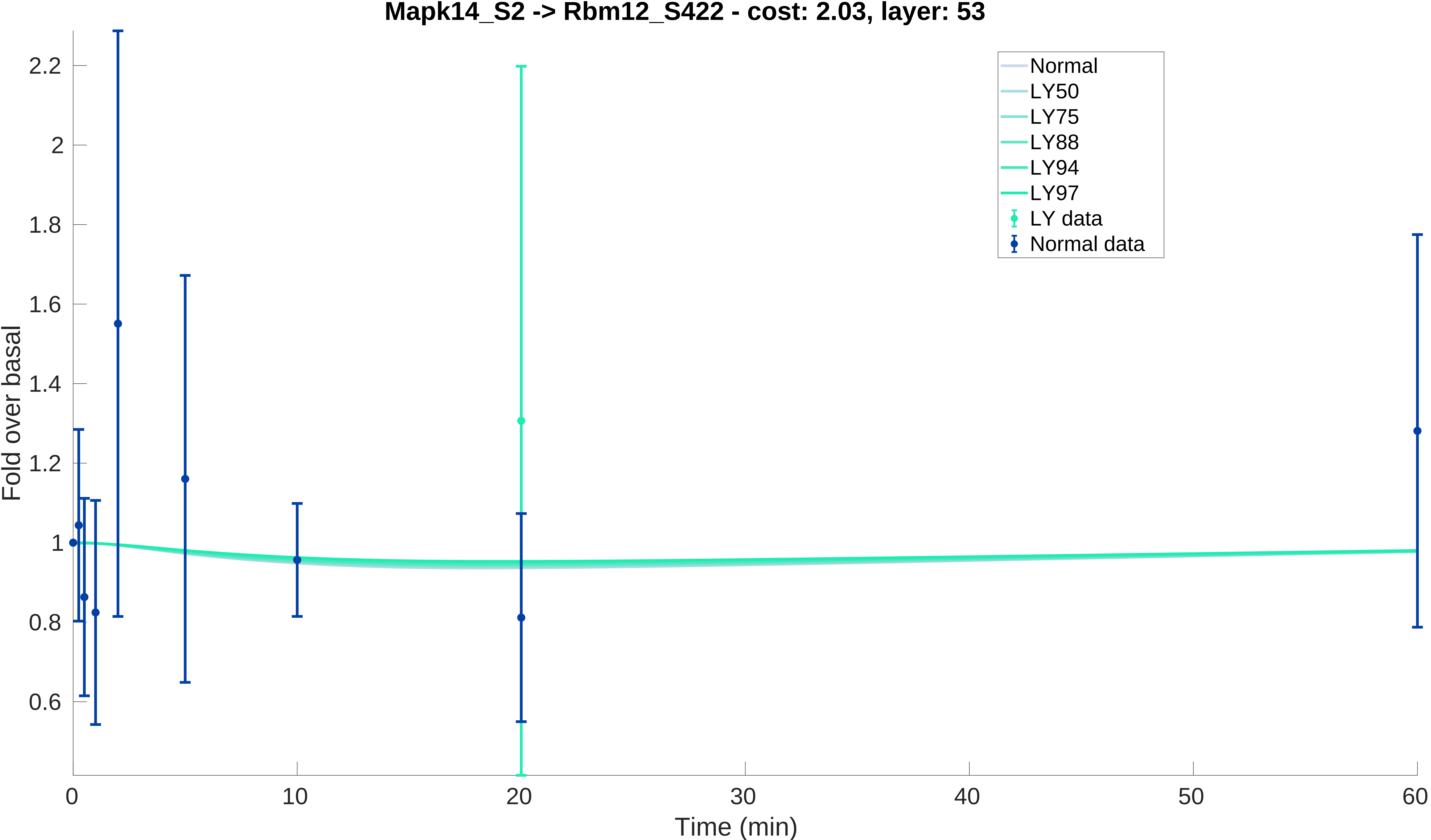

Mapk14_S2 -> Rbm12_S422 - cost: 2.03, layer: 53
Fold over basal
Time (min)
Normal
LY50
LY75
LY88
LY94
LY97
LY data
Normal data
0.6
0.8
1
1.2
1.4
1.6
1.8
2
2.2
0
10
20
30
40
50
60

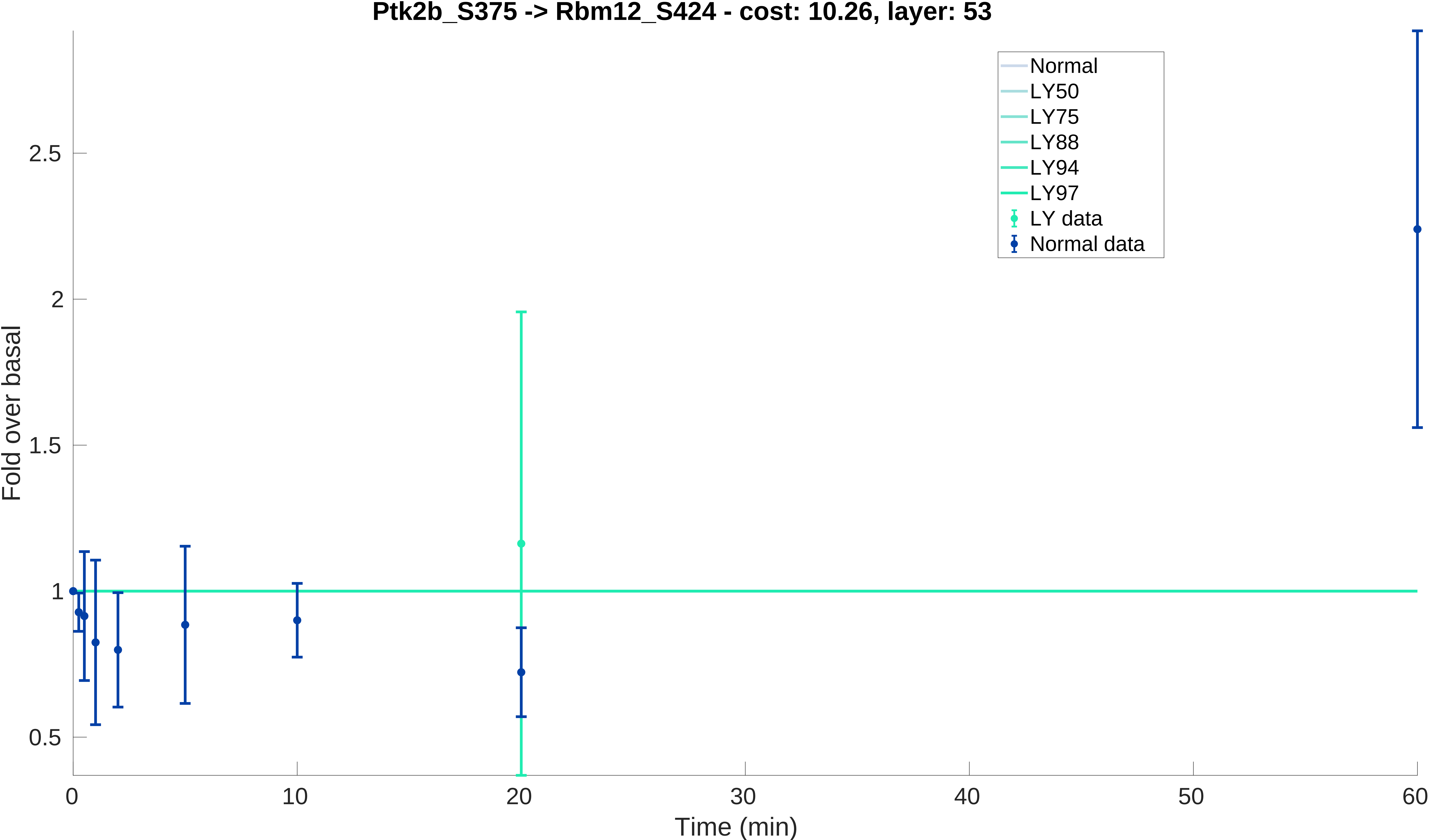

Ptk2b_S375 -> Rbm12_S424 - cost: 10.26, layer: 53
Normal
LY50
LY75
LY88
LY94
LY97
LY data
Normal data
Fold over basal
2.5
2
1.5
1
0.5
0
10
20
30
40
50
60
Time (min)

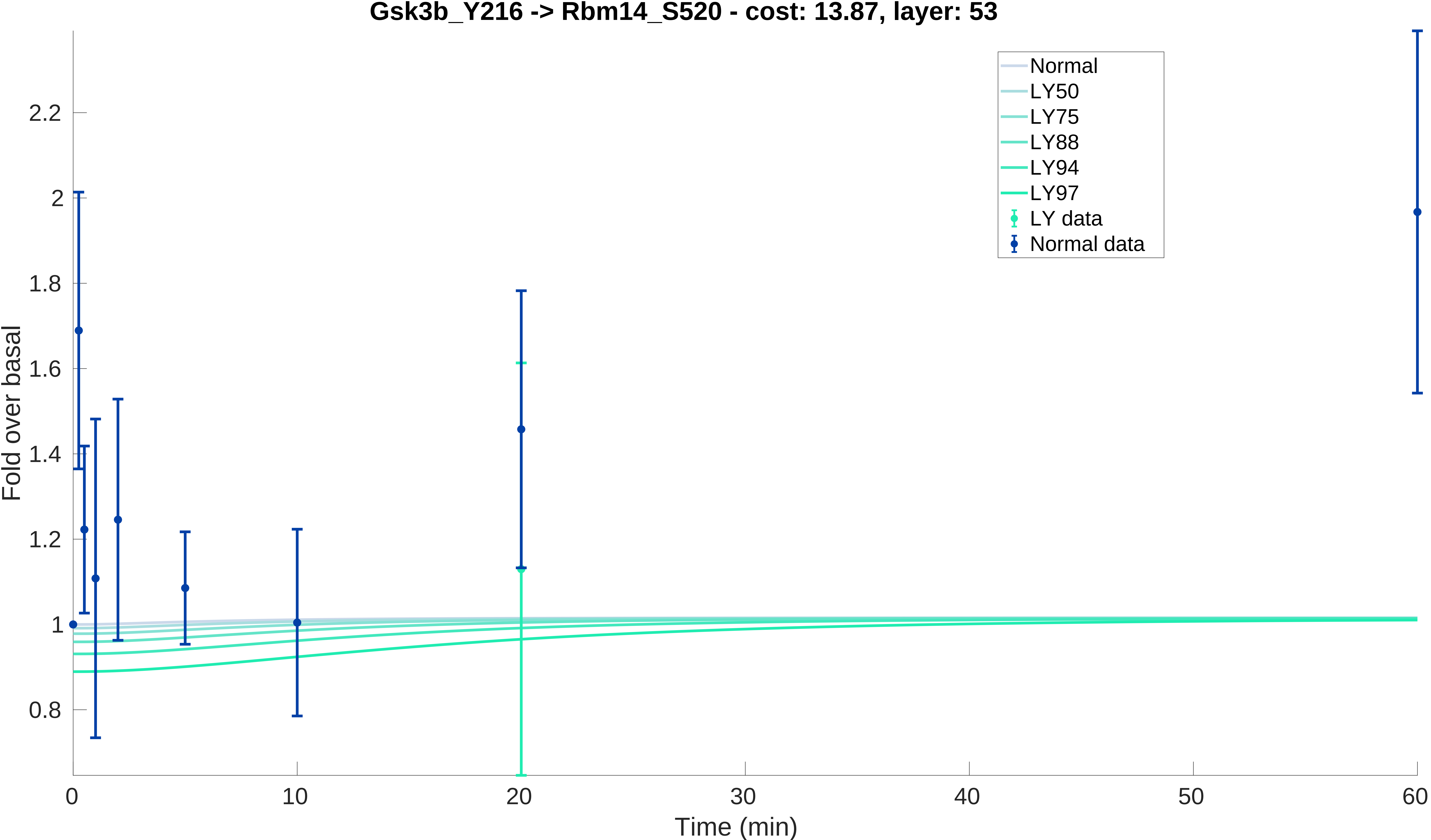

Gsk3b_Y216 -> Rbm14_S520 - cost: 13.87, layer: 53
Normal
LY50
LY75
LY88
LY94
LY97
LY data
Normal data
Fold over basal
0.8
1
1.2
1.4
1.6
1.8
2
2.2
0
10
20
30
40
50
60
Time (min)

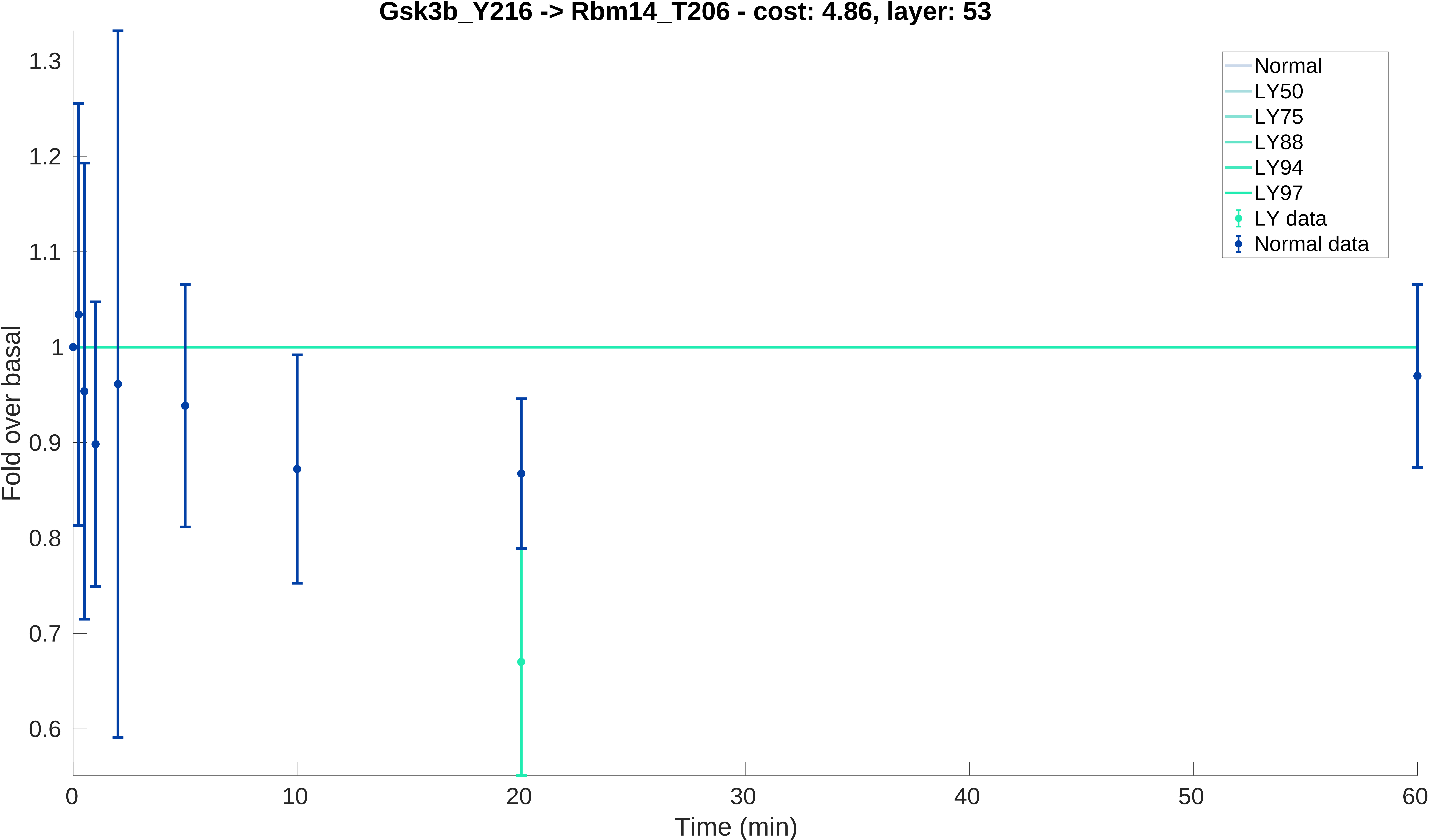

Gsk3b_Y216 -> Rbm14_T206 - cost: 4.86, layer: 53
Fold over basal
Time (min)
1.3
1.2
1.1
1
0.9
0.8
0.7
0.6
0
10
20
30
40
50
60
Normal
LY50
LY75
LY88
LY94
LY97
LY data
Normal data

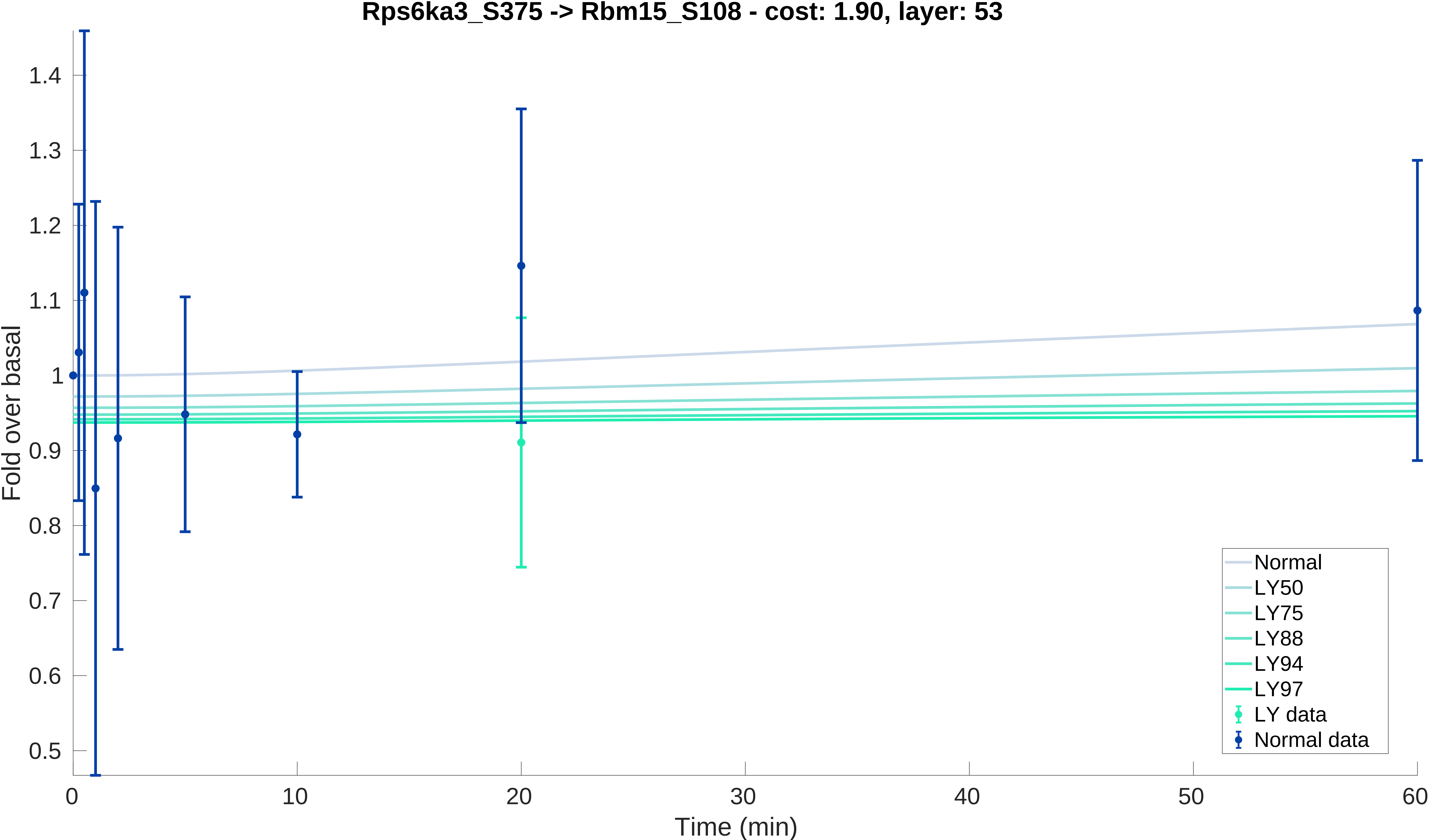

Rps6ka3_S375 -> Rbm15_S108 - cost: 1.90, layer: 53
Fold over basal
Time (min)
1.4
1.3
1.2
1.1
1
0.9
0.8
0.7
0.6
0.5
0
10
20
30
40
50
60
Normal
LY50
LY75
LY88
LY94
LY97
LY data
Normal data

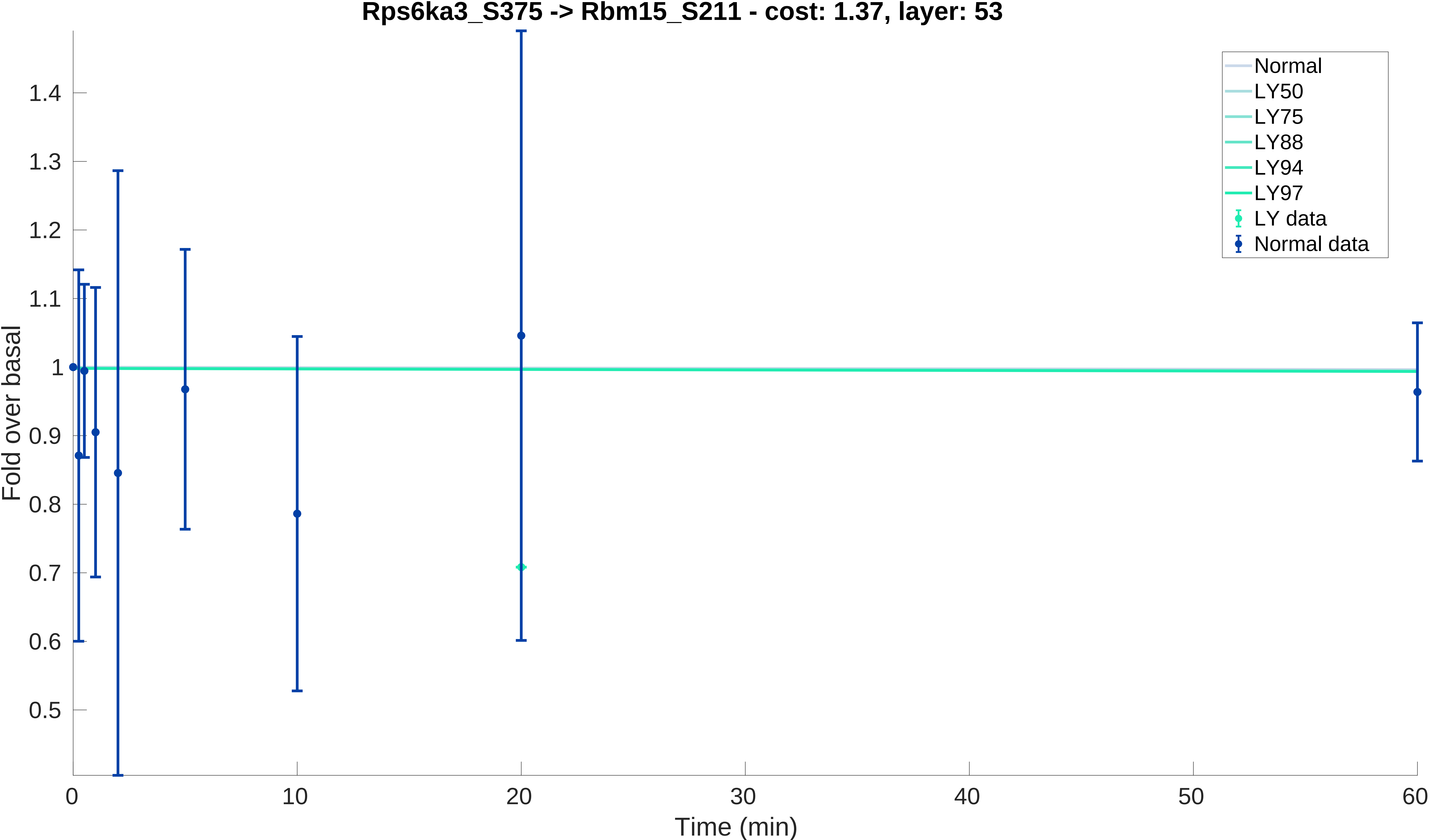

Rps6ka3_S375 -> Rbm15_S211 - cost: 1.37, layer: 53
Normal
LY50
LY75
LY88
LY94
LY97
LY data
Normal data
Fold over basal
1.4
1.3
1.2
1.1
1
0.9
0.8
0.7
0.6
0.5
0
10
20
30
40
50
60
Time (min)

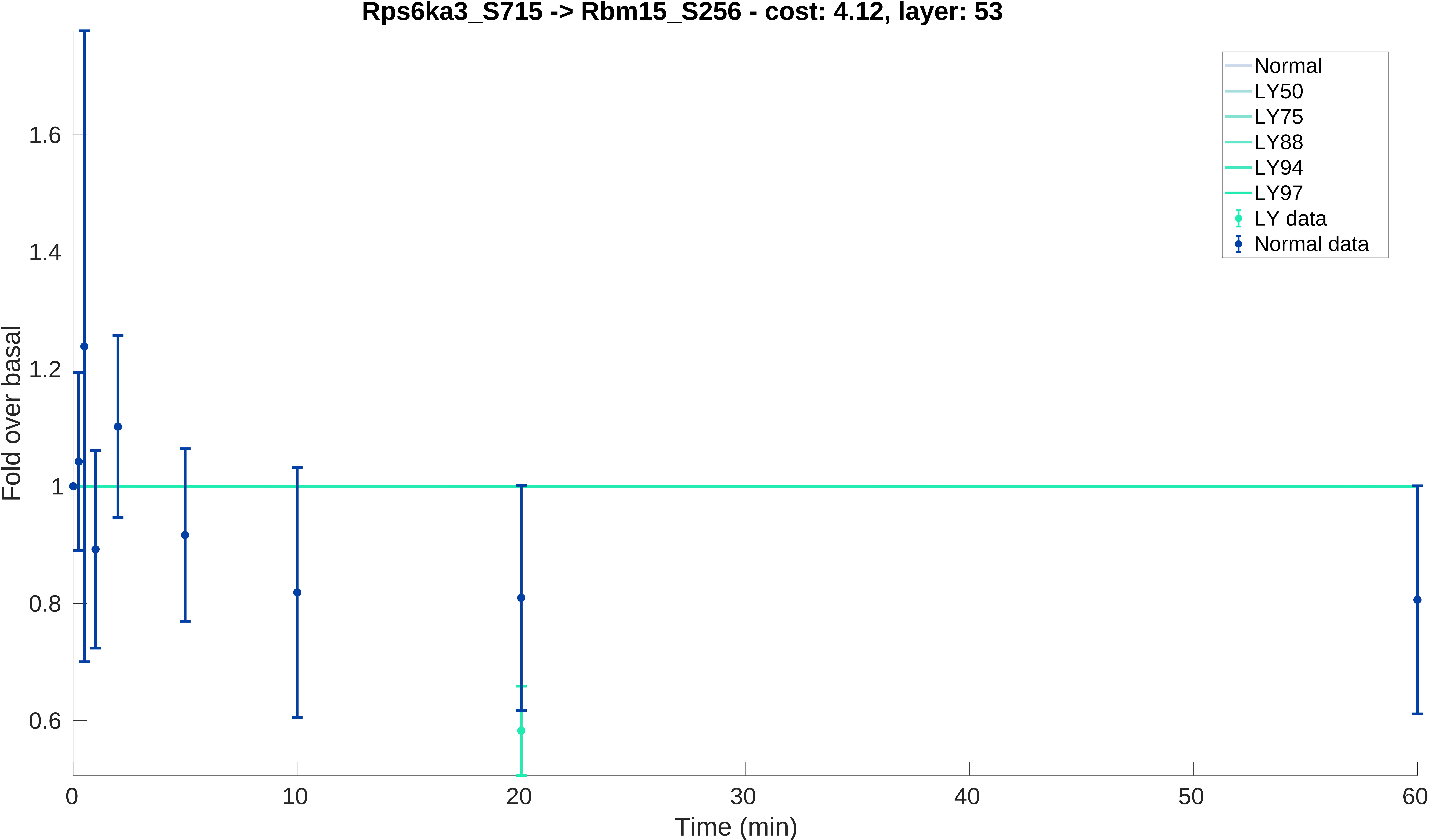
Rps6ka3_S715 -> Rbm15_S256 - cost: 4.12, layer: 53
Normal
LY50
LY75
LY88
LY94
LY97
LY data
Normal data
Fold over basal
1.6
1.4
1.2
1
0.8
0.6
0
10
20
30
40
50
60
Time (min)

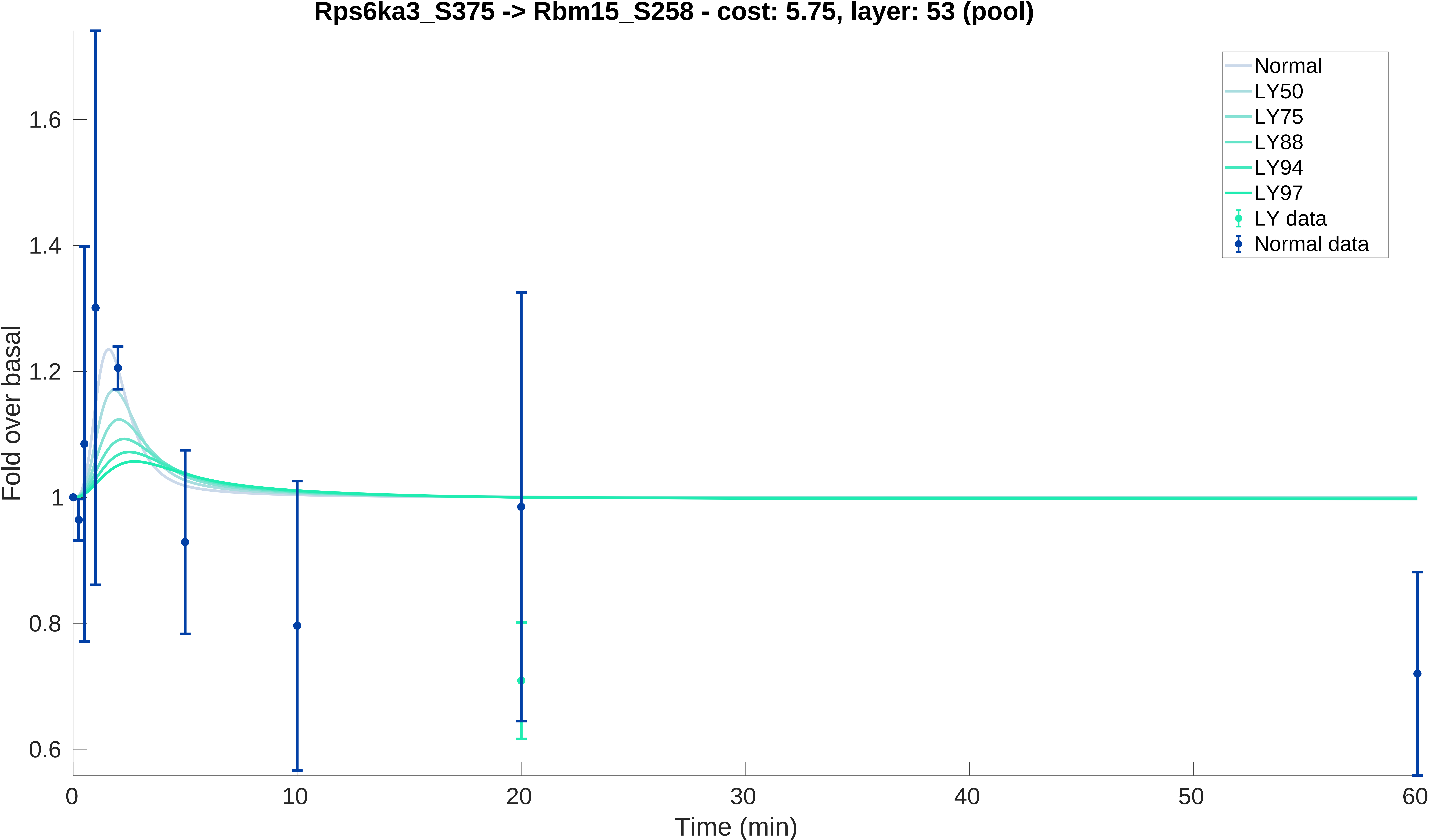

Rps6ka3_S375 -> Rbm15_S258 - cost: 5.75, layer: 53 (pool)
Normal
LY50
LY75
LY88
LY94
LY97
LY data
Normal data
Fold over basal
1.6
1.4
1.2
1
0.8
0.6
0
10
20
30
40
50
60
Time (min)

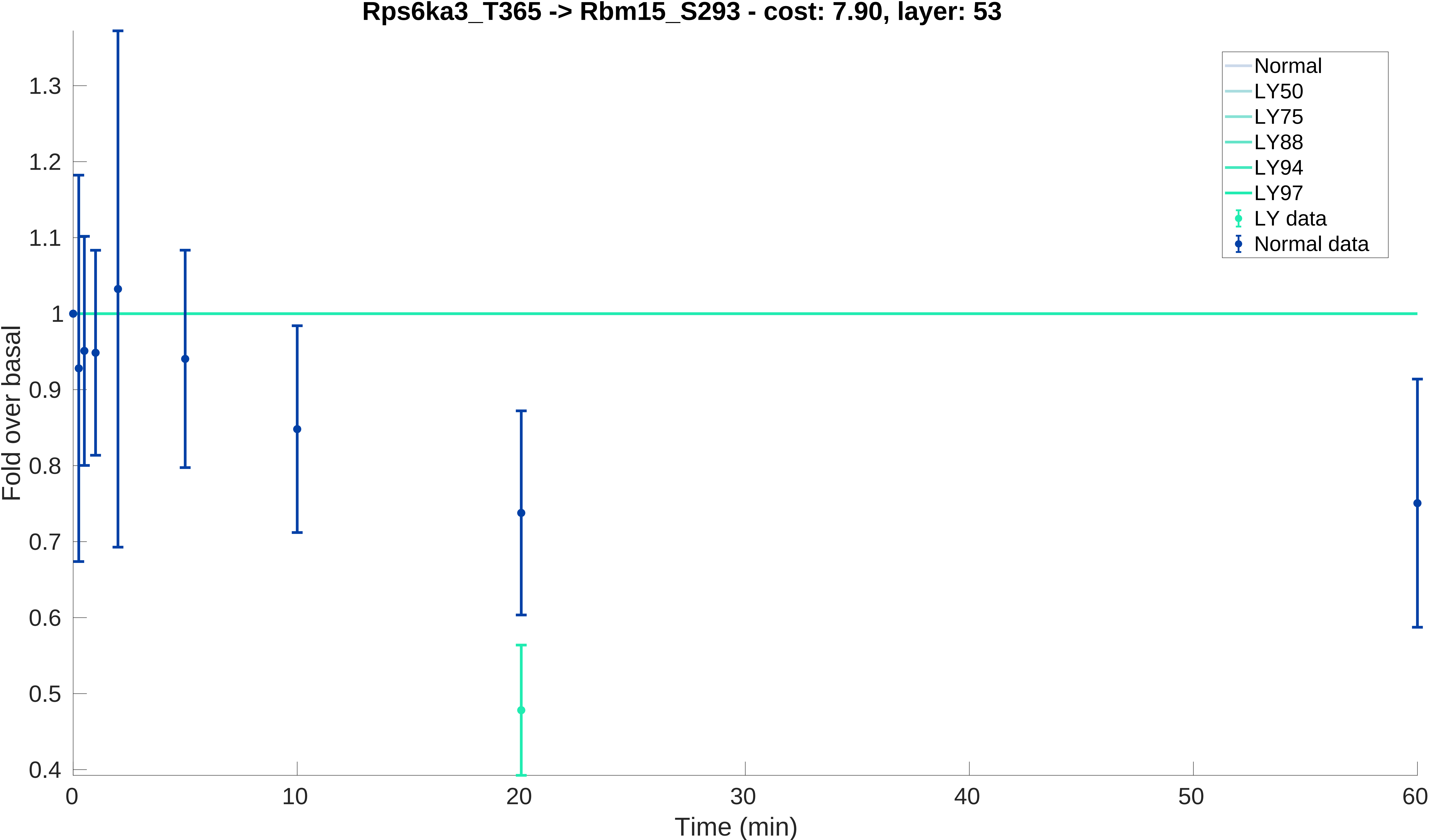

Rps6ka3_T365 -> Rbm15_S293 - cost: 7.90, layer: 53
Fold over basal
1.3
1.2
1.1
1
0.9
0.8
0.7
0.6
0.5
0.4
0
10
20
30
40
50
60
Time (min)
Normal
LY50
LY75
LY88
LY94
LY97
LY data
Normal data

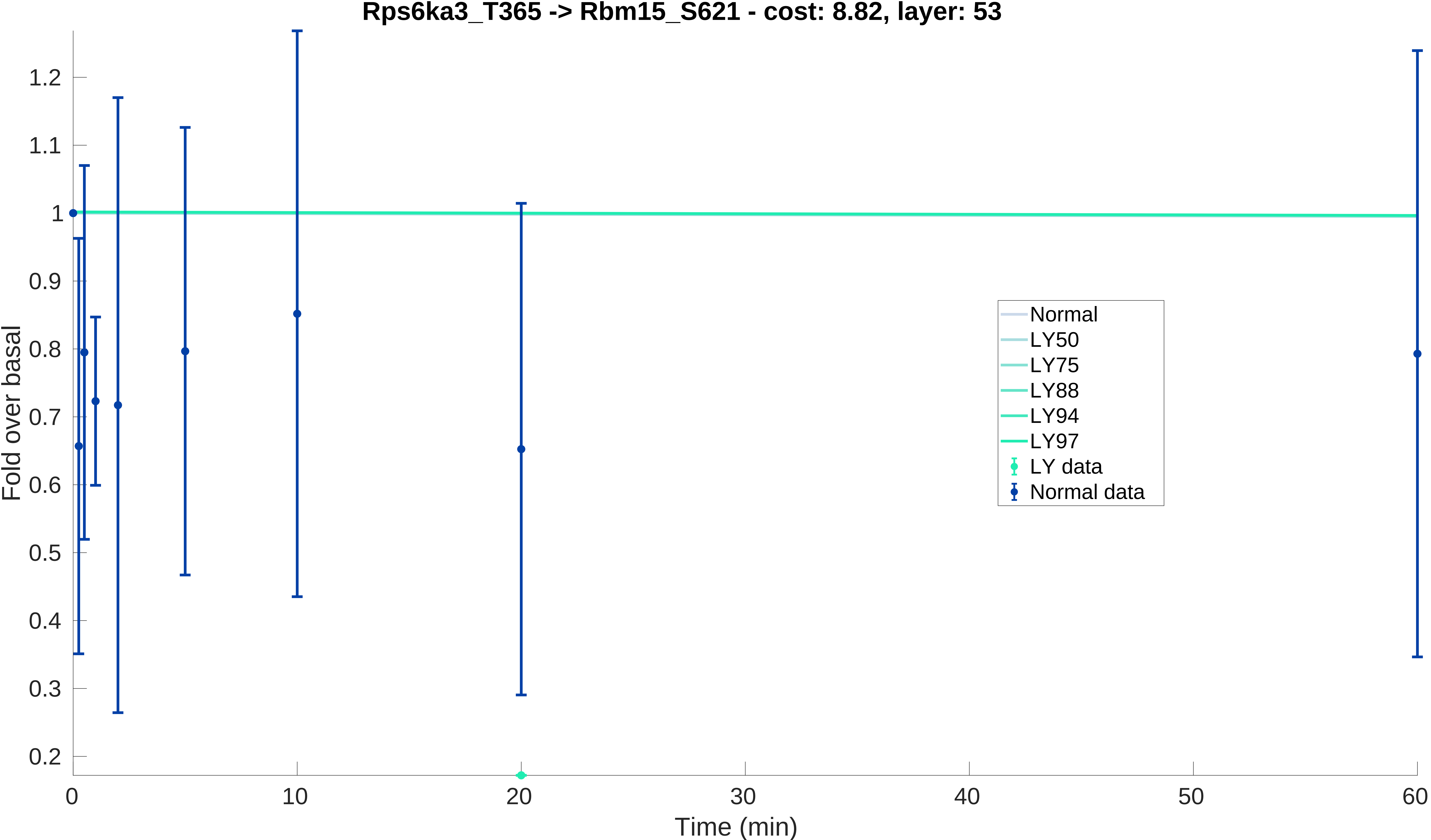

Rps6ka3_T365 -> Rbm15_S621 - cost: 8.82, layer: 53
Fold over basal
Time (min)
0.2
0.3
0.4
0.5
0.6
0.7
0.8
0.9
1
1.1
1.2
0
10
20
30
40
50
60
Normal
LY50
LY75
LY88
LY94
LY97
LY data
Normal data

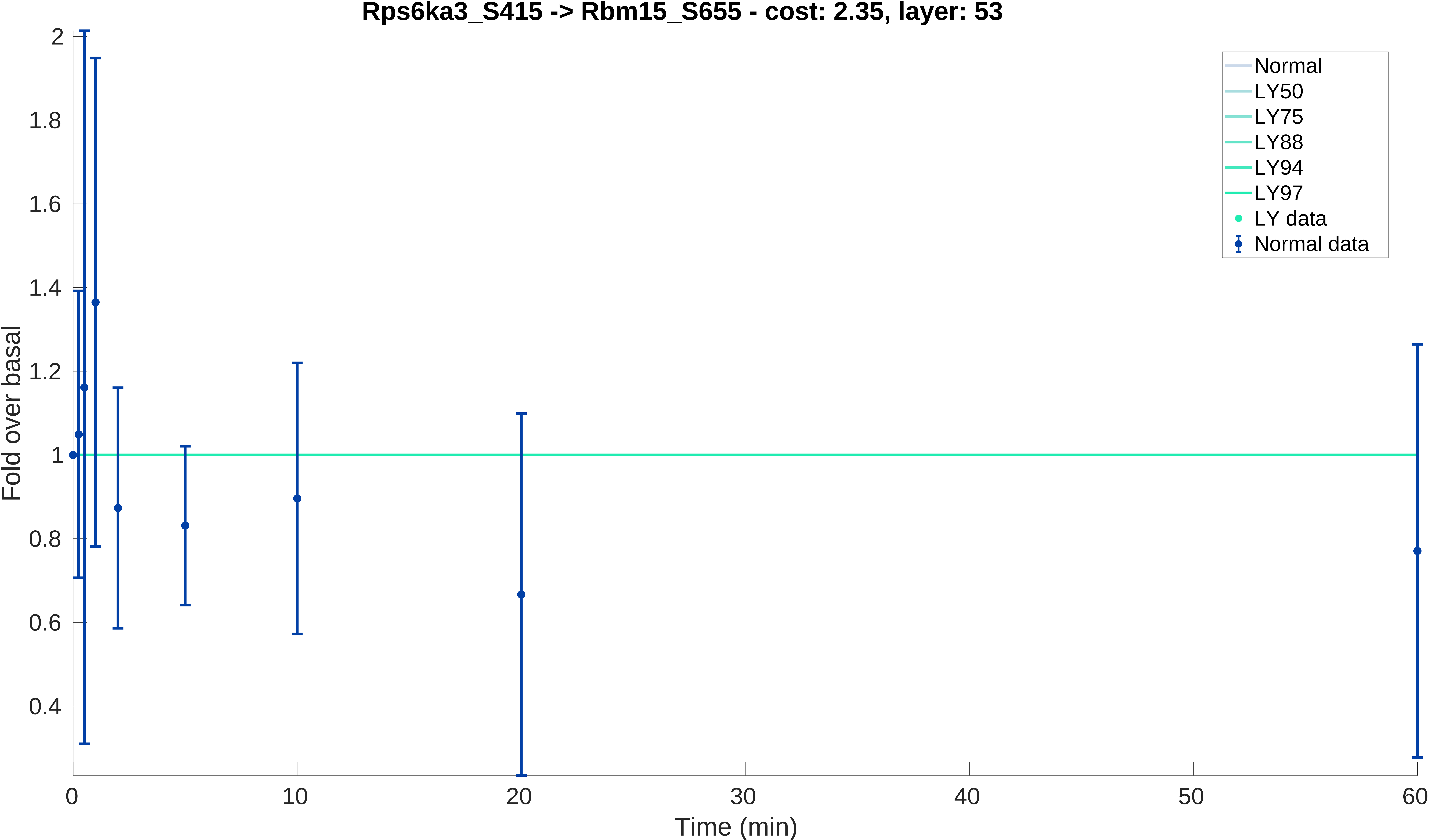

Rps6ka3_S415 -> Rbm15_S655 - cost: 2.35, layer: 53
Fold over basal
Time (min)
0
10
20
30
40
50
60
0.4
0.6
0.8
1
1.2
1.4
1.6
1.8
2
Normal
LY50
LY75
LY88
LY94
LY97
LY data
Normal data

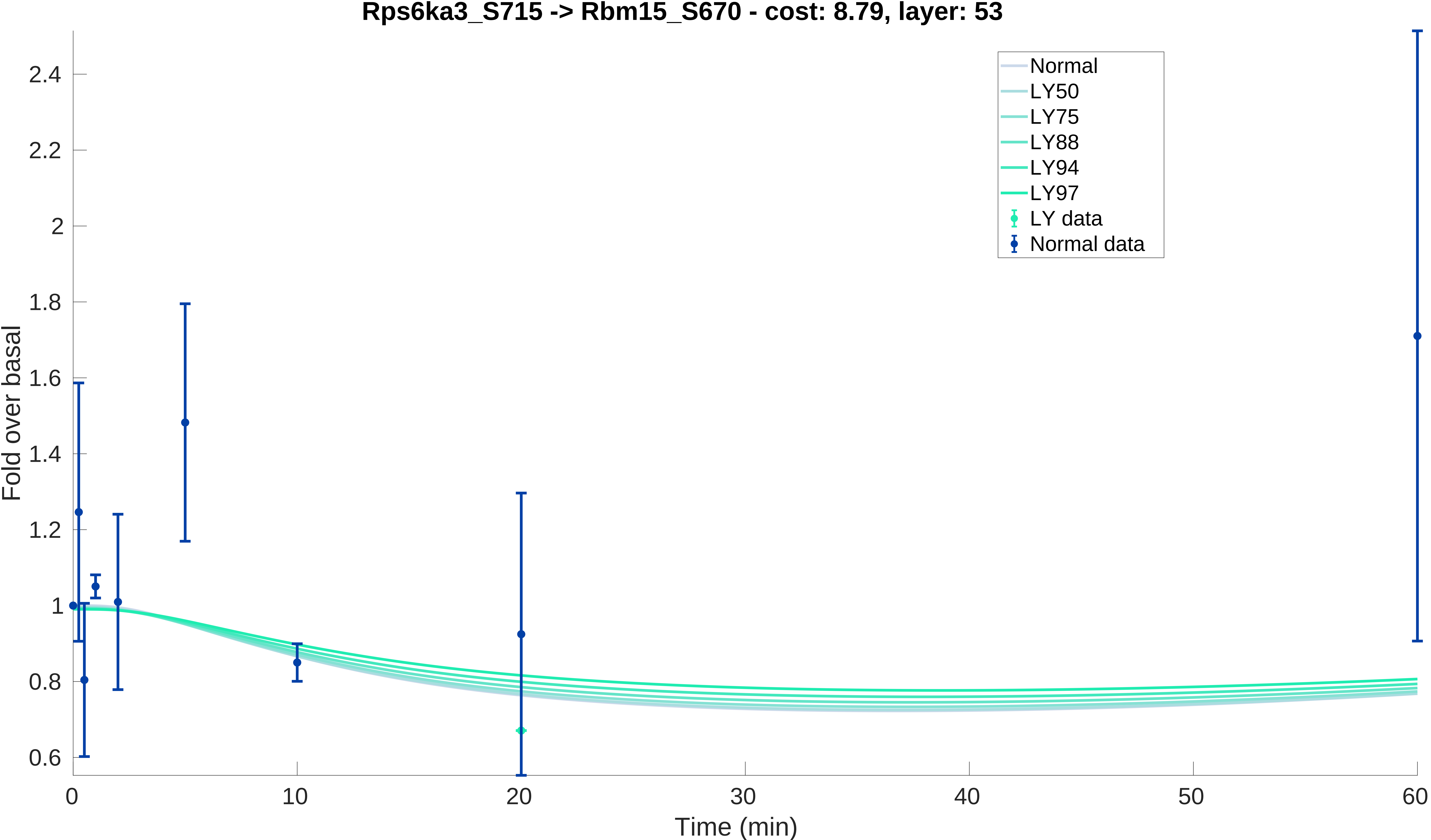

Rps6ka3_S715 -> Rbm15_S670 - cost: 8.79, layer: 53
Fold over basal
Time (min)
Normal
LY50
LY75
LY88
LY94
LY97
LY data
Normal data
0.6
0.8
1
1.2
1.4
1.6
1.8
2
2.2
2.4
0
10
20
30
40
50
60

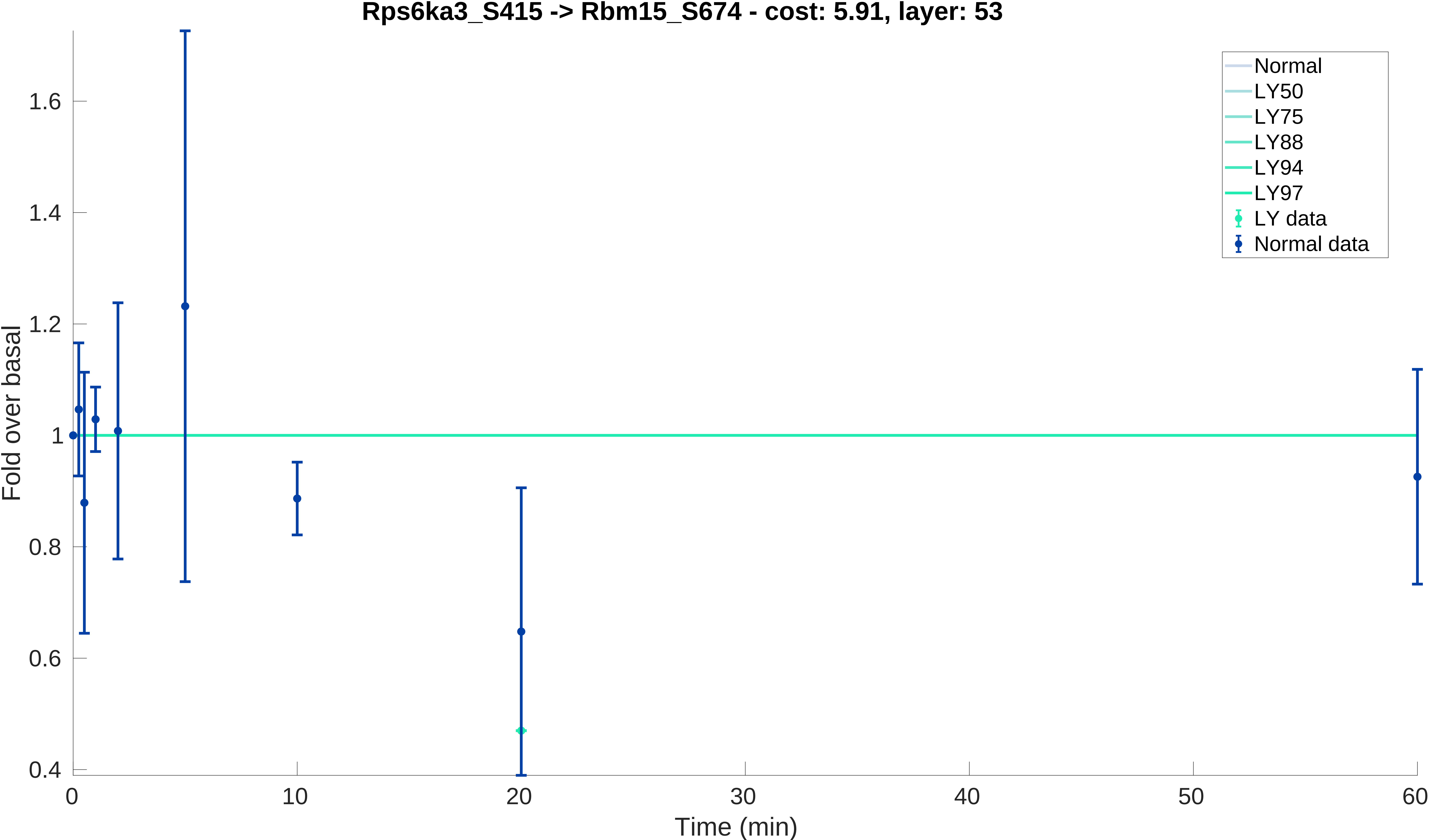
Rps6ka3_S415 -> Rbm15_S674 - cost: 5.91, layer: 53
Fold over basal
Time (min)
0.4
0.6
0.8
1
1.2
1.4
1.6
0
10
20
30
40
50
60
Normal
LY50
LY75
LY88
LY94
LY97
LY data
Normal data

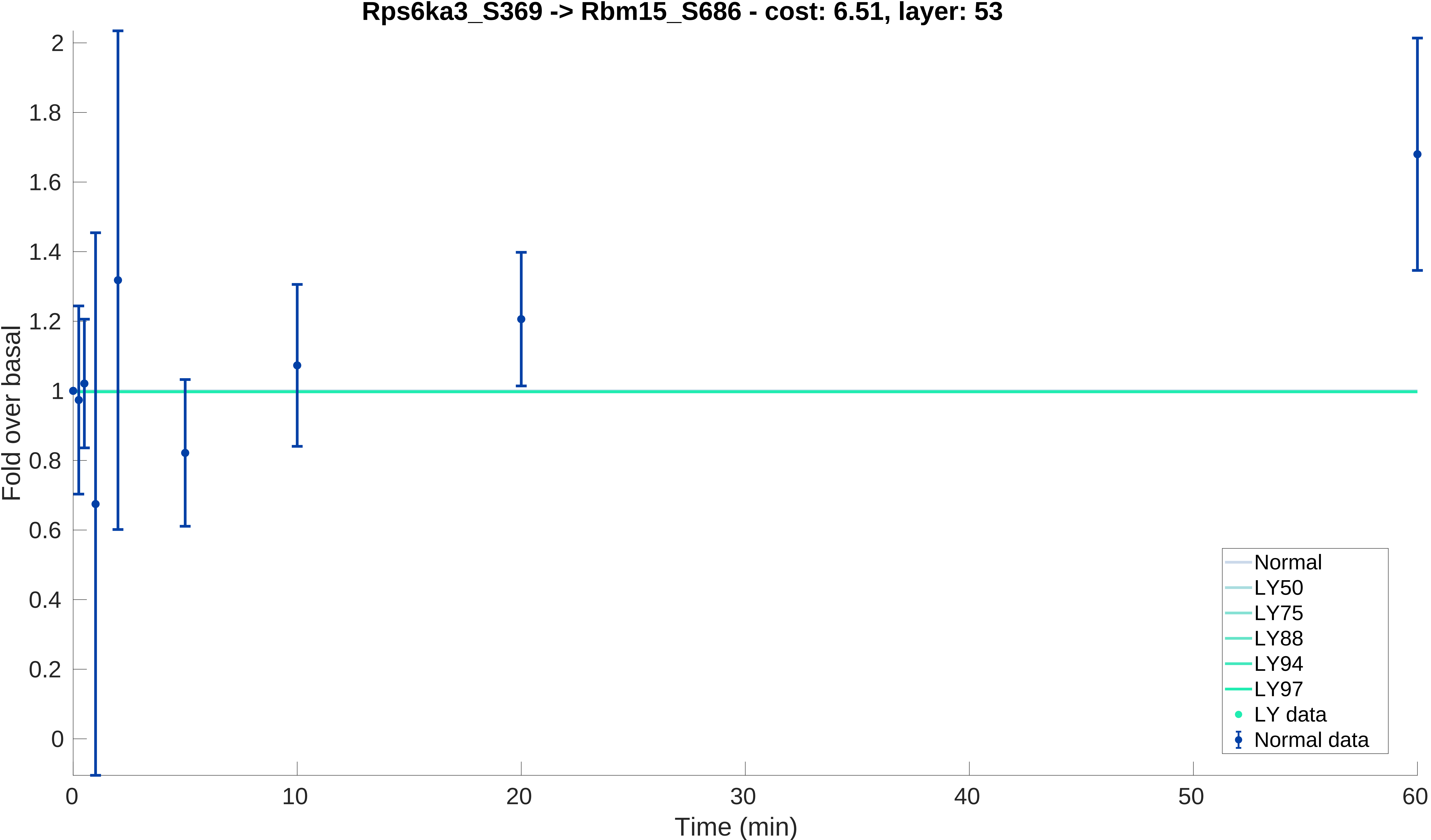

Rps6ka3_S369 -> Rbm15_S686 - cost: 6.51, layer: 53
Fold over basal
Time (min)
0
0.2
0.4
0.6
0.8
1
1.2
1.4
1.6
1.8
2
0
10
20
30
40
50
60
Normal
LY50
LY75
LY88
LY94
LY97
LY data
Normal data

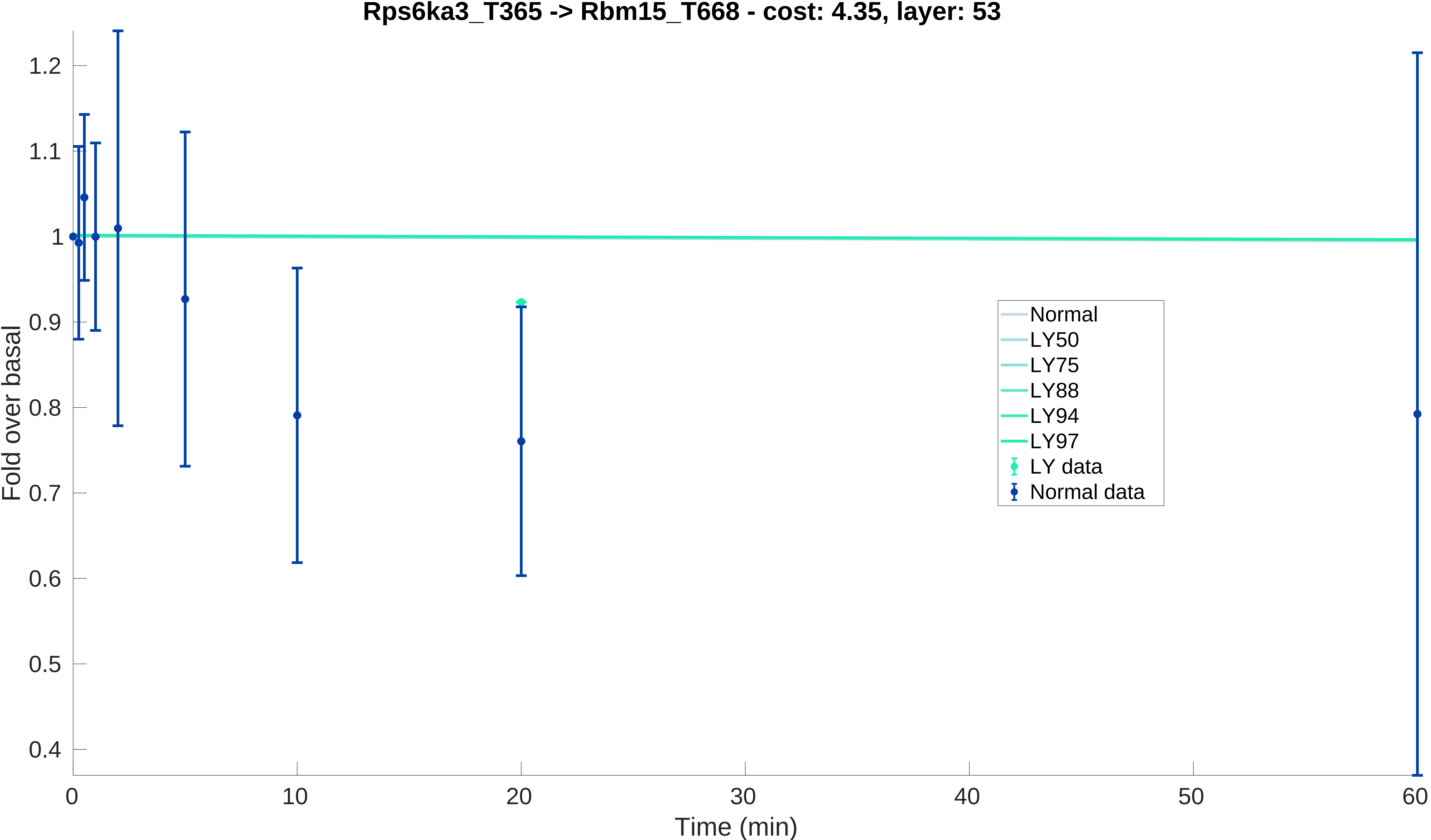

Rps6ka3_T365 -> Rbm15_T668 - cost: 4.35, layer: 53
Fold over basal
1.2
1.1
1
0.9
0.8
0.7
0.6
0.5
0.4
0
10
20
30
40
50
60
Time (min)
Normal
LY50
LY75
LY88
LY94
LY97
LY data
Normal data

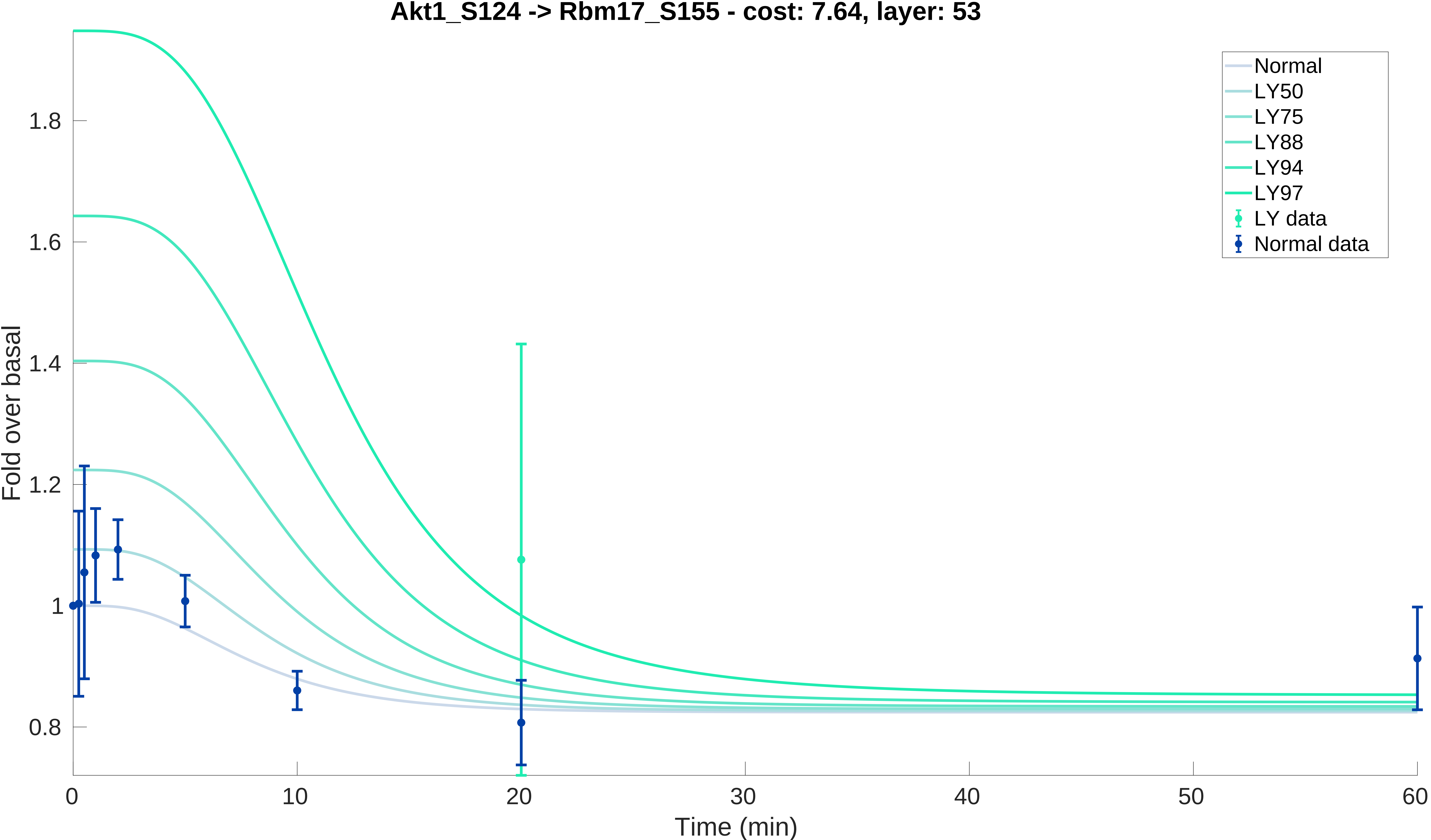

Akt1_S124 -> Rbm17_S155 - cost: 7.64, layer: 53
Fold over basal
Time (min)
0.8
1
1.2
1.4
1.6
1.8
0
10
20
30
40
50
60
Normal
LY50
LY75
LY88
LY94
LY97
LY data
Normal data

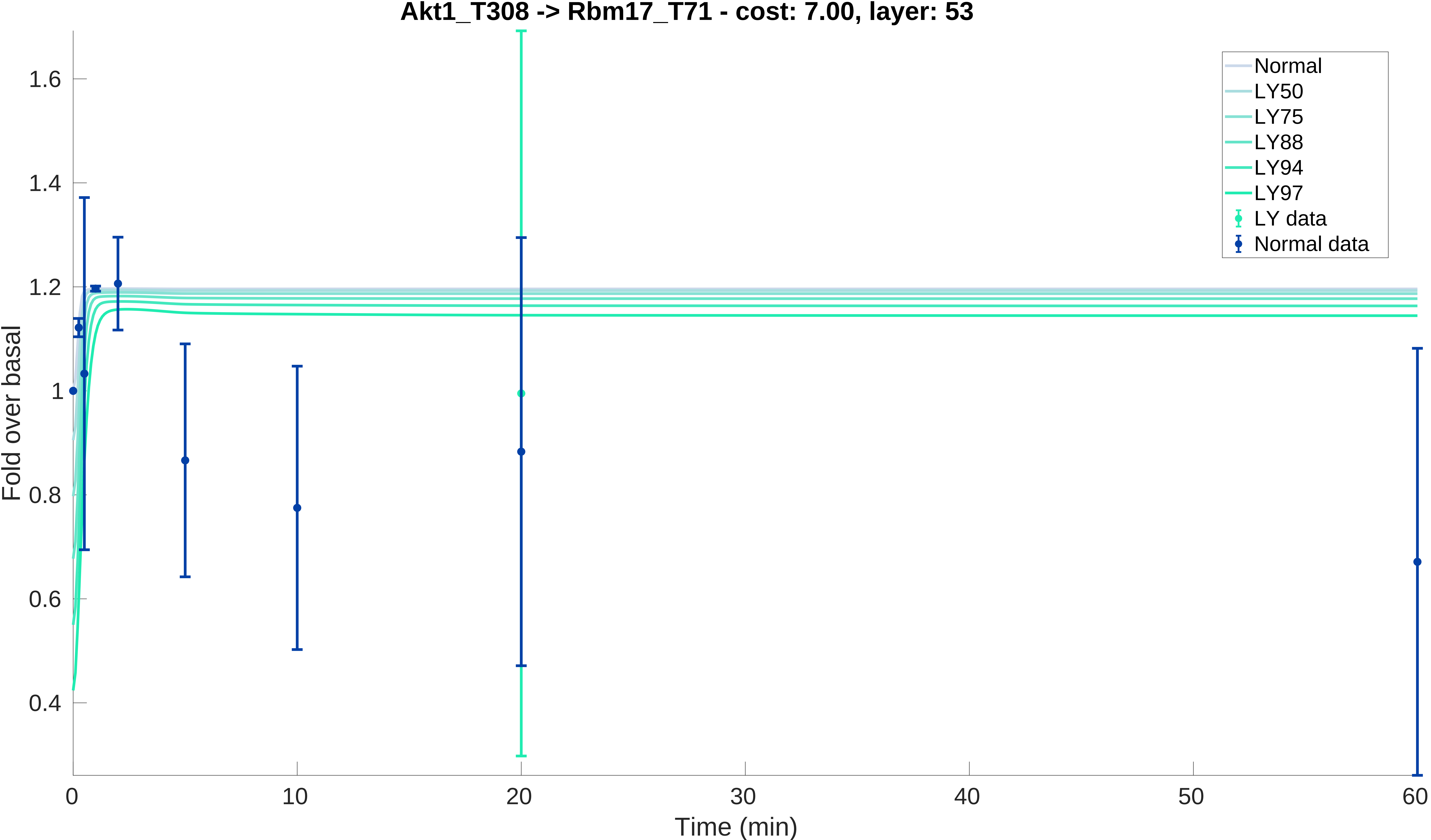
Akt1_T308 -> Rbm17_T71 - cost: 7.00, layer: 53
Fold over basal
Time (min)
0.4
0.6
0.8
1
1.2
1.4
1.6
0
10
20
30
40
50
60
Normal
LY50
LY75
LY88
LY94
LY97
LY data
Normal data

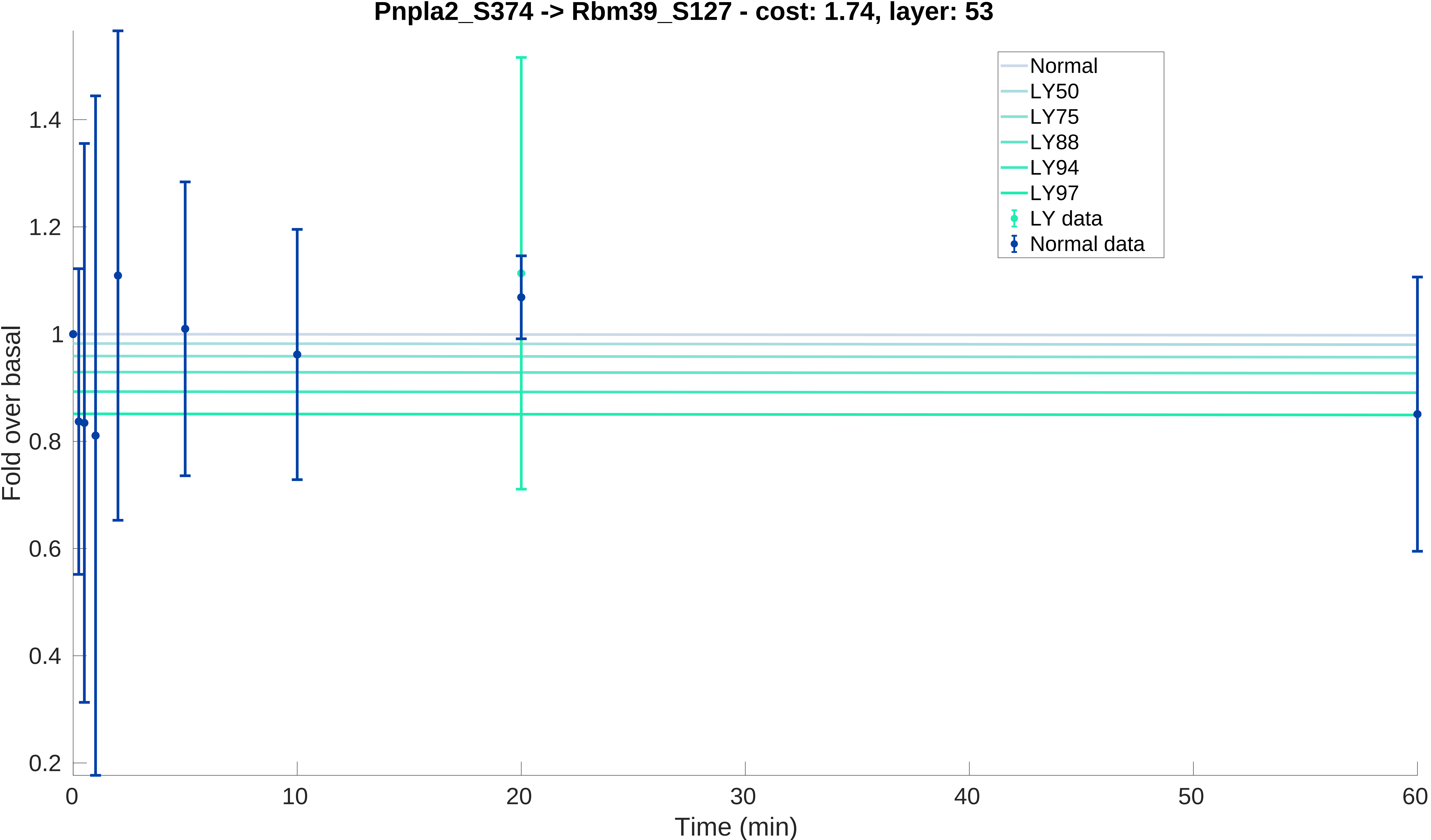

Pnpla2_S374 -> Rbm39_S127 - cost: 1.74, layer: 53
Normal
LY50
LY75
LY88
LY94
LY97
LY data
Normal data
Fold over basal
1.4
1.2
1
0.8
0.6
0.4
0.2
0
10
20
30
40
50
60
Time (min)

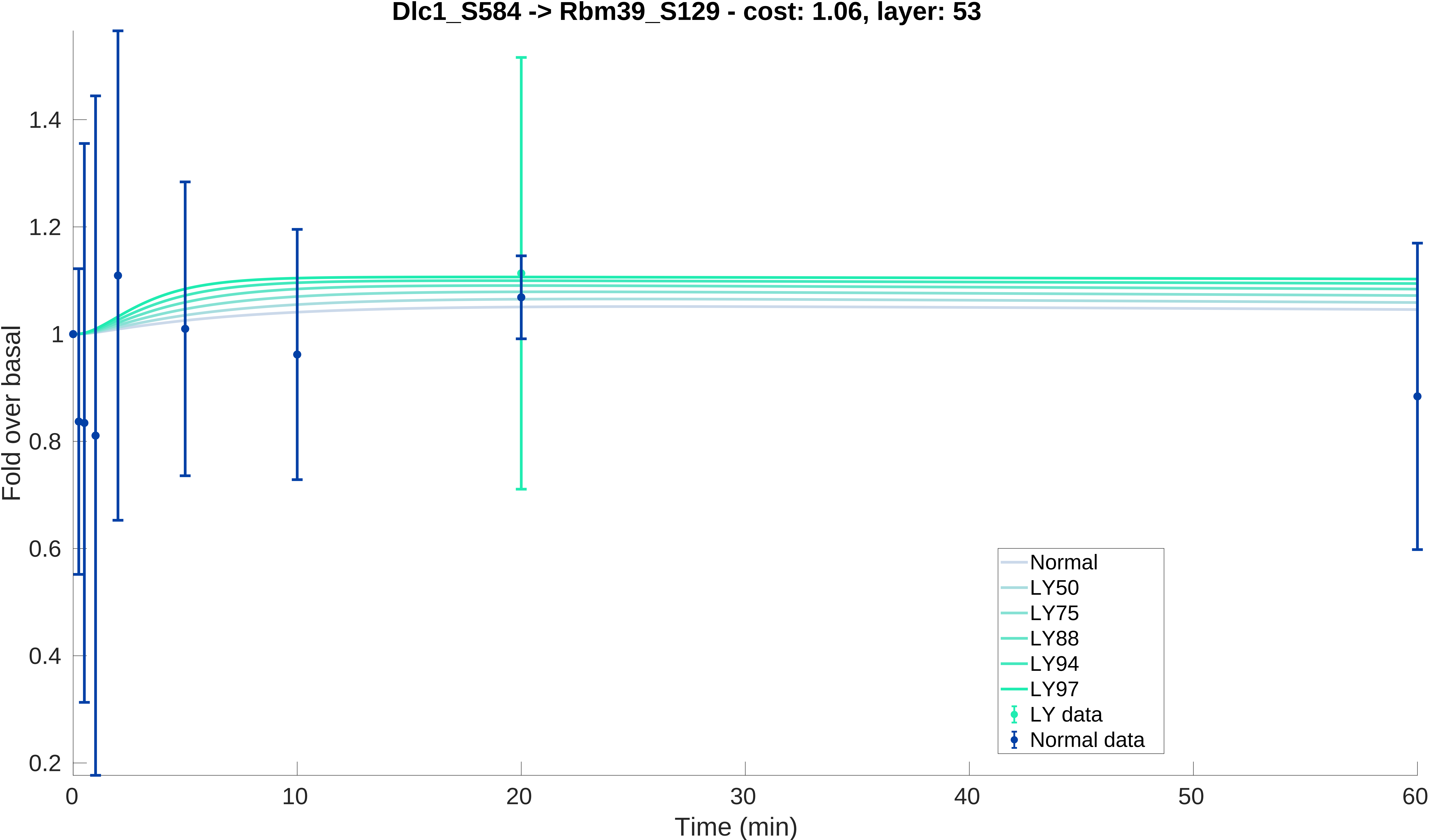

Dlc1_S584 -> Rbm39_S129 - cost: 1.06, layer: 53
Fold over basal
Time (min)
0.2
0.4
0.6
0.8
1
1.2
1.4
0
10
20
30
40
50
60
Normal
LY50
LY75
LY88
LY94
LY97
LY data
Normal data

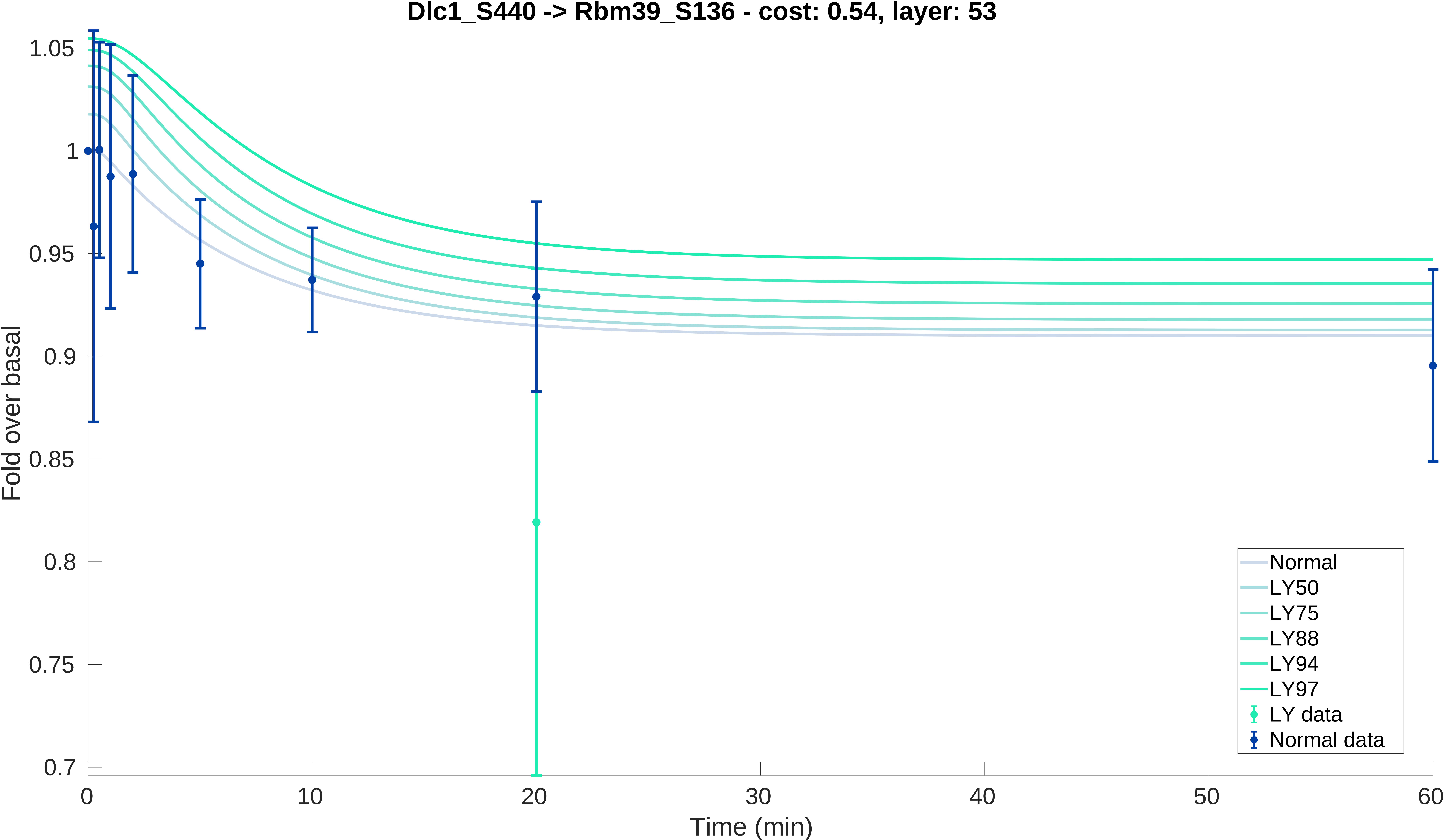

Dlc1_S440 -> Rbm39_S136 - cost: 0.54, layer: 53
Fold over basal
1.05
1
0.95
0.9
0.85
0.8
0.75
0.7
0
10
20
30
40
50
60
Time (min)
Normal
LY50
LY75
LY88
LY94
LY97
LY data
Normal data

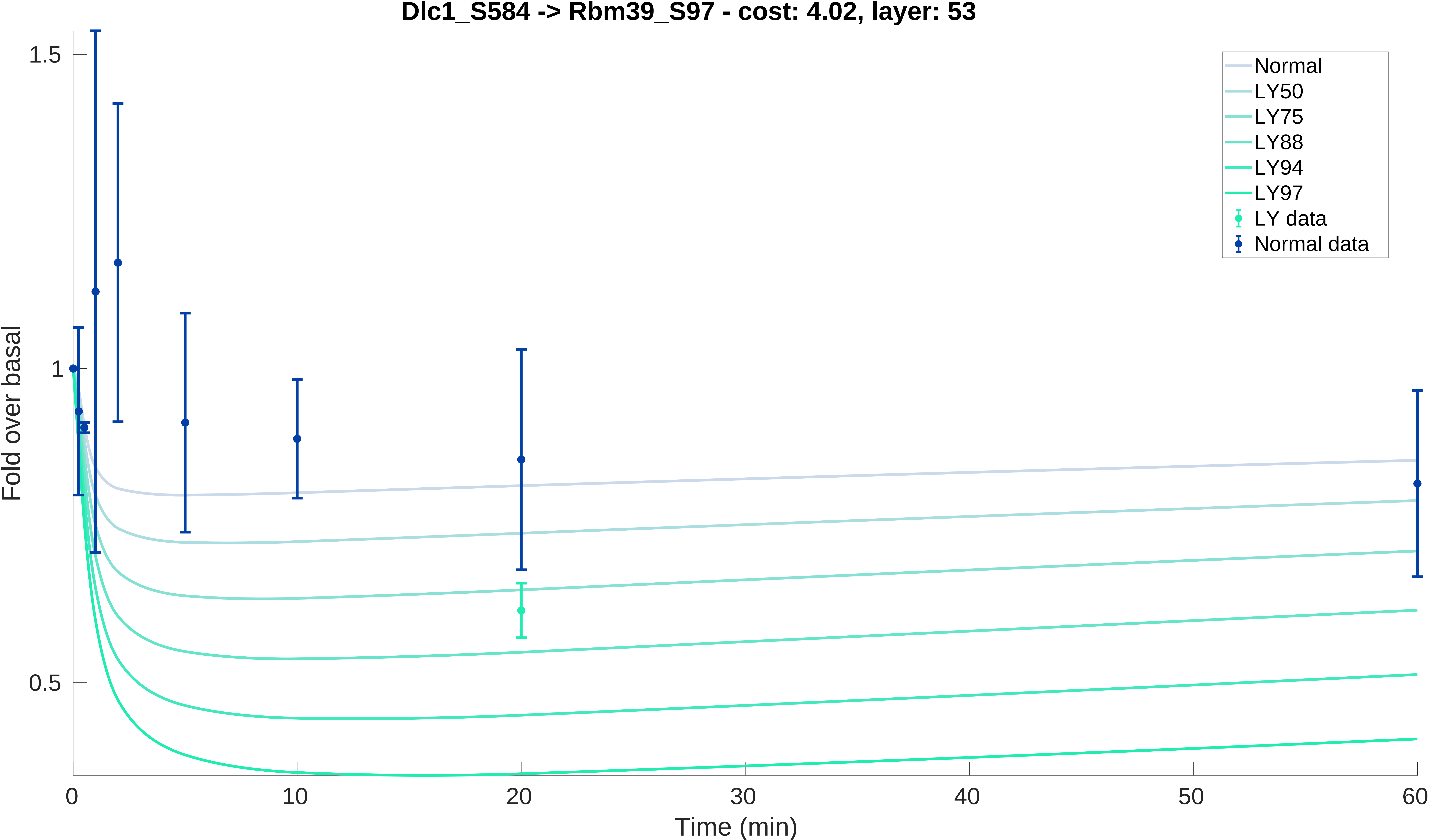

Dlc1_S584 -> Rbm39_S97 - cost: 4.02, layer: 53
Fold over basal
Time (min)
1.5
1
0.5
0
10
20
30
40
50
60
Normal
LY50
LY75
LY88
LY94
LY97
LY data
Normal data

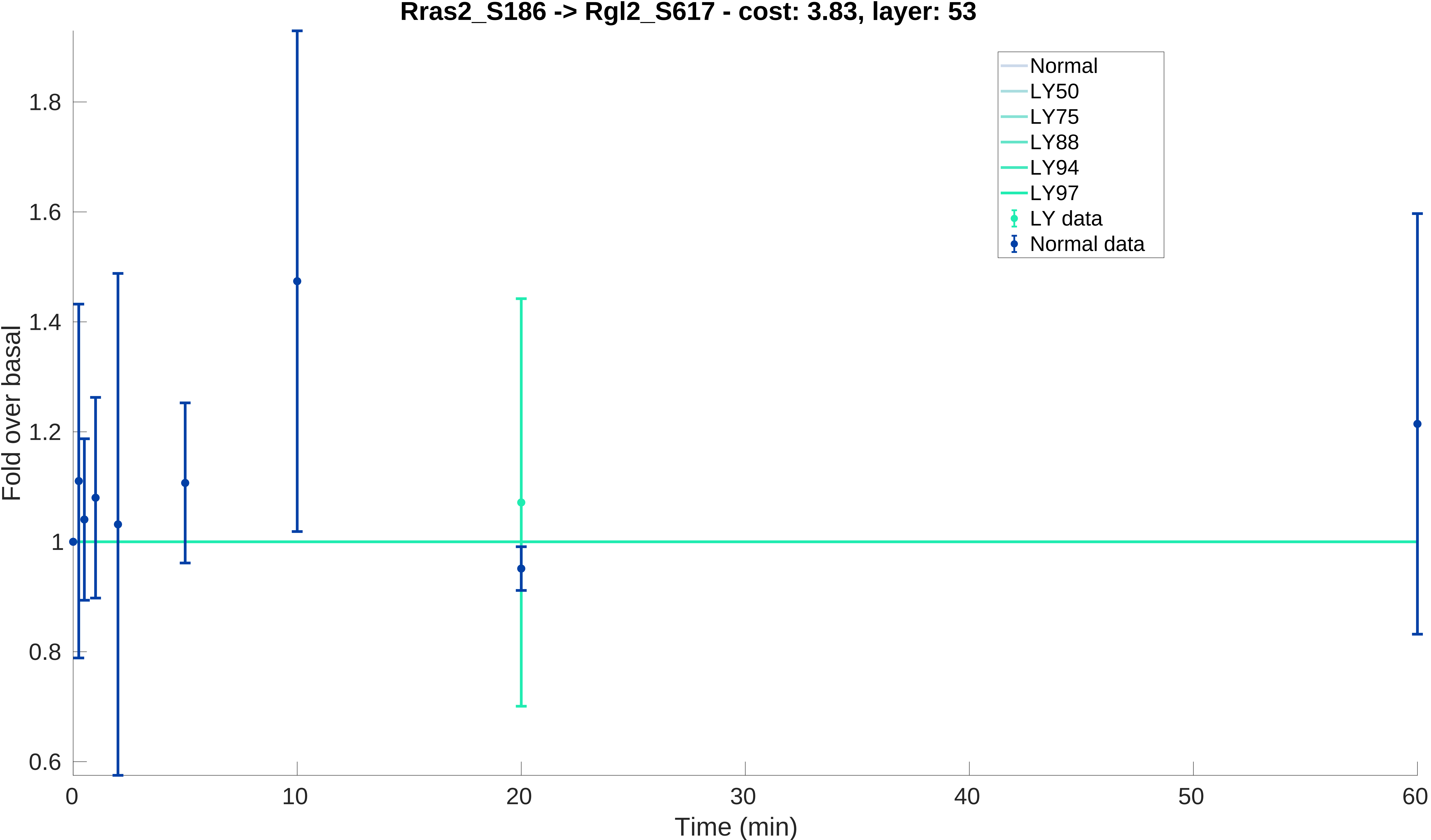
Rras2_S186 -> Rgl2_S617 - cost: 3.83, layer: 53
Normal
LY50
LY75
LY88
LY94
LY97
LY data
Normal data
Fold over basal
1.8
1.6
1.4
1.2
1
0.8
0.6
0
10
20
30
40
50
60
Time (min)

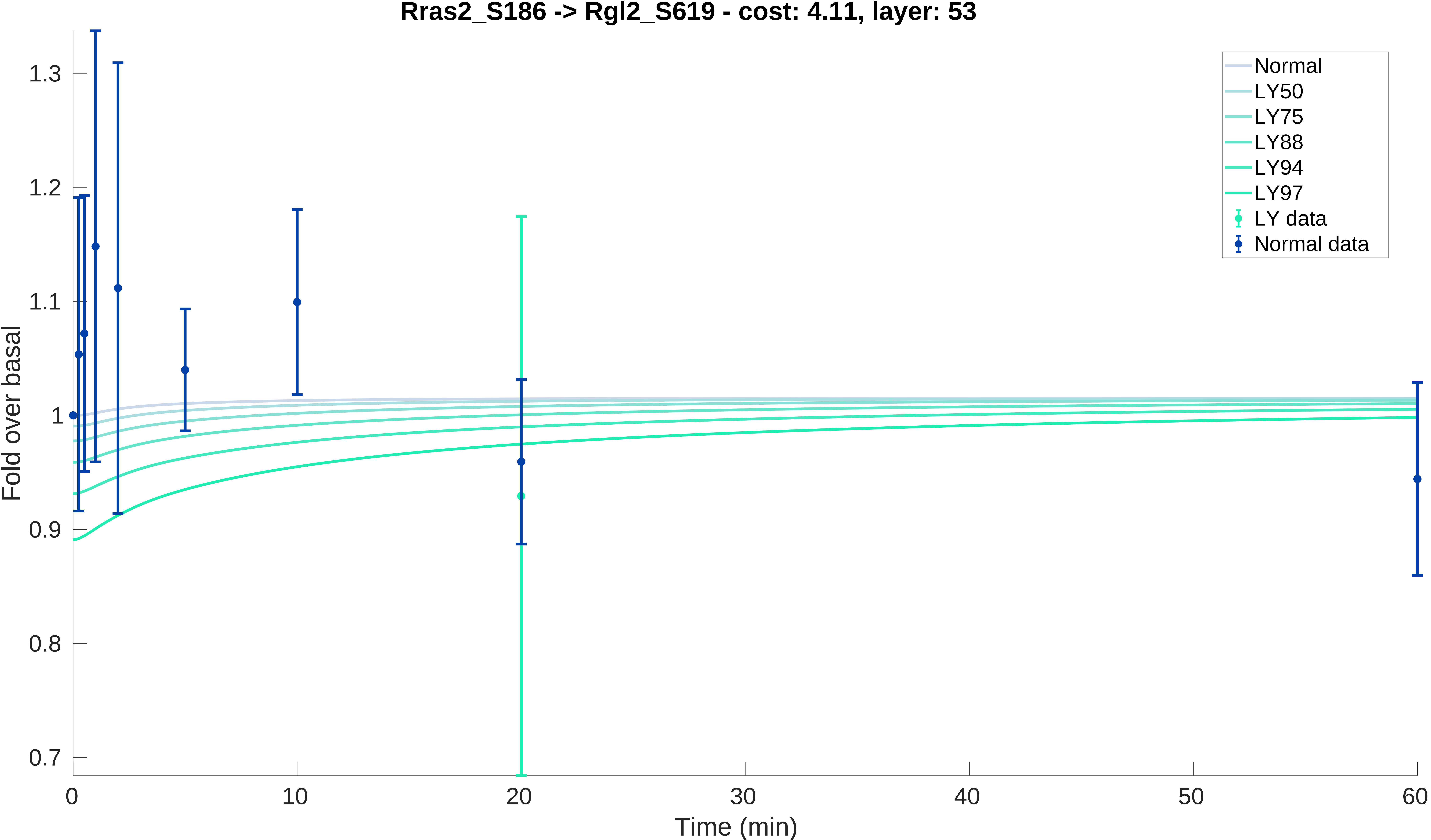

Rras2_S186 -> Rgl2_S619 - cost: 4.11, layer: 53
Normal
LY50
LY75
LY88
LY94
LY97
LY data
Normal data
Fold over basal
1.3
1.2
1.1
1
0.9
0.8
0.7
0
10
20
30
40
50
60
Time (min)

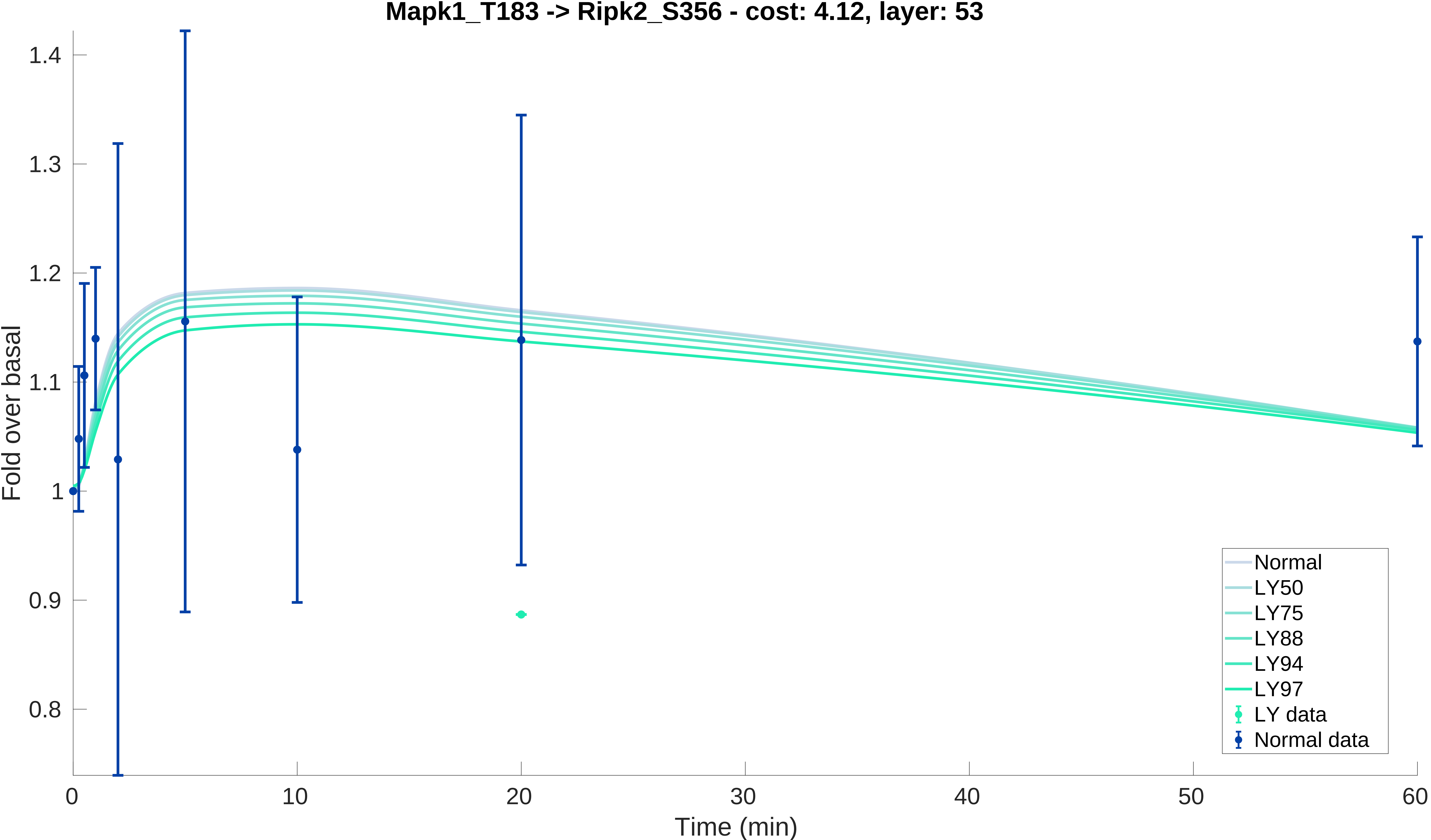

Mapk1_T183 -> Ripk2_S356 - cost: 4.12, layer: 53
Fold over basal
1.4
1.3
1.2
1.1
1
0.9
0.8
0
10
20
30
40
50
60
Time (min)
Normal
LY50
LY75
LY88
LY94
LY97
LY data
Normal data

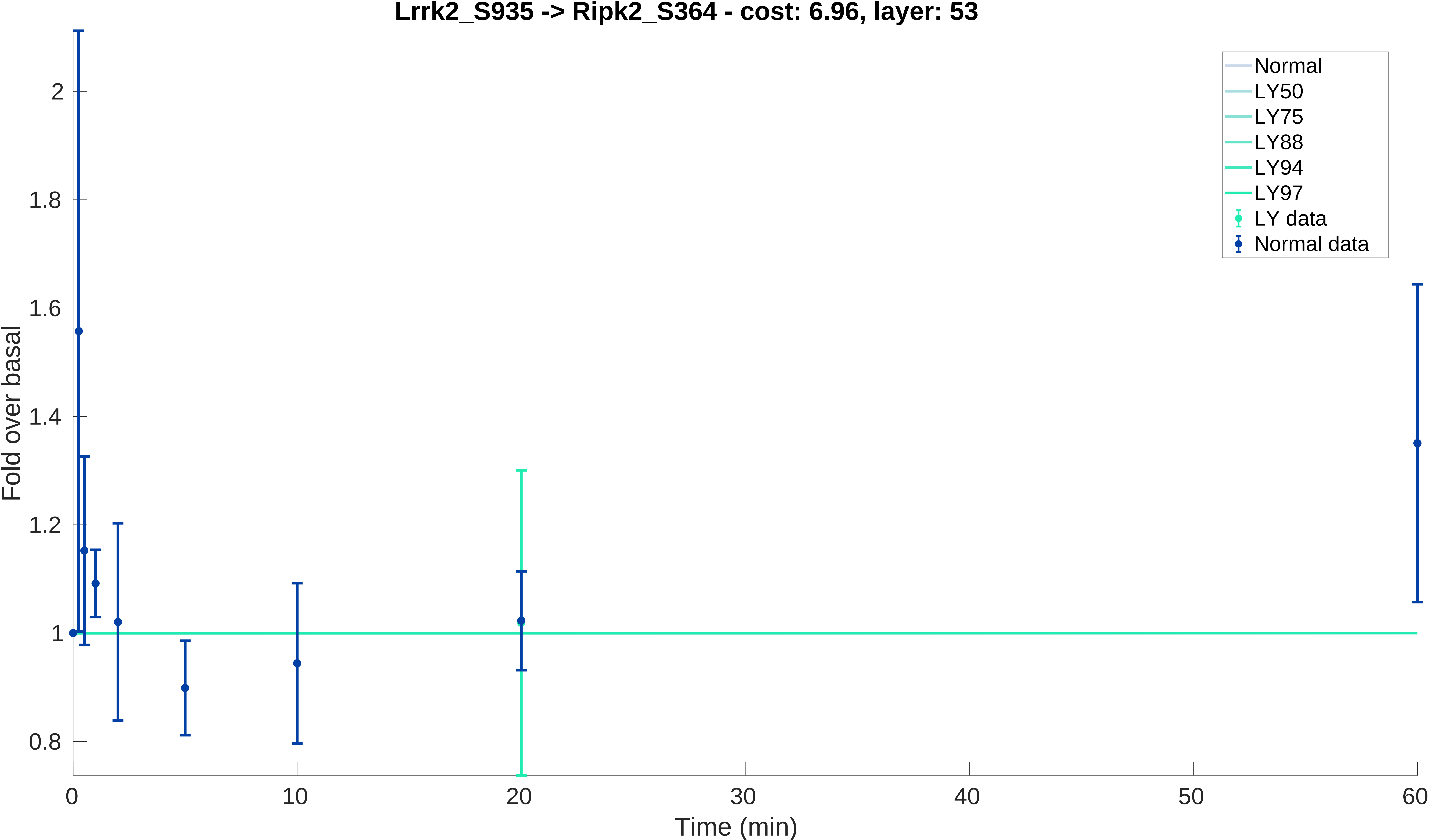

Lrrk2_S935 -> Ripk2_S364 - cost: 6.96, layer: 53
Fold over basal
Time (min)
0.8
1
1.2
1.4
1.6
1.8
2
0
10
20
30
40
50
60
Normal
LY50
LY75
LY88
LY94
LY97
LY data
Normal data

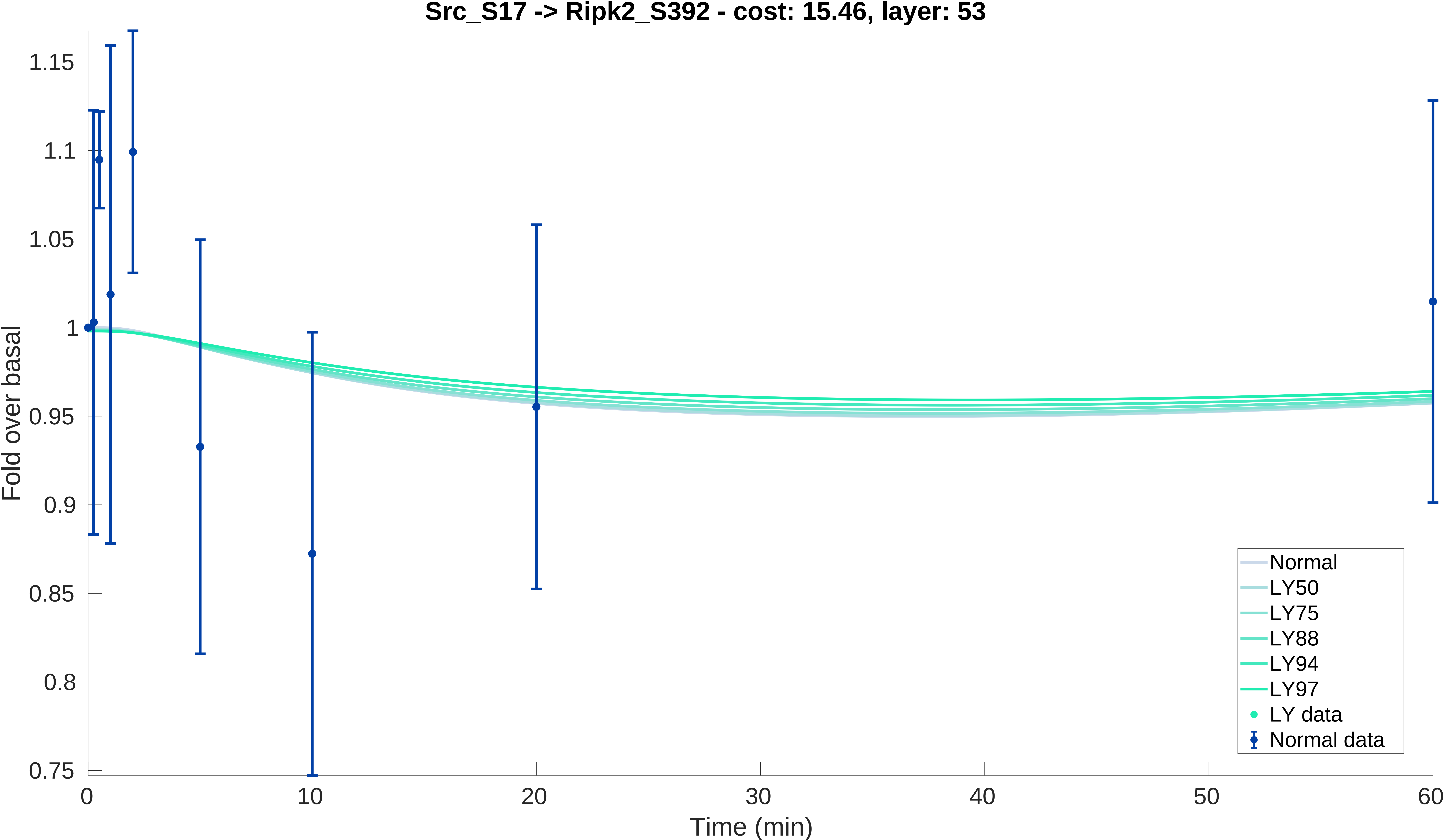

Src_S17 -> Ripk2_S392 - cost: 15.46, layer: 53
Fold over basal
1.15
1.1
1.05
1
0.95
0.9
0.85
0.8
0.75
0
10
20
30
40
50
60
Time (min)
Normal
LY50
LY75
LY88
LY94
LY97
LY data
Normal data

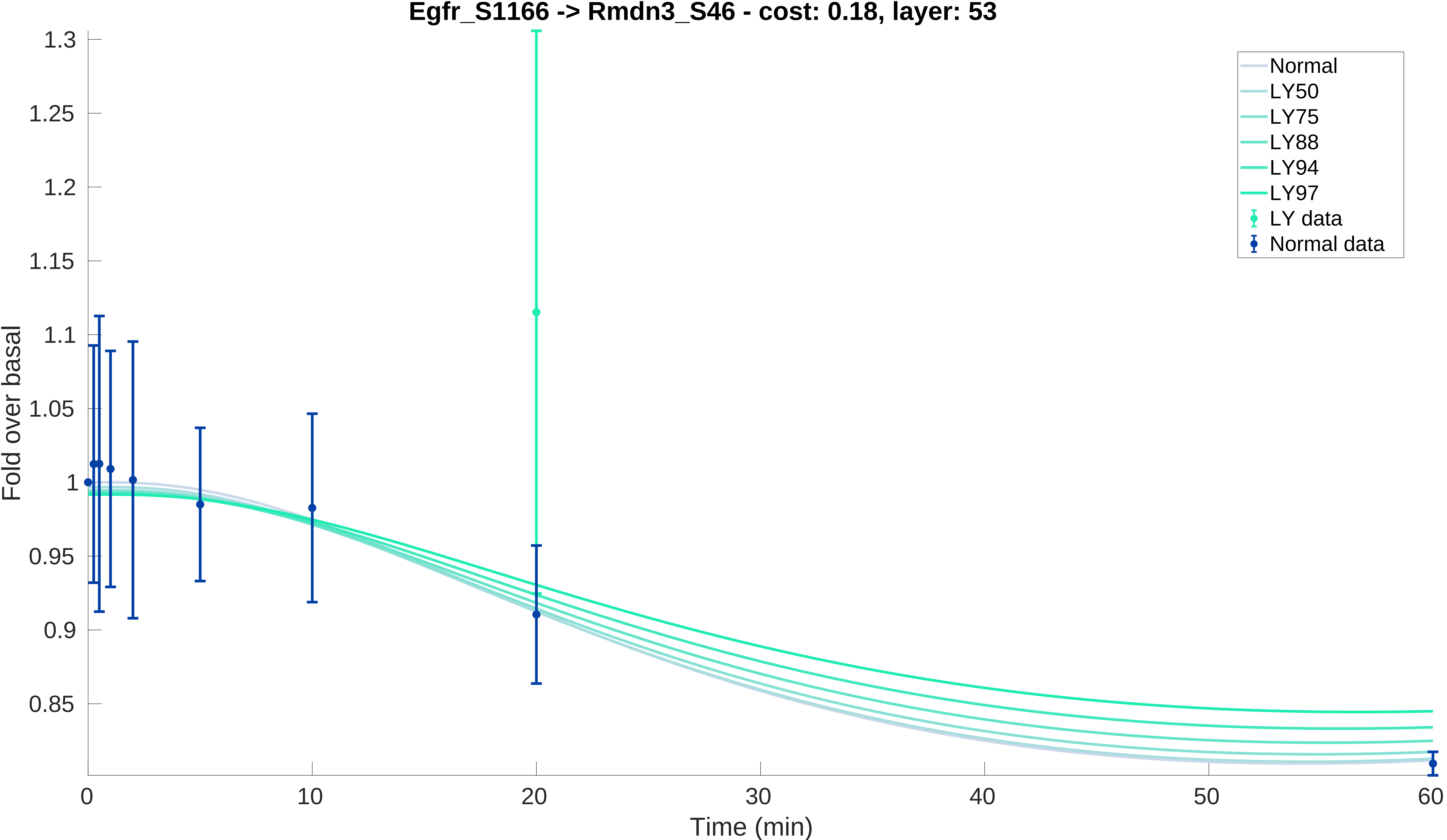

Egfr_S1166 -> Rmdn3_S46 - cost: 0.18, layer: 53
Fold over basal
1.3
1.25
1.2
1.15
1.1
1.05
1
0.95
0.9
0.85
0
10
20
30
40
50
60
Time (min)
Normal
LY50
LY75
LY88
LY94
LY97
LY data
Normal data

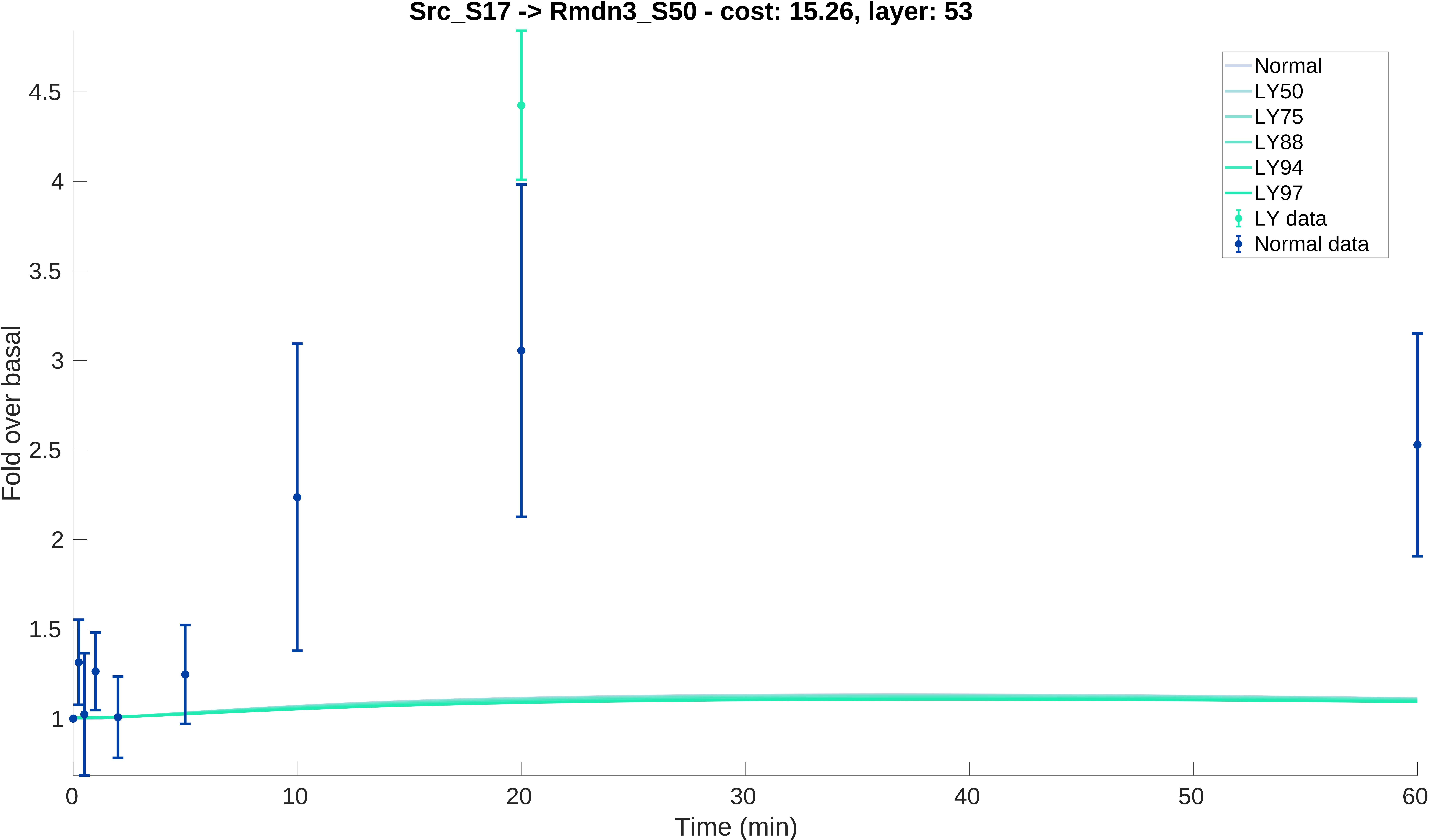
Src_S17 -> Rmdn3_S50 - cost: 15.26, layer: 53
Fold over basal
Time (min)
Normal
LY50
LY75
LY88
LY94
LY97
LY data
Normal data
4.5
4
3.5
3
2.5
2
1.5
1
0
10
20
30
40
50
60

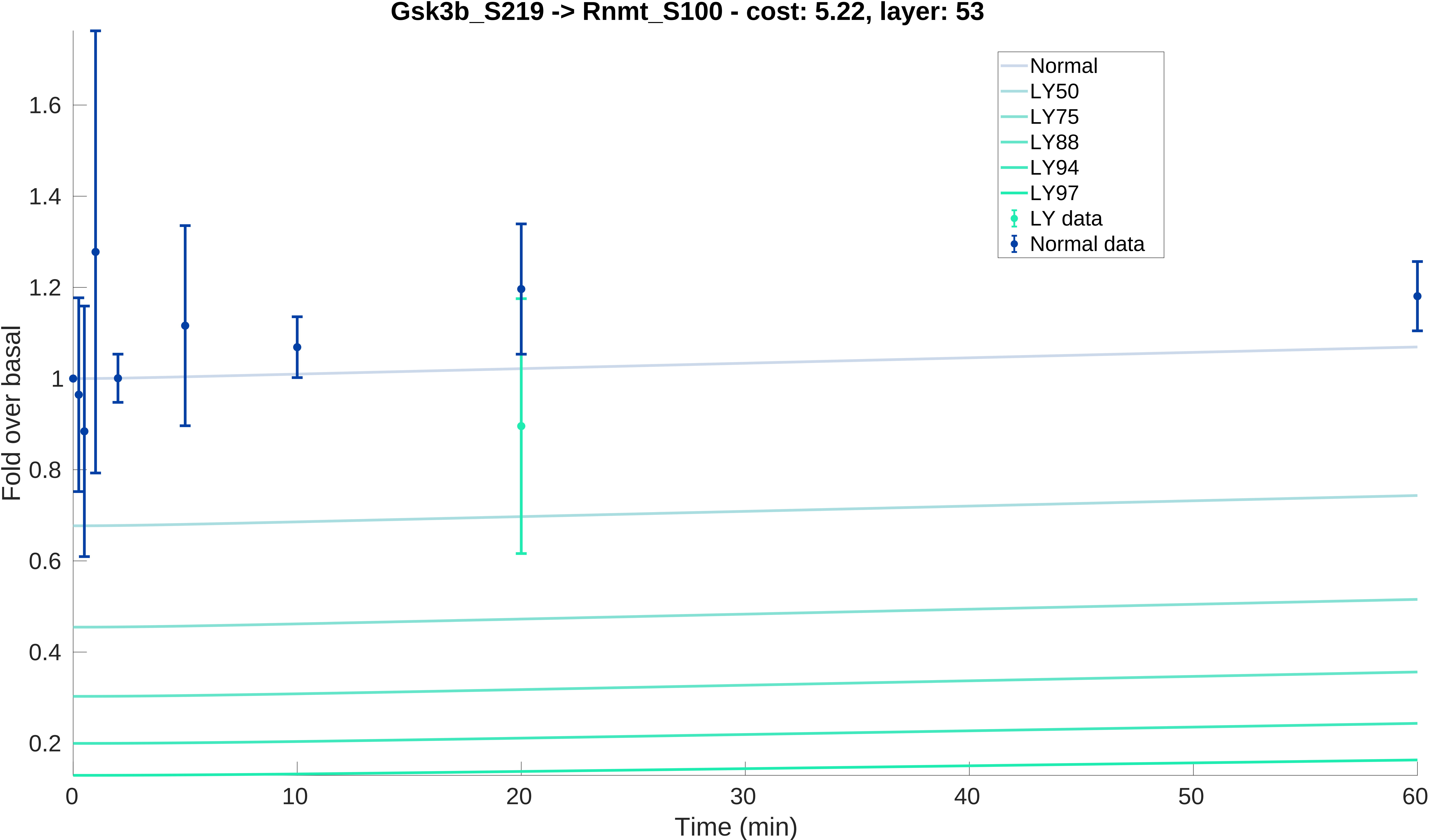

Gsk3b_S219 -> Rnmt_S100 - cost: 5.22, layer: 53
Fold over basal
Time (min)
1.6
1.4
1.2
1
0.8
0.6
0.4
0.2
0
10
20
30
40
50
60
Normal
LY50
LY75
LY88
LY94
LY97
LY data
Normal data

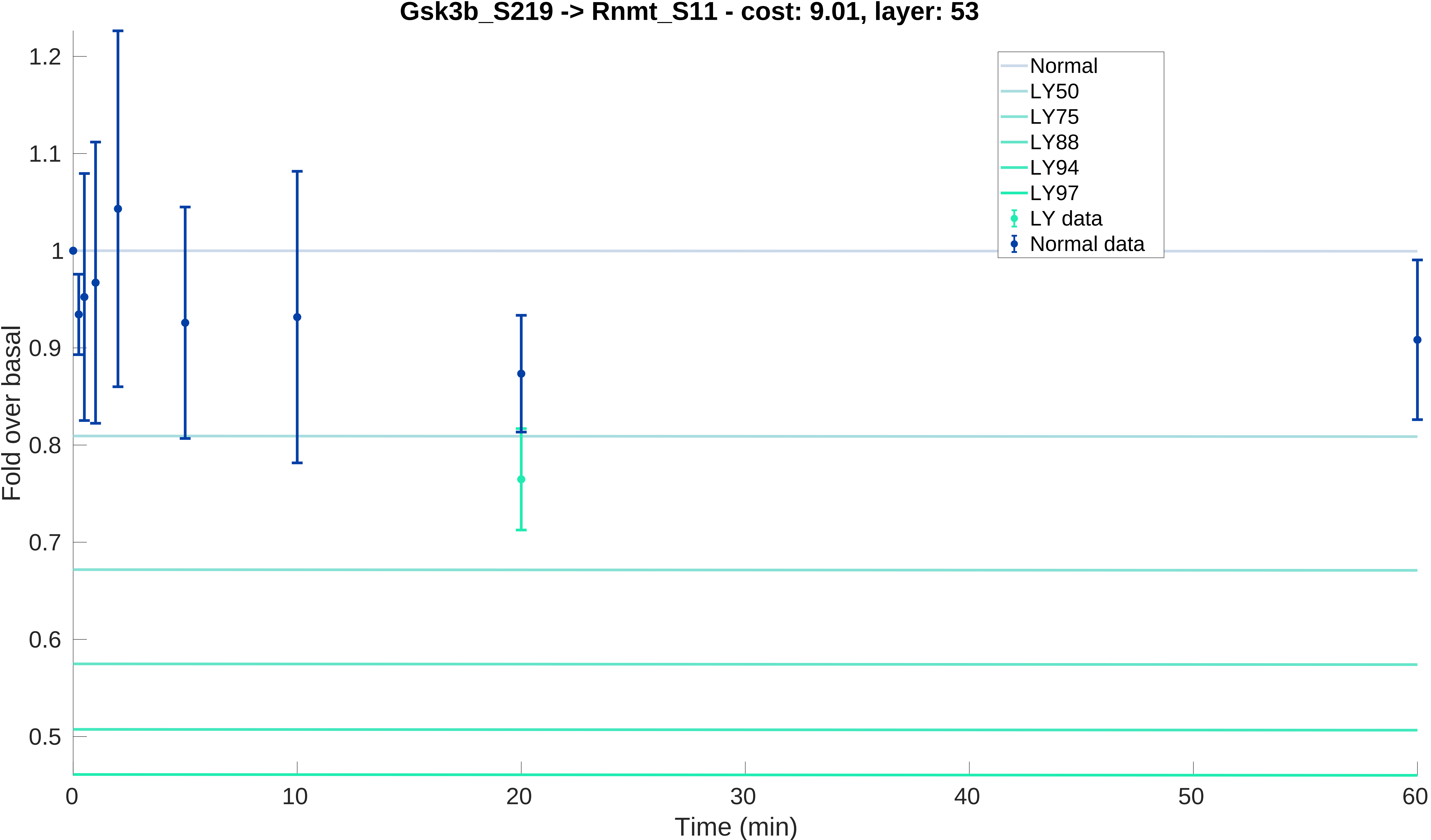

Gsk3b_S219 -> Rnmt_S11 - cost: 9.01, layer: 53
Normal
LY50
LY75
LY88
LY94
LY97
LY data
Normal data
Fold over basal
1.2
1.1
1
0.9
0.8
0.7
0.6
0.5
0
10
20
30
40
50
60
Time (min)

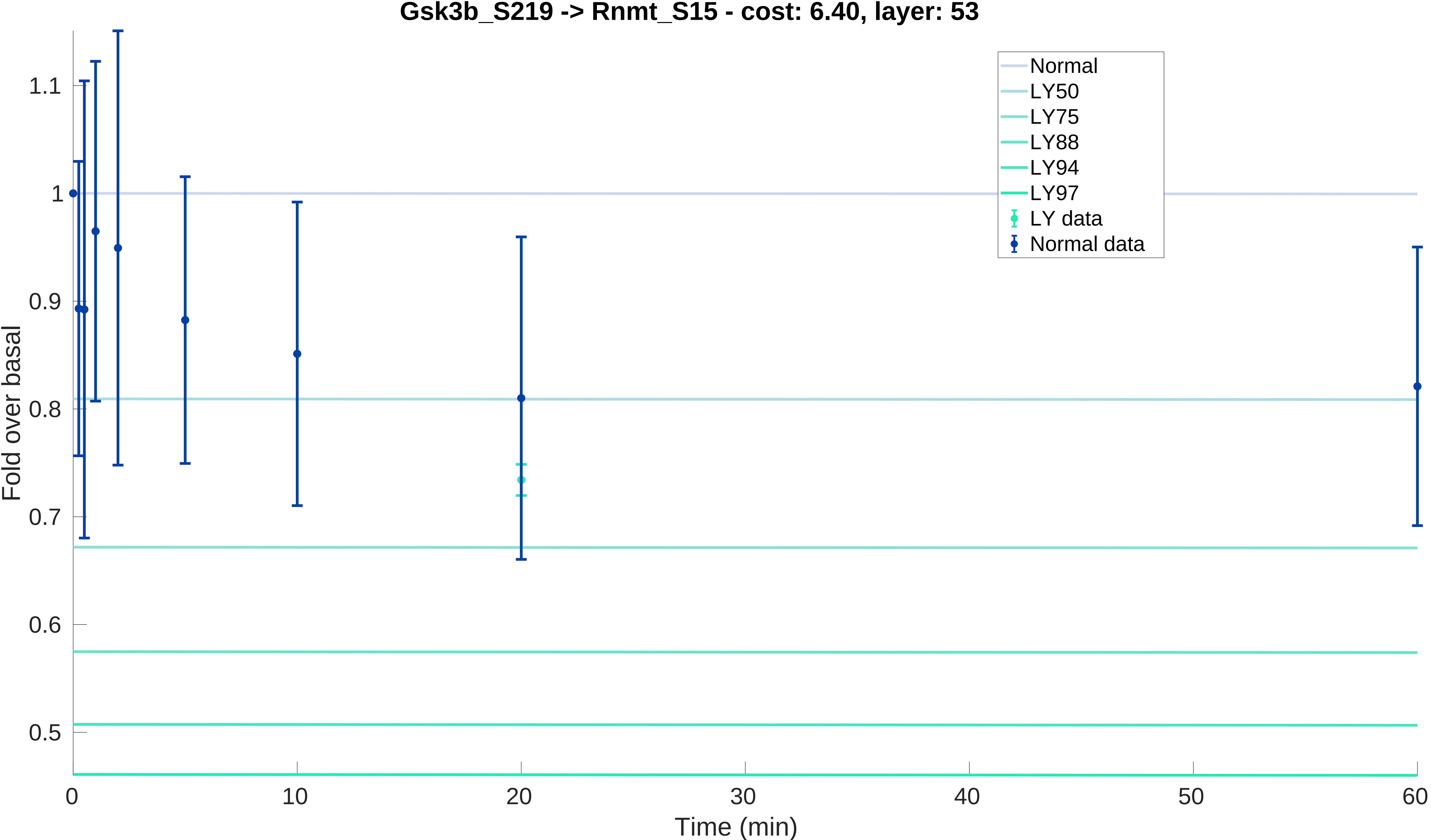

Gsk3b_S219 -> Rnmt_S15 - cost: 6.40, layer: 53
Normal
LY50
LY75
LY88
LY94
LY97
LY data
Normal data
Fold over basal
Time (min)
1.1
1
0.9
0.8
0.7
0.6
0.5
0
10
20
30
40
50
60

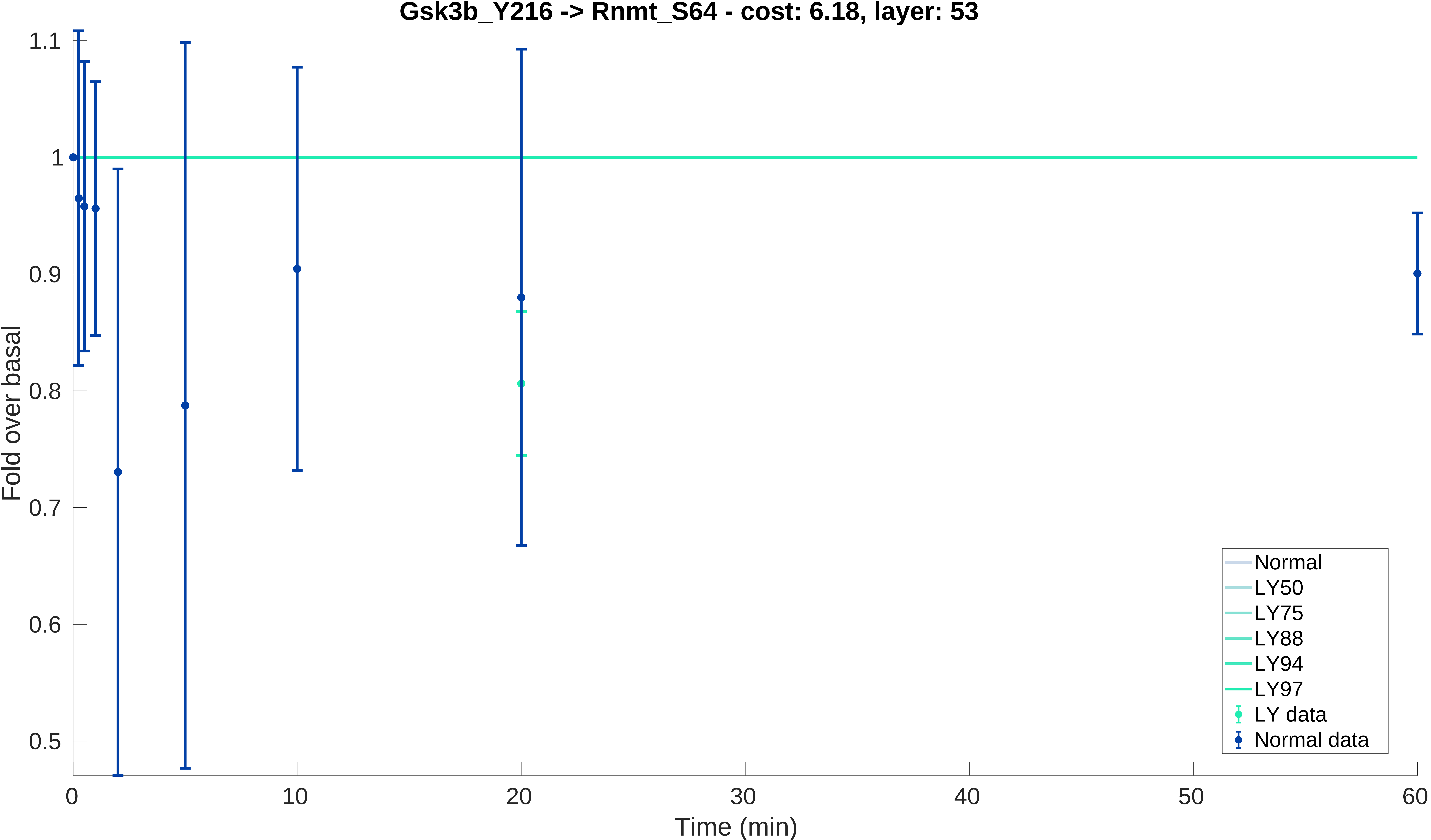
Gsk3b_Y216 -> Rnmt_S64 - cost: 6.18, layer: 53
Fold over basal
1.1
1
0.9
0.8
0.7
0.6
0.5
0
10
20
30
40
50
60
Time (min)
Normal
LY50
LY75
LY88
LY94
LY97
LY data
Normal data

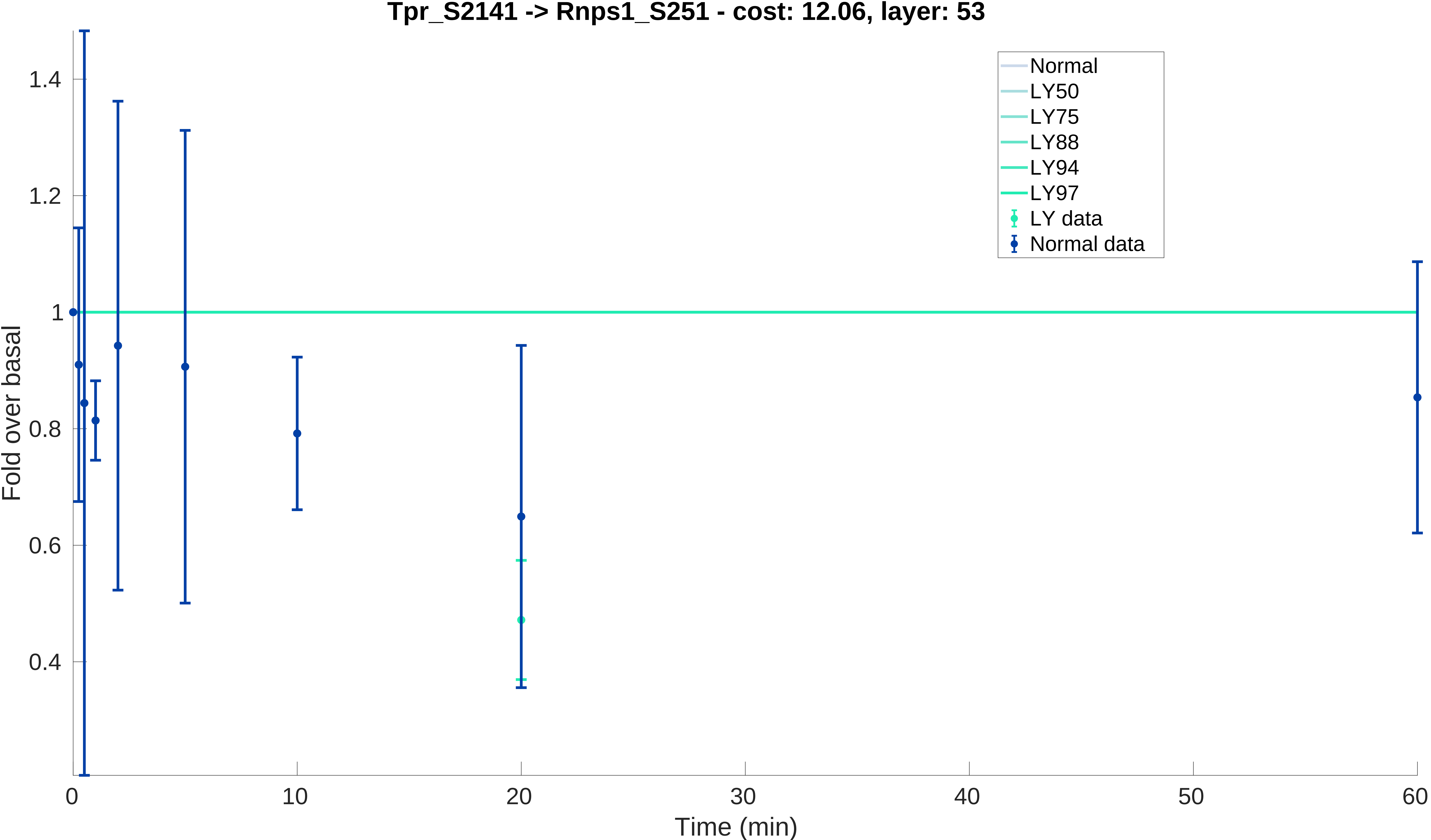

Tpr_S2141 -> Rnps1_S251 - cost: 12.06, layer: 53
Normal
LY50
LY75
LY88
LY94
LY97
LY data
Normal data
Fold over basal
1.4
1.2
1
0.8
0.6
0.4
0
10
20
30
40
50
60
Time (min)

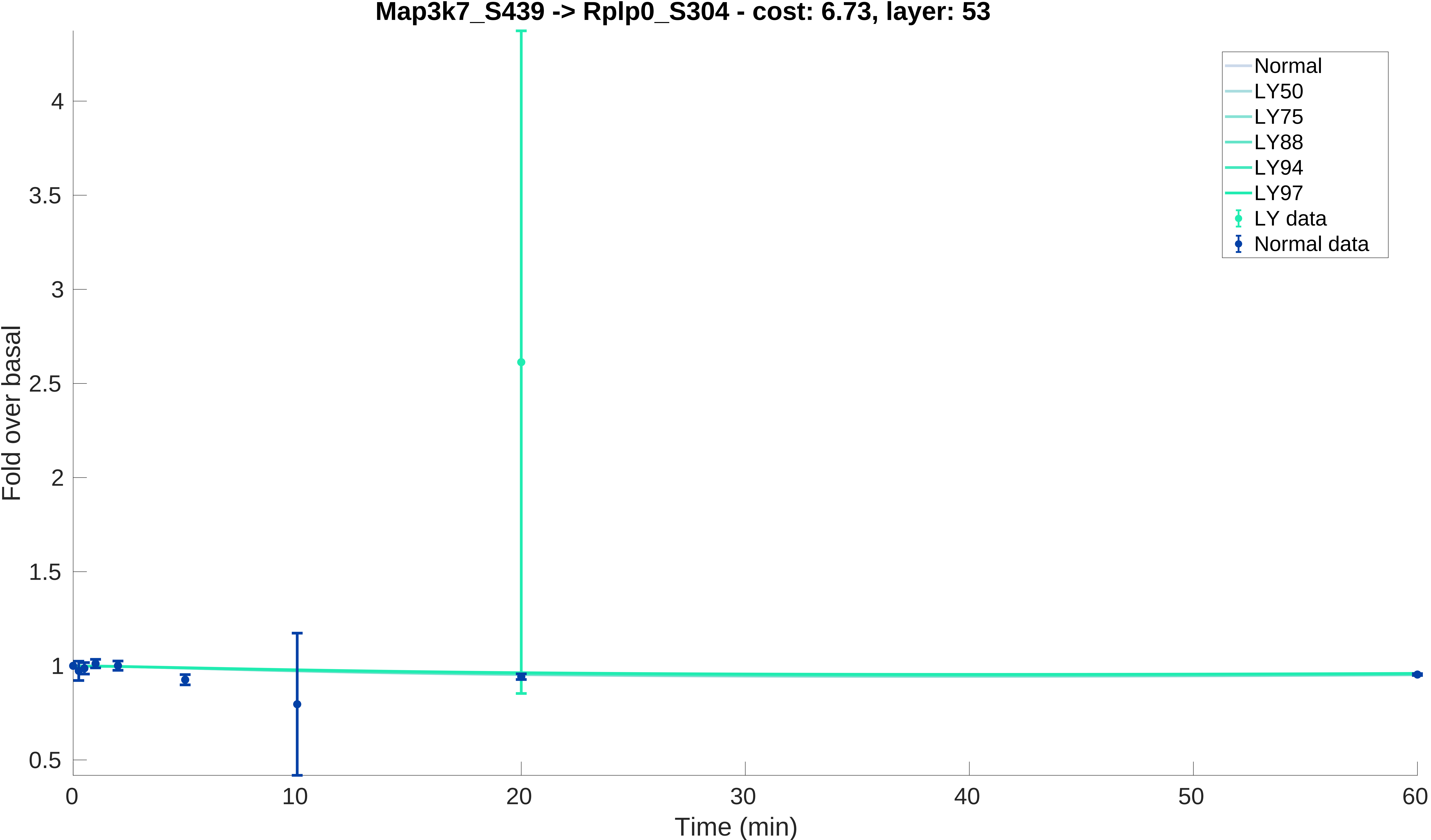

Map3k7_S439 -> Rplp0_S304 - cost: 6.73, layer: 53
Fold over basal
Time (min)
0.5
1
1.5
2
2.5
3
3.5
4
0
10
20
30
40
50
60
Normal
LY50
LY75
LY88
LY94
LY97
LY data
Normal data

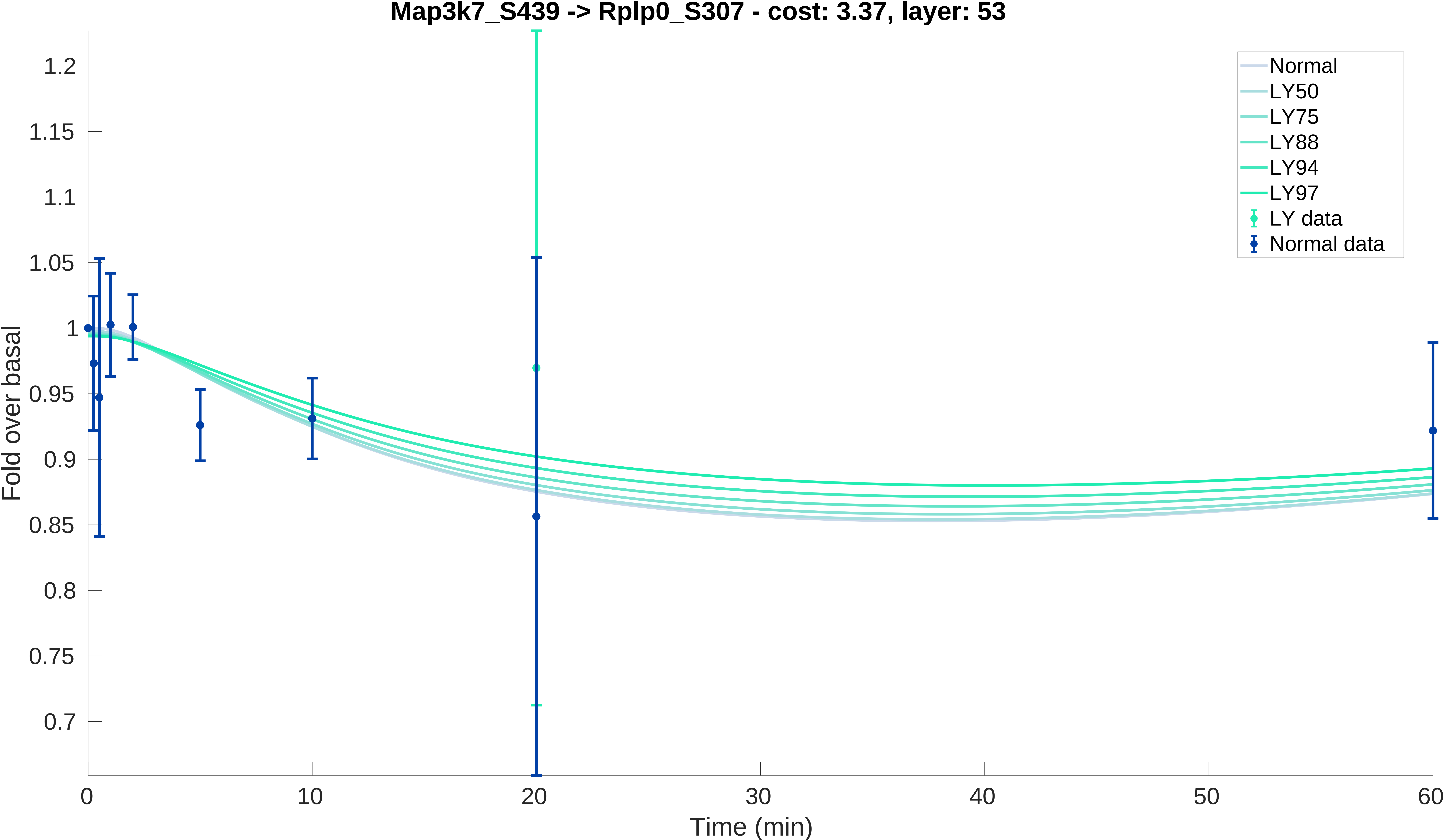

Map3k7_S439 -> Rplp0_S307 - cost: 3.37, layer: 53
Fold over basal
Time (min)
1.2
1.15
1.1
1.05
1
0.95
0.9
0.85
0.8
0.75
0.7
0
10
20
30
40
50
60
Normal
LY50
LY75
LY88
LY94
LY97
LY data
Normal data

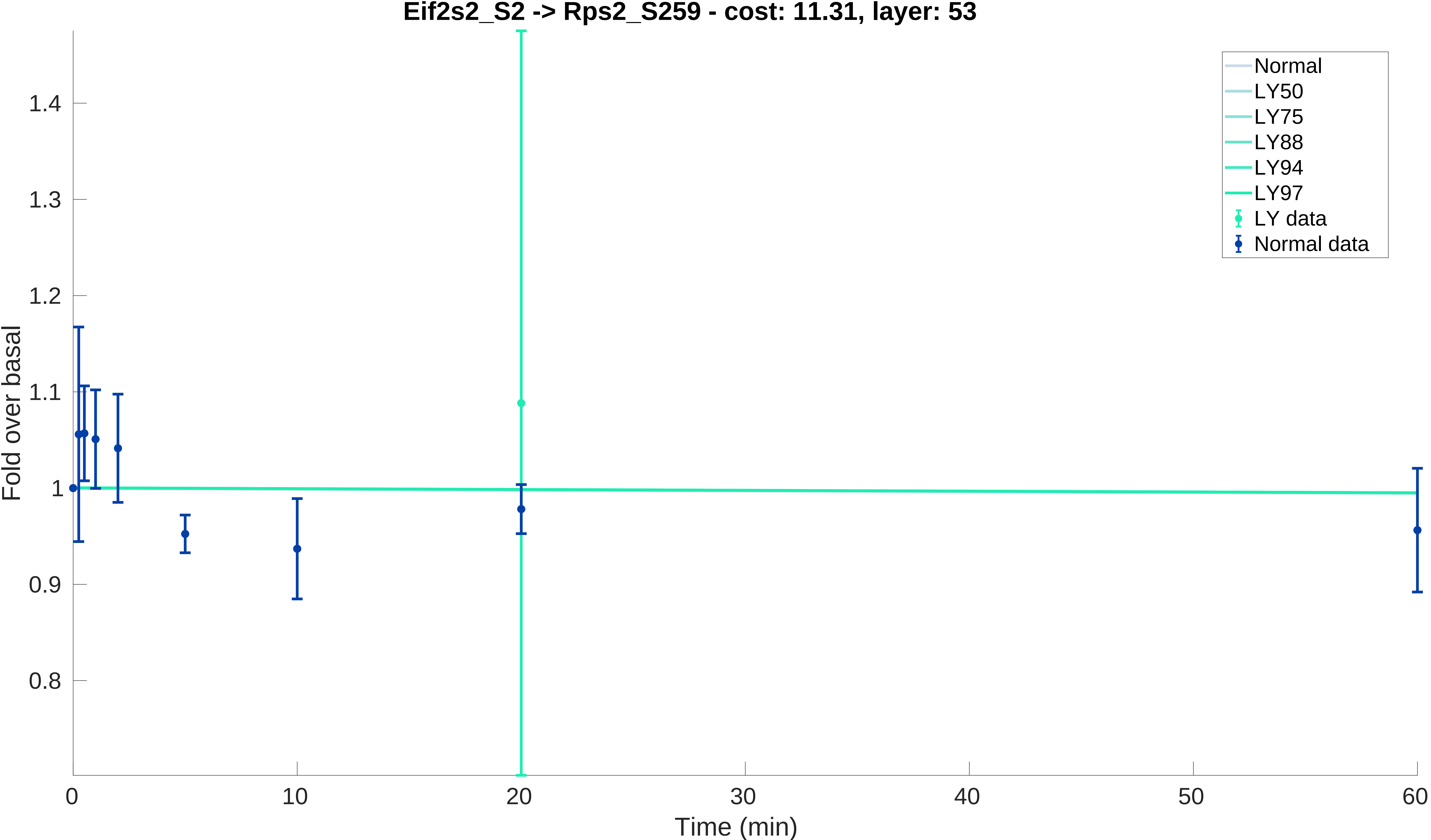

Eif2s2_S2 -> Rps2_S259 - cost: 11.31, layer: 53
Fold over basal
Time (min)
Normal
LY50
LY75
LY88
LY94
LY97
LY data
Normal data
1.4
1.3
1.2
1.1
1
0.9
0.8
0
10
20
30
40
50
60

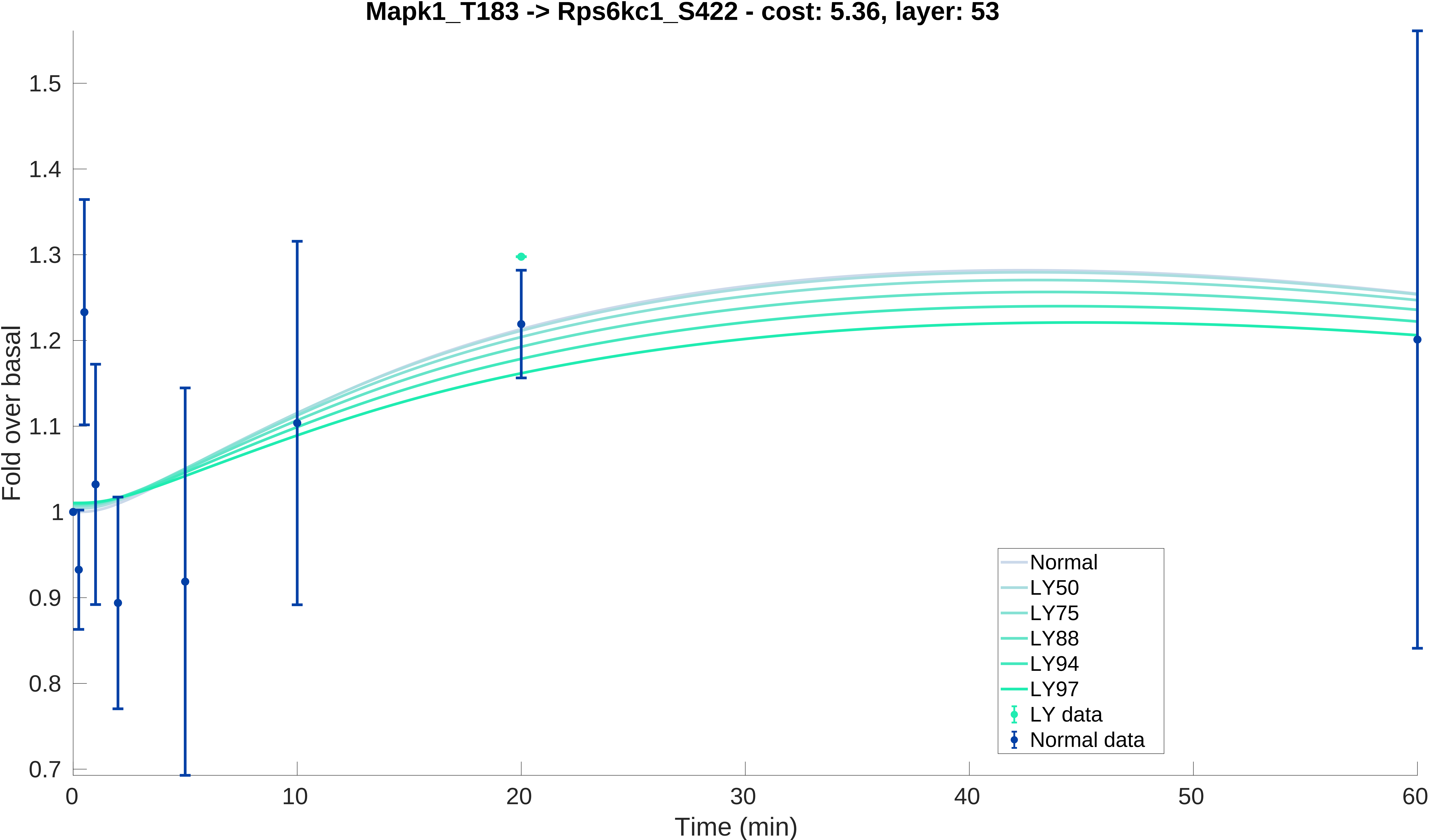
Mapk1_T183 -> Rps6kc1_S422 - cost: 5.36, layer: 53
Fold over basal
Time (min)
0.7
0.8
0.9
1
1.1
1.2
1.3
1.4
1.5
0
10
20
30
40
50
60
Normal
LY50
LY75
LY88
LY94
LY97
LY data
Normal data

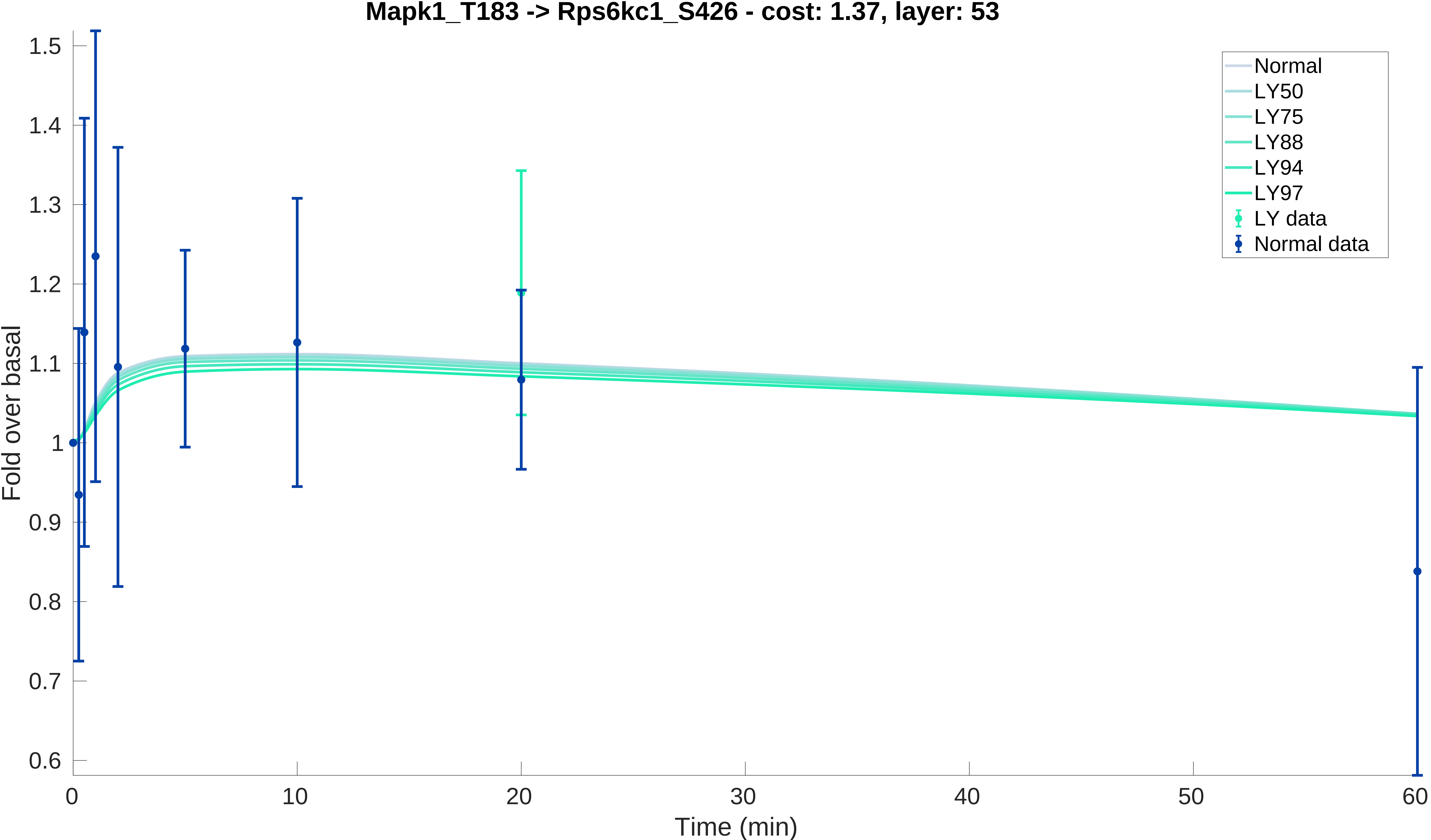

Mapk1_T183 -> Rps6kc1_S426 - cost: 1.37, layer: 53
Fold over basal
Time (min)
Normal
LY50
LY75
LY88
LY94
LY97
LY data
Normal data
0.6
0.7
0.8
0.9
1
1.1
1.2
1.3
1.4
1.5
0
10
20
30
40
50
60

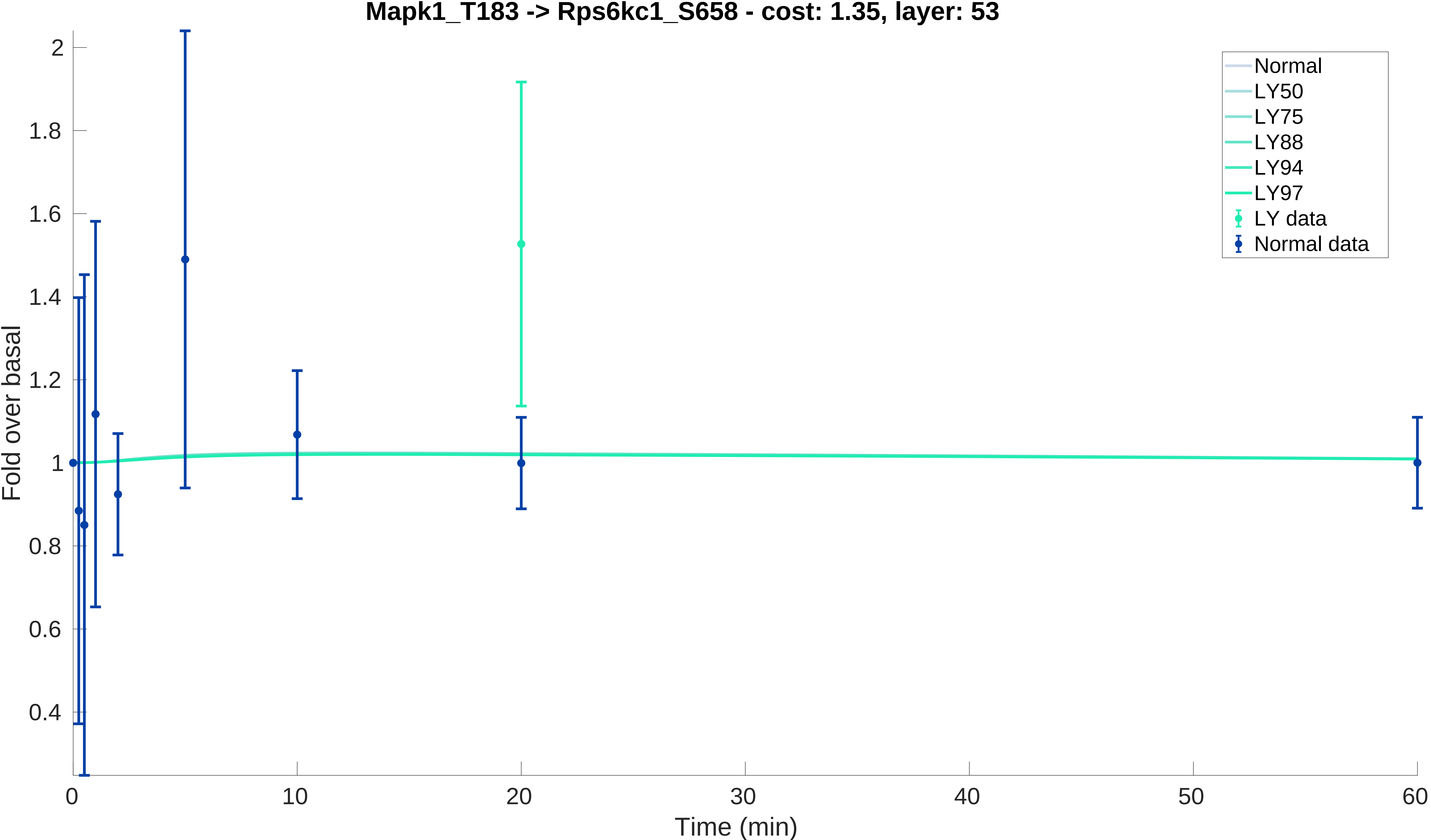
Mapk1_T183 -> Rps6kc1_S658 - cost: 1.35, layer: 53
Fold over basal
Time (min)
2
1.8
1.6
1.4
1.2
1
0.8
0.6
0.4
0
10
20
30
40
50
60
Normal
LY50
LY75
LY88
LY94
LY97
LY data
Normal data

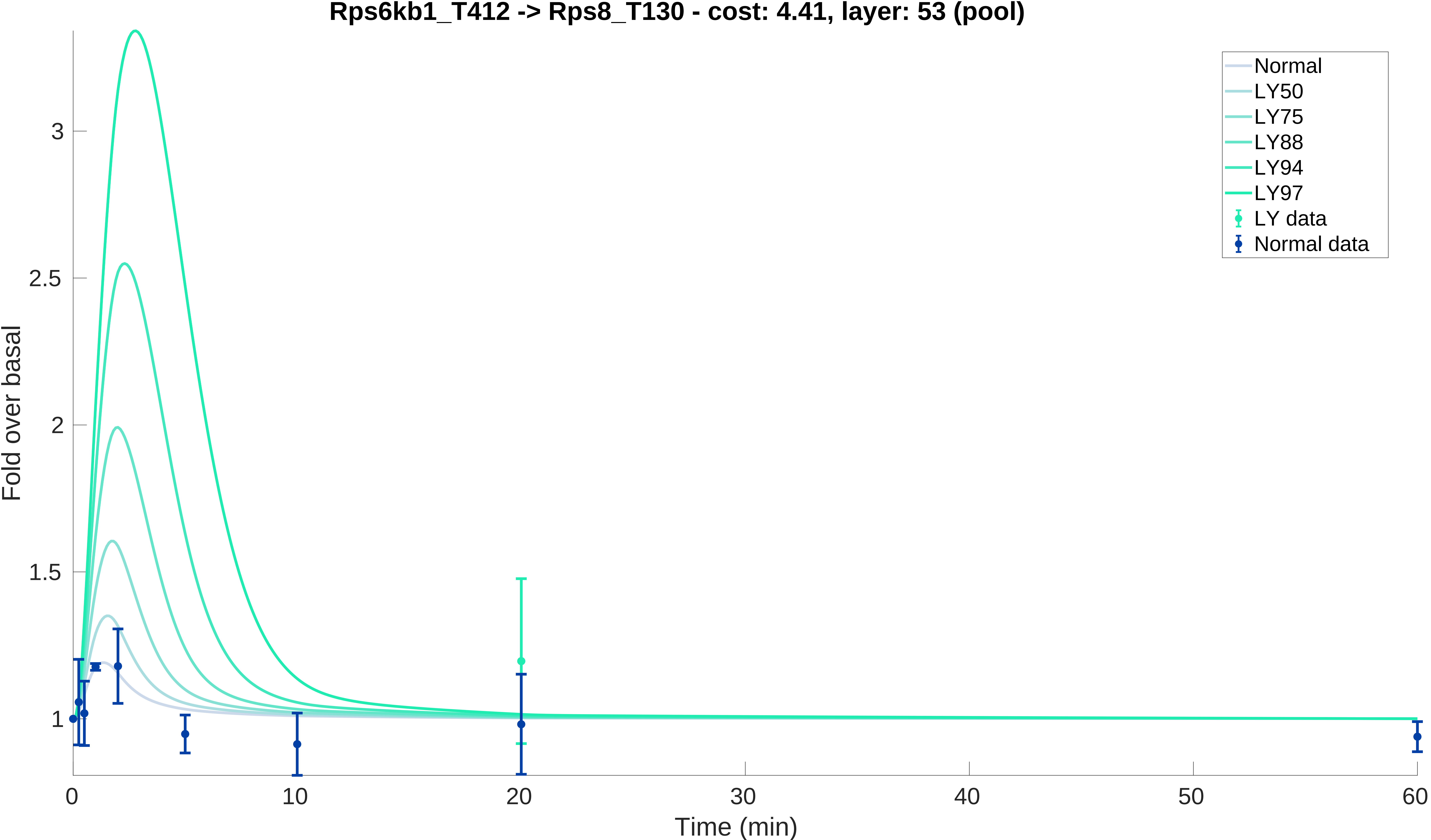

Rps6kb1_T412 -> Rps8_T130 - cost: 4.41, layer: 53 (pool)
Fold over basal
Time (min)
3
2.5
2
1.5
1
0
10
20
30
40
50
60
Normal
LY50
LY75
LY88
LY94
LY97
LY data
Normal data

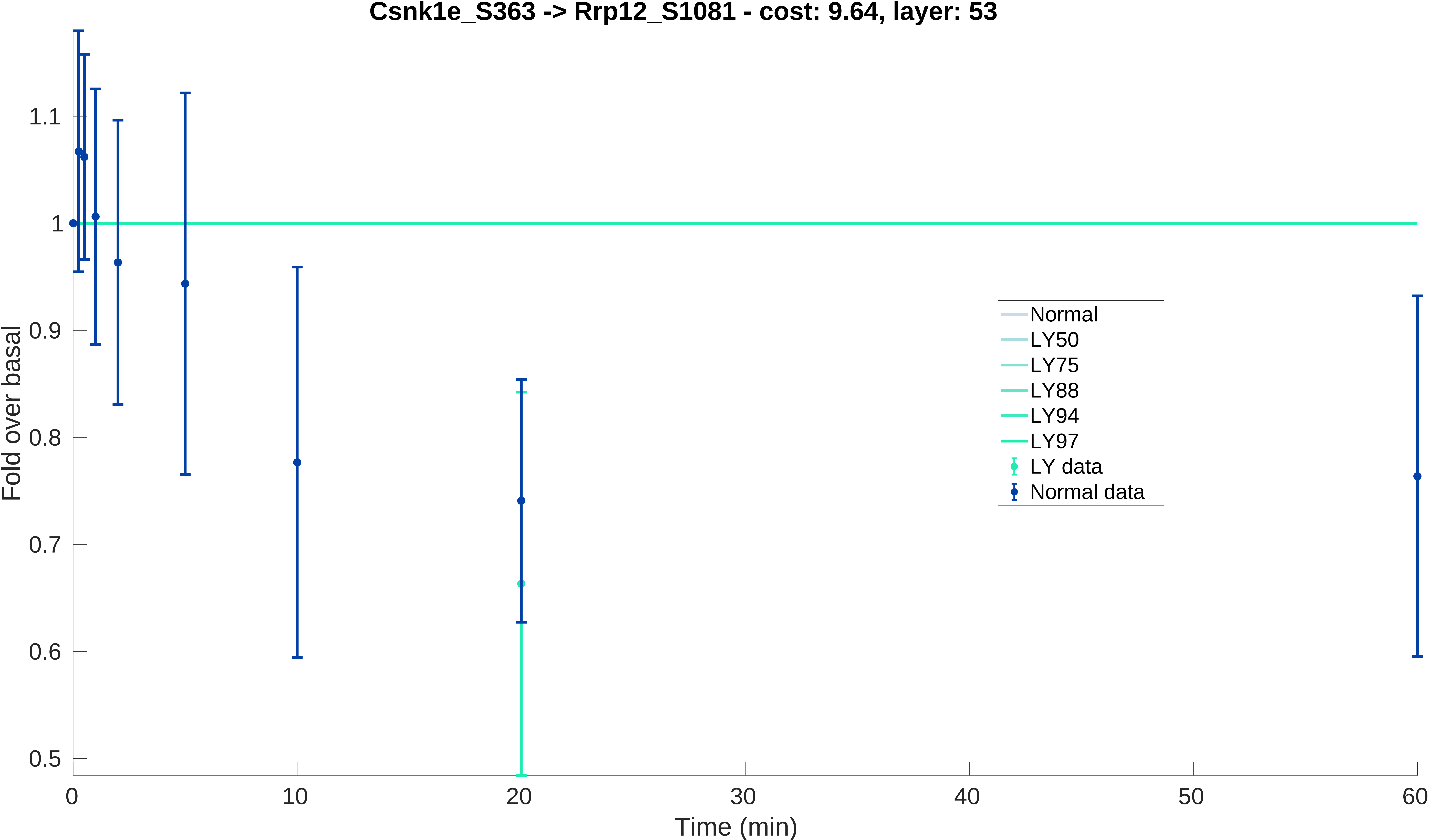
Csnk1e_S363 -> Rrp12_S1081 - cost: 9.64, layer: 53
Fold over basal
1.1
1
0.9
0.8
0.7
0.6
0.5
0
10
20
30
40
50
60
Time (min)
Normal
LY50
LY75
LY88
LY94
LY97
LY data
Normal data

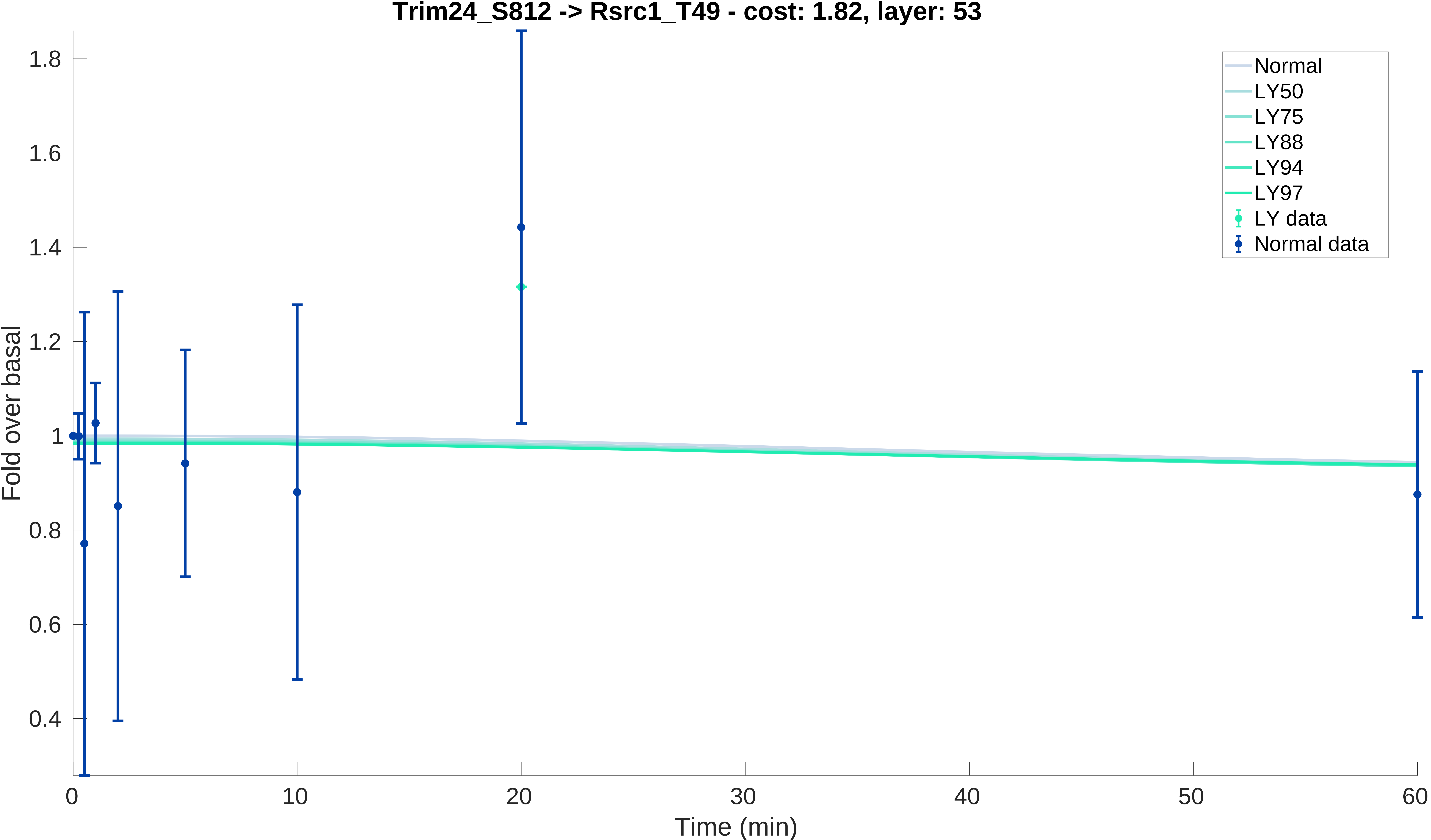

Trim24_S812 -> Rsrc1_T49 - cost: 1.82, layer: 53
Fold over basal
Time (min)
1.8
1.6
1.4
1.2
1
0.8
0.6
0.4
0
10
20
30
40
50
60
Normal
LY50
LY75
LY88
LY94
LY97
LY data
Normal data

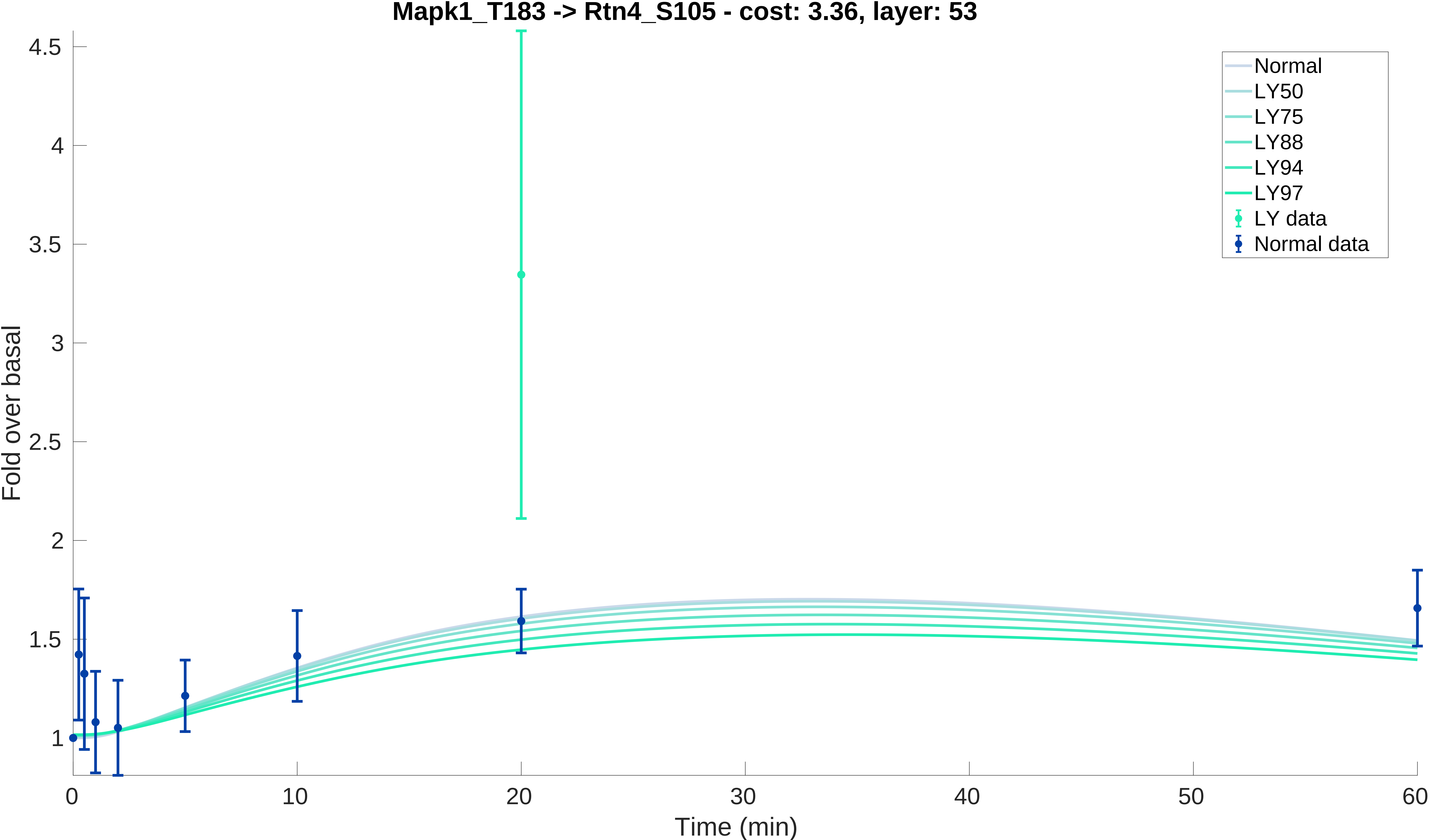
Mapk1_T183 -> Rtn4_S105 - cost: 3.36, layer: 53
Fold over basal
Time (min)
0
10
20
30
40
50
60
1
1.5
2
2.5
3
3.5
4
4.5
Normal
LY50
LY75
LY88
LY94
LY97
LY data
Normal data

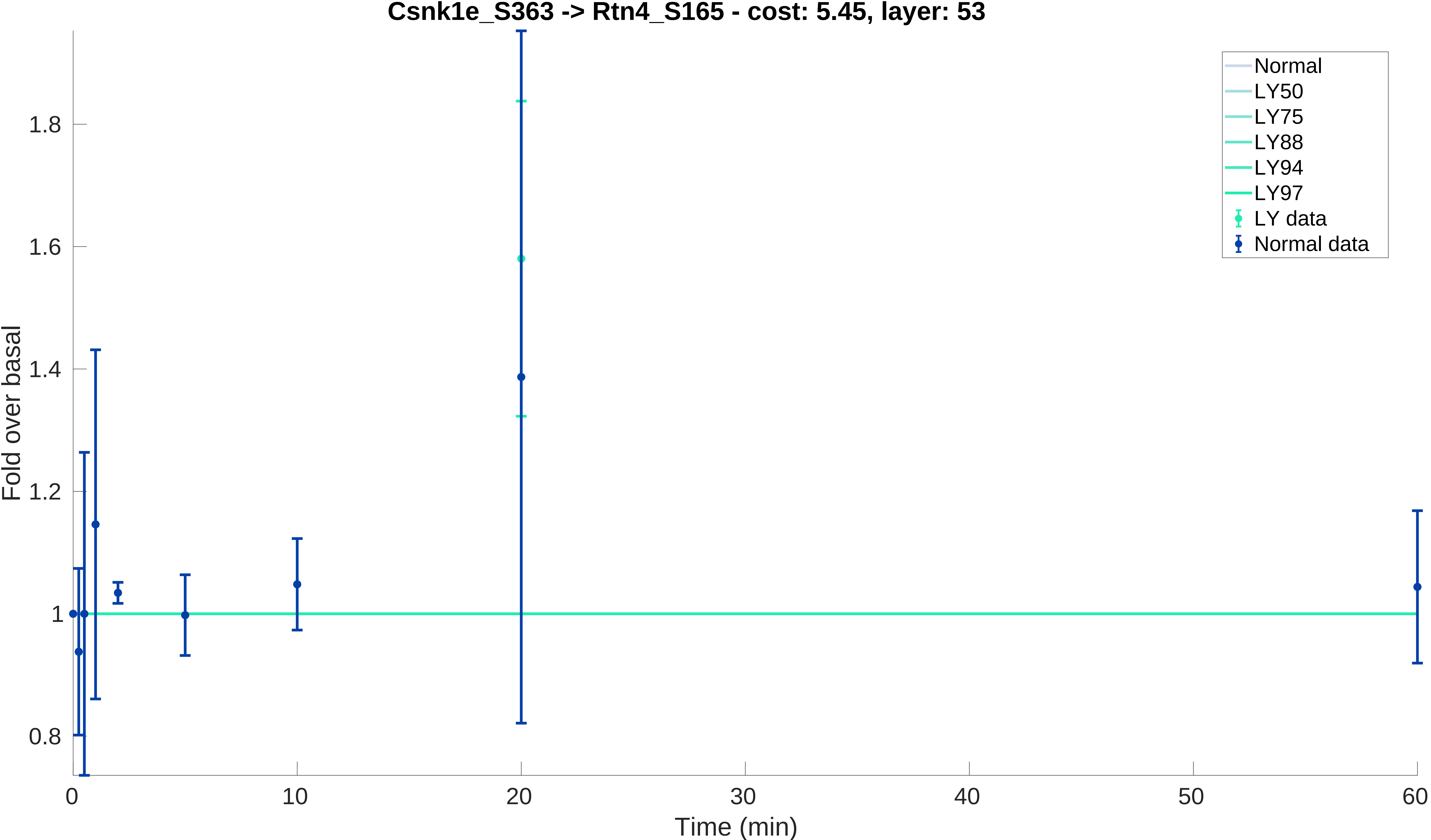
Csnk1e_S363 -> Rtn4_S165 - cost: 5.45, layer: 53
Fold over basal
1.8
1.6
1.4
1.2
1
0.8
0
10
20
30
40
50
60
Time (min)
Normal
LY50
LY75
LY88
LY94
LY97
LY data
Normal data

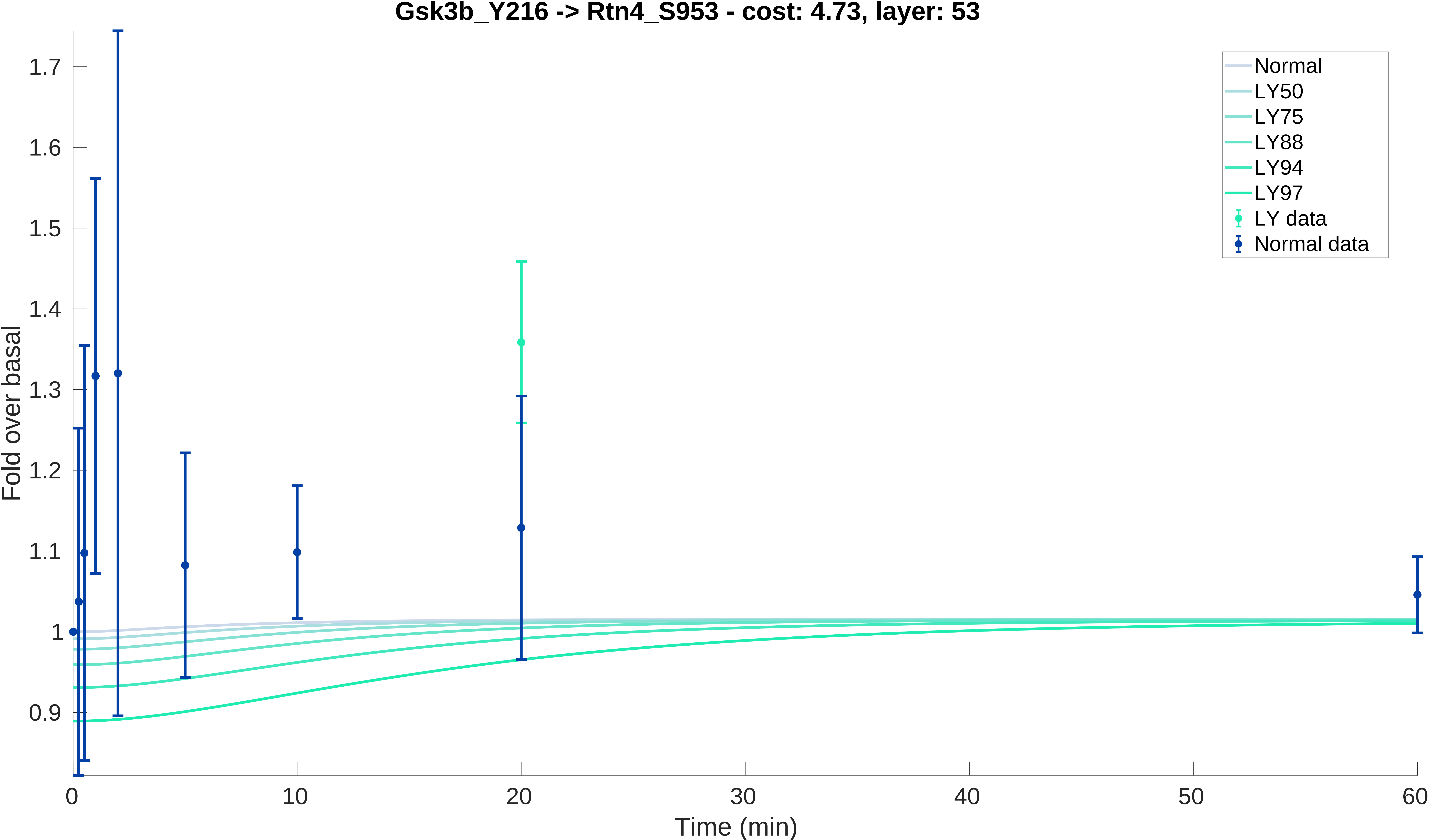

Gsk3b_Y216 -> Rtn4_S953 - cost: 4.73, layer: 53
Fold over basal
Time (min)
0.9
1
1.1
1.2
1.3
1.4
1.5
1.6
1.7
0
10
20
30
40
50
60
Normal
LY50
LY75
LY88
LY94
LY97
LY data
Normal data

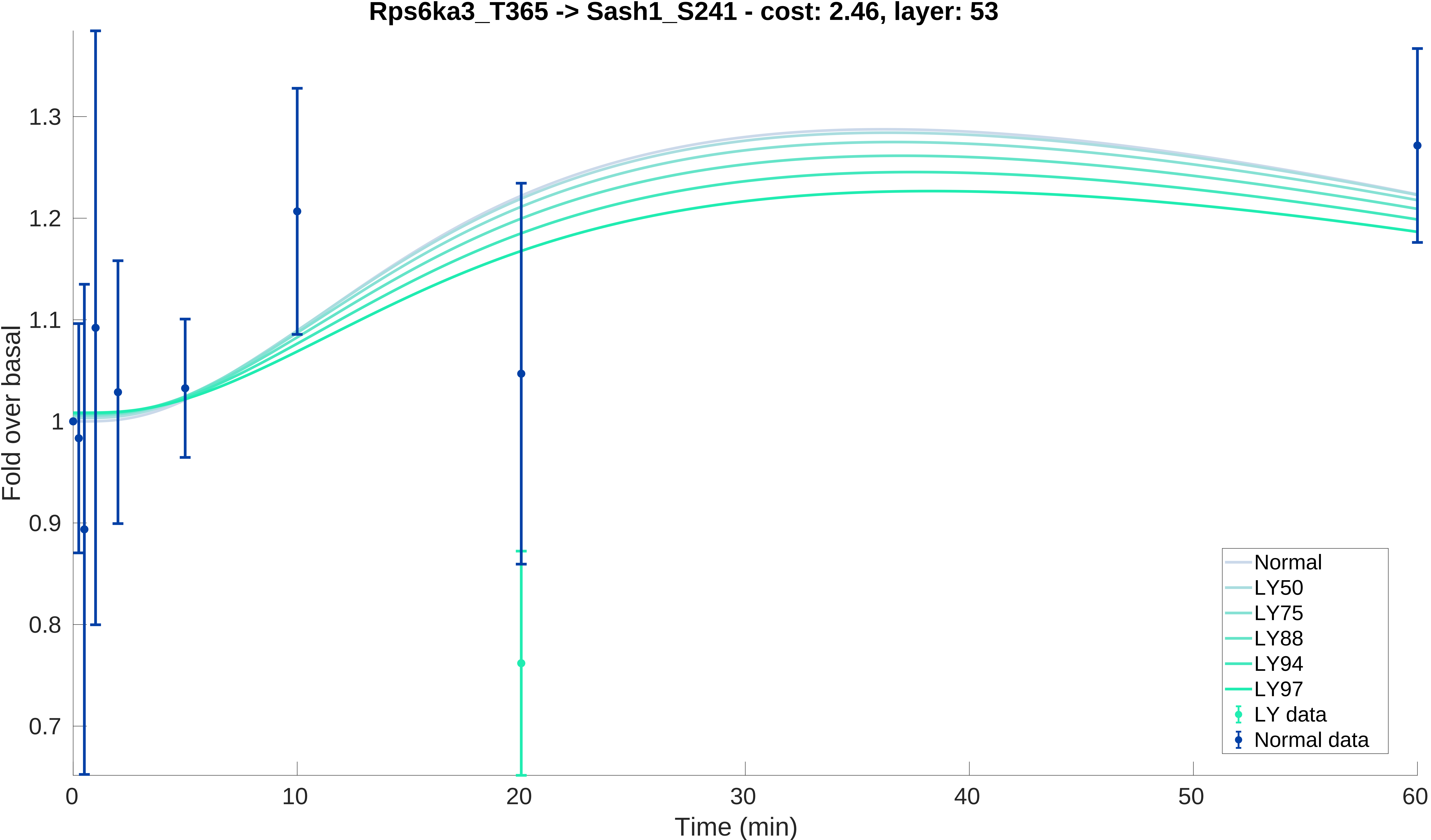

Rps6ka3_T365 -> Sash1_S241 - cost: 2.46, layer: 53
Fold over basal
1.3
1.2
1.1
1
0.9
0.8
0.7
0
10
20
30
40
50
60
Time (min)
Normal
LY50
LY75
LY88
LY94
LY97
LY data
Normal data

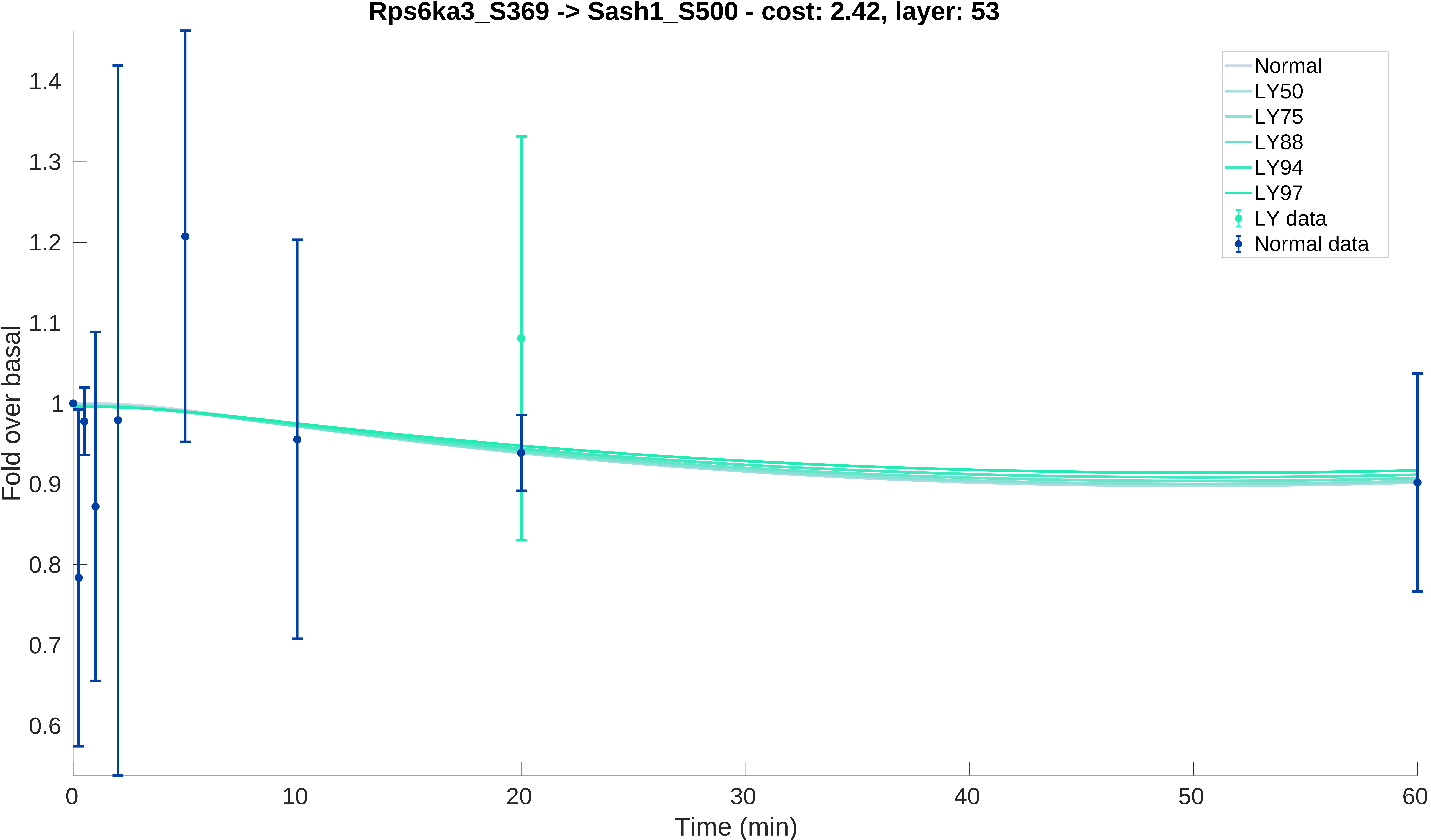

Rps6ka3_S369 -> Sash1_S500 - cost: 2.42, layer: 53
Fold over basal
Time (min)
1.4
1.3
1.2
1.1
1
0.9
0.8
0.7
0.6
0
10
20
30
40
50
60
Normal
LY50
LY75
LY88
LY94
LY97
LY data
Normal data

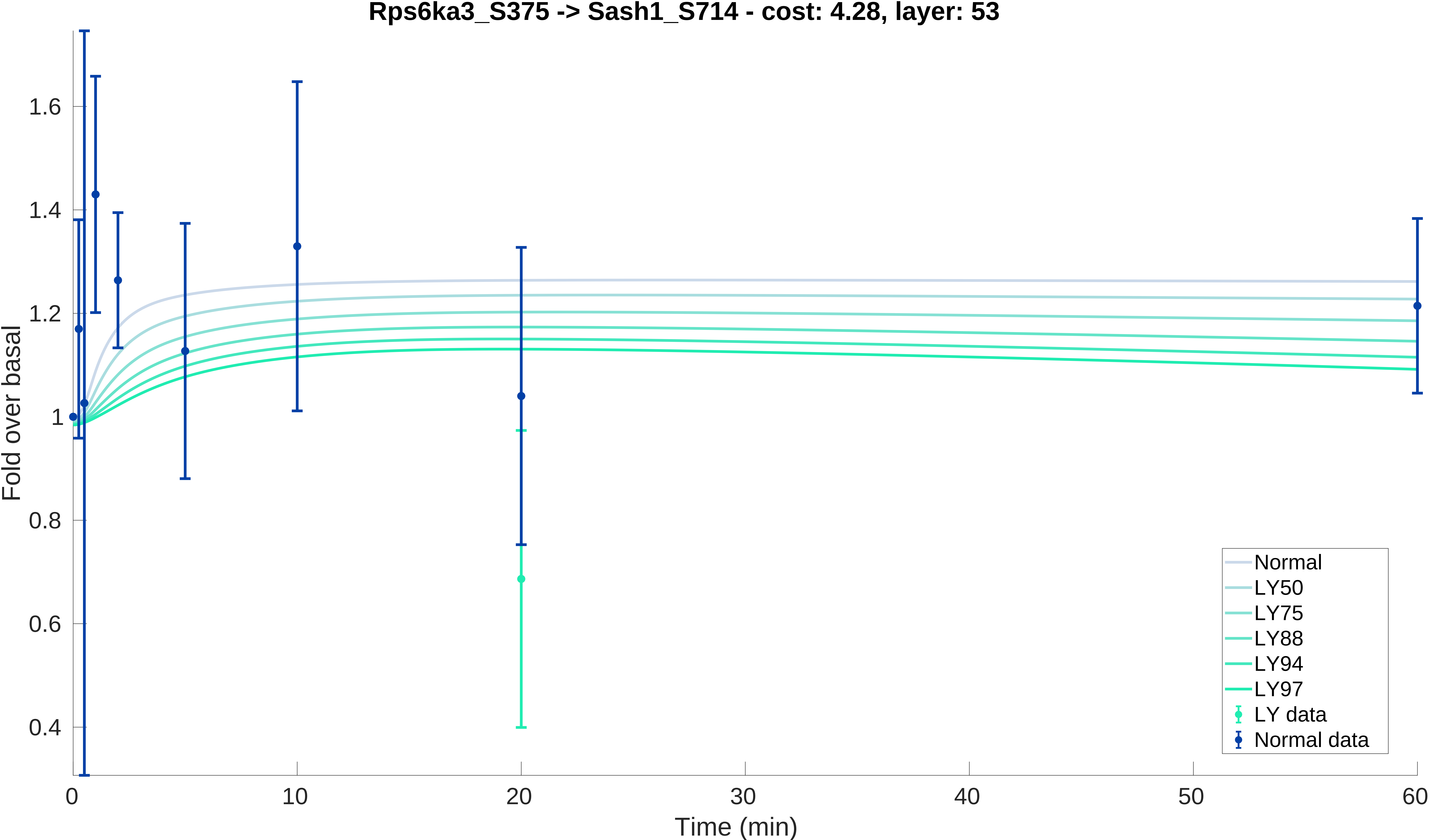

Rps6ka3_S375 -> Sash1_S714 - cost: 4.28, layer: 53
Fold over basal
Time (min)
1.6
1.4
1.2
1
0.8
0.6
0.4
0
10
20
30
40
50
60
Normal
LY50
LY75
LY88
LY94
LY97
LY data
Normal data

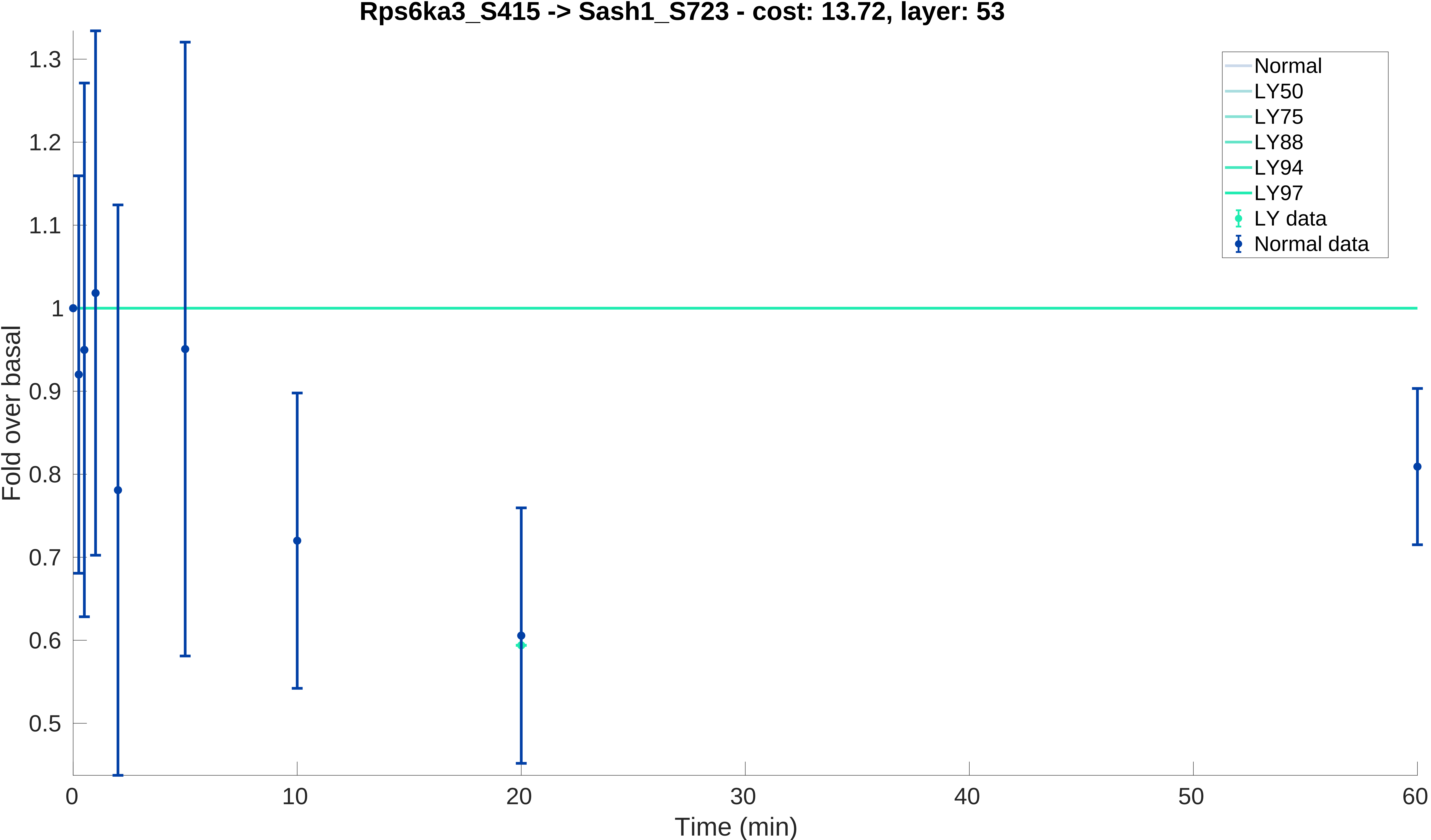

Rps6ka3_S415 -> Sash1_S723 - cost: 13.72, layer: 53
Fold over basal
1.3
1.2
1.1
1
0.9
0.8
0.7
0.6
0.5
0
10
20
30
40
50
60
Time (min)
Normal
LY50
LY75
LY88
LY94
LY97
LY data
Normal data

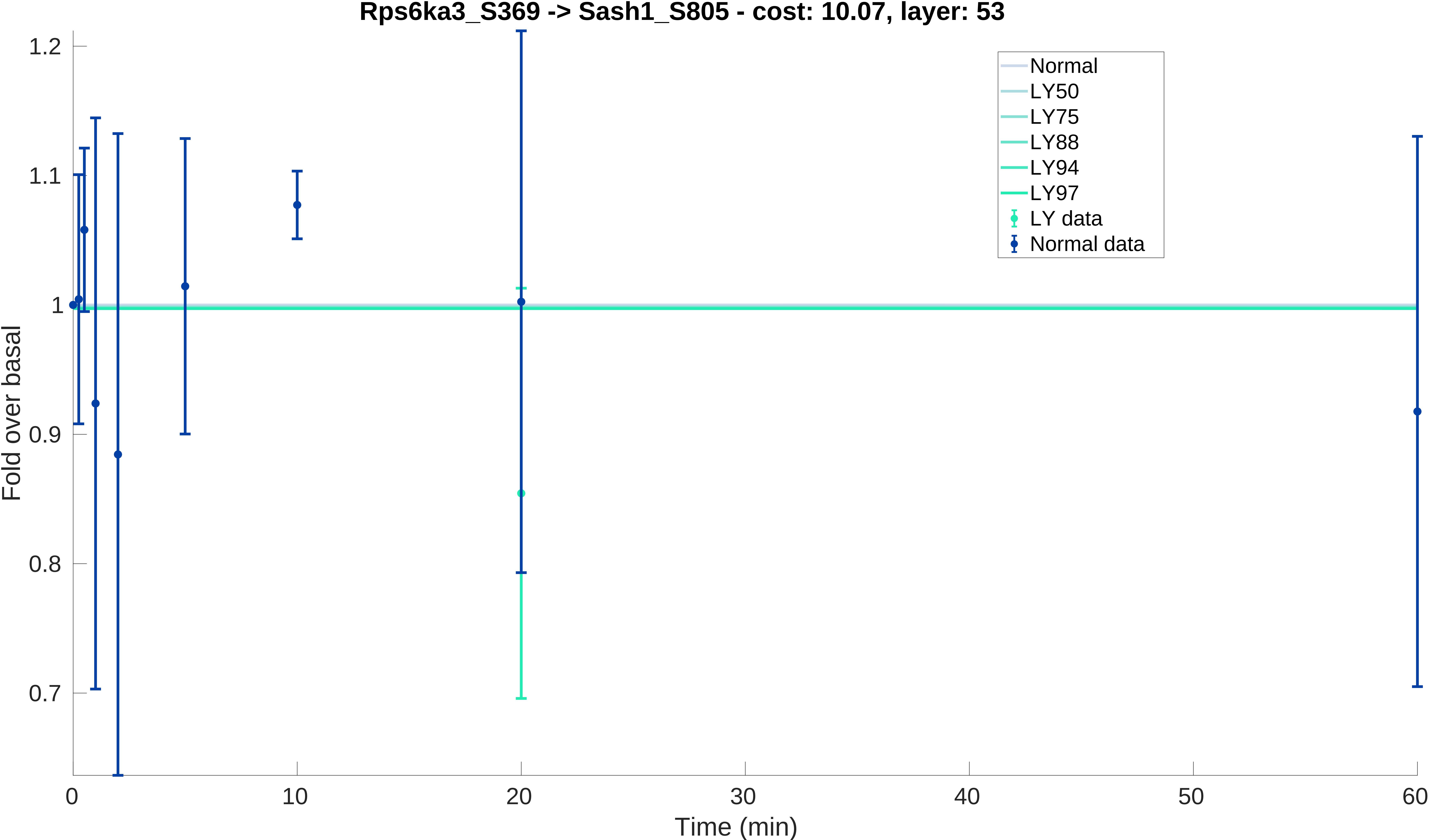

Rps6ka3_S369 -> Sash1_S805 - cost: 10.07, layer: 53
Normal
LY50
LY75
LY88
LY94
LY97
LY data
Normal data
Fold over basal
Time (min)
1.2
1.1
1
0.9
0.8
0.7
0
10
20
30
40
50
60

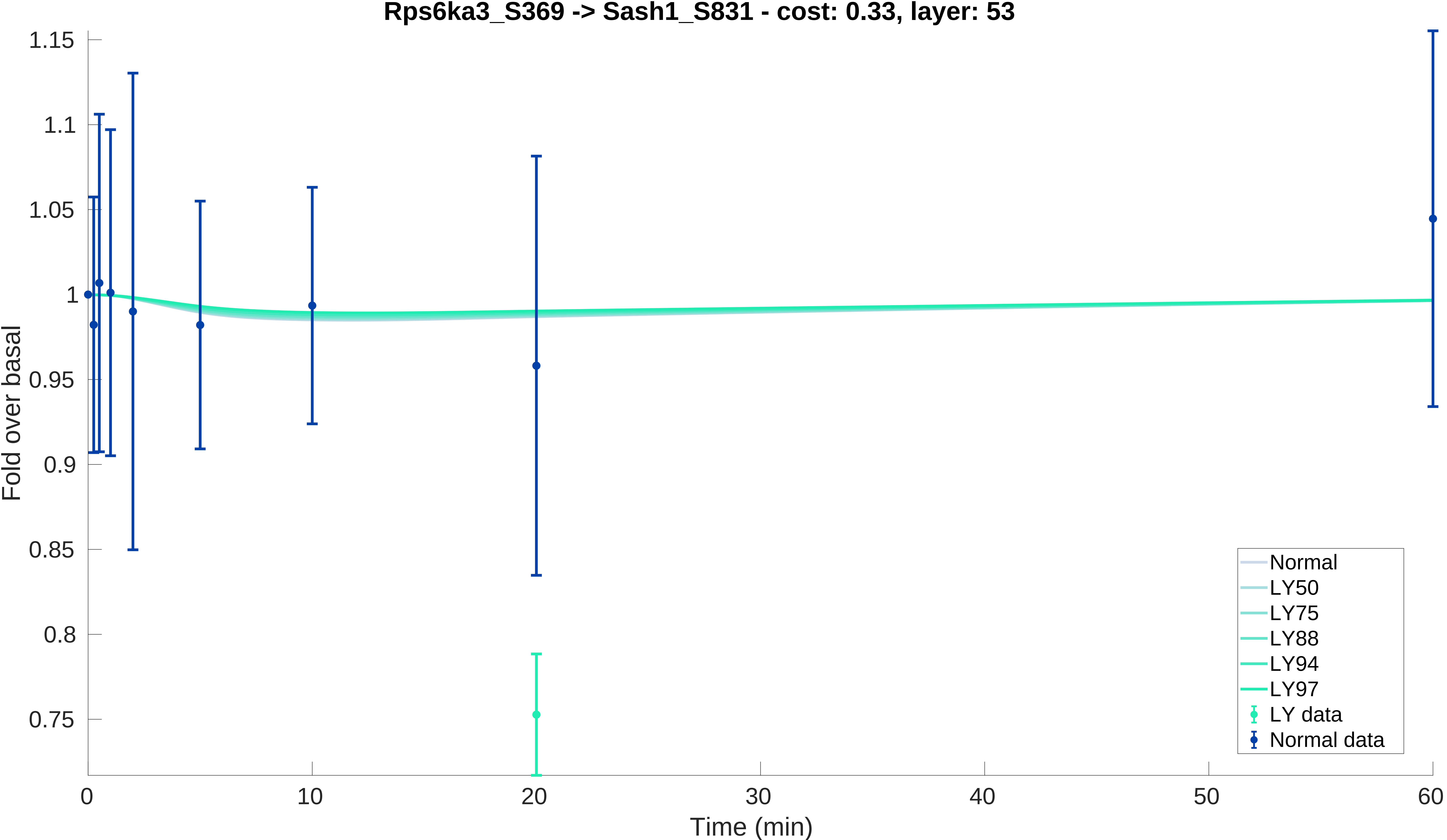

Rps6ka3_S369 -> Sash1_S831 - cost: 0.33, layer: 53
Fold over basal
Time (min)
1.15
1.1
1.05
1
0.95
0.9
0.85
0.8
0.75
0
10
20
30
40
50
60
Normal
LY50
LY75
LY88
LY94
LY97
LY data
Normal data

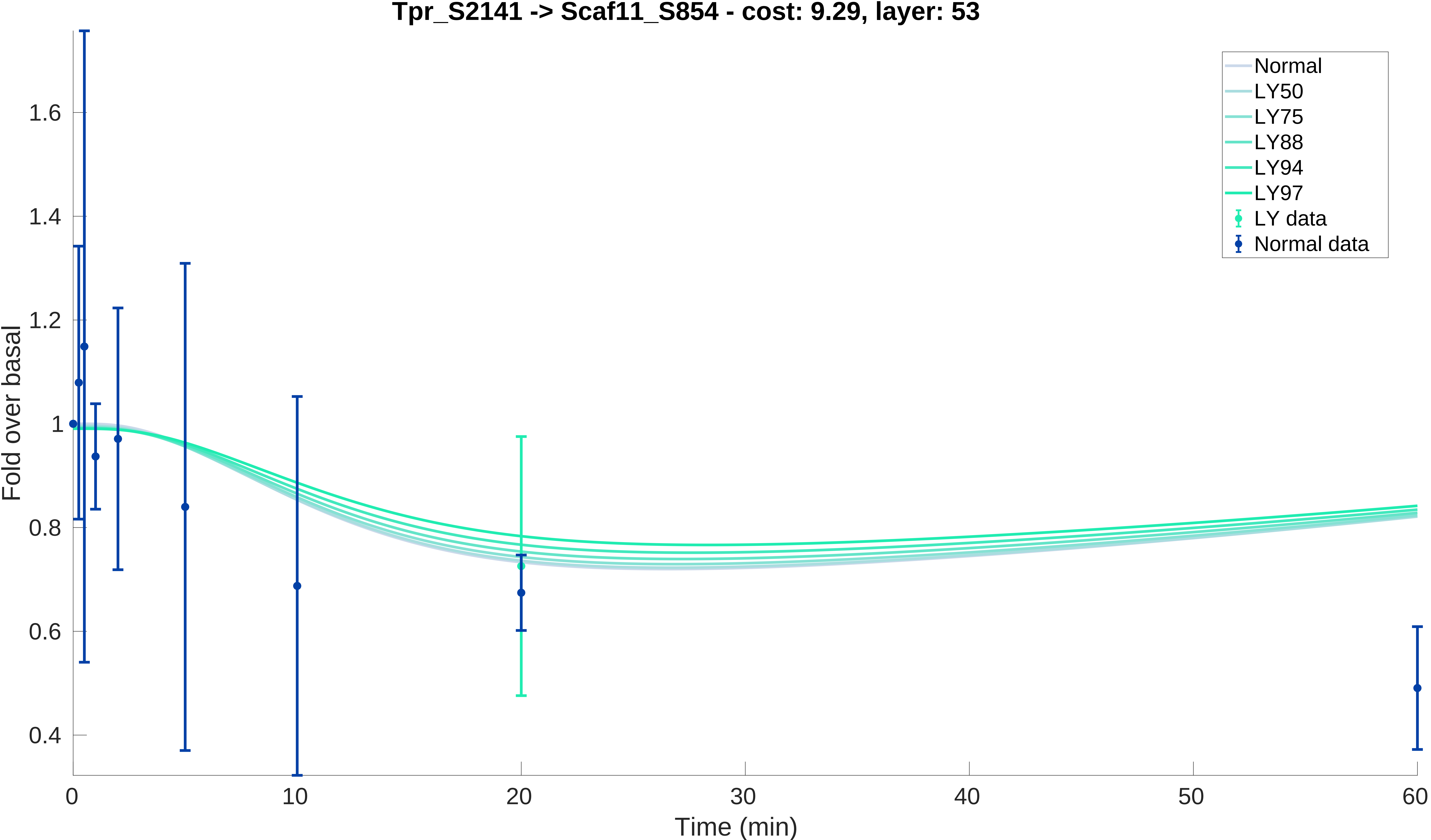

Tpr_S2141 -> Scaf11_S854 - cost: 9.29, layer: 53
Fold over basal
Time (min)
1.6
1.4
1.2
1
0.8
0.6
0.4
0
10
20
30
40
50
60
Normal
LY50
LY75
LY88
LY94
LY97
LY data
Normal data

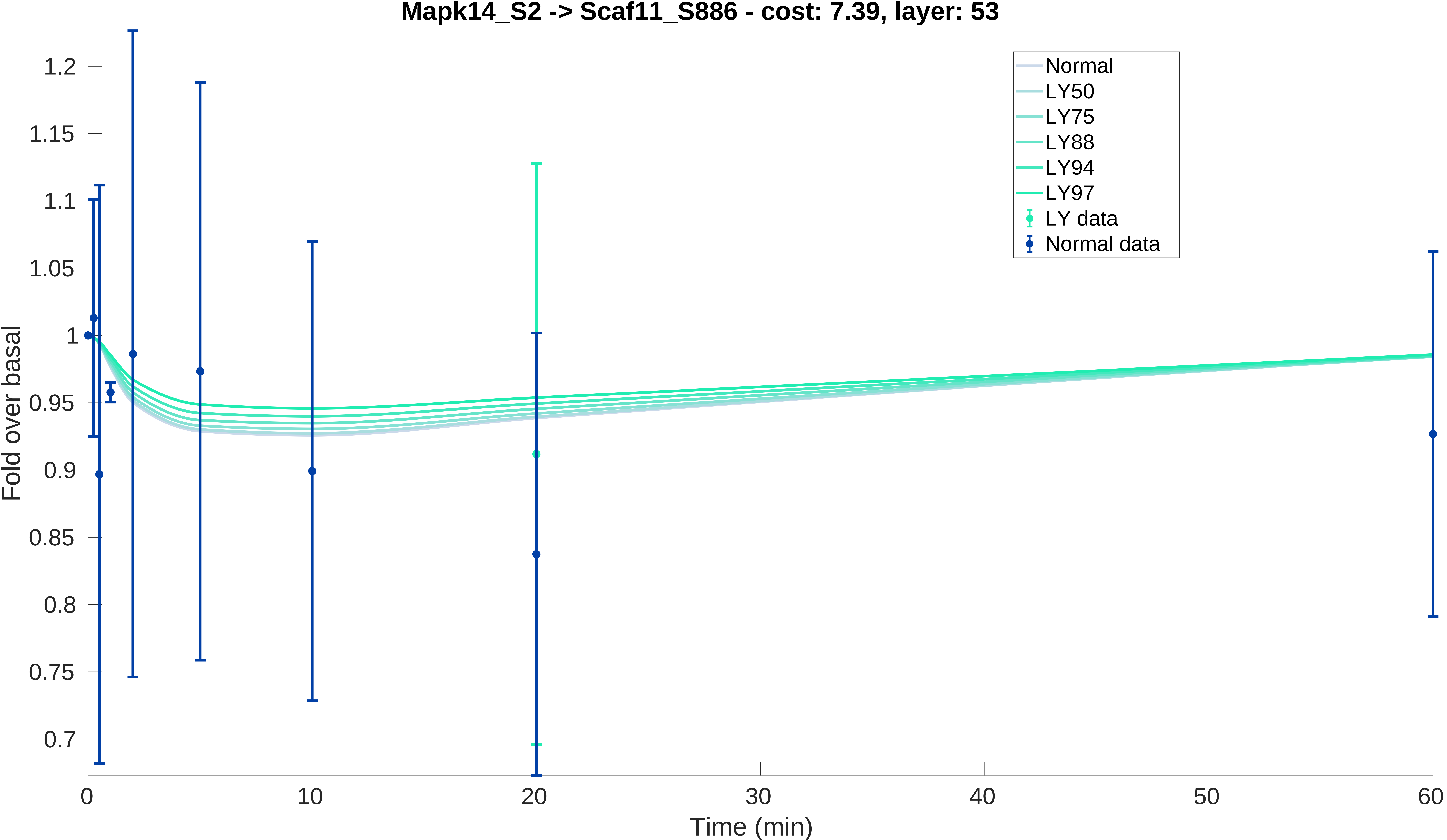
Mapk14_S2 -> Scaf11_S886 - cost: 7.39, layer: 53
Fold over basal
1.2
1.15
1.1
1.05
1
0.95
0.9
0.85
0.8
0.75
0.7
0
10
20
30
40
50
60
Time (min)
Normal
LY50
LY75
LY88
LY94
LY97
LY data
Normal data

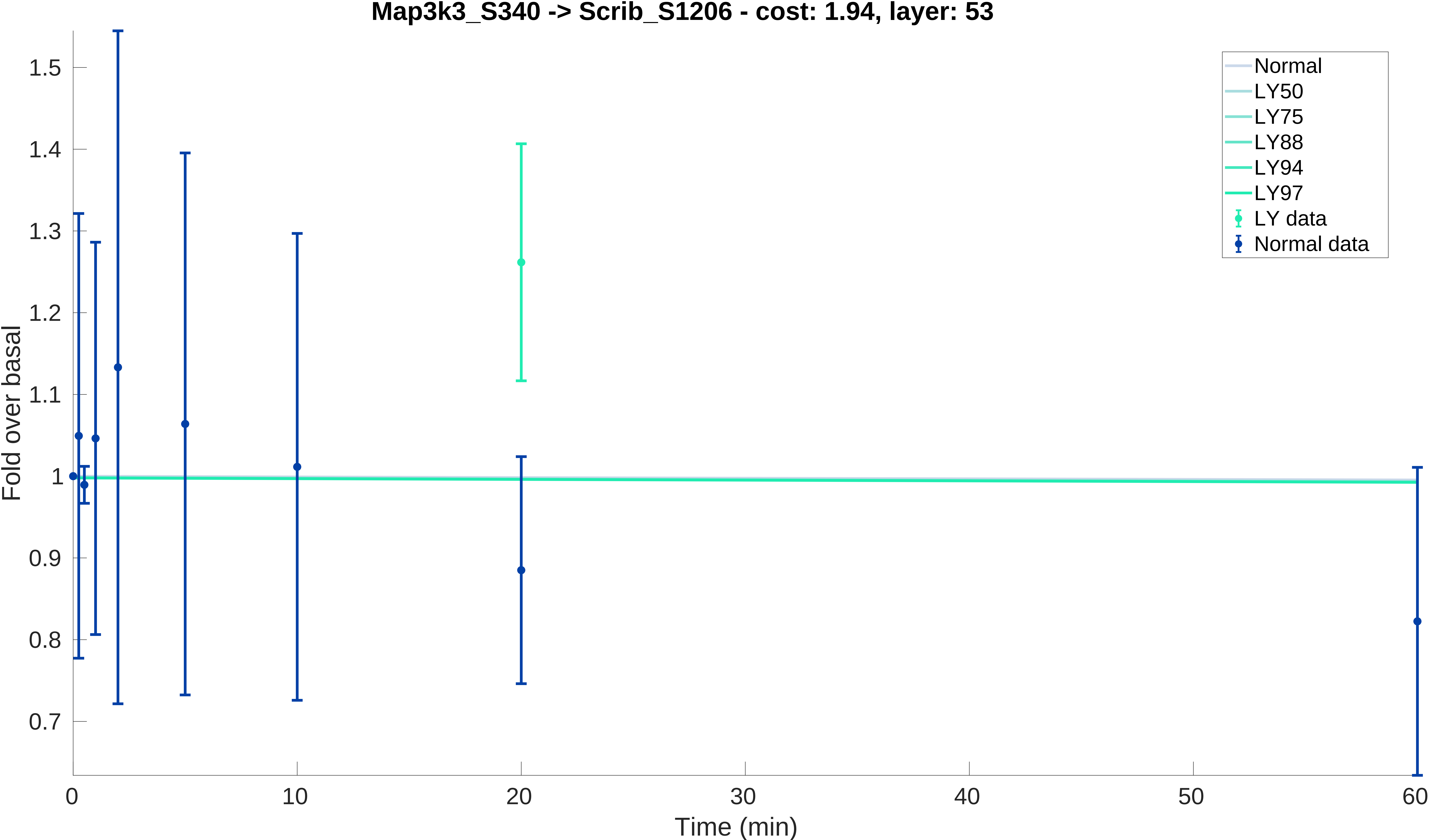

Map3k3_S340 -> Scrib_S1206 - cost: 1.94, layer: 53
Fold over basal
Time (min)
1.5
1.4
1.3
1.2
1.1
1
0.9
0.8
0.7
0
10
20
30
40
50
60
Normal
LY50
LY75
LY88
LY94
LY97
LY data
Normal data

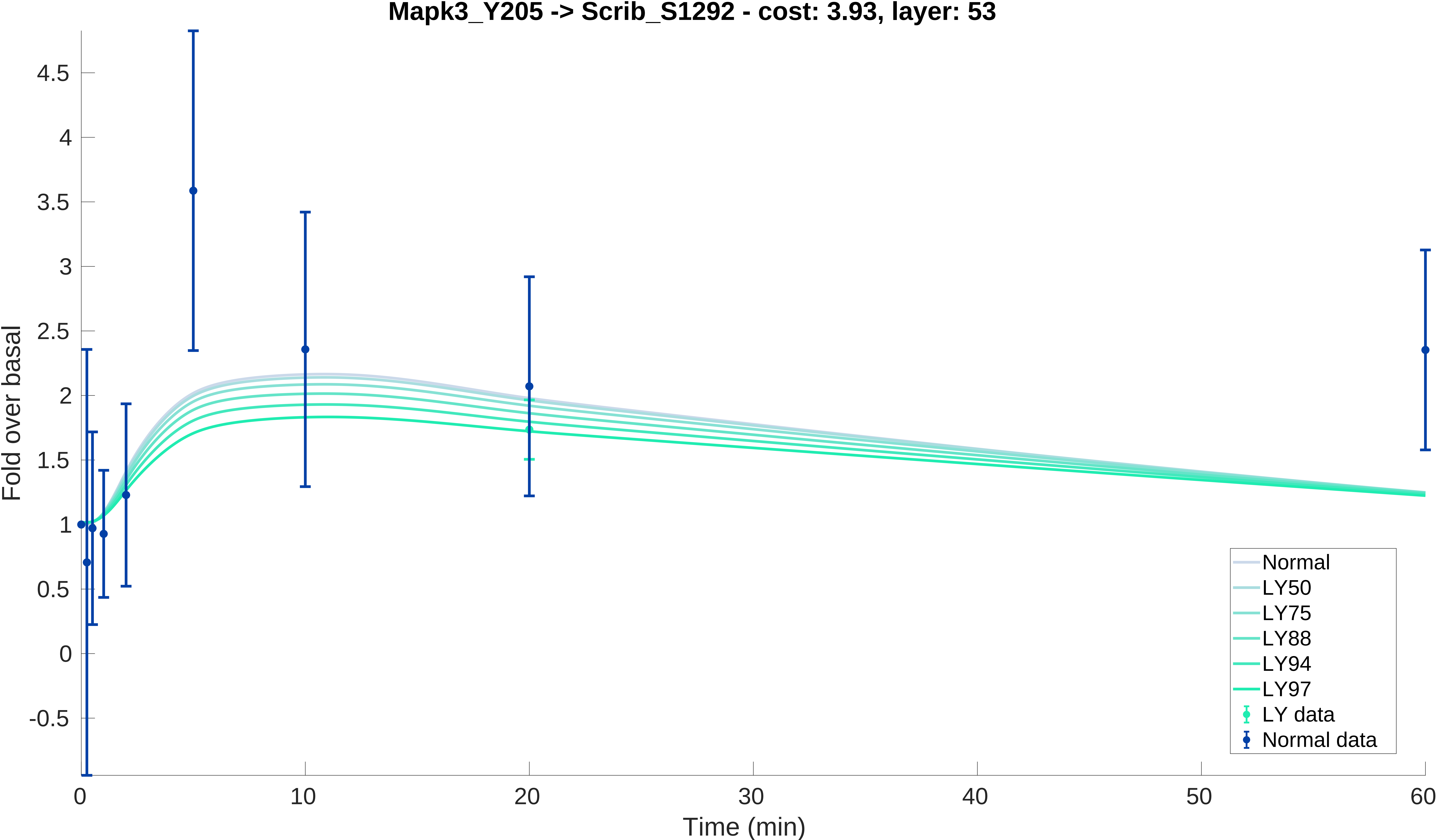

Mapk3_Y205 -> Scrib_S1292 - cost: 3.93, layer: 53
Fold over basal
4.5
4
3.5
3
2.5
2
1.5
1
0.5
0
-0.5
0
10
20
30
40
50
60
Time (min)
Normal
LY50
LY75
LY88
LY94
LY97
LY data
Normal data

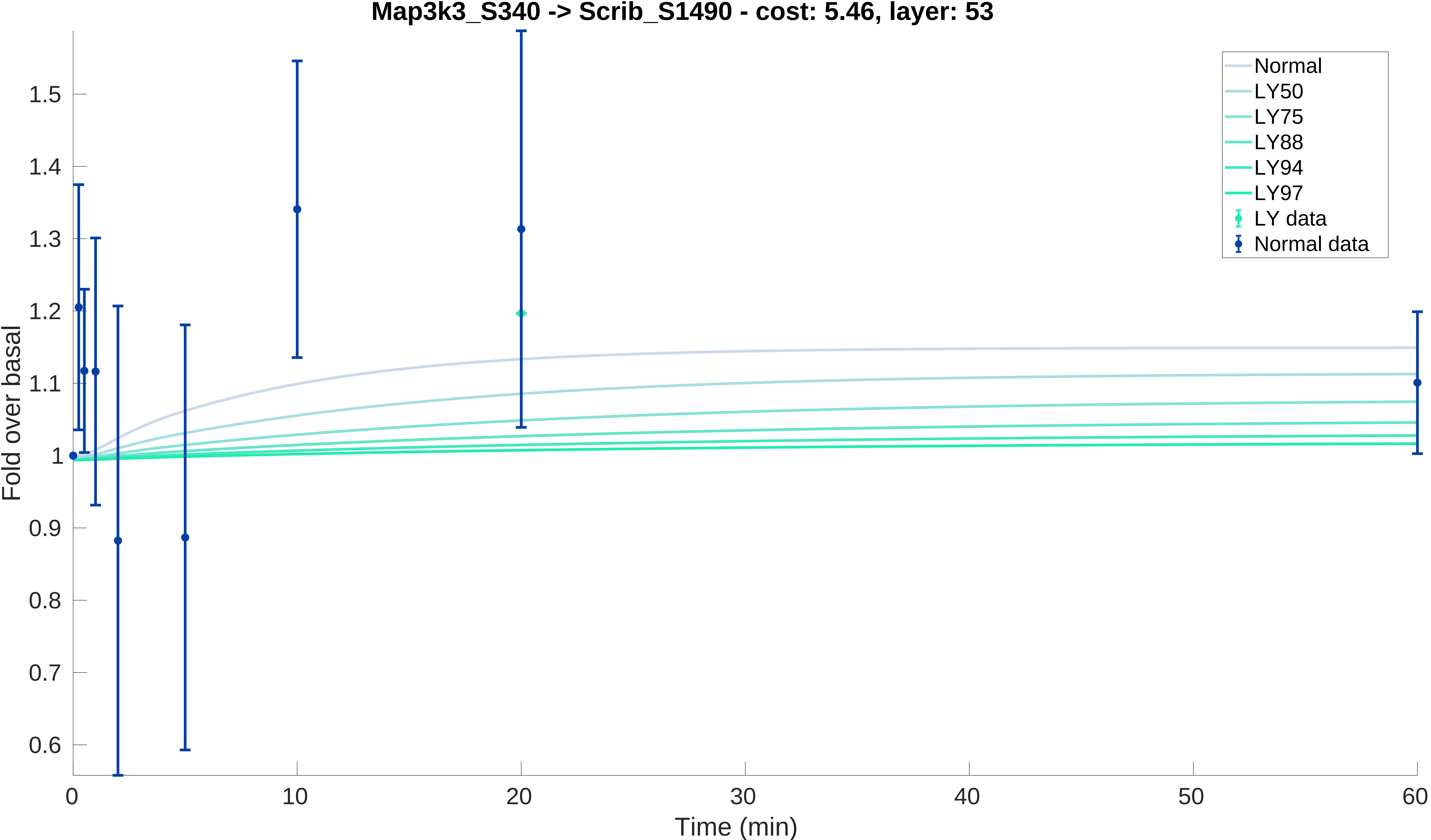

Map3k3_S340 -> Scrib_S1490 - cost: 5.46, layer: 53
Fold over basal
Time (min)
1.5
1.4
1.3
1.2
1.1
1
0.9
0.8
0.7
0.6
0
10
20
30
40
50
60
Normal
LY50
LY75
LY88
LY94
LY97
LY data
Normal data

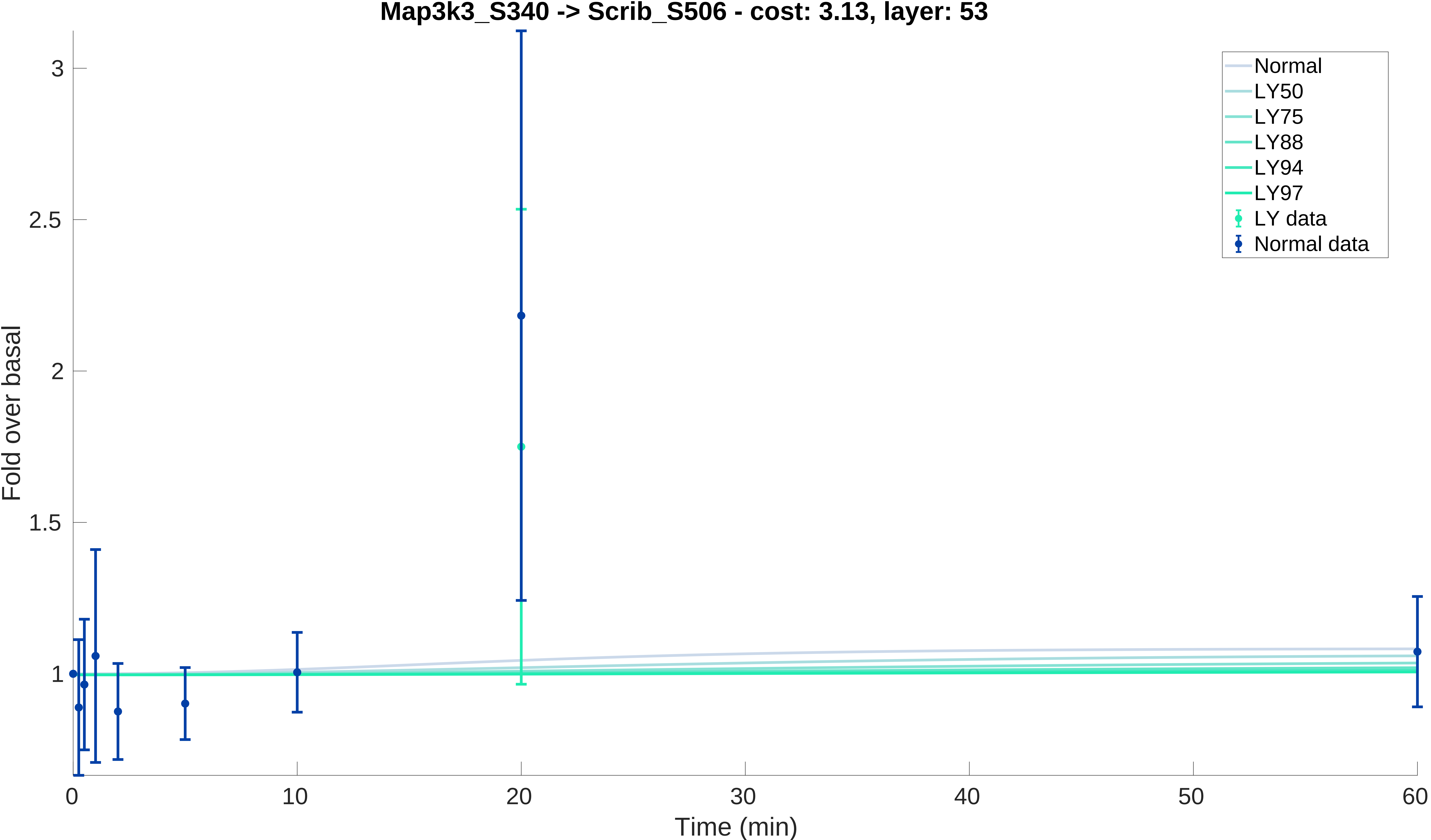

Map3k3_S340 -> Scrib_S506 - cost: 3.13, layer: 53
Fold over basal
Time (min)
0
10
20
30
40
50
60
1
1.5
2
2.5
3
Normal
LY50
LY75
LY88
LY94
LY97
LY data
Normal data

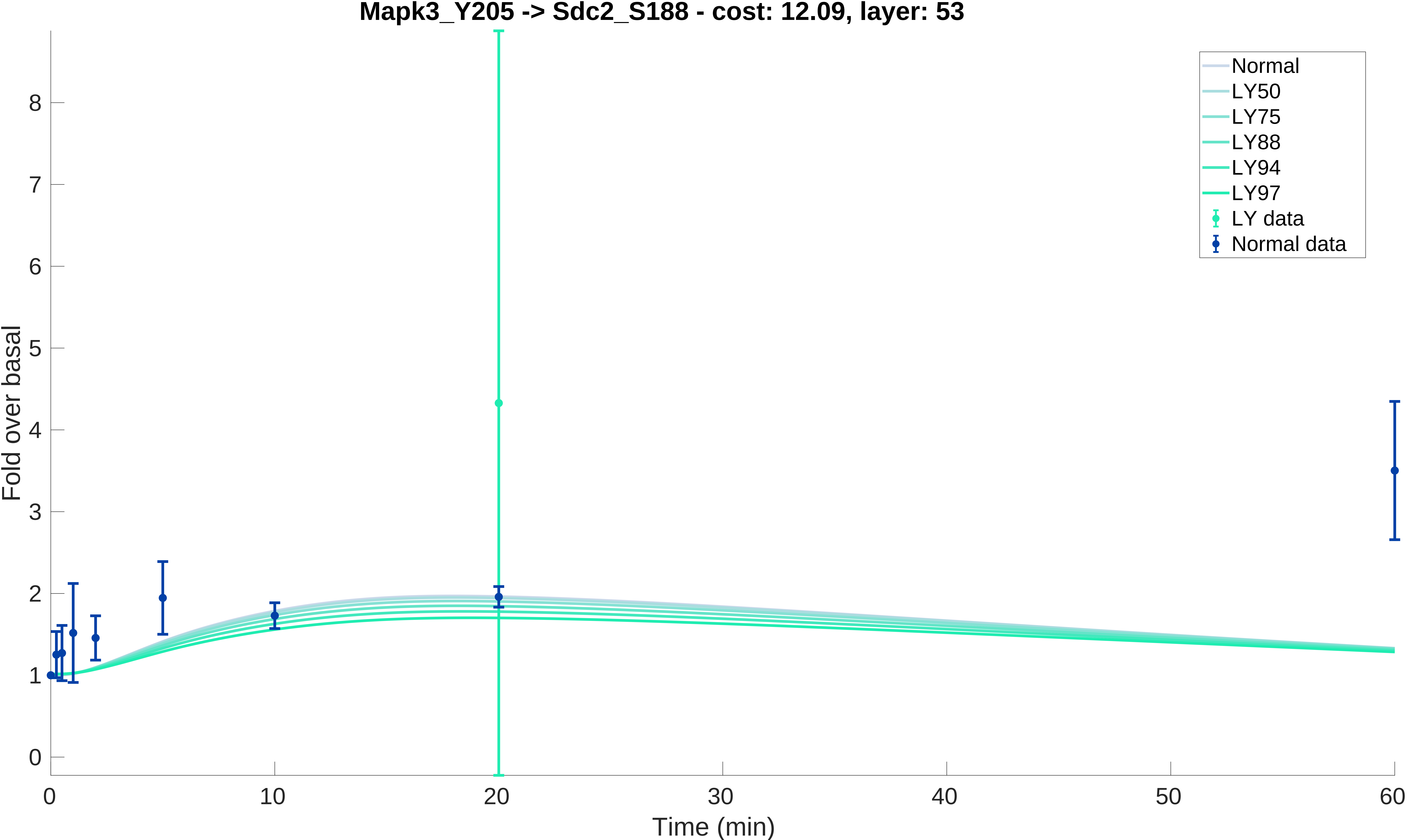

Mapk3_Y205 -> Sdc2_S188 - cost: 12.09, layer: 53
Fold over basal
Time (min)
0
1
2
3
4
5
6
7
8
0
10
20
30
40
50
60
Normal
LY50
LY75
LY88
LY94
LY97
LY data
Normal data

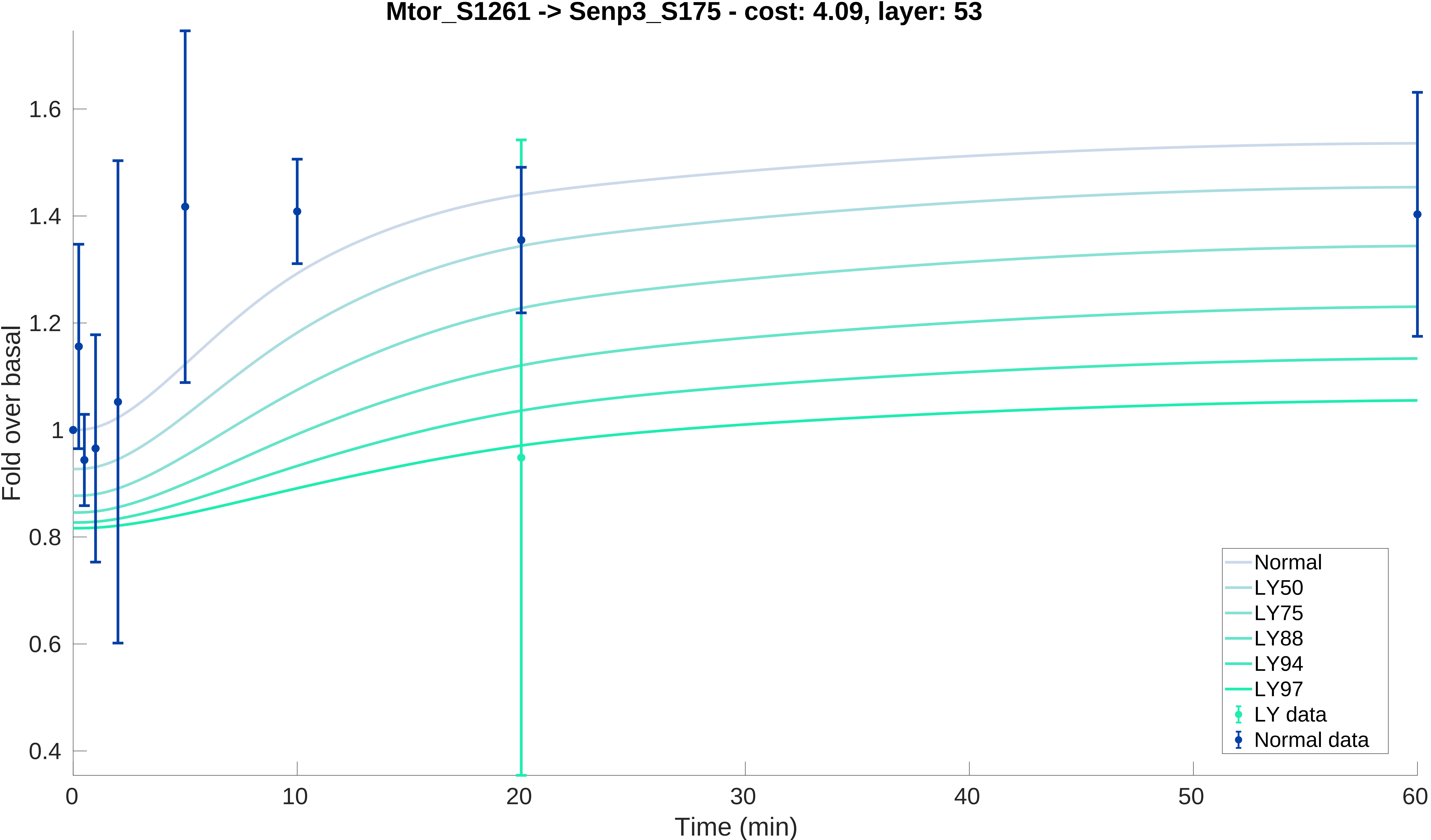

Mtor_S1261 -> Senp3_S175 - cost: 4.09, layer: 53
Fold over basal
1.6
1.4
1.2
1
0.8
0.6
0.4
0
10
20
30
40
50
60
Time (min)
Normal
LY50
LY75
LY88
LY94
LY97
LY data
Normal data

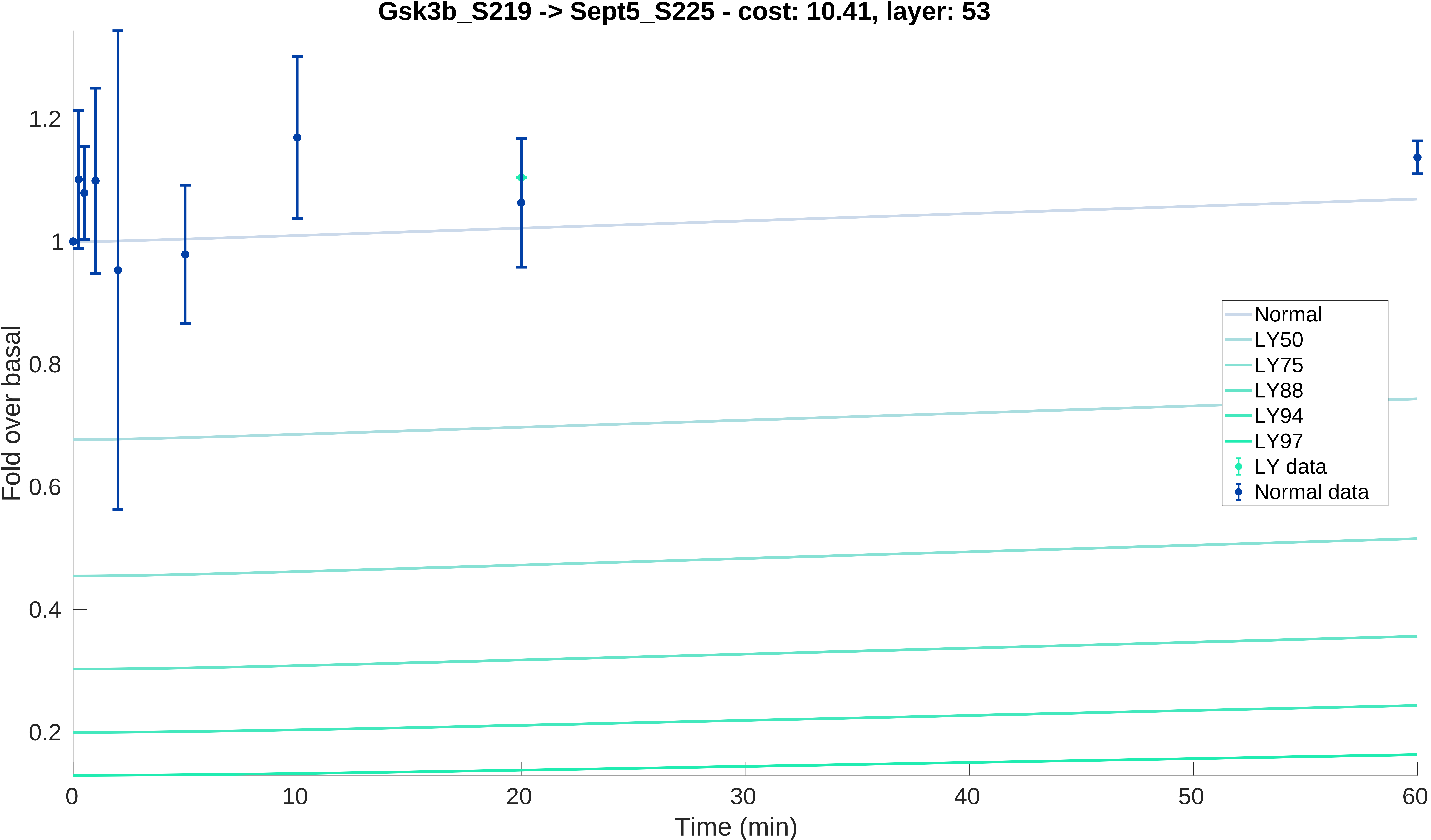

Gsk3b_S219 -> Sept5_S225 - cost: 10.41, layer: 53
Fold over basal
1.2
1
0.8
0.6
0.4
0.2
0
10
20
30
40
50
60
Time (min)
Normal
LY50
LY75
LY88
LY94
LY97
LY data
Normal data

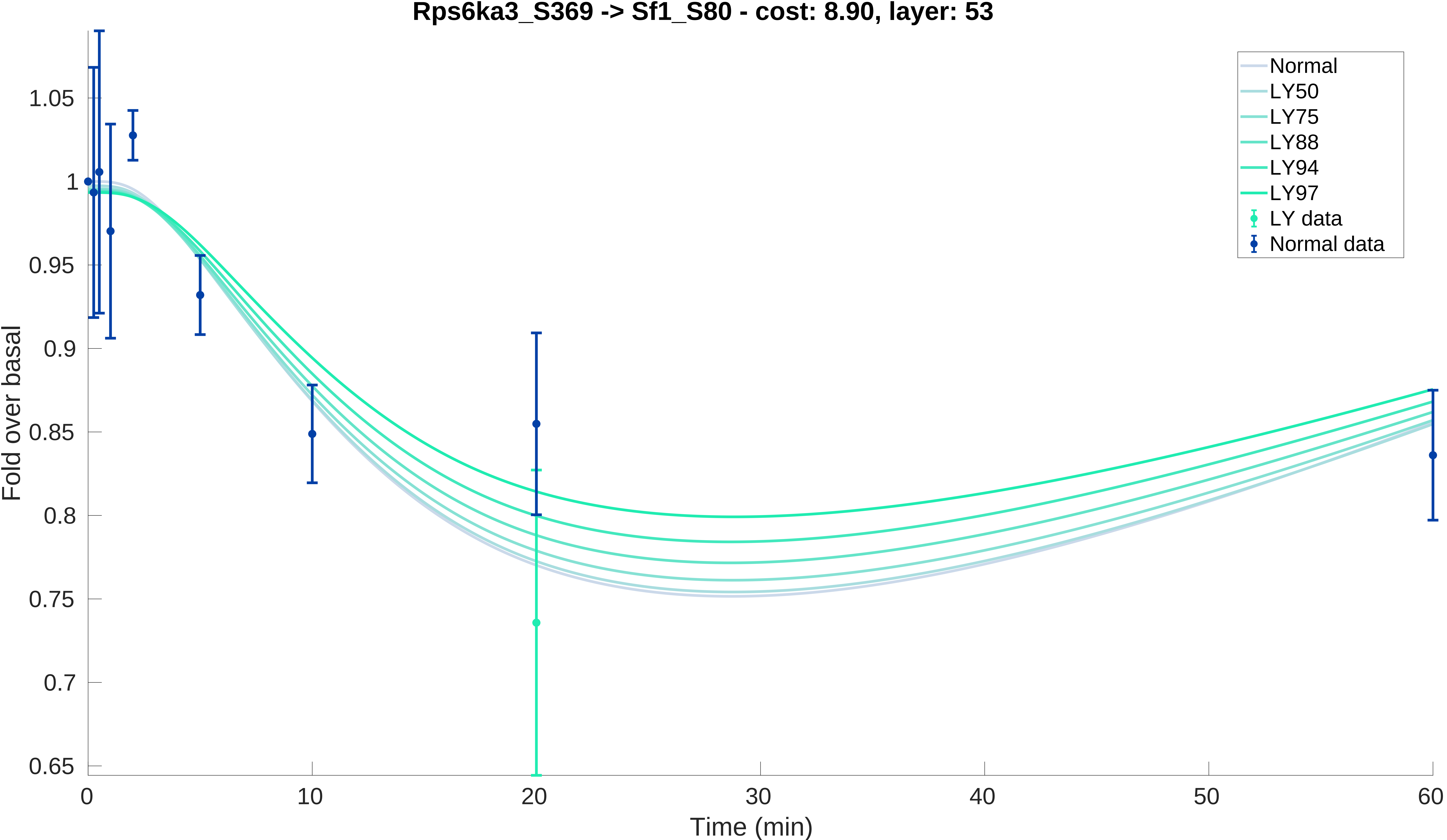
Rps6ka3_S369 -> Sf1_S80 - cost: 8.90, layer: 53
Normal
LY50
LY75
LY88
LY94
LY97
LY data
Normal data
Fold over basal
1.05
1
0.95
0.9
0.85
0.8
0.75
0.7
0.65
0
10
20
30
40
50
60
Time (min)

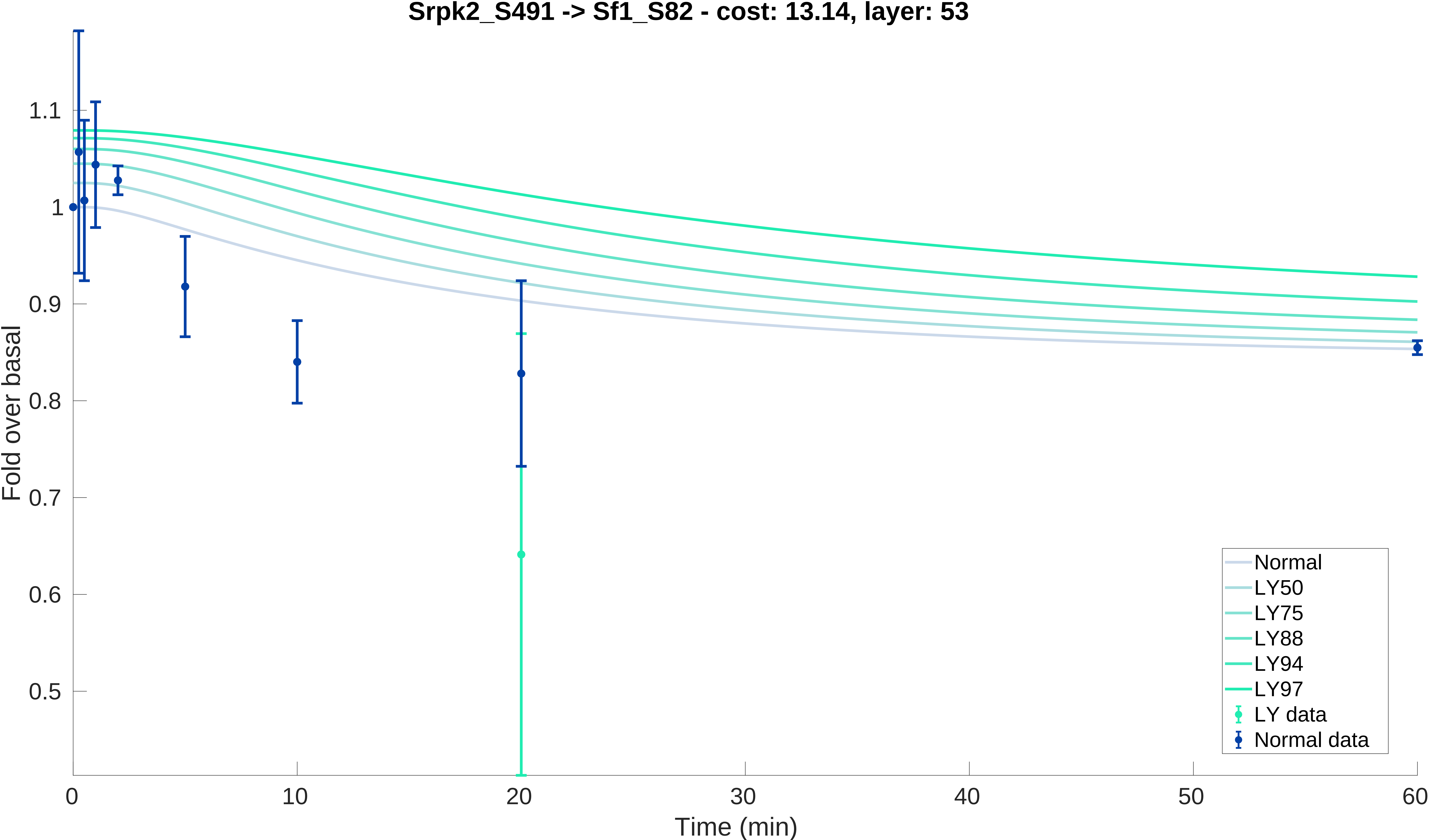

Srpk2_S491 -> Sf1_S82 - cost: 13.14, layer: 53
Fold over basal
1.1
1
0.9
0.8
0.7
0.6
0.5
0
10
20
30
40
50
60
Time (min)
Normal
LY50
LY75
LY88
LY94
LY97
LY data
Normal data

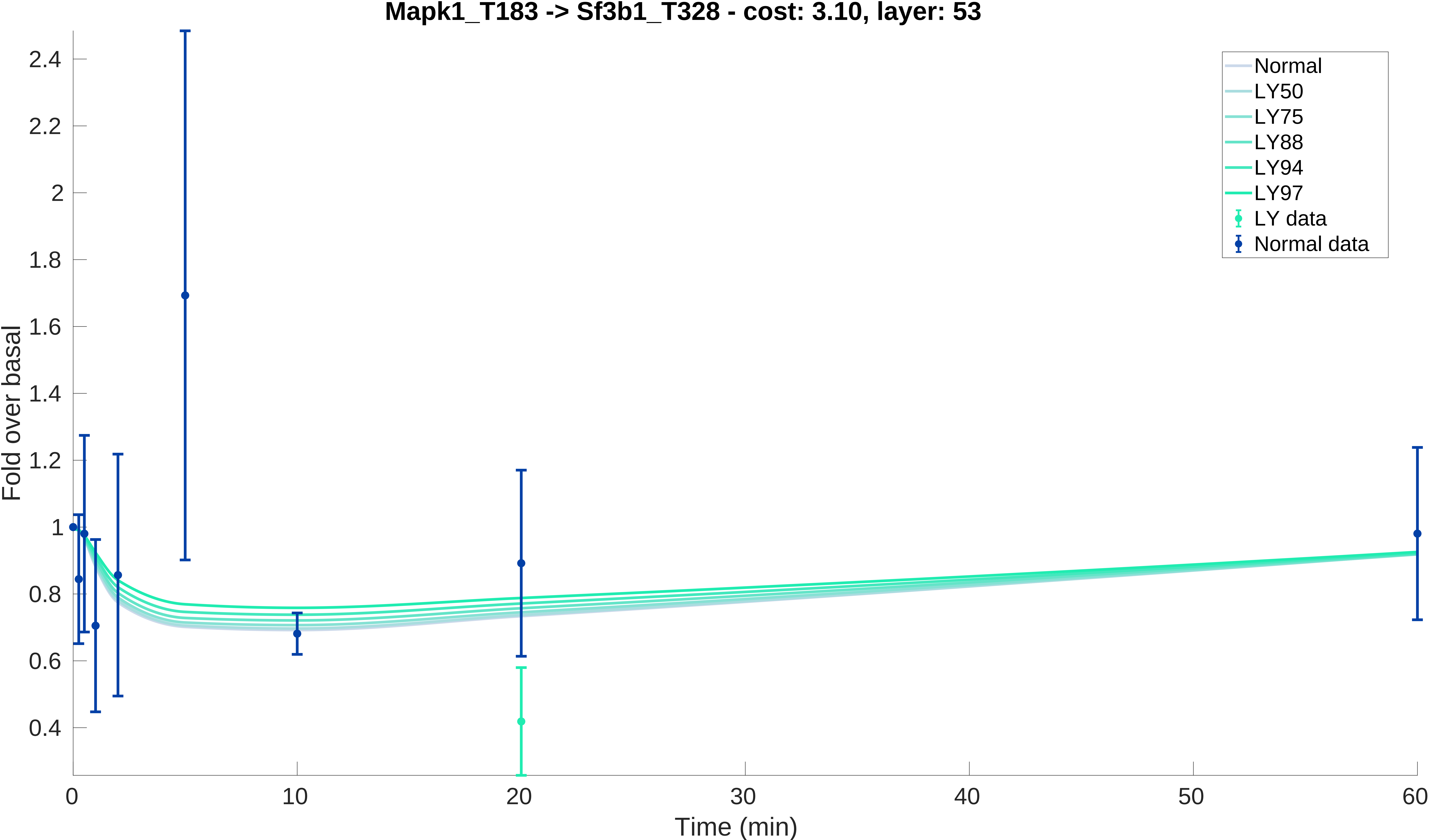

Mapk1_T183 -> Sf3b1_T328 - cost: 3.10, layer: 53
Fold over basal
2.4
2.2
2
1.8
1.6
1.4
1.2
1
0.8
0.6
0.4
0
10
20
30
40
50
60
Time (min)
Normal
LY50
LY75
LY88
LY94
LY97
LY data
Normal data

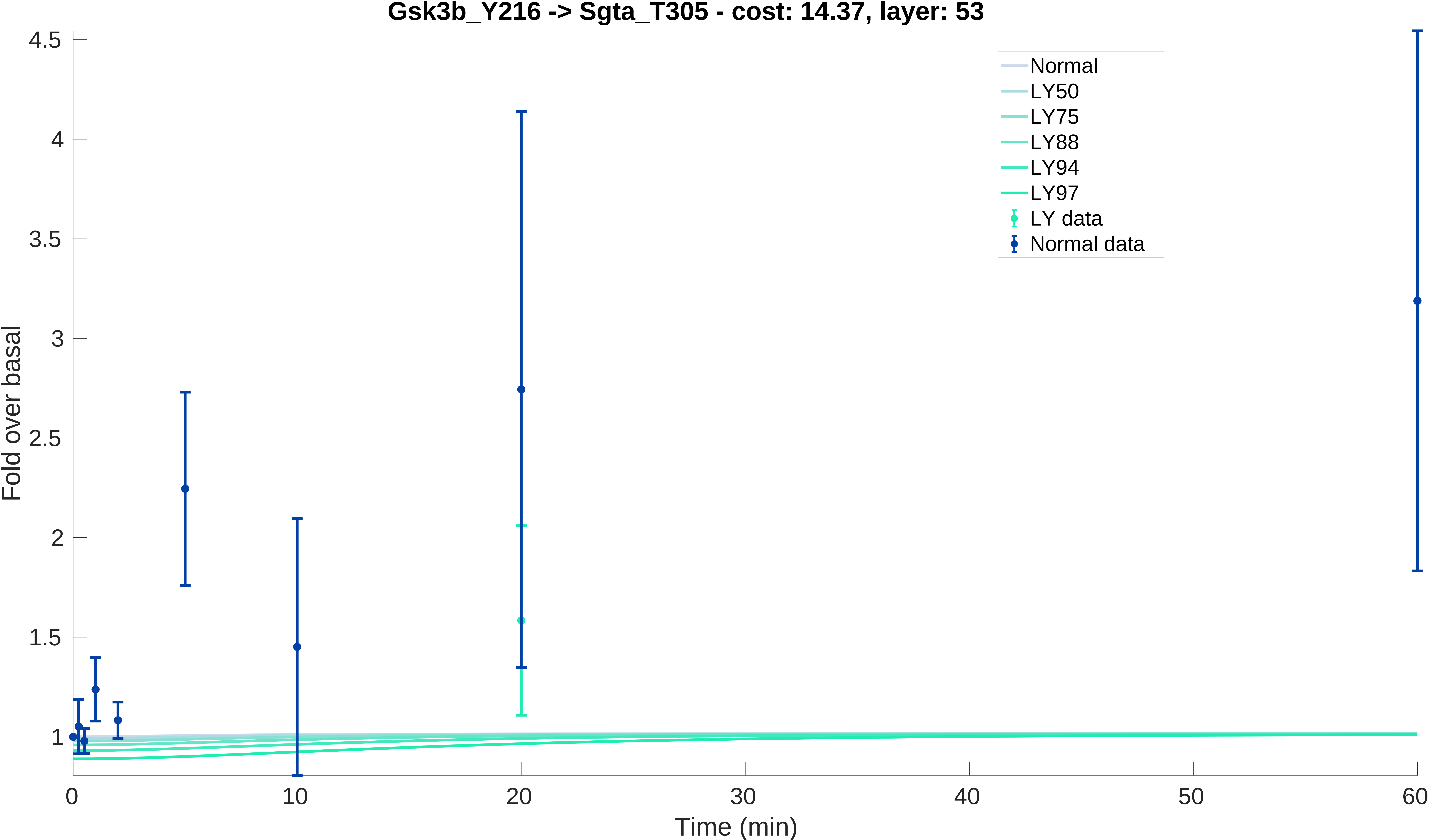

Gsk3b_Y216 -> Sgta_T305 - cost: 14.37, layer: 53
Fold over basal
Time (min)
4.5
4
3.5
3
2.5
2
1.5
1
0
10
20
30
40
50
60
Normal
LY50
LY75
LY88
LY94
LY97
LY data
Normal data

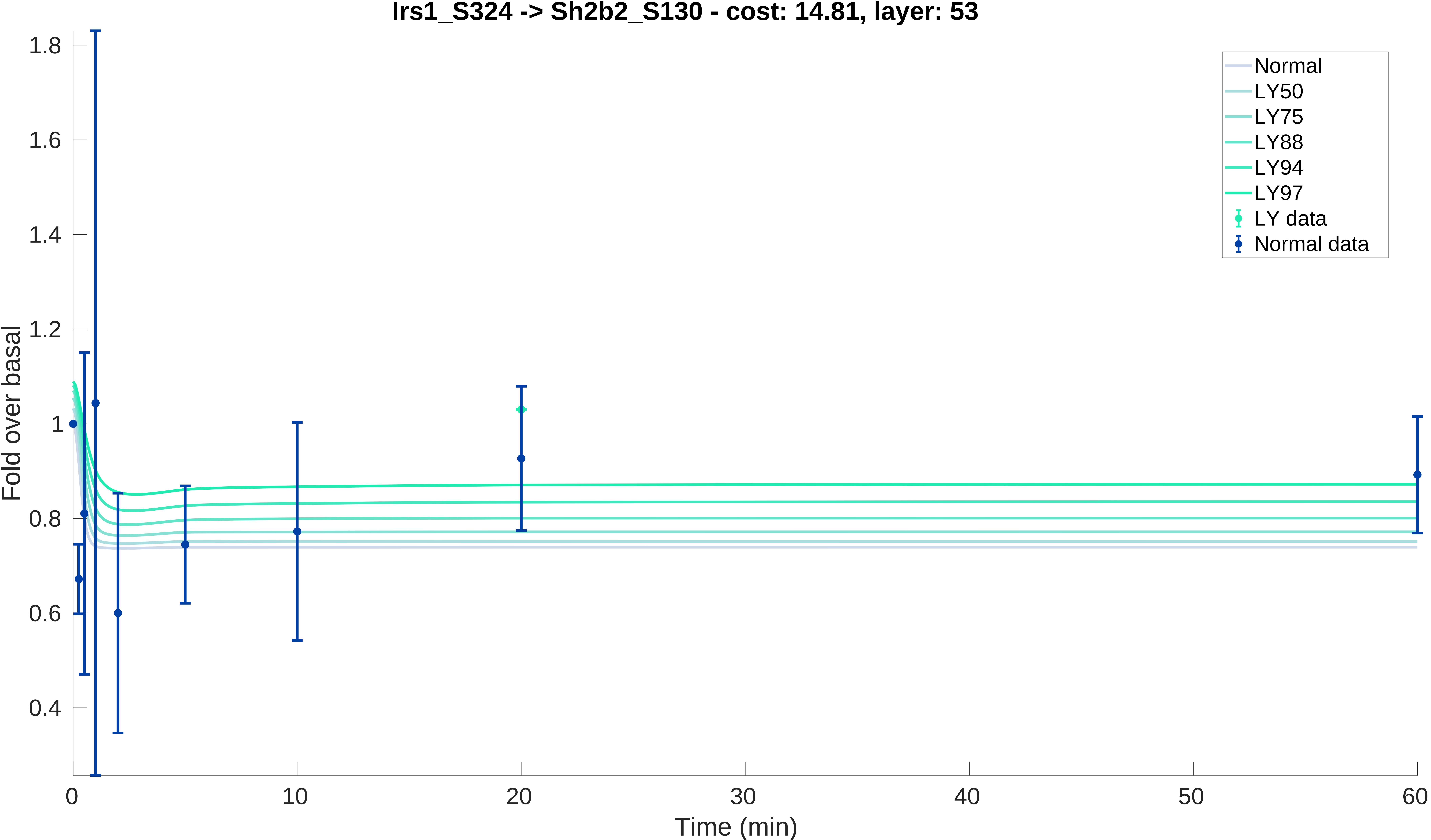
Irs1_S324 -> Sh2b2_S130 - cost: 14.81, layer: 53
Fold over basal
Time (min)
Normal
LY50
LY75
LY88
LY94
LY97
LY data
Normal data
0.4
0.6
0.8
1
1.2
1.4
1.6
1.8
0
10
20
30
40
50
60

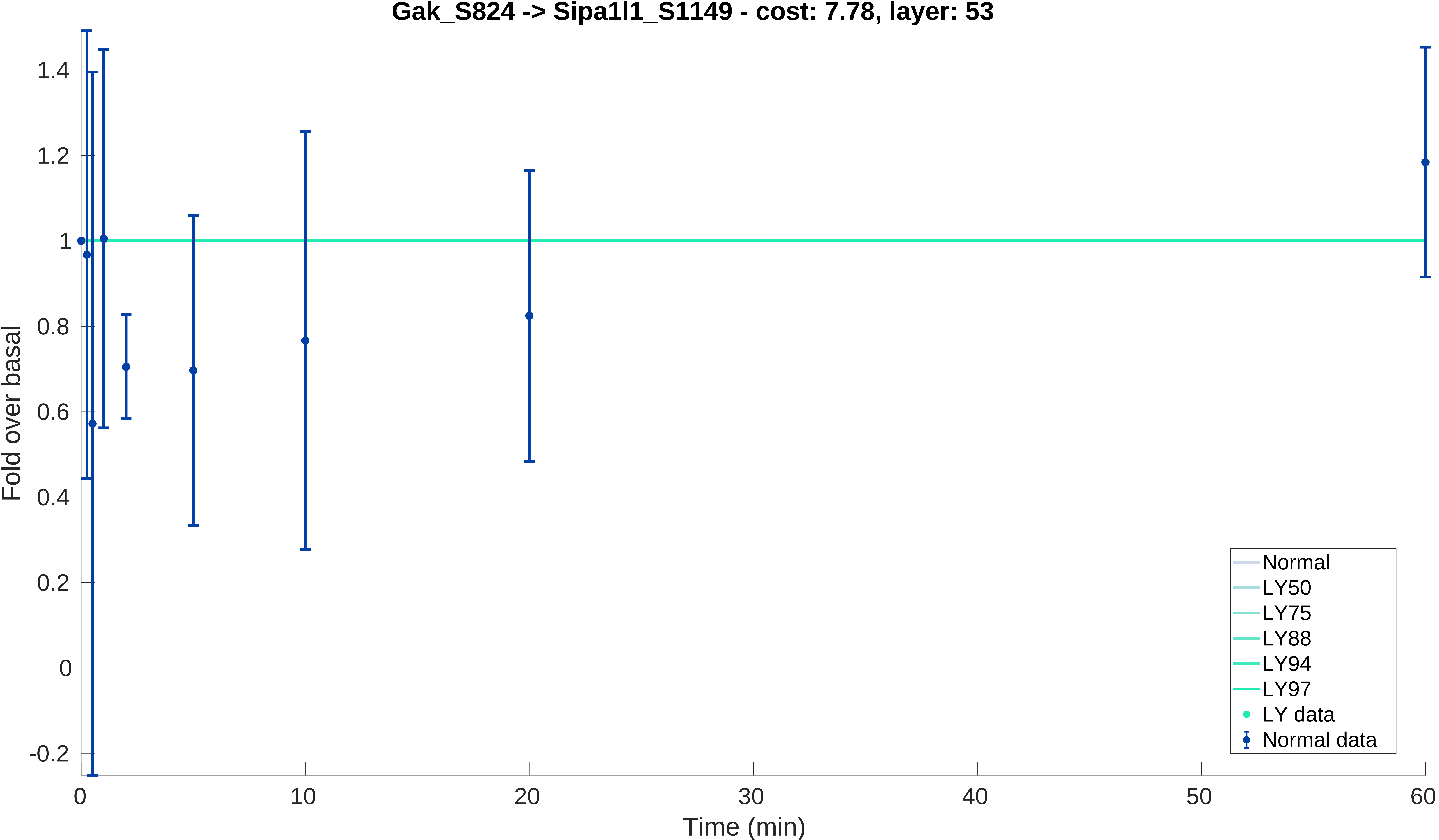

Gak_S824 -> Sipa1l1_S1149 - cost: 7.78, layer: 53
Fold over basal
Time (min)
1.4
1.2
1
0.8
0.6
0.4
0.2
0
-0.2
0
10
20
30
40
50
60
Normal
LY50
LY75
LY88
LY94
LY97
LY data
Normal data

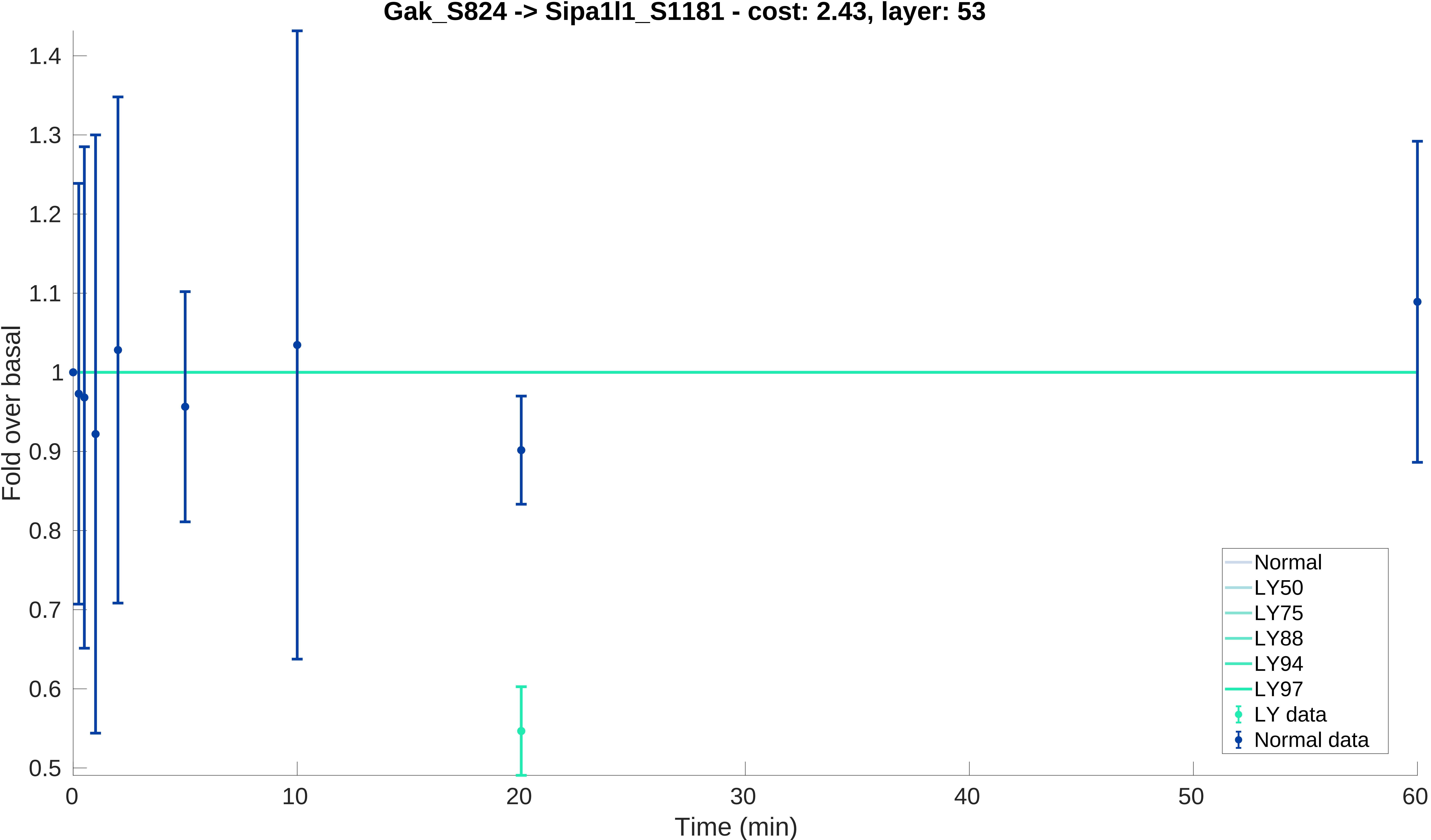
Gak_S824 -> Sipa1l1_S1181 - cost: 2.43, layer: 53
Fold over basal
Time (min)
1.4
1.3
1.2
1.1
1
0.9
0.8
0.7
0.6
0.5
0
10
20
30
40
50
60
Normal
LY50
LY75
LY88
LY94
LY97
LY data
Normal data

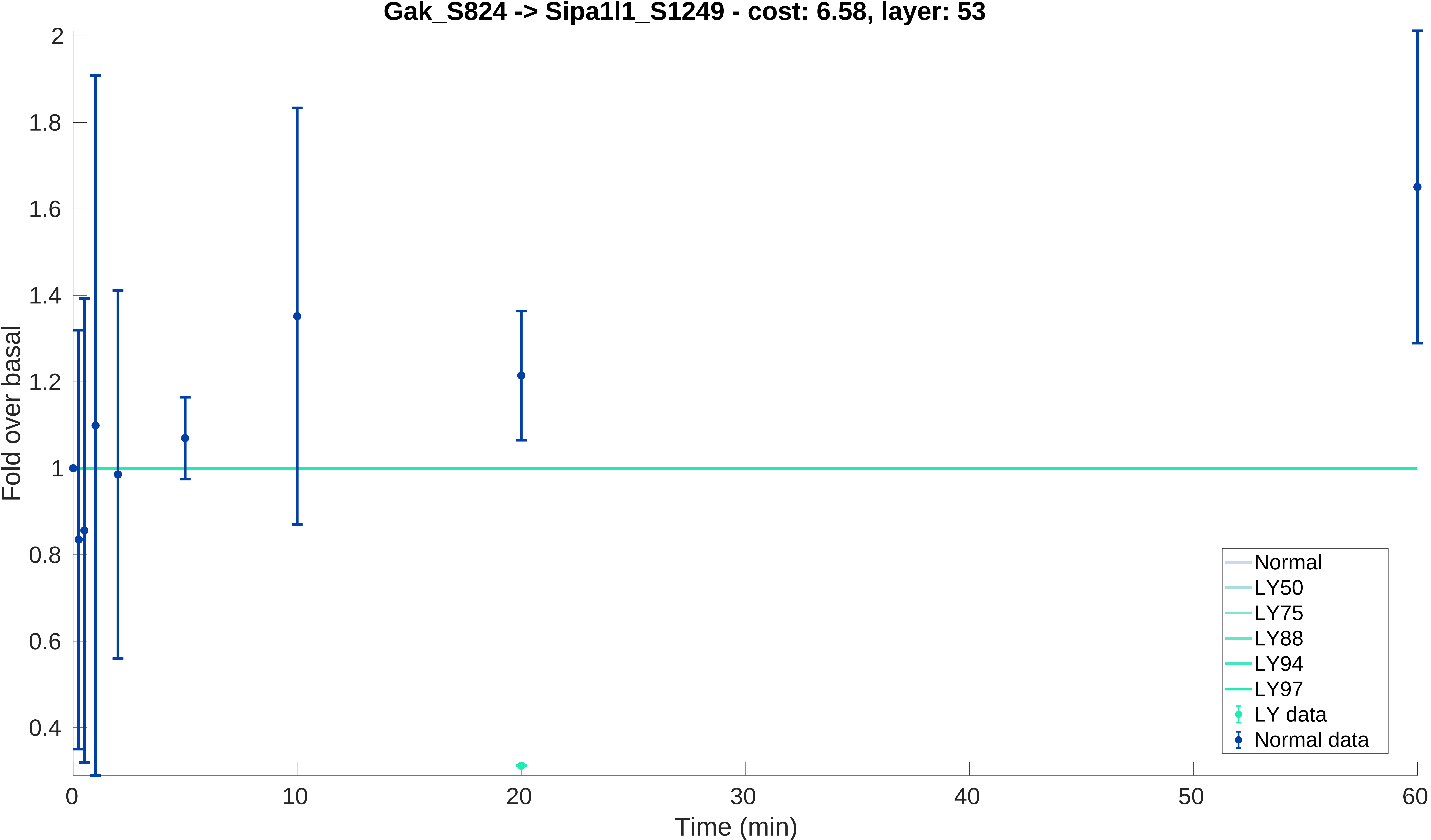
Gak_S824 -> Sipa1l1_S1249 - cost: 6.58, layer: 53
Fold over basal
Time (min)
0.4
0.6
0.8
1
1.2
1.4
1.6
1.8
2
0
10
20
30
40
50
60
Normal
LY50
LY75
LY88
LY94
LY97
LY data
Normal data

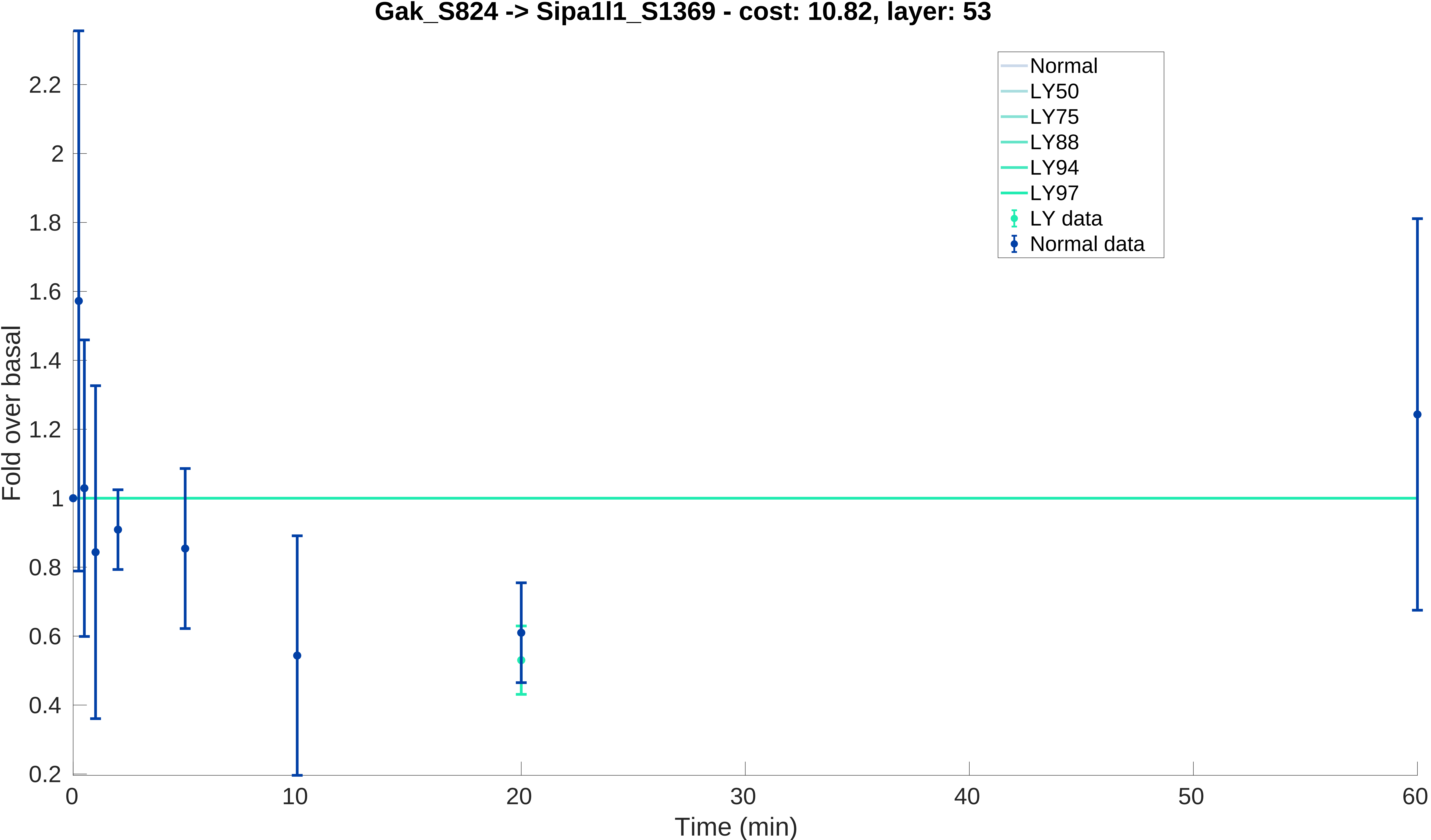
Gak_S824 -> Sipa1l1_S1369 - cost: 10.82, layer: 53
Normal
LY50
LY75
LY88
LY94
LY97
LY data
Normal data
Fold over basal
0.2
0.4
0.6
0.8
1
1.2
1.4
1.6
1.8
2
2.2
0
10
20
30
40
50
60
Time (min)

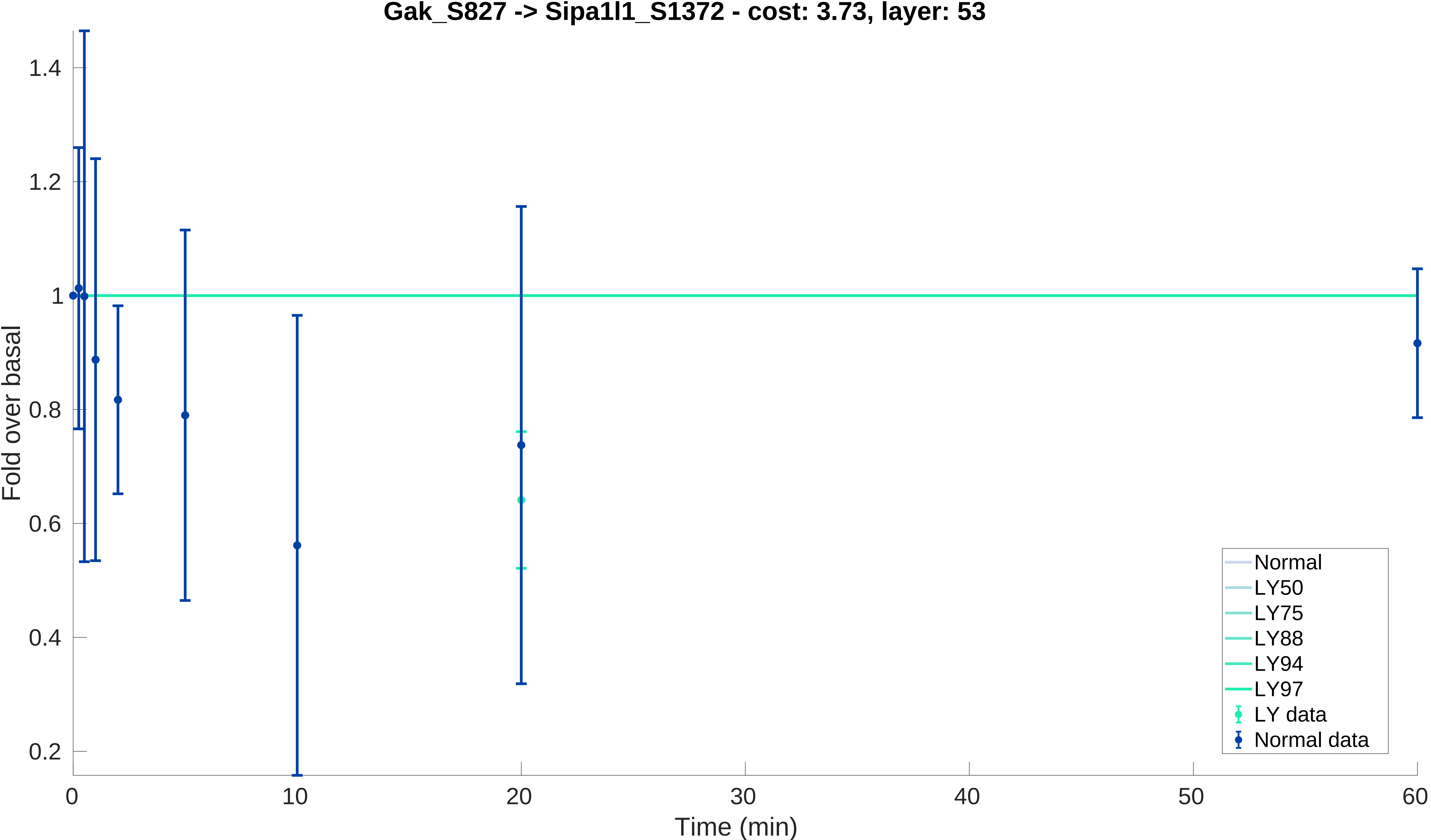

Gak_S827 -> Sipa1l1_S1372 - cost: 3.73, layer: 53
Fold over basal
Time (min)
0.2
0.4
0.6
0.8
1
1.2
1.4
0
10
20
30
40
50
60
Normal
LY50
LY75
LY88
LY94
LY97
LY data
Normal data

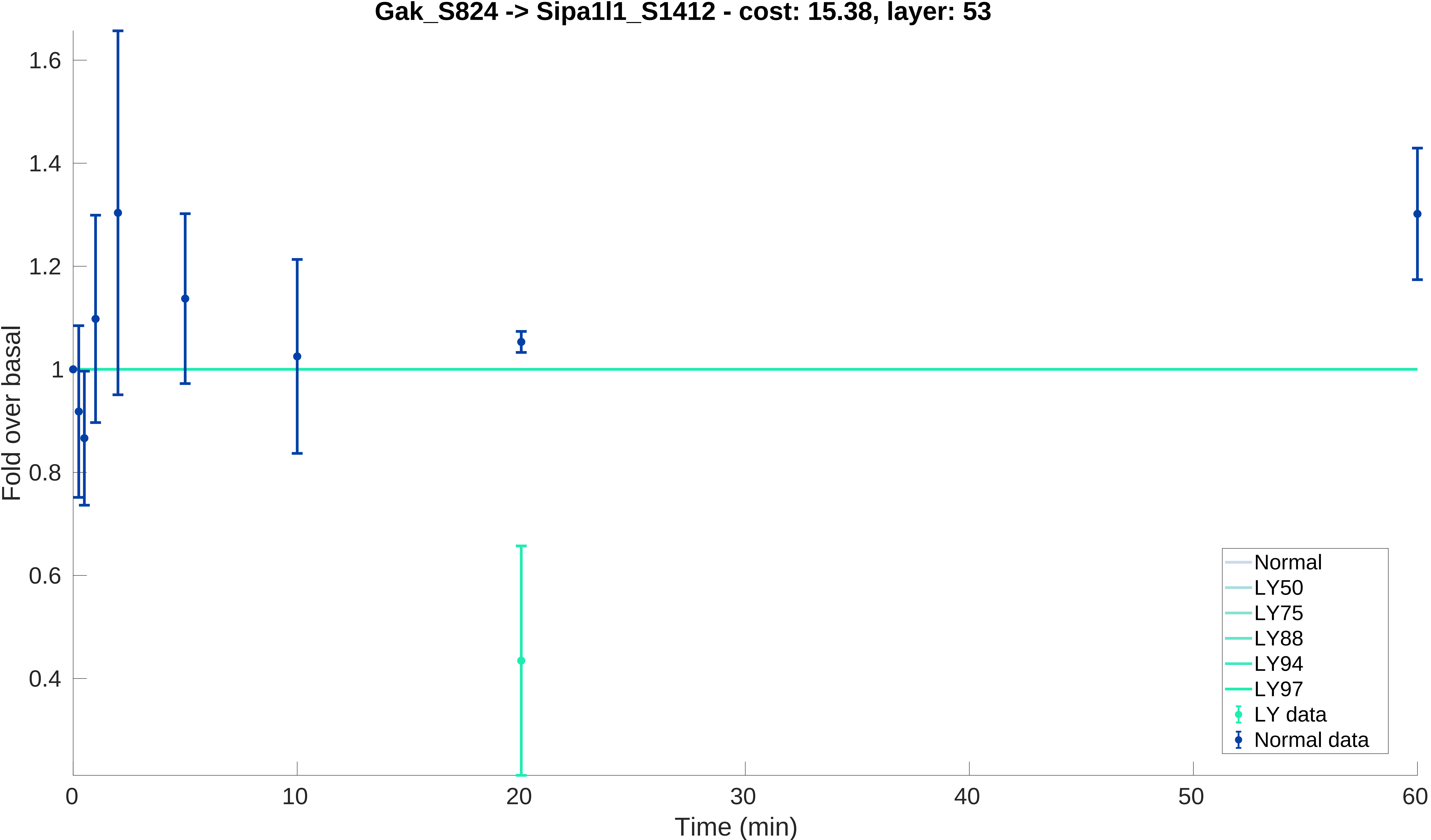
Gak_S824 -> Sipa1l1_S1412 - cost: 15.38, layer: 53
Fold over basal
Time (min)
1.6
1.4
1.2
1
0.8
0.6
0.4
0
10
20
30
40
50
60
Normal
LY50
LY75
LY88
LY94
LY97
LY data
Normal data

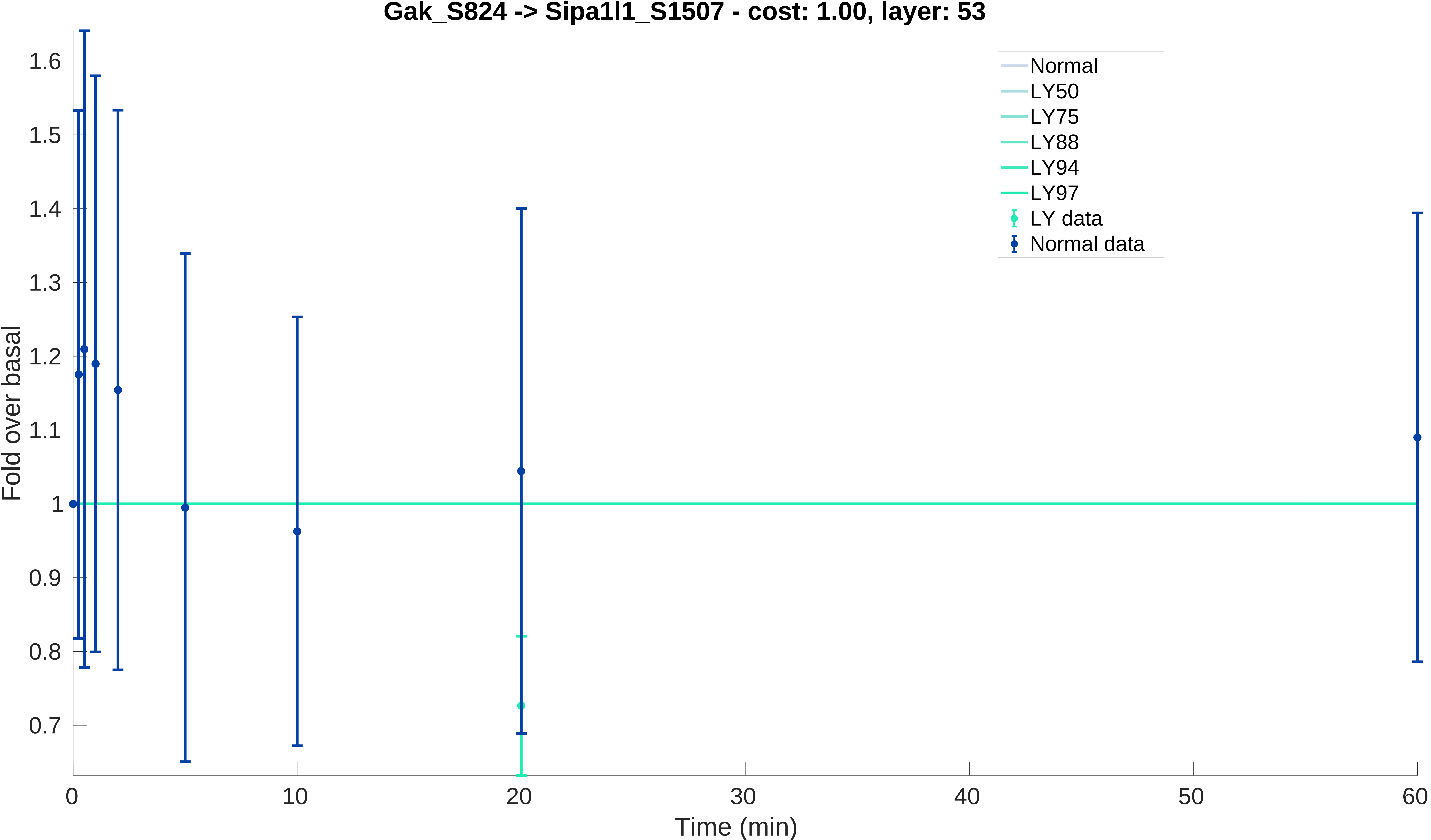
Gak_S824 -> Sipa1l1_S1507 - cost: 1.00, layer: 53
Fold over basal
Time (min)
Normal
LY50
LY75
LY88
LY94
LY97
LY data
Normal data
1.6
1.5
1.4
1.3
1.2
1.1
1
0.9
0.8
0.7
0
10
20
30
40
50
60

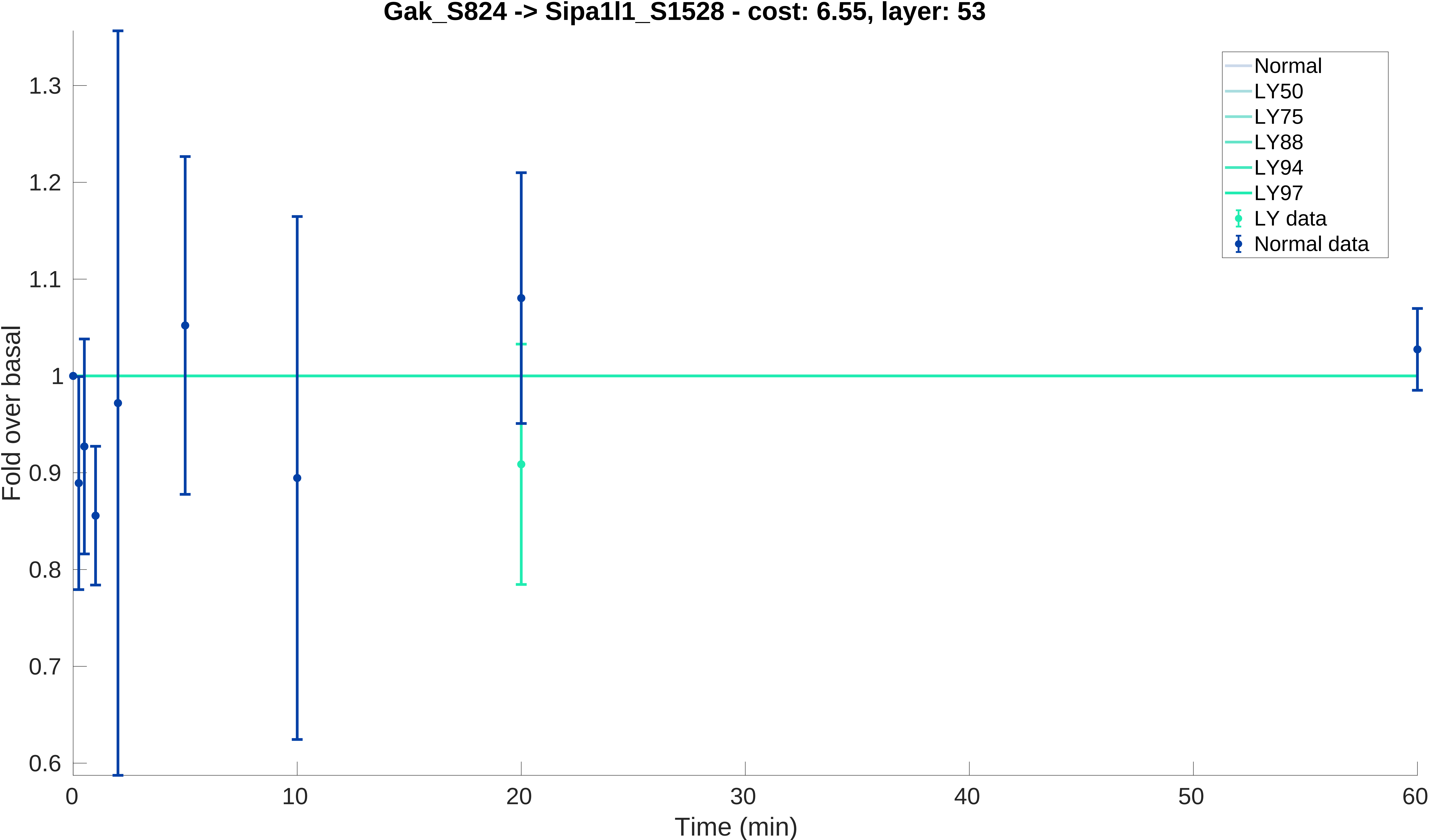

Gak_S824 -> Sipa1l1_S1528 - cost: 6.55, layer: 53
Normal
LY50
LY75
LY88
LY94
LY97
LY data
Normal data
Fold over basal
1.3
1.2
1.1
1
0.9
0.8
0.7
0.6
0
10
20
30
40
50
60
Time (min)

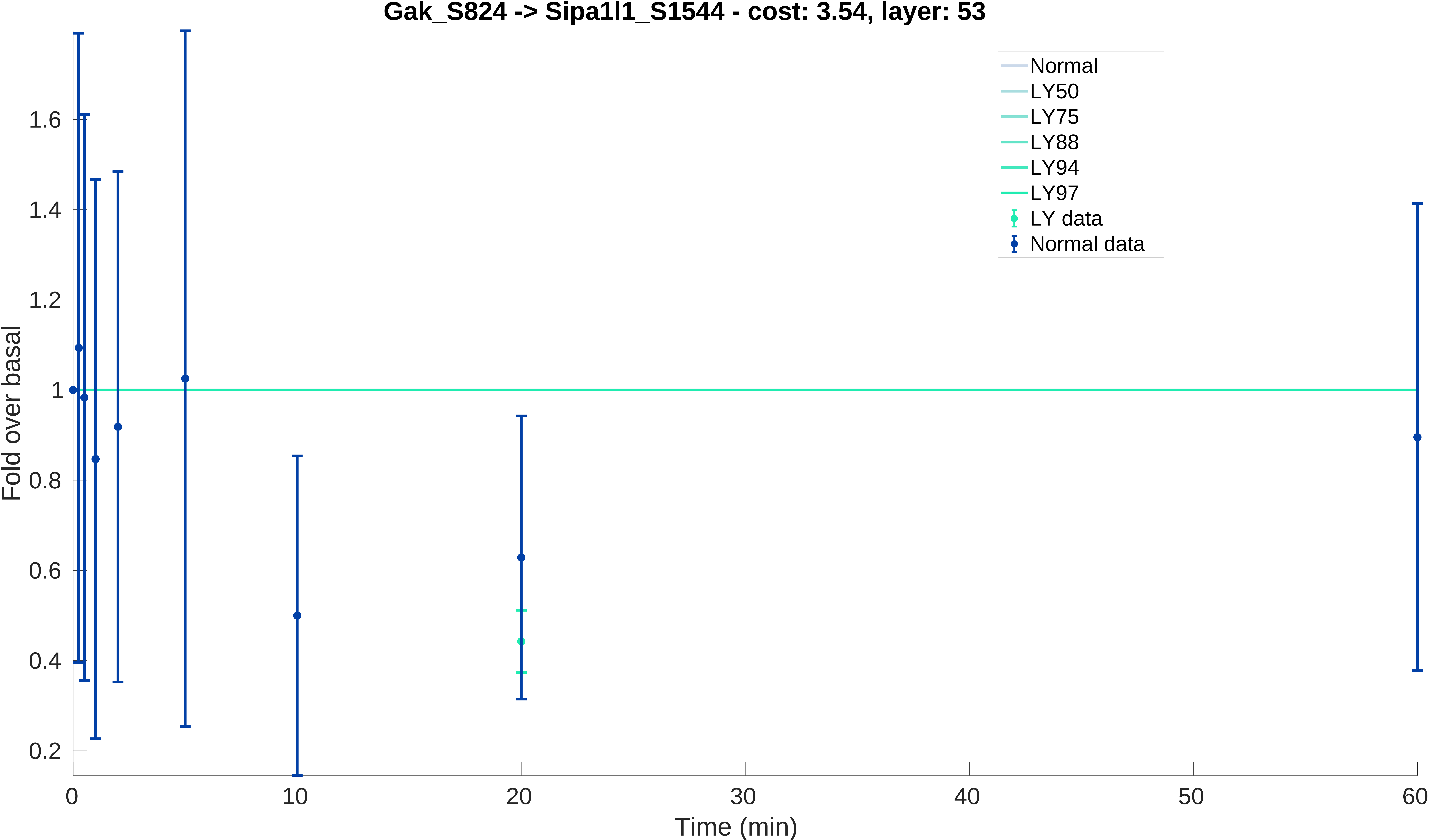

Gak_S824 -> Sipa1l1_S1544 - cost: 3.54, layer: 53
Normal
LY50
LY75
LY88
LY94
LY97
LY data
Normal data
Fold over basal
1.6
1.4
1.2
1
0.8
0.6
0.4
0.2
0
10
20
30
40
50
60
Time (min)

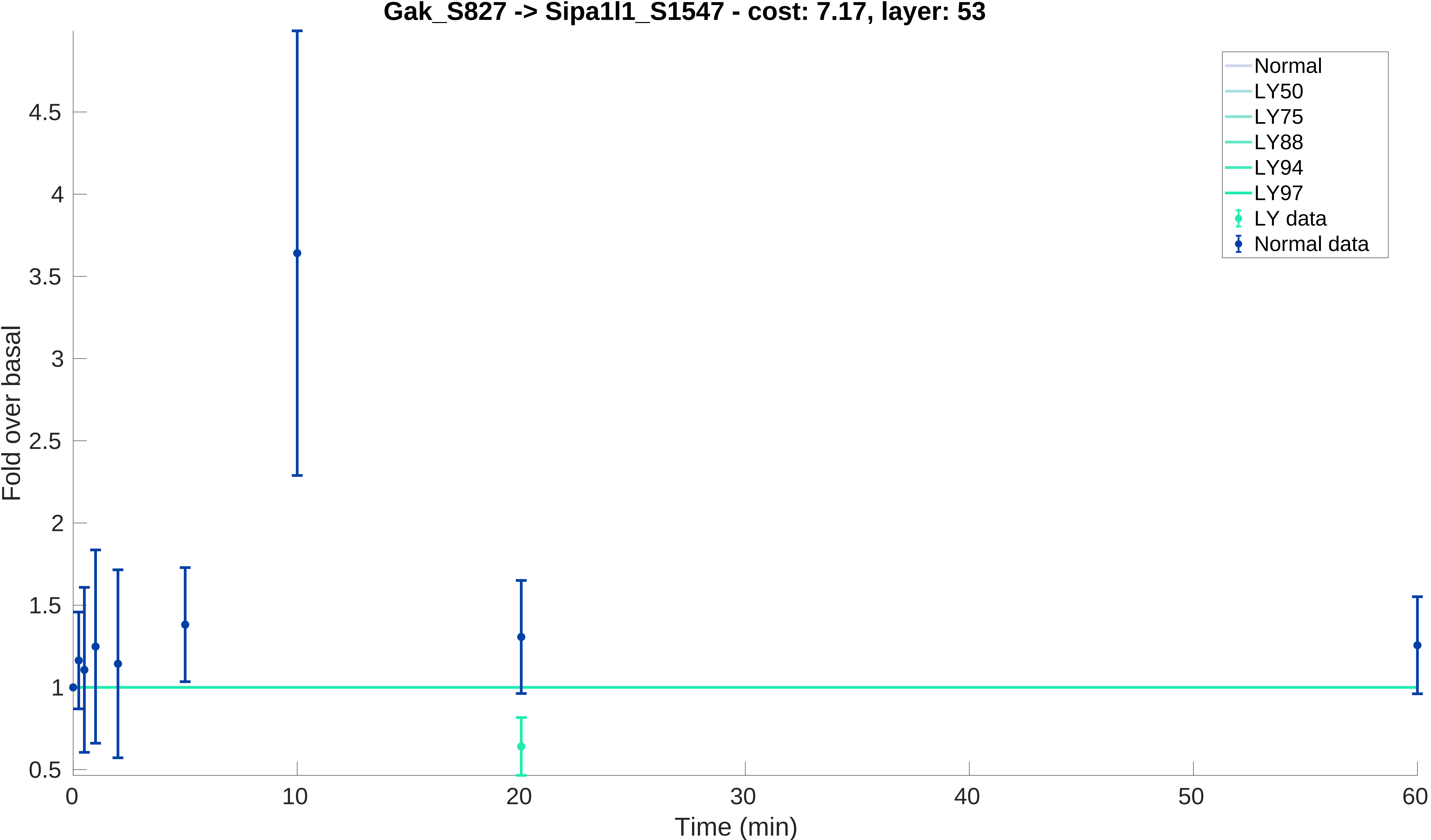

Gak_S827 -> Sipa1l1_S1547 - cost: 7.17, layer: 53
Fold over basal
Time (min)
0.5
1
1.5
2
2.5
3
3.5
4
4.5
0
10
20
30
40
50
60
Normal
LY50
LY75
LY88
LY94
LY97
LY data
Normal data

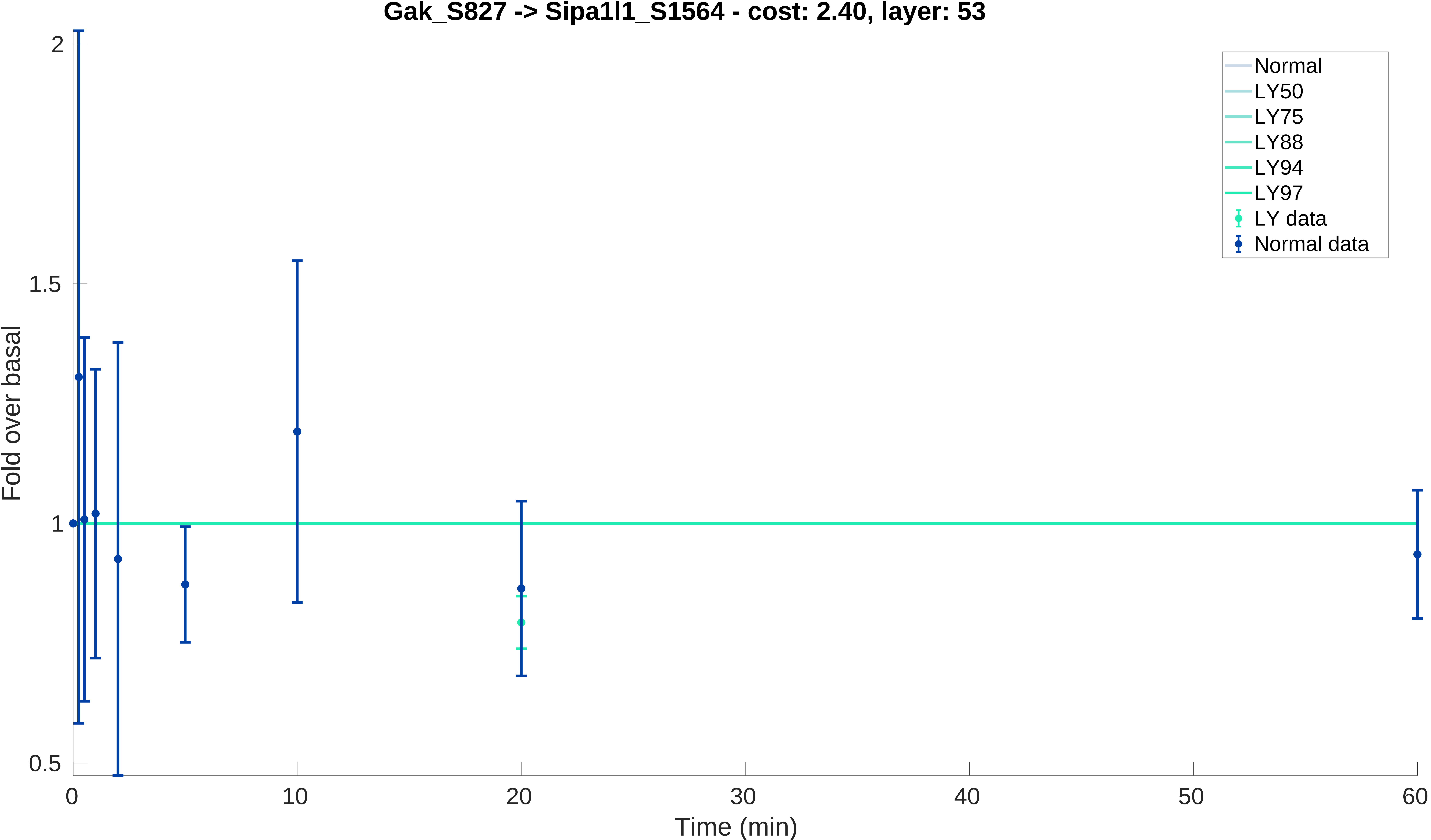
Gak_S827 -> Sipa1l1_S1564 - cost: 2.40, layer: 53
Fold over basal
Time (min)
2
1.5
1
0.5
0
10
20
30
40
50
60
Normal
LY50
LY75
LY88
LY94
LY97
LY data
Normal data

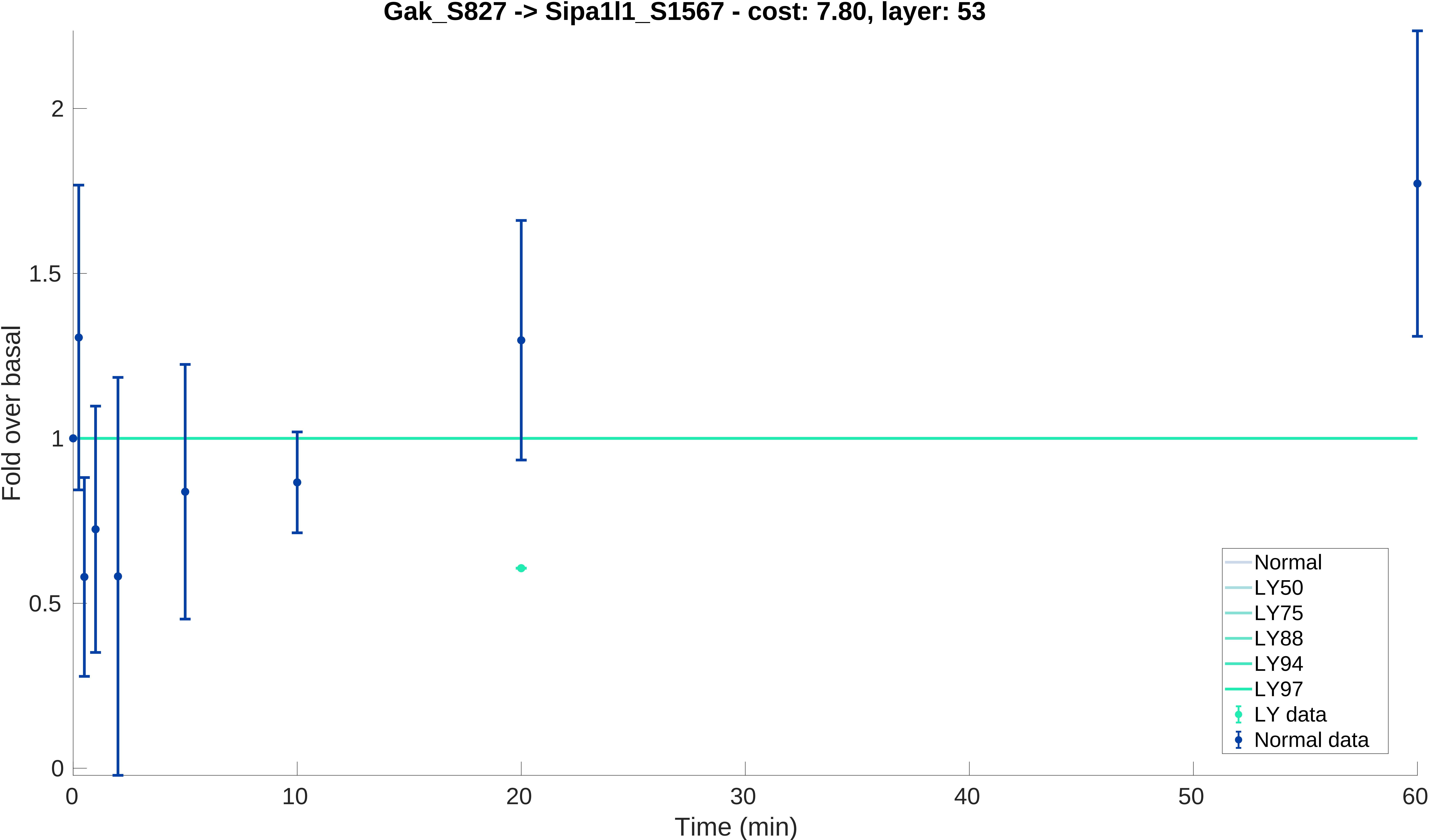

Gak_S827 -> Sipa1l1_S1567 - cost: 7.80, layer: 53
Fold over basal
Time (min)
0
0.5
1
1.5
2
0
10
20
30
40
50
60
Normal
LY50
LY75
LY88
LY94
LY97
LY data
Normal data

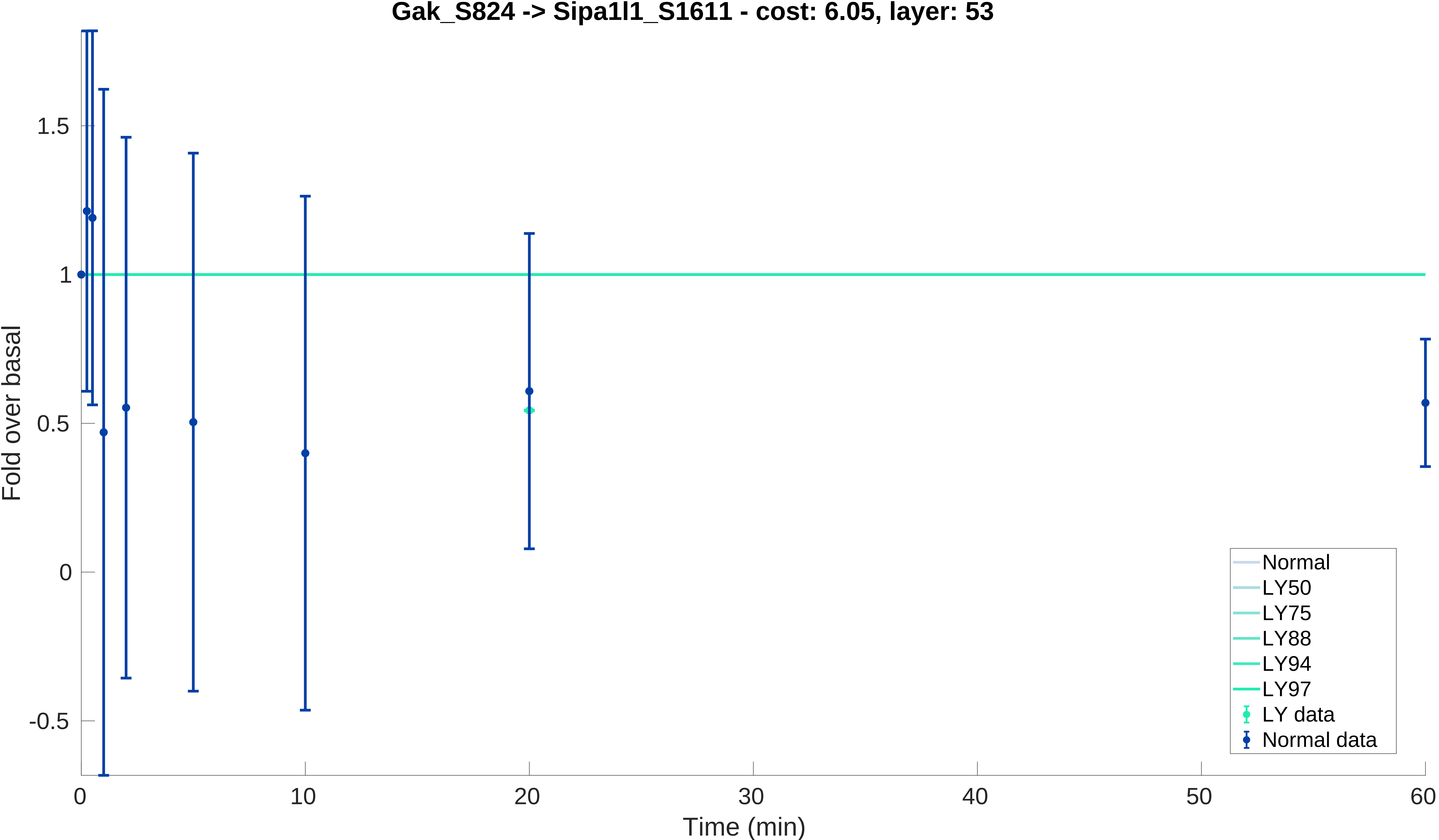

Gak_S824 -> Sipa1l1_S1611 - cost: 6.05, layer: 53
Fold over basal
Time (min)
1.5
1
0.5
0
-0.5
0
10
20
30
40
50
60
Normal
LY50
LY75
LY88
LY94
LY97
LY data
Normal data

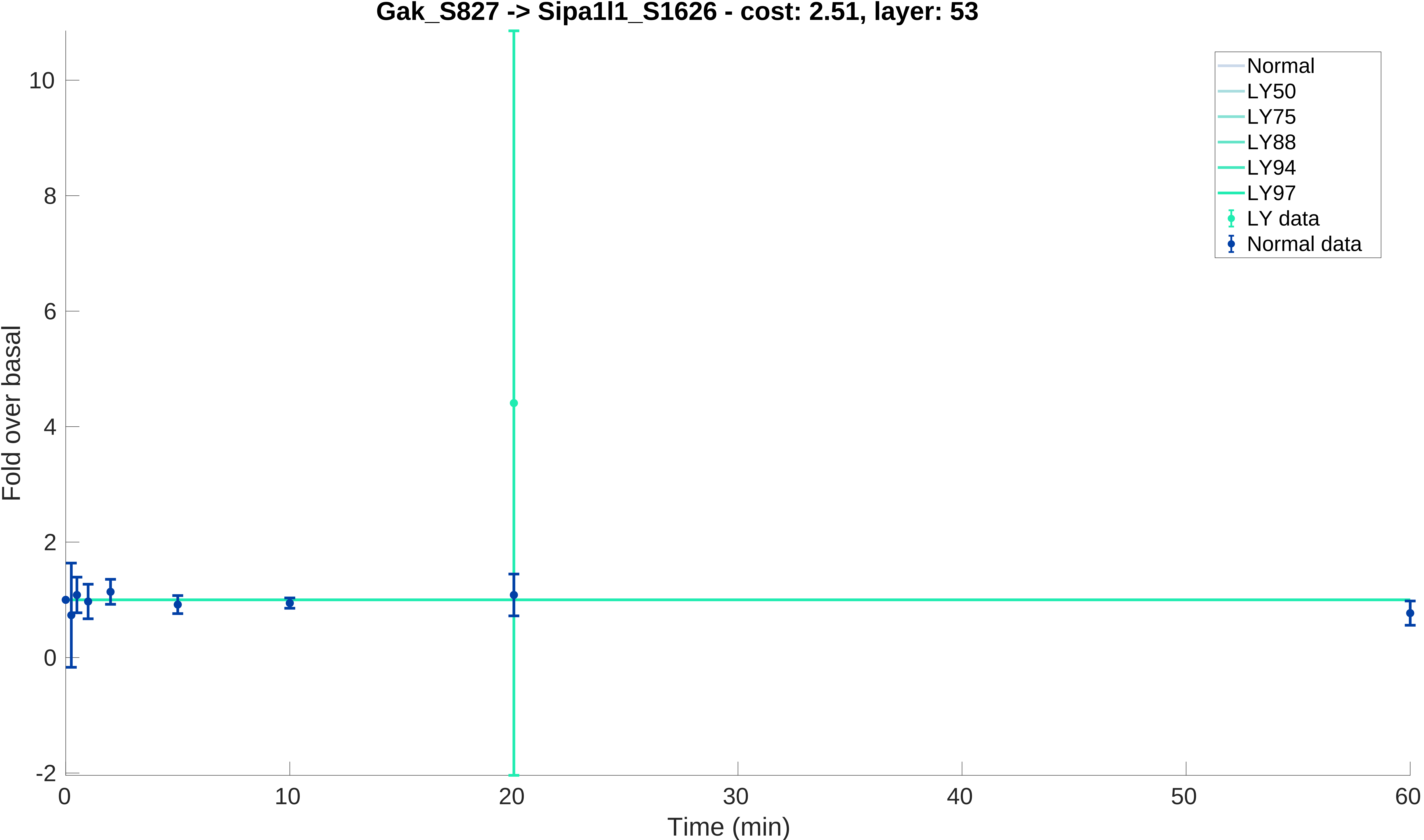
Gak_S827 -> Sipa1l1_S1626 - cost: 2.51, layer: 53
Normal
LY50
LY75
LY88
LY94
LY97
LY data
Normal data
Fold over basal
Time (min)
-2
0
2
4
6
8
10
0
10
20
30
40
50
60

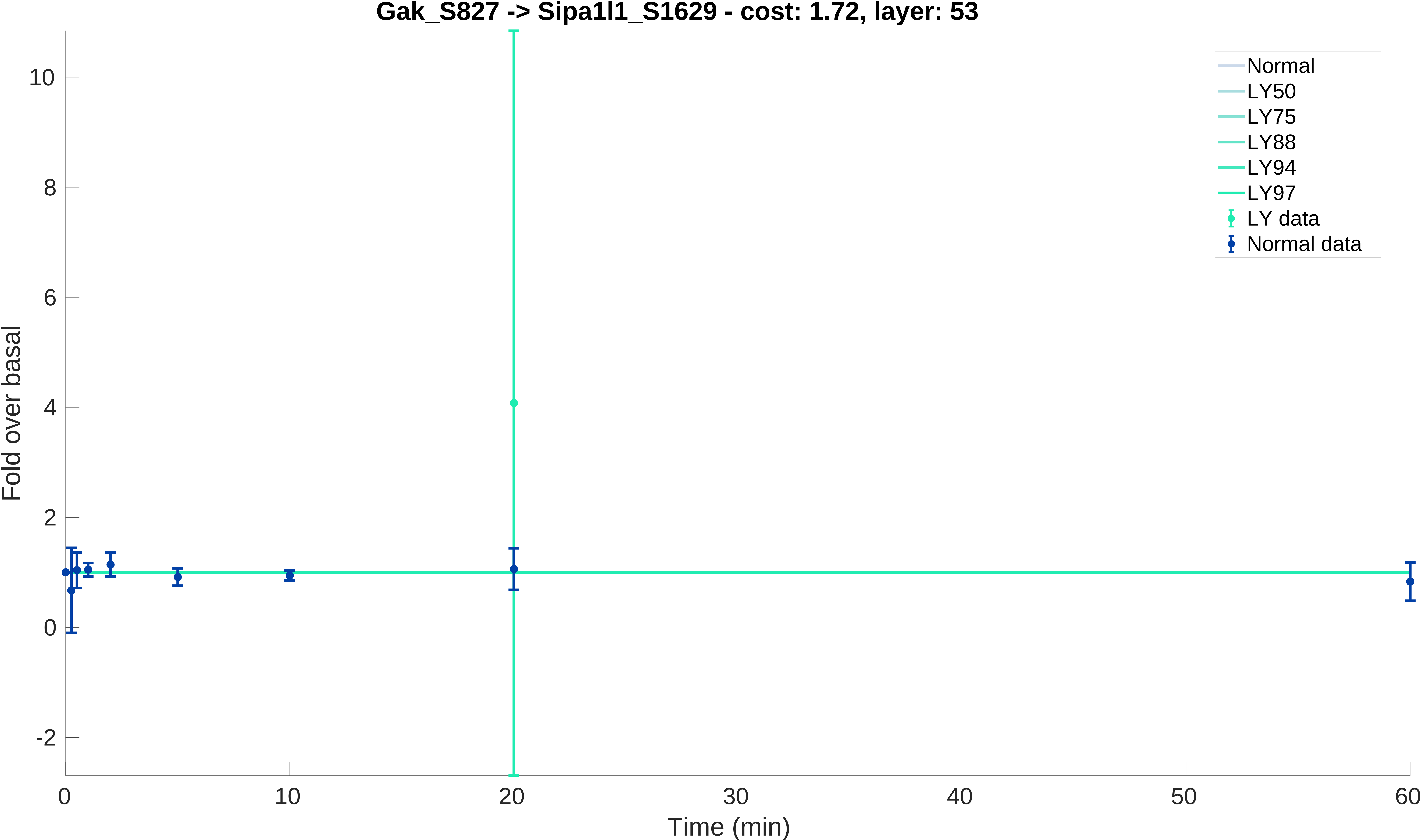

Gak_S827 -> Sipa1l1_S1629 - cost: 1.72, layer: 53
Fold over basal
Time (min)
-2
0
2
4
6
8
10
0
10
20
30
40
50
60
Normal
LY50
LY75
LY88
LY94
LY97
LY data
Normal data

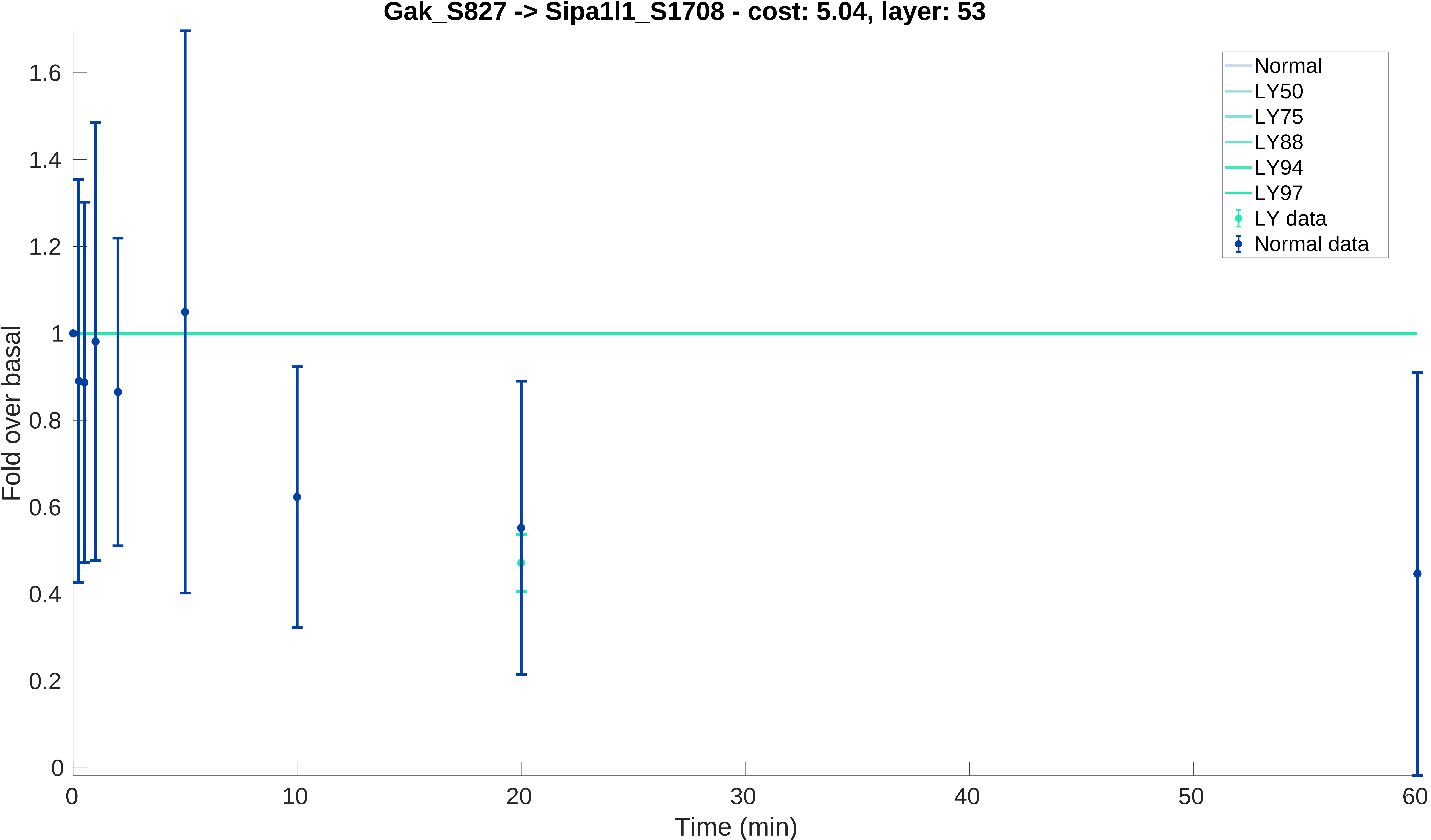

Gak_S827 -> Sipa1l1_S1708 - cost: 5.04, layer: 53
Fold over basal
Time (min)
0
0.2
0.4
0.6
0.8
1
1.2
1.4
1.6
0
10
20
30
40
50
60
Normal
LY50
LY75
LY88
LY94
LY97
LY data
Normal data

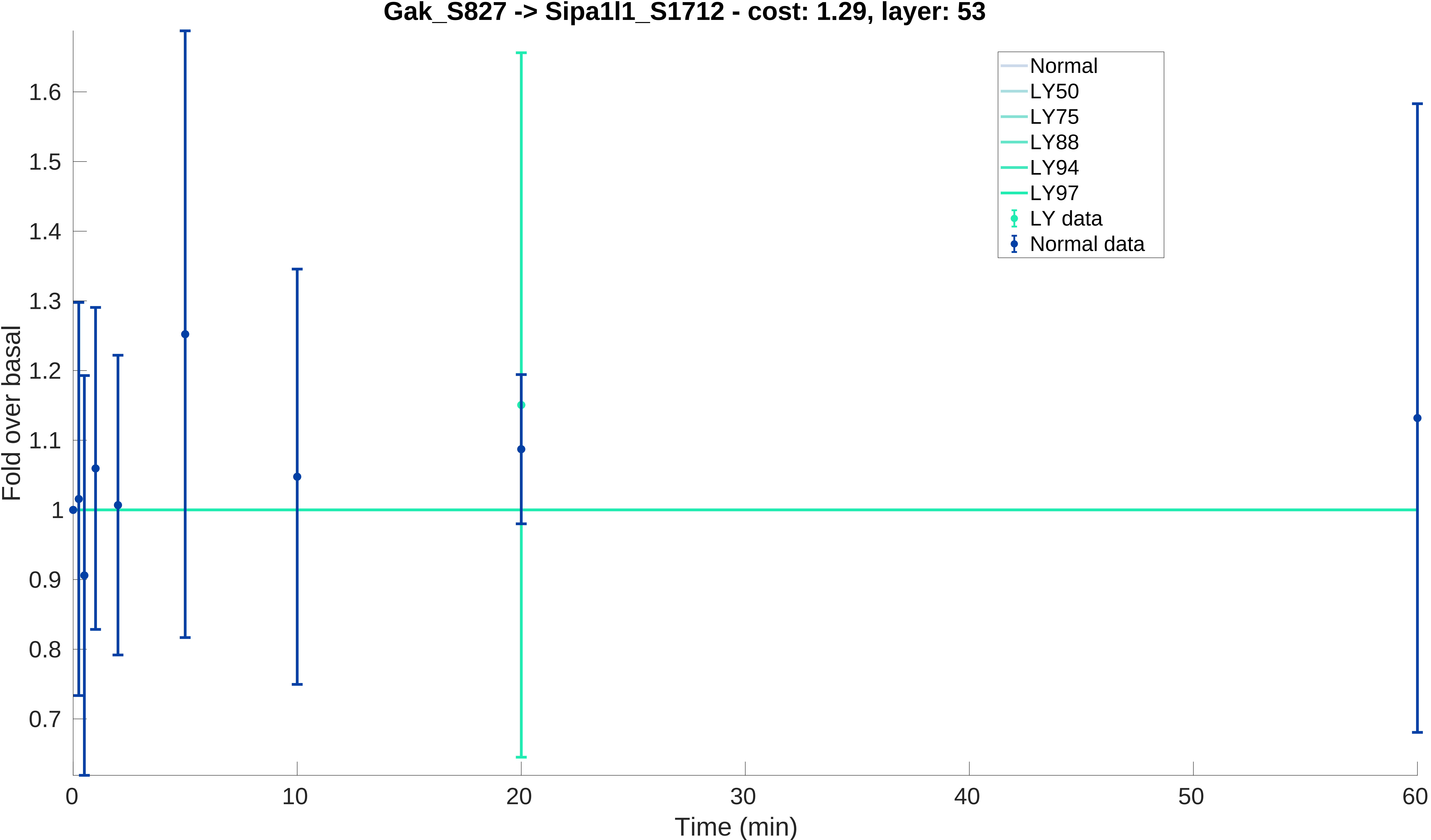

Gak_S827 -> Sipa1l1_S1712 - cost: 1.29, layer: 53
Normal
LY50
LY75
LY88
LY94
LY97
LY data
Normal data
Fold over basal
Time (min)
1.6
1.5
1.4
1.3
1.2
1.1
1
0.9
0.8
0.7
0
10
20
30
40
50
60

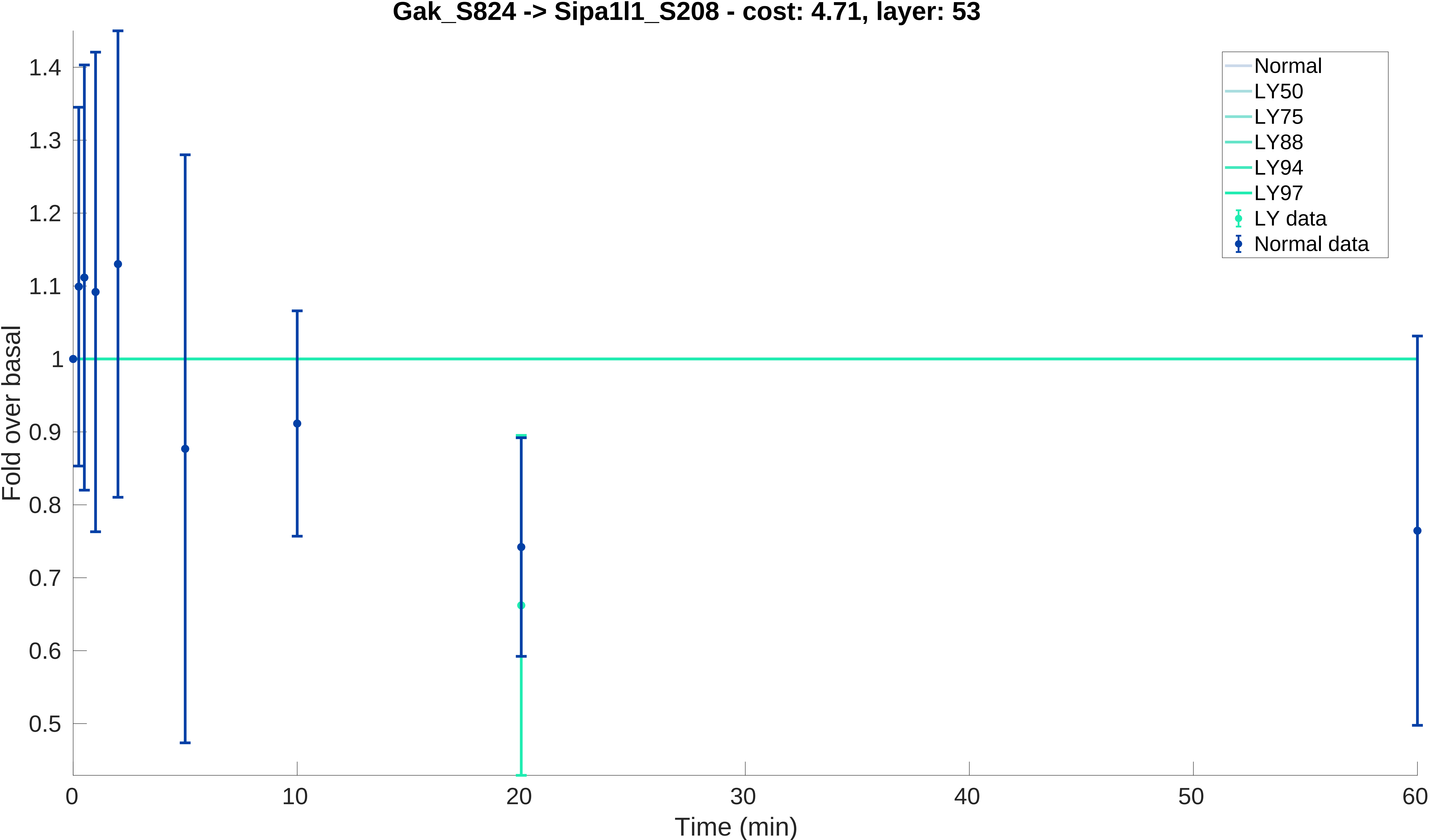
Gak_S824 -> Sipa1l1_S208 - cost: 4.71, layer: 53
Fold over basal
Time (min)
1.4
1.3
1.2
1.1
1
0.9
0.8
0.7
0.6
0.5
0
10
20
30
40
50
60
Normal
LY50
LY75
LY88
LY94
LY97
LY data
Normal data

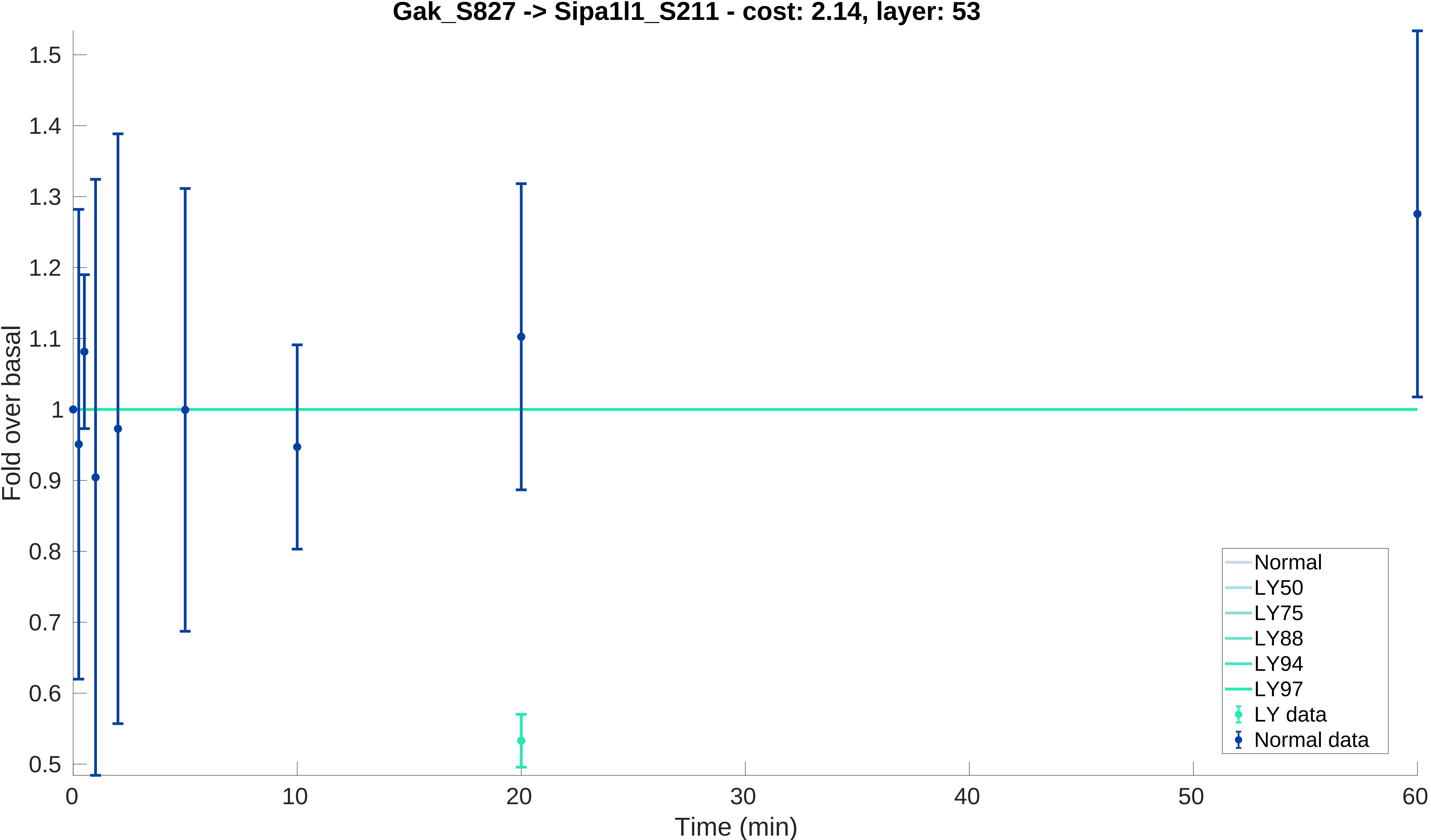

Gak_S827 -> Sipa1l1_S211 - cost: 2.14, layer: 53
Fold over basal
Time (min)
1.5
1.4
1.3
1.2
1.1
1
0.9
0.8
0.7
0.6
0.5
0
10
20
30
40
50
60
Normal
LY50
LY75
LY88
LY94
LY97
LY data
Normal data

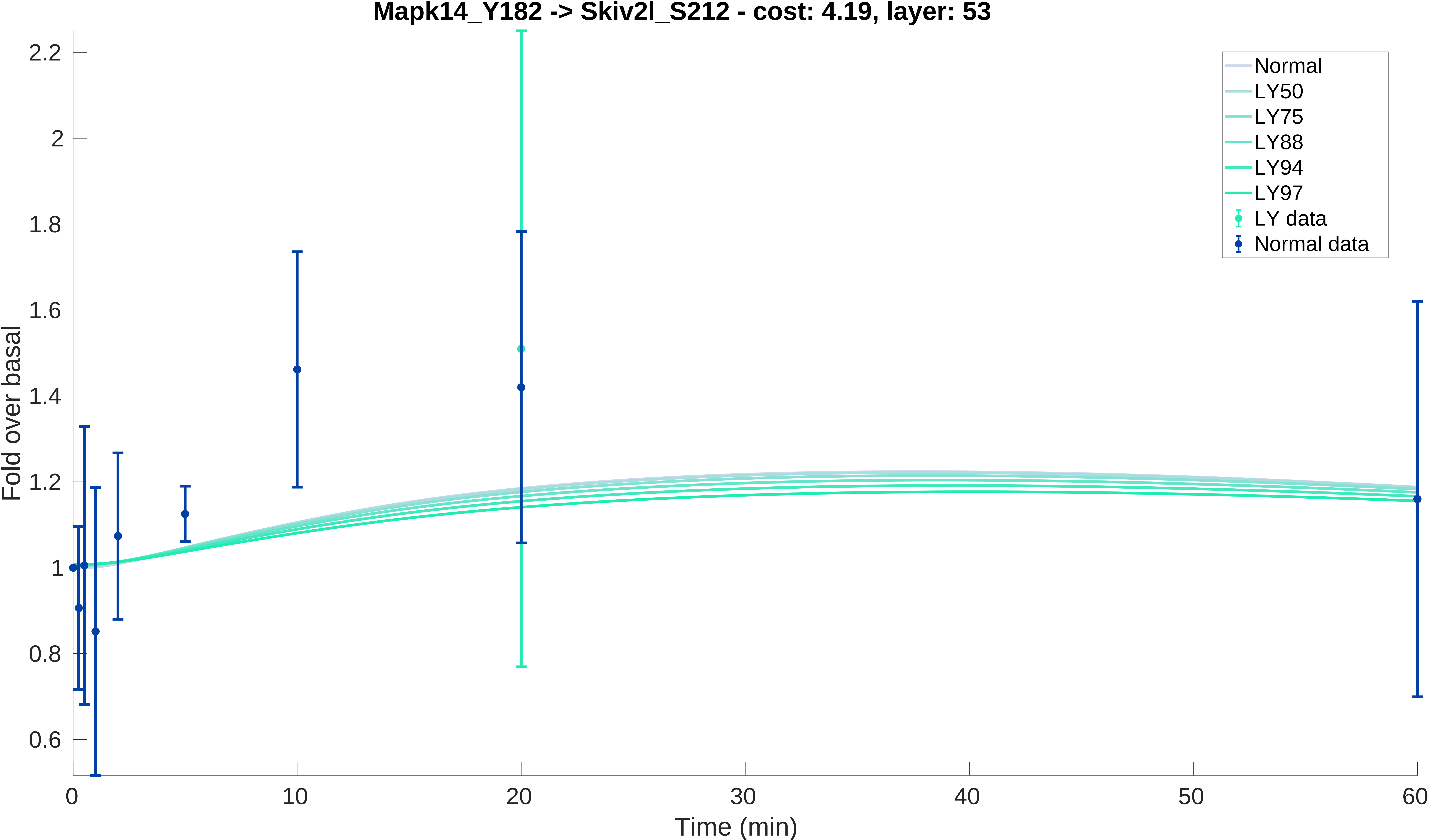

Mapk14_Y182 -> Skiv2l_S212 - cost: 4.19, layer: 53
Fold over basal
Time (min)
Normal
LY50
LY75
LY88
LY94
LY97
LY data
Normal data
2.2
2
1.8
1.6
1.4
1.2
1
0.8
0.6
0
10
20
30
40
50
60

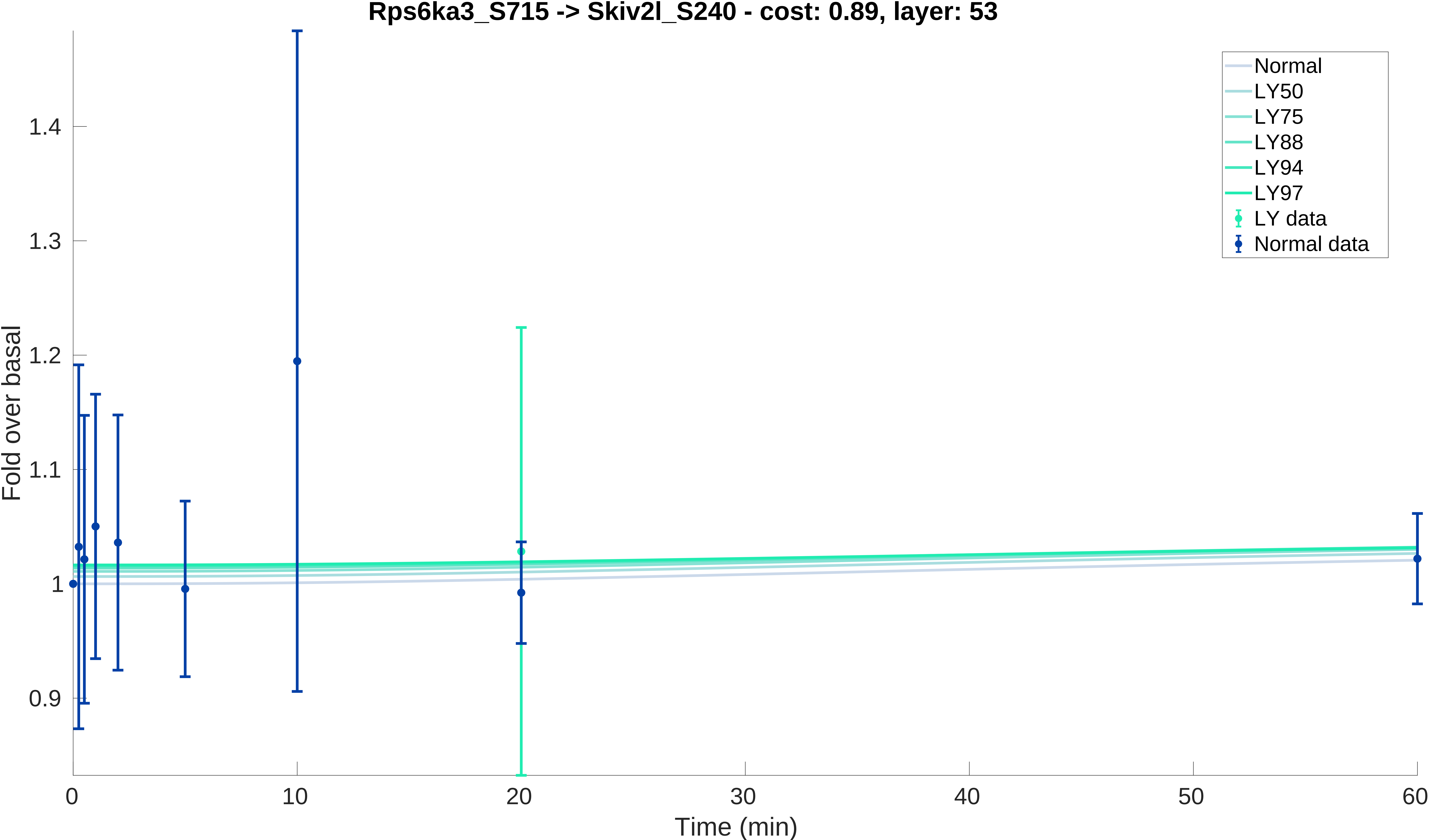

Rps6ka3_S715 -> Skiv2l_S240 - cost: 0.89, layer: 53
Normal
LY50
LY75
LY88
LY94
LY97
LY data
Normal data
Fold over basal
1.4
1.3
1.2
1.1
1
0.9
Time (min)
0
10
20
30
40
50
60

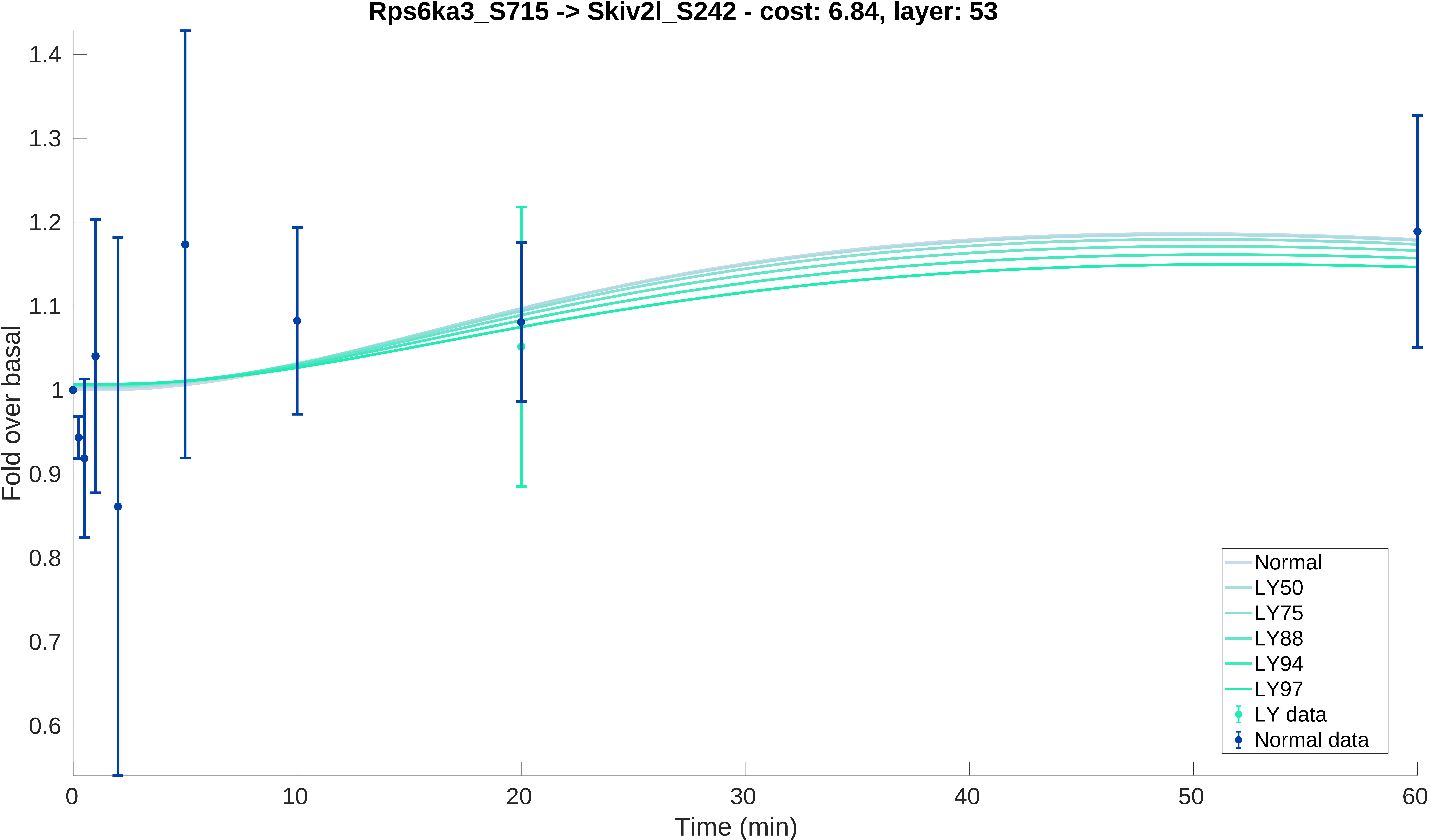

Rps6ka3_S715 -> Skiv2l_S242 - cost: 6.84, layer: 53
Fold over basal
Time (min)
0.6
0.7
0.8
0.9
1
1.1
1.2
1.3
1.4
0
10
20
30
40
50
60
Normal
LY50
LY75
LY88
LY94
LY97
LY data
Normal data

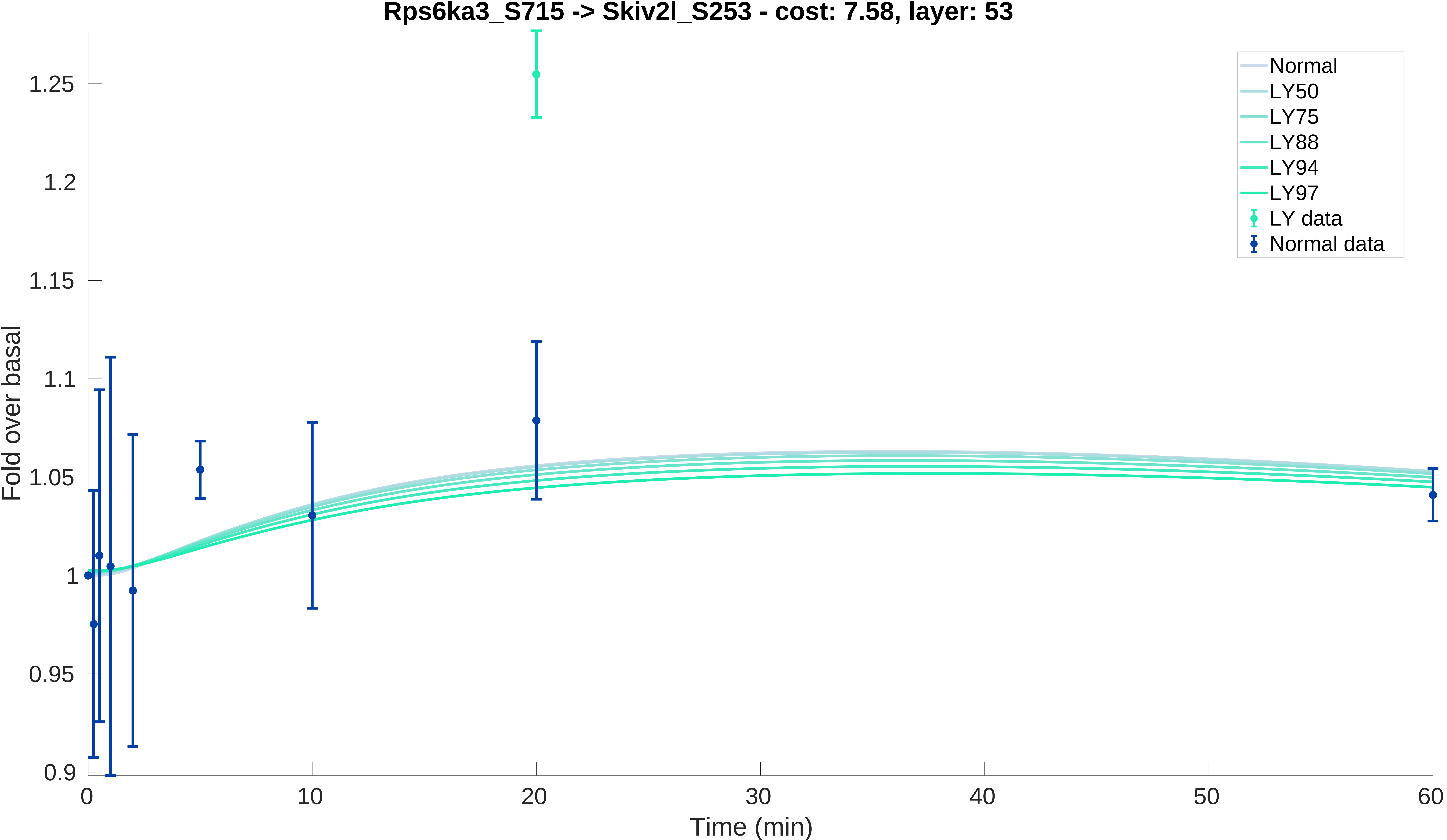
Rps6ka3_S715 -> Skiv2l_S253 - cost: 7.58, layer: 53
Fold over basal
Time (min)
0.9
0.95
1
1.05
1.1
1.15
1.2
1.25
0
10
20
30
40
50
60
Normal
LY50
LY75
LY88
LY94
LY97
LY data
Normal data

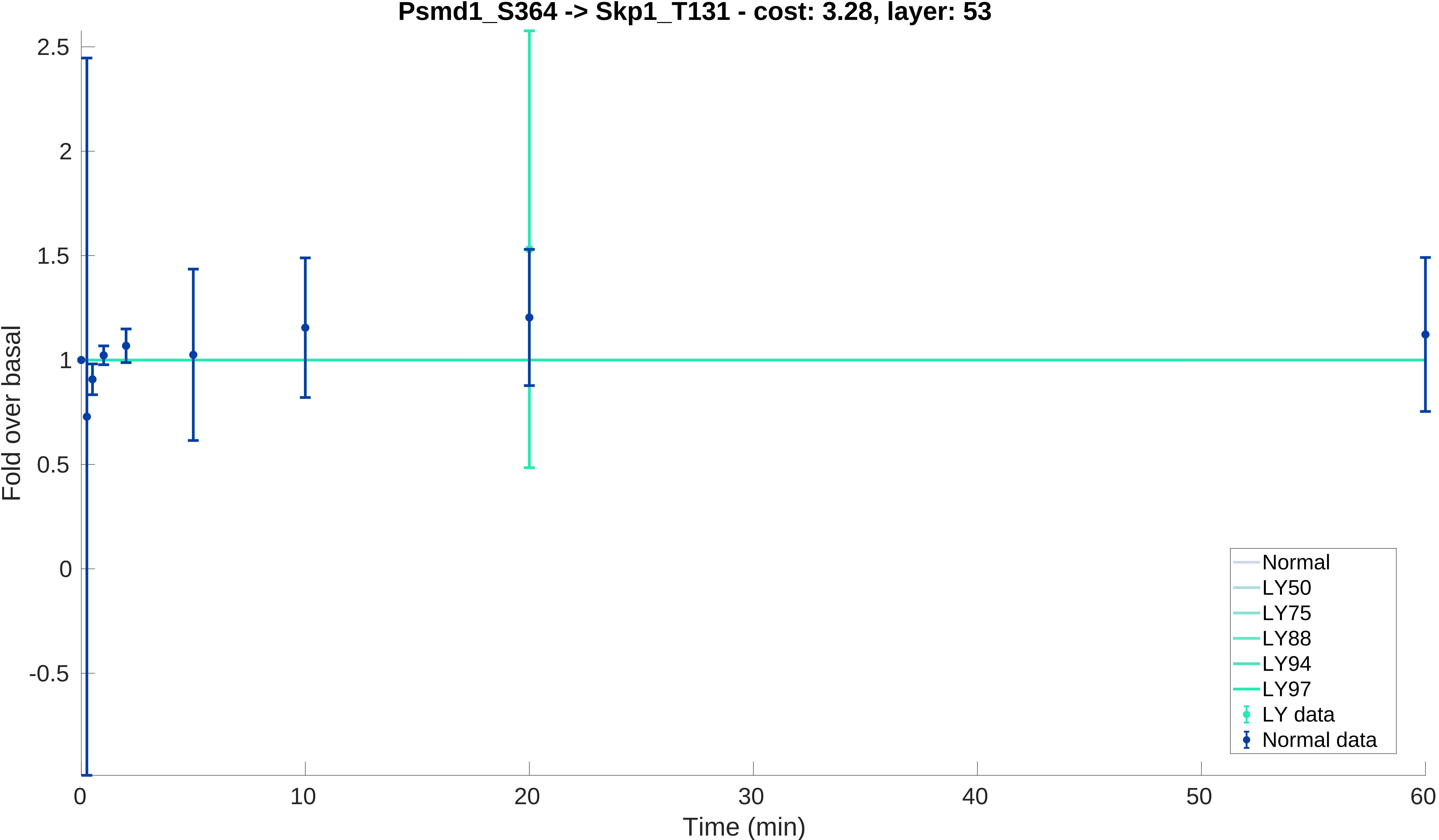

Psmd1_S364 -> Skp1_T131 - cost: 3.28, layer: 53
Fold over basal
2.5
2
1.5
1
0.5
0
-0.5
0
10
20
30
40
50
60
Time (min)
Normal
LY50
LY75
LY88
LY94
LY97
LY data
Normal data

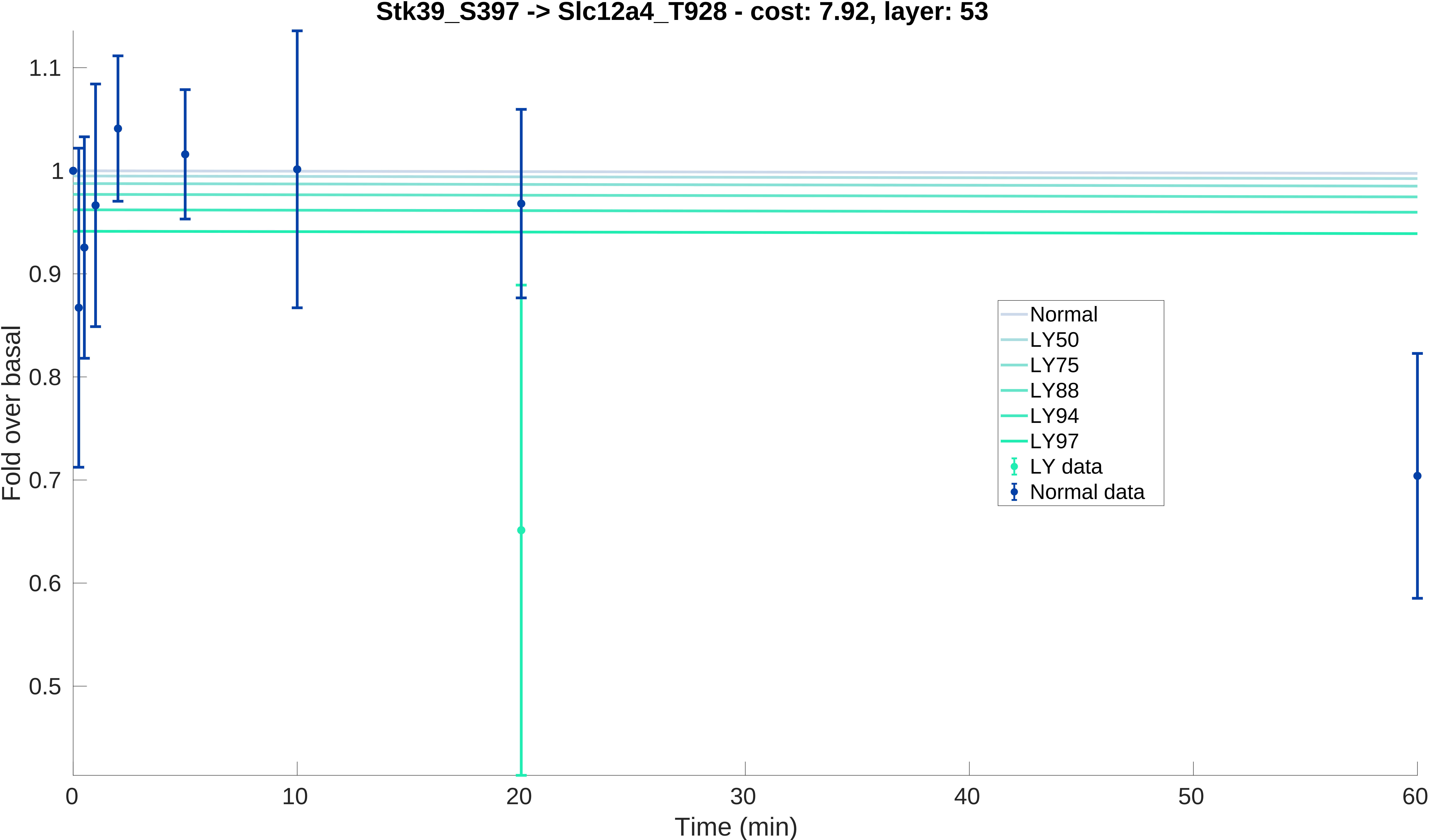

Stk39_S397 -> Slc12a4_T928 - cost: 7.92, layer: 53
Fold over basal
Time (min)
0.5
0.6
0.7
0.8
0.9
1
1.1
0
10
20
30
40
50
60
Normal
LY50
LY75
LY88
LY94
LY97
LY data
Normal data

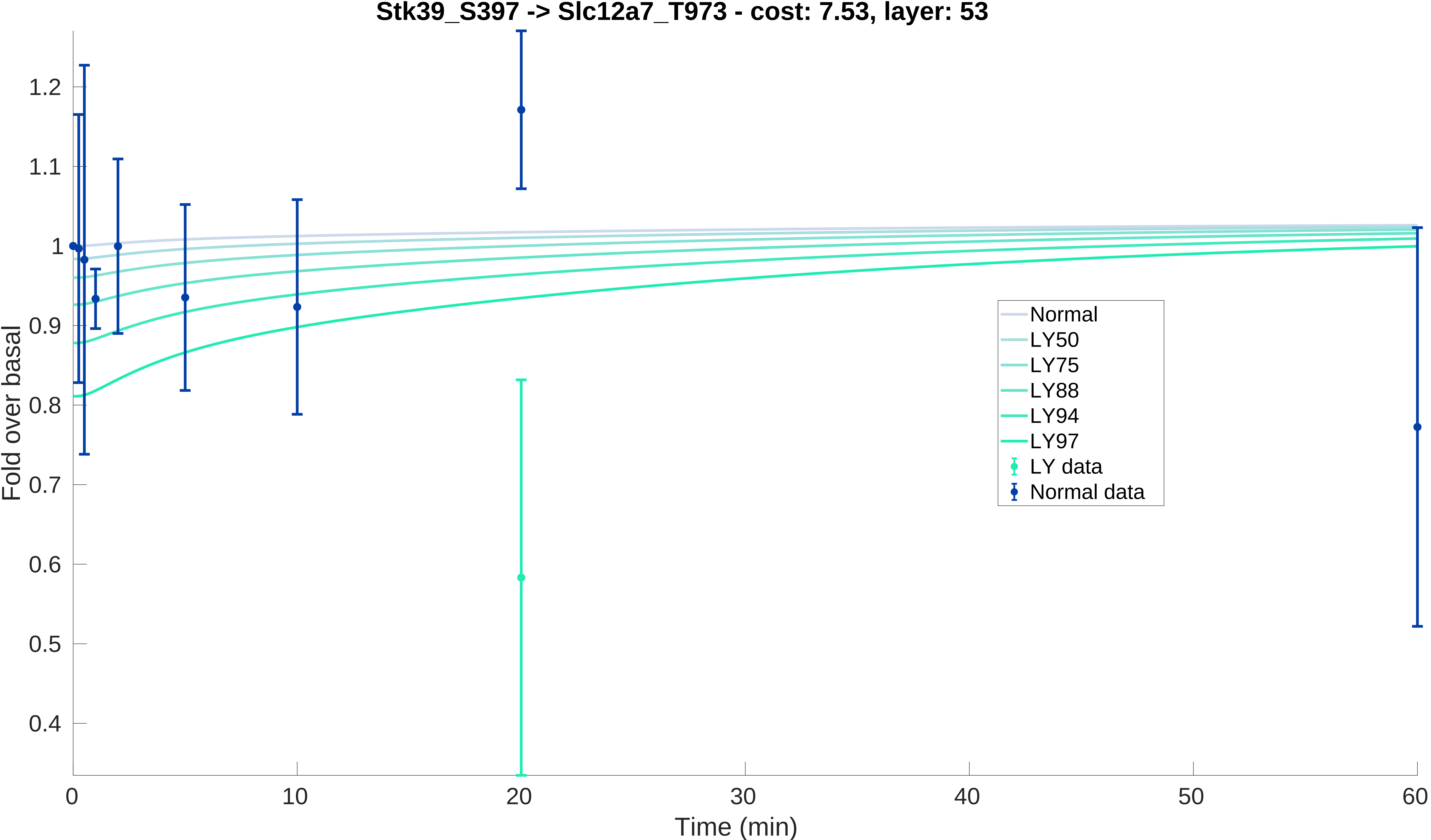

Stk39_S397 -> Slc12a7_T973 - cost: 7.53, layer: 53
Fold over basal
1.2
1.1
1
0.9
0.8
0.7
0.6
0.5
0.4
0
10
20
30
40
50
60
Time (min)
Normal
LY50
LY75
LY88
LY94
LY97
LY data
Normal data

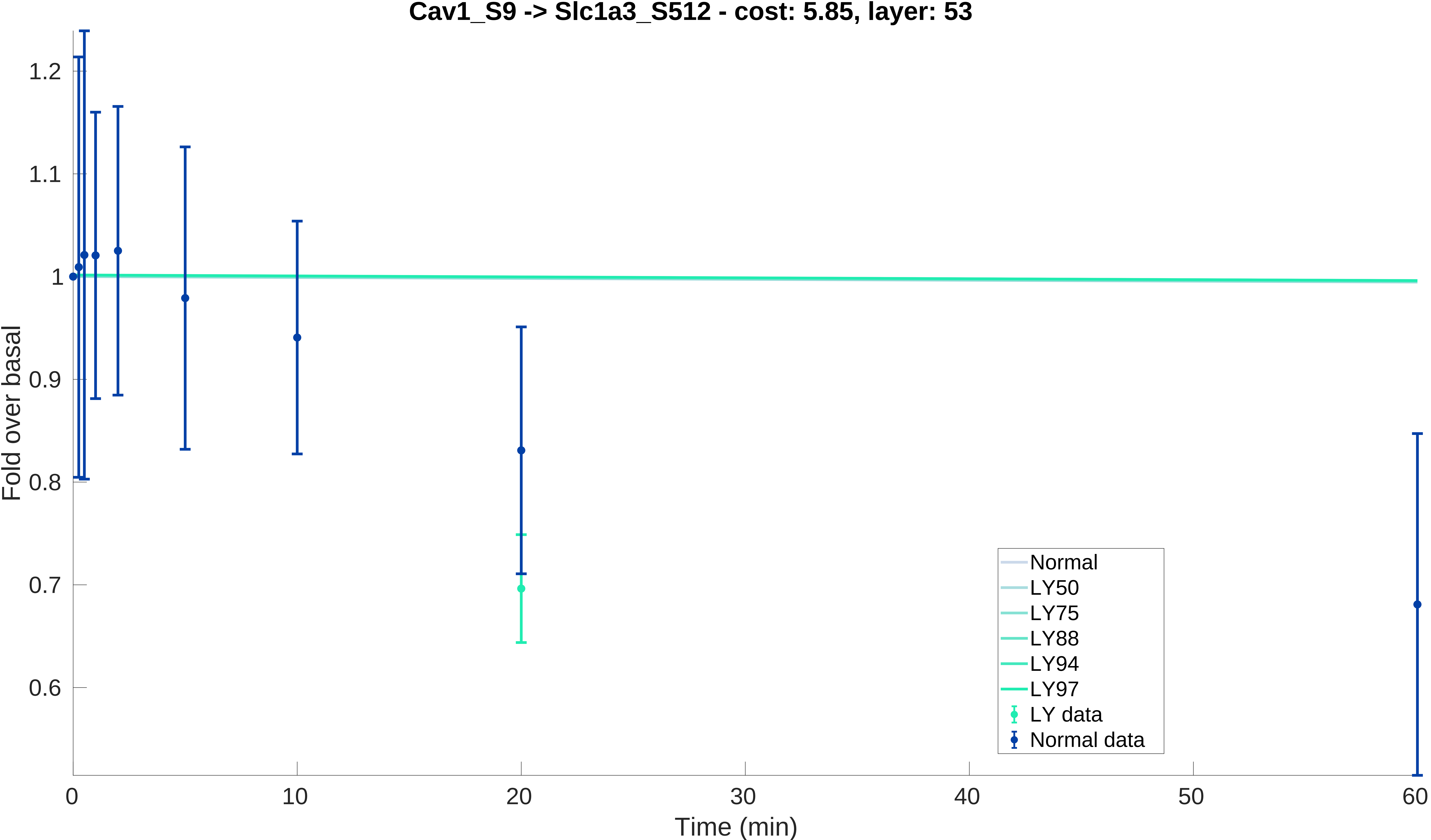

Cav1_S9 -> Slc1a3_S512 - cost: 5.85, layer: 53
Fold over basal
Time (min)
0.6
0.7
0.8
0.9
1
1.1
1.2
0
10
20
30
40
50
60
Normal
LY50
LY75
LY88
LY94
LY97
LY data
Normal data

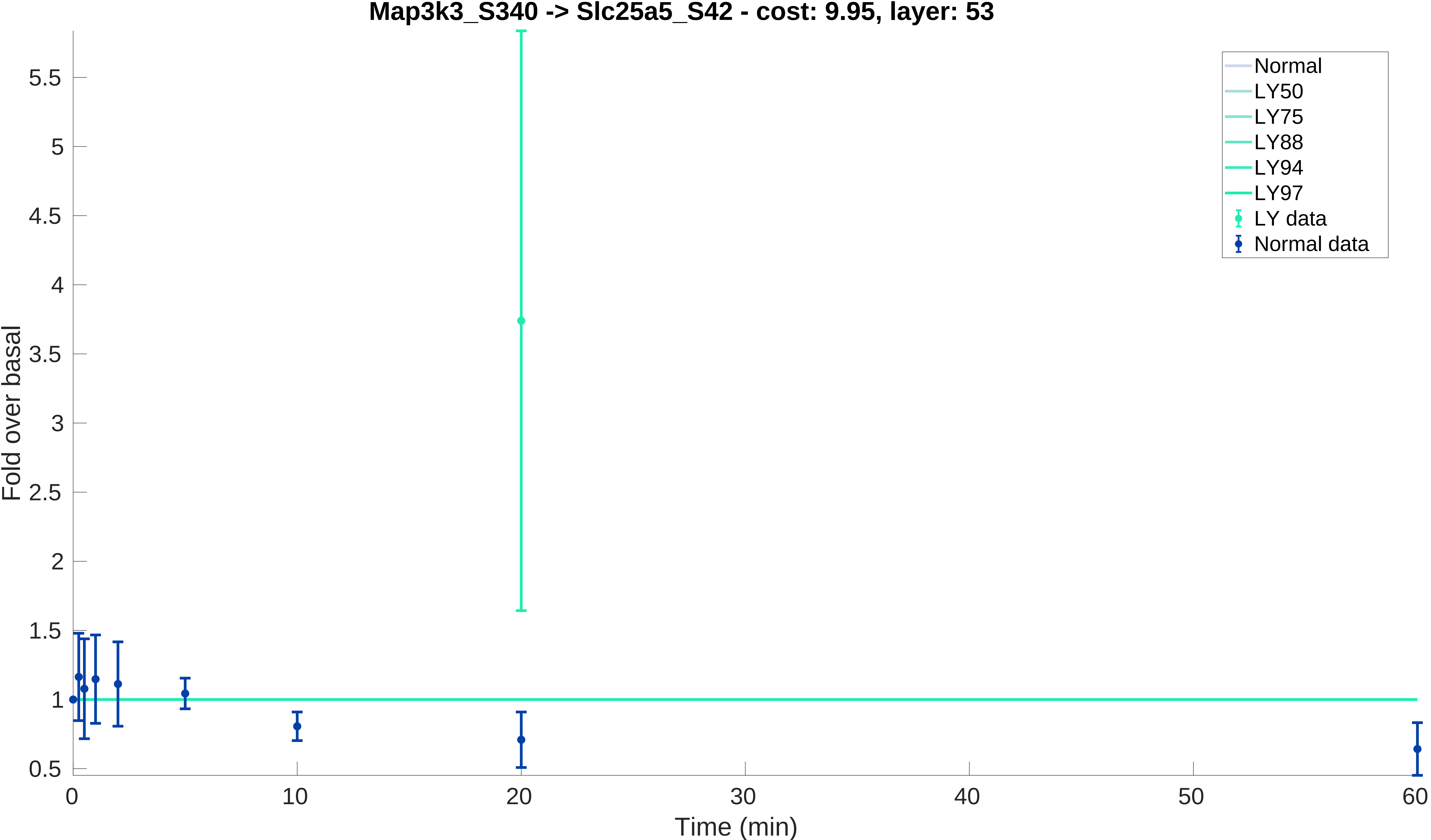
Map3k3_S340 -> Slc25a5_S42 - cost: 9.95, layer: 53
Fold over basal
Time (min)
0.5
1
1.5
2
2.5
3
3.5
4
4.5
5
5.5
0
10
20
30
40
50
60
Normal
LY50
LY75
LY88
LY94
LY97
LY data
Normal data

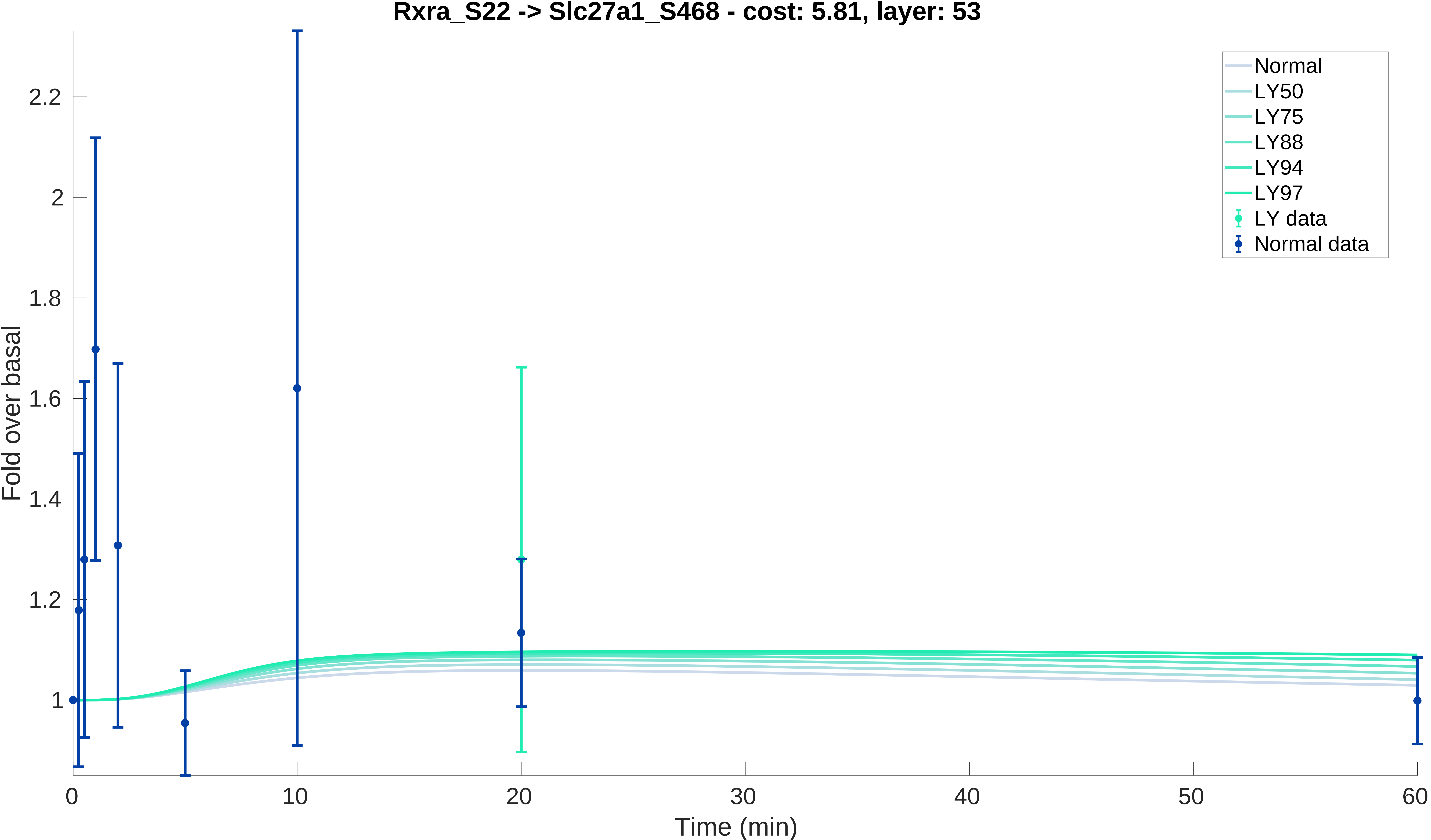
Rxra_S22 -> Slc27a1_S468 - cost: 5.81, layer: 53
Normal
LY50
LY75
LY88
LY94
LY97
LY data
Normal data
Fold over basal
2.2
2
1.8
1.6
1.4
1.2
1
0
10
20
30
40
50
60
Time (min)

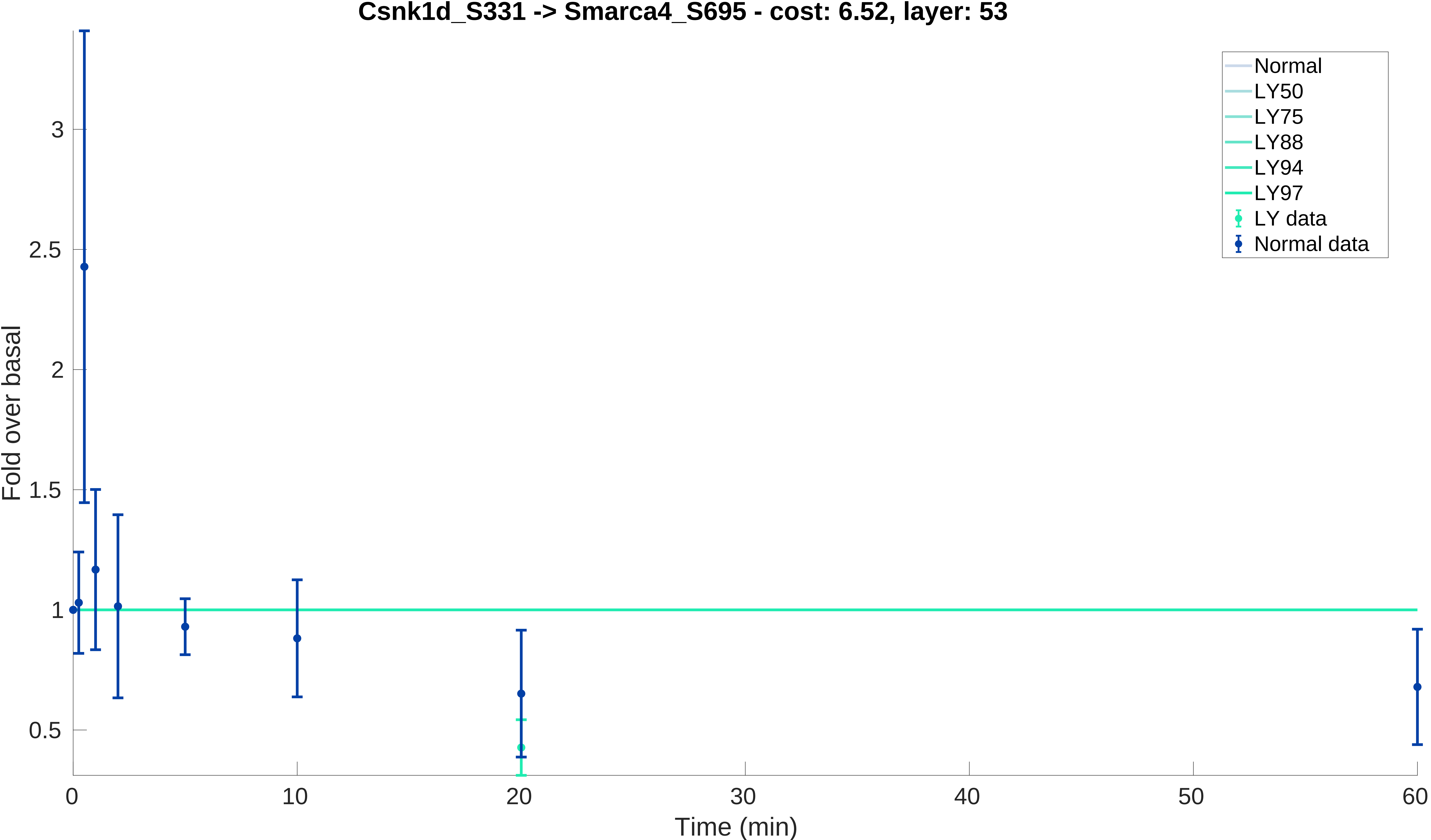

Csnk1d_S331 -> Smarca4_S695 - cost: 6.52, layer: 53
Normal
LY50
LY75
LY88
LY94
LY97
LY data
Normal data
Fold over basal
3
2.5
2
1.5
1
0.5
0
10
20
30
40
50
60
Time (min)

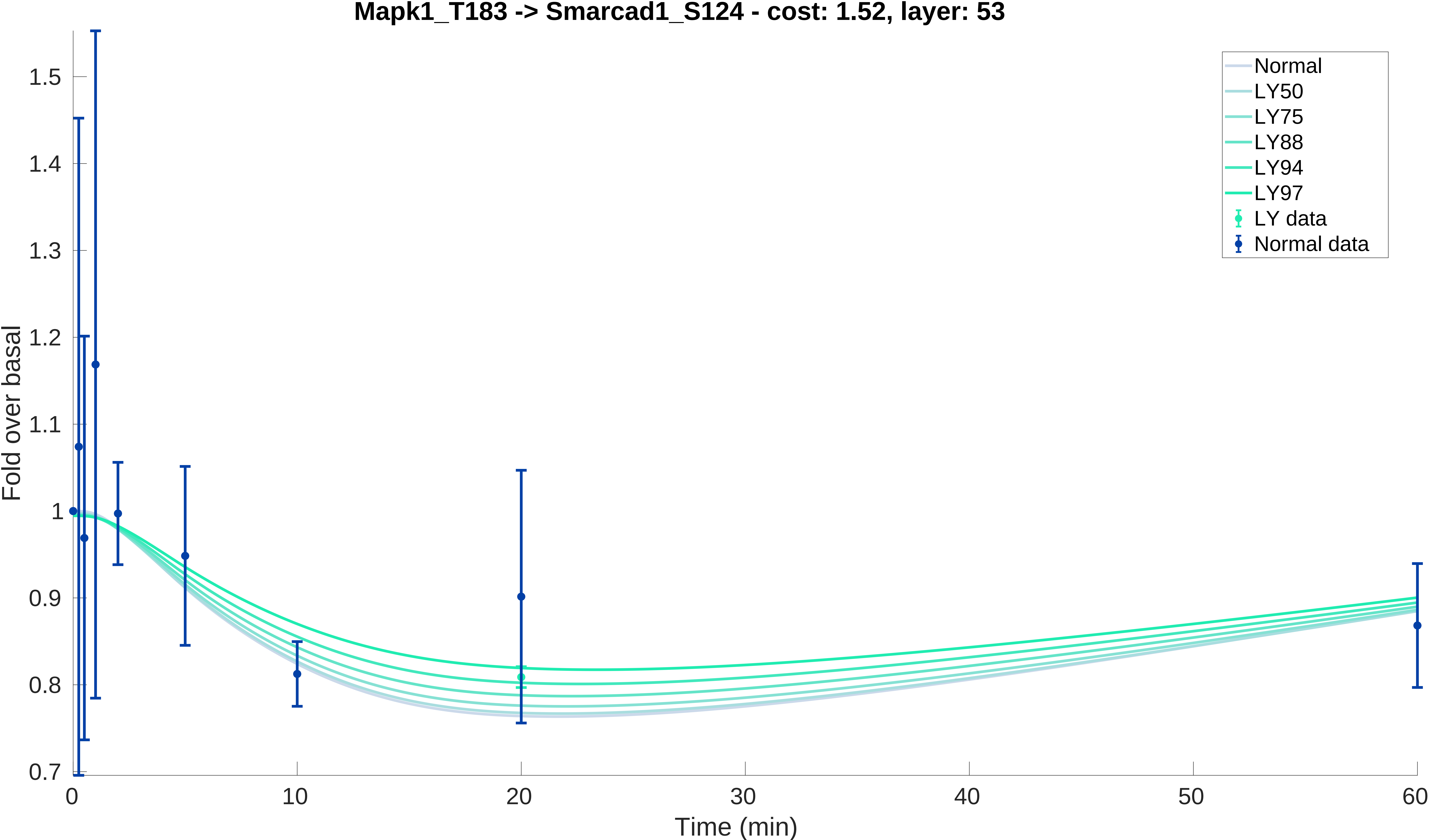

Mapk1_T183 -> Smarcad1_S124 - cost: 1.52, layer: 53
Normal
LY50
LY75
LY88
LY94
LY97
LY data
Normal data
Fold over basal
Time (min)
1.5
1.4
1.3
1.2
1.1
1
0.9
0.8
0.7
0
10
20
30
40
50
60

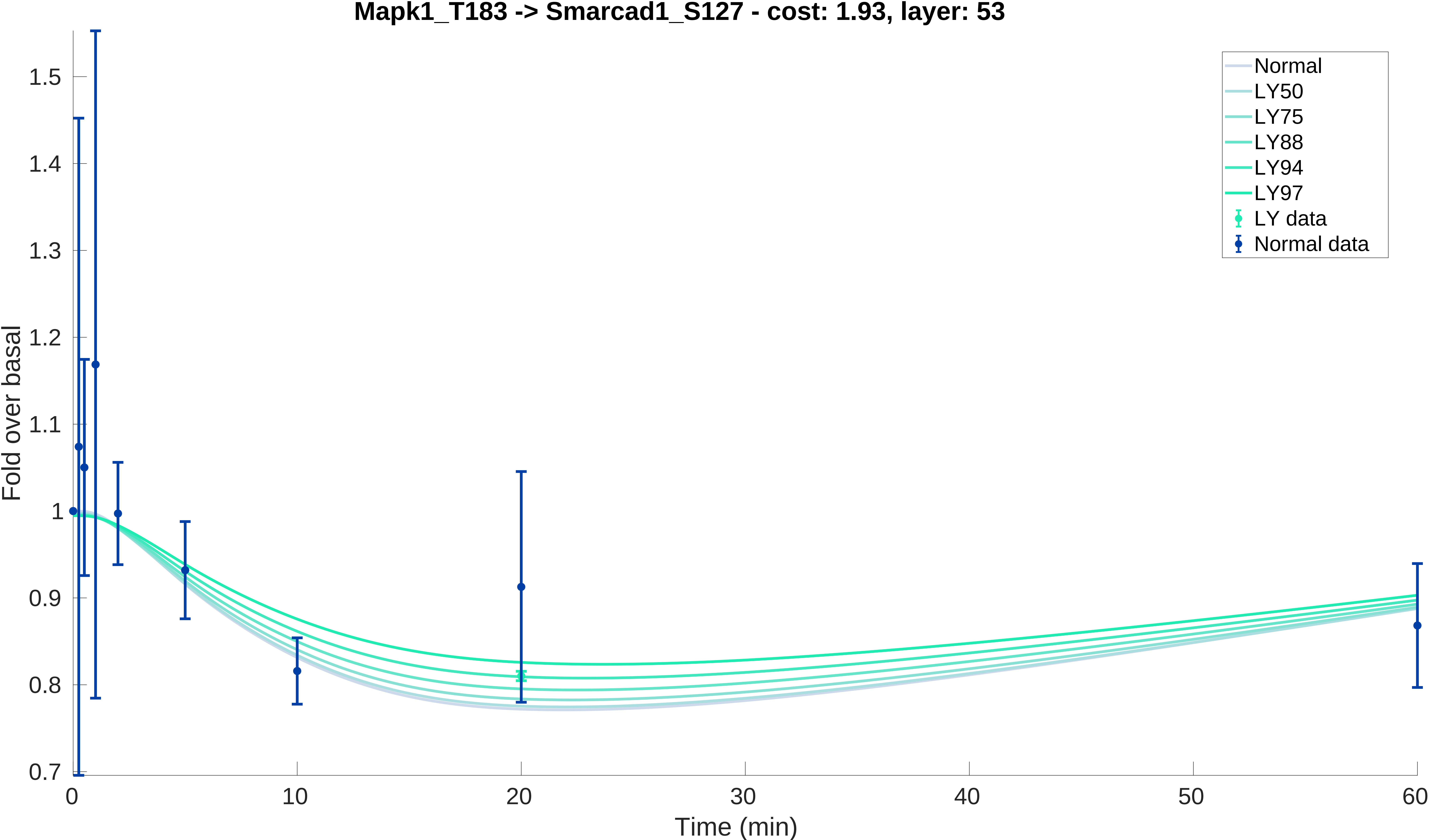
Mapk1_T183 -> Smarcad1_S127 - cost: 1.93, layer: 53
Fold over basal
Time (min)
0.7
0.8
0.9
1
1.1
1.2
1.3
1.4
1.5
0
10
20
30
40
50
60
Normal
LY50
LY75
LY88
LY94
LY97
LY data
Normal data

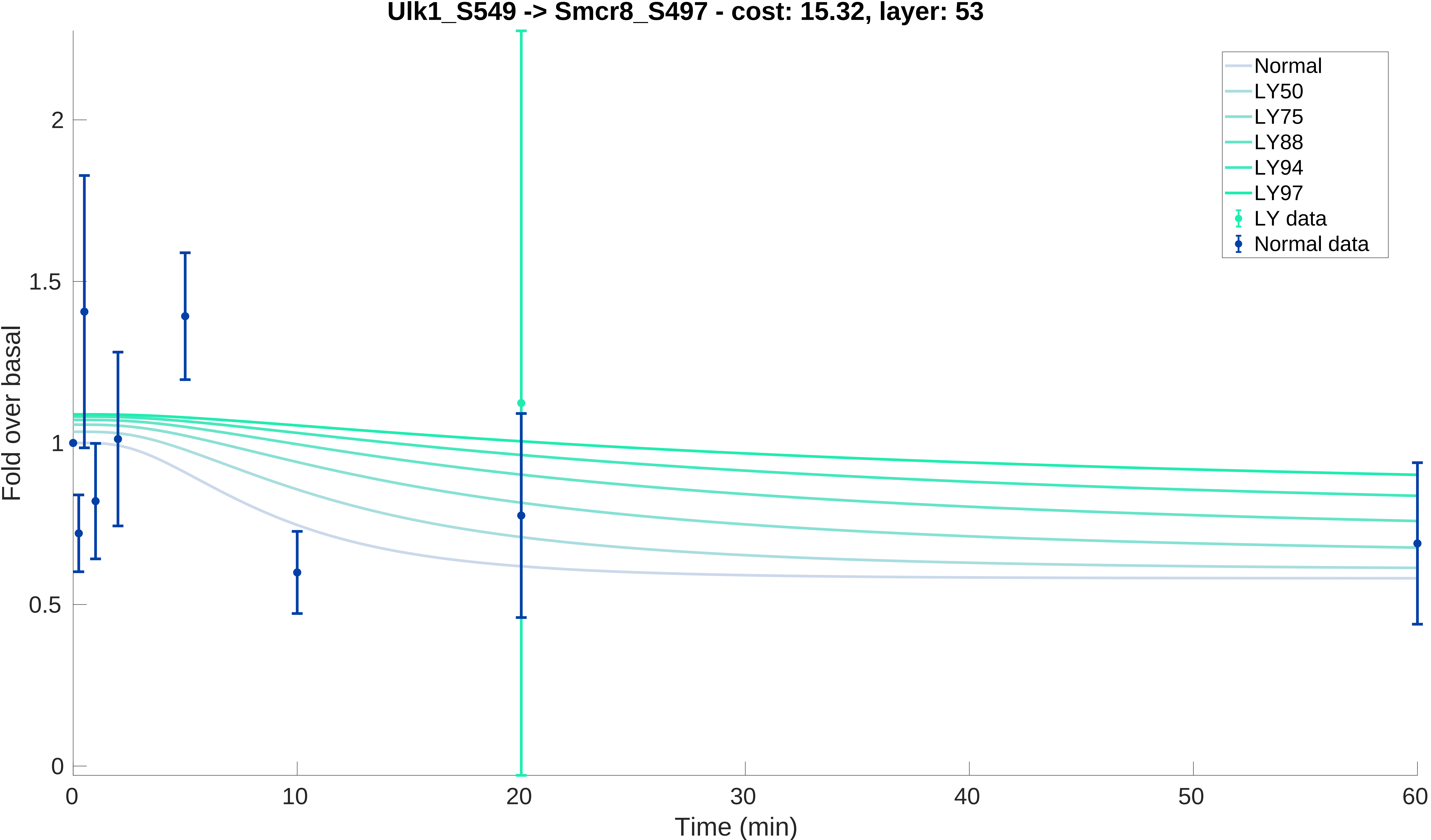

Ulk1_S549 -> Smcr8_S497 - cost: 15.32, layer: 53
Fold over basal
Time (min)
0
0.5
1
1.5
2
0
10
20
30
40
50
60
Normal
LY50
LY75
LY88
LY94
LY97
LY data
Normal data

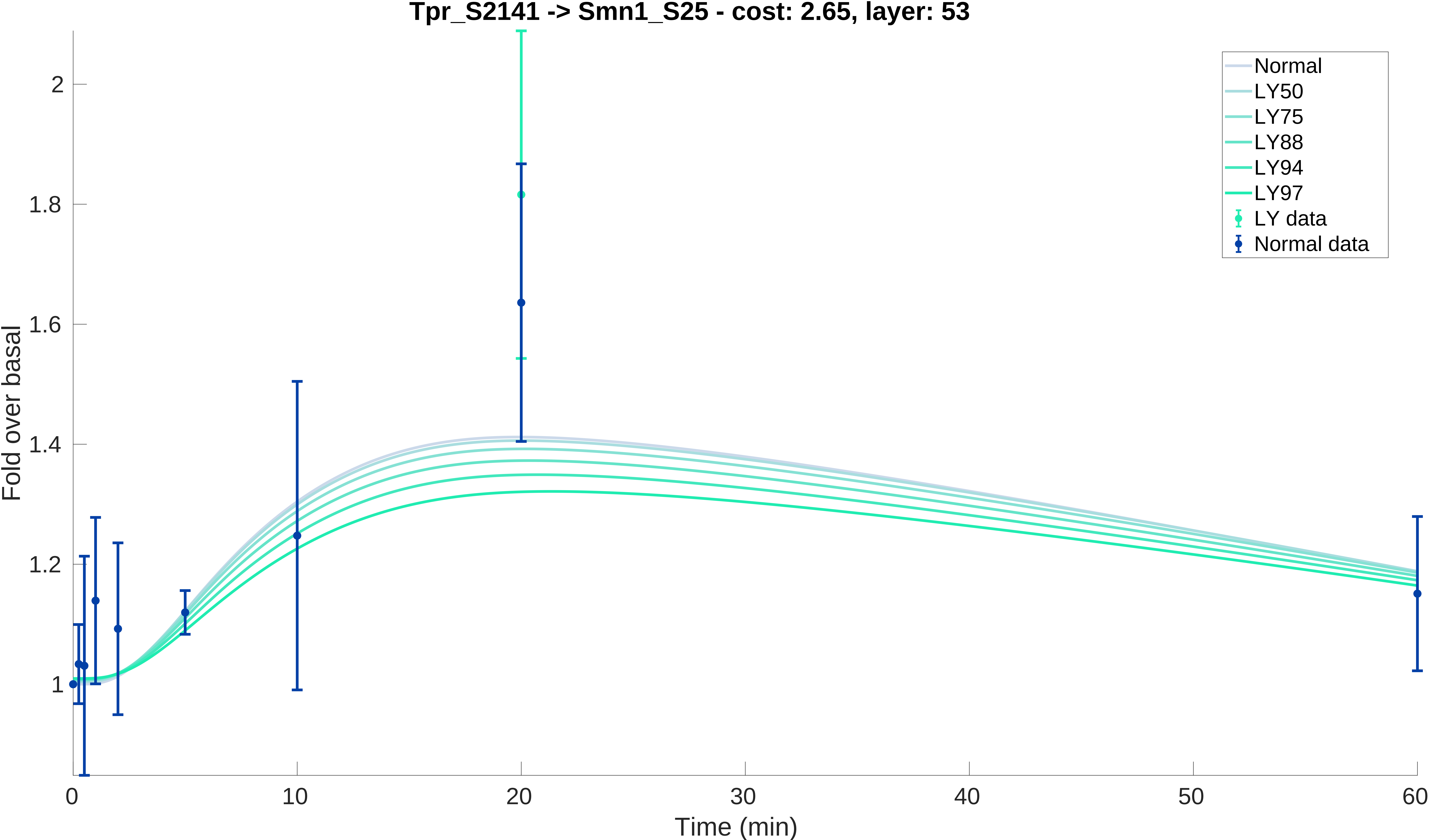
Tpr_S2141 -> Smn1_S25 - cost: 2.65, layer: 53
Fold over basal
Time (min)
Normal
LY50
LY75
LY88
LY94
LY97
LY data
Normal data
0
10
20
30
40
50
60
1
1.2
1.4
1.6
1.8
2

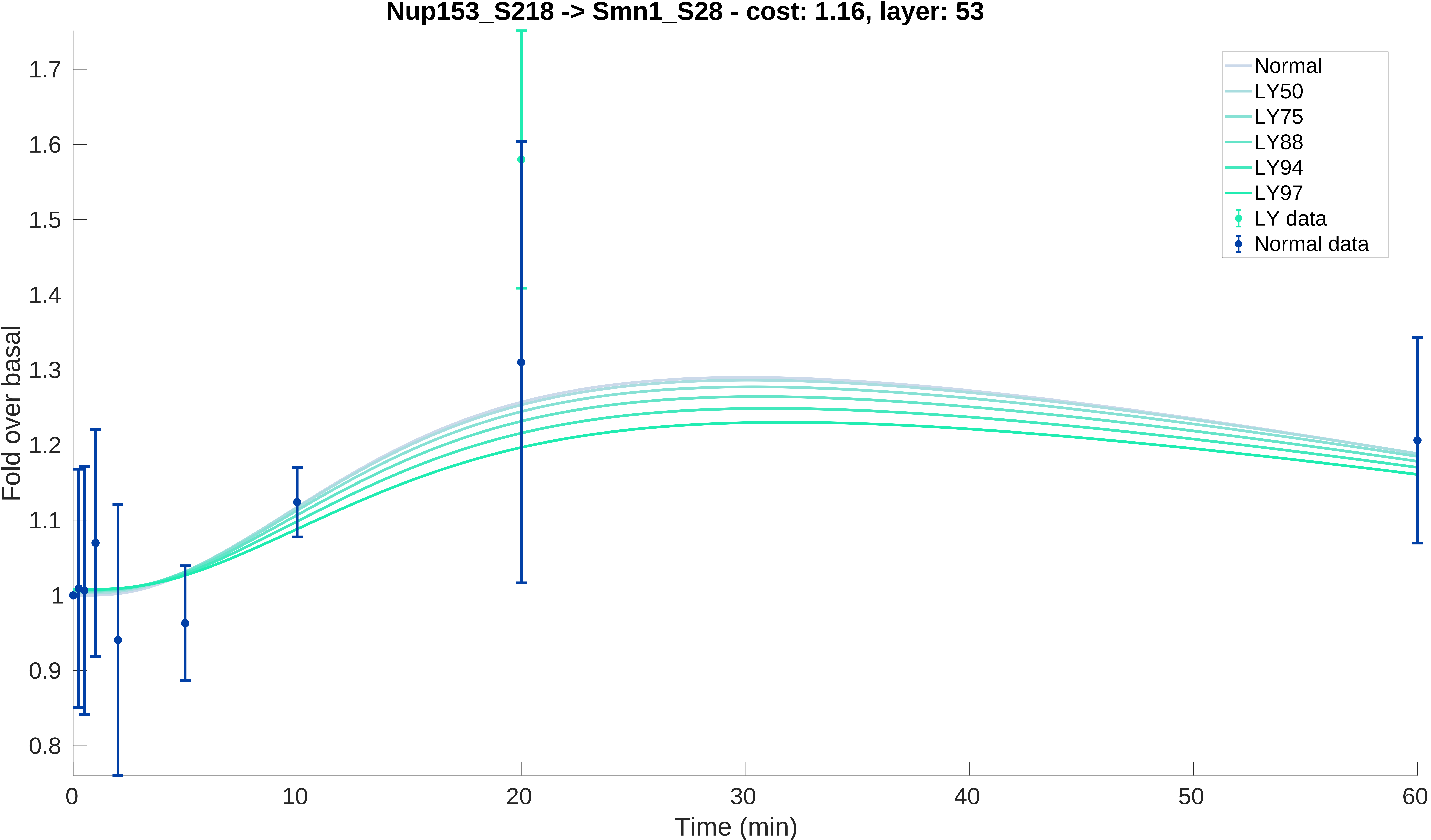

Nup153_S218 -> Smn1_S28 - cost: 1.16, layer: 53
Fold over basal
Time (min)
0.8
0.9
1
1.1
1.2
1.3
1.4
1.5
1.6
1.7
0
10
20
30
40
50
60
Normal
LY50
LY75
LY88
LY94
LY97
LY data
Normal data

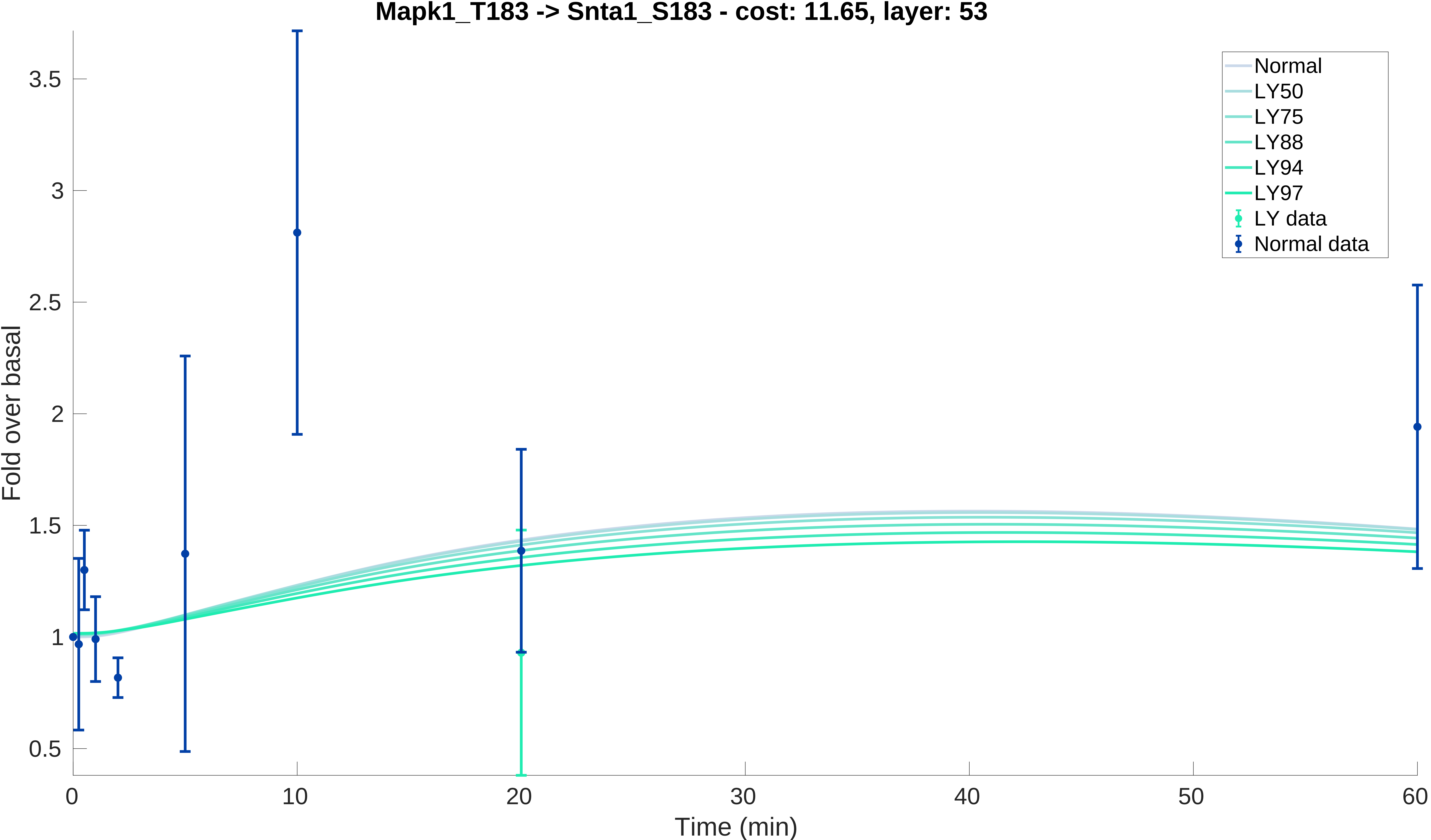

Mapk1_T183 -> Snta1_S183 - cost: 11.65, layer: 53
Fold over basal
Time (min)
Normal
LY50
LY75
LY88
LY94
LY97
LY data
Normal data
0.5
1
1.5
2
2.5
3
3.5
0
10
20
30
40
50
60

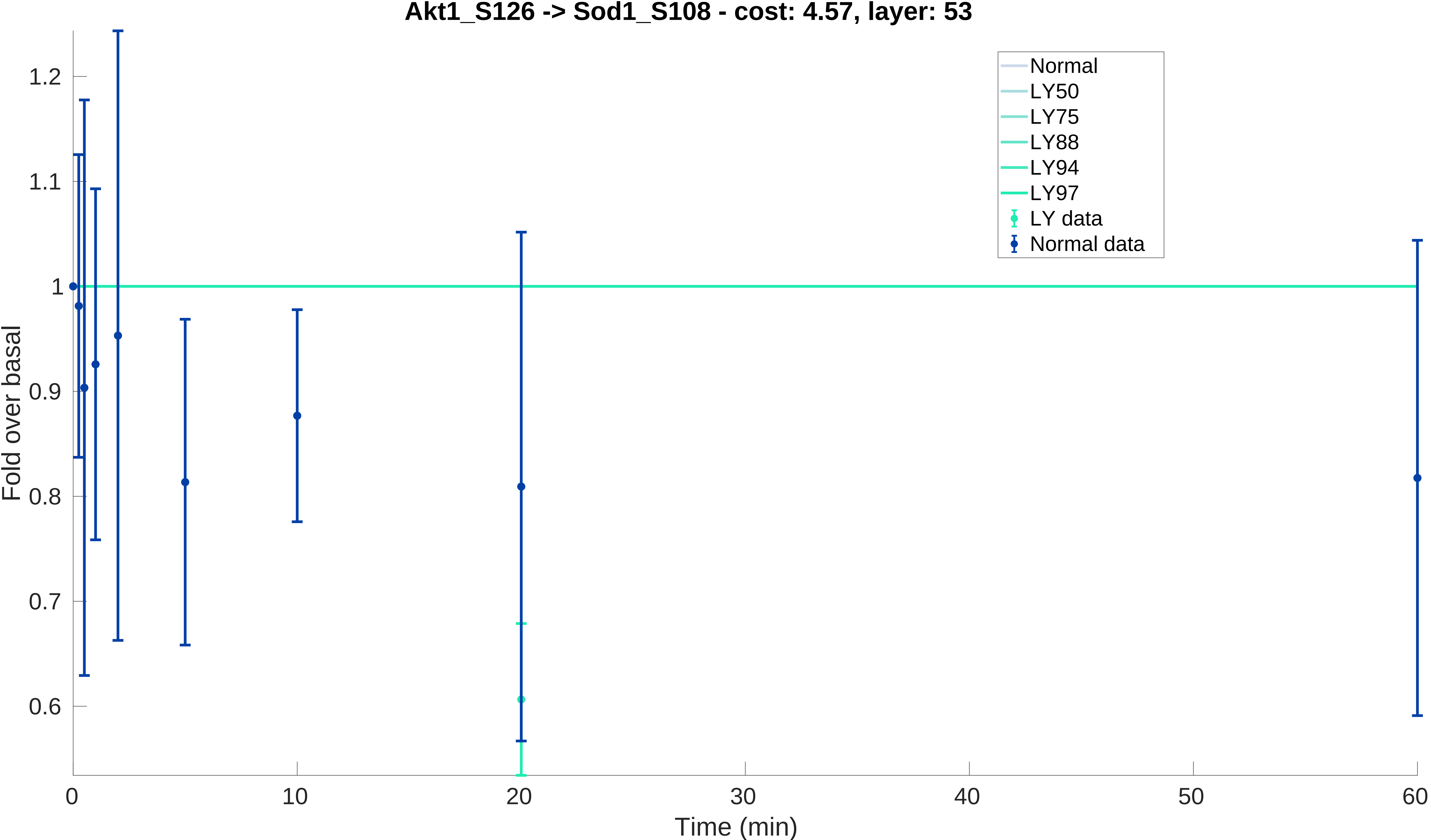
Akt1_S126 -> Sod1_S108 - cost: 4.57, layer: 53
Fold over basal
Time (min)
1.2
1.1
1
0.9
0.8
0.7
0.6
0
10
20
30
40
50
60
Normal
LY50
LY75
LY88
LY94
LY97
LY data
Normal data

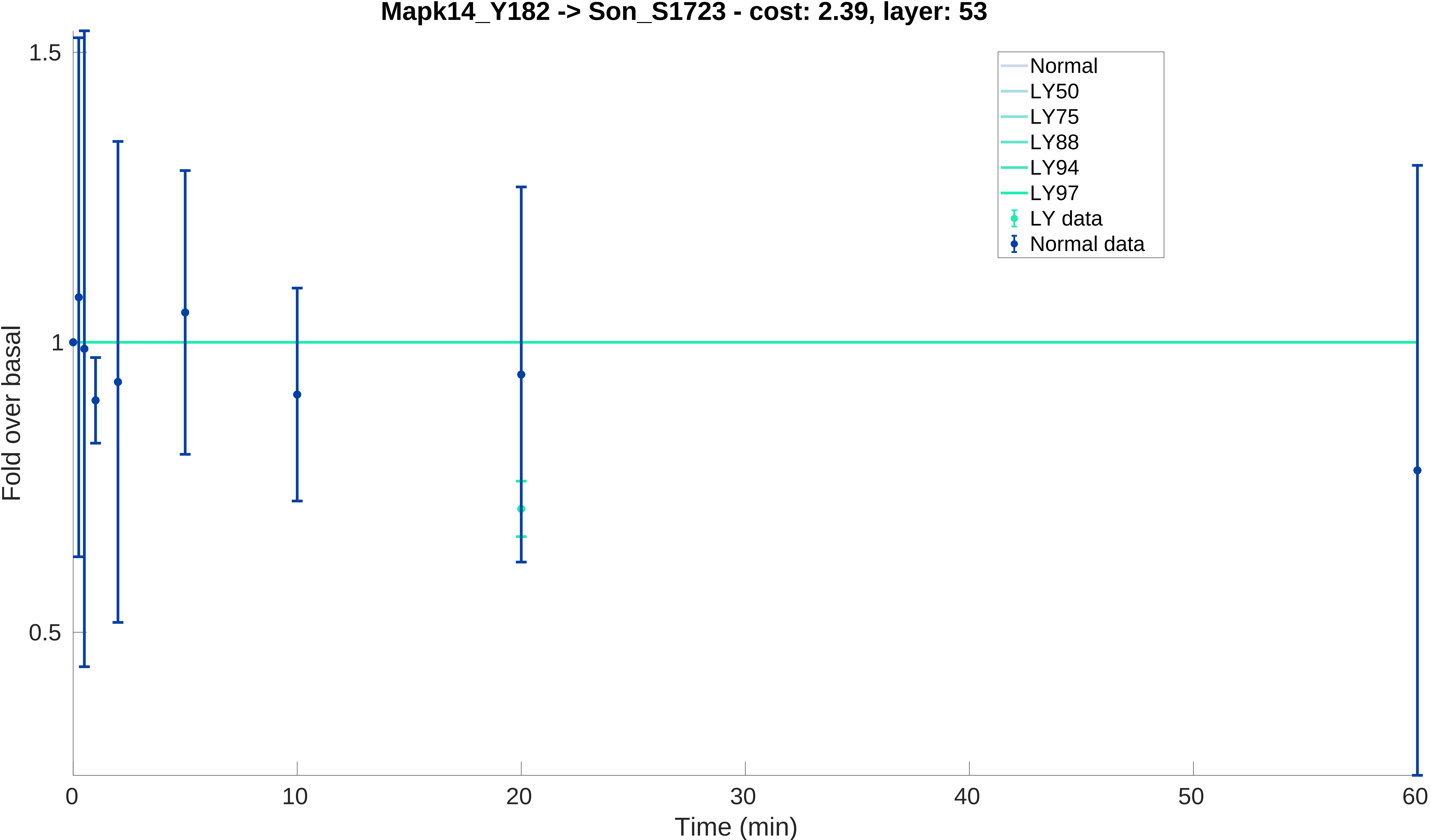

Mapk14_Y182 -> Son_S1723 - cost: 2.39, layer: 53
Normal
LY50
LY75
LY88
LY94
LY97
LY data
Normal data
Fold over basal
1.5
1
0.5
0
10
20
30
40
50
60
Time (min)

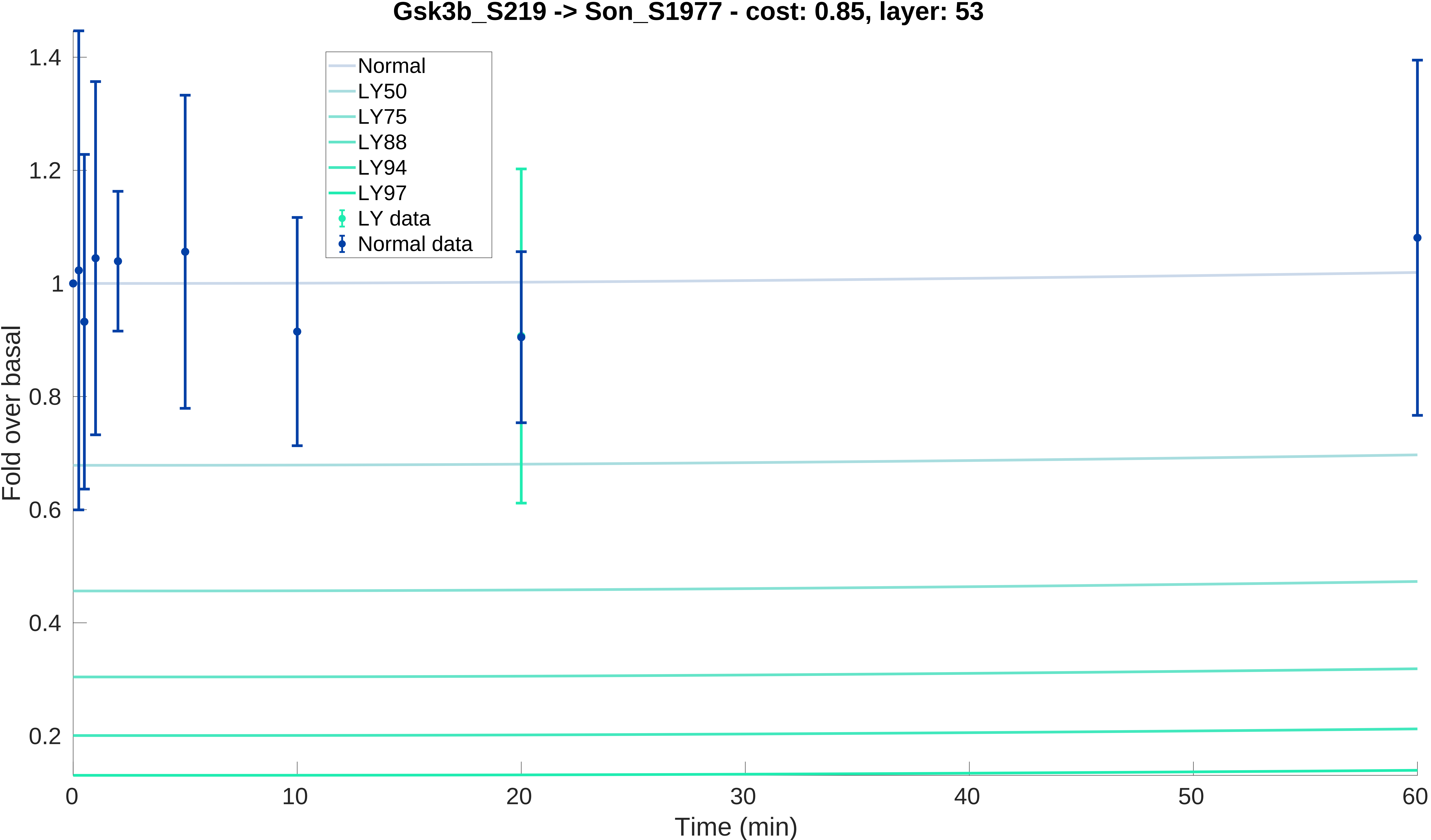
Gsk3b_S219 -> Son_S1977 - cost: 0.85, layer: 53
Normal
LY50
LY75
LY88
LY94
LY97
LY data
Normal data
Fold over basal
Time (min)
1.4
1.2
1
0.8
0.6
0.4
0.2
0
10
20
30
40
50
60

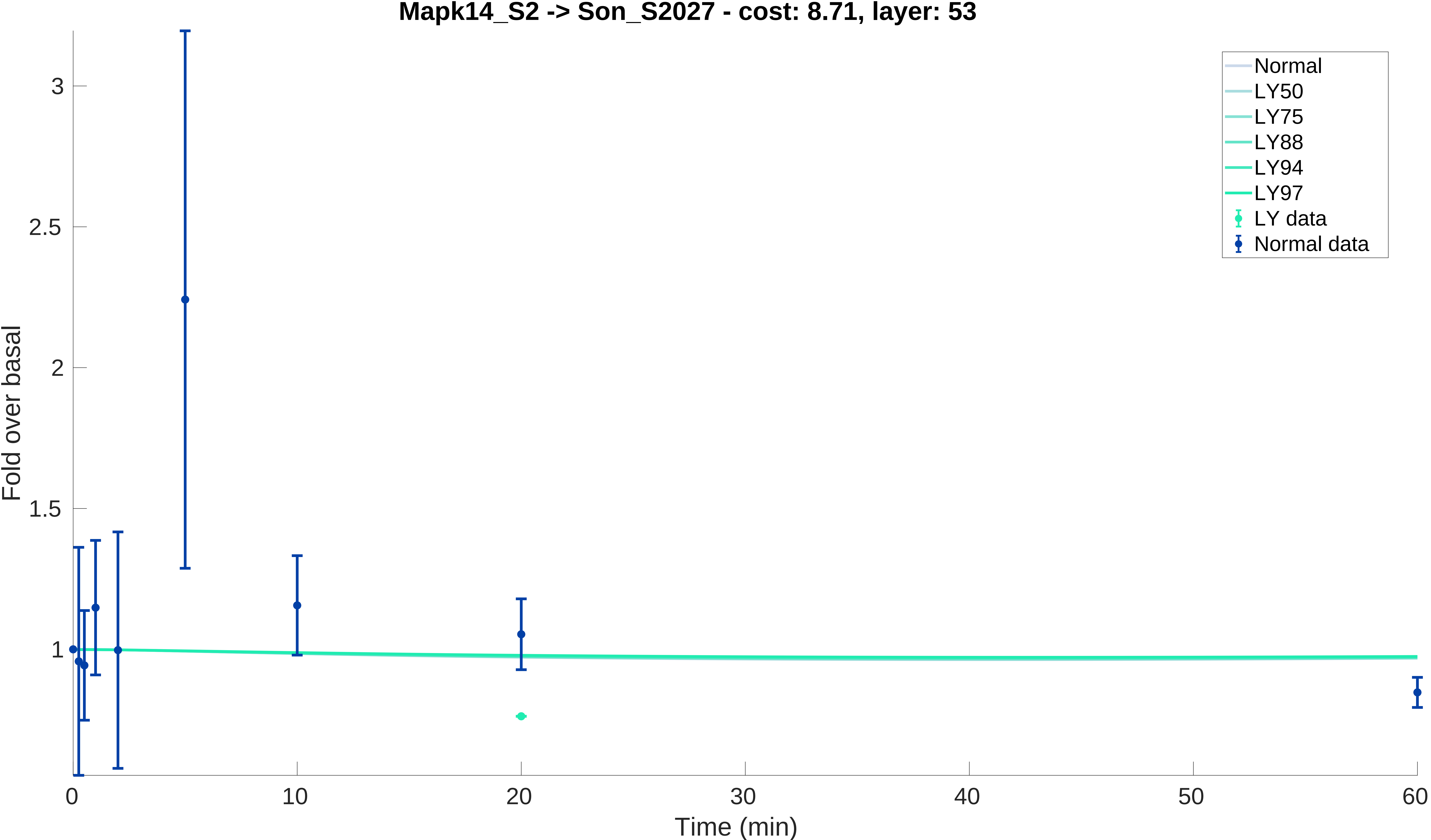

Mapk14_S2 -> Son_S2027 - cost: 8.71, layer: 53
Normal
LY50
LY75
LY88
LY94
LY97
LY data
Normal data
Fold over basal
3
2.5
2
1.5
1
0
10
20
30
40
50
60
Time (min)

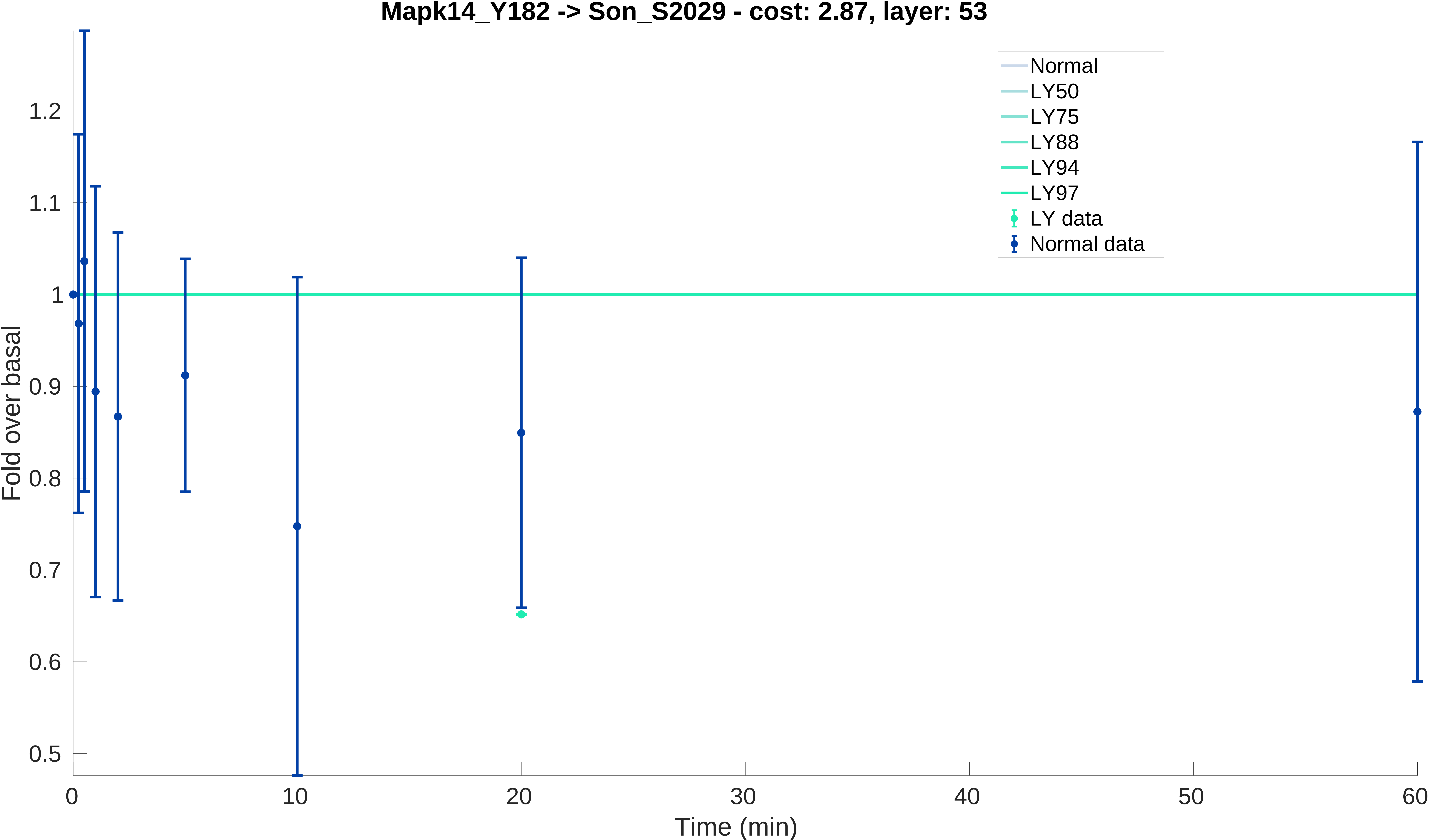

Mapk14_Y182 -> Son_S2029 - cost: 2.87, layer: 53
Normal
LY50
LY75
LY88
LY94
LY97
LY data
Normal data
Fold over basal
1.2
1.1
1
0.9
0.8
0.7
0.6
0.5
0
10
20
30
40
50
60
Time (min)

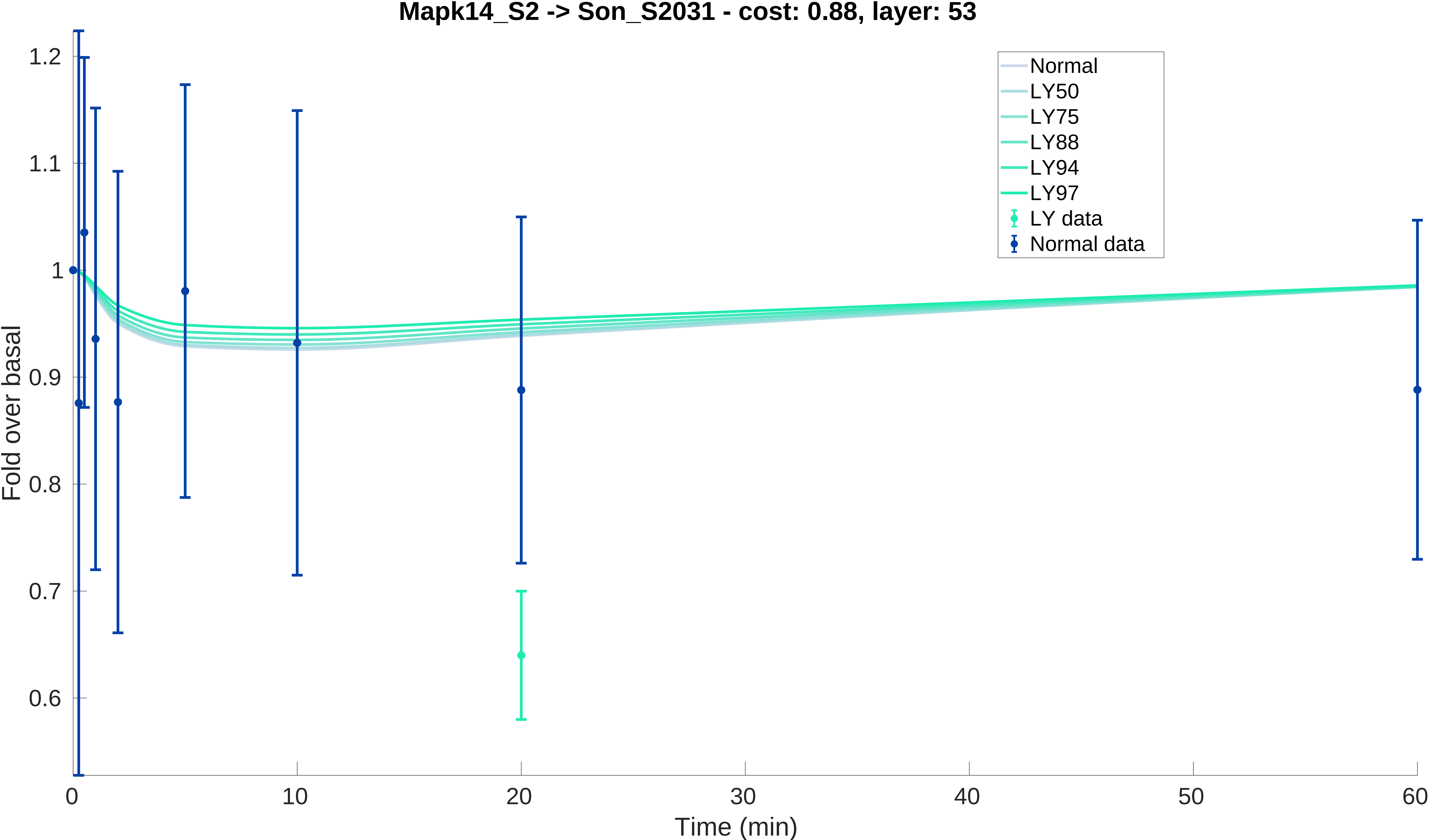
Mapk14_S2 -> Son_S2031 - cost: 0.88, layer: 53
Fold over basal
Time (min)
Normal
LY50
LY75
LY88
LY94
LY97
LY data
Normal data
1.2
1.1
1
0.9
0.8
0.7
0.6
0
10
20
30
40
50
60

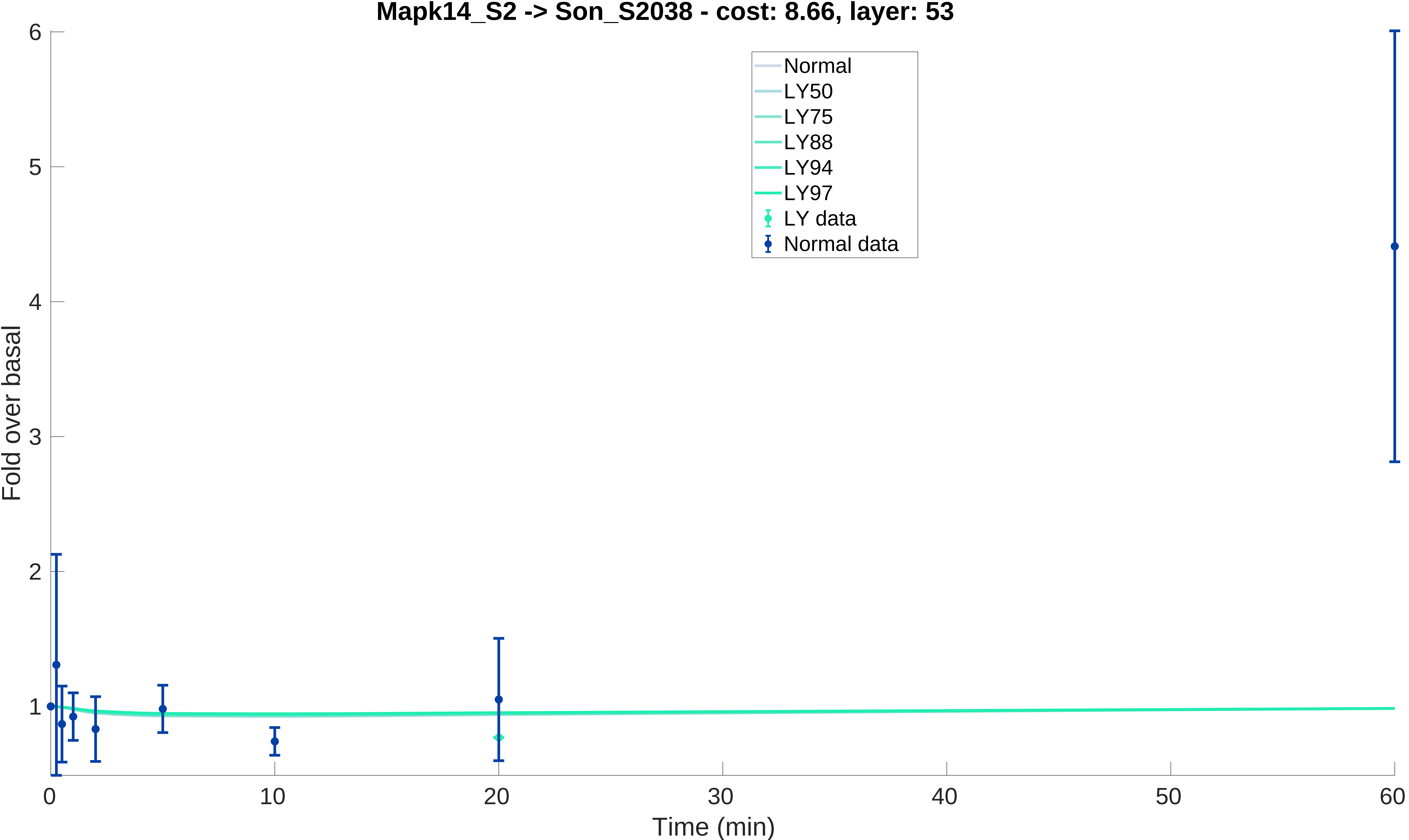
Mapk14_S2 -> Son_S2038 - cost: 8.66, layer: 53
Normal
LY50
LY75
LY88
LY94
LY97
LY data
Normal data
Fold over basal
Time (min)
0
10
20
30
40
50
60
1
2
3
4
5
6

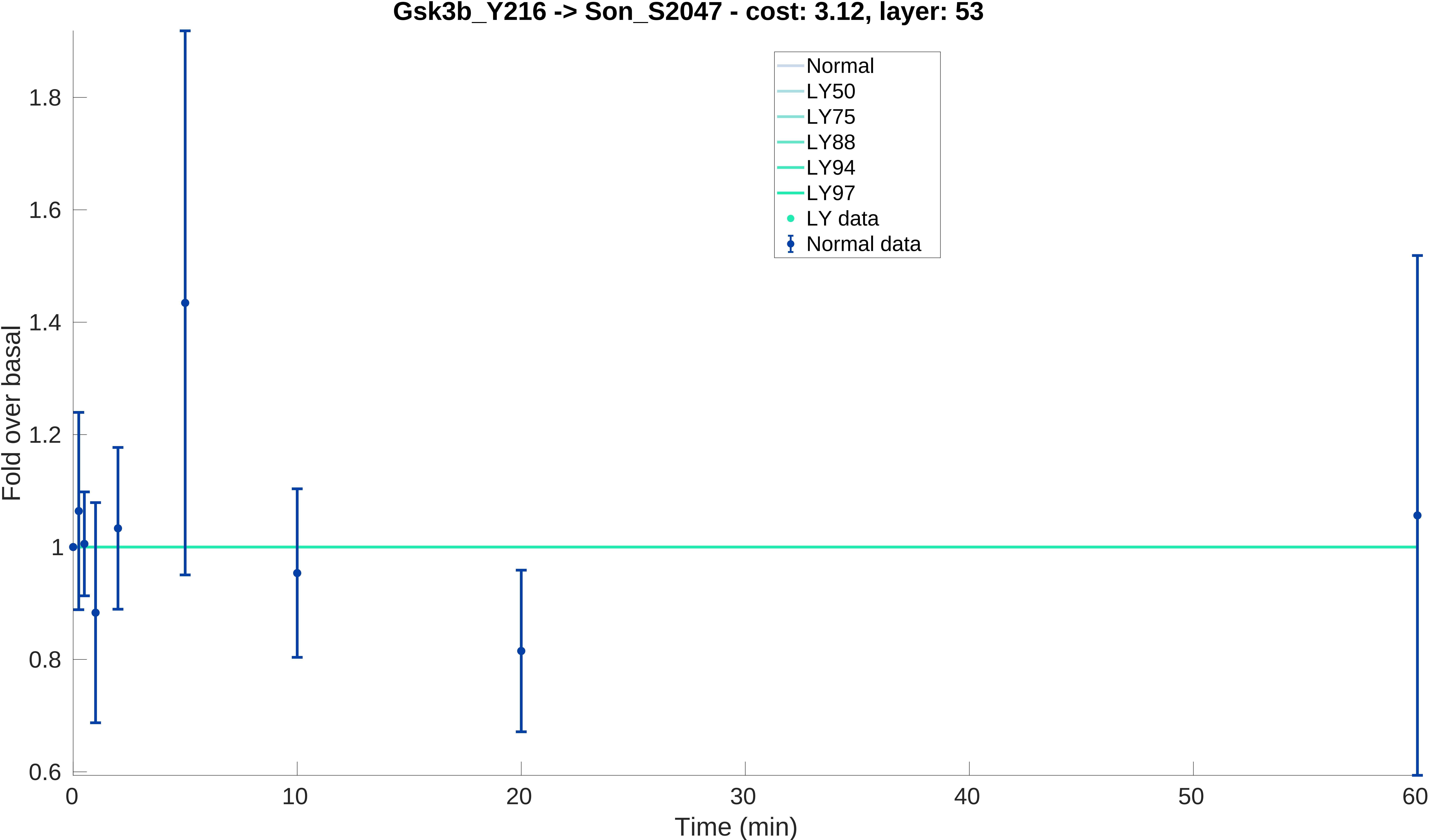

Gsk3b_Y216 -> Son_S2047 - cost: 3.12, layer: 53
Normal
LY50
LY75
LY88
LY94
LY97
LY data
Normal data
Fold over basal
Time (min)
0.6
0.8
1
1.2
1.4
1.6
1.8
0
10
20
30
40
50
60

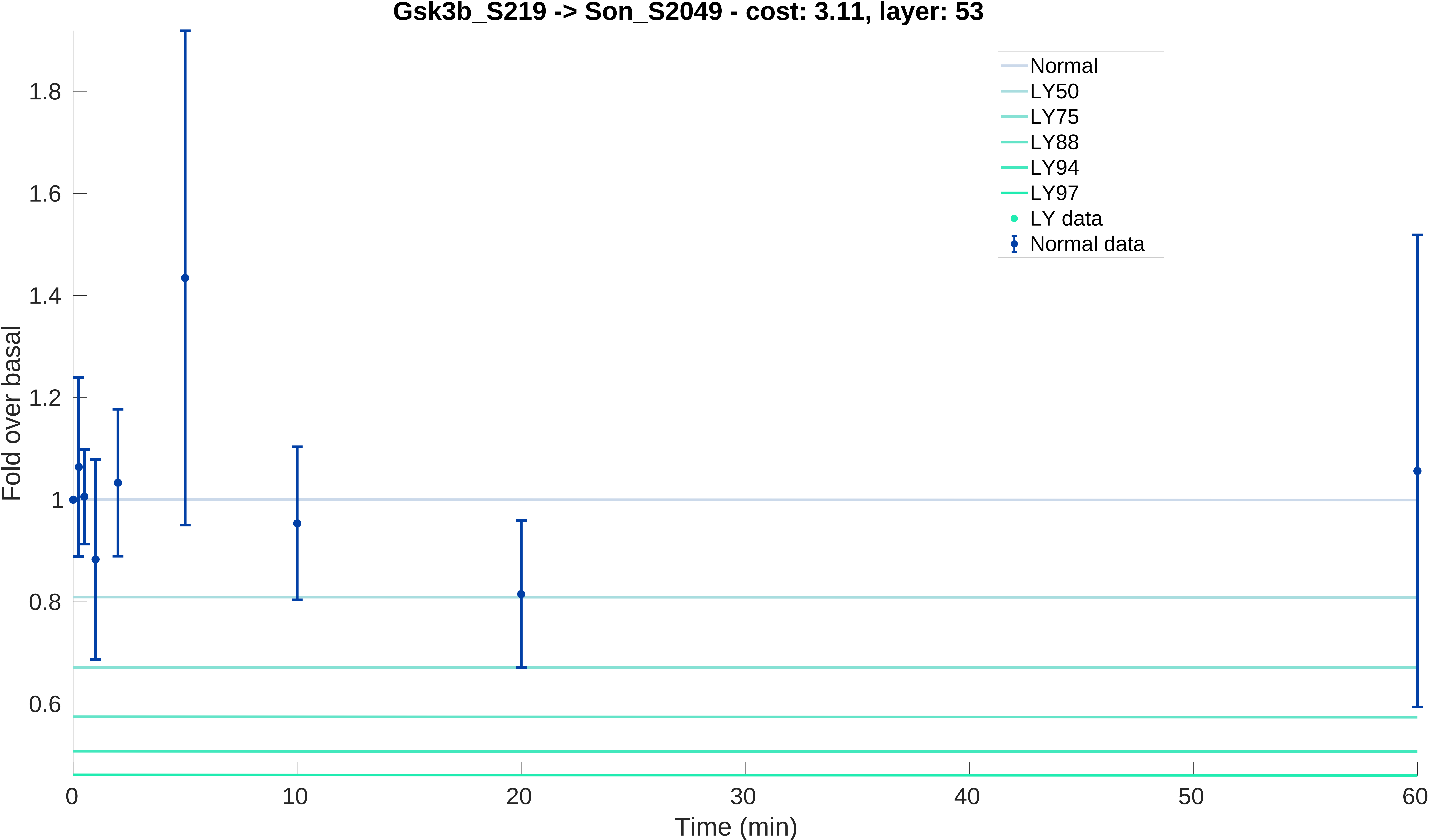

Gsk3b_S219 -> Son_S2049 - cost: 3.11, layer: 53
Normal
LY50
LY75
LY88
LY94
LY97
LY data
Normal data
Fold over basal
1.8
1.6
1.4
1.2
1
0.8
0.6
0
10
20
30
40
50
60
Time (min)

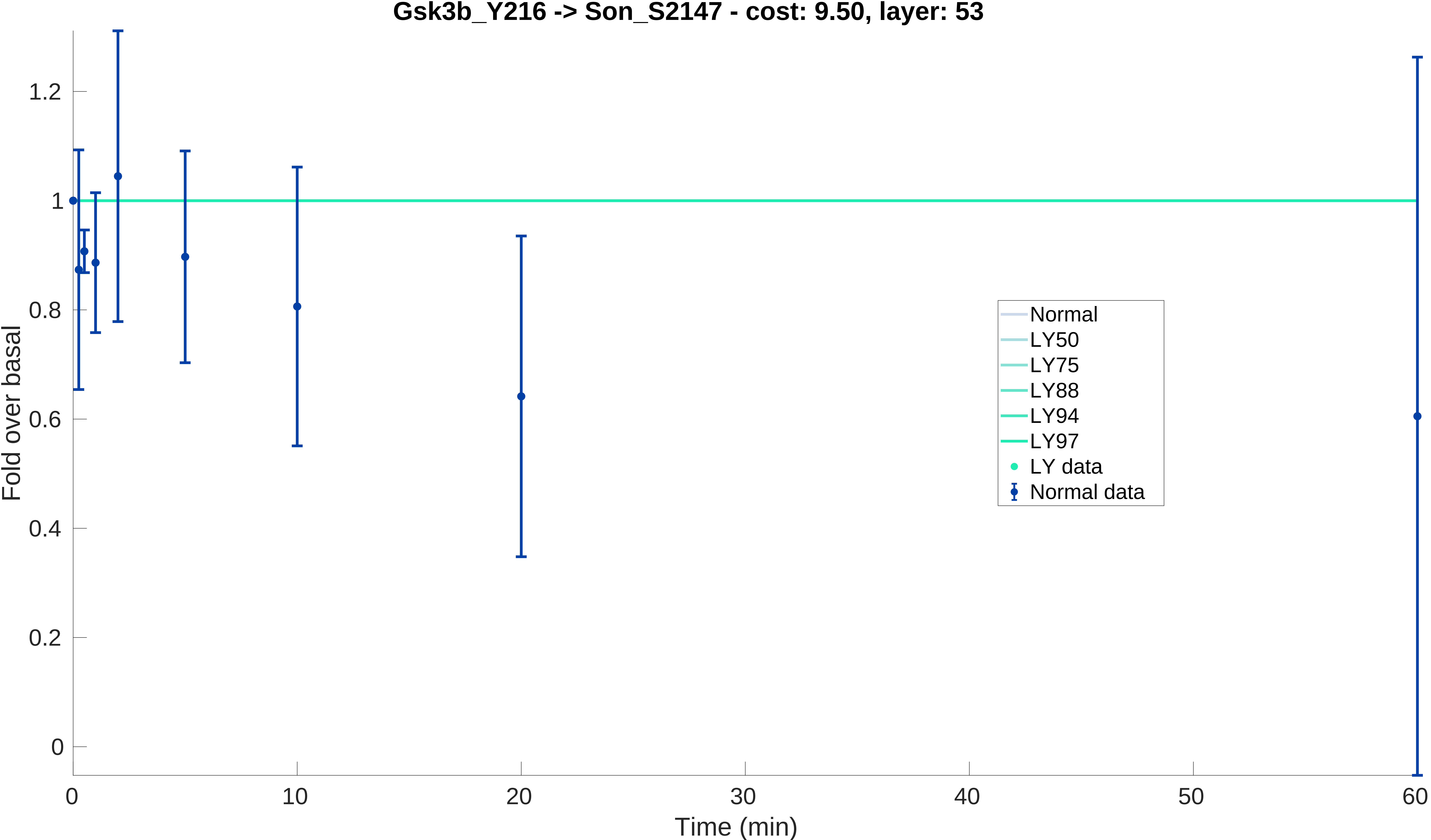
Gsk3b_Y216 -> Son_S2147 - cost: 9.50, layer: 53
Fold over basal
Time (min)
0
0.2
0.4
0.6
0.8
1
1.2
0
10
20
30
40
50
60
Normal
LY50
LY75
LY88
LY94
LY97
LY data
Normal data

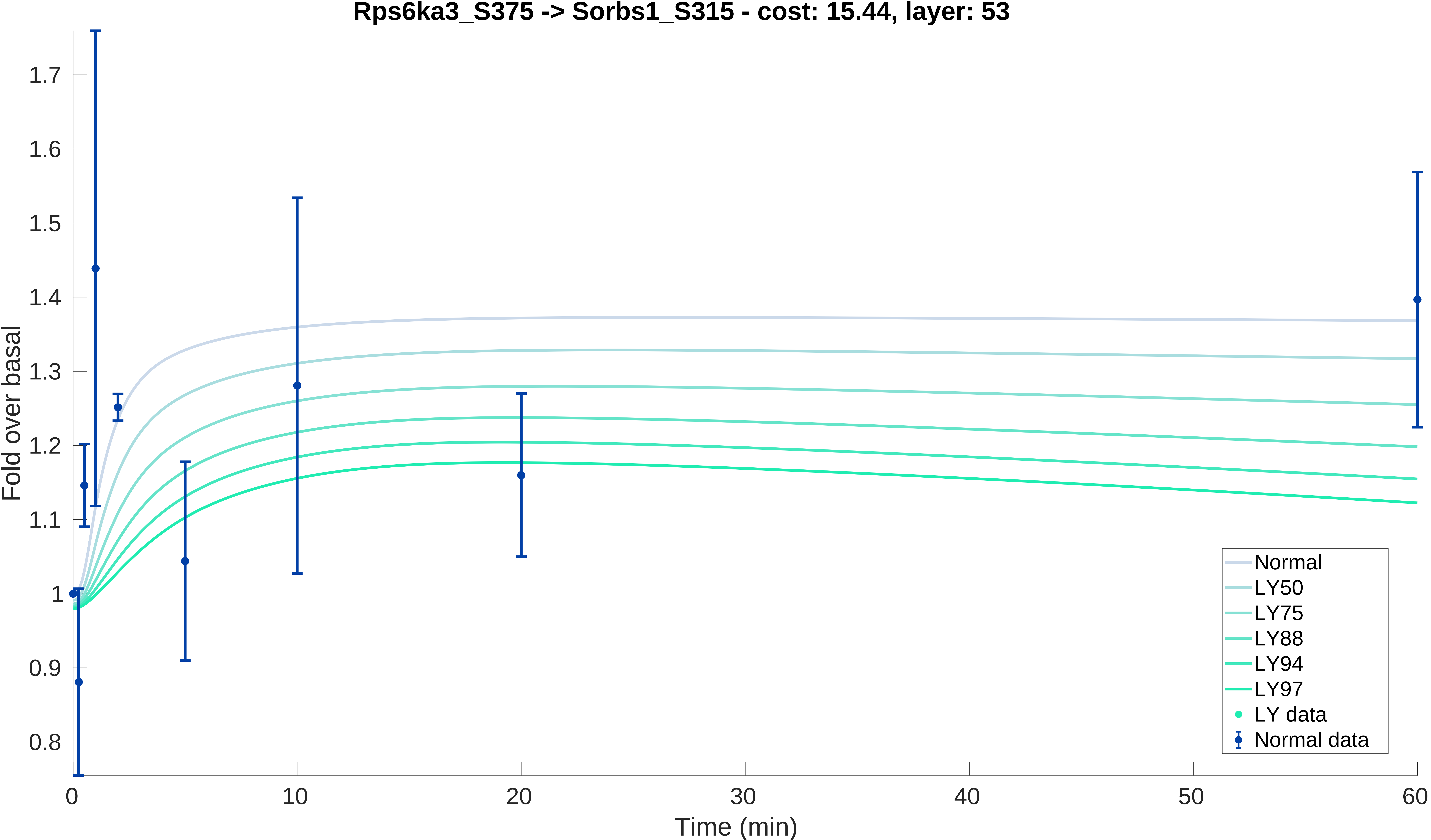
Rps6ka3_S375 -> Sorbs1_S315 - cost: 15.44, layer: 53
Fold over basal
Time (min)
0.8
0.9
1
1.1
1.2
1.3
1.4
1.5
1.6
1.7
0
10
20
30
40
50
60
Normal
LY50
LY75
LY88
LY94
LY97
LY data
Normal data

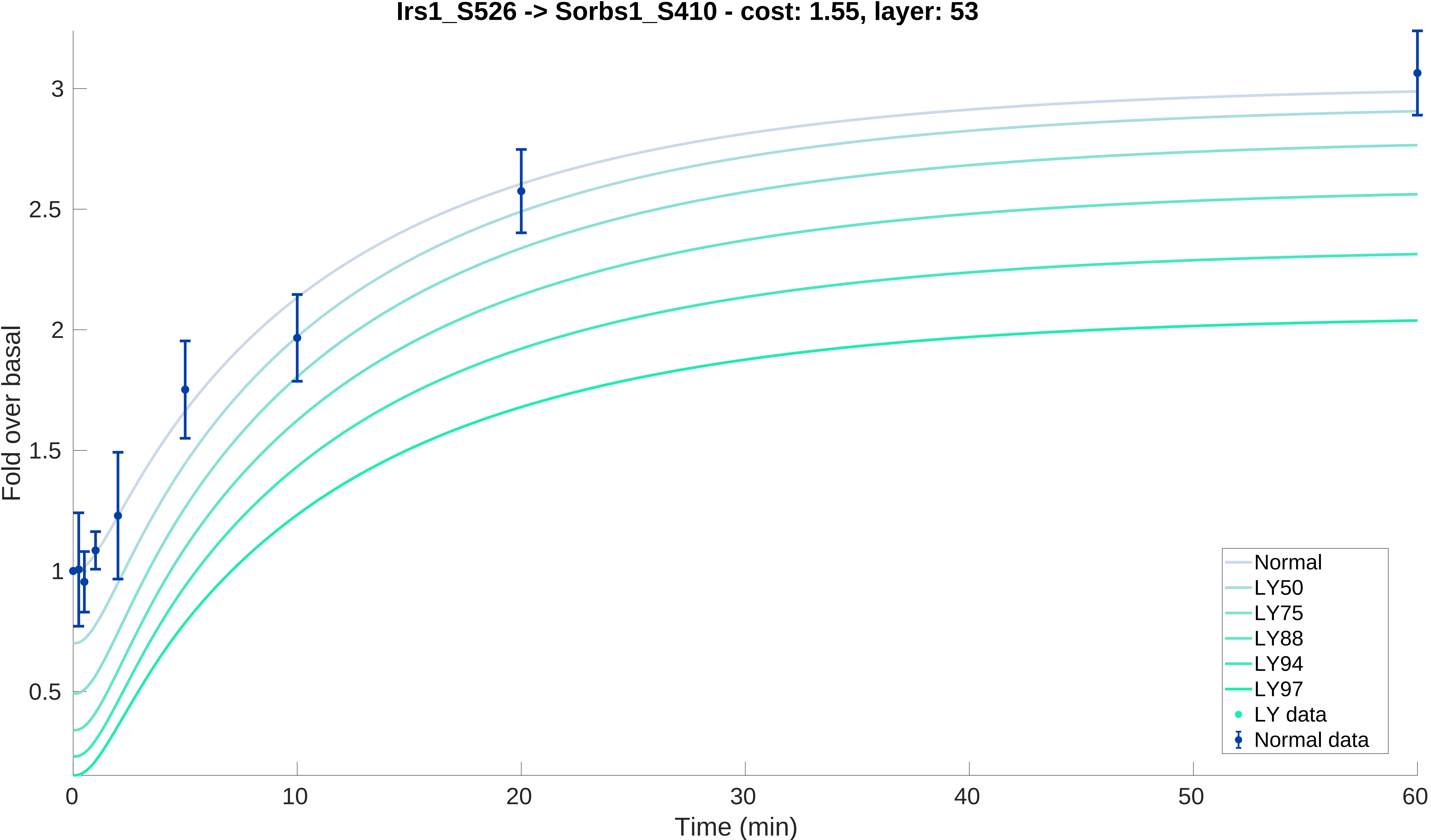

Irs1_S526 -> Sorbs1_S410 - cost: 1.55, layer: 53
Fold over basal
Time (min)
0.5
1
1.5
2
2.5
3
0
10
20
30
40
50
60
Normal
LY50
LY75
LY88
LY94
LY97
LY data
Normal data

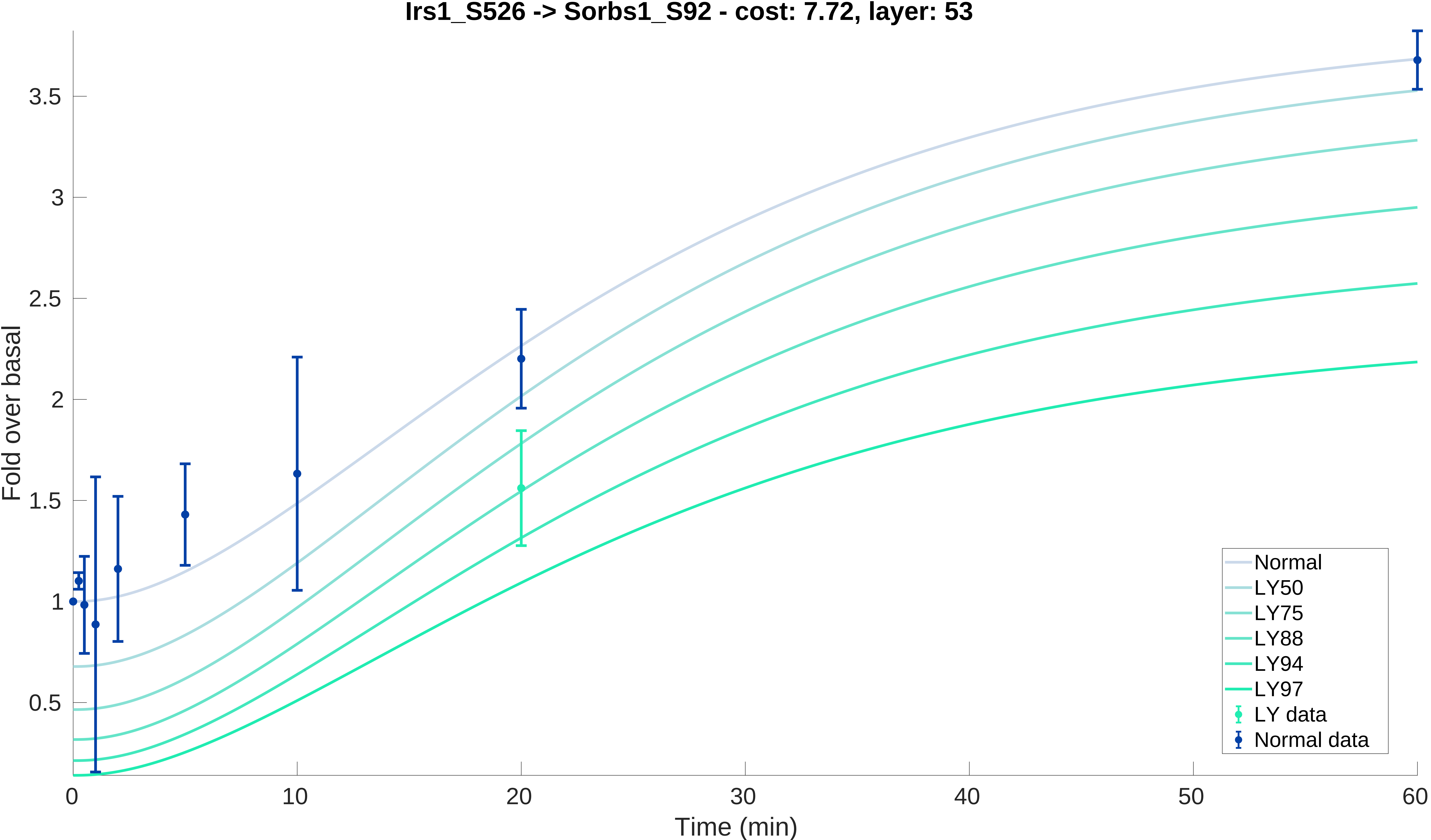

Irs1_S526 -> Sorbs1_S92 - cost: 7.72, layer: 53
Fold over basal
Time (min)
0.5
1
1.5
2
2.5
3
3.5
0
10
20
30
40
50
60
Normal
LY50
LY75
LY88
LY94
LY97
LY data
Normal data

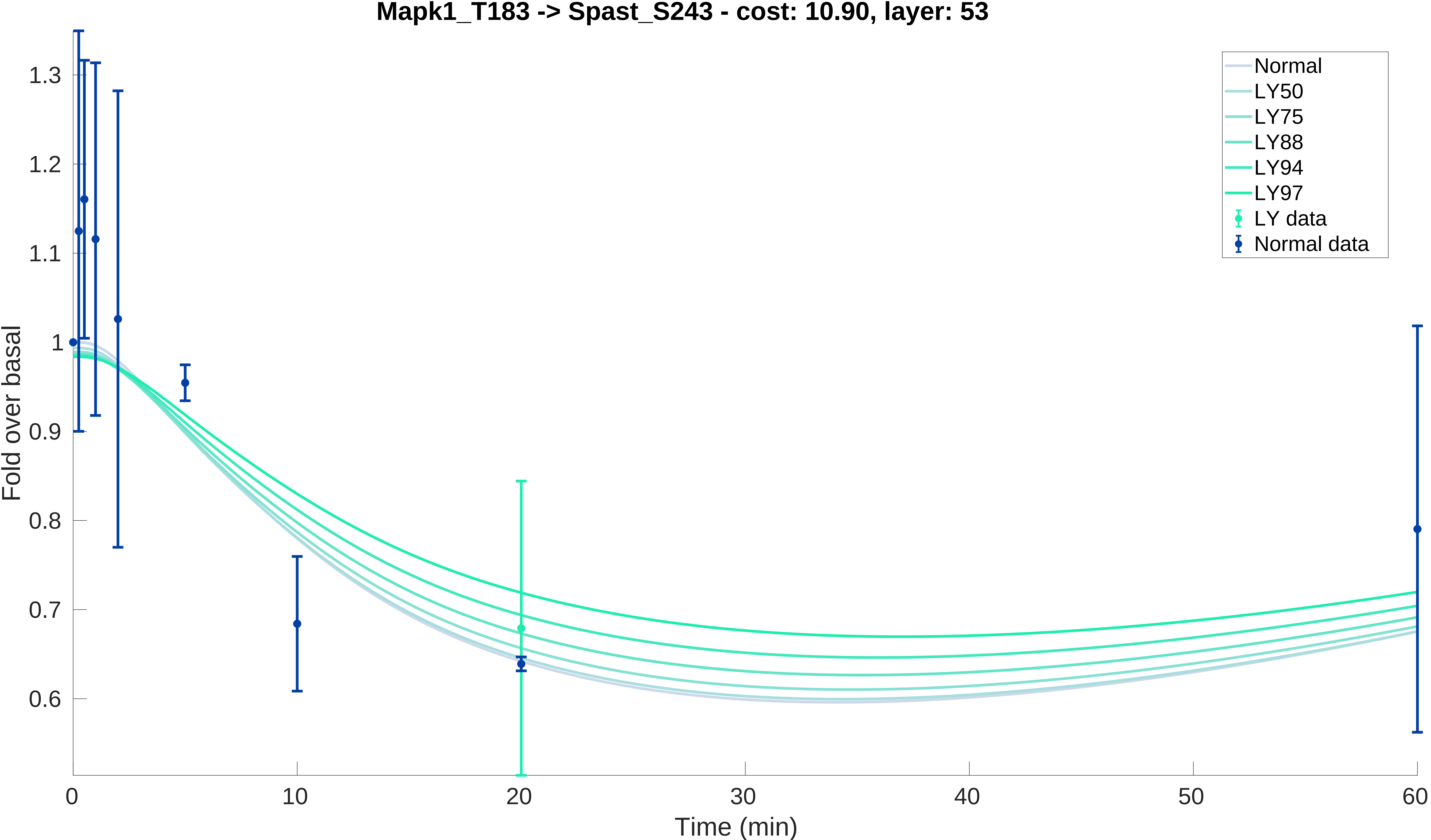

Mapk1_T183 -> Spast_S243 - cost: 10.90, layer: 53
Fold over basal
Time (min)
0
10
20
30
40
50
60
0.6
0.7
0.8
0.9
1
1.1
1.2
1.3
Normal
LY50
LY75
LY88
LY94
LY97
LY data
Normal data

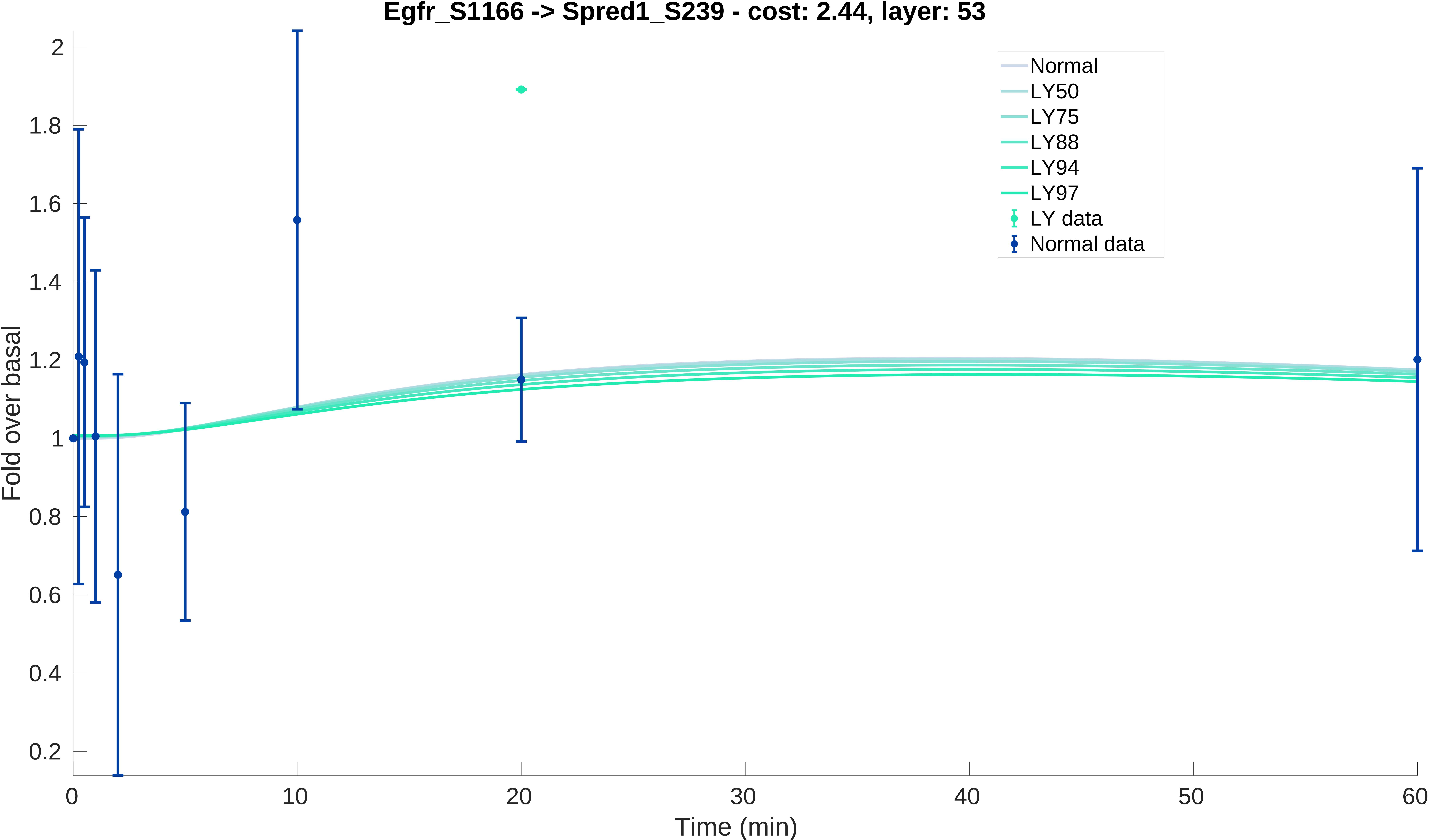

Egfr_S1166 -> Spred1_S239 - cost: 2.44, layer: 53
Fold over basal
Time (min)
0.2
0.4
0.6
0.8
1
1.2
1.4
1.6
1.8
2
0
10
20
30
40
50
60
Normal
LY50
LY75
LY88
LY94
LY97
LY data
Normal data

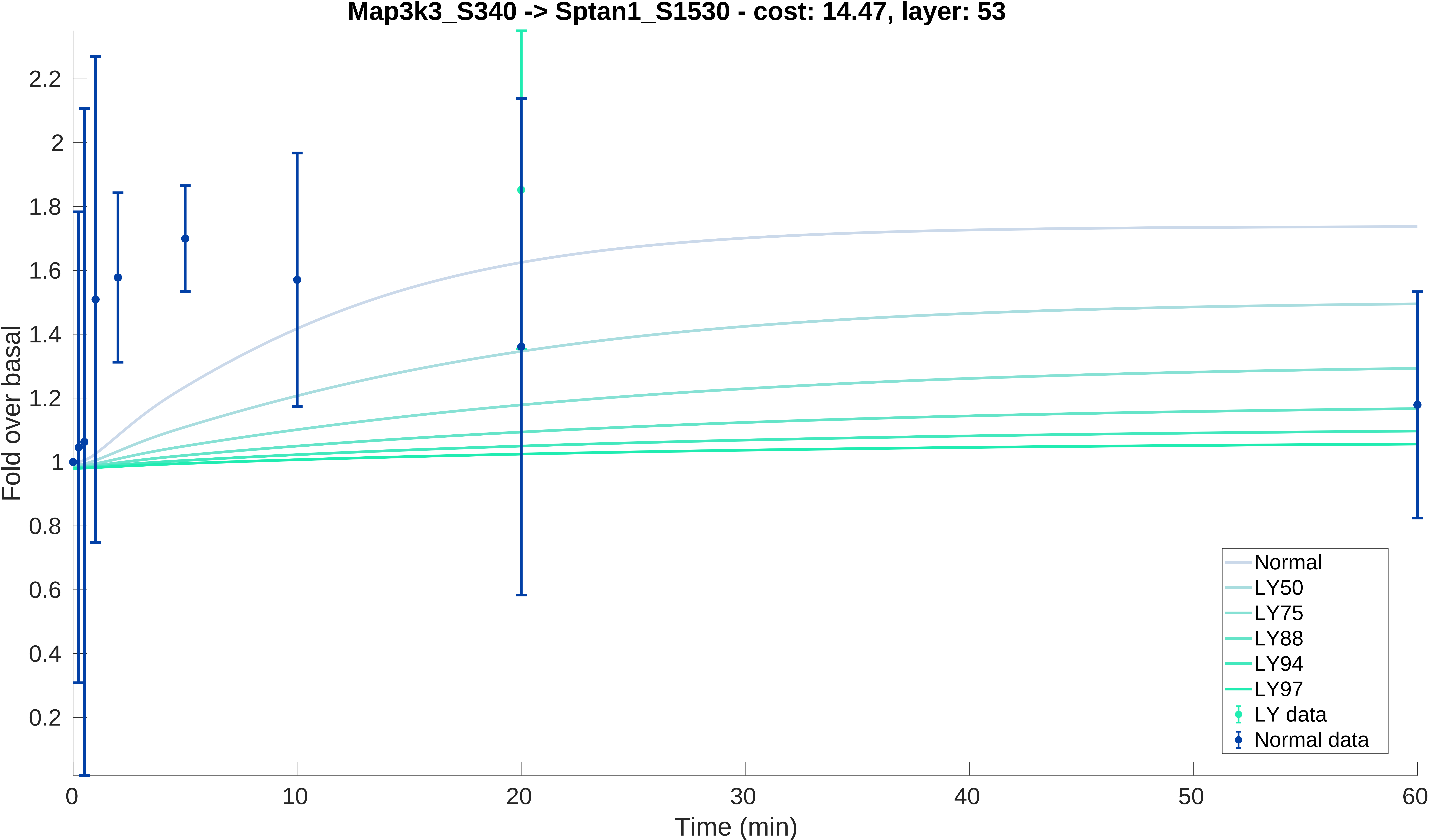

Map3k3_S340 -> Sptan1_S1530 - cost: 14.47, layer: 53
Fold over basal
Time (min)
0.2
0.4
0.6
0.8
1
1.2
1.4
1.6
1.8
2
2.2
0
10
20
30
40
50
60
Normal
LY50
LY75
LY88
LY94
LY97
LY data
Normal data

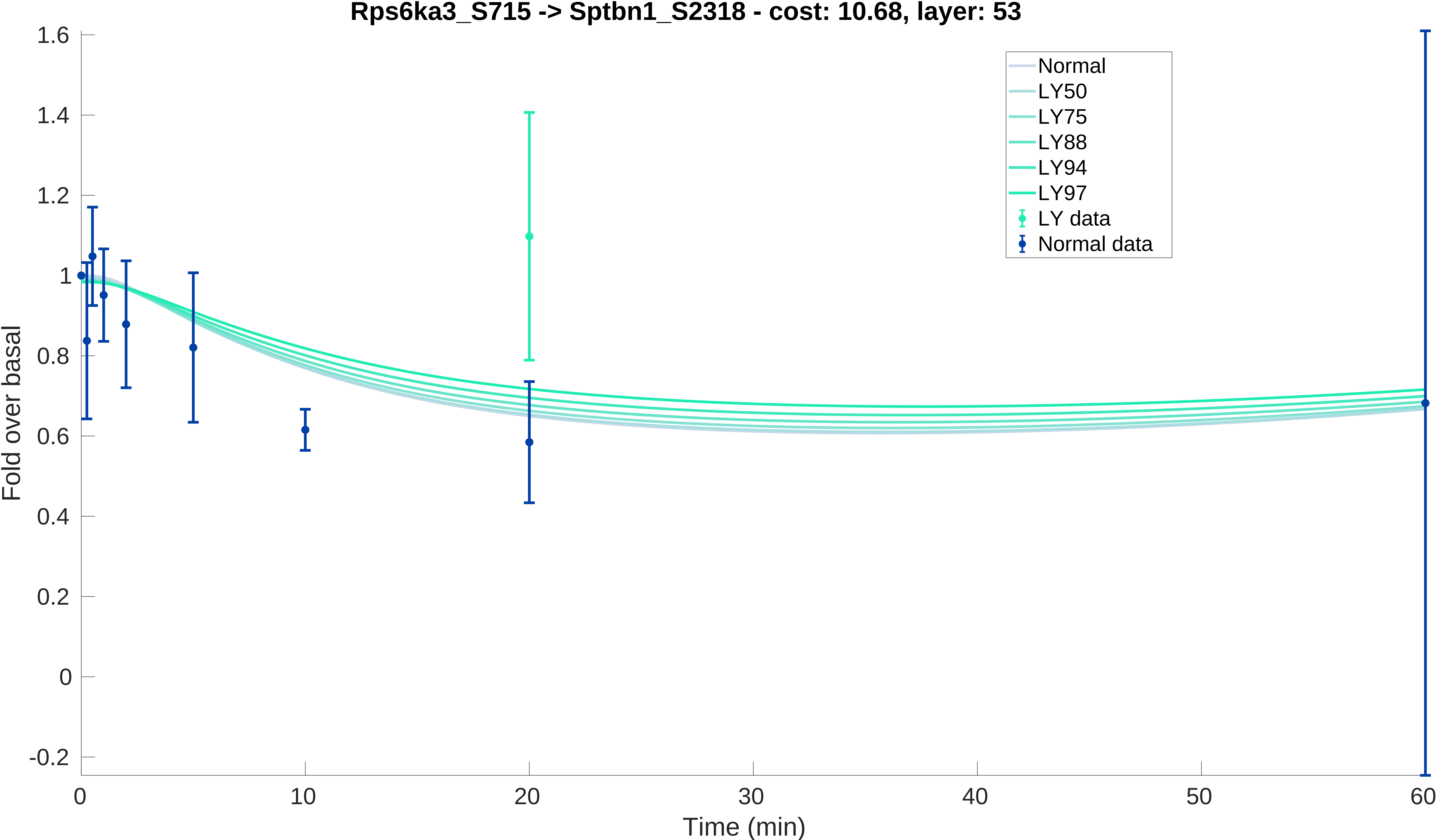

Rps6ka3_S715 -> Sptbn1_S2318 - cost: 10.68, layer: 53
Fold over basal
Time (min)
1.6
1.4
1.2
1
0.8
0.6
0.4
0.2
0
-0.2
0
10
20
30
40
50
60
Normal
LY50
LY75
LY88
LY94
LY97
LY data
Normal data

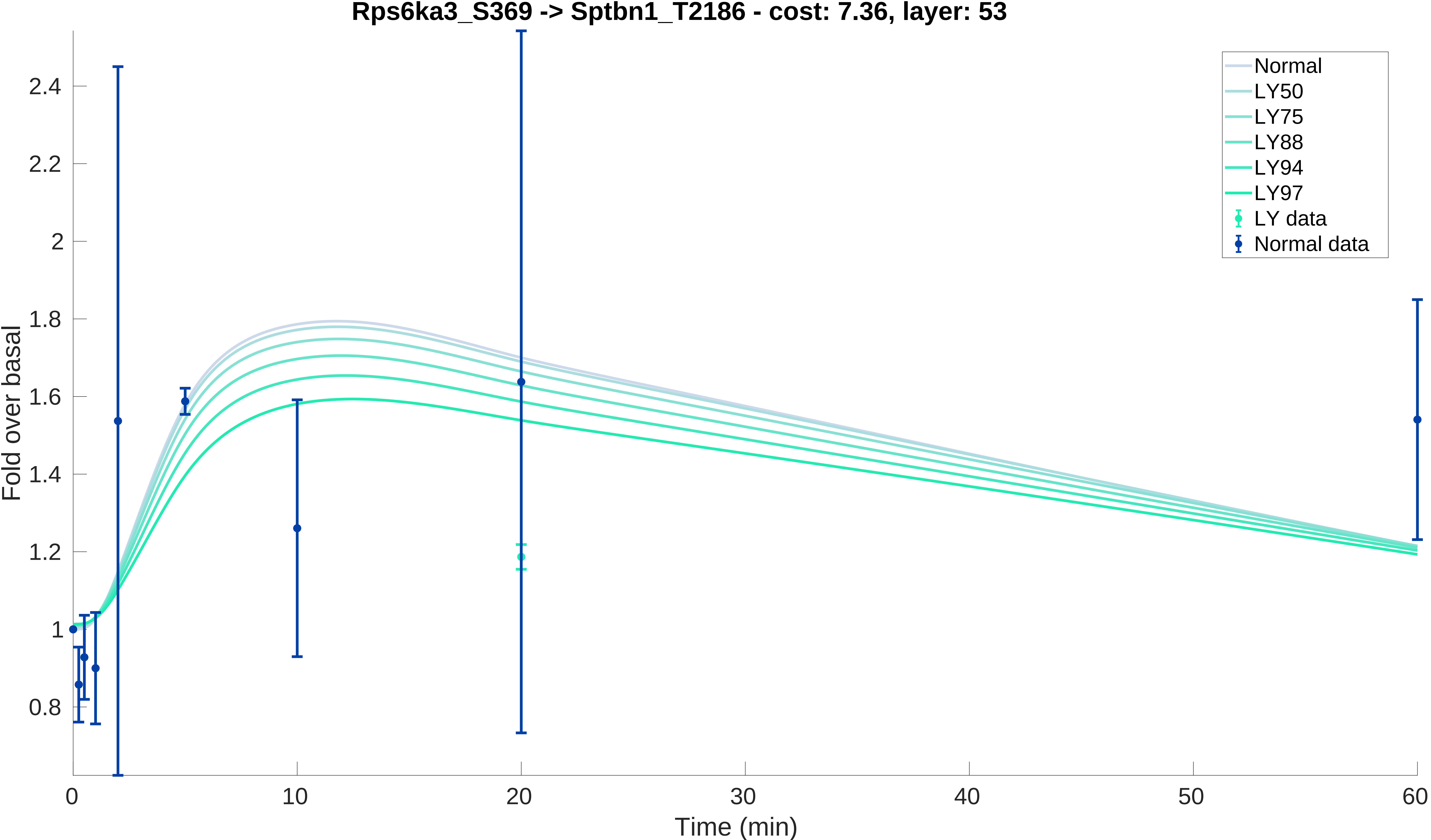

Rps6ka3_S369 -> Sptbn1_T2186 - cost: 7.36, layer: 53
Fold over basal
Time (min)
2.4
2.2
2
1.8
1.6
1.4
1.2
1
0.8
0
10
20
30
40
50
60
Normal
LY50
LY75
LY88
LY94
LY97
LY data
Normal data

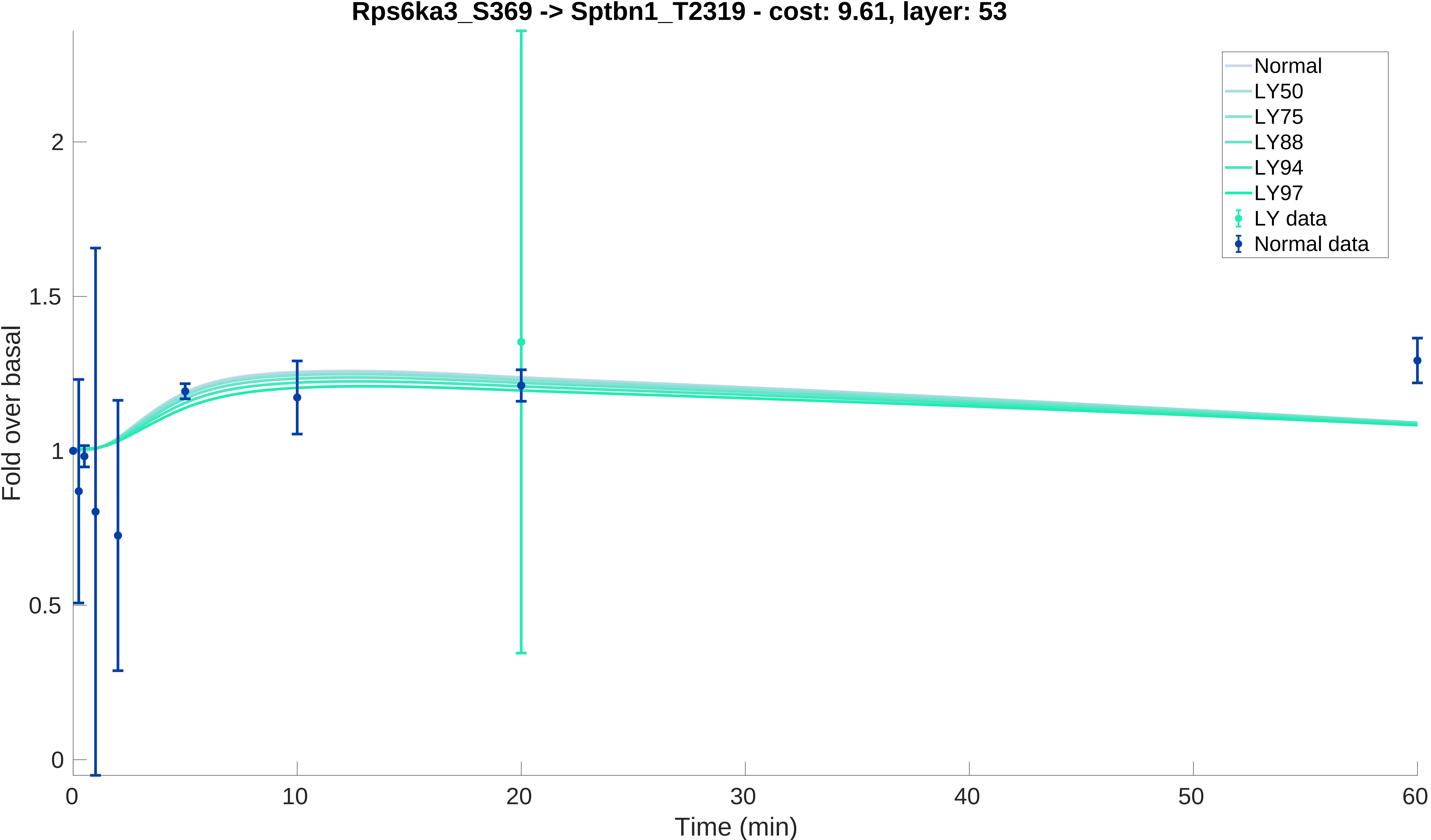

Rps6ka3_S369 -> Sptbn1_T2319 - cost: 9.61, layer: 53
Normal
LY50
LY75
LY88
LY94
LY97
LY data
Normal data
Fold over basal
0
0.5
1
1.5
2
0
10
20
30
40
50
60
Time (min)

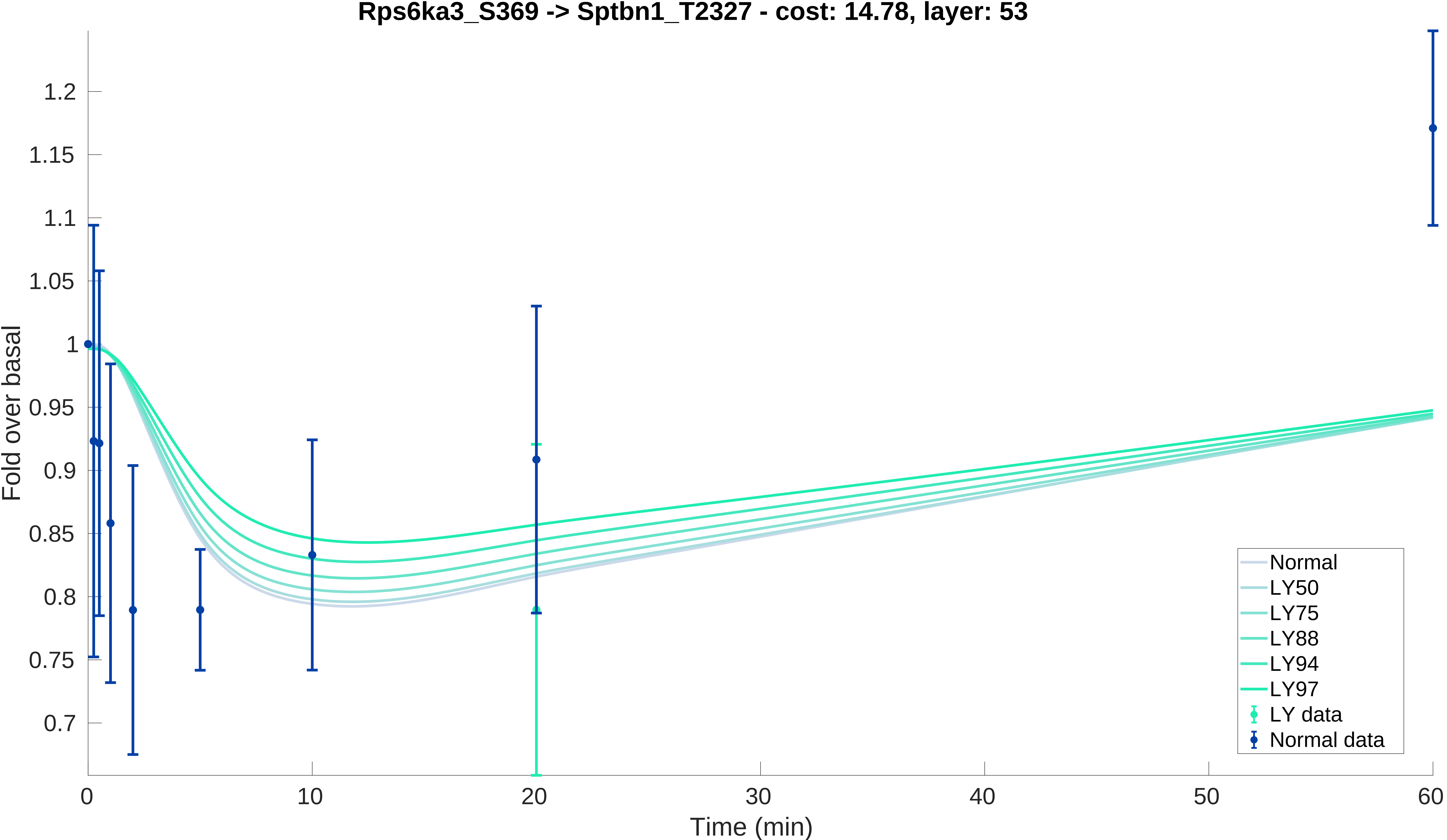

Rps6ka3_S369 -> Sptbn1_T2327 - cost: 14.78, layer: 53
Fold over basal
1.2
1.15
1.1
1.05
1
0.95
0.9
0.85
0.8
0.75
0.7
0
10
20
30
40
50
60
Time (min)
Normal
LY50
LY75
LY88
LY94
LY97
LY data
Normal data

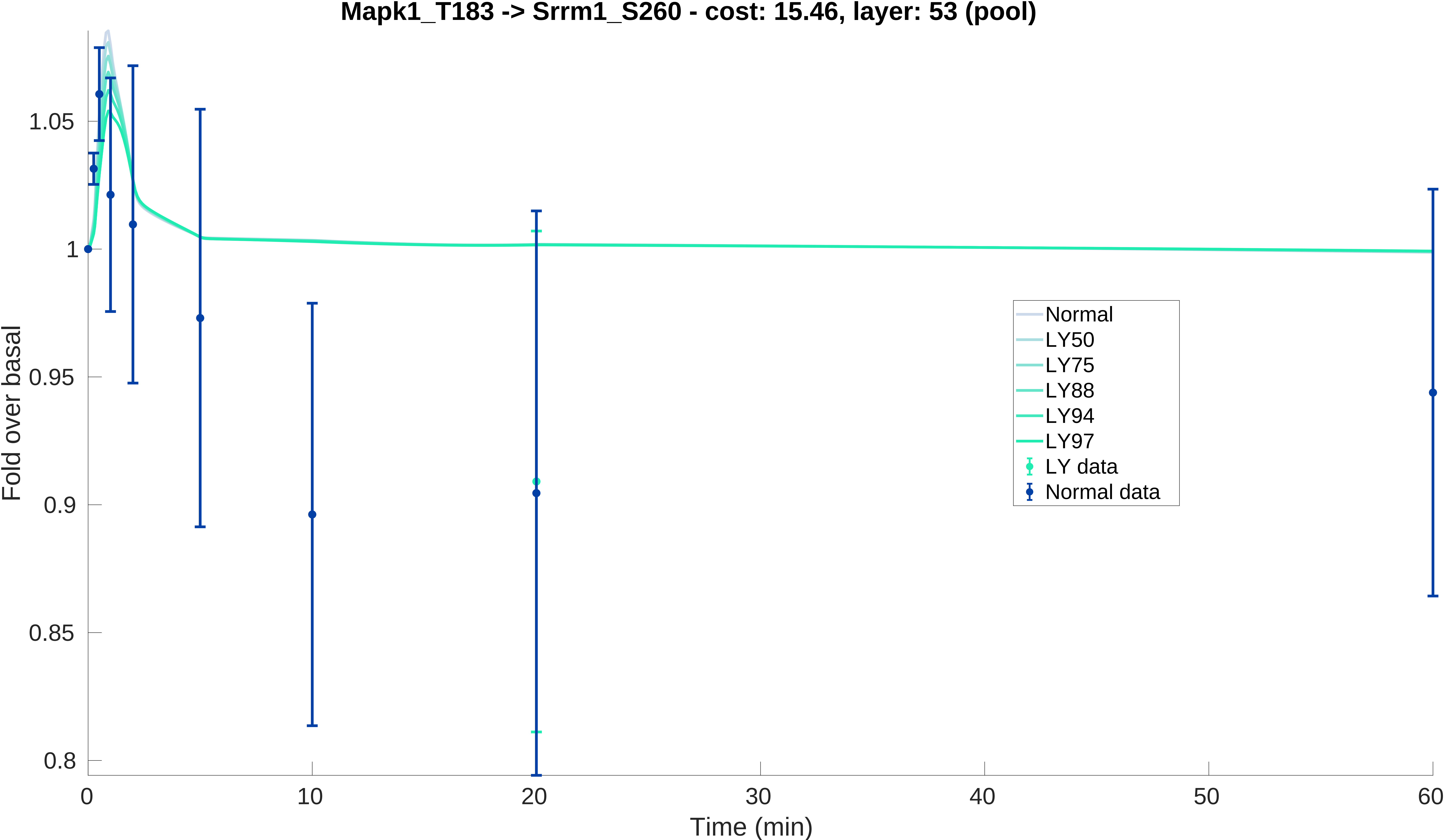
Mapk1_T183 -> Srrm1_S260 - cost: 15.46, layer: 53 (pool)
Fold over basal
1.05
1
0.95
0.9
0.85
0.8
0
10
20
30
40
50
60
Time (min)
Normal
LY50
LY75
LY88
LY94
LY97
LY data
Normal data

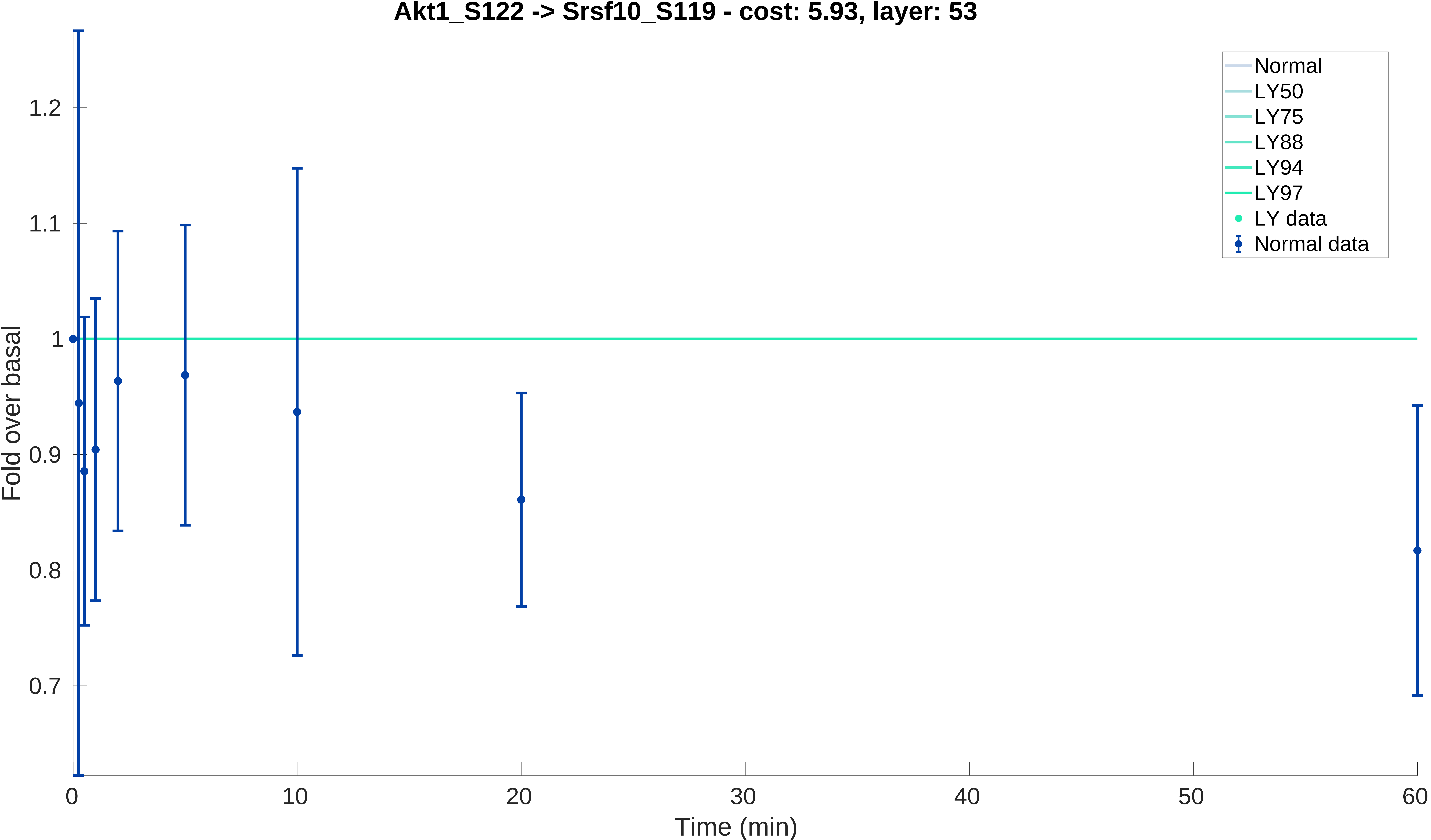
Akt1_S122 -> Srsf10_S119 - cost: 5.93, layer: 53
Fold over basal
Time (min)
0.7
0.8
0.9
1
1.1
1.2
0
10
20
30
40
50
60
Normal
LY50
LY75
LY88
LY94
LY97
LY data
Normal data

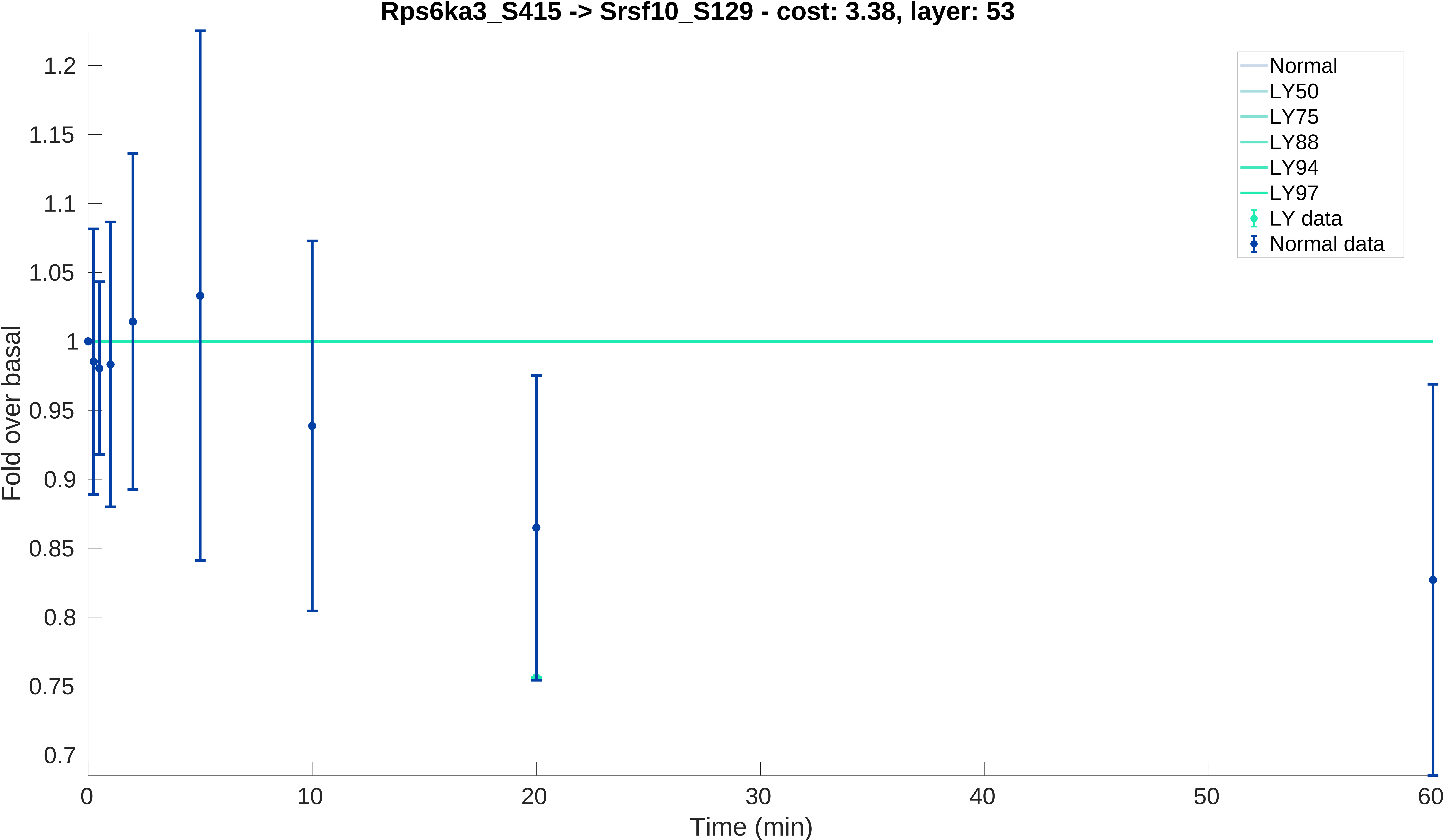

Rps6ka3_S415 -> Srsf10_S129 - cost: 3.38, layer: 53
Fold over basal
Time (min)
1.2
1.15
1.1
1.05
1
0.95
0.9
0.85
0.8
0.75
0.7
0
10
20
30
40
50
60
Normal
LY50
LY75
LY88
LY94
LY97
LY data
Normal data

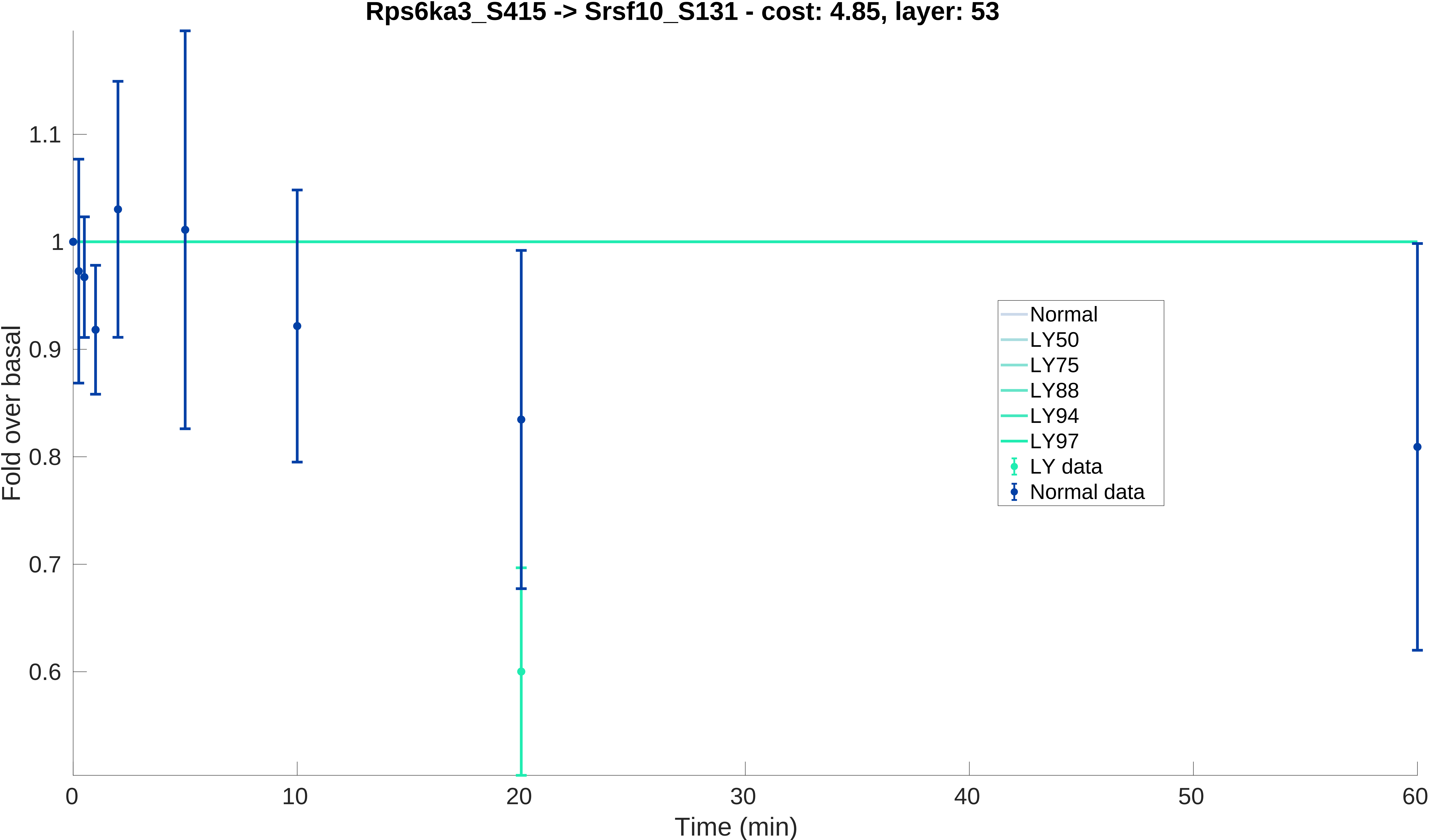
Rps6ka3_S415 -> Srsf10_S131 - cost: 4.85, layer: 53
Fold over basal
Time (min)
0
10
20
30
40
50
60
0.6
0.7
0.8
0.9
1
1.1
Normal
LY50
LY75
LY88
LY94
LY97
LY data
Normal data

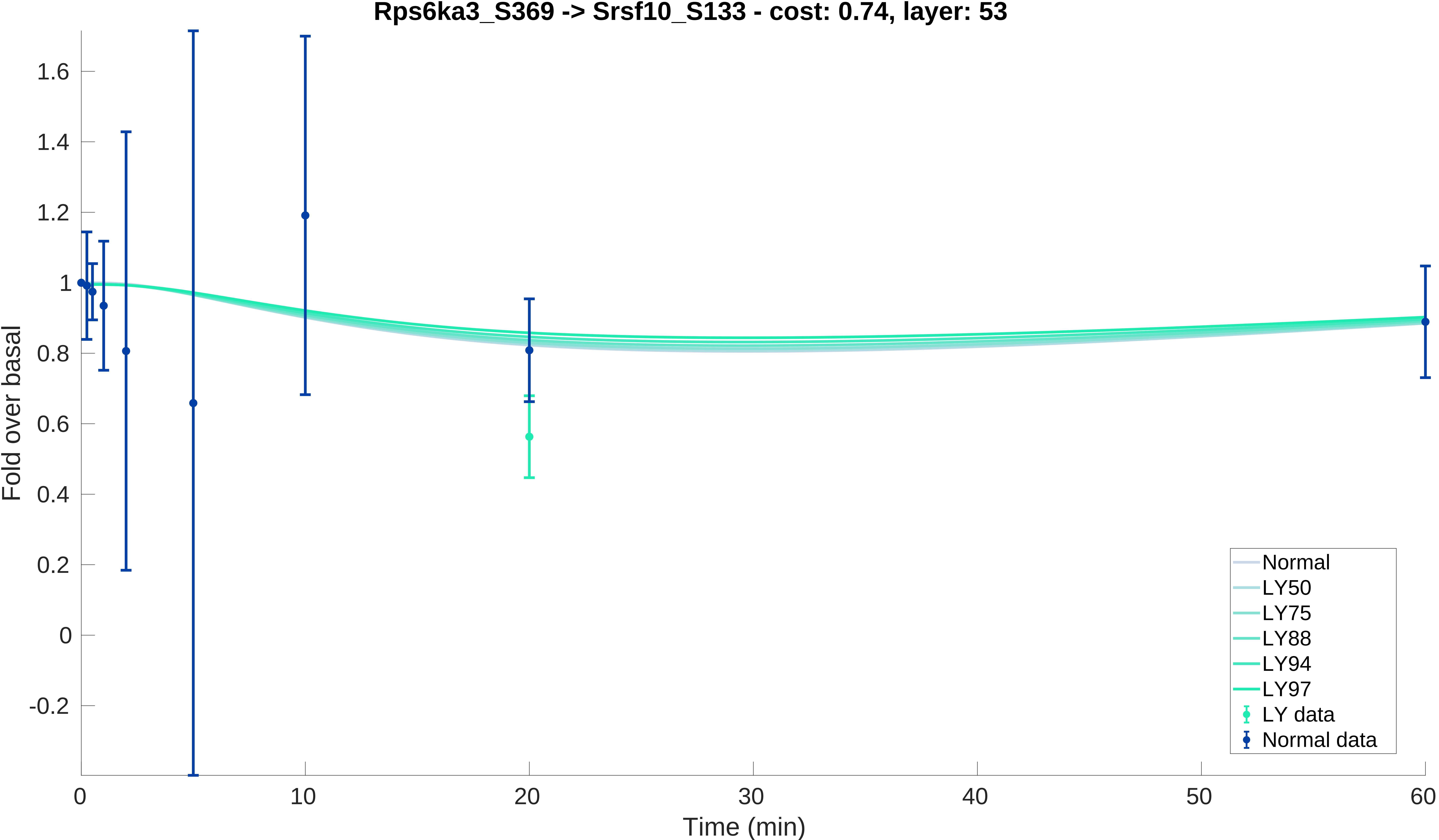

Rps6ka3_S369 -> Srsf10_S133 - cost: 0.74, layer: 53
Fold over basal
1.6
1.4
1.2
1
0.8
0.6
0.4
0.2
0
-0.2
0
10
20
30
40
50
60
Time (min)
Normal
LY50
LY75
LY88
LY94
LY97
LY data
Normal data

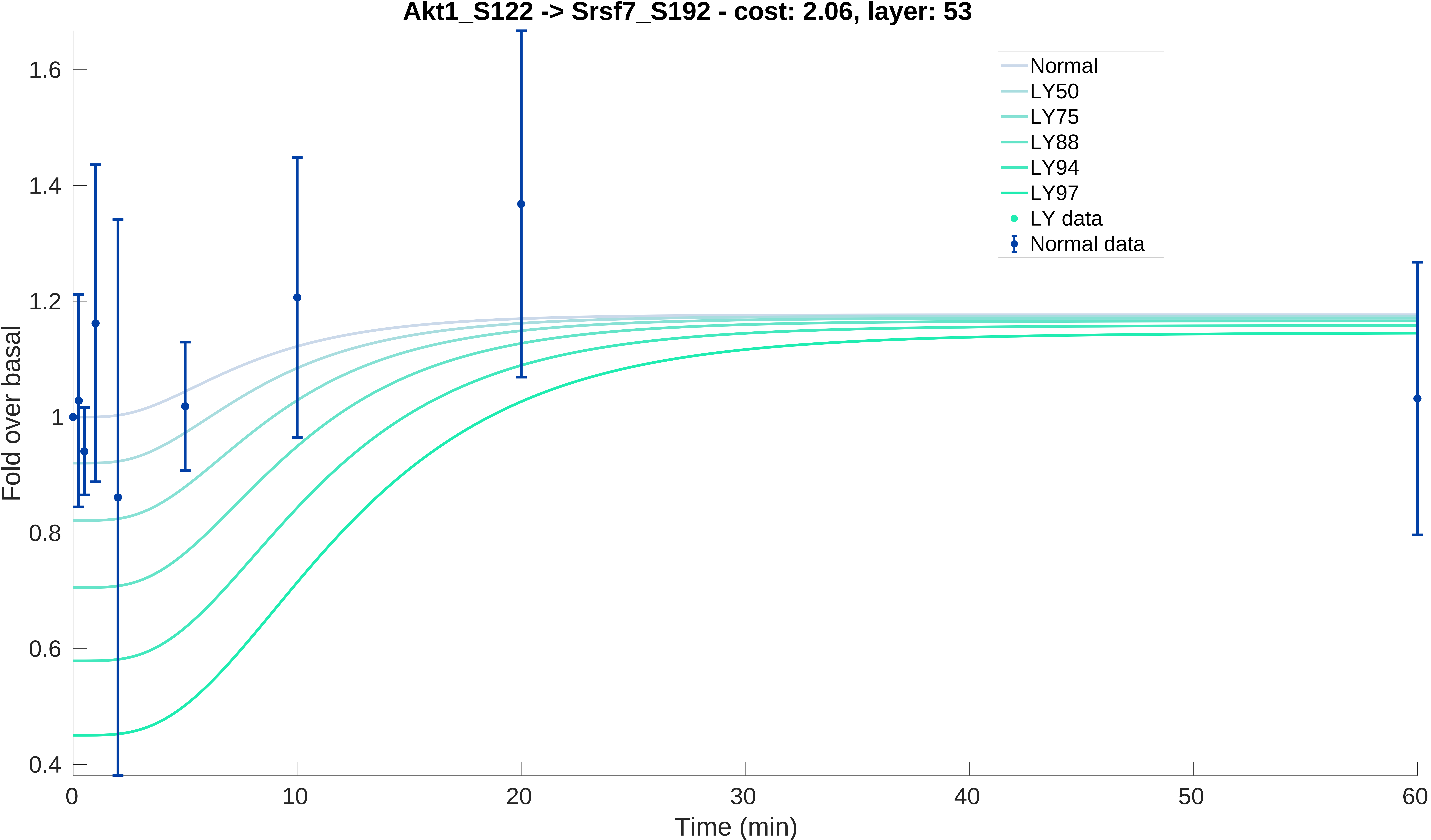

Akt1_S122 -> Srsf7_S192 - cost: 2.06, layer: 53
Normal
LY50
LY75
LY88
LY94
LY97
LY data
Normal data
Fold over basal
Time (min)
0.4
0.6
0.8
1
1.2
1.4
1.6
0
10
20
30
40
50
60

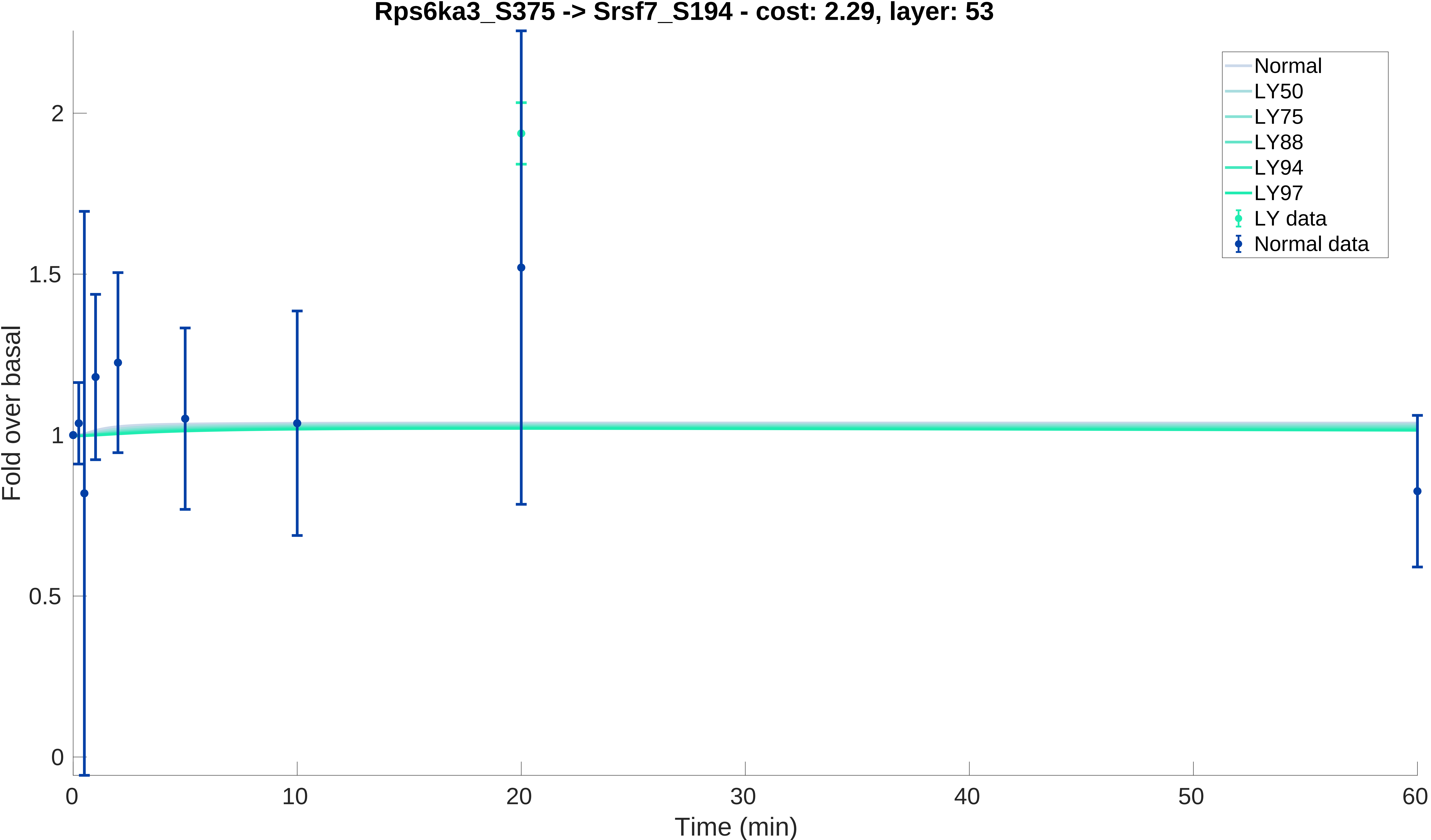

Rps6ka3_S375 -> Srsf7_S194 - cost: 2.29, layer: 53
Fold over basal
Time (min)
0
0.5
1
1.5
2
0
10
20
30
40
50
60
Normal
LY50
LY75
LY88
LY94
LY97
LY data
Normal data

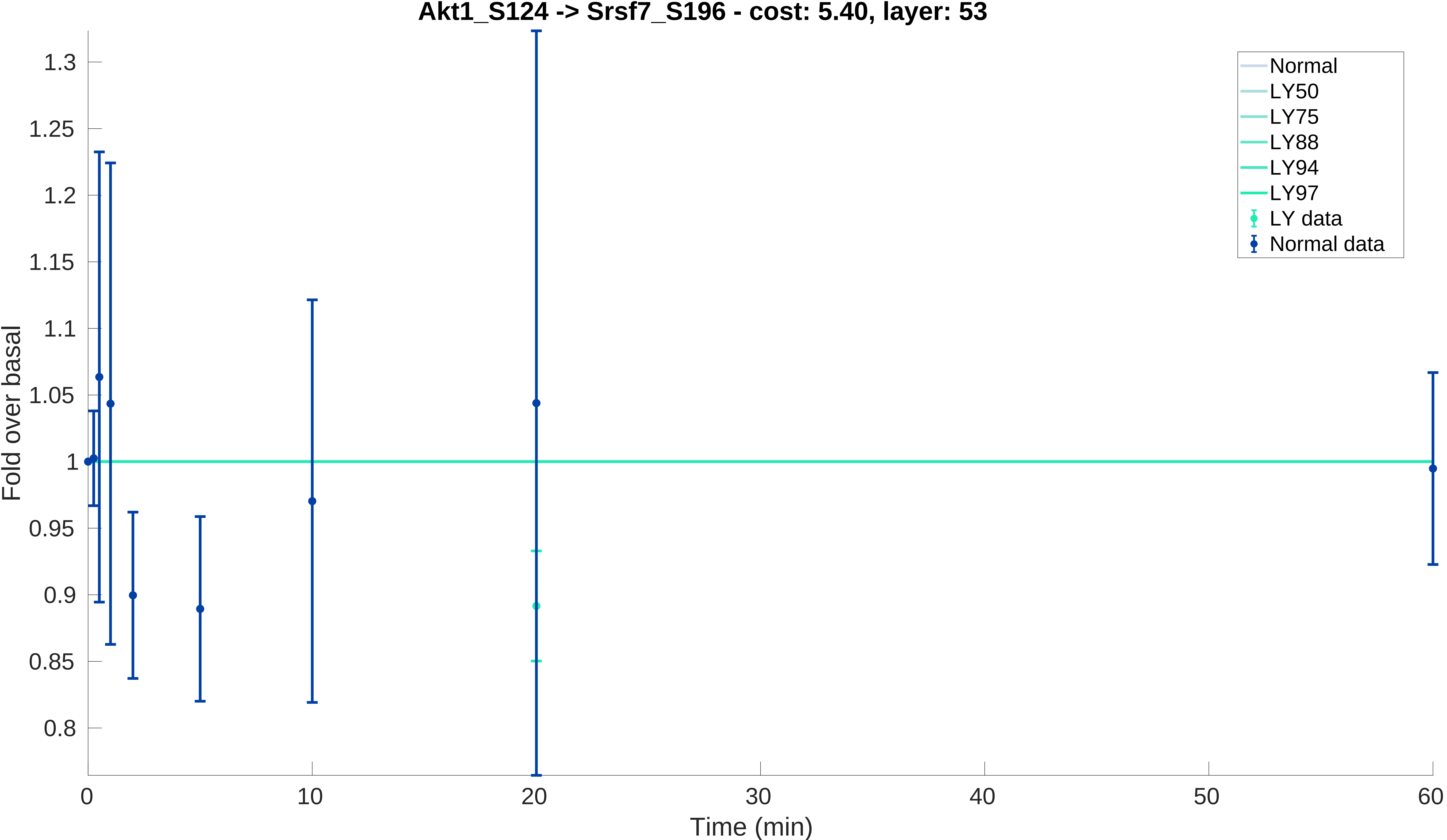

Akt1_S124 -> Srsf7_S196 - cost: 5.40, layer: 53
1.3
1.25
1.2
1.15
1.1
1.05
1
0.95
0.9
0.85
0.8
Fold over basal
0
10
20
30
40
50
60
Time (min)
Normal
LY50
LY75
LY88
LY94
LY97
LY data
Normal data

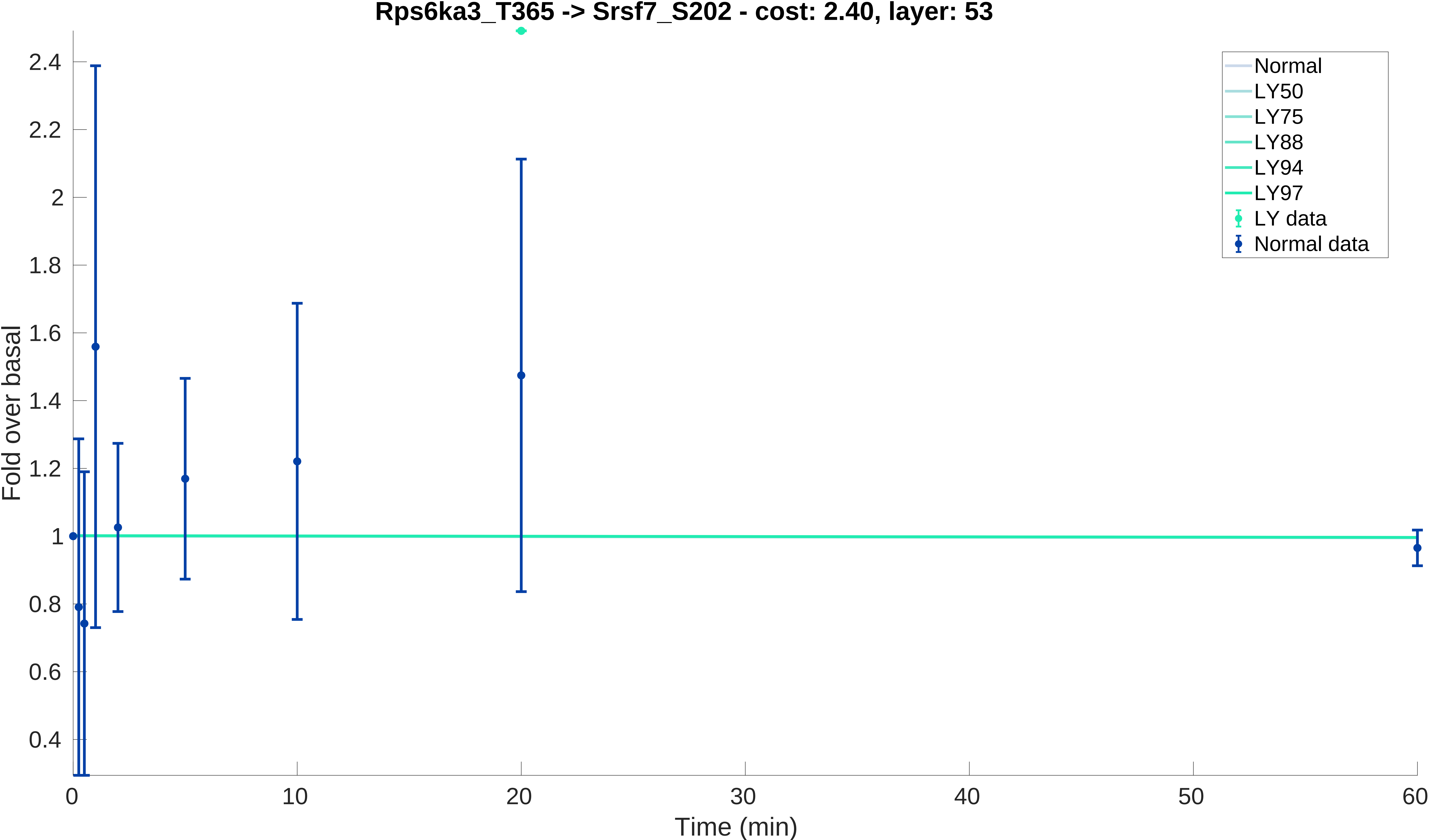

Rps6ka3_T365 -> Srsf7_S202 - cost: 2.40, layer: 53
Fold over basal
Time (min)
0.4
0.6
0.8
1
1.2
1.4
1.6
1.8
2
2.2
2.4
0
10
20
30
40
50
60
Normal
LY50
LY75
LY88
LY94
LY97
LY data
Normal data

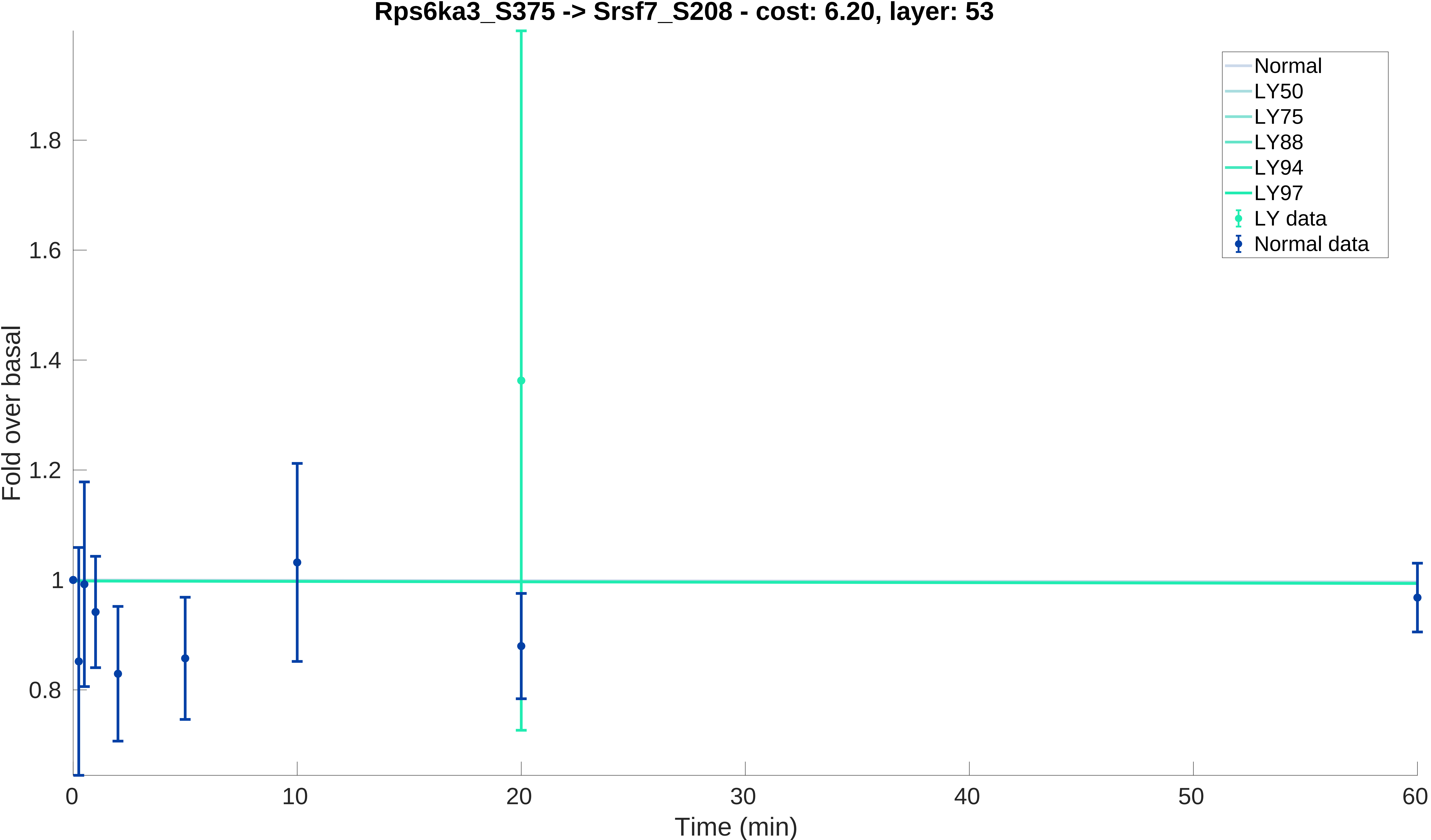

Rps6ka3_S375 -> Srsf7_S208 - cost: 6.20, layer: 53
Fold over basal
Time (min)
0.8
1
1.2
1.4
1.6
1.8
0
10
20
30
40
50
60
Normal
LY50
LY75
LY88
LY94
LY97
LY data
Normal data

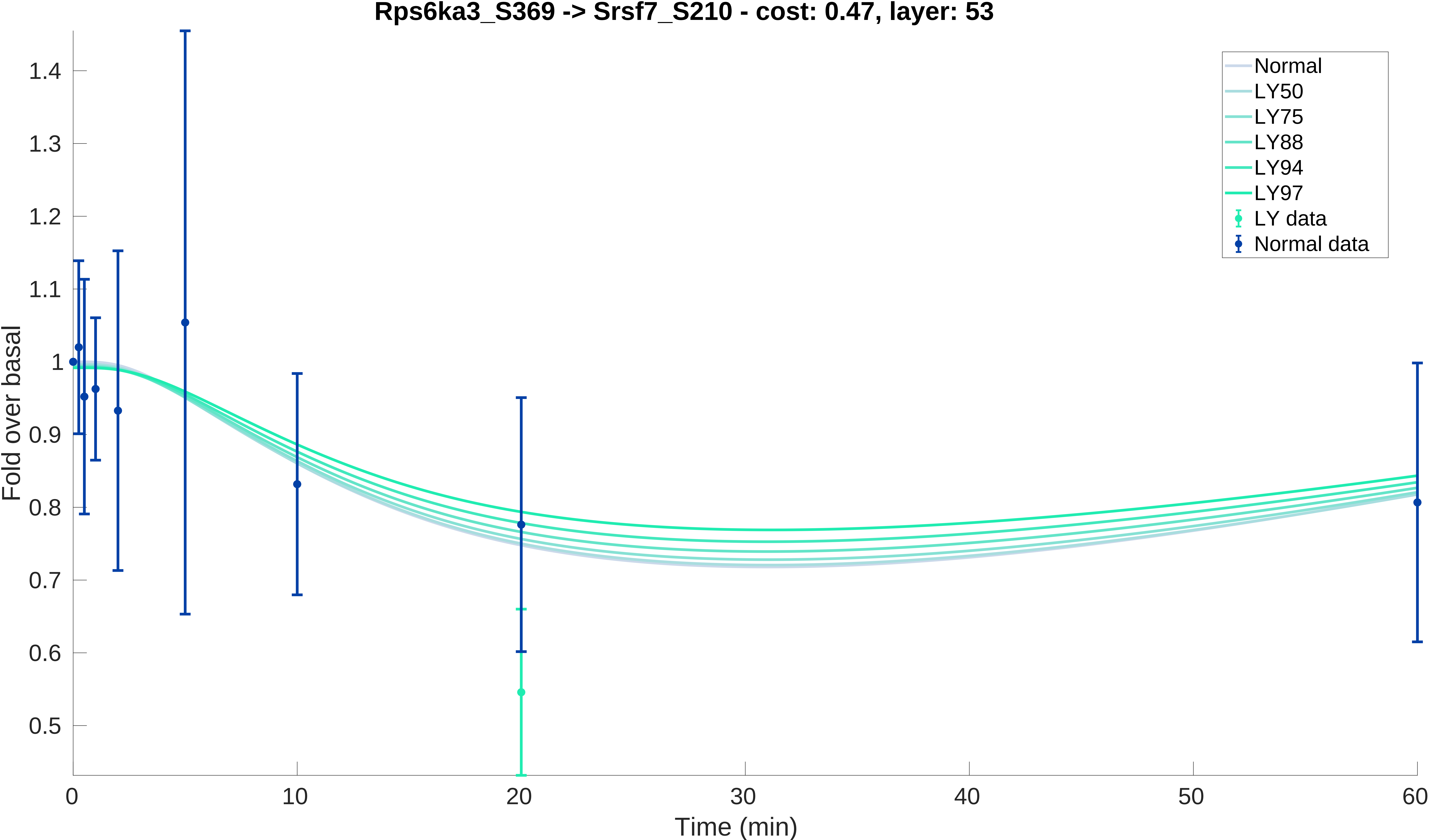

Rps6ka3_S369 -> Srsf7_S210 - cost: 0.47, layer: 53
Fold over basal
1.4
1.3
1.2
1.1
1
0.9
0.8
0.7
0.6
0.5
0
10
20
30
40
50
60
Time (min)
Normal
LY50
LY75
LY88
LY94
LY97
LY data
Normal data

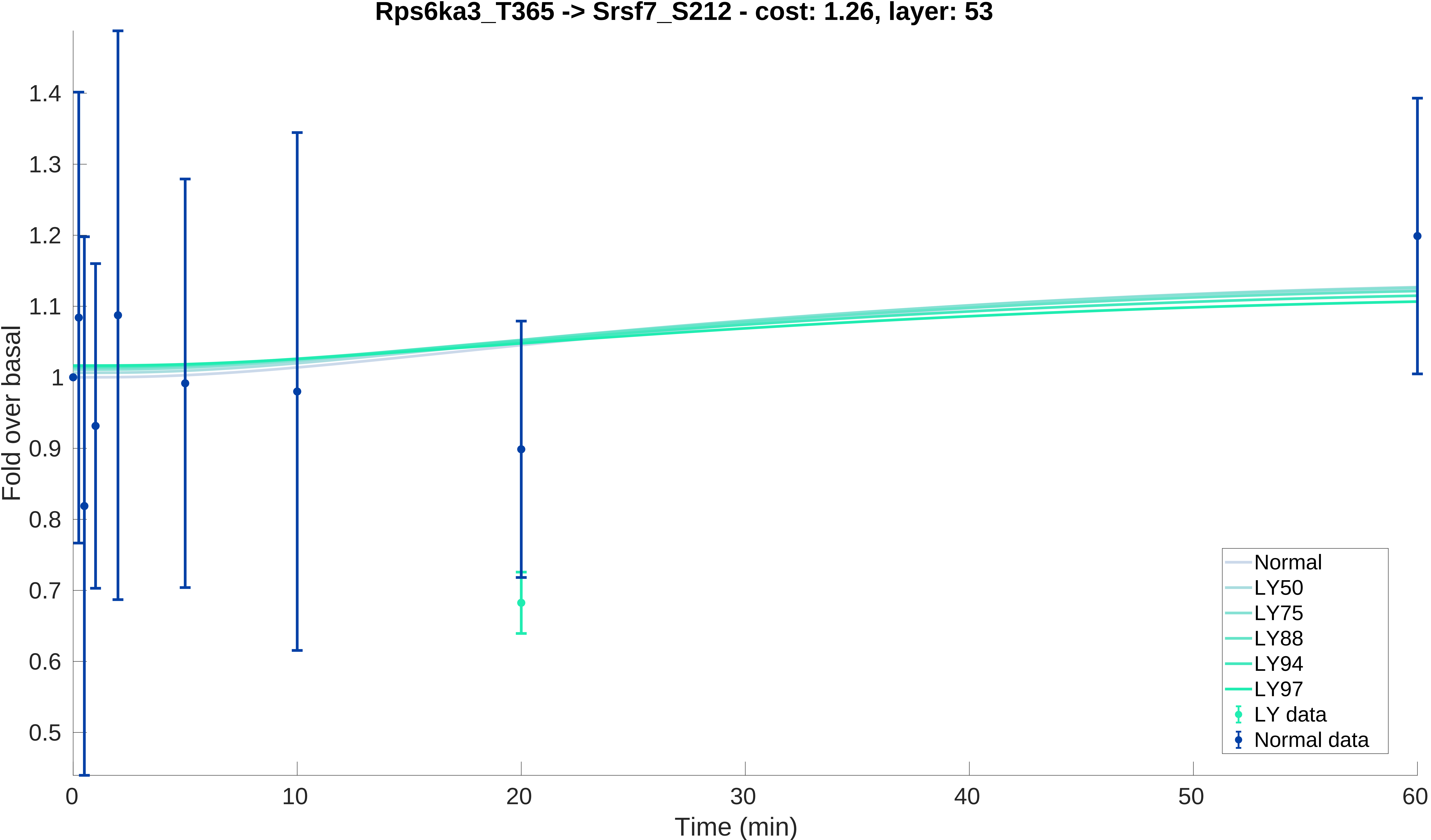

Rps6ka3_T365 -> Srsf7_S212 - cost: 1.26, layer: 53
Fold over basal
Time (min)
1.4
1.3
1.2
1.1
1
0.9
0.8
0.7
0.6
0.5
0
10
20
30
40
50
60
Normal
LY50
LY75
LY88
LY94
LY97
LY data
Normal data

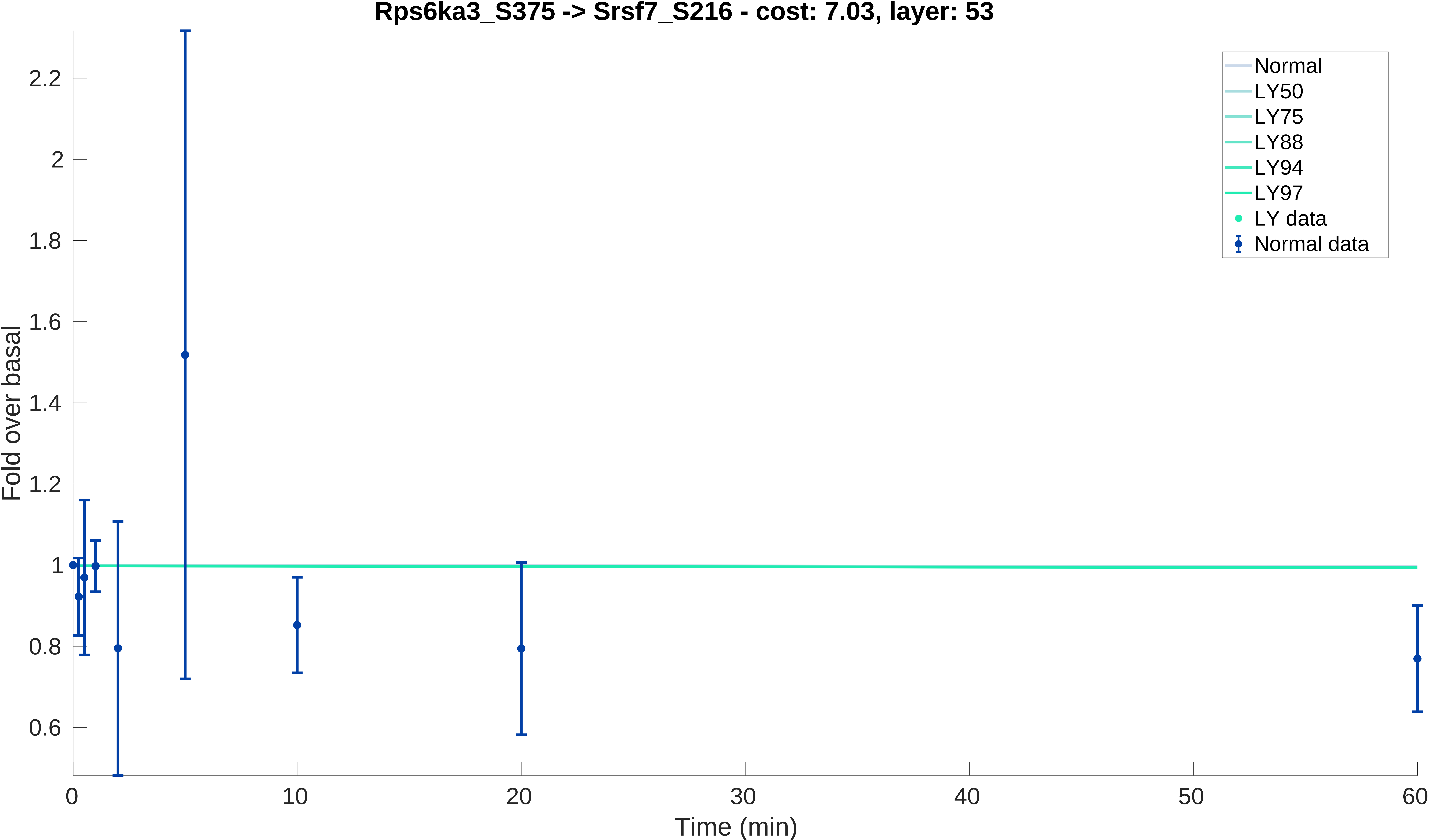

Rps6ka3_S375 -> Srsf7_S216 - cost: 7.03, layer: 53
Fold over basal
Time (min)
0.6
0.8
1
1.2
1.4
1.6
1.8
2
2.2
0
10
20
30
40
50
60
Normal
LY50
LY75
LY88
LY94
LY97
LY data
Normal data

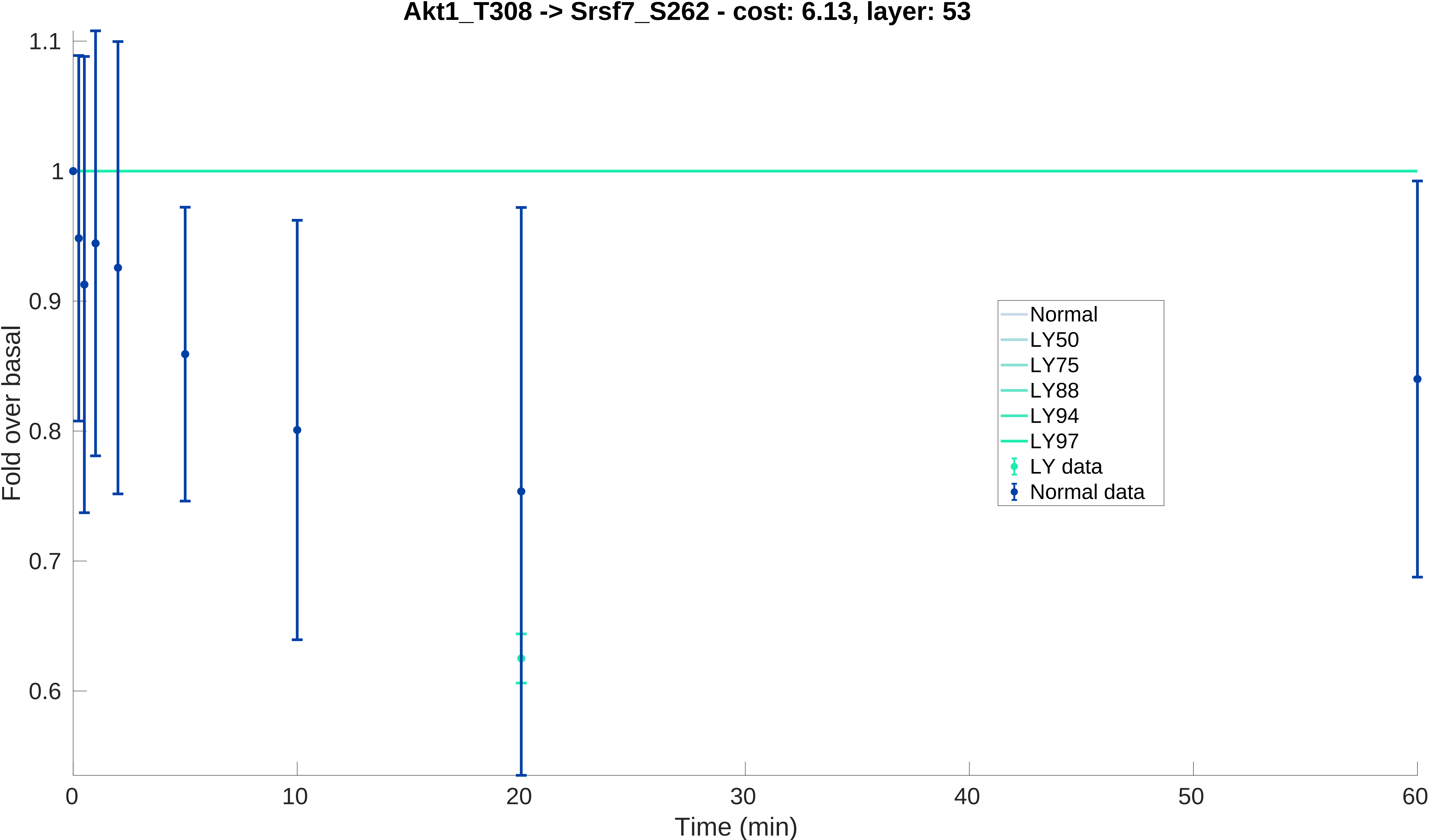

Akt1_T308 -> Srsf7_S262 - cost: 6.13, layer: 53
Fold over basal
1.1
1
0.9
0.8
0.7
0.6
0
10
20
30
40
50
60
Time (min)
Normal
LY50
LY75
LY88
LY94
LY97
LY data
Normal data

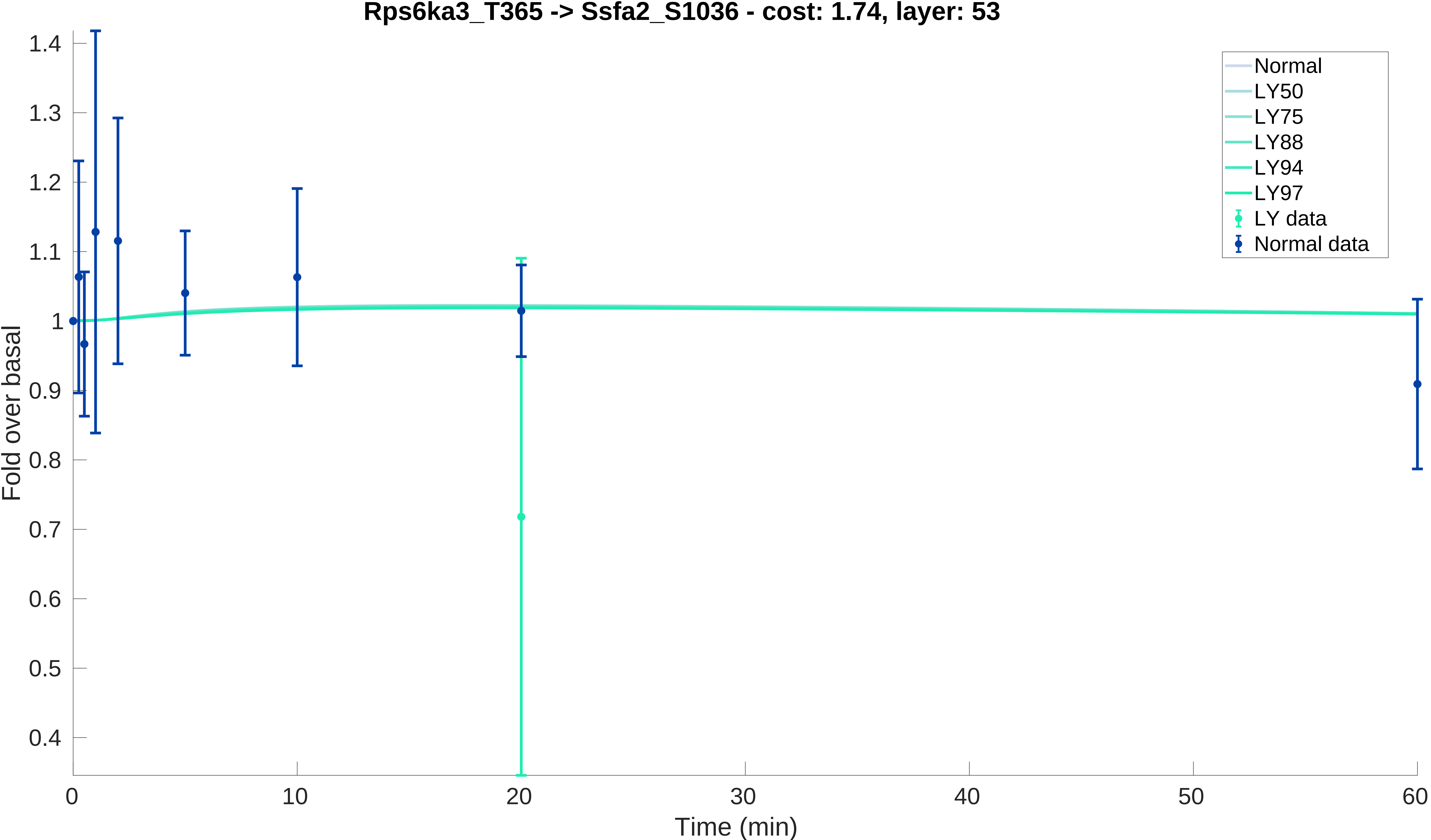

Rps6ka3_T365 -> Ssfa2_S1036 - cost: 1.74, layer: 53
Fold over basal
Time (min)
1.4
1.3
1.2
1.1
1
0.9
0.8
0.7
0.6
0.5
0.4
0
10
20
30
40
50
60
Normal
LY50
LY75
LY88
LY94
LY97
LY data
Normal data

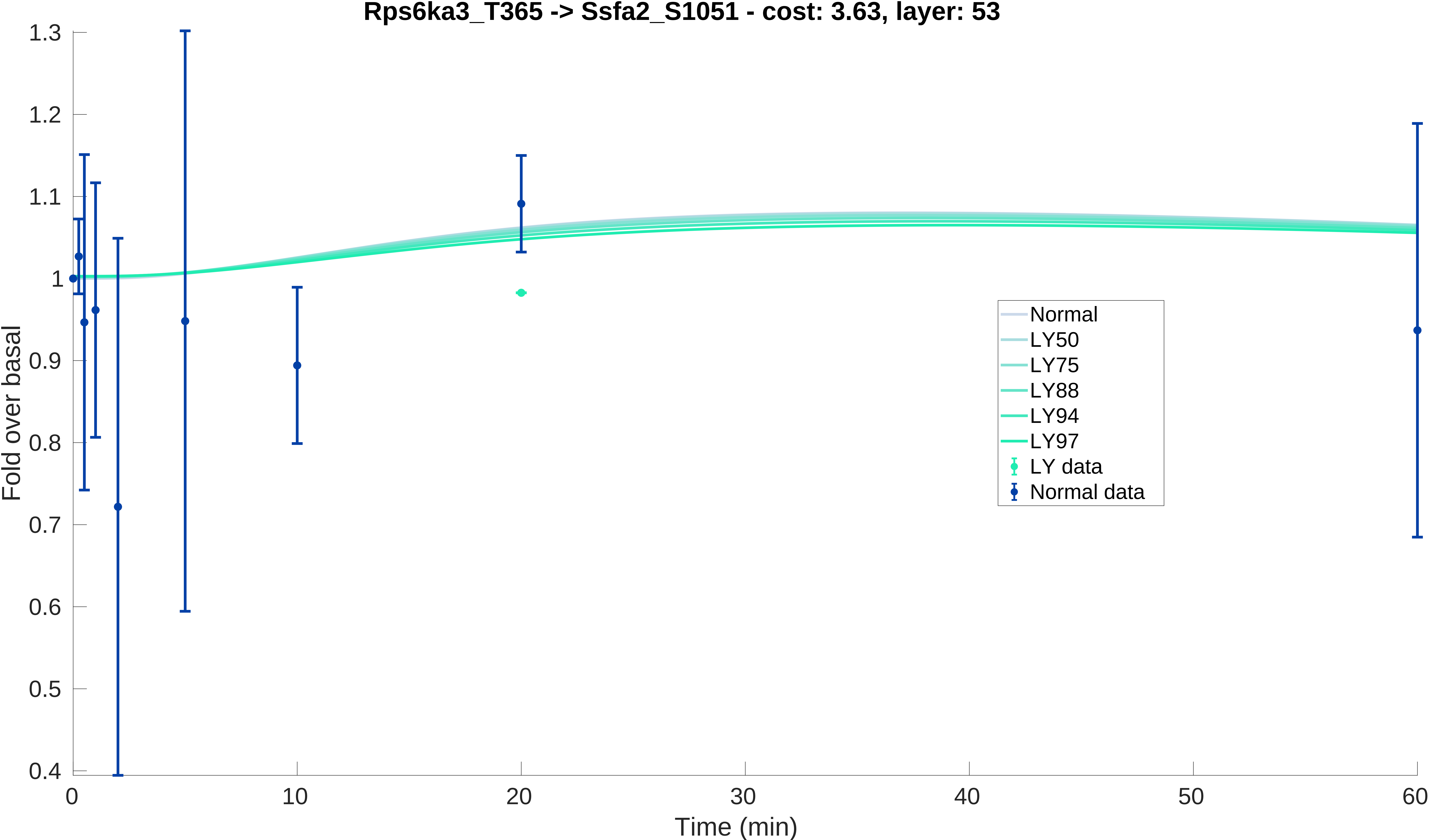

Rps6ka3_T365 -> Ssfa2_S1051 - cost: 3.63, layer: 53
Fold over basal
Time (min)
1.3
1.2
1.1
1
0.9
0.8
0.7
0.6
0.5
0.4
0
10
20
30
40
50
60
Normal
LY50
LY75
LY88
LY94
LY97
LY data
Normal data

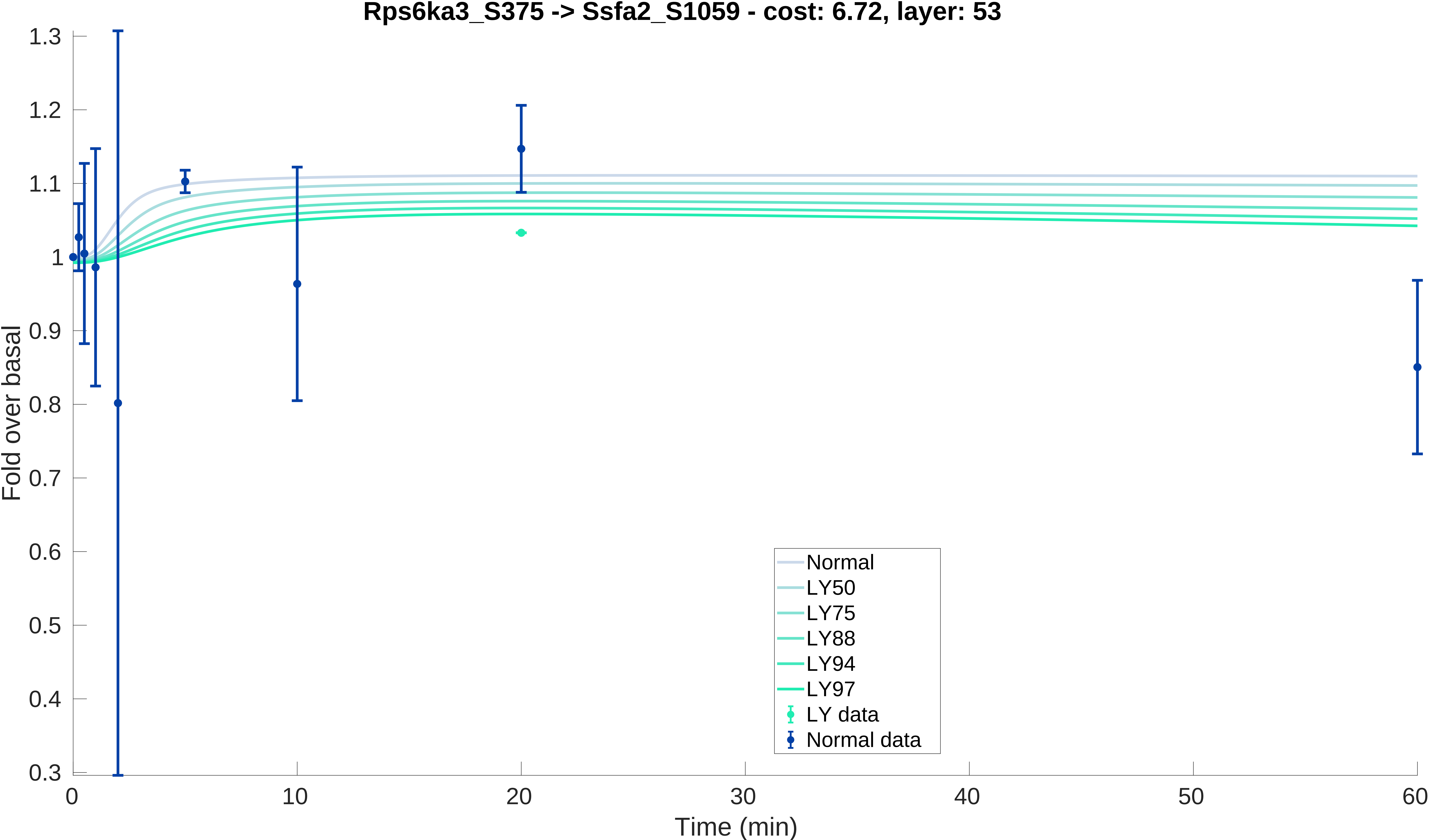

Rps6ka3_S375 -> Ssfa2_S1059 - cost: 6.72, layer: 53
Fold over basal
1.3
1.2
1.1
1
0.9
0.8
0.7
0.6
0.5
0.4
0.3
0
10
20
30
40
50
60
Time (min)
Normal
LY50
LY75
LY88
LY94
LY97
LY data
Normal data

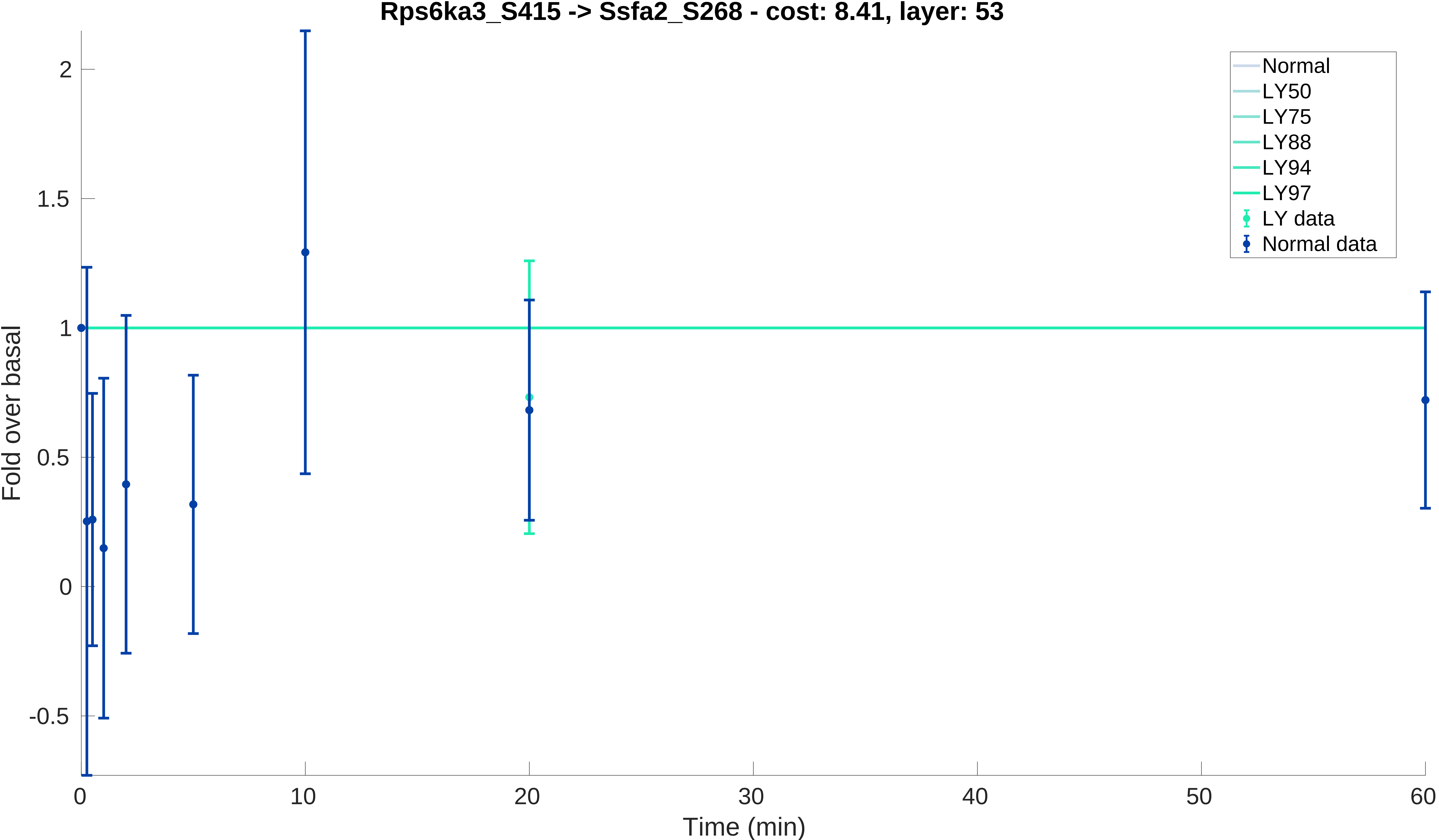

Rps6ka3_S415 -> Ssfa2_S268 - cost: 8.41, layer: 53
Normal
LY50
LY75
LY88
LY94
LY97
LY data
Normal data
Fold over basal
Time (min)
2
1.5
1
0.5
0
-0.5
0
10
20
30
40
50
60

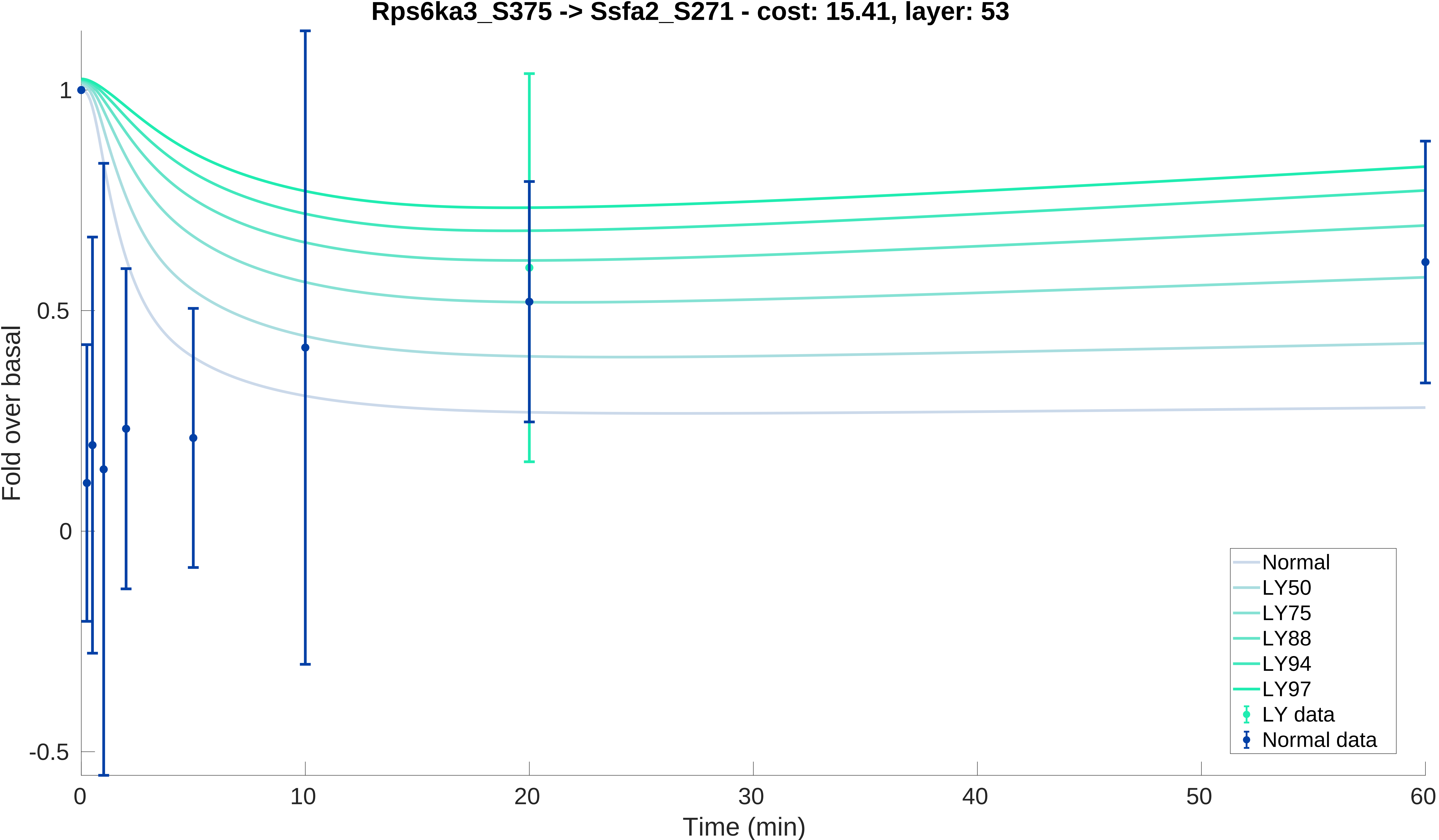
Rps6ka3_S375 -> Ssfa2_S271 - cost: 15.41, layer: 53
Fold over basal
Time (min)
1
0.5
0
-0.5
0
10
20
30
40
50
60
Normal
LY50
LY75
LY88
LY94
LY97
LY data
Normal data

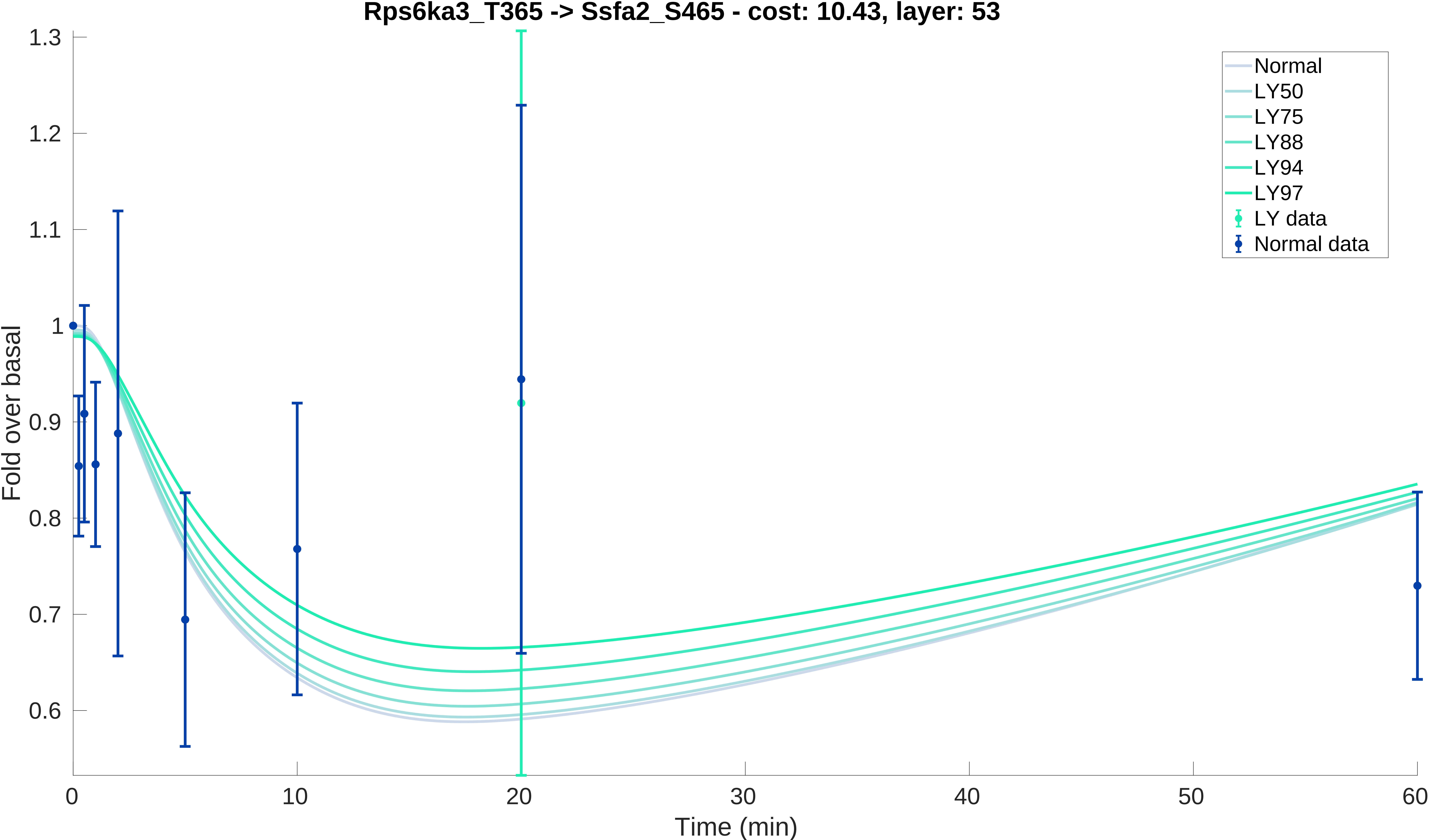

Rps6ka3_T365 -> Ssfa2_S465 - cost: 10.43, layer: 53
Fold over basal
Time (min)
0
10
20
30
40
50
60
0.6
0.7
0.8
0.9
1
1.1
1.2
1.3
Normal
LY50
LY75
LY88
LY94
LY97
LY data
Normal data

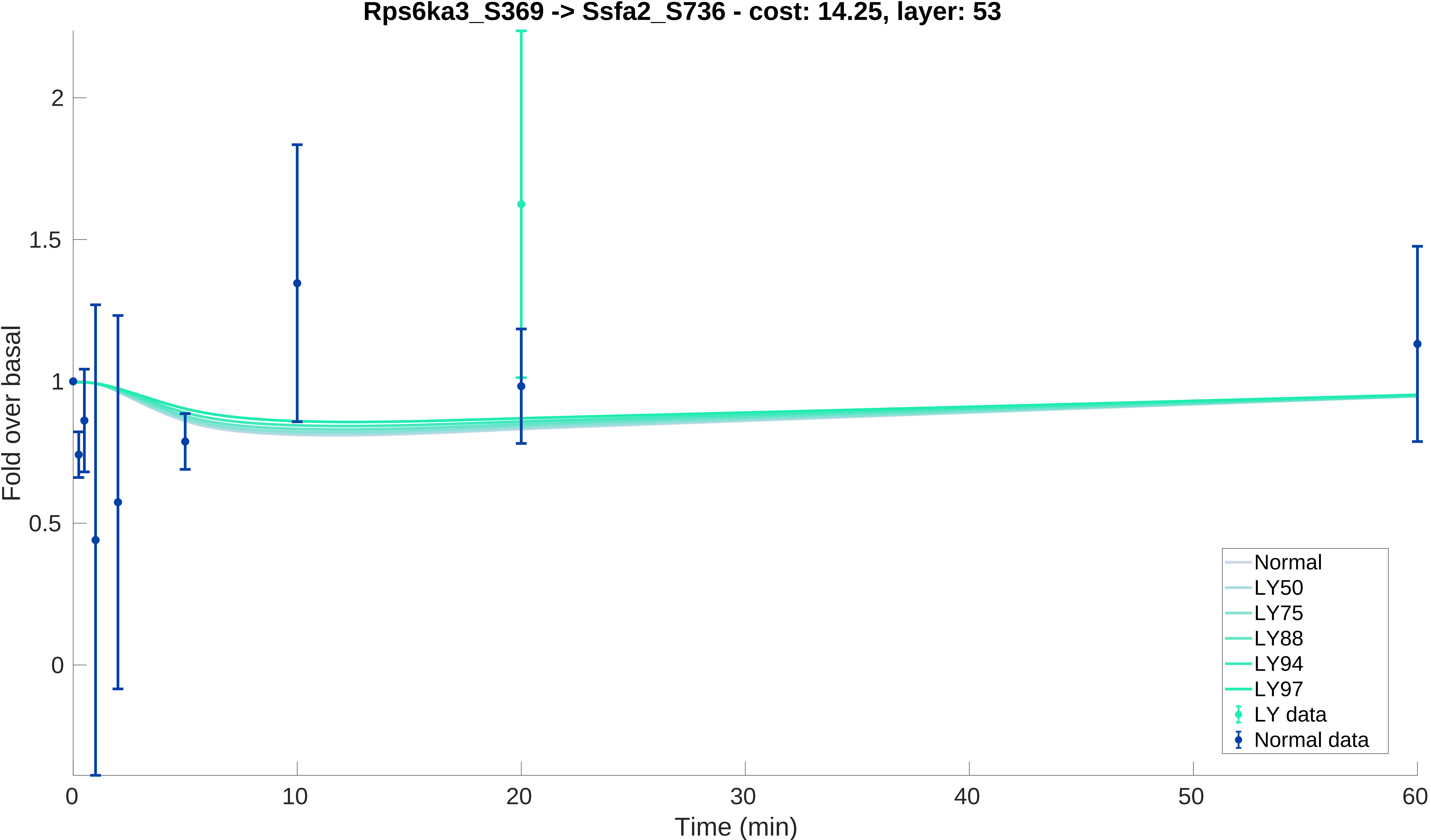

Rps6ka3_S369 -> Ssfa2_S736 - cost: 14.25, layer: 53
Fold over basal
Time (min)
0
0.5
1
1.5
2
0
10
20
30
40
50
60
Normal
LY50
LY75
LY88
LY94
LY97
LY data
Normal data

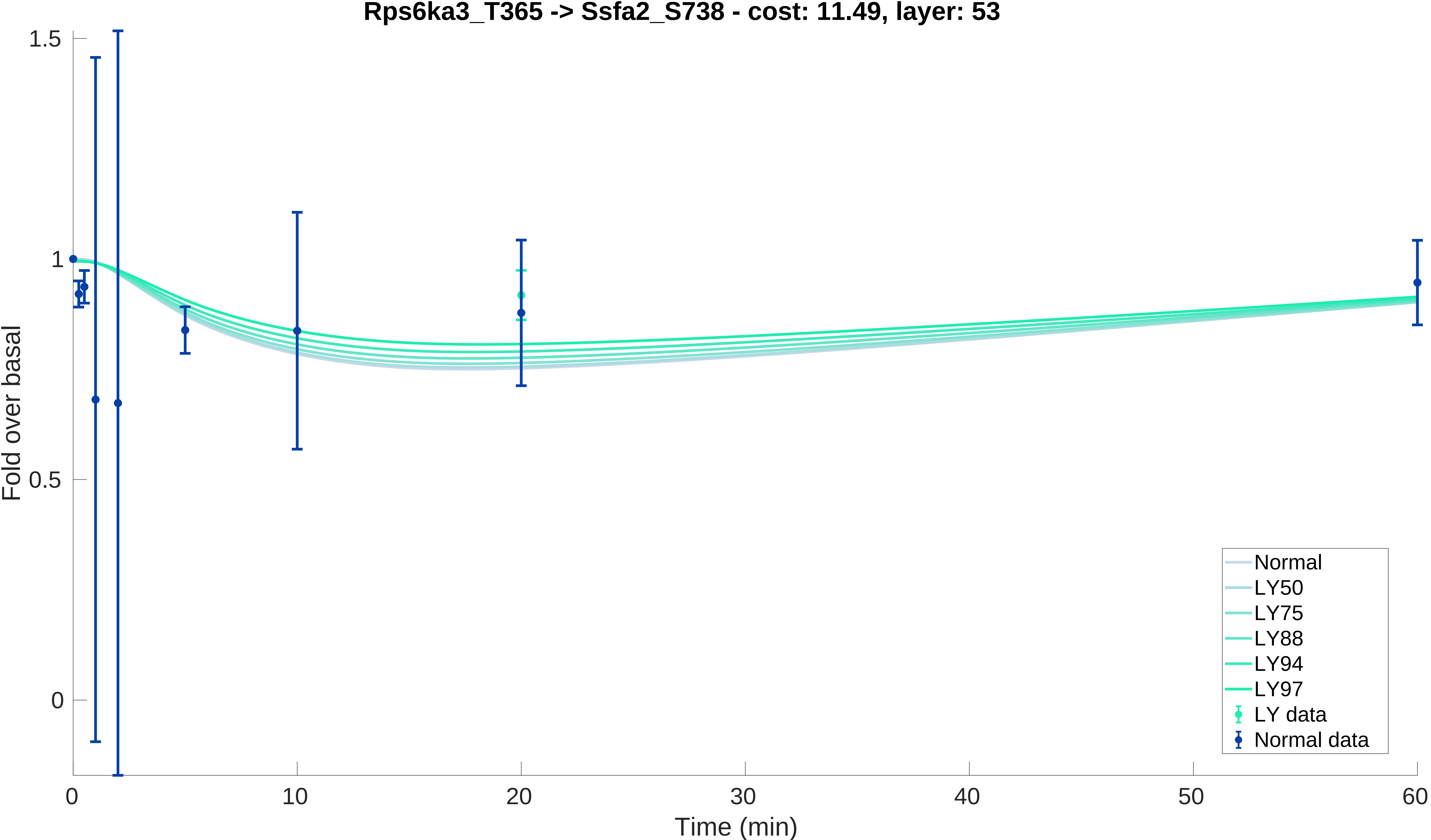

Rps6ka3_T365 -> Ssfa2_S738 - cost: 11.49, layer: 53
Fold over basal
Time (min)
1.5
1
0.5
0
0
10
20
30
40
50
60
Normal
LY50
LY75
LY88
LY94
LY97
LY data
Normal data

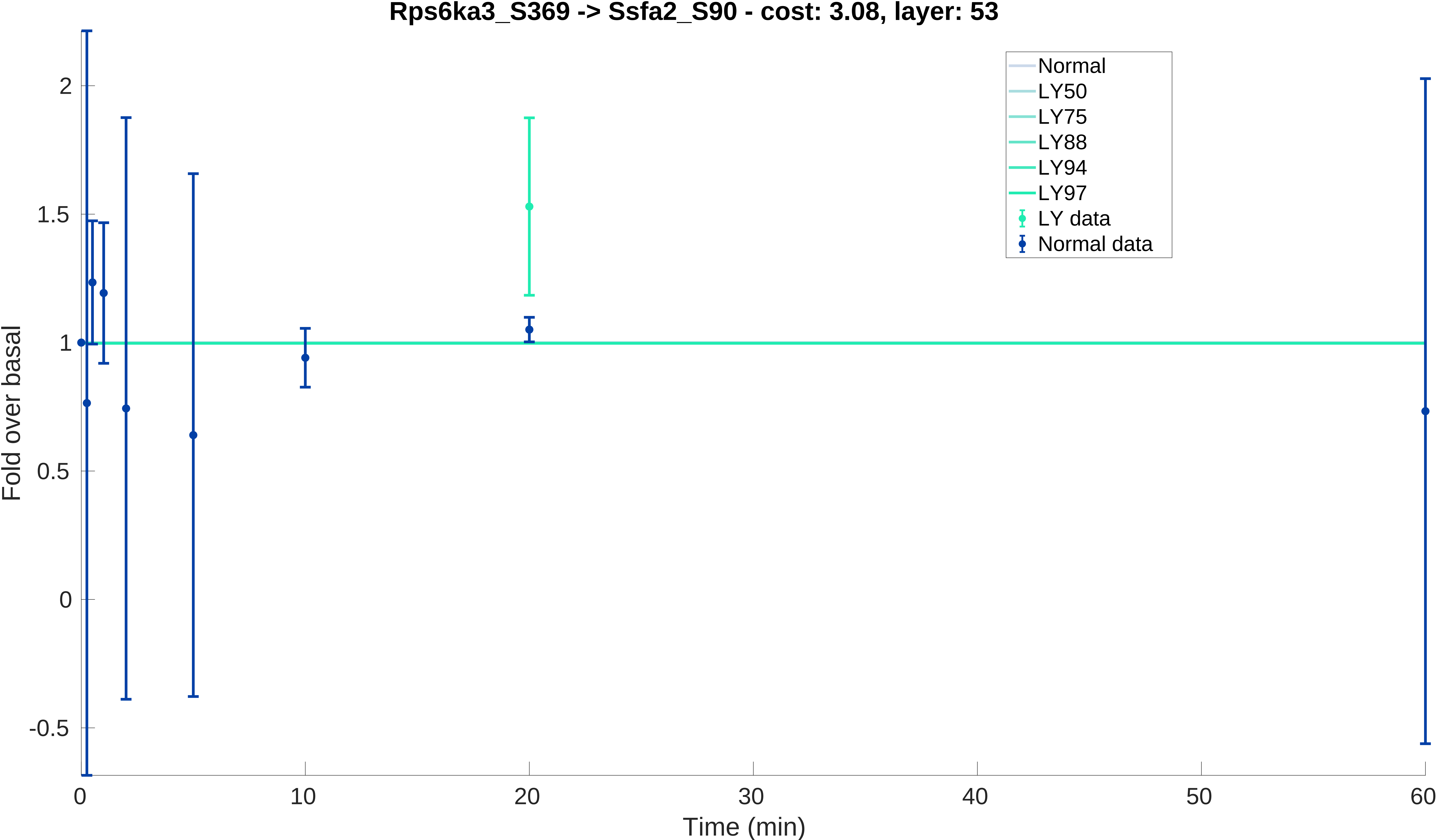

Rps6ka3_S369 -> Ssfa2_S90 - cost: 3.08, layer: 53
Normal
LY50
LY75
LY88
LY94
LY97
LY data
Normal data
Fold over basal
Time (min)
2
1.5
1
0.5
0
-0.5
0
10
20
30
40
50
60

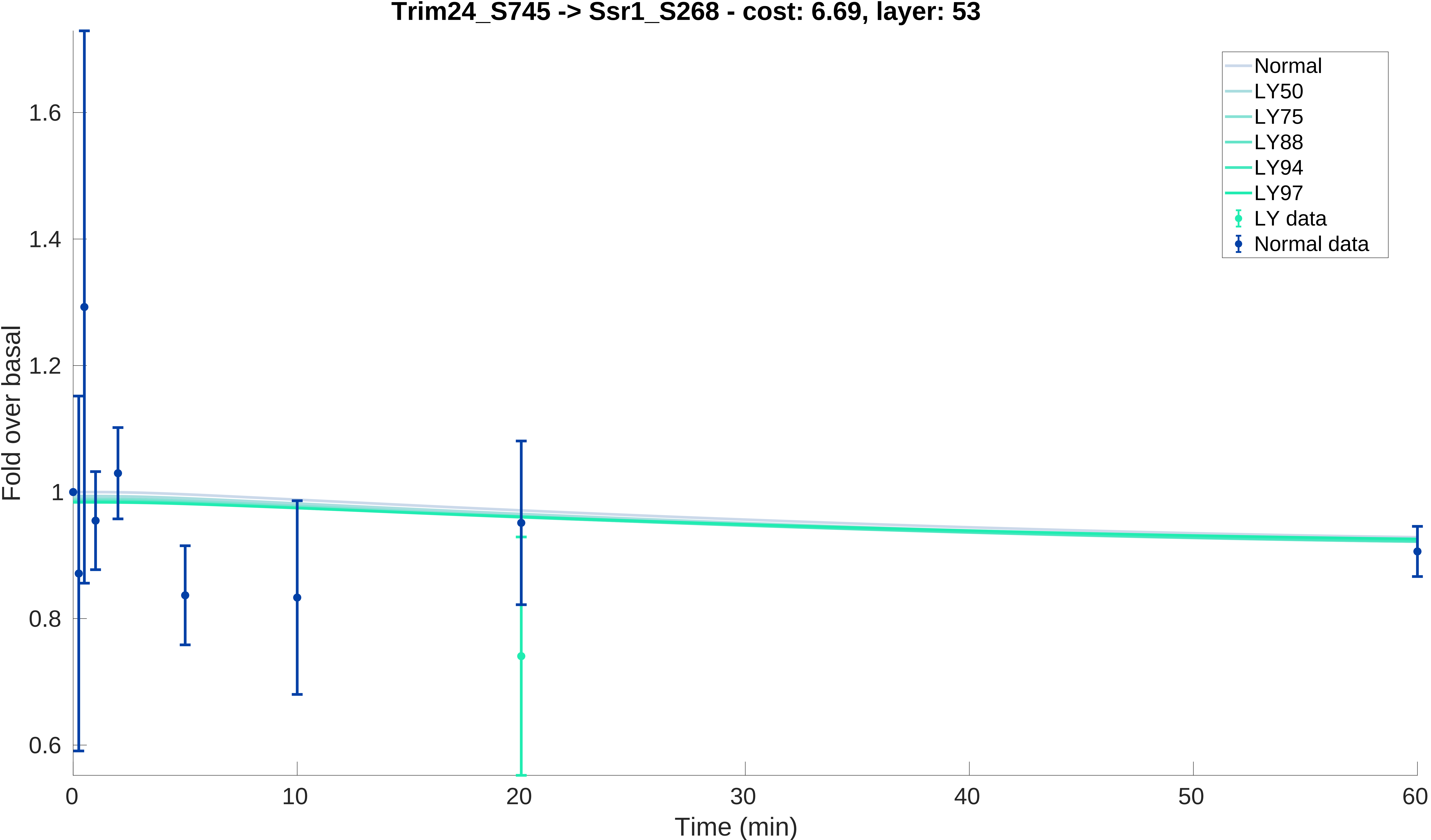

Trim24_S745 -> Ssr1_S268 - cost: 6.69, layer: 53
Fold over basal
Time (min)
1.6
1.4
1.2
1
0.8
0.6
0
10
20
30
40
50
60
Normal
LY50
LY75
LY88
LY94
LY97
LY data
Normal data

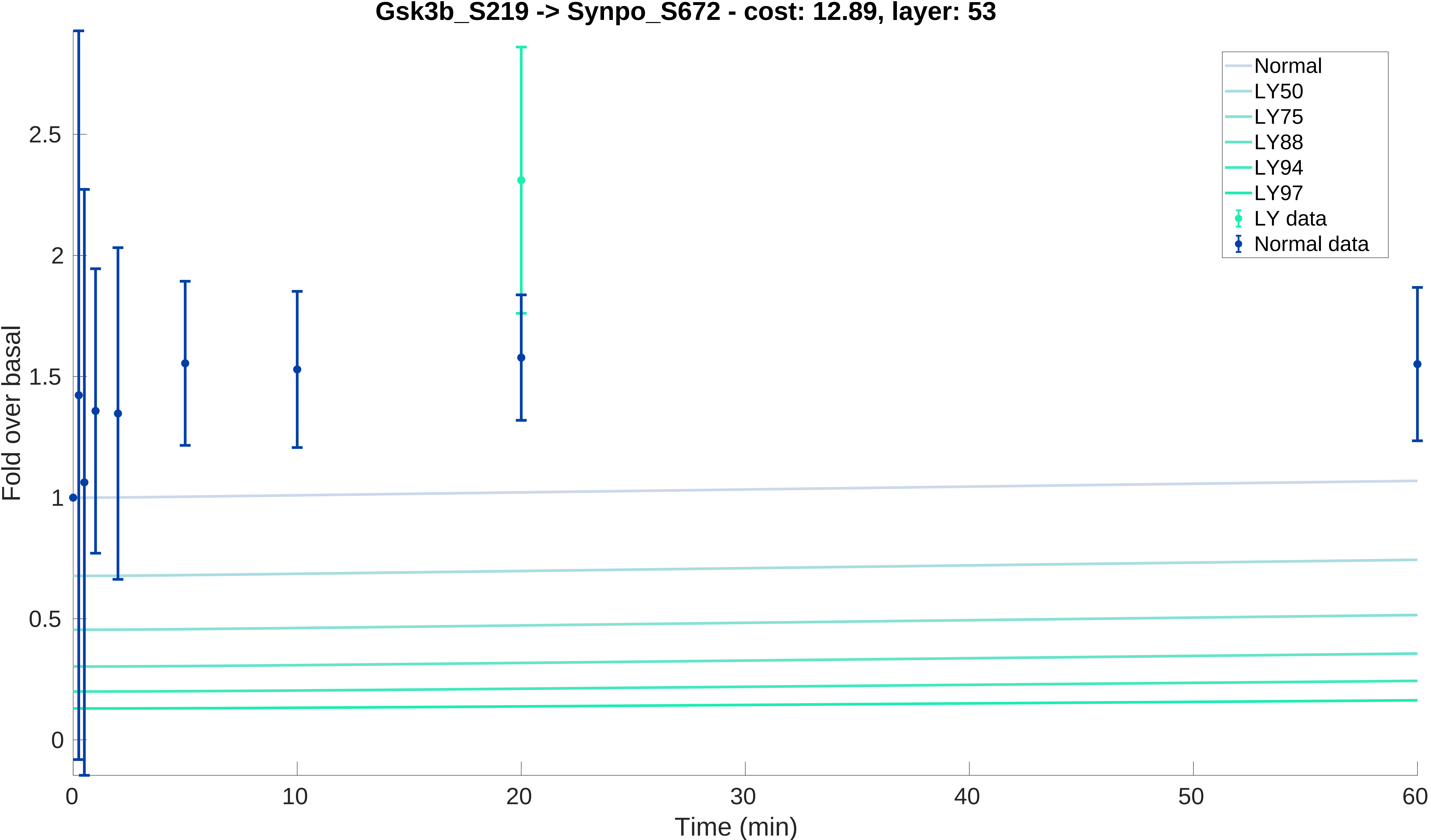

Gsk3b_S219 -> Synpo_S672 - cost: 12.89, layer: 53
Fold over basal
Time (min)
0
0.5
1
1.5
2
2.5
0
10
20
30
40
50
60
Normal
LY50
LY75
LY88
LY94
LY97
LY data
Normal data

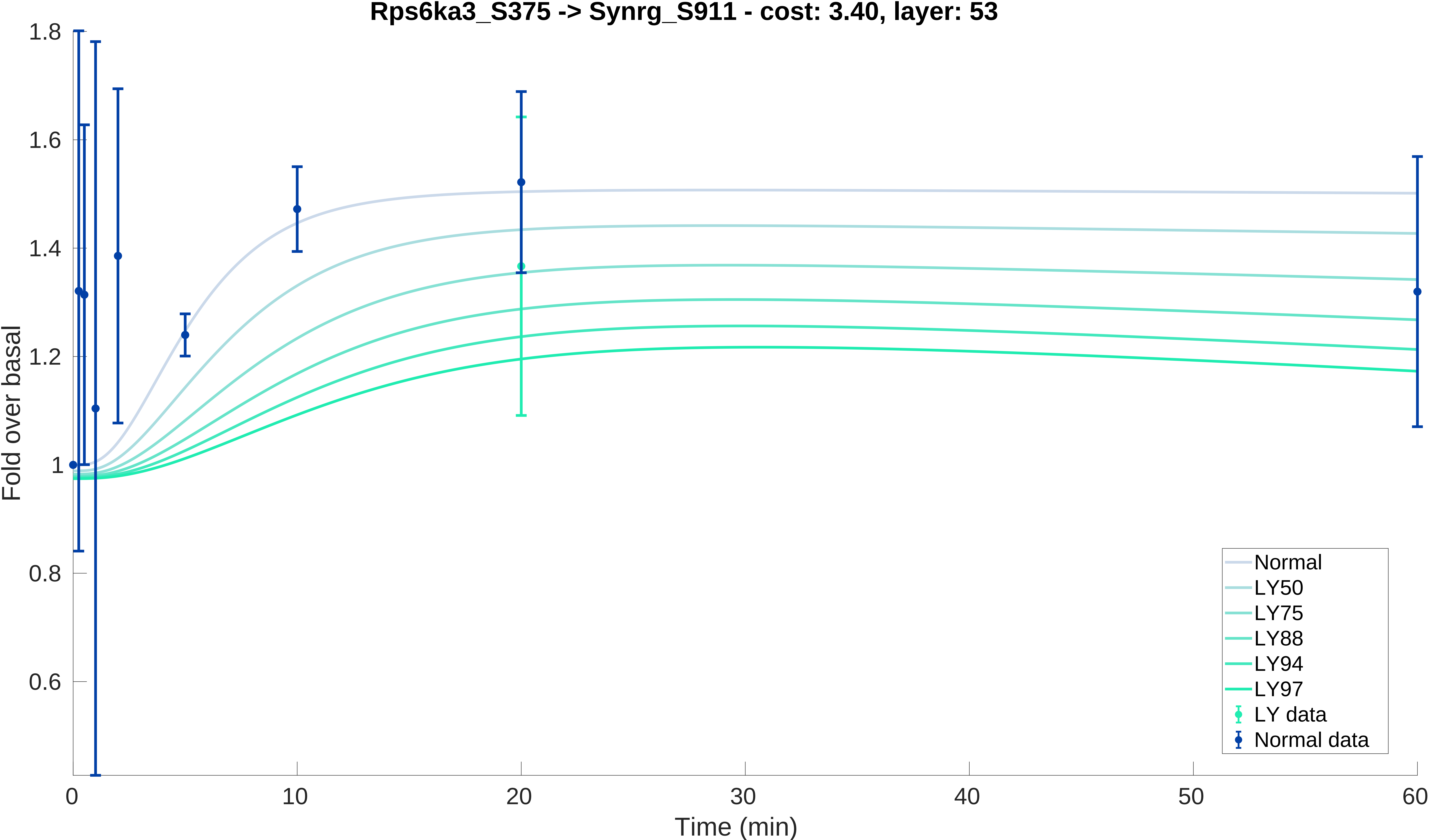
Rps6ka3_S375 -> Synrg_S911 - cost: 3.40, layer: 53
Fold over basal
1.8
1.6
1.4
1.2
1
0.8
0.6
0
10
20
30
40
50
60
Time (min)
Normal
LY50
LY75
LY88
LY94
LY97
LY data
Normal data

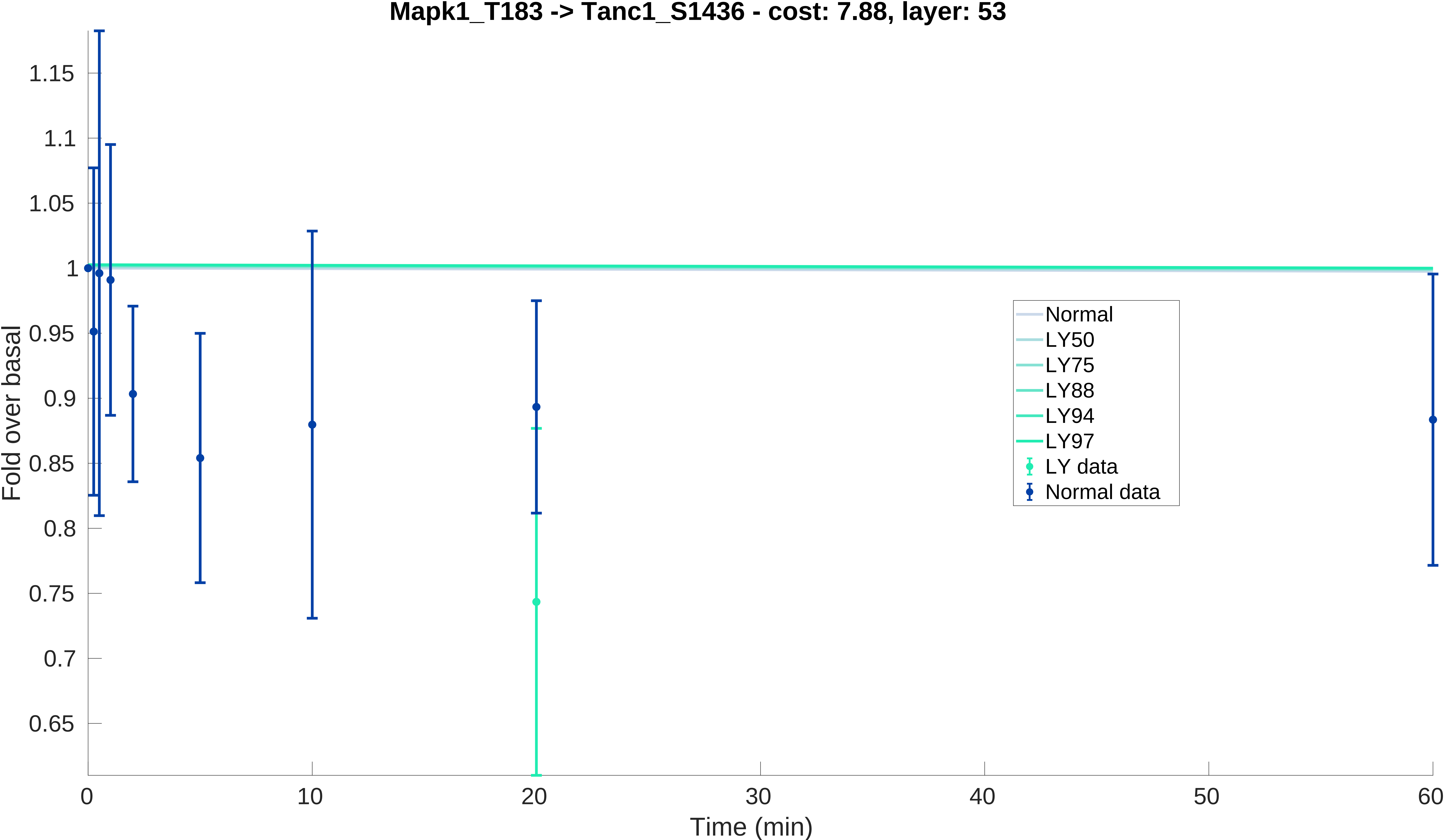

Mapk1_T183 -> Tanc1_S1436 - cost: 7.88, layer: 53
Fold over basal
Time (min)
1.15
1.1
1.05
1
0.95
0.9
0.85
0.8
0.75
0.7
0.65
0
10
20
30
40
50
60
Normal
LY50
LY75
LY88
LY94
LY97
LY data
Normal data

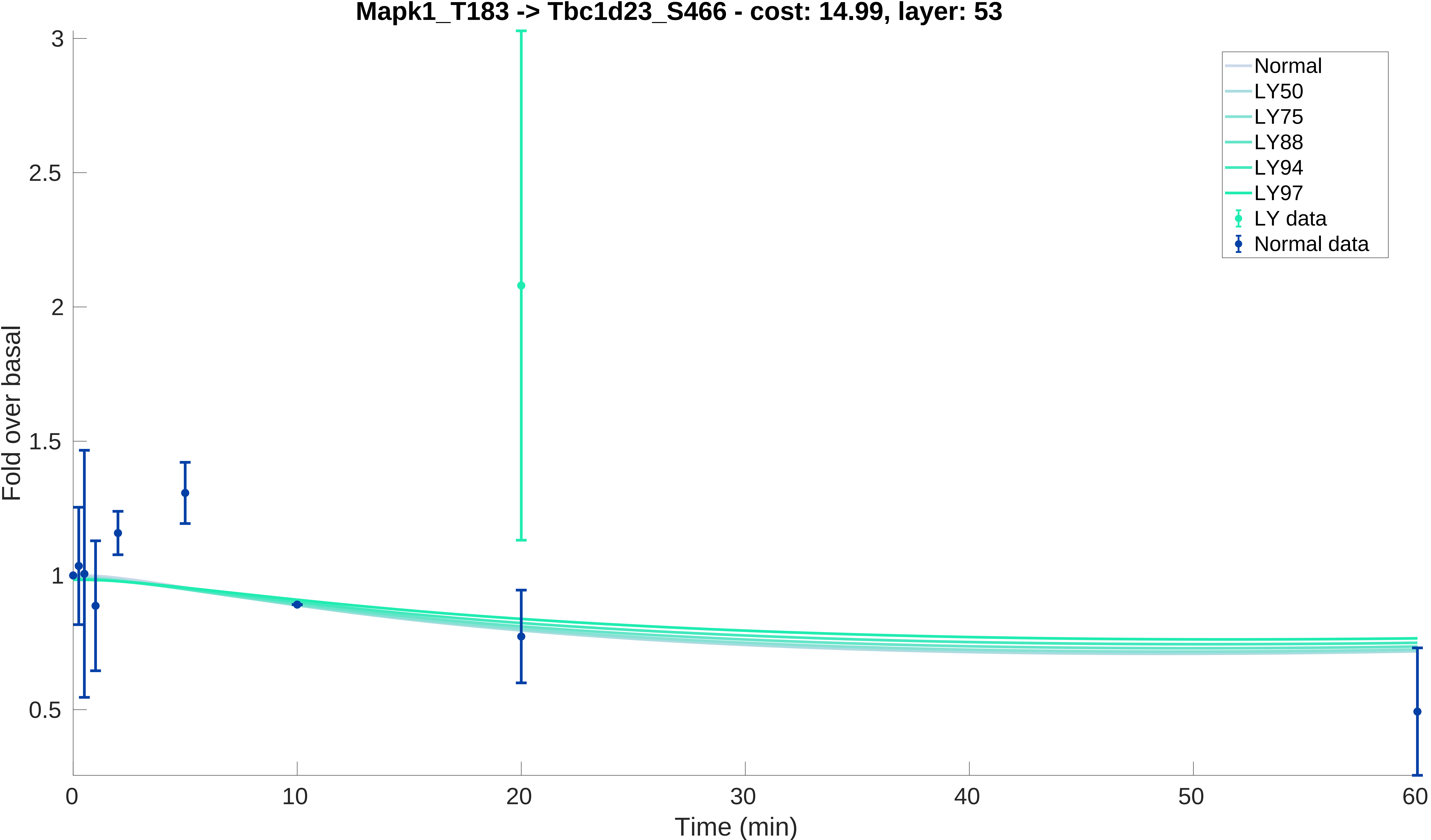

Mapk1_T183 -> Tbc1d23_S466 - cost: 14.99, layer: 53
Fold over basal
Time (min)
3
2.5
2
1.5
1
0.5
0
10
20
30
40
50
60
Normal
LY50
LY75
LY88
LY94
LY97
LY data
Normal data

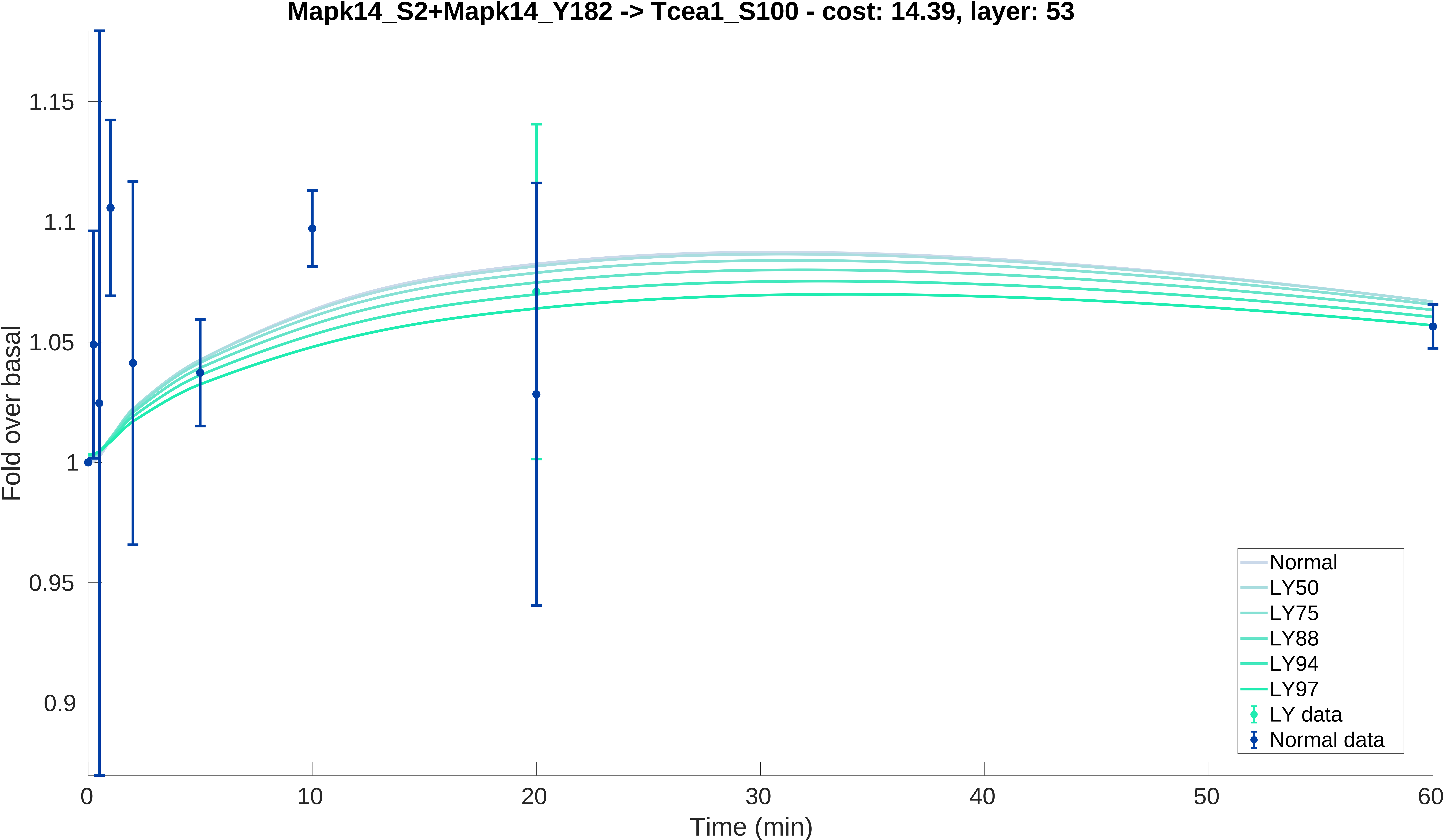
Mapk14_S2+Mapk14_Y182 -> Tcea1_S100 - cost: 14.39, layer: 53
Fold over basal
Time (min)
0.9
0.95
1
1.05
1.1
1.15
0
10
20
30
40
50
60
Normal
LY50
LY75
LY88
LY94
LY97
LY data
Normal data

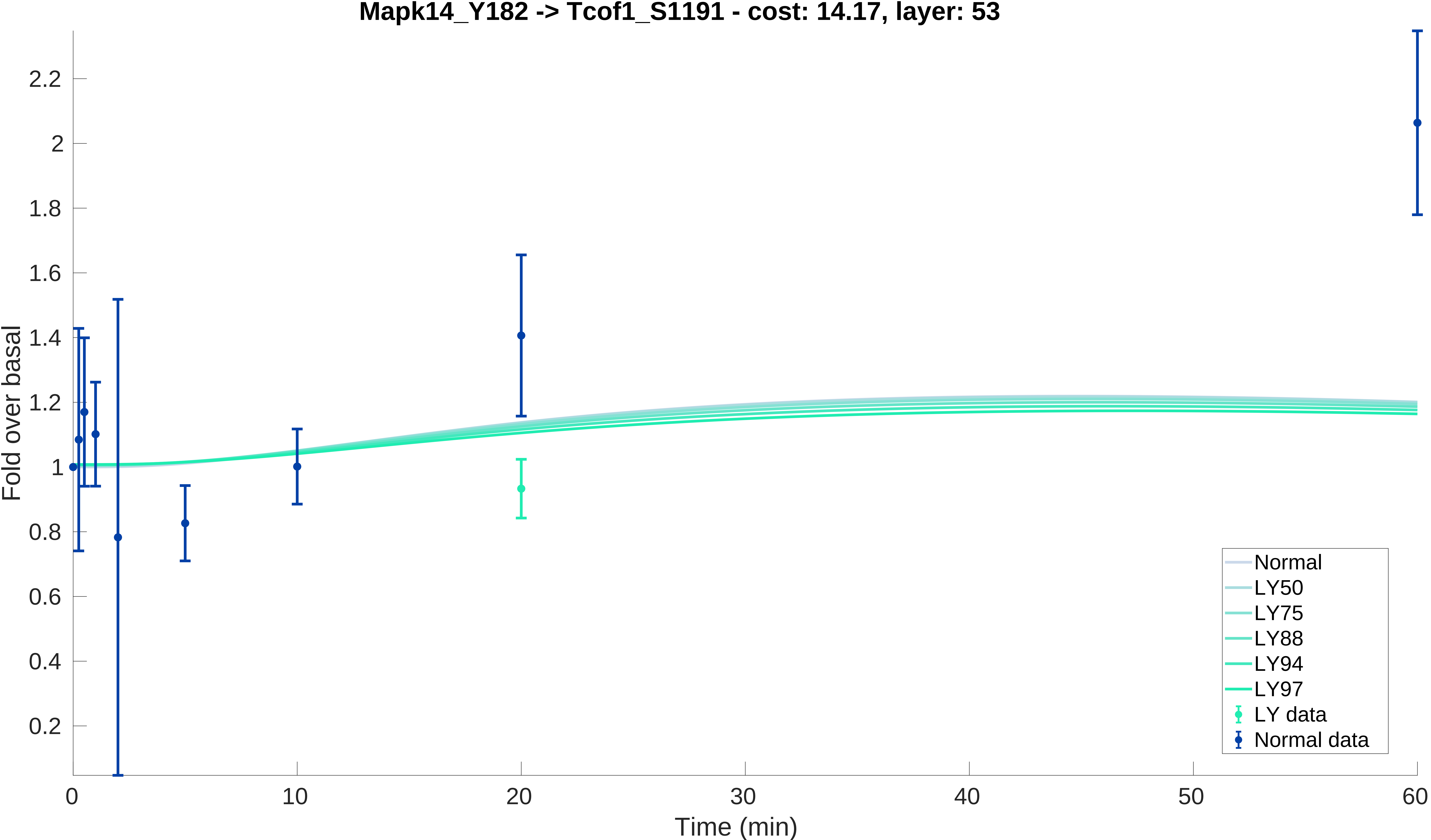

Mapk14_Y182 -> Tcof1_S1191 - cost: 14.17, layer: 53
Fold over basal
Time (min)
2.2
2
1.8
1.6
1.4
1.2
1
0.8
0.6
0.4
0.2
0
10
20
30
40
50
60
Normal
LY50
LY75
LY88
LY94
LY97
LY data
Normal data

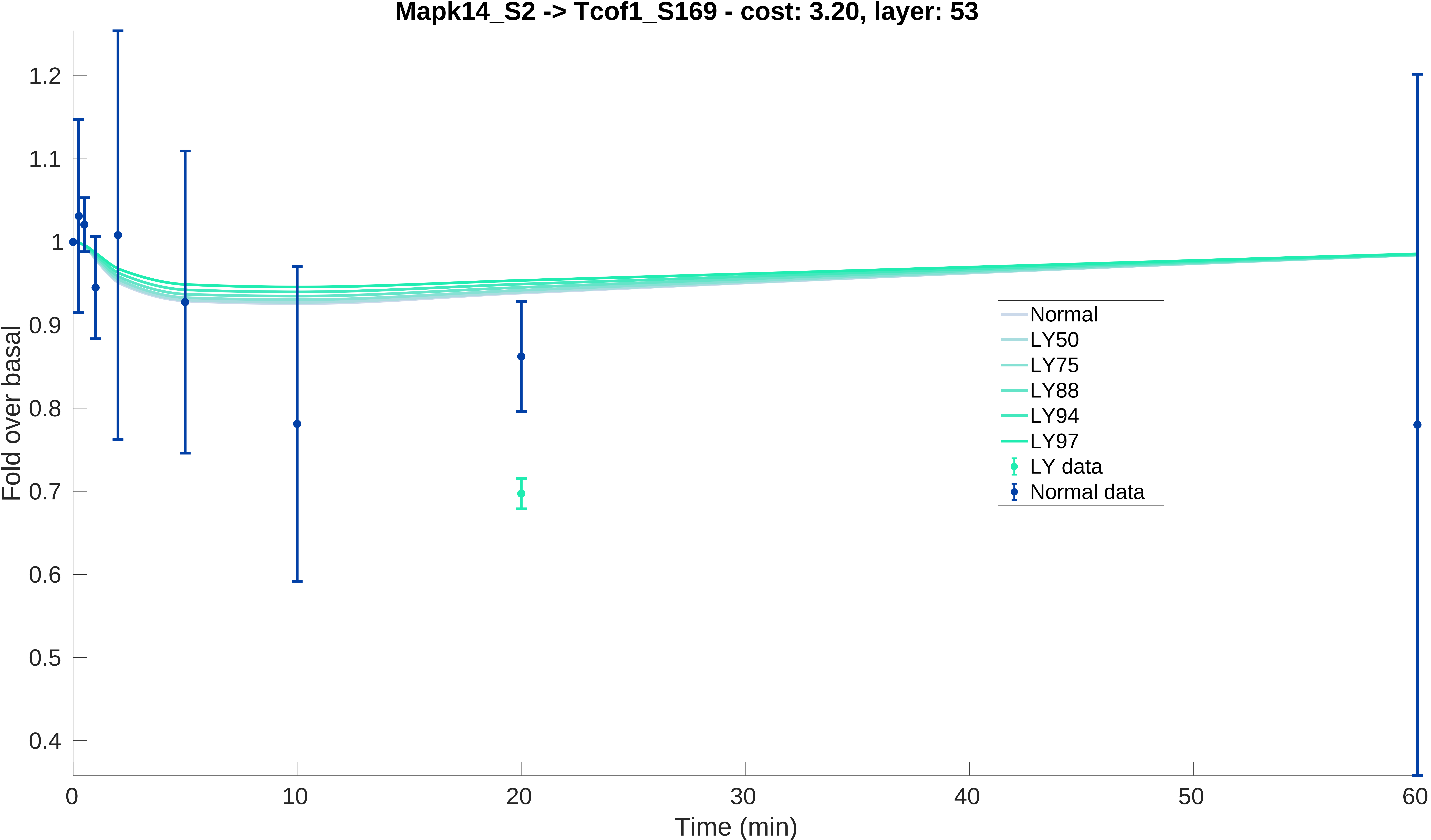

Mapk14_S2 -> Tcof1_S169 - cost: 3.20, layer: 53
Fold over basal
Time (min)
1.2
1.1
1
0.9
0.8
0.7
0.6
0.5
0.4
0
10
20
30
40
50
60
Normal
LY50
LY75
LY88
LY94
LY97
LY data
Normal data

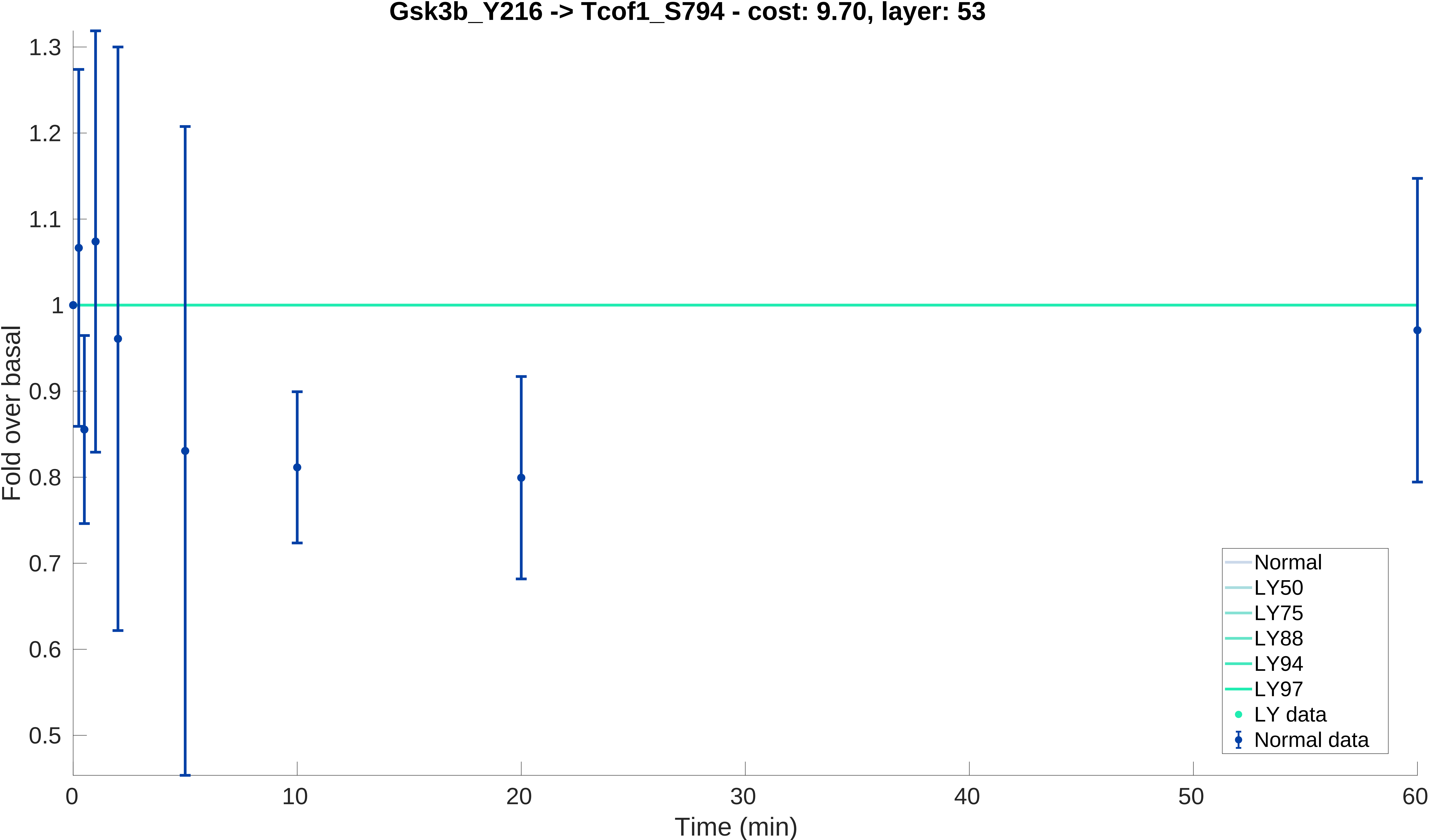
Gsk3b_Y216 -> Tcof1_S794 - cost: 9.70, layer: 53
Fold over basal
1.3
1.2
1.1
1
0.9
0.8
0.7
0.6
0.5
0
10
20
30
40
50
60
Time (min)
Normal
LY50
LY75
LY88
LY94
LY97
LY data
Normal data

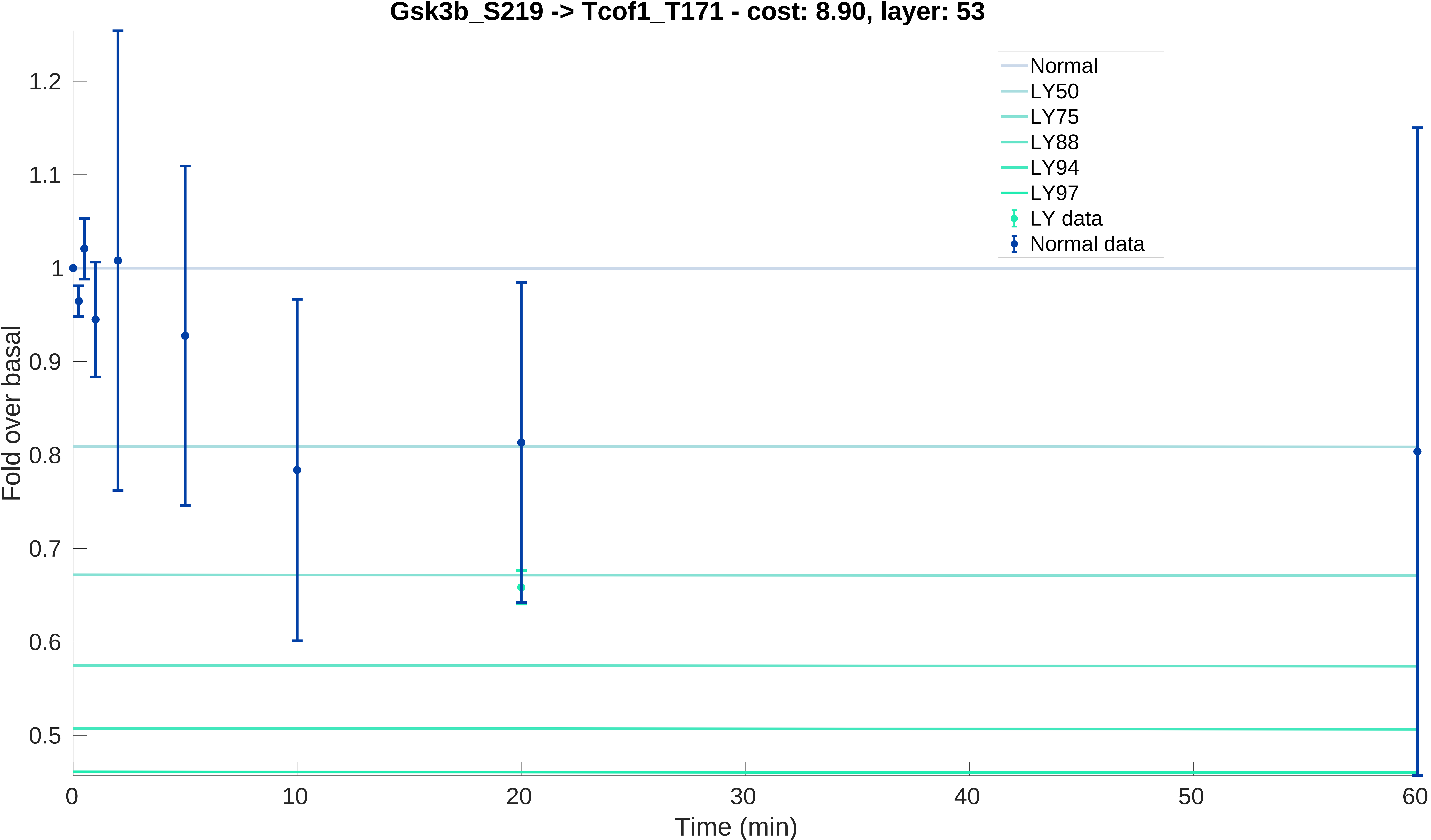
Gsk3b_S219 -> Tcof1_T171 - cost: 8.90, layer: 53
Fold over basal
1.2
1.1
1
0.9
0.8
0.7
0.6
0.5
0
10
20
30
40
50
60
Time (min)
Normal
LY50
LY75
LY88
LY94
LY97
LY data
Normal data

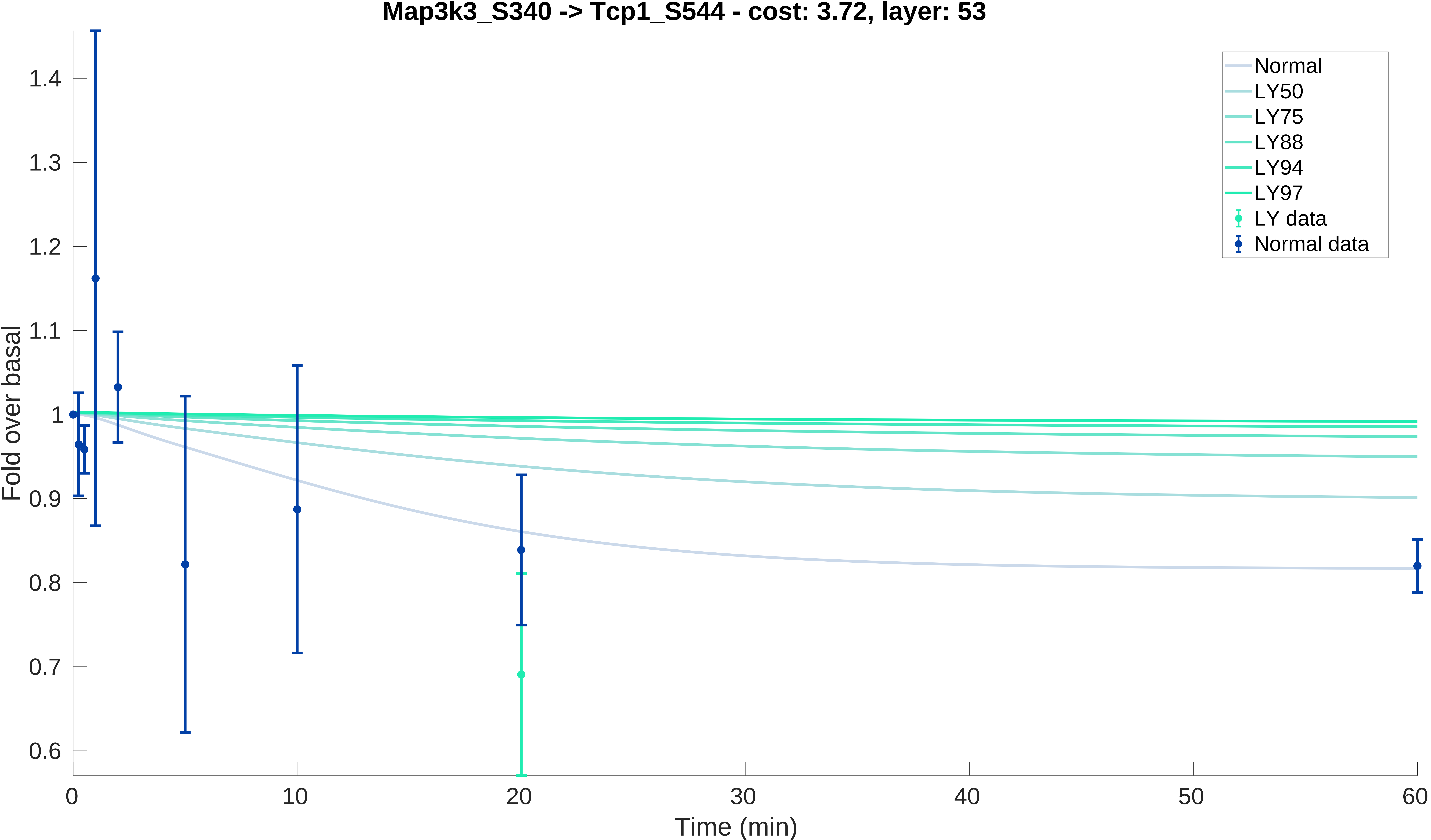

Map3k3_S340 -> Tcp1_S544 - cost: 3.72, layer: 53
Normal
LY50
LY75
LY88
LY94
LY97
LY data
Normal data
Fold over basal
Time (min)
1.4
1.3
1.2
1.1
1
0.9
0.8
0.7
0.6
0
10
20
30
40
50
60

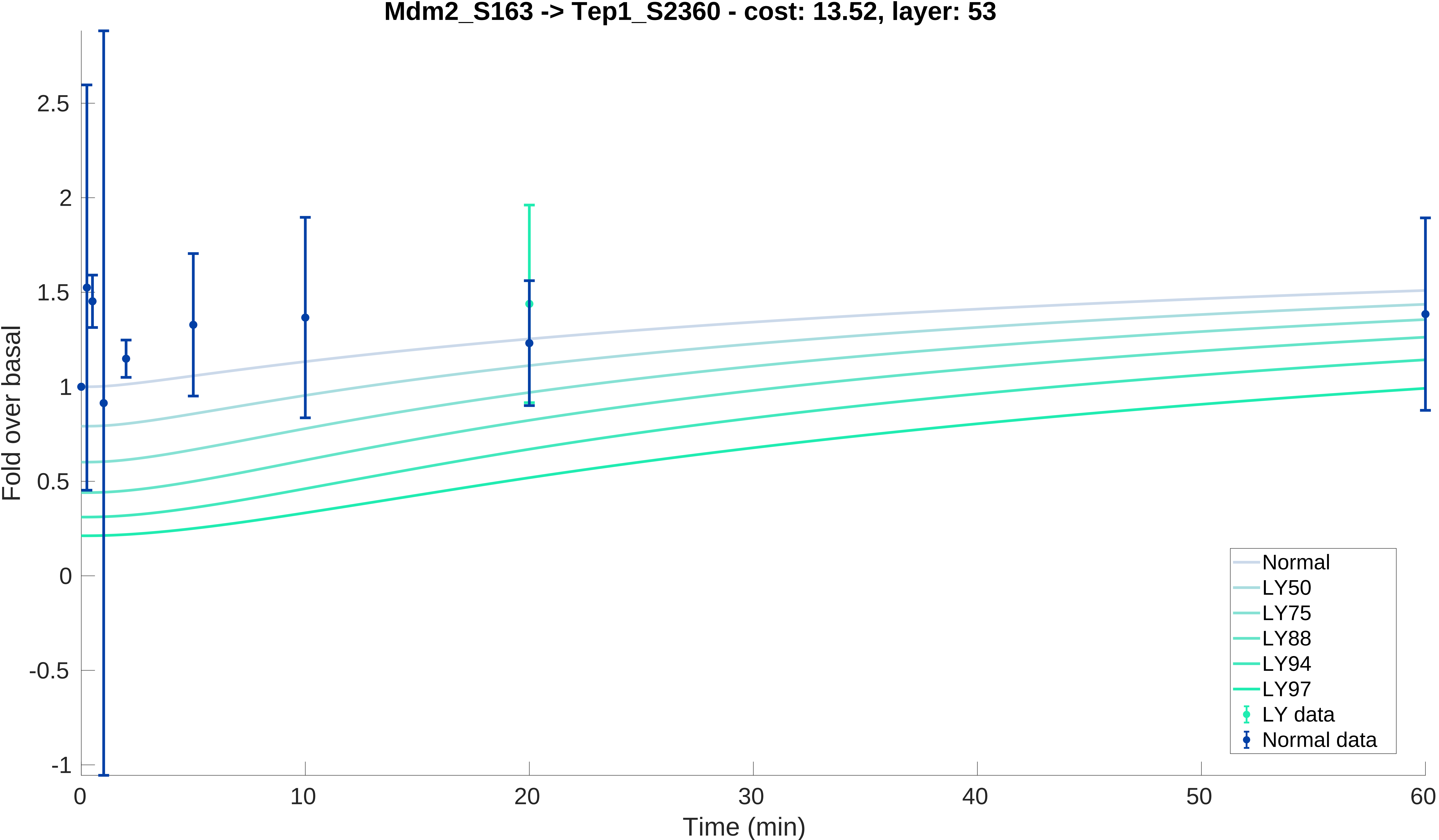

Mdm2_S163 -> Tep1_S2360 - cost: 13.52, layer: 53
Fold over basal
2.5
2
1.5
1
0.5
0
-0.5
-1
0
10
20
30
40
50
60
Time (min)
Normal
LY50
LY75
LY88
LY94
LY97
LY data
Normal data

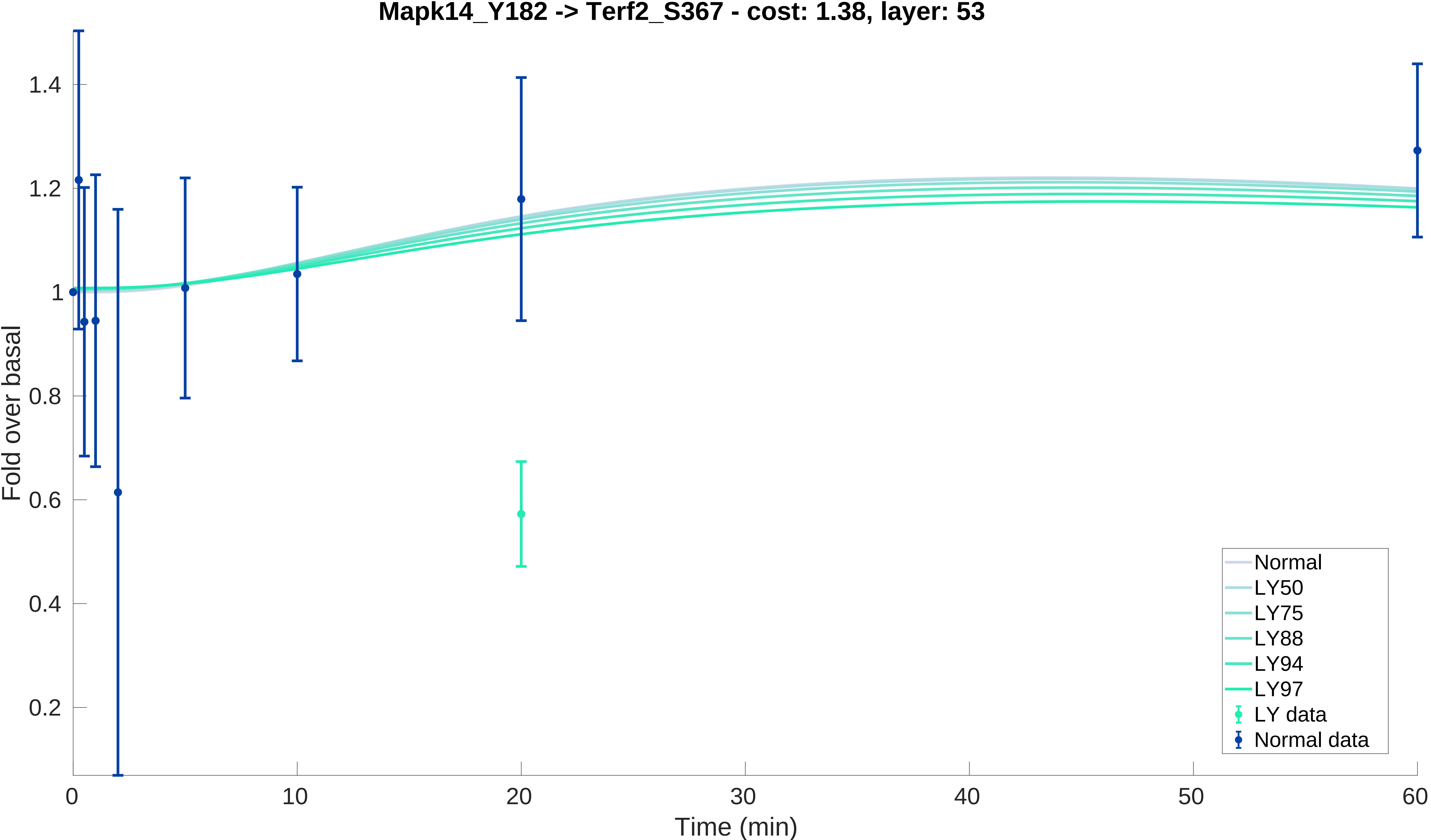

Mapk14_Y182 -> Terf2_S367 - cost: 1.38, layer: 53
Fold over basal
Time (min)
0.2
0.4
0.6
0.8
1
1.2
1.4
0
10
20
30
40
50
60
Normal
LY50
LY75
LY88
LY94
LY97
LY data
Normal data

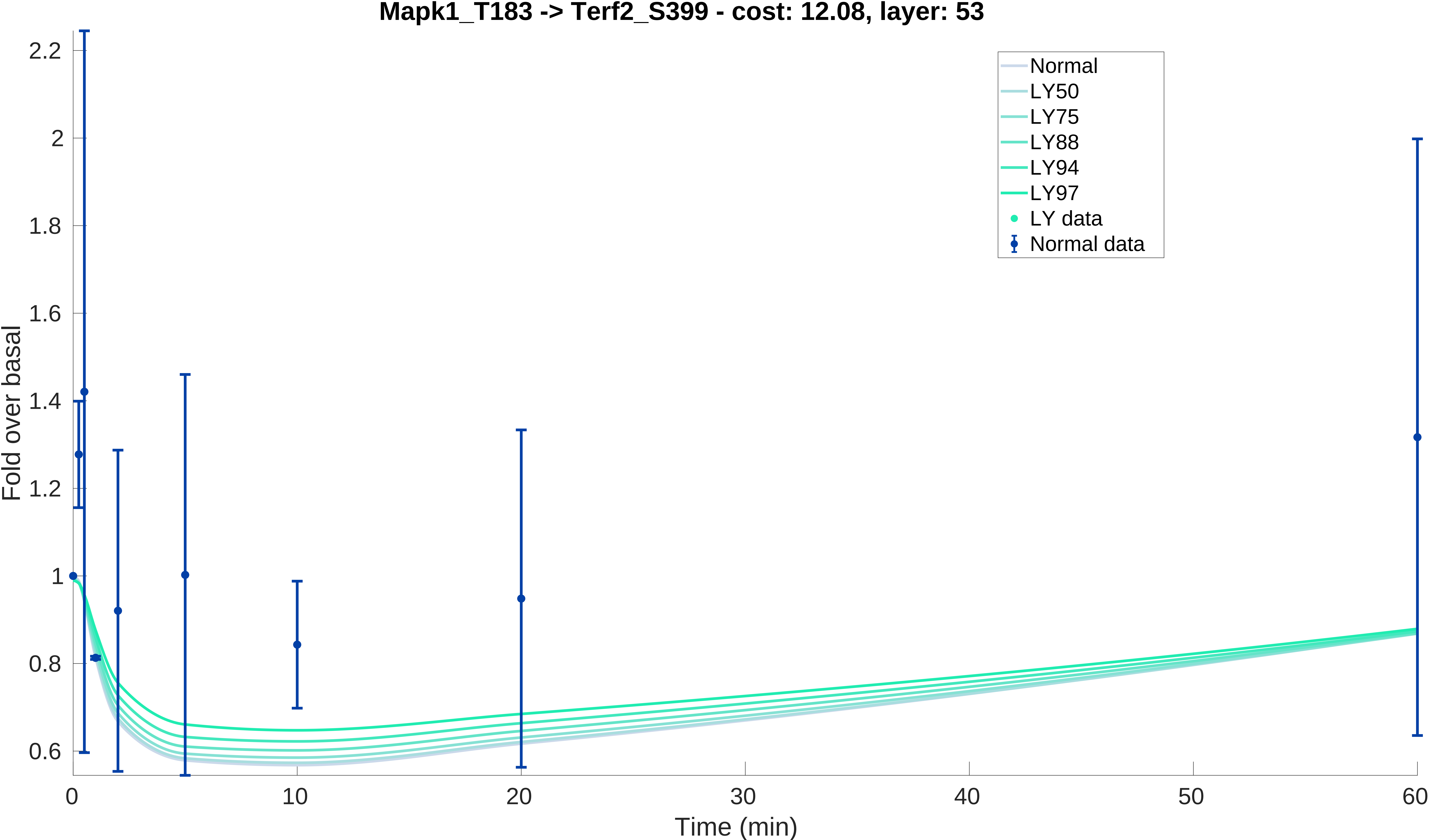

Mapk1_T183 -> Terf2_S399 - cost: 12.08, layer: 53
Normal
LY50
LY75
LY88
LY94
LY97
LY data
Normal data
Fold over basal
Time (min)
0
10
20
30
40
50
60
0.6
0.8
1
1.2
1.4
1.6
1.8
2
2.2

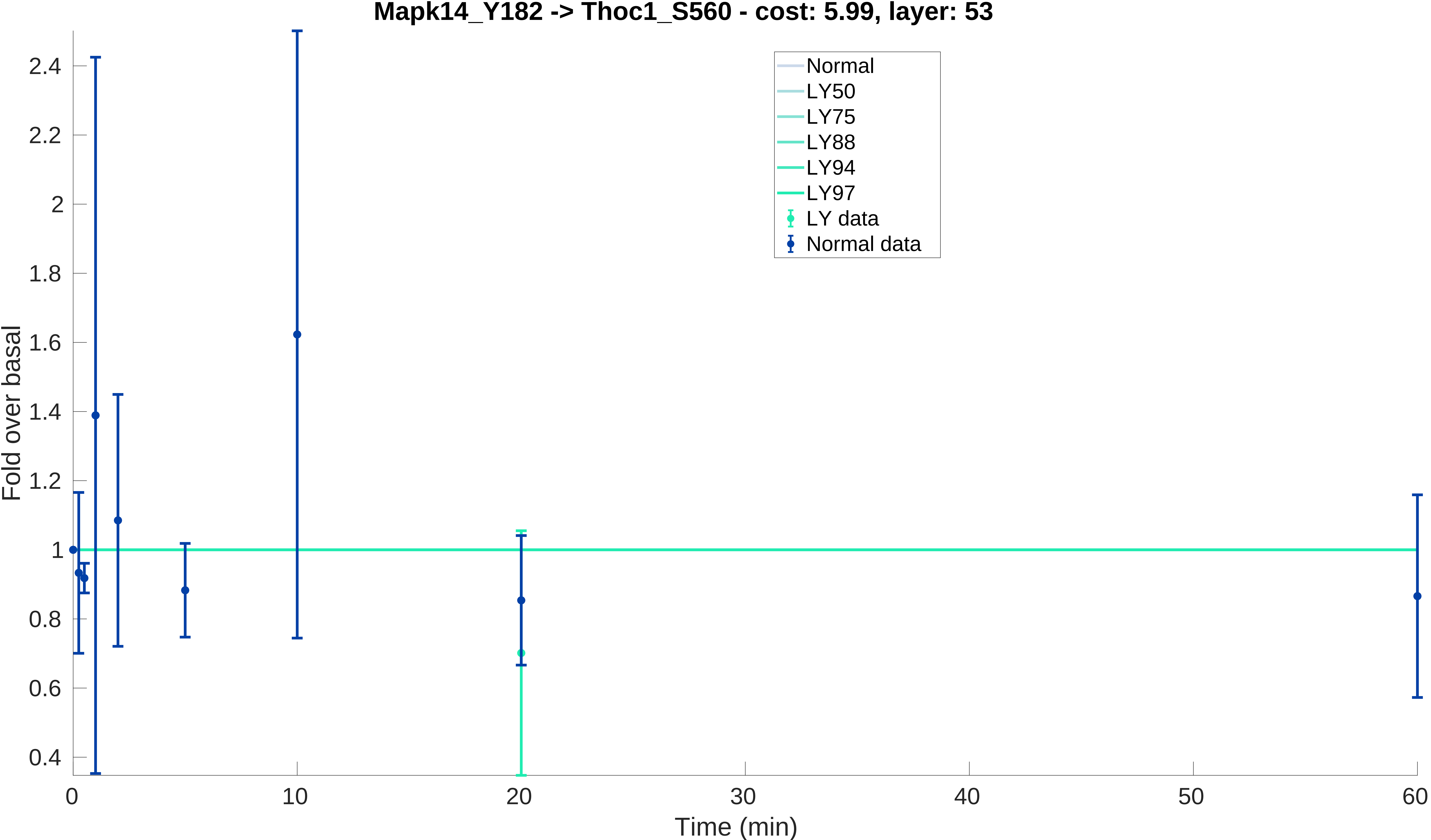

Mapk14_Y182 -> Thoc1_S560 - cost: 5.99, layer: 53
Normal
LY50
LY75
LY88
LY94
LY97
LY data
Normal data
Fold over basal
2.4
2.2
2
1.8
1.6
1.4
1.2
1
0.8
0.6
0.4
0
10
20
30
40
50
60
Time (min)

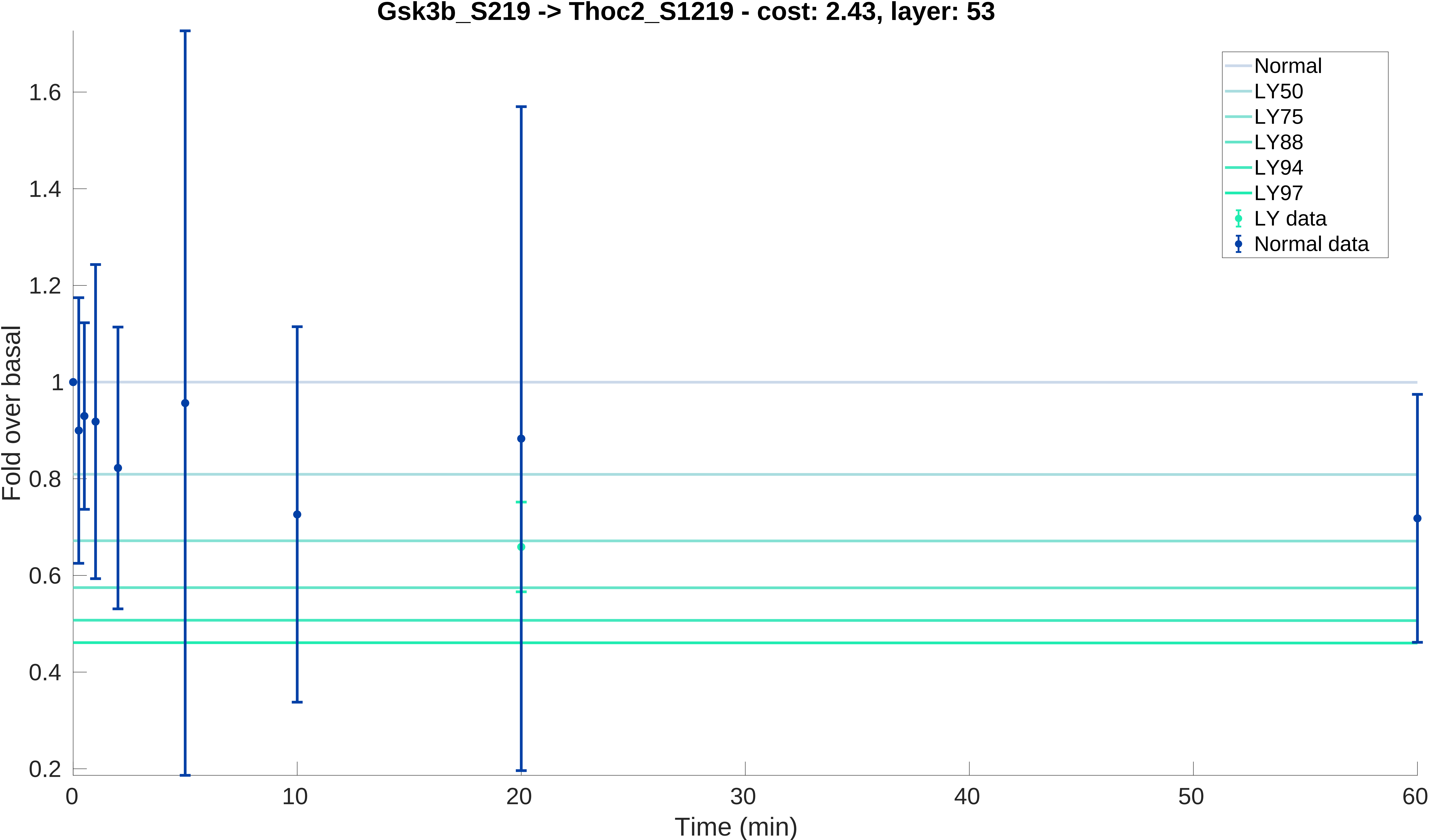
Gsk3b_S219 -> Thoc2_S1219 - cost: 2.43, layer: 53
Fold over basal
Time (min)
1.6
1.4
1.2
1
0.8
0.6
0.4
0.2
0
10
20
30
40
50
60
Normal
LY50
LY75
LY88
LY94
LY97
LY data
Normal data

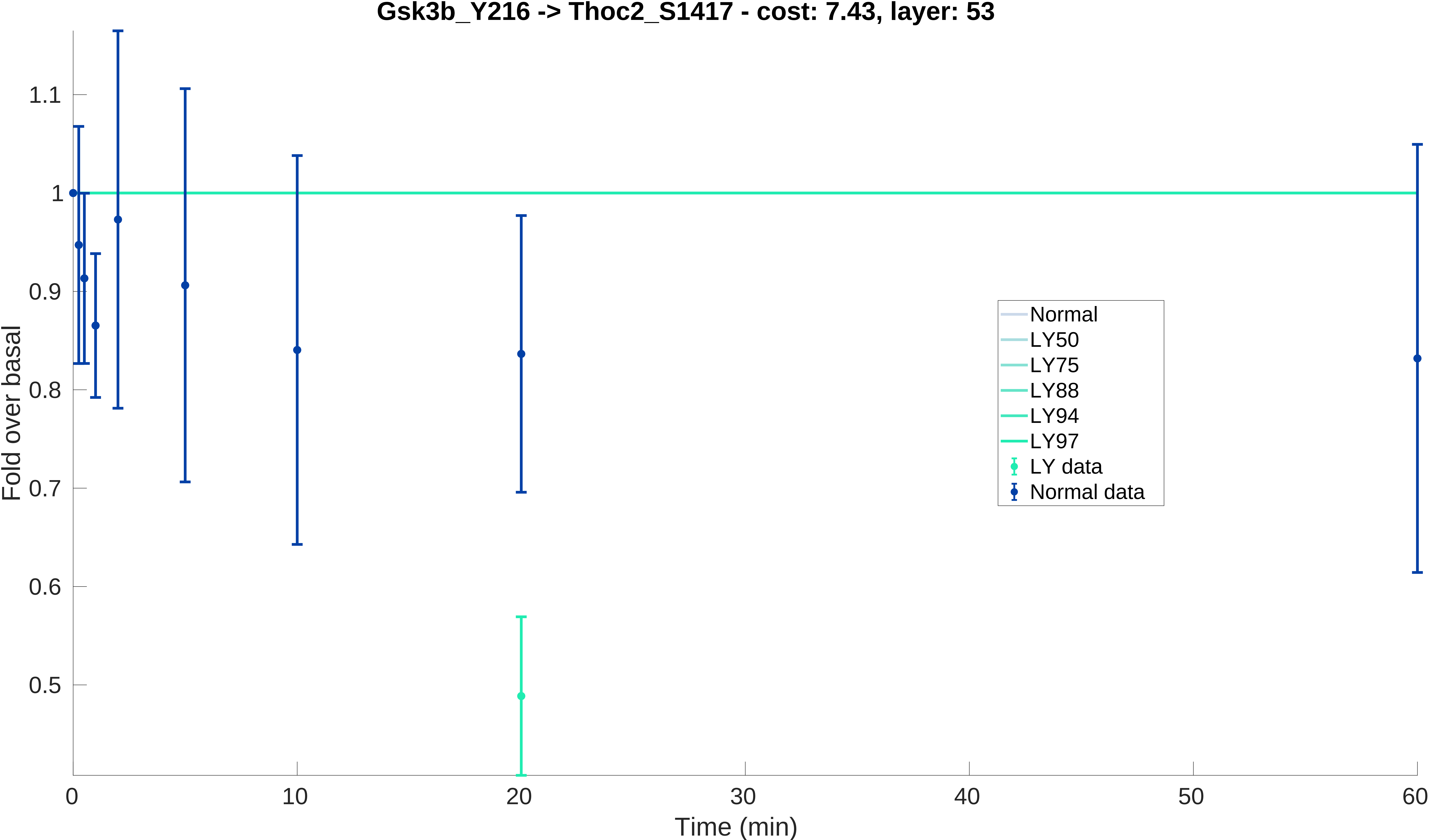

Gsk3b_Y216 -> Thoc2_S1417 - cost: 7.43, layer: 53
Fold over basal
1.1
1
0.9
0.8
0.7
0.6
0.5
0
10
20
30
40
50
60
Time (min)
Normal
LY50
LY75
LY88
LY94
LY97
LY data
Normal data

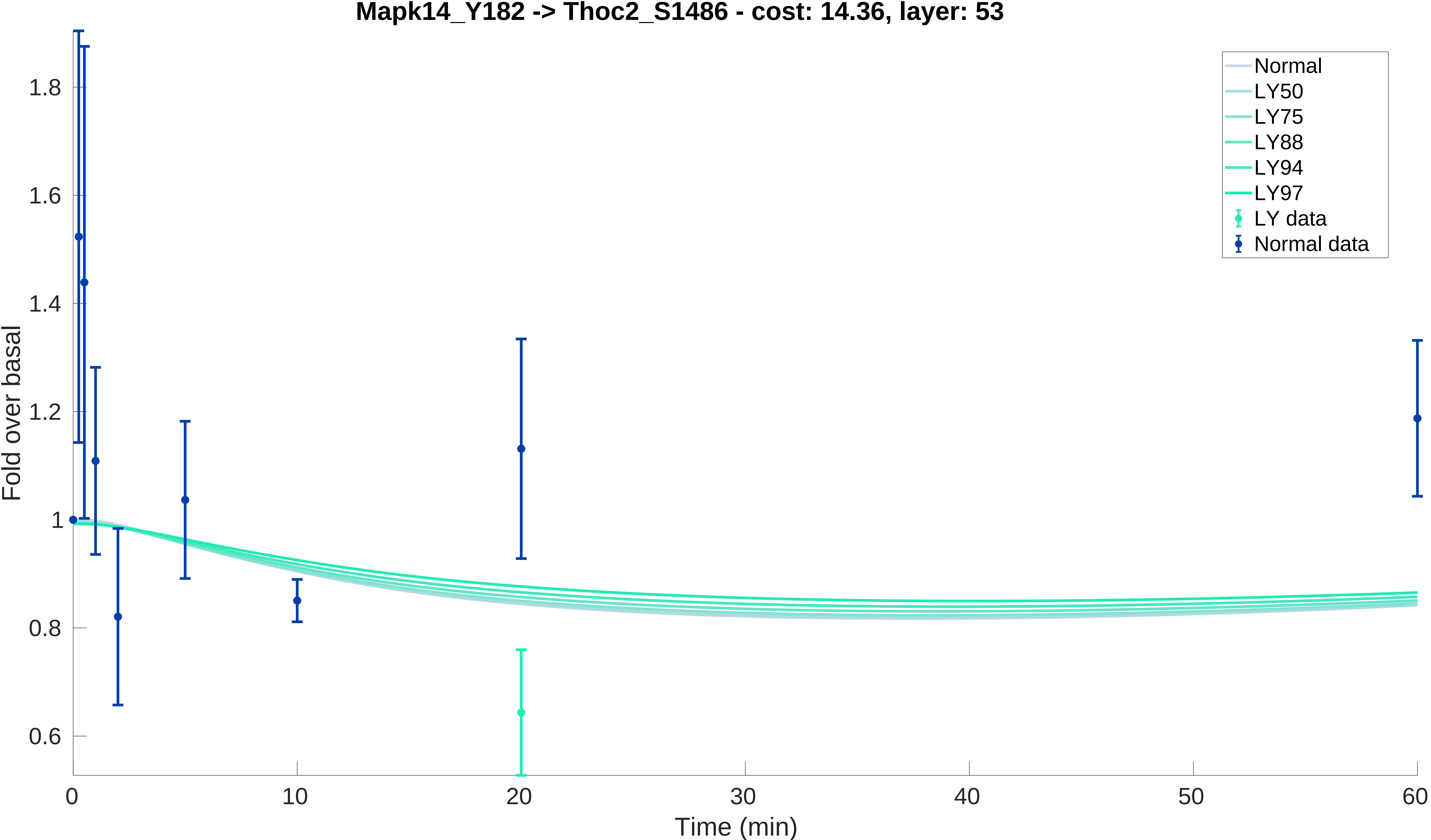

Mapk14_Y182 -> Thoc2_S1486 - cost: 14.36, layer: 53
Fold over basal
Time (min)
0.6
0.8
1
1.2
1.4
1.6
1.8
0
10
20
30
40
50
60
Normal
LY50
LY75
LY88
LY94
LY97
LY data
Normal data

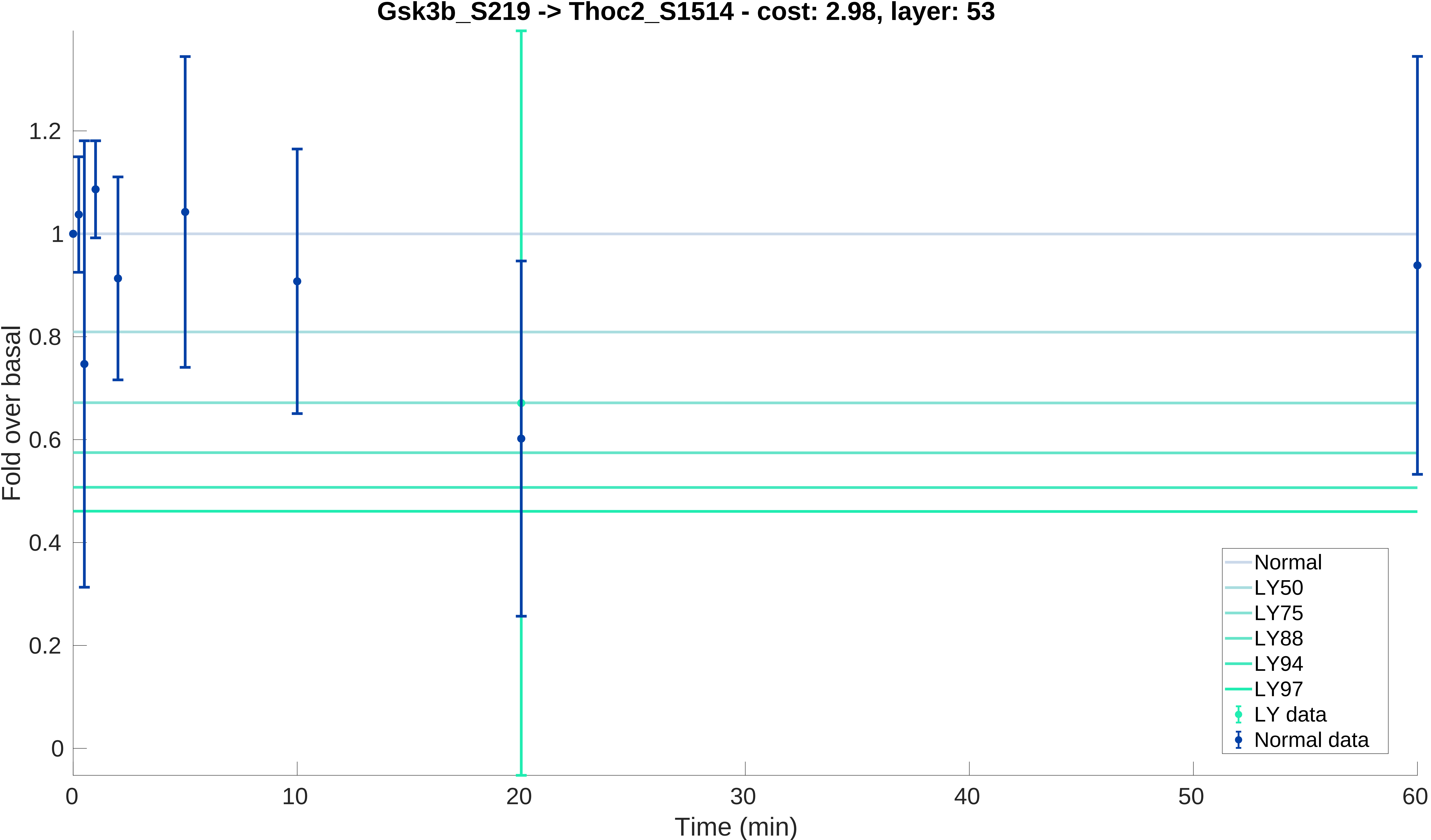

Gsk3b_S219 -> Thoc2_S1514 - cost: 2.98, layer: 53
Fold over basal
Time (min)
0
0.2
0.4
0.6
0.8
1
1.2
0
10
20
30
40
50
60
Normal
LY50
LY75
LY88
LY94
LY97
LY data
Normal data

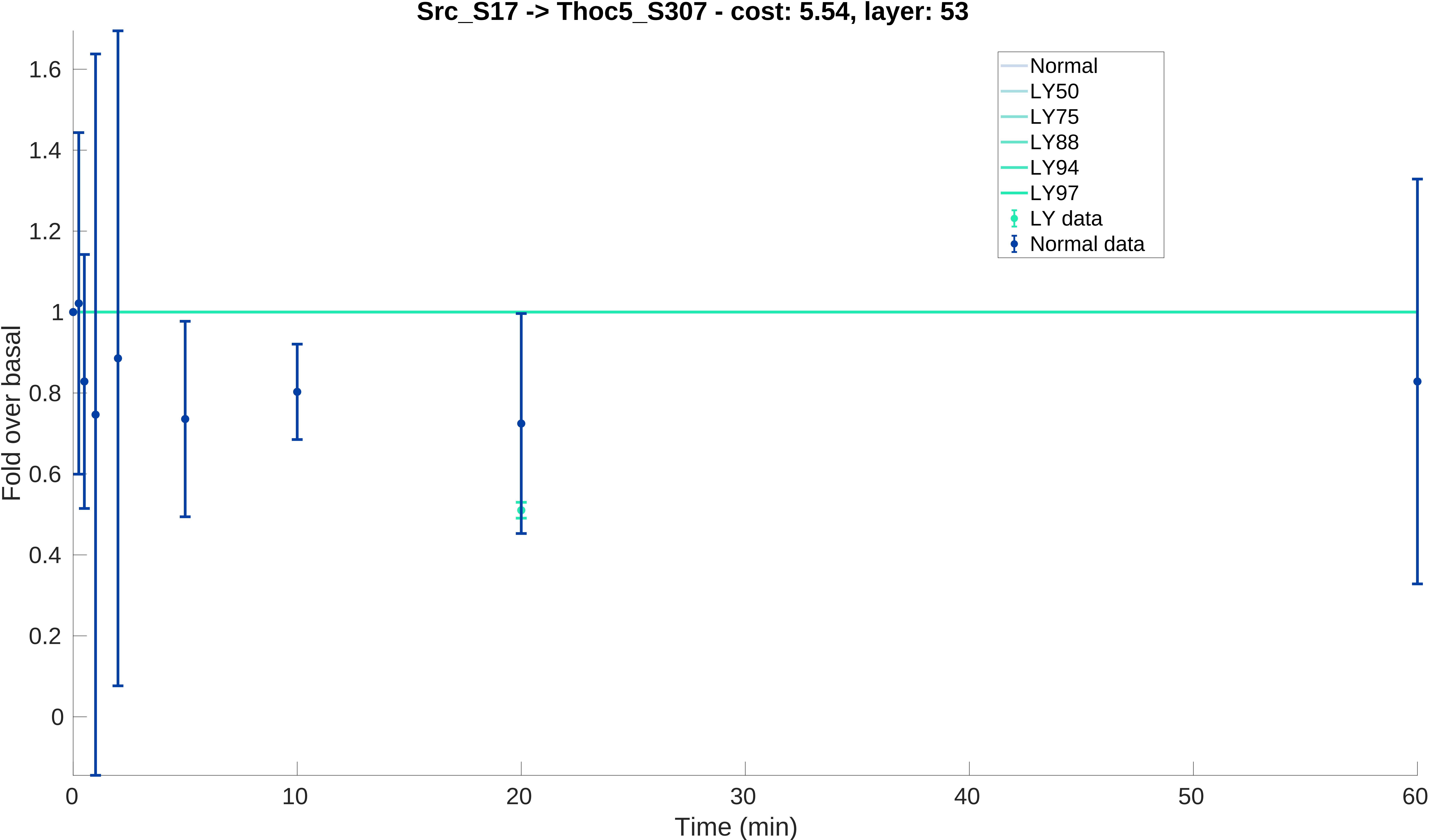

Src_S17 -> Thoc5_S307 - cost: 5.54, layer: 53
Normal
LY50
LY75
LY88
LY94
LY97
LY data
Normal data
Fold over basal
Time (min)
0
0.2
0.4
0.6
0.8
1
1.2
1.4
1.6
0
10
20
30
40
50
60

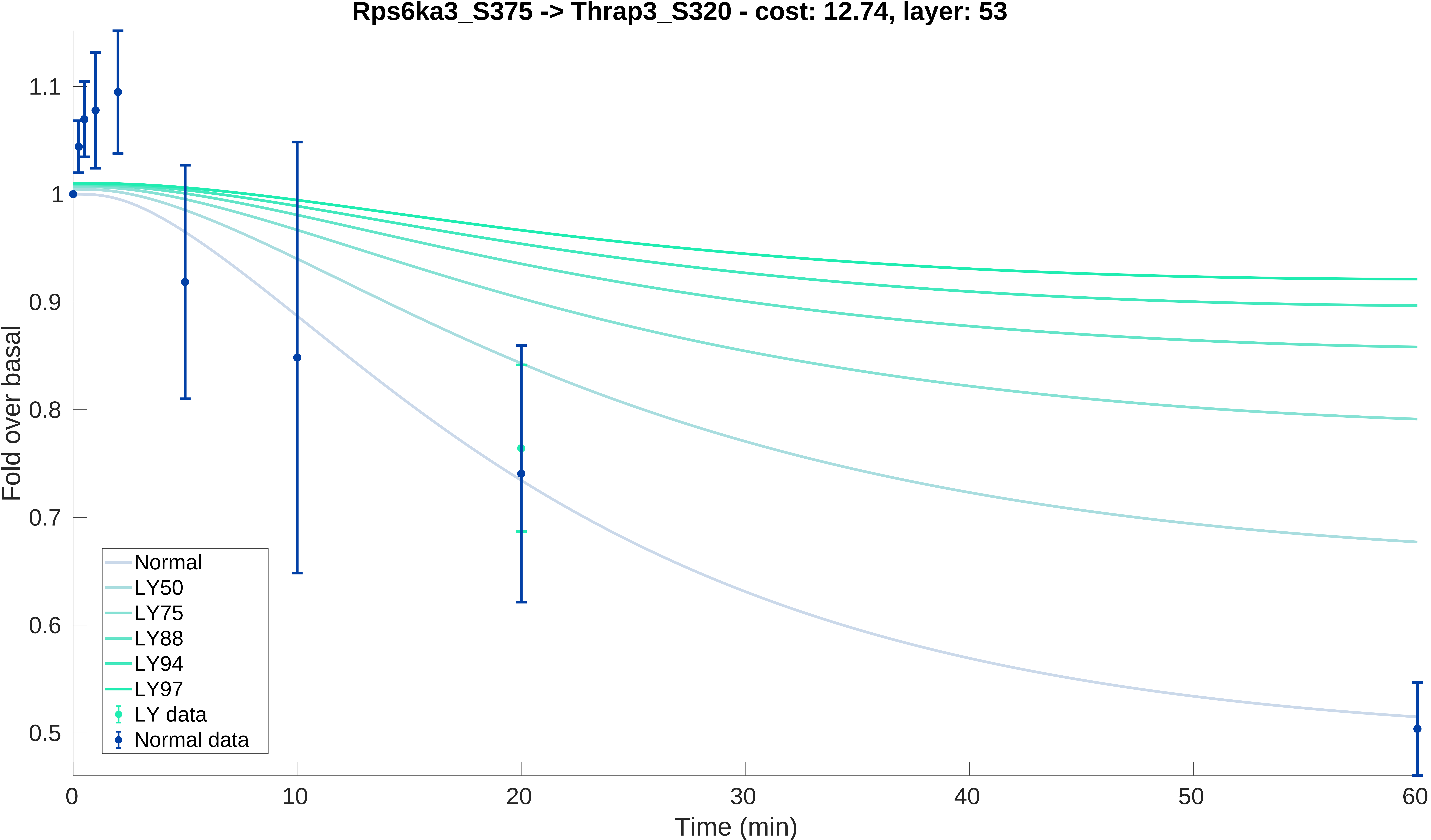

Rps6ka3_S375 -> Thrap3_S320 - cost: 12.74, layer: 53
Fold over basal
Time (min)
0
10
20
30
40
50
60
0.5
0.6
0.7
0.8
0.9
1
1.1
Normal
LY50
LY75
LY88
LY94
LY97
LY data
Normal data

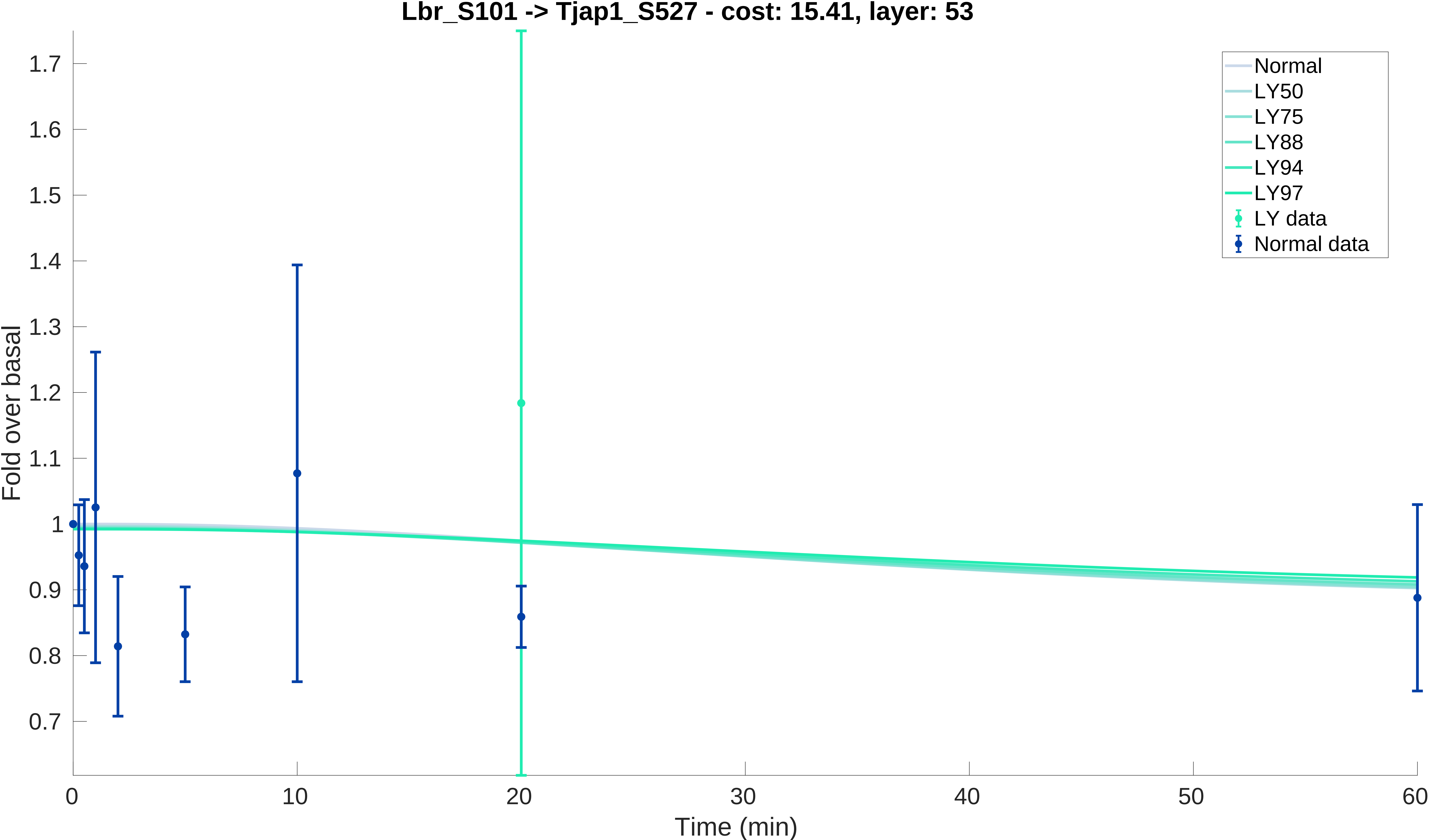
Lbr_S101 -> Tjap1_S527 - cost: 15.41, layer: 53
Fold over basal
Time (min)
0
10
20
30
40
50
60
0.7
0.8
0.9
1
1.1
1.2
1.3
1.4
1.5
1.6
1.7
Normal
LY50
LY75
LY88
LY94
LY97
LY data
Normal data

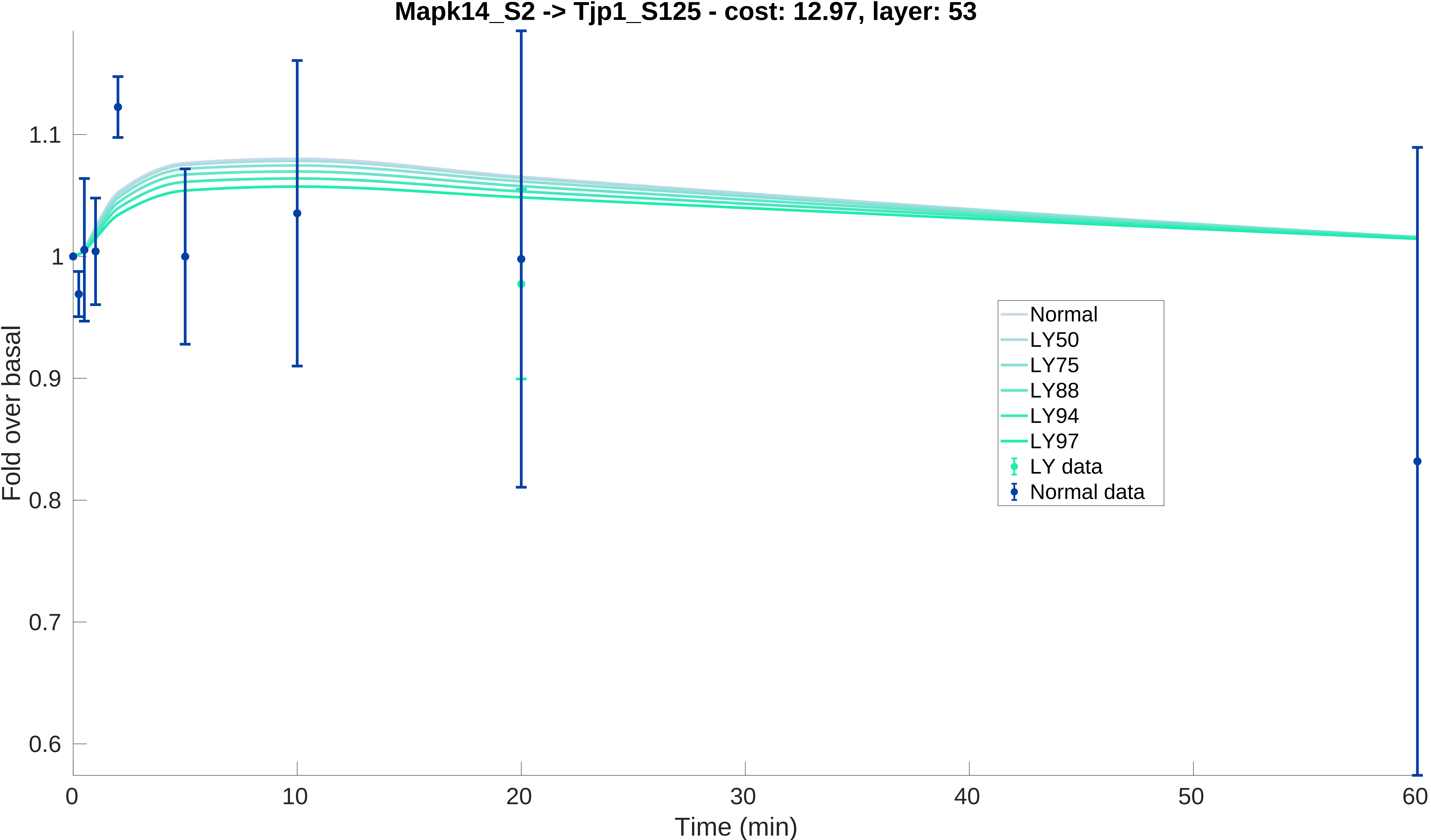

Mapk14_S2 -> Tjp1_S125 - cost: 12.97, layer: 53
Fold over basal
Time (min)
1.1
1
0.9
0.8
0.7
0.6
0
10
20
30
40
50
60
Normal
LY50
LY75
LY88
LY94
LY97
LY data
Normal data

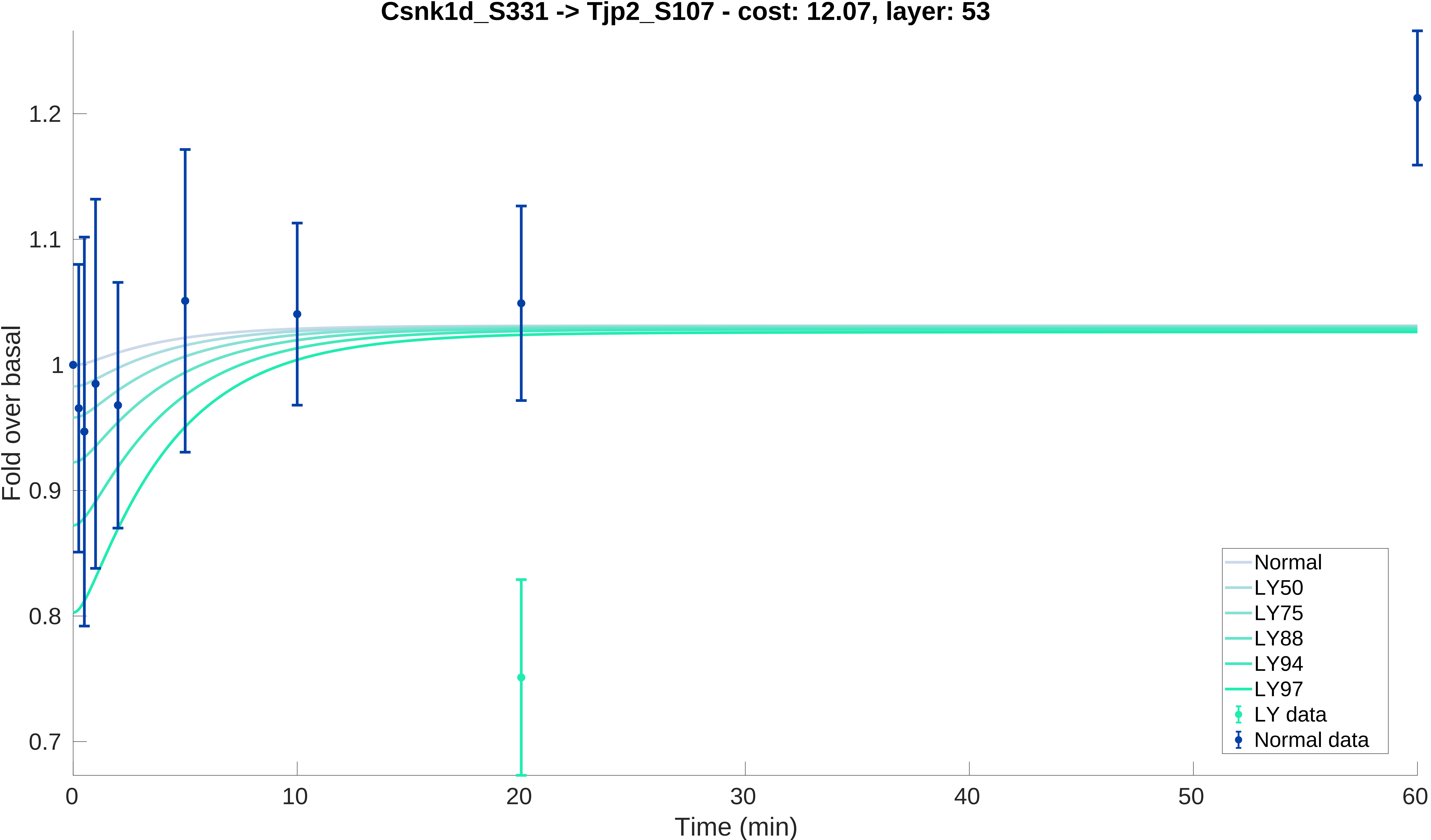

Csnk1d_S331 -> Tjp2_S107 - cost: 12.07, layer: 53
1.2
1.1
1
0.9
0.8
0.7
Fold over basal
0
10
20
30
40
50
60
Time (min)
Normal
LY50
LY75
LY88
LY94
LY97
LY data
Normal data

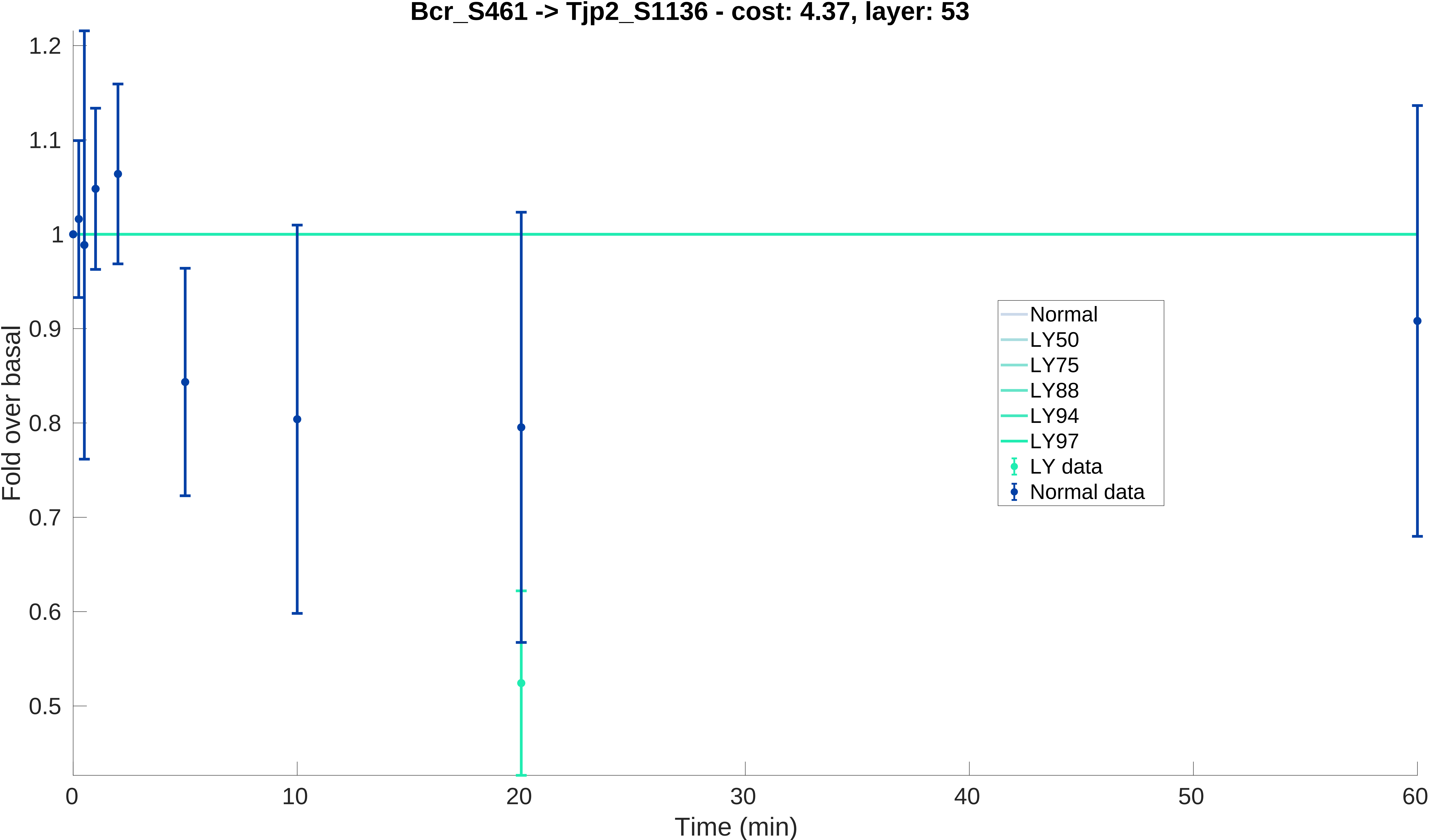

Bcr_S461 -> Tjp2_S1136 - cost: 4.37, layer: 53
Fold over basal
1.2
1.1
1
0.9
0.8
0.7
0.6
0.5
0
10
20
30
40
50
60
Time (min)
Normal
LY50
LY75
LY88
LY94
LY97
LY data
Normal data

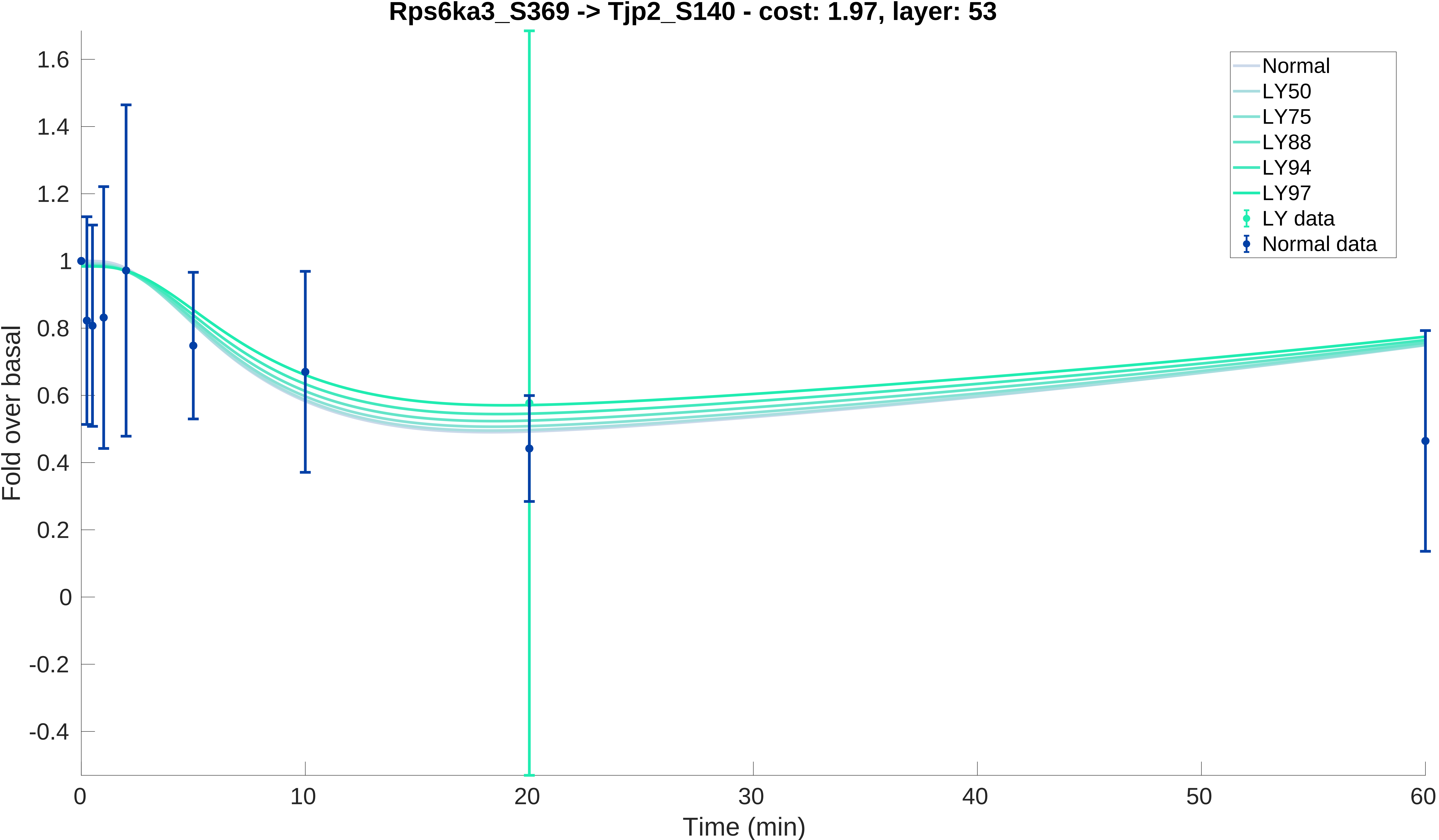
Rps6ka3_S369 -> Tjp2_S140 - cost: 1.97, layer: 53
Fold over basal
Time (min)
1.6
1.4
1.2
1
0.8
0.6
0.4
0.2
0
-0.2
-0.4
0
10
20
30
40
50
60
Normal
LY50
LY75
LY88
LY94
LY97
LY data
Normal data

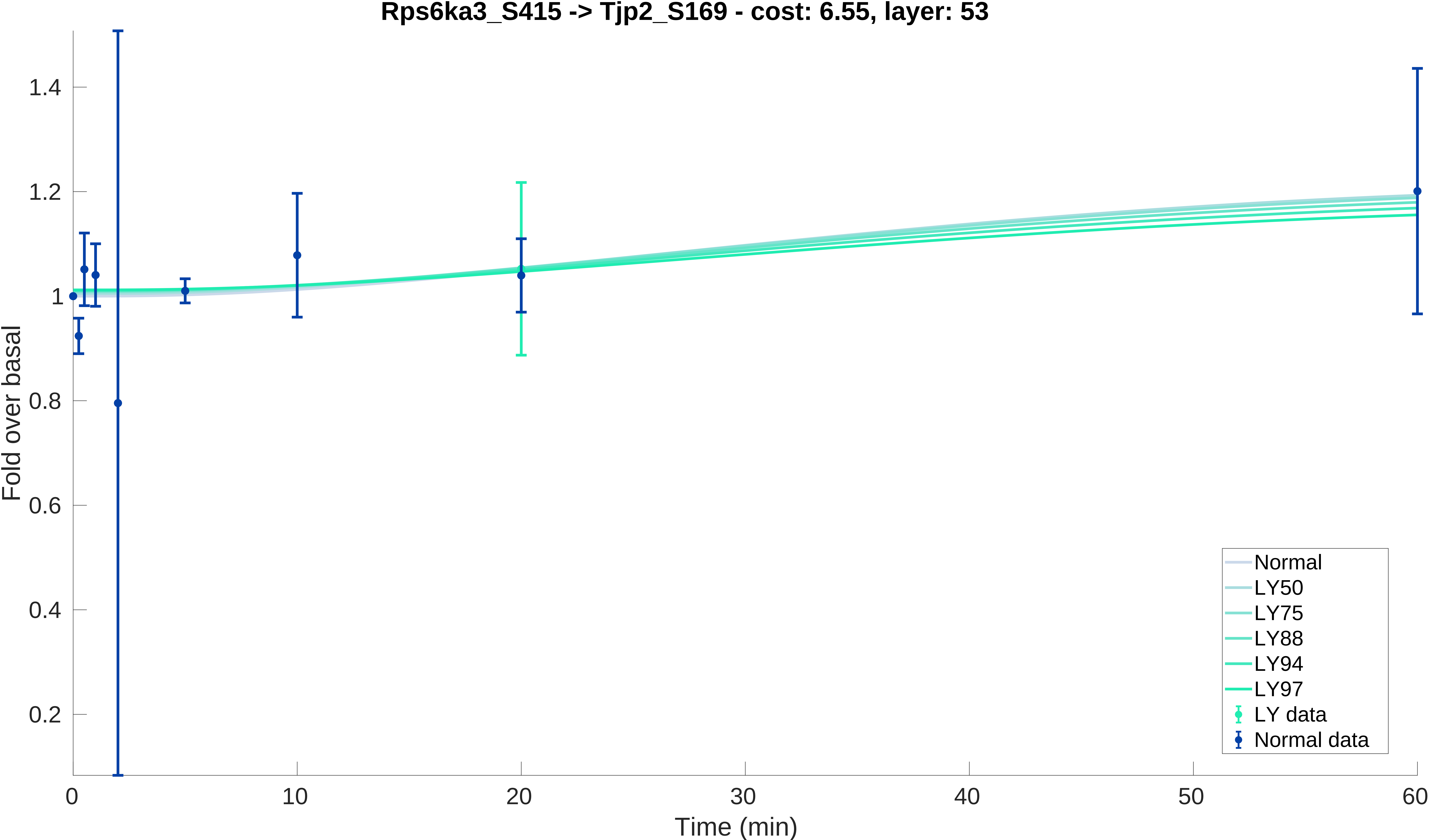

Rps6ka3_S415 -> Tjp2_S169 - cost: 6.55, layer: 53
Fold over basal
Time (min)
0.2
0.4
0.6
0.8
1
1.2
1.4
0
10
20
30
40
50
60
Normal
LY50
LY75
LY88
LY94
LY97
LY data
Normal data

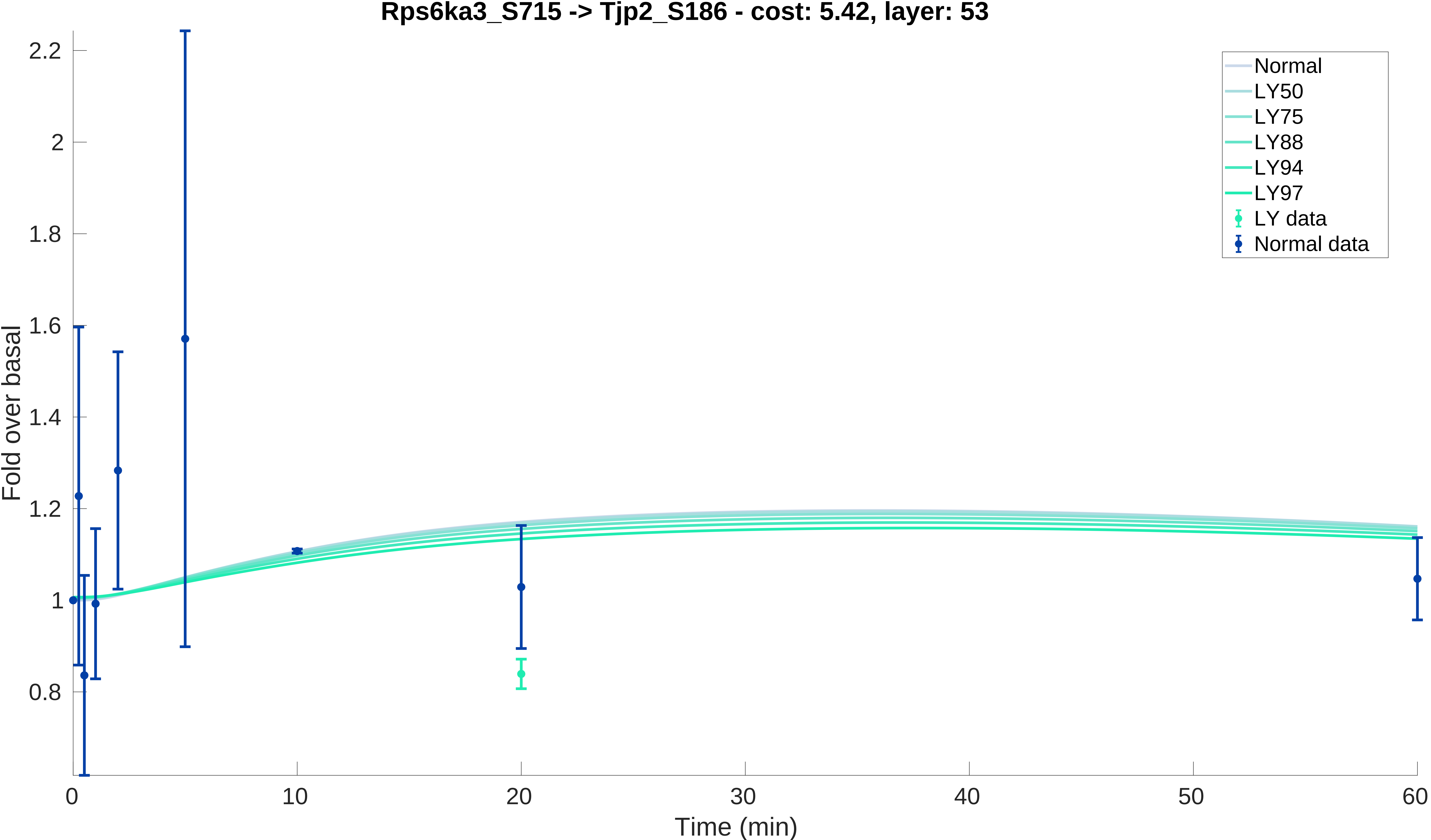

Rps6ka3_S715 -> Tjp2_S186 - cost: 5.42, layer: 53
Normal
LY50
LY75
LY88
LY94
LY97
LY data
Normal data
Fold over basal
2.2
2
1.8
1.6
1.4
1.2
1
0.8
0
10
20
30
40
50
60
Time (min)

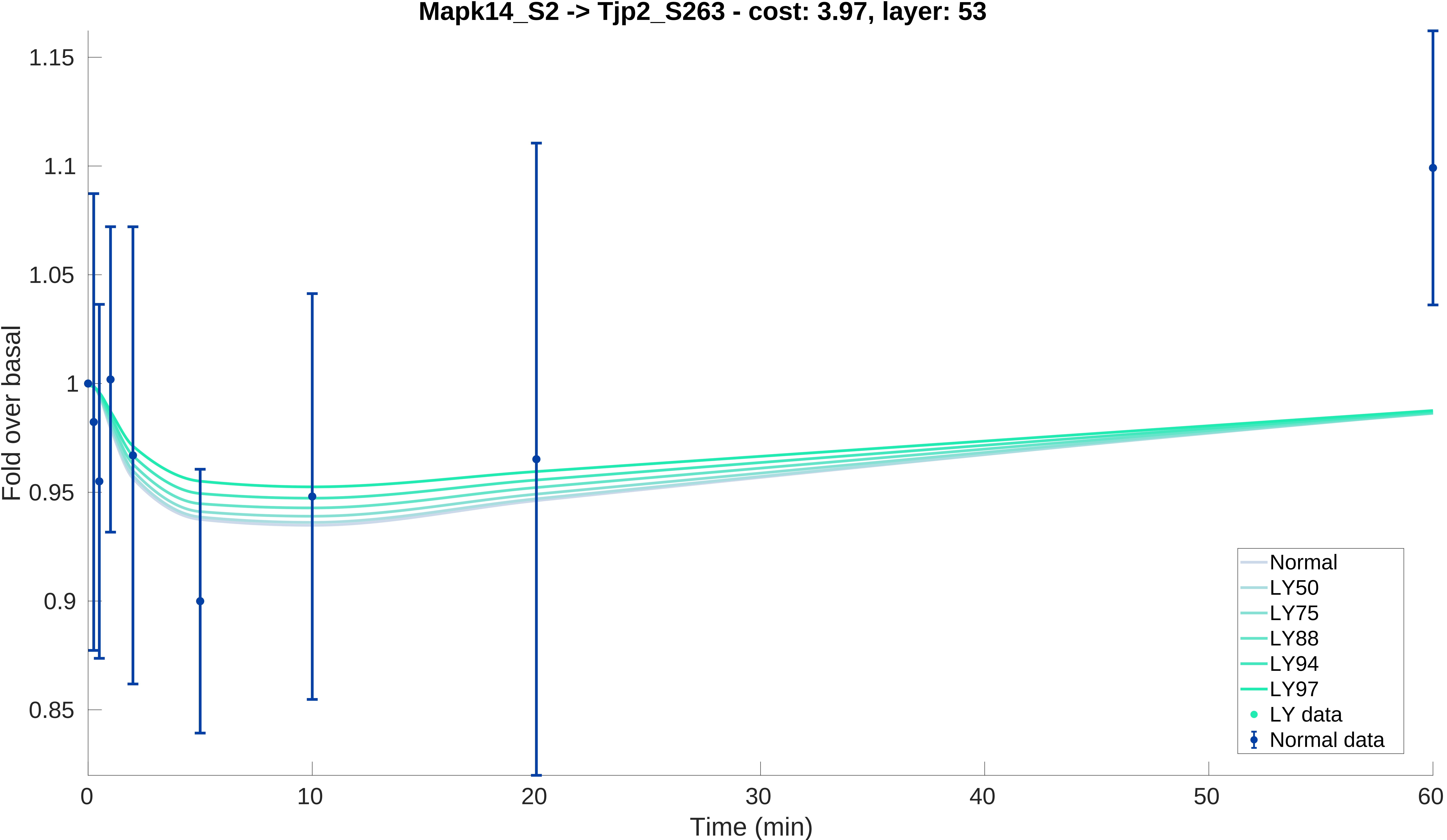
Mapk14_S2 -> Tjp2_S263 - cost: 3.97, layer: 53
Fold over basal
1.15
1.1
1.05
1
0.95
0.9
0.85
0
10
20
30
40
50
60
Time (min)
Normal
LY50
LY75
LY88
LY94
LY97
LY data
Normal data

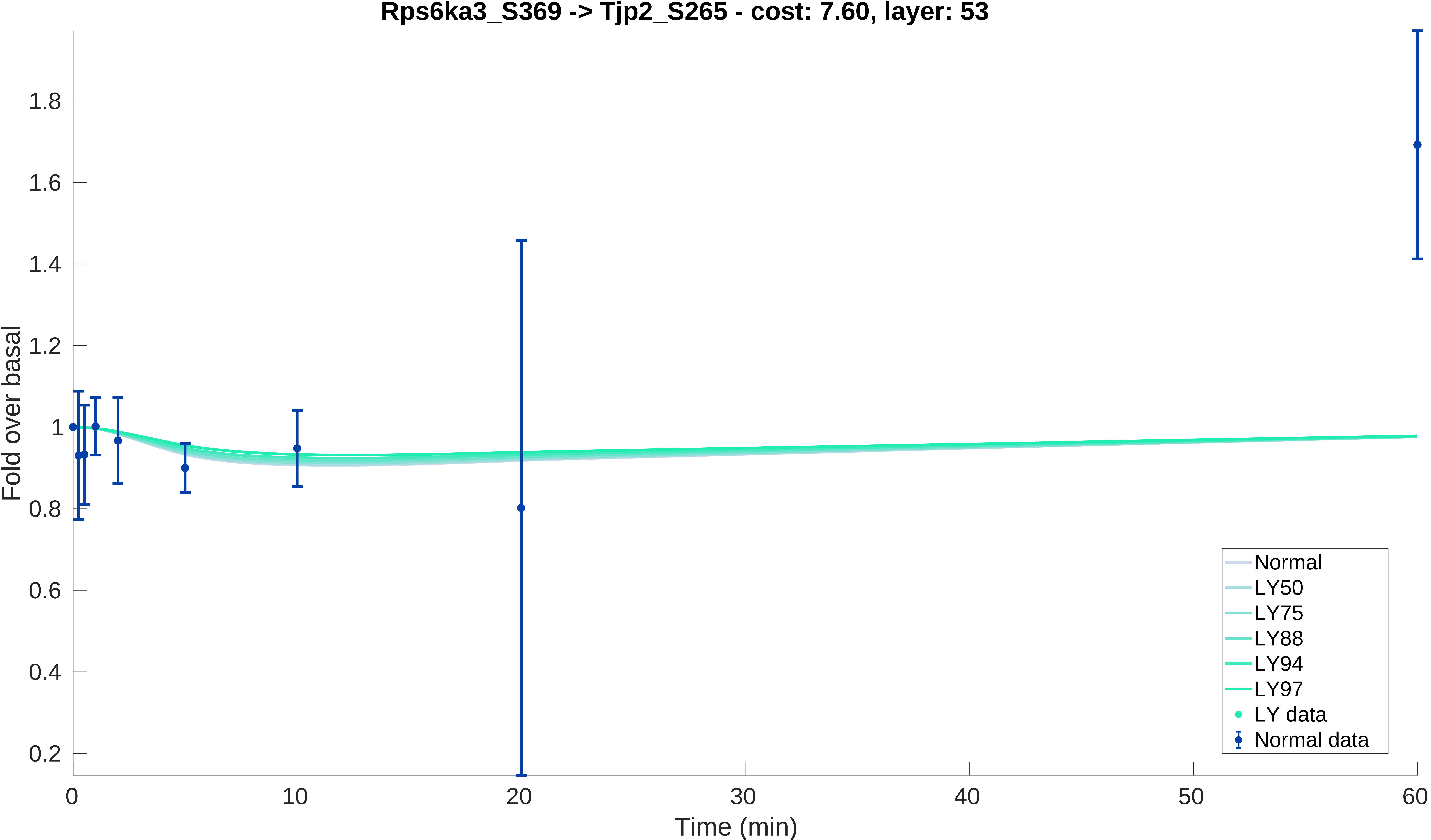
Rps6ka3_S369 -> Tjp2_S265 - cost: 7.60, layer: 53
Fold over basal
Time (min)
1.8
1.6
1.4
1.2
1
0.8
0.6
0.4
0.2
0
10
20
30
40
50
60
Normal
LY50
LY75
LY88
LY94
LY97
LY data
Normal data

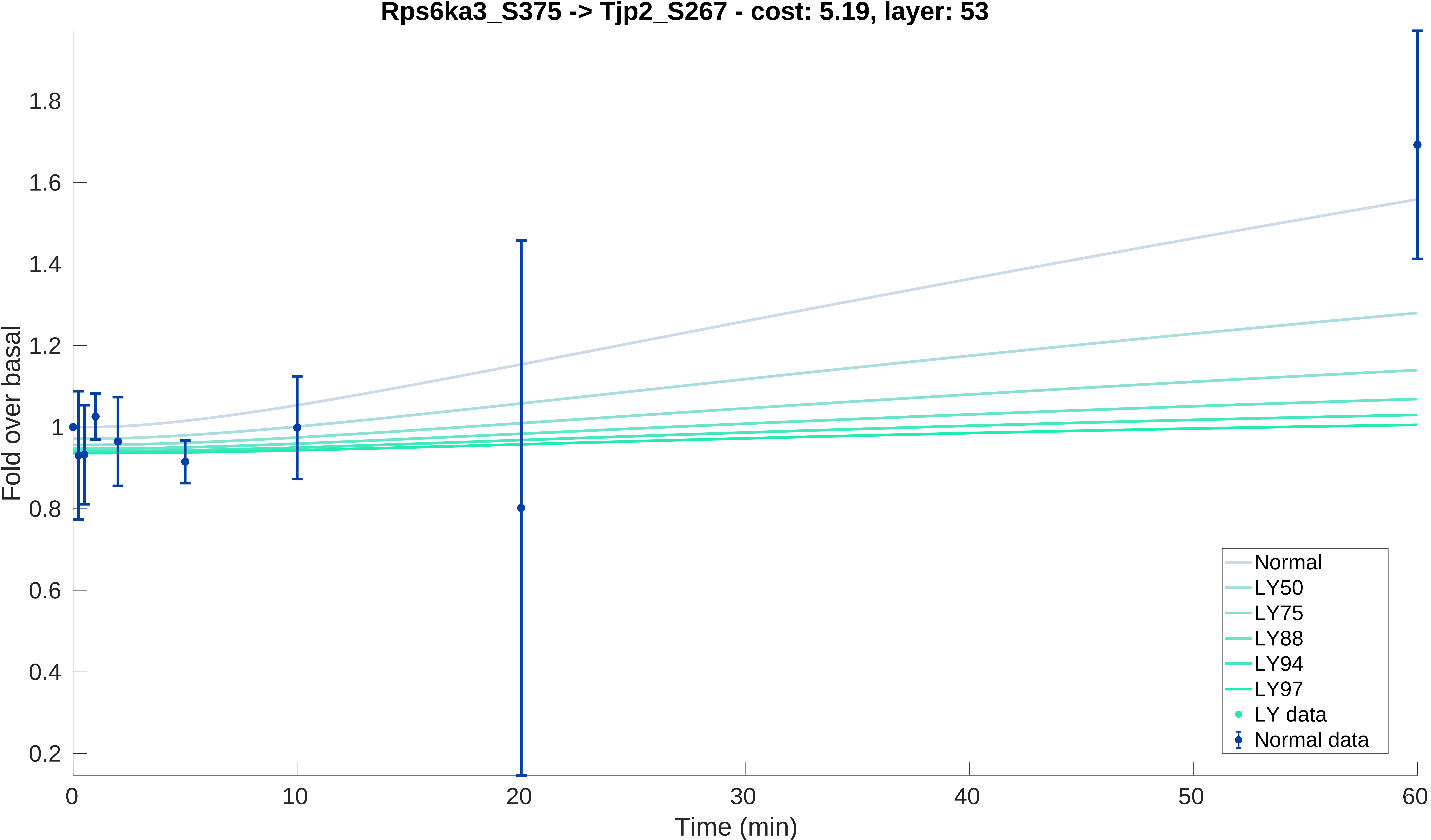

Rps6ka3_S375 -> Tjp2_S267 - cost: 5.19, layer: 53
Fold over basal
1.8
1.6
1.4
1.2
1
0.8
0.6
0.4
0.2
0
10
20
30
40
50
60
Time (min)
Normal
LY50
LY75
LY88
LY94
LY97
LY data
Normal data

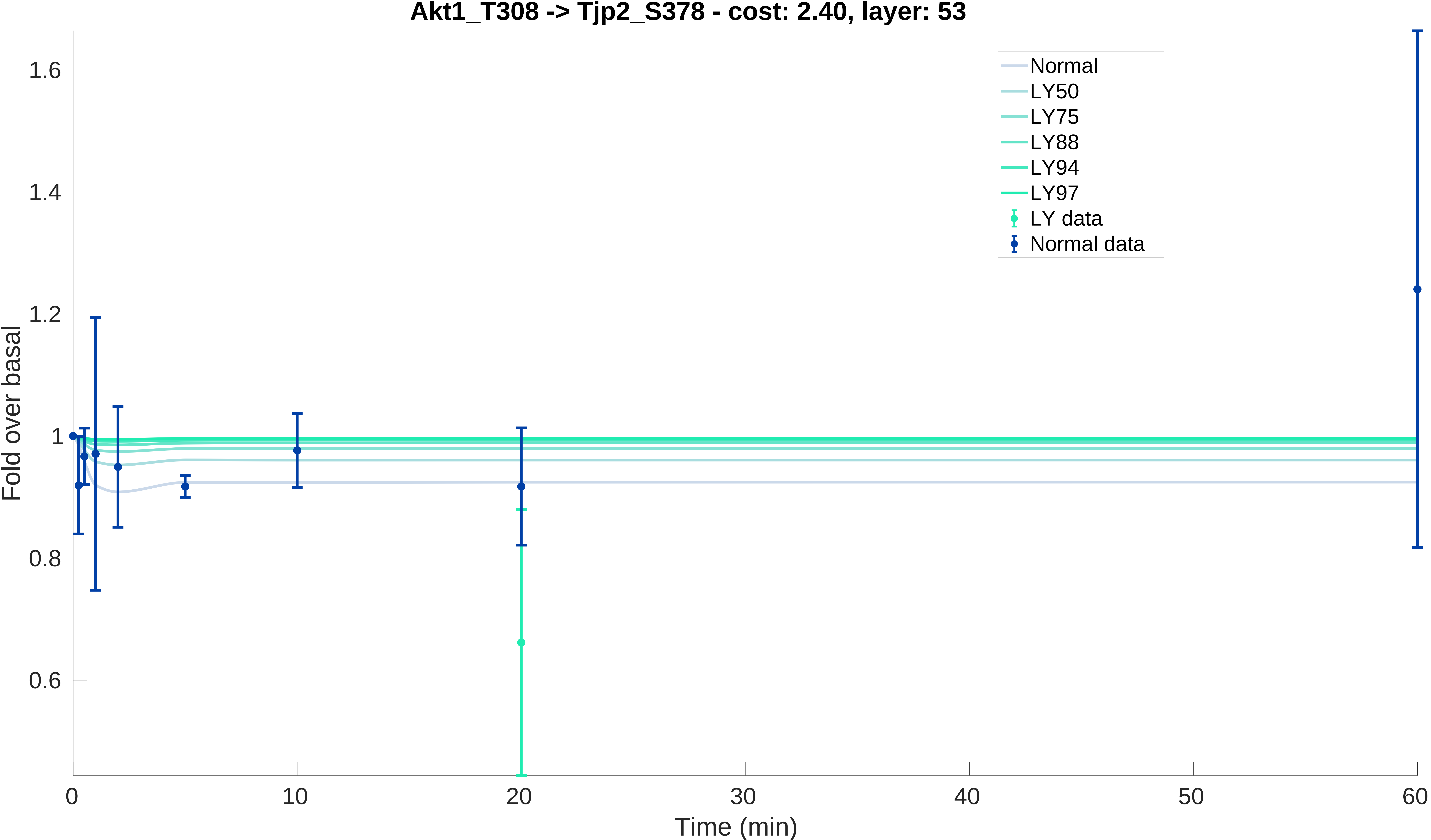

Akt1_T308 -> Tjp2_S378 - cost: 2.40, layer: 53
Fold over basal
Time (min)
Normal
LY50
LY75
LY88
LY94
LY97
LY data
Normal data
1.6
1.4
1.2
1
0.8
0.6
0
10
20
30
40
50
60

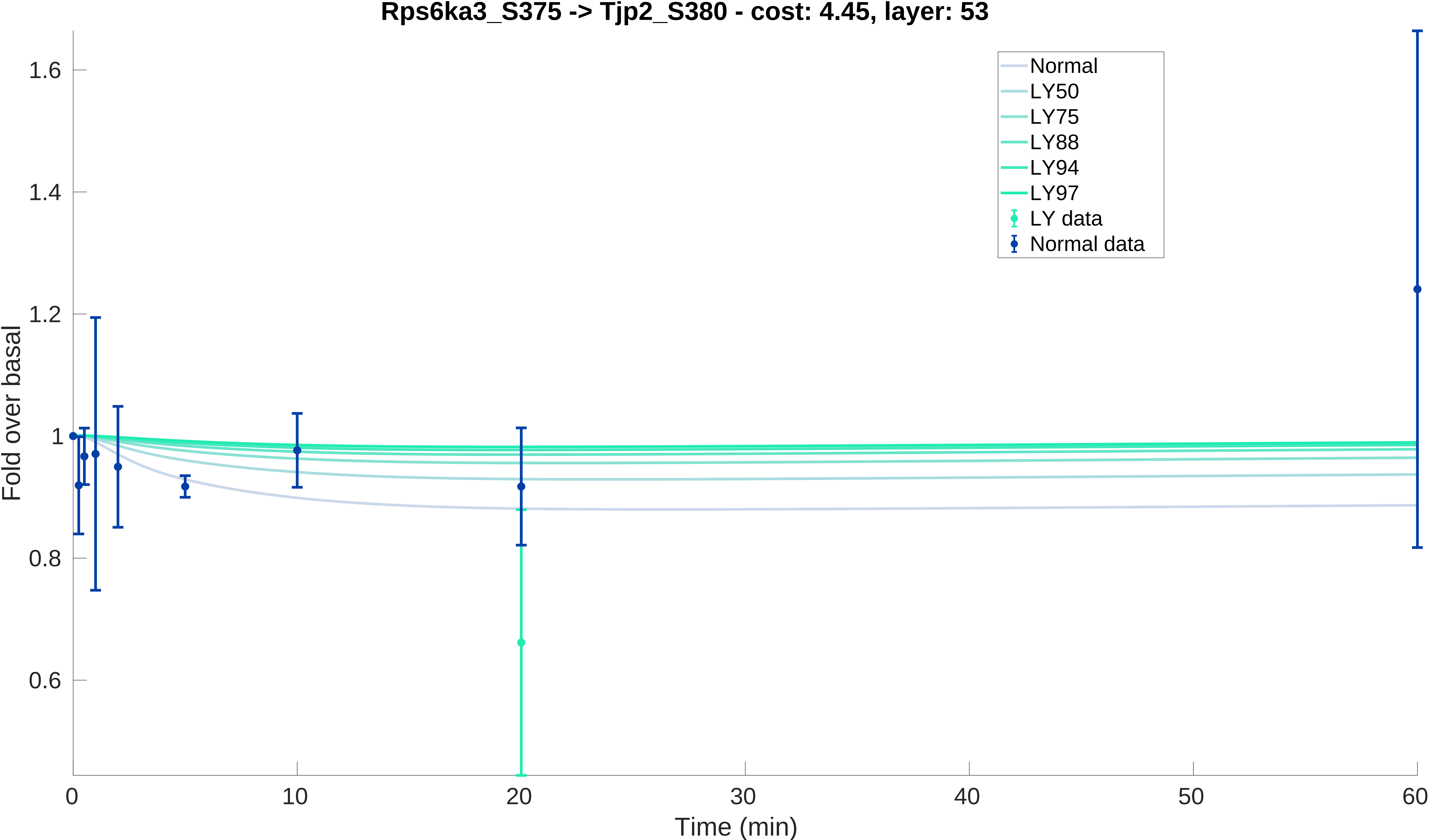

Rps6ka3_S375 -> Tjp2_S380 - cost: 4.45, layer: 53
Normal
LY50
LY75
LY88
LY94
LY97
LY data
Normal data
Fold over basal
1.6
1.4
1.2
1
0.8
0.6
0
10
20
30
40
50
60
Time (min)
